## Supplementary material for "Microbial, dietary insect, and pathogen communities in fresh and decomposing guano of anthropic little brown bat (Myotis lucifugus) maternity colonies": Table S14

| OTU | Size | Taxonomy | Taxon | Taxon Level | Trophic Mode | Guild | Growth Morphology | Trait | Confidence Ranking | Notes | Citation/Source |
|---|---|---|---|---|---|---|---|---|---|---|---|
| Otu00001 | 415107 | k__Fungi(100);p__Ascomycota(100);c__Dothideomycetes(100);o__Capnodiales(100);f__Davidiellaceae(100);f__Davidiellaceae_unclassified(93);f__Davidiellaceae_unclassified(93); | - | - | - | - | - | - | - | Unassigned | - |
| Otu00002 | 246130 | k__Fungi(100);p__Basidiomycota(100);c__Microbotryomycetes(100);o__Sporidiobolales(100);f__Sporidiobolales_family_Incertae_sedis(100);g__Sporobolomyces(100);s__Sporobolomyces_lactosus(100); | Sporobolomyces | 13 | Pathotroph-Saprotroph | \|Fungal Parasite\|-Plant Saprotroph | Yeast | NULL | Probable | As litter_saprotroph (Põlme et al. 2020); As mycoparasite (Põlme et al. 2020); Decay fungal material (Põlme et al. 2020) | [illegible] |
| Otu00003 | 154527 | k__Fungi(100);p__Ascomycota(100);c__Dothideomycetes(100);o__Pleosporales(100);f__Pleosporales_family_Incertae_sedis(77);g__Phoma(68);g__Phoma_unclassified(57); | Phoma | 13 | Pathotroph-Saprotroph-Symbiotroph | Dung Saprotroph-Endophyte-Lichen Parasite-\|Plant Pathogen\|-Plant Saprotroph-Undefined Saprotroph-Wood Saprotroph | Microfungus | Soft Rot | Possible | As foliar_endophyte (Põlme et al. 2020); As litter_saprotroph (Põlme et al. 2020); Decay leaf, fruit, seed, roots, soil (Põlme et al. 2020) | [illegible] |
| Otu00004 | 100581 | k__Fungi(100);p__Ascomycota(100);c__Saccharomycetes(100);o__Saccharomycetales(100);f__Saccharomycetaceae(100);g__Kazachstania(100);s__Kazachstania_intestinalis(100); | Kazachstania | 13 | Saprotroph | \|Undefined Saprotroph\| | Yeast | NULL | Probable | As nectar-tap_saprotroph (Põlme et al. 2020); Decay sugar-rich substrates (Põlme et al. 2020) | [illegible] |
| Otu00005 | 69036 | k__Fungi(100);p__Zygomycota(100);c__Zygomycota_class_Incertae_sedis(100);o__Mucorales(100);f__Mucoraceae(100);g__Mucor(100);s__Mucor_circinelloides_f._lusitanicus(54); | Mucor | 13 | Saprotroph | \|Undefined Saprotroph\| | Microfungus | NULL | Probable | As soil_saprotroph (Põlme et al. 2020); Decay soil (Põlme et al. 2020) | [illegible] |
| Otu00006 | 67850 | k__Plantae(100);p__unclassified_Plantae(100);c__unclassified_Plantae(100);o__unclassified_Plantae(100);f__unclassified_Plantae(100);g__unclassified_Plantae(100);s__Plantae_sp(100); | - | - | - | - | - | - | - | Unassigned | - |
| Otu00007 | 67060 | k__Fungi(100);p__Ascomycota(52);p__Ascomycota_unclassified(52);p__Ascomycota_unclassified(52);p__Ascomycota_unclassified(52);p__Ascomycota_unclassified(52);p__Ascomycota_unclassified(52); | - | - | - | - | - | - | - | Unassigned | - |
| Otu00008 | 56550 | k__Fungi(100);p__Basidiomycota(100);c__Microbotryomycetes(100);o__Sporidiobolales(100);f__Sporidiobolales_family_Incertae_sedis(100);g__Rhodotorula(100);s__Rhodotorula_taiwanensis(87); | Rhodotorula | 13 | Pathotroph-Saprotroph-Symbiotroph | Animal Pathogen-Animal Symbiotroph-Endophyte-Plant Pathogen-\|Undefined Saprotroph\| | Yeast | NULL | Probable | As foliar_endophyte (Põlme et al. 2020); Opportunistic human pathogen | [illegible] |
| Otu00009 | 51420 | k__Fungi(100);p__Ascomycota(100);c__Sordariomycetes(100);o__Hypocreales(100);f__Nectriaceae(100);g__Gibberella(94);s__Gibberella_intricans(94); | Gibberella | 13 | Pathotroph-Saprotroph-Symbiotroph | Endophyte-\|Plant Pathogen\|-Plant Saprotroph-Undefined Saprotroph | Microfungus | Soft Rot | Probable | As foliar_endophyte (Põlme et al. 2020); As litter_saprotroph (Põlme et al. 2020); Causing damping-off disease (Põlme et al. 2020); Decay leaf, fruit, seed, roots, soil, animal material (Põlme et al. 2020) | [illegible] |
| Otu00010 | 44358 | k__Fungi(100);p__Ascomycota(100);c__Dothideomycetes(100);o__Dothideales(100);f__Dothioraceae(100);g__Aureobasidium(100);s__Aureobasidium_pullulans(96); | Aureobasidium | 13 | Pathotroph-Saprotroph-Symbiotroph | Animal Pathogen-Endophyte-Epiphyte-\|Plant Pathogen\|-Plant Saprotroph-Undefined Saprotroph | Dimorphic-Facultative Yeast | NULL | Possible | As foliar_endophyte (Põlme et al. 2020); As litter_saprotroph (Põlme et al. 2020); Causes sooty mold (Põlme et al. 2020); Decay leaf, fruit, seed, roots, animal material, sugar-rich substrates (Põlme et al. 2020) | [illegible] |
| Otu00011 | 40245 | k__Fungi(100);p__Ascomycota(100);c__Dothideomycetes(100);o__Pleosporales(100);f__Pleosporaceae(100);g__Alternaria(83);s__Alternaria_porri(82); | Alternaria | 13 | Pathotroph-Saprotroph-Symbiotroph | Animal Pathogen-Endophyte-\|Plant Pathogen\|-Plant Saprotroph-Wood Saprotroph | Microfungus | Soft Rot | Possible | As foliar_endophyte (Põlme et al. 2020); As litter_saprotroph (Põlme et al. 2020); Decay leaf, fruit, seed (Põlme et al. 2020) | [illegible] |
| Otu00012 | 39843 | k__Fungi(100);p__Ascomycota(100);c__Eurotiomycetes(100);o__Eurotiales(100);f__Trichocomaceae(100);f__Trichocomaceae_unclassified(99);f__Trichocomaceae_unclassified(99); | - | - | - | - | - | - | - | Unassigned | - |
| Otu00013 | 39335 | k__Fungi(100);p__Basidiomycota(100);c__Tremellomycetes(100);o__Tremellales(100);f__Tremellales_family_Incertae_sedis(100);g__Cryptococcus(100);s__Cryptococcus_laurentii(100); | Cryptococcus | 13 | Pathotroph-Saprotroph-Symbiotroph | Animal Pathogen-Endophyte-Epiphyte-\|Undefined Saprotroph\| | Dimorphic Yeast-Yeast | NULL | Possible | As foliar_endophyte (Põlme et al. 2020); Decay leaf, fruit, seed, soil, dung, animal material (Põlme et al. 2020) | [illegible] |
| Otu00014 | 37282 | k__Fungi(100);p__Ascomycota(100);c__Sordariomycetes(97);o__Hypocreales(82);o__Hypocreales_unclassified(82);o__Hypocreales_unclassified(82);o__Hypocreales_unclassified(82); | Hypocreales | 7 | Saprotroph | Undefined Saprotroph | Microfungus | NULL | Possible | NULL | [illegible] |
| Otu00015 | 35560 | k__Fungi(100);p__Ascomycota(100);c__Eurotiomycetes(97);o__Chaetothyriales(95);f__Herpotrichiellaceae(95);g__Rhinocladiella(95);s__Rhinocladiella_sp_VL271(95); | Rhinocladiella | 13 | Pathotroph-Saprotroph-Symbiotroph | Animal Parasite-Endophyte-\|Undefined Saprotroph\|-Wood Saprotroph | Dimorphic-Facultative Yeast | Soft Rot | Probable | As root_endophyte_dark_septate (Põlme et al. 2020); As soil_saprotroph (Põlme et al. 2020); Decay soil, animal material (Põlme et al. 2020) | [illegible] |
| Otu00016 | 34395 | k__Fungi(100);p__Ascomycota(100);c__Sordariomycetes(98);c__Sordariomycetes_unclassified(96);c__Sordariomycetes_unclassified(96);c__Sordariomycetes_unclassified(96);c__Sordariomycetes_unclassified(96); | - | - | - | - | - | - | - | Unassigned | - |
| Otu00017 | 33430 | k__Fungi(100);p__Ascomycota(82);p__Ascomycota_unclassified(82);p__Ascomycota_unclassified(82);p__Ascomycota_unclassified(82);p__Ascomycota_unclassified(82);p__Ascomycota_unclassified(82); | - | - | - | - | - | - | - | Unassigned | - |
| Otu00018 | 28799 | k__Fungi(100);p__Basidiomycota(100);c__Microbotryomycetes(78);o__Sporidiobolales(78);f__Sporidiobolales_family_Incertae_sedis(78);g__Rhodotorula(78);s__Rhodotorula_sp(78); | Rhodotorula | 13 | Pathotroph-Saprotroph-Symbiotroph | Animal Pathogen-Animal Symbiotroph-Endophyte-Plant Pathogen-\|Undefined Saprotroph\| | Yeast | NULL | Probable | As foliar_endophyte (Põlme et al. 2020); Opportunistic human pathogen | [illegible] |
| Otu00019 | 28267 | k__Fungi(100);p__Basidiomycota(100);c__Microbotryomycetes(100);o__Sporidiobolales(100);f__Sporidiobolales_family_Incertae_sedis(100);g__Rhodotorula(100);s__Rhodotorula_mucilaginosa(100); | Rhodotorula | 13 | Pathotroph-Saprotroph-Symbiotroph | Animal Pathogen-Animal Symbiotroph-Endophyte-Plant Pathogen-\|Undefined Saprotroph\| | Yeast | NULL | Probable | As foliar_endophyte (Põlme et al. 2020); Opportunistic human pathogen | [illegible] |
| Otu00020 | 26913 | k__Fungi(100);p__Ascomycota(100);p__Ascomycota_unclassified(97);p__Ascomycota_unclassified(97);p__Ascomycota_unclassified(97);p__Ascomycota_unclassified(97);p__Ascomycota_unclassified(97); | - | - | - | - | - | - | - | Unassigned | - |
| Otu00021 | 26704 | k__Fungi(100);p__Zygomycota(100);c__Zygomycota_class_Incertae_sedis(100);o__Mucorales(100);f__Mucoraceae(100);g__Mucor(100);s__Mucor_genevensis(100); | Mucor | 13 | Saprotroph | \|Undefined Saprotroph\| | Microfungus | NULL | Probable | As soil_saprotroph (Põlme et al. 2020); Decay soil (Põlme et al. 2020) | [illegible] |
| Otu00022 | 20504 | k__Fungi(100);p__Ascomycota(100);c__Sordariomycetes(100);o__Hypocreales(100);f__Hypocreales_family_Incertae_sedis(100);g__Acremonium(100);s__Acremonium_asperulatum(100); | Acremonium | 13 | Pathotroph-Saprotroph-Symbiotroph | Animal Pathogen-Endophyte-Fungal Parasite-Plant Pathogen-\|Undefined Saprotroph\|-Wood Saprotroph | Microfungus | Soft Rot | Possible | As foliar_endophyte (Põlme et al. 2020); culture (Hanada et al. 2010, Del Olmo; Decay leaf, fruit, seed, soil, dung, animal material (Põlme et al. 2020); Endophyte detection method; Ruiz & Arnold 2014); Likely opportunistic human pathogen (Irinyi et al. 2015) | [illegible] |
| Otu00023 | 20295 | k__Fungi(100);p__Ascomycota(100);c__Sordariomycetes(100);o__Hypocreales(100);f__Nectriaceae(100);g__Fusarium(100);s__Fusarium_tricinctum(100); | Fusarium | 13 | Pathotroph-Saprotroph-Symbiotroph | Animal Pathogen-Endophyte-Lichen Parasite-\|Plant Pathogen\|-Plant Saprotroph-Undefined Saprotroph-Wood Saprotroph | Microfungus | Soft Rot | Possible | As foliar_endophyte (Põlme et al. 2020); As litter_saprotroph (Põlme et al. 2020); Causing damping-off disease (Põlme et al. 2020); Decay leaf, fruit, seed, roots, soil, animal material (Põlme et al. 2020); Likely opportunistic human pathogen (Irinyi et al. 2015) | [illegible] |
| Otu00024 | 19758 | k__Fungi(100);p__Basidiomycota(100);c__Wallemiomycetes(100);o__Wallemiales(100);f__Wallemiaceae(100);g__Wallemia(100);s__Wallemia_sebi(100); | Wallemia | 13 | Saprotroph | \|Undefined Saprotroph\| | Microfungus | NULL | Probable | Decay sugar-rich substrates (Põlme et al. 2020) | [illegible] |
| Otu00025 | 19592 | k__Fungi(100);p__Basidiomycota(100);c__Agaricomycetes(100);o__Russulales(100);f__Peniophoraceae(57);g__Peniophora(54);g__Peniophora_unclassified(54); | Peniophora | 13 | Pathotroph-Saprotroph | Plant Pathogen-\|Wood Saprotroph\| | Corticioid | White Rot | Probable | As white_rot (Põlme et al. 2020); As wood_pathogen (Põlme et al. 2020); Decay wood (Põlme et al. 2020) | [illegible] |
| Otu00026 | 19199 | k__Fungi(100);p__Zygomycota(100);c__Zygomycota_class_Incertae_sedis(100);o__Mortierellales(100);f__Mortierellaceae(100);g__unclassified_Mortierellaceae(100);s__Mortierellaceae_sp(100); | Mortierellaceae | 9 | Saprotroph-Symbiotroph | Endophyte-Litter Saprotroph-Soil Saprotroph-Undefined Saprotroph | Microfungus | NULL | Possible | Saprobic in soil and thus frequenctly isolated from soil and material contaminated with soil such as hay or rotten silage and spoil tips (Cannon & Kirk 2007) | [illegible] |
| Otu00027 | 18659 | k__Fungi(100);p__Ascomycota(100);c__Dothideomycetes(100);o__Pleosporales(99);o__Pleosporales_unclassified(67);o__Pleosporales_unclassified(67);o__Pleosporales_unclassified(67); | - | - | - | - | - | - | - | Unassigned | - |
| Otu00028 | 16462 | k__Fungi(100);p__Ascomycota(100);c__Saccharomycetes(100);o__Saccharomycetales(100);f__Saccharomycetaceae(100);g__Saccharomyces(100);s__Saccharomyces_eubayanus(79); | Saccharomyces | 13 | Saprotroph | \|Undefined Saprotroph\| | Yeast | NULL | Probable | As nectar-tap_saprotroph (Põlme et al. 2020); Decay sugar-rich substrates (Põlme et al. 2020) | [illegible] |
| Otu00029 | 15881 | k__Fungi(100);p__Ascomycota(100);c__Dothideomycetes(100);o__Pleosporales(100);f__Lophiostomataceae(100);g__Lophiostoma(100);g__Lophiostoma_unclassified(100); | Lophiostoma | 13 | Saprotroph | Plant Saprotroph-Undefined Saprotroph-\|Wood Saprotroph\| | Microfungus | NULL | Probable | As litter_saprotroph (Põlme et al. 2020); Decay wood, leaf, fruit, seed (Põlme et al. 2020) | [illegible] |
| Otu00030 | 15426 | k__Fungi(100);p__Basidiomycota(100);c__Microbotryomycetes(100);o__Sporidiobolales(100);f__Sporidiobolales_family_Incertae_sedis(100);g__unclassified_Sporidiobolales(64);s__Sporidiobolales_sp(64); | - | - | - | - | - | - | - | Unassigned | - |
| Otu00031 | 15037 | k__Fungi(100);p__Ascomycota(100);c__Sordariomycetes(100);o__Sordariales(100);f__Chaetomiaceae(100);g__Humicola(98);s__Humicola_grisea_var._grisea(92); | Humicola | 13 | Pathotroph-Saprotroph | Plant Pathogen-Undefined Saprotroph-\|Wood Saprotroph\| | Microfungus | Soft Rot | Probable | Decay wood (Põlme et al. 2020) | [illegible] |
| Otu00032 | 14642 | k__Fungi(100);p__Ascomycota(100);c__Sordariomycetes(100);o__Hypocreales(100);f__Nectriaceae(100);g__Fusarium(85);g__Fusarium_unclassified(78); | Fusarium | 13 | Pathotroph-Saprotroph-Symbiotroph | Animal Pathogen-Endophyte-Lichen Parasite-\|Plant Pathogen\|-Plant Saprotroph-Undefined Saprotroph-Wood Saprotroph | Microfungus | Soft Rot | Possible | As foliar_endophyte (Põlme et al. 2020); As litter_saprotroph (Põlme et al. 2020); Causing damping-off disease (Põlme et al. 2020); Decay leaf, fruit, seed, roots, soil, animal material (Põlme et al. 2020); Likely opportunistic human pathogen (Irinyi et al. 2015) | [illegible] |
| Otu00033 | 14542 | k__Plantae(100);p__unclassified_Plantae(100);c__unclassified_Plantae(100);o__unclassified_Plantae(100);f__unclassified_Plantae(100);g__unclassified_Plantae(100);s__Plantae_sp(100); | - | - | - | - | - | - | - | Unassigned | - |
| Otu00034 | 14238 | k__Fungi(100);p__Ascomycota(100);c__Saccharomycetes(100);o__Saccharomycetales(100);f__Saccharomycetaceae(100);g__Lachancea(100);s__Lachancea_thermotolerans(99); | Lachancea | 13 | Pathotroph-Saprotroph | Animal Parasite-Animal Pathogen-\|Undefined Saprotroph\| | Yeast | NULL | Possible | As nectar-tap_saprotroph (Põlme et al. 2020); Decay animal material, sugar-rich substrates (Põlme et al. 2020); Likely opportunistic human pathogen (Irinyi et al. 2015; see L. fermentati) | [illegible] |
| Otu00035 | 13405 | k__Fungi(100);p__Ascomycota(100);c__Sordariomycetes(100);o__Sordariales(100);f__Lasiosphaeriaceae(100);g__Cladorrhinum(100);s__Cladorrhinum_sp(100); | Cladorrhinum | 13 | Pathotroph-Saprotroph-Symbiotroph | Animal Pathogen-Dung Saprotroph-Endophyte-Plant Pathogen-Plant Saprotroph-\|Undefined Saprotroph\|-Wood Saprotroph | Microfungus | Soft Rot | Probable | As foliar_endophyte (Põlme et al. 2020); Decay leaf, fruit, seed (Põlme et al. 2020); Reported in soils (including agricultural) as saprotrophs on dung or plant material, or in roots as endophytes, while some species are associated with human and animal opportunistic diseases (Carmaran et al.2015) | [illegible] |
| Otu00036 | 10393 | k__Fungi(100);p__Ascomycota(100);c__Dothideomycetes(100);o__Dothideales(100);f__Dothideaceae(100);g__Endoconidioma(100);s__Endoconidioma_populi(100); | Endoconidioma | 13 | Saprotroph | Undefined Saprotroph-\|Wood Saprotroph\| | Microfungus | NULL | Probable | Decay: wood (Põlme et al. 2020) | [illegible] |
| Otu00037 | 10211 | k__Fungi(100);p__Ascomycota(100);c__Dothideomycetes(100);o__Pleosporales(100);f__Pleosporaceae(100);g__Curvularia(100);s__Curvularia_gladioli(94); | Curvularia | 13 | Pathotroph-Saprotroph | \|Plant Pathogen\|-Plant Saprotroph | Microfungus | NULL | Probable | As litter_saprotroph (Põlme et al. 2020); Decay wood, leaf, fruit, seed, roots, algal material (Põlme et al. 2020) | [illegible] |
| Otu00038 | 10156 | k__Fungi(100);k__Fungi_unclassified(51);k__Fungi_unclassified(51);k__Fungi_unclassified(51);k__Fungi_unclassified(51);k__Fungi_unclassified(51);k__Fungi_unclassified(51); | - | - | - | - | - | - | - | Unassigned | - |
| Otu00039 | 10063 | k__Fungi(100);p__Ascomycota(100);c__Dothideomycetes(100);o__Pleosporales(100);f__Phaeosphaeriaceae(100);g__unclassified_Phaeosphaeriaceae(99);s__Phaeosphaeriaceae_sp(99); | Phaeosphaeriaceae | 9 | Pathotroph-Saprotroph | Fungal Parasite-Plant Pathogen-Plant Saprotroph | Microfungus | NULL | Probable | Necrotrophic or saprobic on a wide range of plants, especially monocotyledons; a few species parasitic on other fungi (Cannon & Kirk 2007) | [illegible] |
| Otu00040 | 9811 | k__Fungi(100);p__Ascomycota(100);c__Dothideomycetes(100);o__Pleosporales(100);f__Montagnulaceae(100);f__Montagnulaceae_unclassified(74);f__Montagnulaceae_unclassified(74); | Montagnulaceae | 9 | Pathotroph-Saprotroph | Fungal Parasite-Plant Pathogen-Undefined Saprotroph | Microfungus | NULL | Probable | Species in this clade can be saprobic, plant pathogenic, or mycoparasitic (Zhang et al. 2009) | [illegible] |
| Otu00041 | 9683 | k__Fungi(100);p__Zygomycota(100);c__Zygomycota_class_Incertae_sedis(100);o__Mucorales(100);f__Mucoraceae(100);g__Mucor(100);s__Mucor_hiemalis(100); | Mucor | 13 | Saprotroph | \|Undefined Saprotroph\| | Microfungus | NULL | Probable | As soil_saprotroph (Põlme et al. 2020); Decay soil (Põlme et al. 2020) | [illegible] |
| Otu00042 | 9202 | k__Fungi(100);p__Ascomycota(100);c__Saccharomycetes(99);o__Saccharomycetales(99);f__Saccharomycetales_family_Incertae_sedis(96);g__Candida(79);g__Candida_unclassified(67); | Candida | 13 | Pathotroph-Saprotroph-Symbiotroph | Animal Pathogen-Endophyte-Endosymbiont-Epiphyte-\|Undefined Saprotroph\| | Dimorphic Yeast-Yeast | NULL | Probable | As nectar-tap_saprotroph (Põlme et al. 2020); Decay sugar-rich substrates (Põlme et al. 2020); This asexual genus is polyphyletic and contain species that function very differently depend on what they are. Caution should be taken when interpreting guilds for Candida species. | [illegible] |
| Otu00043 | 8691 | k__Fungi(100);p__Basidiomycota(100);c__Microbotryomycetes(100);o__Sporidiobolales(82);f__Sporidiobolales_family_Incertae_sedis(82);g__Rhodotorula(82);s__Rhodotorula_nothofagi(82); | Rhodotorula | 13 | Pathotroph-Saprotroph-Symbiotroph | Animal Pathogen-Animal Symbiotroph-Endophyte-Plant Pathogen-\|Undefined Saprotroph\| | Yeast | NULL | Probable | As foliar_endophyte (Põlme et al. 2020); Opportunistic human pathogen | [illegible] |
| Otu00044 | 8689 | k__Fungi(100);k__Fungi_unclassified(82);k__Fungi_unclassified(82);k__Fungi_unclassified(82);k__Fungi_unclassified(82);k__Fungi_unclassified(82);k__Fungi_unclassified(82); | - | - | - | - | - | - | - | Unassigned | - |
| Otu00045 | 8273 | k__Fungi(100);p__Ascomycota(58);p__Ascomycota_unclassified(58);p__Ascomycota_unclassified(58);p__Ascomycota_unclassified(58);p__Ascomycota_unclassified(58);p__Ascomycota_unclassified(58); | - | - | - | - | - | - | - | Unassigned | - |
| Otu00046 | 8154 | k__Fungi(100);p__Ascomycota(100);c__Taphrinomycetes(100);o__Taphrinales(100);f__Taphrinaceae(100);g__Lalaria(100);s__Lalaria_inositophila(100); | - | - | - | - | - | - | - | Unassigned | - |
| Otu00047 | 7601 | k__Fungi(100);p__Basidiomycota(100);c__Agaricomycetes(100);o__Russulales(100);f__Peniophoraceae(100);g__Peniophora(99);g__Peniophora_unclassified(99); | Peniophora | 13 | Pathotroph-Saprotroph | Plant Pathogen-\|Wood Saprotroph\| | Corticioid | White Rot | Probable | As white_rot (Põlme et al. 2020); As wood_pathogen (Põlme et al. 2020); Decay wood (Põlme et al. 2020) | [illegible] |
| Otu00048 | 7489 | k__Fungi(100);p__Basidiomycota(100);c__Pucciniomycetes(100);o__Pucciniales(100);f__Pucciniaceae(100);g__Puccinia(100);s__Puccinia_bartholomaei(100); | Puccinia | 13 | Pathotroph | \|Plant Pathogen\| | Rust | NULL | Highly Probable | NULL | [illegible] |
| Otu00049 | 7148 | k__Fungi(100);p__Ascomycota(100);c__Pezizomycetes(100);o__Pezizales(100);f__Pyronemataceae(100);g__unclassified_Pyronemataceae(100);s__Pyronemataceae_sp(100); | Pyronemataceae | 9 | Saprotroph-Symbiotroph | Dung Saprotroph-Ectomycorrhizal-Soil Saprotroph-Wood Saprotroph | Gasteroid-Pezizoid | NULL | Possible | Saprobic on soil, dung or rotten wood or mycorrhizal, some species hypogeous, others associated with fire sites (Cannon & Kirk 2007) | [illegible] |
| Otu00050 | 7142 | k__Fungi(100);p__Basidiomycota(100);c__Agaricomycetes(100);o__Polyporales(100);f__Ganodermataceae(96);g__Ganoderma(95);g__Ganoderma_unclassified(95); | Ganoderma | 13 | Pathotroph-Saprotroph | \|Plant Pathogen\|-Wood Saprotroph | Polyporoid | White Rot | Highly Probable | As white_rot (Põlme et al. 2020); As wood_pathogen (Põlme et al. 2020); Decay wood (Põlme et al. 2020); Lignicolous, causing sometimes agressive white rots of timber (Cannon & Kirk 2007) | [illegible] |
| Otu00051 | 6948 | k__Fungi(100);p__Ascomycota(100);c__Dothideomycetes(100);o__Pleosporales(100);f__Sporormiaceae(100);f__Sporormiaceae_unclassified(100);f__Sporormiaceae_unclassified(100); | Sporormiaceae | 9 | Saprotroph | Dung Saprotroph-Plant Saprotroph | Microfungus | NULL | Probable | Saprobic, associated with dung and rotting vegetation (Cannon & Kirk 2007) | [illegible] |
| Otu00052 | 6554 | k__Fungi(100);k__Fungi_unclassified(68);k__Fungi_unclassified(68);k__Fungi_unclassified(68);k__Fungi_unclassified(68);k__Fungi_unclassified(68);k__Fungi_unclassified(68); | - | - | - | - | - | - | - | Unassigned | - |
| Otu00053 | 6512 | k__Fungi(100);p__Basidiomycota(100);c__Agaricomycetes(100);o__Hymenochaetales(100);f__Schizoporaceae(100);g__Schizopora(99);s__Schizopora_flavipora(99); | Schizopora | 13 | Pathotroph-Saprotroph | Plant Pathogen-\|Wood Saprotroph\| | Corticioid | White Rot | Highly Probable | As white_rot (Põlme et al. 2020); As wood_pathogen (Põlme et al. 2020); Decay wood (Põlme et al. 2020) | [illegible] |
| Otu00054 | 5908 | k__Fungi(100);p__Ascomycota(100);c__Dothideomycetes(100);o__Pleosporales(100);o__Pleosporales_unclassified(71);o__Pleosporales_unclassified(71);o__Pleosporales_unclassified(71); | - | - | - | - | - | - | - | Unassigned | - |
| Otu00055 | 5899 | k__Fungi(100);p__Chytridiomycota(100);c__Chytridiomycetes(100);o__Spizellomycetales(100);f__Spizellomycetaceae(100);g__Spizellomyces(99);s__Spizellomyces_acuminatus(94); | Spizellomyces | 13 | Pathotroph-Saprotroph | Plant Pathogen-\|Pollen Saprotroph\|-Undefined Parasite-Undefined Saprotroph | Chytrid-Microfungus | NULL | Probable | Decay pollen (Põlme et al. 2020); Pollen degrader; Common in the soil and also as parasites of a range of soil organisms and plants (Cannon & Kirk 2007) | [illegible] |
| Otu00056 | 5728 | k__Fungi(100);p__Ascomycota(100);c__Saccharomycetes(98);o__Saccharomycetales(98);f__Saccharomycetales_family_Incertae_sedis(92);g__Candida(86);g__Candida_unclassified(69); | Candida | 13 | Pathotroph-Saprotroph-Symbiotroph | Animal Pathogen-Endophyte-Endosymbiont-Epiphyte-\|Undefined Saprotroph\| | Dimorphic Yeast-Yeast | NULL | Probable | As nectar-tap_saprotroph (Põlme et al. 2020); Decay sugar-rich substrates (Põlme et al. 2020); This asexual genus is polyphyletic and contain species that function very differently depend on what they are. Caution should be taken when interpreting guilds for Candida species. | [illegible] |
| Otu00057 | 5606 | k__Fungi(100);k__Fungi_unclassified(60);k__Fungi_unclassified(60);k__Fungi_unclassified(60);k__Fungi_unclassified(60);k__Fungi_unclassified(60);k__Fungi_unclassified(60); | - | - | - | - | - | - | - | Unassigned | - |
| Otu00058 | 5210 | k__Fungi(100);p__Ascomycota(100);c__Sordariomycetes(100);o__Hypocreales(100);f__Nectriaceae(100);g__Fusarium(100);s__Fusarium_acetilerea(100); | Fusarium | 13 | Pathotroph-Saprotroph-Symbiotroph | Animal Pathogen-Endophyte-Lichen Parasite-\|Plant Pathogen\|-Plant Saprotroph-Undefined Saprotroph-Wood Saprotroph | Microfungus | Soft Rot | Possible | As foliar_endophyte (Põlme et al. 2020); As litter_saprotroph (Põlme et al. 2020); Causing damping-off disease (Põlme et al. 2020); Decay leaf, fruit, seed, roots, soil, animal material (Põlme et al. 2020); Likely opportunistic human pathogen (Irinyi et al. 2015) | [illegible] |
| Otu00059 | 4876 | k__Fungi(100);p__Ascomycota(100);c__Eurotiomycetes(100);o__Eurotiales(100);f__Trichocomaceae(100);f__Trichocomaceae_unclassified(79);f__Trichocomaceae_unclassified(79); | - | - | - | - | - | - | - | Unassigned | - |
| Otu00060 | 4590 | k__Fungi(100);p__Basidiomycota(100);c__Tremellomycetes(100);c__Tremellomycetes_unclassified(98);c__Tremellomycetes_unclassified(98);c__Tremellomycetes_unclassified(98);c__Tremellomycetes_unclassified(98); | - | - | - | - | - | - | - | Unassigned | - |
| Otu00061 | 4449 | k__Fungi(100);p__Zygomycota(100);c__Zygomycota_class_Incertae_sedis(100);o__Mucorales(100);o__Mucorales_unclassified(96);o__Mucorales_unclassified(96);o__Mucorales_unclassified(96); | Mucorales | 7 | Saprotroph | Undefined Saprotroph | Microfungus | NULL | Possible | NULL | [illegible] |
| Otu00062 | 4403 | k__Fungi(100);p__Basidiomycota(100);c__Agaricomycetes(100);o__Polyporales(100);f__Meruliaceae(100);g__Irpex(100);s__Irpex_sp_JSL_2009a(100); | Irpex | 13 | Saprotroph | \|Wood Saprotroph\| | Corticioid | White Rot | Highly Probable | As white_rot (Põlme et al. 2020); Decay wood (Põlme et al. 2020); White rot of dead hardwoods, rarely on conifers | [illegible] |
| Otu00063 | 4208 | k__Fungi(100);p__Basidiomycota(100);c__Tremellomycetes(100);o__Filobasidiales(100);f__Filobasidiaceae(100);g__Cryptococcus(100);s__Cryptococcus_friedmannii(82); | Cryptococcus | 13 | Pathotroph-Saprotroph-Symbiotroph | Animal Pathogen-Endophyte-Epiphyte-\|Undefined Saprotroph\| | Dimorphic Yeast-Yeast | NULL | Possible | As foliar_endophyte (Põlme et al. 2020); Decay leaf, fruit, seed, soil, dung, animal material (Põlme et al. 2020) | [illegible] |
| Otu00064 | 4206 | k__Fungi(100);p__Ascomycota(100);c__Dothideomycetes(100);o__Pleosporales(100);f__Sporormiaceae(100);g__Preussia(99);g__Preussia_unclassified(77); | Preussia | 13 | Saprotroph | \|Dung Saprotroph\|-Undefined Saprotroph | Microfungus | NULL | Probable | Decay dung (Põlme et al. 2020) | [illegible] |
| Otu00065 | 4171 | k__Plantae(100);p__unclassified_Plantae(100);c__unclassified_Plantae(100);o__unclassified_Plantae(100);f__unclassified_Plantae(100);g__unclassified_Plantae(100);s__Plantae_sp(100); | - | - | - | - | - | - | - | Unassigned | - |
| Otu00066 | 3945 | k__Fungi(100);p__Basidiomycota(100);c__Agaricomycetes(100);o__Polyporales(100);f__Meripilaceae(100);g__Rigidoporus(94);s__Rigidoporus_sp_JV0511_23(94); | Rigidoporus | 13 | Saprotroph | \|Wood Saprotroph\| | Polyporoid | White Rot | Highly Probable | As white_rot (Põlme et al. 2020); Decay wood (Põlme et al. 2020) | [illegible] |
| Otu00067 | 3900 | k__Fungi(100);p__Basidiomycota(100);c__Agaricomycetes(100);o__Agaricales(100);f__Strophariaceae(100);g__Kuehneromyces(100);s__Kuehneromyces_rostratus(100); | Kuehneromyces | 13 | Saprotroph | Undefined Saprotroph-\|Wood Saprotroph\| | Agaricoid | White Rot | Probable | As white_rot (Põlme et al. 2020); Decay wood (Põlme et al. 2020) | [illegible] |
| Otu00068 | 3859 | k__Fungi(100);p__Basidiomycota(100);c__Agaricomycetes(100);o__Polyporales(100);f__Meruliaceae(100);g__Phlebia(100);s__Phlebia_fuscoatra(100); | Phlebia | 13 | Saprotroph | \|Wood Saprotroph\| | Corticioid | White Rot | Highly Probable | As white_rot (Põlme et al. 2020); Decay wood (Põlme et al. 2020) | [illegible] |
| Otu00069 | 3839 | k__Fungi(100);p__Basidiomycota(100);c__Tremellomycetes(100);o__Tremellales(100);f__Tremellales_family_Incertae_sedis(100);g__Cryptococcus(100);s__Cryptococcus_sp_JCM_11356(93); | Cryptococcus | 13 | Pathotroph-Saprotroph-Symbiotroph | Animal Pathogen-Endophyte-Epiphyte-\|Undefined Saprotroph\| | Dimorphic Yeast-Yeast | NULL | Possible | As foliar_endophyte (Põlme et al. 2020); Decay leaf, fruit, seed, soil, dung, animal material (Põlme et al. 2020) | [illegible] |
| Otu00070 | 3804 | k__Fungi(100);p__Ascomycota(81);p__Ascomycota_unclassified(81);p__Ascomycota_unclassified(81);p__Ascomycota_unclassified(81);p__Ascomycota_unclassified(81);p__Ascomycota_unclassified(81); | - | - | - | - | - | - | - | Unassigned | - |
| Otu00071 | 3472 | k__Fungi(100);p__Ascomycota(100);c__Dothideomycetes(100);o__Pleosporales(100);f__Pleosporales_family_Incertae_sedis(100);g__unclassified_Pleosporales(100);s__Pleosporales_sp(100); | - | - | - | - | - | - | - | Unassigned | - |
| Otu00072 | 3466 | k__Fungi(100);p__Ascomycota(100);c__Eurotiomycetes(100);o__Eurotiales(100);f__Trichocomaceae(100);f__Trichocomaceae_unclassified(100);f__Trichocomaceae_unclassified(100); | - | - | - | - | - | - | - | Unassigned | - |
| Otu00073 | 3444 | k__Fungi(100);p__Ascomycota(99);c__Dothideomycetes(95);o__Dothideomycetes_order_Incertae_sedis(95);f__Pseudeurotiaceae(95);g__Pseudeurotium(78);g__Pseudeurotium_unclassified(76); | Pseudeurotium | 13 | Saprotroph | \|Dung Saprotroph\|-Undefined Saprotroph | Microfungus | NULL | Probable | As soil_saprotroph (Põlme et al. 2020); Decay soil, dung (Põlme et al. 2020) | [illegible] |
| Otu00074 | 3210 | k__Fungi(100);p__Ascomycota(100);c__Dothideomycetes(100);o__Pleosporales(100);f__Phaeosphaeriaceae(100);g__Phaeosphaeria(100);s__Phaeosphaeria_sp(99); | Phaeosphaeria | 13 | Pathotroph-Saprotroph-Symbiotroph | Endophyte-Plant Pathogen-\|Plant Saprotroph\| | Microfungus | NULL | Probable | As litter_saprotroph (Põlme et al. 2020); Decay leaf, fruit, seed (Põlme et al. 2020) | [illegible] |
| Otu00075 | 3168 | k__Fungi(100);p__Ascomycota(100);c__Taphrinomycetes(100);o__Taphrinales(100);f__Taphrinaceae(100);g__Taphrina(58);g__Taphrina_unclassified(58); | Taphrina | 13 | Pathotroph | \|Plant Pathogen\| | Dimorphic-Microfungus | NULL | Probable | NULL | [illegible] |
| Otu00076 | 3129 | k__Plantae(100);p__unclassified_Plantae(100);c__unclassified_Plantae(100);o__unclassified_Plantae(100);f__unclassified_Plantae(100);g__unclassified_Plantae(100);s__Plantae_sp(100); | - | - | - | - | - | - | - | Unassigned | - |
| Otu00077 | 3089 | k__Fungi(100);p__Zygomycota(100);c__Zygomycota_class_Incertae_sedis(100);o__Mucorales(100);f__Umbelopsidaceae(100);g__Umbelopsis(100);s__Umbelopsis_isabellina(98); | Umbelopsis | 13 | Saprotroph | \|Undefined Saprotroph\| | Microfungus | NULL | Probable | As soil_saprotroph (Põlme et al. 2020); Decay roots, soil (Põlme et al. 2020) | [illegible] |
| Otu00078 | 2978 | k__Fungi(100);p__Basidiomycota(100);c__Agaricomycetes(100);o__Russulales(95);o__Russulales_unclassified(95);o__Russulales_unclassified(95);o__Russulales_unclassified(95); | - | - | - | - | - | - | - | Unassigned | - |
| Otu00079 | 2933 | k__Fungi(100);p__Ascomycota(100);p__Ascomycota_unclassified(100);p__Ascomycota_unclassified(100);p__Ascomycota_unclassified(100);p__Ascomycota_unclassified(100);p__Ascomycota_unclassified(100); | - | - | - | - | - | - | - | Unassigned | - |
| Otu00080 | 2902 | k__Fungi(100);p__Basidiomycota(100);c__Agaricomycetes(100);o__Russulales(100);f__Peniophoraceae(100);g__Peniophora(77);g__Peniophora_unclassified(77); | Peniophora | 13 | Pathotroph-Saprotroph | Plant Pathogen-\|Wood Saprotroph\| | Corticioid | White Rot | Probable | As white_rot (Põlme et al. 2020); As wood_pathogen (Põlme et al. 2020); Decay wood (Põlme et al. 2020) | [illegible] |
| Otu00081 | 2883 | k__Fungi(100);p__Basidiomycota(100);c__Microbotryomycetes(100);o__Sporidiobolales(100);f__Sporidiobolales_family_Incertae_sedis(100);g__Rhodosporidium(70);g__Rhodosporidium_unclassified(69); | Rhodosporidium | 13 | Saprotroph | Undefined Saprotroph | Yeast | NULL | Highly Probable | NULL | [illegible] |
| Otu00082 | 2740 | k__Fungi(100);p__Basidiomycota(100);c__Agaricomycetes(100);o__Corticiales(98);f__unclassified_Corticiales(98);g__unclassified_Corticiales(98);s__Corticiales_sp(98); | - | - | - | - | - | - | - | Unassigned | - |
| Otu00083 | 2715 | k__Fungi(100);p__Basidiomycota(100);c__Microbotryomycetes(100);o__Sporidiobolales(100);f__Sporidiobolales_family_Incertae_sedis(79);g__Sporobolomyces(79);g__Sporobolomyces_unclassified(58); | Sporobolomyces | 13 | Pathotroph-Saprotroph | \|Fungal Parasite\|-Plant Saprotroph | Yeast | NULL | Probable | As litter_saprotroph (Põlme et al. 2020); As mycoparasite (Põlme et al. 2020); Decay fungal material (Põlme et al. 2020) | [illegible] |
| Otu00084 | 2660 | k__Fungi(100);p__Basidiomycota(100);c__Agaricomycetes(100);o__Hymenochaetales(100);f__Hymenochaetales_family_Incertae_sedis(100);g__Trichaptum(100);s__Trichaptum_biforme(100); | Trichaptum | 13 | Saprotroph | \|Wood Saprotroph\| | Polyporoid | White Rot | Highly Probable | As white_rot (Põlme et al. 2020); Decay wood (Põlme et al. 2020) | [illegible] |
| Otu00085 | 2618 | k__Fungi(100);p__Basidiomycota(100);c__Agaricomycetes(100);o__Polyporales(100);f__Polyporaceae(100);g__Perenniporia(100);s__Perenniporia_subacida(89); | Perenniporia | 13 | Saprotroph | \|Wood Saprotroph\| | Corticioid-Polyporoid | White Rot | Highly Probable | As white_rot (Põlme et al. 2020); Decay wood (Põlme et al. 2020) | [illegible] |
| Otu00086 | 2590 | k__Fungi(100);p__Ascomycota(100);c__Leotiomycetes(100);o__Helotiales(100);f__Sclerotiniaceae(100);g__Botrytis(90);s__Botrytis_caroliniana(90); | Botrytis | 13 | Pathotroph-Saprotroph | \|Plant Pathogen\|-Plant Saprotroph | Microfungus | NULL | Probable | As litter_saprotroph (Põlme et al. 2020); Decay leaf, fruit, seed (Põlme et al. 2020) | [illegible] |
| Otu00087 | 2576 | k__Fungi(100);p__Ascomycota(100);c__Taphrinomycetes(100);o__Taphrinales(100);f__Taphrinaceae(100);g__Taphrina(100);g__Taphrina_unclassified(100); | Taphrina | 13 | Pathotroph | \|Plant Pathogen\| | Dimorphic-Microfungus | NULL | Probable | NULL | [illegible] |
| Otu00088 | 2531 | k__Fungi(100);p__Ascomycota(100);c__Dothideomycetes(100);o__Capnodiales(100);f__Mycosphaerellaceae(100);f__Mycosphaerellaceae_unclassified(100);f__Mycosphaerellaceae_unclassified(100); | Mycosphaerellaceae | 9 | Pathotroph-Saprotroph | Plant Pathogen-Undefined Saprotroph | Microfungus | NULL | Probable | Biotrophic, necrotrophic or saprobic on various plant tissues (Cannon & Kirk 2007) | [illegible] |
| Otu00089 | 2485 | k__Fungi(100);p__Ascomycota(100);c__Sordariomycetes(100);o__Xylariales(100);f__Amphisphaeriaceae(100);g__Bartalinia(99);s__Bartalinia_robillardoides(99); | Bartalinia | 13 | Saprotroph-Symbiotroph | \|Endophyte\|-Plant Saprotroph-Undefined Saprotroph | Microfungus | NULL | Probable | As foliar_endophyte (Põlme et al. 2020); As litter_saprotroph (Põlme et al. 2020); Decay leaf, fruit, seed (Põlme et al. 2020) | [illegible] |
| Otu00090 | 2436 | k__Fungi(100);p__Basidiomycota(100);c__Tremellomycetes(100);o__Tremellales(100);f__Tremellales_family_Incertae_sedis(100);g__Cryptococcus(100);s__Cryptococcus_sp_CBS_8358(100); | Cryptococcus | 13 | Pathotroph-Saprotroph-Symbiotroph | Animal Pathogen-Endophyte-Epiphyte-\|Undefined Saprotroph\| | Dimorphic Yeast-Yeast | NULL | Possible | As foliar_endophyte (Põlme et al. 2020); Decay leaf, fruit, seed, soil, dung, animal material (Põlme et al. 2020) | [illegible] |
| Otu00091 | 2399 | k__Fungi(100);p__Basidiomycota(100);c__Agaricomycetes(100);o__Polyporales(100);f__Meruliaceae(90);g__Phlebia(90);s__Phlebia_sp_olrim964(90); | Phlebia | 13 | Saprotroph | \|Wood Saprotroph\| | Corticioid | White Rot | Highly Probable | As white_rot (Põlme et al. 2020); Decay wood (Põlme et al. 2020) | [illegible] |
| Otu00092 | 2339 | k__Fungi(100);p__Ascomycota(100);c__Sordariomycetes(100);o__Diaporthales(95);f__Valsaceae(95);f__Valsaceae_unclassified(95);f__Valsaceae_unclassified(95); | Valsaceae | 9 | Pathotroph-Saprotroph-Symbiotroph | Endophyte-Plant Pathogen-Undefined Saprotroph | Microfungus | NULL | Probable | Saprobes, endophytes and weak to virulent parasites of woody plants (Cannon & Kirk 2007) | [illegible] |
| Otu00093 | 2329 | k__Fungi(100);p__Zygomycota(100);c__Zygomycota_class_Incertae_sedis(100);o__Mucorales(100);f__Umbelopsidaceae(100);g__Umbelopsis(100);g__Umbelopsis_unclassified(89); | Umbelopsis | 13 | Saprotroph | \|Undefined Saprotroph\| | Microfungus | NULL | Probable | As soil_saprotroph (Põlme et al. 2020); Decay roots, soil (Põlme et al. 2020) | [illegible] |
| Otu00094 | 2293 | k__Fungi(100);p__Basidiomycota(67);p__Basidiomycota_unclassified(67);p__Basidiomycota_unclassified(67);p__Basidiomycota_unclassified(67);p__Basidiomycota_unclassified(67);p__Basidiomycota_unclassified(67); | - | - | - | - | - | - | - | Unassigned | - |
| Otu00095 | 2276 | k__Fungi(100);p__Ascomycota(100);c__Dothideomycetes(100);o__Pleosporales(100);o__Pleosporales_unclassified(100);o__Pleosporales_unclassified(100);o__Pleosporales_unclassified(100); | - | - | - | - | - | - | - | Unassigned | - |
| Otu00096 | 2202 | k__Fungi(100);p__Ascomycota(100);c__Dothideomycetes(100);o__Pleosporales(100);f__Pleosporales_family_Incertae_sedis(100);g__Pyrenochaeta(100);s__Pyrenochaeta_keratinophila(100); | Pyrenochaeta | 13 | Pathotroph-Saprotroph | Plant Pathogen-Undefined Saprotroph-\|Wood Saprotroph\| | Microfungus | Soft Rot | Highly Probable | Decay wood (Põlme et al. 2020) | [illegible] |
| Otu00097 | 2201 | k__Fungi(100);p__Basidiomycota(100);c__Agaricomycetes(100);o__Boletales(100);f__Suillaceae(100);g__Suillus(100);s__Suillus_placidus(100); | Suillus | 13 | Symbiotroph | \|Ectomycorrhizal\| | Agaricoid-Boletoid | NULL | Highly Probable | Host - Pinaceae (Põlme et al. 2020) | [illegible] |
| Otu00098 | 2191 | k__Fungi(100);p__Basidiomycota(100);c__Agaricomycetes(100);o__Russulales(100);f__Peniophoraceae(100);g__Peniophora(100);g__Peniophora_unclassified(99); | Peniophora | 13 | Pathotroph-Saprotroph | Plant Pathogen-\|Wood Saprotroph\| | Corticioid | White Rot | Probable | As white_rot (Põlme et al. 2020); As wood_pathogen (Põlme et al. 2020); Decay wood (Põlme et al. 2020) | [illegible] |
| Otu00099 | 2157 | k__Fungi(100);p__Basidiomycota(100);c__Exobasidiomycetes(100);o__unclassified_Exobasidiomycetes(100);f__unclassified_Exobasidiomycetes(100);g__unclassified_Exobasidiomycetes(100);s__Exobasidiomycetes_sp(100); | - | - | - | - | - | - | - | Unassigned | - |
| Otu00100 | 1960 | k__Fungi(100);p__Ascomycota(100);c__Leotiomycetes(90);c__Leotiomycetes_unclassified(90);c__Leotiomycetes_unclassified(90);c__Leotiomycetes_unclassified(90);c__Leotiomycetes_unclassified(90); | - | - | - | - | - | - | - | Unassigned | - |
| Otu00101 | 1959 | k__Fungi(100);p__Ascomycota(100);c__Sordariomycetes(100);o__Xylariales(100);f__Amphisphaeriaceae(100);g__Monochaetia(100);s__Monochaetia_sp(72); | Monochaetia | 13 | Pathotroph | \|Plant Pathogen\| | Microfungus | NULL | Probable | NULL | [illegible] |
| Otu00102 | 1956 | k__Fungi(100);p__Ascomycota(100);c__Saccharomycetes(100);o__Saccharomycetales(100);f__Saccharomycetaceae(100);g__Torulaspora(100);s__Torulaspora_delbrueckii(98); | Torulaspora | 13 | Pathotroph-Saprotroph | Animal Parasite-\|Undefined Saprotroph\| | Yeast | NULL | Probable | As nectar-tap_saprotroph (Põlme et al. 2020); Decay animal material, sugar-rich substrates (Põlme et al. 2020) | [illegible] |
| Otu00103 | 1950 | k__Fungi(100);p__Ascomycota(100);c__Sordariomycetes(100);o__Hypocreales(100);f__Cordycipitaceae(100);g__Lecanicillium(80);s__Lecanicillium_muscarium(56); | Lecanicillium | 13 | Pathotroph-Saprotroph-Symbiotroph | \|Animal Parasite\|-Animal Pathogen-Endophyte-Undefined Saprotroph | Microfungus | NULL | Probable | As animal_decomposer (Põlme et al. 2020); As foliar_endophyte (Põlme et al. 2020); Decay animal material (Põlme et al. 2020) | [illegible] |
| Otu00104 | 1937 | k__Fungi(100);p__Ascomycota(100);c__Dothideomycetes(100);o__Capnodiales(100);f__Mycosphaerellaceae(100);g__Pseudocercosporella(100);s__Pseudocercosporella_sp(100); | Pseudocercosporella | 13 | Pathotroph | \|Plant Pathogen\| | Microfungus | NULL | Probable | NULL | [illegible] |
| Otu00105 | 1923 | k__Fungi(100);p__Basidiomycota(100);c__Agaricomycetes(100);o__Polyporales(100);f__Meruliaceae(78);g__Scopuloides(78);s__Scopuloides_hydnoides(78); | Scopuloides | 13 | Saprotroph | Undefined Saprotroph-\|Wood Saprotroph\| | Corticioid | White Rot | Probable | As white_rot (Põlme et al. 2020); Decay wood (Põlme et al. 2020) | [illegible] |
| Otu00106 | 1885 | k__Fungi(100);p__Basidiomycota(100);c__Agaricomycetes(100);o__Polyporales(100);o__Polyporales_unclassified(100);o__Polyporales_unclassified(100);o__Polyporales_unclassified(100); | - | - | - | - | - | - | - | Unassigned | - |
| Otu00107 | 1873 | k__Fungi(100);p__Ascomycota(100);c__Sordariomycetes(100);o__Hypocreales(100);f__Ophiocordycipitaceae(100);f__Ophiocordycipitaceae_unclassified(100);f__Ophiocordycipitaceae_unclassified(100); | Ophiocordycipitaceae | 9 | Pathotroph-Symbiotroph | Endophyte-Insect Pathogen | Microfungus | NULL | Probable | A family of parasitic fungi (Sung et al. 2007) | [illegible] |
| Otu00108 | 1835 | k__Fungi(100);p__Ascomycota(100);c__Sordariomycetes(100);o__Microascales(100);f__Microascaceae(100);g__Microascus(100);s__Microascus_manginii(100); | Microascus | 13 | Pathotroph-Saprotroph-Symbiotroph | Animal Pathogen-Dung Saprotroph-Endophyte-Plant Pathogen-Plant Saprotroph-\|Undefined Saprotroph\| | Microfungus | NULL | Possible | As foliar_endophyte (Põlme et al. 2020); As litter_saprotroph (Põlme et al. 2020); As soil_saprotroph (Põlme et al. 2020); Decay leaf, fruit, seed, soil (Põlme et al. 2020) | [illegible] |
| Otu00109 | 1816 | k__Fungi(100);p__Zygomycota(100);c__Zygomycota_class_Incertae_sedis(100);o__Mucorales(100);f__Mucoraceae(100);g__Mucor(97);g__Mucor_unclassified(56); | Mucor | 13 | Saprotroph | \|Undefined Saprotroph\| | Microfungus | NULL | Probable | As soil_saprotroph (Põlme et al. 2020); Decay soil (Põlme et al. 2020) | [illegible] |
| Otu00110 | 1788 | k__Fungi(100);p__Ascomycota(100);c__Saccharomycetes(100);o__Saccharomycetales(100);f__Saccharomycetaceae(100);g__Kazachstania(100);s__Kazachstania_martiniae(100); | Kazachstania | 13 | Saprotroph | \|Undefined Saprotroph\| | Yeast | NULL | Probable | As nectar-tap_saprotroph (Põlme et al. 2020); Decay sugar-rich substrates (Põlme et al. 2020) | [illegible] |
| Otu00111 | 1783 | k__Fungi(100);p__Ascomycota(100);c__Dothideomycetes(100);o__Pleosporales(100);o__Pleosporales_unclassified(94);o__Pleosporales_unclassified(94);o__Pleosporales_unclassified(94); | - | - | - | - | - | - | - | Unassigned | - |
| Otu00112 | 1759 | k__Fungi(100);p__Basidiomycota(100);c__Tremellomycetes(100);o__Tremellales(99);f__Tremellales_family_Incertae_sedis(99);g__Cryptococcus(99);s__Cryptococcus_magnus(99); | Cryptococcus | 13 | Pathotroph-Saprotroph-Symbiotroph | Animal Pathogen-Endophyte-Epiphyte-\|Undefined Saprotroph\| | Dimorphic Yeast-Yeast | NULL | Possible | As foliar_endophyte (Põlme et al. 2020); Decay leaf, fruit, seed, soil, dung, animal material (Põlme et al. 2020) | [illegible] |
| Otu00113 | 1722 | k__Fungi(100);p__Ascomycota(100);c__Eurotiomycetes(100);c__Eurotiomycetes_unclassified(100);c__Eurotiomycetes_unclassified(100);c__Eurotiomycetes_unclassified(100);c__Eurotiomycetes_unclassified(100); | - | - | - | - | - | - | - | Unassigned | - |
| Otu00114 | 1716 | k__Fungi(100);p__Ascomycota(100);c__Sordariomycetes(100);o__Hypocreales(100);f__Nectriaceae(100);g__Gibberella(100);s__Gibberella_zeae(100); | Gibberella | 13 | Pathotroph-Saprotroph-Symbiotroph | Endophyte-\|Plant Pathogen\|-Plant Saprotroph-Undefined Saprotroph | Microfungus | Soft Rot | Probable | As foliar_endophyte (Põlme et al. 2020); As litter_saprotroph (Põlme et al. 2020); Causing damping-off disease (Põlme et al. 2020); Decay leaf, fruit, seed, roots, soil, animal material (Põlme et al. 2020) | [illegible] |
| Otu00115 | 1660 | k__Fungi(100);p__Basidiomycota(78);p__Basidiomycota_unclassified(78);p__Basidiomycota_unclassified(78);p__Basidiomycota_unclassified(78);p__Basidiomycota_unclassified(78);p__Basidiomycota_unclassified(78); | - | - | - | - | - | - | - | Unassigned | - |
| Otu00116 | 1647 | k__Fungi(100);p__Ascomycota(100);c__Dothideomycetes(100);o__Pleosporales(100);f__Venturiaceae(100);g__Fusicladium(100);s__Fusicladium_caryigenum(100); | Fusicladium | 13 | Pathotroph-Symbiotroph | Endophyte-Lichen Parasite-\|Plant Pathogen\| | Microfungus | NULL | Probable | NULL | [illegible] |
| Otu00117 | 1639 | k__Fungi(100);p__Basidiomycota(100);c__Agaricomycetes(100);o__Polyporales(100);f__Polyporaceae(100);g__Perenniporia(100);s__Perenniporia_sp(100); | Perenniporia | 13 | Saprotroph | \|Wood Saprotroph\| | Corticioid-Polyporoid | White Rot | Highly Probable | As white_rot (Põlme et al. 2020); Decay wood (Põlme et al. 2020) | [illegible] |
| Otu00118 | 1616 | k__Fungi(100);p__Chytridiomycota(100);c__Chytridiomycetes(100);o__Rhizophlyctidales(100);f__Rhizophlyctidaceae(100);g__Rhizophlyctis(100);s__Rhizophlyctis_rosea(100); | Rhizophlyctis | 13 | Pathotroph-Saprotroph | Algal Parasite-\|Plant Saprotroph\|-Undefined Saprotroph | Chytrid-Microfungus | NULL | Probable | As litter_saprotroph (Põlme et al. 2020); Decay leaf, fruit, seed, algal material (Põlme et al. 2020) | [illegible] |
| Otu00119 | 1584 | k__Fungi(100);p__Ascomycota(100);c__Eurotiomycetes(100);o__unclassified_Eurotiomycetes(100);f__unclassified_Eurotiomycetes(100);g__unclassified_Eurotiomycetes(100);s__Eurotiomycetes_sp(100); | - | - | - | - | - | - | - | Unassigned | - |
| Otu00120 | 1577 | k__Fungi(100);p__Ascomycota(100);p__Ascomycota_unclassified(100);p__Ascomycota_unclassified(100);p__Ascomycota_unclassified(100);p__Ascomycota_unclassified(100);p__Ascomycota_unclassified(100); | - | - | - | - | - | - | - | Unassigned | - |
| Otu00121 | 1573 | k__Fungi(100);p__Basidiomycota(100);c__Agaricomycetes(100);o__Polyporales(100);f__Meruliaceae(100);g__Phlebia(100);s__Phlebia_sp_DABAC9(95); | Phlebia | 13 | Saprotroph | \|Wood Saprotroph\| | Corticioid | White Rot | Highly Probable | As white_rot (Põlme et al. 2020); Decay wood (Põlme et al. 2020) | [illegible] |
| Otu00122 | 1482 | k__Fungi(100);p__Ascomycota(100);c__Dothideomycetes(100);o__Pleosporales(100);f__Pleosporales_family_Incertae_sedis(100);g__Ascochyta(100);s__Ascochyta_sp(100); | Ascochyta | 13 | Pathotroph | Lichen Parasite-\|Plant Pathogen\| | Microfungus | NULL | Probable | NULL | [illegible] |
| Otu00123 | 1474 | k__Fungi(100);p__Basidiomycota(100);c__Agaricomycetes(100);o__Polyporales(100);f__Meruliaceae(100);g__Phlebia(100);s__Phlebia_tremellosa(100); | Phlebia | 13 | Saprotroph | \|Wood Saprotroph\| | Corticioid | White Rot | Highly Probable | As white_rot (Põlme et al. 2020); Decay wood (Põlme et al. 2020) | [illegible] |

| | | | | | | | | | | | | |
|---|---|---|---|---|---|---|---|---|---|---|---|---|
| Otu00124 | 1385 | k__Fungi(100);p__Zygomycota(100);c__Zygomycota_class_Incertae_sedis(100);o__Mucorales(100);f__Mucoraceae(100);g__Mucor(100);s__Mucor_sp_JJP_2009a(69); | Mucor | | 13 | Saprotroph | \|Undefined Saprotroph\| | Microfungus | NULL | Probable | As soil_saprotroph (Põlme et al. 2020); Decay soil (Põlme et al. 2020) | [illegible] |
| Otu00125 | 1385 | k__Fungi(100);p__Zygomycota(100);c__Zygomycota_class_Incertae_sedis(100);o__Mucorales(100);f__Mucoraceae(100);g__Mucor(100);g__Mucor_unclassified(63); | Mucor | | 13 | Saprotroph | \|Undefined Saprotroph\| | Microfungus | NULL | Probable | As soil_saprotroph (Põlme et al. 2020); Decay soil (Põlme et al. 2020) | [illegible] |
| Otu00126 | 1360 | k__Fungi(100);p__Ascomycota(100);c__Eurotiomycetes(100);c__Eurotiomycetes_unclassified(99);c__Eurotiomycetes_unclassified(99);c__Eurotiomycetes_unclassified(99);c__Eurotiomycetes_unclassified(99); | - | - | | - | - | - | - | - | Unassigned | - |
| Otu00127 | 1324 | k__Fungi(100);p__Basidiomycota(100);c__Agaricomycetes(100);o__Agaricales(100);f__Psathyrellaceae(100);g__Psathyrella(100);s__Psathyrella_maculata(100); | Psathyrella | | 13 | Saprotroph | Plant Saprotroph-\|Wood Saprotroph\| | Agaricoid | NULL | Probable | As litter_saprotroph (Põlme et al. 2020); Decay wood, leaf, fruit, seed (Põlme et al. 2020) | [illegible] |
| Otu00128 | 1293 | k__Fungi(100);p__Ascomycota(100);c__Dothideomycetes(99);o__Capnodiales(94);o__Capnodiales_unclassified(94);o__Capnodiales_unclassified(94);o__Capnodiales_unclassified(94); | - | - | | - | - | - | - | - | Unassigned | - |
| Otu00129 | 1279 | k__Fungi(100);p__Basidiomycota(100);c__Agaricomycetes(100);o__Russulales(100);f__Stereaceae(100);g__Gloeocystidiellum(100);s__Gloeocystidiellum_sp(99); | Gloeocystidiellum | | 13 | Saprotroph | Undefined Saprotroph-\|Wood Saprotroph\| | Corticioid | White Rot | Probable | As white_rot (Põlme et al. 2020); Decay wood (Põlme et al. 2020) | [illegible] |
| Otu00130 | 1268 | k__Fungi(100);p__Ascomycota(100);c__Dothideomycetes(100);o__Dothideomycetes_order_Incertae_sedis(100);f__Myxotrichaceae(100);g__Oidiodendron(100);s__Oidiodendron_sp_22MS12(100); | Oidiodendron | | 13 | Pathotroph-Saprotroph-Symbiotroph | Endophyte-Ericoid Mycorrhizal-\|Undefined Saprotroph\| | Microfungus | NULL | Probable | As root_endophyte (Põlme et al. 2020); As soil_saprotroph (Põlme et al. 2020); Dark Septate Endophyte; Decay soil (Põlme et al. 2020) | [illegible] |
| Otu00131 | 1266 | k__Fungi(100);p__Basidiomycota(100);c__Agaricomycetes(100);o__Polyporales(100);f__Polyporaceae(100);g__Tyromyces(100);s__Tyromyces_chioneus(100); | Tyromyces | | 13 | Saprotroph | \|Wood Saprotroph\| | Polyporoid | White Rot | Highly Probable | As white_rot (Põlme et al. 2020); Decay wood (Põlme et al. 2020) | [illegible] |
| Otu00132 | 1264 | k__Fungi(100);p__Ascomycota(100);c__Dothideomycetes(100);o__Botryosphaeriales(100);f__Botryosphaeriaceae(100);g__Diplodia(98);g__Diplodia_unclassified(61); | Diplodia | | 13 | Pathotroph-Saprotroph-Symbiotroph | Epiphyte-\|Plant Pathogen\|-Plant Saprotroph-Wood Saprotroph | Microfungus | NULL | Possible | As litter_saprotroph (Põlme et al. 2020); Cause of soft; culture (Carisse et al. 2000); Decay leaf, fruit, seed (Põlme et al. 2020); rot; Epiphyte detection method | [illegible] |
| Otu00133 | 1230 | k__Fungi(100);p__Zygomycota(100);c__Zygomycota_class_Incertae_sedis(100);o__Basidiobolales(100);f__Basidiobolaceae(100);g__Basidiobolus(100);s__Basidiobolus_ranarum(87); | Basidiobolus | | 13 | Pathotroph-Saprotroph-Symbiotroph | Animal Pathogen-Animal Symbiotroph-\|Undefined Saprotroph\| | Microfungus | NULL | Probable | Cause of subcutaneous zygomycosis in humans (Cannon & Kirk 2007); Basidiobolus ranarum is commonly present in decaying fruit and vegetable matter, and as a commensal in the intestinal tract of frogs, toads, and lizards ; Decay leaf, fruit, seed, soil, dung, animal material (Põlme et al. 2020) | [illegible] |
| Otu00134 | 1220 | k__Fungi(100);k__Fungi_unclassified(97);k__Fungi_unclassified(97);k__Fungi_unclassified(97);k__Fungi_unclassified(97);k__Fungi_unclassified(97);k__Fungi_unclassified(97); | - | - | | - | - | - | - | - | Unassigned | - |
| Otu00135 | 1213 | k__Fungi(100);p__Zygomycota(100);c__Zygomycota_class_Incertae_sedis(100);o__Mucorales(100);f__Mucoraceae(100);g__Mucor(100);g__Mucor_unclassified(68); | Mucor | | 13 | Saprotroph | \|Undefined Saprotroph\| | Microfungus | NULL | Probable | As soil_saprotroph (Põlme et al. 2020); Decay soil (Põlme et al. 2020) | [illegible] |
| Otu00136 | 1205 | k__Fungi(100);p__Ascomycota(100);c__Dothideomycetes(100);o__Capnodiales(100);f__Mycosphaerellaceae(100);f__Mycosphaerellaceae_unclassified(95);f__Mycosphaerellaceae_unclassified(95); | Mycosphaerellaceae | | 9 | Pathotroph-Saprotroph | Plant Pathogen-Undefined Saprotroph | Microfungus | NULL | Probable | Biotrophic, necrotrophic or saprobic on various plant tissues (Cannon & Kirk 2007) | [illegible] |
| Otu00137 | 1189 | k__Fungi(100);p__Basidiomycota(100);c__Agaricomycetes(100);o__Russulales(100);f__Stereaceae(100);g__Stereum(100);s__Stereum_sp_Vega483(100); | Stereum | | 13 | Saprotroph | Undefined Saprotroph-\|Wood Saprotroph\| | Polyporoid | White Rot | Probable | As white_rot (Põlme et al. 2020); Decay wood (Põlme et al. 2020) | [illegible] |
| Otu00138 | 1174 | k__Fungi(100);p__Ascomycota(100);c__Dothideomycetes(100);o__Pleosporales(100);f__Cucurbitariaceae(100);g__Pyrenochaetopsis(100);s__Pyrenochaetopsis_leptospora(100); | Pyrenochaetopsis | | 13 | Pathotroph-Saprotroph-Symbiotroph | \|Endophyte\|-Lichen Parasite-Plant Saprotroph-Undefined Saprotroph | Microfungus | NULL | Possible | As foliar_endophyte (Põlme et al. 2020); As litter_saprotroph (Põlme et al. 2020); Decay leaf, fruit, seed (Põlme et al. 2020) | [illegible] |
| Otu00139 | 1171 | k__Fungi(100);p__Ascomycota(100);c__Dothideomycetes(100);o__Pleosporales(100);f__Phaeosphaeriaceae(100);g__unclassified_Phaeosphaeriaceae(99);s__Phaeosphaeriaceae_sp(99); | Phaeosphaeriaceae | | 9 | Pathotroph-Saprotroph | Fungal Parasite-Plant Pathogen-Plant Saprotroph | Microfungus | NULL | Probable | Necrotrophic or saprobic on a wide range of plants, especially monocotyledons; a few species parasitic on other fungi (Cannon & Kirk 2007) | [illegible] |
| Otu00140 | 1163 | k__Fungi(100);p__Ascomycota(100);c__Eurotiomycetes(100);o__Eurotiales(100);f__Trichocomaceae(100);g__Aspergillus(100);s__Aspergillus_cibarius(100); | Aspergillus | | 13 | Pathotroph-Saprotroph-Symbiotroph | Animal Pathogen-Endophyte-Plant Saprotroph-\|Undefined Saprotroph\|-Wood Saprotroph | Microfungus | NULL | Probable | As foliar_endophyte (Põlme et al. 2020); Decay leaf, fruit, seed, soil, dung, animal material (Põlme et al. 2020) | [illegible] |
| Otu00141 | 1156 | k__Fungi(100);p__Ascomycota(100);c__Eurotiomycetes(98);c__Eurotiomycetes_unclassified(91);c__Eurotiomycetes_unclassified(91);c__Eurotiomycetes_unclassified(91);c__Eurotiomycetes_unclassified(91); | - | - | | - | - | - | - | - | Unassigned | - |
| Otu00142 | 1147 | k__Fungi(100);p__Ascomycota(100);c__Taphrinomycetes(100);o__Taphrinales(100);f__Taphrinaceae(100);g__Taphrina(100);s__Taphrina_sp(100); | Taphrina | | 13 | Pathotroph | \|Plant Pathogen\| | Dimorphic-Microfungus | NULL | Probable | NULL | [illegible] |
| Otu00143 | 1142 | k__Fungi(100);p__Ascomycota(90);p__Ascomycota_unclassified(90);p__Ascomycota_unclassified(90);p__Ascomycota_unclassified(90);p__Ascomycota_unclassified(90);p__Ascomycota_unclassified(90); | - | - | | - | - | - | - | - | Unassigned | - |
| Otu00144 | 1137 | k__Fungi(100);p__Ascomycota(100);c__Eurotiomycetes(100);o__Eurotiomycetes_order_Incertae_sedis(100);f__Eurotiomycetes_family_Incertae_sedis(100);g__Sarcinomyces(100);s__Sarcinomyces_sp_SL_2011(100); | Sarcinomyces | | 13 | Saprotroph-Symbiotroph | \|Endophyte\|-Plant Saprotroph | Dimorphic | NULL | Probable | As foliar_endophyte (Põlme et al. 2020); As litter_saprotroph (Põlme et al. 2020); Decay leaf, fruit, seed (Põlme et al. 2020); Host - Pinus_galls_of_Cronartium (Põlme et al. 2020) | [illegible] |
| Otu00145 | 1135 | k__Fungi(100);p__Basidiomycota(100);c__Agaricomycetes(100);o__Russulales(100);f__Peniophoraceae(100);g__Dendrophora(81);s__Dendrophora_sp(81); | Dendrophora | | 13 | Saprotroph | Undefined Saprotroph-\|Wood Saprotroph\| | Corticioid | White Rot | Probable | As white_rot (Põlme et al. 2020); Decay wood (Põlme et al. 2020) | [illegible] |
| Otu00146 | 1094 | k__Fungi(100);p__Ascomycota(100);c__Dothideomycetes(97);o__Pleosporales(97);o__Pleosporales_unclassified(96);o__Pleosporales_unclassified(96);o__Pleosporales_unclassified(96); | - | - | | - | - | - | - | - | Unassigned | - |
| Otu00147 | 1079 | k__Fungi(100);p__Basidiomycota(100);c__Agaricomycetes(100);o__Russulales(100);f__Auriscalpiaceae(100);g__Lentinellus(100);s__Lentinellus_ursinus(100); | Lentinellus | | 13 | Pathotroph-Saprotroph | Plant Pathogen-Undefined Saprotroph-\|Wood Saprotroph\| | Agaricoid | White Rot | Probable | As white_rot (Põlme et al. 2020); As wood_pathogen (Põlme et al. 2020); Decay wood (Põlme et al. 2020) | [illegible] |
| Otu00148 | 1073 | k__Fungi(100);p__Basidiomycota(100);c__Tremellomycetes(100);o__Holtermanniales(89);f__Holtermanniales_family_Incertae_sedis(89);g__Holtermannia(89);s__Holtermannia_corniformis(89); | Holtermannia | | 13 | Pathotroph | \|Animal Parasite\|-Fungal Parasite | Dimorphic | NULL | Probable | As mycoparasite (Põlme et al. 2020); Decay animal material (Põlme et al. 2020) | [illegible] |
| Otu00149 | 1073 | k__Fungi(100);p__Basidiomycota(55);p__Basidiomycota_unclassified(54);p__Basidiomycota_unclassified(54);p__Basidiomycota_unclassified(54);p__Basidiomycota_unclassified(54);p__Basidiomycota_unclassified(54); | - | - | | - | - | - | - | - | Unassigned | - |
| Otu00150 | 1062 | k__Fungi(100);p__Ascomycota(100);c__Dothideomycetes(100);o__Capnodiales(100);f__Capnodiales_family_Incertae_sedis(100);g__Toxicocladosporium(100);s__Toxicocladosporium_sp(73); | Toxicocladosporium | | 13 | Pathotroph-Symbiotroph | Endophyte-\|Plant Pathogen\| | Microfungus | NULL | Probable | As foliar_endophyte (Põlme et al. 2020); Decay leaf, fruit, seed (Põlme et al. 2020); Isolated from leaves of Posoqueria and Eucalyptus camaldulensis (occurring within a single small lesion) as well as moldy paint | [illegible] |
| Otu00151 | 1062 | k__Fungi(100);p__Ascomycota(100);c__Dothideomycetes(100);o__Pleosporales(100);f__Pleosporales_family_Incertae_sedis(100);g__Paraphoma(89);s__Paraphoma_chrysanthemicola(67); | Paraphoma | | 13 | Pathotroph-Saprotroph | \|Plant Pathogen\|-Plant Saprotroph-Undefined Saprotroph-Wood Saprotroph | Microfungus | NULL | Probable | As wood_pathogen (Põlme et al. 2020); Associated with leaf and crown infection of pyrethrum (Moslemi et al. 2017); Decay wood (Põlme et al. 2020) | [illegible] |
| Otu00152 | 1059 | k__Fungi(100);p__Ascomycota(100);c__Dothideomycetes(100);o__Pleosporales(100);f__Leptosphaeriaceae(100);g__Leptosphaeria(100);s__Leptosphaeria_sp(100); | Leptosphaeria | | 13 | Pathotroph-Saprotroph | \|Plant Pathogen\|-Wood Saprotroph | Microfungus | NULL | Probable | As wood_pathogen (Põlme et al. 2020); Decay wood (Põlme et al. 2020) | [illegible] |
| Otu00153 | 1046 | k__Fungi(100);p__Zygomycota(100);c__Zygomycota_class_Incertae_sedis(100);o__Mucorales(100);f__Mucoraceae(100);g__Mucor(100);s__Mucor_sp_JJP_2009a(100); | Mucor | | 13 | Saprotroph | \|Undefined Saprotroph\| | Microfungus | NULL | Probable | As soil_saprotroph (Põlme et al. 2020); Decay soil (Põlme et al. 2020) | [illegible] |
| Otu00154 | 1021 | k__Fungi(100);p__Basidiomycota(100);c__Agaricomycetes(100);o__Polyporales(100);f__Meruliaceae(100);g__Phlebia(100);s__Phlebia_uda(100); | Phlebia | | 13 | Saprotroph | \|Wood Saprotroph\| | Corticioid | White Rot | Highly Probable | As white_rot (Põlme et al. 2020); Decay wood (Põlme et al. 2020) | [illegible] |
| Otu00155 | 1013 | k__Fungi(100);p__Basidiomycota(100);c__Agaricomycetes(100);o__Hymenochaetales(100);f__Schizoporaceae(100);g__Hyphodontia(100);s__Hyphodontia_sp_DLL2011_1(100); | Hyphodontia | | 13 | Saprotroph | Undefined Saprotroph-\|Wood Saprotroph\| | Corticioid | White Rot | Probable | As white_rot (Põlme et al. 2020); Decay wood (Põlme et al. 2020) | [illegible] |
| Otu00156 | 1013 | k__Fungi(100);p__Ascomycota(100);p__Ascomycota_unclassified(98);p__Ascomycota_unclassified(98);p__Ascomycota_unclassified(98);p__Ascomycota_unclassified(98);p__Ascomycota_unclassified(98); | - | - | | - | - | - | - | - | Unassigned | - |
| Otu00157 | 997 | k__Fungi(100);p__Basidiomycota(100);c__Agaricomycetes(100);o__Hymenochaetales(100);f__Schizoporaceae(100);g__Hyphodontia(100);s__Hyphodontia_pallidula(100); | Hyphodontia | | 13 | Saprotroph | Undefined Saprotroph-\|Wood Saprotroph\| | Corticioid | White Rot | Probable | As white_rot (Põlme et al. 2020); Decay wood (Põlme et al. 2020) | [illegible] |
| Otu00158 | 992 | k__Fungi(100);p__Basidiomycota(100);c__Agaricomycetes(100);o__Trechisporales(100);f__Hydnodontaceae(100);g__Sistotremastrum(100);s__Sistotremastrum_guttuliferum(100); | Sistotremastrum | | 13 | Saprotroph | Undefined Saprotroph-\|Wood Saprotroph\| | Corticioid | White Rot | Probable | As white_rot (Põlme et al. 2020); Decay wood (Põlme et al. 2020) | [illegible] |
| Otu00159 | 988 | k__Fungi(100);p__Basidiomycota(100);c__Tremellomycetes(100);o__Filobasidiales(100);f__Filobasidiaceae(100);g__Cryptococcus(100);s__Cryptococcus_oeirensis(95); | Cryptococcus | | 13 | Pathotroph-Saprotroph-Symbiotroph | Animal Pathogen-Endophyte-Epiphyte-\|Undefined Saprotroph\| | Dimorphic Yeast-Yeast | NULL | Possible | As foliar_endophyte (Põlme et al. 2020); Decay leaf, fruit, seed, soil, dung, animal material (Põlme et al. 2020) | [illegible] |
| Otu00160 | 986 | k__Fungi(100);p__Ascomycota(100);c__Eurotiomycetes(100);o__Eurotiales(100);f__Trichocomaceae(100);g__Aspergillus(100);s__Aspergillus_vitricola(98); | Aspergillus | | 13 | Pathotroph-Saprotroph-Symbiotroph | Animal Pathogen-Endophyte-Plant Saprotroph-\|Undefined Saprotroph\|-Wood Saprotroph | Microfungus | NULL | Probable | As foliar_endophyte (Põlme et al. 2020); Decay leaf, fruit, seed, soil, dung, animal material (Põlme et al. 2020) | [illegible] |
| Otu00161 | 982 | k__Fungi(100);p__Basidiomycota(100);c__Agaricomycetes(100);c__Agaricomycetes_unclassified(100);c__Agaricomycetes_unclassified(100);c__Agaricomycetes_unclassified(100);c__Agaricomycetes_unclassified(100); | - | - | | - | - | - | - | - | Unassigned | - |
| Otu00162 | 973 | k__Fungi(100);p__Ascomycota(97);p__Ascomycota_unclassified(97);p__Ascomycota_unclassified(97);p__Ascomycota_unclassified(97);p__Ascomycota_unclassified(97);p__Ascomycota_unclassified(97); | - | - | | - | - | - | - | - | Unassigned | - |
| Otu00163 | 959 | k__Fungi(100);p__Ascomycota(100);c__Dothideomycetes(100);o__Capnodiales(100);f__Davidiellaceae(100);g__Cladosporium(100);s__Cladosporium_sphaerospermum(79); | Cladosporium | | 13 | Pathotroph-Saprotroph-Symbiotroph | Animal Pathogen-Endophyte-Lichen Parasite-Plant Pathogen-\|Plant Saprotroph\|-Wood Saprotroph | Microfungus | NULL | Possible | As foliar_endophyte (Põlme et al. 2020); As litter_saprotroph (Põlme et al. 2020); Decay wood, leaf, fruit, seed, soil, dung, animal material (Põlme et al. 2020) | [illegible] |
| Otu00164 | 941 | k__Fungi(100);p__Ascomycota(100);c__Dothideomycetes(100);o__Pleosporales(100);f__Montagnulaceae(100);f__Montagnulaceae_unclassified(100);f__Montagnulaceae_unclassified(100); | Montagnulaceae | | 9 | Pathotroph-Saprotroph | Fungal Parasite-Plant Pathogen-Undefined Saprotroph | Microfungus | NULL | Probable | Species in this clade can be saprobic, plant pathogenic, or mycoparasitic (Zhang et al. 2009) | [illegible] |
| Otu00165 | 934 | k__Fungi(100);p__Ascomycota(100);c__Leotiomycetes(100);o__Thelebolales(100);f__Thelebolaceae(100);g__Antarctomyces(100);s__Antarctomyces_psychrotrophicus(100); | Antarctomyces | | 13 | Saprotroph | \|Undefined Saprotroph\| | Microfungus | NULL | Probable | As soil_saprotroph (Põlme et al. 2020); Decay soil (Põlme et al. 2020) | [illegible] |
| Otu00166 | 934 | k__Fungi(100);k__Fungi_unclassified(74);k__Fungi_unclassified(74);k__Fungi_unclassified(74);k__Fungi_unclassified(74);k__Fungi_unclassified(74);k__Fungi_unclassified(74); | - | - | | - | - | - | - | - | Unassigned | - |
| Otu00167 | 929 | k__Fungi(100);p__Basidiomycota(100);c__Microbotryomycetes(100);o__Sporidiobolales(100);f__Sporidiobolales_family_Incertae_sedis(100);g__Rhodotorula(100);s__Rhodotorula_slooffiae(99); | Rhodotorula | | 13 | Pathotroph-Saprotroph-Symbiotroph | Animal Pathogen-Animal Symbiotroph-Endophyte-Plant Pathogen-\|Undefined Saprotroph\| | Yeast | NULL | Probable | As foliar_endophyte (Põlme et al. 2020); Opportunistic human pathogen | [illegible] |
| Otu00168 | 922 | k__Fungi(100);p__Ascomycota(100);c__Sordariomycetes(100);o__Hypocreales(100);f__Hypocreaceae(96);g__Hypocrea(87);g__Hypocrea_unclassified(87); | Hypocrea | | 13 | Pathotroph | Endophyte-Fungal Parasite | Microfungus | NULL | Probable | NULL | [illegible] |
| Otu00169 | 921 | k__Fungi(100);p__Zygomycota(100);c__Zygomycota_class_Incertae_sedis(100);o__Mucorales(100);f__Backusellaceae(100);g__Backusella(100);s__Backusella_recurva(100); | Backusella | | 13 | Saprotroph | Dung Saprotroph-\|Undefined Saprotroph\| | Microfungus-Sporophores | NULL | Probable | As soil_saprotroph (Põlme et al. 2020); Decay soil, dung (Põlme et al. 2020) | [illegible] |
| Otu00170 | 915 | k__Fungi(100);p__Basidiomycota(100);c__Agaricomycetes(100);o__Agaricales(100);f__Psathyrellaceae(100);g__Coprinellus(100);s__Coprinellus_micaceus(61); | Coprinellus | | 13 | Saprotroph | \|Undefined Saprotroph\| | Agaricoid | NULL | Probable | As soil_saprotroph (Põlme et al. 2020); Decay soil (Põlme et al. 2020) | [illegible] |
| Otu00171 | 901 | k__Fungi(100);p__Basidiomycota(100);c__Agaricomycetes(100);o__Polyporales(100);f__Polyporales_family_Incertae_sedis(100);g__Phlebiella(100);s__Phlebiella_sp(99); | Phlebiella | | 13 | Saprotroph | Undefined Saprotroph-\|Wood Saprotroph\| | Corticioid | White Rot | Probable | As white_rot (Põlme et al. 2020); Decay wood (Põlme et al. 2020) | [illegible] |
| Otu00172 | 897 | k__Fungi(100);p__Basidiomycota(100);c__Agaricomycetes(100);o__Trechisporales(100);f__Hydnodontaceae(100);g__Trechispora(100);g__Trechispora_unclassified(81); | Trechispora | | 13 | Saprotroph | \|Wood Saprotroph\| | Corticioid | White Rot | Highly Probable | As white_rot (Põlme et al. 2020); Decay wood (Põlme et al. 2020) | [illegible] |
| Otu00173 | 894 | k__Plantae(100);p__unclassified_Plantae(100);c__unclassified_Plantae(100);o__unclassified_Plantae(100);f__unclassified_Plantae(100);g__unclassified_Plantae(100);s__Plantae_sp(100); | - | - | | - | - | - | - | - | Unassigned | - |
| Otu00174 | 875 | k__Fungi(100);p__Ascomycota(100);c__Dothideomycetes(100);o__Pleosporales(100);f__Cucurbitariaceae(100);g__Curreya(100);s__Curreya_sp(96); | Curreya | | 13 | Saprotroph | \|Plant Saprotroph\|-Undefined Saprotroph | Microfungus | NULL | Probable | As litter_saprotroph (Põlme et al. 2020); Decay leaf, fruit, seed (Põlme et al. 2020) | [illegible] |
| Otu00176 | 860 | k__Fungi(100);p__Basidiomycota(100);c__Agaricomycetes(100);o__Agaricales(100);f__Inocybaceae(100);g__Crepidotus(100);s__Crepidotus_applanatus(100); | Crepidotus | | 13 | Saprotroph | \|Wood Saprotroph\| | Agaricoid | NULL | Highly Probable | Decay wood (Põlme et al. 2020) | [illegible] |
| Otu00175 | 860 | k__Fungi(100);p__Ascomycota(93);p__Ascomycota_unclassified(93);p__Ascomycota_unclassified(93);p__Ascomycota_unclassified(93);p__Ascomycota_unclassified(93);p__Ascomycota_unclassified(93); | - | - | | - | - | - | - | - | Unassigned | - |
| Otu00177 | 857 | k__Fungi(100);p__Basidiomycota(100);c__Agaricomycetes(100);o__Polyporales(100);f__Meruliaceae(100);g__Phlebia(100);s__Phlebia_tuberculata(100); | Phlebia | | 13 | Saprotroph | \|Wood Saprotroph\| | Corticioid | White Rot | Highly Probable | As white_rot (Põlme et al. 2020); Decay wood (Põlme et al. 2020) | [illegible] |
| Otu00178 | 829 | k__Fungi(100);p__Ascomycota(100);c__Eurotiomycetes(100);o__Chaetothyriales(100);f__Herpotrichiellaceae(100);g__Rhinocladiella(100);s__Rhinocladiella_sp_04001a(64); | Rhinocladiella | | 13 | Pathotroph-Saprotroph-Symbiotroph | Animal Parasite-Endophyte-\|Undefined Saprotroph\|-Wood Saprotroph | Dimorphic-Facultative Yeast | Soft Rot | Probable | As root_endophyte_dark_septate (Põlme et al. 2020); As soil_saprotroph (Põlme et al. 2020); Decay soil, animal material (Põlme et al. 2020) | [illegible] |
| Otu00179 | 829 | k__Fungi(100);p__Ascomycota(100);c__Dothideomycetes(100);o__Pleosporales(100);o__Pleosporales_unclassified(92);o__Pleosporales_unclassified(92);o__Pleosporales_unclassified(92); | - | - | | - | - | - | - | - | Unassigned | - |
| Otu00180 | 821 | k__Fungi(100);p__Basidiomycota(100);c__Agaricomycetes(100);o__Agaricales(100);f__Strophariaceae(100);g__Hypholoma(100);s__Hypholoma_lateritium(100); | Hypholoma | | 13 | Saprotroph | Undefined Saprotroph-\|Wood Saprotroph\| | Agaricoid | White Rot | Probable | As white_rot (Põlme et al. 2020); Decay wood (Põlme et al. 2020) | [illegible] |
| Otu00181 | 814 | k__Fungi(100);p__Ascomycota(100);c__Saccharomycetes(100);o__Saccharomycetales(100);f__Saccharomycetaceae(100);g__Lachancea(100);s__Lachancea_fermentati(100); | Lachancea | | 13 | Pathotroph-Saprotroph | Animal Parasite-Animal Pathogen-\|Undefined Saprotroph\| | Yeast | NULL | Possible | As nectar-tap_saprotroph (Põlme et al. 2020); Decay animal material, sugar-rich substrates (Põlme et al. 2020); Likely opportunistic human pathogen (Irinyi et al. 2015; see L. fermentati) | [illegible] |
| Otu00182 | 806 | k__Fungi(100);p__Ascomycota(100);c__Sordariomycetes(100);o__Hypocreales(100);f__Hypocreales_family_Incertae_sedis(100);g__Sarocladium(100);s__Sarocladium_strictum(100); | Sarocladium | | 13 | Pathotroph-Saprotroph | \|Plant Pathogen\|-Plant Saprotroph-Undefined Saprotroph | Microfungus | NULL | Probable | As litter_saprotroph (Põlme et al. 2020); Decay leaf, fruit, seed (Põlme et al. 2020) | [illegible] |
| Otu00183 | 793 | k__Fungi(100);p__Basidiomycota(100);c__Tremellomycetes(100);o__Filobasidiales(100);f__Filobasidiaceae(100);f__Filobasidiaceae_unclassified(85);f__Filobasidiaceae_unclassified(85); | - | - | | - | - | - | - | - | Unassigned | - |
| Otu00184 | 785 | k__Fungi(100);p__Ascomycota(100);c__Sordariomycetes(100);o__Xylariales(100);f__Diatrypaceae(100);g__Eutypella(100);s__Eutypella_vitis(99); | Eutypella | | 13 | Pathotroph | \|Plant Pathogen\| | Microfungus | NULL | Probable | NULL | [illegible] |
| Otu00185 | 782 | k__Fungi(100);p__Ascomycota(100);c__Dothideomycetes(100);o__Dothideales(100);f__Dothideaceae(100);g__Endoconidioma(100);s__Endoconidioma_populi(100); | Endoconidioma | | 13 | Saprotroph | Undefined Saprotroph-\|Wood Saprotroph\| | Microfungus | NULL | Probable | Decay: wood (Põlme et al. 2020) | [illegible] |
| Otu00186 | 777 | k__Fungi(100);p__Basidiomycota(100);c__Agaricomycetes(100);o__Polyporales(100);f__Meripilaceae(99);g__Physisporinus(81);g__Physisporinus_unclassified(80); | Physisporinus | | 13 | Saprotroph | \|Wood Saprotroph\| | Corticioid | White Rot | Highly Probable | As white_rot (Põlme et al. 2020); Decay wood (Põlme et al. 2020) | [illegible] |
| Otu00187 | 771 | k__Fungi(100);p__Basidiomycota(100);c__Agaricomycetes(100);o__Trechisporales(100);f__Hydnodontaceae(100);g__Trechispora(100);s__Trechispora_sp(100); | Trechispora | | 13 | Saprotroph | \|Wood Saprotroph\| | Corticioid | White Rot | Highly Probable | As white_rot (Põlme et al. 2020); Decay wood (Põlme et al. 2020) | [illegible] |
| Otu00188 | 764 | k__Fungi(100);p__Basidiomycota(100);c__Agaricomycetes(100);o__Agaricales(100);f__Schizophyllaceae(100);g__Schizophyllum(100);s__Schizophyllum_commune(100); | Schizophyllum | | 13 | Saprotroph | Plant Saprotroph-\|Wood Saprotroph\| | Agaricoid | White Rot | Highly Probable | As litter_saprotroph (Põlme et al. 2020); As white_rot (Põlme et al. 2020); Decay wood, leaf, fruit, seed (Põlme et al. 2020) | [illegible] |
| Otu00189 | 760 | k__Fungi(100);p__Basidiomycota(100);c__Agaricomycetes(100);o__Agaricales(100);f__Marasmiaceae(100);g__Rhodocollybia(100);s__Rhodocollybia_butyracea_f._asema(100); | Rhodocollybia | | 13 | Saprotroph | \|Plant Saprotroph\|-Undefined Saprotroph | Agaricoid | NULL | Probable | As litter_saprotroph (Põlme et al. 2020); Decay leaf, fruit, seed (Põlme et al. 2020) | [illegible] |
| Otu00190 | 752 | k__Fungi(100);p__Ascomycota(100);c__Sordariomycetes(100);o__Sordariales(100);f__Sordariaceae(100);g__Sordaria(99);s__Sordaria_sp_SS_1479(99); | Sordaria | | 13 | Pathotroph-Saprotroph | \|Dung Saprotroph\|-Plant Pathogen-Wood Saprotroph | Microfungus | Soft Rot | Probable | Decay dung (Põlme et al. 2020) | [illegible] |
| Otu00191 | 747 | k__Fungi(100);p__Ascomycota(100);c__Sordariomycetes(100);o__Microascales(100);f__Microascaceae(100);g__unclassified_Microascaceae(100);s__Microascaceae_sp(100); | Microascaceae | | 9 | Pathotroph-Saprotroph-Symbiotroph | Animal Pathogen-Endophyte-Plant Pathogen-Undefined Saprotroph | Microfungus | NULL | Possible | Saprobic in soil or rotting vegetation, some species heat-tolerant and associatated with silage and compost, a few associated with humans and animals (Cannon & Kirk 2007) | [illegible] |
| Otu00192 | 741 | k__Fungi(100);p__Ascomycota(100);c__Leotiomycetes(94);c__Leotiomycetes_unclassified(67);c__Leotiomycetes_unclassified(67);c__Leotiomycetes_unclassified(67);c__Leotiomycetes_unclassified(67); | - | - | | - | - | - | - | - | Unassigned | - |
| Otu00194 | 739 | k__Fungi(100);p__Basidiomycota(100);c__Agaricomycetes(100);o__Polyporales(100);f__Meruliaceae(100);g__Gloeoporus(100);s__Gloeoporus_pannocinctus(100); | Gloeoporus | | 13 | Saprotroph | \|Wood Saprotroph\| | Corticioid-Polyporoid | White Rot | Highly Probable | As white_rot (Põlme et al. 2020); Decay wood (Põlme et al. 2020) | [illegible] |
| Otu00193 | 739 | k__Fungi(100);p__Ascomycota(100);c__Dothideomycetes(100);o__Pleosporales(100);o__Pleosporales_unclassified(100);o__Pleosporales_unclassified(100);o__Pleosporales_unclassified(100); | - | - | | - | - | - | - | - | Unassigned | - |
| Otu00195 | 732 | k__Fungi(100);p__Basidiomycota(100);c__Agaricomycetes(100);o__Hymenochaetales(100);f__Schizoporaceae(100);g__Hyphodontia(96);s__Hyphodontia_radula(91); | Hyphodontia | | 13 | Saprotroph | Undefined Saprotroph-\|Wood Saprotroph\| | Corticioid | White Rot | Probable | As white_rot (Põlme et al. 2020); Decay wood (Põlme et al. 2020) | [illegible] |
| Otu00196 | 729 | k__Fungi(100);p__Ascomycota(98);p__Ascomycota_unclassified(98);p__Ascomycota_unclassified(98);p__Ascomycota_unclassified(98);p__Ascomycota_unclassified(98);p__Ascomycota_unclassified(98); | - | - | | - | - | - | - | - | Unassigned | - |
| Otu00197 | 723 | k__Fungi(100);p__Ascomycota(100);c__Sordariomycetes(100);o__Hypocreales(100);f__Ophiocordycipitaceae(100);g__Purpureocillium(100);s__Purpureocillium_sp(99); | Purpureocillium | | 13 | Pathotroph-Saprotroph | \|Animal Parasite\|-Fungal Parasite-Undefined Saprotroph | Microfungus | NULL | Probable | As animal_decomposer (Põlme et al. 2020); As keratinolytic (Põlme et al. 2020); Decay animal material (Põlme et al. 2020) | [illegible] |
| Otu00198 | 706 | k__Fungi(100);p__Ascomycota(100);c__Dothideomycetes(100);o__Pleosporales(100);f__Pleosporaceae(83);f__Pleosporaceae_unclassified(71);f__Pleosporaceae_unclassified(71); | Pleosporaceae | | 9 | Pathotroph-Saprotroph | Endophyte-Lichen Parasite-Plant Pathogen-Undefined Saprotroph | Microfungus | NULL | Probable | Necrotrophic pathogens and saprobes, especially associated with grasses. Many species are seed-borne (Cannon & Kirk 2007) | [illegible] |
| Otu00199 | 705 | k__Fungi(100);p__Basidiomycota(100);c__Agaricomycetes(100);o__Polyporales(100);f__Meruliaceae(100);g__Steccherinum(100);s__Steccherinum_ochraceum(100); | Steccherinum | | 13 | Saprotroph | Undefined Saprotroph-\|Wood Saprotroph\| | Corticioid | White Rot | Probable | As white_rot (Põlme et al. 2020); Decay wood (Põlme et al. 2020) | [illegible] |
| Otu00200 | 702 | k__Fungi(100);p__Basidiomycota(100);c__Microbotryomycetes(100);o__Sporidiobolales(100);f__Sporidiobolales_family_Incertae_sedis(100);g__Rhodotorula(100);s__Rhodotorula_pinicola(100); | Rhodotorula | | 13 | Pathotroph-Saprotroph-Symbiotroph | Animal Pathogen-Animal Symbiotroph-Endophyte-Plant Pathogen-\|Undefined Saprotroph\| | Yeast | NULL | Probable | As foliar_endophyte (Põlme et al. 2020); Opportunistic human pathogen | [illegible] |
| Otu00201 | 701 | k__Fungi(100);p__Basidiomycota(100);c__Agaricomycetes(100);o__Agaricales(100);f__Inocybaceae(100);g__Crepidotus(100);s__Crepidotus_mollis(100); | Crepidotus | | 13 | Saprotroph | \|Wood Saprotroph\| | Agaricoid | NULL | Highly Probable | Decay wood (Põlme et al. 2020) | [illegible] |
| Otu00202 | 701 | k__Fungi(100);p__Basidiomycota(100);c__Agaricomycetes(100);o__Polyporales(100);f__Phanerochaetaceae(100);g__Antrodiella(100);g__Antrodiella_unclassified(55); | Antrodiella | | 13 | Saprotroph | \|Wood Saprotroph\| | Corticioid-Polyporoid | White Rot | Highly Probable | As white_rot (Põlme et al. 2020); Decay wood (Põlme et al. 2020) | [illegible] |
| Otu00203 | 701 | k__Fungi(100);p__Ascomycota(100);c__Dothideomycetes(100);o__Pleosporales(100);f__Sporormiaceae(100);g__Preussia(100);s__Preussia_sp_GP_2006a(100); | Preussia | | 13 | Saprotroph | \|Dung Saprotroph\|-Undefined Saprotroph | Microfungus | NULL | Probable | Decay dung (Põlme et al. 2020) | [illegible] |
| Otu00204 | 701 | k__Fungi(100);p__Ascomycota(100);c__Sordariomycetes(100);o__Sordariomycetes_order_Incertae_sedis(100);f__Glomerellaceae(100);g__Colletotrichum(100);g__Colletotrichum_unclassified(70); | Colletotrichum | | 13 | Pathotroph-Saprotroph-Symbiotroph | Endophyte-\|Plant Pathogen\|-Plant Saprotroph | Microfungus | NULL | Probable | As foliar_endophyte (Põlme et al. 2020); As litter_saprotroph (Põlme et al. 2020); Decay leaf, fruit, seed (Põlme et al. 2020) | [illegible] |
| Otu00205 | 700 | k__Fungi(100);p__Ascomycota(100);c__Dothideomycetes(100);o__Pleosporales(100);f__Leptosphaeriaceae(94);g__Coniothyrium(85);g__Coniothyrium_unclassified(73); | Coniothyrium | | 13 | Pathotroph-Saprotroph | Fungal Parasite-\|Plant Pathogen\|-Plant Saprotroph-Wood Saprotroph | Microfungus | Soft Rot | Highly Probable | As litter_saprotroph (Põlme et al. 2020); As mycoparasite (Põlme et al. 2020); Decay leaf, fruit, seed, wood (Põlme et al. 2020) | [illegible] |
| Otu00206 | 699 | k__Fungi(100);p__Ascomycota(100);c__Sordariomycetes(100);o__Xylariales(100);f__Xylariaceae(100);g__Daldinia(100);g__Daldinia_unclassified(100); | Daldinia | | 13 | Saprotroph-Symbiotroph | Endophyte-Undefined Saprotroph-\|Wood Saprotroph\| | Daldinioid | NULL | Probable | As foliar_endophyte (Põlme et al. 2020); Decay wood, leaf, fruit, seed (Põlme et al. 2020); Host - angiosperms (Põlme et al. 2020) | [illegible] |
| Otu00207 | 699 | k__Fungi(100);p__Basidiomycota(100);c__Agaricomycetes(100);o__Agaricales(100);o__Agaricales_unclassified(100);o__Agaricales_unclassified(100);o__Agaricales_unclassified(100); | Agaricales | | 7 | Pathotroph-Saprotroph-Symbiotroph | Bryophyte Parasite-Dung Saprotroph-Ectomycorrhizal-Fungal Parasite-Leaf Saprotroph-Plant Parasite-Undefined Saprotroph-Wood Saprotroph | Agaricoid-Gasteroid-Microfungus-Secotioid-Yeast | NULL | Possible | Mushrooms and toadstools, Gill fungi, Agarics... lignicolous, sometimes muscicolous or fungicolous, saprobic, mycorrhizal, rarely parasitic on plants or fungi (Kirk et al. 2008) | [illegible] |
| Otu00208 | 693 | k__Fungi(100);p__Ascomycota(100);c__Dothideomycetes(100);o__Capnodiales(100);f__Capnodiales_family_Incertae_sedis(89);g__Capnobotryella(89);s__Capnobotryella_sp(89); | Capnobotryella | | 13 | Saprotroph | \|Plant Saprotroph\|-Undefined Saprotroph | Microfungus | NULL | Probable | As litter_saprotroph (Põlme et al. 2020); Decay leaf, fruit, seed, soil (Põlme et al. 2020) | [illegible] |
| Otu00209 | 684 | k__Fungi(100);p__Ascomycota(100);c__Sordariomycetes(100);o__Diaporthales(100);f__Gnomoniaceae(100);f__Gnomoniaceae_unclassified(82);f__Gnomoniaceae_unclassified(82); | Gnomoniaceae | | 9 | Pathotroph-Saprotroph-Symbiotroph | Endophyte-Plant Pathogen-Undefined Saprotroph | Microfungus | NULL | Probable | Most species are saprobes or endophytes in leaves and bark, but a few are neurotrophic pathogens (Cannon & Kirk 2007) | [illegible] |
| Otu00210 | 681 | k__Fungi(100);p__Ascomycota(100);p__Ascomycota_unclassified(92);p__Ascomycota_unclassified(92);p__Ascomycota_unclassified(92);p__Ascomycota_unclassified(92);p__Ascomycota_unclassified(92); | - | - | | - | - | - | - | - | Unassigned | - |
| Otu00211 | 680 | k__Fungi(100);p__Ascomycota(100);c__Dothideomycetes(100);o__Capnodiales(100);f__Mycosphaerellaceae(100);f__Mycosphaerellaceae_unclassified(100);f__Mycosphaerellaceae_unclassified(100); | Mycosphaerellaceae | | 9 | Pathotroph-Saprotroph | Plant Pathogen-Undefined Saprotroph | Microfungus | NULL | Probable | Biotrophic, necrotrophic or saprobic on various plant tissues (Cannon & Kirk 2007) | [illegible] |
| Otu00212 | 675 | k__Fungi(100);p__Basidiomycota(100);c__Pucciniomycetes(100);o__Pucciniales(100);f__Melampsoraceae(100);g__Melampsora(100);s__Melampsora_abietis_canadensis(85); | Melampsora | | 13 | Pathotroph | \|Plant Pathogen\| | Rust | NULL | Probable | A prominent group of species produces aecia on conifer needles and telia or leaves of Salicaceae (Cannon & Kirk 2007); Biotrophic, heteroecious or autoecious | [illegible] |
| Otu00213 | 674 | k__Fungi(100);p__Zygomycota(100);c__Zygomycota_class_Incertae_sedis(100);o__Mortierellales(100);f__Mortierellaceae(100);g__Mortierella(100);g__Mortierella_unclassified(87); | Mortierella | | 13 | Saprotroph-Symbiotroph | Endophyte-Plant Saprotroph-\|Undefined Saprotroph\| | Microfungus | NULL | Probable | As chitinolytic (Põlme et al. 2020); As soil_saprotroph (Põlme et al. 2020); Decay roots, soil (Põlme et al. 2020); Saprobic in soil and thus frequently isolated from soil and material contaminated with soil such as hay or rotten silage and spoil tips (Cannon & Kirk 2007) | [illegible] |
| Otu00214 | 673 | k__Fungi(100);p__Ascomycota(100);c__Eurotiomycetes(100);o__Eurotiales(100);f__Trichocomaceae(100);g__Penicillium(100);s__Penicillium_thomii(90); | Penicillium | | 13 | Pathotroph-Saprotroph-Symbiotroph | Animal Parasite-Dung Saprotroph-Endophyte-Fungal Parasite-Plant Pathogen-\|Undefined Saprotroph\|-Wood Saprotroph | Microfungus | Soft Rot | Highly Probable | As foliar_endophyte (Põlme et al. 2020); As mycoparasite (Põlme et al. 2020); Decay soil (Põlme et al. 2020) | [illegible] |
| Otu00215 | 671 | k__Fungi(100);p__Basidiomycota(100);c__Agaricomycetes(100);o__Polyporales(100);f__Polyporaceae(100);g__Tyromyces(100);g__Tyromyces_unclassified(60); | Tyromyces | | 13 | Saprotroph | \|Wood Saprotroph\| | Polyporoid | White Rot | Highly Probable | As white_rot (Põlme et al. 2020); Decay wood (Põlme et al. 2020) | [illegible] |
| Otu00216 | 671 | k__Fungi(100);p__Basidiomycota(100);c__Agaricomycetes(100);o__Boletales(100);f__Suillaceae(100);g__Suillus(100);s__Suillus_intermedius(75); | Suillus | | 13 | Symbiotroph | \|Ectomycorrhizal\| | Agaricoid-Boletoid | NULL | Highly Probable | Host - Pinaceae (Põlme et al. 2020) | [illegible] |
| Otu00217 | 669 | k__Fungi(100);p__Basidiomycota(100);c__Agaricomycetes(100);o__Agaricales(100);f__Pluteaceae(100);g__Pluteus(100);s__Pluteus_hongoi(72); | Pluteus | | 13 | Pathotroph-Saprotroph | Bryophyte Parasite-\|Plant Saprotroph\|-Wood Saprotroph | Agaricoid | White Rot | Probable | As litter_saprotroph (Põlme et al. 2020); As white_rot (Põlme et al. 2020); Decay leaf, fruit, seed (Põlme et al. 2020) | [illegible] |
| Otu00218 | 669 | k__Fungi(100);p__Basidiomycota(100);c__Agaricomycetes(100);o__Polyporales(100);f__Meruliaceae(100);g__Steccherinum(100);s__Steccherinum_sp(100); | Steccherinum | | 13 | Saprotroph | Undefined Saprotroph-\|Wood Saprotroph\| | Corticioid | White Rot | Probable | As white_rot (Põlme et al. 2020); Decay wood (Põlme et al. 2020) | [illegible] |
| Otu00219 | 651 | k__Fungi(100);p__Basidiomycota(100);c__Agaricomycetes(100);o__Polyporales(100);f__Polyporaceae(100);g__Trametes(100);s__Trametes_versicolor(99); | Trametes | | 13 | Pathotroph-Saprotroph | Plant Pathogen-\|Wood Saprotroph\| | Polyporoid | White Rot | Highly Probable | As white_rot (Põlme et al. 2020); As wood_pathogen (Põlme et al. 2020); Decay wood (Põlme et al. 2020) | [illegible] |
| Otu00220 | 641 | k__Fungi(100);p__Ascomycota(100);c__Sordariomycetes(100);c__Sordariomycetes_unclassified(100);c__Sordariomycetes_unclassified(100);c__Sordariomycetes_unclassified(100);c__Sordariomycetes_unclassified(100); | - | - | | - | - | - | - | - | Unassigned | - |
| Otu00221 | 640 | k__Fungi(100);p__Ascomycota(100);c__Saccharomycetes(100);o__Saccharomycetales(100);f__Saccharomycetaceae(100);g__Kluyveromyces(100);s__Kluyveromyces_lactis(99); | Kluyveromyces | | 13 | Saprotroph-Symbiotroph | Endophyte-\|Undefined Saprotroph\| | Yeast | NULL | Probable | Boraginaceae; As foliar_endophyte (Põlme et al. 2020); As nectar-tap_saprotroph (Põlme et al. 2020); Decay sugar-rich substrates (Põlme et al. 2020); Host | [illegible] |
| Otu00222 | 636 | k__Fungi(100);p__Zygomycota(100);c__Zygomycota_class_Incertae_sedis(100);o__Mucorales(100);f__Mucoraceae(100);g__Mucor(100);s__Mucor_mucedo(100); | Mucor | | 13 | Saprotroph | \|Undefined Saprotroph\| | Microfungus | NULL | Probable | As soil_saprotroph (Põlme et al. 2020); Decay soil (Põlme et al. 2020) | [illegible] |
| Otu00224 | 634 | k__Fungi(100);p__Ascomycota(100);c__Lecanoromycetes(100);o__Teloschistales(100);f__Physciaceae(100);g__Physcia(100);s__Physcia_sp(100); | Physcia | | 13 | Symbiotroph | \|Lichenized\| | Thallus | NULL | Highly Probable | NULL | [illegible] |
| Otu00223 | 634 | k__Fungi(100);p__Ascomycota(53);p__Ascomycota_unclassified(52);p__Ascomycota_unclassified(52);p__Ascomycota_unclassified(52);p__Ascomycota_unclassified(52);p__Ascomycota_unclassified(52); | - | - | | - | - | - | - | - | Unassigned | - |
| Otu00225 | 620 | k__Fungi(100);p__Ascomycota(100);c__Taphrinomycetes(100);o__Taphrinales(100);f__Taphrinaceae(100);g__Taphrina(100);s__Taphrina_communis(100); | Taphrina | | 13 | Pathotroph | \|Plant Pathogen\| | Dimorphic-Microfungus | NULL | Probable | NULL | [illegible] |
| Otu00226 | 618 | k__Fungi(100);p__Ascomycota(58);p__Ascomycota_unclassified(58);p__Ascomycota_unclassified(58);p__Ascomycota_unclassified(58);p__Ascomycota_unclassified(58);p__Ascomycota_unclassified(58); | - | - | | - | - | - | - | - | Unassigned | - |
| Otu00227 | 616 | k__Fungi(100);p__Ascomycota(100);c__Sordariomycetes(100);o__Xylariales(100);f__Amphisphaeriaceae(100);g__Pestalotiopsis(100);s__Pestalotiopsis_rosea(69); | Pestalotiopsis | | 13 | Pathotroph | \|Plant Pathogen\| | Microfungus | NULL | Probable | NULL | [illegible] |
| Otu00228 | 610 | k__Fungi(100);p__Basidiomycota(100);c__Agaricomycetes(100);o__Cantharellales(100);f__Hydnaceae(61);g__Sistotrema(61);s__Sistotrema_brinkmannii(61); | Sistotrema | | 13 | Saprotroph-Symbiotroph | Ectomycorrhizal-\|Plant Saprotroph\|-Wood Saprotroph | Corticioid | White Rot | Possible | As litter_saprotroph (Põlme et al. 2020); Decay leaf, fruit, seed (Põlme et al. 2020) | [illegible] |
| Otu00229 | 609 | k__Fungi(100);p__Basidiomycota(100);c__Agaricomycetes(100);o__Agaricales(100);f__Strophariaceae(100);g__Hypholoma(100);s__Hypholoma_subviride(100); | Hypholoma | | 13 | Saprotroph | Undefined Saprotroph-\|Wood Saprotroph\| | Agaricoid | White Rot | Probable | As white_rot (Põlme et al. 2020); Decay wood (Põlme et al. 2020) | [illegible] |
| Otu00230 | 604 | k__Fungi(100);p__Zygomycota(100);c__Zygomycota_class_Incertae_sedis(100);o__Mucorales(100);f__Mucoraceae(100);g__Mucor(100);s__Mucor_hiemalis(100); | Mucor | | 13 | Saprotroph | \|Undefined Saprotroph\| | Microfungus | NULL | Probable | As soil_saprotroph (Põlme et al. 2020); Decay soil (Põlme et al. 2020) | [illegible] |
| Otu00231 | 602 | k__Fungi(100);p__Ascomycota(100);c__Sordariomycetes(100);o__Sordariomycetes_order_Incertae_sedis(100);f__Annulatascaceae(100);g__Rhodoveronaea(100);s__Rhodoveronaea_varioseptata(100); | Rhodoveronaea | | 13 | Saprotroph | \|Undefined Saprotroph\| | Microfungus | NULL | Probable | NULL | [illegible] |
| Otu00232 | 599 | k__Fungi(100);k__Fungi_unclassified(99);k__Fungi_unclassified(99);k__Fungi_unclassified(99);k__Fungi_unclassified(99);k__Fungi_unclassified(99);k__Fungi_unclassified(99); | - | - | | - | - | - | - | - | Unassigned | - |
| Otu00233 | 598 | k__Fungi(100);p__Basidiomycota(100);c__Agaricomycetes(100);o__Agaricales(100);f__Entolomataceae(100);g__Entoloma(100);s__Entoloma_abortivum(100); | Entoloma | | 13 | Pathotroph-Saprotroph-Symbiotroph | Ectomycorrhizal-Fungal Parasite-\|Undefined Saprotroph\|-Undefined Symbiotroph | Agaricoid | NULL | Possible | As soil_saprotroph (Põlme et al. 2020); Decay roots, soil (Põlme et al. 2020) | [illegible] |
| Otu00234 | 598 | k__Fungi(100);p__Ascomycota(100);c__Sordariomycetes(100);o__Xylariales(100);f__Amphisphaeriaceae(100);g__Pestalotiopsis(100);g__Pestalotiopsis_unclassified(100); | Pestalotiopsis | | 13 | Pathotroph | \|Plant Pathogen\| | Microfungus | NULL | Probable | NULL | [illegible] |
| Otu00235 | 596 | k__Fungi(100);p__Ascomycota(100);c__Dothideomycetes(100);o__Pleosporales(100);f__Cucurbitariaceae(100);g__Curreya(87);g__Curreya_unclassified(84); | Curreya | | 13 | Saprotroph | \|Plant Saprotroph\|-Undefined Saprotroph | Microfungus | NULL | Probable | As litter_saprotroph (Põlme et al. 2020); Decay leaf, fruit, seed (Põlme et al. 2020) | [illegible] |
| Otu00236 | 593 | k__Fungi(100);p__Ascomycota(100);c__Taphrinomycetes(100);o__Taphrinales(100);f__Taphrinaceae(100);f__Taphrinaceae_unclassified(100);f__Taphrinaceae_unclassified(100); | Taphrinaceae | | 9 | Pathotroph | Plant Pathogen | Microfungus | NULL | Probable | Biotrophic on plants, usually causing hyperplasia (galls, witches broom) or lesions (Cannon & Kirk 2007) | [illegible] |
| Otu00237 | 592 | k__Fungi(100);p__Ascomycota(100);p__Ascomycota_unclassified(100);p__Ascomycota_unclassified(100);p__Ascomycota_unclassified(100);p__Ascomycota_unclassified(100);p__Ascomycota_unclassified(100); | - | - | | - | - | - | - | - | Unassigned | - |
| Otu00238 | 581 | k__Fungi(100);p__Basidiomycota(100);c__Agaricomycetes(100);o__Polyporales(100);o__Polyporales_unclassified(100);o__Polyporales_unclassified(100);o__Polyporales_unclassified(100); | - | - | | - | - | - | - | - | Unassigned | - |
| Otu00239 | 580 | k__Fungi(100);p__Basidiomycota(100);c__Agaricomycetes(100);o__Agaricales(100);f__Inocybaceae(100);g__Crepidotus(100);s__Crepidotus_crocophyllus(100); | Crepidotus | | 13 | Saprotroph | \|Wood Saprotroph\| | Agaricoid | NULL | Highly Probable | Decay wood (Põlme et al. 2020) | [illegible] |
| Otu00240 | 571 | k__Fungi(100);p__Ascomycota(100);c__Dothideomycetes(100);o__Pleosporales(100);f__Pleosporales_family_Incertae_sedis(100);g__Periconia(100);g__Periconia_unclassified(100); | Periconia | | 13 | Pathotroph-Saprotroph-Symbiotroph | Endophyte-\|Plant Pathogen\|-Wood Saprotroph | Microfungus | Soft Rot | Possible | Asteraceae, Musaceae (Costa et al. 2012); As foliar_endophyte (Põlme et al. 2020); Dark Septate Endophyte, known from plant roots (Mandyam et al. 2010); Endophyte Host ; Decay leaf, fruit, seed (Põlme et al. 2020) | [illegible] |
| Otu00241 | 568 | k__Fungi(100);p__Ascomycota(100);c__Ascomycota_class_Incertae_sedis(97);o__Ascomycota_order_Incertae_sedis(97);f__Ascomycota_family_Incertae_sedis(97);g__Slimacomyces(97);s__Slimacomyces_isiola(97); | Slimacomyces | | 13 | Saprotroph | \|Plant Saprotroph\| | Microfungus | NULL | Probable | As litter_saprotroph (Põlme et al. 2020); Decay leaf, fruit, seed (Põlme et al. 2020) | [illegible] |
| Otu00242 | 566 | k__Fungi(100);p__Ascomycota(100);c__Leotiomycetes(100);o__Helotiales(100);f__Helotiaceae(99);g__Articulospora(99);s__Articulospora_sp(99); | Articulospora | | 13 | Saprotroph | \|Plant Saprotroph\|-Undefined Saprotroph | Microfungus | NULL | Probable | As litter_saprotroph (Põlme et al. 2020); Decay leaf, fruit, seed (Põlme et al. 2020) | [illegible] |
| Otu00243 | 563 | k__Fungi(100);p__Chytridiomycota(100);c__Chytridiomycetes(100);o__Rhizophlyctidales(100);f__Rhizophlyctidaceae(100);g__Rhizophlyctis(100);s__Rhizophlyctis_rosea(100); | Rhizophlyctis | | 13 | Pathotroph-Saprotroph | Algal Parasite-\|Plant Saprotroph\|-Undefined Saprotroph | Chytrid-Microfungus | NULL | Probable | As litter_saprotroph (Põlme et al. 2020); Decay leaf, fruit, seed, algal material (Põlme et al. 2020) | [illegible] |
| Otu00244 | 556 | k__Fungi(100);p__Ascomycota(100);c__Dothideomycetes(100);o__Pleosporales(100);f__Pleosporales_family_Incertae_sedis(99);f__Pleosporales_family_Incertae_sedis_unclassified(97);f__Pleosporales_family_Incertae_sedis_unclassified(97); | - | - | | - | - | - | - | - | Unassigned | - |
| Otu00245 | 551 | k__Fungi(100);p__Ascomycota(100);c__Eurotiomycetes(100);c__Eurotiomycetes_unclassified(99);c__Eurotiomycetes_unclassified(99);c__Eurotiomycetes_unclassified(99);c__Eurotiomycetes_unclassified(99); | - | - | | - | - | - | - | - | Unassigned | - |

| | | | | | | | | | | | |
|---|---|---|---|---|---|---|---|---|---|---|---|
| Otu00246 | 548 | k__Fungi(100);p__Zygomycota(100);c__Zygomycota_class_Incertae_sedis(100);o__Mucorales(100);f__Mucoraceae(100);g__Mucor(100);s__Mucor_sp_JJP_2009a(85); | Mucor | 13 | Saprotroph | \|Undefined Saprotroph\| | Microfungus | NULL | Probable | As soil_saprotroph (Põlme et al. 2020); Decay soil (Põlme et al. 2020) | Tedersoo L, et al. 2014. Science 346:e1256688 ((https://doi.org/10.1126/science.1256688)); Põlme S, et al. 2020. Fungal Diversity 105:1-16 ((https://doi.org/10.1007/s13225-020-00466-2)) |
| Otu00247 | 542 | k__Fungi(100);p__Ascomycota(100);c__Sordariomycetes(100);o__Ophiostomatales(100);f__Ophiostomataceae(100);g__Sporothrix(95);g__Sporothrix_unclassified(95); | Sporothrix | 13 | Pathotroph-Saprotroph-Symbiotroph | Animal Pathogen-Endophyte-\|Plant Pathogen\|-Plant Saprotroph-Undefined Saprotroph-Wood Saprotroph | Microfungus | NULL | Probable | As foliar_endophyte (Põlme et al. 2020); As wood_pathogen (Põlme et al. 2020); culture (Dik et al. 1998); The fungal genus Sporothrix comprises a group of thermodimorphic pathogens that cause skin infections in humans and other mammals. Rose gardenerâ€™s disease (Sporotrichosis) occurs worldwide, with hyperendemic areas in Brazil, China, and South Africa (Rodrigues et al. 2016); The fungus lives throughout the world in soil and on plant matter such as sphagnum moss, rose bushes, and hay (www.cdc.gov); Decay wood (Põlme et al. 2020); Endophyte detection method | Dik AJ, et al. 1998. European Journal of Plant Pathology 104:413-423 ((https://doi.org/10.1023/A:1008025416672)); Rodrigues AM, et al. 2016. PLoS Pathogens 12:e1005638 ((https://doi.org/10.1371/journal.ppat.1005638)); Põlme S, et al. 2020. Fungal Diversity 105:1-16 ((https://doi.org/10.1007/s13225-020-00466-2)); https://www.cdc.gov ((https://www.cdc.gov)) |
| Otu00248 | 533 | k__Fungi(100);p__Ascomycota(100);c__Dothideomycetes(100);o__Dothideales(100);f__Dothioraceae(100);g__Sydowia(100);s__Sydowia_sp(100); | Sydowia | 13 | Pathotroph-Saprotroph | Plant Pathogen-Undefined Saprotroph-\|Wood Saprotroph\| | Microfungus | NULL | Probable | Decay: wood, leaf, fruit, seed (Põlme et al. 2020) | Tedersoo L, et al. 2014. Science 346:e1256688 ((https://doi.org/10.1126/science.1256688)); Põlme S, et al. 2020. Fungal Diversity 105:1-16 ((https://doi.org/10.1007/s13225-020-00466-2)) |
| Otu00249 | 529 | k__Fungi(100);p__Basidiomycota(100);c__Agaricomycetes(100);o__Hymenochaetales(100);f__Hymenochaetales_family_Incertae_sedis(100);g__Trichaptum(100);s__Trichaptum_sp(100); | Trichaptum | 13 | Saprotroph | \|Wood Saprotroph\| | Polyporoid | White Rot | Highly Probable | As white_rot (Põlme et al. 2020); Decay wood (Põlme et al. 2020) | Gilbertson RL, Ryvarden L. 1987-1987. North American Polypores. Fungiflora, Oslo (ISBN: 978-0945345060); Põlme S, et al. 2020. Fungal Diversity 105:1-16 ((https://doi.org/10.1007/s13225-020-00466-2)) |
| Otu00250 | 527 | k__Fungi(100);p__Ascomycota(100);c__Eurotiomycetes(100);o__Mycocaliciales(100);f__Mycocaliciaceae(100);g__Chaenothecopsis(100);s__Chaenothecopsis_epithallina(100); | Chaenothecopsis | 13 | Pathotroph-Saprotroph-Symbiotroph | Lichen Parasite-Lichenized-\|Undefined Saprotroph\| | Calicioid-Thallus | NULL | Possible | As resin_saprotroph (Põlme et al. 2020); licenized taxa; Lichenized and non | Esslinger TL. 2014. North Dakota State University ((http://www.ndsu.edu/pubweb/~esslinge/chcklst/chcklst7.htm)); Põlme S, et al. 2020. Fungal Diversity 105:1-16 ((https://doi.org/10.1007/s13225-020-00466-2)) |
| Otu00251 | 522 | k__Fungi(100);p__Ascomycota(100);c__Eurotiomycetes(100);o__Eurotiales(100);f__Trichocomaceae(100);f__Trichocomaceae_unclassified(100);f__Trichocomaceae_unclassified(100); | - | - | - | - | - | - | - | Unassigned | - |
| Otu00252 | 519 | k__Fungi(100);k__Fungi_unclassified(66);k__Fungi_unclassified(66);k__Fungi_unclassified(66);k__Fungi_unclassified(66);k__Fungi_unclassified(66);k__Fungi_unclassified(66); | - | - | - | - | - | - | - | Unassigned | - |
| Otu00253 | 509 | k__Fungi(100);p__Ascomycota(100);c__Sordariomycetes(100);o__Xylariales(100);f__Xylariaceae(100);g__Annulohypoxylon(100);s__Annulohypoxylon_truncatum(100); | Annulohypoxylon | 13 | Saprotroph-Symbiotroph | Endophyte-Undefined Saprotroph-\|Wood Saprotroph\| | Corticioid-Microfungus-Xylarioid | NULL | Probable | As foliar_endophyte (Põlme et al. 2020); culture (Del Olmo; Decay wood, leaf, fruit, seed (Põlme et al. 2020); Endophyte detection method; Host - angiosperms (Põlme et al. 2020); Ruiz & Arnold 2014) | Del Olmo-Ruiz M, Arnold AE. 2014. Mycologia 106:8-21 ((https://doi.org/10.3852/13-098)); Põlme S, et al. 2020. Fungal Diversity 105:1-16 ((https://doi.org/10.1007/s13225-020-00466-2)) |
| Otu00254 | 507 | k__Fungi(100);p__Basidiomycota(100);c__Agaricomycetes(100);o__Cantharellales(100);f__Ceratobasidiaceae(100);g__unclassified_Ceratobasidiaceae(51);g__unclassified_Ceratobasidiaceae_unclassified(51); | Ceratobasidiaceae | 9 | Pathotroph-Saprotroph-Symbiotroph | Endomycorrhizal-Plant Pathogen-Undefined Saprotroph | Microfungus | NULL | Possible | Best known as plant pathogens, but also saprobic. Species are endomycorrhizal with a wide range of plant roots - especially of Orchidaceae but also even bryophytes (Cannon & Kirk 2007) | Cannon PF, Kirk PM. 2007. Fungal Families of the World. CAB International, Cambridge (ISBN: 978-0851998275) |
| Otu00255 | 500 | k__Fungi(100);p__Ascomycota(100);c__Dothideomycetes(100);o__Pleosporales(100);f__unclassified_Pleosporales(100);g__unclassified_Pleosporales(100);s__Pleosporales_sp(100); | - | - | - | - | - | - | - | Unassigned | - |
| Otu00256 | 499 | k__Fungi(100);p__Basidiomycota(100);c__Agaricomycetes(100);c__Agaricomycetes_unclassified(100);c__Agaricomycetes_unclassified(100);c__Agaricomycetes_unclassified(100);c__Agaricomycetes_unclassified(100); | - | - | - | - | - | - | - | Unassigned | - |
| Otu00257 | 498 | k__Fungi(100);p__Ascomycota(100);c__Sordariomycetes(100);c__Sordariomycetes_unclassified(100);c__Sordariomycetes_unclassified(100);c__Sordariomycetes_unclassified(100);c__Sordariomycetes_unclassified(100); | - | - | - | - | - | - | - | Unassigned | - |
| Otu00258 | 497 | k__Fungi(100);p__Ascomycota(100);c__Dothideomycetes(100);o__Pleosporales(100);f__Montagnulaceae(100);g__Paraphaeosphaeria(82);s__Paraphaeosphaeria_michotii(61); | Paraphaeosphaeria | 13 | Saprotroph | Undefined Saprotroph-\|Wood Saprotroph\| | Microfungus | NULL | Probable | Decay wood (Põlme et al. 2020) | Tedersoo L, et al. 2014. Science 346:e1256688 ((https://doi.org/10.1126/science.1256688)); Põlme S, et al. 2020. Fungal Diversity 105:1-16 ((https://doi.org/10.1007/s13225-020-00466-2)) |
| Otu00259 | 492 | k__Fungi(100);p__Ascomycota(100);c__Eurotiomycetes(100);o__Eurotiales(100);f__Trichocomaceae(100);f__Trichocomaceae_unclassified(100);f__Trichocomaceae_unclassified(100); | - | - | - | - | - | - | - | Unassigned | - |
| Otu00260 | 491 | k__Fungi(100);k__Fungi_unclassified(95);k__Fungi_unclassified(95);k__Fungi_unclassified(95);k__Fungi_unclassified(95);k__Fungi_unclassified(95);k__Fungi_unclassified(95); | - | - | - | - | - | - | - | Unassigned | - |
| Otu00261 | 487 | k__Fungi(100);p__Basidiomycota(100);c__Agaricomycetes(100);o__Agaricales(100);f__Pleurotaceae(100);g__Pleurotus(100);s__Pleurotus_sp(88); | Pleurotus | 13 | Pathotroph-Saprotroph | Endophyte-Nematophagous-Plant Pathogen-\|Wood Saprotroph\| | Agaricoid | White Rot | Probable | As white_rot (Põlme et al. 2020); As wood_pathogen (Põlme et al. 2020); Decay wood, animal material (Põlme et al. 2020); This genus is known to be nematophagous, capturing and consuming nematodes. | James TY, et al. 2006. Nature 443:818-822 ((https://doi.org/10.1038/nature05110)); Costa IPMW, et al. 2012. Mycotaxon 119:e26 ((http://mycotaxon.com/resources/checklists/costa_v119_checklist.pdf)); Tedersoo L, et al. 2014. Science 346:e1256688 ((https://doi.org/10.1126/science.1256688)); Põlme S, et al. 2020. Fungal Diversity 105:1-16 ((https://doi.org/10.1007/s13225-020-00466-2)); The American Phytopathological Society Common Names of Plant Diseases ((http://www.apsnet.org/publications/commonnames/Pages/default.aspx)) |
| Otu00262 | 484 | k__Fungi(100);p__Ascomycota(100);c__Sordariomycetes(100);o__Xylariales(100);f__Xylariaceae(100);g__Camillea(100);s__Camillea_tinctor(82); | Camillea | 13 | Saprotroph-Symbiotroph | Endophyte-Undefined Saprotroph-\|Wood Saprotroph\| | Microfungus | NULL | Probable | As foliar_endophyte (Põlme et al. 2020); Decay wood, leaf, fruit, seed (Põlme et al. 2020); Host - angiosperms (Põlme et al. 2020) | Tedersoo L, et al. 2014. Science 346:e1256688 ((https://doi.org/10.1126/science.1256688)); Põlme S, et al. 2020. Fungal Diversity 105:1-16 ((https://doi.org/10.1007/s13225-020-00466-2)) |
| Otu00263 | 482 | k__Fungi(100);p__Basidiomycota(100);c__Agaricomycetes(100);o__Agaricales(100);f__Psathyrellaceae(100);g__Coprinopsis(100);s__Coprinopsis_strossmayeri(100); | Coprinopsis | 13 | Saprotroph | \|Undefined Saprotroph\| | Agaricoid | NULL | Probable | As soil_saprotroph (Põlme et al. 2020); Decay soil (Põlme et al. 2020) | Tedersoo L, et al. 2014. Science 346:e1256688 ((https://doi.org/10.1126/science.1256688)); Põlme S, et al. 2020. Fungal Diversity 105:1-16 ((https://doi.org/10.1007/s13225-020-00466-2)) |
| Otu00264 | 481 | k__Fungi(100);p__Basidiomycota(100);c__Agaricomycetes(100);o__Polyporales(100);f__Meruliaceae(100);g__Hyphoderma(100);s__Hyphoderma_setigerum(100); | Hyphoderma | 13 | Saprotroph | Undefined Saprotroph-\|Wood Saprotroph\| | Corticioid | White Rot | Probable | As white_rot (Põlme et al. 2020); Decay wood (Põlme et al. 2020) | Tedersoo L, et al. 2014. Science 346:e1256688 ((https://doi.org/10.1126/science.1256688)); Põlme S, et al. 2020. Fungal Diversity 105:1-16 ((https://doi.org/10.1007/s13225-020-00466-2)) |
| Otu00265 | 474 | k__Fungi(100);k__Fungi_unclassified(100);k__Fungi_unclassified(100);k__Fungi_unclassified(100);k__Fungi_unclassified(100);k__Fungi_unclassified(100);k__Fungi_unclassified(100); | - | - | - | - | - | - | - | Unassigned | - |
| Otu00266 | 471 | k__Fungi(100);p__Basidiomycota(100);c__Agaricomycetes(100);o__Polyporales(100);f__Meruliaceae(100);g__Steccherinum(100);s__Steccherinum_sp_MLB_2010(100); | Steccherinum | 13 | Saprotroph | Undefined Saprotroph-\|Wood Saprotroph\| | Corticioid | White Rot | Probable | As white_rot (Põlme et al. 2020); Decay wood (Põlme et al. 2020) | Tedersoo L, et al. 2014. Science 346:e1256688 ((https://doi.org/10.1126/science.1256688)); Põlme S, et al. 2020. Fungal Diversity 105:1-16 ((https://doi.org/10.1007/s13225-020-00466-2)) |
| Otu00267 | 470 | k__Fungi(100);p__Basidiomycota(100);c__Exobasidiomycetes(100);o__Exobasidiales(100);f__Exobasidiaceae(100);g__Exobasidium(100);g__Exobasidium_unclassified(100); | Exobasidium | 13 | Pathotroph | \|Plant Pathogen\| | Leaf Blister-Leaf Gall-Leaf Spot | NULL | Probable | NULL | Tedersoo L, et al. 2014. Science 346:e1256688 ((https://doi.org/10.1126/science.1256688)); Põlme S, et al. 2020. Fungal Diversity 105:1-16 ((https://doi.org/10.1007/s13225-020-00466-2)) |
| Otu00268 | 467 | k__Fungi(100);p__Basidiomycota(100);c__Agaricomycetes(100);o__Cantharellales(100);f__Ceratobasidiaceae(100);f__Ceratobasidiaceae_unclassified(60);f__Ceratobasidiaceae_unclassified(60); | Ceratobasidiaceae | 9 | Pathotroph-Saprotroph-Symbiotroph | Endomycorrhizal-Plant Pathogen-Undefined Saprotroph | Microfungus | NULL | Possible | Best known as plant pathogens, but also saprobic. Species are endomycorrhizal with a wide range of plant roots - especially of Orchidaceae but also even bryophytes (Cannon & Kirk 2007) | Cannon PF, Kirk PM. 2007. Fungal Families of the World. CAB International, Cambridge (ISBN: 978-0851998275) |
| Otu00269 | 462 | k__Fungi(100);p__Ascomycota(100);c__Dothideomycetes(100);o__Pleosporales(100);f__unclassified_Pleosporales(97);g__unclassified_Pleosporales(97);s__Pleosporales_sp(97); | - | - | - | - | - | - | - | Unassigned | - |
| Otu00270 | 462 | k__Plantae(100);p__unclassified_Plantae(100);c__unclassified_Plantae(100);o__unclassified_Plantae(100);f__unclassified_Plantae(100);g__unclassified_Plantae(100);s__Plantae_sp(100); | - | - | - | - | - | - | - | Unassigned | - |
| Otu00271 | 460 | k__Fungi(100);k__Fungi_unclassified(100);k__Fungi_unclassified(100);k__Fungi_unclassified(100);k__Fungi_unclassified(100);k__Fungi_unclassified(100);k__Fungi_unclassified(100); | - | - | - | - | - | - | - | Unassigned | - |
| Otu00272 | 457 | k__Fungi(100);p__Basidiomycota(100);c__Microbotryomycetes(100);o__Sporidiobolales(100);f__Sporidiobolales_family_Incertae_sedis(100);g__Rhodotorula(100);s__Rhodotorula_minuta(100); | Rhodotorula | 13 | Pathotroph-Saprotroph-Symbiotroph | Animal Pathogen-Animal Symbiotroph-Endophyte-Plant Pathogen-\|Undefined Saprotroph\| | Yeast | NULL | Probable | As foliar_endophyte (Põlme et al. 2020); Opportunistic human pathogen | Põlme S, et al. 2020. Fungal Diversity 105:1-16 ((https://doi.org/10.1007/s13225-020-00466-2)); https://www.cdc.gov ((https://www.cdc.gov)) |
| Otu00273 | 456 | k__Fungi(100);p__Ascomycota(100);c__Sordariomycetes(84);c__Sordariomycetes_unclassified(83);c__Sordariomycetes_unclassified(83);c__Sordariomycetes_unclassified(83);c__Sordariomycetes_unclassified(83); | - | - | - | - | - | - | - | Unassigned | - |
| Otu00274 | 453 | k__Fungi(100);p__Basidiomycota(100);c__Exobasidiomycetes(100);o__Exobasidiales(100);f__Exobasidiaceae(100);g__Exobasidium(100);g__Exobasidium_unclassified(100); | Exobasidium | 13 | Pathotroph | \|Plant Pathogen\| | Leaf Blister-Leaf Gall-Leaf Spot | NULL | Probable | NULL | Tedersoo L, et al. 2014. Science 346:e1256688 ((https://doi.org/10.1126/science.1256688)); Põlme S, et al. 2020. Fungal Diversity 105:1-16 ((https://doi.org/10.1007/s13225-020-00466-2)) |
| Otu00275 | 447 | k__Fungi(100);p__Basidiomycota(100);c__Agaricomycetes(100);o__Cantharellales(100);o__Cantharellales_unclassified(95);o__Cantharellales_unclassified(95);o__Cantharellales_unclassified(95); | - | - | - | - | - | - | - | Unassigned | - |
| Otu00276 | 446 | k__Fungi(100);p__Basidiomycota(100);c__Agaricomycetes(100);o__Polyporales(100);f__Fomitopsidaceae(100);g__Piptoporus(100);s__Piptoporus_betulinus(100); | Piptoporus | 13 | Pathotroph-Saprotroph | \|Plant Pathogen\|-Wood Saprotroph | Polyporoid | Brown Rot | Highly Probable | As brown-rot (Põlme et al. 2020); As wood_pathogen (Põlme et al. 2020); Decay wood (Põlme et al. 2020) | Gilbertson RL, Ryvarden L. 1987-1987. North American Polypores. Fungiflora, Oslo (ISBN: 978-0945345060); Põlme S, et al. 2020. Fungal Diversity 105:1-16 ((https://doi.org/10.1007/s13225-020-00466-2)) |
| Otu00277 | 445 | k__Fungi(100);p__Basidiomycota(100);c__Agaricomycetes(100);o__Cantharellales(100);f__Botryobasidiaceae(100);g__Botryobasidium(100);s__Botryobasidium_subcoronatum(100); | Botryobasidium | 13 | Saprotroph | Undefined Saprotroph-\|Wood Saprotroph\| | Corticioid | White Rot | Probable | As white_rot (Põlme et al. 2020); Decay wood (Põlme et al. 2020) | Tedersoo L, et al. 2014. Science 346:e1256688 ((https://doi.org/10.1126/science.1256688)); Põlme S, et al. 2020. Fungal Diversity 105:1-16 ((https://doi.org/10.1007/s13225-020-00466-2)) |
| Otu00278 | 443 | k__Fungi(100);p__Basidiomycota(100);c__Agaricomycetes(100);o__Polyporales(100);f__Meruliaceae(100);g__Steccherinum(100);s__Steccherinum_sp_DLL2011_2(100); | Steccherinum | 13 | Saprotroph | Undefined Saprotroph-\|Wood Saprotroph\| | Corticioid | White Rot | Probable | As white_rot (Põlme et al. 2020); Decay wood (Põlme et al. 2020) | Tedersoo L, et al. 2014. Science 346:e1256688 ((https://doi.org/10.1126/science.1256688)); Põlme S, et al. 2020. Fungal Diversity 105:1-16 ((https://doi.org/10.1007/s13225-020-00466-2)) |
| Otu00279 | 437 | k__Fungi(100);p__Ascomycota(100);c__Dothideomycetes(100);o__Pleosporales(100);f__Lophiostomataceae(100);g__Lophiostoma(100);s__Lophiostoma_sp_DMW2181(100); | Lophiostoma | 13 | Saprotroph | Plant Saprotroph-Undefined Saprotroph-\|Wood Saprotroph\| | Microfungus | NULL | Probable | As litter_saprotroph (Põlme et al. 2020); Decay wood, leaf, fruit, seed (Põlme et al. 2020) | Tedersoo L, et al. 2014. Science 346:e1256688 ((https://doi.org/10.1126/science.1256688)); Põlme S, et al. 2020. Fungal Diversity 105:1-16 ((https://doi.org/10.1007/s13225-020-00466-2)) |
| Otu00280 | 437 | k__Fungi(100);p__Basidiomycota(100);c__Agaricomycetes(100);o__Russulales(100);f__Peniophoraceae(100);g__Peniophora(100);s__Peniophora_sp(99); | Peniophora | 13 | Pathotroph-Saprotroph | Plant Pathogen-\|Wood Saprotroph\| | Corticioid | White Rot | Probable | As white_rot (Põlme et al. 2020); As wood_pathogen (Põlme et al. 2020); Decay wood (Põlme et al. 2020) | Tedersoo L, et al. 2014. Science 346:e1256688 ((https://doi.org/10.1126/science.1256688)); Põlme S, et al. 2020. Fungal Diversity 105:1-16 ((https://doi.org/10.1007/s13225-020-00466-2)) |
| Otu00281 | 436 | k__Fungi(100);p__Zygomycota(100);c__Zygomycota_class_Incertae_sedis(100);o__Mortierellales(100);f__Mortierellaceae(100);g__Mortierella(100);s__Mortierella_gemmifera(100); | Mortierella | 13 | Saprotroph-Symbiotroph | Endophyte-Plant Saprotroph-\|Undefined Saprotroph\| | Microfungus | NULL | Probable | As chitinolytic (Põlme et al. 2020); As soil_saprotroph (Põlme et al. 2020); Decay roots, soil (Põlme et al. 2020); Saprobic in soil and thus frequenctly isolated from soil and material contaminated with soil such as hay or rotten silage and spoil tips (Cannon & Kirk 2007) | Cannon PF, Kirk PM. 2007. Fungal Families of the World. CAB International, Cambridge (ISBN: 978-0851998275); Tedersoo L, et al. 2014. Science 346:e1256688 ((https://doi.org/10.1126/science.1256688)); Purahong W, et al. 2016. Molecular Ecology 25:4059-4074 ((https://doi.org/10.1111/mec.13739)); Põlme S, et al. 2020. Fungal Diversity 105:1-16 ((https://doi.org/10.1007/s13225-020-00466-2)) |
| Otu00282 | 435 | k__Fungi(100);p__Ascomycota(100);c__Saccharomycetes(100);o__Saccharomycetales(100);f__Metschnikowiaceae(100);g__Metschnikowia(100);s__Metschnikowia_pulcherrima(95); | Metschnikowia | 13 | Saprotroph | \|Undefined Saprotroph\| | Yeast | NULL | Probable | As nectar-tap_saprotroph (Põlme et al. 2020); Decay sugar-rich substrates (Põlme et al. 2020) | Põlme S, et al. 2020. Fungal Diversity 105:1-16 ((https://doi.org/10.1007/s13225-020-00466-2)) |
| Otu00283 | 435 | k__Fungi(100);p__Ascomycota(100);c__Eurotiomycetes(100);o__Mycocaliciales(100);f__Mycocaliciaceae(100);g__Mycocalicium(100);s__Mycocalicium_subtile(100); | Mycocalicium | 13 | Pathotroph-Saprotroph-Symbiotroph | Lichen Parasite-Lichenized-\|Undefined Saprotroph\| | Calicioid-Thallus | NULL | Possible | As resin_saprotroph (Põlme et al. 2020); licenized taxa; Lichenized and non | Esslinger TL. 2014. North Dakota State University ((http://www.ndsu.edu/pubweb/~esslinge/chcklst/chcklst7.htm)); Põlme S, et al. 2020. Fungal Diversity 105:1-16 ((https://doi.org/10.1007/s13225-020-00466-2)) |
| Otu00284 | 434 | k__Fungi(100);p__Ascomycota(100);c__Dothideomycetes(100);o__Pleosporales(100);f__Pleosporales_family_Incertae_sedis(100);g__Periconia(100);s__Periconia_sp(100); | Periconia | 13 | Pathotroph-Saprotroph-Symbiotroph | Endophyte-\|Plant Pathogen\|-Wood Saprotroph | Microfungus | Soft Rot | Possible | Asteraceae, Musaceae (Costa et al. 2012); As foliar_endophyte (Põlme et al. 2020); Dark Septate Endophyte, known from plant roots (Mandyam et al. 2010); Endophyte Host ; Decay leaf, fruit, seed (Põlme et al. 2020) | Duncan CG, Eslyn WE. 1966. Mycologia 58:642-645 ((https://doi.org/10.2307/3757045)); Costa IPMW, et al. 2012. Mycotaxon 119:e26 ((http://mycotaxon.com/resources/checklists/costa_v119_checklist.pdf)); Põlme S, et al. 2020. Fungal Diversity 105:1-16 ((https://doi.org/10.1007/s13225-020-00466-2)); Mandyam K, et al. 2012. Mycorrhiza 22:109-119 ((https://doi.org/10.1007/s00572-011-0386-y)); Knapp DG, et al. 2012. PLOS ONE 7:e32570 (( https://doi.org/10.1371/journal.pone.0032570)); The American Phytopathological Society Common Names of Plant Diseases ((http://www.apsnet.org/publications/commonnames/Pages/default.aspx)) |
| Otu00285 | 432 | k__Fungi(100);p__Basidiomycota(100);c__Agaricomycetes(100);o__Cantharellales(100);f__Hydnaceae(100);g__Sistotrema(100);s__Sistotrema_brinkmannii(100); | Sistotrema | 13 | Saprotroph-Symbiotroph | Ectomycorrhizal-\|Plant Saprotroph\|-Wood Saprotroph | Corticioid | White Rot | Possible | As litter_saprotroph (Põlme et al. 2020); Decay leaf, fruit, seed (Põlme et al. 2020) | Gilbertson RL, Ryvarden L. 1987-1987. North American Polypores. Fungiflora, Oslo (ISBN: 978-0945345060); Rinaldi AC, et al. 2008. Fungal Diversity 33:1-45 ((http://www.fungaldiversity.org/fdp/sfdp/33-1.pdf)) (as ECM/not ECM); Tedersoo L, et al. 2010. Mycorrhiza 20:217-263 ((https://doi.org/10.1007/s00572-009-0274-x)) (pro parte); Põlme S, et al. 2020. Fungal Diversity 105:1-16 ((https://doi.org/10.1007/s13225-020-00466-2)); http://mycorrhizas.info/ecmf.html ((http://mycorrhizas.info/ecmf.html)) (as ECM/not ECM). |
| Otu00286 | 425 | k__Fungi(100);p__Basidiomycota(100);c__Agaricomycetes(100);o__Polyporales(100);f__Meruliaceae(100);g__Junghuhnia(100);s__Junghuhnia_nitida(100); | Junghuhnia | 13 | Saprotroph | \|Wood Saprotroph\| | Corticioid | White Rot | Highly Probable | As white_rot (Põlme et al. 2020); Decay wood (Põlme et al. 2020) | Gilbertson RL, Ryvarden L. 1987-1987. North American Polypores. Fungiflora, Oslo (ISBN: 978-0945345060); Põlme S, et al. 2020. Fungal Diversity 105:1-16 ((https://doi.org/10.1007/s13225-020-00466-2)) |
| Otu00287 | 422 | k__Fungi(100);p__Ascomycota(100);c__Eurotiomycetes(100);o__Chaetothyriales(100);f__Herpotrichiellaceae(100);g__Cladophialophora(100);g__Cladophialophora_unclassified(97); | Cladophialophora | 13 | Pathotroph-Saprotroph-Symbiotroph | Animal Pathogen-Endophyte-Fungal Parasite-\|Undefined Saprotroph\| | Dimorphic-Microfungus | NULL | Probable | As mycoparasite (Põlme et al. 2020); As root_endophyte_dark_septate (Põlme et al. 2020); As soil_saprotroph (Põlme et al. 2020); Decay soil (Põlme et al. 2020) | James TY, et al. 2006. Nature 443:818-822 ((https://doi.org/10.1038/nature05110)); Põlme S, et al. 2020. Fungal Diversity 105:1-16 ((https://doi.org/10.1007/s13225-020-00466-2)) |
| Otu00288 | 422 | k__Fungi(100);p__Ascomycota(100);p__Ascomycota_unclassified(100);p__Ascomycota_unclassified(100);p__Ascomycota_unclassified(100);p__Ascomycota_unclassified(100);p__Ascomycota_unclassified(100); | - | - | - | - | - | - | - | Unassigned | - |
| Otu00289 | 420 | k__Fungi(100);p__Ascomycota(100);c__Eurotiomycetes(98);o__Chaetothyriales(95);o__Chaetothyriales_unclassified(95);o__Chaetothyriales_unclassified(95);o__Chaetothyriales_unclassified(95); | - | - | - | - | - | - | - | Unassigned | - |
| Otu00290 | 419 | k__Fungi(100);p__Basidiomycota(100);c__Agaricomycetes(100);o__Hymenochaetales(100);f__Hymenochaetaceae(100);g__Pseudochaete(100);s__Pseudochaete_corrugata(99); | Pseudochaete | 13 | Saprotroph | Undefined Saprotroph-\|Wood Saprotroph\| | Corticioid | White Rot | Probable | As white_rot (Põlme et al. 2020); Decay wood (Põlme et al. 2020) | Tedersoo L, et al. 2014. Science 346:e1256688 ((https://doi.org/10.1126/science.1256688)); Põlme S, et al. 2020. Fungal Diversity 105:1-16 ((https://doi.org/10.1007/s13225-020-00466-2)) |
| Otu00291 | 418 | k__Fungi(100);p__Basidiomycota(100);c__Agaricomycetes(100);o__Trechisporales(100);f__Hydnodontaceae(100);g__Trechispora(100);g__Trechispora_unclassified(100); | Trechispora | 13 | Saprotroph | \|Wood Saprotroph\| | Corticioid | White Rot | Highly Probable | As white_rot (Põlme et al. 2020); Decay wood (Põlme et al. 2020) | Gilbertson RL, Ryvarden L. 1987-1987. North American Polypores. Fungiflora, Oslo (ISBN: 978-0945345060); Rinaldi AC, et al. 2008. Fungal Diversity 33:1-45 ((http://www.fungaldiversity.org/fdp/sfdp/33-1.pdf)) (as Uncertain); Tedersoo L, et al. 2010. Mycorrhiza 20:217-263 ((https://doi.org/10.1007/s00572-009-0274-x)) (as not ECM).; Põlme S, et al. 2020. Fungal Diversity 105:1-16 ((https://doi.org/10.1007/s13225-020-00466-2)) |
| Otu00292 | 415 | k__Fungi(100);p__Basidiomycota(100);c__Agaricomycetes(100);o__Agaricomycetes_order_Incertae_sedis(100);f__Agaricomycetes_family_Incertae_sedis(100);g__Resinicium(100);s__Resinicium_sp(100); | Resinicium | 13 | Saprotroph | Undefined Saprotroph-\|Wood Saprotroph\| | Corticioid | White Rot | Probable | As white_rot (Põlme et al. 2020); Decay wood (Põlme et al. 2020) | Tedersoo L, et al. 2014. Science 346:e1256688 ((https://doi.org/10.1126/science.1256688)); Põlme S, et al. 2020. Fungal Diversity 105:1-16 ((https://doi.org/10.1007/s13225-020-00466-2)) |
| Otu00293 | 415 | k__Fungi(100);p__Basidiomycota(100);c__Agaricomycetes(100);o__Polyporales(100);f__Polyporaceae(100);g__Perenniporia(100);s__Perenniporia_tenuis(58); | Perenniporia | 13 | Saprotroph | \|Wood Saprotroph\| | Corticioid-Polyporoid | White Rot | Highly Probable | As white_rot (Põlme et al. 2020); Decay wood (Põlme et al. 2020) | Gilbertson RL, Ryvarden L. 1987-1987. North American Polypores. Fungiflora, Oslo (ISBN: 978-0945345060); Tedersoo L, et al. 2014. Science 346:e1256688 ((https://doi.org/10.1126/science.1256688)); Põlme S, et al. 2020. Fungal Diversity 105:1-16 ((https://doi.org/10.1007/s13225-020-00466-2)) |
| Otu00294 | 414 | k__Fungi(100);p__Basidiomycota(100);c__Agaricomycetes(100);o__Hymenochaetales(100);f__Hymenochaetaceae(100);g__Botryodontia(100);s__Botryodontia_millavensis(100); | Botryodontia | 13 | Saprotroph | Undefined Saprotroph-\|Wood Saprotroph\| | Corticioid | White Rot | Probable | As white_rot (Põlme et al. 2020); Decay wood (Põlme et al. 2020) | Tedersoo L, et al. 2014. Science 346:e1256688 ((https://doi.org/10.1126/science.1256688)); Põlme S, et al. 2020. Fungal Diversity 105:1-16 ((https://doi.org/10.1007/s13225-020-00466-2)) |
| Otu00295 | 410 | k__Fungi(100);p__Basidiomycota(100);c__Tremellomycetes(100);o__Tremellales(100);f__Tremellales_family_Incertae_sedis(100);g__Bulleromyces(100);s__Bulleromyces_albus(100); | Bulleromyces | 13 | Pathotroph-Saprotroph | Fungal Parasite-\|Undefined Saprotroph\| | Dimorphic | NULL | Probable | As mycoparasite (Põlme et al. 2020) | Põlme S, et al. 2020. Fungal Diversity 105:1-16 ((https://doi.org/10.1007/s13225-020-00466-2)) |
| Otu00296 | 405 | k__Fungi(100);p__Ascomycota(100);c__Sordariomycetes(100);o__Trichosphaeriales(100);f__Trichosphaeriales_family_Incertae_sedis(100);g__unclassified_Trichosphaeriales(100);s__Trichosphaeriales_sp(100); | - | - | - | - | - | - | - | Unassigned | - |
| Otu00298 | 403 | k__Fungi(100);p__Ascomycota(100);c__Eurotiomycetes(100);o__Eurotiales(100);f__Trichocomaceae(100);g__Penicillium(100);g__Penicillium_unclassified(70); | Penicillium | 13 | Pathotroph-Saprotroph-Symbiotroph | Animal Parasite-Dung Saprotroph-Endophyte-Fungal Parasite-Plant Pathogen-\|Undefined Saprotroph\|-Wood Saprotroph | Microfungus | Soft Rot | Highly Probable | As foliar_endophyte (Põlme et al. 2020); As mycoparasite (Põlme et al. 2020); Decay soil (Põlme et al. 2020) | Duncan CG, Eslyn WE. 1966. Mycologia 58:642-664 ((https://doi.org/10.2307/3757045)); Seehann G, et al. 1975. List of Fungi in Soft-Rot Tissue Tests. Institute fur Holzbiologie und Holzschutz, Bundesforschungsanstalt fur Forst - und Holzwirtshaft, Hamburg - Lohbrugge, Germany ((https://www.irg-wp.com/irgdocs/details.php?d6493033-c9ea-40f3-8e3e-5697f32d640d)); Costa IPMW, et al. 2012. Mycotaxon 119:e26 ((http://mycotaxon.com/resources/checklists/costa_v119_checklist.pdf)); Bills GF, et al. 2013. Current Opinion in Microbiology 16:549-565 ((https://doi.org/10.1016/j.mib.2013.08.001)); Tedersoo L, et al. 2014. Science 346:e1256688 ((https://doi.org/10.1126/science.1256688)); Põlme S, et al. 2020. Fungal Diversity 105:1-16 ((https://doi.org/10.1007/s13225-020-00466-2)) |
| Otu00297 | 403 | k__Plantae(100);p__unclassified_Plantae(100);c__unclassified_Plantae(100);o__unclassified_Plantae(100);f__unclassified_Plantae(100);g__unclassified_Plantae(100);s__Plantae_sp(100); | - | - | - | - | - | - | - | Unassigned | - |
| Otu00299 | 400 | k__Fungi(100);p__Ascomycota(100);c__Sordariomycetes(100);o__Xylariales(100);f__Diatrypaceae(100);g__Diatrype(100);s__Diatrype_stigma(91); | Diatrype | 13 | Pathotroph | \|Plant Pathogen\| | Microfungus | NULL | Probable | NULL | Tedersoo L, et al. 2014. Science 346:e1256688 ((https://doi.org/10.1126/science.1256688)); Põlme S, et al. 2020. Fungal Diversity 105:1-16 ((https://doi.org/10.1007/s13225-020-00466-2)) |
| Otu00300 | 397 | k__Fungi(100);p__Basidiomycota(100);c__Tremellomycetes(100);o__Tremellales(100);f__Tremellales_family_Incertae_sedis(100);g__Cryptococcus(100);s__Cryptococcus_sp_QMW_2009a(100); | Cryptococcus | 13 | Pathotroph-Saprotroph-Symbiotroph | Animal Pathogen-Endophyte-Epiphyte-\|Undefined Saprotroph\| | Dimorphic Yeast-Yeast | NULL | Possible | As foliar_endophyte (Põlme et al. 2020); Decay leaf, fruit, seed, soil, dung, animal material (Põlme et al. 2020) | Andrews JH, et al. 1983. Phytopathology 73:228-234 ((https://doi.org/10.1094/Phyto-73-228)); Kurtzman CP, et al. (eds.) 2011. The Yeasts, a Taxonomic Study. Fifth Edition. Vols 1-3. Elsevier, San Diego (ISBN: 9780444521491); Põlme S, et al. 2020. Fungal Diversity 105:1-16 ((https://doi.org/10.1007/s13225-020-00466-2)) |
| Otu00301 | 395 | k__Fungi(100);p__Basidiomycota(97);c__Agaricomycetes(93);c__Agaricomycetes_unclassified(93);c__Agaricomycetes_unclassified(93);c__Agaricomycetes_unclassified(93);c__Agaricomycetes_unclassified(93); | - | - | - | - | - | - | - | Unassigned | - |
| Otu00302 | 392 | k__Fungi(100);p__Basidiomycota(100);c__Agaricomycetes(100);o__Agaricales(100);f__Psathyrellaceae(100);g__Coprinopsis(100);s__Coprinopsis_pachyderma(68); | Coprinopsis | 13 | Saprotroph | \|Undefined Saprotroph\| | Agaricoid | NULL | Probable | As soil_saprotroph (Põlme et al. 2020); Decay soil (Põlme et al. 2020) | Tedersoo L, et al. 2014. Science 346:e1256688 ((https://doi.org/10.1126/science.1256688)); Põlme S, et al. 2020. Fungal Diversity 105:1-16 ((https://doi.org/10.1007/s13225-020-00466-2)) |
| Otu00303 | 389 | k__Fungi(100);p__Basidiomycota(100);c__Agaricomycetes(100);o__Polyporales(100);f__Meripilaceae(100);g__Physisporinus(100);s__Physisporinus_vitreus(100); | Physisporinus | 13 | Saprotroph | \|Wood Saprotroph\| | Corticioid | White Rot | Highly Probable | As white_rot (Põlme et al. 2020); Decay wood (Põlme et al. 2020) | Gilbertson RL, Ryvarden L. 1987-1987. North American Polypores. Fungiflora, Oslo (ISBN: 978-0945345060); Põlme S, et al. 2020. Fungal Diversity 105:1-16 ((https://doi.org/10.1007/s13225-020-00466-2)) |
| Otu00304 | 386 | k__Fungi(100);p__Basidiomycota(100);c__Tremellomycetes(100);o__Tremellales(100);f__Tremellales_family_Incertae_sedis(100);g__Cryptococcus(100);s__Cryptococcus_sp(100); | Cryptococcus | 13 | Pathotroph-Saprotroph-Symbiotroph | Animal Pathogen-Endophyte-Epiphyte-\|Undefined Saprotroph\| | Dimorphic Yeast-Yeast | NULL | Possible | As foliar_endophyte (Põlme et al. 2020); Decay leaf, fruit, seed, soil, dung, animal material (Põlme et al. 2020) | Andrews JH, et al. 1983. Phytopathology 73:228-234 ((https://doi.org/10.1094/Phyto-73-228)); Kurtzman CP, et al. (eds.) 2011. The Yeasts, a Taxonomic Study. Fifth Edition. Vols 1-3. Elsevier, San Diego (ISBN: 9780444521491); Põlme S, et al. 2020. Fungal Diversity 105:1-16 ((https://doi.org/10.1007/s13225-020-00466-2)) |
| Otu00305 | 377 | k__Fungi(100);p__Ascomycota(100);c__Dothideomycetes(100);o__Pleosporales(100);f__Phaeosphaeriaceae(100);g__Stagonospora(54);s__Stagonospora_pseudovitensis(54); | Stagonospora | 13 | Pathotroph | \|Plant Pathogen\| | Microfungus | NULL | Probable | NULL | Tedersoo L, et al. 2014. Science 346:e1256688 ((https://doi.org/10.1126/science.1256688)); Põlme S, et al. 2020. Fungal Diversity 105:1-16 ((https://doi.org/10.1007/s13225-020-00466-2)) |
| Otu00306 | 376 | k__Plantae(100);p__unclassified_Plantae(100);c__unclassified_Plantae(100);o__unclassified_Plantae(100);f__unclassified_Plantae(100);g__unclassified_Plantae(100);s__Plantae_sp(100); | - | - | - | - | - | - | - | Unassigned | - |
| Otu00307 | 375 | k__Fungi(100);p__Ascomycota(100);c__Dothideomycetes(100);o__Pleosporales(100);f__Pleosporaceae(99);f__Pleosporaceae_unclassified(99);f__Pleosporaceae_unclassified(99); | Pleosporaceae | 9 | Pathotroph-Saprotroph | Endophyte-Lichen Parasite-Plant Pathogen-Undefined Saprotroph | Microfungus | NULL | Probable | Necrotrophic pathogens and saprobes, especially associated with grasses. Many species are seed-borne (Cannon & Kirk 2007) | Cannon PF, Kirk PM. 2007. Fungal Families of the World. CAB International, Cambridge (ISBN: 978-0851998275); David AS, et al. 2016. Microbial Ecology 71:912-926 ((https://doi.org/10.1007/s00248-015-0712-6)); Lawrey JD, Diederich P. 2016. http://www.lichenicolous.net (accessed 24 May 2017) ((http://www.lichenicolous.net)) |
| Otu00308 | 371 | k__Fungi(100);p__Basidiomycota(100);c__Agaricomycetes(99);c__Agaricomycetes_unclassified(99);c__Agaricomycetes_unclassified(99);c__Agaricomycetes_unclassified(99);c__Agaricomycetes_unclassified(99); | - | - | - | - | - | - | - | Unassigned | - |
| Otu00310 | 370 | k__Fungi(100);p__Ascomycota(100);c__Dothideomycetes(100);o__Pleosporales(100);f__Didymosphaeriaceae(91);g__Roussoella(91);s__Roussoella_japanensis(91); | Roussoella | 13 | Saprotroph | Undefined Saprotroph-\|Wood Saprotroph\| | Microfungus | NULL | Probable | Decay wood (Põlme et al. 2020) | Tedersoo L, et al. 2014. Science 346:e1256688 ((https://doi.org/10.1126/science.1256688)); Põlme S, et al. 2020. Fungal Diversity 105:1-16 ((https://doi.org/10.1007/s13225-020-00466-2)) |
| Otu00309 | 370 | k__Plantae(100);p__unclassified_Plantae(100);c__unclassified_Plantae(100);o__unclassified_Plantae(100);f__unclassified_Plantae(100);g__unclassified_Plantae(100);s__Plantae_sp(100); | - | - | - | - | - | - | - | Unassigned | - |
| Otu00311 | 366 | k__Plantae(99);p__unclassified_Plantae(99);c__unclassified_Plantae(99);o__unclassified_Plantae(99);f__unclassified_Plantae(99);g__unclassified_Plantae(99);s__Plantae_sp(99); | - | - | - | - | - | - | - | Unassigned | - |
| Otu00312 | 360 | k__Fungi(100);p__Ascomycota(100);p__Ascomycota_unclassified(100);p__Ascomycota_unclassified(100);p__Ascomycota_unclassified(100);p__Ascomycota_unclassified(100);p__Ascomycota_unclassified(100); | - | - | - | - | - | - | - | Unassigned | - |
| Otu00313 | 359 | k__Fungi(100);p__Basidiomycota(100);p__Basidiomycota_unclassified(100);p__Basidiomycota_unclassified(100);p__Basidiomycota_unclassified(100);p__Basidiomycota_unclassified(100);p__Basidiomycota_unclassified(100); | - | - | - | - | - | - | - | Unassigned | - |
| Otu00314 | 357 | k__Fungi(100);p__Basidiomycota(100);c__Agaricomycetes(100);o__Polyporales(100);f__Phanerochaetaceae(100);g__Ceriporia(100);s__Ceriporia_bubalinomarginata(100); | Ceriporia | 13 | Saprotroph | \|Wood Saprotroph\| | Corticioid | White Rot | Highly Probable | As white_rot (Põlme et al. 2020); Decay wood (Põlme et al. 2020) | Tedersoo L, et al. 2014. Science 346:e1256688 ((https://doi.org/10.1126/science.1256688)); Põlme S, et al. 2020. Fungal Diversity 105:1-16 ((https://doi.org/10.1007/s13225-020-00466-2)) |
| Otu00315 | 357 | k__Fungi(100);p__Ascomycota(100);c__Eurotiomycetes(100);o__Eurotiales(100);f__Trichocomaceae(100);g__Penicillium(100);s__Penicillium_bialowiezense(63); | Penicillium | 13 | Pathotroph-Saprotroph-Symbiotroph | Animal Parasite-Dung Saprotroph-Endophyte-Fungal Parasite-Plant Pathogen-\|Undefined Saprotroph\|-Wood Saprotroph | Microfungus | Soft Rot | Highly Probable | As foliar_endophyte (Põlme et al. 2020); As mycoparasite (Põlme et al. 2020); Decay soil (Põlme et al. 2020) | Duncan CG, Eslyn WE. 1966. Mycologia 58:642-664 ((https://doi.org/10.2307/3757045)); Seehann G, et al. 1975. List of Fungi in Soft-Rot Tissue Tests. Institute fur Holzbiologie und Holzschutz, Bundesforschungsanstalt fur Forst - und Holzwirtshaft, Hamburg - Lohbrugge, Germany ((https://www.irg-wp.com/irgdocs/details.php?d6493033-c9ea-40f3-8e3e-5697f32d640d)); Costa IPMW, et al. 2012. Mycotaxon 119:e26 ((http://mycotaxon.com/resources/checklists/costa_v119_checklist.pdf)); Bills GF, et al. 2013. Current Opinion in Microbiology 16:549-565 ((https://doi.org/10.1016/j.mib.2013.08.001)); Tedersoo L, et al. 2014. Science 346:e1256688 ((https://doi.org/10.1126/science.1256688)); Põlme S, et al. 2020. Fungal Diversity 105:1-16 ((https://doi.org/10.1007/s13225-020-00466-2)) |
| Otu00316 | 354 | k__Fungi(100);k__Fungi_unclassified(58);k__Fungi_unclassified(58);k__Fungi_unclassified(58);k__Fungi_unclassified(58);k__Fungi_unclassified(58);k__Fungi_unclassified(58); | - | - | - | - | - | - | - | Unassigned | - |
| Otu00317 | 352 | k__Fungi(100);p__Basidiomycota(100);c__Agaricomycetes(100);o__Polyporales(100);f__Polyporaceae(100);g__Perenniporia(100);s__Perenniporia_sp_DLL2011_1(100); | Perenniporia | 13 | Saprotroph | \|Wood Saprotroph\| | Corticioid-Polyporoid | White Rot | Highly Probable | As white_rot (Põlme et al. 2020); Decay wood (Põlme et al. 2020) | Gilbertson RL, Ryvarden L. 1987-1987. North American Polypores. Fungiflora, Oslo (ISBN: 978-0945345060); Tedersoo L, et al. 2014. Science 346:e1256688 ((https://doi.org/10.1126/science.1256688)); Põlme S, et al. 2020. Fungal Diversity 105:1-16 ((https://doi.org/10.1007/s13225-020-00466-2)) |
| Otu00318 | 352 | k__Fungi(100);k__Fungi_unclassified(52);k__Fungi_unclassified(52);k__Fungi_unclassified(52);k__Fungi_unclassified(52);k__Fungi_unclassified(52);k__Fungi_unclassified(52); | - | - | - | - | - | - | - | Unassigned | - |
| Otu00319 | 348 | k__Fungi(100);p__Ascomycota(100);c__Dothideomycetes(100);o__Botryosphaeriales(100);f__Botryosphaeriaceae(100);g__Thyrostroma(100);s__Thyrostroma_carpophilum(100); | Thyrostroma | 13 | Pathotroph-Saprotroph-Symbiotroph | Endophyte-\|Plant Pathogen\|-Plant Saprotroph | Microfungus | NULL | Probable | As foliar_endophyte (Põlme et al. 2020); As litter_saprotroph (Põlme et al. 2020); Decay wood, leaf, fruit, seed, roots (Põlme et al. 2020) | Tedersoo L, et al. 2014. Science 346:e1256688 ((https://doi.org/10.1126/science.1256688)); Põlme S, et al. 2020. Fungal Diversity 105:1-16 ((https://doi.org/10.1007/s13225-020-00466-2)) |
| Otu00320 | 346 | k__Fungi(100);p__Basidiomycota(99);p__Basidiomycota_unclassified(99);p__Basidiomycota_unclassified(99);p__Basidiomycota_unclassified(99);p__Basidiomycota_unclassified(99);p__Basidiomycota_unclassified(99); | - | - | - | - | - | - | - | Unassigned | - |
| Otu00321 | 344 | k__Fungi(100);p__Basidiomycota(100);c__Agaricomycetes(100);o__Agaricales(100);o__Agaricales_unclassified(100);o__Agaricales_unclassified(100);o__Agaricales_unclassified(100); | Agaricales | 7 | Pathotroph-Saprotroph-Symbiotroph | Bryophyte Parasite-Dung Saprotroph-Ectomycorrhizal-Fungal Parasite-Leaf Saprotroph-Plant Parasite-Undefined Saprotroph-Wood Saprotroph | Agaricoid-Gasteroid-Microfungus-Secotioid-Yeast | NULL | Possible | Mushrooms and toadstools, Gill fungi, Agarics... lignicolous, sometimes muscicolous or fungicolous, saprobic, mycorrhizal, rarely parasitic on plants or fungi (Kirk et al. 2008) | Kirk PM et al. 2008. Dictionary of the Fungi. Tenth Edition. CAB International, Wallingford (ISBN: 978-0851998268) |
| Otu00322 | 344 | k__Fungi(100);p__Basidiomycota(100);c__Agaricomycetes(100);o__Cantharellales(100);o__Cantharellales_unclassified(75);o__Cantharellales_unclassified(75);o__Cantharellales_unclassified(75); | - | - | - | - | - | - | - | Unassigned | - |
| Otu00323 | 342 | k__Fungi(100);p__Ascomycota(100);c__Sordariomycetes(100);o__Hypocreales(100);f__Clavicipitaceae(100);g__Pochonia(100);s__Pochonia_bulbillosa(100); | Pochonia | 13 | Pathotroph-Saprotroph | \|Animal Parasite\|-Animal Pathogen-Undefined Saprotroph | Microfungus | NULL | Probable | As animal_decomposer (Põlme et al. 2020); Decay animal material (Põlme et al. 2020) | Tedersoo L, et al. 2014. Science 346:e1256688 ((https://doi.org/10.1126/science.1256688)); Põlme S, et al. 2020. Fungal Diversity 105:1-16 ((https://doi.org/10.1007/s13225-020-00466-2)) |
| Otu00324 | 340 | k__Fungi(100);p__Basidiomycota(100);c__Tremellomycetes(100);o__Tremellales(100);f__Tremellales_family_Incertae_sedis(100);g__Hannaella(100);s__Hannaella_sp_CMON52(100); | Hannaella | 13 | Pathotroph-Saprotroph | \|Animal Parasite\|-Undefined Saprotroph | Yeast | NULL | Probable | As animal_decomposer (Põlme et al. 2020); Decay animal material (Põlme et al. 2020) | Põlme S, et al. 2020. Fungal Diversity 105:1-16 ((https://doi.org/10.1007/s13225-020-00466-2)) |
| Otu00325 | 338 | k__Fungi(100);p__Ascomycota(100);c__Dothideomycetes(100);o__Capnodiales(100);f__Mycosphaerellaceae(100);g__Ramichloridium(100);s__Ramichloridium_anceps(100); | Ramichloridium | 13 | Pathotroph-Pathotroph-Saprotroph | \|Plant Pathogen\|-Plant Saprotroph | Microfungus | NULL | Probable | As litter_saprotroph (Põlme et al. 2020); Decay leaf, fruit, seed (Põlme et al. 2020) | Tedersoo L, et al. 2014. Science 346:e1256688 ((https://doi.org/10.1126/science.1256688)); Põlme S, et al. 2020. Fungal Diversity 105:1-16 ((https://doi.org/10.1007/s13225-020-00466-2)) |
| Otu00326 | 338 | k__Fungi(100);p__Basidiomycota(100);c__Agaricomycetes(100);o__Agaricales(100);f__Psathyrellaceae(95);g__Coprinellus(95);s__Coprinellus_xanthothrix(95); | Coprinellus | 13 | Saprotroph | \|Undefined Saprotroph\| | Agaricoid | NULL | Probable | As soil_saprotroph (Põlme et al. 2020); Decay soil (Põlme et al. 2020) | Tedersoo L, et al. 2014. Science 346:e1256688 ((https://doi.org/10.1126/science.1256688)); Põlme S, et al. 2020. Fungal Diversity 105:1-16 ((https://doi.org/10.1007/s13225-020-00466-2)) |
| Otu00327 | 336 | k__Fungi(100);p__Ascomycota(100);c__Dothideomycetes(100);o__Capnodiales(98);o__Capnodiales_unclassified(66);o__Capnodiales_unclassified(66);o__Capnodiales_unclassified(66); | - | - | - | - | - | - | - | Unassigned | - |
| Otu00328 | 333 | k__Fungi(100);p__Basidiomycota(100);c__Agaricomycetes(100);o__Polyporales(100);f__Polyporaceae(100);g__Royoporus(100);s__Royoporus_badius(100); | Royoporus | 13 | Saprotroph | Undefined Saprotroph-\|Wood Saprotroph\| | Polyporoid | White Rot | Probable | As white_rot (Põlme et al. 2020); Decay wood (Põlme et al. 2020) | Tedersoo L, et al. 2014. Science 346:e1256688 ((https://doi.org/10.1126/science.1256688)); Põlme S, et al. 2020. Fungal Diversity 105:1-16 ((https://doi.org/10.1007/s13225-020-00466-2)) |
| Otu00330 | 332 | k__Fungi(100);p__Ascomycota(100);c__Sordariomycetes(100);o__Hypocreales(100);f__Nectriaceae(100);g__Fusarium(100);g__Fusarium_unclassified(94); | Fusarium | 13 | Pathotroph-Saprotroph-Symbiotroph | Animal Pathogen-Endophyte-Lichen Parasite-\|Plant Pathogen\|-Plant Saprotroph-Undefined Saprotroph-Wood Saprotroph | Microfungus | Soft Rot | Possible | As foliar_endophyte (Põlme et al. 2020); As litter_saprotroph (Põlme et al. 2020); Causing damping-off disease (Põlme et al. 2020); Decay leaf, fruit, seed, roots, soil, animal material (Põlme et al. 2020); Likely opportunistic human pathogen (Irinyi et al. 2015) | Duncan CG, Eslyn WE. 1966. Mycologia 58:642-664 ((https://doi.org/10.2307/3757045)); Seehann G, et al. 1975. List of Fungi in Soft-Rot Tissue Tests. Institute fur Holzbiologie und Holzschutz, Bundesforschungsanstalt fur Forst - und Holzwirtshaft, Hamburg - Lohbrugge, Germany; Costa IPMW, et al. 2012. Mycotaxon 119:e26 ((http://mycotaxon.com/resources/checklists/costa_v119_checklist.pdf)); Tedersoo L, et al. 2014. Science 346:e1256688 ((https://doi.org/10.1126/science.1256688)); Irinyi L, et al. 2015. Medical Mycology 53:313-337 ((https://doi.org/10.1093/mmy/myv008)); Schroers, et al. 2016. Mycologia 108:806-819 ((https://doi.org/10.3852/15-255)); Põlme S, et al. 2020. Fungal Diversity 105:1-16 ((https://doi.org/10.1007/s13225-020-00466-2)); The American Phytopathological Society Common Names of Plant Diseases ((http://www.apsnet.org/publications/commonnames/Pages/default.aspx)); Lawrey JD, Diederich P. 2016. http://www.lichenicolous.net (accessed 24 May 2017) ((http://www.lichenicolous.net)) |
| Otu00331 | 332 | k__Fungi(100);p__Ascomycota(100);c__Dothideomycetes(100);o__Pleosporales(100);f__Venturiaceae(100);f__Venturiaceae_unclassified(67);f__Venturiaceae_unclassified(67); | Venturiaceae | 9 | Pathotroph-Saprotroph-Symbiotroph | Endophyte-Plant Pathogen-Undefined Saprotroph | Microfungus | NULL | Probable | Mostly endophytes and weak parasites of aerial plant parts, including bark and wood, leaf and fruit tissue; rarely biotrophic. Little is known except for the important pathogens (Cannon & Kirk 2007) | Cannon PF, Kirk PM. 2007. Fungal Families of the World. CAB International, Cambridge (ISBN: 978-0851998275) |
| Otu00329 | 332 | k__Fungi(100);k__Fungi_unclassified(87);k__Fungi_unclassified(87);k__Fungi_unclassified(87);k__Fungi_unclassified(87);k__Fungi_unclassified(87);k__Fungi_unclassified(87); | - | - | - | - | - | - | - | Unassigned | - |
| Otu00332 | 331 | k__Fungi(100);p__Basidiomycota(100);c__Agaricomycetes(100);o__Polyporales(100);f__Meruliaceae(100);g__Bjerkandera(100);s__Bjerkandera_adusta(72); | Bjerkandera | 13 | Pathotroph-Saprotroph-Symbiotroph | Endophyte-Plant Pathogen-\|Wood Saprotroph\| | Polyporoid | White Rot | Highly Probable | As root_endophyte_dark_septate (Põlme et al. 2020); As white_rot (Põlme et al. 2020); As wood_pathogen (Põlme et al. 2020); Decay wood (Põlme et al. 2020) | Gilbertson RL, Ryvarden L. 1987-1987. North American Polypores. Fungiflora, Oslo (ISBN: 978-0945345060); Põlme S, et al. 2020. Fungal Diversity 105:1-16 ((https://doi.org/10.1007/s13225-020-00466-2)) |
| Otu00333 | 331 | k__Fungi(100);p__Basidiomycota(100);c__Agaricomycetes(100);o__Polyporales(100);f__Phanerochaetaceae(100);g__Phanerochaete(100);g__Phanerochaete_unclassified(96); | Phanerochaete | 13 | Saprotroph | \|Wood Saprotroph\| | Corticioid | White Rot | Highly Probable | As white_rot (Põlme et al. 2020); Decay wood (Põlme et al. 2020) | Glenn JK, et al. 1983. Biochemical and Biophysical Research Communications 114:1077-1083 ((https://doi.org/10.1016/0006-291x((83)) 90672-1)); Põlme S, et al. 2020. Fungal Diversity 105:1-16 ((https://doi.org/10.1007/s13225-020-00466-2)) |
| Otu00334 | 330 | k__Fungi(100);p__Basidiomycota(100);c__Agaricomycetes(100);o__Polyporales(100);f__Meruliaceae(100);g__Phlebia(100);s__Phlebia_sp_604(100); | Phlebia | 13 | Saprotroph | \|Wood Saprotroph\| | Corticioid | White Rot | Highly Probable | As white_rot (Põlme et al. 2020); Decay wood (Põlme et al. 2020) | Kirk PM et al. 2008. Dictionary of the Fungi. Tenth Edition. CAB International, Wallingford (ISBN: 978-0851998268); Põlme S, et al. 2020. Fungal Diversity 105:1-16 ((https://doi.org/10.1007/s13225-020-00466-2)) |
| Otu00335 | 327 | k__Fungi(100);p__Ascomycota(100);c__Sordariomycetes(100);c__Sordariomycetes_unclassified(98);c__Sordariomycetes_unclassified(98);c__Sordariomycetes_unclassified(98);c__Sordariomycetes_unclassified(98); | - | - | - | - | - | - | - | Unassigned | - |
| Otu00336 | 326 | k__Fungi(100);p__Basidiomycota(100);c__Agaricomycetes(100);o__Polyporales(100);f__Meripilaceae(100);g__Physisporinus(100);s__Physisporinus_sp(100); | Physisporinus | 13 | Saprotroph | \|Wood Saprotroph\| | Corticioid | White Rot | Highly Probable | As white_rot (Põlme et al. 2020); Decay wood (Põlme et al. 2020) | Gilbertson RL, Ryvarden L. 1987-1987. North American Polypores. Fungiflora, Oslo (ISBN: 978-0945345060); Põlme S, et al. 2020. Fungal Diversity 105:1-16 ((https://doi.org/10.1007/s13225-020-00466-2)) |
| Otu00337 | 325 | k__Fungi(100);p__Ascomycota(100);c__Dothideomycetes(100);o__Pleosporales(100);f__Pleosporales_family_Incertae_sedis(99);f__Pleosporales_family_Incertae_sedis_unclassified(68);f__Pleosporales_family_Incertae_sedis_unclassified(68); | - | - | - | - | - | - | - | Unassigned | - |
| Otu00338 | 323 | k__Fungi(100);p__Basidiomycota(100);c__Agaricomycetes(100);o__Cantharellales(100);f__Clavulinaceae(100);g__Clavulina(100);g__Clavulina_unclassified(100); | Clavulina | 13 | Symbiotroph | \|Ectomycorrhizal\| | Clavarioid | NULL | Highly Probable | NULL | Tedersoo L, et al. 2014. Science 346:e1256688 ((https://doi.org/10.1126/science.1256688)); Põlme S, et al. 2020. Fungal Diversity 105:1-16 ((https://doi.org/10.1007/s13225-020-00466-2)) |
| Otu00339 | 318 | k__Fungi(100);p__Basidiomycota(100);c__Agaricomycetes(100);o__Polyporales(100);f__Meruliaceae(100);g__Phlebia(100);s__Phlebia_subserialis(100); | Phlebia | 13 | Saprotroph | \|Wood Saprotroph\| | Corticioid | White Rot | Highly Probable | As white_rot (Põlme et al. 2020); Decay wood (Põlme et al. 2020) | Kirk PM et al. 2008. Dictionary of the Fungi. Tenth Edition. CAB International, Wallingford (ISBN: 978-0851998268); Põlme S, et al. 2020. Fungal Diversity 105:1-16 ((https://doi.org/10.1007/s13225-020-00466-2)) |
| Otu00340 | 318 | k__Fungi(100);p__Basidiomycota(100);c__Agaricomycetes(100);o__Hymenochaetales(100);f__Schizoporaceae(100);g__Xylodon(78);s__Xylodon_rimosissimus(78); | Xylodon | 13 | Saprotroph | Undefined Saprotroph-\|Wood Saprotroph\| | Corticioid | White Rot | Probable | As white_rot (Põlme et al. 2020); Decay wood (Põlme et al. 2020) | Tedersoo L, et al. 2014. Science 346:e1256688 ((https://doi.org/10.1126/science.1256688)); Põlme S, et al. 2020. Fungal Diversity 105:1-16 ((https://doi.org/10.1007/s13225-020-00466-2)) |
| Otu00341 | 318 | k__Fungi(100);p__Basidiomycota(100);c__Agaricomycetes(100);o__Hymenochaetales(100);f__Hymenochaetaceae(100);g__Inonotus(100);s__Inonotus_sp_CLZ_2013a(100); | Inonotus | 13 | Pathotroph-Saprotroph | Plant Pathogen-\|Wood Saprotroph\| | Polyporoid | White Rot | Highly Probable | As white_rot (Põlme et al. 2020); As wood_pathogen (Põlme et al. 2020); Decay wood (Põlme et al. 2020) | Gilbertson RL, Ryvarden L. 1987-1987. North American Polypores. Fungiflora, Oslo (ISBN: 978-0945345060); Põlme S, et al. 2020. Fungal Diversity 105:1-16 ((https://doi.org/10.1007/s13225-020-00466-2)) |
| Otu00342 | 318 | k__Fungi(100);p__Ascomycota(100);c__Dothideomycetes(100);o__Capnodiales(100);f__Mycosphaerellaceae(100);g__Pseudocercosporella(100);s__Pseudocercosporella_sp(100); | Pseudocercosporella | 13 | Pathotroph | \|Plant Pathogen\| | Microfungus | NULL | Probable | NULL | Tedersoo L, et al. 2014. Science 346:e1256688 ((https://doi.org/10.1126/science.1256688)); Põlme S, et al. 2020. Fungal Diversity 105:1-16 ((https://doi.org/10.1007/s13225-020-00466-2)) |
| Otu00343 | 316 | k__Fungi(100);p__Ascomycota(100);c__Sordariomycetes(100);o__Hypocreales(100);f__Cordycipitaceae(100);g__Simplicillium(100);s__Simplicillium_sp(100); | Simplicillium | 13 | Pathotroph-Saprotroph | \|Animal Parasite\|-Animal Pathogen-Fungal Parasite-Undefined Saprotroph | Microfungus | NULL | Probable | As animal_decomposer (Põlme et al. 2020); As mycoparasite (Põlme et al. 2020); Decay animal material (Põlme et al. 2020) | Tedersoo L, et al. 2014. Science 346:e1256688 ((https://doi.org/10.1126/science.1256688)); Põlme S, et al. 2020. Fungal Diversity 105:1-16 ((https://doi.org/10.1007/s13225-020-00466-2)) |
| Otu00344 | 315 | k__Fungi(100);p__Basidiomycota(100);c__Agaricomycetes(100);o__Atheliales(100);f__Atheliaceae(100);g__Byssocorticium(99);s__Byssocorticium_sp(99); | Byssocorticium | 13 | Saprotroph-Symbiotroph | \|Ectomycorrhizal\|-Undefined Saprotroph | Corticioid | NULL | Highly Probable | NULL | Rinaldi AC, et al. 2008. Fungal Diversity 33:1-45 ((http://www.fungaldiversity.org/fdp/sfdp/33-1.pdf)); Tedersoo L, et al. 2010. Mycorrhiza 20:217-263 ((https://doi.org/10.1007/s00572-009-0274-x)); Taylor DL, et al. 2014. Ecological Monographs 84:3-20 ((https://doi.org/10.1890/12-1693.1)); Põlme S, et al. 2020. Fungal Diversity 105:1-16 ((https://doi.org/10.1007/s13225-020-00466-2)); http://mycorrhizas.info/ecmf.html ((http://mycorrhizas.info/ecmf.html)) |
| Otu00345 | 314 | k__Fungi(100);p__Ascomycota(100);c__Dothideomycetes(100);o__Pleosporales(100);f__Leptosphaeriaceae(93);g__unclassified_Leptosphaeriaceae(67);s__Leptosphaeriaceae_sp(67); | Leptosphaeriaceae | 9 | Pathotroph-Saprotroph | Plant Pathogen-Wood Saprotroph | Microfungus | NULL | Probable | Saprobic or weakly necrotrophic on stems or leaves (Cannon & Kirk 2007) | Cannon PF, Kirk PM. 2007. Fungal Families of the World. CAB International, Cambridge (ISBN: 978-0851998275) |
| Otu00346 | 313 | k__Fungi(100);p__Ascomycota(100);c__Dothideomycetes(100);o__Pleosporales(100);o__Pleosporales_unclassified(99);o__Pleosporales_unclassified(99);o__Pleosporales_unclassified(99); | - | - | - | - | - | - | - | Unassigned | - |
| Otu00347 | 312 | k__Fungi(100);p__Basidiomycota(100);c__Agaricomycetes(100);o__Agaricales(100);f__Tricholomataceae(100);f__Tricholomataceae_unclassified(72);f__Tricholomataceae_unclassified(72); | Tricholomataceae | 9 | Pathotroph-Symbiotroph | Ectomycorrhizal-Fungal Parasite | Agaricoid | NULL | Probable | Tricholoma species are ectomycorrhizal, occurring in coniferous and broadleaved forests (Cannon & Kirk 2007); T. sclerotoideum is parasitic on Helvella spp. | Cannon PF, Kirk PM. 2007. Fungal Families of the World. CAB International, Cambridge (ISBN: 978-0851998275) |
| Otu00348 | 311 | k__Fungi(100);p__Basidiomycota(100);c__Agaricomycetes(100);o__Agaricales(100);f__Strophariaceae(100);g__Pholiota(100);s__Pholiota_squarrosoides(100); | Pholiota | 13 | Pathotroph-Saprotroph | Plant Pathogen-Plant Saprotroph-Undefined Saprotroph-\|Wood Saprotroph\| | Agaricoid | White Rot | Probable | As litter_saprotroph (Põlme et al. 2020); As white_rot (Põlme et al. 2020); As wood_pathogen (Põlme et al. 2020); Decay wood, leaf, fruit, seed (Põlme et al. 2020) | Tedersoo L, et al. 2014. Science 346:e1256688 ((https://doi.org/10.1126/science.1256688)); Põlme S, et al. 2020. Fungal Diversity 105:1-16 ((https://doi.org/10.1007/s13225-020-00466-2)) |
| Otu00349 | 308 | k__Fungi(100);p__Basidiomycota(100);c__Agaricomycetes(100);o__Atheliales(100);f__unclassified_Atheliales(90);g__unclassified_Atheliales(90);s__Atheliales_sp(90); | - | - | - | - | - | - | - | Unassigned | - |
| Otu00350 | 306 | k__Fungi(100);p__Zygomycota(100);c__Zygomycota_class_Incertae_sedis(100);o__Mortierellales(100);f__Mortierellaceae(100);g__Mortierella(100);g__Mortierella_unclassified(100); | Mortierella | 13 | Saprotroph-Symbiotroph | Endophyte-Plant Saprotroph-\|Undefined Saprotroph\| | Microfungus | NULL | Probable | As chitinolytic (Põlme et al. 2020); As soil_saprotroph (Põlme et al. 2020); Decay roots, soil (Põlme et al. 2020); Saprobic in soil and thus frequenctly isolated from soil and material contaminated with soil such as hay or rotten silage and spoil tips (Cannon & Kirk 2007) | Cannon PF, Kirk PM. 2007. Fungal Families of the World. CAB International, Cambridge (ISBN: 978-0851998275); Tedersoo L, et al. 2014. Science 346:e1256688 ((https://doi.org/10.1126/science.1256688)); Purahong W, et al. 2016. Molecular Ecology 25:4059-4074 ((https://doi.org/10.1111/mec.13739)); Põlme S, et al. 2020. Fungal Diversity 105:1-16 ((https://doi.org/10.1007/s13225-020-00466-2)) |
| Otu00351 | 306 | k__Fungi(100);p__Ascomycota(100);c__Sordariomycetes(100);o__Hypocreales(100);f__Hypocreaceae(100);g__Hypocrea(94);s__Hypocrea_atroviridis(89); | Hypocrea | 13 | Pathotroph | Endophyte-Fungal Parasite | Microfungus | NULL | Probable | NULL | Tedersoo L, et al. 2014. Science 346:e1256688 ((https://doi.org/10.1126/science.1256688)); Christian N, et al. 2016. Microbial Ecology 72:621-632 ((https://doi.org/10.1007/s00248-016-0804-y)) |
| Otu00352 | 306 | k__Fungi(100);k__Fungi_unclassified(100);k__Fungi_unclassified(100);k__Fungi_unclassified(100);k__Fungi_unclassified(100);k__Fungi_unclassified(100);k__Fungi_unclassified(100); | - | - | - | - | - | - | - | Unassigned | - |
| Otu00353 | 304 | k__Fungi(100);p__Ascomycota(100);c__Dothideomycetes(100);o__Capnodiales(100);o__Capnodiales_unclassified(100);o__Capnodiales_unclassified(100);o__Capnodiales_unclassified(100); | - | - | - | - | - | - | - | Unassigned | - |
| Otu00355 | 302 | k__Fungi(100);p__Ascomycota(100);c__Dothideomycetes(100);o__Capnodiales(100);f__Mycosphaerellaceae(100);f__Mycosphaerellaceae_unclassified(100);f__Mycosphaerellaceae_unclassified(100); | Mycosphaerellaceae | 9 | Pathotroph-Saprotroph | Plant Pathogen-Undefined Saprotroph | Microfungus | NULL | Probable | Biotrophic, necrotrophic or saprobic on various plant tissues (Cannon & Kirk 2007) | Cannon PF, Kirk PM. 2007. Fungal Families of the World. CAB International, Cambridge (ISBN: 978-0851998275) |
| Otu00354 | 302 | k__Fungi(100);p__Basidiomycota(93);p__Basidiomycota_unclassified(93);p__Basidiomycota_unclassified(93);p__Basidiomycota_unclassified(93);p__Basidiomycota_unclassified(93);p__Basidiomycota_unclassified(93); | - | - | - | - | - | - | - | Unassigned | - |
| Otu00356 | 301 | k__Fungi(100);p__Basidiomycota(100);c__Agaricomycetes(100);o__Polyporales(100);f__Phanerochaetaceae(100);g__Antrodiella(100);s__Antrodiella_americana(98); | Antrodiella | 13 | Saprotroph | \|Wood Saprotroph\| | Corticioid-Polyporoid | White Rot | Highly Probable | As white_rot (Põlme et al. 2020); Decay wood (Põlme et al. 2020) | Gilbertson RL, Ryvarden L. 1987-1987. North American Polypores. Fungiflora, Oslo (ISBN: 978-0945345060); Põlme S, et al. 2020. Fungal Diversity 105:1-16 ((https://doi.org/10.1007/s13225-020-00466-2)) |
| Otu00357 | 301 | k__Fungi(100);p__Basidiomycota(100);c__Agaricomycetes(100);o__Agaricales(100);o__Agaricales_unclassified(100);o__Agaricales_unclassified(100);o__Agaricales_unclassified(100); | Agaricales | 7 | Pathotroph-Saprotroph-Symbiotroph | Bryophyte Parasite-Dung Saprotroph-Ectomycorrhizal-Fungal Parasite-Leaf Saprotroph-Plant Parasite-Undefined Saprotroph-Wood Saprotroph | Agaricoid-Gasteroid-Microfungus-Secotioid-Yeast | NULL | Possible | Mushrooms and toadstools, Gill fungi, Agarics... lignicolous, sometimes muscicolous or fungicolous, saprobic, mycorrhizal, rarely parasitic on plants or fungi (Kirk et al. 2008) | Kirk PM et al. 2008. Dictionary of the Fungi. Tenth Edition. CAB International, Wallingford (ISBN: 978-0851998268) |
| Otu00358 | 301 | k__Fungi(100);p__Basidiomycota(100);c__Exobasidiomycetes(100);o__Exobasidiales(100);f__Exobasidiaceae(100);g__Exobasidium(100);s__Exobasidium_arescens(100); | Exobasidium | 13 | Pathotroph | \|Plant Pathogen\| | Leaf Blister-Leaf Gall-Leaf Spot | NULL | Probable | NULL | Tedersoo L, et al. 2014. Science 346:e1256688 ((https://doi.org/10.1126/science.1256688)); Põlme S, et al. 2020. Fungal Diversity 105:1-16 ((https://doi.org/10.1007/s13225-020-00466-2)) |
| Otu00359 | 297 | k__Fungi(100);p__Ascomycota(100);c__Sordariomycetes(100);o__Xylariales(100);f__Xylariaceae(100);g__Rosellinia(100);s__Rosellinia_sp(100); | Rosellinia | 13 | Pathotroph-Saprotroph | \|Plant Pathogen\|-Undefined Saprotroph-Wood Saprotroph | Microfungus | NULL | Probable | As wood_pathogen (Põlme et al. 2020); Decay wood (Põlme et al. 2020); Host - angiosperms (Põlme et al. 2020) | Tedersoo L, et al. 2014. Science 346:e1256688 ((https://doi.org/10.1126/science.1256688)); Põlme S, et al. 2020. Fungal Diversity 105:1-16 ((https://doi.org/10.1007/s13225-020-00466-2)) |
| Otu00360 | 297 | k__Fungi(100);p__Basidiomycota(100);c__Agaricomycetes(100);o__Polyporales(100);f__Fomitopsidaceae(100);g__Antrodia(100);s__Antrodia_serialis(100); | Antrodia | 13 | Saprotroph | \|Wood Saprotroph\| | Corticioid | Brown Rot | Highly Probable | As brown-rot (Põlme et al. 2020); Decay wood (Põlme et al. 2020) | Gilbertson RL, Ryvarden L. 1987-1987. North American Polypores. Fungiflora, Oslo (ISBN: 978-0945345060); Põlme S, et al. 2020. Fungal Diversity 105:1-16 ((https://doi.org/10.1007/s13225-020-00466-2)) |
| Otu00361 | 296 | k__Fungi(100);p__Basidiomycota(100);c__Agaricomycetes(100);c__Agaricomycetes_unclassified(100);c__Agaricomycetes_unclassified(100);c__Agaricomycetes_unclassified(100);c__Agaricomycetes_unclassified(100); | - | - | - | - | - | - | - | Unassigned | - |
| Otu00362 | 295 | k__Fungi(100);p__Basidiomycota(100);c__Microbotryomycetes(100);o__Sporidiobolales(100);f__Sporidiobolales_family_Incertae_sedis(100);g__Rhodotorula(100);s__Rhodotorula_aurantiaca(100); | Rhodotorula | 13 | Pathotroph-Saprotroph-Symbiotroph | Animal Pathogen-Animal Symbiotroph-Endophyte-Plant Pathogen-\|Undefined Saprotroph\| | Yeast | NULL | Probable | As foliar_endophyte (Põlme et al. 2020); Opportunistic human pathogen | Põlme S, et al. 2020. Fungal Diversity 105:1-16 ((https://doi.org/10.1007/s13225-020-00466-2)); https://www.cdc.gov ((https://www.cdc.gov)) |
| Otu00363 | 293 | k__Fungi(100);p__Ascomycota(100);c__Dothideomycetes(100);o__Capnodiales(100);f__Capnodiales_family_Incertae_sedis(100);g__Capnobotryella(100);s__Capnobotryella_renispora(100); | Capnobotryella | 13 | Saprotroph | \|Plant Saprotroph\|-Undefined Saprotroph | Microfungus | NULL | Probable | As litter_saprotroph (Põlme et al. 2020); Decay leaf, fruit, seed, soil (Põlme et al. 2020) | Tedersoo L, et al. 2014. Science 346:e1256688 ((https://doi.org/10.1126/science.1256688)); Põlme S, et al. 2020. Fungal Diversity 105:1-16 ((https://doi.org/10.1007/s13225-020-00466-2)) |
| Otu00364 | 291 | k__Fungi(100);p__Basidiomycota(100);c__Agaricomycetes(100);o__Polyporales(100);o__Polyporales_unclassified(99);o__Polyporales_unclassified(99);o__Polyporales_unclassified(99); | - | - | - | - | - | - | - | Unassigned | - |
| Otu00365 | 289 | k__Fungi(100);p__Basidiomycota(100);c__Agaricomycetes(100);o__Polyporales(100);f__Meripilaceae(100);g__Rigidoporus(100);s__Rigidoporus_corticola(100); | Rigidoporus | 13 | Saprotroph | \|Wood Saprotroph\| | Polyporoid | White Rot | Highly Probable | As white_rot (Põlme et al. 2020); Decay wood (Põlme et al. 2020) | Gilbertson RL, Ryvarden L. 1987-1987. North American Polypores. Fungiflora, Oslo (ISBN: 978-0945345060); Põlme S, et al. 2020. Fungal Diversity 105:1-16 ((https://doi.org/10.1007/s13225-020-00466-2)) |
| Otu00367 | 286 | k__Fungi(100);p__Basidiomycota(100);c__Agaricomycetes(100);o__Polyporales(100);f__Meruliaceae(100);g__Phlebia(100);s__Phlebia_radiata(100); | Phlebia | 13 | Saprotroph | \|Wood Saprotroph\| | Corticioid | White Rot | Highly Probable | As white_rot (Põlme et al. 2020); Decay wood (Põlme et al. 2020) | Kirk PM et al. 2008. Dictionary of the Fungi. Tenth Edition. CAB International, Wallingford (ISBN: 978-0851998268); Põlme S, et al. 2020. Fungal Diversity 105:1-16 ((https://doi.org/10.1007/s13225-020-00466-2)) |
| Otu00366 | 286 | k__Fungi(100);p__Basidiomycota(100);c__Microbotryomycetes(100);o__Sporidiobolales(100);f__Sporidiobolales_family_Incertae_sedis(100);g__unclassified_Sporidiobolales(100);s__Sporidiobolales_sp(100); | - | - | - | - | - | - | - | Unassigned | - |
| Otu00369 | 285 | k__Fungi(100);p__Basidiomycota(100);c__Agaricomycetes(100);o__Trechisporales(100);f__Hydnodontaceae(100);g__Trechispora(96);g__Trechispora_unclassified(93); | Trechispora | 13 | Saprotroph | \|Wood Saprotroph\| | Corticioid | White Rot | Highly Probable | As white_rot (Põlme et al. 2020); Decay wood (Põlme et al. 2020) | Gilbertson RL, Ryvarden L. 1987-1987. North American Polypores. Fungiflora, Oslo (ISBN: 978-0945345060); Rinaldi AC, et al. 2008. Fungal Diversity 33:1-45 ((http://www.fungaldiversity.org/fdp/sfdp/33-1.pdf)) (as Uncertain); Tedersoo L, et al. 2010. Mycorrhiza 20:217-263 ((https://doi.org/10.1007/s00572-009-0274-x)) (as not ECM).; Põlme S, et al. 2020. Fungal Diversity 105:1-16 ((https://doi.org/10.1007/s13225-020-00466-2)) |
| Otu00368 | 285 | k__Fungi(100);p__Basidiomycota(100);c__Agaricomycetes(100);c__Agaricomycetes_unclassified(99);c__Agaricomycetes_unclassified(99);c__Agaricomycetes_unclassified(99);c__Agaricomycetes_unclassified(99); | - | - | - | - | - | - | - | Unassigned | - |
| Otu00370 | 284 | k__Fungi(100);p__Ascomycota(100);c__Taphrinomycetes(100);o__Taphrinales(100);f__Taphrinaceae(100);g__Lalaria(100);s__Lalaria_inositophila(100); | - | - | - | - | - | - | - | Unassigned | - |
| Otu00371 | 283 | k__Fungi(100);p__Basidiomycota(100);c__Agaricomycetes(100);o__Polyporales(98);o__Polyporales_unclassified(98);o__Polyporales_unclassified(98);o__Polyporales_unclassified(98); | - | - | - | - | - | - | - | Unassigned | - |
| Otu00372 | 281 | k__Fungi(100);p__Basidiomycota(100);c__Agaricomycetes(100);o__Polyporales(100);f__Meruliaceae(100);g__Hyphoderma(100);s__Hyphoderma_puberum(100); | Hyphoderma | 13 | Saprotroph | Undefined Saprotroph-\|Wood Saprotroph\| | Corticioid | White Rot | Probable | As white_rot (Põlme et al. 2020); Decay wood (Põlme et al. 2020) | Tedersoo L, et al. 2014. Science 346:e1256688 ((https://doi.org/10.1126/science.1256688)); Põlme S, et al. 2020. Fungal Diversity 105:1-16 ((https://doi.org/10.1007/s13225-020-00466-2)) |
| Otu00373 | 279 | k__Fungi(100);p__Ascomycota(100);c__Dothideomycetes(100);o__Pleosporales(97);f__Venturiaceae(97);g__Rhizosphaera(97);s__Rhizosphaera_oudemansii(81); | Rhizosphaera | 13 | Pathotroph-Saprotroph | \|Plant Pathogen\|-Plant Saprotroph | Microfungus | NULL | Probable | As litter_saprotroph (Põlme et al. 2020); Decay leaf, fruit, seed (Põlme et al. 2020) | Tedersoo L, et al. 2014. Science 346:e1256688 ((https://doi.org/10.1126/science.1256688)); Põlme S, et al. 2020. Fungal Diversity 105:1-16 ((https://doi.org/10.1007/s13225-020-00466-2)) |
| Otu00375 | 277 | k__Fungi(100);p__Basidiomycota(100);c__Exobasidiomycetes(100);o__Exobasidiales(100);f__Exobasidiaceae(100);g__Exobasidium(100);s__Exobasidium_miyabei(99); | Exobasidium | 13 | Pathotroph | \|Plant Pathogen\| | Leaf Blister-Leaf Gall-Leaf Spot | NULL | Probable | NULL | Tedersoo L, et al. 2014. Science 346:e1256688 ((https://doi.org/10.1126/science.1256688)); Põlme S, et al. 2020. Fungal Diversity 105:1-16 ((https://doi.org/10.1007/s13225-020-00466-2)) |
| Otu00374 | 277 | k__Fungi(100);k__Fungi_unclassified(91);k__Fungi_unclassified(91);k__Fungi_unclassified(91);k__Fungi_unclassified(91);k__Fungi_unclassified(91);k__Fungi_unclassified(91); | - | - | - | - | - | - | - | Unassigned | - |
| Otu00376 | 276 | k__Fungi(100);p__Basidiomycota(100);c__Agaricomycetes(100);o__Polyporales(100);f__Meruliaceae(100);g__Hypochnicium(100);s__Hypochnicium_cremicolor(100); | Hypochnicium | 13 | Saprotroph | Undefined Saprotroph-\|Wood Saprotroph\| | Corticioid | White Rot | Probable | As white_rot (Põlme et al. 2020); Decay wood (Põlme et al. 2020) | Tedersoo L, et al. 2014. Science 346:e1256688 ((https://doi.org/10.1126/science.1256688)); Põlme S, et al. 2020. Fungal Diversity 105:1-16 ((https://doi.org/10.1007/s13225-020-00466-2)) |
| Otu00377 | 275 | k__Fungi(100);p__Ascomycota(100);c__Leotiomycetes(100);o__Helotiales(100);f__Vibrisseaceae(100);g__Phialocephala(100);s__Phialocephala_dimorphospora(91); | Phialocephala | 13 | Saprotroph-Symbiotroph | Endophyte-\|Undefined Saprotroph\| | Microfungus | NULL | Highly Probable | As root_endophyte (Põlme et al. 2020); As root_endophyte_dark_septate (Põlme et al. 2020); As soil_saprotroph (Põlme et al. 2020); Dark Septate Endophyte, known from plant roots.; Decay roots, soil (Põlme et al. 2020) | Newsham KK. 2011. New Phytologist 190:783-793 ((https://doi.org/10.1111/j.1469-8137.2010.03611.x)) (as dark septate root endophytes); Põlme S, et al. 2020. Fungal Diversity 105:1-16 ((https://doi.org/10.1007/s13225-020-00466-2)) |
| Otu00378 | 275 | k__Fungi(100);p__Chytridiomycota(98);c__Chytridiomycetes(98);o__unclassified_Chytridiomycetes(98);f__unclassified_Chytridiomycetes(98);g__unclassified_Chytridiomycetes(98);s__Chytridiomycetes_sp(98); | - | - | - | - | - | - | - | Unassigned | - |
| Otu00379 | 272 | k__Fungi(100);p__Ascomycota(100);c__Saccharomycetes(98);o__Saccharomycetales(98);f__Saccharomycetales_family_Incertae_sedis(95);g__Candida(91);g__Candida_unclassified(64); | Candida | 13 | Pathotroph-Saprotroph-Symbiotroph | Animal Pathogen-Endophyte-Endosymbiont-Epiphyte-\|Undefined Saprotroph\| | Dimorphic Yeast-Yeast | NULL | Probable | As nectar-tap_saprotroph (Põlme et al. 2020); Decay sugar-rich substrates (Põlme et al. 2020); This asexual genus is polyphyletic and contain species that function very differently depend on what they are. Caution should be taken when interpreting guilds for Candida species. | Manolakaki D, et al. 2010. Virulence. 1: 367-375 ((https://doi.org/10.4161/viru.1.5.12796)); Mekbib SB, et al. 2011. Tropical Plant Pathology 36:233-240 ((https://doi.org/10.1590/S1982-56762011000400004)); Põlme S, et al. 2020. Fungal Diversity 105:1-16 ((https://doi.org/10.1007/s13225-020-00466-2)) |
| Otu00380 | 272 | k__Fungi(100);p__Zygomycota(100);c__Zygomycota_class_Incertae_sedis(100);o__Mortierellales(100);f__Mortierellaceae(100);g__Mortierella(100);s__Mortierella_reticulata(97); | Mortierella | 13 | Saprotroph-Symbiotroph | Endophyte-Plant Saprotroph-\|Undefined Saprotroph\| | Microfungus | NULL | Probable | As chitinolytic (Põlme et al. 2020); As soil_saprotroph (Põlme et al. 2020); Decay roots, soil (Põlme et al. 2020); Saprobic in soil and thus frequenctly isolated from soil and material contaminated with soil such as hay or rotten silage and spoil tips (Cannon & Kirk 2007) | Cannon PF, Kirk PM. 2007. Fungal Families of the World. CAB International, Cambridge (ISBN: 978-0851998275); Tedersoo L, et al. 2014. Science 346:e1256688 ((https://doi.org/10.1126/science.1256688)); Purahong W, et al. 2016. Molecular Ecology 25:4059-4074 ((https://doi.org/10.1111/mec.13739)); Põlme S, et al. 2020. Fungal Diversity 105:1-16 ((https://doi.org/10.1007/s13225-020-00466-2)) |
| Otu00381 | 270 | k__Fungi(100);p__Ascomycota(100);c__Saccharomycetes(100);o__Saccharomycetales(100);f__Metschnikowiaceae(100);g__Metschnikowia(100);s__Metschnikowia_pulcherrima(94); | Metschnikowia | 13 | Saprotroph | \|Undefined Saprotroph\| | Yeast | NULL | Probable | As nectar-tap_saprotroph (Põlme et al. 2020); Decay sugar-rich substrates (Põlme et al. 2020) | Põlme S, et al. 2020. Fungal Diversity 105:1-16 ((https://doi.org/10.1007/s13225-020-00466-2)) |
| Otu00383 | 270 | k__Fungi(100);p__Ascomycota(100);c__Dothideomycetes(100);o__Capnodiales(100);f__Capnodiaceae(100);g__Capnodium(100);s__Capnodium_sp(100); | Capnodium | 13 | Pathotroph-Saprotroph | \|Plant Pathogen\|-Plant Saprotroph-Undefined Saprotroph | Microfungus | NULL | Probable | As litter_saprotroph (Põlme et al. 2020); Causes sooty mold (Põlme et al. 2020); Decay leaf, fruit, seed, sugar-rich substrates (Põlme et al. 2020) | Tedersoo L, et al. 2014. Science 346:e1256688 ((https://doi.org/10.1126/science.1256688)); Põlme S, et al. 2020. Fungal Diversity 105:1-16 ((https://doi.org/10.1007/s13225-020-00466-2)) |

| | | | | | | | | | | | | |
|---|---|---|---|---|---|---|---|---|---|---|---|---|
| Otu00382 | 270 | k__Fungi(100);k__Fungi_unclassified(100);k__Fungi_unclassified(100);k__Fungi_unclassified(100);k__Fungi_unclassified(100);k__Fungi_unclassified(100);k__Fungi_unclassified(100); | - | - | | - | - | - | - | - | Unassigned | - |
| Otu00384 | 269 | k__Fungi(100);p__Basidiomycota(98);p__Basidiomycota_unclassified(98);p__Basidiomycota_unclassified(98);p__Basidiomycota_unclassified(98);p__Basidiomycota_unclassified(98);p__Basidiomycota_unclassified(98); | - | - | | - | - | - | - | - | Unassigned | - |
| Otu00386 | 268 | k__Fungi(100);p__Basidiomycota(100);c__Agaricomycetes(100);o__Russulales(100);f__Bondarzewiaceae(100);g__Heterobasidion(100);g__Heterobasidion_unclassified(92); | Heterobasidion | | 13 | Pathotroph-Saprotroph | \|Plant Pathogen\|-Wood Saprotroph | Polyporoid | White Rot | Highly Probable | [illegible] | [illegible] |
| Otu00385 | 268 | k__Fungi(100);p__Basidiomycota(100);p__Basidiomycota_unclassified(100);p__Basidiomycota_unclassified(100);p__Basidiomycota_unclassified(100);p__Basidiomycota_unclassified(100);p__Basidiomycota_unclassified(100); | - | - | | - | - | - | - | - | Unassigned | - |
| Otu00387 | 267 | k__Fungi(100);p__Ascomycota(100);c__Saccharomycetes(100);o__Saccharomycetales(100);f__Metschnikowiaceae(100);g__Metschnikowia(100);s__Metschnikowia_pulcherrima(90); | Metschnikowia | | 13 | Saprotroph | \|Undefined Saprotroph\| | Yeast | NULL | Probable | [illegible] | [illegible] |
| Otu00388 | 265 | k__Fungi(100);p__Basidiomycota(100);c__Agaricomycetes(100);o__Russulales(100);f__Stereaceae(100);g__Gloeocystidiellum(100);s__Gloeocystidiellum_porosum(100); | Gloeocystidiellum | | 13 | Saprotroph | Undefined Saprotroph-\|Wood Saprotroph\| | Corticioid | White Rot | Probable | [illegible] | [illegible] |
| Otu00389 | 265 | k__Fungi(100);p__Ascomycota(80);p__Ascomycota_unclassified(80);p__Ascomycota_unclassified(80);p__Ascomycota_unclassified(80);p__Ascomycota_unclassified(80);p__Ascomycota_unclassified(80); | - | - | | - | - | - | - | - | Unassigned | - |
| Otu00390 | 265 | k__Fungi(100);p__Basidiomycota(100);c__Agaricomycetes(100);c__Agaricomycetes_unclassified(100);c__Agaricomycetes_unclassified(100);c__Agaricomycetes_unclassified(100);c__Agaricomycetes_unclassified(100); | - | - | | - | - | - | - | - | Unassigned | - |
| Otu00391 | 264 | k__Fungi(100);p__Ascomycota(100);c__Dothideomycetes(100);o__Pleosporales(100);f__Pleosporaceae(100);f__Pleosporaceae_unclassified(77);f__Pleosporaceae_unclassified(77); | Pleosporaceae | | 9 | Pathotroph-Saprotroph | Endophyte-Lichen Parasite-Plant Pathogen-Undefined Saprotroph | Microfungus | NULL | Probable | [illegible] | [illegible] |
| Otu00393 | 264 | k__Fungi(100);p__Basidiomycota(100);c__Exobasidiomycetes(100);o__Exobasidiales(100);f__Exobasidiaceae(100);g__Exobasidium(100);s__Exobasidium_yoshinagae(100); | Exobasidium | | 13 | Pathotroph | \|Plant Pathogen\| | Leaf Blister-Leaf Gall-Leaf Spot | NULL | Probable | NULL | [illegible] |
| Otu00392 | 264 | k__Fungi(100);p__Ascomycota(100);p__Ascomycota_unclassified(64);p__Ascomycota_unclassified(64);p__Ascomycota_unclassified(64);p__Ascomycota_unclassified(64);p__Ascomycota_unclassified(64); | - | - | | - | - | - | - | - | Unassigned | - |
| Otu00394 | 263 | k__Fungi(100);k__Fungi_unclassified(100);k__Fungi_unclassified(100);k__Fungi_unclassified(100);k__Fungi_unclassified(100);k__Fungi_unclassified(100);k__Fungi_unclassified(100); | - | - | | - | - | - | - | - | Unassigned | - |
| Otu00395 | 261 | k__Fungi(100);p__Ascomycota(55);p__Ascomycota_unclassified(55);p__Ascomycota_unclassified(55);p__Ascomycota_unclassified(55);p__Ascomycota_unclassified(55);p__Ascomycota_unclassified(55); | - | - | | - | - | - | - | - | Unassigned | - |
| Otu00396 | 260 | k__Fungi(100);p__Ascomycota(100);c__Eurotiomycetes(100);o__Chaetothyriales(100);f__Chaetothyriaceae(96);g__Cyphellophora(96);g__Cyphellophora_unclassified(96); | Cyphellophora | | 13 | Pathotroph-Saprotroph | Animal Parasite-Animal Pathogen-\|Plant Saprotroph\|-Undefined Saprotroph | Dimorphic-Facultative Yeast-Microfungus | NULL | Probable | [illegible] | [illegible] |
| Otu00397 | 259 | k__Fungi(100);p__Zygomycota(100);c__Zygomycota_class_Incertae_sedis(100);o__Mucorales(100);f__Mucoraceae(100);g__Mucor(100);s__Mucor_luteus(100); | Mucor | | 13 | Saprotroph | \|Undefined Saprotroph\| | Microfungus | NULL | Probable | [illegible] | [illegible] |
| Otu00398 | 259 | k__Fungi(100);p__Basidiomycota(100);c__Agaricomycetes(100);o__Auriculariales(100);f__Auriculariales_family_Incertae_sedis(100);g__Basidiodendron(100);s__Basidiodendron_sp(99); | Basidiodendron | | 13 | Saprotroph | Undefined Saprotroph-\|Wood Saprotroph\| | Corticioid | NULL | Probable | [illegible] | [illegible] |
| Otu00399 | 259 | k__Fungi(100);p__Basidiomycota(100);c__Agaricomycetes(100);o__Polyporales(100);f__Polyporaceae(100);g__Trametes(100);s__Trametes_hirsuta(100); | Trametes | | 13 | Pathotroph-Saprotroph | Plant Pathogen-\|Wood Saprotroph\| | Polyporoid | White Rot | Highly Probable | [illegible] | [illegible] |
| Otu00400 | 259 | k__Fungi(100);p__Ascomycota(100);c__Sordariomycetes(100);o__Hypocreales(100);f__Hypocreales_family_Incertae_sedis(100);g__Myrothecium(100);s__Myrothecium_sp(82); | Myrothecium | | 13 | Pathotroph-Saprotroph | Fungal Parasite-\|Plant Saprotroph\|-Undefined Saprotroph | Microfungus | NULL | Probable | [illegible] | [illegible] |
| Otu00401 | 258 | k__Fungi(100);p__Basidiomycota(100);c__Agaricomycetes(100);o__Agaricales(100);f__Hebelomataceae(97);g__unclassified_Hebelomataceae(97);s__Hebelomataceae_sp(97); | - | - | | - | - | - | - | - | Unassigned | - |
| Otu00402 | 257 | k__Fungi(100);k__Fungi_unclassified(58);k__Fungi_unclassified(58);k__Fungi_unclassified(58);k__Fungi_unclassified(58);k__Fungi_unclassified(58);k__Fungi_unclassified(58); | - | - | | - | - | - | - | - | Unassigned | - |
| Otu00403 | 255 | k__Fungi(100);p__Ascomycota(100);c__Saccharomycetes(100);o__Saccharomycetales(100);f__Saccharomycetales_family_Incertae_sedis(100);g__Candida(100);s__Candida_albicans(100); | Candida | | 13 | Pathotroph-Saprotroph-Symbiotroph | Animal Pathogen-Endophyte-Endosymbiont-Epiphyte-\|Undefined Saprotroph\| | Dimorphic Yeast-Yeast | NULL | Probable | [illegible] | [illegible] |
| Otu00405 | 255 | k__Fungi(100);p__Ascomycota(100);c__Dothideomycetes(100);o__Pleosporales(100);f__Phaeosphaeriaceae(100);f__Phaeosphaeriaceae_unclassified(63);f__Phaeosphaeriaceae_unclassified(63); | Phaeosphaeriaceae | | 9 | Pathotroph-Saprotroph | Fungal Parasite-Plant Pathogen-Plant Saprotroph | Microfungus | NULL | Probable | [illegible] | [illegible] |
| Otu00404 | 255 | k__Fungi(100);p__Ascomycota(100);c__Dothideomycetes(100);o__Pleosporales(100);o__Pleosporales_unclassified(100);o__Pleosporales_unclassified(100);o__Pleosporales_unclassified(100); | - | - | | - | - | - | - | - | Unassigned | - |
| Otu00406 | 254 | k__Fungi(100);p__Ascomycota(100);c__Dothideomycetes(100);o__Pleosporales(100);f__Venturiaceae(100);f__Venturiaceae_unclassified(100);f__Venturiaceae_unclassified(100); | Venturiaceae | | 9 | Pathotroph-Saprotroph-Symbiotroph | Endophyte-Plant Pathogen-Undefined Saprotroph | Microfungus | NULL | Probable | [illegible] | [illegible] |
| Otu00407 | 252 | k__Fungi(100);k__Fungi_unclassified(100);k__Fungi_unclassified(100);k__Fungi_unclassified(100);k__Fungi_unclassified(100);k__Fungi_unclassified(100);k__Fungi_unclassified(100); | - | - | | - | - | - | - | - | Unassigned | - |
| Otu00408 | 251 | k__Fungi(100);p__Basidiomycota(100);c__Agaricomycetes(100);o__Polyporales(100);f__Polyporaceae(100);f__Polyporaceae_unclassified(92);f__Polyporaceae_unclassified(92); | Polyporaceae | | 9 | Saprotroph | Wood Saprotroph | Polyporoid | Brown Rot-White | Probable | [illegible] | [illegible] |
| Otu00409 | 251 | k__Fungi(100);p__Basidiomycota(100);c__Agaricomycetes(100);o__Agaricales(100);f__Inocybaceae(100);g__Crepidotus(100);s__Crepidotus_applanatus(100); | Crepidotus | | 13 | Saprotroph | \|Wood Saprotroph\| | Agaricoid | NULL | Highly Probable | [illegible] | [illegible] |
| Otu00410 | 251 | k__Fungi(100);p__Ascomycota(100);p__Ascomycota_unclassified(100);p__Ascomycota_unclassified(100);p__Ascomycota_unclassified(100);p__Ascomycota_unclassified(100);p__Ascomycota_unclassified(100); | - | - | | - | - | - | - | - | Unassigned | - |
| Otu00411 | 251 | k__Fungi(100);k__Fungi_unclassified(58);k__Fungi_unclassified(58);k__Fungi_unclassified(58);k__Fungi_unclassified(58);k__Fungi_unclassified(58);k__Fungi_unclassified(58); | - | - | | - | - | - | - | - | Unassigned | - |
| Otu00412 | 245 | k__Fungi(100);p__Basidiomycota(100);c__Agaricomycetes(100);o__Russulales(100);f__Stereaceae(100);g__Stereum(100);s__Stereum_sp(100); | Stereum | | 13 | Saprotroph | Undefined Saprotroph-\|Wood Saprotroph\| | Polyporoid | White Rot | Probable | [illegible] | [illegible] |
| Otu00413 | 245 | k__Fungi(100);p__Basidiomycota(100);c__Agaricomycetes(100);o__Polyporales(100);f__Phanerochaetaceae(100);g__Hyphodermella(100);s__Hyphodermella_rosae(100); | Hyphodermella | | 13 | Saprotroph | Undefined Saprotroph-\|Wood Saprotroph\| | Corticioid | White Rot | Probable | [illegible] | [illegible] |
| Otu00414 | 243 | k__Fungi(100);p__Basidiomycota(100);c__Agaricomycetes(100);o__Polyporales(100);f__Meruliaceae(100);g__Phlebia(100);s__Phlebia_aurea(100); | Phlebia | | 13 | Saprotroph | \|Wood Saprotroph\| | Corticioid | White Rot | Highly Probable | [illegible] | [illegible] |
| Otu00415 | 242 | k__Fungi(100);p__Basidiomycota(100);c__Microbotryomycetes(100);o__Sporidiobolales(100);f__Sporidiobolales_family_Incertae_sedis(100);g__Rhodotorula(67);g__Rhodotorula_unclassified(67); | Rhodotorula | | 13 | Pathotroph-Saprotroph-Symbiotroph | Animal Pathogen-Animal Symbiotroph-Endophyte-Plant Pathogen-\|Undefined Saprotroph\| | Yeast | NULL | Probable | [illegible] | [illegible] |
| Otu00416 | 242 | k__Fungi(100);p__Basidiomycota(100);c__Agaricomycetes(100);o__Russulales(100);f__Stereaceae(100);g__Gloeocystidiellum(100);s__Gloeocystidiellum_kenyense(100); | Gloeocystidiellum | | 13 | Saprotroph | Undefined Saprotroph-\|Wood Saprotroph\| | Corticioid | White Rot | Probable | [illegible] | [illegible] |
| Otu00417 | 240 | k__Fungi(100);p__Ascomycota(100);c__Sordariomycetes(100);o__Hypocreales(100);f__Hypocreaceae(100);f__Hypocreaceae_unclassified(98);f__Hypocreaceae_unclassified(98); | - | - | | - | - | - | - | - | Unassigned | - |
| Otu00418 | 239 | k__Fungi(100);p__Ascomycota(100);c__Sordariomycetes(100);o__Microascales(100);f__Microascaceae(100);g__unclassified_Microascaceae(100);s__Microascaceae_sp(100); | Microascaceae | | 9 | Pathotroph-Saprotroph-Symbiotroph | Animal Pathogen-Endophyte-Plant Pathogen-Undefined Saprotroph | Microfungus | NULL | Possible | [illegible] | [illegible] |
| Otu00419 | 239 | k__Fungi(100);p__Ascomycota(100);c__Saccharomycetes(100);o__Saccharomycetales(100);f__Metschnikowiaceae(100);g__Metschnikowia(100);s__Metschnikowia_pulcherrima(93); | Metschnikowia | | 13 | Saprotroph | \|Undefined Saprotroph\| | Yeast | NULL | Probable | [illegible] | [illegible] |
| Otu00420 | 239 | k__Fungi(100);p__Ascomycota(100);c__Dothideomycetes(100);o__Pleosporales(100);f__Leptosphaeriaceae(100);g__Leptosphaeria(100);s__Leptosphaeria_sp_MH_2001(100); | Leptosphaeria | | 13 | Pathotroph-Saprotroph | \|Plant Pathogen\|-Wood Saprotroph | Microfungus | NULL | Probable | [illegible] | [illegible] |
| Otu00421 | 239 | k__Fungi(100);p__Basidiomycota(100);c__Agaricomycetes(100);o__Hymenochaetales(100);f__Schizoporaceae(100);g__Hyphodontia(100);s__Hyphodontia_alutacea(100); | Hyphodontia | | 13 | Saprotroph | Undefined Saprotroph-\|Wood Saprotroph\| | Corticioid | White Rot | Probable | [illegible] | [illegible] |
| Otu00422 | 238 | k__Fungi(100);p__Basidiomycota(100);c__Agaricomycetes(100);o__Agaricales(100);f__Marasmiaceae(100);g__Baeospora(100);s__Baeospora_myosura(100); | Baeospora | | 13 | Saprotroph | \|Plant Saprotroph\|-Undefined Saprotroph | Agaricoid | NULL | Probable | [illegible] | [illegible] |
| Otu00424 | 236 | k__Fungi(100);p__Ascomycota(100);c__Lecanoromycetes(100);o__Lecanorales(100);o__Lecanorales_unclassified(98);o__Lecanorales_unclassified(98);o__Lecanorales_unclassified(98); | Lecanorales | | 7 | Pathotroph-Symbiotroph | Lichen Parasite-Lichenized | Microfungus-Thallus | NULL | Probable | [illegible] | [illegible] |
| Otu00423 | 236 | k__Fungi(100);k__Fungi_unclassified(56);k__Fungi_unclassified(56);k__Fungi_unclassified(56);k__Fungi_unclassified(56);k__Fungi_unclassified(56);k__Fungi_unclassified(56); | - | - | | - | - | - | - | - | Unassigned | - |
| Otu00425 | 236 | k__Fungi(100);p__Ascomycota(71);p__Ascomycota_unclassified(71);p__Ascomycota_unclassified(71);p__Ascomycota_unclassified(71);p__Ascomycota_unclassified(71);p__Ascomycota_unclassified(71); | - | - | | - | - | - | - | - | Unassigned | - |
| Otu00426 | 235 | k__Fungi(100);p__Basidiomycota(100);c__Agaricomycetes(100);o__Russulales(97);o__Russulales_unclassified(97);o__Russulales_unclassified(97);o__Russulales_unclassified(97); | - | - | | - | - | - | - | - | Unassigned | - |
| Otu00427 | 234 | k__Fungi(100);p__Ascomycota(100);c__Saccharomycetes(100);o__Saccharomycetales(100);f__Saccharomycetales_family_Incertae_sedis(100);g__Candida(100);s__Candida_sp_AS23486(76); | Candida | | 13 | Pathotroph-Saprotroph-Symbiotroph | Animal Pathogen-Endophyte-Endosymbiont-Epiphyte-\|Undefined Saprotroph\| | Dimorphic Yeast-Yeast | NULL | Probable | [illegible] | [illegible] |
| Otu00428 | 234 | k__Fungi(100);p__Chytridiomycota(100);c__Chytridiomycetes(100);o__Spizellomycetales(100);f__Spizellomycetaceae(100);g__unclassified_Spizellomycetaceae(100);s__Spizellomycetaceae_sp(100); | Spizellomycetaceae | | 9 | Pathotroph | Plant Pathogen-Undefined Parasite-Undefined Saprotroph | Microfungus | NULL | Probable | [illegible] | [illegible] |
| Otu00431 | 233 | k__Fungi(100);p__Ascomycota(100);c__Sordariomycetes(100);o__Diaporthales(100);f__Gnomoniaceae(99);f__Gnomoniaceae_unclassified(97);f__Gnomoniaceae_unclassified(97); | Gnomoniaceae | | 9 | Pathotroph-Saprotroph-Symbiotroph | Endophyte-Plant Pathogen-Undefined Saprotroph | Microfungus | NULL | Probable | [illegible] | [illegible] |
| Otu00429 | 233 | k__Fungi(100);p__Ascomycota(100);c__Dothideomycetes(100);c__Dothideomycetes_unclassified(97);c__Dothideomycetes_unclassified(97);c__Dothideomycetes_unclassified(97);c__Dothideomycetes_unclassified(97); | - | - | | - | - | - | - | - | Unassigned | - |
| Otu00430 | 233 | k__Fungi(100);p__Basidiomycota(100);c__Agaricomycetes(100);o__Polyporales(100);o__Polyporales_unclassified(76);o__Polyporales_unclassified(76);o__Polyporales_unclassified(76); | - | - | | - | - | - | - | - | Unassigned | - |
| Otu00432 | 232 | k__Fungi(100);p__Basidiomycota(100);c__Agaricomycetes(100);o__Polyporales(100);f__Meripilaceae(100);g__Meripilus(100);s__Meripilus_giganteus(100); | Meripilus | | 13 | Pathotroph-Saprotroph | \|Plant Pathogen\|-Wood Saprotroph | Polyporoid | White Rot | Highly Probable | [illegible] | [illegible] |
| Otu00433 | 232 | k__Fungi(100);p__Basidiomycota(100);c__Tremellomycetes(100);o__Filobasidiales(100);f__Filobasidiaceae(100);g__Cryptococcus(100);s__Cryptococcus_wieringae(100); | Cryptococcus | | 13 | Pathotroph-Saprotroph-Symbiotroph | Animal Pathogen-Endophyte-Epiphyte-\|Undefined Saprotroph\| | Dimorphic Yeast-Yeast | NULL | Possible | [illegible] | [illegible] |
| Otu00434 | 232 | k__Fungi(100);p__Basidiomycota(100);c__Agaricomycetes(100);o__Cantharellales(100);o__Cantharellales_unclassified(69);o__Cantharellales_unclassified(69);o__Cantharellales_unclassified(69); | - | - | | - | - | - | - | - | Unassigned | - |
| Otu00437 | 231 | k__Fungi(100);p__Chytridiomycota(100);c__Chytridiomycetes(100);o__Spizellomycetales(100);f__Spizellomycetaceae(100);g__Powellomyces(100);s__Powellomyces_sp(100); | Powellomyces | | 13 | Pathotroph-Saprotroph | Algal Parasite-Plant Pathogen-\|Pollen Saprotroph\| | Chytrid-Microfungus | NULL | Probable | [illegible] | [illegible] |
| Otu00438 | 231 | k__Fungi(100);p__Ascomycota(100);c__Eurotiomycetes(100);o__Onygenales(100);f__Onygenaceae(100);g__Chrysosporium(100);s__Chrysosporium_vallenarense(100); | Chrysosporium | | 13 | Pathotroph-Saprotroph | Plant Pathogen-Undefined Saprotroph-\|Wood Saprotroph\| | Microfungus | Soft Rot | Highly Probable | [illegible] | [illegible] |
| Otu00435 | 231 | k__Fungi(100);p__Basidiomycota(100);c__Agaricomycetes(100);o__Cantharellales(100);o__Cantharellales_unclassified(100);o__Cantharellales_unclassified(100);o__Cantharellales_unclassified(100); | - | - | | - | - | - | - | - | Unassigned | - |
| Otu00436 | 231 | k__Fungi(100);p__Ascomycota(79);p__Ascomycota_unclassified(79);p__Ascomycota_unclassified(79);p__Ascomycota_unclassified(79);p__Ascomycota_unclassified(79);p__Ascomycota_unclassified(79); | - | - | | - | - | - | - | - | Unassigned | - |
| Otu00439 | 227 | k__Fungi(100);k__Fungi_unclassified(68);k__Fungi_unclassified(68);k__Fungi_unclassified(68);k__Fungi_unclassified(68);k__Fungi_unclassified(68);k__Fungi_unclassified(68); | - | - | | - | - | - | - | - | Unassigned | - |
| Otu00440 | 226 | k__Fungi(100);p__Basidiomycota(100);c__Tremellomycetes(100);o__Filobasidiales(53);f__Filobasidiaceae(53);g__Cryptococcus(53);s__Cryptococcus_heimaeyensis(53); | Cryptococcus | | 13 | Pathotroph-Saprotroph-Symbiotroph | Animal Pathogen-Endophyte-Epiphyte-\|Undefined Saprotroph\| | Dimorphic Yeast-Yeast | NULL | Possible | [illegible] | [illegible] |
| Otu00442 | 225 | k__Fungi(100);p__Basidiomycota(100);c__Agaricomycetes(100);o__Agaricales(100);f__Inocybaceae(100);g__Crepidotus(100);s__Crepidotus_aff_alabamensis_PBM2979(100); | Crepidotus | | 13 | Saprotroph | \|Wood Saprotroph\| | Agaricoid | NULL | Highly Probable | [illegible] | [illegible] |
| Otu00441 | 225 | k__Fungi(100);p__Ascomycota(100);c__Eurotiomycetes(93);c__Eurotiomycetes_unclassified(92);c__Eurotiomycetes_unclassified(92);c__Eurotiomycetes_unclassified(92);c__Eurotiomycetes_unclassified(92); | - | - | | - | - | - | - | - | Unassigned | - |
| Otu00444 | 224 | k__Fungi(100);p__Ascomycota(100);c__Dothideomycetes(100);o__Capnodiales(100);f__Mycosphaerellaceae(83);f__Mycosphaerellaceae_unclassified(83);f__Mycosphaerellaceae_unclassified(83); | Mycosphaerellaceae | | 9 | Pathotroph-Saprotroph | Plant Pathogen-Undefined Saprotroph | Microfungus | NULL | Probable | [illegible] | [illegible] |
| Otu00445 | 224 | k__Fungi(100);p__Basidiomycota(100);c__Agaricomycetes(100);o__Polyporales(100);f__Meruliaceae(100);g__Hypochnicium(100);s__Hypochnicium_subrigescens(100); | Hypochnicium | | 13 | Saprotroph | Undefined Saprotroph-\|Wood Saprotroph\| | Corticioid | White Rot | Probable | [illegible] | [illegible] |
| Otu00443 | 224 | k__Fungi(100);p__Basidiomycota(100);c__Agaricomycetes(100);o__Polyporales(100);f__unclassified_Polyporales(99);g__unclassified_Polyporales(99);s__Polyporales_sp(99); | - | - | | - | - | - | - | - | Unassigned | - |
| Otu00446 | 223 | k__Fungi(100);p__Basidiomycota(100);c__Tremellomycetes(100);o__Tremellales(100);f__Tremellales_family_Incertae_sedis(100);g__Cryptococcus(100);s__Cryptococcus_paraflavus(100); | Cryptococcus | | 13 | Pathotroph-Saprotroph-Symbiotroph | Animal Pathogen-Endophyte-Epiphyte-\|Undefined Saprotroph\| | Dimorphic Yeast-Yeast | NULL | Possible | [illegible] | [illegible] |
| Otu00447 | 222 | k__Fungi(100);p__Basidiomycota(100);c__Agaricomycetes(100);o__Agaricales(100);f__Strophariaceae(100);g__Agrocybe(100);s__Agrocybe_erebia(100); | Agrocybe | | 13 | Saprotroph | Dung Saprotroph-\|Undefined Saprotroph\| | Agaricoid | NULL | Probable | [illegible] | [illegible] |
| Otu00448 | 220 | k__Fungi(100);p__Ascomycota(100);c__Dothideomycetes(100);o__Pleosporales(100);o__Pleosporales_unclassified(65);o__Pleosporales_unclassified(65);o__Pleosporales_unclassified(65); | - | - | | - | - | - | - | - | Unassigned | - |
| Otu00449 | 218 | k__Fungi(100);p__Ascomycota(100);c__Sordariomycetes(100);o__Hypocreales(100);f__Clavicipitaceae(100);g__Claviceps(100);s__Claviceps_purpurea_var_phalaridis(100); | Claviceps | | 13 | Pathotroph | \|Plant Pathogen\| | Microfungus | NULL | Probable | NULL | [illegible] |
| Otu00450 | 218 | k__Fungi(100);p__Basidiomycota(100);c__Agaricomycetes(100);o__Polyporales(100);f__Meripilaceae(100);f__Meripilaceae_unclassified(95);f__Meripilaceae_unclassified(95); | Meripilaceae | | 9 | Pathotroph-Saprotroph | Plant Pathogen-Undefined Saprotroph | Polyporoid | NULL | Probable | [illegible] | [illegible] |
| Otu00452 | 217 | k__Fungi(100);p__Ascomycota(100);c__Dothideomycetes(100);o__Dothideomycetes_order_Incertae_sedis(100);f__Pseudeurotiaceae(100);g__unclassified_Pseudeurotiaceae(100);s__Pseudeurotiaceae_sp(100); | Pseudeurotiaceae | | 9 | Saprotroph | Plant Saprotroph-Wood Saprotroph | Microfungus | NULL | Probable | [illegible] | [illegible] |
| Otu00451 | 217 | k__Fungi(100);p__Basidiomycota(96);p__Basidiomycota_unclassified(89);p__Basidiomycota_unclassified(89);p__Basidiomycota_unclassified(89);p__Basidiomycota_unclassified(89);p__Basidiomycota_unclassified(89); | - | - | | - | - | - | - | - | Unassigned | - |
| Otu00453 | 216 | k__Fungi(100);p__Ascomycota(100);c__Sordariomycetes(100);o__Sordariales(100);f__Cephalothecaceae(100);g__Cryptendoxyla(100);s__Cryptendoxyla_hypophloia(100); | Cryptendoxyla | | 13 | Saprotroph | Undefined Saprotroph-\|Wood Saprotroph\| | Microfungus | NULL | Probable | [illegible] | [illegible] |
| Otu00454 | 216 | k__Fungi(100);p__Ascomycota(100);c__Dothideomycetes(100);o__Pleosporales(100);f__Pleosporales_family_Incertae_sedis(99);g__Pseudochaetosphaeronema(99);s__Pseudochaetosphaeronema_larense(99); | Pseudochaetosphaerone | | 13 | Pathotroph-Saprotroph-Symbiotroph | Endophyte-Plant Pathogen-\|Plant Saprotroph\| | Microfungus | NULL | Probable | [illegible] | [illegible] |
| Otu00455 | 216 | k__Fungi(100);p__Ascomycota(100);p__Ascomycota_unclassified(100);p__Ascomycota_unclassified(100);p__Ascomycota_unclassified(100);p__Ascomycota_unclassified(100);p__Ascomycota_unclassified(100); | - | - | | - | - | - | - | - | Unassigned | - |
| Otu00456 | 214 | k__Fungi(100);p__Ascomycota(100);c__Eurotiomycetes(100);o__Chaetothyriales(100);f__Herpotrichiellaceae(100);g__Cladophialophora(100);g__Cladophialophora_unclassified(56); | Cladophialophora | | 13 | Pathotroph-Saprotroph-Symbiotroph | Animal Pathogen-Endophyte-Fungal Parasite-\|Undefined Saprotroph\| | Dimorphic-Microfungus | NULL | Probable | [illegible] | [illegible] |
| Otu00457 | 213 | k__Fungi(100);p__Basidiomycota(100);c__Agaricomycetes(100);o__Polyporales(100);f__Meruliaceae(100);g__Steccherinum(100);s__Steccherinum_sp_DLL2011_2(100); | Steccherinum | | 13 | Saprotroph | Undefined Saprotroph-\|Wood Saprotroph\| | Corticioid | White Rot | Probable | [illegible] | [illegible] |
| Otu00458 | 212 | k__Fungi(100);p__Basidiomycota(100);c__Tremellomycetes(100);o__Cystofilobasidiales(100);f__Cystofilobasidiaceae(100);g__Cystofilobasidium(100);s__Cystofilobasidium_macerans(100); | Cystofilobasidium | | 13 | Saprotroph | \|Plant Saprotroph\| | Yeast | NULL | Possible | [illegible] | [illegible] |
| Otu00459 | 210 | k__Fungi(100);p__Basidiomycota(100);c__Agaricomycetes(100);o__Hymenochaetales(100);f__Schizoporaceae(100);g__Hyphodontia(100);s__Hyphodontia_subalutacea(100); | Hyphodontia | | 13 | Saprotroph | Undefined Saprotroph-\|Wood Saprotroph\| | Corticioid | White Rot | Probable | [illegible] | [illegible] |
| Otu00461 | 209 | k__Fungi(100);p__Basidiomycota(100);c__Agaricomycetes(100);o__Russulales(100);f__Peniophoraceae(100);g__Peniophora(100);s__Peniophora_pini(100); | Peniophora | | 13 | Pathotroph-Saprotroph | Plant Pathogen-\|Wood Saprotroph\| | Corticioid | White Rot | Probable | [illegible] | [illegible] |
| Otu00460 | 209 | k__Fungi(100);p__Basidiomycota(100);c__Agaricostilbomycetes(100);o__unclassified_Agaricostilbomycetes(66);f__unclassified_Agaricostilbomycetes(66);g__unclassified_Agaricostilbomycetes(66);s__Agaricostilbomycetes_sp(66); | - | - | | - | - | - | - | - | Unassigned | - |
| Otu00462 | 208 | k__Fungi(100);p__Ascomycota(100);c__Taphrinomycetes(100);o__Taphrinales(100);o__Taphrinales_unclassified(74);o__Taphrinales_unclassified(74);o__Taphrinales_unclassified(74); | - | - | | - | - | - | - | - | Unassigned | - |
| Otu00463 | 207 | k__Fungi(100);p__Basidiomycota(100);c__Exobasidiomycetes(100);o__Exobasidiales(100);f__Exobasidiaceae(100);g__Exobasidium(100);g__Exobasidium_unclassified(100); | Exobasidium | | 13 | Pathotroph | \|Plant Pathogen\| | Leaf Blister-Leaf Gall-Leaf Spot | NULL | Probable | NULL | [illegible] |
| Otu00464 | 206 | k__Fungi(100);p__Zygomycota(100);c__Zygomycota_class_Incertae_sedis(100);o__Mucorales(100);f__Mucoraceae(100);g__Mucor(100);g__Mucor_unclassified(93); | Mucor | | 13 | Saprotroph | \|Undefined Saprotroph\| | Microfungus | NULL | Probable | [illegible] | [illegible] |
| Otu00466 | 206 | k__Fungi(100);p__Ascomycota(100);c__Sordariomycetes(100);o__Diaporthales(100);f__Valsaceae(100);g__Cytospora(100);s__Cytospora_sp_Fk34(99); | Cytospora | | 13 | Pathotroph-Saprotroph-Symbiotroph | Endophyte-\|Plant Pathogen\|-Plant Saprotroph-Wood Saprotroph | Microfungus | Soft Rot | Probable | [illegible] | [illegible] |
| Otu00465 | 206 | k__Fungi(100);k__Fungi_unclassified(100);k__Fungi_unclassified(100);k__Fungi_unclassified(100);k__Fungi_unclassified(100);k__Fungi_unclassified(100);k__Fungi_unclassified(100); | - | - | | - | - | - | - | - | Unassigned | - |
| Otu00468 | 205 | k__Fungi(100);p__Basidiomycota(100);c__Agaricomycetes(100);o__Polyporales(100);f__Meripilaceae(100);g__Rigidoporus(94);s__Rigidoporus_sp_JV0511_23(94); | Rigidoporus | | 13 | Saprotroph | \|Wood Saprotroph\| | Polyporoid | White Rot | Highly Probable | [illegible] | [illegible] |
| Otu00469 | 205 | k__Fungi(100);p__Basidiomycota(100);c__Agaricomycetes(100);o__Auriculariales(100);f__Auriculariales_family_Incertae_sedis(100);g__Exidia(100);s__Exidia_sp(100); | Exidia | | 13 | Saprotroph-Symbiotroph | Endophyte-Plant Saprotroph-Undefined Saprotroph-\|Wood Saprotroph\| | Tremelloid | NULL | Probable | [illegible] | [illegible] |
| Otu00467 | 205 | k__Fungi(100);p__Basidiomycota(100);c__Agaricomycetes(100);o__Polyporales(100);o__Polyporales_unclassified(100);o__Polyporales_unclassified(100);o__Polyporales_unclassified(100); | - | - | | - | - | - | - | - | Unassigned | - |
| Otu00470 | 204 | k__Fungi(100);p__Zygomycota(100);c__Zygomycota_class_Incertae_sedis(100);o__Mortierellales(100);f__Mortierellaceae(100);g__Mortierella(100);s__Mortierella_rishikesha(60); | Mortierella | | 13 | Saprotroph-Symbiotroph | Endophyte-Plant Saprotroph-\|Undefined Saprotroph\| | Microfungus | NULL | Probable | [illegible] | [illegible] |
| Otu00471 | 204 | k__Fungi(100);p__Ascomycota(100);c__Leotiomycetes(100);o__Leotiales(74);f__Bulgariaceae(74);g__Bulgaria(74);s__Bulgaria_inquinans(74); | Bulgaria | | 13 | Pathotroph-Saprotroph-Symbiotroph | Endophyte-Plant Pathogen-\|Wood Saprotroph\| | Microfungus-Pezizoid | NULL | Probable | [illegible] | [illegible] |
| Otu00472 | 203 | k__Fungi(100);k__Fungi_unclassified(59);k__Fungi_unclassified(59);k__Fungi_unclassified(59);k__Fungi_unclassified(59);k__Fungi_unclassified(59);k__Fungi_unclassified(59); | - | - | | - | - | - | - | - | Unassigned | - |
| Otu00473 | 202 | k__Fungi(100);p__Ascomycota(100);c__Dothideomycetes(100);o__Pleosporales(100);f__Venturiaceae(100);f__Venturiaceae_unclassified(100);f__Venturiaceae_unclassified(100); | Venturiaceae | | 9 | Pathotroph-Saprotroph-Symbiotroph | Endophyte-Plant Pathogen-Undefined Saprotroph | Microfungus | NULL | Probable | [illegible] | [illegible] |
| Otu00474 | 202 | k__Fungi(100);k__Fungi_unclassified(73);k__Fungi_unclassified(73);k__Fungi_unclassified(73);k__Fungi_unclassified(73);k__Fungi_unclassified(73);k__Fungi_unclassified(73); | - | - | | - | - | - | - | - | Unassigned | - |
| Otu00476 | 201 | k__Fungi(100);p__Basidiomycota(100);c__Agaricomycetes(100);o__Hymenochaetales(100);f__Hymenochaetaceae(100);g__Phellinus(100);s__Phellinus_betulinus_subsp_orienticus(100); | Phellinus | | 13 | Pathotroph-Saprotroph | \|Plant Pathogen\|-Wood Saprotroph | Polyporoid | White Rot | Highly Probable | [illegible] | [illegible] |
| Otu00475 | 201 | k__Fungi(100);k__Fungi_unclassified(99);k__Fungi_unclassified(99);k__Fungi_unclassified(99);k__Fungi_unclassified(99);k__Fungi_unclassified(99);k__Fungi_unclassified(99); | - | - | | - | - | - | - | - | Unassigned | - |
| Otu00477 | 199 | k__Fungi(100);p__Basidiomycota(100);c__Cystobasidiomycetes(100);o__Erythrobasidiales(100);f__Erythrobasidiales_family_Incertae_sedis(100);g__Erythrobasidium(100);s__Erythrobasidium_hasegawianum(100); | Erythrobasidium | | 13 | Pathotroph-Saprotroph | \|Fungal Parasite\|-Undefined Saprotroph | Yeast | NULL | Possible | [illegible] | [illegible] |
| Otu00479 | 198 | k__Fungi(100);p__Ascomycota(100);c__Saccharomycetes(100);o__Saccharomycetales(100);f__Saccharomycetales_family_Incertae_sedis(100);g__Candida(100);g__Candida_unclassified(56); | Candida | | 13 | Pathotroph-Saprotroph-Symbiotroph | Animal Pathogen-Endophyte-Endosymbiont-Epiphyte-\|Undefined Saprotroph\| | Dimorphic Yeast-Yeast | NULL | Probable | [illegible] | [illegible] |
| Otu00478 | 198 | k__Fungi(100);k__Fungi_unclassified(100);k__Fungi_unclassified(100);k__Fungi_unclassified(100);k__Fungi_unclassified(100);k__Fungi_unclassified(100);k__Fungi_unclassified(100); | - | - | | - | - | - | - | - | Unassigned | - |
| Otu00480 | 198 | k__Fungi(100);p__Ascomycota(100);c__Dothideomycetes(100);o__Capnodiales(100);o__Capnodiales_unclassified(100);o__Capnodiales_unclassified(100);o__Capnodiales_unclassified(100); | - | - | | - | - | - | - | - | Unassigned | - |
| Otu00481 | 197 | k__Fungi(100);p__Ascomycota(100);c__Saccharomycetes(100);o__Saccharomycetales(100);f__Metschnikowiaceae(100);g__Metschnikowia(100);s__Metschnikowia_pulcherrima(97); | Metschnikowia | | 13 | Saprotroph | \|Undefined Saprotroph\| | Yeast | NULL | Probable | [illegible] | [illegible] |
| Otu00482 | 196 | k__Fungi(100);p__Basidiomycota(100);c__Agaricomycetes(100);o__Agaricales(100);f__Strophariaceae(100);g__Hebeloma(100);g__Hebeloma_unclassified(100); | Hebeloma | | 13 | Symbiotroph | \|Ectomycorrhizal\| | Agaricoid | NULL | Highly Probable | NULL | [illegible] |
| Otu00483 | 196 | k__Plantae(100);p__unclassified_Plantae(100);c__unclassified_Plantae(100);o__unclassified_Plantae(100);f__unclassified_Plantae(100);g__unclassified_Plantae(100);s__Plantae_sp(100); | - | - | | - | - | - | - | - | Unassigned | - |
| Otu00484 | 195 | k__Fungi(100);p__Ascomycota(100);c__Dothideomycetes(100);o__Capnodiales(100);f__Mycosphaerellaceae(100);g__Septoria(100);s__Septoria_sp(100); | Septoria | | 13 | Pathotroph-Saprotroph-Symbiotroph | Endophyte-\|Plant Pathogen\|-Plant Saprotroph | Microfungus | NULL | Probable | [illegible] | [illegible] |
| Otu00486 | 194 | k__Fungi(100);p__Ascomycota(100);c__Dothideomycetes(100);o__Dothideales(100);f__Dothioraceae(100);g__Aureobasidium(100);s__Aureobasidium_pullulans(100); | Aureobasidium | | 13 | Pathotroph-Saprotroph-Symbiotroph | Animal Pathogen-Endophyte-Epiphyte-\|Plant Pathogen\|-Plant Saprotroph-Undefined Saprotroph | Dimorphic-Facultative Yeast | NULL | Possible | [illegible] | [illegible] |
| Otu00485 | 194 | k__Fungi(100);k__Fungi_unclassified(100);k__Fungi_unclassified(100);k__Fungi_unclassified(100);k__Fungi_unclassified(100);k__Fungi_unclassified(100);k__Fungi_unclassified(100); | - | - | | - | - | - | - | - | Unassigned | - |
| Otu00487 | 193 | k__Fungi(100);p__Ascomycota(100);p__Ascomycota_unclassified(100);p__Ascomycota_unclassified(100);p__Ascomycota_unclassified(100);p__Ascomycota_unclassified(100);p__Ascomycota_unclassified(100); | - | - | | - | - | - | - | - | Unassigned | - |
| Otu00488 | 193 | k__Fungi(100);k__Fungi_unclassified(98);k__Fungi_unclassified(98);k__Fungi_unclassified(98);k__Fungi_unclassified(98);k__Fungi_unclassified(98);k__Fungi_unclassified(98); | - | - | | - | - | - | - | - | Unassigned | - |
| Otu00489 | 192 | k__Fungi(100);p__Ascomycota(67);p__Ascomycota_unclassified(67);p__Ascomycota_unclassified(67);p__Ascomycota_unclassified(67);p__Ascomycota_unclassified(67);p__Ascomycota_unclassified(67); | - | - | | - | - | - | - | - | Unassigned | - |
| Otu00490 | 191 | k__Fungi(100);p__Ascomycota(100);c__Saccharomycetes(100);o__Saccharomycetales(100);f__Metschnikowiaceae(100);g__Metschnikowia(100);s__Metschnikowia_pulcherrima(97); | Metschnikowia | | 13 | Saprotroph | \|Undefined Saprotroph\| | Yeast | NULL | Probable | [illegible] | [illegible] |
| Otu00491 | 191 | k__Fungi(100);p__Basidiomycota(100);c__Agaricomycetes(100);o__Polyporales(100);f__Ganodermataceae(100);g__Ganoderma(100);s__Ganoderma_lucidum(100); | Ganoderma | | 13 | Pathotroph-Saprotroph | \|Plant Pathogen\|-Wood Saprotroph | Polyporoid | White Rot | Highly Probable | [illegible] | [illegible] |
| Otu00492 | 191 | k__Fungi(100);p__Ascomycota(100);c__Eurotiomycetes(100);o__Eurotiales(100);f__Trichocomaceae(100);g__Penicillium(100);s__Penicillium_herquei(88); | Penicillium | | 13 | Pathotroph-Saprotroph-Symbiotroph | Animal Parasite-Dung Saprotroph-Endophyte-Fungal Parasite-Plant Pathogen-\|Undefined Saprotroph\|-Wood Saprotroph | Microfungus | Soft Rot | Highly Probable | [illegible] | [illegible] |
| Otu00493 | 191 | k__Fungi(100);p__Basidiomycota(100);c__Agaricomycetes(100);o__Polyporales(100);f__Meruliaceae(100);g__Hyphoderma(100);s__Hyphoderma_setigerum(100); | Hyphoderma | | 13 | Saprotroph | Undefined Saprotroph-\|Wood Saprotroph\| | Corticioid | White Rot | Probable | [illegible] | [illegible] |
| Otu00495 | 191 | k__Fungi(100);p__Basidiomycota(100);c__Agaricomycetes(100);o__Agaricales(100);f__Tricholomataceae(100);g__Lepista(93);s__Lepista_personata(93); | Lepista | | 13 | Saprotroph | \|Plant Saprotroph\|-Undefined Saprotroph | Agaricoid | NULL | Probable | [illegible] | [illegible] |
| Otu00496 | 191 | k__Fungi(100);p__Ascomycota(100);c__Dothideomycetes(100);o__Capnodiales(100);f__Capnodiales_family_Incertae_sedis(100);g__Pseudotaeniolina(100);s__Pseudotaeniolina_globosa(100); | Pseudotaeniolina | | 13 | Saprotroph-Symbiotroph | Endophyte-\|Plant Saprotroph\| | Microfungus | NULL | Probable | [illegible] | [illegible] |
| Otu00497 | 191 | k__Fungi(100);p__Ascomycota(100);c__Dothideomycetes(100);o__Pleosporales(100);f__Lophiostomataceae(100);g__Lophiostoma(100);s__Lophiostoma_cf_cynaroidis_A33(97); | Lophiostoma | | 13 | Saprotroph | Plant Saprotroph-Undefined Saprotroph-\|Wood Saprotroph\| | Microfungus | NULL | Probable | [illegible] | [illegible] |
| Otu00494 | 191 | k__Fungi(100);p__Ascomycota(100);c__Dothideomycetes(100);o__Pleosporales(100);o__Pleosporales_unclassified(100);o__Pleosporales_unclassified(100);o__Pleosporales_unclassified(100); | - | - | | - | - | - | - | - | Unassigned | - |
| Otu00498 | 190 | k__Fungi(100);p__Ascomycota(100);c__Taphrinomycetes(100);o__Taphrinales(100);f__Taphrinaceae(75);g__Taphrina(75);g__Taphrina_unclassified(75); | Taphrina | | 13 | Pathotroph | \|Plant Pathogen\| | Dimorphic-Microfungus | NULL | Probable | NULL | [illegible] |
| Otu00500 | 190 | k__Fungi(100);p__Ascomycota(100);c__Ascomycota_class_Incertae_sedis(100);o__Ascomycota_order_Incertae_sedis(100);f__Ascomycota_family_Incertae_sedis(100);g__Hansfordia(100);s__Hansfordia_pulvinata(100); | Hansfordia | | 13 | Saprotroph | \|Undefined Saprotroph\| | Microfungus | NULL | Probable | NULL | [illegible] |
| Otu00501 | 190 | k__Fungi(100);p__Zygomycota(100);c__Zygomycota_class_Incertae_sedis(100);o__Mucorales(100);f__Cunninghamellaceae(100);g__Absidia(100);s__Absidia_glauca(100); | Absidia | | 13 | Saprotroph | \|Undefined Saprotroph\| | Microfungus | NULL | Probable | [illegible] | [illegible] |
| Otu00502 | 190 | k__Fungi(100);p__Ascomycota(100);c__Dothideomycetes(100);o__Pleosporales(100);f__Pleosporales_family_Incertae_sedis(100);g__Setomelanomma(100);s__Setomelanomma_holmii(98); | Setomelanomma | | 13 | Pathotroph-Saprotroph | \|Plant Pathogen\|-Wood Saprotroph | Microfungus | NULL | Probable | [illegible] | [illegible] |
| Otu00499 | 190 | k__Fungi(100);p__Ascomycota(100);c__Sordariomycetes(100);o__Trichosphaeriales(100);f__Trichosphaeriales_family_Incertae_sedis(100);g__unclassified_Trichosphaeriales(100);s__Trichosphaeriales_sp(100); | - | - | | - | - | - | - | - | Unassigned | - |
| Otu00503 | 189 | k__Fungi(100);p__Ascomycota(100);c__Dothideomycetes(100);o__Pleosporales(100);o__Pleosporales_unclassified(100);o__Pleosporales_unclassified(100);o__Pleosporales_unclassified(100); | - | - | | - | - | - | - | - | Unassigned | - |
| Otu00504 | 188 | k__Fungi(100);p__Basidiomycota(100);c__Microbotryomycetes(100);o__Sporidiobolales(100);f__Sporidiobolales_family_Incertae_sedis(100);g__Rhodosporidium(100);s__Rhodosporidium_paludigenum(100); | Rhodosporidium | | 13 | Saprotroph | Undefined Saprotroph | Yeast | NULL | Highly Probable | NULL | [illegible] |
| Otu00505 | 187 | k__Fungi(100);p__Basidiomycota(100);c__Agaricomycetes(100);o__Agaricales(100);f__Psathyrellaceae(100);g__Psathyrella(100);s__Psathyrella_candolleana(99); | Psathyrella | | 13 | Saprotroph | Plant Saprotroph-\|Wood Saprotroph\| | Agaricoid | NULL | Probable | [illegible] | [illegible] |
| Otu00506 | 186 | k__Fungi(100);k__Fungi_unclassified(75);k__Fungi_unclassified(75);k__Fungi_unclassified(75);k__Fungi_unclassified(75);k__Fungi_unclassified(75);k__Fungi_unclassified(75); | - | - | | - | - | - | - | - | Unassigned | - |
| Otu00507 | 185 | k__Fungi(100);p__Basidiomycota(100);c__Agaricostilbomycetes(100);o__Agaricostilbales(99);f__Agaricostilbaceae(99);g__Bensingtonia(92);s__Bensingtonia_sakaguchii(84); | Bensingtonia | | 13 | Saprotroph | \|Undefined Saprotroph\| | Dimorphic-Facultative Yeast | NULL | Highly Probable | NULL | [illegible] |
| Otu00508 | 185 | k__Fungi(100);p__Ascomycota(100);c__Sordariomycetes(100);o__Sordariomycetes_order_Incertae_sedis(100);f__Plectosphaerellaceae(100);g__unclassified_Plectosphaerellaceae(100);s__Plectosphaerellaceae_sp(100); | Plectosphaerellaceae | | 13 | Pathotroph-Saprotroph-Symbiotroph | Animal Pathogen-Endophyte-Fungal Parasite-Plant Pathogen-Wood Saprotroph | Microfungus | NULL | Possible | NULL | [illegible] |
| Otu00509 | 184 | k__Fungi(100);p__Ascomycota(100);c__Ascomycota_class_Incertae_sedis(100);o__Ascomycota_order_Incertae_sedis(100);f__Ascomycota_family_Incertae_sedis(100);g__Zymoseptoria(100);s__Zymoseptoria_verkleyi(99); | Zymoseptoria | | 13 | Pathotroph-Saprotroph | \|Plant Pathogen\|-Plant Saprotroph | Microfungus | NULL | Probable | [illegible] | [illegible] |
| Otu00510 | 184 | k__Fungi(100);p__Ascomycota(100);c__Sordariomycetes(100);o__Xylariales(100);f__Graphostromataceae(100);g__Graphostroma(100);s__Graphostroma_platystoma(100); | Graphostroma | | 13 | Pathotroph-Saprotroph-Symbiotroph | Endophyte-Plant Pathogen-Undefined Saprotroph-\|Wood Saprotroph\| | Microfungus | NULL | Probable | [illegible] | [illegible] |
| Otu00511 | 182 | k__Fungi(100);p__Basidiomycota(100);c__Tremellomycetes(100);o__Tremellales(100);f__Tremellales_family_Incertae_sedis(100);f__Tremellales_family_Incertae_sedis_unclassified(74);f__Tremellales_family_Incertae_sedis_unclassified(74); | Tremellales | | 7 | Pathotroph-Saprotroph-Symbiotroph | Fungal Parasite-Undefined Saprotroph | Tremelloid-Yeast | NULL | Possible | [illegible] | [illegible] |
| Otu00512 | 181 | k__Fungi(100);p__Ascomycota(100);c__Saccharomycetes(94);o__Saccharomycetales(94);f__Saccharomycetales_family_Incertae_sedis(90);g__Candida(85);g__Candida_unclassified(83); | Candida | | 13 | Pathotroph-Saprotroph-Symbiotroph | Animal Pathogen-Endophyte-Endosymbiont-Epiphyte-\|Undefined Saprotroph\| | Dimorphic Yeast-Yeast | NULL | Probable | [illegible] | [illegible] |
| Otu00514 | 181 | k__Fungi(100);p__Basidiomycota(100);c__Agaricomycetes(100);o__Cantharellales(100);f__Ceratobasidiaceae(100);g__unclassified_Ceratobasidiaceae(59);s__Ceratobasidiaceae_sp(59); | Ceratobasidiaceae | | 9 | Pathotroph-Saprotroph-Symbiotroph | Endomycorrhizal-Plant Pathogen-Undefined Saprotroph | Microfungus | NULL | Possible | [illegible] | [illegible] |
| Otu00515 | 181 | k__Fungi(100);p__Basidiomycota(100);c__Tremellomycetes(100);o__Tremellales(100);f__Tremellales_family_Incertae_sedis(100);g__Dioszegia(100);s__Dioszegia_zsoltii_var._zsoltii(51); | Dioszegia | | 13 | Saprotroph | \|Plant Saprotroph\|-Undefined Saprotroph | Yeast | NULL | Probable | [illegible] | [illegible] |
| Otu00513 | 181 | k__Fungi(100);p__Ascomycota(100);p__Ascomycota_unclassified(100);p__Ascomycota_unclassified(100);p__Ascomycota_unclassified(100);p__Ascomycota_unclassified(100);p__Ascomycota_unclassified(100); | - | - | | - | - | - | - | - | Unassigned | - |
| Otu00516 | 181 | k__Fungi(100);p__Ascomycota(100);c__Taphrinomycetes(100);o__Taphrinales(100);f__unclassified_Taphrinales(76);g__unclassified_Taphrinales(76);s__Taphrinales_sp(76); | - | - | | - | - | - | - | - | Unassigned | - |
| Otu00517 | 179 | k__Fungi(100);k__Fungi_unclassified(73);k__Fungi_unclassified(73);k__Fungi_unclassified(73);k__Fungi_unclassified(73);k__Fungi_unclassified(73);k__Fungi_unclassified(73); | - | - | | - | - | - | - | - | Unassigned | - |
| Otu00519 | 178 | k__Fungi(100);p__Basidiomycota(100);c__Agaricomycetes(100);o__Russulales(100);f__Peniophoraceae(100);g__Peniophora(100);s__Peniophora_sp_J33(100); | Peniophora | | 13 | Pathotroph-Saprotroph | Plant Pathogen-\|Wood Saprotroph\| | Corticioid | White Rot | Probable | [illegible] | [illegible] |
| Otu00518 | 178 | k__Fungi(100);p__Ascomycota(84);c__Leotiomycetes(73);o__Helotiales(72);o__Helotiales_unclassified(72);o__Helotiales_unclassified(72);o__Helotiales_unclassified(72); | - | - | | - | - | - | - | - | Unassigned | - |
| Otu00520 | 178 | k__Fungi(100);p__Basidiomycota(100);c__Agaricomycetes(99);c__Agaricomycetes_unclassified(99);c__Agaricomycetes_unclassified(99);c__Agaricomycetes_unclassified(99);c__Agaricomycetes_unclassified(99); | - | - | | - | - | - | - | - | Unassigned | - |
| Otu00521 | 177 | k__Fungi(100);p__Basidiomycota(100);c__Agaricomycetes(100);o__Cantharellales(97);o__Cantharellales_unclassified(93);o__Cantharellales_unclassified(93);o__Cantharellales_unclassified(93); | - | - | | - | - | - | - | - | Unassigned | - |
| Otu00522 | 176 | k__Fungi(100);p__Ascomycota(100);c__Dothideomycetes(100);o__Capnodiales(100);o__Capnodiales_unclassified(100);o__Capnodiales_unclassified(100);o__Capnodiales_unclassified(100); | - | - | | - | - | - | - | - | Unassigned | - |
| Otu00523 | 175 | k__Fungi(100);p__Basidiomycota(100);c__Agaricomycetes(100);o__Agaricales(100);f__Pleurotaceae(100);g__unclassified_Pleurotaceae(100);s__Pleurotaceae_sp(100); | Pleurotaceae | | 9 | Pathotroph-Saprotroph | Endophyte-Plant Pathogen-Wood Saprotroph | Agaricoid | White Rot | Probable | [illegible] | [illegible] |
| Otu00525 | 174 | k__Fungi(100);p__Basidiomycota(100);c__Agaricomycetes(100);o__Boletales(100);f__Suillaceae(100);g__Suillus(100);s__Suillus_pictus(100); | Suillus | | 13 | Symbiotroph | \|Ectomycorrhizal\| | Agaricoid-Boletoid | NULL | Highly Probable | [illegible] | [illegible] |
| Otu00524 | 174 | k__Fungi(100);p__Ascomycota(100);p__Ascomycota_unclassified(92);p__Ascomycota_unclassified(92);p__Ascomycota_unclassified(92);p__Ascomycota_unclassified(92);p__Ascomycota_unclassified(92); | - | - | | - | - | - | - | - | Unassigned | - |
| Otu00526 | 173 | k__Fungi(100);p__Basidiomycota(100);c__Agaricomycetes(100);o__Agaricales(100);f__Psathyrellaceae(100);g__Psathyrella(100);s__Psathyrella_gossypina(100); | Psathyrella | | 13 | Saprotroph | Plant Saprotroph-\|Wood Saprotroph\| | Agaricoid | NULL | Probable | [illegible] | [illegible] |
| Otu00527 | 173 | k__Fungi(100);p__Ascomycota(90);p__Ascomycota_unclassified(90);p__Ascomycota_unclassified(90);p__Ascomycota_unclassified(90);p__Ascomycota_unclassified(90);p__Ascomycota_unclassified(90); | - | - | | - | - | - | - | - | Unassigned | - |
| Otu00528 | 172 | k__Fungi(100);p__Basidiomycota(100);c__Agaricomycetes(100);o__Auriculariales(100);f__Auriculariales_family_Incertae_sedis(100);g__Exidia(100);s__Exidia_sp(100); | Exidia | | 13 | Saprotroph-Symbiotroph | Endophyte-Plant Saprotroph-Undefined Saprotroph-\|Wood Saprotroph\| | Tremelloid | NULL | Probable | [illegible] | [illegible] |
| Otu00529 | 171 | k__Fungi(100);p__Ascomycota(100);c__Saccharomycetes(100);o__Saccharomycetales(100);f__Metschnikowiaceae(100);g__Metschnikowia(100);s__Metschnikowia_pulcherrima(97); | Metschnikowia | | 13 | Saprotroph | \|Undefined Saprotroph\| | Yeast | NULL | Probable | [illegible] | [illegible] |

| | | | | | | | | | | | | |
|---|---|---|---|---|---|---|---|---|---|---|---|---|
| Otu00530 | 171 | k__Fungi(100);p__Ascomycota(100);c__Eurotiomycetes(100);o__Eurotiales(100);f__Trichocomaceae(100);g__Aspergillus(100);s__Aspergillus_conicus(81); | Aspergillus | | 13 | Pathotroph-Saprotroph-Symbiotroph | Animal Pathogen-Endophyte-Plant Saprotroph-\|Undefined Saprotroph\|-Wood Saprotroph | Microfungus | NULL | Probable | [illegible] | [illegible] |
| Otu00531 | 171 | k__Fungi(100);p__Ascomycota(100);c__Dothideomycetes(100);o__Pleosporales(100);o__Pleosporales_unclassified(100);o__Pleosporales_unclassified(100);o__Pleosporales_unclassified(100); | - | - | | - | - | - | - | - | Unassigned | |
| Otu00532 | 170 | k__Fungi(100);p__Ascomycota(100);c__Dothideomycetes(100);o__Pleosporales(100);f__Lophiostomataceae(100);g__Lophiostoma(100);s__Lophiostoma_sp(100); | Lophiostoma | | 13 | Saprotroph | Plant Saprotroph-Undefined Saprotroph-\|Wood Saprotroph\| | Microfungus | NULL | Probable | [illegible] | [illegible] |
| Otu00533 | 170 | k__Fungi(100);p__Ascomycota(100);p__Ascomycota_unclassified(93);p__Ascomycota_unclassified(93);p__Ascomycota_unclassified(93);p__Ascomycota_unclassified(93);p__Ascomycota_unclassified(93); | - | - | | - | - | - | - | - | Unassigned | |
| Otu00536 | 169 | k__Fungi(100);p__Basidiomycota(100);c__Agaricomycetes(100);o__Agaricales(100);f__Psathyrellaceae(100);g__Psathyrella(100);s__Psathyrella_cernua(72); | Psathyrella | | 13 | Saprotroph | Plant Saprotroph-\|Wood Saprotroph\| | Agaricoid | NULL | Probable | [illegible] | [illegible] |
| Otu00534 | 169 | k__Fungi(100);k__Fungi_unclassified(58);k__Fungi_unclassified(58);k__Fungi_unclassified(58);k__Fungi_unclassified(58);k__Fungi_unclassified(58); | - | - | | - | - | - | - | - | Unassigned | |
| Otu00535 | 169 | k__Fungi(100);p__Ascomycota(100);c__Dothideomycetes(100);o__Pleosporales(100);o__Pleosporales_unclassified(98);o__Pleosporales_unclassified(98);o__Pleosporales_unclassified(98); | - | - | | - | - | - | - | - | Unassigned | |
| Otu00537 | 168 | k__Fungi(100);p__Basidiomycota(100);c__Exobasidiomycetes(100);o__Exobasidiales(100);f__Exobasidiaceae(100);g__Exobasidium(100);s__Exobasidium_sp(100); | Exobasidium | | 13 | Pathotroph | \|Plant Pathogen\| | Leaf Blister-Leaf Gall-Leaf Spot | NULL | Probable | NULL | [illegible] |
| Otu00538 | 167 | k__Fungi(100);p__Ascomycota(100);c__Dothideomycetes(100);o__Pleosporales(100);f__Montagnulaceae(100);g__unclassified_Montagnulaceae(100);s__Montagnulaceae_sp(100); | Montagnulaceae | | 9 | Pathotroph-Saprotroph | Fungal Parasite-Plant Pathogen-Undefined Saprotroph | Microfungus | NULL | Probable | [illegible] | [illegible] |
| Otu00539 | 167 | k__Fungi(100);p__Basidiomycota(100);c__Exobasidiomycetes(100);o__Exobasidiomycetes_order_Incertae_sedis(100);f__Exobasidiomycetes_family_Incertae_sedis(100);g__Meira(100);s__Meira_nashicola(98); | Meira | | 13 | Pathotroph | \|Animal Parasite\|-Animal Pathogen-Plant Pathogen | Dimorphic | NULL | Probable | [illegible] | [illegible] |
| Otu00540 | 166 | k__Fungi(100);p__Ascomycota(100);c__Eurotiomycetes(100);o__Eurotiales(99);f__Trichocomaceae(99);g__Talaromyces(96);g__Talaromyces_unclassified(92); | Talaromyces | | 13 | Pathotroph-Saprotroph | Animal Parasite-\|Undefined Saprotroph\| | Microfungus | NULL | Probable | [illegible] | [illegible] |
| Otu00541 | 166 | k__Fungi(100);p__Basidiomycota(100);c__Agaricomycetes(100);o__Russulales(100);f__Auriscalpiaceae(100);g__Artomyces(100);s__Artomyces_pyxidatus(100); | Artomyces | | 13 | Saprotroph | Undefined Saprotroph-\|Wood Saprotroph\| | Clavarioid | White Rot | Probable | [illegible] | [illegible] |
| Otu00542 | 165 | k__Fungi(100);p__Ascomycota(100);c__Leotiomycetes(100);o__Erysiphales(100);f__Erysiphaceae(100);g__Golovinomyces(100);s__Golovinomyces_magnicellulatus(100); | Golovinomyces | | 13 | Pathotroph | \|Plant Pathogen\| | Microfungus | NULL | Probable | [illegible] | [illegible] |
| Otu00543 | 165 | k__Fungi(100);k__Fungi_unclassified(54);k__Fungi_unclassified(54);k__Fungi_unclassified(54);k__Fungi_unclassified(54);k__Fungi_unclassified(54);k__Fungi_unclassified(54); | - | - | | - | - | - | - | - | Unassigned | |
| Otu00544 | 164 | k__Fungi(100);p__Basidiomycota(100);c__Tremellomycetes(100);o__Tremellales(100);f__Tremellales_family_Incertae_sedis(100);g__Bullera(100);s__Bullera_variabilis(100); | Bullera | | 13 | Saprotroph-Symbiotroph | Endophyte-Epiphyte-Plant Saprotroph-\|Undefined Saprotroph\| | Yeast | NULL | Probable | [illegible] | [illegible] |
| Otu00545 | 164 | k__Fungi(100);p__Basidiomycota(100);c__Exobasidiomycetes(100);o__Exobasidiales(100);f__Exobasidiaceae(100);g__Exobasidium(100);s__Exobasidium_inconspicuum(88); | Exobasidium | | 13 | Pathotroph | \|Plant Pathogen\| | Leaf Blister-Leaf Gall-Leaf Spot | NULL | Probable | NULL | [illegible] |
| Otu00546 | 163 | k__Fungi(100);p__Basidiomycota(100);c__Agaricomycetes(100);o__Auriculariales(100);f__Auriculariales_family_Incertae_sedis(100);g__Heterochaetella(100);s__Heterochaetella_sp(100); | Heterochaetella | | 13 | Saprotroph | Undefined Saprotroph-\|Wood Saprotroph\| | Corticioid | NULL | Probable | [illegible] | [illegible] |
| Otu00548 | 163 | k__Fungi(100);p__Ascomycota(100);c__Sordariomycetes(100);o__Xylariales(100);f__Amphisphaeriaceae(100);f__Amphisphaeriaceae_unclassified(59);f__Amphisphaeriaceae_unclassified(59); | Amphisphaeriaceae | | 9 | Saprotroph | Undefined Saprotroph | Microfungus | NULL | Possible | [illegible] | [illegible] |
| Otu00549 | 163 | k__Fungi(100);p__Ascomycota(100);c__Sordariomycetes(100);o__Xylariales(100);f__Xylariales_family_Incertae_sedis(100);g__Microdochium(99);s__Microdochium_sp_E9329c(99); | Microdochium | | 13 | Pathotroph-Symbiotroph | Endophyte-\|Plant Pathogen\| | Microfungus | NULL | Possible | [illegible] | [illegible] |
| Otu00550 | 163 | k__Fungi(100);p__Ascomycota(100);c__Saccharomycetes(100);o__Saccharomycetales(100);f__Metschnikowiaceae(100);g__Metschnikowia(100);s__Metschnikowia_pulcherrima(99); | Metschnikowia | | 13 | Saprotroph | \|Undefined Saprotroph\| | Yeast | NULL | Probable | [illegible] | [illegible] |
| Otu00547 | 163 | k__Fungi(100);k__Fungi_unclassified(56);k__Fungi_unclassified(56);k__Fungi_unclassified(56);k__Fungi_unclassified(56);k__Fungi_unclassified(56);k__Fungi_unclassified(56); | - | - | | - | - | - | - | - | Unassigned | |
| Otu00551 | 162 | k__Fungi(100);p__Ascomycota(100);c__Saccharomycetes(100);o__Saccharomycetales(100);f__Debaryomycetaceae(100);g__Millerozyma(100);s__Millerozyma_farinosa(100); | Millerozyma | | 13 | Saprotroph | \|Undefined Saprotroph\| | Yeast | NULL | Probable | [illegible] | [illegible] |
| Otu00552 | 162 | k__Fungi(100);p__Ascomycota(94);c__Dothideomycetes(94);o__Pleosporales(92);o__Pleosporales_unclassified(92);o__Pleosporales_unclassified(92);o__Pleosporales_unclassified(92); | - | - | | - | - | - | - | - | Unassigned | |
| Otu00554 | 161 | k__Fungi(100);p__Ascomycota(100);c__Sordariomycetes(100);o__Xylariales(56);f__Xylariales_family_Incertae_sedis(56);g__Monographella(56);s__Monographella_cucumerina(56); | Monographella | | 13 | Pathotroph | \|Plant Pathogen\| | Microfungus | NULL | Probable | NULL | [illegible] |
| Otu00555 | 161 | k__Fungi(100);p__Ascomycota(100);c__Eurotiomycetes(100);o__Chaetothyriales(100);f__Herpotrichiellaceae(100);g__Capronia(87);s__Capronia_sp_CBS_16492(87); | Capronia | | 13 | Pathotroph-Saprotroph-Symbiotroph | Animal Pathogen-Endophyte-Fungal Parasite-\|Undefined Saprotroph\| | Dimorphic-Facultative Yeast | NULL | Highly Probable | [illegible] | [illegible] |
| Otu00553 | 161 | k__Fungi(100);p__Ascomycota(100);p__Ascomycota_unclassified(100);p__Ascomycota_unclassified(100);p__Ascomycota_unclassified(100);p__Ascomycota_unclassified(100);p__Ascomycota_unclassified(100); | - | - | | - | - | - | - | - | Unassigned | |
| Otu00556 | 160 | k__Fungi(100);p__Basidiomycota(100);c__Tremellomycetes(100);o__Trichosporonales(100);f__Trichosporonaceae(100);g__Trichosporon(100);g__Trichosporon_unclassified(100); | Trichosporon | | 13 | Pathotroph-Saprotroph | \|Animal Parasite\|-Animal Pathogen-Undefined Saprotroph | Yeast | NULL | Probable | [illegible] | [illegible] |
| Otu00557 | 159 | k__Fungi(100);p__Ascomycota(100);c__Sordariomycetes(100);o__Diaporthales(100);f__Diaporthales_family_Incertae_sedis(100);g__Ophiognomonia(100);s__Ophiognomonia_rosae(85); | Ophiognomonia | | 13 | Pathotroph | \|Plant Pathogen\| | Microfungus | NULL | Highly Probable | [illegible] | [illegible] |
| Otu00558 | 159 | k__Fungi(100);p__Ascomycota(100);c__Sordariomycetes(100);o__Xylariales(100);f__Xylariaceae(100);g__Lopadostoma(100);g__Lopadostoma_unclassified(95); | Lopadostoma | | 13 | Saprotroph-Symbiotroph | Endophyte-Undefined Saprotroph-\|Wood Saprotroph\| | Microfungus | NULL | Probable | [illegible] | [illegible] |
| Otu00560 | 157 | k__Fungi(100);p__Ascomycota(100);c__Dothideomycetes(100);o__Pleosporales(100);f__Massarinaceae(99);g__unclassified_Massarinaceae(78);s__Massarinaceae_sp(78); | Massarinaceae | | 9 | Pathotroph-Saprotroph | Plant Pathogen-Undefined Saprotroph | Microfungus | NULL | Probable | NULL | [illegible] |
| Otu00559 | 157 | k__Fungi(100);p__Ascomycota(96);p__Ascomycota_unclassified(95);p__Ascomycota_unclassified(95);p__Ascomycota_unclassified(95);p__Ascomycota_unclassified(95);p__Ascomycota_unclassified(95); | - | - | | - | - | - | - | - | Unassigned | |
| Otu00561 | 156 | k__Fungi(100);p__Ascomycota(100);c__Eurotiomycetes(100);o__Eurotiales(100);f__Trichocomaceae(100);g__Penicillium(100);s__Penicillium_charlesii(100); | Penicillium | | 13 | Pathotroph-Saprotroph-Symbiotroph | Animal Parasite-Dung Saprotroph-Endophyte-Fungal Parasite-Plant Pathogen-\|Undefined Saprotroph\|-Wood Saprotroph | Microfungus | Soft Rot | Highly Probable | [illegible] | [illegible] |
| Otu00562 | 155 | k__Fungi(100);p__Basidiomycota(100);c__Agaricomycetes(100);o__Corticiales(100);f__Corticiaceae(100);g__Laetisaria(100);s__Laetisaria_arvalis(100); | Laetisaria | | 13 | Pathotroph-Saprotroph | \|Plant Pathogen\|-Wood Saprotroph | Corticioid | NULL | Probable | [illegible] | [illegible] |
| Otu00565 | 155 | k__Fungi(100);p__Zygomycota(100);c__Zygomycota_class_Incertae_sedis(100);o__Mortierellales(100);f__Mortierellaceae(100);g__Mortierella(100);g__Mortierella_unclassified(57); | Mortierella | | 13 | Saprotroph-Symbiotroph | Endophyte-Plant Saprotroph-\|Undefined Saprotroph\| | Microfungus | NULL | Probable | [illegible] | [illegible] |
| Otu00563 | 155 | k__Fungi(100);p__Basidiomycota(100);p__Basidiomycota_unclassified(100);p__Basidiomycota_unclassified(100);p__Basidiomycota_unclassified(100);p__Basidiomycota_unclassified(100);p__Basidiomycota_unclassified(100); | - | - | | - | - | - | - | - | Unassigned | |
| Otu00564 | 155 | k__Fungi(100);p__Ascomycota(100);c__Eurotiomycetes(100);o__Eurotiales(100);f__Trichocomaceae(100);f__Trichocomaceae_unclassified(100);f__Trichocomaceae_unclassified(100); | - | - | | - | - | - | - | - | Unassigned | |
| Otu00567 | 154 | k__Fungi(100);p__Chytridiomycota(100);c__Chytridiomycetes(100);o__Rhizophlyctidales(100);f__Rhizophlyctidaceae(100);g__Rhizophlyctis(100);s__Rhizophlyctis_rosea(100); | Rhizophlyctis | | 13 | Pathotroph-Saprotroph | Algal Parasite-\|Plant Saprotroph\|-Undefined Saprotroph | Chytrid-Microfungus | NULL | Probable | [illegible] | [illegible] |
| Otu00566 | 154 | k__Fungi(100);p__Basidiomycota(100);c__Agaricomycetes(100);o__Polyporales(100);f__Phanerochaetaceae(96);f__Phanerochaetaceae_unclassified(95);f__Phanerochaetaceae_unclassified(95); | - | - | | - | - | - | - | - | Unassigned | |
| Otu00568 | 153 | k__Fungi(100);p__Basidiomycota(100);c__Agaricomycetes(100);o__Polyporales(100);f__Meruliaceae(100);g__Phlebia(97);g__Phlebia_unclassified(85); | Phlebia | | 13 | Saprotroph | \|Wood Saprotroph\| | Corticioid | White Rot | Highly Probable | [illegible] | [illegible] |
| Otu00571 | 153 | k__Fungi(100);p__Ascomycota(100);c__Dothideomycetes(100);o__Pleosporales(100);f__Leptosphaeriaceae(100);g__unclassified_Leptosphaeriaceae(99);s__Leptosphaeriaceae_sp(99); | Leptosphaeriaceae | | 9 | Pathotroph-Saprotroph | Plant Pathogen-Wood Saprotroph | Microfungus | NULL | Probable | [illegible] | [illegible] |
| Otu00572 | 153 | k__Fungi(100);p__Ascomycota(100);c__Sordariomycetes(100);o__Hypocreales(100);f__Hypocreales_family_Incertae_sedis(100);g__Acremonium(100);s__Acremonium_sp_24_OA_2013(100); | Acremonium | | 13 | Pathotroph-Saprotroph-Symbiotroph | Animal Pathogen-Endophyte-Fungal Parasite-Plant Pathogen-\|Undefined Saprotroph\|-Wood Saprotroph | Microfungus | Soft Rot | Possible | [illegible] | [illegible] |
| Otu00573 | 153 | k__Fungi(100);p__Basidiomycota(100);c__Exobasidiomycetes(100);o__Microstromatales(100);f__Microstromataceae(100);g__Microstroma(100);s__Microstroma_juglandis(100); | Microstroma | | 13 | Pathotroph | \|Plant Pathogen\| | Dimorphic | NULL | Probable | NULL | [illegible] |
| Otu00569 | 153 | k__Fungi(100);k__Fungi_unclassified(60);k__Fungi_unclassified(60);k__Fungi_unclassified(60);k__Fungi_unclassified(60);k__Fungi_unclassified(60);k__Fungi_unclassified(60); | - | - | | - | - | - | - | - | Unassigned | |
| Otu00570 | 153 | k__Fungi(100);p__Ascomycota(100);c__Dothideomycetes(100);o__Pleosporales(100);f__unclassified_Pleosporales(95);g__unclassified_Pleosporales(95);s__Pleosporales_sp(95); | - | - | | - | - | - | - | - | Unassigned | |
| Otu00574 | 152 | k__Fungi(100);p__Ascomycota(100);c__Eurotiomycetes(100);o__Chaetothyriales(100);f__Chaetothyriales_family_Incertae_sedis(100);g__Strelitziana(100);s__Strelitziana_albiziae(100); | Strelitziana | | 13 | Pathotroph | \|Plant Pathogen\| | Microfungus | NULL | Probable | NULL | [illegible] |
| Otu00577 | 152 | k__Fungi(100);p__Basidiomycota(100);c__Agaricomycetes(100);o__Trechisporales(100);f__Hydnodontaceae(100);g__Brevicellicium(100);s__Brevicellicium_olivascens(100); | Brevicellicium | | 13 | Saprotroph | Undefined Saprotroph-\|Wood Saprotroph\| | Corticioid | White Rot | Probable | [illegible] | [illegible] |
| Otu00575 | 152 | k__Fungi(100);k__Fungi_unclassified(99);k__Fungi_unclassified(99);k__Fungi_unclassified(99);k__Fungi_unclassified(99);k__Fungi_unclassified(99);k__Fungi_unclassified(99); | - | - | | - | - | - | - | - | Unassigned | |
| Otu00576 | 152 | k__Fungi(100);p__Ascomycota(95);p__Ascomycota_unclassified(95);p__Ascomycota_unclassified(95);p__Ascomycota_unclassified(95);p__Ascomycota_unclassified(95);p__Ascomycota_unclassified(95); | - | - | | - | - | - | - | - | Unassigned | |
| Otu00579 | 149 | k__Fungi(100);p__Basidiomycota(100);c__Agaricomycetes(100);o__Polyporales(100);f__Polyporaceae(100);g__Skeletocutis(100);s__Skeletocutis_chrysella(90); | Skeletocutis | | 13 | Saprotroph | \|Wood Saprotroph\| | Polyporoid | White Rot | Highly Probable | [illegible] | [illegible] |
| Otu00578 | 149 | k__Fungi(100);p__Ascomycota(83);p__Ascomycota_unclassified(83);p__Ascomycota_unclassified(83);p__Ascomycota_unclassified(83);p__Ascomycota_unclassified(83);p__Ascomycota_unclassified(83); | - | - | | - | - | - | - | - | Unassigned | |
| Otu00581 | 148 | k__Fungi(100);p__Basidiomycota(100);c__Agaricomycetes(100);o__Polyporales(100);f__Phanerochaetaceae(100);g__Phanerochaete(100);s__Phanerochaete_stereoides(94); | Phanerochaete | | 13 | Saprotroph | \|Wood Saprotroph\| | Corticioid | White Rot | Highly Probable | [illegible] | [illegible] |
| Otu00580 | 148 | k__Fungi(100);p__Ascomycota(100);p__Ascomycota_unclassified(100);p__Ascomycota_unclassified(100);p__Ascomycota_unclassified(100);p__Ascomycota_unclassified(100);p__Ascomycota_unclassified(100); | - | - | | - | - | - | - | - | Unassigned | |
| Otu00582 | 147 | k__Fungi(100);p__Basidiomycota(100);c__Agaricomycetes(100);o__Polyporales(100);f__Fomitopsidaceae(100);g__Laetiporus(100);s__Laetiporus_sulphureus(100); | Laetiporus | | 13 | Pathotroph-Saprotroph | \|Plant Pathogen\|-Wood Saprotroph | Polyporoid | Brown Rot | Highly Probable | [illegible] | [illegible] |
| Otu00583 | 147 | k__Fungi(100);p__Ascomycota(100);c__Saccharomycetes(100);o__Saccharomycetales(100);f__Metschnikowiaceae(100);g__Metschnikowia(100);s__Metschnikowia_pulcherrima(99); | Metschnikowia | | 13 | Saprotroph | \|Undefined Saprotroph\| | Yeast | NULL | Probable | [illegible] | [illegible] |
| Otu00584 | 146 | k__Fungi(100);p__Ascomycota(100);c__Saccharomycetes(100);o__Saccharomycetales(100);f__Metschnikowiaceae(100);g__Metschnikowia(100);s__Metschnikowia_pulcherrima(96); | Metschnikowia | | 13 | Saprotroph | \|Undefined Saprotroph\| | Yeast | NULL | Probable | [illegible] | [illegible] |
| Otu00585 | 146 | k__Fungi(100);p__Basidiomycota(100);c__Agaricomycetes(100);o__Polyporales(100);f__Xenasmataceae(100);g__Xenasmatella(100);s__Xenasmatella_sp(100); | Xenasmatella | | 13 | Saprotroph | Undefined Saprotroph-\|Wood Saprotroph\| | Corticioid | White Rot | Probable | [illegible] | [illegible] |
| Otu00586 | 146 | k__Fungi(100);p__Zygomycota(100);c__Zygomycota_class_Incertae_sedis(100);o__Mortierellales(100);f__Mortierellaceae(100);g__Mortierella(100);g__Mortierella_unclassified(100); | Mortierella | | 13 | Saprotroph-Symbiotroph | Endophyte-Plant Saprotroph-\|Undefined Saprotroph\| | Microfungus | NULL | Probable | [illegible] | [illegible] |
| Otu00587 | 146 | k__Fungi(100);k__Fungi_unclassified(57);k__Fungi_unclassified(57);k__Fungi_unclassified(57);k__Fungi_unclassified(57);k__Fungi_unclassified(57);k__Fungi_unclassified(57); | - | - | | - | - | - | - | - | Unassigned | |
| Otu00588 | 145 | k__Fungi(100);p__Basidiomycota(100);c__Agaricomycetes(100);o__Polyporales(100);f__Polyporaceae(100);g__Melanoporia(100);s__Melanoporia_nigra(100); | Melanoporia | | 13 | Saprotroph | \|Wood Saprotroph\| | Polyporoid | White Rot | Highly Probable | [illegible] | [illegible] |
| Otu00589 | 145 | k__Fungi(100);p__Basidiomycota(100);c__Agaricomycetes(100);o__Hymenochaetales(100);f__Schizoporaceae(100);g__Hyphodontia(100);s__Hyphodontia_radula(100); | Hyphodontia | | 13 | Saprotroph | Undefined Saprotroph-\|Wood Saprotroph\| | Corticioid | White Rot | Probable | [illegible] | [illegible] |
| Otu00590 | 145 | k__Fungi(100);p__Ascomycota(100);c__Sordariomycetes(100);o__Xylariales(100);f__Amphisphaeriaceae(99);g__Pestalotiopsis(82);s__Pestalotiopsis_sp_YM1(72); | Pestalotiopsis | | 13 | Pathotroph | \|Plant Pathogen\| | Microfungus | NULL | Probable | NULL | [illegible] |
| Otu00591 | 144 | k__Fungi(100);p__Basidiomycota(100);c__Agaricomycetes(100);o__Agaricales(100);f__Psathyrellaceae(100);g__Coprinopsis(100);s__Coprinopsis_romagnesiana(85); | Coprinopsis | | 13 | Saprotroph | \|Undefined Saprotroph\| | Agaricoid | NULL | Probable | [illegible] | [illegible] |
| Otu00592 | 144 | k__Fungi(100);p__Ascomycota(100);c__Sordariomycetes(100);o__Hypocreales(100);f__Hypocreaceae(100);g__Hypocrea(95);s__Hypocrea_lixii(93); | Hypocrea | | 13 | Pathotroph | Endophyte-Fungal Parasite | Microfungus | NULL | Probable | NULL | [illegible] |
| Otu00595 | 144 | k__Fungi(100);p__Basidiomycota(100);c__Agaricomycetes(100);o__Russulales(100);f__Stereaceae(100);g__Stereum(100);s__Stereum_sp(100); | Stereum | | 13 | Saprotroph | Undefined Saprotroph-\|Wood Saprotroph\| | Polyporoid | White Rot | Probable | [illegible] | [illegible] |
| Otu00593 | 144 | k__Fungi(100);k__Fungi_unclassified(67);k__Fungi_unclassified(67);k__Fungi_unclassified(67);k__Fungi_unclassified(67);k__Fungi_unclassified(67);k__Fungi_unclassified(67); | - | - | | - | - | - | - | - | Unassigned | |
| Otu00594 | 144 | k__Fungi(100);p__Basidiomycota(97);p__Basidiomycota_unclassified(97);p__Basidiomycota_unclassified(97);p__Basidiomycota_unclassified(97);p__Basidiomycota_unclassified(97);p__Basidiomycota_unclassified(97); | - | - | | - | - | - | - | - | Unassigned | |
| Otu00596 | 143 | k__Fungi(100);k__Fungi_unclassified(77);k__Fungi_unclassified(77);k__Fungi_unclassified(77);k__Fungi_unclassified(77);k__Fungi_unclassified(77);k__Fungi_unclassified(77); | - | - | | - | - | - | - | - | Unassigned | |
| Otu00597 | 143 | k__Fungi(100);p__Basidiomycota(100);c__Agaricomycetes(100);o__Russulales(100);o__Russulales_unclassified(81);o__Russulales_unclassified(81);o__Russulales_unclassified(81); | - | - | | - | - | - | - | - | Unassigned | |
| Otu00598 | 142 | k__Fungi(100);p__Ascomycota(100);c__Sordariomycetes(100);o__Sordariales(100);f__Chaetomiaceae(100);g__Chaetomium(100);g__Chaetomium_unclassified(83); | Chaetomium | | 13 | Pathotroph-Saprotroph-Symbiotroph | Animal Pathogen-Dung Saprotroph-Endophyte-Epiphyte-Plant Pathogen-\|Plant Saprotroph\|-Wood Saprotroph | Microfungus | Soft Rot | Possible | [illegible] | [illegible] |
| Otu00599 | 142 | k__Fungi(100);p__Zygomycota(100);c__Zygomycota_class_Incertae_sedis(100);o__Kickxellales(100);f__Kickxellaceae(100);g__Ramicandelaber(100);s__Ramicandelaber_sp(79); | Ramicandelaber | | 13 | Saprotroph | \|Undefined Saprotroph\| | Microfungus | NULL | Probable | [illegible] | [illegible] |
| Otu00600 | 142 | k__Fungi(100);p__Zygomycota(100);c__Zygomycota_class_Incertae_sedis(100);o__Mortierellales(100);f__Mortierellaceae(100);g__Mortierella(100);g__Mortierella_unclassified(100); | Mortierella | | 13 | Saprotroph-Symbiotroph | Endophyte-Plant Saprotroph-\|Undefined Saprotroph\| | Microfungus | NULL | Probable | [illegible] | [illegible] |
| Otu00601 | 142 | k__Fungi(100);p__Ascomycota(100);c__Sordariomycetes(100);o__Hypocreales(100);f__Hypocreales_family_Incertae_sedis(100);g__Myrothecium(100);s__Myrothecium_tongaense(100); | Myrothecium | | 13 | Pathotroph-Saprotroph | Fungal Parasite-\|Plant Saprotroph\|-Undefined Saprotroph | Microfungus | NULL | Probable | [illegible] | [illegible] |
| Otu00602 | 141 | k__Fungi(100);k__Fungi_unclassified(67);k__Fungi_unclassified(67);k__Fungi_unclassified(67);k__Fungi_unclassified(67);k__Fungi_unclassified(67);k__Fungi_unclassified(67); | - | - | | - | - | - | - | - | Unassigned | |
| Otu00603 | 141 | k__Fungi(100);p__Ascomycota(56);p__Ascomycota_unclassified(56);p__Ascomycota_unclassified(56);p__Ascomycota_unclassified(56);p__Ascomycota_unclassified(56);p__Ascomycota_unclassified(56); | - | - | | - | - | - | - | - | Unassigned | |
| Otu00604 | 140 | k__Fungi(100);p__Basidiomycota(100);c__Agaricomycetes(100);o__Trechisporales(100);f__Hydnodontaceae(100);g__Trechispora(100);s__Trechispora_sp(100); | Trechispora | | 13 | Saprotroph | \|Wood Saprotroph\| | Corticioid | White Rot | Highly Probable | [illegible] | [illegible] |
| Otu00605 | 140 | k__Fungi(100);p__Ascomycota(100);c__Sordariomycetes(100);o__Diaporthales(100);f__Diaporthales_family_Incertae_sedis(100);g__Stegonsporium(100);g__Stegonsporium_unclassified(100); | Stegonsporium | | 13 | Saprotroph | \|Wood Saprotroph\| | Microfungus | NULL | Probable | [illegible] | [illegible] |
| Otu00606 | 140 | k__Fungi(100);p__Ascomycota(100);c__Sordariomycetes(100);o__Xylariales(100);f__Diatrypaceae(100);g__unclassified_Diatrypaceae(100);s__Diatrypaceae_sp(100); | Diatrypaceae | | 9 | Pathotroph-Saprotroph | Plant Pathogen-Plant Saprotroph | Microfungus | NULL | Probable | [illegible] | [illegible] |
| Otu00607 | 139 | k__Fungi(100);p__Basidiomycota(100);c__Ustilaginomycetes(100);o__Ustilaginales(100);f__Ustilaginaceae(100);g__unclassified_Ustilaginaceae(100);s__Ustilaginaceae_sp(100); | Ustilaginaceae | | 9 | Pathotroph | Plant Pathogen | Smut | NULL | Probable | [illegible] | [illegible] |
| Otu00610 | 139 | k__Fungi(100);p__Ascomycota(100);c__Sordariomycetes(100);o__Diaporthales(100);f__Diaporthales_family_Incertae_sedis(53);g__Ophiognomonia(53);g__Ophiognomonia_unclassified(52); | Ophiognomonia | | 13 | Pathotroph | \|Plant Pathogen\| | Microfungus | NULL | Highly Probable | [illegible] | [illegible] |
| Otu00611 | 139 | k__Fungi(100);p__Ascomycota(100);c__Saccharomycetes(100);o__Saccharomycetales(100);f__Saccharomycetales_family_Incertae_sedis(100);g__Candida(100);s__Candida_boleticola(83); | Candida | | 13 | Pathotroph-Saprotroph-Symbiotroph | Animal Pathogen-Endophyte-Endosymbiont-Epiphyte-\|Undefined Saprotroph\| | Dimorphic Yeast-Yeast | NULL | Probable | [illegible] | [illegible] |
| Otu00612 | 139 | k__Fungi(100);p__Basidiomycota(100);c__Tremellomycetes(100);o__Cystofilobasidiales(100);f__Cystofilobasidiaceae(100);g__Itersonilia(100);s__Itersonilia_perplexans(100); | Itersonilia | | 13 | Pathotroph | \|Plant Pathogen\|-Plant Saprotroph | Dimorphic-Dimorphic Yeast | NULL | Probable | [illegible] | [illegible] |
| Otu00608 | 139 | k__Fungi(100);p__Basidiomycota(100);p__Basidiomycota_unclassified(100);p__Basidiomycota_unclassified(100);p__Basidiomycota_unclassified(100);p__Basidiomycota_unclassified(100);p__Basidiomycota_unclassified(100); | - | - | | - | - | - | - | - | Unassigned | |
| Otu00609 | 139 | k__Fungi(100);p__Basidiomycota(100);c__Pucciniomycetes(100);c__Pucciniomycetes_unclassified(74);c__Pucciniomycetes_unclassified(74);c__Pucciniomycetes_unclassified(74);c__Pucciniomycetes_unclassified(74); | - | - | | - | - | - | - | - | Unassigned | |
| Otu00613 | 138 | k__Fungi(100);p__Basidiomycota(100);c__Tremellomycetes(100);o__Tremellales(100);f__Tremellales_family_Incertae_sedis(100);g__Dioszegia(100);g__Dioszegia_unclassified(61); | Dioszegia | | 13 | Saprotroph | \|Plant Saprotroph\|-Undefined Saprotroph | Yeast | NULL | Probable | [illegible] | [illegible] |
| Otu00614 | 138 | k__Fungi(100);p__Zygomycota(100);c__Zygomycota_class_Incertae_sedis(100);o__Mortierellales(100);f__Mortierellaceae(100);g__Mortierella(100);s__Mortierella_umbellata(100); | Mortierella | | 13 | Saprotroph-Symbiotroph | Endophyte-Plant Saprotroph-\|Undefined Saprotroph\| | Microfungus | NULL | Probable | [illegible] | [illegible] |
| Otu00615 | 138 | k__Fungi(100);p__Ascomycota(100);c__Saccharomycetes(100);o__Saccharomycetales(100);f__Metschnikowiaceae(100);g__Metschnikowia(100);s__Metschnikowia_pulcherrima(95); | Metschnikowia | | 13 | Saprotroph | \|Undefined Saprotroph\| | Yeast | NULL | Probable | [illegible] | [illegible] |
| Otu00618 | 137 | k__Fungi(100);p__Basidiomycota(100);c__Tremellomycetes(100);o__Tremellales(89);f__Tremellales_family_Incertae_sedis(89);f__Tremellales_family_Incertae_sedis_unclassified(89);f__Tremellales_family_Incertae_sedis_unclassified(89); | Tremellales | | 7 | Pathotroph-Saprotroph-Symbiotroph | Fungal Parasite-Undefined Saprotroph | Tremelloid-Yeast | NULL | Possible | [illegible] | [illegible] |
| Otu00620 | 137 | k__Fungi(100);p__Ascomycota(100);c__Eurotiomycetes(100);o__Chaetothyriales(96);f__Herpotrichiellaceae(96);f__Herpotrichiellaceae_unclassified(96);f__Herpotrichiellaceae_unclassified(96); | Herpotrichiellaceae | | 9 | Pathotroph-Saprotroph | Animal Pathogen-Fungal Parasite-Undefined Saprotroph | Facultative Yeast-Microfungus | NULL | Probable | [illegible] | [illegible] |
| Otu00621 | 137 | k__Fungi(100);p__Basidiomycota(100);c__Agaricomycetes(100);o__Polyporales(100);f__Polyporales_family_Incertae_sedis(100);g__Phlebiella(100);s__Phlebiella_sp(87); | Phlebiella | | 13 | Saprotroph | Undefined Saprotroph-\|Wood Saprotroph\| | Corticioid | White Rot | Probable | [illegible] | [illegible] |
| Otu00616 | 137 | k__Fungi(100);k__Fungi_unclassified(100);k__Fungi_unclassified(100);k__Fungi_unclassified(100);k__Fungi_unclassified(100);k__Fungi_unclassified(100);k__Fungi_unclassified(100); | - | - | | - | - | - | - | - | Unassigned | |
| Otu00617 | 137 | k__Fungi(100);p__Ascomycota(82);p__Ascomycota_unclassified(82);p__Ascomycota_unclassified(82);p__Ascomycota_unclassified(82);p__Ascomycota_unclassified(82);p__Ascomycota_unclassified(82); | - | - | | - | - | - | - | - | Unassigned | |
| Otu00619 | 137 | k__Fungi(100);p__Basidiomycota(100);c__Agaricomycetes(100);c__Agaricomycetes_unclassified(67);c__Agaricomycetes_unclassified(67);c__Agaricomycetes_unclassified(67);c__Agaricomycetes_unclassified(67); | - | - | | - | - | - | - | - | Unassigned | |
| Otu00622 | 136 | k__Fungi(100);p__Ascomycota(100);c__Eurotiomycetes(100);o__Eurotiales(100);f__Trichocomaceae(100);g__Hemicarpenteles(100);s__Hemicarpenteles_thaxteri(100); | Hemicarpenteles | | 13 | Saprotroph | \|Undefined Saprotroph\| | Microfungus | NULL | Probable | NULL | [illegible] |
| Otu00623 | 136 | k__Fungi(100);p__Ascomycota(100);c__Saccharomycetes(100);o__Saccharomycetales(100);f__Metschnikowiaceae(100);g__Metschnikowia(100);s__Metschnikowia_pulcherrima(98); | Metschnikowia | | 13 | Saprotroph | \|Undefined Saprotroph\| | Yeast | NULL | Probable | [illegible] | [illegible] |
| Otu00624 | 136 | k__Fungi(100);p__Ascomycota(100);c__Dothideomycetes(100);o__Capnodiales(100);f__Mycosphaerellaceae(100);g__Mycosphaerella(100);s__Mycosphaerella_sp_AA_2012(98); | Mycosphaerella | | 13 | Pathotroph-Saprotroph | \|Plant Pathogen\|-Plant Saprotroph | Microfungus | NULL | Probable | [illegible] | [illegible] |
| Otu00625 | 135 | k__Fungi(100);p__Ascomycota(100);c__Sordariomycetes(100);o__Coniochaetales(100);f__Coniochaetaceae(100);g__Coniochaeta(78);s__Coniochaeta_sp(76); | Coniochaeta | | 13 | Pathotroph-Saprotroph-Symbiotroph | Animal Pathogen-Dung Saprotroph-Endophyte-Lichen Parasite-Plant Pathogen-\|Undefined Saprotroph\| | Microfungus | NULL | Possible | [illegible] | [illegible] |
| Otu00628 | 134 | k__Fungi(100);p__Ascomycota(100);c__Eurotiomycetes(100);o__Onygenales(100);f__Gymnoascaceae(100);g__unclassified_Gymnoascaceae(95);s__Gymnoascaceae_sp(95); | Gymnoascaceae | | 9 | Saprotroph | Dung Saprotroph-Plant Saprotroph | Microfungus | NULL | Probable | [illegible] | [illegible] |
| Otu00626 | 134 | k__Fungi(100);p__Ascomycota(100);c__Leotiomycetes(100);o__Helotiales(100);f__Helotiales_family_Incertae_sedis(100);f__Helotiales_family_Incertae_sedis_unclassified(65);f__Helotiales_family_Incertae_sedis_unclassified(65); | - | - | | - | - | - | - | - | Unassigned | |
| Otu00627 | 134 | k__Fungi(100);p__Ascomycota(77);p__Ascomycota_unclassified(77);p__Ascomycota_unclassified(77);p__Ascomycota_unclassified(77);p__Ascomycota_unclassified(77);p__Ascomycota_unclassified(77); | - | - | | - | - | - | - | - | Unassigned | |
| Otu00629 | 133 | k__Fungi(100);p__Basidiomycota(100);c__Agaricomycetes(100);o__Boletales(100);o__Boletales_unclassified(58);o__Boletales_unclassified(58);o__Boletales_unclassified(58); | - | - | | - | - | - | - | - | Unassigned | |
| Otu00630 | 132 | k__Fungi(100);p__Ascomycota(100);c__Dothideomycetes(100);o__Capnodiales(95);o__Capnodiales_unclassified(67);o__Capnodiales_unclassified(67);o__Capnodiales_unclassified(67); | - | - | | - | - | - | - | - | Unassigned | |
| Otu00631 | 132 | k__Fungi(100);p__Ascomycota(100);p__Ascomycota_unclassified(100);p__Ascomycota_unclassified(100);p__Ascomycota_unclassified(100);p__Ascomycota_unclassified(100);p__Ascomycota_unclassified(100); | - | - | | - | - | - | - | - | Unassigned | |
| Otu00632 | 132 | k__Fungi(100);k__Fungi_unclassified(60);k__Fungi_unclassified(60);k__Fungi_unclassified(60);k__Fungi_unclassified(60);k__Fungi_unclassified(60);k__Fungi_unclassified(60); | - | - | | - | - | - | - | - | Unassigned | |
| Otu00633 | 132 | k__Fungi(100);p__Ascomycota(51);p__Ascomycota_unclassified(51);p__Ascomycota_unclassified(51);p__Ascomycota_unclassified(51);p__Ascomycota_unclassified(51);p__Ascomycota_unclassified(51); | - | - | | - | - | - | - | - | Unassigned | |
| Otu00634 | 131 | k__Fungi(100);p__Ascomycota(100);c__Sordariomycetes(100);o__Diaporthales(100);f__Gnomoniaceae(100);f__Gnomoniaceae_unclassified(96);f__Gnomoniaceae_unclassified(96); | Gnomoniaceae | | 9 | Pathotroph-Saprotroph-Symbiotroph | Endophyte-Plant Pathogen-Undefined Saprotroph | Microfungus | NULL | Probable | [illegible] | [illegible] |
| Otu00635 | 131 | k__Fungi(100);p__Basidiomycota(100);c__Agaricomycetes(100);o__Agaricales(100);f__Strophariaceae(100);g__Gymnopilus(100);s__Gymnopilus_junonius(100); | Gymnopilus | | 13 | Saprotroph | Plant Saprotroph-\|Wood Saprotroph\| | Agaricoid | White Rot | Probable | [illegible] | [illegible] |
| Otu00636 | 131 | k__Fungi(100);p__Basidiomycota(100);c__Agaricomycetes(100);o__Polyporales(100);f__Fomitopsidaceae(100);g__Antrodia(100);s__Antrodia_oleracea(100); | Antrodia | | 13 | Saprotroph | \|Wood Saprotroph\| | Corticioid | Brown Rot | Highly Probable | [illegible] | [illegible] |
| Otu00637 | 131 | k__Fungi(100);p__Ascomycota(100);c__Sordariomycetes(100);o__Hypocreales(100);f__Nectriaceae(100);g__Cosmospora(90);g__Cosmospora_unclassified(53); | Cosmospora | | 13 | Pathotroph | \|Fungal Parasite\| | Microfungus | NULL | Highly Probable | [illegible] | [illegible] |
| Otu00638 | 131 | k__Fungi(100);p__Ascomycota(100);c__Sordariomycetes(100);o__Hypocreales(100);f__Hypocreaceae(100);f__Hypocreaceae_unclassified(71);f__Hypocreaceae_unclassified(71); | - | - | | - | - | - | - | - | Unassigned | |
| Otu00639 | 130 | k__Fungi(100);p__Ascomycota(100);c__Sordariomycetes(100);o__Xylariales(100);f__Amphisphaeriaceae(100);f__Amphisphaeriaceae_unclassified(84);f__Amphisphaeriaceae_unclassified(84); | Amphisphaeriaceae | | 9 | Saprotroph | Undefined Saprotroph | Microfungus | NULL | Possible | [illegible] | [illegible] |
| Otu00641 | 130 | k__Fungi(100);p__Basidiomycota(100);c__Cystobasidiomycetes(98);o__Erythrobasidiales(98);f__Erythrobasidiales_family_Incertae_sedis(98);g__Erythrobasidium(98);g__Erythrobasidium_unclassified(97); | Erythrobasidium | | 13 | Pathotroph-Saprotroph | \|Fungal Parasite\|-Undefined Saprotroph | Yeast | NULL | Possible | [illegible] | [illegible] |
| Otu00640 | 130 | k__Fungi(100);p__Ascomycota(100);p__Ascomycota_unclassified(100);p__Ascomycota_unclassified(100);p__Ascomycota_unclassified(100);p__Ascomycota_unclassified(100);p__Ascomycota_unclassified(100); | - | - | | - | - | - | - | - | Unassigned | |
| Otu00642 | 130 | k__Fungi(100);p__Basidiomycota(100);c__Agaricomycetes(100);o__Polyporales(100);o__Polyporales_unclassified(100);o__Polyporales_unclassified(100);o__Polyporales_unclassified(100); | - | - | | - | - | - | - | - | Unassigned | |
| Otu00643 | 129 | k__Fungi(100);p__Ascomycota(100);c__Sordariomycetes(100);o__Xylariales(100);f__Xylariaceae(100);f__Xylariaceae_unclassified(94);f__Xylariaceae_unclassified(94); | Xylariaceae | | 9 | Pathotroph-Saprotroph-Symbiotroph | Dung Saprotroph-Endophyte-Plant Pathogen-Undefined Saprotroph | Xylarioid | NULL | Probable | [illegible] | [illegible] |
| Otu00644 | 129 | k__Fungi(100);p__Basidiomycota(100);c__Tremellomycetes(100);o__Tremellales(100);f__Tremellales_family_Incertae_sedis(100);g__Cryptococcus(100);s__Cryptococcus_dimennae(100); | Cryptococcus | | 13 | Pathotroph-Saprotroph-Symbiotroph | Animal Pathogen-Endophyte-Epiphyte-\|Undefined Saprotroph\| | Dimorphic Yeast-Yeast | NULL | Possible | [illegible] | [illegible] |
| Otu00645 | 128 | k__Fungi(100);p__Ascomycota(100);c__Saccharomycetes(100);o__Saccharomycetales(100);f__Metschnikowiaceae(100);g__Metschnikowia(100);s__Metschnikowia_pulcherrima(97); | Metschnikowia | | 13 | Saprotroph | \|Undefined Saprotroph\| | Yeast | NULL | Probable | [illegible] | [illegible] |
| Otu00646 | 128 | k__Fungi(100);p__Ascomycota(100);c__Eurotiomycetes(100);o__Chaetothyriales(100);f__Chaetothyriaceae(100);g__Cyphellophora(100);g__Cyphellophora_unclassified(100); | Cyphellophora | | 13 | Pathotroph-Saprotroph | Animal Parasite-Animal Pathogen-\|Plant Saprotroph\|-Undefined Saprotroph | Dimorphic-Facultative Yeast-Microfungus | NULL | Probable | [illegible] | [illegible] |
| Otu00647 | 127 | k__Fungi(100);p__Ascomycota(100);c__Sordariomycetes(100);o__Hypocreales(100);f__Clavicipitaceae(100);g__Metarhiziopsis(100);s__Metarhiziopsis_microspora(100); | Metarhiziopsis | | 13 | Pathotroph-Saprotroph | \|Animal Parasite\|-Undefined Saprotroph | Microfungus | NULL | Probable | [illegible] | [illegible] |
| Otu00648 | 127 | k__Fungi(100);k__Fungi_unclassified(86);k__Fungi_unclassified(86);k__Fungi_unclassified(86);k__Fungi_unclassified(86);k__Fungi_unclassified(86);k__Fungi_unclassified(86); | - | - | | - | - | - | - | - | Unassigned | |
| Otu00650 | 126 | k__Fungi(100);p__Ascomycota(100);c__Sordariomycetes(100);o__Xylariales(100);f__Xylariaceae(100);g__Camillea(100);g__Camillea_unclassified(95); | Camillea | | 13 | Saprotroph-Symbiotroph | Endophyte-Undefined Saprotroph-\|Wood Saprotroph\| | Microfungus | NULL | Probable | [illegible] | [illegible] |
| Otu00651 | 126 | k__Fungi(100);p__Basidiomycota(100);c__Tremellomycetes(100);o__Tremellales(100);f__Tremellales_family_Incertae_sedis(100);g__Hannaella(98);s__Hannaella_luteola(93); | Hannaella | | 13 | Pathotroph-Saprotroph | \|Animal Parasite\|-Undefined Saprotroph | Yeast | NULL | Probable | [illegible] | [illegible] |
| Otu00649 | 126 | k__Fungi(100);p__Ascomycota(81);p__Ascomycota_unclassified(81);p__Ascomycota_unclassified(81);p__Ascomycota_unclassified(81);p__Ascomycota_unclassified(81);p__Ascomycota_unclassified(81); | - | - | | - | - | - | - | - | Unassigned | |
| Otu00652 | 125 | k__Fungi(100);p__Basidiomycota(100);c__Agaricomycetes(100);o__Agaricales(100);f__Psathyrellaceae(100);g__Lacrymaria(100);s__Lacrymaria_lacrymabunda(100); | Lacrymaria | | 13 | Saprotroph | \|Plant Saprotroph\|-Undefined Saprotroph | Agaricoid | NULL | Probable | [illegible] | [illegible] |
| Otu00653 | 125 | k__Fungi(100);p__Basidiomycota(100);c__Agaricomycetes(100);o__Cantharellales(100);f__Botryobasidiaceae(100);g__Haplotrichum(100);s__Haplotrichum_conspersum(100); | Haplotrichum | | 13 | Saprotroph | \|Wood Saprotroph\| | Corticioid | White Rot | Probable | [illegible] | [illegible] |
| Otu00655 | 124 | k__Fungi(100);p__Basidiomycota(100);c__Agaricomycetes(100);o__Polyporales(100);f__Ganodermataceae(100);g__Ganoderma(100);s__Ganoderma_sp_JM98/1(100); | Ganoderma | | 13 | Pathotroph-Saprotroph | \|Plant Pathogen\|-Wood Saprotroph | Polyporoid | White Rot | Highly Probable | [illegible] | [illegible] |
| Otu00656 | 124 | k__Fungi(100);p__Ascomycota(100);c__Dothideomycetes(100);o__Capnodiales(100);f__Capnodiales_family_Incertae_sedis(100);g__Rachicladosporium(62);g__Rachicladosporium_unclassified(57); | Rachicladosporium | | 13 | Saprotroph | \|Plant Saprotroph\| | Microfungus | NULL | Probable | [illegible] | [illegible] |
| Otu00657 | 124 | k__Fungi(100);p__Basidiomycota(100);c__Tremellomycetes(100);o__Cystofilobasidiales(100);f__Cystofilobasidiaceae(100);g__Udeniomyces(100);s__Udeniomyces_pyricola(100); | Udeniomyces | | 13 | Saprotroph-Symbiotroph | Epiphyte-Plant Saprotroph-\|Undefined Saprotroph\| | Yeast | NULL | Probable | [illegible] | [illegible] |
| Otu00654 | 124 | k__Fungi(100);p__Basidiomycota(94);p__Basidiomycota_unclassified(94);p__Basidiomycota_unclassified(94);p__Basidiomycota_unclassified(94);p__Basidiomycota_unclassified(94);p__Basidiomycota_unclassified(94); | - | - | | - | - | - | - | - | Unassigned | |
| Otu00658 | 123 | k__Fungi(100);p__Ascomycota(100);c__Dothideomycetes(100);o__Pleosporales(100);f__Lophiostomataceae(66);g__Lophiostoma(66);g__Lophiostoma_unclassified(65); | Lophiostoma | | 13 | Saprotroph | Plant Saprotroph-Undefined Saprotroph-\|Wood Saprotroph\| | Microfungus | NULL | Probable | [illegible] | [illegible] |
| Otu00659 | 123 | k__Fungi(100);p__Ascomycota(100);c__Sordariomycetes(100);o__Chaetosphaeriales(100);f__Chaetosphaeriaceae(100);f__Chaetosphaeriaceae_unclassified(100);f__Chaetosphaeriaceae_unclassified(100); | Chaetosphaeriaceae | | 9 | Saprotroph | Plant Saprotroph-Wood Saprotroph | Microfungus | NULL | Probable | [illegible] | [illegible] |
| Otu00661 | 123 | k__Fungi(100);p__Basidiomycota(100);c__Agaricomycetes(100);o__Agaricales(100);f__Psathyrellaceae(100);g__Coprinopsis(100);s__Coprinopsis_spelaiophila(100); | Coprinopsis | | 13 | Saprotroph | \|Undefined Saprotroph\| | Agaricoid | NULL | Probable | [illegible] | [illegible] |
| Otu00660 | 123 | k__Fungi(100);p__Ascomycota(82);p__Ascomycota_unclassified(82);p__Ascomycota_unclassified(82);p__Ascomycota_unclassified(82);p__Ascomycota_unclassified(82);p__Ascomycota_unclassified(82); | - | - | | - | - | - | - | - | Unassigned | |
| Otu00664 | 122 | k__Fungi(100);p__Ascomycota(100);c__Sordariomycetes(100);o__Diaporthales(100);f__Valsaceae(100);g__unclassified_Valsaceae(85);s__Valsaceae_sp(85); | Valsaceae | | 9 | Pathotroph-Saprotroph-Symbiotroph | Endophyte-Plant Pathogen-Undefined Saprotroph | Microfungus | NULL | Probable | [illegible] | [illegible] |
| Otu00665 | 122 | k__Fungi(100);p__Basidiomycota(100);c__Agaricomycetes(100);o__Agaricales(100);f__Psathyrellaceae(100);g__Coprinellus(100);s__Coprinellus_sp(100); | Coprinellus | | 13 | Saprotroph | \|Undefined Saprotroph\| | Agaricoid | NULL | Probable | [illegible] | [illegible] |
| Otu00662 | 122 | k__Fungi(100);p__Ascomycota(59);p__Ascomycota_unclassified(59);p__Ascomycota_unclassified(59);p__Ascomycota_unclassified(59);p__Ascomycota_unclassified(59);p__Ascomycota_unclassified(59); | - | - | | - | - | - | - | - | Unassigned | |
| Otu00663 | 122 | k__Fungi(100);k__Fungi_unclassified(56);k__Fungi_unclassified(56);k__Fungi_unclassified(56);k__Fungi_unclassified(56);k__Fungi_unclassified(56);k__Fungi_unclassified(56); | - | - | | - | - | - | - | - | Unassigned | |
| Otu00666 | 121 | k__Fungi(100);p__Ascomycota(100);c__Sordariomycetes(99);o__Sordariomycetes_order_Incertae_sedis(90);f__Magnaporthaceae(90);g__Mycoleptodiscus(90);g__Mycoleptodiscus_unclassified(90); | Mycoleptodiscus | | 13 | Pathotroph | \|Plant Pathogen\| | Microfungus | NULL | Probable | NULL | [illegible] |
| Otu00667 | 120 | k__Fungi(100);p__Ascomycota(100);c__Dothideomycetes(100);o__Pleosporales(100);f__Pleosporaceae(100);f__Pleosporaceae_unclassified(63);f__Pleosporaceae_unclassified(63); | Pleosporaceae | | 9 | Pathotroph-Saprotroph | Endophyte-Lichen Parasite-Plant Pathogen-Undefined Saprotroph | Microfungus | NULL | Probable | [illegible] | [illegible] |
| Otu00668 | 120 | k__Fungi(100);p__Ascomycota(100);c__Sordariomycetes(100);o__Diaporthales(100);f__Diaporthales_family_Incertae_sedis(100);g__Tubakia(100);s__Tubakia_sp_CBS_129013(81); | Tubakia | | 13 | Pathotroph | \|Plant Pathogen\| | Microfungus | NULL | Probable | NULL | [illegible] |
| Otu00669 | 120 | k__Fungi(100);p__Basidiomycota(100);c__Agaricomycetes(100);o__Cantharellales(100);f__Botryobasidiaceae(100);g__unclassified_Botryobasidiaceae(100);s__Botryobasidiaceae_sp(100); | Botryobasidiaceae | | 9 | Saprotroph | Undefined Saprotroph-Wood Saprotroph | Corticioid | NULL | Probable | [illegible] | [illegible] |
| Otu00671 | 119 | k__Fungi(100);p__Basidiomycota(100);c__Agaricomycetes(100);o__Agaricales(100);o__Agaricales_unclassified(100);o__Agaricales_unclassified(100);o__Agaricales_unclassified(100); | Agaricales | | 7 | Pathotroph-Saprotroph-Symbiotroph | Bryophyte Parasite-Dung Saprotroph-Ectomycorrhizal-Fungal Parasite-Leaf Saprotroph-Plant Parasite-Undefined Saprotroph-Wood Saprotroph | Agaricoid-Gasteroid-Microfungus-Secotioid-Yeast | NULL | Possible | [illegible] | [illegible] |
| Otu00672 | 119 | k__Fungi(100);p__Ascomycota(100);c__Dothideomycetes(100);o__Dothideomycetes_order_Incertae_sedis(100);f__Dothideomycetes_family_Incertae_sedis(100);g__Peltaster(100);s__Peltaster_fructicola(100); | Peltaster | | 13 | Pathotroph | \|Plant Pathogen\| | Microfungus | NULL | Probable | NULL | [illegible] |
| Otu00670 | 119 | k__Fungi(100);k__Fungi_unclassified(68);k__Fungi_unclassified(68);k__Fungi_unclassified(68);k__Fungi_unclassified(68);k__Fungi_unclassified(68);k__Fungi_unclassified(68); | - | - | | - | - | - | - | - | Unassigned | |
| Otu00673 | 119 | k__Fungi(100);k__Fungi_unclassified(99);k__Fungi_unclassified(99);k__Fungi_unclassified(99);k__Fungi_unclassified(99);k__Fungi_unclassified(99);k__Fungi_unclassified(99); | - | - | | - | - | - | - | - | Unassigned | |
| Otu00676 | 118 | k__Fungi(100);p__Ascomycota(100);c__Dothideomycetes(100);o__Capnodiales(100);f__Capnodiales_family_Incertae_sedis(98);g__Rachicladosporium(98);s__Rachicladosporium_sp_HX6(98); | Rachicladosporium | | 13 | Saprotroph | \|Plant Saprotroph\| | Microfungus | NULL | Probable | [illegible] | [illegible] |
| Otu00674 | 118 | k__Fungi(100);k__Fungi_unclassified(57);k__Fungi_unclassified(57);k__Fungi_unclassified(57);k__Fungi_unclassified(57);k__Fungi_unclassified(57);k__Fungi_unclassified(57); | - | - | | - | - | - | - | - | Unassigned | |

| | | | | | | | | | | | | |
|---|---|---|---|---|---|---|---|---|---|---|---|---|
| Otu00675 | 118 | k__Fungi(100);p__Ascomycota(100);c__Leotiomycetes(99);o__Helotiales(99);f__Helotiales_family_Incertae_sedis(99);f__Helotiales_family_Incertae_sedis_unclassified(98);f__Helotiales_family_Incertae_sedis_unclassified(98); | - | - | - | - | - | - | - | - | Unassigned | - |
| Otu00677 | 117 | k__Fungi(100);p__Ascomycota(100);c__Eurotiomycetes(100);o__Chaetothyriales(100);f__Herpotrichiellaceae(100);g__unclassified_Herpotrichiellaceae(100);s__Herpotrichiellaceae_sp(100); | Herpotrichiellaceae | 9 | Pathotroph-Saprotroph | Animal Pathogen-Fungal Parasite-Undefined Saprotroph | Facultative Yeast-Microfungus | NULL | Probable | Mostly saprobic on plants or other fungi, isolated from soil etc., and from medical sources (Cannon & Kirk 2007) | Cannon PF, Kirk PM. 2007. Fungal Families of the World. CAB International, Cambridge (ISBN: 978-0851998275) |
| Otu00678 | 117 | k__Fungi(100);p__Ascomycota(100);c__Sordariomycetes(100);o__Hypocreales(100);f__Nectriaceae(100);f__Nectriaceae_unclassified(83);f__Nectriaceae_unclassified(83); | Nectriaceae | 9 | Pathotroph-Saprotroph-Symbiotroph | Animal Pathogen-Endophyte-Fungal Parasite-Lichen Parasite-Plant Pathogen-Wood Saprotroph | Microfungus | NULL | Possible | Fusarium species are amongst the economically significant plant pathogens (Cannon & Kirk 2007) | Cannon PF, Kirk PM. 2007. Fungal Families of the World. CAB International, Cambridge (ISBN: 978-0851998275); Hirooka Y, et al. 2012. Studies in Mycology 711-210 ((https://doi.org/10.3114/sim0001)); Tedersoo L, et al. 2014. Science 346:e1256688 ((https://doi.org/10.1126/science.1256688)); The American Phytopathological Society Common Names of Plant Diseases ((http://www.apsnet.org/publications/commonnames/Pages/default.aspx)); https://www.cdc.gov ((https://www.cdc.gov)) |
| Otu00679 | 117 | k__Fungi(100);p__Basidiomycota(100);c__Tremellomycetes(100);o__Tremellales(100);f__Tremellales_family_Incertae_sedis(95);f__Tremellales_family_Incertae_sedis_unclassified(94);f__Tremellales_family_Incertae_sedis_unclassified(94); | Tremellales | 7 | Pathotroph-Saprotroph-Symbiotroph | Fungal Parasite-Undefined Saprotroph | Tremelloid-Yeast | NULL | Possible | The order contains both teleomorphic and anamorphic species, most of the latter being yeasts | Sterkenburg E, et al. 2015. New Phytologist 207:1145-1158 ((https://doi.org/10.1111/nph.13426)) |
| Otu00680 | 117 | k__Fungi(100);p__Basidiomycota(100);c__Agaricomycetes(100);o__Agaricomycetes_order_Incertae_sedis(100);f__Agaricomycetes_family_Incertae_sedis(100);g__Resinicium(100);s__Resinicium_bicolor(97); | Resinicium | 13 | Saprotroph | Undefined Saprotroph-\|Wood Saprotroph\| | Corticioid | White Rot | Probable | As white_rot (Põlme et al. 2020); Decay wood (Põlme et al. 2020) | Tedersoo L, et al. 2014. Science 346:e1256688 ((https://doi.org/10.1126/science.1256688)); Põlme S, et al. 2020. Fungal Diversity 105:1-16 ((https://doi.org/10.1007/s13225-020-00466-2)) |
| Otu00681 | 116 | k__Fungi(100);p__Ascomycota(100);c__Saccharomycetes(100);o__Saccharomycetales(100);f__Metschnikowiaceae(100);g__Metschnikowia(100);s__Metschnikowia_pulcherrima(97); | Metschnikowia | 13 | Saprotroph | \|Undefined Saprotroph\| | Yeast | NULL | Probable | As nectar-tap_saprotroph (Põlme et al. 2020); Decay sugar-rich substrates (Põlme et al. 2020) | Põlme S, et al. 2020. Fungal Diversity 105:1-16 ((https://doi.org/10.1007/s13225-020-00466-2)) |
| Otu00682 | 116 | k__Fungi(100);p__Ascomycota(100);c__Dothideomycetes(100);o__Capnodiales(100);f__Teratosphaeriaceae(100);g__Devriesia(100);s__Devriesia_imbrexigena(100); | Devriesia | 13 | Pathotroph-Saprotroph | \|Plant Pathogen\|-Plant Saprotroph | Microfungus | NULL | Probable | As litter_saprotroph (Põlme et al. 2020); Decay leaf, fruit, seed (Põlme et al. 2020) | Tedersoo L, et al. 2014. Science 346:e1256688 ((https://doi.org/10.1126/science.1256688)); Põlme S, et al. 2020. Fungal Diversity 105:1-16 ((https://doi.org/10.1007/s13225-020-00466-2)) |
| Otu00683 | 116 | k__Fungi(100);p__Ascomycota(100);c__Dothideomycetes(100);o__Pleosporales(100);f__Pleosporaceae(100);g__Curvularia(54);g__Curvularia_unclassified(54); | Curvularia | 13 | Pathotroph-Saprotroph | \|Plant Pathogen\|-Plant Saprotroph | Microfungus | NULL | Probable | As litter_saprotroph (Põlme et al. 2020); Decay wood, leaf, fruit, seed, roots, algal material (Põlme et al. 2020) | Tedersoo L, et al. 2014. Science 346:e1256688 ((https://doi.org/10.1126/science.1256688)); Põlme S, et al. 2020. Fungal Diversity 105:1-16 ((https://doi.org/10.1007/s13225-020-00466-2)) |
| Otu00684 | 115 | k__Fungi(100);p__Basidiomycota(100);c__Agaricomycetes(100);o__Agaricales(100);f__Pluteaceae(100);g__Pluteus(100);s__Pluteus_longistriatus(100); | Pluteus | 13 | Pathotroph-Saprotroph | Bryophyte Parasite-\|Plant Saprotroph\|-Wood Saprotroph | Agaricoid | White Rot | Probable | As litter_saprotroph (Põlme et al. 2020); As white_rot (Põlme et al. 2020); Decay leaf, fruit, seed (Põlme et al. 2020) | Grgurinovic CA, Simpson JA. 2001. Fungal Diversity 8:97-127 ((http://www.fungaldiversity.org/fdp/sfdp/FD_8_97-127.pdf)); James TY, et al. 2006. Nature 443:818-822 ((https://doi.org/10.1038/nature05110)); Tedersoo L, et al. 2014. Science 346:e1256688 ((https://doi.org/10.1126/science.1256688)); Põlme S, et al. 2020. Fungal Diversity 105:1-16 ((https://doi.org/10.1007/s13225-020-00466-2)) |
| Otu00686 | 115 | k__Fungi(100);p__Ascomycota(100);c__Dothideomycetes(100);o__Dothideomycetes_order_Incertae_sedis(100);f__Dothideomycetes_family_Incertae_sedis(100);g__Leptospora(100);s__Leptospora_rubella(100); | Leptospora | 13 | Saprotroph | \|Plant Saprotroph\| | Microfungus | NULL | Probable | As litter_saprotroph (Põlme et al. 2020); Decay leaf, fruit, seed (Põlme et al. 2020) | Põlme S, et al. 2020. Fungal Diversity 105:1-16 ((https://doi.org/10.1007/s13225-020-00466-2)) |
| Otu00687 | 115 | k__Fungi(100);p__Ascomycota(100);c__Sordariomycetes(100);o__Xylariales(100);f__Xylariaceae(100);g__Virgaria(100);s__Virgaria_sp(82); | Virgaria | 13 | Saprotroph | Plant Saprotroph-Undefined Saprotroph-\|Wood Saprotroph\| | Microfungus | NULL | Probable | As litter_saprotroph (Põlme et al. 2020); Decay wood, leaf, fruit, seed (Põlme et al. 2020) | Tedersoo L, et al. 2014. Science 346:e1256688 ((https://doi.org/10.1126/science.1256688)); Põlme S, et al. 2020. Fungal Diversity 105:1-16 ((https://doi.org/10.1007/s13225-020-00466-2)) |
| Otu00690 | 115 | k__Fungi(100);p__Ascomycota(100);c__Leotiomycetes(100);o__Leotiomycetes_order_Incertae_sedis(100);f__Leotiomycetes_family_Incertae_sedis(100);g__Sarea(100);s__Sarea_sp(100); | Sarea | 13 | Saprotroph-Symbiotroph | Lichenized-\|Undefined Saprotroph\| | Microfungus-Thallus | NULL | Highly Probable | NULL | Esslinger TL. 2014. North Dakota State University ((http://www.ndsu.edu/pubweb/~esslinge/chcklst/chcklst7.htm)); Põlme S, et al. 2020. Fungal Diversity 105:1-16 ((https://doi.org/10.1007/s13225-020-00466-2)) |
| Otu00691 | 115 | k__Fungi(100);p__Basidiomycota(100);c__Agaricomycetes(100);o__Polyporales(100);f__Polyporaceae(100);f__Polyporaceae_unclassified(94);f__Polyporaceae_unclassified(94); | Polyporaceae | 9 | Saprotroph | Wood Saprotroph | Polyporoid | Brown Rot-White Rot | Probable | Lignicolous, causing brown or white rot, mostly of broadleaved trees (Cannon & Kirk 2007) | Cannon PF, Kirk PM. 2007. Fungal Families of the World. CAB International, Cambridge (ISBN: 978-0851998275) |
| Otu00685 | 115 | k__Fungi(100);p__Ascomycota(100);c__Dothideomycetes(100);o__Capnodiales(100);o__Capnodiales_unclassified(99);o__Capnodiales_unclassified(99);o__Capnodiales_unclassified(99); | - | - | - | - | - | - | - | Unassigned | - |
| Otu00688 | 115 | k__Fungi(100);p__Ascomycota(53);p__Ascomycota_unclassified(53);p__Ascomycota_unclassified(53);p__Ascomycota_unclassified(53);p__Ascomycota_unclassified(53);p__Ascomycota_unclassified(53); | - | - | - | - | - | - | - | Unassigned | - |
| Otu00689 | 115 | k__Fungi(100);p__Ascomycota(58);p__Ascomycota_unclassified(58);p__Ascomycota_unclassified(58);p__Ascomycota_unclassified(58);p__Ascomycota_unclassified(58);p__Ascomycota_unclassified(58); | - | - | - | - | - | - | - | Unassigned | - |
| Otu00694 | 114 | k__Fungi(100);p__Ascomycota(100);c__Saccharomycetes(100);o__Saccharomycetales(100);f__Saccharomycetaceae(100);g__Zygosaccharomyces(100);s__Zygosaccharomyces_bailii(100); | Zygosaccharomyces | 13 | Pathotroph-Saprotroph | Plant Pathogen-\|Undefined Saprotroph\| | Facultative Yeast-Yeast | NULL | Probable | As nectar-tap_saprotroph (Põlme et al. 2020); Decay sugar-rich substrates (Põlme et al. 2020) | Kurtzman CP, et al. (eds.) 2011. The Yeasts, a Taxonomic Study. Fifth Edition. Vols 1-3. Elsevier, San Diego (ISBN: 9780444521491); Põlme S, et al. 2020. Fungal Diversity 105:1-16 ((https://doi.org/10.1007/s13225-020-00466-2)); The American Phytopathological Society Common Names of Plant Diseases ((http://www.apsnet.org/publications/commonnames/Pages/default.aspx)) |
| Otu00695 | 114 | k__Fungi(100);p__Basidiomycota(100);c__Agaricomycetes(100);o__Agaricales(100);f__Tricholomataceae(100);f__Tricholomataceae_unclassified(100);f__Tricholomataceae_unclassified(100); | Tricholomataceae | 9 | Pathotroph-Symbiotroph | Ectomycorrhizal-Fungal Parasite | Agaricoid | NULL | Probable | Tricholoma species are ectomycorrhizal, occurring in coniferous and broadleaved forests (Cannon & Kirk 2007); T. sclerotoideum is parasitic on Helvella spp. | Cannon PF, Kirk PM. 2007. Fungal Families of the World. CAB International, Cambridge (ISBN: 978-0851998275) |
| Otu00696 | 114 | k__Fungi(100);p__Ascomycota(100);c__Dothideomycetes(100);o__Pleosporales(100);f__Leptosphaeriaceae(100);g__Coniothyrium(100);s__Coniothyrium_sidae(100); | Coniothyrium | 13 | Pathotroph-Saprotroph | Fungal Parasite-\|Plant Pathogen\|-Plant Saprotroph-Wood Saprotroph | Microfungus | Soft Rot | Highly Probable | As litter_saprotroph (Põlme et al. 2020); As mycoparasite (Põlme et al. 2020); Decay leaf, fruit, seed, wood (Põlme et al. 2020) | Duncan CG, Eslyn WE. 1966. Mycologia 58:642-664 ((https://doi.org/10.2307/3757045)); Nilsson T. 1973. Studia Forestalia Suecica 104: 1-40 ((https://res.slu.se/id/publ/125502)); Seehann G, et al. 1975. List of Fungi in Soft-Rot Tissue Tests. Institute fur Holzbiologie und Holzschutz, Bundesforschungsanstalt fur Forst - und Holzwirtshaft, Hamburg - Lohbrugge, Germany ((https://www.irg-wp.com/irgdocs/details.php?d6493033-c9ea-40f3-8e3e-5697f32d640d)); Põlme S, et al. 2020. Fungal Diversity 105:1-16 ((https://doi.org/10.1007/s13225-020-00466-2)); The American Phytopathological Society Common Names of Plant Diseases ((https://www.apsnet.org/edcenter/resources/commonnames/Pages/default.aspx)) |
| Otu00698 | 114 | k__Fungi(100);p__unclassified_Fungi(100);c__unclassified_Fungi(100);o__unclassified_Fungi(100);f__unclassified_Fungi(100);g__unclassified_Fungi(100);s__Cystobasidium_pallidum(100); | Cystobasidium | 13 | Pathotroph | \|Fungal Parasite\| | Dimorphic-Facultative Yeast | NULL | Highly Probable | As mycoparasite (Põlme et al. 2020); Decay fungal material (Põlme et al. 2020); Host - Thelebolales (Põlme et al. 2020) | Kurtzman CP, et al. (eds.) 2011. The Yeasts, a Taxonomic Study. Fifth Edition. Vols 1-3. Elsevier, San Diego (ISBN: 9780444521491); Põlme S, et al. 2020. Fungal Diversity 105:1-16 ((https://doi.org/10.1007/s13225-020-00466-2)) |
| Otu00700 | 114 | k__Fungi(100);p__Ascomycota(100);c__Ascomycota_class_Incertae_sedis(100);o__Ascomycota_order_Incertae_sedis(100);f__Ascomycota_family_Incertae_sedis(100);g__Hansfordia(100);s__Hansfordia_pulvinata(100); | Hansfordia | 13 | Saprotroph | \|Undefined Saprotroph\| | Microfungus | NULL | Probable | NULL | Tedersoo L, et al. 2014. Science 346:e1256688 ((https://doi.org/10.1126/science.1256688)); Põlme S, et al. 2020. Fungal Diversity 105:1-16 ((https://doi.org/10.1007/s13225-020-00466-2)) |
| Otu00701 | 114 | k__Fungi(100);p__Ascomycota(100);c__Leotiomycetes(100);o__Helotiales(100);f__Sclerotiniaceae(100);g__Monilinia(100);s__Monilinia_linhartiana(100); | Monilinia | 13 | Pathotroph-Saprotroph | \|Plant Pathogen\|-Plant Saprotroph | Microfungus | NULL | Probable | As litter_saprotroph (Põlme et al. 2020); Decay leaf, fruit, seed (Põlme et al. 2020) | Tedersoo L, et al. 2014. Science 346:e1256688 ((https://doi.org/10.1126/science.1256688)); Põlme S, et al. 2020. Fungal Diversity 105:1-16 ((https://doi.org/10.1007/s13225-020-00466-2)) |
| Otu00702 | 114 | k__Fungi(100);p__Ascomycota(100);c__Dothideomycetes(100);o__Pleosporales(100);f__Phaeosphaeriaceae(96);f__Phaeosphaeriaceae_unclassified(95);f__Phaeosphaeriaceae_unclassified(95); | Phaeosphaeriaceae | 9 | Pathotroph-Saprotroph | Fungal Parasite-Plant Pathogen-Plant Saprotroph | Microfungus | NULL | Probable | Necrotrophic or saprobic on a wide range of plants, especially monocotyledons; a few species parasitic on other fungi (Cannon & Kirk 2007) | Cannon PF, Kirk PM. 2007. Fungal Families of the World. CAB International, Cambridge (ISBN: 978-0851998275) |
| Otu00692 | 114 | k__Fungi(100);k__Fungi_unclassified(89);k__Fungi_unclassified(89);k__Fungi_unclassified(89);k__Fungi_unclassified(89);k__Fungi_unclassified(89); | - | - | - | - | - | - | - | Unassigned | - |
| Otu00693 | 114 | k__Fungi(100);p__Ascomycota(88);p__Ascomycota_unclassified(88);p__Ascomycota_unclassified(88);p__Ascomycota_unclassified(88);p__Ascomycota_unclassified(88);p__Ascomycota_unclassified(88); | - | - | - | - | - | - | - | Unassigned | - |
| Otu00697 | 114 | k__Fungi(100);k__Fungi_unclassified(65);k__Fungi_unclassified(65);k__Fungi_unclassified(65);k__Fungi_unclassified(65);k__Fungi_unclassified(65); | - | - | - | - | - | - | - | Unassigned | - |
| Otu00699 | 114 | k__Fungi(100);p__Ascomycota(92);p__Ascomycota_unclassified(91);p__Ascomycota_unclassified(91);p__Ascomycota_unclassified(91);p__Ascomycota_unclassified(91);p__Ascomycota_unclassified(91); | - | - | - | - | - | - | - | Unassigned | - |
| Otu00703 | 113 | k__Fungi(100);p__Basidiomycota(100);c__Tremellomycetes(100);o__Tremellales(100);f__Tremellales_family_Incertae_sedis(100);g__Filobasidiella(100);s__Filobasidiella_depauperata(100); | Filobasidiella | 13 | Pathotroph-Saprotroph | \|Animal Parasite\|-Fungal Parasite-Undefined Saprotroph | Dimorphic-Facultative Yeast | NULL | Highly Probable | As mycoparasite (Põlme et al. 2020); Decay animal material (Põlme et al. 2020) | Kurtzman CP, et al. (eds.) 2011. The Yeasts, a Taxonomic Study. Fifth Edition. Vols 1-3. Elsevier, San Diego (ISBN: 9780444521491); Põlme S, et al. 2020. Fungal Diversity 105:1-16 ((https://doi.org/10.1007/s13225-020-00466-2)) |
| Otu00704 | 113 | k__Fungi(100);p__Ascomycota(100);c__Saccharomycetes(100);o__Saccharomycetales(100);f__Metschnikowiaceae(100);g__Metschnikowia(100);s__Metschnikowia_pulcherrima(84); | Metschnikowia | 13 | Saprotroph | \|Undefined Saprotroph\| | Yeast | NULL | Probable | As nectar-tap_saprotroph (Põlme et al. 2020); Decay sugar-rich substrates (Põlme et al. 2020) | Põlme S, et al. 2020. Fungal Diversity 105:1-16 ((https://doi.org/10.1007/s13225-020-00466-2)) |
| Otu00705 | 113 | k__Fungi(100);p__Ascomycota(100);c__Dothideomycetes(100);o__Pleosporales(100);f__Lophiostomataceae(55);g__Lophiostoma(55);g__Lophiostoma_unclassified(55); | Lophiostoma | 13 | Saprotroph | Plant Saprotroph-Undefined Saprotroph-\|Wood Saprotroph\| | Microfungus | NULL | Probable | As litter_saprotroph (Põlme et al. 2020); Decay wood, leaf, fruit, seed (Põlme et al. 2020) | Tedersoo L, et al. 2014. Science 346:e1256688 ((https://doi.org/10.1126/science.1256688)); Põlme S, et al. 2020. Fungal Diversity 105:1-16 ((https://doi.org/10.1007/s13225-020-00466-2)) |
| Otu00706 | 113 | k__Fungi(100);p__Ascomycota(100);c__Sordariomycetes(100);o__Diaporthales(100);f__Valsaceae(100);f__Valsaceae_unclassified(70);f__Valsaceae_unclassified(70); | Valsaceae | 9 | Pathotroph-Saprotroph-Symbiotroph | Endophyte-Plant Pathogen-Undefined Saprotroph | Microfungus | NULL | Probable | Saprobes, endophytes and weak to virulent parasites of woody plants (Cannon & Kirk 2007) | Cannon PF, Kirk PM. 2007. Fungal Families of the World. CAB International, Cambridge (ISBN: 978-0851998275) |
| Otu00707 | 113 | k__Fungi(100);p__Ascomycota(95);p__Ascomycota_unclassified(94);p__Ascomycota_unclassified(94);p__Ascomycota_unclassified(94);p__Ascomycota_unclassified(94);p__Ascomycota_unclassified(94); | - | - | - | - | - | - | - | Unassigned | - |
| Otu00708 | 112 | k__Fungi(100);p__Ascomycota(100);c__Sordariomycetes(100);o__Hypocreales(100);o__Hypocreales_unclassified(100);o__Hypocreales_unclassified(100);o__Hypocreales_unclassified(100); | Hypocreales | 7 | Saprotroph | Undefined Saprotroph | Microfungus | NULL | Possible | NULL | Sterkenburg E, et al. 2015. New Phytologist 207:1145-1158 ((https://doi.org/10.1111/nph.13426)) |
| Otu00709 | 112 | k__Fungi(100);p__Basidiomycota(100);c__Agaricomycetes(100);o__Polyporales(100);f__Meruliaceae(100);g__Steccherinum(100);s__Steccherinum_fimbriatum(100); | Steccherinum | 13 | Saprotroph | Undefined Saprotroph-\|Wood Saprotroph\| | Corticioid | White Rot | Probable | As white_rot (Põlme et al. 2020); Decay wood (Põlme et al. 2020) | Tedersoo L, et al. 2014. Science 346:e1256688 ((https://doi.org/10.1126/science.1256688)); Põlme S, et al. 2020. Fungal Diversity 105:1-16 ((https://doi.org/10.1007/s13225-020-00466-2)) |
| Otu00710 | 112 | k__Fungi(100);k__Fungi_unclassified(66);k__Fungi_unclassified(66);k__Fungi_unclassified(66);k__Fungi_unclassified(66);k__Fungi_unclassified(66); | - | - | - | - | - | - | - | Unassigned | - |
| Otu00711 | 111 | k__Fungi(100);p__Ascomycota(100);c__Dothideomycetes(100);o__Dothideomycetes_order_Incertae_sedis(100);f__Myxotrichaceae(100);g__Oidiodendron(100);s__Oidiodendron_maius(100); | Oidiodendron | 13 | Pathotroph-Saprotroph-Symbiotroph | Endophyte-Ericoid Mycorrhizal-\|Undefined Saprotroph\| | Microfungus | NULL | Probable | As root_endophyte (Põlme et al. 2020); As soil_saprotroph (Põlme et al. 2020); Dark Septate Endophyte; Decay soil (Põlme et al. 2020) | Newsham KK. 2011. New Phytologist 190:783-793 ((https://doi.org/10.1111/j.1469-8137.2010.03611.x)) (as dark septate root endophytes); Põlme S, et al. 2020. Fungal Diversity 105:1-16 ((https://doi.org/10.1007/s13225-020-00466-2)) |
| Otu00712 | 111 | k__Fungi(100);p__Ascomycota(100);c__Pezizomycetes(100);o__unclassified_Pezizomycetes(100);f__unclassified_Pezizomycetes(100);g__unclassified_Pezizomycetes(100);s__Pezizomycetes_sp(100); | - | - | - | - | - | - | - | Unassigned | - |
| Otu00713 | 110 | k__Fungi(100);p__Basidiomycota(100);c__Agaricomycetes(100);o__Hymenochaetales(100);f__Hymenochaetaceae(100);g__Inonotus(100);g__Inonotus_unclassified(100); | Inonotus | 13 | Pathotroph-Saprotroph | Plant Pathogen-\|Wood Saprotroph\| | Polyporoid | White Rot | Highly Probable | As white_rot (Põlme et al. 2020); As wood_pathogen (Põlme et al. 2020); Decay wood (Põlme et al. 2020) | Gilbertson RL, Ryvarden L. 1987-1987. North American Polypores. Fungiflora, Oslo (ISBN: 978-0945345060); Põlme S, et al. 2020. Fungal Diversity 105:1-16 ((https://doi.org/10.1007/s13225-020-00466-2)) |
| Otu00715 | 110 | k__Fungi(100);p__Ascomycota(100);c__Dothideomycetes(100);o__Dothideomycetes_order_Incertae_sedis(100);f__Dothideomycetes_family_Incertae_sedis(100);g__Leptospora(100);s__Leptospora_rubella(100); | Leptospora | 13 | Saprotroph | \|Plant Saprotroph\| | Microfungus | NULL | Probable | As litter_saprotroph (Põlme et al. 2020); Decay leaf, fruit, seed (Põlme et al. 2020) | Põlme S, et al. 2020. Fungal Diversity 105:1-16 ((https://doi.org/10.1007/s13225-020-00466-2)) |
| Otu00717 | 110 | k__Fungi(100);p__Ascomycota(100);c__Dothideomycetes(100);o__Capnodiales(100);f__Davidiellaceae(100);g__Cladosporium(100);g__Cladosporium_unclassified(97); | Cladosporium | 13 | Pathotroph-Saprotroph-Symbiotroph | Animal Pathogen-Endophyte-Lichen Parasite-Plant Pathogen-\|Plant Saprotroph\|-Wood Saprotroph | Microfungus | NULL | Possible | As foliar_endophyte (Põlme et al. 2020); As litter_saprotroph (Põlme et al. 2020); Decay wood, leaf, fruit, seed, soil, dung, animal material (Põlme et al. 2020) | Seehann G, et al. 1975. List of Fungi in Soft-Rot Tissue Tests. Institute fur Holzbiologie und Holzschutz, Bundesforschungsanstalt fur Forst - und Holzwirtshaft, Hamburg - Lohbrugge, Germany; Lawrey JD, Diederich P. 2016. http://www.lichenicolous.net (accessed 24 May 2017) ((http://www.lichenicolous.net)); Põlme S, et al. 2020. Fungal Diversity 105:1-16 ((https://doi.org/10.1007/s13225-020-00466-2)); The American Phytopathological Society Common Names of Plant Diseases ((http://www.apsnet.org/publications/commonnames/Pages/default.aspx)) |
| Otu00718 | 110 | k__Fungi(100);p__Basidiomycota(100);c__Agaricomycetes(100);o__Agaricales(100);f__Lyophyllaceae(100);g__Hypsizygus(99);s__Hypsizygus_ulmarius(99); | Hypsizygus | 13 | Saprotroph | Undefined Saprotroph-\|Wood Saprotroph\| | Agaricoid | White Rot | Probable | As white_rot (Põlme et al. 2020); Decay wood (Põlme et al. 2020) | Tedersoo L, et al. 2014. Science 346:e1256688 ((https://doi.org/10.1126/science.1256688)); Põlme S, et al. 2020. Fungal Diversity 105:1-16 ((https://doi.org/10.1007/s13225-020-00466-2)) |
| Otu00714 | 110 | k__Fungi(100);p__Ascomycota(100);c__Dothideomycetes(100);o__Pleosporales(100);f__unclassified_Pleosporales(100);g__unclassified_Pleosporales(100);s__Pleosporales_sp(100); | - | - | - | - | - | - | - | Unassigned | - |
| Otu00716 | 110 | k__Fungi(100);k__Fungi_unclassified(70);k__Fungi_unclassified(70);k__Fungi_unclassified(70);k__Fungi_unclassified(70);k__Fungi_unclassified(70); | - | - | - | - | - | - | - | Unassigned | - |
| Otu00719 | 109 | k__Fungi(100);p__Basidiomycota(100);c__Tremellomycetes(100);o__Tremellales(100);f__Tremellales_family_Incertae_sedis(100);g__Cryptococcus(100);s__Cryptococcus_aff_amylolyticus_AS_22501(100); | Cryptococcus | 13 | Pathotroph-Saprotroph-Symbiotroph | Animal Pathogen-Endophyte-Epiphyte-\|Undefined Saprotroph\| | Dimorphic Yeast-Yeast | NULL | Possible | As foliar_endophyte (Põlme et al. 2020); Decay leaf, fruit, seed, soil, dung, animal material (Põlme et al. 2020) | Andrews JH, et al. 1983. Phytopathology 73:228-234 ((https://doi.org/10.1094/Phyto-73-228)); Kurtzman CP, et al. (eds.) 2011. The Yeasts, a Taxonomic Study. Fifth Edition. Vols 1-3. Elsevier, San Diego (ISBN: 9780444521491); Põlme S, et al. 2020. Fungal Diversity 105:1-16 ((https://doi.org/10.1007/s13225-020-00466-2)) |
| Otu00720 | 109 | k__Fungi(100);p__Ascomycota(100);c__Dothideomycetes(100);o__Pleosporales(100);f__Sporormiaceae(100);g__Preussia(100);s__Preussia_flanaganii(100); | Preussia | 13 | Saprotroph | \|Dung Saprotroph\|-Undefined Saprotroph | Microfungus | NULL | Probable | Decay dung (Põlme et al. 2020) | Tedersoo L, et al. 2014. Science 346:e1256688 ((https://doi.org/10.1126/science.1256688)); Põlme S, et al. 2020. Fungal Diversity 105:1-16 ((https://doi.org/10.1007/s13225-020-00466-2)) |
| Otu00721 | 109 | k__Fungi(100);p__Ascomycota(100);c__Lecanoromycetes(91);o__Teloschistales(59);o__Teloschistales_unclassified(59);o__Teloschistales_unclassified(59);o__Teloschistales_unclassified(59); | - | - | - | - | - | - | - | Unassigned | - |
| Otu00722 | 109 | k__Fungi(100);k__Fungi_unclassified(97);k__Fungi_unclassified(97);k__Fungi_unclassified(97);k__Fungi_unclassified(97);k__Fungi_unclassified(97); | - | - | - | - | - | - | - | Unassigned | - |
| Otu00723 | 108 | k__Fungi(100);p__Basidiomycota(100);c__Agaricomycetes(100);o__Agaricales(100);f__Psathyrellaceae(100);g__Coprinopsis(100);g__Coprinopsis_unclassified(51); | Coprinopsis | 13 | Saprotroph | \|Undefined Saprotroph\| | Agaricoid | NULL | Probable | As soil_saprotroph (Põlme et al. 2020); Decay soil (Põlme et al. 2020) | Tedersoo L, et al. 2014. Science 346:e1256688 ((https://doi.org/10.1126/science.1256688)); Põlme S, et al. 2020. Fungal Diversity 105:1-16 ((https://doi.org/10.1007/s13225-020-00466-2)) |
| Otu00724 | 108 | k__Fungi(100);p__Ascomycota(100);c__Sordariomycetes(100);o__Hypocreales(100);f__Hypocreales_family_Incertae_sedis(100);g__Acremonium(100);s__Acremonium_spinosum(100); | Acremonium | 13 | Pathotroph-Saprotroph-Symbiotroph | Animal Pathogen-Endophyte-Fungal Parasite-Plant Pathogen-\|Undefined Saprotroph\|-Wood Saprotroph | Microfungus | Soft Rot | Possible | As foliar_endophyte (Põlme et al. 2020); culture (Hanada et al. 2010, Del Olmo; Decay leaf, fruit, seed, soil, dung, animal material (Põlme et al. 2020); Endophyte detection method; Ruiz & Arnold 2014); Likely opportunistic human pathogen (Irinyi et al. 2015) | Duncan CG, Eslyn WE. 1966. Mycologia 58:642-645 ((https://doi.org/10.2307/3757045)); Nilsson T. 1973. Studia Forestalia Suecica 104: 1-40 ((https://pub.epsilon.slu.se/5711/)); Worrall JJ, et al. 1997. Mycologia 89:199-219 ((https://doi.org/10.2307/3761073)); Hanada RE, et al. 2010. Fungal Biology 114:901-910 ((https://doi.org/10.1016/j.funbio.2010.08.006)); Del Olmo-Ruiz M, Arnold AE. 2014. Mycologia 106: 8-21 ((https://doi.org/10.3852/13-098)); Irinyi L, et al. 2015. Medical Mycology 53313-337 ((https://doi.org/10.1093/mmy/myv008)); Põlme S, et al. 2020. Fungal Diversity 105:1-16 ((https://doi.org/10.1007/s13225-020-00466-2)) |
| Otu00725 | 108 | k__Fungi(100);p__Basidiomycota(100);c__Cystobasidiomycetes(100);o__Erythrobasidiales(100);f__Erythrobasidiales_family_Incertae_sedis(100);g__Erythrobasidium(100);s__Erythrobasidium_hasegawianum(100); | Erythrobasidium | 13 | Pathotroph-Saprotroph | \|Fungal Parasite\|-Undefined Saprotroph | Yeast | NULL | Possible | As mycoparasite (Põlme et al. 2020); Decay fungal material (Põlme et al. 2020) | Sugiyama J, Hamamoto M. 1998. The Yeasts 4:654-655 ((https://doi.org/10.1016/B978-044481312-1/50086-1)); Põlme S, et al. 2020. Fungal Diversity 105:1-16 ((https://doi.org/10.1007/s13225-020-00466-2)) |
| Otu00726 | 108 | k__Fungi(100);p__Ascomycota(100);c__Lecanoromycetes(100);o__Lecanorales(100);o__Lecanorales_unclassified(100);o__Lecanorales_unclassified(100);o__Lecanorales_unclassified(100); | Lecanorales | 7 | Pathotroph-Symbiotroph | Lichen Parasite-Lichenized | Microfungus-Thallus | NULL | Probable | Lichenized with green algae, occasionally lichenicolous, on rocks, mosses and wood (Cannon & Kirk 2007); ...an order of mostly lichen-forming fungi (Kirk et al. 2008) | Cannon PF, Kirk PM. 2007. Fungal Families of the World. CAB International, Cambridge (ISBN: 978-0851998275); Kirk PM et al. 2008. Dictionary of the Fungi. Tenth Edition. CAB International, Wallingford (ISBN: 978-0851998268); Esslinger TL. 2014. North Dakota State University ((http://www.ndsu.edu/pubweb/~esslinge/chcklst/chcklst7.htm)) |
| Otu00727 | 107 | k__Fungi(100);p__Basidiomycota(100);c__Agaricomycetes(100);o__Polyporales(100);f__Phanerochaetaceae(94);g__Pseudolagarobasidium(94);g__Pseudolagarobasidium_unclassified(91); | Pseudolagarobasidium | 13 | Saprotroph | Undefined Saprotroph-\|Wood Saprotroph\| | Corticioid | White Rot | Probable | As white_rot (Põlme et al. 2020); Decay wood (Põlme et al. 2020) | Tedersoo L, et al. 2014. Science 346:e1256688 ((https://doi.org/10.1126/science.1256688)); Põlme S, et al. 2020. Fungal Diversity 105:1-16 ((https://doi.org/10.1007/s13225-020-00466-2)) |
| Otu00729 | 107 | k__Fungi(100);p__Basidiomycota(100);c__Agaricomycetes(100);o__Polyporales(100);f__Meruliaceae(100);g__Abortiporus(100);s__Abortiporus_biennis(100); | Abortiporus | 13 | Pathotroph-Saprotroph | Plant Pathogen-\|Wood Saprotroph\| | Polyporoid | White Rot | Highly Probable | As root_pathogen (Põlme et al. 2020); As white_rot (Põlme et al. 2020); Decay wood, roots (Põlme et al. 2020) | Gilbertson RL, Ryvarden L. 1987-1987. North American Polypores. Fungiflora, Oslo (ISBN: 978-0945345060); Põlme S, et al. 2020. Fungal Diversity 105:1-16 ((https://doi.org/10.1007/s13225-020-00466-2)) |
| Otu00730 | 107 | k__Fungi(100);p__Basidiomycota(100);c__Agaricomycetes(100);o__Polyporales(100);f__Meruliaceae(100);g__Mycoacia(100);s__Mycoacia_nothofagi_var._nothofagi(100); | Mycoacia | 13 | Saprotroph | Undefined Saprotroph-\|Wood Saprotroph\| | Corticioid | White Rot | Probable | As white_rot (Põlme et al. 2020); Decay wood (Põlme et al. 2020) | Tedersoo L, et al. 2014. Science 346:e1256688 ((https://doi.org/10.1126/science.1256688)); Põlme S, et al. 2020. Fungal Diversity 105:1-16 ((https://doi.org/10.1007/s13225-020-00466-2)) |
| Otu00732 | 107 | k__Fungi(100);p__Basidiomycota(100);c__Agaricomycetes(100);o__Polyporales(100);f__Meruliaceae(100);g__Phlebia(100);s__Phlebia_tuberculata(84); | Phlebia | 13 | Saprotroph | \|Wood Saprotroph\| | Corticioid | White Rot | Highly Probable | As white_rot (Põlme et al. 2020); Decay wood (Põlme et al. 2020) | Kirk PM et al. 2008. Dictionary of the Fungi. Tenth Edition. CAB International, Wallingford (ISBN: 978-0851998268); Põlme S, et al. 2020. Fungal Diversity 105:1-16 ((https://doi.org/10.1007/s13225-020-00466-2)) |
| Otu00728 | 107 | k__Fungi(100);k__Fungi_unclassified(56);k__Fungi_unclassified(56);k__Fungi_unclassified(56);k__Fungi_unclassified(56);k__Fungi_unclassified(56); | - | - | - | - | - | - | - | Unassigned | - |
| Otu00731 | 107 | k__Fungi(100);p__Ascomycota(68);p__Ascomycota_unclassified(68);p__Ascomycota_unclassified(68);p__Ascomycota_unclassified(68);p__Ascomycota_unclassified(68);p__Ascomycota_unclassified(68); | - | - | - | - | - | - | - | Unassigned | - |
| Otu00733 | 105 | k__Fungi(100);p__Basidiomycota(100);c__Microbotryomycetes(100);o__Sporidiobolales(100);f__Sporidiobolales_family_Incertae_sedis(100);g__Sporobolomyces(100);s__Sporobolomyces_lactosus(100); | Sporobolomyces | 13 | Pathotroph-Saprotroph | \|Fungal Parasite\|-Plant Saprotroph | Yeast | NULL | Probable | As litter_saprotroph (Põlme et al. 2020); As mycoparasite (Põlme et al. 2020); Decay fungal material (Põlme et al. 2020) | Tedersoo L, et al. 2014. Science 346:e1256688 ((https://doi.org/10.1126/science.1256688)); Purahong W, et al. 2016. Molecular Ecology 25:4059-4074 ((https://doi.org/10.1111/mec.13739)); Põlme S, et al. 2020. Fungal Diversity 105:1-16 ((https://doi.org/10.1007/s13225-020-00466-2)) |
| Otu00736 | 105 | k__Fungi(100);p__Basidiomycota(100);c__Agaricomycetes(100);o__Polyporales(100);f__Polyporaceae(100);g__Skeletocutis(100);s__Skeletocutis_nivea(100); | Skeletocutis | 13 | Saprotroph | \|Wood Saprotroph\| | Polyporoid | White Rot | Highly Probable | As white_rot (Põlme et al. 2020); Decay wood (Põlme et al. 2020) | Gilbertson RL, Ryvarden L. 1987-1987. North American Polypores. Fungiflora, Oslo (ISBN: 978-0945345060); Põlme S, et al. 2020. Fungal Diversity 105:1-16 ((https://doi.org/10.1007/s13225-020-00466-2)) |
| Otu00737 | 105 | k__Fungi(100);p__Basidiomycota(100);c__Agaricomycetes(100);o__Polyporales(100);f__Phanerochaetaceae(56);g__Phanerochaete(56);s__Phanerochaete_rimosa(56); | Phanerochaete | 13 | Saprotroph | \|Wood Saprotroph\| | Corticioid | White Rot | Highly Probable | As white_rot (Põlme et al. 2020); Decay wood (Põlme et al. 2020) | Glenn JK, et al. 1983. Biochemical and Biophysical Research Communications 114:1077-1083 ((https://doi.org/10.1016/0006-291x((83)) 90672-1)); Põlme S, et al. 2020. Fungal Diversity 105:1-16 ((https://doi.org/10.1007/s13225-020-00466-2)) |
| Otu00738 | 105 | k__Fungi(100);p__Ascomycota(100);c__Saccharomycetes(100);o__Saccharomycetales(100);f__Metschnikowiaceae(100);g__Metschnikowia(100);s__Metschnikowia_pulcherrima(97); | Metschnikowia | 13 | Saprotroph | \|Undefined Saprotroph\| | Yeast | NULL | Probable | As nectar-tap_saprotroph (Põlme et al. 2020); Decay sugar-rich substrates (Põlme et al. 2020) | Põlme S, et al. 2020. Fungal Diversity 105:1-16 ((https://doi.org/10.1007/s13225-020-00466-2)) |
| Otu00734 | 105 | k__Fungi(100);k__Fungi_unclassified(100);k__Fungi_unclassified(100);k__Fungi_unclassified(100);k__Fungi_unclassified(100);k__Fungi_unclassified(100); | - | - | - | - | - | - | - | Unassigned | - |
| Otu00735 | 105 | k__Fungi(100);p__Ascomycota(67);p__Ascomycota_unclassified(67);p__Ascomycota_unclassified(67);p__Ascomycota_unclassified(67);p__Ascomycota_unclassified(67);p__Ascomycota_unclassified(67); | - | - | - | - | - | - | - | Unassigned | - |
| Otu00739 | 104 | k__Fungi(100);p__Ascomycota(100);c__Ascomycota_class_Incertae_sedis(100);o__Ascomycota_order_Incertae_sedis(100);f__Ascomycota_family_Incertae_sedis(100);g__Chalara(100);s__Chalara_sp(100); | Chalara | 13 | Pathotroph-Saprotroph-Symbiotroph | Endophyte-Plant Pathogen-\|Plant Saprotroph\|-Wood Saprotroph | Microfungus | Soft Rot | Probable | As foliar_endophyte (Põlme et al. 2020); As litter_saprotroph (Põlme et al. 2020); As wood_pathogen (Põlme et al. 2020); Decay wood, leaf, fruit, seed (Põlme et al. 2020) | Duncan CG, Eslyn WE. 1966. Mycologia 58:642-645 ((https://doi.org/10.2307/3757045)); Tedersoo L, et al. 2014. Science 346:e1256688 ((https://doi.org/10.1126/science.1256688)); Põlme S, et al. 2020. Fungal Diversity 105:1-16 ((https://doi.org/10.1007/s13225-020-00466-2)) |
| Otu00742 | 104 | k__Fungi(100);p__Ascomycota(100);c__Saccharomycetes(100);o__Saccharomycetales(100);f__Metschnikowiaceae(100);g__Metschnikowia(100);s__Metschnikowia_pulcherrima(97); | Metschnikowia | 13 | Saprotroph | \|Undefined Saprotroph\| | Yeast | NULL | Probable | As nectar-tap_saprotroph (Põlme et al. 2020); Decay sugar-rich substrates (Põlme et al. 2020) | Põlme S, et al. 2020. Fungal Diversity 105:1-16 ((https://doi.org/10.1007/s13225-020-00466-2)) |
| Otu00740 | 104 | k__Fungi(100);p__Basidiomycota(100);c__Agaricomycetes(100);o__Trechisporales(100);f__unclassified_Trechisporales(100);g__unclassified_Trechisporales(100);s__Trechisporales_sp(100); | - | - | - | - | - | - | - | Unassigned | - |
| Otu00741 | 104 | k__Fungi(100);k__Fungi_unclassified(63);k__Fungi_unclassified(63);k__Fungi_unclassified(63);k__Fungi_unclassified(63);k__Fungi_unclassified(63); | - | - | - | - | - | - | - | Unassigned | - |
| Otu00744 | 103 | k__Fungi(100);p__Ascomycota(100);c__Sordariomycetes(100);o__Xylariales(100);f__Xylariaceae(100);g__Daldinia(100);g__Daldinia_unclassified(99); | Daldinia | 13 | Saprotroph-Symbiotroph | Endophyte-Undefined Saprotroph-\|Wood Saprotroph\| | Daldinioid | NULL | Probable | As foliar_endophyte (Põlme et al. 2020); Decay wood, leaf, fruit, seed (Põlme et al. 2020); Host - angiosperms (Põlme et al. 2020) | Tedersoo L, et al. 2014. Science 346:e1256688 ((https://doi.org/10.1126/science.1256688)); Põlme S, et al. 2020. Fungal Diversity 105:1-16 ((https://doi.org/10.1007/s13225-020-00466-2)) |
| Otu00745 | 103 | k__Fungi(100);p__Basidiomycota(100);c__Agaricomycetes(100);o__Agaricales(100);f__Psathyrellaceae(100);g__Psathyrella(100);s__Psathyrella_romagnesii(100); | Psathyrella | 13 | Saprotroph | Plant Saprotroph-\|Wood Saprotroph\| | Agaricoid | NULL | Probable | As litter_saprotroph (Põlme et al. 2020); Decay wood, leaf, fruit, seed (Põlme et al. 2020) | Tedersoo L, et al. 2014. Science 346:e1256688 ((https://doi.org/10.1126/science.1256688)); Põlme S, et al. 2020. Fungal Diversity 105:1-16 ((https://doi.org/10.1007/s13225-020-00466-2)) |
| Otu00749 | 103 | k__Fungi(100);p__Basidiomycota(100);c__Agaricomycetes(100);o__Trechisporales(100);f__Hydnodontaceae(100);g__Trechispora(100);g__Trechispora_unclassified(64); | Trechispora | 13 | Saprotroph | \|Wood Saprotroph\| | Corticioid | White Rot | Highly Probable | As white_rot (Põlme et al. 2020); Decay wood (Põlme et al. 2020) | Gilbertson RL, Ryvarden L. 1987-1987. North American Polypores. Fungiflora, Oslo (ISBN: 978-0945345060); Rinaldi AC, et al. 2008. Fungal Diversity 33:1-45 ((http://www.fungaldiversity.org/fdp/sfdp/33-1.pdf)) (as Uncertain); Tedersoo L, et al. 2010. Mycorrhiza 20:217-263 ((https://doi.org/10.1007/s00572-009-0274-x)) (as not ECM).; Põlme S, et al. 2020. Fungal Diversity 105:1-16 ((https://doi.org/10.1007/s13225-020-00466-2)) |
| Otu00750 | 103 | k__Fungi(100);p__Ascomycota(100);c__Dothideomycetes(100);o__Pleosporales(100);f__Pleosporales_family_Incertae_sedis(100);g__Paraphoma(100);s__Paraphoma_dioscoreae(100); | Paraphoma | 13 | Pathotroph-Saprotroph | \|Plant Pathogen\|-Plant Saprotroph-Undefined Saprotroph-Wood Saprotroph | Microfungus | NULL | Probable | As wood_pathogen (Põlme et al. 2020); Associated with leaf and crown infection of pyrethrum (Moslemi et al. 2017); Decay wood (Põlme et al. 2020) | Quaedvlieg W, et al. 2013. Studies in Mycology 75:307-390 ((https://doi.org/10.3114/sim0017)); Moslemi A, et al. 2017. Plant Pathology 67:124-135 ((https://doi.org/10.1111/ppa.12719)); Põlme S, et al. 2020. Fungal Diversity 105:1-16 ((https://doi.org/10.1007/s13225-020-00466-2)) |
| Otu00743 | 103 | k__Fungi(100);p__Basidiomycota(100);p__Basidiomycota_unclassified(100);p__Basidiomycota_unclassified(100);p__Basidiomycota_unclassified(100);p__Basidiomycota_unclassified(100);p__Basidiomycota_unclassified(100); | - | - | - | - | - | - | - | Unassigned | - |
| Otu00746 | 103 | k__Fungi(100);p__Ascomycota(53);p__Ascomycota_unclassified(53);p__Ascomycota_unclassified(53);p__Ascomycota_unclassified(53);p__Ascomycota_unclassified(53);p__Ascomycota_unclassified(53); | - | - | - | - | - | - | - | Unassigned | - |
| Otu00747 | 103 | k__Fungi(100);p__Ascomycota(73);p__Ascomycota_unclassified(73);p__Ascomycota_unclassified(73);p__Ascomycota_unclassified(73);p__Ascomycota_unclassified(73);p__Ascomycota_unclassified(73); | - | - | - | - | - | - | - | Unassigned | - |
| Otu00748 | 103 | k__Fungi(100);p__Ascomycota(100);p__Ascomycota_unclassified(100);p__Ascomycota_unclassified(100);p__Ascomycota_unclassified(100);p__Ascomycota_unclassified(100);p__Ascomycota_unclassified(100); | - | - | - | - | - | - | - | Unassigned | - |
| Otu00751 | 103 | k__Fungi(100);p__Ascomycota(100);c__Dothideomycetes(100);o__Pleosporales(100);f__unclassified_Pleosporales(71);g__unclassified_Pleosporales(71);s__Pleosporales_sp(71); | - | - | - | - | - | - | - | Unassigned | - |
| Otu00752 | 102 | k__Fungi(100);p__Ascomycota(100);c__Lecanoromycetes(100);o__Lecanorales(100);o__Lecanorales_unclassified(100);o__Lecanorales_unclassified(100);o__Lecanorales_unclassified(100); | Lecanorales | 7 | Pathotroph-Symbiotroph | Lichen Parasite-Lichenized | Microfungus-Thallus | NULL | Probable | Lichenized with green algae, occasionally lichenicolous, on rocks, mosses and wood (Cannon & Kirk 2007); ...an order of mostly lichen-forming fungi (Kirk et al. 2008) | Cannon PF, Kirk PM. 2007. Fungal Families of the World. CAB International, Cambridge (ISBN: 978-0851998275); Kirk PM et al. 2008. Dictionary of the Fungi. Tenth Edition. CAB International, Wallingford (ISBN: 978-0851998268); Esslinger TL. 2014. North Dakota State University ((http://www.ndsu.edu/pubweb/~esslinge/chcklst/chcklst7.htm)) |
| Otu00753 | 102 | k__Fungi(100);p__Ascomycota(100);c__Sordariomycetes(100);o__Sordariomycetes_order_Incertae_sedis(100);f__Magnaporthaceae(100);g__Mycoleptodiscus(100);g__Mycoleptodiscus_unclassified(98); | Mycoleptodiscus | 13 | Pathotroph | \|Plant Pathogen\| | Microfungus | NULL | Probable | NULL | Tedersoo L, et al. 2014. Science 346:e1256688 ((https://doi.org/10.1126/science.1256688)); Põlme S, et al. 2020. Fungal Diversity 105:1-16 ((https://doi.org/10.1007/s13225-020-00466-2)) |
| Otu00754 | 101 | k__Fungi(100);p__Zygomycota(100);c__Zygomycota_class_Incertae_sedis(100);o__Mucorales(100);f__Mucoraceae(100);g__Mucor(100);s__Mucor_sp_XSCG02(100); | Mucor | 13 | Saprotroph | \|Undefined Saprotroph\| | Microfungus | NULL | Probable | As soil_saprotroph (Põlme et al. 2020); Decay soil (Põlme et al. 2020) | Tedersoo L, et al. 2014. Science 346:e1256688 ((https://doi.org/10.1126/science.1256688)); Põlme S, et al. 2020. Fungal Diversity 105:1-16 ((https://doi.org/10.1007/s13225-020-00466-2)) |
| Otu00756 | 101 | k__Fungi(100);p__Ascomycota(100);c__Dothideomycetes(100);o__Capnodiales(100);f__Teratosphaeriaceae(96);g__Devriesia(72);s__Devriesia_americana(54); | Devriesia | 13 | Pathotroph-Saprotroph | \|Plant Pathogen\|-Plant Saprotroph | Microfungus | NULL | Probable | As litter_saprotroph (Põlme et al. 2020); Decay leaf, fruit, seed (Põlme et al. 2020) | Tedersoo L, et al. 2014. Science 346:e1256688 ((https://doi.org/10.1126/science.1256688)); Põlme S, et al. 2020. Fungal Diversity 105:1-16 ((https://doi.org/10.1007/s13225-020-00466-2)) |
| Otu00755 | 101 | k__Fungi(100);p__Ascomycota(100);p__Ascomycota_unclassified(100);p__Ascomycota_unclassified(100);p__Ascomycota_unclassified(100);p__Ascomycota_unclassified(100);p__Ascomycota_unclassified(100); | - | - | - | - | - | - | - | Unassigned | - |
| Otu00757 | 100 | k__Fungi(100);p__Ascomycota(100);c__Dothideomycetes(99);o__Capnodiales(98);f__Teratosphaeriaceae(92);g__Devriesia(88);s__Devriesia_sp(77); | Devriesia | 13 | Pathotroph-Saprotroph | \|Plant Pathogen\|-Plant Saprotroph | Microfungus | NULL | Probable | As litter_saprotroph (Põlme et al. 2020); Decay leaf, fruit, seed (Põlme et al. 2020) | Tedersoo L, et al. 2014. Science 346:e1256688 ((https://doi.org/10.1126/science.1256688)); Põlme S, et al. 2020. Fungal Diversity 105:1-16 ((https://doi.org/10.1007/s13225-020-00466-2)) |
| Otu00760 | 100 | k__Fungi(100);p__Ascomycota(100);c__Saccharomycetes(99);o__Saccharomycetales(99);f__Saccharomycetales_family_Incertae_sedis(94);g__Candida(82);g__Candida_unclassified(56); | Candida | 13 | Pathotroph-Saprotroph-Symbiotroph | Animal Pathogen-Endophyte-Endosymbiont-Epiphyte-\|Undefined Saprotroph\| | Dimorphic Yeast-Yeast | NULL | Probable | As nectar-tap_saprotroph (Põlme et al. 2020); Decay sugar-rich substrates (Põlme et al. 2020); This asexual genus is polyphyletic and contain species that function very differently depend on what they are. Caution should be taken when interpreting guilds for Candida species. | Manolakaki D, et al. 2010. Virulence. 1: 367-375 ((https://doi.org/10.4161/viru.1.5.12796)); Mekbib SB, et al. 2011. Tropical Plant Pathology 36:233-240 ((https://doi.org/10.1590/S1982-56762011000400004)); Põlme S, et al. 2020. Fungal Diversity 105:1-16 ((https://doi.org/10.1007/s13225-020-00466-2)) |
| Otu00761 | 100 | k__Fungi(100);p__Ascomycota(100);c__Sordariomycetes(100);o__Diaporthales(100);f__Diaporthales_family_Incertae_sedis(100);g__Ophiognomonia(100);s__Ophiognomonia_lenticulispora(100); | Ophiognomonia | 13 | Pathotroph | \|Plant Pathogen\| | Microfungus | NULL | Highly Probable | See Marssoniella juglandis, Ophiognomonia leptostyla, and Syn. Gnomonia leptostyla; Causes leaf blotch and spots, also known as walnut anthracnose or walnut black spot (Sogonov et al. 2008) | Sogonov MV, et al. 2008. Studies in Mycology 62:1-79 ((https://www.studiesinmycology.org/sim/Sim62/03_Leaf-inhabiting_genera_of_the_Gnomoniaceae,_Diaporthales.pdf)); Põlme S, et al. 2020. Fungal Diversity 105:1-16 ((https://doi.org/10.1007/s13225-020-00466-2)) |
| Otu00762 | 100 | k__Fungi(100);p__Ascomycota(100);c__Dothideomycetes(100);o__Capnodiales(100);f__Mycosphaerellaceae(100);g__Mycosphaerella(99);s__Mycosphaerella_nyssicola(99); | Mycosphaerella | 13 | Pathotroph-Saprotroph | \|Plant Pathogen\|-Plant Saprotroph | Microfungus | NULL | Probable | As litter_saprotroph (Põlme et al. 2020); Decay leaf, fruit, seed (Põlme et al. 2020) | Tedersoo L, et al. 2014. Science 346:e1256688 ((https://doi.org/10.1126/science.1256688)); Põlme S, et al. 2020. Fungal Diversity 105:1-16 ((https://doi.org/10.1007/s13225-020-00466-2)) |
| Otu00763 | 100 | k__Fungi(100);p__Basidiomycota(100);c__Tremellomycetes(100);o__Tremellales(100);f__Tremellales_family_Incertae_sedis(100);g__Bullera(100);s__Bullera_pseudoalba(100); | Bullera | 13 | Saprotroph-Symbiotroph | Endophyte-Epiphyte-Plant Saprotroph-\|Undefined Saprotroph\| | Yeast | NULL | Probable | As foliar_endophyte (Põlme et al. 2020); Isolated from dead leaves of Oryza sativa, Miscanthus sinensis and Sasa sp. in Japan (Nakase & Suzuki 1986; Syn. Udeniomyces puniceus); Foliar, isolated from leaves of Melostomae candidum (Nakase et al. 2004) | Nakase T, Suzuki M. 1986. Journal of General and Applied Microbiology Tokyo 32:225-240 ((https://doi.org/10.2323/jgam.32.225)); Nakase T, et al. 2004. Systematic and Applied Microbiology 27:558-564 ((https://doi.org/10.1078/0723202041748118)) |
| Otu00758 | 100 | k__Fungi(100);p__Basidiomycota(100);c__Agaricomycetes(100);o__Trechisporales(100);f__unclassified_Trechisporales(100);g__unclassified_Trechisporales(100);s__Trechisporales_sp(100); | - | - | - | - | - | - | - | Unassigned | - |
| Otu00759 | 100 | k__Fungi(100);k__Fungi_unclassified(61);k__Fungi_unclassified(61);k__Fungi_unclassified(61);k__Fungi_unclassified(61);k__Fungi_unclassified(61); | - | - | - | - | - | - | - | Unassigned | - |
| Otu00764 | 99 | k__Fungi(100);p__Ascomycota(100);c__Leotiomycetes(100);o__Helotiales(100);f__Helotiales_family_Incertae_sedis(100);g__Trimmatostroma(100);s__Trimmatostroma_salinum(100); | Trimmatostroma | 13 | Pathotroph-Saprotroph | \|Lichen Parasite\|-Wood Saprotroph | Microfungus | NULL | Probable | Decay fungal material, algal material (Põlme et al. 2020) | Tedersoo L, et al. 2014. Science 346:e1256688 ((https://doi.org/10.1126/science.1256688)); Lawrey JD, Diederich P. 2016. http://www.lichenicolous.net (accessed 24 May 2017) ((http://www.lichenicolous.net)); Põlme S, et al. 2020. Fungal Diversity 105:1-16 ((https://doi.org/10.1007/s13225-020-00466-2)) |
| Otu00765 | 99 | k__Fungi(100);p__Ascomycota(100);c__Eurotiomycetes(100);o__Eurotiales(100);f__Trichocomaceae(100);g__Penicillium(100);g__Penicillium_unclassified(54); | Penicillium | 13 | Pathotroph-Saprotroph-Symbiotroph | Animal Parasite-Dung Saprotroph-Endophyte-Fungal Parasite-Plant Pathogen-\|Undefined Saprotroph\|-Wood Saprotroph | Microfungus | Soft Rot | Highly Probable | As foliar_endophyte (Põlme et al. 2020); As mycoparasite (Põlme et al. 2020); Decay soil (Põlme et al. 2020) | Duncan CG, Eslyn WE. 1966. Mycologia 58:642-664 ((https://doi.org/10.2307/3757045)); Seehann G, et al. 1975. List of Fungi in Soft-Rot Tissue Tests. Institute fur Holzbiologie und Holzschutz, Bundesforschungsanstalt fur Forst - und Holzwirtshaft, Hamburg - Lohbrugge, Germany ((https://www.irg-wp.com/irgdocs/details.php?d6493033-c9ea-40f3-8e3e-5697f32d640d)); Costa IPMW, et al. 2012. Mycotaxon 119:e26 ((http://mycotaxon.com/resources/checklists/costa_v119_checklist.pdf)); Bills GF, et al. 2013. Current Opinion in Microbiology 16: 549-565 ((https://doi.org/10.1016/j.mib.2013.08.001)); Tedersoo L, et al. 2014. Science 346:e1256688 ((https://doi.org/10.1126/science.1256688)); Põlme S, et al. 2020. Fungal Diversity 105:1-16 ((https://doi.org/10.1007/s13225-020-00466-2)) |
| Otu00766 | 99 | k__Fungi(100);p__Basidiomycota(100);c__Agaricomycetes(100);o__Hymenochaetales(100);f__Hymenochaetaceae(100);g__Phellinus(100);s__Phellinus_robiniae(100); | Phellinus | 13 | Pathotroph-Saprotroph | \|Plant Pathogen\|-Wood Saprotroph | Polyporoid | White Rot | Highly Probable | As white_rot (Põlme et al. 2020); As wood_pathogen (Põlme et al. 2020); Decay wood (Põlme et al. 2020) | Gilbertson RL, Ryvarden L. 1987-1987. North American Polypores. Fungiflora, Oslo (ISBN: 978-0945345060); Põlme S, et al. 2020. Fungal Diversity 105:1-16 ((https://doi.org/10.1007/s13225-020-00466-2)) |
| Otu00768 | 99 | k__Fungi(100);p__Basidiomycota(100);c__Tremellomycetes(100);o__Tremellales(100);f__Tremellales_family_Incertae_sedis(100);g__Cryptococcus(100);s__Cryptococcus_sp_CBS_9459(100); | Cryptococcus | 13 | Pathotroph-Saprotroph-Symbiotroph | Animal Pathogen-Endophyte-Epiphyte-\|Undefined Saprotroph\| | Dimorphic Yeast-Yeast | NULL | Possible | As foliar_endophyte (Põlme et al. 2020); Decay leaf, fruit, seed, soil, dung, animal material (Põlme et al. 2020) | Andrews JH, et al. 1983. Phytopathology 73:228-234 ((https://doi.org/10.1094/Phyto-73-228)); Kurtzman CP, et al. (eds.) 2011. The Yeasts, a Taxonomic Study. Fifth Edition. Vols 1-3. Elsevier, San Diego (ISBN: 9780444521491); Põlme S, et al. 2020. Fungal Diversity 105:1-16 ((https://doi.org/10.1007/s13225-020-00466-2)) |
| Otu00769 | 99 | k__Fungi(100);p__Ascomycota(100);c__Dothideomycetes(100);o__Pleosporales(100);f__Lophiostomataceae(100);g__Lophiostoma(100);s__Lophiostoma_sp(100); | Lophiostoma | 13 | Saprotroph | Plant Saprotroph-Undefined Saprotroph-\|Wood Saprotroph\| | Microfungus | NULL | Probable | As litter_saprotroph (Põlme et al. 2020); Decay wood, leaf, fruit, seed (Põlme et al. 2020) | Tedersoo L, et al. 2014. Science 346:e1256688 ((https://doi.org/10.1126/science.1256688)); Põlme S, et al. 2020. Fungal Diversity 105:1-16 ((https://doi.org/10.1007/s13225-020-00466-2)) |
| Otu00767 | 99 | k__Fungi(100);k__Fungi_unclassified(91);k__Fungi_unclassified(91);k__Fungi_unclassified(91);k__Fungi_unclassified(91);k__Fungi_unclassified(91); | - | - | - | - | - | - | - | Unassigned | - |
| Otu00770 | 99 | k__Fungi(100);p__Ascomycota(100);c__Dothideomycetes(100);o__Pleosporales(97);o__Pleosporales_unclassified(97);o__Pleosporales_unclassified(97);o__Pleosporales_unclassified(97); | - | - | - | - | - | - | - | Unassigned | - |
| Otu00771 | 99 | k__Fungi(100);p__Ascomycota(100);p__Ascomycota_unclassified(100);p__Ascomycota_unclassified(100);p__Ascomycota_unclassified(100);p__Ascomycota_unclassified(100);p__Ascomycota_unclassified(100); | - | - | - | - | - | - | - | Unassigned | - |
| Otu00772 | 99 | k__Fungi(100);p__Basidiomycota(100);c__Agaricomycetes(100);o__Polyporales(100);f__Phanerochaetaceae(100);f__Phanerochaetaceae_unclassified(90);f__Phanerochaetaceae_unclassified(90); | - | - | - | - | - | - | - | Unassigned | - |
| Otu00773 | 98 | k__Fungi(100);p__Ascomycota(100);c__Ascomycota_class_Incertae_sedis(100);o__Ascomycota_order_Incertae_sedis(100);f__Ascomycota_family_Incertae_sedis(100);g__Pseudorobillarda(100);s__Pseudorobillarda_sp_MFLUCC12_0422(100); | Pseudorobillarda | 13 | Pathotroph-Saprotroph | Lichen Parasite-\|Wood Saprotroph\| | Microfungus | NULL | Probable | Decay wood (Põlme et al. 2020) | Põlme S, et al. 2020. Fungal Diversity 105:1-16 ((https://doi.org/10.1007/s13225-020-00466-2)) |
| Otu00774 | 98 | k__Fungi(100);p__Ascomycota(100);c__Sordariomycetes(100);o__Hypocreales(100);f__Cordycipitaceae(100);g__Cordyceps(86);s__Cordyceps_bassiana(86); | Cordyceps | 13 | Pathotroph-Symbiotroph | Clavicipitaceous Endophyte-Fungal Parasite-\|Insect Pathogen\| | Cordycepitoid | NULL | Possible | As foliar_endophyte (Põlme et al. 2020); Decay animal material (Põlme et al. 2020) | Samuels GJ. 2007. Self-published; Costa IPMW, et al. 2012. Mycotaxon 119:e26 ((http://mycotaxon.com/resources/checklists/costa_v119_checklist.pdf)); Tedersoo L, et al. 2014. Science 346:e1256688 ((https://doi.org/10.1126/science.1256688)); Põlme S, et al. 2020. Fungal Diversity 105:1-16 ((https://doi.org/10.1007/s13225-020-00466-2)) |
| Otu00775 | 98 | k__Fungi(100);p__Basidiomycota(100);c__Microbotryomycetes(100);o__Sporidiobolales(100);f__Sporidiobolales_family_Incertae_sedis(100);g__Rhodotorula(100);s__Rhodotorula_sp_JCM_11351(100); | Rhodotorula | 13 | Pathotroph-Saprotroph-Symbiotroph | Animal Pathogen-Animal Symbiotroph-Endophyte-Plant Pathogen-\|Undefined Saprotroph\| | Yeast | NULL | Probable | As foliar_endophyte (Põlme et al. 2020); Opportunistic human pathogen | Põlme S, et al. 2020. Fungal Diversity 105:1-16 ((https://doi.org/10.1007/s13225-020-00466-2)); https://www.cdc.gov ((https://www.cdc.gov)) |
| Otu00776 | 98 | k__Fungi(100);k__Fungi_unclassified(71);k__Fungi_unclassified(71);k__Fungi_unclassified(71);k__Fungi_unclassified(71);k__Fungi_unclassified(71); | - | - | - | - | - | - | - | Unassigned | - |
| Otu00778 | 97 | k__Fungi(100);p__Basidiomycota(100);c__Agaricomycetes(100);o__Trechisporales(88);f__Hydnodontaceae(86);g__Trechispora(68);g__Trechispora_unclassified(68); | Trechispora | 13 | Saprotroph | \|Wood Saprotroph\| | Corticioid | White Rot | Highly Probable | As white_rot (Põlme et al. 2020); Decay wood (Põlme et al. 2020) | Gilbertson RL, Ryvarden L. 1987-1987. North American Polypores. Fungiflora, Oslo (ISBN: 978-0945345060); Rinaldi AC, et al. 2008. Fungal Diversity 33:1-45 ((http://www.fungaldiversity.org/fdp/sfdp/33-1.pdf)) (as Uncertain); Tedersoo L, et al. 2010. Mycorrhiza 20:217-263 ((https://doi.org/10.1007/s00572-009-0274-x)) (as not ECM).; Põlme S, et al. 2020. Fungal Diversity 105:1-16 ((https://doi.org/10.1007/s13225-020-00466-2)) |
| Otu00777 | 97 | k__Fungi(100);p__Basidiomycota(100);c__Agaricomycetes(100);o__Polyporales(100);o__Polyporales_unclassified(89);o__Polyporales_unclassified(89);o__Polyporales_unclassified(89); | - | - | - | - | - | - | - | Unassigned | - |
| Otu00779 | 97 | k__Fungi(100);k__Fungi_unclassified(61);k__Fungi_unclassified(61);k__Fungi_unclassified(61);k__Fungi_unclassified(61);k__Fungi_unclassified(61); | - | - | - | - | - | - | - | Unassigned | - |
| Otu00780 | 96 | k__Fungi(100);p__Basidiomycota(100);c__Agaricomycetes(100);o__Hymenochaetales(100);f__Hymenochaetales_family_Incertae_sedis(100);g__Trichaptum(100);s__Trichaptum_biforme(100); | Trichaptum | 13 | Saprotroph | \|Wood Saprotroph\| | Polyporoid | White Rot | Highly Probable | As white_rot (Põlme et al. 2020); Decay wood (Põlme et al. 2020) | Gilbertson RL, Ryvarden L. 1987-1987. North American Polypores. Fungiflora, Oslo (ISBN: 978-0945345060); Põlme S, et al. 2020. Fungal Diversity 105:1-16 ((https://doi.org/10.1007/s13225-020-00466-2)) |
| Otu00784 | 96 | k__Fungi(100);p__Basidiomycota(100);c__Agaricomycetes(100);o__Cantharellales(100);f__Ceratobasidiaceae(100);g__Ceratobasidium(99);g__Ceratobasidium_unclassified(99); | Ceratobasidium | 13 | Pathotroph-Saprotroph-Symbiotroph | Ectomycorrhizal-Endomycorrhizal-Orchid Mycorrhizal-\|Plant Pathogen\|-Plant Saprotroph-Undefined Saprotroph | Corticioid-Microfungus | NULL | Possible | especially of Orchidaceae but also even bryophytes (Cannon & Kirk 2007); As litter_saprotroph (Põlme et al. 2020); Best known as plant pathogens, but also saprobic. Species are endomycorrhizal with a wide range of plant roots ; Decay leaf, fruit, seed, roots, soil (Põlme et al. 2020) | Cannon PF, Kirk PM. 2007. Fungal Families of the World. CAB International, Cambridge (ISBN: 978-0851998275); Tedersoo L, et al. 2014. Science 346:e1256688 ((https://doi.org/10.1126/science.1256688)); Põlme S, et al. 2020. Fungal Diversity 105:1-16 ((https://doi.org/10.1007/s13225-020-00466-2)) |
| Otu00785 | 96 | k__Fungi(100);p__Ascomycota(100);c__Sordariomycetes(100);o__Xylariales(100);f__Xylariaceae(100);f__Xylariaceae_unclassified(100);f__Xylariaceae_unclassified(100); | Xylariaceae | 9 | Pathotroph-Saprotroph-Symbiotroph | Dung Saprotroph-Endophyte-Plant Pathogen-Undefined Saprotroph | Xylarioid | NULL | Probable | Saprobes or weak pathogens, many endophytes, mainly in wood and bark, occasionally in soil and dung (Cannon & Kirk 2007) | Cannon PF, Kirk PM. 2007. Fungal Families of the World. CAB International, Cambridge (ISBN: 978-0851998275) |
| Otu00781 | 96 | k__Fungi(100);p__Basidiomycota(100);p__Basidiomycota_unclassified(86);p__Basidiomycota_unclassified(86);p__Basidiomycota_unclassified(86);p__Basidiomycota_unclassified(86);p__Basidiomycota_unclassified(86); | - | - | - | - | - | - | - | Unassigned | - |
| Otu00782 | 96 | k__Fungi(100);k__Fungi_unclassified(73);k__Fungi_unclassified(73);k__Fungi_unclassified(73);k__Fungi_unclassified(73);k__Fungi_unclassified(73); | - | - | - | - | - | - | - | Unassigned | - |
| Otu00783 | 96 | k__Fungi(100);p__Ascomycota(64);p__Ascomycota_unclassified(64);p__Ascomycota_unclassified(64);p__Ascomycota_unclassified(64);p__Ascomycota_unclassified(64);p__Ascomycota_unclassified(64); | - | - | - | - | - | - | - | Unassigned | - |
| Otu00786 | 95 | k__Fungi(100);p__Ascomycota(100);c__Dothideomycetes(100);o__Pleosporales(100);f__Montagnulaceae(100);f__Montagnulaceae_unclassified(78);f__Montagnulaceae_unclassified(78); | Montagnulaceae | 9 | Pathotroph-Saprotroph | Fungal Parasite-Plant Pathogen-Undefined Saprotroph | Microfungus | NULL | Probable | Species in this clade can be saprobic, plant pathogenic, or mycoparasitic (Zhang et al. 2009) | Zhang Y, et al. 2009. Studies in Mycology 64:85-102 ((https://doi.org/10.3114/sim.2009.64.04)) |
| Otu00788 | 95 | k__Fungi(100);p__Ascomycota(100);c__Saccharomycetes(100);o__Saccharomycetales(100);f__Metschnikowiaceae(100);g__Metschnikowia(100);s__Metschnikowia_pulcherrima(90); | Metschnikowia | 13 | Saprotroph | \|Undefined Saprotroph\| | Yeast | NULL | Probable | As nectar-tap_saprotroph (Põlme et al. 2020); Decay sugar-rich substrates (Põlme et al. 2020) | Põlme S, et al. 2020. Fungal Diversity 105:1-16 ((https://doi.org/10.1007/s13225-020-00466-2)) |
| Otu00789 | 95 | k__Fungi(100);p__Ascomycota(100);c__Sordariomycetes(100);o__Sordariomycetes_order_Incertae_sedis(100);f__Glomerellaceae(100);g__Colletotrichum(100);s__Colletotrichum_anthrisci(100); | Colletotrichum | 13 | Pathotroph-Saprotroph-Symbiotroph | Endophyte-\|Plant Pathogen\|-Plant Saprotroph | Microfungus | NULL | Probable | As foliar_endophyte (Põlme et al. 2020); As litter_saprotroph (Põlme et al. 2020); Decay leaf, fruit, seed (Põlme et al. 2020) | Costa IPMW, et al. 2012. Mycotaxon 119:e26 ((http://mycotaxon.com/resources/checklists/costa_v119_checklist.pdf)); Tedersoo L, et al. 2014. Science 346:e1256688 ((https://doi.org/10.1126/science.1256688)); Põlme S, et al. 2020. Fungal Diversity 105:1-16 ((https://doi.org/10.1007/s13225-020-00466-2)); The American Phytopathological Society Common Names of Plant Diseases ((http://www.apsnet.org/publications/commonnames/Pages/default.aspx)) |
| Otu00791 | 95 | k__Fungi(100);p__Basidiomycota(100);c__Pucciniomycetes(100);o__Septobasidiales(99);f__Septobasidiaceae(99);g__Septobasidium(99);s__Septobasidium_sp_JBO_2010(99); | Septobasidium | 13 | Pathotroph | \|Animal Parasite\|-Animal Pathogen-Arthropod Parasite | Dimorphic-Facultative Yeast | NULL | Probable | Decay animal material (Põlme et al. 2020); Host - scale_insects (Põlme et al. 2020) | Tedersoo L, et al. 2014. Science 346:e1256688 ((https://doi.org/10.1126/science.1256688)); Põlme S, et al. 2020. Fungal Diversity 105:1-16 ((https://doi.org/10.1007/s13225-020-00466-2)) |
| Otu00792 | 95 | k__Fungi(100);p__Basidiomycota(100);c__Agaricomycetes(100);o__Hymenochaetales(100);f__Hymenochaetaceae(100);g__Hymenochaete(100);g__Hymenochaete_unclassified(99); | Hymenochaete | 13 | Pathotroph-Saprotroph | Plant Pathogen-Undefined Saprotroph-\|Wood Saprotroph\| | Corticioid-Polyporoid | White Rot | Probable | As white_rot (Põlme et al. 2020); As wood_pathogen (Põlme et al. 2020); Decay wood (Põlme et al. 2020) | Tedersoo L, et al. 2014. Science 346:e1256688 ((https://doi.org/10.1126/science.1256688)); Põlme S, et al. 2020. Fungal Diversity 105:1-16 ((https://doi.org/10.1007/s13225-020-00466-2)) |
| Otu00787 | 95 | k__Fungi(100);p__Ascomycota(100);c__Dothideomycetes(100);o__Capnodiales(100);o__Capnodiales_unclassified(88);o__Capnodiales_unclassified(88);o__Capnodiales_unclassified(88); | - | - | - | - | - | - | - | Unassigned | - |
| Otu00790 | 95 | k__Fungi(100);p__Ascomycota(100);p__Ascomycota_unclassified(100);p__Ascomycota_unclassified(100);p__Ascomycota_unclassified(100);p__Ascomycota_unclassified(100);p__Ascomycota_unclassified(100); | - | - | - | - | - | - | - | Unassigned | - |
| Otu00793 | 94 | k__Fungi(100);p__Basidiomycota(100);c__Exobasidiomycetes(100);o__Exobasidiomycetes_order_Incertae_sedis(100);f__Exobasidiomycetes_family_Incertae_sedis(100);g__Tilletiopsis(100);s__Tilletiopsis_pallescens(100); | Tilletiopsis | 13 | Pathotroph-Saprotroph | Endophyte-Fungal Parasite-\|Plant Pathogen\|-Plant Saprotroph-Undefined Saprotroph | Microfungus-Yeast | NULL | Probable | As litter_saprotroph (Põlme et al. 2020); As mycoparasite (Põlme et al. 2020); Decay leaf, fruit, seed (Põlme et al. 2020); Isolated from leaves...dead leaves (Nyland 1950) | Nyland G. 1950. Mycologia 42:487-496 ((ttps://doi.org/10.1080/00275514.1950.12017853)); Põlme S, et al. 2020. Fungal Diversity 105:1-16 ((https://doi.org/10.1007/s13225-020-00466-2)) |
| Otu00794 | 94 | k__Fungi(100);p__Ascomycota(100);c__Sordariomycetes(100);o__Xylariales(100);f__Xylariaceae(100);g__Hypoxylon(100);s__Hypoxylon_sp(86); | Hypoxylon | 13 | Saprotroph-Symbiotroph | Endophyte-Undefined Saprotroph-\|Wood Saprotroph\| | Corticioid-Microfungus-Xylarioid | NULL | Probable | As foliar_endophyte (Põlme et al. 2020); Decay wood, leaf, fruit, seed (Põlme et al. 2020); Host - angiosperms (Põlme et al. 2020) | Tedersoo L, et al. 2014. Science 346:e1256688 ((https://doi.org/10.1126/science.1256688)); Põlme S, et al. 2020. Fungal Diversity 105:1-16 ((https://doi.org/10.1007/s13225-020-00466-2)) |
| Otu00795 | 94 | k__Fungi(100);p__Basidiomycota(100);c__Agaricomycetes(100);o__Agaricales(100);f__Psathyrellaceae(100);g__Coprinellus(100);g__Coprinellus_unclassified(100); | Coprinellus | 13 | Saprotroph | \|Undefined Saprotroph\| | Agaricoid | NULL | Probable | As soil_saprotroph (Põlme et al. 2020); Decay soil (Põlme et al. 2020) | Tedersoo L, et al. 2014. Science 346:e1256688 ((https://doi.org/10.1126/science.1256688)); Põlme S, et al. 2020. Fungal Diversity 105:1-16 ((https://doi.org/10.1007/s13225-020-00466-2)) |
| Otu00796 | 94 | k__Fungi(100);p__Basidiomycota(100);c__Agaricomycetes(100);o__Polyporales(100);f__Fomitopsidaceae(100);g__unclassified_Fomitopsidaceae(100);s__Fomitopsidaceae_sp(100); | Fomitopsidaceae | 9 | Pathotroph-Saprotroph | Plant Pathogen-Wood Saprotroph | Polyporoid | Brown Rot | Probable | Most species are parasitic on woody plants, causing brown rots (Cannon & Kirk 2007) | Cannon PF, Kirk PM. 2007. Fungal Families of the World. CAB International, Cambridge (ISBN: 978-0851998275) |
| Otu00797 | 94 | k__Fungi(100);p__Ascomycota(100);c__Sordariomycetes(100);o__Hypocreales(100);f__Nectriaceae(100);g__Gibberella(81);s__Gibberella_intricans(76); | Gibberella | 13 | Pathotroph-Saprotroph-Symbiotroph | Endophyte-\|Plant Pathogen\|-Plant Saprotroph-Undefined Saprotroph | Microfungus | Soft Rot | Probable | As foliar_endophyte (Põlme et al. 2020); As litter_saprotroph (Põlme et al. 2020); Causing damping-off disease (Põlme et al. 2020); Decay leaf, fruit, seed, roots, soil, animal material (Põlme et al. 2020) | Tedersoo L, et al. 2014. Science 346:e1256688 ((https://doi.org/10.1126/science.1256688)); Põlme S, et al. 2020. Fungal Diversity 105:1-16 ((https://doi.org/10.1007/s13225-020-00466-2)) |
| Otu00798 | 94 | k__Fungi(100);p__Ascomycota(100);c__Sordariomycetes(100);o__Ophiostomatales(100);f__Ophiostomataceae(100);f__Ophiostomataceae_unclassified(100);f__Ophiostomataceae_unclassified(100); | Ophiostomataceae | 9 | Pathotroph | Plant Pathogen | Microfungus | Blue-Staining | Probable | At least most species are transmitted via insect vectors, with mycelial colonies developing in bark-boring beetle galleries and spreading through the vascular system; they are highly specific to their beetle vectors. In some cases spores and mycelium are transmitted in specialized organs, termed mycangia (Cannon & Kirk 2007) | Cannon PF, Kirk PM. 2007. Fungal Families of the World. CAB International, Cambridge (ISBN: 978-0851998275); Tedersoo L, et al. 2014. Science 346:e1256688 ((https://doi.org/10.1126/science.1256688)) |
| Otu00799 | 93 | k__Fungi(100);p__Ascomycota(100);c__Sordariomycetes(100);o__Hypocreales(100);f__Nectriaceae(100);g__Fusarium(100);s__Fusarium_fujikuroi(92); | Fusarium | 13 | Pathotroph-Saprotroph-Symbiotroph | Animal Pathogen-Endophyte-Lichen Parasite-\|Plant Pathogen\|-Plant Saprotroph-Undefined Saprotroph-Wood Saprotroph | Microfungus | Soft Rot | Possible | As foliar_endophyte (Põlme et al. 2020); As litter_saprotroph (Põlme et al. 2020); Causing damping-off disease (Põlme et al. 2020); Decay leaf, fruit, seed, roots, soil, animal material (Põlme et al. 2020); Likely opportunistic human pathogen (Irinyi et al. 2015) | Duncan CG, Eslyn WE. 1966. Mycologia 58:642-664 ((https://doi.org/10.2307/3757045)); Seehann G, et al. 1975. List of Fungi in Soft-Rot Tissue Tests. Institute fur Holzbiologie und Holzschutz, Bundesforschungsanstalt fur Forst - und Holzwirtshaft, Hamburg - Lohbrugge, Germany; Costa IPMW, et al. 2012. Mycotaxon 119:e26 ((http://mycotaxon.com/resources/checklists/costa_v119_checklist.pdf)); Tedersoo L, et al. 2014. Science 346:e1256688 ((https://doi.org/10.1126/science.1256688)); Irinyi L, et al. 2015. Medical Mycology 53:313-337 ((https://doi.org/10.1093/mmy/myv008)); Schroers, et al. 2016. Mycologia 108:806-819 ((https://doi.org/10.3852/15-255)); Põlme S, et al. 2020. Fungal Diversity 105:1-16 ((https://doi.org/10.1007/s13225-020-00466-2)); The American Phytopathological Society Common Names of Plant Diseases ((http://www.apsnet.org/publications/commonnames/Pages/default.aspx)); Lawrey JD, Diederich P. 2016. http://www.lichenicolous.net (accessed 24 May 2017) ((http://www.lichenicolous.net)) |
| Otu00801 | 93 | k__Fungi(100);p__Basidiomycota(100);c__Tremellomycetes(100);o__Tremellales(100);f__Tremellales_family_Incertae_sedis(100);g__Bullera(100);s__Bullera_coprosmae(100); | Bullera | 13 | Saprotroph-Symbiotroph | Endophyte-Epiphyte-Plant Saprotroph-\|Undefined Saprotroph\| | Yeast | NULL | Probable | As foliar_endophyte (Põlme et al. 2020); Isolated from dead leaves of Oryza sativa, Miscanthus sinensis and Sasa sp. in Japan (Nakase & Suzuki 1986; Syn. Udeniomyces puniceus); Foliar, isolated from leaves of Melostomae candidum (Nakase et al. 2004) | Nakase T, Suzuki M. 1986. Journal of General and Applied Microbiology Tokyo 32:225-240 ((https://doi.org/10.2323/jgam.32.225)); Nakase T, et al. 2004. Systematic and Applied Microbiology 27:558-564 ((https://doi.org/10.1078/0723202041748118)) |
| Otu00803 | 93 | k__Fungi(100);p__Ascomycota(100);c__Saccharomycetes(100);o__Saccharomycetales(100);f__Metschnikowiaceae(100);g__Metschnikowia(100);s__Metschnikowia_pulcherrima(96); | Metschnikowia | 13 | Saprotroph | \|Undefined Saprotroph\| | Yeast | NULL | Probable | As nectar-tap_saprotroph (Põlme et al. 2020); Decay sugar-rich substrates (Põlme et al. 2020) | Põlme S, et al. 2020. Fungal Diversity 105:1-16 ((https://doi.org/10.1007/s13225-020-00466-2)) |
| Otu00805 | 93 | k__Fungi(100);p__Ascomycota(99);c__Saccharomycetes(99);o__Saccharomycetales(99);f__Metschnikowiaceae(99);g__Metschnikowia(99);s__Metschnikowia_pulcherrima(97); | Metschnikowia | 13 | Saprotroph | \|Undefined Saprotroph\| | Yeast | NULL | Probable | As nectar-tap_saprotroph (Põlme et al. 2020); Decay sugar-rich substrates (Põlme et al. 2020) | Põlme S, et al. 2020. Fungal Diversity 105:1-16 ((https://doi.org/10.1007/s13225-020-00466-2)) |
| Otu00800 | 93 | k__Fungi(100);p__Ascomycota(100);c__Sordariomycetes(100);o__Diaporthales(100);f__Togniniaceae(100);g__unclassified_Togniniaceae(94);s__Togniniaceae_sp(94); | - | - | - | - | - | - | - | Unassigned | - |
| Otu00802 | 93 | k__Fungi(100);p__Ascomycota(100);c__Sordariomycetes(100);o__Hypocreales(100);f__Hypocreaceae(74);f__Hypocreaceae_unclassified(74);f__Hypocreaceae_unclassified(74); | - | - | - | - | - | - | - | Unassigned | - |
| Otu00804 | 93 | k__Fungi(100);k__Fungi_unclassified(57);k__Fungi_unclassified(57);k__Fungi_unclassified(57);k__Fungi_unclassified(57);k__Fungi_unclassified(57); | - | - | - | - | - | - | - | Unassigned | - |
| Otu00806 | 92 | k__Fungi(100);p__Ascomycota(100);c__Dothideomycetes(100);o__Pleosporales(100);f__Massarinaceae(100);g__Massarina(100);s__Massarina_sp_HF12700(100); | Massarina | 13 | Pathotroph-Saprotroph | \|Plant Pathogen\|-Undefined Saprotroph | Microfungus | NULL | Probable | NULL | Tedersoo L, et al. 2014. Science 346:e1256688 ((https://doi.org/10.1126/science.1256688)); Põlme S, et al. 2020. Fungal Diversity 105:1-16 ((https://doi.org/10.1007/s13225-020-00466-2)) |
| Otu00807 | 92 | k__Fungi(100);p__Ascomycota(100);c__Eurotiomycetes(100);o__Chaetothyriales(100);f__Herpotrichiellaceae(100);g__Exophiala(100);s__Exophiala_sp_4_11c(100); | Exophiala | 13 | Pathotroph-Saprotroph-Symbiotroph | \|Animal Parasite\|-Animal Pathogen-Endophyte-Plant Saprotroph-Undefined Saprotroph | Dimorphic-Facultative Yeast | NULL | Probable | As litter_saprotroph (Põlme et al. 2020); As root_endophyte_dark_septate (Põlme et al. 2020); Decay leaf, fruit, seed, animal material (Põlme et al. 2020) | Tedersoo L, et al. 2014. Science 346:e1256688 ((https://doi.org/10.1126/science.1256688)), Nagy LG, et al. 2014. Nature Communications 5:e4471 ((https://doi.org/10.1038/ncomms5471)); Põlme S, et al. 2020. Fungal Diversity 105:1-16 ((https://doi.org/10.1007/s13225-020-00466-2)) |
| Otu00808 | 92 | k__Fungi(100);p__Ascomycota(100);c__Sordariomycetes(100);o__Hypocreales(100);f__Hypocreaceae(100);g__Protocrea(100);s__Protocrea_farinosa(100); | Protocrea | 13 | Pathotroph | \|Fungal Parasite\| | Microfungus | NULL | Probable | As mycoparasite (Põlme et al. 2020); Decay fungal material (Põlme et al. 2020) | Tedersoo L, et al. 2014. Science 346:e1256688 ((https://doi.org/10.1126/science.1256688)); Põlme S, et al. 2020. Fungal Diversity 105:1-16 ((https://doi.org/10.1007/s13225-020-00466-2)) |
| Otu00809 | 92 | k__Fungi(100);p__Ascomycota(100);c__Dothideomycetes(99);o__Pleosporales(99);f__Venturiaceae(99);f__Venturiaceae_unclassified(99);f__Venturiaceae_unclassified(99); | Venturiaceae | 9 | Pathotroph-Saprotroph-Symbiotroph | Endophyte-Plant Pathogen-Undefined Saprotroph | Microfungus | NULL | Probable | Mostly endophytes and weak parasites of aerial plant parts, including bark and wood, leaf and fruit tissue; rarely biotrophic. Little is known except for the important pathogens (Cannon & Kirk 2007) | Cannon PF, Kirk PM. 2007. Fungal Families of the World. CAB International, Cambridge (ISBN: 978-0851998275) |
| Otu00811 | 92 | k__Fungi(100);p__Ascomycota(100);c__Dothideomycetes(100);o__Pleosporales(100);f__Phaeosphaeriaceae(100);g__Ampelomyces(100);s__Ampelomyces_sp_B34(100); | Ampelomyces | 13 | Pathotroph-Saprotroph-Symbiotroph | Endophyte-Fungal Parasite-\|Plant Pathogen\|-Plant Saprotroph-Undefined Saprotroph | Microfungus | NULL | Probable | As foliar_endophyte (Põlme et al. 2020); As litter_saprotroph (Põlme et al. 2020); As mycoparasite (Põlme et al. 2020); culture (Dik et al. 1998); Decay leaf, fruit, seed (Põlme et al. 2020); Endophyte detection method; Host - Erysiphaceae (Põlme et al. 2020) | Dik AJ, et al. 1998. European Journal of Plant Pathology 104:413-423 ((https://doi.org/10.1023/A:1008025416672)); Tedersoo L, et al. 2014. Science 346:e1256688 ((https://doi.org/10.1126/science.1256688)); Põlme S, et al. 2020. Fungal Diversity 105:1-16 ((https://doi.org/10.1007/s13225-020-00466-2)) |
| Otu00810 | 92 | k__Fungi(100);p__Ascomycota(90);p__Ascomycota_unclassified(90);p__Ascomycota_unclassified(90);p__Ascomycota_unclassified(90);p__Ascomycota_unclassified(90);p__Ascomycota_unclassified(90); | - | - | - | - | - | - | - | Unassigned | - |
| Otu00812 | 91 | k__Fungi(100);p__Ascomycota(100);c__Ascomycota_class_Incertae_sedis(95);o__Ascomycota_order_Incertae_sedis(95);f__Ascomycota_family_Incertae_sedis(95);g__Polyscytalum(95);s__Polyscytalum_algarvense(95); | Polyscytalum | 13 | Pathotroph | \|Plant Pathogen\| | Microfungus | NULL | Probable | NULL | Põlme S, et al. 2020. Fungal Diversity 105:1-16 ((https://doi.org/10.1007/s13225-020-00466-2)); The American Phytopathological Society Common Names of Plant Diseases ((http://www.apsnet.org/publications/commonnames/Pages/default.aspx)) |

| | | | | | | | | | | | |
|---|---|---|---|---|---|---|---|---|---|---|---|
| Otu00813 | 91 | k__Fungi(100);p__Basidiomycota(100);c__Agaricomycetes(100);o__Polyporales(100);f__Fomitopsidaceae(100);g__Dacryobolus(100);s__Dacryobolus_sudans(100); | Dacryobolus | 13 | Saprotroph | \|Wood Saprotroph\| | Corticioid | Brown Rot | Probable | As brown-rot (Põlme et al. 2020); Decay wood (Põlme et al. 2020) | Tedersoo L, et al. 2014. Science 346:e1256688 ((https://doi.org/10.1126/science.1256688)); Põlme S, et al. 2020. Fungal Diversity 105:1-16 ((https://doi.org/10.1007/s13225-020-00466-2)) |
| Otu00814 | 91 | k__Fungi(100);p__Ascomycota(100);c__Sordariomycetes(100);o__Microascales(100);f__Microascaceae(100);g__unclassified_Microascaceae(100);s__Microascaceae_sp(100); | Microascaceae | 9 | Pathotroph-Saprotroph-Symbiotroph | Animal Pathogen-Endophyte-Plant Pathogen-Undefined Saprotroph | Microfungus | NULL | Possible | Saprobic in soil or rotting vegetation, some species heat-tolerant and associatated with silage and compost, a few associated with humans and animals (Cannon & Kirk 2007) | Cannon PF, Kirk PM. 2007. Fungal Families of the World. CAB International, Cambridge (ISBN: 978-0851998275) |
| Otu00815 | 91 | k__Fungi(100);p__Basidiomycota(100);c__Agaricomycetes(100);o__Agaricales(100);o__Agaricales_unclassified(100);o__Agaricales_unclassified(100);o__Agaricales_unclassified(100); | Agaricales | 7 | Pathotroph-Saprotroph-Symbiotroph | Bryophyte Parasite-Dung Saprotroph-Ectomycorrhizal-Fungal Parasite-Leaf Saprotroph-Plant Parasite-Undefined Saprotroph-Wood Saprotroph | Agaricoid-Gasteroid-Microfungus-Secotioid-Yeast | NULL | Possible | Mushrooms and toadstools, Gill fungi, Agarics... lignicolous, sometimes muscicolous or fungicolous, saprobic, mycorrhizal, rarely parasitic on plants or fungi (Kirk et al. 2008) | Kirk PM et al. 2008. Dictionary of the Fungi. Tenth Edition. CAB International, Wallingford (ISBN: 978-0851998268) |
| Otu00816 | 91 | k__Fungi(100);p__Basidiomycota(100);c__Agaricomycetes(100);o__Russulales(100);f__Stereaceae(100);g__Gloeocystidiellum(100);s__Gloeocystidiellum_sp(100); | Gloeocystidiellum | 13 | Saprotroph | Undefined Saprotroph-\|Wood Saprotroph\| | Corticioid | White Rot | Probable | As white_rot (Põlme et al. 2020); Decay wood (Põlme et al. 2020) | Tedersoo L, et al. 2014. Science 346:e1256688 ((https://doi.org/10.1126/science.1256688)); Põlme S, et al. 2020. Fungal Diversity 105:1-16 ((https://doi.org/10.1007/s13225-020-00466-2)) |
| Otu00819 | 90 | k__Fungi(100);p__Ascomycota(100);c__Dothideomycetes(100);o__Pleosporales(100);f__Pleosporaceae(100);g__unclassified_Pleosporaceae(59);s__Pleosporaceae_sp(59); | Pleosporaceae | 9 | Pathotroph-Saprotroph | Endophyte-Lichen Parasite-Plant Pathogen-Undefined Saprotroph | Microfungus | NULL | Probable | Necrotrophic pathogens and saprobes, especially associated with grasses. Many species are seed-borne (Cannon & Kirk 2007) | Cannon PF, Kirk PM. 2007. Fungal Families of the World. CAB International, Cambridge (ISBN: 978-0851998275); David AS, et al. 2016. Microbial Ecology 71:912-926 ((https://doi.org/10.1007/s00248-015-0712-6)); Lawrey JD, Diederich P. 2016. http://www.lichenicolous.net (accessed 24 May 2017) ((http://www.lichenicolous.net)) |
| Otu00820 | 90 | k__Fungi(100);p__Ascomycota(100);c__Eurotiomycetes(100);o__Eurotiales(100);f__Trichocomaceae(100);g__Penicillium(100);s__Penicillium_onobense(76); | Penicillium | 13 | Pathotroph-Saprotroph-Symbiotroph | Animal Parasite-Dung Saprotroph-Endophyte-Fungal Parasite-Plant Pathogen-\|Undefined Saprotroph\|-Wood Saprotroph | Microfungus | Soft Rot | Highly Probable | As foliar_endophyte (Põlme et al. 2020); As mycoparasite (Põlme et al. 2020); Decay soil (Põlme et al. 2020) | Duncan CG, Eslyn WE. 1966. Mycologia 58:642-664 ((https://doi.org/10.2307/3757045)); Seehann G, et al. 1975. List of Fungi in Soft-Rot Tissue Tests. Institute fur Holzbiologie und Holzschutz, Bundesforschungsanstalt fur Forst - und Holzwirtshaft, Hamburg - Lohbrugge, Germany ((https://www.irg-wp.com/irgdocs/details.php?d6493033-c9ea-40f3-8e3e-5697f32d640d)); Costa IPMW, et al. 2012. Mycotaxon 119:e26 ((http://mycotaxon.com/resources/checklists/costa_v119_checklist.pdf)); Bills GF, et al. 2013. Current Opinion in Microbiology 16:549-565 ((https://doi.org/10.1016/j.mib.2013.08.001)); Tedersoo L, et al. 2014. Science 346:e1256688 ((https://doi.org/10.1126/science.1256688)); Põlme S, et al. 2020. Fungal Diversity 105:1-16 ((https://doi.org/10.1007/s13225-020-00466-2)) |
| Otu00821 | 90 | k__Fungi(100);p__Basidiomycota(100);c__Agaricomycetes(100);o__Polyporales(100);f__Fomitopsidaceae(100);g__Postia(100);s__Postia_caesia(72); | Postia | 13 | Saprotroph | \|Wood Saprotroph\| | Polyporoid | Brown Rot | Highly Probable | As brown-rot (Põlme et al. 2020); Decay wood (Põlme et al. 2020) | Jordan CR, et al. 1996. Holzforschung - International Journal of the Biology, Chemistry, Physics and Technology of Wood 50:312-318 ((https://doi.org/10.1515/hfsg.1996.50.4.312)); Põlme S, et al. 2020. Fungal Diversity 105:1-16 ((https://doi.org/10.1007/s13225-020-00466-2)) |
| Otu00822 | 90 | k__Fungi(100);p__Ascomycota(100);c__Dothideomycetes(100);o__Pleosporales(100);f__Phaeosphaeriaceae(100);g__Phaeosphaeria(100);s__Phaeosphaeria_triglochinicola(100); | Phaeosphaeria | 13 | Pathotroph-Saprotroph-Symbiotroph | Endophyte-Plant Pathogen-\|Plant Saprotroph\| | Microfungus | NULL | Probable | As litter_saprotroph (Põlme et al. 2020); Decay leaf, fruit, seed (Põlme et al. 2020) | Tedersoo L, et al. 2014. Science 346:e1256688 ((https://doi.org/10.1126/science.1256688));David AS, et al. 2016. Microbial Ecology 71:912-926 ((https://doi.org/10.1007/s00248-015-0712-6)); Põlme S, et al. 2020. Fungal Diversity 105:1-16 ((https://doi.org/10.1007/s13225-020-00466-2)) |
| Otu00817 | 90 | k__Fungi(100);k__Fungi_unclassified(52);k__Fungi_unclassified(52);k__Fungi_unclassified(52);k__Fungi_unclassified(52);k__Fungi_unclassified(52);k__Fungi_unclassified(52); | - | - | - | - | - | - | - | Unassigned | - |
| Otu00818 | 90 | k__Fungi(100);p__Ascomycota(53);p__Ascomycota_unclassified(53);p__Ascomycota_unclassified(53);p__Ascomycota_unclassified(53);p__Ascomycota_unclassified(53);p__Ascomycota_unclassified(53); | - | - | - | - | - | - | - | Unassigned | - |
| Otu00823 | 90 | k__Fungi(100);k__Fungi_unclassified(66);k__Fungi_unclassified(66);k__Fungi_unclassified(66);k__Fungi_unclassified(66);k__Fungi_unclassified(66);k__Fungi_unclassified(66); | - | - | - | - | - | - | - | Unassigned | - |
| Otu00828 | 89 | k__Fungi(100);p__Ascomycota(100);c__Dothideomycetes(100);o__Pleosporales(100);f__Pleosporales_family_Incertae_sedis(100);g__Setomelanomma(100);s__Setomelanomma_sp(68); | Setomelanomma | 13 | Pathotroph-Saprotroph | \|Plant Pathogen\|-Wood Saprotroph | Microfungus | NULL | Probable | As wood_pathogen (Põlme et al. 2020); Decay wood (Põlme et al. 2020) | Põlme S, et al. 2020. Fungal Diversity 105:1-16 ((https://doi.org/10.1007/s13225-020-00466-2)) |
| Otu00829 | 89 | k__Fungi(100);p__Ascomycota(100);c__Dothideomycetes(100);o__Capnodiales(100);f__Mycosphaerellaceae(100);g__Pseudocercospora(100);s__Pseudocercospora_norchiensis(93); | Pseudocercospora | 13 | Pathotroph | \|Plant Pathogen\| | Microfungus | NULL | Probable | NULL | Tedersoo L, et al. 2014. Science 346:e1256688 ((https://doi.org/10.1126/science.1256688)); Põlme S, et al. 2020. Fungal Diversity 105:1-16 ((https://doi.org/10.1007/s13225-020-00466-2)) |
| Otu00830 | 89 | k__Fungi(100);p__Ascomycota(100);c__Sordariomycetes(100);o__Hypocreales(100);f__Hypocreales_family_Incertae_sedis(100);g__Myrothecium(100);s__Myrothecium_sp(100); | Myrothecium | 13 | Pathotroph-Saprotroph | Fungal Parasite-\|Plant Saprotroph\|-Undefined Saprotroph | Microfungus | NULL | Probable | As litter_saprotroph (Põlme et al. 2020); As mycoparasite (Põlme et al. 2020); Decay leaf, fruit, seed (Põlme et al. 2020) | Tedersoo L, et al. 2014. Science 346:e1256688 ((https://doi.org/10.1126/science.1256688)); Põlme S, et al. 2020. Fungal Diversity 105:1-16 ((https://doi.org/10.1007/s13225-020-00466-2)) |
| Otu00831 | 89 | k__Fungi(100);p__Basidiomycota(100);c__Agaricomycetes(100);o__Hymenochaetales(100);f__Schizoporaceae(100);g__Xylodon(100);s__Xylodon_sambuci(100); | Xylodon | 13 | Saprotroph | Undefined Saprotroph-\|Wood Saprotroph\| | Corticioid | White Rot | Probable | As white_rot (Põlme et al. 2020); Decay wood (Põlme et al. 2020) | Tedersoo L, et al. 2014. Science 346:e1256688 ((https://doi.org/10.1126/science.1256688)); Põlme S, et al. 2020. Fungal Diversity 105:1-16 ((https://doi.org/10.1007/s13225-020-00466-2)) |
| Otu00833 | 89 | k__Fungi(100);p__Basidiomycota(100);c__Agaricomycetes(100);o__Trechisporales(100);f__Hydnodontaceae(100);g__Trechispora(100);s__Trechispora_sp(100); | Trechispora | 13 | Saprotroph | \|Wood Saprotroph\| | Corticioid | White Rot | Highly Probable | As white_rot (Põlme et al. 2020); Decay wood (Põlme et al. 2020) | Gilbertson RL, Ryvarden L. 1987-1987. North American Polypores. Fungiflora, Oslo (ISBN: 978-0945345060); Rinaldi AC, et al. 2008. Fungal Diversity 33:1-45 ((http://www.fungaldiversity.org/fdp/sfdp/33-1.pdf)) (as Uncertain); Tedersoo L, et al. 2010. Mycorrhiza 20:217-263 ((https://doi.org/10.1007/s00572-009-0274-x)) (as not ECM).; Põlme S, et al. 2020. Fungal Diversity 105:1-16 ((https://doi.org/10.1007/s13225-020-00466-2)) |
| Otu00824 | 89 | k__Fungi(100);k__Fungi_unclassified(62);k__Fungi_unclassified(62);k__Fungi_unclassified(62);k__Fungi_unclassified(62);k__Fungi_unclassified(62);k__Fungi_unclassified(62); | - | - | - | - | - | - | - | Unassigned | - |
| Otu00825 | 89 | k__Plantae(100);p__unclassified_Plantae(100);c__unclassified_Plantae(100);o__unclassified_Plantae(100);f__unclassified_Plantae(100);g__unclassified_Plantae(100);s__Plantae_sp(100); | - | - | - | - | - | - | - | Unassigned | - |
| Otu00826 | 89 | k__Fungi(100);p__Ascomycota(100);c__Sordariomycetes(100);c__Sordariomycetes_unclassified(100);c__Sordariomycetes_unclassified(100);c__Sordariomycetes_unclassified(100);c__Sordariomycetes_unclassified(100); | - | - | - | - | - | - | - | Unassigned | - |
| Otu00827 | 89 | k__Fungi(100);p__Ascomycota(100);c__Dothideomycetes(100);o__Pleosporales(100);o__Pleosporales_unclassified(59);o__Pleosporales_unclassified(59);o__Pleosporales_unclassified(59); | - | - | - | - | - | - | - | Unassigned | - |
| Otu00832 | 89 | k__Fungi(100);p__Ascomycota(100);c__Sordariomycetes(97);c__Sordariomycetes_unclassified(92);c__Sordariomycetes_unclassified(92);c__Sordariomycetes_unclassified(92);c__Sordariomycetes_unclassified(92); | - | - | - | - | - | - | - | Unassigned | - |
| Otu00834 | 88 | k__Fungi(100);p__Ascomycota(100);c__Sordariomycetes(100);o__Hypocreales(100);f__Hypocreales_family_Incertae_sedis(100);g__Acremonium(100);s__Acremonium_sp_24_OA_2013(100); | Acremonium | 13 | Pathotroph-Saprotroph-Symbiotroph | Animal Pathogen-Endophyte-Fungal Parasite-Plant Pathogen-\|Undefined Saprotroph\|-Wood Saprotroph | Microfungus | Soft Rot | Possible | As foliar_endophyte (Põlme et al. 2020); culture (Hanada et al. 2010, Del Olmo; Decay leaf, fruit, seed, soil, dung, animal material (Põlme et al. 2020); Endophyte detection method; Ruiz & Arnold 2014); Likely opportunistic human pathogen (Irinyi et al. 2015) | Duncan CG, Eslyn WE. 1966. Mycologia 58:642-645 ((https://doi.org/10.2307/3757045)); Nilsson T. 1973. Studia Forestalia Suecica 104:1-40 ((https://pub.epsilon.slu.se/5711/)); Worrall JJ, et al. 1997. Mycologia 89:199-219 ((https://doi.org/10.2307/3761073)); Hanada RE, et al. 2010. Fungal Biology 114:901-910 ((https://doi.org/10.1016/j.funbio.2010.08.006)); Del Olmo-Ruiz M, Arnold AE. 2014. Mycologia 106:8-21 ((https://doi.org/10.3852/13-098)); Irinyi L, et al. 2015. Medical Mycology 53313-337 ((https://doi.org/10.1093/mmy/myv008)); Põlme S, et al. 2020. Fungal Diversity 105:1-16 ((https://doi.org/10.1007/s13225-020-00466-2)) |
| Otu00835 | 88 | k__Fungi(100);p__Ascomycota(100);c__Dothideomycetes(100);o__Capnodiales(100);f__Davidiellaceae(100);g__Cladosporium(100);s__Cladosporium_sp_CBS_55279(100); | Cladosporium | 13 | Pathotroph-Saprotroph-Symbiotroph | Animal Pathogen-Endophyte-Lichen Parasite-Plant Pathogen-\|Plant Saprotroph\|-Wood Saprotroph | Microfungus | NULL | Possible | As foliar_endophyte (Põlme et al. 2020); As litter_saprotroph (Põlme et al. 2020); Decay wood, leaf, fruit, seed, soil, dung, animal material (Põlme et al. 2020) | Seehann G, et al. 1975. List of Fungi in Soft-Rot Tissue Tests. Institute fur Holzbiologie und Holzschutz, Bundesforschungsanstalt fur Forst - und Holzwirtshaft, Hamburg - Lohbrugge, Germany; Lawrey JD, Diederich P. 2016. http://www.lichenicolous.net (accessed 24 May 2017) ((http://www.lichenicolous.net)); Põlme S, et al. 2020. Fungal Diversity 105:1-16 ((https://doi.org/10.1007/s13225-020-00466-2)); The American Phytopathological Society Common Names of Plant Diseases ((http://www.apsnet.org/publications/commonnames/Pages/default.aspx)) |
| Otu00836 | 88 | k__Fungi(100);p__Ascomycota(100);c__Taphrinomycetes(100);o__Taphrinales(100);f__Taphrinaceae(100);g__Taphrina(73);g__Taphrina_unclassified(73); | Taphrina | 13 | Pathotroph | \|Plant Pathogen\| | Dimorphic-Microfungus | NULL | Probable | NULL | Põlme S, et al. 2020. Fungal Diversity 105:1-16 ((https://doi.org/10.1007/s13225-020-00466-2)); The American Phytopathological Society Common Names of Plant Diseases ((http://www.apsnet.org/publications/commonnames/Pages/default.aspx)) |
| Otu00837 | 88 | k__Fungi(100);p__Ascomycota(100);c__Eurotiomycetes(100);c__Eurotiomycetes_unclassified(93);c__Eurotiomycetes_unclassified(93);c__Eurotiomycetes_unclassified(93);c__Eurotiomycetes_unclassified(93); | - | - | - | - | - | - | - | Unassigned | - |
| Otu00839 | 87 | k__Fungi(100);p__Ascomycota(100);c__Dothideomycetes(100);o__Pleosporales(100);f__Sporormiaceae(100);g__Preussia(100);s__Preussia_isomera(100); | Preussia | 13 | Saprotroph | \|Dung Saprotroph\|-Undefined Saprotroph | Microfungus | NULL | Probable | Decay dung (Põlme et al. 2020) | Tedersoo L, et al. 2014. Science 346:e1256688 ((https://doi.org/10.1126/science.1256688)); Põlme S, et al. 2020. Fungal Diversity 105:1-16 ((https://doi.org/10.1007/s13225-020-00466-2)) |
| Otu00840 | 87 | k__Fungi(100);p__Basidiomycota(100);c__Agaricomycetes(100);o__Phallales(100);f__Phallaceae(100);g__Mutinus(100);s__Mutinus_sp_MA_2013(97); | Mutinus | 13 | Saprotroph | Plant Saprotroph-\|Undefined Saprotroph\| | Phalloid | NULL | Probable | stinkhorn; As soil_saprotroph (Põlme et al. 2020); Common name ; Decay soil (Põlme et al. 2020) | Grgurinovic CA, Simpson JA. 2001. Fungal Diversity 8:97-127 ((http://www.fungaldiversity.org/fdp/sfdp/FD_8_97-127.pdf)); Tedersoo L, et al. 2014. Science 346:e1256688 ((https://doi.org/10.1126/science.1256688)); Põlme S, et al. 2020. Fungal Diversity 105:1-16 ((https://doi.org/10.1007/s13225-020-00466-2)) |
| Otu00838 | 87 | k__Fungi(100);p__Basidiomycota(100);c__Agaricomycetes(100);o__Trechisporales(100);f__Hydnodontaceae(100);f__Hydnodontaceae_unclassified(87);f__Hydnodontaceae_unclassified(87); | - | - | - | - | - | - | - | Unassigned | - |
| Otu00841 | 86 | k__Fungi(100);p__Basidiomycota(100);c__Agaricomycetes(100);o__Agaricales(100);f__Pluteaceae(100);g__Pluteus(100);s__Pluteus_petasatus(99); | Pluteus | 13 | Pathotroph-Saprotroph | Bryophyte Parasite-\|Plant Saprotroph\|-Wood Saprotroph | Agaricoid | White Rot | Probable | As litter_saprotroph (Põlme et al. 2020); As white_rot (Põlme et al. 2020); Decay leaf, fruit, seed (Põlme et al. 2020) | Grgurinovic CA, Simpson JA. 2001. Fungal Diversity 8:97-127 ((http://www.fungaldiversity.org/fdp/sfdp/FD_8_97-127.pdf)); James TY, et al. 2006. Nature 443:818-822 ((https://doi.org/10.1038/nature05110)); Tedersoo L, et al. 2014. Science 346:e1256688 ((https://doi.org/10.1126/science.1256688)); Põlme S, et al. 2020. Fungal Diversity 105:1-16 ((https://doi.org/10.1007/s13225-020-00466-2)) |
| Otu00844 | 86 | k__Fungi(100);p__Ascomycota(100);c__Saccharomycetes(100);o__Saccharomycetales(100);f__Metschnikowiaceae(100);g__Metschnikowia(100);s__Metschnikowia_pulcherrima(99); | Metschnikowia | 13 | Saprotroph | \|Undefined Saprotroph\| | Yeast | NULL | Probable | As nectar-tap_saprotroph (Põlme et al. 2020); Decay sugar-rich substrates (Põlme et al. 2020) | Põlme S, et al. 2020. Fungal Diversity 105:1-16 ((https://doi.org/10.1007/s13225-020-00466-2)) |
| Otu00846 | 86 | k__Fungi(100);p__Basidiomycota(100);c__Microbotryomycetes(100);o__Sporidiobolales(100);f__Sporidiobolales_family_Incertae_sedis(100);g__Sporobolomyces(97);g__Sporobolomyces_unclassified(97); | Sporobolomyces | 13 | Pathotroph-Saprotroph | \|Fungal Parasite\|-Plant Saprotroph | Yeast | NULL | Probable | As litter_saprotroph (Põlme et al. 2020); As mycoparasite (Põlme et al. 2020); Decay fungal material (Põlme et al. 2020) | Tedersoo L, et al. 2014. Science 346:e1256688 ((https://doi.org/10.1126/science.1256688)); Purahong W, et al. 2016. Molecular Ecology 25:4059-4074 ((https://doi.org/10.1111/mec.13739)); Põlme S, et al. 2020. Fungal Diversity 105:1-16 ((https://doi.org/10.1007/s13225-020-00466-2)) |
| Otu00842 | 86 | k__Fungi(100);p__Basidiomycota(100);c__Agaricomycetes(100);o__Auriculariales(100);f__Auriculariales_family_Incertae_sedis(100);f__Auriculariales_family_Incertae_sedis_unclassified(76);f__Auriculariales_family_Incertae_sedis_unclassified(76); | - | - | - | - | - | - | - | Unassigned | - |
| Otu00843 | 86 | k__Fungi(100);p__Ascomycota(100);p__Ascomycota_unclassified(100);p__Ascomycota_unclassified(100);p__Ascomycota_unclassified(100);p__Ascomycota_unclassified(100);p__Ascomycota_unclassified(100); | - | - | - | - | - | - | - | Unassigned | - |
| Otu00845 | 86 | k__Fungi(100);p__Basidiomycota(100);c__Agaricomycetes(100);o__Atheliales(100);f__Atheliaceae(100);g__Fibulorhizoctonia(100);s__Fibulorhizoctonia_sp(100); | - | - | - | - | - | - | - | Unassigned | - |
| Otu00847 | 85 | k__Fungi(100);p__Zygomycota(100);c__Zygomycota_class_Incertae_sedis(100);o__Mortierellales(100);f__Mortierellaceae(100);g__Mortierella(100);s__Mortierella_ambigua(100); | Mortierella | 13 | Saprotroph-Symbiotroph | Endophyte-Plant Saprotroph-\|Undefined Saprotroph\| | Microfungus | NULL | Probable | As chitinolytic (Põlme et al. 2020); As soil_saprotroph (Põlme et al. 2020); Decay roots, soil (Põlme et al. 2020); Saprobic in soil and thus frequently isolated from soil and material contaminated with soil such as hay or rotten silage and spoil tips (Cannon & Kirk 2007) | Cannon PF, Kirk PM. 2007. Fungal Families of the World. CAB International, Cambridge (ISBN: 978-0851998275); Tedersoo L, et al. 2014. Science 346:e1256688 ((https://doi.org/10.1126/science.1256688)); Purahong W, et al. 2016. Molecular Ecology 25:4059-4074 ((https://doi.org/10.1111/mec.13739)); Põlme S, et al. 2020. Fungal Diversity 105:1-16 ((https://doi.org/10.1007/s13225-020-00466-2)) |
| Otu00850 | 85 | k__Fungi(100);p__Basidiomycota(100);c__Agaricomycetes(100);o__Polyporales(100);f__Polyporaceae(100);g__unclassified_Polyporaceae(96);s__Polyporaceae_sp(96); | Polyporaceae | 9 | Saprotroph | Wood Saprotroph | Polyporoid | Brown Rot-White | Probable | Lignicolous, causing brown or white rot, mostly of broadleaved trees (Cannon & Kirk 2007) | Cannon PF, Kirk PM. 2007. Fungal Families of the World. CAB International, Cambridge (ISBN: 978-0851998275) |
| Otu00854 | 85 | k__Fungi(100);p__Ascomycota(100);c__Lecanoromycetes(100);o__Lecanorales(100);f__Lecanoraceae(100);g__Lecanora(100);g__Lecanora_unclassified(92); | Lecanora | 13 | Symbiotroph | \|Lichenized\| | Thallus | NULL | Highly Probable | NULL | Esslinger TL. 2014. North Dakota State University ((http://www.ndsu.edu/pubweb/~esslinge/chcklst/chcklst7.htm)); Põlme S, et al. 2020. Fungal Diversity 105:1-16 ((https://doi.org/10.1007/s13225-020-00466-2)) |
| Otu00848 | 85 | k__Fungi(100);k__Fungi_unclassified(62);k__Fungi_unclassified(62);k__Fungi_unclassified(62);k__Fungi_unclassified(62);k__Fungi_unclassified(62);k__Fungi_unclassified(62); | - | - | - | - | - | - | - | Unassigned | - |
| Otu00849 | 85 | k__Fungi(100);p__Ascomycota(85);p__Ascomycota_unclassified(85);p__Ascomycota_unclassified(85);p__Ascomycota_unclassified(85);p__Ascomycota_unclassified(85);p__Ascomycota_unclassified(85); | - | - | - | - | - | - | - | Unassigned | - |
| Otu00851 | 85 | k__Fungi(100);k__Fungi_unclassified(79);k__Fungi_unclassified(79);k__Fungi_unclassified(79);k__Fungi_unclassified(79);k__Fungi_unclassified(79);k__Fungi_unclassified(79); | - | - | - | - | - | - | - | Unassigned | - |
| Otu00852 | 85 | k__Fungi(100);p__Basidiomycota(100);c__Agaricomycetes(100);c__Agaricomycetes_unclassified(100);c__Agaricomycetes_unclassified(100);c__Agaricomycetes_unclassified(100);c__Agaricomycetes_unclassified(100); | - | - | - | - | - | - | - | Unassigned | - |
| Otu00853 | 85 | k__Fungi(100);k__Fungi_unclassified(64);k__Fungi_unclassified(64);k__Fungi_unclassified(64);k__Fungi_unclassified(64);k__Fungi_unclassified(64);k__Fungi_unclassified(64); | - | - | - | - | - | - | - | Unassigned | - |
| Otu00855 | 84 | k__Fungi(100);p__Basidiomycota(100);c__Agaricomycetes(100);o__Atheliales(100);f__Atheliaceae(100);g__Athelia(100);s__Athelia_sp(100); | Athelia | 13 | Pathotroph-Saprotroph-Symbiotroph | Algal Parasite-Epiphyte-\|Lichen Parasite\|-Lichenized-Plant Pathogen-Plant Saprotroph-Wood Saprotroph | Corticioid-Thallus | NULL | Possible | Decay fungal material, algal material (Põlme et al. 2020); Host - algae (Põlme et al. 2020); Lichenized and non; lichenized taxa | Adams GC, Kropp BR. 1996. Mycologia 88:459-472 ((https://doi.org/10.1080/00275514.1996.12026672)); Carisse O, et al. 2000. Phytopathology 90:31-37 ((https://doi.org/10.1094/PHYTO.2000.90.1.31)); Esslinger TL. 2014. North Dakota State University ((http://www.ndsu.edu/pubweb/~esslinge/chcklst/chcklst7.htm)); Põlme S, et al. 2020. Fungal Diversity 105:1-16 ((https://doi.org/10.1007/s13225-020-00466-2)) |
| Otu00856 | 84 | k__Fungi(100);p__Basidiomycota(100);c__Agaricomycetes(100);o__Polyporales(100);f__Meruliaceae(100);g__Steccherinum(100);s__Steccherinum_sp_DLL2011_2(100); | Steccherinum | 13 | Saprotroph | Undefined Saprotroph-\|Wood Saprotroph\| | Corticioid | White Rot | Probable | As white_rot (Põlme et al. 2020); Decay wood (Põlme et al. 2020) | Tedersoo L, et al. 2014. Science 346:e1256688 ((https://doi.org/10.1126/science.1256688)); Põlme S, et al. 2020. Fungal Diversity 105:1-16 ((https://doi.org/10.1007/s13225-020-00466-2)) |
| Otu00858 | 84 | k__Fungi(100);p__Basidiomycota(100);c__Agaricomycetes(100);o__Trechisporales(100);f__Hydnodontaceae(100);g__Trechispora(99);s__Trechispora_sp(80); | Trechispora | 13 | Saprotroph | \|Wood Saprotroph\| | Corticioid | White Rot | Highly Probable | As white_rot (Põlme et al. 2020); Decay wood (Põlme et al. 2020) | Gilbertson RL, Ryvarden L. 1987-1987. North American Polypores. Fungiflora, Oslo (ISBN: 978-0945345060); Rinaldi AC, et al. 2008. Fungal Diversity 33:1-45 ((http://www.fungaldiversity.org/fdp/sfdp/33-1.pdf)) (as Uncertain); Tedersoo L, et al. 2010. Mycorrhiza 20:217-263 ((https://doi.org/10.1007/s00572-009-0274-x)) (as not ECM).; Põlme S, et al. 2020. Fungal Diversity 105:1-16 ((https://doi.org/10.1007/s13225-020-00466-2)) |
| Otu00859 | 84 | k__Fungi(100);p__Zygomycota(100);c__Zygomycota_class_Incertae_sedis(100);o__Mucorales(100);f__Mucoraceae(100);g__Mucor(100);s__Mucor_brunneogriseus(100); | Mucor | 13 | Saprotroph | \|Undefined Saprotroph\| | Microfungus | NULL | Probable | As soil_saprotroph (Põlme et al. 2020); Decay soil (Põlme et al. 2020) | Tedersoo L, et al. 2014. Science 346:e1256688 ((https://doi.org/10.1126/science.1256688)); Põlme S, et al. 2020. Fungal Diversity 105:1-16 ((https://doi.org/10.1007/s13225-020-00466-2)) |
| Otu00862 | 84 | k__Fungi(100);p__Basidiomycota(100);c__Exobasidiomycetes(100);o__Exobasidiales(100);f__Exobasidiaceae(100);g__Exobasidium(100);s__Exobasidium_vaccinii(83); | Exobasidium | 13 | Pathotroph | \|Plant Pathogen\| | Leaf Blister-Leaf Gall-Leaf Spot | NULL | Probable | NULL | Tedersoo L, et al. 2014. Science 346:e1256688 ((https://doi.org/10.1126/science.1256688)); Põlme S, et al. 2020. Fungal Diversity 105:1-16 ((https://doi.org/10.1007/s13225-020-00466-2)) |
| Otu00864 | 84 | k__Fungi(100);p__Ascomycota(100);c__Sordariomycetes(100);o__Sordariomycetes_order_Incertae_sedis(100);f__Annulatascaceae(100);g__Rhodoveronaea(100);s__Rhodoveronaea_varioseptata(100); | Rhodoveronaea | 13 | Saprotroph | \|Undefined Saprotroph\| | Microfungus | NULL | Probable | NULL | Tedersoo L, et al. 2014. Science 346:e1256688 ((https://doi.org/10.1126/science.1256688)); Põlme S, et al. 2020. Fungal Diversity 105:1-16 ((https://doi.org/10.1007/s13225-020-00466-2)) |
| Otu00857 | 84 | k__Fungi(100);k__Fungi_unclassified(87);k__Fungi_unclassified(87);k__Fungi_unclassified(87);k__Fungi_unclassified(87);k__Fungi_unclassified(87);k__Fungi_unclassified(87); | - | - | - | - | - | - | - | Unassigned | - |
| Otu00860 | 84 | k__Fungi(100);p__Ascomycota(72);p__Ascomycota_unclassified(72);p__Ascomycota_unclassified(72);p__Ascomycota_unclassified(72);p__Ascomycota_unclassified(72);p__Ascomycota_unclassified(72); | - | - | - | - | - | - | - | Unassigned | - |
| Otu00861 | 84 | k__Fungi(100);p__Basidiomycota(100);c__Agaricomycetes(98);c__Agaricomycetes_unclassified(98);c__Agaricomycetes_unclassified(98);c__Agaricomycetes_unclassified(98);c__Agaricomycetes_unclassified(98); | - | - | - | - | - | - | - | Unassigned | - |
| Otu00863 | 84 | k__Fungi(100);p__Ascomycota(70);p__Ascomycota_unclassified(70);p__Ascomycota_unclassified(70);p__Ascomycota_unclassified(70);p__Ascomycota_unclassified(70);p__Ascomycota_unclassified(70); | - | - | - | - | - | - | - | Unassigned | - |
| Otu00865 | 83 | k__Fungi(100);p__Zygomycota(100);c__Zygomycota_class_Incertae_sedis(100);o__Mucorales(100);f__Mucoraceae(100);g__Mucor(100);g__Mucor_unclassified(98); | Mucor | 13 | Saprotroph | \|Undefined Saprotroph\| | Microfungus | NULL | Probable | As soil_saprotroph (Põlme et al. 2020); Decay soil (Põlme et al. 2020) | Tedersoo L, et al. 2014. Science 346:e1256688 ((https://doi.org/10.1126/science.1256688)); Põlme S, et al. 2020. Fungal Diversity 105:1-16 ((https://doi.org/10.1007/s13225-020-00466-2)) |
| Otu00867 | 83 | k__Fungi(100);p__Basidiomycota(100);c__Agaricomycetes(100);o__Polyporales(100);f__Polyporaceae(100);g__Sarcoporia(100);s__Sarcoporia_polyspora(100); | Sarcoporia | 13 | Saprotroph | \|Wood Saprotroph\| | Polyporoid | Brown Rot | Probable | As brown-rot (Põlme et al. 2020); Decay wood (Põlme et al. 2020) | Tedersoo L, et al. 2014. Science 346:e1256688 ((https://doi.org/10.1126/science.1256688)); Põlme S, et al. 2020. Fungal Diversity 105:1-16 ((https://doi.org/10.1007/s13225-020-00466-2)) |
| Otu00869 | 83 | k__Fungi(100);p__Basidiomycota(100);c__Agaricomycetes(100);o__Agaricales(100);f__Pleurotaceae(100);g__Pleurotus(100);s__Pleurotus_citrinopileatus(100); | Pleurotus | 13 | Pathotroph-Saprotroph | Endophyte-Nematophagous-Plant Pathogen-\|Wood Saprotroph\| | Agaricoid | White Rot | Probable | As white_rot (Põlme et al. 2020); As wood_pathogen (Põlme et al. 2020); Decay wood, animal material (Põlme et al. 2020); This genus is known to be nematophagous, capturing and consuming nematodes. | James TY, et al. 2006. Nature 443:818-822 ((https://doi.org/10.1038/nature05110)); Costa IPMW, et al. 2012. Mycotaxon 119:e26 ((http://mycotaxon.com/resources/checklists/costa_v119_checklist.pdf)); Tedersoo L, et al. 2014. Science 346:e1256688 ((https://doi.org/10.1126/science.1256688)); Põlme S, et al. 2020. Fungal Diversity 105:1-16 ((https://doi.org/10.1007/s13225-020-00466-2)); The American Phytopathological Society Common Names of Plant Diseases ((http://www.apsnet.org/publications/commonnames/Pages/default.aspx)) |
| Otu00866 | 83 | k__Fungi(100);p__Basidiomycota(100);c__Agaricomycetes(100);c__Agaricomycetes_unclassified(92);c__Agaricomycetes_unclassified(92);c__Agaricomycetes_unclassified(92);c__Agaricomycetes_unclassified(92); | - | - | - | - | - | - | - | Unassigned | - |
| Otu00868 | 83 | k__Fungi(100);p__Ascomycota(78);p__Ascomycota_unclassified(78);p__Ascomycota_unclassified(78);p__Ascomycota_unclassified(78);p__Ascomycota_unclassified(78);p__Ascomycota_unclassified(78); | - | - | - | - | - | - | - | Unassigned | - |
| Otu00870 | 83 | k__Fungi(100);k__Fungi_unclassified(100);k__Fungi_unclassified(100);k__Fungi_unclassified(100);k__Fungi_unclassified(100);k__Fungi_unclassified(100);k__Fungi_unclassified(100); | - | - | - | - | - | - | - | Unassigned | - |
| Otu00871 | 83 | k__Fungi(100);p__Ascomycota(100);p__Ascomycota_unclassified(100);p__Ascomycota_unclassified(100);p__Ascomycota_unclassified(100);p__Ascomycota_unclassified(100);p__Ascomycota_unclassified(100); | - | - | - | - | - | - | - | Unassigned | - |
| Otu00872 | 83 | k__Fungi(100);k__Fungi_unclassified(99);k__Fungi_unclassified(99);k__Fungi_unclassified(99);k__Fungi_unclassified(99);k__Fungi_unclassified(99);k__Fungi_unclassified(99); | - | - | - | - | - | - | - | Unassigned | - |
| Otu00873 | 82 | k__Fungi(100);p__Basidiomycota(100);c__Tremellomycetes(100);o__Tremellales(100);f__Tremellales_family_Incertae_sedis(100);g__Bullera(100);s__Bullera_unica(100); | Bullera | 13 | Saprotroph-Symbiotroph | Endophyte-Epiphyte-Plant Saprotroph-\|Undefined Saprotroph\| | Yeast | NULL | Probable | As foliar_endophyte (Põlme et al. 2020); Isolated from dead leaves of Oryza sativa, Miscanthus sinensis and Sasa sp. in Japan (Nakase & Suzuki 1986; Syn. Udeniomyces puniceus); Foliar, isolated from leaves of Melostomae candidum (Nakase et al. 2004) | Nakase T, Suzuki M. 1986. Journal of General and Applied Microbiology Tokyo 32:225-240 ((https://doi.org/10.2323/jgam.32.225)); Nakase T, et al. 2004. Systematic and Applied Microbiology 27:558-564 ((https://doi.org/10.1078/0723202041748118)) |
| Otu00874 | 82 | k__Fungi(100);p__Ascomycota(100);c__Dothideomycetes(100);o__Pleosporales(100);f__Pleosporales_family_Incertae_sedis(100);g__Phoma(100);s__Phoma_paspali(100); | Phoma | 13 | Pathotroph-Saprotroph-Symbiotroph | Dung Saprotroph-Endophyte-Lichen Parasite-\|Plant Pathogen\|-Plant Saprotroph-Undefined Saprotroph-Wood Saprotroph | Microfungus | Soft Rot | Possible | As foliar_endophyte (Põlme et al. 2020); As litter_saprotroph (Põlme et al. 2020); Decay leaf, fruit, seed, roots, soil (Põlme et al. 2020) | Richardson MJ. 1972. Transactions of the British Mycological Society 58:37-48 ((https://doi.org/10.1016/S0007-1536(72)80069-X)); Seehann G, et al. 1975. List of Fungi in Soft-Rot Tissue Tests. Institute fur Holzbiologie und Holzschutz, Bundesforschungsanstalt fur Forst - und Holzwirtshaft, Hamburg - Lohbrugge, Germany; de Gruyter J, et al. 2010. Mycologia 102:1066-1081 ((https://doi.org/10.3852/09-240)); Costa IPMW, et al. 2012. Mycotaxon 119:e26 ((http://mycotaxon.com/resources/checklists/costa_v119_checklist.pdf)); Tedersoo L, et al. 2014. Science 346:e1256688 ((https://doi.org/10.1126/science.1256688)); Lawrey JD, Diederich P. 2016. http://www.lichenicolous.net (accessed 24 May 2017) ((http://www.lichenicolous.net)); Põlme S, et al. 2020. Fungal Diversity 105:1-16 ((https://doi.org/10.1007/s13225-020-00466-2)) |
| Otu00876 | 82 | k__Fungi(100);p__Basidiomycota(100);c__Agaricomycetes(100);o__Polyporales(100);f__Phanerochaetaceae(100);g__Phanerochaete(100);s__Phanerochaete_sp(83); | Phanerochaete | 13 | Saprotroph | \|Wood Saprotroph\| | Corticioid | White Rot | Highly Probable | As white_rot (Põlme et al. 2020); Decay wood (Põlme et al. 2020) | Glenn JK, et al. 1983. Biochemical and Biophysical Research Communications 114:1077-1083 ((https://doi.org/10.1016/0006-291x((83)) 90672-1)); Põlme S, et al. 2020. Fungal Diversity 105:1-16 ((https://doi.org/10.1007/s13225-020-00466-2)) |
| Otu00877 | 82 | k__Fungi(100);p__Basidiomycota(100);c__Ustilaginomycetes(100);o__Ustilaginales(100);f__Ustilaginaceae(100);g__Ustilago(100);s__Ustilago_filiformis(100); | Ustilago | 13 | Pathotroph | \|Plant Pathogen\| | Smut | NULL | Probable | Host - Poaceae (Põlme et al. 2020) | Tedersoo L, et al. 2014. Science 346:e1256688 ((https://doi.org/10.1126/science.1256688)); Põlme S, et al. 2020. Fungal Diversity 105:1-16 ((https://doi.org/10.1007/s13225-020-00466-2)) |
| Otu00875 | 82 | k__Fungi(100);k__Fungi_unclassified(54);k__Fungi_unclassified(54);k__Fungi_unclassified(54);k__Fungi_unclassified(54);k__Fungi_unclassified(54);k__Fungi_unclassified(54); | - | - | - | - | - | - | - | Unassigned | - |
| Otu00878 | 82 | k__Fungi(100);k__Fungi_unclassified(64);k__Fungi_unclassified(64);k__Fungi_unclassified(64);k__Fungi_unclassified(64);k__Fungi_unclassified(64);k__Fungi_unclassified(64); | - | - | - | - | - | - | - | Unassigned | - |
| Otu00879 | 82 | k__Fungi(100);p__Ascomycota(100);p__Ascomycota_unclassified(100);p__Ascomycota_unclassified(100);p__Ascomycota_unclassified(100);p__Ascomycota_unclassified(100);p__Ascomycota_unclassified(100); | - | - | - | - | - | - | - | Unassigned | - |
| Otu00880 | 81 | k__Fungi(100);p__Zygomycota(100);c__Zygomycota_class_Incertae_sedis(100);o__Mucorales(100);f__Backusellaceae(100);g__Backusella(100);s__Backusella_circina(100); | Backusella | 13 | Saprotroph | Dung Saprotroph-\|Undefined Saprotroph\| | Microfungus-Sporophores | NULL | Probable | As soil_saprotroph (Põlme et al. 2020); Decay soil, dung (Põlme et al. 2020) | Walther G, et al. 2013. Persoonia 30:11-47 ((https://doi.org/10.3767/003158513X665070)); Põlme S, et al. 2020. Fungal Diversity 105:1-16 ((https://doi.org/10.1007/s13225-020-00466-2)) |
| Otu00881 | 81 | k__Fungi(100);p__Ascomycota(71);p__Ascomycota_unclassified(71);p__Ascomycota_unclassified(71);p__Ascomycota_unclassified(71);p__Ascomycota_unclassified(71);p__Ascomycota_unclassified(71); | - | - | - | - | - | - | - | Unassigned | - |
| Otu00882 | 81 | k__Fungi(100);p__Ascomycota(100);c__Sordariomycetes(100);o__Xylariales(100);o__Xylariales_unclassified(92);o__Xylariales_unclassified(92);o__Xylariales_unclassified(92); | - | - | - | - | - | - | - | Unassigned | - |
| Otu00883 | 81 | k__Fungi(100);p__Ascomycota(100);p__Ascomycota_unclassified(98);p__Ascomycota_unclassified(98);p__Ascomycota_unclassified(98);p__Ascomycota_unclassified(98);p__Ascomycota_unclassified(98); | - | - | - | - | - | - | - | Unassigned | - |
| Otu00884 | 81 | k__Fungi(100);p__Ascomycota(100);c__Eurotiomycetes(100);o__Chaetothyriales(100);o__Chaetothyriales_unclassified(72);o__Chaetothyriales_unclassified(72);o__Chaetothyriales_unclassified(72); | - | - | - | - | - | - | - | Unassigned | - |
| Otu00885 | 81 | k__Fungi(100);p__Ascomycota(73);p__Ascomycota_unclassified(73);p__Ascomycota_unclassified(73);p__Ascomycota_unclassified(73);p__Ascomycota_unclassified(73);p__Ascomycota_unclassified(73); | - | - | - | - | - | - | - | Unassigned | - |
| Otu00886 | 80 | k__Fungi(100);p__Ascomycota(100);c__Sordariomycetes(100);o__Xylariales(100);f__Xylariaceae(100);g__Hypoxylon(100);g__Hypoxylon_unclassified(100); | Hypoxylon | 13 | Saprotroph-Saprotroph-Symbiotroph | Endophyte-Undefined Saprotroph-\|Wood Saprotroph\| | Corticioid-Microfungus-Xylarioid | NULL | Probable | As foliar_endophyte (Põlme et al. 2020); Decay wood, leaf, fruit, seed (Põlme et al. 2020); Host - angiosperms (Põlme et al. 2020) | Tedersoo L, et al. 2014. Science 346:e1256688 ((https://doi.org/10.1126/science.1256688)); Põlme S, et al. 2020. Fungal Diversity 105:1-16 ((https://doi.org/10.1007/s13225-020-00466-2)) |
| Otu00887 | 80 | k__Fungi(100);p__Ascomycota(100);c__Eurotiomycetes(100);o__Chaetothyriales(100);f__Herpotrichiellaceae(100);g__Exophiala(100);s__Exophiala_sideris(100); | Exophiala | 13 | Pathotroph-Saprotroph-Symbiotroph | \|Animal Parasite\|-Animal Pathogen-Endophyte-Plant Saprotroph-Undefined Saprotroph | Dimorphic-Facultative Yeast | NULL | Probable | As litter_saprotroph (Põlme et al. 2020); As root_endophyte_dark_septate (Põlme et al. 2020); Decay leaf, fruit, seed, animal material (Põlme et al. 2020) | Tedersoo L, et al. 2014. Science 346:e1256688 ((https://doi.org/10.1126/science.1256688)), Nagy LG, et al. 2014. Nature Communications 5:e4471 ((https://doi.org/10.1038/ncomms5471)); Põlme S, et al. 2020. Fungal Diversity 105:1-16 ((https://doi.org/10.1007/s13225-020-00466-2)) |
| Otu00888 | 80 | k__Fungi(100);p__Basidiomycota(100);c__Agaricomycetes(100);o__Agaricales(100);f__Strophariaceae(100);g__Pholiota(100);s__Pholiota_terrestris(90); | Pholiota | 13 | Pathotroph-Saprotroph | Plant Pathogen-Plant Saprotroph-Undefined Saprotroph-\|Wood Saprotroph\| | Agaricoid | White Rot | Probable | As litter_saprotroph (Põlme et al. 2020); As white_rot (Põlme et al. 2020); As wood_pathogen (Põlme et al. 2020); Decay wood, leaf, fruit, seed (Põlme et al. 2020) | Tedersoo L, et al. 2014. Science 346:e1256688 ((https://doi.org/10.1126/science.1256688)); Põlme S, et al. 2020. Fungal Diversity 105:1-16 ((https://doi.org/10.1007/s13225-020-00466-2)) |
| Otu00889 | 80 | k__Fungi(100);p__Ascomycota(100);c__Dothideomycetes(100);o__Pleosporales(100);f__unclassified_Pleosporales(94);g__unclassified_Pleosporales(94);s__Pleosporales_sp(94); | - | - | - | - | - | - | - | Unassigned | - |
| Otu00890 | 80 | k__Fungi(100);k__Fungi_unclassified(100);k__Fungi_unclassified(100);k__Fungi_unclassified(100);k__Fungi_unclassified(100);k__Fungi_unclassified(100);k__Fungi_unclassified(100); | - | - | - | - | - | - | - | Unassigned | - |
| Otu00891 | 80 | k__Fungi(100);p__Basidiomycota(99);p__Basidiomycota_unclassified(99);p__Basidiomycota_unclassified(99);p__Basidiomycota_unclassified(99);p__Basidiomycota_unclassified(99);p__Basidiomycota_unclassified(99); | - | - | - | - | - | - | - | Unassigned | - |
| Otu00893 | 79 | k__Fungi(100);p__Basidiomycota(100);c__Ustilaginomycetes(100);o__Ustilaginales(100);f__Ustilaginaceae(100);g__Pseudozyma(100);s__Pseudozyma_sp(100); | Pseudozyma | 13 | Pathotroph-Symbiotroph | Endophyte-\|Epiphyte\|-Fungal Parasite | Yeast | NULL | Probable | As foliar_endophyte (Põlme et al. 2020); As mycoparasite (Põlme et al. 2020); Decay leaf, fruit, seed, fungal material (Põlme et al. 2020) | Põlme S, et al. 2020. Fungal Diversity 105:1-16 ((https://doi.org/10.1007/s13225-020-00466-2)) |
| Otu00894 | 79 | k__Fungi(100);p__Ascomycota(100);c__Lecanoromycetes(100);o__Lecanorales(100);o__Lecanorales_unclassified(98);o__Lecanorales_unclassified(98);o__Lecanorales_unclassified(98); | Lecanorales | 7 | Pathotroph-Symbiotroph | Lichen Parasite-Lichenized | Microfungus-Thallus | NULL | Probable | Lichenized with green algae, occasionally lichenicolous, on rocks, mosses and wood (Cannon & Kirk 2007); ...an order of mostly lichen-forming fungi (Kirk et al. 2008) | Cannon PF, Kirk PM. 2007. Fungal Families of the World. CAB International, Cambridge (ISBN: 978-0851998275); Kirk PM et al. 2008. Dictionary of the Fungi. Tenth Edition. CAB International, Wallingford (ISBN: 978-0851998268); Esslinger TL. 2014. North Dakota State University ((http://www.ndsu.edu/pubweb/~esslinge/chcklst/chcklst7.htm)) |
| Otu00895 | 79 | k__Fungi(100);p__Ascomycota(100);c__Eurotiomycetes(100);o__Eurotiales(100);f__Trichocomaceae(100);g__Penicillium(92);g__Penicillium_unclassified(68); | Penicillium | 13 | Pathotroph-Saprotroph-Symbiotroph | Animal Parasite-Dung Saprotroph-Endophyte-Fungal Parasite-Plant Pathogen-\|Undefined Saprotroph\|-Wood Saprotroph | Microfungus | Soft Rot | Highly Probable | As foliar_endophyte (Põlme et al. 2020); As mycoparasite (Põlme et al. 2020); Decay soil (Põlme et al. 2020) | Duncan CG, Eslyn WE. 1966. Mycologia 58:642-664 ((https://doi.org/10.2307/3757045)); Seehann G, et al. 1975. List of Fungi in Soft-Rot Tissue Tests. Institute fur Holzbiologie und Holzschutz, Bundesforschungsanstalt fur Forst - und Holzwirtshaft, Hamburg - Lohbrugge, Germany ((https://www.irg-wp.com/irgdocs/details.php?d6493033-c9ea-40f3-8e3e-5697f32d640d)); Costa IPMW, et al. 2012. Mycotaxon 119:e26 ((http://mycotaxon.com/resources/checklists/costa_v119_checklist.pdf)); Bills GF, et al. 2013. Current Opinion in Microbiology 16:549-565 ((https://doi.org/10.1016/j.mib.2013.08.001)); Tedersoo L, et al. 2014. Science 346:e1256688 ((https://doi.org/10.1126/science.1256688)); Põlme S, et al. 2020. Fungal Diversity 105:1-16 ((https://doi.org/10.1007/s13225-020-00466-2)) |
| Otu00896 | 79 | k__Fungi(100);p__Basidiomycota(100);c__Agaricomycetes(100);o__Polyporales(100);f__Polyporaceae(100);g__Diplomitoporus(100);s__Diplomitoporus_flavescens(100); | Diplomitoporus | 13 | Saprotroph | \|Wood Saprotroph\| | Corticioid-Polyporoid | White Rot | Highly Probable | As white_rot (Põlme et al. 2020); Decay wood (Põlme et al. 2020) | Gilbertson RL, Ryvarden L. 1987-1987. North American Polypores. Fungiflora, Oslo (ISBN: 978-0945345060); Põlme S, et al. 2020. Fungal Diversity 105:1-16 ((https://doi.org/10.1007/s13225-020-00466-2)) |
| Otu00900 | 79 | k__Fungi(100);p__Ascomycota(100);c__Saccharomycetes(100);o__Saccharomycetales(100);f__Metschnikowiaceae(100);g__Metschnikowia(100);s__Metschnikowia_pulcherrima(94); | Metschnikowia | 13 | Saprotroph | \|Undefined Saprotroph\| | Yeast | NULL | Probable | As nectar-tap_saprotroph (Põlme et al. 2020); Decay sugar-rich substrates (Põlme et al. 2020) | Põlme S, et al. 2020. Fungal Diversity 105:1-16 ((https://doi.org/10.1007/s13225-020-00466-2)) |
| Otu00901 | 79 | k__Fungi(100);p__Ascomycota(100);c__Dothideomycetes(100);o__Pleosporales(100);f__Montagnulaceae(100);f__Montagnulaceae_unclassified(88);f__Montagnulaceae_unclassified(88); | Montagnulaceae | 9 | Pathotroph-Saprotroph | Fungal Parasite-Plant Pathogen-Undefined Saprotroph | Microfungus | NULL | Probable | Species in this clade can be saprobic, plant pathogenic, or mycoparasitic (Zhang et al. 2009) | Zhang Y, et al. 2009. Studies in Mycology 64:85-102 ((https://doi.org/10.3114/sim.2009.64.04)) |
| Otu00892 | 79 | k__Fungi(100);p__Ascomycota(100);c__Sordariomycetes(100);o__Trichosphaeriales(100);f__Trichosphaeriales_family_Incertae_sedis(100);g__unclassified_Trichosphaeriales(100);s__Trichosphaeriales_sp(100); | - | - | - | - | - | - | - | Unassigned | - |
| Otu00897 | 79 | k__Fungi(100);k__Fungi_unclassified(74);k__Fungi_unclassified(74);k__Fungi_unclassified(74);k__Fungi_unclassified(74);k__Fungi_unclassified(74);k__Fungi_unclassified(74); | - | - | - | - | - | - | - | Unassigned | - |
| Otu00898 | 79 | k__Fungi(100);k__Fungi_unclassified(97);k__Fungi_unclassified(97);k__Fungi_unclassified(97);k__Fungi_unclassified(97);k__Fungi_unclassified(97);k__Fungi_unclassified(97); | - | - | - | - | - | - | - | Unassigned | - |
| Otu00899 | 79 | k__Fungi(100);k__Fungi_unclassified(74);k__Fungi_unclassified(74);k__Fungi_unclassified(74);k__Fungi_unclassified(74);k__Fungi_unclassified(74);k__Fungi_unclassified(74); | - | - | - | - | - | - | - | Unassigned | - |
| Otu00902 | 78 | k__Fungi(100);p__Ascomycota(100);c__Saccharomycetes(100);o__Saccharomycetales(100);f__Metschnikowiaceae(100);g__Metschnikowia(100);s__Metschnikowia_pulcherrima(98); | Metschnikowia | 13 | Saprotroph | \|Undefined Saprotroph\| | Yeast | NULL | Probable | As nectar-tap_saprotroph (Põlme et al. 2020); Decay sugar-rich substrates (Põlme et al. 2020) | Põlme S, et al. 2020. Fungal Diversity 105:1-16 ((https://doi.org/10.1007/s13225-020-00466-2)) |
| Otu00904 | 78 | k__Fungi(100);p__Basidiomycota(100);c__Agaricomycetes(100);o__Polyporales(100);f__Meruliaceae(100);g__Phlebia(100);s__Phlebia_setulosa(100); | Phlebia | 13 | Saprotroph | \|Wood Saprotroph\| | Corticioid | White Rot | Highly Probable | As white_rot (Põlme et al. 2020); Decay wood (Põlme et al. 2020) | Kirk PM et al. 2008. Dictionary of the Fungi. Tenth Edition. CAB International, Wallingford (ISBN: 978-0851998268); Põlme S, et al. 2020. Fungal Diversity 105:1-16 ((https://doi.org/10.1007/s13225-020-00466-2)) |
| Otu00905 | 78 | k__Fungi(100);p__Basidiomycota(100);c__Pucciniomycetes(100);o__Pucciniales(100);f__Pucciniastraceae(100);g__Melampsoridium(100);s__Melampsoridium_hiratsukanum(100); | Melampsoridium | 13 | Pathotroph | \|Plant Pathogen\| | Rust | NULL | Probable | NULL | Tedersoo L, et al. 2014. Science 346:e1256688 ((https://doi.org/10.1126/science.1256688)); Põlme S, et al. 2020. Fungal Diversity 105:1-16 ((https://doi.org/10.1007/s13225-020-00466-2)) |
| Otu00906 | 78 | k__Fungi(100);p__Ascomycota(100);c__Saccharomycetes(100);o__Saccharomycetales(100);f__Metschnikowiaceae(100);g__Metschnikowia(100);s__Metschnikowia_pulcherrima(95); | Metschnikowia | 13 | Saprotroph | \|Undefined Saprotroph\| | Yeast | NULL | Probable | As nectar-tap_saprotroph (Põlme et al. 2020); Decay sugar-rich substrates (Põlme et al. 2020) | Põlme S, et al. 2020. Fungal Diversity 105:1-16 ((https://doi.org/10.1007/s13225-020-00466-2)) |
| Otu00908 | 78 | k__Fungi(100);p__Ascomycota(100);c__Sordariomycetes(100);o__Hypocreales(100);f__Hypocreales_family_Incertae_sedis(100);g__Acremonium(74);s__Acremonium_sp_24_OA_2013(74); | Acremonium | 13 | Pathotroph-Saprotroph-Symbiotroph | Animal Pathogen-Endophyte-Fungal Parasite-Plant Pathogen-\|Undefined Saprotroph\|-Wood Saprotroph | Microfungus | Soft Rot | Possible | As foliar_endophyte (Põlme et al. 2020); culture (Hanada et al. 2010, Del Olmo; Decay leaf, fruit, seed, soil, dung, animal material (Põlme et al. 2020); Endophyte detection method; Ruiz & Arnold 2014); Likely opportunistic human pathogen (Irinyi et al. 2015) | Duncan CG, Eslyn WE. 1966. Mycologia 58:642-645 ((https://doi.org/10.2307/3757045)); Nilsson T. 1973. Studia Forestalia Suecica 104:1-40 ((https://pub.epsilon.slu.se/5711/)); Worrall JJ, et al. 1997. Mycologia 89:199-219 ((https://doi.org/10.2307/3761073)); Hanada RE, et al. 2010. Fungal Biology 114:901-910 ((https://doi.org/10.1016/j.funbio.2010.08.006)); Del Olmo-Ruiz M, Arnold AE. 2014. Mycologia 106:8-21 ((https://doi.org/10.3852/13-098)); Irinyi L, et al. 2015. Medical Mycology 53313-337 ((https://doi.org/10.1093/mmy/myv008)); Põlme S, et al. 2020. Fungal Diversity 105:1-16 ((https://doi.org/10.1007/s13225-020-00466-2)) |
| Otu00903 | 78 | k__Fungi(100);k__Fungi_unclassified(75);k__Fungi_unclassified(75);k__Fungi_unclassified(75);k__Fungi_unclassified(75);k__Fungi_unclassified(75);k__Fungi_unclassified(75); | - | - | - | - | - | - | - | Unassigned | - |
| Otu00907 | 78 | k__Fungi(100);p__Ascomycota(100);c__Eurotiomycetes(99);o__Chaetothyriales(93);o__Chaetothyriales_unclassified(93);o__Chaetothyriales_unclassified(93);o__Chaetothyriales_unclassified(93); | - | - | - | - | - | - | - | Unassigned | - |
| Otu00909 | 78 | k__Fungi(100);k__Fungi_unclassified(100);k__Fungi_unclassified(100);k__Fungi_unclassified(100);k__Fungi_unclassified(100);k__Fungi_unclassified(100);k__Fungi_unclassified(100); | - | - | - | - | - | - | - | Unassigned | - |
| Otu00910 | 77 | k__Fungi(100);p__Ascomycota(100);c__Saccharomycetes(100);o__Saccharomycetales(100);f__Metschnikowiaceae(100);g__Metschnikowia(100);s__Metschnikowia_pulcherrima(99); | Metschnikowia | 13 | Saprotroph | \|Undefined Saprotroph\| | Yeast | NULL | Probable | As nectar-tap_saprotroph (Põlme et al. 2020); Decay sugar-rich substrates (Põlme et al. 2020) | Põlme S, et al. 2020. Fungal Diversity 105:1-16 ((https://doi.org/10.1007/s13225-020-00466-2)) |
| Otu00911 | 77 | k__Fungi(100);p__Ascomycota(100);c__Eurotiomycetes(100);o__Chaetothyriales(98);f__Herpotrichiellaceae(98);g__unclassified_Herpotrichiellaceae(98);s__Herpotrichiellaceae_sp(98); | Herpotrichiellaceae | 9 | Pathotroph-Saprotroph | Animal Pathogen-Fungal Parasite-Undefined Saprotroph | Facultative Yeast-Microfungus | NULL | Probable | Mostly saprobic on plants or other fungi, isolated from soil etc., and from medical sources (Cannon & Kirk 2007) | Cannon PF, Kirk PM. 2007. Fungal Families of the World. CAB International, Cambridge (ISBN: 978-0851998275) |
| Otu00912 | 77 | k__Fungi(100);p__Basidiomycota(100);c__Agaricomycetes(100);o__Agaricales(100);f__Psathyrellaceae(100);g__Coprinellus(52);g__Coprinellus_unclassified(52); | Coprinellus | 13 | Saprotroph | \|Undefined Saprotroph\| | Agaricoid | NULL | Probable | As soil_saprotroph (Põlme et al. 2020); Decay soil (Põlme et al. 2020) | Tedersoo L, et al. 2014. Science 346:e1256688 ((https://doi.org/10.1126/science.1256688)); Põlme S, et al. 2020. Fungal Diversity 105:1-16 ((https://doi.org/10.1007/s13225-020-00466-2)) |
| Otu00913 | 77 | k__Fungi(100);p__Ascomycota(100);c__Eurotiomycetes(100);o__Chaetothyriales(100);f__Chaetothyriaceae(100);g__Cyphellophora(100);g__Cyphellophora_unclassified(100); | Cyphellophora | 13 | Pathotroph-Saprotroph | Animal Parasite-Animal Pathogen-\|Plant Saprotroph\|-Undefined Saprotroph | Dimorphic-Facultative Yeast-Microfungus | NULL | Probable | As litter_saprotroph (Põlme et al. 2020); Consult the current literature for updated names on this group that included the Black Yeasts; Some species are likely opportunistic human pathogen (Irinyi et al. 2015); Decay leaf, fruit, seed, animal material (Põlme et al. 2020) | Tedersoo L, et al. 2014. Science 346:e1256688 ((https://doi.org/10.1126/science.1256688)); Irinyi L, et al. 2015. Medical Mycology 53:313-337 ((https://doi.org/10.1093/mmy/myv008)); Põlme S, et al. 2020. Fungal Diversity 105:1-16 ((https://doi.org/10.1007/s13225-020-00466-2)) |
| Otu00915 | 76 | k__Fungi(100);p__Ascomycota(100);c__Eurotiomycetes(100);o__Chaetothyriales(100);f__Herpotrichiellaceae(100);g__Capronia(99);s__Capronia_semi-immersa(99); | Capronia | 13 | Pathotroph-Saprotroph-Symbiotroph | Animal Pathogen-Endophyte-Fungal Parasite-\|Undefined Saprotroph\| | Dimorphic-Facultative Yeast | NULL | Highly Probable | As mycoparasite (Põlme et al. 2020); As root_endophyte_dark_septate (Põlme et al. 2020); As soil_saprotroph (Põlme et al. 2020); Dark Septate Endophyte, known from plant roots.; Decay soil (Põlme et al. 2020) | Tedersoo L, et al. 2014. Science 346:e1256688 ((https://doi.org/10.1126/science.1256688)); Põlme S, et al. 2020. Fungal Diversity 105:1-16 ((https://doi.org/10.1007/s13225-020-00466-2)) |
| Otu00916 | 76 | k__Fungi(100);p__Basidiomycota(100);c__Microbotryomycetes(100);o__Sporidiobolales(100);f__Sporidiobolales_family_Incertae_sedis(100);g__Rhodotorula(100);s__Rhodotorula_minuta(100); | Rhodotorula | 13 | Pathotroph-Saprotroph-Symbiotroph | Animal Pathogen-Animal Symbiotroph-Endophyte-Plant Pathogen-\|Undefined Saprotroph\| | Yeast | NULL | Probable | As foliar_endophyte (Põlme et al. 2020); Opportunistic human pathogen | Põlme S, et al. 2020. Fungal Diversity 105:1-16 ((https://doi.org/10.1007/s13225-020-00466-2)); https://www.cdc.gov ((https://www.cdc.gov)) |
| Otu00917 | 76 | k__Fungi(100);p__Zygomycota(100);c__Zygomycota_class_Incertae_sedis(100);o__Mortierellales(100);f__Mortierellaceae(100);g__Mortierella(100);s__Mortierella_humilis(61); | Mortierella | 13 | Saprotroph-Symbiotroph | Endophyte-Plant Saprotroph-\|Undefined Saprotroph\| | Microfungus | NULL | Probable | As chitinolytic (Põlme et al. 2020); As soil_saprotroph (Põlme et al. 2020); Decay roots, soil (Põlme et al. 2020); Saprobic in soil and thus frequently isolated from soil and material contaminated with soil such as hay or rotten silage and spoil tips (Cannon & Kirk 2007) | Cannon PF, Kirk PM. 2007. Fungal Families of the World. CAB International, Cambridge (ISBN: 978-0851998275); Tedersoo L, et al. 2014. Science 346:e1256688 ((https://doi.org/10.1126/science.1256688)); Purahong W, et al. 2016. Molecular Ecology 25:4059-4074 ((https://doi.org/10.1111/mec.13739)); Põlme S, et al. 2020. Fungal Diversity 105:1-16 ((https://doi.org/10.1007/s13225-020-00466-2)) |
| Otu00919 | 76 | k__Fungi(100);p__Ascomycota(100);c__Eurotiomycetes(100);o__Eurotiales(100);f__Trichocomaceae(100);g__Penicillium(100);s__Penicillium_penicillioides(100); | Penicillium | 13 | Pathotroph-Saprotroph-Symbiotroph | Animal Parasite-Dung Saprotroph-Endophyte-Fungal Parasite-Plant Pathogen-\|Undefined Saprotroph\|-Wood Saprotroph | Microfungus | Soft Rot | Highly Probable | As foliar_endophyte (Põlme et al. 2020); As mycoparasite (Põlme et al. 2020); Decay soil (Põlme et al. 2020) | Duncan CG, Eslyn WE. 1966. Mycologia 58:642-664 ((https://doi.org/10.2307/3757045)); Seehann G, et al. 1975. List of Fungi in Soft-Rot Tissue Tests. Institute fur Holzbiologie und Holzschutz, Bundesforschungsanstalt fur Forst - und Holzwirtshaft, Hamburg - Lohbrugge, Germany ((https://www.irg-wp.com/irgdocs/details.php?d6493033-c9ea-40f3-8e3e-5697f32d640d)); Costa IPMW, et al. 2012. Mycotaxon 119:e26 ((http://mycotaxon.com/resources/checklists/costa_v119_checklist.pdf)); Bills GF, et al. 2013. Current Opinion in Microbiology 16:549-565 ((https://doi.org/10.1016/j.mib.2013.08.001)); Tedersoo L, et al. 2014. Science 346:e1256688 ((https://doi.org/10.1126/science.1256688)); Põlme S, et al. 2020. Fungal Diversity 105:1-16 ((https://doi.org/10.1007/s13225-020-00466-2)) |
| Otu00914 | 76 | k__Fungi(100);k__Fungi_unclassified(65);k__Fungi_unclassified(65);k__Fungi_unclassified(65);k__Fungi_unclassified(65);k__Fungi_unclassified(65);k__Fungi_unclassified(65); | - | - | - | - | - | - | - | Unassigned | - |
| Otu00918 | 76 | k__Fungi(100);k__Fungi_unclassified(58);k__Fungi_unclassified(58);k__Fungi_unclassified(58);k__Fungi_unclassified(58);k__Fungi_unclassified(58);k__Fungi_unclassified(58); | - | - | - | - | - | - | - | Unassigned | - |
| Otu00920 | 75 | k__Fungi(100);p__Ascomycota(100);c__Sordariomycetes(100);o__Xylariales(100);f__Xylariaceae(100);g__Xylaria(100);g__Xylaria_unclassified(63); | Xylaria | 13 | Saprotroph-Symbiotroph | Endophyte-Plant Saprotroph-Undefined Saprotroph-\|Wood Saprotroph\| | Xylarioid | Soft Rot | Probable | As foliar_endophyte (Põlme et al. 2020); As litter_saprotroph (Põlme et al. 2020); Decay wood, leaf, fruit, seed (Põlme et al. 2020); Host - angiosperms (Põlme et al. 2020) | Duncan CG, Eslyn WE. 1966. Mycologia 58:642-645 ((https://doi.org/10.2307/3757045)); Costa IPMW, et al. 2012. Mycotaxon 119:e26 ((http://mycotaxon.com/resources/checklists/costa_v119_checklist.pdf)); Põlme S, et al. 2020. Fungal Diversity 105:1-16 ((https://doi.org/10.1007/s13225-020-00466-2)); The American Phytopathological Society Common Names of Plant Diseases ((http://www.apsnet.org/publications/commonnames/Pages/default.aspx)) |
| Otu00922 | 75 | k__Fungi(100);p__Ascomycota(100);c__Sordariomycetes(100);o__Hypocreales(100);f__Clavicipitaceae(100);g__Pochonia(100);s__Pochonia_bulbillosa(100); | Pochonia | 13 | Pathotroph-Saprotroph | \|Animal Parasite\|-Animal Pathogen-Undefined Saprotroph | Microfungus | NULL | Probable | As animal_decomposer (Põlme et al. 2020); Decay animal material (Põlme et al. 2020) | Tedersoo L, et al. 2014. Science 346:e1256688 ((https://doi.org/10.1126/science.1256688)); Põlme S, et al. 2020. Fungal Diversity 105:1-16 ((https://doi.org/10.1007/s13225-020-00466-2)) |
| Otu00924 | 75 | k__Fungi(100);p__Basidiomycota(100);c__Agaricomycetes(100);o__Hymenochaetales(100);f__Hymenochaetaceae(100);g__Tubulicrinis(100);s__Tubulicrinis_sp(100); | Tubulicrinis | 13 | Saprotroph | Undefined Saprotroph-\|Wood Saprotroph\| | Corticioid | White Rot | Probable | As white_rot (Põlme et al. 2020); Decay wood (Põlme et al. 2020) | Tedersoo L, et al. 2014. Science 346:e1256688 ((https://doi.org/10.1126/science.1256688)); Põlme S, et al. 2020. Fungal Diversity 105:1-16 ((https://doi.org/10.1007/s13225-020-00466-2)) |
| Otu00926 | 75 | k__Fungi(100);p__Basidiomycota(100);c__Agaricomycetes(100);o__Agaricales(100);f__Mycenaceae(100);g__Mycena(100);g__Mycena_unclassified(88); | Mycena | 13 | Pathotroph-Saprotroph | Plant Pathogen-\|Plant Saprotroph\|-Undefined Saprotroph-Wood Saprotroph | Agaricoid | NULL | Probable | As litter_saprotroph (Põlme et al. 2020); Decay leaf, fruit, seed (Põlme et al. 2020) | Grgurinovic CA, Simpson JA. 2001. Fungal Diversity 8:97-127 ((http://www.fungaldiversity.org/fdp/sfdp/FD_8_97-127.pdf)); Tedersoo L, et al. 2014. Science 346:e1256688 ((https://doi.org/10.1126/science.1256688)); Põlme S, et al. 2020. Fungal Diversity 105:1-16 ((https://doi.org/10.1007/s13225-020-00466-2)); The American Phytopathological Society Common Names of Plant Diseases ((http://www.apsnet.org/publications/commonnames/Pages/default.aspx)) |
| Otu00928 | 75 | k__Fungi(100);p__Ascomycota(100);c__Sordariomycetes(100);o__Sordariales(100);f__Chaetomiaceae(100);f__Chaetomiaceae_unclassified(76);f__Chaetomiaceae_unclassified(76); | Chaetomiaceae | 9 | Pathotroph-Saprotroph-Symbiotroph | Animal Pathogen-Dung Saprotroph-Endophyte-Epiphyte-Plant Saprotroph-Wood Saprotroph | Microfungus | Soft Rot | Possible | Saprobic, acting as key decay organisms of plant materials and plant-derived commodities (Cannon & Kirk 2007); Epiphyte detection method-culture (Massimo et al. 2015); Endophyte detection method-culture (Busby et al. 2016, David et al. 2016); Likely opportunistic human pathogen (Irinyi et al. 2015) | Seehann G, et al. 1975. List of Fungi in Soft-Rot Tissue Tests. Institute fur Holzbiologie und Holzschutz, Bundesforschungsanstalt fur Forst - und Holzwirtshaft, Hamburg - Lohbrugge, Germany; Cannon PF, Kirk PM. 2007. Fungal Families of the World. CAB International, Cambridge ((https://isbnsearch.org/isbn/9780851998275)); Bills GF, et al. 2013. Current Opinion in Microbiology 16:549-565 ((https://doi.org/10.1016/j.mib.2013.08.001)); Tedersoo L, et al. 2014. Science 346:e1256688 ((https://doi.org/10.1126/science.1256688)); Irinyi L, et al. 2015. Medical Mycology 53:313-337 ((https://doi.org/10.1093/mmy/myv008)); Massimo NC, et al. 2015. Microbial Ecology 70:61-76 ((https://doi.org/10.1007/s00248-014-0563-6)); Busby PE, et al. 2016. New Phytologist 209:1681-1692 ((https://doi.org/10.1111/nph.13742)); David AS, et al. 2016. Microbial Ecology 71:912-926 ((https://doi.org/10.1007/s00248-015-0712-6)) |
| Otu00929 | 75 | k__Fungi(100);p__Ascomycota(100);c__Sordariomycetes(100);o__Diaporthales(100);f__Diaporthaceae(100);f__Diaporthaceae_unclassified(100);f__Diaporthaceae_unclassified(100); | Diaporthaceae | 9 | Pathotroph-Saprotroph-Symbiotroph | Endophyte-Plant Pathogen-Plant Saprotroph | Microfungus | NULL | Probable | Most species are weak pathogens or endophytes in leaf or stem tissue, growing saprobically after the plant dies (Cannon & Kirk 2007) | Cannon PF, Kirk PM. 2007. Fungal Families of the World. CAB International, Cambridge (ISBN: 978-0851998275) |
| Otu00921 | 75 | k__Fungi(100);k__Fungi_unclassified(70);k__Fungi_unclassified(70);k__Fungi_unclassified(70);k__Fungi_unclassified(70);k__Fungi_unclassified(70);k__Fungi_unclassified(70); | - | - | - | - | - | - | - | Unassigned | - |
| Otu00923 | 75 | k__Fungi(100);p__Basidiomycota(100);c__Pucciniomycetes(100);o__Pucciniales(100);o__Pucciniales_unclassified(100);o__Pucciniales_unclassified(100);o__Pucciniales_unclassified(100); | - | - | - | - | - | - | - | Unassigned | - |
| Otu00925 | 75 | k__Fungi(100);p__Ascomycota(76);p__Ascomycota_unclassified(76);p__Ascomycota_unclassified(76);p__Ascomycota_unclassified(76);p__Ascomycota_unclassified(76);p__Ascomycota_unclassified(76); | - | - | - | - | - | - | - | Unassigned | - |
| Otu00927 | 75 | k__Fungi(100);p__Ascomycota(100);c__Dothideomycetes(100);o__Capnodiales(100);o__Capnodiales_unclassified(99);o__Capnodiales_unclassified(99);o__Capnodiales_unclassified(99); | - | - | - | - | - | - | - | Unassigned | - |
| Otu00930 | 75 | k__Fungi(100);k__Fungi_unclassified(66);k__Fungi_unclassified(66);k__Fungi_unclassified(66);k__Fungi_unclassified(66);k__Fungi_unclassified(66);k__Fungi_unclassified(66); | - | - | - | - | - | - | - | Unassigned | - |
| Otu00931 | 74 | k__Fungi(100);p__Ascomycota(100);c__Saccharomycetes(100);o__Saccharomycetales(100);f__Metschnikowiaceae(100);g__Metschnikowia(100);s__Metschnikowia_pulcherrima(91); | Metschnikowia | 13 | Saprotroph | \|Undefined Saprotroph\| | Yeast | NULL | Probable | As nectar-tap_saprotroph (Põlme et al. 2020); Decay sugar-rich substrates (Põlme et al. 2020) | Põlme S, et al. 2020. Fungal Diversity 105:1-16 ((https://doi.org/10.1007/s13225-020-00466-2)) |
| Otu00932 | 74 | k__Fungi(100);p__Ascomycota(100);c__Eurotiomycetes(100);o__Eurotiales(100);f__Trichocomaceae(100);g__Penicillium(100);s__Penicillium_steckii(68); | Penicillium | 13 | Pathotroph-Saprotroph-Symbiotroph | Animal Parasite-Dung Saprotroph-Endophyte-Fungal Parasite-Plant Pathogen-\|Undefined Saprotroph\|-Wood Saprotroph | Microfungus | Soft Rot | Highly Probable | As foliar_endophyte (Põlme et al. 2020); As mycoparasite (Põlme et al. 2020); Decay soil (Põlme et al. 2020) | Duncan CG, Eslyn WE. 1966. Mycologia 58:642-664 ((https://doi.org/10.2307/3757045)); Seehann G, et al. 1975. List of Fungi in Soft-Rot Tissue Tests. Institute fur Holzbiologie und Holzschutz, Bundesforschungsanstalt fur Forst - und Holzwirtshaft, Hamburg - Lohbrugge, Germany ((https://www.irg-wp.com/irgdocs/details.php?d6493033-c9ea-40f3-8e3e-5697f32d640d)); Costa IPMW, et al. 2012. Mycotaxon 119:e26 ((http://mycotaxon.com/resources/checklists/costa_v119_checklist.pdf)); Bills GF, et al. 2013. Current Opinion in Microbiology 16:549-565 ((https://doi.org/10.1016/j.mib.2013.08.001)); Tedersoo L, et al. 2014. Science 346:e1256688 ((https://doi.org/10.1126/science.1256688)); Põlme S, et al. 2020. Fungal Diversity 105:1-16 ((https://doi.org/10.1007/s13225-020-00466-2)) |
| Otu00933 | 74 | k__Fungi(100);p__Basidiomycota(100);c__Agaricomycetes(100);o__Cantharellales(100);f__Clavulinaceae(96);f__Clavulinaceae_unclassified(68);f__Clavulinaceae_unclassified(68); | Clavulinaceae | 9 | Saprotroph-Symbiotroph | Ectomycorrhizal-Lichenized-Wood Saprotroph | Clavarioid | NULL | Possible | Some species form ectomycorrhizas, with fruit bodies in leaf litter and on soil. Others are probably saprobes associated with rotten wood (Cannon & Kirk 2007) | Cannon PF, Kirk PM. 2007. Fungal Families of the World. CAB International, Cambridge ((https://isbnsearch.org/isbn/9780851998275)); Nelsen MP, et al. 2007. American Journal of Botany 94:1289-1296 ((https://doi.org/10.3732/ajb.94.8.1289)) |
| Otu00934 | 74 | k__Fungi(100);p__Basidiomycota(100);c__Microbotryomycetes(76);o__Sporidiobolales(76);f__Sporidiobolales_family_Incertae_sedis(76);g__Rhodotorula(64);g__Rhodotorula_unclassified(64); | Rhodotorula | 13 | Pathotroph-Saprotroph-Symbiotroph | Animal Pathogen-Animal Symbiotroph-Endophyte-Plant Pathogen-\|Undefined Saprotroph\| | Yeast | NULL | Probable | As foliar_endophyte (Põlme et al. 2020); Opportunistic human pathogen | Põlme S, et al. 2020. Fungal Diversity 105:1-16 ((https://doi.org/10.1007/s13225-020-00466-2)); https://www.cdc.gov ((https://www.cdc.gov)) |
| Otu00935 | 74 | k__Fungi(100);p__Ascomycota(100);c__Dothideomycetes(100);o__Pleosporales(100);f__Montagnulaceae(100);g__unclassified_Montagnulaceae(99);s__Montagnulaceae_sp(99); | Montagnulaceae | 9 | Pathotroph-Saprotroph | Fungal Parasite-Plant Pathogen-Undefined Saprotroph | Microfungus | NULL | Probable | Species in this clade can be saprobic, plant pathogenic, or mycoparasitic (Zhang et al. 2009) | Zhang Y, et al. 2009. Studies in Mycology 64:85-102 ((https://doi.org/10.3114/sim.2009.64.04)) |
| Otu00936 | 74 | k__Fungi(100);p__Ascomycota(100);c__Eurotiomycetes(100);o__Mycocaliciales(100);f__Mycocaliciaceae(100);g__Chaenothecopsis(100);s__Chaenothecopsis_sitchensis(100); | Chaenothecopsis | 13 | Pathotroph-Saprotroph-Symbiotroph | Lichen Parasite-Lichenized-\|Undefined Saprotroph\| | Calicioid-Thallus | NULL | Possible | As resin_saprotroph (Põlme et al. 2020); licenized taxa; Lichenized and non | Esslinger TL. 2014. North Dakota State University ((http://www.ndsu.edu/pubweb/~esslinge/chcklst/chcklst7.htm)); Põlme S, et al. 2020. Fungal Diversity 105:1-16 ((https://doi.org/10.1007/s13225-020-00466-2)) |
| Otu00937 | 74 | k__Fungi(100);p__Ascomycota(68);p__Ascomycota_unclassified(68);p__Ascomycota_unclassified(68);p__Ascomycota_unclassified(68);p__Ascomycota_unclassified(68);p__Ascomycota_unclassified(68); | - | - | - | - | - | - | - | Unassigned | - |
| Otu00942 | 73 | k__Fungi(100);p__Basidiomycota(100);c__Agaricomycetes(100);o__Polyporales(100);f__Polyporaceae(100);f__Polyporaceae_unclassified(100);f__Polyporaceae_unclassified(100); | Polyporaceae | 9 | Saprotroph | Wood Saprotroph | Polyporoid | Brown Rot-White | Probable | Lignicolous, causing brown or white rot, mostly of broadleaved trees (Cannon & Kirk 2007) | Cannon PF, Kirk PM. 2007. Fungal Families of the World. CAB International, Cambridge (ISBN: 978-0851998275) |
| Otu00945 | 73 | k__Fungi(100);p__Basidiomycota(100);c__Agaricomycetes(100);o__Hymenochaetales(100);f__Hymenochaetaceae(100);g__Phellinus(100);s__Phellinus_viticola(100); | Phellinus | 13 | Pathotroph-Saprotroph | \|Plant Pathogen\|-Wood Saprotroph | Polyporoid | White Rot | Highly Probable | As white_rot (Põlme et al. 2020); As wood_pathogen (Põlme et al. 2020); Decay wood (Põlme et al. 2020) | Gilbertson RL, Ryvarden L. 1987-1987. North American Polypores. Fungiflora, Oslo (ISBN: 978-0945345060); Põlme S, et al. 2020. Fungal Diversity 105:1-16 ((https://doi.org/10.1007/s13225-020-00466-2)) |
| Otu00938 | 73 | k__Fungi(100);k__Fungi_unclassified(65);k__Fungi_unclassified(65);k__Fungi_unclassified(65);k__Fungi_unclassified(65);k__Fungi_unclassified(65);k__Fungi_unclassified(65); | - | - | - | - | - | - | - | Unassigned | - |
| Otu00939 | 73 | k__Fungi(100);p__Ascomycota(55);p__Ascomycota_unclassified(55);p__Ascomycota_unclassified(55);p__Ascomycota_unclassified(55);p__Ascomycota_unclassified(55);p__Ascomycota_unclassified(55); | - | - | - | - | - | - | - | Unassigned | - |
| Otu00940 | 73 | k__Fungi(100);p__Ascomycota(100);p__Ascomycota_unclassified(100);p__Ascomycota_unclassified(100);p__Ascomycota_unclassified(100);p__Ascomycota_unclassified(100);p__Ascomycota_unclassified(100); | - | - | - | - | - | - | - | Unassigned | - |
| Otu00941 | 73 | k__Fungi(100);k__Fungi_unclassified(57);k__Fungi_unclassified(57);k__Fungi_unclassified(57);k__Fungi_unclassified(57);k__Fungi_unclassified(57);k__Fungi_unclassified(57); | - | - | - | - | - | - | - | Unassigned | - |
| Otu00943 | 73 | k__Fungi(100);p__Ascomycota(100);c__Saccharomycetes(100);o__Saccharomycetales(100);f__Saccharomycetaceae(100);f__Saccharomycetaceae_unclassified(100);f__Saccharomycetaceae_unclassified(100); | - | - | - | - | - | - | - | Unassigned | - |
| Otu00944 | 73 | k__Fungi(100);p__Basidiomycota(100);c__Agaricomycetes(100);o__Sebacinales(100);o__Sebacinales_unclassified(87);o__Sebacinales_unclassified(87);o__Sebacinales_unclassified(87); | - | - | - | - | - | - | - | Unassigned | - |
| Otu00947 | 72 | k__Fungi(100);p__Ascomycota(100);c__Pezizomycetes(100);o__Pezizales(100);f__Ascobolaceae(100);g__unclassified_Ascobolaceae(100);s__Ascobolaceae_sp(100); | Ascobolaceae | 9 | Saprotroph | Dung Saprotroph-Soil Saprotroph-Wood Saprotroph | NULL | NULL | Probable | NULL | Cannon PF, Kirk PM. 2007. Fungal Families of the World. CAB International, Cambridge (ISBN: 978-0851998275) |
| Otu00948 | 72 | k__Fungi(100);p__Basidiomycota(100);c__Tremellomycetes(100);o__Tremellales(100);f__Tremellales_family_Incertae_sedis(100);g__Cryptococcus(100);s__Cryptococcus_laurentii(95); | Cryptococcus | 13 | Pathotroph-Saprotroph-Symbiotroph | Animal Pathogen-Endophyte-Epiphyte-\|Undefined Saprotroph\| | Dimorphic Yeast-Yeast | NULL | Possible | As foliar_endophyte (Põlme et al. 2020); Decay leaf, fruit, seed, soil, dung, animal material (Põlme et al. 2020) | Andrews JH, et al. 1983. Phytopathology 73:228-234 ((https://doi.org/10.1094/Phyto-73-228)); Kurtzman CP, et al. (eds.) 2011. The Yeasts, a Taxonomic Study. Fifth Edition. Vols 1-3. Elsevier, San Diego (ISBN: 9780444521491); Põlme S, et al. 2020. Fungal Diversity 105:1-16 ((https://doi.org/10.1007/s13225-020-00466-2)) |

| | | | | | | | | | | | |
|---|---|---|---|---|---|---|---|---|---|---|---|
| Otu00949 | 72 | k__Fungi(100);p__Ascomycota(100);c__Eurotiomycetes(100);o__Eurotiales(100);f__Trichocomaceae(100);g__Aspergillus(100);s__Aspergillus_sp(96); | Aspergillus | 13 | Pathotroph-Saprotroph-Symbiotroph | Animal Pathogen-Endophyte-Plant Saprotroph-\|Undefined Saprotroph\|-Wood Saprotroph | Microfungus | NULL | Probable | As foliar_endophyte (Põlme et al. 2020); Decay leaf, fruit, seed, soil, dung, animal material (Põlme et al. 2020) | [illegible] |
| Otu00951 | 72 | k__Fungi(100);p__Basidiomycota(100);c__Agaricomycetes(100);o__Boletales(100);f__Serpulaceae(100);g__Serpula(100);s__Serpula_himantioides(100); | Serpula | 13 | Saprotroph | \|Wood Saprotroph\| | Corticioid | Brown Rot | Probable | As brown-rot (Põlme et al. 2020); Decay wood (Põlme et al. 2020); rot fungus that causes very significant economic losses through degradation of wooden parts of buildings (Cannon & Kirk 2007); Serpula lacyrmans (dry rot) is an aggressive brown | [illegible] |
| Otu00953 | 72 | k__Fungi(100);p__Zygomycota(100);c__Zygomycota_class_Incertae_sedis(100);o__Mortierellales(100);f__Mortierellaceae(100);g__Mortierella(100);s__Mortierella_parvispora(100); | Mortierella | 13 | Saprotroph-Symbiotroph | Endophyte-Plant Saprotroph-\|Undefined Saprotroph\| | Microfungus | NULL | Probable | As chitinolytic (Põlme et al. 2020); As soil_saprotroph (Põlme et al. 2020); Decay roots, soil (Põlme et al. 2020); Saprobic in soil and thus frequenctly isolated from soil and material contaminated with soil such as hay or rotten silage and spoil tips (Cannon & Kirk 2007) | [illegible] |
| Otu00954 | 72 | k__Fungi(100);p__Basidiomycota(100);c__Microbotryomycetes(100);o__Sporidiobolales(100);f__Sporidiobolales_family_Incertae_sedis(100);g__Rhodotorula(96);g__Rhodotorula_unclassified(95); | Rhodotorula | 13 | Pathotroph-Saprotroph-Symbiotroph | Animal Pathogen-Animal Symbiotroph-Endophyte-Plant Pathogen-\|Undefined Saprotroph\| | Yeast | NULL | Probable | As foliar_endophyte (Põlme et al. 2020); Opportunistic human pathogen | [illegible] |
| Otu00946 | 72 | k__Fungi(100);p__Ascomycota(87);p__Ascomycota_unclassified(87);p__Ascomycota_unclassified(87);p__Ascomycota_unclassified(87);p__Ascomycota_unclassified(87);p__Ascomycota_unclassified(87); | - | - | - | - | - | - | - | Unassigned | - |
| Otu00950 | 72 | k__Fungi(100);p__Basidiomycota(100);c__Agaricomycetes(100);o__Cantharellales(100);o__Cantharellales_unclassified(100);o__Cantharellales_unclassified(100);o__Cantharellales_unclassified(100); | - | - | - | - | - | - | - | Unassigned | - |
| Otu00952 | 72 | k__Fungi(100);k__Fungi_unclassified(81);k__Fungi_unclassified(81);k__Fungi_unclassified(81);k__Fungi_unclassified(81);k__Fungi_unclassified(81);k__Fungi_unclassified(81); | - | - | - | - | - | - | - | Unassigned | - |
| Otu00956 | 71 | k__Fungi(100);p__Basidiomycota(100);c__Agaricomycetes(100);o__Trechisporales(100);f__Hydnodontaceae(100);g__Trechispora(100);s__Trechispora_sp(100); | Trechispora | 13 | Saprotroph | \|Wood Saprotroph\| | Corticioid | White Rot | Highly Probable | As white_rot (Põlme et al. 2020); Decay wood (Põlme et al. 2020) | [illegible] |
| Otu00958 | 71 | k__Fungi(100);p__Basidiomycota(100);c__Microbotryomycetes(100);o__Sporidiobolales(100);f__Sporidiobolales_family_Incertae_sedis(100);g__Rhodotorula(100);s__Rhodotorula_sp_MJL_2011(96); | Rhodotorula | 13 | Pathotroph-Saprotroph-Symbiotroph | Animal Pathogen-Animal Symbiotroph-Endophyte-Plant Pathogen-\|Undefined Saprotroph\| | Yeast | NULL | Probable | As foliar_endophyte (Põlme et al. 2020); Opportunistic human pathogen | [illegible] |
| Otu00959 | 71 | k__Fungi(100);p__Basidiomycota(100);c__Agaricomycetes(100);o__Agaricales(100);o__Agaricales_unclassified(93);o__Agaricales_unclassified(93);o__Agaricales_unclassified(93); | Agaricales | 7 | Pathotroph-Saprotroph-Symbiotroph | Bryophyte Parasite-Dung Saprotroph-Ectomycorrhizal-Fungal Parasite-Leaf Saprotroph-Plant Parasite-Undefined Saprotroph-Wood Saprotroph | Agaricoid-Gasteroid-Microfungus-Secotioid-Yeast | NULL | Possible | Mushrooms and toadstools, Gill fungi, Agarics... lignicolous, sometimes muscicolous or fungicolous, saprobic, mycorrhizal, rarely parasitic on plants or fungi (Kirk et al. 2008) | [illegible] |
| Otu00955 | 71 | k__Fungi(100);p__Ascomycota(64);p__Ascomycota_unclassified(64);p__Ascomycota_unclassified(64);p__Ascomycota_unclassified(64);p__Ascomycota_unclassified(64);p__Ascomycota_unclassified(64); | - | - | - | - | - | - | - | Unassigned | - |
| Otu00957 | 71 | k__Fungi(100);p__Ascomycota(58);p__Ascomycota_unclassified(58);p__Ascomycota_unclassified(58);p__Ascomycota_unclassified(58);p__Ascomycota_unclassified(58);p__Ascomycota_unclassified(58); | - | - | - | - | - | - | - | Unassigned | - |
| Otu00960 | 71 | k__Fungi(100);p__Ascomycota(100);c__Eurotiomycetes(100);o__Chaetothyriales(100);o__Chaetothyriales_unclassified(99);o__Chaetothyriales_unclassified(99);o__Chaetothyriales_unclassified(99); | - | - | - | - | - | - | - | Unassigned | - |
| Otu00962 | 70 | k__Fungi(100);p__Ascomycota(100);c__Eurotiomycetes(100);o__Chaetothyriales(100);f__Herpotrichiellaceae(100);g__Rhinocladiella(100);s__Rhinocladiella_sp_04001a(56); | Rhinocladiella | 13 | Pathotroph-Saprotroph-Symbiotroph | Animal Parasite-Endophyte-\|Undefined Saprotroph\|-Wood Saprotroph | Dimorphic-Facultative Yeast | Soft Rot | Probable | As root_endophyte_dark_septate (Põlme et al. 2020); As soil_saprotroph (Põlme et al. 2020); Decay soil, animal material (Põlme et al. 2020) | [illegible] |
| Otu00963 | 70 | k__Fungi(100);p__Basidiomycota(100);c__Agaricomycetes(100);o__Polyporales(100);f__Meripilaceae(100);g__Hydnopolyporus(100);s__Hydnopolyporus_fimbriatus(100); | Hydnopolyporus | 13 | Pathotroph-Saprotroph | \|Plant Pathogen\|-Wood Saprotroph | Clavarioid | White Rot | Highly Probable | As white_rot (Põlme et al. 2020); As wood_pathogen (Põlme et al. 2020); Decay wood (Põlme et al. 2020) | [illegible] |
| Otu00964 | 70 | k__Fungi(100);p__Ascomycota(99);c__Saccharomycetes(99);o__Saccharomycetales(99);f__Metschnikowiaceae(99);g__Metschnikowia(99);s__Metschnikowia_pulcherrima(96); | Metschnikowia | 13 | Saprotroph | \|Undefined Saprotroph\| | Yeast | NULL | Probable | As nectar-tap_saprotroph (Põlme et al. 2020); Decay sugar-rich substrates (Põlme et al. 2020) | [illegible] |
| Otu00965 | 70 | k__Fungi(100);p__Zygomycota(100);c__Zygomycota_class_Incertae_sedis(100);o__Mucorales(100);f__Mucoraceae(100);g__Helicostylum(100);s__Helicostylum_pulchrum(100); | Helicostylum | 13 | Saprotroph | \|Undefined Saprotroph\| | Microfungus | NULL | Probable | As soil_saprotroph (Põlme et al. 2020); Decay soil (Põlme et al. 2020) | [illegible] |
| Otu00966 | 70 | k__Fungi(100);p__Zygomycota(100);c__Zygomycota_class_Incertae_sedis(100);o__Mucorales(100);f__Cunninghamellaceae(99);g__Absidia(99);g__Absidia_unclassified(99); | Absidia | 13 | Saprotroph | \|Undefined Saprotroph\| | Microfungus | NULL | Probable | As soil_saprotroph (Põlme et al. 2020); Decay soil (Põlme et al. 2020) | [illegible] |
| Otu00970 | 70 | k__Fungi(100);p__Ascomycota(100);c__Sordariomycetes(100);o__Diaporthales(100);f__Schizoparmaceae(100);g__Pilidiella(100);s__Pilidiella_tibouchinae(75); | Pilidiella | 13 | Pathotroph | \|Plant Pathogen\| | Microfungus | NULL | Probable | NULL | [illegible] |
| Otu00973 | 70 | k__Fungi(100);p__Ascomycota(100);c__Sordariomycetes(100);o__Diaporthales(100);f__Diaporthaceae(100);f__Diaporthaceae_unclassified(100);f__Diaporthaceae_unclassified(100); | Diaporthaceae | 9 | Pathotroph-Saprotroph-Symbiotroph | Endophyte-Plant Pathogen-Plant Saprotroph | Microfungus | NULL | Probable | Most species are weak pathogens or endophytes in leaf or stem tissue, growing saprobically after the plant dies (Cannon & Kirk 2007) | [illegible] |
| Otu00976 | 70 | k__Fungi(100);p__Basidiomycota(100);c__Agaricomycetes(100);o__Polyporales(100);f__Polyporaceae(100);g__Spongipellis(100);s__Spongipellis_pachyodon(100); | Spongipellis | 13 | Pathotroph-Saprotroph | Plant Pathogen-\|Wood Saprotroph\| | Polyporoid | White Rot | Highly Probable | As white_rot (Põlme et al. 2020); As wood_pathogen (Põlme et al. 2020); Decay wood (Põlme et al. 2020) | [illegible] |
| Otu00961 | 70 | k__Fungi(100);k__Fungi_unclassified(100);k__Fungi_unclassified(100);k__Fungi_unclassified(100);k__Fungi_unclassified(100);k__Fungi_unclassified(100);k__Fungi_unclassified(100); | - | - | - | - | - | - | - | Unassigned | - |
| Otu00967 | 70 | k__Fungi(100);p__Basidiomycota(100);c__Agaricomycetes(100);o__Polyporales(100);f__unclassified_Polyporales(90);g__unclassified_Polyporales(90);s__Polyporales_sp(90); | - | - | - | - | - | - | - | Unassigned | - |
| Otu00968 | 70 | k__Fungi(100);p__Ascomycota(100);c__Sordariomycetes(98);c__Sordariomycetes_unclassified(92);c__Sordariomycetes_unclassified(92);c__Sordariomycetes_unclassified(92);c__Sordariomycetes_unclassified(92); | - | - | - | - | - | - | - | Unassigned | - |
| Otu00969 | 70 | k__Fungi(100);k__Fungi_unclassified(58);k__Fungi_unclassified(58);k__Fungi_unclassified(58);k__Fungi_unclassified(58);k__Fungi_unclassified(58);k__Fungi_unclassified(58); | - | - | - | - | - | - | - | Unassigned | - |
| Otu00971 | 70 | k__Fungi(100);k__Fungi_unclassified(63);k__Fungi_unclassified(63);k__Fungi_unclassified(63);k__Fungi_unclassified(63);k__Fungi_unclassified(63);k__Fungi_unclassified(63); | - | - | - | - | - | - | - | Unassigned | - |
| Otu00972 | 70 | k__Fungi(100);k__Fungi_unclassified(55);k__Fungi_unclassified(55);k__Fungi_unclassified(55);k__Fungi_unclassified(55);k__Fungi_unclassified(55);k__Fungi_unclassified(55); | - | - | - | - | - | - | - | Unassigned | - |
| Otu00974 | 70 | k__Fungi(100);p__Ascomycota(100);c__Dothideomycetes(100);o__Pleosporales(100);f__unclassified_Pleosporales(98);g__unclassified_Pleosporales(98);s__Pleosporales_sp(98); | - | - | - | - | - | - | - | Unassigned | - |
| Otu00975 | 70 | k__Fungi(100);p__Basidiomycota(100);c__Agaricomycetes(100);o__Corticiales(100);f__unclassified_Corticiales(100);g__unclassified_Corticiales(100);s__Corticiales_sp(100); | - | - | - | - | - | - | - | Unassigned | - |
| Otu00977 | 69 | k__Fungi(100);p__Basidiomycota(100);c__Agaricomycetes(100);o__Polyporales(100);f__Meruliaceae(100);g__Hypochnicium(100);g__Hypochnicium_unclassified(100); | Hypochnicium | 13 | Saprotroph | Undefined Saprotroph-\|Wood Saprotroph\| | Corticioid | White Rot | Probable | As white_rot (Põlme et al. 2020); Decay wood (Põlme et al. 2020) | [illegible] |
| Otu00978 | 69 | k__Fungi(100);p__Zygomycota(100);c__Zygomycota_class_Incertae_sedis(100);o__Mortierellales(100);f__Mortierellaceae(100);g__Mortierella(100);g__Mortierella_unclassified(100); | Mortierella | 13 | Saprotroph-Symbiotroph | Endophyte-Plant Saprotroph-\|Undefined Saprotroph\| | Microfungus | NULL | Probable | As chitinolytic (Põlme et al. 2020); As soil_saprotroph (Põlme et al. 2020); Decay roots, soil (Põlme et al. 2020); Saprobic in soil and thus frequenctly isolated from soil and material contaminated with soil such as hay or rotten silage and spoil tips (Cannon & Kirk 2007) | [illegible] |
| Otu00979 | 69 | k__Fungi(100);p__Basidiomycota(100);c__Agaricomycetes(100);o__Cantharellales(100);f__Ceratobasidiaceae(100);g__Ceratobasidium(100);s__Ceratobasidium_sp(100); | Ceratobasidium | 13 | Pathotroph-Saprotroph-Symbiotroph | Ectomycorrhizal-Endomycorrhizal-Orchid Mycorrhizal-\|Plant Pathogen\|-Plant Saprotroph-Undefined Saprotroph | Corticioid-Microfungus | NULL | Possible | especially of Orchidaceae but also even bryophytes (Cannon & Kirk 2007); As litter_saprotroph (Põlme et al. 2020); Best known as plant pathogens, but also saprobic. Species are endomycorrhizal with a wide range of plant roots ; Decay leaf, fruit, seed, roots, soil (Põlme et al. 2020) | [illegible] |
| Otu00980 | 69 | k__Fungi(100);p__Basidiomycota(100);c__Agaricomycetes(100);o__Hymenochaetales(100);f__Schizoporaceae(100);g__Hyphodontia(100);s__Hyphodontia_alutaria(100); | Hyphodontia | 13 | Saprotroph | Undefined Saprotroph-\|Wood Saprotroph\| | Corticioid | White Rot | Probable | As white_rot (Põlme et al. 2020); Decay wood (Põlme et al. 2020) | [illegible] |
| Otu00983 | 69 | k__Fungi(100);p__Basidiomycota(100);c__Agaricomycetes(100);o__Auriculariales(100);f__Auriculariales_family_Incertae_sedis(100);g__Protomerulius(100);s__Protomerulius_sp(100); | Protomerulius | 13 | Saprotroph | Undefined Saprotroph-\|Wood Saprotroph\| | Corticioid | White Rot | Probable | As white_rot (Põlme et al. 2020); Decay wood (Põlme et al. 2020) | [illegible] |
| Otu00981 | 69 | k__Fungi(100);p__Basidiomycota(100);c__Agaricomycetes(100);c__Agaricomycetes_unclassified(100);c__Agaricomycetes_unclassified(100);c__Agaricomycetes_unclassified(100);c__Agaricomycetes_unclassified(100); | - | - | - | - | - | - | - | Unassigned | - |
| Otu00982 | 69 | k__Fungi(100);k__Fungi_unclassified(100);k__Fungi_unclassified(100);k__Fungi_unclassified(100);k__Fungi_unclassified(100);k__Fungi_unclassified(100);k__Fungi_unclassified(100); | - | - | - | - | - | - | - | Unassigned | - |
| Otu00984 | 69 | k__Fungi(100);p__Basidiomycota(100);c__Agaricomycetes(100);c__Agaricomycetes_unclassified(87);c__Agaricomycetes_unclassified(87);c__Agaricomycetes_unclassified(87);c__Agaricomycetes_unclassified(87); | - | - | - | - | - | - | - | Unassigned | - |
| Otu00985 | 69 | k__Fungi(100);p__Ascomycota(90);p__Ascomycota_unclassified(90);p__Ascomycota_unclassified(90);p__Ascomycota_unclassified(90);p__Ascomycota_unclassified(90);p__Ascomycota_unclassified(90); | - | - | - | - | - | - | - | Unassigned | - |
| Otu00986 | 69 | k__Fungi(100);p__Ascomycota(100);p__Ascomycota_unclassified(100);p__Ascomycota_unclassified(100);p__Ascomycota_unclassified(100);p__Ascomycota_unclassified(100);p__Ascomycota_unclassified(100); | - | - | - | - | - | - | - | Unassigned | - |
| Otu00987 | 68 | k__Fungi(100);p__Ascomycota(99);c__Saccharomycetes(99);o__Saccharomycetales(99);f__Metschnikowiaceae(99);g__Metschnikowia(99);s__Metschnikowia_pulcherrima(90); | Metschnikowia | 13 | Saprotroph | \|Undefined Saprotroph\| | Yeast | NULL | Probable | As nectar-tap_saprotroph (Põlme et al. 2020); Decay sugar-rich substrates (Põlme et al. 2020) | [illegible] |
| Otu00988 | 68 | k__Fungi(100);p__Ascomycota(100);c__Dothideomycetes(100);o__Capnodiales(100);f__Mycosphaerellaceae(100);f__Mycosphaerellaceae_unclassified(100);f__Mycosphaerellaceae_unclassified(100); | Mycosphaerellaceae | 9 | Pathotroph-Saprotroph | Plant Pathogen-Undefined Saprotroph | Microfungus | NULL | Probable | Biotrophic, necrotrophic or saprobic on various plant tissues (Cannon & Kirk 2007) | [illegible] |
| Otu00989 | 68 | k__Fungi(100);p__Basidiomycota(100);c__Agaricomycetes(100);o__Russulales(100);f__Lachnocladiaceae(100);g__Vararia(100);s__Vararia_sp_PBM2521(100); | Vararia | 13 | Saprotroph | Undefined Saprotroph-\|Wood Saprotroph\| | Corticioid | White Rot | Probable | As white_rot (Põlme et al. 2020); Decay wood (Põlme et al. 2020) | [illegible] |
| Otu00992 | 68 | k__Fungi(100);p__Basidiomycota(100);c__Agaricomycetes(100);o__Polyporales(100);f__Fomitopsidaceae(100);g__Fomitopsis(100);s__Fomitopsis_pinicola(100); | Fomitopsis | 13 | Pathotroph-Saprotroph | Plant Pathogen-\|Wood Saprotroph\| | Polyporoid | Brown Rot | Highly Probable | As brown-rot (Põlme et al. 2020); As wood_pathogen (Põlme et al. 2020); Decay wood (Põlme et al. 2020); Most species are parasitic on woody plants, causing brown rots (Cannon & Kirk 2007) | [illegible] |
| Otu00994 | 68 | k__Fungi(100);p__Basidiomycota(100);c__Tremellomycetes(100);o__Tremellales(100);f__Tremellales_family_Incertae_sedis(100);g__Cryptococcus(100);g__Cryptococcus_unclassified(100); | Cryptococcus | 13 | Pathotroph-Saprotroph-Symbiotroph | Animal Pathogen-Endophyte-Epiphyte-\|Undefined Saprotroph\| | Dimorphic Yeast-Yeast | NULL | Possible | As foliar_endophyte (Põlme et al. 2020); Decay leaf, fruit, seed, soil, dung, animal material (Põlme et al. 2020) | [illegible] |
| Otu00995 | 68 | k__Fungi(100);p__Basidiomycota(100);c__Agaricomycetes(100);o__Hymenochaetales(100);f__Schizoporaceae(100);f__Schizoporaceae_unclassified(100);f__Schizoporaceae_unclassified(100); | Schizoporaceae | 9 | Saprotroph | Wood Saprotroph | Corticioid | White Rot | Probable | Saprobic, causing white rot of standing and fallen wood of coniferous and broadleaved trees, rarely on herbaceous stems (Cannon & Kirk 2007) | [illegible] |
| Otu00997 | 68 | k__Fungi(100);p__Ascomycota(100);c__Saccharomycetes(100);o__Saccharomycetales(100);f__Metschnikowiaceae(100);g__Metschnikowia(100);s__Metschnikowia_pulcherrima(98); | Metschnikowia | 13 | Saprotroph | \|Undefined Saprotroph\| | Yeast | NULL | Probable | As nectar-tap_saprotroph (Põlme et al. 2020); Decay sugar-rich substrates (Põlme et al. 2020) | [illegible] |
| Otu00990 | 68 | k__Fungi(100);p__Ascomycota(100);p__Ascomycota_unclassified(100);p__Ascomycota_unclassified(100);p__Ascomycota_unclassified(100);p__Ascomycota_unclassified(100);p__Ascomycota_unclassified(100); | - | - | - | - | - | - | - | Unassigned | - |
| Otu00991 | 68 | k__Fungi(100);k__Fungi_unclassified(99);k__Fungi_unclassified(99);k__Fungi_unclassified(99);k__Fungi_unclassified(99);k__Fungi_unclassified(99);k__Fungi_unclassified(99); | - | - | - | - | - | - | - | Unassigned | - |
| Otu00993 | 68 | k__Fungi(100);p__Basidiomycota(100);c__Agaricomycetes(100);o__Polyporales(84);o__Polyporales_unclassified(84);o__Polyporales_unclassified(84);o__Polyporales_unclassified(84); | - | - | - | - | - | - | - | Unassigned | - |
| Otu00996 | 68 | k__Fungi(100);p__Ascomycota(100);c__Dothideomycetes(100);o__Pleosporales(100);f__unclassified_Pleosporales(100);g__unclassified_Pleosporales(100);s__Pleosporales_sp(100); | - | - | - | - | - | - | - | Unassigned | - |
| Otu00999 | 67 | k__Fungi(100);p__Ascomycota(100);c__Dothideomycetes(100);o__Pleosporales(100);f__Sporormiaceae(100);g__Preussia(100);g__Preussia_unclassified(87); | Preussia | 13 | Saprotroph | \|Dung Saprotroph\|-Undefined Saprotroph | Microfungus | NULL | Probable | Decay dung (Põlme et al. 2020) | [illegible] |
| Otu01001 | 67 | k__Fungi(100);p__Ascomycota(100);c__Eurotiomycetes(100);o__Chaetothyriales(100);f__Chaetothyriaceae(100);g__Cyphellophora(100);s__Cyphellophora_vermispora(72); | Cyphellophora | 13 | Pathotroph-Saprotroph | Animal Parasite-Animal Pathogen-\|Plant Saprotroph\|-Undefined Saprotroph | Dimorphic-Facultative Yeast-Microfungus | NULL | Probable | As litter_saprotroph (Põlme et al. 2020); Consult the current literature for updated names on this group that included the Black Yeasts; Some species are likely opportunistic human pathogen (Irinyi et al. 2015); Decay leaf, fruit, seed, animal material (Põlme et al. 2020) | [illegible] |
| Otu01003 | 67 | k__Fungi(100);p__Ascomycota(100);c__Sordariomycetes(65);o__Hypocreales(65);f__Nectriaceae(65);g__Cylindrium(65);s__Cylindrium_sp(65); | Cylindrium | 13 | Pathotroph | \|Plant Pathogen\| | Microfungus | NULL | Probable | NULL | [illegible] |
| Otu01004 | 67 | k__Fungi(100);p__Ascomycota(100);c__Dothideomycetes(100);o__Pleosporales(100);f__Cucurbitariaceae(100);g__Pyrenochaetopsis(100);s__Pyrenochaetopsis_pratorum(100); | Pyrenochaetopsis | 13 | Pathotroph-Saprotroph-Symbiotroph | \|Endophyte\|-Lichen Parasite-Plant Saprotroph-Undefined Saprotroph | Microfungus | NULL | Possible | As foliar_endophyte (Põlme et al. 2020); As litter_saprotroph (Põlme et al. 2020); Decay leaf, fruit, seed (Põlme et al. 2020) | [illegible] |
| Otu01005 | 67 | k__Fungi(100);p__Ascomycota(100);c__Dothideomycetes(100);o__Pleosporales(100);f__Pleosporales_family_Incertae_sedis(100);g__Periconia(100);g__Periconia_unclassified(63); | Periconia | 13 | Pathotroph-Saprotroph-Symbiotroph | Endophyte-\|Plant Pathogen\|-Wood Saprotroph | Microfungus | Soft Rot | Possible | Asteraceae, Musaceae (Costa et al. 2012); As foliar_endophyte (Põlme et al. 2020); Dark Septate Endophyte, known from plant roots (Mandyam et al. 2010); Endophyte Host ; Decay leaf, fruit, seed (Põlme et al. 2020) | [illegible] |
| Otu01006 | 67 | k__Fungi(100);p__Ascomycota(100);c__Lecanoromycetes(100);o__Teloschistales(100);f__Physciaceae(100);g__Phaeophyscia(100);g__Phaeophyscia_unclassified(100); | Phaeophyscia | 13 | Symbiotroph | \|Lichenized\| | Thallus | NULL | Highly Probable | NULL | [illegible] |
| Otu01007 | 67 | k__Fungi(100);p__Basidiomycota(100);c__Agaricomycetes(100);o__Russulales(100);f__Stereaceae(100);f__Stereaceae_unclassified(90);f__Stereaceae_unclassified(90); | Stereaceae | 9 | Saprotroph | Undefined Saprotroph-Wood Saprotroph | Corticioid-Polyporoid | NULL | Probable | Lignicolous or terrestrial (in leaf litter); typically saprobic (Cannon & Kirk 2007) | [illegible] |
| Otu00998 | 67 | k__Fungi(100);k__Fungi_unclassified(68);k__Fungi_unclassified(68);k__Fungi_unclassified(68);k__Fungi_unclassified(68);k__Fungi_unclassified(68);k__Fungi_unclassified(68); | - | - | - | - | - | - | - | Unassigned | - |
| Otu01000 | 67 | k__Fungi(100);p__Ascomycota(100);c__Dothideomycetes(100);o__Pleosporales(100);o__Pleosporales_unclassified(93);o__Pleosporales_unclassified(93);o__Pleosporales_unclassified(93); | - | - | - | - | - | - | - | Unassigned | - |
| Otu01002 | 67 | k__Fungi(100);k__Fungi_unclassified(98);k__Fungi_unclassified(98);k__Fungi_unclassified(98);k__Fungi_unclassified(98);k__Fungi_unclassified(98);k__Fungi_unclassified(98); | - | - | - | - | - | - | - | Unassigned | - |
| Otu01008 | 66 | k__Fungi(100);p__Basidiomycota(100);c__Agaricomycetes(100);o__Polyporales(100);f__Polyporaceae(100);g__Sarcoporia(100);s__Sarcoporia_polyspora(100); | Sarcoporia | 13 | Saprotroph | \|Wood Saprotroph\| | Polyporoid | Brown Rot | Probable | As brown-rot (Põlme et al. 2020); Decay wood (Põlme et al. 2020) | [illegible] |
| Otu01009 | 66 | k__Fungi(100);p__Basidiomycota(100);c__Agaricomycetes(100);o__Agaricales(100);f__Strophariaceae(100);g__Agrocybe(100);s__Agrocybe_praecox(100); | Agrocybe | 13 | Saprotroph | Dung Saprotroph-\|Undefined Saprotroph\| | Agaricoid | NULL | Probable | As soil_saprotroph (Põlme et al. 2020); Decay soil (Põlme et al. 2020) | [illegible] |
| Otu01010 | 66 | k__Fungi(100);p__Ascomycota(100);c__Saccharomycetes(100);o__Saccharomycetales(100);f__Metschnikowiaceae(100);g__Metschnikowia(100);s__Metschnikowia_pulcherrima(88); | Metschnikowia | 13 | Saprotroph | \|Undefined Saprotroph\| | Yeast | NULL | Probable | As nectar-tap_saprotroph (Põlme et al. 2020); Decay sugar-rich substrates (Põlme et al. 2020) | [illegible] |
| Otu01011 | 66 | k__Fungi(100);p__Ascomycota(100);c__Sordariomycetes(100);o__Xylariales(100);f__Xylariaceae(100);g__Xylaria(100);s__Xylaria_longipes(96); | Xylaria | 13 | Saprotroph-Symbiotroph | Endophyte-Plant Saprotroph-Undefined Saprotroph-\|Wood Saprotroph\| | Xylarioid | Soft Rot | Probable | As foliar_endophyte (Põlme et al. 2020); As litter_saprotroph (Põlme et al. 2020); Decay wood, leaf, fruit, seed (Põlme et al. 2020); Host - angiosperms (Põlme et al. 2020) | [illegible] |
| Otu01012 | 66 | k__Fungi(100);p__Basidiomycota(100);c__Agaricomycetes(100);o__Cantharellales(100);f__Ceratobasidiaceae(100);f__Ceratobasidiaceae_unclassified(54);f__Ceratobasidiaceae_unclassified(54); | Ceratobasidiaceae | 9 | Pathotroph-Saprotroph-Symbiotroph | Endomycorrhizal-Plant Pathogen-Undefined Saprotroph | Microfungus | NULL | Possible | Best known as plant pathogens, but also saprobic. Species are endomycorrhizal with a wide range of plant roots - especially of Orchidaceae but also even bryophytes (Cannon & Kirk 2007) | [illegible] |
| Otu01013 | 66 | k__Fungi(100);p__Ascomycota(82);p__Ascomycota_unclassified(82);p__Ascomycota_unclassified(82);p__Ascomycota_unclassified(82);p__Ascomycota_unclassified(82);p__Ascomycota_unclassified(82); | - | - | - | - | - | - | - | Unassigned | - |
| Otu01014 | 66 | k__Fungi(100);k__Fungi_unclassified(70);k__Fungi_unclassified(70);k__Fungi_unclassified(70);k__Fungi_unclassified(70);k__Fungi_unclassified(70);k__Fungi_unclassified(70); | - | - | - | - | - | - | - | Unassigned | - |
| Otu01015 | 66 | k__Fungi(100);k__Fungi_unclassified(81);k__Fungi_unclassified(81);k__Fungi_unclassified(81);k__Fungi_unclassified(81);k__Fungi_unclassified(81);k__Fungi_unclassified(81); | - | - | - | - | - | - | - | Unassigned | - |
| Otu01016 | 66 | k__Fungi(100);p__Basidiomycota(100);c__Agaricomycetes(100);o__Cantharellales(100);o__Cantharellales_unclassified(97);o__Cantharellales_unclassified(97);o__Cantharellales_unclassified(97); | - | - | - | - | - | - | - | Unassigned | - |
| Otu01017 | 65 | k__Fungi(100);p__Ascomycota(100);c__Eurotiomycetes(100);o__Eurotiales(100);f__Trichocomaceae(100);g__Penicillium(100);s__Penicillium_cainii(90); | Penicillium | 13 | Pathotroph-Saprotroph-Symbiotroph | Animal Parasite-Dung Saprotroph-Endophyte-Fungal Parasite-Plant Pathogen-\|Undefined Saprotroph\|-Wood Saprotroph | Microfungus | Soft Rot | Highly Probable | As foliar_endophyte (Põlme et al. 2020); As mycoparasite (Põlme et al. 2020); Decay soil (Põlme et al. 2020) | [illegible] |
| Otu01018 | 65 | k__Fungi(100);p__Zygomycota(100);c__Zygomycota_class_Incertae_sedis(100);o__Mucorales(100);f__Mucoraceae(100);g__Mucor(100);g__Mucor_unclassified(100); | Mucor | 13 | Saprotroph | \|Undefined Saprotroph\| | Microfungus | NULL | Probable | As soil_saprotroph (Põlme et al. 2020); Decay soil (Põlme et al. 2020) | [illegible] |
| Otu01019 | 65 | k__Fungi(100);p__Ascomycota(100);c__Sordariomycetes(100);o__Hypocreales(100);o__Hypocreales_unclassified(54);o__Hypocreales_unclassified(54);o__Hypocreales_unclassified(54); | Hypocreales | 7 | Saprotroph | Undefined Saprotroph | Microfungus | NULL | Possible | NULL | [illegible] |
| Otu01020 | 65 | k__Fungi(100);p__Basidiomycota(100);c__Agaricomycetes(100);o__Agaricales(100);o__Agaricales_unclassified(100);o__Agaricales_unclassified(100);o__Agaricales_unclassified(100); | Agaricales | 7 | Pathotroph-Saprotroph-Symbiotroph | Bryophyte Parasite-Dung Saprotroph-Ectomycorrhizal-Fungal Parasite-Leaf Saprotroph-Plant Parasite-Undefined Saprotroph-Wood Saprotroph | Agaricoid-Gasteroid-Microfungus-Secotioid-Yeast | NULL | Possible | Mushrooms and toadstools, Gill fungi, Agarics... lignicolous, sometimes muscicolous or fungicolous, saprobic, mycorrhizal, rarely parasitic on plants or fungi (Kirk et al. 2008) | [illegible] |
| Otu01021 | 65 | k__Fungi(100);p__Basidiomycota(100);c__Agaricomycetes(100);o__Atheliales(100);o__Atheliales_unclassified(61);o__Atheliales_unclassified(61);o__Atheliales_unclassified(61); | - | - | - | - | - | - | - | Unassigned | - |
| Otu01022 | 65 | k__Fungi(100);k__Fungi_unclassified(65);k__Fungi_unclassified(65);k__Fungi_unclassified(65);k__Fungi_unclassified(65);k__Fungi_unclassified(65);k__Fungi_unclassified(65); | - | - | - | - | - | - | - | Unassigned | - |
| Otu01023 | 65 | k__Fungi(100);p__Basidiomycota(100);c__Agaricomycetes(100);o__Trechisporales(100);f__Hydnodontaceae(100);f__Hydnodontaceae_unclassified(94);f__Hydnodontaceae_unclassified(94); | - | - | - | - | - | - | - | Unassigned | - |
| Otu01024 | 65 | k__Fungi(100);p__Basidiomycota(100);p__Basidiomycota_unclassified(100);p__Basidiomycota_unclassified(100);p__Basidiomycota_unclassified(100);p__Basidiomycota_unclassified(100);p__Basidiomycota_unclassified(100); | - | - | - | - | - | - | - | Unassigned | - |
| Otu01025 | 65 | k__Fungi(100);p__Ascomycota(100);c__Dothideomycetes(100);o__Dothideales(100);f__unclassified_Dothideales(56);g__unclassified_Dothideales(56);s__Dothideales_sp(56); | - | - | - | - | - | - | - | Unassigned | - |
| Otu01026 | 65 | k__Fungi(100);k__Fungi_unclassified(97);k__Fungi_unclassified(97);k__Fungi_unclassified(97);k__Fungi_unclassified(97);k__Fungi_unclassified(97);k__Fungi_unclassified(97); | - | - | - | - | - | - | - | Unassigned | - |
| Otu01027 | 65 | k__Fungi(100);p__Ascomycota(99);p__Ascomycota_unclassified(99);p__Ascomycota_unclassified(99);p__Ascomycota_unclassified(99);p__Ascomycota_unclassified(99);p__Ascomycota_unclassified(99); | - | - | - | - | - | - | - | Unassigned | - |
| Otu01028 | 64 | k__Fungi(100);p__Ascomycota(100);c__Dothideomycetes(100);o__Pleosporales(100);f__Sporormiaceae(100);g__Sporormiella(100);s__Sporormiella_sp(100); | Sporormiella | 13 | Saprotroph | Dung Saprotroph-Endophyte-Lichen Parasite-\|Plant Saprotroph\| | Microfungus | NULL | Probable | As litter_saprotroph (Põlme et al. 2020); Decay leaf, fruit, seed (Põlme et al. 2020) | [illegible] |
| Otu01029 | 64 | k__Fungi(100);p__Basidiomycota(100);c__Agaricomycetes(100);o__Polyporales(100);f__Polyporaceae(100);g__unclassified_Polyporaceae(100);s__Polyporaceae_sp(100); | Polyporaceae | 9 | Saprotroph | Wood Saprotroph | Polyporoid | Brown Rot-White Rot | Probable | Lignicolous, causing brown or white rot, mostly of broadleaved trees (Cannon & Kirk 2007) | [illegible] |
| Otu01032 | 64 | k__Fungi(100);p__Basidiomycota(100);c__Agaricomycetes(100);o__Polyporales(100);f__Fomitopsidaceae(100);g__Postia(100);s__Postia_leucomallela(100); | Postia | 13 | Saprotroph | \|Wood Saprotroph\| | Polyporoid | Brown Rot | Highly Probable | As brown-rot (Põlme et al. 2020); Decay wood (Põlme et al. 2020) | [illegible] |
| Otu01035 | 64 | k__Fungi(100);p__Ascomycota(100);c__Sordariomycetes(100);o__Sordariomycetes_order_Incertae_sedis(100);f__Glomerellaceae(100);g__Colletotrichum(97);g__Colletotrichum_unclassified(97); | Colletotrichum | 13 | Pathotroph-Saprotroph-Symbiotroph | Endophyte-\|Plant Pathogen\|-Plant Saprotroph | Microfungus | NULL | Probable | As foliar_endophyte (Põlme et al. 2020); As litter_saprotroph (Põlme et al. 2020); Decay leaf, fruit, seed (Põlme et al. 2020) | [illegible] |
| Otu01036 | 64 | k__Fungi(100);p__Basidiomycota(100);c__Agaricomycetes(100);o__Polyporales(100);f__Phanerochaetaceae(100);g__Ceriporia(100);s__Ceriporia_aurantiocarnescens(100); | Ceriporia | 13 | Saprotroph | \|Wood Saprotroph\| | Corticioid | White Rot | Highly Probable | As white_rot (Põlme et al. 2020); Decay wood (Põlme et al. 2020) | [illegible] |
| Otu01038 | 64 | k__Fungi(100);p__Ascomycota(100);c__Saccharomycetes(100);o__Saccharomycetales(100);f__Metschnikowiaceae(100);g__Metschnikowia(100);s__Metschnikowia_pulcherrima(90); | Metschnikowia | 13 | Saprotroph | \|Undefined Saprotroph\| | Yeast | NULL | Probable | As nectar-tap_saprotroph (Põlme et al. 2020); Decay sugar-rich substrates (Põlme et al. 2020) | [illegible] |
| Otu01030 | 64 | k__Fungi(100);k__Fungi_unclassified(60);k__Fungi_unclassified(60);k__Fungi_unclassified(60);k__Fungi_unclassified(60);k__Fungi_unclassified(60);k__Fungi_unclassified(60); | - | - | - | - | - | - | - | Unassigned | - |
| Otu01031 | 64 | k__Fungi(100);p__Ascomycota(100);c__Eurotiomycetes(99);c__Eurotiomycetes_unclassified(99);c__Eurotiomycetes_unclassified(99);c__Eurotiomycetes_unclassified(99);c__Eurotiomycetes_unclassified(99); | - | - | - | - | - | - | - | Unassigned | - |
| Otu01033 | 64 | k__Fungi(100);p__Basidiomycota(100);c__Tremellomycetes(100);c__Tremellomycetes_unclassified(97);c__Tremellomycetes_unclassified(97);c__Tremellomycetes_unclassified(97);c__Tremellomycetes_unclassified(97); | - | - | - | - | - | - | - | Unassigned | - |
| Otu01034 | 64 | k__Fungi(100);k__Fungi_unclassified(66);k__Fungi_unclassified(66);k__Fungi_unclassified(66);k__Fungi_unclassified(66);k__Fungi_unclassified(66);k__Fungi_unclassified(66); | - | - | - | - | - | - | - | Unassigned | - |
| Otu01037 | 64 | k__Fungi(100);k__Fungi_unclassified(97);k__Fungi_unclassified(97);k__Fungi_unclassified(97);k__Fungi_unclassified(97);k__Fungi_unclassified(97);k__Fungi_unclassified(97); | - | - | - | - | - | - | - | Unassigned | - |
| Otu01039 | 63 | k__Fungi(100);p__Ascomycota(100);c__Saccharomycetes(100);o__Saccharomycetales(100);f__Metschnikowiaceae(100);g__Metschnikowia(100);s__Metschnikowia_pulcherrima(99); | Metschnikowia | 13 | Saprotroph | \|Undefined Saprotroph\| | Yeast | NULL | Probable | As nectar-tap_saprotroph (Põlme et al. 2020); Decay sugar-rich substrates (Põlme et al. 2020) | [illegible] |
| Otu01040 | 63 | k__Fungi(100);p__Ascomycota(100);c__Saccharomycetes(100);o__Saccharomycetales(100);f__Metschnikowiaceae(100);g__Metschnikowia(100);s__Metschnikowia_pulcherrima(91); | Metschnikowia | 13 | Saprotroph | \|Undefined Saprotroph\| | Yeast | NULL | Probable | As nectar-tap_saprotroph (Põlme et al. 2020); Decay sugar-rich substrates (Põlme et al. 2020) | [illegible] |
| Otu01041 | 63 | k__Fungi(100);p__Basidiomycota(100);c__Agaricomycetes(100);o__Cantharellales(100);f__Ceratobasidiaceae(100);g__Ceratobasidium(100);s__Ceratobasidium_sp_Cs162(100); | Ceratobasidium | 13 | Pathotroph-Saprotroph-Symbiotroph | Ectomycorrhizal-Endomycorrhizal-Orchid Mycorrhizal-\|Plant Pathogen\|-Plant Saprotroph-Undefined Saprotroph | Corticioid-Microfungus | NULL | Possible | especially of Orchidaceae but also even bryophytes (Cannon & Kirk 2007); As litter_saprotroph (Põlme et al. 2020); Best known as plant pathogens, but also saprobic. Species are endomycorrhizal with a wide range of plant roots ; Decay leaf, fruit, seed, roots, soil (Põlme et al. 2020) | [illegible] |
| Otu01042 | 63 | k__Fungi(100);p__Ascomycota(100);c__Dothideomycetes(100);o__Pleosporales(100);f__Phaeosphaeriaceae(100);g__Ophiosphaerella(100);g__Ophiosphaerella_unclassified(54); | Ophiosphaerella | 13 | Pathotroph-Saprotroph | \|Plant Pathogen\|-Plant Saprotroph-Undefined Saprotroph | Microfungus | NULL | Probable | As litter_saprotroph (Põlme et al. 2020); Decay leaf, fruit, seed (Põlme et al. 2020) | [illegible] |
| Otu01043 | 63 | k__Fungi(100);p__Ascomycota(100);c__Saccharomycetes(100);o__Saccharomycetales(100);f__Metschnikowiaceae(100);g__Metschnikowia(100);s__Metschnikowia_pulcherrima(99); | Metschnikowia | 13 | Saprotroph | \|Undefined Saprotroph\| | Yeast | NULL | Probable | As nectar-tap_saprotroph (Põlme et al. 2020); Decay sugar-rich substrates (Põlme et al. 2020) | [illegible] |
| Otu01045 | 63 | k__Fungi(100);p__Ascomycota(100);c__Eurotiomycetes(100);o__Mycocaliciales(100);f__Mycocaliciaceae(100);g__Mycocalicium(100);g__Mycocalicium_unclassified(62); | Mycocalicium | 13 | Pathotroph-Saprotroph-Symbiotroph | Lichen Parasite-Lichenized-\|Undefined Saprotroph\| | Calicioid-Thallus | NULL | Possible | As resin_saprotroph (Põlme et al. 2020); licenized taxa; Lichenized and non | [illegible] |
| Otu01046 | 63 | k__Fungi(100);p__Ascomycota(100);c__Leotiomycetes(100);o__Erysiphales(100);f__Erysiphaceae(100);g__Podosphaera(100);s__Podosphaera_sp_DB_2011(100); | Podosphaera | 13 | Pathotroph | \|Plant Pathogen\| | Microfungus | NULL | Probable | Biotrophic, with affected plant tissues damaged but usually not killing outright (Cannon & Kirk 2007); Common name - powdery mildew | [illegible] |
| Otu01044 | 63 | k__Fungi(100);p__Ascomycota(100);c__Dothideomycetes(100);o__Capnodiales(100);o__Capnodiales_unclassified(89);o__Capnodiales_unclassified(89);o__Capnodiales_unclassified(89); | - | - | - | - | - | - | - | Unassigned | - |
| Otu01047 | 63 | k__Fungi(100);k__Fungi_unclassified(72);k__Fungi_unclassified(72);k__Fungi_unclassified(72);k__Fungi_unclassified(72);k__Fungi_unclassified(72);k__Fungi_unclassified(72); | - | - | - | - | - | - | - | Unassigned | - |
| Otu01048 | 62 | k__Fungi(100);p__Basidiomycota(100);c__Ustilaginomycetes(100);o__Ustilaginales(100);f__Ustilaginaceae(100);g__Ustilago(100);s__Ustilago_striiformis(99); | Ustilago | 13 | Pathotroph | \|Plant Pathogen\| | Smut | NULL | Probable | Host - Poaceae (Põlme et al. 2020) | [illegible] |
| Otu01051 | 62 | k__Fungi(100);p__Basidiomycota(100);c__Agaricomycetes(100);o__Phallales(100);f__Phallaceae(100);g__Clathrus(100);s__Clathrus_archeri(100); | Clathrus | 13 | Saprotroph | \|Undefined Saprotroph\| | Clathroid | NULL | Probable | stinkhorn; As soil_saprotroph (Põlme et al. 2020); Common name ; Decay soil (Põlme et al. 2020) | [illegible] |
| Otu01049 | 62 | k__Fungi(100);p__Basidiomycota(100);c__Agaricomycetes(83);c__Agaricomycetes_unclassified(83);c__Agaricomycetes_unclassified(83);c__Agaricomycetes_unclassified(83);c__Agaricomycetes_unclassified(83); | - | - | - | - | - | - | - | Unassigned | - |
| Otu01050 | 62 | k__Fungi(100);p__Ascomycota(67);p__Ascomycota_unclassified(65);p__Ascomycota_unclassified(65);p__Ascomycota_unclassified(65);p__Ascomycota_unclassified(65);p__Ascomycota_unclassified(65); | - | - | - | - | - | - | - | Unassigned | - |
| Otu01052 | 62 | k__Fungi(100);p__Ascomycota(100);p__Ascomycota_unclassified(100);p__Ascomycota_unclassified(100);p__Ascomycota_unclassified(100);p__Ascomycota_unclassified(100);p__Ascomycota_unclassified(100); | - | - | - | - | - | - | - | Unassigned | - |
| Otu01053 | 62 | k__Fungi(100);p__Ascomycota(80);p__Ascomycota_unclassified(80);p__Ascomycota_unclassified(80);p__Ascomycota_unclassified(80);p__Ascomycota_unclassified(80);p__Ascomycota_unclassified(80); | - | - | - | - | - | - | - | Unassigned | - |
| Otu01054 | 62 | k__Fungi(100);k__Fungi_unclassified(76);k__Fungi_unclassified(76);k__Fungi_unclassified(76);k__Fungi_unclassified(76);k__Fungi_unclassified(76);k__Fungi_unclassified(76); | - | - | - | - | - | - | - | Unassigned | - |
| Otu01055 | 62 | k__Fungi(100);p__Basidiomycota(100);c__Agaricomycetes(100);c__Agaricomycetes_unclassified(96);c__Agaricomycetes_unclassified(96);c__Agaricomycetes_unclassified(96);c__Agaricomycetes_unclassified(96); | - | - | - | - | - | - | - | Unassigned | - |
| Otu01056 | 61 | k__Fungi(100);p__Ascomycota(100);c__Dothideomycetes(100);o__Capnodiales(100);f__Davidiellaceae(100);g__Cladosporium(100);s__Cladosporium_halotolerans(96); | Cladosporium | 13 | Pathotroph-Saprotroph-Symbiotroph | Animal Pathogen-Endophyte-Lichen Parasite-Plant Pathogen-\|Plant Saprotroph\|-Wood Saprotroph | Microfungus | NULL | Possible | As foliar_endophyte (Põlme et al. 2020); As litter_saprotroph (Põlme et al. 2020); Decay wood, leaf, fruit, seed, soil, dung, animal material (Põlme et al. 2020) | [illegible] |
| Otu01058 | 61 | k__Fungi(100);p__Ascomycota(100);c__Saccharomycetes(100);o__Saccharomycetales(100);f__Metschnikowiaceae(100);g__Metschnikowia(100);s__Metschnikowia_pulcherrima(97); | Metschnikowia | 13 | Saprotroph | \|Undefined Saprotroph\| | Yeast | NULL | Probable | As nectar-tap_saprotroph (Põlme et al. 2020); Decay sugar-rich substrates (Põlme et al. 2020) | [illegible] |
| Otu01059 | 61 | k__Fungi(100);p__Ascomycota(100);c__Lecanoromycetes(100);o__Lecanorales(100);o__Lecanorales_unclassified(87);o__Lecanorales_unclassified(87);o__Lecanorales_unclassified(87); | Lecanorales | 7 | Pathotroph-Symbiotroph | Lichen Parasite-Lichenized | Microfungus-Thallus | NULL | Probable | Lichenized with green algae, occasionally lichenicolous, on rocks, mosses and wood (Cannon & Kirk 2007); ...an order of mostly lichen-forming fungi (Kirk et al. 2008) | [illegible] |
| Otu01061 | 61 | k__Fungi(100);p__Ascomycota(100);c__Sordariomycetes(100);o__Xylariales(100);f__Xylariaceae(100);g__Nemania(100);s__Nemania_sp(100); | Nemania | 13 | Saprotroph | Undefined Saprotroph-\|Wood Saprotroph\| | Microfungus | NULL | Probable | Decay wood (Põlme et al. 2020) | [illegible] |
| Otu01062 | 61 | k__Fungi(100);p__Chytridiomycota(96);c__Chytridiomycetes(96);o__Spizellomycetales(96);f__Spizellomycetaceae(96);f__Spizellomycetaceae_unclassified(96);f__Spizellomycetaceae_unclassified(96); | Spizellomycetaceae | 9 | Pathotroph | Plant Pathogen-Undefined Parasite-Undefined Saprotroph | Microfungus | NULL | Probable | Common in the soil and also as parasites of a range of soil organisms and plants (Cannon & Kirk 2007) | [illegible] |
| Otu01063 | 61 | k__Fungi(100);p__Basidiomycota(100);c__Agaricomycetes(100);o__Agaricales(100);f__Entolomataceae(100);g__Clitopilus(100);s__Clitopilus_hobsonii(78); | Clitopilus | 13 | Pathotroph-Saprotroph | Fungal Parasite-\|Plant Saprotroph\|-Undefined Saprotroph | Agaricoid | NULL | Probable | As litter_saprotroph (Põlme et al. 2020); As mycoparasite (Põlme et al. 2020); Decay leaf, fruit, seed (Põlme et al. 2020) | [illegible] |
| Otu01064 | 61 | k__Fungi(100);p__Ascomycota(100);c__Sordariomycetes(100);o__Xylariales(100);f__Graphostromataceae(100);g__Graphostroma(100);s__Graphostroma_platystoma(100); | Graphostroma | 13 | Pathotroph-Saprotroph-Symbiotroph | Endophyte-Plant Pathogen-Undefined Saprotroph-\|Wood Saprotroph\| | Microfungus | NULL | Probable | As foliar_endophyte (Põlme et al. 2020); As wood_pathogen (Põlme et al. 2020); Decay wood, leaf, fruit, seed (Põlme et al. 2020); Host - angiosperms (Põlme et al. 2020) | [illegible] |
| Otu01065 | 61 | k__Fungi(100);p__Ascomycota(100);c__Sordariomycetes(100);o__Xylariales(100);f__Xylariaceae(100);g__Hypoxylon(100);s__Hypoxylon_sp(100); | Hypoxylon | 13 | Saprotroph-Saprotroph-Symbiotroph | Endophyte-Undefined Saprotroph-\|Wood Saprotroph\| | Corticioid-Microfungus-Xylarioid | NULL | Probable | As foliar_endophyte (Põlme et al. 2020); Decay wood, leaf, fruit, seed (Põlme et al. 2020); Host - angiosperms (Põlme et al. 2020) | [illegible] |
| Otu01067 | 61 | k__Fungi(100);p__Basidiomycota(100);c__Agaricomycetes(100);o__Hymenochaetales(100);f__Hymenochaetaceae(100);g__Hymenochaete(100);g__Hymenochaete_unclassified(100); | Hymenochaete | 13 | Pathotroph-Saprotroph | Plant Pathogen-Undefined Saprotroph-\|Wood Saprotroph\| | Corticioid-Polyporoid | White Rot | Probable | As white_rot (Põlme et al. 2020); As wood_pathogen (Põlme et al. 2020); Decay wood (Põlme et al. 2020) | [illegible] |
| Otu01068 | 61 | k__Fungi(100);p__Basidiomycota(100);c__Agaricomycetes(100);o__Polyporales(100);f__Polyporaceae(100);g__Polyporus(100);s__Polyporus_sp(100); | Polyporus | 13 | Pathotroph-Saprotroph | Plant Pathogen-\|Wood Saprotroph\| | Polyporoid | White Rot | Highly Probable | As white_rot (Põlme et al. 2020); As wood_pathogen (Põlme et al. 2020); Decay wood (Põlme et al. 2020) | [illegible] |
| Otu01057 | 61 | k__Fungi(100);p__Ascomycota(100);p__Ascomycota_unclassified(100);p__Ascomycota_unclassified(100);p__Ascomycota_unclassified(100);p__Ascomycota_unclassified(100);p__Ascomycota_unclassified(100); | - | - | - | - | - | - | - | Unassigned | - |
| Otu01060 | 61 | k__Fungi(100);p__Ascomycota(100);c__Dothideomycetes(100);o__Capnodiales(100);o__Capnodiales_unclassified(100);o__Capnodiales_unclassified(100);o__Capnodiales_unclassified(100); | - | - | - | - | - | - | - | Unassigned | - |
| Otu01066 | 61 | k__Fungi(100);p__Basidiomycota(100);p__Basidiomycota_unclassified(100);p__Basidiomycota_unclassified(100);p__Basidiomycota_unclassified(100);p__Basidiomycota_unclassified(100);p__Basidiomycota_unclassified(100); | - | - | - | - | - | - | - | Unassigned | - |
| Otu01069 | 60 | k__Fungi(100);p__Basidiomycota(100);c__Agaricomycetes(100);o__Agaricales(100);f__Hygrophoraceae(100);g__Ampulloclitocybe(100);s__Ampulloclitocybe_clavipes(100); | Ampulloclitocybe | 13 | Saprotroph | \|Plant Saprotroph\|-Undefined Saprotroph | Agaricoid | NULL | Probable | As litter_saprotroph (Põlme et al. 2020); Decay leaf, fruit, seed (Põlme et al. 2020) | [illegible] |
| Otu01072 | 60 | k__Fungi(100);p__Basidiomycota(100);c__Agaricomycetes(100);o__Agaricales(100);f__Mycenaceae(100);g__Mycena(100);s__Mycena_zephirus(64); | Mycena | 13 | Pathotroph-Saprotroph | Plant Pathogen-\|Plant Saprotroph\|-Undefined Saprotroph-Wood Saprotroph | Agaricoid | NULL | Probable | As litter_saprotroph (Põlme et al. 2020); Decay leaf, fruit, seed (Põlme et al. 2020) | [illegible] |
| Otu01074 | 60 | k__Fungi(100);p__Ascomycota(100);c__Dothideomycetes(100);o__Capnodiales(100);f__Teratosphaeriaceae(100);g__Devriesia(100);s__Devriesia_sp(100); | Devriesia | 13 | Pathotroph-Saprotroph | \|Plant Pathogen\|-Plant Saprotroph | Microfungus | NULL | Probable | As litter_saprotroph (Põlme et al. 2020); Decay leaf, fruit, seed (Põlme et al. 2020) | [illegible] |
| Otu01076 | 60 | k__Fungi(100);p__Ascomycota(100);c__Sordariomycetes(85);o__Hypocreales(85);f__Cordycipitaceae(85);g__Isaria(85);s__Isaria_fumosorosea(80); | Isaria | 13 | Pathotroph-Saprotroph | \|Animal Parasite\|-Animal Pathogen-Undefined Saprotroph | Microfungus | NULL | Probable | As animal_decomposer (Põlme et al. 2020); Decay animal material (Põlme et al. 2020) | [illegible] |
| Otu01078 | 60 | k__Fungi(100);p__Basidiomycota(100);c__Agaricomycetes(100);o__Polyporales(100);f__Meruliaceae(100);g__Crustoderma(100);s__Crustoderma_flavescens(100); | Crustoderma | 13 | Saprotroph | \|Wood Saprotroph\| | Corticioid | Brown Rot | Probable | As brown-rot (Põlme et al. 2020); Decay wood (Põlme et al. 2020) | [illegible] |
| Otu01081 | 60 | k__Fungi(100);p__Ascomycota(100);c__Sordariomycetes(100);o__Xylariales(100);f__Xylariaceae(100);g__Hypoxylon(100);s__Hypoxylon_howeanum(100); | Hypoxylon | 13 | Saprotroph-Saprotroph-Symbiotroph | Endophyte-Undefined Saprotroph-\|Wood Saprotroph\| | Corticioid-Microfungus-Xylarioid | NULL | Probable | As foliar_endophyte (Põlme et al. 2020); Decay wood, leaf, fruit, seed (Põlme et al. 2020); Host - angiosperms (Põlme et al. 2020) | [illegible] |
| Otu01082 | 60 | k__Fungi(100);p__Ascomycota(100);c__Pezizomycetes(100);o__Pezizales(100);f__Sarcosomataceae(100);g__Galiella(100);s__Galiella_sp(100); | Galiella | 13 | Saprotroph | Undefined Saprotroph-\|Wood Saprotroph\| | Microfungus | NULL | Probable | Decay wood (Põlme et al. 2020) | [illegible] |
| Otu01070 | 60 | k__Fungi(100);p__Ascomycota(74);p__Ascomycota_unclassified(74);p__Ascomycota_unclassified(74);p__Ascomycota_unclassified(74);p__Ascomycota_unclassified(74);p__Ascomycota_unclassified(74); | - | - | - | - | - | - | - | Unassigned | - |
| Otu01071 | 60 | k__Fungi(100);p__Ascomycota(100);p__Ascomycota_unclassified(100);p__Ascomycota_unclassified(100);p__Ascomycota_unclassified(100);p__Ascomycota_unclassified(100);p__Ascomycota_unclassified(100); | - | - | - | - | - | - | - | Unassigned | - |
| Otu01073 | 60 | k__Fungi(100);p__Ascomycota(100);p__Ascomycota_unclassified(100);p__Ascomycota_unclassified(100);p__Ascomycota_unclassified(100);p__Ascomycota_unclassified(100);p__Ascomycota_unclassified(100); | - | - | - | - | - | - | - | Unassigned | - |
| Otu01075 | 60 | k__Fungi(100);p__Ascomycota(84);p__Ascomycota_unclassified(84);p__Ascomycota_unclassified(84);p__Ascomycota_unclassified(84);p__Ascomycota_unclassified(84);p__Ascomycota_unclassified(84); | - | - | - | - | - | - | - | Unassigned | - |
| Otu01077 | 60 | k__Fungi(100);p__Basidiomycota(100);c__Agaricomycetes(100);c__Agaricomycetes_unclassified(52);c__Agaricomycetes_unclassified(52);c__Agaricomycetes_unclassified(52);c__Agaricomycetes_unclassified(52); | - | - | - | - | - | - | - | Unassigned | - |
| Otu01079 | 60 | k__Fungi(100);p__Ascomycota(100);c__Dothideomycetes(100);o__Capnodiales(99);o__Capnodiales_unclassified(99);o__Capnodiales_unclassified(99);o__Capnodiales_unclassified(99); | - | - | - | - | - | - | - | Unassigned | - |
| Otu01080 | 60 | k__Fungi(100);p__Ascomycota(100);c__Sordariomycetes(100);c__Sordariomycetes_unclassified(100);c__Sordariomycetes_unclassified(100);c__Sordariomycetes_unclassified(100);c__Sordariomycetes_unclassified(100); | - | - | - | - | - | - | - | Unassigned | - |
| Otu01083 | 59 | k__Fungi(100);p__Ascomycota(100);c__Lecanoromycetes(100);o__Lecanorales(100);f__Parmeliaceae(100);g__Parmotrema(100);s__Parmotrema_hypoleucinum(90); | Parmotrema | 13 | Symbiotroph | \|Lichenized\| | Thallus | NULL | Highly Probable | NULL | [illegible] |
| Otu01087 | 59 | k__Fungi(100);p__Basidiomycota(100);c__Agaricomycetes(100);o__Russulales(100);f__Lachnocladiaceae(70);f__Lachnocladiaceae_unclassified(70);f__Lachnocladiaceae_unclassified(70); | Lachnocladiaceae | 9 | Saprotroph | Wood Saprotroph | Coraloid-Corticioid | NULL | Probable | On decaying coniferous or broadleaved wood, sometimes on herbaceous stems, presumably saprobic (Cannon & Kirk 2007) | [illegible] |
| Otu01088 | 59 | k__Fungi(100);p__Basidiomycota(100);c__Agaricomycetes(100);o__Agaricales(100);f__Psathyrellaceae(100);g__unclassified_Psathyrellaceae(63);s__Psathyrellaceae_sp(63); | Psathyrellaceae | 9 | Saprotroph | Dung Saprotroph-Plant Saprotroph-Wood Saprotroph | Agaricoid | NULL | Probable | Saprobic, in grass, on dung, rotten wood and similar substrata (Cannon & Kirk 2007) | [illegible] |
| Otu01089 | 59 | k__Fungi(100);p__Basidiomycota(100);c__Microbotryomycetes(100);o__Leucosporidiales(100);f__Leucosporidiaceae(100);g__Leucosporidiella(95);g__Leucosporidiella_unclassified(94); | Leucosporidiella | 13 | Saprotroph | Soil Saprotroph-Undefined Saprotroph | Yeast | NULL | Probable | A yeast from fruit (Barnett & Buhagiar 1971; Syn. Candida-Leucosporidium-Rhodotorula-Torulopsis-Vanrijia spp.);Isolated from permafrost soil (Golubev 1998; Syn. Leucosporidium/Rhodotorula spp.) | [illegible] |
| Otu01090 | 59 | k__Fungi(100);p__Basidiomycota(100);c__Tremellomycetes(100);o__Tremellales(100);f__Tremellales_family_Incertae_sedis(100);g__Cryptococcus(100);s__Cryptococcus_aff_amylolyticus_AS_22501(90); | Cryptococcus | 13 | Pathotroph-Saprotroph-Symbiotroph | Animal Pathogen-Endophyte-Epiphyte-\|Undefined Saprotroph\| | Dimorphic Yeast-Yeast | NULL | Possible | As foliar_endophyte (Põlme et al. 2020); Decay leaf, fruit, seed, soil, dung, animal material (Põlme et al. 2020) | [illegible] |
| Otu01091 | 59 | k__Fungi(100);p__Ascomycota(100);c__Saccharomycetes(100);o__Saccharomycetales(100);f__Metschnikowiaceae(100);g__Metschnikowia(100);s__Metschnikowia_pulcherrima(97); | Metschnikowia | 13 | Saprotroph | \|Undefined Saprotroph\| | Yeast | NULL | Probable | As nectar-tap_saprotroph (Põlme et al. 2020); Decay sugar-rich substrates (Põlme et al. 2020) | [illegible] |
| Otu01092 | 59 | k__Fungi(100);p__Zygomycota(100);c__Zygomycota_class_Incertae_sedis(100);o__Mortierellales(100);f__Mortierellaceae(100);g__Mortierella(100);g__Mortierella_unclassified(100); | Mortierella | 13 | Saprotroph-Symbiotroph | Endophyte-Plant Saprotroph-\|Undefined Saprotroph\| | Microfungus | NULL | Probable | As chitinolytic (Põlme et al. 2020); As soil_saprotroph (Põlme et al. 2020); Decay roots, soil (Põlme et al. 2020); Saprobic in soil and thus frequenctly isolated from soil and material contaminated with soil such as hay or rotten silage and spoil tips (Cannon & Kirk 2007) | [illegible] |
| Otu01093 | 59 | k__Fungi(100);p__Basidiomycota(100);c__Agaricomycetes(100);o__Polyporales(100);f__Meruliaceae(100);g__Hypochnicium(100);s__Hypochnicium_sp_DLL2011_2(100); | Hypochnicium | 13 | Saprotroph | Undefined Saprotroph-\|Wood Saprotroph\| | Corticioid | White Rot | Probable | As white_rot (Põlme et al. 2020); Decay wood (Põlme et al. 2020) | [illegible] |
| Otu01094 | 59 | k__Fungi(100);p__Basidiomycota(100);c__Agaricomycetes(100);o__Russulales(100);f__Stereaceae(100);g__Stereum(100);s__Stereum_subtomentosum(100); | Stereum | 13 | Saprotroph | Undefined Saprotroph-\|Wood Saprotroph\| | Polyporoid | White Rot | Probable | As white_rot (Põlme et al. 2020); Decay wood (Põlme et al. 2020) | [illegible] |
| Otu01095 | 59 | k__Fungi(100);p__Ascomycota(100);c__Lecanoromycetes(100);o__Lecanorales(100);f__Parmeliaceae(100);g__Flavoparmelia(100);s__Flavoparmelia_caperata(100); | Flavoparmelia | 13 | Symbiotroph | \|Lichenized\| | Thallus | NULL | Highly Probable | NULL | [illegible] |

| | | | | | | | | | | | |
|---|---|---|---|---|---|---|---|---|---|---|---|
| Otu01097 | 59 | k__Fungi(100);p__Basidiomycota(100);c__Tremellomycetes(100);o__Filobasidiales(100);f__Filobasidiaceae(100);g__Cryptococcus(100);g__Cryptococcus_unclassified(100); | Cryptococcus | 13 | Pathotroph-Saprotroph-Symbiotroph | Animal Pathogen-Endophyte-Epiphyte-Undefined Saprotroph | Dimorphic Yeast-Yeast | NULL | Possible | As foliar_endophyte (Põlme et al. 2020); Decay leaf, fruit, seed, soil, dung, animal material (Põlme et al. 2020) | Andrews JH, et al. 1983. Phytopathology 73:228-234 ((https://doi.org/10.1094/Phyto-73-228)); Kurtzman CP, et al. (eds.) 2011. The Yeasts, a Taxonomic Study. Fifth Edition. Vols 1-3. Elsevier, San Diego (ISBN: 9780444521491); Põlme S, et al. 2020. Fungal Diversity 105:1-16 ((https://doi.org/10.1007/s13225-020-00466-2)) |
| Otu01098 | 59 | k__Fungi(100);p__Ascomycota(100);c__Eurotiomycetes(100);o__Eurotiales(100);f__Trichocomaceae(100);g__Talaromyces(97);g__Talaromyces_unclassified(89); | Talaromyces | 13 | Pathotroph-Saprotroph | Animal Parasite-\|Undefined Saprotroph\| | Microfungus | NULL | Probable | Decay leaf, fruit, seed, soil, animal material (Põlme et al. 2020) | Tedersoo L, et al. 2014. Science 346:e1256688 ((https://doi.org/10.1126/science.1256688)); Põlme S, et al. 2020. Fungal Diversity 105:1-16 ((https://doi.org/10.1007/s13225-020-00466-2)) |
| Otu01099 | 59 | k__Fungi(100);p__Ascomycota(100);c__Saccharomycetes(100);o__Saccharomycetales(100);f__Saccharomycetaceae(100);g__Kazachstania(100);s__Kazachstania_intestinalis(100); | Kazachstania | 13 | Saprotroph | \|Undefined Saprotroph\| | Yeast | NULL | Probable | As nectar-tap_saprotroph (Põlme et al. 2020); Decay sugar-rich substrates (Põlme et al. 2020) | Põlme S, et al. 2020. Fungal Diversity 105:1-16 ((https://doi.org/10.1007/s13225-020-00466-2)) |
| Otu01084 | 59 | k__Fungi(100);k__Fungi_unclassified(95);k__Fungi_unclassified(95);k__Fungi_unclassified(95);k__Fungi_unclassified(95);k__Fungi_unclassified(95);k__Fungi_unclassified(95); | - | - | - | - | - | - | - | Unassigned | - |
| Otu01085 | 59 | k__Fungi(100);k__Fungi_unclassified(55);k__Fungi_unclassified(55);k__Fungi_unclassified(55);k__Fungi_unclassified(55);k__Fungi_unclassified(55);k__Fungi_unclassified(55); | - | - | - | - | - | - | - | Unassigned | - |
| Otu01086 | 59 | k__Fungi(100);k__Fungi_unclassified(62);k__Fungi_unclassified(62);k__Fungi_unclassified(62);k__Fungi_unclassified(62);k__Fungi_unclassified(62);k__Fungi_unclassified(62); | - | - | - | - | - | - | - | Unassigned | - |
| Otu01096 | 59 | k__Fungi(100);p__Ascomycota(53);p__Ascomycota_unclassified(53);p__Ascomycota_unclassified(53);p__Ascomycota_unclassified(53);p__Ascomycota_unclassified(53);p__Ascomycota_unclassified(53); | - | - | - | - | - | - | - | Unassigned | - |
| Otu01102 | 58 | k__Fungi(100);p__Basidiomycota(100);c__Agaricomycetes(100);o__Polyporales(100);f__Meruliaceae(100);g__Hyphoderma(100);s__Hyphoderma_setigerum(100); | Hyphoderma | 13 | Saprotroph | Undefined Saprotroph-\|Wood Saprotroph\| | Corticioid | White Rot | Probable | As white_rot (Põlme et al. 2020); Decay wood (Põlme et al. 2020) | Tedersoo L, et al. 2014. Science 346:e1256688 ((https://doi.org/10.1126/science.1256688)); Põlme S, et al. 2020. Fungal Diversity 105:1-16 ((https://doi.org/10.1007/s13225-020-00466-2)) |
| Otu01103 | 58 | k__Fungi(100);p__Basidiomycota(100);c__Agaricomycetes(100);o__Hymenochaetales(100);f__Hymenochaetaceae(100);g__Phellinus(100);s__Phellinus_ferruginosus(100); | Phellinus | 13 | Pathotroph-Saprotroph | \|Plant Pathogen\|-Wood Saprotroph | Polyporoid | White Rot | Highly Probable | As white_rot (Põlme et al. 2020); As wood_pathogen (Põlme et al. 2020); Decay wood (Põlme et al. 2020) | Gilbertson RL, Ryvarden L. 1987-1987. North American Polypores. Fungiflora, Oslo (ISBN: 978-0945345060); Põlme S, et al. 2020. Fungal Diversity 105:1-16 ((https://doi.org/10.1007/s13225-020-00466-2)) |
| Otu01105 | 58 | k__Fungi(100);p__Ascomycota(100);c__Saccharomycetes(100);o__Saccharomycetales(100);f__unclassified_Saccharomycetales(95);g__unclassified_Saccharomycetales(95);s__Saccharomycetales_sp(95); | Saccharomycetales | 7 | Saprotroph | Undefined Saprotroph | Yeast | NULL | Possible | NULL | Sterkenburg E, et al. 2015. New Phytologist 207:1145-1158 ((https://doi.org/10.1111/nph.13426)) |
| Otu01106 | 58 | k__Fungi(100);p__Basidiomycota(100);c__Agaricomycetes(100);o__Hymenochaetales(100);f__Hymenochaetaceae(100);g__Fuscoporia(100);s__Fuscoporia_sp_JV0509/78(100); | Fuscoporia | 13 | Saprotroph | Undefined Saprotroph-\|Wood Saprotroph\| | Polyporoid | White Rot | Probable | As white_rot (Põlme et al. 2020); Decay wood (Põlme et al. 2020) | Tedersoo L, et al. 2014. Science 346:e1256688 ((https://doi.org/10.1126/science.1256688)); Põlme S, et al. 2020. Fungal Diversity 105:1-16 ((https://doi.org/10.1007/s13225-020-00466-2)) |
| Otu01107 | 58 | k__Fungi(100);p__Basidiomycota(100);c__Agaricomycetes(100);o__Trechisporales(100);f__Hydnodontaceae(100);g__Trechispora(100);s__Trechispora_sp_1_MPM_2013(95); | Trechispora | 13 | Saprotroph | \|Wood Saprotroph\| | Corticioid | White Rot | Highly Probable | As white_rot (Põlme et al. 2020); Decay wood (Põlme et al. 2020) | Gilbertson RL, Ryvarden L. 1987-1987. North American Polypores. Fungiflora, Oslo (ISBN: 978-0945345060); Rinaldi AC, et al. 2008. Fungal Diversity 33:1-45 ((http://www.fungaldiversity.org/fdp/sfdp/33-1.pdf)) (as Uncertain); Tedersoo L, et al. 2010. Mycorrhiza 20:217-263 ((https://doi.org/10.1007/s00572-009-0274-x)) (as not ECM).; Põlme S, et al. 2020. Fungal Diversity 105:1-16 ((https://doi.org/10.1007/s13225-020-00466-2)) |
| Otu01108 | 58 | k__Fungi(100);p__Basidiomycota(100);c__Agaricomycetes(100);o__Boletales(100);f__Suillaceae(100);g__Suillus(100);s__Suillus_grevillei(97); | Suillus | 13 | Symbiotroph | \|Ectomycorrhizal\| | Agaricoid-Boletoid | NULL | Highly Probable | Host - Pinaceae (Põlme et al. 2020) | Rinaldi AC, et al. 2008. Fungal Diversity 33:1-45 ((http://www.fungaldiversity.org/fdp/sfdp/33-1.pdf)); Tedersoo L, et al. 2010. Mycorrhiza 20:217-263 ((https://doi.org/10.1007/s00572-009-0274-x)); Põlme S, et al. 2020. Fungal Diversity 105:1-16 ((https://doi.org/10.1007/s13225-020-00466-2)); http://mycorrhizas.info/ecmf.html ((http://mycorrhizas.info/ecmf.html)) |
| Otu01109 | 58 | k__Fungi(100);p__Ascomycota(100);c__Dothideomycetes(100);o__Pleosporales(100);f__Leptosphaeriaceae(100);g__Plenodomus(100);s__Plenodomus_sp(100); | Plenodomus | 13 | Pathotroph-Saprotroph | \|Plant Pathogen\|-Plant Saprotroph-Undefined Saprotroph | Microfungus | NULL | Probable | As litter_saprotroph (Põlme et al. 2020); Decay leaf, fruit, seed (Põlme et al. 2020) | Tedersoo L, et al. 2014. Science 346:e1256688 ((https://doi.org/10.1126/science.1256688)); Põlme S, et al. 2020. Fungal Diversity 105:1-16 ((https://doi.org/10.1007/s13225-020-00466-2)) |
| Otu01111 | 58 | k__Fungi(100);p__Basidiomycota(100);c__Agaricomycetes(99);o__Agaricales(97);o__Agaricales_unclassified(97);o__Agaricales_unclassified(97);o__Agaricales_unclassified(97); | Agaricales | 7 | Pathotroph-Saprotroph-Symbiotroph | Bryophyte Parasite-Dung Saprotroph-Ectomycorrhizal-Fungal Parasite-Leaf Saprotroph-Plant Parasite-Undefined Saprotroph-Wood Saprotroph | Agaricoid-Gasteroid-Microfungus-Secotioid-Yeast | NULL | Possible | Mushrooms and toadstools, Gill fungi, Agarics... lignicolous, sometimes muscicolous or fungicolous, saprobic, mycorrhizal, rarely parasitic on plants or fungi (Kirk et al. 2008) | Kirk PM et al. 2008. Dictionary of the Fungi. Tenth Edition. CAB International, Wallingford (ISBN: 978-0851998268) |
| Otu01100 | 58 | k__Fungi(100);p__Ascomycota(100);c__Dothideomycetes(100);c__Dothideomycetes_unclassified(95);c__Dothideomycetes_unclassified(95);c__Dothideomycetes_unclassified(95);c__Dothideomycetes_unclassified(95); | - | - | - | - | - | - | - | Unassigned | - |
| Otu01101 | 58 | k__Fungi(100);k__Fungi_unclassified(59);k__Fungi_unclassified(59);k__Fungi_unclassified(59);k__Fungi_unclassified(59);k__Fungi_unclassified(59);k__Fungi_unclassified(59); | - | - | - | - | - | - | - | Unassigned | - |
| Otu01104 | 58 | k__Fungi(100);k__Fungi_unclassified(56);k__Fungi_unclassified(56);k__Fungi_unclassified(56);k__Fungi_unclassified(56);k__Fungi_unclassified(56);k__Fungi_unclassified(56); | - | - | - | - | - | - | - | Unassigned | - |
| Otu01110 | 58 | k__Fungi(100);p__Ascomycota(94);p__Ascomycota_unclassified(94);p__Ascomycota_unclassified(94);p__Ascomycota_unclassified(94);p__Ascomycota_unclassified(94);p__Ascomycota_unclassified(94); | - | - | - | - | - | - | - | Unassigned | - |
| Otu01112 | 57 | k__Fungi(100);p__Basidiomycota(100);c__Agaricomycetes(100);o__Agaricales(100);f__Mycenaceae(100);g__Mycena(100);g__Mycena_unclassified(53); | Mycena | 13 | Pathotroph-Saprotroph | Plant Pathogen-\|Plant Saprotroph\|-Undefined Saprotroph-Wood Saprotroph | Agaricoid | NULL | Probable | As litter_saprotroph (Põlme et al. 2020); Decay leaf, fruit, seed (Põlme et al. 2020) | Grgurinovic CA, Simpson JA. 2001. Fungal Diversity 8:97-127 ((http://www.fungaldiversity.org/fdp/sfdp/FD_8_97-127.pdf)); Tedersoo L, et al. 2014. Science 346:e1256688 ((https://doi.org/10.1126/science.1256688)); Põlme S, et al. 2020. Fungal Diversity 105:1-16 ((https://doi.org/10.1007/s13225-020-00466-2)); The American Phytopathological Society Common Names of Plant Diseases ((http://www.apsnet.org/publications/commonnames/Pages/default.aspx)) |
| Otu01113 | 57 | k__Fungi(100);p__Ascomycota(100);c__Sordariomycetes(100);o__Hypocreales(100);f__Hypocreales_family_Incertae_sedis(100);g__Acremonium(100);s__Acremonium_rutilum(100); | Acremonium | 13 | Pathotroph-Saprotroph-Symbiotroph | Animal Pathogen-Endophyte-Fungal Parasite-Plant Pathogen-\|Undefined Saprotroph\|-Wood Saprotroph | Microfungus | Soft Rot | Possible | As foliar_endophyte (Põlme et al. 2020); culture (Hanada et al. 2010, Del Olmo; Decay leaf, fruit, seed, soil, dung, animal material (Põlme et al. 2020); Endophyte detection method; Ruiz & Arnold 2014); Likely opportunistic human pathogen (Irinyi et al. 2015) | Duncan CG, Eslyn WE. 1966. Mycologia 58:642-645 ((https://doi.org/10.2307/3757045)); Nilsson T. 1973. Studia Forestalia Suecica 104: 1-40 ((https://pub.epsilon.slu.se/5711/)); Worrall JJ, et al. 1997. Mycologia 89:199-219 ((https://doi.org/10.2307/3761073)); Hanada RE, et al. 2010. Fungal Biology 114:901-910 ((https://doi.org/10.1016/j.funbio.2010.08.006)); Del Olmo-Ruiz M, Arnold AE. 2014. Mycologia 106: 8-21 ((https://doi.org/10.3852/13-098)); Irinyi L, et al. 2015. Medical Mycology 53313-337 ((https://doi.org/10.1093/mmy/myv008)); Põlme S, et al. 2020. Fungal Diversity 105:1-16 ((https://doi.org/10.1007/s13225-020-00466-2)) |
| Otu01115 | 57 | k__Fungi(100);p__Ascomycota(100);c__Dothideomycetes(100);o__Pleosporales(100);f__Cucurbitariaceae(100);g__Curreya(100);g__Curreya_unclassified(76); | Curreya | 13 | Saprotroph | \|Plant Saprotroph\|-Undefined Saprotroph | Microfungus | NULL | Probable | As litter_saprotroph (Põlme et al. 2020); Decay leaf, fruit, seed (Põlme et al. 2020) | Tedersoo L, et al. 2014. Science 346:e1256688 ((https://doi.org/10.1126/science.1256688)); Põlme S, et al. 2020. Fungal Diversity 105:1-16 ((https://doi.org/10.1007/s13225-020-00466-2)) |
| Otu01117 | 57 | k__Fungi(100);p__Ascomycota(100);c__Saccharomycetes(100);o__Saccharomycetales(100);f__Metschnikowiaceae(100);g__Metschnikowia(100);s__Metschnikowia_pulcherrima(97); | Metschnikowia | 13 | Saprotroph | \|Undefined Saprotroph\| | Yeast | NULL | Probable | As nectar-tap_saprotroph (Põlme et al. 2020); Decay sugar-rich substrates (Põlme et al. 2020) | Põlme S, et al. 2020. Fungal Diversity 105:1-16 ((https://doi.org/10.1007/s13225-020-00466-2)) |
| Otu01118 | 57 | k__Fungi(100);p__Basidiomycota(100);c__Agaricomycetes(100);o__Russulales(100);f__Stereaceae(100);g__Stereum(81);g__Stereum_unclassified(51); | Stereum | 13 | Saprotroph | Undefined Saprotroph-\|Wood Saprotroph\| | Polyporoid | White Rot | Probable | As white_rot (Põlme et al. 2020); Decay wood (Põlme et al. 2020) | Tedersoo L, et al. 2014. Science 346:e1256688 ((https://doi.org/10.1126/science.1256688)); Põlme S, et al. 2020. Fungal Diversity 105:1-16 ((https://doi.org/10.1007/s13225-020-00466-2)) |
| Otu01119 | 57 | k__Fungi(100);p__Basidiomycota(100);c__Agaricomycetes(100);o__Agaricales(100);f__Strophariaceae(100);f__Strophariaceae_unclassified(100);f__Strophariaceae_unclassified(100); | Strophariaceae | 9 | Pathotroph-Saprotroph | Dung Saprotroph-Plant Parasite-Soil Saprotroph-Undefined Saprotroph-Wood Saprotroph | Agaricoid | NULL | Probable | Very variable ecology; Pholiota species are lignicolous and can act as weak parasites while most other taxa are either saprobes in grassy habitats or are found in dung. Species are not ectomycorrhizal (Cannon & Kirk 2007) | Cannon PF, Kirk PM. 2007. Fungal Families of the World. CAB International, Cambridge (ISBN: 978-0851998275) |
| Otu01120 | 57 | k__Fungi(100);p__Basidiomycota(100);c__Agaricomycetes(100);o__Polyporales(100);f__Phanerochaetaceae(100);g__Phanerochaete(100);s__Phanerochaete_sp_DLL2011_1(100); | Phanerochaete | 13 | Saprotroph | \|Wood Saprotroph\| | Corticioid | White Rot | Highly Probable | As white_rot (Põlme et al. 2020); Decay wood (Põlme et al. 2020) | Glenn JK, et al. 1983. Biochemical and Biophysical Research Communications 114:1077-1083 ((https://doi.org/10.1016/0006-291x((83)) 90672-1)); Põlme S, et al. 2020. Fungal Diversity 105:1-16 ((https://doi.org/10.1007/s13225-020-00466-2)) |
| Otu01121 | 57 | k__Fungi(100);p__Basidiomycota(100);c__Tremellomycetes(100);o__Filobasidiales(100);f__Filobasidiaceae(100);g__Cryptococcus(90);g__Cryptococcus_unclassified(90); | Cryptococcus | 13 | Pathotroph-Saprotroph-Symbiotroph | Animal Pathogen-Endophyte-Epiphyte-\|Undefined Saprotroph\| | Dimorphic Yeast-Yeast | NULL | Possible | As foliar_endophyte (Põlme et al. 2020); Decay leaf, fruit, seed, soil, dung, animal material (Põlme et al. 2020) | Andrews JH, et al. 1983. Phytopathology 73:228-234 ((https://doi.org/10.1094/Phyto-73-228)); Kurtzman CP, et al. (eds.) 2011. The Yeasts, a Taxonomic Study. Fifth Edition. Vols 1-3. Elsevier, San Diego (ISBN: 9780444521491); Põlme S, et al. 2020. Fungal Diversity 105:1-16 ((https://doi.org/10.1007/s13225-020-00466-2)) |
| Otu01122 | 57 | k__Fungi(100);p__Ascomycota(100);c__Saccharomycetes(100);o__Saccharomycetales(100);f__Metschnikowiaceae(100);g__Metschnikowia(100);s__Metschnikowia_pulcherrima(99); | Metschnikowia | 13 | Saprotroph | \|Undefined Saprotroph\| | Yeast | NULL | Probable | As nectar-tap_saprotroph (Põlme et al. 2020); Decay sugar-rich substrates (Põlme et al. 2020) | Põlme S, et al. 2020. Fungal Diversity 105:1-16 ((https://doi.org/10.1007/s13225-020-00466-2)) |
| Otu01114 | 57 | k__Fungi(100);p__Ascomycota(51);p__Ascomycota_unclassified(51);p__Ascomycota_unclassified(51);p__Ascomycota_unclassified(51);p__Ascomycota_unclassified(51);p__Ascomycota_unclassified(51); | - | - | - | - | - | - | - | Unassigned | - |
| Otu01116 | 57 | k__Fungi(100);k__Fungi_unclassified(55);k__Fungi_unclassified(55);k__Fungi_unclassified(55);k__Fungi_unclassified(55);k__Fungi_unclassified(55);k__Fungi_unclassified(55); | - | - | - | - | - | - | - | Unassigned | - |
| Otu01123 | 57 | k__Fungi(100);p__Ascomycota(100);p__Ascomycota_unclassified(100);p__Ascomycota_unclassified(100);p__Ascomycota_unclassified(100);p__Ascomycota_unclassified(100);p__Ascomycota_unclassified(100); | - | - | - | - | - | - | - | Unassigned | - |
| Otu01128 | 56 | k__Fungi(100);p__Basidiomycota(100);c__Agaricomycetes(100);o__Cantharellales(100);f__Hydnaceae(100);g__Sistotrema(100);s__Sistotrema_sp(100); | Sistotrema | 13 | Saprotroph-Symbiotroph | Ectomycorrhizal-\|Plant Saprotroph\|-Wood Saprotroph | Corticioid | White Rot | Possible | As litter_saprotroph (Põlme et al. 2020); Decay leaf, fruit, seed (Põlme et al. 2020) | Gilbertson RL, Ryvarden L. 1987-1987. North American Polypores. Fungiflora, Oslo (ISBN: 978-0945345060); Rinaldi AC, et al. 2008. Fungal Diversity 33:1-45 ((http://www.fungaldiversity.org/fdp/sfdp/33-1.pdf)) (as ECM/not ECM); Tedersoo L, et al. 2010. Mycorrhiza 20:217-263 ((https://doi.org/10.1007/s00572-009-0274-x)) (pro parte); Põlme S, et al. 2020. Fungal Diversity 105:1-16 ((https://doi.org/10.1007/s13225-020-00466-2)); http://mycorrhizas.info/ecmf.html ((http://mycorrhizas.info/ecmf.html)) (as ECM/not ECM). |
| Otu01129 | 56 | k__Fungi(100);p__Ascomycota(100);c__Sordariomycetes(100);o__Diaporthales(100);f__Diaporthaceae(100);f__Diaporthaceae_unclassified(86);f__Diaporthaceae_unclassified(86); | Diaporthaceae | 9 | Pathotroph-Saprotroph-Symbiotroph | Endophyte-Plant Pathogen-Plant Saprotroph | Microfungus | NULL | Probable | Most species are weak pathogens or endophytes in leaf or stem tissue, growing saprobically after the plant dies (Cannon & Kirk 2007) | Cannon PF, Kirk PM. 2007. Fungal Families of the World. CAB International, Cambridge (ISBN: 978-0851998275) |
| Otu01132 | 56 | k__Fungi(100);p__Ascomycota(100);c__Sordariomycetes(100);o__Coniochaetales(100);f__Coniochaetaceae(100);g__Coniochaeta(86);s__Coniochaeta_sp(86); | Coniochaeta | 13 | Pathotroph-Saprotroph-Symbiotroph | Animal Pathogen-Dung Saprotroph-Endophyte-Lichen Parasite-Plant Pathogen-\|Undefined Saprotroph\| | Microfungus | NULL | Possible | As foliar_endophyte (Põlme et al. 2020); Decay leaf, fruit, seed, soil, dung, animal material (Põlme et al. 2020) | Bell A. 1983. Dung Fungi: An Illustrated Guide to Coprophilous Fungi in New Zealand. Victoria University Press, Wellington (ISBN: 978-0864730015); Damm U, et al. 2010. Persoonia 24:60-80 ((http://doi.org/10.3767/003158510X500705)); Costa IPMW, et al. 2012. Mycotaxon 119:e26 ((http://mycotaxon.com/resources/checklists/costa_v119_checklist.pdf)); Tedersoo L, et al. 2014. Science 346: 1256688 ((https://doi.org/:10.1126/science.1256688)); Irinyi L, et al. 2015. Medical Mycology 53:313-337 ((https://doi.org/10.1093/mmy/myv008)); Lawrey JD, Diederich P. 2016. http://www.lichenicolous.net (accessed 24 May 2017) ((http://www.lichenicolous.net)); Põlme S, et al. 2020. Fungal Diversity 105:1-16 ((https://doi.org/10.1007/s13225-020-00466-2)) |
| Otu01135 | 56 | k__Fungi(100);p__Basidiomycota(100);c__Tremellomycetes(100);o__Tremellales(100);f__Tremellales_family_Incertae_sedis(100);g__Bandoniozyma(100);s__Bandoniozyma_tunnelae(95); | Bandoniozyma | 13 | Saprotroph | \|Undefined Saprotroph\| | Dimorphic-Facultative Yeast | NULL | Probable | NULL | Tedersoo L, et al. 2014. Science 346:e1256688 ((https://doi.org/10.1126/science.1256688)); Põlme S, et al. 2020. Fungal Diversity 105:1-16 ((https://doi.org/10.1007/s13225-020-00466-2)) |
| Otu01136 | 56 | k__Fungi(100);p__Basidiomycota(100);c__Agaricomycetes(100);o__Thelephorales(100);f__Thelephoraceae(100);g__Tomentella(99);s__Tomentella_sublilacina(99); | Tomentella | 13 | Symbiotroph | \|Ectomycorrhizal\| | Corticioid | NULL | Highly Probable | NULL | Rinaldi AC, et al. 2008. Fungal Diversity 33:1-45 ((http://www.fungaldiversity.org/fdp/sfdp/33-1.pdf)); Tedersoo L, et al. 2010. Mycorrhiza 20:217-263 ((https://doi.org/10.1007/s00572-009-0274-x)); Põlme S, et al. 2020. Fungal Diversity 105:1-16 ((https://doi.org/10.1007/s13225-020-00466-2)); http://mycorrhizas.info/ecmf.html ((http://mycorrhizas.info/ecmf.html)) |
| Otu01137 | 56 | k__Fungi(100);p__Ascomycota(100);c__Ascomycota_class_Incertae_sedis(100);o__Ascomycota_order_Incertae_sedis(100);f__Ascomycota_family_Incertae_sedis(100);g__Rhinocladium(100);s__Rhinocladium_lesnei(100); | Rhinocladium | 13 | Pathotroph | \|Plant Pathogen\| | Microfungus | NULL | Probable | NULL | Põlme S, et al. 2020. Fungal Diversity 105:1-16 ((https://doi.org/10.1007/s13225-020-00466-2)); The American Phytopathological Society Common Names of Plant Diseases ((http://www.apsnet.org/publications/commonnames/Pages/default.aspx)) |
| Otu01139 | 56 | k__Fungi(100);p__Basidiomycota(100);c__Agaricomycetes(100);o__Polyporales(100);f__Phanerochaetaceae(100);g__Ceriporia(100);g__Ceriporia_unclassified(100); | Ceriporia | 13 | Saprotroph | \|Wood Saprotroph\| | Corticioid | White Rot | Highly Probable | As white_rot (Põlme et al. 2020); Decay wood (Põlme et al. 2020) | Tedersoo L, et al. 2014. Science 346:e1256688 ((https://doi.org/10.1126/science.1256688)); Põlme S, et al. 2020. Fungal Diversity 105:1-16 ((https://doi.org/10.1007/s13225-020-00466-2)) |
| Otu01124 | 56 | k__Fungi(100);p__Basidiomycota(100);c__Agaricomycetes(100);c__Agaricomycetes_unclassified(100);c__Agaricomycetes_unclassified(100);c__Agaricomycetes_unclassified(100);c__Agaricomycetes_unclassified(100); | - | - | - | - | - | - | - | Unassigned | - |
| Otu01125 | 56 | k__Fungi(100);p__Ascomycota(74);p__Ascomycota_unclassified(74);p__Ascomycota_unclassified(74);p__Ascomycota_unclassified(74);p__Ascomycota_unclassified(74);p__Ascomycota_unclassified(74); | - | - | - | - | - | - | - | Unassigned | - |
| Otu01126 | 56 | k__Fungi(100);p__Basidiomycota(100);c__Agaricomycetes(100);c__Agaricomycetes_unclassified(100);c__Agaricomycetes_unclassified(100);c__Agaricomycetes_unclassified(100);c__Agaricomycetes_unclassified(100); | - | - | - | - | - | - | - | Unassigned | - |
| Otu01127 | 56 | k__Fungi(100);p__Ascomycota(100);c__Eurotiomycetes(83);c__Eurotiomycetes_unclassified(83);c__Eurotiomycetes_unclassified(83);c__Eurotiomycetes_unclassified(83);c__Eurotiomycetes_unclassified(83); | - | - | - | - | - | - | - | Unassigned | - |
| Otu01130 | 56 | k__Fungi(100);k__Fungi_unclassified(95);k__Fungi_unclassified(95);k__Fungi_unclassified(95);k__Fungi_unclassified(95);k__Fungi_unclassified(95);k__Fungi_unclassified(95); | - | - | - | - | - | - | - | Unassigned | - |
| Otu01131 | 56 | k__Fungi(100);k__Fungi_unclassified(67);k__Fungi_unclassified(67);k__Fungi_unclassified(67);k__Fungi_unclassified(67);k__Fungi_unclassified(67);k__Fungi_unclassified(67); | - | - | - | - | - | - | - | Unassigned | - |
| Otu01133 | 56 | k__Fungi(100);k__Fungi_unclassified(61);k__Fungi_unclassified(61);k__Fungi_unclassified(61);k__Fungi_unclassified(61);k__Fungi_unclassified(61);k__Fungi_unclassified(61); | - | - | - | - | - | - | - | Unassigned | - |
| Otu01134 | 56 | k__Fungi(100);k__Fungi_unclassified(83);k__Fungi_unclassified(83);k__Fungi_unclassified(83);k__Fungi_unclassified(83);k__Fungi_unclassified(83);k__Fungi_unclassified(83); | - | - | - | - | - | - | - | Unassigned | - |
| Otu01138 | 56 | k__Fungi(100);p__Ascomycota(100);c__Sordariomycetes(100);c__Sordariomycetes_unclassified(86);c__Sordariomycetes_unclassified(86);c__Sordariomycetes_unclassified(86);c__Sordariomycetes_unclassified(86); | - | - | - | - | - | - | - | Unassigned | - |
| Otu01140 | 55 | k__Fungi(100);p__Ascomycota(100);c__Sordariomycetes(100);o__Xylariales(100);f__Xylariaceae(100);g__Hypoxylon(100);s__Hypoxylon_perforatum(100); | Hypoxylon | 13 | Saprotroph-Saprotroph-Symbiotroph | Endophyte-Undefined Saprotroph-\|Wood Saprotroph\| | Corticioid-Microfungus-Xylarioid | NULL | Probable | As foliar_endophyte (Põlme et al. 2020); Decay wood, leaf, fruit, seed (Põlme et al. 2020); Host - angiosperms (Põlme et al. 2020) | Tedersoo L, et al. 2014. Science 346:e1256688 ((https://doi.org/10.1126/science.1256688)); Põlme S, et al. 2020. Fungal Diversity 105:1-16 ((https://doi.org/10.1007/s13225-020-00466-2)) |
| Otu01141 | 55 | k__Fungi(100);p__Ascomycota(100);c__Saccharomycetes(100);o__Saccharomycetales(100);f__Metschnikowiaceae(100);g__Metschnikowia(100);s__Metschnikowia_pulcherrima(95); | Metschnikowia | 13 | Saprotroph | \|Undefined Saprotroph\| | Yeast | NULL | Probable | As nectar-tap_saprotroph (Põlme et al. 2020); Decay sugar-rich substrates (Põlme et al. 2020) | Põlme S, et al. 2020. Fungal Diversity 105:1-16 ((https://doi.org/10.1007/s13225-020-00466-2)) |
| Otu01144 | 55 | k__Fungi(100);p__Ascomycota(100);c__Saccharomycetes(100);o__Saccharomycetales(100);f__Metschnikowiaceae(100);g__Metschnikowia(100);s__Metschnikowia_pulcherrima(97); | Metschnikowia | 13 | Saprotroph | \|Undefined Saprotroph\| | Yeast | NULL | Probable | As nectar-tap_saprotroph (Põlme et al. 2020); Decay sugar-rich substrates (Põlme et al. 2020) | Põlme S, et al. 2020. Fungal Diversity 105:1-16 ((https://doi.org/10.1007/s13225-020-00466-2)) |
| Otu01146 | 55 | k__Fungi(100);p__Ascomycota(100);c__Sordariomycetes(100);o__Calosphaeriales(100);f__Calosphaeriaceae(100);g__Jattaea(100);s__Jattaea_prunicola(100); | Jattaea | 13 | Saprotroph | \|Undefined Saprotroph\| | Microfungus | NULL | Probable | NULL | Tedersoo L, et al. 2014. Science 346:e1256688 ((https://doi.org/10.1126/science.1256688)); Põlme S, et al. 2020. Fungal Diversity 105:1-16 ((https://doi.org/10.1007/s13225-020-00466-2)) |
| Otu01142 | 55 | k__Fungi(100);p__Ascomycota(100);p__Ascomycota_unclassified(100);p__Ascomycota_unclassified(100);p__Ascomycota_unclassified(100);p__Ascomycota_unclassified(100);p__Ascomycota_unclassified(100); | - | - | - | - | - | - | - | Unassigned | - |
| Otu01143 | 55 | k__Fungi(100);p__Ascomycota(82);p__Ascomycota_unclassified(82);p__Ascomycota_unclassified(82);p__Ascomycota_unclassified(82);p__Ascomycota_unclassified(82);p__Ascomycota_unclassified(82); | - | - | - | - | - | - | - | Unassigned | - |
| Otu01145 | 55 | k__Fungi(100);p__Ascomycota(79);p__Ascomycota_unclassified(79);p__Ascomycota_unclassified(79);p__Ascomycota_unclassified(79);p__Ascomycota_unclassified(79);p__Ascomycota_unclassified(79); | - | - | - | - | - | - | - | Unassigned | - |
| Otu01147 | 55 | k__Fungi(100);k__Fungi_unclassified(68);k__Fungi_unclassified(68);k__Fungi_unclassified(68);k__Fungi_unclassified(68);k__Fungi_unclassified(68);k__Fungi_unclassified(68); | - | - | - | - | - | - | - | Unassigned | - |
| Otu01148 | 55 | k__Fungi(100);k__Fungi_unclassified(55);k__Fungi_unclassified(55);k__Fungi_unclassified(55);k__Fungi_unclassified(55);k__Fungi_unclassified(55);k__Fungi_unclassified(55); | - | - | - | - | - | - | - | Unassigned | - |
| Otu01149 | 55 | k__Fungi(100);p__Ascomycota(100);c__Dothideomycetes(100);o__Pleosporales(99);o__Pleosporales_unclassified(99);o__Pleosporales_unclassified(99);o__Pleosporales_unclassified(99); | - | - | - | - | - | - | - | Unassigned | - |
| Otu01150 | 54 | k__Fungi(100);p__Basidiomycota(100);c__Agaricomycetes(100);o__Agaricales(100);f__Strophariaceae(100);g__Pholiota(100);s__Pholiota_adiposa(100); | Pholiota | 13 | Pathotroph-Saprotroph | Plant Pathogen-Plant Saprotroph-Undefined Saprotroph-\|Wood Saprotroph\| | Agaricoid | White Rot | Probable | As litter_saprotroph (Põlme et al. 2020); As white_rot (Põlme et al. 2020); As wood_pathogen (Põlme et al. 2020); Decay wood, leaf, fruit, seed (Põlme et al. 2020) | Tedersoo L, et al. 2014. Science 346:e1256688 ((https://doi.org/10.1126/science.1256688)); Põlme S, et al. 2020. Fungal Diversity 105:1-16 ((https://doi.org/10.1007/s13225-020-00466-2)) |
| Otu01154 | 54 | k__Fungi(100);p__Ascomycota(100);c__Saccharomycetes(100);o__Saccharomycetales(100);f__Metschnikowiaceae(100);g__Metschnikowia(100);g__Metschnikowia_unclassified(73); | Metschnikowia | 13 | Saprotroph | \|Undefined Saprotroph\| | Yeast | NULL | Probable | As nectar-tap_saprotroph (Põlme et al. 2020); Decay sugar-rich substrates (Põlme et al. 2020) | Põlme S, et al. 2020. Fungal Diversity 105:1-16 ((https://doi.org/10.1007/s13225-020-00466-2)) |
| Otu01156 | 54 | k__Fungi(100);p__Ascomycota(100);c__Sordariomycetes(56);o__Sordariales(56);f__Cephalothecaceae(56);g__unclassified_Cephalothecaceae(56);s__Cephalothecaceae_sp(56); | Cephalothecaceae | 9 | Saprotroph | Fungal Parasite-Wood Saprotroph | Microfungus | NULL | Probable | Typically saprobic, fruiting on very rotten wood or other fungi (Cannon & Kirk 2007) | Cannon PF, Kirk PM. 2007. Fungal Families of the World. CAB International, Cambridge (ISBN: 978-0851998275) |
| Otu01157 | 54 | k__Fungi(100);p__Ascomycota(100);c__Saccharomycetes(97);o__Saccharomycetales(97);f__Saccharomycetales_family_Incertae_sedis(93);g__Candida(89);g__Candida_unclassified(73); | Candida | 13 | Pathotroph-Saprotroph-Symbiotroph | Animal Pathogen-Endophyte-Endosymbiont-Epiphyte-\|Undefined Saprotroph\| | Dimorphic Yeast-Yeast | NULL | Probable | As nectar-tap_saprotroph (Põlme et al. 2020); Decay sugar-rich substrates (Põlme et al. 2020); This asexual genus is polyphyletic and contain species that function very differently depend on what they are. Caution should be taken when interpreting guilds for Candida species. | Manolakaki D, et al. 2010. Virulence. 1: 367-375 ((https://doi.org/10.4161/viru.1.5.12796)); Mekbib SB, et al. 2011. Tropical Plant Pathology 36:233-240 ((https://doi.org/10.1590/S1982-56762011000400004)); Põlme S, et al. 2020. Fungal Diversity 105:1-16 ((https://doi.org/10.1007/s13225-020-00466-2)) |
| Otu01158 | 54 | k__Fungi(100);p__Basidiomycota(100);c__Tremellomycetes(100);o__Tremellales(100);f__Tremellales_family_Incertae_sedis(100);g__Fibulobasidium(54);s__Fibulobasidium_inconspicuum(54); | Fibulobasidium | 13 | Saprotroph | Undefined Saprotroph-\|Wood Saprotroph\| | Dimorphic-Facultative Yeast | NULL | Highly Probable | Decay wood (Põlme et al. 2020) | Kurtzman CP, et al. (eds.) 2011. The Yeasts, a Taxonomic Study. Fifth Edition. Vols 1-3. Elsevier, San Diego (ISBN: 9780444521491); Põlme S, et al. 2020. Fungal Diversity 105:1-16 ((https://doi.org/10.1007/s13225-020-00466-2)) |
| Otu01160 | 54 | k__Fungi(100);p__Ascomycota(100);c__Sordariomycetes(100);o__Xylariales(100);f__Xylariaceae(100);g__Nemania(100);g__Nemania_unclassified(97); | Nemania | 13 | Saprotroph | Undefined Saprotroph-\|Wood Saprotroph\| | Microfungus | NULL | Probable | Decay wood (Põlme et al. 2020) | Tedersoo L, et al. 2014. Science 346:e1256688 ((https://doi.org/10.1126/science.1256688)); Põlme S, et al. 2020. Fungal Diversity 105:1-16 ((https://doi.org/10.1007/s13225-020-00466-2)) |
| Otu01161 | 54 | k__Fungi(100);p__Ascomycota(100);c__Leotiomycetes(100);o__Erysiphales(100);f__Erysiphaceae(100);g__Erysiphe(100);s__Erysiphe_sp(63); | Erysiphe | 13 | Pathotroph | \|Plant Pathogen\| | Microfungus | NULL | Probable | Biotrophic, with affected plant tissues damaged but usually not killing outright (Cannon & Kirk 2007); Common name - powdery mildew | Tedersoo L, et al. 2014. Science 346:e1256688 ((https://doi.org/10.1126/science.1256688)); Põlme S, et al. 2020. Fungal Diversity 105:1-16 ((https://doi.org/10.1007/s13225-020-00466-2)) |
| Otu01162 | 54 | k__Fungi(100);p__Basidiomycota(100);c__Agaricomycetes(100);o__Polyporales(100);f__Polyporaceae(100);g__Skeletocutis(100);g__Skeletocutis_unclassified(100); | Skeletocutis | 13 | Saprotroph | \|Wood Saprotroph\| | Polyporoid | White Rot | Highly Probable | As white_rot (Põlme et al. 2020); Decay wood (Põlme et al. 2020) | Gilbertson RL, Ryvarden L. 1987-1987. North American Polypores. Fungiflora, Oslo (ISBN: 978-0945345060); Põlme S, et al. 2020. Fungal Diversity 105:1-16 ((https://doi.org/10.1007/s13225-020-00466-2)) |
| Otu01151 | 54 | k__Fungi(100);p__Ascomycota(93);p__Ascomycota_unclassified(93);p__Ascomycota_unclassified(93);p__Ascomycota_unclassified(93);p__Ascomycota_unclassified(93);p__Ascomycota_unclassified(93); | - | - | - | - | - | - | - | Unassigned | - |
| Otu01152 | 54 | k__Fungi(100);p__Ascomycota(100);c__Leotiomycetes(95);c__Leotiomycetes_unclassified(95);c__Leotiomycetes_unclassified(95);c__Leotiomycetes_unclassified(95);c__Leotiomycetes_unclassified(95); | - | - | - | - | - | - | - | Unassigned | - |
| Otu01153 | 54 | k__Fungi(100);p__Basidiomycota(100);c__Agaricomycetes(100);c__Agaricomycetes_unclassified(100);c__Agaricomycetes_unclassified(100);c__Agaricomycetes_unclassified(100);c__Agaricomycetes_unclassified(100); | - | - | - | - | - | - | - | Unassigned | - |
| Otu01155 | 54 | k__Fungi(100);p__Ascomycota(73);p__Ascomycota_unclassified(73);p__Ascomycota_unclassified(73);p__Ascomycota_unclassified(73);p__Ascomycota_unclassified(73);p__Ascomycota_unclassified(73); | - | - | - | - | - | - | - | Unassigned | - |
| Otu01159 | 54 | k__Fungi(100);p__Ascomycota(100);c__Dothideomycetes(100);o__Pleosporales(100);o__Pleosporales_unclassified(67);o__Pleosporales_unclassified(67);o__Pleosporales_unclassified(67); | - | - | - | - | - | - | - | Unassigned | - |
| Otu01164 | 53 | k__Fungi(100);p__Basidiomycota(100);c__Agaricomycetes(100);o__Cantharellales(100);f__Ceratobasidiaceae(100);g__Ceratobasidium(78);g__Ceratobasidium_unclassified(78); | Ceratobasidium | 13 | Pathotroph-Saprotroph-Symbiotroph | Ectomycorrhizal-Endomycorrhizal-Orchid Mycorrhizal-\|Plant Pathogen\|-Plant Saprotroph-Undefined Saprotroph | Corticioid-Microfungus | NULL | Possible | especially of Orchidaceae but also even bryophytes (Cannon & Kirk 2007); As litter_saprotroph (Põlme et al. 2020); Best known as plant pathogens, but also saprobic. Species are endomycorrhizal with a wide range of plant roots ; Decay leaf, fruit, seed, roots, soil (Põlme et al. 2020) | Cannon PF, Kirk PM. 2007. Fungal Families of the World. CAB International, Cambridge (ISBN: 978-0851998275); Tedersoo L, et al. 2014. Science 346:e1256688 ((https://doi.org/10.1126/science.1256688)); Põlme S, et al. 2020. Fungal Diversity 105:1-16 ((https://doi.org/10.1007/s13225-020-00466-2)) |
| Otu01165 | 53 | k__Fungi(100);p__Basidiomycota(100);c__Exobasidiomycetes(100);o__Exobasidiales(100);f__Exobasidiaceae(100);g__Exobasidium(100);g__Exobasidium_unclassified(99); | Exobasidium | 13 | Pathotroph | \|Plant Pathogen\| | Leaf Blister-Leaf Gall-Leaf Spot | NULL | Probable | NULL | Tedersoo L, et al. 2014. Science 346:e1256688 ((https://doi.org/10.1126/science.1256688)); Põlme S, et al. 2020. Fungal Diversity 105:1-16 ((https://doi.org/10.1007/s13225-020-00466-2)) |
| Otu01166 | 53 | k__Fungi(100);p__Ascomycota(100);c__Sordariomycetes(100);o__Hypocreales(100);f__Ophiocordycipitaceae(100);f__Ophiocordycipitaceae_unclassified(97);f__Ophiocordycipitaceae_unclassified(97); | Ophiocordycipitaceae | 9 | Pathotroph-Symbiotroph | Endophyte-Insect Pathogen | Microfungus | NULL | Probable | A family of parasitic fungi (Sung et al. 2007) | Sung GH, et al. 2007. Studies in Mycology 57:5-59 ((https://doi.org/10.3114/sim.2007.57.01)); Costa IPMW, et al. 2012. Mycotaxon 119: e26 ((http://mycotaxon.com/resources/checklists/costa_v119_checklist.pdf)) |
| Otu01169 | 53 | k__Fungi(100);p__Basidiomycota(100);c__Agaricomycetes(100);o__Agaricales(97);o__Agaricales_unclassified(97);o__Agaricales_unclassified(97);o__Agaricales_unclassified(97); | Agaricales | 7 | Pathotroph-Saprotroph-Symbiotroph | Bryophyte Parasite-Dung Saprotroph-Ectomycorrhizal-Fungal Parasite-Leaf Saprotroph-Plant Parasite-Undefined Saprotroph-Wood Saprotroph | Agaricoid-Gasteroid-Microfungus-Secotioid-Yeast | NULL | Possible | Mushrooms and toadstools, Gill fungi, Agarics... lignicolous, sometimes muscicolous or fungicolous, saprobic, mycorrhizal, rarely parasitic on plants or fungi (Kirk et al. 2008) | Kirk PM et al. 2008. Dictionary of the Fungi. Tenth Edition. CAB International, Wallingford (ISBN: 978-0851998268) |
| Otu01170 | 53 | k__Fungi(100);p__Basidiomycota(100);c__Agaricomycetes(100);o__Polyporales(100);f__Fomitopsidaceae(100);g__Antrodia(100);s__Antrodia_xantha(100); | Antrodia | 13 | Saprotroph | \|Wood Saprotroph\| | Corticioid | Brown Rot | Highly Probable | As brown-rot (Põlme et al. 2020); Decay wood (Põlme et al. 2020) | Gilbertson RL, Ryvarden L. 1987-1987. North American Polypores. Fungiflora, Oslo (ISBN: 978-0945345060); Põlme S, et al. 2020. Fungal Diversity 105:1-16 ((https://doi.org/10.1007/s13225-020-00466-2)) |
| Otu01173 | 53 | k__Fungi(100);p__Ascomycota(100);c__Lecanoromycetes(100);o__Lecanorales(100);o__Lecanorales_unclassified(100);o__Lecanorales_unclassified(100);o__Lecanorales_unclassified(100); | Lecanorales | 7 | Pathotroph-Symbiotroph | Lichen Parasite-Lichenized | Microfungus-Thallus | NULL | Probable | Lichenized with green algae, occasionally lichenicolous, on rocks, mosses and wood (Cannon & Kirk 2007); ...an order of mostly lichen-forming fungi (Kirk et al. 2008) | Cannon PF, Kirk PM. 2007. Fungal Families of the World. CAB International, Cambridge (ISBN: 978-0851998275); Kirk PM et al. 2008. Dictionary of the Fungi. Tenth Edition. CAB International, Wallingford (ISBN: 978-0851998268); Esslinger TL. 2014. North Dakota State University ((http://www.ndsu.edu/pubweb/~esslinge/chcklst/chcklst7.htm)) |
| Otu01163 | 53 | k__Fungi(100);p__Basidiomycota(99);c__Agaricomycetes(99);c__Agaricomycetes_unclassified(99);c__Agaricomycetes_unclassified(99);c__Agaricomycetes_unclassified(99);c__Agaricomycetes_unclassified(99); | - | - | - | - | - | - | - | Unassigned | - |
| Otu01167 | 53 | k__Fungi(100);p__Ascomycota(51);p__Ascomycota_unclassified(51);p__Ascomycota_unclassified(51);p__Ascomycota_unclassified(51);p__Ascomycota_unclassified(51);p__Ascomycota_unclassified(51); | - | - | - | - | - | - | - | Unassigned | - |
| Otu01168 | 53 | k__Fungi(100);k__Fungi_unclassified(57);k__Fungi_unclassified(57);k__Fungi_unclassified(57);k__Fungi_unclassified(57);k__Fungi_unclassified(57);k__Fungi_unclassified(57); | - | - | - | - | - | - | - | Unassigned | - |
| Otu01171 | 53 | k__Fungi(100);k__Fungi_unclassified(67);k__Fungi_unclassified(67);k__Fungi_unclassified(67);k__Fungi_unclassified(67);k__Fungi_unclassified(67);k__Fungi_unclassified(67); | - | - | - | - | - | - | - | Unassigned | - |
| Otu01172 | 53 | k__Fungi(100);p__Ascomycota(85);p__Ascomycota_unclassified(85);p__Ascomycota_unclassified(85);p__Ascomycota_unclassified(85);p__Ascomycota_unclassified(85);p__Ascomycota_unclassified(85); | - | - | - | - | - | - | - | Unassigned | - |
| Otu01175 | 52 | k__Fungi(100);p__Ascomycota(100);c__Saccharomycetes(100);o__Saccharomycetales(100);f__Metschnikowiaceae(100);g__Metschnikowia(100);s__Metschnikowia_pulcherrima(97); | Metschnikowia | 13 | Saprotroph | \|Undefined Saprotroph\| | Yeast | NULL | Probable | As nectar-tap_saprotroph (Põlme et al. 2020); Decay sugar-rich substrates (Põlme et al. 2020) | Põlme S, et al. 2020. Fungal Diversity 105:1-16 ((https://doi.org/10.1007/s13225-020-00466-2)) |
| Otu01176 | 52 | k__Fungi(100);p__Ascomycota(100);c__Saccharomycetes(99);o__Saccharomycetales(99);f__Saccharomycetales_family_Incertae_sedis(95);g__Candida(91);g__Candida_unclassified(75); | Candida | 13 | Pathotroph-Saprotroph-Symbiotroph | Animal Pathogen-Endophyte-Endosymbiont-Epiphyte-\|Undefined Saprotroph\| | Dimorphic Yeast-Yeast | NULL | Probable | As nectar-tap_saprotroph (Põlme et al. 2020); Decay sugar-rich substrates (Põlme et al. 2020); This asexual genus is polyphyletic and contain species that function very differently depend on what they are. Caution should be taken when interpreting guilds for Candida species. | Manolakaki D, et al. 2010. Virulence. 1: 367-375 ((https://doi.org/10.4161/viru.1.5.12796)); Mekbib SB, et al. 2011. Tropical Plant Pathology 36:233-240 ((https://doi.org/10.1590/S1982-56762011000400004)); Põlme S, et al. 2020. Fungal Diversity 105:1-16 ((https://doi.org/10.1007/s13225-020-00466-2)) |
| Otu01178 | 52 | k__Fungi(100);p__Ascomycota(100);c__Dothideomycetes(100);o__Pleosporales(100);f__Pleosporales_family_Incertae_sedis(77);g__Phoma(77);s__Phoma_cladoniicola(56); | Phoma | 13 | Pathotroph-Saprotroph-Symbiotroph | Dung Saprotroph-Endophyte-Lichen Parasite-\|Plant Pathogen\|-Plant Saprotroph-Undefined Saprotroph-Wood Saprotroph | Microfungus | Soft Rot | Possible | As foliar_endophyte (Põlme et al. 2020); As litter_saprotroph (Põlme et al. 2020); Decay leaf, fruit, seed, roots, soil (Põlme et al. 2020) | Richardson MJ. 1972. Transactions of the British Mycological Society 58:37-48 ((https://doi.org/10.1016/S0007-1536(72)80069-X)); Seehann G, et al. 1975. List of Fungi in Soft-Rot Tissue Tests. Institute fur Holzbiologie und Holzschutz, Bundesforschungsanstalt fur Forst - und Holzwirtshaft, Hamburg - Lohbrugge, Germany; de Gruyter J, et al. 2010. Mycologia 102:1066-1081 ((https://doi.org/10.3852/09-240)); Costa IPMW, et al. 2012. Mycotaxon 119:e26 ((http://mycotaxon.com/resources/checklists/costa_v119_checklist.pdf)); Tedersoo L, et al. 2014. Science 346:e1256688 ((https://doi.org/10.1126/science.1256688)); Lawrey JD, Diederich P. 2016. http://www.lichenicolous.net (accessed 24 May 2017) ((http://www.lichenicolous.net)); Põlme S, et al. 2020. Fungal Diversity 105:1-16 ((https://doi.org/10.1007/s13225-020-00466-2)) |
| Otu01181 | 52 | k__Fungi(100);p__Ascomycota(100);c__Dothideomycetes(97);o__Capnodiales(97);f__Mycosphaerellaceae(97);f__Mycosphaerellaceae_unclassified(97);f__Mycosphaerellaceae_unclassified(97); | Mycosphaerellaceae | 9 | Pathotroph-Saprotroph | Plant Pathogen-Undefined Saprotroph | Microfungus | NULL | Probable | Biotrophic, necrotrophic or saprobic on various plant tissues (Cannon & Kirk 2007) | Cannon PF, Kirk PM. 2007. Fungal Families of the World. CAB International, Cambridge (ISBN: 978-0851998275) |
| Otu01182 | 52 | k__Fungi(100);p__Ascomycota(100);c__Sordariomycetes(100);o__Sordariomycetes_order_Incertae_sedis(100);f__Sordariomycetes_family_Incertae_sedis(100);g__Myrmecridium(100);s__Myrmecridium_schulzeri(66); | Myrmecridium | 13 | Saprotroph | \|Undefined Saprotroph\| | Microfungus | NULL | Probable | As soil_saprotroph (Põlme et al. 2020); Decay soil (Põlme et al. 2020) | Tedersoo L, et al. 2014. Science 346:e1256688 ((https://doi.org/10.1126/science.1256688)); Põlme S, et al. 2020. Fungal Diversity 105:1-16 ((https://doi.org/10.1007/s13225-020-00466-2)) |
| Otu01183 | 52 | k__Fungi(100);p__Basidiomycota(100);c__Agaricomycetes(100);o__Hymenochaetales(100);f__Hymenochaetaceae(100);g__Tubulicrinis(100);s__Tubulicrinis_sp(100); | Tubulicrinis | 13 | Saprotroph | Undefined Saprotroph-\|Wood Saprotroph\| | Corticioid | White Rot | Probable | As white_rot (Põlme et al. 2020); Decay wood (Põlme et al. 2020) | Tedersoo L, et al. 2014. Science 346:e1256688 ((https://doi.org/10.1126/science.1256688)); Põlme S, et al. 2020. Fungal Diversity 105:1-16 ((https://doi.org/10.1007/s13225-020-00466-2)) |
| Otu01185 | 52 | k__Fungi(100);p__Ascomycota(100);c__Sordariomycetes(100);o__Sordariomycetes_order_Incertae_sedis(100);f__Annulatascaceae(100);g__Rhodoveronaea(100);s__Rhodoveronaea_varioseptata(100); | Rhodoveronaea | 13 | Saprotroph | \|Undefined Saprotroph\| | Microfungus | NULL | Probable | NULL | Tedersoo L, et al. 2014. Science 346:e1256688 ((https://doi.org/10.1126/science.1256688)); Põlme S, et al. 2020. Fungal Diversity 105:1-16 ((https://doi.org/10.1007/s13225-020-00466-2)) |
| Otu01187 | 52 | k__Fungi(100);p__Ascomycota(100);c__Saccharomycetes(100);o__Saccharomycetales(100);f__Metschnikowiaceae(100);g__Metschnikowia(100);s__Metschnikowia_pulcherrima(97); | Metschnikowia | 13 | Saprotroph | \|Undefined Saprotroph\| | Yeast | NULL | Probable | As nectar-tap_saprotroph (Põlme et al. 2020); Decay sugar-rich substrates (Põlme et al. 2020) | Põlme S, et al. 2020. Fungal Diversity 105:1-16 ((https://doi.org/10.1007/s13225-020-00466-2)) |
| Otu01174 | 52 | k__Fungi(100);p__Ascomycota(100);c__Pezizomycetes(93);o__Pezizales(93);o__Pezizales_unclassified(93);o__Pezizales_unclassified(93);o__Pezizales_unclassified(93); | - | - | - | - | - | - | - | Unassigned | - |
| Otu01177 | 52 | k__Fungi(100);k__Fungi_unclassified(66);k__Fungi_unclassified(66);k__Fungi_unclassified(66);k__Fungi_unclassified(66);k__Fungi_unclassified(66);k__Fungi_unclassified(66); | - | - | - | - | - | - | - | Unassigned | - |
| Otu01179 | 52 | k__Fungi(100);p__Ascomycota(100);c__Dothideomycetes(100);c__Dothideomycetes_unclassified(100);c__Dothideomycetes_unclassified(100);c__Dothideomycetes_unclassified(100);c__Dothideomycetes_unclassified(100); | - | - | - | - | - | - | - | Unassigned | - |
| Otu01180 | 52 | k__Fungi(100);p__Ascomycota(58);p__Ascomycota_unclassified(58);p__Ascomycota_unclassified(58);p__Ascomycota_unclassified(58);p__Ascomycota_unclassified(58);p__Ascomycota_unclassified(58); | - | - | - | - | - | - | - | Unassigned | - |
| Otu01184 | 52 | k__Fungi(100);p__Ascomycota(56);p__Ascomycota_unclassified(56);p__Ascomycota_unclassified(56);p__Ascomycota_unclassified(56);p__Ascomycota_unclassified(56);p__Ascomycota_unclassified(56); | - | - | - | - | - | - | - | Unassigned | - |
| Otu01186 | 52 | k__Fungi(100);p__Basidiomycota(100);c__Pucciniomycetes(100);o__Pucciniales(100);f__Pucciniastraceae(100);f__Pucciniastraceae_unclassified(99);f__Pucciniastraceae_unclassified(99); | - | - | - | - | - | - | - | Unassigned | - |
| Otu01188 | 52 | k__Fungi(100);p__Basidiomycota(100);c__Agaricomycetes(100);o__Polyporales(100);f__Phanerochaetaceae(100);g__unclassified_Phanerochaetaceae(100);s__Phanerochaetaceae_sp(100); | - | - | - | - | - | - | - | Unassigned | - |
| Otu01189 | 51 | k__Fungi(100);p__Basidiomycota(100);c__Agaricomycetes(100);o__Cantharellales(100);f__Hydnaceae(100);g__Sistotrema(99);s__Sistotrema_sp(53); | Sistotrema | 13 | Saprotroph-Symbiotroph | Ectomycorrhizal-\|Plant Saprotroph\|-Wood Saprotroph | Corticioid | White Rot | Possible | As litter_saprotroph (Põlme et al. 2020); Decay leaf, fruit, seed (Põlme et al. 2020) | Gilbertson RL, Ryvarden L. 1987-1987. North American Polypores. Fungiflora, Oslo (ISBN: 978-0945345060); Rinaldi AC, et al. 2008. Fungal Diversity 33:1-45 ((http://www.fungaldiversity.org/fdp/sfdp/33-1.pdf)) (as ECM/not ECM); Tedersoo L, et al. 2010. Mycorrhiza 20:217-263 ((https://doi.org/10.1007/s00572-009-0274-x)) (pro parte); Põlme S, et al. 2020. Fungal Diversity 105:1-16 ((https://doi.org/10.1007/s13225-020-00466-2)); http://mycorrhizas.info/ecmf.html ((http://mycorrhizas.info/ecmf.html)) (as ECM/not ECM). |
| Otu01190 | 51 | k__Fungi(100);p__Ascomycota(100);c__Sordariomycetes(100);o__Xylariales(100);f__Xylariaceae(100);f__Xylariaceae_unclassified(73);f__Xylariaceae_unclassified(73); | Xylariaceae | 9 | Pathotroph-Saprotroph-Symbiotroph | Dung Saprotroph-Endophyte-Plant Pathogen-Undefined Saprotroph | Xylarioid | NULL | Probable | Saprobes or weak pathogens, many endophytes, mainly in wood and bark, occasionally in soil and dung (Cannon & Kirk 2007) | Cannon PF, Kirk PM. 2007. Fungal Families of the World. CAB International, Cambridge (ISBN: 978-0851998275) |
| Otu01192 | 51 | k__Fungi(100);p__Basidiomycota(100);c__Tremellomycetes(100);o__Filobasidiales(100);f__Filobasidiaceae(100);g__Cryptococcus(100);s__Cryptococcus_flavus(100); | Cryptococcus | 13 | Pathotroph-Saprotroph-Symbiotroph | Animal Pathogen-Endophyte-Epiphyte-\|Undefined Saprotroph\| | Dimorphic Yeast-Yeast | NULL | Possible | As foliar_endophyte (Põlme et al. 2020); Decay leaf, fruit, seed, soil, dung, animal material (Põlme et al. 2020) | Andrews JH, et al. 1983. Phytopathology 73:228-234 ((https://doi.org/10.1094/Phyto-73-228)); Kurtzman CP, et al. (eds.) 2011. The Yeasts, a Taxonomic Study. Fifth Edition. Vols 1-3. Elsevier, San Diego (ISBN: 9780444521491); Põlme S, et al. 2020. Fungal Diversity 105:1-16 ((https://doi.org/10.1007/s13225-020-00466-2)) |
| Otu01193 | 51 | k__Fungi(100);p__Basidiomycota(100);c__Agaricomycetes(100);o__Agaricales(100);f__Entolomataceae(100);g__Entoloma(100);s__Entoloma_sp_AJ108(100); | Entoloma | 13 | Pathotroph-Saprotroph-Symbiotroph | Ectomycorrhizal-Fungal Parasite-\|Undefined Saprotroph\|-Undefined Symbiotroph | Agaricoid | NULL | Possible | As soil_saprotroph (Põlme et al. 2020); Decay roots, soil (Põlme et al. 2020) | Rinaldi AC, et al. 2008. Fungal Diversity 33:1-45 ((http://www.fungaldiversity.org/fdp/sfdp/33-1.pdf)) (as ECM/not ECM); Tedersoo L, et al. 2010. Mycorrhiza 20:217-263 ((https://doi.org/10.1007/s00572-009-0274-x; Põlme S, et al. 2020. Fungal Diversity 105:1-16 ((https://doi.org/10.1007/s13225-020-00466-2)); sensu stricto)); http://mycorrhizas.info/ecmf.html ((http://mycorrhizas.info/ecmf.html)) ((as ECM/not ECM)) |
| Otu01194 | 51 | k__Fungi(100);p__Ascomycota(100);c__Leotiomycetes(100);o__Helotiales(100);f__Helotiales_family_Incertae_sedis(100);g__Leptodontidium(100);g__Leptodontidium_unclassified(51); | Leptodontidium | 13 | Saprotroph-Symbiotroph | Endophyte-Ericoid Mycorrhizal-\|Plant Saprotroph\| | Microfungus | NULL | Highly Probable | As litter_saprotroph (Põlme et al. 2020); As root_endophyte_dark_septate (Põlme et al. 2020); Dark Septate Endophyte, known from plant roots.; Decay leaf, fruit, seed (Põlme et al. 2020) | Jumpponen A, Trappe J. 1998. New Phytologist 140:295-310 ((https://doi.org/10.1046/j.1469-8137.1998.00265.x)) (as DSE); Taylor DL, et al. 2014. Ecological Monographs 8:43-20 ((https://esajournals.onlinelibrary.wiley.com/doi/10.1890/12-1693.1)); Põlme S, et al. 2020. Fungal Diversity 105:1-16 ((https://doi.org/10.1007/s13225-020-00466-2)) |
| Otu01197 | 51 | k__Fungi(100);p__Basidiomycota(100);c__Agaricomycetes(100);o__Agaricales(100);f__Psathyrellaceae(100);g__Coprinopsis(100);s__Coprinopsis_calospora(100); | Coprinopsis | 13 | Saprotroph | \|Undefined Saprotroph\| | Agaricoid | NULL | Probable | As soil_saprotroph (Põlme et al. 2020); Decay soil (Põlme et al. 2020) | Tedersoo L, et al. 2014. Science 346:e1256688 ((https://doi.org/10.1126/science.1256688)); Põlme S, et al. 2020. Fungal Diversity 105:1-16 ((https://doi.org/10.1007/s13225-020-00466-2)) |
| Otu01198 | 51 | k__Fungi(100);p__Ascomycota(100);c__Saccharomycetes(100);o__Saccharomycetales(100);f__Metschnikowiaceae(100);g__Metschnikowia(100);s__Metschnikowia_pulcherrima(95); | Metschnikowia | 13 | Saprotroph | \|Undefined Saprotroph\| | Yeast | NULL | Probable | As nectar-tap_saprotroph (Põlme et al. 2020); Decay sugar-rich substrates (Põlme et al. 2020) | Põlme S, et al. 2020. Fungal Diversity 105:1-16 ((https://doi.org/10.1007/s13225-020-00466-2)) |
| Otu01203 | 51 | k__Fungi(100);p__Ascomycota(100);c__Sordariomycetes(100);o__Diaporthales(100);f__Valsaceae(100);f__Valsaceae_unclassified(99);f__Valsaceae_unclassified(99); | Valsaceae | 9 | Pathotroph-Saprotroph-Symbiotroph | Endophyte-Plant Pathogen-Undefined Saprotroph | Microfungus | NULL | Probable | Saprobes, endophytes and weak to virulent parasites of woody plants (Cannon & Kirk 2007) | Cannon PF, Kirk PM. 2007. Fungal Families of the World. CAB International, Cambridge (ISBN: 978-0851998275) |
| Otu01191 | 51 | k__Fungi(100);k__Fungi_unclassified(65);k__Fungi_unclassified(65);k__Fungi_unclassified(65);k__Fungi_unclassified(65);k__Fungi_unclassified(65);k__Fungi_unclassified(65); | - | - | - | - | - | - | - | Unassigned | - |
| Otu01195 | 51 | k__Fungi(100);k__Fungi_unclassified(61);k__Fungi_unclassified(61);k__Fungi_unclassified(61);k__Fungi_unclassified(61);k__Fungi_unclassified(61);k__Fungi_unclassified(61); | - | - | - | - | - | - | - | Unassigned | - |
| Otu01196 | 51 | k__Fungi(100);k__Fungi_unclassified(75);k__Fungi_unclassified(75);k__Fungi_unclassified(75);k__Fungi_unclassified(75);k__Fungi_unclassified(75);k__Fungi_unclassified(75); | - | - | - | - | - | - | - | Unassigned | - |
| Otu01199 | 51 | k__Fungi(100);p__Ascomycota(100);c__Dothideomycetes(100);o__Pleosporales(100);o__Pleosporales_unclassified(100);o__Pleosporales_unclassified(100);o__Pleosporales_unclassified(100); | - | - | - | - | - | - | - | Unassigned | - |
| Otu01200 | 51 | k__Fungi(100);p__Ascomycota(95);p__Ascomycota_unclassified(95);p__Ascomycota_unclassified(95);p__Ascomycota_unclassified(95);p__Ascomycota_unclassified(95);p__Ascomycota_unclassified(95); | - | - | - | - | - | - | - | Unassigned | - |
| Otu01201 | 51 | k__Fungi(100);k__Fungi_unclassified(79);k__Fungi_unclassified(79);k__Fungi_unclassified(79);k__Fungi_unclassified(79);k__Fungi_unclassified(79);k__Fungi_unclassified(79); | - | - | - | - | - | - | - | Unassigned | - |
| Otu01202 | 51 | k__Fungi(100);k__Fungi_unclassified(69);k__Fungi_unclassified(69);k__Fungi_unclassified(69);k__Fungi_unclassified(69);k__Fungi_unclassified(69);k__Fungi_unclassified(69); | - | - | - | - | - | - | - | Unassigned | - |
| Otu01207 | 50 | k__Fungi(100);p__Zygomycota(100);c__Zygomycota_class_Incertae_sedis(100);o__Mortierellales(100);f__Mortierellaceae(100);g__Mortierella(100);g__Mortierella_unclassified(98); | Mortierella | 13 | Saprotroph-Symbiotroph | Endophyte-Plant Saprotroph-\|Undefined Saprotroph\| | Microfungus | NULL | Probable | As chitinolytic (Põlme et al. 2020); As soil_saprotroph (Põlme et al. 2020); Decay roots, soil (Põlme et al. 2020); Saprobic in soil and thus frequenctly isolated from soil and material contaminated with soil such as hay or rotten silage and spoil tips (Cannon & Kirk 2007) | Cannon PF, Kirk PM. 2007. Fungal Families of the World. CAB International, Cambridge (ISBN: 978-0851998275); Tedersoo L, et al. 2014. Science 346:e1256688 ((https://doi.org/10.1126/science.1256688)); Purahong W, et al. 2016. Molecular Ecology 25:4059-4074 ((https://doi.org/10.1111/mec.13739)); Põlme S, et al. 2020. Fungal Diversity 105:1-16 ((https://doi.org/10.1007/s13225-020-00466-2)) |
| Otu01208 | 50 | k__Fungi(100);p__Basidiomycota(100);c__Agaricomycetes(100);o__Boletales(100);f__Boletaceae(96);g__Boletus(92);g__Boletus_unclassified(92); | Boletus | 13 | Pathotroph-Symbiotroph | \|Ectomycorrhizal\|-Fungal Parasite | Agaricoid-Boletoid | NULL | Highly Probable | NULL | Rinaldi AC, et al. 2008. Fungal Diversity 33:1-45 ((http://www.fungaldiversity.org/fdp/sfdp/33-1.pdf)); Tedersoo L, et al. 2010. Mycorrhiza 20:217-263 ((https://doi.org/10.1007/s00572-009-0274-x)); Põlme S, et al. 2020. Fungal Diversity 105:1-16 ((https://doi.org/10.1007/s13225-020-00466-2)); http://mycorrhizas.info/ecmf.html ((http://mycorrhizas.info/ecmf.html)); List compiled by John Plischke III ((https://goodmushroom.weebly.com/uploads/5/7/1/7/57176535/fungi_on_fungi_pdf_lq.pdf)) |
| Otu01211 | 50 | k__Fungi(100);p__Ascomycota(100);c__Saccharomycetes(100);o__Saccharomycetales(100);f__Metschnikowiaceae(100);g__Metschnikowia(100);s__Metschnikowia_pulcherrima(92); | Metschnikowia | 13 | Saprotroph | \|Undefined Saprotroph\| | Yeast | NULL | Probable | As nectar-tap_saprotroph (Põlme et al. 2020); Decay sugar-rich substrates (Põlme et al. 2020) | Põlme S, et al. 2020. Fungal Diversity 105:1-16 ((https://doi.org/10.1007/s13225-020-00466-2)) |
| Otu01212 | 50 | k__Fungi(100);p__Basidiomycota(100);c__Agaricomycetes(100);o__Agaricales(100);f__Psathyrellaceae(100);g__Coprinopsis(100);s__Coprinopsis_radiata(86); | Coprinopsis | 13 | Saprotroph | \|Undefined Saprotroph\| | Agaricoid | NULL | Probable | As soil_saprotroph (Põlme et al. 2020); Decay soil (Põlme et al. 2020) | Tedersoo L, et al. 2014. Science 346:e1256688 ((https://doi.org/10.1126/science.1256688)); Põlme S, et al. 2020. Fungal Diversity 105:1-16 ((https://doi.org/10.1007/s13225-020-00466-2)) |
| Otu01214 | 50 | k__Fungi(100);p__Ascomycota(100);c__Sordariomycetes(100);o__Xylariales(100);f__Xylariaceae(100);g__Hypoxylon(100);s__Hypoxylon_cercidicola(100); | Hypoxylon | 13 | Saprotroph-Saprotroph-Symbiotroph | Endophyte-Undefined Saprotroph-\|Wood Saprotroph\| | Corticioid-Microfungus-Xylarioid | NULL | Probable | As foliar_endophyte (Põlme et al. 2020); Decay wood, leaf, fruit, seed (Põlme et al. 2020); Host - angiosperms (Põlme et al. 2020) | Tedersoo L, et al. 2014. Science 346:e1256688 ((https://doi.org/10.1126/science.1256688)); Põlme S, et al. 2020. Fungal Diversity 105:1-16 ((https://doi.org/10.1007/s13225-020-00466-2)) |
| Otu01215 | 50 | k__Fungi(100);p__Basidiomycota(100);c__Agaricomycetes(100);o__Trechisporales(100);f__Hydnodontaceae(100);g__Trechispora(100);s__Trechispora_sp_DLL2009_177(100); | Trechispora | 13 | Saprotroph | \|Wood Saprotroph\| | Corticioid | White Rot | Highly Probable | As white_rot (Põlme et al. 2020); Decay wood (Põlme et al. 2020) | Gilbertson RL, Ryvarden L. 1987-1987. North American Polypores. Fungiflora, Oslo (ISBN: 978-0945345060); Rinaldi AC, et al. 2008. Fungal Diversity 33:1-45 ((http://www.fungaldiversity.org/fdp/sfdp/33-1.pdf)) (as Uncertain); Tedersoo L, et al. 2010. Mycorrhiza 20:217-263 ((https://doi.org/10.1007/s00572-009-0274-x)) (as not ECM).; Põlme S, et al. 2020. Fungal Diversity 105:1-16 ((https://doi.org/10.1007/s13225-020-00466-2)) |
| Otu01219 | 50 | k__Fungi(100);p__Basidiomycota(100);c__Tremellomycetes(100);o__Tremellales(100);f__Tremellales_family_Incertae_sedis(100);g__Tremella(100);s__Tremella_sp_BEA_2010(100); | Tremella | 13 | Pathotroph | \|Fungal Parasite\|-Lichen Parasite | Dimorphic-Facultative Yeast-Microfungus-Tremelloid | NULL | Probable | As mycoparasite (Põlme et al. 2020); Decay fungal material (Põlme et al. 2020) | Tedersoo L, et al. 2014. Science 346:e1256688 ((https://doi.org/10.1126/science.1256688)); Põlme S, et al. 2020. Fungal Diversity 105:1-16 ((https://doi.org/10.1007/s13225-020-00466-2)) |
| Otu01221 | 50 | k__Fungi(100);p__Basidiomycota(100);c__Agaricomycetes(100);o__Polyporales(100);f__Phanerochaetaceae(100);g__Rhizochaete(100);s__Rhizochaete_filamentosa(100); | Rhizochaete | 13 | Saprotroph | Undefined Saprotroph-\|Wood Saprotroph\| | Corticioid | White Rot | Probable | As white_rot (Põlme et al. 2020); Decay wood (Põlme et al. 2020) | Tedersoo L, et al. 2014. Science 346:e1256688 ((https://doi.org/10.1126/science.1256688)); Põlme S, et al. 2020. Fungal Diversity 105:1-16 ((https://doi.org/10.1007/s13225-020-00466-2)) |
| Otu01227 | 50 | k__Fungi(100);p__Basidiomycota(100);c__Agaricomycetes(100);o__Polyporales(100);f__Polyporaceae(100);g__Fomes(100);s__Fomes_fomentarius(100); | Fomes | 13 | Pathotroph-Saprotroph | Plant Pathogen-\|Wood Saprotroph\| | Polyporoid | White Rot | Highly Probable | As white_rot (Põlme et al. 2020); As wood_pathogen (Põlme et al. 2020); Decay wood (Põlme et al. 2020) | Gilbertson RL, Ryvarden L. 1987-1987. North American Polypores. Fungiflora, Oslo (ISBN: 978-0945345060); Põlme S, et al. 2020. Fungal Diversity 105:1-16 ((https://doi.org/10.1007/s13225-020-00466-2)) |
| Otu01204 | 50 | k__Fungi(100);k__Fungi_unclassified(72);k__Fungi_unclassified(72);k__Fungi_unclassified(72);k__Fungi_unclassified(72);k__Fungi_unclassified(72);k__Fungi_unclassified(72); | - | - | - | - | - | - | - | Unassigned | - |
| Otu01205 | 50 | k__Fungi(100);p__Ascomycota(100);c__Dothideomycetes(100);o__Capnodiales(100);f__Capnodiales_family_Incertae_sedis(96);f__Capnodiales_family_Incertae_sedis_unclassified(96);f__Capnodiales_family_Incertae_sedis_unclassified(96); | - | - | - | - | - | - | - | Unassigned | - |
| Otu01206 | 50 | k__Fungi(100);p__Basidiomycota(100);c__Agaricomycetes(100);o__Auriculariales(100);f__Auriculariales_family_Incertae_sedis(100);g__unclassified_Auriculariales(100);s__Auriculariales_sp(100); | - | - | - | - | - | - | - | Unassigned | - |
| Otu01209 | 50 | k__Fungi(100);p__Ascomycota(100);p__Ascomycota_unclassified(58);p__Ascomycota_unclassified(58);p__Ascomycota_unclassified(58);p__Ascomycota_unclassified(58);p__Ascomycota_unclassified(58); | - | - | - | - | - | - | - | Unassigned | - |
| Otu01210 | 50 | k__Fungi(100);p__Basidiomycota(100);c__Agaricomycetes(100);c__Agaricomycetes_unclassified(100);c__Agaricomycetes_unclassified(100);c__Agaricomycetes_unclassified(100);c__Agaricomycetes_unclassified(100); | - | - | - | - | - | - | - | Unassigned | - |
| Otu01213 | 50 | k__Fungi(100);k__Fungi_unclassified(61);k__Fungi_unclassified(61);k__Fungi_unclassified(61);k__Fungi_unclassified(61);k__Fungi_unclassified(61);k__Fungi_unclassified(61); | - | - | - | - | - | - | - | Unassigned | - |
| Otu01216 | 50 | k__Fungi(100);p__Ascomycota(100);c__Dothideomycetes(100);o__Pleosporales(100);o__Pleosporales_unclassified(98);o__Pleosporales_unclassified(98);o__Pleosporales_unclassified(98); | - | - | - | - | - | - | - | Unassigned | - |
| Otu01217 | 50 | k__Fungi(100);p__Ascomycota(100);c__Sordariomycetes(100);c__Sordariomycetes_unclassified(100);c__Sordariomycetes_unclassified(100);c__Sordariomycetes_unclassified(100);c__Sordariomycetes_unclassified(100); | - | - | - | - | - | - | - | Unassigned | - |
| Otu01218 | 50 | k__Fungi(100);p__Basidiomycota(100);p__Basidiomycota_unclassified(100);p__Basidiomycota_unclassified(100);p__Basidiomycota_unclassified(100);p__Basidiomycota_unclassified(100);p__Basidiomycota_unclassified(100); | - | - | - | - | - | - | - | Unassigned | - |
| Otu01220 | 50 | k__Fungi(100);p__Ascomycota(100);c__Dothideomycetes(100);c__Dothideomycetes_unclassified(100);c__Dothideomycetes_unclassified(100);c__Dothideomycetes_unclassified(100);c__Dothideomycetes_unclassified(100); | - | - | - | - | - | - | - | Unassigned | - |
| Otu01222 | 50 | k__Fungi(100);p__Ascomycota(100);c__Dothideomycetes(100);o__Dothideales(100);f__Dothioraceae(100);g__unclassified_Dothioraceae(100);s__Dothioraceae_sp(100); | - | - | - | - | - | - | - | Unassigned | - |
| Otu01223 | 50 | k__Fungi(100);k__Fungi_unclassified(94);k__Fungi_unclassified(94);k__Fungi_unclassified(94);k__Fungi_unclassified(94);k__Fungi_unclassified(94);k__Fungi_unclassified(94); | - | - | - | - | - | - | - | Unassigned | - |
| Otu01224 | 50 | k__Fungi(100);p__Ascomycota(100);p__Ascomycota_unclassified(94);p__Ascomycota_unclassified(94);p__Ascomycota_unclassified(94);p__Ascomycota_unclassified(94);p__Ascomycota_unclassified(94); | - | - | - | - | - | - | - | Unassigned | - |
| Otu01225 | 50 | k__Fungi(100);k__Fungi_unclassified(62);k__Fungi_unclassified(62);k__Fungi_unclassified(62);k__Fungi_unclassified(62);k__Fungi_unclassified(62);k__Fungi_unclassified(62); | - | - | - | - | - | - | - | Unassigned | - |
| Otu01226 | 50 | k__Fungi(100);k__Fungi_unclassified(66);k__Fungi_unclassified(66);k__Fungi_unclassified(66);k__Fungi_unclassified(66);k__Fungi_unclassified(66);k__Fungi_unclassified(66); | - | - | - | - | - | - | - | Unassigned | - |
| Otu01228 | 49 | k__Fungi(100);p__Ascomycota(100);c__Saccharomycetes(100);o__Saccharomycetales(100);f__Metschnikowiaceae(100);g__Metschnikowia(100);s__Metschnikowia_pulcherrima(90); | Metschnikowia | 13 | Saprotroph | \|Undefined Saprotroph\| | Yeast | NULL | Probable | As nectar-tap_saprotroph (Põlme et al. 2020); Decay sugar-rich substrates (Põlme et al. 2020) | Põlme S, et al. 2020. Fungal Diversity 105:1-16 ((https://doi.org/10.1007/s13225-020-00466-2)) |
| Otu01229 | 49 | k__Fungi(100);p__Basidiomycota(100);c__Exobasidiomycetes(100);o__Exobasidiales(100);f__Exobasidiaceae(100);g__Exobasidium(100);g__Exobasidium_unclassified(68); | Exobasidium | 13 | Pathotroph | \|Plant Pathogen\| | Leaf Blister-Leaf Gall-Leaf Spot | NULL | Probable | NULL | Tedersoo L, et al. 2014. Science 346:e1256688 ((https://doi.org/10.1126/science.1256688)); Põlme S, et al. 2020. Fungal Diversity 105:1-16 ((https://doi.org/10.1007/s13225-020-00466-2)) |
| Otu01230 | 49 | k__Fungi(100);p__Ascomycota(100);c__Ascomycota_class_Incertae_sedis(100);o__Ascomycota_order_Incertae_sedis(100);f__Ascomycota_family_Incertae_sedis(100);g__Cryptostroma(100);s__Cryptostroma_corticale(100); | Cryptostroma | 13 | Pathotroph-Saprotroph | \|Plant Pathogen\|-Wood Saprotroph | Microfungus | NULL | Possible | As wood_pathogen (Põlme et al. 2020); Decay wood (Põlme et al. 2020); Host - Acer (Põlme et al. 2020) | Gregory PH, Waller S.1951. Transactions of the British Mycological Society 34:579-597 ((https://doi.org/10.1016/S0007-1536((51))80043-3)); Põlme S, et al. 2020. Fungal Diversity 105:1-16 ((https://doi.org/10.1007/s13225-020-00466-2)) |
| Otu01231 | 49 | k__Fungi(100);p__Ascomycota(100);c__Dothideomycetes(100);o__Pleosporales(100);f__Pleosporaceae(100);f__Pleosporaceae_unclassified(78);f__Pleosporaceae_unclassified(78); | Pleosporaceae | 9 | Pathotroph-Saprotroph | Endophyte-Lichen Parasite-Plant Pathogen-Undefined Saprotroph | Microfungus | NULL | Probable | Necrotrophic pathogens and saprobes, especially associated with grasses. Many species are seed-borne (Cannon & Kirk 2007) | Cannon PF, Kirk PM. 2007. Fungal Families of the World. CAB International, Cambridge (ISBN: 978-0851998275); David AS, et al. 2016. Microbial Ecology 71:912-926 ((https://doi.org/10.1007/s00248-015-0712-6)); Lawrey JD, Diederich P. 2016. http://www.lichenicolous.net (accessed 24 May 2017) ((http://www.lichenicolous.net)) |
| Otu01233 | 49 | k__Fungi(100);p__Ascomycota(100);c__Saccharomycetes(100);o__Saccharomycetales(100);f__Metschnikowiaceae(100);g__Metschnikowia(100);s__Metschnikowia_pulcherrima(98); | Metschnikowia | 13 | Saprotroph | \|Undefined Saprotroph\| | Yeast | NULL | Probable | As nectar-tap_saprotroph (Põlme et al. 2020); Decay sugar-rich substrates (Põlme et al. 2020) | Põlme S, et al. 2020. Fungal Diversity 105:1-16 ((https://doi.org/10.1007/s13225-020-00466-2)) |
| Otu01234 | 49 | k__Fungi(100);p__Basidiomycota(100);c__Agaricomycetes(100);o__Polyporales(100);f__Meruliaceae(100);g__Junghuhnia(100);s__Junghuhnia_luteoalba(100); | Junghuhnia | 13 | Saprotroph | \|Wood Saprotroph\| | Corticioid | White Rot | Highly Probable | As white_rot (Põlme et al. 2020); Decay wood (Põlme et al. 2020) | Gilbertson RL, Ryvarden L. 1987-1987. North American Polypores. Fungiflora, Oslo (ISBN: 978-0945345060); Põlme S, et al. 2020. Fungal Diversity 105:1-16 ((https://doi.org/10.1007/s13225-020-00466-2)) |
| Otu01235 | 49 | k__Fungi(100);p__Ascomycota(100);c__Saccharomycetes(100);o__Saccharomycetales(100);f__Metschnikowiaceae(100);g__Metschnikowia(100);g__Metschnikowia_unclassified(78); | Metschnikowia | 13 | Saprotroph | \|Undefined Saprotroph\| | Yeast | NULL | Probable | As nectar-tap_saprotroph (Põlme et al. 2020); Decay sugar-rich substrates (Põlme et al. 2020) | Põlme S, et al. 2020. Fungal Diversity 105:1-16 ((https://doi.org/10.1007/s13225-020-00466-2)) |
| Otu01236 | 49 | k__Fungi(100);p__Basidiomycota(100);c__Exobasidiomycetes(100);o__Exobasidiales(100);f__Exobasidiaceae(100);g__Exobasidium(100);g__Exobasidium_unclassified(86); | Exobasidium | 13 | Pathotroph | \|Plant Pathogen\| | Leaf Blister-Leaf Gall-Leaf Spot | NULL | Probable | NULL | Tedersoo L, et al. 2014. Science 346:e1256688 ((https://doi.org/10.1126/science.1256688)); Põlme S, et al. 2020. Fungal Diversity 105:1-16 ((https://doi.org/10.1007/s13225-020-00466-2)) |
| Otu01237 | 49 | k__Fungi(100);p__Ascomycota(100);c__Dothideomycetes(100);o__Botryosphaeriales(100);f__Botryosphaeriaceae(100);g__Dothiorella(100);s__Dothiorella_sp(100); | Dothiorella | 13 | Pathotroph-Saprotroph-Symbiotroph | Endophyte-\|Plant Pathogen\|-Plant Saprotroph | Microfungus | NULL | Probable | As foliar_endophyte (Põlme et al. 2020); As litter_saprotroph (Põlme et al. 2020); Decay leaf, fruit, seed (Põlme et al. 2020) | Tedersoo L, et al. 2014. Science 346:e1256688 ((https://doi.org/10.1126/science.1256688)); Põlme S, et al. 2020. Fungal Diversity 105:1-16 ((https://doi.org/10.1007/s13225-020-00466-2)) |
| Otu01240 | 49 | k__Fungi(100);p__Ascomycota(100);c__Dothideomycetes(100);o__Pleosporales(100);f__Didymosphaeriaceae(100);g__Roussoella(100);s__Roussoella_japanensis(100); | Roussoella | 13 | Saprotroph | Undefined Saprotroph-\|Wood Saprotroph\| | Microfungus | NULL | Probable | Decay wood (Põlme et al. 2020) | Tedersoo L, et al. 2014. Science 346:e1256688 ((https://doi.org/10.1126/science.1256688)); Põlme S, et al. 2020. Fungal Diversity 105:1-16 ((https://doi.org/10.1007/s13225-020-00466-2)) |
| Otu01241 | 49 | k__Fungi(100);p__Basidiomycota(100);c__Agaricomycetes(100);o__Polyporales(100);f__Fomitopsidaceae(100);g__Antrodia(100);s__Antrodia_sp_Lohmus_521_(TU)(100); | Antrodia | 13 | Saprotroph | \|Wood Saprotroph\| | Corticioid | Brown Rot | Highly Probable | As brown-rot (Põlme et al. 2020); Decay wood (Põlme et al. 2020) | Gilbertson RL, Ryvarden L. 1987-1987. North American Polypores. Fungiflora, Oslo (ISBN: 978-0945345060); Põlme S, et al. 2020. Fungal Diversity 105:1-16 ((https://doi.org/10.1007/s13225-020-00466-2)) |
| Otu01232 | 49 | k__Fungi(100);p__Ascomycota(100);c__Leotiomycetes(98);o__Helotiales(98);o__Helotiales_unclassified(98);o__Helotiales_unclassified(98);o__Helotiales_unclassified(98); | - | - | - | - | - | - | - | Unassigned | - |
| Otu01238 | 49 | k__Fungi(100);k__Fungi_unclassified(98);k__Fungi_unclassified(98);k__Fungi_unclassified(98);k__Fungi_unclassified(98);k__Fungi_unclassified(98);k__Fungi_unclassified(98); | - | - | - | - | - | - | - | Unassigned | - |
| Otu01239 | 49 | k__Fungi(100);p__Basidiomycota(100);c__Agaricomycetes(100);c__Agaricomycetes_unclassified(100);c__Agaricomycetes_unclassified(100);c__Agaricomycetes_unclassified(100);c__Agaricomycetes_unclassified(100); | - | - | - | - | - | - | - | Unassigned | - |
| Otu01242 | 49 | k__Fungi(100);p__Basidiomycota(100);p__Basidiomycota_unclassified(96);p__Basidiomycota_unclassified(96);p__Basidiomycota_unclassified(96);p__Basidiomycota_unclassified(96);p__Basidiomycota_unclassified(96); | - | - | - | - | - | - | - | Unassigned | - |

| | | | | | | | | | | | |
|---|---|---|---|---|---|---|---|---|---|---|---|
| Otu01245 | 48 | k__Fungi(100);p__Basidiomycota(100);c__Agaricomycetes(100);o__Polyporales(100);f__Polyporaceae(100);g__Spongipellis(100);s__Spongipellis_delectans(100); | Spongipellis | 13 | Pathotroph-Saprotroph | Plant Pathogen-\|Wood Saprotroph\| | Polyporoid | White Rot | Highly Probable | As white_rot (Põlme et al. 2020); As wood_pathogen (Põlme et al. 2020); Decay wood (Põlme et al. 2020) | Gilbertson RL, Ryvarden L. 1987-1987. North American Polypores. Fungiflora, Oslo (ISBN: 978-0945345060); Põlme S, et al. 2020. Fungal Diversity 105:1-16 ((https://doi.org/10.1007/s13225-020-00466-2)) |
| Otu01248 | 48 | k__Fungi(100);p__Basidiomycota(100);c__Agaricomycetes(100);o__Polyporales(100);f__Phanerochaetaceae(100);g__Ceriporiopsis(100);s__Ceriporiopsis_gilvescens(100); | Ceriporiopsis | 13 | Saprotroph | \|Wood Saprotroph\| | Corticioid | White Rot | Highly Probable | As white_rot (Põlme et al. 2020); Decay wood (Põlme et al. 2020) | Gilbertson RL, Ryvarden L. 1987-1987. North American Polypores. Fungiflora, Oslo (ISBN: 978-0945345060); Tedersoo L, et al. 2014. Science 346:e1256688 ((https://doi.org/10.1126/science.1256688)); Põlme S, et al. 2020. Fungal Diversity 105:1-16 ((https://doi.org/10.1007/s13225-020-00466-2)) |
| Otu01249 | 48 | k__Fungi(100);p__Ascomycota(100);c__Sordariomycetes(100);o__Xylariales(100);f__Diatrypaceae(100);g__Eutypa(90);g__Eutypa_unclassified(65); | Eutypa | 13 | Pathotroph | \|Plant Pathogen\| | Microfungus | NULL | Probable | NULL | Tedersoo L, et al. 2014. Science 346:e1256688 ((https://doi.org/10.1126/science.1256688)); Põlme S, et al. 2020. Fungal Diversity 105:1-16 ((https://doi.org/10.1007/s13225-020-00466-2)) |
| Otu01252 | 48 | k__Fungi(100);p__Basidiomycota(100);c__Agaricomycetes(100);o__Polyporales(100);f__Meruliaceae(100);g__Phlebia(100);s__Phlebia_sp_olrim964(100); | Phlebia | 13 | Saprotroph | \|Wood Saprotroph\| | Corticioid | White Rot | Highly Probable | As white_rot (Põlme et al. 2020); Decay wood (Põlme et al. 2020) | Kirk PM et al. 2008. Dictionary of the Fungi. Tenth Edition. CAB International, Wallingford (ISBN: 978-0851998268); Põlme S, et al. 2020. Fungal Diversity 105:1-16 ((https://doi.org/10.1007/s13225-020-00466-2)) |
| Otu01253 | 48 | k__Fungi(100);p__Ascomycota(100);c__Dothideomycetes(100);o__Pleosporales(100);f__Leptosphaeriaceae(100);g__Leptosphaeria(98);g__Leptosphaeria_unclassified(98); | Leptosphaeria | 13 | Pathotroph-Saprotroph | \|Plant Pathogen\|-Wood Saprotroph | Microfungus | NULL | Probable | As wood_pathogen (Põlme et al. 2020); Decay wood (Põlme et al. 2020) | Tedersoo L, et al. 2014. Science 346:e1256688 ((https://doi.org/10.1126/science.1256688)); Põlme S, et al. 2020. Fungal Diversity 105:1-16 ((https://doi.org/10.1007/s13225-020-00466-2)) |
| Otu01254 | 48 | k__Fungi(100);p__Basidiomycota(100);c__Agaricomycetes(100);o__Russulales(100);f__Auriscalpiaceae(100);g__Lentinellus(100);s__Lentinellus_flabelliformis(53); | Lentinellus | 13 | Pathotroph-Saprotroph | Plant Pathogen-Undefined Saprotroph-\|Wood Saprotroph\| | Agaricoid | White Rot | Probable | As white_rot (Põlme et al. 2020); As wood_pathogen (Põlme et al. 2020); Decay wood (Põlme et al. 2020) | Tedersoo L, et al. 2014. Science 346:e1256688 ((https://doi.org/10.1126/science.1256688)); Põlme S, et al. 2020. Fungal Diversity 105:1-16 ((https://doi.org/10.1007/s13225-020-00466-2)) |
| Otu01256 | 48 | k__Fungi(100);p__Ascomycota(100);c__Sordariomycetes(100);o__Hypocreales(100);o__Hypocreales_unclassified(96);o__Hypocreales_unclassified(96);o__Hypocreales_unclassified(96); | Hypocreales | 7 | Saprotroph | Undefined Saprotroph | Microfungus | NULL | Possible | NULL | Sterkenburg E, et al. 2015. New Phytologist 207:1145-1158 ((https://doi.org/10.1111/nph.13426)) |
| Otu01258 | 48 | k__Fungi(100);p__Basidiomycota(100);c__Agaricomycetes(100);o__Agaricales(100);f__Agaricaceae(100);g__Lycoperdon(98);g__Lycoperdon_unclassified(69); | Lycoperdon | 13 | Saprotroph | \|Plant Saprotroph\|-Undefined Saprotroph | Gasteroid | NULL | Probable | puffball; As litter_saprotroph (Põlme et al. 2020); Common name ; Decay leaf, fruit, seed (Põlme et al. 2020) | Grgurinovic CA, Simpson JA. 2001. Fungal Diversity 8:97-127 ((http://www.fungaldiversity.org/fdp/sfdp/FD_8_97-127.pdf)); Tedersoo L, et al. 2014. Science 346:e1256688 ((https://doi.org/10.1126/science.1256688)); Põlme S, et al. 2020. Fungal Diversity 105:1-16 ((https://doi.org/10.1007/s13225-020-00466-2)) |
| Otu01259 | 48 | k__Fungi(100);p__Ascomycota(100);c__Leotiomycetes(100);o__Helotiales(100);f__Dermateaceae(90);g__Pezicula(90);s__Pezicula_alnicola(90); | Pezicula | 13 | Pathotroph-Saprotroph-Symbiotroph | \|Endophyte\|-Plant Pathogen-Plant Saprotroph-Undefined Saprotroph | Microfungus | NULL | Probable | As litter_saprotroph (Põlme et al. 2020); As root_endophyte (Põlme et al. 2020); As soil_saprotroph (Põlme et al. 2020); Decay soil (Põlme et al. 2020); Most species in Pezicula are only known as saprobes or harmless endophytes, and occasionally as weak plant pathogens. | Tedersoo L, et al. 2014. Science 346:e1256688 ((https://doi.org/10.1126/science.1256688)); Chen C, et al. 2015. Fungal Biology 120: 1291-1322 ((https://doi.org/10.1016/j.funbio.2015.09.013)); Põlme S, et al. 2020. Fungal Diversity 105:1-16 ((https://doi.org/10.1007/s13225-020-00466-2)) |
| Otu01243 | 48 | k__Fungi(100);k__Fungi_unclassified(100);k__Fungi_unclassified(100);k__Fungi_unclassified(100);k__Fungi_unclassified(100);k__Fungi_unclassified(100);k__Fungi_unclassified(100); | - | - | - | - | - | - | - | Unassigned | - |
| Otu01244 | 48 | k__Fungi(100);p__Ascomycota(88);p__Ascomycota_unclassified(88);p__Ascomycota_unclassified(88);p__Ascomycota_unclassified(88);p__Ascomycota_unclassified(88);p__Ascomycota_unclassified(88); | - | - | - | - | - | - | - | Unassigned | - |
| Otu01246 | 48 | k__Fungi(100);k__Fungi_unclassified(55);k__Fungi_unclassified(55);k__Fungi_unclassified(55);k__Fungi_unclassified(55);k__Fungi_unclassified(55);k__Fungi_unclassified(55); | - | - | - | - | - | - | - | Unassigned | - |
| Otu01247 | 48 | k__Fungi(100);p__Ascomycota(57);p__Ascomycota_unclassified(57);p__Ascomycota_unclassified(57);p__Ascomycota_unclassified(57);p__Ascomycota_unclassified(57);p__Ascomycota_unclassified(57); | - | - | - | - | - | - | - | Unassigned | - |
| Otu01250 | 48 | k__Fungi(100);k__Fungi_unclassified(86);k__Fungi_unclassified(86);k__Fungi_unclassified(86);k__Fungi_unclassified(86);k__Fungi_unclassified(86);k__Fungi_unclassified(86); | - | - | - | - | - | - | - | Unassigned | - |
| Otu01251 | 48 | k__Fungi(100);k__Fungi_unclassified(53);k__Fungi_unclassified(53);k__Fungi_unclassified(53);k__Fungi_unclassified(53);k__Fungi_unclassified(53);k__Fungi_unclassified(53); | - | - | - | - | - | - | - | Unassigned | - |
| Otu01255 | 48 | k__Fungi(100);k__Fungi_unclassified(67);k__Fungi_unclassified(67);k__Fungi_unclassified(67);k__Fungi_unclassified(67);k__Fungi_unclassified(67);k__Fungi_unclassified(67); | - | - | - | - | - | - | - | Unassigned | - |
| Otu01257 | 48 | k__Fungi(100);p__Ascomycota(100);c__Leotiomycetes(96);o__Helotiales(94);o__Helotiales_unclassified(90);o__Helotiales_unclassified(90);o__Helotiales_unclassified(90); | - | - | - | - | - | - | - | Unassigned | - |
| Otu01261 | 47 | k__Fungi(100);p__Ascomycota(100);c__Sordariomycetes(100);o__Hypocreales(100);f__Ophiocordycipitaceae(100);g__Elaphocordyceps(100);s__Elaphocordyceps_sp_20124794d(88); | Elaphocordyceps | 13 | Pathotroph | Fungal Parasite | Secotioid-Spathuloid | NULL | Highly Probable | NULL | Samuels GJ. 2007. Self-published; Sung G-H, et al. 2007. Studies in Mycology 57:5-59 ((https://www.studiesinmycology.org/index.php/issue/59-studies-in-mycology-no-57)); List compiled by John Plischke III ((https://goodmushroom.weebly.com/uploads/5/7/1/7/57176535/fungi_on_fungi_pdf_lq.pdf)) |
| Otu01262 | 47 | k__Fungi(100);p__Basidiomycota(100);c__Tremellomycetes(100);o__Cystofilobasidiales(100);f__Cystofilobasidiaceae(100);g__Guehomyces(100);s__Guehomyces_pullulans(100); | Guehomyces | 13 | Saprotroph | Undefined Saprotroph | Yeast | NULL | Probable | Isolated from an experimental brewery (Fell & Scorzetti 2004; Syn. Basidiotrichosporon-Monilia-Oidium-Oospora-Tausonia-Trichosporon sp.) | Fell JW, Scorzetti G. 2004. International Journal of Systematic and Evolutionary Microbiology 54:995-998 ((https://doi.org/10.1099/ijs.0.03017-0)) |
| Otu01265 | 47 | k__Fungi(100);p__Basidiomycota(100);c__Agaricomycetes(100);o__Polyporales(100);f__Phanerochaetaceae(100);g__Byssomerulius(100);s__Byssomerulius_sp(100); | Byssomerulius | 13 | Saprotroph | Undefined Saprotroph-\|Wood Saprotroph\| | Corticioid | White Rot | Probable | As white_rot (Põlme et al. 2020); Decay wood (Põlme et al. 2020) | Tedersoo L, et al. 2014. Science 346:e1256688 ((https://doi.org/10.1126/science.1256688)); Põlme S, et al. 2020. Fungal Diversity 105:1-16 ((https://doi.org/10.1007/s13225-020-00466-2)) |
| Otu01266 | 47 | k__Fungi(100);p__Basidiomycota(100);c__Pucciniomycetes(100);o__Pucciniales(100);f__Phragmidiaceae(100);g__Kuehneola(100);s__Kuehneola_uredinis(100); | Kuehneola | 13 | Pathotroph | \|Plant Pathogen\| | Rust | NULL | Probable | NULL | Tedersoo L, et al. 2014. Science 346:e1256688 ((https://doi.org/10.1126/science.1256688)); Põlme S, et al. 2020. Fungal Diversity 105:1-16 ((https://doi.org/10.1007/s13225-020-00466-2)) |
| Otu01268 | 47 | k__Fungi(100);p__Ascomycota(100);c__Saccharomycetes(100);o__Saccharomycetales(100);f__Metschnikowiaceae(100);g__Metschnikowia(100);s__Metschnikowia_pulcherrima(96); | Metschnikowia | 13 | Saprotroph | \|Undefined Saprotroph\| | Yeast | NULL | Probable | As nectar-tap_saprotroph (Põlme et al. 2020); Decay sugar-rich substrates (Põlme et al. 2020) | Põlme S, et al. 2020. Fungal Diversity 105:1-16 ((https://doi.org/10.1007/s13225-020-00466-2)) |
| Otu01269 | 47 | k__Fungi(100);p__Basidiomycota(100);c__Agaricomycetes(100);o__Trechisporales(100);f__Hydnodontaceae(100);g__Trechispora(100);s__Trechispora_sp(100); | Trechispora | 13 | Saprotroph | \|Wood Saprotroph\| | Corticioid | White Rot | Highly Probable | As white_rot (Põlme et al. 2020); Decay wood (Põlme et al. 2020) | Gilbertson RL, Ryvarden L. 1987-1987. North American Polypores. Fungiflora, Oslo (ISBN: 978-0945345060); Rinaldi AC, et al. 2008. Fungal Diversity 33:1-45 ((http://www.fungaldiversity.org/fdp/sfdp/33-1.pdf)) (as Uncertain); Tedersoo L, et al. 2010. Mycorrhiza 20:217-263 ((https://doi.org/10.1007/s00572-009-0274-x)) (as not ECM).; Põlme S, et al. 2020. Fungal Diversity 105:1-16 ((https://doi.org/10.1007/s13225-020-00466-2)) |
| Otu01270 | 47 | k__Fungi(100);p__Ascomycota(100);c__Saccharomycetes(100);o__Saccharomycetales(100);f__Metschnikowiaceae(100);g__Metschnikowia(100);s__Metschnikowia_pulcherrima(100); | Metschnikowia | 13 | Saprotroph | \|Undefined Saprotroph\| | Yeast | NULL | Probable | As nectar-tap_saprotroph (Põlme et al. 2020); Decay sugar-rich substrates (Põlme et al. 2020) | Põlme S, et al. 2020. Fungal Diversity 105:1-16 ((https://doi.org/10.1007/s13225-020-00466-2)) |
| Otu01272 | 47 | k__Fungi(100);p__Basidiomycota(100);c__Agaricomycetes(100);o__Polyporales(100);f__Phanerochaetaceae(100);g__Phanerochaete(100);s__Phanerochaete_sp_DLL2011_1(98); | Phanerochaete | 13 | Saprotroph | \|Wood Saprotroph\| | Corticioid | White Rot | Highly Probable | As white_rot (Põlme et al. 2020); Decay wood (Põlme et al. 2020) | Glenn JK, et al. 1983. Biochemical and Biophysical Research Communications 114:1077-1083 ((https://doi.org/10.1016/0006-291x((83)) 90672-1)); Põlme S, et al. 2020. Fungal Diversity 105:1-16 ((https://doi.org/10.1007/s13225-020-00466-2)) |
| Otu01274 | 47 | k__Fungi(100);p__Ascomycota(100);c__Ascomycota_class_Incertae_sedis(100);o__Ascomycota_order_Incertae_sedis(100);f__Ascomycota_family_Incertae_sedis(100);g__Torula(100);s__Torula_sp(100); | Torula | 13 | Saprotroph-Symbiotroph | \|Endophyte\|-Plant Saprotroph-Wood Saprotroph | Facultative Yeast-Microfungus | Soft Rot | Probable | As foliar_endophyte (Põlme et al. 2020); Decay wood, leaf, fruit, seed (Põlme et al. 2020); Yeast can be propagated for long periods in wood sugar solutions (Fink & Lechner 1936) | Fink H, Lechner R. 1936. Biochemische Zeitschrift 286:83-90 ((https://archive.org/details/sim_biochemische-zeitschrift-beitraege-zur-chemischen_1936_286)); Crous PW, et al. 2015. IMA Fungus 6: 163-198 ((https://doi.org/10.5598/imafungus.2015.06.01.11)); Põlme S, et al. 2020. Fungal Diversity 105:1-16 ((https://doi.org/10.1007/s13225-020-00466-2)) |
| Otu01275 | 47 | k__Fungi(100);p__Ascomycota(100);c__Ascomycota_class_Incertae_sedis(100);o__Ascomycota_order_Incertae_sedis(100);f__Ascomycota_family_Incertae_sedis(100);g__Ochroconis(100);s__Ochroconis_constricta(100); | Ochroconis | 13 | Pathotroph-Saprotroph | \|Animal Parasite\|-Plant Pathogen-Undefined Saprotroph | Microfungus | NULL | Probable | Decay leaf, fruit, seed, animal material (Põlme et al. 2020) | Tedersoo L, et al. 2014. Science 346:e1256688 ((https://doi.org/10.1126/science.1256688)); Põlme S, et al. 2020. Fungal Diversity 105:1-16 ((https://doi.org/10.1007/s13225-020-00466-2)) |
| Otu01278 | 47 | k__Fungi(100);p__Ascomycota(100);c__Saccharomycetes(100);o__Saccharomycetales(100);f__Metschnikowiaceae(100);g__Metschnikowia(100);s__Metschnikowia_pulcherrima(94); | Metschnikowia | 13 | Saprotroph | \|Undefined Saprotroph\| | Yeast | NULL | Probable | As nectar-tap_saprotroph (Põlme et al. 2020); Decay sugar-rich substrates (Põlme et al. 2020) | Põlme S, et al. 2020. Fungal Diversity 105:1-16 ((https://doi.org/10.1007/s13225-020-00466-2)) |
| Otu01279 | 47 | k__Fungi(100);p__Ascomycota(100);c__Sordariomycetes(100);o__Hypocreales(100);f__Ophiocordycipitaceae(100);g__Hirsutella(100);s__Hirsutella_aphidis(100); | Hirsutella | 13 | Pathotroph-Saprotroph | \|Animal Parasite\|-Animal Pathogen-Undefined Saprotroph | Microfungus | NULL | Probable | As animal_decomposer (Põlme et al. 2020); Decay animal material (Põlme et al. 2020) | Tedersoo L, et al. 2014. Science 346:e1256688 ((https://doi.org/10.1126/science.1256688)); Põlme S, et al. 2020. Fungal Diversity 105:1-16 ((https://doi.org/10.1007/s13225-020-00466-2)) |
| Otu01280 | 47 | k__Fungi(100);p__Basidiomycota(100);c__Agaricomycetes(100);o__Agaricales(100);f__Psathyrellaceae(100);g__Coprinopsis(100);s__Coprinopsis_sp_JKR_2010(73); | Coprinopsis | 13 | Saprotroph | \|Undefined Saprotroph\| | Agaricoid | NULL | Probable | As soil_saprotroph (Põlme et al. 2020); Decay soil (Põlme et al. 2020) | Tedersoo L, et al. 2014. Science 346:e1256688 ((https://doi.org/10.1126/science.1256688)); Põlme S, et al. 2020. Fungal Diversity 105:1-16 ((https://doi.org/10.1007/s13225-020-00466-2)) |
| Otu01281 | 47 | k__Fungi(100);p__Ascomycota(98);c__Saccharomycetes(98);o__Saccharomycetales(98);f__Metschnikowiaceae(98);g__Metschnikowia(98);s__Metschnikowia_pulcherrima(96); | Metschnikowia | 13 | Saprotroph | \|Undefined Saprotroph\| | Yeast | NULL | Probable | As nectar-tap_saprotroph (Põlme et al. 2020); Decay sugar-rich substrates (Põlme et al. 2020) | Põlme S, et al. 2020. Fungal Diversity 105:1-16 ((https://doi.org/10.1007/s13225-020-00466-2)) |
| Otu01260 | 47 | k__Fungi(100);p__Ascomycota(100);c__Dothideomycetes(98);o__Pleosporales(98);o__Pleosporales_unclassified(98);o__Pleosporales_unclassified(98);o__Pleosporales_unclassified(98); | - | - | - | - | - | - | - | Unassigned | - |
| Otu01263 | 47 | k__Fungi(100);p__Basidiomycota(100);c__Agaricomycetes(100);o__Polyporales(100);o__Polyporales_unclassified(100);o__Polyporales_unclassified(100);o__Polyporales_unclassified(100); | - | - | - | - | - | - | - | Unassigned | - |
| Otu01264 | 47 | k__Fungi(100);p__Ascomycota(100);c__Dothideomycetes(100);o__Pleosporales(100);o__Pleosporales_unclassified(73);o__Pleosporales_unclassified(73);o__Pleosporales_unclassified(73); | - | - | - | - | - | - | - | Unassigned | - |
| Otu01267 | 47 | k__Fungi(100);k__Fungi_unclassified(56);k__Fungi_unclassified(56);k__Fungi_unclassified(56);k__Fungi_unclassified(56);k__Fungi_unclassified(56);k__Fungi_unclassified(56); | - | - | - | - | - | - | - | Unassigned | - |
| Otu01271 | 47 | k__Fungi(100);k__Fungi_unclassified(98);k__Fungi_unclassified(98);k__Fungi_unclassified(98);k__Fungi_unclassified(98);k__Fungi_unclassified(98);k__Fungi_unclassified(98); | - | - | - | - | - | - | - | Unassigned | - |
| Otu01273 | 47 | k__Fungi(100);k__Fungi_unclassified(58);k__Fungi_unclassified(58);k__Fungi_unclassified(58);k__Fungi_unclassified(58);k__Fungi_unclassified(58);k__Fungi_unclassified(58); | - | - | - | - | - | - | - | Unassigned | - |
| Otu01276 | 47 | k__Fungi(100);p__Ascomycota(60);p__Ascomycota_unclassified(60);p__Ascomycota_unclassified(60);p__Ascomycota_unclassified(60);p__Ascomycota_unclassified(60);p__Ascomycota_unclassified(60); | - | - | - | - | - | - | - | Unassigned | - |
| Otu01277 | 47 | k__Fungi(100);p__Ascomycota(100);p__Ascomycota_unclassified(100);p__Ascomycota_unclassified(100);p__Ascomycota_unclassified(100);p__Ascomycota_unclassified(100);p__Ascomycota_unclassified(100); | - | - | - | - | - | - | - | Unassigned | - |
| Otu01283 | 46 | k__Fungi(100);p__Basidiomycota(100);c__Agaricomycetes(100);o__Trechisporales(100);f__Hydnodontaceae(100);g__Trechispora(100);s__Trechispora_sp(100); | Trechispora | 13 | Saprotroph | \|Wood Saprotroph\| | Corticioid | White Rot | Highly Probable | As white_rot (Põlme et al. 2020); Decay wood (Põlme et al. 2020) | Gilbertson RL, Ryvarden L. 1987-1987. North American Polypores. Fungiflora, Oslo (ISBN: 978-0945345060); Rinaldi AC, et al. 2008. Fungal Diversity 33:1-45 ((http://www.fungaldiversity.org/fdp/sfdp/33-1.pdf)) (as Uncertain); Tedersoo L, et al. 2010. Mycorrhiza 20:217-263 ((https://doi.org/10.1007/s00572-009-0274-x)) (as not ECM).; Põlme S, et al. 2020. Fungal Diversity 105:1-16 ((https://doi.org/10.1007/s13225-020-00466-2)) |
| Otu01284 | 46 | k__Fungi(100);p__Ascomycota(100);c__Ascomycota_class_Incertae_sedis(100);o__Ascomycota_order_Incertae_sedis(100);f__Ascomycota_family_Incertae_sedis(100);g__Tumularia(100);s__Tumularia_sp(100); | Tumularia | 13 | Saprotroph | \|Plant Saprotroph\|-Undefined Saprotroph | Microfungus | NULL | Probable | As litter_saprotroph (Põlme et al. 2020); Decay leaf, fruit, seed (Põlme et al. 2020) | Tedersoo L, et al. 2014. Science 346:e1256688 ((https://doi.org/10.1126/science.1256688)); Põlme S, et al. 2020. Fungal Diversity 105:1-16 ((https://doi.org/10.1007/s13225-020-00466-2)) |
| Otu01285 | 46 | k__Fungi(100);p__Basidiomycota(100);c__Agaricomycetes(100);o__Agaricomycetes_order_Incertae_sedis(100);f__Agaricomycetes_family_Incertae_sedis(100);g__Resinicium(100);s__Resinicium_rimulosum(100); | Resinicium | 13 | Saprotroph | Undefined Saprotroph-\|Wood Saprotroph\| | Corticioid | White Rot | Probable | As white_rot (Põlme et al. 2020); Decay wood (Põlme et al. 2020) | Tedersoo L, et al. 2014. Science 346:e1256688 ((https://doi.org/10.1126/science.1256688)); Põlme S, et al. 2020. Fungal Diversity 105:1-16 ((https://doi.org/10.1007/s13225-020-00466-2)) |
| Otu01290 | 46 | k__Fungi(100);p__Ascomycota(100);c__Saccharomycetes(100);o__Saccharomycetales(100);f__Saccharomycetales_family_Incertae_sedis(100);g__Debaryomyces(100);g__Debaryomyces_unclassified(72); | Debaryomyces | 13 | Saprotroph | \|Undefined Saprotroph\| | Yeast | NULL | Highly Probable | As nectar-tap_saprotroph (Põlme et al. 2020); Decay sugar-rich substrates (Põlme et al. 2020) | Kurtzman CP, et al. (eds.) 2011. The Yeasts, a Taxonomic Study. Fifth Edition. Vols 1-3. Elsevier, San Diego (ISBN: 9780444521491); Põlme S, et al. 2020. Fungal Diversity 105:1-16 ((https://doi.org/10.1007/s13225-020-00466-2)) |
| Otu01294 | 46 | k__Fungi(100);p__Basidiomycota(100);c__Agaricomycetes(100);o__Russulales(100);f__Hericiaceae(100);g__Dentipellis(100);s__Dentipellis_fragilis(100); | Dentipellis | 13 | Saprotroph | Undefined Saprotroph-\|Wood Saprotroph\| | Corticioid-Hydnoid | White Rot | Probable | As white_rot (Põlme et al. 2020); Decay wood (Põlme et al. 2020) | Tedersoo L, et al. 2014. Science 346:e1256688 ((https://doi.org/10.1126/science.1256688)); Põlme S, et al. 2020. Fungal Diversity 105:1-16 ((https://doi.org/10.1007/s13225-020-00466-2)) |
| Otu01282 | 46 | k__Fungi(100);p__Ascomycota(100);c__Sordariomycetes(100);o__Diaporthales(100);o__Diaporthales_unclassified(70);o__Diaporthales_unclassified(70);o__Diaporthales_unclassified(70); | - | - | - | - | - | - | - | Unassigned | - |
| Otu01286 | 46 | k__Fungi(100);p__Ascomycota(100);p__Ascomycota_unclassified(100);p__Ascomycota_unclassified(100);p__Ascomycota_unclassified(100);p__Ascomycota_unclassified(100);p__Ascomycota_unclassified(100); | - | - | - | - | - | - | - | Unassigned | - |
| Otu01287 | 46 | k__Fungi(100);k__Fungi_unclassified(77);k__Fungi_unclassified(77);k__Fungi_unclassified(77);k__Fungi_unclassified(77);k__Fungi_unclassified(77);k__Fungi_unclassified(77); | - | - | - | - | - | - | - | Unassigned | - |
| Otu01288 | 46 | k__Fungi(100);p__Basidiomycota(100);c__Agaricomycetes(100);c__Agaricomycetes_unclassified(100);c__Agaricomycetes_unclassified(100);c__Agaricomycetes_unclassified(100);c__Agaricomycetes_unclassified(100); | - | - | - | - | - | - | - | Unassigned | - |
| Otu01289 | 46 | k__Fungi(100);k__Fungi_unclassified(66);k__Fungi_unclassified(66);k__Fungi_unclassified(66);k__Fungi_unclassified(66);k__Fungi_unclassified(66);k__Fungi_unclassified(66); | - | - | - | - | - | - | - | Unassigned | - |
| Otu01291 | 46 | k__Fungi(100);k__Fungi_unclassified(64);k__Fungi_unclassified(64);k__Fungi_unclassified(64);k__Fungi_unclassified(64);k__Fungi_unclassified(64);k__Fungi_unclassified(64); | - | - | - | - | - | - | - | Unassigned | - |
| Otu01292 | 46 | k__Fungi(100);p__Ascomycota(100);c__Dothideomycetes(100);c__Dothideomycetes_unclassified(70);c__Dothideomycetes_unclassified(70);c__Dothideomycetes_unclassified(70);c__Dothideomycetes_unclassified(70); | - | - | - | - | - | - | - | Unassigned | - |
| Otu01293 | 46 | k__Fungi(100);k__Fungi_unclassified(98);k__Fungi_unclassified(98);k__Fungi_unclassified(98);k__Fungi_unclassified(98);k__Fungi_unclassified(98);k__Fungi_unclassified(98); | - | - | - | - | - | - | - | Unassigned | - |
| Otu01295 | 45 | k__Fungi(100);p__Basidiomycota(100);c__Exobasidiomycetes(100);o__Exobasidiales(100);f__Exobasidiaceae(100);g__Exobasidium(100);s__Exobasidium_sp(100); | Exobasidium | 13 | Pathotroph | \|Plant Pathogen\| | Leaf Blister-Leaf Gall-Leaf Spot | NULL | Probable | NULL | Tedersoo L, et al. 2014. Science 346:e1256688 ((https://doi.org/10.1126/science.1256688)); Põlme S, et al. 2020. Fungal Diversity 105:1-16 ((https://doi.org/10.1007/s13225-020-00466-2)) |
| Otu01296 | 45 | k__Fungi(100);p__Ascomycota(100);c__Dothideomycetes(100);o__Pleosporales(100);f__Phaeosphaeriaceae(100);f__Phaeosphaeriaceae_unclassified(80);f__Phaeosphaeriaceae_unclassified(80); | Phaeosphaeriaceae | 9 | Pathotroph-Saprotroph | Fungal Parasite-Plant Pathogen-Plant Saprotroph | Microfungus | NULL | Probable | Necrotrophic or saprobic on a wide range of plants, especially monocotyledons; a few species parasitic on other fungi (Cannon & Kirk 2007) | Cannon PF, Kirk PM. 2007. Fungal Families of the World. CAB International, Cambridge (ISBN: 978-0851998275) |
| Otu01297 | 45 | k__Fungi(100);p__Basidiomycota(100);c__Agaricomycetes(100);o__Agaricales(100);f__Strophariaceae(100);g__Pholiota(100);s__Pholiota_lenta(87); | Pholiota | 13 | Pathotroph-Saprotroph | Plant Pathogen-Plant Saprotroph-Undefined Saprotroph-\|Wood Saprotroph\| | Agaricoid | White Rot | Probable | As litter_saprotroph (Põlme et al. 2020); As white_rot (Põlme et al. 2020); As wood_pathogen (Põlme et al. 2020); Decay wood, leaf, fruit, seed (Põlme et al. 2020) | Tedersoo L, et al. 2014. Science 346:e1256688 ((https://doi.org/10.1126/science.1256688)); Põlme S, et al. 2020. Fungal Diversity 105:1-16 ((https://doi.org/10.1007/s13225-020-00466-2)) |
| Otu01300 | 45 | k__Fungi(100);p__Zygomycota(100);c__Zygomycota_class_Incertae_sedis(100);o__Mortierellales(100);f__Mortierellaceae(100);g__Mortierella(100);s__Mortierella_histoplasmatoides(89); | Mortierella | 13 | Saprotroph-Symbiotroph | Endophyte-Plant Saprotroph-\|Undefined Saprotroph\| | Microfungus | NULL | Probable | As chitinolytic (Põlme et al. 2020); As soil_saprotroph (Põlme et al. 2020); Decay roots, soil (Põlme et al. 2020); Saprobic in soil and thus frequently isolated from soil and material contaminated with soil such as hay or rotten silage and spoil tips (Cannon & Kirk 2007) | Cannon PF, Kirk PM. 2007. Fungal Families of the World. CAB International, Cambridge (ISBN: 978-0851998275); Tedersoo L, et al. 2014. Science 346:e1256688 ((https://doi.org/10.1126/science.1256688)); Purahong W, et al. 2016. Molecular Ecology 25:4059-4074 ((https://doi.org/10.1111/mec.13739)); Põlme S, et al. 2020. Fungal Diversity 105:1-16 ((https://doi.org/10.1007/s13225-020-00466-2)) |
| Otu01301 | 45 | k__Fungi(100);p__Ascomycota(100);c__Eurotiomycetes(100);o__Eurotiales(100);f__Trichocomaceae(98);g__Aspergillus(98);g__Aspergillus_unclassified(65); | Aspergillus | 13 | Pathotroph-Saprotroph-Symbiotroph | Animal Pathogen-Endophyte-Plant Saprotroph-\|Undefined Saprotroph\|-Wood Saprotroph | Microfungus | NULL | Probable | As foliar_endophyte (Põlme et al. 2020); Decay leaf, fruit, seed, soil, dung, animal material (Põlme et al. 2020) | Seehann G, et al. 1975. List of Fungi in Soft-Rot Tissue Tests. Institute fur Holzbiologie und Holzschutz, Bundesforschungsanstalt fur Forst - und Holzwirtshaft, Hamburg - Lohbrugge, Germany; Tedersoo L, et al. 2014. Science 346:e1256688 ((https://doi.org/10.1126/science.1256688)); Irinyi L, et al. 2015. Medical Mycology 53:313-337 ((https://doi.org/10.1093/mmy/myv008)); Põlme S, et al. 2020. Fungal Diversity 105:1-16 ((https://doi.org/10.1007/s13225-020-00466-2)); Massimo NC, et al. 2015. Microbial Ecology 70:61-76 ((https://doi.org/10.1007/s00248-014-0563-6)); The American Phytopathological Society Common Names of Plant Diseases ((http://www.apsnet.org/publications/commonnames/Pages/default.aspx)); https://www.cdc.gov ((https://www.cdc.gov)) |
| Otu01303 | 45 | k__Fungi(100);p__Ascomycota(100);c__Saccharomycetes(100);o__Saccharomycetales(100);f__Metschnikowiaceae(100);g__Metschnikowia(100);s__Metschnikowia_pulcherrima(96); | Metschnikowia | 13 | Saprotroph | \|Undefined Saprotroph\| | Yeast | NULL | Probable | As nectar-tap_saprotroph (Põlme et al. 2020); Decay sugar-rich substrates (Põlme et al. 2020) | Põlme S, et al. 2020. Fungal Diversity 105:1-16 ((https://doi.org/10.1007/s13225-020-00466-2)) |
| Otu01304 | 45 | k__Fungi(100);p__Ascomycota(100);c__Sordariomycetes(100);o__Hypocreales(100);o__Hypocreales_unclassified(61);o__Hypocreales_unclassified(61);o__Hypocreales_unclassified(61); | Hypocreales | 7 | Saprotroph | Undefined Saprotroph | Microfungus | NULL | Possible | NULL | Sterkenburg E, et al. 2015. New Phytologist 207:1145-1158 ((https://doi.org/10.1111/nph.13426)) |
| Otu01306 | 45 | k__Fungi(100);p__Basidiomycota(100);c__Agaricomycetes(100);o__Russulales(100);f__Peniophoraceae(100);g__Peniophora(100);s__Peniophora_incarnata(100); | Peniophora | 13 | Pathotroph-Saprotroph | Plant Pathogen-\|Wood Saprotroph\| | Corticioid | White Rot | Probable | As white_rot (Põlme et al. 2020); As wood_pathogen (Põlme et al. 2020); Decay wood (Põlme et al. 2020) | Tedersoo L, et al. 2014. Science 346:e1256688 ((https://doi.org/10.1126/science.1256688)); Põlme S, et al. 2020. Fungal Diversity 105:1-16 ((https://doi.org/10.1007/s13225-020-00466-2)) |
| Otu01307 | 45 | k__Fungi(100);p__Ascomycota(100);c__Sordariomycetes(100);o__Sordariales(100);f__Lasiosphaeriaceae(100);g__Podospora(100);s__Podospora_glutinans(100); | Podospora | 13 | Saprotroph-Symbiotroph | \|Dung Saprotroph\|-Endophyte-Plant Saprotroph-Undefined Saprotroph | Microfungus | NULL | Possible | As foliar_endophyte (Põlme et al. 2020); As litter_saprotroph (Põlme et al. 2020); culture (David et al. 2016); Decay leaf, fruit, seed, dung (Põlme et al. 2020); Endophyte detection method | Bell A. 1983. Dung Fungi: An Illustrated Guide to Coprophilous Fungi in New Zealand. Victoria University Press, Wellington (ISBN: 978-0864730015); Tedersoo L, et al. 2014. Science 346:e1256688 ((https://doi.org/10.1126/science.1256688)); David AS, et al. 2016. Microbial Ecology 71:912-926 ((https://doi.org/10.1007/s00248-015-0712-6)); Põlme S, et al. 2020. Fungal Diversity 105:1-16 ((https://doi.org/10.1007/s13225-020-00466-2)) |
| Otu01308 | 45 | k__Fungi(100);p__Ascomycota(100);c__Lecanoromycetes(100);o__Candelariales(100);f__Candelariaceae(100);f__Candelariaceae_unclassified(83);f__Candelariaceae_unclassified(83); | Candelariaceae | 9 | Symbiotroph | Lichenized | Thallus | NULL | Highly Probable | Lichenized with green algae, usually associated with nitrogen-rich substrata (Cannon & Kirk 2007) | Cannon PF, Kirk PM. 2007. Fungal Families of the World. CAB International, Cambridge (ISBN: 978-0851998275) |
| Otu01309 | 45 | k__Fungi(100);p__Zygomycota(100);c__Zygomycota_class_Incertae_sedis(100);o__Mucorales(100);f__Umbelopsidaceae(100);g__Umbelopsis(100);s__Umbelopsis_ramanniana(100); | Umbelopsis | 13 | Saprotroph | \|Undefined Saprotroph\| | Microfungus | NULL | Probable | As soil_saprotroph (Põlme et al. 2020); Decay roots, soil (Põlme et al. 2020) | Tedersoo L, et al. 2014. Science 346:e1256688 ((https://doi.org/10.1126/science.1256688)); Põlme S, et al. 2020. Fungal Diversity 105:1-16 ((https://doi.org/10.1007/s13225-020-00466-2)) |
| Otu01311 | 45 | k__Fungi(100);p__Basidiomycota(100);c__Agaricomycetes(100);o__Corticiales(100);f__Corticiaceae(100);g__Corticium(100);s__Lyomyces_sp_8_MPM_2012(100); | Lyomyces | 13 | Saprotroph | \|Wood Saprotroph\| | Corticioid | White Rot | Probable | As white_rot (Põlme et al. 2020); Decay wood (Põlme et al. 2020) | Põlme S, et al. 2020. Fungal Diversity 105:1-16 ((https://doi.org/10.1007/s13225-020-00466-2)) |
| Otu01312 | 45 | k__Fungi(100);p__Basidiomycota(100);c__Microbotryomycetes(100);o__Sporidiobolales(100);f__Sporidiobolales_family_Incertae_sedis(100);g__Sporobolomyces(100);s__Sporobolomyces_coprosmae(100); | Sporobolomyces | 13 | Pathotroph-Saprotroph | \|Fungal Parasite\|-Plant Saprotroph | Yeast | NULL | Probable | As litter_saprotroph (Põlme et al. 2020); As mycoparasite (Põlme et al. 2020); Decay fungal material (Põlme et al. 2020) | Tedersoo L, et al. 2014. Science 346:e1256688 ((https://doi.org/10.1126/science.1256688)); Purahong W, et al. 2016. Molecular Ecology 25:4059-4074 ((https://doi.org/10.1111/mec.13739)); Põlme S, et al. 2020. Fungal Diversity 105:1-16 ((https://doi.org/10.1007/s13225-020-00466-2)) |
| Otu01313 | 45 | k__Fungi(100);p__Basidiomycota(100);c__Agaricomycetes(100);o__Polyporales(96);f__Meruliaceae(94);g__Hyphoderma(94);g__Hyphoderma_unclassified(94); | Hyphoderma | 13 | Saprotroph | Undefined Saprotroph-\|Wood Saprotroph\| | Corticioid | White Rot | Probable | As white_rot (Põlme et al. 2020); Decay wood (Põlme et al. 2020) | Tedersoo L, et al. 2014. Science 346:e1256688 ((https://doi.org/10.1126/science.1256688)); Põlme S, et al. 2020. Fungal Diversity 105:1-16 ((https://doi.org/10.1007/s13225-020-00466-2)) |
| Otu01298 | 45 | k__Fungi(100);p__Ascomycota(100);p__Ascomycota_unclassified(92);p__Ascomycota_unclassified(92);p__Ascomycota_unclassified(92);p__Ascomycota_unclassified(92);p__Ascomycota_unclassified(92); | - | - | - | - | - | - | - | Unassigned | - |
| Otu01299 | 45 | k__Fungi(100);p__Ascomycota(100);c__Dothideomycetes(100);c__Dothideomycetes_unclassified(100);c__Dothideomycetes_unclassified(100);c__Dothideomycetes_unclassified(100);c__Dothideomycetes_unclassified(100); | - | - | - | - | - | - | - | Unassigned | - |
| Otu01302 | 45 | k__Fungi(100);p__Basidiomycota(100);c__Agaricostilbomycetes(100);c__Agaricostilbomycetes_unclassified(85);c__Agaricostilbomycetes_unclassified(85);c__Agaricostilbomycetes_unclassified(85);c__Agaricostilbomycetes_unclassified(85); | - | - | - | - | - | - | - | Unassigned | - |
| Otu01305 | 45 | k__Fungi(100);p__Ascomycota(100);c__Dothideomycetes(100);o__Pleosporales(100);f__Pleosporales_family_Incertae_sedis(100);f__Pleosporales_family_Incertae_sedis_unclassified(80);f__Pleosporales_family_Incertae_sedis_unclassified(80); | - | - | - | - | - | - | - | Unassigned | - |
| Otu01310 | 45 | k__Fungi(100);p__Ascomycota(100);c__Dothideomycetes(87);c__Dothideomycetes_unclassified(87);c__Dothideomycetes_unclassified(87);c__Dothideomycetes_unclassified(87);c__Dothideomycetes_unclassified(87); | - | - | - | - | - | - | - | Unassigned | - |
| Otu01314 | 44 | k__Fungi(100);p__Ascomycota(100);c__Dothideomycetes(100);o__Pleosporales(100);f__Lophiostomataceae(100);g__Lophiostoma(100);g__Lophiostoma_unclassified(87); | Lophiostoma | 13 | Saprotroph | Plant Saprotroph-Undefined Saprotroph-\|Wood Saprotroph\| | Microfungus | NULL | Probable | As litter_saprotroph (Põlme et al. 2020); Decay wood, leaf, fruit, seed (Põlme et al. 2020) | Tedersoo L, et al. 2014. Science 346:e1256688 ((https://doi.org/10.1126/science.1256688)); Põlme S, et al. 2020. Fungal Diversity 105:1-16 ((https://doi.org/10.1007/s13225-020-00466-2)) |
| Otu01316 | 44 | k__Fungi(100);p__Basidiomycota(100);c__Agaricomycetes(100);o__Trechisporales(100);f__Hydnodontaceae(100);g__Trechispora(100);g__Trechispora_unclassified(94); | Trechispora | 13 | Saprotroph | \|Wood Saprotroph\| | Corticioid | White Rot | Highly Probable | As white_rot (Põlme et al. 2020); Decay wood (Põlme et al. 2020) | Gilbertson RL, Ryvarden L. 1987-1987. North American Polypores. Fungiflora, Oslo (ISBN: 978-0945345060); Rinaldi AC, et al. 2008. Fungal Diversity 33:1-45 ((http://www.fungaldiversity.org/fdp/sfdp/33-1.pdf)) (as Uncertain); Tedersoo L, et al. 2010. Mycorrhiza 20:217-263 ((https://doi.org/10.1007/s00572-009-0274-x)) (as not ECM).; Põlme S, et al. 2020. Fungal Diversity 105:1-16 ((https://doi.org/10.1007/s13225-020-00466-2)) |
| Otu01317 | 44 | k__Fungi(100);p__Basidiomycota(100);c__Agaricomycetes(100);o__Agaricales(100);f__Psathyrellaceae(96);f__Psathyrellaceae_unclassified(53);f__Psathyrellaceae_unclassified(53); | Psathyrellaceae | 9 | Saprotroph | Dung Saprotroph-Plant Saprotroph-Wood Saprotroph | Agaricoid | NULL | Probable | Saprobic, in grass, on dung, rotten wood and similar substrata (Cannon & Kirk 2007) | Cannon PF, Kirk PM. 2007. Fungal Families of the World. CAB International, Cambridge (ISBN: 978-0851998275) |
| Otu01318 | 44 | k__Fungi(100);p__Basidiomycota(100);c__Agaricomycetes(100);o__Corticiales(100);f__Corticiaceae(100);g__Punctularia(100);s__Punctularia_sp_XL_A10(100); | Punctularia | 13 | Pathotroph-Saprotroph | Plant Pathogen-Undefined Saprotroph-\|Wood Saprotroph\| | Corticioid | White Rot | Probable | As white_rot (Põlme et al. 2020); As wood_pathogen (Põlme et al. 2020); Decay wood (Põlme et al. 2020) | Tedersoo L, et al. 2014. Science 346:e1256688 ((https://doi.org/10.1126/science.1256688)); Põlme S, et al. 2020. Fungal Diversity 105:1-16 ((https://doi.org/10.1007/s13225-020-00466-2)) |
| Otu01322 | 44 | k__Fungi(100);p__Ascomycota(100);c__Leotiomycetes(100);o__Helotiales(100);f__Helotiaceae(100);g__Hymenoscyphus(100);s__Hymenoscyphus_macroguttatus(100); | Hymenoscyphus | 13 | Pathotroph-Saprotroph-Symbiotroph | Bryophyte Parasite-Ectomycorrhizal-Ericoid Mycorrhizal-Plant Pathogen-\|Plant Saprotroph\|-Undefined Saprotroph | Microfungus | NULL | Possible | As litter_saprotroph (Põlme et al. 2020); Dark Septate Microfungus; Syn. Pezoloma among other (see e.g., Pezoloma ericae); Decay leaf, fruit, seed (Põlme et al. 2020) | Baral HO, Krieglsteiner L. 2006. Acta Mycologica 41:11-20 ((https://doi.org/10.5586/am.2006.003)) (as a bryophyte parasite); Huhtinen S, et al. 2010. Nova Hedwigia 90:413-431 ((https://doi.org/10.1127/0029-5035/2010/0090-0413)); Bent E, et al. 2011. Applied and Environmental Microbiology 77:3351-3359 ((https://doi.org/10.1128/AEM.02575-10)) (as ectomycorrhizae); Tedersoo L, et al. 2014. Science 346:1256688 ((https://doi.org/10.1126/science.1256688)) (as saprotroph); Põlme S, et al. 2020. Fungal Diversity 105:1-16 ((https://doi.org/10.1007/s13225-020-00466-2)) |
| Otu01323 | 44 | k__Fungi(100);p__Ascomycota(100);c__Sordariomycetes(100);o__Coniochaetales(100);f__Coniochaetaceae(100);g__Coniochaeta(80);s__Coniochaeta_sp(80); | Coniochaeta | 13 | Pathotroph-Saprotroph-Symbiotroph | Animal Pathogen-Dung Saprotroph-Endophyte-Lichen Parasite-Plant Pathogen-\|Undefined Saprotroph\| | Microfungus | NULL | Possible | As foliar_endophyte (Põlme et al. 2020); Decay leaf, fruit, seed, soil, dung, animal material (Põlme et al. 2020) | Bell A. 1983. Dung Fungi: An Illustrated Guide to Coprophilous Fungi in New Zealand. Victoria University Press, Wellington (ISBN: 978-0864730015); Damm U, et al. 2010. Persoonia 24:60-80 ((http://doi.org/10.3767/003158510X500705)); Costa IPMW, et al. 2012. Mycotaxon 119:e26 ((http://mycotaxon.com/resources/checklists/costa_v119_checklist.pdf)); Tedersoo L, et al. 2014. Science 346:1256688 ((https://doi.org/:10.1126/science.1256688)); Irinyi L, et al. 2015. Medical Mycology 53:313-337 ((https://doi.org/10.1093/mmy/myv008)); Lawrey JD, Diederich P. 2016. http://www.lichenicolous.net (accessed 24 May 2017) ((http://www.lichenicolous.net)); Põlme S, et al. 2020. Fungal Diversity 105:1-16 ((https://doi.org/10.1007/s13225-020-00466-2)) |
| Otu01324 | 44 | k__Fungi(100);p__Ascomycota(100);c__Eurotiomycetes(100);o__Chaetothyriales(100);f__Herpotrichiellaceae(100);g__Exophiala(100);s__Exophiala_xenobiotica(100); | Exophiala | 13 | Pathotroph-Saprotroph-Symbiotroph | \|Animal Parasite\|-Animal Pathogen-Endophyte-Plant Saprotroph-Undefined Saprotroph | Dimorphic-Facultative Yeast | NULL | Probable | As litter_saprotroph (Põlme et al. 2020); As root_endophyte_dark_septate (Põlme et al. 2020); Decay leaf, fruit, seed, animal material (Põlme et al. 2020) | Tedersoo L, et al. 2014. Science 346:e1256688 ((https://doi.org/10.1126/science.1256688)), Nagy LG, et al. 2014. Nature Communications 5:e4471 ((https://doi.org/10.1038/ncomms5471)); Põlme S, et al. 2020. Fungal Diversity 105:1-16 ((https://doi.org/10.1007/s13225-020-00466-2)) |
| Otu01331 | 44 | k__Fungi(100);p__Ascomycota(100);c__Eurotiomycetes(100);o__Chaetothyriales(100);f__Herpotrichiellaceae(100);g__Exophiala(100);g__Exophiala_unclassified(94); | Exophiala | 13 | Pathotroph-Saprotroph-Symbiotroph | \|Animal Parasite\|-Animal Pathogen-Endophyte-Plant Saprotroph-Undefined Saprotroph | Dimorphic-Facultative Yeast | NULL | Probable | As litter_saprotroph (Põlme et al. 2020); As root_endophyte_dark_septate (Põlme et al. 2020); Decay leaf, fruit, seed, animal material (Põlme et al. 2020) | Tedersoo L, et al. 2014. Science 346:e1256688 ((https://doi.org/10.1126/science.1256688)), Nagy LG, et al. 2014. Nature Communications 5:e4471 ((https://doi.org/10.1038/ncomms5471)); Põlme S, et al. 2020. Fungal Diversity 105:1-16 ((https://doi.org/10.1007/s13225-020-00466-2)) |
| Otu01332 | 44 | k__Fungi(100);p__Basidiomycota(100);c__Agaricomycetes(100);o__Agaricales(100);f__Mycenaceae(100);g__Panellus(100);s__Panellus_stipticus(100); | Panellus | 13 | Saprotroph | \|Wood Saprotroph\| | Agaricoid | White Rot | Probable | As white_rot (Põlme et al. 2020); Decay wood (Põlme et al. 2020) | Grgurinovic CA, Simpson JA. 2001. Fungal Diversity 8:97-127 ((http://www.fungaldiversity.org/fdp/sfdp/FD_8_97-127.pdf)); Tedersoo L, et al. 2014. Science 346:e1256688 ((https://doi.org/10.1126/science.1256688)); Põlme S, et al. 2020. Fungal Diversity 105:1-16 ((https://doi.org/10.1007/s13225-020-00466-2)) |
| Otu01333 | 44 | k__Fungi(100);p__Basidiomycota(100);c__Tremellomycetes(100);o__Tremellales(100);f__Tremellales_family_Incertae_sedis(100);g__Tremella(100);s__Tremella_globispora(100); | Tremella | 13 | Pathotroph | \|Fungal Parasite\|-Lichen Parasite | Dimorphic-Facultative Yeast-Microfungus-Tremelloid | NULL | Probable | As mycoparasite (Põlme et al. 2020); Decay fungal material (Põlme et al. 2020) | Tedersoo L, et al. 2014. Science 346:e1256688 ((https://doi.org/10.1126/science.1256688)); Põlme S, et al. 2020. Fungal Diversity 105:1-16 ((https://doi.org/10.1007/s13225-020-00466-2)) |
| Otu01334 | 44 | k__Fungi(100);p__Basidiomycota(100);c__Agaricomycetes(100);o__Agaricales(100);f__Mycenaceae(100);g__Mycena(100);s__Mycena_inclinata(100); | Mycena | 13 | Pathotroph-Saprotroph | Plant Pathogen-\|Plant Saprotroph\|-Undefined Saprotroph-Wood Saprotroph | Agaricoid | NULL | Probable | As litter_saprotroph (Põlme et al. 2020); Decay leaf, fruit, seed (Põlme et al. 2020) | Grgurinovic CA, Simpson JA. 2001. Fungal Diversity 8:97-127 ((http://www.fungaldiversity.org/fdp/sfdp/FD_8_97-127.pdf)); Tedersoo L, et al. 2014. Science 346:e1256688 ((https://doi.org/10.1126/science.1256688)); Põlme S, et al. 2020. Fungal Diversity 105:1-16 ((https://doi.org/10.1007/s13225-020-00466-2)); The American Phytopathological Society Common Names of Plant Diseases ((http://www.apsnet.org/publications/commonnames/Pages/default.aspx)) |
| Otu01335 | 44 | k__Fungi(100);p__Basidiomycota(100);c__Agaricostilbomycetes(100);o__Agaricostilbales(100);f__Agaricostilbaceae(100);g__Bensingtonia(100);g__Bensingtonia_unclassified(100); | Bensingtonia | 13 | Saprotroph | \|Undefined Saprotroph\| | Dimorphic-Facultative Yeast | NULL | Highly Probable | NULL | Kurtzman CP, et al. (eds.) 2011. The Yeasts, a Taxonomic Study. Fifth Edition. Vols 1-3. Elsevier, San Diego (ISBN: 9780444521491); Põlme S, et al. 2020. Fungal Diversity 105:1-16 ((https://doi.org/10.1007/s13225-020-00466-2)) |
| Otu01336 | 44 | k__Fungi(100);p__Basidiomycota(100);c__Agaricomycetes(100);o__Hymenochaetales(100);f__Schizoporaceae(100);f__Schizoporaceae_unclassified(94);f__Schizoporaceae_unclassified(94); | Schizoporaceae | 9 | Saprotroph | Wood Saprotroph | Corticioid | White Rot | Probable | Saprobic, causing white rot of standing and fallen wood of coniferous and broadleaved trees, rarely on herbaceous stems (Cannon & Kirk 2007) | Cannon PF, Kirk PM. 2007. Fungal Families of the World. CAB International, Cambridge (ISBN: 978-0851998275) |
| Otu01315 | 44 | k__Fungi(100);p__Ascomycota(100);p__Ascomycota_unclassified(75);p__Ascomycota_unclassified(75);p__Ascomycota_unclassified(75);p__Ascomycota_unclassified(75);p__Ascomycota_unclassified(75); | - | - | - | - | - | - | - | Unassigned | - |
| Otu01319 | 44 | k__Fungi(100);k__Fungi_unclassified(98);k__Fungi_unclassified(98);k__Fungi_unclassified(98);k__Fungi_unclassified(98);k__Fungi_unclassified(98);k__Fungi_unclassified(98); | - | - | - | - | - | - | - | Unassigned | - |
| Otu01320 | 44 | k__Fungi(100);p__Ascomycota(100);c__Dothideomycetes(100);o__Pleosporales(100);o__Pleosporales_unclassified(62);o__Pleosporales_unclassified(62);o__Pleosporales_unclassified(62); | - | - | - | - | - | - | - | Unassigned | - |
| Otu01321 | 44 | k__Fungi(100);p__Ascomycota(100);c__Dothideomycetes(96);o__Capnodiales(78);o__Capnodiales_unclassified(78);o__Capnodiales_unclassified(78);o__Capnodiales_unclassified(78); | - | - | - | - | - | - | - | Unassigned | - |
| Otu01325 | 44 | k__Fungi(100);k__Fungi_unclassified(75);k__Fungi_unclassified(75);k__Fungi_unclassified(75);k__Fungi_unclassified(75);k__Fungi_unclassified(75);k__Fungi_unclassified(75); | - | - | - | - | - | - | - | Unassigned | - |
| Otu01326 | 44 | k__Fungi(100);k__Fungi_unclassified(66);k__Fungi_unclassified(66);k__Fungi_unclassified(66);k__Fungi_unclassified(66);k__Fungi_unclassified(66);k__Fungi_unclassified(66); | - | - | - | - | - | - | - | Unassigned | - |
| Otu01327 | 44 | k__Fungi(100);k__Fungi_unclassified(53);k__Fungi_unclassified(53);k__Fungi_unclassified(53);k__Fungi_unclassified(53);k__Fungi_unclassified(53);k__Fungi_unclassified(53); | - | - | - | - | - | - | - | Unassigned | - |
| Otu01328 | 44 | k__Fungi(100);p__Ascomycota(100);p__Ascomycota_unclassified(100);p__Ascomycota_unclassified(100);p__Ascomycota_unclassified(100);p__Ascomycota_unclassified(100);p__Ascomycota_unclassified(100); | - | - | - | - | - | - | - | Unassigned | - |
| Otu01329 | 44 | k__Fungi(100);p__Ascomycota(62);p__Ascomycota_unclassified(62);p__Ascomycota_unclassified(62);p__Ascomycota_unclassified(62);p__Ascomycota_unclassified(62);p__Ascomycota_unclassified(62); | - | - | - | - | - | - | - | Unassigned | - |
| Otu01330 | 44 | k__Fungi(100);p__Ascomycota(100);c__Dothideomycetes(100);c__Dothideomycetes_unclassified(100);c__Dothideomycetes_unclassified(100);c__Dothideomycetes_unclassified(100);c__Dothideomycetes_unclassified(100); | - | - | - | - | - | - | - | Unassigned | - |
| Otu01337 | 44 | k__Fungi(100);k__Fungi_unclassified(80);k__Fungi_unclassified(80);k__Fungi_unclassified(80);k__Fungi_unclassified(80);k__Fungi_unclassified(80);k__Fungi_unclassified(80); | - | - | - | - | - | - | - | Unassigned | - |
| Otu01340 | 43 | k__Fungi(100);p__Ascomycota(100);c__Leotiomycetes(100);o__Helotiales(100);f__Hyaloscyphaceae(100);f__Hyaloscyphaceae_unclassified(100);f__Hyaloscyphaceae_unclassified(100); | Hyaloscyphaceae | 9 | Saprotroph | Plant Saprotroph-Wood Saprotroph | Microfungus | NULL | Probable | Saprobic on woody and herbaceous material (Cannon & Kirk 2007) | Cannon PF, Kirk PM. 2007. Fungal Families of the World. CAB International, Cambridge (ISBN: 978-0851998275) |
| Otu01341 | 43 | k__Fungi(100);p__Basidiomycota(100);c__Agaricomycetes(100);o__Russulales(100);f__Stereaceae(100);g__Gloeocystidiellum(100);s__Gloeocystidiellum_porosum(100); | Gloeocystidiellum | 13 | Saprotroph | Undefined Saprotroph-\|Wood Saprotroph\| | Corticioid | White Rot | Probable | As white_rot (Põlme et al. 2020); Decay wood (Põlme et al. 2020) | Tedersoo L, et al. 2014. Science 346:e1256688 ((https://doi.org/10.1126/science.1256688)); Põlme S, et al. 2020. Fungal Diversity 105:1-16 ((https://doi.org/10.1007/s13225-020-00466-2)) |
| Otu01342 | 43 | k__Fungi(100);p__Ascomycota(100);c__Ascomycota_class_Incertae_sedis(100);o__Ascomycota_order_Incertae_sedis(100);f__Ascomycota_family_Incertae_sedis(100);g__Pseudoveronaea(100);s__Pseudoveronaea_ellipsoidea(100); | Pseudoveronaea | 13 | Saprotroph | \|Plant Saprotroph\| | Microfungus | NULL | Probable | As litter_saprotroph (Põlme et al. 2020); Decay leaf, fruit, seed (Põlme et al. 2020) | Põlme S, et al. 2020. Fungal Diversity 105:1-16 ((https://doi.org/10.1007/s13225-020-00466-2)) |
| Otu01343 | 43 | k__Fungi(100);p__Ascomycota(100);c__Saccharomycetes(100);o__Saccharomycetales(100);f__Metschnikowiaceae(100);g__Metschnikowia(100);s__Metschnikowia_pulcherrima(91); | Metschnikowia | 13 | Saprotroph | \|Undefined Saprotroph\| | Yeast | NULL | Probable | As nectar-tap_saprotroph (Põlme et al. 2020); Decay sugar-rich substrates (Põlme et al. 2020) | Põlme S, et al. 2020. Fungal Diversity 105:1-16 ((https://doi.org/10.1007/s13225-020-00466-2)) |
| Otu01346 | 43 | k__Fungi(100);p__Basidiomycota(100);c__Agaricomycetes(100);o__Gomphales(100);f__Gomphaceae(100);g__Ramaria(100);s__Ramaria_stricta(100); | Ramaria | 13 | Saprotroph-Symbiotroph | \|Ectomycorrhizal\|-Plant Saprotroph | Clavarioid | NULL | Possible | As litter_saprotroph (Põlme et al. 2020) | Rinaldi AC, et al. 2008. Fungal Diversity 33:1-45 ((http://www.fungaldiversity.org/fdp/sfdp/33-1.pdf)) (as ECM/not ECM); Tedersoo L, et al. 2010. Mycorrhiza 20:217-263 ((10.1007/s00572-009-0274-x)) (pro parte); Põlme S, et al. 2020. Fungal Diversity 105:1-16 ((https://doi.org/10.1007/s13225-020-00466-2)); http://mycorrhizas.info/ecmf.html ((http://mycorrhizas.info/ecmf.html)) (as ECM) |
| Otu01347 | 43 | k__Fungi(100);p__Ascomycota(100);c__Saccharomycetes(100);o__Saccharomycetales(100);f__Metschnikowiaceae(100);g__Metschnikowia(100);s__Metschnikowia_pulcherrima(98); | Metschnikowia | 13 | Saprotroph | \|Undefined Saprotroph\| | Yeast | NULL | Probable | As nectar-tap_saprotroph (Põlme et al. 2020); Decay sugar-rich substrates (Põlme et al. 2020) | Põlme S, et al. 2020. Fungal Diversity 105:1-16 ((https://doi.org/10.1007/s13225-020-00466-2)) |
| Otu01348 | 43 | k__Fungi(100);p__Basidiomycota(100);c__Agaricomycetes(100);o__Russulales(100);f__Peniophoraceae(100);f__Peniophoraceae_unclassified(100);f__Peniophoraceae_unclassified(100); | Peniophoraceae | 9 | Pathotroph-Saprotroph | Plant Pathogen-Wood Saprotroph | Corticioid | White Rot | Probable | Most species are saprobic, causing rots of standing and fallen wood. Many species are adapted to xeric conditions, being able to grow on rapidly drying dead attached branches (Cannon & Kirk 2007) | Cannon PF, Kirk PM. 2007. Fungal Families of the World. CAB International, Cambridge (ISBN: 978-0851998275) |
| Otu01349 | 43 | k__Fungi(100);p__Ascomycota(100);c__Saccharomycetes(100);o__Saccharomycetales(100);f__Metschnikowiaceae(100);g__Metschnikowia(100);s__Metschnikowia_pulcherrima(98); | Metschnikowia | 13 | Saprotroph | \|Undefined Saprotroph\| | Yeast | NULL | Probable | As nectar-tap_saprotroph (Põlme et al. 2020); Decay sugar-rich substrates (Põlme et al. 2020) | Põlme S, et al. 2020. Fungal Diversity 105:1-16 ((https://doi.org/10.1007/s13225-020-00466-2)) |
| Otu01352 | 43 | k__Fungi(100);p__Ascomycota(100);c__Saccharomycetes(100);o__Saccharomycetales(100);f__Metschnikowiaceae(100);g__Metschnikowia(100);s__Metschnikowia_pulcherrima(91); | Metschnikowia | 13 | Saprotroph | \|Undefined Saprotroph\| | Yeast | NULL | Probable | As nectar-tap_saprotroph (Põlme et al. 2020); Decay sugar-rich substrates (Põlme et al. 2020) | Põlme S, et al. 2020. Fungal Diversity 105:1-16 ((https://doi.org/10.1007/s13225-020-00466-2)) |
| Otu01357 | 43 | k__Fungi(100);p__Basidiomycota(100);c__Agaricomycetes(100);o__Polyporales(100);f__Meruliaceae(100);g__Hyphoderma(100);s__Hyphoderma_setigerum(100); | Hyphoderma | 13 | Saprotroph | Undefined Saprotroph-\|Wood Saprotroph\| | Corticioid | White Rot | Probable | As white_rot (Põlme et al. 2020); Decay wood (Põlme et al. 2020) | Tedersoo L, et al. 2014. Science 346:e1256688 ((https://doi.org/10.1126/science.1256688)); Põlme S, et al. 2020. Fungal Diversity 105:1-16 ((https://doi.org/10.1007/s13225-020-00466-2)) |
| Otu01358 | 43 | k__Fungi(100);p__Ascomycota(100);c__Saccharomycetes(100);o__Saccharomycetales(100);f__Metschnikowiaceae(100);g__Metschnikowia(100);s__Metschnikowia_pulcherrima(98); | Metschnikowia | 13 | Saprotroph | \|Undefined Saprotroph\| | Yeast | NULL | Probable | As nectar-tap_saprotroph (Põlme et al. 2020); Decay sugar-rich substrates (Põlme et al. 2020) | Põlme S, et al. 2020. Fungal Diversity 105:1-16 ((https://doi.org/10.1007/s13225-020-00466-2)) |
| Otu01359 | 43 | k__Fungi(100);p__Ascomycota(100);c__Saccharomycetes(100);o__Saccharomycetales(100);f__Metschnikowiaceae(100);g__Metschnikowia(100);s__Metschnikowia_pulcherrima(96); | Metschnikowia | 13 | Saprotroph | \|Undefined Saprotroph\| | Yeast | NULL | Probable | As nectar-tap_saprotroph (Põlme et al. 2020); Decay sugar-rich substrates (Põlme et al. 2020) | Põlme S, et al. 2020. Fungal Diversity 105:1-16 ((https://doi.org/10.1007/s13225-020-00466-2)) |
| Otu01338 | 43 | k__Fungi(100);p__Ascomycota(63);p__Ascomycota_unclassified(63);p__Ascomycota_unclassified(63);p__Ascomycota_unclassified(63);p__Ascomycota_unclassified(63);p__Ascomycota_unclassified(63); | - | - | - | - | - | - | - | Unassigned | - |
| Otu01339 | 43 | k__Fungi(100);p__Ascomycota(84);p__Ascomycota_unclassified(84);p__Ascomycota_unclassified(84);p__Ascomycota_unclassified(84);p__Ascomycota_unclassified(84);p__Ascomycota_unclassified(84); | - | - | - | - | - | - | - | Unassigned | - |
| Otu01344 | 43 | k__Fungi(100);p__Basidiomycota(100);p__Basidiomycota_unclassified(52);p__Basidiomycota_unclassified(52);p__Basidiomycota_unclassified(52);p__Basidiomycota_unclassified(52);p__Basidiomycota_unclassified(52); | - | - | - | - | - | - | - | Unassigned | - |
| Otu01345 | 43 | k__Fungi(100);p__Ascomycota(100);c__Sordariomycetes(100);o__Sordariales(96);o__Sordariales_unclassified(87);o__Sordariales_unclassified(87);o__Sordariales_unclassified(87); | - | - | - | - | - | - | - | Unassigned | - |
| Otu01350 | 43 | k__Fungi(100);p__Ascomycota(100);c__Eurotiomycetes(100);o__Eurotiales(96);f__Trichocomaceae(96);f__Trichocomaceae_unclassified(96);f__Trichocomaceae_unclassified(96); | - | - | - | - | - | - | - | Unassigned | - |
| Otu01351 | 43 | k__Fungi(100);p__Basidiomycota(100);c__Tremellomycetes(100);o__Cystofilobasidiales(100);f__unclassified_Cystofilobasidiales(100);g__unclassified_Cystofilobasidiales(100);s__Cystofilobasidiales_sp(100); | - | - | - | - | - | - | - | Unassigned | - |
| Otu01353 | 43 | k__Fungi(100);p__Ascomycota(100);p__Ascomycota_unclassified(70);p__Ascomycota_unclassified(70);p__Ascomycota_unclassified(70);p__Ascomycota_unclassified(70);p__Ascomycota_unclassified(70); | - | - | - | - | - | - | - | Unassigned | - |
| Otu01354 | 43 | k__Fungi(100);k__Fungi_unclassified(73);k__Fungi_unclassified(73);k__Fungi_unclassified(73);k__Fungi_unclassified(73);k__Fungi_unclassified(73);k__Fungi_unclassified(73); | - | - | - | - | - | - | - | Unassigned | - |
| Otu01355 | 43 | k__Fungi(100);p__Ascomycota(100);p__Ascomycota_unclassified(100);p__Ascomycota_unclassified(100);p__Ascomycota_unclassified(100);p__Ascomycota_unclassified(100);p__Ascomycota_unclassified(100); | - | - | - | - | - | - | - | Unassigned | - |
| Otu01356 | 43 | k__Fungi(100);k__Fungi_unclassified(70);k__Fungi_unclassified(70);k__Fungi_unclassified(70);k__Fungi_unclassified(70);k__Fungi_unclassified(70);k__Fungi_unclassified(70); | - | - | - | - | - | - | - | Unassigned | - |
| Otu01362 | 42 | k__Fungi(100);p__Ascomycota(100);c__Saccharomycetes(100);o__Saccharomycetales(100);f__Metschnikowiaceae(100);g__Metschnikowia(100);s__Metschnikowia_pulcherrima(93); | Metschnikowia | 13 | Saprotroph | \|Undefined Saprotroph\| | Yeast | NULL | Probable | As nectar-tap_saprotroph (Põlme et al. 2020); Decay sugar-rich substrates (Põlme et al. 2020) | Põlme S, et al. 2020. Fungal Diversity 105:1-16 ((https://doi.org/10.1007/s13225-020-00466-2)) |
| Otu01363 | 42 | k__Fungi(100);p__Basidiomycota(100);c__Agaricomycetes(100);o__Polyporales(100);f__Meripilaceae(100);g__Grifola(100);s__Grifola_frondosa(100); | Grifola | 13 | Pathotroph-Saprotroph | Plant Pathogen-\|Wood Saprotroph\| | Polyporoid | White Rot | Highly Probable | As white_rot (Põlme et al. 2020); As wood_pathogen (Põlme et al. 2020); Decay wood (Põlme et al. 2020) | Gilbertson RL, Ryvarden L. 1987-1987. North American Polypores. Fungiflora, Oslo (ISBN: 978-0945345060); Põlme S, et al. 2020. Fungal Diversity 105:1-16 ((https://doi.org/10.1007/s13225-020-00466-2)) |
| Otu01365 | 42 | k__Fungi(100);p__Ascomycota(100);c__Saccharomycetes(100);o__Saccharomycetales(100);f__Metschnikowiaceae(100);g__Metschnikowia(100);s__Metschnikowia_pulcherrima(98); | Metschnikowia | 13 | Saprotroph | \|Undefined Saprotroph\| | Yeast | NULL | Probable | As nectar-tap_saprotroph (Põlme et al. 2020); Decay sugar-rich substrates (Põlme et al. 2020) | Põlme S, et al. 2020. Fungal Diversity 105:1-16 ((https://doi.org/10.1007/s13225-020-00466-2)) |
| Otu01368 | 42 | k__Fungi(100);p__Basidiomycota(100);c__Agaricomycetes(100);o__Agaricales(100);f__Bolbitiaceae(100);g__Conocybe(100);s__Conocybe_apala(100); | Conocybe | 13 | Saprotroph | Dung Saprotroph-Plant Saprotroph-\|Undefined Saprotroph\| | Agaricoid | NULL | Probable | As litter_saprotroph (Põlme et al. 2020); As soil_saprotroph (Põlme et al. 2020); Decay soil (Põlme et al. 2020) | Tedersoo L, et al. 2014. Science 346:e1256688 ((https://doi.org/10.1126/science.1256688)); Põlme S, et al. 2020. Fungal Diversity 105:1-16 ((https://doi.org/10.1007/s13225-020-00466-2)) |
| Otu01369 | 42 | k__Fungi(100);p__Ascomycota(100);c__Eurotiomycetes(100);o__Chaetothyriales(100);f__Chaetothyriaceae(100);g__Cyphellophora(100);s__Cyphellophora_europaea(84); | Cyphellophora | 13 | Pathotroph-Saprotroph | Animal Parasite-Animal Pathogen-\|Plant Saprotroph\|-Undefined Saprotroph | Dimorphic-Facultative Yeast-Microfungus | NULL | Probable | As litter_saprotroph (Põlme et al. 2020); Consult the current literature for updated names on this group that included the Black Yeasts; Some species are likely opportunistic human pathogen (Irinyi et al. 2015); Decay leaf, fruit, seed, animal material (Põlme et al. 2020) | Tedersoo L, et al. 2014. Science 346:e1256688 ((https://doi.org/10.1126/science.1256688)); Irinyi L, et al. 2015. Medical Mycology 53:313-337 ((https://doi.org/10.1093/mmy/myv008)); Põlme S, et al. 2020. Fungal Diversity 105:1-16 ((https://doi.org/10.1007/s13225-020-00466-2)) |
| Otu01371 | 42 | k__Fungi(100);p__Basidiomycota(100);c__Agaricomycetes(100);o__Agaricales(100);f__Cortinariaceae(100);g__Cortinarius(100);s__Cortinarius_leiocastaneus(100); | Cortinarius | 13 | Symbiotroph | \|Ectomycorrhizal\| | Agaricoid | NULL | Highly Probable | NULL | Rinaldi AC, et al. 2008. Fungal Diversity 33:1-45 ((http://www.fungaldiversity.org/fdp/sfdp/33-1.pdf)); Tedersoo L, et al. 2010. Mycorrhiza 20:217-263 ((https://doi.org/10.1007/s00572-009-0274-x)); Põlme S, et al. 2020. Fungal Diversity 105:1-16 ((https://doi.org/10.1007/s13225-020-00466-2)); http://mycorrhizas.info/ecmf.html ((http://mycorrhizas.info/ecmf.html)) |
| Otu01373 | 42 | k__Fungi(100);p__Basidiomycota(100);c__Agaricomycetes(100);o__Polyporales(93);f__Meruliaceae(93);g__Hyphoderma(93);g__Hyphoderma_unclassified(93); | Hyphoderma | 13 | Saprotroph | Undefined Saprotroph-\|Wood Saprotroph\| | Corticioid | White Rot | Probable | As white_rot (Põlme et al. 2020); Decay wood (Põlme et al. 2020) | Tedersoo L, et al. 2014. Science 346:e1256688 ((https://doi.org/10.1126/science.1256688)); Põlme S, et al. 2020. Fungal Diversity 105:1-16 ((https://doi.org/10.1007/s13225-020-00466-2)) |
| Otu01375 | 42 | k__Fungi(100);p__Basidiomycota(100);c__Agaricomycetes(100);o__Boletales(100);f__Tapinellaceae(100);g__Pseudomerulius(100);s__Pseudomerulius_curtisii(100); | Pseudomerulius | 13 | Saprotroph | \|Wood Saprotroph\| | Corticioid | Brown Rot | Probable | As brown-rot (Põlme et al. 2020); Decay wood (Põlme et al. 2020) | Tedersoo L, et al. 2014. Science 346:e1256688 ((https://doi.org/10.1126/science.1256688)); Põlme S, et al. 2020. Fungal Diversity 105:1-16 ((https://doi.org/10.1007/s13225-020-00466-2)) |
| Otu01376 | 42 | k__Fungi(100);p__Ascomycota(100);c__Eurotiomycetes(100);o__Eurotiales(100);f__Trichocomaceae(100);g__Aspergillus(100);s__Aspergillus_penicillioides(96); | Aspergillus | 13 | Pathotroph-Saprotroph-Symbiotroph | Animal Pathogen-Endophyte-Plant Saprotroph-\|Undefined Saprotroph\|-Wood Saprotroph | Microfungus | NULL | Probable | As foliar_endophyte (Põlme et al. 2020); Decay leaf, fruit, seed, soil, dung, animal material (Põlme et al. 2020) | Seehann G, et al. 1975. List of Fungi in Soft-Rot Tissue Tests. Institute fur Holzbiologie und Holzschutz, Bundesforschungsanstalt fur Forst - und Holzwirtshaft, Hamburg - Lohbrugge, Germany; Tedersoo L, et al. 2014. Science 346:e1256688 ((https://doi.org/10.1126/science.1256688)); Irinyi L, et al. 2015. Medical Mycology 53:313-337 ((https://doi.org/10.1093/mmy/myv008)); Põlme S, et al. 2020. Fungal Diversity 105:1-16 ((https://doi.org/10.1007/s13225-020-00466-2)); Massimo NC, et al. 2015. Microbial Ecology 70:61-76 ((https://doi.org/10.1007/s00248-014-0563-6)); The American Phytopathological Society Common Names of Plant Diseases ((http://www.apsnet.org/publications/commonnames/Pages/default.aspx)); https://www.cdc.gov ((https://www.cdc.gov)) |
| Otu01377 | 42 | k__Fungi(100);p__Basidiomycota(100);c__Microbotryomycetes(100);o__Sporidiobolales(100);f__Sporidiobolales_family_Incertae_sedis(100);g__Rhodotorula(100);s__Rhodotorula_mucilaginosa(100); | Rhodotorula | 13 | Pathotroph-Saprotroph-Symbiotroph | Animal Pathogen-Animal Symbiotroph-Endophyte-Plant Pathogen-\|Undefined Saprotroph\| | Yeast | NULL | Probable | As foliar_endophyte (Põlme et al. 2020); Opportunistic human pathogen | Põlme S, et al. 2020. Fungal Diversity 105:1-16 ((https://doi.org/10.1007/s13225-020-00466-2)); https://www.cdc.gov ((https://www.cdc.gov)) |
| Otu01378 | 42 | k__Fungi(100);p__Basidiomycota(100);c__Agaricomycetes(100);o__Hymenochaetales(100);f__Schizoporaceae(100);g__Hyphodontia(100);s__Hyphodontia_palmae(100); | Hyphodontia | 13 | Saprotroph | Undefined Saprotroph-\|Wood Saprotroph\| | Corticioid | White Rot | Probable | As white_rot (Põlme et al. 2020); Decay wood (Põlme et al. 2020) | Tedersoo L, et al. 2014. Science 346:e1256688 ((https://doi.org/10.1126/science.1256688)); Põlme S, et al. 2020. Fungal Diversity 105:1-16 ((https://doi.org/10.1007/s13225-020-00466-2)) |
| Otu01380 | 42 | k__Fungi(100);p__Basidiomycota(100);c__Microbotryomycetes(100);o__Sporidiobolales(100);f__Sporidiobolales_family_Incertae_sedis(100);g__Sporobolomyces(100);s__Sporobolomyces_lactosus(100); | Sporobolomyces | 13 | Pathotroph-Saprotroph | \|Fungal Parasite\|-Plant Saprotroph | Yeast | NULL | Probable | As litter_saprotroph (Põlme et al. 2020); As mycoparasite (Põlme et al. 2020); Decay fungal material (Põlme et al. 2020) | Tedersoo L, et al. 2014. Science 346:e1256688 ((https://doi.org/10.1126/science.1256688)); Purahong W, et al. 2016. Molecular Ecology 25:4059-4074 ((https://doi.org/10.1111/mec.13739)); Põlme S, et al. 2020. Fungal Diversity 105:1-16 ((https://doi.org/10.1007/s13225-020-00466-2)) |
| Otu01381 | 42 | k__Fungi(100);p__Ascomycota(100);c__Saccharomycetes(100);o__Saccharomycetales(100);f__Metschnikowiaceae(100);g__Metschnikowia(100);s__Metschnikowia_pulcherrima(93); | Metschnikowia | 13 | Saprotroph | \|Undefined Saprotroph\| | Yeast | NULL | Probable | As nectar-tap_saprotroph (Põlme et al. 2020); Decay sugar-rich substrates (Põlme et al. 2020) | Põlme S, et al. 2020. Fungal Diversity 105:1-16 ((https://doi.org/10.1007/s13225-020-00466-2)) |
| Otu01360 | 42 | k__Fungi(100);p__Basidiomycota(100);c__Agaricomycetes(100);o__Polyporales(100);f__Phanerochaetaceae(98);f__Phanerochaetaceae_unclassified(96);f__Phanerochaetaceae_unclassified(96); | - | - | - | - | - | - | - | Unassigned | - |
| Otu01361 | 42 | k__Fungi(100);p__Ascomycota(100);c__Eurotiomycetes(96);c__Eurotiomycetes_unclassified(96);c__Eurotiomycetes_unclassified(96);c__Eurotiomycetes_unclassified(96);c__Eurotiomycetes_unclassified(96); | - | - | - | - | - | - | - | Unassigned | - |
| Otu01364 | 42 | k__Fungi(100);p__Ascomycota(100);c__Dothideomycetes(100);o__Pleosporales(100);o__Pleosporales_unclassified(98);o__Pleosporales_unclassified(98);o__Pleosporales_unclassified(98); | - | - | - | - | - | - | - | Unassigned | - |
| Otu01366 | 42 | k__Fungi(100);k__Fungi_unclassified(67);k__Fungi_unclassified(67);k__Fungi_unclassified(67);k__Fungi_unclassified(67);k__Fungi_unclassified(67);k__Fungi_unclassified(67); | - | - | - | - | - | - | - | Unassigned | - |
| Otu01367 | 42 | k__Fungi(100);p__Ascomycota(100);c__Leotiomycetes(100);o__Rhytismatales(98);o__Rhytismatales_unclassified(98);o__Rhytismatales_unclassified(98);o__Rhytismatales_unclassified(98); | - | - | - | - | - | - | - | Unassigned | - |
| Otu01370 | 42 | k__Fungi(100);k__Fungi_unclassified(60);k__Fungi_unclassified(60);k__Fungi_unclassified(60);k__Fungi_unclassified(60);k__Fungi_unclassified(60);k__Fungi_unclassified(60); | - | - | - | - | - | - | - | Unassigned | - |
| Otu01372 | 42 | k__Fungi(100);p__Basidiomycota(100);c__Agaricomycetes(100);c__Agaricomycetes_unclassified(100);c__Agaricomycetes_unclassified(100);c__Agaricomycetes_unclassified(100);c__Agaricomycetes_unclassified(100); | - | - | - | - | - | - | - | Unassigned | - |
| Otu01374 | 42 | k__Fungi(100);k__Fungi_unclassified(100);k__Fungi_unclassified(100);k__Fungi_unclassified(100);k__Fungi_unclassified(100);k__Fungi_unclassified(100);k__Fungi_unclassified(100); | - | - | - | - | - | - | - | Unassigned | - |
| Otu01379 | 42 | k__Fungi(100);k__Fungi_unclassified(67);k__Fungi_unclassified(67);k__Fungi_unclassified(67);k__Fungi_unclassified(67);k__Fungi_unclassified(67);k__Fungi_unclassified(67); | - | - | - | - | - | - | - | Unassigned | - |
| Otu01383 | 41 | k__Fungi(100);p__Basidiomycota(100);c__Microbotryomycetes(100);o__Sporidiobolales(100);f__Sporidiobolales_family_Incertae_sedis(100);g__Sporobolomyces(100);s__Sporobolomyces_lactosus(100); | Sporobolomyces | 13 | Pathotroph-Saprotroph | \|Fungal Parasite\|-Plant Saprotroph | Yeast | NULL | Probable | As litter_saprotroph (Põlme et al. 2020); As mycoparasite (Põlme et al. 2020); Decay fungal material (Põlme et al. 2020) | Tedersoo L, et al. 2014. Science 346:e1256688 ((https://doi.org/10.1126/science.1256688)); Purahong W, et al. 2016. Molecular Ecology 25:4059-4074 ((https://doi.org/10.1111/mec.13739)); Põlme S, et al. 2020. Fungal Diversity 105:1-16 ((https://doi.org/10.1007/s13225-020-00466-2)) |
| Otu01385 | 41 | k__Fungi(100);p__Ascomycota(100);c__Eurotiomycetes(100);o__Onygenales(100);f__Gymnoascaceae(100);g__Gymnoascus(100);s__Gymnoascus_sp(100); | Gymnoascus | 13 | Saprotroph | \|Undefined Saprotroph\| | Microfungus | NULL | Probable | As soil_saprotroph (Põlme et al. 2020); Decay soil (Põlme et al. 2020) | Tedersoo L, et al. 2014. Science 346:e1256688 ((https://doi.org/10.1126/science.1256688)); Põlme S, et al. 2020. Fungal Diversity 105:1-16 ((https://doi.org/10.1007/s13225-020-00466-2)) |
| Otu01386 | 41 | k__Fungi(100);p__Basidiomycota(100);c__Agaricomycetes(100);o__Agaricales(100);f__Physalacriaceae(100);g__Cylindrobasidium(100);s__Cylindrobasidium_evolvens(100); | Cylindrobasidium | 13 | Saprotroph | Undefined Saprotroph-\|Wood Saprotroph\| | Corticioid | White Rot | Possible | As white_rot (Põlme et al. 2020); Decay wood (Põlme et al. 2020) | Jülich W. 1974. Persoonia 8:59-98 ((http://www.cybertruffle.org.uk/cyberliber/61056/0008/001/0072.htm)); Põlme S, et al. 2020. Fungal Diversity 105:1-16 ((https://doi.org/10.1007/s13225-020-00466-2)) |
| Otu01387 | 41 | k__Fungi(100);p__Basidiomycota(100);c__Microbotryomycetes(100);o__Sporidiobolales(100);f__Sporidiobolales_family_Incertae_sedis(100);g__Rhodotorula(100);s__Rhodotorula_taiwanensis(100); | Rhodotorula | 13 | Pathotroph-Saprotroph-Symbiotroph | Animal Pathogen-Animal Symbiotroph-Endophyte-Plant Pathogen-\|Undefined Saprotroph\| | Yeast | NULL | Probable | As foliar_endophyte (Põlme et al. 2020); Opportunistic human pathogen | Põlme S, et al. 2020. Fungal Diversity 105:1-16 ((https://doi.org/10.1007/s13225-020-00466-2)); https://www.cdc.gov ((https://www.cdc.gov)) |
| Otu01390 | 41 | k__Fungi(100);p__Basidiomycota(100);c__Tremellomycetes(100);o__Cystofilobasidiales(100);f__Cystofilobasidiales_family_Incertae_sedis(100);g__Syzygospora(100);s__Syzygospora_bachmannii(100); | Syzygospora | 13 | Pathotroph | \|Fungal Parasite\|-Lichen Parasite | Dimorphic-Tremelloid | NULL | Highly Probable | As mycoparasite (Põlme et al. 2020); Decay fungal material (Põlme et al. 2020) | Põlme S, et al. 2020. Fungal Diversity 105:1-16 ((https://doi.org/10.1007/s13225-020-00466-2)); List compiled by John Plischke III ((https://goodmushroom.weebly.com/uploads/5/7/1/7/57176535/fungi_on_fungi_pdf_lq.pdf)) |
| Otu01391 | 41 | k__Fungi(100);p__Basidiomycota(100);c__Agaricomycetes(100);o__Polyporales(100);f__Meruliaceae(100);g__Steccherinum(100);s__Steccherinum_ochraceum(98); | Steccherinum | 13 | Saprotroph | Undefined Saprotroph-\|Wood Saprotroph\| | Corticioid | White Rot | Probable | As white_rot (Põlme et al. 2020); Decay wood (Põlme et al. 2020) | Tedersoo L, et al. 2014. Science 346:e1256688 ((https://doi.org/10.1126/science.1256688)); Põlme S, et al. 2020. Fungal Diversity 105:1-16 ((https://doi.org/10.1007/s13225-020-00466-2)) |
| Otu01392 | 41 | k__Fungi(100);p__Basidiomycota(100);c__Agaricomycetes(100);o__Russulales(88);f__Peniophoraceae(83);g__Gloiothele(83);s__Gloiothele_citrina(83); | Gloiothele | 13 | Saprotroph | Undefined Saprotroph-\|Wood Saprotroph\| | Corticioid | White Rot | Probable | As white_rot (Põlme et al. 2020); Decay wood (Põlme et al. 2020) | Tedersoo L, et al. 2014. Science 346:e1256688 ((https://doi.org/10.1126/science.1256688)); Põlme S, et al. 2020. Fungal Diversity 105:1-16 ((https://doi.org/10.1007/s13225-020-00466-2)) |

| | | | | | | | | | | | |
|---|---|---|---|---|---|---|---|---|---|---|---|
| Otu01394 | 41 | k__Fungi(100);p__Basidiomycota(100);c__Agaricomycetes(100);o__Agaricales(100);o__Agaricales_unclassified(100);o__Agaricales_unclassified(100);o__Agaricales_unclassified(100); | Agaricales | 7 | Pathotroph-Saprotroph-Symbiotroph | Bryophyte Parasite-Dung Saprotroph-Ectomycorrhizal-Fungal Parasite-Leaf Saprotroph-Plant Parasite-Undefined Saprotroph-Wood Saprotroph | Agaricoid-Gasteroid-Microfungus-Secotioid-Yeast | NULL | Possible | Mushrooms and toadstools, Gill fungi, Agarics... lignicolous, sometimes muscicolous or fungicolous, saprobic, mycorrhizal, rarely parasitic on plants or fungi (Kirk et al. 2008) | [illegible] |
| Otu01382 | 41 | k__Fungi(100);k__Fungi_unclassified(59);k__Fungi_unclassified(59);k__Fungi_unclassified(59);k__Fungi_unclassified(59);k__Fungi_unclassified(59);k__Fungi_unclassified(59); | - | - | - | - | - | - | - | Unassigned | - |
| Otu01384 | 41 | k__Fungi(100);p__Ascomycota(100);c__Dothideomycetes(100);o__Capnodiales(100);o__Capnodiales_unclassified(86);o__Capnodiales_unclassified(86);o__Capnodiales_unclassified(86); | - | - | - | - | - | - | - | Unassigned | - |
| Otu01388 | 41 | k__Fungi(100);p__Ascomycota(100);c__Dothideomycetes(100);o__Pleosporales(100);o__Pleosporales_unclassified(100);o__Pleosporales_unclassified(100);o__Pleosporales_unclassified(100); | - | - | - | - | - | - | - | Unassigned | - |
| Otu01389 | 41 | k__Fungi(100);p__Basidiomycota(100);c__Agaricomycetes(100);o__Polyporales(98);o__Polyporales_unclassified(98);o__Polyporales_unclassified(98);o__Polyporales_unclassified(98); | - | - | - | - | - | - | - | Unassigned | - |
| Otu01393 | 41 | k__Fungi(100);p__Basidiomycota(100);c__Microbotryomycetes(100);o__Sporidiobolales(100);f__Sporidiobolales_family_Incertae_sedis(100);g__unclassified_Sporidiobolales(57);s__Sporidiobolales_sp(57); | - | - | - | - | - | - | - | Unassigned | - |
| Otu01395 | 40 | k__Fungi(100);p__Basidiomycota(100);c__Agaricomycetes(100);o__Agaricales(70);o__Agaricales_unclassified(70);o__Agaricales_unclassified(70);o__Agaricales_unclassified(70); | Agaricales | 7 | Pathotroph-Saprotroph-Symbiotroph | Bryophyte Parasite-Dung Saprotroph-Ectomycorrhizal-Fungal Parasite-Leaf Saprotroph-Plant Parasite-Undefined Saprotroph-Wood Saprotroph | Agaricoid-Gasteroid-Microfungus-Secotioid-Yeast | NULL | Possible | Mushrooms and toadstools, Gill fungi, Agarics... lignicolous, sometimes muscicolous or fungicolous, saprobic, mycorrhizal, rarely parasitic on plants or fungi (Kirk et al. 2008) | [illegible] |
| Otu01396 | 40 | k__Fungi(100);p__Ascomycota(100);c__Sordariomycetes(100);o__Xylariales(100);f__Diatrypaceae(100);g__Eutypella(100);s__Eutypella_scoparia(98); | Eutypella | 13 | Pathotroph | \|Plant Pathogen\| | Microfungus | NULL | Probable | NULL | [illegible] |
| Otu01398 | 40 | k__Fungi(100);p__Zygomycota(100);c__Zygomycota_class_Incertae_sedis(100);o__Mucorales(100);f__Mucoraceae(100);g__Mucor(100);g__Mucor_unclassified(100); | Mucor | 13 | Saprotroph | \|Undefined Saprotroph\| | Microfungus | NULL | Probable | As soil_saprotroph (Põlme et al. 2020); Decay soil (Põlme et al. 2020) | [illegible] |
| Otu01399 | 40 | k__Fungi(100);p__Basidiomycota(100);c__Agaricomycetes(100);o__Agaricales(100);f__Marasmiaceae(100);g__Calyptella(100);s__Calyptella_capula(100); | Calyptella | 13 | Pathotroph-Saprotroph | Plant Pathogen-\|Plant Saprotroph\|-Undefined Saprotroph-Wood Saprotroph | Agaricoid | NULL | Probable | As litter_saprotroph (Põlme et al. 2020); Decay wood, leaf, fruit, seed (Põlme et al. 2020) | [illegible] |
| Otu01401 | 40 | k__Fungi(100);p__Basidiomycota(100);c__Agaricomycetes(100);o__Agaricales(100);f__Psathyrellaceae(100);g__Coprinopsis(100);s__Coprinopsis_sp(100); | Coprinopsis | 13 | Saprotroph | \|Undefined Saprotroph\| | Agaricoid | NULL | Probable | As soil_saprotroph (Põlme et al. 2020); Decay soil (Põlme et al. 2020) | [illegible] |
| Otu01403 | 40 | k__Fungi(100);p__Ascomycota(100);c__Dothideomycetes(100);o__Capnodiales(100);f__Mycosphaerellaceae(100);g__Uwebraunia(100);s__Uwebraunia_dekkeri(53); | Uwebraunia | 13 | Saprotroph | \|Plant Saprotroph\|-Undefined Saprotroph | Microfungus | NULL | Probable | As litter_saprotroph (Põlme et al. 2020); Decay leaf, fruit, seed (Põlme et al. 2020) | [illegible] |
| Otu01404 | 40 | k__Fungi(100);p__Basidiomycota(100);c__Tremellomycetes(100);o__Tremellales(100);f__Tremellales_family_Incertae_sedis(100);g__Cryptococcus(90);s__Cryptococcus_dimennae(90); | Cryptococcus | 13 | Pathotroph-Saprotroph-Symbiotroph | Animal Pathogen-Endophyte-Epiphyte-\|Undefined Saprotroph\| | Dimorphic Yeast-Yeast | NULL | Possible | As foliar_endophyte (Põlme et al. 2020); Decay leaf, fruit, seed, soil, dung, animal material (Põlme et al. 2020) | [illegible] |
| Otu01405 | 40 | k__Fungi(100);p__Basidiomycota(100);c__Microbotryomycetes(100);o__Leucosporidiales(100);f__Leucosporidiaceae(100);g__Leucosporidium(100);s__Leucosporidium_golubevii(100); | Leucosporidium | 13 | Saprotroph | \|Undefined Saprotroph\| | Yeast | NULL | Probable | A yeast from fruit (Barnett & Buhagiar 1971; Syn. Candida; As soil_saprotroph (Põlme et al. 2020); Decay soil (Põlme et al. 2020); Leucosporidiella; Rhodotorula; Torulopsis; Vanrijia spp.);Isolated from permafrost soil (Golubev 1998; Syn. Leucosporidium/Rhodotorula spp.) | [illegible] |
| Otu01408 | 40 | k__Fungi(100);p__Ascomycota(100);c__Sordariomycetes(100);o__Microascales(100);f__Halosphaeriaceae(100);g__unclassified_Halosphaeriaceae(100);s__Halosphaeriaceae_sp(100); | Halosphaeriaceae | 9 | Saprotroph | Undefined Saprotroph-Wood Saprotroph | Microfungus | NULL | Probable | Found in aquatic habitats, mostly in marine environments; saprobic on submerged or intertidal wood, occasionally associated with sand grains and similar substrata (Cannon & Kirk 2007) | [illegible] |
| Otu01410 | 40 | k__Fungi(100);p__Basidiomycota(100);c__Agaricomycetes(100);o__Trechisporales(100);f__Hydnodontaceae(100);g__Trechispora(100);s__Trechispora_sp(100); | Trechispora | 13 | Saprotroph | \|Wood Saprotroph\| | Corticioid | White Rot | Highly Probable | As white_rot (Põlme et al. 2020); Decay wood (Põlme et al. 2020) | [illegible] |
| Otu01411 | 40 | k__Fungi(100);p__Basidiomycota(100);c__Agaricomycetes(100);o__Boletales(100);f__Boletaceae(100);f__Boletaceae_unclassified(95);f__Boletaceae_unclassified(95); | Boletaceae | 9 | Pathotroph-Saprotroph-Symbiotroph | Ectomycorrhizal-Fungal Parasite-Plant Saprotroph-Wood Saprotroph | Boletoid | NULL | Probable | Most species form ectomycorrhizal associations with gymnosperm or angiosperm trees. Some are saprobes on decaying wood and leaf litter, a few are parasitic on other fungi (Cannon & Kirk 2007) | [illegible] |
| Otu01412 | 40 | k__Fungi(100);p__Ascomycota(100);c__Dothideomycetes(100);o__Myriangiales(100);f__Elsinoaceae(100);g__Elsinoe(93);g__Elsinoe_unclassified(93); | Elsinoe | 13 | Pathotroph-Saprotroph | \|Plant Pathogen\|-Plant Saprotroph | Microfungus | NULL | Probable | As litter_saprotroph (Põlme et al. 2020); Decay leaf, fruit, seed (Põlme et al. 2020) | [illegible] |
| Otu01413 | 40 | k__Fungi(100);p__Ascomycota(100);c__Ascomycota_class_Incertae_sedis(100);o__Ascomycota_order_Incertae_sedis(100);f__Ascomycota_family_Incertae_sedis(100);g__Phialemoniopsis(100);s__Phialemoniopsis_ocularis(83); | Phialemoniopsis | 13 | Pathotroph-Saprotroph | \|Animal Parasite\|-Undefined Saprotroph | Microfungus | NULL | Probable | As animal_decomposer (Põlme et al. 2020); Decay animal material (Põlme et al. 2020) | [illegible] |
| Otu01414 | 40 | k__Fungi(100);p__Ascomycota(100);c__Saccharomycetes(100);o__Saccharomycetales(100);f__Metschnikowiaceae(100);g__Metschnikowia(100);s__Metschnikowia_pulcherrima(83); | Metschnikowia | 13 | Saprotroph | \|Undefined Saprotroph\| | Yeast | NULL | Probable | As nectar-tap_saprotroph (Põlme et al. 2020); Decay sugar-rich substrates (Põlme et al. 2020) | [illegible] |
| Otu01397 | 40 | k__Fungi(100);p__Basidiomycota(100);c__Agaricomycetes(100);o__Polyporales(100);f__Phanerochaetaceae(95);f__Phanerochaetaceae_unclassified(95);f__Phanerochaetaceae_unclassified(95); | - | - | - | - | - | - | - | Unassigned | - |
| Otu01400 | 40 | k__Fungi(100);p__Ascomycota(100);c__Dothideomycetes(88);c__Dothideomycetes_unclassified(63);c__Dothideomycetes_unclassified(63);c__Dothideomycetes_unclassified(63);c__Dothideomycetes_unclassified(63); | - | - | - | - | - | - | - | Unassigned | - |
| Otu01402 | 40 | k__Fungi(100);k__Fungi_unclassified(78);k__Fungi_unclassified(78);k__Fungi_unclassified(78);k__Fungi_unclassified(78);k__Fungi_unclassified(78);k__Fungi_unclassified(78); | - | - | - | - | - | - | - | Unassigned | - |
| Otu01406 | 40 | k__Fungi(100);k__Fungi_unclassified(65);k__Fungi_unclassified(65);k__Fungi_unclassified(65);k__Fungi_unclassified(65);k__Fungi_unclassified(65);k__Fungi_unclassified(65); | - | - | - | - | - | - | - | Unassigned | - |
| Otu01407 | 40 | k__Fungi(100);p__Ascomycota(80);p__Ascomycota_unclassified(80);p__Ascomycota_unclassified(80);p__Ascomycota_unclassified(80);p__Ascomycota_unclassified(80);p__Ascomycota_unclassified(80); | - | - | - | - | - | - | - | Unassigned | - |
| Otu01409 | 40 | k__Fungi(100);p__Basidiomycota(100);c__Agaricomycetes(100);c__Agaricomycetes_unclassified(100);c__Agaricomycetes_unclassified(100);c__Agaricomycetes_unclassified(100);c__Agaricomycetes_unclassified(100); | - | - | - | - | - | - | - | Unassigned | - |
| Otu01415 | 40 | k__Fungi(100);k__Fungi_unclassified(55);k__Fungi_unclassified(55);k__Fungi_unclassified(55);k__Fungi_unclassified(55);k__Fungi_unclassified(55);k__Fungi_unclassified(55); | - | - | - | - | - | - | - | Unassigned | - |
| Otu01416 | 40 | k__Fungi(100);p__Basidiomycota(100);c__Atractiellomycetes(100);o__Atractiellales(100);f__unclassified_Atractiellales(100);g__unclassified_Atractiellales(100);s__Atractiellales_sp(100); | - | - | - | - | - | - | - | Unassigned | - |
| Otu01417 | 40 | k__Fungi(100);p__Ascomycota(100);p__Ascomycota_unclassified(100);p__Ascomycota_unclassified(100);p__Ascomycota_unclassified(100);p__Ascomycota_unclassified(100);p__Ascomycota_unclassified(100); | - | - | - | - | - | - | - | Unassigned | - |
| Otu01418 | 39 | k__Fungi(100);p__Ascomycota(100);c__Saccharomycetes(100);o__Saccharomycetales(100);f__Metschnikowiaceae(100);g__Metschnikowia(100);g__Metschnikowia_unclassified(90); | Metschnikowia | 13 | Saprotroph | \|Undefined Saprotroph\| | Yeast | NULL | Probable | As nectar-tap_saprotroph (Põlme et al. 2020); Decay sugar-rich substrates (Põlme et al. 2020) | [illegible] |
| Otu01421 | 39 | k__Fungi(100);p__Ascomycota(100);c__Taphrinomycetes(100);o__Taphrinales(100);f__Taphrinaceae(100);g__Taphrina(100);s__Taphrina_confusa(95); | Taphrina | 13 | Pathotroph | \|Plant Pathogen\| | Dimorphic-Microfungus | NULL | Probable | NULL | [illegible] |
| Otu01426 | 39 | k__Fungi(100);p__Ascomycota(100);c__Dothideomycetes(100);o__Capnodiales(100);f__Mycosphaerellaceae(100);g__Pseudocercosporella(100);s__Pseudocercosporella_sp_RH22(100); | Pseudocercosporella | 13 | Pathotroph | \|Plant Pathogen\| | Microfungus | NULL | Probable | NULL | [illegible] |
| Otu01427 | 39 | k__Fungi(100);p__Ascomycota(100);c__Saccharomycetes(100);o__Saccharomycetales(100);f__Metschnikowiaceae(100);g__Metschnikowia(100);s__Metschnikowia_pulcherrima(93); | Metschnikowia | 13 | Saprotroph | \|Undefined Saprotroph\| | Yeast | NULL | Probable | As nectar-tap_saprotroph (Põlme et al. 2020); Decay sugar-rich substrates (Põlme et al. 2020) | [illegible] |
| Otu01431 | 39 | k__Fungi(100);p__Ascomycota(100);c__Saccharomycetes(98);o__Saccharomycetales(98);f__Saccharomycetales_family_Incertae_sedis(90);g__Candida(80);g__Candida_unclassified(67); | Candida | 13 | Pathotroph-Saprotroph-Symbiotroph | Animal Pathogen-Endophyte-Endosymbiont-Epiphyte-\|Undefined Saprotroph\| | Dimorphic Yeast-Yeast | NULL | Probable | As nectar-tap_saprotroph (Põlme et al. 2020); Decay sugar-rich substrates (Põlme et al. 2020); This asexual genus is polyphyletic and contain species that function very differently depend on what they are. Caution should be taken when interpreting guilds for Candida species. | [illegible] |
| Otu01433 | 39 | k__Fungi(100);p__Basidiomycota(100);c__Agaricomycetes(100);o__Polyporales(100);f__Phanerochaetaceae(85);g__Ceriporia(70);g__Ceriporia_unclassified(70); | Ceriporia | 13 | Saprotroph | \|Wood Saprotroph\| | Corticioid | White Rot | Highly Probable | As white_rot (Põlme et al. 2020); Decay wood (Põlme et al. 2020) | [illegible] |
| Otu01435 | 39 | k__Fungi(100);p__Ascomycota(100);c__Sordariomycetes(100);o__Hypocreales(100);o__Hypocreales_unclassified(100);o__Hypocreales_unclassified(100);o__Hypocreales_unclassified(100); | Hypocreales | 7 | Saprotroph | Undefined Saprotroph | Microfungus | NULL | Possible | NULL | [illegible] |
| Otu01437 | 39 | k__Fungi(100);p__Basidiomycota(100);c__Agaricomycetes(100);o__Agaricales(100);f__Marasmiaceae(100);g__Megacollybia(100);s__Megacollybia_rodmanii(100); | Megacollybia | 13 | Saprotroph | \|Plant Saprotroph\|-Undefined Saprotroph-Wood Saprotroph | Agaricoid | White Rot | Probable | As litter_saprotroph (Põlme et al. 2020); As white_rot (Põlme et al. 2020); Decay wood, leaf, fruit, seed (Põlme et al. 2020) | [illegible] |
| Otu01438 | 39 | k__Fungi(100);p__Basidiomycota(100);c__Tremellomycetes(100);o__Tremellales(100);f__Tremellales_family_Incertae_sedis(100);f__Tremellales_family_Incertae_sedis_unclassified(100);f__Tremellales_family_Incertae_sedis_unclassified(100); | Tremellales | 7 | Pathotroph-Saprotroph-Symbiotroph | Fungal Parasite-Undefined Saprotroph | Tremelloid-Yeast | NULL | Possible | The order contains both teleomorphic and anamorphic species, most of the latter being yeasts | [illegible] |
| Otu01441 | 39 | k__Fungi(100);p__Ascomycota(100);c__Saccharomycetes(100);o__Saccharomycetales(100);f__Metschnikowiaceae(100);g__Metschnikowia(100);s__Metschnikowia_pulcherrima(93); | Metschnikowia | 13 | Saprotroph | \|Undefined Saprotroph\| | Yeast | NULL | Probable | As nectar-tap_saprotroph (Põlme et al. 2020); Decay sugar-rich substrates (Põlme et al. 2020) | [illegible] |
| Otu01445 | 39 | k__Fungi(100);p__Basidiomycota(100);c__Agaricomycetes(100);o__Polyporales(100);f__Phanerochaetaceae(100);g__Byssomerulius(100);s__Byssomerulius_sp(100); | Byssomerulius | 13 | Saprotroph | Undefined Saprotroph-\|Wood Saprotroph\| | Corticioid | White Rot | Probable | As white_rot (Põlme et al. 2020); Decay wood (Põlme et al. 2020) | [illegible] |
| Otu01419 | 39 | k__Fungi(100);p__Basidiomycota(100);c__Agaricomycetes(100);c__Agaricomycetes_unclassified(100);c__Agaricomycetes_unclassified(100);c__Agaricomycetes_unclassified(100);c__Agaricomycetes_unclassified(100); | - | - | - | - | - | - | - | Unassigned | - |
| Otu01420 | 39 | k__Fungi(100);p__Basidiomycota(100);c__Agaricomycetes(100);c__Agaricomycetes_unclassified(100);c__Agaricomycetes_unclassified(100);c__Agaricomycetes_unclassified(100);c__Agaricomycetes_unclassified(100); | - | - | - | - | - | - | - | Unassigned | - |
| Otu01422 | 39 | k__Fungi(100);p__Ascomycota(100);p__Ascomycota_unclassified(100);p__Ascomycota_unclassified(100);p__Ascomycota_unclassified(100);p__Ascomycota_unclassified(100);p__Ascomycota_unclassified(100); | - | - | - | - | - | - | - | Unassigned | - |
| Otu01423 | 39 | k__Fungi(100);k__Fungi_unclassified(54);k__Fungi_unclassified(54);k__Fungi_unclassified(54);k__Fungi_unclassified(54);k__Fungi_unclassified(54);k__Fungi_unclassified(54); | - | - | - | - | - | - | - | Unassigned | - |
| Otu01424 | 39 | k__Fungi(100);p__Ascomycota(75);p__Ascomycota_unclassified(75);p__Ascomycota_unclassified(75);p__Ascomycota_unclassified(75);p__Ascomycota_unclassified(75);p__Ascomycota_unclassified(75); | - | - | - | - | - | - | - | Unassigned | - |
| Otu01425 | 39 | k__Fungi(100);p__Ascomycota(100);c__Dothideomycetes(100);o__Pleosporales(100);o__Pleosporales_unclassified(100);o__Pleosporales_unclassified(100);o__Pleosporales_unclassified(100); | - | - | - | - | - | - | - | Unassigned | - |
| Otu01428 | 39 | k__Fungi(100);p__Basidiomycota(100);p__Basidiomycota_unclassified(100);p__Basidiomycota_unclassified(100);p__Basidiomycota_unclassified(100);p__Basidiomycota_unclassified(100);p__Basidiomycota_unclassified(100); | - | - | - | - | - | - | - | Unassigned | - |
| Otu01429 | 39 | k__Fungi(100);k__Fungi_unclassified(70);k__Fungi_unclassified(70);k__Fungi_unclassified(70);k__Fungi_unclassified(70);k__Fungi_unclassified(70);k__Fungi_unclassified(70); | - | - | - | - | - | - | - | Unassigned | - |
| Otu01430 | 39 | k__Fungi(100);p__Ascomycota(72);p__Ascomycota_unclassified(72);p__Ascomycota_unclassified(72);p__Ascomycota_unclassified(72);p__Ascomycota_unclassified(72);p__Ascomycota_unclassified(72); | - | - | - | - | - | - | - | Unassigned | - |
| Otu01432 | 39 | k__Fungi(100);k__Fungi_unclassified(65);k__Fungi_unclassified(65);k__Fungi_unclassified(65);k__Fungi_unclassified(65);k__Fungi_unclassified(65);k__Fungi_unclassified(65); | - | - | - | - | - | - | - | Unassigned | - |
| Otu01434 | 39 | k__Fungi(100);k__Fungi_unclassified(100);k__Fungi_unclassified(100);k__Fungi_unclassified(100);k__Fungi_unclassified(100);k__Fungi_unclassified(100);k__Fungi_unclassified(100); | - | - | - | - | - | - | - | Unassigned | - |
| Otu01436 | 39 | k__Fungi(100);k__Fungi_unclassified(54);k__Fungi_unclassified(54);k__Fungi_unclassified(54);k__Fungi_unclassified(54);k__Fungi_unclassified(54);k__Fungi_unclassified(54); | - | - | - | - | - | - | - | Unassigned | - |
| Otu01439 | 39 | k__Fungi(100);p__Ascomycota(98);p__Ascomycota_unclassified(95);p__Ascomycota_unclassified(95);p__Ascomycota_unclassified(95);p__Ascomycota_unclassified(95);p__Ascomycota_unclassified(95); | - | - | - | - | - | - | - | Unassigned | - |
| Otu01440 | 39 | k__Fungi(100);p__Ascomycota(80);p__Ascomycota_unclassified(80);p__Ascomycota_unclassified(80);p__Ascomycota_unclassified(80);p__Ascomycota_unclassified(80);p__Ascomycota_unclassified(80); | - | - | - | - | - | - | - | Unassigned | - |
| Otu01442 | 39 | k__Fungi(100);p__Ascomycota(80);p__Ascomycota_unclassified(80);p__Ascomycota_unclassified(80);p__Ascomycota_unclassified(80);p__Ascomycota_unclassified(80);p__Ascomycota_unclassified(80); | - | - | - | - | - | - | - | Unassigned | - |
| Otu01443 | 39 | k__Fungi(100);k__Fungi_unclassified(88);k__Fungi_unclassified(88);k__Fungi_unclassified(88);k__Fungi_unclassified(88);k__Fungi_unclassified(88);k__Fungi_unclassified(88); | - | - | - | - | - | - | - | Unassigned | - |
| Otu01444 | 39 | k__Fungi(100);k__Fungi_unclassified(100);k__Fungi_unclassified(100);k__Fungi_unclassified(100);k__Fungi_unclassified(100);k__Fungi_unclassified(100);k__Fungi_unclassified(100); | - | - | - | - | - | - | - | Unassigned | - |
| Otu01446 | 39 | k__Fungi(100);k__Fungi_unclassified(54);k__Fungi_unclassified(54);k__Fungi_unclassified(54);k__Fungi_unclassified(54);k__Fungi_unclassified(54);k__Fungi_unclassified(54); | - | - | - | - | - | - | - | Unassigned | - |
| Otu01447 | 38 | k__Fungi(100);p__Ascomycota(100);c__Dothideomycetes(100);o__Pleosporales(100);f__Phaeosphaeriaceae(100);g__Neostagonospora(95);s__Neostagonospora_caricis(95); | Neostagonospora | 13 | Saprotroph | \|Plant Saprotroph\|-Wood Saprotroph | Microfungus | NULL | Probable | As litter_saprotroph (Põlme et al. 2020); Decay wood, leaf, fruit, seed (Põlme et al. 2020) | [illegible] |
| Otu01452 | 38 | k__Fungi(100);p__Ascomycota(100);c__Dothideomycetes(100);o__Pleosporales(100);f__Venturiaceae(100);g__Fusicladium(100);s__Fusicladium_amoenum(98); | Fusicladium | 13 | Pathotroph-Symbiotroph | Endophyte-Lichen Parasite-\|Plant Pathogen\| | Microfungus | NULL | Probable | NULL | [illegible] |
| Otu01455 | 38 | k__Fungi(100);p__Basidiomycota(100);c__Agaricomycetes(100);o__Cantharellales(100);f__Ceratobasidiaceae(100);g__Rhizoctonia(100);s__Rhizoctonia_sp(100); | Rhizoctonia | 13 | Pathotroph-Saprotroph-Symbiotroph | Ectomycorrhizal-Orchid Mycorrhizal-\|Plant Pathogen\|-Plant Saprotroph-Wood Saprotroph | Corticioid-Microfungus | Soft Rot | Probable | As litter_saprotroph (Põlme et al. 2020); Decay leaf, fruit, seed, roots, soil (Põlme et al. 2020); Only symbiotic (perhaps leaning parasitic) in special cases with orchids, otherwise typically a plant pathogen. | [illegible] |
| Otu01456 | 38 | k__Fungi(100);p__Basidiomycota(100);c__Microbotryomycetes(100);o__Sporidiobolales(100);f__Sporidiobolales_family_Incertae_sedis(100);g__Sporobolomyces(100);s__Sporobolomyces_symmetricus(100); | Sporobolomyces | 13 | Pathotroph-Saprotroph | \|Fungal Parasite\|-Plant Saprotroph | Yeast | NULL | Probable | As litter_saprotroph (Põlme et al. 2020); As mycoparasite (Põlme et al. 2020); Decay fungal material (Põlme et al. 2020) | [illegible] |
| Otu01457 | 38 | k__Fungi(100);p__Ascomycota(100);c__Saccharomycetes(100);o__Saccharomycetales(100);f__Metschnikowiaceae(100);g__Metschnikowia(100);s__Metschnikowia_pulcherrima(100); | Metschnikowia | 13 | Saprotroph | \|Undefined Saprotroph\| | Yeast | NULL | Probable | As nectar-tap_saprotroph (Põlme et al. 2020); Decay sugar-rich substrates (Põlme et al. 2020) | [illegible] |
| Otu01459 | 38 | k__Fungi(100);p__Ascomycota(100);c__Leotiomycetes(100);o__Helotiales(100);f__Vibrisseaceae(98);g__Acephala(95);s__Acephala_sp_1(95); | Acephala | 13 | Saprotroph-Symbiotroph | Ectomycorrhizal-Endophyte-\|Undefined Saprotroph\| | Microfungus | NULL | Possible | As root_endophyte_dark_septate (Põlme et al. 2020); As soil_saprotroph (Põlme et al. 2020); Decay soil (Põlme et al. 2020) | [illegible] |
| Otu01460 | 38 | k__Fungi(100);p__Ascomycota(100);c__Dothideomycetes(100);o__Pleosporales(100);f__Pleosporales_family_Incertae_sedis(82);g__Massariosphaeria(79);s__Massariosphaeria_sp_FAEII23a(79); | Massariosphaeria | 13 | Saprotroph | Undefined Saprotroph-\|Wood Saprotroph\| | Microfungus | NULL | Probable | Decay wood (Põlme et al. 2020) | [illegible] |
| Otu01462 | 38 | k__Fungi(100);p__Basidiomycota(100);c__Agaricomycetes(100);o__Russulales(100);f__Bondarzewiaceae(100);g__Bondarzewia(100);s__Bondarzewia_sp_MB_2014(100); | Bondarzewia | 13 | Saprotroph | \|Wood Saprotroph\| | Polyporoid | White Rot | Highly Probable | As white_rot (Põlme et al. 2020); Decay wood (Põlme et al. 2020) | [illegible] |
| Otu01448 | 38 | k__Fungi(100);k__Fungi_unclassified(100);k__Fungi_unclassified(100);k__Fungi_unclassified(100);k__Fungi_unclassified(100);k__Fungi_unclassified(100);k__Fungi_unclassified(100); | - | - | - | - | - | - | - | Unassigned | - |
| Otu01449 | 38 | k__Fungi(100);k__Fungi_unclassified(66);k__Fungi_unclassified(66);k__Fungi_unclassified(66);k__Fungi_unclassified(66);k__Fungi_unclassified(66);k__Fungi_unclassified(66); | - | - | - | - | - | - | - | Unassigned | - |
| Otu01450 | 38 | k__Fungi(100);p__Basidiomycota(100);p__Basidiomycota_unclassified(100);p__Basidiomycota_unclassified(100);p__Basidiomycota_unclassified(100);p__Basidiomycota_unclassified(100);p__Basidiomycota_unclassified(100); | - | - | - | - | - | - | - | Unassigned | - |
| Otu01451 | 38 | k__Fungi(100);p__Ascomycota(79);p__Ascomycota_unclassified(79);p__Ascomycota_unclassified(79);p__Ascomycota_unclassified(79);p__Ascomycota_unclassified(79);p__Ascomycota_unclassified(79); | - | - | - | - | - | - | - | Unassigned | - |
| Otu01453 | 38 | k__Fungi(100);k__Fungi_unclassified(74);k__Fungi_unclassified(74);k__Fungi_unclassified(74);k__Fungi_unclassified(74);k__Fungi_unclassified(74);k__Fungi_unclassified(74); | - | - | - | - | - | - | - | Unassigned | - |
| Otu01454 | 38 | k__Fungi(100);p__Ascomycota(85);p__Ascomycota_unclassified(85);p__Ascomycota_unclassified(85);p__Ascomycota_unclassified(85);p__Ascomycota_unclassified(85);p__Ascomycota_unclassified(85); | - | - | - | - | - | - | - | Unassigned | - |
| Otu01458 | 38 | k__Fungi(100);k__Fungi_unclassified(82);k__Fungi_unclassified(82);k__Fungi_unclassified(82);k__Fungi_unclassified(82);k__Fungi_unclassified(82);k__Fungi_unclassified(82); | - | - | - | - | - | - | - | Unassigned | - |
| Otu01461 | 38 | k__Fungi(100);p__Basidiomycota(100);c__Agaricomycetes(100);o__Cantharellales(100);o__Cantharellales_unclassified(93);o__Cantharellales_unclassified(93);o__Cantharellales_unclassified(93); | - | - | - | - | - | - | - | Unassigned | - |
| Otu01463 | 37 | k__Fungi(100);p__Ascomycota(100);c__Saccharomycetes(100);o__Saccharomycetales(100);f__Metschnikowiaceae(100);g__Metschnikowia(100);s__Metschnikowia_pulcherrima(95); | Metschnikowia | 13 | Saprotroph | \|Undefined Saprotroph\| | Yeast | NULL | Probable | As nectar-tap_saprotroph (Põlme et al. 2020); Decay sugar-rich substrates (Põlme et al. 2020) | [illegible] |
| Otu01464 | 37 | k__Fungi(100);p__Basidiomycota(100);c__Agaricomycetes(100);o__Polyporales(100);f__Meruliaceae(100);g__Hypochnicium(100);s__Hypochnicium_subrigescens(100); | Hypochnicium | 13 | Saprotroph | Undefined Saprotroph-\|Wood Saprotroph\| | Corticioid | White Rot | Probable | As white_rot (Põlme et al. 2020); Decay wood (Põlme et al. 2020) | [illegible] |
| Otu01465 | 37 | k__Fungi(100);p__Basidiomycota(100);c__Agaricomycetes(100);o__Polyporales(100);f__Meruliaceae(100);g__Hyphoderma(100);s__Hyphoderma_praetermissum(100); | Hyphoderma | 13 | Saprotroph | Undefined Saprotroph-\|Wood Saprotroph\| | Corticioid | White Rot | Probable | As white_rot (Põlme et al. 2020); Decay wood (Põlme et al. 2020) | [illegible] |
| Otu01470 | 37 | k__Fungi(100);p__Basidiomycota(100);c__Tremellomycetes(100);o__Tremellales(100);f__Tremellales_family_Incertae_sedis(100);g__Hannaella(95);s__Hannaella_sp(82); | Hannaella | 13 | Pathotroph-Saprotroph | \|Animal Parasite\|-Undefined Saprotroph | Yeast | NULL | Probable | As animal_decomposer (Põlme et al. 2020); Decay animal material (Põlme et al. 2020) | [illegible] |
| Otu01471 | 37 | k__Fungi(100);p__Ascomycota(60);c__Sordariomycetes(60);o__Xylariales(60);f__Amphisphaeriaceae(60);g__Pestalotiopsis(60);s__Pestalotiopsis_saprophytica(55); | Pestalotiopsis | 13 | Pathotroph | \|Plant Pathogen\| | Microfungus | NULL | Probable | NULL | [illegible] |
| Otu01474 | 37 | k__Fungi(100);p__Ascomycota(100);c__Saccharomycetes(100);o__Saccharomycetales(100);f__Metschnikowiaceae(100);g__Metschnikowia(100);s__Metschnikowia_pulcherrima(92); | Metschnikowia | 13 | Saprotroph | \|Undefined Saprotroph\| | Yeast | NULL | Probable | As nectar-tap_saprotroph (Põlme et al. 2020); Decay sugar-rich substrates (Põlme et al. 2020) | [illegible] |
| Otu01476 | 37 | k__Fungi(100);p__Basidiomycota(100);c__Agaricomycetes(100);o__Cantharellales(100);f__Botryobasidiaceae(100);g__unclassified_Botryobasidiaceae(100);s__Botryobasidiaceae_sp(100); | Botryobasidiaceae | 9 | Saprotroph | Undefined Saprotroph-Wood Saprotroph | Corticioid | NULL | Probable | Saprobic, typically on rotten wood (Cannon & Kirk 2007) | [illegible] |
| Otu01477 | 37 | k__Fungi(100);p__Basidiomycota(100);c__Agaricomycetes(100);o__Agaricales(100);f__Agaricales_family_Incertae_sedis(100);g__unclassified_Agaricales(100);s__Agaricales_sp(100); | Agaricales | 7 | Pathotroph-Saprotroph-Symbiotroph | Bryophyte Parasite-Dung Saprotroph-Ectomycorrhizal-Fungal Parasite-Leaf Saprotroph-Plant Parasite-Undefined Saprotroph-Wood Saprotroph | Agaricoid-Gasteroid-Microfungus-Secotioid-Yeast | NULL | Possible | Mushrooms and toadstools, Gill fungi, Agarics... lignicolous, sometimes muscicolous or fungicolous, saprobic, mycorrhizal, rarely parasitic on plants or fungi (Kirk et al. 2008) | [illegible] |
| Otu01480 | 37 | k__Fungi(100);p__Ascomycota(100);c__Sordariomycetes(100);o__Hypocreales(100);f__Hypocreales_family_Incertae_sedis(100);g__Acremonium(100);g__Acremonium_unclassified(100); | Acremonium | 13 | Pathotroph-Saprotroph-Symbiotroph | Animal Pathogen-Endophyte-Fungal Parasite-Plant Pathogen-\|Undefined Saprotroph\|-Wood Saprotroph | Microfungus | Soft Rot | Possible | As foliar_endophyte (Põlme et al. 2020); culture (Hanada et al. 2010, Del Olmo; Decay leaf, fruit, seed, soil, dung, animal material (Põlme et al. 2020); Endophyte detection method; Ruiz & Arnold 2014); Likely opportunistic human pathogen (Irinyi et al. 2015) | [illegible] |
| Otu01481 | 37 | k__Fungi(100);p__Basidiomycota(100);c__Agaricomycetes(100);o__Agaricales(100);o__Agaricales_unclassified(100);o__Agaricales_unclassified(100);o__Agaricales_unclassified(100); | Agaricales | 7 | Pathotroph-Saprotroph-Symbiotroph | Bryophyte Parasite-Dung Saprotroph-Ectomycorrhizal-Fungal Parasite-Leaf Saprotroph-Plant Parasite-Undefined Saprotroph-Wood Saprotroph | Agaricoid-Gasteroid-Microfungus-Secotioid-Yeast | NULL | Possible | Mushrooms and toadstools, Gill fungi, Agarics... lignicolous, sometimes muscicolous or fungicolous, saprobic, mycorrhizal, rarely parasitic on plants or fungi (Kirk et al. 2008) | [illegible] |
| Otu01482 | 37 | k__Fungi(100);p__Ascomycota(100);c__Leotiomycetes(100);o__Helotiales(100);f__Helotiales_family_Incertae_sedis(100);g__Leptodontidium(100);s__Leptodontidium_elatius(82); | Leptodontidium | 13 | Saprotroph-Symbiotroph | Endophyte-Ericoid Mycorrhizal-\|Plant Saprotroph\| | Microfungus | NULL | Highly Probable | As litter_saprotroph (Põlme et al. 2020); As root_endophyte_dark_septate (Põlme et al. 2020); Dark Septate Endophyte, known from plant roots.; Decay leaf, fruit, seed (Põlme et al. 2020) | [illegible] |
| Otu01484 | 37 | k__Fungi(100);p__Ascomycota(100);c__Dothideomycetes(100);o__Pleosporales(100);f__Pleosporales_family_Incertae_sedis(100);g__Pyrenochaeta(100);g__Pyrenochaeta_unclassified(100); | Pyrenochaeta | 13 | Pathotroph-Saprotroph | Plant Pathogen-Undefined Saprotroph-\|Wood Saprotroph\| | Microfungus | Soft Rot | Highly Probable | Decay wood (Põlme et al. 2020) | [illegible] |
| Otu01466 | 37 | k__Fungi(100);k__Fungi_unclassified(63);k__Fungi_unclassified(63);k__Fungi_unclassified(63);k__Fungi_unclassified(63);k__Fungi_unclassified(63);k__Fungi_unclassified(63); | - | - | - | - | - | - | - | Unassigned | - |
| Otu01467 | 37 | k__Fungi(100);p__Ascomycota(100);c__Dothideomycetes(100);o__Pleosporales(100);o__Pleosporales_unclassified(100);o__Pleosporales_unclassified(100);o__Pleosporales_unclassified(100); | - | - | - | - | - | - | - | Unassigned | - |
| Otu01468 | 37 | k__Fungi(100);k__Fungi_unclassified(55);k__Fungi_unclassified(55);k__Fungi_unclassified(55);k__Fungi_unclassified(55);k__Fungi_unclassified(55);k__Fungi_unclassified(55); | - | - | - | - | - | - | - | Unassigned | - |
| Otu01469 | 37 | k__Fungi(100);k__Fungi_unclassified(79);k__Fungi_unclassified(79);k__Fungi_unclassified(79);k__Fungi_unclassified(79);k__Fungi_unclassified(79);k__Fungi_unclassified(79); | - | - | - | - | - | - | - | Unassigned | - |
| Otu01472 | 37 | k__Fungi(100);k__Fungi_unclassified(82);k__Fungi_unclassified(82);k__Fungi_unclassified(82);k__Fungi_unclassified(82);k__Fungi_unclassified(82);k__Fungi_unclassified(82); | - | - | - | - | - | - | - | Unassigned | - |
| Otu01473 | 37 | k__Fungi(100);k__Fungi_unclassified(73);k__Fungi_unclassified(73);k__Fungi_unclassified(73);k__Fungi_unclassified(73);k__Fungi_unclassified(73);k__Fungi_unclassified(73); | - | - | - | - | - | - | - | Unassigned | - |
| Otu01475 | 37 | k__Fungi(100);p__Basidiomycota(100);p__Basidiomycota_unclassified(95);p__Basidiomycota_unclassified(95);p__Basidiomycota_unclassified(95);p__Basidiomycota_unclassified(95);p__Basidiomycota_unclassified(95); | - | - | - | - | - | - | - | Unassigned | - |
| Otu01478 | 37 | k__Fungi(100);p__Basidiomycota(100);c__Agaricomycetes(73);c__Agaricomycetes_unclassified(73);c__Agaricomycetes_unclassified(73);c__Agaricomycetes_unclassified(73);c__Agaricomycetes_unclassified(73); | - | - | - | - | - | - | - | Unassigned | - |
| Otu01479 | 37 | k__Fungi(100);p__Ascomycota(90);p__Ascomycota_unclassified(90);p__Ascomycota_unclassified(90);p__Ascomycota_unclassified(90);p__Ascomycota_unclassified(90);p__Ascomycota_unclassified(90); | - | - | - | - | - | - | - | Unassigned | - |
| Otu01483 | 37 | k__Fungi(100);p__Ascomycota(65);p__Ascomycota_unclassified(65);p__Ascomycota_unclassified(65);p__Ascomycota_unclassified(65);p__Ascomycota_unclassified(65);p__Ascomycota_unclassified(65); | - | - | - | - | - | - | - | Unassigned | - |
| Otu01485 | 37 | k__Fungi(100);p__Ascomycota(100);c__Dothideomycetes(100);c__Dothideomycetes_unclassified(100);c__Dothideomycetes_unclassified(100);c__Dothideomycetes_unclassified(100);c__Dothideomycetes_unclassified(100); | - | - | - | - | - | - | - | Unassigned | - |
| Otu01486 | 36 | k__Fungi(100);p__Ascomycota(100);c__Sordariomycetes(100);o__Sordariomycetes_order_Incertae_sedis(100);f__Apiosporaceae(100);g__Arthrinium(100);s__Arthrinium_marii(78); | Arthrinium | 13 | Pathotroph-Saprotroph-Symbiotroph | Dung Saprotroph-Endophyte-\|Plant Pathogen\|-Undefined Saprotroph | Microfungus | NULL | Probable | As foliar_endophyte (Põlme et al. 2020); Decay leaf, fruit, seed (Põlme et al. 2020) | [illegible] |
| Otu01487 | 36 | k__Fungi(100);p__Ascomycota(100);c__Sordariomycetes(100);o__Calosphaeriales(98);f__Calosphaeriaceae(98);g__Jattaea(98);g__Jattaea_unclassified(98); | Jattaea | 13 | Saprotroph | \|Undefined Saprotroph\| | Microfungus | NULL | Probable | NULL | [illegible] |
| Otu01489 | 36 | k__Fungi(100);p__Ascomycota(100);c__Saccharomycetes(100);o__Saccharomycetales(100);f__Metschnikowiaceae(100);g__Metschnikowia(100);g__Metschnikowia_unclassified(87); | Metschnikowia | 13 | Saprotroph | \|Undefined Saprotroph\| | Yeast | NULL | Probable | As nectar-tap_saprotroph (Põlme et al. 2020); Decay sugar-rich substrates (Põlme et al. 2020) | [illegible] |
| Otu01493 | 36 | k__Fungi(100);p__Basidiomycota(100);c__Agaricomycetes(100);o__Agaricales(100);f__Lyophyllaceae(100);g__Ossicaulis(100);s__Ossicaulis_lignatilis(100); | Ossicaulis | 13 | Saprotroph | \|Wood Saprotroph\| | Agaricoid | Brown Rot | Probable | As brown-rot (Põlme et al. 2020); Decay wood (Põlme et al. 2020) | [illegible] |
| Otu01494 | 36 | k__Fungi(100);p__Ascomycota(89);c__Dothideomycetes(81);o__Dothideomycetes_order_Incertae_sedis(81);f__Pseudeurotiaceae(81);g__Pseudeurotium(70);g__Pseudeurotium_unclassified(64); | Pseudeurotium | 13 | Saprotroph | \|Dung Saprotroph\|-Undefined Saprotroph | Microfungus | NULL | Probable | As soil_saprotroph (Põlme et al. 2020); Decay soil, dung (Põlme et al. 2020) | [illegible] |
| Otu01496 | 36 | k__Fungi(100);p__Basidiomycota(100);c__Agaricomycetes(100);o__Agaricales(100);f__Strophariaceae(100);g__Pholiota(100);s__Pholiota_conissans(100); | Pholiota | 13 | Pathotroph-Saprotroph | Plant Pathogen-Plant Saprotroph-Undefined Saprotroph-\|Wood Saprotroph\| | Agaricoid | White Rot | Probable | As litter_saprotroph (Põlme et al. 2020); As white_rot (Põlme et al. 2020); As wood_pathogen (Põlme et al. 2020); Decay wood, leaf, fruit, seed (Põlme et al. 2020) | [illegible] |
| Otu01497 | 36 | k__Fungi(100);p__Ascomycota(100);c__Dothideomycetes(100);o__Pleosporales(100);f__Venturiaceae(100);g__Venturia(59);g__Venturia_unclassified(59); | Venturia | 13 | Pathotroph-Pathotroph-Saprotroph | \|Plant Pathogen\|-Plant Saprotroph | Microfungus | NULL | Probable | As litter_saprotroph (Põlme et al. 2020); Decay leaf, fruit, seed (Põlme et al. 2020) | [illegible] |
| Otu01498 | 36 | k__Fungi(100);p__Ascomycota(100);c__Saccharomycetes(100);o__Saccharomycetales(100);f__Metschnikowiaceae(100);g__Metschnikowia(100);s__Metschnikowia_pulcherrima(89); | Metschnikowia | 13 | Saprotroph | \|Undefined Saprotroph\| | Yeast | NULL | Probable | As nectar-tap_saprotroph (Põlme et al. 2020); Decay sugar-rich substrates (Põlme et al. 2020) | [illegible] |
| Otu01499 | 36 | k__Fungi(100);p__Ascomycota(100);c__Saccharomycetes(100);o__Saccharomycetales(100);f__Metschnikowiaceae(100);g__Metschnikowia(100);s__Metschnikowia_pulcherrima(92); | Metschnikowia | 13 | Saprotroph | \|Undefined Saprotroph\| | Yeast | NULL | Probable | As nectar-tap_saprotroph (Põlme et al. 2020); Decay sugar-rich substrates (Põlme et al. 2020) | [illegible] |
| Otu01500 | 36 | k__Fungi(100);p__Ascomycota(100);c__Dothideomycetes(100);o__Capnodiales(100);f__Mycosphaerellaceae(100);f__Mycosphaerellaceae_unclassified(70);f__Mycosphaerellaceae_unclassified(70); | Mycosphaerellaceae | 9 | Pathotroph-Saprotroph | Plant Pathogen-Undefined Saprotroph | Microfungus | NULL | Probable | Biotrophic, necrotrophic or saprobic on various plant tissues (Cannon & Kirk 2007) | [illegible] |
| Otu01503 | 36 | k__Fungi(100);p__Basidiomycota(100);c__Agaricomycetes(100);o__Polyporales(100);f__Polyporaceae(100);g__Hapalopilus(100);s__Hapalopilus_rutilans(100); | Hapalopilus | 13 | Saprotroph | \|Wood Saprotroph\| | Polyporoid | White Rot | Highly Probable | As white_rot (Põlme et al. 2020); Decay wood (Põlme et al. 2020) | [illegible] |
| Otu01504 | 36 | k__Fungi(100);p__Ascomycota(100);c__Eurotiomycetes(100);o__Eurotiales(100);f__Trichocomaceae(100);g__Penicillium(100);s__Penicillium_melinii(59); | Penicillium | 13 | Pathotroph-Saprotroph-Symbiotroph | Animal Parasite-Dung Saprotroph-Endophyte-Fungal Parasite-Plant Pathogen-\|Undefined Saprotroph\|-Wood Saprotroph | Microfungus | Soft Rot | Highly Probable | As foliar_endophyte (Põlme et al. 2020); As mycoparasite (Põlme et al. 2020); Decay soil (Põlme et al. 2020) | [illegible] |
| Otu01505 | 36 | k__Fungi(100);p__Ascomycota(100);c__Saccharomycetes(100);o__Saccharomycetales(100);f__Metschnikowiaceae(100);g__Metschnikowia(100);s__Metschnikowia_pulcherrima(89); | Metschnikowia | 13 | Saprotroph | \|Undefined Saprotroph\| | Yeast | NULL | Probable | As nectar-tap_saprotroph (Põlme et al. 2020); Decay sugar-rich substrates (Põlme et al. 2020) | [illegible] |
| Otu01488 | 36 | k__Fungi(100);p__Ascomycota(100);c__Dothideomycetes(100);o__Pleosporales(100);o__Pleosporales_unclassified(87);o__Pleosporales_unclassified(87);o__Pleosporales_unclassified(87); | - | - | - | - | - | - | - | Unassigned | - |
| Otu01490 | 36 | k__Fungi(100);k__Fungi_unclassified(53);k__Fungi_unclassified(53);k__Fungi_unclassified(53);k__Fungi_unclassified(53);k__Fungi_unclassified(53);k__Fungi_unclassified(53); | - | - | - | - | - | - | - | Unassigned | - |
| Otu01491 | 36 | k__Fungi(100);k__Fungi_unclassified(73);k__Fungi_unclassified(73);k__Fungi_unclassified(73);k__Fungi_unclassified(73);k__Fungi_unclassified(73);k__Fungi_unclassified(73); | - | - | - | - | - | - | - | Unassigned | - |
| Otu01492 | 36 | k__Fungi(100);p__Ascomycota(84);p__Ascomycota_unclassified(84);p__Ascomycota_unclassified(84);p__Ascomycota_unclassified(84);p__Ascomycota_unclassified(84);p__Ascomycota_unclassified(84); | - | - | - | - | - | - | - | Unassigned | - |
| Otu01495 | 36 | k__Fungi(100);p__Basidiomycota(100);c__Agaricomycetes(100);o__Polyporales(100);o__Polyporales_unclassified(98);o__Polyporales_unclassified(98);o__Polyporales_unclassified(98); | - | - | - | - | - | - | - | Unassigned | - |
| Otu01501 | 36 | k__Fungi(100);p__Ascomycota(100);c__Leotiomycetes(92);o__Helotiales(89);o__Helotiales_unclassified(89);o__Helotiales_unclassified(89);o__Helotiales_unclassified(89); | - | - | - | - | - | - | - | Unassigned | - |
| Otu01502 | 36 | k__Fungi(100);p__Basidiomycota(100);c__Agaricomycetes(100);o__Cantharellales(100);o__Cantharellales_unclassified(100);o__Cantharellales_unclassified(100);o__Cantharellales_unclassified(100); | - | - | - | - | - | - | - | Unassigned | - |
| Otu01507 | 35 | k__Fungi(100);p__Basidiomycota(100);c__Agaricomycetes(100);o__Polyporales(100);f__Fomitopsidaceae(100);g__Fibroporia(100);s__Fibroporia_radiculosa(100); | Fibroporia | 13 | Saprotroph | \|Wood Saprotroph\| | Corticioid | Brown Rot | Probable | As brown-rot (Põlme et al. 2020); Decay wood (Põlme et al. 2020) | [illegible] |
| Otu01509 | 35 | k__Fungi(100);p__Basidiomycota(100);c__Agaricomycetes(100);o__Agaricales(100);f__Cortinariaceae(100);g__Cortinarius(100);g__Cortinarius_unclassified(95); | Cortinarius | 13 | Symbiotroph | \|Ectomycorrhizal\| | Agaricoid | NULL | Highly Probable | NULL | [illegible] |
| Otu01511 | 35 | k__Fungi(100);p__Basidiomycota(100);c__Agaricomycetes(100);o__Agaricales(100);o__Agaricales_unclassified(100);o__Agaricales_unclassified(100);o__Agaricales_unclassified(100); | Agaricales | 7 | Pathotroph-Saprotroph-Symbiotroph | Bryophyte Parasite-Dung Saprotroph-Ectomycorrhizal-Fungal Parasite-Leaf Saprotroph-Plant Parasite-Undefined Saprotroph-Wood Saprotroph | Agaricoid-Gasteroid-Microfungus-Secotioid-Yeast | NULL | Possible | Mushrooms and toadstools, Gill fungi, Agarics... lignicolous, sometimes muscicolous or fungicolous, saprobic, mycorrhizal, rarely parasitic on plants or fungi (Kirk et al. 2008) | [illegible] |
| Otu01513 | 35 | k__Fungi(100);p__Ascomycota(100);c__Saccharomycetes(100);o__Saccharomycetales(100);f__Metschnikowiaceae(100);g__Metschnikowia(100);s__Metschnikowia_pulcherrima(100); | Metschnikowia | 13 | Saprotroph | \|Undefined Saprotroph\| | Yeast | NULL | Probable | As nectar-tap_saprotroph (Põlme et al. 2020); Decay sugar-rich substrates (Põlme et al. 2020) | [illegible] |
| Otu01514 | 35 | k__Fungi(100);p__Ascomycota(100);c__Saccharomycetes(100);o__Saccharomycetales(100);f__Metschnikowiaceae(100);g__Metschnikowia(100);s__Metschnikowia_pulcherrima(86); | Metschnikowia | 13 | Saprotroph | \|Undefined Saprotroph\| | Yeast | NULL | Probable | As nectar-tap_saprotroph (Põlme et al. 2020); Decay sugar-rich substrates (Põlme et al. 2020) | [illegible] |
| Otu01516 | 35 | k__Fungi(100);p__Ascomycota(100);c__Dothideomycetes(100);o__Pleosporales(100);f__Lophiostomataceae(69);g__Lophiostoma(69);s__Lophiostoma_cf_cynaroidis_A33(69); | Lophiostoma | 13 | Saprotroph | Plant Saprotroph-Undefined Saprotroph-\|Wood Saprotroph\| | Microfungus | NULL | Probable | As litter_saprotroph (Põlme et al. 2020); Decay wood, leaf, fruit, seed (Põlme et al. 2020) | [illegible] |
| Otu01518 | 35 | k__Fungi(100);p__Ascomycota(100);c__Saccharomycetes(100);o__Saccharomycetales(100);f__Metschnikowiaceae(100);g__Metschnikowia(100);s__Metschnikowia_pulcherrima(100); | Metschnikowia | 13 | Saprotroph | \|Undefined Saprotroph\| | Yeast | NULL | Probable | As nectar-tap_saprotroph (Põlme et al. 2020); Decay sugar-rich substrates (Põlme et al. 2020) | [illegible] |
| Otu01520 | 35 | k__Fungi(100);p__Zygomycota(100);c__Zygomycota_class_Incertae_sedis(100);o__Mortierellales(100);f__Mortierellaceae(100);g__Mortierella(100);s__Mortierella_schmuckeri(61); | Mortierella | 13 | Saprotroph-Symbiotroph | Endophyte-Plant Saprotroph-\|Undefined Saprotroph\| | Microfungus | NULL | Probable | As chitinolytic (Põlme et al. 2020); As soil_saprotroph (Põlme et al. 2020); Decay roots, soil (Põlme et al. 2020); Saprobic in soil and thus frequenctly isolated from soil and material contaminated with soil such as hay or rotten silage and spoil tips (Cannon & Kirk 2007) | [illegible] |
| Otu01521 | 35 | k__Fungi(100);p__Ascomycota(100);c__Saccharomycetes(100);o__Saccharomycetales(100);f__Metschnikowiaceae(100);g__Metschnikowia(100);s__Metschnikowia_pulcherrima(100); | Metschnikowia | 13 | Saprotroph | \|Undefined Saprotroph\| | Yeast | NULL | Probable | As nectar-tap_saprotroph (Põlme et al. 2020); Decay sugar-rich substrates (Põlme et al. 2020) | [illegible] |
| Otu01522 | 35 | k__Fungi(100);p__Ascomycota(100);c__Saccharomycetes(100);o__Saccharomycetales(100);f__Metschnikowiaceae(100);g__Metschnikowia(100);s__Metschnikowia_pulcherrima(95); | Metschnikowia | 13 | Saprotroph | \|Undefined Saprotroph\| | Yeast | NULL | Probable | As nectar-tap_saprotroph (Põlme et al. 2020); Decay sugar-rich substrates (Põlme et al. 2020) | [illegible] |
| Otu01523 | 35 | k__Fungi(100);p__Basidiomycota(100);c__Agaricomycetes(100);o__Cantharellales(98);f__Hydnaceae(98);g__Sistotrema(98);s__Sistotrema_sp(98); | Sistotrema | 13 | Saprotroph-Symbiotroph | Ectomycorrhizal-\|Plant Saprotroph\|-Wood Saprotroph | Corticioid | White Rot | Possible | As litter_saprotroph (Põlme et al. 2020); Decay leaf, fruit, seed (Põlme et al. 2020) | [illegible] |
| Otu01524 | 35 | k__Fungi(100);p__Ascomycota(100);c__Saccharomycetes(100);o__Saccharomycetales(100);f__Metschnikowiaceae(100);g__Metschnikowia(100);s__Metschnikowia_pulcherrima(92); | Metschnikowia | 13 | Saprotroph | \|Undefined Saprotroph\| | Yeast | NULL | Probable | As nectar-tap_saprotroph (Põlme et al. 2020); Decay sugar-rich substrates (Põlme et al. 2020) | [illegible] |
| Otu01526 | 35 | k__Fungi(100);p__Ascomycota(100);c__Saccharomycetes(100);o__Saccharomycetales(100);f__Metschnikowiaceae(100);g__Metschnikowia(100);s__Metschnikowia_pulcherrima(95); | Metschnikowia | 13 | Saprotroph | \|Undefined Saprotroph\| | Yeast | NULL | Probable | As nectar-tap_saprotroph (Põlme et al. 2020); Decay sugar-rich substrates (Põlme et al. 2020) | [illegible] |
| Otu01527 | 35 | k__Fungi(100);p__Ascomycota(100);c__Saccharomycetes(100);o__Saccharomycetales(100);f__Metschnikowiaceae(100);g__Metschnikowia(100);s__Metschnikowia_pulcherrima(92); | Metschnikowia | 13 | Saprotroph | \|Undefined Saprotroph\| | Yeast | NULL | Probable | As nectar-tap_saprotroph (Põlme et al. 2020); Decay sugar-rich substrates (Põlme et al. 2020) | [illegible] |
| Otu01529 | 35 | k__Fungi(100);p__Ascomycota(100);c__Saccharomycetes(100);o__Saccharomycetales(100);f__Metschnikowiaceae(100);g__Metschnikowia(100);g__Metschnikowia_unclassified(63); | Metschnikowia | 13 | Saprotroph | \|Undefined Saprotroph\| | Yeast | NULL | Probable | As nectar-tap_saprotroph (Põlme et al. 2020); Decay sugar-rich substrates (Põlme et al. 2020) | [illegible] |
| Otu01533 | 35 | k__Fungi(100);p__Ascomycota(98);c__Saccharomycetes(98);o__Saccharomycetales(98);f__Metschnikowiaceae(98);g__Metschnikowia(98);s__Metschnikowia_pulcherrima(98); | Metschnikowia | 13 | Saprotroph | \|Undefined Saprotroph\| | Yeast | NULL | Probable | As nectar-tap_saprotroph (Põlme et al. 2020); Decay sugar-rich substrates (Põlme et al. 2020) | [illegible] |
| Otu01535 | 35 | k__Fungi(100);p__Ascomycota(100);c__Saccharomycetes(100);o__Saccharomycetales(100);f__Metschnikowiaceae(100);g__Metschnikowia(100);s__Metschnikowia_pulcherrima(92); | Metschnikowia | 13 | Saprotroph | \|Undefined Saprotroph\| | Yeast | NULL | Probable | As nectar-tap_saprotroph (Põlme et al. 2020); Decay sugar-rich substrates (Põlme et al. 2020) | [illegible] |
| Otu01536 | 35 | k__Fungi(100);p__Ascomycota(100);c__Ascomycota_class_Incertae_sedis(100);o__Ascomycota_order_Incertae_sedis(100);f__Ascomycota_family_Incertae_sedis(100);g__Microcyclospora(100);s__Microcyclospora_quercina(95); | Microcyclospora | 13 | Pathotroph | \|Plant Pathogen\| | Microfungus | NULL | Probable | NULL | [illegible] |
| Otu01506 | 35 | k__Fungi(100);k__Fungi_unclassified(100);k__Fungi_unclassified(100);k__Fungi_unclassified(100);k__Fungi_unclassified(100);k__Fungi_unclassified(100);k__Fungi_unclassified(100); | - | - | - | - | - | - | - | Unassigned | - |
| Otu01508 | 35 | k__Fungi(100);p__Ascomycota(100);c__Sordariomycetes(100);c__Sordariomycetes_unclassified(100);c__Sordariomycetes_unclassified(100);c__Sordariomycetes_unclassified(100);c__Sordariomycetes_unclassified(100); | - | - | - | - | - | - | - | Unassigned | - |
| Otu01510 | 35 | k__Fungi(100);p__Ascomycota(52);p__Ascomycota_unclassified(52);p__Ascomycota_unclassified(52);p__Ascomycota_unclassified(52);p__Ascomycota_unclassified(52);p__Ascomycota_unclassified(52); | - | - | - | - | - | - | - | Unassigned | - |
| Otu01512 | 35 | k__Fungi(100);p__Ascomycota(58);p__Ascomycota_unclassified(58);p__Ascomycota_unclassified(58);p__Ascomycota_unclassified(58);p__Ascomycota_unclassified(58);p__Ascomycota_unclassified(58); | - | - | - | - | - | - | - | Unassigned | - |
| Otu01515 | 35 | k__Fungi(100);k__Fungi_unclassified(63);k__Fungi_unclassified(63);k__Fungi_unclassified(63);k__Fungi_unclassified(63);k__Fungi_unclassified(63);k__Fungi_unclassified(63); | - | - | - | - | - | - | - | Unassigned | - |
| Otu01517 | 35 | k__Fungi(100);k__Fungi_unclassified(100);k__Fungi_unclassified(100);k__Fungi_unclassified(100);k__Fungi_unclassified(100);k__Fungi_unclassified(100);k__Fungi_unclassified(100); | - | - | - | - | - | - | - | Unassigned | - |
| Otu01519 | 35 | k__Fungi(100);p__Ascomycota(100);c__Lecanoromycetes(100);o__Teloschistales(86);o__Teloschistales_unclassified(83);o__Teloschistales_unclassified(83);o__Teloschistales_unclassified(83); | - | - | - | - | - | - | - | Unassigned | - |
| Otu01525 | 35 | k__Fungi(100);p__Basidiomycota(100);c__Agaricomycetes(100);c__Agaricomycetes_unclassified(100);c__Agaricomycetes_unclassified(100);c__Agaricomycetes_unclassified(100);c__Agaricomycetes_unclassified(100); | - | - | - | - | - | - | - | Unassigned | - |
| Otu01528 | 35 | k__Fungi(100);k__Fungi_unclassified(52);k__Fungi_unclassified(52);k__Fungi_unclassified(52);k__Fungi_unclassified(52);k__Fungi_unclassified(52);k__Fungi_unclassified(52); | - | - | - | - | - | - | - | Unassigned | - |
| Otu01530 | 35 | k__Fungi(100);k__Fungi_unclassified(52);k__Fungi_unclassified(52);k__Fungi_unclassified(52);k__Fungi_unclassified(52);k__Fungi_unclassified(52);k__Fungi_unclassified(52); | - | - | - | - | - | - | - | Unassigned | - |
| Otu01531 | 35 | k__Fungi(100);k__Fungi_unclassified(58);k__Fungi_unclassified(58);k__Fungi_unclassified(58);k__Fungi_unclassified(58);k__Fungi_unclassified(58);k__Fungi_unclassified(58); | - | - | - | - | - | - | - | Unassigned | - |
| Otu01532 | 35 | k__Fungi(100);k__Fungi_unclassified(83);k__Fungi_unclassified(83);k__Fungi_unclassified(83);k__Fungi_unclassified(83);k__Fungi_unclassified(83);k__Fungi_unclassified(83); | - | - | - | - | - | - | - | Unassigned | - |
| Otu01534 | 35 | k__Plantae(98);p__unclassified_Plantae(98);c__unclassified_Plantae(98);o__unclassified_Plantae(98);f__unclassified_Plantae(98);g__unclassified_Plantae(98);s__Plantae_sp(98); | - | - | - | - | - | - | - | Unassigned | - |
| Otu01537 | 34 | k__Fungi(100);p__Ascomycota(100);c__Dothideomycetes(100);o__Capnodiales(100);f__Teratosphaeriaceae(100);g__Devriesia(100);s__Devriesia_sp(100); | Devriesia | 13 | Pathotroph-Saprotroph | \|Plant Pathogen\|-Plant Saprotroph | Microfungus | NULL | Probable | As litter_saprotroph (Põlme et al. 2020); Decay leaf, fruit, seed (Põlme et al. 2020) | [illegible] |
| Otu01540 | 34 | k__Fungi(100);p__Ascomycota(100);c__Dothideomycetes(100);o__Capnodiales(100);f__Mycosphaerellaceae(100);f__Mycosphaerellaceae_unclassified(100);f__Mycosphaerellaceae_unclassified(100); | Mycosphaerellaceae | 9 | Pathotroph-Saprotroph | Plant Pathogen-Undefined Saprotroph | Microfungus | NULL | Probable | Biotrophic, necrotrophic or saprobic on various plant tissues (Cannon & Kirk 2007) | [illegible] |
| Otu01541 | 34 | k__Fungi(100);p__Basidiomycota(100);c__Agaricomycetes(100);o__Atheliales(100);f__Atheliaceae(100);g__Piloderma(100);g__Piloderma_unclassified(77); | Piloderma | 13 | Symbiotroph | \|Ectomycorrhizal\| | Corticioid | NULL | Highly Probable | NULL | [illegible] |
| Otu01542 | 34 | k__Fungi(100);p__Basidiomycota(100);c__Tremellomycetes(100);o__Tremellales(100);f__Tremellales_family_Incertae_sedis(100);g__Cryptococcus(100);s__Cryptococcus_sp(100); | Cryptococcus | 13 | Pathotroph-Saprotroph-Symbiotroph | Animal Pathogen-Endophyte-Epiphyte-\|Undefined Saprotroph\| | Dimorphic Yeast-Yeast | NULL | Possible | As foliar_endophyte (Põlme et al. 2020); Decay leaf, fruit, seed, soil, dung, animal material (Põlme et al. 2020) | [illegible] |
| Otu01543 | 34 | k__Fungi(100);p__Ascomycota(100);c__Eurotiomycetes(100);o__Chaetothyriales(100);f__Herpotrichiellaceae(100);g__Capronia(100);s__Capronia_sp_94006a(62); | Capronia | 13 | Pathotroph-Saprotroph-Symbiotroph | Animal Pathogen-Endophyte-Fungal Parasite-\|Undefined Saprotroph\| | Dimorphic-Facultative Yeast | NULL | Highly Probable | As mycoparasite (Põlme et al. 2020); As root_endophyte_dark_septate (Põlme et al. 2020); As soil_saprotroph (Põlme et al. 2020); Dark Septate Endophyte, known from plant roots.; Decay soil (Põlme et al. 2020) | [illegible] |
| Otu01545 | 34 | k__Fungi(100);p__Ascomycota(100);c__Dothideomycetes(100);o__Capnodiales(100);f__Mycosphaerellaceae(100);f__Mycosphaerellaceae_unclassified(71);f__Mycosphaerellaceae_unclassified(71); | Mycosphaerellaceae | 9 | Pathotroph-Saprotroph | Plant Pathogen-Undefined Saprotroph | Microfungus | NULL | Probable | Biotrophic, necrotrophic or saprobic on various plant tissues (Cannon & Kirk 2007) | [illegible] |
| Otu01546 | 34 | k__Fungi(100);p__Ascomycota(100);c__Sordariomycetes(100);o__Hypocreales(100);f__Bionectriaceae(100);g__Clonostachys(74);s__Clonostachys_rosea_f._catenulata(68); | Clonostachys | 13 | Pathotroph-Saprotroph | Fungal Parasite-Plant Pathogen-Plant Saprotroph-\|Wood Saprotroph\| | Microfungus | NULL | Probable | As litter_saprotroph (Põlme et al. 2020); As mycoparasite (Põlme et al. 2020); Decay wood, leaf, fruit, seed, fungal material, animal material (Põlme et al. 2020) | [illegible] |
| Otu01547 | 34 | k__Fungi(100);p__Ascomycota(100);c__Sordariomycetes(100);o__Xylariales(100);f__Amphisphaeriaceae(98);f__Amphisphaeriaceae_unclassified(98);f__Amphisphaeriaceae_unclassified(98); | Amphisphaeriaceae | 9 | Saprotroph | Undefined Saprotroph | Microfungus | NULL | Possible | Saprobes or weak parasites of leaves, stems, etc., often appearing as secondary colonizers following damage to plants by other agents (Cannon & Kirk 2007) | [illegible] |
| Otu01549 | 34 | k__Fungi(100);p__Zygomycota(100);c__Zygomycota_class_Incertae_sedis(100);o__Mortierellales(100);f__Mortierellaceae(100);g__Mortierella(100);s__Mortierella_amoeboidea(83); | Mortierella | 13 | Saprotroph-Symbiotroph | Endophyte-Plant Saprotroph-\|Undefined Saprotroph\| | Microfungus | NULL | Probable | As chitinolytic (Põlme et al. 2020); As soil_saprotroph (Põlme et al. 2020); Decay roots, soil (Põlme et al. 2020); Saprobic in soil and thus frequenctly isolated from soil and material contaminated with soil such as hay or rotten silage and spoil tips (Cannon & Kirk 2007) | [illegible] |
| Otu01550 | 34 | k__Fungi(100);p__Zygomycota(100);c__Zygomycota_class_Incertae_sedis(100);o__Mortierellales(100);f__Mortierellaceae(100);g__Mortierella(100);s__Mortierella_dichotoma(100); | Mortierella | 13 | Saprotroph-Symbiotroph | Endophyte-Plant Saprotroph-\|Undefined Saprotroph\| | Microfungus | NULL | Probable | As chitinolytic (Põlme et al. 2020); As soil_saprotroph (Põlme et al. 2020); Decay roots, soil (Põlme et al. 2020); Saprobic in soil and thus frequenctly isolated from soil and material contaminated with soil such as hay or rotten silage and spoil tips (Cannon & Kirk 2007) | [illegible] |
| Otu01551 | 34 | k__Fungi(100);p__Ascomycota(100);c__Leotiomycetes(100);o__Leotiomycetes_order_Incertae_sedis(100);f__Leotiomycetes_family_Incertae_sedis(100);g__Sarea(100);s__Sarea_resinae(100); | Sarea | 13 | Saprotroph-Symbiotroph | Lichenized-\|Undefined Saprotroph\| | Microfungus-Thallus | NULL | Highly Probable | NULL | [illegible] |
| Otu01552 | 34 | k__Fungi(100);p__Basidiomycota(100);c__Agaricomycetes(100);o__Polyporales(100);f__Polyporaceae(100);g__Rhodonia(100);s__Rhodonia_placenta(100); | Rhodonia | 13 | Saprotroph | \|Wood Saprotroph\| | Corticioid | Brown Rot | Probable | As brown-rot (Põlme et al. 2020); Decay wood (Põlme et al. 2020) | [illegible] |
| Otu01554 | 34 | k__Fungi(100);p__Basidiomycota(100);c__Agaricomycetes(100);o__Trechisporales(100);f__Hydnodontaceae(100);g__Trechispora(98);g__Trechispora_unclassified(98); | Trechispora | 13 | Saprotroph | \|Wood Saprotroph\| | Corticioid | White Rot | Highly Probable | As white_rot (Põlme et al. 2020); Decay wood (Põlme et al. 2020) | [illegible] |

| | | | | | | | | | | | |
|---|---|---|---|---|---|---|---|---|---|---|---|
| Otu01555 | 34 | k__Fungi(100);p__Zygomycota(100);c__Zygomycota_class_Incertae_sedis(100);o__Mucorales(100);f__Cunninghamellaceae(100);g__Absidia(100);s__Absidia_caerulea(100); | Absidia | 13 | Saprotroph | \|Undefined Saprotroph\| | Microfungus | NULL | Probable | As soil_saprotroph (Põlme et al. 2020); Decay soil (Põlme et al. 2020) | Tedersoo L, et al. 2014. Science 346:e1256688 ((https://doi.org/10.1126/science.1256688)); Põlme S, et al. 2020. Fungal Diversity 105:1-16 ((https://doi.org/10.1007/s13225-020-00466-2)) |
| Otu01556 | 34 | k__Fungi(100);p__Ascomycota(100);c__Taphrinomycetes(100);o__Taphrinales(100);f__Taphrinaceae(100);g__Taphrina(100);s__Taphrina_betulina(100); | Taphrina | 13 | Pathotroph | \|Plant Pathogen\| | Dimorphic-Microfungus | NULL | Probable | NULL | Põlme S, et al. 2020. Fungal Diversity 105:1-16 ((https://doi.org/10.1007/s13225-020-00466-2)); The American Phytopathological Society Common Names of Plant Diseases ((http://www.apsnet.org/publications/commonnames/Pages/default.aspx)) |
| Otu01559 | 34 | k__Fungi(100);p__Ascomycota(100);c__Dothideomycetes(100);o__Pleosporales(100);f__Phaeosphaeriaceae(100);g__Phaeosphaeria(100);s__Phaeosphaeria_juncicola(100); | Phaeosphaeria | 13 | Pathotroph-Saprotroph-Symbiotroph | Endophyte-Plant Pathogen-\|Plant Saprotroph\| | Microfungus | NULL | Probable | As litter_saprotroph (Põlme et al. 2020); Decay leaf, fruit, seed (Põlme et al. 2020) | Tedersoo L, et al. 2014. Science 346:e1256688 ((https://doi.org/10.1126/science.1256688));David AS, et al. 2016. Microbial Ecology 71: 912-926 ((https://doi.org/10.1007/s00248-015-0712-6)); Põlme S, et al. 2020. Fungal Diversity 105:1-16 ((https://doi.org/10.1007/s13225-020-00466-2)) |
| Otu01538 | 34 | k__Fungi(100);k__Fungi_unclassified(59);k__Fungi_unclassified(59);k__Fungi_unclassified(59);k__Fungi_unclassified(59);k__Fungi_unclassified(59);k__Fungi_unclassified(59); | - | - | - | - | - | - | - | Unassigned | - |
| Otu01539 | 34 | k__Fungi(100);k__Fungi_unclassified(68);k__Fungi_unclassified(68);k__Fungi_unclassified(68);k__Fungi_unclassified(68);k__Fungi_unclassified(68);k__Fungi_unclassified(68); | - | - | - | - | - | - | - | Unassigned | - |
| Otu01544 | 34 | k__Fungi(100);k__Fungi_unclassified(71);k__Fungi_unclassified(71);k__Fungi_unclassified(71);k__Fungi_unclassified(71);k__Fungi_unclassified(71);k__Fungi_unclassified(71); | - | - | - | - | - | - | - | Unassigned | - |
| Otu01548 | 34 | k__Fungi(100);p__Basidiomycota(100);p__Basidiomycota_unclassified(100);p__Basidiomycota_unclassified(100);p__Basidiomycota_unclassified(100);p__Basidiomycota_unclassified(100);p__Basidiomycota_unclassified(100); | - | - | - | - | - | - | - | Unassigned | - |
| Otu01553 | 34 | k__Fungi(100);k__Fungi_unclassified(83);k__Fungi_unclassified(83);k__Fungi_unclassified(83);k__Fungi_unclassified(83);k__Fungi_unclassified(83);k__Fungi_unclassified(83); | - | - | - | - | - | - | - | Unassigned | - |
| Otu01557 | 34 | k__Fungi(100);p__Ascomycota(100);p__Ascomycota_unclassified(100);p__Ascomycota_unclassified(100);p__Ascomycota_unclassified(100);p__Ascomycota_unclassified(100);p__Ascomycota_unclassified(100); | - | - | - | - | - | - | - | Unassigned | - |
| Otu01558 | 34 | k__Fungi(100);p__Ascomycota(100);p__Ascomycota_unclassified(100);p__Ascomycota_unclassified(100);p__Ascomycota_unclassified(100);p__Ascomycota_unclassified(100);p__Ascomycota_unclassified(100); | - | - | - | - | - | - | - | Unassigned | - |
| Otu01562 | 33 | k__Fungi(100);p__Ascomycota(100);c__Eurotiomycetes(100);o__Chaetothyriales(100);f__Herpotrichiellaceae(100);g__Capronia(94);s__Capronia_sp_CBS_16492(94); | Capronia | 13 | Pathotroph-Saprotroph-Symbiotroph | Animal Pathogen-Endophyte-Fungal Parasite-\|Undefined Saprotroph\| | Dimorphic-Facultative Yeast | NULL | Highly Probable | As mycoparasite (Põlme et al. 2020); As root_endophyte_dark_septate (Põlme et al. 2020); As soil_saprotroph (Põlme et al. 2020); Dark Septate Endophyte, known from plant roots.; Decay soil (Põlme et al. 2020) | Tedersoo L, et al. 2014. Science 346:e1256688 ((https://doi.org/10.1126/science.1256688)); Põlme S, et al. 2020. Fungal Diversity 105:1-16 ((https://doi.org/10.1007/s13225-020-00466-2)) |
| Otu01564 | 33 | k__Fungi(100);p__Ascomycota(100);c__Dothideomycetes(100);o__Pleosporales(100);f__Venturiaceae(100);g__Rhizosphaera(100);s__Rhizosphaera_sp(100); | Rhizosphaera | 13 | Pathotroph-Saprotroph | \|Plant Pathogen\|-Plant Saprotroph | Microfungus | NULL | Probable | As litter_saprotroph (Põlme et al. 2020); Decay leaf, fruit, seed (Põlme et al. 2020) | Tedersoo L, et al. 2014. Science 346:e1256688 ((https://doi.org/10.1126/science.1256688)); Põlme S, et al. 2020. Fungal Diversity 105:1-16 ((https://doi.org/10.1007/s13225-020-00466-2)) |
| Otu01567 | 33 | k__Fungi(100);p__Basidiomycota(100);c__Agaricomycetes(100);o__Polyporales(100);f__Fomitopsidaceae(100);g__Postia(100);s__Postia_stiptica(100); | Postia | 13 | Saprotroph | \|Wood Saprotroph\| | Polyporoid | Brown Rot | Highly Probable | As brown-rot (Põlme et al. 2020); Decay wood (Põlme et al. 2020) | Jordan CR, et al. 1996. Holzforschung - International Journal of the Biology, Chemistry, Physics and Technology of Wood 50:312-318 ((https://doi.org/10.1515/hfsg.1996.50.4.312)); Põlme S, et al. 2020. Fungal Diversity 105:1-16 ((https://doi.org/10.1007/s13225-020-00466-2)) |
| Otu01571 | 33 | k__Fungi(100);p__Basidiomycota(100);c__Microbotryomycetes(100);o__Sporidiobolales(100);f__Sporidiobolales_family_Incertae_sedis(100);g__Rhodotorula(100);g__Rhodotorula_unclassified(100); | Rhodotorula | 13 | Pathotroph-Saprotroph-Symbiotroph | Animal Pathogen-Animal Symbiotroph-Endophyte-Plant Pathogen-\|Undefined Saprotroph\| | Yeast | NULL | Probable | As foliar_endophyte (Põlme et al. 2020); Opportunistic human pathogen | Põlme S, et al. 2020. Fungal Diversity 105:1-16 ((https://doi.org/10.1007/s13225-020-00466-2)); https://www.cdc.gov ((https://www.cdc.gov)) |
| Otu01574 | 33 | k__Fungi(100);p__Ascomycota(100);c__Saccharomycetes(100);o__Saccharomycetales(100);f__Metschnikowiaceae(100);g__Metschnikowia(100);s__Metschnikowia_pulcherrima(82); | Metschnikowia | 13 | Saprotroph | \|Undefined Saprotroph\| | Yeast | NULL | Probable | As nectar-tap_saprotroph (Põlme et al. 2020); Decay sugar-rich substrates (Põlme et al. 2020) | Põlme S, et al. 2020. Fungal Diversity 105:1-16 ((https://doi.org/10.1007/s13225-020-00466-2)) |
| Otu01576 | 33 | k__Fungi(100);p__Ascomycota(100);c__Saccharomycetes(100);o__Saccharomycetales(100);f__Metschnikowiaceae(100);g__Metschnikowia(100);g__Metschnikowia_unclassified(58); | Metschnikowia | 13 | Saprotroph | \|Undefined Saprotroph\| | Yeast | NULL | Probable | As nectar-tap_saprotroph (Põlme et al. 2020); Decay sugar-rich substrates (Põlme et al. 2020) | Põlme S, et al. 2020. Fungal Diversity 105:1-16 ((https://doi.org/10.1007/s13225-020-00466-2)) |
| Otu01582 | 33 | k__Fungi(100);p__Basidiomycota(100);c__Agaricomycetes(100);o__Agaricales(100);f__Inocybaceae(100);g__Flammulaster(100);s__Flammulaster_limulatus_var._limulatus(100); | Flammulaster | 13 | Saprotroph | \|Plant Saprotroph\|-Undefined Saprotroph-Wood Saprotroph | Agaricoid | NULL | Probable | As litter_saprotroph (Põlme et al. 2020); Decay leaf, fruit, seed (Põlme et al. 2020) | Tedersoo L, et al. 2014. Science 346:e1256688 ((https://doi.org/10.1126/science.1256688)); Põlme S, et al. 2020. Fungal Diversity 105:1-16 ((https://doi.org/10.1007/s13225-020-00466-2)) |
| Otu01560 | 33 | k__Fungi(100);k__Fungi_unclassified(73);k__Fungi_unclassified(73);k__Fungi_unclassified(73);k__Fungi_unclassified(73);k__Fungi_unclassified(73);k__Fungi_unclassified(73); | - | - | - | - | - | - | - | Unassigned | - |
| Otu01561 | 33 | k__Fungi(100);p__Ascomycota(67);p__Ascomycota_unclassified(67);p__Ascomycota_unclassified(67);p__Ascomycota_unclassified(67);p__Ascomycota_unclassified(67);p__Ascomycota_unclassified(67); | - | - | - | - | - | - | - | Unassigned | - |
| Otu01563 | 33 | k__Fungi(100);k__Fungi_unclassified(100);k__Fungi_unclassified(100);k__Fungi_unclassified(100);k__Fungi_unclassified(100);k__Fungi_unclassified(100);k__Fungi_unclassified(100); | - | - | - | - | - | - | - | Unassigned | - |
| Otu01565 | 33 | k__Fungi(100);k__Fungi_unclassified(64);k__Fungi_unclassified(64);k__Fungi_unclassified(64);k__Fungi_unclassified(64);k__Fungi_unclassified(64);k__Fungi_unclassified(64); | - | - | - | - | - | - | - | Unassigned | - |
| Otu01566 | 33 | k__Fungi(100);p__Ascomycota(100);p__Ascomycota_unclassified(100);p__Ascomycota_unclassified(100);p__Ascomycota_unclassified(100);p__Ascomycota_unclassified(100);p__Ascomycota_unclassified(100); | - | - | - | - | - | - | - | Unassigned | - |
| Otu01568 | 33 | k__Fungi(100);k__Fungi_unclassified(55);k__Fungi_unclassified(55);k__Fungi_unclassified(55);k__Fungi_unclassified(55);k__Fungi_unclassified(55);k__Fungi_unclassified(55); | - | - | - | - | - | - | - | Unassigned | - |
| Otu01569 | 33 | k__Fungi(100);p__Ascomycota(100);c__Eurotiomycetes(70);o__Eurotiales(70);f__Elaphomycetaceae(70);g__unclassified_Elaphomycetaceae(70);s__Elaphomycetaceae_sp(70); | - | - | - | - | - | - | - | Unassigned | - |
| Otu01570 | 33 | k__Fungi(100);p__Basidiomycota(100);c__Agaricomycetes(100);o__unclassified_Agaricomycetes(85);f__unclassified_Agaricomycetes(85);g__unclassified_Agaricomycetes(85);s__Agaricomycetes_sp(85); | - | - | - | - | - | - | - | Unassigned | - |
| Otu01572 | 33 | k__Fungi(100);p__Ascomycota(100);c__Dothideomycetes(100);c__Dothideomycetes_unclassified(88);c__Dothideomycetes_unclassified(88);c__Dothideomycetes_unclassified(88);c__Dothideomycetes_unclassified(88); | - | - | - | - | - | - | - | Unassigned | - |
| Otu01573 | 33 | k__Fungi(100);k__Fungi_unclassified(67);k__Fungi_unclassified(67);k__Fungi_unclassified(67);k__Fungi_unclassified(67);k__Fungi_unclassified(67);k__Fungi_unclassified(67); | - | - | - | - | - | - | - | Unassigned | - |
| Otu01575 | 33 | k__Fungi(100);k__Fungi_unclassified(73);k__Fungi_unclassified(73);k__Fungi_unclassified(73);k__Fungi_unclassified(73);k__Fungi_unclassified(73);k__Fungi_unclassified(73); | - | - | - | - | - | - | - | Unassigned | - |
| Otu01577 | 33 | k__Plantae(97);p__unclassified_Plantae(97);c__unclassified_Plantae(97);o__unclassified_Plantae(97);f__unclassified_Plantae(97);g__unclassified_Plantae(97);s__Plantae_sp(97); | - | - | - | - | - | - | - | Unassigned | - |
| Otu01578 | 33 | k__Fungi(100);k__Fungi_unclassified(67);k__Fungi_unclassified(67);k__Fungi_unclassified(67);k__Fungi_unclassified(67);k__Fungi_unclassified(67);k__Fungi_unclassified(67); | - | - | - | - | - | - | - | Unassigned | - |
| Otu01579 | 33 | k__Fungi(100);k__Fungi_unclassified(61);k__Fungi_unclassified(61);k__Fungi_unclassified(61);k__Fungi_unclassified(61);k__Fungi_unclassified(61);k__Fungi_unclassified(61); | - | - | - | - | - | - | - | Unassigned | - |
| Otu01580 | 33 | k__Fungi(100);p__Basidiomycota(100);c__Agaricomycetes(100);c__Agaricomycetes_unclassified(82);c__Agaricomycetes_unclassified(82);c__Agaricomycetes_unclassified(82);c__Agaricomycetes_unclassified(82); | - | - | - | - | - | - | - | Unassigned | - |
| Otu01581 | 33 | k__Fungi(100);p__Ascomycota(100);p__Ascomycota_unclassified(100);p__Ascomycota_unclassified(100);p__Ascomycota_unclassified(100);p__Ascomycota_unclassified(100);p__Ascomycota_unclassified(100); | - | - | - | - | - | - | - | Unassigned | - |
| Otu01583 | 33 | k__Fungi(100);k__Fungi_unclassified(67);k__Fungi_unclassified(67);k__Fungi_unclassified(67);k__Fungi_unclassified(67);k__Fungi_unclassified(67);k__Fungi_unclassified(67); | - | - | - | - | - | - | - | Unassigned | - |
| Otu01586 | 32 | k__Fungi(100);p__Ascomycota(100);c__Dothideomycetes(100);o__Capnodiales(100);f__Mycosphaerellaceae(100);g__Mycosphaerella(100);s__Mycosphaerella_latebrosa(100); | Mycosphaerella | 13 | Pathotroph-Saprotroph | \|Plant Pathogen\|-Plant Saprotroph | Microfungus | NULL | Probable | As litter_saprotroph (Põlme et al. 2020); Decay leaf, fruit, seed (Põlme et al. 2020) | Tedersoo L, et al. 2014. Science 346:e1256688 ((https://doi.org/10.1126/science.1256688)); Põlme S, et al. 2020. Fungal Diversity 105:1-16 ((https://doi.org/10.1007/s13225-020-00466-2)) |
| Otu01589 | 32 | k__Fungi(100);p__Ascomycota(100);c__Saccharomycetes(100);o__Saccharomycetales(100);f__Metschnikowiaceae(100);g__Metschnikowia(100);s__Metschnikowia_pulcherrima(100); | Metschnikowia | 13 | Saprotroph | \|Undefined Saprotroph\| | Yeast | NULL | Probable | As nectar-tap_saprotroph (Põlme et al. 2020); Decay sugar-rich substrates (Põlme et al. 2020) | Põlme S, et al. 2020. Fungal Diversity 105:1-16 ((https://doi.org/10.1007/s13225-020-00466-2)) |
| Otu01594 | 32 | k__Fungi(100);p__Ascomycota(100);c__Leotiomycetes(100);o__Erysiphales(100);f__Erysiphaceae(100);g__Golovinomyces(100);s__Golovinomyces_sp(100); | Golovinomyces | 13 | Pathotroph | \|Plant Pathogen\| | Microfungus | NULL | Probable | Common name - powdery mildew | Tedersoo L, et al. 2014. Science 346:e1256688 ((https://doi.org/10.1126/science.1256688)); Põlme S, et al. 2020. Fungal Diversity 105:1-16 ((https://doi.org/10.1007/s13225-020-00466-2)) |
| Otu01597 | 32 | k__Fungi(100);p__Basidiomycota(100);c__Exobasidiomycetes(100);o__Microstromatales(100);f__Microstromataceae(100);g__Microstroma(100);s__Microstroma_juglandis(100); | Microstroma | 13 | Pathotroph | \|Plant Pathogen\| | Dimorphic | NULL | Probable | NULL | Tedersoo L, et al. 2014. Science 346:e1256688 ((https://doi.org/10.1126/science.1256688)); Põlme S, et al. 2020. Fungal Diversity 105:1-16 ((https://doi.org/10.1007/s13225-020-00466-2)) |
| Otu01598 | 32 | k__Fungi(100);p__Basidiomycota(100);c__Exobasidiomycetes(100);o__Exobasidiales(100);f__Exobasidiaceae(100);g__Exobasidium(100);s__Exobasidium_woronichinii(100); | Exobasidium | 13 | Pathotroph | \|Plant Pathogen\| | Leaf Blister-Leaf Gall-Leaf Spot | NULL | Probable | NULL | Tedersoo L, et al. 2014. Science 346:e1256688 ((https://doi.org/10.1126/science.1256688)); Põlme S, et al. 2020. Fungal Diversity 105:1-16 ((https://doi.org/10.1007/s13225-020-00466-2)) |
| Otu01601 | 32 | k__Fungi(100);p__Basidiomycota(100);c__Microbotryomycetes(100);o__Sporidiobolales(100);f__Sporidiobolales_family_Incertae_sedis(100);g__Sporobolomyces(100);s__Sporobolomyces_sp_CBS_10200(100); | Sporobolomyces | 13 | Pathotroph-Saprotroph | \|Fungal Parasite\|-Plant Saprotroph | Yeast | NULL | Probable | As litter_saprotroph (Põlme et al. 2020); As mycoparasite (Põlme et al. 2020); Decay fungal material (Põlme et al. 2020) | Tedersoo L, et al. 2014. Science 346:e1256688 ((https://doi.org/10.1126/science.1256688)); Purahong W, et al. 2016. Molecular Ecology 25:4059-4074 ((https://doi.org/10.1111/mec.13739)); Põlme S, et al. 2020. Fungal Diversity 105:1-16 ((https://doi.org/10.1007/s13225-020-00466-2)) |
| Otu01609 | 32 | k__Fungi(100);p__Basidiomycota(100);c__Tremellomycetes(100);o__Tremellales(100);f__Tremellales_family_Incertae_sedis(100);g__Hannaella(100);s__Hannaella_oryzae(100); | Hannaella | 13 | Pathotroph-Saprotroph | \|Animal Parasite\|-Undefined Saprotroph | Yeast | NULL | Probable | As animal_decomposer (Põlme et al. 2020); Decay animal material (Põlme et al. 2020) | Põlme S, et al. 2020. Fungal Diversity 105:1-16 ((https://doi.org/10.1007/s13225-020-00466-2)) |
| Otu01610 | 32 | k__Fungi(100);p__Ascomycota(100);c__Eurotiomycetes(100);o__Mycocaliciales(100);f__Mycocaliciaceae(100);g__Chaenothecopsis(100);s__Chaenothecopsis_tibellii(100); | Chaenothecopsis | 13 | Pathotroph-Saprotroph-Symbiotroph | Lichen Parasite-Lichenized-\|Undefined Saprotroph\| | Calicioid-Thallus | NULL | Possible | As resin_saprotroph (Põlme et al. 2020); licenized taxa; Lichenized and non | Esslinger TL. 2014. North Dakota State University ((http://www.ndsu.edu/pubweb/~esslinge/chcklst/chcklst7.htm)); Põlme S, et al. 2020. Fungal Diversity 105:1-16 ((https://doi.org/10.1007/s13225-020-00466-2)) |
| Otu01611 | 32 | k__Fungi(100);p__Ascomycota(100);c__Lecanoromycetes(100);o__Lecanorales(100);f__Parmeliaceae(100);g__Punctelia(100);s__Punctelia_rudecta(94); | Punctelia | 13 | Symbiotroph | \|Lichenized\| | Thallus | NULL | Highly Probable | NULL | Esslinger TL. 2014. North Dakota State University ((http://www.ndsu.edu/pubweb/~esslinge/chcklst/chcklst7.htm)); Põlme S, et al. 2020. Fungal Diversity 105:1-16 ((https://doi.org/10.1007/s13225-020-00466-2)) |
| Otu01612 | 32 | k__Fungi(100);p__Ascomycota(100);c__Lecanoromycetes(100);o__Lecanorales(97);o__Lecanorales_unclassified(97);o__Lecanorales_unclassified(97);o__Lecanorales_unclassified(97); | Lecanorales | 7 | Pathotroph-Symbiotroph | Lichen Parasite-Lichenized | Microfungus-Thallus | NULL | Probable | Lichenized with green algae, occasionally lichenicolous, on rocks, mosses and wood (Cannon & Kirk 2007); ...an order of mostly lichen-forming fungi (Kirk et al. 2008) | Cannon PF, Kirk PM. 2007. Fungal Families of the World. CAB International, Cambridge (ISBN: 978-0851998275); Kirk PM et al. 2008. Dictionary of the Fungi. Tenth Edition. CAB International, Wallingford (ISBN: 978-0851998268); Esslinger TL. 2014. North Dakota State University ((http://www.ndsu.edu/pubweb/~esslinge/chcklst/chcklst7.htm)) |
| Otu01614 | 32 | k__Fungi(100);p__Ascomycota(100);c__Saccharomycetes(100);o__Saccharomycetales(100);f__Metschnikowiaceae(100);g__Metschnikowia(100);s__Metschnikowia_pulcherrima(97); | Metschnikowia | 13 | Saprotroph | \|Undefined Saprotroph\| | Yeast | NULL | Probable | As nectar-tap_saprotroph (Põlme et al. 2020); Decay sugar-rich substrates (Põlme et al. 2020) | Põlme S, et al. 2020. Fungal Diversity 105:1-16 ((https://doi.org/10.1007/s13225-020-00466-2)) |
| Otu01616 | 32 | k__Fungi(100);p__Ascomycota(100);c__Saccharomycetes(100);o__Saccharomycetales(100);f__Metschnikowiaceae(100);g__Metschnikowia(100);g__Metschnikowia_unclassified(82); | Metschnikowia | 13 | Saprotroph | \|Undefined Saprotroph\| | Yeast | NULL | Probable | As nectar-tap_saprotroph (Põlme et al. 2020); Decay sugar-rich substrates (Põlme et al. 2020) | Põlme S, et al. 2020. Fungal Diversity 105:1-16 ((https://doi.org/10.1007/s13225-020-00466-2)) |
| Otu01617 | 32 | k__Fungi(100);p__Ascomycota(100);c__Sordariomycetes(100);o__Sordariales(100);f__Chaetomiaceae(100);g__Chaetomium(100);s__Chaetomium_caprinum(91); | Chaetomium | 13 | Pathotroph-Saprotroph-Symbiotroph | Animal Pathogen-Dung Saprotroph-Endophyte-Epiphyte-Plant Pathogen-\|Plant Saprotroph\|-Wood Saprotroph | Microfungus | Soft Rot | Possible | As foliar_endophyte (Põlme et al. 2020); As litter_saprotroph (Põlme et al. 2020); culture (Busby et al. 2016, David et al. 2016); Likely opportunistic human pathogen (Irinyi et al. 2015); culture (Massimo et al. 2015); Endophyte detection method; Decay wood, leaf, fruit, seed, soil, dung (Põlme et al. 2020); derived commodities (Cannon & Kirk 2007); Epiphyte detection method; Saprobic, acting as key decay organisms of plant materials and plant | Seehann G, et al. 1975. List of Fungi in Soft-Rot Tissue Tests. Institute fur Holzbiologie und Holzschutz, Bundesforschungsanstalt fur Forst - und Holzwirtshaft, Hamburg - Lohbrugge, Germany; Cannon PF, Kirk PM. 2007. Fungal Families of the World. CAB International, Cambridge ((https://isbnsearch.org/isbn/9780851998275)); Bills GF, et al. 2013. Current Opinion in Microbiology 16:549-565 ((https://doi.org/10.1016/j.mib.2013.08.001)); Tedersoo L, et al. 2014. Science 346:e1256688 ((https://doi.org/10.1126/science.1256688)); Irinyi L, et al. 2015. Medical Mycology 53:313-337 ((https://doi.org/10.1093/mmy/myv008)); Busby PE, et al. 2016. New Phytologist 209:1681-1692 ((https://doi.org/10.1111/nph.13742)); Põlme S, et al. 2020. Fungal Diversity 105:1-16 ((https://doi.org/10.1007/s13225-020-00466-2)) |
| Otu01618 | 32 | k__Fungi(100);p__Ascomycota(100);c__Dothideomycetes(100);o__Pleosporales(100);f__Phaeosphaeriaceae(100);g__unclassified_Phaeosphaeriaceae(66);s__Phaeosphaeriaceae_sp(66); | Phaeosphaeriaceae | 9 | Pathotroph-Saprotroph | Fungal Parasite-Plant Pathogen-Plant Saprotroph | Microfungus | NULL | Probable | Necrotrophic or saprobic on a wide range of plants, especially monocotyledons; a few species parasitic on other fungi (Cannon & Kirk 2007) | Cannon PF, Kirk PM. 2007. Fungal Families of the World. CAB International, Cambridge (ISBN: 978-0851998275) |
| Otu01619 | 32 | k__Fungi(100);p__Ascomycota(100);c__Leotiomycetes(100);o__Leotiomycetes_order_Incertae_sedis(100);f__Leotiomycetes_family_Incertae_sedis(100);g__Sarea(100);s__Sarea_resinae(100); | Sarea | 13 | Saprotroph-Symbiotroph | Lichenized-\|Undefined Saprotroph\| | Microfungus-Thallus | NULL | Highly Probable | NULL | Esslinger TL. 2014. North Dakota State University ((http://www.ndsu.edu/pubweb/~esslinge/chcklst/chcklst7.htm)); Põlme S, et al. 2020. Fungal Diversity 105:1-16 ((https://doi.org/10.1007/s13225-020-00466-2)) |
| Otu01620 | 32 | k__Fungi(100);p__Basidiomycota(100);c__Tremellomycetes(100);o__Tremellales(91);f__Tremellales_family_Incertae_sedis(88);f__Tremellales_family_Incertae_sedis_unclassified(88);f__Tremellales_family_Incertae_sedis_unclassified(88); | Tremellales | 7 | Pathotroph-Saprotroph-Symbiotroph | Fungal Parasite-Undefined Saprotroph | Tremelloid-Yeast | NULL | Possible | The order contains both teleomorphic and anamorphic species, most of the latter being yeasts | Sterkenburg E, et al. 2015. New Phytologist 207:1145-1158 ((https://doi.org/10.1111/nph.13426)) |
| Otu01584 | 32 | k__Fungi(100);p__Ascomycota(85);p__Ascomycota_unclassified(85);p__Ascomycota_unclassified(85);p__Ascomycota_unclassified(85);p__Ascomycota_unclassified(85);p__Ascomycota_unclassified(85); | - | - | - | - | - | - | - | Unassigned | - |
| Otu01585 | 32 | k__Fungi(100);k__Fungi_unclassified(69);k__Fungi_unclassified(69);k__Fungi_unclassified(69);k__Fungi_unclassified(69);k__Fungi_unclassified(69);k__Fungi_unclassified(69); | - | - | - | - | - | - | - | Unassigned | - |
| Otu01587 | 32 | k__Fungi(100);k__Fungi_unclassified(57);k__Fungi_unclassified(57);k__Fungi_unclassified(57);k__Fungi_unclassified(57);k__Fungi_unclassified(57);k__Fungi_unclassified(57); | - | - | - | - | - | - | - | Unassigned | - |
| Otu01588 | 32 | k__Fungi(100);p__Ascomycota(82);p__Ascomycota_unclassified(82);p__Ascomycota_unclassified(82);p__Ascomycota_unclassified(82);p__Ascomycota_unclassified(82);p__Ascomycota_unclassified(82); | - | - | - | - | - | - | - | Unassigned | - |
| Otu01590 | 32 | k__Fungi(100);p__Ascomycota(100);c__Leotiomycetes(88);o__Helotiales(72);o__Helotiales_unclassified(72);o__Helotiales_unclassified(72);o__Helotiales_unclassified(72); | - | - | - | - | - | - | - | Unassigned | - |
| Otu01591 | 32 | k__Fungi(100);p__Ascomycota(60);p__Ascomycota_unclassified(60);p__Ascomycota_unclassified(60);p__Ascomycota_unclassified(60);p__Ascomycota_unclassified(60);p__Ascomycota_unclassified(60); | - | - | - | - | - | - | - | Unassigned | - |
| Otu01592 | 32 | k__Fungi(100);p__Ascomycota(100);c__Dothideomycetes(100);o__Capnodiales(97);o__Capnodiales_unclassified(54);o__Capnodiales_unclassified(54);o__Capnodiales_unclassified(54); | - | - | - | - | - | - | - | Unassigned | - |
| Otu01593 | 32 | k__Fungi(100);p__Ascomycota(85);p__Ascomycota_unclassified(85);p__Ascomycota_unclassified(85);p__Ascomycota_unclassified(85);p__Ascomycota_unclassified(85);p__Ascomycota_unclassified(85); | - | - | - | - | - | - | - | Unassigned | - |
| Otu01595 | 32 | k__Fungi(100);p__Ascomycota(100);c__Leotiomycetes(100);o__Helotiales(79);o__Helotiales_unclassified(79);o__Helotiales_unclassified(79);o__Helotiales_unclassified(79); | - | - | - | - | - | - | - | Unassigned | - |
| Otu01596 | 32 | k__Fungi(100);p__Ascomycota(79);p__Ascomycota_unclassified(79);p__Ascomycota_unclassified(79);p__Ascomycota_unclassified(79);p__Ascomycota_unclassified(79);p__Ascomycota_unclassified(79); | - | - | - | - | - | - | - | Unassigned | - |
| Otu01599 | 32 | k__Fungi(100);k__Fungi_unclassified(57);k__Fungi_unclassified(57);k__Fungi_unclassified(57);k__Fungi_unclassified(57);k__Fungi_unclassified(57);k__Fungi_unclassified(57); | - | - | - | - | - | - | - | Unassigned | - |
| Otu01600 | 32 | k__Fungi(100);k__Fungi_unclassified(60);k__Fungi_unclassified(60);k__Fungi_unclassified(60);k__Fungi_unclassified(60);k__Fungi_unclassified(60);k__Fungi_unclassified(60); | - | - | - | - | - | - | - | Unassigned | - |
| Otu01602 | 32 | k__Fungi(100);k__Fungi_unclassified(75);k__Fungi_unclassified(75);k__Fungi_unclassified(75);k__Fungi_unclassified(75);k__Fungi_unclassified(75);k__Fungi_unclassified(75); | - | - | - | - | - | - | - | Unassigned | - |
| Otu01603 | 32 | k__Fungi(100);k__Fungi_unclassified(57);k__Fungi_unclassified(57);k__Fungi_unclassified(57);k__Fungi_unclassified(57);k__Fungi_unclassified(57);k__Fungi_unclassified(57); | - | - | - | - | - | - | - | Unassigned | - |
| Otu01604 | 32 | k__Fungi(100);k__Fungi_unclassified(94);k__Fungi_unclassified(94);k__Fungi_unclassified(94);k__Fungi_unclassified(94);k__Fungi_unclassified(94);k__Fungi_unclassified(94); | - | - | - | - | - | - | - | Unassigned | - |
| Otu01605 | 32 | k__Fungi(100);p__Ascomycota(88);p__Ascomycota_unclassified(88);p__Ascomycota_unclassified(88);p__Ascomycota_unclassified(88);p__Ascomycota_unclassified(88);p__Ascomycota_unclassified(88); | - | - | - | - | - | - | - | Unassigned | - |
| Otu01606 | 32 | k__Fungi(100);p__Ascomycota(82);p__Ascomycota_unclassified(82);p__Ascomycota_unclassified(82);p__Ascomycota_unclassified(82);p__Ascomycota_unclassified(82);p__Ascomycota_unclassified(82); | - | - | - | - | - | - | - | Unassigned | - |
| Otu01607 | 32 | k__Fungi(100);k__Fungi_unclassified(75);k__Fungi_unclassified(75);k__Fungi_unclassified(75);k__Fungi_unclassified(75);k__Fungi_unclassified(75);k__Fungi_unclassified(75); | - | - | - | - | - | - | - | Unassigned | - |
| Otu01608 | 32 | k__Fungi(100);p__Ascomycota(54);p__Ascomycota_unclassified(54);p__Ascomycota_unclassified(54);p__Ascomycota_unclassified(54);p__Ascomycota_unclassified(54);p__Ascomycota_unclassified(54); | - | - | - | - | - | - | - | Unassigned | - |
| Otu01613 | 32 | k__Fungi(100);k__Fungi_unclassified(60);k__Fungi_unclassified(60);k__Fungi_unclassified(60);k__Fungi_unclassified(60);k__Fungi_unclassified(60);k__Fungi_unclassified(60); | - | - | - | - | - | - | - | Unassigned | - |
| Otu01615 | 32 | k__Fungi(100);p__Ascomycota(100);c__Dothideomycetes(100);c__Dothideomycetes_unclassified(100);c__Dothideomycetes_unclassified(100);c__Dothideomycetes_unclassified(100);c__Dothideomycetes_unclassified(100); | - | - | - | - | - | - | - | Unassigned | - |
| Otu01621 | 32 | k__Fungi(100);p__Ascomycota(100);c__Leotiomycetes(100);o__Helotiales(100);o__Helotiales_unclassified(100);o__Helotiales_unclassified(100);o__Helotiales_unclassified(100); | - | - | - | - | - | - | - | Unassigned | - |
| Otu01622 | 32 | k__Fungi(100);p__Basidiomycota(100);c__Agaricomycetes(100);o__Russulales(100);o__Russulales_unclassified(100);o__Russulales_unclassified(100);o__Russulales_unclassified(100); | - | - | - | - | - | - | - | Unassigned | - |
| Otu01625 | 31 | k__Fungi(100);p__Basidiomycota(100);c__Agaricomycetes(100);o__Hymenochaetales(100);f__Schizoporaceae(100);g__Hyphodontia(100);s__Hyphodontia_alutacea(100); | Hyphodontia | 13 | Saprotroph | Undefined Saprotroph-\|Wood Saprotroph\| | Corticioid | White Rot | Probable | As white_rot (Põlme et al. 2020); Decay wood (Põlme et al. 2020) | Tedersoo L, et al. 2014. Science 346:e1256688 ((https://doi.org/10.1126/science.1256688)); Põlme S, et al. 2020. Fungal Diversity 105:1-16 ((https://doi.org/10.1007/s13225-020-00466-2)) |
| Otu01627 | 31 | k__Fungi(100);p__Basidiomycota(100);c__Microbotryomycetes(100);o__Sporidiobolales(100);f__Sporidiobolales_family_Incertae_sedis(100);g__Rhodotorula(100);s__Rhodotorula_oryzicola(100); | Rhodotorula | 13 | Pathotroph-Saprotroph-Symbiotroph | Animal Pathogen-Animal Symbiotroph-Endophyte-Plant Pathogen-\|Undefined Saprotroph\| | Yeast | NULL | Probable | As foliar_endophyte (Põlme et al. 2020); Opportunistic human pathogen | Põlme S, et al. 2020. Fungal Diversity 105:1-16 ((https://doi.org/10.1007/s13225-020-00466-2)); https://www.cdc.gov ((https://www.cdc.gov)) |
| Otu01628 | 31 | k__Fungi(100);p__Ascomycota(100);c__Saccharomycetes(100);o__Saccharomycetales(100);f__Saccharomycetales_family_Incertae_sedis(100);g__Candida(100);g__Candida_unclassified(59); | Candida | 13 | Pathotroph-Saprotroph-Symbiotroph | Animal Pathogen-Endophyte-Endosymbiont-Epiphyte-\|Undefined Saprotroph\| | Dimorphic Yeast-Yeast | NULL | Probable | As nectar-tap_saprotroph (Põlme et al. 2020); Decay sugar-rich substrates (Põlme et al. 2020); This asexual genus is polyphyletic and contain species that function very differently depend on what they are. Caution should be taken when interpreting guilds for Candida species. | Manolakaki D, et al. 2010. Virulence. 1: 367-375 ((https://doi.org/10.4161/viru.1.5.12796)); Mekbib SB, et al. 2011. Tropical Plant Pathology 36:233-240 ((https://doi.org/10.1590/S1982-56762011000400004)); Põlme S, et al. 2020. Fungal Diversity 105:1-16 ((https://doi.org/10.1007/s13225-020-00466-2)) |
| Otu01631 | 31 | k__Fungi(100);p__Basidiomycota(100);c__Agaricomycetes(100);o__Agaricales(100);f__Strophariaceae(100);g__Gymnopilus(94);g__Gymnopilus_unclassified(94); | Gymnopilus | 13 | Saprotroph | Plant Saprotroph-\|Wood Saprotroph\| | Agaricoid | White Rot | Probable | As litter_saprotroph (Põlme et al. 2020); As white_rot (Põlme et al. 2020); Decay wood (Põlme et al. 2020) | Tedersoo L, et al. 2014. Science 346:e1256688 ((https://doi.org/10.1126/science.1256688)); Põlme S, et al. 2020. Fungal Diversity 105:1-16 ((https://doi.org/10.1007/s13225-020-00466-2)) |
| Otu01632 | 31 | k__Fungi(100);p__Ascomycota(100);c__Leotiomycetes(100);o__Helotiales(100);f__Helotiaceae(97);g__Crocicreas(81);s__Crocicreas_sp(81); | Crocicreas | 13 | Saprotroph | \|Plant Saprotroph\|-Undefined Saprotroph | Microfungus-Pezizoid | NULL | Probable | As litter_saprotroph (Põlme et al. 2020); Decay leaf, fruit, seed (Põlme et al. 2020) | Carpenter SE, 1983. Memoirs of The New York Botanical Garden 33:1-290 (ISBN: 978-0893272302); Põlme S, et al. 2020. Fungal Diversity 105:1-16 ((https://doi.org/10.1007/s13225-020-00466-2)) |
| Otu01634 | 31 | k__Fungi(100);p__Basidiomycota(100);c__Agaricomycetes(100);o__Cantharellales(100);f__Ceratobasidiaceae(100);f__Ceratobasidiaceae_unclassified(100);f__Ceratobasidiaceae_unclassified(100); | Ceratobasidiaceae | 9 | Pathotroph-Saprotroph-Symbiotroph | Endomycorrhizal-Plant Pathogen-Undefined Saprotroph | Microfungus | NULL | Possible | Best known as plant pathogens, but also saprobic. Species are endomycorrhizal with a wide range of plant roots - especially of Orchidaceae but also even bryophytes (Cannon & Kirk 2007) | Cannon PF, Kirk PM. 2007. Fungal Families of the World. CAB International, Cambridge (ISBN: 978-0851998275) |
| Otu01635 | 31 | k__Fungi(100);p__Ascomycota(100);c__Saccharomycetes(100);o__Saccharomycetales(100);f__Metschnikowiaceae(100);g__Metschnikowia(100);s__Metschnikowia_pulcherrima(84); | Metschnikowia | 13 | Saprotroph | \|Undefined Saprotroph\| | Yeast | NULL | Probable | As nectar-tap_saprotroph (Põlme et al. 2020); Decay sugar-rich substrates (Põlme et al. 2020) | Põlme S, et al. 2020. Fungal Diversity 105:1-16 ((https://doi.org/10.1007/s13225-020-00466-2)) |
| Otu01636 | 31 | k__Fungi(100);p__Basidiomycota(100);c__Agaricomycetes(100);o__Hymenochaetales(100);f__Hymenochaetaceae(100);g__Phellinus(100);s__Phellinus_sp_LWZ_2012b(100); | Phellinus | 13 | Pathotroph-Saprotroph | \|Plant Pathogen\|-Wood Saprotroph | Polyporoid | White Rot | Highly Probable | As white_rot (Põlme et al. 2020); As wood_pathogen (Põlme et al. 2020); Decay wood (Põlme et al. 2020) | Gilbertson RL, Ryvarden L. 1987-1987. North American Polypores. Fungiflora, Oslo (ISBN: 978-0945345060); Põlme S, et al. 2020. Fungal Diversity 105:1-16 ((https://doi.org/10.1007/s13225-020-00466-2)) |
| Otu01642 | 31 | k__Fungi(100);p__Ascomycota(100);c__Saccharomycetes(100);o__Saccharomycetales(100);f__Metschnikowiaceae(100);g__Metschnikowia(100);g__Metschnikowia_unclassified(84); | Metschnikowia | 13 | Saprotroph | \|Undefined Saprotroph\| | Yeast | NULL | Probable | As nectar-tap_saprotroph (Põlme et al. 2020); Decay sugar-rich substrates (Põlme et al. 2020) | Põlme S, et al. 2020. Fungal Diversity 105:1-16 ((https://doi.org/10.1007/s13225-020-00466-2)) |
| Otu01644 | 31 | k__Fungi(100);p__Basidiomycota(100);c__Agaricomycetes(100);o__Polyporales(100);f__Meruliaceae(100);g__Hyphoderma(100);s__Hyphoderma_guttuliferum(100); | Hyphoderma | 13 | Saprotroph | Undefined Saprotroph-\|Wood Saprotroph\| | Corticioid | White Rot | Probable | As white_rot (Põlme et al. 2020); Decay wood (Põlme et al. 2020) | Tedersoo L, et al. 2014. Science 346:e1256688 ((https://doi.org/10.1126/science.1256688)); Põlme S, et al. 2020. Fungal Diversity 105:1-16 ((https://doi.org/10.1007/s13225-020-00466-2)) |
| Otu01645 | 31 | k__Fungi(100);p__Ascomycota(100);c__Saccharomycetes(100);o__Saccharomycetales(100);f__Saccharomycetales_family_Incertae_sedis(100);g__Candida(100);g__Candida_unclassified(100); | Candida | 13 | Pathotroph-Saprotroph-Symbiotroph | Animal Pathogen-Endophyte-Endosymbiont-Epiphyte-\|Undefined Saprotroph\| | Dimorphic Yeast-Yeast | NULL | Probable | As nectar-tap_saprotroph (Põlme et al. 2020); Decay sugar-rich substrates (Põlme et al. 2020); This asexual genus is polyphyletic and contain species that function very differently depend on what they are. Caution should be taken when interpreting guilds for Candida species. | Manolakaki D, et al. 2010. Virulence. 1: 367-375 ((https://doi.org/10.4161/viru.1.5.12796)); Mekbib SB, et al. 2011. Tropical Plant Pathology 36:233-240 ((https://doi.org/10.1590/S1982-56762011000400004)); Põlme S, et al. 2020. Fungal Diversity 105:1-16 ((https://doi.org/10.1007/s13225-020-00466-2)) |
| Otu01646 | 31 | k__Fungi(100);p__Basidiomycota(100);c__Microbotryomycetes(100);o__Sporidiobolales(100);f__Sporidiobolales_family_Incertae_sedis(100);g__Sporobolomyces(100);s__Sporobolomyces_linderae(100); | Sporobolomyces | 13 | Pathotroph-Saprotroph | \|Fungal Parasite\|-Plant Saprotroph | Yeast | NULL | Probable | As litter_saprotroph (Põlme et al. 2020); As mycoparasite (Põlme et al. 2020); Decay fungal material (Põlme et al. 2020) | Tedersoo L, et al. 2014. Science 346:e1256688 ((https://doi.org/10.1126/science.1256688)); Purahong W, et al. 2016. Molecular Ecology 25:4059-4074 ((https://doi.org/10.1111/mec.13739)); Põlme S, et al. 2020. Fungal Diversity 105:1-16 ((https://doi.org/10.1007/s13225-020-00466-2)) |
| Otu01647 | 31 | k__Fungi(100);p__Ascomycota(100);c__Sordariomycetes(100);o__Hypocreales(100);f__Hypocreales_family_Incertae_sedis(100);g__Myrothecium(100);s__Myrothecium_sp_TA26_35(100); | Myrothecium | 13 | Pathotroph-Saprotroph | Fungal Parasite-\|Plant Saprotroph\|-Undefined Saprotroph | Microfungus | NULL | Probable | As litter_saprotroph (Põlme et al. 2020); As mycoparasite (Põlme et al. 2020); Decay leaf, fruit, seed (Põlme et al. 2020) | Tedersoo L, et al. 2014. Science 346:e1256688 ((https://doi.org/10.1126/science.1256688)); Põlme S, et al. 2020. Fungal Diversity 105:1-16 ((https://doi.org/10.1007/s13225-020-00466-2)) |
| Otu01648 | 31 | k__Fungi(100);p__Ascomycota(100);c__Saccharomycetes(100);o__Saccharomycetales(100);f__Metschnikowiaceae(100);g__Metschnikowia(100);g__Metschnikowia_unclassified(75); | Metschnikowia | 13 | Saprotroph | \|Undefined Saprotroph\| | Yeast | NULL | Probable | As nectar-tap_saprotroph (Põlme et al. 2020); Decay sugar-rich substrates (Põlme et al. 2020) | Põlme S, et al. 2020. Fungal Diversity 105:1-16 ((https://doi.org/10.1007/s13225-020-00466-2)) |
| Otu01649 | 31 | k__Fungi(100);p__Ascomycota(100);c__Eurotiomycetes(100);o__Eurotiales(100);f__Trichocomaceae(100);g__Paecilomyces(100);s__Paecilomyces_lilacinus(100); | Paecilomyces | 13 | Pathotroph-Saprotroph-Symbiotroph | Clavicipitaceous \|Endophyte\|-Plant Pathogen-Plant Saprotroph-Undefined Saprotroph-Wood Saprotroph | Microfungus | Soft Rot | Probable | As foliar_endophyte (Põlme et al. 2020); As litter_saprotroph (Põlme et al. 2020); culture (Lopez et al. 2014); Decay leaf, fruit, seed, animal material (Põlme et al. 2020); Endophyte detection method | Seehann G, et al. 1975. List of Fungi in Soft-Rot Tissue Tests. Institute fur Holzbiologie und Holzschutz, Bundesforschungsanstalt fur Forst - und Holzwirtshaft, Hamburg - Lohbrugge, Germany; Costa IPMW, et al. 2012. Mycotaxon 119:e26 ((http://mycotaxon.com/resources/checklists/costa_v119_checklist.pdf)); Tedersoo L, et al. 2014. Science 346:e1256688 ((https://doi.org/10.1126/science.1256688)); Põlme S, et al. 2020. Fungal Diversity 105:1-16 ((https://doi.org/10.1007/s13225-020-00466-2)); The American Phytopathological Society Common Names of Plant Diseases ((http://www.apsnet.org/publications/commonnames/Pages/default.aspx)); Lopez DC, et al. 2014. Plos One 9:e103891 ((https://doi.org/10.1371/journal.pone.0103891)) |
| Otu01651 | 31 | k__Fungi(100);p__Ascomycota(100);c__Dothideomycetes(100);o__Capnodiales(100);f__Mycosphaerellaceae(100);g__Pseudocercospora(100);g__Pseudocercospora_unclassified(100); | Pseudocercospora | 13 | Pathotroph | \|Plant Pathogen\| | Microfungus | NULL | Probable | NULL | Tedersoo L, et al. 2014. Science 346:e1256688 ((https://doi.org/10.1126/science.1256688)); Põlme S, et al. 2020. Fungal Diversity 105:1-16 ((https://doi.org/10.1007/s13225-020-00466-2)) |
| Otu01653 | 31 | k__Fungi(100);p__Ascomycota(100);c__Leotiomycetes(100);o__Helotiales(100);f__Helotiaceae(100);g__Hymenoscyphus(100);g__Hymenoscyphus_unclassified(88); | Hymenoscyphus | 13 | Pathotroph-Saprotroph-Symbiotroph | Bryophyte Parasite-Ectomycorrhizal-Ericoid Mycorrhizal-Plant Pathogen-\|Plant Saprotroph\|-Undefined Saprotroph | Microfungus | NULL | Possible | As litter_saprotroph (Põlme et al. 2020); Dark Septate Microfungus; Syn. Pezoloma among other (see e.g., Pezoloma ericae); Decay leaf, fruit, seed (Põlme et al. 2020) | Baral HO, Krieglsteiner L. 2006. Acta Mycologica 41:11-20 ((https://doi.org/10.5586/am.2006.003)) (as a bryophyte parasite); Huhtinen S, et al. 2010. Nova Hedwigia 90:413-431 ((https://doi.org/10.1127/0029-5035/2010/0090-0413)); Bent E, et al. 2011. Applied and Environmental Microbiology 77:3351-3359 ((https://doi.org/10.1128/AEM.02575-10)) (as ectomycorrhizae); Tedersoo L, et al. 2014. Science 346:1256688 ((https://doi.org/10.1126/science.1256688)) (as saprotroph); Põlme S, et al. 2020. Fungal Diversity 105:1-16 ((https://doi.org/10.1007/s13225-020-00466-2)) |
| Otu01655 | 31 | k__Fungi(100);p__Ascomycota(100);c__Saccharomycetes(100);o__Saccharomycetales(100);f__Metschnikowiaceae(100);g__Metschnikowia(100);s__Metschnikowia_pulcherrima(100); | Metschnikowia | 13 | Saprotroph | \|Undefined Saprotroph\| | Yeast | NULL | Probable | As nectar-tap_saprotroph (Põlme et al. 2020); Decay sugar-rich substrates (Põlme et al. 2020) | Põlme S, et al. 2020. Fungal Diversity 105:1-16 ((https://doi.org/10.1007/s13225-020-00466-2)) |
| Otu01660 | 31 | k__Fungi(100);p__Basidiomycota(100);c__Agaricomycetes(100);o__Polyporales(100);f__Phanerochaetaceae(100);g__Ceriporia(100);s__Ceriporia_alachuana(100); | Ceriporia | 13 | Saprotroph | \|Wood Saprotroph\| | Corticioid | White Rot | Highly Probable | As white_rot (Põlme et al. 2020); Decay wood (Põlme et al. 2020) | Tedersoo L, et al. 2014. Science 346:e1256688 ((https://doi.org/10.1126/science.1256688)); Põlme S, et al. 2020. Fungal Diversity 105:1-16 ((https://doi.org/10.1007/s13225-020-00466-2)) |
| Otu01664 | 31 | k__Fungi(100);p__Ascomycota(100);c__Eurotiomycetes(100);o__Chaetothyriales(100);f__Herpotrichiellaceae(100);g__Capronia(100);s__Capronia_sp_MUCL_39977(100); | Capronia | 13 | Pathotroph-Saprotroph-Symbiotroph | Animal Pathogen-Endophyte-Fungal Parasite-\|Undefined Saprotroph\| | Dimorphic-Facultative Yeast | NULL | Highly Probable | As mycoparasite (Põlme et al. 2020); As root_endophyte_dark_septate (Põlme et al. 2020); As soil_saprotroph (Põlme et al. 2020); Dark Septate Endophyte, known from plant roots.; Decay soil (Põlme et al. 2020) | Tedersoo L, et al. 2014. Science 346:e1256688 ((https://doi.org/10.1126/science.1256688)); Põlme S, et al. 2020. Fungal Diversity 105:1-16 ((https://doi.org/10.1007/s13225-020-00466-2)) |
| Otu01665 | 31 | k__Fungi(100);p__Ascomycota(100);c__Saccharomycetes(100);o__Saccharomycetales(100);f__Saccharomycetaceae(100);g__Kazachstania(100);s__Kazachstania_intestinalis(100); | Kazachstania | 13 | Saprotroph | \|Undefined Saprotroph\| | Yeast | NULL | Probable | As nectar-tap_saprotroph (Põlme et al. 2020); Decay sugar-rich substrates (Põlme et al. 2020) | Põlme S, et al. 2020. Fungal Diversity 105:1-16 ((https://doi.org/10.1007/s13225-020-00466-2)) |
| Otu01666 | 31 | k__Fungi(100);p__Basidiomycota(100);c__Agaricomycetes(100);o__Polyporales(100);f__Fomitopsidaceae(100);g__Fomitopsis(100);s__Fomitopsis_spraguei(100); | Fomitopsis | 13 | Pathotroph-Saprotroph | Plant Pathogen-\|Wood Saprotroph\| | Polyporoid | Brown Rot | Highly Probable | As brown-rot (Põlme et al. 2020); As wood_pathogen (Põlme et al. 2020); Decay wood (Põlme et al. 2020); Most species are parasitic on woody plants, causing brown rots (Cannon & Kirk 2007) | Gilbertson RL, Ryvarden L. 1987-1987. North American Polypores. Fungiflora, Oslo (ISBN: 978-0945345060); Cannon PF, Kirk PM. 2007. Fungal Families of the World. CAB International, Cambridge (ISBN: 978-0851998275); Põlme S, et al. 2020. Fungal Diversity 105:1-16 ((https://doi.org/10.1007/s13225-020-00466-2)) |
| Otu01623 | 31 | k__Fungi(100);p__Ascomycota(81);p__Ascomycota_unclassified(81);p__Ascomycota_unclassified(81);p__Ascomycota_unclassified(81);p__Ascomycota_unclassified(81);p__Ascomycota_unclassified(81); | - | - | - | - | - | - | - | Unassigned | - |
| Otu01624 | 31 | k__Fungi(100);k__Fungi_unclassified(100);k__Fungi_unclassified(100);k__Fungi_unclassified(100);k__Fungi_unclassified(100);k__Fungi_unclassified(100);k__Fungi_unclassified(100); | - | - | - | - | - | - | - | Unassigned | - |
| Otu01626 | 31 | k__Fungi(100);p__Ascomycota(100);p__Ascomycota_unclassified(52);p__Ascomycota_unclassified(52);p__Ascomycota_unclassified(52);p__Ascomycota_unclassified(52);p__Ascomycota_unclassified(52); | - | - | - | - | - | - | - | Unassigned | - |
| Otu01629 | 31 | k__Fungi(100);k__Fungi_unclassified(52);k__Fungi_unclassified(52);k__Fungi_unclassified(52);k__Fungi_unclassified(52);k__Fungi_unclassified(52);k__Fungi_unclassified(52); | - | - | - | - | - | - | - | Unassigned | - |
| Otu01630 | 31 | k__Fungi(100);k__Fungi_unclassified(55);k__Fungi_unclassified(55);k__Fungi_unclassified(55);k__Fungi_unclassified(55);k__Fungi_unclassified(55);k__Fungi_unclassified(55); | - | - | - | - | - | - | - | Unassigned | - |
| Otu01633 | 31 | k__Fungi(100);k__Fungi_unclassified(55);k__Fungi_unclassified(55);k__Fungi_unclassified(55);k__Fungi_unclassified(55);k__Fungi_unclassified(55);k__Fungi_unclassified(55); | - | - | - | - | - | - | - | Unassigned | - |
| Otu01637 | 31 | k__Fungi(100);p__Ascomycota(52);p__Ascomycota_unclassified(52);p__Ascomycota_unclassified(52);p__Ascomycota_unclassified(52);p__Ascomycota_unclassified(52);p__Ascomycota_unclassified(52); | - | - | - | - | - | - | - | Unassigned | - |
| Otu01638 | 31 | k__Fungi(100);k__Fungi_unclassified(52);k__Fungi_unclassified(52);k__Fungi_unclassified(52);k__Fungi_unclassified(52);k__Fungi_unclassified(52);k__Fungi_unclassified(52); | - | - | - | - | - | - | - | Unassigned | - |
| Otu01639 | 31 | k__Fungi(100);p__Ascomycota(84);p__Ascomycota_unclassified(84);p__Ascomycota_unclassified(84);p__Ascomycota_unclassified(84);p__Ascomycota_unclassified(84);p__Ascomycota_unclassified(84); | - | - | - | - | - | - | - | Unassigned | - |
| Otu01640 | 31 | k__Fungi(100);k__Fungi_unclassified(52);k__Fungi_unclassified(52);k__Fungi_unclassified(52);k__Fungi_unclassified(52);k__Fungi_unclassified(52);k__Fungi_unclassified(52); | - | - | - | - | - | - | - | Unassigned | - |
| Otu01641 | 31 | k__Fungi(100);p__Basidiomycota(100);c__Agaricomycetes(100);o__Boletales(100);o__Boletales_unclassified(94);o__Boletales_unclassified(94);o__Boletales_unclassified(94); | - | - | - | - | - | - | - | Unassigned | - |
| Otu01643 | 31 | k__Fungi(100);k__Fungi_unclassified(100);k__Fungi_unclassified(100);k__Fungi_unclassified(100);k__Fungi_unclassified(100);k__Fungi_unclassified(100);k__Fungi_unclassified(100); | - | - | - | - | - | - | - | Unassigned | - |
| Otu01650 | 31 | k__Fungi(100);k__Fungi_unclassified(75);k__Fungi_unclassified(75);k__Fungi_unclassified(75);k__Fungi_unclassified(75);k__Fungi_unclassified(75);k__Fungi_unclassified(75); | - | - | - | - | - | - | - | Unassigned | - |
| Otu01652 | 31 | k__Fungi(100);p__Ascomycota(88);p__Ascomycota_unclassified(88);p__Ascomycota_unclassified(88);p__Ascomycota_unclassified(88);p__Ascomycota_unclassified(88);p__Ascomycota_unclassified(88); | - | - | - | - | - | - | - | Unassigned | - |
| Otu01654 | 31 | k__Fungi(100);p__Ascomycota(100);c__Sordariomycetes(100);c__Sordariomycetes_unclassified(97);c__Sordariomycetes_unclassified(97);c__Sordariomycetes_unclassified(97);c__Sordariomycetes_unclassified(97); | - | - | - | - | - | - | - | Unassigned | - |
| Otu01656 | 31 | k__Fungi(100);p__Ascomycota(55);p__Ascomycota_unclassified(55);p__Ascomycota_unclassified(55);p__Ascomycota_unclassified(55);p__Ascomycota_unclassified(55);p__Ascomycota_unclassified(55); | - | - | - | - | - | - | - | Unassigned | - |
| Otu01657 | 31 | k__Fungi(100);k__Fungi_unclassified(88);k__Fungi_unclassified(88);k__Fungi_unclassified(88);k__Fungi_unclassified(88);k__Fungi_unclassified(88);k__Fungi_unclassified(88); | - | - | - | - | - | - | - | Unassigned | - |
| Otu01658 | 31 | k__Fungi(100);k__Fungi_unclassified(59);k__Fungi_unclassified(59);k__Fungi_unclassified(59);k__Fungi_unclassified(59);k__Fungi_unclassified(59);k__Fungi_unclassified(59); | - | - | - | - | - | - | - | Unassigned | - |
| Otu01659 | 31 | k__Fungi(100);p__Ascomycota(100);p__Ascomycota_unclassified(55);p__Ascomycota_unclassified(55);p__Ascomycota_unclassified(55);p__Ascomycota_unclassified(55);p__Ascomycota_unclassified(55); | - | - | - | - | - | - | - | Unassigned | - |
| Otu01661 | 31 | k__Fungi(100);p__Ascomycota(62);p__Ascomycota_unclassified(62);p__Ascomycota_unclassified(62);p__Ascomycota_unclassified(62);p__Ascomycota_unclassified(62);p__Ascomycota_unclassified(62); | - | - | - | - | - | - | - | Unassigned | - |
| Otu01662 | 31 | k__Fungi(100);p__Ascomycota(100);p__Ascomycota_unclassified(100);p__Ascomycota_unclassified(100);p__Ascomycota_unclassified(100);p__Ascomycota_unclassified(100);p__Ascomycota_unclassified(100); | - | - | - | - | - | - | - | Unassigned | - |
| Otu01663 | 31 | k__Fungi(100);p__Ascomycota(100);c__Dothideomycetes(100);o__Pleosporales(100);o__Pleosporales_unclassified(71);o__Pleosporales_unclassified(71);o__Pleosporales_unclassified(71); | - | - | - | - | - | - | - | Unassigned | - |
| Otu01667 | 30 | k__Fungi(100);p__Basidiomycota(100);c__Microbotryomycetes(100);o__Sporidiobolales(100);f__Sporidiobolales_family_Incertae_sedis(100);g__Sporobolomyces(100);s__Sporobolomyces_lactosus(100); | Sporobolomyces | 13 | Pathotroph-Saprotroph | \|Fungal Parasite\|-Plant Saprotroph | Yeast | NULL | Probable | As litter_saprotroph (Põlme et al. 2020); As mycoparasite (Põlme et al. 2020); Decay fungal material (Põlme et al. 2020) | Tedersoo L, et al. 2014. Science 346:e1256688 ((https://doi.org/10.1126/science.1256688)); Purahong W, et al. 2016. Molecular Ecology 25:4059-4074 ((https://doi.org/10.1111/mec.13739)); Põlme S, et al. 2020. Fungal Diversity 105:1-16 ((https://doi.org/10.1007/s13225-020-00466-2)) |
| Otu01668 | 30 | k__Fungi(100);p__Basidiomycota(100);c__Agaricomycetes(100);o__Agaricales(100);o__Agaricales_unclassified(100);o__Agaricales_unclassified(100);o__Agaricales_unclassified(100); | Agaricales | 7 | Pathotroph-Saprotroph-Symbiotroph | Bryophyte Parasite-Dung Saprotroph-Ectomycorrhizal-Fungal Parasite-Leaf Saprotroph-Plant Parasite-Undefined Saprotroph-Wood Saprotroph | Agaricoid-Gasteroid-Microfungus-Secotioid-Yeast | NULL | Possible | Mushrooms and toadstools, Gill fungi, Agarics... lignicolous, sometimes muscicolous or fungicolous, saprobic, mycorrhizal, rarely parasitic on plants or fungi (Kirk et al. 2008) | Kirk PM et al. 2008. Dictionary of the Fungi. Tenth Edition. CAB International, Wallingford (ISBN: 978-0851998268) |
| Otu01669 | 30 | k__Fungi(100);p__Ascomycota(100);c__Saccharomycetes(100);o__Saccharomycetales(100);f__Metschnikowiaceae(100);g__Metschnikowia(100);g__Metschnikowia_unclassified(80); | Metschnikowia | 13 | Saprotroph | \|Undefined Saprotroph\| | Yeast | NULL | Probable | As nectar-tap_saprotroph (Põlme et al. 2020); Decay sugar-rich substrates (Põlme et al. 2020) | Põlme S, et al. 2020. Fungal Diversity 105:1-16 ((https://doi.org/10.1007/s13225-020-00466-2)) |
| Otu01670 | 30 | k__Fungi(100);p__Ascomycota(100);c__Saccharomycetes(100);o__Saccharomycetales(100);f__Metschnikowiaceae(100);g__Metschnikowia(100);s__Metschnikowia_pulcherrima(90); | Metschnikowia | 13 | Saprotroph | \|Undefined Saprotroph\| | Yeast | NULL | Probable | As nectar-tap_saprotroph (Põlme et al. 2020); Decay sugar-rich substrates (Põlme et al. 2020) | Põlme S, et al. 2020. Fungal Diversity 105:1-16 ((https://doi.org/10.1007/s13225-020-00466-2)) |
| Otu01672 | 30 | k__Fungi(100);p__Zygomycota(100);c__Zygomycota_class_Incertae_sedis(100);o__Mortierellales(100);f__Mortierellaceae(100);g__Mortierella(100);s__Mortierella_capitata(100); | Mortierella | 13 | Saprotroph-Symbiotroph | Endophyte-Plant Saprotroph-\|Undefined Saprotroph\| | Microfungus | NULL | Probable | As chitinolytic (Põlme et al. 2020); As soil_saprotroph (Põlme et al. 2020); Decay roots, soil (Põlme et al. 2020); Saprobic in soil and thus frequently isolated from soil and material contaminated with soil such as hay or rotten silage and spoil tips (Cannon & Kirk 2007) | Cannon PF, Kirk PM. 2007. Fungal Families of the World. CAB International, Cambridge (ISBN: 978-0851998275); Tedersoo L, et al. 2014. Science 346:e1256688 ((https://doi.org/10.1126/science.1256688)); Purahong W, et al. 2016. Molecular Ecology 25:4059-4074 ((https://doi.org/10.1111/mec.13739)); Põlme S, et al. 2020. Fungal Diversity 105:1-16 ((https://doi.org/10.1007/s13225-020-00466-2)) |
| Otu01673 | 30 | k__Fungi(100);p__Ascomycota(100);c__Lecanoromycetes(100);o__Lecanorales(100);f__Cladoniaceae(100);g__Cladonia(100);g__Cladonia_unclassified(97); | Cladonia | 13 | Symbiotroph | \|Lichenized\| | Thallus | NULL | Highly Probable | NULL | James TY, et al. 2006. Nature 443:818-822 ((https://doi.org/10.1038/nature05110)); Põlme S, et al. 2020. Fungal Diversity 105:1-16 ((https://doi.org/10.1007/s13225-020-00466-2)) |
| Otu01674 | 30 | k__Fungi(100);p__Ascomycota(100);c__Sordariomycetes(100);o__Coniochaetales(100);f__Coniochaetaceae(100);f__Coniochaetaceae_unclassified(94);f__Coniochaetaceae_unclassified(94); | Coniochaetaceae | 9 | Pathotroph-Saprotroph-Symbiotroph | Animal Pathogen-Dung Saprotroph-Endophyte-Lichen Parasite-Plant Pathogen-Undefined Saprotroph | Microfungus | NULL | Possible | Saprobic on wood and bark (especially in dry habitats), coprophilous or isolated from soil and similar substrata (Cannon & Kirk 2007) | Cannon PF, Kirk PM. 2007. Fungal Families of the World. CAB International, Cambridge (ISBN: 978-0851998275); Damm U, et al. 2010. Persoonia 24:60-80 ((http://doi.org/10.3767/003158510X500705)); Irinyi L, et al. 2015. Medical Mycology 53:313-337 ((https://doi.org/10.1093/mmy/myv008)); Lawrey JD, Diederich P. 2016. http://www.lichenicolous.net (accessed 24 May 2017) ((http://www.lichenicolous.net)) |
| Otu01676 | 30 | k__Fungi(100);p__Basidiomycota(100);c__Agaricomycetes(100);o__Cantharellales(97);f__Clavulinaceae(97);g__Clavulina(97);g__Clavulina_unclassified(97); | Clavulina | 13 | Symbiotroph | \|Ectomycorrhizal\| | Clavarioid | NULL | Highly Probable | NULL | Tedersoo L, et al. 2014. Science 346:e1256688 ((https://doi.org/10.1126/science.1256688)); Põlme S, et al. 2020. Fungal Diversity 105:1-16 ((https://doi.org/10.1007/s13225-020-00466-2)) |
| Otu01679 | 30 | k__Fungi(100);p__Ascomycota(100);c__Saccharomycetes(100);o__Saccharomycetales(100);f__Metschnikowiaceae(100);g__Metschnikowia(100);s__Metschnikowia_viticola(100); | Metschnikowia | 13 | Saprotroph | \|Undefined Saprotroph\| | Yeast | NULL | Probable | As nectar-tap_saprotroph (Põlme et al. 2020); Decay sugar-rich substrates (Põlme et al. 2020) | Põlme S, et al. 2020. Fungal Diversity 105:1-16 ((https://doi.org/10.1007/s13225-020-00466-2)) |
| Otu01682 | 30 | k__Fungi(100);p__Ascomycota(100);c__Saccharomycetes(100);o__Saccharomycetales(100);f__Metschnikowiaceae(100);g__Metschnikowia(100);g__Metschnikowia_unclassified(84); | Metschnikowia | 13 | Saprotroph | \|Undefined Saprotroph\| | Yeast | NULL | Probable | As nectar-tap_saprotroph (Põlme et al. 2020); Decay sugar-rich substrates (Põlme et al. 2020) | Põlme S, et al. 2020. Fungal Diversity 105:1-16 ((https://doi.org/10.1007/s13225-020-00466-2)) |
| Otu01684 | 30 | k__Fungi(100);p__Ascomycota(100);c__Sordariomycetes(100);o__Hypocreales(100);f__Ophiocordycipitaceae(100);g__Chaunopycnis(100);s__Chaunopycnis_pustulata(100); | Chaunopycnis | 13 | Pathotroph | Fungal Parasite | NULL | NULL | Probable | NULL | Tedersoo L, et al. 2014. Science 346:e1256688 ((https://doi.org/10.1126/science.1256688)) |
| Otu01685 | 30 | k__Fungi(100);p__Ascomycota(100);c__Leotiomycetes(100);o__Helotiales(100);f__Helotiales_family_Incertae_sedis(100);g__Scytalidium(100);s__Scytalidium_cuboideum(100); | Scytalidium | 13 | Pathotroph-Saprotroph | Animal Parasite-Fungal Parasite-Plant Pathogen-Plant Saprotroph-\|Wood Saprotroph\| | Microfungus | Soft Rot | Probable | As litter_saprotroph (Põlme et al. 2020); As mycoparasite (Põlme et al. 2020); Decay wood, leaf, fruit, seed, soil, animal material (Põlme et al. 2020) | Nilsson T. 1973. Studia Forestalia Suecica 104:1-40 ((https://pub.epsilon.slu.se/5711/)); Costa IPMW, et al. 2012. Mycotaxon 119:e26 ((http://mycotaxon.com/resources/checklists/costa_v119_checklist.pdf)); Tedersoo L, et al. 2014. Science 346:e1256688 ((https://doi.org/10.1126/science.1256688)); Põlme S, et al. 2020. Fungal Diversity 105:1-16 ((https://doi.org/10.1007/s13225-020-00466-2)); The American Phytopathological Society Common Names of Plant Diseases ((http://www.apsnet.org/publications/commonnames/Pages/default.aspx)) |
| Otu01686 | 30 | k__Fungi(100);p__Zygomycota(100);c__Zygomycota_class_Incertae_sedis(100);o__Mucorales(100);f__Cunninghamellaceae(100);g__Cunninghamella(100);s__Cunninghamella_elegans(100); | Cunninghamella | 13 | Saprotroph | \|Undefined Saprotroph\| | Microfungus | NULL | Probable | As soil_saprotroph (Põlme et al. 2020); Decay soil (Põlme et al. 2020) | Tedersoo L, et al. 2014. Science 346:e1256688 ((https://doi.org/10.1126/science.1256688)); Põlme S, et al. 2020. Fungal Diversity 105:1-16 ((https://doi.org/10.1007/s13225-020-00466-2)) |
| Otu01688 | 30 | k__Fungi(100);p__Ascomycota(100);c__Sordariomycetes(100);o__Hypocreales(100);f__Cordycipitaceae(100);g__Simplicillium(100);g__Simplicillium_unclassified(77); | Simplicillium | 13 | Pathotroph-Saprotroph | \|Animal Parasite\|-Animal Pathogen-Fungal Parasite-Undefined Saprotroph | Microfungus | NULL | Probable | As animal_decomposer (Põlme et al. 2020); As mycoparasite (Põlme et al. 2020); Decay animal material (Põlme et al. 2020) | Tedersoo L, et al. 2014. Science 346:e1256688 ((https://doi.org/10.1126/science.1256688)); Põlme S, et al. 2020. Fungal Diversity 105:1-16 ((https://doi.org/10.1007/s13225-020-00466-2)) |
| Otu01689 | 30 | k__Fungi(100);p__Chytridiomycota(100);c__Chytridiomycetes(100);o__Spizellomycetales(100);f__Spizellomycetaceae(100);g__Powellomyces(100);s__Powellomyces_sp_DAOM_226231(100); | Powellomyces | 13 | Pathotroph-Saprotroph | Algal Parasite-Plant Pathogen-\|Pollen Saprotroph\| | Chytrid-Microfungus | NULL | Probable | Decay pollen (Põlme et al. 2020) | Tedersoo L, et al. 2014. Science 346:e1256688 ((https://doi.org/10.1126/science.1256688)); Põlme S, et al. 2020. Fungal Diversity 105:1-16 ((https://doi.org/10.1007/s13225-020-00466-2)) |
| Otu01691 | 30 | k__Fungi(100);p__Basidiomycota(100);c__Tremellomycetes(100);o__Cystofilobasidiales(100);f__Cystofilobasidiaceae(100);g__Udeniomyces(100);s__Udeniomyces_megalosporus(100); | Udeniomyces | 13 | Saprotroph-Symbiotroph | Epiphyte-Plant Saprotroph-\|Undefined Saprotroph\| | Yeast | NULL | Probable | Isolated from the pear phylloplane (Stadelmann 1975); Isolated from dead leaves of Oryza sativa, Miscanthus sinensis and Sasa sp. in Japan (Nakase & Suzuki 1986; Bullera spp.) | Stadelmann F. 1975. Antonie van Leeuwenhoek 41:575-582 ((https://doi.org/10.1007/BF02565099)); Nakase T, Suzuki M. 1986. Journal of General and Applied Microbiology Tokyo 32:225-240 ((https://doi.org/10.2323/jgam.32.225)); Põlme S, et al. 2020. Fungal Diversity 105:1-16 ((https://doi.org/10.1007/s13225-020-00466-2)) |
| Otu01692 | 30 | k__Fungi(100);p__Ascomycota(100);c__Sordariomycetes(100);o__Xylariales(100);f__Xylariaceae(100);g__Hypoxylon(100);s__Hypoxylon_fragiforme(100); | Hypoxylon | 13 | Saprotroph-Symbiotroph | Endophyte-Undefined Saprotroph-\|Wood Saprotroph\| | Corticioid-Microfungus-Xylarioid | NULL | Probable | As foliar_endophyte (Põlme et al. 2020); Decay wood, leaf, fruit, seed (Põlme et al. 2020); Host - angiosperms (Põlme et al. 2020) | Tedersoo L, et al. 2014. Science 346:e1256688 ((https://doi.org/10.1126/science.1256688)); Põlme S, et al. 2020. Fungal Diversity 105:1-16 ((https://doi.org/10.1007/s13225-020-00466-2)) |
| Otu01693 | 30 | k__Fungi(100);p__Ascomycota(100);c__Saccharomycetes(100);o__Saccharomycetales(100);f__Metschnikowiaceae(100);g__Metschnikowia(100);s__Metschnikowia_pulcherrima(100); | Metschnikowia | 13 | Saprotroph | \|Undefined Saprotroph\| | Yeast | NULL | Probable | As nectar-tap_saprotroph (Põlme et al. 2020); Decay sugar-rich substrates (Põlme et al. 2020) | Põlme S, et al. 2020. Fungal Diversity 105:1-16 ((https://doi.org/10.1007/s13225-020-00466-2)) |
| Otu01695 | 30 | k__Fungi(100);p__Ascomycota(100);c__Dothideomycetes(100);o__Pleosporales(100);f__Phaeosphaeriaceae(67);f__Phaeosphaeriaceae_unclassified(67);f__Phaeosphaeriaceae_unclassified(67); | Phaeosphaeriaceae | 9 | Pathotroph-Saprotroph | Fungal Parasite-Plant Pathogen-Plant Saprotroph | Microfungus | NULL | Probable | Necrotrophic or saprobic on a wide range of plants, especially monocotyledons; a few species parasitic on other fungi (Cannon & Kirk 2007) | Cannon PF, Kirk PM. 2007. Fungal Families of the World. CAB International, Cambridge (ISBN: 978-0851998275) |
| Otu01696 | 30 | k__Fungi(100);p__Ascomycota(100);c__Sordariomycetes(100);o__Ophiostomatales(100);f__Ophiostomataceae(100);g__Sporothrix(100);s__Sporothrix_sp_FMR_9334(100); | Sporothrix | 13 | Pathotroph-Saprotroph-Symbiotroph | Animal Pathogen-Endophyte-\|Plant Pathogen\|-Plant Saprotroph-Undefined Saprotroph-Wood Saprotroph | Microfungus | NULL | Probable | As foliar_endophyte (Põlme et al. 2020); As wood_pathogen (Põlme et al. 2020); culture (Dik et al. 1998); The fungal genus Sporothrix comprises a group of thermodimorphic pathogens that cause skin infections in humans and other mammals. Rose gardenerâ€™s disease (Sporotrichosis) occurs worldwide, with hyperendemic areas in Brazil, China, and South Africa (Rodrigues et al. 2016); The fungus lives throughout the world in soil and on plant matter such as sphagnum moss, rose bushes, and hay (www.cdc.gov); Decay wood (Põlme et al. 2020); Endophyte detection method | Dik AJ, et al. 1998. European Journal of Plant Pathology 104:413-423 ((https://doi.org/10.1023/A:1008025416672)); Rodrigues AM, et al. 2016. PLoS Pathogens 12:e1005638 ((https://doi.org/10.1371/journal.ppat.1005638)); Põlme S, et al. 2020. Fungal Diversity 105:1-16 ((https://doi.org/10.1007/s13225-020-00466-2)); https://www.cdc.gov ((https://www.cdc.gov)) |
| Otu01671 | 30 | k__Fungi(100);p__Ascomycota(74);p__Ascomycota_unclassified(74);p__Ascomycota_unclassified(74);p__Ascomycota_unclassified(74);p__Ascomycota_unclassified(74);p__Ascomycota_unclassified(74); | - | - | - | - | - | - | - | Unassigned | - |
| Otu01675 | 30 | k__Fungi(100);p__Ascomycota(97);p__Ascomycota_unclassified(90);p__Ascomycota_unclassified(90);p__Ascomycota_unclassified(90);p__Ascomycota_unclassified(90);p__Ascomycota_unclassified(90); | - | - | - | - | - | - | - | Unassigned | - |
| Otu01677 | 30 | k__Fungi(100);k__Fungi_unclassified(67);k__Fungi_unclassified(67);k__Fungi_unclassified(67);k__Fungi_unclassified(67);k__Fungi_unclassified(67);k__Fungi_unclassified(67); | - | - | - | - | - | - | - | Unassigned | - |
| Otu01678 | 30 | k__Fungi(100);k__Fungi_unclassified(80);k__Fungi_unclassified(80);k__Fungi_unclassified(80);k__Fungi_unclassified(80);k__Fungi_unclassified(80);k__Fungi_unclassified(80); | - | - | - | - | - | - | - | Unassigned | - |
| Otu01680 | 30 | k__Fungi(100);p__Basidiomycota(100);c__Agaricomycetes(100);c__Agaricomycetes_unclassified(100);c__Agaricomycetes_unclassified(100);c__Agaricomycetes_unclassified(100);c__Agaricomycetes_unclassified(100); | - | - | - | - | - | - | - | Unassigned | - |
| Otu01681 | 30 | k__Fungi(100);p__Ascomycota(100);p__Ascomycota_unclassified(87);p__Ascomycota_unclassified(87);p__Ascomycota_unclassified(87);p__Ascomycota_unclassified(87);p__Ascomycota_unclassified(87); | - | - | - | - | - | - | - | Unassigned | - |
| Otu01683 | 30 | k__Fungi(100);k__Fungi_unclassified(97);k__Fungi_unclassified(97);k__Fungi_unclassified(97);k__Fungi_unclassified(97);k__Fungi_unclassified(97);k__Fungi_unclassified(97); | - | - | - | - | - | - | - | Unassigned | - |
| Otu01687 | 30 | k__Fungi(100);k__Fungi_unclassified(100);k__Fungi_unclassified(100);k__Fungi_unclassified(100);k__Fungi_unclassified(100);k__Fungi_unclassified(100);k__Fungi_unclassified(100); | - | - | - | - | - | - | - | Unassigned | - |
| Otu01690 | 30 | k__Fungi(100);p__Ascomycota(90);p__Ascomycota_unclassified(90);p__Ascomycota_unclassified(90);p__Ascomycota_unclassified(90);p__Ascomycota_unclassified(90);p__Ascomycota_unclassified(90); | - | - | - | - | - | - | - | Unassigned | - |
| Otu01694 | 30 | k__Fungi(100);p__Basidiomycota(100);c__Agaricomycetes(100);c__Agaricomycetes_unclassified(100);c__Agaricomycetes_unclassified(100);c__Agaricomycetes_unclassified(100);c__Agaricomycetes_unclassified(100); | - | - | - | - | - | - | - | Unassigned | - |
| Otu01697 | 30 | k__Fungi(100);p__Ascomycota(74);p__Ascomycota_unclassified(74);p__Ascomycota_unclassified(74);p__Ascomycota_unclassified(74);p__Ascomycota_unclassified(74);p__Ascomycota_unclassified(74); | - | - | - | - | - | - | - | Unassigned | - |
| Otu01700 | 29 | k__Fungi(100);p__Ascomycota(100);c__Saccharomycetes(100);o__Saccharomycetales(100);f__Metschnikowiaceae(100);g__Metschnikowia(100);s__Metschnikowia_pulcherrima(100); | Metschnikowia | 13 | Saprotroph | \|Undefined Saprotroph\| | Yeast | NULL | Probable | As nectar-tap_saprotroph (Põlme et al. 2020); Decay sugar-rich substrates (Põlme et al. 2020) | Põlme S, et al. 2020. Fungal Diversity 105:1-16 ((https://doi.org/10.1007/s13225-020-00466-2)) |
| Otu01702 | 29 | k__Fungi(100);p__Basidiomycota(100);c__Agaricomycetes(100);o__Cantharellales(100);f__Ceratobasidiaceae(100);f__Ceratobasidiaceae_unclassified(52);f__Ceratobasidiaceae_unclassified(52); | Ceratobasidiaceae | 9 | Pathotroph-Saprotroph-Symbiotroph | Endomycorrhizal-Plant Pathogen-Undefined Saprotroph | Microfungus | NULL | Possible | Best known as plant pathogens, but also saprobic. Species are endomycorrhizal with a wide range of plant roots - especially of Orchidaceae but also even bryophytes (Cannon & Kirk 2007) | Cannon PF, Kirk PM. 2007. Fungal Families of the World. CAB International, Cambridge (ISBN: 978-0851998275) |
| Otu01704 | 29 | k__Fungi(100);p__Ascomycota(100);c__Saccharomycetes(100);o__Saccharomycetales(100);f__Metschnikowiaceae(100);g__Metschnikowia(100);s__Metschnikowia_pulcherrima(90); | Metschnikowia | 13 | Saprotroph | \|Undefined Saprotroph\| | Yeast | NULL | Probable | As nectar-tap_saprotroph (Põlme et al. 2020); Decay sugar-rich substrates (Põlme et al. 2020) | Põlme S, et al. 2020. Fungal Diversity 105:1-16 ((https://doi.org/10.1007/s13225-020-00466-2)) |
| Otu01705 | 29 | k__Fungi(100);p__Ascomycota(100);c__Saccharomycetes(100);o__Saccharomycetales(100);f__Metschnikowiaceae(100);g__Metschnikowia(100);s__Metschnikowia_pulcherrima(87); | Metschnikowia | 13 | Saprotroph | \|Undefined Saprotroph\| | Yeast | NULL | Probable | As nectar-tap_saprotroph (Põlme et al. 2020); Decay sugar-rich substrates (Põlme et al. 2020) | Põlme S, et al. 2020. Fungal Diversity 105:1-16 ((https://doi.org/10.1007/s13225-020-00466-2)) |
| Otu01706 | 29 | k__Fungi(100);p__Ascomycota(100);c__Saccharomycetes(100);o__Saccharomycetales(100);f__Metschnikowiaceae(100);g__Metschnikowia(100);g__Metschnikowia_unclassified(90); | Metschnikowia | 13 | Saprotroph | \|Undefined Saprotroph\| | Yeast | NULL | Probable | As nectar-tap_saprotroph (Põlme et al. 2020); Decay sugar-rich substrates (Põlme et al. 2020) | Põlme S, et al. 2020. Fungal Diversity 105:1-16 ((https://doi.org/10.1007/s13225-020-00466-2)) |
| Otu01710 | 29 | k__Fungi(100);p__Basidiomycota(100);c__Agaricomycetes(100);o__Polyporales(100);f__Ganodermataceae(100);g__Ganoderma(100);s__Ganoderma_meredithae(100); | Ganoderma | 13 | Pathotroph-Saprotroph | \|Plant Pathogen\|-Wood Saprotroph | Polyporoid | White Rot | Highly Probable | As white_rot (Põlme et al. 2020); As wood_pathogen (Põlme et al. 2020); Decay wood (Põlme et al. 2020); Lignicolous, causing sometimes agressive white rots of timber (Cannon & Kirk 2007) | Gilbertson RL, Ryvarden L. 1987-1987. North American Polypores. Fungiflora, Oslo (ISBN: 978-0945345060); Cannon PF, Kirk PM. 2007. Fungal Families of the World. CAB International, Cambridge (ISBN: 978-0851998275); Põlme S, et al. 2020. Fungal Diversity 105:1-16 ((https://doi.org/10.1007/s13225-020-00466-2)) |
| Otu01713 | 29 | k__Fungi(100);p__Basidiomycota(100);c__Agaricomycetes(100);o__Agaricales(100);o__Agaricales_unclassified(100);o__Agaricales_unclassified(100);o__Agaricales_unclassified(100); | Agaricales | 7 | Pathotroph-Saprotroph-Symbiotroph | Bryophyte Parasite-Dung Saprotroph-Ectomycorrhizal-Fungal Parasite-Leaf Saprotroph-Plant Parasite-Undefined Saprotroph-Wood Saprotroph | Agaricoid-Gasteroid-Microfungus-Secotioid-Yeast | NULL | Possible | Mushrooms and toadstools, Gill fungi, Agarics... lignicolous, sometimes muscicolous or fungicolous, saprobic, mycorrhizal, rarely parasitic on plants or fungi (Kirk et al. 2008) | Kirk PM et al. 2008. Dictionary of the Fungi. Tenth Edition. CAB International, Wallingford (ISBN: 978-0851998268) |
| Otu01714 | 29 | k__Fungi(100);p__Basidiomycota(100);c__Agaricomycetes(100);o__Agaricales(100);f__Psathyrellaceae(100);g__Psathyrella(100);g__Psathyrella_unclassified(100); | Psathyrella | 13 | Saprotroph | Plant Saprotroph-\|Wood Saprotroph\| | Agaricoid | NULL | Probable | As litter_saprotroph (Põlme et al. 2020); Decay wood, leaf, fruit, seed (Põlme et al. 2020) | Tedersoo L, et al. 2014. Science 346:e1256688 ((https://doi.org/10.1126/science.1256688)); Põlme S, et al. 2020. Fungal Diversity 105:1-16 ((https://doi.org/10.1007/s13225-020-00466-2)) |
| Otu01715 | 29 | k__Fungi(100);p__Basidiomycota(100);c__Agaricomycetes(100);o__Polyporales(100);f__Phanerochaetaceae(100);g__Phanerochaete(100);s__Phanerochaete_sp(56); | Phanerochaete | 13 | Saprotroph | \|Wood Saprotroph\| | Corticioid | White Rot | Highly Probable | As white_rot (Põlme et al. 2020); Decay wood (Põlme et al. 2020) | Glenn JK, et al. 1983. Biochemical and Biophysical Research Communications 114:1077-1083 ((https://doi.org/10.1016/0006-291x((83)) 90672-1)); Põlme S, et al. 2020. Fungal Diversity 105:1-16 ((https://doi.org/10.1007/s13225-020-00466-2)) |
| Otu01716 | 29 | k__Fungi(100);p__Basidiomycota(100);c__Agaricomycetes(100);o__Cantharellales(100);f__Hydnaceae(100);g__Sistotrema(100);g__Sistotrema_unclassified(100); | Sistotrema | 13 | Saprotroph-Symbiotroph | Ectomycorrhizal-\|Plant Saprotroph\|-Wood Saprotroph | Corticioid | White Rot | Possible | As litter_saprotroph (Põlme et al. 2020); Decay leaf, fruit, seed (Põlme et al. 2020) | Gilbertson RL, Ryvarden L. 1987-1987. North American Polypores. Fungiflora, Oslo (ISBN: 978-0945345060); Rinaldi AC, et al. 2008. Fungal Diversity 33:1-45 ((http://www.fungaldiversity.org/fdp/sfdp/33-1.pdf)) (as ECM/not ECM); Tedersoo L, et al. 2010. Mycorrhiza 20: 217-263 ((https://doi.org/10.1007/s00572-009-0274-x)) (pro parte); Põlme S, et al. 2020. Fungal Diversity 105:1-16 ((https://doi.org/10.1007/s13225-020-00466-2)); http://mycorrhizas.info/ecmf.html ((http://mycorrhizas.info/ecmf.html)) (as ECM/not ECM). |
| Otu01717 | 29 | k__Fungi(100);p__Ascomycota(100);c__Saccharomycetes(100);o__Saccharomycetales(100);f__Metschnikowiaceae(100);g__Metschnikowia(100);s__Metschnikowia_pulcherrima(97); | Metschnikowia | 13 | Saprotroph | \|Undefined Saprotroph\| | Yeast | NULL | Probable | As nectar-tap_saprotroph (Põlme et al. 2020); Decay sugar-rich substrates (Põlme et al. 2020) | Põlme S, et al. 2020. Fungal Diversity 105:1-16 ((https://doi.org/10.1007/s13225-020-00466-2)) |
| Otu01718 | 29 | k__Fungi(100);p__Ascomycota(97);c__Leotiomycetes(97);o__Helotiales(97);f__Rutstroemiaceae(97);g__Rutstroemia(73);g__Rutstroemia_unclassified(73); | Rutstroemia | 13 | Saprotroph | \|Plant Saprotroph\| | Microfungus | NULL | Probable | As litter_saprotroph (Põlme et al. 2020); Decay leaf, fruit, seed (Põlme et al. 2020); Essentially saprobic on various plant parts, especially seeds and fruits, though initial infection may take place in living tissue (Cannon & Kirk 2007) | Cannon PF, Kirk PM. 2007. Fungal Families of the World. CAB International, Cambridge (ISBN: 978-0851998275); Tedersoo L, et al. 2014. Science 346:e1256688 ((https://doi.org/10.1126/science.1256688)); Põlme S, et al. 2020. Fungal Diversity 105:1-16 ((https://doi.org/10.1007/s13225-020-00466-2)) |

| | | | | | | | | | | | |
|---|---|---|---|---|---|---|---|---|---|---|---|
| Otu01719 | 29 | k__Fungi(100);p__Basidiomycota(100);c__Microbotryomycetes(100);o__Sporidiobolales(100);f__Sporidiobolales_family_Incertae_sedis(100);g__Rhodotorula(100);s__Rhodotorula_unclassified(94); | Rhodotorula | 13 | Pathotroph-Saprotroph-Symbiotroph | Animal Pathogen-Animal Symbiotroph-Endophyte-Plant Pathogen-(Undefined Saprotroph) | Yeast | NULL | Probable | As foliar_endophyte (Põlme et al. 2020); Opportunistic human pathogen | Põlme S, et al. 2020. Fungal Diversity 105:1-16 (https://doi.org/10.1007/s13225-020-00466-2); https://www.cdc.gov (https://www.cdc.gov/) |
| Otu01720 | 29 | k__Fungi(100);p__Basidiomycota(100);c__Agaricomycetes(100);o__Cantharellales(100);f__Ceratobasidiaceae(100);g__Ceratobasidium(100);s__Ceratobasidium_unclassified(100); | Ceratobasidium | 13 | Pathotroph-Saprotroph-Symbiotroph | Ectomycorrhizal-Endomycorrhizal-Orchid Mycorrhizal-(Plant Pathogen)-Plant Saprotroph-Undefined Saprotroph | Corticioid-Microfungus | NULL | Possible | especially of Orchidaceae but also even bryophytes (Cannon & Kirk 2007); As litter_saprotroph (Põlme et al. 2020); Best known as plant pathogens, but also saprobic. Species are endomycorrhizal with a wide range of plant roots ; Decay leaf, fruit, seed, roots, soil (Põlme et al. 2020) | Cannon PF, Kirk PM. 2007. Fungal Families of the World. CAB International, Cambridge (ISBN: 978-0851998275); Tedersoo L, et al. 2014. Science 346:e1256688 (https://doi.org/10.1126/science.1256688); Põlme S, et al. 2020. Fungal Diversity 105:1-16 (https://doi.org/10.1007/s13225-020-00466-2) |
| Otu01722 | 29 | k__Fungi(100);p__Ascomycota(100);c__Saccharomycetes(100);o__Saccharomycetales(100);f__Metschnikowiaceae(100);g__Metschnikowia(100);s__Metschnikowia_pulcherrima(97); | Metschnikowia | 13 | Saprotroph | (Undefined Saprotroph) | Yeast | NULL | Probable | As nectar-tap_saprotroph (Põlme et al. 2020); Decay sugar-rich substrates (Põlme et al. 2020) | Põlme S, et al. 2020. Fungal Diversity 105:1-16 (https://doi.org/10.1007/s13225-020-00466-2) |
| Otu01724 | 29 | k__Fungi(100);p__Basidiomycota(100);c__Agaricomycetes(100);o__Agaricales(100);f__Psathyrellaceae(100);g__Psathyrella(100);s__Psathyrella_squamosa(100); | Psathyrella | 13 | Saprotroph | Plant Saprotroph-(Wood Saprotroph) | Agaricoid | NULL | Probable | As litter_saprotroph (Põlme et al. 2020); Decay wood, leaf, fruit, seed (Põlme et al. 2020) | Tedersoo L, et al. 2014. Science 346:e1256688 (https://doi.org/10.1126/science.1256688); Põlme S, et al. 2020. Fungal Diversity 105:1-16 (https://doi.org/10.1007/s13225-020-00466-2) |
| Otu01725 | 29 | k__Fungi(100);p__Basidiomycota(100);c__Agaricomycetes(100);o__Cantharellales(100);f__Clavulinaceae(73);f__Clavulinaceae_unclassified(63); | Clavulinaceae | 9 | Saprotroph-Symbiotroph | Ectomycorrhizal-Lichenized-Wood Saprotroph | Clavarioid | NULL | Probable | Some species form ectomycorrhizas, with fruit bodies in leaf litter and on soil. Others are probably saprotes associated with rotten wood (Cannon & Kirk 2007) | Cannon PF, Kirk PM. 2007. Fungal Families of the World. CAB International, Cambridge (https://isbnsearch.org/isbn/9780851998275); Nelsen MP, et al. 2007. American Journal of Botany 94:1289-1296 (https://doi.org/10.3732/ajb.94.8.1289) |
| Otu01732 | 29 | k__Fungi(100);p__Ascomycota(100);c__Sordariomycetes(100);o__Xylariales(100);f__Xylariaceae(100);g__Hypoxylon(100);s__Hypoxylon_submonticulosum(100); | Hypoxylon | 13 | Saprotroph-Symbiotroph | Endophyte-Undefined Saprotroph-(Wood Saprotroph) | Corticioid-Microfungus-Xylarioid | NULL | Probable | As foliar_endophyte (Põlme et al. 2020); Decay wood, leaf, fruit, seed (Põlme et al. 2020); Host - angiosperms (Põlme et al. 2020) | Tedersoo L, et al. 2014. Science 346:e1256688 (https://doi.org/10.1126/science.1256688); Põlme S, et al. 2020. Fungal Diversity 105:1-16 (https://doi.org/10.1007/s13225-020-00466-2) |
| Otu01735 | 29 | k__Fungi(100);p__Basidiomycota(100);c__Agaricomycetes(100);o__Polyporales(100);f__Phanerochaetaceae(100);g__Hyphodermella(100);s__Hyphodermella_unclassified(100); | Hyphodermella | 13 | Saprotroph | Undefined Saprotroph-(Wood Saprotroph) | Corticioid | White Rot | Probable | As white_rot (Põlme et al. 2020); Decay wood (Põlme et al. 2020) | Tedersoo L, et al. 2014. Science 346:e1256688 (https://doi.org/10.1126/science.1256688); Põlme S, et al. 2020. Fungal Diversity 105:1-16 (https://doi.org/10.1007/s13225-020-00466-2) |
| Otu01736 | 29 | k__Fungi(100);p__Ascomycota(100);c__Saccharomycetes(100);o__Saccharomycetales(100);f__Metschnikowiaceae(100);g__Metschnikowia(100);s__Metschnikowia_pulcherrima(97); | Metschnikowia | 13 | Saprotroph | (Undefined Saprotroph) | Yeast | NULL | Probable | As nectar-tap_saprotroph (Põlme et al. 2020); Decay sugar-rich substrates (Põlme et al. 2020) | Põlme S, et al. 2020. Fungal Diversity 105:1-16 (https://doi.org/10.1007/s13225-020-00466-2) |
| Otu01737 | 29 | k__Fungi(100);p__Basidiomycota(100);c__Microbotryomycetes(69);o__Sporidiobolales(69);f__Sporidiobolales_family_Incertae_sedis(69);g__Rhodotorula(69);s__Rhodotorula_sp(69); | Rhodotorula | 13 | Pathotroph-Saprotroph-Symbiotroph | Animal Pathogen-Animal Symbiotroph-Endophyte-Plant Pathogen-(Undefined Saprotroph) | Yeast | NULL | Probable | As foliar_endophyte (Põlme et al. 2020); Opportunistic human pathogen | Põlme S, et al. 2020. Fungal Diversity 105:1-16 (https://doi.org/10.1007/s13225-020-00466-2); https://www.cdc.gov (https://www.cdc.gov/) |
| Otu01740 | 29 | k__Fungi(100);p__Ascomycota(100);c__Sordariomycetes(100);o__Hypocreales(100);f__Nectriaceae(100);g__Nectria(90);s__Nectria_sp_strim170(90); | Nectria | 13 | Pathotroph-Saprotroph-Symbiotroph | Animal Pathogen-Endophyte-Fungal Parasite-Lichen Parasite-(Plant Pathogen)-Plant Saprotroph-Wood Saprotroph | Microfungus | NULL | Possible | As foliar_endophyte (Põlme et al. 2020); As litter_saprotroph (Põlme et al. 2020); Decay wood, leaf, fruit, seed, roots, soil (Põlme et al. 2020) | Hirooka Y, et al. 2012. Studies in Mycology 71:1-210 (https://doi.org/10.3114/sim0001); Tedersoo L, et al. 2014. Science 346:e1256688 (https://doi.org/10.1126/science.1256688); Põlme S, et al. 2020. Fungal Diversity 105:1-16 (https://doi.org/10.1007/s13225-020-00466-2) |
| Otu01698 | 29 | k__Fungi(100);k__Fungi_unclassified(66);k__Fungi_unclassified(66);k__Fungi_unclassified(66);k__Fungi_unclassified(66);k__Fungi_unclassified(66);k__Fungi_unclassified(66); | - | - | - | - | - | - | - | Unassigned | - |
| Otu01699 | 29 | k__Fungi(100);p__Basidiomycota(100);c__Pucciniomycetes(100);o__Pucciniales(100);o__Pucciniales_unclassified(100);o__Pucciniales_unclassified(100);o__Pucciniales_unclassified(100); | - | - | - | - | - | - | - | Unassigned | - |
| Otu01701 | 29 | k__Fungi(100);k__Fungi_unclassified(83);k__Fungi_unclassified(83);k__Fungi_unclassified(83);k__Fungi_unclassified(83);k__Fungi_unclassified(83);k__Fungi_unclassified(83); | - | - | - | - | - | - | - | Unassigned | - |
| Otu01703 | 29 | k__Fungi(100);p__Ascomycota(52);p__Ascomycota_unclassified(52);p__Ascomycota_unclassified(52);p__Ascomycota_unclassified(52);p__Ascomycota_unclassified(52);p__Ascomycota_unclassified(52); | - | - | - | - | - | - | - | Unassigned | - |
| Otu01707 | 29 | k__Fungi(100);p__Ascomycota(59);p__Ascomycota_unclassified(59);p__Ascomycota_unclassified(59);p__Ascomycota_unclassified(59);p__Ascomycota_unclassified(59);p__Ascomycota_unclassified(59); | - | - | - | - | - | - | - | Unassigned | - |
| Otu01708 | 29 | k__Fungi(100);p__Basidiomycota(100);c__Agaricomycetes(100);o__Trechisporales(100);f__Hydnodontaceae(100);f__Hydnodontaceae_unclassified(66);f__Hydnodontaceae_unclassified(66); | - | - | - | - | - | - | - | Unassigned | - |
| Otu01709 | 29 | k__Fungi(100);p__Ascomycota(69);p__Ascomycota_unclassified(69);p__Ascomycota_unclassified(69);p__Ascomycota_unclassified(69);p__Ascomycota_unclassified(69);p__Ascomycota_unclassified(69); | - | - | - | - | - | - | - | Unassigned | - |
| Otu01711 | 29 | k__Fungi(100);k__Fungi_unclassified(73);k__Fungi_unclassified(73);k__Fungi_unclassified(73);k__Fungi_unclassified(73);k__Fungi_unclassified(73);k__Fungi_unclassified(73); | - | - | - | - | - | - | - | Unassigned | - |
| Otu01712 | 29 | k__Fungi(100);p__Ascomycota(52);p__Ascomycota_unclassified(52);p__Ascomycota_unclassified(52);p__Ascomycota_unclassified(52);p__Ascomycota_unclassified(52);p__Ascomycota_unclassified(52); | - | - | - | - | - | - | - | Unassigned | - |
| Otu01721 | 29 | k__Fungi(100);k__Fungi_unclassified(56);k__Fungi_unclassified(56);k__Fungi_unclassified(56);k__Fungi_unclassified(56);k__Fungi_unclassified(56);k__Fungi_unclassified(56); | - | - | - | - | - | - | - | Unassigned | - |
| Otu01723 | 29 | k__Fungi(100);p__Basidiomycota(100);p__Basidiomycota_unclassified(69);p__Basidiomycota_unclassified(69);p__Basidiomycota_unclassified(69);p__Basidiomycota_unclassified(69);p__Basidiomycota_unclassified(69); | - | - | - | - | - | - | - | Unassigned | - |
| Otu01726 | 29 | k__Fungi(100);p__Ascomycota(100);c__Dothideomycetes(100);o__Pleosporales(90);o__Pleosporales_unclassified(90);o__Pleosporales_unclassified(90);o__Pleosporales_unclassified(90); | - | - | - | - | - | - | - | Unassigned | - |
| Otu01727 | 29 | k__Fungi(100);k__Fungi_unclassified(59);k__Fungi_unclassified(59);k__Fungi_unclassified(59);k__Fungi_unclassified(59);k__Fungi_unclassified(59);k__Fungi_unclassified(59); | - | - | - | - | - | - | - | Unassigned | - |
| Otu01728 | 29 | k__Fungi(100);p__Ascomycota(100);p__Ascomycota_unclassified(100);p__Ascomycota_unclassified(100);p__Ascomycota_unclassified(100);p__Ascomycota_unclassified(100);p__Ascomycota_unclassified(100); | - | - | - | - | - | - | - | Unassigned | - |
| Otu01729 | 29 | k__Fungi(100);p__Ascomycota(100);c__Eurotiomycetes(100);c__Eurotiomycetes_unclassified(66);c__Eurotiomycetes_unclassified(66);c__Eurotiomycetes_unclassified(66);c__Eurotiomycetes_unclassified(66); | - | - | - | - | - | - | - | Unassigned | - |
| Otu01730 | 29 | k__Fungi(100);k__Fungi_unclassified(94);k__Fungi_unclassified(94);k__Fungi_unclassified(94);k__Fungi_unclassified(94);k__Fungi_unclassified(94);k__Fungi_unclassified(94); | - | - | - | - | - | - | - | Unassigned | - |
| Otu01731 | 29 | k__Fungi(100);k__Fungi_unclassified(69);k__Fungi_unclassified(69);k__Fungi_unclassified(69);k__Fungi_unclassified(69);k__Fungi_unclassified(69);k__Fungi_unclassified(69); | - | - | - | - | - | - | - | Unassigned | - |
| Otu01733 | 29 | k__Fungi(100);p__Basidiomycota(100);c__Agaricomycetes(100);c__Agaricomycetes_unclassified(100);c__Agaricomycetes_unclassified(100);c__Agaricomycetes_unclassified(100);c__Agaricomycetes_unclassified(100); | - | - | - | - | - | - | - | Unassigned | - |
| Otu01734 | 29 | k__Fungi(100);k__Fungi_unclassified(87);k__Fungi_unclassified(87);k__Fungi_unclassified(87);k__Fungi_unclassified(87);k__Fungi_unclassified(87);k__Fungi_unclassified(87); | - | - | - | - | - | - | - | Unassigned | - |
| Otu01738 | 29 | k__Fungi(100);k__Fungi_unclassified(100);k__Fungi_unclassified(100);k__Fungi_unclassified(100);k__Fungi_unclassified(100);k__Fungi_unclassified(100);k__Fungi_unclassified(100); | - | - | - | - | - | - | - | Unassigned | - |
| Otu01739 | 29 | k__Fungi(100);p__Ascomycota(100);c__Sordariomycetes(100);c__Sordariomycetes_unclassified(56);c__Sordariomycetes_unclassified(56);c__Sordariomycetes_unclassified(56);c__Sordariomycetes_unclassified(56); | - | - | - | - | - | - | - | Unassigned | - |
| Otu01741 | 28 | k__Fungi(100);p__Ascomycota(100);c__Leotiomycetes(100);o__Helotiales(100);f__Hyaloscyphaceae(100);g__Arachnopeziza(100);s__Arachnopeziza_sp(75); | Arachnopeziza | 13 | Saprotroph | Undefined Saprotroph-(Wood Saprotroph) | Microfungus-Pezizoid | NULL | Probable | Decay wood (Põlme et al. 2020) | Tedersoo L, et al. 2014. Science 346:e1256688 (https://doi.org/10.1126/science.1256688); Põlme S, et al. 2020. Fungal Diversity 105:1-16 (https://doi.org/10.1007/s13225-020-00466-2) |
| Otu01743 | 28 | k__Fungi(100);p__Ascomycota(100);c__Saccharomycetes(100);o__Saccharomycetales(100);f__Metschnikowiaceae(100);g__Metschnikowia(100);s__Metschnikowia_pulcherrima(100); | Metschnikowia | 13 | Saprotroph | (Undefined Saprotroph) | Yeast | NULL | Probable | As nectar-tap_saprotroph (Põlme et al. 2020); Decay sugar-rich substrates (Põlme et al. 2020) | Põlme S, et al. 2020. Fungal Diversity 105:1-16 (https://doi.org/10.1007/s13225-020-00466-2) |
| Otu01745 | 28 | k__Fungi(100);p__Ascomycota(100);c__Saccharomycetes(100);o__Saccharomycetales(100);f__Metschnikowiaceae(100);g__Metschnikowia(100);s__Metschnikowia_pulcherrima(90); | Metschnikowia | 13 | Saprotroph | (Undefined Saprotroph) | Yeast | NULL | Probable | As nectar-tap_saprotroph (Põlme et al. 2020); Decay sugar-rich substrates (Põlme et al. 2020) | Põlme S, et al. 2020. Fungal Diversity 105:1-16 (https://doi.org/10.1007/s13225-020-00466-2) |
| Otu01747 | 28 | k__Fungi(100);p__Ascomycota(100);c__Sordariomycetes(100);o__Diaporthales(100);f__Gnomoniaceae(100);g__Cryptodiaporthe(100);s__Cryptodiaporthe_galericulata(100); | Cryptodiaporthe | 13 | Pathotroph | (Plant Pathogen) | Microfungus | NULL | Probable | Cause of canker diseases (Redlin & Rossman 1991) | Redlin SC, Rossman AY. 1991. Mycologia 83:200-209 (https://doi.org/10.2307/3759935); Tedersoo L, et al. 2014. Science 346:e1256688 (https://doi.org/10.1126/science.1256688); Põlme S, et al. 2020. Fungal Diversity 105:1-16 (https://doi.org/10.1007/s13225-020-00466-2) |
| Otu01750 | 28 | k__Fungi(100);p__Ascomycota(100);c__Sordariomycetes(100);o__Xylariales(100);f__Xylariaceae(100);g__Kretzschmaria(100);s__Kretzschmaria_sp(86); | Kretzschmaria | 13 | Pathotroph-Saprotroph | Plant Pathogen-Undefined Saprotroph-(Wood Saprotroph) | Microfungus | NULL | Probable | As wood_pathogen (Põlme et al. 2020); Decay wood (Põlme et al. 2020) | Tedersoo L, et al. 2014. Science 346:e1256688 (https://doi.org/10.1126/science.1256688); Põlme S, et al. 2020. Fungal Diversity 105:1-16 (https://doi.org/10.1007/s13225-020-00466-2) |
| Otu01751 | 28 | k__Fungi(100);p__Ascomycota(100);c__Sordariomycetes(100);o__Xylariales(100);f__Amphisphaeriaceae(100);g__Seimatosporium(58);s__Seimatosporium_unclassified(58); | Seimatosporium | 13 | Pathotroph | (Plant Pathogen) | Microfungus | NULL | Probable | Some species causing leaf spot disease (www.apsnet.org) | Tedersoo L, et al. 2014. Science 346:e1256688 (https://doi.org/10.1126/science.1256688); Põlme S, et al. 2020. Fungal Diversity 105:1-16 (https://doi.org/10.1007/s13225-020-00466-2); The American Phytopathological Society Common Names of Plant Diseases (http://www.apsnet.org/publications/commonnames/Pages/default.aspx) |
| Otu01754 | 28 | k__Fungi(100);p__Ascomycota(100);c__Saccharomycetes(100);o__Saccharomycetales(100);f__Metschnikowiaceae(100);g__Metschnikowia(100);s__Metschnikowia_pulcherrima(90); | Metschnikowia | 13 | Saprotroph | (Undefined Saprotroph) | Yeast | NULL | Probable | As nectar-tap_saprotroph (Põlme et al. 2020); Decay sugar-rich substrates (Põlme et al. 2020) | Põlme S, et al. 2020. Fungal Diversity 105:1-16 (https://doi.org/10.1007/s13225-020-00466-2) |
| Otu01756 | 28 | k__Fungi(100);p__Ascomycota(100);c__Saccharomycetes(100);o__Saccharomycetales(100);f__Metschnikowiaceae(100);g__Metschnikowia(100);s__Metschnikowia_pulcherrima(97); | Metschnikowia | 13 | Saprotroph | (Undefined Saprotroph) | Yeast | NULL | Probable | As nectar-tap_saprotroph (Põlme et al. 2020); Decay sugar-rich substrates (Põlme et al. 2020) | Põlme S, et al. 2020. Fungal Diversity 105:1-16 (https://doi.org/10.1007/s13225-020-00466-2) |
| Otu01757 | 28 | k__Fungi(100);p__Ascomycota(100);c__Saccharomycetes(100);o__Saccharomycetales(100);f__Metschnikowiaceae(100);g__Metschnikowia(100);s__Metschnikowia_pulcherrima(93); | Metschnikowia | 13 | Saprotroph | (Undefined Saprotroph) | Yeast | NULL | Probable | As nectar-tap_saprotroph (Põlme et al. 2020); Decay sugar-rich substrates (Põlme et al. 2020) | Põlme S, et al. 2020. Fungal Diversity 105:1-16 (https://doi.org/10.1007/s13225-020-00466-2) |
| Otu01760 | 28 | k__Fungi(100);p__Ascomycota(100);c__Sordariomycetes(100);o__Xylariales(100);f__Diatrypaceae(100);g__Eutypella(100);s__Eutypella_cerviculata(100); | Eutypella | 13 | Pathotroph | (Plant Pathogen) | Microfungus | NULL | Probable | NULL | Tedersoo L, et al. 2014. Science 346:e1256688 (https://doi.org/10.1126/science.1256688); Põlme S, et al. 2020. Fungal Diversity 105:1-16 (https://doi.org/10.1007/s13225-020-00466-2) |
| Otu01762 | 28 | k__Fungi(100);p__Ascomycota(100);c__Dothideomycetes(100);o__Pleosporales(100);f__Venturiaceae(100);f__Venturiaceae_unclassified(97);f__Venturiaceae_unclassified(97); | Venturiaceae | 9 | Pathotroph-Saprotroph-Symbiotroph | Endophyte-Plant Pathogen-Undefined Saprotroph | Microfungus | NULL | Probable | Mostly endophytes and weak parasites of aerial plant parts, including bark and wood, leaf and fruit tissue; rarely biotrophic. Little is known except for the important pathogens (Cannon & Kirk 2007) | Cannon PF, Kirk PM. 2007. Fungal Families of the World. CAB International, Cambridge (ISBN: 978-0851998275) |
| Otu01763 | 28 | k__Fungi(100);p__Ascomycota(100);c__Sordariomycetes(97);o__Hypocreales(97);o__Hypocreales_unclassified(97);o__Hypocreales_unclassified(97);o__Hypocreales_unclassified(97); | Hypocreales | 7 | Saprotroph | Undefined Saprotroph | Microfungus | NULL | Possible | NULL | Sterkenburg E, et al. 2015. New Phytologist 207:1145-1158 (https://doi.org/10.1111/nph.13426) |
| Otu01765 | 28 | k__Fungi(100);p__Ascomycota(100);c__Sordariomycetes(100);o__Diaporthales(100);f__Gnomoniaceae(100);g__Cryptodiaporthe(100);s__Cryptodiaporthe_aceina(100); | Cryptodiaporthe | 13 | Pathotroph | (Plant Pathogen) | Microfungus | NULL | Probable | Cause of canker diseases (Redlin & Rossman 1991) | Redlin SC, Rossman AY. 1991. Mycologia 83:200-209 (https://doi.org/10.2307/3759935); Tedersoo L, et al. 2014. Science 346:e1256688 (https://doi.org/10.1126/science.1256688); Põlme S, et al. 2020. Fungal Diversity 105:1-16 (https://doi.org/10.1007/s13225-020-00466-2) |
| Otu01768 | 28 | k__Fungi(100);p__Basidiomycota(100);c__Agaricomycetes(100);o__Agaricales(100);f__Strophariaceae(100);g__Gymnopilus(100);s__Gymnopilus_penetrans(100); | Gymnopilus | 13 | Saprotroph | Plant Saprotroph-(Wood Saprotroph) | Agaricoid | White Rot | Probable | As litter_saprotroph (Põlme et al. 2020); As white_rot (Põlme et al. 2020); Decay wood (Põlme et al. 2020) | Tedersoo L, et al. 2014. Science 346:e1256688 (https://doi.org/10.1126/science.1256688); Põlme S, et al. 2020. Fungal Diversity 105:1-16 (https://doi.org/10.1007/s13225-020-00466-2) |
| Otu01770 | 28 | k__Fungi(100);p__Ascomycota(100);c__Saccharomycetes(100);o__Saccharomycetales(100);f__Metschnikowiaceae(100);g__Metschnikowia(100);s__Metschnikowia_pulcherrima(93); | Metschnikowia | 13 | Saprotroph | (Undefined Saprotroph) | Yeast | NULL | Probable | As nectar-tap_saprotroph (Põlme et al. 2020); Decay sugar-rich substrates (Põlme et al. 2020) | Põlme S, et al. 2020. Fungal Diversity 105:1-16 (https://doi.org/10.1007/s13225-020-00466-2) |
| Otu01771 | 28 | k__Fungi(100);p__Ascomycota(100);c__Dothideomycetes(100);o__Capnodiales(100);f__Mycosphaerellaceae(90);f__Mycosphaerellaceae_unclassified(90);f__Mycosphaerellaceae_unclassified(90); | Mycosphaerellaceae | 9 | Pathotroph-Saprotroph | Plant Pathogen-Undefined Saprotroph | Microfungus | NULL | Probable | Biotrophic, necrotrophic or saprobic on various plant tissues (Cannon & Kirk 2007) | Cannon PF, Kirk PM. 2007. Fungal Families of the World. CAB International, Cambridge (ISBN: 978-0851998275) |
| Otu01772 | 28 | k__Fungi(100);p__Ascomycota(100);c__Saccharomycetes(100);o__Saccharomycetales(100);f__Metschnikowiaceae(100);g__Metschnikowia(100);s__Metschnikowia_pulcherrima(93); | Metschnikowia | 13 | Saprotroph | (Undefined Saprotroph) | Yeast | NULL | Probable | As nectar-tap_saprotroph (Põlme et al. 2020); Decay sugar-rich substrates (Põlme et al. 2020) | Põlme S, et al. 2020. Fungal Diversity 105:1-16 (https://doi.org/10.1007/s13225-020-00466-2) |
| Otu01777 | 28 | k__Fungi(100);p__Basidiomycota(100);c__Agaricomycetes(100);o__Agaricales(100);f__Mycenaceae(100);g__Mycena(100);s__Mycena_zephirus(83); | Mycena | 13 | Pathotroph-Saprotroph | Plant Pathogen-(Plant Saprotroph)-Undefined Saprotroph-Wood Saprotroph | Agaricoid | NULL | Probable | As litter_saprotroph (Põlme et al. 2020); Decay leaf, fruit, seed (Põlme et al. 2020) | Grgurinovic CA, Simpson JA. 2001. Fungal Diversity 8:97-127 (http://www.fungaldiversity.org/fdp/sfdp/FD_8_97-127.pdf); Tedersoo L, et al. 2014. Science 346:e1256688 (https://doi.org/10.1126/science.1256688); Põlme S, et al. 2020. Fungal Diversity 105:1-16 (https://doi.org/10.1007/s13225-020-00466-2); The American Phytopathological Society Common Names of Plant Diseases (http://www.apsnet.org/publications/commonnames/Pages/default.aspx) |
| Otu01780 | 28 | k__Fungi(100);p__Basidiomycota(100);c__Agaricomycetes(100);o__Polyporales(100);f__Fomitopsidaceae(100);g__Postia(100);s__Postia_fragilis(97); | Postia | 13 | Saprotroph | (Wood Saprotroph) | Polyporoid | Brown Rot | Highly Probable | As brown-rot (Põlme et al. 2020); Decay wood (Põlme et al. 2020) | Jordan CR, et al. 1996. Holzforschung - International Journal of the Biology, Chemistry, Physics and Technology of Wood 50:312-318 (https://doi.org/10.1515/hfsg.1996.50.4.312)); Põlme S, et al. 2020. Fungal Diversity 105:1-16 (https://doi.org/10.1007/s13225-020-00466-2) |
| Otu01781 | 28 | k__Fungi(100);p__Basidiomycota(100);c__Agaricomycetes(100);o__Boletales(100);f__Tapinellaceae(100);g__Pseudomerulius(100);s__Pseudomerulius_aureus(100); | Pseudomerulius | 13 | Saprotroph | (Wood Saprotroph) | Corticioid | Brown Rot | Probable | As brown-rot (Põlme et al. 2020); Decay wood (Põlme et al. 2020) | Tedersoo L, et al. 2014. Science 346:e1256688 (https://doi.org/10.1126/science.1256688); Põlme S, et al. 2020. Fungal Diversity 105:1-16 (https://doi.org/10.1007/s13225-020-00466-2) |
| Otu01742 | 28 | k__Fungi(100);p__Ascomycota(58);p__Ascomycota_unclassified(58);p__Ascomycota_unclassified(58);p__Ascomycota_unclassified(58);p__Ascomycota_unclassified(58);p__Ascomycota_unclassified(58); | - | - | - | - | - | - | - | Unassigned | - |
| Otu01744 | 28 | k__Fungi(100);p__Ascomycota(54);p__Ascomycota_unclassified(54);p__Ascomycota_unclassified(54);p__Ascomycota_unclassified(54);p__Ascomycota_unclassified(54);p__Ascomycota_unclassified(54); | - | - | - | - | - | - | - | Unassigned | - |
| Otu01746 | 28 | k__Fungi(100);p__Basidiomycota(100);c__Agaricomycetes(58);c__Agaricomycetes_unclassified(58);c__Agaricomycetes_unclassified(58);c__Agaricomycetes_unclassified(58);c__Agaricomycetes_unclassified(58); | - | - | - | - | - | - | - | Unassigned | - |
| Otu01748 | 28 | k__Fungi(100);p__Ascomycota(100);p__Ascomycota_unclassified(86);p__Ascomycota_unclassified(86);p__Ascomycota_unclassified(86);p__Ascomycota_unclassified(86);p__Ascomycota_unclassified(86); | - | - | - | - | - | - | - | Unassigned | - |
| Otu01749 | 28 | k__Fungi(100);p__Ascomycota(100);c__Eurotiomycetes(100);c__Eurotiomycetes_unclassified(100);c__Eurotiomycetes_unclassified(100);c__Eurotiomycetes_unclassified(100);c__Eurotiomycetes_unclassified(100); | - | - | - | - | - | - | - | Unassigned | - |
| Otu01752 | 28 | k__Fungi(100);p__Basidiomycota(100);c__Pucciniomycetes(97);o__Pucciniales(86);o__Pucciniales_unclassified(86);o__Pucciniales_unclassified(86);o__Pucciniales_unclassified(86); | - | - | - | - | - | - | - | Unassigned | - |
| Otu01753 | 28 | k__Fungi(100);p__Basidiomycota(100);c__Agaricomycetes(100);o__Polyporales(100);o__Polyporales_unclassified(100);o__Polyporales_unclassified(100);o__Polyporales_unclassified(100); | - | - | - | - | - | - | - | Unassigned | - |
| Otu01755 | 28 | k__Fungi(100);k__Fungi_unclassified(58);k__Fungi_unclassified(58);k__Fungi_unclassified(58);k__Fungi_unclassified(58);k__Fungi_unclassified(58);k__Fungi_unclassified(58); | - | - | - | - | - | - | - | Unassigned | - |
| Otu01758 | 28 | k__Fungi(100);p__Ascomycota(100);p__Ascomycota_unclassified(75);p__Ascomycota_unclassified(75);p__Ascomycota_unclassified(75);p__Ascomycota_unclassified(75);p__Ascomycota_unclassified(75); | - | - | - | - | - | - | - | Unassigned | - |
| Otu01759 | 28 | k__Fungi(100);p__Ascomycota(100);c__Sordariomycetes(100);c__Sordariomycetes_unclassified(100);c__Sordariomycetes_unclassified(100);c__Sordariomycetes_unclassified(100);c__Sordariomycetes_unclassified(100); | - | - | - | - | - | - | - | Unassigned | - |
| Otu01761 | 28 | k__Fungi(100);k__Fungi_unclassified(61);k__Fungi_unclassified(61);k__Fungi_unclassified(61);k__Fungi_unclassified(61);k__Fungi_unclassified(61);k__Fungi_unclassified(61); | - | - | - | - | - | - | - | Unassigned | - |
| Otu01764 | 28 | k__Fungi(100);k__Fungi_unclassified(72);k__Fungi_unclassified(72);k__Fungi_unclassified(72);k__Fungi_unclassified(72);k__Fungi_unclassified(72);k__Fungi_unclassified(72); | - | - | - | - | - | - | - | Unassigned | - |
| Otu01766 | 28 | k__Fungi(100);p__Ascomycota(100);c__Sordariomycetes(100);c__Sordariomycetes_unclassified(100);c__Sordariomycetes_unclassified(100);c__Sordariomycetes_unclassified(100);c__Sordariomycetes_unclassified(100); | - | - | - | - | - | - | - | Unassigned | - |
| Otu01767 | 28 | k__Fungi(100);k__Fungi_unclassified(75);k__Fungi_unclassified(75);k__Fungi_unclassified(75);k__Fungi_unclassified(75);k__Fungi_unclassified(75);k__Fungi_unclassified(75); | - | - | - | - | - | - | - | Unassigned | - |
| Otu01769 | 28 | k__Fungi(100);p__Ascomycota(100);p__Ascomycota_unclassified(100);p__Ascomycota_unclassified(100);p__Ascomycota_unclassified(100);p__Ascomycota_unclassified(100);p__Ascomycota_unclassified(100); | - | - | - | - | - | - | - | Unassigned | - |
| Otu01773 | 28 | k__Fungi(100);p__Ascomycota(65);p__Ascomycota_unclassified(65);p__Ascomycota_unclassified(65);p__Ascomycota_unclassified(65);p__Ascomycota_unclassified(65);p__Ascomycota_unclassified(65); | - | - | - | - | - | - | - | Unassigned | - |
| Otu01774 | 28 | k__Fungi(100);k__Fungi_unclassified(97);k__Fungi_unclassified(97);k__Fungi_unclassified(97);k__Fungi_unclassified(97);k__Fungi_unclassified(97);k__Fungi_unclassified(97); | - | - | - | - | - | - | - | Unassigned | - |
| Otu01775 | 28 | k__Fungi(100);p__Ascomycota(58);p__Ascomycota_unclassified(58);p__Ascomycota_unclassified(58);p__Ascomycota_unclassified(58);p__Ascomycota_unclassified(58);p__Ascomycota_unclassified(58); | - | - | - | - | - | - | - | Unassigned | - |
| Otu01776 | 28 | k__Fungi(100);p__Ascomycota(100);c__Dothideomycetes(100);o__Capnodiales(100);o__Capnodiales_unclassified(100);o__Capnodiales_unclassified(100);o__Capnodiales_unclassified(100); | - | - | - | - | - | - | - | Unassigned | - |
| Otu01778 | 28 | k__Fungi(100);k__Fungi_unclassified(100);k__Fungi_unclassified(100);k__Fungi_unclassified(100);k__Fungi_unclassified(100);k__Fungi_unclassified(100);k__Fungi_unclassified(100); | - | - | - | - | - | - | - | Unassigned | - |
| Otu01779 | 28 | k__Fungi(100);k__Fungi_unclassified(58);k__Fungi_unclassified(58);k__Fungi_unclassified(58);k__Fungi_unclassified(58);k__Fungi_unclassified(58);k__Fungi_unclassified(58); | - | - | - | - | - | - | - | Unassigned | - |
| Otu01782 | 27 | k__Fungi(100);p__Ascomycota(100);c__Saccharomycetes(100);o__Saccharomycetales(100);f__Metschnikowiaceae(100);g__Metschnikowia(100);s__Metschnikowia_unclassified(89); | Metschnikowia | 13 | Saprotroph | (Undefined Saprotroph) | Yeast | NULL | Probable | As nectar-tap_saprotroph (Põlme et al. 2020); Decay sugar-rich substrates (Põlme et al. 2020) | Põlme S, et al. 2020. Fungal Diversity 105:1-16 (https://doi.org/10.1007/s13225-020-00466-2) |
| Otu01784 | 27 | k__Fungi(100);p__Ascomycota(100);c__Dothideomycetes(100);o__Pleosporales(100);f__Pleosporales_family_Incertae_sedis(100);g__Ascochyta(100);s__Ascochyta_sp(100); | Ascochyta | 13 | Pathotroph | Lichen Parasite-(Plant Pathogen) | Microfungus | NULL | Probable | NULL | Tedersoo L, et al. 2014. Science 346:e1256688 (https://doi.org/10.1126/science.1256688); Lawrey JD, Diederich P. 2016. http://www.lichenicolous.net (accessed 24 May 2017) (http://www.lichenicolous.net/); Põlme S, et al. 2020. Fungal Diversity 105:1-16 (https://doi.org/10.1007/s13225-020-00466-2); The American Phytopathological Society Common Names of Plant Diseases (https://www.apsnet.org/edcenter/resources/commonnames/Pages/default.aspx) |
| Otu01785 | 27 | k__Fungi(100);p__Ascomycota(100);c__Dothideomycetes(100);o__Capnodiales(100);f__Mycosphaerellaceae(100);f__Mycosphaerellaceae_unclassified(100);f__Mycosphaerellaceae_unclassified(100); | Mycosphaerellaceae | 9 | Pathotroph-Saprotroph | Plant Pathogen-Undefined Saprotroph | Microfungus | NULL | Probable | Biotrophic, necrotrophic or saprobic on various plant tissues (Cannon & Kirk 2007) | Cannon PF, Kirk PM. 2007. Fungal Families of the World. CAB International, Cambridge (ISBN: 978-0851998275) |
| Otu01786 | 27 | k__Fungi(100);p__Ascomycota(100);c__Orbiliomycetes(100);o__Orbiliales(100);f__Orbiliaceae(100);g__Dactylella(100);s__Dactylella_oviparasitica(100); | Dactylella | 13 | Saprotroph | Plant Saprotroph-Undefined Saprotroph-(Wood Saprotroph) | Microfungus | NULL | Probable | As litter_saprotroph (Põlme et al. 2020); Decay wood, leaf, fruit, seed, soil, animal material (Põlme et al. 2020) | Tedersoo L, et al. 2014. Science 346:e1256688 (https://doi.org/10.1126/science.1256688); Põlme S, et al. 2020. Fungal Diversity 105:1-16 (https://doi.org/10.1007/s13225-020-00466-2) |
| Otu01787 | 27 | k__Fungi(100);p__Ascomycota(100);c__Saccharomycetes(100);o__Saccharomycetales(100);f__Metschnikowiaceae(100);g__Metschnikowia(100);s__Metschnikowia_pulcherrima(93); | Metschnikowia | 13 | Saprotroph | (Undefined Saprotroph) | Yeast | NULL | Probable | As nectar-tap_saprotroph (Põlme et al. 2020); Decay sugar-rich substrates (Põlme et al. 2020) | Põlme S, et al. 2020. Fungal Diversity 105:1-16 (https://doi.org/10.1007/s13225-020-00466-2) |
| Otu01788 | 27 | k__Fungi(100);p__Ascomycota(100);c__Saccharomycetes(100);o__Saccharomycetales(100);f__Metschnikowiaceae(100);g__Metschnikowia(100);s__Metschnikowia_pulcherrima(100); | Metschnikowia | 13 | Saprotroph | (Undefined Saprotroph) | Yeast | NULL | Probable | As nectar-tap_saprotroph (Põlme et al. 2020); Decay sugar-rich substrates (Põlme et al. 2020) | Põlme S, et al. 2020. Fungal Diversity 105:1-16 (https://doi.org/10.1007/s13225-020-00466-2) |
| Otu01791 | 27 | k__Fungi(100);p__Basidiomycota(100);c__Microbotryomycetes(100);o__Microbotryales(100);f__Microbotryaceae(100);g__Microbotryum(100);s__Microbotryum_anomalum(100); | Microbotryum | 13 | Pathotroph | (Plant Pathogen) | Smut | NULL | Probable | NULL | Tedersoo L, et al. 2014. Science 346:e1256688 (https://doi.org/10.1126/science.1256688); Põlme S, et al. 2020. Fungal Diversity 105:1-16 (https://doi.org/10.1007/s13225-020-00466-2) |
| Otu01793 | 27 | k__Fungi(100);p__Ascomycota(100);c__Saccharomycetes(100);o__Saccharomycetales(100);f__Metschnikowiaceae(100);g__Metschnikowia(100);s__Metschnikowia_pulcherrima(100); | Metschnikowia | 13 | Saprotroph | (Undefined Saprotroph) | Yeast | NULL | Probable | As nectar-tap_saprotroph (Põlme et al. 2020); Decay sugar-rich substrates (Põlme et al. 2020) | Põlme S, et al. 2020. Fungal Diversity 105:1-16 (https://doi.org/10.1007/s13225-020-00466-2) |
| Otu01798 | 27 | k__Fungi(100);p__Basidiomycota(100);c__Agaricomycetes(100);o__Hymenochaetales(100);f__Schizoporaceae(100);f__Schizoporaceae_unclassified(100);f__Schizoporaceae_unclassified(100); | Schizoporaceae | 9 | Saprotroph | Wood Saprotroph | Corticioid | White Rot | Probable | Saprobic, causing white rot of standing and fallen wood of coniferous and broadleaved trees, rarely on herbaceous stems (Cannon & Kirk 2007) | Cannon PF, Kirk PM. 2007. Fungal Families of the World. CAB International, Cambridge (ISBN: 978-0851998275) |
| Otu01800 | 27 | k__Fungi(100);p__Ascomycota(100);c__Saccharomycetes(100);o__Saccharomycetales(100);f__Metschnikowiaceae(100);g__Metschnikowia(100);s__Metschnikowia_pulcherrima(75); | Metschnikowia | 13 | Saprotroph | (Undefined Saprotroph) | Yeast | NULL | Probable | As nectar-tap_saprotroph (Põlme et al. 2020); Decay sugar-rich substrates (Põlme et al. 2020) | Põlme S, et al. 2020. Fungal Diversity 105:1-16 (https://doi.org/10.1007/s13225-020-00466-2) |
| Otu01801 | 27 | k__Fungi(100);p__Ascomycota(100);c__Saccharomycetes(100);o__Saccharomycetales(100);f__Metschnikowiaceae(100);g__Metschnikowia(100);s__Metschnikowia_pulcherrima(97); | Metschnikowia | 13 | Saprotroph | (Undefined Saprotroph) | Yeast | NULL | Probable | As nectar-tap_saprotroph (Põlme et al. 2020); Decay sugar-rich substrates (Põlme et al. 2020) | Põlme S, et al. 2020. Fungal Diversity 105:1-16 (https://doi.org/10.1007/s13225-020-00466-2) |
| Otu01802 | 27 | k__Fungi(100);p__Basidiomycota(100);c__Exobasidiomycetes(100);o__Microstromatales(100);f__Microstromataceae(100);g__Microstroma(100);s__Microstroma_juglandis(100); | Microstroma | 13 | Pathotroph | (Plant Pathogen) | Dimorphic | NULL | Probable | NULL | Tedersoo L, et al. 2014. Science 346:e1256688 (https://doi.org/10.1126/science.1256688); Põlme S, et al. 2020. Fungal Diversity 105:1-16 (https://doi.org/10.1007/s13225-020-00466-2) |
| Otu01805 | 27 | k__Fungi(100);p__Basidiomycota(100);c__Agaricomycetes(100);o__Agaricales(100);f__Inocybaceae(97);g__Crepidotus(86);s__Crepidotus_unclassified(86); | Crepidotus | 13 | Saprotroph | (Wood Saprotroph) | Agaricoid | NULL | Highly Probable | Decay wood (Põlme et al. 2020) | Grgurinovic CA, Simpson JA. 2001. Fungal Diversity 8:97-127 (http://www.fungaldiversity.org/fdp/sfdp/FD_8_97-127.pdf); Aime MC, et al. 2005. American Journal of Botany 92:74-82 (https://doi.org/10.3732/ajb.92.1.74); Tedersoo L, et al. 2014. Science 346:e1256688 (https://doi.org/10.1126/science.1256688); Põlme S, et al. 2020. Fungal Diversity 105:1-16 (https://doi.org/10.1007/s13225-020-00466-2) |
| Otu01806 | 27 | k__Fungi(100);p__Ascomycota(100);c__Saccharomycetes(100);o__Saccharomycetales(100);f__Metschnikowiaceae(100);g__Metschnikowia(100);s__Metschnikowia_pulcherrima(100); | Metschnikowia | 13 | Saprotroph | (Undefined Saprotroph) | Yeast | NULL | Probable | As nectar-tap_saprotroph (Põlme et al. 2020); Decay sugar-rich substrates (Põlme et al. 2020) | Põlme S, et al. 2020. Fungal Diversity 105:1-16 (https://doi.org/10.1007/s13225-020-00466-2) |
| Otu01807 | 27 | k__Fungi(100);p__Ascomycota(100);c__Saccharomycetes(100);o__Saccharomycetales(100);f__Metschnikowiaceae(100);g__Metschnikowia(100);s__Metschnikowia_pulcherrima(89); | Metschnikowia | 13 | Saprotroph | (Undefined Saprotroph) | Yeast | NULL | Probable | As nectar-tap_saprotroph (Põlme et al. 2020); Decay sugar-rich substrates (Põlme et al. 2020) | Põlme S, et al. 2020. Fungal Diversity 105:1-16 (https://doi.org/10.1007/s13225-020-00466-2) |
| Otu01809 | 27 | k__Fungi(100);p__Basidiomycota(100);c__Agaricomycetes(100);o__Polyporales(100);f__Polyporaceae(100);g__Pycnoporus(100);s__Pycnoporus_cinnabarinus(100); | Pycnoporus | 13 | Saprotroph | (Wood Saprotroph) | Polyporoid | White Rot | Highly Probable | As white_rot (Põlme et al. 2020); Decay wood (Põlme et al. 2020) | Gilbertson RL, Ryvarden L. 1987-1987. North American Polypores. Fungiflora, Oslo (ISBN: 978-0945345060); Põlme S, et al. 2020. Fungal Diversity 105:1-16 (https://doi.org/10.1007/s13225-020-00466-2) |
| Otu01811 | 27 | k__Fungi(100);p__Ascomycota(100);c__Sordariomycetes(100);o__Hypocreales(100);f__Nectriaceae(82);f__Nectriaceae_unclassified(82);f__Nectriaceae_unclassified(82); | Nectriaceae | 9 | Pathotroph-Saprotroph-Symbiotroph | Animal Pathogen-Endophyte-Fungal Parasite-Lichen Parasite-Plant Pathogen-Wood Saprotroph | Microfungus | NULL | Possible | Fusarium species are amongst the economically significant plant pathogens (Cannon & Kirk 2007) | Cannon PF, Kirk PM. 2007. Fungal Families of the World. CAB International, Cambridge (ISBN: 978-0851998275); Hirooka Y, et al. 2012. Studies in Mycology 71:1-210 (https://doi.org/10.3114/sim0001); Tedersoo L, et al. 2014. Science 346:e1256688 (https://doi.org/10.1126/science.1256688); The American Phytopathological Society Common Names of Plant Diseases (http://www.apsnet.org/publications/commonnames/Pages/default.aspx); https://www.cdc.gov (https://www.cdc.gov/) |
| Otu01812 | 27 | k__Fungi(100);p__Basidiomycota(100);c__Agaricomycetes(100);o__Russulales(100);f__Russulaceae(100);g__Lactarius(100);s__Lactarius_mammosus(100); | Lactarius | 13 | Symbiotroph | (Ectomycorrhizal) | Agaricoid | NULL | Highly Probable | milk; cap; Common name | Rinaldi AC, et al. 2008. Fungal Diversity 33:1-45 (http://www.fungaldiversity.org/fdp/sfdp/33-1.pdf)); Tedersoo L, et al. 2010. Mycorrhiza 20:217-263 (https://doi.org/10.1007/s00572-009-0274-x)); Põlme S, et al. 2020. Fungal Diversity 105:1-16 (https://doi.org/10.1007/s13225-020-00466-2)); http://mycorrhizas.info/ecmf.html (http://mycorrhizas.info/ecmf.html) |
| Otu01813 | 27 | k__Fungi(100);p__Basidiomycota(100);c__Agaricomycetes(100);o__Agaricales(100);f__Pluteaceae(100);g__Pluteus(100);s__Pluteus_unclassified(100); | Pluteus | 13 | Pathotroph-Saprotroph | Bryophyte Parasite-(Plant Saprotroph)-Wood Saprotroph | Agaricoid | White Rot | Probable | As litter_saprotroph (Põlme et al. 2020); As white_rot (Põlme et al. 2020); Decay leaf, fruit, seed (Põlme et al. 2020) | Grgurinovic CA, Simpson JA. 2001. Fungal Diversity 8:97-127 (http://www.fungaldiversity.org/fdp/sfdp/FD_8_97-127.pdf); James TY, et al. 2006. Nature 443:818-822 (https://doi.org/10.1038/nature05110); Tedersoo L, et al. 2014. Science 346:e1256688 (https://doi.org/10.1126/science.1256688); Põlme S, et al. 2020. Fungal Diversity 105:1-16 (https://doi.org/10.1007/s13225-020-00466-2) |
| Otu01815 | 27 | k__Fungi(100);p__Basidiomycota(100);c__Agaricomycetes(100);o__Boletales(100);f__Boletinellaceae(100);g__Boletinellus(100);s__Boletinellus_sp(100); | Boletinellus | 13 | Saprotroph-Symbiotroph | Animal Associated Biotroph-Root Associated Biotroph | Boletoid | Brown Rot | Probable | Some species used to be known as Gyrodon. | Tedersoo L, et al. 2014. Science 346:e1256688 (https://doi.org/10.1126/science.1256688) |
| Otu01816 | 27 | k__Fungi(100);p__Basidiomycota(100);c__Agaricomycetes(100);o__Cantharellales(100);f__Ceratobasidiaceae(100);g__Ceratobasidium(100);s__Ceratobasidium_unclassified(56); | Ceratobasidium | 13 | Pathotroph-Saprotroph-Symbiotroph | Ectomycorrhizal-Endomycorrhizal-Orchid Mycorrhizal-(Plant Pathogen)-Plant Saprotroph-Undefined Saprotroph | Corticioid-Microfungus | NULL | Possible | especially of Orchidaceae but also even bryophytes (Cannon & Kirk 2007); As litter_saprotroph (Põlme et al. 2020); Best known as plant pathogens, but also saprobic. Species are endomycorrhizal with a wide range of plant roots ; Decay leaf, fruit, seed, roots, soil (Põlme et al. 2020) | Cannon PF, Kirk PM. 2007. Fungal Families of the World. CAB International, Cambridge (ISBN: 978-0851998275); Tedersoo L, et al. 2014. Science 346:e1256688 (https://doi.org/10.1126/science.1256688); Põlme S, et al. 2020. Fungal Diversity 105:1-16 (https://doi.org/10.1007/s13225-020-00466-2) |
| Otu01817 | 27 | k__Fungi(100);p__Ascomycota(100);c__Sordariomycetes(100);o__Sordariales(100);f__Chaetomiaceae(100);g__Chaetomium(97);s__Chaetomium_unclassified(93); | Chaetomium | 13 | Pathotroph-Saprotroph-Symbiotroph | Animal Pathogen-Dung Saprotroph-Endophyte-Epiphyte-Plant Pathogen-(Plant Saprotroph)-Wood Saprotroph | Microfungus | Soft Rot | Possible | As foliar_endophyte (Põlme et al. 2020); As litter_saprotroph (Põlme et al. 2020); culture (Busby et al. 2016, David et al. 2016); Likely opportunistic human pathogen (Irinyi et al. 2015); culture (Massimo et al. 2015); Endophyte detection method; Decay wood, leaf, fruit, seed, soil, dung (Põlme et al. 2020); derived commodities (Cannon & Kirk 2007); Epiphyte detection method; Saprobic, acting as key decay organisms of plant materials and plant | Seehann G, et al. 1975. List of Fungi in Soft-Rot Tissue Tests. Institute für Holzbiologie und Holzschutz, Bundesforschungsanstalt für Forst - und Holzwirtschaft, Hamburg - Lohbrugge, Germany; Cannon PF, Kirk PM. 2007. Fungal Families of the World. CAB International, Cambridge (https://isbnsearch.org/isbn/9780851998275); Bills GF, et al. 2013. Current Opinion in Microbiology 16:549-565 (https://doi.org/10.1016/j.mib.2013.08.001); Tedersoo L, et al. 2014. Science 346:e1256688 (https://doi.org/10.1126/science.1256688); Irinyi L, et al. 2015. Medical Mycology 53:313-337 (https://doi.org/10.1093/mmy/myv008); Busby PE, et al. 2016. New Phytologist 209:1681-1692 (https://doi.org/10.1111/nph.13742); Põlme S, et al. 2020. Fungal Diversity 105:1-16 (https://doi.org/10.1007/s13225-020-00466-2) |
| Otu01818 | 27 | k__Fungi(100);p__Ascomycota(100);c__Saccharomycetes(100);o__Saccharomycetales(100);f__Metschnikowiaceae(100);g__Metschnikowia(100);s__Metschnikowia_pulcherrima(97); | Metschnikowia | 13 | Saprotroph | (Undefined Saprotroph) | Yeast | NULL | Probable | As nectar-tap_saprotroph (Põlme et al. 2020); Decay sugar-rich substrates (Põlme et al. 2020) | Põlme S, et al. 2020. Fungal Diversity 105:1-16 (https://doi.org/10.1007/s13225-020-00466-2) |
| Otu01819 | 27 | k__Fungi(100);p__Basidiomycota(100);c__Agaricomycetes(100);o__Agaricales(100);f__Agaricaceae(100);g__Agaricus(100);s__Agaricus_sp_TL2321(100); | Agaricus | 13 | Saprotroph | (Undefined Saprotroph) | Agaricoid-Gasteroid-Secotioid | NULL | Probable | As soil_saprotroph (Põlme et al. 2020); Decay soil (Põlme et al. 2020) | Tedersoo L, et al. 2014. Science 346:e1256688 (https://doi.org/10.1126/science.1256688); Põlme S, et al. 2020. Fungal Diversity 105:1-16 (https://doi.org/10.1007/s13225-020-00466-2) |
| Otu01821 | 27 | k__Fungi(100);p__Ascomycota(100);c__Sordariomycetes(100);o__Sordariomycetes_order_Incertae_sedis(100);f__Sordariomycetes_family_Incertae_sedis(100);g__Myrmecridium(100);s__Myrmecridium_sp(86); | Myrmecridium | 13 | Saprotroph | (Undefined Saprotroph) | Microfungus | NULL | Probable | As soil_saprotroph (Põlme et al. 2020); Decay soil (Põlme et al. 2020) | Tedersoo L, et al. 2014. Science 346:e1256688 (https://doi.org/10.1126/science.1256688); Põlme S, et al. 2020. Fungal Diversity 105:1-16 (https://doi.org/10.1007/s13225-020-00466-2) |
| Otu01822 | 27 | k__Fungi(100);p__Ascomycota(100);c__Dothideomycetes(100);o__Capnodiales(100);f__Mycosphaerellaceae(97);g__Mycosphaerella(97);s__Mycosphaerella_nyssicola(97); | Mycosphaerella | 13 | Pathotroph-Saprotroph | (Plant Pathogen)-Plant Saprotroph | Microfungus | NULL | Probable | As litter_saprotroph (Põlme et al. 2020); Decay leaf, fruit, seed (Põlme et al. 2020) | Tedersoo L, et al. 2014. Science 346:e1256688 (https://doi.org/10.1126/science.1256688); Põlme S, et al. 2020. Fungal Diversity 105:1-16 (https://doi.org/10.1007/s13225-020-00466-2) |
| Otu01827 | 27 | k__Fungi(100);p__Basidiomycota(100);c__Agaricomycetes(100);o__Polyporales(100);f__Polyporaceae(100);g__Panus(63);s__Panus_unclassified(63); | Panus | 13 | Saprotroph | Undefined Saprotroph-(Wood Saprotroph) | Agaricoid | White Rot | Probable | As white_rot (Põlme et al. 2020); Decay wood (Põlme et al. 2020) | Tedersoo L, et al. 2014. Science 346:e1256688 (https://doi.org/10.1126/science.1256688); Põlme S, et al. 2020. Fungal Diversity 105:1-16 (https://doi.org/10.1007/s13225-020-00466-2) |
| Otu01828 | 27 | k__Fungi(100);p__Ascomycota(100);c__Saccharomycetes(100);o__Saccharomycetales(100);f__Saccharomycetales_family_Incertae_sedis(100);g__Candida(100);s__Candida_sp_AS23486(89); | Candida | 13 | Pathotroph-Saprotroph-Symbiotroph | Animal Pathogen-Endophyte-Endosymbiont-Epiphyte-(Undefined Saprotroph) | Dimorphic Yeast-Yeast | NULL | Probable | As nectar-tap_saprotroph (Põlme et al. 2020); Decay sugar-rich substrates (Põlme et al. 2020). This asexual genus is polyphyletic and contain species that function very differently depend on what they are. Caution should be taken when interpreting guilds for Candida species. | Manolakaki D, et al. 2010. Virulence. 1:367-375 (https://doi.org/10.4161/viru.1.5.12796)); Mekbib SB, et al. 2011. Tropical Plant Pathology 36:233-240 (https://doi.org/10.1590/S1982-56762011000400004)); Põlme S, et al. 2020. Fungal Diversity 105:1-16 (https://doi.org/10.1007/s13225-020-00466-2) |
| Otu01830 | 27 | k__Fungi(100);p__Ascomycota(100);c__Sordariomycetes(100);o__Hypocreales(100);f__Hypocreales_family_Incertae_sedis(78);g__Acremonium(78);s__Acremonium_cavaraeanum(78); | Acremonium | 13 | Pathotroph-Saprotroph-Symbiotroph | Animal Pathogen-Endophyte-Fungal Parasite-Plant Pathogen-(Undefined Saprotroph)-Wood Saprotroph | Microfungus | Soft Rot | Possible | As foliar_endophyte (Põlme et al. 2020); culture (Hanada et al. 2010, Del Olmo; Decay leaf, fruit, seed, soil, dung, animal material (Põlme et al. 2020); Endophyte detection method, Ruiz & Arnold 2014); Likely opportunistic human pathogen (Irinyi et al. 2015) | Duncan CG, Eslyn WE. 1966. Mycologia 58:642-645 (https://doi.org/10.2307/3757045)); Nilsson T. 1973. Studia Forestalia Suecica 104: 1-40 (https://pub.epsilon.slu.se/5711/)); Worrall JJ, et al. 1997. Mycologia 89:199-219 (https://doi.org/10.2307/3761073)); Hanada RE, et al. 2010. Fungal Biology 114:901-910 (https://doi.org/10.1016/j.funbio.2010.08.006)); Del Olmo-Ruiz M, Arnold AE. 2014. Mycologia 106: 8-21 (https://doi.org/10.3852/13-066)); Irinyi L, et al. 2015. Medical Mycology 53:313-337 (https://doi.org/10.1093/mmy/myv008)); Põlme S, et al. 2020. Fungal Diversity 105:1-16 (https://doi.org/10.1007/s13225-020-00466-2) |
| Otu01831 | 27 | k__Fungi(100);p__Ascomycota(100);c__Eurotiomycetes(86);o__Chaetothyriales(86);f__Herpotrichiellaceae(86);g__unclassified_Herpotrichiellaceae(60);s__Herpotrichiellaceae_sp(60); | Herpotrichiellaceae | 9 | Pathotroph-Saprotroph | Animal Pathogen-Undefined Saprotroph | Facultative Yeast-Microfungus | NULL | Probable | Mostly saprobic on plants or other fungi, isolated from soil etc., and from medical sources (Cannon & Kirk 2007) | Cannon PF, Kirk PM. 2007. Fungal Families of the World. CAB International, Cambridge (ISBN: 978-0851998275) |
| Otu01832 | 27 | k__Fungi(100);p__Basidiomycota(100);c__Agaricomycetes(100);o__Agaricales(100);f__Strophariaceae(100);g__Stropharia(100);s__Stropharia_rugosoannulata(100); | Stropharia | 13 | Saprotroph | Dung Saprotroph-Plant Parasite-(Plant Saprotroph)-Undefined Saprotroph-Wood Saprotroph | Agaricoid | NULL | Probable | As litter_saprotroph (Põlme et al. 2020); Decay leaf, fruit, seed (Põlme et al. 2020); Very variable ecology; Pholiota species are lignicolous and can act as weak parasites while most other taxa are either saprobes in grassy habitats or are found in dung. Species are not ectomycorrhizal (Cannon & Kirk 2007) | Cannon PF, Kirk PM. 2007. Fungal Families of the World. CAB International, Cambridge (ISBN: 978-0851998275); Tedersoo L, et al. 2014. Science 346:e1256688 (https://doi.org/10.1126/science.1256688); Põlme S, et al. 2020. Fungal Diversity 105:1-16 (https://doi.org/10.1007/s13225-020-00466-2) |
| Otu01783 | 27 | k__Fungi(100);p__Ascomycota(100);c__Dothideomycetes(100);o__Pleosporales(100);o__Pleosporales_unclassified(100);o__Pleosporales_unclassified(100);o__Pleosporales_unclassified(100); | - | - | - | - | - | - | - | Unassigned | - |
| Otu01789 | 27 | k__Fungi(100);p__Ascomycota(100);c__Dothideomycetes(100);o__Pleosporales(100);f__Pleosporales_family_Incertae_sedis(52);f__Pleosporales_family_Incertae_sedis_unclassified(52);f__Pleosporales_family_Incertae_sedis_unclassified(52); | - | - | - | - | - | - | - | Unassigned | - |
| Otu01790 | 27 | k__Fungi(100);k__Fungi_unclassified(86);k__Fungi_unclassified(86);k__Fungi_unclassified(86);k__Fungi_unclassified(86);k__Fungi_unclassified(86);k__Fungi_unclassified(86); | - | - | - | - | - | - | - | Unassigned | - |
| Otu01792 | 27 | k__Fungi(100);p__Ascomycota(100);c__Dothideomycetes(100);o__Pleosporales(100);f__unclassified_Pleosporales(52);g__unclassified_Pleosporales(52);s__Pleosporales_sp(52); | - | - | - | - | - | - | - | Unassigned | - |
| Otu01794 | 27 | k__Fungi(100);k__Fungi_unclassified(63);k__Fungi_unclassified(63);k__Fungi_unclassified(63);k__Fungi_unclassified(63);k__Fungi_unclassified(63);k__Fungi_unclassified(63); | - | - | - | - | - | - | - | Unassigned | - |
| Otu01795 | 27 | k__Fungi(100);k__Fungi_unclassified(78);k__Fungi_unclassified(78);k__Fungi_unclassified(78);k__Fungi_unclassified(78);k__Fungi_unclassified(78);k__Fungi_unclassified(78); | - | - | - | - | - | - | - | Unassigned | - |
| Otu01796 | 27 | k__Fungi(100);p__Ascomycota(56);p__Ascomycota_unclassified(56);p__Ascomycota_unclassified(56);p__Ascomycota_unclassified(56);p__Ascomycota_unclassified(56);p__Ascomycota_unclassified(56); | - | - | - | - | - | - | - | Unassigned | - |
| Otu01797 | 27 | k__Fungi(100);p__Ascomycota(78);p__Ascomycota_unclassified(78);p__Ascomycota_unclassified(78);p__Ascomycota_unclassified(78);p__Ascomycota_unclassified(78);p__Ascomycota_unclassified(78); | - | - | - | - | - | - | - | Unassigned | - |
| Otu01799 | 27 | k__Fungi(100);p__Ascomycota(100);p__Ascomycota_unclassified(97);p__Ascomycota_unclassified(97);p__Ascomycota_unclassified(97);p__Ascomycota_unclassified(97);p__Ascomycota_unclassified(97); | - | - | - | - | - | - | - | Unassigned | - |
| Otu01803 | 27 | k__Fungi(100);k__Fungi_unclassified(75);k__Fungi_unclassified(75);k__Fungi_unclassified(75);k__Fungi_unclassified(75);k__Fungi_unclassified(75);k__Fungi_unclassified(75); | - | - | - | - | - | - | - | Unassigned | - |
| Otu01804 | 27 | k__Fungi(100);p__Ascomycota(100);p__Ascomycota_unclassified(100);p__Ascomycota_unclassified(100);p__Ascomycota_unclassified(100);p__Ascomycota_unclassified(100);p__Ascomycota_unclassified(100); | - | - | - | - | - | - | - | Unassigned | - |
| Otu01808 | 27 | k__Fungi(100);p__Basidiomycota(100);c__Agaricomycetes(100);o__Cantharellales(100);f__unclassified_Cantharellales(52);g__unclassified_Cantharellales(52);s__Cantharellales_sp(52); | - | - | - | - | - | - | - | Unassigned | - |
| Otu01810 | 27 | k__Fungi(100);p__Ascomycota(100);c__Leotiomycetes(100);c__Leotiomycetes_unclassified(78);c__Leotiomycetes_unclassified(78);c__Leotiomycetes_unclassified(78);c__Leotiomycetes_unclassified(78); | - | - | - | - | - | - | - | Unassigned | - |
| Otu01814 | 27 | k__Fungi(100);p__Ascomycota(82);p__Ascomycota_unclassified(82);p__Ascomycota_unclassified(82);p__Ascomycota_unclassified(82);p__Ascomycota_unclassified(82);p__Ascomycota_unclassified(82); | - | - | - | - | - | - | - | Unassigned | - |
| Otu01820 | 27 | k__Fungi(100);p__Ascomycota(100);p__Ascomycota_unclassified(100);p__Ascomycota_unclassified(100);p__Ascomycota_unclassified(100);p__Ascomycota_unclassified(100);p__Ascomycota_unclassified(100); | - | - | - | - | - | - | - | Unassigned | - |
| Otu01823 | 27 | k__Fungi(100);p__Basidiomycota(100);c__Agaricomycetes(100);o__Polyporales(100);o__Polyporales_unclassified(89);o__Polyporales_unclassified(89);o__Polyporales_unclassified(89); | - | - | - | - | - | - | - | Unassigned | - |
| Otu01824 | 27 | k__Fungi(100);k__Fungi_unclassified(63);k__Fungi_unclassified(63);k__Fungi_unclassified(63);k__Fungi_unclassified(63);k__Fungi_unclassified(63);k__Fungi_unclassified(63); | - | - | - | - | - | - | - | Unassigned | - |
| Otu01825 | 27 | k__Fungi(100);p__Ascomycota(56);p__Ascomycota_unclassified(56);p__Ascomycota_unclassified(56);p__Ascomycota_unclassified(56);p__Ascomycota_unclassified(56);p__Ascomycota_unclassified(56); | - | - | - | - | - | - | - | Unassigned | - |
| Otu01826 | 27 | k__Fungi(100);p__Ascomycota(71);p__Ascomycota_unclassified(71);p__Ascomycota_unclassified(71);p__Ascomycota_unclassified(71);p__Ascomycota_unclassified(71);p__Ascomycota_unclassified(71); | - | - | - | - | - | - | - | Unassigned | - |
| Otu01829 | 27 | k__Fungi(100);p__Ascomycota(63);p__Ascomycota_unclassified(63);p__Ascomycota_unclassified(63);p__Ascomycota_unclassified(63);p__Ascomycota_unclassified(63);p__Ascomycota_unclassified(63); | - | - | - | - | - | - | - | Unassigned | - |
| Otu01833 | 26 | k__Fungi(100);p__Ascomycota(100);c__Saccharomycetes(100);o__Saccharomycetales(100);f__Metschnikowiaceae(100);g__Metschnikowia(100);s__Metschnikowia_pulcherrima(70); | Metschnikowia | 13 | Saprotroph | (Undefined Saprotroph) | Yeast | NULL | Probable | As nectar-tap_saprotroph (Põlme et al. 2020); Decay sugar-rich substrates (Põlme et al. 2020) | Põlme S, et al. 2020. Fungal Diversity 105:1-16 (https://doi.org/10.1007/s13225-020-00466-2) |
| Otu01834 | 26 | k__Fungi(100);p__Ascomycota(100);c__Saccharomycetes(100);o__Saccharomycetales(100);f__Metschnikowiaceae(100);g__Metschnikowia(100);s__Metschnikowia_pulcherrima(97); | Metschnikowia | 13 | Saprotroph | (Undefined Saprotroph) | Yeast | NULL | Probable | As nectar-tap_saprotroph (Põlme et al. 2020); Decay sugar-rich substrates (Põlme et al. 2020) | Põlme S, et al. 2020. Fungal Diversity 105:1-16 (https://doi.org/10.1007/s13225-020-00466-2) |
| Otu01836 | 26 | k__Fungi(100);p__Ascomycota(100);c__Saccharomycetes(100);o__Saccharomycetales(100);f__Metschnikowiaceae(100);g__Metschnikowia(100);s__Metschnikowia_unclassified(66); | Metschnikowia | 13 | Saprotroph | (Undefined Saprotroph) | Yeast | NULL | Probable | As nectar-tap_saprotroph (Põlme et al. 2020); Decay sugar-rich substrates (Põlme et al. 2020) | Põlme S, et al. 2020. Fungal Diversity 105:1-16 (https://doi.org/10.1007/s13225-020-00466-2) |
| Otu01839 | 26 | k__Fungi(100);p__Ascomycota(100);c__Saccharomycetes(100);o__Saccharomycetales(100);f__Metschnikowiaceae(100);g__Metschnikowia(100);s__Metschnikowia_pulcherrima(93); | Metschnikowia | 13 | Saprotroph | (Undefined Saprotroph) | Yeast | NULL | Probable | As nectar-tap_saprotroph (Põlme et al. 2020); Decay sugar-rich substrates (Põlme et al. 2020) | Põlme S, et al. 2020. Fungal Diversity 105:1-16 (https://doi.org/10.1007/s13225-020-00466-2) |
| Otu01840 | 26 | k__Fungi(100);p__Basidiomycota(100);c__Agaricomycetes(100);o__Agaricales(100);f__Psathyrellaceae(100);g__Psathyrella(100);s__Psathyrella_spadicea(97); | Psathyrella | 13 | Saprotroph | Plant Saprotroph-(Wood Saprotroph) | Agaricoid | NULL | Probable | As litter_saprotroph (Põlme et al. 2020); Decay wood, leaf, fruit, seed (Põlme et al. 2020) | Tedersoo L, et al. 2014. Science 346:e1256688 (https://doi.org/10.1126/science.1256688); Põlme S, et al. 2020. Fungal Diversity 105:1-16 (https://doi.org/10.1007/s13225-020-00466-2) |
| Otu01841 | 26 | k__Fungi(100);p__Ascomycota(100);c__Saccharomycetes(100);o__Saccharomycetales(100);f__Metschnikowiaceae(100);g__Metschnikowia(100);s__Metschnikowia_pulcherrima(100); | Metschnikowia | 13 | Saprotroph | (Undefined Saprotroph) | Yeast | NULL | Probable | As nectar-tap_saprotroph (Põlme et al. 2020); Decay sugar-rich substrates (Põlme et al. 2020) | Põlme S, et al. 2020. Fungal Diversity 105:1-16 (https://doi.org/10.1007/s13225-020-00466-2) |
| Otu01844 | 26 | k__Fungi(100);p__Ascomycota(100);c__Dothideomycetes(100);o__Pleosporales(100);f__Phaeosphaeriaceae(100);g__Sclerostagonospora(100);s__Sclerostagonospora_sp(100); | Sclerostagonospora | 13 | Pathotroph-Saprotroph | Plant Pathogen-(Plant Saprotroph)-Undefined Saprotroph | Microfungus | NULL | Probable | As litter_saprotroph (Põlme et al. 2020); Decay leaf, fruit, seed (Põlme et al. 2020) | Tedersoo L, et al. 2014. Science 346:e1256688 (https://doi.org/10.1126/science.1256688); Põlme S, et al. 2020. Fungal Diversity 105:1-16 (https://doi.org/10.1007/s13225-020-00466-2) |
| Otu01845 | 26 | k__Fungi(100);p__Ascomycota(100);c__Saccharomycetes(100);o__Saccharomycetales(100);f__Metschnikowiaceae(100);g__Metschnikowia(100);s__Metschnikowia_pulcherrima(93); | Metschnikowia | 13 | Saprotroph | (Undefined Saprotroph) | Yeast | NULL | Probable | As nectar-tap_saprotroph (Põlme et al. 2020); Decay sugar-rich substrates (Põlme et al. 2020) | Põlme S, et al. 2020. Fungal Diversity 105:1-16 (https://doi.org/10.1007/s13225-020-00466-2) |
| Otu01846 | 26 | k__Fungi(100);p__Zygomycota(100);c__Zygomycota_class_Incertae_sedis(100);o__Mortierellales(100);f__Mortierellaceae(100);g__Mortierella(100);s__Mortierella_turficola(70); | Mortierella | 13 | Saprotroph-Symbiotroph | Endophyte-Plant Saprotroph-(Undefined Saprotroph) | Microfungus | NULL | Probable | As chitinolytic (Põlme et al. 2020); As soil_saprotroph (Põlme et al. 2020); Decay roots, soil (Põlme et al. 2020); Saprobic in soil and thus frequently isolated from soil and material contaminated with soil such as hay or rotten silage and spoil tips (Cannon & Kirk 2007) | Cannon PF, Kirk PM. 2007. Fungal Families of the World. CAB International, Cambridge (ISBN: 978-0851998275); Tedersoo L, et al. 2014. Science 346:e1256688 (https://doi.org/10.1126/science.1256688); Purahong W, et al. 2016. Molecular Ecology 25:4059-4074 (https://doi.org/10.1111/mec.13739); Põlme S, et al. 2020. Fungal Diversity 105:1-16 (https://doi.org/10.1007/s13225-020-00466-2) |
| Otu01848 | 26 | k__Fungi(100);p__Ascomycota(100);c__Saccharomycetes(97);o__Saccharomycetales(97);f__Saccharomycetales_family_Incertae_sedis(97);g__Candida(81);s__Candida_unclassified(58); | Candida | 13 | Pathotroph-Saprotroph-Symbiotroph | Animal Pathogen-Endophyte-Endosymbiont-Epiphyte-(Undefined Saprotroph) | Dimorphic Yeast-Yeast | NULL | Probable | As nectar-tap_saprotroph (Põlme et al. 2020); Decay sugar-rich substrates (Põlme et al. 2020). This asexual genus is polyphyletic and contain species that function very differently depend on what they are. Caution should be taken when interpreting guilds for Candida species. | Manolakaki D, et al. 2010. Virulence. 1:367-375 (https://doi.org/10.4161/viru.1.5.12796)); Mekbib SB, et al. 2011. Tropical Plant Pathology 36:233-240 (https://doi.org/10.1590/S1982-56762011000400004)); Põlme S, et al. 2020. Fungal Diversity 105:1-16 (https://doi.org/10.1007/s13225-020-00466-2) |
| Otu01849 | 26 | k__Fungi(100);p__Basidiomycota(100);c__Agaricomycetes(100);o__Thelephorales(100);f__Thelephoraceae(100);g__Tomentella(100);s__Tomentella_sp(62); | Tomentella | 13 | Symbiotroph | (Ectomycorrhizal) | Corticioid | NULL | Highly Probable | NULL | Rinaldi AC, et al. 2008. Fungal Diversity 33:1-45 (http://www.fungaldiversity.org/fdp/sfdp/33-1.pdf)); Tedersoo L, et al. 2010. Mycorrhiza 20:217-263 (https://doi.org/10.1007/s00572-009-0274-x)); Põlme S, et al. 2020. Fungal Diversity 105:1-16 (https://doi.org/10.1007/s13225-020-00466-2)); http://mycorrhizas.info/ecmf.html (http://mycorrhizas.info/ecmf.html) |
| Otu01850 | 26 | k__Fungi(100);p__Basidiomycota(100);c__Exobasidiomycetes(100);o__Exobasidiales(100);f__Exobasidiaceae(100);g__Exobasidium(100);s__Exobasidium_japonicum(77); | Exobasidium | 13 | Pathotroph | (Plant Pathogen) | Leaf Blister-Leaf Gall-Leaf Spot | NULL | Probable | NULL | Tedersoo L, et al. 2014. Science 346:e1256688 (https://doi.org/10.1126/science.1256688); Põlme S, et al. 2020. Fungal Diversity 105:1-16 (https://doi.org/10.1007/s13225-020-00466-2) |
| Otu01853 | 26 | k__Fungi(100);p__Basidiomycota(100);c__Tremellomycetes(100);o__Trichosporonales(100);f__Trichosporonaceae(100);g__Trichosporon(100);s__Trichosporon_cutaneum(100); | Trichosporon | 13 | Pathotroph-Saprotroph | (Animal Parasite)-Animal Pathogen-Undefined Saprotroph | Yeast | NULL | Probable | As animal_decomposer (Põlme et al. 2020); Decay animal material (Põlme et al. 2020); Likely opportunistic human pathogen (Irinyi et al. 2015) | Irinyi L, et al. 2015. Medical Mycology 53:313-337 (https://doi.org/10.1093/mmy/myv008); Põlme S, et al. 2020. Fungal Diversity 105:1-16 (https://doi.org/10.1007/s13225-020-00466-2); https://www.cdc.gov (https://www.cdc.gov/) |
| Otu01855 | 26 | k__Fungi(100);p__Ascomycota(100);c__Lecanoromycetes(100);o__Teloschistales(100);f__Physciaceae(100);g__Physcia(100);s__Physcia_aipolia(100); | Physcia | 13 | Symbiotroph | (Lichenized) | Thallus | NULL | Highly Probable | NULL | Esslinger TL. 2014. North Dakota State University (http://www.ndsu.edu/pubweb/~esslinge/chcklst/chcklst7.htm)); Põlme S, et al. 2020. Fungal Diversity 105:1-16 (https://doi.org/10.1007/s13225-020-00466-2) |
| Otu01858 | 26 | k__Fungi(100);p__Ascomycota(100);c__Saccharomycetes(100);o__Saccharomycetales(100);f__Metschnikowiaceae(100);g__Metschnikowia(100);s__Metschnikowia_pulcherrima(97); | Metschnikowia | 13 | Saprotroph | (Undefined Saprotroph) | Yeast | NULL | Probable | As nectar-tap_saprotroph (Põlme et al. 2020); Decay sugar-rich substrates (Põlme et al. 2020) | Põlme S, et al. 2020. Fungal Diversity 105:1-16 (https://doi.org/10.1007/s13225-020-00466-2) |
| Otu01860 | 26 | k__Fungi(100);p__Basidiomycota(100);c__Agaricomycetes(100);o__Agaricales(100);f__Agaricaceae(100);g__Lycoperdon(100);s__Lycoperdon_pyriforme(100); | Lycoperdon | 13 | Saprotroph | (Plant Saprotroph)-Undefined Saprotroph | Gasteroid | NULL | Probable | puffball; As litter_saprotroph (Põlme et al. 2020); Common name ; Decay leaf, fruit, seed (Põlme et al. 2020) | Grgurinovic CA, Simpson JA. 2001. Fungal Diversity 8:97-127 (http://www.fungaldiversity.org/fdp/sfdp/FD_8_97-127.pdf); Tedersoo L, et al. 2014. Science 346:e1256688 (https://doi.org/10.1126/science.1256688); Põlme S, et al. 2020. Fungal Diversity 105:1-16 (https://doi.org/10.1007/s13225-020-00466-2) |
| Otu01861 | 26 | k__Fungi(100);p__Basidiomycota(100);c__Agaricomycetes(100);o__Hymenochaetales(100);f__Schizoporaceae(100);g__Schizopora(100);s__Schizopora_unclassified(100); | Schizopora | 13 | Pathotroph-Saprotroph | Plant Pathogen-(Wood Saprotroph) | Corticioid | White Rot | Highly Probable | As white_rot (Põlme et al. 2020); As wood_pathogen (Põlme et al. 2020); Decay wood (Põlme et al. 2020) | Gilbertson RL, Ryvarden L. 1987-1987. North American Polypores. Fungiflora, Oslo (ISBN: 978-0945345060); Põlme S, et al. 2020. Fungal Diversity 105:1-16 (https://doi.org/10.1007/s13225-020-00466-2) |
| Otu01862 | 26 | k__Fungi(100);p__Ascomycota(100);c__Eurotiomycetes(100);o__Chaetothyriales(100);f__Herpotrichiellaceae(100);g__Exophiala(100);s__Exophiala_unclassified(100); | Exophiala | 13 | Pathotroph-Saprotroph-Symbiotroph | (Animal Parasite)-Animal Pathogen-Endophyte-Plant Saprotroph-Undefined Saprotroph | Dimorphic-Facultative Yeast | NULL | Probable | As litter_saprotroph (Põlme et al. 2020); As root_endophyte_dark_septate (Põlme et al. 2020); Decay leaf, fruit, seed, animal material (Põlme et al. 2020) | Tedersoo L, et al. 2014. Science 346:e1256688 (https://doi.org/10.1126/science.1256688); Nagy LG, et al. 2014. Nature Communications 5:e4471 (https://doi.org/10.1038/ncomms5471)); Põlme S, et al. 2020. Fungal Diversity 105:1-16 (https://doi.org/10.1007/s13225-020-00466-2) |
| Otu01865 | 26 | k__Fungi(100);p__Ascomycota(100);c__Sordariomycetes(100);o__Ophiostomatales(100);f__Ophiostomataceae(100);g__Ophiostoma(100);s__Ophiostoma_auroreae(93); | Ophiostoma | 13 | Pathotroph-Saprotroph | (Plant Pathogen)-Wood Saprotroph | Microfungus | Blue-Staining | Probable | As wood_pathogen (Põlme et al. 2020); At least most species are transmitted via insect vectors, with mycelial colonies developing in bark, boring beetle galleries and spreading through the vascular system; they are highly specific to their beetle vectors. In some cases spores and mycelium are transmitted in specialized organs, termed mycangia (Cannon & Kirk 2007); Decay wood (Põlme et al. 2020) | Cannon PF, Kirk PM. 2007. Fungal Families of the World. CAB International, Cambridge (ISBN: 978-0851998275); Tedersoo L, et al. 2014. Science 346:e1256688 (https://doi.org/10.1126/science.1256688); Põlme S, et al. 2020. Fungal Diversity 105:1-16 (https://doi.org/10.1007/s13225-020-00466-2) |
| Otu01866 | 26 | k__Fungi(100);p__Ascomycota(100);c__Saccharomycetes(100);o__Saccharomycetales(100);f__Metschnikowiaceae(100);g__Metschnikowia(100);s__Metschnikowia_pulcherrima(93); | Metschnikowia | 13 | Saprotroph | (Undefined Saprotroph) | Yeast | NULL | Probable | As nectar-tap_saprotroph (Põlme et al. 2020); Decay sugar-rich substrates (Põlme et al. 2020) | Põlme S, et al. 2020. Fungal Diversity 105:1-16 (https://doi.org/10.1007/s13225-020-00466-2) |
| Otu01868 | 26 | k__Fungi(100);p__Ascomycota(100);c__Lecanoromycetes(100);o__Teloschistales(100);f__Teloschistaceae(100);g__Caloplaca(100);s__Caloplaca_unclassified(93); | Caloplaca | 13 | Symbiotroph | (Lichenized) | Thallus | NULL | Highly Probable | Commonly called firedot lichens | Esslinger TL. 2014. North Dakota State University (http://www.ndsu.edu/pubweb/~esslinge/chcklst/chcklst7.htm)); Põlme S, et al. 2020. Fungal Diversity 105:1-16 (https://doi.org/10.1007/s13225-020-00466-2) |
| Otu01869 | 26 | k__Fungi(100);p__Ascomycota(100);c__Dothideomycetes(100);o__Pleosporales(100);f__Pleosporales_family_Incertae_sedis(100);g__Massarosphaeria(100);s__Massarosphaeria_sp_FAEI23a(100); | Massarosphaeria | 13 | Saprotroph | Undefined Saprotroph-(Wood Saprotroph) | Microfungus | NULL | Probable | Decay wood (Põlme et al. 2020) | Tedersoo L, et al. 2014. Science 346:e1256688 (https://doi.org/10.1126/science.1256688); Põlme S, et al. 2020. Fungal Diversity 105:1-16 (https://doi.org/10.1007/s13225-020-00466-2) |
| Otu01870 | 26 | k__Fungi(100);p__Basidiomycota(100);c__Agaricomycetes(100);o__Polyporales(100);f__Fomitopsidaceae(100);g__Postia(100);s__Postia_lateritia(100); | Postia | 13 | Saprotroph | (Wood Saprotroph) | Polyporoid | Brown Rot | Highly Probable | As brown-rot (Põlme et al. 2020); Decay wood (Põlme et al. 2020) | Jordan CR, et al. 1996. Holzforschung - International Journal of the Biology, Chemistry, Physics and Technology of Wood 50:312-318 (https://doi.org/10.1515/hfsg.1996.50.4.312)); Põlme S, et al. 2020. Fungal Diversity 105:1-16 (https://doi.org/10.1007/s13225-020-00466-2) |
| Otu01872 | 26 | k__Fungi(100);p__Ascomycota(100);c__Dothideomycetes(100);o__Pleosporales(100);f__Tubeufiaceae(100);g__Tubeufia(100);s__Tubeufia_cerea(100); | Tubeufia | 13 | Saprotroph | Undefined Saprotroph-(Wood Saprotroph) | Microfungus | NULL | Probable | Decay wood (Põlme et al. 2020) | Tedersoo L, et al. 2014. Science 346:e1256688 (https://doi.org/10.1126/science.1256688); Põlme S, et al. 2020. Fungal Diversity 105:1-16 (https://doi.org/10.1007/s13225-020-00466-2) |
| Otu01873 | 26 | k__Fungi(100);p__Basidiomycota(100);c__Agaricomycetes(100);o__Thelephorales(100);f__Thelephoraceae(100);g__Tomentella(100);s__Tomentella_unclassified(85); | Tomentella | 13 | Symbiotroph | (Ectomycorrhizal) | Corticioid | NULL | Highly Probable | NULL | Rinaldi AC, et al. 2008. Fungal Diversity 33:1-45 (http://www.fungaldiversity.org/fdp/sfdp/33-1.pdf)); Tedersoo L, et al. 2010. Mycorrhiza 20:217-263 (https://doi.org/10.1007/s00572-009-0274-x)); Põlme S, et al. 2020. Fungal Diversity 105:1-16 (https://doi.org/10.1007/s13225-020-00466-2)); http://mycorrhizas.info/ecmf.html (http://mycorrhizas.info/ecmf.html) |
| Otu01874 | 26 | k__Fungi(100);p__Basidiomycota(100);c__Agaricomycetes(100);o__Hymenochaetales(100);f__Hymenochaetaceae(100);g__Phellinopsis(100);s__Phellinopsis_resupinata(100); | Phellinopsis | 13 | Pathotroph-Saprotroph | Plant Pathogen-Undefined Saprotroph-(Wood Saprotroph) | Polyporoid | White Rot | Probable | As white_rot (Põlme et al. 2020); As wood_pathogen (Põlme et al. 2020); Decay wood (Põlme et al. 2020) | Tedersoo L, et al. 2014. Science 346:e1256688 (https://doi.org/10.1126/science.1256688); Põlme S, et al. 2020. Fungal Diversity 105:1-16 (https://doi.org/10.1007/s13225-020-00466-2) |
| Otu01876 | 26 | k__Fungi(100);p__Ascomycota(100);c__Saccharomycetes(100);o__Saccharomycetales(100);f__Metschnikowiaceae(100);g__Metschnikowia(100);s__Metschnikowia_pulcherrima(93); | Metschnikowia | 13 | Saprotroph | (Undefined Saprotroph) | Yeast | NULL | Probable | As nectar-tap_saprotroph (Põlme et al. 2020); Decay sugar-rich substrates (Põlme et al. 2020) | Põlme S, et al. 2020. Fungal Diversity 105:1-16 (https://doi.org/10.1007/s13225-020-00466-2) |
| Otu01879 | 26 | k__Fungi(100);p__Ascomycota(100);c__Sordariomycetes(100);o__Hypocreales(100);f__Cordycipitaceae(100);g__Simplicillium(100);s__Simplicillium_lamellicola(100); | Simplicillium | 13 | Pathotroph-Saprotroph | (Animal Parasite)-Animal Pathogen-Fungal Parasite-Undefined Saprotroph | Microfungus | NULL | Probable | As animal_decomposer (Põlme et al. 2020); As mycoparasite (Põlme et al. 2020); Decay animal material (Põlme et al. 2020) | Tedersoo L, et al. 2014. Science 346:e1256688 (https://doi.org/10.1126/science.1256688); Põlme S, et al. 2020. Fungal Diversity 105:1-16 (https://doi.org/10.1007/s13225-020-00466-2) |
| Otu01835 | 26 | k__Fungi(100);p__Ascomycota(100);c__Sordariomycetes(100);o__Xylariales(100);o__Xylariales_unclassified(93);o__Xylariales_unclassified(93);o__Xylariales_unclassified(93); | - | - | - | - | - | - | - | Unassigned | - |
| Otu01837 | 26 | k__Fungi(100);k__Fungi_unclassified(58);k__Fungi_unclassified(58);k__Fungi_unclassified(58);k__Fungi_unclassified(58);k__Fungi_unclassified(58);k__Fungi_unclassified(58); | - | - | - | - | - | - | - | Unassigned | - |
| Otu01838 | 26 | k__Fungi(100);p__Ascomycota(100);c__Taphrinomycetes(100);o__Taphrinales(100);f__Taphrinaceae(100);g__Lalaria(100);s__Lalaria_inositophila(100); | - | - | - | - | - | - | - | Unassigned | - |
| Otu01842 | 26 | k__Fungi(100);p__Ascomycota(100);p__Ascomycota_unclassified(100);p__Ascomycota_unclassified(100);p__Ascomycota_unclassified(100);p__Ascomycota_unclassified(100);p__Ascomycota_unclassified(100); | - | - | - | - | - | - | - | Unassigned | - |
| Otu01843 | 26 | k__Fungi(100);k__Fungi_unclassified(74);k__Fungi_unclassified(74);k__Fungi_unclassified(74);k__Fungi_unclassified(74);k__Fungi_unclassified(74);k__Fungi_unclassified(74); | - | - | - | - | - | - | - | Unassigned | - |

| | | | | | | | | | | | |
|---|---|---|---|---|---|---|---|---|---|---|---|
| Otu01847 | 26 | k__Fungi(100);p__Ascomycota(58);p__Ascomycota_unclassified(58);p__Ascomycota_unclassified(58);p__Ascomycota_unclassified(58);p__Ascomycota_unclassified(58);p__Ascomycota_unclassified(58); | - | - | - | - | - | - | - | Unassigned | - |
| Otu01851 | 26 | k__Fungi(100);p__Ascomycota(100);c__Dothideomycetes(66);c__Dothideomycetes_unclassified(66);c__Dothideomycetes_unclassified(66);c__Dothideomycetes_unclassified(66);c__Dothideomycetes_unclassified(66); | - | - | - | - | - | - | - | Unassigned | - |
| Otu01852 | 26 | k__Fungi(100);k__Fungi_unclassified(66);k__Fungi_unclassified(66);k__Fungi_unclassified(66);k__Fungi_unclassified(66);k__Fungi_unclassified(66);k__Fungi_unclassified(66); | - | - | - | - | - | - | - | Unassigned | - |
| Otu01854 | 26 | k__Fungi(100);p__Ascomycota(54);p__Ascomycota_unclassified(54);p__Ascomycota_unclassified(54);p__Ascomycota_unclassified(54);p__Ascomycota_unclassified(54);p__Ascomycota_unclassified(54); | - | - | - | - | - | - | - | Unassigned | - |
| Otu01856 | 26 | k__Fungi(100);p__Ascomycota(66);p__Ascomycota_unclassified(66);p__Ascomycota_unclassified(66);p__Ascomycota_unclassified(66);p__Ascomycota_unclassified(66);p__Ascomycota_unclassified(66); | - | - | - | - | - | - | - | Unassigned | - |
| Otu01857 | 26 | k__Fungi(100);k__Fungi_unclassified(54);k__Fungi_unclassified(54);k__Fungi_unclassified(54);k__Fungi_unclassified(54);k__Fungi_unclassified(54);k__Fungi_unclassified(54); | - | - | - | - | - | - | - | Unassigned | - |
| Otu01859 | 26 | k__Fungi(100);p__Ascomycota(93);p__Ascomycota_unclassified(93);p__Ascomycota_unclassified(93);p__Ascomycota_unclassified(93);p__Ascomycota_unclassified(93);p__Ascomycota_unclassified(93); | - | - | - | - | - | - | - | Unassigned | - |
| Otu01863 | 26 | k__Plantae(100);p__unclassified_Plantae(100);c__unclassified_Plantae(100);o__unclassified_Plantae(100);f__unclassified_Plantae(100);g__unclassified_Plantae(100);s__Plantae_sp(100); | - | - | - | - | - | - | - | Unassigned | - |
| Otu01864 | 26 | k__Fungi(100);p__Ascomycota(93);p__Ascomycota_unclassified(93);p__Ascomycota_unclassified(93);p__Ascomycota_unclassified(93);p__Ascomycota_unclassified(93);p__Ascomycota_unclassified(93); | - | - | - | - | - | - | - | Unassigned | - |
| Otu01867 | 26 | k__Fungi(100);k__Fungi_unclassified(66);k__Fungi_unclassified(66);k__Fungi_unclassified(66);k__Fungi_unclassified(66);k__Fungi_unclassified(66);k__Fungi_unclassified(66); | - | - | - | - | - | - | - | Unassigned | - |
| Otu01871 | 26 | k__Fungi(100);p__Ascomycota(89);p__Ascomycota_unclassified(89);p__Ascomycota_unclassified(89);p__Ascomycota_unclassified(89);p__Ascomycota_unclassified(89);p__Ascomycota_unclassified(89); | - | - | - | - | - | - | - | Unassigned | - |
| Otu01875 | 26 | k__Fungi(100);k__Fungi_unclassified(54);k__Fungi_unclassified(54);k__Fungi_unclassified(54);k__Fungi_unclassified(54);k__Fungi_unclassified(54);k__Fungi_unclassified(54); | - | - | - | - | - | - | - | Unassigned | - |
| Otu01877 | 26 | k__Fungi(100);k__Fungi_unclassified(100);k__Fungi_unclassified(100);k__Fungi_unclassified(100);k__Fungi_unclassified(100);k__Fungi_unclassified(100);k__Fungi_unclassified(100); | - | - | - | - | - | - | - | Unassigned | - |
| Otu01878 | 26 | k__Fungi(100);k__Fungi_unclassified(77);k__Fungi_unclassified(77);k__Fungi_unclassified(77);k__Fungi_unclassified(77);k__Fungi_unclassified(77);k__Fungi_unclassified(77); | - | - | - | - | - | - | - | Unassigned | - |
| Otu01880 | 26 | k__Fungi(100);k__Fungi_unclassified(100);k__Fungi_unclassified(100);k__Fungi_unclassified(100);k__Fungi_unclassified(100);k__Fungi_unclassified(100);k__Fungi_unclassified(100); | - | - | - | - | - | - | - | Unassigned | - |
| Otu01881 | 26 | k__Fungi(100);k__Fungi_unclassified(93);k__Fungi_unclassified(93);k__Fungi_unclassified(93);k__Fungi_unclassified(93);k__Fungi_unclassified(93);k__Fungi_unclassified(93); | - | - | - | - | - | - | - | Unassigned | - |
| Otu01882 | 25 | k__Fungi(100);p__Basidiomycota(100);c__Agaricomycetes(100);o__Thelephorales(100);f__Thelephoraceae(100);g__Tomentella(100);g__Tomentella_unclassified(64); | Tomentella | 13 | Symbiotroph | \|Ectomycorrhizal\| | Corticioid | NULL | Highly Probable | NULL | Rinaldi AC, et al. 2008. Fungal Diversity 33:1-45 ((http://www.fungaldiversity.org/fdp/sfdp/33-1.pdf)); Tedersoo L, et al. 2010. Mycorrhiza 20:217-263 ((https://doi.org/10.1007/s00572-009-0274-x)); Põlme S, et al. 2020. Fungal Diversity 105:1-16 ((https://doi.org/10.1007/s13225-020-00466-2)); http://mycorrhizas.info/ecmf.html ((http://mycorrhizas.info/ecmf.html)) |
| Otu01883 | 25 | k__Fungi(100);p__Ascomycota(100);c__Saccharomycetes(100);o__Saccharomycetales(100);f__Saccharomycetaceae(100);g__Kazachstania(100);s__Kazachstania_intestinalis(100); | Kazachstania | 13 | Saprotroph | \|Undefined Saprotroph\| | Yeast | NULL | Probable | As nectar-tap_saprotroph (Põlme et al. 2020); Decay sugar-rich substrates (Põlme et al. 2020) | Põlme S, et al. 2020. Fungal Diversity 105:1-16 ((https://doi.org/10.1007/s13225-020-00466-2)) |
| Otu01885 | 25 | k__Fungi(100);p__Ascomycota(100);c__Saccharomycetes(100);o__Saccharomycetales(100);f__Metschnikowiaceae(100);g__Metschnikowia(100);s__Metschnikowia_pulcherrima(96); | Metschnikowia | 13 | Saprotroph | \|Undefined Saprotroph\| | Yeast | NULL | Probable | As nectar-tap_saprotroph (Põlme et al. 2020); Decay sugar-rich substrates (Põlme et al. 2020) | Põlme S, et al. 2020. Fungal Diversity 105:1-16 ((https://doi.org/10.1007/s13225-020-00466-2)) |
| Otu01889 | 25 | k__Fungi(100);p__Ascomycota(100);c__Saccharomycetes(100);o__Saccharomycetales(100);f__Metschnikowiaceae(100);g__Metschnikowia(100);g__Metschnikowia_unclassified(92); | Metschnikowia | 13 | Saprotroph | \|Undefined Saprotroph\| | Yeast | NULL | Probable | As nectar-tap_saprotroph (Põlme et al. 2020); Decay sugar-rich substrates (Põlme et al. 2020) | Põlme S, et al. 2020. Fungal Diversity 105:1-16 ((https://doi.org/10.1007/s13225-020-00466-2)) |
| Otu01890 | 25 | k__Fungi(100);p__Ascomycota(100);c__Saccharomycetes(100);o__Saccharomycetales(100);f__Metschnikowiaceae(100);g__Metschnikowia(100);s__Metschnikowia_pulcherrima(96); | Metschnikowia | 13 | Saprotroph | \|Undefined Saprotroph\| | Yeast | NULL | Probable | As nectar-tap_saprotroph (Põlme et al. 2020); Decay sugar-rich substrates (Põlme et al. 2020) | Põlme S, et al. 2020. Fungal Diversity 105:1-16 ((https://doi.org/10.1007/s13225-020-00466-2)) |
| Otu01892 | 25 | k__Fungi(100);p__Basidiomycota(100);c__Agaricomycetes(100);o__Agaricales(100);f__Entolomataceae(100);g__unclassified_Entolomataceae(100);s__Entolomataceae_sp(100); | Entolomataceae | 9 | Pathotroph-Saprotroph-Symbiotroph | Ectomycorrhizal-Fungal Parasite-Soil Saprotroph-Undefined Saprotroph | Agaricoid | NULL | Possible | Saprobic or ectomycorrhizal, in both broadleaved and coniferous forests; a few species associated with other substrata such as dung and mosses (Cannon & Kirk 2007) | Cannon PF, Kirk PM. 2007. Fungal Families of the World. CAB International, Cambridge (ISBN: 978-0851998275); List compiled by John Plischke III ((https://goodmushroom.weebly.com/uploads/5/7/1/7/57176535/fungi_on_fungi_pdf_lq.pdf)) |
| Otu01894 | 25 | k__Fungi(100);p__Ascomycota(100);c__Dothideomycetes(100);o__Pleosporales(100);f__Venturiaceae(100);g__Venturia(100);s__Venturia_macularis(61); | Venturia | 13 | Pathotroph-Pathotroph-Saprotroph | \|Plant Pathogen\|-Plant Saprotroph | Microfungus | NULL | Probable | As litter_saprotroph (Põlme et al. 2020); Decay leaf, fruit, seed (Põlme et al. 2020) | Tedersoo L, et al. 2014. Science 346:e1256688 ((https://doi.org/10.1126/science.1256688)); Põlme S, et al. 2020. Fungal Diversity 105:1-16 ((https://doi.org/10.1007/s13225-020-00466-2)) |
| Otu01895 | 25 | k__Fungi(100);p__Ascomycota(100);c__Sordariomycetes(100);o__Xylariales(100);f__Xylariaceae(100);g__Hypoxylon(61);g__Hypoxylon_unclassified(61); | Hypoxylon | 13 | Saprotroph-Saprotroph-Symbiotroph | Endophyte-Undefined Saprotroph-\|Wood Saprotroph\| | Corticioid-Microfungus-Xylarioid | NULL | Probable | As foliar_endophyte (Põlme et al. 2020); Decay wood, leaf, fruit, seed (Põlme et al. 2020); Host - angiosperms (Põlme et al. 2020) | Tedersoo L, et al. 2014. Science 346:e1256688 ((https://doi.org/10.1126/science.1256688)); Põlme S, et al. 2020. Fungal Diversity 105:1-16 ((https://doi.org/10.1007/s13225-020-00466-2)) |
| Otu01896 | 25 | k__Fungi(100);p__Ascomycota(100);c__Sordariomycetes(100);o__Xylariales(100);f__Xylariaceae(100);g__Xylaria(56);g__Xylaria_unclassified(56); | Xylaria | 13 | Saprotroph-Symbiotroph | Endophyte-Plant Saprotroph-Undefined Saprotroph-\|Wood Saprotroph\| | Xylarioid | Soft Rot | Probable | As foliar_endophyte (Põlme et al. 2020); As litter_saprotroph (Põlme et al. 2020); Decay wood, leaf, fruit, seed (Põlme et al. 2020); Host - angiosperms (Põlme et al. 2020) | Duncan CG, Eslyn WE. 1966. Mycologia 58:642-645 ((https://doi.org/10.2307/3757045)); Costa IPMW, et al. 2012. Mycotaxon 119:e26 ((http://mycotaxon.com/resources/checklists/costa_v119_checklist.pdf)); Põlme S, et al. 2020. Fungal Diversity 105:1-16 ((https://doi.org/10.1007/s13225-020-00466-2)); The American Phytopathological Society Common Names of Plant Diseases ((http://www.apsnet.org/publications/commonnames/Pages/default.aspx)) |
| Otu01897 | 25 | k__Fungi(100);p__Basidiomycota(100);c__Cystobasidiomycetes(72);o__Cystobasidiales(72);f__Cystobasidiaceae(72);g__Occultifur(72);g__Occultifur_unclassified(64); | Occultifur | 13 | Pathotroph-Saprotroph | \|Fungal Parasite\|-Plant Saprotroph-Undefined Saprotroph | Dimorphic-Yeast | NULL | Probable | As litter_saprotroph (Põlme et al. 2020); As mycoparasite (Põlme et al. 2020); Decay leaf, fruit, seed, fungal material (Põlme et al. 2020) | Gomes FCO, et al. 2015. Antonie van Leeuwenhoek 107:597-611 ((https://doi.org/10.1007/s10482-014-0356-4)); Khunnamwong P, et al. 2017. International Journal of Systematic and Evolutionary Microbiology 67:2628-2633 ((https://doi.org/10.1099/ijsem.0.001988)); Põlme S, et al. 2020. Fungal Diversity 105:1-16 ((https://doi.org/10.1007/s13225-020-00466-2)) |
| Otu01899 | 25 | k__Fungi(100);p__Ascomycota(100);c__Saccharomycetes(100);o__Saccharomycetales(100);f__Metschnikowiaceae(100);g__Metschnikowia(100);s__Metschnikowia_pulcherrima(96); | Metschnikowia | 13 | Saprotroph | \|Undefined Saprotroph\| | Yeast | NULL | Probable | As nectar-tap_saprotroph (Põlme et al. 2020); Decay sugar-rich substrates (Põlme et al. 2020) | Põlme S, et al. 2020. Fungal Diversity 105:1-16 ((https://doi.org/10.1007/s13225-020-00466-2)) |
| Otu01905 | 25 | k__Fungi(100);p__Basidiomycota(100);c__Agaricomycetes(100);o__Polyporales(100);f__Fomitopsidaceae(100);g__Dacryobolus(100);s__Dacryobolus_karstenii(100); | Dacryobolus | 13 | Saprotroph | \|Wood Saprotroph\| | Corticioid | Brown Rot | Probable | As brown-rot (Põlme et al. 2020); Decay wood (Põlme et al. 2020) | Tedersoo L, et al. 2014. Science 346:e1256688 ((https://doi.org/10.1126/science.1256688)); Põlme S, et al. 2020. Fungal Diversity 105:1-16 ((https://doi.org/10.1007/s13225-020-00466-2)) |
| Otu01906 | 25 | k__Fungi(100);p__Ascomycota(100);c__Saccharomycetes(100);o__Saccharomycetales(100);f__Metschnikowiaceae(100);g__Metschnikowia(100);s__Metschnikowia_pulcherrima(92); | Metschnikowia | 13 | Saprotroph | \|Undefined Saprotroph\| | Yeast | NULL | Probable | As nectar-tap_saprotroph (Põlme et al. 2020); Decay sugar-rich substrates (Põlme et al. 2020) | Põlme S, et al. 2020. Fungal Diversity 105:1-16 ((https://doi.org/10.1007/s13225-020-00466-2)) |
| Otu01908 | 25 | k__Fungi(100);p__Basidiomycota(100);c__Agaricomycetes(100);o__Polyporales(100);f__Polyporaceae(100);g__Cinereomyces(100);s__Cinereomyces_lindbladii(100); | Cinereomyces | 13 | Saprotroph | Undefined Saprotroph-\|Wood Saprotroph\| | Corticioid | White Rot | Probable | As white_rot (Põlme et al. 2020); Decay wood (Põlme et al. 2020) | Tedersoo L, et al. 2014. Science 346:e1256688 ((https://doi.org/10.1126/science.1256688)); Põlme S, et al. 2020. Fungal Diversity 105:1-16 ((https://doi.org/10.1007/s13225-020-00466-2)) |
| Otu01914 | 25 | k__Fungi(100);p__Ascomycota(100);c__Saccharomycetes(100);o__Saccharomycetales(100);f__Metschnikowiaceae(100);g__Metschnikowia(100);s__Metschnikowia_pulcherrima(100); | Metschnikowia | 13 | Saprotroph | \|Undefined Saprotroph\| | Yeast | NULL | Probable | As nectar-tap_saprotroph (Põlme et al. 2020); Decay sugar-rich substrates (Põlme et al. 2020) | Põlme S, et al. 2020. Fungal Diversity 105:1-16 ((https://doi.org/10.1007/s13225-020-00466-2)) |
| Otu01916 | 25 | k__Fungi(100);p__Ascomycota(100);c__Saccharomycetes(100);o__Saccharomycetales(100);f__Metschnikowiaceae(100);g__Metschnikowia(100);s__Metschnikowia_pulcherrima(96); | Metschnikowia | 13 | Saprotroph | \|Undefined Saprotroph\| | Yeast | NULL | Probable | As nectar-tap_saprotroph (Põlme et al. 2020); Decay sugar-rich substrates (Põlme et al. 2020) | Põlme S, et al. 2020. Fungal Diversity 105:1-16 ((https://doi.org/10.1007/s13225-020-00466-2)) |
| Otu01918 | 25 | k__Fungi(100);p__Ascomycota(100);c__Taphrinomycetes(100);o__Taphrinales(100);f__Taphrinaceae(100);g__Taphrina(100);s__Taphrina_alni(100); | Taphrina | 13 | Pathotroph | \|Plant Pathogen\| | Dimorphic-Microfungus | NULL | Probable | NULL | Põlme S, et al. 2020. Fungal Diversity 105:1-16 ((https://doi.org/10.1007/s13225-020-00466-2)); The American Phytopathological Society Common Names of Plant Diseases ((http://www.apsnet.org/publications/commonnames/Pages/default.aspx)) |
| Otu01919 | 25 | k__Fungi(100);p__Ascomycota(100);c__Dothideomycetes(100);o__Dothideales(100);f__Dothioraceae(100);g__Aureobasidium(100);s__Aureobasidium_pullulans_var._subglaciale(88); | Aureobasidium | 13 | Pathotroph-Saprotroph-Symbiotroph | Animal Pathogen-Endophyte-Epiphyte-\|Plant Pathogen\|-Plant Saprotroph-Undefined Saprotroph | Dimorphic-Facultative Yeast | NULL | Possible | As foliar_endophyte (Põlme et al. 2020); As litter_saprotroph (Põlme et al. 2020); Causes sooty mold (Põlme et al. 2020); Decay leaf, fruit, seed, roots, animal material, sugar-rich substrates (Põlme et al. 2020) | Costa IPMW, et al. 2012. Mycotaxon 119:e26 ((http://mycotaxon.com/resources/checklists/costa_v119_checklist.pdf)); Tedersoo L, et al. 2014. Science 346:e1256688 ((https://doi.org/10.1126/science.1256688)); Irinyia L, et al. 2016. Fungal Biology 120:125-136 ((https://doi.org/10.1016/j.funbio.2015.04.007)); Põlme S, et al. 2020. Fungal Diversity 105:1-16 ((https://doi.org/10.1007/s13225-020-00466-2)) |
| Otu01920 | 25 | k__Fungi(100);p__Basidiomycota(100);c__Agaricomycetes(100);o__Hymenochaetales(100);f__Schizoporaceae(100);f__Schizoporaceae_unclassified(100);f__Schizoporaceae_unclassified(100); | Schizoporaceae | 9 | Saprotroph | Wood Saprotroph | Corticioid | White Rot | Probable | Saprobic, causing white rot of standing and fallen wood of coniferous and broadleaved trees, rarely on herbaceous stems (Cannon & Kirk 2007) | Cannon PF, Kirk PM. 2007. Fungal Families of the World. CAB International, Cambridge (ISBN: 978-0851998275) |
| Otu01921 | 25 | k__Fungi(100);p__Ascomycota(100);c__Saccharomycetes(100);o__Saccharomycetales(100);f__Metschnikowiaceae(100);g__Metschnikowia(100);s__Metschnikowia_pulcherrima(84); | Metschnikowia | 13 | Saprotroph | \|Undefined Saprotroph\| | Yeast | NULL | Probable | As nectar-tap_saprotroph (Põlme et al. 2020); Decay sugar-rich substrates (Põlme et al. 2020) | Põlme S, et al. 2020. Fungal Diversity 105:1-16 ((https://doi.org/10.1007/s13225-020-00466-2)) |
| Otu01923 | 25 | k__Fungi(100);p__Ascomycota(100);c__Saccharomycetes(100);o__Saccharomycetales(100);f__Metschnikowiaceae(100);g__Metschnikowia(100);s__Metschnikowia_pulcherrima(96); | Metschnikowia | 13 | Saprotroph | \|Undefined Saprotroph\| | Yeast | NULL | Probable | As nectar-tap_saprotroph (Põlme et al. 2020); Decay sugar-rich substrates (Põlme et al. 2020) | Põlme S, et al. 2020. Fungal Diversity 105:1-16 ((https://doi.org/10.1007/s13225-020-00466-2)) |
| Otu01925 | 25 | k__Fungi(100);p__Basidiomycota(100);c__Agaricomycetes(100);o__Trechisporales(100);f__Hydnodontaceae(100);g__Trechispora(100);s__Trechispora_sp(100); | Trechispora | 13 | Saprotroph | \|Wood Saprotroph\| | Corticioid | White Rot | Highly Probable | As white_rot (Põlme et al. 2020); Decay wood (Põlme et al. 2020) | Gilbertson RL, Ryvarden L. 1987-1987. North American Polypores. Fungiflora, Oslo (ISBN: 978-0945345060); Rinaldi AC, et al. 2008. Fungal Diversity 33:1-45 ((http://www.fungaldiversity.org/fdp/sfdp/33-1.pdf)) (as Uncertain); Tedersoo L, et al. 2010. Mycorrhiza 20:217-263 ((https://doi.org/10.1007/s00572-009-0274-x)) (as not ECM).; Põlme S, et al. 2020. Fungal Diversity 105:1-16 ((https://doi.org/10.1007/s13225-020-00466-2)) |
| Otu01926 | 25 | k__Fungi(100);p__Ascomycota(100);c__Leotiomycetes(100);o__Helotiales(100);f__Helotiales_family_Incertae_sedis(100);g__Tetracladium(100);s__Tetracladium_maxilliforme(96); | Tetracladium | 13 | Saprotroph | \|Plant Saprotroph\|-Undefined Saprotroph | Microfungus | NULL | Probable | As litter_saprotroph (Põlme et al. 2020); Decay leaf, fruit, seed (Põlme et al. 2020) | Tedersoo L, et al. 2014. Science 346:e1256688 ((https://doi.org/10.1126/science.1256688)); Põlme S, et al. 2020. Fungal Diversity 105:1-16 ((https://doi.org/10.1007/s13225-020-00466-2)) |
| Otu01884 | 25 | k__Fungi(100);p__Basidiomycota(100);c__Microbotryomycetes(61);o__Sporidiobolales(61);f__Sporidiobolales_family_Incertae_sedis(61);f__Sporidiobolales_family_Incertae_sedis_unclassified(61);f__Sporidiobolales_family_Incertae_sedis_unclassified(61); | - | - | - | - | - | - | - | Unassigned | - |
| Otu01886 | 25 | k__Fungi(100);p__Ascomycota(80);p__Ascomycota_unclassified(80);p__Ascomycota_unclassified(80);p__Ascomycota_unclassified(80);p__Ascomycota_unclassified(80);p__Ascomycota_unclassified(80); | - | - | - | - | - | - | - | Unassigned | - |
| Otu01887 | 25 | k__Fungi(100);k__Fungi_unclassified(52);k__Fungi_unclassified(52);k__Fungi_unclassified(52);k__Fungi_unclassified(52);k__Fungi_unclassified(52);k__Fungi_unclassified(52); | - | - | - | - | - | - | - | Unassigned | - |
| Otu01888 | 25 | k__Fungi(100);k__Fungi_unclassified(84);k__Fungi_unclassified(84);k__Fungi_unclassified(84);k__Fungi_unclassified(84);k__Fungi_unclassified(84);k__Fungi_unclassified(84); | - | - | - | - | - | - | - | Unassigned | - |
| Otu01891 | 25 | k__Fungi(100);k__Fungi_unclassified(96);k__Fungi_unclassified(96);k__Fungi_unclassified(96);k__Fungi_unclassified(96);k__Fungi_unclassified(96);k__Fungi_unclassified(96); | - | - | - | - | - | - | - | Unassigned | - |
| Otu01893 | 25 | k__Fungi(100);p__Ascomycota(100);c__Dothideomycetes(100);o__Pleosporales(100);o__Pleosporales_unclassified(92);o__Pleosporales_unclassified(92);o__Pleosporales_unclassified(92); | - | - | - | - | - | - | - | Unassigned | - |
| Otu01898 | 25 | k__Fungi(100);p__Ascomycota(80);p__Ascomycota_unclassified(80);p__Ascomycota_unclassified(80);p__Ascomycota_unclassified(80);p__Ascomycota_unclassified(80);p__Ascomycota_unclassified(80); | - | - | - | - | - | - | - | Unassigned | - |
| Otu01900 | 25 | k__Fungi(100);k__Fungi_unclassified(56);k__Fungi_unclassified(56);k__Fungi_unclassified(56);k__Fungi_unclassified(56);k__Fungi_unclassified(56);k__Fungi_unclassified(56); | - | - | - | - | - | - | - | Unassigned | - |
| Otu01901 | 25 | k__Fungi(100);p__Ascomycota(72);p__Ascomycota_unclassified(72);p__Ascomycota_unclassified(72);p__Ascomycota_unclassified(72);p__Ascomycota_unclassified(72);p__Ascomycota_unclassified(72); | - | - | - | - | - | - | - | Unassigned | - |
| Otu01902 | 25 | k__Fungi(100);k__Fungi_unclassified(61);k__Fungi_unclassified(61);k__Fungi_unclassified(61);k__Fungi_unclassified(61);k__Fungi_unclassified(61);k__Fungi_unclassified(61); | - | - | - | - | - | - | - | Unassigned | - |
| Otu01903 | 25 | k__Fungi(100);k__Fungi_unclassified(64);k__Fungi_unclassified(64);k__Fungi_unclassified(64);k__Fungi_unclassified(64);k__Fungi_unclassified(64);k__Fungi_unclassified(64); | - | - | - | - | - | - | - | Unassigned | - |
| Otu01904 | 25 | k__Fungi(100);k__Fungi_unclassified(61);k__Fungi_unclassified(61);k__Fungi_unclassified(61);k__Fungi_unclassified(61);k__Fungi_unclassified(61);k__Fungi_unclassified(61); | - | - | - | - | - | - | - | Unassigned | - |
| Otu01907 | 25 | k__Fungi(100);p__Ascomycota(68);p__Ascomycota_unclassified(68);p__Ascomycota_unclassified(68);p__Ascomycota_unclassified(68);p__Ascomycota_unclassified(68);p__Ascomycota_unclassified(68); | - | - | - | - | - | - | - | Unassigned | - |
| Otu01909 | 25 | k__Fungi(100);p__Ascomycota(56);p__Ascomycota_unclassified(56);p__Ascomycota_unclassified(56);p__Ascomycota_unclassified(56);p__Ascomycota_unclassified(56);p__Ascomycota_unclassified(56); | - | - | - | - | - | - | - | Unassigned | - |
| Otu01910 | 25 | k__Fungi(100);p__Ascomycota(100);c__Leotiomycetes(100);o__unclassified_Leotiomycetes(100);f__unclassified_Leotiomycetes(100);g__unclassified_Leotiomycetes(100);s__Leotiomycetes_sp(100); | - | - | - | - | - | - | - | Unassigned | - |
| Otu01911 | 25 | k__Fungi(100);p__Ascomycota(100);c__Leotiomycetes(100);c__Leotiomycetes_unclassified(96);c__Leotiomycetes_unclassified(96);c__Leotiomycetes_unclassified(96);c__Leotiomycetes_unclassified(96); | - | - | - | - | - | - | - | Unassigned | - |
| Otu01912 | 25 | k__Fungi(100);k__Fungi_unclassified(88);k__Fungi_unclassified(88);k__Fungi_unclassified(88);k__Fungi_unclassified(88);k__Fungi_unclassified(88);k__Fungi_unclassified(88); | - | - | - | - | - | - | - | Unassigned | - |
| Otu01913 | 25 | k__Fungi(100);k__Fungi_unclassified(64);k__Fungi_unclassified(64);k__Fungi_unclassified(64);k__Fungi_unclassified(64);k__Fungi_unclassified(64);k__Fungi_unclassified(64); | - | - | - | - | - | - | - | Unassigned | - |
| Otu01915 | 25 | k__Fungi(100);p__Ascomycota(68);p__Ascomycota_unclassified(68);p__Ascomycota_unclassified(68);p__Ascomycota_unclassified(68);p__Ascomycota_unclassified(68);p__Ascomycota_unclassified(68); | - | - | - | - | - | - | - | Unassigned | - |
| Otu01917 | 25 | k__Fungi(100);p__Ascomycota(100);p__Ascomycota_unclassified(100);p__Ascomycota_unclassified(100);p__Ascomycota_unclassified(100);p__Ascomycota_unclassified(100);p__Ascomycota_unclassified(100); | - | - | - | - | - | - | - | Unassigned | - |
| Otu01922 | 25 | k__Fungi(100);k__Fungi_unclassified(100);k__Fungi_unclassified(100);k__Fungi_unclassified(100);k__Fungi_unclassified(100);k__Fungi_unclassified(100);k__Fungi_unclassified(100); | - | - | - | - | - | - | - | Unassigned | - |
| Otu01924 | 25 | k__Fungi(100);k__Fungi_unclassified(100);k__Fungi_unclassified(100);k__Fungi_unclassified(100);k__Fungi_unclassified(100);k__Fungi_unclassified(100);k__Fungi_unclassified(100); | - | - | - | - | - | - | - | Unassigned | - |
| Otu01927 | 25 | k__Fungi(100);p__Ascomycota(100);c__Leotiomycetes(100);c__Leotiomycetes_unclassified(100);c__Leotiomycetes_unclassified(100);c__Leotiomycetes_unclassified(100);c__Leotiomycetes_unclassified(100); | - | - | - | - | - | - | - | Unassigned | - |
| Otu01929 | 24 | k__Fungi(100);p__Ascomycota(100);c__Dothideomycetes(100);o__Dothideomycetes_order_Incertae_sedis(100);f__Myxotrichaceae(100);g__Oidiodendron(100);g__Oidiodendron_unclassified(100); | Oidiodendron | 13 | Pathotroph-Saprotroph-Symbiotroph | Endophyte-Ericoid Mycorrhizal-\|Undefined Saprotroph\| | Microfungus | NULL | Probable | As root_endophyte (Põlme et al. 2020); As soil_saprotroph (Põlme et al. 2020); Dark Septate Endophyte; Decay soil (Põlme et al. 2020) | Newsham KK. 2011. New Phytologist 190:783-793 ((https://doi.org/10.1111/j.1469-8137.2010.03611.x)) (as dark septate root endophytes); Põlme S, et al. 2020. Fungal Diversity 105:1-16 ((https://doi.org/10.1007/s13225-020-00466-2)) |
| Otu01930 | 24 | k__Fungi(100);p__Ascomycota(100);c__Sordariomycetes(100);o__Sordariomycetes_order_Incertae_sedis(100);f__Sordariomycetes_family_Incertae_sedis(100);g__Eucasphaeria(100);s__Eucasphaeria_capensis(100); | Eucasphaeria | 13 | Pathotroph-Saprotroph | \|Plant Pathogen\|-Plant Saprotroph-Undefined Saprotroph | Microfungus | NULL | Probable | As litter_saprotroph (Põlme et al. 2020); Decay leaf, fruit, seed (Põlme et al. 2020) | Tedersoo L, et al. 2014. Science 346:e1256688 ((https://doi.org/10.1126/science.1256688)); Põlme S, et al. 2020. Fungal Diversity 105:1-16 ((https://doi.org/10.1007/s13225-020-00466-2)) |
| Otu01931 | 24 | k__Fungi(100);p__Ascomycota(100);c__Eurotiomycetes(100);o__Chaetothyriales(55);f__Herpotrichiellaceae(55);f__Herpotrichiellaceae_unclassified(55);f__Herpotrichiellaceae_unclassified(55); | Herpotrichiellaceae | 9 | Pathotroph-Saprotroph | Animal Pathogen-Fungal Parasite-Undefined Saprotroph | Facultative Yeast-Microfungus | NULL | Probable | Mostly saprobic on plants or other fungi, isolated from soil etc., and from medical sources (Cannon & Kirk 2007) | Cannon PF, Kirk PM. 2007. Fungal Families of the World. CAB International, Cambridge (ISBN: 978-0851998275) |
| Otu01935 | 24 | k__Fungi(100);p__Basidiomycota(100);c__Tremellomycetes(100);o__Tremellales(100);f__Tremellales_family_Incertae_sedis(100);f__Tremellales_family_Incertae_sedis_unclassified(67);f__Tremellales_family_Incertae_sedis_unclassified(67); | Tremellales | 7 | Pathotroph-Saprotroph-Symbiotroph | Fungal Parasite-Undefined Saprotroph | Tremelloid-Yeast | NULL | Possible | The order contains both teleomorphic and anamorphic species, most of the latter being yeasts | Sterkenburg E, et al. 2015. New Phytologist 207:1145-1158 ((https://doi.org/10.1111/nph.13426)) |
| Otu01936 | 24 | k__Fungi(100);p__Basidiomycota(100);c__Agaricomycetes(100);o__Polyporales(100);f__Fomitopsidaceae(100);g__Phaeolus(100);s__Phaeolus_schweinitzii(100); | Phaeolus | 13 | Pathotroph-Saprotroph | Plant Pathogen-\|Wood Saprotroph\| | Polyporoid | Brown Rot | Highly Probable | As brown-rot (Põlme et al. 2020); As root_pathogen (Põlme et al. 2020); Decay wood, roots (Põlme et al. 2020) | Gilbertson RL, Ryvarden L. 1987-1987. North American Polypores. Fungiflora, Oslo (ISBN: 978-0945345060); Põlme S, et al. 2020. Fungal Diversity 105:1-16 ((https://doi.org/10.1007/s13225-020-00466-2)) |
| Otu01937 | 24 | k__Fungi(100);p__Basidiomycota(100);c__Agaricomycetes(100);o__Agaricales(100);f__Strophariaceae(100);g__Pholiota(100);s__Pholiota_sp_TENN61700(100); | Pholiota | 13 | Pathotroph-Saprotroph | Plant Pathogen-Plant Saprotroph-Undefined Saprotroph-\|Wood Saprotroph\| | Agaricoid | White Rot | Probable | As litter_saprotroph (Põlme et al. 2020); As white_rot (Põlme et al. 2020); As wood_pathogen (Põlme et al. 2020); Decay wood, leaf, fruit, seed (Põlme et al. 2020) | Tedersoo L, et al. 2014. Science 346:e1256688 ((https://doi.org/10.1126/science.1256688)); Põlme S, et al. 2020. Fungal Diversity 105:1-16 ((https://doi.org/10.1007/s13225-020-00466-2)) |
| Otu01938 | 24 | k__Fungi(100);p__Basidiomycota(100);c__Agaricomycetes(100);o__Agaricales(100);f__Entolomataceae(100);g__unclassified_Entolomataceae(100);s__Entolomataceae_sp(100); | Entolomataceae | 9 | Pathotroph-Saprotroph-Symbiotroph | Ectomycorrhizal-Fungal Parasite-Soil Saprotroph-Undefined Saprotroph | Agaricoid | NULL | Possible | Saprobic or ectomycorrhizal, in both broadleaved and coniferous forests; a few species associated with other substrata such as dung and mosses (Cannon & Kirk 2007) | Cannon PF, Kirk PM. 2007. Fungal Families of the World. CAB International, Cambridge (ISBN: 978-0851998275); List compiled by John Plischke III ((https://goodmushroom.weebly.com/uploads/5/7/1/7/57176535/fungi_on_fungi_pdf_lq.pdf)) |
| Otu01945 | 24 | k__Fungi(100);p__Ascomycota(100);c__Eurotiomycetes(100);o__Eurotiales(100);f__Trichocomaceae(100);g__Penicillium(100);g__Penicillium_unclassified(100); | Penicillium | 13 | Pathotroph-Saprotroph-Symbiotroph | Animal Parasite-Dung Saprotroph-Endophyte-Fungal Parasite-Plant Pathogen-\|Undefined Saprotroph\|-Wood Saprotroph | Microfungus | Soft Rot | Highly Probable | As foliar_endophyte (Põlme et al. 2020); As mycoparasite (Põlme et al. 2020); Decay soil (Põlme et al. 2020) | Duncan CG, Eslyn WE. 1966. Mycologia 58:642-664 ((https://doi.org/10.2307/3757045)); Seehann G, et al. 1975. List of Fungi in Soft-Rot Tissue Tests. Institute fur Holzbiologie und Holzschutz, Bundesforschungsanstalt fur Forst - und Holzwirtshaft, Hamburg - Lohbrugge, Germany ((https://www.irg-wp.com/irgdocs/details.php?d6493033-c9ea-40f3-8e3e-5697f32d640d)); Costa IPMW, et al. 2012. Mycotaxon 119:e26 ((http://mycotaxon.com/resources/checklists/costa_v119_checklist.pdf)); Bills GF, et al. 2013. Current Opinion in Microbiology 16:549-565 ((https://doi.org/10.1016/j.mib.2013.08.001)); Tedersoo L, et al. 2014. Science 346:e1256688 ((https://doi.org/10.1126/science.1256688)); Põlme S, et al. 2020. Fungal Diversity 105:1-16 ((https://doi.org/10.1007/s13225-020-00466-2)) |
| Otu01947 | 24 | k__Fungi(100);p__Ascomycota(100);c__Saccharomycetes(100);o__Saccharomycetales(100);f__Metschnikowiaceae(100);g__Metschnikowia(100);s__Metschnikowia_pulcherrima(100); | Metschnikowia | 13 | Saprotroph | \|Undefined Saprotroph\| | Yeast | NULL | Probable | As nectar-tap_saprotroph (Põlme et al. 2020); Decay sugar-rich substrates (Põlme et al. 2020) | Põlme S, et al. 2020. Fungal Diversity 105:1-16 ((https://doi.org/10.1007/s13225-020-00466-2)) |
| Otu01948 | 24 | k__Fungi(100);p__Basidiomycota(100);c__Agaricomycetes(100);o__Atheliales(100);f__Atheliaceae(100);g__Amphinema(100);s__Amphinema_sp_6_UK_2011(100); | Amphinema | 13 | Symbiotroph | \|Ectomycorrhizal\| | Corticioid | NULL | Highly Probable | NULL | Rinaldi AC, et al. 2008. Fungal Diversity 33:1-45 ((http://www.fungaldiversity.org/fdp/sfdp/33-1.pdf)); Tedersoo L, et al. 2010. Mycorrhiza 20:217-263 ((https://doi.org/10.1007/s00572-009-0274-x)); Põlme S, et al. 2020. Fungal Diversity 105:1-16 ((https://doi.org/10.1007/s13225-020-00466-2)); http://mycorrhizas.info/ecmf.html ((http://mycorrhizas.info/ecmf.html)) |
| Otu01949 | 24 | k__Fungi(100);p__Basidiomycota(100);c__Agaricomycetes(100);o__Cantharellales(100);f__Botryobasidiaceae(100);g__unclassified_Botryobasidiaceae(96);s__Botryobasidiaceae_sp(96); | Botryobasidiaceae | 9 | Saprotroph | Undefined Saprotroph-Wood Saprotroph | Corticioid | NULL | Probable | Saprobic, typically on rotten wood (Cannon & Kirk 2007) | Cannon PF, Kirk PM. 2007. Fungal Families of the World. CAB International, Cambridge (ISBN: 978-0851998275) |
| Otu01950 | 24 | k__Fungi(100);p__Ascomycota(100);c__Saccharomycetes(100);o__Saccharomycetales(100);f__Metschnikowiaceae(100);g__Metschnikowia(100);s__Metschnikowia_pulcherrima(92); | Metschnikowia | 13 | Saprotroph | \|Undefined Saprotroph\| | Yeast | NULL | Probable | As nectar-tap_saprotroph (Põlme et al. 2020); Decay sugar-rich substrates (Põlme et al. 2020) | Põlme S, et al. 2020. Fungal Diversity 105:1-16 ((https://doi.org/10.1007/s13225-020-00466-2)) |
| Otu01952 | 24 | k__Fungi(100);p__Ascomycota(100);c__Saccharomycetes(100);o__Saccharomycetales(100);f__Metschnikowiaceae(100);g__Metschnikowia(100);s__Metschnikowia_pulcherrima(96); | Metschnikowia | 13 | Saprotroph | \|Undefined Saprotroph\| | Yeast | NULL | Probable | As nectar-tap_saprotroph (Põlme et al. 2020); Decay sugar-rich substrates (Põlme et al. 2020) | Põlme S, et al. 2020. Fungal Diversity 105:1-16 ((https://doi.org/10.1007/s13225-020-00466-2)) |
| Otu01954 | 24 | k__Fungi(100);p__Ascomycota(100);c__Saccharomycetes(100);o__Saccharomycetales(100);f__Metschnikowiaceae(100);g__Metschnikowia(100);s__Metschnikowia_pulcherrima(96); | Metschnikowia | 13 | Saprotroph | \|Undefined Saprotroph\| | Yeast | NULL | Probable | As nectar-tap_saprotroph (Põlme et al. 2020); Decay sugar-rich substrates (Põlme et al. 2020) | Põlme S, et al. 2020. Fungal Diversity 105:1-16 ((https://doi.org/10.1007/s13225-020-00466-2)) |
| Otu01956 | 24 | k__Fungi(100);p__Ascomycota(100);c__Saccharomycetes(100);o__Saccharomycetales(100);f__Metschnikowiaceae(100);g__Metschnikowia(100);s__Metschnikowia_pulcherrima(92); | Metschnikowia | 13 | Saprotroph | \|Undefined Saprotroph\| | Yeast | NULL | Probable | As nectar-tap_saprotroph (Põlme et al. 2020); Decay sugar-rich substrates (Põlme et al. 2020) | Põlme S, et al. 2020. Fungal Diversity 105:1-16 ((https://doi.org/10.1007/s13225-020-00466-2)) |
| Otu01957 | 24 | k__Fungi(100);p__Ascomycota(100);c__Dothideomycetes(100);o__Capnodiales(100);f__Teratosphaeriaceae(55);g__Catenulostroma(55);s__Catenulostroma_hermanusense(55); | Catenulostroma | 13 | Pathotroph | \|Plant Pathogen\| | Microfungus | NULL | Probable | NULL | Tedersoo L, et al. 2014. Science 346:e1256688 ((https://doi.org/10.1126/science.1256688)); Põlme S, et al. 2020. Fungal Diversity 105:1-16 ((https://doi.org/10.1007/s13225-020-00466-2)) |
| Otu01958 | 24 | k__Fungi(100);p__Basidiomycota(100);c__Agaricomycetes(100);o__Polyporales(100);f__Polyporaceae(100);g__Coriolopsis(100);g__Coriolopsis_unclassified(100); | Coriolopsis | 13 | Saprotroph | \|Wood Saprotroph\| | Polyporoid | White Rot | Highly Probable | As white_rot (Põlme et al. 2020); Decay wood (Põlme et al. 2020) | Gilbertson RL, Ryvarden L. 1987-1987. North American Polypores. Fungiflora, Oslo (ISBN: 978-0945345060); Põlme S, et al. 2020. Fungal Diversity 105:1-16 ((https://doi.org/10.1007/s13225-020-00466-2)) |
| Otu01959 | 24 | k__Fungi(100);p__Ascomycota(100);c__Eurotiomycetes(100);o__Eurotiales(100);f__Trichocomaceae(100);g__Aspergillus(100);g__Aspergillus_unclassified(100); | Aspergillus | 13 | Pathotroph-Saprotroph-Symbiotroph | Animal Pathogen-Endophyte-Plant Saprotroph-\|Undefined Saprotroph\|-Wood Saprotroph | Microfungus | NULL | Probable | As foliar_endophyte (Põlme et al. 2020); Decay leaf, fruit, seed, soil, dung, animal material (Põlme et al. 2020) | Seehann G, et al. 1975. List of Fungi in Soft-Rot Tissue Tests. Institute fur Holzbiologie und Holzschutz, Bundesforschungsanstalt fur Forst - und Holzwirtshaft, Hamburg - Lohbrugge, Germany; Tedersoo L, et al. 2014. Science 346:e1256688 ((https://doi.org/10.1126/science.1256688)); Irinyi L, et al. 2015. Medical Mycology 53:313-337 ((https://doi.org/10.1093/mmy/myv008)); Põlme S, et al. 2020. Fungal Diversity 105:1-16 ((https://doi.org/10.1007/s13225-020-00466-2)); Massimo NC, et al. 2015. Microbial Ecology 70:61-76 ((https://doi.org/10.1007/s00248-014-0563-6)); The American Phytopathological Society Common Names of Plant Diseases ((http://www.apsnet.org/publications/commonnames/Pages/default.aspx)); https://www.cdc.gov ((https://www.cdc.gov)) |
| Otu01960 | 24 | k__Fungi(100);p__Ascomycota(100);c__Sordariomycetes(100);o__Hypocreales(100);f__Hypocreaceae(100);g__Hypomyces(100);s__Hypomyces_aurantius(100); | Hypomyces | 13 | Pathotroph | \|Fungal Parasite\| | Microfungus | NULL | Highly Probable | As mycoparasite (Põlme et al. 2020); Decay fungal material (Põlme et al. 2020); Sexual form with asexual forms spread across multiple genera | Põlme S, et al. 2020. Fungal Diversity 105:1-16 ((https://doi.org/10.1007/s13225-020-00466-2)); List compiled by John Plischke III ((https://goodmushroom.weebly.com/uploads/5/7/1/7/57176535/fungi_on_fungi_pdf_lq.pdf)) |
| Otu01962 | 24 | k__Fungi(100);p__Ascomycota(100);c__Saccharomycetes(100);o__Saccharomycetales(100);f__Metschnikowiaceae(100);g__Metschnikowia(100);s__Metschnikowia_pulcherrima(100); | Metschnikowia | 13 | Saprotroph | \|Undefined Saprotroph\| | Yeast | NULL | Probable | As nectar-tap_saprotroph (Põlme et al. 2020); Decay sugar-rich substrates (Põlme et al. 2020) | Põlme S, et al. 2020. Fungal Diversity 105:1-16 ((https://doi.org/10.1007/s13225-020-00466-2)) |
| Otu01963 | 24 | k__Fungi(100);p__Ascomycota(100);c__Dothideomycetes(100);o__Capnodiales(100);f__Teratosphaeriaceae(100);g__Devriesia(100);s__Devriesia_sp(100); | Devriesia | 13 | Pathotroph-Saprotroph | \|Plant Pathogen\|-Plant Saprotroph | Microfungus | NULL | Probable | As litter_saprotroph (Põlme et al. 2020); Decay leaf, fruit, seed (Põlme et al. 2020) | Tedersoo L, et al. 2014. Science 346:e1256688 ((https://doi.org/10.1126/science.1256688)); Põlme S, et al. 2020. Fungal Diversity 105:1-16 ((https://doi.org/10.1007/s13225-020-00466-2)) |
| Otu01965 | 24 | k__Fungi(100);p__Basidiomycota(100);c__Agaricomycetes(100);o__Polyporales(100);f__Fomitopsidaceae(100);g__Fibroporia(100);s__Fibroporia_vaillantii(100); | Fibroporia | 13 | Saprotroph | \|Wood Saprotroph\| | Corticioid | Brown Rot | Probable | As brown-rot (Põlme et al. 2020); Decay wood (Põlme et al. 2020) | Tedersoo L, et al. 2014. Science 346:e1256688 ((https://doi.org/10.1126/science.1256688)); Põlme S, et al. 2020. Fungal Diversity 105:1-16 ((https://doi.org/10.1007/s13225-020-00466-2)) |
| Otu01966 | 24 | k__Fungi(100);p__Basidiomycota(100);c__Agaricomycetes(100);o__Polyporales(100);f__Ganodermataceae(100);g__Ganoderma(100);s__Ganoderma_resinaceum(100); | Ganoderma | 13 | Pathotroph-Saprotroph | \|Plant Pathogen\|-Wood Saprotroph | Polyporoid | White Rot | Highly Probable | As white_rot (Põlme et al. 2020); As wood_pathogen (Põlme et al. 2020); Decay wood (Põlme et al. 2020); Lignicolous, causing sometimes agressive white rots of timber (Cannon & Kirk 2007) | Gilbertson RL, Ryvarden L. 1987-1987. North American Polypores. Fungiflora, Oslo (ISBN: 978-0945345060); Cannon PF, Kirk PM. 2007. Fungal Families of the World. CAB International, Cambridge (ISBN: 978-0851998275); Põlme S, et al. 2020. Fungal Diversity 105:1-16 ((https://doi.org/10.1007/s13225-020-00466-2)) |
| Otu01968 | 24 | k__Fungi(100);p__Ascomycota(100);c__Eurotiomycetes(100);o__Chaetothyriales(100);f__Herpotrichiellaceae(100);f__Herpotrichiellaceae_unclassified(96);f__Herpotrichiellaceae_unclassified(96); | Herpotrichiellaceae | 9 | Pathotroph-Saprotroph | Animal Pathogen-Fungal Parasite-Undefined Saprotroph | Facultative Yeast-Microfungus | NULL | Probable | Mostly saprobic on plants or other fungi, isolated from soil etc., and from medical sources (Cannon & Kirk 2007) | Cannon PF, Kirk PM. 2007. Fungal Families of the World. CAB International, Cambridge (ISBN: 978-0851998275) |
| Otu01969 | 24 | k__Fungi(100);p__Basidiomycota(100);c__Tremellomycetes(100);o__Tremellales(100);f__Tremellales_family_Incertae_sedis(100);g__Cryptococcus(100);s__Cryptococcus_aff_amylolyticus_AS_22501(100); | Cryptococcus | 13 | Pathotroph-Saprotroph-Symbiotroph | Animal Pathogen-Endophyte-Epiphyte-\|Undefined Saprotroph\| | Dimorphic Yeast-Yeast | NULL | Possible | As foliar_endophyte (Põlme et al. 2020); Decay leaf, fruit, seed, soil, dung, animal material (Põlme et al. 2020) | Andrews JH, et al. 1983. Phytopathology 73:228-234 ((https://doi.org/10.1094/Phyto-73-228)); Kurtzman CP, et al. (eds.) 2011. The Yeasts, a Taxonomic Study. Fifth Edition. Vols 1-3. Elsevier, San Diego (ISBN: 9780444521491); Põlme S, et al. 2020. Fungal Diversity 105:1-16 ((https://doi.org/10.1007/s13225-020-00466-2)) |
| Otu01970 | 24 | k__Fungi(100);p__Ascomycota(100);c__Saccharomycetes(100);o__Saccharomycetales(100);f__Metschnikowiaceae(100);g__Metschnikowia(100);s__Metschnikowia_pulcherrima(100); | Metschnikowia | 13 | Saprotroph | \|Undefined Saprotroph\| | Yeast | NULL | Probable | As nectar-tap_saprotroph (Põlme et al. 2020); Decay sugar-rich substrates (Põlme et al. 2020) | Põlme S, et al. 2020. Fungal Diversity 105:1-16 ((https://doi.org/10.1007/s13225-020-00466-2)) |
| Otu01972 | 24 | k__Fungi(100);p__Ascomycota(100);c__Saccharomycetes(100);o__Saccharomycetales(100);f__Metschnikowiaceae(100);g__Metschnikowia(100);g__Metschnikowia_unclassified(88); | Metschnikowia | 13 | Saprotroph | \|Undefined Saprotroph\| | Yeast | NULL | Probable | As nectar-tap_saprotroph (Põlme et al. 2020); Decay sugar-rich substrates (Põlme et al. 2020) | Põlme S, et al. 2020. Fungal Diversity 105:1-16 ((https://doi.org/10.1007/s13225-020-00466-2)) |
| Otu01973 | 24 | k__Fungi(100);p__Ascomycota(100);c__Saccharomycetes(100);o__Saccharomycetales(100);f__Metschnikowiaceae(100);g__Metschnikowia(100);s__Metschnikowia_pulcherrima(96); | Metschnikowia | 13 | Saprotroph | \|Undefined Saprotroph\| | Yeast | NULL | Probable | As nectar-tap_saprotroph (Põlme et al. 2020); Decay sugar-rich substrates (Põlme et al. 2020) | Põlme S, et al. 2020. Fungal Diversity 105:1-16 ((https://doi.org/10.1007/s13225-020-00466-2)) |
| Otu01928 | 24 | k__Fungi(100);p__Ascomycota(100);p__Ascomycota_unclassified(100);p__Ascomycota_unclassified(100);p__Ascomycota_unclassified(100);p__Ascomycota_unclassified(100);p__Ascomycota_unclassified(100); | - | - | - | - | - | - | - | Unassigned | - |
| Otu01932 | 24 | k__Fungi(100);p__Basidiomycota(100);c__Microbotryomycetes(100);o__Sporidiobolales(100);f__Sporidiobolales_family_Incertae_sedis(100);f__Sporidiobolales_family_Incertae_sedis_unclassified(63);f__Sporidiobolales_family_Incertae_sedis_unclassified(63); | - | - | - | - | - | - | - | Unassigned | - |
| Otu01933 | 24 | k__Fungi(100);k__Fungi_unclassified(84);k__Fungi_unclassified(84);k__Fungi_unclassified(84);k__Fungi_unclassified(84);k__Fungi_unclassified(84);k__Fungi_unclassified(84); | - | - | - | - | - | - | - | Unassigned | - |
| Otu01934 | 24 | k__Fungi(100);k__Fungi_unclassified(80);k__Fungi_unclassified(80);k__Fungi_unclassified(80);k__Fungi_unclassified(80);k__Fungi_unclassified(80);k__Fungi_unclassified(80); | - | - | - | - | - | - | - | Unassigned | - |
| Otu01939 | 24 | k__Fungi(100);k__Fungi_unclassified(55);k__Fungi_unclassified(55);k__Fungi_unclassified(55);k__Fungi_unclassified(55);k__Fungi_unclassified(55);k__Fungi_unclassified(55); | - | - | - | - | - | - | - | Unassigned | - |
| Otu01940 | 24 | k__Fungi(100);k__Fungi_unclassified(100);k__Fungi_unclassified(100);k__Fungi_unclassified(100);k__Fungi_unclassified(100);k__Fungi_unclassified(100);k__Fungi_unclassified(100); | - | - | - | - | - | - | - | Unassigned | - |
| Otu01941 | 24 | k__Fungi(100);p__Ascomycota(75);p__Ascomycota_unclassified(75);p__Ascomycota_unclassified(75);p__Ascomycota_unclassified(75);p__Ascomycota_unclassified(75);p__Ascomycota_unclassified(75); | - | - | - | - | - | - | - | Unassigned | - |
| Otu01942 | 24 | k__Fungi(100);k__Fungi_unclassified(55);k__Fungi_unclassified(55);k__Fungi_unclassified(55);k__Fungi_unclassified(55);k__Fungi_unclassified(55);k__Fungi_unclassified(55); | - | - | - | - | - | - | - | Unassigned | - |
| Otu01943 | 24 | k__Fungi(100);p__Ascomycota(100);p__Ascomycota_unclassified(96);p__Ascomycota_unclassified(96);p__Ascomycota_unclassified(96);p__Ascomycota_unclassified(96);p__Ascomycota_unclassified(96); | - | - | - | - | - | - | - | Unassigned | - |
| Otu01944 | 24 | k__Fungi(100);k__Fungi_unclassified(75);k__Fungi_unclassified(75);k__Fungi_unclassified(75);k__Fungi_unclassified(75);k__Fungi_unclassified(75);k__Fungi_unclassified(75); | - | - | - | - | - | - | - | Unassigned | - |
| Otu01946 | 24 | k__Fungi(100);p__Ascomycota(100);p__Ascomycota_unclassified(100);p__Ascomycota_unclassified(100);p__Ascomycota_unclassified(100);p__Ascomycota_unclassified(100);p__Ascomycota_unclassified(100); | - | - | - | - | - | - | - | Unassigned | - |
| Otu01951 | 24 | k__Fungi(100);p__Ascomycota(100);c__Dothideomycetes(100);o__Capnodiales(100);o__Capnodiales_unclassified(100);o__Capnodiales_unclassified(100);o__Capnodiales_unclassified(100); | - | - | - | - | - | - | - | Unassigned | - |
| Otu01953 | 24 | k__Fungi(100);p__Ascomycota(63);p__Ascomycota_unclassified(63);p__Ascomycota_unclassified(63);p__Ascomycota_unclassified(63);p__Ascomycota_unclassified(63);p__Ascomycota_unclassified(63); | - | - | - | - | - | - | - | Unassigned | - |
| Otu01955 | 24 | k__Fungi(100);p__Ascomycota(67);p__Ascomycota_unclassified(67);p__Ascomycota_unclassified(67);p__Ascomycota_unclassified(67);p__Ascomycota_unclassified(67);p__Ascomycota_unclassified(67); | - | - | - | - | - | - | - | Unassigned | - |
| Otu01961 | 24 | k__Fungi(100);k__Fungi_unclassified(71);k__Fungi_unclassified(71);k__Fungi_unclassified(71);k__Fungi_unclassified(71);k__Fungi_unclassified(71);k__Fungi_unclassified(71); | - | - | - | - | - | - | - | Unassigned | - |
| Otu01964 | 24 | k__Fungi(100);p__Basidiomycota(100);c__Tremellomycetes(67);c__Tremellomycetes_unclassified(67);c__Tremellomycetes_unclassified(67);c__Tremellomycetes_unclassified(67);c__Tremellomycetes_unclassified(67); | - | - | - | - | - | - | - | Unassigned | - |
| Otu01967 | 24 | k__Fungi(100);k__Fungi_unclassified(92);k__Fungi_unclassified(92);k__Fungi_unclassified(92);k__Fungi_unclassified(92);k__Fungi_unclassified(92);k__Fungi_unclassified(92); | - | - | - | - | - | - | - | Unassigned | - |
| Otu01971 | 24 | k__Fungi(100);p__Basidiomycota(100);c__Agaricomycetes(100);c__Agaricomycetes_unclassified(55);c__Agaricomycetes_unclassified(55);c__Agaricomycetes_unclassified(55);c__Agaricomycetes_unclassified(55); | - | - | - | - | - | - | - | Unassigned | - |
| Otu01975 | 23 | k__Fungi(100);p__Ascomycota(100);c__Saccharomycetes(100);o__Saccharomycetales(100);f__Metschnikowiaceae(100);g__Metschnikowia(100);s__Metschnikowia_pulcherrima(92); | Metschnikowia | 13 | Saprotroph | \|Undefined Saprotroph\| | Yeast | NULL | Probable | As nectar-tap_saprotroph (Põlme et al. 2020); Decay sugar-rich substrates (Põlme et al. 2020) | Põlme S, et al. 2020. Fungal Diversity 105:1-16 ((https://doi.org/10.1007/s13225-020-00466-2)) |
| Otu01976 | 23 | k__Fungi(100);p__Ascomycota(100);c__Saccharomycetes(100);o__Saccharomycetales(100);f__Metschnikowiaceae(100);g__Metschnikowia(100);s__Metschnikowia_pulcherrima(92); | Metschnikowia | 13 | Saprotroph | \|Undefined Saprotroph\| | Yeast | NULL | Probable | As nectar-tap_saprotroph (Põlme et al. 2020); Decay sugar-rich substrates (Põlme et al. 2020) | Põlme S, et al. 2020. Fungal Diversity 105:1-16 ((https://doi.org/10.1007/s13225-020-00466-2)) |
| Otu01977 | 23 | k__Fungi(100);p__Ascomycota(96);c__Saccharomycetes(96);o__Saccharomycetales(96);f__Metschnikowiaceae(96);g__Metschnikowia(96);s__Metschnikowia_pulcherrima(96); | Metschnikowia | 13 | Saprotroph | \|Undefined Saprotroph\| | Yeast | NULL | Probable | As nectar-tap_saprotroph (Põlme et al. 2020); Decay sugar-rich substrates (Põlme et al. 2020) | Põlme S, et al. 2020. Fungal Diversity 105:1-16 ((https://doi.org/10.1007/s13225-020-00466-2)) |
| Otu01978 | 23 | k__Fungi(100);p__Ascomycota(100);c__Sordariomycetes(100);o__Xylariales(100);f__Xylariaceae(100);f__Xylariaceae_unclassified(83);f__Xylariaceae_unclassified(83); | Xylariaceae | 9 | Pathotroph-Saprotroph-Symbiotroph | Dung Saprotroph-Endophyte-Plant Pathogen-Undefined Saprotroph | Xylarioid | NULL | Probable | Saprobes or weak pathogens, many endophytes, mainly in wood and bark, occasionally in soil and dung (Cannon & Kirk 2007) | Cannon PF, Kirk PM. 2007. Fungal Families of the World. CAB International, Cambridge (ISBN: 978-0851998275) |
| Otu01979 | 23 | k__Fungi(100);p__Ascomycota(100);c__Saccharomycetes(100);o__Saccharomycetales(100);f__Metschnikowiaceae(100);g__Metschnikowia(100);s__Metschnikowia_pulcherrima(87); | Metschnikowia | 13 | Saprotroph | \|Undefined Saprotroph\| | Yeast | NULL | Probable | As nectar-tap_saprotroph (Põlme et al. 2020); Decay sugar-rich substrates (Põlme et al. 2020) | Põlme S, et al. 2020. Fungal Diversity 105:1-16 ((https://doi.org/10.1007/s13225-020-00466-2)) |
| Otu01981 | 23 | k__Fungi(100);p__Ascomycota(100);c__Saccharomycetes(100);o__Saccharomycetales(100);f__Metschnikowiaceae(100);g__Metschnikowia(100);s__Metschnikowia_pulcherrima(100); | Metschnikowia | 13 | Saprotroph | \|Undefined Saprotroph\| | Yeast | NULL | Probable | As nectar-tap_saprotroph (Põlme et al. 2020); Decay sugar-rich substrates (Põlme et al. 2020) | Põlme S, et al. 2020. Fungal Diversity 105:1-16 ((https://doi.org/10.1007/s13225-020-00466-2)) |
| Otu01982 | 23 | k__Fungi(100);p__Basidiomycota(100);c__Microbotryomycetes(100);o__Sporidiobolales(100);f__Sporidiobolales_family_Incertae_sedis(100);g__Sporobolomyces(100);s__Sporobolomyces_lactosus(100); | Sporobolomyces | 13 | Pathotroph-Saprotroph | \|Fungal Parasite\|-Plant Saprotroph | Yeast | NULL | Probable | As litter_saprotroph (Põlme et al. 2020); As mycoparasite (Põlme et al. 2020); Decay fungal material (Põlme et al. 2020) | Tedersoo L, et al. 2014. Science 346:e1256688 ((https://doi.org/10.1126/science.1256688)); Purahong W, et al. 2016. Molecular Ecology 25:4059-4074 ((https://doi.org/10.1111/mec.13739)); Põlme S, et al. 2020. Fungal Diversity 105:1-16 ((https://doi.org/10.1007/s13225-020-00466-2)) |
| Otu01984 | 23 | k__Fungi(100);p__Ascomycota(100);c__Sordariomycetes(100);o__Hypocreales(100);f__Ophiocordycipitaceae(100);g__Elaphocordyceps(100);g__Elaphocordyceps_unclassified(100); | Elaphocordyceps | 13 | Pathotroph | Fungal Parasite | Secotioid-Spathuloid | NULL | Highly Probable | NULL | Samuels GJ. 2007. Self-published; Sung G-H, et al. 2007. Studies in Mycology 57:5-59 ((https://www.studiesinmycology.org/index.php/issue/59-studies-in-mycology-no-57)); List compiled by John Plischke III ((https://goodmushroom.weebly.com/uploads/5/7/1/7/57176535/fungi_on_fungi_pdf_lq.pdf)) |
| Otu01985 | 23 | k__Fungi(100);p__Ascomycota(100);c__Dothideomycetes(100);o__Pleosporales(100);f__Venturiaceae(100);g__Venturia(96);g__Venturia_unclassified(96); | Venturia | 13 | Pathotroph-Pathotroph-Saprotroph | \|Plant Pathogen\|-Plant Saprotroph | Microfungus | NULL | Probable | As litter_saprotroph (Põlme et al. 2020); Decay leaf, fruit, seed (Põlme et al. 2020) | Tedersoo L, et al. 2014. Science 346:e1256688 ((https://doi.org/10.1126/science.1256688)); Põlme S, et al. 2020. Fungal Diversity 105:1-16 ((https://doi.org/10.1007/s13225-020-00466-2)) |
| Otu01987 | 23 | k__Fungi(100);p__Ascomycota(100);c__Saccharomycetes(100);o__Saccharomycetales(100);f__Metschnikowiaceae(100);g__Metschnikowia(100);s__Metschnikowia_pulcherrima(100); | Metschnikowia | 13 | Saprotroph | \|Undefined Saprotroph\| | Yeast | NULL | Probable | As nectar-tap_saprotroph (Põlme et al. 2020); Decay sugar-rich substrates (Põlme et al. 2020) | Põlme S, et al. 2020. Fungal Diversity 105:1-16 ((https://doi.org/10.1007/s13225-020-00466-2)) |
| Otu01990 | 23 | k__Fungi(100);p__Basidiomycota(96);c__Microbotryomycetes(96);o__Sporidiobolales(96);f__Sporidiobolales_family_Incertae_sedis(96);g__Sporobolomyces(96);s__Sporobolomyces_lactosus(96); | Sporobolomyces | 13 | Pathotroph-Saprotroph | \|Fungal Parasite\|-Plant Saprotroph | Yeast | NULL | Probable | As litter_saprotroph (Põlme et al. 2020); As mycoparasite (Põlme et al. 2020); Decay fungal material (Põlme et al. 2020) | Tedersoo L, et al. 2014. Science 346:e1256688 ((https://doi.org/10.1126/science.1256688)); Purahong W, et al. 2016. Molecular Ecology 25:4059-4074 ((https://doi.org/10.1111/mec.13739)); Põlme S, et al. 2020. Fungal Diversity 105:1-16 ((https://doi.org/10.1007/s13225-020-00466-2)) |
| Otu01991 | 23 | k__Fungi(100);p__Ascomycota(100);c__Saccharomycetes(100);o__Saccharomycetales(100);f__Metschnikowiaceae(100);g__Metschnikowia(100);s__Metschnikowia_pulcherrima(92); | Metschnikowia | 13 | Saprotroph | \|Undefined Saprotroph\| | Yeast | NULL | Probable | As nectar-tap_saprotroph (Põlme et al. 2020); Decay sugar-rich substrates (Põlme et al. 2020) | Põlme S, et al. 2020. Fungal Diversity 105:1-16 ((https://doi.org/10.1007/s13225-020-00466-2)) |
| Otu01997 | 23 | k__Fungi(100);p__Ascomycota(100);c__Saccharomycetes(100);o__Saccharomycetales(100);f__Metschnikowiaceae(100);g__Metschnikowia(100);s__Metschnikowia_pulcherrima(100); | Metschnikowia | 13 | Saprotroph | \|Undefined Saprotroph\| | Yeast | NULL | Probable | As nectar-tap_saprotroph (Põlme et al. 2020); Decay sugar-rich substrates (Põlme et al. 2020) | Põlme S, et al. 2020. Fungal Diversity 105:1-16 ((https://doi.org/10.1007/s13225-020-00466-2)) |
| Otu01998 | 23 | k__Fungi(100);p__Ascomycota(100);c__Saccharomycetes(100);o__Saccharomycetales(100);f__Metschnikowiaceae(100);g__Metschnikowia(100);s__Metschnikowia_pulcherrima(92); | Metschnikowia | 13 | Saprotroph | \|Undefined Saprotroph\| | Yeast | NULL | Probable | As nectar-tap_saprotroph (Põlme et al. 2020); Decay sugar-rich substrates (Põlme et al. 2020) | Põlme S, et al. 2020. Fungal Diversity 105:1-16 ((https://doi.org/10.1007/s13225-020-00466-2)) |
| Otu01999 | 23 | k__Fungi(100);p__Basidiomycota(100);c__Tremellomycetes(100);o__Tremellales(100);f__unclassified_Tremellales(100);g__unclassified_Tremellales(100);s__Tremellales_sp(100); | Tremellales | 7 | Pathotroph-Saprotroph-Symbiotroph | Fungal Parasite-Undefined Saprotroph | Tremelloid-Yeast | NULL | Possible | The order contains both teleomorphic and anamorphic species, most of the latter being yeasts | Sterkenburg E, et al. 2015. New Phytologist 207:1145-1158 ((https://doi.org/10.1111/nph.13426)) |
| Otu02000 | 23 | k__Fungi(100);p__Ascomycota(100);c__Eurotiomycetes(100);o__Chaetothyriales(100);f__Herpotrichiellaceae(100);g__Cladophialophora(100);g__Cladophialophora_unclassified(74); | Cladophialophora | 13 | Pathotroph-Saprotroph-Symbiotroph | Animal Pathogen-Endophyte-Fungal Parasite-\|Undefined Saprotroph\| | Dimorphic-Microfungus | NULL | Probable | As mycoparasite (Põlme et al. 2020); As root_endophyte_dark_septate (Põlme et al. 2020); As soil_saprotroph (Põlme et al. 2020); Decay soil (Põlme et al. 2020) | James TY, et al. 2006. Nature 443:818-822 ((https://doi.org/10.1038/nature05110)); Põlme S, et al. 2020. Fungal Diversity 105:1-16 ((https://doi.org/10.1007/s13225-020-00466-2)) |
| Otu02001 | 23 | k__Fungi(100);p__Basidiomycota(100);c__Microbotryomycetes(100);o__Sporidiobolales(100);f__Sporidiobolales_family_Incertae_sedis(100);g__Rhodotorula(100);s__Rhodotorula_taiwanensis(100); | Rhodotorula | 13 | Pathotroph-Saprotroph-Symbiotroph | Animal Pathogen-Animal Symbiotroph-Endophyte-Plant Pathogen-\|Undefined Saprotroph\| | Yeast | NULL | Probable | As foliar_endophyte (Põlme et al. 2020); Opportunistic human pathogen | Põlme S, et al. 2020. Fungal Diversity 105:1-16 ((https://doi.org/10.1007/s13225-020-00466-2)); https://www.cdc.gov ((https://www.cdc.gov)) |
| Otu02003 | 23 | k__Fungi(100);p__Ascomycota(100);c__Dothideomycetes(100);o__Capnodiales(100);f__Davidiellaceae(100);g__Cladosporium(100);s__Cladosporium_sp(100); | Cladosporium | 13 | Pathotroph-Saprotroph-Symbiotroph | Animal Pathogen-Endophyte-Lichen Parasite-Plant Pathogen-\|Plant Saprotroph\|-Wood Saprotroph | Microfungus | NULL | Possible | As foliar_endophyte (Põlme et al. 2020); As litter_saprotroph (Põlme et al. 2020); Decay wood, leaf, fruit, seed, soil, dung, animal material (Põlme et al. 2020) | Seehann G, et al. 1975. List of Fungi in Soft-Rot Tissue Tests. Institute fur Holzbiologie und Holzschutz, Bundesforschungsanstalt fur Forst - und Holzwirtshaft, Hamburg - Lohbrugge, Germany; Lawrey JD, Diederich P. 2016. http://www.lichenicolous.net (accessed 24 May 2017) ((http://www.lichenicolous.net)); Põlme S, et al. 2020. Fungal Diversity 105:1-16 ((https://doi.org/10.1007/s13225-020-00466-2)); The American Phytopathological Society Common Names of Plant Diseases ((http://www.apsnet.org/publications/commonnames/Pages/default.aspx)) |
| Otu02004 | 23 | k__Fungi(100);p__Basidiomycota(100);c__Exobasidiomycetes(100);o__Exobasidiales(100);f__Exobasidiaceae(100);g__Exobasidium(100);g__Exobasidium_unclassified(100); | Exobasidium | 13 | Pathotroph | \|Plant Pathogen\| | Leaf Blister-Leaf Gall-Leaf Spot | NULL | Probable | NULL | Tedersoo L, et al. 2014. Science 346:e1256688 ((https://doi.org/10.1126/science.1256688)); Põlme S, et al. 2020. Fungal Diversity 105:1-16 ((https://doi.org/10.1007/s13225-020-00466-2)) |
| Otu02005 | 23 | k__Fungi(100);p__Ascomycota(100);c__Saccharomycetes(100);o__Saccharomycetales(100);f__Metschnikowiaceae(100);g__Metschnikowia(100);s__Metschnikowia_pulcherrima(96); | Metschnikowia | 13 | Saprotroph | \|Undefined Saprotroph\| | Yeast | NULL | Probable | As nectar-tap_saprotroph (Põlme et al. 2020); Decay sugar-rich substrates (Põlme et al. 2020) | Põlme S, et al. 2020. Fungal Diversity 105:1-16 ((https://doi.org/10.1007/s13225-020-00466-2)) |
| Otu02007 | 23 | k__Fungi(100);p__Ascomycota(100);c__Saccharomycetes(100);o__Saccharomycetales(100);f__Metschnikowiaceae(100);g__Metschnikowia(100);s__Metschnikowia_pulcherrima(79); | Metschnikowia | 13 | Saprotroph | \|Undefined Saprotroph\| | Yeast | NULL | Probable | As nectar-tap_saprotroph (Põlme et al. 2020); Decay sugar-rich substrates (Põlme et al. 2020) | Põlme S, et al. 2020. Fungal Diversity 105:1-16 ((https://doi.org/10.1007/s13225-020-00466-2)) |
| Otu02008 | 23 | k__Fungi(100);p__Ascomycota(100);c__Saccharomycetes(100);o__Saccharomycetales(100);o__Saccharomycetales_unclassified(100);o__Saccharomycetales_unclassified(100);o__Saccharomycetales_unclassified(100); | Saccharomycetales | 7 | Saprotroph | Undefined Saprotroph | Yeast | NULL | Possible | NULL | Sterkenburg E, et al. 2015. New Phytologist 207:1145-1158 ((https://doi.org/10.1111/nph.13426)) |
| Otu02009 | 23 | k__Fungi(100);p__Basidiomycota(100);c__Agaricomycetes(100);o__Hymenochaetales(100);f__Schizoporaceae(100);g__Schizopora(100);s__Schizopora_paradoxa(100); | Schizopora | 13 | Pathotroph-Saprotroph | Plant Pathogen-\|Wood Saprotroph\| | Corticioid | White Rot | Highly Probable | As white_rot (Põlme et al. 2020); As wood_pathogen (Põlme et al. 2020); Decay wood (Põlme et al. 2020) | Gilbertson RL, Ryvarden L. 1987-1987. North American Polypores. Fungiflora, Oslo (ISBN: 978-0945345060); Põlme S, et al. 2020. Fungal Diversity 105:1-16 ((https://doi.org/10.1007/s13225-020-00466-2)) |
| Otu02012 | 23 | k__Fungi(100);p__Basidiomycota(100);c__Agaricomycetes(100);o__Trechisporales(100);f__Hydnodontaceae(100);g__Trechispora(100);s__Trechispora_sp(100); | Trechispora | 13 | Saprotroph | \|Wood Saprotroph\| | Corticioid | White Rot | Highly Probable | As white_rot (Põlme et al. 2020); Decay wood (Põlme et al. 2020) | Gilbertson RL, Ryvarden L. 1987-1987. North American Polypores. Fungiflora, Oslo (ISBN: 978-0945345060); Rinaldi AC, et al. 2008. Fungal Diversity 33:1-45 ((http://www.fungaldiversity.org/fdp/sfdp/33-1.pdf)) (as Uncertain); Tedersoo L, et al. 2010. Mycorrhiza 20:217-263 ((https://doi.org/10.1007/s00572-009-0274-x)) (as not ECM).; Põlme S, et al. 2020. Fungal Diversity 105:1-16 ((https://doi.org/10.1007/s13225-020-00466-2)) |
| Otu02014 | 23 | k__Fungi(100);p__Ascomycota(100);c__Dothideomycetes(100);o__Capnodiales(100);f__Mycosphaerellaceae(100);g__Zasmidium(100);g__Zasmidium_unclassified(87); | Zasmidium | 13 | Pathotroph-Saprotroph | \|Plant Pathogen\|-Plant Saprotroph-Undefined Saprotroph | Microfungus | NULL | Probable | As litter_saprotroph (Põlme et al. 2020); Decay leaf, fruit, seed (Põlme et al. 2020) | Tedersoo L, et al. 2014. Science 346:e1256688 ((https://doi.org/10.1126/science.1256688)); Põlme S, et al. 2020. Fungal Diversity 105:1-16 ((https://doi.org/10.1007/s13225-020-00466-2)) |
| Otu02015 | 23 | k__Fungi(100);p__Basidiomycota(100);c__Agaricomycetes(100);o__Polyporales(100);f__Polyporales_family_Incertae_sedis(100);g__Phlebiella(100);s__Phlebiella_christiansenii(100); | Phlebiella | 13 | Saprotroph | Undefined Saprotroph-\|Wood Saprotroph\| | Corticioid | White Rot | Probable | As white_rot (Põlme et al. 2020); Decay wood (Põlme et al. 2020) | Tedersoo L, et al. 2014. Science 346:e1256688 ((https://doi.org/10.1126/science.1256688)); Põlme S, et al. 2020. Fungal Diversity 105:1-16 ((https://doi.org/10.1007/s13225-020-00466-2)) |
| Otu02017 | 23 | k__Fungi(100);p__Basidiomycota(100);c__Agaricomycetes(100);o__Agaricales(100);f__Psathyrellaceae(100);g__Coprinopsis(100);s__Coprinopsis_pseudofriesii(100); | Coprinopsis | 13 | Saprotroph | \|Undefined Saprotroph\| | Agaricoid | NULL | Probable | As soil_saprotroph (Põlme et al. 2020); Decay soil (Põlme et al. 2020) | Tedersoo L, et al. 2014. Science 346:e1256688 ((https://doi.org/10.1126/science.1256688)); Põlme S, et al. 2020. Fungal Diversity 105:1-16 ((https://doi.org/10.1007/s13225-020-00466-2)) |
| Otu02018 | 23 | k__Fungi(100);p__Basidiomycota(100);c__Agaricomycetes(100);o__Polyporales(100);f__Meruliaceae(100);g__Phlebia(100);s__Phlebia_sp(100); | Phlebia | 13 | Saprotroph | \|Wood Saprotroph\| | Corticioid | White Rot | Highly Probable | As white_rot (Põlme et al. 2020); Decay wood (Põlme et al. 2020) | Kirk PM et al. 2008. Dictionary of the Fungi. Tenth Edition. CAB International, Wallingford (ISBN: 978-0851998268); Põlme S, et al. 2020. Fungal Diversity 105:1-16 ((https://doi.org/10.1007/s13225-020-00466-2)) |
| Otu02022 | 23 | k__Fungi(100);p__Basidiomycota(100);c__Tremellomycetes(100);o__Tremellales(100);f__Tremellales_family_Incertae_sedis(100);g__Cryptococcus(100);g__Cryptococcus_unclassified(100); | Cryptococcus | 13 | Pathotroph-Saprotroph-Symbiotroph | Animal Pathogen-Endophyte-Epiphyte-\|Undefined Saprotroph\| | Dimorphic Yeast-Yeast | NULL | Possible | As foliar_endophyte (Põlme et al. 2020); Decay leaf, fruit, seed, soil, dung, animal material (Põlme et al. 2020) | Andrews JH, et al. 1983. Phytopathology 73:228-234 ((https://doi.org/10.1094/Phyto-73-228)); Kurtzman CP, et al. (eds.) 2011. The Yeasts, a Taxonomic Study. Fifth Edition. Vols 1-3. Elsevier, San Diego (ISBN: 9780444521491); Põlme S, et al. 2020. Fungal Diversity 105:1-16 ((https://doi.org/10.1007/s13225-020-00466-2)) |
| Otu02023 | 23 | k__Fungi(100);p__Ascomycota(100);c__Saccharomycetes(100);o__Saccharomycetales(100);f__Metschnikowiaceae(100);g__Metschnikowia(100);s__Metschnikowia_pulcherrima(92); | Metschnikowia | 13 | Saprotroph | \|Undefined Saprotroph\| | Yeast | NULL | Probable | As nectar-tap_saprotroph (Põlme et al. 2020); Decay sugar-rich substrates (Põlme et al. 2020) | Põlme S, et al. 2020. Fungal Diversity 105:1-16 ((https://doi.org/10.1007/s13225-020-00466-2)) |
| Otu02024 | 23 | k__Fungi(100);p__Basidiomycota(100);c__Tremellomycetes(100);o__Tremellales(100);f__Tremellales_family_Incertae_sedis(100);g__Fibulobasidium(100);s__Fibulobasidium_inconspicuum(100); | Fibulobasidium | 13 | Saprotroph | Undefined Saprotroph-\|Wood Saprotroph\| | Dimorphic-Facultative Yeast | NULL | Highly Probable | Decay wood (Põlme et al. 2020) | Kurtzman CP, et al. (eds.) 2011. The Yeasts, a Taxonomic Study. Fifth Edition. Vols 1-3. Elsevier, San Diego (ISBN: 9780444521491); Põlme S, et al. 2020. Fungal Diversity 105:1-16 ((https://doi.org/10.1007/s13225-020-00466-2)) |
| Otu02025 | 23 | k__Fungi(100);p__Basidiomycota(100);c__Tremellomycetes(100);o__Tremellales(100);f__Tremellales_family_Incertae_sedis(100);g__Cryptococcus(100);s__Cryptococcus_dimennae(100); | Cryptococcus | 13 | Pathotroph-Saprotroph-Symbiotroph | Animal Pathogen-Endophyte-Epiphyte-\|Undefined Saprotroph\| | Dimorphic Yeast-Yeast | NULL | Possible | As foliar_endophyte (Põlme et al. 2020); Decay leaf, fruit, seed, soil, dung, animal material (Põlme et al. 2020) | Andrews JH, et al. 1983. Phytopathology 73:228-234 ((https://doi.org/10.1094/Phyto-73-228)); Kurtzman CP, et al. (eds.) 2011. The Yeasts, a Taxonomic Study. Fifth Edition. Vols 1-3. Elsevier, San Diego (ISBN: 9780444521491); Põlme S, et al. 2020. Fungal Diversity 105:1-16 ((https://doi.org/10.1007/s13225-020-00466-2)) |
| Otu02026 | 23 | k__Fungi(100);p__Ascomycota(100);c__Leotiomycetes(100);o__Helotiales(100);f__Helotiales_family_Incertae_sedis(100);g__Xenopolyscytalum(100);s__Xenopolyscytalum_sp(100); | Xenopolyscytalum | 13 | Saprotroph-Symbiotroph | Endophyte-\|Plant Saprotroph\|-Undefined Saprotroph | Microfungus | NULL | Probable | As foliar_endophyte (Põlme et al. 2020); As litter_saprotroph (Põlme et al. 2020); Decay leaf, fruit, seed (Põlme et al. 2020) | Tedersoo L, et al. 2014. Science 346:e1256688 ((https://doi.org/10.1126/science.1256688)); Põlme S, et al. 2020. Fungal Diversity 105:1-16 ((https://doi.org/10.1007/s13225-020-00466-2)) |
| Otu02028 | 23 | k__Fungi(100);p__Basidiomycota(100);c__Tremellomycetes(100);o__Tremellales(87);f__Tremellales_family_Incertae_sedis(87);g__Cryptococcus(87);s__Cryptococcus_aerius(87); | Cryptococcus | 13 | Pathotroph-Saprotroph-Symbiotroph | Animal Pathogen-Endophyte-Epiphyte-\|Undefined Saprotroph\| | Dimorphic Yeast-Yeast | NULL | Possible | As foliar_endophyte (Põlme et al. 2020); Decay leaf, fruit, seed, soil, dung, animal material (Põlme et al. 2020) | Andrews JH, et al. 1983. Phytopathology 73:228-234 ((https://doi.org/10.1094/Phyto-73-228)); Kurtzman CP, et al. (eds.) 2011. The Yeasts, a Taxonomic Study. Fifth Edition. Vols 1-3. Elsevier, San Diego (ISBN: 9780444521491); Põlme S, et al. 2020. Fungal Diversity 105:1-16 ((https://doi.org/10.1007/s13225-020-00466-2)) |
| Otu02031 | 23 | k__Fungi(100);p__Chytridiomycota(100);c__Chytridiomycetes(100);o__Spizellomycetales(100);f__Spizellomycetaceae(100);f__Spizellomycetaceae_unclassified(100);f__Spizellomycetaceae_unclassified(100); | Spizellomycetaceae | 9 | Pathotroph | Plant Pathogen-Undefined Parasite-Undefined Saprotroph | Microfungus | NULL | Probable | Common in the soil and also as parasites of a range of soil organisms and plants (Cannon & Kirk 2007) | Cannon PF, Kirk PM. 2007. Fungal Families of the World. CAB International, Cambridge (ISBN: 978-0851998275) |
| Otu02032 | 23 | k__Fungi(100);p__Ascomycota(100);c__Saccharomycetes(100);o__Saccharomycetales(100);f__Metschnikowiaceae(100);g__Metschnikowia(100);s__Metschnikowia_pulcherrima(100); | Metschnikowia | 13 | Saprotroph | \|Undefined Saprotroph\| | Yeast | NULL | Probable | As nectar-tap_saprotroph (Põlme et al. 2020); Decay sugar-rich substrates (Põlme et al. 2020) | Põlme S, et al. 2020. Fungal Diversity 105:1-16 ((https://doi.org/10.1007/s13225-020-00466-2)) |
| Otu02033 | 23 | k__Fungi(100);p__Ascomycota(100);c__Saccharomycetes(100);o__Saccharomycetales(100);f__Metschnikowiaceae(100);g__Metschnikowia(100);s__Metschnikowia_pulcherrima(100); | Metschnikowia | 13 | Saprotroph | \|Undefined Saprotroph\| | Yeast | NULL | Probable | As nectar-tap_saprotroph (Põlme et al. 2020); Decay sugar-rich substrates (Põlme et al. 2020) | Põlme S, et al. 2020. Fungal Diversity 105:1-16 ((https://doi.org/10.1007/s13225-020-00466-2)) |
| Otu02034 | 23 | k__Fungi(100);p__Basidiomycota(100);c__Agaricomycetes(100);o__Trechisporales(100);f__Hydnodontaceae(100);g__Trechispora(100);s__Trechispora_sp(100); | Trechispora | 13 | Saprotroph | \|Wood Saprotroph\| | Corticioid | White Rot | Highly Probable | As white_rot (Põlme et al. 2020); Decay wood (Põlme et al. 2020) | Gilbertson RL, Ryvarden L. 1987-1987. North American Polypores. Fungiflora, Oslo (ISBN: 978-0945345060); Rinaldi AC, et al. 2008. Fungal Diversity 33:1-45 ((http://www.fungaldiversity.org/fdp/sfdp/33-1.pdf)) (as Uncertain); Tedersoo L, et al. 2010. Mycorrhiza 20:217-263 ((https://doi.org/10.1007/s00572-009-0274-x)) (as not ECM).; Põlme S, et al. 2020. Fungal Diversity 105:1-16 ((https://doi.org/10.1007/s13225-020-00466-2)) |
| Otu01974 | 23 | k__Fungi(100);k__Fungi_unclassified(74);k__Fungi_unclassified(74);k__Fungi_unclassified(74);k__Fungi_unclassified(74);k__Fungi_unclassified(74);k__Fungi_unclassified(74); | - | - | - | - | - | - | - | Unassigned | - |
| Otu01980 | 23 | k__Fungi(100);k__Fungi_unclassified(100);k__Fungi_unclassified(100);k__Fungi_unclassified(100);k__Fungi_unclassified(100);k__Fungi_unclassified(100);k__Fungi_unclassified(100); | - | - | - | - | - | - | - | Unassigned | - |
| Otu01983 | 23 | k__Fungi(100);p__Ascomycota(100);c__Dothideomycetes(100);o__Pleosporales(100);o__Pleosporales_unclassified(96);o__Pleosporales_unclassified(96);o__Pleosporales_unclassified(96); | - | - | - | - | - | - | - | Unassigned | - |
| Otu01986 | 23 | k__Fungi(100);p__Ascomycota(100);c__Lecanoromycetes(100);c__Lecanoromycetes_unclassified(61);c__Lecanoromycetes_unclassified(61);c__Lecanoromycetes_unclassified(61);c__Lecanoromycetes_unclassified(61); | - | - | - | - | - | - | - | Unassigned | - |
| Otu01988 | 23 | k__Fungi(100);p__Ascomycota(100);p__Ascomycota_unclassified(87);p__Ascomycota_unclassified(87);p__Ascomycota_unclassified(87);p__Ascomycota_unclassified(87);p__Ascomycota_unclassified(87); | - | - | - | - | - | - | - | Unassigned | - |
| Otu01989 | 23 | k__Fungi(100);p__Basidiomycota(100);p__Basidiomycota_unclassified(100);p__Basidiomycota_unclassified(100);p__Basidiomycota_unclassified(100);p__Basidiomycota_unclassified(100);p__Basidiomycota_unclassified(100); | - | - | - | - | - | - | - | Unassigned | - |
| Otu01992 | 23 | k__Fungi(100);p__Basidiomycota(100);c__Agaricomycetes(100);o__Cantharellales(100);f__unclassified_Cantharellales(83);g__unclassified_Cantharellales(83);s__Cantharellales_sp(83); | - | - | - | - | - | - | - | Unassigned | - |
| Otu01993 | 23 | k__Fungi(100);p__Ascomycota(100);c__Sordariomycetes(100);o__Diaporthales(100);o__Diaporthales_unclassified(96);o__Diaporthales_unclassified(96);o__Diaporthales_unclassified(96); | - | - | - | - | - | - | - | Unassigned | - |
| Otu01994 | 23 | k__Fungi(100);p__Ascomycota(100);c__Dothideomycetes(100);o__Pleosporales(96);o__Pleosporales_unclassified(96);o__Pleosporales_unclassified(96);o__Pleosporales_unclassified(96); | - | - | - | - | - | - | - | Unassigned | - |
| Otu01995 | 23 | k__Fungi(100);p__Ascomycota(100);c__Sordariomycetes(100);o__Sordariales(100);o__Sordariales_unclassified(87);o__Sordariales_unclassified(87);o__Sordariales_unclassified(87); | - | - | - | - | - | - | - | Unassigned | - |
| Otu01996 | 23 | k__Fungi(100);p__Ascomycota(70);p__Ascomycota_unclassified(70);p__Ascomycota_unclassified(70);p__Ascomycota_unclassified(70);p__Ascomycota_unclassified(70);p__Ascomycota_unclassified(70); | - | - | - | - | - | - | - | Unassigned | - |
| Otu02002 | 23 | k__Fungi(100);k__Fungi_unclassified(61);k__Fungi_unclassified(61);k__Fungi_unclassified(61);k__Fungi_unclassified(61);k__Fungi_unclassified(61);k__Fungi_unclassified(61); | - | - | - | - | - | - | - | Unassigned | - |
| Otu02006 | 23 | k__Fungi(100);p__Ascomycota(100);p__Ascomycota_unclassified(100);p__Ascomycota_unclassified(100);p__Ascomycota_unclassified(100);p__Ascomycota_unclassified(100);p__Ascomycota_unclassified(100); | - | - | - | - | - | - | - | Unassigned | - |
| Otu02010 | 23 | k__Fungi(100);p__Basidiomycota(100);c__Agaricomycetes(100);c__Agaricomycetes_unclassified(100);c__Agaricomycetes_unclassified(100);c__Agaricomycetes_unclassified(100);c__Agaricomycetes_unclassified(100); | - | - | - | - | - | - | - | Unassigned | - |
| Otu02011 | 23 | k__Fungi(100);k__Fungi_unclassified(61);k__Fungi_unclassified(61);k__Fungi_unclassified(61);k__Fungi_unclassified(61);k__Fungi_unclassified(61);k__Fungi_unclassified(61); | - | - | - | - | - | - | - | Unassigned | - |
| Otu02013 | 23 | k__Fungi(100);k__Fungi_unclassified(83);k__Fungi_unclassified(83);k__Fungi_unclassified(83);k__Fungi_unclassified(83);k__Fungi_unclassified(83);k__Fungi_unclassified(83); | - | - | - | - | - | - | - | Unassigned | - |
| Otu02016 | 23 | k__Fungi(100);k__Fungi_unclassified(57);k__Fungi_unclassified(57);k__Fungi_unclassified(57);k__Fungi_unclassified(57);k__Fungi_unclassified(57);k__Fungi_unclassified(57); | - | - | - | - | - | - | - | Unassigned | - |

| | | | | | | | | | | | |
|---|---|---|---|---|---|---|---|---|---|---|---|
| Otu02019 | 23 | k__Fungi(100);p__Ascomycota(100);c__Dothideomycetes(100);o__Capnodiales(100);o__Capnodiales_unclassified(100);o__Capnodiales_unclassified(100);o__Capnodiales_unclassified(100); | - | - | - | - | - | - | - | Unassigned | |
| Otu02020 | 23 | k__Fungi(100);k__Fungi_unclassified(100);k__Fungi_unclassified(100);k__Fungi_unclassified(100);k__Fungi_unclassified(100);k__Fungi_unclassified(100);k__Fungi_unclassified(100); | - | - | - | - | - | - | - | Unassigned | |
| Otu02021 | 23 | k__Fungi(100);k__Fungi_unclassified(87);k__Fungi_unclassified(87);k__Fungi_unclassified(87);k__Fungi_unclassified(87);k__Fungi_unclassified(87);k__Fungi_unclassified(87); | - | - | - | - | - | - | - | Unassigned | |
| Otu02027 | 23 | k__Fungi(100);p__Basidiomycota(100);p__Basidiomycota_unclassified(100);p__Basidiomycota_unclassified(100);p__Basidiomycota_unclassified(100);p__Basidiomycota_unclassified(100);p__Basidiomycota_unclassified(100); | - | - | - | - | - | - | - | Unassigned | |
| Otu02029 | 23 | k__Fungi(100);p__Ascomycota(66);p__Ascomycota_unclassified(66);p__Ascomycota_unclassified(66);p__Ascomycota_unclassified(66);p__Ascomycota_unclassified(66);p__Ascomycota_unclassified(66); | - | - | - | - | - | - | - | Unassigned | |
| Otu02030 | 23 | k__Fungi(100);p__Ascomycota(96);p__Ascomycota_unclassified(96);p__Ascomycota_unclassified(96);p__Ascomycota_unclassified(96);p__Ascomycota_unclassified(96);p__Ascomycota_unclassified(96); | - | - | - | - | - | - | - | Unassigned | |
| Otu02035 | 23 | k__Fungi(100);p__Ascomycota(79);p__Ascomycota_unclassified(79);p__Ascomycota_unclassified(79);p__Ascomycota_unclassified(79);p__Ascomycota_unclassified(79);p__Ascomycota_unclassified(79); | - | - | - | - | - | - | - | Unassigned | |
| Otu02036 | 23 | k__Fungi(100);k__Fungi_unclassified(79);k__Fungi_unclassified(79);k__Fungi_unclassified(79);k__Fungi_unclassified(79);k__Fungi_unclassified(79);k__Fungi_unclassified(79); | - | - | - | - | - | - | - | Unassigned | |
| Otu02038 | 22 | k__Fungi(100);p__Ascomycota(100);c__Eurotiomycetes(100);o__Verrucariales(100);f__Verrucariaceae(100);f__Verrucariaceae_unclassified(96);f__Verrucariaceae_unclassified(96); | Verrucariaceae | 9 | Symbiotroph | Lichenized | Thallus | NULL | Highly Probable | [illegible] | [illegible] |
| Otu02040 | 22 | k__Fungi(100);p__Basidiomycota(100);c__Agaricomycetes(100);o__Polyporales(100);f__Fomitopsidaceae(100);g__Postia(100);s__Postia_guttulata(100); | Postia | 13 | Saprotroph | \|Wood Saprotroph\| | Polyporoid | Brown Rot | Highly Probable | [illegible] | [illegible] |
| Otu02042 | 22 | k__Fungi(100);p__Basidiomycota(100);c__Agaricomycetes(100);o__Agaricales(100);f__Strophariaceae(100);g__Gymnopilus(100);s__Gymnopilus_cf_flavidellus_Murphy_2452(100); | Gymnopilus | 13 | Saprotroph | Plant Saprotroph-\|Wood Saprotroph\| | Agaricoid | White Rot | Probable | [illegible] | [illegible] |
| Otu02044 | 22 | k__Fungi(100);p__Ascomycota(100);c__Sordariomycetes(100);o__Hypocreales(100);f__Hypocreales_family_Incertae_sedis(100);g__Acremonium(100);s__Acremonium_rutilum(100); | Acremonium | 13 | Pathotroph-Saprotroph-Symbiotroph | Animal Pathogen-Endophyte-Fungal Parasite-Plant Pathogen-\|Undefined Saprotroph\|-Wood Saprotroph | Microfungus | Soft Rot | Possible | [illegible] | [illegible] |
| Otu02046 | 22 | k__Fungi(100);p__Basidiomycota(100);c__Agaricomycetes(100);o__Polyporales(100);f__Meruliaceae(100);f__Meruliaceae_unclassified(78);f__Meruliaceae_unclassified(78); | Meruliaceae | 9 | Pathotroph-Saprotroph | Plant Pathogen-Undefined Saprotroph | Corticioid-Polyporoid | NULL | Probable | [illegible] | [illegible] |
| Otu02047 | 22 | k__Fungi(100);p__Basidiomycota(100);c__Agaricomycetes(100);o__Polyporales(100);f__Phanerochaetaceae(100);g__Rhizochaete(100);s__Rhizochaete_radicata(100); | Rhizochaete | 13 | Saprotroph | Undefined Saprotroph-\|Wood Saprotroph\| | Corticioid | White Rot | Probable | [illegible] | [illegible] |
| Otu02048 | 22 | k__Fungi(100);p__Ascomycota(100);c__Sordariomycetes(100);o__Xylariales(100);f__Amphisphaeriaceae(100);g__Adisciso(100);s__Adisciso_yakushimense(91); | Adisciso | 13 | Pathotroph | \|Plant Pathogen\| | Microfungus | NULL | Probable | NULL | [illegible] |
| Otu02049 | 22 | k__Fungi(100);p__Ascomycota(100);c__Dothideomycetes(100);o__Pleosporales(100);f__Leptosphaeriaceae(100);g__Coniothyrium(100);s__Coniothyrium_sp_JK27(100); | Coniothyrium | 13 | Pathotroph-Saprotroph | Fungal Parasite-\|Plant Pathogen\|-Plant Saprotroph-Wood Saprotroph | Microfungus | Soft Rot | Highly Probable | [illegible] | [illegible] |
| Otu02050 | 22 | k__Fungi(100);p__Basidiomycota(100);c__Agaricomycetes(100);o__Polyporales(100);f__Xenasmataceae(100);g__Xenasmatella(100);s__Xenasmatella_sp(100); | Xenasmatella | 13 | Saprotroph | Undefined Saprotroph-\|Wood Saprotroph\| | Corticioid | White Rot | Probable | [illegible] | [illegible] |
| Otu02051 | 22 | k__Fungi(100);p__Ascomycota(100);c__Saccharomycetes(100);o__Saccharomycetales(100);f__Metschnikowiaceae(100);g__Metschnikowia(100);g__Metschnikowia_unclassified(96); | Metschnikowia | 13 | Saprotroph | \|Undefined Saprotroph\| | Yeast | NULL | Probable | [illegible] | [illegible] |
| Otu02052 | 22 | k__Fungi(100);p__Ascomycota(100);c__Saccharomycetes(100);o__Saccharomycetales(100);f__Metschnikowiaceae(100);g__Metschnikowia(100);s__Metschnikowia_pulcherrima(82); | Metschnikowia | 13 | Saprotroph | \|Undefined Saprotroph\| | Yeast | NULL | Probable | [illegible] | [illegible] |
| Otu02053 | 22 | k__Fungi(100);p__Basidiomycota(100);c__Agaricomycetes(100);o__Agaricales(100);f__Strophariaceae(100);g__Psilocybe(100);g__Psilocybe_unclassified(91); | Psilocybe | 13 | Saprotroph | \|Plant Saprotroph\|-Undefined Saprotroph | Agaricoid | Poisonous | Probable | [illegible] | [illegible] |
| Otu02056 | 22 | k__Fungi(100);p__Ascomycota(100);c__Dothideomycetes(100);o__Pleosporales(100);f__Didymosphaeriaceae(91);g__Roussoella(91);g__Roussoella_unclassified(87); | Roussoella | 13 | Saprotroph | Undefined Saprotroph-\|Wood Saprotroph\| | Microfungus | NULL | Probable | [illegible] | [illegible] |
| Otu02058 | 22 | k__Fungi(100);p__Ascomycota(100);c__Dothideomycetes(100);o__Myriangiales(100);f__Myriangiales_family_Incertae_sedis(100);g__Endosporium(100);s__Endosporium_aviarium(100); | Endosporium | 13 | Saprotroph | \|Plant Saprotroph\|-Undefined Saprotroph | Microfungus | NULL | Probable | [illegible] | [illegible] |
| Otu02059 | 22 | k__Fungi(100);p__Ascomycota(100);c__Saccharomycetes(100);o__Saccharomycetales(100);f__Metschnikowiaceae(100);g__Metschnikowia(100);s__Metschnikowia_pulcherrima(87); | Metschnikowia | 13 | Saprotroph | \|Undefined Saprotroph\| | Yeast | NULL | Probable | [illegible] | [illegible] |
| Otu02061 | 22 | k__Fungi(100);p__Ascomycota(100);c__Eurotiomycetes(100);o__Eurotiales(100);f__Trichocomaceae(100);g__Aspergillus(87);s__Aspergillus_fumigatus(78); | Aspergillus | 13 | Pathotroph-Saprotroph-Symbiotroph | Animal Pathogen-Endophyte-Plant Saprotroph-\|Undefined Saprotroph\|-Wood Saprotroph | Microfungus | NULL | Probable | [illegible] | [illegible] |
| Otu02063 | 22 | k__Fungi(100);p__Ascomycota(100);c__Sordariomycetes(100);o__Chaetosphaeriales(100);f__Chaetosphaeriaceae(100);g__Chaetosphaeria(100);s__Chaetosphaeria_sp(96); | Chaetosphaeria | 13 | Saprotroph-Symbiotroph | Endophyte-\|Plant Saprotroph\|-Wood Saprotroph | Microfungus | NULL | Probable | [illegible] | [illegible] |
| Otu02064 | 22 | k__Fungi(100);p__Ascomycota(100);c__Saccharomycetes(100);o__Saccharomycetales(100);f__Metschnikowiaceae(100);g__Metschnikowia(100);s__Metschnikowia_pulcherrima(96); | Metschnikowia | 13 | Saprotroph | \|Undefined Saprotroph\| | Yeast | NULL | Probable | [illegible] | [illegible] |
| Otu02065 | 22 | k__Fungi(100);p__Basidiomycota(100);c__Microbotryomycetes(100);o__Sporidiobolales(100);f__Sporidiobolales_family_Incertae_sedis(100);g__Sporobolomyces(100);s__Sporobolomyces_ogasawarensis(96); | Sporobolomyces | 13 | Pathotroph-Saprotroph | \|Fungal Parasite\|-Plant Saprotroph | Yeast | NULL | Probable | [illegible] | [illegible] |
| Otu02067 | 22 | k__Fungi(100);p__Basidiomycota(100);c__Agaricomycetes(100);o__Agaricales(100);f__Mycenaceae(100);g__Xeromphalina(100);s__Xeromphalina_sp(100); | Xeromphalina | 13 | Saprotroph | Undefined Saprotroph-\|Wood Saprotroph\| | Agaricoid | White Rot | Probable | [illegible] | [illegible] |
| Otu02068 | 22 | k__Fungi(100);p__Ascomycota(100);c__Saccharomycetes(100);o__Saccharomycetales(100);f__Metschnikowiaceae(100);g__Metschnikowia(100);s__Metschnikowia_pulcherrima(91); | Metschnikowia | 13 | Saprotroph | \|Undefined Saprotroph\| | Yeast | NULL | Probable | [illegible] | [illegible] |
| Otu02073 | 22 | k__Fungi(100);p__Ascomycota(100);c__Sordariomycetes(100);o__Chaetosphaeriales(100);f__Chaetosphaeriaceae(100);g__Chloridium(100);s__Chloridium_virescens_var_chlamydosporum(100); | Chloridium | 13 | Pathotroph-Saprotroph-Symbiotroph | Ectomycorrhizal-Endophyte-Plant Pathogen-\|Plant Saprotroph\|-Wood Saprotroph | Microfungus | NULL | Possible | [illegible] | [illegible] |
| Otu02077 | 22 | k__Fungi(100);p__Ascomycota(100);c__Saccharomycetes(100);o__Saccharomycetales(100);f__Metschnikowiaceae(100);g__Metschnikowia(100);g__Metschnikowia_unclassified(96); | Metschnikowia | 13 | Saprotroph | \|Undefined Saprotroph\| | Yeast | NULL | Probable | [illegible] | [illegible] |
| Otu02080 | 22 | k__Fungi(100);p__Basidiomycota(100);c__Agaricomycetes(100);o__Polyporales(100);f__Fomitopsidaceae(100);g__Climacocystis(100);s__Climacocystis_borealis(100); | Climacocystis | 13 | Saprotroph | \|Wood Saprotroph\| | Polyporoid | White Rot | Highly Probable | [illegible] | [illegible] |
| Otu02081 | 22 | k__Fungi(100);p__Basidiomycota(100);c__Agaricomycetes(100);o__Russulales(100);f__Hericiaceae(100);g__Hericium(100);s__Hericium_erinaceum(100); | Hericium | 13 | Saprotroph-Pathotroph-Saprotroph | Plant Pathogen-Undefined Saprotroph-\|Wood Saprotroph\| | Hydnoid-Polyporoid | White Rot | Probable | [illegible] | [illegible] |
| Otu02082 | 22 | k__Fungi(100);p__Basidiomycota(100);c__Agaricomycetes(100);o__Polyporales(100);f__Meruliaceae(100);g__Hyphoderma(100);g__Hyphoderma_unclassified(100); | Hyphoderma | 13 | Saprotroph | Undefined Saprotroph-\|Wood Saprotroph\| | Corticioid | White Rot | Probable | [illegible] | [illegible] |
| Otu02083 | 22 | k__Fungi(100);p__Ascomycota(100);c__Saccharomycetes(100);o__Saccharomycetales(100);f__Saccharomycetales_family_Incertae_sedis(100);g__Candida(82);g__Candida_unclassified(69); | Candida | 13 | Pathotroph-Saprotroph-Symbiotroph | Animal Pathogen-Endophyte-Endosymbiont-Epiphyte-\|Undefined Saprotroph\| | Dimorphic Yeast-Yeast | NULL | Probable | [illegible] | [illegible] |
| Otu02084 | 22 | k__Fungi(100);p__Basidiomycota(100);c__Agaricostilbomycetes(100);o__Agaricostilbales(100);f__Agaricostilbaceae(100);g__Bensingtonia(100);s__Bensingtonia_yuccicola(100); | Bensingtonia | 13 | Saprotroph | \|Undefined Saprotroph\| | Dimorphic-Facultative Yeast | NULL | Highly Probable | NULL | [illegible] |
| Otu02086 | 22 | k__Fungi(100);p__Basidiomycota(100);c__Agaricomycetes(100);o__Polyporales(100);f__Meruliaceae(100);g__Phlebia(96);g__Phlebia_unclassified(96); | Phlebia | 13 | Saprotroph | \|Wood Saprotroph\| | Corticioid | White Rot | Highly Probable | [illegible] | [illegible] |
| Otu02087 | 22 | k__Fungi(100);p__Basidiomycota(100);c__Agaricomycetes(100);o__Cantharellales(100);f__Botryobasidiaceae(100);g__Botryobasidium(100);s__Botryobasidium_subcoronatum(100); | Botryobasidium | 13 | Saprotroph | Undefined Saprotroph-\|Wood Saprotroph\| | Corticioid | White Rot | Probable | [illegible] | [illegible] |
| Otu02093 | 22 | k__Fungi(100);p__Ascomycota(100);c__Sordariomycetes(100);o__Hypocreales(100);f__unclassified_Hypocreales(87);g__unclassified_Hypocreales(87);s__Hypocreales_sp(87); | Hypocreales | 7 | Saprotroph | Undefined Saprotroph | Microfungus | NULL | Possible | NULL | [illegible] |
| Otu02094 | 22 | k__Fungi(100);p__Basidiomycota(100);c__Agaricomycetes(100);o__Agaricales(100);f__Psathyrellaceae(100);g__Psathyrella(100);s__Psathyrella_impexa(96); | Psathyrella | 13 | Saprotroph | Plant Saprotroph-\|Wood Saprotroph\| | Agaricoid | NULL | Probable | [illegible] | [illegible] |
| Otu02097 | 22 | k__Fungi(100);p__Basidiomycota(100);c__Agaricomycetes(100);o__Agaricales(100);f__Cortinariaceae(100);g__Cortinarius(100);g__Cortinarius_unclassified(100); | Cortinarius | 13 | Symbiotroph | \|Ectomycorrhizal\| | Agaricoid | NULL | Highly Probable | NULL | [illegible] |
| Otu02099 | 22 | k__Fungi(100);p__Basidiomycota(100);c__Microbotryomycetes(100);o__Sporidiobolales(100);f__Sporidiobolales_family_Incertae_sedis(100);g__Sporobolomyces(100);s__Sporobolomyces_lactosus(100); | Sporobolomyces | 13 | Pathotroph-Saprotroph | \|Fungal Parasite\|-Plant Saprotroph | Yeast | NULL | Probable | [illegible] | [illegible] |
| Otu02100 | 22 | k__Fungi(100);p__Ascomycota(100);c__Saccharomycetes(100);o__Saccharomycetales(100);f__Metschnikowiaceae(100);g__Metschnikowia(100);s__Metschnikowia_pulcherrima(96); | Metschnikowia | 13 | Saprotroph | \|Undefined Saprotroph\| | Yeast | NULL | Probable | [illegible] | [illegible] |
| Otu02101 | 22 | k__Fungi(100);p__Basidiomycota(100);c__Tremellomycetes(100);o__Trichosporonales(100);f__Trichosporonaceae(100);g__Trichosporon(100);s__Trichosporon_dulcitum(55); | Trichosporon | 13 | Pathotroph-Saprotroph | \|Animal Parasite\|-Animal Pathogen-Undefined Saprotroph | Yeast | NULL | Probable | [illegible] | [illegible] |
| Otu02102 | 22 | k__Fungi(100);p__Basidiomycota(100);c__Agaricomycetes(100);o__Boletales(100);f__Sclerodermataceae(100);g__Scleroderma(100);s__Scleroderma_cepa(100); | Scleroderma | 13 | Symbiotroph | \|Ectomycorrhizal\| | Gasteroid | NULL | Highly Probable | puffball; Common name | [illegible] |
| Otu02037 | 22 | k__Fungi(100);k__Fungi_unclassified(69);k__Fungi_unclassified(69);k__Fungi_unclassified(69);k__Fungi_unclassified(69);k__Fungi_unclassified(69);k__Fungi_unclassified(69); | - | - | - | - | - | - | - | Unassigned | |
| Otu02039 | 22 | k__Fungi(100);k__Fungi_unclassified(64);k__Fungi_unclassified(64);k__Fungi_unclassified(64);k__Fungi_unclassified(64);k__Fungi_unclassified(64);k__Fungi_unclassified(64); | - | - | - | - | - | - | - | Unassigned | |
| Otu02041 | 22 | k__Fungi(100);p__Ascomycota(96);p__Ascomycota_unclassified(96);p__Ascomycota_unclassified(96);p__Ascomycota_unclassified(96);p__Ascomycota_unclassified(96);p__Ascomycota_unclassified(96); | - | - | - | - | - | - | - | Unassigned | |
| Otu02043 | 22 | k__Fungi(100);p__Ascomycota(73);p__Ascomycota_unclassified(73);p__Ascomycota_unclassified(73);p__Ascomycota_unclassified(73);p__Ascomycota_unclassified(73);p__Ascomycota_unclassified(73); | - | - | - | - | - | - | - | Unassigned | |
| Otu02045 | 22 | k__Fungi(100);k__Fungi_unclassified(69);k__Fungi_unclassified(69);k__Fungi_unclassified(69);k__Fungi_unclassified(69);k__Fungi_unclassified(69);k__Fungi_unclassified(69); | - | - | - | - | - | - | - | Unassigned | |
| Otu02054 | 22 | k__Fungi(100);p__Ascomycota(55);p__Ascomycota_unclassified(55);p__Ascomycota_unclassified(55);p__Ascomycota_unclassified(55);p__Ascomycota_unclassified(55);p__Ascomycota_unclassified(55); | - | - | - | - | - | - | - | Unassigned | |
| Otu02055 | 22 | k__Fungi(100);k__Fungi_unclassified(100);k__Fungi_unclassified(100);k__Fungi_unclassified(100);k__Fungi_unclassified(100);k__Fungi_unclassified(100);k__Fungi_unclassified(100); | - | - | - | - | - | - | - | Unassigned | |
| Otu02057 | 22 | k__Fungi(100);p__Basidiomycota(100);p__Basidiomycota_unclassified(100);p__Basidiomycota_unclassified(100);p__Basidiomycota_unclassified(100);p__Basidiomycota_unclassified(100);p__Basidiomycota_unclassified(100); | - | - | - | - | - | - | - | Unassigned | |
| Otu02060 | 22 | k__Fungi(100);p__Ascomycota(73);p__Ascomycota_unclassified(73);p__Ascomycota_unclassified(73);p__Ascomycota_unclassified(73);p__Ascomycota_unclassified(73);p__Ascomycota_unclassified(73); | - | - | - | - | - | - | - | Unassigned | |
| Otu02062 | 22 | k__Fungi(100);p__Basidiomycota(100);c__Microbotryomycetes(100);o__Sporidiobolales(100);f__Sporidiobolales_family_Incertae_sedis(100);f__Sporidiobolales_family_Incertae_sedis_unclassified(55);f__Sporidiobolales_family_Incertae_sedis_unclassified(55); | - | - | - | - | - | - | - | Unassigned | |
| Otu02066 | 22 | k__Fungi(100);p__Basidiomycota(100);c__Agaricomycetes(100);o__Trechisporales(100);f__Hydnodontaceae(100);f__Hydnodontaceae_unclassified(82);f__Hydnodontaceae_unclassified(82); | - | - | - | - | - | - | - | Unassigned | |
| Otu02069 | 22 | k__Fungi(100);k__Fungi_unclassified(64);k__Fungi_unclassified(64);k__Fungi_unclassified(64);k__Fungi_unclassified(64);k__Fungi_unclassified(64);k__Fungi_unclassified(64); | - | - | - | - | - | - | - | Unassigned | |
| Otu02070 | 22 | k__Fungi(100);k__Fungi_unclassified(73);k__Fungi_unclassified(73);k__Fungi_unclassified(73);k__Fungi_unclassified(73);k__Fungi_unclassified(73);k__Fungi_unclassified(73); | - | - | - | - | - | - | - | Unassigned | |
| Otu02071 | 22 | k__Fungi(100);k__Fungi_unclassified(78);k__Fungi_unclassified(78);k__Fungi_unclassified(78);k__Fungi_unclassified(78);k__Fungi_unclassified(78);k__Fungi_unclassified(78); | - | - | - | - | - | - | - | Unassigned | |
| Otu02072 | 22 | k__Fungi(100);k__Fungi_unclassified(96);k__Fungi_unclassified(96);k__Fungi_unclassified(96);k__Fungi_unclassified(96);k__Fungi_unclassified(96);k__Fungi_unclassified(96); | - | - | - | - | - | - | - | Unassigned | |
| Otu02074 | 22 | k__Fungi(100);k__Fungi_unclassified(100);k__Fungi_unclassified(100);k__Fungi_unclassified(100);k__Fungi_unclassified(100);k__Fungi_unclassified(100);k__Fungi_unclassified(100); | - | - | - | - | - | - | - | Unassigned | |
| Otu02075 | 22 | k__Fungi(100);k__Fungi_unclassified(100);k__Fungi_unclassified(100);k__Fungi_unclassified(100);k__Fungi_unclassified(100);k__Fungi_unclassified(100);k__Fungi_unclassified(100); | - | - | - | - | - | - | - | Unassigned | |
| Otu02076 | 22 | k__Fungi(100);p__Ascomycota(55);p__Ascomycota_unclassified(55);p__Ascomycota_unclassified(55);p__Ascomycota_unclassified(55);p__Ascomycota_unclassified(55);p__Ascomycota_unclassified(55); | - | - | - | - | - | - | - | Unassigned | |
| Otu02078 | 22 | k__Fungi(100);p__Ascomycota(100);p__Ascomycota_unclassified(100);p__Ascomycota_unclassified(100);p__Ascomycota_unclassified(100);p__Ascomycota_unclassified(100);p__Ascomycota_unclassified(100); | - | - | - | - | - | - | - | Unassigned | |
| Otu02079 | 22 | k__Fungi(100);p__Basidiomycota(100);c__Agaricomycetes(100);c__Agaricomycetes_unclassified(100);c__Agaricomycetes_unclassified(100);c__Agaricomycetes_unclassified(100);c__Agaricomycetes_unclassified(100); | - | - | - | - | - | - | - | Unassigned | |
| Otu02085 | 22 | k__Fungi(100);k__Fungi_unclassified(78);k__Fungi_unclassified(78);k__Fungi_unclassified(78);k__Fungi_unclassified(78);k__Fungi_unclassified(78);k__Fungi_unclassified(78); | - | - | - | - | - | - | - | Unassigned | |
| Otu02088 | 22 | k__Fungi(100);p__Basidiomycota(100);c__Agaricomycetes(100);o__Polyporales(100);f__Phanerochaetaceae(96);f__Phanerochaetaceae_unclassified(91);f__Phanerochaetaceae_unclassified(91); | - | - | - | - | - | - | - | Unassigned | |
| Otu02089 | 22 | k__Fungi(100);k__Fungi_unclassified(69);k__Fungi_unclassified(69);k__Fungi_unclassified(69);k__Fungi_unclassified(69);k__Fungi_unclassified(69);k__Fungi_unclassified(69); | - | - | - | - | - | - | - | Unassigned | |
| Otu02090 | 22 | k__Fungi(100);k__Fungi_unclassified(100);k__Fungi_unclassified(100);k__Fungi_unclassified(100);k__Fungi_unclassified(100);k__Fungi_unclassified(100);k__Fungi_unclassified(100); | - | - | - | - | - | - | - | Unassigned | |
| Otu02091 | 22 | k__Fungi(100);k__Fungi_unclassified(78);k__Fungi_unclassified(78);k__Fungi_unclassified(78);k__Fungi_unclassified(78);k__Fungi_unclassified(78);k__Fungi_unclassified(78); | - | - | - | - | - | - | - | Unassigned | |
| Otu02092 | 22 | k__Fungi(100);k__Fungi_unclassified(91);k__Fungi_unclassified(91);k__Fungi_unclassified(91);k__Fungi_unclassified(91);k__Fungi_unclassified(91);k__Fungi_unclassified(91); | - | - | - | - | - | - | - | Unassigned | |
| Otu02095 | 22 | k__Fungi(100);p__Ascomycota(96);p__Ascomycota_unclassified(96);p__Ascomycota_unclassified(96);p__Ascomycota_unclassified(96);p__Ascomycota_unclassified(96);p__Ascomycota_unclassified(96); | - | - | - | - | - | - | - | Unassigned | |
| Otu02096 | 22 | k__Fungi(100);p__Ascomycota(100);c__Dothideomycetes(91);c__Dothideomycetes_unclassified(78);c__Dothideomycetes_unclassified(78);c__Dothideomycetes_unclassified(78);c__Dothideomycetes_unclassified(78); | - | - | - | - | - | - | - | Unassigned | |
| Otu02098 | 22 | k__Fungi(100);k__Fungi_unclassified(100);k__Fungi_unclassified(100);k__Fungi_unclassified(100);k__Fungi_unclassified(100);k__Fungi_unclassified(100);k__Fungi_unclassified(100); | - | - | - | - | - | - | - | Unassigned | |
| Otu02104 | 21 | k__Fungi(100);p__Ascomycota(100);c__Leotiomycetes(100);o__Helotiales(100);f__Sclerotiniaceae(100);g__Amphobotrys(100);s__Amphobotrys_ricini(100); | Amphobotrys | 13 | Pathotroph | \|Plant Pathogen\| | Microfungus | NULL | Probable | NULL | [illegible] |
| Otu02106 | 21 | k__Fungi(100);p__Ascomycota(100);c__Saccharomycetes(100);o__Saccharomycetales(100);f__Metschnikowiaceae(100);g__Metschnikowia(100);s__Metschnikowia_pulcherrima(91); | Metschnikowia | 13 | Saprotroph | \|Undefined Saprotroph\| | Yeast | NULL | Probable | [illegible] | [illegible] |
| Otu02107 | 21 | k__Fungi(100);p__Ascomycota(100);c__Saccharomycetes(100);o__Saccharomycetales(100);f__Metschnikowiaceae(100);g__Metschnikowia(100);s__Metschnikowia_pulcherrima(96); | Metschnikowia | 13 | Saprotroph | \|Undefined Saprotroph\| | Yeast | NULL | Probable | [illegible] | [illegible] |
| Otu02108 | 21 | k__Fungi(100);p__Basidiomycota(100);c__Agaricomycetes(100);o__Hymenochaetales(100);f__Schizoporaceae(100);g__Basidioradulum(100);s__Basidioradulum_radula(100); | Basidioradulum | 13 | Saprotroph | Undefined Saprotroph-\|Wood Saprotroph\| | Corticioid | White Rot | Probable | [illegible] | [illegible] |
| Otu02109 | 21 | k__Fungi(100);p__Basidiomycota(100);c__Agaricomycetes(100);o__Cantharellales(100);f__Ceratobasidiaceae(100);g__Ceratobasidium(58);g__Ceratobasidium_unclassified(58); | Ceratobasidium | 13 | Pathotroph-Saprotroph-Symbiotroph | Ectomycorrhizal-Endomycorrhizal-Orchid Mycorrhizal-\|Plant Pathogen\|-Plant Saprotroph-Undefined Saprotroph | Corticioid-Microfungus | NULL | Possible | [illegible] | [illegible] |
| Otu02110 | 21 | k__Fungi(100);p__Basidiomycota(100);c__Agaricomycetes(100);o__Agaricales(100);f__Entolomataceae(100);g__Entoloma(100);s__Entoloma_sp_AJ108(100); | Entoloma | 13 | Pathotroph-Saprotroph-Symbiotroph | Ectomycorrhizal-Fungal Parasite-\|Undefined Saprotroph\|-Undefined Symbiotroph | Agaricoid | NULL | Possible | [illegible] | [illegible] |
| Otu02111 | 21 | k__Fungi(100);p__Ascomycota(100);c__Saccharomycetes(100);o__Saccharomycetales(100);f__Saccharomycetales_family_Incertae_sedis(100);g__Candida(100);s__Candida_quercitrusa(100); | Candida | 13 | Pathotroph-Saprotroph-Symbiotroph | Animal Pathogen-Endophyte-Endosymbiont-Epiphyte-\|Undefined Saprotroph\| | Dimorphic Yeast-Yeast | NULL | Probable | [illegible] | [illegible] |
| Otu02113 | 21 | k__Fungi(100);p__Ascomycota(100);c__Saccharomycetes(100);o__Saccharomycetales(100);f__Metschnikowiaceae(100);g__Metschnikowia(100);s__Metschnikowia_pulcherrima(100); | Metschnikowia | 13 | Saprotroph | \|Undefined Saprotroph\| | Yeast | NULL | Probable | [illegible] | [illegible] |
| Otu02114 | 21 | k__Fungi(100);p__Ascomycota(100);c__Sordariomycetes(100);o__Hypocreales(100);f__Hypocreales_family_Incertae_sedis(100);g__Gliomastix(91);s__Gliomastix_cerealis(91); | Gliomastix | 13 | Saprotroph | \|Undefined Saprotroph\| | Microfungus | NULL | Probable | NULL | [illegible] |
| Otu02117 | 21 | k__Fungi(100);p__Ascomycota(100);c__Saccharomycetes(100);o__Saccharomycetales(100);f__Metschnikowiaceae(100);g__Metschnikowia(100);s__Metschnikowia_pulcherrima(86); | Metschnikowia | 13 | Saprotroph | \|Undefined Saprotroph\| | Yeast | NULL | Probable | [illegible] | [illegible] |
| Otu02120 | 21 | k__Fungi(100);p__Basidiomycota(100);c__Agaricomycetes(100);o__Agaricales(100);f__Psathyrellaceae(100);g__Parasola(100);g__Parasola_unclassified(100); | Parasola | 13 | Saprotroph | \|Undefined Saprotroph\| | Agaricoid | NULL | Probable | [illegible] | [illegible] |
| Otu02121 | 21 | k__Fungi(100);p__Basidiomycota(100);c__Agaricomycetes(100);o__Polyporales(100);f__Meruliaceae(100);g__Hypochnicium(100);g__Hypochnicium_unclassified(91); | Hypochnicium | 13 | Saprotroph | Undefined Saprotroph-\|Wood Saprotroph\| | Corticioid | White Rot | Probable | [illegible] | [illegible] |
| Otu02123 | 21 | k__Fungi(100);p__Ascomycota(100);c__Leotiomycetes(100);o__Helotiales(100);f__Hyaloscyphaceae(58);f__Hyaloscyphaceae_unclassified(58);f__Hyaloscyphaceae_unclassified(58); | Hyaloscyphaceae | 9 | Saprotroph | Plant Saprotroph-Wood Saprotroph | Microfungus | NULL | Probable | [illegible] | [illegible] |
| Otu02124 | 21 | k__Fungi(100);p__Basidiomycota(100);c__Exobasidiomycetes(100);o__Tilletiales(100);f__Tilletiaceae(100);g__Tilletia(100);g__Tilletia_unclassified(62); | Tilletia | 13 | Pathotroph | \|Plant Pathogen\| | Smut | NULL | Probable | [illegible] | [illegible] |
| Otu02126 | 21 | k__Fungi(100);p__Ascomycota(86);c__Saccharomycetes(86);o__Saccharomycetales(86);f__Metschnikowiaceae(86);g__Metschnikowia(86);s__Metschnikowia_pulcherrima(81); | Metschnikowia | 13 | Saprotroph | \|Undefined Saprotroph\| | Yeast | NULL | Probable | [illegible] | [illegible] |
| Otu02132 | 21 | k__Fungi(100);p__Ascomycota(100);c__Dothideomycetes(100);o__Pleosporales(100);f__Massarinaceae(100);g__Massarina(100);s__Massarina_sp_HF12700(100); | Massarina | 13 | Pathotroph-Saprotroph | \|Plant Pathogen\|-Undefined Saprotroph | Microfungus | NULL | Probable | NULL | [illegible] |
| Otu02134 | 21 | k__Fungi(100);p__Ascomycota(100);c__Sordariomycetes(100);o__Microascales(100);f__Halosphaeriaceae(100);g__unclassified_Halosphaeriaceae(100);s__Halosphaeriaceae_sp(100); | Halosphaeriaceae | 9 | Saprotroph | Undefined Saprotroph-Wood Saprotroph | Microfungus | NULL | Probable | [illegible] | [illegible] |
| Otu02135 | 21 | k__Fungi(100);p__Ascomycota(100);c__Eurotiomycetes(100);o__Eurotiales(100);f__Trichocomaceae(100);g__Penicillium(100);s__Penicillium_adametzoides(100); | Penicillium | 13 | Pathotroph-Saprotroph-Symbiotroph | Animal Parasite-Dung Saprotroph-Endophyte-Fungal Parasite-Plant Pathogen-\|Undefined Saprotroph\|-Wood Saprotroph | Microfungus | Soft Rot | Highly Probable | [illegible] | [illegible] |
| Otu02139 | 21 | k__Fungi(100);p__Ascomycota(100);c__Leotiomycetes(100);o__Rhytismatales(100);f__Rhytismataceae(100);g__Lophodermium(100);s__Lophodermium_sp(100); | Lophodermium | 13 | Pathotroph-Saprotroph-Symbiotroph | Endophyte-\|Plant Pathogen\|-Plant Saprotroph | Microfungus | NULL | Probable | [illegible] | [illegible] |
| Otu02141 | 21 | k__Fungi(100);p__Basidiomycota(100);c__Tremellomycetes(100);o__Tremellales(100);f__Tremellales_family_Incertae_sedis(100);g__Cryptococcus(100);s__Cryptococcus_laurentii(100); | Cryptococcus | 13 | Pathotroph-Saprotroph-Symbiotroph | Animal Pathogen-Endophyte-Epiphyte-\|Undefined Saprotroph\| | Dimorphic Yeast-Yeast | NULL | Possible | [illegible] | [illegible] |
| Otu02142 | 21 | k__Fungi(100);p__Basidiomycota(100);c__Agaricomycetes(100);o__Agaricales(100);f__Entolomataceae(100);f__Entolomataceae_unclassified(53);f__Entolomataceae_unclassified(53); | Entolomataceae | 9 | Pathotroph-Saprotroph-Symbiotroph | Ectomycorrhizal-Fungal Parasite-Soil Saprotroph-Undefined Saprotroph | Agaricoid | NULL | Possible | [illegible] | [illegible] |
| Otu02143 | 21 | k__Fungi(100);p__Ascomycota(100);c__Ascomycota_class_Incertae_sedis(100);o__Ascomycota_order_Incertae_sedis(100);f__Ascomycota_family_Incertae_sedis(100);g__Microcyclospora(100);s__Microcyclospora_tardicrescens(100); | Microcyclospora | 13 | Pathotroph | \|Plant Pathogen\| | Microfungus | NULL | Probable | NULL | [illegible] |
| Otu02145 | 21 | k__Fungi(100);p__Basidiomycota(100);c__Agaricomycetes(100);o__Trechisporales(100);f__Hydnodontaceae(100);g__Trechispora(100);g__Trechispora_unclassified(100); | Trechispora | 13 | Saprotroph | \|Wood Saprotroph\| | Corticioid | White Rot | Highly Probable | [illegible] | [illegible] |
| Otu02146 | 21 | k__Fungi(100);p__Ascomycota(100);c__Sordariomycetes(100);o__Diaporthales(100);f__Gnomoniaceae(100);g__Amphiporthe(100);s__Amphiporthe_sp(100); | Amphiporthe | 13 | Saprotroph | \|Undefined Saprotroph\| | Microfungus | NULL | Probable | NULL | [illegible] |
| Otu02148 | 21 | k__Fungi(100);p__Basidiomycota(100);c__Agaricomycetes(100);o__Trechisporales(100);f__Hydnodontaceae(100);g__Trechispora(100);s__Trechispora_sp(100); | Trechispora | 13 | Saprotroph | \|Wood Saprotroph\| | Corticioid | White Rot | Highly Probable | [illegible] | [illegible] |
| Otu02149 | 21 | k__Fungi(100);p__Basidiomycota(100);c__Tremellomycetes(100);o__Filobasidiales(100);f__Filobasidiaceae(100);g__Cryptococcus(100);s__Cryptococcus_skinneri(100); | Cryptococcus | 13 | Pathotroph-Saprotroph-Symbiotroph | Animal Pathogen-Endophyte-Epiphyte-\|Undefined Saprotroph\| | Dimorphic Yeast-Yeast | NULL | Possible | [illegible] | [illegible] |
| Otu02150 | 21 | k__Fungi(100);p__Basidiomycota(100);c__Microbotryomycetes(100);o__Sporidiobolales(100);f__Sporidiobolales_family_Incertae_sedis(100);g__Rhodotorula(100);g__Rhodotorula_unclassified(100); | Rhodotorula | 13 | Pathotroph-Saprotroph-Symbiotroph | Animal Pathogen-Animal Symbiotroph-Endophyte-Plant Pathogen-\|Undefined Saprotroph\| | Yeast | NULL | Probable | [illegible] | [illegible] |
| Otu02151 | 21 | k__Fungi(100);p__Ascomycota(100);c__Saccharomycetes(100);o__Saccharomycetales(100);f__Metschnikowiaceae(100);g__Metschnikowia(100);s__Metschnikowia_pulcherrima(96); | Metschnikowia | 13 | Saprotroph | \|Undefined Saprotroph\| | Yeast | NULL | Probable | [illegible] | [illegible] |
| Otu02153 | 21 | k__Fungi(100);p__Basidiomycota(100);c__Agaricomycetes(100);o__Agaricales(100);f__Psathyrellaceae(100);g__Coprinopsis(100);s__Coprinopsis_lagopus(62); | Coprinopsis | 13 | Saprotroph | \|Undefined Saprotroph\| | Agaricoid | NULL | Probable | [illegible] | [illegible] |
| Otu02154 | 21 | k__Fungi(100);p__Basidiomycota(100);c__Microbotryomycetes(100);o__Sporidiobolales(100);f__Sporidiobolales_family_Incertae_sedis(100);g__Sporobolomyces(100);s__Sporobolomyces_sasicola(100); | Sporobolomyces | 13 | Pathotroph-Saprotroph | \|Fungal Parasite\|-Plant Saprotroph | Yeast | NULL | Probable | [illegible] | [illegible] |
| Otu02155 | 21 | k__Fungi(100);p__Ascomycota(100);c__Saccharomycetes(100);o__Saccharomycetales(100);f__Saccharomycetaceae(100);g__Saccharomyces(100);g__Saccharomyces_unclassified(53); | Saccharomyces | 13 | Saprotroph | \|Undefined Saprotroph\| | Yeast | NULL | Probable | [illegible] | [illegible] |
| Otu02157 | 21 | k__Fungi(100);p__Ascomycota(100);c__Eurotiomycetes(100);o__Eurotiales(100);o__Eurotiales_unclassified(100);o__Eurotiales_unclassified(100);o__Eurotiales_unclassified(100); | Eurotiales | 7 | Saprotroph | Undefined Saprotroph | Microfungus | NULL | Possible | NULL | [illegible] |
| Otu02158 | 21 | k__Fungi(100);p__Ascomycota(96);c__Saccharomycetes(96);o__Saccharomycetales(96);f__Saccharomycetales_family_Incertae_sedis(91);g__Candida(91);g__Candida_unclassified(72); | Candida | 13 | Pathotroph-Saprotroph-Symbiotroph | Animal Pathogen-Endophyte-Endosymbiont-Epiphyte-\|Undefined Saprotroph\| | Dimorphic Yeast-Yeast | NULL | Probable | [illegible] | [illegible] |
| Otu02103 | 21 | k__Fungi(100);p__Ascomycota(100);p__Ascomycota_unclassified(91);p__Ascomycota_unclassified(91);p__Ascomycota_unclassified(91);p__Ascomycota_unclassified(91);p__Ascomycota_unclassified(91); | - | - | - | - | - | - | - | Unassigned | |
| Otu02105 | 21 | k__Fungi(100);p__Ascomycota(100);p__Ascomycota_unclassified(100);p__Ascomycota_unclassified(100);p__Ascomycota_unclassified(100);p__Ascomycota_unclassified(100);p__Ascomycota_unclassified(100); | - | - | - | - | - | - | - | Unassigned | |
| Otu02112 | 21 | k__Fungi(100);k__Fungi_unclassified(100);k__Fungi_unclassified(100);k__Fungi_unclassified(100);k__Fungi_unclassified(100);k__Fungi_unclassified(100);k__Fungi_unclassified(100); | - | - | - | - | - | - | - | Unassigned | |
| Otu02115 | 21 | k__Fungi(100);k__Fungi_unclassified(72);k__Fungi_unclassified(72);k__Fungi_unclassified(72);k__Fungi_unclassified(72);k__Fungi_unclassified(72);k__Fungi_unclassified(72); | - | - | - | - | - | - | - | Unassigned | |
| Otu02116 | 21 | k__Fungi(100);p__Ascomycota(62);p__Ascomycota_unclassified(62);p__Ascomycota_unclassified(62);p__Ascomycota_unclassified(62);p__Ascomycota_unclassified(62);p__Ascomycota_unclassified(62); | - | - | - | - | - | - | - | Unassigned | |
| Otu02118 | 21 | k__Fungi(100);p__Ascomycota(81);p__Ascomycota_unclassified(81);p__Ascomycota_unclassified(81);p__Ascomycota_unclassified(81);p__Ascomycota_unclassified(81);p__Ascomycota_unclassified(81); | - | - | - | - | - | - | - | Unassigned | |
| Otu02119 | 21 | k__Fungi(100);k__Fungi_unclassified(62);k__Fungi_unclassified(62);k__Fungi_unclassified(62);k__Fungi_unclassified(62);k__Fungi_unclassified(62);k__Fungi_unclassified(62); | - | - | - | - | - | - | - | Unassigned | |
| Otu02122 | 21 | k__Fungi(100);k__Fungi_unclassified(72);k__Fungi_unclassified(72);k__Fungi_unclassified(72);k__Fungi_unclassified(72);k__Fungi_unclassified(72);k__Fungi_unclassified(72); | - | - | - | - | - | - | - | Unassigned | |
| Otu02125 | 21 | k__Fungi(100);k__Fungi_unclassified(100);k__Fungi_unclassified(100);k__Fungi_unclassified(100);k__Fungi_unclassified(100);k__Fungi_unclassified(100);k__Fungi_unclassified(100); | - | - | - | - | - | - | - | Unassigned | |
| Otu02127 | 21 | k__Fungi(100);p__Basidiomycota(100);p__Basidiomycota_unclassified(81);p__Basidiomycota_unclassified(81);p__Basidiomycota_unclassified(81);p__Basidiomycota_unclassified(81);p__Basidiomycota_unclassified(81); | - | - | - | - | - | - | - | Unassigned | |
| Otu02128 | 21 | k__Fungi(100);p__Ascomycota(58);p__Ascomycota_unclassified(58);p__Ascomycota_unclassified(58);p__Ascomycota_unclassified(58);p__Ascomycota_unclassified(58);p__Ascomycota_unclassified(58); | - | - | - | - | - | - | - | Unassigned | |
| Otu02129 | 21 | k__Fungi(100);p__Ascomycota(100);p__Ascomycota_unclassified(100);p__Ascomycota_unclassified(100);p__Ascomycota_unclassified(100);p__Ascomycota_unclassified(100);p__Ascomycota_unclassified(100); | - | - | - | - | - | - | - | Unassigned | |
| Otu02130 | 21 | k__Fungi(100);k__Fungi_unclassified(86);k__Fungi_unclassified(86);k__Fungi_unclassified(86);k__Fungi_unclassified(86);k__Fungi_unclassified(86);k__Fungi_unclassified(86); | - | - | - | - | - | - | - | Unassigned | |
| Otu02131 | 21 | k__Fungi(100);p__Ascomycota(58);p__Ascomycota_unclassified(58);p__Ascomycota_unclassified(58);p__Ascomycota_unclassified(58);p__Ascomycota_unclassified(58);p__Ascomycota_unclassified(58); | - | - | - | - | - | - | - | Unassigned | |
| Otu02133 | 21 | k__Fungi(100);p__Basidiomycota(100);c__Agaricomycetes(100);o__Auriculariales(100);o__Auriculariales_unclassified(96);o__Auriculariales_unclassified(96);o__Auriculariales_unclassified(96); | - | - | - | - | - | - | - | Unassigned | |
| Otu02136 | 21 | k__Fungi(100);p__Ascomycota(81);p__Ascomycota_unclassified(81);p__Ascomycota_unclassified(81);p__Ascomycota_unclassified(81);p__Ascomycota_unclassified(81);p__Ascomycota_unclassified(81); | - | - | - | - | - | - | - | Unassigned | |
| Otu02137 | 21 | k__Fungi(100);p__Ascomycota(58);p__Ascomycota_unclassified(58);p__Ascomycota_unclassified(58);p__Ascomycota_unclassified(58);p__Ascomycota_unclassified(58);p__Ascomycota_unclassified(58); | - | - | - | - | - | - | - | Unassigned | |
| Otu02138 | 21 | k__Fungi(100);k__Fungi_unclassified(67);k__Fungi_unclassified(67);k__Fungi_unclassified(67);k__Fungi_unclassified(67);k__Fungi_unclassified(67);k__Fungi_unclassified(67); | - | - | - | - | - | - | - | Unassigned | |
| Otu02140 | 21 | k__Fungi(100);k__Fungi_unclassified(58);k__Fungi_unclassified(58);k__Fungi_unclassified(58);k__Fungi_unclassified(58);k__Fungi_unclassified(58);k__Fungi_unclassified(58); | - | - | - | - | - | - | - | Unassigned | |
| Otu02144 | 21 | k__Fungi(100);p__Basidiomycota(100);c__Agaricomycetes(100);o__Boletales(96);o__Boletales_unclassified(96);o__Boletales_unclassified(96);o__Boletales_unclassified(96); | - | - | - | - | - | - | - | Unassigned | |
| Otu02147 | 21 | k__Fungi(100);p__Ascomycota(67);p__Ascomycota_unclassified(67);p__Ascomycota_unclassified(67);p__Ascomycota_unclassified(67);p__Ascomycota_unclassified(67);p__Ascomycota_unclassified(67); | - | - | - | - | - | - | - | Unassigned | |
| Otu02152 | 21 | k__Fungi(100);k__Fungi_unclassified(53);k__Fungi_unclassified(53);k__Fungi_unclassified(53);k__Fungi_unclassified(53);k__Fungi_unclassified(53);k__Fungi_unclassified(53); | - | - | - | - | - | - | - | Unassigned | |
| Otu02156 | 21 | k__Fungi(100);p__Ascomycota(86);p__Ascomycota_unclassified(86);p__Ascomycota_unclassified(86);p__Ascomycota_unclassified(86);p__Ascomycota_unclassified(86);p__Ascomycota_unclassified(86); | - | - | - | - | - | - | - | Unassigned | |
| Otu02160 | 20 | k__Fungi(100);p__Ascomycota(100);c__Pezizomycetes(100);o__Pezizales(100);f__Pyronemataceae(100);g__Geopyxis(100);s__Geopyxis_sp(100); | Geopyxis | 13 | Saprotroph-Symbiotroph | Ectomycorrhizal-Plant Saprotroph-\|Undefined Saprotroph\| | Pezizoid | NULL | Possible | [illegible] | [illegible] |
| Otu02162 | 20 | k__Fungi(100);p__Ascomycota(100);c__Saccharomycetes(100);o__Saccharomycetales(100);f__Metschnikowiaceae(100);g__Metschnikowia(100);s__Metschnikowia_pulcherrima(80); | Metschnikowia | 13 | Saprotroph | \|Undefined Saprotroph\| | Yeast | NULL | Probable | [illegible] | [illegible] |
| Otu02165 | 20 | k__Fungi(100);p__Ascomycota(100);c__Saccharomycetes(100);o__Saccharomycetales(100);f__Metschnikowiaceae(100);g__Metschnikowia(100);s__Metschnikowia_pulcherrima(100); | Metschnikowia | 13 | Saprotroph | \|Undefined Saprotroph\| | Yeast | NULL | Probable | [illegible] | [illegible] |
| Otu02166 | 20 | k__Fungi(100);p__Basidiomycota(100);c__Agaricomycetes(100);o__Sebacinales(85);f__Sebacinaceae(75);g__Craterocolla(75);s__Craterocolla_cerasi(75); | Craterocolla | 13 | Saprotroph | \|Plant Saprotroph\|-Undefined Saprotroph | Corticioid | NULL | Probable | [illegible] | [illegible] |
| Otu02174 | 20 | k__Fungi(100);p__Ascomycota(100);c__Saccharomycetes(100);o__Saccharomycetales(100);f__Metschnikowiaceae(100);g__Metschnikowia(100);s__Metschnikowia_pulcherrima(95); | Metschnikowia | 13 | Saprotroph | \|Undefined Saprotroph\| | Yeast | NULL | Probable | [illegible] | [illegible] |
| Otu02176 | 20 | k__Fungi(100);p__Ascomycota(100);c__Ascomycota_class_Incertae_sedis(100);o__Ascomycota_order_Incertae_sedis(100);f__Ascomycota_family_Incertae_sedis(100);g__Chalara(100);s__Chalara_microspora(100); | Chalara | 13 | Pathotroph-Saprotroph-Symbiotroph | Endophyte-Plant Pathogen-\|Plant Saprotroph\|-Wood Saprotroph | Microfungus | Soft Rot | Probable | [illegible] | [illegible] |
| Otu02177 | 20 | k__Fungi(100);p__Basidiomycota(100);c__Agaricomycetes(100);o__Agaricales(100);f__Psathyrellaceae(100);f__Psathyrellaceae_unclassified(70);f__Psathyrellaceae_unclassified(70); | Psathyrellaceae | 9 | Saprotroph | Dung Saprotroph-Plant Saprotroph-Wood Saprotroph | Agaricoid | NULL | Probable | [illegible] | [illegible] |
| Otu02179 | 20 | k__Fungi(100);p__Zygomycota(100);c__Zygomycota_class_Incertae_sedis(100);o__Mortierellales(100);f__Mortierellaceae(100);g__Mortierella(100);g__Mortierella_unclassified(100); | Mortierella | 13 | Saprotroph-Symbiotroph | Endophyte-Plant Saprotroph-\|Undefined Saprotroph\| | Microfungus | NULL | Probable | [illegible] | [illegible] |
| Otu02180 | 20 | k__Fungi(100);p__Basidiomycota(100);c__Agaricomycetes(100);o__Trechisporales(100);f__Hydnodontaceae(100);g__Sistotremastrum(100);s__Sistotremastrum_guttuliferum(100); | Sistotremastrum | 13 | Saprotroph | Undefined Saprotroph-\|Wood Saprotroph\| | Corticioid | White Rot | Probable | [illegible] | [illegible] |
| Otu02181 | 20 | k__Fungi(100);p__Basidiomycota(100);c__Agaricomycetes(100);o__Agaricales(100);f__Inocybaceae(100);g__Crepidotus(100);s__Crepidotus_sphaerosporus(100); | Crepidotus | 13 | Saprotroph | \|Wood Saprotroph\| | Agaricoid | NULL | Highly Probable | [illegible] | [illegible] |
| Otu02185 | 20 | k__Fungi(100);p__Ascomycota(100);c__Dothideomycetes(100);o__Capnodiales(100);f__Mycosphaerellaceae(100);g__Pseudocercosporella(100);s__Pseudocercosporella_sp(100); | Pseudocercosporella | 13 | Pathotroph | \|Plant Pathogen\| | Microfungus | NULL | Probable | NULL | [illegible] |
| Otu02187 | 20 | k__Fungi(100);p__Ascomycota(100);c__Saccharomycetes(100);o__Saccharomycetales(100);f__Metschnikowiaceae(100);g__Metschnikowia(100);g__Metschnikowia_unclassified(80); | Metschnikowia | 13 | Saprotroph | \|Undefined Saprotroph\| | Yeast | NULL | Probable | [illegible] | [illegible] |
| Otu02188 | 20 | k__Fungi(100);p__Ascomycota(100);c__Saccharomycetes(100);o__Saccharomycetales(100);f__Metschnikowiaceae(100);g__Metschnikowia(100);s__Metschnikowia_pulcherrima(95); | Metschnikowia | 13 | Saprotroph | \|Undefined Saprotroph\| | Yeast | NULL | Probable | [illegible] | [illegible] |
| Otu02189 | 20 | k__Fungi(100);p__Basidiomycota(100);c__Agaricomycetes(100);o__Polyporales(100);f__Polyporaceae(100);g__Datronia(100);s__Datronia_scutellata(75); | Datronia | 13 | Saprotroph | \|Wood Saprotroph\| | Corticioid | White Rot | Highly Probable | [illegible] | [illegible] |
| Otu02190 | 20 | k__Fungi(100);p__Basidiomycota(100);c__Agaricomycetes(100);o__Russulales(100);f__Stereaceae(100);f__Stereaceae_unclassified(100);f__Stereaceae_unclassified(100); | Stereaceae | 9 | Saprotroph | Undefined Saprotroph-Wood Saprotroph | Corticioid-Polyporoid | NULL | Probable | [illegible] | [illegible] |
| Otu02194 | 20 | k__Fungi(100);p__Basidiomycota(100);c__Agaricomycetes(100);o__Hymenochaetales(100);f__Schizoporaceae(100);g__Hyphodontia(100);s__Hyphodontia_pallidula(100); | Hyphodontia | 13 | Saprotroph | Undefined Saprotroph-\|Wood Saprotroph\| | Corticioid | White Rot | Probable | [illegible] | [illegible] |
| Otu02195 | 20 | k__Fungi(100);p__Basidiomycota(100);c__Agaricomycetes(100);o__Polyporales(100);f__Phanerochaetaceae(100);g__Hyphodermella(95);g__Hyphodermella_unclassified(95); | Hyphodermella | 13 | Saprotroph | Undefined Saprotroph-\|Wood Saprotroph\| | Corticioid | White Rot | Probable | [illegible] | [illegible] |

| | | | | | | | | | | | |
|---|---|---|---|---|---|---|---|---|---|---|---|
| Otu02197 | 20 | k__Fungi(100);p__Basidiomycota(100);c__Microbotryomycetes(100);o__Sporidiobolales(100);f__Sporidiobolales_family_Incertae_sedis(100);g__Rhodotorula(100);g__Rhodotorula_unclassified(95); | Rhodotorula | 13 | Pathotroph-Saprotroph-Symbiotroph | Animal Pathogen-Animal Symbiotroph-Endophyte-Plant Pathogen-\|Undefined Saprotroph\| | Yeast | NULL | Probable | As foliar_endophyte (Põlme et al. 2020); Opportunistic human pathogen | Põlme S, et al. 2020. Fungal Diversity 105:1-16 ((https://doi.org/10.1007/s13225-020-00466-2)); https://www.cdc.gov ((https://www.cdc.gov)) |
| Otu02198 | 20 | k__Fungi(100);p__Ascomycota(100);c__Sordariomycetes(100);o__Hypocreales(100);f__Hypocreales_family_Incertae_sedis(100);g__Acremonium(100);s__Acremonium_sp_274_OA_2013(100); | Acremonium | 13 | Pathotroph-Saprotroph-Symbiotroph | Animal Pathogen-Endophyte-Fungal Parasite-Plant Pathogen-\|Undefined Saprotroph\|-Wood Saprotroph | Microfungus | Soft Rot | Possible | As foliar_endophyte (Põlme et al. 2020); culture (Hanada et al. 2010, Del Olmo; Decay leaf, fruit, seed, soil, dung, animal material (Põlme et al. 2020); Endophyte detection method; Ruiz & Arnold 2014); Likely opportunistic human pathogen (Irinyi et al. 2015) | Duncan CG, Eslyn WE. 1966. Mycologia 58:642-645 ((https://doi.org/10.2307/3757045)); Nilsson T. 1973. Studia Forestalia Suecica 104: 1-40 ((https://pub.epsilon.slu.se/5711/)); Worrall JJ, et al. 1997. Mycologia 89:199-219 ((https://doi.org/10.2307/3761073)); Hanada RE, et al. 2010. Fungal Biology 114:901-910 ((https://doi.org/10.1016/j.funbio.2010.08.006)); Del Olmo-Ruiz M, Arnold AE. 2014. Mycologia 106: 8-21 ((https://doi.org/10.3852/13-098)); Irinyi L, et al. 2015. Medical Mycology 53313-337 ((https://doi.org/10.1093/mmy/myv008)); Põlme S, et al. 2020. Fungal Diversity 105:1-16 ((https://doi.org/10.1007/s13225-020-00466-2)) |
| Otu02200 | 20 | k__Fungi(100);p__Basidiomycota(100);c__Agaricomycetes(100);o__Russulales(100);f__Russulaceae(100);g__Russula(100);s__Russula_badia(100); | Russula | 13 | Symbiotroph | \|Ectomycorrhizal\| | Agaricoid | NULL | Highly Probable | NULL | Rinaldi AC, et al. 2008. Fungal Diversity 33:1-45 ((http://www.fungaldiversity.org/fdp/sfdp/33-1.pdf)); Tedersoo L, et al. 2010. Mycorrhiza 20:217-263 ((https://doi.org/10.1007/s00572-009-0274-x)); Põlme S, et al. 2020. Fungal Diversity 105:1-16 ((https://doi.org/10.1007/s13225-020-00466-2)); http://mycorrhizas.info/ecmf.html ((http://mycorrhizas.info/ecmf.html)) |
| Otu02202 | 20 | k__Fungi(100);p__Ascomycota(100);c__Dothideomycetes(95);o__Dothideomycetes_order_Incertae_sedis(65);f__Dothideomycetes_family_Incertae_sedis(65);g__Phaeosclera(65);s__Phaeosclera_sp(65); | Phaeosclera | 13 | Saprotroph | \|Plant Saprotroph\| | Microfungus | NULL | Probable | As litter_saprotroph (Põlme et al. 2020); Decay leaf, fruit, seed (Põlme et al. 2020) | Põlme S, et al. 2020. Fungal Diversity 105:1-16 ((https://doi.org/10.1007/s13225-020-00466-2)) |
| Otu02203 | 20 | k__Fungi(100);p__Ascomycota(100);c__Dothideomycetes(100);o__Dothideomycetes_order_Incertae_sedis(100);f__Myxotrichaceae(100);g__Oidiodendron(100);g__Oidiodendron_unclassified(100); | Oidiodendron | 13 | Pathotroph-Saprotroph-Symbiotroph | Endophyte-Ericoid Mycorrhizal-\|Undefined Saprotroph\| | Microfungus | NULL | Probable | As root_endophyte (Põlme et al. 2020); As soil_saprotroph (Põlme et al. 2020); Dark Septate Endophyte; Decay soil (Põlme et al. 2020) | Newsham KK. 2011. New Phytologist 190:783-793 ((https://doi.org/10.1111/j.1469-8137.2010.03611.x)) (as dark septate root endophytes); Põlme S, et al. 2020. Fungal Diversity 105:1-16 ((https://doi.org/10.1007/s13225-020-00466-2)) |
| Otu02204 | 20 | k__Fungi(100);p__Ascomycota(100);c__Pezizomycetes(100);o__Pezizales(100);f__Pezizaceae(100);g__Peziza(100);s__Peziza_howsei(65); | Peziza | 13 | Saprotroph-Symbiotroph | Dung Saprotroph-Ectomycorrhizal-Endophyte-Plant Saprotroph-\|Undefined Saprotroph\| | Pezizoid | NULL | Possible | As foliar_endophyte (Põlme et al. 2020); As soil_saprotroph (Põlme et al. 2020); Decay soil (Põlme et al. 2020); Saprobic or ectomycorrhizal, on or in soil, leafmould etc., some species associated with dung or fire sites (Cannon & Kirk 2007) | Cannon PF, Kirk PM. 2007. Fungal Families of the World. CAB International, Cambridge (ISBN: 978-0851998275); Rinaldi AC, et al. 2008. Fungal Diversity 33:1-45 ((http://www.fungaldiversity.org/fdp/sfdp/33-1.pdf)); Tedersoo L, et al. 2010. Mycorrhiza 20:217-263 ((https://doi.org/10.1007/s00572-009-0274-x)) (pro parte); Põlme S, et al. 2020. Fungal Diversity 105:1-16 ((https://doi.org/10.1007/s13225-020-00466-2)) |
| Otu02205 | 20 | k__Fungi(100);p__Basidiomycota(100);c__Agaricomycetes(100);o__Cantharellales(100);f__Ceratobasidiaceae(100);g__unclassified_Ceratobasidiaceae(100);s__Ceratobasidiaceae_sp(100); | Ceratobasidiaceae | 9 | Pathotroph-Saprotroph-Symbiotroph | Endomycorrhizal-Plant Pathogen-Undefined Saprotroph | Microfungus | NULL | Possible | Best known as plant pathogens, but also saprobic. Species are endomycorrhizal with a wide range of plant roots - especially of Orchidaceae but also even bryophytes (Cannon & Kirk 2007) | Cannon PF, Kirk PM. 2007. Fungal Families of the World. CAB International, Cambridge (ISBN: 978-0851998275) |
| Otu02206 | 20 | k__Fungi(100);p__Basidiomycota(100);c__Agaricomycetes(100);o__Polyporales(100);f__Fomitopsidaceae(100);g__Postia(100);s__Postia_sericeomollis(100); | Postia | 13 | Saprotroph | \|Wood Saprotroph\| | Polyporoid | Brown Rot | Highly Probable | As brown-rot (Põlme et al. 2020); Decay wood (Põlme et al. 2020) | Jordan CR, et al. 1996. Holzforschung - International Journal of the Biology, Chemistry, Physics and Technology of Wood 50:312-318 ((https://doi.org/10.1515/hfsg.1996.50.4.312)); Põlme S, et al. 2020. Fungal Diversity 105:1-16 ((https://doi.org/10.1007/s13225-020-00466-2)) |
| Otu02208 | 20 | k__Fungi(100);p__Ascomycota(100);c__Dothideomycetes(100);o__Capnodiales(100);f__Teratosphaeriaceae(100);g__Devriesia(100);s__Devriesia_sp(100); | Devriesia | 13 | Pathotroph-Saprotroph | \|Plant Pathogen\|-Plant Saprotroph | Microfungus | NULL | Probable | As litter_saprotroph (Põlme et al. 2020); Decay leaf, fruit, seed (Põlme et al. 2020) | Tedersoo L, et al. 2014. Science 346:e1256688 ((https://doi.org/10.1126/science.1256688)); Põlme S, et al. 2020. Fungal Diversity 105:1-16 ((https://doi.org/10.1007/s13225-020-00466-2)) |
| Otu02210 | 20 | k__Fungi(100);p__Ascomycota(100);c__Saccharomycetes(100);o__Saccharomycetales(100);f__Metschnikowiaceae(100);g__Metschnikowia(100);s__Metschnikowia_pulcherrima(95); | Metschnikowia | 13 | Saprotroph | \|Undefined Saprotroph\| | Yeast | NULL | Probable | As nectar-tap_saprotroph (Põlme et al. 2020); Decay sugar-rich substrates (Põlme et al. 2020) | Põlme S, et al. 2020. Fungal Diversity 105:1-16 ((https://doi.org/10.1007/s13225-020-00466-2)) |
| Otu02214 | 20 | k__Fungi(100);p__Ascomycota(100);c__Dothideomycetes(100);o__Capnodiales(100);f__Teratosphaeriaceae(100);g__Teratosphaeria(100);s__Teratosphaeria_knoxdavesii(100); | Teratosphaeria | 13 | Pathotroph-Saprotroph | \|Plant Pathogen\|-Plant Saprotroph-Wood Saprotroph | Microfungus | NULL | Probable | As litter_saprotroph (Põlme et al. 2020); As wood_pathogen (Põlme et al. 2020); Decay wood, leaf, fruit, seed (Põlme et al. 2020) | Yamazaki A, et al. 2010. Mycoscience 51:443-455 ((https://www.jstage.jst.go.jp/article/mycosci/51/6/51_MYC51443/_pdf)); Tedersoo L, et al. 2014. Science 346:e1256688 ((https://doi.org/10.1126/science.1256688)); Põlme S, et al. 2020. Fungal Diversity 105:1-16 ((https://doi.org/10.1007/s13225-020-00466-2)) |
| Otu02216 | 20 | k__Fungi(100);p__Basidiomycota(100);c__Agaricomycetes(100);o__Thelephorales(100);f__Thelephoraceae(100);f__Thelephoraceae_unclassified(100);f__Thelephoraceae_unclassified(100); | Thelephoraceae | 9 | Saprotroph-Symbiotroph | Ectomycorrhizal-Undefined Saprotroph | Clavarioid | NULL | Probable | On the ground, on soil, a few on wood. At least some speices ectomycorrhizal (Cannon & Kirk 2007) | Cannon PF, Kirk PM. 2007. Fungal Families of the World. CAB International, Cambridge (ISBN: 978-0851998275) |
| Otu02219 | 20 | k__Fungi(100);p__Basidiomycota(100);c__Agaricomycetes(100);o__Agaricales(100);f__Psathyrellaceae(100);g__Coprinopsis(100);s__Coprinopsis_sp_CCG_2012(100); | Coprinopsis | 13 | Saprotroph | \|Undefined Saprotroph\| | Agaricoid | NULL | Probable | As soil_saprotroph (Põlme et al. 2020); Decay soil (Põlme et al. 2020) | Tedersoo L, et al. 2014. Science 346:e1256688 ((https://doi.org/10.1126/science.1256688)); Põlme S, et al. 2020. Fungal Diversity 105:1-16 ((https://doi.org/10.1007/s13225-020-00466-2)) |
| Otu02223 | 20 | k__Fungi(100);p__Basidiomycota(100);c__Exobasidiomycetes(100);o__Exobasidiales(100);f__Exobasidiaceae(100);g__Exobasidium(100);s__Exobasidium_formosanum(100); | Exobasidium | 13 | Pathotroph | \|Plant Pathogen\| | Leaf Blister-Leaf Gall-Leaf Spot | NULL | Probable | NULL | Tedersoo L, et al. 2014. Science 346:e1256688 ((https://doi.org/10.1126/science.1256688)); Põlme S, et al. 2020. Fungal Diversity 105:1-16 ((https://doi.org/10.1007/s13225-020-00466-2)) |
| Otu02225 | 20 | k__Fungi(100);p__Zygomycota(100);c__Zygomycota_class_Incertae_sedis(100);o__Mortierellales(100);f__Mortierellaceae(100);g__Mortierella(100);g__Mortierella_unclassified(100); | Mortierella | 13 | Saprotroph-Symbiotroph | Endophyte-Plant Saprotroph-\|Undefined Saprotroph\| | Microfungus | NULL | Probable | As chitinolytic (Põlme et al. 2020); As soil_saprotroph (Põlme et al. 2020); Decay roots, soil (Põlme et al. 2020); Saprobic in soil and thus frequently isolated from soil and material contaminated with soil such as hay or rotten silage and spoil tips (Cannon & Kirk 2007) | Cannon PF, Kirk PM. 2007. Fungal Families of the World. CAB International, Cambridge (ISBN: 978-0851998275); Tedersoo L, et al. 2014. Science 346:e1256688 ((https://doi.org/10.1126/science.1256688)); Purahong W, et al. 2016. Molecular Ecology 25:4059-4074 ((https://doi.org/10.1111/mec.13739)); Põlme S, et al. 2020. Fungal Diversity 105:1-16 ((https://doi.org/10.1007/s13225-020-00466-2)) |
| Otu02226 | 20 | k__Fungi(100);p__Basidiomycota(100);c__Agaricomycetes(100);o__Cantharellales(100);f__Hydnaceae(100);g__Sistotrema(100);s__Sistotrema_sp(100); | Sistotrema | 13 | Saprotroph-Symbiotroph | Ectomycorrhizal-\|Plant Saprotroph\|-Wood Saprotroph | Corticioid | White Rot | Possible | As litter_saprotroph (Põlme et al. 2020); Decay leaf, fruit, seed (Põlme et al. 2020) | Gilbertson RL, Ryvarden L. 1987-1987. North American Polypores. Fungiflora, Oslo (ISBN: 978-0945345060); Rinaldi AC, et al. 2008. Fungal Diversity 33:1-45 ((http://www.fungaldiversity.org/fdp/sfdp/33-1.pdf)) (as ECM/not ECM); Tedersoo L, et al. 2010. Mycorrhiza 20: 217-263 ((https://doi.org/10.1007/s00572-009-0274-x)) (pro parte); Põlme S, et al. 2020. Fungal Diversity 105:1-16 ((https://doi.org/10.1007/s13225-020-00466-2)); http://mycorrhizas.info/ecmf.html ((http://mycorrhizas.info/ecmf.html)) (as ECM/not ECM). |
| Otu02228 | 20 | k__Fungi(100);p__Ascomycota(100);c__Saccharomycetes(100);o__Saccharomycetales(100);f__Metschnikowiaceae(100);g__Metschnikowia(100);s__Metschnikowia_pulcherrima(100); | Metschnikowia | 13 | Saprotroph | \|Undefined Saprotroph\| | Yeast | NULL | Probable | As nectar-tap_saprotroph (Põlme et al. 2020); Decay sugar-rich substrates (Põlme et al. 2020) | Põlme S, et al. 2020. Fungal Diversity 105:1-16 ((https://doi.org/10.1007/s13225-020-00466-2)) |
| Otu02231 | 20 | k__Fungi(100);p__Ascomycota(100);c__Saccharomycetes(100);o__Saccharomycetales(100);f__Metschnikowiaceae(100);g__Metschnikowia(100);s__Metschnikowia_pulcherrima(80); | Metschnikowia | 13 | Saprotroph | \|Undefined Saprotroph\| | Yeast | NULL | Probable | As nectar-tap_saprotroph (Põlme et al. 2020); Decay sugar-rich substrates (Põlme et al. 2020) | Põlme S, et al. 2020. Fungal Diversity 105:1-16 ((https://doi.org/10.1007/s13225-020-00466-2)) |
| Otu02234 | 20 | k__Fungi(100);p__Ascomycota(100);c__Lecanoromycetes(100);o__Lecanorales(100);f__Parmeliaceae(100);g__Parmelia(100);s__Parmelia_sulcata(95); | Parmelia | 13 | Symbiotroph | \|Lichenized\| | Thallus | NULL | Highly Probable | NULL | Esslinger TL. 2014. North Dakota State University ((http://www.ndsu.edu/pubweb/~esslinge/chcklst/chcklst7.htm)); Põlme S, et al. 2020. Fungal Diversity 105:1-16 ((https://doi.org/10.1007/s13225-020-00466-2)) |
| Otu02236 | 20 | k__Fungi(100);p__Ascomycota(100);c__Dothideomycetes(100);o__Pleosporales(100);f__Massarinaceae(100);g__Helminthosporium(100);s__Helminthosporium_sp_E9213f(100); | Helminthosporium | 13 | Pathotroph | Fungal Parasite-\|Plant Pathogen\| | Microfungus | NULL | Probable | NULL | Samuels GJ. 2007. Self-published; Tedersoo L, et al. 2014. Science 346:e1256688 ((https://doi.org/10.1126/science.12566)); Põlme S, et al. 2020. Fungal Diversity 105:1-16 ((https://doi.org/10.1007/s13225-020-00466-2)) |
| Otu02159 | 20 | k__Fungi(100);k__Fungi_unclassified(90);k__Fungi_unclassified(90);k__Fungi_unclassified(90);k__Fungi_unclassified(90);k__Fungi_unclassified(90);k__Fungi_unclassified(90); | - | - | - | - | - | - | - | Unassigned | - |
| Otu02161 | 20 | k__Fungi(100);p__Ascomycota(65);p__Ascomycota_unclassified(65);p__Ascomycota_unclassified(65);p__Ascomycota_unclassified(65);p__Ascomycota_unclassified(65);p__Ascomycota_unclassified(65); | - | - | - | - | - | - | - | Unassigned | - |
| Otu02163 | 20 | k__Fungi(100);k__Fungi_unclassified(100);k__Fungi_unclassified(100);k__Fungi_unclassified(100);k__Fungi_unclassified(100);k__Fungi_unclassified(100);k__Fungi_unclassified(100); | - | - | - | - | - | - | - | Unassigned | - |
| Otu02164 | 20 | k__Fungi(100);k__Fungi_unclassified(61);k__Fungi_unclassified(61);k__Fungi_unclassified(61);k__Fungi_unclassified(61);k__Fungi_unclassified(61);k__Fungi_unclassified(61); | - | - | - | - | - | - | - | Unassigned | - |
| Otu02167 | 20 | k__Fungi(100);p__Ascomycota(100);c__Dothideomycetes(100);c__Dothideomycetes_unclassified(80);c__Dothideomycetes_unclassified(80);c__Dothideomycetes_unclassified(80);c__Dothideomycetes_unclassified(80); | - | - | - | - | - | - | - | Unassigned | - |
| Otu02168 | 20 | k__Fungi(100);p__Ascomycota(65);p__Ascomycota_unclassified(65);p__Ascomycota_unclassified(65);p__Ascomycota_unclassified(65);p__Ascomycota_unclassified(65);p__Ascomycota_unclassified(65); | - | - | - | - | - | - | - | Unassigned | - |
| Otu02169 | 20 | k__Fungi(100);k__Fungi_unclassified(80);k__Fungi_unclassified(80);k__Fungi_unclassified(80);k__Fungi_unclassified(80);k__Fungi_unclassified(80);k__Fungi_unclassified(80); | - | - | - | - | - | - | - | Unassigned | - |
| Otu02170 | 20 | k__Fungi(100);p__Ascomycota(100);c__Sordariomycetes(100);c__Sordariomycetes_unclassified(100);c__Sordariomycetes_unclassified(100);c__Sordariomycetes_unclassified(100);c__Sordariomycetes_unclassified(100); | - | - | - | - | - | - | - | Unassigned | - |
| Otu02171 | 20 | k__Fungi(100);k__Fungi_unclassified(100);k__Fungi_unclassified(100);k__Fungi_unclassified(100);k__Fungi_unclassified(100);k__Fungi_unclassified(100);k__Fungi_unclassified(100); | - | - | - | - | - | - | - | Unassigned | - |
| Otu02172 | 20 | k__Fungi(100);k__Fungi_unclassified(65);k__Fungi_unclassified(65);k__Fungi_unclassified(65);k__Fungi_unclassified(65);k__Fungi_unclassified(65);k__Fungi_unclassified(65); | - | - | - | - | - | - | - | Unassigned | - |
| Otu02173 | 20 | k__Fungi(100);p__Basidiomycota(100);c__Exobasidiomycetes(100);o__unclassified_Exobasidiomycetes(95);f__unclassified_Exobasidiomycetes(95);g__unclassified_Exobasidiomycetes(95);s__Exobasidiomycetes_sp(95); | - | - | - | - | - | - | - | Unassigned | - |
| Otu02175 | 20 | k__Fungi(100);p__Ascomycota(100);p__Ascomycota_unclassified(100);p__Ascomycota_unclassified(100);p__Ascomycota_unclassified(100);p__Ascomycota_unclassified(100);p__Ascomycota_unclassified(100); | - | - | - | - | - | - | - | Unassigned | - |
| Otu02178 | 20 | k__Fungi(100);k__Fungi_unclassified(61);k__Fungi_unclassified(61);k__Fungi_unclassified(61);k__Fungi_unclassified(61);k__Fungi_unclassified(61);k__Fungi_unclassified(61); | - | - | - | - | - | - | - | Unassigned | - |
| Otu02182 | 20 | k__Fungi(100);p__Basidiomycota(100);c__Agaricomycetes(100);c__Agaricomycetes_unclassified(100);c__Agaricomycetes_unclassified(100);c__Agaricomycetes_unclassified(100);c__Agaricomycetes_unclassified(100); | - | - | - | - | - | - | - | Unassigned | - |
| Otu02183 | 20 | k__Fungi(100);k__Fungi_unclassified(61);k__Fungi_unclassified(61);k__Fungi_unclassified(61);k__Fungi_unclassified(61);k__Fungi_unclassified(61);k__Fungi_unclassified(61); | - | - | - | - | - | - | - | Unassigned | - |
| Otu02184 | 20 | k__Fungi(100);k__Fungi_unclassified(80);k__Fungi_unclassified(80);k__Fungi_unclassified(80);k__Fungi_unclassified(80);k__Fungi_unclassified(80);k__Fungi_unclassified(80); | - | - | - | - | - | - | - | Unassigned | - |
| Otu02186 | 20 | k__Fungi(100);k__Fungi_unclassified(100);k__Fungi_unclassified(100);k__Fungi_unclassified(100);k__Fungi_unclassified(100);k__Fungi_unclassified(100);k__Fungi_unclassified(100); | - | - | - | - | - | - | - | Unassigned | - |
| Otu02191 | 20 | k__Fungi(100);p__Ascomycota(100);c__Sordariomycetes(100);o__Diaporthales(100);o__Diaporthales_unclassified(100);o__Diaporthales_unclassified(100);o__Diaporthales_unclassified(100); | - | - | - | - | - | - | - | Unassigned | - |
| Otu02192 | 20 | k__Fungi(100);p__Ascomycota(100);c__Sordariomycetes(100);c__Sordariomycetes_unclassified(100);c__Sordariomycetes_unclassified(100);c__Sordariomycetes_unclassified(100);c__Sordariomycetes_unclassified(100); | - | - | - | - | - | - | - | Unassigned | - |
| Otu02193 | 20 | k__Fungi(100);p__Basidiomycota(100);c__Agaricomycetes(75);c__Agaricomycetes_unclassified(75);c__Agaricomycetes_unclassified(75);c__Agaricomycetes_unclassified(75);c__Agaricomycetes_unclassified(75); | - | - | - | - | - | - | - | Unassigned | - |
| Otu02196 | 20 | k__Fungi(100);p__Ascomycota(100);p__Ascomycota_unclassified(85);p__Ascomycota_unclassified(85);p__Ascomycota_unclassified(85);p__Ascomycota_unclassified(85);p__Ascomycota_unclassified(85); | - | - | - | - | - | - | - | Unassigned | - |
| Otu02199 | 20 | k__Plantae(100);p__unclassified_Plantae(100);c__unclassified_Plantae(100);o__unclassified_Plantae(100);f__unclassified_Plantae(100);g__unclassified_Plantae(100);s__Plantae_sp(100); | - | - | - | - | - | - | - | Unassigned | - |
| Otu02201 | 20 | k__Fungi(100);p__Ascomycota(100);c__Dothideomycetes(100);o__Pleosporales(100);o__Pleosporales_unclassified(100);o__Pleosporales_unclassified(100);o__Pleosporales_unclassified(100); | - | - | - | - | - | - | - | Unassigned | - |
| Otu02207 | 20 | k__Fungi(100);p__Ascomycota(100);c__Sordariomycetes(100);o__Xylariales(80);o__Xylariales_unclassified(80);o__Xylariales_unclassified(80);o__Xylariales_unclassified(80); | - | - | - | - | - | - | - | Unassigned | - |
| Otu02209 | 20 | k__Fungi(100);k__Fungi_unclassified(55);k__Fungi_unclassified(55);k__Fungi_unclassified(55);k__Fungi_unclassified(55);k__Fungi_unclassified(55);k__Fungi_unclassified(55); | - | - | - | - | - | - | - | Unassigned | - |
| Otu02211 | 20 | k__Fungi(100);p__Ascomycota(100);c__Eurotiomycetes(100);o__Eurotiales(100);f__Trichocomaceae(100);g__unclassified_Trichocomaceae(90);s__Trichocomaceae_sp(90); | - | - | - | - | - | - | - | Unassigned | - |
| Otu02212 | 20 | k__Fungi(100);p__Ascomycota(100);p__Ascomycota_unclassified(100);p__Ascomycota_unclassified(100);p__Ascomycota_unclassified(100);p__Ascomycota_unclassified(100);p__Ascomycota_unclassified(100); | - | - | - | - | - | - | - | Unassigned | - |
| Otu02213 | 20 | k__Fungi(100);p__Basidiomycota(55);p__Basidiomycota_unclassified(55);p__Basidiomycota_unclassified(55);p__Basidiomycota_unclassified(55);p__Basidiomycota_unclassified(55);p__Basidiomycota_unclassified(55); | - | - | - | - | - | - | - | Unassigned | - |
| Otu02215 | 20 | k__Fungi(100);p__Basidiomycota(100);c__Microbotryomycetes(80);o__Sporidiobolales(80);f__Sporidiobolales_family_Incertae_sedis(80);f__Sporidiobolales_family_Incertae_sedis_unclassified(65);f__Sporidiobolales_family_Incertae_sedis_unclassified(65); | - | - | - | - | - | - | - | Unassigned | - |
| Otu02217 | 20 | k__Fungi(100);p__Basidiomycota(100);c__Agaricomycetes(100);c__Agaricomycetes_unclassified(100);c__Agaricomycetes_unclassified(100);c__Agaricomycetes_unclassified(100);c__Agaricomycetes_unclassified(100); | - | - | - | - | - | - | - | Unassigned | - |
| Otu02218 | 20 | k__Fungi(100);p__Ascomycota(100);c__Leotiomycetes(100);o__Helotiales(100);o__Helotiales_unclassified(75);o__Helotiales_unclassified(75);o__Helotiales_unclassified(75); | - | - | - | - | - | - | - | Unassigned | - |
| Otu02220 | 20 | k__Fungi(100);k__Fungi_unclassified(100);k__Fungi_unclassified(100);k__Fungi_unclassified(100);k__Fungi_unclassified(100);k__Fungi_unclassified(100);k__Fungi_unclassified(100); | - | - | - | - | - | - | - | Unassigned | - |
| Otu02221 | 20 | k__Fungi(100);p__Ascomycota(100);p__Ascomycota_unclassified(100);p__Ascomycota_unclassified(100);p__Ascomycota_unclassified(100);p__Ascomycota_unclassified(100);p__Ascomycota_unclassified(100); | - | - | - | - | - | - | - | Unassigned | - |
| Otu02222 | 20 | k__Fungi(100);p__Ascomycota(61);p__Ascomycota_unclassified(61);p__Ascomycota_unclassified(61);p__Ascomycota_unclassified(61);p__Ascomycota_unclassified(61);p__Ascomycota_unclassified(61); | - | - | - | - | - | - | - | Unassigned | - |
| Otu02224 | 20 | k__Fungi(100);k__Fungi_unclassified(55);k__Fungi_unclassified(55);k__Fungi_unclassified(55);k__Fungi_unclassified(55);k__Fungi_unclassified(55);k__Fungi_unclassified(55); | - | - | - | - | - | - | - | Unassigned | - |
| Otu02227 | 20 | k__Fungi(100);p__Ascomycota(100);c__Dothideomycetes(100);o__Pleosporales(100);f__unclassified_Pleosporales(80);g__unclassified_Pleosporales(80);s__Pleosporales_sp(80); | - | - | - | - | - | - | - | Unassigned | - |
| Otu02229 | 20 | k__Fungi(100);p__Ascomycota(70);p__Ascomycota_unclassified(70);p__Ascomycota_unclassified(70);p__Ascomycota_unclassified(70);p__Ascomycota_unclassified(70);p__Ascomycota_unclassified(70); | - | - | - | - | - | - | - | Unassigned | - |
| Otu02230 | 20 | k__Fungi(100);p__Ascomycota(100);p__Ascomycota_unclassified(100);p__Ascomycota_unclassified(100);p__Ascomycota_unclassified(100);p__Ascomycota_unclassified(100);p__Ascomycota_unclassified(100); | - | - | - | - | - | - | - | Unassigned | - |
| Otu02232 | 20 | k__Fungi(100);p__Ascomycota(100);p__Ascomycota_unclassified(100);p__Ascomycota_unclassified(100);p__Ascomycota_unclassified(100);p__Ascomycota_unclassified(100);p__Ascomycota_unclassified(100); | - | - | - | - | - | - | - | Unassigned | - |
| Otu02233 | 20 | k__Fungi(100);p__Ascomycota(100);c__Sordariomycetes(100);c__Sordariomycetes_unclassified(100);c__Sordariomycetes_unclassified(100);c__Sordariomycetes_unclassified(100);c__Sordariomycetes_unclassified(100); | - | - | - | - | - | - | - | Unassigned | - |
| Otu02235 | 20 | k__Fungi(100);p__Ascomycota(85);p__Ascomycota_unclassified(85);p__Ascomycota_unclassified(85);p__Ascomycota_unclassified(85);p__Ascomycota_unclassified(85);p__Ascomycota_unclassified(85); | - | - | - | - | - | - | - | Unassigned | - |
| Otu02237 | 20 | k__Fungi(100);p__Ascomycota(100);c__Sordariomycetes(100);c__Sordariomycetes_unclassified(100);c__Sordariomycetes_unclassified(100);c__Sordariomycetes_unclassified(100);c__Sordariomycetes_unclassified(100); | - | - | - | - | - | - | - | Unassigned | - |
| Otu02243 | 19 | k__Fungi(100);p__Ascomycota(100);c__Saccharomycetes(100);o__Saccharomycetales(100);f__Metschnikowiaceae(100);g__Metschnikowia(100);s__Metschnikowia_pulcherrima(95); | Metschnikowia | 13 | Saprotroph | \|Undefined Saprotroph\| | Yeast | NULL | Probable | As nectar-tap_saprotroph (Põlme et al. 2020); Decay sugar-rich substrates (Põlme et al. 2020) | Põlme S, et al. 2020. Fungal Diversity 105:1-16 ((https://doi.org/10.1007/s13225-020-00466-2)) |
| Otu02244 | 19 | k__Fungi(100);p__Zygomycota(100);c__Zygomycota_class_Incertae_sedis(100);o__Mucorales(100);f__Mucoraceae(100);g__Mucor(100);g__Mucor_unclassified(95); | Mucor | 13 | Saprotroph | \|Undefined Saprotroph\| | Microfungus | NULL | Probable | As soil_saprotroph (Põlme et al. 2020); Decay soil (Põlme et al. 2020) | Tedersoo L, et al. 2014. Science 346:e1256688 ((https://doi.org/10.1126/science.1256688)); Põlme S, et al. 2020. Fungal Diversity 105:1-16 ((https://doi.org/10.1007/s13225-020-00466-2)) |
| Otu02245 | 19 | k__Fungi(100);p__Basidiomycota(100);c__Agaricomycetes(100);o__Agaricales(100);f__Psathyrellaceae(100);g__Coprinellus(100);s__Coprinellus_disseminatus(100); | Coprinellus | 13 | Saprotroph | \|Undefined Saprotroph\| | Agaricoid | NULL | Probable | As soil_saprotroph (Põlme et al. 2020); Decay soil (Põlme et al. 2020) | Tedersoo L, et al. 2014. Science 346:e1256688 ((https://doi.org/10.1126/science.1256688)); Põlme S, et al. 2020. Fungal Diversity 105:1-16 ((https://doi.org/10.1007/s13225-020-00466-2)) |
| Otu02246 | 19 | k__Fungi(100);p__Ascomycota(100);c__Saccharomycetes(100);o__Saccharomycetales(100);f__Metschnikowiaceae(100);g__Metschnikowia(100);g__Metschnikowia_unclassified(53); | Metschnikowia | 13 | Saprotroph | \|Undefined Saprotroph\| | Yeast | NULL | Probable | As nectar-tap_saprotroph (Põlme et al. 2020); Decay sugar-rich substrates (Põlme et al. 2020) | Põlme S, et al. 2020. Fungal Diversity 105:1-16 ((https://doi.org/10.1007/s13225-020-00466-2)) |
| Otu02247 | 19 | k__Fungi(100);p__Ascomycota(100);c__Saccharomycetes(100);o__Saccharomycetales(100);f__Metschnikowiaceae(100);g__Metschnikowia(100);g__Metschnikowia_unclassified(85); | Metschnikowia | 13 | Saprotroph | \|Undefined Saprotroph\| | Yeast | NULL | Probable | As nectar-tap_saprotroph (Põlme et al. 2020); Decay sugar-rich substrates (Põlme et al. 2020) | Põlme S, et al. 2020. Fungal Diversity 105:1-16 ((https://doi.org/10.1007/s13225-020-00466-2)) |
| Otu02248 | 19 | k__Fungi(100);p__Ascomycota(100);c__Saccharomycetes(100);o__Saccharomycetales(100);f__Metschnikowiaceae(100);g__Metschnikowia(100);s__Metschnikowia_pulcherrima(95); | Metschnikowia | 13 | Saprotroph | \|Undefined Saprotroph\| | Yeast | NULL | Probable | As nectar-tap_saprotroph (Põlme et al. 2020); Decay sugar-rich substrates (Põlme et al. 2020) | Põlme S, et al. 2020. Fungal Diversity 105:1-16 ((https://doi.org/10.1007/s13225-020-00466-2)) |
| Otu02249 | 19 | k__Fungi(100);p__Ascomycota(100);c__Saccharomycetes(100);o__Saccharomycetales(100);f__Metschnikowiaceae(100);g__Metschnikowia(100);s__Metschnikowia_pulcherrima(100); | Metschnikowia | 13 | Saprotroph | \|Undefined Saprotroph\| | Yeast | NULL | Probable | As nectar-tap_saprotroph (Põlme et al. 2020); Decay sugar-rich substrates (Põlme et al. 2020) | Põlme S, et al. 2020. Fungal Diversity 105:1-16 ((https://doi.org/10.1007/s13225-020-00466-2)) |
| Otu02252 | 19 | k__Fungi(100);p__Ascomycota(100);c__Saccharomycetes(100);o__Saccharomycetales(100);f__Metschnikowiaceae(100);g__Metschnikowia(100);g__Metschnikowia_unclassified(79); | Metschnikowia | 13 | Saprotroph | \|Undefined Saprotroph\| | Yeast | NULL | Probable | As nectar-tap_saprotroph (Põlme et al. 2020); Decay sugar-rich substrates (Põlme et al. 2020) | Põlme S, et al. 2020. Fungal Diversity 105:1-16 ((https://doi.org/10.1007/s13225-020-00466-2)) |
| Otu02255 | 19 | k__Fungi(100);p__Basidiomycota(100);c__Microbotryomycetes(100);o__Sporidiobolales(100);f__Sporidiobolales_family_Incertae_sedis(100);g__Sporobolomyces(100);s__Sporobolomyces_sp_TY_223(100); | Sporobolomyces | 13 | Pathotroph-Saprotroph | \|Fungal Parasite\|-Plant Saprotroph | Yeast | NULL | Probable | As litter_saprotroph (Põlme et al. 2020); As mycoparasite (Põlme et al. 2020); Decay fungal material (Põlme et al. 2020) | Tedersoo L, et al. 2014. Science 346:e1256688 ((https://doi.org/10.1126/science.1256688)); Purahong W, et al. 2016. Molecular Ecology 25:4059-4074 ((https://doi.org/10.1111/mec.13739)); Põlme S, et al. 2020. Fungal Diversity 105:1-16 ((https://doi.org/10.1007/s13225-020-00466-2)) |
| Otu02256 | 19 | k__Fungi(100);p__Basidiomycota(100);c__Agaricomycetes(100);o__Russulales(100);f__Auriscalpiaceae(100);g__Lentinellus(100);s__Lentinellus_cochleatus(100); | Lentinellus | 13 | Pathotroph-Saprotroph | Plant Pathogen-Undefined Saprotroph-\|Wood Saprotroph\| | Agaricoid | White Rot | Probable | As white_rot (Põlme et al. 2020); As wood_pathogen (Põlme et al. 2020); Decay wood (Põlme et al. 2020) | Tedersoo L, et al. 2014. Science 346:e1256688 ((https://doi.org/10.1126/science.1256688)); Põlme S, et al. 2020. Fungal Diversity 105:1-16 ((https://doi.org/10.1007/s13225-020-00466-2)) |
| Otu02261 | 19 | k__Fungi(100);p__Ascomycota(100);c__Dothideomycetes(100);o__Capnodiales(100);f__Teratosphaeriaceae(100);g__Devriesia(100);s__Devriesia_sp(100); | Devriesia | 13 | Pathotroph-Saprotroph | \|Plant Pathogen\|-Plant Saprotroph | Microfungus | NULL | Probable | As litter_saprotroph (Põlme et al. 2020); Decay leaf, fruit, seed (Põlme et al. 2020) | Tedersoo L, et al. 2014. Science 346:e1256688 ((https://doi.org/10.1126/science.1256688)); Põlme S, et al. 2020. Fungal Diversity 105:1-16 ((https://doi.org/10.1007/s13225-020-00466-2)) |
| Otu02262 | 19 | k__Fungi(100);p__Ascomycota(100);c__Saccharomycetes(100);o__Saccharomycetales(100);f__Metschnikowiaceae(100);g__Metschnikowia(100);s__Metschnikowia_pulcherrima(95); | Metschnikowia | 13 | Saprotroph | \|Undefined Saprotroph\| | Yeast | NULL | Probable | As nectar-tap_saprotroph (Põlme et al. 2020); Decay sugar-rich substrates (Põlme et al. 2020) | Põlme S, et al. 2020. Fungal Diversity 105:1-16 ((https://doi.org/10.1007/s13225-020-00466-2)) |
| Otu02265 | 19 | k__Fungi(100);p__Ascomycota(100);c__Saccharomycetes(100);o__Saccharomycetales(100);f__Metschnikowiaceae(100);g__Metschnikowia(100);s__Metschnikowia_pulcherrima(100); | Metschnikowia | 13 | Saprotroph | \|Undefined Saprotroph\| | Yeast | NULL | Probable | As nectar-tap_saprotroph (Põlme et al. 2020); Decay sugar-rich substrates (Põlme et al. 2020) | Põlme S, et al. 2020. Fungal Diversity 105:1-16 ((https://doi.org/10.1007/s13225-020-00466-2)) |
| Otu02266 | 19 | k__Fungi(100);p__Ascomycota(100);c__Sordariomycetes(100);o__Hypocreales(100);f__Hypocreales_family_Incertae_sedis(100);g__Acremonium(100);s__Acremonium_sp(100); | Acremonium | 13 | Pathotroph-Saprotroph-Symbiotroph | Animal Pathogen-Endophyte-Fungal Parasite-Plant Pathogen-\|Undefined Saprotroph\|-Wood Saprotroph | Microfungus | Soft Rot | Possible | As foliar_endophyte (Põlme et al. 2020); culture (Hanada et al. 2010, Del Olmo; Decay leaf, fruit, seed, soil, dung, animal material (Põlme et al. 2020); Endophyte detection method; Ruiz & Arnold 2014); Likely opportunistic human pathogen (Irinyi et al. 2015) | Duncan CG, Eslyn WE. 1966. Mycologia 58:642-645 ((https://doi.org/10.2307/3757045)); Nilsson T. 1973. Studia Forestalia Suecica 104: 1-40 ((https://pub.epsilon.slu.se/5711/)); Worrall JJ, et al. 1997. Mycologia 89:199-219 ((https://doi.org/10.2307/3761073)); Hanada RE, et al. 2010. Fungal Biology 114:901-910 ((https://doi.org/10.1016/j.funbio.2010.08.006)); Del Olmo-Ruiz M, Arnold AE. 2014. Mycologia 106: 8-21 ((https://doi.org/10.3852/13-098)); Irinyi L, et al. 2015. Medical Mycology 53313-337 ((https://doi.org/10.1093/mmy/myv008)); Põlme S, et al. 2020. Fungal Diversity 105:1-16 ((https://doi.org/10.1007/s13225-020-00466-2)) |
| Otu02276 | 19 | k__Fungi(100);p__Ascomycota(100);c__Saccharomycetes(100);o__Saccharomycetales(100);f__Metschnikowiaceae(100);g__Metschnikowia(100);s__Metschnikowia_pulcherrima(95); | Metschnikowia | 13 | Saprotroph | \|Undefined Saprotroph\| | Yeast | NULL | Probable | As nectar-tap_saprotroph (Põlme et al. 2020); Decay sugar-rich substrates (Põlme et al. 2020) | Põlme S, et al. 2020. Fungal Diversity 105:1-16 ((https://doi.org/10.1007/s13225-020-00466-2)) |
| Otu02277 | 19 | k__Fungi(100);p__Ascomycota(100);c__Sordariomycetes(100);o__Hypocreales(100);o__Hypocreales_unclassified(100);o__Hypocreales_unclassified(100);o__Hypocreales_unclassified(100); | Hypocreales | 7 | Saprotroph | Undefined Saprotroph | Microfungus | NULL | Possible | NULL | Sterkenburg E, et al. 2015. New Phytologist 207:1145-1158 ((https://doi.org/10.1111/nph.13426)) |
| Otu02278 | 19 | k__Fungi(100);p__Ascomycota(100);c__Sordariomycetes(100);o__Hypocreales(100);f__Hypocreales_family_Incertae_sedis(100);g__Acremonium(100);g__Acremonium_unclassified(100); | Acremonium | 13 | Pathotroph-Saprotroph-Symbiotroph | Animal Pathogen-Endophyte-Fungal Parasite-Plant Pathogen-\|Undefined Saprotroph\|-Wood Saprotroph | Microfungus | Soft Rot | Possible | As foliar_endophyte (Põlme et al. 2020); culture (Hanada et al. 2010, Del Olmo; Decay leaf, fruit, seed, soil, dung, animal material (Põlme et al. 2020); Endophyte detection method; Ruiz & Arnold 2014); Likely opportunistic human pathogen (Irinyi et al. 2015) | Duncan CG, Eslyn WE. 1966. Mycologia 58:642-645 ((https://doi.org/10.2307/3757045)); Nilsson T. 1973. Studia Forestalia Suecica 104: 1-40 ((https://pub.epsilon.slu.se/5711/)); Worrall JJ, et al. 1997. Mycologia 89:199-219 ((https://doi.org/10.2307/3761073)); Hanada RE, et al. 2010. Fungal Biology 114:901-910 ((https://doi.org/10.1016/j.funbio.2010.08.006)); Del Olmo-Ruiz M, Arnold AE. 2014. Mycologia 106: 8-21 ((https://doi.org/10.3852/13-098)); Irinyi L, et al. 2015. Medical Mycology 53313-337 ((https://doi.org/10.1093/mmy/myv008)); Põlme S, et al. 2020. Fungal Diversity 105:1-16 ((https://doi.org/10.1007/s13225-020-00466-2)) |
| Otu02281 | 19 | k__Fungi(100);p__Ascomycota(100);c__Sordariomycetes(100);o__Hypocreales(100);o__Hypocreales_unclassified(90);o__Hypocreales_unclassified(90);o__Hypocreales_unclassified(90); | Hypocreales | 7 | Saprotroph | Undefined Saprotroph | Microfungus | NULL | Possible | NULL | Sterkenburg E, et al. 2015. New Phytologist 207:1145-1158 ((https://doi.org/10.1111/nph.13426)) |
| Otu02285 | 19 | k__Fungi(100);p__Ascomycota(100);c__Sordariomycetes(100);o__Hypocreales(100);f__Nectriaceae(100);g__Fusarium(100);s__Fusarium_sp_OTU_011(100); | Fusarium | 13 | Pathotroph-Saprotroph-Symbiotroph | Animal Pathogen-Endophyte-Lichen Parasite-\|Plant Pathogen\|-Plant Saprotroph-Undefined Saprotroph-Wood Saprotroph | Microfungus | Soft Rot | Possible | As foliar_endophyte (Põlme et al. 2020); As litter_saprotroph (Põlme et al. 2020); Causing damping-off disease (Põlme et al. 2020); Decay leaf, fruit, seed, roots, soil, animal material (Põlme et al. 2020); Likely opportunistic human pathogen (Irinyi et al. 2015) | Duncan CG, Eslyn WE. 1966. Mycologia 58:642-664 ((https://doi.org/10.2307/3757045)); Seehann G, et al. 1975. List of Fungi in Soft-Rot Tissue Tests. Institute fur Holzbiologie und Holzschutz, Bundesforschungsanstalt fur Forst - und Holzwirtshaft, Hamburg - Lohbrugge, Germany; Costa IPMW, et al. 2012. Mycotaxon 119:e26 ((http://mycotaxon.com/resources/checklists/costa_v119_checklist.pdf)); Tedersoo L, et al. 2014. Science 346:e1256688 ((https://doi.org/10.1126/science.1256688)); Irinyi L, et al. 2015. Medical Mycology 53:313-337 ((https://doi.org/10.1093/mmy/myv008)); Schroers, et al. 2016. Mycologia 108:806-819 ((https://doi.org/10.3852/15-255)); Põlme S, et al. 2020. Fungal Diversity 105:1-16 ((https://doi.org/10.1007/s13225-020-00466-2)); The American Phytopathological Society Common Names of Plant Diseases ((http://www.apsnet.org/publications/commonnames/Pages/default.aspx)); Lawrey JD, Diederich P. 2016. http://www.lichenicolous.net (accessed 24 May 2017) ((http://www.lichenicolous.net)) |
| Otu02289 | 19 | k__Fungi(100);p__Basidiomycota(100);c__Tremellomycetes(100);o__Tremellales(100);f__Tremellales_family_Incertae_sedis(100);g__Cryptococcus(100);s__Cryptococcus_pinus(100); | Cryptococcus | 13 | Pathotroph-Saprotroph-Symbiotroph | Animal Pathogen-Endophyte-Epiphyte-\|Undefined Saprotroph\| | Dimorphic Yeast-Yeast | NULL | Possible | As foliar_endophyte (Põlme et al. 2020); Decay leaf, fruit, seed, soil, dung, animal material (Põlme et al. 2020) | Andrews JH, et al. 1983. Phytopathology 73:228-234 ((https://doi.org/10.1094/Phyto-73-228)); Kurtzman CP, et al. (eds.) 2011. The Yeasts, a Taxonomic Study. Fifth Edition. Vols 1-3. Elsevier, San Diego (ISBN: 9780444521491); Põlme S, et al. 2020. Fungal Diversity 105:1-16 ((https://doi.org/10.1007/s13225-020-00466-2)) |
| Otu02290 | 19 | k__Fungi(100);p__Ascomycota(100);c__Sordariomycetes(100);o__Hypocreales(100);o__Hypocreales_unclassified(79);o__Hypocreales_unclassified(79);o__Hypocreales_unclassified(79); | Hypocreales | 7 | Saprotroph | Undefined Saprotroph | Microfungus | NULL | Possible | NULL | Sterkenburg E, et al. 2015. New Phytologist 207:1145-1158 ((https://doi.org/10.1111/nph.13426)) |
| Otu02291 | 19 | k__Fungi(100);p__Ascomycota(100);c__Saccharomycetes(100);o__Saccharomycetales(100);f__Metschnikowiaceae(100);g__Metschnikowia(100);s__Metschnikowia_pulcherrima(100); | Metschnikowia | 13 | Saprotroph | \|Undefined Saprotroph\| | Yeast | NULL | Probable | As nectar-tap_saprotroph (Põlme et al. 2020); Decay sugar-rich substrates (Põlme et al. 2020) | Põlme S, et al. 2020. Fungal Diversity 105:1-16 ((https://doi.org/10.1007/s13225-020-00466-2)) |
| Otu02294 | 19 | k__Fungi(100);p__Ascomycota(100);c__Sordariomycetes(100);o__Coniochaetales(100);f__Coniochaetaceae(100);f__Coniochaetaceae_unclassified(53);f__Coniochaetaceae_unclassified(53); | Coniochaetaceae | 9 | Pathotroph-Saprotroph-Symbiotroph | Animal Pathogen-Dung Saprotroph-Endophyte-Lichen Parasite-Plant Pathogen-Undefined Saprotroph | Microfungus | NULL | Possible | Saprobic on wood and bark (especially in dry habitats), coprophilous or isolated from soil and similar substrata (Cannon & Kirk 2007) | Cannon PF, Kirk PM. 2007. Fungal Families of the World. CAB International, Cambridge (ISBN: 978-0851998275); Damm U, et al. 2010. Persoonia 24:60-80 ((http://doi.org/10.3767/003158510X500705)); Irinyi L, et al. 2015. Medical Mycology 53:313-337 ((https://doi.org/10.1093/mmy/myv008)); Lawrey JD, Diederich P. 2016. http://www.lichenicolous.net (accessed 24 May 2017) ((http://www.lichenicolous.net)) |
| Otu02295 | 19 | k__Fungi(100);p__Basidiomycota(100);c__Agaricomycetes(100);o__Polyporales(100);f__Phanerochaetaceae(100);g__Phanerochaete(100);s__Phanerochaete_sordida(100); | Phanerochaete | 13 | Saprotroph | \|Wood Saprotroph\| | Corticioid | White Rot | Highly Probable | As white_rot (Põlme et al. 2020); Decay wood (Põlme et al. 2020) | Glenn JK, et al. 1983. Biochemical and Biophysical Research Communications 114:1077-1083 ((https://doi.org/10.1016/0006-291x((83)) 90672-1)); Põlme S, et al. 2020. Fungal Diversity 105:1-16 ((https://doi.org/10.1007/s13225-020-00466-2)) |
| Otu02296 | 19 | k__Fungi(100);p__Ascomycota(100);c__Saccharomycetes(100);o__Saccharomycetales(100);f__Metschnikowiaceae(100);g__Metschnikowia(100);s__Metschnikowia_pulcherrima(95); | Metschnikowia | 13 | Saprotroph | \|Undefined Saprotroph\| | Yeast | NULL | Probable | As nectar-tap_saprotroph (Põlme et al. 2020); Decay sugar-rich substrates (Põlme et al. 2020) | Põlme S, et al. 2020. Fungal Diversity 105:1-16 ((https://doi.org/10.1007/s13225-020-00466-2)) |
| Otu02297 | 19 | k__Fungi(100);p__Ascomycota(100);c__Saccharomycetes(100);o__Saccharomycetales(100);f__Metschnikowiaceae(100);g__Metschnikowia(100);s__Metschnikowia_pulcherrima(100); | Metschnikowia | 13 | Saprotroph | \|Undefined Saprotroph\| | Yeast | NULL | Probable | As nectar-tap_saprotroph (Põlme et al. 2020); Decay sugar-rich substrates (Põlme et al. 2020) | Põlme S, et al. 2020. Fungal Diversity 105:1-16 ((https://doi.org/10.1007/s13225-020-00466-2)) |
| Otu02298 | 19 | k__Fungi(100);p__Ascomycota(100);c__Saccharomycetes(100);o__Saccharomycetales(100);f__Saccharomycopsidaceae(100);g__Saccharomycopsis(100);s__Saccharomycopsis_microspora(100); | Saccharomycopsis | 13 | Pathotroph-Saprotroph | Animal Parasite-\|Undefined Saprotroph\| | Yeast | NULL | Probable | As nectar-tap_saprotroph (Põlme et al. 2020); Decay animal material, sugar-rich substrates (Põlme et al. 2020) | Põlme S, et al. 2020. Fungal Diversity 105:1-16 ((https://doi.org/10.1007/s13225-020-00466-2)) |
| Otu02299 | 19 | k__Fungi(100);p__Ascomycota(100);c__Saccharomycetes(100);o__Saccharomycetales(100);f__Metschnikowiaceae(100);g__Metschnikowia(100);s__Metschnikowia_pulcherrima(95); | Metschnikowia | 13 | Saprotroph | \|Undefined Saprotroph\| | Yeast | NULL | Probable | As nectar-tap_saprotroph (Põlme et al. 2020); Decay sugar-rich substrates (Põlme et al. 2020) | Põlme S, et al. 2020. Fungal Diversity 105:1-16 ((https://doi.org/10.1007/s13225-020-00466-2)) |
| Otu02301 | 19 | k__Fungi(100);p__Ascomycota(100);c__Dothideomycetes(100);o__Capnodiales(100);f__Mycosphaerellaceae(100);f__Mycosphaerellaceae_unclassified(100);f__Mycosphaerellaceae_unclassified(100); | Mycosphaerellaceae | 9 | Pathotroph-Saprotroph | Plant Pathogen-Undefined Saprotroph | Microfungus | NULL | Probable | Biotrophic, necrotrophic or saprobic on various plant tissues (Cannon & Kirk 2007) | Cannon PF, Kirk PM. 2007. Fungal Families of the World. CAB International, Cambridge (ISBN: 978-0851998275) |
| Otu02302 | 19 | k__Fungi(100);p__Ascomycota(100);c__Dothideomycetes(100);o__Pleosporales(100);f__Massarinaceae(100);g__Massarina(100);s__Massarina_sp_HF12700(100); | Massarina | 13 | Pathotroph-Saprotroph | \|Plant Pathogen\|-Undefined Saprotroph | Microfungus | NULL | Probable | NULL | Tedersoo L, et al. 2014. Science 346:e1256688 ((https://doi.org/10.1126/science.1256688)); Põlme S, et al. 2020. Fungal Diversity 105:1-16 ((https://doi.org/10.1007/s13225-020-00466-2)) |
| Otu02303 | 19 | k__Fungi(100);p__Ascomycota(100);c__Sordariomycetes(100);o__Hypocreales(100);f__Nectriaceae(100);f__Nectriaceae_unclassified(53);f__Nectriaceae_unclassified(53); | Nectriaceae | 9 | Pathotroph-Saprotroph-Symbiotroph | Animal Pathogen-Endophyte-Fungal Parasite-Lichen Parasite-Plant Pathogen-Wood Saprotroph | Microfungus | NULL | Possible | Fusarium species are amongst the economically significant plant pathogens (Cannon & Kirk 2007) | Cannon PF, Kirk PM. 2007. Fungal Families of the World. CAB International, Cambridge (ISBN: 978-0851998275); Hirooka Y, et al. 2012. Studies in Mycology 711-210 ((https://doi.org/10.3114/sim0001)); Tedersoo L, et al. 2014. Science 346:e1256688 ((https://doi.org/10.1126/science.1256688)); The American Phytopathological Society Common Names of Plant Diseases ((http://www.apsnet.org/publications/commonnames/Pages/default.aspx)); https://www.cdc.gov ((https://www.cdc.gov)) |
| Otu02306 | 19 | k__Fungi(100);p__Ascomycota(100);c__Saccharomycetes(100);o__Saccharomycetales(100);f__Saccharomycetales_family_Incertae_sedis(100);g__Candida(95);g__Candida_unclassified(90); | Candida | 13 | Pathotroph-Saprotroph-Symbiotroph | Animal Pathogen-Endophyte-Endosymbiont-Epiphyte-\|Undefined Saprotroph\| | Dimorphic Yeast-Yeast | NULL | Probable | As nectar-tap_saprotroph (Põlme et al. 2020); Decay sugar-rich substrates (Põlme et al. 2020); This asexual genus is polyphyletic and contain species that function very differently depend on what they are. Caution should be taken when interpreting guilds for Candida species. | Manolakaki D, et al. 2010. Virulence. 1: 367-375 ((https://doi.org/10.4161/viru.1.5.12796)); Mekbib SB, et al. 2011. Tropical Plant Pathology 36:233-240 ((https://doi.org/10.1590/S1982-56762011000400004)); Põlme S, et al. 2020. Fungal Diversity 105:1-16 ((https://doi.org/10.1007/s13225-020-00466-2)) |
| Otu02307 | 19 | k__Fungi(100);p__Ascomycota(100);c__Saccharomycetes(100);o__Saccharomycetales(100);f__Metschnikowiaceae(100);g__Metschnikowia(100);g__Metschnikowia_unclassified(95); | Metschnikowia | 13 | Saprotroph | \|Undefined Saprotroph\| | Yeast | NULL | Probable | As nectar-tap_saprotroph (Põlme et al. 2020); Decay sugar-rich substrates (Põlme et al. 2020) | Põlme S, et al. 2020. Fungal Diversity 105:1-16 ((https://doi.org/10.1007/s13225-020-00466-2)) |
| Otu02238 | 19 | k__Fungi(100);p__Basidiomycota(100);c__Agaricomycetes(100);o__Polyporales(100);f__Phanerochaetaceae(100);f__Phanerochaetaceae_unclassified(100);f__Phanerochaetaceae_unclassified(100); | - | - | - | - | - | - | - | Unassigned | - |
| Otu02239 | 19 | k__Fungi(100);p__Ascomycota(85);p__Ascomycota_unclassified(85);p__Ascomycota_unclassified(85);p__Ascomycota_unclassified(85);p__Ascomycota_unclassified(85);p__Ascomycota_unclassified(85); | - | - | - | - | - | - | - | Unassigned | - |
| Otu02240 | 19 | k__Fungi(100);k__Fungi_unclassified(58);k__Fungi_unclassified(58);k__Fungi_unclassified(58);k__Fungi_unclassified(58);k__Fungi_unclassified(58);k__Fungi_unclassified(58); | - | - | - | - | - | - | - | Unassigned | - |
| Otu02241 | 19 | k__Fungi(100);p__Ascomycota(58);p__Ascomycota_unclassified(58);p__Ascomycota_unclassified(58);p__Ascomycota_unclassified(58);p__Ascomycota_unclassified(58);p__Ascomycota_unclassified(58); | - | - | - | - | - | - | - | Unassigned | - |
| Otu02242 | 19 | k__Fungi(100);p__Basidiomycota(100);c__Agaricomycetes(100);c__Agaricomycetes_unclassified(100);c__Agaricomycetes_unclassified(100);c__Agaricomycetes_unclassified(100);c__Agaricomycetes_unclassified(100); | - | - | - | - | - | - | - | Unassigned | - |
| Otu02250 | 19 | k__Fungi(100);p__Ascomycota(58);p__Ascomycota_unclassified(58);p__Ascomycota_unclassified(58);p__Ascomycota_unclassified(58);p__Ascomycota_unclassified(58);p__Ascomycota_unclassified(58); | - | - | - | - | - | - | - | Unassigned | - |
| Otu02251 | 19 | k__Fungi(100);p__Ascomycota(95);p__Ascomycota_unclassified(90);p__Ascomycota_unclassified(90);p__Ascomycota_unclassified(90);p__Ascomycota_unclassified(90);p__Ascomycota_unclassified(90); | - | - | - | - | - | - | - | Unassigned | - |
| Otu02253 | 19 | k__Fungi(100);p__Ascomycota(100);c__Dothideomycetes(95);o__Pleosporales(95);o__Pleosporales_unclassified(95);o__Pleosporales_unclassified(95);o__Pleosporales_unclassified(95); | - | - | - | - | - | - | - | Unassigned | - |
| Otu02254 | 19 | k__Fungi(100);p__Basidiomycota(100);c__Microbotryomycetes(100);o__Sporidiobolales(100);f__Sporidiobolales_family_Incertae_sedis(100);f__Sporidiobolales_family_Incertae_sedis_unclassified(64);f__Sporidiobolales_family_Incertae_sedis_unclassified(64); | - | - | - | - | - | - | - | Unassigned | - |
| Otu02257 | 19 | k__Fungi(100);k__Fungi_unclassified(85);k__Fungi_unclassified(85);k__Fungi_unclassified(85);k__Fungi_unclassified(85);k__Fungi_unclassified(85);k__Fungi_unclassified(85); | - | - | - | - | - | - | - | Unassigned | - |
| Otu02258 | 19 | k__Fungi(100);p__Ascomycota(95);p__Ascomycota_unclassified(95);p__Ascomycota_unclassified(95);p__Ascomycota_unclassified(95);p__Ascomycota_unclassified(95);p__Ascomycota_unclassified(95); | - | - | - | - | - | - | - | Unassigned | - |
| Otu02259 | 19 | k__Fungi(100);p__Ascomycota(64);p__Ascomycota_unclassified(64);p__Ascomycota_unclassified(64);p__Ascomycota_unclassified(64);p__Ascomycota_unclassified(64);p__Ascomycota_unclassified(64); | - | - | - | - | - | - | - | Unassigned | - |
| Otu02260 | 19 | k__Fungi(100);p__Basidiomycota(100);c__Tremellomycetes(100);o__Filobasidiales(100);f__Filobasidiaceae(100);f__Filobasidiaceae_unclassified(100);f__Filobasidiaceae_unclassified(100); | - | - | - | - | - | - | - | Unassigned | - |
| Otu02263 | 19 | k__Fungi(100);k__Fungi_unclassified(74);k__Fungi_unclassified(74);k__Fungi_unclassified(74);k__Fungi_unclassified(74);k__Fungi_unclassified(74);k__Fungi_unclassified(74); | - | - | - | - | - | - | - | Unassigned | - |
| Otu02264 | 19 | k__Fungi(100);p__Ascomycota(100);p__Ascomycota_unclassified(100);p__Ascomycota_unclassified(100);p__Ascomycota_unclassified(100);p__Ascomycota_unclassified(100);p__Ascomycota_unclassified(100); | - | - | - | - | - | - | - | Unassigned | - |
| Otu02267 | 19 | k__Fungi(100);k__Fungi_unclassified(53);k__Fungi_unclassified(53);k__Fungi_unclassified(53);k__Fungi_unclassified(53);k__Fungi_unclassified(53);k__Fungi_unclassified(53); | - | - | - | - | - | - | - | Unassigned | - |
| Otu02268 | 19 | k__Fungi(100);p__Basidiomycota(100);c__Agaricomycetes(100);o__Cantharellales(100);o__Cantharellales_unclassified(69);o__Cantharellales_unclassified(69);o__Cantharellales_unclassified(69); | - | - | - | - | - | - | - | Unassigned | - |
| Otu02269 | 19 | k__Fungi(100);p__Ascomycota(100);p__Ascomycota_unclassified(79);p__Ascomycota_unclassified(79);p__Ascomycota_unclassified(79);p__Ascomycota_unclassified(79);p__Ascomycota_unclassified(79); | - | - | - | - | - | - | - | Unassigned | - |
| Otu02270 | 19 | k__Fungi(100);p__Ascomycota(95);p__Ascomycota_unclassified(95);p__Ascomycota_unclassified(95);p__Ascomycota_unclassified(95);p__Ascomycota_unclassified(95);p__Ascomycota_unclassified(95); | - | - | - | - | - | - | - | Unassigned | - |
| Otu02271 | 19 | k__Fungi(100);p__Basidiomycota(100);c__Agaricomycetes(100);c__Agaricomycetes_unclassified(100);c__Agaricomycetes_unclassified(100);c__Agaricomycetes_unclassified(100);c__Agaricomycetes_unclassified(100); | - | - | - | - | - | - | - | Unassigned | - |
| Otu02272 | 19 | k__Fungi(100);p__Ascomycota(100);p__Ascomycota_unclassified(100);p__Ascomycota_unclassified(100);p__Ascomycota_unclassified(100);p__Ascomycota_unclassified(100);p__Ascomycota_unclassified(100); | - | - | - | - | - | - | - | Unassigned | - |
| Otu02273 | 19 | k__Fungi(100);p__Ascomycota(69);p__Ascomycota_unclassified(69);p__Ascomycota_unclassified(69);p__Ascomycota_unclassified(69);p__Ascomycota_unclassified(69);p__Ascomycota_unclassified(69); | - | - | - | - | - | - | - | Unassigned | - |
| Otu02274 | 19 | k__Fungi(100);p__Ascomycota(58);p__Ascomycota_unclassified(58);p__Ascomycota_unclassified(58);p__Ascomycota_unclassified(58);p__Ascomycota_unclassified(58);p__Ascomycota_unclassified(58); | - | - | - | - | - | - | - | Unassigned | - |
| Otu02275 | 19 | k__Fungi(100);p__Ascomycota(100);p__Ascomycota_unclassified(100);p__Ascomycota_unclassified(100);p__Ascomycota_unclassified(100);p__Ascomycota_unclassified(100);p__Ascomycota_unclassified(100); | - | - | - | - | - | - | - | Unassigned | - |
| Otu02279 | 19 | k__Fungi(100);p__Ascomycota(85);p__Ascomycota_unclassified(85);p__Ascomycota_unclassified(85);p__Ascomycota_unclassified(85);p__Ascomycota_unclassified(85);p__Ascomycota_unclassified(85); | - | - | - | - | - | - | - | Unassigned | - |
| Otu02280 | 19 | k__Fungi(100);p__Ascomycota(90);p__Ascomycota_unclassified(90);p__Ascomycota_unclassified(90);p__Ascomycota_unclassified(90);p__Ascomycota_unclassified(90);p__Ascomycota_unclassified(90); | - | - | - | - | - | - | - | Unassigned | - |
| Otu02282 | 19 | k__Fungi(100);k__Fungi_unclassified(100);k__Fungi_unclassified(100);k__Fungi_unclassified(100);k__Fungi_unclassified(100);k__Fungi_unclassified(100);k__Fungi_unclassified(100); | - | - | - | - | - | - | - | Unassigned | - |
| Otu02283 | 19 | k__Fungi(100);k__Fungi_unclassified(58);k__Fungi_unclassified(58);k__Fungi_unclassified(58);k__Fungi_unclassified(58);k__Fungi_unclassified(58);k__Fungi_unclassified(58); | - | - | - | - | - | - | - | Unassigned | - |
| Otu02284 | 19 | k__Fungi(100);p__Ascomycota(100);p__Ascomycota_unclassified(100);p__Ascomycota_unclassified(100);p__Ascomycota_unclassified(100);p__Ascomycota_unclassified(100);p__Ascomycota_unclassified(100); | - | - | - | - | - | - | - | Unassigned | - |
| Otu02286 | 19 | k__Fungi(100);p__Basidiomycota(100);c__Agaricomycetes(100);c__Agaricomycetes_unclassified(100);c__Agaricomycetes_unclassified(100);c__Agaricomycetes_unclassified(100);c__Agaricomycetes_unclassified(100); | - | - | - | - | - | - | - | Unassigned | - |
| Otu02287 | 19 | k__Fungi(100);k__Fungi_unclassified(58);k__Fungi_unclassified(58);k__Fungi_unclassified(58);k__Fungi_unclassified(58);k__Fungi_unclassified(58);k__Fungi_unclassified(58); | - | - | - | - | - | - | - | Unassigned | - |
| Otu02288 | 19 | k__Fungi(100);k__Fungi_unclassified(90);k__Fungi_unclassified(90);k__Fungi_unclassified(90);k__Fungi_unclassified(90);k__Fungi_unclassified(90);k__Fungi_unclassified(90); | - | - | - | - | - | - | - | Unassigned | - |
| Otu02292 | 19 | k__Fungi(100);p__Ascomycota(64);p__Ascomycota_unclassified(64);p__Ascomycota_unclassified(64);p__Ascomycota_unclassified(64);p__Ascomycota_unclassified(64);p__Ascomycota_unclassified(64); | - | - | - | - | - | - | - | Unassigned | - |
| Otu02293 | 19 | k__Fungi(100);k__Fungi_unclassified(90);k__Fungi_unclassified(90);k__Fungi_unclassified(90);k__Fungi_unclassified(90);k__Fungi_unclassified(90);k__Fungi_unclassified(90); | - | - | - | - | - | - | - | Unassigned | - |
| Otu02300 | 19 | k__Fungi(100);p__Ascomycota(100);p__Ascomycota_unclassified(74);p__Ascomycota_unclassified(74);p__Ascomycota_unclassified(74);p__Ascomycota_unclassified(74);p__Ascomycota_unclassified(74); | - | - | - | - | - | - | - | Unassigned | - |
| Otu02304 | 19 | k__Fungi(100);k__Fungi_unclassified(100);k__Fungi_unclassified(100);k__Fungi_unclassified(100);k__Fungi_unclassified(100);k__Fungi_unclassified(100);k__Fungi_unclassified(100); | - | - | - | - | - | - | - | Unassigned | - |
| Otu02305 | 19 | k__Fungi(100);p__Ascomycota(79);p__Ascomycota_unclassified(79);p__Ascomycota_unclassified(79);p__Ascomycota_unclassified(79);p__Ascomycota_unclassified(79);p__Ascomycota_unclassified(79); | - | - | - | - | - | - | - | Unassigned | - |
| Otu02308 | 19 | k__Fungi(100);k__Fungi_unclassified(100);k__Fungi_unclassified(100);k__Fungi_unclassified(100);k__Fungi_unclassified(100);k__Fungi_unclassified(100);k__Fungi_unclassified(100); | - | - | - | - | - | - | - | Unassigned | - |
| Otu02309 | 19 | k__Fungi(100);p__Ascomycota(79);p__Ascomycota_unclassified(79);p__Ascomycota_unclassified(79);p__Ascomycota_unclassified(79);p__Ascomycota_unclassified(79);p__Ascomycota_unclassified(79); | - | - | - | - | - | - | - | Unassigned | - |
| Otu02310 | 19 | k__Fungi(100);p__Basidiomycota(100);c__Agaricomycetes(100);o__Polyporales(100);o__Polyporales_unclassified(95);o__Polyporales_unclassified(95);o__Polyporales_unclassified(95); | - | - | - | - | - | - | - | Unassigned | - |
| Otu02311 | 19 | k__Fungi(100);p__Ascomycota(64);p__Ascomycota_unclassified(64);p__Ascomycota_unclassified(64);p__Ascomycota_unclassified(64);p__Ascomycota_unclassified(64);p__Ascomycota_unclassified(64); | - | - | - | - | - | - | - | Unassigned | - |
| Otu02312 | 18 | k__Fungi(100);p__Basidiomycota(89);c__Microbotryomycetes(89);o__Sporidiobolales(89);f__Sporidiobolales_family_Incertae_sedis(89);g__Sporobolomyces(89);s__Sporobolomyces_lactosus(89); | Sporobolomyces | 13 | Pathotroph-Saprotroph | \|Fungal Parasite\|-Plant Saprotroph | Yeast | NULL | Probable | As litter_saprotroph (Põlme et al. 2020); As mycoparasite (Põlme et al. 2020); Decay fungal material (Põlme et al. 2020) | Tedersoo L, et al. 2014. Science 346:e1256688 ((https://doi.org/10.1126/science.1256688)); Purahong W, et al. 2016. Molecular Ecology 25:4059-4074 ((https://doi.org/10.1111/mec.13739)); Põlme S, et al. 2020. Fungal Diversity 105:1-16 ((https://doi.org/10.1007/s13225-020-00466-2)) |
| Otu02313 | 18 | k__Fungi(100);p__Basidiomycota(100);c__Agaricomycetes(100);o__Agaricales(100);f__Strophariaceae(100);g__Psilocybe(100);g__Psilocybe_unclassified(100); | Psilocybe | 13 | Saprotroph | \|Plant Saprotroph\|-Undefined Saprotroph | Agaricoid | Poisonous | Probable | As litter_saprotroph (Põlme et al. 2020); bluing clade; Decay leaf, fruit, seed (Põlme et al. 2020); Typified by P. semilanceata (Norvell 2007); Psilocybe = blue staining hallucinogenic clade and Deconica = non | Norvell L. 2007. Taxon. 59: 291-293 ((https://doi.org/10.1002/tax.591029)); Tedersoo L, et al. 2014. Science 346:e1256688 ((https://doi.org/10.1126/science.1256688)); Põlme S, et al. 2020. Fungal Diversity 105:1-16 ((https://doi.org/10.1007/s13225-020-00466-2)) |
| Otu02316 | 18 | k__Fungi(100);p__Basidiomycota(100);c__Agaricomycetes(100);o__Agaricales(100);f__Cortinariaceae(100);g__Cortinarius(100);g__Cortinarius_unclassified(89); | Cortinarius | 13 | Symbiotroph | \|Ectomycorrhizal\| | Agaricoid | NULL | Highly Probable | NULL | Rinaldi AC, et al. 2008. Fungal Diversity 33:1-45 ((http://www.fungaldiversity.org/fdp/sfdp/33-1.pdf)); Tedersoo L, et al. 2010. Mycorrhiza 20:217-263 ((https://doi.org/10.1007/s00572-009-0274-x)); Põlme S, et al. 2020. Fungal Diversity 105:1-16 ((https://doi.org/10.1007/s13225-020-00466-2)); http://mycorrhizas.info/ecmf.html ((http://mycorrhizas.info/ecmf.html)) |
| Otu02319 | 18 | k__Fungi(100);p__Ascomycota(100);c__Saccharomycetes(100);o__Saccharomycetales(100);f__Metschnikowiaceae(100);g__Metschnikowia(100);s__Metschnikowia_pulcherrima(95); | Metschnikowia | 13 | Saprotroph | \|Undefined Saprotroph\| | Yeast | NULL | Probable | As nectar-tap_saprotroph (Põlme et al. 2020); Decay sugar-rich substrates (Põlme et al. 2020) | Põlme S, et al. 2020. Fungal Diversity 105:1-16 ((https://doi.org/10.1007/s13225-020-00466-2)) |
| Otu02320 | 18 | k__Fungi(100);p__Ascomycota(100);c__Saccharomycetes(100);o__Saccharomycetales(100);f__Metschnikowiaceae(100);g__Metschnikowia(100);s__Metschnikowia_pulcherrima(100); | Metschnikowia | 13 | Saprotroph | \|Undefined Saprotroph\| | Yeast | NULL | Probable | As nectar-tap_saprotroph (Põlme et al. 2020); Decay sugar-rich substrates (Põlme et al. 2020) | Põlme S, et al. 2020. Fungal Diversity 105:1-16 ((https://doi.org/10.1007/s13225-020-00466-2)) |
| Otu02321 | 18 | k__Fungi(100);p__Basidiomycota(100);c__Pucciniomycetes(100);o__Septobasidiales(100);f__Septobasidiaceae(100);g__Septobasidium(100);s__Septobasidium_sp_JBO_2010i(100); | Septobasidium | 13 | Pathotroph | \|Animal Parasite\|-Animal Pathogen-Arthropod Parasite | Dimorphic-Facultative Yeast | NULL | Probable | Decay animal material (Põlme et al. 2020); Host - scale_insects (Põlme et al. 2020) | Tedersoo L, et al. 2014. Science 346:e1256688 ((https://doi.org/10.1126/science.1256688)); Põlme S, et al. 2020. Fungal Diversity 105:1-16 ((https://doi.org/10.1007/s13225-020-00466-2)) |
| Otu02324 | 18 | k__Fungi(100);p__Ascomycota(100);c__Taphrinomycetes(100);o__Taphrinales(100);f__Taphrinaceae(100);g__Taphrina(100);g__Taphrina_unclassified(100); | Taphrina | 13 | Pathotroph | \|Plant Pathogen\| | Dimorphic-Microfungus | NULL | Probable | NULL | Põlme S, et al. 2020. Fungal Diversity 105:1-16 ((https://doi.org/10.1007/s13225-020-00466-2)); The American Phytopathological Society Common Names of Plant Diseases ((http://www.apsnet.org/publications/commonnames/Pages/default.aspx)) |
| Otu02327 | 18 | k__Fungi(100);p__Basidiomycota(100);c__Agaricostilbomycetes(100);o__Agaricostilbales(100);f__Chionosphaeraceae(100);g__Chionosphaera(73);g__Chionosphaera_unclassified(73); | Chionosphaera | 20 | Pathotroph-Saprotroph | Lichen Parasite-\|Plant Saprotroph\| | Dimorphic-Microfungus | NULL | Probable | As litter_saprotroph (Põlme et al. 2020); Decay leaf, fruit, seed (Põlme et al. 2020); Lichen parasite on various lichens (Kirschner 2001) | Kirschner R. 2001. Mycological Research 1051403-1408 ((https://doi.org/10.1017/S0953756201005019)); Lawrey JD, Diederich P. 2016. http://www.lichenicolous.net (accessed 24 May 2017) ((http://www.lichenicolous.net)); Põlme S, et al. 2020. Fungal Diversity 105:1-16 ((https://doi.org/10.1007/s13225-020-00466-2)) |
| Otu02328 | 18 | k__Fungi(100);p__Ascomycota(100);c__Saccharomycetes(100);o__Saccharomycetales(100);f__Metschnikowiaceae(100);g__Metschnikowia(100);s__Metschnikowia_pulcherrima(100); | Metschnikowia | 13 | Saprotroph | \|Undefined Saprotroph\| | Yeast | NULL | Probable | As nectar-tap_saprotroph (Põlme et al. 2020); Decay sugar-rich substrates (Põlme et al. 2020) | Põlme S, et al. 2020. Fungal Diversity 105:1-16 ((https://doi.org/10.1007/s13225-020-00466-2)) |
| Otu02332 | 18 | k__Fungi(100);p__Basidiomycota(100);c__Agaricomycetes(100);o__Agaricales(100);f__Physalacriaceae(100);g__Cylindrobasidium(100);s__Cylindrobasidium_sp_DLL2011_1(100); | Cylindrobasidium | 13 | Saprotroph | Undefined Saprotroph-\|Wood Saprotroph\| | Corticioid | White Rot | Possible | As white_rot (Põlme et al. 2020); Decay wood (Põlme et al. 2020) | Jülich W. 1974. Persoonia 8:59-98 ((http://www.cybertruffle.org.uk/cyberliber/61056/0008/001/0072.htm)); Põlme S, et al. 2020. Fungal Diversity 105:1-16 ((https://doi.org/10.1007/s13225-020-00466-2)) |
| Otu02333 | 18 | k__Fungi(100);p__Basidiomycota(100);c__Microbotryomycetes(78);o__Sporidiobolales(78);f__Sporidiobolales_family_Incertae_sedis(78);g__Rhodotorula(78);s__Rhodotorula_oryzicola(78); | Rhodotorula | 13 | Pathotroph-Saprotroph-Symbiotroph | Animal Pathogen-Animal Symbiotroph-Endophyte-Plant Pathogen-\|Undefined Saprotroph\| | Yeast | NULL | Probable | As foliar_endophyte (Põlme et al. 2020); Opportunistic human pathogen | Põlme S, et al. 2020. Fungal Diversity 105:1-16 ((https://doi.org/10.1007/s13225-020-00466-2)); https://www.cdc.gov ((https://www.cdc.gov)) |
| Otu02335 | 18 | k__Fungi(100);p__Zygomycota(100);c__Zygomycota_class_Incertae_sedis(100);o__Kickxellales(100);f__Kickxellaceae(100);g__Ramicandelaber(100);g__Ramicandelaber_unclassified(100); | Ramicandelaber | 13 | Saprotroph | \|Undefined Saprotroph\| | Microfungus | NULL | Probable | As soil_saprotroph (Põlme et al. 2020); Decay soil (Põlme et al. 2020) | Tedersoo L, et al. 2014. Science 346:e1256688 ((https://doi.org/10.1126/science.1256688)); Põlme S, et al. 2020. Fungal Diversity 105:1-16 ((https://doi.org/10.1007/s13225-020-00466-2)) |
| Otu02336 | 18 | k__Fungi(100);p__Ascomycota(100);c__Saccharomycetes(100);o__Saccharomycetales(100);f__Metschnikowiaceae(100);g__Metschnikowia(100);s__Metschnikowia_pulcherrima(100); | Metschnikowia | 13 | Saprotroph | \|Undefined Saprotroph\| | Yeast | NULL | Probable | As nectar-tap_saprotroph (Põlme et al. 2020); Decay sugar-rich substrates (Põlme et al. 2020) | Põlme S, et al. 2020. Fungal Diversity 105:1-16 ((https://doi.org/10.1007/s13225-020-00466-2)) |
| Otu02337 | 18 | k__Fungi(100);p__Ascomycota(100);c__Ascomycota_class_Incertae_sedis(100);o__Ascomycota_order_Incertae_sedis(100);f__Ascomycota_family_Incertae_sedis(100);g__Veronaea(100);g__Veronaea_unclassified(100); | Veronaea | 13 | Pathotroph-Saprotroph | Dung Saprotroph-\|Plant Pathogen\|-Plant Saprotroph | Dimorphic | NULL | Probable | As litter_saprotroph (Põlme et al. 2020); Decay leaf, fruit, seed (Põlme et al. 2020) | Põlme S, et al. 2020. Fungal Diversity 105:1-16 ((https://doi.org/10.1007/s13225-020-00466-2)); The American Phytopathological Society Common Names of Plant Diseases ((http://www.apsnet.org/publications/commonnames/Pages/default.aspx)) |
| Otu02342 | 18 | k__Fungi(100);p__Basidiomycota(100);c__Agaricomycetes(100);o__Agaricales(100);o__Agaricales_unclassified(100);o__Agaricales_unclassified(100);o__Agaricales_unclassified(100); | Agaricales | 7 | Pathotroph-Saprotroph-Symbiotroph | Bryophyte Parasite-Dung Saprotroph-Ectomycorrhizal-Fungal Parasite-Leaf Saprotroph-Plant Parasite-Undefined Saprotroph-Wood Saprotroph | Agaricoid-Gasteroid-Microfungus-Secotioid-Yeast | NULL | Possible | Mushrooms and toadstools, Gill fungi, Agarics... lignicolous, sometimes muscicolous or fungicolous, saprobic, mycorrhizal, rarely parasitic on plants or fungi (Kirk et al. 2008) | Kirk PM et al. 2008. Dictionary of the Fungi. Tenth Edition. CAB International, Wallingford (ISBN: 978-0851998268) |
| Otu02344 | 18 | k__Fungi(100);p__Ascomycota(100);c__Saccharomycetes(100);o__Saccharomycetales(100);f__Metschnikowiaceae(100);g__Metschnikowia(100);s__Metschnikowia_pulcherrima(95); | Metschnikowia | 13 | Saprotroph | \|Undefined Saprotroph\| | Yeast | NULL | Probable | As nectar-tap_saprotroph (Põlme et al. 2020); Decay sugar-rich substrates (Põlme et al. 2020) | Põlme S, et al. 2020. Fungal Diversity 105:1-16 ((https://doi.org/10.1007/s13225-020-00466-2)) |
| Otu02346 | 18 | k__Fungi(100);p__Ascomycota(100);c__Sordariomycetes(100);o__Diaporthales(100);f__Gnomoniaceae(100);g__Gnomonia(100);s__Gnomonia_sp(100); | Gnomonia | 13 | Pathotroph-Saprotroph-Symbiotroph | \|Endophyte\|-Plant Pathogen-Plant Saprotroph-Undefined Saprotroph | Microfungus | NULL | Probable | As foliar_endophyte (Põlme et al. 2020); As litter_saprotroph (Põlme et al. 2020); Decay leaf, fruit, seed (Põlme et al. 2020); Most species are saprobes or endophytes in leaves and bark, but a few are neurotrophic pathogens (Cannon & Kirk 2007) | Cannon PF, Kirk PM. 2007. Fungal Families of the World. CAB International, Cambridge (ISBN: 978-0851998275); Tedersoo L, et al. 2014. Science 346:e1256688 ((https://doi.org/10.1126/science.1256688)); Põlme S, et al. 2020. Fungal Diversity 105:1-16 ((https://doi.org/10.1007/s13225-020-00466-2)) |
| Otu02347 | 18 | k__Fungi(100);p__Ascomycota(100);c__Saccharomycetes(100);o__Saccharomycetales(100);f__Metschnikowiaceae(100);g__Metschnikowia(100);s__Metschnikowia_pulcherrima(95); | Metschnikowia | 13 | Saprotroph | \|Undefined Saprotroph\| | Yeast | NULL | Probable | As nectar-tap_saprotroph (Põlme et al. 2020); Decay sugar-rich substrates (Põlme et al. 2020) | Põlme S, et al. 2020. Fungal Diversity 105:1-16 ((https://doi.org/10.1007/s13225-020-00466-2)) |
| Otu02349 | 18 | k__Fungi(100);p__Ascomycota(100);c__Eurotiomycetes(100);o__Chaetothyriales(100);f__Herpotrichiellaceae(100);f__Herpotrichiellaceae_unclassified(100);f__Herpotrichiellaceae_unclassified(100); | Herpotrichiellaceae | 9 | Pathotroph-Saprotroph | Animal Pathogen-Fungal Parasite-Undefined Saprotroph | Facultative Yeast-Microfungus | NULL | Probable | Mostly saprobic on plants or other fungi, isolated from soil etc., and from medical sources (Cannon & Kirk 2007) | Cannon PF, Kirk PM. 2007. Fungal Families of the World. CAB International, Cambridge (ISBN: 978-0851998275) |
| Otu02352 | 18 | k__Fungi(100);p__Basidiomycota(100);c__Agaricomycetes(100);o__Sebacinales(100);f__Sebacinaceae(100);g__Craterocolla(100);s__Craterocolla_cerasi(100); | Craterocolla | 13 | Saprotroph | \|Plant Saprotroph\|-Undefined Saprotroph | Corticioid | NULL | Probable | As litter_saprotroph (Põlme et al. 2020); Decay leaf, fruit, seed (Põlme et al. 2020) | Tedersoo L, et al. 2014. Science 346:e1256688 ((https://doi.org/10.1126/science.1256688)); Põlme S, et al. 2020. Fungal Diversity 105:1-16 ((https://doi.org/10.1007/s13225-020-00466-2)) |
| Otu02353 | 18 | k__Fungi(100);p__Ascomycota(100);c__Dothideomycetes(100);o__Pleosporales(100);f__Venturiaceae(100);g__Fusicladium(100);s__Fusicladium_caryigenum(100); | Fusicladium | 13 | Pathotroph-Symbiotroph | Endophyte-Lichen Parasite-\|Plant Pathogen\| | Microfungus | NULL | Probable | NULL | Tedersoo L, et al. 2014. Science 346:e1256688 ((https://doi.org/10.1126/science.1256688)); Põlme S, et al. 2020. Fungal Diversity 105:1-16 ((https://doi.org/10.1007/s13225-020-00466-2)) |
| Otu02356 | 18 | k__Fungi(100);p__Basidiomycota(100);c__Agaricomycetes(100);o__Polyporales(100);f__Fomitopsidaceae(100);g__Postia(100);s__Postia_alni(100); | Postia | 13 | Saprotroph | \|Wood Saprotroph\| | Polyporoid | Brown Rot | Highly Probable | As brown-rot (Põlme et al. 2020); Decay wood (Põlme et al. 2020) | Jordan CR, et al. 1996. Holzforschung - International Journal of the Biology, Chemistry, Physics and Technology of Wood 50:312-318 ((https://doi.org/10.1515/hfsg.1996.50.4.312)); Põlme S, et al. 2020. Fungal Diversity 105:1-16 ((https://doi.org/10.1007/s13225-020-00466-2)) |
| Otu02357 | 18 | k__Fungi(100);p__Ascomycota(100);c__Lecanoromycetes(100);o__Teloschistales(100);f__Physciaceae(100);g__Buellia(100);s__Buellia_griseovirens(100); | Buellia | 13 | Symbiotroph | \|Lichenized\| | Thallus | NULL | Highly Probable | NULL | Esslinger TL. 2014. North Dakota State University ((http://www.ndsu.edu/pubweb/~esslinge/chcklst/chcklst7.htm)); Põlme S, et al. 2020. Fungal Diversity 105:1-16 ((https://doi.org/10.1007/s13225-020-00466-2)) |
| Otu02358 | 18 | k__Fungi(100);p__Basidiomycota(100);c__Agaricomycetes(100);o__Agaricales(100);f__Pluteaceae(100);g__Pluteus(100);s__Pluteus_seticeps(100); | Pluteus | 13 | Pathotroph-Saprotroph | Bryophyte Parasite-\|Plant Saprotroph\|-Wood Saprotroph | Agaricoid | White Rot | Probable | As litter_saprotroph (Põlme et al. 2020); As white_rot (Põlme et al. 2020); Decay leaf, fruit, seed (Põlme et al. 2020) | Grgurinovic CA, Simpson JA. 2001. Fungal Diversity 8:97-127 ((http://www.fungaldiversity.org/fdp/sfdp/FD_8_97-127.pdf)); James TY, et al. 2006. Nature 443:818-822 ((https://doi.org/10.1038/nature05110)); Tedersoo L, et al. 2014. Science 346:e1256688 ((https://doi.org/10.1126/science.1256688)); Põlme S, et al. 2020. Fungal Diversity 105:1-16 ((https://doi.org/10.1007/s13225-020-00466-2)) |
| Otu02360 | 18 | k__Fungi(100);p__Zygomycota(100);c__Zygomycota_class_Incertae_sedis(100);o__Mortierellales(100);f__Mortierellaceae(100);g__Mortierella(100);s__Mortierella_sp(67); | Mortierella | 13 | Saprotroph-Symbiotroph | Endophyte-Plant Saprotroph-\|Undefined Saprotroph\| | Microfungus | NULL | Probable | As chitinolytic (Põlme et al. 2020); As soil_saprotroph (Põlme et al. 2020); Decay roots, soil (Põlme et al. 2020); Saprobic in soil and thus frequently isolated from soil and material contaminated with soil such as hay or rotten silage and spoil tips (Cannon & Kirk 2007) | Cannon PF, Kirk PM. 2007. Fungal Families of the World. CAB International, Cambridge (ISBN: 978-0851998275); Tedersoo L, et al. 2014. Science 346:e1256688 ((https://doi.org/10.1126/science.1256688)); Purahong W, et al. 2016. Molecular Ecology 25:4059-4074 ((https://doi.org/10.1111/mec.13739)); Põlme S, et al. 2020. Fungal Diversity 105:1-16 ((https://doi.org/10.1007/s13225-020-00466-2)) |

| | | | | | | | | | | | |
|---|---|---|---|---|---|---|---|---|---|---|---|
| Otu02361 | 18 | k__Fungi(100);p__Basidiomycota(100);c__Agaricomycetes(100);o__Corticiales(100);f__Corticiaceae(100);g__Corticium(100);s__Lyomyces_sp_8_MPM_2012(100); | Lyomyces | 13 | Saprotroph | \|Wood Saprotroph\| | Corticioid | White Rot | Probable | [illegible] | [illegible] |
| Otu02363 | 18 | k__Fungi(100);p__Basidiomycota(100);c__Agaricomycetes(100);o__Polyporales(100);f__Fomitopsidaceae(100);g__Laetiporus(100);s__Laetiporus_cincinnatus(100); | Laetiporus | 13 | Pathotroph-Saprotroph | \|Plant Pathogen\|-Wood Saprotroph | Polyporoid | Brown Rot | Highly Probable | [illegible] | [illegible] |
| Otu02365 | 18 | k__Fungi(100);p__Ascomycota(100);c__Dothideomycetes(100);o__Capnodiales(100);f__Mycosphaerellaceae(100);f__Mycosphaerellaceae_unclassified(95);f__Mycosphaerellaceae_unclassified(95); | Mycosphaerellaceae | 9 | Pathotroph-Saprotroph | Plant Pathogen-Undefined Saprotroph | Microfungus | NULL | Probable | [illegible] | [illegible] |
| Otu02366 | 18 | k__Fungi(100);p__Ascomycota(100);c__Saccharomycetes(100);o__Saccharomycetales(100);f__Metschnikowiaceae(100);g__Metschnikowia(100);s__Metschnikowia_pulcherrima(95); | Metschnikowia | 13 | Saprotroph | \|Undefined Saprotroph\| | Yeast | NULL | Probable | [illegible] | [illegible] |
| Otu02368 | 18 | k__Fungi(100);p__Basidiomycota(100);c__Agaricomycetes(100);o__Agaricales(100);o__Agaricales_unclassified(100);o__Agaricales_unclassified(100);o__Agaricales_unclassified(100); | Agaricales | 7 | Pathotroph-Saprotroph-Symbiotroph | Bryophyte Parasite-Dung Saprotroph-Ectomycorrhizal-Fungal Parasite-Leaf Saprotroph-Plant Parasite-Undefined Saprotroph-Wood Saprotroph | Agaricoid-Gasteroid-Microfungus-Secotioid-Yeast | NULL | Possible | [illegible] | [illegible] |
| Otu02370 | 18 | k__Fungi(100);p__Ascomycota(100);c__Saccharomycetes(100);o__Saccharomycetales(100);f__Metschnikowiaceae(100);g__Metschnikowia(100);s__Metschnikowia_pulcherrima(100); | Metschnikowia | 13 | Saprotroph | \|Undefined Saprotroph\| | Yeast | NULL | Probable | [illegible] | [illegible] |
| Otu02373 | 18 | k__Fungi(100);p__Ascomycota(100);c__Eurotiomycetes(100);o__Chaetothyriales(100);f__Herpotrichiellaceae(100);g__Exophiala(100);s__Exophiala_equina(100); | Exophiala | 13 | Pathotroph-Saprotroph-Symbiotroph | \|Animal Parasite\|-Animal Pathogen-Endophyte-Plant Saprotroph-Undefined Saprotroph | Dimorphic-Facultative Yeast | NULL | Probable | [illegible] | [illegible] |
| Otu02374 | 18 | k__Fungi(100);p__Ascomycota(100);c__Saccharomycetes(95);o__Saccharomycetales(95);f__Saccharomycetales_family_Incertae_sedis(78);g__Candida(73);g__Candida_unclassified(56); | Candida | 13 | Pathotroph-Saprotroph-Symbiotroph | Animal Pathogen-Endophyte-Endosymbiont-Epiphyte-\|Undefined Saprotroph\| | Dimorphic Yeast-Yeast | NULL | Probable | [illegible] | [illegible] |
| Otu02376 | 18 | k__Fungi(100);p__Basidiomycota(100);c__Tremellomycetes(100);o__Tremellales(100);f__Tremellales_family_Incertae_sedis(100);g__Cryptococcus(100);s__Cryptococcus_nemorosus(100); | Cryptococcus | 13 | Pathotroph-Saprotroph-Symbiotroph | Animal Pathogen-Endophyte-Epiphyte-\|Undefined Saprotroph\| | Dimorphic Yeast-Yeast | NULL | Possible | [illegible] | [illegible] |
| Otu02382 | 18 | k__Fungi(100);p__Ascomycota(100);c__Dothideomycetes(100);o__Pleosporales(100);f__Phaeosphaeriaceae(100);f__Phaeosphaeriaceae_unclassified(100);f__Phaeosphaeriaceae_unclassified(100); | Phaeosphaeriaceae | 9 | Pathotroph-Saprotroph | Fungal Parasite-Plant Pathogen-Plant Saprotroph | Microfungus | NULL | Probable | [illegible] | [illegible] |
| Otu02385 | 18 | k__Fungi(100);p__Ascomycota(100);c__Ascomycota_class_Incertae_sedis(100);o__Ascomycota_order_Incertae_sedis(100);f__Ascomycota_family_Incertae_sedis(100);g__Hansfordia(100);s__Hansfordia_sp_XC_2012(100); | Hansfordia | 13 | Saprotroph | \|Undefined Saprotroph\| | Microfungus | NULL | Probable | NULL | [illegible] |
| Otu02386 | 18 | k__Fungi(100);p__Ascomycota(100);c__Sordariomycetes(100);o__Xylariales(100);f__Amphisphaeriaceae(100);f__Amphisphaeriaceae_unclassified(100);f__Amphisphaeriaceae_unclassified(100); | Amphisphaeriaceae | 9 | Saprotroph | Undefined Saprotroph | Microfungus | NULL | Possible | [illegible] | [illegible] |
| Otu02314 | 18 | k__Fungi(100);p__Basidiomycota(100);c__Agaricomycetes(56);c__Agaricomycetes_unclassified(56);c__Agaricomycetes_unclassified(56);c__Agaricomycetes_unclassified(56);c__Agaricomycetes_unclassified(56); | - | - | - | - | - | - | - | Unassigned | - |
| Otu02315 | 18 | k__Fungi(100);p__Ascomycota(56);p__Ascomycota_unclassified(56);p__Ascomycota_unclassified(56);p__Ascomycota_unclassified(56);p__Ascomycota_unclassified(56);p__Ascomycota_unclassified(56); | - | - | - | - | - | - | - | Unassigned | - |
| Otu02317 | 18 | k__Fungi(100);p__Ascomycota(100);p__Ascomycota_unclassified(100);p__Ascomycota_unclassified(100);p__Ascomycota_unclassified(100);p__Ascomycota_unclassified(100);p__Ascomycota_unclassified(100); | - | - | - | - | - | - | - | Unassigned | - |
| Otu02318 | 18 | k__Fungi(100);k__Fungi_unclassified(56);k__Fungi_unclassified(56);k__Fungi_unclassified(56);k__Fungi_unclassified(56);k__Fungi_unclassified(56);k__Fungi_unclassified(56); | - | - | - | - | - | - | - | Unassigned | - |
| Otu02322 | 18 | k__Fungi(100);k__Fungi_unclassified(67);k__Fungi_unclassified(67);k__Fungi_unclassified(67);k__Fungi_unclassified(67);k__Fungi_unclassified(67);k__Fungi_unclassified(67); | - | - | - | - | - | - | - | Unassigned | - |
| Otu02323 | 18 | k__Fungi(100);k__Fungi_unclassified(62);k__Fungi_unclassified(62);k__Fungi_unclassified(62);k__Fungi_unclassified(62);k__Fungi_unclassified(62);k__Fungi_unclassified(62); | - | - | - | - | - | - | - | Unassigned | - |
| Otu02325 | 18 | k__Fungi(100);k__Fungi_unclassified(95);k__Fungi_unclassified(95);k__Fungi_unclassified(95);k__Fungi_unclassified(95);k__Fungi_unclassified(95);k__Fungi_unclassified(95); | - | - | - | - | - | - | - | Unassigned | - |
| Otu02326 | 18 | k__Fungi(100);p__Ascomycota(95);p__Ascomycota_unclassified(95);p__Ascomycota_unclassified(95);p__Ascomycota_unclassified(95);p__Ascomycota_unclassified(95);p__Ascomycota_unclassified(95); | - | - | - | - | - | - | - | Unassigned | - |
| Otu02329 | 18 | k__Fungi(100);p__Ascomycota(100);p__Ascomycota_unclassified(100);p__Ascomycota_unclassified(100);p__Ascomycota_unclassified(100);p__Ascomycota_unclassified(100);p__Ascomycota_unclassified(100); | - | - | - | - | - | - | - | Unassigned | - |
| Otu02330 | 18 | k__Fungi(100);p__Ascomycota(100);p__Ascomycota_unclassified(100);p__Ascomycota_unclassified(100);p__Ascomycota_unclassified(100);p__Ascomycota_unclassified(100);p__Ascomycota_unclassified(100); | - | - | - | - | - | - | - | Unassigned | - |
| Otu02331 | 18 | k__Fungi(100);k__Fungi_unclassified(100);k__Fungi_unclassified(100);k__Fungi_unclassified(100);k__Fungi_unclassified(100);k__Fungi_unclassified(100);k__Fungi_unclassified(100); | - | - | - | - | - | - | - | Unassigned | - |
| Otu02334 | 18 | k__Fungi(100);k__Fungi_unclassified(100);k__Fungi_unclassified(100);k__Fungi_unclassified(100);k__Fungi_unclassified(100);k__Fungi_unclassified(100);k__Fungi_unclassified(100); | - | - | - | - | - | - | - | Unassigned | - |
| Otu02338 | 18 | k__Fungi(100);p__Basidiomycota(84);c__Agaricomycetes(78);c__Agaricomycetes_unclassified(78);c__Agaricomycetes_unclassified(78);c__Agaricomycetes_unclassified(78);c__Agaricomycetes_unclassified(78); | - | - | - | - | - | - | - | Unassigned | - |
| Otu02339 | 18 | k__Fungi(100);p__Ascomycota(73);p__Ascomycota_unclassified(73);p__Ascomycota_unclassified(73);p__Ascomycota_unclassified(73);p__Ascomycota_unclassified(73);p__Ascomycota_unclassified(73); | - | - | - | - | - | - | - | Unassigned | - |
| Otu02340 | 18 | k__Fungi(100);k__Fungi_unclassified(100);k__Fungi_unclassified(100);k__Fungi_unclassified(100);k__Fungi_unclassified(100);k__Fungi_unclassified(100);k__Fungi_unclassified(100); | - | - | - | - | - | - | - | Unassigned | - |
| Otu02341 | 18 | k__Fungi(100);k__Fungi_unclassified(100);k__Fungi_unclassified(100);k__Fungi_unclassified(100);k__Fungi_unclassified(100);k__Fungi_unclassified(100);k__Fungi_unclassified(100); | - | - | - | - | - | - | - | Unassigned | - |
| Otu02343 | 18 | k__Fungi(100);p__Ascomycota(100);p__Ascomycota_unclassified(100);p__Ascomycota_unclassified(100);p__Ascomycota_unclassified(100);p__Ascomycota_unclassified(100);p__Ascomycota_unclassified(100); | - | - | - | - | - | - | - | Unassigned | - |
| Otu02345 | 18 | k__Fungi(100);k__Fungi_unclassified(78);k__Fungi_unclassified(78);k__Fungi_unclassified(78);k__Fungi_unclassified(78);k__Fungi_unclassified(78);k__Fungi_unclassified(78); | - | - | - | - | - | - | - | Unassigned | - |
| Otu02348 | 18 | k__Fungi(100);p__Ascomycota(67);p__Ascomycota_unclassified(67);p__Ascomycota_unclassified(67);p__Ascomycota_unclassified(67);p__Ascomycota_unclassified(67);p__Ascomycota_unclassified(67); | - | - | - | - | - | - | - | Unassigned | - |
| Otu02350 | 18 | k__Fungi(100);p__Ascomycota(89);p__Ascomycota_unclassified(89);p__Ascomycota_unclassified(89);p__Ascomycota_unclassified(89);p__Ascomycota_unclassified(89);p__Ascomycota_unclassified(89); | - | - | - | - | - | - | - | Unassigned | - |
| Otu02351 | 18 | k__Fungi(100);p__Ascomycota(84);p__Ascomycota_unclassified(84);p__Ascomycota_unclassified(84);p__Ascomycota_unclassified(84);p__Ascomycota_unclassified(84);p__Ascomycota_unclassified(84); | - | - | - | - | - | - | - | Unassigned | - |
| Otu02354 | 18 | k__Fungi(100);k__Fungi_unclassified(100);k__Fungi_unclassified(100);k__Fungi_unclassified(100);k__Fungi_unclassified(100);k__Fungi_unclassified(100);k__Fungi_unclassified(100); | - | - | - | - | - | - | - | Unassigned | - |
| Otu02355 | 18 | k__Fungi(100);p__Ascomycota(62);p__Ascomycota_unclassified(62);p__Ascomycota_unclassified(62);p__Ascomycota_unclassified(62);p__Ascomycota_unclassified(62);p__Ascomycota_unclassified(62); | - | - | - | - | - | - | - | Unassigned | - |
| Otu02359 | 18 | k__Fungi(100);p__Basidiomycota(100);c__Microbotryomycetes(100);o__Sporidiobolales(100);f__Sporidiobolales_family_Incertae_sedis(100);g__unclassified_Sporidiobolales(56);s__Sporidiobolales_sp(56); | - | - | - | - | - | - | - | Unassigned | - |
| Otu02362 | 18 | k__Fungi(100);p__Ascomycota(67);p__Ascomycota_unclassified(67);p__Ascomycota_unclassified(67);p__Ascomycota_unclassified(67);p__Ascomycota_unclassified(67);p__Ascomycota_unclassified(67); | - | - | - | - | - | - | - | Unassigned | - |
| Otu02364 | 18 | k__Fungi(100);k__Fungi_unclassified(78);k__Fungi_unclassified(78);k__Fungi_unclassified(78);k__Fungi_unclassified(78);k__Fungi_unclassified(78);k__Fungi_unclassified(78); | - | - | - | - | - | - | - | Unassigned | - |
| Otu02367 | 18 | k__Fungi(100);k__Fungi_unclassified(78);k__Fungi_unclassified(78);k__Fungi_unclassified(78);k__Fungi_unclassified(78);k__Fungi_unclassified(78);k__Fungi_unclassified(78); | - | - | - | - | - | - | - | Unassigned | - |
| Otu02369 | 18 | k__Fungi(100);p__Basidiomycota(100);c__Agaricomycetes(100);o__Polyporales(100);o__Polyporales_unclassified(100);o__Polyporales_unclassified(100);o__Polyporales_unclassified(100); | - | - | - | - | - | - | - | Unassigned | - |
| Otu02371 | 18 | k__Fungi(100);k__Fungi_unclassified(95);k__Fungi_unclassified(95);k__Fungi_unclassified(95);k__Fungi_unclassified(95);k__Fungi_unclassified(95);k__Fungi_unclassified(95); | - | - | - | - | - | - | - | Unassigned | - |
| Otu02372 | 18 | k__Plantae(100);p__unclassified_Plantae(100);c__unclassified_Plantae(100);o__unclassified_Plantae(100);f__unclassified_Plantae(100);g__unclassified_Plantae(100);s__Plantae_sp(100); | - | - | - | - | - | - | - | Unassigned | - |
| Otu02375 | 18 | k__Fungi(100);k__Fungi_unclassified(100);k__Fungi_unclassified(100);k__Fungi_unclassified(100);k__Fungi_unclassified(100);k__Fungi_unclassified(100);k__Fungi_unclassified(100); | - | - | - | - | - | - | - | Unassigned | - |
| Otu02377 | 18 | k__Fungi(100);p__Ascomycota(100);c__Leotiomycetes(95);o__Helotiales(78);o__Helotiales_unclassified(78);o__Helotiales_unclassified(78);o__Helotiales_unclassified(78); | - | - | - | - | - | - | - | Unassigned | - |
| Otu02378 | 18 | k__Fungi(100);k__Fungi_unclassified(67);k__Fungi_unclassified(67);k__Fungi_unclassified(67);k__Fungi_unclassified(67);k__Fungi_unclassified(67);k__Fungi_unclassified(67); | - | - | - | - | - | - | - | Unassigned | - |
| Otu02379 | 18 | k__Fungi(100);k__Fungi_unclassified(100);k__Fungi_unclassified(100);k__Fungi_unclassified(100);k__Fungi_unclassified(100);k__Fungi_unclassified(100);k__Fungi_unclassified(100); | - | - | - | - | - | - | - | Unassigned | - |
| Otu02380 | 18 | k__Fungi(100);p__Ascomycota(100);c__Dothideomycetes(78);o__Pleosporales(78);o__Pleosporales_unclassified(78);o__Pleosporales_unclassified(78);o__Pleosporales_unclassified(78); | - | - | - | - | - | - | - | Unassigned | - |
| Otu02381 | 18 | k__Fungi(100);k__Fungi_unclassified(73);k__Fungi_unclassified(73);k__Fungi_unclassified(73);k__Fungi_unclassified(73);k__Fungi_unclassified(73);k__Fungi_unclassified(73); | - | - | - | - | - | - | - | Unassigned | - |
| Otu02383 | 18 | k__Fungi(100);p__Ascomycota(78);p__Ascomycota_unclassified(78);p__Ascomycota_unclassified(78);p__Ascomycota_unclassified(78);p__Ascomycota_unclassified(78);p__Ascomycota_unclassified(78); | - | - | - | - | - | - | - | Unassigned | - |
| Otu02384 | 18 | k__Fungi(100);k__Fungi_unclassified(100);k__Fungi_unclassified(100);k__Fungi_unclassified(100);k__Fungi_unclassified(100);k__Fungi_unclassified(100);k__Fungi_unclassified(100); | - | - | - | - | - | - | - | Unassigned | - |
| Otu02387 | 18 | k__Fungi(100);p__Ascomycota(100);c__Dothideomycetes(100);o__Pleosporales(100);o__Pleosporales_unclassified(100);o__Pleosporales_unclassified(100);o__Pleosporales_unclassified(100); | - | - | - | - | - | - | - | Unassigned | - |
| Otu02388 | 17 | k__Fungi(100);p__Basidiomycota(100);c__Agaricomycetes(100);o__Phallales(100);f__Phallaceae(100);g__Phallus(100);s__Phallus_impudicus(100); | Phallus | 13 | Saprotroph | Plant Saprotroph-\|Undefined Saprotroph\|-Wood Saprotroph | Phalloid | White Rot | Probable | [illegible] | [illegible] |
| Otu02389 | 17 | k__Fungi(100);p__Basidiomycota(100);c__Agaricomycetes(100);o__Agaricales(100);f__Cortinariaceae(100);g__Cortinarius(100);g__Cortinarius_unclassified(100); | Cortinarius | 13 | Symbiotroph | \|Ectomycorrhizal\| | Agaricoid | NULL | Highly Probable | NULL | [illegible] |
| Otu02390 | 17 | k__Fungi(100);p__Ascomycota(100);c__Saccharomycetes(100);o__Saccharomycetales(100);f__Metschnikowiaceae(100);g__Metschnikowia(100);s__Metschnikowia_pulcherrima(89); | Metschnikowia | 13 | Saprotroph | \|Undefined Saprotroph\| | Yeast | NULL | Probable | [illegible] | [illegible] |
| Otu02391 | 17 | k__Fungi(100);p__Ascomycota(100);c__Saccharomycetes(100);o__Saccharomycetales(100);f__Metschnikowiaceae(100);g__Metschnikowia(100);s__Metschnikowia_pulcherrima(83); | Metschnikowia | 13 | Saprotroph | \|Undefined Saprotroph\| | Yeast | NULL | Probable | [illegible] | [illegible] |
| Otu02392 | 17 | k__Fungi(100);p__Ascomycota(100);c__Dothideomycetes(100);o__Capnodiales(100);f__Mycosphaerellaceae(100);g__Mycosphaerella(100);s__Mycosphaerella_sumatrensis(100); | Mycosphaerella | 13 | Pathotroph-Saprotroph | \|Plant Pathogen\|-Plant Saprotroph | Microfungus | NULL | Probable | [illegible] | [illegible] |
| Otu02393 | 17 | k__Fungi(100);p__Ascomycota(100);c__Saccharomycetes(100);o__Saccharomycetales(100);f__Metschnikowiaceae(100);g__Metschnikowia(100);s__Metschnikowia_pulcherrima(83); | Metschnikowia | 13 | Saprotroph | \|Undefined Saprotroph\| | Yeast | NULL | Probable | [illegible] | [illegible] |
| Otu02399 | 17 | k__Fungi(100);p__Ascomycota(100);c__Saccharomycetes(100);o__Saccharomycetales(100);f__Saccharomycetales_family_Incertae_sedis(100);g__Candida(100);s__Candida_sp_AS23486(59); | Candida | 13 | Pathotroph-Saprotroph-Symbiotroph | Animal Pathogen-Endophyte-Endosymbiont-Epiphyte-\|Undefined Saprotroph\| | Dimorphic Yeast-Yeast | NULL | Probable | [illegible] | [illegible] |
| Otu02401 | 17 | k__Fungi(100);p__Ascomycota(100);c__Saccharomycetes(100);o__Saccharomycetales(100);f__Metschnikowiaceae(100);g__Metschnikowia(100);s__Metschnikowia_pulcherrima(100); | Metschnikowia | 13 | Saprotroph | \|Undefined Saprotroph\| | Yeast | NULL | Probable | [illegible] | [illegible] |
| Otu02402 | 17 | k__Fungi(100);p__Ascomycota(100);c__Saccharomycetes(100);o__Saccharomycetales(100);f__Metschnikowiaceae(100);g__Metschnikowia(100);s__Metschnikowia_pulcherrima(95); | Metschnikowia | 13 | Saprotroph | \|Undefined Saprotroph\| | Yeast | NULL | Probable | [illegible] | [illegible] |
| Otu02406 | 17 | k__Fungi(100);p__Ascomycota(100);c__Saccharomycetes(100);o__Saccharomycetales(100);f__Metschnikowiaceae(100);g__Metschnikowia(100);s__Metschnikowia_pulcherrima(95); | Metschnikowia | 13 | Saprotroph | \|Undefined Saprotroph\| | Yeast | NULL | Probable | [illegible] | [illegible] |
| Otu02407 | 17 | k__Fungi(100);p__Ascomycota(100);c__Saccharomycetes(100);o__Saccharomycetales(100);f__Metschnikowiaceae(100);g__Metschnikowia(100);s__Metschnikowia_pulcherrima(100); | Metschnikowia | 13 | Saprotroph | \|Undefined Saprotroph\| | Yeast | NULL | Probable | [illegible] | [illegible] |
| Otu02409 | 17 | k__Fungi(100);p__Ascomycota(100);c__Leotiomycetes(100);o__Helotiales(100);f__Rutstroemiaceae(100);g__unclassified_Rutstroemiaceae(100);s__Rutstroemiaceae_sp(100); | Rutstroemiaceae | 9 | Saprotroph | Plant Saprotroph | Microfungus | NULL | Probable | [illegible] | [illegible] |
| Otu02410 | 17 | k__Fungi(100);p__Ascomycota(100);c__Saccharomycetes(100);o__Saccharomycetales(100);f__Metschnikowiaceae(100);g__Metschnikowia(100);g__Metschnikowia_unclassified(89); | Metschnikowia | 13 | Saprotroph | \|Undefined Saprotroph\| | Yeast | NULL | Probable | [illegible] | [illegible] |
| Otu02413 | 17 | k__Fungi(100);p__Ascomycota(100);c__Saccharomycetes(100);o__Saccharomycetales(100);f__Metschnikowiaceae(100);g__Metschnikowia(100);s__Metschnikowia_pulcherrima(100); | Metschnikowia | 13 | Saprotroph | \|Undefined Saprotroph\| | Yeast | NULL | Probable | [illegible] | [illegible] |
| Otu02414 | 17 | k__Fungi(100);p__Ascomycota(100);c__Leotiomycetes(100);o__Helotiales(100);f__Dermateaceae(83);g__Mollisia(83);g__Mollisia_unclassified(65); | Mollisia | 13 | Pathotroph-Saprotroph-Symbiotroph | Endophyte-\|Plant Pathogen\|-Plant Saprotroph | Microfungus-Pezizoid | Soft Rot | Probable | [illegible] | [illegible] |
| Otu02416 | 17 | k__Fungi(100);p__Ascomycota(100);c__Dothideomycetes(100);o__Dothideomycetes_order_Incertae_sedis(100);f__Myxotrichaceae(100);g__Oidiodendron(100);s__Oidiodendron_truncatum(100); | Oidiodendron | 13 | Pathotroph-Saprotroph-Symbiotroph | Endophyte-Ericoid Mycorrhizal-\|Undefined Saprotroph\| | Microfungus | NULL | Probable | [illegible] | [illegible] |
| Otu02417 | 17 | k__Fungi(100);p__Ascomycota(100);c__Saccharomycetes(100);o__Saccharomycetales(100);f__Metschnikowiaceae(100);g__Metschnikowia(100);s__Metschnikowia_pulcherrima(95); | Metschnikowia | 13 | Saprotroph | \|Undefined Saprotroph\| | Yeast | NULL | Probable | [illegible] | [illegible] |
| Otu02421 | 17 | k__Fungi(100);p__Ascomycota(100);c__Sordariomycetes(100);o__Sordariomycetes_order_Incertae_sedis(100);f__Plectosphaerellaceae(100);g__Gibellulopsis(100);s__Gibellulopsis_sp_YH_2012(100); | Gibellulopsis | 13 | Pathotroph-Saprotroph | \|Plant Pathogen\|-Plant Saprotroph | Microfungus | NULL | Probable | [illegible] | [illegible] |
| Otu02422 | 17 | k__Fungi(100);p__Ascomycota(100);c__Leotiomycetes(100);o__Helotiales(100);f__Helotiales_family_Incertae_sedis(100);g__Pilidium(100);s__Pilidium_concavum(83); | Pilidium | 13 | Pathotroph | \|Plant Pathogen\| | Microfungus | NULL | Probable | NULL | [illegible] |
| Otu02424 | 17 | k__Fungi(100);p__Ascomycota(100);c__Eurotiomycetes(100);o__Eurotiales(100);f__Trichocomaceae(100);g__Aspergillus(100);s__Aspergillus_candidus(100); | Aspergillus | 13 | Pathotroph-Saprotroph-Symbiotroph | Animal Pathogen-Endophyte-Plant Saprotroph-\|Undefined Saprotroph\|-Wood Saprotroph | Microfungus | NULL | Probable | [illegible] | [illegible] |
| Otu02426 | 17 | k__Fungi(100);p__Ascomycota(100);c__Leotiomycetes(100);o__Helotiales(100);f__Vibrisseaceae(100);g__Phialocephala(100);s__Phialocephala_fusca(100); | Phialocephala | 13 | Saprotroph-Symbiotroph | Endophyte-\|Undefined Saprotroph\| | Microfungus | NULL | Highly Probable | [illegible] | [illegible] |
| Otu02427 | 17 | k__Fungi(100);p__Basidiomycota(100);c__Tremellomycetes(100);o__Tremellales(100);f__Tremellales_family_Incertae_sedis(100);f__Tremellales_family_Incertae_sedis_unclassified(100);f__Tremellales_family_Incertae_sedis_unclassified(100); | Tremellales | 7 | Pathotroph-Saprotroph-Symbiotroph | Fungal Parasite-Undefined Saprotroph | Tremelloid-Yeast | NULL | Possible | [illegible] | [illegible] |
| Otu02429 | 17 | k__Fungi(100);p__Ascomycota(100);c__Eurotiomycetes(100);o__Chaetothyriales(100);f__Herpotrichiellaceae(100);g__Phaeomoniella(100);s__Phaeomoniella_prunicola(100); | Phaeomoniella | 13 | Pathotroph | \|Plant Pathogen\| | Dimorphic-Microfungus; Facultative Yeast (Tedersoo et | NULL | Highly Probable | [illegible] | [illegible] |
| Otu02430 | 17 | k__Fungi(100);p__Zygomycota(100);c__Zygomycota_class_Incertae_sedis(100);o__Mortierellales(100);f__Mortierellaceae(100);g__Mortierella(100);s__Mortierella_globulifera(100); | Mortierella | 13 | Saprotroph-Symbiotroph | Endophyte-Plant Saprotroph-\|Undefined Saprotroph\| | Microfungus | NULL | Probable | [illegible] | [illegible] |
| Otu02434 | 17 | k__Fungi(100);p__Ascomycota(100);c__Eurotiomycetes(100);o__Chaetothyriales(100);f__Herpotrichiellaceae(100);g__Phaeomoniella(100);s__Phaeomoniella_niveniae(100); | Phaeomoniella | 13 | Pathotroph | \|Plant Pathogen\| | Dimorphic-Microfungus; Facultative Yeast (Tedersoo et | NULL | Highly Probable | [illegible] | [illegible] |
| Otu02435 | 17 | k__Fungi(100);p__Ascomycota(100);c__Sordariomycetes(100);o__Hypocreales(100);f__Hypocreales_family_Incertae_sedis(100);g__Acremonium(100);s__Acremonium_pilosum(100); | Acremonium | 13 | Pathotroph-Saprotroph-Symbiotroph | Animal Pathogen-Endophyte-Fungal Parasite-Plant Pathogen-\|Undefined Saprotroph\|-Wood Saprotroph | Microfungus | Soft Rot | Possible | [illegible] | [illegible] |
| Otu02436 | 17 | k__Fungi(100);p__Ascomycota(100);c__Dothideomycetes(100);o__Pleosporales(100);f__Massarinaceae(100);g__Saccharicola(100);s__Saccharicola_sp(100); | Saccharicola | 13 | Saprotroph | Undefined Saprotroph | NULL | NULL | Probable | NULL | [illegible] |
| Otu02440 | 17 | k__Fungi(100);p__Basidiomycota(100);c__Agaricomycetes(100);o__Boletales(100);f__Tapinellaceae(100);g__Tapinella(100);s__Tapinella_atrotomentosa(100); | Tapinella | 13 | Saprotroph | \|Wood Saprotroph\| | Agaricoid | Brown Rot | Probable | [illegible] | [illegible] |
| Otu02441 | 17 | k__Fungi(100);p__Ascomycota(100);c__Eurotiomycetes(100);o__Eurotiales(100);f__Trichocomaceae(100);g__Paecilomyces(100);g__Paecilomyces_unclassified(59); | Paecilomyces | 13 | Pathotroph-Saprotroph-Symbiotroph | Clavicipitaceous \|Endophyte\|-Plant Pathogen-Plant Saprotroph-Undefined Saprotroph-Wood Saprotroph | Microfungus | Soft Rot | Probable | [illegible] | [illegible] |
| Otu02442 | 17 | k__Fungi(100);p__Basidiomycota(100);c__Agaricomycetes(100);o__Russulales(100);f__Peniophoraceae(100);g__Peniophora(100);g__Peniophora_unclassified(95); | Peniophora | 13 | Pathotroph-Saprotroph | Plant Pathogen-\|Wood Saprotroph\| | Corticioid | White Rot | Probable | [illegible] | [illegible] |
| Otu02447 | 17 | k__Fungi(100);p__Ascomycota(100);c__Sordariomycetes(100);o__Diaporthales(100);f__Melanconidaceae(100);g__Melanconiella(100);s__Melanconiella_ellisii(100); | Melanconiella | 13 | Saprotroph | Undefined Saprotroph-\|Wood Saprotroph\| | Microfungus | NULL | Probable | [illegible] | [illegible] |
| Otu02448 | 17 | k__Fungi(100);p__Ascomycota(100);c__Eurotiomycetes(100);o__Eurotiales(100);f__Trichocomaceae(100);g__Penicillium(100);s__Penicillium_sp(100); | Penicillium | 13 | Pathotroph-Saprotroph-Symbiotroph | Animal Parasite-Dung Saprotroph-Endophyte-Fungal Parasite-Plant Pathogen-\|Undefined Saprotroph\|-Wood Saprotroph | Microfungus | Soft Rot | Highly Probable | [illegible] | [illegible] |
| Otu02450 | 17 | k__Fungi(100);p__Ascomycota(100);c__Eurotiomycetes(100);o__Onygenales(100);f__Gymnoascaceae(100);g__Gymnoascus(100);s__Gymnoascus_sp_22MS07(100); | Gymnoascus | 13 | Saprotroph | \|Undefined Saprotroph\| | Microfungus | NULL | Probable | [illegible] | [illegible] |
| Otu02452 | 17 | k__Fungi(100);p__Ascomycota(100);c__Sordariomycetes(100);o__Diaporthales(100);f__Togniniaceae(100);g__Phaeoacremonium(100);s__Phaeoacremonium_tuscanum(100); | Phaeoacremonium | 13 | Pathotroph | \|Plant Pathogen\| | Microfungus | NULL | Probable | NULL | [illegible] |
| Otu02453 | 17 | k__Fungi(100);p__Basidiomycota(100);c__Pucciniomycetes(100);o__Pucciniales(100);f__Pucciniaceae(100);g__Gymnosporangium(100);s__Gymnosporangium_juniperi_virginianae(100); | Gymnosporangium | 13 | Pathotroph | \|Plant Pathogen\| | Rust-Telial Horns | NULL | Probable | NULL | [illegible] |
| Otu02454 | 17 | k__Fungi(100);p__Ascomycota(100);c__Saccharomycetes(100);o__Saccharomycetales(100);f__Metschnikowiaceae(100);g__Metschnikowia(100);s__Metschnikowia_pulcherrima(100); | Metschnikowia | 13 | Saprotroph | \|Undefined Saprotroph\| | Yeast | NULL | Probable | [illegible] | [illegible] |
| Otu02455 | 17 | k__Fungi(100);p__Basidiomycota(100);c__Agaricomycetes(100);o__Agaricales(100);f__Cortinariaceae(100);g__Cortinarius(100);s__Cortinarius_helvelloides(100); | Cortinarius | 13 | Symbiotroph | \|Ectomycorrhizal\| | Agaricoid | NULL | Highly Probable | NULL | [illegible] |
| Otu02458 | 17 | k__Fungi(100);p__Basidiomycota(100);c__Agaricomycetes(100);o__Polyporales(100);f__Meruliaceae(100);g__Crustoderma(100);s__Crustoderma_corneum(100); | Crustoderma | 13 | Saprotroph | \|Wood Saprotroph\| | Corticioid | Brown Rot | Probable | [illegible] | [illegible] |
| Otu02459 | 17 | k__Fungi(100);p__Ascomycota(100);c__Sordariomycetes(100);o__Microascales(100);f__Microascaceae(100);g__Microascus(100);s__Microascus_manginii(100); | Microascus | 13 | Pathotroph-Saprotroph-Symbiotroph | Animal Pathogen-Dung Saprotroph-Endophyte-Plant Pathogen-Plant Saprotroph-\|Undefined Saprotroph\| | Microfungus | NULL | Possible | [illegible] | [illegible] |
| Otu02460 | 17 | k__Fungi(100);p__Ascomycota(100);c__Sordariomycetes(100);o__Sordariomycetes_order_Incertae_sedis(100);f__Plectosphaerellaceae(100);g__Lectera(100);s__Lectera_longa(100); | Lectera | 13 | Pathotroph | \|Plant Pathogen\| | Microfungus | NULL | Probable | NULL | [illegible] |
| Otu02462 | 17 | k__Fungi(100);p__Ascomycota(100);c__Eurotiomycetes(100);o__Chaetothyriales(100);f__Herpotrichiellaceae(100);g__Phaeomoniella(100);g__Phaeomoniella_unclassified(100); | Phaeomoniella | 13 | Pathotroph | \|Plant Pathogen\| | Dimorphic-Microfungus; Facultative Yeast (Tedersoo et | NULL | Highly Probable | [illegible] | [illegible] |
| Otu02465 | 17 | k__Fungi(100);p__Ascomycota(100);c__Eurotiomycetes(100);o__Eurotiales(100);f__Trichocomaceae(100);g__Penicillium(100);s__Penicillium_oxalicum(89); | Penicillium | 13 | Pathotroph-Saprotroph-Symbiotroph | Animal Parasite-Dung Saprotroph-Endophyte-Fungal Parasite-Plant Pathogen-\|Undefined Saprotroph\|-Wood Saprotroph | Microfungus | Soft Rot | Highly Probable | [illegible] | [illegible] |
| Otu02466 | 17 | k__Fungi(100);p__Basidiomycota(100);c__Agaricomycetes(100);o__Corticiales(100);f__Corticiaceae(100);g__unclassified_Corticiaceae(100);s__Corticiaceae_sp(100); | Corticiaceae | 9 | Pathotroph-Saprotroph | Lichen Parasite-Plant Pathogen-Wood Saprotroph | Corticioid-Microfungus | NULL | Possible | [illegible] | [illegible] |
| Otu02468 | 17 | k__Fungi(100);p__Basidiomycota(100);c__Microbotryomycetes(100);o__Sporidiobolales(100);f__Sporidiobolales_family_Incertae_sedis(100);g__Rhodotorula(100);g__Rhodotorula_unclassified(100); | Rhodotorula | 13 | Pathotroph-Saprotroph-Symbiotroph | Animal Pathogen-Animal Symbiotroph-Endophyte-Plant Pathogen-\|Undefined Saprotroph\| | Yeast | NULL | Probable | [illegible] | [illegible] |
| Otu02469 | 17 | k__Fungi(100);p__Ascomycota(100);c__Saccharomycetes(100);o__Saccharomycetales(100);f__Metschnikowiaceae(100);g__Metschnikowia(100);s__Metschnikowia_pulcherrima(100); | Metschnikowia | 13 | Saprotroph | \|Undefined Saprotroph\| | Yeast | NULL | Probable | [illegible] | [illegible] |
| Otu02472 | 17 | k__Fungi(100);p__Basidiomycota(100);c__Microbotryomycetes(100);o__Sporidiobolales(100);f__Sporidiobolales_family_Incertae_sedis(100);g__Rhodotorula(100);s__Rhodotorula_sp_JCM_11351(100); | Rhodotorula | 13 | Pathotroph-Saprotroph-Symbiotroph | Animal Pathogen-Animal Symbiotroph-Endophyte-Plant Pathogen-\|Undefined Saprotroph\| | Yeast | NULL | Probable | [illegible] | [illegible] |
| Otu02473 | 17 | k__Fungi(100);p__Basidiomycota(100);c__Microbotryomycetes(100);o__Sporidiobolales(100);f__Sporidiobolales_family_Incertae_sedis(100);g__Sporobolomyces(100);s__Sporobolomyces_sasicola(100); | Sporobolomyces | 13 | Pathotroph-Saprotroph | \|Fungal Parasite\|-Plant Saprotroph | Yeast | NULL | Probable | [illegible] | [illegible] |
| Otu02394 | 17 | k__Fungi(100);p__Ascomycota(100);p__Ascomycota_unclassified(100);p__Ascomycota_unclassified(100);p__Ascomycota_unclassified(100);p__Ascomycota_unclassified(100);p__Ascomycota_unclassified(100); | - | - | - | - | - | - | - | Unassigned | - |
| Otu02395 | 17 | k__Fungi(100);p__Ascomycota(100);c__Saccharomycetes(100);o__Saccharomycetales(100);f__Saccharomycetaceae(100);g__unclassified_Saccharomycetaceae(100);s__Saccharomycetaceae_sp(100); | - | - | - | - | - | - | - | Unassigned | - |
| Otu02396 | 17 | k__Fungi(100);p__Ascomycota(100);p__Ascomycota_unclassified(95);p__Ascomycota_unclassified(95);p__Ascomycota_unclassified(95);p__Ascomycota_unclassified(95);p__Ascomycota_unclassified(95); | - | - | - | - | - | - | - | Unassigned | - |
| Otu02397 | 17 | k__Fungi(100);k__Fungi_unclassified(100);k__Fungi_unclassified(100);k__Fungi_unclassified(100);k__Fungi_unclassified(100);k__Fungi_unclassified(100);k__Fungi_unclassified(100); | - | - | - | - | - | - | - | Unassigned | - |
| Otu02398 | 17 | k__Fungi(100);p__Ascomycota(53);p__Ascomycota_unclassified(53);p__Ascomycota_unclassified(53);p__Ascomycota_unclassified(53);p__Ascomycota_unclassified(53);p__Ascomycota_unclassified(53); | - | - | - | - | - | - | - | Unassigned | - |
| Otu02400 | 17 | k__Fungi(100);p__Ascomycota(59);p__Ascomycota_unclassified(59);p__Ascomycota_unclassified(59);p__Ascomycota_unclassified(59);p__Ascomycota_unclassified(59);p__Ascomycota_unclassified(59); | - | - | - | - | - | - | - | Unassigned | - |
| Otu02403 | 17 | k__Fungi(100);p__Ascomycota(71);p__Ascomycota_unclassified(71);p__Ascomycota_unclassified(71);p__Ascomycota_unclassified(71);p__Ascomycota_unclassified(71);p__Ascomycota_unclassified(71); | - | - | - | - | - | - | - | Unassigned | - |
| Otu02404 | 17 | k__Fungi(100);p__Ascomycota(100);c__Sordariomycetes(100);o__Hypocreales(100);f__Hypocreaceae(100);f__Hypocreaceae_unclassified(100);f__Hypocreaceae_unclassified(100); | - | - | - | - | - | - | - | Unassigned | - |
| Otu02405 | 17 | k__Fungi(100);k__Fungi_unclassified(65);k__Fungi_unclassified(65);k__Fungi_unclassified(65);k__Fungi_unclassified(65);k__Fungi_unclassified(65);k__Fungi_unclassified(65); | - | - | - | - | - | - | - | Unassigned | - |
| Otu02408 | 17 | k__Fungi(100);k__Fungi_unclassified(65);k__Fungi_unclassified(65);k__Fungi_unclassified(65);k__Fungi_unclassified(65);k__Fungi_unclassified(65);k__Fungi_unclassified(65); | - | - | - | - | - | - | - | Unassigned | - |
| Otu02411 | 17 | k__Fungi(100);p__Ascomycota(65);p__Ascomycota_unclassified(65);p__Ascomycota_unclassified(65);p__Ascomycota_unclassified(65);p__Ascomycota_unclassified(65);p__Ascomycota_unclassified(65); | - | - | - | - | - | - | - | Unassigned | - |
| Otu02412 | 17 | k__Fungi(100);p__Basidiomycota(83);p__Basidiomycota_unclassified(83);p__Basidiomycota_unclassified(83);p__Basidiomycota_unclassified(83);p__Basidiomycota_unclassified(83);p__Basidiomycota_unclassified(83); | - | - | - | - | - | - | - | Unassigned | - |
| Otu02415 | 17 | k__Fungi(100);k__Fungi_unclassified(65);k__Fungi_unclassified(65);k__Fungi_unclassified(65);k__Fungi_unclassified(65);k__Fungi_unclassified(65);k__Fungi_unclassified(65); | - | - | - | - | - | - | - | Unassigned | - |
| Otu02418 | 17 | k__Fungi(100);p__Basidiomycota(100);c__Agaricomycetes(100);c__Agaricomycetes_unclassified(100);c__Agaricomycetes_unclassified(100);c__Agaricomycetes_unclassified(100);c__Agaricomycetes_unclassified(100); | - | - | - | - | - | - | - | Unassigned | - |
| Otu02419 | 17 | k__Fungi(100);p__Basidiomycota(100);c__Microbotryomycetes(100);o__Sporidiobolales(100);f__Sporidiobolales_family_Incertae_sedis(100);f__Sporidiobolales_family_Incertae_sedis_unclassified(53);f__Sporidiobolales_family_Incertae_sedis_unclassified(53); | - | - | - | - | - | - | - | Unassigned | - |
| Otu02420 | 17 | k__Fungi(100);k__Fungi_unclassified(100);k__Fungi_unclassified(100);k__Fungi_unclassified(100);k__Fungi_unclassified(100);k__Fungi_unclassified(100);k__Fungi_unclassified(100); | - | - | - | - | - | - | - | Unassigned | - |
| Otu02423 | 17 | k__Fungi(100);p__Ascomycota(71);p__Ascomycota_unclassified(71);p__Ascomycota_unclassified(71);p__Ascomycota_unclassified(71);p__Ascomycota_unclassified(71);p__Ascomycota_unclassified(71); | - | - | - | - | - | - | - | Unassigned | - |
| Otu02425 | 17 | k__Fungi(100);k__Fungi_unclassified(71);k__Fungi_unclassified(71);k__Fungi_unclassified(71);k__Fungi_unclassified(71);k__Fungi_unclassified(71);k__Fungi_unclassified(71); | - | - | - | - | - | - | - | Unassigned | - |
| Otu02428 | 17 | k__Fungi(100);k__Fungi_unclassified(77);k__Fungi_unclassified(77);k__Fungi_unclassified(77);k__Fungi_unclassified(77);k__Fungi_unclassified(77);k__Fungi_unclassified(77); | - | - | - | - | - | - | - | Unassigned | - |
| Otu02431 | 17 | k__Fungi(100);p__Basidiomycota(100);p__Basidiomycota_unclassified(100);p__Basidiomycota_unclassified(100);p__Basidiomycota_unclassified(100);p__Basidiomycota_unclassified(100);p__Basidiomycota_unclassified(100); | - | - | - | - | - | - | - | Unassigned | - |
| Otu02432 | 17 | k__Fungi(100);p__Ascomycota(100);c__Sordariomycetes(100);o__Diaporthales(100);o__Diaporthales_unclassified(100);o__Diaporthales_unclassified(100);o__Diaporthales_unclassified(100); | - | - | - | - | - | - | - | Unassigned | - |
| Otu02433 | 17 | k__Fungi(100);k__Fungi_unclassified(71);k__Fungi_unclassified(71);k__Fungi_unclassified(71);k__Fungi_unclassified(71);k__Fungi_unclassified(71);k__Fungi_unclassified(71); | - | - | - | - | - | - | - | Unassigned | - |
| Otu02437 | 17 | k__Fungi(100);p__Ascomycota(53);p__Ascomycota_unclassified(53);p__Ascomycota_unclassified(53);p__Ascomycota_unclassified(53);p__Ascomycota_unclassified(53);p__Ascomycota_unclassified(53); | - | - | - | - | - | - | - | Unassigned | - |
| Otu02438 | 17 | k__Fungi(100);k__Fungi_unclassified(59);k__Fungi_unclassified(59);k__Fungi_unclassified(59);k__Fungi_unclassified(59);k__Fungi_unclassified(59);k__Fungi_unclassified(59); | - | - | - | - | - | - | - | Unassigned | - |
| Otu02439 | 17 | k__Fungi(100);p__Ascomycota(100);p__Ascomycota_unclassified(95);p__Ascomycota_unclassified(95);p__Ascomycota_unclassified(95);p__Ascomycota_unclassified(95);p__Ascomycota_unclassified(95); | - | - | - | - | - | - | - | Unassigned | - |
| Otu02443 | 17 | k__Fungi(100);p__Ascomycota(65);p__Ascomycota_unclassified(65);p__Ascomycota_unclassified(65);p__Ascomycota_unclassified(65);p__Ascomycota_unclassified(65);p__Ascomycota_unclassified(65); | - | - | - | - | - | - | - | Unassigned | - |
| Otu02444 | 17 | k__Fungi(100);p__Ascomycota(100);p__Ascomycota_unclassified(100);p__Ascomycota_unclassified(100);p__Ascomycota_unclassified(100);p__Ascomycota_unclassified(100);p__Ascomycota_unclassified(100); | - | - | - | - | - | - | - | Unassigned | - |
| Otu02445 | 17 | k__Fungi(100);k__Fungi_unclassified(100);k__Fungi_unclassified(100);k__Fungi_unclassified(100);k__Fungi_unclassified(100);k__Fungi_unclassified(100);k__Fungi_unclassified(100); | - | - | - | - | - | - | - | Unassigned | - |
| Otu02446 | 17 | k__Fungi(100);k__Fungi_unclassified(89);k__Fungi_unclassified(89);k__Fungi_unclassified(89);k__Fungi_unclassified(89);k__Fungi_unclassified(89);k__Fungi_unclassified(89); | - | - | - | - | - | - | - | Unassigned | - |
| Otu02449 | 17 | k__Fungi(100);p__Ascomycota(100);p__Ascomycota_unclassified(100);p__Ascomycota_unclassified(100);p__Ascomycota_unclassified(100);p__Ascomycota_unclassified(100);p__Ascomycota_unclassified(100); | - | - | - | - | - | - | - | Unassigned | - |
| Otu02451 | 17 | k__Fungi(100);p__Ascomycota(100);c__Dothideomycetes(100);o__Pleosporales(100);o__Pleosporales_unclassified(100);o__Pleosporales_unclassified(100);o__Pleosporales_unclassified(100); | - | - | - | - | - | - | - | Unassigned | - |
| Otu02456 | 17 | k__Fungi(100);p__Basidiomycota(100);c__Agaricomycetes(100);c__Agaricomycetes_unclassified(100);c__Agaricomycetes_unclassified(100);c__Agaricomycetes_unclassified(100);c__Agaricomycetes_unclassified(100); | - | - | - | - | - | - | - | Unassigned | - |
| Otu02457 | 17 | k__Fungi(100);k__Fungi_unclassified(83);k__Fungi_unclassified(83);k__Fungi_unclassified(83);k__Fungi_unclassified(83);k__Fungi_unclassified(83);k__Fungi_unclassified(83); | - | - | - | - | - | - | - | Unassigned | - |
| Otu02461 | 17 | k__Fungi(100);p__Basidiomycota(100);p__Basidiomycota_unclassified(100);p__Basidiomycota_unclassified(100);p__Basidiomycota_unclassified(100);p__Basidiomycota_unclassified(100);p__Basidiomycota_unclassified(100); | - | - | - | - | - | - | - | Unassigned | - |
| Otu02463 | 17 | k__Fungi(100);p__Ascomycota(100);c__Dothideomycetes(100);c__Dothideomycetes_unclassified(100);c__Dothideomycetes_unclassified(100);c__Dothideomycetes_unclassified(100);c__Dothideomycetes_unclassified(100); | - | - | - | - | - | - | - | Unassigned | - |
| Otu02464 | 17 | k__Fungi(100);p__Ascomycota(100);p__Ascomycota_unclassified(100);p__Ascomycota_unclassified(100);p__Ascomycota_unclassified(100);p__Ascomycota_unclassified(100);p__Ascomycota_unclassified(100); | - | - | - | - | - | - | - | Unassigned | - |
| Otu02467 | 17 | k__Fungi(100);p__Ascomycota(100);c__Dothideomycetes(100);o__Capnodiales(100);o__Capnodiales_unclassified(100);o__Capnodiales_unclassified(100);o__Capnodiales_unclassified(100); | - | - | - | - | - | - | - | Unassigned | - |
| Otu02470 | 17 | k__Fungi(100);p__Ascomycota(100);c__Sordariomycetes(100);o__Diaporthales(100);o__Diaporthales_unclassified(100);o__Diaporthales_unclassified(100);o__Diaporthales_unclassified(100); | - | - | - | - | - | - | - | Unassigned | - |
| Otu02471 | 17 | k__Fungi(100);p__Ascomycota(100);c__Sordariomycetes(100);c__Sordariomycetes_unclassified(100);c__Sordariomycetes_unclassified(100);c__Sordariomycetes_unclassified(100);c__Sordariomycetes_unclassified(100); | - | - | - | - | - | - | - | Unassigned | - |
| Otu02474 | 16 | k__Fungi(100);p__Ascomycota(100);c__Dothideomycetes(100);o__Myriangiales(100);f__Elsinoaceae(100);f__Elsinoaceae_unclassified(94);f__Elsinoaceae_unclassified(94); | Elsinoaceae | 9 | Pathotroph | Plant Pathogen | Microfungus | NULL | Probable | [illegible] | [illegible] |
| Otu02478 | 16 | k__Fungi(100);p__Ascomycota(100);c__Saccharomycetes(100);o__Saccharomycetales(100);f__Metschnikowiaceae(100);g__Metschnikowia(100);s__Metschnikowia_pulcherrima(63); | Metschnikowia | 13 | Saprotroph | \|Undefined Saprotroph\| | Yeast | NULL | Probable | [illegible] | [illegible] |
| Otu02480 | 16 | k__Fungi(100);p__Basidiomycota(100);c__Wallemiomycetes(100);o__Wallemiales(100);f__Wallemiaceae(100);g__Wallemia(100);s__Wallemia_sp_SJ_2014(100); | Wallemia | 13 | Saprotroph | \|Undefined Saprotroph\| | Microfungus | NULL | Probable | [illegible] | [illegible] |
| Otu02483 | 16 | k__Fungi(100);p__Ascomycota(100);c__Saccharomycetes(100);o__Saccharomycetales(100);f__Metschnikowiaceae(100);g__Metschnikowia(100);g__Metschnikowia_unclassified(82); | Metschnikowia | 13 | Saprotroph | \|Undefined Saprotroph\| | Yeast | NULL | Probable | [illegible] | [illegible] |
| Otu02487 | 16 | k__Fungi(100);p__Basidiomycota(100);c__Agaricomycetes(100);o__Agaricales(100);f__Entolomataceae(100);g__Clitopilus(100);s__Clitopilus_cystidiatus(100); | Clitopilus | 13 | Pathotroph-Saprotroph | Fungal Parasite-\|Plant Saprotroph\|-Undefined Saprotroph | Agaricoid | NULL | Probable | [illegible] | [illegible] |
| Otu02490 | 16 | k__Fungi(100);p__Zygomycota(100);c__Zygomycota_class_Incertae_sedis(100);o__Mucorales(100);f__Mucoraceae(100);g__Mucor(94);g__Mucor_unclassified(94); | Mucor | 13 | Saprotroph | \|Undefined Saprotroph\| | Microfungus | NULL | Probable | [illegible] | [illegible] |
| Otu02494 | 16 | k__Fungi(100);p__Basidiomycota(100);c__Agaricomycetes(100);o__Boletales(100);f__Boletaceae(100);g__Boletus(100);g__Boletus_unclassified(100); | Boletus | 13 | Pathotroph-Symbiotroph | \|Ectomycorrhizal\|-Fungal Parasite | Agaricoid-Boletoid | NULL | Highly Probable | NULL | [illegible] |
| Otu02496 | 16 | k__Fungi(100);p__Basidiomycota(100);c__Agaricomycetes(100);o__Thelephorales(100);f__Thelephoraceae(100);g__Tomentella(100);g__Tomentella_unclassified(100); | Tomentella | 13 | Symbiotroph | \|Ectomycorrhizal\| | Corticioid | NULL | Highly Probable | NULL | [illegible] |
| Otu02498 | 16 | k__Fungi(100);p__Ascomycota(100);c__Saccharomycetes(100);o__Saccharomycetales(100);f__Metschnikowiaceae(100);g__Metschnikowia(100);g__Metschnikowia_unclassified(75); | Metschnikowia | 13 | Saprotroph | \|Undefined Saprotroph\| | Yeast | NULL | Probable | [illegible] | [illegible] |
| Otu02501 | 16 | k__Fungi(100);p__Ascomycota(100);c__Saccharomycetes(100);o__Saccharomycetales(100);f__Metschnikowiaceae(100);g__Metschnikowia(100);g__Metschnikowia_unclassified(75); | Metschnikowia | 13 | Saprotroph | \|Undefined Saprotroph\| | Yeast | NULL | Probable | [illegible] | [illegible] |
| Otu02502 | 16 | k__Fungi(100);p__Basidiomycota(100);c__Agaricomycetes(100);o__Boletales(100);f__Paxillaceae(100);g__Paxillus(100);s__Paxillus_involutus(100); | Paxillus | 13 | Symbiotroph | \|Ectomycorrhizal\| | Agaricoid | NULL | Highly Probable | NULL | [illegible] |
| Otu02507 | 16 | k__Fungi(100);p__Ascomycota(100);c__Saccharomycetes(100);o__Saccharomycetales(100);f__Metschnikowiaceae(100);g__Metschnikowia(100);s__Metschnikowia_pulcherrima(88); | Metschnikowia | 13 | Saprotroph | \|Undefined Saprotroph\| | Yeast | NULL | Probable | [illegible] | [illegible] |
| Otu02508 | 16 | k__Fungi(100);p__Ascomycota(100);c__Saccharomycetes(100);o__Saccharomycetales(100);f__Metschnikowiaceae(100);g__Metschnikowia(100);s__Metschnikowia_pulcherrima(88); | Metschnikowia | 13 | Saprotroph | \|Undefined Saprotroph\| | Yeast | NULL | Probable | [illegible] | [illegible] |
| Otu02509 | 16 | k__Fungi(100);p__Ascomycota(100);c__Saccharomycetes(100);o__Saccharomycetales(100);f__Saccharomycetaceae(100);g__Kazachstania(100);s__Kazachstania_intestinalis(100); | Kazachstania | 13 | Saprotroph | \|Undefined Saprotroph\| | Yeast | NULL | Probable | [illegible] | [illegible] |
| Otu02513 | 16 | k__Fungi(100);p__Ascomycota(100);c__Pezizomycetes(100);o__Pezizales(100);f__Pyronemataceae(100);g__Geopyxis(100);s__Geopyxis_sp(100); | Geopyxis | 13 | Saprotroph-Symbiotroph | Ectomycorrhizal-Plant Saprotroph-\|Undefined Saprotroph\| | Pezizoid | NULL | Possible | [illegible] | [illegible] |
| Otu02515 | 16 | k__Fungi(100);p__Ascomycota(100);c__Saccharomycetes(100);o__Saccharomycetales(100);f__Metschnikowiaceae(100);g__Metschnikowia(100);s__Metschnikowia_pulcherrima(94); | Metschnikowia | 13 | Saprotroph | \|Undefined Saprotroph\| | Yeast | NULL | Probable | [illegible] | [illegible] |
| Otu02517 | 16 | k__Fungi(100);p__Basidiomycota(100);c__Ustilaginomycetes(100);o__Ustilaginales(100);f__Ustilaginaceae(100);g__unclassified_Ustilaginaceae(88);s__Ustilaginaceae_sp(88); | Ustilaginaceae | 9 | Pathotroph | Plant Pathogen | Smut | NULL | Probable | [illegible] | [illegible] |
| Otu02519 | 16 | k__Fungi(100);p__Ascomycota(100);c__Saccharomycetes(100);o__Saccharomycetales(100);f__Metschnikowiaceae(100);g__Metschnikowia(100);s__Metschnikowia_pulcherrima(100); | Metschnikowia | 13 | Saprotroph | \|Undefined Saprotroph\| | Yeast | NULL | Probable | [illegible] | [illegible] |
| Otu02520 | 16 | k__Fungi(100);p__Ascomycota(100);c__Eurotiomycetes(82);o__Chaetothyriales(82);f__Herpotrichiellaceae(82);f__Herpotrichiellaceae_unclassified(82);f__Herpotrichiellaceae_unclassified(82); | Herpotrichiellaceae | 9 | Pathotroph-Saprotroph | Animal Pathogen-Fungal Parasite-Undefined Saprotroph | Facultative Yeast-Microfungus | NULL | Probable | [illegible] | [illegible] |
| Otu02524 | 16 | k__Fungi(100);p__Basidiomycota(100);c__Agaricomycetes(100);o__Agaricales(100);f__Strophariaceae(100);g__Galerina(100);s__Galerina_unicolor_(Fr.)_Singer(100); | Galerina | 13 | Pathotroph-Saprotroph | Bryophyte Parasite-Plant Pathogen-Plant Saprotroph-Undefined Saprotroph-\|Wood Saprotroph\| | Agaricoid | White Rot | Probable | [illegible] | [illegible] |

| | | | | | | | | | | | |
|---|---|---|---|---|---|---|---|---|---|---|---|
| Otu02525 | 16 | k__Fungi(100);p__Ascomycota(100);c__Dothideomycetes(100);o__Capnodiales(100);f__Capnodiales_family_Incertae_sedis(100);g__Rachicladosporium(69);g__Rachicladosporium_unclassified(69); | Rachicladosporium | 13 | Saprotroph | \|Plant Saprotroph\| | Microfungus | NULL | Probable | As litter_saprotroph (Põlme et al. 2020); Decay leaf, fruit, seed (Põlme et al. 2020) | [illegible] |
| Otu02526 | 16 | k__Fungi(100);p__Basidiomycota(100);c__Pucciniomycetes(100);o__Pucciniales(100);f__Cronartiaceae(100);g__unclassified_Cronartiaceae(75);s__Cronartiaceae_sp(75); | Cronartiaceae | 9 | Pathotroph | Plant Pathogen | Microfungus | NULL | Probable | Common Name - Blister Rust, Rust Fungus; Most species are heteroecious, with the aecial stages forming on stems and cones of Pinus species and uredinia and telia on a range of dicotyledonous plants (Cannon & Kirk 2007) | [illegible] |
| Otu02527 | 16 | k__Fungi(100);p__Basidiomycota(100);c__Tremellomycetes(100);o__Tremellales(100);f__Tremellales_family_Incertae_sedis(100);g__Kockovaella(94);s__Kockovaella_schimae(57); | Kockovaella | 13 | Saprotroph-Symbiotroph | Endophyte-\|Epiphyte\|-Undefined Saprotroph | Yeast | NULL | Probable | As foliar_endophyte (Põlme et al. 2020); Decay leaf, fruit, seed (Põlme et al. 2020); Foliar, isolated from a plant sample collected in a tropical rain forest in Thailand. | [illegible] |
| Otu02528 | 16 | k__Fungi(100);p__Basidiomycota(100);c__Tremellomycetes(100);o__Tremellales(100);f__Tremellales_family_Incertae_sedis(100);g__Fibulobasidium(75);s__Fibulobasidium_inconspicuum(75); | Fibulobasidium | 13 | Saprotroph | Undefined Saprotroph-\|Wood Saprotroph\| | Dimorphic-Facultative Yeast | NULL | Highly Probable | Decay wood (Põlme et al. 2020) | [illegible] |
| Otu02529 | 16 | k__Fungi(100);p__Ascomycota(100);c__Lecanoromycetes(100);o__Lecanorales(100);o__Lecanorales_unclassified(100);o__Lecanorales_unclassified(100);o__Lecanorales_unclassified(100); | Lecanorales | 7 | Pathotroph-Symbiotroph | Lichen Parasite-Lichenized | Microfungus-Thallus | NULL | Probable | Lichenized with green algae, occasionally lichenicolous, on rocks, mosses and wood (Cannon & Kirk 2007); ...an order of mostly lichen-forming fungi (Kirk et al. 2008) | [illegible] |
| Otu02532 | 16 | k__Fungi(100);p__Basidiomycota(100);c__Agaricomycetes(100);o__Trechisporales(100);f__Hydnodontaceae(100);g__Trechispora(100);g__Trechispora_unclassified(100); | Trechispora | 13 | Saprotroph | \|Wood Saprotroph\| | Corticioid | White Rot | Highly Probable | As white_rot (Põlme et al. 2020); Decay wood (Põlme et al. 2020) | [illegible] |
| Otu02536 | 16 | k__Fungi(100);p__Ascomycota(100);c__Saccharomycetes(100);o__Saccharomycetales(100);f__Metschnikowiaceae(100);g__Metschnikowia(100);s__Metschnikowia_pulcherrima(100); | Metschnikowia | 13 | Saprotroph | \|Undefined Saprotroph\| | Yeast | NULL | Probable | As nectar-tap_saprotroph (Põlme et al. 2020); Decay sugar-rich substrates (Põlme et al. 2020) | [illegible] |
| Otu02537 | 16 | k__Fungi(100);p__Ascomycota(100);c__Sordariomycetes(100);o__Hypocreales(100);o__Hypocreales_unclassified(88);o__Hypocreales_unclassified(88);o__Hypocreales_unclassified(88); | Hypocreales | 7 | Saprotroph | Undefined Saprotroph | Microfungus | NULL | Possible | NULL | [illegible] |
| Otu02539 | 16 | k__Fungi(100);p__Ascomycota(100);c__Saccharomycetes(100);o__Saccharomycetales(100);f__Metschnikowiaceae(100);g__Metschnikowia(100);s__Metschnikowia_pulcherrima(100); | Metschnikowia | 13 | Saprotroph | \|Undefined Saprotroph\| | Yeast | NULL | Probable | As nectar-tap_saprotroph (Põlme et al. 2020); Decay sugar-rich substrates (Põlme et al. 2020) | [illegible] |
| Otu02542 | 16 | k__Fungi(100);p__Ascomycota(100);c__Saccharomycetes(100);o__Saccharomycetales(100);f__Metschnikowiaceae(100);g__Metschnikowia(100);s__Metschnikowia_pulcherrima(100); | Metschnikowia | 13 | Saprotroph | \|Undefined Saprotroph\| | Yeast | NULL | Probable | As nectar-tap_saprotroph (Põlme et al. 2020); Decay sugar-rich substrates (Põlme et al. 2020) | [illegible] |
| Otu02543 | 16 | k__Fungi(100);p__Ascomycota(100);c__Sordariomycetes(100);o__Ophiostomatales(88);f__Ophiostomataceae(88);f__Ophiostomataceae_unclassified(88);f__Ophiostomataceae_unclassified(88); | Ophiostomataceae | 9 | Pathotroph | Plant Pathogen | Microfungus | Blue-Staining | Probable | At least most species are transmitted via insect vectors, with mycelial colonies developing in bark-boring beetle galleries and spreading through the vascular system; they are highly specific to their beetle vectors. In some cases spores and mycelium are transmitted in specialized organs, termed mycangia (Cannon & Kirk 2007) | [illegible] |
| Otu02544 | 16 | k__Fungi(100);p__Ascomycota(100);c__Eurotiomycetes(100);o__Eurotiales(100);f__Eurotiales_family_Incertae_sedis(100);g__Thermomyces(100);s__Thermomyces_lanuginosus(100); | Thermomyces | 13 | Pathotroph-Saprotroph-Symbiotroph | Endophyte-Plant Pathogen-Plant Saprotroph-\|Undefined Saprotroph\| | Microfungus | NULL | Possible | Cannabaceae, Moraceae, Oleaceae; As foliar_endophyte (Põlme et al. 2020); As litter_saprotroph (Põlme et al. 2020); As soil_saprotroph (Põlme et al. 2020); Decay leaf, fruit, seed, soil (Põlme et al. 2020); Host | [illegible] |
| Otu02545 | 16 | k__Fungi(100);p__Basidiomycota(100);c__Microbotryomycetes(100);o__Sporidiobolales(100);f__Sporidiobolales_family_Incertae_sedis(100);g__Sporobolomyces(100);s__Sporobolomyces_lactosus(100); | Sporobolomyces | 13 | Pathotroph-Saprotroph | \|Fungal Parasite\|-Plant Saprotroph | Yeast | NULL | Probable | As litter_saprotroph (Põlme et al. 2020); As mycoparasite (Põlme et al. 2020); Decay fungal material (Põlme et al. 2020) | [illegible] |
| Otu02547 | 16 | k__Fungi(100);p__Basidiomycota(100);c__Agaricomycetes(100);o__Agaricales(100);f__Strophariaceae(100);g__Stropharia(100);s__Stropharia_rugosoannulata(100); | Stropharia | 13 | Saprotroph | Dung Saprotroph-Plant Parasite-\|Plant Saprotroph\|-Undefined Saprotroph-Wood Saprotroph | Agaricoid | NULL | Probable | As litter_saprotroph (Põlme et al. 2020); Decay leaf, fruit, seed (Põlme et al. 2020); Very variable ecology; Pholiota species are lignicolous and can act as weak parasites while most other taxa are either saprobes in grassy habitats or are found in dung. Species are not ectomycorrhizal (Cannon & Kirk 2007) | [illegible] |
| Otu02549 | 16 | k__Fungi(100);p__Ascomycota(100);c__Saccharomycetes(100);o__Saccharomycetales(100);f__Metschnikowiaceae(100);g__Metschnikowia(100);s__Metschnikowia_pulcherrima(94); | Metschnikowia | 13 | Saprotroph | \|Undefined Saprotroph\| | Yeast | NULL | Probable | As nectar-tap_saprotroph (Põlme et al. 2020); Decay sugar-rich substrates (Põlme et al. 2020) | [illegible] |
| Otu02551 | 16 | k__Fungi(100);p__Basidiomycota(100);c__Agaricomycetes(100);o__Polyporales(100);f__Meruliaceae(100);g__Steccherinum(100);s__Steccherinum_cf_ciliolatum(100); | Steccherinum | 13 | Saprotroph | Undefined Saprotroph-\|Wood Saprotroph\| | Corticioid | White Rot | Probable | As white_rot (Põlme et al. 2020); Decay wood (Põlme et al. 2020) | [illegible] |
| Otu02554 | 16 | k__Fungi(100);p__Basidiomycota(100);c__Exobasidiomycetes(100);o__Exobasidiales(100);f__Exobasidiaceae(100);g__Exobasidium(100);s__Exobasidium_yoshinagae(100); | Exobasidium | 13 | Pathotroph | \|Plant Pathogen\| | Leaf Blister-Leaf Gall-Leaf Spot | NULL | Probable | NULL | [illegible] |
| Otu02555 | 16 | k__Fungi(100);p__Ascomycota(100);c__Eurotiomycetes(100);o__Chaetothyriales(100);f__Herpotrichiellaceae(100);f__Herpotrichiellaceae_unclassified(100);f__Herpotrichiellaceae_unclassified(100); | Herpotrichiellaceae | 9 | Pathotroph-Saprotroph | Animal Pathogen-Fungal Parasite-Undefined Saprotroph | Facultative Yeast-Microfungus | NULL | Probable | Mostly saprobic on plants or other fungi, isolated from soil etc., and from medical sources (Cannon & Kirk 2007) | [illegible] |
| Otu02557 | 16 | k__Fungi(100);p__Basidiomycota(100);c__Agaricomycetes(100);o__Agaricales(100);f__Tricholomataceae(100);g__Melanoleuca(100);s__Melanoleuca_strictipes(100); | Melanoleuca | 13 | Saprotroph | Plant Saprotroph-\|Undefined Saprotroph\| | Agaricoid | NULL | Probable | As litter_saprotroph (Põlme et al. 2020); As soil_saprotroph (Põlme et al. 2020); Decay leaf, fruit, seed, soil (Põlme et al. 2020) | [illegible] |
| Otu02559 | 16 | k__Fungi(100);p__Basidiomycota(100);c__Agaricomycetes(100);o__Polyporales(100);f__Phanerochaetaceae(100);g__Phlebiopsis(100);s__Phlebiopsis_gigantea(100); | Phlebiopsis | 13 | Saprotroph | Undefined Saprotroph-\|Wood Saprotroph\| | Corticioid | White Rot | Probable | As white_rot (Põlme et al. 2020); Decay wood (Põlme et al. 2020) | [illegible] |
| Otu02560 | 16 | k__Fungi(100);p__Basidiomycota(100);c__Agaricomycetes(100);o__Agaricales(100);f__Marasmiaceae(100);g__Marasmiellus(100);s__Marasmiellus_opacus(100); | Marasmiellus | 13 | Saprotroph | \|Plant Saprotroph\|-Undefined Saprotroph-Wood Saprotroph | Agaricoid | NULL | Probable | As litter_saprotroph (Põlme et al. 2020); Decay leaf, fruit, seed (Põlme et al. 2020) | [illegible] |
| Otu02561 | 16 | k__Fungi(100);p__Basidiomycota(100);c__Agaricomycetes(100);o__Agaricales(100);f__Physalacriaceae(100);g__Cylindrobasidium(100);s__Cylindrobasidium_evolvens(100); | Cylindrobasidium | 13 | Saprotroph | Undefined Saprotroph-\|Wood Saprotroph\| | Corticioid | White Rot | Possible | As white_rot (Põlme et al. 2020); Decay wood (Põlme et al. 2020) | [illegible] |
| Otu02563 | 16 | k__Fungi(100);p__Basidiomycota(100);c__Agaricomycetes(100);o__Russulales(100);f__Stereaceae(100);g__unclassified_Stereaceae(100);s__Stereaceae_sp(100); | Stereaceae | 9 | Saprotroph | Undefined Saprotroph-Wood Saprotroph | Corticioid-Polyporoid | NULL | Probable | Lignicolous or terrestrial (in leaf litter); typically saprobic (Cannon & Kirk 2007) | [illegible] |
| Otu02566 | 16 | k__Fungi(100);p__Chytridiomycota(100);c__Chytridiomycetes(100);o__Spizellomycetales(100);f__Spizellomycetaceae(100);f__Spizellomycetaceae_unclassified(100);f__Spizellomycetaceae_unclassified(100); | Spizellomycetaceae | 9 | Pathotroph | Plant Pathogen-Undefined Parasite-Undefined Saprotroph | Microfungus | NULL | Probable | Common in the soil and also as parasites of a range of soil organisms and plants (Cannon & Kirk 2007) | [illegible] |
| Otu02570 | 16 | k__Fungi(100);p__Basidiomycota(100);c__Agaricomycetes(100);o__Trechisporales(100);f__Hydnodontaceae(100);g__Trechispora(100);s__Trechispora_sp(100); | Trechispora | 13 | Saprotroph | \|Wood Saprotroph\| | Corticioid | White Rot | Highly Probable | As white_rot (Põlme et al. 2020); Decay wood (Põlme et al. 2020) | [illegible] |
| Otu02571 | 16 | k__Fungi(100);p__Ascomycota(100);c__Saccharomycetes(100);o__Saccharomycetales(100);f__Saccharomycetales_family_Incertae_sedis(100);g__Candida(100);s__Candida_stellata(100); | Candida | 13 | Pathotroph-Saprotroph-Symbiotroph | Animal Pathogen-Endophyte-Endosymbiont-Epiphyte-\|Undefined Saprotroph\| | Dimorphic Yeast-Yeast | NULL | Probable | As nectar-tap_saprotroph (Põlme et al. 2020); Decay sugar-rich substrates (Põlme et al. 2020); This asexual genus is polyphyletic and contain species that function very differently depend on what they are. Caution should be taken when interpreting guilds for Candida species. | [illegible] |
| Otu02574 | 16 | k__Fungi(100);p__Ascomycota(100);c__Saccharomycetes(100);o__Saccharomycetales(100);f__Metschnikowiaceae(100);g__Metschnikowia(100);s__Metschnikowia_pulcherrima(88); | Metschnikowia | 13 | Saprotroph | \|Undefined Saprotroph\| | Yeast | NULL | Probable | As nectar-tap_saprotroph (Põlme et al. 2020); Decay sugar-rich substrates (Põlme et al. 2020) | [illegible] |
| Otu02575 | 16 | k__Fungi(100);p__Ascomycota(100);c__Eurotiomycetes(100);o__Onygenales(100);f__Ajellomycetaceae(100);g__Spiromastix(100);g__Spiromastix_unclassified(100); | Spiromastix | 13 | Pathotroph-Saprotroph | Animal Pathogen-Dung Saprotroph-\|Undefined Saprotroph\| | Microfungus | NULL | Probable | As soil_saprotroph (Põlme et al. 2020); Decay soil (Põlme et al. 2020) | [illegible] |
| Otu02475 | 16 | k__Fungi(100);p__Ascomycota(100);p__Ascomycota_unclassified(100);p__Ascomycota_unclassified(100);p__Ascomycota_unclassified(100);p__Ascomycota_unclassified(100);p__Ascomycota_unclassified(100); | - | - | - | - | - | - | - | Unassigned | - |
| Otu02476 | 16 | k__Fungi(100);p__Ascomycota(100);p__Ascomycota_unclassified(100);p__Ascomycota_unclassified(100);p__Ascomycota_unclassified(100);p__Ascomycota_unclassified(100);p__Ascomycota_unclassified(100); | - | - | - | - | - | - | - | Unassigned | - |
| Otu02477 | 16 | k__Fungi(100);p__Ascomycota(57);p__Ascomycota_unclassified(57);p__Ascomycota_unclassified(57);p__Ascomycota_unclassified(57);p__Ascomycota_unclassified(57);p__Ascomycota_unclassified(57); | - | - | - | - | - | - | - | Unassigned | - |
| Otu02479 | 16 | k__Fungi(100);p__Ascomycota(100);c__Sordariomycetes(100);c__Sordariomycetes_unclassified(100);c__Sordariomycetes_unclassified(100);c__Sordariomycetes_unclassified(100);c__Sordariomycetes_unclassified(100); | - | - | - | - | - | - | - | Unassigned | - |
| Otu02481 | 16 | k__Fungi(100);p__Ascomycota(100);p__Ascomycota_unclassified(94);p__Ascomycota_unclassified(94);p__Ascomycota_unclassified(94);p__Ascomycota_unclassified(94);p__Ascomycota_unclassified(94); | - | - | - | - | - | - | - | Unassigned | - |
| Otu02482 | 16 | k__Fungi(100);p__Ascomycota(100);c__Dothideomycetes(100);o__Capnodiales(94);o__Capnodiales_unclassified(94);o__Capnodiales_unclassified(94);o__Capnodiales_unclassified(94); | - | - | - | - | - | - | - | Unassigned | - |
| Otu02484 | 16 | k__Fungi(100);p__Basidiomycota(100);p__Basidiomycota_unclassified(82);p__Basidiomycota_unclassified(82);p__Basidiomycota_unclassified(82);p__Basidiomycota_unclassified(82);p__Basidiomycota_unclassified(82); | - | - | - | - | - | - | - | Unassigned | - |
| Otu02485 | 16 | k__Fungi(100);p__Ascomycota(75);p__Ascomycota_unclassified(75);p__Ascomycota_unclassified(75);p__Ascomycota_unclassified(75);p__Ascomycota_unclassified(75);p__Ascomycota_unclassified(75); | - | - | - | - | - | - | - | Unassigned | - |
| Otu02486 | 16 | k__Fungi(100);k__Fungi_unclassified(57);k__Fungi_unclassified(57);k__Fungi_unclassified(57);k__Fungi_unclassified(57);k__Fungi_unclassified(57);k__Fungi_unclassified(57); | - | - | - | - | - | - | - | Unassigned | - |
| Otu02488 | 16 | k__Fungi(100);k__Fungi_unclassified(69);k__Fungi_unclassified(69);k__Fungi_unclassified(69);k__Fungi_unclassified(69);k__Fungi_unclassified(69);k__Fungi_unclassified(69); | - | - | - | - | - | - | - | Unassigned | - |
| Otu02489 | 16 | k__Fungi(100);k__Fungi_unclassified(100);k__Fungi_unclassified(100);k__Fungi_unclassified(100);k__Fungi_unclassified(100);k__Fungi_unclassified(100);k__Fungi_unclassified(100); | - | - | - | - | - | - | - | Unassigned | - |
| Otu02491 | 16 | k__Fungi(100);k__Fungi_unclassified(69);k__Fungi_unclassified(69);k__Fungi_unclassified(69);k__Fungi_unclassified(69);k__Fungi_unclassified(69);k__Fungi_unclassified(69); | - | - | - | - | - | - | - | Unassigned | - |
| Otu02492 | 16 | k__Fungi(100);k__Fungi_unclassified(69);k__Fungi_unclassified(69);k__Fungi_unclassified(69);k__Fungi_unclassified(69);k__Fungi_unclassified(69);k__Fungi_unclassified(69); | - | - | - | - | - | - | - | Unassigned | - |
| Otu02493 | 16 | k__Fungi(100);k__Fungi_unclassified(100);k__Fungi_unclassified(100);k__Fungi_unclassified(100);k__Fungi_unclassified(100);k__Fungi_unclassified(100);k__Fungi_unclassified(100); | - | - | - | - | - | - | - | Unassigned | - |
| Otu02495 | 16 | k__Fungi(100);p__Ascomycota(100);c__Eurotiomycetes(100);o__Chaetothyriales(100);o__Chaetothyriales_unclassified(100);o__Chaetothyriales_unclassified(100);o__Chaetothyriales_unclassified(100); | - | - | - | - | - | - | - | Unassigned | - |
| Otu02497 | 16 | k__Fungi(100);p__Ascomycota(100);p__Ascomycota_unclassified(100);p__Ascomycota_unclassified(100);p__Ascomycota_unclassified(100);p__Ascomycota_unclassified(100);p__Ascomycota_unclassified(100); | - | - | - | - | - | - | - | Unassigned | - |
| Otu02499 | 16 | k__Fungi(100);p__Basidiomycota(100);c__Agaricomycetes(100);o__Sebacinales(100);o__Sebacinales_unclassified(100);o__Sebacinales_unclassified(100);o__Sebacinales_unclassified(100); | - | - | - | - | - | - | - | Unassigned | - |
| Otu02500 | 16 | k__Fungi(100);k__Fungi_unclassified(75);k__Fungi_unclassified(75);k__Fungi_unclassified(75);k__Fungi_unclassified(75);k__Fungi_unclassified(75);k__Fungi_unclassified(75); | - | - | - | - | - | - | - | Unassigned | - |
| Otu02503 | 16 | k__Fungi(100);p__Basidiomycota(100);c__Agaricomycetes(100);c__Agaricomycetes_unclassified(100);c__Agaricomycetes_unclassified(100);c__Agaricomycetes_unclassified(100);c__Agaricomycetes_unclassified(100); | - | - | - | - | - | - | - | Unassigned | - |
| Otu02504 | 16 | k__Fungi(100);k__Fungi_unclassified(57);k__Fungi_unclassified(57);k__Fungi_unclassified(57);k__Fungi_unclassified(57);k__Fungi_unclassified(57);k__Fungi_unclassified(57); | - | - | - | - | - | - | - | Unassigned | - |
| Otu02505 | 16 | k__Fungi(100);k__Fungi_unclassified(88);k__Fungi_unclassified(88);k__Fungi_unclassified(88);k__Fungi_unclassified(88);k__Fungi_unclassified(88);k__Fungi_unclassified(88); | - | - | - | - | - | - | - | Unassigned | - |
| Otu02506 | 16 | k__Fungi(100);p__Basidiomycota(100);c__Agaricomycetes(100);c__Agaricomycetes_unclassified(100);c__Agaricomycetes_unclassified(100);c__Agaricomycetes_unclassified(100);c__Agaricomycetes_unclassified(100); | - | - | - | - | - | - | - | Unassigned | - |
| Otu02510 | 16 | k__Fungi(100);p__Ascomycota(100);p__Ascomycota_unclassified(100);p__Ascomycota_unclassified(100);p__Ascomycota_unclassified(100);p__Ascomycota_unclassified(100);p__Ascomycota_unclassified(100); | - | - | - | - | - | - | - | Unassigned | - |
| Otu02511 | 16 | k__Fungi(100);p__Ascomycota(100);c__Dothideomycetes(100);o__Pleosporales(100);f__unclassified_Pleosporales(63);g__unclassified_Pleosporales(63);s__Pleosporales_sp(63); | - | - | - | - | - | - | - | Unassigned | - |
| Otu02512 | 16 | k__Fungi(100);p__Basidiomycota(100);c__Microbotryomycetes(100);o__Sporidiobolales(100);f__Sporidiobolales_family_Incertae_sedis(100);f__Sporidiobolales_family_Incertae_sedis_unclassified(57);f__Sporidiobolales_family_Incertae_sedis_unclassified(57); | - | - | - | - | - | - | - | Unassigned | - |
| Otu02514 | 16 | k__Fungi(100);k__Fungi_unclassified(69);k__Fungi_unclassified(69);k__Fungi_unclassified(69);k__Fungi_unclassified(69);k__Fungi_unclassified(69);k__Fungi_unclassified(69); | - | - | - | - | - | - | - | Unassigned | - |
| Otu02516 | 16 | k__Fungi(100);k__Fungi_unclassified(88);k__Fungi_unclassified(88);k__Fungi_unclassified(88);k__Fungi_unclassified(88);k__Fungi_unclassified(88);k__Fungi_unclassified(88); | - | - | - | - | - | - | - | Unassigned | - |
| Otu02518 | 16 | k__Fungi(100);k__Fungi_unclassified(69);k__Fungi_unclassified(69);k__Fungi_unclassified(69);k__Fungi_unclassified(69);k__Fungi_unclassified(69);k__Fungi_unclassified(69); | - | - | - | - | - | - | - | Unassigned | - |
| Otu02521 | 16 | k__Fungi(100);p__Ascomycota(100);c__Dothideomycetes(100);o__Pleosporales(100);o__Pleosporales_unclassified(100);o__Pleosporales_unclassified(100);o__Pleosporales_unclassified(100); | - | - | - | - | - | - | - | Unassigned | - |
| Otu02522 | 16 | k__Fungi(100);p__Basidiomycota(100);c__Agaricomycetes(100);o__Polyporales(100);o__Polyporales_unclassified(75);o__Polyporales_unclassified(75);o__Polyporales_unclassified(75); | - | - | - | - | - | - | - | Unassigned | - |
| Otu02523 | 16 | k__Fungi(100);p__Ascomycota(100);p__Ascomycota_unclassified(57);p__Ascomycota_unclassified(57);p__Ascomycota_unclassified(57);p__Ascomycota_unclassified(57);p__Ascomycota_unclassified(57); | - | - | - | - | - | - | - | Unassigned | - |
| Otu02530 | 16 | k__Fungi(100);p__Basidiomycota(100);c__Agaricomycetes(100);o__Russulales(100);o__Russulales_unclassified(100);o__Russulales_unclassified(100);o__Russulales_unclassified(100); | - | - | - | - | - | - | - | Unassigned | - |
| Otu02531 | 16 | k__Fungi(100);k__Fungi_unclassified(69);k__Fungi_unclassified(69);k__Fungi_unclassified(69);k__Fungi_unclassified(69);k__Fungi_unclassified(69);k__Fungi_unclassified(69); | - | - | - | - | - | - | - | Unassigned | - |
| Otu02533 | 16 | k__Fungi(100);k__Fungi_unclassified(63);k__Fungi_unclassified(63);k__Fungi_unclassified(63);k__Fungi_unclassified(63);k__Fungi_unclassified(63);k__Fungi_unclassified(63); | - | - | - | - | - | - | - | Unassigned | - |
| Otu02534 | 16 | k__Fungi(100);p__Basidiomycota(100);c__Agaricomycetes(100);o__Auriculariales(100);f__unclassified_Auriculariales(100);g__unclassified_Auriculariales(100);s__Auriculariales_sp(100); | - | - | - | - | - | - | - | Unassigned | - |
| Otu02535 | 16 | k__Fungi(100);k__Fungi_unclassified(75);k__Fungi_unclassified(75);k__Fungi_unclassified(75);k__Fungi_unclassified(75);k__Fungi_unclassified(75);k__Fungi_unclassified(75); | - | - | - | - | - | - | - | Unassigned | - |
| Otu02538 | 16 | k__Fungi(100);k__Fungi_unclassified(82);k__Fungi_unclassified(82);k__Fungi_unclassified(82);k__Fungi_unclassified(82);k__Fungi_unclassified(82);k__Fungi_unclassified(82); | - | - | - | - | - | - | - | Unassigned | - |
| Otu02540 | 16 | k__Fungi(100);k__Fungi_unclassified(69);k__Fungi_unclassified(69);k__Fungi_unclassified(69);k__Fungi_unclassified(69);k__Fungi_unclassified(69);k__Fungi_unclassified(69); | - | - | - | - | - | - | - | Unassigned | - |
| Otu02541 | 16 | k__Fungi(100);p__Ascomycota(100);p__Ascomycota_unclassified(94);p__Ascomycota_unclassified(94);p__Ascomycota_unclassified(94);p__Ascomycota_unclassified(94);p__Ascomycota_unclassified(94); | - | - | - | - | - | - | - | Unassigned | - |
| Otu02546 | 16 | k__Fungi(100);k__Fungi_unclassified(69);k__Fungi_unclassified(69);k__Fungi_unclassified(69);k__Fungi_unclassified(69);k__Fungi_unclassified(69);k__Fungi_unclassified(69); | - | - | - | - | - | - | - | Unassigned | - |
| Otu02548 | 16 | k__Fungi(100);p__Ascomycota(57);p__Ascomycota_unclassified(57);p__Ascomycota_unclassified(57);p__Ascomycota_unclassified(57);p__Ascomycota_unclassified(57);p__Ascomycota_unclassified(57); | - | - | - | - | - | - | - | Unassigned | - |
| Otu02550 | 16 | k__Fungi(100);p__Ascomycota(100);c__Taphrinomycetes(88);o__Taphrinales(88);o__Taphrinales_unclassified(69);o__Taphrinales_unclassified(69);o__Taphrinales_unclassified(69); | - | - | - | - | - | - | - | Unassigned | - |
| Otu02552 | 16 | k__Fungi(100);p__Ascomycota(57);p__Ascomycota_unclassified(57);p__Ascomycota_unclassified(57);p__Ascomycota_unclassified(57);p__Ascomycota_unclassified(57);p__Ascomycota_unclassified(57); | - | - | - | - | - | - | - | Unassigned | - |
| Otu02553 | 16 | k__Fungi(100);p__Basidiomycota(100);c__Microbotryomycetes(100);o__Sporidiobolales(57);f__Sporidiobolales_family_Incertae_sedis(57);f__Sporidiobolales_family_Incertae_sedis_unclassified(57);f__Sporidiobolales_family_Incertae_sedis_unclassified(57); | - | - | - | - | - | - | - | Unassigned | - |
| Otu02556 | 16 | k__Fungi(100);p__Ascomycota(100);c__Dothideomycetes(100);o__Pleosporales(100);f__unclassified_Pleosporales(94);g__unclassified_Pleosporales(94);s__Pleosporales_sp(94); | - | - | - | - | - | - | - | Unassigned | - |
| Otu02558 | 16 | k__Fungi(100);p__Ascomycota(100);p__Ascomycota_unclassified(100);p__Ascomycota_unclassified(100);p__Ascomycota_unclassified(100);p__Ascomycota_unclassified(100);p__Ascomycota_unclassified(100); | - | - | - | - | - | - | - | Unassigned | - |
| Otu02562 | 16 | k__Fungi(100);k__Fungi_unclassified(75);k__Fungi_unclassified(75);k__Fungi_unclassified(75);k__Fungi_unclassified(75);k__Fungi_unclassified(75);k__Fungi_unclassified(75); | - | - | - | - | - | - | - | Unassigned | - |
| Otu02564 | 16 | k__Fungi(100);p__Ascomycota(100);c__Sordariomycetes(100);c__Sordariomycetes_unclassified(100);c__Sordariomycetes_unclassified(100);c__Sordariomycetes_unclassified(100);c__Sordariomycetes_unclassified(100); | - | - | - | - | - | - | - | Unassigned | - |
| Otu02565 | 16 | k__Fungi(100);p__Ascomycota(100);c__Dothideomycetes(100);o__Pleosporales(100);o__Pleosporales_unclassified(94);o__Pleosporales_unclassified(94);o__Pleosporales_unclassified(94); | - | - | - | - | - | - | - | Unassigned | - |
| Otu02567 | 16 | k__Fungi(100);k__Fungi_unclassified(94);k__Fungi_unclassified(94);k__Fungi_unclassified(94);k__Fungi_unclassified(94);k__Fungi_unclassified(94);k__Fungi_unclassified(94); | - | - | - | - | - | - | - | Unassigned | - |
| Otu02568 | 16 | k__Fungi(100);p__Ascomycota(100);c__Eurotiomycetes(100);o__Eurotiales(100);f__Trichocomaceae(100);f__Trichocomaceae_unclassified(100);f__Trichocomaceae_unclassified(100); | - | - | - | - | - | - | - | Unassigned | - |
| Otu02569 | 16 | k__Fungi(100);k__Fungi_unclassified(100);k__Fungi_unclassified(100);k__Fungi_unclassified(100);k__Fungi_unclassified(100);k__Fungi_unclassified(100);k__Fungi_unclassified(100); | - | - | - | - | - | - | - | Unassigned | - |
| Otu02572 | 16 | k__Fungi(100);p__Ascomycota(63);p__Ascomycota_unclassified(63);p__Ascomycota_unclassified(63);p__Ascomycota_unclassified(63);p__Ascomycota_unclassified(63);p__Ascomycota_unclassified(63); | - | - | - | - | - | - | - | Unassigned | - |
| Otu02573 | 16 | k__Fungi(100);p__Basidiomycota(100);c__Agaricomycetes(100);c__Agaricomycetes_unclassified(100);c__Agaricomycetes_unclassified(100);c__Agaricomycetes_unclassified(100);c__Agaricomycetes_unclassified(100); | - | - | - | - | - | - | - | Unassigned | - |
| Otu02576 | 16 | k__Fungi(100);k__Fungi_unclassified(69);k__Fungi_unclassified(69);k__Fungi_unclassified(69);k__Fungi_unclassified(69);k__Fungi_unclassified(69);k__Fungi_unclassified(69); | - | - | - | - | - | - | - | Unassigned | - |
| Otu02577 | 16 | k__Fungi(100);p__Ascomycota(100);c__Sordariomycetes(100);c__Sordariomycetes_unclassified(100);c__Sordariomycetes_unclassified(100);c__Sordariomycetes_unclassified(100);c__Sordariomycetes_unclassified(100); | - | - | - | - | - | - | - | Unassigned | - |
| Otu02578 | 16 | k__Fungi(100);p__Ascomycota(100);p__Ascomycota_unclassified(94);p__Ascomycota_unclassified(94);p__Ascomycota_unclassified(94);p__Ascomycota_unclassified(94);p__Ascomycota_unclassified(94); | - | - | - | - | - | - | - | Unassigned | - |
| Otu02579 | 15 | k__Fungi(100);p__Basidiomycota(100);c__Agaricomycetes(100);o__Russulales(100);f__Peniophoraceae(100);f__Peniophoraceae_unclassified(100);f__Peniophoraceae_unclassified(100); | Peniophoraceae | 9 | Pathotroph-Saprotroph | Plant Pathogen-Wood Saprotroph | Corticioid | White Rot | Probable | Most species are saprobic, causing rots of standing and fallen wood. Many species are adapted to xeric conditions, being able to grow on rapidly drying dead attached branches (Cannon & Kirk 2007) | [illegible] |
| Otu02580 | 15 | k__Fungi(100);p__Basidiomycota(100);c__Agaricomycetes(100);o__Agaricales(100);f__Mycenaceae(100);g__Mycena(100);s__Mycena_sanguinolenta(100); | Mycena | 13 | Pathotroph-Saprotroph | Plant Pathogen-\|Plant Saprotroph\|-Undefined Saprotroph-Wood Saprotroph | Agaricoid | NULL | Probable | As litter_saprotroph (Põlme et al. 2020); Decay leaf, fruit, seed (Põlme et al. 2020) | [illegible] |
| Otu02582 | 15 | k__Fungi(100);p__Basidiomycota(100);c__Microbotryomycetes(100);o__Sporidiobolales(100);f__Sporidiobolales_family_Incertae_sedis(100);g__Sporobolomyces(100);s__Sporobolomyces_aff_jilinensis_MCA_3774(100); | Sporobolomyces | 13 | Pathotroph-Saprotroph | \|Fungal Parasite\|-Plant Saprotroph | Yeast | NULL | Probable | As litter_saprotroph (Põlme et al. 2020); As mycoparasite (Põlme et al. 2020); Decay fungal material (Põlme et al. 2020) | [illegible] |
| Otu02583 | 15 | k__Fungi(100);p__Ascomycota(100);c__Sordariomycetes(100);o__Hypocreales(54);o__Hypocreales_unclassified(54);o__Hypocreales_unclassified(54);o__Hypocreales_unclassified(54); | Hypocreales | 7 | Saprotroph | Undefined Saprotroph | Microfungus | NULL | Possible | NULL | [illegible] |
| Otu02585 | 15 | k__Fungi(100);p__Basidiomycota(100);c__Agaricomycetes(100);o__Agaricales(100);f__Marasmiaceae(100);g__Marasmius(100);s__Marasmius_rotula(100); | Marasmius | 13 | Pathotroph-Saprotroph-Symbiotroph | Endophyte-Plant Pathogen-\|Plant Saprotroph\|-Undefined Saprotroph | Agaricoid | NULL | Probable | As litter_saprotroph (Põlme et al. 2020); Decay leaf, fruit, seed (Põlme et al. 2020); Often one of the most prominent fungal components of tropical forests, playing a major role in nutrient cycling (Cannon & Kirk 2007) | [illegible] |
| Otu02588 | 15 | k__Fungi(100);p__Ascomycota(100);c__Eurotiomycetes(100);o__Chaetothyriales(100);f__Herpotrichiellaceae(100);f__Herpotrichiellaceae_unclassified(100);f__Herpotrichiellaceae_unclassified(100); | Herpotrichiellaceae | 9 | Pathotroph-Saprotroph | Animal Pathogen-Fungal Parasite-Undefined Saprotroph | Facultative Yeast-Microfungus | NULL | Probable | Mostly saprobic on plants or other fungi, isolated from soil etc., and from medical sources (Cannon & Kirk 2007) | [illegible] |
| Otu02589 | 15 | k__Fungi(100);p__Ascomycota(100);c__Saccharomycetes(100);o__Saccharomycetales(100);f__Metschnikowiaceae(100);g__Metschnikowia(100);s__Metschnikowia_pulcherrima(100); | Metschnikowia | 13 | Saprotroph | \|Undefined Saprotroph\| | Yeast | NULL | Probable | As nectar-tap_saprotroph (Põlme et al. 2020); Decay sugar-rich substrates (Põlme et al. 2020) | [illegible] |
| Otu02590 | 15 | k__Fungi(100);p__Ascomycota(100);c__Ascomycota_class_Incertae_sedis(94);o__Ascomycota_order_Incertae_sedis(94);f__Ascomycota_family_Incertae_sedis(94);g__Filosporella(94);g__Filosporella_unclassified(94); | Filosporella | 13 | Saprotroph | \|Plant Saprotroph\| | Microfungus | NULL | Probable | As litter_saprotroph (Põlme et al. 2020); Decay leaf, fruit, seed (Põlme et al. 2020) | [illegible] |
| Otu02591 | 15 | k__Fungi(100);p__Ascomycota(100);c__Saccharomycetes(100);o__Saccharomycetales(100);f__Metschnikowiaceae(100);g__Metschnikowia(100);s__Metschnikowia_pulcherrima(100); | Metschnikowia | 13 | Saprotroph | \|Undefined Saprotroph\| | Yeast | NULL | Probable | As nectar-tap_saprotroph (Põlme et al. 2020); Decay sugar-rich substrates (Põlme et al. 2020) | [illegible] |
| Otu02594 | 15 | k__Fungi(100);p__Ascomycota(100);c__Dothideomycetes(100);o__Pleosporales(100);f__Pleosporaceae(100);g__Drechslera(100);s__Drechslera_poae(100); | Drechslera | 13 | Pathotroph | \|Plant Pathogen\| | Microfungus | NULL | Probable | NULL | [illegible] |
| Otu02596 | 15 | k__Fungi(100);p__Basidiomycota(100);c__Agaricomycetes(100);o__Russulales(100);f__Russulaceae(100);g__Russula(100);s__Russula_sp_FH_HES04F1SY(100); | Russula | 13 | Symbiotroph | \|Ectomycorrhizal\| | Agaricoid | NULL | Highly Probable | NULL | [illegible] |
| Otu02597 | 15 | k__Fungi(100);p__Ascomycota(100);c__Saccharomycetes(100);o__Saccharomycetales(100);f__Metschnikowiaceae(100);g__Metschnikowia(100);s__Metschnikowia_pulcherrima(87); | Metschnikowia | 13 | Saprotroph | \|Undefined Saprotroph\| | Yeast | NULL | Probable | As nectar-tap_saprotroph (Põlme et al. 2020); Decay sugar-rich substrates (Põlme et al. 2020) | [illegible] |
| Otu02598 | 15 | k__Fungi(100);p__Basidiomycota(100);c__Agaricomycetes(100);o__Boletales(100);f__Paxillaceae(100);g__Hydnomerulius(100);s__Hydnomerulius_pinastri(100); | Hydnomerulius | 13 | Saprotroph | \|Wood Saprotroph\| | Corticioid | Brown Rot | Probable | As brown-rot (Põlme et al. 2020); Decay wood (Põlme et al. 2020) | [illegible] |
| Otu02599 | 15 | k__Fungi(100);p__Ascomycota(100);c__Ascomycota_class_Incertae_sedis(100);o__Ascomycota_order_Incertae_sedis(100);f__Ascomycota_family_Incertae_sedis(100);g__Cordana(100);s__Cordana_ellipsoidea(100); | Cordana | 13 | Pathotroph-Saprotroph | \|Plant Pathogen\| | Microfungus | NULL | Probable | NULL | [illegible] |
| Otu02601 | 15 | k__Fungi(100);p__Ascomycota(100);c__Dothideomycetes(100);o__Capnodiales(100);f__Mycosphaerellaceae(100);g__Mycosphaerella(100);s__Mycosphaerella_nyssicola(100); | Mycosphaerella | 13 | Pathotroph-Saprotroph | \|Plant Pathogen\|-Plant Saprotroph | Microfungus | NULL | Probable | As litter_saprotroph (Põlme et al. 2020); Decay leaf, fruit, seed (Põlme et al. 2020) | [illegible] |
| Otu02604 | 15 | k__Fungi(100);p__Basidiomycota(100);c__Agaricomycetes(100);o__Boletales(100);f__Hygrophoropsidaceae(100);g__Leucogyrophana(100);s__Leucogyrophana_mollusca(100); | Leucogyrophana | 13 | Pathotroph-Saprotroph | Lichen Parasite-\|Wood Saprotroph\| | Corticioid-Microfungus | Brown Rot | Probable | As brown-rot (Põlme et al. 2020); Decay wood (Põlme et al. 2020); Syn. Penttilamyces | [illegible] |
| Otu02605 | 15 | k__Fungi(100);p__Ascomycota(100);c__Lecanoromycetes(100);o__Teloschistales(100);f__Caliciaceae(100);g__Calicium(100);s__Calicium_glaucellum(100); | Calicium | 13 | Symbiotroph | \|Lichenized\| | Calicioid-Thallus | NULL | Highly Probable | NULL | [illegible] |
| Otu02606 | 15 | k__Fungi(100);p__Ascomycota(100);c__Leotiomycetes(100);o__Rhytismatales(100);f__Rhytismataceae(100);f__Rhytismataceae_unclassified(80);f__Rhytismataceae_unclassified(80); | Rhytismataceae | 9 | Pathotroph-Saprotroph-Symbiotroph | Endophyte-Leaf Saprotroph-Plant Pathogen | Microfungus | NULL | Probable | Biotrophic, necrotrophic or saprobic on leaves and bark, often initially as endophytes, causing tar spots, premature leaf fall and occasionally witches broom symptoms (Cannon & Kirk 2007) | [illegible] |
| Otu02608 | 15 | k__Fungi(100);p__Ascomycota(100);c__Dothideomycetes(100);o__Pleosporales(100);f__Pleosporaceae(100);g__Dendryphion(100);s__Dendryphion_europaeum(100); | Dendryphion | 13 | Pathotroph-Saprotroph-Symbiotroph | Endophyte-\|Plant Pathogen\|-Plant Saprotroph | Microfungus | NULL | Probable | As foliar_endophyte (Põlme et al. 2020); As litter_saprotroph (Põlme et al. 2020); Decay leaf, fruit, seed (Põlme et al. 2020) | [illegible] |
| Otu02614 | 15 | k__Fungi(100);p__Ascomycota(100);c__Saccharomycetes(100);o__Saccharomycetales(100);f__Metschnikowiaceae(100);g__Metschnikowia(100);s__Metschnikowia_pulcherrima(74); | Metschnikowia | 13 | Saprotroph | \|Undefined Saprotroph\| | Yeast | NULL | Probable | As nectar-tap_saprotroph (Põlme et al. 2020); Decay sugar-rich substrates (Põlme et al. 2020) | [illegible] |
| Otu02617 | 15 | k__Fungi(100);p__Ascomycota(100);c__Leotiomycetes(100);o__Helotiales(100);f__Helotiales_family_Incertae_sedis(100);g__Tetracladium(100);s__Tetracladium_marchalianum(94); | Tetracladium | 13 | Saprotroph | \|Plant Saprotroph\|-Undefined Saprotroph | Microfungus | NULL | Probable | As litter_saprotroph (Põlme et al. 2020); Decay leaf, fruit, seed (Põlme et al. 2020) | [illegible] |
| Otu02620 | 15 | k__Fungi(100);p__Basidiomycota(100);c__Microbotryomycetes(100);o__Sporidiobolales(100);f__Sporidiobolales_family_Incertae_sedis(100);g__Rhodotorula(100);s__Rhodotorula_sp_JCM_11351(100); | Rhodotorula | 13 | Pathotroph-Saprotroph-Symbiotroph | Animal Pathogen-Animal Symbiotroph-Endophyte-Plant Pathogen-\|Undefined Saprotroph\| | Yeast | NULL | Probable | As foliar_endophyte (Põlme et al. 2020); Opportunistic human pathogen | [illegible] |
| Otu02623 | 15 | k__Fungi(100);p__Ascomycota(100);c__Dothideomycetes(100);o__Capnodiales(100);f__Mycosphaerellaceae(100);g__Mycosphaerella(100);s__Mycosphaerella_dearnessii(100); | Mycosphaerella | 13 | Pathotroph-Saprotroph | \|Plant Pathogen\|-Plant Saprotroph | Microfungus | NULL | Probable | As litter_saprotroph (Põlme et al. 2020); Decay leaf, fruit, seed (Põlme et al. 2020) | [illegible] |
| Otu02625 | 15 | k__Fungi(100);p__Basidiomycota(100);c__Agaricomycetes(100);o__Agaricales(100);f__Psathyrellaceae(100);g__Parasola(100);s__Parasola_schroeterii(100); | Parasola | 13 | Saprotroph | \|Undefined Saprotroph\| | Agaricoid | NULL | Probable | As soil_saprotroph (Põlme et al. 2020); Decay soil (Põlme et al. 2020) | [illegible] |
| Otu02626 | 15 | k__Fungi(100);p__Basidiomycota(100);c__Agaricomycetes(100);o__Cantharellales(100);f__Botryobasidiaceae(100);g__Botryobasidium(100);s__Botryobasidium_subcoronatum(100); | Botryobasidium | 13 | Saprotroph | Undefined Saprotroph-\|Wood Saprotroph\| | Corticioid | White Rot | Probable | As white_rot (Põlme et al. 2020); Decay wood (Põlme et al. 2020) | [illegible] |
| Otu02630 | 15 | k__Fungi(100);p__Ascomycota(100);c__Saccharomycetes(100);o__Saccharomycetales(100);f__Metschnikowiaceae(100);g__Metschnikowia(100);s__Metschnikowia_pulcherrima(94); | Metschnikowia | 13 | Saprotroph | \|Undefined Saprotroph\| | Yeast | NULL | Probable | As nectar-tap_saprotroph (Põlme et al. 2020); Decay sugar-rich substrates (Põlme et al. 2020) | [illegible] |
| Otu02631 | 15 | k__Fungi(100);p__Basidiomycota(100);c__Agaricomycetes(100);o__Polyporales(100);f__Fomitopsidaceae(100);g__Antrodia(100);s__Antrodia_serialis(100); | Antrodia | 13 | Saprotroph | \|Wood Saprotroph\| | Corticioid | Brown Rot | Highly Probable | As brown-rot (Põlme et al. 2020); Decay wood (Põlme et al. 2020) | [illegible] |
| Otu02632 | 15 | k__Fungi(100);p__Ascomycota(100);c__Leotiomycetes(100);o__Helotiales(100);f__Helotiaceae(100);g__Articulospora(100);s__Articulospora_sp(100); | Articulospora | 13 | Saprotroph | \|Plant Saprotroph\|-Undefined Saprotroph | Microfungus | NULL | Probable | As litter_saprotroph (Põlme et al. 2020); Decay leaf, fruit, seed (Põlme et al. 2020) | [illegible] |
| Otu02635 | 15 | k__Fungi(100);p__Zygomycota(100);c__Zygomycota_class_Incertae_sedis(100);o__Mortierellales(100);f__Mortierellaceae(100);g__Mortierella(100);s__Mortierella_zychae(67); | Mortierella | 13 | Saprotroph-Symbiotroph | Endophyte-Plant Saprotroph-\|Undefined Saprotroph\| | Microfungus | NULL | Probable | As chitinolytic (Põlme et al. 2020); As soil_saprotroph (Põlme et al. 2020); Decay roots, soil (Põlme et al. 2020); Saprobic in soil and thus frequently isolated from soil and material contaminated with soil such as hay or rotten silage and spoil tips (Cannon & Kirk 2007) | [illegible] |
| Otu02637 | 15 | k__Fungi(100);p__Basidiomycota(100);c__Agaricomycetes(100);o__Agaricales(100);f__Tricholomataceae(100);g__Clitocybe(100);s__Clitocybe_subditopoda(100); | Clitocybe | 13 | Pathotroph-Saprotroph | \|Plant Saprotroph\| | Agaricoid | NULL | Probable | As litter_saprotroph (Põlme et al. 2020); Decay leaf, fruit, seed (Põlme et al. 2020) | [illegible] |
| Otu02639 | 15 | k__Fungi(100);p__Ascomycota(100);c__Saccharomycetes(94);o__Saccharomycetales(94);f__Saccharomycetales_family_Incertae_sedis(87);g__Candida(80);g__Candida_unclassified(74); | Candida | 13 | Pathotroph-Saprotroph-Symbiotroph | Animal Pathogen-Endophyte-Endosymbiont-Epiphyte-\|Undefined Saprotroph\| | Dimorphic Yeast-Yeast | NULL | Probable | As nectar-tap_saprotroph (Põlme et al. 2020); Decay sugar-rich substrates (Põlme et al. 2020); This asexual genus is polyphyletic and contain species that function very differently depend on what they are. Caution should be taken when interpreting guilds for Candida species. | [illegible] |
| Otu02640 | 15 | k__Fungi(100);p__Ascomycota(100);c__Saccharomycetes(100);o__Saccharomycetales(100);f__Metschnikowiaceae(100);g__Metschnikowia(100);g__Metschnikowia_unclassified(80); | Metschnikowia | 13 | Saprotroph | \|Undefined Saprotroph\| | Yeast | NULL | Probable | As nectar-tap_saprotroph (Põlme et al. 2020); Decay sugar-rich substrates (Põlme et al. 2020) | [illegible] |
| Otu02642 | 15 | k__Fungi(100);p__Basidiomycota(100);c__Agaricomycetes(100);o__Thelephorales(100);f__Thelephoraceae(100);g__Tomentella(61);g__Tomentella_unclassified(61); | Tomentella | 13 | Symbiotroph | \|Ectomycorrhizal\| | Corticioid | NULL | Highly Probable | NULL | [illegible] |
| Otu02643 | 15 | k__Fungi(100);p__Basidiomycota(100);c__Agaricomycetes(100);o__Agaricales(100);f__Bolbitiaceae(100);g__Bolbitius(100);s__Bolbitius_variicolor(100); | Bolbitius | 13 | Saprotroph | \|Dung Saprotroph\|-Plant Saprotroph | Agaricoid | NULL | Probable | As litter_saprotroph (Põlme et al. 2020); Decay leaf, fruit, seed, dung (Põlme et al. 2020) | [illegible] |
| Otu02644 | 15 | k__Fungi(100);p__Ascomycota(100);c__Sordariomycetes(100);o__Hypocreales(100);o__Hypocreales_unclassified(100);o__Hypocreales_unclassified(100);o__Hypocreales_unclassified(100); | Hypocreales | 7 | Saprotroph | Undefined Saprotroph | Microfungus | NULL | Possible | NULL | [illegible] |
| Otu02645 | 15 | k__Fungi(100);p__Ascomycota(100);c__Saccharomycetes(100);o__Saccharomycetales(100);f__Metschnikowiaceae(100);g__Metschnikowia(100);s__Metschnikowia_pulcherrima(87); | Metschnikowia | 13 | Saprotroph | \|Undefined Saprotroph\| | Yeast | NULL | Probable | As nectar-tap_saprotroph (Põlme et al. 2020); Decay sugar-rich substrates (Põlme et al. 2020) | [illegible] |
| Otu02646 | 15 | k__Fungi(100);p__Ascomycota(100);c__Eurotiomycetes(100);o__Eurotiales(100);f__Trichocomaceae(100);g__Talaromyces(100);s__Talaromyces_variabilis(100); | Talaromyces | 13 | Pathotroph-Saprotroph | Animal Parasite-\|Undefined Saprotroph\| | Microfungus | NULL | Probable | Decay leaf, fruit, seed, soil, animal material (Põlme et al. 2020) | [illegible] |
| Otu02647 | 15 | k__Fungi(100);p__Basidiomycota(100);c__Agaricomycetes(100);o__Hymenochaetales(100);f__Schizoporaceae(67);g__unclassified_Schizoporaceae(67);s__Schizoporaceae_sp(67); | Schizoporaceae | 9 | Saprotroph | Wood Saprotroph | Corticioid | White Rot | Probable | Saprobic, causing white rot of standing and fallen wood of coniferous and broadleaved trees, rarely on herbaceous stems (Cannon & Kirk 2007) | [illegible] |
| Otu02650 | 15 | k__Fungi(100);p__Ascomycota(100);c__Sordariomycetes(100);o__Xylariales(100);f__Xylariaceae(100);f__Xylariaceae_unclassified(100);f__Xylariaceae_unclassified(100); | Xylariaceae | 9 | Pathotroph-Saprotroph-Symbiotroph | Dung Saprotroph-Endophyte-Plant Pathogen-Undefined Saprotroph | Xylarioid | NULL | Probable | Saprobes or weak pathogens, many endophytes, mainly in wood and bark, occasionally in soil and dung (Cannon & Kirk 2007) | [illegible] |
| Otu02652 | 15 | k__Fungi(100);p__Ascomycota(100);c__Dothideomycetes(100);o__Capnodiales(100);f__Mycosphaerellaceae(100);f__Mycosphaerellaceae_unclassified(100);f__Mycosphaerellaceae_unclassified(100); | Mycosphaerellaceae | 9 | Pathotroph-Saprotroph | Plant Pathogen-Undefined Saprotroph | Microfungus | NULL | Probable | Biotrophic, necrotrophic or saprobic on various plant tissues (Cannon & Kirk 2007) | [illegible] |
| Otu02653 | 15 | k__Fungi(100);p__Ascomycota(100);c__Leotiomycetes(87);o__Leotiomycetes_order_Incertae_sedis(87);f__Leotiomycetes_family_Incertae_sedis(87);g__Sarea(87);g__Sarea_unclassified(87); | Sarea | 13 | Saprotroph-Symbiotroph | Lichenized-\|Undefined Saprotroph\| | Microfungus-Thallus | NULL | Highly Probable | NULL | [illegible] |
| Otu02655 | 15 | k__Fungi(100);p__Basidiomycota(100);c__Microbotryomycetes(100);o__Sporidiobolales(100);f__Sporidiobolales_family_Incertae_sedis(100);g__Sporobolomyces(100);s__Sporobolomyces_lactosus(100); | Sporobolomyces | 13 | Pathotroph-Saprotroph | \|Fungal Parasite\|-Plant Saprotroph | Yeast | NULL | Probable | As litter_saprotroph (Põlme et al. 2020); As mycoparasite (Põlme et al. 2020); Decay fungal material (Põlme et al. 2020) | [illegible] |
| Otu02657 | 15 | k__Fungi(100);p__Ascomycota(100);c__Sordariomycetes(100);o__Hypocreales(100);f__Hypocreales_family_Incertae_sedis(100);g__Acremonium(100);s__Acremonium_stromaticum(100); | Acremonium | 13 | Pathotroph-Saprotroph-Symbiotroph | Animal Pathogen-Endophyte-Fungal Parasite-Plant Pathogen-\|Undefined Saprotroph\|-Wood Saprotroph | Microfungus | Soft Rot | Possible | As foliar_endophyte (Põlme et al. 2020); culture (Hanada et al. 2010, Del Olmo; Decay leaf, fruit, seed, soil, dung, animal material (Põlme et al. 2020); Endophyte detection method; Ruiz & Arnold 2014); Likely opportunistic human pathogen (Irinyi et al. 2015) | [illegible] |
| Otu02658 | 15 | k__Fungi(100);p__Basidiomycota(100);c__Pucciniomycetes(100);o__Pucciniales(100);f__Cronartiaceae(87);f__Cronartiaceae_unclassified(87);f__Cronartiaceae_unclassified(87); | Cronartiaceae | 9 | Pathotroph | Plant Pathogen | Microfungus | NULL | Probable | Common Name - Blister Rust, Rust Fungus; Most species are heteroecious, with the aecial stages forming on stems and cones of Pinus species and uredinia and telia on a range of dicotyledonous plants (Cannon & Kirk 2007) | [illegible] |
| Otu02660 | 15 | k__Fungi(100);p__Ascomycota(100);c__Eurotiomycetes(100);o__Eurotiales(100);f__Trichocomaceae(100);g__Paecilomyces(100);s__Paecilomyces_sp_CZ_2011a(100); | Paecilomyces | 13 | Pathotroph-Saprotroph-Symbiotroph | Clavicipitaceous \|Endophyte\|-Plant Pathogen-Plant Saprotroph-Undefined Saprotroph-Wood Saprotroph | Microfungus | Soft Rot | Probable | As foliar_endophyte (Põlme et al. 2020); As litter_saprotroph (Põlme et al. 2020); culture (Lopez et al. 2014); Decay leaf, fruit, seed, animal material (Põlme et al. 2020); Endophyte detection method | [illegible] |
| Otu02661 | 15 | k__Fungi(100);p__Ascomycota(100);c__Saccharomycetes(100);o__Saccharomycetales(100);f__Metschnikowiaceae(100);g__Metschnikowia(100);s__Metschnikowia_pulcherrima(100); | Metschnikowia | 13 | Saprotroph | \|Undefined Saprotroph\| | Yeast | NULL | Probable | As nectar-tap_saprotroph (Põlme et al. 2020); Decay sugar-rich substrates (Põlme et al. 2020) | [illegible] |
| Otu02662 | 15 | k__Fungi(100);p__Ascomycota(100);c__Sordariomycetes(100);o__Microascales(100);f__Microascaceae(100);g__unclassified_Microascaceae(100);s__Microascaceae_sp(100); | Microascaceae | 9 | Pathotroph-Saprotroph-Symbiotroph | Animal Pathogen-Endophyte-Plant Pathogen-Undefined Saprotroph | Microfungus | NULL | Possible | Saprobic in soil or rotting vegetation, some species heat-tolerant and associated with silage and compost, a few associated with humans and animals (Cannon & Kirk 2007) | [illegible] |
| Otu02663 | 15 | k__Fungi(100);p__Ascomycota(100);c__Saccharomycetes(100);o__Saccharomycetales(100);f__Metschnikowiaceae(100);g__Metschnikowia(100);s__Metschnikowia_pulcherrima(100); | Metschnikowia | 13 | Saprotroph | \|Undefined Saprotroph\| | Yeast | NULL | Probable | As nectar-tap_saprotroph (Põlme et al. 2020); Decay sugar-rich substrates (Põlme et al. 2020) | [illegible] |
| Otu02664 | 15 | k__Fungi(100);p__Basidiomycota(100);c__Agaricomycetes(100);o__Agaricales(100);f__Psathyrellaceae(100);g__Psathyrella(100);g__Psathyrella_unclassified(100); | Psathyrella | 13 | Saprotroph | Plant Saprotroph-\|Wood Saprotroph\| | Agaricoid | NULL | Probable | As litter_saprotroph (Põlme et al. 2020); Decay wood, leaf, fruit, seed (Põlme et al. 2020) | [illegible] |
| Otu02665 | 15 | k__Fungi(100);p__Ascomycota(100);c__Leotiomycetes(100);o__Helotiales(100);f__Helotiales_family_Incertae_sedis(100);g__Pyrenopeziza(100);s__axenic_grass_root_isolate_(cf_Pyrenopeziza_revincta)(100); | Pyrenopeziza | 13 | Pathotroph-Saprotroph | \|Plant Pathogen\|-Plant Saprotroph | Microfungus | NULL | Probable | As litter_saprotroph (Põlme et al. 2020); Decay leaf, fruit, seed (Põlme et al. 2020) | [illegible] |
| Otu02666 | 15 | k__Fungi(100);p__Ascomycota(100);c__Saccharomycetes(100);o__Saccharomycetales(100);f__Metschnikowiaceae(100);g__Metschnikowia(100);s__Metschnikowia_pulcherrima(87); | Metschnikowia | 13 | Saprotroph | \|Undefined Saprotroph\| | Yeast | NULL | Probable | As nectar-tap_saprotroph (Põlme et al. 2020); Decay sugar-rich substrates (Põlme et al. 2020) | [illegible] |
| Otu02670 | 15 | k__Fungi(100);p__Basidiomycota(100);c__Agaricomycetes(100);o__Agaricales(100);f__Pterulaceae(100);g__Aphanobasidium(100);s__Aphanobasidium_pseudotsugae(100); | Aphanobasidium | 13 | Saprotroph | Undefined Saprotroph-\|Wood Saprotroph\| | Corticioid | White Rot | Probable | As white_rot (Põlme et al. 2020); Decay wood (Põlme et al. 2020) | [illegible] |
| Otu02672 | 15 | k__Fungi(100);p__Basidiomycota(100);c__Tremellomycetes(100);o__Tremellales(100);f__Tremellales_family_Incertae_sedis(100);g__Tremella(100);g__Tremella_unclassified(100); | Tremella | 13 | Pathotroph | \|Fungal Parasite\|-Lichen Parasite | Dimorphic-Facultative Yeast-Microfungus-Tremelloid | NULL | Probable | As mycoparasite (Põlme et al. 2020); Decay fungal material (Põlme et al. 2020) | [illegible] |
| Otu02677 | 15 | k__Fungi(100);p__Basidiomycota(100);c__Agaricomycetes(100);o__Agaricales(100);f__Psathyrellaceae(100);g__Coprinopsis(100);s__Coprinopsis_semitalis(100); | Coprinopsis | 13 | Saprotroph | \|Undefined Saprotroph\| | Agaricoid | NULL | Probable | As soil_saprotroph (Põlme et al. 2020); Decay soil (Põlme et al. 2020) | [illegible] |
| Otu02678 | 15 | k__Fungi(100);p__Ascomycota(100);c__Saccharomycetes(100);o__Saccharomycetales(100);f__Metschnikowiaceae(100);g__Metschnikowia(100);s__Metschnikowia_pulcherrima(100); | Metschnikowia | 13 | Saprotroph | \|Undefined Saprotroph\| | Yeast | NULL | Probable | As nectar-tap_saprotroph (Põlme et al. 2020); Decay sugar-rich substrates (Põlme et al. 2020) | [illegible] |
| Otu02679 | 15 | k__Fungi(100);p__Ascomycota(100);c__Saccharomycetes(100);o__Saccharomycetales(100);f__Metschnikowiaceae(100);g__Metschnikowia(100);s__Metschnikowia_pulcherrima(94); | Metschnikowia | 13 | Saprotroph | \|Undefined Saprotroph\| | Yeast | NULL | Probable | As nectar-tap_saprotroph (Põlme et al. 2020); Decay sugar-rich substrates (Põlme et al. 2020) | [illegible] |
| Otu02681 | 15 | k__Fungi(100);p__Ascomycota(100);c__Saccharomycetes(100);o__Saccharomycetales(100);f__Metschnikowiaceae(100);g__Metschnikowia(100);s__Metschnikowia_pulcherrima(100); | Metschnikowia | 13 | Saprotroph | \|Undefined Saprotroph\| | Yeast | NULL | Probable | As nectar-tap_saprotroph (Põlme et al. 2020); Decay sugar-rich substrates (Põlme et al. 2020) | [illegible] |
| Otu02684 | 15 | k__Fungi(100);p__Ascomycota(100);c__Sordariomycetes(100);o__Hypocreales(100);o__Hypocreales_unclassified(54);o__Hypocreales_unclassified(54);o__Hypocreales_unclassified(54); | Hypocreales | 7 | Saprotroph | Undefined Saprotroph | Microfungus | NULL | Possible | NULL | [illegible] |
| Otu02686 | 15 | k__Fungi(100);p__Basidiomycota(100);c__Agaricomycetes(100);o__Hymenochaetales(100);f__Hymenochaetaceae(100);g__Phellinus(100);s__Phellinus_contiguus(100); | Phellinus | 13 | Pathotroph-Saprotroph | \|Plant Pathogen\|-Wood Saprotroph | Polyporoid | White Rot | Highly Probable | As white_rot (Põlme et al. 2020); As wood_pathogen (Põlme et al. 2020); Decay wood (Põlme et al. 2020) | [illegible] |
| Otu02687 | 15 | k__Fungi(100);p__Basidiomycota(100);c__Microbotryomycetes(100);o__Sporidiobolales(100);f__Sporidiobolales_family_Incertae_sedis(100);g__Sporobolomyces(100);s__Sporobolomyces_lactosus(100); | Sporobolomyces | 13 | Pathotroph-Saprotroph | \|Fungal Parasite\|-Plant Saprotroph | Yeast | NULL | Probable | As litter_saprotroph (Põlme et al. 2020); As mycoparasite (Põlme et al. 2020); Decay fungal material (Põlme et al. 2020) | [illegible] |
| Otu02689 | 15 | k__Fungi(100);p__Ascomycota(100);c__Saccharomycetes(100);o__Saccharomycetales(100);f__Metschnikowiaceae(100);g__Metschnikowia(100);s__Metschnikowia_pulcherrima(94); | Metschnikowia | 13 | Saprotroph | \|Undefined Saprotroph\| | Yeast | NULL | Probable | As nectar-tap_saprotroph (Põlme et al. 2020); Decay sugar-rich substrates (Põlme et al. 2020) | [illegible] |
| Otu02690 | 15 | k__Fungi(100);p__Ascomycota(100);c__Saccharomycetes(100);o__Saccharomycetales(100);f__Metschnikowiaceae(100);g__Metschnikowia(100);g__Metschnikowia_unclassified(67); | Metschnikowia | 13 | Saprotroph | \|Undefined Saprotroph\| | Yeast | NULL | Probable | As nectar-tap_saprotroph (Põlme et al. 2020); Decay sugar-rich substrates (Põlme et al. 2020) | [illegible] |
| Otu02692 | 15 | k__Fungi(100);p__Basidiomycota(100);c__Agaricomycetes(100);o__Polyporales(100);f__Meruliaceae(100);g__Junghuhnia(87);g__Junghuhnia_unclassified(67); | Junghuhnia | 13 | Saprotroph | \|Wood Saprotroph\| | Corticioid | White Rot | Highly Probable | As white_rot (Põlme et al. 2020); Decay wood (Põlme et al. 2020) | [illegible] |
| Otu02697 | 15 | k__Fungi(100);p__Basidiomycota(100);c__Agaricomycetes(100);o__Atheliales(100);f__Atheliaceae(100);g__Piloderma(100);s__Piloderma_fallax(100); | Piloderma | 13 | Symbiotroph | \|Ectomycorrhizal\| | Corticioid | NULL | Highly Probable | NULL | [illegible] |
| Otu02699 | 15 | k__Fungi(100);p__Ascomycota(100);c__Dothideomycetes(100);o__Capnodiales(100);f__Mycosphaerellaceae(100);g__Passalora(94);s__Passalora_californica(94); | Passalora | 13 | Pathotroph | \|Plant Pathogen\| | Microfungus | NULL | Probable | NULL | [illegible] |
| Otu02702 | 15 | k__Fungi(100);p__Ascomycota(100);c__Leotiomycetes(67);o__Helotiales(67);f__Helotiales_family_Incertae_sedis(67);g__Infundichalara(67);s__Infundichalara_sp(67); | Infundichalara | 13 | Saprotroph | \|Plant Saprotroph\|-Undefined Saprotroph | Microfungus | NULL | Probable | As litter_saprotroph (Põlme et al. 2020); Decay leaf, fruit, seed (Põlme et al. 2020); Host - Pinus (Põlme et al. 2020) | [illegible] |
| Otu02703 | 15 | k__Fungi(100);p__Ascomycota(100);c__Sordariomycetes(100);o__Hypocreales(100);f__Nectriaceae(100);f__Nectriaceae_unclassified(100);f__Nectriaceae_unclassified(100); | Nectriaceae | 9 | Pathotroph-Saprotroph-Symbiotroph | Animal Pathogen-Endophyte-Fungal Parasite-Lichen Parasite-Plant Pathogen-Wood Saprotroph | Microfungus | NULL | Possible | Fusarium species are amongst the economically significant plant pathogens (Cannon & Kirk 2007) | [illegible] |
| Otu02704 | 15 | k__Fungi(100);p__Ascomycota(100);c__Saccharomycetes(100);o__Saccharomycetales(100);f__Metschnikowiaceae(100);g__Metschnikowia(100);s__Metschnikowia_pulcherrima(94); | Metschnikowia | 13 | Saprotroph | \|Undefined Saprotroph\| | Yeast | NULL | Probable | As nectar-tap_saprotroph (Põlme et al. 2020); Decay sugar-rich substrates (Põlme et al. 2020) | [illegible] |

| | | | | | | | | | | | |
|---|---|---|---|---|---|---|---|---|---|---|---|
| Otu02708 | 15 | k__Fungi(100);p__Basidiomycota(100);c__Agaricomycetes(100);o__Cantharellales(100);f__Ceratobasidiaceae(100);g__Rhizoctonia(100);s__Rhizoctonia_sp(100); | Rhizoctonia | 13 | Pathotroph-Saprotroph-Symbiotroph | Ectomycorrhizal-Orchid Mycorrhizal-\|Plant Pathogen\|-Plant Saprotroph-Wood Saprotroph | Corticioid-Microfungus | Soft Rot | Probable | As litter_saprotroph (Põlme et al. 2020); Decay leaf, fruit, seed, roots, soil (Põlme et al. 2020); Only symbiotic (perhaps leaning parasitic) in special cases with orchids, otherwise typically a plant pathogen. | Seehann G, et al. 1975. List of Fungi in Soft-Rot Tissue Tests. Institute fur Holzbiologie und Holzschutz, Bundesforschungsanstalt fur Forst - und Holzwirtschaft, Hamburg - Lohbrugge, Germany; Costa IPMW, et al. 2012. Mycotaxon 119:e26 ((http://mycotaxon.com/resources/checklists/costa_v119_checklist.pdf)); Tedersoo L, et al. 2014. Science 346:e1256688 ((https://doi.org/10.1126/science.1256688)); Põlme S, et al. 2020. Fungal Diversity 105:1-16 ((https://doi.org/10.1007/s13225-020-00466-2)); List compiled by Sara Branco with input from Nhu Nguyen and Else Vellinga; The American Phytopathological Society Common Names of Plant Diseases ((http://www.apsnet.org/publications/commonnames/Pages/default.aspx)) |
| Otu02581 | 15 | k__Fungi(100);k__Fungi_unclassified(80);k__Fungi_unclassified(80);k__Fungi_unclassified(80);k__Fungi_unclassified(80);k__Fungi_unclassified(80);k__Fungi_unclassified(80); | - | - | - | - | - | - | - | Unassigned | [illegible] |
| Otu02584 | 15 | k__Fungi(100);p__Ascomycota(80);p__Ascomycota_unclassified(80);p__Ascomycota_unclassified(80);p__Ascomycota_unclassified(80);p__Ascomycota_unclassified(80);p__Ascomycota_unclassified(80); | - | - | - | - | - | - | - | Unassigned | [illegible] |
| Otu02586 | 15 | k__Fungi(100);p__Ascomycota(87);p__Ascomycota_unclassified(87);p__Ascomycota_unclassified(87);p__Ascomycota_unclassified(87);p__Ascomycota_unclassified(87);p__Ascomycota_unclassified(87); | - | - | - | - | - | - | - | Unassigned | [illegible] |
| Otu02587 | 15 | k__Fungi(100);p__Ascomycota(80);p__Ascomycota_unclassified(80);p__Ascomycota_unclassified(80);p__Ascomycota_unclassified(80);p__Ascomycota_unclassified(80);p__Ascomycota_unclassified(80); | - | - | - | - | - | - | - | Unassigned | [illegible] |
| Otu02592 | 15 | k__Fungi(100);p__Ascomycota(100);p__Ascomycota_unclassified(100);p__Ascomycota_unclassified(100);p__Ascomycota_unclassified(100);p__Ascomycota_unclassified(100);p__Ascomycota_unclassified(100); | - | - | - | - | - | - | - | Unassigned | [illegible] |
| Otu02593 | 15 | k__Fungi(100);p__Ascomycota(100);p__Ascomycota_unclassified(100);p__Ascomycota_unclassified(100);p__Ascomycota_unclassified(100);p__Ascomycota_unclassified(100);p__Ascomycota_unclassified(100); | - | - | - | - | - | - | - | Unassigned | [illegible] |
| Otu02595 | 15 | k__Fungi(100);p__Ascomycota(74);p__Ascomycota_unclassified(74);p__Ascomycota_unclassified(74);p__Ascomycota_unclassified(74);p__Ascomycota_unclassified(74);p__Ascomycota_unclassified(74); | - | - | - | - | - | - | - | Unassigned | [illegible] |
| Otu02600 | 15 | k__Fungi(100);p__Basidiomycota(100);c__Agaricomycetes(100);o__Russulales(100);o__Russulales_unclassified(100);o__Russulales_unclassified(100);o__Russulales_unclassified(100); | - | - | - | - | - | - | - | Unassigned | [illegible] |
| Otu02602 | 15 | k__Fungi(100);k__Fungi_unclassified(54);k__Fungi_unclassified(54);k__Fungi_unclassified(54);k__Fungi_unclassified(54);k__Fungi_unclassified(54);k__Fungi_unclassified(54); | - | - | - | - | - | - | - | Unassigned | [illegible] |
| Otu02603 | 15 | k__Fungi(100);k__Fungi_unclassified(100);k__Fungi_unclassified(100);k__Fungi_unclassified(100);k__Fungi_unclassified(100);k__Fungi_unclassified(100);k__Fungi_unclassified(100); | - | - | - | - | - | - | - | Unassigned | [illegible] |
| Otu02607 | 15 | k__Fungi(100);p__Basidiomycota(100);c__Agaricomycetes(100);c__Agaricomycetes_unclassified(100);c__Agaricomycetes_unclassified(100);c__Agaricomycetes_unclassified(100);c__Agaricomycetes_unclassified(100); | - | - | - | - | - | - | - | Unassigned | [illegible] |
| Otu02609 | 15 | k__Fungi(100);k__Fungi_unclassified(54);k__Fungi_unclassified(54);k__Fungi_unclassified(54);k__Fungi_unclassified(54);k__Fungi_unclassified(54);k__Fungi_unclassified(54); | - | - | - | - | - | - | - | Unassigned | [illegible] |
| Otu02610 | 15 | k__Fungi(100);k__Fungi_unclassified(100);k__Fungi_unclassified(100);k__Fungi_unclassified(100);k__Fungi_unclassified(100);k__Fungi_unclassified(100);k__Fungi_unclassified(100); | - | - | - | - | - | - | - | Unassigned | [illegible] |
| Otu02611 | 15 | k__Fungi(100);p__Ascomycota(100);c__Sordariomycetes(100);c__Sordariomycetes_unclassified(100);c__Sordariomycetes_unclassified(100);c__Sordariomycetes_unclassified(100);c__Sordariomycetes_unclassified(100); | - | - | - | - | - | - | - | Unassigned | [illegible] |
| Otu02612 | 15 | k__Fungi(100);p__Ascomycota(61);p__Ascomycota_unclassified(61);p__Ascomycota_unclassified(61);p__Ascomycota_unclassified(61);p__Ascomycota_unclassified(61);p__Ascomycota_unclassified(61); | - | - | - | - | - | - | - | Unassigned | [illegible] |
| Otu02613 | 15 | k__Fungi(100);k__Fungi_unclassified(54);k__Fungi_unclassified(54);k__Fungi_unclassified(54);k__Fungi_unclassified(54);k__Fungi_unclassified(54);k__Fungi_unclassified(54); | - | - | - | - | - | - | - | Unassigned | [illegible] |
| Otu02615 | 15 | k__Fungi(100);k__Fungi_unclassified(100);k__Fungi_unclassified(100);k__Fungi_unclassified(100);k__Fungi_unclassified(100);k__Fungi_unclassified(100);k__Fungi_unclassified(100); | - | - | - | - | - | - | - | Unassigned | [illegible] |
| Otu02616 | 15 | k__Fungi(100);p__Ascomycota(100);p__Ascomycota_unclassified(100);p__Ascomycota_unclassified(100);p__Ascomycota_unclassified(100);p__Ascomycota_unclassified(100);p__Ascomycota_unclassified(100); | - | - | - | - | - | - | - | Unassigned | [illegible] |
| Otu02618 | 15 | k__Fungi(100);k__Fungi_unclassified(54);k__Fungi_unclassified(54);k__Fungi_unclassified(54);k__Fungi_unclassified(54);k__Fungi_unclassified(54);k__Fungi_unclassified(54); | - | - | - | - | - | - | - | Unassigned | [illegible] |
| Otu02619 | 15 | k__Fungi(100);k__Fungi_unclassified(61);k__Fungi_unclassified(61);k__Fungi_unclassified(61);k__Fungi_unclassified(61);k__Fungi_unclassified(61);k__Fungi_unclassified(61); | - | - | - | - | - | - | - | Unassigned | [illegible] |
| Otu02621 | 15 | k__Fungi(100);p__Ascomycota(100);p__Ascomycota_unclassified(100);p__Ascomycota_unclassified(100);p__Ascomycota_unclassified(100);p__Ascomycota_unclassified(100);p__Ascomycota_unclassified(100); | - | - | - | - | - | - | - | Unassigned | [illegible] |
| Otu02622 | 15 | k__Fungi(100);p__Basidiomycota(100);p__Basidiomycota_unclassified(100);p__Basidiomycota_unclassified(100);p__Basidiomycota_unclassified(100);p__Basidiomycota_unclassified(100);p__Basidiomycota_unclassified(100); | - | - | - | - | - | - | - | Unassigned | [illegible] |
| Otu02624 | 15 | k__Fungi(100);p__Ascomycota(100);p__Ascomycota_unclassified(80);p__Ascomycota_unclassified(80);p__Ascomycota_unclassified(80);p__Ascomycota_unclassified(80);p__Ascomycota_unclassified(80); | - | - | - | - | - | - | - | Unassigned | [illegible] |
| Otu02627 | 15 | k__Fungi(100);k__Fungi_unclassified(74);k__Fungi_unclassified(74);k__Fungi_unclassified(74);k__Fungi_unclassified(74);k__Fungi_unclassified(74);k__Fungi_unclassified(74); | - | - | - | - | - | - | - | Unassigned | [illegible] |
| Otu02628 | 15 | k__Fungi(100);p__Ascomycota(100);c__Dothideomycetes(100);o__Capnodiales(100);o__Capnodiales_unclassified(100);o__Capnodiales_unclassified(100);o__Capnodiales_unclassified(100); | - | - | - | - | - | - | - | Unassigned | [illegible] |
| Otu02629 | 15 | k__Fungi(100);k__Fungi_unclassified(61);k__Fungi_unclassified(61);k__Fungi_unclassified(61);k__Fungi_unclassified(61);k__Fungi_unclassified(61);k__Fungi_unclassified(61); | - | - | - | - | - | - | - | Unassigned | [illegible] |
| Otu02633 | 15 | k__Fungi(100);k__Fungi_unclassified(54);k__Fungi_unclassified(54);k__Fungi_unclassified(54);k__Fungi_unclassified(54);k__Fungi_unclassified(54);k__Fungi_unclassified(54); | - | - | - | - | - | - | - | Unassigned | - |
| Otu02634 | 15 | k__Fungi(100);p__Ascomycota(54);p__Ascomycota_unclassified(54);p__Ascomycota_unclassified(54);p__Ascomycota_unclassified(54);p__Ascomycota_unclassified(54);p__Ascomycota_unclassified(54); | - | - | - | - | - | - | - | Unassigned | - |
| Otu02636 | 15 | k__Fungi(100);p__Ascomycota(67);p__Ascomycota_unclassified(67);p__Ascomycota_unclassified(67);p__Ascomycota_unclassified(67);p__Ascomycota_unclassified(67);p__Ascomycota_unclassified(67); | - | - | - | - | - | - | - | Unassigned | - |
| Otu02638 | 15 | k__Fungi(100);p__Ascomycota(80);p__Ascomycota_unclassified(80);p__Ascomycota_unclassified(80);p__Ascomycota_unclassified(80);p__Ascomycota_unclassified(80);p__Ascomycota_unclassified(80); | - | - | - | - | - | - | - | Unassigned | - |
| Otu02641 | 15 | k__Fungi(100);p__Ascomycota(54);p__Ascomycota_unclassified(54);p__Ascomycota_unclassified(54);p__Ascomycota_unclassified(54);p__Ascomycota_unclassified(54);p__Ascomycota_unclassified(54); | - | - | - | - | - | - | - | Unassigned | - |
| Otu02648 | 15 | k__Fungi(100);k__Fungi_unclassified(54);k__Fungi_unclassified(54);k__Fungi_unclassified(54);k__Fungi_unclassified(54);k__Fungi_unclassified(54);k__Fungi_unclassified(54); | - | - | - | - | - | - | - | Unassigned | - |
| Otu02649 | 15 | k__Fungi(100);p__Ascomycota(100);c__Dothideomycetes(100);o__Pleosporales(100);o__Pleosporales_unclassified(67);o__Pleosporales_unclassified(67);o__Pleosporales_unclassified(67); | - | - | - | - | - | - | - | Unassigned | - |
| Otu02651 | 15 | k__Fungi(100);p__Ascomycota(67);p__Ascomycota_unclassified(67);p__Ascomycota_unclassified(67);p__Ascomycota_unclassified(67);p__Ascomycota_unclassified(67);p__Ascomycota_unclassified(67); | - | - | - | - | - | - | - | Unassigned | - |
| Otu02654 | 15 | k__Fungi(100);p__Ascomycota(100);c__Dothideomycetes(100);o__Pleosporales(100);o__Pleosporales_unclassified(100);o__Pleosporales_unclassified(100);o__Pleosporales_unclassified(100); | - | - | - | - | - | - | - | Unassigned | - |
| Otu02656 | 15 | k__Plantae(100);p__unclassified_Plantae(100);c__unclassified_Plantae(100);o__unclassified_Plantae(100);f__unclassified_Plantae(100);g__unclassified_Plantae(100);s__Plantae_sp(100); | - | - | - | - | - | - | - | Unassigned | - |
| Otu02659 | 15 | k__Fungi(100);k__Fungi_unclassified(100);k__Fungi_unclassified(100);k__Fungi_unclassified(100);k__Fungi_unclassified(100);k__Fungi_unclassified(100);k__Fungi_unclassified(100); | - | - | - | - | - | - | - | Unassigned | - |
| Otu02667 | 15 | k__Fungi(100);k__Fungi_unclassified(100);k__Fungi_unclassified(100);k__Fungi_unclassified(100);k__Fungi_unclassified(100);k__Fungi_unclassified(100);k__Fungi_unclassified(100); | - | - | - | - | - | - | - | Unassigned | - |
| Otu02668 | 15 | k__Fungi(100);k__Fungi_unclassified(61);k__Fungi_unclassified(61);k__Fungi_unclassified(61);k__Fungi_unclassified(61);k__Fungi_unclassified(61);k__Fungi_unclassified(61); | - | - | - | - | - | - | - | Unassigned | - |
| Otu02669 | 15 | k__Fungi(100);k__Fungi_unclassified(67);k__Fungi_unclassified(67);k__Fungi_unclassified(67);k__Fungi_unclassified(67);k__Fungi_unclassified(67);k__Fungi_unclassified(67); | - | - | - | - | - | - | - | Unassigned | - |
| Otu02671 | 15 | k__Fungi(100);p__Ascomycota(100);c__Lecanoromycetes(80);c__Lecanoromycetes_unclassified(80);c__Lecanoromycetes_unclassified(80);c__Lecanoromycetes_unclassified(80);c__Lecanoromycetes_unclassified(80); | - | - | - | - | - | - | - | Unassigned | - |
| Otu02673 | 15 | k__Fungi(100);k__Fungi_unclassified(54);k__Fungi_unclassified(54);k__Fungi_unclassified(54);k__Fungi_unclassified(54);k__Fungi_unclassified(54);k__Fungi_unclassified(54); | - | - | - | - | - | - | - | Unassigned | - |
| Otu02674 | 15 | k__Fungi(100);p__Ascomycota(100);p__Ascomycota_unclassified(100);p__Ascomycota_unclassified(100);p__Ascomycota_unclassified(100);p__Ascomycota_unclassified(100);p__Ascomycota_unclassified(100); | - | - | - | - | - | - | - | Unassigned | - |
| Otu02675 | 15 | k__Fungi(100);k__Fungi_unclassified(67);k__Fungi_unclassified(67);k__Fungi_unclassified(67);k__Fungi_unclassified(67);k__Fungi_unclassified(67);k__Fungi_unclassified(67); | - | - | - | - | - | - | - | Unassigned | - |
| Otu02676 | 15 | k__Fungi(100);p__Ascomycota(100);c__Leotiomycetes(100);o__Helotiales(100);o__Helotiales_unclassified(74);o__Helotiales_unclassified(74);o__Helotiales_unclassified(74); | - | - | - | - | - | - | - | Unassigned | - |
| Otu02680 | 15 | k__Fungi(100);p__Basidiomycota(100);c__Atractiellomycetes(100);o__Atractiellales(100);f__unclassified_Atractiellales(100);g__unclassified_Atractiellales(100);s__Atractiellales_sp(100); | - | - | - | - | - | - | - | Unassigned | - |
| Otu02682 | 15 | k__Plantae(100);p__unclassified_Plantae(100);c__unclassified_Plantae(100);o__unclassified_Plantae(100);f__unclassified_Plantae(100);g__unclassified_Plantae(100);s__Plantae_sp(100); | - | - | - | - | - | - | - | Unassigned | - |
| Otu02683 | 15 | k__Fungi(100);p__Ascomycota(61);p__Ascomycota_unclassified(61);p__Ascomycota_unclassified(61);p__Ascomycota_unclassified(61);p__Ascomycota_unclassified(61);p__Ascomycota_unclassified(61); | - | - | - | - | - | - | - | Unassigned | - |
| Otu02685 | 15 | k__Fungi(100);k__Fungi_unclassified(61);k__Fungi_unclassified(61);k__Fungi_unclassified(61);k__Fungi_unclassified(61);k__Fungi_unclassified(61);k__Fungi_unclassified(61); | - | - | - | - | - | - | - | Unassigned | - |
| Otu02688 | 15 | k__Fungi(100);p__Ascomycota(80);p__Ascomycota_unclassified(80);p__Ascomycota_unclassified(80);p__Ascomycota_unclassified(80);p__Ascomycota_unclassified(80);p__Ascomycota_unclassified(80); | - | - | - | - | - | - | - | Unassigned | - |
| Otu02691 | 15 | k__Fungi(100);p__Basidiomycota(100);c__Agaricostilbomycetes(100);c__Agaricostilbomycetes_unclassified(100);c__Agaricostilbomycetes_unclassified(100);c__Agaricostilbomycetes_unclassified(100);c__Agaricostilbomycetes_unclassified(100); | - | - | - | - | - | - | - | Unassigned | - |
| Otu02693 | 15 | k__Fungi(100);k__Fungi_unclassified(74);k__Fungi_unclassified(74);k__Fungi_unclassified(74);k__Fungi_unclassified(74);k__Fungi_unclassified(74);k__Fungi_unclassified(74); | - | - | - | - | - | - | - | Unassigned | - |
| Otu02694 | 15 | k__Fungi(100);p__Ascomycota(100);c__Dothideomycetes(100);o__Pleosporales(100);f__unclassified_Pleosporales(80);g__unclassified_Pleosporales(80);s__Pleosporales_sp(80); | - | - | - | - | - | - | - | Unassigned | - |
| Otu02695 | 15 | k__Fungi(100);k__Fungi_unclassified(67);k__Fungi_unclassified(67);k__Fungi_unclassified(67);k__Fungi_unclassified(67);k__Fungi_unclassified(67);k__Fungi_unclassified(67); | - | - | - | - | - | - | - | Unassigned | - |
| Otu02696 | 15 | k__Fungi(100);p__Ascomycota(100);p__Ascomycota_unclassified(94);p__Ascomycota_unclassified(94);p__Ascomycota_unclassified(94);p__Ascomycota_unclassified(94);p__Ascomycota_unclassified(94); | - | - | - | - | - | - | - | Unassigned | - |
| Otu02698 | 15 | k__Fungi(100);p__Ascomycota(61);p__Ascomycota_unclassified(61);p__Ascomycota_unclassified(61);p__Ascomycota_unclassified(61);p__Ascomycota_unclassified(61);p__Ascomycota_unclassified(61); | - | - | - | - | - | - | - | Unassigned | - |
| Otu02700 | 15 | k__Fungi(100);p__Basidiomycota(100);c__Agaricomycetes(100);c__Agaricomycetes_unclassified(94);c__Agaricomycetes_unclassified(94);c__Agaricomycetes_unclassified(94);c__Agaricomycetes_unclassified(94); | - | - | - | - | - | - | - | Unassigned | - |
| Otu02701 | 15 | k__Fungi(100);p__Basidiomycota(100);c__Agaricomycetes(100);o__Polyporales(100);o__Polyporales_unclassified(100);o__Polyporales_unclassified(100);o__Polyporales_unclassified(100); | - | - | - | - | - | - | - | Unassigned | - |
| Otu02705 | 15 | k__Fungi(100);k__Fungi_unclassified(67);k__Fungi_unclassified(67);k__Fungi_unclassified(67);k__Fungi_unclassified(67);k__Fungi_unclassified(67);k__Fungi_unclassified(67); | - | - | - | - | - | - | - | Unassigned | - |
| Otu02706 | 15 | k__Fungi(100);p__Ascomycota(67);p__Ascomycota_unclassified(67);p__Ascomycota_unclassified(67);p__Ascomycota_unclassified(67);p__Ascomycota_unclassified(67);p__Ascomycota_unclassified(67); | - | - | - | - | - | - | - | Unassigned | - |
| Otu02707 | 15 | k__Fungi(100);k__Fungi_unclassified(87);k__Fungi_unclassified(87);k__Fungi_unclassified(87);k__Fungi_unclassified(87);k__Fungi_unclassified(87);k__Fungi_unclassified(87); | - | - | - | - | - | - | - | Unassigned | - |
| Otu02709 | 14 | k__Fungi(100);p__Ascomycota(100);c__Saccharomycetes(100);o__Saccharomycetales(100);f__Metschnikowiaceae(100);g__Metschnikowia(100);s__Metschnikowia_pulcherrima(93); | Metschnikowia | 13 | Saprotroph | \|Undefined Saprotroph\| | Yeast | NULL | Probable | As nectar-tap_saprotroph (Põlme et al. 2020); Decay sugar-rich substrates (Põlme et al. 2020) | Põlme S, et al. 2020. Fungal Diversity 105:1-16 ((https://doi.org/10.1007/s13225-020-00466-2)) |
| Otu02711 | 14 | k__Fungi(100);p__Ascomycota(100);c__Eurotiomycetes(100);o__Chaetothyriales(100);f__Herpotrichiellaceae(100);g__unclassified_Herpotrichiellaceae(93);s__Herpotrichiellaceae_sp(93); | Herpotrichiellaceae | 9 | Pathotroph-Saprotroph | Animal Pathogen-Fungal Parasite-Undefined Saprotroph | Facultative Yeast-Microfungus | NULL | Probable | Mostly saprobic on plants or other fungi, isolated from soil etc., and from medical sources (Cannon & Kirk 2007) | Cannon PF, Kirk PM. 2007. Fungal Families of the World. CAB International, Cambridge (ISBN: 978-0851998275) |
| Otu02716 | 14 | k__Fungi(100);p__Ascomycota(100);c__Sordariomycetes(100);o__Hypocreales(100);f__Hypocreales_family_Incertae_sedis(100);g__Myrothecium(100);s__Myrothecium_sp_E2(100); | Myrothecium | 13 | Pathotroph-Saprotroph | Fungal Parasite-\|Plant Saprotroph\|-Undefined Saprotroph | Microfungus | NULL | Probable | As litter_saprotroph (Põlme et al. 2020); As mycoparasite (Põlme et al. 2020); Decay leaf, fruit, seed (Põlme et al. 2020) | Tedersoo L, et al. 2014. Science 346:e1256688 ((https://doi.org/10.1126/science.1256688)); Põlme S, et al. 2020. Fungal Diversity 105:1-16 ((https://doi.org/10.1007/s13225-020-00466-2)) |
| Otu02717 | 14 | k__Fungi(100);p__Basidiomycota(100);c__Microbotryomycetes(100);o__Sporidiobolales(100);f__Sporidiobolales_family_Incertae_sedis(100);g__Sporobolomyces(100);s__Sporobolomyces_lactosus(100); | Sporobolomyces | 13 | Pathotroph-Saprotroph | \|Fungal Parasite\|-Plant Saprotroph | Yeast | NULL | Probable | As litter_saprotroph (Põlme et al. 2020); As mycoparasite (Põlme et al. 2020); Decay fungal material (Põlme et al. 2020) | Tedersoo L, et al. 2014. Science 346:e1256688 ((https://doi.org/10.1126/science.1256688)); Purahong W, et al. 2016. Molecular Ecology 25:4059-4074 ((https://doi.org/10.1111/mec.13739)); Põlme S, et al. 2020. Fungal Diversity 105:1-16 ((https://doi.org/10.1007/s13225-020-00466-2)) |
| Otu02719 | 14 | k__Fungi(100);p__Basidiomycota(100);c__Agaricomycetes(100);o__Agaricales(100);f__Inocybaceae(100);g__Inocybe(100);s__Inocybe_grammata(100); | Inocybe | 13 | Symbiotroph | \|Ectomycorrhizal\| | Agaricoid | NULL | Highly Probable | Sensu stricto (Matheny 2009). | Rinaldi AC, et al. 2008. Fungal Diversity 33:1-45 ((http://www.fungaldiversity.org/fdp/sfdp/33-1.pdf)); Matheny PB. 2009. McIlvainea 18:11-21 ((https://namyco.org/docs/3_Matheny.pdf)); Tedersoo L, et al. 2010. Mycorrhiza 20:217-263 ((https://doi.org/10.1007/s00572-009-0274-x)); Põlme S, et al. 2020. Fungal Diversity 105:1-16 ((https://doi.org/10.1007/s13225-020-00466-2)); http://mycorrhizas.info/ecmf.html ((http://mycorrhizas.info/ecmf.html)) |
| Otu02722 | 14 | k__Fungi(100);p__Basidiomycota(100);c__Agaricomycetes(100);o__Agaricales(100);f__Marasmiaceae(100);g__Marasmius(100);s__Marasmius_scorodonius(100); | Marasmius | 13 | Pathotroph-Saprotroph-Symbiotroph | Endophyte-Plant Pathogen-\|Plant Saprotroph\|-Undefined Saprotroph | Agaricoid | NULL | Probable | As litter_saprotroph (Põlme et al. 2020); Decay leaf, fruit, seed (Põlme et al. 2020); Often one of the most prominent fungal components of tropical forests, playing a major role in nutrient cycling (Cannon & Kirk 2007) | Cannon PF, Kirk PM. 2007. Fungal Families of the World. CAB International, Cambridge (ISBN: 978-0851998275); Tedersoo L, et al. 2014. Science 346:e1256688 ((https://doi.org/10.1126/science.1256688)); David AS, et al. 2016. Microbial Ecology 71:912-926 ((https://doi.org/10.1007/s00248-015-0712-6)); Põlme S, et al. 2020. Fungal Diversity 105:1-16 ((https://doi.org/10.1007/s13225-020-00466-2)) |
| Otu02723 | 14 | k__Fungi(100);p__Ascomycota(100);c__Sordariomycetes(100);o__Hypocreales(100);f__Hypocreaceae(100);g__Hypocrea(79);g__Hypocrea_unclassified(79); | Hypocrea | 13 | Pathotroph | Endophyte-Fungal Parasite | Microfungus | NULL | Probable | NULL | Tedersoo L, et al. 2014. Science 346:e1256688 ((https://doi.org/10.1126/science.1256688)); Christian N, et al. 2016. Microbial Ecology 72:621-632 ((https://doi.org/10.1007/s00248-016-0804-y)) |
| Otu02726 | 14 | k__Fungi(100);p__Ascomycota(100);c__Saccharomycetes(100);o__Saccharomycetales(100);f__Metschnikowiaceae(100);g__Metschnikowia(100);s__Metschnikowia_pulcherrima(86); | Metschnikowia | 13 | Saprotroph | \|Undefined Saprotroph\| | Yeast | NULL | Probable | As nectar-tap_saprotroph (Põlme et al. 2020); Decay sugar-rich substrates (Põlme et al. 2020) | Põlme S, et al. 2020. Fungal Diversity 105:1-16 ((https://doi.org/10.1007/s13225-020-00466-2)) |
| Otu02729 | 14 | k__Fungi(100);p__Ascomycota(100);c__Leotiomycetes(100);o__Helotiales(100);f__Helotiales_family_Incertae_sedis(100);g__Phacidiella(100);s__Phacidiella_eucalypti(100); | Phacidiella | 13 | Pathotroph | \|Plant Pathogen\| | Microfungus | NULL | Highly Probable | Canker; see Allantophomopsiella pseudotsugae | Crous PW, et al. 2014. IMA Fungus 5:173-193 ((https://doi.org/10.5598/imafungus.2014.05.02.02)); Põlme S, et al. 2020. Fungal Diversity 105:1-16 ((https://doi.org/10.1007/s13225-020-00466-2)) |
| Otu02730 | 14 | k__Fungi(100);p__Ascomycota(100);c__Leotiomycetes(100);o__Helotiales(100);f__Helotiaceae(100);g__Hymenoscyphus(100);s__Hymenoscyphus_fructigenus(100); | Hymenoscyphus | 13 | Pathotroph-Saprotroph-Symbiotroph | Bryophyte Parasite-Ectomycorrhizal-Ericoid Mycorrhizal-Plant Pathogen-\|Plant Saprotroph\|-Undefined Saprotroph | Microfungus | NULL | Possible | As litter_saprotroph (Põlme et al. 2020); Dark Septate Microfungus; Syn. Pezoloma among other (see e.g., Pezoloma ericae); Decay leaf, fruit, seed (Põlme et al. 2020) | Baral HO, Krieglsteiner L. 2006. Acta Mycologica 41:11-20 ((https://doi.org/10.5586/am.2006.003)) (as a bryophyte parasite); Huhtinen S, et al. 2010. Nova Hedwigia 90:413-431 ((https://doi.org/10.1127/0029-5035/2010/0090-0413)); Bent E, et al. 2011. Applied and Environmental Microbiology 77:3351-3359 ((https://doi.org/10.1128/AEM.02575-10)) (as ectomycorrhizae); Tedersoo L, et al. 2014. Science 346:1256688 ((https://doi.org/10.1126/science.1256688)) (as saprotroph); Põlme S, et al. 2020. Fungal Diversity 105:1-16 ((https://doi.org/10.1007/s13225-020-00466-2)) |
| Otu02731 | 14 | k__Fungi(100);p__Basidiomycota(100);c__Agaricomycetes(100);o__Agaricales(100);f__Cortinariaceae(100);g__Cortinarius(100);s__Cortinarius_illitus(100); | Cortinarius | 13 | Symbiotroph | \|Ectomycorrhizal\| | Agaricoid | NULL | Highly Probable | NULL | Rinaldi AC, et al. 2008. Fungal Diversity 33:1-45 ((http://www.fungaldiversity.org/fdp/sfdp/33-1.pdf)); Tedersoo L, et al. 2010. Mycorrhiza 20:217-263 ((https://doi.org/10.1007/s00572-009-0274-x)); Põlme S, et al. 2020. Fungal Diversity 105:1-16 ((https://doi.org/10.1007/s13225-020-00466-2)); http://mycorrhizas.info/ecmf.html ((http://mycorrhizas.info/ecmf.html)) |
| Otu02732 | 14 | k__Fungi(100);p__Basidiomycota(100);c__Agaricomycetes(100);o__Agaricales(100);f__Cortinariaceae(100);g__Cortinarius(100);g__Cortinarius_unclassified(100); | Cortinarius | 13 | Symbiotroph | \|Ectomycorrhizal\| | Agaricoid | NULL | Highly Probable | NULL | Rinaldi AC, et al. 2008. Fungal Diversity 33:1-45 ((http://www.fungaldiversity.org/fdp/sfdp/33-1.pdf)); Tedersoo L, et al. 2010. Mycorrhiza 20:217-263 ((https://doi.org/10.1007/s00572-009-0274-x)); Põlme S, et al. 2020. Fungal Diversity 105:1-16 ((https://doi.org/10.1007/s13225-020-00466-2)); http://mycorrhizas.info/ecmf.html ((http://mycorrhizas.info/ecmf.html)) |
| Otu02734 | 14 | k__Fungi(100);p__Ascomycota(100);c__Sordariomycetes(100);o__Xylariales(100);f__Xylariaceae(100);g__Virgaria(100);s__Virgaria_sp(100); | Virgaria | 13 | Saprotroph | Plant Saprotroph-Undefined Saprotroph-\|Wood Saprotroph\| | Microfungus | NULL | Probable | As litter_saprotroph (Põlme et al. 2020); Decay wood, leaf, fruit, seed (Põlme et al. 2020) | Tedersoo L, et al. 2014. Science 346:e1256688 ((https://doi.org/10.1126/science.1256688)); Põlme S, et al. 2020. Fungal Diversity 105:1-16 ((https://doi.org/10.1007/s13225-020-00466-2)) |
| Otu02737 | 14 | k__Fungi(100);p__Basidiomycota(100);c__Microbotryomycetes(100);o__Sporidiobolales(100);f__Sporidiobolales_family_Incertae_sedis(100);g__Rhodotorula(100);s__Rhodotorula_mucilaginosa(100); | Rhodotorula | 13 | Pathotroph-Saprotroph-Symbiotroph | Animal Pathogen-Animal Symbiotroph-Endophyte-Plant Pathogen-\|Undefined Saprotroph\| | Yeast | NULL | Probable | As foliar_endophyte (Põlme et al. 2020); Opportunistic human pathogen | Põlme S, et al. 2020. Fungal Diversity 105:1-16 ((https://doi.org/10.1007/s13225-020-00466-2)); https://www.cdc.gov ((https://www.cdc.gov)) |
| Otu02739 | 14 | k__Fungi(100);p__Ascomycota(100);c__Saccharomycetes(100);o__Saccharomycetales(100);f__Metschnikowiaceae(100);g__Metschnikowia(100);s__Metschnikowia_pulcherrima(93); | Metschnikowia | 13 | Saprotroph | \|Undefined Saprotroph\| | Yeast | NULL | Probable | As nectar-tap_saprotroph (Põlme et al. 2020); Decay sugar-rich substrates (Põlme et al. 2020) | Põlme S, et al. 2020. Fungal Diversity 105:1-16 ((https://doi.org/10.1007/s13225-020-00466-2)) |
| Otu02742 | 14 | k__Fungi(100);p__Basidiomycota(100);c__Agaricomycetes(100);o__Trechisporales(100);f__Hydnodontaceae(100);g__Trechispora(100);g__Trechispora_unclassified(100); | Trechispora | 13 | Saprotroph | \|Wood Saprotroph\| | Corticioid | White Rot | Highly Probable | As white_rot (Põlme et al. 2020); Decay wood (Põlme et al. 2020) | Gilbertson RL, Ryvarden L. 1987-1987. North American Polypores. Fungiflora, Oslo (ISBN: 978-0945345060); Rinaldi AC, et al. 2008. Fungal Diversity 33:1-45 ((http://www.fungaldiversity.org/fdp/sfdp/33-1.pdf)) (as Uncertain); Tedersoo L, et al. 2010. Mycorrhiza 20:217-263 ((https://doi.org/10.1007/s00572-009-0274-x)) (as not ECM).; Põlme S, et al. 2020. Fungal Diversity 105:1-16 ((https://doi.org/10.1007/s13225-020-00466-2)) |
| Otu02743 | 14 | k__Fungi(100);p__Basidiomycota(100);c__Tremellomycetes(100);o__Tremellales(100);f__Tremellales_family_Incertae_sedis(100);f__Tremellales_family_Incertae_sedis_unclassified(100);f__Tremellales_family_Incertae_sedis_unclassified(100); | Tremellales | 7 | Pathotroph-Saprotroph-Symbiotroph | Fungal Parasite-Undefined Saprotroph | Tremelloid-Yeast | NULL | Possible | The order contains both teleomorphic and anamorphic species, most of the latter being yeasts | Sterkenburg E, et al. 2015. New Phytologist 207:1145-1158 ((https://doi.org/10.1111/nph.13426)) |
| Otu02745 | 14 | k__Fungi(100);p__Ascomycota(100);c__Saccharomycetes(100);o__Saccharomycetales(100);f__Metschnikowiaceae(100);g__Metschnikowia(100);s__Metschnikowia_pulcherrima(93); | Metschnikowia | 13 | Saprotroph | \|Undefined Saprotroph\| | Yeast | NULL | Probable | As nectar-tap_saprotroph (Põlme et al. 2020); Decay sugar-rich substrates (Põlme et al. 2020) | Põlme S, et al. 2020. Fungal Diversity 105:1-16 ((https://doi.org/10.1007/s13225-020-00466-2)) |
| Otu02747 | 14 | k__Fungi(100);p__Zygomycota(100);c__Zygomycota_class_Incertae_sedis(100);o__Mortierellales(100);f__Mortierellaceae(100);g__Mortierella(100);g__Mortierella_unclassified(100); | Mortierella | 13 | Saprotroph-Symbiotroph | Endophyte-Plant Saprotroph-\|Undefined Saprotroph\| | Microfungus | NULL | Probable | As chitinolytic (Põlme et al. 2020); As soil_saprotroph (Põlme et al. 2020); Decay roots, soil (Põlme et al. 2020); Saprobic in soil and thus frequently isolated from soil and material contaminated with soil such as hay or rotten silage and spoil tips (Cannon & Kirk 2007) | Cannon PF, Kirk PM. 2007. Fungal Families of the World. CAB International, Cambridge (ISBN: 978-0851998275); Tedersoo L, et al. 2014. Science 346:e1256688 ((https://doi.org/10.1126/science.1256688)); Purahong W, et al. 2016. Molecular Ecology 25:4059-4074 ((https://doi.org/10.1111/mec.13739)); Põlme S, et al. 2020. Fungal Diversity 105:1-16 ((https://doi.org/10.1007/s13225-020-00466-2)) |
| Otu02749 | 14 | k__Fungi(100);p__Ascomycota(100);c__Saccharomycetes(100);o__Saccharomycetales(100);f__Metschnikowiaceae(100);g__Metschnikowia(100);s__Metschnikowia_pulcherrima(93); | Metschnikowia | 13 | Saprotroph | \|Undefined Saprotroph\| | Yeast | NULL | Probable | As nectar-tap_saprotroph (Põlme et al. 2020); Decay sugar-rich substrates (Põlme et al. 2020) | Põlme S, et al. 2020. Fungal Diversity 105:1-16 ((https://doi.org/10.1007/s13225-020-00466-2)) |
| Otu02750 | 14 | k__Fungi(100);p__Ascomycota(100);c__Saccharomycetes(100);o__Saccharomycetales(100);f__Saccharomycetales_family_Incertae_sedis(100);g__Candida(100);g__Candida_unclassified(100); | Candida | 13 | Pathotroph-Saprotroph-Symbiotroph | Animal Pathogen-Endophyte-Endosymbiont-Epiphyte-\|Undefined Saprotroph\| | Dimorphic Yeast-Yeast | NULL | Probable | As nectar-tap_saprotroph (Põlme et al. 2020); Decay sugar-rich substrates (Põlme et al. 2020); This asexual genus is polyphyletic and contain species that function very differently depend on what they are. Caution should be taken when interpreting guilds for Candida species. | Manolakaki D, et al. 2010. Virulence. 1: 367-375 ((https://doi.org/10.4161/viru.1.5.12796)); Mekbib SB, et al. 2011. Tropical Plant Pathology 36:233-240 ((https://doi.org/10.1590/S1982-56762011000400004)); Põlme S, et al. 2020. Fungal Diversity 105:1-16 ((https://doi.org/10.1007/s13225-020-00466-2)) |
| Otu02751 | 14 | k__Fungi(100);p__Ascomycota(100);c__Saccharomycetes(86);o__Saccharomycetales(86);f__Saccharomycetales_family_Incertae_sedis(79);g__Candida(79);g__Candida_unclassified(58); | Candida | 13 | Pathotroph-Saprotroph-Symbiotroph | Animal Pathogen-Endophyte-Endosymbiont-Epiphyte-\|Undefined Saprotroph\| | Dimorphic Yeast-Yeast | NULL | Probable | As nectar-tap_saprotroph (Põlme et al. 2020); Decay sugar-rich substrates (Põlme et al. 2020); This asexual genus is polyphyletic and contain species that function very differently depend on what they are. Caution should be taken when interpreting guilds for Candida species. | Manolakaki D, et al. 2010. Virulence. 1: 367-375 ((https://doi.org/10.4161/viru.1.5.12796)); Mekbib SB, et al. 2011. Tropical Plant Pathology 36:233-240 ((https://doi.org/10.1590/S1982-56762011000400004)); Põlme S, et al. 2020. Fungal Diversity 105:1-16 ((https://doi.org/10.1007/s13225-020-00466-2)) |
| Otu02752 | 14 | k__Fungi(100);p__Basidiomycota(100);c__Tremellomycetes(100);o__Filobasidiales(100);f__Filobasidiaceae(100);g__Cryptococcus(100);s__Cryptococcus_cylindricus(100); | Cryptococcus | 13 | Pathotroph-Saprotroph-Symbiotroph | Animal Pathogen-Endophyte-Epiphyte-\|Undefined Saprotroph\| | Dimorphic Yeast-Yeast | NULL | Possible | As foliar_endophyte (Põlme et al. 2020); Decay leaf, fruit, seed, soil, dung, animal material (Põlme et al. 2020) | Andrews JH, et al. 1983. Phytopathology 73:228-234 ((https://doi.org/10.1094/Phyto-73-228)); Kurtzman CP, et al. (eds.) 2011. The Yeasts, a Taxonomic Study. Fifth Edition. Vols 1-3. Elsevier, San Diego (ISBN: 9780444521491); Põlme S, et al. 2020. Fungal Diversity 105:1-16 ((https://doi.org/10.1007/s13225-020-00466-2)) |
| Otu02757 | 14 | k__Fungi(100);p__Ascomycota(100);c__Sordariomycetes(100);o__Hypocreales(100);f__unclassified_Hypocreales(100);g__unclassified_Hypocreales(100);s__Hypocreales_sp(100); | Hypocreales | 7 | Saprotroph | Undefined Saprotroph | Microfungus | NULL | Possible | NULL | Sterkenburg E, et al. 2015. New Phytologist 207:1145-1158 ((https://doi.org/10.1111/nph.13426)) |
| Otu02758 | 14 | k__Fungi(100);p__Basidiomycota(100);c__Agaricomycetes(100);o__Trechisporales(100);f__Hydnodontaceae(100);g__Trechispora(100);s__Trechispora_sp(100); | Trechispora | 13 | Saprotroph | \|Wood Saprotroph\| | Corticioid | White Rot | Highly Probable | As white_rot (Põlme et al. 2020); Decay wood (Põlme et al. 2020) | Gilbertson RL, Ryvarden L. 1987-1987. North American Polypores. Fungiflora, Oslo (ISBN: 978-0945345060); Rinaldi AC, et al. 2008. Fungal Diversity 33:1-45 ((http://www.fungaldiversity.org/fdp/sfdp/33-1.pdf)) (as Uncertain); Tedersoo L, et al. 2010. Mycorrhiza 20:217-263 ((https://doi.org/10.1007/s00572-009-0274-x)) (as not ECM).; Põlme S, et al. 2020. Fungal Diversity 105:1-16 ((https://doi.org/10.1007/s13225-020-00466-2)) |
| Otu02759 | 14 | k__Fungi(100);p__Ascomycota(100);c__Eurotiomycetes(100);o__Chaetothyriales(100);f__Herpotrichiellaceae(100);f__Herpotrichiellaceae_unclassified(100);f__Herpotrichiellaceae_unclassified(100); | Herpotrichiellaceae | 9 | Pathotroph-Saprotroph | Animal Pathogen-Fungal Parasite-Undefined Saprotroph | Facultative Yeast-Microfungus | NULL | Probable | Mostly saprobic on plants or other fungi, isolated from soil etc., and from medical sources (Cannon & Kirk 2007) | Cannon PF, Kirk PM. 2007. Fungal Families of the World. CAB International, Cambridge (ISBN: 978-0851998275) |
| Otu02760 | 14 | k__Fungi(100);p__Basidiomycota(100);c__Pucciniomycetes(100);o__Pucciniales(100);f__Pucciniaceae(100);g__Puccinia(100);g__Puccinia_unclassified(100); | Puccinia | 13 | Pathotroph | \|Plant Pathogen\| | Rust | NULL | Highly Probable | NULL | Tedersoo L, et al. 2014. Science 346:e1256688 ((https://doi.org/10.1126/science.1256688)); Põlme S, et al. 2020. Fungal Diversity 105:1-16 ((https://doi.org/10.1007/s13225-020-00466-2)) |
| Otu02761 | 14 | k__Fungi(100);p__Basidiomycota(100);c__Microbotryomycetes(100);o__Microbotryales(100);f__Ustilentylomataceae(100);g__Microbotryozyma(100);s__Microbotryozyma_collariae(100); | Microbotryozyma | 13 | Pathotroph-Saprotroph | \|Animal Parasite\|-Undefined Saprotroph | Yeast | NULL | Probable | As animal_decomposer (Põlme et al. 2020); Decay animal material (Põlme et al. 2020) | Põlme S, et al. 2020. Fungal Diversity 105:1-16 ((https://doi.org/10.1007/s13225-020-00466-2)) |
| Otu02764 | 14 | k__Fungi(100);p__Basidiomycota(100);c__Tremellomycetes(100);o__Tremellales(100);f__Tremellales_family_Incertae_sedis(100);f__Tremellales_family_Incertae_sedis_unclassified(100);f__Tremellales_family_Incertae_sedis_unclassified(100); | Tremellales | 7 | Pathotroph-Saprotroph-Symbiotroph | Fungal Parasite-Undefined Saprotroph | Tremelloid-Yeast | NULL | Possible | The order contains both teleomorphic and anamorphic species, most of the latter being yeasts | Sterkenburg E, et al. 2015. New Phytologist 207:1145-1158 ((https://doi.org/10.1111/nph.13426)) |
| Otu02765 | 14 | k__Fungi(100);p__Basidiomycota(100);c__Tremellomycetes(100);o__Tremellales(100);f__Tremellales_family_Incertae_sedis(100);g__Kockovaella(79);g__Kockovaella_unclassified(79); | Kockovaella | 13 | Saprotroph-Symbiotroph | Endophyte-\|Epiphyte\|-Undefined Saprotroph | Yeast | NULL | Probable | As foliar_endophyte (Põlme et al. 2020); Decay leaf, fruit, seed (Põlme et al. 2020); Foliar, isolated from a plant sample collected in a tropical rain forest in Thailand. | Fungsin B, et al. 2002. International Journal of Systematic and Evolutionary Microbiology 52:281-284 ((https://doi.org/10.1099/00207713-52-1-281)); Põlme S, et al. 2020. Fungal Diversity 105:1-16 ((https://doi.org/10.1007/s13225-020-00466-2)) |
| Otu02767 | 14 | k__Fungi(100);p__Zygomycota(100);c__Zygomycota_class_Incertae_sedis(100);o__Mortierellales(100);f__Mortierellaceae(100);g__Mortierella(100);g__Mortierella_unclassified(58); | Mortierella | 13 | Saprotroph-Symbiotroph | Endophyte-Plant Saprotroph-\|Undefined Saprotroph\| | Microfungus | NULL | Probable | As chitinolytic (Põlme et al. 2020); As soil_saprotroph (Põlme et al. 2020); Decay roots, soil (Põlme et al. 2020); Saprobic in soil and thus frequently isolated from soil and material contaminated with soil such as hay or rotten silage and spoil tips (Cannon & Kirk 2007) | Cannon PF, Kirk PM. 2007. Fungal Families of the World. CAB International, Cambridge (ISBN: 978-0851998275); Tedersoo L, et al. 2014. Science 346:e1256688 ((https://doi.org/10.1126/science.1256688)); Purahong W, et al. 2016. Molecular Ecology 25:4059-4074 ((https://doi.org/10.1111/mec.13739)); Põlme S, et al. 2020. Fungal Diversity 105:1-16 ((https://doi.org/10.1007/s13225-020-00466-2)) |
| Otu02768 | 14 | k__Fungi(100);p__Basidiomycota(100);c__Agaricomycetes(100);o__Boletales(100);f__Boletaceae(100);g__Xerocomus(100);s__Xerocomus_perplexus(100); | Xerocomus | 13 | Symbiotroph | \|Ectomycorrhizal\| | Agaricoid-Boletoid | NULL | Highly Probable | NULL | Tedersoo L, et al. 2010. Mycorrhiza 20:217-263 ((https://doi.org/10.1007/s00572-009-0274-x)); Põlme S, et al. 2020. Fungal Diversity 105:1-16 ((https://doi.org/10.1007/s13225-020-00466-2)); http://mycorrhizas.info/ecmf.html ((http://mycorrhizas.info/ecmf.html)) |
| Otu02769 | 14 | k__Fungi(100);p__Ascomycota(100);c__Dothideomycetes(100);o__Pleosporales(100);f__Cucurbitariaceae(100);g__Curreya(100);g__Curreya_unclassified(93); | Curreya | 13 | Saprotroph | \|Plant Saprotroph\|-Undefined Saprotroph | Microfungus | NULL | Probable | As litter_saprotroph (Põlme et al. 2020); Decay leaf, fruit, seed (Põlme et al. 2020) | Tedersoo L, et al. 2014. Science 346:e1256688 ((https://doi.org/10.1126/science.1256688)); Põlme S, et al. 2020. Fungal Diversity 105:1-16 ((https://doi.org/10.1007/s13225-020-00466-2)) |
| Otu02770 | 14 | k__Fungi(100);p__Ascomycota(100);c__Dothideomycetes(100);o__Capnodiales(100);f__Mycosphaerellaceae(100);f__Mycosphaerellaceae_unclassified(100);f__Mycosphaerellaceae_unclassified(100); | Mycosphaerellaceae | 9 | Pathotroph-Saprotroph | Plant Pathogen-Undefined Saprotroph | Microfungus | NULL | Probable | Biotrophic, necrotrophic or saprobic on various plant tissues (Cannon & Kirk 2007) | Cannon PF, Kirk PM. 2007. Fungal Families of the World. CAB International, Cambridge (ISBN: 978-0851998275) |
| Otu02771 | 14 | k__Fungi(100);p__Basidiomycota(100);c__Agaricomycetes(100);o__Russulales(100);f__Stereaceae(100);g__unclassified_Stereaceae(72);s__Stereaceae_sp(72); | Stereaceae | 9 | Saprotroph | Undefined Saprotroph-Wood Saprotroph | Corticioid-Polyporoid | NULL | Probable | Lignicolous or terrestrial (in leaf litter); typically saprobic (Cannon & Kirk 2007) | Cannon PF, Kirk PM. 2007. Fungal Families of the World. CAB International, Cambridge (ISBN: 978-0851998275) |
| Otu02772 | 14 | k__Fungi(100);p__Ascomycota(100);c__Saccharomycetes(100);o__Saccharomycetales(100);f__Metschnikowiaceae(100);g__Metschnikowia(100);s__Metschnikowia_pulcherrima(100); | Metschnikowia | 13 | Saprotroph | \|Undefined Saprotroph\| | Yeast | NULL | Probable | As nectar-tap_saprotroph (Põlme et al. 2020); Decay sugar-rich substrates (Põlme et al. 2020) | Põlme S, et al. 2020. Fungal Diversity 105:1-16 ((https://doi.org/10.1007/s13225-020-00466-2)) |
| Otu02775 | 14 | k__Fungi(100);p__Basidiomycota(100);c__Agaricomycetes(100);o__Cantharellales(100);f__Clavulinaceae(100);g__Clavulina(100);s__Clavulina_sp(100); | Clavulina | 13 | Symbiotroph | \|Ectomycorrhizal\| | Clavarioid | NULL | Highly Probable | NULL | Tedersoo L, et al. 2014. Science 346:e1256688 ((https://doi.org/10.1126/science.1256688)); Põlme S, et al. 2020. Fungal Diversity 105:1-16 ((https://doi.org/10.1007/s13225-020-00466-2)) |
| Otu02776 | 14 | k__Fungi(100);p__Basidiomycota(100);c__Agaricomycetes(100);o__Thelephorales(100);f__Thelephoraceae(100);f__Thelephoraceae_unclassified(93);f__Thelephoraceae_unclassified(93); | Thelephoraceae | 9 | Saprotroph-Symbiotroph | Ectomycorrhizal-Undefined Saprotroph | Clavarioid | NULL | Probable | On the ground, on soil, a few on wood. At least some speices ectomycorrhizal (Cannon & Kirk 2007) | Cannon PF, Kirk PM. 2007. Fungal Families of the World. CAB International, Cambridge (ISBN: 978-0851998275) |
| Otu02781 | 14 | k__Fungi(100);p__Basidiomycota(100);c__Agaricomycetes(100);o__Agaricales(100);f__Strophariaceae(100);g__Gymnopilus(100);s__Gymnopilus_aeruginosus(100); | Gymnopilus | 13 | Saprotroph | Plant Saprotroph-\|Wood Saprotroph\| | Agaricoid | White Rot | Probable | As litter_saprotroph (Põlme et al. 2020); As white_rot (Põlme et al. 2020); Decay wood (Põlme et al. 2020) | Tedersoo L, et al. 2014. Science 346:e1256688 ((https://doi.org/10.1126/science.1256688)); Põlme S, et al. 2020. Fungal Diversity 105:1-16 ((https://doi.org/10.1007/s13225-020-00466-2)) |
| Otu02783 | 14 | k__Fungi(100);p__Basidiomycota(100);c__Agaricomycetes(100);o__Agaricales(100);f__Psathyrellaceae(100);g__Coprinellus(100);s__Coprinellus_sp_PB_2000_1(100); | Coprinellus | 13 | Saprotroph | \|Undefined Saprotroph\| | Agaricoid | NULL | Probable | As soil_saprotroph (Põlme et al. 2020); Decay soil (Põlme et al. 2020) | Tedersoo L, et al. 2014. Science 346:e1256688 ((https://doi.org/10.1126/science.1256688)); Põlme S, et al. 2020. Fungal Diversity 105:1-16 ((https://doi.org/10.1007/s13225-020-00466-2)) |
| Otu02784 | 14 | k__Fungi(100);p__Basidiomycota(100);c__Pucciniomycetes(100);o__Pucciniales(100);f__Cronartiaceae(79);f__Cronartiaceae_unclassified(79);f__Cronartiaceae_unclassified(79); | Cronartiaceae | 9 | Pathotroph | Plant Pathogen | Microfungus | NULL | Probable | Common Name - Blister Rust, Rust Fungus; Most species are heteroecious, with the aecial stages forming on stems and cones of Pinus species and uredinia and telia on a range of dicotyledonous plants (Cannon & Kirk 2007) | Cannon PF, Kirk PM. 2007. Fungal Families of the World. CAB International, Cambridge (ISBN: 978-0851998275) |
| Otu02786 | 14 | k__Fungi(100);p__Ascomycota(100);c__Dothideomycetes(100);o__Capnodiales(100);f__Mycosphaerellaceae(100);g__Ramularia(79);s__Ramularia_cynarae(72); | Ramularia | 13 | Pathotroph | \|Plant Pathogen\| | Microfungus | NULL | Probable | NULL | Tedersoo L, et al. 2014. Science 346:e1256688 ((https://doi.org/10.1126/science.1256688)); Põlme S, et al. 2020. Fungal Diversity 105:1-16 ((https://doi.org/10.1007/s13225-020-00466-2)) |
| Otu02789 | 14 | k__Fungi(100);p__Ascomycota(100);c__Saccharomycetes(100);o__Saccharomycetales(100);f__Saccharomycetales_family_Incertae_sedis(100);g__Candida(100);s__Candida_stellata(93); | Candida | 13 | Pathotroph-Saprotroph-Symbiotroph | Animal Pathogen-Endophyte-Endosymbiont-Epiphyte-\|Undefined Saprotroph\| | Dimorphic Yeast-Yeast | NULL | Probable | As nectar-tap_saprotroph (Põlme et al. 2020); Decay sugar-rich substrates (Põlme et al. 2020); This asexual genus is polyphyletic and contain species that function very differently depend on what they are. Caution should be taken when interpreting guilds for Candida species. | Manolakaki D, et al. 2010. Virulence. 1: 367-375 ((https://doi.org/10.4161/viru.1.5.12796)); Mekbib SB, et al. 2011. Tropical Plant Pathology 36:233-240 ((https://doi.org/10.1590/S1982-56762011000400004)); Põlme S, et al. 2020. Fungal Diversity 105:1-16 ((https://doi.org/10.1007/s13225-020-00466-2)) |
| Otu02790 | 14 | k__Fungi(100);p__Ascomycota(100);c__Saccharomycetes(100);o__Saccharomycetales(100);f__Saccharomycetaceae(100);g__Kazachstania(100);s__Kazachstania_intestinalis(100); | Kazachstania | 13 | Saprotroph | \|Undefined Saprotroph\| | Yeast | NULL | Probable | As nectar-tap_saprotroph (Põlme et al. 2020); Decay sugar-rich substrates (Põlme et al. 2020) | Põlme S, et al. 2020. Fungal Diversity 105:1-16 ((https://doi.org/10.1007/s13225-020-00466-2)) |
| Otu02791 | 14 | k__Fungi(100);p__Ascomycota(100);c__Saccharomycetes(100);o__Saccharomycetales(100);f__Metschnikowiaceae(100);g__Metschnikowia(100);g__Metschnikowia_unclassified(86); | Metschnikowia | 13 | Saprotroph | \|Undefined Saprotroph\| | Yeast | NULL | Probable | As nectar-tap_saprotroph (Põlme et al. 2020); Decay sugar-rich substrates (Põlme et al. 2020) | Põlme S, et al. 2020. Fungal Diversity 105:1-16 ((https://doi.org/10.1007/s13225-020-00466-2)) |
| Otu02792 | 14 | k__Fungi(100);p__Ascomycota(100);c__Dothideomycetes(100);o__Capnodiales(100);f__Teratosphaeriaceae(100);g__Teratosphaeria(100);g__Teratosphaeria_unclassified(100); | Teratosphaeria | 13 | Pathotroph-Saprotroph | \|Plant Pathogen\|-Plant Saprotroph-Wood Saprotroph | Microfungus | NULL | Probable | As litter_saprotroph (Põlme et al. 2020); As wood_pathogen (Põlme et al. 2020); Decay wood, leaf, fruit, seed (Põlme et al. 2020) | Yamazaki A, et al. 2010. Mycoscience 51:443-455 ((https://www.jstage.jst.go.jp/article/mycosci/51/6/51_MYC51443/_pdf)); Tedersoo L, et al. 2014. Science 346:e1256688 ((https://doi.org/10.1126/science.1256688)); Põlme S, et al. 2020. Fungal Diversity 105:1-16 ((https://doi.org/10.1007/s13225-020-00466-2)) |
| Otu02794 | 14 | k__Fungi(100);p__Basidiomycota(100);c__Agaricostilbomycetes(100);o__Agaricostilbales(100);f__Agaricostilbaceae(93);f__Agaricostilbaceae_unclassified(93);f__Agaricostilbaceae_unclassified(93); | Agaricostilbaceae | 9 | Saprotroph | Undefined Saprotroph | NULL | NULL | Possible | Probably saprobic, usually associated with dead palm tissues (Cannon & Kirk 2007) | Cannon PF, Kirk PM. 2007. Fungal Families of the World. CAB International, Cambridge (ISBN: 978-0851998275) |
| Otu02796 | 14 | k__Fungi(100);p__Ascomycota(100);c__Leotiomycetes(100);o__Helotiales(100);f__Vibrisseaceae(100);g__Phialocephala(100);s__Phialocephala_urceolata(100); | Phialocephala | 13 | Saprotroph-Symbiotroph | Endophyte-\|Undefined Saprotroph\| | Microfungus | NULL | Highly Probable | As root_endophyte (Põlme et al. 2020); As root_endophyte_dark_septate (Põlme et al. 2020); As soil_saprotroph (Põlme et al. 2020); Dark Septate Endophyte, known from plant roots.; Decay roots, soil (Põlme et al. 2020) | Newsham KK. 2011. New Phytologist 190:783-793 ((https://doi.org/10.1111/j.1469-8137.2010.03611.x)) (as dark septate root endophytes); Põlme S, et al. 2020. Fungal Diversity 105:1-16 ((https://doi.org/10.1007/s13225-020-00466-2)) |
| Otu02797 | 14 | k__Fungi(100);p__Ascomycota(100);c__Saccharomycetes(100);o__Saccharomycetales(100);f__Metschnikowiaceae(100);g__Metschnikowia(100);s__Metschnikowia_viticola(100); | Metschnikowia | 13 | Saprotroph | \|Undefined Saprotroph\| | Yeast | NULL | Probable | As nectar-tap_saprotroph (Põlme et al. 2020); Decay sugar-rich substrates (Põlme et al. 2020) | Põlme S, et al. 2020. Fungal Diversity 105:1-16 ((https://doi.org/10.1007/s13225-020-00466-2)) |
| Otu02800 | 14 | k__Fungi(100);p__Ascomycota(100);c__Saccharomycetes(100);o__Saccharomycetales(100);f__Metschnikowiaceae(100);g__Metschnikowia(100);s__Metschnikowia_pulcherrima(100); | Metschnikowia | 13 | Saprotroph | \|Undefined Saprotroph\| | Yeast | NULL | Probable | As nectar-tap_saprotroph (Põlme et al. 2020); Decay sugar-rich substrates (Põlme et al. 2020) | Põlme S, et al. 2020. Fungal Diversity 105:1-16 ((https://doi.org/10.1007/s13225-020-00466-2)) |
| Otu02803 | 14 | k__Fungi(100);p__Ascomycota(100);c__Saccharomycetes(86);o__Saccharomycetales(86);f__Saccharomycetales_family_Incertae_sedis(79);g__Candida(58);g__Candida_unclassified(58); | Candida | 13 | Pathotroph-Saprotroph-Symbiotroph | Animal Pathogen-Endophyte-Endosymbiont-Epiphyte-\|Undefined Saprotroph\| | Dimorphic Yeast-Yeast | NULL | Probable | As nectar-tap_saprotroph (Põlme et al. 2020); Decay sugar-rich substrates (Põlme et al. 2020); This asexual genus is polyphyletic and contain species that function very differently depend on what they are. Caution should be taken when interpreting guilds for Candida species. | Manolakaki D, et al. 2010. Virulence. 1: 367-375 ((https://doi.org/10.4161/viru.1.5.12796)); Mekbib SB, et al. 2011. Tropical Plant Pathology 36:233-240 ((https://doi.org/10.1590/S1982-56762011000400004)); Põlme S, et al. 2020. Fungal Diversity 105:1-16 ((https://doi.org/10.1007/s13225-020-00466-2)) |
| Otu02805 | 14 | k__Fungi(100);p__Ascomycota(100);c__Saccharomycetes(100);o__Saccharomycetales(100);f__Metschnikowiaceae(100);g__Metschnikowia(100);s__Metschnikowia_pulcherrima(100); | Metschnikowia | 13 | Saprotroph | \|Undefined Saprotroph\| | Yeast | NULL | Probable | As nectar-tap_saprotroph (Põlme et al. 2020); Decay sugar-rich substrates (Põlme et al. 2020) | Põlme S, et al. 2020. Fungal Diversity 105:1-16 ((https://doi.org/10.1007/s13225-020-00466-2)) |
| Otu02806 | 14 | k__Fungi(100);p__Basidiomycota(100);c__Agaricomycetes(100);o__Agaricales(100);f__Cortinariaceae(100);g__Cortinarius(100);g__Cortinarius_unclassified(100); | Cortinarius | 13 | Symbiotroph | \|Ectomycorrhizal\| | Agaricoid | NULL | Highly Probable | NULL | Rinaldi AC, et al. 2008. Fungal Diversity 33:1-45 ((http://www.fungaldiversity.org/fdp/sfdp/33-1.pdf)); Tedersoo L, et al. 2010. Mycorrhiza 20:217-263 ((https://doi.org/10.1007/s00572-009-0274-x)); Põlme S, et al. 2020. Fungal Diversity 105:1-16 ((https://doi.org/10.1007/s13225-020-00466-2)); http://mycorrhizas.info/ecmf.html ((http://mycorrhizas.info/ecmf.html)) |
| Otu02807 | 14 | k__Fungi(100);p__Ascomycota(100);c__Saccharomycetes(100);o__Saccharomycetales(100);f__Metschnikowiaceae(100);g__Metschnikowia(100);s__Metschnikowia_pulcherrima(100); | Metschnikowia | 13 | Saprotroph | \|Undefined Saprotroph\| | Yeast | NULL | Probable | As nectar-tap_saprotroph (Põlme et al. 2020); Decay sugar-rich substrates (Põlme et al. 2020) | Põlme S, et al. 2020. Fungal Diversity 105:1-16 ((https://doi.org/10.1007/s13225-020-00466-2)) |
| Otu02808 | 14 | k__Fungi(100);p__Ascomycota(100);c__Dothideomycetes(100);o__Capnodiales(100);f__Mycosphaerellaceae(100);g__Dissoconium(100);s__Dissoconium_eucalypti(72); | Dissoconium | 13 | Pathotroph-Saprotroph | Plant Pathogen-\|Plant Saprotroph\|-Undefined Saprotroph | Microfungus | NULL | Probable | As litter_saprotroph (Põlme et al. 2020); Decay leaf, fruit, seed (Põlme et al. 2020) | Tedersoo L, et al. 2014. Science 346:e1256688 ((https://doi.org/10.1126/science.1256688)); Põlme S, et al. 2020. Fungal Diversity 105:1-16 ((https://doi.org/10.1007/s13225-020-00466-2)) |
| Otu02809 | 14 | k__Fungi(100);p__Ascomycota(100);c__Saccharomycetes(93);o__Saccharomycetales(93);f__Saccharomycetales_family_Incertae_sedis(65);f__Saccharomycetales_family_Incertae_sedis_unclassified(58);f__Saccharomycetales_family_Incertae_sedis_unclassified(58); | Saccharomycetales | 7 | Saprotroph | Undefined Saprotroph | Yeast | NULL | Possible | NULL | Sterkenburg E, et al. 2015. New Phytologist 207:1145-1158 ((https://doi.org/10.1111/nph.13426)) |
| Otu02810 | 14 | k__Fungi(100);p__Ascomycota(100);c__Sordariomycetes(100);o__Xylariales(100);f__Xylariales_family_Incertae_sedis(100);g__Monographella(100);s__Monographella_albescens(100); | Monographella | 13 | Pathotroph | \|Plant Pathogen\| | Microfungus | NULL | Probable | NULL | Tedersoo L, et al. 2014. Science 346:e1256688 ((https://doi.org/10.1126/science.1256688)); Põlme S, et al. 2020. Fungal Diversity 105:1-16 ((https://doi.org/10.1007/s13225-020-00466-2)) |
| Otu02811 | 14 | k__Fungi(100);p__Basidiomycota(100);c__Agaricomycetes(100);o__Polyporales(100);f__Phanerochaetaceae(100);g__Ceriporiopsis(100);s__Ceriporiopsis_balaenae(100); | Ceriporiopsis | 13 | Saprotroph | \|Wood Saprotroph\| | Corticioid | White Rot | Highly Probable | As white_rot (Põlme et al. 2020); Decay wood (Põlme et al. 2020) | Gilbertson RL, Ryvarden L. 1987-1987. North American Polypores. Fungiflora, Oslo (ISBN: 978-0945345060); Tedersoo L, et al. 2014. Science 346:e1256688 ((https://doi.org/10.1126/science.1256688)); Põlme S, et al. 2020. Fungal Diversity 105:1-16 ((https://doi.org/10.1007/s13225-020-00466-2)) |
| Otu02813 | 14 | k__Fungi(100);p__Ascomycota(100);c__Saccharomycetes(100);o__Saccharomycetales(100);f__Metschnikowiaceae(100);g__Metschnikowia(100);s__Metschnikowia_pulcherrima(72); | Metschnikowia | 13 | Saprotroph | \|Undefined Saprotroph\| | Yeast | NULL | Probable | As nectar-tap_saprotroph (Põlme et al. 2020); Decay sugar-rich substrates (Põlme et al. 2020) | Põlme S, et al. 2020. Fungal Diversity 105:1-16 ((https://doi.org/10.1007/s13225-020-00466-2)) |
| Otu02815 | 14 | k__Fungi(100);p__Ascomycota(100);c__Dothideomycetes(100);o__Capnodiales(100);f__Teratosphaeriaceae(79);f__Teratosphaeriaceae_unclassified(79);f__Teratosphaeriaceae_unclassified(79); | Teratosphaeriaceae | 9 | Pathotroph-Saprotroph | Animal Pathogen-Plant Pathogen-Undefined Saprotroph | NULL | NULL | Probable | Includes numerous saprobic, extremophilic, human opportunistic, and plant pathogenic fungi (Quaedvlieg et al. 2014) | Quaedvlieg W, et al. 2014. Persoonia 3:31-40 ((https://doi.org/10.3767/003158514X681981)) |
| Otu02817 | 14 | k__Fungi(100);p__Ascomycota(100);c__Saccharomycetes(100);o__Saccharomycetales(100);f__Metschnikowiaceae(100);g__Metschnikowia(100);s__Metschnikowia_pulcherrima(100); | Metschnikowia | 13 | Saprotroph | \|Undefined Saprotroph\| | Yeast | NULL | Probable | As nectar-tap_saprotroph (Põlme et al. 2020); Decay sugar-rich substrates (Põlme et al. 2020) | Põlme S, et al. 2020. Fungal Diversity 105:1-16 ((https://doi.org/10.1007/s13225-020-00466-2)) |
| Otu02823 | 14 | k__Fungi(100);p__Basidiomycota(100);c__Agaricomycetes(100);o__Trechisporales(100);f__Hydnodontaceae(100);g__Trechispora(100);s__Trechispora_sp(100); | Trechispora | 13 | Saprotroph | \|Wood Saprotroph\| | Corticioid | White Rot | Highly Probable | As white_rot (Põlme et al. 2020); Decay wood (Põlme et al. 2020) | Gilbertson RL, Ryvarden L. 1987-1987. North American Polypores. Fungiflora, Oslo (ISBN: 978-0945345060); Rinaldi AC, et al. 2008. Fungal Diversity 33:1-45 ((http://www.fungaldiversity.org/fdp/sfdp/33-1.pdf)) (as Uncertain); Tedersoo L, et al. 2010. Mycorrhiza 20:217-263 ((https://doi.org/10.1007/s00572-009-0274-x)) (as not ECM).; Põlme S, et al. 2020. Fungal Diversity 105:1-16 ((https://doi.org/10.1007/s13225-020-00466-2)) |
| Otu02825 | 14 | k__Fungi(100);p__Basidiomycota(100);c__Microbotryomycetes(100);o__Sporidiobolales(100);f__Sporidiobolales_family_Incertae_sedis(100);g__Sporobolomyces(93);g__Sporobolomyces_unclassified(93); | Sporobolomyces | 13 | Pathotroph-Saprotroph | \|Fungal Parasite\|-Plant Saprotroph | Yeast | NULL | Probable | As litter_saprotroph (Põlme et al. 2020); As mycoparasite (Põlme et al. 2020); Decay fungal material (Põlme et al. 2020) | Tedersoo L, et al. 2014. Science 346:e1256688 ((https://doi.org/10.1126/science.1256688)); Purahong W, et al. 2016. Molecular Ecology 25:4059-4074 ((https://doi.org/10.1111/mec.13739)); Põlme S, et al. 2020. Fungal Diversity 105:1-16 ((https://doi.org/10.1007/s13225-020-00466-2)) |
| Otu02826 | 14 | k__Fungi(100);p__Ascomycota(100);c__Saccharomycetes(100);o__Saccharomycetales(100);f__Metschnikowiaceae(100);g__Metschnikowia(100);s__Metschnikowia_pulcherrima(100); | Metschnikowia | 13 | Saprotroph | \|Undefined Saprotroph\| | Yeast | NULL | Probable | As nectar-tap_saprotroph (Põlme et al. 2020); Decay sugar-rich substrates (Põlme et al. 2020) | Põlme S, et al. 2020. Fungal Diversity 105:1-16 ((https://doi.org/10.1007/s13225-020-00466-2)) |
| Otu02829 | 14 | k__Fungi(100);p__Ascomycota(100);c__Saccharomycetes(100);o__Saccharomycetales(100);f__Metschnikowiaceae(100);g__Metschnikowia(100);s__Metschnikowia_pulcherrima(93); | Metschnikowia | 13 | Saprotroph | \|Undefined Saprotroph\| | Yeast | NULL | Probable | As nectar-tap_saprotroph (Põlme et al. 2020); Decay sugar-rich substrates (Põlme et al. 2020) | Põlme S, et al. 2020. Fungal Diversity 105:1-16 ((https://doi.org/10.1007/s13225-020-00466-2)) |
| Otu02831 | 14 | k__Fungi(100);p__Ascomycota(100);c__Saccharomycetes(100);o__Saccharomycetales(100);f__Saccharomycetaceae(100);g__Kazachstania(100);s__Kazachstania_intestinalis(100); | Kazachstania | 13 | Saprotroph | \|Undefined Saprotroph\| | Yeast | NULL | Probable | As nectar-tap_saprotroph (Põlme et al. 2020); Decay sugar-rich substrates (Põlme et al. 2020) | Põlme S, et al. 2020. Fungal Diversity 105:1-16 ((https://doi.org/10.1007/s13225-020-00466-2)) |
| Otu02834 | 14 | k__Fungi(100);p__Ascomycota(100);c__Dothideomycetes(100);o__Pleosporales(100);f__Pleosporaceae(100);g__Bipolaris(100);s__Bipolaris_sp_Bp_zj_2006a(100); | Bipolaris | 13 | Pathotroph-Saprotroph | \|Plant Pathogen\|-Plant Saprotroph | Microfungus | NULL | Probable | As litter_saprotroph (Põlme et al. 2020); Decay leaf, fruit, seed (Põlme et al. 2020) | Tedersoo L, et al. 2014. Science 346:e1256688 ((https://doi.org/10.1126/science.1256688)); Põlme S, et al. 2020. Fungal Diversity 105:1-16 ((https://doi.org/10.1007/s13225-020-00466-2)) |
| Otu02835 | 14 | k__Fungi(100);p__Basidiomycota(100);c__Agaricomycetes(100);o__Agaricales(100);f__Agaricaceae(100);f__Agaricaceae_unclassified(93);f__Agaricaceae_unclassified(93); | Agaricaceae | 9 | Saprotroph | Undefined Saprotroph | Agaricoid-Gasteroid-Secotioid | NULL | Probable | Primarily saprobes in grassland and woodland situations (Cannon & Kirk 2007) | Cannon PF, Kirk PM. 2007. Fungal Families of the World. CAB International, Cambridge (ISBN: 978-0851998275) |
| Otu02836 | 14 | k__Fungi(100);p__Ascomycota(100);c__Saccharomycetes(100);o__Saccharomycetales(100);f__Metschnikowiaceae(100);g__Metschnikowia(100);g__Metschnikowia_unclassified(93); | Metschnikowia | 13 | Saprotroph | \|Undefined Saprotroph\| | Yeast | NULL | Probable | As nectar-tap_saprotroph (Põlme et al. 2020); Decay sugar-rich substrates (Põlme et al. 2020) | Põlme S, et al. 2020. Fungal Diversity 105:1-16 ((https://doi.org/10.1007/s13225-020-00466-2)) |
| Otu02838 | 14 | k__Fungi(100);p__Ascomycota(100);c__Eurotiomycetes(100);o__Chaetothyriales(100);f__Herpotrichiellaceae(100);g__Cladophialophora(100);g__Cladophialophora_unclassified(86); | Cladophialophora | 13 | Pathotroph-Saprotroph-Symbiotroph | Animal Pathogen-Endophyte-Fungal Parasite-\|Undefined Saprotroph\| | Dimorphic-Microfungus | NULL | Probable | As mycoparasite (Põlme et al. 2020); As root_endophyte_dark_septate (Põlme et al. 2020); As soil_saprotroph (Põlme et al. 2020); Decay soil (Põlme et al. 2020) | James TY, et al. 2006. Nature 443:818-822 ((https://doi.org/10.1038/nature05110)); Põlme S, et al. 2020. Fungal Diversity 105:1-16 ((https://doi.org/10.1007/s13225-020-00466-2)) |
| Otu02839 | 14 | k__Fungi(100);p__Basidiomycota(100);c__Agaricomycetes(100);o__Polyporales(100);f__Meruliaceae(100);g__Hyphoderma(100);s__Hyphoderma_subtestaceum(100); | Hyphoderma | 13 | Saprotroph | Undefined Saprotroph-\|Wood Saprotroph\| | Corticioid | White Rot | Probable | As white_rot (Põlme et al. 2020); Decay wood (Põlme et al. 2020) | Tedersoo L, et al. 2014. Science 346:e1256688 ((https://doi.org/10.1126/science.1256688)); Põlme S, et al. 2020. Fungal Diversity 105:1-16 ((https://doi.org/10.1007/s13225-020-00466-2)) |
| Otu02843 | 14 | k__Fungi(100);p__Ascomycota(100);c__Saccharomycetes(100);o__Saccharomycetales(100);f__Saccharomycetales_family_Incertae_sedis(100);g__Candida(100);g__Candida_unclassified(65); | Candida | 13 | Pathotroph-Saprotroph-Symbiotroph | Animal Pathogen-Endophyte-Endosymbiont-Epiphyte-\|Undefined Saprotroph\| | Dimorphic Yeast-Yeast | NULL | Probable | As nectar-tap_saprotroph (Põlme et al. 2020); Decay sugar-rich substrates (Põlme et al. 2020); This asexual genus is polyphyletic and contain species that function very differently depend on what they are. Caution should be taken when interpreting guilds for Candida species. | Manolakaki D, et al. 2010. Virulence. 1: 367-375 ((https://doi.org/10.4161/viru.1.5.12796)); Mekbib SB, et al. 2011. Tropical Plant Pathology 36:233-240 ((https://doi.org/10.1590/S1982-56762011000400004)); Põlme S, et al. 2020. Fungal Diversity 105:1-16 ((https://doi.org/10.1007/s13225-020-00466-2)) |
| Otu02846 | 14 | k__Fungi(100);p__Zygomycota(100);c__Zygomycota_class_Incertae_sedis(100);o__Mortierellales(100);f__Mortierellaceae(100);g__Mortierella(100);s__Mortierella_strangulata(100); | Mortierella | 13 | Saprotroph-Symbiotroph | Endophyte-Plant Saprotroph-\|Undefined Saprotroph\| | Microfungus | NULL | Probable | As chitinolytic (Põlme et al. 2020); As soil_saprotroph (Põlme et al. 2020); Decay roots, soil (Põlme et al. 2020); Saprobic in soil and thus frequently isolated from soil and material contaminated with soil such as hay or rotten silage and spoil tips (Cannon & Kirk 2007) | Cannon PF, Kirk PM. 2007. Fungal Families of the World. CAB International, Cambridge (ISBN: 978-0851998275); Tedersoo L, et al. 2014. Science 346:e1256688 ((https://doi.org/10.1126/science.1256688)); Purahong W, et al. 2016. Molecular Ecology 25:4059-4074 ((https://doi.org/10.1111/mec.13739)); Põlme S, et al. 2020. Fungal Diversity 105:1-16 ((https://doi.org/10.1007/s13225-020-00466-2)) |
| Otu02710 | 14 | k__Fungi(100);k__Fungi_unclassified(100);k__Fungi_unclassified(100);k__Fungi_unclassified(100);k__Fungi_unclassified(100);k__Fungi_unclassified(100);k__Fungi_unclassified(100); | - | - | - | - | - | - | - | Unassigned | - |
| Otu02712 | 14 | k__Fungi(100);k__Fungi_unclassified(79);k__Fungi_unclassified(79);k__Fungi_unclassified(79);k__Fungi_unclassified(79);k__Fungi_unclassified(79);k__Fungi_unclassified(79); | - | - | - | - | - | - | - | Unassigned | - |
| Otu02713 | 14 | k__Fungi(100);p__Ascomycota(100);p__Ascomycota_unclassified(100);p__Ascomycota_unclassified(100);p__Ascomycota_unclassified(100);p__Ascomycota_unclassified(100);p__Ascomycota_unclassified(100); | - | - | - | - | - | - | - | Unassigned | - |
| Otu02714 | 14 | k__Fungi(100);p__Ascomycota(86);p__Ascomycota_unclassified(86);p__Ascomycota_unclassified(86);p__Ascomycota_unclassified(86);p__Ascomycota_unclassified(86);p__Ascomycota_unclassified(86); | - | - | - | - | - | - | - | Unassigned | - |
| Otu02715 | 14 | k__Fungi(100);k__Fungi_unclassified(58);k__Fungi_unclassified(58);k__Fungi_unclassified(58);k__Fungi_unclassified(58);k__Fungi_unclassified(58);k__Fungi_unclassified(58); | - | - | - | - | - | - | - | Unassigned | - |
| Otu02718 | 14 | k__Fungi(100);k__Fungi_unclassified(58);k__Fungi_unclassified(58);k__Fungi_unclassified(58);k__Fungi_unclassified(58);k__Fungi_unclassified(58);k__Fungi_unclassified(58); | - | - | - | - | - | - | - | Unassigned | - |
| Otu02720 | 14 | k__Fungi(100);k__Fungi_unclassified(79);k__Fungi_unclassified(79);k__Fungi_unclassified(79);k__Fungi_unclassified(79);k__Fungi_unclassified(79);k__Fungi_unclassified(79); | - | - | - | - | - | - | - | Unassigned | - |
| Otu02721 | 14 | k__Fungi(100);k__Fungi_unclassified(79);k__Fungi_unclassified(79);k__Fungi_unclassified(79);k__Fungi_unclassified(79);k__Fungi_unclassified(79);k__Fungi_unclassified(79); | - | - | - | - | - | - | - | Unassigned | - |
| Otu02724 | 14 | k__Fungi(100);p__Ascomycota(100);c__Leotiomycetes(86);o__Helotiales(86);o__Helotiales_unclassified(86);o__Helotiales_unclassified(86);o__Helotiales_unclassified(86); | - | - | - | - | - | - | - | Unassigned | - |
| Otu02725 | 14 | k__Fungi(100);k__Fungi_unclassified(65);k__Fungi_unclassified(65);k__Fungi_unclassified(65);k__Fungi_unclassified(65);k__Fungi_unclassified(65);k__Fungi_unclassified(65); | - | - | - | - | - | - | - | Unassigned | - |
| Otu02727 | 14 | k__Fungi(100);p__Ascomycota(100);c__Eurotiomycetes(100);c__Eurotiomycetes_unclassified(65);c__Eurotiomycetes_unclassified(65);c__Eurotiomycetes_unclassified(65);c__Eurotiomycetes_unclassified(65); | - | - | - | - | - | - | - | Unassigned | - |
| Otu02728 | 14 | k__Fungi(100);k__Fungi_unclassified(100);k__Fungi_unclassified(100);k__Fungi_unclassified(100);k__Fungi_unclassified(100);k__Fungi_unclassified(100);k__Fungi_unclassified(100); | - | - | - | - | - | - | - | Unassigned | - |
| Otu02733 | 14 | k__Fungi(100);p__Ascomycota(100);p__Ascomycota_unclassified(79);p__Ascomycota_unclassified(79);p__Ascomycota_unclassified(79);p__Ascomycota_unclassified(79);p__Ascomycota_unclassified(79); | - | - | - | - | - | - | - | Unassigned | - |
| Otu02735 | 14 | k__Fungi(100);p__Ascomycota(100);p__Ascomycota_unclassified(100);p__Ascomycota_unclassified(100);p__Ascomycota_unclassified(100);p__Ascomycota_unclassified(100);p__Ascomycota_unclassified(100); | - | - | - | - | - | - | - | Unassigned | - |
| Otu02736 | 14 | k__Fungi(100);p__Ascomycota(79);p__Ascomycota_unclassified(79);p__Ascomycota_unclassified(79);p__Ascomycota_unclassified(79);p__Ascomycota_unclassified(79);p__Ascomycota_unclassified(79); | - | - | - | - | - | - | - | Unassigned | - |
| Otu02738 | 14 | k__Fungi(100);p__Ascomycota(100);c__Sordariomycetes(100);c__Sordariomycetes_unclassified(79);c__Sordariomycetes_unclassified(79);c__Sordariomycetes_unclassified(79);c__Sordariomycetes_unclassified(79); | - | - | - | - | - | - | - | Unassigned | - |
| Otu02740 | 14 | k__Fungi(100);k__Fungi_unclassified(72);k__Fungi_unclassified(72);k__Fungi_unclassified(72);k__Fungi_unclassified(72);k__Fungi_unclassified(72);k__Fungi_unclassified(72); | - | - | - | - | - | - | - | Unassigned | - |
| Otu02741 | 14 | k__Fungi(100);p__Ascomycota(100);c__Sordariomycetes(100);c__Sordariomycetes_unclassified(100);c__Sordariomycetes_unclassified(100);c__Sordariomycetes_unclassified(100);c__Sordariomycetes_unclassified(100); | - | - | - | - | - | - | - | Unassigned | - |
| Otu02744 | 14 | k__Fungi(100);p__Ascomycota(58);p__Ascomycota_unclassified(58);p__Ascomycota_unclassified(58);p__Ascomycota_unclassified(58);p__Ascomycota_unclassified(58);p__Ascomycota_unclassified(58); | - | - | - | - | - | - | - | Unassigned | - |
| Otu02746 | 14 | k__Fungi(100);p__Ascomycota(100);c__Eurotiomycetes(79);o__Chaetothyriales(79);o__Chaetothyriales_unclassified(79);o__Chaetothyriales_unclassified(79);o__Chaetothyriales_unclassified(79); | - | - | - | - | - | - | - | Unassigned | - |
| Otu02748 | 14 | k__Fungi(100);k__Fungi_unclassified(100);k__Fungi_unclassified(100);k__Fungi_unclassified(100);k__Fungi_unclassified(100);k__Fungi_unclassified(100);k__Fungi_unclassified(100); | - | - | - | - | - | - | - | Unassigned | - |
| Otu02753 | 14 | k__Fungi(100);p__Basidiomycota(100);c__Agaricostilbomycetes(100);c__Agaricostilbomycetes_unclassified(100);c__Agaricostilbomycetes_unclassified(100);c__Agaricostilbomycetes_unclassified(100);c__Agaricostilbomycetes_unclassified(100); | - | - | - | - | - | - | - | Unassigned | - |
| Otu02754 | 14 | k__Fungi(100);p__Chytridiomycota(100);c__Chytridiomycetes(100);c__Chytridiomycetes_unclassified(100);c__Chytridiomycetes_unclassified(100);c__Chytridiomycetes_unclassified(100);c__Chytridiomycetes_unclassified(100); | - | - | - | - | - | - | - | Unassigned | - |
| Otu02755 | 14 | k__Fungi(100);p__Basidiomycota(100);c__Atractiellomycetes(100);o__Atractiellales(100);f__unclassified_Atractiellales(100);g__unclassified_Atractiellales(100);s__Atractiellales_sp(100); | - | - | - | - | - | - | - | Unassigned | - |
| Otu02756 | 14 | k__Fungi(100);p__Basidiomycota(100);c__Agaricomycetes(100);c__Agaricomycetes_unclassified(100);c__Agaricomycetes_unclassified(100);c__Agaricomycetes_unclassified(100);c__Agaricomycetes_unclassified(100); | - | - | - | - | - | - | - | Unassigned | - |
| Otu02762 | 14 | k__Fungi(100);p__Ascomycota(93);p__Ascomycota_unclassified(93);p__Ascomycota_unclassified(93);p__Ascomycota_unclassified(93);p__Ascomycota_unclassified(93);p__Ascomycota_unclassified(93); | - | - | - | - | - | - | - | Unassigned | - |
| Otu02763 | 14 | k__Fungi(100);p__Ascomycota(86);p__Ascomycota_unclassified(86);p__Ascomycota_unclassified(86);p__Ascomycota_unclassified(86);p__Ascomycota_unclassified(86);p__Ascomycota_unclassified(86); | - | - | - | - | - | - | - | Unassigned | - |
| Otu02766 | 14 | k__Fungi(100);p__Ascomycota(100);c__Sordariomycetes(100);c__Sordariomycetes_unclassified(100);c__Sordariomycetes_unclassified(100);c__Sordariomycetes_unclassified(100);c__Sordariomycetes_unclassified(100); | - | - | - | - | - | - | - | Unassigned | - |
| Otu02773 | 14 | k__Fungi(100);k__Fungi_unclassified(58);k__Fungi_unclassified(58);k__Fungi_unclassified(58);k__Fungi_unclassified(58);k__Fungi_unclassified(58);k__Fungi_unclassified(58); | - | - | - | - | - | - | - | Unassigned | - |
| Otu02774 | 14 | k__Fungi(100);k__Fungi_unclassified(58);k__Fungi_unclassified(58);k__Fungi_unclassified(58);k__Fungi_unclassified(58);k__Fungi_unclassified(58);k__Fungi_unclassified(58); | - | - | - | - | - | - | - | Unassigned | - |
| Otu02777 | 14 | k__Fungi(100);p__Ascomycota(100);p__Ascomycota_unclassified(100);p__Ascomycota_unclassified(100);p__Ascomycota_unclassified(100);p__Ascomycota_unclassified(100);p__Ascomycota_unclassified(100); | - | - | - | - | - | - | - | Unassigned | - |

| | | | | | | | | | | | |
|---|---|---|---|---|---|---|---|---|---|---|---|
| Otu02778 | 14 | k__Fungi(100);p__Basidiomycota(100);c__Agaricomycetes(100);c__Agaricomycetes_unclassified(100);c__Agaricomycetes_unclassified(100);c__Agaricomycetes_unclassified(100);c__Agaricomycetes_unclassified(100); | - | - | - | - | - | - | - | Unassigned | - |
| Otu02779 | 14 | k__Fungi(100);p__Ascomycota(100);c__Lecanoromycetes(100);c__Lecanoromycetes_unclassified(79);c__Lecanoromycetes_unclassified(79);c__Lecanoromycetes_unclassified(79);c__Lecanoromycetes_unclassified(79); | - | - | - | - | - | - | - | Unassigned | - |
| Otu02780 | 14 | k__Fungi(100);p__Ascomycota(100);p__Ascomycota_unclassified(100);p__Ascomycota_unclassified(100);p__Ascomycota_unclassified(100);p__Ascomycota_unclassified(100);p__Ascomycota_unclassified(100); | - | - | - | - | - | - | - | Unassigned | - |
| Otu02782 | 14 | k__Fungi(100);p__Ascomycota(100);p__Ascomycota_unclassified(100);p__Ascomycota_unclassified(100);p__Ascomycota_unclassified(100);p__Ascomycota_unclassified(100);p__Ascomycota_unclassified(100); | - | - | - | - | - | - | - | Unassigned | - |
| Otu02785 | 14 | k__Fungi(100);p__Ascomycota(65);p__Ascomycota_unclassified(65);p__Ascomycota_unclassified(65);p__Ascomycota_unclassified(65);p__Ascomycota_unclassified(65);p__Ascomycota_unclassified(65); | - | - | - | - | - | - | - | Unassigned | - |
| Otu02787 | 14 | k__Fungi(100);k__Fungi_unclassified(58);k__Fungi_unclassified(58);k__Fungi_unclassified(58);k__Fungi_unclassified(58);k__Fungi_unclassified(58);k__Fungi_unclassified(58); | - | - | - | - | - | - | - | Unassigned | - |
| Otu02788 | 14 | k__Fungi(100);k__Fungi_unclassified(100);k__Fungi_unclassified(100);k__Fungi_unclassified(100);k__Fungi_unclassified(100);k__Fungi_unclassified(100);k__Fungi_unclassified(100); | - | - | - | - | - | - | - | Unassigned | - |
| Otu02793 | 14 | k__Fungi(100);k__Fungi_unclassified(100);k__Fungi_unclassified(100);k__Fungi_unclassified(100);k__Fungi_unclassified(100);k__Fungi_unclassified(100);k__Fungi_unclassified(100); | - | - | - | - | - | - | - | Unassigned | - |
| Otu02795 | 14 | k__Fungi(100);p__Ascomycota(100);c__Dothideomycetes(100);o__Pleosporales(100);o__Pleosporales_unclassified(100);o__Pleosporales_unclassified(100);o__Pleosporales_unclassified(100); | - | - | - | - | - | - | - | Unassigned | - |
| Otu02798 | 14 | k__Fungi(100);p__Ascomycota(100);c__Sordariomycetes(100);o__Diaporthales(100);o__Diaporthales_unclassified(100);o__Diaporthales_unclassified(100);o__Diaporthales_unclassified(100); | - | - | - | - | - | - | - | Unassigned | - |
| Otu02799 | 14 | k__Fungi(100);p__Ascomycota(72);p__Ascomycota_unclassified(72);p__Ascomycota_unclassified(72);p__Ascomycota_unclassified(72);p__Ascomycota_unclassified(72);p__Ascomycota_unclassified(72); | - | - | - | - | - | - | - | Unassigned | - |
| Otu02801 | 14 | k__Fungi(100);p__Ascomycota(100);p__Ascomycota_unclassified(100);p__Ascomycota_unclassified(100);p__Ascomycota_unclassified(100);p__Ascomycota_unclassified(100);p__Ascomycota_unclassified(100); | - | - | - | - | - | - | - | Unassigned | - |
| Otu02802 | 14 | k__Fungi(100);k__Fungi_unclassified(65);k__Fungi_unclassified(65);k__Fungi_unclassified(65);k__Fungi_unclassified(65);k__Fungi_unclassified(65);k__Fungi_unclassified(65); | - | - | - | - | - | - | - | Unassigned | - |
| Otu02804 | 14 | k__Fungi(100);p__Ascomycota(65);p__Ascomycota_unclassified(65);p__Ascomycota_unclassified(65);p__Ascomycota_unclassified(65);p__Ascomycota_unclassified(65);p__Ascomycota_unclassified(65); | - | - | - | - | - | - | - | Unassigned | - |
| Otu02812 | 14 | k__Fungi(100);p__Basidiomycota(100);c__Agaricomycetes(100);c__Agaricomycetes_unclassified(100);c__Agaricomycetes_unclassified(100);c__Agaricomycetes_unclassified(100);c__Agaricomycetes_unclassified(100); | - | - | - | - | - | - | - | Unassigned | - |
| Otu02814 | 14 | k__Fungi(100);p__Ascomycota(100);p__Ascomycota_unclassified(100);p__Ascomycota_unclassified(100);p__Ascomycota_unclassified(100);p__Ascomycota_unclassified(100);p__Ascomycota_unclassified(100); | - | - | - | - | - | - | - | Unassigned | - |
| Otu02816 | 14 | k__Fungi(100);k__Fungi_unclassified(93);k__Fungi_unclassified(93);k__Fungi_unclassified(93);k__Fungi_unclassified(93);k__Fungi_unclassified(93);k__Fungi_unclassified(93); | - | - | - | - | - | - | - | Unassigned | - |
| Otu02818 | 14 | k__Fungi(100);p__Ascomycota(86);p__Ascomycota_unclassified(86);p__Ascomycota_unclassified(86);p__Ascomycota_unclassified(86);p__Ascomycota_unclassified(86);p__Ascomycota_unclassified(86); | - | - | - | - | - | - | - | Unassigned | - |
| Otu02819 | 14 | k__Fungi(100);p__Basidiomycota(100);c__Agaricomycetes(100);c__Agaricomycetes_unclassified(100);c__Agaricomycetes_unclassified(100);c__Agaricomycetes_unclassified(100);c__Agaricomycetes_unclassified(100); | - | - | - | - | - | - | - | Unassigned | - |
| Otu02820 | 14 | k__Fungi(100);p__Ascomycota(100);c__Dothideomycetes(100);o__Hysteriales(100);f__unclassified_Hysteriales(100);g__unclassified_Hysteriales(100);s__Hysteriales_sp(100); | - | - | - | - | - | - | - | Unassigned | - |
| Otu02821 | 14 | k__Fungi(100);k__Fungi_unclassified(93);k__Fungi_unclassified(93);k__Fungi_unclassified(93);k__Fungi_unclassified(93);k__Fungi_unclassified(93);k__Fungi_unclassified(93); | - | - | - | - | - | - | - | Unassigned | - |
| Otu02822 | 14 | k__Fungi(100);k__Fungi_unclassified(65);k__Fungi_unclassified(65);k__Fungi_unclassified(65);k__Fungi_unclassified(65);k__Fungi_unclassified(65);k__Fungi_unclassified(65); | - | - | - | - | - | - | - | Unassigned | - |
| Otu02824 | 14 | k__Fungi(100);k__Fungi_unclassified(79);k__Fungi_unclassified(79);k__Fungi_unclassified(79);k__Fungi_unclassified(79);k__Fungi_unclassified(79);k__Fungi_unclassified(79); | - | - | - | - | - | - | - | Unassigned | - |
| Otu02827 | 14 | k__Fungi(100);p__Basidiomycota(100);c__Tremellomycetes(100);c__Tremellomycetes_unclassified(79);c__Tremellomycetes_unclassified(79);c__Tremellomycetes_unclassified(79);c__Tremellomycetes_unclassified(79); | - | - | - | - | - | - | - | Unassigned | - |
| Otu02828 | 14 | k__Fungi(100);k__Fungi_unclassified(100);k__Fungi_unclassified(100);k__Fungi_unclassified(100);k__Fungi_unclassified(100);k__Fungi_unclassified(100);k__Fungi_unclassified(100); | - | - | - | - | - | - | - | Unassigned | - |
| Otu02830 | 14 | k__Fungi(100);k__Fungi_unclassified(72);k__Fungi_unclassified(72);k__Fungi_unclassified(72);k__Fungi_unclassified(72);k__Fungi_unclassified(72);k__Fungi_unclassified(72); | - | - | - | - | - | - | - | Unassigned | - |
| Otu02832 | 14 | k__Fungi(100);k__Fungi_unclassified(100);k__Fungi_unclassified(100);k__Fungi_unclassified(100);k__Fungi_unclassified(100);k__Fungi_unclassified(100);k__Fungi_unclassified(100); | - | - | - | - | - | - | - | Unassigned | - |
| Otu02833 | 14 | k__Fungi(100);p__Basidiomycota(100);c__Agaricomycetes(100);c__Agaricomycetes_unclassified(86);c__Agaricomycetes_unclassified(86);c__Agaricomycetes_unclassified(86);c__Agaricomycetes_unclassified(86); | - | - | - | - | - | - | - | Unassigned | - |
| Otu02837 | 14 | k__Fungi(100);p__Ascomycota(100);p__Ascomycota_unclassified(100);p__Ascomycota_unclassified(100);p__Ascomycota_unclassified(100);p__Ascomycota_unclassified(100);p__Ascomycota_unclassified(100); | - | - | - | - | - | - | - | Unassigned | - |
| Otu02840 | 14 | k__Fungi(100);k__Fungi_unclassified(65);k__Fungi_unclassified(65);k__Fungi_unclassified(65);k__Fungi_unclassified(65);k__Fungi_unclassified(65);k__Fungi_unclassified(65); | - | - | - | - | - | - | - | Unassigned | - |
| Otu02841 | 14 | k__Fungi(100);p__Ascomycota(100);c__Eurotiomycetes(100);o__Eurotiales(100);f__Elaphomycetaceae(100);g__unclassified_Elaphomycetaceae(100);s__Elaphomycetaceae_sp(100); | - | - | - | - | - | - | - | Unassigned | - |
| Otu02842 | 14 | k__Fungi(100);k__Fungi_unclassified(100);k__Fungi_unclassified(100);k__Fungi_unclassified(100);k__Fungi_unclassified(100);k__Fungi_unclassified(100);k__Fungi_unclassified(100); | - | - | - | - | - | - | - | Unassigned | - |
| Otu02844 | 14 | k__Fungi(100);k__Fungi_unclassified(100);k__Fungi_unclassified(100);k__Fungi_unclassified(100);k__Fungi_unclassified(100);k__Fungi_unclassified(100);k__Fungi_unclassified(100); | - | - | - | - | - | - | - | Unassigned | - |
| Otu02845 | 14 | k__Fungi(100);p__Basidiomycota(100);c__Tremellomycetes(100);o__Tremellales(100);f__unclassified_Tremellales(100);g__unclassified_Tremellales(100);s__uncultured_Tremellales(100); | - | - | - | - | - | - | - | Unassigned | - |
| Otu02847 | 14 | k__Fungi(100);p__Ascomycota(65);p__Ascomycota_unclassified(65);p__Ascomycota_unclassified(65);p__Ascomycota_unclassified(65);p__Ascomycota_unclassified(65);p__Ascomycota_unclassified(65); | - | - | - | - | - | - | - | Unassigned | - |
| Otu02848 | 14 | k__Fungi(100);p__Ascomycota(58);p__Ascomycota_unclassified(58);p__Ascomycota_unclassified(58);p__Ascomycota_unclassified(58);p__Ascomycota_unclassified(58);p__Ascomycota_unclassified(58); | - | - | - | - | - | - | - | Unassigned | - |
| Otu02849 | 14 | k__Fungi(100);p__Basidiomycota(100);p__Basidiomycota_unclassified(100);p__Basidiomycota_unclassified(100);p__Basidiomycota_unclassified(100);p__Basidiomycota_unclassified(100);p__Basidiomycota_unclassified(100); | - | - | - | - | - | - | - | Unassigned | - |
| Otu02850 | 14 | k__Fungi(100);p__Ascomycota(100);c__Dothideomycetes(100);o__Pleosporales(100);o__Pleosporales_unclassified(100);o__Pleosporales_unclassified(100);o__Pleosporales_unclassified(100); | - | - | - | - | - | - | - | Unassigned | - |
| Otu02851 | 14 | k__Fungi(100);k__Fungi_unclassified(65);k__Fungi_unclassified(65);k__Fungi_unclassified(65);k__Fungi_unclassified(65);k__Fungi_unclassified(65);k__Fungi_unclassified(65); | - | - | - | - | - | - | - | Unassigned | - |
| Otu02852 | 14 | k__Fungi(100);k__Fungi_unclassified(65);k__Fungi_unclassified(65);k__Fungi_unclassified(65);k__Fungi_unclassified(65);k__Fungi_unclassified(65);k__Fungi_unclassified(65); | - | - | - | - | - | - | - | Unassigned | - |
| Otu02853 | 13 | k__Fungi(100);p__Basidiomycota(100);c__Agaricomycetes(100);o__Agaricales(100);f__Entolomataceae(100);g__Clitopilus(100);s__Clitopilus_sp_1_T_778(100); | Clitopilus | 13 | Pathotroph-Saprotroph | Fungal Parasite-\|Plant Saprotroph\|-Undefined Saprotroph | Agaricoid | NULL | Probable | [illegible] | [illegible] |
| Otu02855 | 13 | k__Fungi(100);p__Ascomycota(100);c__Saccharomycetes(100);o__Saccharomycetales(100);f__Metschnikowiaceae(100);g__Metschnikowia(100);s__Metschnikowia_pulcherrima(100); | Metschnikowia | 13 | Saprotroph | \|Undefined Saprotroph\| | Yeast | NULL | Probable | [illegible] | [illegible] |
| Otu02856 | 13 | k__Fungi(100);p__Ascomycota(100);c__Saccharomycetes(100);o__Saccharomycetales(100);f__Metschnikowiaceae(100);g__Metschnikowia(100);s__Metschnikowia_pulcherrima(100); | Metschnikowia | 13 | Saprotroph | \|Undefined Saprotroph\| | Yeast | NULL | Probable | [illegible] | [illegible] |
| Otu02857 | 13 | k__Fungi(100);p__Basidiomycota(100);c__Agaricomycetes(100);o__Agaricales(100);f__Agaricaceae(100);g__Coprinus(100);s__Coprinopsis_cordispora(100); | Coprinopsis | 13 | Saprotroph | \|Undefined Saprotroph\| | Agaricoid | NULL | Probable | [illegible] | [illegible] |
| Otu02860 | 13 | k__Fungi(100);p__Basidiomycota(100);c__Agaricomycetes(100);o__Cantharellales(100);f__Clavulinaceae(100);f__Clavulinaceae_unclassified(100);f__Clavulinaceae_unclassified(100); | Clavulinaceae | 9 | Saprotroph-Symbiotroph | Ectomycorrhizal-Lichenized-Wood Saprotroph | Clavarioid | NULL | Possible | [illegible] | [illegible] |
| Otu02861 | 13 | k__Fungi(100);p__Ascomycota(100);c__Dothideomycetes(100);o__Capnodiales(100);f__Teratosphaeriaceae(93);f__Teratosphaeriaceae_unclassified(93);f__Teratosphaeriaceae_unclassified(93); | Teratosphaeriaceae | 9 | Pathotroph-Saprotroph | Animal Pathogen-Plant Pathogen-Undefined Saprotroph | NULL | NULL | Probable | [illegible] | [illegible] |
| Otu02865 | 13 | k__Fungi(100);p__Basidiomycota(100);c__Agaricomycetes(100);o__Polyporales(100);f__Fomitopsidaceae(100);g__Pilatoporus(100);s__Pilatoporus_bondartsevae(100); | Pilatoporus | 13 | Saprotroph | \|Wood Saprotroph\| | Polyporoid | Brown Rot | Probable | [illegible] | [illegible] |
| Otu02867 | 13 | k__Fungi(100);p__Ascomycota(100);c__Saccharomycetes(100);o__Saccharomycetales(100);f__Metschnikowiaceae(100);g__Metschnikowia(100);s__Metschnikowia_pulcherrima(85); | Metschnikowia | 13 | Saprotroph | \|Undefined Saprotroph\| | Yeast | NULL | Probable | [illegible] | [illegible] |
| Otu02868 | 13 | k__Fungi(100);p__Ascomycota(100);c__Sordariomycetes(100);o__Hypocreales(100);f__Clavicipitaceae(100);g__Metarhizium(100);s__Metarhizium_flavoviride_var_pemphigi(85); | Metarhizium | 13 | Pathotroph-Saprotroph | \|Animal Parasite\|-Animal Pathogen-Undefined Saprotroph | Microfungus | NULL | Probable | [illegible] | [illegible] |
| Otu02869 | 13 | k__Fungi(100);p__Ascomycota(100);c__Sordariomycetes(100);o__Hypocreales(100);o__Hypocreales_unclassified(93);o__Hypocreales_unclassified(93);o__Hypocreales_unclassified(93); | Hypocreales | 7 | Saprotroph | Undefined Saprotroph | Microfungus | NULL | Possible | NULL | [illegible] |
| Otu02872 | 13 | k__Fungi(100);p__Ascomycota(100);c__Eurotiomycetes(100);o__Eurotiales(100);f__Trichocomaceae(100);g__Aspergillus(100);s__Aspergillus_subversicolor(100); | Aspergillus | 13 | Pathotroph-Saprotroph-Symbiotroph | Animal Pathogen-Endophyte-Plant Saprotroph-\|Undefined Saprotroph\|-Wood Saprotroph | Microfungus | NULL | Probable | [illegible] | [illegible] |
| Otu02875 | 13 | k__Fungi(100);p__Ascomycota(100);c__Eurotiomycetes(100);o__Eurotiales(100);f__Trichocomaceae(100);g__Paecilomyces(100);s__Paecilomyces_lilacinus(100); | Paecilomyces | 13 | Pathotroph-Saprotroph-Symbiotroph | Clavicipitaceous \|Endophyte\|-Plant Pathogen-Plant Saprotroph-Undefined Saprotroph-Wood Saprotroph | Microfungus | Soft Rot | Probable | [illegible] | [illegible] |
| Otu02876 | 13 | k__Fungi(100);p__Ascomycota(100);c__Saccharomycetes(100);o__Saccharomycetales(100);f__Metschnikowiaceae(100);g__Metschnikowia(100);s__Metschnikowia_pulcherrima(62); | Metschnikowia | 13 | Saprotroph | \|Undefined Saprotroph\| | Yeast | NULL | Probable | [illegible] | [illegible] |
| Otu02879 | 13 | k__Fungi(100);p__Ascomycota(100);c__Dothideomycetes(100);o__Capnodiales(100);f__Mycosphaerellaceae(100);g__Pseudocercosporella(100);s__Pseudocercosporella_sp_RH22(100); | Pseudocercosporella | 13 | Pathotroph | \|Plant Pathogen\| | Microfungus | NULL | Probable | NULL | [illegible] |
| Otu02881 | 13 | k__Fungi(100);p__Basidiomycota(100);c__Ustilaginomycetes(100);o__Ustilaginales(100);f__Anthracoideaceae(100);g__Schizonella(100);s__Schizonella_melanogramma(100); | Schizonella | 13 | Pathotroph | \|Plant Pathogen\| | Smut | NULL | Probable | [illegible] | [illegible] |
| Otu02885 | 13 | k__Fungi(100);p__Basidiomycota(100);c__Tremellomycetes(100);o__Tremellales(100);f__Tremellales_family_Incertae_sedis(100);f__Tremellales_family_Incertae_sedis_unclassified(100);f__Tremellales_family_Incertae_sedis_unclassified(100); | Tremellales | 7 | Pathotroph-Saprotroph-Symbiotroph | Fungal Parasite-Undefined Saprotroph | Tremelloid-Yeast | NULL | Possible | [illegible] | [illegible] |
| Otu02886 | 13 | k__Fungi(100);p__Zygomycota(100);c__Zygomycota_class_Incertae_sedis(100);o__Mucorales(100);f__Mucoraceae(100);g__Mucor(100);s__Mucor_flavus(70); | Mucor | 13 | Saprotroph | \|Undefined Saprotroph\| | Microfungus | NULL | Probable | [illegible] | [illegible] |
| Otu02890 | 13 | k__Fungi(100);p__Basidiomycota(100);c__Exobasidiomycetes(100);o__Exobasidiales(100);f__Exobasidiaceae(100);g__Exobasidium(100);s__Exobasidium_yoshinagae(100); | Exobasidium | 13 | Pathotroph | \|Plant Pathogen\| | Leaf Blister-Leaf Gall-Leaf Spot | NULL | Probable | NULL | [illegible] |
| Otu02892 | 13 | k__Fungi(100);p__Ascomycota(100);c__Saccharomycetes(100);o__Saccharomycetales(100);f__Metschnikowiaceae(100);g__Metschnikowia(100);s__Metschnikowia_pulcherrima(100); | Metschnikowia | 13 | Saprotroph | \|Undefined Saprotroph\| | Yeast | NULL | Probable | [illegible] | [illegible] |
| Otu02895 | 13 | k__Fungi(100);p__Basidiomycota(100);c__Ustilaginomycetes(100);o__Urocystidales(100);f__Urocystidaceae(100);g__Urocystis(100);s__Urocystis_agropyri(100); | Urocystis | 13 | Pathotroph | \|Plant Pathogen\| | Smut | NULL | Probable | NULL | [illegible] |
| Otu02898 | 13 | k__Fungi(100);p__Basidiomycota(100);c__Agaricomycetes(100);o__Agaricales(100);f__Physalacriaceae(100);g__Flammulina(100);s__Flammulina_velutipes(100); | Flammulina | 13 | Saprotroph | Undefined Saprotroph-\|Wood Saprotroph\| | Agaricoid | White Rot | Probable | [illegible] | [illegible] |
| Otu02903 | 13 | k__Fungi(100);p__Basidiomycota(100);c__Agaricomycetes(100);o__Agaricales(100);f__Strophariaceae(100);g__Pholiota(100);s__Pholiota_spumosa(100); | Pholiota | 13 | Pathotroph-Saprotroph | Plant Pathogen-Plant Saprotroph-Undefined Saprotroph-\|Wood Saprotroph\| | Agaricoid | White Rot | Probable | [illegible] | [illegible] |
| Otu02904 | 13 | k__Fungi(100);p__Ascomycota(100);c__Sordariomycetes(100);o__Sordariales(100);f__Chaetomiaceae(100);g__Chaetomium(70);g__Chaetomium_unclassified(70); | Chaetomium | 13 | Pathotroph-Saprotroph-Symbiotroph | Animal Pathogen-Dung Saprotroph-Endophyte-Epiphyte-Plant Pathogen-\|Plant Saprotroph\|-Wood Saprotroph | Microfungus | Soft Rot | Possible | [illegible] | [illegible] |
| Otu02905 | 13 | k__Fungi(100);p__Ascomycota(100);c__Saccharomycetes(100);o__Saccharomycetales(100);f__Metschnikowiaceae(100);g__Metschnikowia(100);g__Metschnikowia_unclassified(93); | Metschnikowia | 13 | Saprotroph | \|Undefined Saprotroph\| | Yeast | NULL | Probable | [illegible] | [illegible] |
| Otu02907 | 13 | k__Fungi(100);p__Ascomycota(100);c__Saccharomycetes(100);o__Saccharomycetales(100);f__unclassified_Saccharomycetales(85);g__unclassified_Saccharomycetales(85);s__Saccharomycetales_sp(85); | Saccharomycetales | 7 | Saprotroph | Undefined Saprotroph | Yeast | NULL | Possible | NULL | [illegible] |
| Otu02908 | 13 | k__Fungi(100);p__Ascomycota(100);c__Dothideomycetes(100);o__Capnodiales(100);f__Capnodiales_family_Incertae_sedis(100);g__Hispidoconidioma(100);s__Hispidoconidioma_sp(100); | Hispidoconidioma | 13 | Saprotroph | \|Wood Saprotroph\| | Microfungus | NULL | Probable | [illegible] | [illegible] |
| Otu02909 | 13 | k__Fungi(100);p__Basidiomycota(100);c__Tremellomycetes(100);o__Tremellales(100);f__Tremellales_family_Incertae_sedis(100);g__Hannaella(100);s__Hannaella_zeae(100); | Hannaella | 13 | Pathotroph-Saprotroph | \|Animal Parasite\|-Undefined Saprotroph | Yeast | NULL | Probable | [illegible] | [illegible] |
| Otu02913 | 13 | k__Fungi(100);p__Ascomycota(100);c__Dothideomycetes(100);o__Botryosphaeriales(100);f__Botryosphaeriaceae(100);g__Phaeobotryon(77);s__Phaeobotryon_cupressi(77); | Phaeobotryon | 13 | Pathotroph-Saprotroph-Symbiotroph | Endophyte-\|Plant Pathogen\|-Plant Saprotroph | Microfungus | NULL | Probable | [illegible] | [illegible] |
| Otu02915 | 13 | k__Fungi(100);p__Ascomycota(100);c__Eurotiomycetes(100);o__Chaetothyriales(100);f__Herpotrichiellaceae(100);g__Phaeococcomyces(100);s__Phaeococcomyces_catenatus(100); | Phaeococcomyces | 13 | Saprotroph | \|Undefined Saprotroph\| | Dimorphic-Facultative Yeast | NULL | Probable | NULL | [illegible] |
| Otu02916 | 13 | k__Fungi(100);p__Ascomycota(100);c__Dothideomycetes(100);o__Capnodiales(100);f__Teratosphaeriaceae(100);g__Teratosphaeria(100);s__Teratosphaeria_knoxdavesii(100); | Teratosphaeria | 13 | Pathotroph-Saprotroph | \|Plant Pathogen\|-Plant Saprotroph-Wood Saprotroph | Microfungus | NULL | Probable | [illegible] | [illegible] |
| Otu02917 | 13 | k__Fungi(100);p__Ascomycota(100);c__Sordariomycetes(100);o__Sordariales(100);f__Chaetomiaceae(100);g__Chaetomium(100);g__Chaetomium_unclassified(100); | Chaetomium | 13 | Pathotroph-Saprotroph-Symbiotroph | Animal Pathogen-Dung Saprotroph-Endophyte-Epiphyte-Plant Pathogen-\|Plant Saprotroph\|-Wood Saprotroph | Microfungus | Soft Rot | Possible | [illegible] | [illegible] |
| Otu02919 | 13 | k__Fungi(100);p__Basidiomycota(100);c__Agaricomycetes(100);o__Agaricales(100);f__Cortinariaceae(100);g__Cortinarius(100);g__Cortinarius_unclassified(100); | Cortinarius | 13 | Symbiotroph | \|Ectomycorrhizal\| | Agaricoid | NULL | Highly Probable | NULL | [illegible] |
| Otu02920 | 13 | k__Fungi(100);p__Ascomycota(100);c__Sordariomycetes(100);o__Xylariales(100);f__Diatrypaceae(100);g__Diatrype(100);g__Diatrype_unclassified(100); | Diatrype | 13 | Pathotroph | \|Plant Pathogen\| | Microfungus | NULL | Probable | NULL | [illegible] |
| Otu02922 | 13 | k__Fungi(100);p__Basidiomycota(100);c__Agaricomycetes(100);o__Hymenochaetales(100);f__Hymenochaetaceae(100);g__Hymenochaete(100);g__Hymenochaete_unclassified(100); | Hymenochaete | 13 | Pathotroph-Saprotroph | Plant Pathogen-Undefined Saprotroph-\|Wood Saprotroph\| | Corticioid-Polyporoid | White Rot | Probable | [illegible] | [illegible] |
| Otu02925 | 13 | k__Fungi(100);p__Ascomycota(100);c__Saccharomycetes(100);o__Saccharomycetales(100);f__Metschnikowiaceae(100);g__Metschnikowia(100);s__Metschnikowia_pulcherrima(93); | Metschnikowia | 13 | Saprotroph | \|Undefined Saprotroph\| | Yeast | NULL | Probable | [illegible] | [illegible] |
| Otu02929 | 13 | k__Fungi(100);p__Ascomycota(100);c__Saccharomycetes(100);o__Saccharomycetales(100);f__Metschnikowiaceae(100);g__Metschnikowia(100);s__Metschnikowia_pulcherrima(100); | Metschnikowia | 13 | Saprotroph | \|Undefined Saprotroph\| | Yeast | NULL | Probable | [illegible] | [illegible] |
| Otu02930 | 13 | k__Fungi(100);p__Ascomycota(100);c__Sordariomycetes(100);o__Hypocreales(100);f__Clavicipitaceae(100);g__unclassified_Clavicipitaceae(100);s__Clavicipitaceae_sp(100); | Clavicipitaceae | 9 | Pathotroph-Symbiotroph | Endophyte-Epiphyte-Fungal Parasite-Insect Parasite | Microfungus | NULL | Possible | [illegible] | [illegible] |
| Otu02931 | 13 | k__Fungi(100);p__Ascomycota(100);c__Eurotiomycetes(100);o__Eurotiales(100);f__Trichocomaceae(100);g__Penicillium(100);s__Penicillium_sumatrense(100); | Penicillium | 13 | Pathotroph-Saprotroph-Symbiotroph | Animal Parasite-Dung Saprotroph-Endophyte-Fungal Parasite-Plant Pathogen-\|Undefined Saprotroph\|-Wood Saprotroph | Microfungus | Soft Rot | Highly Probable | [illegible] | [illegible] |
| Otu02933 | 13 | k__Fungi(100);p__Ascomycota(100);c__Saccharomycetes(100);o__Saccharomycetales(100);f__Metschnikowiaceae(100);g__Metschnikowia(100);s__Metschnikowia_pulcherrima(93); | Metschnikowia | 13 | Saprotroph | \|Undefined Saprotroph\| | Yeast | NULL | Probable | [illegible] | [illegible] |
| Otu02935 | 13 | k__Fungi(100);p__Ascomycota(100);c__Saccharomycetes(100);o__Saccharomycetales(100);f__Metschnikowiaceae(100);g__Metschnikowia(100);s__Metschnikowia_pulcherrima(100); | Metschnikowia | 13 | Saprotroph | \|Undefined Saprotroph\| | Yeast | NULL | Probable | [illegible] | [illegible] |
| Otu02938 | 13 | k__Fungi(100);p__Ascomycota(100);c__Eurotiomycetes(100);o__Verrucariales(100);f__Verrucariales_family_Incertae_sedis(100);g__Agonimia(100);s__Agonimia_sp_BGK_2011a(100); | Agonimia | 13 | Symbiotroph | \|Lichenized\| | Microfungus-Thallus | NULL | Highly Probable | NULL | [illegible] |
| Otu02939 | 13 | k__Fungi(100);p__Ascomycota(100);c__Leotiomycetes(100);o__Helotiales(100);f__Vibrisseaceae(62);g__Phialocephala(54);g__Phialocephala_unclassified(54); | Phialocephala | 13 | Saprotroph-Symbiotroph | Endophyte-\|Undefined Saprotroph\| | Microfungus | NULL | Highly Probable | [illegible] | [illegible] |
| Otu02941 | 13 | k__Fungi(100);p__Ascomycota(100);c__Saccharomycetes(100);o__Saccharomycetales(100);f__Metschnikowiaceae(100);g__Metschnikowia(100);s__Metschnikowia_pulcherrima(85); | Metschnikowia | 13 | Saprotroph | \|Undefined Saprotroph\| | Yeast | NULL | Probable | [illegible] | [illegible] |
| Otu02942 | 13 | k__Fungi(100);p__Ascomycota(100);c__Saccharomycetes(100);o__Saccharomycetales(100);f__Metschnikowiaceae(100);g__Metschnikowia(100);s__Metschnikowia_pulcherrima(62); | Metschnikowia | 13 | Saprotroph | \|Undefined Saprotroph\| | Yeast | NULL | Probable | [illegible] | [illegible] |
| Otu02943 | 13 | k__Fungi(100);p__Ascomycota(100);c__Sordariomycetes(100);o__Microascales(100);f__Halosphaeriaceae(100);g__unclassified_Halosphaeriaceae(100);s__Halosphaeriaceae_sp(100); | Halosphaeriaceae | 9 | Saprotroph | Undefined Saprotroph-Wood Saprotroph | Microfungus | NULL | Probable | [illegible] | [illegible] |
| Otu02944 | 13 | k__Fungi(100);p__Ascomycota(100);c__Sordariomycetes(100);o__Xylariales(100);f__Diatrypaceae(100);f__Diatrypaceae_unclassified(100);f__Diatrypaceae_unclassified(100); | Diatrypaceae | 9 | Pathotroph-Saprotroph | Plant Pathogen-Plant Saprotroph | Microfungus | NULL | Probable | [illegible] | [illegible] |
| Otu02946 | 13 | k__Fungi(100);p__Ascomycota(100);c__Saccharomycetes(100);o__Saccharomycetales(100);f__Metschnikowiaceae(100);g__Metschnikowia(100);g__Metschnikowia_unclassified(85); | Metschnikowia | 13 | Saprotroph | \|Undefined Saprotroph\| | Yeast | NULL | Probable | [illegible] | [illegible] |
| Otu02947 | 13 | k__Fungi(100);p__Basidiomycota(100);c__Agaricomycetes(100);o__Polyporales(100);f__Meruliaceae(100);g__Hypochnicium(100);s__Hypochnicium_geogenium(100); | Hypochnicium | 13 | Saprotroph | Undefined Saprotroph-\|Wood Saprotroph\| | Corticioid | White Rot | Probable | [illegible] | [illegible] |
| Otu02948 | 13 | k__Fungi(100);p__Ascomycota(100);c__Sordariomycetes(100);o__Xylariales(100);f__Graphostromataceae(100);g__Graphostroma(100);s__Graphostroma_platystoma(100); | Graphostroma | 13 | Pathotroph-Saprotroph-Symbiotroph | Endophyte-Plant Pathogen-Undefined Saprotroph-\|Wood Saprotroph\| | Microfungus | NULL | Probable | [illegible] | [illegible] |
| Otu02949 | 13 | k__Fungi(100);p__Ascomycota(100);c__Dothideomycetes(100);o__Capnodiales(100);f__Mycosphaerellaceae(100);g__Mycosphaerella(93);s__Mycosphaerella_nyssicola(77); | Mycosphaerella | 13 | Pathotroph-Saprotroph | \|Plant Pathogen\|-Plant Saprotroph | Microfungus | NULL | Probable | [illegible] | [illegible] |
| Otu02951 | 13 | k__Fungi(100);p__Ascomycota(100);c__Saccharomycetes(100);o__Saccharomycetales(100);f__Metschnikowiaceae(100);g__Metschnikowia(100);s__Metschnikowia_pulcherrima(100); | Metschnikowia | 13 | Saprotroph | \|Undefined Saprotroph\| | Yeast | NULL | Probable | [illegible] | [illegible] |
| Otu02953 | 13 | k__Fungi(100);p__Ascomycota(100);c__Saccharomycetes(100);o__Saccharomycetales(100);f__Saccharomycetales_family_Incertae_sedis(100);g__Candida(100);s__Candida_mycetangii(100); | Candida | 13 | Pathotroph-Saprotroph-Symbiotroph | Animal Pathogen-Endophyte-Endosymbiont-Epiphyte-\|Undefined Saprotroph\| | Dimorphic Yeast-Yeast | NULL | Probable | [illegible] | [illegible] |
| Otu02955 | 13 | k__Fungi(100);p__Basidiomycota(100);c__Agaricomycetes(100);o__Agaricales(100);f__Entolomataceae(100);g__Clitopilus(100);s__Clitopilus_cystidiatus(100); | Clitopilus | 13 | Pathotroph-Saprotroph | Fungal Parasite-\|Plant Saprotroph\|-Undefined Saprotroph | Agaricoid | NULL | Probable | [illegible] | [illegible] |
| Otu02956 | 13 | k__Fungi(100);p__Basidiomycota(100);c__Agaricomycetes(100);o__Trechisporales(100);f__Hydnodontaceae(100);g__Trechispora(100);g__Trechispora_unclassified(100); | Trechispora | 13 | Saprotroph | \|Wood Saprotroph\| | Corticioid | White Rot | Highly Probable | [illegible] | [illegible] |
| Otu02958 | 13 | k__Fungi(100);p__Ascomycota(100);c__Saccharomycetes(100);o__Saccharomycetales(100);f__Metschnikowiaceae(100);g__Metschnikowia(100);s__Metschnikowia_pulcherrima(85); | Metschnikowia | 13 | Saprotroph | \|Undefined Saprotroph\| | Yeast | NULL | Probable | [illegible] | [illegible] |
| Otu02960 | 13 | k__Fungi(100);p__Ascomycota(100);c__Sordariomycetes(100);o__Lulworthiales(100);f__Lulworthiaceae(100);g__Zalerion(100);s__Zalerion_arboricola(100); | Zalerion | 13 | Pathotroph-Saprotroph | Plant Pathogen-\|Plant Saprotroph\|-Undefined Saprotroph | Microfungus | NULL | Probable | [illegible] | [illegible] |
| Otu02961 | 13 | k__Fungi(100);p__Ascomycota(100);c__Leotiomycetes(100);o__Helotiales(100);f__Hyaloscyphaceae(100);g__Cistella(100);s__Cistella_sp_KUS_F52527(100); | Cistella | 13 | Saprotroph-Symbiotroph | Endophyte-Plant Saprotroph-Undefined Saprotroph-\|Wood Saprotroph\| | Microfungus | NULL | Probable | [illegible] | [illegible] |
| Otu02965 | 13 | k__Fungi(100);p__Basidiomycota(100);c__Microbotryomycetes(100);o__Sporidiobolales(100);f__Sporidiobolales_family_Incertae_sedis(100);g__Sporobolomyces(100);s__Sporobolomyces_lactosus(100); | Sporobolomyces | 13 | Pathotroph-Saprotroph | \|Fungal Parasite\|-Plant Saprotroph | Yeast | NULL | Probable | [illegible] | [illegible] |
| Otu02966 | 13 | k__Fungi(100);p__Ascomycota(100);c__Eurotiomycetes(100);o__Eurotiales(100);o__Eurotiales_unclassified(100);o__Eurotiales_unclassified(100);o__Eurotiales_unclassified(100); | Eurotiales | 7 | Saprotroph | Undefined Saprotroph | Microfungus | NULL | Possible | NULL | [illegible] |
| Otu02969 | 13 | k__Fungi(100);p__Basidiomycota(100);c__Agaricomycetes(100);o__Agaricales(100);f__Marasmiaceae(100);g__Rhodocollybia(100);s__Rhodocollybia_maculata(100); | Rhodocollybia | 13 | Saprotroph | \|Plant Saprotroph\|-Undefined Saprotroph | Agaricoid | NULL | Probable | [illegible] | [illegible] |
| Otu02970 | 13 | k__Fungi(100);p__Basidiomycota(100);c__Pucciniomycetes(100);o__Septobasidiales(100);f__Septobasidiaceae(100);g__Septobasidium(100);s__Septobasidium_sp_JBO_2010i(100); | Septobasidium | 13 | Pathotroph | \|Animal Parasite\|-Animal Pathogen-Arthropod Parasite | Dimorphic-Facultative Yeast | NULL | Probable | [illegible] | [illegible] |
| Otu02971 | 13 | k__Fungi(100);p__Basidiomycota(100);c__Agaricomycetes(100);o__Agaricales(100);f__Psathyrellaceae(100);g__Psathyrella(100);s__Psathyrella_typhae(100); | Psathyrella | 13 | Saprotroph | Plant Saprotroph-\|Wood Saprotroph\| | Agaricoid | NULL | Probable | [illegible] | [illegible] |
| Otu02972 | 13 | k__Fungi(100);p__Basidiomycota(100);c__Tremellomycetes(100);o__Tremellales(100);f__Tremellales_family_Incertae_sedis(100);g__Cryptococcus(100);s__Cryptococcus_aff_gilvescens_IMUFRJ_51978(100); | Cryptococcus | 13 | Pathotroph-Saprotroph-Symbiotroph | Animal Pathogen-Endophyte-Epiphyte-\|Undefined Saprotroph\| | Dimorphic Yeast-Yeast | NULL | Possible | [illegible] | [illegible] |
| Otu02973 | 13 | k__Fungi(100);p__Basidiomycota(100);c__Agaricomycetes(100);o__Agaricales(100);f__Marasmiaceae(100);g__Omphalotus(100);s__Omphalotus_illudens(100); | Omphalotus | 13 | Saprotroph | \|Wood Saprotroph\| | Agaricoid | White Rot | Probable | [illegible] | [illegible] |
| Otu02975 | 13 | k__Fungi(100);p__Basidiomycota(100);c__Agaricomycetes(100);o__Cantharellales(100);f__Ceratobasidiaceae(100);g__Thanatephorus(100);s__Thanatephorus_cucumeris(100); | Thanatephorus | 13 | Pathotroph-Saprotroph | \|Plant Pathogen\|-Plant Saprotroph | Corticioid | NULL | Probable | [illegible] | [illegible] |
| Otu02977 | 13 | k__Fungi(100);p__Ascomycota(100);c__Sordariomycetes(100);o__Sordariomycetes_order_Incertae_sedis(100);f__Annulatascaceae(100);g__Conlarium(100);s__Conlarium_duplumascospora(100); | Conlarium | 13 | Saprotroph | Undefined Saprotroph-\|Wood Saprotroph\| | Microfungus | NULL | Probable | [illegible] | [illegible] |
| Otu02978 | 13 | k__Fungi(100);p__Ascomycota(100);c__Saccharomycetes(100);o__Saccharomycetales(100);f__Metschnikowiaceae(100);g__Metschnikowia(100);s__Metschnikowia_pulcherrima(100); | Metschnikowia | 13 | Saprotroph | \|Undefined Saprotroph\| | Yeast | NULL | Probable | [illegible] | [illegible] |
| Otu02980 | 13 | k__Fungi(100);p__Basidiomycota(100);c__Tremellomycetes(100);o__Tremellales(100);f__Tremellales_family_Incertae_sedis(100);g__Cryptococcus(100);s__Cryptococcus_fagi(100); | Cryptococcus | 13 | Pathotroph-Saprotroph-Symbiotroph | Animal Pathogen-Endophyte-Epiphyte-\|Undefined Saprotroph\| | Dimorphic Yeast-Yeast | NULL | Possible | [illegible] | [illegible] |
| Otu02981 | 13 | k__Fungi(100);p__Basidiomycota(100);c__Agaricomycetes(100);o__Agaricales(100);f__Strophariaceae(100);g__Hypholoma(100);s__Hypholoma_sp(100); | Hypholoma | 13 | Saprotroph | Undefined Saprotroph-\|Wood Saprotroph\| | Agaricoid | White Rot | Probable | [illegible] | [illegible] |
| Otu02982 | 13 | k__Fungi(100);p__Ascomycota(100);c__Saccharomycetes(100);o__Saccharomycetales(100);f__Metschnikowiaceae(100);g__Metschnikowia(100);s__Metschnikowia_pulcherrima(93); | Metschnikowia | 13 | Saprotroph | \|Undefined Saprotroph\| | Yeast | NULL | Probable | [illegible] | [illegible] |
| Otu02983 | 13 | k__Fungi(100);p__Ascomycota(100);c__Lecanoromycetes(100);o__Lecanorales(100);f__Cladoniaceae(100);g__Cladonia(100);s__Cladonia_novochlorophaea(100); | Cladonia | 13 | Symbiotroph | \|Lichenized\| | Thallus | NULL | Highly Probable | NULL | [illegible] |
| Otu02985 | 13 | k__Fungi(100);p__Basidiomycota(100);c__Agaricostilbomycetes(100);o__Agaricostilbales(100);f__Agaricostilbaceae(100);g__Bensingtonia(100);g__Bensingtonia_unclassified(100); | Bensingtonia | 13 | Saprotroph | \|Undefined Saprotroph\| | Dimorphic-Facultative Yeast | NULL | Highly Probable | NULL | [illegible] |
| Otu02986 | 13 | k__Fungi(100);p__Basidiomycota(100);c__Agaricomycetes(100);o__Hymenochaetales(100);f__Schizoporaceae(100);g__Hyphodontia(100);s__Hyphodontia_radula(85); | Hyphodontia | 13 | Saprotroph | Undefined Saprotroph-\|Wood Saprotroph\| | Corticioid | White Rot | Probable | [illegible] | [illegible] |
| Otu02987 | 13 | k__Fungi(100);p__Ascomycota(100);c__Saccharomycetes(100);o__Saccharomycetales(100);f__Saccharomycetales_family_Incertae_sedis(100);g__Candida(85);g__Candida_unclassified(62); | Candida | 13 | Pathotroph-Saprotroph-Symbiotroph | Animal Pathogen-Endophyte-Endosymbiont-Epiphyte-\|Undefined Saprotroph\| | Dimorphic Yeast-Yeast | NULL | Probable | [illegible] | [illegible] |
| Otu02989 | 13 | k__Fungi(100);p__Ascomycota(100);c__Saccharomycetes(100);o__Saccharomycetales(100);f__Metschnikowiaceae(100);g__Metschnikowia(100);g__Metschnikowia_unclassified(85); | Metschnikowia | 13 | Saprotroph | \|Undefined Saprotroph\| | Yeast | NULL | Probable | [illegible] | [illegible] |
| Otu02990 | 13 | k__Fungi(100);p__Ascomycota(100);c__Dothideomycetes(85);o__Dothideomycetes_order_Incertae_sedis(85);f__Pseudeurotiaceae(85);g__Pseudogymnoascus(85);g__Pseudogymnoascus_unclassified(70); | Pseudogymnoascus | 13 | Pathotroph-Saprotroph-Symbiotroph | Animal Pathogen-\|Undefined Saprotroph\| | Microfungus | NULL | Possible | [illegible] | [illegible] |
| Otu02992 | 13 | k__Fungi(100);p__Ascomycota(100);c__Saccharomycetes(100);o__Saccharomycetales(100);f__Metschnikowiaceae(100);g__Metschnikowia(100);s__Metschnikowia_pulcherrima(100); | Metschnikowia | 13 | Saprotroph | \|Undefined Saprotroph\| | Yeast | NULL | Probable | [illegible] | [illegible] |
| Otu02993 | 13 | k__Fungi(100);p__Basidiomycota(100);c__Tremellomycetes(100);o__Tremellales(100);f__Tremellales_family_Incertae_sedis(100);g__Auriculibuller(100);s__Auriculibuller_fuscus(100); | Auriculibuller | 13 | Saprotroph | \|Undefined Saprotroph\| | Yeast | NULL | Probable | NULL | [illegible] |
| Otu02996 | 13 | k__Fungi(100);p__Ascomycota(100);c__Saccharomycetes(100);o__Saccharomycetales(100);f__Metschnikowiaceae(100);g__Metschnikowia(100);s__Metschnikowia_pulcherrima(85); | Metschnikowia | 13 | Saprotroph | \|Undefined Saprotroph\| | Yeast | NULL | Probable | [illegible] | [illegible] |
| Otu02997 | 13 | k__Fungi(100);p__Zygomycota(100);c__Zygomycota_class_Incertae_sedis(100);o__Mortierellales(100);f__Mortierellaceae(100);g__Mortierella(100);s__Mortierella_fimbricystis(100); | Mortierella | 13 | Saprotroph-Symbiotroph | Endophyte-Plant Saprotroph-\|Undefined Saprotroph\| | Microfungus | NULL | Probable | [illegible] | [illegible] |
| Otu02998 | 13 | k__Fungi(100);p__Ascomycota(100);c__Saccharomycetes(100);o__Saccharomycetales(100);f__Metschnikowiaceae(100);g__Metschnikowia(100);s__Metschnikowia_pulcherrima(100); | Metschnikowia | 13 | Saprotroph | \|Undefined Saprotroph\| | Yeast | NULL | Probable | [illegible] | [illegible] |
| Otu03003 | 13 | k__Fungi(100);p__Ascomycota(100);c__Sordariomycetes(100);o__Sordariomycetes_order_Incertae_sedis(100);f__Annulatascaceae(100);g__Rhodoveronaea(100);s__Rhodoveronaea_varioseptata(100); | Rhodoveronaea | 13 | Saprotroph | \|Undefined Saprotroph\| | Microfungus | NULL | Probable | NULL | [illegible] |
| Otu03006 | 13 | k__Fungi(100);p__Basidiomycota(100);c__Agaricomycetes(100);o__Trechisporales(100);f__Hydnodontaceae(100);g__Trechispora(100);g__Trechispora_unclassified(100); | Trechispora | 13 | Saprotroph | \|Wood Saprotroph\| | Corticioid | White Rot | Highly Probable | [illegible] | [illegible] |
| Otu03007 | 13 | k__Fungi(100);p__Basidiomycota(100);c__Agaricomycetes(100);o__Russulales(100);f__Peniophoraceae(100);g__Gloiothele(100);s__Gloiothele_citrina(100); | Gloiothele | 13 | Saprotroph | Undefined Saprotroph-\|Wood Saprotroph\| | Corticioid | White Rot | Probable | [illegible] | [illegible] |
| Otu03010 | 13 | k__Fungi(100);p__Chytridiomycota(70);c__Chytridiomycetes(70);o__Rhizophlyctidales(70);f__Rhizophlyctidaceae(70);g__Rhizophlyctis(70);s__Rhizophlyctis_rosea(70); | Rhizophlyctis | 13 | Pathotroph-Saprotroph | Algal Parasite-\|Plant Saprotroph\|-Undefined Saprotroph | Chytrid-Microfungus | NULL | Probable | [illegible] | [illegible] |
| Otu03012 | 13 | k__Fungi(100);p__Ascomycota(100);c__Eurotiomycetes(100);o__Eurotiales(100);f__Trichocomaceae(100);g__Aspergillus(100);s__Aspergillus_sclerotiorum(77); | Aspergillus | 13 | Pathotroph-Saprotroph-Symbiotroph | Animal Pathogen-Endophyte-Plant Saprotroph-\|Undefined Saprotroph\|-Wood Saprotroph | Microfungus | NULL | Probable | [illegible] | [illegible] |
| Otu03013 | 13 | k__Fungi(100);p__Ascomycota(100);c__Saccharomycetes(100);o__Saccharomycetales(100);f__Metschnikowiaceae(100);g__Metschnikowia(100);g__Metschnikowia_unclassified(85); | Metschnikowia | 13 | Saprotroph | \|Undefined Saprotroph\| | Yeast | NULL | Probable | [illegible] | [illegible] |
| Otu03014 | 13 | k__Fungi(100);p__Basidiomycota(100);c__Agaricomycetes(100);o__Polyporales(100);f__Meruliaceae(100);g__Hyphoderma(100);s__Hyphoderma_setigerum(100); | Hyphoderma | 13 | Saprotroph | Undefined Saprotroph-\|Wood Saprotroph\| | Corticioid | White Rot | Probable | [illegible] | [illegible] |
| Otu03015 | 13 | k__Fungi(100);p__Ascomycota(100);c__Saccharomycetes(100);o__Saccharomycetales(100);f__Metschnikowiaceae(100);g__Metschnikowia(100);s__Metschnikowia_pulcherrima(100); | Metschnikowia | 13 | Saprotroph | \|Undefined Saprotroph\| | Yeast | NULL | Probable | [illegible] | [illegible] |
| Otu03016 | 13 | k__Fungi(100);p__Basidiomycota(100);c__Agaricomycetes(100);o__Polyporales(100);f__Meruliaceae(100);g__Hypochnicium(100);s__Hypochnicium_lundellii(100); | Hypochnicium | 13 | Saprotroph | Undefined Saprotroph-\|Wood Saprotroph\| | Corticioid | White Rot | Probable | [illegible] | [illegible] |
| Otu03019 | 13 | k__Fungi(100);p__Ascomycota(100);c__Saccharomycetes(100);o__Saccharomycetales(100);f__Metschnikowiaceae(100);g__Metschnikowia(100);s__Metschnikowia_pulcherrima(93); | Metschnikowia | 13 | Saprotroph | \|Undefined Saprotroph\| | Yeast | NULL | Probable | [illegible] | [illegible] |
| Otu02854 | 13 | k__Fungi(100);p__Basidiomycota(100);c__Agaricomycetes(100);o__Sebacinales(100);o__Sebacinales_unclassified(100);o__Sebacinales_unclassified(100);o__Sebacinales_unclassified(100); | - | - | - | - | - | - | - | Unassigned | - |
| Otu02858 | 13 | k__Fungi(100);p__Ascomycota(54);p__Ascomycota_unclassified(54);p__Ascomycota_unclassified(54);p__Ascomycota_unclassified(54);p__Ascomycota_unclassified(54);p__Ascomycota_unclassified(54); | - | - | - | - | - | - | - | Unassigned | - |
| Otu02859 | 13 | k__Fungi(100);k__Fungi_unclassified(100);k__Fungi_unclassified(100);k__Fungi_unclassified(100);k__Fungi_unclassified(100);k__Fungi_unclassified(100);k__Fungi_unclassified(100); | - | - | - | - | - | - | - | Unassigned | - |
| Otu02862 | 13 | k__Fungi(100);p__Ascomycota(70);p__Ascomycota_unclassified(70);p__Ascomycota_unclassified(70);p__Ascomycota_unclassified(70);p__Ascomycota_unclassified(70);p__Ascomycota_unclassified(70); | - | - | - | - | - | - | - | Unassigned | - |
| Otu02863 | 13 | k__Fungi(100);k__Fungi_unclassified(77);k__Fungi_unclassified(77);k__Fungi_unclassified(77);k__Fungi_unclassified(77);k__Fungi_unclassified(77);k__Fungi_unclassified(77); | - | - | - | - | - | - | - | Unassigned | - |
| Otu02864 | 13 | k__Fungi(100);k__Fungi_unclassified(77);k__Fungi_unclassified(77);k__Fungi_unclassified(77);k__Fungi_unclassified(77);k__Fungi_unclassified(77);k__Fungi_unclassified(77); | - | - | - | - | - | - | - | Unassigned | - |
| Otu02866 | 13 | k__Fungi(100);p__Ascomycota(62);p__Ascomycota_unclassified(62);p__Ascomycota_unclassified(62);p__Ascomycota_unclassified(62);p__Ascomycota_unclassified(62);p__Ascomycota_unclassified(62); | - | - | - | - | - | - | - | Unassigned | - |
| Otu02870 | 13 | k__Fungi(100);p__Basidiomycota(100);c__Agaricomycetes(100);c__Agaricomycetes_unclassified(100);c__Agaricomycetes_unclassified(100);c__Agaricomycetes_unclassified(100);c__Agaricomycetes_unclassified(100); | - | - | - | - | - | - | - | Unassigned | - |
| Otu02871 | 13 | k__Fungi(100);k__Fungi_unclassified(62);k__Fungi_unclassified(62);k__Fungi_unclassified(62);k__Fungi_unclassified(62);k__Fungi_unclassified(62);k__Fungi_unclassified(62); | - | - | - | - | - | - | - | Unassigned | - |
| Otu02873 | 13 | k__Fungi(100);p__Ascomycota(100);c__Sordariomycetes(100);c__Sordariomycetes_unclassified(100);c__Sordariomycetes_unclassified(100);c__Sordariomycetes_unclassified(100);c__Sordariomycetes_unclassified(100); | - | - | - | - | - | - | - | Unassigned | - |
| Otu02874 | 13 | k__Fungi(100);k__Fungi_unclassified(93);k__Fungi_unclassified(93);k__Fungi_unclassified(93);k__Fungi_unclassified(93);k__Fungi_unclassified(93);k__Fungi_unclassified(93); | - | - | - | - | - | - | - | Unassigned | - |
| Otu02877 | 13 | k__Fungi(100);p__Ascomycota(100);c__Sordariomycetes(100);c__Sordariomycetes_unclassified(100);c__Sordariomycetes_unclassified(100);c__Sordariomycetes_unclassified(100);c__Sordariomycetes_unclassified(100); | - | - | - | - | - | - | - | Unassigned | - |
| Otu02878 | 13 | k__Fungi(100);p__Ascomycota(100);p__Ascomycota_unclassified(100);p__Ascomycota_unclassified(100);p__Ascomycota_unclassified(100);p__Ascomycota_unclassified(100);p__Ascomycota_unclassified(100); | - | - | - | - | - | - | - | Unassigned | - |
| Otu02880 | 13 | k__Fungi(100);k__Fungi_unclassified(54);k__Fungi_unclassified(54);k__Fungi_unclassified(54);k__Fungi_unclassified(54);k__Fungi_unclassified(54);k__Fungi_unclassified(54); | - | - | - | - | - | - | - | Unassigned | - |
| Otu02882 | 13 | k__Fungi(100);k__Fungi_unclassified(70);k__Fungi_unclassified(70);k__Fungi_unclassified(70);k__Fungi_unclassified(70);k__Fungi_unclassified(70);k__Fungi_unclassified(70); | - | - | - | - | - | - | - | Unassigned | - |
| Otu02883 | 13 | k__Fungi(100);p__Ascomycota(54);p__Ascomycota_unclassified(54);p__Ascomycota_unclassified(54);p__Ascomycota_unclassified(54);p__Ascomycota_unclassified(54);p__Ascomycota_unclassified(54); | - | - | - | - | - | - | - | Unassigned | - |
| Otu02884 | 13 | k__Fungi(100);p__Ascomycota(100);p__Ascomycota_unclassified(70);p__Ascomycota_unclassified(70);p__Ascomycota_unclassified(70);p__Ascomycota_unclassified(70);p__Ascomycota_unclassified(70); | - | - | - | - | - | - | - | Unassigned | - |
| Otu02887 | 13 | k__Fungi(100);k__Fungi_unclassified(85);k__Fungi_unclassified(85);k__Fungi_unclassified(85);k__Fungi_unclassified(85);k__Fungi_unclassified(85);k__Fungi_unclassified(85); | - | - | - | - | - | - | - | Unassigned | - |
| Otu02888 | 13 | k__Fungi(100);p__Ascomycota(54);p__Ascomycota_unclassified(54);p__Ascomycota_unclassified(54);p__Ascomycota_unclassified(54);p__Ascomycota_unclassified(54);p__Ascomycota_unclassified(54); | - | - | - | - | - | - | - | Unassigned | - |
| Otu02889 | 13 | k__Fungi(100);p__Ascomycota(77);p__Ascomycota_unclassified(77);p__Ascomycota_unclassified(77);p__Ascomycota_unclassified(77);p__Ascomycota_unclassified(77);p__Ascomycota_unclassified(77); | - | - | - | - | - | - | - | Unassigned | - |

| | | | | | | | | | | | |
|---|---|---|---|---|---|---|---|---|---|---|---|
| Otu02891 | 13 | k__Fungi(100);p__Ascomycota(77);p__Ascomycota_unclassified(77);p__Ascomycota_unclassified(77);p__Ascomycota_unclassified(77);p__Ascomycota_unclassified(77);p__Ascomycota_unclassified(77); | - | - | - | - | - | - | - | Unassigned | - |
| Otu02893 | 13 | k__Fungi(100);k__Fungi_unclassified(62);k__Fungi_unclassified(62);k__Fungi_unclassified(62);k__Fungi_unclassified(62);k__Fungi_unclassified(62);k__Fungi_unclassified(62); | - | - | - | - | - | - | - | Unassigned | - |
| Otu02894 | 13 | k__Fungi(100);p__Ascomycota(77);p__Ascomycota_unclassified(77);p__Ascomycota_unclassified(77);p__Ascomycota_unclassified(77);p__Ascomycota_unclassified(77);p__Ascomycota_unclassified(77); | - | - | - | - | - | - | - | Unassigned | - |
| Otu02896 | 13 | k__Fungi(100);p__Ascomycota(93);p__Ascomycota_unclassified(93);p__Ascomycota_unclassified(93);p__Ascomycota_unclassified(93);p__Ascomycota_unclassified(93);p__Ascomycota_unclassified(93); | - | - | - | - | - | - | - | Unassigned | - |
| Otu02897 | 13 | k__Fungi(100);p__Ascomycota(100);p__Ascomycota_unclassified(100);p__Ascomycota_unclassified(100);p__Ascomycota_unclassified(100);p__Ascomycota_unclassified(100);p__Ascomycota_unclassified(100); | - | - | - | - | - | - | - | Unassigned | - |
| Otu02899 | 13 | k__Fungi(100);p__Ascomycota(62);p__Ascomycota_unclassified(62);p__Ascomycota_unclassified(62);p__Ascomycota_unclassified(62);p__Ascomycota_unclassified(62);p__Ascomycota_unclassified(62); | - | - | - | - | - | - | - | Unassigned | - |
| Otu02900 | 13 | k__Fungi(100);k__Fungi_unclassified(62);k__Fungi_unclassified(62);k__Fungi_unclassified(62);k__Fungi_unclassified(62);k__Fungi_unclassified(62);k__Fungi_unclassified(62); | - | - | - | - | - | - | - | Unassigned | - |
| Otu02901 | 13 | k__Fungi(100);p__Basidiomycota(100);c__Ustilaginomycetes(100);o__Ustilaginales(100);f__Ustilaginales_family_Incertae_sedis(100);g__Farysizyma(100);s__Farysizyma_itapuensis(100); | - | - | - | - | - | - | - | Unassigned | - |
| Otu02902 | 13 | k__Fungi(100);p__Ascomycota(62);p__Ascomycota_unclassified(62);p__Ascomycota_unclassified(62);p__Ascomycota_unclassified(62);p__Ascomycota_unclassified(62);p__Ascomycota_unclassified(62); | - | - | - | - | - | - | - | Unassigned | - |
| Otu02906 | 13 | k__Fungi(100);k__Fungi_unclassified(70);k__Fungi_unclassified(70);k__Fungi_unclassified(70);k__Fungi_unclassified(70);k__Fungi_unclassified(70);k__Fungi_unclassified(70); | - | - | - | - | - | - | - | Unassigned | - |
| Otu02910 | 13 | k__Fungi(100);p__Basidiomycota(100);c__Agaricomycetes(100);c__Agaricomycetes_unclassified(100);c__Agaricomycetes_unclassified(100);c__Agaricomycetes_unclassified(100);c__Agaricomycetes_unclassified(100); | - | - | - | - | - | - | - | Unassigned | - |
| Otu02911 | 13 | k__Fungi(100);p__Ascomycota(54);p__Ascomycota_unclassified(54);p__Ascomycota_unclassified(54);p__Ascomycota_unclassified(54);p__Ascomycota_unclassified(54);p__Ascomycota_unclassified(54); | - | - | - | - | - | - | - | Unassigned | - |
| Otu02912 | 13 | k__Fungi(100);p__Basidiomycota(100);p__Basidiomycota_unclassified(100);p__Basidiomycota_unclassified(100);p__Basidiomycota_unclassified(100);p__Basidiomycota_unclassified(100);p__Basidiomycota_unclassified(100); | - | - | - | - | - | - | - | Unassigned | - |
| Otu02914 | 13 | k__Fungi(100);p__Ascomycota(100);p__Ascomycota_unclassified(100);p__Ascomycota_unclassified(100);p__Ascomycota_unclassified(100);p__Ascomycota_unclassified(100);p__Ascomycota_unclassified(100); | - | - | - | - | - | - | - | Unassigned | - |
| Otu02918 | 13 | k__Fungi(100);k__Fungi_unclassified(62);k__Fungi_unclassified(62);k__Fungi_unclassified(62);k__Fungi_unclassified(62);k__Fungi_unclassified(62);k__Fungi_unclassified(62); | - | - | - | - | - | - | - | Unassigned | - |
| Otu02921 | 13 | k__Fungi(100);p__Ascomycota(100);c__Sordariomycetes(93);c__Sordariomycetes_unclassified(93);c__Sordariomycetes_unclassified(93);c__Sordariomycetes_unclassified(93);c__Sordariomycetes_unclassified(93); | - | - | - | - | - | - | - | Unassigned | - |
| Otu02923 | 13 | k__Fungi(100);p__Ascomycota(100);p__Ascomycota_unclassified(100);p__Ascomycota_unclassified(100);p__Ascomycota_unclassified(100);p__Ascomycota_unclassified(100);p__Ascomycota_unclassified(100); | - | - | - | - | - | - | - | Unassigned | - |
| Otu02924 | 13 | k__Fungi(100);p__Ascomycota(100);c__Ascomycota_class_Incertae_sedis(100);o__Ascomycota_order_Incertae_sedis(100);f__Ascomycota_family_Incertae_sedis(100);f__Ascomycota_family_Incertae_sedis_unclassified(100);f__Ascomycota_family_Incertae_sedis_unclassified(100); | - | - | - | - | - | - | - | Unassigned | - |
| Otu02926 | 13 | k__Fungi(100);k__Fungi_unclassified(62);k__Fungi_unclassified(62);k__Fungi_unclassified(62);k__Fungi_unclassified(62);k__Fungi_unclassified(62);k__Fungi_unclassified(62); | - | - | - | - | - | - | - | Unassigned | - |
| Otu02927 | 13 | k__Fungi(100);p__Ascomycota(100);p__Ascomycota_unclassified(100);p__Ascomycota_unclassified(100);p__Ascomycota_unclassified(100);p__Ascomycota_unclassified(100);p__Ascomycota_unclassified(100); | - | - | - | - | - | - | - | Unassigned | - |
| Otu02928 | 13 | k__Fungi(100);p__Ascomycota(70);p__Ascomycota_unclassified(70);p__Ascomycota_unclassified(70);p__Ascomycota_unclassified(70);p__Ascomycota_unclassified(70);p__Ascomycota_unclassified(70); | - | - | - | - | - | - | - | Unassigned | - |
| Otu02932 | 13 | k__Fungi(100);k__Fungi_unclassified(100);k__Fungi_unclassified(100);k__Fungi_unclassified(100);k__Fungi_unclassified(100);k__Fungi_unclassified(100);k__Fungi_unclassified(100); | - | - | - | - | - | - | - | Unassigned | - |
| Otu02934 | 13 | k__Fungi(100);k__Fungi_unclassified(62);k__Fungi_unclassified(62);k__Fungi_unclassified(62);k__Fungi_unclassified(62);k__Fungi_unclassified(62);k__Fungi_unclassified(62); | - | - | - | - | - | - | - | Unassigned | - |
| Otu02936 | 13 | k__Fungi(100);p__Ascomycota(100);c__Lecanoromycetes(100);o__Teloschistales(85);o__Teloschistales_unclassified(85);o__Teloschistales_unclassified(85);o__Teloschistales_unclassified(85); | - | - | - | - | - | - | - | Unassigned | - |
| Otu02937 | 13 | k__Fungi(100);k__Fungi_unclassified(54);k__Fungi_unclassified(54);k__Fungi_unclassified(54);k__Fungi_unclassified(54);k__Fungi_unclassified(54);k__Fungi_unclassified(54); | - | - | - | - | - | - | - | Unassigned | - |
| Otu02940 | 13 | k__Fungi(100);k__Fungi_unclassified(62);k__Fungi_unclassified(62);k__Fungi_unclassified(62);k__Fungi_unclassified(62);k__Fungi_unclassified(62);k__Fungi_unclassified(62); | - | - | - | - | - | - | - | Unassigned | - |
| Otu02945 | 13 | k__Fungi(100);k__Fungi_unclassified(100);k__Fungi_unclassified(100);k__Fungi_unclassified(100);k__Fungi_unclassified(100);k__Fungi_unclassified(100);k__Fungi_unclassified(100); | - | - | - | - | - | - | - | Unassigned | - |
| Otu02950 | 13 | k__Fungi(100);p__Ascomycota(100);c__Dothideomycetes(100);c__Dothideomycetes_unclassified(100);c__Dothideomycetes_unclassified(100);c__Dothideomycetes_unclassified(100);c__Dothideomycetes_unclassified(100); | - | - | - | - | - | - | - | Unassigned | - |
| Otu02952 | 13 | k__Fungi(100);p__Ascomycota(100);p__Ascomycota_unclassified(100);p__Ascomycota_unclassified(100);p__Ascomycota_unclassified(100);p__Ascomycota_unclassified(100);p__Ascomycota_unclassified(100); | - | - | - | - | - | - | - | Unassigned | - |
| Otu02954 | 13 | k__Fungi(100);k__Fungi_unclassified(62);k__Fungi_unclassified(62);k__Fungi_unclassified(62);k__Fungi_unclassified(62);k__Fungi_unclassified(62);k__Fungi_unclassified(62); | - | - | - | - | - | - | - | Unassigned | - |
| Otu02957 | 13 | k__Fungi(100);p__Basidiomycota(100);c__Agaricomycetes(93);c__Agaricomycetes_unclassified(93);c__Agaricomycetes_unclassified(93);c__Agaricomycetes_unclassified(93);c__Agaricomycetes_unclassified(93); | - | - | - | - | - | - | - | Unassigned | - |
| Otu02959 | 13 | k__Fungi(100);k__Fungi_unclassified(77);k__Fungi_unclassified(77);k__Fungi_unclassified(77);k__Fungi_unclassified(77);k__Fungi_unclassified(77);k__Fungi_unclassified(77); | - | - | - | - | - | - | - | Unassigned | - |
| Otu02962 | 13 | k__Fungi(100);k__Fungi_unclassified(100);k__Fungi_unclassified(100);k__Fungi_unclassified(100);k__Fungi_unclassified(100);k__Fungi_unclassified(100);k__Fungi_unclassified(100); | - | - | - | - | - | - | - | Unassigned | - |
| Otu02963 | 13 | k__Fungi(100);p__Ascomycota(54);p__Ascomycota_unclassified(54);p__Ascomycota_unclassified(54);p__Ascomycota_unclassified(54);p__Ascomycota_unclassified(54);p__Ascomycota_unclassified(54); | - | - | - | - | - | - | - | Unassigned | - |
| Otu02964 | 13 | k__Fungi(100);k__Fungi_unclassified(70);k__Fungi_unclassified(70);k__Fungi_unclassified(70);k__Fungi_unclassified(70);k__Fungi_unclassified(70);k__Fungi_unclassified(70); | - | - | - | - | - | - | - | Unassigned | - |
| Otu02967 | 13 | k__Fungi(100);p__Ascomycota(100);p__Ascomycota_unclassified(100);p__Ascomycota_unclassified(100);p__Ascomycota_unclassified(100);p__Ascomycota_unclassified(100);p__Ascomycota_unclassified(100); | - | - | - | - | - | - | - | Unassigned | - |
| Otu02968 | 13 | k__Fungi(100);p__Ascomycota(100);p__Ascomycota_unclassified(100);p__Ascomycota_unclassified(100);p__Ascomycota_unclassified(100);p__Ascomycota_unclassified(100);p__Ascomycota_unclassified(100); | - | - | - | - | - | - | - | Unassigned | - |
| Otu02974 | 13 | k__Fungi(100);k__Fungi_unclassified(62);k__Fungi_unclassified(62);k__Fungi_unclassified(62);k__Fungi_unclassified(62);k__Fungi_unclassified(62);k__Fungi_unclassified(62); | - | - | - | - | - | - | - | Unassigned | - |
| Otu02976 | 13 | k__Fungi(100);p__Ascomycota(100);p__Ascomycota_unclassified(100);p__Ascomycota_unclassified(100);p__Ascomycota_unclassified(100);p__Ascomycota_unclassified(100);p__Ascomycota_unclassified(100); | - | - | - | - | - | - | - | Unassigned | - |
| Otu02979 | 13 | k__Fungi(100);k__Fungi_unclassified(70);k__Fungi_unclassified(70);k__Fungi_unclassified(70);k__Fungi_unclassified(70);k__Fungi_unclassified(70);k__Fungi_unclassified(70); | - | - | - | - | - | - | - | Unassigned | - |
| Otu02984 | 13 | k__Fungi(100);p__Ascomycota(100);p__Ascomycota_unclassified(85);p__Ascomycota_unclassified(85);p__Ascomycota_unclassified(85);p__Ascomycota_unclassified(85);p__Ascomycota_unclassified(85); | - | - | - | - | - | - | - | Unassigned | - |
| Otu02988 | 13 | k__Fungi(100);p__Basidiomycota(100);c__Agaricomycetes(100);o__Polyporales(100);o__Polyporales_unclassified(93);o__Polyporales_unclassified(93);o__Polyporales_unclassified(93); | - | - | - | - | - | - | - | Unassigned | - |
| Otu02991 | 13 | k__Fungi(100);p__Ascomycota(100);c__Leotiomycetes(100);o__Helotiales(100);o__Helotiales_unclassified(100);o__Helotiales_unclassified(100);o__Helotiales_unclassified(100); | - | - | - | - | - | - | - | Unassigned | - |
| Otu02994 | 13 | k__Fungi(100);p__Ascomycota(100);c__Sordariomycetes(100);c__Sordariomycetes_unclassified(100);c__Sordariomycetes_unclassified(100);c__Sordariomycetes_unclassified(100);c__Sordariomycetes_unclassified(100); | - | - | - | - | - | - | - | Unassigned | - |
| Otu02995 | 13 | k__Fungi(100);p__Ascomycota(100);c__Dothideomycetes(100);o__Pleosporales(100);o__Pleosporales_unclassified(100);o__Pleosporales_unclassified(100);o__Pleosporales_unclassified(100); | - | - | - | - | - | - | - | Unassigned | - |
| Otu02999 | 13 | k__Fungi(100);p__Ascomycota(100);c__Leotiomycetes(100);o__Helotiales(100);o__Helotiales_unclassified(54);o__Helotiales_unclassified(54);o__Helotiales_unclassified(54); | - | - | - | - | - | - | - | Unassigned | - |
| Otu03000 | 13 | k__Fungi(100);p__Basidiomycota(100);c__Atractiellomycetes(100);o__Atractiellales(100);f__unclassified_Atractiellales(100);g__unclassified_Atractiellales(100);s__Atractiellales_sp(100); | - | - | - | - | - | - | - | Unassigned | - |
| Otu03001 | 13 | k__Fungi(100);k__Fungi_unclassified(54);k__Fungi_unclassified(54);k__Fungi_unclassified(54);k__Fungi_unclassified(54);k__Fungi_unclassified(54);k__Fungi_unclassified(54); | - | - | - | - | - | - | - | Unassigned | - |
| Otu03002 | 13 | k__Fungi(100);k__Fungi_unclassified(54);k__Fungi_unclassified(54);k__Fungi_unclassified(54);k__Fungi_unclassified(54);k__Fungi_unclassified(54);k__Fungi_unclassified(54); | - | - | - | - | - | - | - | Unassigned | - |
| Otu03004 | 13 | k__Plantae(100);p__unclassified_Plantae(100);c__unclassified_Plantae(100);o__unclassified_Plantae(100);f__unclassified_Plantae(100);g__unclassified_Plantae(100);s__Plantae_sp(100); | - | - | - | - | - | - | - | Unassigned | - |
| Otu03005 | 13 | k__Fungi(100);k__Fungi_unclassified(62);k__Fungi_unclassified(62);k__Fungi_unclassified(62);k__Fungi_unclassified(62);k__Fungi_unclassified(62);k__Fungi_unclassified(62); | - | - | - | - | - | - | - | Unassigned | - |
| Otu03008 | 13 | k__Fungi(100);p__Ascomycota(100);p__Ascomycota_unclassified(100);p__Ascomycota_unclassified(100);p__Ascomycota_unclassified(100);p__Ascomycota_unclassified(100);p__Ascomycota_unclassified(100); | - | - | - | - | - | - | - | Unassigned | - |
| Otu03009 | 13 | k__Fungi(100);k__Fungi_unclassified(77);k__Fungi_unclassified(77);k__Fungi_unclassified(77);k__Fungi_unclassified(77);k__Fungi_unclassified(77);k__Fungi_unclassified(77); | - | - | - | - | - | - | - | Unassigned | - |
| Otu03011 | 13 | k__Fungi(100);p__Basidiomycota(100);c__Agaricomycetes(100);c__Agaricomycetes_unclassified(62);c__Agaricomycetes_unclassified(62);c__Agaricomycetes_unclassified(62);c__Agaricomycetes_unclassified(62); | - | - | - | - | - | - | - | Unassigned | - |
| Otu03017 | 13 | k__Fungi(100);k__Fungi_unclassified(70);k__Fungi_unclassified(70);k__Fungi_unclassified(70);k__Fungi_unclassified(70);k__Fungi_unclassified(70);k__Fungi_unclassified(70); | - | - | - | - | - | - | - | Unassigned | - |
| Otu03018 | 13 | k__Fungi(100);k__Fungi_unclassified(62);k__Fungi_unclassified(62);k__Fungi_unclassified(62);k__Fungi_unclassified(62);k__Fungi_unclassified(62);k__Fungi_unclassified(62); | - | - | - | - | - | - | - | Unassigned | - |
| Otu03020 | 13 | k__Fungi(100);p__Basidiomycota(100);c__Agaricomycetes(100);c__Agaricomycetes_unclassified(100);c__Agaricomycetes_unclassified(100);c__Agaricomycetes_unclassified(100);c__Agaricomycetes_unclassified(100); | - | - | - | - | - | - | - | Unassigned | - |
| Otu03021 | 13 | k__Fungi(100);p__Ascomycota(54);p__Ascomycota_unclassified(54);p__Ascomycota_unclassified(54);p__Ascomycota_unclassified(54);p__Ascomycota_unclassified(54);p__Ascomycota_unclassified(54); | - | - | - | - | - | - | - | Unassigned | - |
| Otu03022 | 13 | k__Fungi(100);k__Fungi_unclassified(77);k__Fungi_unclassified(77);k__Fungi_unclassified(77);k__Fungi_unclassified(77);k__Fungi_unclassified(77);k__Fungi_unclassified(77); | - | - | - | - | - | - | - | Unassigned | - |
| Otu03025 | 12 | k__Fungi(100);p__Ascomycota(100);c__Saccharomycetes(100);o__Saccharomycetales(100);f__Metschnikowiaceae(100);g__Metschnikowia(100);s__Metschnikowia_pulcherrima(92); | Metschnikowia | 13 | Saprotroph | \|Undefined Saprotroph\| | Yeast | NULL | Probable | As nectar-tap_saprotroph (Põlme et al. 2020); Decay sugar-rich substrates (Põlme et al. 2020) | Põlme S, et al. 2020. Fungal Diversity 105:1-16 ((https://doi.org/10.1007/s13225-020-00466-2)) |
| Otu03026 | 12 | k__Fungi(100);p__Ascomycota(100);c__Sordariomycetes(100);o__Hypocreales(100);f__Nectriaceae(100);g__Microcera(92);s__Microcera_rubra(92); | Microcera | 13 | Pathotroph-Saprotroph | \|Animal Parasite\|-Undefined Saprotroph | Microfungus | NULL | Probable | As animal_decomposer (Põlme et al. 2020); Decay animal material (Põlme et al. 2020) | Põlme S, et al. 2020. Fungal Diversity 105:1-16 ((https://doi.org/10.1007/s13225-020-00466-2)) |
| Otu03032 | 12 | k__Fungi(100);p__Ascomycota(100);c__Saccharomycetes(100);o__Saccharomycetales(100);f__Metschnikowiaceae(100);g__Metschnikowia(100);s__Metschnikowia_pulcherrima(100); | Metschnikowia | 13 | Saprotroph | \|Undefined Saprotroph\| | Yeast | NULL | Probable | As nectar-tap_saprotroph (Põlme et al. 2020); Decay sugar-rich substrates (Põlme et al. 2020) | Põlme S, et al. 2020. Fungal Diversity 105:1-16 ((https://doi.org/10.1007/s13225-020-00466-2)) |
| Otu03037 | 12 | k__Fungi(100);p__Basidiomycota(100);c__Agaricostilbomycetes(100);o__Agaricostilbales(100);f__Chionosphaeraceae(100);g__Kurtzmanomyces(100);g__Kurtzmanomyces_unclassified(100); | Kurtzmanomyces | 13 | Saprotroph | \|Undefined Saprotroph\| | Dimorphic | NULL | Probable | NULL | Põlme S, et al. 2020. Fungal Diversity 105:1-16 ((https://doi.org/10.1007/s13225-020-00466-2)) |
| Otu03041 | 12 | k__Fungi(100);p__Ascomycota(100);c__Saccharomycetes(100);o__Saccharomycetales(100);f__Metschnikowiaceae(100);g__Metschnikowia(100);s__Metschnikowia_pulcherrima(67); | Metschnikowia | 13 | Saprotroph | \|Undefined Saprotroph\| | Yeast | NULL | Probable | As nectar-tap_saprotroph (Põlme et al. 2020); Decay sugar-rich substrates (Põlme et al. 2020) | Põlme S, et al. 2020. Fungal Diversity 105:1-16 ((https://doi.org/10.1007/s13225-020-00466-2)) |
| Otu03044 | 12 | k__Fungi(100);p__Basidiomycota(100);c__Agaricomycetes(100);o__Thelephorales(100);f__Thelephoraceae(100);g__Tomentella(100);g__Tomentella_unclassified(100); | Tomentella | 13 | Symbiotroph | \|Ectomycorrhizal\| | Corticioid | NULL | Highly Probable | NULL | Rinaldi AC, et al. 2008. Fungal Diversity 33:1-45 ((http://www.fungaldiversity.org/fdp/sfdp/33-1.pdf)); Tedersoo L, et al. 2010. Mycorrhiza 20:217-263 ((https://doi.org/10.1007/s00572-009-0274-x)); Põlme S, et al. 2020. Fungal Diversity 105:1-16 ((https://doi.org/10.1007/s13225-020-00466-2)); http://mycorrhizas.info/ecmf.html ((http://mycorrhizas.info/ecmf.html)) |
| Otu03046 | 12 | k__Fungi(100);p__Ascomycota(100);c__Saccharomycetes(100);o__Saccharomycetales(100);f__Metschnikowiaceae(100);g__Metschnikowia(100);s__Metschnikowia_pulcherrima(100); | Metschnikowia | 13 | Saprotroph | \|Undefined Saprotroph\| | Yeast | NULL | Probable | As nectar-tap_saprotroph (Põlme et al. 2020); Decay sugar-rich substrates (Põlme et al. 2020) | Põlme S, et al. 2020. Fungal Diversity 105:1-16 ((https://doi.org/10.1007/s13225-020-00466-2)) |
| Otu03049 | 12 | k__Fungi(100);p__Ascomycota(100);c__Saccharomycetes(100);o__Saccharomycetales(100);f__Metschnikowiaceae(100);g__Metschnikowia(100);s__Metschnikowia_pulcherrima(100); | Metschnikowia | 13 | Saprotroph | \|Undefined Saprotroph\| | Yeast | NULL | Probable | As nectar-tap_saprotroph (Põlme et al. 2020); Decay sugar-rich substrates (Põlme et al. 2020) | Põlme S, et al. 2020. Fungal Diversity 105:1-16 ((https://doi.org/10.1007/s13225-020-00466-2)) |
| Otu03050 | 12 | k__Fungi(100);p__Ascomycota(100);c__Saccharomycetes(100);o__Saccharomycetales(100);f__Metschnikowiaceae(100);g__Metschnikowia(100);s__Metschnikowia_pulcherrima(100); | Metschnikowia | 13 | Saprotroph | \|Undefined Saprotroph\| | Yeast | NULL | Probable | As nectar-tap_saprotroph (Põlme et al. 2020); Decay sugar-rich substrates (Põlme et al. 2020) | Põlme S, et al. 2020. Fungal Diversity 105:1-16 ((https://doi.org/10.1007/s13225-020-00466-2)) |
| Otu03051 | 12 | k__Fungi(100);p__Ascomycota(100);c__Saccharomycetes(100);o__Saccharomycetales(100);f__Metschnikowiaceae(100);g__Metschnikowia(100);g__Metschnikowia_unclassified(67); | Metschnikowia | 13 | Saprotroph | \|Undefined Saprotroph\| | Yeast | NULL | Probable | As nectar-tap_saprotroph (Põlme et al. 2020); Decay sugar-rich substrates (Põlme et al. 2020) | Põlme S, et al. 2020. Fungal Diversity 105:1-16 ((https://doi.org/10.1007/s13225-020-00466-2)) |
| Otu03052 | 12 | k__Fungi(100);p__Ascomycota(100);c__Saccharomycetes(100);o__Saccharomycetales(100);f__Metschnikowiaceae(100);g__Metschnikowia(100);s__Metschnikowia_pulcherrima(84); | Metschnikowia | 13 | Saprotroph | \|Undefined Saprotroph\| | Yeast | NULL | Probable | As nectar-tap_saprotroph (Põlme et al. 2020); Decay sugar-rich substrates (Põlme et al. 2020) | Põlme S, et al. 2020. Fungal Diversity 105:1-16 ((https://doi.org/10.1007/s13225-020-00466-2)) |
| Otu03054 | 12 | k__Fungi(100);p__Basidiomycota(100);c__Agaricomycetes(100);o__Agaricales(100);f__Tricholomataceae(100);g__Clitocybe(67);g__Clitocybe_unclassified(67); | Clitocybe | 13 | Pathotroph-Saprotroph | \|Plant Saprotroph\| | Agaricoid | NULL | Probable | As litter_saprotroph (Põlme et al. 2020); Decay leaf, fruit, seed (Põlme et al. 2020) | Tedersoo L, et al. 2014. Science 346:e1256688 ((https://doi.org/10.1126/science.1256688)); Põlme S, et al. 2020. Fungal Diversity 105:1-16 ((https://doi.org/10.1007/s13225-020-00466-2)) |
| Otu03055 | 12 | k__Fungi(100);p__Ascomycota(100);c__Sordariomycetes(100);o__Hypocreales(100);f__Hypocreales_family_Incertae_sedis(100);g__Acremonium(59);g__Acremonium_unclassified(59); | Acremonium | 13 | Pathotroph-Saprotroph-Symbiotroph | Animal Pathogen-Endophyte-Fungal Parasite-Plant Pathogen-\|Undefined Saprotroph\|-Wood Saprotroph | Microfungus | Soft Rot | Possible | As foliar_endophyte (Põlme et al. 2020); culture (Hanada et al. 2010, Del Olmo; Decay leaf, fruit, seed, soil, dung, animal material (Põlme et al. 2020); Endophyte detection method; Ruiz & Arnold 2014); Likely opportunistic human pathogen (Irinyi et al. 2015) | Duncan CG, Eslyn WE. 1966. Mycologia 58:642-645 ((https://doi.org/10.2307/3757045)); Nilsson T. 1973. Studia Forestalia Suecica 104: 1-40 ((https://pub.epsilon.slu.se/5711/)); Worrall JJ, et al. 1997. Mycologia 89:199-219 ((https://doi.org/10.2307/3761073)); Hanada RE, et al. 2010. Fungal Biology 114:901-910 ((https://doi.org/10.1016/j.funbio.2010.08.006)); Del Olmo-Ruiz M, Arnold AE. 2014. Mycologia 106: 8-21 ((https://doi.org/10.3852/13-098)); Irinyi L, et al. 2015. Medical Mycology 53313-337 ((https://doi.org/10.1093/mmy/myv008)); Põlme S, et al. 2020. Fungal Diversity 105:1-16 ((https://doi.org/10.1007/s13225-020-00466-2)) |
| Otu03059 | 12 | k__Fungi(100);p__Basidiomycota(100);c__Agaricomycetes(100);o__Thelephorales(100);f__Thelephoraceae(100);g__Tomentella(100);g__Tomentella_unclassified(75); | Tomentella | 13 | Symbiotroph | \|Ectomycorrhizal\| | Corticioid | NULL | Highly Probable | NULL | Rinaldi AC, et al. 2008. Fungal Diversity 33:1-45 ((http://www.fungaldiversity.org/fdp/sfdp/33-1.pdf)); Tedersoo L, et al. 2010. Mycorrhiza 20:217-263 ((https://doi.org/10.1007/s00572-009-0274-x)); Põlme S, et al. 2020. Fungal Diversity 105:1-16 ((https://doi.org/10.1007/s13225-020-00466-2)); http://mycorrhizas.info/ecmf.html ((http://mycorrhizas.info/ecmf.html)) |
| Otu03060 | 12 | k__Fungi(100);p__Basidiomycota(100);c__Agaricomycetes(100);o__Agaricales(100);f__Marasmiaceae(100);g__Gymnopus(100);s__Gymnopus_luxurians(100); | Gymnopus | 13 | Saprotroph | \|Plant Saprotroph\| | Agaricoid | NULL | Probable | As litter_saprotroph (Põlme et al. 2020); Decay leaf, fruit, seed (Põlme et al. 2020) | Põlme S, et al. 2020. Fungal Diversity 105:1-16 ((https://doi.org/10.1007/s13225-020-00466-2)) |
| Otu03064 | 12 | k__Fungi(100);p__Zygomycota(100);c__Zygomycota_class_Incertae_sedis(100);o__Mortierellales(100);f__Mortierellaceae(100);g__Mortierella(100);g__Mortierella_unclassified(100); | Mortierella | 13 | Saprotroph-Symbiotroph | Endophyte-Plant Saprotroph-\|Undefined Saprotroph\| | Microfungus | NULL | Probable | As chitinolytic (Põlme et al. 2020); As soil_saprotroph (Põlme et al. 2020); Decay roots, soil (Põlme et al. 2020); Saprobic in soil and thus frequenctly isolated from soil and material contaminated with soil such as hay or rotten silage and spoil tips (Cannon & Kirk 2007) | Cannon PF, Kirk PM. 2007. Fungal Families of the World. CAB International, Cambridge (ISBN: 978-0851998275); Tedersoo L, et al. 2014. Science 346:e1256688 ((https://doi.org/10.1126/science.1256688)); Purahong W, et al. 2016. Molecular Ecology 25:4059-4074 ((https://doi.org/10.1111/mec.13739)); Põlme S, et al. 2020. Fungal Diversity 105:1-16 ((https://doi.org/10.1007/s13225-020-00466-2)) |
| Otu03068 | 12 | k__Fungi(100);p__Ascomycota(100);c__Saccharomycetes(100);o__Saccharomycetales(100);f__Trichomonascaceae(100);g__Blastobotrys(100);s__Blastobotrys_nivea(100); | Blastobotrys | 13 | Saprotroph | \|Undefined Saprotroph\| | Yeast | Its unusual metal | Highly Probable | As nectar-tap_saprotroph (Põlme et al. 2020); Decay sugar-rich substrates (Põlme et al. 2020); In the course of identifying alternative microorganisms for biotechnological interest, A. adeninivorans has already proved its strengthened competitiveness as a promising cell factory for many more applications. | Kunze G, et al. 2014. Biotechnology for Biofuels 7:66 ((https://doi.org/10.1186/1754-6834-7-66)); Põlme S, et al. 2020. Fungal Diversity 105:1-16 ((https://doi.org/10.1007/s13225-020-00466-2)) |
| Otu03069 | 12 | k__Fungi(100);p__Ascomycota(100);c__Saccharomycetes(100);o__Saccharomycetales(100);f__Metschnikowiaceae(100);g__Metschnikowia(100);s__Metschnikowia_pulcherrima(92); | Metschnikowia | 13 | Saprotroph | \|Undefined Saprotroph\| | Yeast | NULL | Probable | As nectar-tap_saprotroph (Põlme et al. 2020); Decay sugar-rich substrates (Põlme et al. 2020) | Põlme S, et al. 2020. Fungal Diversity 105:1-16 ((https://doi.org/10.1007/s13225-020-00466-2)) |
| Otu03070 | 12 | k__Fungi(100);p__Basidiomycota(100);c__Agaricomycetes(100);o__Agaricales(100);f__Amanitaceae(100);g__Amanita(100);s__Amanita_rubescens(100); | Amanita | 13 | Saprotroph-Symbiotroph | \|Ectomycorrhizal\|-Undefined Saprotroph | Agaricoid | NULL | Probable | Some members of the genus are deadly poisonous | Rinaldi AC, et al. 2008. Fungal Diversity 33:1-45 ((http://www.fungaldiversity.org/fdp/sfdp/33-1.pdf)); Tedersoo L, et al. 2010. Mycorrhiza 20:217-263 ((https://doi.org/10.1007/s00572-009-0274-x)); Wolfe BE, et al. 2012. PLoS ONE 7:e39597 ((https://doi.org/10.1371/journal.pone.0039597)); Põlme S, et al. 2020. Fungal Diversity 105:1-16 ((https://doi.org/10.1007/s13225-020-00466-2)); http://mycorrhizas.info/ecmf.html ((http://mycorrhizas.info/ecmf.html)); http://en.wikipedia.org/wiki/List_of_deadly_fungi ((http://en.wikipedia.org/wiki/List_of_deadly_fungi)) |
| Otu03071 | 12 | k__Fungi(100);p__Zygomycota(100);c__Zygomycota_class_Incertae_sedis(100);o__Mucorales(100);f__Mucoraceae(100);g__Mucor(100);g__Mucor_unclassified(100); | Mucor | 13 | Saprotroph | \|Undefined Saprotroph\| | Microfungus | NULL | Probable | As soil_saprotroph (Põlme et al. 2020); Decay soil (Põlme et al. 2020) | Tedersoo L, et al. 2014. Science 346:e1256688 ((https://doi.org/10.1126/science.1256688)); Põlme S, et al. 2020. Fungal Diversity 105:1-16 ((https://doi.org/10.1007/s13225-020-00466-2)) |
| Otu03073 | 12 | k__Fungi(100);p__Zygomycota(100);c__Zygomycota_class_Incertae_sedis(100);o__Mortierellales(100);f__Mortierellaceae(100);g__Mortierella(100);g__Mortierella_unclassified(100); | Mortierella | 13 | Saprotroph-Symbiotroph | Endophyte-Plant Saprotroph-\|Undefined Saprotroph\| | Microfungus | NULL | Probable | As chitinolytic (Põlme et al. 2020); As soil_saprotroph (Põlme et al. 2020); Decay roots, soil (Põlme et al. 2020); Saprobic in soil and thus frequenctly isolated from soil and material contaminated with soil such as hay or rotten silage and spoil tips (Cannon & Kirk 2007) | Cannon PF, Kirk PM. 2007. Fungal Families of the World. CAB International, Cambridge (ISBN: 978-0851998275); Tedersoo L, et al. 2014. Science 346:e1256688 ((https://doi.org/10.1126/science.1256688)); Purahong W, et al. 2016. Molecular Ecology 25:4059-4074 ((https://doi.org/10.1111/mec.13739)); Põlme S, et al. 2020. Fungal Diversity 105:1-16 ((https://doi.org/10.1007/s13225-020-00466-2)) |
| Otu03074 | 12 | k__Fungi(100);p__Ascomycota(100);c__Saccharomycetes(100);o__Saccharomycetales(100);f__Metschnikowiaceae(100);g__Metschnikowia(100);g__Metschnikowia_unclassified(75); | Metschnikowia | 13 | Saprotroph | \|Undefined Saprotroph\| | Yeast | NULL | Probable | As nectar-tap_saprotroph (Põlme et al. 2020); Decay sugar-rich substrates (Põlme et al. 2020) | Põlme S, et al. 2020. Fungal Diversity 105:1-16 ((https://doi.org/10.1007/s13225-020-00466-2)) |
| Otu03076 | 12 | k__Fungi(100);p__Basidiomycota(100);c__Microbotryomycetes(100);o__Sporidiobolales(100);f__Sporidiobolales_family_Incertae_sedis(100);g__Sporobolomyces(100);s__Sporobolomyces_lactosus(100); | Sporobolomyces | 13 | Pathotroph-Saprotroph | \|Fungal Parasite\|-Plant Saprotroph | Yeast | NULL | Probable | As litter_saprotroph (Põlme et al. 2020); As mycoparasite (Põlme et al. 2020); Decay fungal material (Põlme et al. 2020) | Tedersoo L, et al. 2014. Science 346:e1256688 ((https://doi.org/10.1126/science.1256688)); Purahong W, et al. 2016. Molecular Ecology 25:4059-4074 ((https://doi.org/10.1111/mec.13739)); Põlme S, et al. 2020. Fungal Diversity 105:1-16 ((https://doi.org/10.1007/s13225-020-00466-2)) |
| Otu03077 | 12 | k__Fungi(100);p__Ascomycota(100);c__Dothideomycetes(100);o__Pleosporales(100);f__Phaeosphaeriaceae(100);g__Phaeosphaeria(100);s__Phaeosphaeria_triglochinicola(100); | Phaeosphaeria | 13 | Pathotroph-Saprotroph-Symbiotroph | Endophyte-Plant Pathogen-\|Plant Saprotroph\| | Microfungus | NULL | Probable | As litter_saprotroph (Põlme et al. 2020); Decay leaf, fruit, seed (Põlme et al. 2020) | Tedersoo L, et al. 2014. Science 346:e1256688 ((https://doi.org/10.1126/science.1256688));David AS, et al. 2016. Microbial Ecology 71: 912-926 ((https://doi.org/10.1007/s00248-015-0712-6)); Põlme S, et al. 2020. Fungal Diversity 105:1-16 ((https://doi.org/10.1007/s13225-020-00466-2)) |
| Otu03081 | 12 | k__Fungi(100);p__Ascomycota(100);c__Saccharomycetes(100);o__Saccharomycetales(100);f__Metschnikowiaceae(100);g__Metschnikowia(100);s__Metschnikowia_pulcherrima(92); | Metschnikowia | 13 | Saprotroph | \|Undefined Saprotroph\| | Yeast | NULL | Probable | As nectar-tap_saprotroph (Põlme et al. 2020); Decay sugar-rich substrates (Põlme et al. 2020) | Põlme S, et al. 2020. Fungal Diversity 105:1-16 ((https://doi.org/10.1007/s13225-020-00466-2)) |
| Otu03082 | 12 | k__Fungi(100);p__Ascomycota(100);c__Saccharomycetes(100);o__Saccharomycetales(100);f__Metschnikowiaceae(100);g__Metschnikowia(100);s__Metschnikowia_pulcherrima(84); | Metschnikowia | 13 | Saprotroph | \|Undefined Saprotroph\| | Yeast | NULL | Probable | As nectar-tap_saprotroph (Põlme et al. 2020); Decay sugar-rich substrates (Põlme et al. 2020) | Põlme S, et al. 2020. Fungal Diversity 105:1-16 ((https://doi.org/10.1007/s13225-020-00466-2)) |
| Otu03085 | 12 | k__Fungi(100);p__Ascomycota(100);c__Eurotiomycetes(100);o__Chaetothyriales(100);f__Herpotrichiellaceae(100);g__Phaeomoniella(100);s__Phaeomoniella_sp_Z94(100); | Phaeomoniella | 13 | Pathotroph | \|Plant Pathogen\| | Dimorphic-Microfungus; Facultative Yeast (Tedersoo et | NULL | Highly Probable | Pathotroph isolated from necrosis in Prunus wood (Damm et al. 2010) | Damm U, et al. 2010. Persoonia 24:60-80 ((http://doi.org/10.3767/003158510X500705)); Tedersoo L, et al. 2014. Science 346:e1256688 ((https://doi.org/10.1126/science.1256688)); Põlme S, et al. 2020. Fungal Diversity 105:1-16 ((https://doi.org/10.1007/s13225-020-00466-2)) |
| Otu03086 | 12 | k__Fungi(100);p__Ascomycota(100);c__Saccharomycetes(100);o__Saccharomycetales(100);f__Metschnikowiaceae(100);g__Metschnikowia(100);s__Metschnikowia_pulcherrima(100); | Metschnikowia | 13 | Saprotroph | \|Undefined Saprotroph\| | Yeast | NULL | Probable | As nectar-tap_saprotroph (Põlme et al. 2020); Decay sugar-rich substrates (Põlme et al. 2020) | Põlme S, et al. 2020. Fungal Diversity 105:1-16 ((https://doi.org/10.1007/s13225-020-00466-2)) |
| Otu03087 | 12 | k__Fungi(100);p__Ascomycota(100);c__Saccharomycetes(100);o__Saccharomycetales(100);f__Metschnikowiaceae(100);g__Metschnikowia(100);s__Metschnikowia_pulcherrima(92); | Metschnikowia | 13 | Saprotroph | \|Undefined Saprotroph\| | Yeast | NULL | Probable | As nectar-tap_saprotroph (Põlme et al. 2020); Decay sugar-rich substrates (Põlme et al. 2020) | Põlme S, et al. 2020. Fungal Diversity 105:1-16 ((https://doi.org/10.1007/s13225-020-00466-2)) |
| Otu03088 | 12 | k__Fungi(100);p__Ascomycota(100);c__Sordariomycetes(100);o__Chaetosphaeriales(100);f__Chaetosphaeriaceae(100);g__Chaetosphaeria(100);s__Chaetosphaeria_sp(100); | Chaetosphaeria | 13 | Saprotroph-Symbiotroph | Endophyte-\|Plant Saprotroph\|-Wood Saprotroph | Microfungus | NULL | Probable | As foliar_endophyte (Põlme et al. 2020); As litter_saprotroph (Põlme et al. 2020); Decay wood, leaf, fruit, seed (Põlme et al. 2020) | Réblová M, et al. 2006. Mycological Research 110:104-109 ((https://doi.org/10.1016/j.mycres.2005.09.003)); Tedersoo L, et al. 2014. Science 346:e1256688 ((https://doi.org/10.1126/science.1256688)); Põlme S, et al. 2020. Fungal Diversity 105:1-16 ((https://doi.org/10.1007/s13225-020-00466-2)) |
| Otu03089 | 12 | k__Fungi(100);p__Ascomycota(100);c__Dothideomycetes(100);o__Pleosporales(100);f__Venturiaceae(100);g__Venturia(100);g__Venturia_unclassified(100); | Venturia | 13 | Pathotroph-Pathotroph-Saprotroph | \|Plant Pathogen\|-Plant Saprotroph | Microfungus | NULL | Probable | As litter_saprotroph (Põlme et al. 2020); Decay leaf, fruit, seed (Põlme et al. 2020) | Tedersoo L, et al. 2014. Science 346:e1256688 ((https://doi.org/10.1126/science.1256688)); Põlme S, et al. 2020. Fungal Diversity 105:1-16 ((https://doi.org/10.1007/s13225-020-00466-2)) |
| Otu03090 | 12 | k__Fungi(100);p__Ascomycota(100);c__Dothideomycetes(100);o__Capnodiales(100);f__Teratosphaeriaceae(59);f__Teratosphaeriaceae_unclassified(59);f__Teratosphaeriaceae_unclassified(59); | Teratosphaeriaceae | 9 | Pathotroph-Saprotroph | Animal Pathogen-Plant Pathogen-Undefined Saprotroph | NULL | NULL | Probable | Includes numerous saprobic, extremophilic, human opportunistic, and plant pathogenic fungi (Quaedvlieg et al. 2014) | Quaedvlieg W, et al. 2014. Persoonia 3:31-40 ((https://doi.org/10.3767/003158514X681981)) |
| Otu03093 | 12 | k__Fungi(100);p__Basidiomycota(100);c__Agaricomycetes(100);o__Agaricales(100);f__Cortinariaceae(100);g__Cortinarius(100);s__Cortinarius_scaurus_var._herpeticus(100); | Cortinarius | 13 | Symbiotroph | \|Ectomycorrhizal\| | Agaricoid | NULL | Highly Probable | NULL | Rinaldi AC, et al. 2008. Fungal Diversity 33:1-45 ((http://www.fungaldiversity.org/fdp/sfdp/33-1.pdf)); Tedersoo L, et al. 2010. Mycorrhiza 20:217-263 ((https://doi.org/10.1007/s00572-009-0274-x)); Põlme S, et al. 2020. Fungal Diversity 105:1-16 ((https://doi.org/10.1007/s13225-020-00466-2)); http://mycorrhizas.info/ecmf.html ((http://mycorrhizas.info/ecmf.html)) |
| Otu03094 | 12 | k__Fungi(100);p__Basidiomycota(100);c__Agaricomycetes(100);o__Cantharellales(100);f__Clavulinaceae(92);g__Clavulina(92);g__Clavulina_unclassified(92); | Clavulina | 13 | Symbiotroph | \|Ectomycorrhizal\| | Clavarioid | NULL | Highly Probable | NULL | Tedersoo L, et al. 2014. Science 346:e1256688 ((https://doi.org/10.1126/science.1256688)); Põlme S, et al. 2020. Fungal Diversity 105:1-16 ((https://doi.org/10.1007/s13225-020-00466-2)) |
| Otu03097 | 12 | k__Fungi(100);p__Basidiomycota(100);c__Agaricomycetes(100);o__Corticiales(100);f__Corticiaceae(100);g__Marchandiomyces(100);s__Marchandiomyces_lignicola(100); | Marchandiomyces | 13 | Pathotroph | \|Lichen Parasite\| | Corticioid-Microfungus | NULL | Probable | Decay fungal material, algal material (Põlme et al. 2020) | Tedersoo L, et al. 2014. Science 346:e1256688 ((https://doi.org/10.1126/science.1256688)); Lawrey JD, Diederich P. 2016. http://www.lichenicolous.net (accessed 24 May 2017) ((http://www.lichenicolous.net)); Põlme S, et al. 2020. Fungal Diversity 105:1-16 ((https://doi.org/10.1007/s13225-020-00466-2)) |
| Otu03098 | 12 | k__Fungi(100);p__Basidiomycota(100);c__Agaricomycetes(100);o__Polyporales(100);f__Polyporaceae(100);f__Polyporaceae_unclassified(92);f__Polyporaceae_unclassified(92); | Polyporaceae | 9 | Saprotroph | Wood Saprotroph | Polyporoid | Brown Rot-White | Probable | Lignicolous, causing brown or white rot, mostly of broadleaved trees (Cannon & Kirk 2007) | Cannon PF, Kirk PM. 2007. Fungal Families of the World. CAB International, Cambridge (ISBN: 978-0851998275) |
| Otu03100 | 12 | k__Fungi(100);p__Ascomycota(100);c__Pezizomycetes(100);o__Pezizales(100);f__Tuberaceae(100);g__Tuber(100);g__Tuber_unclassified(100); | Tuber | 13 | Symbiotroph | \|Ectomycorrhizal\| | Gasteroid | Hypogeous | Highly Probable | true truffle; Common name | Rinaldi AC, et al. 2008. Fungal Diversity 33:1-45 ((http://www.fungaldiversity.org/fdp/sfdp/33-1.pdf)); Tedersoo L, et al. 2010. Mycorrhiza 20:217-263 ((https://doi.org/10.1007/s00572-009-0274-x)); Põlme S, et al. 2020. Fungal Diversity 105:1-16 ((https://doi.org/10.1007/s13225-020-00466-2)); http://mycorrhizas.info/ecmf.html ((http://mycorrhizas.info/ecmf.html)) |
| Otu03102 | 12 | k__Fungi(100);p__Ascomycota(100);c__Saccharomycetes(100);o__Saccharomycetales(100);f__Saccharomycodaceae(100);g__Hanseniaspora(100);s__Hanseniaspora_vineae(100); | Hanseniaspora | 13 | Pathotroph-Saprotroph | Animal Parasite-\|Undefined Saprotroph\| | Yeast | NULL | Probable | As nectar-tap_saprotroph (Põlme et al. 2020); Decay animal material, sugar-rich substrates (Põlme et al. 2020) | Põlme S, et al. 2020. Fungal Diversity 105:1-16 ((https://doi.org/10.1007/s13225-020-00466-2)) |
| Otu03107 | 12 | k__Fungi(100);p__Ascomycota(100);c__Eurotiomycetes(100);o__Chaetothyriales(100);f__Herpotrichiellaceae(100);g__Cladophialophora(100);g__Cladophialophora_unclassified(84); | Cladophialophora | 13 | Pathotroph-Saprotroph-Symbiotroph | Animal Pathogen-Endophyte-Fungal Parasite-\|Undefined Saprotroph\| | Dimorphic-Microfungus | NULL | Probable | As mycoparasite (Põlme et al. 2020); As root_endophyte_dark_septate (Põlme et al. 2020); As soil_saprotroph (Põlme et al. 2020); Decay soil (Põlme et al. 2020) | James TY, et al. 2006. Nature 443:818-822 ((https://doi.org/10.1038/nature05110)); Põlme S, et al. 2020. Fungal Diversity 105:1-16 ((https://doi.org/10.1007/s13225-020-00466-2)) |
| Otu03110 | 12 | k__Fungi(100);p__Zygomycota(100);c__Zygomycota_class_Incertae_sedis(100);o__Mucorales(100);f__Phycomycetaceae(100);g__Phycomyces(100);s__Phycomyces_blakesleeanus(100); | Phycomyces | 13 | Saprotroph | \|Dung Saprotroph\|-Undefined Saprotroph | Microfungus | NULL | Probable | As soil_saprotroph (Põlme et al. 2020); Decay dung, soil (Põlme et al. 2020) | Tedersoo L, et al. 2014. Science 346:e1256688 ((https://doi.org/10.1126/science.1256688)); Põlme S, et al. 2020. Fungal Diversity 105:1-16 ((https://doi.org/10.1007/s13225-020-00466-2)) |
| Otu03112 | 12 | k__Fungi(100);p__Ascomycota(100);c__Saccharomycetes(100);o__Saccharomycetales(100);f__Metschnikowiaceae(100);g__Metschnikowia(100);s__Metschnikowia_pulcherrima(100); | Metschnikowia | 13 | Saprotroph | \|Undefined Saprotroph\| | Yeast | NULL | Probable | As nectar-tap_saprotroph (Põlme et al. 2020); Decay sugar-rich substrates (Põlme et al. 2020) | Põlme S, et al. 2020. Fungal Diversity 105:1-16 ((https://doi.org/10.1007/s13225-020-00466-2)) |
| Otu03113 | 12 | k__Fungi(100);p__Ascomycota(100);c__Saccharomycetes(100);o__Saccharomycetales(100);f__Metschnikowiaceae(100);g__Metschnikowia(100);g__Metschnikowia_unclassified(92); | Metschnikowia | 13 | Saprotroph | \|Undefined Saprotroph\| | Yeast | NULL | Probable | As nectar-tap_saprotroph (Põlme et al. 2020); Decay sugar-rich substrates (Põlme et al. 2020) | Põlme S, et al. 2020. Fungal Diversity 105:1-16 ((https://doi.org/10.1007/s13225-020-00466-2)) |
| Otu03114 | 12 | k__Fungi(100);p__Ascomycota(100);c__Leotiomycetes(100);o__Helotiales(100);f__Helotiaceae(100);g__Neobulgaria(100);s__Neobulgaria_sp(100); | Neobulgaria | 13 | Saprotroph | Undefined Saprotroph-\|Wood Saprotroph\| | Microfungus | NULL | Probable | Decay wood (Põlme et al. 2020) | Tedersoo L, et al. 2014. Science 346:e1256688 ((https://doi.org/10.1126/science.1256688)); Põlme S, et al. 2020. Fungal Diversity 105:1-16 ((https://doi.org/10.1007/s13225-020-00466-2)) |
| Otu03117 | 12 | k__Fungi(100);p__Basidiomycota(100);c__Agaricomycetes(100);o__Agaricales(100);f__Strophariaceae(100);g__Galerina(100);g__Galerina_unclassified(100); | Galerina | 13 | Pathotroph-Saprotroph | Bryophyte Parasite-Plant Pathogen-Plant Saprotroph-Undefined Saprotroph-\|Wood Saprotroph\| | Agaricoid | White Rot | Probable | As litter_saprotroph (Põlme et al. 2020); Decay wood, leaf, fruit, seed (Põlme et al. 2020) | Tedersoo L, et al. 2014. Science 346:e1256688 ((https://doi.org/10.1126/science.1256688)); Põlme S, et al. 2020. Fungal Diversity 105:1-16 ((https://doi.org/10.1007/s13225-020-00466-2)) |
| Otu03118 | 12 | k__Fungi(100);p__Ascomycota(100);c__Saccharomycetes(100);o__Saccharomycetales(100);f__Metschnikowiaceae(100);g__Metschnikowia(100);g__Metschnikowia_unclassified(75); | Metschnikowia | 13 | Saprotroph | \|Undefined Saprotroph\| | Yeast | NULL | Probable | As nectar-tap_saprotroph (Põlme et al. 2020); Decay sugar-rich substrates (Põlme et al. 2020) | Põlme S, et al. 2020. Fungal Diversity 105:1-16 ((https://doi.org/10.1007/s13225-020-00466-2)) |
| Otu03119 | 12 | k__Fungi(100);p__Basidiomycota(100);c__Agaricomycetes(100);o__Agaricales(100);f__Cortinariaceae(100);g__Cortinarius(100);g__Cortinarius_unclassified(84); | Cortinarius | 13 | Symbiotroph | \|Ectomycorrhizal\| | Agaricoid | NULL | Highly Probable | NULL | Rinaldi AC, et al. 2008. Fungal Diversity 33:1-45 ((http://www.fungaldiversity.org/fdp/sfdp/33-1.pdf)); Tedersoo L, et al. 2010. Mycorrhiza 20:217-263 ((https://doi.org/10.1007/s00572-009-0274-x)); Põlme S, et al. 2020. Fungal Diversity 105:1-16 ((https://doi.org/10.1007/s13225-020-00466-2)); http://mycorrhizas.info/ecmf.html ((http://mycorrhizas.info/ecmf.html)) |
| Otu03120 | 12 | k__Fungi(100);p__Ascomycota(100);c__Saccharomycetes(100);o__Saccharomycetales(100);f__Metschnikowiaceae(100);g__Metschnikowia(100);s__Metschnikowia_pulcherrima(84); | Metschnikowia | 13 | Saprotroph | \|Undefined Saprotroph\| | Yeast | NULL | Probable | As nectar-tap_saprotroph (Põlme et al. 2020); Decay sugar-rich substrates (Põlme et al. 2020) | Põlme S, et al. 2020. Fungal Diversity 105:1-16 ((https://doi.org/10.1007/s13225-020-00466-2)) |
| Otu03121 | 12 | k__Fungi(100);p__Basidiomycota(100);c__Agaricomycetes(100);o__Russulales(100);f__Peniophoraceae(100);g__Peniophora(100);g__Peniophora_unclassified(100); | Peniophora | 13 | Pathotroph-Saprotroph | Plant Pathogen-\|Wood Saprotroph\| | Corticioid | White Rot | Probable | As white_rot (Põlme et al. 2020); As wood_pathogen (Põlme et al. 2020); Decay wood (Põlme et al. 2020) | Tedersoo L, et al. 2014. Science 346:e1256688 ((https://doi.org/10.1126/science.1256688)); Põlme S, et al. 2020. Fungal Diversity 105:1-16 ((https://doi.org/10.1007/s13225-020-00466-2)) |
| Otu03122 | 12 | k__Fungi(100);p__Ascomycota(100);c__Saccharomycetes(100);o__Saccharomycetales(100);f__Metschnikowiaceae(100);g__Metschnikowia(100);g__Metschnikowia_unclassified(59); | Metschnikowia | 13 | Saprotroph | \|Undefined Saprotroph\| | Yeast | NULL | Probable | As nectar-tap_saprotroph (Põlme et al. 2020); Decay sugar-rich substrates (Põlme et al. 2020) | Põlme S, et al. 2020. Fungal Diversity 105:1-16 ((https://doi.org/10.1007/s13225-020-00466-2)) |
| Otu03124 | 12 | k__Fungi(100);p__Ascomycota(100);c__Sordariomycetes(100);o__Hypocreales(100);f__Cordycipitaceae(100);g__Simplicillium(100);s__Simplicillium_minatense(75); | Simplicillium | 13 | Pathotroph-Saprotroph | \|Animal Parasite\|-Animal Pathogen-Fungal Parasite-Undefined Saprotroph | Microfungus | NULL | Probable | As animal_decomposer (Põlme et al. 2020); As mycoparasite (Põlme et al. 2020); Decay animal material (Põlme et al. 2020) | Tedersoo L, et al. 2014. Science 346:e1256688 ((https://doi.org/10.1126/science.1256688)); Põlme S, et al. 2020. Fungal Diversity 105:1-16 ((https://doi.org/10.1007/s13225-020-00466-2)) |
| Otu03126 | 12 | k__Fungi(100);p__Ascomycota(100);c__Dothideomycetes(100);o__Pleosporales(100);f__Tubeufiaceae(100);g__Helicoma(100);s__Helicoma_conicodentatum(100); | Helicoma | 13 | Saprotroph | Undefined Saprotroph-\|Wood Saprotroph\| | Microfungus | NULL | Probable | Decay wood (Põlme et al. 2020) | Tedersoo L, et al. 2014. Science 346:e1256688 ((https://doi.org/10.1126/science.1256688)); Põlme S, et al. 2020. Fungal Diversity 105:1-16 ((https://doi.org/10.1007/s13225-020-00466-2)) |
| Otu03127 | 12 | k__Fungi(100);p__Ascomycota(100);c__Pezizomycetes(100);o__Pezizales(100);f__Sarcosomataceae(100);g__Urnula(100);s__Urnula_craterium(100); | Urnula | 13 | Pathotroph-Saprotroph | \|Plant Pathogen\|-Plant Saprotroph-Undefined Saprotroph | Microfungus | NULL | Probable | As litter_saprotroph (Põlme et al. 2020); Decay leaf, fruit, seed (Põlme et al. 2020) | Tedersoo L, et al. 2014. Science 346:e1256688 ((https://doi.org/10.1126/science.1256688)); Põlme S, et al. 2020. Fungal Diversity 105:1-16 ((https://doi.org/10.1007/s13225-020-00466-2)) |
| Otu03132 | 12 | k__Fungi(100);p__Basidiomycota(100);c__Microbotryomycetes(100);o__Sporidiobolales(100);f__Sporidiobolales_family_Incertae_sedis(100);g__Sporobolomyces(100);s__Sporobolomyces_lactosus(100); | Sporobolomyces | 13 | Pathotroph-Saprotroph | \|Fungal Parasite\|-Plant Saprotroph | Yeast | NULL | Probable | As litter_saprotroph (Põlme et al. 2020); As mycoparasite (Põlme et al. 2020); Decay fungal material (Põlme et al. 2020) | Tedersoo L, et al. 2014. Science 346:e1256688 ((https://doi.org/10.1126/science.1256688)); Purahong W, et al. 2016. Molecular Ecology 25:4059-4074 ((https://doi.org/10.1111/mec.13739)); Põlme S, et al. 2020. Fungal Diversity 105:1-16 ((https://doi.org/10.1007/s13225-020-00466-2)) |
| Otu03133 | 12 | k__Fungi(100);p__Ascomycota(100);c__Leotiomycetes(100);o__Helotiales(100);f__Helotiaceae(100);g__Hymenoscyphus(100);g__Hymenoscyphus_unclassified(100); | Hymenoscyphus | 13 | Pathotroph-Saprotroph-Symbiotroph | Bryophyte Parasite-Ectomycorrhizal-Ericoid Mycorrhizal-Plant Pathogen-\|Plant Saprotroph\|-Undefined Saprotroph | Microfungus | NULL | Possible | As litter_saprotroph (Põlme et al. 2020); Dark Septate Microfungus; Syn. Pezoloma among other (see e.g., Pezoloma ericae); Decay leaf, fruit, seed (Põlme et al. 2020) | Baral HO, Krieglsteiner L. 2006. Acta Mycologica 41:11-20 ((https://doi.org/10.5586/am.2006.003)) (as a bryophyte parasite); Huhtinen S, et al. 2010. Nova Hedwigia 90:413-431 ((https://doi.org/10.1127/0029-5035/2010/0090-0413)); Bent E, et al. 2011. Applied and Environmental Microbiology 77:3351-3359 ((https://doi.org/10.1128/AEM.02575-10)) (as ectomycorrhizae); Tedersoo L, et al. 2014. Science 346:1256688 ((https://doi.org/10.1126/science.1256688)) (as saprotroph); Põlme S, et al. 2020. Fungal Diversity 105:1-16 ((https://doi.org/10.1007/s13225-020-00466-2)) |
| Otu03136 | 12 | k__Fungi(100);p__Basidiomycota(100);c__Agaricomycetes(100);o__Agaricales(100);f__Agaricaceae(100);f__Agaricaceae_unclassified(100);f__Agaricaceae_unclassified(100); | Agaricaceae | 9 | Saprotroph | Undefined Saprotroph | Agaricoid-Gasteroid-Secotioid | NULL | Probable | Primarily saprobes in grassland and woodland situations (Cannon & Kirk 2007) | Cannon PF, Kirk PM. 2007. Fungal Families of the World. CAB International, Cambridge (ISBN: 978-0851998275) |
| Otu03138 | 12 | k__Fungi(100);p__Basidiomycota(100);c__Agaricomycetes(100);o__Hymenochaetales(100);f__Schizoporaceae(100);g__Xylodon(100);s__Xylodon_nespori(100); | Xylodon | 13 | Saprotroph | Undefined Saprotroph-\|Wood Saprotroph\| | Corticioid | White Rot | Probable | As white_rot (Põlme et al. 2020); Decay wood (Põlme et al. 2020) | Tedersoo L, et al. 2014. Science 346:e1256688 ((https://doi.org/10.1126/science.1256688)); Põlme S, et al. 2020. Fungal Diversity 105:1-16 ((https://doi.org/10.1007/s13225-020-00466-2)) |
| Otu03141 | 12 | k__Fungi(100);p__Ascomycota(100);c__Saccharomycetes(100);o__Saccharomycetales(100);f__Metschnikowiaceae(100);g__Metschnikowia(100);s__Metschnikowia_pulcherrima(92); | Metschnikowia | 13 | Saprotroph | \|Undefined Saprotroph\| | Yeast | NULL | Probable | As nectar-tap_saprotroph (Põlme et al. 2020); Decay sugar-rich substrates (Põlme et al. 2020) | Põlme S, et al. 2020. Fungal Diversity 105:1-16 ((https://doi.org/10.1007/s13225-020-00466-2)) |
| Otu03144 | 12 | k__Fungi(100);p__Basidiomycota(100);c__Agaricomycetes(100);o__Cantharellales(100);f__Hydnaceae(100);g__Sistotrema(100);s__Sistotrema_muscicola(100); | Sistotrema | 13 | Saprotroph-Symbiotroph | Ectomycorrhizal-\|Plant Saprotroph\|-Wood Saprotroph | Corticioid | White Rot | Possible | As litter_saprotroph (Põlme et al. 2020); Decay leaf, fruit, seed (Põlme et al. 2020) | Gilbertson RL, Ryvarden L. 1987-1987. North American Polypores. Fungiflora, Oslo (ISBN: 978-0945345060); Rinaldi AC, et al. 2008. Fungal Diversity 33:1-45 ((http://www.fungaldiversity.org/fdp/sfdp/33-1.pdf)) (as ECM/not ECM); Tedersoo L, et al. 2010. Mycorrhiza 20: 217-263 ((https://doi.org/10.1007/s00572-009-0274-x)) (pro parte); Põlme S, et al. 2020. Fungal Diversity 105:1-16 ((https://doi.org/10.1007/s13225-020-00466-2)); http://mycorrhizas.info/ecmf.html ((http://mycorrhizas.info/ecmf.html)) (as ECM/not ECM). |
| Otu03147 | 12 | k__Fungi(100);p__Ascomycota(100);c__Saccharomycetes(100);o__Saccharomycetales(100);f__Metschnikowiaceae(100);g__Metschnikowia(100);g__Metschnikowia_unclassified(75); | Metschnikowia | 13 | Saprotroph | \|Undefined Saprotroph\| | Yeast | NULL | Probable | As nectar-tap_saprotroph (Põlme et al. 2020); Decay sugar-rich substrates (Põlme et al. 2020) | Põlme S, et al. 2020. Fungal Diversity 105:1-16 ((https://doi.org/10.1007/s13225-020-00466-2)) |
| Otu03148 | 12 | k__Fungi(100);p__Ascomycota(100);c__Saccharomycetes(100);o__Saccharomycetales(100);f__Metschnikowiaceae(100);g__Metschnikowia(100);s__Metschnikowia_pulcherrima(84); | Metschnikowia | 13 | Saprotroph | \|Undefined Saprotroph\| | Yeast | NULL | Probable | As nectar-tap_saprotroph (Põlme et al. 2020); Decay sugar-rich substrates (Põlme et al. 2020) | Põlme S, et al. 2020. Fungal Diversity 105:1-16 ((https://doi.org/10.1007/s13225-020-00466-2)) |
| Otu03149 | 12 | k__Fungi(100);p__Ascomycota(100);c__Saccharomycetes(100);o__Saccharomycetales(100);f__Metschnikowiaceae(100);g__Metschnikowia(100);s__Metschnikowia_pulcherrima(84); | Metschnikowia | 13 | Saprotroph | \|Undefined Saprotroph\| | Yeast | NULL | Probable | As nectar-tap_saprotroph (Põlme et al. 2020); Decay sugar-rich substrates (Põlme et al. 2020) | Põlme S, et al. 2020. Fungal Diversity 105:1-16 ((https://doi.org/10.1007/s13225-020-00466-2)) |
| Otu03151 | 12 | k__Fungi(100);p__Basidiomycota(100);c__Agaricomycetes(100);o__Auriculariales(100);f__Auriculariales_family_Incertae_sedis(100);g__Heterochaetella(84);s__Heterochaetella_sp(84); | Heterochaetella | 13 | Saprotroph | Undefined Saprotroph-\|Wood Saprotroph\| | Corticioid | NULL | Probable | Decay wood (Põlme et al. 2020) | Tedersoo L, et al. 2014. Science 346:e1256688 ((https://doi.org/10.1126/science.1256688)); Põlme S, et al. 2020. Fungal Diversity 105:1-16 ((https://doi.org/10.1007/s13225-020-00466-2)) |
| Otu03152 | 12 | k__Fungi(100);p__Ascomycota(100);c__Saccharomycetes(100);o__Saccharomycetales(100);f__Metschnikowiaceae(100);g__Metschnikowia(100);g__Metschnikowia_unclassified(84); | Metschnikowia | 13 | Saprotroph | \|Undefined Saprotroph\| | Yeast | NULL | Probable | As nectar-tap_saprotroph (Põlme et al. 2020); Decay sugar-rich substrates (Põlme et al. 2020) | Põlme S, et al. 2020. Fungal Diversity 105:1-16 ((https://doi.org/10.1007/s13225-020-00466-2)) |
| Otu03153 | 12 | k__Fungi(100);p__Ascomycota(100);c__Dothideomycetes(100);o__Pleosporales(100);f__Venturiaceae(100);f__Venturiaceae_unclassified(84);f__Venturiaceae_unclassified(84); | Venturiaceae | 9 | Pathotroph-Saprotroph-Symbiotroph | Endophyte-Plant Pathogen-Undefined Saprotroph | Microfungus | NULL | Probable | Mostly endophytes and weak parasites of aerial plant parts, including bark and wood, leaf and fruit tissue; rarely biotrophic. Little is known except for the important pathogens (Cannon & Kirk 2007) | Cannon PF, Kirk PM. 2007. Fungal Families of the World. CAB International, Cambridge (ISBN: 978-0851998275) |
| Otu03154 | 12 | k__Fungi(100);p__Ascomycota(100);c__Saccharomycetes(100);o__Saccharomycetales(100);f__Saccharomycetales_family_Incertae_sedis(92);g__Candida(92);g__Candida_unclassified(59); | Candida | 13 | Pathotroph-Saprotroph-Symbiotroph | Animal Pathogen-Endophyte-Endosymbiont-Epiphyte-\|Undefined Saprotroph\| | Dimorphic Yeast-Yeast | NULL | Probable | As nectar-tap_saprotroph (Põlme et al. 2020); Decay sugar-rich substrates (Põlme et al. 2020); This asexual genus is polyphyletic and contain species that function very differently depend on what they are. Caution should be taken when interpreting guilds for Candida species. | Manolakaki D, et al. 2010. Virulence. 1: 367-375 ((https://doi.org/10.4161/viru.1.5.12796)); Mekbib SB, et al. 2011. Tropical Plant Pathology 36:233-240 ((https://doi.org/10.1590/S1982-56762011000400004)); Põlme S, et al. 2020. Fungal Diversity 105:1-16 ((https://doi.org/10.1007/s13225-020-00466-2)) |
| Otu03155 | 12 | k__Fungi(100);p__Ascomycota(100);c__Leotiomycetes(100);o__Helotiales(100);f__Vibrisseaceae(100);f__Vibrisseaceae_unclassified(84);f__Vibrisseaceae_unclassified(84); | Vibrisseaceae | 9 | Saprotroph-Symbiotroph | Endophyte-Undefined Saprotroph-Wood Saprotroph | Microfungus | NULL | Possible | Saprobic, usually on wood and bark immersed in fresh water. Some species may be root endophytes (Cannon & Kirk 2007) | Cannon PF, Kirk PM. 2007. Fungal Families of the World. CAB International, Cambridge (ISBN: 978-0851998275) |
| Otu03156 | 12 | k__Fungi(100);p__Ascomycota(100);c__Saccharomycetes(100);o__Saccharomycetales(100);f__Metschnikowiaceae(100);g__Metschnikowia(100);s__Metschnikowia_pulcherrima(92); | Metschnikowia | 13 | Saprotroph | \|Undefined Saprotroph\| | Yeast | NULL | Probable | As nectar-tap_saprotroph (Põlme et al. 2020); Decay sugar-rich substrates (Põlme et al. 2020) | Põlme S, et al. 2020. Fungal Diversity 105:1-16 ((https://doi.org/10.1007/s13225-020-00466-2)) |
| Otu03158 | 12 | k__Fungi(100);p__Basidiomycota(100);c__Agaricomycetes(100);o__Agaricales(100);f__Agaricaceae(100);g__Cystolepiota(100);g__Cystolepiota_unclassified(100); | Cystolepiota | 13 | Saprotroph | \|Undefined Saprotroph\| | Agaricoid | NULL | Probable | As soil_saprotroph (Põlme et al. 2020); Decay soil (Põlme et al. 2020) | Tedersoo L, et al. 2014. Science 346:e1256688 ((https://doi.org/10.1126/science.1256688)); Põlme S, et al. 2020. Fungal Diversity 105:1-16 ((https://doi.org/10.1007/s13225-020-00466-2)) |
| Otu03159 | 12 | k__Fungi(100);p__Ascomycota(92);c__Saccharomycetes(92);o__Saccharomycetales(92);f__Metschnikowiaceae(92);g__Metschnikowia(92);s__Metschnikowia_pulcherrima(92); | Metschnikowia | 13 | Saprotroph | \|Undefined Saprotroph\| | Yeast | NULL | Probable | As nectar-tap_saprotroph (Põlme et al. 2020); Decay sugar-rich substrates (Põlme et al. 2020) | Põlme S, et al. 2020. Fungal Diversity 105:1-16 ((https://doi.org/10.1007/s13225-020-00466-2)) |
| Otu03160 | 12 | k__Fungi(100);p__Ascomycota(100);c__Saccharomycetes(100);o__Saccharomycetales(100);f__Metschnikowiaceae(100);g__Metschnikowia(100);s__Metschnikowia_pulcherrima(92); | Metschnikowia | 13 | Saprotroph | \|Undefined Saprotroph\| | Yeast | NULL | Probable | As nectar-tap_saprotroph (Põlme et al. 2020); Decay sugar-rich substrates (Põlme et al. 2020) | Põlme S, et al. 2020. Fungal Diversity 105:1-16 ((https://doi.org/10.1007/s13225-020-00466-2)) |
| Otu03023 | 12 | k__Fungi(100);k__Fungi_unclassified(59);k__Fungi_unclassified(59);k__Fungi_unclassified(59);k__Fungi_unclassified(59);k__Fungi_unclassified(59);k__Fungi_unclassified(59); | - | - | - | - | - | - | - | Unassigned | - |
| Otu03024 | 12 | k__Fungi(100);k__Fungi_unclassified(75);k__Fungi_unclassified(75);k__Fungi_unclassified(75);k__Fungi_unclassified(75);k__Fungi_unclassified(75);k__Fungi_unclassified(75); | - | - | - | - | - | - | - | Unassigned | - |
| Otu03027 | 12 | k__Fungi(100);p__Ascomycota(100);c__Sordariomycetes(100);c__Sordariomycetes_unclassified(100);c__Sordariomycetes_unclassified(100);c__Sordariomycetes_unclassified(100);c__Sordariomycetes_unclassified(100); | - | - | - | - | - | - | - | Unassigned | - |
| Otu03028 | 12 | k__Fungi(100);k__Fungi_unclassified(100);k__Fungi_unclassified(100);k__Fungi_unclassified(100);k__Fungi_unclassified(100);k__Fungi_unclassified(100);k__Fungi_unclassified(100); | - | - | - | - | - | - | - | Unassigned | - |
| Otu03029 | 12 | k__Fungi(100);k__Fungi_unclassified(100);k__Fungi_unclassified(100);k__Fungi_unclassified(100);k__Fungi_unclassified(100);k__Fungi_unclassified(100);k__Fungi_unclassified(100); | - | - | - | - | - | - | - | Unassigned | - |
| Otu03030 | 12 | k__Fungi(100);k__Fungi_unclassified(59);k__Fungi_unclassified(59);k__Fungi_unclassified(59);k__Fungi_unclassified(59);k__Fungi_unclassified(59);k__Fungi_unclassified(59); | - | - | - | - | - | - | - | Unassigned | - |
| Otu03031 | 12 | k__Fungi(100);p__Ascomycota(100);c__Leotiomycetes(100);c__Leotiomycetes_unclassified(100);c__Leotiomycetes_unclassified(100);c__Leotiomycetes_unclassified(100);c__Leotiomycetes_unclassified(100); | - | - | - | - | - | - | - | Unassigned | - |
| Otu03033 | 12 | k__Fungi(100);k__Fungi_unclassified(59);k__Fungi_unclassified(59);k__Fungi_unclassified(59);k__Fungi_unclassified(59);k__Fungi_unclassified(59);k__Fungi_unclassified(59); | - | - | - | - | - | - | - | Unassigned | - |
| Otu03034 | 12 | k__Fungi(100);p__Ascomycota(84);p__Ascomycota_unclassified(84);p__Ascomycota_unclassified(84);p__Ascomycota_unclassified(84);p__Ascomycota_unclassified(84);p__Ascomycota_unclassified(84); | - | - | - | - | - | - | - | Unassigned | - |
| Otu03035 | 12 | k__Fungi(100);k__Fungi_unclassified(67);k__Fungi_unclassified(67);k__Fungi_unclassified(67);k__Fungi_unclassified(67);k__Fungi_unclassified(67);k__Fungi_unclassified(67); | - | - | - | - | - | - | - | Unassigned | - |
| Otu03036 | 12 | k__Fungi(100);k__Fungi_unclassified(67);k__Fungi_unclassified(67);k__Fungi_unclassified(67);k__Fungi_unclassified(67);k__Fungi_unclassified(67);k__Fungi_unclassified(67); | - | - | - | - | - | - | - | Unassigned | - |
| Otu03038 | 12 | k__Fungi(100);k__Fungi_unclassified(67);k__Fungi_unclassified(67);k__Fungi_unclassified(67);k__Fungi_unclassified(67);k__Fungi_unclassified(67);k__Fungi_unclassified(67); | - | - | - | - | - | - | - | Unassigned | - |
| Otu03039 | 12 | k__Fungi(100);p__Ascomycota(67);p__Ascomycota_unclassified(67);p__Ascomycota_unclassified(67);p__Ascomycota_unclassified(67);p__Ascomycota_unclassified(67);p__Ascomycota_unclassified(67); | - | - | - | - | - | - | - | Unassigned | - |
| Otu03040 | 12 | k__Fungi(100);p__Ascomycota(100);p__Ascomycota_unclassified(100);p__Ascomycota_unclassified(100);p__Ascomycota_unclassified(100);p__Ascomycota_unclassified(100);p__Ascomycota_unclassified(100); | - | - | - | - | - | - | - | Unassigned | - |
| Otu03042 | 12 | k__Fungi(100);k__Fungi_unclassified(75);k__Fungi_unclassified(75);k__Fungi_unclassified(75);k__Fungi_unclassified(75);k__Fungi_unclassified(75);k__Fungi_unclassified(75); | - | - | - | - | - | - | - | Unassigned | - |
| Otu03043 | 12 | k__Fungi(100);k__Fungi_unclassified(92);k__Fungi_unclassified(92);k__Fungi_unclassified(92);k__Fungi_unclassified(92);k__Fungi_unclassified(92);k__Fungi_unclassified(92); | - | - | - | - | - | - | - | Unassigned | - |
| Otu03045 | 12 | k__Fungi(100);p__Ascomycota(100);p__Ascomycota_unclassified(100);p__Ascomycota_unclassified(100);p__Ascomycota_unclassified(100);p__Ascomycota_unclassified(100);p__Ascomycota_unclassified(100); | - | - | - | - | - | - | - | Unassigned | - |
| Otu03047 | 12 | k__Fungi(100);k__Fungi_unclassified(75);k__Fungi_unclassified(75);k__Fungi_unclassified(75);k__Fungi_unclassified(75);k__Fungi_unclassified(75);k__Fungi_unclassified(75); | - | - | - | - | - | - | - | Unassigned | - |
| Otu03048 | 12 | k__Fungi(100);p__Ascomycota(100);c__Eurotiomycetes(100);c__Eurotiomycetes_unclassified(100);c__Eurotiomycetes_unclassified(100);c__Eurotiomycetes_unclassified(100);c__Eurotiomycetes_unclassified(100); | - | - | - | - | - | - | - | Unassigned | - |
| Otu03053 | 12 | k__Fungi(100);p__Ascomycota(100);c__Sordariomycetes(100);c__Sordariomycetes_unclassified(92);c__Sordariomycetes_unclassified(92);c__Sordariomycetes_unclassified(92);c__Sordariomycetes_unclassified(92); | - | - | - | - | - | - | - | Unassigned | - |
| Otu03056 | 12 | k__Fungi(100);p__Basidiomycota(100);c__Agaricomycetes(100);c__Agaricomycetes_unclassified(59);c__Agaricomycetes_unclassified(59);c__Agaricomycetes_unclassified(59);c__Agaricomycetes_unclassified(59); | - | - | - | - | - | - | - | Unassigned | - |
| Otu03057 | 12 | k__Fungi(100);k__Fungi_unclassified(100);k__Fungi_unclassified(100);k__Fungi_unclassified(100);k__Fungi_unclassified(100);k__Fungi_unclassified(100);k__Fungi_unclassified(100); | - | - | - | - | - | - | - | Unassigned | - |
| Otu03058 | 12 | k__Fungi(100);k__Fungi_unclassified(84);k__Fungi_unclassified(84);k__Fungi_unclassified(84);k__Fungi_unclassified(84);k__Fungi_unclassified(84);k__Fungi_unclassified(84); | - | - | - | - | - | - | - | Unassigned | - |
| Otu03061 | 12 | k__Fungi(100);p__Basidiomycota(84);p__Basidiomycota_unclassified(84);p__Basidiomycota_unclassified(84);p__Basidiomycota_unclassified(84);p__Basidiomycota_unclassified(84);p__Basidiomycota_unclassified(84); | - | - | - | - | - | - | - | Unassigned | - |
| Otu03062 | 12 | k__Fungi(100);p__Ascomycota(75);p__Ascomycota_unclassified(75);p__Ascomycota_unclassified(75);p__Ascomycota_unclassified(75);p__Ascomycota_unclassified(75);p__Ascomycota_unclassified(75); | - | - | - | - | - | - | - | Unassigned | - |
| Otu03063 | 12 | k__Fungi(100);p__Ascomycota(100);p__Ascomycota_unclassified(100);p__Ascomycota_unclassified(100);p__Ascomycota_unclassified(100);p__Ascomycota_unclassified(100);p__Ascomycota_unclassified(100); | - | - | - | - | - | - | - | Unassigned | - |
| Otu03065 | 12 | k__Fungi(100);p__Ascomycota(100);c__Dothideomycetes(100);o__Capnodiales(100);o__Capnodiales_unclassified(100);o__Capnodiales_unclassified(100);o__Capnodiales_unclassified(100); | - | - | - | - | - | - | - | Unassigned | - |
| Otu03066 | 12 | k__Fungi(100);p__Ascomycota(100);p__Ascomycota_unclassified(100);p__Ascomycota_unclassified(100);p__Ascomycota_unclassified(100);p__Ascomycota_unclassified(100);p__Ascomycota_unclassified(100); | - | - | - | - | - | - | - | Unassigned | - |
| Otu03067 | 12 | k__Fungi(100);p__Ascomycota(100);c__Sordariomycetes(100);c__Sordariomycetes_unclassified(100);c__Sordariomycetes_unclassified(100);c__Sordariomycetes_unclassified(100);c__Sordariomycetes_unclassified(100); | - | - | - | - | - | - | - | Unassigned | - |
| Otu03072 | 12 | k__Fungi(100);k__Fungi_unclassified(100);k__Fungi_unclassified(100);k__Fungi_unclassified(100);k__Fungi_unclassified(100);k__Fungi_unclassified(100);k__Fungi_unclassified(100); | - | - | - | - | - | - | - | Unassigned | - |
| Otu03075 | 12 | k__Fungi(100);p__Ascomycota(100);c__Dothideomycetes(100);o__Pleosporales(100);o__Pleosporales_unclassified(92);o__Pleosporales_unclassified(92);o__Pleosporales_unclassified(92); | - | - | - | - | - | - | - | Unassigned | - |
| Otu03078 | 12 | k__Fungi(100);p__Ascomycota(100);c__Dothideomycetes(100);o__Capnodiales(100);o__Capnodiales_unclassified(100);o__Capnodiales_unclassified(100);o__Capnodiales_unclassified(100); | - | - | - | - | - | - | - | Unassigned | - |
| Otu03079 | 12 | k__Fungi(100);k__Fungi_unclassified(100);k__Fungi_unclassified(100);k__Fungi_unclassified(100);k__Fungi_unclassified(100);k__Fungi_unclassified(100);k__Fungi_unclassified(100); | - | - | - | - | - | - | - | Unassigned | - |
| Otu03080 | 12 | k__Fungi(100);p__Basidiomycota(100);c__Agaricomycetes(100);c__Agaricomycetes_unclassified(100);c__Agaricomycetes_unclassified(100);c__Agaricomycetes_unclassified(100);c__Agaricomycetes_unclassified(100); | - | - | - | - | - | - | - | Unassigned | - |
| Otu03083 | 12 | k__Fungi(100);p__Ascomycota(100);c__Eurotiomycetes(100);o__Chaetothyriales(100);o__Chaetothyriales_unclassified(100);o__Chaetothyriales_unclassified(100);o__Chaetothyriales_unclassified(100); | - | - | - | - | - | - | - | Unassigned | - |
| Otu03084 | 12 | k__Fungi(100);k__Fungi_unclassified(67);k__Fungi_unclassified(67);k__Fungi_unclassified(67);k__Fungi_unclassified(67);k__Fungi_unclassified(67);k__Fungi_unclassified(67); | - | - | - | - | - | - | - | Unassigned | - |
| Otu03091 | 12 | k__Fungi(100);p__Ascomycota(92);p__Ascomycota_unclassified(92);p__Ascomycota_unclassified(92);p__Ascomycota_unclassified(92);p__Ascomycota_unclassified(92);p__Ascomycota_unclassified(92); | - | - | - | - | - | - | - | Unassigned | - |
| Otu03092 | 12 | k__Fungi(100);p__Ascomycota(100);p__Ascomycota_unclassified(100);p__Ascomycota_unclassified(100);p__Ascomycota_unclassified(100);p__Ascomycota_unclassified(100);p__Ascomycota_unclassified(100); | - | - | - | - | - | - | - | Unassigned | - |
| Otu03095 | 12 | k__Fungi(100);k__Fungi_unclassified(59);k__Fungi_unclassified(59);k__Fungi_unclassified(59);k__Fungi_unclassified(59);k__Fungi_unclassified(59);k__Fungi_unclassified(59); | - | - | - | - | - | - | - | Unassigned | - |
| Otu03096 | 12 | k__Fungi(100);k__Fungi_unclassified(67);k__Fungi_unclassified(67);k__Fungi_unclassified(67);k__Fungi_unclassified(67);k__Fungi_unclassified(67);k__Fungi_unclassified(67); | - | - | - | - | - | - | - | Unassigned | - |
| Otu03099 | 12 | k__Fungi(100);k__Fungi_unclassified(100);k__Fungi_unclassified(100);k__Fungi_unclassified(100);k__Fungi_unclassified(100);k__Fungi_unclassified(100);k__Fungi_unclassified(100); | - | - | - | - | - | - | - | Unassigned | - |
| Otu03101 | 12 | k__Fungi(100);p__Ascomycota(92);p__Ascomycota_unclassified(92);p__Ascomycota_unclassified(92);p__Ascomycota_unclassified(92);p__Ascomycota_unclassified(92);p__Ascomycota_unclassified(92); | - | - | - | - | - | - | - | Unassigned | - |
| Otu03103 | 12 | k__Fungi(100);p__Ascomycota(75);p__Ascomycota_unclassified(75);p__Ascomycota_unclassified(75);p__Ascomycota_unclassified(75);p__Ascomycota_unclassified(75);p__Ascomycota_unclassified(75); | - | - | - | - | - | - | - | Unassigned | - |
| Otu03104 | 12 | k__Fungi(100);k__Fungi_unclassified(67);k__Fungi_unclassified(67);k__Fungi_unclassified(67);k__Fungi_unclassified(67);k__Fungi_unclassified(67);k__Fungi_unclassified(67); | - | - | - | - | - | - | - | Unassigned | - |
| Otu03105 | 12 | k__Fungi(100);k__Fungi_unclassified(100);k__Fungi_unclassified(100);k__Fungi_unclassified(100);k__Fungi_unclassified(100);k__Fungi_unclassified(100);k__Fungi_unclassified(100); | - | - | - | - | - | - | - | Unassigned | - |
| Otu03106 | 12 | k__Fungi(100);p__Ascomycota(84);p__Ascomycota_unclassified(84);p__Ascomycota_unclassified(84);p__Ascomycota_unclassified(84);p__Ascomycota_unclassified(84);p__Ascomycota_unclassified(84); | - | - | - | - | - | - | - | Unassigned | - |
| Otu03108 | 12 | k__Fungi(100);p__Ascomycota(100);p__Ascomycota_unclassified(75);p__Ascomycota_unclassified(75);p__Ascomycota_unclassified(75);p__Ascomycota_unclassified(75);p__Ascomycota_unclassified(75); | - | - | - | - | - | - | - | Unassigned | - |

| | | | | | | | | | | | |
|---|---|---|---|---|---|---|---|---|---|---|---|
| Otu03109 | 12 | k__Fungi(100);p__Ascomycota(84);p__Ascomycota_unclassified(84);p__Ascomycota_unclassified(84);p__Ascomycota_unclassified(84);p__Ascomycota_unclassified(84);p__Ascomycota_unclassified(84); | - | - | - | - | - | - | - | Unassigned | - |
| Otu03111 | 12 | k__Fungi(100);p__Ascomycota(100);c__Sordariomycetes(100);c__Sordariomycetes_unclassified(100);c__Sordariomycetes_unclassified(100);c__Sordariomycetes_unclassified(100);c__Sordariomycetes_unclassified(100); | - | - | - | - | - | - | - | Unassigned | - |
| Otu03115 | 12 | k__Fungi(100);k__Fungi_unclassified(100);k__Fungi_unclassified(100);k__Fungi_unclassified(100);k__Fungi_unclassified(100);k__Fungi_unclassified(100);k__Fungi_unclassified(100); | - | - | - | - | - | - | - | Unassigned | - |
| Otu03116 | 12 | k__Fungi(100);k__Fungi_unclassified(75);k__Fungi_unclassified(75);k__Fungi_unclassified(75);k__Fungi_unclassified(75);k__Fungi_unclassified(75);k__Fungi_unclassified(75); | - | - | - | - | - | - | - | Unassigned | - |
| Otu03123 | 12 | k__Fungi(100);k__Fungi_unclassified(59);k__Fungi_unclassified(59);k__Fungi_unclassified(59);k__Fungi_unclassified(59);k__Fungi_unclassified(59);k__Fungi_unclassified(59); | - | - | - | - | - | - | - | Unassigned | - |
| Otu03125 | 12 | k__Fungi(100);p__Ascomycota(100);c__Dothideomycetes(100);o__Capnodiales(84);o__Capnodiales_unclassified(84);o__Capnodiales_unclassified(84);o__Capnodiales_unclassified(84); | - | - | - | - | - | - | - | Unassigned | - |
| Otu03128 | 12 | k__Fungi(100);p__Basidiomycota(100);c__Agaricomycetes(100);o__Polyporales(100);o__Polyporales_unclassified(100);o__Polyporales_unclassified(100);o__Polyporales_unclassified(100); | - | - | - | - | - | - | - | Unassigned | - |
| Otu03129 | 12 | k__Fungi(100);k__Fungi_unclassified(92);k__Fungi_unclassified(92);k__Fungi_unclassified(92);k__Fungi_unclassified(92);k__Fungi_unclassified(92);k__Fungi_unclassified(92); | - | - | - | - | - | - | - | Unassigned | - |
| Otu03130 | 12 | k__Fungi(100);k__Fungi_unclassified(100);k__Fungi_unclassified(100);k__Fungi_unclassified(100);k__Fungi_unclassified(100);k__Fungi_unclassified(100);k__Fungi_unclassified(100); | - | - | - | - | - | - | - | Unassigned | - |
| Otu03131 | 12 | k__Fungi(100);k__Fungi_unclassified(67);k__Fungi_unclassified(67);k__Fungi_unclassified(67);k__Fungi_unclassified(67);k__Fungi_unclassified(67);k__Fungi_unclassified(67); | - | - | - | - | - | - | - | Unassigned | - |
| Otu03134 | 12 | k__Fungi(100);p__Basidiomycota(100);c__Agaricomycetes(100);c__Agaricomycetes_unclassified(100);c__Agaricomycetes_unclassified(100);c__Agaricomycetes_unclassified(100);c__Agaricomycetes_unclassified(100); | - | - | - | - | - | - | - | Unassigned | - |
| Otu03135 | 12 | k__Fungi(100);p__Ascomycota(75);p__Ascomycota_unclassified(75);p__Ascomycota_unclassified(75);p__Ascomycota_unclassified(75);p__Ascomycota_unclassified(75);p__Ascomycota_unclassified(75); | - | - | - | - | - | - | - | Unassigned | - |
| Otu03137 | 12 | k__Fungi(100);k__Fungi_unclassified(59);k__Fungi_unclassified(59);k__Fungi_unclassified(59);k__Fungi_unclassified(59);k__Fungi_unclassified(59);k__Fungi_unclassified(59); | - | - | - | - | - | - | - | Unassigned | - |
| Otu03139 | 12 | k__Fungi(100);p__Basidiomycota(100);p__Basidiomycota_unclassified(100);p__Basidiomycota_unclassified(100);p__Basidiomycota_unclassified(100);p__Basidiomycota_unclassified(100);p__Basidiomycota_unclassified(100); | - | - | - | - | - | - | - | Unassigned | - |
| Otu03140 | 12 | k__Fungi(100);p__Ascomycota(100);c__Dothideomycetes(100);o__Pleosporales(100);o__Pleosporales_unclassified(100);o__Pleosporales_unclassified(100);o__Pleosporales_unclassified(100); | - | - | - | - | - | - | - | Unassigned | - |
| Otu03142 | 12 | k__Fungi(100);k__Fungi_unclassified(59);k__Fungi_unclassified(59);k__Fungi_unclassified(59);k__Fungi_unclassified(59);k__Fungi_unclassified(59);k__Fungi_unclassified(59); | - | - | - | - | - | - | - | Unassigned | - |
| Otu03143 | 12 | k__Fungi(100);k__Fungi_unclassified(100);k__Fungi_unclassified(100);k__Fungi_unclassified(100);k__Fungi_unclassified(100);k__Fungi_unclassified(100);k__Fungi_unclassified(100); | - | - | - | - | - | - | - | Unassigned | - |
| Otu03145 | 12 | k__Fungi(100);p__Ascomycota(100);p__Ascomycota_unclassified(84);p__Ascomycota_unclassified(84);p__Ascomycota_unclassified(84);p__Ascomycota_unclassified(84);p__Ascomycota_unclassified(84); | - | - | - | - | - | - | - | Unassigned | - |
| Otu03146 | 12 | k__Fungi(100);k__Fungi_unclassified(67);k__Fungi_unclassified(67);k__Fungi_unclassified(67);k__Fungi_unclassified(67);k__Fungi_unclassified(67);k__Fungi_unclassified(67); | - | - | - | - | - | - | - | Unassigned | - |
| Otu03150 | 12 | k__Fungi(100);k__Fungi_unclassified(75);k__Fungi_unclassified(75);k__Fungi_unclassified(75);k__Fungi_unclassified(75);k__Fungi_unclassified(75);k__Fungi_unclassified(75); | - | - | - | - | - | - | - | Unassigned | - |
| Otu03157 | 12 | k__Fungi(100);k__Fungi_unclassified(75);k__Fungi_unclassified(75);k__Fungi_unclassified(75);k__Fungi_unclassified(75);k__Fungi_unclassified(75);k__Fungi_unclassified(75); | - | - | - | - | - | - | - | Unassigned | - |
| Otu03161 | 11 | k__Fungi(100);p__Ascomycota(100);c__Saccharomycetes(100);o__Saccharomycetales(100);f__Metschnikowiaceae(100);g__Metschnikowia(100);s__Metschnikowia_pulcherrima(100); | Metschnikowia | 13 | Saprotroph | \|Undefined Saprotroph\| | Yeast | NULL | Probable | [illegible] | [illegible] |
| Otu03171 | 11 | k__Fungi(100);p__Ascomycota(100);c__Sordariomycetes(100);o__Hypocreales(100);o__Hypocreales_unclassified(100);o__Hypocreales_unclassified(100);o__Hypocreales_unclassified(100); | Hypocreales | 7 | Saprotroph | Undefined Saprotroph | Microfungus | NULL | Possible | NULL | [illegible] |
| Otu03172 | 11 | k__Fungi(100);p__Ascomycota(100);c__Saccharomycetes(100);o__Saccharomycetales(100);f__Metschnikowiaceae(100);g__Metschnikowia(100);s__Metschnikowia_pulcherrima(100); | Metschnikowia | 13 | Saprotroph | \|Undefined Saprotroph\| | Yeast | NULL | Probable | [illegible] | [illegible] |
| Otu03173 | 11 | k__Fungi(100);p__Basidiomycota(100);c__Agaricomycetes(100);o__Agaricales(100);f__Marasmiaceae(100);g__Gymnopus(100);s__Gymnopus_subnudus(100); | Gymnopus | 13 | Saprotroph | \|Plant Saprotroph\| | Agaricoid | NULL | Probable | [illegible] | [illegible] |
| Otu03176 | 11 | k__Fungi(100);p__Ascomycota(100);c__Sordariomycetes(100);o__Hypocreales(100);f__Hypocreaceae(100);g__Trichoderma(82);g__Trichoderma_unclassified(73); | Trichoderma | 13 | Pathotroph-Saprotroph-Symbiotroph | Animal Pathogen-Endophyte-Epiphyte-\|Fungal Parasite\|-Plant Pathogen-Wood Saprotroph | Microfungus | Soft Rot | Probable | [illegible] | [illegible] |
| Otu03177 | 11 | k__Fungi(100);p__Ascomycota(100);c__Saccharomycetes(100);o__Saccharomycetales(100);f__Metschnikowiaceae(100);g__Metschnikowia(100);s__Metschnikowia_pulcherrima(91); | Metschnikowia | 13 | Saprotroph | \|Undefined Saprotroph\| | Yeast | NULL | Probable | [illegible] | [illegible] |
| Otu03180 | 11 | k__Fungi(100);p__Ascomycota(100);c__Saccharomycetes(100);o__Saccharomycetales(100);f__Metschnikowiaceae(100);g__Metschnikowia(100);s__Metschnikowia_pulcherrima(100); | Metschnikowia | 13 | Saprotroph | \|Undefined Saprotroph\| | Yeast | NULL | Probable | [illegible] | [illegible] |
| Otu03182 | 11 | k__Fungi(100);p__Ascomycota(100);c__Saccharomycetes(100);o__Saccharomycetales(100);f__Metschnikowiaceae(100);g__Metschnikowia(100);s__Metschnikowia_pulcherrima(100); | Metschnikowia | 13 | Saprotroph | \|Undefined Saprotroph\| | Yeast | NULL | Probable | [illegible] | [illegible] |
| Otu03183 | 11 | k__Fungi(100);p__Ascomycota(100);c__Saccharomycetes(100);o__Saccharomycetales(100);f__Metschnikowiaceae(100);g__Metschnikowia(100);s__Metschnikowia_pulcherrima(100); | Metschnikowia | 13 | Saprotroph | \|Undefined Saprotroph\| | Yeast | NULL | Probable | [illegible] | [illegible] |
| Otu03185 | 11 | k__Fungi(100);p__Ascomycota(100);c__Leotiomycetes(100);o__Leotiales(82);f__Leotiaceae(64);g__Alatospora(64);g__Alatospora_unclassified(64); | Alatospora | 13 | Saprotroph | \|Plant Saprotroph\|-Undefined Saprotroph | Microfungus | NULL | Probable | [illegible] | [illegible] |
| Otu03186 | 11 | k__Fungi(100);p__Ascomycota(100);c__Saccharomycetes(100);o__Saccharomycetales(100);f__Saccharomycetaceae(100);g__Kazachstania(100);s__Kazachstania_intestinalis(100); | Kazachstania | 13 | Saprotroph | \|Undefined Saprotroph\| | Yeast | NULL | Probable | [illegible] | [illegible] |
| Otu03187 | 11 | k__Fungi(100);p__Ascomycota(100);c__Saccharomycetes(100);o__Saccharomycetales(100);f__Metschnikowiaceae(100);g__Metschnikowia(100);g__Metschnikowia_unclassified(91); | Metschnikowia | 13 | Saprotroph | \|Undefined Saprotroph\| | Yeast | NULL | Probable | [illegible] | [illegible] |
| Otu03188 | 11 | k__Fungi(100);p__Ascomycota(100);c__Saccharomycetes(100);o__Saccharomycetales(100);f__Metschnikowiaceae(100);g__Metschnikowia(100);g__Metschnikowia_unclassified(82); | Metschnikowia | 13 | Saprotroph | \|Undefined Saprotroph\| | Yeast | NULL | Probable | [illegible] | [illegible] |
| Otu03190 | 11 | k__Fungi(100);p__Ascomycota(100);c__Saccharomycetes(100);o__Saccharomycetales(100);f__Metschnikowiaceae(100);g__Metschnikowia(100);s__Metschnikowia_pulcherrima(64); | Metschnikowia | 13 | Saprotroph | \|Undefined Saprotroph\| | Yeast | NULL | Probable | [illegible] | [illegible] |
| Otu03191 | 11 | k__Fungi(100);p__Zygomycota(100);c__Zygomycota_class_Incertae_sedis(100);o__Mortierellales(100);f__Mortierellaceae(100);g__Mortierella(100);s__Mortierella_horticola(64); | Mortierella | 13 | Saprotroph-Symbiotroph | Endophyte-Plant Saprotroph-\|Undefined Saprotroph\| | Microfungus | NULL | Probable | [illegible] | [illegible] |
| Otu03192 | 11 | k__Fungi(100);p__Ascomycota(100);c__Saccharomycetes(100);o__Saccharomycetales(100);f__Metschnikowiaceae(100);g__Metschnikowia(100);s__Metschnikowia_pulcherrima(64); | Metschnikowia | 13 | Saprotroph | \|Undefined Saprotroph\| | Yeast | NULL | Probable | [illegible] | [illegible] |
| Otu03193 | 11 | k__Fungi(100);p__Ascomycota(100);c__Saccharomycetes(100);o__Saccharomycetales(100);f__Metschnikowiaceae(100);g__Metschnikowia(100);s__Metschnikowia_pulcherrima(100); | Metschnikowia | 13 | Saprotroph | \|Undefined Saprotroph\| | Yeast | NULL | Probable | [illegible] | [illegible] |
| Otu03195 | 11 | k__Fungi(100);p__Ascomycota(100);c__Saccharomycetes(100);o__Saccharomycetales(100);f__Metschnikowiaceae(100);g__Metschnikowia(100);g__Metschnikowia_unclassified(91); | Metschnikowia | 13 | Saprotroph | \|Undefined Saprotroph\| | Yeast | NULL | Probable | [illegible] | [illegible] |
| Otu03197 | 11 | k__Fungi(100);p__Ascomycota(100);c__Saccharomycetes(100);o__Saccharomycetales(100);f__Metschnikowiaceae(100);g__Metschnikowia(100);s__Metschnikowia_pulcherrima(91); | Metschnikowia | 13 | Saprotroph | \|Undefined Saprotroph\| | Yeast | NULL | Probable | [illegible] | [illegible] |
| Otu03200 | 11 | k__Fungi(100);p__Ascomycota(100);c__Saccharomycetes(100);o__Saccharomycetales(100);f__Metschnikowiaceae(100);g__Metschnikowia(100);s__Metschnikowia_pulcherrima(91); | Metschnikowia | 13 | Saprotroph | \|Undefined Saprotroph\| | Yeast | NULL | Probable | [illegible] | [illegible] |
| Otu03201 | 11 | k__Fungi(100);p__Ascomycota(100);c__Leotiomycetes(100);o__Helotiales(100);f__Hyaloscyphaceae(100);g__Haplographium(100);s__Haplographium_delicatum(100); | Haplographium | 13 | Saprotroph | \|Undefined Saprotroph\| | Microfungus | NULL | Probable | NULL | [illegible] |
| Otu03203 | 11 | k__Fungi(100);p__Ascomycota(100);c__Saccharomycetes(100);o__Saccharomycetales(100);f__Metschnikowiaceae(100);g__Metschnikowia(100);g__Metschnikowia_unclassified(82); | Metschnikowia | 13 | Saprotroph | \|Undefined Saprotroph\| | Yeast | NULL | Probable | [illegible] | [illegible] |
| Otu03204 | 11 | k__Fungi(100);p__Ascomycota(100);c__Saccharomycetes(100);o__Saccharomycetales(100);f__Metschnikowiaceae(100);g__Metschnikowia(100);g__Metschnikowia_unclassified(64); | Metschnikowia | 13 | Saprotroph | \|Undefined Saprotroph\| | Yeast | NULL | Probable | [illegible] | [illegible] |
| Otu03207 | 11 | k__Fungi(100);p__Basidiomycota(100);c__Agaricomycetes(100);o__Agaricales(100);f__Entolomataceae(100);g__Clitopilus(100);s__Clitopilus_scyphoides(73); | Clitopilus | 13 | Pathotroph-Saprotroph | Fungal Parasite-\|Plant Saprotroph\|-Undefined Saprotroph | Agaricoid | NULL | Probable | [illegible] | [illegible] |
| Otu03208 | 11 | k__Fungi(100);p__Basidiomycota(100);c__Tremellomycetes(100);o__Tremellales(100);f__Tremellales_family_Incertae_sedis(100);g__Cryptococcus(100);g__Cryptococcus_unclassified(100); | Cryptococcus | 13 | Pathotroph-Saprotroph-Symbiotroph | Animal Pathogen-Endophyte-Epiphyte-\|Undefined Saprotroph\| | Dimorphic Yeast-Yeast | NULL | Possible | [illegible] | [illegible] |
| Otu03209 | 11 | k__Fungi(100);p__Ascomycota(100);c__Saccharomycetes(100);o__Saccharomycetales(100);f__Metschnikowiaceae(100);g__Metschnikowia(100);s__Metschnikowia_pulcherrima(73); | Metschnikowia | 13 | Saprotroph | \|Undefined Saprotroph\| | Yeast | NULL | Probable | [illegible] | [illegible] |
| Otu03210 | 11 | k__Fungi(100);p__Basidiomycota(100);c__Agaricomycetes(100);o__Agaricales(100);f__Strophariaceae(100);g__Psilocybe(100);s__Psilocybe_cf_subviscida/crobula_X_45(100); | Psilocybe | 13 | Saprotroph | \|Plant Saprotroph\|-Undefined Saprotroph | Agaricoid | Poisonous | Probable | [illegible] | [illegible] |
| Otu03211 | 11 | k__Fungi(100);p__Basidiomycota(100);c__Agaricomycetes(100);o__Agaricales(100);f__Marasmiaceae(100);g__Mycetinis(100);s__Mycetinis_alliaceus(100); | Mycetinis | 13 | Saprotroph | \|Plant Saprotroph\|-Undefined Saprotroph | Agaricoid | NULL | Probable | [illegible] | [illegible] |
| Otu03212 | 11 | k__Fungi(100);p__Basidiomycota(100);c__Agaricomycetes(100);o__Trechisporales(100);f__Hydnodontaceae(100);g__Trechispora(100);s__Trechispora_sp_DLL2011_4(100); | Trechispora | 13 | Saprotroph | \|Wood Saprotroph\| | Corticioid | White Rot | Highly Probable | [illegible] | [illegible] |
| Otu03213 | 11 | k__Fungi(100);p__Ascomycota(100);c__Saccharomycetes(100);o__Saccharomycetales(100);f__Metschnikowiaceae(100);g__Metschnikowia(100);s__Metschnikowia_pulcherrima(55); | Metschnikowia | 13 | Saprotroph | \|Undefined Saprotroph\| | Yeast | NULL | Probable | [illegible] | [illegible] |
| Otu03214 | 11 | k__Fungi(100);p__Basidiomycota(100);c__Agaricomycetes(100);o__Agaricales(100);f__Amanitaceae(100);g__Amanita(100);s__Amanita_muscaria(100); | Amanita | 13 | Saprotroph-Symbiotroph | \|Ectomycorrhizal\|-Undefined Saprotroph | Agaricoid | NULL | Probable | [illegible] | [illegible] |
| Otu03215 | 11 | k__Fungi(100);p__Basidiomycota(100);c__Agaricomycetes(100);o__Agaricales(100);o__Agaricales_unclassified(64);o__Agaricales_unclassified(64);o__Agaricales_unclassified(64); | Agaricales | 7 | Pathotroph-Saprotroph-Symbiotroph | Bryophyte Parasite-Dung Saprotroph-Ectomycorrhizal-Fungal Parasite-Leaf Saprotroph-Plant Parasite-Undefined Saprotroph-Wood Saprotroph | Agaricoid-Gasteroid-Microfungus-Secotioid-Yeast | NULL | Possible | [illegible] | [illegible] |
| Otu03216 | 11 | k__Fungi(100);p__Basidiomycota(100);c__Agaricomycetes(100);o__Trechisporales(100);f__Hydnodontaceae(100);g__Trechispora(100);g__Trechispora_unclassified(64); | Trechispora | 13 | Saprotroph | \|Wood Saprotroph\| | Corticioid | White Rot | Highly Probable | [illegible] | [illegible] |
| Otu03224 | 11 | k__Fungi(100);p__Basidiomycota(100);c__Tremellomycetes(100);o__Tremellales(100);f__Tremellales_family_Incertae_sedis(100);g__Fellomyces(91);g__Fellomyces_unclassified(91); | Fellomyces | 13 | Pathotroph-Saprotroph | Fungal Parasite-\|Undefined Saprotroph\| | Yeast | NULL | Probable | [illegible] | [illegible] |
| Otu03225 | 11 | k__Fungi(100);p__Basidiomycota(100);c__Agaricomycetes(100);o__Cantharellales(100);f__Clavulinaceae(100);f__Clavulinaceae_unclassified(73);f__Clavulinaceae_unclassified(73); | Clavulinaceae | 9 | Saprotroph-Symbiotroph | Ectomycorrhizal-Lichenized-Wood Saprotroph | Clavarioid | NULL | Possible | [illegible] | [illegible] |
| Otu03226 | 11 | k__Fungi(100);p__Basidiomycota(73);c__Microbotryomycetes(73);o__Sporidiobolales(73);f__Sporidiobolales_family_Incertae_sedis(73);g__Sporobolomyces(73);s__Sporobolomyces_lactosus(73); | Sporobolomyces | 13 | Pathotroph-Saprotroph | \|Fungal Parasite\|-Plant Saprotroph | Yeast | NULL | Probable | [illegible] | [illegible] |
| Otu03227 | 11 | k__Fungi(100);p__Basidiomycota(100);c__Agaricomycetes(100);o__Agaricales(100);f__Mycenaceae(100);g__Mycena(100);g__Mycena_unclassified(100); | Mycena | 13 | Pathotroph-Saprotroph | Plant Pathogen-\|Plant Saprotroph\|-Undefined Saprotroph-Wood Saprotroph | Agaricoid | NULL | Probable | [illegible] | [illegible] |
| Otu03228 | 11 | k__Fungi(100);p__Basidiomycota(100);c__Agaricomycetes(100);o__Trechisporales(100);f__Hydnodontaceae(100);g__Sistotremastrum(100);s__Sistotremastrum_guttuliferum(100); | Sistotremastrum | 13 | Saprotroph | Undefined Saprotroph-\|Wood Saprotroph\| | Corticioid | White Rot | Probable | [illegible] | [illegible] |
| Otu03229 | 11 | k__Fungi(100);p__Ascomycota(100);c__Lecanoromycetes(100);o__Lecanorales(100);f__Lecanorales_family_Incertae_sedis(100);g__Leprocaulon(91);g__Leprocaulon_unclassified(91); | Leprocaulon | 13 | Symbiotroph | \|Lichenized\| | Thallus | NULL | Highly Probable | NULL | [illegible] |
| Otu03230 | 11 | k__Fungi(100);p__Ascomycota(100);c__Saccharomycetes(100);o__Saccharomycetales(100);f__Metschnikowiaceae(100);g__Metschnikowia(100);s__Metschnikowia_pulcherrima(100); | Metschnikowia | 13 | Saprotroph | \|Undefined Saprotroph\| | Yeast | NULL | Probable | [illegible] | [illegible] |
| Otu03231 | 11 | k__Fungi(100);p__Basidiomycota(100);c__Pucciniomycetes(100);o__Septobasidiales(100);f__Septobasidiaceae(100);g__Septobasidium(100);s__Septobasidium_sp_JBO_2010i(100); | Septobasidium | 13 | Pathotroph | \|Animal Parasite\|-Animal Pathogen-Arthropod Parasite | Dimorphic-Facultative Yeast | NULL | Probable | [illegible] | [illegible] |
| Otu03232 | 11 | k__Fungi(100);p__Basidiomycota(100);c__Exobasidiomycetes(100);o__Microstromatales(100);f__Quambalariaceae(100);g__Quambalaria(100);s__Quambalaria_cyanescens(100); | Quambalaria | 13 | Pathotroph | \|Plant Pathogen\| | Smut | NULL | Probable | NULL | [illegible] |
| Otu03233 | 11 | k__Fungi(100);p__Ascomycota(100);c__Saccharomycetes(100);o__Saccharomycetales(100);f__Metschnikowiaceae(100);g__Metschnikowia(100);s__Metschnikowia_pulcherrima(100); | Metschnikowia | 13 | Saprotroph | \|Undefined Saprotroph\| | Yeast | NULL | Probable | [illegible] | [illegible] |
| Otu03237 | 11 | k__Fungi(100);p__Basidiomycota(100);c__Agaricomycetes(100);o__Agaricales(100);f__Tricholomataceae(100);g__Tricholoma(100);s__Tricholoma_fulvum(73); | Tricholoma | 13 | Pathotroph-Symbiotroph | \|Ectomycorrhizal\|-Fungal Parasite | Agaricoid | NULL | Highly Probable | [illegible] | [illegible] |
| Otu03238 | 11 | k__Fungi(100);p__Ascomycota(100);c__Dothideomycetes(100);o__Pleosporales(100);f__Pleosporales_family_Incertae_sedis(100);g__Berkleasmium(100);s__Berkleasmium_sp(100); | Berkleasmium | 13 | Saprotroph | \|Wood Saprotroph\| | Microfungus | NULL | Possible | [illegible] | [illegible] |
| Otu03239 | 11 | k__Fungi(100);p__Basidiomycota(100);c__Agaricomycetes(100);o__Polyporales(100);f__Polyporaceae(100);g__Amyloporia(100);s__Amyloporia_sinuosa(100); | Amyloporia | 13 | Saprotroph | \|Wood Saprotroph\| | Corticioid | Brown Rot | Probable | [illegible] | [illegible] |
| Otu03240 | 11 | k__Fungi(100);p__Basidiomycota(100);c__Agaricomycetes(100);o__Boletales(100);f__Boletaceae(100);g__Boletus(100);g__Boletus_unclassified(100); | Boletus | 13 | Pathotroph-Symbiotroph | \|Ectomycorrhizal\|-Fungal Parasite | Agaricoid-Boletoid | NULL | Highly Probable | NULL | [illegible] |
| Otu03241 | 11 | k__Fungi(100);p__Ascomycota(100);c__Saccharomycetes(100);o__Saccharomycetales(100);f__Metschnikowiaceae(100);g__Metschnikowia(100);s__Metschnikowia_pulcherrima(91); | Metschnikowia | 13 | Saprotroph | \|Undefined Saprotroph\| | Yeast | NULL | Probable | [illegible] | [illegible] |
| Otu03242 | 11 | k__Fungi(100);p__Basidiomycota(100);c__Agaricomycetes(100);o__Cantharellales(100);f__Ceratobasidiaceae(100);g__Ceratobasidium(100);s__Ceratobasidium_sp(100); | Ceratobasidium | 13 | Pathotroph-Saprotroph-Symbiotroph | Ectomycorrhizal-Endomycorrhizal-Orchid Mycorrhizal-\|Plant Pathogen\|-Plant Saprotroph-Undefined Saprotroph | Corticioid-Microfungus | NULL | Possible | [illegible] | [illegible] |
| Otu03244 | 11 | k__Fungi(100);p__Basidiomycota(91);c__Microbotryomycetes(91);o__Sporidiobolales(91);f__Sporidiobolales_family_Incertae_sedis(91);g__Sporobolomyces(91);s__Sporobolomyces_lactosus(91); | Sporobolomyces | 13 | Pathotroph-Saprotroph | \|Fungal Parasite\|-Plant Saprotroph | Yeast | NULL | Probable | [illegible] | [illegible] |
| Otu03246 | 11 | k__Fungi(100);p__Ascomycota(100);c__Leotiomycetes(100);o__Helotiales(100);f__Helotiaceae(91);g__Hymenoscyphus(91);g__Hymenoscyphus_unclassified(91); | Hymenoscyphus | 13 | Pathotroph-Saprotroph-Symbiotroph | Bryophyte Parasite-Ectomycorrhizal-Ericoid Mycorrhizal-Plant Pathogen-\|Plant Saprotroph\|-Undefined Saprotroph | Microfungus | NULL | Possible | [illegible] | [illegible] |
| Otu03247 | 11 | k__Fungi(100);p__Ascomycota(100);c__Saccharomycetes(91);o__Saccharomycetales(91);f__Saccharomycetales_family_Incertae_sedis(91);g__Candida(91);g__Candida_unclassified(91); | Candida | 13 | Pathotroph-Saprotroph-Symbiotroph | Animal Pathogen-Endophyte-Endosymbiont-Epiphyte-\|Undefined Saprotroph\| | Dimorphic Yeast-Yeast | NULL | Probable | [illegible] | [illegible] |
| Otu03248 | 11 | k__Fungi(100);p__Basidiomycota(100);c__Agaricomycetes(100);o__Corticiales(100);f__Corticiaceae(100);g__Waitea(100);g__Waitea_unclassified(100); | Waitea | 13 | Pathotroph-Saprotroph | \|Plant Pathogen\|-Plant Saprotroph | Corticioid | NULL | Probable | [illegible] | [illegible] |
| Otu03249 | 11 | k__Fungi(100);p__Basidiomycota(100);c__Agaricomycetes(100);o__Agaricales(100);f__Strophariaceae(100);g__Galerina(100);s__Galerina_triscopa_f_telamonioides(100); | Galerina | 13 | Pathotroph-Saprotroph | Bryophyte Parasite-Plant Pathogen-Plant Saprotroph-Undefined Saprotroph-\|Wood Saprotroph\| | Agaricoid | White Rot | Probable | [illegible] | [illegible] |
| Otu03250 | 11 | k__Fungi(100);p__Ascomycota(100);c__Eurotiomycetes(100);o__Chaetothyriales(100);f__Chaetothyriaceae(100);g__Cyphellophora(100);s__Cyphellophora_sp_CBS_11294(100); | Cyphellophora | 13 | Pathotroph-Saprotroph | Animal Parasite-Animal Pathogen-\|Plant Saprotroph\|-Undefined Saprotroph | Dimorphic-Facultative Yeast-Microfungus | NULL | Probable | [illegible] | [illegible] |
| Otu03251 | 11 | k__Fungi(100);p__Ascomycota(100);c__Saccharomycetes(100);o__Saccharomycetales(100);f__Metschnikowiaceae(100);g__Metschnikowia(100);s__Metschnikowia_pulcherrima(100); | Metschnikowia | 13 | Saprotroph | \|Undefined Saprotroph\| | Yeast | NULL | Probable | [illegible] | [illegible] |
| Otu03252 | 11 | k__Fungi(100);p__Basidiomycota(100);c__Microbotryomycetes(100);o__Sporidiobolales(100);f__Sporidiobolales_family_Incertae_sedis(100);g__Rhodotorula(100);s__Rhodotorula_sp(100); | Rhodotorula | 13 | Pathotroph-Saprotroph-Symbiotroph | Animal Pathogen-Animal Symbiotroph-Endophyte-Plant Pathogen-\|Undefined Saprotroph\| | Yeast | NULL | Probable | [illegible] | [illegible] |
| Otu03253 | 11 | k__Fungi(100);p__Basidiomycota(100);c__Agaricomycetes(100);o__Trechisporales(100);f__Hydnodontaceae(100);g__Trechispora(100);s__Trechispora_sp(100); | Trechispora | 13 | Saprotroph | \|Wood Saprotroph\| | Corticioid | White Rot | Highly Probable | [illegible] | [illegible] |
| Otu03254 | 11 | k__Fungi(100);p__Basidiomycota(100);c__Agaricomycetes(100);o__Agaricales(100);f__Cortinariaceae(100);g__Cortinarius(100);s__Cortinarius_glaucopus(55); | Cortinarius | 13 | Symbiotroph | \|Ectomycorrhizal\| | Agaricoid | NULL | Highly Probable | NULL | [illegible] |
| Otu03255 | 11 | k__Fungi(100);p__Basidiomycota(100);c__Agaricomycetes(100);o__Hymenochaetales(100);f__Hymenochaetaceae(100);g__Fuscoporia(100);s__Fuscoporia_gilva(100); | Fuscoporia | 13 | Saprotroph | Undefined Saprotroph-\|Wood Saprotroph\| | Polyporoid | White Rot | Probable | [illegible] | [illegible] |
| Otu03257 | 11 | k__Fungi(100);p__Basidiomycota(100);c__Agaricomycetes(100);o__Agaricales(100);o__Agaricales_unclassified(100);o__Agaricales_unclassified(100);o__Agaricales_unclassified(100); | Agaricales | 7 | Pathotroph-Saprotroph-Symbiotroph | Bryophyte Parasite-Dung Saprotroph-Ectomycorrhizal-Fungal Parasite-Leaf Saprotroph-Plant Parasite-Undefined Saprotroph-Wood Saprotroph | Agaricoid-Gasteroid-Microfungus-Secotioid-Yeast | NULL | Possible | [illegible] | [illegible] |
| Otu03259 | 11 | k__Fungi(100);p__Ascomycota(100);c__Leotiomycetes(100);o__Helotiales(100);f__Rutstroemiaceae(100);g__Lambertella(100);s__Lambertella_corni_maris(100); | Lambertella | 13 | Pathotroph-Saprotroph | Plant Pathogen-\|Plant Saprotroph\|-Undefined Saprotroph | Microfungus | NULL | Probable | [illegible] | [illegible] |
| Otu03260 | 11 | k__Fungi(100);p__Ascomycota(100);c__Saccharomycetes(100);o__Saccharomycetales(100);f__Metschnikowiaceae(100);g__Metschnikowia(100);s__Metschnikowia_pulcherrima(91); | Metschnikowia | 13 | Saprotroph | \|Undefined Saprotroph\| | Yeast | NULL | Probable | [illegible] | [illegible] |
| Otu03261 | 11 | k__Fungi(100);p__Ascomycota(100);c__Pezizomycetes(100);o__Pezizales(100);f__Pyronemataceae(100);g__unclassified_Pyronemataceae(100);s__Pyronemataceae_sp(100); | Pyronemataceae | 9 | Saprotroph-Symbiotroph | Dung Saprotroph-Ectomycorrhizal-Soil Saprotroph-Wood Saprotroph | Gasteroid-Pezizoid | NULL | Possible | [illegible] | [illegible] |
| Otu03262 | 11 | k__Fungi(100);p__Ascomycota(100);c__Eurotiomycetes(100);o__Chaetothyriales(100);f__Herpotrichiellaceae(100);f__Herpotrichiellaceae_unclassified(100);f__Herpotrichiellaceae_unclassified(100); | Herpotrichiellaceae | 9 | Pathotroph-Saprotroph | Animal Pathogen-Fungal Parasite-Undefined Saprotroph | Facultative Yeast-Microfungus | NULL | Probable | [illegible] | [illegible] |
| Otu03264 | 11 | k__Fungi(100);p__Basidiomycota(100);c__Exobasidiomycetes(100);o__Exobasidiales(100);f__Exobasidiaceae(100);g__Exobasidium(100);s__Exobasidium_gracile(100); | Exobasidium | 13 | Pathotroph | \|Plant Pathogen\| | Leaf Blister-Leaf Gall-Leaf Spot | NULL | Probable | NULL | [illegible] |
| Otu03265 | 11 | k__Fungi(100);p__Chytridiomycota(100);c__Chytridiomycetes(100);o__Spizellomycetales(100);f__Spizellomycetaceae(100);g__Spizellomyces(100);g__Spizellomyces_unclassified(100); | Spizellomyces | 13 | Pathotroph-Saprotroph | Plant Pathogen-\|Pollen Saprotroph\|-Undefined Parasite-Undefined Saprotroph | Chytrid-Microfungus | NULL | Probable | [illegible] | [illegible] |
| Otu03266 | 11 | k__Fungi(100);p__Basidiomycota(100);c__Agaricomycetes(100);o__Polyporales(100);f__Fomitopsidaceae(100);g__Daedalea(100);s__Daedalea_quercina(100); | Daedalea | 13 | Saprotroph | \|Wood Saprotroph\| | Polyporoid | Brown Rot | Highly Probable | [illegible] | [illegible] |
| Otu03267 | 11 | k__Fungi(100);p__Ascomycota(100);c__Eurotiomycetes(100);o__Onygenales(100);f__Gymnoascaceae(100);g__Gymnoascus(100);s__Gymnoascus_sp(100); | Gymnoascus | 13 | Saprotroph | \|Undefined Saprotroph\| | Microfungus | NULL | Probable | [illegible] | [illegible] |
| Otu03268 | 11 | k__Fungi(100);p__Basidiomycota(100);c__Agaricomycetes(100);o__Agaricales(100);f__Cortinariaceae(100);g__Cortinarius(100);g__Cortinarius_unclassified(100); | Cortinarius | 13 | Symbiotroph | \|Ectomycorrhizal\| | Agaricoid | NULL | Highly Probable | NULL | [illegible] |
| Otu03270 | 11 | k__Fungi(100);p__Ascomycota(100);c__Sordariomycetes(100);o__Microascales(100);f__Microascaceae(100);g__Microascus(100);s__Microascus_sp_r249(100); | Microascus | 13 | Pathotroph-Saprotroph-Symbiotroph | Animal Pathogen-Dung Saprotroph-Endophyte-Plant Pathogen-Plant Saprotroph-\|Undefined Saprotroph\| | Microfungus | NULL | Possible | [illegible] | [illegible] |
| Otu03271 | 11 | k__Fungi(100);p__Ascomycota(100);c__Sordariomycetes(100);o__Xylariales(100);f__Xylariaceae(100);g__Hypoxylon(100);s__Hypoxylon_howeanum(100); | Hypoxylon | 13 | Saprotroph-Saprotroph-Symbiotroph | Endophyte-Undefined Saprotroph-\|Wood Saprotroph\| | Corticioid-Microfungus-Xylarioid | NULL | Probable | [illegible] | [illegible] |
| Otu03275 | 11 | k__Fungi(100);p__Basidiomycota(100);c__Tremellomycetes(100);o__Trichosporonales(100);f__Trichosporonaceae(100);g__Trichosporon(100);s__Trichosporon_smithiae(55); | Trichosporon | 13 | Pathotroph-Saprotroph | \|Animal Parasite\|-Animal Pathogen-Undefined Saprotroph | Yeast | NULL | Probable | [illegible] | [illegible] |
| Otu03277 | 11 | k__Fungi(100);p__Ascomycota(100);c__Saccharomycetes(100);o__Saccharomycetales(100);f__Metschnikowiaceae(100);g__Metschnikowia(100);s__Metschnikowia_pulcherrima(100); | Metschnikowia | 13 | Saprotroph | \|Undefined Saprotroph\| | Yeast | NULL | Probable | [illegible] | [illegible] |
| Otu03278 | 11 | k__Fungi(100);p__Ascomycota(100);c__Orbiliomycetes(100);o__Orbiliales(100);f__Orbiliaceae(100);g__Drechslerella(100);s__Drechslerella_sp_ZY_2011a(100); | Drechslerella | 13 | Pathotroph-Saprotroph | \|Animal Parasite\|-Nematophagous-Wood Saprotroph | Microfungus | Nematode-trappi | Probable | [illegible] | [illegible] |
| Otu03280 | 11 | k__Fungi(100);p__Ascomycota(100);c__Saccharomycetes(100);o__Saccharomycetales(100);f__Pichiaceae(100);g__Ogataea(100);s__Ogataea_paradorogensis(100); | Ogataea | 13 | Saprotroph | \|Undefined Saprotroph\| | Yeast | NULL | Probable | [illegible] | [illegible] |
| Otu03281 | 11 | k__Fungi(100);p__Ascomycota(100);c__Leotiomycetes(100);o__Helotiales(100);f__Helotiales_family_Incertae_sedis(100);g__Mitrula(100);s__Mitrula_paludosa(100); | Mitrula | 13 | Saprotroph-Pathotroph-Saprotroph | Plant Pathogen-\|Plant Saprotroph\|-Wood Saprotroph | Microfungus | Soft Rot | Highly Probable | [illegible] | [illegible] |
| Otu03282 | 11 | k__Fungi(100);p__Ascomycota(100);c__Saccharomycetes(100);o__Saccharomycetales(100);f__Metschnikowiaceae(100);g__Metschnikowia(100);s__Metschnikowia_pulcherrima(100); | Metschnikowia | 13 | Saprotroph | \|Undefined Saprotroph\| | Yeast | NULL | Probable | [illegible] | [illegible] |
| Otu03284 | 11 | k__Fungi(100);p__Ascomycota(100);c__Sordariomycetes(100);o__Diaporthales(100);f__Gnomoniaceae(100);g__Gnomonia(100);g__Gnomonia_unclassified(100); | Gnomonia | 13 | Pathotroph-Saprotroph-Symbiotroph | \|Endophyte\|-Plant Pathogen-Plant Saprotroph-Undefined Saprotroph | Microfungus | NULL | Probable | [illegible] | [illegible] |
| Otu03286 | 11 | k__Fungi(100);p__Basidiomycota(100);c__Microbotryomycetes(100);o__Sporidiobolales(100);f__Sporidiobolales_family_Incertae_sedis(100);g__Sporobolomyces(100);s__Sporobolomyces_lactosus(100); | Sporobolomyces | 13 | Pathotroph-Saprotroph | \|Fungal Parasite\|-Plant Saprotroph | Yeast | NULL | Probable | [illegible] | [illegible] |
| Otu03287 | 11 | k__Fungi(100);p__Basidiomycota(100);c__Agaricomycetes(100);o__Cantharellales(100);f__Ceratobasidiaceae(100);f__Ceratobasidiaceae_unclassified(100);f__Ceratobasidiaceae_unclassified(100); | Ceratobasidiaceae | 9 | Pathotroph-Saprotroph-Symbiotroph | Endomycorrhizal-Plant Pathogen-Undefined Saprotroph | Microfungus | NULL | Possible | [illegible] | [illegible] |
| Otu03288 | 11 | k__Fungi(100);p__Ascomycota(100);c__Saccharomycetes(100);o__Saccharomycetales(100);f__Metschnikowiaceae(100);g__Metschnikowia(100);s__Metschnikowia_pulcherrima(100); | Metschnikowia | 13 | Saprotroph | \|Undefined Saprotroph\| | Yeast | NULL | Probable | [illegible] | [illegible] |
| Otu03289 | 11 | k__Fungi(100);p__Ascomycota(100);c__Saccharomycetes(100);o__Saccharomycetales(100);f__Metschnikowiaceae(100);g__Metschnikowia(100);s__Metschnikowia_pulcherrima(91); | Metschnikowia | 13 | Saprotroph | \|Undefined Saprotroph\| | Yeast | NULL | Probable | [illegible] | [illegible] |
| Otu03293 | 11 | k__Fungi(100);p__Ascomycota(100);c__Sordariomycetes(100);o__Lulworthiales(100);f__Lulworthiaceae(100);g__Zalerion(100);s__Zalerion_arboricola(100); | Zalerion | 13 | Pathotroph-Saprotroph | Plant Pathogen-\|Plant Saprotroph\|-Undefined Saprotroph | Microfungus | NULL | Probable | [illegible] | [illegible] |
| Otu03294 | 11 | k__Fungi(100);p__Basidiomycota(100);c__Agaricomycetes(100);o__Agaricales(100);f__Inocybaceae(100);g__Inocybe(100);g__Inocybe_unclassified(100); | Inocybe | 13 | Symbiotroph | \|Ectomycorrhizal\| | Agaricoid | NULL | Highly Probable | [illegible] | [illegible] |
| Otu03296 | 11 | k__Fungi(100);p__Ascomycota(100);c__Saccharomycetes(100);o__Saccharomycetales(100);f__Metschnikowiaceae(100);g__Metschnikowia(100);s__Metschnikowia_pulcherrima(91); | Metschnikowia | 13 | Saprotroph | \|Undefined Saprotroph\| | Yeast | NULL | Probable | [illegible] | [illegible] |
| Otu03297 | 11 | k__Fungi(100);p__Ascomycota(100);c__Saccharomycetes(100);o__Saccharomycetales(100);f__Metschnikowiaceae(100);g__Metschnikowia(100);s__Metschnikowia_pulcherrima(91); | Metschnikowia | 13 | Saprotroph | \|Undefined Saprotroph\| | Yeast | NULL | Probable | [illegible] | [illegible] |
| Otu03301 | 11 | k__Fungi(100);p__Basidiomycota(100);c__Tremellomycetes(100);o__Tremellales(100);f__Tremellales_family_Incertae_sedis(100);f__Tremellales_family_Incertae_sedis_unclassified(100);f__Tremellales_family_Incertae_sedis_unclassified(100); | Tremellales | 7 | Pathotroph-Saprotroph-Symbiotroph | Fungal Parasite-Undefined Saprotroph | Tremelloid-Yeast | NULL | Possible | [illegible] | [illegible] |
| Otu03302 | 11 | k__Fungi(100);p__Basidiomycota(100);c__Agaricomycetes(100);o__Russulales(100);f__Stereaceae(100);g__Gloeocystidiellum(100);s__Gloeocystidiellum_porosum(100); | Gloeocystidiellum | 13 | Saprotroph | Undefined Saprotroph-\|Wood Saprotroph\| | Corticioid | White Rot | Probable | [illegible] | [illegible] |
| Otu03305 | 11 | k__Fungi(100);p__Basidiomycota(100);c__Agaricomycetes(100);o__Atheliales(100);f__Atheliaceae(100);g__Amphinema(100);s__Amphinema_sp(100); | Amphinema | 13 | Symbiotroph | \|Ectomycorrhizal\| | Corticioid | NULL | Highly Probable | NULL | [illegible] |
| Otu03306 | 11 | k__Fungi(100);p__Ascomycota(100);c__Saccharomycetes(100);o__Saccharomycetales(100);f__Metschnikowiaceae(100);g__Metschnikowia(100);s__Metschnikowia_pulcherrima(100); | Metschnikowia | 13 | Saprotroph | \|Undefined Saprotroph\| | Yeast | NULL | Probable | [illegible] | [illegible] |
| Otu03307 | 11 | k__Fungi(100);p__Ascomycota(100);c__Saccharomycetes(100);o__Saccharomycetales(100);f__Metschnikowiaceae(100);g__Metschnikowia(100);g__Metschnikowia_unclassified(82); | Metschnikowia | 13 | Saprotroph | \|Undefined Saprotroph\| | Yeast | NULL | Probable | [illegible] | [illegible] |
| Otu03308 | 11 | k__Fungi(100);p__Ascomycota(100);c__Saccharomycetes(100);o__Saccharomycetales(100);f__Metschnikowiaceae(100);g__Metschnikowia(100);s__Metschnikowia_pulcherrima(91); | Metschnikowia | 13 | Saprotroph | \|Undefined Saprotroph\| | Yeast | NULL | Probable | [illegible] | [illegible] |
| Otu03311 | 11 | k__Fungi(100);p__Basidiomycota(100);c__Agaricomycetes(100);o__Agaricales(100);f__Marasmiaceae(100);g__Gymnopus(100);s__Gymnopus_ceraceicola(100); | Gymnopus | 13 | Saprotroph | \|Plant Saprotroph\| | Agaricoid | NULL | Probable | [illegible] | [illegible] |
| Otu03314 | 11 | k__Fungi(100);p__Basidiomycota(100);c__Agaricomycetes(100);o__Russulales(100);f__Russulaceae(100);g__Russula(100);s__Russula_sp_r_07045(100); | Russula | 13 | Symbiotroph | \|Ectomycorrhizal\| | Agaricoid | NULL | Highly Probable | NULL | [illegible] |
| Otu03316 | 11 | k__Fungi(100);p__Ascomycota(100);c__Saccharomycetes(100);o__Saccharomycetales(100);f__Metschnikowiaceae(100);g__Metschnikowia(100);s__Metschnikowia_pulcherrima(100); | Metschnikowia | 13 | Saprotroph | \|Undefined Saprotroph\| | Yeast | NULL | Probable | [illegible] | [illegible] |
| Otu03319 | 11 | k__Fungi(100);p__Ascomycota(100);c__Sordariomycetes(100);o__Hypocreales(100);f__Hypocreales_family_Incertae_sedis(100);g__Trichothecium(100);s__Trichothecium_ovalisporum(100); | Trichothecium | 13 | Pathotroph-Saprotroph | Fungal Parasite-\|Plant Pathogen\|-Plant Saprotroph | Microfungus | NULL | Probable | [illegible] | [illegible] |
| Otu03321 | 11 | k__Fungi(100);p__Ascomycota(100);c__Sordariomycetes(100);o__Hypocreales(100);f__Nectriaceae(100);g__Neonectria(73);g__Neonectria_unclassified(73); | Neonectria | 13 | Pathotroph | \|Plant Pathogen\| | Microfungus | NULL | Probable | NULL | [illegible] |
| Otu03322 | 11 | k__Fungi(100);p__Basidiomycota(100);c__Agaricomycetes(100);o__Agaricales(100);f__Inocybaceae(100);g__Inocybe(100);g__Inocybe_unclassified(100); | Inocybe | 13 | Symbiotroph | \|Ectomycorrhizal\| | Agaricoid | NULL | Highly Probable | [illegible] | [illegible] |
| Otu03323 | 11 | k__Fungi(100);p__Ascomycota(100);c__Saccharomycetes(100);o__Saccharomycetales(100);f__Metschnikowiaceae(100);g__Metschnikowia(100);s__Metschnikowia_pulcherrima(91); | Metschnikowia | 13 | Saprotroph | \|Undefined Saprotroph\| | Yeast | NULL | Probable | [illegible] | [illegible] |
| Otu03328 | 11 | k__Fungi(100);p__Ascomycota(100);c__Saccharomycetes(100);o__Saccharomycetales(100);f__Metschnikowiaceae(100);g__Metschnikowia(100);s__Metschnikowia_pulcherrima(73); | Metschnikowia | 13 | Saprotroph | \|Undefined Saprotroph\| | Yeast | NULL | Probable | [illegible] | [illegible] |
| Otu03329 | 11 | k__Fungi(100);p__Basidiomycota(100);c__Agaricomycetes(100);o__Hymenochaetales(100);f__Hymenochaetaceae(100);g__Inonotus(100);s__Inonotus_obliquus(100); | Inonotus | 13 | Pathotroph-Saprotroph | Plant Pathogen-\|Wood Saprotroph\| | Polyporoid | White Rot | Highly Probable | [illegible] | [illegible] |
| Otu03330 | 11 | k__Fungi(100);p__Ascomycota(100);c__Eurotiomycetes(100);o__Chaetothyriales(100);f__Herpotrichiellaceae(100);g__Exophiala(100);s__Exophiala_sp_CPC_12171(100); | Exophiala | 13 | Pathotroph-Saprotroph-Symbiotroph | \|Animal Parasite\|-Animal Pathogen-Endophyte-Plant Saprotroph-Undefined Saprotroph | Dimorphic-Facultative Yeast | NULL | Probable | [illegible] | [illegible] |
| Otu03331 | 11 | k__Fungi(100);p__Basidiomycota(100);c__Agaricomycetes(100);o__Agaricales(100);f__Agaricaceae(100);g__Coprinus(100);s__Coprinus_phaeopunctatus(100); | Coprinus | 13 | Pathotroph-Saprotroph | \|Dung Saprotroph\|-Plant Pathogen-Plant Saprotroph | Agaricoid | NULL | Highly Probable | [illegible] | [illegible] |
| Otu03333 | 11 | k__Fungi(100);p__Ascomycota(100);c__Sordariomycetes(100);o__Hypocreales(100);f__Clavicipitaceae(100);g__Metarhizium(100);g__Metarhizium_unclassified(55); | Metarhizium | 13 | Pathotroph-Saprotroph | \|Animal Parasite\|-Animal Pathogen-Undefined Saprotroph | Microfungus | NULL | Probable | [illegible] | [illegible] |
| Otu03334 | 11 | k__Fungi(100);p__Basidiomycota(100);c__Agaricomycetes(100);o__Polyporales(100);f__Fomitopsidaceae(100);g__Postia(100);s__Postia_balsamea(100); | Postia | 13 | Saprotroph | \|Wood Saprotroph\| | Polyporoid | Brown Rot | Highly Probable | [illegible] | [illegible] |
| Otu03335 | 11 | k__Fungi(100);p__Ascomycota(100);c__Eurotiomycetes(100);o__Onygenales(100);f__Onygenaceae(100);g__Chrysosporium(100);s__Chrysosporium_carmichaelii(100); | Chrysosporium | 13 | Pathotroph-Saprotroph | Plant Pathogen-Undefined Saprotroph-\|Wood Saprotroph\| | Microfungus | Soft Rot | Highly Probable | [illegible] | [illegible] |
| Otu03336 | 11 | k__Fungi(100);p__Basidiomycota(100);c__Agaricomycetes(100);o__Russulales(100);f__Stephanosporaceae(100);g__Cristinia(100);s__Cristinia_helvetica(100); | Cristinia | 13 | Saprotroph | Undefined Saprotroph-\|Wood Saprotroph\| | Corticioid | White Rot | Probable | [illegible] | [illegible] |
| Otu03337 | 11 | k__Fungi(100);p__Ascomycota(100);c__Dothideomycetes(100);o__Pleosporales(100);f__Sporormiaceae(100);g__Preussia(100);g__Preussia_unclassified(100); | Preussia | 13 | Saprotroph | \|Dung Saprotroph\|-Undefined Saprotroph | Microfungus | NULL | Probable | [illegible] | [illegible] |
| Otu03340 | 11 | k__Fungi(100);p__Ascomycota(100);c__Sordariomycetes(100);o__Hypocreales(100);f__Hypocreaceae(100);g__Trichoderma(100);g__Trichoderma_unclassified(100); | Trichoderma | 13 | Pathotroph-Saprotroph-Symbiotroph | Animal Pathogen-Endophyte-Epiphyte-\|Fungal Parasite\|-Plant Pathogen-Wood Saprotroph | Microfungus | Soft Rot | Probable | [illegible] | [illegible] |
| Otu03344 | 11 | k__Fungi(100);p__Basidiomycota(100);c__Agaricomycetes(100);o__Geastrales(100);f__Geastraceae(100);g__Geastrum(100);g__Geastrum_unclassified(100); | Geastrum | 13 | Saprotroph | \|Plant Saprotroph\|-Undefined Saprotroph | Gasteroid | NULL | Probable | [illegible] | [illegible] |
| Otu03346 | 11 | k__Fungi(100);p__Ascomycota(100);c__Sordariomycetes(100);o__Hypocreales(100);o__Hypocreales_unclassified(91);o__Hypocreales_unclassified(91);o__Hypocreales_unclassified(91); | Hypocreales | 7 | Saprotroph | Undefined Saprotroph | Microfungus | NULL | Possible | NULL | [illegible] |
| Otu03348 | 11 | k__Fungi(100);p__Ascomycota(100);c__Saccharomycetes(100);o__Saccharomycetales(100);f__Metschnikowiaceae(100);g__Metschnikowia(100);s__Metschnikowia_pulcherrima(73); | Metschnikowia | 13 | Saprotroph | \|Undefined Saprotroph\| | Yeast | NULL | Probable | [illegible] | [illegible] |
| Otu03349 | 11 | k__Fungi(100);p__Ascomycota(100);c__Sordariomycetes(100);o__Diaporthales(100);f__Gnomoniaceae(100);g__Gnomoniopsis(55);g__Gnomoniopsis_unclassified(55); | Gnomoniopsis | 13 | Saprotroph-Symbiotroph | \|Endophyte\|-Plant Saprotroph | Microfungus | NULL | Probable | [illegible] | [illegible] |
| Otu03162 | 11 | k__Fungi(100);p__Ascomycota(64);p__Ascomycota_unclassified(64);p__Ascomycota_unclassified(64);p__Ascomycota_unclassified(64);p__Ascomycota_unclassified(64);p__Ascomycota_unclassified(64); | - | - | - | - | - | - | - | Unassigned | - |

| | | | | | | | | | | | |
|---|---|---|---|---|---|---|---|---|---|---|---|
| Otu03163 | 11 | k__Fungi(100);p__Ascomycota(100);c__Dothideomycetes(100);o__Pleosporales(100);o__Pleosporales_unclassified(55);o__Pleosporales_unclassified(55);o__Pleosporales_unclassified(55); | - | - | - | - | - | - | - | Unassigned | - |
| Otu03164 | 11 | k__Fungi(100);p__Ascomycota(82);p__Ascomycota_unclassified(82);p__Ascomycota_unclassified(82);p__Ascomycota_unclassified(82);p__Ascomycota_unclassified(82);p__Ascomycota_unclassified(82); | - | - | - | - | - | - | - | Unassigned | - |
| Otu03165 | 11 | k__Fungi(100);k__Fungi_unclassified(100);k__Fungi_unclassified(100);k__Fungi_unclassified(100);k__Fungi_unclassified(100);k__Fungi_unclassified(100);k__Fungi_unclassified(100); | - | - | - | - | - | - | - | Unassigned | - |
| Otu03166 | 11 | k__Fungi(100);k__Fungi_unclassified(100);k__Fungi_unclassified(100);k__Fungi_unclassified(100);k__Fungi_unclassified(100);k__Fungi_unclassified(100);k__Fungi_unclassified(100); | - | - | - | - | - | - | - | Unassigned | - |
| Otu03167 | 11 | k__Fungi(100);p__Ascomycota(73);p__Ascomycota_unclassified(73);p__Ascomycota_unclassified(73);p__Ascomycota_unclassified(73);p__Ascomycota_unclassified(73);p__Ascomycota_unclassified(73); | - | - | - | - | - | - | - | Unassigned | - |
| Otu03168 | 11 | k__Fungi(100);p__Ascomycota(64);p__Ascomycota_unclassified(64);p__Ascomycota_unclassified(64);p__Ascomycota_unclassified(64);p__Ascomycota_unclassified(64);p__Ascomycota_unclassified(64); | - | - | - | - | - | - | - | Unassigned | - |
| Otu03169 | 11 | k__Fungi(100);k__Fungi_unclassified(64);k__Fungi_unclassified(64);k__Fungi_unclassified(64);k__Fungi_unclassified(64);k__Fungi_unclassified(64);k__Fungi_unclassified(64); | - | - | - | - | - | - | - | Unassigned | - |
| Otu03170 | 11 | k__Fungi(100);k__Fungi_unclassified(64);k__Fungi_unclassified(64);k__Fungi_unclassified(64);k__Fungi_unclassified(64);k__Fungi_unclassified(64);k__Fungi_unclassified(64); | - | - | - | - | - | - | - | Unassigned | - |
| Otu03174 | 11 | k__Fungi(100);k__Fungi_unclassified(73);k__Fungi_unclassified(73);k__Fungi_unclassified(73);k__Fungi_unclassified(73);k__Fungi_unclassified(73);k__Fungi_unclassified(73); | - | - | - | - | - | - | - | Unassigned | - |
| Otu03175 | 11 | k__Fungi(100);k__Fungi_unclassified(82);k__Fungi_unclassified(82);k__Fungi_unclassified(82);k__Fungi_unclassified(82);k__Fungi_unclassified(82);k__Fungi_unclassified(82); | - | - | - | - | - | - | - | Unassigned | - |
| Otu03178 | 11 | k__Fungi(100);p__Ascomycota(73);p__Ascomycota_unclassified(73);p__Ascomycota_unclassified(73);p__Ascomycota_unclassified(73);p__Ascomycota_unclassified(73);p__Ascomycota_unclassified(73); | - | - | - | - | - | - | - | Unassigned | - |
| Otu03179 | 11 | k__Fungi(100);k__Fungi_unclassified(100);k__Fungi_unclassified(100);k__Fungi_unclassified(100);k__Fungi_unclassified(100);k__Fungi_unclassified(100);k__Fungi_unclassified(100); | - | - | - | - | - | - | - | Unassigned | - |
| Otu03181 | 11 | k__Fungi(100);k__Fungi_unclassified(55);k__Fungi_unclassified(55);k__Fungi_unclassified(55);k__Fungi_unclassified(55);k__Fungi_unclassified(55);k__Fungi_unclassified(55); | - | - | - | - | - | - | - | Unassigned | - |
| Otu03184 | 11 | k__Fungi(100);k__Fungi_unclassified(73);k__Fungi_unclassified(73);k__Fungi_unclassified(73);k__Fungi_unclassified(73);k__Fungi_unclassified(73);k__Fungi_unclassified(73); | - | - | - | - | - | - | - | Unassigned | - |
| Otu03189 | 11 | k__Fungi(100);p__Ascomycota(100);p__Ascomycota_unclassified(100);p__Ascomycota_unclassified(100);p__Ascomycota_unclassified(100);p__Ascomycota_unclassified(100);p__Ascomycota_unclassified(100); | - | - | - | - | - | - | - | Unassigned | - |
| Otu03194 | 11 | k__Fungi(100);k__Fungi_unclassified(73);k__Fungi_unclassified(73);k__Fungi_unclassified(73);k__Fungi_unclassified(73);k__Fungi_unclassified(73);k__Fungi_unclassified(73); | - | - | - | - | - | - | - | Unassigned | - |
| Otu03196 | 11 | k__Fungi(100);p__Ascomycota(100);p__Ascomycota_unclassified(100);p__Ascomycota_unclassified(100);p__Ascomycota_unclassified(100);p__Ascomycota_unclassified(100);p__Ascomycota_unclassified(100); | - | - | - | - | - | - | - | Unassigned | - |
| Otu03198 | 11 | k__Fungi(100);p__Ascomycota(82);p__Ascomycota_unclassified(82);p__Ascomycota_unclassified(82);p__Ascomycota_unclassified(82);p__Ascomycota_unclassified(82);p__Ascomycota_unclassified(82); | - | - | - | - | - | - | - | Unassigned | - |
| Otu03199 | 11 | k__Fungi(100);p__Ascomycota(55);p__Ascomycota_unclassified(55);p__Ascomycota_unclassified(55);p__Ascomycota_unclassified(55);p__Ascomycota_unclassified(55);p__Ascomycota_unclassified(55); | - | - | - | - | - | - | - | Unassigned | - |
| Otu03202 | 11 | k__Fungi(100);p__Ascomycota(100);c__Dothideomycetes(100);o__Capnodiales(100);f__Davidiellaceae(100);f__Davidiellaceae_unclassified(91);f__Davidiellaceae_unclassified(91); | - | - | - | - | - | - | - | Unassigned | - |
| Otu03205 | 11 | k__Fungi(100);k__Fungi_unclassified(82);k__Fungi_unclassified(82);k__Fungi_unclassified(82);k__Fungi_unclassified(82);k__Fungi_unclassified(82);k__Fungi_unclassified(82); | - | - | - | - | - | - | - | Unassigned | - |
| Otu03206 | 11 | k__Fungi(100);p__Ascomycota(100);c__Sordariomycetes(100);c__Sordariomycetes_unclassified(100);c__Sordariomycetes_unclassified(100);c__Sordariomycetes_unclassified(100);c__Sordariomycetes_unclassified(100); | - | - | - | - | - | - | - | Unassigned | - |
| Otu03217 | 11 | k__Fungi(100);k__Fungi_unclassified(64);k__Fungi_unclassified(64);k__Fungi_unclassified(64);k__Fungi_unclassified(64);k__Fungi_unclassified(64);k__Fungi_unclassified(64); | - | - | - | - | - | - | - | Unassigned | - |
| Otu03218 | 11 | k__Fungi(100);p__Basidiomycota(100);c__Agaricomycetes(100);c__Agaricomycetes_unclassified(100);c__Agaricomycetes_unclassified(100);c__Agaricomycetes_unclassified(100);c__Agaricomycetes_unclassified(100); | - | - | - | - | - | - | - | Unassigned | - |
| Otu03219 | 11 | k__Fungi(100);k__Fungi_unclassified(55);k__Fungi_unclassified(55);k__Fungi_unclassified(55);k__Fungi_unclassified(55);k__Fungi_unclassified(55);k__Fungi_unclassified(55); | - | - | - | - | - | - | - | Unassigned | - |
| Otu03220 | 11 | k__Fungi(100);k__Fungi_unclassified(73);k__Fungi_unclassified(73);k__Fungi_unclassified(73);k__Fungi_unclassified(73);k__Fungi_unclassified(73);k__Fungi_unclassified(73); | - | - | - | - | - | - | - | Unassigned | - |
| Otu03221 | 11 | k__Fungi(100);p__Ascomycota(73);p__Ascomycota_unclassified(73);p__Ascomycota_unclassified(73);p__Ascomycota_unclassified(73);p__Ascomycota_unclassified(73);p__Ascomycota_unclassified(73); | - | - | - | - | - | - | - | Unassigned | - |
| Otu03222 | 11 | k__Fungi(100);p__Ascomycota(91);p__Ascomycota_unclassified(91);p__Ascomycota_unclassified(91);p__Ascomycota_unclassified(91);p__Ascomycota_unclassified(91);p__Ascomycota_unclassified(91); | - | - | - | - | - | - | - | Unassigned | - |
| Otu03223 | 11 | k__Fungi(100);p__Ascomycota(100);c__Dothideomycetes(91);o__Capnodiales(82);o__Capnodiales_unclassified(73);o__Capnodiales_unclassified(73);o__Capnodiales_unclassified(73); | - | - | - | - | - | - | - | Unassigned | - |
| Otu03234 | 11 | k__Fungi(100);k__Fungi_unclassified(100);k__Fungi_unclassified(100);k__Fungi_unclassified(100);k__Fungi_unclassified(100);k__Fungi_unclassified(100);k__Fungi_unclassified(100); | - | - | - | - | - | - | - | Unassigned | - |
| Otu03235 | 11 | k__Fungi(100);k__Fungi_unclassified(55);k__Fungi_unclassified(55);k__Fungi_unclassified(55);k__Fungi_unclassified(55);k__Fungi_unclassified(55);k__Fungi_unclassified(55); | - | - | - | - | - | - | - | Unassigned | - |
| Otu03236 | 11 | k__Fungi(100);p__Basidiomycota(100);p__Basidiomycota_unclassified(100);p__Basidiomycota_unclassified(100);p__Basidiomycota_unclassified(100);p__Basidiomycota_unclassified(100);p__Basidiomycota_unclassified(100); | - | - | - | - | - | - | - | Unassigned | - |
| Otu03243 | 11 | k__Fungi(100);k__Fungi_unclassified(100);k__Fungi_unclassified(100);k__Fungi_unclassified(100);k__Fungi_unclassified(100);k__Fungi_unclassified(100);k__Fungi_unclassified(100); | - | - | - | - | - | - | - | Unassigned | - |
| Otu03245 | 11 | k__Fungi(100);p__Basidiomycota(100);c__Agaricostilbomycetes(100);c__Agaricostilbomycetes_unclassified(100);c__Agaricostilbomycetes_unclassified(100);c__Agaricostilbomycetes_unclassified(100);c__Agaricostilbomycetes_unclassified(100); | - | - | - | - | - | - | - | Unassigned | - |
| Otu03256 | 11 | k__Fungi(100);p__Basidiomycota(100);p__Basidiomycota_unclassified(100);p__Basidiomycota_unclassified(100);p__Basidiomycota_unclassified(100);p__Basidiomycota_unclassified(100);p__Basidiomycota_unclassified(100); | - | - | - | - | - | - | - | Unassigned | - |
| Otu03258 | 11 | k__Fungi(100);p__Basidiomycota(100);c__Agaricomycetes(100);c__Agaricomycetes_unclassified(100);c__Agaricomycetes_unclassified(100);c__Agaricomycetes_unclassified(100);c__Agaricomycetes_unclassified(100); | - | - | - | - | - | - | - | Unassigned | - |
| Otu03263 | 11 | k__Fungi(100);k__Fungi_unclassified(55);k__Fungi_unclassified(55);k__Fungi_unclassified(55);k__Fungi_unclassified(55);k__Fungi_unclassified(55);k__Fungi_unclassified(55); | - | - | - | - | - | - | - | Unassigned | - |
| Otu03269 | 11 | k__Fungi(100);k__Fungi_unclassified(100);k__Fungi_unclassified(100);k__Fungi_unclassified(100);k__Fungi_unclassified(100);k__Fungi_unclassified(100);k__Fungi_unclassified(100); | - | - | - | - | - | - | - | Unassigned | - |
| Otu03272 | 11 | k__Fungi(100);k__Fungi_unclassified(55);k__Fungi_unclassified(55);k__Fungi_unclassified(55);k__Fungi_unclassified(55);k__Fungi_unclassified(55);k__Fungi_unclassified(55); | - | - | - | - | - | - | - | Unassigned | - |
| Otu03273 | 11 | k__Fungi(100);p__Ascomycota(55);p__Ascomycota_unclassified(55);p__Ascomycota_unclassified(55);p__Ascomycota_unclassified(55);p__Ascomycota_unclassified(55);p__Ascomycota_unclassified(55); | - | - | - | - | - | - | - | Unassigned | - |
| Otu03274 | 11 | k__Fungi(100);p__Ascomycota(100);c__Eurotiomycetes(100);o__Eurotiales(100);f__Trichocomaceae(100);f__Trichocomaceae_unclassified(100);f__Trichocomaceae_unclassified(100); | - | - | - | - | - | - | - | Unassigned | - |
| Otu03276 | 11 | k__Fungi(100);k__Fungi_unclassified(82);k__Fungi_unclassified(82);k__Fungi_unclassified(82);k__Fungi_unclassified(82);k__Fungi_unclassified(82);k__Fungi_unclassified(82); | - | - | - | - | - | - | - | Unassigned | - |
| Otu03279 | 11 | k__Fungi(100);k__Fungi_unclassified(73);k__Fungi_unclassified(73);k__Fungi_unclassified(73);k__Fungi_unclassified(73);k__Fungi_unclassified(73);k__Fungi_unclassified(73); | - | - | - | - | - | - | - | Unassigned | - |
| Otu03283 | 11 | k__Fungi(100);p__Ascomycota(100);p__Ascomycota_unclassified(100);p__Ascomycota_unclassified(100);p__Ascomycota_unclassified(100);p__Ascomycota_unclassified(100);p__Ascomycota_unclassified(100); | - | - | - | - | - | - | - | Unassigned | - |
| Otu03285 | 11 | k__Fungi(100);k__Fungi_unclassified(100);k__Fungi_unclassified(100);k__Fungi_unclassified(100);k__Fungi_unclassified(100);k__Fungi_unclassified(100);k__Fungi_unclassified(100); | - | - | - | - | - | - | - | Unassigned | - |
| Otu03290 | 11 | k__Fungi(100);k__Fungi_unclassified(64);k__Fungi_unclassified(64);k__Fungi_unclassified(64);k__Fungi_unclassified(64);k__Fungi_unclassified(64);k__Fungi_unclassified(64); | - | - | - | - | - | - | - | Unassigned | - |
| Otu03291 | 11 | k__Fungi(100);p__Ascomycota(64);p__Ascomycota_unclassified(64);p__Ascomycota_unclassified(64);p__Ascomycota_unclassified(64);p__Ascomycota_unclassified(64);p__Ascomycota_unclassified(64); | - | - | - | - | - | - | - | Unassigned | - |
| Otu03292 | 11 | k__Fungi(100);k__Fungi_unclassified(64);k__Fungi_unclassified(64);k__Fungi_unclassified(64);k__Fungi_unclassified(64);k__Fungi_unclassified(64);k__Fungi_unclassified(64); | - | - | - | - | - | - | - | Unassigned | - |
| Otu03295 | 11 | k__Fungi(100);p__Ascomycota(100);p__Ascomycota_unclassified(100);p__Ascomycota_unclassified(100);p__Ascomycota_unclassified(100);p__Ascomycota_unclassified(100);p__Ascomycota_unclassified(100); | - | - | - | - | - | - | - | Unassigned | - |
| Otu03298 | 11 | k__Fungi(100);k__Fungi_unclassified(73);k__Fungi_unclassified(73);k__Fungi_unclassified(73);k__Fungi_unclassified(73);k__Fungi_unclassified(73);k__Fungi_unclassified(73); | - | - | - | - | - | - | - | Unassigned | - |
| Otu03299 | 11 | k__Fungi(100);k__Fungi_unclassified(64);k__Fungi_unclassified(64);k__Fungi_unclassified(64);k__Fungi_unclassified(64);k__Fungi_unclassified(64);k__Fungi_unclassified(64); | - | - | - | - | - | - | - | Unassigned | - |
| Otu03300 | 11 | k__Fungi(100);p__Ascomycota(100);c__Sordariomycetes(100);c__Sordariomycetes_unclassified(100);c__Sordariomycetes_unclassified(100);c__Sordariomycetes_unclassified(100);c__Sordariomycetes_unclassified(100); | - | - | - | - | - | - | - | Unassigned | - |
| Otu03303 | 11 | k__Fungi(100);p__Ascomycota(100);c__Dothideomycetes(91);o__Capnodiales(91);o__Capnodiales_unclassified(91);o__Capnodiales_unclassified(91);o__Capnodiales_unclassified(91); | - | - | - | - | - | - | - | Unassigned | - |
| Otu03304 | 11 | k__Fungi(100);p__Ascomycota(82);p__Ascomycota_unclassified(82);p__Ascomycota_unclassified(82);p__Ascomycota_unclassified(82);p__Ascomycota_unclassified(82);p__Ascomycota_unclassified(82); | - | - | - | - | - | - | - | Unassigned | - |
| Otu03309 | 11 | k__Fungi(100);p__Basidiomycota(100);c__Agaricomycetes(100);o__Polyporales(100);o__Polyporales_unclassified(100);o__Polyporales_unclassified(100);o__Polyporales_unclassified(100); | - | - | - | - | - | - | - | Unassigned | - |
| Otu03310 | 11 | k__Fungi(100);k__Fungi_unclassified(82);k__Fungi_unclassified(82);k__Fungi_unclassified(82);k__Fungi_unclassified(82);k__Fungi_unclassified(82);k__Fungi_unclassified(82); | - | - | - | - | - | - | - | Unassigned | - |
| Otu03312 | 11 | k__Fungi(100);p__Basidiomycota(100);c__Agaricomycetes(100);o__Auriculariales(100);f__unclassified_Auriculariales(100);g__unclassified_Auriculariales(100);s__Auriculariales_sp(100); | - | - | - | - | - | - | - | Unassigned | - |
| Otu03313 | 11 | k__Fungi(100);k__Fungi_unclassified(64);k__Fungi_unclassified(64);k__Fungi_unclassified(64);k__Fungi_unclassified(64);k__Fungi_unclassified(64);k__Fungi_unclassified(64); | - | - | - | - | - | - | - | Unassigned | - |
| Otu03315 | 11 | k__Fungi(100);p__Ascomycota(100);c__Leotiomycetes(100);o__Helotiales(100);o__Helotiales_unclassified(73);o__Helotiales_unclassified(73);o__Helotiales_unclassified(73); | - | - | - | - | - | - | - | Unassigned | - |
| Otu03317 | 11 | k__Fungi(100);p__Ascomycota(100);c__Dothideomycetes(100);o__Pleosporales(100);o__Pleosporales_unclassified(100);o__Pleosporales_unclassified(100);o__Pleosporales_unclassified(100); | - | - | - | - | - | - | - | Unassigned | - |
| Otu03318 | 11 | k__Fungi(100);p__Ascomycota(91);p__Ascomycota_unclassified(91);p__Ascomycota_unclassified(91);p__Ascomycota_unclassified(91);p__Ascomycota_unclassified(91);p__Ascomycota_unclassified(91); | - | - | - | - | - | - | - | Unassigned | - |
| Otu03320 | 11 | k__Fungi(100);p__Ascomycota(73);p__Ascomycota_unclassified(73);p__Ascomycota_unclassified(73);p__Ascomycota_unclassified(73);p__Ascomycota_unclassified(73);p__Ascomycota_unclassified(73); | - | - | - | - | - | - | - | Unassigned | - |
| Otu03324 | 11 | k__Fungi(100);k__Fungi_unclassified(100);k__Fungi_unclassified(100);k__Fungi_unclassified(100);k__Fungi_unclassified(100);k__Fungi_unclassified(100);k__Fungi_unclassified(100); | - | - | - | - | - | - | - | Unassigned | - |
| Otu03325 | 11 | k__Fungi(100);p__Ascomycota(100);c__Sordariomycetes(100);c__Sordariomycetes_unclassified(100);c__Sordariomycetes_unclassified(100);c__Sordariomycetes_unclassified(100);c__Sordariomycetes_unclassified(100); | - | - | - | - | - | - | - | Unassigned | - |
| Otu03326 | 11 | k__Fungi(100);k__Fungi_unclassified(91);k__Fungi_unclassified(91);k__Fungi_unclassified(91);k__Fungi_unclassified(91);k__Fungi_unclassified(91);k__Fungi_unclassified(91); | - | - | - | - | - | - | - | Unassigned | - |
| Otu03327 | 11 | k__Fungi(100);p__Ascomycota(100);c__Sordariomycetes(100);o__Hypocreales(100);f__Hypocreaceae(91);f__Hypocreaceae_unclassified(91);f__Hypocreaceae_unclassified(91); | - | - | - | - | - | - | - | Unassigned | - |
| Otu03332 | 11 | k__Fungi(100);k__Fungi_unclassified(64);k__Fungi_unclassified(64);k__Fungi_unclassified(64);k__Fungi_unclassified(64);k__Fungi_unclassified(64);k__Fungi_unclassified(64); | - | - | - | - | - | - | - | Unassigned | - |
| Otu03338 | 11 | k__Fungi(100);p__Ascomycota(100);p__Ascomycota_unclassified(100);p__Ascomycota_unclassified(100);p__Ascomycota_unclassified(100);p__Ascomycota_unclassified(100);p__Ascomycota_unclassified(100); | - | - | - | - | - | - | - | Unassigned | - |
| Otu03339 | 11 | k__Fungi(100);p__Ascomycota(100);p__Ascomycota_unclassified(100);p__Ascomycota_unclassified(100);p__Ascomycota_unclassified(100);p__Ascomycota_unclassified(100);p__Ascomycota_unclassified(100); | - | - | - | - | - | - | - | Unassigned | - |
| Otu03341 | 11 | k__Fungi(100);k__Fungi_unclassified(73);k__Fungi_unclassified(73);k__Fungi_unclassified(73);k__Fungi_unclassified(73);k__Fungi_unclassified(73);k__Fungi_unclassified(73); | - | - | - | - | - | - | - | Unassigned | - |
| Otu03342 | 11 | k__Fungi(100);p__Basidiomycota(100);c__Agaricomycetes(100);o__Cantharellales(100);o__Cantharellales_unclassified(100);o__Cantharellales_unclassified(100);o__Cantharellales_unclassified(100); | - | - | - | - | - | - | - | Unassigned | - |
| Otu03343 | 11 | k__Fungi(100);k__Fungi_unclassified(91);k__Fungi_unclassified(91);k__Fungi_unclassified(91);k__Fungi_unclassified(91);k__Fungi_unclassified(91);k__Fungi_unclassified(91); | - | - | - | - | - | - | - | Unassigned | - |
| Otu03345 | 11 | k__Fungi(100);p__Ascomycota(100);p__Ascomycota_unclassified(100);p__Ascomycota_unclassified(100);p__Ascomycota_unclassified(100);p__Ascomycota_unclassified(100);p__Ascomycota_unclassified(100); | - | - | - | - | - | - | - | Unassigned | - |
| Otu03347 | 11 | k__Fungi(100);k__Fungi_unclassified(100);k__Fungi_unclassified(100);k__Fungi_unclassified(100);k__Fungi_unclassified(100);k__Fungi_unclassified(100);k__Fungi_unclassified(100); | - | - | - | - | - | - | - | Unassigned | - |
| Otu03350 | 11 | k__Fungi(100);k__Fungi_unclassified(64);k__Fungi_unclassified(64);k__Fungi_unclassified(64);k__Fungi_unclassified(64);k__Fungi_unclassified(64);k__Fungi_unclassified(64); | - | - | - | - | - | - | - | Unassigned | - |
| Otu03351 | 10 | k__Fungi(100);p__Ascomycota(100);c__Saccharomycetes(100);o__Saccharomycetales(100);f__Metschnikowiaceae(100);g__Metschnikowia(100);s__Metschnikowia_pulcherrima(100); | Metschnikowia | 13 | Saprotroph | \|Undefined Saprotroph\| | Yeast | NULL | Probable | As nectar-tap_saprotroph (Põlme et al. 2020); Decay sugar-rich substrates (Põlme et al. 2020) | Põlme S, et al. 2020. Fungal Diversity 105:1-16 ((https://doi.org/10.1007/s13225-020-00466-2)) |
| Otu03353 | 10 | k__Fungi(100);p__Ascomycota(100);c__Sordariomycetes(100);o__Microascales(100);f__Microascaceae(100);g__unclassified_Microascaceae(100);s__Microascaceae_sp(100); | Microascaceae | 9 | Pathotroph-Saprotroph-Symbiotroph | Animal Pathogen-Endophyte-Plant Pathogen-Undefined Saprotroph | Microfungus | NULL | Possible | Saprobic in soil or rotting vegetation, some species heat-tolerant and associatated with silage and compost, a few associated with humans and animals (Cannon & Kirk 2007) | Cannon PF, Kirk PM. 2007. Fungal Families of the World. CAB International, Cambridge (ISBN: 978-0851998275) |
| Otu03355 | 10 | k__Fungi(100);p__Basidiomycota(100);c__Wallemiomycetes(100);o__Wallemiales(100);f__Wallemiaceae(100);g__Wallemia(100);s__Wallemia_sebi(100); | Wallemia | 13 | Saprotroph | \|Undefined Saprotroph\| | Microfungus | NULL | Probable | Decay sugar-rich substrates (Põlme et al. 2020) | Tedersoo L, et al. 2014. Science 346:e1256688 ((https://doi.org/10.1126/science.1256688)); Põlme S, et al. 2020. Fungal Diversity 105:1-16 ((https://doi.org/10.1007/s13225-020-00466-2)) |
| Otu03356 | 10 | k__Fungi(100);p__Ascomycota(100);c__Dothideomycetes(100);o__Capnodiales(100);f__Mycosphaerellaceae(100);f__Mycosphaerellaceae_unclassified(100);f__Mycosphaerellaceae_unclassified(100); | Mycosphaerellaceae | 9 | Pathotroph-Saprotroph | Plant Pathogen-Undefined Saprotroph | Microfungus | NULL | Probable | Biotrophic, necrotrophic or saprobic on various plant tissues (Cannon & Kirk 2007) | Cannon PF, Kirk PM. 2007. Fungal Families of the World. CAB International, Cambridge (ISBN: 978-0851998275) |
| Otu03359 | 10 | k__Fungi(100);p__Basidiomycota(100);c__Agaricomycetes(100);o__Agaricales(100);f__Pleurotaceae(100);f__Pleurotaceae_unclassified(70);f__Pleurotaceae_unclassified(70); | Pleurotaceae | 9 | Pathotroph-Saprotroph | Endophyte-Plant Pathogen-Wood Saprotroph | Agaricoid | White Rot | Probable | Sapwood rot on Pistacia; Saprobic on woody substrata, more rarely associated with roots of herbaceous plants. Many species are known to derive nutrition from nematodes, the hyphae with adhesive knobs or toxigenic secretory appendages (Cannon & Kirk 2007) | Cannon PF, Kirk PM. 2007. Fungal Families of the World. CAB International, Cambridge (ISBN: 978-0851998275); Costa IPMW, et al. 2012. Mycotaxon 119:e26 ((http://mycotaxon.com/resources/checklists/costa_v119_checklist.pdf)); The American Phytopathological Society Common Names of Plant Diseases ((http://www.apsnet.org/publications/commonnames/Pages/default.aspx)) |
| Otu03362 | 10 | k__Fungi(100);p__Ascomycota(100);c__Eurotiomycetes(100);o__Eurotiales(100);f__Trichocomaceae(100);g__Paecilomyces(100);s__Paecilomyces_marquandii(100); | Paecilomyces | 13 | Pathotroph-Saprotroph-Symbiotroph | Clavicipitaceous \|Endophyte\|-Plant Pathogen-Plant Saprotroph-Undefined Saprotroph-Wood Saprotroph | Microfungus | Soft Rot | Probable | As foliar_endophyte (Põlme et al. 2020); As litter_saprotroph (Põlme et al. 2020); culture (Lopez et al. 2014); Decay leaf, fruit, seed, animal material (Põlme et al. 2020); Endophyte detection method | Seehann G, et al. 1975. List of Fungi in Soft-Rot Tissue Tests. Institute fur Holzbiologie und Holzschutz, Bundesforschungsanstalt fur Forst - und Holzwirtshaft, Hamburg - Lohbrugge, Germany; Costa IPMW, et al. 2012. Mycotaxon 119:e26 ((http://mycotaxon.com/resources/checklists/costa_v119_checklist.pdf)); Tedersoo L, et al. 2014. Science 346:e1256688 ((https://doi.org/10.1126/science.1256688)); Põlme S, et al. 2020. Fungal Diversity 105:1-16 ((https://doi.org/10.1007/s13225-020-00466-2)); The American Phytopathological Society Common Names of Plant Diseases ((http://www.apsnet.org/publications/commonnames/Pages/default.aspx)); Lopez DC, et al. 2014. Plos One 9:e103891 ((https://doi.org/10.1371/journal.pone.0103891)) |
| Otu03364 | 10 | k__Fungi(100);p__Ascomycota(100);c__Saccharomycetes(100);o__Saccharomycetales(100);f__Saccharomycetales_family_Incertae_sedis(90);g__Candida(90);g__Candida_unclassified(70); | Candida | 13 | Pathotroph-Saprotroph-Symbiotroph | Animal Pathogen-Endophyte-Endosymbiont-Epiphyte-\|Undefined Saprotroph\| | Dimorphic Yeast-Yeast | NULL | Probable | As nectar-tap_saprotroph (Põlme et al. 2020); Decay sugar-rich substrates (Põlme et al. 2020); This asexual genus is polyphyletic and contain species that function very differently depend on what they are. Caution should be taken when interpreting guilds for Candida species. | Manolakaki D, et al. 2010. Virulence. 1: 367-375 ((https://doi.org/10.4161/viru.1.5.12796)); Mekbib SB, et al. 2011. Tropical Plant Pathology 36:233-240 ((https://doi.org/10.1590/S1982-56762011000400004)); Põlme S, et al. 2020. Fungal Diversity 105:1-16 ((https://doi.org/10.1007/s13225-020-00466-2)) |
| Otu03366 | 10 | k__Fungi(100);p__Ascomycota(100);c__Eurotiomycetes(100);o__Eurotiales(100);f__Trichocomaceae(100);g__Eurotium(100);s__Eurotium_halophilicum(100); | Eurotium | 13 | Saprotroph-Symbiotroph | Endophyte-\|Plant Saprotroph\|-Undefined Saprotroph | Microfungus | NULL | Probable | As foliar_endophyte (Põlme et al. 2020); As litter_saprotroph (Põlme et al. 2020); Decay leaf, fruit, seed (Põlme et al. 2020) | Tedersoo L, et al. 2014. Science 346:e1256688 ((https://doi.org/10.1126/science.1256688)); Massimo NC, et al. 2015. Microbial Ecology 70:61-76 ((https://doi.org/10.1007/s00248-014-0563-6)); Põlme S, et al. 2020. Fungal Diversity 105:1-16 ((https://doi.org/10.1007/s13225-020-00466-2)) |
| Otu03367 | 10 | k__Fungi(100);p__Basidiomycota(100);c__Agaricomycetes(100);o__Thelephorales(100);f__Thelephoraceae(100);g__Tomentella(100);g__Tomentella_unclassified(100); | Tomentella | 13 | Symbiotroph | \|Ectomycorrhizal\| | Corticioid | NULL | Highly Probable | NULL | Rinaldi AC, et al. 2008. Fungal Diversity 33:1-45 ((http://www.fungaldiversity.org/fdp/sfdp/33-1.pdf)); Tedersoo L, et al. 2010. Mycorrhiza 20:217-263 ((https://doi.org/10.1007/s00572-009-0274-x)); Põlme S, et al. 2020. Fungal Diversity 105:1-16 ((https://doi.org/10.1007/s13225-020-00466-2)); http://mycorrhizas.info/ecmf.html ((http://mycorrhizas.info/ecmf.html)) |
| Otu03368 | 10 | k__Fungi(100);p__Ascomycota(100);c__Saccharomycetes(100);o__Saccharomycetales(100);f__Metschnikowiaceae(100);g__Metschnikowia(100);s__Metschnikowia_pulcherrima(90); | Metschnikowia | 13 | Saprotroph | \|Undefined Saprotroph\| | Yeast | NULL | Probable | As nectar-tap_saprotroph (Põlme et al. 2020); Decay sugar-rich substrates (Põlme et al. 2020) | Põlme S, et al. 2020. Fungal Diversity 105:1-16 ((https://doi.org/10.1007/s13225-020-00466-2)) |
| Otu03369 | 10 | k__Fungi(100);p__Ascomycota(100);c__Sordariomycetes(100);o__Hypocreales(100);o__Hypocreales_unclassified(100);o__Hypocreales_unclassified(100);o__Hypocreales_unclassified(100); | Hypocreales | 7 | Saprotroph | Undefined Saprotroph | Microfungus | NULL | Possible | NULL | Sterkenburg E, et al. 2015. New Phytologist 207:1145-1158 ((https://doi.org/10.1111/nph.13426)) |
| Otu03371 | 10 | k__Fungi(100);p__Ascomycota(100);c__Saccharomycetes(100);o__Saccharomycetales(100);f__Metschnikowiaceae(100);g__Metschnikowia(100);s__Metschnikowia_pulcherrima(100); | Metschnikowia | 13 | Saprotroph | \|Undefined Saprotroph\| | Yeast | NULL | Probable | As nectar-tap_saprotroph (Põlme et al. 2020); Decay sugar-rich substrates (Põlme et al. 2020) | Põlme S, et al. 2020. Fungal Diversity 105:1-16 ((https://doi.org/10.1007/s13225-020-00466-2)) |
| Otu03373 | 10 | k__Fungi(100);p__Ascomycota(100);c__Saccharomycetes(100);o__Saccharomycetales(100);f__Saccharomycetales_family_Incertae_sedis(70);f__Saccharomycetales_family_Incertae_sedis_unclassified(61);f__Saccharomycetales_family_Incertae_sedis_unclassified(61); | Saccharomycetales | 7 | Saprotroph | Undefined Saprotroph | Yeast | NULL | Possible | NULL | Sterkenburg E, et al. 2015. New Phytologist 207:1145-1158 ((https://doi.org/10.1111/nph.13426)) |
| Otu03374 | 10 | k__Fungi(100);p__Ascomycota(100);c__Saccharomycetes(100);o__Saccharomycetales(100);f__Metschnikowiaceae(100);g__Metschnikowia(100);s__Metschnikowia_pulcherrima(100); | Metschnikowia | 13 | Saprotroph | \|Undefined Saprotroph\| | Yeast | NULL | Probable | As nectar-tap_saprotroph (Põlme et al. 2020); Decay sugar-rich substrates (Põlme et al. 2020) | Põlme S, et al. 2020. Fungal Diversity 105:1-16 ((https://doi.org/10.1007/s13225-020-00466-2)) |
| Otu03379 | 10 | k__Fungi(100);p__Basidiomycota(100);c__Microbotryomycetes(100);o__Sporidiobolales(100);f__Sporidiobolales_family_Incertae_sedis(100);g__Rhodotorula(100);s__Rhodotorula_mucilaginosa(100); | Rhodotorula | 13 | Pathotroph-Saprotroph-Symbiotroph | Animal Pathogen-Animal Symbiotroph-Endophyte-Plant Pathogen-\|Undefined Saprotroph\| | Yeast | NULL | Probable | As foliar_endophyte (Põlme et al. 2020); Opportunistic human pathogen | Põlme S, et al. 2020. Fungal Diversity 105:1-16 ((https://doi.org/10.1007/s13225-020-00466-2)); https://www.cdc.gov ((https://www.cdc.gov)) |
| Otu03380 | 10 | k__Fungi(100);p__Ascomycota(100);c__Eurotiomycetes(100);o__Chaetothyriales(100);f__Herpotrichiellaceae(100);g__Cladophialophora(100);g__Cladophialophora_unclassified(100); | Cladophialophora | 13 | Pathotroph-Saprotroph-Symbiotroph | Animal Pathogen-Endophyte-Fungal Parasite-\|Undefined Saprotroph\| | Dimorphic-Microfungus | NULL | Probable | As mycoparasite (Põlme et al. 2020); As root_endophyte_dark_septate (Põlme et al. 2020); As soil_saprotroph (Põlme et al. 2020); Decay soil (Põlme et al. 2020) | James TY, et al. 2006. Nature 443:818-822 ((https://doi.org/10.1038/nature05110)); Põlme S, et al. 2020. Fungal Diversity 105:1-16 ((https://doi.org/10.1007/s13225-020-00466-2)) |
| Otu03382 | 10 | k__Fungi(100);p__Ascomycota(100);c__Eurotiomycetes(100);o__Chaetothyriales(100);f__Herpotrichiellaceae(100);g__Exophiala(70);g__Exophiala_unclassified(70); | Exophiala | 13 | Pathotroph-Saprotroph-Symbiotroph | \|Animal Parasite\|-Animal Pathogen-Endophyte-Plant Saprotroph-Undefined Saprotroph | Dimorphic-Facultative Yeast | NULL | Probable | As litter_saprotroph (Põlme et al. 2020); As root_endophyte_dark_septate (Põlme et al. 2020); Decay leaf, fruit, seed, animal material (Põlme et al. 2020) | Tedersoo L, et al. 2014. Science 346:e1256688 ((https://doi.org/10.1126/science.1256688)); Nagy LG, et al. 2014. Nature Communications 5:e4471 ((https://doi.org/10.1038/ncomms5471)); Põlme S, et al. 2020. Fungal Diversity 105:1-16 ((https://doi.org/10.1007/s13225-020-00466-2)) |
| Otu03383 | 10 | k__Fungi(100);p__Ascomycota(100);c__Eurotiomycetes(100);o__Mycocaliciales(100);f__Mycocaliciaceae(100);g__Chaenothecopsis(100);s__Chaenothecopsis_tibellii(100); | Chaenothecopsis | 13 | Pathotroph-Saprotroph-Symbiotroph | Lichen Parasite-Lichenized-\|Undefined Saprotroph\| | Calicioid-Thallus | NULL | Possible | As resin_saprotroph (Põlme et al. 2020); licenized taxa; Lichenized and non | Esslinger TL. 2014. North Dakota State University ((http://www.ndsu.edu/pubweb/~esslinge/chcklst/chcklst7.htm)); Põlme S, et al. 2020. Fungal Diversity 105:1-16 ((https://doi.org/10.1007/s13225-020-00466-2)) |
| Otu03384 | 10 | k__Fungi(100);p__Basidiomycota(100);c__Agaricomycetes(100);o__Agaricales(100);f__Niaceae(100);g__Merismodes(100);s__Merismodes_fasciculata(100); | Merismodes | 13 | Saprotroph | Undefined Saprotroph-\|Wood Saprotroph\| | Cyphelloid | NULL | Probable | Decay wood (Põlme et al. 2020) | Tedersoo L, et al. 2014. Science 346:e1256688 ((https://doi.org/10.1126/science.1256688)); Põlme S, et al. 2020. Fungal Diversity 105:1-16 ((https://doi.org/10.1007/s13225-020-00466-2)) |
| Otu03386 | 10 | k__Fungi(100);p__Ascomycota(100);c__Dothideomycetes(100);o__Capnodiales(100);f__Teratosphaeriaceae(100);g__Devriesia(100);s__Devriesia_sp(90); | Devriesia | 13 | Pathotroph-Saprotroph | \|Plant Pathogen\|-Plant Saprotroph | Microfungus | NULL | Probable | As litter_saprotroph (Põlme et al. 2020); Decay leaf, fruit, seed (Põlme et al. 2020) | Tedersoo L, et al. 2014. Science 346:e1256688 ((https://doi.org/10.1126/science.1256688)); Põlme S, et al. 2020. Fungal Diversity 105:1-16 ((https://doi.org/10.1007/s13225-020-00466-2)) |
| Otu03388 | 10 | k__Fungi(100);p__Ascomycota(100);c__Eurotiomycetes(100);o__Chaetothyriales(100);f__Herpotrichiellaceae(100);g__Phaeococcomyces(100);s__Phaeococcomyces_sp(100); | Phaeococcomyces | 13 | Saprotroph | \|Undefined Saprotroph\| | Dimorphic-Facultative Yeast | NULL | Probable | NULL | Tedersoo L, et al. 2014. Science 346:e1256688 ((https://doi.org/10.1126/science.1256688)); Põlme S, et al. 2020. Fungal Diversity 105:1-16 ((https://doi.org/10.1007/s13225-020-00466-2)) |
| Otu03391 | 10 | k__Fungi(100);p__Ascomycota(100);c__Leotiomycetes(100);o__Helotiales(100);f__Vibrisseaceae(100);g__Phialocephala(90);g__Phialocephala_unclassified(90); | Phialocephala | 13 | Saprotroph-Symbiotroph | Endophyte-\|Undefined Saprotroph\| | Microfungus | NULL | Highly Probable | As root_endophyte (Põlme et al. 2020); As root_endophyte_dark_septate (Põlme et al. 2020); As soil_saprotroph (Põlme et al. 2020); Dark Septate Endophyte, known from plant roots.; Decay roots, soil (Põlme et al. 2020) | Newsham KK. 2011. New Phytologist 190:783-793 ((https://doi.org/10.1111/j.1469-8137.2010.03611.x)) (as dark septate root endophytes); Põlme S, et al. 2020. Fungal Diversity 105:1-16 ((https://doi.org/10.1007/s13225-020-00466-2)) |
| Otu03392 | 10 | k__Fungi(100);p__Basidiomycota(100);c__Agaricomycetes(100);o__Russulales(100);f__Russulaceae(100);g__Russula(100);s__Russula_compacta(100); | Russula | 13 | Symbiotroph | \|Ectomycorrhizal\| | Agaricoid | NULL | Highly Probable | NULL | Rinaldi AC, et al. 2008. Fungal Diversity 33:1-45 ((http://www.fungaldiversity.org/fdp/sfdp/33-1.pdf)); Tedersoo L, et al. 2010. Mycorrhiza 20:217-263 ((https://doi.org/10.1007/s00572-009-0274-x)); Põlme S, et al. 2020. Fungal Diversity 105:1-16 ((https://doi.org/10.1007/s13225-020-00466-2)); http://mycorrhizas.info/ecmf.html ((http://mycorrhizas.info/ecmf.html)) |
| Otu03395 | 10 | k__Fungi(100);p__Ascomycota(100);c__Saccharomycetes(100);o__Saccharomycetales(100);f__Metschnikowiaceae(100);g__Metschnikowia(100);s__Metschnikowia_pulcherrima(90); | Metschnikowia | 13 | Saprotroph | \|Undefined Saprotroph\| | Yeast | NULL | Probable | As nectar-tap_saprotroph (Põlme et al. 2020); Decay sugar-rich substrates (Põlme et al. 2020) | Põlme S, et al. 2020. Fungal Diversity 105:1-16 ((https://doi.org/10.1007/s13225-020-00466-2)) |
| Otu03397 | 10 | k__Fungi(100);p__Ascomycota(100);c__Saccharomycetes(100);o__Saccharomycetales(100);f__Metschnikowiaceae(100);g__Metschnikowia(100);s__Metschnikowia_pulcherrima(100); | Metschnikowia | 13 | Saprotroph | \|Undefined Saprotroph\| | Yeast | NULL | Probable | As nectar-tap_saprotroph (Põlme et al. 2020); Decay sugar-rich substrates (Põlme et al. 2020) | Põlme S, et al. 2020. Fungal Diversity 105:1-16 ((https://doi.org/10.1007/s13225-020-00466-2)) |
| Otu03399 | 10 | k__Fungi(100);p__Basidiomycota(100);c__Pucciniomycetes(100);o__Pucciniales(100);f__Pucciniastraceae(100);g__Thekopsora(70);g__Thekopsora_unclassified(70); | Thekopsora | 13 | Pathotroph | \|Plant Pathogen\| | Rust | NULL | Probable | NULL | Tedersoo L, et al. 2014. Science 346:e1256688 ((https://doi.org/10.1126/science.1256688)); Põlme S, et al. 2020. Fungal Diversity 105:1-16 ((https://doi.org/10.1007/s13225-020-00466-2)) |
| Otu03401 | 10 | k__Fungi(100);p__Ascomycota(100);c__Saccharomycetes(100);o__Saccharomycetales(100);f__Metschnikowiaceae(100);g__Metschnikowia(100);g__Metschnikowia_unclassified(61); | Metschnikowia | 13 | Saprotroph | \|Undefined Saprotroph\| | Yeast | NULL | Probable | As nectar-tap_saprotroph (Põlme et al. 2020); Decay sugar-rich substrates (Põlme et al. 2020) | Põlme S, et al. 2020. Fungal Diversity 105:1-16 ((https://doi.org/10.1007/s13225-020-00466-2)) |
| Otu03404 | 10 | k__Fungi(100);p__Ascomycota(100);c__Saccharomycetes(100);o__Saccharomycetales(100);f__Metschnikowiaceae(100);g__Metschnikowia(100);s__Metschnikowia_pulcherrima(100); | Metschnikowia | 13 | Saprotroph | \|Undefined Saprotroph\| | Yeast | NULL | Probable | As nectar-tap_saprotroph (Põlme et al. 2020); Decay sugar-rich substrates (Põlme et al. 2020) | Põlme S, et al. 2020. Fungal Diversity 105:1-16 ((https://doi.org/10.1007/s13225-020-00466-2)) |
| Otu03405 | 10 | k__Fungi(100);p__Basidiomycota(100);c__Agaricostilbomycetes(100);o__Agaricostilbales(100);f__Chionosphaeraceae(100);g__Chionosphaera(80);s__Chionosphaera_cuniculicola(80); | Chionosphaera | 20 | Pathotroph-Saprotroph | Lichen Parasite-\|Plant Saprotroph\| | Dimorphic-Microfungus | NULL | Probable | As litter_saprotroph (Põlme et al. 2020); Decay leaf, fruit, seed (Põlme et al. 2020); Lichen parasite on various lichens (Kirschner 2001) | Kirschner R. 2001. Mycological Research 1051403-1408 ((https://doi.org/10.1017/S0953756201005019)); Lawrey JD, Diederich P. 2016. http://www.lichenicolous.net (accessed 24 May 2017) ((http://www.lichenicolous.net)); Põlme S, et al. 2020. Fungal Diversity 105:1-16 ((https://doi.org/10.1007/s13225-020-00466-2)) |
| Otu03406 | 10 | k__Fungi(100);p__Ascomycota(100);c__Sordariomycetes(100);o__Hypocreales(100);f__Hypocreaceae(100);g__Hypocrea(100);s__Hypocrea_sp(80); | Hypocrea | 13 | Pathotroph | Endophyte-Fungal Parasite | Microfungus | NULL | Probable | NULL | Tedersoo L, et al. 2014. Science 346:e1256688 ((https://doi.org/10.1126/science.1256688)); Christian N, et al. 2016. Microbial Ecology 72:621-632 ((https://doi.org/10.1007/s00248-016-0804-y)) |
| Otu03407 | 10 | k__Fungi(100);p__Basidiomycota(100);c__Agaricomycetes(100);o__Cantharellales(100);f__Tulasnellaceae(100);g__Tulasnella(100);s__Tulasnella_sp_184(100); | Tulasnella | 13 | Saprotroph-Symbiotroph | Ectomycorrhizal-\|Plant Saprotroph\|-Undefined Saprotroph | Corticioid | NULL | Possible | As litter_saprotroph (Põlme et al. 2020); Decay leaf, fruit, seed (Põlme et al. 2020) | Rinaldi AC, et al. 2008. Fungal Diversity 33:1-45 ((http://www.fungaldiversity.org/fdp/sfdp/33-1.pdf)); Tedersoo L, et al. 2010. Mycorrhiza 20:217-263 ((https://doi.org/10.1007/s00572-009-0274-x)) (pro parte); Põlme S, et al. 2020. Fungal Diversity 105:1-16 ((https://doi.org/10.1007/s13225-020-00466-2)) |
| Otu03408 | 10 | k__Fungi(100);p__Ascomycota(100);c__Saccharomycetes(100);o__Saccharomycetales(100);f__Saccharomycetales_family_Incertae_sedis(90);g__Candida(80);g__Candida_unclassified(80); | Candida | 13 | Pathotroph-Saprotroph-Symbiotroph | Animal Pathogen-Endophyte-Endosymbiont-Epiphyte-\|Undefined Saprotroph\| | Dimorphic Yeast-Yeast | NULL | Probable | As nectar-tap_saprotroph (Põlme et al. 2020); Decay sugar-rich substrates (Põlme et al. 2020); This asexual genus is polyphyletic and contain species that function very differently depend on what they are. Caution should be taken when interpreting guilds for Candida species. | Manolakaki D, et al. 2010. Virulence. 1: 367-375 ((https://doi.org/10.4161/viru.1.5.12796)); Mekbib SB, et al. 2011. Tropical Plant Pathology 36:233-240 ((https://doi.org/10.1590/S1982-56762011000400004)); Põlme S, et al. 2020. Fungal Diversity 105:1-16 ((https://doi.org/10.1007/s13225-020-00466-2)) |
| Otu03411 | 10 | k__Fungi(100);p__Basidiomycota(100);c__Agaricomycetes(100);o__Polyporales(100);f__Phanerochaetaceae(100);g__Ceriporia(100);g__Ceriporia_unclassified(100); | Ceriporia | 13 | Saprotroph | \|Wood Saprotroph\| | Corticioid | White Rot | Highly Probable | As white_rot (Põlme et al. 2020); Decay wood (Põlme et al. 2020) | Tedersoo L, et al. 2014. Science 346:e1256688 ((https://doi.org/10.1126/science.1256688)); Põlme S, et al. 2020. Fungal Diversity 105:1-16 ((https://doi.org/10.1007/s13225-020-00466-2)) |
| Otu03412 | 10 | k__Fungi(100);p__Ascomycota(100);c__Saccharomycetes(100);o__Saccharomycetales(100);f__Metschnikowiaceae(100);g__Metschnikowia(100);s__Metschnikowia_pulcherrima(100); | Metschnikowia | 13 | Saprotroph | \|Undefined Saprotroph\| | Yeast | NULL | Probable | As nectar-tap_saprotroph (Põlme et al. 2020); Decay sugar-rich substrates (Põlme et al. 2020) | Põlme S, et al. 2020. Fungal Diversity 105:1-16 ((https://doi.org/10.1007/s13225-020-00466-2)) |
| Otu03415 | 10 | k__Fungi(100);p__Ascomycota(100);c__Lecanoromycetes(100);o__Lecanorales(100);f__Ramalinaceae(100);g__Bacidina(100);s__Bacidina_sulphurella(100); | Bacidina | 13 | Symbiotroph | \|Lichenized\| | Thallus | NULL | Highly Probable | NULL | Esslinger TL. 2014. North Dakota State University ((http://www.ndsu.edu/pubweb/~esslinge/chcklst/chcklst7.htm)); Põlme S, et al. 2020. Fungal Diversity 105:1-16 ((https://doi.org/10.1007/s13225-020-00466-2)) |
| Otu03418 | 10 | k__Fungi(100);p__Ascomycota(100);c__Saccharomycetes(100);o__Saccharomycetales(100);f__Saccharomycetales_family_Incertae_sedis(80);g__Candida(80);g__Candida_unclassified(70); | Candida | 13 | Pathotroph-Saprotroph-Symbiotroph | Animal Pathogen-Endophyte-Endosymbiont-Epiphyte-\|Undefined Saprotroph\| | Dimorphic Yeast-Yeast | NULL | Probable | As nectar-tap_saprotroph (Põlme et al. 2020); Decay sugar-rich substrates (Põlme et al. 2020); This asexual genus is polyphyletic and contain species that function very differently depend on what they are. Caution should be taken when interpreting guilds for Candida species. | Manolakaki D, et al. 2010. Virulence. 1: 367-375 ((https://doi.org/10.4161/viru.1.5.12796)); Mekbib SB, et al. 2011. Tropical Plant Pathology 36:233-240 ((https://doi.org/10.1590/S1982-56762011000400004)); Põlme S, et al. 2020. Fungal Diversity 105:1-16 ((https://doi.org/10.1007/s13225-020-00466-2)) |
| Otu03420 | 10 | k__Fungi(100);p__Basidiomycota(100);c__Agaricomycetes(100);o__Polyporales(100);f__Fomitopsidaceae(100);g__Ischnoderma(100);s__Ischnoderma_sp(100); | Ischnoderma | 13 | Saprotroph | \|Wood Saprotroph\| | Polyporoid | White Rot | Highly Probable | As white_rot (Põlme et al. 2020); Decay wood (Põlme et al. 2020) | Gilbertson RL, Ryvarden L. 1987-1987. North American Polypores. Fungiflora, Oslo (ISBN: 978-0945345060); Põlme S, et al. 2020. Fungal Diversity 105:1-16 ((https://doi.org/10.1007/s13225-020-00466-2)) |
| Otu03422 | 10 | k__Fungi(100);p__Basidiomycota(100);c__Agaricomycetes(100);o__Agaricales(100);f__Cortinariaceae(100);g__Cortinarius(100);s__Cortinarius_lucorum(100); | Cortinarius | 13 | Symbiotroph | \|Ectomycorrhizal\| | Agaricoid | NULL | Highly Probable | NULL | Rinaldi AC, et al. 2008. Fungal Diversity 33:1-45 ((http://www.fungaldiversity.org/fdp/sfdp/33-1.pdf)); Tedersoo L, et al. 2010. Mycorrhiza 20:217-263 ((https://doi.org/10.1007/s00572-009-0274-x)); Põlme S, et al. 2020. Fungal Diversity 105:1-16 ((https://doi.org/10.1007/s13225-020-00466-2)); http://mycorrhizas.info/ecmf.html ((http://mycorrhizas.info/ecmf.html)) |
| Otu03423 | 10 | k__Fungi(100);p__Ascomycota(100);c__Saccharomycetes(100);o__Saccharomycetales(100);f__Metschnikowiaceae(100);g__Metschnikowia(100);g__Metschnikowia_unclassified(70); | Metschnikowia | 13 | Saprotroph | \|Undefined Saprotroph\| | Yeast | NULL | Probable | As nectar-tap_saprotroph (Põlme et al. 2020); Decay sugar-rich substrates (Põlme et al. 2020) | Põlme S, et al. 2020. Fungal Diversity 105:1-16 ((https://doi.org/10.1007/s13225-020-00466-2)) |
| Otu03424 | 10 | k__Fungi(100);p__Ascomycota(100);c__Sordariomycetes(100);o__Hypocreales(100);f__Nectriaceae(100);g__Gibberella(100);g__Gibberella_unclassified(100); | Gibberella | 13 | Pathotroph-Saprotroph-Symbiotroph | Endophyte-\|Plant Pathogen\|-Plant Saprotroph-Undefined Saprotroph | Microfungus | Soft Rot | Probable | As foliar_endophyte (Põlme et al. 2020); As litter_saprotroph (Põlme et al. 2020); Causing damping-off disease (Põlme et al. 2020); Decay leaf, fruit, seed, roots, soil, animal material (Põlme et al. 2020) | Tedersoo L, et al. 2014. Science 346:e1256688 ((https://doi.org/10.1126/science.1256688)); Põlme S, et al. 2020. Fungal Diversity 105:1-16 ((https://doi.org/10.1007/s13225-020-00466-2)) |
| Otu03430 | 10 | k__Fungi(100);p__Zygomycota(100);c__Zygomycota_class_Incertae_sedis(100);o__Mucorales(100);f__Mucoraceae(100);g__Mucor(100);s__Mucor_sp(100); | Mucor | 13 | Saprotroph | \|Undefined Saprotroph\| | Microfungus | NULL | Probable | As soil_saprotroph (Põlme et al. 2020); Decay soil (Põlme et al. 2020) | Tedersoo L, et al. 2014. Science 346:e1256688 ((https://doi.org/10.1126/science.1256688)); Põlme S, et al. 2020. Fungal Diversity 105:1-16 ((https://doi.org/10.1007/s13225-020-00466-2)) |
| Otu03431 | 10 | k__Fungi(100);p__Zygomycota(100);c__Zygomycota_class_Incertae_sedis(100);o__Mucorales(100);f__Choanephoraceae(100);g__Blakeslea(61);s__Blakeslea_trispora(61); | Blakeslea | 13 | Pathotroph-Saprotroph | Plant Pathogen-\|Undefined Saprotroph\| | Microfungus | NULL | Probable | As soil_saprotroph (Põlme et al. 2020); Decay leaf, fruit, seed, soil (Põlme et al. 2020) | Tedersoo L, et al. 2014. Science 346:e1256688 ((https://doi.org/10.1126/science.1256688)); Põlme S, et al. 2020. Fungal Diversity 105:1-16 ((https://doi.org/10.1007/s13225-020-00466-2)) |
| Otu03432 | 10 | k__Fungi(100);p__Basidiomycota(100);c__Agaricomycetes(100);o__Agaricales(100);f__Psathyrellaceae(100);g__Parasola(100);g__Parasola_unclassified(61); | Parasola | 13 | Saprotroph | \|Undefined Saprotroph\| | Agaricoid | NULL | Probable | As soil_saprotroph (Põlme et al. 2020); Decay soil (Põlme et al. 2020) | Tedersoo L, et al. 2014. Science 346:e1256688 ((https://doi.org/10.1126/science.1256688)); Põlme S, et al. 2020. Fungal Diversity 105:1-16 ((https://doi.org/10.1007/s13225-020-00466-2)) |
| Otu03433 | 10 | k__Fungi(100);p__Ascomycota(100);c__Lecanoromycetes(100);o__Lecanorales(100);o__Lecanorales_unclassified(100);o__Lecanorales_unclassified(100);o__Lecanorales_unclassified(100); | Lecanorales | 7 | Pathotroph-Symbiotroph | Lichen Parasite-Lichenized | Microfungus-Thallus | NULL | Probable | Lichenized with green algae, occasionally lichenicolous, on rocks, mosses and wood (Cannon & Kirk 2007); ...an order of mostly lichen-forming fungi (Kirk et al. 2008) | Cannon PF, Kirk PM. 2007. Fungal Families of the World. CAB International, Cambridge (ISBN: 978-0851998275); Kirk PM et al. 2008. Dictionary of the Fungi. Tenth Edition. CAB International, Wallingford (ISBN: 978-0851998268); Esslinger TL. 2014. North Dakota State University ((http://www.ndsu.edu/pubweb/~esslinge/chcklst/chcklst7.htm)) |
| Otu03434 | 10 | k__Fungi(100);p__Basidiomycota(100);c__Tremellomycetes(100);o__Tremellales(90);f__Tremellales_family_Incertae_sedis(90);f__Tremellales_family_Incertae_sedis_unclassified(90);f__Tremellales_family_Incertae_sedis_unclassified(90); | Tremellales | 7 | Pathotroph-Saprotroph-Symbiotroph | Fungal Parasite-Undefined Saprotroph | Tremelloid-Yeast | NULL | Possible | The order contains both teleomorphic and anamorphic species, most of the latter being yeasts | Sterkenburg E, et al. 2015. New Phytologist 207:1145-1158 ((https://doi.org/10.1111/nph.13426)) |
| Otu03435 | 10 | k__Fungi(100);p__Ascomycota(100);c__Sordariomycetes(100);o__Hypocreales(100);f__Hypocreales_family_Incertae_sedis(100);g__Sarocladium(100);s__Sarocladium_sp(100); | Sarocladium | 13 | Pathotroph-Saprotroph | \|Plant Pathogen\|-Plant Saprotroph-Undefined Saprotroph | Microfungus | NULL | Probable | As litter_saprotroph (Põlme et al. 2020); Decay leaf, fruit, seed (Põlme et al. 2020) | Tedersoo L, et al. 2014. Science 346:e1256688 ((https://doi.org/10.1126/science.1256688)); Põlme S, et al. 2020. Fungal Diversity 105:1-16 ((https://doi.org/10.1007/s13225-020-00466-2)) |
| Otu03436 | 10 | k__Fungi(100);p__Ascomycota(100);c__Eurotiomycetes(100);o__Chaetothyriales(100);f__Herpotrichiellaceae(90);g__unclassified_Herpotrichiellaceae(90);s__Herpotrichiellaceae_sp(90); | Herpotrichiellaceae | 9 | Pathotroph-Saprotroph | Animal Pathogen-Fungal Parasite-Undefined Saprotroph | Facultative Yeast-Microfungus | NULL | Probable | Mostly saprobic on plants or other fungi, isolated from soil etc., and from medical sources (Cannon & Kirk 2007) | Cannon PF, Kirk PM. 2007. Fungal Families of the World. CAB International, Cambridge (ISBN: 978-0851998275) |
| Otu03438 | 10 | k__Fungi(100);p__Basidiomycota(100);c__Tremellomycetes(100);o__Tremellales(100);f__Tremellales_family_Incertae_sedis(100);g__Hannaella(100);g__Hannaella_unclassified(80); | Hannaella | 13 | Pathotroph-Saprotroph | \|Animal Parasite\|-Undefined Saprotroph | Yeast | NULL | Probable | As animal_decomposer (Põlme et al. 2020); Decay animal material (Põlme et al. 2020) | Põlme S, et al. 2020. Fungal Diversity 105:1-16 ((https://doi.org/10.1007/s13225-020-00466-2)) |
| Otu03440 | 10 | k__Fungi(100);p__Ascomycota(100);c__Dothideomycetes(100);o__Capnodiales(100);f__Mycosphaerellaceae(90);f__Mycosphaerellaceae_unclassified(90);f__Mycosphaerellaceae_unclassified(90); | Mycosphaerellaceae | 9 | Pathotroph-Saprotroph | Plant Pathogen-Undefined Saprotroph | Microfungus | NULL | Probable | Biotrophic, necrotrophic or saprobic on various plant tissues (Cannon & Kirk 2007) | Cannon PF, Kirk PM. 2007. Fungal Families of the World. CAB International, Cambridge (ISBN: 978-0851998275) |
| Otu03441 | 10 | k__Fungi(100);p__Ascomycota(100);c__Dothideomycetes(100);o__Pleosporales(100);f__Massarinaceae(100);g__Massarina(100);s__Massarina_sp_HF12700(100); | Massarina | 13 | Pathotroph-Saprotroph | \|Plant Pathogen\|-Undefined Saprotroph | Microfungus | NULL | Probable | NULL | Tedersoo L, et al. 2014. Science 346:e1256688 ((https://doi.org/10.1126/science.1256688)); Põlme S, et al. 2020. Fungal Diversity 105:1-16 ((https://doi.org/10.1007/s13225-020-00466-2)) |
| Otu03445 | 10 | k__Fungi(100);p__Ascomycota(100);c__Saccharomycetes(100);o__Saccharomycetales(100);f__Metschnikowiaceae(100);g__Metschnikowia(100);s__Metschnikowia_pulcherrima(70); | Metschnikowia | 13 | Saprotroph | \|Undefined Saprotroph\| | Yeast | NULL | Probable | As nectar-tap_saprotroph (Põlme et al. 2020); Decay sugar-rich substrates (Põlme et al. 2020) | Põlme S, et al. 2020. Fungal Diversity 105:1-16 ((https://doi.org/10.1007/s13225-020-00466-2)) |
| Otu03446 | 10 | k__Fungi(100);p__Ascomycota(100);c__Leotiomycetes(100);o__Helotiales(100);f__Hyaloscyphaceae(100);g__Lachnum(100);g__Lachnum_unclassified(100); | Lachnum | 13 | Saprotroph-Symbiotroph | Endophyte-\|Plant Saprotroph\|-Undefined Saprotroph-Wood Saprotroph | Microfungus | NULL | Probable | As litter_saprotroph (Põlme et al. 2020); As root_endophyte (Põlme et al. 2020); Decay wood, leaf, fruit, seed (Põlme et al. 2020) | Tedersoo L, et al. 2014. Science 346:e1256688 ((https://doi.org/10.1126/science.1256688)); Põlme S, et al. 2020. Fungal Diversity 105:1-16 ((https://doi.org/10.1007/s13225-020-00466-2)) |
| Otu03448 | 10 | k__Fungi(100);p__Ascomycota(100);c__Leotiomycetes(100);o__Helotiales(100);f__Vibrisseaceae(90);g__Phialocephala(90);s__Phialocephala_scopiformis(80); | Phialocephala | 13 | Saprotroph-Symbiotroph | Endophyte-\|Undefined Saprotroph\| | Microfungus | NULL | Highly Probable | As root_endophyte (Põlme et al. 2020); As root_endophyte_dark_septate (Põlme et al. 2020); As soil_saprotroph (Põlme et al. 2020); Dark Septate Endophyte, known from plant roots.; Decay roots, soil (Põlme et al. 2020) | Newsham KK. 2011. New Phytologist 190:783-793 ((https://doi.org/10.1111/j.1469-8137.2010.03611.x)) (as dark septate root endophytes); Põlme S, et al. 2020. Fungal Diversity 105:1-16 ((https://doi.org/10.1007/s13225-020-00466-2)) |
| Otu03450 | 10 | k__Fungi(100);p__Ascomycota(100);c__Dothideomycetes(100);o__Pleosporales(100);f__Lophiostomataceae(100);g__Lophiostoma(100);g__Lophiostoma_unclassified(100); | Lophiostoma | 13 | Saprotroph | Plant Saprotroph-Undefined Saprotroph-\|Wood Saprotroph\| | Microfungus | NULL | Probable | As litter_saprotroph (Põlme et al. 2020); Decay wood, leaf, fruit, seed (Põlme et al. 2020) | Tedersoo L, et al. 2014. Science 346:e1256688 ((https://doi.org/10.1126/science.1256688)); Põlme S, et al. 2020. Fungal Diversity 105:1-16 ((https://doi.org/10.1007/s13225-020-00466-2)) |
| Otu03452 | 10 | k__Fungi(100);p__Basidiomycota(100);c__Agaricostilbomycetes(100);o__Agaricostilbales(100);f__Agaricostilbaceae(100);f__Agaricostilbaceae_unclassified(100);f__Agaricostilbaceae_unclassified(100); | Agaricostilbaceae | 9 | Saprotroph | Undefined Saprotroph | NULL | NULL | Possible | Probably saprobic, usually associated with dead palm tissues (Cannon & Kirk 2007) | Cannon PF, Kirk PM. 2007. Fungal Families of the World. CAB International, Cambridge (ISBN: 978-0851998275) |
| Otu03453 | 10 | k__Fungi(100);p__Ascomycota(100);c__Sordariomycetes(100);o__Hypocreales(100);f__Hypocreales_family_Incertae_sedis(100);g__Myrothecium(100);s__Myrothecium_cinctum(100); | Myrothecium | 13 | Pathotroph-Saprotroph | Fungal Parasite-\|Plant Saprotroph\|-Undefined Saprotroph | Microfungus | NULL | Probable | As litter_saprotroph (Põlme et al. 2020); As mycoparasite (Põlme et al. 2020); Decay leaf, fruit, seed (Põlme et al. 2020) | Tedersoo L, et al. 2014. Science 346:e1256688 ((https://doi.org/10.1126/science.1256688)); Põlme S, et al. 2020. Fungal Diversity 105:1-16 ((https://doi.org/10.1007/s13225-020-00466-2)) |
| Otu03456 | 10 | k__Fungi(100);p__Basidiomycota(100);c__Agaricomycetes(100);o__Atheliales(100);f__Atheliaceae(100);g__Byssocorticium(100);s__Byssocorticium_sp(100); | Byssocorticium | 13 | Saprotroph-Symbiotroph | \|Ectomycorrhizal\|-Undefined Saprotroph | Corticioid | NULL | Highly Probable | NULL | Rinaldi AC, et al. 2008. Fungal Diversity 33:1-45 ((http://www.fungaldiversity.org/fdp/sfdp/33-1.pdf)); Tedersoo L, et al. 2010. Mycorrhiza 20:217-263 ((https://doi.org/10.1007/s00572-009-0274-x)); Taylor DL, et al. 2014. Ecological Monographs 84:3-20 ((https://doi.org/10.1890/12-1693.1)); Põlme S, et al. 2020. Fungal Diversity 105:1-16 ((https://doi.org/10.1007/s13225-020-00466-2)); http://mycorrhizas.info/ecmf.html ((http://mycorrhizas.info/ecmf.html)) |
| Otu03458 | 10 | k__Fungi(100);p__Ascomycota(100);c__Leotiomycetes(100);o__Helotiales(100);f__Hyaloscyphaceae(100);f__Hyaloscyphaceae_unclassified(70);f__Hyaloscyphaceae_unclassified(70); | Hyaloscyphaceae | 9 | Saprotroph | Plant Saprotroph-Wood Saprotroph | Microfungus | NULL | Probable | Saprobic on woody and herbaceous material (Cannon & Kirk 2007) | Cannon PF, Kirk PM. 2007. Fungal Families of the World. CAB International, Cambridge (ISBN: 978-0851998275) |
| Otu03460 | 10 | k__Fungi(100);p__Ascomycota(100);c__Saccharomycetes(100);o__Saccharomycetales(100);f__Metschnikowiaceae(100);g__Metschnikowia(100);g__Metschnikowia_unclassified(61); | Metschnikowia | 13 | Saprotroph | \|Undefined Saprotroph\| | Yeast | NULL | Probable | As nectar-tap_saprotroph (Põlme et al. 2020); Decay sugar-rich substrates (Põlme et al. 2020) | Põlme S, et al. 2020. Fungal Diversity 105:1-16 ((https://doi.org/10.1007/s13225-020-00466-2)) |
| Otu03462 | 10 | k__Fungi(100);p__Basidiomycota(100);c__Agaricomycetes(100);o__Russulales(100);f__Lachnocladiaceae(100);g__Scytinostroma(100);s__Scytinostroma_sp_DLL2011_1(100); | Scytinostroma | 13 | Saprotroph | Undefined Saprotroph-\|Wood Saprotroph\| | Corticioid | White Rot | Probable | As white_rot (Põlme et al. 2020); Decay wood (Põlme et al. 2020) | Tedersoo L, et al. 2014. Science 346:e1256688 ((https://doi.org/10.1126/science.1256688)); Põlme S, et al. 2020. Fungal Diversity 105:1-16 ((https://doi.org/10.1007/s13225-020-00466-2)) |
| Otu03464 | 10 | k__Fungi(100);p__Ascomycota(100);c__Sordariomycetes(100);o__Hypocreales(100);f__Nectriaceae(100);g__Fusarium(70);g__Fusarium_unclassified(61); | Fusarium | 13 | Pathotroph-Saprotroph-Symbiotroph | Animal Pathogen-Endophyte-Lichen Parasite-\|Plant Pathogen\|-Plant Saprotroph-Undefined Saprotroph-Wood Saprotroph | Microfungus | Soft Rot | Possible | As foliar_endophyte (Põlme et al. 2020); As litter_saprotroph (Põlme et al. 2020); Causing damping-off disease (Põlme et al. 2020); Decay leaf, fruit, seed, roots, soil, animal material (Põlme et al. 2020); Likely opportunistic human pathogen (Irinyi et al. 2015) | Duncan CG, Eslyn WE. 1966. Mycologia 58:642-664 ((https://doi.org/10.2307/3757045)); Seehann G, et al. 1975. List of Fungi in Soft-Rot Tissue Tests. Institute fur Holzbiologie und Holzschutz, Bundesforschungsanstalt fur Forst - und Holzwirtshaft, Hamburg - Lohbrugge, Germany; Costa IPMW, et al. 2012. Mycotaxon 119:e26 ((http://mycotaxon.com/resources/checklists/costa_v119_checklist.pdf)); Tedersoo L, et al. 2014. Science 346:e1256688 ((https://doi.org/10.1126/science.1256688)); Irinyi L, et al. 2015. Medical Mycology 53:313-337 ((https://doi.org/10.1093/mmy/myv008)); Schroers, et al. 2016. Mycologia 108:806-819 ((https://doi.org/10.3852/15-255)); Põlme S, et al. 2020. Fungal Diversity 105:1-16 ((https://doi.org/10.1007/s13225-020-00466-2)); The American Phytopathological Society Common Names of Plant Diseases ((http://www.apsnet.org/publications/commonnames/Pages/default.aspx)); Lawrey JD, Diederich P. 2016. http://www.lichenicolous.net (accessed 24 May 2017) ((http://www.lichenicolous.net)) |
| Otu03465 | 10 | k__Fungi(100);p__Basidiomycota(100);c__Tremellomycetes(100);o__Filobasidiales(100);f__Filobasidiaceae(100);g__Cryptococcus(100);s__Cryptococcus_sp(100); | Cryptococcus | 13 | Pathotroph-Saprotroph-Symbiotroph | Animal Pathogen-Endophyte-Epiphyte-\|Undefined Saprotroph\| | Dimorphic Yeast-Yeast | NULL | Possible | As foliar_endophyte (Põlme et al. 2020); Decay leaf, fruit, seed, soil, dung, animal material (Põlme et al. 2020) | Andrews JH, et al. 1983. Phytopathology 73:228-234 ((https://doi.org/10.1094/Phyto-73-228)); Kurtzman CP, et al. (eds.) 2011. The Yeasts, a Taxonomic Study. Fifth Edition. Vols 1-3. Elsevier, San Diego (ISBN: 9780444521491); Põlme S, et al. 2020. Fungal Diversity 105:1-16 ((https://doi.org/10.1007/s13225-020-00466-2)) |
| Otu03466 | 10 | k__Fungi(100);p__Ascomycota(100);c__Sordariomycetes(100);o__Sordariomycetes_order_Incertae_sedis(100);f__Sordariomycetes_family_Incertae_sedis(100);g__Myrmecridium(100);s__Myrmecridium_schulzeri(80); | Myrmecridium | 13 | Saprotroph | \|Undefined Saprotroph\| | Microfungus | NULL | Probable | As soil_saprotroph (Põlme et al. 2020); Decay soil (Põlme et al. 2020) | Tedersoo L, et al. 2014. Science 346:e1256688 ((https://doi.org/10.1126/science.1256688)); Põlme S, et al. 2020. Fungal Diversity 105:1-16 ((https://doi.org/10.1007/s13225-020-00466-2)) |
| Otu03468 | 10 | k__Fungi(100);p__Ascomycota(100);c__Dothideomycetes(100);o__Pleosporales(100);f__Venturiaceae(100);g__Fusicladium(100);s__Fusicladium_caryigenum(100); | Fusicladium | 13 | Pathotroph-Symbiotroph | Endophyte-Lichen Parasite-\|Plant Pathogen\| | Microfungus | NULL | Probable | NULL | Tedersoo L, et al. 2014. Science 346:e1256688 ((https://doi.org/10.1126/science.1256688)); Põlme S, et al. 2020. Fungal Diversity 105:1-16 ((https://doi.org/10.1007/s13225-020-00466-2)) |
| Otu03469 | 10 | k__Fungi(100);p__Basidiomycota(100);c__Wallemiomycetes(100);o__Wallemiales(100);f__Wallemiaceae(100);g__Wallemia(100);s__Wallemia_muriae(100); | Wallemia | 13 | Saprotroph | \|Undefined Saprotroph\| | Microfungus | NULL | Probable | Decay sugar-rich substrates (Põlme et al. 2020) | Tedersoo L, et al. 2014. Science 346:e1256688 ((https://doi.org/10.1126/science.1256688)); Põlme S, et al. 2020. Fungal Diversity 105:1-16 ((https://doi.org/10.1007/s13225-020-00466-2)) |
| Otu03471 | 10 | k__Fungi(100);p__Ascomycota(100);c__Saccharomycetes(100);o__Saccharomycetales(100);f__Metschnikowiaceae(100);g__Metschnikowia(100);s__Metschnikowia_pulcherrima(90); | Metschnikowia | 13 | Saprotroph | \|Undefined Saprotroph\| | Yeast | NULL | Probable | As nectar-tap_saprotroph (Põlme et al. 2020); Decay sugar-rich substrates (Põlme et al. 2020) | Põlme S, et al. 2020. Fungal Diversity 105:1-16 ((https://doi.org/10.1007/s13225-020-00466-2)) |
| Otu03473 | 10 | k__Fungi(100);p__Basidiomycota(100);c__Agaricomycetes(100);o__Agaricales(100);f__Strophariaceae(100);g__Galerina(100);g__Galerina_unclassified(100); | Galerina | 13 | Pathotroph-Saprotroph | Bryophyte Parasite-Plant Pathogen-Plant Saprotroph-Undefined Saprotroph-\|Wood Saprotroph\| | Agaricoid | White Rot | Probable | As litter_saprotroph (Põlme et al. 2020); Decay wood, leaf, fruit, seed (Põlme et al. 2020) | Tedersoo L, et al. 2014. Science 346:e1256688 ((https://doi.org/10.1126/science.1256688)); Põlme S, et al. 2020. Fungal Diversity 105:1-16 ((https://doi.org/10.1007/s13225-020-00466-2)) |
| Otu03475 | 10 | k__Fungi(100);p__Ascomycota(100);c__Saccharomycetes(100);o__Saccharomycetales(100);f__Metschnikowiaceae(100);g__Metschnikowia(100);s__Metschnikowia_pulcherrima(80); | Metschnikowia | 13 | Saprotroph | \|Undefined Saprotroph\| | Yeast | NULL | Probable | As nectar-tap_saprotroph (Põlme et al. 2020); Decay sugar-rich substrates (Põlme et al. 2020) | Põlme S, et al. 2020. Fungal Diversity 105:1-16 ((https://doi.org/10.1007/s13225-020-00466-2)) |
| Otu03477 | 10 | k__Fungi(100);p__Ascomycota(100);c__Pezizomycetes(100);o__Pezizales(100);f__Pezizaceae(100);g__unclassified_Pezizaceae(100);s__Pezizaceae_sp(100); | Pezizaceae | 9 | Saprotroph-Symbiotroph | Dung Saprotroph-Ectomycorrhizal-Litter Saprotroph-Undefined Saprotroph | Pezizoid | NULL | Possible | Saprobic or ectomycorrhizal, on or in soil, leafmould etc., some species associated with dung or fire sites (Cannon & Kirk 2007) | Cannon PF, Kirk PM. 2007. Fungal Families of the World. CAB International, Cambridge (ISBN: 978-0851998275) |
| Otu03479 | 10 | k__Fungi(100);p__Ascomycota(100);c__Eurotiomycetes(100);o__Chaetothyriales(100);f__Herpotrichiellaceae(100);g__Cladophialophora(100);s__Cladophialophora_sp_20104730d(100); | Cladophialophora | 13 | Pathotroph-Saprotroph-Symbiotroph | Animal Pathogen-Endophyte-Fungal Parasite-\|Undefined Saprotroph\| | Dimorphic-Microfungus | NULL | Probable | As mycoparasite (Põlme et al. 2020); As root_endophyte_dark_septate (Põlme et al. 2020); As soil_saprotroph (Põlme et al. 2020); Decay soil (Põlme et al. 2020) | James TY, et al. 2006. Nature 443:818-822 ((https://doi.org/10.1038/nature05110)); Põlme S, et al. 2020. Fungal Diversity 105:1-16 ((https://doi.org/10.1007/s13225-020-00466-2)) |
| Otu03482 | 10 | k__Fungi(100);p__Ascomycota(100);c__Saccharomycetes(100);o__Saccharomycetales(100);f__Metschnikowiaceae(100);g__Metschnikowia(100);s__Metschnikowia_pulcherrima(100); | Metschnikowia | 13 | Saprotroph | \|Undefined Saprotroph\| | Yeast | NULL | Probable | As nectar-tap_saprotroph (Põlme et al. 2020); Decay sugar-rich substrates (Põlme et al. 2020) | Põlme S, et al. 2020. Fungal Diversity 105:1-16 ((https://doi.org/10.1007/s13225-020-00466-2)) |
| Otu03483 | 10 | k__Fungi(100);p__Basidiomycota(100);c__Agaricomycetes(100);o__Polyporales(100);f__Phanerochaetaceae(100);g__Phanerochaete(100);s__Phanerochaete_chrysorhiza(100); | Phanerochaete | 13 | Saprotroph | \|Wood Saprotroph\| | Corticioid | White Rot | Highly Probable | As white_rot (Põlme et al. 2020); Decay wood (Põlme et al. 2020) | Glenn JK, et al. 1983. Biochemical and Biophysical Research Communications 114:1077-1083 ((https://doi.org/10.1016/0006-291x((83)) 90672-1)); Põlme S, et al. 2020. Fungal Diversity 105:1-16 ((https://doi.org/10.1007/s13225-020-00466-2)) |
| Otu03487 | 10 | k__Fungi(100);p__Ascomycota(100);c__Saccharomycetes(100);o__Saccharomycetales(100);f__Metschnikowiaceae(100);g__Metschnikowia(100);s__Metschnikowia_pulcherrima(100); | Metschnikowia | 13 | Saprotroph | \|Undefined Saprotroph\| | Yeast | NULL | Probable | As nectar-tap_saprotroph (Põlme et al. 2020); Decay sugar-rich substrates (Põlme et al. 2020) | Põlme S, et al. 2020. Fungal Diversity 105:1-16 ((https://doi.org/10.1007/s13225-020-00466-2)) |
| Otu03488 | 10 | k__Fungi(100);p__Ascomycota(100);c__Saccharomycetes(100);o__Saccharomycetales(100);f__Metschnikowiaceae(100);g__Metschnikowia(100);g__Metschnikowia_unclassified(80); | Metschnikowia | 13 | Saprotroph | \|Undefined Saprotroph\| | Yeast | NULL | Probable | As nectar-tap_saprotroph (Põlme et al. 2020); Decay sugar-rich substrates (Põlme et al. 2020) | Põlme S, et al. 2020. Fungal Diversity 105:1-16 ((https://doi.org/10.1007/s13225-020-00466-2)) |
| Otu03492 | 10 | k__Fungi(100);p__Ascomycota(100);c__Saccharomycetes(100);o__Saccharomycetales(100);f__Metschnikowiaceae(100);g__Metschnikowia(100);s__Metschnikowia_pulcherrima(90); | Metschnikowia | 13 | Saprotroph | \|Undefined Saprotroph\| | Yeast | NULL | Probable | As nectar-tap_saprotroph (Põlme et al. 2020); Decay sugar-rich substrates (Põlme et al. 2020) | Põlme S, et al. 2020. Fungal Diversity 105:1-16 ((https://doi.org/10.1007/s13225-020-00466-2)) |
| Otu03493 | 10 | k__Fungi(100);p__Ascomycota(100);c__Saccharomycetes(100);o__Saccharomycetales(100);f__Metschnikowiaceae(100);g__Metschnikowia(100);s__Metschnikowia_pulcherrima(90); | Metschnikowia | 13 | Saprotroph | \|Undefined Saprotroph\| | Yeast | NULL | Probable | As nectar-tap_saprotroph (Põlme et al. 2020); Decay sugar-rich substrates (Põlme et al. 2020) | Põlme S, et al. 2020. Fungal Diversity 105:1-16 ((https://doi.org/10.1007/s13225-020-00466-2)) |
| Otu03494 | 10 | k__Fungi(100);p__Basidiomycota(100);c__Agaricomycetes(100);o__Trechisporales(100);f__Hydnodontaceae(100);g__Trechispora(100);s__Trechispora_sp_1_MPM_2013(61); | Trechispora | 13 | Saprotroph | \|Wood Saprotroph\| | Corticioid | White Rot | Highly Probable | As white_rot (Põlme et al. 2020); Decay wood (Põlme et al. 2020) | Gilbertson RL, Ryvarden L. 1987-1987. North American Polypores. Fungiflora, Oslo (ISBN: 978-0945345060); Rinaldi AC, et al. 2008. Fungal Diversity 33:1-45 ((http://www.fungaldiversity.org/fdp/sfdp/33-1.pdf)) (as Uncertain); Tedersoo L, et al. 2010. Mycorrhiza 20:217-263 ((https://doi.org/10.1007/s00572-009-0274-x)) (as not ECM).; Põlme S, et al. 2020. Fungal Diversity 105:1-16 ((https://doi.org/10.1007/s13225-020-00466-2)) |
| Otu03495 | 10 | k__Fungi(100);p__Ascomycota(100);c__Saccharomycetes(100);o__Saccharomycetales(100);f__Metschnikowiaceae(100);g__Metschnikowia(100);s__Metschnikowia_pulcherrima(100); | Metschnikowia | 13 | Saprotroph | \|Undefined Saprotroph\| | Yeast | NULL | Probable | As nectar-tap_saprotroph (Põlme et al. 2020); Decay sugar-rich substrates (Põlme et al. 2020) | Põlme S, et al. 2020. Fungal Diversity 105:1-16 ((https://doi.org/10.1007/s13225-020-00466-2)) |
| Otu03496 | 10 | k__Fungi(100);p__Basidiomycota(100);c__Agaricomycetes(100);o__Trechisporales(100);f__Hydnodontaceae(100);g__Trechispora(100);s__Trechispora_sp(100); | Trechispora | 13 | Saprotroph | \|Wood Saprotroph\| | Corticioid | White Rot | Highly Probable | As white_rot (Põlme et al. 2020); Decay wood (Põlme et al. 2020) | Gilbertson RL, Ryvarden L. 1987-1987. North American Polypores. Fungiflora, Oslo (ISBN: 978-0945345060); Rinaldi AC, et al. 2008. Fungal Diversity 33:1-45 ((http://www.fungaldiversity.org/fdp/sfdp/33-1.pdf)) (as Uncertain); Tedersoo L, et al. 2010. Mycorrhiza 20:217-263 ((https://doi.org/10.1007/s00572-009-0274-x)) (as not ECM).; Põlme S, et al. 2020. Fungal Diversity 105:1-16 ((https://doi.org/10.1007/s13225-020-00466-2)) |
| Otu03499 | 10 | k__Fungi(100);p__Basidiomycota(100);c__Exobasidiomycetes(100);o__Exobasidiales(100);f__Exobasidiaceae(100);g__Exobasidium(100);g__Exobasidium_unclassified(100); | Exobasidium | 13 | Pathotroph | \|Plant Pathogen\| | Leaf Blister-Leaf Gall-Leaf Spot | NULL | Probable | NULL | Tedersoo L, et al. 2014. Science 346:e1256688 ((https://doi.org/10.1126/science.1256688)); Põlme S, et al. 2020. Fungal Diversity 105:1-16 ((https://doi.org/10.1007/s13225-020-00466-2)) |
| Otu03500 | 10 | k__Fungi(100);p__Basidiomycota(100);c__Microbotryomycetes(100);o__Sporidiobolales(100);f__Sporidiobolales_family_Incertae_sedis(100);g__Sporobolomyces(100);s__Sporobolomyces_griseoflavus(100); | Sporobolomyces | 13 | Pathotroph-Saprotroph | \|Fungal Parasite\|-Plant Saprotroph | Yeast | NULL | Probable | As litter_saprotroph (Põlme et al. 2020); As mycoparasite (Põlme et al. 2020); Decay fungal material (Põlme et al. 2020) | Tedersoo L, et al. 2014. Science 346:e1256688 ((https://doi.org/10.1126/science.1256688)); Purahong W, et al. 2016. Molecular Ecology 25:4059-4074 ((https://doi.org/10.1111/mec.13739)); Põlme S, et al. 2020. Fungal Diversity 105:1-16 ((https://doi.org/10.1007/s13225-020-00466-2)) |
| Otu03502 | 10 | k__Fungi(100);p__Basidiomycota(100);c__Agaricomycetes(100);o__Russulales(100);f__Stephanosporaceae(100);g__Cristinia(100);s__Cristinia_helvetica(100); | Cristinia | 13 | Saprotroph | Undefined Saprotroph-\|Wood Saprotroph\| | Corticioid | White Rot | Probable | As white_rot (Põlme et al. 2020); Decay wood (Põlme et al. 2020) | Tedersoo L, et al. 2014. Science 346:e1256688 ((https://doi.org/10.1126/science.1256688)); Põlme S, et al. 2020. Fungal Diversity 105:1-16 ((https://doi.org/10.1007/s13225-020-00466-2)) |
| Otu03503 | 10 | k__Fungi(100);p__Ascomycota(100);c__Eurotiomycetes(100);o__Onygenales(100);f__Ajellomycetaceae(100);g__Spiromastix(100);s__Spiromastix_sp_F19(100); | Spiromastix | 13 | Pathotroph-Saprotroph | Animal Pathogen-Dung Saprotroph-\|Undefined Saprotroph\| | Microfungus | NULL | Probable | As soil_saprotroph (Põlme et al. 2020); Decay soil (Põlme et al. 2020) | Tedersoo L, et al. 2014. Science 346:e1256688 ((https://doi.org/10.1126/science.1256688)); Põlme S, et al. 2020. Fungal Diversity 105:1-16 ((https://doi.org/10.1007/s13225-020-00466-2)) |
| Otu03504 | 10 | k__Fungi(100);p__Ascomycota(100);c__Saccharomycetes(100);o__Saccharomycetales(100);f__Metschnikowiaceae(100);g__Metschnikowia(100);s__Metschnikowia_pulcherrima(100); | Metschnikowia | 13 | Saprotroph | \|Undefined Saprotroph\| | Yeast | NULL | Probable | As nectar-tap_saprotroph (Põlme et al. 2020); Decay sugar-rich substrates (Põlme et al. 2020) | Põlme S, et al. 2020. Fungal Diversity 105:1-16 ((https://doi.org/10.1007/s13225-020-00466-2)) |
| Otu03505 | 10 | k__Fungi(100);p__Ascomycota(100);c__Sordariomycetes(100);o__Hypocreales(100);f__Ophiocordycipitaceae(100);f__Ophiocordycipitaceae_unclassified(100);f__Ophiocordycipitaceae_unclassified(100); | Ophiocordycipitaceae | 9 | Pathotroph-Symbiotroph | Endophyte-Insect Pathogen | Microfungus | NULL | Probable | A family of parasitic fungi (Sung et al. 2007) | Sung GH, et al. 2007. Studies in Mycology 57:5-59 ((https://doi.org/10.3114/sim.2007.57.01)); Costa IPMW, et al. 2012. Mycotaxon 119:e26 ((http://mycotaxon.com/resources/checklists/costa_v119_checklist.pdf)) |

| | | | | | | | | | | | |
|---|---|---|---|---|---|---|---|---|---|---|---|
| Otu03508 | 10 | k__Fungi(100);p__Ascomycota(100);c__Sordariomycetes(100);o__Microascales(100);f__Microascaceae(100);f__Microascaceae_unclassified(100);f__Microascaceae_unclassified(100); | Microascaceae | 9 | Pathotroph-Saprotroph-Symbiotroph | Animal Pathogen-Endophyte-Plant Pathogen-Undefined Saprotroph | Microfungus | NULL | Possible | Saprobic in soil or rotting vegetation, some species heat-tolerant and associatated with silage and compost, a few associated with humans and animals (Cannon & Kirk 2007) | Cannon PF, Kirk PM. 2007. Fungal Families of the World. CAB International, Cambridge (ISBN: 978-0851998275) |
| Otu03510 | 10 | k__Fungi(100);p__Basidiomycota(100);c__Agaricomycetes(100);o__Russulales(100);f__Stereaceae(100);f__Stereaceae_unclassified(100);f__Stereaceae_unclassified(100); | Stereaceae | 9 | Saprotroph | Undefined Saprotroph-Wood Saprotroph | Corticioid-Polyporoid | NULL | Probable | Lignicolous or terrestrial (in leaf litter); typically saprobic (Cannon & Kirk 2007) | Cannon PF, Kirk PM. 2007. Fungal Families of the World. CAB International, Cambridge (ISBN: 978-0851998275) |
| Otu03511 | 10 | k__Fungi(100);p__Basidiomycota(100);c__Agaricomycetes(100);o__Russulales(100);f__Stereaceae(100);g__Stereum(100);s__Stereum_sp_4_RG_2014(100); | Stereum | 13 | Saprotroph | Undefined Saprotroph-\|Wood Saprotroph\| | Polyporoid | White Rot | Probable | As white_rot (Põlme et al. 2020); Decay wood (Põlme et al. 2020) | Tedersoo L, et al. 2014. Science 346:e1256688 ((https://doi.org/10.1126/science.1256688)); Põlme S, et al. 2020. Fungal Diversity 105:1-16 ((https://doi.org/10.1007/s13225-020-00466-2)) |
| Otu03512 | 10 | k__Fungi(100);p__Ascomycota(100);c__Saccharomycetes(100);o__Saccharomycetales(100);f__Metschnikowiaceae(100);g__Metschnikowia(100);s__Metschnikowia_pulcherrima(90); | Metschnikowia | 13 | Saprotroph | \|Undefined Saprotroph\| | Yeast | NULL | Probable | As nectar-tap_saprotroph (Põlme et al. 2020); Decay sugar-rich substrates (Põlme et al. 2020) | Põlme S, et al. 2020. Fungal Diversity 105:1-16 ((https://doi.org/10.1007/s13225-020-00466-2)) |
| Otu03514 | 10 | k__Fungi(100);p__Basidiomycota(100);c__Agaricomycetes(100);o__Cantharellales(100);f__Hydnaceae(100);g__Sistotrema(100);g__Sistotrema_unclassified(100); | Sistotrema | 13 | Saprotroph-Symbiotroph | Ectomycorrhizal-\|Plant Saprotroph\|-Wood Saprotroph | Corticioid | White Rot | Possible | As litter_saprotroph (Põlme et al. 2020); Decay leaf, fruit, seed (Põlme et al. 2020) | Gilbertson RL, Ryvarden L. 1987-1987. North American Polypores. Fungiflora, Oslo (ISBN: 978-0945345060); Rinaldi AC, et al. 2008. Fungal Diversity 33:1-45 ((http://www.fungaldiversity.org/fdp/sfdp/33-1.pdf)) (as ECM/not ECM); Tedersoo L, et al. 2010. Mycorrhiza 20: 217-263 ((https://doi.org/10.1007/s00572-009-0274-x)) (pro parte); Põlme S, et al. 2020. Fungal Diversity 105:1-16 ((https://doi.org/10.1007/s13225-020-00466-2)); http://mycorrhizas.info/ecmf.html ((http://mycorrhizas.info/ecmf.html)) (as ECM/not ECM). |
| Otu03515 | 10 | k__Fungi(100);p__Basidiomycota(100);c__Agaricomycetes(100);o__Agaricales(100);f__Psathyrellaceae(100);g__Coprinopsis(100);s__Coprinopsis_marcescibilis(100); | Coprinopsis | 13 | Saprotroph | \|Undefined Saprotroph\| | Agaricoid | NULL | Probable | As soil_saprotroph (Põlme et al. 2020); Decay soil (Põlme et al. 2020) | Tedersoo L, et al. 2014. Science 346:e1256688 ((https://doi.org/10.1126/science.1256688)); Põlme S, et al. 2020. Fungal Diversity 105:1-16 ((https://doi.org/10.1007/s13225-020-00466-2)) |
| Otu03516 | 10 | k__Fungi(100);p__Ascomycota(100);c__Saccharomycetes(100);o__Saccharomycetales(100);f__Metschnikowiaceae(100);g__Metschnikowia(100);s__Metschnikowia_pulcherrima(80); | Metschnikowia | 13 | Saprotroph | \|Undefined Saprotroph\| | Yeast | NULL | Probable | As nectar-tap_saprotroph (Põlme et al. 2020); Decay sugar-rich substrates (Põlme et al. 2020) | Põlme S, et al. 2020. Fungal Diversity 105:1-16 ((https://doi.org/10.1007/s13225-020-00466-2)) |
| Otu03517 | 10 | k__Fungi(100);p__Basidiomycota(100);c__Agaricomycetes(100);o__Cantharellales(100);f__Ceratobasidiaceae(100);g__Ceratobasidium(100);g__Ceratobasidium_unclassified(100); | Ceratobasidium | 13 | Pathotroph-Saprotroph-Symbiotroph | Ectomycorrhizal-Endomycorrhizal-Orchid Mycorrhizal-\|Plant Pathogen\|-Plant Saprotroph-Undefined Saprotroph | Corticioid-Microfungus | NULL | Possible | especially of Orchidaceae but also even bryophytes (Cannon & Kirk 2007); As litter_saprotroph (Põlme et al. 2020); Best known as plant pathogens, but also saprobic. Species are endomycorrhizal with a wide range of plant roots ; Decay leaf, fruit, seed, roots, soil (Põlme et al. 2020) | Cannon PF, Kirk PM. 2007. Fungal Families of the World. CAB International, Cambridge (ISBN: 978-0851998275); Tedersoo L, et al. 2014. Science 346:e1256688 ((https://doi.org/10.1126/science.1256688)); Põlme S, et al. 2020. Fungal Diversity 105:1-16 ((https://doi.org/10.1007/s13225-020-00466-2)) |
| Otu03519 | 10 | k__Fungi(100);p__Ascomycota(100);c__Saccharomycetes(100);o__Saccharomycetales(100);f__Metschnikowiaceae(100);g__Metschnikowia(100);s__Metschnikowia_pulcherrima(90); | Metschnikowia | 13 | Saprotroph | \|Undefined Saprotroph\| | Yeast | NULL | Probable | As nectar-tap_saprotroph (Põlme et al. 2020); Decay sugar-rich substrates (Põlme et al. 2020) | Põlme S, et al. 2020. Fungal Diversity 105:1-16 ((https://doi.org/10.1007/s13225-020-00466-2)) |
| Otu03520 | 10 | k__Fungi(100);p__Ascomycota(100);c__Saccharomycetes(100);o__Saccharomycetales(100);f__Metschnikowiaceae(100);g__Metschnikowia(100);s__Metschnikowia_pulcherrima(90); | Metschnikowia | 13 | Saprotroph | \|Undefined Saprotroph\| | Yeast | NULL | Probable | As nectar-tap_saprotroph (Põlme et al. 2020); Decay sugar-rich substrates (Põlme et al. 2020) | Põlme S, et al. 2020. Fungal Diversity 105:1-16 ((https://doi.org/10.1007/s13225-020-00466-2)) |
| Otu03521 | 10 | k__Fungi(100);p__Ascomycota(100);c__Saccharomycetes(100);o__Saccharomycetales(100);f__Metschnikowiaceae(100);g__Metschnikowia(100);s__Metschnikowia_pulcherrima(100); | Metschnikowia | 13 | Saprotroph | \|Undefined Saprotroph\| | Yeast | NULL | Probable | As nectar-tap_saprotroph (Põlme et al. 2020); Decay sugar-rich substrates (Põlme et al. 2020) | Põlme S, et al. 2020. Fungal Diversity 105:1-16 ((https://doi.org/10.1007/s13225-020-00466-2)) |
| Otu03522 | 10 | k__Fungi(100);p__Zygomycota(100);c__Zygomycota_class_Incertae_sedis(100);o__Mortierellales(100);f__Mortierellaceae(100);g__Mortierella(100);g__Mortierella_unclassified(100); | Mortierella | 13 | Saprotroph-Symbiotroph | Endophyte-Plant Saprotroph-\|Undefined Saprotroph\| | Microfungus | NULL | Probable | As chitinolytic (Põlme et al. 2020); As soil_saprotroph (Põlme et al. 2020); Decay roots, soil (Põlme et al. 2020); Saprobic in soil and thus frequently isolated from soil and material contaminated with soil such as hay or rotten silage and spoil tips (Cannon & Kirk 2007) | Cannon PF, Kirk PM. 2007. Fungal Families of the World. CAB International, Cambridge (ISBN: 978-0851998275); Tedersoo L, et al. 2014. Science 346:e1256688 ((https://doi.org/10.1126/science.1256688)); Purahong W, et al. 2016. Molecular Ecology 25:4059-4074 ((https://doi.org/10.1111/mec.13739)); Põlme S, et al. 2020. Fungal Diversity 105:1-16 ((https://doi.org/10.1007/s13225-020-00466-2)) |
| Otu03523 | 10 | k__Fungi(100);p__Ascomycota(100);c__Saccharomycetes(100);o__Saccharomycetales(100);f__Metschnikowiaceae(100);g__Metschnikowia(100);s__Metschnikowia_pulcherrima(90); | Metschnikowia | 13 | Saprotroph | \|Undefined Saprotroph\| | Yeast | NULL | Probable | As nectar-tap_saprotroph (Põlme et al. 2020); Decay sugar-rich substrates (Põlme et al. 2020) | Põlme S, et al. 2020. Fungal Diversity 105:1-16 ((https://doi.org/10.1007/s13225-020-00466-2)) |
| Otu03525 | 10 | k__Fungi(100);p__Ascomycota(100);c__Saccharomycetes(100);o__Saccharomycetales(100);f__Metschnikowiaceae(100);g__Metschnikowia(100);s__Metschnikowia_pulcherrima(100); | Metschnikowia | 13 | Saprotroph | \|Undefined Saprotroph\| | Yeast | NULL | Probable | As nectar-tap_saprotroph (Põlme et al. 2020); Decay sugar-rich substrates (Põlme et al. 2020) | Põlme S, et al. 2020. Fungal Diversity 105:1-16 ((https://doi.org/10.1007/s13225-020-00466-2)) |
| Otu03526 | 10 | k__Fungi(100);p__Ascomycota(100);c__Saccharomycetes(100);o__Saccharomycetales(100);f__Saccharomycetales_family_Incertae_sedis(100);g__Candida(100);g__Candida_unclassified(70); | Candida | 13 | Pathotroph-Saprotroph-Symbiotroph | Animal Pathogen-Endophyte-Endosymbiont-Epiphyte-\|Undefined Saprotroph\| | Dimorphic Yeast-Yeast | NULL | Probable | As nectar-tap_saprotroph (Põlme et al. 2020); Decay sugar-rich substrates (Põlme et al. 2020); This asexual genus is polyphyletic and contain species that function very differently depend on what they are. Caution should be taken when interpreting guilds for Candida species. | Manolakaki D, et al. 2010. Virulence. 1: 367-375 ((https://doi.org/10.4161/viru.1.5.12796)); Mekbib SB, et al. 2011. Tropical Plant Pathology 36:233-240 ((https://doi.org/10.1590/S1982-56762011000400004)); Põlme S, et al. 2020. Fungal Diversity 105:1-16 ((https://doi.org/10.1007/s13225-020-00466-2)) |
| Otu03527 | 10 | k__Fungi(100);p__Ascomycota(100);c__Saccharomycetes(100);o__Saccharomycetales(100);f__Metschnikowiaceae(100);g__Metschnikowia(100);s__Metschnikowia_pulcherrima(80); | Metschnikowia | 13 | Saprotroph | \|Undefined Saprotroph\| | Yeast | NULL | Probable | As nectar-tap_saprotroph (Põlme et al. 2020); Decay sugar-rich substrates (Põlme et al. 2020) | Põlme S, et al. 2020. Fungal Diversity 105:1-16 ((https://doi.org/10.1007/s13225-020-00466-2)) |
| Otu03528 | 10 | k__Fungi(100);p__Ascomycota(100);c__Sordariomycetes(100);o__Calosphaeriales(100);f__Calosphaeriaceae(100);g__Jattaea(100);s__Jattaea_prunicola(100); | Jattaea | 13 | Saprotroph | \|Undefined Saprotroph\| | Microfungus | NULL | Probable | NULL | Tedersoo L, et al. 2014. Science 346:e1256688 ((https://doi.org/10.1126/science.1256688)); Põlme S, et al. 2020. Fungal Diversity 105:1-16 ((https://doi.org/10.1007/s13225-020-00466-2)) |
| Otu03529 | 10 | k__Fungi(100);p__Ascomycota(100);c__Saccharomycetes(100);o__Saccharomycetales(100);f__Metschnikowiaceae(100);g__Metschnikowia(100);s__Metschnikowia_pulcherrima(100); | Metschnikowia | 13 | Saprotroph | \|Undefined Saprotroph\| | Yeast | NULL | Probable | As nectar-tap_saprotroph (Põlme et al. 2020); Decay sugar-rich substrates (Põlme et al. 2020) | Põlme S, et al. 2020. Fungal Diversity 105:1-16 ((https://doi.org/10.1007/s13225-020-00466-2)) |
| Otu03534 | 10 | k__Fungi(100);p__Basidiomycota(100);c__Ustilaginomycetes(100);o__Ustilaginales(100);f__Ustilaginaceae(100);g__unclassified_Ustilaginaceae(90);s__Ustilaginaceae_sp(90); | Ustilaginaceae | 9 | Pathotroph | Plant Pathogen | Smut | NULL | Probable | Biotrophic in living tissue of Gramineae and Cyperaceae (Cannon & Kirk 2007) | Cannon PF, Kirk PM. 2007. Fungal Families of the World. CAB International, Cambridge (ISBN: 978-0851998275) |
| Otu03537 | 10 | k__Fungi(100);p__Ascomycota(100);c__Saccharomycetes(100);o__Saccharomycetales(100);f__Metschnikowiaceae(100);g__Metschnikowia(100);s__Metschnikowia_pulcherrima(100); | Metschnikowia | 13 | Saprotroph | \|Undefined Saprotroph\| | Yeast | NULL | Probable | As nectar-tap_saprotroph (Põlme et al. 2020); Decay sugar-rich substrates (Põlme et al. 2020) | Põlme S, et al. 2020. Fungal Diversity 105:1-16 ((https://doi.org/10.1007/s13225-020-00466-2)) |
| Otu03539 | 10 | k__Fungi(100);p__Ascomycota(100);c__Saccharomycetes(100);o__Saccharomycetales(100);f__Metschnikowiaceae(100);g__Metschnikowia(100);s__Metschnikowia_pulcherrima(90); | Metschnikowia | 13 | Saprotroph | \|Undefined Saprotroph\| | Yeast | NULL | Probable | As nectar-tap_saprotroph (Põlme et al. 2020); Decay sugar-rich substrates (Põlme et al. 2020) | Põlme S, et al. 2020. Fungal Diversity 105:1-16 ((https://doi.org/10.1007/s13225-020-00466-2)) |
| Otu03540 | 10 | k__Fungi(100);p__Ascomycota(100);c__Saccharomycetes(100);o__Saccharomycetales(100);f__Metschnikowiaceae(100);g__Metschnikowia(100);g__Metschnikowia_unclassified(61); | Metschnikowia | 13 | Saprotroph | \|Undefined Saprotroph\| | Yeast | NULL | Probable | As nectar-tap_saprotroph (Põlme et al. 2020); Decay sugar-rich substrates (Põlme et al. 2020) | Põlme S, et al. 2020. Fungal Diversity 105:1-16 ((https://doi.org/10.1007/s13225-020-00466-2)) |
| Otu03542 | 10 | k__Fungi(100);p__Ascomycota(100);c__Saccharomycetes(100);o__Saccharomycetales(100);f__Metschnikowiaceae(100);g__Metschnikowia(100);s__Metschnikowia_pulcherrima(90); | Metschnikowia | 13 | Saprotroph | \|Undefined Saprotroph\| | Yeast | NULL | Probable | As nectar-tap_saprotroph (Põlme et al. 2020); Decay sugar-rich substrates (Põlme et al. 2020) | Põlme S, et al. 2020. Fungal Diversity 105:1-16 ((https://doi.org/10.1007/s13225-020-00466-2)) |
| Otu03544 | 10 | k__Fungi(100);p__Ascomycota(100);c__Sordariomycetes(100);o__Hypocreales(100);f__Nectriaceae(70);f__Nectriaceae_unclassified(70);f__Nectriaceae_unclassified(70); | Nectriaceae | 9 | Pathotroph-Saprotroph-Symbiotroph | Animal Pathogen-Endophyte-Fungal Parasite-Lichen Parasite-Plant Pathogen-Wood Saprotroph | Microfungus | NULL | Possible | Fusarium species are amongst the economically significant plant pathogens (Cannon & Kirk 2007) | Cannon PF, Kirk PM. 2007. Fungal Families of the World. CAB International, Cambridge (ISBN: 978-0851998275); Hirooka Y, et al. 2012. Studies in Mycology 711-210 ((https://doi.org/10.3114/sim0001)); Tedersoo L, et al. 2014. Science 346:e1256688 ((https://doi.org/10.1126/science.1256688)); The American Phytopathological Society Common Names of Plant Diseases ((http://www.apsnet.org/publications/commonnames/Pages/default.aspx)); https://www.cdc.gov ((https://www.cdc.gov)) |
| Otu03545 | 10 | k__Fungi(100);p__Ascomycota(100);c__Saccharomycetes(100);o__Saccharomycetales(100);f__Metschnikowiaceae(100);g__Metschnikowia(100);s__Metschnikowia_pulcherrima(100); | Metschnikowia | 13 | Saprotroph | \|Undefined Saprotroph\| | Yeast | NULL | Probable | As nectar-tap_saprotroph (Põlme et al. 2020); Decay sugar-rich substrates (Põlme et al. 2020) | Põlme S, et al. 2020. Fungal Diversity 105:1-16 ((https://doi.org/10.1007/s13225-020-00466-2)) |
| Otu03546 | 10 | k__Fungi(100);p__Ascomycota(100);c__Saccharomycetes(100);o__Saccharomycetales(100);f__Metschnikowiaceae(100);g__Metschnikowia(100);g__Metschnikowia_unclassified(100); | Metschnikowia | 13 | Saprotroph | \|Undefined Saprotroph\| | Yeast | NULL | Probable | As nectar-tap_saprotroph (Põlme et al. 2020); Decay sugar-rich substrates (Põlme et al. 2020) | Põlme S, et al. 2020. Fungal Diversity 105:1-16 ((https://doi.org/10.1007/s13225-020-00466-2)) |
| Otu03547 | 10 | k__Fungi(100);p__Basidiomycota(100);c__Agaricomycetes(100);o__Trechisporales(100);f__Hydnodontaceae(100);g__Trechispora(100);s__Trechispora_sp(61); | Trechispora | 13 | Saprotroph | \|Wood Saprotroph\| | Corticioid | White Rot | Highly Probable | As white_rot (Põlme et al. 2020); Decay wood (Põlme et al. 2020) | Gilbertson RL, Ryvarden L. 1987-1987. North American Polypores. Fungiflora, Oslo (ISBN: 978-0945345060); Rinaldi AC, et al. 2008. Fungal Diversity 33:1-45 ((http://www.fungaldiversity.org/fdp/sfdp/33-1.pdf)) (as Uncertain); Tedersoo L, et al. 2010. Mycorrhiza 20:217-263 ((https://doi.org/10.1007/s00572-009-0274-x)) (as not ECM).; Põlme S, et al. 2020. Fungal Diversity 105:1-16 ((https://doi.org/10.1007/s13225-020-00466-2)) |
| Otu03548 | 10 | k__Fungi(100);p__Ascomycota(100);c__Saccharomycetes(100);o__Saccharomycetales(100);f__Metschnikowiaceae(100);g__Metschnikowia(100);g__Metschnikowia_unclassified(100); | Metschnikowia | 13 | Saprotroph | \|Undefined Saprotroph\| | Yeast | NULL | Probable | As nectar-tap_saprotroph (Põlme et al. 2020); Decay sugar-rich substrates (Põlme et al. 2020) | Põlme S, et al. 2020. Fungal Diversity 105:1-16 ((https://doi.org/10.1007/s13225-020-00466-2)) |
| Otu03549 | 10 | k__Fungi(100);p__Ascomycota(100);c__Saccharomycetes(100);o__Saccharomycetales(100);f__Metschnikowiaceae(100);g__Metschnikowia(100);s__Metschnikowia_pulcherrima(100); | Metschnikowia | 13 | Saprotroph | \|Undefined Saprotroph\| | Yeast | NULL | Probable | As nectar-tap_saprotroph (Põlme et al. 2020); Decay sugar-rich substrates (Põlme et al. 2020) | Põlme S, et al. 2020. Fungal Diversity 105:1-16 ((https://doi.org/10.1007/s13225-020-00466-2)) |
| Otu03550 | 10 | k__Fungi(100);p__Ascomycota(100);c__Sordariomycetes(90);o__Chaetosphaeriales(80);f__Chaetosphaeriaceae(80);g__unclassified_Chaetosphaeriaceae(70);s__Chaetosphaeriaceae_sp(70); | Chaetosphaeriaceae | 9 | Saprotroph | Plant Saprotroph-Wood Saprotroph | Microfungus | NULL | Probable | Saprobes associated with decaying woody and herbaceous plant material, including timber and timber products in damp buildings (Cannon & Kirk 2007) | Cannon PF, Kirk PM. 2007. Fungal Families of the World. CAB International, Cambridge (ISBN: 978-0851998275) |
| Otu03555 | 10 | k__Fungi(100);p__Ascomycota(100);c__Eurotiomycetes(100);o__Chaetothyriales(100);f__Herpotrichiellaceae(100);g__Phaeomoniella(100);s__Phaeomoniella_sp(100); | Phaeomoniella | 13 | Pathotroph | \|Plant Pathogen\| | Dimorphic-Microfungus; Facultative Yeast (Tedersoo et | NULL | Highly Probable | Pathotroph isolated from necrosis in Prunus wood (Damm et al. 2010) | Damm U, et al. 2010. Persoonia 24:60-80 ((http://doi.org/10.3767/003158510X500705)); Tedersoo L, et al. 2014. Science 346:e1256688 ((https://doi.org/10.1126/science.1256688)); Põlme S, et al. 2020. Fungal Diversity 105:1-16 ((https://doi.org/10.1007/s13225-020-00466-2)) |
| Otu03556 | 10 | k__Fungi(100);p__Ascomycota(100);c__Saccharomycetes(100);o__Saccharomycetales(100);f__Saccharomycetales_family_Incertae_sedis(90);g__Candida(90);g__Candida_unclassified(70); | Candida | 13 | Pathotroph-Saprotroph-Symbiotroph | Animal Pathogen-Endophyte-Endosymbiont-Epiphyte-\|Undefined Saprotroph\| | Dimorphic Yeast-Yeast | NULL | Probable | As nectar-tap_saprotroph (Põlme et al. 2020); Decay sugar-rich substrates (Põlme et al. 2020); This asexual genus is polyphyletic and contain species that function very differently depend on what they are. Caution should be taken when interpreting guilds for Candida species. | Manolakaki D, et al. 2010. Virulence. 1: 367-375 ((https://doi.org/10.4161/viru.1.5.12796)); Mekbib SB, et al. 2011. Tropical Plant Pathology 36:233-240 ((https://doi.org/10.1590/S1982-56762011000400004)); Põlme S, et al. 2020. Fungal Diversity 105:1-16 ((https://doi.org/10.1007/s13225-020-00466-2)) |
| Otu03559 | 10 | k__Fungi(100);p__Ascomycota(100);c__Saccharomycetes(100);o__Saccharomycetales(100);f__Metschnikowiaceae(100);g__Metschnikowia(100);s__Metschnikowia_pulcherrima(100); | Metschnikowia | 13 | Saprotroph | \|Undefined Saprotroph\| | Yeast | NULL | Probable | As nectar-tap_saprotroph (Põlme et al. 2020); Decay sugar-rich substrates (Põlme et al. 2020) | Põlme S, et al. 2020. Fungal Diversity 105:1-16 ((https://doi.org/10.1007/s13225-020-00466-2)) |
| Otu03561 | 10 | k__Fungi(100);p__Ascomycota(100);c__Leotiomycetes(100);o__Helotiales(100);f__Dermateaceae(100);f__Dermateaceae_unclassified(100);f__Dermateaceae_unclassified(100); | Dermateaceae | 9 | Pathotroph-Saprotroph | Endophyte-\|Plant Pathogen\|-Plant Saprotroph-Wood Saprotroph | Microfungus-Pezizoid | NULL | Possible | As foliar_endophyte (Põlme et al. 2020); As litter_saprotroph (Põlme et al. 2020); Dark Septate Endophyte, known from plant roots.; Decay leaf, fruit, seed (Põlme et al. 2020); Saprobic or parasitic on herbaceous and woody material (Cannon & Kirk 2007) | Cannon PF, Kirk PM. 2007. Fungal Families of the World. CAB International, Cambridge (ISBN: 978-0851998275); Põlme S, et al. 2020. Fungal Diversity 105:1-16 ((https://doi.org/10.1007/s13225-020-00466-2)) |
| Otu03564 | 10 | k__Fungi(100);p__Ascomycota(100);c__Saccharomycetes(100);o__Saccharomycetales(100);f__Metschnikowiaceae(100);g__Metschnikowia(100);s__Metschnikowia_pulcherrima(100); | Metschnikowia | 13 | Saprotroph | \|Undefined Saprotroph\| | Yeast | NULL | Probable | As nectar-tap_saprotroph (Põlme et al. 2020); Decay sugar-rich substrates (Põlme et al. 2020) | Põlme S, et al. 2020. Fungal Diversity 105:1-16 ((https://doi.org/10.1007/s13225-020-00466-2)) |
| Otu03568 | 10 | k__Fungi(100);p__Ascomycota(100);c__Sordariomycetes(100);o__Hypocreales(100);o__Hypocreales_unclassified(100);o__Hypocreales_unclassified(100);o__Hypocreales_unclassified(100); | Hypocreales | 7 | Saprotroph | Undefined Saprotroph | Microfungus | NULL | Possible | NULL | Sterkenburg E, et al. 2015. New Phytologist 207:1145-1158 ((https://doi.org/10.1111/nph.13426)) |
| Otu03569 | 10 | k__Fungi(100);p__Ascomycota(100);c__Sordariomycetes(100);o__Hypocreales(100);f__Hypocreales_family_Incertae_sedis(100);f__Hypocreales_family_Incertae_sedis_unclassified(100);f__Hypocreales_family_Incertae_sedis_unclassified(100); | Hypocreales | 7 | Saprotroph | Undefined Saprotroph | Microfungus | NULL | Possible | NULL | Sterkenburg E, et al. 2015. New Phytologist 207:1145-1158 ((https://doi.org/10.1111/nph.13426)) |
| Otu03573 | 10 | k__Fungi(100);p__Ascomycota(100);c__Saccharomycetes(100);o__Saccharomycetales(100);f__Metschnikowiaceae(100);g__Metschnikowia(100);s__Metschnikowia_pulcherrima(80); | Metschnikowia | 13 | Saprotroph | \|Undefined Saprotroph\| | Yeast | NULL | Probable | As nectar-tap_saprotroph (Põlme et al. 2020); Decay sugar-rich substrates (Põlme et al. 2020) | Põlme S, et al. 2020. Fungal Diversity 105:1-16 ((https://doi.org/10.1007/s13225-020-00466-2)) |
| Otu03574 | 10 | k__Fungi(100);p__Ascomycota(100);c__Saccharomycetes(100);o__Saccharomycetales(100);f__Metschnikowiaceae(100);g__Metschnikowia(100);s__Metschnikowia_pulcherrima(90); | Metschnikowia | 13 | Saprotroph | \|Undefined Saprotroph\| | Yeast | NULL | Probable | As nectar-tap_saprotroph (Põlme et al. 2020); Decay sugar-rich substrates (Põlme et al. 2020) | Põlme S, et al. 2020. Fungal Diversity 105:1-16 ((https://doi.org/10.1007/s13225-020-00466-2)) |
| Otu03576 | 10 | k__Fungi(100);p__Zygomycota(100);c__Zygomycota_class_Incertae_sedis(100);o__Mortierellales(100);f__Mortierellaceae(100);g__Mortierella(100);g__Mortierella_unclassified(100); | Mortierella | 13 | Saprotroph-Symbiotroph | Endophyte-Plant Saprotroph-\|Undefined Saprotroph\| | Microfungus | NULL | Probable | As chitinolytic (Põlme et al. 2020); As soil_saprotroph (Põlme et al. 2020); Decay roots, soil (Põlme et al. 2020); Saprobic in soil and thus frequently isolated from soil and material contaminated with soil such as hay or rotten silage and spoil tips (Cannon & Kirk 2007) | Cannon PF, Kirk PM. 2007. Fungal Families of the World. CAB International, Cambridge (ISBN: 978-0851998275); Tedersoo L, et al. 2014. Science 346:e1256688 ((https://doi.org/10.1126/science.1256688)); Purahong W, et al. 2016. Molecular Ecology 25:4059-4074 ((https://doi.org/10.1111/mec.13739)); Põlme S, et al. 2020. Fungal Diversity 105:1-16 ((https://doi.org/10.1007/s13225-020-00466-2)) |
| Otu03577 | 10 | k__Fungi(100);p__Ascomycota(100);c__Saccharomycetes(100);o__Saccharomycetales(100);f__Metschnikowiaceae(100);g__Metschnikowia(100);g__Metschnikowia_unclassified(80); | Metschnikowia | 13 | Saprotroph | \|Undefined Saprotroph\| | Yeast | NULL | Probable | As nectar-tap_saprotroph (Põlme et al. 2020); Decay sugar-rich substrates (Põlme et al. 2020) | Põlme S, et al. 2020. Fungal Diversity 105:1-16 ((https://doi.org/10.1007/s13225-020-00466-2)) |
| Otu03578 | 10 | k__Fungi(100);p__Basidiomycota(100);c__Agaricomycetes(100);o__Agaricales(100);f__Cortinariaceae(100);g__Cortinarius(100);g__Cortinarius_unclassified(100); | Cortinarius | 13 | Symbiotroph | \|Ectomycorrhizal\| | Agaricoid | NULL | Highly Probable | NULL | Rinaldi AC, et al. 2008. Fungal Diversity 33:1-45 ((http://www.fungaldiversity.org/fdp/sfdp/33-1.pdf)); Tedersoo L, et al. 2010. Mycorrhiza 20:217-263 ((https://doi.org/10.1007/s00572-009-0274-x)); Põlme S, et al. 2020. Fungal Diversity 105:1-16 ((https://doi.org/10.1007/s13225-020-00466-2)); http://mycorrhizas.info/ecmf.html ((http://mycorrhizas.info/ecmf.html)) |
| Otu03580 | 10 | k__Fungi(100);p__Ascomycota(100);c__Sordariomycetes(100);o__Diaporthales(100);f__Melanconidaceae(100);g__Melanconiella(100);s__Melanconiella_ellisii(100); | Melanconiella | 13 | Saprotroph | Undefined Saprotroph-\|Wood Saprotroph\| | Microfungus | NULL | Probable | Decay wood (Põlme et al. 2020) | Tedersoo L, et al. 2014. Science 346:e1256688 ((https://doi.org/10.1126/science.1256688)); Põlme S, et al. 2020. Fungal Diversity 105:1-16 ((https://doi.org/10.1007/s13225-020-00466-2)) |
| Otu03352 | 10 | k__Fungi(100);p__Ascomycota(100);c__Dothideomycetes(100);c__Dothideomycetes_unclassified(100);c__Dothideomycetes_unclassified(100);c__Dothideomycetes_unclassified(100);c__Dothideomycetes_unclassified(100); | - | - | - | - | - | - | - | Unassigned | - |
| Otu03354 | 10 | k__Fungi(100);k__Fungi_unclassified(90);k__Fungi_unclassified(90);k__Fungi_unclassified(90);k__Fungi_unclassified(90);k__Fungi_unclassified(90);k__Fungi_unclassified(90); | - | - | - | - | - | - | - | Unassigned | - |
| Otu03357 | 10 | k__Fungi(100);k__Fungi_unclassified(100);k__Fungi_unclassified(100);k__Fungi_unclassified(100);k__Fungi_unclassified(100);k__Fungi_unclassified(100);k__Fungi_unclassified(100); | - | - | - | - | - | - | - | Unassigned | - |
| Otu03358 | 10 | k__Fungi(100);k__Fungi_unclassified(100);k__Fungi_unclassified(100);k__Fungi_unclassified(100);k__Fungi_unclassified(100);k__Fungi_unclassified(100);k__Fungi_unclassified(100); | - | - | - | - | - | - | - | Unassigned | - |
| Otu03360 | 10 | k__Fungi(100);k__Fungi_unclassified(100);k__Fungi_unclassified(100);k__Fungi_unclassified(100);k__Fungi_unclassified(100);k__Fungi_unclassified(100);k__Fungi_unclassified(100); | - | - | - | - | - | - | - | Unassigned | - |
| Otu03361 | 10 | k__Fungi(100);p__Ascomycota(100);c__Dothideomycetes(90);c__Dothideomycetes_unclassified(70);c__Dothideomycetes_unclassified(70);c__Dothideomycetes_unclassified(70);c__Dothideomycetes_unclassified(70); | - | - | - | - | - | - | - | Unassigned | - |
| Otu03363 | 10 | k__Fungi(100);p__Ascomycota(100);p__Ascomycota_unclassified(100);p__Ascomycota_unclassified(100);p__Ascomycota_unclassified(100);p__Ascomycota_unclassified(100);p__Ascomycota_unclassified(100); | - | - | - | - | - | - | - | Unassigned | - |
| Otu03365 | 10 | k__Fungi(100);p__Ascomycota(90);p__Ascomycota_unclassified(90);p__Ascomycota_unclassified(90);p__Ascomycota_unclassified(90);p__Ascomycota_unclassified(90);p__Ascomycota_unclassified(90); | - | - | - | - | - | - | - | Unassigned | - |
| Otu03370 | 10 | k__Fungi(100);p__Ascomycota(100);p__Ascomycota_unclassified(100);p__Ascomycota_unclassified(100);p__Ascomycota_unclassified(100);p__Ascomycota_unclassified(100);p__Ascomycota_unclassified(100); | - | - | - | - | - | - | - | Unassigned | - |
| Otu03372 | 10 | k__Fungi(100);p__Ascomycota(70);p__Ascomycota_unclassified(70);p__Ascomycota_unclassified(70);p__Ascomycota_unclassified(70);p__Ascomycota_unclassified(70);p__Ascomycota_unclassified(70); | - | - | - | - | - | - | - | Unassigned | - |
| Otu03375 | 10 | k__Fungi(100);p__Ascomycota(100);p__Ascomycota_unclassified(100);p__Ascomycota_unclassified(100);p__Ascomycota_unclassified(100);p__Ascomycota_unclassified(100);p__Ascomycota_unclassified(100); | - | - | - | - | - | - | - | Unassigned | - |
| Otu03376 | 10 | k__Fungi(100);p__Ascomycota(61);p__Ascomycota_unclassified(61);p__Ascomycota_unclassified(61);p__Ascomycota_unclassified(61);p__Ascomycota_unclassified(61);p__Ascomycota_unclassified(61); | - | - | - | - | - | - | - | Unassigned | - |
| Otu03377 | 10 | k__Fungi(100);k__Fungi_unclassified(61);k__Fungi_unclassified(61);k__Fungi_unclassified(61);k__Fungi_unclassified(61);k__Fungi_unclassified(61);k__Fungi_unclassified(61); | - | - | - | - | - | - | - | Unassigned | - |
| Otu03378 | 10 | k__Fungi(100);p__Basidiomycota(100);c__Agaricomycetes(100);c__Agaricomycetes_unclassified(90);c__Agaricomycetes_unclassified(90);c__Agaricomycetes_unclassified(90);c__Agaricomycetes_unclassified(90); | - | - | - | - | - | - | - | Unassigned | - |
| Otu03381 | 10 | k__Fungi(100);p__Ascomycota(100);p__Ascomycota_unclassified(100);p__Ascomycota_unclassified(100);p__Ascomycota_unclassified(100);p__Ascomycota_unclassified(100);p__Ascomycota_unclassified(100); | - | - | - | - | - | - | - | Unassigned | - |
| Otu03385 | 10 | k__Fungi(100);p__Basidiomycota(100);c__Agaricomycetes(100);c__Agaricomycetes_unclassified(100);c__Agaricomycetes_unclassified(100);c__Agaricomycetes_unclassified(100);c__Agaricomycetes_unclassified(100); | - | - | - | - | - | - | - | Unassigned | - |
| Otu03387 | 10 | k__Fungi(100);k__Fungi_unclassified(80);k__Fungi_unclassified(80);k__Fungi_unclassified(80);k__Fungi_unclassified(80);k__Fungi_unclassified(80);k__Fungi_unclassified(80); | - | - | - | - | - | - | - | Unassigned | - |
| Otu03389 | 10 | k__Fungi(100);p__Ascomycota(61);p__Ascomycota_unclassified(61);p__Ascomycota_unclassified(61);p__Ascomycota_unclassified(61);p__Ascomycota_unclassified(61);p__Ascomycota_unclassified(61); | - | - | - | - | - | - | - | Unassigned | - |
| Otu03390 | 10 | k__Fungi(100);k__Fungi_unclassified(100);k__Fungi_unclassified(100);k__Fungi_unclassified(100);k__Fungi_unclassified(100);k__Fungi_unclassified(100);k__Fungi_unclassified(100); | - | - | - | - | - | - | - | Unassigned | - |
| Otu03393 | 10 | k__Fungi(100);k__Fungi_unclassified(100);k__Fungi_unclassified(100);k__Fungi_unclassified(100);k__Fungi_unclassified(100);k__Fungi_unclassified(100);k__Fungi_unclassified(100); | - | - | - | - | - | - | - | Unassigned | - |
| Otu03394 | 10 | k__Fungi(100);p__Ascomycota(100);c__Sordariomycetes(100);c__Sordariomycetes_unclassified(100);c__Sordariomycetes_unclassified(100);c__Sordariomycetes_unclassified(100);c__Sordariomycetes_unclassified(100); | - | - | - | - | - | - | - | Unassigned | - |
| Otu03396 | 10 | k__Fungi(100);p__Ascomycota(70);p__Ascomycota_unclassified(70);p__Ascomycota_unclassified(70);p__Ascomycota_unclassified(70);p__Ascomycota_unclassified(70);p__Ascomycota_unclassified(70); | - | - | - | - | - | - | - | Unassigned | - |
| Otu03398 | 10 | k__Fungi(100);p__Ascomycota(70);p__Ascomycota_unclassified(70);p__Ascomycota_unclassified(70);p__Ascomycota_unclassified(70);p__Ascomycota_unclassified(70);p__Ascomycota_unclassified(70); | - | - | - | - | - | - | - | Unassigned | - |
| Otu03400 | 10 | k__Fungi(100);k__Fungi_unclassified(100);k__Fungi_unclassified(100);k__Fungi_unclassified(100);k__Fungi_unclassified(100);k__Fungi_unclassified(100);k__Fungi_unclassified(100); | - | - | - | - | - | - | - | Unassigned | - |
| Otu03402 | 10 | k__Fungi(100);p__Basidiomycota(100);c__Agaricomycetes(100);c__Agaricomycetes_unclassified(100);c__Agaricomycetes_unclassified(100);c__Agaricomycetes_unclassified(100);c__Agaricomycetes_unclassified(100); | - | - | - | - | - | - | - | Unassigned | - |
| Otu03403 | 10 | k__Fungi(100);k__Fungi_unclassified(90);k__Fungi_unclassified(90);k__Fungi_unclassified(90);k__Fungi_unclassified(90);k__Fungi_unclassified(90);k__Fungi_unclassified(90); | - | - | - | - | - | - | - | Unassigned | - |
| Otu03409 | 10 | k__Fungi(100);p__Ascomycota(90);p__Ascomycota_unclassified(90);p__Ascomycota_unclassified(90);p__Ascomycota_unclassified(90);p__Ascomycota_unclassified(90);p__Ascomycota_unclassified(90); | - | - | - | - | - | - | - | Unassigned | - |
| Otu03410 | 10 | k__Fungi(100);p__Ascomycota(100);p__Ascomycota_unclassified(100);p__Ascomycota_unclassified(100);p__Ascomycota_unclassified(100);p__Ascomycota_unclassified(100);p__Ascomycota_unclassified(100); | - | - | - | - | - | - | - | Unassigned | - |
| Otu03413 | 10 | k__Fungi(100);p__Basidiomycota(100);p__Basidiomycota_unclassified(100);p__Basidiomycota_unclassified(100);p__Basidiomycota_unclassified(100);p__Basidiomycota_unclassified(100);p__Basidiomycota_unclassified(100); | - | - | - | - | - | - | - | Unassigned | - |
| Otu03414 | 10 | k__Fungi(100);k__Fungi_unclassified(100);k__Fungi_unclassified(100);k__Fungi_unclassified(100);k__Fungi_unclassified(100);k__Fungi_unclassified(100);k__Fungi_unclassified(100); | - | - | - | - | - | - | - | Unassigned | - |
| Otu03416 | 10 | k__Fungi(100);p__Ascomycota(100);c__Dothideomycetes(100);o__Capnodiales(70);o__Capnodiales_unclassified(70);o__Capnodiales_unclassified(70);o__Capnodiales_unclassified(70); | - | - | - | - | - | - | - | Unassigned | - |
| Otu03417 | 10 | k__Fungi(100);k__Fungi_unclassified(61);k__Fungi_unclassified(61);k__Fungi_unclassified(61);k__Fungi_unclassified(61);k__Fungi_unclassified(61);k__Fungi_unclassified(61); | - | - | - | - | - | - | - | Unassigned | - |
| Otu03419 | 10 | k__Fungi(100);k__Fungi_unclassified(80);k__Fungi_unclassified(80);k__Fungi_unclassified(80);k__Fungi_unclassified(80);k__Fungi_unclassified(80);k__Fungi_unclassified(80); | - | - | - | - | - | - | - | Unassigned | - |
| Otu03421 | 10 | k__Fungi(100);p__Ascomycota(100);c__Sordariomycetes(100);o__Sordariales(100);o__Sordariales_unclassified(100);o__Sordariales_unclassified(100);o__Sordariales_unclassified(100); | - | - | - | - | - | - | - | Unassigned | - |
| Otu03425 | 10 | k__Fungi(100);k__Fungi_unclassified(100);k__Fungi_unclassified(100);k__Fungi_unclassified(100);k__Fungi_unclassified(100);k__Fungi_unclassified(100);k__Fungi_unclassified(100); | - | - | - | - | - | - | - | Unassigned | - |
| Otu03426 | 10 | k__Fungi(100);k__Fungi_unclassified(61);k__Fungi_unclassified(61);k__Fungi_unclassified(61);k__Fungi_unclassified(61);k__Fungi_unclassified(61);k__Fungi_unclassified(61); | - | - | - | - | - | - | - | Unassigned | - |
| Otu03427 | 10 | k__Fungi(100);k__Fungi_unclassified(100);k__Fungi_unclassified(100);k__Fungi_unclassified(100);k__Fungi_unclassified(100);k__Fungi_unclassified(100);k__Fungi_unclassified(100); | - | - | - | - | - | - | - | Unassigned | - |
| Otu03428 | 10 | k__Fungi(100);k__Fungi_unclassified(80);k__Fungi_unclassified(80);k__Fungi_unclassified(80);k__Fungi_unclassified(80);k__Fungi_unclassified(80);k__Fungi_unclassified(80); | - | - | - | - | - | - | - | Unassigned | - |
| Otu03429 | 10 | k__Fungi(100);k__Fungi_unclassified(70);k__Fungi_unclassified(70);k__Fungi_unclassified(70);k__Fungi_unclassified(70);k__Fungi_unclassified(70);k__Fungi_unclassified(70); | - | - | - | - | - | - | - | Unassigned | - |
| Otu03437 | 10 | k__Fungi(100);p__Ascomycota(80);p__Ascomycota_unclassified(80);p__Ascomycota_unclassified(80);p__Ascomycota_unclassified(80);p__Ascomycota_unclassified(80);p__Ascomycota_unclassified(80); | - | - | - | - | - | - | - | Unassigned | - |
| Otu03439 | 10 | k__Fungi(100);p__Ascomycota(80);p__Ascomycota_unclassified(80);p__Ascomycota_unclassified(80);p__Ascomycota_unclassified(80);p__Ascomycota_unclassified(80);p__Ascomycota_unclassified(80); | - | - | - | - | - | - | - | Unassigned | - |
| Otu03442 | 10 | k__Fungi(100);p__Chytridiomycota(70);c__Chytridiomycetes(70);c__Chytridiomycetes_unclassified(70);c__Chytridiomycetes_unclassified(70);c__Chytridiomycetes_unclassified(70);c__Chytridiomycetes_unclassified(70); | - | - | - | - | - | - | - | Unassigned | - |
| Otu03443 | 10 | k__Fungi(100);k__Fungi_unclassified(61);k__Fungi_unclassified(61);k__Fungi_unclassified(61);k__Fungi_unclassified(61);k__Fungi_unclassified(61);k__Fungi_unclassified(61); | - | - | - | - | - | - | - | Unassigned | - |
| Otu03444 | 10 | k__Fungi(100);p__Ascomycota(80);p__Ascomycota_unclassified(80);p__Ascomycota_unclassified(80);p__Ascomycota_unclassified(80);p__Ascomycota_unclassified(80);p__Ascomycota_unclassified(80); | - | - | - | - | - | - | - | Unassigned | - |
| Otu03447 | 10 | k__Fungi(100);k__Fungi_unclassified(100);k__Fungi_unclassified(100);k__Fungi_unclassified(100);k__Fungi_unclassified(100);k__Fungi_unclassified(100);k__Fungi_unclassified(100); | - | - | - | - | - | - | - | Unassigned | - |
| Otu03449 | 10 | k__Fungi(100);p__Ascomycota(100);p__Ascomycota_unclassified(100);p__Ascomycota_unclassified(100);p__Ascomycota_unclassified(100);p__Ascomycota_unclassified(100);p__Ascomycota_unclassified(100); | - | - | - | - | - | - | - | Unassigned | - |
| Otu03451 | 10 | k__Fungi(100);p__Ascomycota(70);p__Ascomycota_unclassified(70);p__Ascomycota_unclassified(70);p__Ascomycota_unclassified(70);p__Ascomycota_unclassified(70);p__Ascomycota_unclassified(70); | - | - | - | - | - | - | - | Unassigned | - |
| Otu03454 | 10 | k__Fungi(100);p__Ascomycota(61);p__Ascomycota_unclassified(61);p__Ascomycota_unclassified(61);p__Ascomycota_unclassified(61);p__Ascomycota_unclassified(61);p__Ascomycota_unclassified(61); | - | - | - | - | - | - | - | Unassigned | - |
| Otu03455 | 10 | k__Fungi(100);k__Fungi_unclassified(100);k__Fungi_unclassified(100);k__Fungi_unclassified(100);k__Fungi_unclassified(100);k__Fungi_unclassified(100);k__Fungi_unclassified(100); | - | - | - | - | - | - | - | Unassigned | - |
| Otu03457 | 10 | k__Fungi(100);p__Ascomycota(61);p__Ascomycota_unclassified(61);p__Ascomycota_unclassified(61);p__Ascomycota_unclassified(61);p__Ascomycota_unclassified(61);p__Ascomycota_unclassified(61); | - | - | - | - | - | - | - | Unassigned | - |
| Otu03459 | 10 | k__Fungi(100);p__Ascomycota(100);c__Dothideomycetes(100);o__Pleosporales(100);o__Pleosporales_unclassified(80);o__Pleosporales_unclassified(80);o__Pleosporales_unclassified(80); | - | - | - | - | - | - | - | Unassigned | - |
| Otu03461 | 10 | k__Fungi(100);p__Ascomycota(100);p__Ascomycota_unclassified(100);p__Ascomycota_unclassified(100);p__Ascomycota_unclassified(100);p__Ascomycota_unclassified(100);p__Ascomycota_unclassified(100); | - | - | - | - | - | - | - | Unassigned | - |
| Otu03463 | 10 | k__Fungi(100);p__Ascomycota(61);p__Ascomycota_unclassified(61);p__Ascomycota_unclassified(61);p__Ascomycota_unclassified(61);p__Ascomycota_unclassified(61);p__Ascomycota_unclassified(61); | - | - | - | - | - | - | - | Unassigned | - |
| Otu03467 | 10 | k__Fungi(100);p__Ascomycota(100);c__Dothideomycetes(100);o__Pleosporales(100);o__Pleosporales_unclassified(100);o__Pleosporales_unclassified(100);o__Pleosporales_unclassified(100); | - | - | - | - | - | - | - | Unassigned | - |
| Otu03470 | 10 | k__Fungi(100);k__Fungi_unclassified(70);k__Fungi_unclassified(70);k__Fungi_unclassified(70);k__Fungi_unclassified(70);k__Fungi_unclassified(70);k__Fungi_unclassified(70); | - | - | - | - | - | - | - | Unassigned | - |
| Otu03472 | 10 | k__Fungi(100);k__Fungi_unclassified(100);k__Fungi_unclassified(100);k__Fungi_unclassified(100);k__Fungi_unclassified(100);k__Fungi_unclassified(100);k__Fungi_unclassified(100); | - | - | - | - | - | - | - | Unassigned | - |
| Otu03474 | 10 | k__Fungi(100);p__Ascomycota(100);p__Ascomycota_unclassified(90);p__Ascomycota_unclassified(90);p__Ascomycota_unclassified(90);p__Ascomycota_unclassified(90);p__Ascomycota_unclassified(90); | - | - | - | - | - | - | - | Unassigned | - |
| Otu03476 | 10 | k__Fungi(100);p__Ascomycota(61);p__Ascomycota_unclassified(61);p__Ascomycota_unclassified(61);p__Ascomycota_unclassified(61);p__Ascomycota_unclassified(61);p__Ascomycota_unclassified(61); | - | - | - | - | - | - | - | Unassigned | - |
| Otu03478 | 10 | k__Fungi(100);p__Ascomycota(100);c__Lecanoromycetes(70);c__Lecanoromycetes_unclassified(70);c__Lecanoromycetes_unclassified(70);c__Lecanoromycetes_unclassified(70);c__Lecanoromycetes_unclassified(70); | - | - | - | - | - | - | - | Unassigned | - |
| Otu03480 | 10 | k__Fungi(100);k__Fungi_unclassified(80);k__Fungi_unclassified(80);k__Fungi_unclassified(80);k__Fungi_unclassified(80);k__Fungi_unclassified(80);k__Fungi_unclassified(80); | - | - | - | - | - | - | - | Unassigned | - |
| Otu03481 | 10 | k__Fungi(100);p__Ascomycota(61);p__Ascomycota_unclassified(61);p__Ascomycota_unclassified(61);p__Ascomycota_unclassified(61);p__Ascomycota_unclassified(61);p__Ascomycota_unclassified(61); | - | - | - | - | - | - | - | Unassigned | - |
| Otu03484 | 10 | k__Plantae(100);p__unclassified_Plantae(100);c__unclassified_Plantae(100);o__unclassified_Plantae(100);f__unclassified_Plantae(100);g__unclassified_Plantae(100);s__Plantae_sp(100); | - | - | - | - | - | - | - | Unassigned | - |
| Otu03485 | 10 | k__Fungi(100);k__Fungi_unclassified(70);k__Fungi_unclassified(70);k__Fungi_unclassified(70);k__Fungi_unclassified(70);k__Fungi_unclassified(70);k__Fungi_unclassified(70); | - | - | - | - | - | - | - | Unassigned | - |
| Otu03486 | 10 | k__Fungi(100);p__Ascomycota(100);p__Ascomycota_unclassified(100);p__Ascomycota_unclassified(100);p__Ascomycota_unclassified(100);p__Ascomycota_unclassified(100);p__Ascomycota_unclassified(100); | - | - | - | - | - | - | - | Unassigned | - |
| Otu03489 | 10 | k__Fungi(100);k__Fungi_unclassified(100);k__Fungi_unclassified(100);k__Fungi_unclassified(100);k__Fungi_unclassified(100);k__Fungi_unclassified(100);k__Fungi_unclassified(100); | - | - | - | - | - | - | - | Unassigned | - |
| Otu03490 | 10 | k__Fungi(100);p__Ascomycota(100);p__Ascomycota_unclassified(100);p__Ascomycota_unclassified(100);p__Ascomycota_unclassified(100);p__Ascomycota_unclassified(100);p__Ascomycota_unclassified(100); | - | - | - | - | - | - | - | Unassigned | - |
| Otu03491 | 10 | k__Fungi(100);p__Basidiomycota(100);c__Agaricomycetes(100);c__Agaricomycetes_unclassified(100);c__Agaricomycetes_unclassified(100);c__Agaricomycetes_unclassified(100);c__Agaricomycetes_unclassified(100); | - | - | - | - | - | - | - | Unassigned | - |
| Otu03497 | 10 | k__Fungi(100);k__Fungi_unclassified(100);k__Fungi_unclassified(100);k__Fungi_unclassified(100);k__Fungi_unclassified(100);k__Fungi_unclassified(100);k__Fungi_unclassified(100); | - | - | - | - | - | - | - | Unassigned | - |
| Otu03498 | 10 | k__Fungi(100);k__Fungi_unclassified(80);k__Fungi_unclassified(80);k__Fungi_unclassified(80);k__Fungi_unclassified(80);k__Fungi_unclassified(80);k__Fungi_unclassified(80); | - | - | - | - | - | - | - | Unassigned | - |
| Otu03501 | 10 | k__Fungi(100);p__Basidiomycota(100);p__Basidiomycota_unclassified(100);p__Basidiomycota_unclassified(100);p__Basidiomycota_unclassified(100);p__Basidiomycota_unclassified(100);p__Basidiomycota_unclassified(100); | - | - | - | - | - | - | - | Unassigned | - |
| Otu03506 | 10 | k__Fungi(100);k__Fungi_unclassified(100);k__Fungi_unclassified(100);k__Fungi_unclassified(100);k__Fungi_unclassified(100);k__Fungi_unclassified(100);k__Fungi_unclassified(100); | - | - | - | - | - | - | - | Unassigned | - |
| Otu03507 | 10 | k__Fungi(100);k__Fungi_unclassified(90);k__Fungi_unclassified(90);k__Fungi_unclassified(90);k__Fungi_unclassified(90);k__Fungi_unclassified(90);k__Fungi_unclassified(90); | - | - | - | - | - | - | - | Unassigned | - |
| Otu03509 | 10 | k__Fungi(100);k__Fungi_unclassified(100);k__Fungi_unclassified(100);k__Fungi_unclassified(100);k__Fungi_unclassified(100);k__Fungi_unclassified(100);k__Fungi_unclassified(100); | - | - | - | - | - | - | - | Unassigned | - |
| Otu03513 | 10 | k__Fungi(100);k__Fungi_unclassified(61);k__Fungi_unclassified(61);k__Fungi_unclassified(61);k__Fungi_unclassified(61);k__Fungi_unclassified(61);k__Fungi_unclassified(61); | - | - | - | - | - | - | - | Unassigned | - |
| Otu03518 | 10 | k__Fungi(100);k__Fungi_unclassified(100);k__Fungi_unclassified(100);k__Fungi_unclassified(100);k__Fungi_unclassified(100);k__Fungi_unclassified(100);k__Fungi_unclassified(100); | - | - | - | - | - | - | - | Unassigned | - |
| Otu03524 | 10 | k__Fungi(100);k__Fungi_unclassified(70);k__Fungi_unclassified(70);k__Fungi_unclassified(70);k__Fungi_unclassified(70);k__Fungi_unclassified(70);k__Fungi_unclassified(70); | - | - | - | - | - | - | - | Unassigned | - |
| Otu03530 | 10 | k__Fungi(100);k__Fungi_unclassified(80);k__Fungi_unclassified(80);k__Fungi_unclassified(80);k__Fungi_unclassified(80);k__Fungi_unclassified(80);k__Fungi_unclassified(80); | - | - | - | - | - | - | - | Unassigned | - |
| Otu03531 | 10 | k__Fungi(100);k__Fungi_unclassified(70);k__Fungi_unclassified(70);k__Fungi_unclassified(70);k__Fungi_unclassified(70);k__Fungi_unclassified(70);k__Fungi_unclassified(70); | - | - | - | - | - | - | - | Unassigned | - |
| Otu03532 | 10 | k__Fungi(100);p__Basidiomycota(100);c__Agaricomycetes(100);o__Russulales(100);o__Russulales_unclassified(100);o__Russulales_unclassified(100);o__Russulales_unclassified(100); | - | - | - | - | - | - | - | Unassigned | - |
| Otu03533 | 10 | k__Fungi(100);p__Ascomycota(61);p__Ascomycota_unclassified(61);p__Ascomycota_unclassified(61);p__Ascomycota_unclassified(61);p__Ascomycota_unclassified(61);p__Ascomycota_unclassified(61); | - | - | - | - | - | - | - | Unassigned | - |
| Otu03535 | 10 | k__Fungi(100);k__Fungi_unclassified(61);k__Fungi_unclassified(61);k__Fungi_unclassified(61);k__Fungi_unclassified(61);k__Fungi_unclassified(61);k__Fungi_unclassified(61); | - | - | - | - | - | - | - | Unassigned | - |
| Otu03536 | 10 | k__Fungi(100);k__Fungi_unclassified(80);k__Fungi_unclassified(80);k__Fungi_unclassified(80);k__Fungi_unclassified(80);k__Fungi_unclassified(80);k__Fungi_unclassified(80); | - | - | - | - | - | - | - | Unassigned | - |
| Otu03538 | 10 | k__Fungi(100);k__Fungi_unclassified(90);k__Fungi_unclassified(90);k__Fungi_unclassified(90);k__Fungi_unclassified(90);k__Fungi_unclassified(90);k__Fungi_unclassified(90); | - | - | - | - | - | - | - | Unassigned | - |
| Otu03541 | 10 | k__Fungi(100);k__Fungi_unclassified(80);k__Fungi_unclassified(80);k__Fungi_unclassified(80);k__Fungi_unclassified(80);k__Fungi_unclassified(80);k__Fungi_unclassified(80); | - | - | - | - | - | - | - | Unassigned | - |
| Otu03543 | 10 | k__Fungi(100);k__Fungi_unclassified(70);k__Fungi_unclassified(70);k__Fungi_unclassified(70);k__Fungi_unclassified(70);k__Fungi_unclassified(70);k__Fungi_unclassified(70); | - | - | - | - | - | - | - | Unassigned | - |
| Otu03551 | 10 | k__Fungi(100);k__Fungi_unclassified(61);k__Fungi_unclassified(61);k__Fungi_unclassified(61);k__Fungi_unclassified(61);k__Fungi_unclassified(61);k__Fungi_unclassified(61); | - | - | - | - | - | - | - | Unassigned | - |
| Otu03552 | 10 | k__Fungi(100);k__Fungi_unclassified(61);k__Fungi_unclassified(61);k__Fungi_unclassified(61);k__Fungi_unclassified(61);k__Fungi_unclassified(61);k__Fungi_unclassified(61); | - | - | - | - | - | - | - | Unassigned | - |
| Otu03553 | 10 | k__Fungi(100);p__Basidiomycota(100);p__Basidiomycota_unclassified(90);p__Basidiomycota_unclassified(90);p__Basidiomycota_unclassified(90);p__Basidiomycota_unclassified(90);p__Basidiomycota_unclassified(90); | - | - | - | - | - | - | - | Unassigned | - |
| Otu03554 | 10 | k__Fungi(100);k__Fungi_unclassified(61);k__Fungi_unclassified(61);k__Fungi_unclassified(61);k__Fungi_unclassified(61);k__Fungi_unclassified(61);k__Fungi_unclassified(61); | - | - | - | - | - | - | - | Unassigned | - |
| Otu03557 | 10 | k__Fungi(100);k__Fungi_unclassified(61);k__Fungi_unclassified(61);k__Fungi_unclassified(61);k__Fungi_unclassified(61);k__Fungi_unclassified(61);k__Fungi_unclassified(61); | - | - | - | - | - | - | - | Unassigned | - |
| Otu03558 | 10 | k__Fungi(100);p__Ascomycota(100);c__Dothideomycetes(100);o__Myriangiales(100);f__unclassified_Myriangiales(100);g__unclassified_Myriangiales(100);s__Myriangiales_sp(100); | - | - | - | - | - | - | - | Unassigned | - |
| Otu03560 | 10 | k__Fungi(100);k__Fungi_unclassified(100);k__Fungi_unclassified(100);k__Fungi_unclassified(100);k__Fungi_unclassified(100);k__Fungi_unclassified(100);k__Fungi_unclassified(100); | - | - | - | - | - | - | - | Unassigned | - |
| Otu03562 | 10 | k__Fungi(100);p__Ascomycota(100);c__Dothideomycetes(100);o__Pleosporales(100);o__Pleosporales_unclassified(100);o__Pleosporales_unclassified(100);o__Pleosporales_unclassified(100); | - | - | - | - | - | - | - | Unassigned | - |
| Otu03563 | 10 | k__Fungi(100);p__Basidiomycota(100);p__Basidiomycota_unclassified(100);p__Basidiomycota_unclassified(100);p__Basidiomycota_unclassified(100);p__Basidiomycota_unclassified(100);p__Basidiomycota_unclassified(100); | - | - | - | - | - | - | - | Unassigned | - |
| Otu03565 | 10 | k__Fungi(100);k__Fungi_unclassified(80);k__Fungi_unclassified(80);k__Fungi_unclassified(80);k__Fungi_unclassified(80);k__Fungi_unclassified(80);k__Fungi_unclassified(80); | - | - | - | - | - | - | - | Unassigned | - |
| Otu03566 | 10 | k__Plantae(100);p__unclassified_Plantae(100);c__unclassified_Plantae(100);o__unclassified_Plantae(100);f__unclassified_Plantae(100);g__unclassified_Plantae(100);s__Plantae_sp(100); | - | - | - | - | - | - | - | Unassigned | - |
| Otu03567 | 10 | k__Fungi(100);p__Ascomycota(61);p__Ascomycota_unclassified(61);p__Ascomycota_unclassified(61);p__Ascomycota_unclassified(61);p__Ascomycota_unclassified(61);p__Ascomycota_unclassified(61); | - | - | - | - | - | - | - | Unassigned | - |
| Otu03570 | 10 | k__Fungi(100);k__Fungi_unclassified(100);k__Fungi_unclassified(100);k__Fungi_unclassified(100);k__Fungi_unclassified(100);k__Fungi_unclassified(100);k__Fungi_unclassified(100); | - | - | - | - | - | - | - | Unassigned | - |
| Otu03571 | 10 | k__Fungi(100);k__Fungi_unclassified(61);k__Fungi_unclassified(61);k__Fungi_unclassified(61);k__Fungi_unclassified(61);k__Fungi_unclassified(61);k__Fungi_unclassified(61); | - | - | - | - | - | - | - | Unassigned | - |
| Otu03572 | 10 | k__Fungi(100);k__Fungi_unclassified(100);k__Fungi_unclassified(100);k__Fungi_unclassified(100);k__Fungi_unclassified(100);k__Fungi_unclassified(100);k__Fungi_unclassified(100); | - | - | - | - | - | - | - | Unassigned | - |
| Otu03575 | 10 | k__Fungi(100);p__Ascomycota(80);p__Ascomycota_unclassified(80);p__Ascomycota_unclassified(80);p__Ascomycota_unclassified(80);p__Ascomycota_unclassified(80);p__Ascomycota_unclassified(80); | - | - | - | - | - | - | - | Unassigned | - |
| Otu03579 | 10 | k__Fungi(100);p__Basidiomycota(100);c__Agaricomycetes(100);o__Polyporales(100);o__Polyporales_unclassified(100);o__Polyporales_unclassified(100);o__Polyporales_unclassified(100); | - | - | - | - | - | - | - | Unassigned | - |
| Otu03581 | 10 | k__Fungi(100);p__Ascomycota(100);p__Ascomycota_unclassified(100);p__Ascomycota_unclassified(100);p__Ascomycota_unclassified(100);p__Ascomycota_unclassified(100);p__Ascomycota_unclassified(100); | - | - | - | - | - | - | - | Unassigned | - |
| Otu03582 | 9 | k__Fungi(100);p__Basidiomycota(100);c__Agaricomycetes(100);o__Agaricales(100);f__Psathyrellaceae(100);g__Coprinopsis(100);s__Coprinopsis_stercorea(100); | Coprinopsis | 13 | Saprotroph | \|Undefined Saprotroph\| | Agaricoid | NULL | Probable | As soil_saprotroph (Põlme et al. 2020); Decay soil (Põlme et al. 2020) | Tedersoo L, et al. 2014. Science 346:e1256688 ((https://doi.org/10.1126/science.1256688)); Põlme S, et al. 2020. Fungal Diversity 105:1-16 ((https://doi.org/10.1007/s13225-020-00466-2)) |
| Otu03584 | 9 | k__Fungi(100);p__Ascomycota(100);c__Saccharomycetes(100);o__Saccharomycetales(100);f__Metschnikowiaceae(100);g__Metschnikowia(100);s__Metschnikowia_pulcherrima(56); | Metschnikowia | 13 | Saprotroph | \|Undefined Saprotroph\| | Yeast | NULL | Probable | As nectar-tap_saprotroph (Põlme et al. 2020); Decay sugar-rich substrates (Põlme et al. 2020) | Põlme S, et al. 2020. Fungal Diversity 105:1-16 ((https://doi.org/10.1007/s13225-020-00466-2)) |
| Otu03585 | 9 | k__Fungi(100);p__Basidiomycota(100);c__Agaricomycetes(100);o__Agaricales(100);f__Agaricaceae(100);g__Agaricus(100);s__Agaricus_sp_0747(67); | Agaricus | 13 | Saprotroph | \|Undefined Saprotroph\| | Agaricoid-Gasteroid-Secotioid | NULL | Probable | As soil_saprotroph (Põlme et al. 2020); Decay soil (Põlme et al. 2020) | Tedersoo L, et al. 2014. Science 346:e1256688 ((https://doi.org/10.1126/science.1256688)); Põlme S, et al. 2020. Fungal Diversity 105:1-16 ((https://doi.org/10.1007/s13225-020-00466-2)) |
| Otu03586 | 9 | k__Fungi(100);p__Basidiomycota(100);c__Agaricomycetes(100);o__Agaricales(100);f__Entolomataceae(100);g__Entoloma(100);g__Entoloma_unclassified(100); | Entoloma | 13 | Pathotroph-Saprotroph-Symbiotroph | Ectomycorrhizal-Fungal Parasite-\|Undefined Saprotroph\|-Undefined Symbiotroph | Agaricoid | NULL | Possible | As soil_saprotroph (Põlme et al. 2020); Decay roots, soil (Põlme et al. 2020) | Rinaldi AC, et al. 2008. Fungal Diversity 33:1-45 ((http://www.fungaldiversity.org/fdp/sfdp/33-1.pdf)) (as ECM/not ECM); Tedersoo L, et al. 2010. Mycorrhiza 20:217-263 ((https://doi.org/10.1007/s00572-009-0274-x; Põlme S, et al. 2020. Fungal Diversity 105:1-16 ((https://doi.org/10.1007/s13225-020-00466-2)); sensu stricto)); http://mycorrhizas.info/ecmf.html ((http://mycorrhizas.info/ecmf.html)) ((as ECM/not ECM)) |
| Otu03587 | 9 | k__Fungi(100);p__Ascomycota(100);c__Saccharomycetes(100);o__Saccharomycetales(100);f__Saccharomycetales_family_Incertae_sedis(100);g__Candida(67);g__Candida_unclassified(56); | Candida | 13 | Pathotroph-Saprotroph-Symbiotroph | Animal Pathogen-Endophyte-Endosymbiont-Epiphyte-\|Undefined Saprotroph\| | Dimorphic Yeast-Yeast | NULL | Probable | As nectar-tap_saprotroph (Põlme et al. 2020); Decay sugar-rich substrates (Põlme et al. 2020); This asexual genus is polyphyletic and contain species that function very differently depend on what they are. Caution should be taken when interpreting guilds for Candida species. | Manolakaki D, et al. 2010. Virulence. 1: 367-375 ((https://doi.org/10.4161/viru.1.5.12796)); Mekbib SB, et al. 2011. Tropical Plant Pathology 36:233-240 ((https://doi.org/10.1590/S1982-56762011000400004)); Põlme S, et al. 2020. Fungal Diversity 105:1-16 ((https://doi.org/10.1007/s13225-020-00466-2)) |
| Otu03588 | 9 | k__Fungi(100);p__Basidiomycota(100);c__Agaricomycetes(100);o__Polyporales(100);f__Phanerochaetaceae(100);g__Antrodiella(100);s__Antrodiella_beschidica(100); | Antrodiella | 13 | Saprotroph | \|Wood Saprotroph\| | Corticioid-Polyporoid | White Rot | Highly Probable | As white_rot (Põlme et al. 2020); Decay wood (Põlme et al. 2020) | Gilbertson RL, Ryvarden L. 1987-1987. North American Polypores. Fungiflora, Oslo (ISBN: 978-0945345060); Põlme S, et al. 2020. Fungal Diversity 105:1-16 ((https://doi.org/10.1007/s13225-020-00466-2)) |
| Otu03589 | 9 | k__Fungi(100);p__Ascomycota(100);c__Eurotiomycetes(100);o__Chaetothyriales(100);f__Herpotrichiellaceae(100);g__Cladophialophora(100);g__Cladophialophora_unclassified(67); | Cladophialophora | 13 | Pathotroph-Saprotroph-Symbiotroph | Animal Pathogen-Endophyte-Fungal Parasite-\|Undefined Saprotroph\| | Dimorphic-Microfungus | NULL | Probable | As mycoparasite (Põlme et al. 2020); As root_endophyte_dark_septate (Põlme et al. 2020); As soil_saprotroph (Põlme et al. 2020); Decay soil (Põlme et al. 2020) | James TY, et al. 2006. Nature 443:818-822 ((https://doi.org/10.1038/nature05110)); Põlme S, et al. 2020. Fungal Diversity 105:1-16 ((https://doi.org/10.1007/s13225-020-00466-2)) |
| Otu03590 | 9 | k__Fungi(100);p__Zygomycota(100);c__Zygomycota_class_Incertae_sedis(100);o__Mucorales(100);f__Lentamycetaceae(100);g__Lentamyces(100);s__Lentamyces_parricida(100); | Lentamyces | 13 | Pathotroph-Symbiotroph | Animal Symbiotroph-\|Fungal Parasite\| | Microfungus | NULL | Probable | Animal gut fungus; As mycoparasite (Põlme et al. 2020); Decay fungal material (Põlme et al. 2020) | Tedersoo L, et al. 2014. Science 346:e1256688 ((https://doi.org/10.1126/science.1256688)); Põlme S, et al. 2020. Fungal Diversity 105:1-16 ((https://doi.org/10.1007/s13225-020-00466-2)) |
| Otu03594 | 9 | k__Fungi(100);p__Basidiomycota(100);c__Agaricomycetes(100);o__Agaricales(100);f__Inocybaceae(100);g__Inocybe(100);s__Inocybe_lilacina(100); | Inocybe | 13 | Symbiotroph | \|Ectomycorrhizal\| | Agaricoid | NULL | Highly Probable | Sensu stricto (Matheny 2009). | Rinaldi AC, et al. 2008. Fungal Diversity 33:1-45 ((http://www.fungaldiversity.org/fdp/sfdp/33-1.pdf)); Matheny PB. 2009. McIlvainea 18:11-21 ((https://namyco.org/docs/3_Matheny.pdf)); Tedersoo L, et al. 2010. Mycorrhiza 20:217-263 ((https://doi.org/10.1007/s00572-009-0274-x)); Põlme S, et al. 2020. Fungal Diversity 105:1-16 ((https://doi.org/10.1007/s13225-020-00466-2)); http://mycorrhizas.info/ecmf.html ((http://mycorrhizas.info/ecmf.html)) |
| Otu03597 | 9 | k__Fungi(100);p__Ascomycota(100);c__Saccharomycetes(100);o__Saccharomycetales(100);f__Metschnikowiaceae(100);g__Metschnikowia(100);s__Metschnikowia_pulcherrima(67); | Metschnikowia | 13 | Saprotroph | \|Undefined Saprotroph\| | Yeast | NULL | Probable | As nectar-tap_saprotroph (Põlme et al. 2020); Decay sugar-rich substrates (Põlme et al. 2020) | Põlme S, et al. 2020. Fungal Diversity 105:1-16 ((https://doi.org/10.1007/s13225-020-00466-2)) |
| Otu03598 | 9 | k__Fungi(100);p__Ascomycota(100);c__Eurotiomycetes(100);o__Onygenales(100);f__Onygenaceae(100);f__Onygenaceae_unclassified(67);f__Onygenaceae_unclassified(67); | Onygenaceae | 9 | Saprotroph | Dung Saprotroph-Soil Saprotroph | Microfungus | NULL | Probable | Generally saprobic and keratinophilic, in soil or coprophilous, occasionally found on hair or horn (Cannon & Kirk 2007) | Cannon PF, Kirk PM. 2007. Fungal Families of the World. CAB International, Cambridge (ISBN: 978-0851998275) |
| Otu03599 | 9 | k__Fungi(100);p__Ascomycota(100);c__Saccharomycetes(100);o__Saccharomycetales(100);f__Metschnikowiaceae(100);g__Metschnikowia(100);s__Metschnikowia_pulcherrima(78); | Metschnikowia | 13 | Saprotroph | \|Undefined Saprotroph\| | Yeast | NULL | Probable | As nectar-tap_saprotroph (Põlme et al. 2020); Decay sugar-rich substrates (Põlme et al. 2020) | Põlme S, et al. 2020. Fungal Diversity 105:1-16 ((https://doi.org/10.1007/s13225-020-00466-2)) |
| Otu03601 | 9 | k__Fungi(100);p__Ascomycota(100);c__Saccharomycetes(89);o__Saccharomycetales(89);f__Saccharomycetales_family_Incertae_sedis(89);g__Candida(89);g__Candida_unclassified(78); | Candida | 13 | Pathotroph-Saprotroph-Symbiotroph | Animal Pathogen-Endophyte-Endosymbiont-Epiphyte-\|Undefined Saprotroph\| | Dimorphic Yeast-Yeast | NULL | Probable | As nectar-tap_saprotroph (Põlme et al. 2020); Decay sugar-rich substrates (Põlme et al. 2020); This asexual genus is polyphyletic and contain species that function very differently depend on what they are. Caution should be taken when interpreting guilds for Candida species. | Manolakaki D, et al. 2010. Virulence. 1: 367-375 ((https://doi.org/10.4161/viru.1.5.12796)); Mekbib SB, et al. 2011. Tropical Plant Pathology 36:233-240 ((https://doi.org/10.1590/S1982-56762011000400004)); Põlme S, et al. 2020. Fungal Diversity 105:1-16 ((https://doi.org/10.1007/s13225-020-00466-2)) |
| Otu03602 | 9 | k__Fungi(100);p__Ascomycota(100);c__Saccharomycetes(100);o__Saccharomycetales(100);f__Metschnikowiaceae(100);g__Metschnikowia(100);s__Metschnikowia_pulcherrima(100); | Metschnikowia | 13 | Saprotroph | \|Undefined Saprotroph\| | Yeast | NULL | Probable | As nectar-tap_saprotroph (Põlme et al. 2020); Decay sugar-rich substrates (Põlme et al. 2020) | Põlme S, et al. 2020. Fungal Diversity 105:1-16 ((https://doi.org/10.1007/s13225-020-00466-2)) |
| Otu03603 | 9 | k__Fungi(100);p__Ascomycota(100);c__Saccharomycetes(100);o__Saccharomycetales(100);f__Metschnikowiaceae(100);g__Metschnikowia(100);s__Metschnikowia_pulcherrima(100); | Metschnikowia | 13 | Saprotroph | \|Undefined Saprotroph\| | Yeast | NULL | Probable | As nectar-tap_saprotroph (Põlme et al. 2020); Decay sugar-rich substrates (Põlme et al. 2020) | Põlme S, et al. 2020. Fungal Diversity 105:1-16 ((https://doi.org/10.1007/s13225-020-00466-2)) |
| Otu03604 | 9 | k__Fungi(100);p__Ascomycota(100);c__Sordariomycetes(100);o__Microascales(100);f__Halosphaeriaceae(100);g__unclassified_Halosphaeriaceae(100);s__Halosphaeriaceae_sp(100); | Halosphaeriaceae | 9 | Saprotroph | Undefined Saprotroph-Wood Saprotroph | Microfungus | NULL | Probable | Found in aquatic habitats, mostly in marine environments; saprobic on submerged or intertidal wood, occasionally associated with sand grains and similar substrata (Cannon & Kirk 2007) | Cannon PF, Kirk PM. 2007. Fungal Families of the World. CAB International, Cambridge (ISBN: 978-0851998275) |
| Otu03608 | 9 | k__Fungi(100);p__Basidiomycota(100);c__Tremellomycetes(100);o__Tremellales(100);f__Tremellales_family_Incertae_sedis(100);f__Tremellales_family_Incertae_sedis_unclassified(100);f__Tremellales_family_Incertae_sedis_unclassified(100); | Tremellales | 7 | Pathotroph-Saprotroph-Symbiotroph | Fungal Parasite-Undefined Saprotroph | Tremelloid-Yeast | NULL | Possible | The order contains both teleomorphic and anamorphic species, most of the latter being yeasts | Sterkenburg E, et al. 2015. New Phytologist 207:1145-1158 ((https://doi.org/10.1111/nph.13426)) |
| Otu03609 | 9 | k__Fungi(100);p__Ascomycota(100);c__Saccharomycetes(100);o__Saccharomycetales(100);f__Metschnikowiaceae(100);g__Metschnikowia(100);s__Metschnikowia_pulcherrima(89); | Metschnikowia | 13 | Saprotroph | \|Undefined Saprotroph\| | Yeast | NULL | Probable | As nectar-tap_saprotroph (Põlme et al. 2020); Decay sugar-rich substrates (Põlme et al. 2020) | Põlme S, et al. 2020. Fungal Diversity 105:1-16 ((https://doi.org/10.1007/s13225-020-00466-2)) |
| Otu03611 | 9 | k__Fungi(100);p__Ascomycota(100);c__Saccharomycetes(100);o__Saccharomycetales(100);f__Metschnikowiaceae(100);g__Metschnikowia(100);s__Metschnikowia_pulcherrima(89); | Metschnikowia | 13 | Saprotroph | \|Undefined Saprotroph\| | Yeast | NULL | Probable | As nectar-tap_saprotroph (Põlme et al. 2020); Decay sugar-rich substrates (Põlme et al. 2020) | Põlme S, et al. 2020. Fungal Diversity 105:1-16 ((https://doi.org/10.1007/s13225-020-00466-2)) |
| Otu03612 | 9 | k__Fungi(100);p__Ascomycota(100);c__Saccharomycetes(100);o__Saccharomycetales(100);f__Saccharomycetaceae(100);g__Kazachstania(100);s__Kazachstania_intestinalis(100); | Kazachstania | 13 | Saprotroph | \|Undefined Saprotroph\| | Yeast | NULL | Probable | As nectar-tap_saprotroph (Põlme et al. 2020); Decay sugar-rich substrates (Põlme et al. 2020) | Põlme S, et al. 2020. Fungal Diversity 105:1-16 ((https://doi.org/10.1007/s13225-020-00466-2)) |
| Otu03613 | 9 | k__Fungi(100);p__Basidiomycota(100);c__Agaricomycetes(100);o__Agaricales(100);f__Strophariaceae(100);f__Strophariaceae_unclassified(100);f__Strophariaceae_unclassified(100); | Strophariaceae | 9 | Pathotroph-Saprotroph | Dung Saprotroph-Plant Parasite-Soil Saprotroph-Undefined Saprotroph-Wood Saprotroph | Agaricoid | NULL | Probable | Very variable ecology; Pholiota species are lignicolous and can act as weak parasites while most other taxa are either sparobes in grassy habitats or are found in dung. Species are not ectomycorrhizal (Cannon & Kirk 2007) | Cannon PF, Kirk PM. 2007. Fungal Families of the World. CAB International, Cambridge (ISBN: 978-0851998275) |
| Otu03614 | 9 | k__Fungi(100);p__Ascomycota(100);c__Saccharomycetes(100);o__Saccharomycetales(100);f__Saccharomycetales_family_Incertae_sedis(100);g__Candida(100);g__Candida_unclassified(56); | Candida | 13 | Pathotroph-Saprotroph-Symbiotroph | Animal Pathogen-Endophyte-Endosymbiont-Epiphyte-\|Undefined Saprotroph\| | Dimorphic Yeast-Yeast | NULL | Probable | As nectar-tap_saprotroph (Põlme et al. 2020); Decay sugar-rich substrates (Põlme et al. 2020); This asexual genus is polyphyletic and contain species that function very differently depend on what they are. Caution should be taken when interpreting guilds for Candida species. | Manolakaki D, et al. 2010. Virulence. 1: 367-375 ((https://doi.org/10.4161/viru.1.5.12796)); Mekbib SB, et al. 2011. Tropical Plant Pathology 36:233-240 ((https://doi.org/10.1590/S1982-56762011000400004)); Põlme S, et al. 2020. Fungal Diversity 105:1-16 ((https://doi.org/10.1007/s13225-020-00466-2)) |
| Otu03617 | 9 | k__Fungi(100);p__Zygomycota(100);c__Zygomycota_class_Incertae_sedis(100);o__Mortierellales(100);f__Mortierellaceae(100);g__Mortierella(100);s__Mortierella_sarnyensis(100); | Mortierella | 13 | Saprotroph-Symbiotroph | Endophyte-Plant Saprotroph-\|Undefined Saprotroph\| | Microfungus | NULL | Probable | As chitinolytic (Põlme et al. 2020); As soil_saprotroph (Põlme et al. 2020); Decay roots, soil (Põlme et al. 2020); Saprobic in soil and thus frequently isolated from soil and material contaminated with soil such as hay or rotten silage and spoil tips (Cannon & Kirk 2007) | Cannon PF, Kirk PM. 2007. Fungal Families of the World. CAB International, Cambridge (ISBN: 978-0851998275); Tedersoo L, et al. 2014. Science 346:e1256688 ((https://doi.org/10.1126/science.1256688)); Purahong W, et al. 2016. Molecular Ecology 25:4059-4074 ((https://doi.org/10.1111/mec.13739)); Põlme S, et al. 2020. Fungal Diversity 105:1-16 ((https://doi.org/10.1007/s13225-020-00466-2)) |
| Otu03623 | 9 | k__Fungi(100);p__Ascomycota(100);c__Saccharomycetes(100);o__Saccharomycetales(100);f__Metschnikowiaceae(100);g__Metschnikowia(100);s__Metschnikowia_pulcherrima(56); | Metschnikowia | 13 | Saprotroph | \|Undefined Saprotroph\| | Yeast | NULL | Probable | As nectar-tap_saprotroph (Põlme et al. 2020); Decay sugar-rich substrates (Põlme et al. 2020) | Põlme S, et al. 2020. Fungal Diversity 105:1-16 ((https://doi.org/10.1007/s13225-020-00466-2)) |
| Otu03627 | 9 | k__Fungi(100);p__Ascomycota(100);c__Eurotiomycetes(100);o__Eurotiales(100);f__Trichocomaceae(100);g__Penicillium(100);s__Penicillium_chrysogenum(100); | Penicillium | 13 | Pathotroph-Saprotroph-Symbiotroph | Animal Parasite-Dung Saprotroph-Endophyte-Fungal Parasite-Plant Pathogen-\|Undefined Saprotroph\|-Wood Saprotroph | Microfungus | Soft Rot | Highly Probable | As foliar_endophyte (Põlme et al. 2020); As mycoparasite (Põlme et al. 2020); Decay soil (Põlme et al. 2020) | Duncan CG, Eslyn WE. 1966. Mycologia 58:642-664 ((https://doi.org/10.2307/3757045)); Seehann G, et al. 1975. List of Fungi in Soft-Rot Tissue Tests. Institute fur Holzbiologie und Holzschutz, Bundesforschungsanstalt fur Forst - und Holzwirtshaft, Hamburg - Lohbrugge, Germany ((https://www.irg-wp.com/irgdocs/details.php?d6493033-c9ea-40f3-8e3e-5697f32d640d)); Costa IPMW, et al. 2012. Mycotaxon 119:e26 ((http://mycotaxon.com/resources/checklists/costa_v119_checklist.pdf)); Bills GF, et al. 2013. Current Opinion in Microbiology 16: 549-565 ((https://doi.org/10.1016/j.mib.2013.08.001)); Tedersoo L, et al. 2014. Science 346:e1256688 ((https://doi.org/10.1126/science.1256688)); Põlme S, et al. 2020. Fungal Diversity 105:1-16 ((https://doi.org/10.1007/s13225-020-00466-2)) |
| Otu03628 | 9 | k__Fungi(100);p__Ascomycota(100);c__Saccharomycetes(100);o__Saccharomycetales(100);f__Saccharomycetales_family_Incertae_sedis(89);g__Candida(78);g__Candida_unclassified(67); | Candida | 13 | Pathotroph-Saprotroph-Symbiotroph | Animal Pathogen-Endophyte-Endosymbiont-Epiphyte-\|Undefined Saprotroph\| | Dimorphic Yeast-Yeast | NULL | Probable | As nectar-tap_saprotroph (Põlme et al. 2020); Decay sugar-rich substrates (Põlme et al. 2020); This asexual genus is polyphyletic and contain species that function very differently depend on what they are. Caution should be taken when interpreting guilds for Candida species. | Manolakaki D, et al. 2010. Virulence. 1: 367-375 ((https://doi.org/10.4161/viru.1.5.12796)); Mekbib SB, et al. 2011. Tropical Plant Pathology 36:233-240 ((https://doi.org/10.1590/S1982-56762011000400004)); Põlme S, et al. 2020. Fungal Diversity 105:1-16 ((https://doi.org/10.1007/s13225-020-00466-2)) |
| Otu03629 | 9 | k__Fungi(100);p__Ascomycota(100);c__Saccharomycetes(100);o__Saccharomycetales(100);f__Metschnikowiaceae(100);g__Metschnikowia(100);s__Metschnikowia_pulcherrima(100); | Metschnikowia | 13 | Saprotroph | \|Undefined Saprotroph\| | Yeast | NULL | Probable | As nectar-tap_saprotroph (Põlme et al. 2020); Decay sugar-rich substrates (Põlme et al. 2020) | Põlme S, et al. 2020. Fungal Diversity 105:1-16 ((https://doi.org/10.1007/s13225-020-00466-2)) |
| Otu03630 | 9 | k__Fungi(100);p__Zygomycota(100);c__Zygomycota_class_Incertae_sedis(100);o__Mucorales(100);f__Mucoraceae(100);g__Mucor(100);s__Mucor_sp_JJP_2009a(100); | Mucor | 13 | Saprotroph | \|Undefined Saprotroph\| | Microfungus | NULL | Probable | As soil_saprotroph (Põlme et al. 2020); Decay soil (Põlme et al. 2020) | Tedersoo L, et al. 2014. Science 346:e1256688 ((https://doi.org/10.1126/science.1256688)); Põlme S, et al. 2020. Fungal Diversity 105:1-16 ((https://doi.org/10.1007/s13225-020-00466-2)) |
| Otu03632 | 9 | k__Fungi(100);p__Ascomycota(100);c__Saccharomycetes(100);o__Saccharomycetales(100);f__Metschnikowiaceae(100);g__Metschnikowia(100);s__Metschnikowia_pulcherrima(56); | Metschnikowia | 13 | Saprotroph | \|Undefined Saprotroph\| | Yeast | NULL | Probable | As nectar-tap_saprotroph (Põlme et al. 2020); Decay sugar-rich substrates (Põlme et al. 2020) | Põlme S, et al. 2020. Fungal Diversity 105:1-16 ((https://doi.org/10.1007/s13225-020-00466-2)) |

| | | | | | | | | | | | |
|---|---|---|---|---|---|---|---|---|---|---|---|
| Otu03633 | 9 | k__Fungi(100);p__Ascomycota(100);c__Eurotiomycetes(100);o__Eurotiales(100);f__Trichocomaceae(100);g__Penicillium(100);s__Penicillium_coprobium(100); | Penicillium | 13 | Pathotroph-Saprotroph-Symbiotroph | Animal Parasite-Dung Saprotroph-Endophyte-Fungal Parasite-Plant Pathogen-\|Undefined Saprotroph\|-Wood Saprotroph | Microfungus | Soft Rot | Highly Probable | As foliar_endophyte (Põlme et al. 2020); As mycoparasite (Põlme et al. 2020); Decay soil (Põlme et al. 2020) | Duncan CG, Eslyn WE. 1966. Mycologia 58:642-664 ((https://doi.org/10.2307/3757045)); Seehann G, et al. 1975. List of Fungi in Soft-Rot Tissue Tests. Institute fur Holzbiologie und Holzschutz, Bundesforschungsanstalt fur Forst - und Holzwirtshaft, Hamburg - Lohbrugge, Germany ((https://www.irg-wp.com/irgdocs/details.php?d6493033-c9ea-40f3-8e3e-5697f32d640d)); Costa IPMW, et al. 2012. Mycotaxon 119:e26 ((http://mycotaxon.com/resources/checklists/costa_v119_checklist.pdf)); Bills GF, et al. 2013. Current Opinion in Microbiology 16:549-565 ((https://doi.org/10.1016/j.mib.2013.08.001)); Tedersoo L, et al. 2014. Science 346:e1256688 ((https://doi.org/10.1126/science.1256688)); Põlme S, et al. 2020. Fungal Diversity 105:1-16 ((https://doi.org/10.1007/s13225-020-00466-2)) |
| Otu03635 | 9 | k__Fungi(100);p__Ascomycota(100);c__Saccharomycetes(100);o__Saccharomycetales(100);f__Saccharomycopsidaceae(100);g__Saccharomycopsis(100);s__Saccharomycopsis_javanensis(100); | Saccharomycopsis | 13 | Pathotroph-Saprotroph | Animal Parasite-\|Undefined Saprotroph\| | Yeast | NULL | Probable | As nectar-tap_saprotroph (Põlme et al. 2020); Decay animal material, sugar-rich substrates (Põlme et al. 2020) | Põlme S, et al. 2020. Fungal Diversity 105:1-16 ((https://doi.org/10.1007/s13225-020-00466-2)) |
| Otu03636 | 9 | k__Fungi(100);p__Basidiomycota(100);c__Agaricomycetes(100);o__Agaricales(100);f__Lyophyllaceae(100);f__Lyophyllaceae_unclassified(100);f__Lyophyllaceae_unclassified(100); | Lyophyllaceae | 9 | Pathotroph-Saprotroph-Symbiotroph | Ectomycorrhizal-Fungal Pathogen-Undefined Saprotroph | Agaricoid | NULL | Possible | NULL | Kirk PM et al. 2008. Dictionary of the Fungi. Tenth Edition. CAB International, Wallingford (ISBN: 978-0851998268) |
| Otu03638 | 9 | k__Fungi(100);p__Basidiomycota(100);c__Agaricomycetes(100);o__Polyporales(100);f__Polyporaceae(100);g__Tyromyces(100);s__Tyromyces_chioneus(100); | Tyromyces | 13 | Saprotroph | \|Wood Saprotroph\| | Polyporoid | White Rot | Highly Probable | As white_rot (Põlme et al. 2020); Decay wood (Põlme et al. 2020) | Gilbertson RL, Ryvarden L. 1987-1987. North American Polypores. Fungiflora, Oslo (ISBN: 978-0945345060); Põlme S, et al. 2020. Fungal Diversity 105:1-16 ((https://doi.org/10.1007/s13225-020-00466-2)) |
| Otu03639 | 9 | k__Fungi(100);p__Ascomycota(100);c__Sordariomycetes(100);o__Xylariales(100);f__Xylariaceae(100);g__Hypoxylon(100);s__Hypoxylon_howeanum(100); | Hypoxylon | 13 | Saprotroph-Saprotroph-Symbiotroph | Endophyte-Undefined Saprotroph-\|Wood Saprotroph\| | Corticioid-Microfungus-Xylarioid | NULL | Probable | As foliar_endophyte (Põlme et al. 2020); Decay wood, leaf, fruit, seed (Põlme et al. 2020); Host - angiosperms (Põlme et al. 2020) | Tedersoo L, et al. 2014. Science 346:e1256688 ((https://doi.org/10.1126/science.1256688)); Põlme S, et al. 2020. Fungal Diversity 105:1-16 ((https://doi.org/10.1007/s13225-020-00466-2)) |
| Otu03640 | 9 | k__Fungi(100);p__Ascomycota(100);c__Ascomycota_class_Incertae_sedis(100);o__Ascomycota_order_Incertae_sedis(100);f__Ascomycota_family_Incertae_sedis(100);g__Slimacomyces(100);s__Slimacomyces_isiola(100); | Slimacomyces | 13 | Saprotroph | \|Plant Saprotroph\| | Microfungus | NULL | Probable | As litter_saprotroph (Põlme et al. 2020); Decay leaf, fruit, seed (Põlme et al. 2020) | Põlme S, et al. 2020. Fungal Diversity 105:1-16 ((https://doi.org/10.1007/s13225-020-00466-2)) |
| Otu03641 | 9 | k__Fungi(100);p__Basidiomycota(100);c__Exobasidiomycetes(100);o__Exobasidiomycetes_order_Incertae_sedis(100);f__Exobasidiomycetes_family_Incertae_sedis(100);g__Meira(100);s__Meira_sp_ANTCW08_165(100); | Meira | 13 | Pathotroph | \|Animal Parasite\|-Animal Pathogen-Plant Pathogen | Dimorphic | NULL | Probable | An anamorphic fungus isolated from cadavers of the citrus rust mite (P. oleivora) infesting grapefruit (C. paradisi) leaves (Boekhout et al. 2003); Plant pathogen (Põlme et al. 2020); Similar to the yeast-like fungi classified in the Ustilaginales, such as Pseudozyma species (Boekhout et al. 2003) | Boekhout T, et al. 2003. International Journal of Systematic and Evolutionary Microbiology 53:1655-1664 ((https://doi.org/10.1099/ijs.0.02434-0)); Põlme S, et al. 2020. Fungal Diversity 105:1-16 ((https://doi.org/10.1007/s13225-020-00466-2)) |
| Otu03642 | 9 | k__Fungi(100);p__Basidiomycota(100);c__Agaricomycetes(100);o__Polyporales(100);f__Meruliaceae(100);g__Phlebia(100);s__Phlebia_sp(100); | Phlebia | 13 | Saprotroph | \|Wood Saprotroph\| | Corticioid | White Rot | Highly Probable | As white_rot (Põlme et al. 2020); Decay wood (Põlme et al. 2020) | Kirk PM et al. 2008. Dictionary of the Fungi. Tenth Edition. CAB International, Wallingford (ISBN: 978-0851998268); Põlme S, et al. 2020. Fungal Diversity 105:1-16 ((https://doi.org/10.1007/s13225-020-00466-2)) |
| Otu03644 | 9 | k__Fungi(100);p__Basidiomycota(100);c__Microbotryomycetes(100);o__Sporidiobolales(100);f__Sporidiobolales_family_Incertae_sedis(100);g__Rhodotorula(56);s__Rhodotorula_graminis(56); | Rhodotorula | 13 | Pathotroph-Saprotroph-Symbiotroph | Animal Pathogen-Animal Symbiotroph-Endophyte-Plant Pathogen-\|Undefined Saprotroph\| | Yeast | NULL | Probable | As foliar_endophyte (Põlme et al. 2020); Opportunistic human pathogen | Põlme S, et al. 2020. Fungal Diversity 105:1-16 ((https://doi.org/10.1007/s13225-020-00466-2)); https://www.cdc.gov ((https://www.cdc.gov)) |
| Otu03645 | 9 | k__Fungi(100);p__Ascomycota(100);c__Leotiomycetes(100);o__Geoglossales(100);f__Geoglossaceae(100);g__Geoglossum(100);s__Geoglossum_umbratile(100); | Geoglossum | 13 | Saprotroph-Symbiotroph | \|Undefined Saprotroph\|-Undefined Symbiotroph | Clavarioid-Spathuloid | NULL | Probable | As soil_saprotroph (Põlme et al. 2020); Decay roots, soil (Põlme et al. 2020) | Tedersoo L, et al. 2014. Science 346:e1256688 ((https://doi.org/10.1126/science.1256688)); Põlme S, et al. 2020. Fungal Diversity 105:1-16 ((https://doi.org/10.1007/s13225-020-00466-2)) |
| Otu03647 | 9 | k__Fungi(100);p__Basidiomycota(100);c__Agaricomycetes(100);o__Agaricales(100);f__Marasmiaceae(100);g__Marasmius(100);g__Marasmius_unclassified(100); | Marasmius | 13 | Pathotroph-Saprotroph-Symbiotroph | Endophyte-Plant Pathogen-\|Plant Saprotroph\|-Undefined Saprotroph | Agaricoid | NULL | Probable | As litter_saprotroph (Põlme et al. 2020); Decay leaf, fruit, seed (Põlme et al. 2020); Often one of the most prominent fungal components of tropical forests, playing a major role in nutrient cycling (Cannon & Kirk 2007) | Cannon PF, Kirk PM. 2007. Fungal Families of the World. CAB International, Cambridge (ISBN: 978-0851998275); Tedersoo L, et al. 2014. Science 346:e1256688 ((https://doi.org/10.1126/science.1256688)); David AS, et al. 2016. Microbial Ecology 71:912-926 ((https://doi.org/10.1007/s00248-015-0712-6)); Põlme S, et al. 2020. Fungal Diversity 105:1-16 ((https://doi.org/10.1007/s13225-020-00466-2)) |
| Otu03650 | 9 | k__Fungi(100);p__Ascomycota(100);c__Leotiomycetes(100);o__Helotiales(100);f__Dermateaceae(100);g__Phaeomollisia(100);s__Phaeomollisia_piceae(100); | Phaeomollisia | 13 | Saprotroph-Symbiotroph | \|Endophyte\|-Undefined Saprotroph-Wood Saprotroph | Microfungus | NULL | Probable | As foliar_endophyte (Põlme et al. 2020); Decay wood, leaf, fruit, seed (Põlme et al. 2020) | Tedersoo L, et al. 2014. Science 346:e1256688 ((https://doi.org/10.1126/science.1256688)); Põlme S, et al. 2020. Fungal Diversity 105:1-16 ((https://doi.org/10.1007/s13225-020-00466-2)) |
| Otu03653 | 9 | k__Fungi(100);p__Basidiomycota(100);c__Microbotryomycetes(100);o__Sporidiobolales(100);f__Sporidiobolales_family_Incertae_sedis(100);g__Rhodotorula(100);g__Rhodotorula_unclassified(100); | Rhodotorula | 13 | Pathotroph-Saprotroph-Symbiotroph | Animal Pathogen-Animal Symbiotroph-Endophyte-Plant Pathogen-\|Undefined Saprotroph\| | Yeast | NULL | Probable | As foliar_endophyte (Põlme et al. 2020); Opportunistic human pathogen | Põlme S, et al. 2020. Fungal Diversity 105:1-16 ((https://doi.org/10.1007/s13225-020-00466-2)); https://www.cdc.gov ((https://www.cdc.gov)) |
| Otu03657 | 9 | k__Fungi(100);p__Basidiomycota(100);c__Agaricomycetes(100);o__Agaricales(100);f__Inocybaceae(100);g__Inocybe(100);s__Inocybe_impexa(78); | Inocybe | 13 | Symbiotroph | \|Ectomycorrhizal\| | Agaricoid | NULL | Highly Probable | Sensu stricto (Matheny 2009). | Rinaldi AC, et al. 2008. Fungal Diversity 33:1-45 ((http://www.fungaldiversity.org/fdp/sfdp/33-1.pdf)); Matheny PB. 2009. McIlvainea 18:11-21 ((https://namyco.org/docs/3_Matheny.pdf)); Tedersoo L, et al. 2010. Mycorrhiza 20:217-263 ((https://doi.org/10.1007/s00572-009-0274-x)); Põlme S, et al. 2020. Fungal Diversity 105:1-16 ((https://doi.org/10.1007/s13225-020-00466-2)); http://mycorrhizas.info/ecmf.html ((http://mycorrhizas.info/ecmf.html)) |
| Otu03660 | 9 | k__Fungi(100);p__Ascomycota(100);c__Saccharomycetes(89);o__Saccharomycetales(89);f__Saccharomycetales_family_Incertae_sedis(78);g__Candida(78);g__Candida_unclassified(78); | Candida | 13 | Pathotroph-Saprotroph-Symbiotroph | Animal Pathogen-Endophyte-Endosymbiont-Epiphyte-\|Undefined Saprotroph\| | Dimorphic Yeast-Yeast | NULL | Probable | As nectar-tap_saprotroph (Põlme et al. 2020); Decay sugar-rich substrates (Põlme et al. 2020); This asexual genus is polyphyletic and contain species that function very differently depend on what they are. Caution should be taken when interpreting guilds for Candida species. | Manolakaki D, et al. 2010. Virulence. 1: 367-375 ((https://doi.org/10.4161/viru.1.5.12796)); Mekbib SB, et al. 2011. Tropical Plant Pathology 36:233-240 ((https://doi.org/10.1590/S1982-56762011000400004)); Põlme S, et al. 2020. Fungal Diversity 105:1-16 ((https://doi.org/10.1007/s13225-020-00466-2)) |
| Otu03663 | 9 | k__Fungi(100);p__Ascomycota(100);c__Saccharomycetes(100);o__Saccharomycetales(100);f__Metschnikowiaceae(100);g__Metschnikowia(100);g__Metschnikowia_unclassified(78); | Metschnikowia | 13 | Saprotroph | \|Undefined Saprotroph\| | Yeast | NULL | Probable | As nectar-tap_saprotroph (Põlme et al. 2020); Decay sugar-rich substrates (Põlme et al. 2020) | Põlme S, et al. 2020. Fungal Diversity 105:1-16 ((https://doi.org/10.1007/s13225-020-00466-2)) |
| Otu03664 | 9 | k__Fungi(100);p__Ascomycota(100);c__Sordariomycetes(100);o__Sordariales(100);f__Lasiosphaeriaceae(100);g__Lasiosphaeris(67);g__Lasiosphaeris_unclassified(67); | Lasiosphaeris | 13 | Saprotroph | \|Undefined Saprotroph\| | Microfungus | NULL | Probable | NULL | Tedersoo L, et al. 2014. Science 346:e1256688 ((https://doi.org/10.1126/science.1256688)); Põlme S, et al. 2020. Fungal Diversity 105:1-16 ((https://doi.org/10.1007/s13225-020-00466-2)) |
| Otu03666 | 9 | k__Fungi(100);p__Ascomycota(100);c__Sordariomycetes(100);o__Microascales(78);f__Halosphaeriaceae(78);g__unclassified_Halosphaeriaceae(78);s__Halosphaeriaceae_sp(78); | Halosphaeriaceae | 9 | Saprotroph | Undefined Saprotroph-Wood Saprotroph | Microfungus | NULL | Probable | Found in aquatic habitats, mostly in marine environments; saprobic on submerged or intertidal wood, occasionally associated with sand grains and similar substrata (Cannon & Kirk 2007) | Cannon PF, Kirk PM. 2007. Fungal Families of the World. CAB International, Cambridge (ISBN: 978-0851998275) |
| Otu03667 | 9 | k__Fungi(100);p__Ascomycota(100);c__Saccharomycetes(100);o__Saccharomycetales(100);f__Metschnikowiaceae(100);g__Metschnikowia(100);s__Metschnikowia_pulcherrima(56); | Metschnikowia | 13 | Saprotroph | \|Undefined Saprotroph\| | Yeast | NULL | Probable | As nectar-tap_saprotroph (Põlme et al. 2020); Decay sugar-rich substrates (Põlme et al. 2020) | Põlme S, et al. 2020. Fungal Diversity 105:1-16 ((https://doi.org/10.1007/s13225-020-00466-2)) |
| Otu03671 | 9 | k__Fungi(100);p__Ascomycota(100);c__Saccharomycetes(100);o__Saccharomycetales(100);f__Metschnikowiaceae(100);g__Metschnikowia(100);s__Metschnikowia_pulcherrima(89); | Metschnikowia | 13 | Saprotroph | \|Undefined Saprotroph\| | Yeast | NULL | Probable | As nectar-tap_saprotroph (Põlme et al. 2020); Decay sugar-rich substrates (Põlme et al. 2020) | Põlme S, et al. 2020. Fungal Diversity 105:1-16 ((https://doi.org/10.1007/s13225-020-00466-2)) |
| Otu03672 | 9 | k__Fungi(100);p__Ascomycota(100);c__Saccharomycetes(100);o__Saccharomycetales(100);f__Metschnikowiaceae(100);g__Metschnikowia(100);s__Metschnikowia_pulcherrima(67); | Metschnikowia | 13 | Saprotroph | \|Undefined Saprotroph\| | Yeast | NULL | Probable | As nectar-tap_saprotroph (Põlme et al. 2020); Decay sugar-rich substrates (Põlme et al. 2020) | Põlme S, et al. 2020. Fungal Diversity 105:1-16 ((https://doi.org/10.1007/s13225-020-00466-2)) |
| Otu03673 | 9 | k__Fungi(100);p__Ascomycota(100);c__Dothideomycetes(100);o__Capnodiales(100);f__Mycosphaerellaceae(100);f__Mycosphaerellaceae_unclassified(100);f__Mycosphaerellaceae_unclassified(100); | Mycosphaerellaceae | 9 | Pathotroph-Saprotroph | Plant Pathogen-Undefined Saprotroph | Microfungus | NULL | Probable | Biotrophic, necrotrophic or saprobic on various plant tissues (Cannon & Kirk 2007) | Cannon PF, Kirk PM. 2007. Fungal Families of the World. CAB International, Cambridge (ISBN: 978-0851998275) |
| Otu03674 | 9 | k__Fungi(100);p__Ascomycota(100);c__Saccharomycetes(100);o__Saccharomycetales(100);f__Metschnikowiaceae(100);g__Metschnikowia(100);s__Metschnikowia_pulcherrima(67); | Metschnikowia | 13 | Saprotroph | \|Undefined Saprotroph\| | Yeast | NULL | Probable | As nectar-tap_saprotroph (Põlme et al. 2020); Decay sugar-rich substrates (Põlme et al. 2020) | Põlme S, et al. 2020. Fungal Diversity 105:1-16 ((https://doi.org/10.1007/s13225-020-00466-2)) |
| Otu03675 | 9 | k__Fungi(100);p__Basidiomycota(100);c__Agaricomycetes(100);o__Agaricales(100);f__Strophariaceae(100);g__Pholiota(100);s__Pholiota_sp_Di267_2(100); | Pholiota | 13 | Pathotroph-Saprotroph | Plant Pathogen-Plant Saprotroph-Undefined Saprotroph-\|Wood Saprotroph\| | Agaricoid | White Rot | Probable | As litter_saprotroph (Põlme et al. 2020); As white_rot (Põlme et al. 2020); As wood_pathogen (Põlme et al. 2020); Decay wood, leaf, fruit, seed (Põlme et al. 2020) | Tedersoo L, et al. 2014. Science 346:e1256688 ((https://doi.org/10.1126/science.1256688)); Põlme S, et al. 2020. Fungal Diversity 105:1-16 ((https://doi.org/10.1007/s13225-020-00466-2)) |
| Otu03677 | 9 | k__Fungi(100);p__Ascomycota(100);c__Saccharomycetes(100);o__Saccharomycetales(100);f__Metschnikowiaceae(100);g__Metschnikowia(100);s__Metschnikowia_pulcherrima(100); | Metschnikowia | 13 | Saprotroph | \|Undefined Saprotroph\| | Yeast | NULL | Probable | As nectar-tap_saprotroph (Põlme et al. 2020); Decay sugar-rich substrates (Põlme et al. 2020) | Põlme S, et al. 2020. Fungal Diversity 105:1-16 ((https://doi.org/10.1007/s13225-020-00466-2)) |
| Otu03681 | 9 | k__Fungi(100);p__Basidiomycota(100);c__Agaricomycetes(100);o__Trechisporales(100);f__Hydnodontaceae(100);g__Trechispora(100);g__Trechispora_unclassified(100); | Trechispora | 13 | Saprotroph | \|Wood Saprotroph\| | Corticioid | White Rot | Highly Probable | As white_rot (Põlme et al. 2020); Decay wood (Põlme et al. 2020) | Gilbertson RL, Ryvarden L. 1987-1987. North American Polypores. Fungiflora, Oslo (ISBN: 978-0945345060); Rinaldi AC, et al. 2008. Fungal Diversity 33:1-45 ((http://www.fungaldiversity.org/fdp/sfdp/33-1.pdf)) (as Uncertain); Tedersoo L, et al. 2010. Mycorrhiza 20:217-263 ((https://doi.org/10.1007/s00572-009-0274-x)) (as not ECM).; Põlme S, et al. 2020. Fungal Diversity 105:1-16 ((https://doi.org/10.1007/s13225-020-00466-2)) |
| Otu03688 | 9 | k__Fungi(100);p__Ascomycota(100);c__Saccharomycetes(100);o__Saccharomycetales(100);f__Metschnikowiaceae(100);g__Metschnikowia(100);s__Metschnikowia_pulcherrima(89); | Metschnikowia | 13 | Saprotroph | \|Undefined Saprotroph\| | Yeast | NULL | Probable | As nectar-tap_saprotroph (Põlme et al. 2020); Decay sugar-rich substrates (Põlme et al. 2020) | Põlme S, et al. 2020. Fungal Diversity 105:1-16 ((https://doi.org/10.1007/s13225-020-00466-2)) |
| Otu03690 | 9 | k__Fungi(100);p__Ascomycota(100);c__Leotiomycetes(100);o__Helotiales(100);f__Rutstroemiaceae(100);f__Rutstroemiaceae_unclassified(100);f__Rutstroemiaceae_unclassified(100); | Rutstroemiaceae | 9 | Saprotroph | Plant Saprotroph | Microfungus | NULL | Probable | Essentially saprobic on various plant parts, especially seeds and fruits, though initial infection may take place in living tissue (Cannon & Kirk 2007) | Cannon PF, Kirk PM. 2007. Fungal Families of the World. CAB International, Cambridge (ISBN: 978-0851998275) |
| Otu03691 | 9 | k__Fungi(100);p__Ascomycota(100);c__Saccharomycetes(100);o__Saccharomycetales(100);f__Metschnikowiaceae(100);g__Metschnikowia(100);s__Metschnikowia_pulcherrima(100); | Metschnikowia | 13 | Saprotroph | \|Undefined Saprotroph\| | Yeast | NULL | Probable | As nectar-tap_saprotroph (Põlme et al. 2020); Decay sugar-rich substrates (Põlme et al. 2020) | Põlme S, et al. 2020. Fungal Diversity 105:1-16 ((https://doi.org/10.1007/s13225-020-00466-2)) |
| Otu03693 | 9 | k__Fungi(100);p__Ascomycota(100);c__Dothideomycetes(100);o__Dothideomycetes_order_Incertae_sedis(100);f__Myxotrichaceae(100);g__Oidiodendron(100);s__Oidiodendron_echinulatum(78); | Oidiodendron | 13 | Pathotroph-Saprotroph-Symbiotroph | Endophyte-Ericoid Mycorrhizal-\|Undefined Saprotroph\| | Microfungus | NULL | Probable | As root_endophyte (Põlme et al. 2020); As soil_saprotroph (Põlme et al. 2020); Dark Septate Endophyte; Decay soil (Põlme et al. 2020) | Newsham KK. 2011. New Phytologist 190:783-793 ((https://doi.org/10.1111/j.1469-8137.2010.03611.x)) (as dark septate root endophytes); Põlme S, et al. 2020. Fungal Diversity 105:1-16 ((https://doi.org/10.1007/s13225-020-00466-2)) |
| Otu03695 | 9 | k__Fungi(100);p__Ascomycota(100);c__Saccharomycetes(100);o__Saccharomycetales(100);f__Metschnikowiaceae(100);g__Metschnikowia(100);s__Metschnikowia_pulcherrima(100); | Metschnikowia | 13 | Saprotroph | \|Undefined Saprotroph\| | Yeast | NULL | Probable | As nectar-tap_saprotroph (Põlme et al. 2020); Decay sugar-rich substrates (Põlme et al. 2020) | Põlme S, et al. 2020. Fungal Diversity 105:1-16 ((https://doi.org/10.1007/s13225-020-00466-2)) |
| Otu03696 | 9 | k__Fungi(100);p__Zygomycota(100);c__Zygomycota_class_Incertae_sedis(100);o__Mucorales(100);f__Mucoraceae(100);g__Mucor(100);g__Mucor_unclassified(100); | Mucor | 13 | Saprotroph | \|Undefined Saprotroph\| | Microfungus | NULL | Probable | As soil_saprotroph (Põlme et al. 2020); Decay soil (Põlme et al. 2020) | Tedersoo L, et al. 2014. Science 346:e1256688 ((https://doi.org/10.1126/science.1256688)); Põlme S, et al. 2020. Fungal Diversity 105:1-16 ((https://doi.org/10.1007/s13225-020-00466-2)) |
| Otu03699 | 9 | k__Fungi(100);p__Ascomycota(100);c__Saccharomycetes(100);o__Saccharomycetales(100);f__Metschnikowiaceae(100);g__Metschnikowia(100);g__Metschnikowia_unclassified(89); | Metschnikowia | 13 | Saprotroph | \|Undefined Saprotroph\| | Yeast | NULL | Probable | As nectar-tap_saprotroph (Põlme et al. 2020); Decay sugar-rich substrates (Põlme et al. 2020) | Põlme S, et al. 2020. Fungal Diversity 105:1-16 ((https://doi.org/10.1007/s13225-020-00466-2)) |
| Otu03700 | 9 | k__Fungi(100);p__Ascomycota(100);c__Saccharomycetes(100);o__Saccharomycetales(100);f__Metschnikowiaceae(100);g__Metschnikowia(100);s__Metschnikowia_pulcherrima(78); | Metschnikowia | 13 | Saprotroph | \|Undefined Saprotroph\| | Yeast | NULL | Probable | As nectar-tap_saprotroph (Põlme et al. 2020); Decay sugar-rich substrates (Põlme et al. 2020) | Põlme S, et al. 2020. Fungal Diversity 105:1-16 ((https://doi.org/10.1007/s13225-020-00466-2)) |
| Otu03702 | 9 | k__Fungi(100);p__Ascomycota(100);c__Saccharomycetes(100);o__Saccharomycetales(100);f__Metschnikowiaceae(100);g__Metschnikowia(100);s__Metschnikowia_pulcherrima(89); | Metschnikowia | 13 | Saprotroph | \|Undefined Saprotroph\| | Yeast | NULL | Probable | As nectar-tap_saprotroph (Põlme et al. 2020); Decay sugar-rich substrates (Põlme et al. 2020) | Põlme S, et al. 2020. Fungal Diversity 105:1-16 ((https://doi.org/10.1007/s13225-020-00466-2)) |
| Otu03704 | 9 | k__Fungi(100);p__Basidiomycota(100);c__Agaricomycetes(100);o__Polyporales(100);f__Phanerochaetaceae(100);g__Ceriporia(100);s__Ceriporia_purpurea(100); | Ceriporia | 13 | Saprotroph | \|Wood Saprotroph\| | Corticioid | White Rot | Highly Probable | As white_rot (Põlme et al. 2020); Decay wood (Põlme et al. 2020) | Tedersoo L, et al. 2014. Science 346:e1256688 ((https://doi.org/10.1126/science.1256688)); Põlme S, et al. 2020. Fungal Diversity 105:1-16 ((https://doi.org/10.1007/s13225-020-00466-2)) |
| Otu03708 | 9 | k__Fungi(100);p__Ascomycota(100);c__Dothideomycetes(100);o__Capnodiales(100);f__Mycosphaerellaceae(100);f__Mycosphaerellaceae_unclassified(100);f__Mycosphaerellaceae_unclassified(100); | Mycosphaerellaceae | 9 | Pathotroph-Saprotroph | Plant Pathogen-Undefined Saprotroph | Microfungus | NULL | Probable | Biotrophic, necrotrophic or saprobic on various plant tissues (Cannon & Kirk 2007) | Cannon PF, Kirk PM. 2007. Fungal Families of the World. CAB International, Cambridge (ISBN: 978-0851998275) |
| Otu03710 | 9 | k__Fungi(100);p__Ascomycota(100);c__Saccharomycetes(100);o__Saccharomycetales(100);f__Metschnikowiaceae(100);g__Metschnikowia(100);s__Metschnikowia_pulcherrima(100); | Metschnikowia | 13 | Saprotroph | \|Undefined Saprotroph\| | Yeast | NULL | Probable | As nectar-tap_saprotroph (Põlme et al. 2020); Decay sugar-rich substrates (Põlme et al. 2020) | Põlme S, et al. 2020. Fungal Diversity 105:1-16 ((https://doi.org/10.1007/s13225-020-00466-2)) |
| Otu03711 | 9 | k__Fungi(100);p__Ascomycota(100);c__Ascomycota_class_Incertae_sedis(100);o__Ascomycota_order_Incertae_sedis(100);f__Ascomycota_family_Incertae_sedis(100);g__Chalara(100);s__Chalara_holubovae(100); | Chalara | 13 | Pathotroph-Saprotroph-Symbiotroph | Endophyte-Plant Pathogen-\|Plant Saprotroph\|-Wood Saprotroph | Microfungus | Soft Rot | Probable | As foliar_endophyte (Põlme et al. 2020); As litter_saprotroph (Põlme et al. 2020); As wood_pathogen (Põlme et al. 2020); Decay wood, leaf, fruit, seed (Põlme et al. 2020) | Duncan CG, Eslyn WE. 1966. Mycologia 58:642-645 ((https://doi.org/10.2307/3757045)); Tedersoo L, et al. 2014. Science 346:e1256688 ((https://doi.org/10.1126/science.1256688)); Põlme S, et al. 2020. Fungal Diversity 105:1-16 ((https://doi.org/10.1007/s13225-020-00466-2)) |
| Otu03712 | 9 | k__Fungi(100);p__Ascomycota(100);c__Saccharomycetes(100);o__Saccharomycetales(100);f__Metschnikowiaceae(100);g__Metschnikowia(100);s__Metschnikowia_pulcherrima(100); | Metschnikowia | 13 | Saprotroph | \|Undefined Saprotroph\| | Yeast | NULL | Probable | As nectar-tap_saprotroph (Põlme et al. 2020); Decay sugar-rich substrates (Põlme et al. 2020) | Põlme S, et al. 2020. Fungal Diversity 105:1-16 ((https://doi.org/10.1007/s13225-020-00466-2)) |
| Otu03716 | 9 | k__Fungi(100);p__Ascomycota(100);c__Eurotiomycetes(100);o__Chaetothyriales(100);f__Herpotrichiellaceae(100);g__Cladophialophora(67);g__Cladophialophora_unclassified(67); | Cladophialophora | 13 | Pathotroph-Saprotroph-Symbiotroph | Animal Pathogen-Endophyte-Fungal Parasite-\|Undefined Saprotroph\| | Dimorphic-Microfungus | NULL | Probable | As mycoparasite (Põlme et al. 2020); As root_endophyte_dark_septate (Põlme et al. 2020); As soil_saprotroph (Põlme et al. 2020); Decay soil (Põlme et al. 2020) | James TY, et al. 2006. Nature 443:818-822 ((https://doi.org/10.1038/nature05110)); Põlme S, et al. 2020. Fungal Diversity 105:1-16 ((https://doi.org/10.1007/s13225-020-00466-2)) |
| Otu03718 | 9 | k__Fungi(100);p__Ascomycota(100);c__Saccharomycetes(100);o__Saccharomycetales(100);f__Metschnikowiaceae(100);g__Metschnikowia(100);s__Metschnikowia_pulcherrima(78); | Metschnikowia | 13 | Saprotroph | \|Undefined Saprotroph\| | Yeast | NULL | Probable | As nectar-tap_saprotroph (Põlme et al. 2020); Decay sugar-rich substrates (Põlme et al. 2020) | Põlme S, et al. 2020. Fungal Diversity 105:1-16 ((https://doi.org/10.1007/s13225-020-00466-2)) |
| Otu03719 | 9 | k__Fungi(100);p__Ascomycota(100);c__Sordariomycetes(100);o__Diaporthales(100);f__Diaporthaceae(100);f__Diaporthaceae_unclassified(56);f__Diaporthaceae_unclassified(56); | Diaporthaceae | 9 | Pathotroph-Saprotroph-Symbiotroph | Endophyte-Plant Pathogen-Plant Saprotroph | Microfungus | NULL | Probable | Most species are weak pathogens or endophytes in leaf or stem tissue, growing saprobically after the plant dies (Cannon & Kirk 2007) | Cannon PF, Kirk PM. 2007. Fungal Families of the World. CAB International, Cambridge (ISBN: 978-0851998275) |
| Otu03720 | 9 | k__Fungi(100);p__Ascomycota(100);c__Eurotiomycetes(100);o__Chaetothyriales(100);f__Chaetothyriaceae(100);g__Cyphellophora(100);g__Cyphellophora_unclassified(100); | Cyphellophora | 13 | Pathotroph-Saprotroph | Animal Parasite-Animal Pathogen-\|Plant Saprotroph\|-Undefined Saprotroph | Dimorphic-Facultative Yeast-Microfungus | NULL | Probable | As litter_saprotroph (Põlme et al. 2020); Consult the current literature for updated names on this group that included the Black Yeasts; Some species are likely opportunistic human pathogen (Irinyi et al. 2015); Decay leaf, fruit, seed, animal material (Põlme et al. 2020) | Tedersoo L, et al. 2014. Science 346:e1256688 ((https://doi.org/10.1126/science.1256688)); Irinyi L, et al. 2015. Medical Mycology 53:313-337 ((https://doi.org/10.1093/mmy/myv008)); Põlme S, et al. 2020. Fungal Diversity 105:1-16 ((https://doi.org/10.1007/s13225-020-00466-2)) |
| Otu03721 | 9 | k__Fungi(100);p__Basidiomycota(100);c__Agaricomycetes(100);o__Agaricales(100);f__Pluteaceae(100);g__Volvariella(100);g__Volvariella_unclassified(100); | Volvariella | 13 | Saprotroph | \|Plant Saprotroph\|-Undefined Saprotroph-Wood Saprotroph | Agaricoid | White Rot | Highly Probable | As litter_saprotroph (Põlme et al. 2020); As white_rot (Põlme et al. 2020); Decay leaf, fruit, seed (Põlme et al. 2020) | Tedersoo L, et al. 2014. Science 346:e1256688 ((https://doi.org/10.1126/science.1256688)); Põlme S, et al. 2020. Fungal Diversity 105:1-16 ((https://doi.org/10.1007/s13225-020-00466-2)) |
| Otu03722 | 9 | k__Fungi(100);p__Ascomycota(100);c__Sordariomycetes(100);o__Hypocreales(100);f__Clavicipitaceae(100);f__Clavicipitaceae_unclassified(89);f__Clavicipitaceae_unclassified(89); | Clavicipitaceae | 9 | Pathotroph-Symbiotroph | Endophyte-Epiphyte-Fungal Parasite-Insect Parasite | Microfungus | NULL | Possible | Parasitic, endophytic or epiphytic on Insecta, other fungi or Gamineae (Cannon & Kirk 2007) | Cannon PF, Kirk PM. 2007. Fungal Families of the World. CAB International, Cambridge (ISBN: 978-0851998275) |
| Otu03724 | 9 | k__Fungi(100);p__Ascomycota(100);c__Sordariomycetes(100);o__Hypocreales(100);f__Hypocreales_family_Incertae_sedis(100);g__Acremonium(100);s__Acremonium_cavaraeanum(100); | Acremonium | 13 | Pathotroph-Saprotroph-Symbiotroph | Animal Pathogen-Endophyte-Fungal Parasite-Plant Pathogen-\|Undefined Saprotroph\|-Wood Saprotroph | Microfungus | Soft Rot | Possible | As foliar_endophyte (Põlme et al. 2020); culture (Hanada et al. 2010, Del Olmo; Decay leaf, fruit, seed, soil, dung, animal material (Põlme et al. 2020); Endophyte detection method; Ruiz & Arnold 2014); Likely opportunistic human pathogen (Irinyi et al. 2015) | Duncan CG, Eslyn WE. 1966. Mycologia 58:642-645 ((https://doi.org/10.2307/3757045)); Nilsson T. 1973. Studia Forestalia Suecica 104: 1-40 ((https://pub.epsilon.slu.se/5711/)); Worrall JJ, et al. 1997. Mycologia 89:199-219 ((https://doi.org/10.2307/3761073)); Hanada RE, et al. 2010. Fungal Biology 114:901-910 ((https://doi.org/10.1016/j.funbio.2010.08.006)); Del Olmo-Ruiz M, Arnold AE. 2014. Mycologia 106: 8-21 ((https://doi.org/10.3852/13-098)); Irinyi L, et al. 2015. Medical Mycology 53313-337 ((https://doi.org/10.1093/mmy/myv008)); Põlme S, et al. 2020. Fungal Diversity 105:1-16 ((https://doi.org/10.1007/s13225-020-00466-2)) |
| Otu03725 | 9 | k__Fungi(100);p__Ascomycota(100);c__Saccharomycetes(100);o__Saccharomycetales(100);f__Metschnikowiaceae(100);g__Metschnikowia(100);g__Metschnikowia_unclassified(89); | Metschnikowia | 13 | Saprotroph | \|Undefined Saprotroph\| | Yeast | NULL | Probable | As nectar-tap_saprotroph (Põlme et al. 2020); Decay sugar-rich substrates (Põlme et al. 2020) | Põlme S, et al. 2020. Fungal Diversity 105:1-16 ((https://doi.org/10.1007/s13225-020-00466-2)) |
| Otu03726 | 9 | k__Fungi(100);p__Ascomycota(100);c__Saccharomycetes(100);o__Saccharomycetales(100);f__Metschnikowiaceae(100);g__Metschnikowia(100);s__Metschnikowia_pulcherrima(89); | Metschnikowia | 13 | Saprotroph | \|Undefined Saprotroph\| | Yeast | NULL | Probable | As nectar-tap_saprotroph (Põlme et al. 2020); Decay sugar-rich substrates (Põlme et al. 2020) | Põlme S, et al. 2020. Fungal Diversity 105:1-16 ((https://doi.org/10.1007/s13225-020-00466-2)) |
| Otu03728 | 9 | k__Fungi(100);p__Ascomycota(100);c__Saccharomycetes(100);o__Saccharomycetales(100);f__Metschnikowiaceae(100);g__Metschnikowia(100);s__Metschnikowia_pulcherrima(89); | Metschnikowia | 13 | Saprotroph | \|Undefined Saprotroph\| | Yeast | NULL | Probable | As nectar-tap_saprotroph (Põlme et al. 2020); Decay sugar-rich substrates (Põlme et al. 2020) | Põlme S, et al. 2020. Fungal Diversity 105:1-16 ((https://doi.org/10.1007/s13225-020-00466-2)) |
| Otu03729 | 9 | k__Fungi(100);p__Ascomycota(100);c__Leotiomycetes(100);o__Helotiales(100);f__Vibrisseaceae(89);g__Phialocephala(89);s__Phialocephala_sp_CM16s1(67); | Phialocephala | 13 | Saprotroph-Symbiotroph | Endophyte-\|Undefined Saprotroph\| | Microfungus | NULL | Highly Probable | As root_endophyte (Põlme et al. 2020); As root_endophyte_dark_septate (Põlme et al. 2020); As soil_saprotroph (Põlme et al. 2020); Dark Septate Endophyte, known from plant roots.; Decay roots, soil (Põlme et al. 2020) | Newsham KK. 2011. New Phytologist 190:783-793 ((https://doi.org/10.1111/j.1469-8137.2010.03611.x)) (as dark septate root endophytes); Põlme S, et al. 2020. Fungal Diversity 105:1-16 ((https://doi.org/10.1007/s13225-020-00466-2)) |
| Otu03732 | 9 | k__Fungi(100);p__Ascomycota(100);c__Saccharomycetes(100);o__Saccharomycetales(100);f__Metschnikowiaceae(100);g__Metschnikowia(100);s__Metschnikowia_pulcherrima(100); | Metschnikowia | 13 | Saprotroph | \|Undefined Saprotroph\| | Yeast | NULL | Probable | As nectar-tap_saprotroph (Põlme et al. 2020); Decay sugar-rich substrates (Põlme et al. 2020) | Põlme S, et al. 2020. Fungal Diversity 105:1-16 ((https://doi.org/10.1007/s13225-020-00466-2)) |
| Otu03733 | 9 | k__Fungi(100);p__Ascomycota(100);c__Saccharomycetes(100);o__Saccharomycetales(100);f__Metschnikowiaceae(100);g__Metschnikowia(100);g__Metschnikowia_unclassified(89); | Metschnikowia | 13 | Saprotroph | \|Undefined Saprotroph\| | Yeast | NULL | Probable | As nectar-tap_saprotroph (Põlme et al. 2020); Decay sugar-rich substrates (Põlme et al. 2020) | Põlme S, et al. 2020. Fungal Diversity 105:1-16 ((https://doi.org/10.1007/s13225-020-00466-2)) |
| Otu03734 | 9 | k__Fungi(100);p__Basidiomycota(100);c__Tremellomycetes(100);o__Tremellales(100);f__Tremellales_family_Incertae_sedis(100);g__Cryptococcus(67);s__Cryptococcus_sp_SJ4L02(67); | Cryptococcus | 13 | Pathotroph-Saprotroph-Symbiotroph | Animal Pathogen-Endophyte-Epiphyte-\|Undefined Saprotroph\| | Dimorphic Yeast-Yeast | NULL | Possible | As foliar_endophyte (Põlme et al. 2020); Decay leaf, fruit, seed, soil, dung, animal material (Põlme et al. 2020) | Andrews JH, et al. 1983. Phytopathology 73:228-234 ((https://doi.org/10.1094/Phyto-73-228)); Kurtzman CP, et al. (eds.) 2011. The Yeasts, a Taxonomic Study. Fifth Edition. Vols 1-3. Elsevier, San Diego (ISBN: 9780444521491); Põlme S, et al. 2020. Fungal Diversity 105:1-16 ((https://doi.org/10.1007/s13225-020-00466-2)) |
| Otu03736 | 9 | k__Fungi(100);p__Ascomycota(100);c__Dothideomycetes(100);o__Pleosporales(100);f__Phaeosphaeriaceae(89);g__unclassified_Phaeosphaeriaceae(56);s__Phaeosphaeriaceae_sp(56); | Phaeosphaeriaceae | 9 | Pathotroph-Saprotroph | Fungal Parasite-Plant Pathogen-Plant Saprotroph | Microfungus | NULL | Probable | Necrotrophic or saprobic on a wide range of plants, especially monocotyledons; a few species parasitic on other fungi (Cannon & Kirk 2007) | Cannon PF, Kirk PM. 2007. Fungal Families of the World. CAB International, Cambridge (ISBN: 978-0851998275) |
| Otu03737 | 9 | k__Fungi(100);p__Ascomycota(100);c__Saccharomycetes(100);o__Saccharomycetales(100);f__Metschnikowiaceae(100);g__Metschnikowia(100);s__Metschnikowia_pulcherrima(56); | Metschnikowia | 13 | Saprotroph | \|Undefined Saprotroph\| | Yeast | NULL | Probable | As nectar-tap_saprotroph (Põlme et al. 2020); Decay sugar-rich substrates (Põlme et al. 2020) | Põlme S, et al. 2020. Fungal Diversity 105:1-16 ((https://doi.org/10.1007/s13225-020-00466-2)) |
| Otu03738 | 9 | k__Fungi(100);p__Ascomycota(100);c__Saccharomycetes(100);o__Saccharomycetales(100);f__Saccharomycetales_family_Incertae_sedis(89);g__Candida(89);g__Candida_unclassified(56); | Candida | 13 | Pathotroph-Saprotroph-Symbiotroph | Animal Pathogen-Endophyte-Endosymbiont-Epiphyte-\|Undefined Saprotroph\| | Dimorphic Yeast-Yeast | NULL | Probable | As nectar-tap_saprotroph (Põlme et al. 2020); Decay sugar-rich substrates (Põlme et al. 2020); This asexual genus is polyphyletic and contain species that function very differently depend on what they are. Caution should be taken when interpreting guilds for Candida species. | Manolakaki D, et al. 2010. Virulence. 1: 367-375 ((https://doi.org/10.4161/viru.1.5.12796)); Mekbib SB, et al. 2011. Tropical Plant Pathology 36:233-240 ((https://doi.org/10.1590/S1982-56762011000400004)); Põlme S, et al. 2020. Fungal Diversity 105:1-16 ((https://doi.org/10.1007/s13225-020-00466-2)) |
| Otu03739 | 9 | k__Fungi(100);p__Basidiomycota(100);c__Microbotryomycetes(56);o__Sporidiobolales(56);f__Sporidiobolales_family_Incertae_sedis(56);g__Rhodotorula(56);s__Rhodotorula_sp(56); | Rhodotorula | 13 | Pathotroph-Saprotroph-Symbiotroph | Animal Pathogen-Animal Symbiotroph-Endophyte-Plant Pathogen-\|Undefined Saprotroph\| | Yeast | NULL | Probable | As foliar_endophyte (Põlme et al. 2020); Opportunistic human pathogen | Põlme S, et al. 2020. Fungal Diversity 105:1-16 ((https://doi.org/10.1007/s13225-020-00466-2)); https://www.cdc.gov ((https://www.cdc.gov)) |
| Otu03741 | 9 | k__Fungi(100);p__Ascomycota(100);c__Saccharomycetes(100);o__Saccharomycetales(100);f__Metschnikowiaceae(100);g__Metschnikowia(100);s__Metschnikowia_pulcherrima(56); | Metschnikowia | 13 | Saprotroph | \|Undefined Saprotroph\| | Yeast | NULL | Probable | As nectar-tap_saprotroph (Põlme et al. 2020); Decay sugar-rich substrates (Põlme et al. 2020) | Põlme S, et al. 2020. Fungal Diversity 105:1-16 ((https://doi.org/10.1007/s13225-020-00466-2)) |
| Otu03745 | 9 | k__Fungi(100);p__Ascomycota(100);c__Lecanoromycetes(100);o__Lecanorales(100);f__Parmeliaceae(100);f__Parmeliaceae_unclassified(56);f__Parmeliaceae_unclassified(56); | Parmeliaceae | 9 | Symbiotroph | Lichenized | Thallus | NULL | Highly Probable | Lichenized with green algae, mostly growing on tree bark and rocks (Cannon & Kirk 2007) | Cannon PF, Kirk PM. 2007. Fungal Families of the World. CAB International, Cambridge (ISBN: 978-0851998275) |
| Otu03746 | 9 | k__Fungi(100);p__Ascomycota(100);c__Sordariomycetes(100);o__Boliniales(67);f__Boliniaceae(67);f__Boliniaceae_unclassified(67);f__Boliniaceae_unclassified(67); | Boliniaceae | 9 | Saprotroph | Wood Saprotroph | Microfungus | NULL | Probable | Saprobic on and in wood and bark (Cannon & Kirk 2007) | Cannon PF, Kirk PM. 2007. Fungal Families of the World. CAB International, Cambridge (ISBN: 978-0851998275) |
| Otu03757 | 9 | k__Fungi(100);p__Ascomycota(100);c__Saccharomycetes(100);o__Saccharomycetales(100);f__Metschnikowiaceae(100);g__Metschnikowia(100);s__Metschnikowia_pulcherrima(89); | Metschnikowia | 13 | Saprotroph | \|Undefined Saprotroph\| | Yeast | NULL | Probable | As nectar-tap_saprotroph (Põlme et al. 2020); Decay sugar-rich substrates (Põlme et al. 2020) | Põlme S, et al. 2020. Fungal Diversity 105:1-16 ((https://doi.org/10.1007/s13225-020-00466-2)) |
| Otu03760 | 9 | k__Fungi(100);p__Ascomycota(100);c__Sordariomycetes(100);o__Sordariales(100);f__Chaetomiaceae(100);g__Humicola(100);s__Humicola_grisea_var._grisea(100); | Humicola | 13 | Pathotroph-Saprotroph | Plant Pathogen-Undefined Saprotroph-\|Wood Saprotroph\| | Microfungus | Soft Rot | Probable | Decay wood (Põlme et al. 2020) | Seehann G, et al. 1975. List of Fungi in Soft-Rot Tissue Tests. Institute fur Holzbiologie und Holzschutz, Bundesforschungsanstalt fur Forst - und Holzwirtshaft, Hamburg - Lohbrugge, Germany; Costa IPMW, et al. 2012. Mycotaxon 119:e26 ((http://mycotaxon.com/resources/checklists/costa_v119_checklist.pdf)); Tedersoo L, et al. 2014. Science 346:e1256688 ((https://doi.org/10.1126/science.1256688)); Põlme S, et al. 2020. Fungal Diversity 105:1-16 ((https://doi.org/10.1007/s13225-020-00466-2)) |
| Otu03761 | 9 | k__Fungi(100);p__Ascomycota(100);c__Eurotiomycetes(100);o__Chaetothyriales(100);f__Chaetothyriaceae(100);g__Ceramothyrium(100);s__Ceramothyrium_carniolicum(100); | Ceramothyrium | 13 | Pathotroph-Saprotroph-Symbiotroph | Endophyte-Plant Pathogen-\|Plant Saprotroph\|-Undefined Saprotroph | Microfungus | NULL | Probable | As foliar_endophyte (Põlme et al. 2020); As litter_saprotroph (Põlme et al. 2020); Causes sooty mold (Põlme et al. 2020); Decay leaf, fruit, seed, sugar-rich substrates (Põlme et al. 2020) | Tedersoo L, et al. 2014. Science 346:e1256688 ((https://doi.org/10.1126/science.1256688)); Põlme S, et al. 2020. Fungal Diversity 105:1-16 ((https://doi.org/10.1007/s13225-020-00466-2)) |
| Otu03762 | 9 | k__Fungi(100);p__Ascomycota(100);c__Saccharomycetes(100);o__Saccharomycetales(100);f__Metschnikowiaceae(100);g__Metschnikowia(100);s__Metschnikowia_pulcherrima(78); | Metschnikowia | 13 | Saprotroph | \|Undefined Saprotroph\| | Yeast | NULL | Probable | As nectar-tap_saprotroph (Põlme et al. 2020); Decay sugar-rich substrates (Põlme et al. 2020) | Põlme S, et al. 2020. Fungal Diversity 105:1-16 ((https://doi.org/10.1007/s13225-020-00466-2)) |
| Otu03766 | 9 | k__Fungi(100);p__Ascomycota(100);c__Sordariomycetes(100);o__Diaporthales(100);f__Diaporthales_family_Incertae_sedis(100);g__Ophiognomonia(67);s__Ophiognomonia_sp_BPI_877592(67); | Ophiognomonia | 13 | Pathotroph | \|Plant Pathogen\| | Microfungus | NULL | Highly Probable | See Marssoniella juglandis, Ophiognomonia leptostyla, and Syn. Gnomonia leptostyla; Causes leaf blotch and spots, also known as walnut anthracnose or walnut black spot (Sogonov et al. 2008) | Sogonov MV, et al. 2008. Studies in Mycology 62:1-79 ((https://www.studiesinmycology.org/sim/Sim62/03_Leaf-inhabiting_genera_of_the_Gnomoniaceae,_Diaporthales.pdf)); Põlme S, et al. 2020. Fungal Diversity 105:1-16 ((https://doi.org/10.1007/s13225-020-00466-2)) |
| Otu03770 | 9 | k__Fungi(100);p__Ascomycota(100);c__Saccharomycetes(100);o__Saccharomycetales(100);f__Metschnikowiaceae(100);g__Metschnikowia(100);s__Metschnikowia_pulcherrima(67); | Metschnikowia | 13 | Saprotroph | \|Undefined Saprotroph\| | Yeast | NULL | Probable | As nectar-tap_saprotroph (Põlme et al. 2020); Decay sugar-rich substrates (Põlme et al. 2020) | Põlme S, et al. 2020. Fungal Diversity 105:1-16 ((https://doi.org/10.1007/s13225-020-00466-2)) |
| Otu03771 | 9 | k__Fungi(100);p__Basidiomycota(100);c__Agaricomycetes(100);o__Agaricales(100);o__Agaricales_unclassified(100);o__Agaricales_unclassified(100);o__Agaricales_unclassified(100); | Agaricales | 7 | Pathotroph-Saprotroph-Symbiotroph | Bryophyte Parasite-Dung Saprotroph-Ectomycorrhizal-Fungal Parasite-Leaf Saprotroph-Plant Parasite-Undefined Saprotroph-Wood Saprotroph | Agaricoid-Gasteroid-Microfungus-Secotioid-Yeast | NULL | Possible | Mushrooms and toadstools, Gill fungi, Agarics... lignicolous, sometimes muscicolous or fungicolous, saprobic, mycorrhizal, rarely parasitic on plants or fungi (Kirk et al. 2008) | Kirk PM et al. 2008. Dictionary of the Fungi. Tenth Edition. CAB International, Wallingford (ISBN: 978-0851998268) |
| Otu03772 | 9 | k__Fungi(100);p__Ascomycota(100);c__Saccharomycetes(100);o__Saccharomycetales(100);f__Metschnikowiaceae(100);g__Metschnikowia(100);g__Metschnikowia_unclassified(67); | Metschnikowia | 13 | Saprotroph | \|Undefined Saprotroph\| | Yeast | NULL | Probable | As nectar-tap_saprotroph (Põlme et al. 2020); Decay sugar-rich substrates (Põlme et al. 2020) | Põlme S, et al. 2020. Fungal Diversity 105:1-16 ((https://doi.org/10.1007/s13225-020-00466-2)) |
| Otu03774 | 9 | k__Fungi(100);p__Basidiomycota(100);c__Agaricomycetes(100);o__Atheliales(100);f__Atheliaceae(100);g__Amphinema(100);s__Amphinema_byssoides(100); | Amphinema | 13 | Symbiotroph | \|Ectomycorrhizal\| | Corticioid | NULL | Highly Probable | NULL | Rinaldi AC, et al. 2008. Fungal Diversity 33:1-45 ((http://www.fungaldiversity.org/fdp/sfdp/33-1.pdf)); Tedersoo L, et al. 2010. Mycorrhiza 20:217-263 ((https://doi.org/10.1007/s00572-009-0274-x)); Põlme S, et al. 2020. Fungal Diversity 105:1-16 ((https://doi.org/10.1007/s13225-020-00466-2)); http://mycorrhizas.info/ecmf.html ((http://mycorrhizas.info/ecmf.html)) |
| Otu03775 | 9 | k__Fungi(100);p__Ascomycota(100);c__Saccharomycetes(100);o__Saccharomycetales(100);f__Metschnikowiaceae(100);g__Metschnikowia(100);s__Metschnikowia_pulcherrima(78); | Metschnikowia | 13 | Saprotroph | \|Undefined Saprotroph\| | Yeast | NULL | Probable | As nectar-tap_saprotroph (Põlme et al. 2020); Decay sugar-rich substrates (Põlme et al. 2020) | Põlme S, et al. 2020. Fungal Diversity 105:1-16 ((https://doi.org/10.1007/s13225-020-00466-2)) |
| Otu03779 | 9 | k__Fungi(100);p__Basidiomycota(100);c__Agaricomycetes(100);o__Polyporales(100);f__Phanerochaetaceae(100);g__Byssomerulius(100);s__Byssomerulius_sp(100); | Byssomerulius | 13 | Saprotroph | Undefined Saprotroph-\|Wood Saprotroph\| | Corticioid | White Rot | Probable | As white_rot (Põlme et al. 2020); Decay wood (Põlme et al. 2020) | Tedersoo L, et al. 2014. Science 346:e1256688 ((https://doi.org/10.1126/science.1256688)); Põlme S, et al. 2020. Fungal Diversity 105:1-16 ((https://doi.org/10.1007/s13225-020-00466-2)) |
| Otu03780 | 9 | k__Fungi(100);p__Ascomycota(100);c__Leotiomycetes(100);o__Helotiales(100);f__Hyaloscyphaceae(100);g__Lachnum(56);g__Lachnum_unclassified(56); | Lachnum | 13 | Saprotroph-Symbiotroph | Endophyte-\|Plant Saprotroph\|-Undefined Saprotroph-Wood Saprotroph | Microfungus | NULL | Probable | As litter_saprotroph (Põlme et al. 2020); As root_endophyte (Põlme et al. 2020); Decay wood, leaf, fruit, seed (Põlme et al. 2020) | Tedersoo L, et al. 2014. Science 346:e1256688 ((https://doi.org/10.1126/science.1256688)); Põlme S, et al. 2020. Fungal Diversity 105:1-16 ((https://doi.org/10.1007/s13225-020-00466-2)) |
| Otu03785 | 9 | k__Fungi(100);p__Ascomycota(100);c__Saccharomycetes(100);o__Saccharomycetales(100);f__Metschnikowiaceae(100);g__Metschnikowia(100);s__Metschnikowia_pulcherrima(100); | Metschnikowia | 13 | Saprotroph | \|Undefined Saprotroph\| | Yeast | NULL | Probable | As nectar-tap_saprotroph (Põlme et al. 2020); Decay sugar-rich substrates (Põlme et al. 2020) | Põlme S, et al. 2020. Fungal Diversity 105:1-16 ((https://doi.org/10.1007/s13225-020-00466-2)) |
| Otu03787 | 9 | k__Fungi(100);p__Ascomycota(100);c__Saccharomycetes(100);o__Saccharomycetales(100);f__Metschnikowiaceae(100);g__Metschnikowia(100);s__Metschnikowia_pulcherrima(67); | Metschnikowia | 13 | Saprotroph | \|Undefined Saprotroph\| | Yeast | NULL | Probable | As nectar-tap_saprotroph (Põlme et al. 2020); Decay sugar-rich substrates (Põlme et al. 2020) | Põlme S, et al. 2020. Fungal Diversity 105:1-16 ((https://doi.org/10.1007/s13225-020-00466-2)) |
| Otu03790 | 9 | k__Fungi(100);p__Ascomycota(100);c__Dothideomycetes(100);o__Dothideomycetes_order_Incertae_sedis(100);f__Myxotrichaceae(100);g__Oidiodendron(100);g__Oidiodendron_unclassified(56); | Oidiodendron | 13 | Pathotroph-Saprotroph-Symbiotroph | Endophyte-Ericoid Mycorrhizal-\|Undefined Saprotroph\| | Microfungus | NULL | Probable | As root_endophyte (Põlme et al. 2020); As soil_saprotroph (Põlme et al. 2020); Dark Septate Endophyte; Decay soil (Põlme et al. 2020) | Newsham KK. 2011. New Phytologist 190:783-793 ((https://doi.org/10.1111/j.1469-8137.2010.03611.x)) (as dark septate root endophytes); Põlme S, et al. 2020. Fungal Diversity 105:1-16 ((https://doi.org/10.1007/s13225-020-00466-2)) |
| Otu03793 | 9 | k__Fungi(100);p__Basidiomycota(100);c__Agaricomycetes(100);o__Agaricales(100);f__Physalacriaceae(100);f__Physalacriaceae_unclassified(100);f__Physalacriaceae_unclassified(100); | Physalacriaceae | 9 | Pathotroph-Saprotroph | Algal Parasite--Leaf Saprotroph-Wood Saprotroph | Agaricoid-Microfungus | NULL | Probable | Presumably saprobic, found on decaying leaves and wood. One genus is a parasite of marine algae (Cannon & Kirk 2007) | Cannon PF, Kirk PM. 2007. Fungal Families of the World. CAB International, Cambridge (ISBN: 978-0851998275) |
| Otu03794 | 9 | k__Fungi(100);p__Ascomycota(100);c__Saccharomycetes(100);o__Saccharomycetales(100);f__Metschnikowiaceae(100);g__Metschnikowia(100);s__Metschnikowia_pulcherrima(78); | Metschnikowia | 13 | Saprotroph | \|Undefined Saprotroph\| | Yeast | NULL | Probable | As nectar-tap_saprotroph (Põlme et al. 2020); Decay sugar-rich substrates (Põlme et al. 2020) | Põlme S, et al. 2020. Fungal Diversity 105:1-16 ((https://doi.org/10.1007/s13225-020-00466-2)) |
| Otu03796 | 9 | k__Fungi(100);p__Ascomycota(100);c__Saccharomycetes(100);o__Saccharomycetales(100);f__Metschnikowiaceae(100);g__Metschnikowia(100);s__Metschnikowia_pulcherrima(78); | Metschnikowia | 13 | Saprotroph | \|Undefined Saprotroph\| | Yeast | NULL | Probable | As nectar-tap_saprotroph (Põlme et al. 2020); Decay sugar-rich substrates (Põlme et al. 2020) | Põlme S, et al. 2020. Fungal Diversity 105:1-16 ((https://doi.org/10.1007/s13225-020-00466-2)) |
| Otu03797 | 9 | k__Fungi(100);p__Ascomycota(100);c__Saccharomycetes(100);o__Saccharomycetales(100);f__Saccharomycetales_family_Incertae_sedis(100);g__Candida(100);g__Candida_unclassified(89); | Candida | 13 | Pathotroph-Saprotroph-Symbiotroph | Animal Pathogen-Endophyte-Endosymbiont-Epiphyte-\|Undefined Saprotroph\| | Dimorphic Yeast-Yeast | NULL | Probable | As nectar-tap_saprotroph (Põlme et al. 2020); Decay sugar-rich substrates (Põlme et al. 2020); This asexual genus is polyphyletic and contain species that function very differently depend on what they are. Caution should be taken when interpreting guilds for Candida species. | Manolakaki D, et al. 2010. Virulence. 1: 367-375 ((https://doi.org/10.4161/viru.1.5.12796)); Mekbib SB, et al. 2011. Tropical Plant Pathology 36:233-240 ((https://doi.org/10.1590/S1982-56762011000400004)); Põlme S, et al. 2020. Fungal Diversity 105:1-16 ((https://doi.org/10.1007/s13225-020-00466-2)) |
| Otu03798 | 9 | k__Fungi(100);p__Chytridiomycota(78);c__Chytridiomycetes(78);o__Spizellomycetales(78);f__Spizellomycetaceae(78);f__Spizellomycetaceae_unclassified(78);f__Spizellomycetaceae_unclassified(78); | Spizellomycetaceae | 9 | Pathotroph | Plant Pathogen-Undefined Parasite-Undefined Saprotroph | Microfungus | NULL | Probable | Common in the soil and also as parasites of a range of soil organisms and plants (Cannon & Kirk 2007) | Cannon PF, Kirk PM. 2007. Fungal Families of the World. CAB International, Cambridge (ISBN: 978-0851998275) |
| Otu03799 | 9 | k__Fungi(100);p__Ascomycota(100);c__Saccharomycetes(100);o__Saccharomycetales(100);f__Metschnikowiaceae(100);g__Metschnikowia(100);g__Metschnikowia_unclassified(56); | Metschnikowia | 13 | Saprotroph | \|Undefined Saprotroph\| | Yeast | NULL | Probable | As nectar-tap_saprotroph (Põlme et al. 2020); Decay sugar-rich substrates (Põlme et al. 2020) | Põlme S, et al. 2020. Fungal Diversity 105:1-16 ((https://doi.org/10.1007/s13225-020-00466-2)) |
| Otu03800 | 9 | k__Fungi(100);p__Basidiomycota(100);c__Basidiomycota_class_Incertae_sedis(100);o__Basidiomycota_order_Incertae_sedis(100);f__Basidiomycota_family_Incertae_sedis(100);g__Frantisekia(100);s__Frantisekia_mentschulensis(100); | Frantisekia | 13 | Saprotroph | Undefined Saprotroph-\|Wood Saprotroph\| | Polyporoid | White Rot | Probable | As white_rot (Põlme et al. 2020); Decay wood (Põlme et al. 2020) | Tedersoo L, et al. 2014. Science 346:e1256688 ((https://doi.org/10.1126/science.1256688)); Põlme S, et al. 2020. Fungal Diversity 105:1-16 ((https://doi.org/10.1007/s13225-020-00466-2)) |
| Otu03802 | 9 | k__Fungi(100);p__Ascomycota(100);c__Saccharomycetes(100);o__Saccharomycetales(100);f__Metschnikowiaceae(100);g__Metschnikowia(100);s__Metschnikowia_pulcherrima(100); | Metschnikowia | 13 | Saprotroph | \|Undefined Saprotroph\| | Yeast | NULL | Probable | As nectar-tap_saprotroph (Põlme et al. 2020); Decay sugar-rich substrates (Põlme et al. 2020) | Põlme S, et al. 2020. Fungal Diversity 105:1-16 ((https://doi.org/10.1007/s13225-020-00466-2)) |
| Otu03806 | 9 | k__Fungi(100);p__Ascomycota(100);c__Saccharomycetes(100);o__Saccharomycetales(100);f__Metschnikowiaceae(100);g__Metschnikowia(100);g__Metschnikowia_unclassified(100); | Metschnikowia | 13 | Saprotroph | \|Undefined Saprotroph\| | Yeast | NULL | Probable | As nectar-tap_saprotroph (Põlme et al. 2020); Decay sugar-rich substrates (Põlme et al. 2020) | Põlme S, et al. 2020. Fungal Diversity 105:1-16 ((https://doi.org/10.1007/s13225-020-00466-2)) |
| Otu03811 | 9 | k__Fungi(100);p__Ascomycota(100);c__Saccharomycetes(100);o__Saccharomycetales(100);f__Metschnikowiaceae(100);g__Metschnikowia(100);s__Metschnikowia_pulcherrima(100); | Metschnikowia | 13 | Saprotroph | \|Undefined Saprotroph\| | Yeast | NULL | Probable | As nectar-tap_saprotroph (Põlme et al. 2020); Decay sugar-rich substrates (Põlme et al. 2020) | Põlme S, et al. 2020. Fungal Diversity 105:1-16 ((https://doi.org/10.1007/s13225-020-00466-2)) |
| Otu03812 | 9 | k__Fungi(100);p__Ascomycota(100);c__Sordariomycetes(100);o__Xylariales(100);f__Xylariaceae(100);g__Hypoxylon(100);s__Hypoxylon_cercidicola(100); | Hypoxylon | 13 | Saprotroph-Saprotroph-Symbiotroph | Endophyte-Undefined Saprotroph-\|Wood Saprotroph\| | Corticioid-Microfungus-Xylarioid | NULL | Probable | As foliar_endophyte (Põlme et al. 2020); Decay wood, leaf, fruit, seed (Põlme et al. 2020); Host - angiosperms (Põlme et al. 2020) | Tedersoo L, et al. 2014. Science 346:e1256688 ((https://doi.org/10.1126/science.1256688)); Põlme S, et al. 2020. Fungal Diversity 105:1-16 ((https://doi.org/10.1007/s13225-020-00466-2)) |
| Otu03814 | 9 | k__Fungi(100);p__Ascomycota(100);c__Pezizomycetes(100);o__Pezizales(100);f__Chorioactidaceae(100);g__Desmazierella(100);s__Desmazierella_acicola(100); | Desmazierella | 13 | Saprotroph | \|Plant Saprotroph\|-Undefined Saprotroph | Microfungus | NULL | Probable | As litter_saprotroph (Põlme et al. 2020); Decay leaf, fruit, seed (Põlme et al. 2020) | Tedersoo L, et al. 2014. Science 346:e1256688 ((https://doi.org/10.1126/science.1256688)); Põlme S, et al. 2020. Fungal Diversity 105:1-16 ((https://doi.org/10.1007/s13225-020-00466-2)) |
| Otu03818 | 9 | k__Fungi(100);p__Ascomycota(100);c__Saccharomycetes(100);o__Saccharomycetales(100);f__Metschnikowiaceae(100);g__Metschnikowia(100);s__Metschnikowia_pulcherrima(89); | Metschnikowia | 13 | Saprotroph | \|Undefined Saprotroph\| | Yeast | NULL | Probable | As nectar-tap_saprotroph (Põlme et al. 2020); Decay sugar-rich substrates (Põlme et al. 2020) | Põlme S, et al. 2020. Fungal Diversity 105:1-16 ((https://doi.org/10.1007/s13225-020-00466-2)) |
| Otu03823 | 9 | k__Fungi(100);p__Basidiomycota(100);c__Agaricomycetes(100);o__Atheliales(100);f__Atheliaceae(100);f__Atheliaceae_unclassified(100);f__Atheliaceae_unclassified(100); | Atheliaceae | 9 | Pathotroph-Saprotroph-Symbiotroph | Ectomycorrhizal-Lichen Parasite-Lichenized-Plant Pathogen | Corticioid | NULL | Possible | Many species form ectomycorrhizal associations, often occupying a dominant position in mycorrhizal communities. Other species are pathogens of many different plants and even lichens, and one species has even been suggested to form a lichen association with cyanobacteria. Yet more species are sparse and active decomposers, and a few are aquatic (Cannon & Kirk 2007) | Cannon PF, Kirk PM. 2007. Fungal Families of the World. CAB International, Cambridge (ISBN: 978-0851998275) |
| Otu03824 | 9 | k__Fungi(100);p__Ascomycota(100);c__Saccharomycetes(100);o__Saccharomycetales(100);f__Metschnikowiaceae(100);g__Metschnikowia(100);s__Metschnikowia_pulcherrima(100); | Metschnikowia | 13 | Saprotroph | \|Undefined Saprotroph\| | Yeast | NULL | Probable | As nectar-tap_saprotroph (Põlme et al. 2020); Decay sugar-rich substrates (Põlme et al. 2020) | Põlme S, et al. 2020. Fungal Diversity 105:1-16 ((https://doi.org/10.1007/s13225-020-00466-2)) |
| Otu03826 | 9 | k__Fungi(100);p__Ascomycota(100);c__Saccharomycetes(100);o__Saccharomycetales(100);f__Metschnikowiaceae(100);g__Metschnikowia(100);s__Metschnikowia_pulcherrima(100); | Metschnikowia | 13 | Saprotroph | \|Undefined Saprotroph\| | Yeast | NULL | Probable | As nectar-tap_saprotroph (Põlme et al. 2020); Decay sugar-rich substrates (Põlme et al. 2020) | Põlme S, et al. 2020. Fungal Diversity 105:1-16 ((https://doi.org/10.1007/s13225-020-00466-2)) |
| Otu03829 | 9 | k__Fungi(100);p__Basidiomycota(100);c__Agaricomycetes(100);o__Atheliales(100);f__Atheliaceae(100);g__Amphinema(100);s__Amphinema_sp(100); | Amphinema | 13 | Symbiotroph | \|Ectomycorrhizal\| | Corticioid | NULL | Highly Probable | NULL | Rinaldi AC, et al. 2008. Fungal Diversity 33:1-45 ((http://www.fungaldiversity.org/fdp/sfdp/33-1.pdf)); Tedersoo L, et al. 2010. Mycorrhiza 20:217-263 ((https://doi.org/10.1007/s00572-009-0274-x)); Põlme S, et al. 2020. Fungal Diversity 105:1-16 ((https://doi.org/10.1007/s13225-020-00466-2)); http://mycorrhizas.info/ecmf.html ((http://mycorrhizas.info/ecmf.html)) |
| Otu03830 | 9 | k__Fungi(100);p__Basidiomycota(100);c__Agaricomycetes(100);o__Agaricales(100);o__Agaricales_unclassified(100);o__Agaricales_unclassified(100);o__Agaricales_unclassified(100); | Agaricales | 7 | Pathotroph-Saprotroph-Symbiotroph | Bryophyte Parasite-Dung Saprotroph-Ectomycorrhizal-Fungal Parasite-Leaf Saprotroph-Plant Parasite-Undefined Saprotroph-Wood Saprotroph | Agaricoid-Gasteroid-Microfungus-Secotioid-Yeast | NULL | Possible | Mushrooms and toadstools, Gill fungi, Agarics... lignicolous, sometimes muscicolous or fungicolous, saprobic, mycorrhizal, rarely parasitic on plants or fungi (Kirk et al. 2008) | Kirk PM et al. 2008. Dictionary of the Fungi. Tenth Edition. CAB International, Wallingford (ISBN: 978-0851998268) |
| Otu03831 | 9 | k__Fungi(100);p__Ascomycota(100);c__Dothideomycetes(100);o__Pleosporales(100);f__Venturiaceae(100);g__Fusicladium(100);s__Fusicladium_caryigenum(100); | Fusicladium | 13 | Pathotroph-Symbiotroph | Endophyte-Lichen Parasite-\|Plant Pathogen\| | Microfungus | NULL | Probable | NULL | Tedersoo L, et al. 2014. Science 346:e1256688 ((https://doi.org/10.1126/science.1256688)); Põlme S, et al. 2020. Fungal Diversity 105:1-16 ((https://doi.org/10.1007/s13225-020-00466-2)) |
| Otu03832 | 9 | k__Fungi(100);p__Ascomycota(100);c__Saccharomycetes(100);o__Saccharomycetales(100);f__Metschnikowiaceae(100);g__Metschnikowia(100);g__Metschnikowia_unclassified(56); | Metschnikowia | 13 | Saprotroph | \|Undefined Saprotroph\| | Yeast | NULL | Probable | As nectar-tap_saprotroph (Põlme et al. 2020); Decay sugar-rich substrates (Põlme et al. 2020) | Põlme S, et al. 2020. Fungal Diversity 105:1-16 ((https://doi.org/10.1007/s13225-020-00466-2)) |
| Otu03833 | 9 | k__Fungi(100);p__Ascomycota(100);c__Saccharomycetes(100);o__Saccharomycetales(100);f__Metschnikowiaceae(100);g__Metschnikowia(100);s__Metschnikowia_pulcherrima(100); | Metschnikowia | 13 | Saprotroph | \|Undefined Saprotroph\| | Yeast | NULL | Probable | As nectar-tap_saprotroph (Põlme et al. 2020); Decay sugar-rich substrates (Põlme et al. 2020) | Põlme S, et al. 2020. Fungal Diversity 105:1-16 ((https://doi.org/10.1007/s13225-020-00466-2)) |
| Otu03836 | 9 | k__Fungi(100);p__Ascomycota(100);c__Leotiomycetes(100);o__Helotiales(100);f__Hyaloscyphaceae(100);g__Proliferodiscus(100);s__Proliferodiscus_sp_TNS_F17436(100); | Proliferodiscus | 13 | Saprotroph | Undefined Saprotroph-\|Wood Saprotroph\| | Microfungus | NULL | Probable | Decay wood (Põlme et al. 2020) | Tedersoo L, et al. 2014. Science 346:e1256688 ((https://doi.org/10.1126/science.1256688)); Põlme S, et al. 2020. Fungal Diversity 105:1-16 ((https://doi.org/10.1007/s13225-020-00466-2)) |
| Otu03838 | 9 | k__Fungi(100);p__Ascomycota(100);c__Eurotiomycetes(100);o__Chaetothyriales(100);f__Herpotrichiellaceae(100);g__Phaeomoniella(100);g__Phaeomoniella_unclassified(100); | Phaeomoniella | 13 | Pathotroph | \|Plant Pathogen\| | Dimorphic-Microfungus; Facultative Yeast (Tedersoo et | NULL | Highly Probable | Pathotroph isolated from necrosis in Prunus wood (Damm et al. 2010) | Damm U, et al. 2010. Persoonia 24:60-80 ((http://doi.org/10.3767/003158510X500705)); Tedersoo L, et al. 2014. Science 346:e1256688 ((https://doi.org/10.1126/science.1256688)); Põlme S, et al. 2020. Fungal Diversity 105:1-16 ((https://doi.org/10.1007/s13225-020-00466-2)) |
| Otu03839 | 9 | k__Fungi(100);p__Ascomycota(100);c__Sordariomycetes(100);o__Sordariales(100);f__Lasiosphaeriaceae(100);g__Zopfiella(100);g__Zopfiella_unclassified(100); | Zopfiella | 13 | Saprotroph | \|Dung Saprotroph\| | Microfungus | NULL | Highly Probable | Decay dung (Põlme et al. 2020) | Bills GF, et al. 2013. Current Opinion in Microbiology 16:549-565 ((https://doi.org/10.1016/j.mib.2013.08.001)); Tedersoo L, et al. 2014. Science 346:e1256688 ((https://doi.org/10.1126/science.1256688)); Põlme S, et al. 2020. Fungal Diversity 105:1-16 ((https://doi.org/10.1007/s13225-020-00466-2)) |
| Otu03840 | 9 | k__Fungi(100);p__Ascomycota(100);c__Saccharomycetes(100);o__Saccharomycetales(100);f__Metschnikowiaceae(100);g__Metschnikowia(100);g__Metschnikowia_unclassified(56); | Metschnikowia | 13 | Saprotroph | \|Undefined Saprotroph\| | Yeast | NULL | Probable | As nectar-tap_saprotroph (Põlme et al. 2020); Decay sugar-rich substrates (Põlme et al. 2020) | Põlme S, et al. 2020. Fungal Diversity 105:1-16 ((https://doi.org/10.1007/s13225-020-00466-2)) |
| Otu03841 | 9 | k__Fungi(100);p__Basidiomycota(100);c__Tremellomycetes(100);o__Cystofilobasidiales(100);f__Cystofilobasidiaceae(100);g__Cystofilobasidium(100);s__Cystofilobasidium_infirmominiatum(100); | Cystofilobasidium | 13 | Saprotroph | \|Plant Saprotroph\| | Yeast | NULL | Possible | As litter_saprotroph (Põlme et al. 2020); Decay leaf, fruit, seed (Põlme et al. 2020) | Oberwinkler F, et al. 1983. Systematic and Applied Microbiology 41:14-122 ((https://doi.org/10.1016/S0723-2020((83))80039-3)) |
| Otu03842 | 9 | k__Fungi(100);p__Ascomycota(100);c__Eurotiomycetes(100);o__Chaetothyriales(100);f__Herpotrichiellaceae(100);g__Capronia(100);s__Capronia_pilosella(100); | Capronia | 13 | Pathotroph-Saprotroph-Symbiotroph | Animal Pathogen-Endophyte-Fungal Parasite-\|Undefined Saprotroph\| | Dimorphic-Facultative Yeast | NULL | Highly Probable | As mycoparasite (Põlme et al. 2020); As root_endophyte_dark_septate (Põlme et al. 2020); As soil_saprotroph (Põlme et al. 2020); Dark Septate Endophyte, known from plant roots.; Decay soil (Põlme et al. 2020) | Tedersoo L, et al. 2014. Science 346:e1256688 ((https://doi.org/10.1126/science.1256688)); Põlme S, et al. 2020. Fungal Diversity 105:1-16 ((https://doi.org/10.1007/s13225-020-00466-2)) |
| Otu03846 | 9 | k__Fungi(100);p__Basidiomycota(100);c__Exobasidiomycetes(100);o__Microstromatales(100);f__Microstromataceae(100);g__Microstroma(100);s__Microstroma_juglandis(100); | Microstroma | 13 | Pathotroph | \|Plant Pathogen\| | Dimorphic | NULL | Probable | NULL | Tedersoo L, et al. 2014. Science 346:e1256688 ((https://doi.org/10.1126/science.1256688)); Põlme S, et al. 2020. Fungal Diversity 105:1-16 ((https://doi.org/10.1007/s13225-020-00466-2)) |
| Otu03848 | 9 | k__Fungi(100);p__Zygomycota(100);c__Zygomycota_class_Incertae_sedis(100);o__Mortierellales(100);f__Mortierellaceae(100);g__Mortierella(100);s__Mortierella_sp(100); | Mortierella | 13 | Saprotroph-Symbiotroph | Endophyte-Plant Saprotroph-\|Undefined Saprotroph\| | Microfungus | NULL | Probable | As chitinolytic (Põlme et al. 2020); As soil_saprotroph (Põlme et al. 2020); Decay roots, soil (Põlme et al. 2020); Saprobic in soil and thus frequenctly isolated from soil and material contaminated with soil such as hay or rotten silage and spoil tips (Cannon & Kirk 2007) | Cannon PF, Kirk PM. 2007. Fungal Families of the World. CAB International, Cambridge (ISBN: 978-0851998275); Tedersoo L, et al. 2014. Science 346:e1256688 ((https://doi.org/10.1126/science.1256688)); Purahong W, et al. 2016. Molecular Ecology 25:4059-4074 ((https://doi.org/10.1111/mec.13739)); Põlme S, et al. 2020. Fungal Diversity 105:1-16 ((https://doi.org/10.1007/s13225-020-00466-2)) |
| Otu03849 | 9 | k__Fungi(100);p__Ascomycota(100);c__Saccharomycetes(100);o__Saccharomycetales(100);f__Metschnikowiaceae(100);g__Metschnikowia(100);s__Metschnikowia_pulcherrima(67); | Metschnikowia | 13 | Saprotroph | \|Undefined Saprotroph\| | Yeast | NULL | Probable | As nectar-tap_saprotroph (Põlme et al. 2020); Decay sugar-rich substrates (Põlme et al. 2020) | Põlme S, et al. 2020. Fungal Diversity 105:1-16 ((https://doi.org/10.1007/s13225-020-00466-2)) |
| Otu03850 | 9 | k__Fungi(100);p__Basidiomycota(100);c__Microbotryomycetes(100);o__Sporidiobolales(100);f__Sporidiobolales_family_Incertae_sedis(100);g__Rhodotorula(100);g__Rhodotorula_unclassified(100); | Rhodotorula | 13 | Pathotroph-Saprotroph-Symbiotroph | Animal Pathogen-Animal Symbiotroph-Endophyte-Plant Pathogen-\|Undefined Saprotroph\| | Yeast | NULL | Probable | As foliar_endophyte (Põlme et al. 2020); Opportunistic human pathogen | Põlme S, et al. 2020. Fungal Diversity 105:1-16 ((https://doi.org/10.1007/s13225-020-00466-2)); https://www.cdc.gov ((https://www.cdc.gov)) |
| Otu03851 | 9 | k__Fungi(100);p__Ascomycota(100);c__Saccharomycetes(100);o__Saccharomycetales(100);f__Metschnikowiaceae(100);g__Metschnikowia(100);g__Metschnikowia_unclassified(78); | Metschnikowia | 13 | Saprotroph | \|Undefined Saprotroph\| | Yeast | NULL | Probable | As nectar-tap_saprotroph (Põlme et al. 2020); Decay sugar-rich substrates (Põlme et al. 2020) | Põlme S, et al. 2020. Fungal Diversity 105:1-16 ((https://doi.org/10.1007/s13225-020-00466-2)) |
| Otu03854 | 9 | k__Fungi(100);p__Basidiomycota(100);c__Microbotryomycetes(100);o__Sporidiobolales(100);f__Sporidiobolales_family_Incertae_sedis(100);g__Rhodotorula(100);s__Rhodotorula_mucilaginosa(78); | Rhodotorula | 13 | Pathotroph-Saprotroph-Symbiotroph | Animal Pathogen-Animal Symbiotroph-Endophyte-Plant Pathogen-\|Undefined Saprotroph\| | Yeast | NULL | Probable | As foliar_endophyte (Põlme et al. 2020); Opportunistic human pathogen | Põlme S, et al. 2020. Fungal Diversity 105:1-16 ((https://doi.org/10.1007/s13225-020-00466-2)); https://www.cdc.gov ((https://www.cdc.gov)) |
| Otu03862 | 9 | k__Fungi(100);p__Ascomycota(100);c__Dothideomycetes(100);o__Capnodiales(100);f__Davidiellaceae(100);g__Cladosporium(100);g__Cladosporium_unclassified(100); | Cladosporium | 13 | Pathotroph-Saprotroph-Symbiotroph | Animal Pathogen-Endophyte-Lichen Parasite-Plant Pathogen-\|Plant Saprotroph\|-Wood Saprotroph | Microfungus | NULL | Possible | As foliar_endophyte (Põlme et al. 2020); As litter_saprotroph (Põlme et al. 2020); Decay wood, leaf, fruit, seed, soil, dung, animal material (Põlme et al. 2020) | Seehann G, et al. 1975. List of Fungi in Soft-Rot Tissue Tests. Institute fur Holzbiologie und Holzschutz, Bundesforschungsanstalt fur Forst - und Holzwirtshaft, Hamburg - Lohbrugge, Germany; Lawrey JD, Diederich P. 2016. http://www.lichenicolous.net (accessed 24 May 2017) ((http://www.lichenicolous.net)); Põlme S, et al. 2020. Fungal Diversity 105:1-16 ((https://doi.org/10.1007/s13225-020-00466-2)); The American Phytopathological Society Common Names of Plant Diseases ((http://www.apsnet.org/publications/commonnames/Pages/default.aspx)) |
| Otu03583 | 9 | k__Fungi(100);p__Ascomycota(56);p__Ascomycota_unclassified(56);p__Ascomycota_unclassified(56);p__Ascomycota_unclassified(56);p__Ascomycota_unclassified(56);p__Ascomycota_unclassified(56); | - | - | - | - | - | - | - | Unassigned | - |
| Otu03591 | 9 | k__Fungi(100);p__Ascomycota(56);p__Ascomycota_unclassified(56);p__Ascomycota_unclassified(56);p__Ascomycota_unclassified(56);p__Ascomycota_unclassified(56);p__Ascomycota_unclassified(56); | - | - | - | - | - | - | - | Unassigned | - |
| Otu03592 | 9 | k__Fungi(100);p__Ascomycota(78);p__Ascomycota_unclassified(78);p__Ascomycota_unclassified(78);p__Ascomycota_unclassified(78);p__Ascomycota_unclassified(78);p__Ascomycota_unclassified(78); | - | - | - | - | - | - | - | Unassigned | - |
| Otu03593 | 9 | k__Fungi(100);p__Basidiomycota(100);c__Agaricomycetes(100);c__Agaricomycetes_unclassified(100);c__Agaricomycetes_unclassified(100);c__Agaricomycetes_unclassified(100);c__Agaricomycetes_unclassified(100); | - | - | - | - | - | - | - | Unassigned | - |
| Otu03595 | 9 | k__Fungi(100);p__Ascomycota(56);p__Ascomycota_unclassified(56);p__Ascomycota_unclassified(56);p__Ascomycota_unclassified(56);p__Ascomycota_unclassified(56);p__Ascomycota_unclassified(56); | - | - | - | - | - | - | - | Unassigned | - |
| Otu03596 | 9 | k__Fungi(100);k__Fungi_unclassified(56);k__Fungi_unclassified(56);k__Fungi_unclassified(56);k__Fungi_unclassified(56);k__Fungi_unclassified(56);k__Fungi_unclassified(56); | - | - | - | - | - | - | - | Unassigned | - |
| Otu03600 | 9 | k__Fungi(100);p__Ascomycota(100);p__Ascomycota_unclassified(100);p__Ascomycota_unclassified(100);p__Ascomycota_unclassified(100);p__Ascomycota_unclassified(100);p__Ascomycota_unclassified(100); | - | - | - | - | - | - | - | Unassigned | - |
| Otu03605 | 9 | k__Fungi(100);p__Ascomycota(89);p__Ascomycota_unclassified(89);p__Ascomycota_unclassified(89);p__Ascomycota_unclassified(89);p__Ascomycota_unclassified(89);p__Ascomycota_unclassified(89); | - | - | - | - | - | - | - | Unassigned | - |
| Otu03606 | 9 | k__Fungi(100);p__Ascomycota(100);c__Dothideomycetes(100);o__Pleosporales(100);o__Pleosporales_unclassified(100);o__Pleosporales_unclassified(100);o__Pleosporales_unclassified(100); | - | - | - | - | - | - | - | Unassigned | - |
| Otu03607 | 9 | k__Fungi(100);k__Fungi_unclassified(56);k__Fungi_unclassified(56);k__Fungi_unclassified(56);k__Fungi_unclassified(56);k__Fungi_unclassified(56);k__Fungi_unclassified(56); | - | - | - | - | - | - | - | Unassigned | - |
| Otu03610 | 9 | k__Fungi(100);p__Ascomycota(56);p__Ascomycota_unclassified(56);p__Ascomycota_unclassified(56);p__Ascomycota_unclassified(56);p__Ascomycota_unclassified(56);p__Ascomycota_unclassified(56); | - | - | - | - | - | - | - | Unassigned | - |
| Otu03615 | 9 | k__Fungi(100);k__Fungi_unclassified(78);k__Fungi_unclassified(78);k__Fungi_unclassified(78);k__Fungi_unclassified(78);k__Fungi_unclassified(78);k__Fungi_unclassified(78); | - | - | - | - | - | - | - | Unassigned | - |
| Otu03616 | 9 | k__Fungi(100);p__Ascomycota(89);p__Ascomycota_unclassified(89);p__Ascomycota_unclassified(89);p__Ascomycota_unclassified(89);p__Ascomycota_unclassified(89);p__Ascomycota_unclassified(89); | - | - | - | - | - | - | - | Unassigned | - |
| Otu03618 | 9 | k__Fungi(100);k__Fungi_unclassified(89);k__Fungi_unclassified(89);k__Fungi_unclassified(89);k__Fungi_unclassified(89);k__Fungi_unclassified(89);k__Fungi_unclassified(89); | - | - | - | - | - | - | - | Unassigned | - |
| Otu03619 | 9 | k__Fungi(100);p__Ascomycota(100);c__Dothideomycetes(89);o__Capnodiales(56);o__Capnodiales_unclassified(56);o__Capnodiales_unclassified(56);o__Capnodiales_unclassified(56); | - | - | - | - | - | - | - | Unassigned | - |
| Otu03620 | 9 | k__Fungi(100);p__Basidiomycota(100);c__Agaricomycetes(100);c__Agaricomycetes_unclassified(100);c__Agaricomycetes_unclassified(100);c__Agaricomycetes_unclassified(100);c__Agaricomycetes_unclassified(100); | - | - | - | - | - | - | - | Unassigned | - |
| Otu03621 | 9 | k__Fungi(100);p__Basidiomycota(89);c__Agaricomycetes(89);c__Agaricomycetes_unclassified(89);c__Agaricomycetes_unclassified(89);c__Agaricomycetes_unclassified(89);c__Agaricomycetes_unclassified(89); | - | - | - | - | - | - | - | Unassigned | - |
| Otu03622 | 9 | k__Fungi(100);p__Ascomycota(67);p__Ascomycota_unclassified(67);p__Ascomycota_unclassified(67);p__Ascomycota_unclassified(67);p__Ascomycota_unclassified(67);p__Ascomycota_unclassified(67); | - | - | - | - | - | - | - | Unassigned | - |
| Otu03624 | 9 | k__Fungi(100);p__Ascomycota(67);p__Ascomycota_unclassified(67);p__Ascomycota_unclassified(67);p__Ascomycota_unclassified(67);p__Ascomycota_unclassified(67);p__Ascomycota_unclassified(67); | - | - | - | - | - | - | - | Unassigned | - |
| Otu03625 | 9 | k__Fungi(100);k__Fungi_unclassified(100);k__Fungi_unclassified(100);k__Fungi_unclassified(100);k__Fungi_unclassified(100);k__Fungi_unclassified(100);k__Fungi_unclassified(100); | - | - | - | - | - | - | - | Unassigned | - |
| Otu03626 | 9 | k__Fungi(100);k__Fungi_unclassified(56);k__Fungi_unclassified(56);k__Fungi_unclassified(56);k__Fungi_unclassified(56);k__Fungi_unclassified(56);k__Fungi_unclassified(56); | - | - | - | - | - | - | - | Unassigned | - |
| Otu03631 | 9 | k__Fungi(100);p__Basidiomycota(100);p__Basidiomycota_unclassified(100);p__Basidiomycota_unclassified(100);p__Basidiomycota_unclassified(100);p__Basidiomycota_unclassified(100);p__Basidiomycota_unclassified(100); | - | - | - | - | - | - | - | Unassigned | - |
| Otu03634 | 9 | k__Fungi(100);p__Ascomycota(100);c__Sordariomycetes(100);c__Sordariomycetes_unclassified(100);c__Sordariomycetes_unclassified(100);c__Sordariomycetes_unclassified(100);c__Sordariomycetes_unclassified(100); | - | - | - | - | - | - | - | Unassigned | - |
| Otu03637 | 9 | k__Fungi(100);p__Ascomycota(100);p__Ascomycota_unclassified(100);p__Ascomycota_unclassified(100);p__Ascomycota_unclassified(100);p__Ascomycota_unclassified(100);p__Ascomycota_unclassified(100); | - | - | - | - | - | - | - | Unassigned | - |
| Otu03643 | 9 | k__Fungi(100);p__Ascomycota(56);p__Ascomycota_unclassified(56);p__Ascomycota_unclassified(56);p__Ascomycota_unclassified(56);p__Ascomycota_unclassified(56);p__Ascomycota_unclassified(56); | - | - | - | - | - | - | - | Unassigned | - |
| Otu03646 | 9 | k__Fungi(100);k__Fungi_unclassified(100);k__Fungi_unclassified(100);k__Fungi_unclassified(100);k__Fungi_unclassified(100);k__Fungi_unclassified(100);k__Fungi_unclassified(100); | - | - | - | - | - | - | - | Unassigned | - |
| Otu03648 | 9 | k__Fungi(100);p__Ascomycota(100);p__Ascomycota_unclassified(100);p__Ascomycota_unclassified(100);p__Ascomycota_unclassified(100);p__Ascomycota_unclassified(100);p__Ascomycota_unclassified(100); | - | - | - | - | - | - | - | Unassigned | - |
| Otu03649 | 9 | k__Fungi(100);k__Fungi_unclassified(56);k__Fungi_unclassified(56);k__Fungi_unclassified(56);k__Fungi_unclassified(56);k__Fungi_unclassified(56);k__Fungi_unclassified(56); | - | - | - | - | - | - | - | Unassigned | - |
| Otu03651 | 9 | k__Fungi(100);p__Ascomycota(100);c__Sordariomycetes(100);o__Xylariales(100);f__Xylariales_family_Incertae_sedis(100);f__Xylariales_family_Incertae_sedis_unclassified(100);f__Xylariales_family_Incertae_sedis_unclassified(100); | - | - | - | - | - | - | - | Unassigned | - |
| Otu03652 | 9 | k__Fungi(100);p__Ascomycota(89);p__Ascomycota_unclassified(89);p__Ascomycota_unclassified(89);p__Ascomycota_unclassified(89);p__Ascomycota_unclassified(89);p__Ascomycota_unclassified(89); | - | - | - | - | - | - | - | Unassigned | - |
| Otu03654 | 9 | k__Fungi(100);p__Ascomycota(100);p__Ascomycota_unclassified(100);p__Ascomycota_unclassified(100);p__Ascomycota_unclassified(100);p__Ascomycota_unclassified(100);p__Ascomycota_unclassified(100); | - | - | - | - | - | - | - | Unassigned | - |
| Otu03655 | 9 | k__Fungi(100);p__Ascomycota(100);c__Leotiomycetes(100);o__Helotiales(100);o__Helotiales_unclassified(100);o__Helotiales_unclassified(100);o__Helotiales_unclassified(100); | - | - | - | - | - | - | - | Unassigned | - |
| Otu03656 | 9 | k__Plantae(100);p__unclassified_Plantae(100);c__unclassified_Plantae(100);o__unclassified_Plantae(100);f__unclassified_Plantae(100);g__unclassified_Plantae(100);s__Plantae_sp(100); | - | - | - | - | - | - | - | Unassigned | - |
| Otu03658 | 9 | k__Fungi(100);p__Ascomycota(100);c__Dothideomycetes(100);o__Dothideales(100);o__Dothideales_unclassified(100);o__Dothideales_unclassified(100);o__Dothideales_unclassified(100); | - | - | - | - | - | - | - | Unassigned | - |
| Otu03659 | 9 | k__Fungi(100);p__Ascomycota(100);p__Ascomycota_unclassified(100);p__Ascomycota_unclassified(100);p__Ascomycota_unclassified(100);p__Ascomycota_unclassified(100);p__Ascomycota_unclassified(100); | - | - | - | - | - | - | - | Unassigned | - |
| Otu03661 | 9 | k__Fungi(100);p__Ascomycota(100);c__Dothideomycetes(100);o__Capnodiales(100);f__Davidiellaceae(100);f__Davidiellaceae_unclassified(100);f__Davidiellaceae_unclassified(100); | - | - | - | - | - | - | - | Unassigned | - |
| Otu03662 | 9 | k__Fungi(100);k__Fungi_unclassified(78);k__Fungi_unclassified(78);k__Fungi_unclassified(78);k__Fungi_unclassified(78);k__Fungi_unclassified(78);k__Fungi_unclassified(78); | - | - | - | - | - | - | - | Unassigned | - |
| Otu03665 | 9 | k__Fungi(100);p__Ascomycota(100);c__Dothideomycetes(100);o__Pleosporales(100);f__unclassified_Pleosporales(78);g__unclassified_Pleosporales(78);s__Pleosporales_sp(78); | - | - | - | - | - | - | - | Unassigned | - |
| Otu03668 | 9 | k__Fungi(100);p__Ascomycota(100);c__Dothideomycetes(100);o__Capnodiales(89);o__Capnodiales_unclassified(89);o__Capnodiales_unclassified(89);o__Capnodiales_unclassified(89); | - | - | - | - | - | - | - | Unassigned | - |
| Otu03669 | 9 | k__Fungi(100);p__Basidiomycota(100);c__Microbotryomycetes(100);o__Sporidiobolales(100);f__Sporidiobolales_family_Incertae_sedis(100);f__Sporidiobolales_family_Incertae_sedis_unclassified(89);f__Sporidiobolales_family_Incertae_sedis_unclassified(89); | - | - | - | - | - | - | - | Unassigned | - |
| Otu03670 | 9 | k__Fungi(100);k__Fungi_unclassified(78);k__Fungi_unclassified(78);k__Fungi_unclassified(78);k__Fungi_unclassified(78);k__Fungi_unclassified(78);k__Fungi_unclassified(78); | - | - | - | - | - | - | - | Unassigned | - |
| Otu03676 | 9 | k__Fungi(100);p__Ascomycota(100);c__Lecanoromycetes(100);o__Teloschistales(56);o__Teloschistales_unclassified(56);o__Teloschistales_unclassified(56);o__Teloschistales_unclassified(56); | - | - | - | - | - | - | - | Unassigned | - |
| Otu03678 | 9 | k__Fungi(100);k__Fungi_unclassified(89);k__Fungi_unclassified(89);k__Fungi_unclassified(89);k__Fungi_unclassified(89);k__Fungi_unclassified(89);k__Fungi_unclassified(89); | - | - | - | - | - | - | - | Unassigned | - |
| Otu03679 | 9 | k__Fungi(100);p__Ascomycota(100);c__Dothideomycetes(100);o__Pleosporales(100);o__Pleosporales_unclassified(100);o__Pleosporales_unclassified(100);o__Pleosporales_unclassified(100); | - | - | - | - | - | - | - | Unassigned | - |
| Otu03680 | 9 | k__Fungi(100);p__Ascomycota(100);c__Dothideomycetes(100);o__Pleosporales(100);f__unclassified_Pleosporales(100);g__unclassified_Pleosporales(100);s__Pleosporales_sp(100); | - | - | - | - | - | - | - | Unassigned | - |
| Otu03682 | 9 | k__Fungi(100);p__Ascomycota(100);c__Dothideomycetes(100);c__Dothideomycetes_unclassified(100);c__Dothideomycetes_unclassified(100);c__Dothideomycetes_unclassified(100);c__Dothideomycetes_unclassified(100); | - | - | - | - | - | - | - | Unassigned | - |
| Otu03683 | 9 | k__Fungi(100);p__Basidiomycota(100);c__Agaricomycetes(100);o__Thelephorales(100);o__Thelephorales_unclassified(100);o__Thelephorales_unclassified(100);o__Thelephorales_unclassified(100); | - | - | - | - | - | - | - | Unassigned | - |
| Otu03684 | 9 | k__Fungi(100);k__Fungi_unclassified(89);k__Fungi_unclassified(89);k__Fungi_unclassified(89);k__Fungi_unclassified(89);k__Fungi_unclassified(89);k__Fungi_unclassified(89); | - | - | - | - | - | - | - | Unassigned | - |
| Otu03685 | 9 | k__Fungi(100);p__Ascomycota(100);c__Leotiomycetes(89);c__Leotiomycetes_unclassified(89);c__Leotiomycetes_unclassified(89);c__Leotiomycetes_unclassified(89);c__Leotiomycetes_unclassified(89); | - | - | - | - | - | - | - | Unassigned | - |
| Otu03686 | 9 | k__Plantae(100);p__unclassified_Plantae(100);c__unclassified_Plantae(100);o__unclassified_Plantae(100);f__unclassified_Plantae(100);g__unclassified_Plantae(100);s__Plantae_sp(100); | - | - | - | - | - | - | - | Unassigned | - |
| Otu03687 | 9 | k__Fungi(100);k__Fungi_unclassified(100);k__Fungi_unclassified(100);k__Fungi_unclassified(100);k__Fungi_unclassified(100);k__Fungi_unclassified(100);k__Fungi_unclassified(100); | - | - | - | - | - | - | - | Unassigned | - |
| Otu03689 | 9 | k__Fungi(100);p__Ascomycota(100);c__Dothideomycetes(100);o__Pleosporales(100);o__Pleosporales_unclassified(78);o__Pleosporales_unclassified(78);o__Pleosporales_unclassified(78); | - | - | - | - | - | - | - | Unassigned | - |

| | | | | | | | | | | | |
|---|---|---|---|---|---|---|---|---|---|---|---|
| Otu03692 | 9 | k__Fungi(100);k__Fungi_unclassified(67);k__Fungi_unclassified(67);k__Fungi_unclassified(67);k__Fungi_unclassified(67);k__Fungi_unclassified(67); | - | - | - | - | - | - | - | Unassigned | - |
| Otu03694 | 9 | k__Fungi(100);p__Ascomycota(67);p__Ascomycota_unclassified(67);p__Ascomycota_unclassified(67);p__Ascomycota_unclassified(67);p__Ascomycota_unclassified(67);p__Ascomycota_unclassified(67); | - | - | - | - | - | - | - | Unassigned | - |
| Otu03697 | 9 | k__Fungi(100);p__Ascomycota(67);p__Ascomycota_unclassified(67);p__Ascomycota_unclassified(67);p__Ascomycota_unclassified(67);p__Ascomycota_unclassified(67);p__Ascomycota_unclassified(67); | - | - | - | - | - | - | - | Unassigned | - |
| Otu03698 | 9 | k__Fungi(100);p__Basidiomycota(100);c__Agaricomycetes(100);o__Trechisporales(100);f__Hydnodontaceae(56);f__Hydnodontaceae_unclassified(56);f__Hydnodontaceae_unclassified(56); | - | - | - | - | - | - | - | Unassigned | - |
| Otu03701 | 9 | k__Fungi(100);p__Ascomycota(100);c__Lecanoromycetes(100);o__Teloschistales(78);o__Teloschistales_unclassified(78);o__Teloschistales_unclassified(78);o__Teloschistales_unclassified(78); | - | - | - | - | - | - | - | Unassigned | - |
| Otu03703 | 9 | k__Fungi(100);p__Ascomycota(67);p__Ascomycota_unclassified(67);p__Ascomycota_unclassified(67);p__Ascomycota_unclassified(67);p__Ascomycota_unclassified(67);p__Ascomycota_unclassified(67); | - | - | - | - | - | - | - | Unassigned | - |
| Otu03705 | 9 | k__Fungi(100);k__Fungi_unclassified(100);k__Fungi_unclassified(100);k__Fungi_unclassified(100);k__Fungi_unclassified(100);k__Fungi_unclassified(100);k__Fungi_unclassified(100); | - | - | - | - | - | - | - | Unassigned | - |
| Otu03706 | 9 | k__Fungi(100);k__Fungi_unclassified(67);k__Fungi_unclassified(67);k__Fungi_unclassified(67);k__Fungi_unclassified(67);k__Fungi_unclassified(67);k__Fungi_unclassified(67); | - | - | - | - | - | - | - | Unassigned | - |
| Otu03707 | 9 | k__Fungi(100);k__Fungi_unclassified(100);k__Fungi_unclassified(100);k__Fungi_unclassified(100);k__Fungi_unclassified(100);k__Fungi_unclassified(100);k__Fungi_unclassified(100); | - | - | - | - | - | - | - | Unassigned | - |
| Otu03709 | 9 | k__Fungi(100);k__Fungi_unclassified(100);k__Fungi_unclassified(100);k__Fungi_unclassified(100);k__Fungi_unclassified(100);k__Fungi_unclassified(100);k__Fungi_unclassified(100); | - | - | - | - | - | - | - | Unassigned | - |
| Otu03713 | 9 | k__Fungi(100);p__Ascomycota(56);p__Ascomycota_unclassified(56);p__Ascomycota_unclassified(56);p__Ascomycota_unclassified(56);p__Ascomycota_unclassified(56);p__Ascomycota_unclassified(56); | - | - | - | - | - | - | - | Unassigned | - |
| Otu03714 | 9 | k__Fungi(100);k__Fungi_unclassified(78);k__Fungi_unclassified(78);k__Fungi_unclassified(78);k__Fungi_unclassified(78);k__Fungi_unclassified(78);k__Fungi_unclassified(78); | - | - | - | - | - | - | - | Unassigned | - |
| Otu03715 | 9 | k__Fungi(100);k__Fungi_unclassified(56);k__Fungi_unclassified(56);k__Fungi_unclassified(56);k__Fungi_unclassified(56);k__Fungi_unclassified(56);k__Fungi_unclassified(56); | - | - | - | - | - | - | - | Unassigned | - |
| Otu03717 | 9 | k__Fungi(100);k__Fungi_unclassified(56);k__Fungi_unclassified(56);k__Fungi_unclassified(56);k__Fungi_unclassified(56);k__Fungi_unclassified(56);k__Fungi_unclassified(56); | - | - | - | - | - | - | - | Unassigned | - |
| Otu03723 | 9 | k__Fungi(100);p__Ascomycota(89);p__Ascomycota_unclassified(89);p__Ascomycota_unclassified(89);p__Ascomycota_unclassified(89);p__Ascomycota_unclassified(89);p__Ascomycota_unclassified(89); | - | - | - | - | - | - | - | Unassigned | - |
| Otu03727 | 9 | k__Fungi(100);k__Fungi_unclassified(100);k__Fungi_unclassified(100);k__Fungi_unclassified(100);k__Fungi_unclassified(100);k__Fungi_unclassified(100);k__Fungi_unclassified(100); | - | - | - | - | - | - | - | Unassigned | - |
| Otu03730 | 9 | k__Fungi(100);p__Ascomycota(56);p__Ascomycota_unclassified(56);p__Ascomycota_unclassified(56);p__Ascomycota_unclassified(56);p__Ascomycota_unclassified(56);p__Ascomycota_unclassified(56); | - | - | - | - | - | - | - | Unassigned | - |
| Otu03731 | 9 | k__Fungi(100);k__Fungi_unclassified(67);k__Fungi_unclassified(67);k__Fungi_unclassified(67);k__Fungi_unclassified(67);k__Fungi_unclassified(67);k__Fungi_unclassified(67); | - | - | - | - | - | - | - | Unassigned | - |
| Otu03735 | 9 | k__Fungi(100);p__Ascomycota(100);c__Dothideomycetes(100);o__Pleosporales(100);o__Pleosporales_unclassified(78);o__Pleosporales_unclassified(78);o__Pleosporales_unclassified(78); | - | - | - | - | - | - | - | Unassigned | - |
| Otu03740 | 9 | k__Fungi(100);p__Ascomycota(100);p__Ascomycota_unclassified(100);p__Ascomycota_unclassified(100);p__Ascomycota_unclassified(100);p__Ascomycota_unclassified(100);p__Ascomycota_unclassified(100); | - | - | - | - | - | - | - | Unassigned | - |
| Otu03742 | 9 | k__Fungi(100);k__Fungi_unclassified(78);k__Fungi_unclassified(78);k__Fungi_unclassified(78);k__Fungi_unclassified(78);k__Fungi_unclassified(78);k__Fungi_unclassified(78); | - | - | - | - | - | - | - | Unassigned | - |
| Otu03743 | 9 | k__Fungi(100);k__Fungi_unclassified(56);k__Fungi_unclassified(56);k__Fungi_unclassified(56);k__Fungi_unclassified(56);k__Fungi_unclassified(56);k__Fungi_unclassified(56); | - | - | - | - | - | - | - | Unassigned | - |
| Otu03744 | 9 | k__Fungi(100);k__Fungi_unclassified(56);k__Fungi_unclassified(56);k__Fungi_unclassified(56);k__Fungi_unclassified(56);k__Fungi_unclassified(56);k__Fungi_unclassified(56); | - | - | - | - | - | - | - | Unassigned | - |
| Otu03747 | 9 | k__Fungi(100);p__Basidiomycota(100);c__Agaricomycetes(100);o__Polyporales(100);o__Polyporales_unclassified(67);o__Polyporales_unclassified(67);o__Polyporales_unclassified(67); | - | - | - | - | - | - | - | Unassigned | - |
| Otu03748 | 9 | k__Fungi(100);p__Ascomycota(67);p__Ascomycota_unclassified(67);p__Ascomycota_unclassified(67);p__Ascomycota_unclassified(67);p__Ascomycota_unclassified(67);p__Ascomycota_unclassified(67); | - | - | - | - | - | - | - | Unassigned | - |
| Otu03749 | 9 | k__Fungi(100);k__Fungi_unclassified(56);k__Fungi_unclassified(56);k__Fungi_unclassified(56);k__Fungi_unclassified(56);k__Fungi_unclassified(56);k__Fungi_unclassified(56); | - | - | - | - | - | - | - | Unassigned | - |
| Otu03750 | 9 | k__Fungi(100);k__Fungi_unclassified(100);k__Fungi_unclassified(100);k__Fungi_unclassified(100);k__Fungi_unclassified(100);k__Fungi_unclassified(100);k__Fungi_unclassified(100); | - | - | - | - | - | - | - | Unassigned | - |
| Otu03751 | 9 | k__Fungi(100);p__Ascomycota(56);p__Ascomycota_unclassified(56);p__Ascomycota_unclassified(56);p__Ascomycota_unclassified(56);p__Ascomycota_unclassified(56);p__Ascomycota_unclassified(56); | - | - | - | - | - | - | - | Unassigned | - |
| Otu03752 | 9 | k__Fungi(100);p__Ascomycota(89);p__Ascomycota_unclassified(89);p__Ascomycota_unclassified(89);p__Ascomycota_unclassified(89);p__Ascomycota_unclassified(89);p__Ascomycota_unclassified(89); | - | - | - | - | - | - | - | Unassigned | - |
| Otu03753 | 9 | k__Fungi(100);p__Basidiomycota(100);c__Agaricomycetes(100);o__Gomphales(100);o__Gomphales_unclassified(78);o__Gomphales_unclassified(78);o__Gomphales_unclassified(78); | - | - | - | - | - | - | - | Unassigned | - |
| Otu03754 | 9 | k__Fungi(100);p__Ascomycota(100);p__Ascomycota_unclassified(100);p__Ascomycota_unclassified(100);p__Ascomycota_unclassified(100);p__Ascomycota_unclassified(100);p__Ascomycota_unclassified(100); | - | - | - | - | - | - | - | Unassigned | - |
| Otu03755 | 9 | k__Fungi(100);k__Fungi_unclassified(67);k__Fungi_unclassified(67);k__Fungi_unclassified(67);k__Fungi_unclassified(67);k__Fungi_unclassified(67);k__Fungi_unclassified(67); | - | - | - | - | - | - | - | Unassigned | - |
| Otu03756 | 9 | k__Fungi(100);p__Ascomycota(100);p__Ascomycota_unclassified(100);p__Ascomycota_unclassified(100);p__Ascomycota_unclassified(100);p__Ascomycota_unclassified(100);p__Ascomycota_unclassified(100); | - | - | - | - | - | - | - | Unassigned | - |
| Otu03758 | 9 | k__Fungi(100);k__Fungi_unclassified(56);k__Fungi_unclassified(56);k__Fungi_unclassified(56);k__Fungi_unclassified(56);k__Fungi_unclassified(56);k__Fungi_unclassified(56); | - | - | - | - | - | - | - | Unassigned | - |
| Otu03759 | 9 | k__Fungi(100);k__Fungi_unclassified(56);k__Fungi_unclassified(56);k__Fungi_unclassified(56);k__Fungi_unclassified(56);k__Fungi_unclassified(56);k__Fungi_unclassified(56); | - | - | - | - | - | - | - | Unassigned | - |
| Otu03763 | 9 | k__Fungi(100);p__Ascomycota(78);p__Ascomycota_unclassified(78);p__Ascomycota_unclassified(78);p__Ascomycota_unclassified(78);p__Ascomycota_unclassified(78);p__Ascomycota_unclassified(78); | - | - | - | - | - | - | - | Unassigned | - |
| Otu03764 | 9 | k__Fungi(100);p__Ascomycota(100);p__Ascomycota_unclassified(100);p__Ascomycota_unclassified(100);p__Ascomycota_unclassified(100);p__Ascomycota_unclassified(100);p__Ascomycota_unclassified(100); | - | - | - | - | - | - | - | Unassigned | - |
| Otu03765 | 9 | k__Fungi(100);k__Fungi_unclassified(78);k__Fungi_unclassified(78);k__Fungi_unclassified(78);k__Fungi_unclassified(78);k__Fungi_unclassified(78);k__Fungi_unclassified(78); | - | - | - | - | - | - | - | Unassigned | - |
| Otu03767 | 9 | k__Fungi(100);p__Ascomycota(100);p__Ascomycota_unclassified(100);p__Ascomycota_unclassified(100);p__Ascomycota_unclassified(100);p__Ascomycota_unclassified(100);p__Ascomycota_unclassified(100); | - | - | - | - | - | - | - | Unassigned | - |
| Otu03768 | 9 | k__Fungi(100);p__Ascomycota(56);p__Ascomycota_unclassified(56);p__Ascomycota_unclassified(56);p__Ascomycota_unclassified(56);p__Ascomycota_unclassified(56);p__Ascomycota_unclassified(56); | - | - | - | - | - | - | - | Unassigned | - |
| Otu03769 | 9 | k__Fungi(100);k__Fungi_unclassified(78);k__Fungi_unclassified(78);k__Fungi_unclassified(78);k__Fungi_unclassified(78);k__Fungi_unclassified(78);k__Fungi_unclassified(78); | - | - | - | - | - | - | - | Unassigned | - |
| Otu03773 | 9 | k__Fungi(100);p__Ascomycota(89);p__Ascomycota_unclassified(89);p__Ascomycota_unclassified(89);p__Ascomycota_unclassified(89);p__Ascomycota_unclassified(89);p__Ascomycota_unclassified(89); | - | - | - | - | - | - | - | Unassigned | - |
| Otu03776 | 9 | k__Fungi(100);p__Ascomycota(100);c__Leotiomycetes(78);c__Leotiomycetes_unclassified(67);c__Leotiomycetes_unclassified(67);c__Leotiomycetes_unclassified(67);c__Leotiomycetes_unclassified(67); | - | - | - | - | - | - | - | Unassigned | - |
| Otu03777 | 9 | k__Fungi(100);p__Ascomycota(100);c__Dothideomycetes(100);o__Pleosporales(100);o__Pleosporales_unclassified(67);o__Pleosporales_unclassified(67);o__Pleosporales_unclassified(67); | - | - | - | - | - | - | - | Unassigned | - |
| Otu03778 | 9 | k__Fungi(100);p__Ascomycota(89);p__Ascomycota_unclassified(89);p__Ascomycota_unclassified(89);p__Ascomycota_unclassified(89);p__Ascomycota_unclassified(89);p__Ascomycota_unclassified(89); | - | - | - | - | - | - | - | Unassigned | - |
| Otu03781 | 9 | k__Fungi(100);p__Ascomycota(100);p__Ascomycota_unclassified(100);p__Ascomycota_unclassified(100);p__Ascomycota_unclassified(100);p__Ascomycota_unclassified(100);p__Ascomycota_unclassified(100); | - | - | - | - | - | - | - | Unassigned | - |
| Otu03782 | 9 | k__Fungi(100);p__Ascomycota(100);p__Ascomycota_unclassified(100);p__Ascomycota_unclassified(100);p__Ascomycota_unclassified(100);p__Ascomycota_unclassified(100);p__Ascomycota_unclassified(100); | - | - | - | - | - | - | - | Unassigned | - |
| Otu03783 | 9 | k__Fungi(100);p__Ascomycota(100);c__Leotiomycetes(100);o__Helotiales(100);o__Helotiales_unclassified(67);o__Helotiales_unclassified(67);o__Helotiales_unclassified(67); | - | - | - | - | - | - | - | Unassigned | - |
| Otu03784 | 9 | k__Fungi(100);p__Basidiomycota(100);c__Agaricomycetes(100);o__Polyporales(100);o__Polyporales_unclassified(100);o__Polyporales_unclassified(100);o__Polyporales_unclassified(100); | - | - | - | - | - | - | - | Unassigned | - |
| Otu03786 | 9 | k__Fungi(100);k__Fungi_unclassified(67);k__Fungi_unclassified(67);k__Fungi_unclassified(67);k__Fungi_unclassified(67);k__Fungi_unclassified(67);k__Fungi_unclassified(67); | - | - | - | - | - | - | - | Unassigned | - |
| Otu03788 | 9 | k__Fungi(100);p__Ascomycota(78);p__Ascomycota_unclassified(78);p__Ascomycota_unclassified(78);p__Ascomycota_unclassified(78);p__Ascomycota_unclassified(78);p__Ascomycota_unclassified(78); | - | - | - | - | - | - | - | Unassigned | - |
| Otu03789 | 9 | k__Fungi(100);p__Ascomycota(56);p__Ascomycota_unclassified(56);p__Ascomycota_unclassified(56);p__Ascomycota_unclassified(56);p__Ascomycota_unclassified(56);p__Ascomycota_unclassified(56); | - | - | - | - | - | - | - | Unassigned | - |
| Otu03791 | 9 | k__Fungi(100);p__Ascomycota(100);p__Ascomycota_unclassified(67);p__Ascomycota_unclassified(67);p__Ascomycota_unclassified(67);p__Ascomycota_unclassified(67);p__Ascomycota_unclassified(67); | - | - | - | - | - | - | - | Unassigned | - |
| Otu03792 | 9 | k__Fungi(100);p__Ascomycota(89);p__Ascomycota_unclassified(89);p__Ascomycota_unclassified(89);p__Ascomycota_unclassified(89);p__Ascomycota_unclassified(89);p__Ascomycota_unclassified(89); | - | - | - | - | - | - | - | Unassigned | - |
| Otu03795 | 9 | k__Fungi(100);k__Fungi_unclassified(89);k__Fungi_unclassified(89);k__Fungi_unclassified(89);k__Fungi_unclassified(89);k__Fungi_unclassified(89);k__Fungi_unclassified(89); | - | - | - | - | - | - | - | Unassigned | - |
| Otu03801 | 9 | k__Fungi(100);p__Ascomycota(56);p__Ascomycota_unclassified(56);p__Ascomycota_unclassified(56);p__Ascomycota_unclassified(56);p__Ascomycota_unclassified(56);p__Ascomycota_unclassified(56); | - | - | - | - | - | - | - | Unassigned | - |
| Otu03803 | 9 | k__Fungi(100);p__Basidiomycota(100);c__Agaricomycetes(100);c__Agaricomycetes_unclassified(100);c__Agaricomycetes_unclassified(100);c__Agaricomycetes_unclassified(100);c__Agaricomycetes_unclassified(100); | - | - | - | - | - | - | - | Unassigned | - |
| Otu03804 | 9 | k__Fungi(100);p__Ascomycota(78);p__Ascomycota_unclassified(78);p__Ascomycota_unclassified(78);p__Ascomycota_unclassified(78);p__Ascomycota_unclassified(78);p__Ascomycota_unclassified(78); | - | - | - | - | - | - | - | Unassigned | - |
| Otu03805 | 9 | k__Fungi(100);p__Basidiomycota(100);c__Agaricomycetes(100);c__Agaricomycetes_unclassified(100);c__Agaricomycetes_unclassified(100);c__Agaricomycetes_unclassified(100);c__Agaricomycetes_unclassified(100); | - | - | - | - | - | - | - | Unassigned | - |
| Otu03807 | 9 | k__Fungi(100);k__Fungi_unclassified(89);k__Fungi_unclassified(89);k__Fungi_unclassified(89);k__Fungi_unclassified(89);k__Fungi_unclassified(89);k__Fungi_unclassified(89); | - | - | - | - | - | - | - | Unassigned | - |
| Otu03808 | 9 | k__Fungi(100);p__Ascomycota(78);p__Ascomycota_unclassified(78);p__Ascomycota_unclassified(78);p__Ascomycota_unclassified(78);p__Ascomycota_unclassified(78);p__Ascomycota_unclassified(78); | - | - | - | - | - | - | - | Unassigned | - |
| Otu03809 | 9 | k__Fungi(100);k__Fungi_unclassified(100);k__Fungi_unclassified(100);k__Fungi_unclassified(100);k__Fungi_unclassified(100);k__Fungi_unclassified(100);k__Fungi_unclassified(100); | - | - | - | - | - | - | - | Unassigned | - |
| Otu03810 | 9 | k__Fungi(100);p__Ascomycota(100);p__Ascomycota_unclassified(100);p__Ascomycota_unclassified(100);p__Ascomycota_unclassified(100);p__Ascomycota_unclassified(100);p__Ascomycota_unclassified(100); | - | - | - | - | - | - | - | Unassigned | - |
| Otu03813 | 9 | k__Fungi(100);p__Ascomycota(100);c__Dothideomycetes(100);c__Dothideomycetes_unclassified(100);c__Dothideomycetes_unclassified(100);c__Dothideomycetes_unclassified(100);c__Dothideomycetes_unclassified(100); | - | - | - | - | - | - | - | Unassigned | - |
| Otu03815 | 9 | k__Fungi(100);k__Fungi_unclassified(78);k__Fungi_unclassified(78);k__Fungi_unclassified(78);k__Fungi_unclassified(78);k__Fungi_unclassified(78);k__Fungi_unclassified(78); | - | - | - | - | - | - | - | Unassigned | - |
| Otu03816 | 9 | k__Fungi(100);p__Ascomycota(78);p__Ascomycota_unclassified(78);p__Ascomycota_unclassified(78);p__Ascomycota_unclassified(78);p__Ascomycota_unclassified(78);p__Ascomycota_unclassified(78); | - | - | - | - | - | - | - | Unassigned | - |
| Otu03817 | 9 | k__Fungi(100);k__Fungi_unclassified(67);k__Fungi_unclassified(67);k__Fungi_unclassified(67);k__Fungi_unclassified(67);k__Fungi_unclassified(67);k__Fungi_unclassified(67); | - | - | - | - | - | - | - | Unassigned | - |
| Otu03819 | 9 | k__Fungi(100);p__Ascomycota(100);p__Ascomycota_unclassified(100);p__Ascomycota_unclassified(100);p__Ascomycota_unclassified(100);p__Ascomycota_unclassified(100);p__Ascomycota_unclassified(100); | - | - | - | - | - | - | - | Unassigned | - |
| Otu03820 | 9 | k__Fungi(100);p__Ascomycota(89);p__Ascomycota_unclassified(89);p__Ascomycota_unclassified(89);p__Ascomycota_unclassified(89);p__Ascomycota_unclassified(89);p__Ascomycota_unclassified(89); | - | - | - | - | - | - | - | Unassigned | - |
| Otu03821 | 9 | k__Fungi(100);p__Basidiomycota(100);c__Agaricomycetes(100);o__Corticiales(100);f__unclassified_Corticiales(100);g__unclassified_Corticiales(100);s__Corticiales_sp(100); | - | - | - | - | - | - | - | Unassigned | - |
| Otu03822 | 9 | k__Fungi(100);p__Ascomycota(78);p__Ascomycota_unclassified(78);p__Ascomycota_unclassified(78);p__Ascomycota_unclassified(78);p__Ascomycota_unclassified(78);p__Ascomycota_unclassified(78); | - | - | - | - | - | - | - | Unassigned | - |
| Otu03825 | 9 | k__Fungi(100);p__Basidiomycota(100);c__Agaricomycetes(100);o__Cantharellales(100);o__Cantharellales_unclassified(56);o__Cantharellales_unclassified(56);o__Cantharellales_unclassified(56); | - | - | - | - | - | - | - | Unassigned | - |
| Otu03827 | 9 | k__Fungi(100);p__Basidiomycota(100);c__Agaricomycetes(100);c__Agaricomycetes_unclassified(100);c__Agaricomycetes_unclassified(100);c__Agaricomycetes_unclassified(100);c__Agaricomycetes_unclassified(100); | - | - | - | - | - | - | - | Unassigned | - |
| Otu03828 | 9 | k__Fungi(100);p__Ascomycota(100);c__Leotiomycetes(100);c__Leotiomycetes_unclassified(100);c__Leotiomycetes_unclassified(100);c__Leotiomycetes_unclassified(100);c__Leotiomycetes_unclassified(100); | - | - | - | - | - | - | - | Unassigned | - |
| Otu03834 | 9 | k__Fungi(100);p__Ascomycota(100);p__Ascomycota_unclassified(100);p__Ascomycota_unclassified(100);p__Ascomycota_unclassified(100);p__Ascomycota_unclassified(100);p__Ascomycota_unclassified(100); | - | - | - | - | - | - | - | Unassigned | - |
| Otu03835 | 9 | k__Fungi(100);k__Fungi_unclassified(100);k__Fungi_unclassified(100);k__Fungi_unclassified(100);k__Fungi_unclassified(100);k__Fungi_unclassified(100);k__Fungi_unclassified(100); | - | - | - | - | - | - | - | Unassigned | - |
| Otu03837 | 9 | k__Fungi(100);p__Ascomycota(78);p__Ascomycota_unclassified(78);p__Ascomycota_unclassified(78);p__Ascomycota_unclassified(78);p__Ascomycota_unclassified(78);p__Ascomycota_unclassified(78); | - | - | - | - | - | - | - | Unassigned | - |
| Otu03843 | 9 | k__Fungi(100);k__Fungi_unclassified(89);k__Fungi_unclassified(89);k__Fungi_unclassified(89);k__Fungi_unclassified(89);k__Fungi_unclassified(89);k__Fungi_unclassified(89); | - | - | - | - | - | - | - | Unassigned | - |
| Otu03844 | 9 | k__Fungi(100);k__Fungi_unclassified(67);k__Fungi_unclassified(67);k__Fungi_unclassified(67);k__Fungi_unclassified(67);k__Fungi_unclassified(67);k__Fungi_unclassified(67); | - | - | - | - | - | - | - | Unassigned | - |
| Otu03845 | 9 | k__Fungi(100);p__Ascomycota(100);p__Ascomycota_unclassified(78);p__Ascomycota_unclassified(78);p__Ascomycota_unclassified(78);p__Ascomycota_unclassified(78);p__Ascomycota_unclassified(78); | - | - | - | - | - | - | - | Unassigned | - |
| Otu03847 | 9 | k__Fungi(100);k__Fungi_unclassified(78);k__Fungi_unclassified(78);k__Fungi_unclassified(78);k__Fungi_unclassified(78);k__Fungi_unclassified(78);k__Fungi_unclassified(78); | - | - | - | - | - | - | - | Unassigned | - |
| Otu03852 | 9 | k__Fungi(100);p__Basidiomycota(100);c__Agaricomycetes(100);c__Agaricomycetes_unclassified(100);c__Agaricomycetes_unclassified(100);c__Agaricomycetes_unclassified(100);c__Agaricomycetes_unclassified(100); | - | - | - | - | - | - | - | Unassigned | - |
| Otu03853 | 9 | k__Fungi(100);k__Fungi_unclassified(67);k__Fungi_unclassified(67);k__Fungi_unclassified(67);k__Fungi_unclassified(67);k__Fungi_unclassified(67);k__Fungi_unclassified(67); | - | - | - | - | - | - | - | Unassigned | - |
| Otu03855 | 9 | k__Fungi(100);k__Fungi_unclassified(100);k__Fungi_unclassified(100);k__Fungi_unclassified(100);k__Fungi_unclassified(100);k__Fungi_unclassified(100);k__Fungi_unclassified(100); | - | - | - | - | - | - | - | Unassigned | - |
| Otu03856 | 9 | k__Fungi(100);p__Ascomycota(100);c__Dothideomycetes(100);o__Pleosporales(100);o__Pleosporales_unclassified(100);o__Pleosporales_unclassified(100);o__Pleosporales_unclassified(100); | - | - | - | - | - | - | - | Unassigned | - |
| Otu03857 | 9 | k__Fungi(100);p__Ascomycota(100);c__Dothideomycetes(100);o__Pleosporales(100);o__Pleosporales_unclassified(100);o__Pleosporales_unclassified(100);o__Pleosporales_unclassified(100); | - | - | - | - | - | - | - | Unassigned | - |
| Otu03858 | 9 | k__Fungi(100);k__Fungi_unclassified(100);k__Fungi_unclassified(100);k__Fungi_unclassified(100);k__Fungi_unclassified(100);k__Fungi_unclassified(100);k__Fungi_unclassified(100); | - | - | - | - | - | - | - | Unassigned | - |
| Otu03859 | 9 | k__Fungi(100);k__Fungi_unclassified(56);k__Fungi_unclassified(56);k__Fungi_unclassified(56);k__Fungi_unclassified(56);k__Fungi_unclassified(56);k__Fungi_unclassified(56); | - | - | - | - | - | - | - | Unassigned | - |
| Otu03860 | 9 | k__Fungi(100);p__Basidiomycota(100);c__Agaricomycetes(100);c__Agaricomycetes_unclassified(78);c__Agaricomycetes_unclassified(78);c__Agaricomycetes_unclassified(78);c__Agaricomycetes_unclassified(78); | - | - | - | - | - | - | - | Unassigned | - |
| Otu03861 | 9 | k__Fungi(100);p__Ascomycota(100);p__Ascomycota_unclassified(89);p__Ascomycota_unclassified(89);p__Ascomycota_unclassified(89);p__Ascomycota_unclassified(89);p__Ascomycota_unclassified(89); | - | - | - | - | - | - | - | Unassigned | - |
| Otu03863 | 8 | k__Fungi(100);p__Ascomycota(100);c__Dothideomycetes(100);o__Pleosporales(100);f__Venturiaceae(100);f__Venturiaceae_unclassified(100);f__Venturiaceae_unclassified(100); | Venturiaceae | 9 | Pathotroph-Saprotroph-Symbiotroph | Endophyte-Plant Pathogen-Undefined Saprotroph | Microfungus | NULL | Probable | Mostly endophytes and weak parasites of aerial plant parts, including bark and wood, leaf and fruit tissue; rarely biotrophic. Little is known except for the important pathogens (Cannon & Kirk 2007) | Cannon PF, Kirk PM. 2007. Fungal Families of the World. CAB International, Cambridge (ISBN: 978-0851998275) |
| Otu03864 | 8 | k__Fungi(100);p__Ascomycota(100);c__Saccharomycetes(100);o__Saccharomycetales(100);f__Metschnikowiaceae(100);g__Metschnikowia(100);s__Metschnikowia_pulcherrima(100); | Metschnikowia | 13 | Saprotroph | \|Undefined Saprotroph\| | Yeast | NULL | Probable | As nectar-tap_saprotroph (Põlme et al. 2020); Decay sugar-rich substrates (Põlme et al. 2020) | Põlme S, et al. 2020. Fungal Diversity 105:1-16 ((https://doi.org/10.1007/s13225-020-00466-2)) |
| Otu03866 | 8 | k__Fungi(100);p__Ascomycota(100);c__Saccharomycetes(100);o__Saccharomycetales(100);f__Metschnikowiaceae(100);g__Metschnikowia(100);s__Metschnikowia_pulcherrima(100); | Metschnikowia | 13 | Saprotroph | \|Undefined Saprotroph\| | Yeast | NULL | Probable | As nectar-tap_saprotroph (Põlme et al. 2020); Decay sugar-rich substrates (Põlme et al. 2020) | Põlme S, et al. 2020. Fungal Diversity 105:1-16 ((https://doi.org/10.1007/s13225-020-00466-2)) |
| Otu03868 | 8 | k__Fungi(100);p__Ascomycota(100);c__Saccharomycetes(100);o__Saccharomycetales(100);f__Metschnikowiaceae(100);g__Metschnikowia(100);g__Metschnikowia_unclassified(75); | Metschnikowia | 13 | Saprotroph | \|Undefined Saprotroph\| | Yeast | NULL | Probable | As nectar-tap_saprotroph (Põlme et al. 2020); Decay sugar-rich substrates (Põlme et al. 2020) | Põlme S, et al. 2020. Fungal Diversity 105:1-16 ((https://doi.org/10.1007/s13225-020-00466-2)) |
| Otu03869 | 8 | k__Fungi(100);p__Ascomycota(100);c__Saccharomycetes(100);o__Saccharomycetales(100);f__Metschnikowiaceae(100);g__Metschnikowia(100);g__Metschnikowia_unclassified(63); | Metschnikowia | 13 | Saprotroph | \|Undefined Saprotroph\| | Yeast | NULL | Probable | As nectar-tap_saprotroph (Põlme et al. 2020); Decay sugar-rich substrates (Põlme et al. 2020) | Põlme S, et al. 2020. Fungal Diversity 105:1-16 ((https://doi.org/10.1007/s13225-020-00466-2)) |
| Otu03870 | 8 | k__Fungi(100);p__Ascomycota(100);c__Saccharomycetes(100);o__Saccharomycetales(100);f__Metschnikowiaceae(100);g__Metschnikowia(100);s__Metschnikowia_pulcherrima(100); | Metschnikowia | 13 | Saprotroph | \|Undefined Saprotroph\| | Yeast | NULL | Probable | As nectar-tap_saprotroph (Põlme et al. 2020); Decay sugar-rich substrates (Põlme et al. 2020) | Põlme S, et al. 2020. Fungal Diversity 105:1-16 ((https://doi.org/10.1007/s13225-020-00466-2)) |
| Otu03871 | 8 | k__Fungi(100);p__Ascomycota(100);c__Saccharomycetes(100);o__Saccharomycetales(100);f__Metschnikowiaceae(100);g__Metschnikowia(100);g__Metschnikowia_unclassified(100); | Metschnikowia | 13 | Saprotroph | \|Undefined Saprotroph\| | Yeast | NULL | Probable | As nectar-tap_saprotroph (Põlme et al. 2020); Decay sugar-rich substrates (Põlme et al. 2020) | Põlme S, et al. 2020. Fungal Diversity 105:1-16 ((https://doi.org/10.1007/s13225-020-00466-2)) |
| Otu03872 | 8 | k__Fungi(100);p__Ascomycota(100);c__Saccharomycetes(100);o__Saccharomycetales(100);f__Metschnikowiaceae(100);g__Metschnikowia(100);s__Metschnikowia_pulcherrima(100); | Metschnikowia | 13 | Saprotroph | \|Undefined Saprotroph\| | Yeast | NULL | Probable | As nectar-tap_saprotroph (Põlme et al. 2020); Decay sugar-rich substrates (Põlme et al. 2020) | Põlme S, et al. 2020. Fungal Diversity 105:1-16 ((https://doi.org/10.1007/s13225-020-00466-2)) |
| Otu03873 | 8 | k__Fungi(100);p__Ascomycota(100);c__Saccharomycetes(100);o__Saccharomycetales(100);f__Metschnikowiaceae(100);g__Metschnikowia(100);s__Metschnikowia_pulcherrima(100); | Metschnikowia | 13 | Saprotroph | \|Undefined Saprotroph\| | Yeast | NULL | Probable | As nectar-tap_saprotroph (Põlme et al. 2020); Decay sugar-rich substrates (Põlme et al. 2020) | Põlme S, et al. 2020. Fungal Diversity 105:1-16 ((https://doi.org/10.1007/s13225-020-00466-2)) |
| Otu03874 | 8 | k__Fungi(100);p__Ascomycota(100);c__Saccharomycetes(100);o__Saccharomycetales(100);f__Metschnikowiaceae(100);g__Metschnikowia(100);g__Metschnikowia_unclassified(100); | Metschnikowia | 13 | Saprotroph | \|Undefined Saprotroph\| | Yeast | NULL | Probable | As nectar-tap_saprotroph (Põlme et al. 2020); Decay sugar-rich substrates (Põlme et al. 2020) | Põlme S, et al. 2020. Fungal Diversity 105:1-16 ((https://doi.org/10.1007/s13225-020-00466-2)) |
| Otu03877 | 8 | k__Fungi(100);p__Ascomycota(100);c__Saccharomycetes(100);o__Saccharomycetales(100);f__Metschnikowiaceae(100);g__Metschnikowia(100);s__Metschnikowia_pulcherrima(100); | Metschnikowia | 13 | Saprotroph | \|Undefined Saprotroph\| | Yeast | NULL | Probable | As nectar-tap_saprotroph (Põlme et al. 2020); Decay sugar-rich substrates (Põlme et al. 2020) | Põlme S, et al. 2020. Fungal Diversity 105:1-16 ((https://doi.org/10.1007/s13225-020-00466-2)) |
| Otu03879 | 8 | k__Fungi(100);p__Ascomycota(100);c__Saccharomycetes(100);o__Saccharomycetales(100);f__Metschnikowiaceae(100);g__Metschnikowia(100);s__Metschnikowia_pulcherrima(100); | Metschnikowia | 13 | Saprotroph | \|Undefined Saprotroph\| | Yeast | NULL | Probable | As nectar-tap_saprotroph (Põlme et al. 2020); Decay sugar-rich substrates (Põlme et al. 2020) | Põlme S, et al. 2020. Fungal Diversity 105:1-16 ((https://doi.org/10.1007/s13225-020-00466-2)) |
| Otu03880 | 8 | k__Fungi(100);p__Ascomycota(100);c__Saccharomycetes(100);o__Saccharomycetales(100);f__Metschnikowiaceae(100);g__Metschnikowia(100);g__Metschnikowia_unclassified(88); | Metschnikowia | 13 | Saprotroph | \|Undefined Saprotroph\| | Yeast | NULL | Probable | As nectar-tap_saprotroph (Põlme et al. 2020); Decay sugar-rich substrates (Põlme et al. 2020) | Põlme S, et al. 2020. Fungal Diversity 105:1-16 ((https://doi.org/10.1007/s13225-020-00466-2)) |
| Otu03883 | 8 | k__Fungi(100);p__Ascomycota(100);c__Saccharomycetes(100);o__Saccharomycetales(100);f__Metschnikowiaceae(100);g__Metschnikowia(100);s__Metschnikowia_pulcherrima(100); | Metschnikowia | 13 | Saprotroph | \|Undefined Saprotroph\| | Yeast | NULL | Probable | As nectar-tap_saprotroph (Põlme et al. 2020); Decay sugar-rich substrates (Põlme et al. 2020) | Põlme S, et al. 2020. Fungal Diversity 105:1-16 ((https://doi.org/10.1007/s13225-020-00466-2)) |
| Otu03885 | 8 | k__Fungi(100);p__Ascomycota(100);c__Saccharomycetes(100);o__Saccharomycetales(100);f__Metschnikowiaceae(100);g__Metschnikowia(100);s__Metschnikowia_pulcherrima(75); | Metschnikowia | 13 | Saprotroph | \|Undefined Saprotroph\| | Yeast | NULL | Probable | As nectar-tap_saprotroph (Põlme et al. 2020); Decay sugar-rich substrates (Põlme et al. 2020) | Põlme S, et al. 2020. Fungal Diversity 105:1-16 ((https://doi.org/10.1007/s13225-020-00466-2)) |
| Otu03887 | 8 | k__Fungi(100);p__Ascomycota(100);c__Saccharomycetes(100);o__Saccharomycetales(100);f__Saccharomycetaceae(100);g__Kazachstania(100);s__Kazachstania_intestinalis(100); | Kazachstania | 13 | Saprotroph | \|Undefined Saprotroph\| | Yeast | NULL | Probable | As nectar-tap_saprotroph (Põlme et al. 2020); Decay sugar-rich substrates (Põlme et al. 2020) | Põlme S, et al. 2020. Fungal Diversity 105:1-16 ((https://doi.org/10.1007/s13225-020-00466-2)) |
| Otu03888 | 8 | k__Fungi(100);p__Ascomycota(100);c__Saccharomycetes(100);o__Saccharomycetales(100);f__Metschnikowiaceae(100);g__Metschnikowia(100);g__Metschnikowia_unclassified(100); | Metschnikowia | 13 | Saprotroph | \|Undefined Saprotroph\| | Yeast | NULL | Probable | As nectar-tap_saprotroph (Põlme et al. 2020); Decay sugar-rich substrates (Põlme et al. 2020) | Põlme S, et al. 2020. Fungal Diversity 105:1-16 ((https://doi.org/10.1007/s13225-020-00466-2)) |
| Otu03892 | 8 | k__Fungi(100);p__Ascomycota(100);c__Saccharomycetes(100);o__Saccharomycetales(100);f__Metschnikowiaceae(100);g__Metschnikowia(100);s__Metschnikowia_pulcherrima(75); | Metschnikowia | 13 | Saprotroph | \|Undefined Saprotroph\| | Yeast | NULL | Probable | As nectar-tap_saprotroph (Põlme et al. 2020); Decay sugar-rich substrates (Põlme et al. 2020) | Põlme S, et al. 2020. Fungal Diversity 105:1-16 ((https://doi.org/10.1007/s13225-020-00466-2)) |
| Otu03896 | 8 | k__Fungi(100);p__Ascomycota(100);c__Saccharomycetes(100);o__Saccharomycetales(100);f__Metschnikowiaceae(100);g__Metschnikowia(100);s__Metschnikowia_pulcherrima(88); | Metschnikowia | 13 | Saprotroph | \|Undefined Saprotroph\| | Yeast | NULL | Probable | As nectar-tap_saprotroph (Põlme et al. 2020); Decay sugar-rich substrates (Põlme et al. 2020) | Põlme S, et al. 2020. Fungal Diversity 105:1-16 ((https://doi.org/10.1007/s13225-020-00466-2)) |
| Otu03897 | 8 | k__Fungi(100);p__Ascomycota(100);c__Saccharomycetes(100);o__Saccharomycetales(100);f__Metschnikowiaceae(100);g__Metschnikowia(100);s__Metschnikowia_pulcherrima(75); | Metschnikowia | 13 | Saprotroph | \|Undefined Saprotroph\| | Yeast | NULL | Probable | As nectar-tap_saprotroph (Põlme et al. 2020); Decay sugar-rich substrates (Põlme et al. 2020) | Põlme S, et al. 2020. Fungal Diversity 105:1-16 ((https://doi.org/10.1007/s13225-020-00466-2)) |
| Otu03898 | 8 | k__Fungi(100);p__Ascomycota(100);c__Saccharomycetes(100);o__Saccharomycetales(100);f__Metschnikowiaceae(100);g__Metschnikowia(100);s__Metschnikowia_pulcherrima(75); | Metschnikowia | 13 | Saprotroph | \|Undefined Saprotroph\| | Yeast | NULL | Probable | As nectar-tap_saprotroph (Põlme et al. 2020); Decay sugar-rich substrates (Põlme et al. 2020) | Põlme S, et al. 2020. Fungal Diversity 105:1-16 ((https://doi.org/10.1007/s13225-020-00466-2)) |
| Otu03899 | 8 | k__Fungi(100);p__Ascomycota(100);c__Saccharomycetes(100);o__Saccharomycetales(100);f__Metschnikowiaceae(100);g__Metschnikowia(100);g__Metschnikowia_unclassified(88); | Metschnikowia | 13 | Saprotroph | \|Undefined Saprotroph\| | Yeast | NULL | Probable | As nectar-tap_saprotroph (Põlme et al. 2020); Decay sugar-rich substrates (Põlme et al. 2020) | Põlme S, et al. 2020. Fungal Diversity 105:1-16 ((https://doi.org/10.1007/s13225-020-00466-2)) |
| Otu03910 | 8 | k__Fungi(100);p__Ascomycota(100);c__Saccharomycetes(100);o__Saccharomycetales(100);f__Metschnikowiaceae(100);g__Metschnikowia(100);s__Metschnikowia_pulcherrima(100); | Metschnikowia | 13 | Saprotroph | \|Undefined Saprotroph\| | Yeast | NULL | Probable | As nectar-tap_saprotroph (Põlme et al. 2020); Decay sugar-rich substrates (Põlme et al. 2020) | Põlme S, et al. 2020. Fungal Diversity 105:1-16 ((https://doi.org/10.1007/s13225-020-00466-2)) |
| Otu03914 | 8 | k__Fungi(100);p__Ascomycota(100);c__Saccharomycetes(100);o__Saccharomycetales(100);f__Metschnikowiaceae(100);g__Metschnikowia(100);s__Metschnikowia_pulcherrima(100); | Metschnikowia | 13 | Saprotroph | \|Undefined Saprotroph\| | Yeast | NULL | Probable | As nectar-tap_saprotroph (Põlme et al. 2020); Decay sugar-rich substrates (Põlme et al. 2020) | Põlme S, et al. 2020. Fungal Diversity 105:1-16 ((https://doi.org/10.1007/s13225-020-00466-2)) |
| Otu03919 | 8 | k__Fungi(100);p__Ascomycota(100);c__Saccharomycetes(100);o__Saccharomycetales(100);f__Metschnikowiaceae(100);g__Metschnikowia(100);s__Metschnikowia_pulcherrima(88); | Metschnikowia | 13 | Saprotroph | \|Undefined Saprotroph\| | Yeast | NULL | Probable | As nectar-tap_saprotroph (Põlme et al. 2020); Decay sugar-rich substrates (Põlme et al. 2020) | Põlme S, et al. 2020. Fungal Diversity 105:1-16 ((https://doi.org/10.1007/s13225-020-00466-2)) |
| Otu03921 | 8 | k__Fungi(100);p__Ascomycota(100);c__Sordariomycetes(100);o__Chaetosphaeriales(75);f__Chaetosphaeriaceae(75);g__unclassified_Chaetosphaeriaceae(75);s__Chaetosphaeriaceae_sp(75); | Chaetosphaeriaceae | 9 | Saprotroph | Plant Saprotroph-Wood Saprotroph | Microfungus | NULL | Probable | Saprobes associated with decaying woody and herbaceous plant material, including timber and timber products in damp buildings (Cannon & Kirk 2007) | Cannon PF, Kirk PM. 2007. Fungal Families of the World. CAB International, Cambridge (ISBN: 978-0851998275) |
| Otu03922 | 8 | k__Fungi(100);p__Ascomycota(100);c__Saccharomycetes(100);o__Saccharomycetales(100);f__Metschnikowiaceae(100);g__Metschnikowia(100);s__Metschnikowia_pulcherrima(100); | Metschnikowia | 13 | Saprotroph | \|Undefined Saprotroph\| | Yeast | NULL | Probable | As nectar-tap_saprotroph (Põlme et al. 2020); Decay sugar-rich substrates (Põlme et al. 2020) | Põlme S, et al. 2020. Fungal Diversity 105:1-16 ((https://doi.org/10.1007/s13225-020-00466-2)) |
| Otu03923 | 8 | k__Fungi(100);p__Ascomycota(100);c__Sordariomycetes(100);o__Sordariomycetes_order_Incertae_sedis(100);f__Magnaporthaceae(100);f__Magnaporthaceae_unclassified(75);f__Magnaporthaceae_unclassified(75); | Magnaporthaceae | 9 | Pathotroph | Plant Pathogen | Microfungus | NULL | Probable | Species in this family cause rice blast and take-all; Usually necrotrophic on roots (Cannon & Kirk 2007) | Cannon PF, Kirk PM. 2007. Fungal Families of the World. CAB International, Cambridge (ISBN: 978-0851998275) |
| Otu03924 | 8 | k__Fungi(100);p__Ascomycota(100);c__Saccharomycetes(100);o__Saccharomycetales(100);f__Metschnikowiaceae(100);g__Metschnikowia(100);g__Metschnikowia_unclassified(100); | Metschnikowia | 13 | Saprotroph | \|Undefined Saprotroph\| | Yeast | NULL | Probable | As nectar-tap_saprotroph (Põlme et al. 2020); Decay sugar-rich substrates (Põlme et al. 2020) | Põlme S, et al. 2020. Fungal Diversity 105:1-16 ((https://doi.org/10.1007/s13225-020-00466-2)) |
| Otu03927 | 8 | k__Fungi(100);p__Ascomycota(100);c__Saccharomycetes(100);o__Saccharomycetales(100);f__Metschnikowiaceae(100);g__Metschnikowia(100);s__Metschnikowia_pulcherrima(100); | Metschnikowia | 13 | Saprotroph | \|Undefined Saprotroph\| | Yeast | NULL | Probable | As nectar-tap_saprotroph (Põlme et al. 2020); Decay sugar-rich substrates (Põlme et al. 2020) | Põlme S, et al. 2020. Fungal Diversity 105:1-16 ((https://doi.org/10.1007/s13225-020-00466-2)) |
| Otu03933 | 8 | k__Fungi(100);p__Ascomycota(100);c__Saccharomycetes(100);o__Saccharomycetales(100);f__Metschnikowiaceae(100);g__Metschnikowia(100);s__Metschnikowia_pulcherrima(100); | Metschnikowia | 13 | Saprotroph | \|Undefined Saprotroph\| | Yeast | NULL | Probable | As nectar-tap_saprotroph (Põlme et al. 2020); Decay sugar-rich substrates (Põlme et al. 2020) | Põlme S, et al. 2020. Fungal Diversity 105:1-16 ((https://doi.org/10.1007/s13225-020-00466-2)) |
| Otu03936 | 8 | k__Fungi(100);p__Ascomycota(100);c__Saccharomycetes(100);o__Saccharomycetales(100);f__Saccharomycetales_family_Incertae_sedis(88);g__Candida(88);g__Candida_unclassified(63); | Candida | 13 | Pathotroph-Saprotroph-Symbiotroph | Animal Pathogen-Endophyte-Endosymbiont-Epiphyte-\|Undefined Saprotroph\| | Dimorphic Yeast-Yeast | NULL | Probable | As nectar-tap_saprotroph (Põlme et al. 2020); Decay sugar-rich substrates (Põlme et al. 2020); This asexual genus is polyphyletic and contain species that function very differently depend on what they are. Caution should be taken when interpreting guilds for Candida species. | Manolakaki D, et al. 2010. Virulence. 1: 367-375 ((https://doi.org/10.4161/viru.1.5.12796)); Mekbib SB, et al. 2011. Tropical Plant Pathology 36:233-240 ((https://doi.org/10.1590/S1982-56762011000400004)); Põlme S, et al. 2020. Fungal Diversity 105:1-16 ((https://doi.org/10.1007/s13225-020-00466-2)) |
| Otu03938 | 8 | k__Fungi(100);p__Ascomycota(100);c__Lecanoromycetes(100);o__Lecanorales(100);f__Lecanoraceae(100);f__Lecanoraceae_unclassified(100);f__Lecanoraceae_unclassified(100); | Lecanoraceae | 9 | Pathotroph-Symbiotroph | Lichen Parasite-Lichenized | Microfungus-Thallus | NULL | Probable | Lichenized with green algae, occasionally lichenicolous, on rocks, mosses and wood (Cannon & Kirk 2007) | Cannon PF, Kirk PM. 2007. Fungal Families of the World. CAB International, Cambridge (ISBN: 978-0851998275) |
| Otu03943 | 8 | k__Fungi(100);p__Ascomycota(100);c__Saccharomycetes(100);o__Saccharomycetales(100);f__Metschnikowiaceae(100);g__Metschnikowia(100);g__Metschnikowia_unclassified(88); | Metschnikowia | 13 | Saprotroph | \|Undefined Saprotroph\| | Yeast | NULL | Probable | As nectar-tap_saprotroph (Põlme et al. 2020); Decay sugar-rich substrates (Põlme et al. 2020) | Põlme S, et al. 2020. Fungal Diversity 105:1-16 ((https://doi.org/10.1007/s13225-020-00466-2)) |
| Otu03945 | 8 | k__Fungi(100);p__Ascomycota(100);c__Saccharomycetes(100);o__Saccharomycetales(100);f__Metschnikowiaceae(100);g__Metschnikowia(100);s__Metschnikowia_pulcherrima(75); | Metschnikowia | 13 | Saprotroph | \|Undefined Saprotroph\| | Yeast | NULL | Probable | As nectar-tap_saprotroph (Põlme et al. 2020); Decay sugar-rich substrates (Põlme et al. 2020) | Põlme S, et al. 2020. Fungal Diversity 105:1-16 ((https://doi.org/10.1007/s13225-020-00466-2)) |
| Otu03948 | 8 | k__Fungi(100);p__Ascomycota(100);c__Saccharomycetes(100);o__Saccharomycetales(100);f__Metschnikowiaceae(100);g__Metschnikowia(100);s__Metschnikowia_pulcherrima(88); | Metschnikowia | 13 | Saprotroph | \|Undefined Saprotroph\| | Yeast | NULL | Probable | As nectar-tap_saprotroph (Põlme et al. 2020); Decay sugar-rich substrates (Põlme et al. 2020) | Põlme S, et al. 2020. Fungal Diversity 105:1-16 ((https://doi.org/10.1007/s13225-020-00466-2)) |
| Otu03950 | 8 | k__Fungi(100);p__Ascomycota(100);c__Saccharomycetes(100);o__Saccharomycetales(100);f__Metschnikowiaceae(100);g__Metschnikowia(100);s__Metschnikowia_pulcherrima(100); | Metschnikowia | 13 | Saprotroph | \|Undefined Saprotroph\| | Yeast | NULL | Probable | As nectar-tap_saprotroph (Põlme et al. 2020); Decay sugar-rich substrates (Põlme et al. 2020) | Põlme S, et al. 2020. Fungal Diversity 105:1-16 ((https://doi.org/10.1007/s13225-020-00466-2)) |
| Otu03952 | 8 | k__Fungi(100);p__Ascomycota(100);c__Saccharomycetes(100);o__Saccharomycetales(100);f__Metschnikowiaceae(100);g__Metschnikowia(100);s__Metschnikowia_pulcherrima(100); | Metschnikowia | 13 | Saprotroph | \|Undefined Saprotroph\| | Yeast | NULL | Probable | As nectar-tap_saprotroph (Põlme et al. 2020); Decay sugar-rich substrates (Põlme et al. 2020) | Põlme S, et al. 2020. Fungal Diversity 105:1-16 ((https://doi.org/10.1007/s13225-020-00466-2)) |
| Otu03954 | 8 | k__Fungi(100);p__Ascomycota(100);c__Taphrinomycetes(100);o__Taphrinales(100);f__Taphrinaceae(100);g__Taphrina(63);g__Taphrina_unclassified(63); | Taphrina | 13 | Pathotroph | \|Plant Pathogen\| | Dimorphic-Microfungus | NULL | Probable | NULL | Põlme S, et al. 2020. Fungal Diversity 105:1-16 ((https://doi.org/10.1007/s13225-020-00466-2)); The American Phytopathological Society Common Names of Plant Diseases ((http://www.apsnet.org/publications/commonnames/Pages/default.aspx)) |
| Otu03955 | 8 | k__Fungi(100);p__Ascomycota(100);c__Sordariomycetes(100);o__Hypocreales(100);o__Hypocreales_unclassified(88);o__Hypocreales_unclassified(88);o__Hypocreales_unclassified(88); | Hypocreales | 7 | Saprotroph | Undefined Saprotroph | Microfungus | NULL | Possible | NULL | Sterkenburg E, et al. 2015. New Phytologist 207:1145-1158 ((https://doi.org/10.1111/nph.13426)) |
| Otu03956 | 8 | k__Fungi(100);p__Ascomycota(100);c__Saccharomycetes(100);o__Saccharomycetales(100);f__Metschnikowiaceae(100);g__Metschnikowia(100);g__Metschnikowia_unclassified(100); | Metschnikowia | 13 | Saprotroph | \|Undefined Saprotroph\| | Yeast | NULL | Probable | As nectar-tap_saprotroph (Põlme et al. 2020); Decay sugar-rich substrates (Põlme et al. 2020) | Põlme S, et al. 2020. Fungal Diversity 105:1-16 ((https://doi.org/10.1007/s13225-020-00466-2)) |
| Otu03957 | 8 | k__Fungi(100);p__Basidiomycota(100);c__Microbotryomycetes(100);o__Sporidiobolales(100);f__Sporidiobolales_family_Incertae_sedis(100);g__Sporobolomyces(100);s__Sporobolomyces_lactosus(100); | Sporobolomyces | 13 | Pathotroph-Saprotroph | \|Fungal Parasite\|-Plant Saprotroph | Yeast | NULL | Probable | As litter_saprotroph (Põlme et al. 2020); As mycoparasite (Põlme et al. 2020); Decay fungal material (Põlme et al. 2020) | Tedersoo L, et al. 2014. Science 346:e1256688 ((https://doi.org/10.1126/science.1256688)); Purahong W, et al. 2016. Molecular Ecology 25:4059-4074 ((https://doi.org/10.1111/mec.13739)); Põlme S, et al. 2020. Fungal Diversity 105:1-16 ((https://doi.org/10.1007/s13225-020-00466-2)) |
| Otu03958 | 8 | k__Fungi(100);p__Ascomycota(100);c__Sordariomycetes(100);o__Ophiostomatales(100);f__Ophiostomataceae(100);f__Ophiostomataceae_unclassified(100);f__Ophiostomataceae_unclassified(100); | Ophiostomataceae | 9 | Pathotroph | Plant Pathogen | Microfungus | Blue-Staining | Probable | At least most species are transmitted via insect vectors, with mycelial colonies developing in bark-boring beetle galleries and spreading through the vascular system; they are highly specific to their beetle vectors. In some cases spores and mycelium are transmitted in specialized organs, termed mycangia (Cannon & Kirk 2007) | Cannon PF, Kirk PM. 2007. Fungal Families of the World. CAB International, Cambridge (ISBN: 978-0851998275); Tedersoo L, et al. 2014. Science 346:e1256688 ((https://doi.org/10.1126/science.1256688)) |
| Otu03962 | 8 | k__Fungi(100);p__Ascomycota(100);c__Saccharomycetes(100);o__Saccharomycetales(100);f__Metschnikowiaceae(100);g__Metschnikowia(100);s__Metschnikowia_pulcherrima(100); | Metschnikowia | 13 | Saprotroph | \|Undefined Saprotroph\| | Yeast | NULL | Probable | As nectar-tap_saprotroph (Põlme et al. 2020); Decay sugar-rich substrates (Põlme et al. 2020) | Põlme S, et al. 2020. Fungal Diversity 105:1-16 ((https://doi.org/10.1007/s13225-020-00466-2)) |
| Otu03964 | 8 | k__Fungi(100);p__Ascomycota(100);c__Saccharomycetes(100);o__Saccharomycetales(100);f__Metschnikowiaceae(100);g__Metschnikowia(100);s__Metschnikowia_pulcherrima(100); | Metschnikowia | 13 | Saprotroph | \|Undefined Saprotroph\| | Yeast | NULL | Probable | As nectar-tap_saprotroph (Põlme et al. 2020); Decay sugar-rich substrates (Põlme et al. 2020) | Põlme S, et al. 2020. Fungal Diversity 105:1-16 ((https://doi.org/10.1007/s13225-020-00466-2)) |
| Otu03965 | 8 | k__Fungi(100);p__Ascomycota(100);c__Saccharomycetes(100);o__Saccharomycetales(100);f__Metschnikowiaceae(100);g__Metschnikowia(100);s__Metschnikowia_pulcherrima(75); | Metschnikowia | 13 | Saprotroph | \|Undefined Saprotroph\| | Yeast | NULL | Probable | As nectar-tap_saprotroph (Põlme et al. 2020); Decay sugar-rich substrates (Põlme et al. 2020) | Põlme S, et al. 2020. Fungal Diversity 105:1-16 ((https://doi.org/10.1007/s13225-020-00466-2)) |
| Otu03966 | 8 | k__Fungi(100);p__Ascomycota(100);c__Saccharomycetes(100);o__Saccharomycetales(100);f__Metschnikowiaceae(100);g__Metschnikowia(100);s__Metschnikowia_pulcherrima(100); | Metschnikowia | 13 | Saprotroph | \|Undefined Saprotroph\| | Yeast | NULL | Probable | As nectar-tap_saprotroph (Põlme et al. 2020); Decay sugar-rich substrates (Põlme et al. 2020) | Põlme S, et al. 2020. Fungal Diversity 105:1-16 ((https://doi.org/10.1007/s13225-020-00466-2)) |
| Otu03967 | 8 | k__Fungi(100);p__Ascomycota(100);c__Saccharomycetes(100);o__Saccharomycetales(100);f__Saccharomycetales_family_Incertae_sedis(100);f__Saccharomycetales_family_Incertae_sedis_unclassified(100);f__Saccharomycetales_family_Incertae_sedis_unclassified(100); | Saccharomycetales | 7 | Saprotroph | Undefined Saprotroph | Yeast | NULL | Possible | NULL | Sterkenburg E, et al. 2015. New Phytologist 207:1145-1158 ((https://doi.org/10.1111/nph.13426)) |
| Otu03968 | 8 | k__Fungi(100);p__Ascomycota(100);c__Saccharomycetes(100);o__Saccharomycetales(100);f__Metschnikowiaceae(100);g__Metschnikowia(100);g__Metschnikowia_unclassified(63); | Metschnikowia | 13 | Saprotroph | \|Undefined Saprotroph\| | Yeast | NULL | Probable | As nectar-tap_saprotroph (Põlme et al. 2020); Decay sugar-rich substrates (Põlme et al. 2020) | Põlme S, et al. 2020. Fungal Diversity 105:1-16 ((https://doi.org/10.1007/s13225-020-00466-2)) |
| Otu03975 | 8 | k__Fungi(100);p__Ascomycota(100);c__Saccharomycetes(100);o__Saccharomycetales(100);f__Metschnikowiaceae(100);g__Metschnikowia(100);s__Metschnikowia_pulcherrima(100); | Metschnikowia | 13 | Saprotroph | \|Undefined Saprotroph\| | Yeast | NULL | Probable | As nectar-tap_saprotroph (Põlme et al. 2020); Decay sugar-rich substrates (Põlme et al. 2020) | Põlme S, et al. 2020. Fungal Diversity 105:1-16 ((https://doi.org/10.1007/s13225-020-00466-2)) |
| Otu03977 | 8 | k__Fungi(100);p__Ascomycota(100);c__Saccharomycetes(100);o__Saccharomycetales(100);f__Metschnikowiaceae(100);g__Metschnikowia(100);s__Metschnikowia_pulcherrima(75); | Metschnikowia | 13 | Saprotroph | \|Undefined Saprotroph\| | Yeast | NULL | Probable | As nectar-tap_saprotroph (Põlme et al. 2020); Decay sugar-rich substrates (Põlme et al. 2020) | Põlme S, et al. 2020. Fungal Diversity 105:1-16 ((https://doi.org/10.1007/s13225-020-00466-2)) |
| Otu03979 | 8 | k__Fungi(100);p__Ascomycota(100);c__Saccharomycetes(100);o__Saccharomycetales(100);f__Metschnikowiaceae(100);g__Metschnikowia(100);s__Metschnikowia_pulcherrima(75); | Metschnikowia | 13 | Saprotroph | \|Undefined Saprotroph\| | Yeast | NULL | Probable | As nectar-tap_saprotroph (Põlme et al. 2020); Decay sugar-rich substrates (Põlme et al. 2020) | Põlme S, et al. 2020. Fungal Diversity 105:1-16 ((https://doi.org/10.1007/s13225-020-00466-2)) |
| Otu03980 | 8 | k__Fungi(100);p__Ascomycota(100);c__Saccharomycetes(100);o__Saccharomycetales(100);f__Metschnikowiaceae(100);g__Metschnikowia(100);s__Metschnikowia_pulcherrima(100); | Metschnikowia | 13 | Saprotroph | \|Undefined Saprotroph\| | Yeast | NULL | Probable | As nectar-tap_saprotroph (Põlme et al. 2020); Decay sugar-rich substrates (Põlme et al. 2020) | Põlme S, et al. 2020. Fungal Diversity 105:1-16 ((https://doi.org/10.1007/s13225-020-00466-2)) |
| Otu03981 | 8 | k__Fungi(100);p__Ascomycota(100);c__Saccharomycetes(100);o__Saccharomycetales(100);f__Metschnikowiaceae(100);g__Metschnikowia(100);s__Metschnikowia_pulcherrima(63); | Metschnikowia | 13 | Saprotroph | \|Undefined Saprotroph\| | Yeast | NULL | Probable | As nectar-tap_saprotroph (Põlme et al. 2020); Decay sugar-rich substrates (Põlme et al. 2020) | Põlme S, et al. 2020. Fungal Diversity 105:1-16 ((https://doi.org/10.1007/s13225-020-00466-2)) |
| Otu03982 | 8 | k__Fungi(100);p__Ascomycota(100);c__Saccharomycetes(100);o__Saccharomycetales(100);f__Metschnikowiaceae(100);g__Metschnikowia(100);g__Metschnikowia_unclassified(100); | Metschnikowia | 13 | Saprotroph | \|Undefined Saprotroph\| | Yeast | NULL | Probable | As nectar-tap_saprotroph (Põlme et al. 2020); Decay sugar-rich substrates (Põlme et al. 2020) | Põlme S, et al. 2020. Fungal Diversity 105:1-16 ((https://doi.org/10.1007/s13225-020-00466-2)) |
| Otu03983 | 8 | k__Fungi(100);p__Ascomycota(100);c__Saccharomycetes(100);o__Saccharomycetales(100);f__Metschnikowiaceae(100);g__Metschnikowia(100);s__Metschnikowia_pulcherrima(88); | Metschnikowia | 13 | Saprotroph | \|Undefined Saprotroph\| | Yeast | NULL | Probable | As nectar-tap_saprotroph (Põlme et al. 2020); Decay sugar-rich substrates (Põlme et al. 2020) | Põlme S, et al. 2020. Fungal Diversity 105:1-16 ((https://doi.org/10.1007/s13225-020-00466-2)) |
| Otu03984 | 8 | k__Fungi(100);p__Basidiomycota(100);c__Microbotryomycetes(100);o__Sporidiobolales(100);f__Sporidiobolales_family_Incertae_sedis(100);g__Sporobolomyces(100);s__Sporobolomyces_lactosus(100); | Sporobolomyces | 13 | Pathotroph-Saprotroph | \|Fungal Parasite\|-Plant Saprotroph | Yeast | NULL | Probable | As litter_saprotroph (Põlme et al. 2020); As mycoparasite (Põlme et al. 2020); Decay fungal material (Põlme et al. 2020) | Tedersoo L, et al. 2014. Science 346:e1256688 ((https://doi.org/10.1126/science.1256688)); Purahong W, et al. 2016. Molecular Ecology 25:4059-4074 ((https://doi.org/10.1111/mec.13739)); Põlme S, et al. 2020. Fungal Diversity 105:1-16 ((https://doi.org/10.1007/s13225-020-00466-2)) |
| Otu03986 | 8 | k__Fungi(100);p__Basidiomycota(100);c__Agaricomycetes(100);o__Agaricales(100);f__Inocybaceae(100);g__Crepidotus(100);s__Crepidotus_sphaerosporus(100); | Crepidotus | 13 | Saprotroph | \|Wood Saprotroph\| | Agaricoid | NULL | Highly Probable | Decay wood (Põlme et al. 2020) | Grgurinovic CA, Simpson JA. 2001. Fungal Diversity 8:97-127 ((http://www.fungaldiversity.org/fdp/sfdp/FD_8_97-127.pdf)); Aime MC, et al. 2005. American Journal of Botany 92:74-82 ((https://doi.org/10.3732/ajb.92.1.74)); Tedersoo L, et al. 2014. Science 346:e1256688 ((https://doi.org/10.1126/science.1256688)); Põlme S, et al. 2020. Fungal Diversity 105:1-16 ((https://doi.org/10.1007/s13225-020-00466-2)) |
| Otu03988 | 8 | k__Fungi(100);p__Ascomycota(100);c__Saccharomycetes(100);o__Saccharomycetales(100);f__Saccharomycetales_family_Incertae_sedis(88);g__Candida(75);g__Candida_unclassified(75); | Candida | 13 | Pathotroph-Saprotroph-Symbiotroph | Animal Pathogen-Endophyte-Endosymbiont-Epiphyte-\|Undefined Saprotroph\| | Dimorphic Yeast-Yeast | NULL | Probable | As nectar-tap_saprotroph (Põlme et al. 2020); Decay sugar-rich substrates (Põlme et al. 2020); This asexual genus is polyphyletic and contain species that function very differently depend on what they are. Caution should be taken when interpreting guilds for Candida species. | Manolakaki D, et al. 2010. Virulence. 1: 367-375 ((https://doi.org/10.4161/viru.1.5.12796)); Mekbib SB, et al. 2011. Tropical Plant Pathology 36:233-240 ((https://doi.org/10.1590/S1982-56762011000400004)); Põlme S, et al. 2020. Fungal Diversity 105:1-16 ((https://doi.org/10.1007/s13225-020-00466-2)) |
| Otu03989 | 8 | k__Fungi(100);p__Basidiomycota(100);c__Agaricomycetes(100);o__Agaricales(100);f__Inocybaceae(100);g__Inocybe(100);s__Inocybe_curvipes(100); | Inocybe | 13 | Symbiotroph | \|Ectomycorrhizal\| | Agaricoid | NULL | Highly Probable | Sensu stricto (Matheny 2009). | Rinaldi AC, et al. 2008. Fungal Diversity 33:1-45 ((http://www.fungaldiversity.org/fdp/sfdp/33-1.pdf)); Matheny PB. 2009. McIlvainea 18:11-21 ((https://namyco.org/docs/3_Matheny.pdf)); Tedersoo L, et al. 2010. Mycorrhiza 20:217-263 ((https://doi.org/10.1007/s00572-009-0274-x)); Põlme S, et al. 2020. Fungal Diversity 105:1-16 ((https://doi.org/10.1007/s13225-020-00466-2)); http://mycorrhizas.info/ecmf.html ((http://mycorrhizas.info/ecmf.html)) |
| Otu03990 | 8 | k__Fungi(100);p__Ascomycota(100);c__Dothideomycetes(100);o__Pleosporales(100);f__Pleosporaceae(100);g__Curvularia(100);g__Curvularia_unclassified(75); | Curvularia | 13 | Pathotroph-Saprotroph | \|Plant Pathogen\|-Plant Saprotroph | Microfungus | NULL | Probable | As litter_saprotroph (Põlme et al. 2020); Decay wood, leaf, fruit, seed, roots, algal material (Põlme et al. 2020) | Tedersoo L, et al. 2014. Science 346:e1256688 ((https://doi.org/10.1126/science.1256688)); Põlme S, et al. 2020. Fungal Diversity 105:1-16 ((https://doi.org/10.1007/s13225-020-00466-2)) |
| Otu03991 | 8 | k__Fungi(100);p__Ascomycota(100);c__Saccharomycetes(100);o__Saccharomycetales(100);f__Metschnikowiaceae(100);g__Metschnikowia(100);s__Metschnikowia_pulcherrima(100); | Metschnikowia | 13 | Saprotroph | \|Undefined Saprotroph\| | Yeast | NULL | Probable | As nectar-tap_saprotroph (Põlme et al. 2020); Decay sugar-rich substrates (Põlme et al. 2020) | Põlme S, et al. 2020. Fungal Diversity 105:1-16 ((https://doi.org/10.1007/s13225-020-00466-2)) |
| Otu03995 | 8 | k__Fungi(100);p__Basidiomycota(100);c__Agaricomycetes(100);o__Agaricales(100);f__Strophariaceae(100);g__Pholiota(100);g__Pholiota_unclassified(100); | Pholiota | 13 | Pathotroph-Saprotroph | Plant Pathogen-Plant Saprotroph-Undefined Saprotroph-\|Wood Saprotroph\| | Agaricoid | White Rot | Probable | As litter_saprotroph (Põlme et al. 2020); As white_rot (Põlme et al. 2020); As wood_pathogen (Põlme et al. 2020); Decay wood, leaf, fruit, seed (Põlme et al. 2020) | Tedersoo L, et al. 2014. Science 346:e1256688 ((https://doi.org/10.1126/science.1256688)); Põlme S, et al. 2020. Fungal Diversity 105:1-16 ((https://doi.org/10.1007/s13225-020-00466-2)) |
| Otu03996 | 8 | k__Fungi(100);p__Ascomycota(100);c__Dothideomycetes(100);o__Pleosporales(100);f__Leptosphaeriaceae(100);g__Subplenodomus(100);s__Subplenodomus_apiicola(100); | Subplenodomus | 13 | Pathotroph-Saprotroph | \|Plant Pathogen\|-Plant Saprotroph-Undefined Saprotroph | Microfungus | NULL | Probable | As litter_saprotroph (Põlme et al. 2020); Decay leaf, fruit, seed (Põlme et al. 2020) | Tedersoo L, et al. 2014. Science 346:e1256688 ((https://doi.org/10.1126/science.1256688)); Põlme S, et al. 2020. Fungal Diversity 105:1-16 ((https://doi.org/10.1007/s13225-020-00466-2)) |
| Otu03997 | 8 | k__Fungi(100);p__Ascomycota(100);c__Sordariomycetes(100);o__Xylariales(100);f__Xylariaceae(100);f__Xylariaceae_unclassified(100);f__Xylariaceae_unclassified(100); | Xylariaceae | 9 | Pathotroph-Saprotroph-Symbiotroph | Dung Saprotroph-Endophyte-Plant Pathogen-Undefined Saprotroph | Xylarioid | NULL | Probable | Saprobes or weak pathogens, many endophytes, mainly in wood and bark, occasionally in soil and dung (Cannon & Kirk 2007) | Cannon PF, Kirk PM. 2007. Fungal Families of the World. CAB International, Cambridge (ISBN: 978-0851998275) |
| Otu03999 | 8 | k__Fungi(100);p__Ascomycota(100);c__Dothideomycetes(100);o__Pleosporales(100);f__Massarinaceae(88);g__Saccharicola(88);s__Saccharicola_sp(88); | Saccharicola | 13 | Saprotroph | Undefined Saprotroph | NULL | NULL | Probable | NULL | Tedersoo L, et al. 2014. Science 346:e1256688 ((https://doi.org/10.1126/science.1256688)) |
| Otu04002 | 8 | k__Fungi(100);p__Basidiomycota(100);c__Agaricomycetes(100);o__Polyporales(100);f__Fomitopsidaceae(100);g__Postia(100);s__Postia_tephroleuca(100); | Postia | 13 | Saprotroph | \|Wood Saprotroph\| | Polyporoid | Brown Rot | Highly Probable | As brown-rot (Põlme et al. 2020); Decay wood (Põlme et al. 2020) | Jordan CR, et al. 1996. Holzforschung - International Journal of the Biology, Chemistry, Physics and Technology of Wood 50:312-318 ((https://doi.org/10.1515/hfsg.1996.50.4.312)); Põlme S, et al. 2020. Fungal Diversity 105:1-16 ((https://doi.org/10.1007/s13225-020-00466-2)) |
| Otu04003 | 8 | k__Fungi(100);p__Ascomycota(100);c__Saccharomycetes(100);o__Saccharomycetales(100);f__Metschnikowiaceae(100);g__Metschnikowia(100);s__Metschnikowia_pulcherrima(100); | Metschnikowia | 13 | Saprotroph | \|Undefined Saprotroph\| | Yeast | NULL | Probable | As nectar-tap_saprotroph (Põlme et al. 2020); Decay sugar-rich substrates (Põlme et al. 2020) | Põlme S, et al. 2020. Fungal Diversity 105:1-16 ((https://doi.org/10.1007/s13225-020-00466-2)) |
| Otu04007 | 8 | k__Fungi(100);p__Basidiomycota(100);c__Agaricomycetes(100);o__Hymenochaetales(100);f__Schizoporaceae(100);g__Hyphodontia(100);s__Hyphodontia_subalutacea(100); | Hyphodontia | 13 | Saprotroph | Undefined Saprotroph-\|Wood Saprotroph\| | Corticioid | White Rot | Probable | As white_rot (Põlme et al. 2020); Decay wood (Põlme et al. 2020) | Tedersoo L, et al. 2014. Science 346:e1256688 ((https://doi.org/10.1126/science.1256688)); Põlme S, et al. 2020. Fungal Diversity 105:1-16 ((https://doi.org/10.1007/s13225-020-00466-2)) |
| Otu04008 | 8 | k__Fungi(100);p__Basidiomycota(88);c__Agaricomycetes(88);o__Agaricales(88);o__Agaricales_unclassified(88);o__Agaricales_unclassified(88);o__Agaricales_unclassified(88); | Agaricales | 7 | Pathotroph-Saprotroph-Symbiotroph | Bryophyte Parasite-Dung Saprotroph-Ectomycorrhizal-Fungal Parasite-Leaf Saprotroph-Plant Parasite-Undefined Saprotroph-Wood Saprotroph | Agaricoid-Gasteroid-Microfungus-Secotioid-Yeast | NULL | Possible | Mushrooms and toadstools, Gill fungi, Agarics... lignicolous, sometimes muscicolous or fungicolous, saprobic, mycorrhizal, rarely parasitic on plants or fungi (Kirk et al. 2008) | Kirk PM et al. 2008. Dictionary of the Fungi. Tenth Edition. CAB International, Wallingford (ISBN: 978-0851998268) |
| Otu04009 | 8 | k__Fungi(100);p__Ascomycota(100);c__Saccharomycetes(100);o__Saccharomycetales(100);f__Metschnikowiaceae(100);g__Metschnikowia(100);s__Metschnikowia_pulcherrima(88); | Metschnikowia | 13 | Saprotroph | \|Undefined Saprotroph\| | Yeast | NULL | Probable | As nectar-tap_saprotroph (Põlme et al. 2020); Decay sugar-rich substrates (Põlme et al. 2020) | Põlme S, et al. 2020. Fungal Diversity 105:1-16 ((https://doi.org/10.1007/s13225-020-00466-2)) |
| Otu04010 | 8 | k__Fungi(100);p__Ascomycota(100);c__Eurotiomycetes(100);o__Chaetothyriales(100);f__Chaetothyriaceae(100);g__Cyphellophora(100);s__Cyphellophora_reptans(100); | Cyphellophora | 13 | Pathotroph-Saprotroph | Animal Parasite-Animal Pathogen-\|Plant Saprotroph\|-Undefined Saprotroph | Dimorphic-Facultative Yeast-Microfungus | NULL | Probable | As litter_saprotroph (Põlme et al. 2020); Consult the current literature for updated names on this group that included the Black Yeasts; Some species are likely opportunistic human pathogen (Irinyi et al. 2015); Decay leaf, fruit, seed, animal material (Põlme et al. 2020) | Tedersoo L, et al. 2014. Science 346:e1256688 ((https://doi.org/10.1126/science.1256688)); Irinyi L, et al. 2015. Medical Mycology 53:313-337 ((https://doi.org/10.1093/mmy/myv008)); Põlme S, et al. 2020. Fungal Diversity 105:1-16 ((https://doi.org/10.1007/s13225-020-00466-2)) |
| Otu04013 | 8 | k__Fungi(100);p__Ascomycota(100);c__Saccharomycetes(100);o__Saccharomycetales(100);f__Metschnikowiaceae(100);g__Metschnikowia(100);s__Metschnikowia_pulcherrima(100); | Metschnikowia | 13 | Saprotroph | \|Undefined Saprotroph\| | Yeast | NULL | Probable | As nectar-tap_saprotroph (Põlme et al. 2020); Decay sugar-rich substrates (Põlme et al. 2020) | Põlme S, et al. 2020. Fungal Diversity 105:1-16 ((https://doi.org/10.1007/s13225-020-00466-2)) |
| Otu04014 | 8 | k__Fungi(100);p__Basidiomycota(100);c__Agaricomycetes(100);o__Polyporales(100);f__Polyporaceae(100);g__Perenniporia(100);g__Perenniporia_unclassified(100); | Perenniporia | 13 | Saprotroph | \|Wood Saprotroph\| | Corticioid-Polyporoid | White Rot | Highly Probable | As white_rot (Põlme et al. 2020); Decay wood (Põlme et al. 2020) | Gilbertson RL, Ryvarden L. 1987-1987. North American Polypores. Fungiflora, Oslo (ISBN: 978-0945345060); Tedersoo L, et al. 2014. Science 346:e1256688 ((https://doi.org/10.1126/science.1256688)); Põlme S, et al. 2020. Fungal Diversity 105:1-16 ((https://doi.org/10.1007/s13225-020-00466-2)) |
| Otu04016 | 8 | k__Fungi(100);p__Basidiomycota(100);c__Microbotryomycetes(100);o__Sporidiobolales(100);f__Sporidiobolales_family_Incertae_sedis(100);g__Rhodotorula(100);s__Rhodotorula_taiwanensis(100); | Rhodotorula | 13 | Pathotroph-Saprotroph-Symbiotroph | Animal Pathogen-Animal Symbiotroph-Endophyte-Plant Pathogen-\|Undefined Saprotroph\| | Yeast | NULL | Probable | As foliar_endophyte (Põlme et al. 2020); Opportunistic human pathogen | Põlme S, et al. 2020. Fungal Diversity 105:1-16 ((https://doi.org/10.1007/s13225-020-00466-2)); https://www.cdc.gov ((https://www.cdc.gov)) |
| Otu04020 | 8 | k__Fungi(100);p__Ascomycota(100);c__Sordariomycetes(100);o__Hypocreales(100);o__Hypocreales_unclassified(100);o__Hypocreales_unclassified(100);o__Hypocreales_unclassified(100); | Hypocreales | 7 | Saprotroph | Undefined Saprotroph | Microfungus | NULL | Possible | NULL | Sterkenburg E, et al. 2015. New Phytologist 207:1145-1158 ((https://doi.org/10.1111/nph.13426)) |
| Otu04024 | 8 | k__Fungi(100);p__Ascomycota(100);c__Saccharomycetes(100);o__Saccharomycetales(100);f__Metschnikowiaceae(100);g__Metschnikowia(100);s__Metschnikowia_pulcherrima(75); | Metschnikowia | 13 | Saprotroph | \|Undefined Saprotroph\| | Yeast | NULL | Probable | As nectar-tap_saprotroph (Põlme et al. 2020); Decay sugar-rich substrates (Põlme et al. 2020) | Põlme S, et al. 2020. Fungal Diversity 105:1-16 ((https://doi.org/10.1007/s13225-020-00466-2)) |
| Otu04026 | 8 | k__Fungi(100);p__Basidiomycota(100);c__Agaricomycetes(100);o__Russulales(100);f__Russulaceae(100);g__Russula(100);s__Russula_compacta(100); | Russula | 13 | Symbiotroph | \|Ectomycorrhizal\| | Agaricoid | NULL | Highly Probable | NULL | Rinaldi AC, et al. 2008. Fungal Diversity 33:1-45 ((http://www.fungaldiversity.org/fdp/sfdp/33-1.pdf)); Tedersoo L, et al. 2010. Mycorrhiza 20:217-263 ((https://doi.org/10.1007/s00572-009-0274-x)); Põlme S, et al. 2020. Fungal Diversity 105:1-16 ((https://doi.org/10.1007/s13225-020-00466-2)); http://mycorrhizas.info/ecmf.html ((http://mycorrhizas.info/ecmf.html)) |
| Otu04028 | 8 | k__Fungi(100);p__Basidiomycota(100);c__Agaricomycetes(100);o__Agaricales(100);f__Marasmiaceae(100);g__Micromphale(100);s__Micromphale_perforans(100); | Micromphale | 13 | Saprotroph | \|Plant Saprotroph\|-Undefined Saprotroph | Agaricoid | NULL | Probable | As litter_saprotroph (Põlme et al. 2020); Decay leaf, fruit, seed (Põlme et al. 2020) | Tedersoo L, et al. 2014. Science 346:e1256688 ((https://doi.org/10.1126/science.1256688)); Põlme S, et al. 2020. Fungal Diversity 105:1-16 ((https://doi.org/10.1007/s13225-020-00466-2)) |
| Otu04032 | 8 | k__Fungi(100);p__Basidiomycota(100);c__Agaricomycetes(100);o__Polyporales(88);f__Meruliaceae(88);g__Hyphoderma(88);g__Hyphoderma_unclassified(88); | Hyphoderma | 13 | Saprotroph | Undefined Saprotroph-\|Wood Saprotroph\| | Corticioid | White Rot | Probable | As white_rot (Põlme et al. 2020); Decay wood (Põlme et al. 2020) | Tedersoo L, et al. 2014. Science 346:e1256688 ((https://doi.org/10.1126/science.1256688)); Põlme S, et al. 2020. Fungal Diversity 105:1-16 ((https://doi.org/10.1007/s13225-020-00466-2)) |
| Otu04033 | 8 | k__Fungi(100);p__Basidiomycota(100);c__Agaricomycetes(100);o__Agaricales(100);f__Pluteaceae(100);g__Pluteus(100);s__Pluteus_sp_UC_1861231(100); | Pluteus | 13 | Pathotroph-Saprotroph | Bryophyte Parasite-\|Plant Saprotroph\|-Wood Saprotroph | Agaricoid | White Rot | Probable | As litter_saprotroph (Põlme et al. 2020); As white_rot (Põlme et al. 2020); Decay leaf, fruit, seed (Põlme et al. 2020) | Grgurinovic CA, Simpson JA. 2001. Fungal Diversity 8:97-127 ((http://www.fungaldiversity.org/fdp/sfdp/FD_8_97-127.pdf)); James TY, et al. 2006. Nature 443:818-822 ((https://doi.org/10.1038/nature05110)); Tedersoo L, et al. 2014. Science 346:e1256688 ((https://doi.org/10.1126/science.1256688)); Põlme S, et al. 2020. Fungal Diversity 105:1-16 ((https://doi.org/10.1007/s13225-020-00466-2)) |
| Otu04036 | 8 | k__Fungi(100);p__Ascomycota(100);c__Saccharomycetes(100);o__Saccharomycetales(100);f__Saccharomycetales_family_Incertae_sedis(100);g__Candida(100);g__Candida_unclassified(88); | Candida | 13 | Pathotroph-Saprotroph-Symbiotroph | Animal Pathogen-Endophyte-Endosymbiont-Epiphyte-\|Undefined Saprotroph\| | Dimorphic Yeast-Yeast | NULL | Probable | As nectar-tap_saprotroph (Põlme et al. 2020); Decay sugar-rich substrates (Põlme et al. 2020); This asexual genus is polyphyletic and contain species that function very differently depend on what they are. Caution should be taken when interpreting guilds for Candida species. | Manolakaki D, et al. 2010. Virulence. 1: 367-375 ((https://doi.org/10.4161/viru.1.5.12796)); Mekbib SB, et al. 2011. Tropical Plant Pathology 36:233-240 ((https://doi.org/10.1590/S1982-56762011000400004)); Põlme S, et al. 2020. Fungal Diversity 105:1-16 ((https://doi.org/10.1007/s13225-020-00466-2)) |
| Otu04038 | 8 | k__Fungi(100);p__Ascomycota(100);c__Saccharomycetes(100);o__Saccharomycetales(100);f__Saccharomycetales_family_Incertae_sedis(75);g__Candida(75);g__Candida_unclassified(63); | Candida | 13 | Pathotroph-Saprotroph-Symbiotroph | Animal Pathogen-Endophyte-Endosymbiont-Epiphyte-\|Undefined Saprotroph\| | Dimorphic Yeast-Yeast | NULL | Probable | As nectar-tap_saprotroph (Põlme et al. 2020); Decay sugar-rich substrates (Põlme et al. 2020); This asexual genus is polyphyletic and contain species that function very differently depend on what they are. Caution should be taken when interpreting guilds for Candida species. | Manolakaki D, et al. 2010. Virulence. 1: 367-375 ((https://doi.org/10.4161/viru.1.5.12796)); Mekbib SB, et al. 2011. Tropical Plant Pathology 36:233-240 ((https://doi.org/10.1590/S1982-56762011000400004)); Põlme S, et al. 2020. Fungal Diversity 105:1-16 ((https://doi.org/10.1007/s13225-020-00466-2)) |
| Otu04040 | 8 | k__Fungi(100);p__Ascomycota(100);c__Sordariomycetes(100);o__Xylariales(100);f__Xylariaceae(100);g__Hypoxylon(75);g__Hypoxylon_unclassified(75); | Hypoxylon | 13 | Saprotroph-Saprotroph-Symbiotroph | Endophyte-Undefined Saprotroph-\|Wood Saprotroph\| | Corticioid-Microfungus-Xylarioid | NULL | Probable | As foliar_endophyte (Põlme et al. 2020); Decay wood, leaf, fruit, seed (Põlme et al. 2020); Host - angiosperms (Põlme et al. 2020) | Tedersoo L, et al. 2014. Science 346:e1256688 ((https://doi.org/10.1126/science.1256688)); Põlme S, et al. 2020. Fungal Diversity 105:1-16 ((https://doi.org/10.1007/s13225-020-00466-2)) |
| Otu04042 | 8 | k__Fungi(100);p__Ascomycota(100);c__Sordariomycetes(100);o__Diaporthales(100);f__Valsaceae(100);f__Valsaceae_unclassified(100);f__Valsaceae_unclassified(100); | Valsaceae | 9 | Pathotroph-Saprotroph-Symbiotroph | Endophyte-Plant Pathogen-Undefined Saprotroph | Microfungus | NULL | Probable | Saprobes, endophytes and weak to virulent parasites of woody plants (Cannon & Kirk 2007) | Cannon PF, Kirk PM. 2007. Fungal Families of the World. CAB International, Cambridge (ISBN: 978-0851998275) |
| Otu04043 | 8 | k__Fungi(100);p__Ascomycota(100);c__Eurotiomycetes(100);o__Ascosphaerales(100);f__Ascosphaeraceae(100);g__Ascosphaera(100);s__Ascosphaera_aggregata(100); | Ascosphaera | 13 | Pathotroph-Saprotroph | \|Animal Parasite\|-Animal Pathogen-Undefined Saprotroph | Microfungus | NULL | Probable | As animal_decomposer (Põlme et al. 2020); Decay animal material (Põlme et al. 2020); Host - Apis (Põlme et al. 2020) | Tedersoo L, et al. 2014. Science 346:e1256688 ((https://doi.org/10.1126/science.1256688)); Põlme S, et al. 2020. Fungal Diversity 105:1-16 ((https://doi.org/10.1007/s13225-020-00466-2)) |
| Otu04044 | 8 | k__Fungi(100);p__Ascomycota(100);c__Sordariomycetes(100);o__Hypocreales(100);o__Hypocreales_unclassified(100);o__Hypocreales_unclassified(100);o__Hypocreales_unclassified(100); | Hypocreales | 7 | Saprotroph | Undefined Saprotroph | Microfungus | NULL | Possible | NULL | Sterkenburg E, et al. 2015. New Phytologist 207:1145-1158 ((https://doi.org/10.1111/nph.13426)) |
| Otu04045 | 8 | k__Fungi(100);p__Ascomycota(100);c__Taphrinomycetes(100);o__Taphrinales(100);f__Taphrinaceae(100);g__Taphrina(100);s__Taphrina_sp(100); | Taphrina | 13 | Pathotroph | \|Plant Pathogen\| | Dimorphic-Microfungus | NULL | Probable | NULL | Põlme S, et al. 2020. Fungal Diversity 105:1-16 ((https://doi.org/10.1007/s13225-020-00466-2)); The American Phytopathological Society Common Names of Plant Diseases ((http://www.apsnet.org/publications/commonnames/Pages/default.aspx)) |
| Otu04046 | 8 | k__Fungi(100);p__Basidiomycota(100);c__Agaricomycetes(100);o__Cantharellales(100);f__Ceratobasidiaceae(100);f__Ceratobasidiaceae_unclassified(100);f__Ceratobasidiaceae_unclassified(100); | Ceratobasidiaceae | 9 | Pathotroph-Saprotroph-Symbiotroph | Endomycorrhizal-Plant Pathogen-Undefined Saprotroph | Microfungus | NULL | Possible | Best known as plant pathogens, but also saprobic. Species are endomycorrhizal with a wide range of plant roots - especially of Orchidaceae but also even bryophytes (Cannon & Kirk 2007) | Cannon PF, Kirk PM. 2007. Fungal Families of the World. CAB International, Cambridge (ISBN: 978-0851998275) |
| Otu04047 | 8 | k__Fungi(100);p__Ascomycota(100);c__Saccharomycetes(100);o__Saccharomycetales(100);f__Saccharomycetaceae(100);g__Williopsis(100);s__Williopsis_californica_var_dimennae(100); | Williopsis | 13 | Saprotroph-Symbiotroph | Endophyte-Plant Saprotroph-\|Undefined Saprotroph\| | Microfungus-Yeast | NULL | Possible | As foliar_endophyte (Põlme et al. 2020); As litter_saprotroph (Põlme et al. 2020); As soil_saprotroph (Põlme et al. 2020); culture (Del Olmo; Decay leaf, fruit, seed, soil (Põlme et al. 2020); Endophyte detection method; Ruiz & Arnold 2014) | Del Olmo-Ruiz M, Arnold AE. 2014. Mycologia 106:8-21 ((https://doi.org/10.3852/13-098)); Põlme S, et al. 2020. Fungal Diversity 105:1-16 ((https://doi.org/10.1007/s13225-020-00466-2)) |
| Otu04048 | 8 | k__Fungi(100);p__Ascomycota(100);c__Eurotiomycetes(100);o__Chaetothyriales(100);f__Herpotrichiellaceae(100);f__Herpotrichiellaceae_unclassified(100);f__Herpotrichiellaceae_unclassified(100); | Herpotrichiellaceae | 9 | Pathotroph-Saprotroph | Animal Pathogen-Fungal Parasite-Undefined Saprotroph | Facultative Yeast-Microfungus | NULL | Probable | Mostly saprobic on plants or other fungi, isolated from soil etc., and from medical sources (Cannon & Kirk 2007) | Cannon PF, Kirk PM. 2007. Fungal Families of the World. CAB International, Cambridge (ISBN: 978-0851998275) |
| Otu04049 | 8 | k__Fungi(100);p__Ascomycota(100);c__Dothideomycetes(100);o__Pleosporales(100);f__Tubeufiaceae(100);g__Acanthostigma(100);s__Acanthostigma_scopulum(100); | Acanthostigma | 13 | Saprotroph | Undefined Saprotroph-\|Wood Saprotroph\| | Microfungus | NULL | Probable | Decay wood (Põlme et al. 2020) | Tedersoo L, et al. 2014. Science 346:e1256688 ((https://doi.org/10.1126/science.1256688)); Põlme S, et al. 2020. Fungal Diversity 105:1-16 ((https://doi.org/10.1007/s13225-020-00466-2)) |
| Otu04051 | 8 | k__Fungi(100);p__Ascomycota(100);c__Pezizomycetes(100);o__Pezizales(100);f__Sarcosomataceae(100);g__Galiella(100);s__Galiella_sp(100); | Galiella | 13 | Saprotroph | Undefined Saprotroph-\|Wood Saprotroph\| | Microfungus | NULL | Probable | Decay wood (Põlme et al. 2020) | Tedersoo L, et al. 2014. Science 346:e1256688 ((https://doi.org/10.1126/science.1256688)); Põlme S, et al. 2020. Fungal Diversity 105:1-16 ((https://doi.org/10.1007/s13225-020-00466-2)) |
| Otu04053 | 8 | k__Fungi(100);p__Basidiomycota(100);c__Agaricomycetes(100);o__Agaricales(100);f__Pluteaceae(100);g__Pluteus(100);s__Pluteus_brunneidiscus(100); | Pluteus | 13 | Pathotroph-Saprotroph | Bryophyte Parasite-\|Plant Saprotroph\|-Wood Saprotroph | Agaricoid | White Rot | Probable | As litter_saprotroph (Põlme et al. 2020); As white_rot (Põlme et al. 2020); Decay leaf, fruit, seed (Põlme et al. 2020) | Grgurinovic CA, Simpson JA. 2001. Fungal Diversity 8:97-127 ((http://www.fungaldiversity.org/fdp/sfdp/FD_8_97-127.pdf)); James TY, et al. 2006. Nature 443:818-822 ((https://doi.org/10.1038/nature05110)); Tedersoo L, et al. 2014. Science 346:e1256688 ((https://doi.org/10.1126/science.1256688)); Põlme S, et al. 2020. Fungal Diversity 105:1-16 ((https://doi.org/10.1007/s13225-020-00466-2)) |
| Otu04054 | 8 | k__Fungi(100);p__Ascomycota(100);c__Dothideomycetes(100);o__Pleosporales(100);f__Lophiostomataceae(88);g__Lophiostoma(88);s__Lophiostoma_cf_cynaroidis_A33(88); | Lophiostoma | 13 | Saprotroph | Plant Saprotroph-Undefined Saprotroph-\|Wood Saprotroph\| | Microfungus | NULL | Probable | As litter_saprotroph (Põlme et al. 2020); Decay wood, leaf, fruit, seed (Põlme et al. 2020) | Tedersoo L, et al. 2014. Science 346:e1256688 ((https://doi.org/10.1126/science.1256688)); Põlme S, et al. 2020. Fungal Diversity 105:1-16 ((https://doi.org/10.1007/s13225-020-00466-2)) |

| | | | | | | | | | | | | |
|---|---|---|---|---|---|---|---|---|---|---|---|---|
| Otu04055 | 8 | k__Fungi(100);p__Basidiomycota(100);c__Agaricomycetes(100);o__Atheliales(100);f__Atheliaceae(100);g__Byssocorticium(100);s__Byssocorticium_sp(100); | Byssocorticium | | 13 | Saprotroph-Symbiotroph | [illegible] | [illegible] | [illegible] | [illegible] | [illegible] | [illegible] |
| Otu04056 | 8 | k__Fungi(100);p__Ascomycota(100);c__Sordariomycetes(100);o__Hypocreales(100);f__Hypocreales_family_Incertae_sedis(100);g__Myrothecium(100);g__Myrothecium_unclassified(100); | Myrothecium | | 13 | Pathotroph-Saprotroph | [illegible] | [illegible] | [illegible] | [illegible] | [illegible] | [illegible] |
| Otu04057 | 8 | k__Fungi(100);p__Ascomycota(100);c__Saccharomycetes(100);o__Saccharomycetales(100);f__Metschnikowiaceae(100);g__Metschnikowia(100);s__Metschnikowia_pulcherrima(88); | Metschnikowia | | 13 | Saprotroph | [illegible] | [illegible] | [illegible] | [illegible] | [illegible] | [illegible] |
| Otu04058 | 8 | k__Fungi(100);p__Ascomycota(100);c__Saccharomycetes(100);o__Saccharomycetales(100);f__Metschnikowiaceae(100);g__Metschnikowia(100);g__Metschnikowia_unclassified(63); | Metschnikowia | | 13 | Saprotroph | [illegible] | [illegible] | [illegible] | [illegible] | [illegible] | [illegible] |
| Otu04060 | 8 | k__Fungi(100);p__Ascomycota(100);c__Taphrinomycetes(100);o__Taphrinales(100);f__Taphrinaceae(100);g__Taphrina(100);g__Taphrina_unclassified(100); | Taphrina | | 13 | Pathotroph | [illegible] | [illegible] | [illegible] | [illegible] | [illegible] | [illegible] |
| Otu04063 | 8 | k__Fungi(100);p__Ascomycota(100);c__Sordariomycetes(100);o__Hypocreales(100);f__Cordycipitaceae(100);g__Beauveria(100);g__Beauveria_unclassified(100); | Beauveria | | 13 | Pathotroph-Saprotroph-Symbiotroph | [illegible] | [illegible] | [illegible] | [illegible] | [illegible] | [illegible] |
| Otu04067 | 8 | k__Fungi(100);p__Ascomycota(100);c__Pezizomycetes(100);o__Pezizales(100);f__Pyronemataceae(100);g__Wilcoxina(100);s__Wilcoxina_sp(100); | Wilcoxina | | 13 | Symbiotroph | [illegible] | [illegible] | [illegible] | [illegible] | [illegible] | [illegible] |
| Otu04068 | 8 | k__Fungi(100);p__Zygomycota(100);c__Zygomycota_class_Incertae_sedis(100);o__Mortierellales(100);f__Mortierellaceae(100);g__Mortierella(100);s__Mortierella_bisporalis(100); | Mortierella | | 13 | Saprotroph-Symbiotroph | [illegible] | [illegible] | [illegible] | [illegible] | [illegible] | [illegible] |
| Otu04069 | 8 | k__Fungi(100);p__Ascomycota(100);c__Saccharomycetes(100);o__Saccharomycetales(100);f__Metschnikowiaceae(100);g__Metschnikowia(100);s__Metschnikowia_pulcherrima(100); | Metschnikowia | | 13 | Saprotroph | [illegible] | [illegible] | [illegible] | [illegible] | [illegible] | [illegible] |
| Otu04070 | 8 | k__Fungi(100);p__Basidiomycota(100);c__Microbotryomycetes(100);o__Sporidiobolales(100);f__Sporidiobolales_family_Incertae_sedis(100);g__Sporobolomyces(100);s__Sporobolomyces_lactosus(100); | Sporobolomyces | | 13 | Pathotroph-Saprotroph | [illegible] | [illegible] | [illegible] | [illegible] | [illegible] | [illegible] |
| Otu04071 | 8 | k__Fungi(100);p__Ascomycota(100);c__Eurotiomycetes(100);o__Eurotiales(100);f__Trichocomaceae(100);g__Penicillium(88);g__Penicillium_unclassified(88); | Penicillium | | 13 | Pathotroph-Saprotroph-Symbiotroph | [illegible] | [illegible] | [illegible] | [illegible] | [illegible] | [illegible] |
| Otu04073 | 8 | k__Fungi(100);p__Ascomycota(100);c__Saccharomycetes(100);o__Saccharomycetales(100);f__Metschnikowiaceae(100);g__Metschnikowia(100);s__Metschnikowia_pulcherrima(100); | Metschnikowia | | 13 | Saprotroph | [illegible] | [illegible] | [illegible] | [illegible] | [illegible] | [illegible] |
| Otu04075 | 8 | k__Fungi(100);p__Ascomycota(100);c__Saccharomycetes(100);o__Saccharomycetales(100);f__Metschnikowiaceae(100);g__Metschnikowia(100);s__Metschnikowia_pulcherrima(100); | Metschnikowia | | 13 | Saprotroph | [illegible] | [illegible] | [illegible] | [illegible] | [illegible] | [illegible] |
| Otu04076 | 8 | k__Fungi(100);p__Basidiomycota(100);c__Agaricomycetes(100);o__Polyporales(100);f__Phanerochaetaceae(100);g__Ceriporiopsis(100);s__Ceriporiopsis_resinascens(100); | Ceriporiopsis | | 13 | Saprotroph | [illegible] | [illegible] | [illegible] | [illegible] | [illegible] | [illegible] |
| Otu04077 | 8 | k__Fungi(100);p__Ascomycota(100);c__Saccharomycetes(100);o__Saccharomycetales(100);f__Metschnikowiaceae(100);g__Metschnikowia(100);s__Metschnikowia_pulcherrima(63); | Metschnikowia | | 13 | Saprotroph | [illegible] | [illegible] | [illegible] | [illegible] | [illegible] | [illegible] |
| Otu04079 | 8 | k__Fungi(100);p__Ascomycota(100);c__Sordariomycetes(100);o__Xylariales(100);f__Xylariaceae(100);f__Xylariaceae_unclassified(100);f__Xylariaceae_unclassified(100); | Xylariaceae | | 9 | Pathotroph-Saprotroph-Symbiotroph | [illegible] | [illegible] | [illegible] | [illegible] | [illegible] | [illegible] |
| Otu04080 | 8 | k__Fungi(100);p__Ascomycota(100);c__Dothideomycetes(100);o__Pleosporales(100);f__Venturiaceae(100);g__Fusicladium(100);s__Fusicladium_caryigenum(100); | Fusicladium | | 13 | Pathotroph-Symbiotroph | [illegible] | [illegible] | [illegible] | [illegible] | [illegible] | [illegible] |
| Otu04082 | 8 | k__Fungi(100);p__Ascomycota(100);c__Ascomycota_class_Incertae_sedis(100);o__Ascomycota_order_Incertae_sedis(100);f__Ascomycota_family_Incertae_sedis(100);g__Slimacomyces(100);s__Slimacomyces_isiola(100); | Slimacomyces | | 13 | Saprotroph | [illegible] | [illegible] | [illegible] | [illegible] | [illegible] | [illegible] |
| Otu04086 | 8 | k__Fungi(100);p__Ascomycota(100);c__Saccharomycetes(100);o__Saccharomycetales(100);f__Saccharomycetales_family_Incertae_sedis(100);g__Candida(100);g__Candida_unclassified(75); | Candida | | 13 | Pathotroph-Saprotroph-Symbiotroph | [illegible] | [illegible] | [illegible] | [illegible] | [illegible] | [illegible] |
| Otu04092 | 8 | k__Fungi(100);p__Basidiomycota(100);c__Agaricomycetes(100);o__Polyporales(100);f__Polyporaceae(100);g__unclassified_Polyporaceae(100);s__Polyporaceae_sp(100); | Polyporaceae | | 9 | Saprotroph | [illegible] | [illegible] | [illegible] | [illegible] | [illegible] | [illegible] |
| Otu04096 | 8 | k__Fungi(100);p__Basidiomycota(100);c__Agaricomycetes(100);o__Agaricales(100);f__Inocybaceae(100);g__Crepidotus(100);s__Crepidotus_crocophyllus(100); | Crepidotus | | 13 | Saprotroph | [illegible] | [illegible] | [illegible] | [illegible] | [illegible] | [illegible] |
| Otu04097 | 8 | k__Fungi(100);p__Ascomycota(100);c__Saccharomycetes(100);o__Saccharomycetales(100);f__Saccharomycetales_family_Incertae_sedis(100);g__Candida(75);g__Candida_unclassified(75); | Candida | | 13 | Pathotroph-Saprotroph-Symbiotroph | [illegible] | [illegible] | [illegible] | [illegible] | [illegible] | [illegible] |
| Otu04098 | 8 | k__Fungi(100);p__Ascomycota(100);c__Dothideomycetes(100);o__Dothideomycetes_order_Incertae_sedis(100);f__Pseudeurotiaceae(100);g__unclassified_Pseudeurotiaceae(100);s__Pseudeurotiaceae_sp(100); | Pseudeurotiaceae | | 9 | Saprotroph | [illegible] | [illegible] | [illegible] | [illegible] | [illegible] | [illegible] |
| Otu04099 | 8 | k__Fungi(100);p__Ascomycota(100);c__Saccharomycetes(100);o__Saccharomycetales(100);f__Metschnikowiaceae(100);g__Metschnikowia(100);s__Metschnikowia_pulcherrima(88); | Metschnikowia | | 13 | Saprotroph | [illegible] | [illegible] | [illegible] | [illegible] | [illegible] | [illegible] |
| Otu04100 | 8 | k__Fungi(100);p__Basidiomycota(100);c__Agaricomycetes(100);o__Agaricales(100);f__Cortinariaceae(100);g__Cortinarius(100);g__Cortinarius_unclassified(100); | Cortinarius | | 13 | Symbiotroph | [illegible] | [illegible] | [illegible] | [illegible] | [illegible] | [illegible] |
| Otu04102 | 8 | k__Fungi(100);p__Basidiomycota(100);c__Agaricomycetes(100);o__Boletales(100);f__Boletaceae(100);g__Boletus(100);s__Boletus_sp(100); | Boletus | | 13 | Pathotroph-Symbiotroph | [illegible] | [illegible] | [illegible] | [illegible] | [illegible] | [illegible] |
| Otu04104 | 8 | k__Fungi(100);p__Basidiomycota(100);c__Agaricomycetes(100);o__Boletales(100);f__Boletaceae(100);g__Boletus(100);s__Boletus_cf_subvelutipes_MES244(100); | Boletus | | 13 | Pathotroph-Symbiotroph | [illegible] | [illegible] | [illegible] | [illegible] | [illegible] | [illegible] |
| Otu04105 | 8 | k__Fungi(100);p__Basidiomycota(100);c__Agaricomycetes(100);o__Trechisporales(100);f__Hydnodontaceae(100);g__Brevicellicium(75);s__Brevicellicium_olivascens(75); | Brevicellicium | | 13 | Saprotroph | [illegible] | [illegible] | [illegible] | [illegible] | [illegible] | [illegible] |
| Otu04108 | 8 | k__Fungi(100);p__Basidiomycota(100);c__Pucciniomycetes(100);o__Septobasidiales(100);f__Septobasidiaceae(100);g__Septobasidium(100);s__Septobasidium_sp_JBO_2010i(100); | Septobasidium | | 13 | Pathotroph | [illegible] | [illegible] | [illegible] | [illegible] | [illegible] | [illegible] |
| Otu04110 | 8 | k__Fungi(100);p__Basidiomycota(100);c__Agaricomycetes(100);o__Agaricales(100);f__Inocybaceae(100);g__Crepidotus(100);s__Crepidotus_mollis(100); | Crepidotus | | 13 | Saprotroph | [illegible] | [illegible] | [illegible] | [illegible] | [illegible] | [illegible] |
| Otu04112 | 8 | k__Fungi(100);p__Ascomycota(100);c__Saccharomycetes(100);o__Saccharomycetales(100);f__Metschnikowiaceae(100);g__Metschnikowia(100);s__Metschnikowia_pulcherrima(88); | Metschnikowia | | 13 | Saprotroph | [illegible] | [illegible] | [illegible] | [illegible] | [illegible] | [illegible] |
| Otu04113 | 8 | k__Fungi(100);p__Ascomycota(100);c__Dothideomycetes(100);o__Capnodiales(100);f__Teratosphaeriaceae(100);g__Devriesia(100);s__Devriesia_sp(100); | Devriesia | | 13 | Pathotroph-Saprotroph | [illegible] | [illegible] | [illegible] | [illegible] | [illegible] | [illegible] |
| Otu04114 | 8 | k__Fungi(100);p__Ascomycota(100);c__Eurotiomycetes(100);o__Chaetothyriales(100);f__Herpotrichiellaceae(100);g__Phaeomoniella(100);s__Phaeomoniella_niveniae(63); | Phaeomoniella | | 13 | Pathotroph | [illegible] | [illegible] | [illegible] | [illegible] | [illegible] | [illegible] |
| Otu04116 | 8 | k__Fungi(100);p__Basidiomycota(100);c__Tremellomycetes(100);o__Tremellales(100);f__Tremellales_family_Incertae_sedis(100);g__Cryptococcus(100);s__Cryptococcus_sp_D162_2(100); | Cryptococcus | | 13 | Pathotroph-Saprotroph-Symbiotroph | [illegible] | [illegible] | [illegible] | [illegible] | [illegible] | [illegible] |
| Otu04117 | 8 | k__Fungi(100);p__Basidiomycota(100);c__Tremellomycetes(100);o__Tremellales(100);f__Tremellales_family_Incertae_sedis(100);g__Cryptococcus(100);s__Cryptococcus_podzolicus(100); | Cryptococcus | | 13 | Pathotroph-Saprotroph-Symbiotroph | [illegible] | [illegible] | [illegible] | [illegible] | [illegible] | [illegible] |
| Otu04118 | 8 | k__Fungi(100);p__Basidiomycota(100);c__Agaricomycetes(100);o__Cantharellales(100);f__Ceratobasidiaceae(100);f__Ceratobasidiaceae_unclassified(63);f__Ceratobasidiaceae_unclassified(63); | Ceratobasidiaceae | | 9 | Pathotroph-Saprotroph-Symbiotroph | [illegible] | [illegible] | [illegible] | [illegible] | [illegible] | [illegible] |
| Otu04121 | 8 | k__Fungi(100);p__Basidiomycota(100);c__Agaricomycetes(100);o__Boletales(100);f__Boletaceae(100);g__unclassified_Boletaceae(100);s__Boletaceae_sp(100); | Boletaceae | | 9 | Pathotroph-Saprotroph-Symbiotroph | [illegible] | [illegible] | [illegible] | [illegible] | [illegible] | [illegible] |
| Otu04123 | 8 | k__Fungi(100);p__Ascomycota(100);c__Sordariomycetes(100);o__Xylariales(100);f__Diatrypaceae(100);g__Cryptosphaeria(100);s__Cryptosphaeria_sp_NSW05PO(100); | Cryptosphaeria | | 13 | Pathotroph | [illegible] | [illegible] | [illegible] | [illegible] | [illegible] | [illegible] |
| Otu04124 | 8 | k__Fungi(100);p__Basidiomycota(100);c__Agaricomycetes(100);o__Auriculariales(100);f__Auriculariales_family_Incertae_sedis(100);g__Auricularia(100);s__Auricularia_delicata(100); | Auricularia | | 13 | Saprotroph | [illegible] | [illegible] | [illegible] | [illegible] | [illegible] | [illegible] |
| Otu04129 | 8 | k__Fungi(100);p__Basidiomycota(100);c__Agaricomycetes(100);o__Polyporales(100);f__Polyporaceae(100);g__Wolfiporia(100);s__Wolfiporia_dilatohypha(100); | Wolfiporia | | 13 | Saprotroph | [illegible] | [illegible] | [illegible] | [illegible] | [illegible] | [illegible] |
| Otu04130 | 8 | k__Fungi(100);p__Ascomycota(100);c__Saccharomycetes(100);o__Saccharomycetales(100);f__Metschnikowiaceae(100);g__Metschnikowia(100);s__Metschnikowia_pulcherrima(100); | Metschnikowia | | 13 | Saprotroph | [illegible] | [illegible] | [illegible] | [illegible] | [illegible] | [illegible] |
| Otu04131 | 8 | k__Fungi(100);p__Ascomycota(100);c__Sordariomycetes(100);o__Xylariales(100);f__Xylariaceae(100);g__Hypoxylon(100);s__Hypoxylon_sp(100); | Hypoxylon | | 13 | Saprotroph-Saprotroph-Symbiotroph | [illegible] | [illegible] | [illegible] | [illegible] | [illegible] | [illegible] |
| Otu04132 | 8 | k__Fungi(100);p__Ascomycota(100);c__Sordariomycetes(100);o__Hypocreales(100);f__Hypocreaceae(100);g__Hypocrea(100);g__Hypocrea_unclassified(100); | Hypocrea | | 13 | Pathotroph | [illegible] | [illegible] | [illegible] | [illegible] | [illegible] | [illegible] |
| Otu04133 | 8 | k__Fungi(100);p__Basidiomycota(100);c__Microbotryomycetes(100);o__Microbotryales(100);f__Microbotryaceae(100);g__Sphacelotheca(100);s__Sphacelotheca_destruens(100); | Sphacelotheca | | 13 | Pathotroph | [illegible] | [illegible] | [illegible] | [illegible] | [illegible] | [illegible] |
| Otu04135 | 8 | k__Fungi(100);p__Basidiomycota(100);c__Tremellomycetes(100);o__Tremellales(100);f__Tremellales_family_Incertae_sedis(100);g__Kockovaella(100);g__Kockovaella_unclassified(100); | Kockovaella | | 13 | Saprotroph-Symbiotroph | [illegible] | [illegible] | [illegible] | [illegible] | [illegible] | [illegible] |
| Otu04137 | 8 | k__Fungi(100);p__Basidiomycota(100);c__Agaricomycetes(100);o__Agaricales(100);o__Agaricales_unclassified(100);o__Agaricales_unclassified(100);o__Agaricales_unclassified(100); | Agaricales | | 7 | Pathotroph-Saprotroph-Symbiotroph | [illegible] | [illegible] | [illegible] | [illegible] | [illegible] | [illegible] |
| Otu04138 | 8 | k__Fungi(100);p__Ascomycota(100);c__Sordariomycetes(100);o__Hypocreales(100);f__Nectriaceae(100);g__unclassified_Nectriaceae(100);s__Nectriaceae_sp(100); | Nectriaceae | | 9 | Pathotroph-Saprotroph-Symbiotroph | [illegible] | [illegible] | [illegible] | [illegible] | [illegible] | [illegible] |
| Otu04140 | 8 | k__Fungi(100);p__Basidiomycota(100);c__Agaricomycetes(100);o__Agaricales(100);o__Agaricales_unclassified(100);o__Agaricales_unclassified(100);o__Agaricales_unclassified(100); | Agaricales | | 7 | Pathotroph-Saprotroph-Symbiotroph | [illegible] | [illegible] | [illegible] | [illegible] | [illegible] | [illegible] |
| Otu04144 | 8 | k__Fungi(100);p__Basidiomycota(100);c__Cystobasidiomycetes(100);o__Erythrobasidiales(100);f__Erythrobasidiales_family_Incertae_sedis(100);g__Erythrobasidium(100);s__Erythrobasidium_sp(100); | Erythrobasidium | | 13 | Pathotroph-Saprotroph | [illegible] | [illegible] | [illegible] | [illegible] | [illegible] | [illegible] |
| Otu04145 | 8 | k__Fungi(100);p__Basidiomycota(100);c__Tremellomycetes(100);o__Cystofilobasidiales(100);f__Cystofilobasidiales_family_Incertae_sedis(100);g__Syzygospora(100);s__Syzygospora_sp(100); | Syzygospora | | 13 | Pathotroph | [illegible] | [illegible] | [illegible] | [illegible] | [illegible] | [illegible] |
| Otu04146 | 8 | k__Fungi(100);p__Basidiomycota(100);c__Agaricomycetes(100);o__Agaricales(100);f__Marasmiaceae(100);f__Marasmiaceae_unclassified(100);f__Marasmiaceae_unclassified(100); | Marasmiaceae | | 9 | Pathotroph-Saprotroph-Symbiotroph | [illegible] | [illegible] | [illegible] | [illegible] | [illegible] | [illegible] |
| Otu04147 | 8 | k__Fungi(100);p__Ascomycota(100);c__Ascomycota_class_Incertae_sedis(100);o__Ascomycota_order_Incertae_sedis(100);f__Ascomycota_family_Incertae_sedis(100);g__Veronaea(100);g__Veronaea_unclassified(100); | Veronaea | | 13 | Pathotroph-Saprotroph | [illegible] | [illegible] | [illegible] | [illegible] | [illegible] | [illegible] |
| Otu04148 | 8 | k__Fungi(100);p__Ascomycota(100);c__Sordariomycetes(100);o__Xylariales(100);f__Xylariaceae(100);g__unclassified_Xylariaceae(100);s__Xylariaceae_sp(100); | Xylariaceae | | 9 | Pathotroph-Saprotroph-Symbiotroph | [illegible] | [illegible] | [illegible] | [illegible] | [illegible] | [illegible] |
| Otu04149 | 8 | k__Fungi(100);p__Ascomycota(100);c__Eurotiomycetes(100);o__Chaetothyriales(100);f__Herpotrichiellaceae(100);f__Herpotrichiellaceae_unclassified(100);f__Herpotrichiellaceae_unclassified(100); | Herpotrichiellaceae | | 9 | Pathotroph-Saprotroph | [illegible] | [illegible] | [illegible] | [illegible] | [illegible] | [illegible] |
| Otu04150 | 8 | k__Fungi(100);p__Basidiomycota(100);c__Agaricomycetes(100);o__Boletales(100);f__Gyroporaceae(100);g__Gyroporus(100);s__Gyroporus_sp(100); | Gyroporus | | 13 | Symbiotroph | [illegible] | [illegible] | [illegible] | [illegible] | [illegible] | [illegible] |
| Otu04151 | 8 | k__Fungi(100);p__Basidiomycota(100);c__Agaricomycetes(100);o__Agaricomycetes_order_Incertae_sedis(100);f__Agaricomycetes_family_Incertae_sedis(100);g__Peniophorella(100);s__Peniophorella_pertenuis(100); | Peniophorella | | 13 | Saprotroph | [illegible] | [illegible] | [illegible] | [illegible] | [illegible] | [illegible] |
| Otu04152 | 8 | k__Fungi(100);p__Ascomycota(100);c__Eurotiomycetes(100);o__Chaetothyriales(100);f__Herpotrichiellaceae(100);g__Phaeomoniella(100);s__Phaeomoniella_prunicola(100); | Phaeomoniella | | 13 | Pathotroph | [illegible] | [illegible] | [illegible] | [illegible] | [illegible] | [illegible] |
| Otu04153 | 8 | k__Fungi(100);p__Zygomycota(100);c__Zygomycota_class_Incertae_sedis(100);o__Mortierellales(100);f__Mortierellaceae(100);g__Mortierella(100);g__Mortierella_unclassified(100); | Mortierella | | 13 | Saprotroph-Symbiotroph | [illegible] | [illegible] | [illegible] | [illegible] | [illegible] | [illegible] |
| Otu04154 | 8 | k__Fungi(100);p__Ascomycota(100);c__Dothideomycetes(100);o__Capnodiales(100);f__Mycosphaerellaceae(100);f__Mycosphaerellaceae_unclassified(63);f__Mycosphaerellaceae_unclassified(63); | Mycosphaerellaceae | | 9 | Pathotroph-Saprotroph | [illegible] | [illegible] | [illegible] | [illegible] | [illegible] | [illegible] |
| Otu04155 | 8 | k__Fungi(100);p__Ascomycota(100);c__Sordariomycetes(100);o__Xylariales(100);f__Xylariaceae(100);g__Xylaria(100);s__Xylaria_sp_MUCL_51697(100); | Xylaria | | 13 | Saprotroph-Symbiotroph | [illegible] | [illegible] | [illegible] | [illegible] | [illegible] | [illegible] |
| Otu04157 | 8 | k__Fungi(100);p__Ascomycota(100);c__Dothideomycetes(100);o__Capnodiales(100);f__Davidiellaceae(100);g__Cladosporium(100);g__Cladosporium_unclassified(100); | Cladosporium | | 13 | Pathotroph-Saprotroph-Symbiotroph | [illegible] | [illegible] | [illegible] | [illegible] | [illegible] | [illegible] |
| Otu04159 | 8 | k__Fungi(100);p__Ascomycota(100);c__Sordariomycetes(100);o__Hypocreales(100);f__Hypocreales_family_Incertae_sedis(100);g__Gliomastix(100);s__Gliomastix_tumulicola(100); | Gliomastix | | 13 | Saprotroph | [illegible] | [illegible] | [illegible] | [illegible] | [illegible] | [illegible] |
| Otu04172 | 8 | k__Fungi(100);p__Ascomycota(100);c__Sordariomycetes(100);o__Hypocreales(100);f__Hypocreales_family_Incertae_sedis(100);g__Acremonium(100);s__Acremonium_cavaraeanum(100); | Acremonium | | 13 | Pathotroph-Saprotroph-Symbiotroph | [illegible] | [illegible] | [illegible] | [illegible] | [illegible] | [illegible] |
| Otu04173 | 8 | k__Fungi(100);p__Basidiomycota(100);c__Agaricomycetes(100);o__Agaricales(100);f__Strophariaceae(100);g__Hebeloma(100);s__Hebeloma_helodes(75); | Hebeloma | | 13 | Symbiotroph | [illegible] | [illegible] | [illegible] | [illegible] | [illegible] | [illegible] |
| Otu04174 | 8 | k__Fungi(100);p__Ascomycota(100);c__Saccharomycetes(100);o__Saccharomycetales(100);f__Metschnikowiaceae(100);g__Metschnikowia(100);s__Metschnikowia_pulcherrima(88); | Metschnikowia | | 13 | Saprotroph | [illegible] | [illegible] | [illegible] | [illegible] | [illegible] | [illegible] |
| Otu04175 | 8 | k__Fungi(100);p__Ascomycota(100);c__Sordariomycetes(100);o__Microascales(100);f__Halosphaeriaceae(100);g__unclassified_Halosphaeriaceae(100);s__Halosphaeriaceae_sp(100); | Halosphaeriaceae | | 9 | Saprotroph | [illegible] | [illegible] | [illegible] | [illegible] | [illegible] | [illegible] |
| Otu04177 | 8 | k__Fungi(100);p__Ascomycota(100);c__Saccharomycetes(100);o__Saccharomycetales(100);f__Metschnikowiaceae(100);g__Metschnikowia(100);s__Metschnikowia_pulcherrima(100); | Metschnikowia | | 13 | Saprotroph | [illegible] | [illegible] | [illegible] | [illegible] | [illegible] | [illegible] |
| Otu04178 | 8 | k__Fungi(100);p__Basidiomycota(100);c__Agaricomycetes(100);o__Polyporales(100);f__Phanerochaetaceae(100);g__Ceriporia(100);g__Ceriporia_unclassified(100); | Ceriporia | | 13 | Saprotroph | [illegible] | [illegible] | [illegible] | [illegible] | [illegible] | [illegible] |
| Otu04183 | 8 | k__Fungi(100);p__Ascomycota(100);c__Saccharomycetes(100);o__Saccharomycetales(100);f__Metschnikowiaceae(100);g__Metschnikowia(100);s__Metschnikowia_pulcherrima(100); | Metschnikowia | | 13 | Saprotroph | [illegible] | [illegible] | [illegible] | [illegible] | [illegible] | [illegible] |
| Otu04184 | 8 | k__Fungi(100);p__Basidiomycota(100);c__Agaricomycetes(100);o__Russulales(100);f__Stereaceae(100);g__Gloeocystidiellum(100);s__Gloeocystidiellum_sp(100); | Gloeocystidiellum | | 13 | Saprotroph | [illegible] | [illegible] | [illegible] | [illegible] | [illegible] | [illegible] |
| Otu04188 | 8 | k__Fungi(100);p__Ascomycota(100);c__Saccharomycetes(100);o__Saccharomycetales(100);f__Metschnikowiaceae(100);g__Metschnikowia(100);s__Metschnikowia_pulcherrima(75); | Metschnikowia | | 13 | Saprotroph | [illegible] | [illegible] | [illegible] | [illegible] | [illegible] | [illegible] |
| Otu04189 | 8 | k__Fungi(100);p__Ascomycota(100);c__Saccharomycetes(100);o__Saccharomycetales(100);f__Metschnikowiaceae(100);g__Metschnikowia(100);s__Metschnikowia_pulcherrima(100); | Metschnikowia | | 13 | Saprotroph | [illegible] | [illegible] | [illegible] | [illegible] | [illegible] | [illegible] |
| Otu04191 | 8 | k__Fungi(100);p__Basidiomycota(100);c__Agaricomycetes(100);o__Agaricales(100);f__Cortinariaceae(100);g__Cortinarius(100);g__Cortinarius_unclassified(100); | Cortinarius | | 13 | Symbiotroph | [illegible] | [illegible] | [illegible] | [illegible] | [illegible] | [illegible] |
| Otu04192 | 8 | k__Fungi(100);p__Basidiomycota(100);c__Agaricomycetes(100);o__Agaricales(100);f__Psathyrellaceae(100);g__Psathyrella(100);g__Psathyrella_unclassified(100); | Psathyrella | | 13 | Saprotroph | [illegible] | [illegible] | [illegible] | [illegible] | [illegible] | [illegible] |
| Otu04193 | 8 | k__Fungi(100);p__Ascomycota(100);c__Eurotiomycetes(100);o__Eurotiales(100);f__Trichocomaceae(100);g__Penicillium(100);g__Penicillium_unclassified(100); | Penicillium | | 13 | Pathotroph-Saprotroph-Symbiotroph | [illegible] | [illegible] | [illegible] | [illegible] | [illegible] | [illegible] |
| Otu04194 | 8 | k__Fungi(100);p__Basidiomycota(100);c__Agaricomycetes(100);o__Agaricales(100);f__Agaricaceae(100);g__Coprinus(100);s__Coprinus_foetidellus(100); | Coprinus | | 13 | Pathotroph-Saprotroph | [illegible] | [illegible] | [illegible] | [illegible] | [illegible] | [illegible] |
| Otu04196 | 8 | k__Fungi(100);p__Ascomycota(100);c__Saccharomycetes(100);o__Saccharomycetales(100);f__Metschnikowiaceae(100);g__Metschnikowia(100);s__Metschnikowia_pulcherrima(63); | Metschnikowia | | 13 | Saprotroph | [illegible] | [illegible] | [illegible] | [illegible] | [illegible] | [illegible] |
| Otu04197 | 8 | k__Fungi(100);p__Basidiomycota(100);c__Agaricomycetes(100);o__Agaricales(100);f__Mycenaceae(100);g__Mycena(100);s__Mycena_plumipes(100); | Mycena | | 13 | Pathotroph-Saprotroph | [illegible] | [illegible] | [illegible] | [illegible] | [illegible] | [illegible] |
| Otu04198 | 8 | k__Fungi(100);p__Basidiomycota(100);c__Agaricomycetes(100);o__Polyporales(100);f__Meruliaceae(100);g__Steccherinum(100);s__Steccherinum_cf_ciliolatum(100); | Steccherinum | | 13 | Saprotroph | [illegible] | [illegible] | [illegible] | [illegible] | [illegible] | [illegible] |
| Otu04199 | 8 | k__Fungi(100);p__Ascomycota(100);c__Saccharomycetes(100);o__Saccharomycetales(100);f__Metschnikowiaceae(100);g__Metschnikowia(100);s__Metschnikowia_pulcherrima(100); | Metschnikowia | | 13 | Saprotroph | [illegible] | [illegible] | [illegible] | [illegible] | [illegible] | [illegible] |
| Otu04200 | 8 | k__Fungi(100);p__Ascomycota(100);c__Lecanoromycetes(100);o__Ostropales(100);f__Stictidaceae(100);g__Cryptodiscus(100);s__Cryptodiscus_pallidus(100); | Cryptodiscus | | 13 | Pathotroph-Saprotroph | [illegible] | [illegible] | [illegible] | [illegible] | [illegible] | [illegible] |
| Otu04202 | 8 | k__Fungi(100);p__Ascomycota(100);c__Saccharomycetes(100);o__Saccharomycetales(100);f__Metschnikowiaceae(100);g__Metschnikowia(100);s__Metschnikowia_pulcherrima(88); | Metschnikowia | | 13 | Saprotroph | [illegible] | [illegible] | [illegible] | [illegible] | [illegible] | [illegible] |
| Otu04205 | 8 | k__Fungi(100);p__Basidiomycota(100);c__Tremellomycetes(100);o__Tremellales(100);f__Tremellales_family_Incertae_sedis(100);f__Tremellales_family_Incertae_sedis_unclassified(100);f__Tremellales_family_Incertae_sedis_unclassified(100); | Tremellales | | 7 | Pathotroph-Saprotroph-Symbiotroph | [illegible] | [illegible] | [illegible] | [illegible] | [illegible] | [illegible] |
| Otu04207 | 8 | k__Fungi(100);p__Ascomycota(100);c__Dothideomycetes(100);o__Pleosporales(100);f__Cucurbitariaceae(100);g__Curreya(88);g__Curreya_unclassified(88); | Curreya | | 13 | Saprotroph | [illegible] | [illegible] | [illegible] | [illegible] | [illegible] | [illegible] |
| Otu04209 | 8 | k__Fungi(100);p__Ascomycota(100);c__Dothideomycetes(100);o__Capnodiales(100);f__Mycosphaerellaceae(100);f__Mycosphaerellaceae_unclassified(100);f__Mycosphaerellaceae_unclassified(100); | Mycosphaerellaceae | | 9 | Pathotroph-Saprotroph | [illegible] | [illegible] | [illegible] | [illegible] | [illegible] | [illegible] |
| Otu04210 | 8 | k__Fungi(100);p__Ascomycota(100);c__Sordariomycetes(100);o__Hypocreales(100);o__Hypocreales_unclassified(100);o__Hypocreales_unclassified(100);o__Hypocreales_unclassified(100); | Hypocreales | | 7 | Saprotroph | [illegible] | [illegible] | [illegible] | [illegible] | [illegible] | [illegible] |
| Otu04213 | 8 | k__Fungi(100);p__Ascomycota(100);c__Dothideomycetes(100);o__Pleosporales(100);f__Venturiaceae(100);g__Venturia(75);g__Venturia_unclassified(63); | Venturia | | 13 | Pathotroph-Pathotroph-Saprotroph | [illegible] | [illegible] | [illegible] | [illegible] | [illegible] | [illegible] |
| Otu04214 | 8 | k__Fungi(100);p__Basidiomycota(100);c__Agaricomycetes(100);o__Agaricales(100);f__Fistulinaceae(100);g__Porodisculus(100);s__Porodisculus_pendulus(100); | Porodisculus | | 13 | Pathotroph-Saprotroph | [illegible] | [illegible] | [illegible] | [illegible] | [illegible] | [illegible] |
| Otu04216 | 8 | k__Fungi(100);p__Ascomycota(100);c__Leotiomycetes(100);o__Helotiales(100);f__Helotiales_family_Incertae_sedis(100);g__Ascocoryne(100);s__Ascocoryne_sp(100); | Ascocoryne | | 13 | Saprotroph | [illegible] | [illegible] | [illegible] | [illegible] | [illegible] | [illegible] |
| Otu04218 | 8 | k__Fungi(100);p__Ascomycota(100);c__Saccharomycetes(100);o__Saccharomycetales(100);f__Metschnikowiaceae(100);g__Metschnikowia(100);s__Metschnikowia_pulcherrima(100); | Metschnikowia | | 13 | Saprotroph | [illegible] | [illegible] | [illegible] | [illegible] | [illegible] | [illegible] |
| Otu04219 | 8 | k__Fungi(100);p__Ascomycota(100);c__Sordariomycetes(100);o__Hypocreales(100);f__Hypocreales_family_Incertae_sedis(100);g__Myrothecium(100);s__Myrothecium_inundatum(100); | Myrothecium | | 13 | Pathotroph-Saprotroph | [illegible] | [illegible] | [illegible] | [illegible] | [illegible] | [illegible] |
| Otu03865 | 8 | k__Fungi(100);p__Ascomycota(88);p__Ascomycota_unclassified(88);p__Ascomycota_unclassified(88);p__Ascomycota_unclassified(88);p__Ascomycota_unclassified(88);p__Ascomycota_unclassified(88); | - | - | - | - | - | - | - | - | Unassigned | |
| Otu03867 | 8 | k__Fungi(100);k__Fungi_unclassified(75);k__Fungi_unclassified(75);k__Fungi_unclassified(75);k__Fungi_unclassified(75);k__Fungi_unclassified(75);k__Fungi_unclassified(75); | - | - | - | - | - | - | - | - | Unassigned | |
| Otu03875 | 8 | k__Fungi(100);p__Ascomycota(100);p__Ascomycota_unclassified(100);p__Ascomycota_unclassified(100);p__Ascomycota_unclassified(100);p__Ascomycota_unclassified(100);p__Ascomycota_unclassified(100); | - | - | - | - | - | - | - | - | Unassigned | |
| Otu03876 | 8 | k__Fungi(100);p__Ascomycota(75);p__Ascomycota_unclassified(75);p__Ascomycota_unclassified(75);p__Ascomycota_unclassified(75);p__Ascomycota_unclassified(75);p__Ascomycota_unclassified(75); | - | - | - | - | - | - | - | - | Unassigned | |
| Otu03878 | 8 | k__Fungi(100);k__Fungi_unclassified(63);k__Fungi_unclassified(63);k__Fungi_unclassified(63);k__Fungi_unclassified(63);k__Fungi_unclassified(63);k__Fungi_unclassified(63); | - | - | - | - | - | - | - | - | Unassigned | |
| Otu03881 | 8 | k__Fungi(100);p__Ascomycota(63);p__Ascomycota_unclassified(63);p__Ascomycota_unclassified(63);p__Ascomycota_unclassified(63);p__Ascomycota_unclassified(63);p__Ascomycota_unclassified(63); | - | - | - | - | - | - | - | - | Unassigned | |
| Otu03882 | 8 | k__Fungi(100);k__Fungi_unclassified(63);k__Fungi_unclassified(63);k__Fungi_unclassified(63);k__Fungi_unclassified(63);k__Fungi_unclassified(63);k__Fungi_unclassified(63); | - | - | - | - | - | - | - | - | Unassigned | |
| Otu03884 | 8 | k__Fungi(100);k__Fungi_unclassified(100);k__Fungi_unclassified(100);k__Fungi_unclassified(100);k__Fungi_unclassified(100);k__Fungi_unclassified(100);k__Fungi_unclassified(100); | - | - | - | - | - | - | - | - | Unassigned | |
| Otu03886 | 8 | k__Fungi(100);k__Fungi_unclassified(75);k__Fungi_unclassified(75);k__Fungi_unclassified(75);k__Fungi_unclassified(75);k__Fungi_unclassified(75);k__Fungi_unclassified(75); | - | - | - | - | - | - | - | - | Unassigned | |
| Otu03889 | 8 | k__Fungi(100);k__Fungi_unclassified(88);k__Fungi_unclassified(88);k__Fungi_unclassified(88);k__Fungi_unclassified(88);k__Fungi_unclassified(88);k__Fungi_unclassified(88); | - | - | - | - | - | - | - | - | Unassigned | |
| Otu03890 | 8 | k__Fungi(100);k__Fungi_unclassified(100);k__Fungi_unclassified(100);k__Fungi_unclassified(100);k__Fungi_unclassified(100);k__Fungi_unclassified(100);k__Fungi_unclassified(100); | - | - | - | - | - | - | - | - | Unassigned | |
| Otu03891 | 8 | k__Fungi(100);k__Fungi_unclassified(100);k__Fungi_unclassified(100);k__Fungi_unclassified(100);k__Fungi_unclassified(100);k__Fungi_unclassified(100);k__Fungi_unclassified(100); | - | - | - | - | - | - | - | - | Unassigned | |
| Otu03893 | 8 | k__Fungi(100);k__Fungi_unclassified(88);k__Fungi_unclassified(88);k__Fungi_unclassified(88);k__Fungi_unclassified(88);k__Fungi_unclassified(88);k__Fungi_unclassified(88); | - | - | - | - | - | - | - | - | Unassigned | |
| Otu03894 | 8 | k__Fungi(100);p__Ascomycota(100);p__Ascomycota_unclassified(100);p__Ascomycota_unclassified(100);p__Ascomycota_unclassified(100);p__Ascomycota_unclassified(100);p__Ascomycota_unclassified(100); | - | - | - | - | - | - | - | - | Unassigned | |
| Otu03895 | 8 | k__Fungi(100);k__Fungi_unclassified(100);k__Fungi_unclassified(100);k__Fungi_unclassified(100);k__Fungi_unclassified(100);k__Fungi_unclassified(100);k__Fungi_unclassified(100); | - | - | - | - | - | - | - | - | Unassigned | |
| Otu03900 | 8 | k__Fungi(100);k__Fungi_unclassified(100);k__Fungi_unclassified(100);k__Fungi_unclassified(100);k__Fungi_unclassified(100);k__Fungi_unclassified(100);k__Fungi_unclassified(100); | - | - | - | - | - | - | - | - | Unassigned | |
| Otu03901 | 8 | k__Fungi(100);p__Ascomycota(63);p__Ascomycota_unclassified(63);p__Ascomycota_unclassified(63);p__Ascomycota_unclassified(63);p__Ascomycota_unclassified(63);p__Ascomycota_unclassified(63); | - | - | - | - | - | - | - | - | Unassigned | |
| Otu03902 | 8 | k__Fungi(100);k__Fungi_unclassified(75);k__Fungi_unclassified(75);k__Fungi_unclassified(75);k__Fungi_unclassified(75);k__Fungi_unclassified(75);k__Fungi_unclassified(75); | - | - | - | - | - | - | - | - | Unassigned | |
| Otu03903 | 8 | k__Fungi(100);p__Ascomycota(88);p__Ascomycota_unclassified(88);p__Ascomycota_unclassified(88);p__Ascomycota_unclassified(88);p__Ascomycota_unclassified(88);p__Ascomycota_unclassified(88); | - | - | - | - | - | - | - | - | Unassigned | |
| Otu03904 | 8 | k__Fungi(100);k__Fungi_unclassified(75);k__Fungi_unclassified(75);k__Fungi_unclassified(75);k__Fungi_unclassified(75);k__Fungi_unclassified(75);k__Fungi_unclassified(75); | - | - | - | - | - | - | - | - | Unassigned | |
| Otu03905 | 8 | k__Fungi(100);k__Fungi_unclassified(88);k__Fungi_unclassified(88);k__Fungi_unclassified(88);k__Fungi_unclassified(88);k__Fungi_unclassified(88);k__Fungi_unclassified(88); | - | - | - | - | - | - | - | - | Unassigned | |
| Otu03906 | 8 | k__Fungi(100);k__Fungi_unclassified(75);k__Fungi_unclassified(75);k__Fungi_unclassified(75);k__Fungi_unclassified(75);k__Fungi_unclassified(75);k__Fungi_unclassified(75); | - | - | - | - | - | - | - | - | Unassigned | |
| Otu03907 | 8 | k__Fungi(100);p__Ascomycota(63);p__Ascomycota_unclassified(63);p__Ascomycota_unclassified(63);p__Ascomycota_unclassified(63);p__Ascomycota_unclassified(63);p__Ascomycota_unclassified(63); | - | - | - | - | - | - | - | - | Unassigned | |
| Otu03908 | 8 | k__Fungi(100);k__Fungi_unclassified(75);k__Fungi_unclassified(75);k__Fungi_unclassified(75);k__Fungi_unclassified(75);k__Fungi_unclassified(75);k__Fungi_unclassified(75); | - | - | - | - | - | - | - | - | Unassigned | |
| Otu03909 | 8 | k__Fungi(100);p__Ascomycota(63);p__Ascomycota_unclassified(63);p__Ascomycota_unclassified(63);p__Ascomycota_unclassified(63);p__Ascomycota_unclassified(63);p__Ascomycota_unclassified(63); | - | - | - | - | - | - | - | - | Unassigned | |
| Otu03911 | 8 | k__Fungi(100);p__Ascomycota(63);p__Ascomycota_unclassified(63);p__Ascomycota_unclassified(63);p__Ascomycota_unclassified(63);p__Ascomycota_unclassified(63);p__Ascomycota_unclassified(63); | - | - | - | - | - | - | - | - | Unassigned | |
| Otu03912 | 8 | k__Fungi(100);p__Ascomycota(75);p__Ascomycota_unclassified(75);p__Ascomycota_unclassified(75);p__Ascomycota_unclassified(75);p__Ascomycota_unclassified(75);p__Ascomycota_unclassified(75); | - | - | - | - | - | - | - | - | Unassigned | |
| Otu03913 | 8 | k__Fungi(100);p__Ascomycota(75);p__Ascomycota_unclassified(75);p__Ascomycota_unclassified(75);p__Ascomycota_unclassified(75);p__Ascomycota_unclassified(75);p__Ascomycota_unclassified(75); | - | - | - | - | - | - | - | - | Unassigned | |
| Otu03915 | 8 | k__Fungi(100);k__Fungi_unclassified(100);k__Fungi_unclassified(100);k__Fungi_unclassified(100);k__Fungi_unclassified(100);k__Fungi_unclassified(100);k__Fungi_unclassified(100); | - | - | - | - | - | - | - | - | Unassigned | |
| Otu03916 | 8 | k__Fungi(100);k__Fungi_unclassified(100);k__Fungi_unclassified(100);k__Fungi_unclassified(100);k__Fungi_unclassified(100);k__Fungi_unclassified(100);k__Fungi_unclassified(100); | - | - | - | - | - | - | - | - | Unassigned | |
| Otu03917 | 8 | k__Fungi(100);p__Ascomycota(100);c__Dothideomycetes(100);o__Capnodiales(100);o__Capnodiales_unclassified(100);o__Capnodiales_unclassified(100);o__Capnodiales_unclassified(100); | - | - | - | - | - | - | - | - | Unassigned | |
| Otu03918 | 8 | k__Fungi(100);p__Ascomycota(88);p__Ascomycota_unclassified(88);p__Ascomycota_unclassified(88);p__Ascomycota_unclassified(88);p__Ascomycota_unclassified(88);p__Ascomycota_unclassified(88); | - | - | - | - | - | - | - | - | Unassigned | |
| Otu03920 | 8 | k__Fungi(100);p__Ascomycota(75);p__Ascomycota_unclassified(75);p__Ascomycota_unclassified(75);p__Ascomycota_unclassified(75);p__Ascomycota_unclassified(75);p__Ascomycota_unclassified(75); | - | - | - | - | - | - | - | - | Unassigned | |
| Otu03925 | 8 | k__Plantae(100);p__unclassified_Plantae(100);c__unclassified_Plantae(100);o__unclassified_Plantae(100);f__unclassified_Plantae(100);g__unclassified_Plantae(100);s__Plantae_sp(100); | - | - | - | - | - | - | - | - | Unassigned | |
| Otu03926 | 8 | k__Fungi(100);k__Fungi_unclassified(100);k__Fungi_unclassified(100);k__Fungi_unclassified(100);k__Fungi_unclassified(100);k__Fungi_unclassified(100);k__Fungi_unclassified(100); | - | - | - | - | - | - | - | - | Unassigned | |
| Otu03928 | 8 | k__Fungi(100);p__Ascomycota(75);p__Ascomycota_unclassified(75);p__Ascomycota_unclassified(75);p__Ascomycota_unclassified(75);p__Ascomycota_unclassified(75);p__Ascomycota_unclassified(75); | - | - | - | - | - | - | - | - | Unassigned | |
| Otu03929 | 8 | k__Fungi(100);p__Ascomycota(63);p__Ascomycota_unclassified(63);p__Ascomycota_unclassified(63);p__Ascomycota_unclassified(63);p__Ascomycota_unclassified(63);p__Ascomycota_unclassified(63); | - | - | - | - | - | - | - | - | Unassigned | |
| Otu03930 | 8 | k__Fungi(100);k__Fungi_unclassified(100);k__Fungi_unclassified(100);k__Fungi_unclassified(100);k__Fungi_unclassified(100);k__Fungi_unclassified(100);k__Fungi_unclassified(100); | - | - | - | - | - | - | - | - | Unassigned | |
| Otu03931 | 8 | k__Fungi(100);p__Basidiomycota(100);p__Basidiomycota_unclassified(100);p__Basidiomycota_unclassified(100);p__Basidiomycota_unclassified(100);p__Basidiomycota_unclassified(100);p__Basidiomycota_unclassified(100); | - | - | - | - | - | - | - | - | Unassigned | |
| Otu03932 | 8 | k__Fungi(100);p__Ascomycota(75);p__Ascomycota_unclassified(75);p__Ascomycota_unclassified(75);p__Ascomycota_unclassified(75);p__Ascomycota_unclassified(75);p__Ascomycota_unclassified(75); | - | - | - | - | - | - | - | - | Unassigned | |
| Otu03934 | 8 | k__Fungi(100);k__Fungi_unclassified(63);k__Fungi_unclassified(63);k__Fungi_unclassified(63);k__Fungi_unclassified(63);k__Fungi_unclassified(63);k__Fungi_unclassified(63); | - | - | - | - | - | - | - | - | Unassigned | |
| Otu03935 | 8 | k__Fungi(100);p__Ascomycota(100);p__Ascomycota_unclassified(100);p__Ascomycota_unclassified(100);p__Ascomycota_unclassified(100);p__Ascomycota_unclassified(100);p__Ascomycota_unclassified(100); | - | - | - | - | - | - | - | - | Unassigned | |
| Otu03937 | 8 | k__Fungi(100);p__Ascomycota(88);p__Ascomycota_unclassified(88);p__Ascomycota_unclassified(88);p__Ascomycota_unclassified(88);p__Ascomycota_unclassified(88);p__Ascomycota_unclassified(88); | - | - | - | - | - | - | - | - | Unassigned | |
| Otu03939 | 8 | k__Fungi(100);k__Fungi_unclassified(63);k__Fungi_unclassified(63);k__Fungi_unclassified(63);k__Fungi_unclassified(63);k__Fungi_unclassified(63);k__Fungi_unclassified(63); | - | - | - | - | - | - | - | - | Unassigned | |
| Otu03940 | 8 | k__Fungi(100);p__Ascomycota(100);p__Ascomycota_unclassified(100);p__Ascomycota_unclassified(100);p__Ascomycota_unclassified(100);p__Ascomycota_unclassified(100);p__Ascomycota_unclassified(100); | - | - | - | - | - | - | - | - | Unassigned | |
| Otu03941 | 8 | k__Fungi(100);k__Fungi_unclassified(100);k__Fungi_unclassified(100);k__Fungi_unclassified(100);k__Fungi_unclassified(100);k__Fungi_unclassified(100);k__Fungi_unclassified(100); | - | - | - | - | - | - | - | - | Unassigned | |
| Otu03942 | 8 | k__Fungi(100);p__Ascomycota(100);p__Ascomycota_unclassified(100);p__Ascomycota_unclassified(100);p__Ascomycota_unclassified(100);p__Ascomycota_unclassified(100);p__Ascomycota_unclassified(100); | - | - | - | - | - | - | - | - | Unassigned | |
| Otu03944 | 8 | k__Fungi(100);k__Fungi_unclassified(100);k__Fungi_unclassified(100);k__Fungi_unclassified(100);k__Fungi_unclassified(100);k__Fungi_unclassified(100);k__Fungi_unclassified(100); | - | - | - | - | - | - | - | - | Unassigned | |
| Otu03946 | 8 | k__Fungi(100);p__Ascomycota(100);p__Ascomycota_unclassified(88);p__Ascomycota_unclassified(88);p__Ascomycota_unclassified(88);p__Ascomycota_unclassified(88);p__Ascomycota_unclassified(88); | - | - | - | - | - | - | - | - | Unassigned | |
| Otu03947 | 8 | k__Fungi(100);k__Fungi_unclassified(100);k__Fungi_unclassified(100);k__Fungi_unclassified(100);k__Fungi_unclassified(100);k__Fungi_unclassified(100);k__Fungi_unclassified(100); | - | - | - | - | - | - | - | - | Unassigned | |
| Otu03949 | 8 | k__Fungi(100);k__Fungi_unclassified(100);k__Fungi_unclassified(100);k__Fungi_unclassified(100);k__Fungi_unclassified(100);k__Fungi_unclassified(100);k__Fungi_unclassified(100); | - | - | - | - | - | - | - | - | Unassigned | |
| Otu03951 | 8 | k__Fungi(100);p__Ascomycota(75);p__Ascomycota_unclassified(75);p__Ascomycota_unclassified(75);p__Ascomycota_unclassified(75);p__Ascomycota_unclassified(75);p__Ascomycota_unclassified(75); | - | - | - | - | - | - | - | - | Unassigned | |
| Otu03953 | 8 | k__Fungi(100);p__Ascomycota(100);c__Lecanoromycetes(88);c__Lecanoromycetes_unclassified(63);c__Lecanoromycetes_unclassified(63);c__Lecanoromycetes_unclassified(63);c__Lecanoromycetes_unclassified(63); | - | - | - | - | - | - | - | - | Unassigned | |
| Otu03959 | 8 | k__Fungi(100);k__Fungi_unclassified(63);k__Fungi_unclassified(63);k__Fungi_unclassified(63);k__Fungi_unclassified(63);k__Fungi_unclassified(63);k__Fungi_unclassified(63); | - | - | - | - | - | - | - | - | Unassigned | |
| Otu03960 | 8 | k__Fungi(100);k__Fungi_unclassified(75);k__Fungi_unclassified(75);k__Fungi_unclassified(75);k__Fungi_unclassified(75);k__Fungi_unclassified(75);k__Fungi_unclassified(75); | - | - | - | - | - | - | - | - | Unassigned | |

| | | | | | | | | | | | |
|---|---|---|---|---|---|---|---|---|---|---|---|
| Otu03961 | 8 | k__Fungi(100);p__Ascomycota(63);p__Ascomycota_unclassified(63);p__Ascomycota_unclassified(63);p__Ascomycota_unclassified(63);p__Ascomycota_unclassified(63);p__Ascomycota_unclassified(63); | - | - | - | - | - | - | - | Unassigned | - |
| Otu03963 | 8 | k__Fungi(100);k__Fungi_unclassified(63);k__Fungi_unclassified(63);k__Fungi_unclassified(63);k__Fungi_unclassified(63);k__Fungi_unclassified(63);k__Fungi_unclassified(63); | - | - | - | - | - | - | - | Unassigned | - |
| Otu03969 | 8 | k__Fungi(100);p__Ascomycota(63);p__Ascomycota_unclassified(63);p__Ascomycota_unclassified(63);p__Ascomycota_unclassified(63);p__Ascomycota_unclassified(63);p__Ascomycota_unclassified(63); | - | - | - | - | - | - | - | Unassigned | - |
| Otu03970 | 8 | k__Fungi(100);p__Ascomycota(63);p__Ascomycota_unclassified(63);p__Ascomycota_unclassified(63);p__Ascomycota_unclassified(63);p__Ascomycota_unclassified(63);p__Ascomycota_unclassified(63); | - | - | - | - | - | - | - | Unassigned | - |
| Otu03971 | 8 | k__Fungi(100);k__Fungi_unclassified(63);k__Fungi_unclassified(63);k__Fungi_unclassified(63);k__Fungi_unclassified(63);k__Fungi_unclassified(63);k__Fungi_unclassified(63); | - | - | - | - | - | - | - | Unassigned | - |
| Otu03972 | 8 | k__Fungi(100);k__Fungi_unclassified(100);k__Fungi_unclassified(100);k__Fungi_unclassified(100);k__Fungi_unclassified(100);k__Fungi_unclassified(100);k__Fungi_unclassified(100); | - | - | - | - | - | - | - | Unassigned | - |
| Otu03973 | 8 | k__Fungi(100);k__Fungi_unclassified(75);k__Fungi_unclassified(75);k__Fungi_unclassified(75);k__Fungi_unclassified(75);k__Fungi_unclassified(75);k__Fungi_unclassified(75); | - | - | - | - | - | - | - | Unassigned | - |
| Otu03974 | 8 | k__Fungi(100);p__Ascomycota(63);p__Ascomycota_unclassified(63);p__Ascomycota_unclassified(63);p__Ascomycota_unclassified(63);p__Ascomycota_unclassified(63);p__Ascomycota_unclassified(63); | - | - | - | - | - | - | - | Unassigned | - |
| Otu03976 | 8 | k__Fungi(100);k__Fungi_unclassified(75);k__Fungi_unclassified(75);k__Fungi_unclassified(75);k__Fungi_unclassified(75);k__Fungi_unclassified(75);k__Fungi_unclassified(75); | - | - | - | - | - | - | - | Unassigned | - |
| Otu03978 | 8 | k__Fungi(100);k__Fungi_unclassified(100);k__Fungi_unclassified(100);k__Fungi_unclassified(100);k__Fungi_unclassified(100);k__Fungi_unclassified(100);k__Fungi_unclassified(100); | - | - | - | - | - | - | - | Unassigned | - |
| Otu03985 | 8 | k__Fungi(100);k__Fungi_unclassified(75);k__Fungi_unclassified(75);k__Fungi_unclassified(75);k__Fungi_unclassified(75);k__Fungi_unclassified(75);k__Fungi_unclassified(75); | - | - | - | - | - | - | - | Unassigned | - |
| Otu03987 | 8 | k__Fungi(100);k__Fungi_unclassified(100);k__Fungi_unclassified(100);k__Fungi_unclassified(100);k__Fungi_unclassified(100);k__Fungi_unclassified(100);k__Fungi_unclassified(100); | - | - | - | - | - | - | - | Unassigned | - |
| Otu03992 | 8 | k__Fungi(100);p__Ascomycota(100);p__Ascomycota_unclassified(100);p__Ascomycota_unclassified(100);p__Ascomycota_unclassified(100);p__Ascomycota_unclassified(100);p__Ascomycota_unclassified(100); | - | - | - | - | - | - | - | Unassigned | - |
| Otu03993 | 8 | k__Fungi(100);p__Ascomycota(75);p__Ascomycota_unclassified(75);p__Ascomycota_unclassified(75);p__Ascomycota_unclassified(75);p__Ascomycota_unclassified(75);p__Ascomycota_unclassified(75); | - | - | - | - | - | - | - | Unassigned | - |
| Otu03994 | 8 | k__Fungi(100);p__Ascomycota(75);p__Ascomycota_unclassified(75);p__Ascomycota_unclassified(75);p__Ascomycota_unclassified(75);p__Ascomycota_unclassified(75);p__Ascomycota_unclassified(75); | - | - | - | - | - | - | - | Unassigned | - |
| Otu03998 | 8 | k__Fungi(100);p__Ascomycota(100);p__Ascomycota_unclassified(88);p__Ascomycota_unclassified(88);p__Ascomycota_unclassified(88);p__Ascomycota_unclassified(88);p__Ascomycota_unclassified(88); | - | - | - | - | - | - | - | Unassigned | - |
| Otu04000 | 8 | k__Fungi(100);k__Fungi_unclassified(100);k__Fungi_unclassified(100);k__Fungi_unclassified(100);k__Fungi_unclassified(100);k__Fungi_unclassified(100);k__Fungi_unclassified(100); | - | - | - | - | - | - | - | Unassigned | - |
| Otu04001 | 8 | k__Fungi(100);k__Fungi_unclassified(88);k__Fungi_unclassified(88);k__Fungi_unclassified(88);k__Fungi_unclassified(88);k__Fungi_unclassified(88);k__Fungi_unclassified(88); | - | - | - | - | - | - | - | Unassigned | - |
| Otu04004 | 8 | k__Fungi(100);p__Ascomycota(75);p__Ascomycota_unclassified(75);p__Ascomycota_unclassified(75);p__Ascomycota_unclassified(75);p__Ascomycota_unclassified(75);p__Ascomycota_unclassified(75); | - | - | - | - | - | - | - | Unassigned | - |
| Otu04005 | 8 | k__Fungi(100);p__Ascomycota(75);p__Ascomycota_unclassified(75);p__Ascomycota_unclassified(75);p__Ascomycota_unclassified(75);p__Ascomycota_unclassified(75);p__Ascomycota_unclassified(75); | - | - | - | - | - | - | - | Unassigned | - |
| Otu04006 | 8 | k__Fungi(100);p__Basidiomycota(100);p__Basidiomycota_unclassified(100);p__Basidiomycota_unclassified(100);p__Basidiomycota_unclassified(100);p__Basidiomycota_unclassified(100);p__Basidiomycota_unclassified(100); | - | - | - | - | - | - | - | Unassigned | - |
| Otu04011 | 8 | k__Fungi(100);k__Fungi_unclassified(100);k__Fungi_unclassified(100);k__Fungi_unclassified(100);k__Fungi_unclassified(100);k__Fungi_unclassified(100);k__Fungi_unclassified(100); | - | - | - | - | - | - | - | Unassigned | - |
| Otu04012 | 8 | k__Fungi(100);p__Ascomycota(100);p__Ascomycota_unclassified(88);p__Ascomycota_unclassified(88);p__Ascomycota_unclassified(88);p__Ascomycota_unclassified(88);p__Ascomycota_unclassified(88); | - | - | - | - | - | - | - | Unassigned | - |
| Otu04015 | 8 | k__Fungi(100);p__Basidiomycota(100);c__Agaricomycetes(100);o__Trechisporales(100);f__unclassified_Trechisporales(100);g__unclassified_Trechisporales(100);s__Trechisporales_sp(100); | - | - | - | - | - | - | - | Unassigned | - |
| Otu04017 | 8 | k__Fungi(100);p__Ascomycota(100);c__Leotiomycetes(88);c__Leotiomycetes_unclassified(88);c__Leotiomycetes_unclassified(88);c__Leotiomycetes_unclassified(88);c__Leotiomycetes_unclassified(88); | - | - | - | - | - | - | - | Unassigned | - |
| Otu04018 | 8 | k__Fungi(100);p__Ascomycota(100);c__Dothideomycetes(88);c__Dothideomycetes_unclassified(63);c__Dothideomycetes_unclassified(63);c__Dothideomycetes_unclassified(63);c__Dothideomycetes_unclassified(63); | - | - | - | - | - | - | - | Unassigned | - |
| Otu04019 | 8 | k__Fungi(100);p__Ascomycota(100);c__Dothideomycetes(100);o__Pleosporales(100);f__Pleosporales_family_Incertae_sedis(88);f__Pleosporales_family_Incertae_sedis_unclassified(88);f__Pleosporales_family_Incertae_sedis_unclassified(88); | - | - | - | - | - | - | - | Unassigned | - |
| Otu04021 | 8 | k__Fungi(100);p__Basidiomycota(100);c__Agaricomycetes(100);o__Polyporales(100);f__Phanerochaetaceae(100);f__Phanerochaetaceae_unclassified(100);f__Phanerochaetaceae_unclassified(100); | - | - | - | - | - | - | - | Unassigned | - |
| Otu04022 | 8 | k__Fungi(100);p__unclassified_Fungi(100);c__unclassified_Fungi(100);o__unclassified_Fungi(100);f__unclassified_Fungi(100);g__unclassified_Fungi(100);s__uncultured_fungus(100); | - | - | - | - | - | - | - | Unassigned | - |
| Otu04023 | 8 | k__Fungi(100);p__Basidiomycota(100);c__Agaricomycetes(100);o__Cantharellales(100);o__Cantharellales_unclassified(100);o__Cantharellales_unclassified(100);o__Cantharellales_unclassified(100); | - | - | - | - | - | - | - | Unassigned | - |
| Otu04025 | 8 | k__Fungi(100);p__Basidiomycota(100);c__Microbotryomycetes(100);o__Sporidiobolales(100);f__Sporidiobolales_family_Incertae_sedis(100);f__Sporidiobolales_family_Incertae_sedis_unclassified(100);f__Sporidiobolales_family_Incertae_sedis_unclassified(100); | - | - | - | - | - | - | - | Unassigned | - |
| Otu04027 | 8 | k__Fungi(100);k__Fungi_unclassified(88);k__Fungi_unclassified(88);k__Fungi_unclassified(88);k__Fungi_unclassified(88);k__Fungi_unclassified(88);k__Fungi_unclassified(88); | - | - | - | - | - | - | - | Unassigned | - |
| Otu04029 | 8 | k__Fungi(100);k__Fungi_unclassified(100);k__Fungi_unclassified(100);k__Fungi_unclassified(100);k__Fungi_unclassified(100);k__Fungi_unclassified(100);k__Fungi_unclassified(100); | - | - | - | - | - | - | - | Unassigned | - |
| Otu04030 | 8 | k__Fungi(100);k__Fungi_unclassified(63);k__Fungi_unclassified(63);k__Fungi_unclassified(63);k__Fungi_unclassified(63);k__Fungi_unclassified(63);k__Fungi_unclassified(63); | - | - | - | - | - | - | - | Unassigned | - |
| Otu04031 | 8 | k__Fungi(100);k__Fungi_unclassified(88);k__Fungi_unclassified(88);k__Fungi_unclassified(88);k__Fungi_unclassified(88);k__Fungi_unclassified(88);k__Fungi_unclassified(88); | - | - | - | - | - | - | - | Unassigned | - |
| Otu04034 | 8 | k__Fungi(100);p__Basidiomycota(100);c__Agaricomycetes(100);c__Agaricomycetes_unclassified(100);c__Agaricomycetes_unclassified(100);c__Agaricomycetes_unclassified(100);c__Agaricomycetes_unclassified(100); | - | - | - | - | - | - | - | Unassigned | - |
| Otu04035 | 8 | k__Fungi(100);k__Fungi_unclassified(63);k__Fungi_unclassified(63);k__Fungi_unclassified(63);k__Fungi_unclassified(63);k__Fungi_unclassified(63);k__Fungi_unclassified(63); | - | - | - | - | - | - | - | Unassigned | - |
| Otu04037 | 8 | k__Fungi(100);p__Ascomycota(100);c__Lecanoromycetes(100);c__Lecanoromycetes_unclassified(100);c__Lecanoromycetes_unclassified(100);c__Lecanoromycetes_unclassified(100);c__Lecanoromycetes_unclassified(100); | - | - | - | - | - | - | - | Unassigned | - |
| Otu04039 | 8 | k__Fungi(100);p__Ascomycota(100);c__Eurotiomycetes(100);o__Chaetothyriales(100);o__Chaetothyriales_unclassified(100);o__Chaetothyriales_unclassified(100);o__Chaetothyriales_unclassified(100); | - | - | - | - | - | - | - | Unassigned | - |
| Otu04041 | 8 | k__Fungi(100);p__Ascomycota(100);p__Ascomycota_unclassified(100);p__Ascomycota_unclassified(100);p__Ascomycota_unclassified(100);p__Ascomycota_unclassified(100);p__Ascomycota_unclassified(100); | - | - | - | - | - | - | - | Unassigned | - |
| Otu04050 | 8 | k__Fungi(100);p__Ascomycota(100);c__Pezizomycetes(100);o__Pezizales(100);o__Pezizales_unclassified(100);o__Pezizales_unclassified(100);o__Pezizales_unclassified(100); | - | - | - | - | - | - | - | Unassigned | - |
| Otu04052 | 8 | k__Fungi(100);p__Ascomycota(75);p__Ascomycota_unclassified(75);p__Ascomycota_unclassified(75);p__Ascomycota_unclassified(75);p__Ascomycota_unclassified(75);p__Ascomycota_unclassified(75); | - | - | - | - | - | - | - | Unassigned | - |
| Otu04059 | 8 | k__Fungi(100);k__Fungi_unclassified(75);k__Fungi_unclassified(75);k__Fungi_unclassified(75);k__Fungi_unclassified(75);k__Fungi_unclassified(75);k__Fungi_unclassified(75); | - | - | - | - | - | - | - | Unassigned | - |
| Otu04061 | 8 | k__Fungi(100);p__Ascomycota(100);p__Ascomycota_unclassified(88);p__Ascomycota_unclassified(88);p__Ascomycota_unclassified(88);p__Ascomycota_unclassified(88);p__Ascomycota_unclassified(88); | - | - | - | - | - | - | - | Unassigned | - |
| Otu04062 | 8 | k__Fungi(100);p__Ascomycota(75);p__Ascomycota_unclassified(75);p__Ascomycota_unclassified(75);p__Ascomycota_unclassified(75);p__Ascomycota_unclassified(75);p__Ascomycota_unclassified(75); | - | - | - | - | - | - | - | Unassigned | - |
| Otu04064 | 8 | k__Fungi(100);k__Fungi_unclassified(75);k__Fungi_unclassified(75);k__Fungi_unclassified(75);k__Fungi_unclassified(75);k__Fungi_unclassified(75);k__Fungi_unclassified(75); | - | - | - | - | - | - | - | Unassigned | - |
| Otu04065 | 8 | k__Fungi(100);k__Fungi_unclassified(88);k__Fungi_unclassified(88);k__Fungi_unclassified(88);k__Fungi_unclassified(88);k__Fungi_unclassified(88);k__Fungi_unclassified(88); | - | - | - | - | - | - | - | Unassigned | - |
| Otu04066 | 8 | k__Fungi(100);k__Fungi_unclassified(75);k__Fungi_unclassified(75);k__Fungi_unclassified(75);k__Fungi_unclassified(75);k__Fungi_unclassified(75);k__Fungi_unclassified(75); | - | - | - | - | - | - | - | Unassigned | - |
| Otu04072 | 8 | k__Fungi(100);k__Fungi_unclassified(100);k__Fungi_unclassified(100);k__Fungi_unclassified(100);k__Fungi_unclassified(100);k__Fungi_unclassified(100);k__Fungi_unclassified(100); | - | - | - | - | - | - | - | Unassigned | - |
| Otu04074 | 8 | k__Fungi(100);p__Ascomycota(100);c__Dothideomycetes(100);o__Pleosporales(88);o__Pleosporales_unclassified(88);o__Pleosporales_unclassified(88);o__Pleosporales_unclassified(88); | - | - | - | - | - | - | - | Unassigned | - |
| Otu04078 | 8 | k__Fungi(100);p__Ascomycota(63);p__Ascomycota_unclassified(63);p__Ascomycota_unclassified(63);p__Ascomycota_unclassified(63);p__Ascomycota_unclassified(63);p__Ascomycota_unclassified(63); | - | - | - | - | - | - | - | Unassigned | - |
| Otu04081 | 8 | k__Fungi(100);k__Fungi_unclassified(75);k__Fungi_unclassified(75);k__Fungi_unclassified(75);k__Fungi_unclassified(75);k__Fungi_unclassified(75);k__Fungi_unclassified(75); | - | - | - | - | - | - | - | Unassigned | - |
| Otu04083 | 8 | k__Fungi(100);p__Ascomycota(100);p__Ascomycota_unclassified(100);p__Ascomycota_unclassified(100);p__Ascomycota_unclassified(100);p__Ascomycota_unclassified(100);p__Ascomycota_unclassified(100); | - | - | - | - | - | - | - | Unassigned | - |
| Otu04084 | 8 | k__Fungi(100);p__Ascomycota(100);c__Leotiomycetes(100);o__Helotiales(100);o__Helotiales_unclassified(100);o__Helotiales_unclassified(100);o__Helotiales_unclassified(100); | - | - | - | - | - | - | - | Unassigned | - |
| Otu04085 | 8 | k__Fungi(100);k__Fungi_unclassified(63);k__Fungi_unclassified(63);k__Fungi_unclassified(63);k__Fungi_unclassified(63);k__Fungi_unclassified(63);k__Fungi_unclassified(63); | - | - | - | - | - | - | - | Unassigned | - |
| Otu04087 | 8 | k__Fungi(100);k__Fungi_unclassified(63);k__Fungi_unclassified(63);k__Fungi_unclassified(63);k__Fungi_unclassified(63);k__Fungi_unclassified(63);k__Fungi_unclassified(63); | - | - | - | - | - | - | - | Unassigned | - |
| Otu04088 | 8 | k__Fungi(100);p__Basidiomycota(100);c__Agaricomycetes(100);c__Agaricomycetes_unclassified(100);c__Agaricomycetes_unclassified(100);c__Agaricomycetes_unclassified(100);c__Agaricomycetes_unclassified(100); | - | - | - | - | - | - | - | Unassigned | - |
| Otu04089 | 8 | k__Fungi(100);k__Fungi_unclassified(100);k__Fungi_unclassified(100);k__Fungi_unclassified(100);k__Fungi_unclassified(100);k__Fungi_unclassified(100);k__Fungi_unclassified(100); | - | - | - | - | - | - | - | Unassigned | - |
| Otu04090 | 8 | k__Fungi(100);p__Ascomycota(63);p__Ascomycota_unclassified(63);p__Ascomycota_unclassified(63);p__Ascomycota_unclassified(63);p__Ascomycota_unclassified(63);p__Ascomycota_unclassified(63); | - | - | - | - | - | - | - | Unassigned | - |
| Otu04091 | 8 | k__Fungi(100);k__Fungi_unclassified(75);k__Fungi_unclassified(75);k__Fungi_unclassified(75);k__Fungi_unclassified(75);k__Fungi_unclassified(75);k__Fungi_unclassified(75); | - | - | - | - | - | - | - | Unassigned | - |
| Otu04093 | 8 | k__Fungi(100);p__Ascomycota(75);p__Ascomycota_unclassified(75);p__Ascomycota_unclassified(75);p__Ascomycota_unclassified(75);p__Ascomycota_unclassified(75);p__Ascomycota_unclassified(75); | - | - | - | - | - | - | - | Unassigned | - |
| Otu04094 | 8 | k__Fungi(100);p__Ascomycota(100);p__Ascomycota_unclassified(100);p__Ascomycota_unclassified(100);p__Ascomycota_unclassified(100);p__Ascomycota_unclassified(100);p__Ascomycota_unclassified(100); | - | - | - | - | - | - | - | Unassigned | - |
| Otu04095 | 8 | k__Fungi(100);k__Fungi_unclassified(100);k__Fungi_unclassified(100);k__Fungi_unclassified(100);k__Fungi_unclassified(100);k__Fungi_unclassified(100);k__Fungi_unclassified(100); | - | - | - | - | - | - | - | Unassigned | - |
| Otu04101 | 8 | k__Fungi(100);p__Basidiomycota(100);c__Agaricostilbomycetes(100);o__Agaricostilbales(100);o__Agaricostilbales_unclassified(100);o__Agaricostilbales_unclassified(100);o__Agaricostilbales_unclassified(100); | - | - | - | - | - | - | - | Unassigned | - |
| Otu04103 | 8 | k__Fungi(100);p__Basidiomycota(100);c__Agaricomycetes(100);c__Agaricomycetes_unclassified(100);c__Agaricomycetes_unclassified(100);c__Agaricomycetes_unclassified(100);c__Agaricomycetes_unclassified(100); | - | - | - | - | - | - | - | Unassigned | - |
| Otu04106 | 8 | k__Fungi(100);k__Fungi_unclassified(75);k__Fungi_unclassified(75);k__Fungi_unclassified(75);k__Fungi_unclassified(75);k__Fungi_unclassified(75);k__Fungi_unclassified(75); | - | - | - | - | - | - | - | Unassigned | - |
| Otu04107 | 8 | k__Fungi(100);k__Fungi_unclassified(63);k__Fungi_unclassified(63);k__Fungi_unclassified(63);k__Fungi_unclassified(63);k__Fungi_unclassified(63);k__Fungi_unclassified(63); | - | - | - | - | - | - | - | Unassigned | - |
| Otu04109 | 8 | k__Fungi(100);k__Fungi_unclassified(75);k__Fungi_unclassified(75);k__Fungi_unclassified(75);k__Fungi_unclassified(75);k__Fungi_unclassified(75);k__Fungi_unclassified(75); | - | - | - | - | - | - | - | Unassigned | - |
| Otu04111 | 8 | k__Fungi(100);k__Fungi_unclassified(75);k__Fungi_unclassified(75);k__Fungi_unclassified(75);k__Fungi_unclassified(75);k__Fungi_unclassified(75);k__Fungi_unclassified(75); | - | - | - | - | - | - | - | Unassigned | - |
| Otu04115 | 8 | k__Fungi(100);p__Basidiomycota(100);c__Agaricomycetes(100);o__Trechisporales(100);f__unclassified_Trechisporales(100);g__unclassified_Trechisporales(100);s__Trechisporales_sp(100); | - | - | - | - | - | - | - | Unassigned | - |
| Otu04119 | 8 | k__Fungi(100);p__Basidiomycota(100);c__Agaricomycetes(100);c__Agaricomycetes_unclassified(100);c__Agaricomycetes_unclassified(100);c__Agaricomycetes_unclassified(100);c__Agaricomycetes_unclassified(100); | - | - | - | - | - | - | - | Unassigned | - |
| Otu04120 | 8 | k__Fungi(100);p__Basidiomycota(100);p__Basidiomycota_unclassified(88);p__Basidiomycota_unclassified(88);p__Basidiomycota_unclassified(88);p__Basidiomycota_unclassified(88);p__Basidiomycota_unclassified(88); | - | - | - | - | - | - | - | Unassigned | - |
| Otu04122 | 8 | k__Fungi(100);p__Ascomycota(75);p__Ascomycota_unclassified(75);p__Ascomycota_unclassified(75);p__Ascomycota_unclassified(75);p__Ascomycota_unclassified(75);p__Ascomycota_unclassified(75); | - | - | - | - | - | - | - | Unassigned | - |
| Otu04125 | 8 | k__Fungi(100);k__Fungi_unclassified(75);k__Fungi_unclassified(75);k__Fungi_unclassified(75);k__Fungi_unclassified(75);k__Fungi_unclassified(75);k__Fungi_unclassified(75); | - | - | - | - | - | - | - | Unassigned | - |
| Otu04126 | 8 | k__Fungi(100);p__Ascomycota(100);c__Dothideomycetes(100);o__Dothideales(100);f__Dothioraceae(100);f__Dothioraceae_unclassified(100);f__Dothioraceae_unclassified(100); | - | - | - | - | - | - | - | Unassigned | - |
| Otu04127 | 8 | k__Fungi(100);k__Fungi_unclassified(100);k__Fungi_unclassified(100);k__Fungi_unclassified(100);k__Fungi_unclassified(100);k__Fungi_unclassified(100);k__Fungi_unclassified(100); | - | - | - | - | - | - | - | Unassigned | - |
| Otu04128 | 8 | k__Fungi(100);k__Fungi_unclassified(100);k__Fungi_unclassified(100);k__Fungi_unclassified(100);k__Fungi_unclassified(100);k__Fungi_unclassified(100);k__Fungi_unclassified(100); | - | - | - | - | - | - | - | Unassigned | - |
| Otu04134 | 8 | k__Fungi(100);p__Ascomycota(100);c__Eurotiomycetes(100);o__Onygenales(100);o__Onygenales_unclassified(100);o__Onygenales_unclassified(100);o__Onygenales_unclassified(100); | - | - | - | - | - | - | - | Unassigned | - |
| Otu04136 | 8 | k__Fungi(100);p__Ascomycota(100);p__Ascomycota_unclassified(100);p__Ascomycota_unclassified(100);p__Ascomycota_unclassified(100);p__Ascomycota_unclassified(100);p__Ascomycota_unclassified(100); | - | - | - | - | - | - | - | Unassigned | - |
| Otu04139 | 8 | k__Fungi(100);p__Ascomycota(100);c__Dothideomycetes(100);o__Capnodiales(100);o__Capnodiales_unclassified(88);o__Capnodiales_unclassified(88);o__Capnodiales_unclassified(88); | - | - | - | - | - | - | - | Unassigned | - |
| Otu04141 | 8 | k__Fungi(100);p__Basidiomycota(100);c__Pucciniomycetes(100);o__Pucciniales(100);o__Pucciniales_unclassified(100);o__Pucciniales_unclassified(100);o__Pucciniales_unclassified(100); | - | - | - | - | - | - | - | Unassigned | - |
| Otu04142 | 8 | k__Fungi(100);p__Ascomycota(63);p__Ascomycota_unclassified(63);p__Ascomycota_unclassified(63);p__Ascomycota_unclassified(63);p__Ascomycota_unclassified(63);p__Ascomycota_unclassified(63); | - | - | - | - | - | - | - | Unassigned | - |
| Otu04143 | 8 | k__Fungi(100);k__Fungi_unclassified(100);k__Fungi_unclassified(100);k__Fungi_unclassified(100);k__Fungi_unclassified(100);k__Fungi_unclassified(100);k__Fungi_unclassified(100); | - | - | - | - | - | - | - | Unassigned | - |
| Otu04156 | 8 | k__Fungi(100);k__Fungi_unclassified(100);k__Fungi_unclassified(100);k__Fungi_unclassified(100);k__Fungi_unclassified(100);k__Fungi_unclassified(100);k__Fungi_unclassified(100); | - | - | - | - | - | - | - | Unassigned | - |
| Otu04158 | 8 | k__Fungi(100);p__Basidiomycota(100);c__Agaricomycetes(100);c__Agaricomycetes_unclassified(100);c__Agaricomycetes_unclassified(100);c__Agaricomycetes_unclassified(100);c__Agaricomycetes_unclassified(100); | - | - | - | - | - | - | - | Unassigned | - |
| Otu04160 | 8 | k__Fungi(100);k__Fungi_unclassified(100);k__Fungi_unclassified(100);k__Fungi_unclassified(100);k__Fungi_unclassified(100);k__Fungi_unclassified(100);k__Fungi_unclassified(100); | - | - | - | - | - | - | - | Unassigned | - |
| Otu04161 | 8 | k__Fungi(100);k__Fungi_unclassified(100);k__Fungi_unclassified(100);k__Fungi_unclassified(100);k__Fungi_unclassified(100);k__Fungi_unclassified(100);k__Fungi_unclassified(100); | - | - | - | - | - | - | - | Unassigned | - |
| Otu04162 | 8 | k__Fungi(100);p__Ascomycota(100);c__Dothideomycetes(100);o__Pleosporales(100);o__Pleosporales_unclassified(100);o__Pleosporales_unclassified(100);o__Pleosporales_unclassified(100); | - | - | - | - | - | - | - | Unassigned | - |
| Otu04163 | 8 | k__Fungi(100);p__Basidiomycota(100);c__Agaricomycetes(100);c__Agaricomycetes_unclassified(100);c__Agaricomycetes_unclassified(100);c__Agaricomycetes_unclassified(100);c__Agaricomycetes_unclassified(100); | - | - | - | - | - | - | - | Unassigned | - |
| Otu04164 | 8 | k__Fungi(100);k__Fungi_unclassified(88);k__Fungi_unclassified(88);k__Fungi_unclassified(88);k__Fungi_unclassified(88);k__Fungi_unclassified(88);k__Fungi_unclassified(88); | - | - | - | - | - | - | - | Unassigned | - |
| Otu04165 | 8 | k__Fungi(100);p__Ascomycota(100);c__Sordariomycetes(100);o__Hypocreales(100);f__Hypocreaceae(100);f__Hypocreaceae_unclassified(100);f__Hypocreaceae_unclassified(100); | - | - | - | - | - | - | - | Unassigned | - |
| Otu04166 | 8 | k__Fungi(100);p__Ascomycota(100);p__Ascomycota_unclassified(100);p__Ascomycota_unclassified(100);p__Ascomycota_unclassified(100);p__Ascomycota_unclassified(100);p__Ascomycota_unclassified(100); | - | - | - | - | - | - | - | Unassigned | - |
| Otu04167 | 8 | k__Fungi(100);k__Fungi_unclassified(75);k__Fungi_unclassified(75);k__Fungi_unclassified(75);k__Fungi_unclassified(75);k__Fungi_unclassified(75);k__Fungi_unclassified(75); | - | - | - | - | - | - | - | Unassigned | - |
| Otu04168 | 8 | k__Fungi(100);p__Ascomycota(100);c__Eurotiomycetes(100);o__Chaetothyriales(100);o__Chaetothyriales_unclassified(63);o__Chaetothyriales_unclassified(63);o__Chaetothyriales_unclassified(63); | - | - | - | - | - | - | - | Unassigned | - |
| Otu04169 | 8 | k__Fungi(100);p__Ascomycota(100);c__Dothideomycetes(100);c__Dothideomycetes_unclassified(88);c__Dothideomycetes_unclassified(88);c__Dothideomycetes_unclassified(88);c__Dothideomycetes_unclassified(88); | - | - | - | - | - | - | - | Unassigned | - |
| Otu04170 | 8 | k__Fungi(100);k__Fungi_unclassified(100);k__Fungi_unclassified(100);k__Fungi_unclassified(100);k__Fungi_unclassified(100);k__Fungi_unclassified(100);k__Fungi_unclassified(100); | - | - | - | - | - | - | - | Unassigned | - |
| Otu04171 | 8 | k__Fungi(100);p__Ascomycota(100);p__Ascomycota_unclassified(100);p__Ascomycota_unclassified(100);p__Ascomycota_unclassified(100);p__Ascomycota_unclassified(100);p__Ascomycota_unclassified(100); | - | - | - | - | - | - | - | Unassigned | - |
| Otu04176 | 8 | k__Fungi(100);p__Ascomycota(100);c__Dothideomycetes(100);o__Pleosporales(100);o__Pleosporales_unclassified(100);o__Pleosporales_unclassified(100);o__Pleosporales_unclassified(100); | - | - | - | - | - | - | - | Unassigned | - |
| Otu04179 | 8 | k__Fungi(100);p__Basidiomycota(100);c__Agaricomycetes(100);c__Agaricomycetes_unclassified(100);c__Agaricomycetes_unclassified(100);c__Agaricomycetes_unclassified(100);c__Agaricomycetes_unclassified(100); | - | - | - | - | - | - | - | Unassigned | - |
| Otu04180 | 8 | k__Fungi(100);p__Ascomycota(63);p__Ascomycota_unclassified(63);p__Ascomycota_unclassified(63);p__Ascomycota_unclassified(63);p__Ascomycota_unclassified(63);p__Ascomycota_unclassified(63); | - | - | - | - | - | - | - | Unassigned | - |
| Otu04181 | 8 | k__Fungi(100);p__Ascomycota(100);p__Ascomycota_unclassified(100);p__Ascomycota_unclassified(100);p__Ascomycota_unclassified(100);p__Ascomycota_unclassified(100);p__Ascomycota_unclassified(100); | - | - | - | - | - | - | - | Unassigned | - |
| Otu04182 | 8 | k__Fungi(100);p__Ascomycota(100);c__Leotiomycetes(100);c__Leotiomycetes_unclassified(100);c__Leotiomycetes_unclassified(100);c__Leotiomycetes_unclassified(100);c__Leotiomycetes_unclassified(100); | - | - | - | - | - | - | - | Unassigned | - |
| Otu04185 | 8 | k__Fungi(100);p__Basidiomycota(100);p__Basidiomycota_unclassified(100);p__Basidiomycota_unclassified(100);p__Basidiomycota_unclassified(100);p__Basidiomycota_unclassified(100);p__Basidiomycota_unclassified(100); | - | - | - | - | - | - | - | Unassigned | - |
| Otu04186 | 8 | k__Fungi(100);p__Ascomycota(75);p__Ascomycota_unclassified(75);p__Ascomycota_unclassified(75);p__Ascomycota_unclassified(75);p__Ascomycota_unclassified(75);p__Ascomycota_unclassified(75); | - | - | - | - | - | - | - | Unassigned | - |
| Otu04187 | 8 | k__Fungi(100);k__Fungi_unclassified(100);k__Fungi_unclassified(100);k__Fungi_unclassified(100);k__Fungi_unclassified(100);k__Fungi_unclassified(100);k__Fungi_unclassified(100); | - | - | - | - | - | - | - | Unassigned | - |
| Otu04190 | 8 | k__Fungi(100);p__Ascomycota(100);p__Ascomycota_unclassified(100);p__Ascomycota_unclassified(100);p__Ascomycota_unclassified(100);p__Ascomycota_unclassified(100);p__Ascomycota_unclassified(100); | - | - | - | - | - | - | - | Unassigned | - |
| Otu04195 | 8 | k__Fungi(100);p__Basidiomycota(100);p__Basidiomycota_unclassified(88);p__Basidiomycota_unclassified(88);p__Basidiomycota_unclassified(88);p__Basidiomycota_unclassified(88);p__Basidiomycota_unclassified(88); | - | - | - | - | - | - | - | Unassigned | - |
| Otu04201 | 8 | k__Fungi(100);p__Ascomycota(100);c__Dothideomycetes(100);o__Myriangiales(100);o__Myriangiales_unclassified(100);o__Myriangiales_unclassified(100);o__Myriangiales_unclassified(100); | - | - | - | - | - | - | - | Unassigned | - |
| Otu04203 | 8 | k__Fungi(100);k__Fungi_unclassified(100);k__Fungi_unclassified(100);k__Fungi_unclassified(100);k__Fungi_unclassified(100);k__Fungi_unclassified(100);k__Fungi_unclassified(100); | - | - | - | - | - | - | - | Unassigned | - |
| Otu04204 | 8 | k__Fungi(100);p__Basidiomycota(100);c__Agaricomycetes(100);c__Agaricomycetes_unclassified(100);c__Agaricomycetes_unclassified(100);c__Agaricomycetes_unclassified(100);c__Agaricomycetes_unclassified(100); | - | - | - | - | - | - | - | Unassigned | - |
| Otu04206 | 8 | k__Fungi(100);p__Ascomycota(100);p__Ascomycota_unclassified(100);p__Ascomycota_unclassified(100);p__Ascomycota_unclassified(100);p__Ascomycota_unclassified(100);p__Ascomycota_unclassified(100); | - | - | - | - | - | - | - | Unassigned | - |
| Otu04208 | 8 | k__Fungi(100);p__Ascomycota(100);c__Sordariomycetes(100);o__Sordariales(75);o__Sordariales_unclassified(75);o__Sordariales_unclassified(75);o__Sordariales_unclassified(75); | - | - | - | - | - | - | - | Unassigned | - |
| Otu04211 | 8 | k__Fungi(100);k__Fungi_unclassified(100);k__Fungi_unclassified(100);k__Fungi_unclassified(100);k__Fungi_unclassified(100);k__Fungi_unclassified(100);k__Fungi_unclassified(100); | - | - | - | - | - | - | - | Unassigned | - |
| Otu04212 | 8 | k__Fungi(100);p__Basidiomycota(100);c__Agaricomycetes(100);c__Agaricomycetes_unclassified(100);c__Agaricomycetes_unclassified(100);c__Agaricomycetes_unclassified(100);c__Agaricomycetes_unclassified(100); | - | - | - | - | - | - | - | Unassigned | - |
| Otu04215 | 8 | k__Fungi(100);p__Ascomycota(63);p__Ascomycota_unclassified(63);p__Ascomycota_unclassified(63);p__Ascomycota_unclassified(63);p__Ascomycota_unclassified(63);p__Ascomycota_unclassified(63); | - | - | - | - | - | - | - | Unassigned | - |
| Otu04217 | 8 | k__Fungi(100);p__Basidiomycota(100);c__Agaricomycetes(100);c__Agaricomycetes_unclassified(100);c__Agaricomycetes_unclassified(100);c__Agaricomycetes_unclassified(100);c__Agaricomycetes_unclassified(100); | - | - | - | - | - | - | - | Unassigned | - |
| Otu04221 | 7 | k__Fungi(100);p__Ascomycota(100);c__Dothideomycetes(100);o__Pleosporales(100);f__Leptosphaeriaceae(100);f__Leptosphaeriaceae_unclassified(100);f__Leptosphaeriaceae_unclassified(100); | Leptosphaeriaceae | 9 | Pathotroph-Saprotroph | Plant Pathogen-Wood Saprotroph | Microfungus | NULL | Probable | Saprobic or weakly necrotrophic on stems or leaves (Cannon & Kirk 2007) | Cannon PF, Kirk PM. 2007. Fungal Families of the World. CAB International, Cambridge (ISBN: 978-0851998275) |
| Otu04227 | 7 | k__Fungi(100);p__Basidiomycota(100);c__Agaricomycetes(100);o__Agaricales(100);f__Agaricaceae(100);g__Agaricus(100);s__Agaricus_campestris(58); | Agaricus | 13 | Saprotroph | \|Undefined Saprotroph\| | Agaricoid-Gasteroid-Secotioid | NULL | Probable | As soil_saprotroph (Põlme et al. 2020); Decay soil (Põlme et al. 2020) | Tedersoo L, et al. 2014. Science 346:e1256688 ((https://doi.org/10.1126/science.1256688)); Põlme S, et al. 2020. Fungal Diversity 105:1-16 ((https://doi.org/10.1007/s13225-020-00466-2)) |
| Otu04228 | 7 | k__Fungi(100);p__Ascomycota(100);c__Saccharomycetes(100);o__Saccharomycetales(100);f__Metschnikowiaceae(100);g__Metschnikowia(100);s__Metschnikowia_pulcherrima(100); | Metschnikowia | 13 | Saprotroph | \|Undefined Saprotroph\| | Yeast | NULL | Probable | As nectar-tap_saprotroph (Põlme et al. 2020); Decay sugar-rich substrates (Põlme et al. 2020) | Põlme S, et al. 2020. Fungal Diversity 105:1-16 ((https://doi.org/10.1007/s13225-020-00466-2)) |
| Otu04231 | 7 | k__Fungi(100);p__Basidiomycota(100);c__Agaricomycetes(100);o__Agaricales(100);o__Agaricales_unclassified(100);o__Agaricales_unclassified(100);o__Agaricales_unclassified(100); | Agaricales | 7 | Pathotroph-Saprotroph-Symbiotroph | Bryophyte Parasite-Dung Saprotroph-Ectomycorrhizal-Fungal Parasite-Leaf Saprotroph-Plant Parasite-Undefined Saprotroph-Wood Saprotroph | Agaricoid-Gasteroid-Microfungus-Secotioid-Yeast | NULL | Possible | Mushrooms and toadstools, Gill fungi, Agarics... lignicolous, sometimes muscicolous or fungicolous, saprobic, mycorrhizal, rarely parasitic on plants or fungi (Kirk et al. 2008) | Kirk PM et al. 2008. Dictionary of the Fungi. Tenth Edition. CAB International, Wallingford (ISBN: 978-0851998268) |
| Otu04232 | 7 | k__Fungi(100);p__Ascomycota(100);c__Saccharomycetes(100);o__Saccharomycetales(100);f__Metschnikowiaceae(100);g__Metschnikowia(100);s__Metschnikowia_pulcherrima(100); | Metschnikowia | 13 | Saprotroph | \|Undefined Saprotroph\| | Yeast | NULL | Probable | As nectar-tap_saprotroph (Põlme et al. 2020); Decay sugar-rich substrates (Põlme et al. 2020) | Põlme S, et al. 2020. Fungal Diversity 105:1-16 ((https://doi.org/10.1007/s13225-020-00466-2)) |
| Otu04233 | 7 | k__Fungi(100);p__Ascomycota(100);c__Dothideomycetes(100);o__Capnodiales(100);f__Teratosphaeriaceae(100);g__Devriesia(100);s__Devriesia_americana(100); | Devriesia | 13 | Pathotroph-Saprotroph | \|Plant Pathogen\|-Plant Saprotroph | Microfungus | NULL | Probable | As litter_saprotroph (Põlme et al. 2020); Decay leaf, fruit, seed (Põlme et al. 2020) | Tedersoo L, et al. 2014. Science 346:e1256688 ((https://doi.org/10.1126/science.1256688)); Põlme S, et al. 2020. Fungal Diversity 105:1-16 ((https://doi.org/10.1007/s13225-020-00466-2)) |
| Otu04236 | 7 | k__Fungi(100);p__Ascomycota(100);c__Saccharomycetes(100);o__Saccharomycetales(100);f__Metschnikowiaceae(100);g__Metschnikowia(100);s__Metschnikowia_pulcherrima(100); | Metschnikowia | 13 | Saprotroph | \|Undefined Saprotroph\| | Yeast | NULL | Probable | As nectar-tap_saprotroph (Põlme et al. 2020); Decay sugar-rich substrates (Põlme et al. 2020) | Põlme S, et al. 2020. Fungal Diversity 105:1-16 ((https://doi.org/10.1007/s13225-020-00466-2)) |
| Otu04239 | 7 | k__Fungi(100);p__Ascomycota(100);c__Saccharomycetes(100);o__Saccharomycetales(100);f__Saccharomycetales_family_Incertae_sedis(100);g__Candida(58);g__Candida_unclassified(58); | Candida | 13 | Pathotroph-Saprotroph-Symbiotroph | Animal Pathogen-Endophyte-Endosymbiont-Epiphyte-\|Undefined Saprotroph\| | Dimorphic Yeast-Yeast | NULL | Probable | As nectar-tap_saprotroph (Põlme et al. 2020); Decay sugar-rich substrates (Põlme et al. 2020); This asexual genus is polyphyletic and contain species that function very differently depend on what they are. Caution should be taken when interpreting guilds for Candida species. | Manolakaki D, et al. 2010. Virulence. 1: 367-375 ((https://doi.org/10.4161/viru.1.5.12796)); Mekbib SB, et al. 2011. Tropical Plant Pathology 36:233-240 ((https://doi.org/10.1590/S1982-56762011000400004)); Põlme S, et al. 2020. Fungal Diversity 105:1-16 ((https://doi.org/10.1007/s13225-020-00466-2)) |
| Otu04245 | 7 | k__Fungi(100);p__Ascomycota(100);c__Saccharomycetes(100);o__Saccharomycetales(100);f__Metschnikowiaceae(100);g__Metschnikowia(100);g__Metschnikowia_unclassified(58); | Metschnikowia | 13 | Saprotroph | \|Undefined Saprotroph\| | Yeast | NULL | Probable | As nectar-tap_saprotroph (Põlme et al. 2020); Decay sugar-rich substrates (Põlme et al. 2020) | Põlme S, et al. 2020. Fungal Diversity 105:1-16 ((https://doi.org/10.1007/s13225-020-00466-2)) |
| Otu04246 | 7 | k__Fungi(100);p__Ascomycota(100);c__Saccharomycetes(100);o__Saccharomycetales(100);f__Saccharomycetales_family_Incertae_sedis(86);g__Candida(72);g__Candida_unclassified(58); | Candida | 13 | Pathotroph-Saprotroph-Symbiotroph | Animal Pathogen-Endophyte-Endosymbiont-Epiphyte-\|Undefined Saprotroph\| | Dimorphic Yeast-Yeast | NULL | Probable | As nectar-tap_saprotroph (Põlme et al. 2020); Decay sugar-rich substrates (Põlme et al. 2020); This asexual genus is polyphyletic and contain species that function very differently depend on what they are. Caution should be taken when interpreting guilds for Candida species. | Manolakaki D, et al. 2010. Virulence. 1: 367-375 ((https://doi.org/10.4161/viru.1.5.12796)); Mekbib SB, et al. 2011. Tropical Plant Pathology 36:233-240 ((https://doi.org/10.1590/S1982-56762011000400004)); Põlme S, et al. 2020. Fungal Diversity 105:1-16 ((https://doi.org/10.1007/s13225-020-00466-2)) |
| Otu04247 | 7 | k__Fungi(100);p__Ascomycota(100);c__Saccharomycetes(100);o__Saccharomycetales(100);f__Metschnikowiaceae(100);g__Metschnikowia(100);g__Metschnikowia_unclassified(72); | Metschnikowia | 13 | Saprotroph | \|Undefined Saprotroph\| | Yeast | NULL | Probable | As nectar-tap_saprotroph (Põlme et al. 2020); Decay sugar-rich substrates (Põlme et al. 2020) | Põlme S, et al. 2020. Fungal Diversity 105:1-16 ((https://doi.org/10.1007/s13225-020-00466-2)) |
| Otu04248 | 7 | k__Fungi(100);p__Ascomycota(100);c__Sordariomycetes(100);o__Calosphaeriales(100);f__Calosphaeriaceae(100);g__Jattaea(100);s__Jattaea_prunicola(100); | Jattaea | 13 | Saprotroph | \|Undefined Saprotroph\| | Microfungus | NULL | Probable | NULL | Tedersoo L, et al. 2014. Science 346:e1256688 ((https://doi.org/10.1126/science.1256688)); Põlme S, et al. 2020. Fungal Diversity 105:1-16 ((https://doi.org/10.1007/s13225-020-00466-2)) |
| Otu04250 | 7 | k__Fungi(100);p__Ascomycota(100);c__Sordariomycetes(100);o__Xylariales(100);f__Xylariaceae(100);g__Nodulisporium(100);s__Nodulisporium_sp(100); | Nodulisporium | 13 | Saprotroph | \|Undefined Saprotroph\| | Microfungus | NULL | Probable | NULL | Tedersoo L, et al. 2014. Science 346:e1256688 ((https://doi.org/10.1126/science.1256688)); Põlme S, et al. 2020. Fungal Diversity 105:1-16 ((https://doi.org/10.1007/s13225-020-00466-2)) |
| Otu04251 | 7 | k__Fungi(100);p__Basidiomycota(100);c__Agaricomycetes(100);o__Hymenochaetales(100);f__Hymenochaetaceae(100);f__Hymenochaetaceae_unclassified(100);f__Hymenochaetaceae_unclassified(100); | Hymenochaetaceae | 9 | Saprotroph-Symbiotroph | Ectomycorrhizal-Wood Saprotroph | Corticioid-Polyporoid | NULL | Probable | Lignicolous, causing white rots, rarely ectomycorrhizal (Cannon & Kirk 2007) | Cannon PF, Kirk PM. 2007. Fungal Families of the World. CAB International, Cambridge (ISBN: 978-0851998275) |
| Otu04254 | 7 | k__Fungi(100);p__Ascomycota(100);c__Saccharomycetes(100);o__Saccharomycetales(100);f__Metschnikowiaceae(100);g__Metschnikowia(100);s__Metschnikowia_pulcherrima(86); | Metschnikowia | 13 | Saprotroph | \|Undefined Saprotroph\| | Yeast | NULL | Probable | As nectar-tap_saprotroph (Põlme et al. 2020); Decay sugar-rich substrates (Põlme et al. 2020) | Põlme S, et al. 2020. Fungal Diversity 105:1-16 ((https://doi.org/10.1007/s13225-020-00466-2)) |
| Otu04255 | 7 | k__Fungi(100);p__Basidiomycota(100);c__Agaricomycetes(100);o__Agaricales(100);f__Typhulaceae(100);g__Typhula(100);s__Typhula_variabilis(100); | Typhula | 13 | Pathotroph-Saprotroph | \|Plant Pathogen\|-Plant Saprotroph | Clavarioid | NULL | Probable | As litter_saprotroph (Põlme et al. 2020); Decay leaf, fruit, seed (Põlme et al. 2020) | Tedersoo L, et al. 2014. Science 346:e1256688 ((https://doi.org/10.1126/science.1256688)); Põlme S, et al. 2020. Fungal Diversity 105:1-16 ((https://doi.org/10.1007/s13225-020-00466-2)) |
| Otu04257 | 7 | k__Fungi(100);p__Ascomycota(100);c__Sordariomycetes(100);o__Sordariales(100);f__Cephalothecaceae(100);g__Phialemonium(100);s__Phialemonium_atrogriseum(100); | Phialemonium | 13 | Pathotroph-Saprotroph | \|Animal Parasite\|-Undefined Saprotroph | Microfungus | NULL | Probable | As animal_decomposer (Põlme et al. 2020); Decay animal material (Põlme et al. 2020) | Tedersoo L, et al. 2014. Science 346:e1256688 ((https://doi.org/10.1126/science.1256688)); Põlme S, et al. 2020. Fungal Diversity 105:1-16 ((https://doi.org/10.1007/s13225-020-00466-2)) |
| Otu04261 | 7 | k__Fungi(100);p__Ascomycota(100);c__Saccharomycetes(100);o__Saccharomycetales(100);f__Metschnikowiaceae(100);g__Metschnikowia(100);s__Metschnikowia_pulcherrima(86); | Metschnikowia | 13 | Saprotroph | \|Undefined Saprotroph\| | Yeast | NULL | Probable | As nectar-tap_saprotroph (Põlme et al. 2020); Decay sugar-rich substrates (Põlme et al. 2020) | Põlme S, et al. 2020. Fungal Diversity 105:1-16 ((https://doi.org/10.1007/s13225-020-00466-2)) |
| Otu04264 | 7 | k__Fungi(100);p__Ascomycota(100);c__Leotiomycetes(100);o__Helotiales(100);f__Hyaloscyphaceae(100);g__Lachnum(100);g__Lachnum_unclassified(86); | Lachnum | 13 | Saprotroph-Symbiotroph | Endophyte-\|Plant Saprotroph\|-Undefined Saprotroph-Wood Saprotroph | Microfungus | NULL | Probable | As litter_saprotroph (Põlme et al. 2020); As root_endophyte (Põlme et al. 2020); Decay wood, leaf, fruit, seed (Põlme et al. 2020) | Tedersoo L, et al. 2014. Science 346:e1256688 ((https://doi.org/10.1126/science.1256688)); Põlme S, et al. 2020. Fungal Diversity 105:1-16 ((https://doi.org/10.1007/s13225-020-00466-2)) |
| Otu04265 | 7 | k__Fungi(100);p__Ascomycota(100);c__Sordariomycetes(72);o__Hypocreales(58);o__Hypocreales_unclassified(58);o__Hypocreales_unclassified(58);o__Hypocreales_unclassified(58); | Hypocreales | 7 | Saprotroph | Undefined Saprotroph | Microfungus | NULL | Possible | NULL | Sterkenburg E, et al. 2015. New Phytologist 207:1145-1158 ((https://doi.org/10.1111/nph.13426)) |
| Otu04268 | 7 | k__Fungi(100);p__Ascomycota(100);c__Saccharomycetes(100);o__Saccharomycetales(100);f__Metschnikowiaceae(100);g__Metschnikowia(100);g__Metschnikowia_unclassified(100); | Metschnikowia | 13 | Saprotroph | \|Undefined Saprotroph\| | Yeast | NULL | Probable | As nectar-tap_saprotroph (Põlme et al. 2020); Decay sugar-rich substrates (Põlme et al. 2020) | Põlme S, et al. 2020. Fungal Diversity 105:1-16 ((https://doi.org/10.1007/s13225-020-00466-2)) |
| Otu04272 | 7 | k__Fungi(100);p__Basidiomycota(100);c__Agaricomycetes(100);o__Agaricales(100);f__Pleurotaceae(100);g__unclassified_Pleurotaceae(100);s__Pleurotaceae_sp(100); | Pleurotaceae | 9 | Pathotroph-Saprotroph | Endophyte-Plant Pathogen-Wood Saprotroph | Agaricoid | White Rot | Probable | Sapwood rot on Pistacia; Saprobic on woody substrata, more rarely associated with roots of herbaceous plants. Many species are known to derive nutrition from nematodes, the hyphae with adhesive knobs or toxigenic secretory appendages (Cannon & Kirk 2007) | Cannon PF, Kirk PM. 2007. Fungal Families of the World. CAB International, Cambridge (ISBN: 978-0851998275); Costa IPMW, et al. 2012. Mycotaxon 119:e26 ((http://mycotaxon.com/resources/checklists/costa_v119_checklist.pdf)); The American Phytopathological Society Common Names of Plant Diseases ((http://www.apsnet.org/publications/commonnames/Pages/default.aspx)) |
| Otu04273 | 7 | k__Fungi(100);p__Ascomycota(100);c__Dothideomycetes(100);o__Dothideales(100);f__Dothioraceae(100);g__Pringsheimia(100);s__Pringsheimia_sp(100); | Pringsheimia | 13 | Saprotroph | Undefined Saprotroph-\|Wood Saprotroph\| | Microfungus | NULL | Probable | Decay: wood (Põlme et al. 2020) | Tedersoo L, et al. 2014. Science 346:e1256688 ((https://doi.org/10.1126/science.1256688)); Põlme S, et al. 2020. Fungal Diversity 105:1-16 ((https://doi.org/10.1007/s13225-020-00466-2)) |
| Otu04274 | 7 | k__Fungi(100);p__Basidiomycota(100);c__Agaricomycetes(100);o__Agaricales(100);f__Mycenaceae(100);g__Mycena(100);g__Mycena_unclassified(100); | Mycena | 13 | Pathotroph-Saprotroph | Plant Pathogen-\|Plant Saprotroph\|-Undefined Saprotroph-Wood Saprotroph | Agaricoid | NULL | Probable | As litter_saprotroph (Põlme et al. 2020); Decay leaf, fruit, seed (Põlme et al. 2020) | Grgurinovic CA, Simpson JA. 2001. Fungal Diversity 8:97-127 ((http://www.fungaldiversity.org/fdp/sfdp/FD_8_97-127.pdf)); Tedersoo L, et al. 2014. Science 346:e1256688 ((https://doi.org/10.1126/science.1256688)); Põlme S, et al. 2020. Fungal Diversity 105:1-16 ((https://doi.org/10.1007/s13225-020-00466-2)); The American Phytopathological Society Common Names of Plant Diseases ((http://www.apsnet.org/publications/commonnames/Pages/default.aspx)) |
| Otu04275 | 7 | k__Fungi(100);p__Ascomycota(100);c__Saccharomycetes(100);o__Saccharomycetales(100);f__Metschnikowiaceae(100);g__Metschnikowia(100);s__Metschnikowia_pulcherrima(100); | Metschnikowia | 13 | Saprotroph | \|Undefined Saprotroph\| | Yeast | NULL | Probable | As nectar-tap_saprotroph (Põlme et al. 2020); Decay sugar-rich substrates (Põlme et al. 2020) | Põlme S, et al. 2020. Fungal Diversity 105:1-16 ((https://doi.org/10.1007/s13225-020-00466-2)) |
| Otu04278 | 7 | k__Fungi(100);p__Ascomycota(100);c__Leotiomycetes(100);o__Helotiales(100);f__Vibrisseaceae(100);g__Phialocephala(100);s__Phialocephala_humicola(100); | Phialocephala | 13 | Saprotroph-Symbiotroph | Endophyte-\|Undefined Saprotroph\| | Microfungus | NULL | Highly Probable | As root_endophyte (Põlme et al. 2020); As root_endophyte_dark_septate (Põlme et al. 2020); As soil_saprotroph (Põlme et al. 2020); Dark Septate Endophyte, known from plant roots.; Decay roots, soil (Põlme et al. 2020) | Newsham KK. 2011. New Phytologist 190:783-793 ((https://doi.org/10.1111/j.1469-8137.2010.03611.x)) (as dark septate root endophytes); Põlme S, et al. 2020. Fungal Diversity 105:1-16 ((https://doi.org/10.1007/s13225-020-00466-2)) |
| Otu04279 | 7 | k__Fungi(100);p__Basidiomycota(100);c__Agaricomycetes(100);o__Agaricales(100);f__Entolomataceae(100);g__Entoloma(100);g__Entoloma_unclassified(86); | Entoloma | 13 | Pathotroph-Saprotroph-Symbiotroph | Ectomycorrhizal-Fungal Parasite-\|Undefined Saprotroph\|-Undefined Symbiotroph | Agaricoid | NULL | Possible | As soil_saprotroph (Põlme et al. 2020); Decay roots, soil (Põlme et al. 2020) | Rinaldi AC, et al. 2008. Fungal Diversity 33:1-45 ((http://www.fungaldiversity.org/fdp/sfdp/33-1.pdf)) (as ECM/not ECM); Tedersoo L, et al. 2010. Mycorrhiza 20:217-263 ((https://doi.org/10.1007/s00572-009-0274-x; Põlme S, et al. 2020. Fungal Diversity 105:1-16 ((https://doi.org/10.1007/s13225-020-00466-2)); sensu stricto)); http://mycorrhizas.info/ecmf.html ((http://mycorrhizas.info/ecmf.html)) ((as ECM/not ECM)) |
| Otu04281 | 7 | k__Fungi(100);p__Ascomycota(100);c__Saccharomycetes(100);o__Saccharomycetales(100);f__Metschnikowiaceae(100);g__Metschnikowia(100);g__Metschnikowia_unclassified(72); | Metschnikowia | 13 | Saprotroph | \|Undefined Saprotroph\| | Yeast | NULL | Probable | As nectar-tap_saprotroph (Põlme et al. 2020); Decay sugar-rich substrates (Põlme et al. 2020) | Põlme S, et al. 2020. Fungal Diversity 105:1-16 ((https://doi.org/10.1007/s13225-020-00466-2)) |
| Otu04285 | 7 | k__Fungi(100);p__Ascomycota(100);c__Saccharomycetes(100);o__Saccharomycetales(100);f__Metschnikowiaceae(100);g__Metschnikowia(100);s__Metschnikowia_pulcherrima(100); | Metschnikowia | 13 | Saprotroph | \|Undefined Saprotroph\| | Yeast | NULL | Probable | As nectar-tap_saprotroph (Põlme et al. 2020); Decay sugar-rich substrates (Põlme et al. 2020) | Põlme S, et al. 2020. Fungal Diversity 105:1-16 ((https://doi.org/10.1007/s13225-020-00466-2)) |
| Otu04290 | 7 | k__Fungi(100);p__Ascomycota(100);c__Sordariomycetes(100);o__Xylariales(100);f__Xylariaceae(100);g__Hypoxylon(100);s__Hypoxylon_rubiginosum(100); | Hypoxylon | 13 | Saprotroph-Saprotroph-Symbiotroph | Endophyte-Undefined Saprotroph-\|Wood Saprotroph\| | Corticioid-Microfungus-Xylarioid | NULL | Probable | As foliar_endophyte (Põlme et al. 2020); Decay wood, leaf, fruit, seed (Põlme et al. 2020); Host - angiosperms (Põlme et al. 2020) | Tedersoo L, et al. 2014. Science 346:e1256688 ((https://doi.org/10.1126/science.1256688)); Põlme S, et al. 2020. Fungal Diversity 105:1-16 ((https://doi.org/10.1007/s13225-020-00466-2)) |
| Otu04291 | 7 | k__Fungi(100);p__Ascomycota(100);c__Eurotiomycetes(100);o__Chaetothyriales(100);f__Herpotrichiellaceae(100);g__Phaeomoniella(100);s__Phaeomoniella_sp(58); | Phaeomoniella | 13 | Pathotroph | \|Plant Pathogen\| | Dimorphic-Microfungus; Facultative Yeast (Tedersoo et | NULL | Highly Probable | Pathotroph isolated from necrosis in Prunus wood (Damm et al. 2010) | Damm U, et al. 2010. Persoonia 24:60-80 ((http://doi.org/10.3767/003158510X500705)); Tedersoo L, et al. 2014. Science 346:e1256688 ((https://doi.org/10.1126/science.1256688)); Põlme S, et al. 2020. Fungal Diversity 105:1-16 ((https://doi.org/10.1007/s13225-020-00466-2)) |
| Otu04294 | 7 | k__Fungi(100);p__Ascomycota(100);c__Saccharomycetes(100);o__Saccharomycetales(100);f__Metschnikowiaceae(100);g__Metschnikowia(100);s__Metschnikowia_pulcherrima(100); | Metschnikowia | 13 | Saprotroph | \|Undefined Saprotroph\| | Yeast | NULL | Probable | As nectar-tap_saprotroph (Põlme et al. 2020); Decay sugar-rich substrates (Põlme et al. 2020) | Põlme S, et al. 2020. Fungal Diversity 105:1-16 ((https://doi.org/10.1007/s13225-020-00466-2)) |
| Otu04295 | 7 | k__Fungi(100);p__Ascomycota(100);c__Saccharomycetes(100);o__Saccharomycetales(100);f__Metschnikowiaceae(100);g__Metschnikowia(100);s__Metschnikowia_pulcherrima(86); | Metschnikowia | 13 | Saprotroph | \|Undefined Saprotroph\| | Yeast | NULL | Probable | As nectar-tap_saprotroph (Põlme et al. 2020); Decay sugar-rich substrates (Põlme et al. 2020) | Põlme S, et al. 2020. Fungal Diversity 105:1-16 ((https://doi.org/10.1007/s13225-020-00466-2)) |
| Otu04296 | 7 | k__Fungi(100);p__Ascomycota(100);c__Saccharomycetes(100);o__Saccharomycetales(100);f__Metschnikowiaceae(100);g__Metschnikowia(100);s__Metschnikowia_pulcherrima(100); | Metschnikowia | 13 | Saprotroph | \|Undefined Saprotroph\| | Yeast | NULL | Probable | As nectar-tap_saprotroph (Põlme et al. 2020); Decay sugar-rich substrates (Põlme et al. 2020) | Põlme S, et al. 2020. Fungal Diversity 105:1-16 ((https://doi.org/10.1007/s13225-020-00466-2)) |
| Otu04300 | 7 | k__Fungi(100);p__Ascomycota(100);c__Saccharomycetes(100);o__Saccharomycetales(100);f__Metschnikowiaceae(100);g__Metschnikowia(100);s__Metschnikowia_pulcherrima(100); | Metschnikowia | 13 | Saprotroph | \|Undefined Saprotroph\| | Yeast | NULL | Probable | As nectar-tap_saprotroph (Põlme et al. 2020); Decay sugar-rich substrates (Põlme et al. 2020) | Põlme S, et al. 2020. Fungal Diversity 105:1-16 ((https://doi.org/10.1007/s13225-020-00466-2)) |
| Otu04301 | 7 | k__Fungi(100);p__Ascomycota(100);c__Saccharomycetes(100);o__Saccharomycetales(100);f__Metschnikowiaceae(100);g__Metschnikowia(100);s__Metschnikowia_pulcherrima(58); | Metschnikowia | 13 | Saprotroph | \|Undefined Saprotroph\| | Yeast | NULL | Probable | As nectar-tap_saprotroph (Põlme et al. 2020); Decay sugar-rich substrates (Põlme et al. 2020) | Põlme S, et al. 2020. Fungal Diversity 105:1-16 ((https://doi.org/10.1007/s13225-020-00466-2)) |
| Otu04303 | 7 | k__Fungi(100);p__Basidiomycota(100);c__Agaricomycetes(100);o__Polyporales(100);f__Meruliaceae(100);g__Steccherinum(86);g__Steccherinum_unclassified(86); | Steccherinum | 13 | Saprotroph | Undefined Saprotroph-\|Wood Saprotroph\| | Corticioid | White Rot | Probable | As white_rot (Põlme et al. 2020); Decay wood (Põlme et al. 2020) | Tedersoo L, et al. 2014. Science 346:e1256688 ((https://doi.org/10.1126/science.1256688)); Põlme S, et al. 2020. Fungal Diversity 105:1-16 ((https://doi.org/10.1007/s13225-020-00466-2)) |
| Otu04305 | 7 | k__Fungi(100);p__Ascomycota(100);c__Saccharomycetes(100);o__Saccharomycetales(100);f__Metschnikowiaceae(100);g__Metschnikowia(100);s__Metschnikowia_pulcherrima(86); | Metschnikowia | 13 | Saprotroph | \|Undefined Saprotroph\| | Yeast | NULL | Probable | As nectar-tap_saprotroph (Põlme et al. 2020); Decay sugar-rich substrates (Põlme et al. 2020) | Põlme S, et al. 2020. Fungal Diversity 105:1-16 ((https://doi.org/10.1007/s13225-020-00466-2)) |
| Otu04306 | 7 | k__Fungi(100);p__Ascomycota(100);c__Pezizomycetes(100);o__Pezizales(100);f__Sarcosomataceae(100);f__Sarcosomataceae_unclassified(100);f__Sarcosomataceae_unclassified(100); | Sarcosomataceae | 9 | Saprotroph | Wood Saprotroph | Pezizoid | NULL | Probable | Saprobic on rotten, sometimes buried, wood (Cannon & Kirk 2007) | Cannon PF, Kirk PM. 2007. Fungal Families of the World. CAB International, Cambridge (ISBN: 978-0851998275) |
| Otu04307 | 7 | k__Fungi(100);p__Basidiomycota(100);c__Exobasidiomycetes(100);o__Exobasidiales(100);f__Exobasidiaceae(100);g__Exobasidium(100);g__Exobasidium_unclassified(100); | Exobasidium | 13 | Pathotroph | \|Plant Pathogen\| | Leaf Blister-Leaf Gall-Leaf Spot | NULL | Probable | NULL | Tedersoo L, et al. 2014. Science 346:e1256688 ((https://doi.org/10.1126/science.1256688)); Põlme S, et al. 2020. Fungal Diversity 105:1-16 ((https://doi.org/10.1007/s13225-020-00466-2)) |
| Otu04308 | 7 | k__Fungi(100);p__Ascomycota(100);c__Saccharomycetes(100);o__Saccharomycetales(100);f__Metschnikowiaceae(100);g__Metschnikowia(100);s__Metschnikowia_pulcherrima(100); | Metschnikowia | 13 | Saprotroph | \|Undefined Saprotroph\| | Yeast | NULL | Probable | As nectar-tap_saprotroph (Põlme et al. 2020); Decay sugar-rich substrates (Põlme et al. 2020) | Põlme S, et al. 2020. Fungal Diversity 105:1-16 ((https://doi.org/10.1007/s13225-020-00466-2)) |
| Otu04313 | 7 | k__Fungi(100);p__Ascomycota(100);c__Saccharomycetes(100);o__Saccharomycetales(100);f__Metschnikowiaceae(100);g__Metschnikowia(100);s__Metschnikowia_pulcherrima(100); | Metschnikowia | 13 | Saprotroph | \|Undefined Saprotroph\| | Yeast | NULL | Probable | As nectar-tap_saprotroph (Põlme et al. 2020); Decay sugar-rich substrates (Põlme et al. 2020) | Põlme S, et al. 2020. Fungal Diversity 105:1-16 ((https://doi.org/10.1007/s13225-020-00466-2)) |
| Otu04314 | 7 | k__Fungi(100);p__Basidiomycota(100);c__Agaricostilbomycetes(100);o__Agaricostilbales(100);f__Chionosphaeraceae(100);g__Kurtzmanomyces(100);g__Kurtzmanomyces_unclassified(100); | Kurtzmanomyces | 13 | Saprotroph | \|Undefined Saprotroph\| | Dimorphic | NULL | Probable | NULL | Põlme S, et al. 2020. Fungal Diversity 105:1-16 ((https://doi.org/10.1007/s13225-020-00466-2)) |
| Otu04315 | 7 | k__Fungi(100);p__Ascomycota(100);c__Saccharomycetes(100);o__Saccharomycetales(100);f__Metschnikowiaceae(100);g__Metschnikowia(100);s__Metschnikowia_pulcherrima(100); | Metschnikowia | 13 | Saprotroph | \|Undefined Saprotroph\| | Yeast | NULL | Probable | As nectar-tap_saprotroph (Põlme et al. 2020); Decay sugar-rich substrates (Põlme et al. 2020) | Põlme S, et al. 2020. Fungal Diversity 105:1-16 ((https://doi.org/10.1007/s13225-020-00466-2)) |
| Otu04317 | 7 | k__Fungi(100);p__Ascomycota(100);c__Saccharomycetes(100);o__Saccharomycetales(100);f__Metschnikowiaceae(100);g__Metschnikowia(100);s__Metschnikowia_pulcherrima(72); | Metschnikowia | 13 | Saprotroph | \|Undefined Saprotroph\| | Yeast | NULL | Probable | As nectar-tap_saprotroph (Põlme et al. 2020); Decay sugar-rich substrates (Põlme et al. 2020) | Põlme S, et al. 2020. Fungal Diversity 105:1-16 ((https://doi.org/10.1007/s13225-020-00466-2)) |
| Otu04318 | 7 | k__Fungi(100);p__Ascomycota(100);c__Saccharomycetes(100);o__Saccharomycetales(100);f__Metschnikowiaceae(100);g__Metschnikowia(100);g__Metschnikowia_unclassified(58); | Metschnikowia | 13 | Saprotroph | \|Undefined Saprotroph\| | Yeast | NULL | Probable | As nectar-tap_saprotroph (Põlme et al. 2020); Decay sugar-rich substrates (Põlme et al. 2020) | Põlme S, et al. 2020. Fungal Diversity 105:1-16 ((https://doi.org/10.1007/s13225-020-00466-2)) |
| Otu04319 | 7 | k__Fungi(100);p__Ascomycota(100);c__Sordariomycetes(100);o__Melanosporales(100);f__Ceratostomataceae(100);g__Harzia(100);s__Harzia_acremonioides(100); | Harzia | 13 | Saprotroph | Plant Saprotroph-Undefined Saprotroph-\|Wood Saprotroph\| | Microfungus | NULL | Probable | As litter_saprotroph (Põlme et al. 2020); Decay wood, leaf, fruit, seed (Põlme et al. 2020) | Tedersoo L, et al. 2014. Science 346:e1256688 ((https://doi.org/10.1126/science.1256688)); Põlme S, et al. 2020. Fungal Diversity 105:1-16 ((https://doi.org/10.1007/s13225-020-00466-2)) |
| Otu04322 | 7 | k__Fungi(100);p__Basidiomycota(100);c__Agaricomycetes(100);o__Agaricales(100);o__Agaricales_unclassified(100);o__Agaricales_unclassified(100);o__Agaricales_unclassified(100); | Agaricales | 7 | Pathotroph-Saprotroph-Symbiotroph | Bryophyte Parasite-Dung Saprotroph-Ectomycorrhizal-Fungal Parasite-Leaf Saprotroph-Plant Parasite-Undefined Saprotroph-Wood Saprotroph | Agaricoid-Gasteroid-Microfungus-Secotioid-Yeast | NULL | Possible | Mushrooms and toadstools, Gill fungi, Agarics... lignicolous, sometimes muscicolous or fungicolous, saprobic, mycorrhizal, rarely parasitic on plants or fungi (Kirk et al. 2008) | Kirk PM et al. 2008. Dictionary of the Fungi. Tenth Edition. CAB International, Wallingford (ISBN: 978-0851998268) |
| Otu04323 | 7 | k__Fungi(100);p__Basidiomycota(100);c__Agaricomycetes(100);o__Agaricales(100);f__Inocybaceae(100);g__Inocybe(100);g__Inocybe_unclassified(100); | Inocybe | 13 | Symbiotroph | \|Ectomycorrhizal\| | Agaricoid | NULL | Highly Probable | Sensu stricto (Matheny 2009). | Rinaldi AC, et al. 2008. Fungal Diversity 33:1-45 ((http://www.fungaldiversity.org/fdp/sfdp/33-1.pdf)); Matheny PB. 2009. McIlvainea 18:11-21 ((https://namyco.org/docs/3_Matheny.pdf)); Tedersoo L, et al. 2010. Mycorrhiza 20:217-263 ((https://doi.org/10.1007/s00572-009-0274-x)); Põlme S, et al. 2020. Fungal Diversity 105:1-16 ((https://doi.org/10.1007/s13225-020-00466-2)); http://mycorrhizas.info/ecmf.html ((http://mycorrhizas.info/ecmf.html)) |
| Otu04324 | 7 | k__Fungi(100);p__Ascomycota(100);c__Saccharomycetes(100);o__Saccharomycetales(100);f__Saccharomycetales_family_Incertae_sedis(100);g__Candida(86);g__Candida_unclassified(72); | Candida | 13 | Pathotroph-Saprotroph-Symbiotroph | Animal Pathogen-Endophyte-Endosymbiont-Epiphyte-\|Undefined Saprotroph\| | Dimorphic Yeast-Yeast | NULL | Probable | As nectar-tap_saprotroph (Põlme et al. 2020); Decay sugar-rich substrates (Põlme et al. 2020); This asexual genus is polyphyletic and contain species that function very differently depend on what they are. Caution should be taken when interpreting guilds for Candida species. | Manolakaki D, et al. 2010. Virulence. 1: 367-375 ((https://doi.org/10.4161/viru.1.5.12796)); Mekbib SB, et al. 2011. Tropical Plant Pathology 36:233-240 ((https://doi.org/10.1590/S1982-56762011000400004)); Põlme S, et al. 2020. Fungal Diversity 105:1-16 ((https://doi.org/10.1007/s13225-020-00466-2)) |
| Otu04328 | 7 | k__Fungi(100);p__Ascomycota(100);c__Leotiomycetes(100);o__Helotiales(100);f__Helotiaceae(86);g__Tricladium(86);s__Tricladium_splendens(86); | Tricladium | 13 | Saprotroph | \|Plant Saprotroph\|-Undefined Saprotroph | Microfungus | NULL | Probable | As litter_saprotroph (Põlme et al. 2020); Decay leaf, fruit, seed (Põlme et al. 2020) | Tedersoo L, et al. 2014. Science 346:e1256688 ((https://doi.org/10.1126/science.1256688)); Põlme S, et al. 2020. Fungal Diversity 105:1-16 ((https://doi.org/10.1007/s13225-020-00466-2)) |
| Otu04329 | 7 | k__Fungi(100);p__Ascomycota(100);c__Leotiomycetes(100);o__Helotiales(100);f__Rutstroemiaceae(100);f__Rutstroemiaceae_unclassified(100);f__Rutstroemiaceae_unclassified(100); | Rutstroemiaceae | 9 | Saprotroph | Plant Saprotroph | Microfungus | NULL | Probable | Essentially saprobic on various plant parts, especially seeds and fruits, though initial infection may take place in living tissue (Cannon & Kirk 2007) | Cannon PF, Kirk PM. 2007. Fungal Families of the World. CAB International, Cambridge (ISBN: 978-0851998275) |
| Otu04330 | 7 | k__Fungi(100);p__Ascomycota(100);c__Saccharomycetes(72);o__Saccharomycetales(72);o__Saccharomycetales_unclassified(72);o__Saccharomycetales_unclassified(72);o__Saccharomycetales_unclassified(72); | Saccharomycetales | 7 | Saprotroph | Undefined Saprotroph | Yeast | NULL | Possible | NULL | Sterkenburg E, et al. 2015. New Phytologist 207:1145-1158 ((https://doi.org/10.1111/nph.13426)) |
| Otu04332 | 7 | k__Fungi(100);p__Zygomycota(100);c__Zygomycota_class_Incertae_sedis(100);o__Mucorales(100);f__Mucoraceae(100);g__Mucor(100);g__Mucor_unclassified(100); | Mucor | 13 | Saprotroph | \|Undefined Saprotroph\| | Microfungus | NULL | Probable | As soil_saprotroph (Põlme et al. 2020); Decay soil (Põlme et al. 2020) | Tedersoo L, et al. 2014. Science 346:e1256688 ((https://doi.org/10.1126/science.1256688)); Põlme S, et al. 2020. Fungal Diversity 105:1-16 ((https://doi.org/10.1007/s13225-020-00466-2)) |
| Otu04334 | 7 | k__Fungi(100);p__Ascomycota(100);c__Saccharomycetes(100);o__Saccharomycetales(100);f__Metschnikowiaceae(100);g__Metschnikowia(100);s__Metschnikowia_pulcherrima(58); | Metschnikowia | 13 | Saprotroph | \|Undefined Saprotroph\| | Yeast | NULL | Probable | As nectar-tap_saprotroph (Põlme et al. 2020); Decay sugar-rich substrates (Põlme et al. 2020) | Põlme S, et al. 2020. Fungal Diversity 105:1-16 ((https://doi.org/10.1007/s13225-020-00466-2)) |
| Otu04335 | 7 | k__Fungi(100);p__Ascomycota(100);c__Dothideomycetes(100);o__Pleosporales(100);f__Pleosporales_family_Incertae_sedis(100);g__Letendraea(100);s__Letendraea_sp(100); | Letendraea | 13 | Pathotroph | \|Fungal Parasite\| | Microfungus | NULL | Probable | As mycoparasite (Põlme et al. 2020); Decay fungal material (Põlme et al. 2020) | Põlme S, et al. 2020. Fungal Diversity 105:1-16 ((https://doi.org/10.1007/s13225-020-00466-2)) |
| Otu04337 | 7 | k__Fungi(100);p__Basidiomycota(100);c__Agaricomycetes(100);o__Russulales(100);f__Auriscalpiaceae(100);g__Lentinellus(100);g__Lentinellus_unclassified(100); | Lentinellus | 13 | Pathotroph-Saprotroph | Plant Pathogen-Undefined Saprotroph-\|Wood Saprotroph\| | Agaricoid | White Rot | Probable | As white_rot (Põlme et al. 2020); As wood_pathogen (Põlme et al. 2020); Decay wood (Põlme et al. 2020) | Tedersoo L, et al. 2014. Science 346:e1256688 ((https://doi.org/10.1126/science.1256688)); Põlme S, et al. 2020. Fungal Diversity 105:1-16 ((https://doi.org/10.1007/s13225-020-00466-2)) |
| Otu04339 | 7 | k__Fungi(100);p__Ascomycota(100);c__Saccharomycetes(100);o__Saccharomycetales(100);f__Metschnikowiaceae(100);g__Metschnikowia(100);s__Metschnikowia_pulcherrima(100); | Metschnikowia | 13 | Saprotroph | \|Undefined Saprotroph\| | Yeast | NULL | Probable | As nectar-tap_saprotroph (Põlme et al. 2020); Decay sugar-rich substrates (Põlme et al. 2020) | Põlme S, et al. 2020. Fungal Diversity 105:1-16 ((https://doi.org/10.1007/s13225-020-00466-2)) |
| Otu04340 | 7 | k__Fungi(100);p__Basidiomycota(100);c__Agaricomycetes(100);o__Hymenochaetales(100);f__Hymenochaetaceae(100);g__Inonotus(100);s__Inonotus_hispidus(100); | Inonotus | 13 | Pathotroph-Saprotroph | Plant Pathogen-\|Wood Saprotroph\| | Polyporoid | White Rot | Highly Probable | As white_rot (Põlme et al. 2020); As wood_pathogen (Põlme et al. 2020); Decay wood (Põlme et al. 2020) | Gilbertson RL, Ryvarden L. 1987-1987. North American Polypores. Fungiflora, Oslo (ISBN: 978-0945345060); Põlme S, et al. 2020. Fungal Diversity 105:1-16 ((https://doi.org/10.1007/s13225-020-00466-2)) |
| Otu04344 | 7 | k__Fungi(100);p__Ascomycota(100);c__Dothideomycetes(100);o__Botryosphaeriales(100);f__Botryosphaeriales_family_Incertae_sedis(100);g__Camarosporium(100);s__Camarosporium_sp(100); | Camarosporium | 13 | Pathotroph-Saprotroph-Symbiotroph | \|Endophyte\|-Plant Pathogen-Plant Saprotroph | Microfungus | NULL | Possible | Chenipodiaceae; As foliar_endophyte (Põlme et al. 2020); As litter_saprotroph (Põlme et al. 2020); culture (Del Olmo; Decay leaf, fruit, seed (Põlme et al. 2020); Endophyte detection method; Ruiz & Arnold 2014); Endophyte host | Costa IPMW, et al. 2012. Mycotaxon 119:e26 ((http://mycotaxon.com/resources/checklists/costa_v119_checklist.pdf)); Del Olmo-Ruiz M, Arnold AE. 2014. Mycologia 106:8-21 ((https://doi.org/10.3852/13-098)); Põlme S, et al. 2020. Fungal Diversity 105:1-16 ((https://doi.org/10.1007/s13225-020-00466-2)); The American Phytopathological Society Common Names of Plant Diseases ((http://www.apsnet.org/publications/commonnames/Pages/default.aspx)) |
| Otu04345 | 7 | k__Fungi(100);p__Ascomycota(100);c__Saccharomycetes(100);o__Saccharomycetales(100);f__Metschnikowiaceae(100);g__Metschnikowia(100);s__Metschnikowia_pulcherrima(100); | Metschnikowia | 13 | Saprotroph | \|Undefined Saprotroph\| | Yeast | NULL | Probable | As nectar-tap_saprotroph (Põlme et al. 2020); Decay sugar-rich substrates (Põlme et al. 2020) | Põlme S, et al. 2020. Fungal Diversity 105:1-16 ((https://doi.org/10.1007/s13225-020-00466-2)) |
| Otu04346 | 7 | k__Fungi(100);p__Ascomycota(100);c__Saccharomycetes(100);o__Saccharomycetales(100);f__Metschnikowiaceae(100);g__Metschnikowia(100);s__Metschnikowia_pulcherrima(86); | Metschnikowia | 13 | Saprotroph | \|Undefined Saprotroph\| | Yeast | NULL | Probable | As nectar-tap_saprotroph (Põlme et al. 2020); Decay sugar-rich substrates (Põlme et al. 2020) | Põlme S, et al. 2020. Fungal Diversity 105:1-16 ((https://doi.org/10.1007/s13225-020-00466-2)) |
| Otu04349 | 7 | k__Fungi(100);p__Basidiomycota(100);c__Agaricomycetes(100);o__Agaricales(100);f__Marasmiaceae(100);g__Gymnopus(100);s__Gymnopus_iocephalus(86); | Gymnopus | 13 | Saprotroph | \|Plant Saprotroph\| | Agaricoid | NULL | Probable | As litter_saprotroph (Põlme et al. 2020); Decay leaf, fruit, seed (Põlme et al. 2020) | Põlme S, et al. 2020. Fungal Diversity 105:1-16 ((https://doi.org/10.1007/s13225-020-00466-2)) |
| Otu04350 | 7 | k__Fungi(100);p__Basidiomycota(100);c__Agaricomycetes(100);o__Russulales(100);f__Auriscalpiaceae(100);g__Lentinellus(100);s__Lentinellus_castoreus(100); | Lentinellus | 13 | Pathotroph-Saprotroph | Plant Pathogen-Undefined Saprotroph-\|Wood Saprotroph\| | Agaricoid | White Rot | Probable | As white_rot (Põlme et al. 2020); As wood_pathogen (Põlme et al. 2020); Decay wood (Põlme et al. 2020) | Tedersoo L, et al. 2014. Science 346:e1256688 ((https://doi.org/10.1126/science.1256688)); Põlme S, et al. 2020. Fungal Diversity 105:1-16 ((https://doi.org/10.1007/s13225-020-00466-2)) |
| Otu04354 | 7 | k__Fungi(100);p__Ascomycota(100);c__Saccharomycetes(100);o__Saccharomycetales(100);f__Metschnikowiaceae(100);g__Metschnikowia(100);s__Metschnikowia_pulcherrima(86); | Metschnikowia | 13 | Saprotroph | \|Undefined Saprotroph\| | Yeast | NULL | Probable | As nectar-tap_saprotroph (Põlme et al. 2020); Decay sugar-rich substrates (Põlme et al. 2020) | Põlme S, et al. 2020. Fungal Diversity 105:1-16 ((https://doi.org/10.1007/s13225-020-00466-2)) |
| Otu04359 | 7 | k__Fungi(100);p__Ascomycota(100);c__Saccharomycetes(100);o__Saccharomycetales(100);f__Metschnikowiaceae(100);g__Metschnikowia(100);g__Metschnikowia_unclassified(72); | Metschnikowia | 13 | Saprotroph | \|Undefined Saprotroph\| | Yeast | NULL | Probable | As nectar-tap_saprotroph (Põlme et al. 2020); Decay sugar-rich substrates (Põlme et al. 2020) | Põlme S, et al. 2020. Fungal Diversity 105:1-16 ((https://doi.org/10.1007/s13225-020-00466-2)) |
| Otu04360 | 7 | k__Fungi(100);p__Ascomycota(100);c__Saccharomycetes(100);o__Saccharomycetales(100);f__Metschnikowiaceae(100);g__Metschnikowia(100);s__Metschnikowia_pulcherrima(86); | Metschnikowia | 13 | Saprotroph | \|Undefined Saprotroph\| | Yeast | NULL | Probable | As nectar-tap_saprotroph (Põlme et al. 2020); Decay sugar-rich substrates (Põlme et al. 2020) | Põlme S, et al. 2020. Fungal Diversity 105:1-16 ((https://doi.org/10.1007/s13225-020-00466-2)) |
| Otu04361 | 7 | k__Fungi(100);p__Basidiomycota(100);c__Agaricomycetes(100);o__Agaricales(100);o__Agaricales_unclassified(100);o__Agaricales_unclassified(100);o__Agaricales_unclassified(100); | Agaricales | 7 | Pathotroph-Saprotroph-Symbiotroph | Bryophyte Parasite-Dung Saprotroph-Ectomycorrhizal-Fungal Parasite-Leaf Saprotroph-Plant Parasite-Undefined Saprotroph-Wood Saprotroph | Agaricoid-Gasteroid-Microfungus-Secotioid-Yeast | NULL | Possible | Mushrooms and toadstools, Gill fungi, Agarics... lignicolous, sometimes muscicolous or fungicolous, saprobic, mycorrhizal, rarely parasitic on plants or fungi (Kirk et al. 2008) | Kirk PM et al. 2008. Dictionary of the Fungi. Tenth Edition. CAB International, Wallingford (ISBN: 978-0851998268) |
| Otu04362 | 7 | k__Fungi(100);p__Ascomycota(100);c__Saccharomycetes(100);o__Saccharomycetales(100);f__Metschnikowiaceae(100);g__Metschnikowia(100);g__Metschnikowia_unclassified(86); | Metschnikowia | 13 | Saprotroph | \|Undefined Saprotroph\| | Yeast | NULL | Probable | As nectar-tap_saprotroph (Põlme et al. 2020); Decay sugar-rich substrates (Põlme et al. 2020) | Põlme S, et al. 2020. Fungal Diversity 105:1-16 ((https://doi.org/10.1007/s13225-020-00466-2)) |
| Otu04364 | 7 | k__Fungi(100);p__Ascomycota(100);c__Sordariomycetes(100);o__Hypocreales(100);f__Hypocreales_family_Incertae_sedis(100);g__Stachybotrys(100);s__Stachybotrys_sp(100); | Stachybotrys | 13 | Pathotroph-Saprotroph | Plant Pathogen-Plant Saprotroph-\|Wood Saprotroph\| | Microfungus | Soft Rot | Highly Probable | As litter_saprotroph (Põlme et al. 2020); Decay wood, leaf, fruit, seed (Põlme et al. 2020) | Seehann G, et al. 1975. List of Fungi in Soft-Rot Tissue Tests. Institute fur Holzbiologie und Holzschutz, Bundesforschungsanstalt fur Forst - und Holzwirtshaft, Hamburg - Lohbrugge, Germany; Costa IPMW, et al. 2012. Mycotaxon 119:e26 ((http://mycotaxon.com/resources/checklists/costa_v119_checklist.pdf)); Tedersoo L, et al. 2014. Science 346:e1256688 ((https://doi.org/10.1126/science.1256688)); Põlme S, et al. 2020. Fungal Diversity 105:1-16 ((https://doi.org/10.1007/s13225-020-00466-2)); The American Phytopathological Society Common Names of Plant Diseases ((http://www.apsnet.org/publications/commonnames/Pages/default.aspx)) |

| | | | | | | | | | | | |
|---|---|---|---|---|---|---|---|---|---|---|---|
| Otu04366 | 7 | k__Fungi(100);p__Ascomycota(100);c__Lecanoromycetes(100);o__Ostropales(100);f__Stictidaceae(100);f__Stictidaceae_unclassified(100);f__Stictidaceae_unclassified(100); | Stictidaceae | 9 | Symbiotroph | Lichenized-Wood Saprotroph | Microfungus-Thallus | NULL | Probable | [illegible] | [illegible] |
| Otu04367 | 7 | k__Fungi(100);p__Ascomycota(100);c__Dothideomycetes(100);o__Capnodiales(100);f__Mycosphaerellaceae(100);f__Mycosphaerellaceae_unclassified(58);f__Mycosphaerellaceae_unclassified(58); | Mycosphaerellaceae | 9 | Pathotroph-Saprotroph | Plant Pathogen-Undefined Saprotroph | Microfungus | NULL | Probable | [illegible] | [illegible] |
| Otu04368 | 7 | k__Fungi(100);p__Ascomycota(100);c__Eurotiomycetes(100);o__Chaetothyriales(100);f__Herpotrichiellaceae(100);f__Herpotrichiellaceae_unclassified(100);f__Herpotrichiellaceae_unclassified(100); | Herpotrichiellaceae | 9 | Pathotroph-Saprotroph | Animal Pathogen-Fungal Parasite-Undefined Saprotroph | Facultative Yeast-Microfungus | NULL | Probable | [illegible] | [illegible] |
| Otu04369 | 7 | k__Fungi(100);p__Ascomycota(100);c__Eurotiomycetes(100);o__Chaetothyriales(100);f__Herpotrichiellaceae(100);g__Exophiala(100);g__Exophiala_unclassified(100); | Exophiala | 13 | Pathotroph-Saprotroph-Symbiotroph | \|Animal Parasite\|-Animal Pathogen-Endophyte-Plant Saprotroph-Undefined Saprotroph | Dimorphic-Facultative Yeast | NULL | Probable | [illegible] | [illegible] |
| Otu04370 | 7 | k__Fungi(100);p__Basidiomycota(100);c__Agaricomycetes(100);o__Agaricales(100);o__Agaricales_unclassified(100);o__Agaricales_unclassified(100);o__Agaricales_unclassified(100); | Agaricales | 7 | Pathotroph-Saprotroph-Symbiotroph | Bryophyte Parasite-Dung Saprotroph-Ectomycorrhizal-Fungal Parasite-Leaf Saprotroph-Plant Parasite-Undefined Saprotroph-Wood Saprotroph | Agaricoid-Gasteroid-Microfungus-Secotioid-Yeast | NULL | Possible | [illegible] | [illegible] |
| Otu04371 | 7 | k__Fungi(100);p__Basidiomycota(100);c__Agaricomycetes(100);o__Agaricales(100);o__Agaricales_unclassified(100);o__Agaricales_unclassified(100);o__Agaricales_unclassified(100); | Agaricales | 7 | Pathotroph-Saprotroph-Symbiotroph | Bryophyte Parasite-Dung Saprotroph-Ectomycorrhizal-Fungal Parasite-Leaf Saprotroph-Plant Parasite-Undefined Saprotroph-Wood Saprotroph | Agaricoid-Gasteroid-Microfungus-Secotioid-Yeast | NULL | Possible | [illegible] | [illegible] |
| Otu04372 | 7 | k__Fungi(100);p__Basidiomycota(100);c__Agaricomycetes(100);o__Thelephorales(100);f__Bankeraceae(100);g__Phellodon(100);s__Phellodon_confluens(100); | Phellodon | 13 | Symbiotroph | \|Ectomycorrhizal\| | Agaricoid-Hydnoid | NULL | Highly Probable | NULL | [illegible] |
| Otu04373 | 7 | k__Fungi(100);p__Basidiomycota(100);c__Agaricomycetes(100);o__Agaricales(100);f__Inocybaceae(100);g__Inocybe(100);s__Inocybe_phaeocomis(100); | Inocybe | 13 | Symbiotroph | \|Ectomycorrhizal\| | Agaricoid | NULL | Highly Probable | [illegible] | [illegible] |
| Otu04374 | 7 | k__Fungi(100);p__Ascomycota(100);c__Saccharomycetes(100);o__Saccharomycetales(100);f__Metschnikowiaceae(100);g__Metschnikowia(100);s__Metschnikowia_pulcherrima(86); | Metschnikowia | 13 | Saprotroph | \|Undefined Saprotroph\| | Yeast | NULL | Probable | [illegible] | [illegible] |
| Otu04375 | 7 | k__Fungi(100);p__Ascomycota(100);c__Eurotiomycetes(100);o__Chaetothyriales(100);f__Herpotrichiellaceae(100);g__Rhinocladiella(100);s__Rhinocladiella_sp_VL271(100); | Rhinocladiella | 13 | Pathotroph-Saprotroph-Symbiotroph | Animal Parasite-Endophyte-\|Undefined Saprotroph\|-Wood Saprotroph | Dimorphic-Facultative Yeast | Soft Rot | Probable | [illegible] | [illegible] |
| Otu04377 | 7 | k__Fungi(100);p__Basidiomycota(100);c__Agaricomycetes(100);o__Agaricales(100);f__Entolomataceae(100);g__unclassified_Entolomataceae(100);s__Entolomataceae_sp(100); | Entolomataceae | 9 | Pathotroph-Saprotroph-Symbiotroph | Ectomycorrhizal-Fungal Parasite-Soil Saprotroph-Undefined Saprotroph | Agaricoid | NULL | Possible | [illegible] | [illegible] |
| Otu04379 | 7 | k__Fungi(100);p__Ascomycota(100);c__Lecanoromycetes(100);o__Lecanorales(100);o__Lecanorales_unclassified(100);o__Lecanorales_unclassified(100);o__Lecanorales_unclassified(100); | Lecanorales | 7 | Pathotroph-Symbiotroph | Lichen Parasite-Lichenized | Microfungus-Thallus | NULL | Probable | [illegible] | [illegible] |
| Otu04380 | 7 | k__Fungi(100);p__Ascomycota(100);c__Saccharomycetes(100);o__Saccharomycetales(100);f__Metschnikowiaceae(100);g__Metschnikowia(100);s__Metschnikowia_pulcherrima(100); | Metschnikowia | 13 | Saprotroph | \|Undefined Saprotroph\| | Yeast | NULL | Probable | [illegible] | [illegible] |
| Otu04382 | 7 | k__Fungi(100);p__Ascomycota(100);c__Dothideomycetes(100);o__Pleosporales(100);f__Phaeosphaeriaceae(100);g__Neostagonospora(100);s__Neostagonospora_caricis(100); | Neostagonospora | 13 | Saprotroph | \|Plant Saprotroph\|-Wood Saprotroph | Microfungus | NULL | Probable | [illegible] | [illegible] |
| Otu04387 | 7 | k__Fungi(100);p__Basidiomycota(100);c__Agaricomycetes(100);o__Atheliales(100);f__Atheliaceae(100);g__Athelopsis(100);s__Athelopsis_lembospora(100); | Athelopsis | 13 | Saprotroph | Undefined Saprotroph-\|Wood Saprotroph\| | Corticioid | White Rot | Probable | [illegible] | [illegible] |
| Otu04389 | 7 | k__Fungi(100);p__Basidiomycota(100);c__Tremellomycetes(100);o__Tremellales(100);f__Tremellales_family_Incertae_sedis(100);g__Cryptococcus(86);s__Cryptococcus_sp_S5_2(86); | Cryptococcus | 13 | Pathotroph-Saprotroph-Symbiotroph | Animal Pathogen-Endophyte-Epiphyte-\|Undefined Saprotroph\| | Dimorphic Yeast-Yeast | NULL | Possible | [illegible] | [illegible] |
| Otu04390 | 7 | k__Fungi(100);p__Ascomycota(100);c__Saccharomycetes(100);o__Saccharomycetales(100);f__Metschnikowiaceae(100);g__Metschnikowia(100);s__Metschnikowia_pulcherrima(86); | Metschnikowia | 13 | Saprotroph | \|Undefined Saprotroph\| | Yeast | NULL | Probable | [illegible] | [illegible] |
| Otu04391 | 7 | k__Fungi(100);p__Basidiomycota(100);c__Agaricomycetes(100);o__Polyporales(100);f__Polyporaceae(100);g__Polyporus(100);s__Polyporus_umbellatus(100); | Polyporus | 13 | Pathotroph-Saprotroph | Plant Pathogen-\|Wood Saprotroph\| | Polyporoid | White Rot | Highly Probable | [illegible] | [illegible] |
| Otu04392 | 7 | k__Fungi(100);p__Basidiomycota(100);c__Agaricomycetes(100);o__Polyporales(100);f__Polyporaceae(100);g__Datronia(100);s__Datronia_mollis(100); | Datronia | 13 | Saprotroph | \|Wood Saprotroph\| | Corticioid | White Rot | Highly Probable | [illegible] | [illegible] |
| Otu04393 | 7 | k__Fungi(100);p__Ascomycota(100);c__Saccharomycetes(100);o__Saccharomycetales(100);f__Metschnikowiaceae(100);g__Metschnikowia(100);s__Metschnikowia_pulcherrima(72); | Metschnikowia | 13 | Saprotroph | \|Undefined Saprotroph\| | Yeast | NULL | Probable | [illegible] | [illegible] |
| Otu04394 | 7 | k__Fungi(100);p__Basidiomycota(100);c__Agaricomycetes(100);o__Boletales(100);f__Boletaceae(100);g__Boletus(100);g__Boletus_unclassified(100); | Boletus | 13 | Pathotroph-Symbiotroph | \|Ectomycorrhizal\|-Fungal Parasite | Agaricoid-Boletoid | NULL | Highly Probable | NULL | [illegible] |
| Otu04400 | 7 | k__Fungi(100);p__Ascomycota(100);c__Saccharomycetes(100);o__Saccharomycetales(100);f__Metschnikowiaceae(100);g__Metschnikowia(100);s__Metschnikowia_pulcherrima(100); | Metschnikowia | 13 | Saprotroph | \|Undefined Saprotroph\| | Yeast | NULL | Probable | [illegible] | [illegible] |
| Otu04404 | 7 | k__Fungi(100);p__Ascomycota(100);c__Saccharomycetes(100);o__Saccharomycetales(100);f__Metschnikowiaceae(100);g__Metschnikowia(100);s__Metschnikowia_pulcherrima(58); | Metschnikowia | 13 | Saprotroph | \|Undefined Saprotroph\| | Yeast | NULL | Probable | [illegible] | [illegible] |
| Otu04406 | 7 | k__Fungi(100);p__Ascomycota(100);c__Saccharomycetes(100);o__Saccharomycetales(100);f__Saccharomycetales_family_Incertae_sedis(100);g__Candida(100);g__Candida_unclassified(72); | Candida | 13 | Pathotroph-Saprotroph-Symbiotroph | Animal Pathogen-Endophyte-Endosymbiont-Epiphyte-\|Undefined Saprotroph\| | Dimorphic Yeast-Yeast | NULL | Probable | [illegible] | [illegible] |
| Otu04407 | 7 | k__Fungi(100);p__Ascomycota(100);c__Saccharomycetes(100);o__Saccharomycetales(100);f__Metschnikowiaceae(100);g__Metschnikowia(100);s__Metschnikowia_pulcherrima(86); | Metschnikowia | 13 | Saprotroph | \|Undefined Saprotroph\| | Yeast | NULL | Probable | [illegible] | [illegible] |
| Otu04408 | 7 | k__Fungi(100);p__Basidiomycota(100);c__Microbotryomycetes(100);o__Sporidiobolales(100);f__Sporidiobolales_family_Incertae_sedis(100);g__Rhodotorula(100);s__Rhodotorula_oryzicola(100); | Rhodotorula | 13 | Pathotroph-Saprotroph-Symbiotroph | Animal Pathogen-Animal Symbiotroph-Endophyte-Plant Pathogen-\|Undefined Saprotroph\| | Yeast | NULL | Probable | [illegible] | [illegible] |
| Otu04410 | 7 | k__Fungi(100);p__Ascomycota(100);c__Eurotiomycetes(100);o__Chaetothyriales(100);f__Herpotrichiellaceae(100);g__Cladophialophora(100);g__Cladophialophora_unclassified(100); | Cladophialophora | 13 | Pathotroph-Saprotroph-Symbiotroph | Animal Pathogen-Endophyte-Fungal Parasite-\|Undefined Saprotroph\| | Dimorphic-Microfungus | NULL | Probable | [illegible] | [illegible] |
| Otu04413 | 7 | k__Fungi(100);p__Chytridiomycota(100);c__Chytridiomycetes(100);o__Spizellomycetales(100);f__Spizellomycetaceae(100);g__Spizellomyces(100);g__Spizellomyces_unclassified(100); | Spizellomyces | 13 | Pathotroph-Saprotroph | Plant Pathogen-\|Pollen Saprotroph\|-Undefined Parasite-Undefined Saprotroph | Chytrid-Microfungus | NULL | Probable | [illegible] | [illegible] |
| Otu04414 | 7 | k__Fungi(100);p__Basidiomycota(100);c__Agaricomycetes(100);o__Agaricales(100);f__Lyophyllaceae(100);g__Lyophyllum(100);s__Lyophyllum_decastes(100); | Lyophyllum | 13 | Saprotroph-Symbiotroph | \|Ectomycorrhizal\|-Undefined Saprotroph | Agaricoid | NULL | Possible | [illegible] | [illegible] |
| Otu04416 | 7 | k__Fungi(100);p__Basidiomycota(100);c__Tremellomycetes(100);o__Tremellales(100);f__Sirobasidiaceae(100);g__Sirobasidium(100);s__Sirobasidium_intermedium(100); | Sirobasidium | 13 | Pathotroph | \|Fungal Parasite\| | Dimorphic | NULL | Probable | [illegible] | [illegible] |
| Otu04421 | 7 | k__Fungi(100);p__Basidiomycota(100);c__Agaricomycetes(100);o__Russulales(100);f__Stereaceae(100);g__Gloeocystidiellum(100);s__Gloeocystidiellum_porosum(100); | Gloeocystidiellum | 13 | Saprotroph | Undefined Saprotroph-\|Wood Saprotroph\| | Corticioid | White Rot | Probable | [illegible] | [illegible] |
| Otu04423 | 7 | k__Fungi(100);p__Basidiomycota(100);c__Agaricomycetes(100);o__Thelephorales(100);f__Thelephoraceae(100);g__Tomentella(100);g__Tomentella_unclassified(58); | Tomentella | 13 | Symbiotroph | \|Ectomycorrhizal\| | Corticioid | NULL | Highly Probable | NULL | [illegible] |
| Otu04426 | 7 | k__Fungi(100);p__Basidiomycota(100);c__Pucciniomycetes(100);o__Pucciniales(100);f__Pucciniaceae(100);g__Puccinia(100);s__Puccinia_coronata_var._avenae(58); | Puccinia | 13 | Pathotroph | \|Plant Pathogen\| | Rust | NULL | Highly Probable | NULL | [illegible] |
| Otu04429 | 7 | k__Fungi(100);p__Basidiomycota(100);c__Agaricomycetes(100);o__Agaricales(100);o__Agaricales_unclassified(100);o__Agaricales_unclassified(100);o__Agaricales_unclassified(100); | Agaricales | 7 | Pathotroph-Saprotroph-Symbiotroph | Bryophyte Parasite-Dung Saprotroph-Ectomycorrhizal-Fungal Parasite-Leaf Saprotroph-Plant Parasite-Undefined Saprotroph-Wood Saprotroph | Agaricoid-Gasteroid-Microfungus-Secotioid-Yeast | NULL | Possible | [illegible] | [illegible] |
| Otu04430 | 7 | k__Fungi(100);p__Basidiomycota(100);c__Agaricomycetes(100);o__Agaricales(100);f__Psathyrellaceae(100);g__Coprinopsis(100);s__Coprinopsis_sp_CCG_2012(100); | Coprinopsis | 13 | Saprotroph | \|Undefined Saprotroph\| | Agaricoid | NULL | Probable | [illegible] | [illegible] |
| Otu04431 | 7 | k__Fungi(100);p__Ascomycota(100);c__Saccharomycetes(100);o__Saccharomycetales(100);f__Metschnikowiaceae(100);g__Metschnikowia(100);s__Metschnikowia_pulcherrima(86); | Metschnikowia | 13 | Saprotroph | \|Undefined Saprotroph\| | Yeast | NULL | Probable | [illegible] | [illegible] |
| Otu04433 | 7 | k__Fungi(100);p__Ascomycota(100);c__Saccharomycetes(100);o__Saccharomycetales(100);f__Metschnikowiaceae(100);g__Metschnikowia(100);s__Metschnikowia_pulcherrima(100); | Metschnikowia | 13 | Saprotroph | \|Undefined Saprotroph\| | Yeast | NULL | Probable | [illegible] | [illegible] |
| Otu04439 | 7 | k__Fungi(100);p__Basidiomycota(100);c__Agaricomycetes(100);o__Trechisporales(100);f__Hydnodontaceae(100);g__Trechispora(100);g__Trechispora_unclassified(100); | Trechispora | 13 | Saprotroph | \|Wood Saprotroph\| | Corticioid | White Rot | Highly Probable | [illegible] | [illegible] |
| Otu04443 | 7 | k__Fungi(100);p__Ascomycota(100);c__Saccharomycetes(100);o__Saccharomycetales(100);f__Metschnikowiaceae(100);g__Metschnikowia(100);s__Metschnikowia_pulcherrima(100); | Metschnikowia | 13 | Saprotroph | \|Undefined Saprotroph\| | Yeast | NULL | Probable | [illegible] | [illegible] |
| Otu04446 | 7 | k__Fungi(100);p__Ascomycota(100);c__Saccharomycetes(100);o__Saccharomycetales(100);f__Metschnikowiaceae(100);g__Metschnikowia(100);s__Metschnikowia_pulcherrima(86); | Metschnikowia | 13 | Saprotroph | \|Undefined Saprotroph\| | Yeast | NULL | Probable | [illegible] | [illegible] |
| Otu04448 | 7 | k__Fungi(100);p__Ascomycota(100);c__Sordariomycetes(100);o__Lulworthiales(100);f__Lulworthiaceae(100);g__Zalerion(100);s__Zalerion_arboricola(100); | Zalerion | 13 | Pathotroph-Saprotroph | Plant Pathogen-\|Plant Saprotroph\|-Undefined Saprotroph | Microfungus | NULL | Probable | [illegible] | [illegible] |
| Otu04449 | 7 | k__Fungi(100);p__Ascomycota(100);c__Saccharomycetes(100);o__Saccharomycetales(100);f__Metschnikowiaceae(100);g__Metschnikowia(100);s__Metschnikowia_pulcherrima(86); | Metschnikowia | 13 | Saprotroph | \|Undefined Saprotroph\| | Yeast | NULL | Probable | [illegible] | [illegible] |
| Otu04451 | 7 | k__Fungi(100);p__Ascomycota(100);c__Saccharomycetes(100);o__Saccharomycetales(100);f__Metschnikowiaceae(100);g__Metschnikowia(100);s__Metschnikowia_pulcherrima(100); | Metschnikowia | 13 | Saprotroph | \|Undefined Saprotroph\| | Yeast | NULL | Probable | [illegible] | [illegible] |
| Otu04452 | 7 | k__Fungi(100);p__Basidiomycota(100);c__Tremellomycetes(100);o__Tremellales(100);f__Tremellales_family_Incertae_sedis(100);g__Tremella(100);s__Tremella_nivalis(72); | Tremella | 13 | Pathotroph | \|Fungal Parasite\|-Lichen Parasite | Dimorphic-Facultative Yeast-Microfungus-Tremelloid | NULL | Probable | [illegible] | [illegible] |
| Otu04453 | 7 | k__Fungi(100);p__Basidiomycota(100);c__Agaricostilbomycetes(100);o__Agaricostilbales(100);f__Agaricostilbaceae(100);g__Sterigmatomyces(100);s__Sterigmatomyces_halophilus(100); | Sterigmatomyces | 13 | Saprotroph | \|Undefined Saprotroph\| | Dimorphic | NULL | Probable | NULL | [illegible] |
| Otu04457 | 7 | k__Fungi(100);p__Ascomycota(100);c__Saccharomycetes(100);o__Saccharomycetales(100);f__Metschnikowiaceae(100);g__Metschnikowia(100);g__Metschnikowia_unclassified(86); | Metschnikowia | 13 | Saprotroph | \|Undefined Saprotroph\| | Yeast | NULL | Probable | [illegible] | [illegible] |
| Otu04459 | 7 | k__Fungi(100);p__Ascomycota(100);c__Sordariomycetes(100);o__Coniochaetales(100);f__Coniochaetales_family_Incertae_sedis(100);g__Pseudobotrytis(100);s__Pseudobotrytis_terrestris(100); | Pseudobotrytis | 13 | Pathotroph-Saprotroph-Symbiotroph | Endophyte-Plant Pathogen-Plant Saprotroph-\|Undefined Saprotroph\| | Microfungus | NULL | Probable | [illegible] | [illegible] |
| Otu04460 | 7 | k__Fungi(100);p__Zygomycota(100);c__Zygomycota_class_Incertae_sedis(100);o__Mortierellales(100);f__Mortierellaceae(100);g__Mortierella(100);g__Mortierella_unclassified(100); | Mortierella | 13 | Saprotroph-Symbiotroph | Endophyte-Plant Saprotroph-\|Undefined Saprotroph\| | Microfungus | NULL | Probable | [illegible] | [illegible] |
| Otu04461 | 7 | k__Fungi(100);p__Ascomycota(100);c__Saccharomycetes(100);o__Saccharomycetales(100);f__Metschnikowiaceae(100);g__Metschnikowia(100);s__Metschnikowia_pulcherrima(100); | Metschnikowia | 13 | Saprotroph | \|Undefined Saprotroph\| | Yeast | NULL | Probable | [illegible] | [illegible] |
| Otu04463 | 7 | k__Fungi(100);p__Ascomycota(100);c__Dothideomycetes(100);o__Pleosporales(86);f__Montagnulaceae(86);g__Paraphaeosphaeria(86);g__Paraphaeosphaeria_unclassified(86); | Paraphaeosphaeria | 13 | Saprotroph | Undefined Saprotroph-\|Wood Saprotroph\| | Microfungus | NULL | Probable | [illegible] | [illegible] |
| Otu04464 | 7 | k__Fungi(100);p__Basidiomycota(100);c__Basidiomycota_class_Incertae_sedis(100);o__Malasseziales(100);f__Malasseziales_family_Incertae_sedis(100);g__Malassezia(100);g__Malassezia_unclassified(100); | Malassezia | 13 | Pathotroph-Saprotroph | Animal Pathogen-\|Undefined Saprotroph\| | Facultative Yeast-Yeast | NULL | Probable | [illegible] | [illegible] |
| Otu04465 | 7 | k__Fungi(100);p__Ascomycota(100);c__Saccharomycetes(100);o__Saccharomycetales(100);f__Metschnikowiaceae(100);g__Metschnikowia(100);g__Metschnikowia_unclassified(58); | Metschnikowia | 13 | Saprotroph | \|Undefined Saprotroph\| | Yeast | NULL | Probable | [illegible] | [illegible] |
| Otu04466 | 7 | k__Fungi(100);p__Ascomycota(100);c__Saccharomycetes(100);o__Saccharomycetales(100);f__Metschnikowiaceae(100);g__Metschnikowia(100);s__Metschnikowia_pulcherrima(100); | Metschnikowia | 13 | Saprotroph | \|Undefined Saprotroph\| | Yeast | NULL | Probable | [illegible] | [illegible] |
| Otu04467 | 7 | k__Fungi(100);p__Basidiomycota(100);c__Microbotryomycetes(100);o__Sporidiobolales(100);f__Sporidiobolales_family_Incertae_sedis(100);g__Rhodotorula(100);s__Rhodotorula_hordea(100); | Rhodotorula | 13 | Pathotroph-Saprotroph-Symbiotroph | Animal Pathogen-Animal Symbiotroph-Endophyte-Plant Pathogen-\|Undefined Saprotroph\| | Yeast | NULL | Probable | [illegible] | [illegible] |
| Otu04469 | 7 | k__Fungi(100);p__Ascomycota(100);c__Leotiomycetes(100);o__Helotiales(100);f__Hyaloscyphaceae(100);g__Lachnum(100);g__Lachnum_unclassified(100); | Lachnum | 13 | Saprotroph-Symbiotroph | Endophyte-\|Plant Saprotroph\|-Undefined Saprotroph-Wood Saprotroph | Microfungus | NULL | Probable | [illegible] | [illegible] |
| Otu04471 | 7 | k__Fungi(100);p__Ascomycota(100);c__Dothideomycetes(100);o__Capnodiales(100);f__Mycosphaerellaceae(86);f__Mycosphaerellaceae_unclassified(86);f__Mycosphaerellaceae_unclassified(86); | Mycosphaerellaceae | 9 | Pathotroph-Saprotroph | Plant Pathogen-Undefined Saprotroph | Microfungus | NULL | Probable | [illegible] | [illegible] |
| Otu04474 | 7 | k__Fungi(100);p__Basidiomycota(100);c__Wallemiomycetes(100);o__Wallemiales(100);f__Wallemiaceae(100);g__Wallemia(100);s__Wallemia_muriae(100); | Wallemia | 13 | Saprotroph | \|Undefined Saprotroph\| | Microfungus | NULL | Probable | [illegible] | [illegible] |
| Otu04475 | 7 | k__Fungi(100);p__Basidiomycota(100);c__Agaricomycetes(100);o__Agaricales(100);f__Psathyrellaceae(100);g__Coprinopsis(100);g__Coprinopsis_unclassified(100); | Coprinopsis | 13 | Saprotroph | \|Undefined Saprotroph\| | Agaricoid | NULL | Probable | [illegible] | [illegible] |
| Otu04477 | 7 | k__Fungi(100);p__Basidiomycota(100);c__Agaricomycetes(100);o__Polyporales(100);f__Polyporaceae(100);g__Trametes(58);g__Trametes_unclassified(58); | Trametes | 13 | Pathotroph-Saprotroph | Plant Pathogen-\|Wood Saprotroph\| | Polyporoid | White Rot | Highly Probable | [illegible] | [illegible] |
| Otu04479 | 7 | k__Fungi(100);p__Ascomycota(100);c__Saccharomycetes(100);o__Saccharomycetales(100);f__Metschnikowiaceae(100);g__Metschnikowia(100);s__Metschnikowia_pulcherrima(100); | Metschnikowia | 13 | Saprotroph | \|Undefined Saprotroph\| | Yeast | NULL | Probable | [illegible] | [illegible] |
| Otu04480 | 7 | k__Fungi(100);p__Ascomycota(100);c__Saccharomycetes(100);o__Saccharomycetales(100);f__Saccharomycetales_family_Incertae_sedis(100);g__Candida(86);g__Candida_unclassified(58); | Candida | 13 | Pathotroph-Saprotroph-Symbiotroph | Animal Pathogen-Endophyte-Endosymbiont-Epiphyte-\|Undefined Saprotroph\| | Dimorphic Yeast-Yeast | NULL | Probable | [illegible] | [illegible] |
| Otu04482 | 7 | k__Fungi(100);p__Basidiomycota(100);c__Agaricomycetes(100);o__Hymenochaetales(100);f__Schizoporaceae(100);g__Hyphodontia(100);s__Hyphodontia_abieticola(100); | Hyphodontia | 13 | Saprotroph | Undefined Saprotroph-\|Wood Saprotroph\| | Corticioid | White Rot | Probable | [illegible] | [illegible] |
| Otu04483 | 7 | k__Fungi(100);p__Ascomycota(100);c__Saccharomycetes(100);o__Saccharomycetales(100);f__Metschnikowiaceae(100);g__Metschnikowia(100);s__Metschnikowia_pulcherrima(58); | Metschnikowia | 13 | Saprotroph | \|Undefined Saprotroph\| | Yeast | NULL | Probable | [illegible] | [illegible] |
| Otu04485 | 7 | k__Fungi(100);p__Basidiomycota(100);c__Agaricomycetes(100);o__Agaricales(100);f__Agaricaceae(86);f__Agaricaceae_unclassified(86);f__Agaricaceae_unclassified(86); | Agaricaceae | 9 | Saprotroph | Undefined Saprotroph | Agaricoid-Gasteroid-Secotioid | NULL | Probable | [illegible] | [illegible] |
| Otu04487 | 7 | k__Fungi(100);p__Basidiomycota(100);c__Agaricomycetes(100);o__Agaricales(100);f__Psathyrellaceae(100);g__Coprinellus(72);g__Coprinellus_unclassified(72); | Coprinellus | 13 | Saprotroph | \|Undefined Saprotroph\| | Agaricoid | NULL | Probable | [illegible] | [illegible] |
| Otu04488 | 7 | k__Fungi(100);p__Basidiomycota(100);c__Agaricomycetes(100);o__Agaricales(100);f__Psathyrellaceae(100);g__Psathyrella(100);s__Psathyrella_maculata(100); | Psathyrella | 13 | Saprotroph | Plant Saprotroph-\|Wood Saprotroph\| | Agaricoid | NULL | Probable | [illegible] | [illegible] |
| Otu04489 | 7 | k__Fungi(100);p__Ascomycota(100);c__Saccharomycetes(100);o__Saccharomycetales(100);f__Metschnikowiaceae(100);g__Metschnikowia(100);s__Metschnikowia_pulcherrima(86); | Metschnikowia | 13 | Saprotroph | \|Undefined Saprotroph\| | Yeast | NULL | Probable | [illegible] | [illegible] |
| Otu04492 | 7 | k__Fungi(100);p__Basidiomycota(100);c__Agaricomycetes(100);o__Polyporales(100);f__Phanerochaetaceae(100);g__Ceriporia(72);g__Ceriporia_unclassified(72); | Ceriporia | 13 | Saprotroph | \|Wood Saprotroph\| | Corticioid | White Rot | Highly Probable | [illegible] | [illegible] |
| Otu04493 | 7 | k__Fungi(100);p__Basidiomycota(100);c__Exobasidiomycetes(100);o__Exobasidiales(100);f__Exobasidiaceae(100);g__Exobasidium(100);s__Exobasidium_arescens(58); | Exobasidium | 13 | Pathotroph | \|Plant Pathogen\| | Leaf Blister-Leaf Gall-Leaf Spot | NULL | Probable | NULL | [illegible] |
| Otu04501 | 7 | k__Fungi(100);p__Ascomycota(100);c__Saccharomycetes(100);o__Saccharomycetales(100);f__Saccharomycetales_family_Incertae_sedis(100);g__Candida(100);s__Candida_cretensis(58); | Candida | 13 | Pathotroph-Saprotroph-Symbiotroph | Animal Pathogen-Endophyte-Endosymbiont-Epiphyte-\|Undefined Saprotroph\| | Dimorphic Yeast-Yeast | NULL | Probable | [illegible] | [illegible] |
| Otu04503 | 7 | k__Fungi(100);p__Ascomycota(100);c__Saccharomycetes(100);o__Saccharomycetales(100);f__Saccharomycetaceae(100);g__Kazachstania(100);s__Kazachstania_intestinalis(100); | Kazachstania | 13 | Saprotroph | \|Undefined Saprotroph\| | Yeast | NULL | Probable | [illegible] | [illegible] |
| Otu04504 | 7 | k__Fungi(100);p__Ascomycota(100);c__Saccharomycetes(100);o__Saccharomycetales(100);f__Saccharomycetales_family_Incertae_sedis(100);g__Candida(100);g__Candida_unclassified(86); | Candida | 13 | Pathotroph-Saprotroph-Symbiotroph | Animal Pathogen-Endophyte-Endosymbiont-Epiphyte-\|Undefined Saprotroph\| | Dimorphic Yeast-Yeast | NULL | Probable | [illegible] | [illegible] |
| Otu04505 | 7 | k__Fungi(100);p__Ascomycota(100);c__Saccharomycetes(100);o__Saccharomycetales(100);f__Metschnikowiaceae(100);g__Metschnikowia(100);s__Metschnikowia_pulcherrima(100); | Metschnikowia | 13 | Saprotroph | \|Undefined Saprotroph\| | Yeast | NULL | Probable | [illegible] | [illegible] |
| Otu04509 | 7 | k__Fungi(100);p__Ascomycota(100);c__Saccharomycetes(100);o__Saccharomycetales(100);f__Metschnikowiaceae(100);g__Metschnikowia(100);g__Metschnikowia_unclassified(100); | Metschnikowia | 13 | Saprotroph | \|Undefined Saprotroph\| | Yeast | NULL | Probable | [illegible] | [illegible] |
| Otu04510 | 7 | k__Fungi(100);p__Ascomycota(100);c__Saccharomycetes(100);o__Saccharomycetales(100);f__Metschnikowiaceae(100);g__Metschnikowia(100);s__Metschnikowia_pulcherrima(100); | Metschnikowia | 13 | Saprotroph | \|Undefined Saprotroph\| | Yeast | NULL | Probable | [illegible] | [illegible] |
| Otu04512 | 7 | k__Fungi(100);p__Ascomycota(100);c__Dothideomycetes(100);o__Pleosporales(100);f__Montagnulaceae(100);f__Montagnulaceae_unclassified(100);f__Montagnulaceae_unclassified(100); | Montagnulaceae | 9 | Pathotroph-Saprotroph | Fungal Parasite-Plant Pathogen-Undefined Saprotroph | Microfungus | NULL | Probable | [illegible] | [illegible] |
| Otu04513 | 7 | k__Fungi(100);p__Ascomycota(100);c__Saccharomycetes(100);o__Saccharomycetales(100);f__Metschnikowiaceae(100);g__Metschnikowia(100);s__Metschnikowia_pulcherrima(72); | Metschnikowia | 13 | Saprotroph | \|Undefined Saprotroph\| | Yeast | NULL | Probable | [illegible] | [illegible] |
| Otu04517 | 7 | k__Fungi(100);p__Basidiomycota(100);c__Microbotryomycetes(100);o__Sporidiobolales(100);f__Sporidiobolales_family_Incertae_sedis(100);g__Sporobolomyces(100);s__Sporobolomyces_lactosus(100); | Sporobolomyces | 13 | Pathotroph-Saprotroph | \|Fungal Parasite\|-Plant Saprotroph | Yeast | NULL | Probable | [illegible] | [illegible] |
| Otu04518 | 7 | k__Fungi(100);p__Ascomycota(100);c__Saccharomycetes(100);o__Saccharomycetales(100);f__Metschnikowiaceae(100);g__Metschnikowia(100);s__Metschnikowia_pulcherrima(86); | Metschnikowia | 13 | Saprotroph | \|Undefined Saprotroph\| | Yeast | NULL | Probable | [illegible] | [illegible] |
| Otu04523 | 7 | k__Fungi(100);p__Ascomycota(100);c__Saccharomycetes(100);o__Saccharomycetales(100);f__Metschnikowiaceae(100);g__Metschnikowia(100);s__Metschnikowia_pulcherrima(72); | Metschnikowia | 13 | Saprotroph | \|Undefined Saprotroph\| | Yeast | NULL | Probable | [illegible] | [illegible] |
| Otu04524 | 7 | k__Fungi(100);p__Ascomycota(100);c__Leotiomycetes(100);o__Helotiales(100);f__Hyaloscyphaceae(100);f__Hyaloscyphaceae_unclassified(100);f__Hyaloscyphaceae_unclassified(100); | Hyaloscyphaceae | 9 | Saprotroph | Plant Saprotroph-Wood Saprotroph | Microfungus | NULL | Probable | [illegible] | [illegible] |
| Otu04527 | 7 | k__Fungi(100);p__Ascomycota(100);c__Saccharomycetes(100);o__Saccharomycetales(100);f__Metschnikowiaceae(100);g__Metschnikowia(100);g__Metschnikowia_unclassified(72); | Metschnikowia | 13 | Saprotroph | \|Undefined Saprotroph\| | Yeast | NULL | Probable | [illegible] | [illegible] |
| Otu04528 | 7 | k__Fungi(100);p__Ascomycota(100);c__Sordariomycetes(86);o__Hypocreales(86);o__Hypocreales_unclassified(72);o__Hypocreales_unclassified(72);o__Hypocreales_unclassified(72); | Hypocreales | 7 | Saprotroph | Undefined Saprotroph | Microfungus | NULL | Possible | NULL | [illegible] |
| Otu04529 | 7 | k__Fungi(100);p__Ascomycota(100);c__Saccharomycetes(100);o__Saccharomycetales(100);f__Metschnikowiaceae(100);g__Metschnikowia(100);s__Metschnikowia_pulcherrima(100); | Metschnikowia | 13 | Saprotroph | \|Undefined Saprotroph\| | Yeast | NULL | Probable | [illegible] | [illegible] |
| Otu04531 | 7 | k__Fungi(100);p__Ascomycota(100);c__Saccharomycetes(100);o__Saccharomycetales(100);f__Metschnikowiaceae(100);g__Metschnikowia(100);g__Metschnikowia_unclassified(58); | Metschnikowia | 13 | Saprotroph | \|Undefined Saprotroph\| | Yeast | NULL | Probable | [illegible] | [illegible] |
| Otu04532 | 7 | k__Fungi(100);p__Zygomycota(100);c__Zygomycota_class_Incertae_sedis(100);o__Mucorales(100);f__Mucoraceae(100);g__Mucor(100);g__Mucor_unclassified(100); | Mucor | 13 | Saprotroph | \|Undefined Saprotroph\| | Microfungus | NULL | Probable | [illegible] | [illegible] |
| Otu04534 | 7 | k__Fungi(100);p__Ascomycota(100);c__Saccharomycetes(100);o__Saccharomycetales(100);f__Metschnikowiaceae(100);g__Metschnikowia(100);s__Metschnikowia_pulcherrima(72); | Metschnikowia | 13 | Saprotroph | \|Undefined Saprotroph\| | Yeast | NULL | Probable | [illegible] | [illegible] |
| Otu04536 | 7 | k__Fungi(100);p__Basidiomycota(100);c__Agaricomycetes(100);o__Agaricales(100);f__Cortinariaceae(100);g__Cortinarius(100);s__Cortinarius_scaurus_var._herpeticus(100); | Cortinarius | 13 | Symbiotroph | \|Ectomycorrhizal\| | Agaricoid | NULL | Highly Probable | NULL | [illegible] |
| Otu04537 | 7 | k__Fungi(100);p__Ascomycota(100);c__Saccharomycetes(100);o__Saccharomycetales(100);f__Metschnikowiaceae(100);g__Metschnikowia(100);s__Metschnikowia_pulcherrima(86); | Metschnikowia | 13 | Saprotroph | \|Undefined Saprotroph\| | Yeast | NULL | Probable | [illegible] | [illegible] |
| Otu04539 | 7 | k__Fungi(100);p__Ascomycota(100);c__Sordariomycetes(100);o__Coniochaetales(100);f__Coniochaetaceae(100);g__Coniochaeta(72);s__Coniochaeta_sp(72); | Coniochaeta | 13 | Pathotroph-Saprotroph-Symbiotroph | Animal Pathogen-Dung Saprotroph-Endophyte-Lichen Parasite-Plant Pathogen-\|Undefined Saprotroph\| | Microfungus | NULL | Possible | [illegible] | [illegible] |
| Otu04540 | 7 | k__Fungi(100);p__Basidiomycota(100);c__Agaricomycetes(100);o__Agaricales(100);f__Inocybaceae(100);g__Tubaria(100);s__Tubaria_conspersa(100); | Tubaria | 13 | Saprotroph | \|Plant Saprotroph\|-Wood Saprotroph | Agaricoid | NULL | Probable | [illegible] | [illegible] |
| Otu04541 | 7 | k__Fungi(100);p__Ascomycota(100);c__Sordariomycetes(100);o__Xylariales(100);f__Xylariaceae(100);g__Hypoxylon(100);g__Hypoxylon_unclassified(100); | Hypoxylon | 13 | Saprotroph-Saprotroph-Symbiotroph | Endophyte-Undefined Saprotroph-\|Wood Saprotroph\| | Corticioid-Microfungus-Xylarioid | NULL | Probable | [illegible] | [illegible] |
| Otu04545 | 7 | k__Fungi(100);p__Ascomycota(100);c__Leotiomycetes(100);o__Helotiales(100);f__Hyaloscyphaceae(100);g__Lachnum(100);s__Lachnum_virgineum(100); | Lachnum | 13 | Saprotroph-Symbiotroph | Endophyte-\|Plant Saprotroph\|-Undefined Saprotroph-Wood Saprotroph | Microfungus | NULL | Probable | [illegible] | [illegible] |
| Otu04550 | 7 | k__Fungi(100);p__Ascomycota(100);c__Saccharomycetes(100);o__Saccharomycetales(100);f__Metschnikowiaceae(100);g__Metschnikowia(100);g__Metschnikowia_unclassified(86); | Metschnikowia | 13 | Saprotroph | \|Undefined Saprotroph\| | Yeast | NULL | Probable | [illegible] | [illegible] |
| Otu04551 | 7 | k__Fungi(100);p__Ascomycota(100);c__Saccharomycetes(100);o__Saccharomycetales(100);f__Metschnikowiaceae(100);g__Metschnikowia(100);g__Metschnikowia_unclassified(58); | Metschnikowia | 13 | Saprotroph | \|Undefined Saprotroph\| | Yeast | NULL | Probable | [illegible] | [illegible] |
| Otu04552 | 7 | k__Fungi(100);p__Ascomycota(100);c__Saccharomycetes(100);o__Saccharomycetales(100);f__Saccharomycetales_family_Incertae_sedis(58);g__Candida(58);g__Candida_unclassified(58); | Candida | 13 | Pathotroph-Saprotroph-Symbiotroph | Animal Pathogen-Endophyte-Endosymbiont-Epiphyte-\|Undefined Saprotroph\| | Dimorphic Yeast-Yeast | NULL | Probable | [illegible] | [illegible] |
| Otu04555 | 7 | k__Fungi(100);p__Ascomycota(100);c__Saccharomycetes(100);o__Saccharomycetales(100);f__Metschnikowiaceae(100);g__Metschnikowia(100);s__Metschnikowia_pulcherrima(58); | Metschnikowia | 13 | Saprotroph | \|Undefined Saprotroph\| | Yeast | NULL | Probable | [illegible] | [illegible] |
| Otu04556 | 7 | k__Fungi(100);p__Ascomycota(100);c__Saccharomycetes(100);o__Saccharomycetales(100);f__Metschnikowiaceae(100);g__Metschnikowia(100);s__Metschnikowia_pulcherrima(100); | Metschnikowia | 13 | Saprotroph | \|Undefined Saprotroph\| | Yeast | NULL | Probable | [illegible] | [illegible] |
| Otu04560 | 7 | k__Fungi(100);p__Ascomycota(100);c__Saccharomycetes(100);o__Saccharomycetales(100);f__Metschnikowiaceae(100);g__Metschnikowia(100);s__Metschnikowia_pulcherrima(100); | Metschnikowia | 13 | Saprotroph | \|Undefined Saprotroph\| | Yeast | NULL | Probable | [illegible] | [illegible] |
| Otu04561 | 7 | k__Fungi(100);p__Ascomycota(100);c__Saccharomycetes(100);o__Saccharomycetales(100);f__Metschnikowiaceae(100);g__Metschnikowia(100);s__Metschnikowia_pulcherrima(72); | Metschnikowia | 13 | Saprotroph | \|Undefined Saprotroph\| | Yeast | NULL | Probable | [illegible] | [illegible] |
| Otu04562 | 7 | k__Fungi(100);p__Ascomycota(100);c__Saccharomycetes(100);o__Saccharomycetales(100);f__Metschnikowiaceae(100);g__Metschnikowia(100);s__Metschnikowia_pulcherrima(72); | Metschnikowia | 13 | Saprotroph | \|Undefined Saprotroph\| | Yeast | NULL | Probable | [illegible] | [illegible] |
| Otu04563 | 7 | k__Fungi(100);p__Ascomycota(100);c__Saccharomycetes(100);o__Saccharomycetales(100);f__Metschnikowiaceae(100);g__Metschnikowia(100);s__Metschnikowia_pulcherrima(86); | Metschnikowia | 13 | Saprotroph | \|Undefined Saprotroph\| | Yeast | NULL | Probable | [illegible] | [illegible] |
| Otu04564 | 7 | k__Fungi(100);p__Ascomycota(100);c__Saccharomycetes(100);o__Saccharomycetales(100);f__Metschnikowiaceae(100);g__Metschnikowia(100);g__Metschnikowia_unclassified(86); | Metschnikowia | 13 | Saprotroph | \|Undefined Saprotroph\| | Yeast | NULL | Probable | [illegible] | [illegible] |
| Otu04566 | 7 | k__Fungi(100);p__Ascomycota(100);c__Saccharomycetes(100);o__Saccharomycetales(100);f__Metschnikowiaceae(100);g__Metschnikowia(100);s__Metschnikowia_viticola(100); | Metschnikowia | 13 | Saprotroph | \|Undefined Saprotroph\| | Yeast | NULL | Probable | [illegible] | [illegible] |
| Otu04568 | 7 | k__Fungi(100);p__Ascomycota(100);c__Saccharomycetes(100);o__Saccharomycetales(100);f__Metschnikowiaceae(100);g__Metschnikowia(100);s__Metschnikowia_pulcherrima(72); | Metschnikowia | 13 | Saprotroph | \|Undefined Saprotroph\| | Yeast | NULL | Probable | [illegible] | [illegible] |
| Otu04569 | 7 | k__Fungi(100);p__Ascomycota(100);c__Saccharomycetes(100);o__Saccharomycetales(100);f__Metschnikowiaceae(100);g__Metschnikowia(100);g__Metschnikowia_unclassified(72); | Metschnikowia | 13 | Saprotroph | \|Undefined Saprotroph\| | Yeast | NULL | Probable | [illegible] | [illegible] |
| Otu04570 | 7 | k__Fungi(100);p__Ascomycota(100);c__Saccharomycetes(100);o__Saccharomycetales(100);f__Metschnikowiaceae(100);g__Metschnikowia(100);s__Metschnikowia_pulcherrima(58); | Metschnikowia | 13 | Saprotroph | \|Undefined Saprotroph\| | Yeast | NULL | Probable | [illegible] | [illegible] |
| Otu04571 | 7 | k__Fungi(100);p__Ascomycota(100);c__Saccharomycetes(100);o__Saccharomycetales(100);f__Metschnikowiaceae(100);g__Metschnikowia(100);g__Metschnikowia_unclassified(72); | Metschnikowia | 13 | Saprotroph | \|Undefined Saprotroph\| | Yeast | NULL | Probable | [illegible] | [illegible] |
| Otu04573 | 7 | k__Fungi(100);p__Ascomycota(100);c__Saccharomycetes(100);o__Saccharomycetales(100);f__Metschnikowiaceae(100);g__Metschnikowia(100);g__Metschnikowia_unclassified(72); | Metschnikowia | 13 | Saprotroph | \|Undefined Saprotroph\| | Yeast | NULL | Probable | [illegible] | [illegible] |
| Otu04574 | 7 | k__Fungi(100);p__Ascomycota(100);c__Dothideomycetes(100);o__Capnodiales(100);f__Mycosphaerellaceae(100);g__Pseudocercosporella(72);s__Pseudocercosporella_chaenomelis(72); | Pseudocercosporella | 13 | Pathotroph | \|Plant Pathogen\| | Microfungus | NULL | Probable | NULL | [illegible] |
| Otu04578 | 7 | k__Fungi(100);p__Ascomycota(100);c__Eurotiomycetes(100);o__Eurotiales(100);f__Trichocomaceae(100);g__Penicillium(100);s__Penicillium_decumbens(58); | Penicillium | 13 | Pathotroph-Saprotroph-Symbiotroph | Animal Parasite-Dung Saprotroph-Endophyte-Fungal Parasite-Plant Pathogen-\|Undefined Saprotroph\|-Wood Saprotroph | Microfungus | Soft Rot | Highly Probable | [illegible] | [illegible] |
| Otu04580 | 7 | k__Fungi(100);p__Basidiomycota(100);c__Microbotryomycetes(100);o__Sporidiobolales(100);f__Sporidiobolales_family_Incertae_sedis(100);g__Rhodotorula(100);s__Rhodotorula_taiwanensis(100); | Rhodotorula | 13 | Pathotroph-Saprotroph-Symbiotroph | Animal Pathogen-Animal Symbiotroph-Endophyte-Plant Pathogen-\|Undefined Saprotroph\| | Yeast | NULL | Probable | [illegible] | [illegible] |
| Otu04582 | 7 | k__Fungi(100);p__Ascomycota(100);c__Saccharomycetes(100);o__Saccharomycetales(100);f__Metschnikowiaceae(100);g__Metschnikowia(100);g__Metschnikowia_unclassified(86); | Metschnikowia | 13 | Saprotroph | \|Undefined Saprotroph\| | Yeast | NULL | Probable | [illegible] | [illegible] |
| Otu04584 | 7 | k__Fungi(100);p__Ascomycota(100);c__Saccharomycetes(100);o__Saccharomycetales(100);f__Metschnikowiaceae(100);g__Metschnikowia(100);s__Metschnikowia_pulcherrima(100); | Metschnikowia | 13 | Saprotroph | \|Undefined Saprotroph\| | Yeast | NULL | Probable | [illegible] | [illegible] |
| Otu04585 | 7 | k__Fungi(100);p__Ascomycota(100);c__Saccharomycetes(100);o__Saccharomycetales(100);f__Saccharomycetales_family_Incertae_sedis(100);g__Candida(100);g__Candida_unclassified(72); | Candida | 13 | Pathotroph-Saprotroph-Symbiotroph | Animal Pathogen-Endophyte-Endosymbiont-Epiphyte-\|Undefined Saprotroph\| | Dimorphic Yeast-Yeast | NULL | Probable | [illegible] | [illegible] |
| Otu04586 | 7 | k__Fungi(100);p__Ascomycota(100);c__Saccharomycetes(100);o__Saccharomycetales(100);f__Metschnikowiaceae(100);g__Metschnikowia(100);s__Metschnikowia_pulcherrima(100); | Metschnikowia | 13 | Saprotroph | \|Undefined Saprotroph\| | Yeast | NULL | Probable | [illegible] | [illegible] |
| Otu04587 | 7 | k__Fungi(100);p__Ascomycota(100);c__Sordariomycetes(100);o__Chaetosphaeriales(100);f__Chaetosphaeriaceae(100);g__Chloridium(100);s__Chloridium_virescens_var_chlamydosporum(100); | Chloridium | 13 | Pathotroph-Saprotroph-Symbiotroph | Ectomycorrhizal-Endophyte-Plant Pathogen-\|Plant Saprotroph\|-Wood Saprotroph | Microfungus | NULL | Possible | [illegible] | [illegible] |
| Otu04588 | 7 | k__Fungi(100);p__Ascomycota(100);c__Saccharomycetes(72);o__Saccharomycetales(72);f__Saccharomycetales_family_Incertae_sedis(72);g__Candida(72);g__Candida_unclassified(72); | Candida | 13 | Pathotroph-Saprotroph-Symbiotroph | Animal Pathogen-Endophyte-Endosymbiont-Epiphyte-\|Undefined Saprotroph\| | Dimorphic Yeast-Yeast | NULL | Probable | [illegible] | [illegible] |
| Otu04592 | 7 | k__Fungi(100);p__Ascomycota(100);c__Sordariomycetes(100);o__Hypocreales(72);o__Hypocreales_unclassified(72);o__Hypocreales_unclassified(72);o__Hypocreales_unclassified(72); | Hypocreales | 7 | Saprotroph | Undefined Saprotroph | Microfungus | NULL | Possible | NULL | [illegible] |
| Otu04594 | 7 | k__Fungi(100);p__Ascomycota(100);c__Dothideomycetes(100);o__Capnodiales(100);f__Capnodiales_family_Incertae_sedis(100);g__Phaeotheca(100);s__Phaeotheca_sp(100); | Phaeotheca | 13 | Pathotroph-Saprotroph | \|Plant Pathogen\|-Plant Saprotroph | Microfungus | NULL | Probable | [illegible] | [illegible] |
| Otu04595 | 7 | k__Fungi(100);p__Basidiomycota(100);c__Agaricomycetes(100);o__Agaricales(100);f__Psathyrellaceae(100);g__Coprinellus(100);s__Coprinellus_xanthothrix(100); | Coprinellus | 13 | Saprotroph | \|Undefined Saprotroph\| | Agaricoid | NULL | Probable | [illegible] | [illegible] |
| Otu04596 | 7 | k__Fungi(100);p__Ascomycota(100);c__Saccharomycetes(100);o__Saccharomycetales(100);f__Metschnikowiaceae(100);g__Metschnikowia(100);s__Metschnikowia_pulcherrima(100); | Metschnikowia | 13 | Saprotroph | \|Undefined Saprotroph\| | Yeast | NULL | Probable | [illegible] | [illegible] |
| Otu04600 | 7 | k__Fungi(100);p__Ascomycota(100);c__Saccharomycetes(100);o__Saccharomycetales(100);f__Metschnikowiaceae(100);g__Metschnikowia(100);s__Metschnikowia_pulcherrima(86); | Metschnikowia | 13 | Saprotroph | \|Undefined Saprotroph\| | Yeast | NULL | Probable | [illegible] | [illegible] |
| Otu04602 | 7 | k__Fungi(100);p__Basidiomycota(100);c__Agaricomycetes(100);o__Agaricales(100);f__Strophariaceae(100);g__Kuehneromyces(100);s__Kuehneromyces_lignicola(100); | Kuehneromyces | 13 | Saprotroph | Undefined Saprotroph-\|Wood Saprotroph\| | Agaricoid | White Rot | Probable | [illegible] | [illegible] |
| Otu04603 | 7 | k__Fungi(100);p__Basidiomycota(100);c__Agaricomycetes(100);o__Phallales(100);f__Phallaceae(100);g__Phallus(86);s__Phallus_sp_1_LL_2013(86); | Phallus | 13 | Saprotroph | Plant Saprotroph-\|Undefined Saprotroph\|-Wood Saprotroph | Phalloid | White Rot | Probable | [illegible] | [illegible] |
| Otu04604 | 7 | k__Fungi(100);p__Ascomycota(100);c__Leotiomycetes(100);o__Helotiales(100);f__Vibrisseaceae(100);g__Phialocephala(100);s__Phialocephala_sp_CM16s1(100); | Phialocephala | 13 | Saprotroph-Symbiotroph | Endophyte-\|Undefined Saprotroph\| | Microfungus | NULL | Highly Probable | [illegible] | [illegible] |
| Otu04607 | 7 | k__Fungi(100);p__Ascomycota(100);c__Sordariomycetes(100);o__Hypocreales(100);f__Hypocreaceae(100);g__Protocrea(100);s__Protocrea_farinosa(100); | Protocrea | 13 | Pathotroph | \|Fungal Parasite\| | Microfungus | NULL | Probable | [illegible] | [illegible] |
| Otu04609 | 7 | k__Fungi(100);p__Ascomycota(100);c__Saccharomycetes(100);o__Saccharomycetales(100);f__Metschnikowiaceae(100);g__Metschnikowia(100);s__Metschnikowia_pulcherrima(72); | Metschnikowia | 13 | Saprotroph | \|Undefined Saprotroph\| | Yeast | NULL | Probable | [illegible] | [illegible] |
| Otu04610 | 7 | k__Fungi(100);p__Basidiomycota(100);c__Agaricomycetes(100);o__Trechisporales(100);f__Hydnodontaceae(100);g__Trechispora(100);g__Trechispora_unclassified(100); | Trechispora | 13 | Saprotroph | \|Wood Saprotroph\| | Corticioid | White Rot | Highly Probable | [illegible] | [illegible] |
| Otu04611 | 7 | k__Fungi(100);p__Ascomycota(100);c__Saccharomycetes(100);o__Saccharomycetales(100);f__Metschnikowiaceae(100);g__Metschnikowia(100);s__Metschnikowia_pulcherrima(86); | Metschnikowia | 13 | Saprotroph | \|Undefined Saprotroph\| | Yeast | NULL | Probable | [illegible] | [illegible] |
| Otu04220 | 7 | k__Fungi(100);p__Basidiomycota(58);p__Basidiomycota_unclassified(58);p__Basidiomycota_unclassified(58);p__Basidiomycota_unclassified(58);p__Basidiomycota_unclassified(58);p__Basidiomycota_unclassified(58); | - | - | - | - | - | - | - | Unassigned | |
| Otu04222 | 7 | k__Fungi(100);p__Ascomycota(72);p__Ascomycota_unclassified(72);p__Ascomycota_unclassified(72);p__Ascomycota_unclassified(72);p__Ascomycota_unclassified(72);p__Ascomycota_unclassified(72); | - | - | - | - | - | - | - | Unassigned | |
| Otu04223 | 7 | k__Fungi(100);p__Ascomycota(100);p__Ascomycota_unclassified(100);p__Ascomycota_unclassified(100);p__Ascomycota_unclassified(100);p__Ascomycota_unclassified(100);p__Ascomycota_unclassified(100); | - | - | - | - | - | - | - | Unassigned | |
| Otu04224 | 7 | k__Fungi(100);k__Fungi_unclassified(58);k__Fungi_unclassified(58);k__Fungi_unclassified(58);k__Fungi_unclassified(58);k__Fungi_unclassified(58);k__Fungi_unclassified(58); | - | - | - | - | - | - | - | Unassigned | |
| Otu04225 | 7 | k__Fungi(100);p__Ascomycota(72);p__Ascomycota_unclassified(72);p__Ascomycota_unclassified(72);p__Ascomycota_unclassified(72);p__Ascomycota_unclassified(72);p__Ascomycota_unclassified(72); | - | - | - | - | - | - | - | Unassigned | |
| Otu04226 | 7 | k__Fungi(100);k__Fungi_unclassified(72);k__Fungi_unclassified(72);k__Fungi_unclassified(72);k__Fungi_unclassified(72);k__Fungi_unclassified(72);k__Fungi_unclassified(72); | - | - | - | - | - | - | - | Unassigned | |
| Otu04229 | 7 | k__Fungi(100);k__Fungi_unclassified(58);k__Fungi_unclassified(58);k__Fungi_unclassified(58);k__Fungi_unclassified(58);k__Fungi_unclassified(58);k__Fungi_unclassified(58); | - | - | - | - | - | - | - | Unassigned | |
| Otu04230 | 7 | k__Fungi(100);p__Zygomycota(100);c__Zygomycota_class_Incertae_sedis(100);o__Mucorales(100);f__Mucoraceae(100);g__unclassified_Mucoraceae(100);s__Mucoraceae_sp(100); | - | - | - | - | - | - | - | Unassigned | |
| Otu04234 | 7 | k__Fungi(100);p__Basidiomycota(100);p__Basidiomycota_unclassified(100);p__Basidiomycota_unclassified(100);p__Basidiomycota_unclassified(100);p__Basidiomycota_unclassified(100);p__Basidiomycota_unclassified(100); | - | - | - | - | - | - | - | Unassigned | |
| Otu04235 | 7 | k__Fungi(100);k__Fungi_unclassified(100);k__Fungi_unclassified(100);k__Fungi_unclassified(100);k__Fungi_unclassified(100);k__Fungi_unclassified(100);k__Fungi_unclassified(100); | - | - | - | - | - | - | - | Unassigned | |
| Otu04237 | 7 | k__Fungi(100);k__Fungi_unclassified(72);k__Fungi_unclassified(72);k__Fungi_unclassified(72);k__Fungi_unclassified(72);k__Fungi_unclassified(72);k__Fungi_unclassified(72); | - | - | - | - | - | - | - | Unassigned | |
| Otu04238 | 7 | k__Fungi(100);k__Fungi_unclassified(72);k__Fungi_unclassified(72);k__Fungi_unclassified(72);k__Fungi_unclassified(72);k__Fungi_unclassified(72);k__Fungi_unclassified(72); | - | - | - | - | - | - | - | Unassigned | |
| Otu04240 | 7 | k__Plantae(100);p__unclassified_Plantae(100);c__unclassified_Plantae(100);o__unclassified_Plantae(100);f__unclassified_Plantae(100);g__unclassified_Plantae(100);s__Plantae_sp(100); | - | - | - | - | - | - | - | Unassigned | |
| Otu04241 | 7 | k__Fungi(100);p__Ascomycota(100);c__Dothideomycetes(100);o__Pleosporales(100);o__Pleosporales_unclassified(100);o__Pleosporales_unclassified(100);o__Pleosporales_unclassified(100); | - | - | - | - | - | - | - | Unassigned | |
| Otu04242 | 7 | k__Plantae(100);p__unclassified_Plantae(100);c__unclassified_Plantae(100);o__unclassified_Plantae(100);f__unclassified_Plantae(100);g__unclassified_Plantae(100);s__Plantae_sp(100); | - | - | - | - | - | - | - | Unassigned | |
| Otu04243 | 7 | k__Fungi(100);k__Fungi_unclassified(72);k__Fungi_unclassified(72);k__Fungi_unclassified(72);k__Fungi_unclassified(72);k__Fungi_unclassified(72);k__Fungi_unclassified(72); | - | - | - | - | - | - | - | Unassigned | |

| | | | | | | | | | | | |
|---|---|---|---|---|---|---|---|---|---|---|---|
| Otu04244 | 7 | k__Fungi(100);k__Fungi_unclassified(86);k__Fungi_unclassified(86);k__Fungi_unclassified(86);k__Fungi_unclassified(86);k__Fungi_unclassified(86);k__Fungi_unclassified(86); | - | - | - | - | - | - | - | Unassigned | - |
| Otu04249 | 7 | k__Fungi(100);p__Basidiomycota(100);c__Agaricomycetes(100);c__Agaricomycetes_unclassified(72);c__Agaricomycetes_unclassified(72);c__Agaricomycetes_unclassified(72);c__Agaricomycetes_unclassified(72); | - | - | - | - | - | - | - | Unassigned | - |
| Otu04252 | 7 | k__Fungi(100);p__Basidiomycota(100);p__Basidiomycota_unclassified(58);p__Basidiomycota_unclassified(58);p__Basidiomycota_unclassified(58);p__Basidiomycota_unclassified(58);p__Basidiomycota_unclassified(58); | - | - | - | - | - | - | - | Unassigned | - |
| Otu04253 | 7 | k__Fungi(100);k__Fungi_unclassified(72);k__Fungi_unclassified(72);k__Fungi_unclassified(72);k__Fungi_unclassified(72);k__Fungi_unclassified(72);k__Fungi_unclassified(72); | - | - | - | - | - | - | - | Unassigned | - |
| Otu04256 | 7 | k__Fungi(100);k__Fungi_unclassified(58);k__Fungi_unclassified(58);k__Fungi_unclassified(58);k__Fungi_unclassified(58);k__Fungi_unclassified(58);k__Fungi_unclassified(58); | - | - | - | - | - | - | - | Unassigned | - |
| Otu04258 | 7 | k__Fungi(100);k__Fungi_unclassified(72);k__Fungi_unclassified(72);k__Fungi_unclassified(72);k__Fungi_unclassified(72);k__Fungi_unclassified(72);k__Fungi_unclassified(72); | - | - | - | - | - | - | - | Unassigned | - |
| Otu04259 | 7 | k__Fungi(100);k__Fungi_unclassified(72);k__Fungi_unclassified(72);k__Fungi_unclassified(72);k__Fungi_unclassified(72);k__Fungi_unclassified(72);k__Fungi_unclassified(72); | - | - | - | - | - | - | - | Unassigned | - |
| Otu04260 | 7 | k__Fungi(100);k__Fungi_unclassified(58);k__Fungi_unclassified(58);k__Fungi_unclassified(58);k__Fungi_unclassified(58);k__Fungi_unclassified(58);k__Fungi_unclassified(58); | - | - | - | - | - | - | - | Unassigned | - |
| Otu04262 | 7 | k__Fungi(100);p__Ascomycota(58);p__Ascomycota_unclassified(58);p__Ascomycota_unclassified(58);p__Ascomycota_unclassified(58);p__Ascomycota_unclassified(58);p__Ascomycota_unclassified(58); | - | - | - | - | - | - | - | Unassigned | - |
| Otu04263 | 7 | k__Fungi(100);k__Fungi_unclassified(100);k__Fungi_unclassified(100);k__Fungi_unclassified(100);k__Fungi_unclassified(100);k__Fungi_unclassified(100);k__Fungi_unclassified(100); | - | - | - | - | - | - | - | Unassigned | - |
| Otu04266 | 7 | k__Fungi(100);k__Fungi_unclassified(86);k__Fungi_unclassified(86);k__Fungi_unclassified(86);k__Fungi_unclassified(86);k__Fungi_unclassified(86);k__Fungi_unclassified(86); | - | - | - | - | - | - | - | Unassigned | - |
| Otu04267 | 7 | k__Fungi(100);p__Basidiomycota(100);c__Tremellomycetes(100);c__Tremellomycetes_unclassified(86);c__Tremellomycetes_unclassified(86);c__Tremellomycetes_unclassified(86);c__Tremellomycetes_unclassified(86); | - | - | - | - | - | - | - | Unassigned | - |
| Otu04269 | 7 | k__Fungi(100);k__Fungi_unclassified(58);k__Fungi_unclassified(58);k__Fungi_unclassified(58);k__Fungi_unclassified(58);k__Fungi_unclassified(58);k__Fungi_unclassified(58); | - | - | - | - | - | - | - | Unassigned | - |
| Otu04270 | 7 | k__Fungi(100);p__Ascomycota(100);c__Dothideomycetes(100);o__Venturiales(72);o__Venturiales_unclassified(72);o__Venturiales_unclassified(72);o__Venturiales_unclassified(72); | - | - | - | - | - | - | - | Unassigned | - |
| Otu04271 | 7 | k__Fungi(100);k__Fungi_unclassified(58);k__Fungi_unclassified(58);k__Fungi_unclassified(58);k__Fungi_unclassified(58);k__Fungi_unclassified(58);k__Fungi_unclassified(58); | - | - | - | - | - | - | - | Unassigned | - |
| Otu04276 | 7 | k__Fungi(100);p__Ascomycota(72);p__Ascomycota_unclassified(72);p__Ascomycota_unclassified(72);p__Ascomycota_unclassified(72);p__Ascomycota_unclassified(72);p__Ascomycota_unclassified(72); | - | - | - | - | - | - | - | Unassigned | - |
| Otu04277 | 7 | k__Fungi(100);p__Ascomycota(72);p__Ascomycota_unclassified(72);p__Ascomycota_unclassified(72);p__Ascomycota_unclassified(72);p__Ascomycota_unclassified(72);p__Ascomycota_unclassified(72); | - | - | - | - | - | - | - | Unassigned | - |
| Otu04280 | 7 | k__Fungi(100);k__Fungi_unclassified(58);k__Fungi_unclassified(58);k__Fungi_unclassified(58);k__Fungi_unclassified(58);k__Fungi_unclassified(58);k__Fungi_unclassified(58); | - | - | - | - | - | - | - | Unassigned | - |
| Otu04282 | 7 | k__Fungi(100);p__Basidiomycota(100);c__Agaricomycetes(100);o__Polyporales(100);f__unclassified_Polyporales(100);g__unclassified_Polyporales(100);s__Polyporales_sp(100); | - | - | - | - | - | - | - | Unassigned | - |
| Otu04283 | 7 | k__Fungi(100);k__Fungi_unclassified(58);k__Fungi_unclassified(58);k__Fungi_unclassified(58);k__Fungi_unclassified(58);k__Fungi_unclassified(58);k__Fungi_unclassified(58); | - | - | - | - | - | - | - | Unassigned | - |
| Otu04284 | 7 | k__Fungi(100);k__Fungi_unclassified(100);k__Fungi_unclassified(100);k__Fungi_unclassified(100);k__Fungi_unclassified(100);k__Fungi_unclassified(100);k__Fungi_unclassified(100); | - | - | - | - | - | - | - | Unassigned | - |
| Otu04286 | 7 | k__Fungi(100);p__Ascomycota(100);c__Sordariomycetes(100);c__Sordariomycetes_unclassified(100);c__Sordariomycetes_unclassified(100);c__Sordariomycetes_unclassified(100);c__Sordariomycetes_unclassified(100); | - | - | - | - | - | - | - | Unassigned | - |
| Otu04287 | 7 | k__Fungi(100);k__Fungi_unclassified(100);k__Fungi_unclassified(100);k__Fungi_unclassified(100);k__Fungi_unclassified(100);k__Fungi_unclassified(100);k__Fungi_unclassified(100); | - | - | - | - | - | - | - | Unassigned | - |
| Otu04288 | 7 | k__Fungi(100);p__Ascomycota(86);p__Ascomycota_unclassified(86);p__Ascomycota_unclassified(86);p__Ascomycota_unclassified(86);p__Ascomycota_unclassified(86);p__Ascomycota_unclassified(86); | - | - | - | - | - | - | - | Unassigned | - |
| Otu04289 | 7 | k__Fungi(100);k__Fungi_unclassified(58);k__Fungi_unclassified(58);k__Fungi_unclassified(58);k__Fungi_unclassified(58);k__Fungi_unclassified(58);k__Fungi_unclassified(58); | - | - | - | - | - | - | - | Unassigned | - |
| Otu04292 | 7 | k__Fungi(100);k__Fungi_unclassified(58);k__Fungi_unclassified(58);k__Fungi_unclassified(58);k__Fungi_unclassified(58);k__Fungi_unclassified(58);k__Fungi_unclassified(58); | - | - | - | - | - | - | - | Unassigned | - |
| Otu04293 | 7 | k__Fungi(100);p__Ascomycota(58);p__Ascomycota_unclassified(58);p__Ascomycota_unclassified(58);p__Ascomycota_unclassified(58);p__Ascomycota_unclassified(58);p__Ascomycota_unclassified(58); | - | - | - | - | - | - | - | Unassigned | - |
| Otu04297 | 7 | k__Fungi(100);k__Fungi_unclassified(58);k__Fungi_unclassified(58);k__Fungi_unclassified(58);k__Fungi_unclassified(58);k__Fungi_unclassified(58);k__Fungi_unclassified(58); | - | - | - | - | - | - | - | Unassigned | - |
| Otu04298 | 7 | k__Fungi(100);p__Ascomycota(86);p__Ascomycota_unclassified(72);p__Ascomycota_unclassified(72);p__Ascomycota_unclassified(72);p__Ascomycota_unclassified(72);p__Ascomycota_unclassified(72); | - | - | - | - | - | - | - | Unassigned | - |
| Otu04299 | 7 | k__Fungi(100);k__Fungi_unclassified(86);k__Fungi_unclassified(86);k__Fungi_unclassified(86);k__Fungi_unclassified(86);k__Fungi_unclassified(86);k__Fungi_unclassified(86); | - | - | - | - | - | - | - | Unassigned | - |
| Otu04302 | 7 | k__Fungi(100);k__Fungi_unclassified(86);k__Fungi_unclassified(86);k__Fungi_unclassified(86);k__Fungi_unclassified(86);k__Fungi_unclassified(86);k__Fungi_unclassified(86); | - | - | - | - | - | - | - | Unassigned | - |
| Otu04304 | 7 | k__Fungi(100);p__Ascomycota(72);p__Ascomycota_unclassified(72);p__Ascomycota_unclassified(72);p__Ascomycota_unclassified(72);p__Ascomycota_unclassified(72);p__Ascomycota_unclassified(72); | - | - | - | - | - | - | - | Unassigned | - |
| Otu04309 | 7 | k__Fungi(100);p__Ascomycota(58);p__Ascomycota_unclassified(58);p__Ascomycota_unclassified(58);p__Ascomycota_unclassified(58);p__Ascomycota_unclassified(58);p__Ascomycota_unclassified(58); | - | - | - | - | - | - | - | Unassigned | - |
| Otu04310 | 7 | k__Fungi(100);k__Fungi_unclassified(86);k__Fungi_unclassified(86);k__Fungi_unclassified(86);k__Fungi_unclassified(86);k__Fungi_unclassified(86);k__Fungi_unclassified(86); | - | - | - | - | - | - | - | Unassigned | - |
| Otu04311 | 7 | k__Fungi(100);p__Ascomycota(100);c__Sordariomycetes(100);o__Xylariales(100);o__Xylariales_unclassified(100);o__Xylariales_unclassified(100);o__Xylariales_unclassified(100); | - | - | - | - | - | - | - | Unassigned | - |
| Otu04312 | 7 | k__Fungi(100);p__Basidiomycota(100);c__Agaricomycetes(100);c__Agaricomycetes_unclassified(100);c__Agaricomycetes_unclassified(100);c__Agaricomycetes_unclassified(100);c__Agaricomycetes_unclassified(100); | - | - | - | - | - | - | - | Unassigned | - |
| Otu04316 | 7 | k__Fungi(100);p__Ascomycota(86);p__Ascomycota_unclassified(86);p__Ascomycota_unclassified(86);p__Ascomycota_unclassified(86);p__Ascomycota_unclassified(86);p__Ascomycota_unclassified(86); | - | - | - | - | - | - | - | Unassigned | - |
| Otu04320 | 7 | k__Fungi(100);k__Fungi_unclassified(100);k__Fungi_unclassified(100);k__Fungi_unclassified(100);k__Fungi_unclassified(100);k__Fungi_unclassified(100);k__Fungi_unclassified(100); | - | - | - | - | - | - | - | Unassigned | - |
| Otu04321 | 7 | k__Fungi(100);p__Ascomycota(100);p__Ascomycota_unclassified(100);p__Ascomycota_unclassified(100);p__Ascomycota_unclassified(100);p__Ascomycota_unclassified(100);p__Ascomycota_unclassified(100); | - | - | - | - | - | - | - | Unassigned | - |
| Otu04325 | 7 | k__Fungi(100);p__Ascomycota(100);c__Dothideomycetes(100);c__Dothideomycetes_unclassified(86);c__Dothideomycetes_unclassified(86);c__Dothideomycetes_unclassified(86);c__Dothideomycetes_unclassified(86); | - | - | - | - | - | - | - | Unassigned | - |
| Otu04326 | 7 | k__Fungi(100);k__Fungi_unclassified(100);k__Fungi_unclassified(100);k__Fungi_unclassified(100);k__Fungi_unclassified(100);k__Fungi_unclassified(100);k__Fungi_unclassified(100); | - | - | - | - | - | - | - | Unassigned | - |
| Otu04327 | 7 | k__Fungi(100);k__Fungi_unclassified(72);k__Fungi_unclassified(72);k__Fungi_unclassified(72);k__Fungi_unclassified(72);k__Fungi_unclassified(72);k__Fungi_unclassified(72); | - | - | - | - | - | - | - | Unassigned | - |
| Otu04331 | 7 | k__Fungi(100);k__Fungi_unclassified(100);k__Fungi_unclassified(100);k__Fungi_unclassified(100);k__Fungi_unclassified(100);k__Fungi_unclassified(100);k__Fungi_unclassified(100); | - | - | - | - | - | - | - | Unassigned | - |
| Otu04333 | 7 | k__Fungi(100);p__Ascomycota(100);c__Sordariomycetes(100);c__Sordariomycetes_unclassified(100);c__Sordariomycetes_unclassified(100);c__Sordariomycetes_unclassified(100);c__Sordariomycetes_unclassified(100); | - | - | - | - | - | - | - | Unassigned | - |
| Otu04336 | 7 | k__Fungi(100);p__Ascomycota(100);p__Ascomycota_unclassified(100);p__Ascomycota_unclassified(100);p__Ascomycota_unclassified(100);p__Ascomycota_unclassified(100);p__Ascomycota_unclassified(100); | - | - | - | - | - | - | - | Unassigned | - |
| Otu04338 | 7 | k__Fungi(100);p__Basidiomycota(100);c__Agaricomycetes(100);c__Agaricomycetes_unclassified(100);c__Agaricomycetes_unclassified(100);c__Agaricomycetes_unclassified(100);c__Agaricomycetes_unclassified(100); | - | - | - | - | - | - | - | Unassigned | - |
| Otu04341 | 7 | k__Fungi(100);p__Basidiomycota(100);p__Basidiomycota_unclassified(100);p__Basidiomycota_unclassified(100);p__Basidiomycota_unclassified(100);p__Basidiomycota_unclassified(100);p__Basidiomycota_unclassified(100); | - | - | - | - | - | - | - | Unassigned | - |
| Otu04342 | 7 | k__Fungi(100);k__Fungi_unclassified(58);k__Fungi_unclassified(58);k__Fungi_unclassified(58);k__Fungi_unclassified(58);k__Fungi_unclassified(58);k__Fungi_unclassified(58); | - | - | - | - | - | - | - | Unassigned | - |
| Otu04343 | 7 | k__Fungi(100);k__Fungi_unclassified(100);k__Fungi_unclassified(100);k__Fungi_unclassified(100);k__Fungi_unclassified(100);k__Fungi_unclassified(100);k__Fungi_unclassified(100); | - | - | - | - | - | - | - | Unassigned | - |
| Otu04347 | 7 | k__Fungi(100);p__Ascomycota(72);p__Ascomycota_unclassified(72);p__Ascomycota_unclassified(72);p__Ascomycota_unclassified(72);p__Ascomycota_unclassified(72);p__Ascomycota_unclassified(72); | - | - | - | - | - | - | - | Unassigned | - |
| Otu04348 | 7 | k__Fungi(100);k__Fungi_unclassified(100);k__Fungi_unclassified(100);k__Fungi_unclassified(100);k__Fungi_unclassified(100);k__Fungi_unclassified(100);k__Fungi_unclassified(100); | - | - | - | - | - | - | - | Unassigned | - |
| Otu04351 | 7 | k__Fungi(100);k__Fungi_unclassified(58);k__Fungi_unclassified(58);k__Fungi_unclassified(58);k__Fungi_unclassified(58);k__Fungi_unclassified(58);k__Fungi_unclassified(58); | - | - | - | - | - | - | - | Unassigned | - |
| Otu04352 | 7 | k__Fungi(100);p__Ascomycota(72);p__Ascomycota_unclassified(72);p__Ascomycota_unclassified(72);p__Ascomycota_unclassified(72);p__Ascomycota_unclassified(72);p__Ascomycota_unclassified(72); | - | - | - | - | - | - | - | Unassigned | - |
| Otu04353 | 7 | k__Fungi(100);p__Ascomycota(72);p__Ascomycota_unclassified(72);p__Ascomycota_unclassified(72);p__Ascomycota_unclassified(72);p__Ascomycota_unclassified(72);p__Ascomycota_unclassified(72); | - | - | - | - | - | - | - | Unassigned | - |
| Otu04355 | 7 | k__Fungi(100);k__Fungi_unclassified(86);k__Fungi_unclassified(86);k__Fungi_unclassified(86);k__Fungi_unclassified(86);k__Fungi_unclassified(86);k__Fungi_unclassified(86); | - | - | - | - | - | - | - | Unassigned | - |
| Otu04356 | 7 | k__Fungi(100);p__Basidiomycota(100);c__Agaricomycetes(100);c__Agaricomycetes_unclassified(100);c__Agaricomycetes_unclassified(100);c__Agaricomycetes_unclassified(100);c__Agaricomycetes_unclassified(100); | - | - | - | - | - | - | - | Unassigned | - |
| Otu04357 | 7 | k__Fungi(100);p__Basidiomycota(100);c__Agaricomycetes(100);o__Cantharellales(100);o__Cantharellales_unclassified(86);o__Cantharellales_unclassified(86);o__Cantharellales_unclassified(86); | - | - | - | - | - | - | - | Unassigned | - |
| Otu04358 | 7 | k__Fungi(100);p__Ascomycota(86);p__Ascomycota_unclassified(86);p__Ascomycota_unclassified(86);p__Ascomycota_unclassified(86);p__Ascomycota_unclassified(86);p__Ascomycota_unclassified(86); | - | - | - | - | - | - | - | Unassigned | - |
| Otu04363 | 7 | k__Fungi(100);p__Basidiomycota(100);p__Basidiomycota_unclassified(100);p__Basidiomycota_unclassified(100);p__Basidiomycota_unclassified(100);p__Basidiomycota_unclassified(100);p__Basidiomycota_unclassified(100); | - | - | - | - | - | - | - | Unassigned | - |
| Otu04365 | 7 | k__Fungi(100);k__Fungi_unclassified(72);k__Fungi_unclassified(72);k__Fungi_unclassified(72);k__Fungi_unclassified(72);k__Fungi_unclassified(72);k__Fungi_unclassified(72); | - | - | - | - | - | - | - | Unassigned | - |
| Otu04376 | 7 | k__Fungi(100);p__Basidiomycota(100);c__Tremellomycetes(100);o__Cystofilobasidiales(100);o__Cystofilobasidiales_unclassified(100);o__Cystofilobasidiales_unclassified(100);o__Cystofilobasidiales_unclassified(100); | - | - | - | - | - | - | - | Unassigned | - |
| Otu04378 | 7 | k__Fungi(100);k__Fungi_unclassified(100);k__Fungi_unclassified(100);k__Fungi_unclassified(100);k__Fungi_unclassified(100);k__Fungi_unclassified(100);k__Fungi_unclassified(100); | - | - | - | - | - | - | - | Unassigned | - |
| Otu04381 | 7 | k__Fungi(100);p__Ascomycota(86);p__Ascomycota_unclassified(86);p__Ascomycota_unclassified(86);p__Ascomycota_unclassified(86);p__Ascomycota_unclassified(86);p__Ascomycota_unclassified(86); | - | - | - | - | - | - | - | Unassigned | - |
| Otu04383 | 7 | k__Fungi(100);p__Ascomycota(86);p__Ascomycota_unclassified(86);p__Ascomycota_unclassified(86);p__Ascomycota_unclassified(86);p__Ascomycota_unclassified(86);p__Ascomycota_unclassified(86); | - | - | - | - | - | - | - | Unassigned | - |
| Otu04384 | 7 | k__Fungi(100);p__Ascomycota(100);c__Leotiomycetes(100);o__Helotiales(72);o__Helotiales_unclassified(72);o__Helotiales_unclassified(72);o__Helotiales_unclassified(72); | - | - | - | - | - | - | - | Unassigned | - |
| Otu04385 | 7 | k__Fungi(100);k__Fungi_unclassified(100);k__Fungi_unclassified(100);k__Fungi_unclassified(100);k__Fungi_unclassified(100);k__Fungi_unclassified(100);k__Fungi_unclassified(100); | - | - | - | - | - | - | - | Unassigned | - |
| Otu04386 | 7 | k__Fungi(100);p__Ascomycota(58);p__Ascomycota_unclassified(58);p__Ascomycota_unclassified(58);p__Ascomycota_unclassified(58);p__Ascomycota_unclassified(58);p__Ascomycota_unclassified(58); | - | - | - | - | - | - | - | Unassigned | - |
| Otu04388 | 7 | k__Fungi(100);p__Ascomycota(72);p__Ascomycota_unclassified(72);p__Ascomycota_unclassified(72);p__Ascomycota_unclassified(72);p__Ascomycota_unclassified(72);p__Ascomycota_unclassified(72); | - | - | - | - | - | - | - | Unassigned | - |
| Otu04395 | 7 | k__Fungi(100);p__Basidiomycota(100);c__Agaricomycetes(100);c__Agaricomycetes_unclassified(100);c__Agaricomycetes_unclassified(100);c__Agaricomycetes_unclassified(100);c__Agaricomycetes_unclassified(100); | - | - | - | - | - | - | - | Unassigned | - |
| Otu04396 | 7 | k__Fungi(100);p__Basidiomycota(100);c__Exobasidiomycetes(100);o__unclassified_Exobasidiomycetes(100);f__unclassified_Exobasidiomycetes(100);g__unclassified_Exobasidiomycetes(100);s__Exobasidiomycetes_sp(100); | - | - | - | - | - | - | - | Unassigned | - |
| Otu04397 | 7 | k__Fungi(100);p__Basidiomycota(100);c__Agaricomycetes(100);o__Russulales(100);o__Russulales_unclassified(100);o__Russulales_unclassified(100);o__Russulales_unclassified(100); | - | - | - | - | - | - | - | Unassigned | - |
| Otu04398 | 7 | k__Fungi(100);k__Fungi_unclassified(100);k__Fungi_unclassified(100);k__Fungi_unclassified(100);k__Fungi_unclassified(100);k__Fungi_unclassified(100);k__Fungi_unclassified(100); | - | - | - | - | - | - | - | Unassigned | - |
| Otu04399 | 7 | k__Fungi(100);p__Basidiomycota(100);c__Agaricomycetes(100);o__Atheliales(100);f__unclassified_Atheliales(100);g__unclassified_Atheliales(100);s__Atheliales_sp(100); | - | - | - | - | - | - | - | Unassigned | - |
| Otu04401 | 7 | k__Fungi(100);p__Basidiomycota(100);c__Agaricomycetes(100);o__Trechisporales(100);o__Trechisporales_unclassified(86);o__Trechisporales_unclassified(86);o__Trechisporales_unclassified(86); | - | - | - | - | - | - | - | Unassigned | - |
| Otu04402 | 7 | k__Fungi(100);k__Fungi_unclassified(58);k__Fungi_unclassified(58);k__Fungi_unclassified(58);k__Fungi_unclassified(58);k__Fungi_unclassified(58);k__Fungi_unclassified(58); | - | - | - | - | - | - | - | Unassigned | - |
| Otu04403 | 7 | k__Fungi(100);p__Basidiomycota(100);c__Agaricomycetes(100);c__Agaricomycetes_unclassified(100);c__Agaricomycetes_unclassified(100);c__Agaricomycetes_unclassified(100);c__Agaricomycetes_unclassified(100); | - | - | - | - | - | - | - | Unassigned | - |
| Otu04405 | 7 | k__Fungi(100);p__Ascomycota(86);p__Ascomycota_unclassified(86);p__Ascomycota_unclassified(86);p__Ascomycota_unclassified(86);p__Ascomycota_unclassified(86);p__Ascomycota_unclassified(86); | - | - | - | - | - | - | - | Unassigned | - |
| Otu04409 | 7 | k__Fungi(100);p__Ascomycota(100);p__Ascomycota_unclassified(100);p__Ascomycota_unclassified(100);p__Ascomycota_unclassified(100);p__Ascomycota_unclassified(100);p__Ascomycota_unclassified(100); | - | - | - | - | - | - | - | Unassigned | - |
| Otu04411 | 7 | k__Fungi(100);k__Fungi_unclassified(86);k__Fungi_unclassified(86);k__Fungi_unclassified(86);k__Fungi_unclassified(86);k__Fungi_unclassified(86);k__Fungi_unclassified(86); | - | - | - | - | - | - | - | Unassigned | - |
| Otu04412 | 7 | k__Fungi(100);p__Ascomycota(100);c__Dothideomycetes(100);o__Pleosporales(86);o__Pleosporales_unclassified(86);o__Pleosporales_unclassified(86);o__Pleosporales_unclassified(86); | - | - | - | - | - | - | - | Unassigned | - |
| Otu04415 | 7 | k__Fungi(100);k__Fungi_unclassified(72);k__Fungi_unclassified(72);k__Fungi_unclassified(72);k__Fungi_unclassified(72);k__Fungi_unclassified(72);k__Fungi_unclassified(72); | - | - | - | - | - | - | - | Unassigned | - |
| Otu04417 | 7 | k__Fungi(100);k__Fungi_unclassified(58);k__Fungi_unclassified(58);k__Fungi_unclassified(58);k__Fungi_unclassified(58);k__Fungi_unclassified(58);k__Fungi_unclassified(58); | - | - | - | - | - | - | - | Unassigned | - |
| Otu04418 | 7 | k__Fungi(100);p__Ascomycota(100);c__Sordariomycetes(100);c__Sordariomycetes_unclassified(58);c__Sordariomycetes_unclassified(58);c__Sordariomycetes_unclassified(58);c__Sordariomycetes_unclassified(58); | - | - | - | - | - | - | - | Unassigned | - |
| Otu04419 | 7 | k__Fungi(100);p__Basidiomycota(100);c__Agaricomycetes(100);o__Cantharellales(100);f__Hydnaceae(100);f__Hydnaceae_unclassified(100);f__Hydnaceae_unclassified(100); | - | - | - | - | - | - | - | Unassigned | - |
| Otu04420 | 7 | k__Fungi(100);p__Ascomycota(100);p__Ascomycota_unclassified(100);p__Ascomycota_unclassified(100);p__Ascomycota_unclassified(100);p__Ascomycota_unclassified(100);p__Ascomycota_unclassified(100); | - | - | - | - | - | - | - | Unassigned | - |
| Otu04422 | 7 | k__Fungi(100);k__Fungi_unclassified(72);k__Fungi_unclassified(72);k__Fungi_unclassified(72);k__Fungi_unclassified(72);k__Fungi_unclassified(72);k__Fungi_unclassified(72); | - | - | - | - | - | - | - | Unassigned | - |
| Otu04424 | 7 | k__Fungi(100);p__Basidiomycota(100);c__Agaricomycetes(100);c__Agaricomycetes_unclassified(100);c__Agaricomycetes_unclassified(100);c__Agaricomycetes_unclassified(100);c__Agaricomycetes_unclassified(100); | - | - | - | - | - | - | - | Unassigned | - |
| Otu04425 | 7 | k__Fungi(100);k__Fungi_unclassified(72);k__Fungi_unclassified(72);k__Fungi_unclassified(72);k__Fungi_unclassified(72);k__Fungi_unclassified(72);k__Fungi_unclassified(72); | - | - | - | - | - | - | - | Unassigned | - |
| Otu04427 | 7 | k__Fungi(100);p__Ascomycota(86);p__Ascomycota_unclassified(86);p__Ascomycota_unclassified(86);p__Ascomycota_unclassified(86);p__Ascomycota_unclassified(86);p__Ascomycota_unclassified(86); | - | - | - | - | - | - | - | Unassigned | - |
| Otu04428 | 7 | k__Fungi(100);p__Basidiomycota(100);c__Microbotryomycetes(100);o__Sporidiobolales(100);f__Sporidiobolales_family_Incertae_sedis(100);f__Sporidiobolales_family_Incertae_sedis_unclassified(58);f__Sporidiobolales_family_Incertae_sedis_unclassified(58); | - | - | - | - | - | - | - | Unassigned | - |
| Otu04432 | 7 | k__Fungi(100);k__Fungi_unclassified(72);k__Fungi_unclassified(72);k__Fungi_unclassified(72);k__Fungi_unclassified(72);k__Fungi_unclassified(72);k__Fungi_unclassified(72); | - | - | - | - | - | - | - | Unassigned | - |
| Otu04434 | 7 | k__Fungi(100);p__Ascomycota(58);p__Ascomycota_unclassified(58);p__Ascomycota_unclassified(58);p__Ascomycota_unclassified(58);p__Ascomycota_unclassified(58);p__Ascomycota_unclassified(58); | - | - | - | - | - | - | - | Unassigned | - |
| Otu04435 | 7 | k__Fungi(100);k__Fungi_unclassified(100);k__Fungi_unclassified(100);k__Fungi_unclassified(100);k__Fungi_unclassified(100);k__Fungi_unclassified(100);k__Fungi_unclassified(100); | - | - | - | - | - | - | - | Unassigned | - |
| Otu04436 | 7 | k__Fungi(100);k__Fungi_unclassified(86);k__Fungi_unclassified(86);k__Fungi_unclassified(86);k__Fungi_unclassified(86);k__Fungi_unclassified(86);k__Fungi_unclassified(86); | - | - | - | - | - | - | - | Unassigned | - |
| Otu04437 | 7 | k__Fungi(100);p__Ascomycota(58);p__Ascomycota_unclassified(58);p__Ascomycota_unclassified(58);p__Ascomycota_unclassified(58);p__Ascomycota_unclassified(58);p__Ascomycota_unclassified(58); | - | - | - | - | - | - | - | Unassigned | - |
| Otu04438 | 7 | k__Fungi(100);k__Fungi_unclassified(72);k__Fungi_unclassified(72);k__Fungi_unclassified(72);k__Fungi_unclassified(72);k__Fungi_unclassified(72);k__Fungi_unclassified(72); | - | - | - | - | - | - | - | Unassigned | - |
| Otu04440 | 7 | k__Fungi(100);p__Ascomycota(72);p__Ascomycota_unclassified(72);p__Ascomycota_unclassified(72);p__Ascomycota_unclassified(72);p__Ascomycota_unclassified(72);p__Ascomycota_unclassified(72); | - | - | - | - | - | - | - | Unassigned | - |
| Otu04441 | 7 | k__Fungi(100);p__Ascomycota(100);p__Ascomycota_unclassified(100);p__Ascomycota_unclassified(100);p__Ascomycota_unclassified(100);p__Ascomycota_unclassified(100);p__Ascomycota_unclassified(100); | - | - | - | - | - | - | - | Unassigned | - |
| Otu04442 | 7 | k__Fungi(100);p__Basidiomycota(100);c__Agaricomycetes(100);c__Agaricomycetes_unclassified(100);c__Agaricomycetes_unclassified(100);c__Agaricomycetes_unclassified(100);c__Agaricomycetes_unclassified(100); | - | - | - | - | - | - | - | Unassigned | - |
| Otu04444 | 7 | k__Fungi(100);p__Ascomycota(100);p__Ascomycota_unclassified(100);p__Ascomycota_unclassified(100);p__Ascomycota_unclassified(100);p__Ascomycota_unclassified(100);p__Ascomycota_unclassified(100); | - | - | - | - | - | - | - | Unassigned | - |
| Otu04445 | 7 | k__Fungi(100);k__Fungi_unclassified(86);k__Fungi_unclassified(86);k__Fungi_unclassified(86);k__Fungi_unclassified(86);k__Fungi_unclassified(86);k__Fungi_unclassified(86); | - | - | - | - | - | - | - | Unassigned | - |
| Otu04447 | 7 | k__Fungi(100);p__Ascomycota(58);p__Ascomycota_unclassified(58);p__Ascomycota_unclassified(58);p__Ascomycota_unclassified(58);p__Ascomycota_unclassified(58);p__Ascomycota_unclassified(58); | - | - | - | - | - | - | - | Unassigned | - |
| Otu04450 | 7 | k__Fungi(100);p__Ascomycota(100);p__Ascomycota_unclassified(100);p__Ascomycota_unclassified(100);p__Ascomycota_unclassified(100);p__Ascomycota_unclassified(100);p__Ascomycota_unclassified(100); | - | - | - | - | - | - | - | Unassigned | - |
| Otu04454 | 7 | k__Fungi(100);p__Basidiomycota(100);c__Agaricomycetes(100);c__Agaricomycetes_unclassified(100);c__Agaricomycetes_unclassified(100);c__Agaricomycetes_unclassified(100);c__Agaricomycetes_unclassified(100); | - | - | - | - | - | - | - | Unassigned | - |
| Otu04455 | 7 | k__Fungi(100);p__Basidiomycota(100);p__Basidiomycota_unclassified(100);p__Basidiomycota_unclassified(100);p__Basidiomycota_unclassified(100);p__Basidiomycota_unclassified(100);p__Basidiomycota_unclassified(100); | - | - | - | - | - | - | - | Unassigned | - |
| Otu04456 | 7 | k__Fungi(100);k__Fungi_unclassified(100);k__Fungi_unclassified(100);k__Fungi_unclassified(100);k__Fungi_unclassified(100);k__Fungi_unclassified(100);k__Fungi_unclassified(100); | - | - | - | - | - | - | - | Unassigned | - |
| Otu04458 | 7 | k__Fungi(100);k__Fungi_unclassified(100);k__Fungi_unclassified(100);k__Fungi_unclassified(100);k__Fungi_unclassified(100);k__Fungi_unclassified(100);k__Fungi_unclassified(100); | - | - | - | - | - | - | - | Unassigned | - |
| Otu04462 | 7 | k__Fungi(100);p__Chytridiomycota(58);p__Chytridiomycota_unclassified(58);p__Chytridiomycota_unclassified(58);p__Chytridiomycota_unclassified(58);p__Chytridiomycota_unclassified(58);p__Chytridiomycota_unclassified(58); | - | - | - | - | - | - | - | Unassigned | - |
| Otu04468 | 7 | k__Fungi(100);k__Fungi_unclassified(100);k__Fungi_unclassified(100);k__Fungi_unclassified(100);k__Fungi_unclassified(100);k__Fungi_unclassified(100);k__Fungi_unclassified(100); | - | - | - | - | - | - | - | Unassigned | - |
| Otu04470 | 7 | k__Fungi(100);p__Basidiomycota(100);p__Basidiomycota_unclassified(100);p__Basidiomycota_unclassified(100);p__Basidiomycota_unclassified(100);p__Basidiomycota_unclassified(100);p__Basidiomycota_unclassified(100); | - | - | - | - | - | - | - | Unassigned | - |
| Otu04472 | 7 | k__Fungi(100);p__Ascomycota(100);c__Eurotiomycetes(100);o__Eurotiales(100);f__Trichocomaceae(100);f__Trichocomaceae_unclassified(86);f__Trichocomaceae_unclassified(86); | - | - | - | - | - | - | - | Unassigned | - |
| Otu04473 | 7 | k__Fungi(100);p__Ascomycota(100);p__Ascomycota_unclassified(100);p__Ascomycota_unclassified(100);p__Ascomycota_unclassified(100);p__Ascomycota_unclassified(100);p__Ascomycota_unclassified(100); | - | - | - | - | - | - | - | Unassigned | - |
| Otu04476 | 7 | k__Fungi(100);k__Fungi_unclassified(72);k__Fungi_unclassified(72);k__Fungi_unclassified(72);k__Fungi_unclassified(72);k__Fungi_unclassified(72);k__Fungi_unclassified(72); | - | - | - | - | - | - | - | Unassigned | - |
| Otu04478 | 7 | k__Fungi(100);p__Basidiomycota(100);c__Agaricomycetes(100);o__Polyporales(100);o__Polyporales_unclassified(100);o__Polyporales_unclassified(100);o__Polyporales_unclassified(100); | - | - | - | - | - | - | - | Unassigned | - |
| Otu04481 | 7 | k__Fungi(100);k__Fungi_unclassified(58);k__Fungi_unclassified(58);k__Fungi_unclassified(58);k__Fungi_unclassified(58);k__Fungi_unclassified(58);k__Fungi_unclassified(58); | - | - | - | - | - | - | - | Unassigned | - |
| Otu04484 | 7 | k__Fungi(100);p__Ascomycota(100);p__Ascomycota_unclassified(100);p__Ascomycota_unclassified(100);p__Ascomycota_unclassified(100);p__Ascomycota_unclassified(100);p__Ascomycota_unclassified(100); | - | - | - | - | - | - | - | Unassigned | - |
| Otu04486 | 7 | k__Fungi(100);p__Ascomycota(58);p__Ascomycota_unclassified(58);p__Ascomycota_unclassified(58);p__Ascomycota_unclassified(58);p__Ascomycota_unclassified(58);p__Ascomycota_unclassified(58); | - | - | - | - | - | - | - | Unassigned | - |
| Otu04490 | 7 | k__Fungi(100);k__Fungi_unclassified(72);k__Fungi_unclassified(72);k__Fungi_unclassified(72);k__Fungi_unclassified(72);k__Fungi_unclassified(72);k__Fungi_unclassified(72); | - | - | - | - | - | - | - | Unassigned | - |
| Otu04491 | 7 | k__Fungi(100);k__Fungi_unclassified(100);k__Fungi_unclassified(100);k__Fungi_unclassified(100);k__Fungi_unclassified(100);k__Fungi_unclassified(100);k__Fungi_unclassified(100); | - | - | - | - | - | - | - | Unassigned | - |
| Otu04494 | 7 | k__Fungi(100);p__Ascomycota(100);c__Ascomycota_class_Incertae_sedis(58);o__Ascomycota_order_Incertae_sedis(58);f__Ascomycota_family_Incertae_sedis(58);f__Ascomycota_family_Incertae_sedis_unclassified(58);f__Ascomycota_family_Incertae_sedis_unclassified(58); | - | - | - | - | - | - | - | Unassigned | - |
| Otu04495 | 7 | k__Fungi(100);p__Ascomycota(100);c__Leotiomycetes(100);c__Leotiomycetes_unclassified(100);c__Leotiomycetes_unclassified(100);c__Leotiomycetes_unclassified(100);c__Leotiomycetes_unclassified(100); | - | - | - | - | - | - | - | Unassigned | - |
| Otu04496 | 7 | k__Fungi(100);p__Basidiomycota(100);c__Agaricomycetes(100);c__Agaricomycetes_unclassified(100);c__Agaricomycetes_unclassified(100);c__Agaricomycetes_unclassified(100);c__Agaricomycetes_unclassified(100); | - | - | - | - | - | - | - | Unassigned | - |
| Otu04497 | 7 | k__Fungi(100);k__Fungi_unclassified(100);k__Fungi_unclassified(100);k__Fungi_unclassified(100);k__Fungi_unclassified(100);k__Fungi_unclassified(100);k__Fungi_unclassified(100); | - | - | - | - | - | - | - | Unassigned | - |
| Otu04498 | 7 | k__Fungi(100);p__Ascomycota(100);c__Dothideomycetes(100);o__Capnodiales(100);o__Capnodiales_unclassified(86);o__Capnodiales_unclassified(86);o__Capnodiales_unclassified(86); | - | - | - | - | - | - | - | Unassigned | - |
| Otu04499 | 7 | k__Fungi(100);p__Ascomycota(86);p__Ascomycota_unclassified(86);p__Ascomycota_unclassified(86);p__Ascomycota_unclassified(86);p__Ascomycota_unclassified(86);p__Ascomycota_unclassified(86); | - | - | - | - | - | - | - | Unassigned | - |
| Otu04500 | 7 | k__Fungi(100);p__Ascomycota(100);p__Ascomycota_unclassified(100);p__Ascomycota_unclassified(100);p__Ascomycota_unclassified(100);p__Ascomycota_unclassified(100);p__Ascomycota_unclassified(100); | - | - | - | - | - | - | - | Unassigned | - |
| Otu04502 | 7 | k__Fungi(100);p__Ascomycota(100);c__Leotiomycetes(86);o__Helotiales(86);o__Helotiales_unclassified(86);o__Helotiales_unclassified(86);o__Helotiales_unclassified(86); | - | - | - | - | - | - | - | Unassigned | - |
| Otu04506 | 7 | k__Fungi(100);p__Ascomycota(72);p__Ascomycota_unclassified(72);p__Ascomycota_unclassified(72);p__Ascomycota_unclassified(72);p__Ascomycota_unclassified(72);p__Ascomycota_unclassified(72); | - | - | - | - | - | - | - | Unassigned | - |
| Otu04507 | 7 | k__Fungi(100);p__Ascomycota(100);c__Taphrinomycetes(100);o__Taphrinales(100);f__Taphrinaceae(100);g__Lalaria(100);s__Lalaria_inositophila(100); | - | - | - | - | - | - | - | Unassigned | - |
| Otu04508 | 7 | k__Fungi(100);p__Ascomycota(86);p__Ascomycota_unclassified(86);p__Ascomycota_unclassified(86);p__Ascomycota_unclassified(86);p__Ascomycota_unclassified(86);p__Ascomycota_unclassified(86); | - | - | - | - | - | - | - | Unassigned | - |
| Otu04511 | 7 | k__Fungi(100);k__Fungi_unclassified(72);k__Fungi_unclassified(72);k__Fungi_unclassified(72);k__Fungi_unclassified(72);k__Fungi_unclassified(72);k__Fungi_unclassified(72); | - | - | - | - | - | - | - | Unassigned | - |
| Otu04514 | 7 | k__Fungi(100);p__Ascomycota(72);p__Ascomycota_unclassified(72);p__Ascomycota_unclassified(72);p__Ascomycota_unclassified(72);p__Ascomycota_unclassified(72);p__Ascomycota_unclassified(72); | - | - | - | - | - | - | - | Unassigned | - |
| Otu04515 | 7 | k__Fungi(100);p__Ascomycota(86);p__Ascomycota_unclassified(86);p__Ascomycota_unclassified(86);p__Ascomycota_unclassified(86);p__Ascomycota_unclassified(86);p__Ascomycota_unclassified(86); | - | - | - | - | - | - | - | Unassigned | - |
| Otu04516 | 7 | k__Fungi(100);p__Ascomycota(86);p__Ascomycota_unclassified(86);p__Ascomycota_unclassified(86);p__Ascomycota_unclassified(86);p__Ascomycota_unclassified(86);p__Ascomycota_unclassified(86); | - | - | - | - | - | - | - | Unassigned | - |
| Otu04519 | 7 | k__Fungi(100);p__Ascomycota(86);p__Ascomycota_unclassified(86);p__Ascomycota_unclassified(86);p__Ascomycota_unclassified(86);p__Ascomycota_unclassified(86);p__Ascomycota_unclassified(86); | - | - | - | - | - | - | - | Unassigned | - |
| Otu04520 | 7 | k__Fungi(100);p__Ascomycota(86);p__Ascomycota_unclassified(86);p__Ascomycota_unclassified(86);p__Ascomycota_unclassified(86);p__Ascomycota_unclassified(86);p__Ascomycota_unclassified(86); | - | - | - | - | - | - | - | Unassigned | - |
| Otu04521 | 7 | k__Fungi(100);p__Ascomycota(100);c__Dothideomycetes(100);o__Pleosporales(100);o__Pleosporales_unclassified(72);o__Pleosporales_unclassified(72);o__Pleosporales_unclassified(72); | - | - | - | - | - | - | - | Unassigned | - |
| Otu04522 | 7 | k__Fungi(100);k__Fungi_unclassified(86);k__Fungi_unclassified(86);k__Fungi_unclassified(86);k__Fungi_unclassified(86);k__Fungi_unclassified(86);k__Fungi_unclassified(86); | - | - | - | - | - | - | - | Unassigned | - |
| Otu04525 | 7 | k__Plantae(100);p__unclassified_Plantae(100);c__unclassified_Plantae(100);o__unclassified_Plantae(100);f__unclassified_Plantae(100);g__unclassified_Plantae(100);s__Plantae_sp(100); | - | - | - | - | - | - | - | Unassigned | - |
| Otu04526 | 7 | k__Fungi(100);p__Ascomycota(100);p__Ascomycota_unclassified(100);p__Ascomycota_unclassified(100);p__Ascomycota_unclassified(100);p__Ascomycota_unclassified(100);p__Ascomycota_unclassified(100); | - | - | - | - | - | - | - | Unassigned | - |
| Otu04530 | 7 | k__Fungi(100);k__Fungi_unclassified(72);k__Fungi_unclassified(72);k__Fungi_unclassified(72);k__Fungi_unclassified(72);k__Fungi_unclassified(72);k__Fungi_unclassified(72); | - | - | - | - | - | - | - | Unassigned | - |
| Otu04533 | 7 | k__Fungi(100);k__Fungi_unclassified(72);k__Fungi_unclassified(72);k__Fungi_unclassified(72);k__Fungi_unclassified(72);k__Fungi_unclassified(72);k__Fungi_unclassified(72); | - | - | - | - | - | - | - | Unassigned | - |
| Otu04535 | 7 | k__Fungi(100);p__Ascomycota(100);c__Dothideomycetes(100);o__Pleosporales(100);o__Pleosporales_unclassified(100);o__Pleosporales_unclassified(100);o__Pleosporales_unclassified(100); | - | - | - | - | - | - | - | Unassigned | - |
| Otu04538 | 7 | k__Fungi(100);p__Ascomycota(58);p__Ascomycota_unclassified(58);p__Ascomycota_unclassified(58);p__Ascomycota_unclassified(58);p__Ascomycota_unclassified(58);p__Ascomycota_unclassified(58); | - | - | - | - | - | - | - | Unassigned | - |
| Otu04542 | 7 | k__Fungi(100);k__Fungi_unclassified(58);k__Fungi_unclassified(58);k__Fungi_unclassified(58);k__Fungi_unclassified(58);k__Fungi_unclassified(58);k__Fungi_unclassified(58); | - | - | - | - | - | - | - | Unassigned | - |
| Otu04543 | 7 | k__Fungi(100);p__Ascomycota(100);p__Ascomycota_unclassified(100);p__Ascomycota_unclassified(100);p__Ascomycota_unclassified(100);p__Ascomycota_unclassified(100);p__Ascomycota_unclassified(100); | - | - | - | - | - | - | - | Unassigned | - |
| Otu04544 | 7 | k__Fungi(100);p__Basidiomycota(100);c__Tremellomycetes(100);o__Filobasidiales(100);f__Filobasidiaceae(100);f__Filobasidiaceae_unclassified(58);f__Filobasidiaceae_unclassified(58); | - | - | - | - | - | - | - | Unassigned | - |
| Otu04546 | 7 | k__Fungi(100);p__Ascomycota(100);c__Eurotiomycetes(72);c__Eurotiomycetes_unclassified(58);c__Eurotiomycetes_unclassified(58);c__Eurotiomycetes_unclassified(58);c__Eurotiomycetes_unclassified(58); | - | - | - | - | - | - | - | Unassigned | - |
| Otu04547 | 7 | k__Fungi(100);p__Ascomycota(86);p__Ascomycota_unclassified(86);p__Ascomycota_unclassified(86);p__Ascomycota_unclassified(86);p__Ascomycota_unclassified(86);p__Ascomycota_unclassified(86); | - | - | - | - | - | - | - | Unassigned | - |
| Otu04548 | 7 | k__Fungi(100);p__Basidiomycota(100);c__Agaricomycetes(72);c__Agaricomycetes_unclassified(72);c__Agaricomycetes_unclassified(72);c__Agaricomycetes_unclassified(72);c__Agaricomycetes_unclassified(72); | - | - | - | - | - | - | - | Unassigned | - |
| Otu04549 | 7 | k__Fungi(100);k__Fungi_unclassified(58);k__Fungi_unclassified(58);k__Fungi_unclassified(58);k__Fungi_unclassified(58);k__Fungi_unclassified(58);k__Fungi_unclassified(58); | - | - | - | - | - | - | - | Unassigned | - |
| Otu04553 | 7 | k__Fungi(100);p__Ascomycota(100);c__Sordariomycetes(100);c__Sordariomycetes_unclassified(100);c__Sordariomycetes_unclassified(100);c__Sordariomycetes_unclassified(100);c__Sordariomycetes_unclassified(100); | - | - | - | - | - | - | - | Unassigned | - |
| Otu04554 | 7 | k__Fungi(100);k__Fungi_unclassified(100);k__Fungi_unclassified(100);k__Fungi_unclassified(100);k__Fungi_unclassified(100);k__Fungi_unclassified(100);k__Fungi_unclassified(100); | - | - | - | - | - | - | - | Unassigned | - |
| Otu04557 | 7 | k__Fungi(100);k__Fungi_unclassified(72);k__Fungi_unclassified(72);k__Fungi_unclassified(72);k__Fungi_unclassified(72);k__Fungi_unclassified(72);k__Fungi_unclassified(72); | - | - | - | - | - | - | - | Unassigned | - |
| Otu04558 | 7 | k__Fungi(100);k__Fungi_unclassified(86);k__Fungi_unclassified(86);k__Fungi_unclassified(86);k__Fungi_unclassified(86);k__Fungi_unclassified(86);k__Fungi_unclassified(86); | - | - | - | - | - | - | - | Unassigned | - |
| Otu04559 | 7 | k__Fungi(100);k__Fungi_unclassified(86);k__Fungi_unclassified(86);k__Fungi_unclassified(86);k__Fungi_unclassified(86);k__Fungi_unclassified(86);k__Fungi_unclassified(86); | - | - | - | - | - | - | - | Unassigned | - |
| Otu04565 | 7 | k__Fungi(100);p__Ascomycota(100);c__Sordariomycetes(100);c__Sordariomycetes_unclassified(100);c__Sordariomycetes_unclassified(100);c__Sordariomycetes_unclassified(100);c__Sordariomycetes_unclassified(100); | - | - | - | - | - | - | - | Unassigned | - |
| Otu04567 | 7 | k__Fungi(100);k__Fungi_unclassified(72);k__Fungi_unclassified(72);k__Fungi_unclassified(72);k__Fungi_unclassified(72);k__Fungi_unclassified(72);k__Fungi_unclassified(72); | - | - | - | - | - | - | - | Unassigned | - |
| Otu04572 | 7 | k__Fungi(100);p__Ascomycota(100);c__Dothideomycetes(100);o__Pleosporales(100);o__Pleosporales_unclassified(86);o__Pleosporales_unclassified(86);o__Pleosporales_unclassified(86); | - | - | - | - | - | - | - | Unassigned | - |
| Otu04575 | 7 | k__Fungi(100);p__Ascomycota(100);p__Ascomycota_unclassified(100);p__Ascomycota_unclassified(100);p__Ascomycota_unclassified(100);p__Ascomycota_unclassified(100);p__Ascomycota_unclassified(100); | - | - | - | - | - | - | - | Unassigned | - |
| Otu04576 | 7 | k__Fungi(100);p__Ascomycota(72);p__Ascomycota_unclassified(72);p__Ascomycota_unclassified(72);p__Ascomycota_unclassified(72);p__Ascomycota_unclassified(72);p__Ascomycota_unclassified(72); | - | - | - | - | - | - | - | Unassigned | - |
| Otu04577 | 7 | k__Fungi(100);p__Ascomycota(58);p__Ascomycota_unclassified(58);p__Ascomycota_unclassified(58);p__Ascomycota_unclassified(58);p__Ascomycota_unclassified(58);p__Ascomycota_unclassified(58); | - | - | - | - | - | - | - | Unassigned | - |
| Otu04579 | 7 | k__Fungi(100);p__Ascomycota(100);c__Sordariomycetes(100);c__Sordariomycetes_unclassified(58);c__Sordariomycetes_unclassified(58);c__Sordariomycetes_unclassified(58);c__Sordariomycetes_unclassified(58); | - | - | - | - | - | - | - | Unassigned | - |
| Otu04581 | 7 | k__Fungi(100);p__Ascomycota(100);p__Ascomycota_unclassified(100);p__Ascomycota_unclassified(100);p__Ascomycota_unclassified(100);p__Ascomycota_unclassified(100);p__Ascomycota_unclassified(100); | - | - | - | - | - | - | - | Unassigned | - |
| Otu04583 | 7 | k__Fungi(100);k__Fungi_unclassified(58);k__Fungi_unclassified(58);k__Fungi_unclassified(58);k__Fungi_unclassified(58);k__Fungi_unclassified(58);k__Fungi_unclassified(58); | - | - | - | - | - | - | - | Unassigned | - |
| Otu04589 | 7 | k__Fungi(100);k__Fungi_unclassified(58);k__Fungi_unclassified(58);k__Fungi_unclassified(58);k__Fungi_unclassified(58);k__Fungi_unclassified(58);k__Fungi_unclassified(58); | - | - | - | - | - | - | - | Unassigned | - |
| Otu04590 | 7 | k__Fungi(100);p__Ascomycota(72);p__Ascomycota_unclassified(72);p__Ascomycota_unclassified(72);p__Ascomycota_unclassified(72);p__Ascomycota_unclassified(72);p__Ascomycota_unclassified(72); | - | - | - | - | - | - | - | Unassigned | - |
| Otu04591 | 7 | k__Fungi(100);p__Ascomycota(100);p__Ascomycota_unclassified(72);p__Ascomycota_unclassified(72);p__Ascomycota_unclassified(72);p__Ascomycota_unclassified(72);p__Ascomycota_unclassified(72); | - | - | - | - | - | - | - | Unassigned | - |
| Otu04593 | 7 | k__Fungi(100);p__Ascomycota(86);p__Ascomycota_unclassified(86);p__Ascomycota_unclassified(86);p__Ascomycota_unclassified(86);p__Ascomycota_unclassified(86);p__Ascomycota_unclassified(86); | - | - | - | - | - | - | - | Unassigned | - |
| Otu04597 | 7 | k__Fungi(100);p__Rozellomycota(100);c__unclassified_Rozellomycota(100);o__unclassified_Rozellomycota(100);f__unclassified_Rozellomycota(100);g__unclassified_Rozellomycota(100);s__Rozellomycota_sp(100); | - | - | - | - | - | - | - | Unassigned | - |
| Otu04598 | 7 | k__Fungi(100);p__Ascomycota(58);p__Ascomycota_unclassified(58);p__Ascomycota_unclassified(58);p__Ascomycota_unclassified(58);p__Ascomycota_unclassified(58);p__Ascomycota_unclassified(58); | - | - | - | - | - | - | - | Unassigned | - |
| Otu04599 | 7 | k__Fungi(100);p__Ascomycota(100);p__Ascomycota_unclassified(100);p__Ascomycota_unclassified(100);p__Ascomycota_unclassified(100);p__Ascomycota_unclassified(100);p__Ascomycota_unclassified(100); | - | - | - | - | - | - | - | Unassigned | - |
| Otu04601 | 7 | k__Fungi(100);p__Basidiomycota(100);c__Microbotryomycetes(100);o__Sporidiobolales(100);f__Sporidiobolales_family_Incertae_sedis(100);g__unclassified_Sporidiobolales(72);s__Sporidiobolales_sp(72); | - | - | - | - | - | - | - | Unassigned | - |
| Otu04605 | 7 | k__Fungi(100);k__Fungi_unclassified(86);k__Fungi_unclassified(86);k__Fungi_unclassified(86);k__Fungi_unclassified(86);k__Fungi_unclassified(86);k__Fungi_unclassified(86); | - | - | - | - | - | - | - | Unassigned | - |
| Otu04606 | 7 | k__Fungi(100);p__Ascomycota(86);p__Ascomycota_unclassified(86);p__Ascomycota_unclassified(86);p__Ascomycota_unclassified(86);p__Ascomycota_unclassified(86);p__Ascomycota_unclassified(86); | - | - | - | - | - | - | - | Unassigned | - |
| Otu04608 | 7 | k__Fungi(100);p__Ascomycota(72);p__Ascomycota_unclassified(72);p__Ascomycota_unclassified(72);p__Ascomycota_unclassified(72);p__Ascomycota_unclassified(72);p__Ascomycota_unclassified(72); | - | - | - | - | - | - | - | Unassigned | - |
| Otu04612 | 7 | k__Fungi(100);p__Ascomycota(100);p__Ascomycota_unclassified(100);p__Ascomycota_unclassified(100);p__Ascomycota_unclassified(100);p__Ascomycota_unclassified(100);p__Ascomycota_unclassified(100); | - | - | - | - | - | - | - | Unassigned | - |
| Otu04613 | 7 | k__Fungi(100);k__Fungi_unclassified(72);k__Fungi_unclassified(72);k__Fungi_unclassified(72);k__Fungi_unclassified(72);k__Fungi_unclassified(72);k__Fungi_unclassified(72); | - | - | - | - | - | - | - | Unassigned | - |
| Otu04616 | 6 | k__Fungi(100);p__Basidiomycota(100);c__Agaricomycetes(100);o__Agaricales(100);f__Entolomataceae(100);g__Entoloma(100);g__Entoloma_unclassified(100); | Entoloma | 13 | Pathotroph-Saprotroph-Symbiotroph | Ectomycorrhizal-Fungal Parasite-\|Undefined Saprotroph\|-Undefined Symbiotroph | Agaricoid | NULL | Possible | As soil_saprotroph (Põlme et al. 2020); Decay roots, soil (Põlme et al. 2020) | Rinaldi AC, et al. 2008. Fungal Diversity 33:1-45 ((http://www.fungaldiversity.org/fdp/sfdp/33-1.pdf)) (as ECM/not ECM); Tedersoo L, et al. 2010. Mycorrhiza 20:217-263 ((https://doi.org/10.1007/s00572-009-0274-x; Põlme S, et al. 2020. Fungal Diversity 105:1-16 ((https://doi.org/10.1007/s13225-020-00466-2)); sensu stricto)); http://mycorrhizas.info/ecmf.html ((http://mycorrhizas.info/ecmf.html)) ((as ECM/not ECM)) |
| Otu04618 | 6 | k__Fungi(100);p__Ascomycota(100);c__Saccharomycetes(100);o__Saccharomycetales(100);f__Metschnikowiaceae(100);g__Metschnikowia(100);s__Metschnikowia_pulcherrima(67); | Metschnikowia | 13 | Saprotroph | \|Undefined Saprotroph\| | Yeast | NULL | Probable | As nectar-tap_saprotroph (Põlme et al. 2020); Decay sugar-rich substrates (Põlme et al. 2020) | Põlme S, et al. 2020. Fungal Diversity 105:1-16 ((https://doi.org/10.1007/s13225-020-00466-2)) |
| Otu04619 | 6 | k__Fungi(100);p__Ascomycota(100);c__Saccharomycetes(100);o__Saccharomycetales(100);f__Metschnikowiaceae(100);g__Metschnikowia(100);s__Metschnikowia_pulcherrima(100); | Metschnikowia | 13 | Saprotroph | \|Undefined Saprotroph\| | Yeast | NULL | Probable | As nectar-tap_saprotroph (Põlme et al. 2020); Decay sugar-rich substrates (Põlme et al. 2020) | Põlme S, et al. 2020. Fungal Diversity 105:1-16 ((https://doi.org/10.1007/s13225-020-00466-2)) |
| Otu04620 | 6 | k__Fungi(100);p__Ascomycota(100);c__Dothideomycetes(100);o__Dothideomycetes_order_Incertae_sedis(100);f__Eremomycetaceae(100);g__Arthrographis(100);g__Arthrographis_unclassified(84); | Arthrographis | 13 | Saprotroph | \|Undefined Saprotroph\| | Microfungus | NULL | Probable | As animal_decomposer (Põlme et al. 2020); Decay animal material (Põlme et al. 2020) | Tedersoo L, et al. 2014. Science 346:e1256688 ((https://doi.org/10.1126/science.1256688)); Põlme S, et al. 2020. Fungal Diversity 105:1-16 ((https://doi.org/10.1007/s13225-020-00466-2)) |
| Otu04622 | 6 | k__Fungi(100);p__Ascomycota(100);c__Saccharomycetes(100);o__Saccharomycetales(100);f__Metschnikowiaceae(100);g__Metschnikowia(100);s__Metschnikowia_pulcherrima(100); | Metschnikowia | 13 | Saprotroph | \|Undefined Saprotroph\| | Yeast | NULL | Probable | As nectar-tap_saprotroph (Põlme et al. 2020); Decay sugar-rich substrates (Põlme et al. 2020) | Põlme S, et al. 2020. Fungal Diversity 105:1-16 ((https://doi.org/10.1007/s13225-020-00466-2)) |
| Otu04623 | 6 | k__Fungi(100);p__Basidiomycota(100);c__Agaricomycetes(100);o__Corticiales(100);f__Corticiaceae(100);f__Corticiaceae_unclassified(100);f__Corticiaceae_unclassified(100); | Corticiaceae | 9 | Pathotroph-Saprotroph | Lichen Parasite-Plant Pathogen-Wood Saprotroph | Corticioid-Microfungus | NULL | Possible | Saprobes or pathogens on a wide range of mainly woody substrata; a few species are lichenicolous and some are grass pathogens (Cannon & Kirk 2007) | Cannon PF, Kirk PM. 2007. Fungal Families of the World. CAB International, Cambridge (ISBN: 978-0851998275) |
| Otu04624 | 6 | k__Fungi(100);p__Basidiomycota(100);c__Agaricomycetes(100);o__Thelephorales(100);f__Thelephoraceae(100);g__Tomentellopsis(100);s__Tomentellopsis_sp_DLL2011_1(100); | Tomentellopsis | 13 | Symbiotroph | \|Ectomycorrhizal\| | Corticioid | NULL | Highly Probable | NULL | Rinaldi AC, et al. 2008. Fungal Diversity 33:1-45 ((http://www.fungaldiversity.org/fdp/sfdp/33-1.pdf)); Tedersoo L, et al. 2010. Mycorrhiza 20:217-263 ((https://doi.org/10.1007/s00572-009-0274-x)); Põlme S, et al. 2020. Fungal Diversity 105:1-16 ((https://doi.org/10.1007/s13225-020-00466-2)); http://mycorrhizas.info/ecmf.html ((http://mycorrhizas.info/ecmf.html)) |
| Otu04626 | 6 | k__Fungi(100);p__Ascomycota(100);c__Saccharomycetes(100);o__Saccharomycetales(100);f__Metschnikowiaceae(100);g__Metschnikowia(100);g__Metschnikowia_unclassified(84); | Metschnikowia | 13 | Saprotroph | \|Undefined Saprotroph\| | Yeast | NULL | Probable | As nectar-tap_saprotroph (Põlme et al. 2020); Decay sugar-rich substrates (Põlme et al. 2020) | Põlme S, et al. 2020. Fungal Diversity 105:1-16 ((https://doi.org/10.1007/s13225-020-00466-2)) |
| Otu04627 | 6 | k__Fungi(100);p__Ascomycota(100);c__Dothideomycetes(100);o__Pleosporales(100);f__Pleosporaceae(100);g__Alternaria(100);s__Alternaria_porri(100); | Alternaria | 13 | Pathotroph-Saprotroph-Symbiotroph | Animal Pathogen-Endophyte-\|Plant Pathogen\|-Plant Saprotroph-Wood Saprotroph | Microfungus | Soft Rot | Possible | As foliar_endophyte (Põlme et al. 2020); As litter_saprotroph (Põlme et al. 2020); Decay leaf, fruit, seed (Põlme et al. 2020) | Duncan CG, Eslyn WE. 1966. Mycologia 58:642-664 ((https://doi.org/10.2307/3757045)); Seehann G, et al. 1975. List of Fungi in Soft-Rot Tissue Tests. Institute fur Holzbiologie und Holzschutz, Bundesforschungsanstalt fur Forst - und Holzwirtshaft, Hamburg - Lohbrugge, Germany ((https://www.irg-wp.com/irgdocs/details.php?d6493033-c9ea-40f3-8e3e-5697f32d640d)); Tedersoo L, et al. 2014. Science 346:e1256688 ((https://doi.org/10.1126/science.1256688)); Põlme S, et al. 2020. Fungal Diversity 105:1-16 ((https://doi.org/10.1007/s13225-020-00466-2)) |
| Otu04631 | 6 | k__Fungi(100);p__Ascomycota(100);c__Saccharomycetes(100);o__Saccharomycetales(100);f__Saccharomycetaceae(100);g__Kazachstania(100);s__Kazachstania_intestinalis(100); | Kazachstania | 13 | Saprotroph | \|Undefined Saprotroph\| | Yeast | NULL | Probable | As nectar-tap_saprotroph (Põlme et al. 2020); Decay sugar-rich substrates (Põlme et al. 2020) | Põlme S, et al. 2020. Fungal Diversity 105:1-16 ((https://doi.org/10.1007/s13225-020-00466-2)) |
| Otu04633 | 6 | k__Fungi(100);p__Ascomycota(100);c__Saccharomycetes(100);o__Saccharomycetales(100);f__Metschnikowiaceae(100);g__Metschnikowia(100);g__Metschnikowia_unclassified(100); | Metschnikowia | 13 | Saprotroph | \|Undefined Saprotroph\| | Yeast | NULL | Probable | As nectar-tap_saprotroph (Põlme et al. 2020); Decay sugar-rich substrates (Põlme et al. 2020) | Põlme S, et al. 2020. Fungal Diversity 105:1-16 ((https://doi.org/10.1007/s13225-020-00466-2)) |
| Otu04637 | 6 | k__Fungi(100);p__Ascomycota(100);c__Saccharomycetes(100);o__Saccharomycetales(100);f__Metschnikowiaceae(100);g__Metschnikowia(100);s__Metschnikowia_pulcherrima(100); | Metschnikowia | 13 | Saprotroph | \|Undefined Saprotroph\| | Yeast | NULL | Probable | As nectar-tap_saprotroph (Põlme et al. 2020); Decay sugar-rich substrates (Põlme et al. 2020) | Põlme S, et al. 2020. Fungal Diversity 105:1-16 ((https://doi.org/10.1007/s13225-020-00466-2)) |
| Otu04640 | 6 | k__Fungi(100);p__Ascomycota(100);c__Eurotiomycetes(100);o__Chaetothyriales(100);f__Herpotrichiellaceae(100);g__Exophiala(100);s__Exophiala_opportunistica(100); | Exophiala | 13 | Pathotroph-Saprotroph-Symbiotroph | \|Animal Parasite\|-Animal Pathogen-Endophyte-Plant Saprotroph-Undefined Saprotroph | Dimorphic-Facultative Yeast | NULL | Probable | As litter_saprotroph (Põlme et al. 2020); As root_endophyte_dark_septate (Põlme et al. 2020); Decay leaf, fruit, seed, animal material (Põlme et al. 2020) | Tedersoo L, et al. 2014. Science 346:e1256688 ((https://doi.org/10.1126/science.1256688)), Nagy LG, et al. 2014. Nature Communications 5:e4471 ((https://doi.org/10.1038/ncomms5471)); Põlme S, et al. 2020. Fungal Diversity 105:1-16 ((https://doi.org/10.1007/s13225-020-00466-2)) |
| Otu04641 | 6 | k__Fungi(100);p__Basidiomycota(100);c__Microbotryomycetes(100);o__Sporidiobolales(100);f__Sporidiobolales_family_Incertae_sedis(100);g__Sporobolomyces(100);s__Sporobolomyces_lactosus(100); | Sporobolomyces | 13 | Pathotroph-Saprotroph | \|Fungal Parasite\|-Plant Saprotroph | Yeast | NULL | Probable | As litter_saprotroph (Põlme et al. 2020); As mycoparasite (Põlme et al. 2020); Decay fungal material (Põlme et al. 2020) | Tedersoo L, et al. 2014. Science 346:e1256688 ((https://doi.org/10.1126/science.1256688)); Purahong W, et al. 2016. Molecular Ecology 25:4059-4074 ((https://doi.org/10.1111/mec.13739)); Põlme S, et al. 2020. Fungal Diversity 105:1-16 ((https://doi.org/10.1007/s13225-020-00466-2)) |
| Otu04645 | 6 | k__Fungi(100);p__Ascomycota(100);c__Saccharomycetes(100);o__Saccharomycetales(100);f__Metschnikowiaceae(100);g__Metschnikowia(100);s__Metschnikowia_pulcherrima(100); | Metschnikowia | 13 | Saprotroph | \|Undefined Saprotroph\| | Yeast | NULL | Probable | As nectar-tap_saprotroph (Põlme et al. 2020); Decay sugar-rich substrates (Põlme et al. 2020) | Põlme S, et al. 2020. Fungal Diversity 105:1-16 ((https://doi.org/10.1007/s13225-020-00466-2)) |
| Otu04646 | 6 | k__Fungi(100);p__Ascomycota(100);c__Saccharomycetes(100);o__Saccharomycetales(100);f__Metschnikowiaceae(100);g__Metschnikowia(100);g__Metschnikowia_unclassified(67); | Metschnikowia | 13 | Saprotroph | \|Undefined Saprotroph\| | Yeast | NULL | Probable | As nectar-tap_saprotroph (Põlme et al. 2020); Decay sugar-rich substrates (Põlme et al. 2020) | Põlme S, et al. 2020. Fungal Diversity 105:1-16 ((https://doi.org/10.1007/s13225-020-00466-2)) |
| Otu04647 | 6 | k__Fungi(100);p__Basidiomycota(100);c__Pucciniomycetes(100);o__Septobasidiales(100);f__Septobasidiaceae(100);g__Septobasidium(100);s__Septobasidium_sp_JBO_2010i(100); | Septobasidium | 13 | Pathotroph | \|Animal Parasite\|-Animal Pathogen-Arthropod Parasite | Dimorphic-Facultative Yeast | NULL | Probable | Decay animal material (Põlme et al. 2020); Host - scale_insects (Põlme et al. 2020) | Tedersoo L, et al. 2014. Science 346:e1256688 ((https://doi.org/10.1126/science.1256688)); Põlme S, et al. 2020. Fungal Diversity 105:1-16 ((https://doi.org/10.1007/s13225-020-00466-2)) |
| Otu04649 | 6 | k__Fungi(100);p__Ascomycota(100);c__Dothideomycetes(100);o__Capnodiales(100);f__Davidiellaceae(100);g__Cladosporium(100);s__Cladosporium_sphaerospermum(100); | Cladosporium | 13 | Pathotroph-Saprotroph-Symbiotroph | Animal Pathogen-Endophyte-Lichen Parasite-Plant Pathogen-\|Plant Saprotroph\|-Wood Saprotroph | Microfungus | NULL | Possible | As foliar_endophyte (Põlme et al. 2020); As litter_saprotroph (Põlme et al. 2020); Decay wood, leaf, fruit, seed, soil, dung, animal material (Põlme et al. 2020) | Seehann G, et al. 1975. List of Fungi in Soft-Rot Tissue Tests. Institute fur Holzbiologie und Holzschutz, Bundesforschungsanstalt fur Forst - und Holzwirtshaft, Hamburg - Lohbrugge, Germany; Lawrey JD, Diederich P. 2016. http://www.lichenicolous.net (accessed 24 May 2017) ((http://www.lichenicolous.net)); Põlme S, et al. 2020. Fungal Diversity 105:1-16 ((https://doi.org/10.1007/s13225-020-00466-2)); The American Phytopathological Society Common Names of Plant Diseases ((http://www.apsnet.org/publications/commonnames/Pages/default.aspx)) |
| Otu04651 | 6 | k__Fungi(100);p__Basidiomycota(100);c__Agaricomycetes(100);o__Agaricales(100);f__Agaricales_family_Incertae_sedis(100);g__Plicaturopsis(100);s__Plicaturopsis_crispa(100); | Plicaturopsis | 13 | Saprotroph | \|Wood Saprotroph\| | Corticioid-Polyporoid | White Rot | Probable | As white_rot (Põlme et al. 2020); Decay wood (Põlme et al. 2020) | Tedersoo L, et al. 2014. Science 346:e1256688 ((https://doi.org/10.1126/science.1256688)); Põlme S, et al. 2020. Fungal Diversity 105:1-16 ((https://doi.org/10.1007/s13225-020-00466-2)) |
| Otu04653 | 6 | k__Fungi(100);p__Ascomycota(100);c__Saccharomycetes(100);o__Saccharomycetales(100);f__Metschnikowiaceae(100);g__Metschnikowia(100);s__Metschnikowia_pulcherrima(67); | Metschnikowia | 13 | Saprotroph | \|Undefined Saprotroph\| | Yeast | NULL | Probable | As nectar-tap_saprotroph (Põlme et al. 2020); Decay sugar-rich substrates (Põlme et al. 2020) | Põlme S, et al. 2020. Fungal Diversity 105:1-16 ((https://doi.org/10.1007/s13225-020-00466-2)) |
| Otu04654 | 6 | k__Fungi(100);p__Basidiomycota(100);c__Agaricomycetes(100);o__Agaricales(100);f__Bolbitiaceae(100);g__Bolbitius(100);s__Bolbitius_psittacinus(100); | Bolbitius | 13 | Saprotroph | \|Dung Saprotroph\|-Plant Saprotroph | Agaricoid | NULL | Probable | As litter_saprotroph (Põlme et al. 2020); Decay leaf, fruit, seed, dung (Põlme et al. 2020) | Tedersoo L, et al. 2014. Science 346:e1256688 ((https://doi.org/10.1126/science.1256688)); Põlme S, et al. 2020. Fungal Diversity 105:1-16 ((https://doi.org/10.1007/s13225-020-00466-2)) |
| Otu04655 | 6 | k__Fungi(100);p__Ascomycota(100);c__Saccharomycetes(100);o__Saccharomycetales(100);f__Metschnikowiaceae(100);g__Metschnikowia(100);s__Metschnikowia_pulcherrima(100); | Metschnikowia | 13 | Saprotroph | \|Undefined Saprotroph\| | Yeast | NULL | Probable | As nectar-tap_saprotroph (Põlme et al. 2020); Decay sugar-rich substrates (Põlme et al. 2020) | Põlme S, et al. 2020. Fungal Diversity 105:1-16 ((https://doi.org/10.1007/s13225-020-00466-2)) |

| | | | | | | |
|---|---|---|---|---|---|---|
| Otu04656 | 6 | k__Fungi(100);p__Basidiomycota(100);c__Microbotryomycetes(100);o__Sporidiobolales(100);f__Sporidiobolales_family_Incertae_sedis(100);g__Sporobolomyces(100);s__Sporobolomyces_lactosus(100); | Sporobolomyces | 13 | Pathotroph-Saprotroph | [illegible] |
| Otu04658 | 6 | k__Fungi(100);p__Basidiomycota(100);c__Agaricomycetes(100);o__Agaricales(100);f__Cortinariaceae(100);g__Cortinarius(100);g__Cortinarius_unclassified(100); | Cortinarius | 13 | Symbiotroph | [illegible] |
| Otu04659 | 6 | k__Fungi(100);p__Basidiomycota(100);c__Agaricomycetes(100);o__Agaricales(100);f__Cortinariaceae(100);g__Cortinarius(100);g__Cortinarius_unclassified(100); | Cortinarius | 13 | Symbiotroph | [illegible] |
| Otu04660 | 6 | k__Fungi(100);p__Ascomycota(100);c__Saccharomycetes(100);o__Saccharomycetales(100);f__Metschnikowiaceae(100);g__Metschnikowia(100);s__Metschnikowia_pulcherrima(100); | Metschnikowia | 13 | Saprotroph | [illegible] |
| Otu04663 | 6 | k__Fungi(100);p__Basidiomycota(100);c__Agaricomycetes(100);o__Agaricales(100);o__Agaricales_unclassified(100);o__Agaricales_unclassified(100);o__Agaricales_unclassified(100); | Agaricales | 7 | Pathotroph-Saprotroph-Symbiotroph | [illegible] |
| Otu04665 | 6 | k__Fungi(100);p__Ascomycota(100);c__Sordariomycetes(100);o__Lulworthiales(100);f__Lulworthiaceae(100);g__Zalerion(100);s__Zalerion_arboricola(100); | Zalerion | 13 | Pathotroph-Saprotroph | [illegible] |
| Otu04666 | 6 | k__Fungi(100);p__Ascomycota(100);c__Saccharomycetes(100);o__Saccharomycetales(100);f__Metschnikowiaceae(100);g__Metschnikowia(100);s__Metschnikowia_pulcherrima(100); | Metschnikowia | 13 | Saprotroph | [illegible] |
| Otu04671 | 6 | k__Fungi(100);p__Ascomycota(100);c__Leotiomycetes(100);o__Helotiales(100);f__Vibrisseaceae(100);f__Vibrisseaceae_unclassified(100);f__Vibrisseaceae_unclassified(100); | Vibrisseaceae | 9 | Saprotroph-Symbiotroph | [illegible] |
| Otu04673 | 6 | k__Fungi(100);p__Ascomycota(100);c__Sordariomycetes(100);o__Hypocreales(100);o__Hypocreales_unclassified(100);o__Hypocreales_unclassified(100);o__Hypocreales_unclassified(100); | Hypocreales | 7 | Saprotroph | [illegible] |
| Otu04674 | 6 | k__Fungi(100);p__Ascomycota(100);c__Sordariomycetes(100);o__Hypocreales(100);f__Clavicipitaceae(100);g__Pochonia(100);s__Pochonia_chlamydosporia(100); | Pochonia | 13 | Pathotroph-Saprotroph | [illegible] |
| Otu04677 | 6 | k__Fungi(100);p__Ascomycota(100);c__Sordariomycetes(100);o__Diaporthales(100);f__Melanconidaceae(100);g__Melanconiella(100);s__Melanconiella_elegans(100); | Melanconiella | 13 | Saprotroph | [illegible] |
| Otu04678 | 6 | k__Fungi(100);p__Ascomycota(100);c__Sordariomycetes(100);o__Hypocreales(100);f__Nectriaceae(100);g__Gibberella(84);g__Gibberella_unclassified(84); | Gibberella | 13 | Pathotroph-Saprotroph-Symbiotroph | [illegible] |
| Otu04679 | 6 | k__Fungi(100);p__Ascomycota(100);c__Saccharomycetes(100);o__Saccharomycetales(100);f__Metschnikowiaceae(100);g__Metschnikowia(100);s__Metschnikowia_pulcherrima(84); | Metschnikowia | 13 | Saprotroph | [illegible] |
| Otu04681 | 6 | k__Fungi(100);p__Ascomycota(100);c__Dothideomycetes(100);o__Capnodiales(100);f__Capnodiales_family_Incertae_sedis(100);g__Rachicladosporium(67);g__Rachicladosporium_unclassified(67); | Rachicladosporium | 13 | Saprotroph | [illegible] |
| Otu04682 | 6 | k__Fungi(100);p__Ascomycota(100);c__Saccharomycetes(100);o__Saccharomycetales(100);f__Metschnikowiaceae(100);g__Metschnikowia(100);s__Metschnikowia_pulcherrima(100); | Metschnikowia | 13 | Saprotroph | [illegible] |
| Otu04684 | 6 | k__Fungi(100);p__Ascomycota(100);c__Eurotiomycetes(100);o__Chaetothyriales(100);f__Herpotrichiellaceae(100);f__Herpotrichiellaceae_unclassified(100);f__Herpotrichiellaceae_unclassified(100); | Herpotrichiellaceae | 9 | Pathotroph-Saprotroph | [illegible] |
| Otu04687 | 6 | k__Fungi(100);p__Ascomycota(100);c__Sordariomycetes(100);o__Hypocreales(100);f__Hypocreales_family_Incertae_sedis(100);f__Hypocreales_family_Incertae_sedis_unclassified(84);f__Hypocreales_family_Incertae_sedis_unclassified(84); | Hypocreales | 7 | Saprotroph | [illegible] |
| Otu04689 | 6 | k__Fungi(100);p__Ascomycota(100);c__Dothideomycetes(100);o__Capnodiales(100);f__Mycosphaerellaceae(100);f__Mycosphaerellaceae_unclassified(84);f__Mycosphaerellaceae_unclassified(84); | Mycosphaerellaceae | 9 | Pathotroph-Saprotroph | [illegible] |
| Otu04690 | 6 | k__Fungi(100);p__Ascomycota(100);c__Saccharomycetes(100);o__Saccharomycetales(100);f__Metschnikowiaceae(100);g__Metschnikowia(100);s__Metschnikowia_pulcherrima(100); | Metschnikowia | 13 | Saprotroph | [illegible] |
| Otu04691 | 6 | k__Fungi(100);p__Ascomycota(100);c__Saccharomycetes(100);o__Saccharomycetales(100);f__Metschnikowiaceae(100);g__Metschnikowia(100);s__Metschnikowia_pulcherrima(84); | Metschnikowia | 13 | Saprotroph | [illegible] |
| Otu04692 | 6 | k__Fungi(100);p__Basidiomycota(100);c__Agaricomycetes(100);o__Hymenochaetales(100);f__Schizoporaceae(100);g__Schizopora(84);s__Schizopora_paradoxa(84); | Schizopora | 13 | Pathotroph-Saprotroph | [illegible] |
| Otu04694 | 6 | k__Fungi(100);p__Basidiomycota(100);c__Tremellomycetes(100);o__Filobasidiales(100);f__Filobasidiaceae(100);g__Cryptococcus(100);s__Cryptococcus_heimaeyensis(100); | Cryptococcus | 13 | Pathotroph-Saprotroph-Symbiotroph | [illegible] |
| Otu04695 | 6 | k__Fungi(100);p__Ascomycota(100);c__Saccharomycetes(100);o__Saccharomycetales(100);f__Metschnikowiaceae(100);g__Metschnikowia(100);s__Metschnikowia_pulcherrima(100); | Metschnikowia | 13 | Saprotroph | [illegible] |
| Otu04696 | 6 | k__Fungi(100);p__Ascomycota(100);c__Saccharomycetes(100);o__Saccharomycetales(100);f__Metschnikowiaceae(100);g__Metschnikowia(100);s__Metschnikowia_pulcherrima(100); | Metschnikowia | 13 | Saprotroph | [illegible] |
| Otu04697 | 6 | k__Fungi(100);p__Ascomycota(100);c__Saccharomycetes(100);o__Saccharomycetales(100);f__Metschnikowiaceae(100);g__Metschnikowia(100);s__Metschnikowia_pulcherrima(84); | Metschnikowia | 13 | Saprotroph | [illegible] |
| Otu04698 | 6 | k__Fungi(100);p__Basidiomycota(100);c__Agaricomycetes(100);o__Agaricales(100);o__Agaricales_unclassified(100);o__Agaricales_unclassified(100);o__Agaricales_unclassified(100); | Agaricales | 7 | Pathotroph-Saprotroph-Symbiotroph | [illegible] |
| Otu04699 | 6 | k__Fungi(100);p__Basidiomycota(100);c__Agaricomycetes(100);o__Agaricales(100);f__Pluteaceae(100);g__Pluteus(100);g__Pluteus_unclassified(100); | Pluteus | 13 | Pathotroph-Saprotroph | [illegible] |
| Otu04701 | 6 | k__Fungi(100);p__Ascomycota(100);c__Eurotiomycetes(100);o__Arachnomycetales(84);f__Arachnomycetaceae(84);g__Arachnomyces(84);s__Arachnomyces_nodosetosus(84); | Arachnomyces | 13 | Pathotroph-Saprotroph | [illegible] |
| Otu04702 | 6 | k__Fungi(100);p__Ascomycota(100);c__Eurotiomycetes(100);o__Eurotiales(100);f__Trichocomaceae(100);g__Aspergillus(100);s__Aspergillus_flavus(100); | Aspergillus | 13 | Pathotroph-Saprotroph-Symbiotroph | [illegible] |
| Otu04706 | 6 | k__Fungi(100);p__Ascomycota(100);c__Sordariomycetes(100);o__Hypocreales(100);f__Nectriaceae(100);g__Gibberella(100);s__Gibberella_intricans(100); | Gibberella | 13 | Pathotroph-Saprotroph-Symbiotroph | [illegible] |
| Otu04707 | 6 | k__Fungi(100);p__Basidiomycota(100);c__Agaricomycetes(100);o__Thelephorales(100);f__Thelephoraceae(100);g__Tomentella(100);s__Tomentella_coerulea(100); | Tomentella | 13 | Symbiotroph | [illegible] |
| Otu04709 | 6 | k__Fungi(100);p__Ascomycota(100);c__Saccharomycetes(100);o__Saccharomycetales(100);f__Metschnikowiaceae(100);g__Metschnikowia(100);g__Metschnikowia_unclassified(67); | Metschnikowia | 13 | Saprotroph | [illegible] |
| Otu04712 | 6 | k__Fungi(100);p__Basidiomycota(100);c__Cystobasidiomycetes(100);o__Erythrobasidiales(100);f__Erythrobasidiales_family_Incertae_sedis(100);g__Bannoa(100);s__Bannoa_hahajimensis(100); | Bannoa | 13 | Saprotroph | [illegible] |
| Otu04713 | 6 | k__Fungi(100);p__Ascomycota(100);c__Leotiomycetes(100);o__Helotiales(100);f__Hyaloscyphaceae(100);g__Lachnum(100);s__Lachnum_sp_1_KO_2013(67); | Lachnum | 13 | Saprotroph-Symbiotroph | [illegible] |
| Otu04716 | 6 | k__Fungi(100);p__Ascomycota(100);c__Dothideomycetes(100);o__Capnodiales(100);f__Mycosphaerellaceae(100);f__Mycosphaerellaceae_unclassified(100);f__Mycosphaerellaceae_unclassified(100); | Mycosphaerellaceae | 9 | Pathotroph-Saprotroph | [illegible] |
| Otu04718 | 6 | k__Fungi(100);p__Basidiomycota(100);c__Agaricomycetes(100);o__Boletales(100);f__Suillaceae(100);g__Suillus(100);g__Suillus_unclassified(100); | Suillus | 13 | Symbiotroph | [illegible] |
| Otu04719 | 6 | k__Fungi(100);p__Basidiomycota(100);c__Agaricomycetes(100);o__Hymenochaetales(100);f__Schizoporaceae(100);g__Hyphodontia(100);g__Hyphodontia_unclassified(100); | Hyphodontia | 13 | Saprotroph | [illegible] |
| Otu04721 | 6 | k__Fungi(100);p__Ascomycota(100);c__Sordariomycetes(100);o__Sordariales(100);f__Cephalothecaceae(100);g__Phialemonium(100);s__Phialemonium_atrogriseum(100); | Phialemonium | 13 | Pathotroph-Saprotroph | [illegible] |
| Otu04722 | 6 | k__Fungi(100);p__Ascomycota(100);c__Saccharomycetes(100);o__Saccharomycetales(100);f__Metschnikowiaceae(100);g__Metschnikowia(100);s__Metschnikowia_pulcherrima(100); | Metschnikowia | 13 | Saprotroph | [illegible] |
| Otu04724 | 6 | k__Fungi(100);p__Basidiomycota(100);c__Microbotryomycetes(100);o__Sporidiobolales(100);f__Sporidiobolales_family_Incertae_sedis(100);g__Rhodotorula(100);g__Rhodotorula_unclassified(67); | Rhodotorula | 13 | Pathotroph-Saprotroph-Symbiotroph | [illegible] |
| Otu04725 | 6 | k__Fungi(100);p__Ascomycota(84);c__Saccharomycetes(84);o__Saccharomycetales(84);f__Metschnikowiaceae(84);g__Metschnikowia(84);s__Metschnikowia_pulcherrima(67); | Metschnikowia | 13 | Saprotroph | [illegible] |
| Otu04726 | 6 | k__Fungi(100);p__Chytridiomycota(100);c__Chytridiomycetes(100);o__Spizellomycetales(100);f__Spizellomycetaceae(100);g__Powellomyces(100);g__Powellomyces_unclassified(100); | Powellomyces | 13 | Pathotroph-Saprotroph | [illegible] |
| Otu04727 | 6 | k__Fungi(100);p__Basidiomycota(100);c__Microbotryomycetes(100);o__Sporidiobolales(100);f__Sporidiobolales_family_Incertae_sedis(100);g__Sporobolomyces(100);s__Sporobolomyces_lactosus(100); | Sporobolomyces | 13 | Pathotroph-Saprotroph | [illegible] |
| Otu04729 | 6 | k__Fungi(100);p__Ascomycota(100);c__Saccharomycetes(100);o__Saccharomycetales(100);f__Metschnikowiaceae(100);g__Metschnikowia(100);s__Metschnikowia_pulcherrima(100); | Metschnikowia | 13 | Saprotroph | [illegible] |
| Otu04733 | 6 | k__Fungi(100);p__Zygomycota(100);c__Zygomycota_class_Incertae_sedis(100);o__Mortierellales(100);f__Mortierellaceae(100);g__Mortierella(100);s__Mortierella_sp(100); | Mortierella | 13 | Saprotroph-Symbiotroph | [illegible] |
| Otu04734 | 6 | k__Fungi(100);p__Ascomycota(100);c__Saccharomycetes(100);o__Saccharomycetales(100);f__Metschnikowiaceae(100);g__Metschnikowia(100);s__Metschnikowia_pulcherrima(84); | Metschnikowia | 13 | Saprotroph | [illegible] |
| Otu04735 | 6 | k__Fungi(100);p__Ascomycota(100);c__Saccharomycetes(100);o__Saccharomycetales(100);f__Metschnikowiaceae(100);g__Metschnikowia(100);s__Metschnikowia_pulcherrima(84); | Metschnikowia | 13 | Saprotroph | [illegible] |
| Otu04737 | 6 | k__Fungi(100);p__Ascomycota(100);c__Saccharomycetes(100);o__Saccharomycetales(100);f__Metschnikowiaceae(100);g__Metschnikowia(100);s__Metschnikowia_pulcherrima(100); | Metschnikowia | 13 | Saprotroph | [illegible] |
| Otu04739 | 6 | k__Fungi(100);p__Basidiomycota(100);c__Microbotryomycetes(100);o__Sporidiobolales(100);f__Sporidiobolales_family_Incertae_sedis(100);g__Sporobolomyces(100);s__Sporobolomyces_lactosus(100); | Sporobolomyces | 13 | Pathotroph-Saprotroph | [illegible] |
| Otu04741 | 6 | k__Fungi(100);p__Ascomycota(100);c__Saccharomycetes(100);o__Saccharomycetales(100);f__Metschnikowiaceae(100);g__Metschnikowia(100);s__Metschnikowia_pulcherrima(100); | Metschnikowia | 13 | Saprotroph | [illegible] |
| Otu04743 | 6 | k__Fungi(100);p__Ascomycota(100);c__Leotiomycetes(100);o__Helotiales(100);f__Helotiales_family_Incertae_sedis(100);g__Hyphodiscus(100);s__Hyphodiscus_hymeniophilus(100); | Hyphodiscus | 13 | Pathotroph-Saprotroph | [illegible] |
| Otu04744 | 6 | k__Fungi(100);p__Ascomycota(100);c__Sordariomycetes(100);o__Hypocreales(100);f__Nectriaceae(100);f__Nectriaceae_unclassified(100);f__Nectriaceae_unclassified(100); | Nectriaceae | 9 | Pathotroph-Saprotroph-Symbiotroph | [illegible] |
| Otu04746 | 6 | k__Fungi(100);p__Basidiomycota(100);c__Pucciniomycetes(100);o__Pucciniales(100);f__Pucciniaceae(100);f__Pucciniaceae_unclassified(100);f__Pucciniaceae_unclassified(100); | Pucciniaceae | 9 | Pathotroph | [illegible] |
| Otu04747 | 6 | k__Fungi(100);p__Ascomycota(100);c__Saccharomycetes(100);o__Saccharomycetales(100);f__Pichiaceae(100);g__Ogataea(100);g__Ogataea_unclassified(100); | Ogataea | 13 | Saprotroph | [illegible] |
| Otu04748 | 6 | k__Fungi(100);p__Ascomycota(100);c__Saccharomycetes(100);o__Saccharomycetales(100);f__Saccharomycetales_family_Incertae_sedis(100);g__Candida(100);g__Candida_unclassified(100); | Candida | 13 | Pathotroph-Saprotroph-Symbiotroph | [illegible] |
| Otu04749 | 6 | k__Fungi(100);p__Ascomycota(100);c__Saccharomycetes(100);o__Saccharomycetales(100);f__Metschnikowiaceae(100);g__Metschnikowia(100);s__Metschnikowia_pulcherrima(100); | Metschnikowia | 13 | Saprotroph | [illegible] |
| Otu04751 | 6 | k__Fungi(100);p__Ascomycota(100);c__Saccharomycetes(100);o__Saccharomycetales(100);f__Metschnikowiaceae(100);g__Metschnikowia(100);g__Metschnikowia_unclassified(67); | Metschnikowia | 13 | Saprotroph | [illegible] |
| Otu04753 | 6 | k__Fungi(100);p__Ascomycota(100);c__Saccharomycetes(100);o__Saccharomycetales(100);f__Metschnikowiaceae(100);g__Metschnikowia(100);g__Metschnikowia_unclassified(100); | Metschnikowia | 13 | Saprotroph | [illegible] |
| Otu04757 | 6 | k__Fungi(100);p__Ascomycota(100);c__Saccharomycetes(100);o__Saccharomycetales(100);f__Metschnikowiaceae(100);g__Metschnikowia(100);g__Metschnikowia_unclassified(84); | Metschnikowia | 13 | Saprotroph | [illegible] |
| Otu04759 | 6 | k__Fungi(100);p__Basidiomycota(100);c__Agaricomycetes(100);o__Auriculariales(100);f__Auriculariales_family_Incertae_sedis(100);g__Basidiodendron(100);g__Basidiodendron_unclassified(100); | Basidiodendron | 13 | Saprotroph | [illegible] |
| Otu04761 | 6 | k__Fungi(100);p__Basidiomycota(100);c__Microbotryomycetes(100);o__Sporidiobolales(100);f__Sporidiobolales_family_Incertae_sedis(100);g__Rhodotorula(100);g__Rhodotorula_unclassified(100); | Rhodotorula | 13 | Pathotroph-Saprotroph-Symbiotroph | [illegible] |
| Otu04762 | 6 | k__Fungi(100);p__Basidiomycota(100);c__Agaricomycetes(100);o__Russulales(100);f__Bondarzewiaceae(100);g__Gloiodon(100);s__Gloiodon_strigosus(100); | Gloiodon | 13 | Saprotroph | [illegible] |
| Otu04763 | 6 | k__Fungi(100);p__Basidiomycota(100);c__Agaricomycetes(100);o__Thelephorales(100);f__Thelephoraceae(100);f__Thelephoraceae_unclassified(100);f__Thelephoraceae_unclassified(100); | Thelephoraceae | 9 | Saprotroph-Symbiotroph | [illegible] |
| Otu04764 | 6 | k__Fungi(100);p__Basidiomycota(100);c__Tremellomycetes(100);o__Tremellales(100);f__Tremellales_family_Incertae_sedis(100);f__Tremellales_family_Incertae_sedis_unclassified(100);f__Tremellales_family_Incertae_sedis_unclassified(100); | Tremellales | 7 | Pathotroph-Saprotroph-Symbiotroph | [illegible] |
| Otu04766 | 6 | k__Fungi(100);p__Ascomycota(100);c__Saccharomycetes(100);o__Saccharomycetales(100);f__Metschnikowiaceae(100);g__Metschnikowia(100);g__Metschnikowia_unclassified(100); | Metschnikowia | 13 | Saprotroph | [illegible] |
| Otu04768 | 6 | k__Fungi(100);p__Basidiomycota(100);c__Agaricomycetes(100);o__Agaricales(100);f__Entolomataceae(100);g__Entoloma(100);s__Entoloma_sp_AJ108(100); | Entoloma | 13 | Pathotroph-Saprotroph-Symbiotroph | [illegible] |
| Otu04770 | 6 | k__Fungi(100);p__Basidiomycota(100);c__Agaricomycetes(100);o__Hymenochaetales(100);f__Hymenochaetaceae(100);g__Hymenochaete(100);g__Hymenochaete_unclassified(100); | Hymenochaete | 13 | Pathotroph-Saprotroph | [illegible] |
| Otu04772 | 6 | k__Fungi(100);p__Ascomycota(100);c__Schizosaccharomycetes(100);o__Schizosaccharomycetales(100);f__Schizosaccharomycetaceae(100);g__Schizosaccharomyces(100);s__Schizosaccharomyces_cryophilus(100); | Schizosaccharomyces | 13 | Saprotroph | [illegible] |
| Otu04773 | 6 | k__Fungi(100);p__Ascomycota(100);c__Saccharomycetes(100);o__Saccharomycetales(100);f__Saccharomycetales_family_Incertae_sedis(100);g__Candida(84);g__Candida_unclassified(84); | Candida | 13 | Pathotroph-Saprotroph-Symbiotroph | [illegible] |
| Otu04774 | 6 | k__Fungi(100);p__Ascomycota(100);c__Dothideomycetes(100);o__Capnodiales(100);f__Teratosphaeriaceae(100);g__Devriesia(100);s__Devriesia_fraseriae(100); | Devriesia | 13 | Pathotroph-Saprotroph | [illegible] |
| Otu04777 | 6 | k__Fungi(100);p__Ascomycota(100);c__Saccharomycetes(100);o__Saccharomycetales(100);f__Metschnikowiaceae(100);g__Metschnikowia(100);s__Metschnikowia_pulcherrima(84); | Metschnikowia | 13 | Saprotroph | [illegible] |
| Otu04778 | 6 | k__Fungi(100);p__Basidiomycota(100);c__Agaricomycetes(100);o__Hymenochaetales(100);f__Hymenochaetaceae(100);g__Inonotus(100);g__Inonotus_unclassified(100); | Inonotus | 13 | Pathotroph-Saprotroph | [illegible] |
| Otu04783 | 6 | k__Fungi(100);p__Ascomycota(100);c__Saccharomycetes(100);o__Saccharomycetales(100);f__Metschnikowiaceae(100);g__Metschnikowia(100);s__Metschnikowia_pulcherrima(100); | Metschnikowia | 13 | Saprotroph | [illegible] |
| Otu04784 | 6 | k__Fungi(100);p__Ascomycota(100);c__Eurotiomycetes(100);o__Eurotiales(100);f__Trichocomaceae(100);g__Penicillium(100);g__Penicillium_unclassified(100); | Penicillium | 13 | Pathotroph-Saprotroph-Symbiotroph | [illegible] |
| Otu04786 | 6 | k__Fungi(100);p__Ascomycota(100);c__Lecanoromycetes(100);o__Lecanorales(100);f__Cladoniaceae(100);g__Cladonia(100);g__Cladonia_unclassified(100); | Cladonia | 13 | Symbiotroph | [illegible] |
| Otu04788 | 6 | k__Fungi(100);p__Basidiomycota(100);c__Microbotryomycetes(100);o__Sporidiobolales(100);f__Sporidiobolales_family_Incertae_sedis(100);g__Sporobolomyces(100);s__Sporobolomyces_lactosus(100); | Sporobolomyces | 13 | Pathotroph-Saprotroph | [illegible] |
| Otu04791 | 6 | k__Fungi(100);p__Ascomycota(100);c__Sordariomycetes(100);o__Xylariales(100);f__Xylariaceae(100);g__Nemania(100);s__Nemania_aenea(100); | Nemania | 13 | Saprotroph | [illegible] |
| Otu04792 | 6 | k__Fungi(100);p__Ascomycota(100);c__Leotiomycetes(100);o__Leotiales(100);f__Bulgariaceae(100);g__Bulgaria(100);s__Bulgaria_inquinans(100); | Bulgaria | 13 | Pathotroph-Saprotroph-Symbiotroph | [illegible] |
| Otu04794 | 6 | k__Fungi(100);p__Ascomycota(100);c__Saccharomycetes(100);o__Saccharomycetales(100);f__Metschnikowiaceae(100);g__Metschnikowia(100);s__Metschnikowia_pulcherrima(100); | Metschnikowia | 13 | Saprotroph | [illegible] |
| Otu04799 | 6 | k__Fungi(100);p__Ascomycota(100);c__Saccharomycetes(84);o__Saccharomycetales(84);f__Saccharomycetales_family_Incertae_sedis(84);g__Candida(84);g__Candida_unclassified(67); | Candida | 13 | Pathotroph-Saprotroph-Symbiotroph | [illegible] |
| Otu04800 | 6 | k__Fungi(100);p__Ascomycota(100);c__Saccharomycetes(100);o__Saccharomycetales(100);f__Metschnikowiaceae(100);g__Metschnikowia(100);s__Metschnikowia_pulcherrima(100); | Metschnikowia | 13 | Saprotroph | [illegible] |
| Otu04802 | 6 | k__Fungi(100);p__Ascomycota(100);c__Saccharomycetes(100);o__Saccharomycetales(100);f__Metschnikowiaceae(100);g__Metschnikowia(100);g__Metschnikowia_unclassified(84); | Metschnikowia | 13 | Saprotroph | [illegible] |
| Otu04805 | 6 | k__Fungi(100);p__Ascomycota(100);c__Saccharomycetes(100);o__Saccharomycetales(100);f__Metschnikowiaceae(100);g__Metschnikowia(100);s__Metschnikowia_pulcherrima(100); | Metschnikowia | 13 | Saprotroph | [illegible] |
| Otu04806 | 6 | k__Fungi(100);p__Ascomycota(100);c__Saccharomycetes(100);o__Saccharomycetales(100);f__Metschnikowiaceae(100);g__Metschnikowia(100);s__Metschnikowia_pulcherrima(100); | Metschnikowia | 13 | Saprotroph | [illegible] |
| Otu04807 | 6 | k__Fungi(100);p__Ascomycota(100);c__Sordariomycetes(100);o__Hypocreales(100);o__Hypocreales_unclassified(100);o__Hypocreales_unclassified(100);o__Hypocreales_unclassified(100); | Hypocreales | 7 | Saprotroph | [illegible] |
| Otu04811 | 6 | k__Fungi(100);p__Ascomycota(100);c__Dothideomycetes(100);o__Capnodiales(100);f__Mycosphaerellaceae(100);g__Mycosphaerella(100);s__Mycosphaerella_nyssicola(100); | Mycosphaerella | 13 | Pathotroph-Saprotroph | [illegible] |
| Otu04813 | 6 | k__Fungi(100);p__Ascomycota(100);c__Saccharomycetes(100);o__Saccharomycetales(100);f__Metschnikowiaceae(100);g__Metschnikowia(100);g__Metschnikowia_unclassified(84); | Metschnikowia | 13 | Saprotroph | [illegible] |
| Otu04814 | 6 | k__Fungi(100);p__Ascomycota(100);c__Saccharomycetes(100);o__Saccharomycetales(100);f__Metschnikowiaceae(100);g__Metschnikowia(100);s__Metschnikowia_pulcherrima(67); | Metschnikowia | 13 | Saprotroph | [illegible] |
| Otu04816 | 6 | k__Fungi(100);p__Basidiomycota(100);c__Agaricomycetes(100);o__Polyporales(100);f__Meruliaceae(100);g__Hypochnicium(100);g__Hypochnicium_unclassified(100); | Hypochnicium | 13 | Saprotroph | [illegible] |
| Otu04817 | 6 | k__Fungi(100);p__Basidiomycota(100);c__Agaricomycetes(100);o__Hymenochaetales(100);f__Schizoporaceae(100);g__Hyphodontia(100);s__Hyphodontia_floccosa(100); | Hyphodontia | 13 | Saprotroph | [illegible] |
| Otu04819 | 6 | k__Fungi(100);p__Ascomycota(100);c__Saccharomycetes(100);o__Saccharomycetales(100);f__Metschnikowiaceae(100);g__Metschnikowia(100);g__Metschnikowia_unclassified(84); | Metschnikowia | 13 | Saprotroph | [illegible] |
| Otu04821 | 6 | k__Fungi(100);p__Ascomycota(100);c__Saccharomycetes(100);o__Saccharomycetales(100);f__Metschnikowiaceae(100);g__Metschnikowia(100);s__Metschnikowia_pulcherrima(84); | Metschnikowia | 13 | Saprotroph | [illegible] |
| Otu04822 | 6 | k__Fungi(100);p__Basidiomycota(100);c__Agaricomycetes(100);o__Agaricales(100);o__Agaricales_unclassified(100);o__Agaricales_unclassified(100);o__Agaricales_unclassified(100); | Agaricales | 7 | Pathotroph-Saprotroph-Symbiotroph | [illegible] |
| Otu04824 | 6 | k__Fungi(100);p__Basidiomycota(100);c__Agaricomycetes(100);o__Trechisporales(100);f__Hydnodontaceae(100);g__Trechispora(100);g__Trechispora_unclassified(100); | Trechispora | 13 | Saprotroph | [illegible] |
| Otu04825 | 6 | k__Fungi(100);p__Basidiomycota(100);c__Agaricomycetes(100);o__Agaricales(100);f__Inocybaceae(100);g__Inocybe(100);s__Inocybe_sindonia(100); | Inocybe | 13 | Symbiotroph | [illegible] |
| Otu04827 | 6 | k__Fungi(100);p__Zygomycota(100);c__Zygomycota_class_Incertae_sedis(100);o__Mucorales(100);f__Cunninghamellaceae(100);g__Gongronella(100);s__Gongronella_sp_11_KF_2011a(100); | Gongronella | 13 | Saprotroph | [illegible] |
| Otu04829 | 6 | k__Fungi(100);p__Basidiomycota(100);c__Agaricomycetes(100);o__Agaricales(100);f__Inocybaceae(100);g__Inocybe(100);g__Inocybe_unclassified(100); | Inocybe | 13 | Symbiotroph | [illegible] |
| Otu04830 | 6 | k__Fungi(100);p__Basidiomycota(100);c__Agaricomycetes(100);o__Agaricales(100);f__Inocybaceae(100);g__unclassified_Inocybaceae(100);s__Inocybaceae_sp(100); | Inocybaceae | 9 | Symbiotroph | [illegible] |
| Otu04832 | 6 | k__Fungi(100);p__Ascomycota(100);c__Saccharomycetes(100);o__Saccharomycetales(100);f__Metschnikowiaceae(100);g__Metschnikowia(100);s__Metschnikowia_pulcherrima(67); | Metschnikowia | 13 | Saprotroph | [illegible] |
| Otu04833 | 6 | k__Fungi(100);p__Basidiomycota(100);c__Agaricomycetes(100);o__Agaricales(100);f__Agaricaceae(100);g__Agaricus(100);g__Agaricus_unclassified(100); | Agaricus | 13 | Saprotroph | [illegible] |
| Otu04834 | 6 | k__Fungi(100);p__Ascomycota(100);c__Saccharomycetes(100);o__Saccharomycetales(100);f__Metschnikowiaceae(100);g__Metschnikowia(100);s__Metschnikowia_pulcherrima(84); | Metschnikowia | 13 | Saprotroph | [illegible] |
| Otu04836 | 6 | k__Fungi(100);p__Ascomycota(100);c__Saccharomycetes(100);o__Saccharomycetales(100);f__Metschnikowiaceae(100);g__Metschnikowia(100);g__Metschnikowia_unclassified(100); | Metschnikowia | 13 | Saprotroph | [illegible] |
| Otu04838 | 6 | k__Fungi(100);p__Ascomycota(100);c__Dothideomycetes(100);o__Pleosporales(100);f__Sporormiaceae(100);g__Preussia(100);s__Preussia_sp_2_JAS_2013(100); | Preussia | 13 | Saprotroph | [illegible] |
| Otu04839 | 6 | k__Fungi(100);p__Ascomycota(100);c__Dothideomycetes(100);o__Dothideomycetes_order_Incertae_sedis(100);f__Myxotrichaceae(100);g__Oidiodendron(100);g__Oidiodendron_unclassified(67); | Oidiodendron | 13 | Pathotroph-Saprotroph-Symbiotroph | [illegible] |
| Otu04843 | 6 | k__Fungi(100);p__Ascomycota(100);c__Saccharomycetes(100);o__Saccharomycetales(100);f__Metschnikowiaceae(100);g__Metschnikowia(100);g__Metschnikowia_unclassified(84); | Metschnikowia | 13 | Saprotroph | [illegible] |
| Otu04845 | 6 | k__Fungi(100);p__Ascomycota(100);c__Eurotiomycetes(100);o__Chaetothyriales(100);f__Chaetothyriales_family_Incertae_sedis(100);g__Strelitziana(100);s__Strelitziana_mali(100); | Strelitziana | 13 | Pathotroph | [illegible] |
| Otu04846 | 6 | k__Fungi(100);p__Ascomycota(100);c__Sordariomycetes(100);o__Diaporthales(100);f__Gnomoniaceae(84);f__Gnomoniaceae_unclassified(84);f__Gnomoniaceae_unclassified(84); | Gnomoniaceae | 9 | Pathotroph-Saprotroph-Symbiotroph | [illegible] |
| Otu04847 | 6 | k__Fungi(100);p__Ascomycota(100);c__Saccharomycetes(100);o__Saccharomycetales(100);f__Metschnikowiaceae(100);g__Metschnikowia(100);s__Metschnikowia_pulcherrima(67); | Metschnikowia | 13 | Saprotroph | [illegible] |
| Otu04849 | 6 | k__Fungi(100);p__Ascomycota(100);c__Eurotiomycetes(100);o__Chaetothyriales(100);f__Herpotrichiellaceae(100);g__Rhinocladiella(100);s__Rhinocladiella_sp_VL271(100); | Rhinocladiella | 13 | Pathotroph-Saprotroph-Symbiotroph | [illegible] |
| Otu04853 | 6 | k__Fungi(100);p__Ascomycota(100);c__Saccharomycetes(100);o__Saccharomycetales(100);f__Metschnikowiaceae(100);g__Metschnikowia(100);s__Metschnikowia_pulcherrima(84); | Metschnikowia | 13 | Saprotroph | [illegible] |
| Otu04854 | 6 | k__Fungi(100);p__Ascomycota(100);c__Saccharomycetes(100);o__Saccharomycetales(100);f__Metschnikowiaceae(100);g__Metschnikowia(100);g__Metschnikowia_unclassified(67); | Metschnikowia | 13 | Saprotroph | [illegible] |
| Otu04855 | 6 | k__Fungi(100);p__Ascomycota(100);c__Saccharomycetes(100);o__Saccharomycetales(100);f__Metschnikowiaceae(100);g__Metschnikowia(100);s__Metschnikowia_pulcherrima(84); | Metschnikowia | 13 | Saprotroph | [illegible] |
| Otu04858 | 6 | k__Fungi(100);p__Ascomycota(100);c__Saccharomycetes(100);o__Saccharomycetales(100);f__Metschnikowiaceae(100);g__Metschnikowia(100);g__Metschnikowia_unclassified(100); | Metschnikowia | 13 | Saprotroph | [illegible] |
| Otu04859 | 6 | k__Fungi(100);p__Ascomycota(100);c__Leotiomycetes(100);o__Helotiales(100);f__Hyaloscyphaceae(100);g__Lachnum(100);s__Lachnum_rachidicola(100); | Lachnum | 13 | Saprotroph-Symbiotroph | [illegible] |
| Otu04860 | 6 | k__Fungi(100);p__Ascomycota(100);c__Saccharomycetes(100);o__Saccharomycetales(100);f__Metschnikowiaceae(100);g__Metschnikowia(100);s__Metschnikowia_pulcherrima(84); | Metschnikowia | 13 | Saprotroph | [illegible] |
| Otu04861 | 6 | k__Fungi(100);p__Ascomycota(100);c__Saccharomycetes(100);o__Saccharomycetales(100);f__Saccharomycetaceae(100);g__Kazachstania(100);s__Kazachstania_intestinalis(100); | Kazachstania | 13 | Saprotroph | [illegible] |
| Otu04862 | 6 | k__Fungi(100);p__Ascomycota(100);c__Saccharomycetes(100);o__Saccharomycetales(100);f__Metschnikowiaceae(100);g__Metschnikowia(100);s__Metschnikowia_pulcherrima(100); | Metschnikowia | 13 | Saprotroph | [illegible] |
| Otu04864 | 6 | k__Fungi(100);p__Basidiomycota(100);c__Agaricomycetes(100);o__Agaricales(100);f__Amanitaceae(100);g__Amanita(100);s__Amanita_sp(100); | Amanita | 13 | Saprotroph-Symbiotroph | [illegible] |
| Otu04865 | 6 | k__Fungi(100);p__Basidiomycota(100);c__Agaricomycetes(100);o__Gomphales(100);f__Gomphaceae(100);g__Ramaria(100);s__Ramaria_stricta(100); | Ramaria | 13 | Saprotroph-Symbiotroph | [illegible] |
| Otu04866 | 6 | k__Fungi(100);p__Ascomycota(100);c__Saccharomycetes(100);o__Saccharomycetales(100);f__Metschnikowiaceae(100);g__Metschnikowia(100);g__Metschnikowia_unclassified(100); | Metschnikowia | 13 | Saprotroph | [illegible] |
| Otu04867 | 6 | k__Fungi(100);p__Basidiomycota(100);c__Agaricomycetes(100);o__Agaricales(100);f__Inocybaceae(100);g__Inocybe(100);s__Inocybe_sp(100); | Inocybe | 13 | Symbiotroph | [illegible] |
| Otu04868 | 6 | k__Fungi(100);p__Basidiomycota(100);c__Microbotryomycetes(100);o__Sporidiobolales(100);f__Sporidiobolales_family_Incertae_sedis(100);g__Sporobolomyces(100);s__Sporobolomyces_lactosus(100); | Sporobolomyces | 13 | Pathotroph-Saprotroph | [illegible] |
| Otu04871 | 6 | k__Fungi(100);p__Ascomycota(100);c__Lecanoromycetes(100);o__Ostropales(100);f__Stictidaceae(67);f__Stictidaceae_unclassified(67);f__Stictidaceae_unclassified(67); | Stictidaceae | 9 | Symbiotroph | [illegible] |
| Otu04875 | 6 | k__Fungi(100);p__Ascomycota(100);c__Saccharomycetes(100);o__Saccharomycetales(100);f__Metschnikowiaceae(100);g__Metschnikowia(100);g__Metschnikowia_unclassified(84); | Metschnikowia | 13 | Saprotroph | [illegible] |
| Otu04876 | 6 | k__Fungi(100);p__Ascomycota(100);c__Saccharomycetes(100);o__Saccharomycetales(100);f__Metschnikowiaceae(100);g__Metschnikowia(100);g__Metschnikowia_unclassified(100); | Metschnikowia | 13 | Saprotroph | [illegible] |
| Otu04877 | 6 | k__Fungi(100);p__Ascomycota(100);c__Dothideomycetes(100);o__Pleosporales(100);f__Massarinaceae(100);g__Massarina(67);s__Massarina_eburnea(67); | Massarina | 13 | Pathotroph-Saprotroph | [illegible] |
| Otu04878 | 6 | k__Fungi(100);p__Ascomycota(100);c__Saccharomycetes(100);o__Saccharomycetales(100);f__Metschnikowiaceae(100);g__Metschnikowia(100);s__Metschnikowia_pulcherrima(84); | Metschnikowia | 13 | Saprotroph | [illegible] |
| Otu04879 | 6 | k__Fungi(100);p__Ascomycota(100);c__Ascomycota_class_Incertae_sedis(100);o__Ascomycota_order_Incertae_sedis(100);f__Ascomycota_family_Incertae_sedis(100);g__Wojnowicia(100);s__Wojnowicia_viburni(100); | Wojnowicia | 13 | Saprotroph | [illegible] |
| Otu04880 | 6 | k__Fungi(100);p__Ascomycota(100);c__Dothideomycetes(100);o__Botryosphaeriales(100);f__Botryosphaeriaceae(100);g__Neofusicoccum(100);s__Neofusicoccum_parvum(100); | Neofusicoccum | 13 | Pathotroph-Saprotroph-Symbiotroph | [illegible] |
| Otu04885 | 6 | k__Fungi(100);p__Ascomycota(100);c__Saccharomycetes(100);o__Saccharomycetales(100);f__Metschnikowiaceae(100);g__Metschnikowia(100);g__Metschnikowia_unclassified(100); | Metschnikowia | 13 | Saprotroph | [illegible] |
| Otu04886 | 6 | k__Fungi(100);p__Ascomycota(100);c__Ascomycota_class_Incertae_sedis(100);o__Ascomycota_order_Incertae_sedis(100);f__Ascomycota_family_Incertae_sedis(100);g__Knufia(100);s__Knufia_cryptophialidica(100); | Knufia | 13 | Pathotroph-Saprotroph | [illegible] |
| Otu04888 | 6 | k__Fungi(100);p__Basidiomycota(100);c__Agaricomycetes(100);o__Agaricales(100);f__Agaricaceae(100);g__Cystolepiota(84);g__Cystolepiota_unclassified(84); | Cystolepiota | 13 | Saprotroph | [illegible] |
| Otu04889 | 6 | k__Fungi(100);p__Basidiomycota(100);c__Agaricomycetes(100);o__Agaricales(100);o__Agaricales_unclassified(100);o__Agaricales_unclassified(100);o__Agaricales_unclassified(100); | Agaricales | 7 | Pathotroph-Saprotroph-Symbiotroph | [illegible] |
| Otu04890 | 6 | k__Fungi(100);p__Basidiomycota(100);c__Agaricomycetes(100);o__Agaricales(100);f__Psathyrellaceae(100);g__unclassified_Psathyrellaceae(100);s__Psathyrellaceae_sp(100); | Psathyrellaceae | 9 | Saprotroph | [illegible] |
| Otu04892 | 6 | k__Fungi(100);p__Ascomycota(100);c__Saccharomycetes(100);o__Saccharomycetales(100);f__Metschnikowiaceae(100);g__Metschnikowia(100);s__Metschnikowia_pulcherrima(84); | Metschnikowia | 13 | Saprotroph | [illegible] |
| Otu04896 | 6 | k__Fungi(100);p__Basidiomycota(100);c__Agaricomycetes(100);o__Agaricales(100);f__Bolbitiaceae(100);g__Conocybe(100);g__Conocybe_unclassified(100); | Conocybe | 13 | Saprotroph | [illegible] |
| Otu04897 | 6 | k__Fungi(100);p__Ascomycota(100);c__Saccharomycetes(100);o__Saccharomycetales(100);f__Metschnikowiaceae(100);g__Metschnikowia(100);s__Metschnikowia_pulcherrima(100); | Metschnikowia | 13 | Saprotroph | [illegible] |
| Otu04898 | 6 | k__Fungi(100);p__Basidiomycota(100);c__Agaricomycetes(100);o__Agaricales(100);f__Psathyrellaceae(100);g__Coprinopsis(100);s__Coprinopsis_clastophylla(100); | Coprinopsis | 13 | Saprotroph | [illegible] |
| Otu04899 | 6 | k__Fungi(100);p__Basidiomycota(100);c__Microbotryomycetes(100);o__Sporidiobolales(100);f__Sporidiobolales_family_Incertae_sedis(100);g__Sporobolomyces(100);s__Sporobolomyces_inositophilus(100); | Sporobolomyces | 13 | Pathotroph-Saprotroph | [illegible] |

| | | | | | | | | | | | |
|---|---|---|---|---|---|---|---|---|---|---|---|
| Otu04901 | 6 | k__Fungi(100);p__Ascomycota(100);c__Dothideomycetes(100);o__Dothideomycetes_order_Incertae_sedis(100);f__Myxotrichaceae(100);g__Oidiodendron(100);s__Oidiodendron_myxotrichoides(100); | Oidiodendron | 13 | Pathotroph-Saprotroph-Symbiotroph | Endophyte-Ericoid Mycorrhizal-\|Undefined Saprotroph\| | Microfungus | NULL | Probable | As root_endophyte (Põlme et al. 2020); As soil_saprotroph (Põlme et al. 2020); Dark Septate Endophyte; Decay soil (Põlme et al. 2020) | Newsham KK. 2011. New Phytologist 190:783-793 ((https://doi.org/10.1111/j.1469-8137.2010.03611.x)) (as dark septate root endophytes); Põlme S, et al. 2020. Fungal Diversity 105:1-16 ((https://doi.org/10.1007/s13225-020-00466-2)) |
| Otu04903 | 6 | k__Fungi(100);p__Chytridiomycota(100);c__Chytridiomycetes(100);o__Rhizophydiales(100);f__Rhizophydiaceae(100);g__Rhizophydium(100);s__Rhizophydium_sp(100); | Rhizophydium | 13 | Pathotroph-Pathotroph-Saprotroph | \|Algal Parasite\|-Plant Pathogen-Plant Saprotroph-Protist Parasite-Undefined Saprotroph | Chytrid-Microfungus | NULL | Probable | As animal_decomposer (Põlme et al. 2020); As keratinolytic (Põlme et al. 2020); As litter_saprotroph (Põlme et al. 2020); Decay leaf, fruit, seed, algal material, fungal material, protist material (Põlme et al. 2020) | Põlme S, et al. 2020. Fungal Diversity 105:1-16 ((https://doi.org/10.1007/s13225-020-00466-2)); The American Phytopathological Society Common Names of Plant Diseases ((http://www.apsnet.org/publications/commonnames/Pages/default.aspx)) |
| Otu04904 | 6 | k__Fungi(100);p__Ascomycota(100);c__Saccharomycetes(100);o__Saccharomycetales(100);f__Metschnikowiaceae(100);g__Metschnikowia(100);s__Metschnikowia_pulcherrima(84); | Metschnikowia | 13 | Saprotroph | \|Undefined Saprotroph\| | Yeast | NULL | Probable | As nectar-tap_saprotroph (Põlme et al. 2020); Decay sugar-rich substrates (Põlme et al. 2020) | Põlme S, et al. 2020. Fungal Diversity 105:1-16 ((https://doi.org/10.1007/s13225-020-00466-2)) |
| Otu04908 | 6 | k__Fungi(100);p__Ascomycota(100);c__Eurotiomycetes(100);o__Chaetothyriales(100);f__Herpotrichiellaceae(100);f__Herpotrichiellaceae_unclassified(100);f__Herpotrichiellaceae_unclassified(100); | Herpotrichiellaceae | 9 | Pathotroph-Saprotroph | Animal Pathogen-Fungal Parasite-Undefined Saprotroph | Facultative Yeast-Microfungus | NULL | Probable | Mostly saprobic on plants or other fungi, isolated from soil etc., and from medical sources (Cannon & Kirk 2007) | Cannon PF, Kirk PM. 2007. Fungal Families of the World. CAB International, Cambridge (ISBN: 978-0851998275) |
| Otu04909 | 6 | k__Fungi(100);p__Ascomycota(100);c__Eurotiomycetes(100);o__Chaetothyriales(100);f__Herpotrichiellaceae(100);g__Cladophialophora(100);g__Cladophialophora_unclassified(100); | Cladophialophora | 13 | Pathotroph-Saprotroph-Symbiotroph | Animal Pathogen-Endophyte-Fungal Parasite-\|Undefined Saprotroph\| | Dimorphic-Microfungus | NULL | Probable | As mycoparasite (Põlme et al. 2020); As root_endophyte_dark_septate (Põlme et al. 2020); As soil_saprotroph (Põlme et al. 2020); Decay soil (Põlme et al. 2020) | James TY, et al. 2006. Nature 443:818-822 ((https://doi.org/10.1038/nature05110)); Põlme S, et al. 2020. Fungal Diversity 105:1-16 ((https://doi.org/10.1007/s13225-020-00466-2)) |
| Otu04912 | 6 | k__Fungi(100);p__Ascomycota(100);c__Saccharomycetes(100);o__Saccharomycetales(100);f__Trichomonascaceae(100);g__Blastobotrys(100);s__Blastobotrys_proliferans(100); | Blastobotrys | 13 | Saprotroph | \|Undefined Saprotroph\| | Yeast | Its unusual metal | Highly Probable | As nectar-tap_saprotroph (Põlme et al. 2020); Decay sugar-rich substrates (Põlme et al. 2020); In the course of identifying alternative microorganisms for biotechnological interest, A. adeninivorans has already proved its strengthened competitiveness as a promising cell factory for many more applications. | Kunze G, et al. 2014. Biotechnology for Biofuels 7:66 ((https://doi.org/10.1186/1754-6834-7-66)); Põlme S, et al. 2020. Fungal Diversity 105:1-16 ((https://doi.org/10.1007/s13225-020-00466-2)) |
| Otu04913 | 6 | k__Fungi(100);p__Ascomycota(100);c__Saccharomycetes(100);o__Saccharomycetales(100);f__Metschnikowiaceae(100);g__Metschnikowia(100);s__Metschnikowia_pulcherrima(100); | Metschnikowia | 13 | Saprotroph | \|Undefined Saprotroph\| | Yeast | NULL | Probable | As nectar-tap_saprotroph (Põlme et al. 2020); Decay sugar-rich substrates (Põlme et al. 2020) | Põlme S, et al. 2020. Fungal Diversity 105:1-16 ((https://doi.org/10.1007/s13225-020-00466-2)) |
| Otu04914 | 6 | k__Fungi(100);p__Ascomycota(100);c__Saccharomycetes(100);o__Saccharomycetales(100);f__Metschnikowiaceae(100);g__Metschnikowia(100);s__Metschnikowia_pulcherrima(84); | Metschnikowia | 13 | Saprotroph | \|Undefined Saprotroph\| | Yeast | NULL | Probable | As nectar-tap_saprotroph (Põlme et al. 2020); Decay sugar-rich substrates (Põlme et al. 2020) | Põlme S, et al. 2020. Fungal Diversity 105:1-16 ((https://doi.org/10.1007/s13225-020-00466-2)) |
| Otu04916 | 6 | k__Fungi(100);p__Ascomycota(100);c__Saccharomycetes(100);o__Saccharomycetales(100);f__Metschnikowiaceae(100);g__Metschnikowia(100);s__Metschnikowia_pulcherrima(84); | Metschnikowia | 13 | Saprotroph | \|Undefined Saprotroph\| | Yeast | NULL | Probable | As nectar-tap_saprotroph (Põlme et al. 2020); Decay sugar-rich substrates (Põlme et al. 2020) | Põlme S, et al. 2020. Fungal Diversity 105:1-16 ((https://doi.org/10.1007/s13225-020-00466-2)) |
| Otu04917 | 6 | k__Fungi(100);p__Basidiomycota(100);c__Tremellomycetes(100);o__Tremellales(100);f__Tremellales_family_Incertae_sedis(100);g__Filobasidiella(100);s__Filobasidiella_depauperata(100); | Filobasidiella | 13 | Pathotroph-Saprotroph | \|Animal Parasite\|-Fungal Parasite-Undefined Saprotroph | Dimorphic-Facultative Yeast | NULL | Highly Probable | As mycoparasite (Põlme et al. 2020); Decay animal material (Põlme et al. 2020) | Kurtzman CP, et al. (eds.) 2011. The Yeasts, a Taxonomic Study. Fifth Edition. Vols 1-3. Elsevier, San Diego (ISBN: 9780444521491); Põlme S, et al. 2020. Fungal Diversity 105:1-16 ((https://doi.org/10.1007/s13225-020-00466-2)) |
| Otu04919 | 6 | k__Fungi(100);p__Ascomycota(100);c__Saccharomycetes(100);o__Saccharomycetales(100);f__Metschnikowiaceae(100);g__Metschnikowia(100);g__Metschnikowia_unclassified(84); | Metschnikowia | 13 | Saprotroph | \|Undefined Saprotroph\| | Yeast | NULL | Probable | As nectar-tap_saprotroph (Põlme et al. 2020); Decay sugar-rich substrates (Põlme et al. 2020) | Põlme S, et al. 2020. Fungal Diversity 105:1-16 ((https://doi.org/10.1007/s13225-020-00466-2)) |
| Otu04921 | 6 | k__Fungi(100);p__Ascomycota(100);c__Saccharomycetes(100);o__Saccharomycetales(100);f__Metschnikowiaceae(100);g__Metschnikowia(100);g__Metschnikowia_unclassified(84); | Metschnikowia | 13 | Saprotroph | \|Undefined Saprotroph\| | Yeast | NULL | Probable | As nectar-tap_saprotroph (Põlme et al. 2020); Decay sugar-rich substrates (Põlme et al. 2020) | Põlme S, et al. 2020. Fungal Diversity 105:1-16 ((https://doi.org/10.1007/s13225-020-00466-2)) |
| Otu04922 | 6 | k__Fungi(100);p__Ascomycota(100);c__Saccharomycetes(100);o__Saccharomycetales(100);f__Metschnikowiaceae(100);g__Metschnikowia(100);s__Metschnikowia_pulcherrima(100); | Metschnikowia | 13 | Saprotroph | \|Undefined Saprotroph\| | Yeast | NULL | Probable | As nectar-tap_saprotroph (Põlme et al. 2020); Decay sugar-rich substrates (Põlme et al. 2020) | Põlme S, et al. 2020. Fungal Diversity 105:1-16 ((https://doi.org/10.1007/s13225-020-00466-2)) |
| Otu04924 | 6 | k__Fungi(100);p__Basidiomycota(100);c__Agaricomycetes(100);o__Agaricales(100);f__Agaricaceae(100);g__Cystodermella(100);s__Cystodermella_granulosa_var._granulosa(100); | Cystodermella | 13 | Saprotroph | \|Plant Saprotroph\|-Undefined Saprotroph | Agaricoid | NULL | Probable | As litter_saprotroph (Põlme et al. 2020); Decay leaf, fruit, seed (Põlme et al. 2020) | Tedersoo L, et al. 2014. Science 346:e1256688 ((https://doi.org/10.1126/science.1256688)); Põlme S, et al. 2020. Fungal Diversity 105:1-16 ((https://doi.org/10.1007/s13225-020-00466-2)) |
| Otu04926 | 6 | k__Fungi(100);p__Ascomycota(100);c__Saccharomycetes(100);o__Saccharomycetales(100);f__Metschnikowiaceae(100);g__Metschnikowia(100);s__Metschnikowia_pulcherrima(84); | Metschnikowia | 13 | Saprotroph | \|Undefined Saprotroph\| | Yeast | NULL | Probable | As nectar-tap_saprotroph (Põlme et al. 2020); Decay sugar-rich substrates (Põlme et al. 2020) | Põlme S, et al. 2020. Fungal Diversity 105:1-16 ((https://doi.org/10.1007/s13225-020-00466-2)) |
| Otu04928 | 6 | k__Fungi(100);p__Ascomycota(100);c__Dothideomycetes(100);o__Pleosporales(100);f__Lentitheciaceae(84);g__Setoseptoria(84);s__Setoseptoria_phragmitis(84); | Setoseptoria | 13 | Pathotroph-Saprotroph | \|Plant Pathogen\|-Plant Saprotroph | Microfungus | NULL | Probable | As litter_saprotroph (Põlme et al. 2020); As wood_pathogen (Põlme et al. 2020); Decay wood, leaf, fruit, seed (Põlme et al. 2020) | Põlme S, et al. 2020. Fungal Diversity 105:1-16 ((https://doi.org/10.1007/s13225-020-00466-2)) |
| Otu04929 | 6 | k__Fungi(100);p__Ascomycota(100);c__Saccharomycetes(84);o__Saccharomycetales(84);f__Saccharomycetales_family_Incertae_sedis(84);g__Candida(84);g__Candida_unclassified(67); | Candida | 13 | Pathotroph-Saprotroph-Symbiotroph | Animal Pathogen-Endophyte-Endosymbiont-Epiphyte-\|Undefined Saprotroph\| | Dimorphic Yeast-Yeast | NULL | Probable | As nectar-tap_saprotroph (Põlme et al. 2020); Decay sugar-rich substrates (Põlme et al. 2020); This asexual genus is polyphyletic and contain species that function very differently depend on what they are. Caution should be taken when interpreting guilds for Candida species. | Manolakaki D, et al. 2010. Virulence. 1: 367-375 ((https://doi.org/10.4161/viru.1.5.12796)); Mekbib SB, et al. 2011. Tropical Plant Pathology 36:233-240 ((https://doi.org/10.1590/S1982-56762011000400004)); Põlme S, et al. 2020. Fungal Diversity 105:1-16 ((https://doi.org/10.1007/s13225-020-00466-2)) |
| Otu04930 | 6 | k__Fungi(100);p__Ascomycota(100);c__Dothideomycetes(100);o__Pleosporales(100);f__Phaeosphaeriaceae(100);g__Phaeosphaeria(100);s__Phaeosphaeria_triglochinicola(100); | Phaeosphaeria | 13 | Pathotroph-Saprotroph-Symbiotroph | Endophyte-Plant Pathogen-\|Plant Saprotroph\| | Microfungus | NULL | Probable | As litter_saprotroph (Põlme et al. 2020); Decay leaf, fruit, seed (Põlme et al. 2020) | Tedersoo L, et al. 2014. Science 346:e1256688 ((https://doi.org/10.1126/science.1256688));David AS, et al. 2016. Microbial Ecology 71:912-926 ((https://doi.org/10.1007/s00248-015-0712-6)); Põlme S, et al. 2020. Fungal Diversity 105:1-16 ((https://doi.org/10.1007/s13225-020-00466-2)) |
| Otu04933 | 6 | k__Fungi(100);p__Basidiomycota(100);c__Agaricomycetes(100);o__Polyporales(100);f__Polyporaceae(100);f__Polyporaceae_unclassified(100);f__Polyporaceae_unclassified(100); | Polyporaceae | 9 | Saprotroph | Wood Saprotroph | Polyporoid | Brown Rot-White | Probable | Lignicolous, causing brown or white rot, mostly of broadleaved trees (Cannon & Kirk 2007) | Cannon PF, Kirk PM. 2007. Fungal Families of the World. CAB International, Cambridge (ISBN: 978-0851998275) |
| Otu04937 | 6 | k__Fungi(100);p__Ascomycota(100);c__Ascomycota_class_Incertae_sedis(100);o__Ascomycota_order_Incertae_sedis(100);f__Ascomycota_family_Incertae_sedis(100);g__Zymoseptoria(100);s__Zymoseptoria_verkleyi(100); | Zymoseptoria | 13 | Pathotroph-Saprotroph | \|Plant Pathogen\|-Plant Saprotroph | Microfungus | NULL | Probable | As litter_saprotroph (Põlme et al. 2020); Decay leaf, fruit, seed (Põlme et al. 2020) | Põlme S, et al. 2020. Fungal Diversity 105:1-16 ((https://doi.org/10.1007/s13225-020-00466-2)) |
| Otu04938 | 6 | k__Fungi(100);p__Basidiomycota(100);c__Agaricomycetes(100);o__Agaricales(100);o__Agaricales_unclassified(100);o__Agaricales_unclassified(100);o__Agaricales_unclassified(100); | Agaricales | 7 | Pathotroph-Saprotroph-Symbiotroph | Bryophyte Parasite-Dung Saprotroph-Ectomycorrhizal-Fungal Parasite-Leaf Saprotroph-Plant Parasite-Undefined Saprotroph-Wood Saprotroph | Agaricoid-Gasteroid-Microfungus-Secotioid-Yeast | NULL | Possible | Mushrooms and toadstools, Gill fungi, Agarics... lignicolous, sometimes muscicolous or fungicolous, saprobic, mycorrhizal, rarely parasitic on plants or fungi (Kirk et al. 2008) | Kirk PM et al. 2008. Dictionary of the Fungi. Tenth Edition. CAB International, Wallingford (ISBN: 978-0851998268) |
| Otu04940 | 6 | k__Fungi(100);p__Basidiomycota(100);c__Agaricomycetes(100);o__Agaricales(100);f__Tricholomataceae(100);g__Leucopaxillus(100);s__Leucopaxillus_giganteus(100); | Leucopaxillus | 13 | Pathotroph-Saprotroph | Plant Pathogen-\|Plant Saprotroph\|-Undefined Saprotroph | Agaricoid | NULL | Probable | As litter_saprotroph (Põlme et al. 2020); Decay leaf, fruit, seed (Põlme et al. 2020) | Tedersoo L, et al. 2014. Science 346:e1256688 ((https://doi.org/10.1126/science.1256688)); Põlme S, et al. 2020. Fungal Diversity 105:1-16 ((https://doi.org/10.1007/s13225-020-00466-2)) |
| Otu04941 | 6 | k__Fungi(100);p__Ascomycota(100);c__Saccharomycetes(100);o__Saccharomycetales(100);f__Saccharomycetales_family_Incertae_sedis(100);g__Candida(100);g__Candida_unclassified(67); | Candida | 13 | Pathotroph-Saprotroph-Symbiotroph | Animal Pathogen-Endophyte-Endosymbiont-Epiphyte-\|Undefined Saprotroph\| | Dimorphic Yeast-Yeast | NULL | Probable | As nectar-tap_saprotroph (Põlme et al. 2020); Decay sugar-rich substrates (Põlme et al. 2020); This asexual genus is polyphyletic and contain species that function very differently depend on what they are. Caution should be taken when interpreting guilds for Candida species. | Manolakaki D, et al. 2010. Virulence. 1: 367-375 ((https://doi.org/10.4161/viru.1.5.12796)); Mekbib SB, et al. 2011. Tropical Plant Pathology 36:233-240 ((https://doi.org/10.1590/S1982-56762011000400004)); Põlme S, et al. 2020. Fungal Diversity 105:1-16 ((https://doi.org/10.1007/s13225-020-00466-2)) |
| Otu04942 | 6 | k__Fungi(100);p__Ascomycota(100);c__Saccharomycetes(100);o__Saccharomycetales(100);f__Metschnikowiaceae(100);g__Metschnikowia(100);g__Metschnikowia_unclassified(100); | Metschnikowia | 13 | Saprotroph | \|Undefined Saprotroph\| | Yeast | NULL | Probable | As nectar-tap_saprotroph (Põlme et al. 2020); Decay sugar-rich substrates (Põlme et al. 2020) | Põlme S, et al. 2020. Fungal Diversity 105:1-16 ((https://doi.org/10.1007/s13225-020-00466-2)) |
| Otu04944 | 6 | k__Fungi(100);p__Basidiomycota(100);c__Agaricomycetes(100);o__Agaricales(100);f__Entolomataceae(100);g__Clitopilus(100);g__Clitopilus_unclassified(100); | Clitopilus | 13 | Pathotroph-Saprotroph | Fungal Parasite-\|Plant Saprotroph\|-Undefined Saprotroph | Agaricoid | NULL | Probable | As litter_saprotroph (Põlme et al. 2020); As mycoparasite (Põlme et al. 2020); Decay leaf, fruit, seed (Põlme et al. 2020) | Tedersoo L, et al. 2014. Science 346:e1256688 ((https://doi.org/10.1126/science.1256688)); Põlme S, et al. 2020. Fungal Diversity 105:1-16 ((https://doi.org/10.1007/s13225-020-00466-2)) |
| Otu04947 | 6 | k__Fungi(100);p__Basidiomycota(100);c__Agaricomycetes(100);o__Agaricales(100);f__Bolbitiaceae(100);g__Conocybe(100);s__Conocybe_hausknechtii(100); | Conocybe | 13 | Saprotroph | Dung Saprotroph-Plant Saprotroph-\|Undefined Saprotroph\| | Agaricoid | NULL | Probable | As litter_saprotroph (Põlme et al. 2020); As soil_saprotroph (Põlme et al. 2020); Decay soil (Põlme et al. 2020) | Tedersoo L, et al. 2014. Science 346:e1256688 ((https://doi.org/10.1126/science.1256688)); Põlme S, et al. 2020. Fungal Diversity 105:1-16 ((https://doi.org/10.1007/s13225-020-00466-2)) |
| Otu04949 | 6 | k__Fungi(100);p__Ascomycota(100);c__Saccharomycetes(100);o__Saccharomycetales(100);f__Saccharomycetales_family_Incertae_sedis(84);g__Candida(84);g__Candida_unclassified(67); | Candida | 13 | Pathotroph-Saprotroph-Symbiotroph | Animal Pathogen-Endophyte-Endosymbiont-Epiphyte-\|Undefined Saprotroph\| | Dimorphic Yeast-Yeast | NULL | Probable | As nectar-tap_saprotroph (Põlme et al. 2020); Decay sugar-rich substrates (Põlme et al. 2020); This asexual genus is polyphyletic and contain species that function very differently depend on what they are. Caution should be taken when interpreting guilds for Candida species. | Manolakaki D, et al. 2010. Virulence. 1: 367-375 ((https://doi.org/10.4161/viru.1.5.12796)); Mekbib SB, et al. 2011. Tropical Plant Pathology 36:233-240 ((https://doi.org/10.1590/S1982-56762011000400004)); Põlme S, et al. 2020. Fungal Diversity 105:1-16 ((https://doi.org/10.1007/s13225-020-00466-2)) |
| Otu04951 | 6 | k__Fungi(100);p__Ascomycota(100);c__Saccharomycetes(100);o__Saccharomycetales(100);f__Metschnikowiaceae(100);g__Metschnikowia(100);s__Metschnikowia_pulcherrima(100); | Metschnikowia | 13 | Saprotroph | \|Undefined Saprotroph\| | Yeast | NULL | Probable | As nectar-tap_saprotroph (Põlme et al. 2020); Decay sugar-rich substrates (Põlme et al. 2020) | Põlme S, et al. 2020. Fungal Diversity 105:1-16 ((https://doi.org/10.1007/s13225-020-00466-2)) |
| Otu04952 | 6 | k__Fungi(100);p__Ascomycota(100);c__Saccharomycetes(100);o__Saccharomycetales(100);f__Metschnikowiaceae(100);g__Metschnikowia(100);s__Metschnikowia_pulcherrima(67); | Metschnikowia | 13 | Saprotroph | \|Undefined Saprotroph\| | Yeast | NULL | Probable | As nectar-tap_saprotroph (Põlme et al. 2020); Decay sugar-rich substrates (Põlme et al. 2020) | Põlme S, et al. 2020. Fungal Diversity 105:1-16 ((https://doi.org/10.1007/s13225-020-00466-2)) |
| Otu04955 | 6 | k__Fungi(100);p__Basidiomycota(100);c__Agaricomycetes(100);o__Agaricales(100);f__Marasmiaceae(100);g__Tetrapyrgos(100);g__Tetrapyrgos_unclassified(100); | Tetrapyrgos | 13 | Saprotroph | \|Plant Saprotroph\|-Undefined Saprotroph | Agaricoid | NULL | Probable | As litter_saprotroph (Põlme et al. 2020); Decay leaf, fruit, seed (Põlme et al. 2020) | Tedersoo L, et al. 2014. Science 346:e1256688 ((https://doi.org/10.1126/science.1256688)); Põlme S, et al. 2020. Fungal Diversity 105:1-16 ((https://doi.org/10.1007/s13225-020-00466-2)) |
| Otu04956 | 6 | k__Fungi(100);p__Ascomycota(100);c__Saccharomycetes(100);o__Saccharomycetales(100);f__Metschnikowiaceae(100);g__Metschnikowia(100);s__Metschnikowia_pulcherrima(100); | Metschnikowia | 13 | Saprotroph | \|Undefined Saprotroph\| | Yeast | NULL | Probable | As nectar-tap_saprotroph (Põlme et al. 2020); Decay sugar-rich substrates (Põlme et al. 2020) | Põlme S, et al. 2020. Fungal Diversity 105:1-16 ((https://doi.org/10.1007/s13225-020-00466-2)) |
| Otu04958 | 6 | k__Fungi(100);p__Ascomycota(100);c__Lecanoromycetes(100);o__Lecanorales(67);o__Lecanorales_unclassified(67);o__Lecanorales_unclassified(67);o__Lecanorales_unclassified(67); | Lecanorales | 7 | Pathotroph-Symbiotroph | Lichen Parasite-Lichenized | Microfungus-Thallus | NULL | Probable | Lichenized with green algae, occasionally lichenicolous, on rocks, mosses and wood (Cannon & Kirk 2007); ...an order of mostly lichen-forming fungi (Kirk et al. 2008) | Cannon PF, Kirk PM. 2007. Fungal Families of the World. CAB International, Cambridge (ISBN: 978-0851998275); Kirk PM et al. 2008. Dictionary of the Fungi. Tenth Edition. CAB International, Wallingford (ISBN: 978-0851998268); Esslinger TL. 2014. North Dakota State University ((http://www.ndsu.edu/pubweb/~esslinge/chcklst/chcklst7.htm)) |
| Otu04960 | 6 | k__Fungi(100);p__Ascomycota(100);c__Saccharomycetes(100);o__Saccharomycetales(100);f__Metschnikowiaceae(100);g__Metschnikowia(100);s__Metschnikowia_pulcherrima(67); | Metschnikowia | 13 | Saprotroph | \|Undefined Saprotroph\| | Yeast | NULL | Probable | As nectar-tap_saprotroph (Põlme et al. 2020); Decay sugar-rich substrates (Põlme et al. 2020) | Põlme S, et al. 2020. Fungal Diversity 105:1-16 ((https://doi.org/10.1007/s13225-020-00466-2)) |
| Otu04961 | 6 | k__Fungi(100);p__Basidiomycota(100);c__Agaricomycetes(100);o__Russulales(100);f__Russulaceae(100);g__Lactifluus(100);s__Lactifluus_volemus(100); | Lactifluus | 13 | Symbiotroph | \|Ectomycorrhizal\| | Agaricoid | NULL | Highly Probable | NULL | Põlme S, et al. 2020. Fungal Diversity 105:1-16 ((https://doi.org/10.1007/s13225-020-00466-2)); List compiled by Sara Branco with input from Nhu Nguyen and Else Vellinga |
| Otu04962 | 6 | k__Fungi(100);p__Ascomycota(100);c__Saccharomycetes(100);o__Saccharomycetales(100);f__Metschnikowiaceae(100);g__Metschnikowia(100);s__Metschnikowia_pulcherrima(67); | Metschnikowia | 13 | Saprotroph | \|Undefined Saprotroph\| | Yeast | NULL | Probable | As nectar-tap_saprotroph (Põlme et al. 2020); Decay sugar-rich substrates (Põlme et al. 2020) | Põlme S, et al. 2020. Fungal Diversity 105:1-16 ((https://doi.org/10.1007/s13225-020-00466-2)) |
| Otu04963 | 6 | k__Fungi(100);p__Ascomycota(100);c__Saccharomycetes(100);o__Saccharomycetales(100);f__Metschnikowiaceae(100);g__Metschnikowia(100);s__Metschnikowia_pulcherrima(67); | Metschnikowia | 13 | Saprotroph | \|Undefined Saprotroph\| | Yeast | NULL | Probable | As nectar-tap_saprotroph (Põlme et al. 2020); Decay sugar-rich substrates (Põlme et al. 2020) | Põlme S, et al. 2020. Fungal Diversity 105:1-16 ((https://doi.org/10.1007/s13225-020-00466-2)) |
| Otu04966 | 6 | k__Fungi(100);p__Ascomycota(100);c__Saccharomycetes(100);o__Saccharomycetales(100);f__Metschnikowiaceae(100);g__Metschnikowia(100);s__Metschnikowia_pulcherrima(100); | Metschnikowia | 13 | Saprotroph | \|Undefined Saprotroph\| | Yeast | NULL | Probable | As nectar-tap_saprotroph (Põlme et al. 2020); Decay sugar-rich substrates (Põlme et al. 2020) | Põlme S, et al. 2020. Fungal Diversity 105:1-16 ((https://doi.org/10.1007/s13225-020-00466-2)) |
| Otu04967 | 6 | k__Fungi(100);p__Ascomycota(100);c__Saccharomycetes(100);o__Saccharomycetales(100);f__Metschnikowiaceae(100);g__Metschnikowia(100);g__Metschnikowia_unclassified(100); | Metschnikowia | 13 | Saprotroph | \|Undefined Saprotroph\| | Yeast | NULL | Probable | As nectar-tap_saprotroph (Põlme et al. 2020); Decay sugar-rich substrates (Põlme et al. 2020) | Põlme S, et al. 2020. Fungal Diversity 105:1-16 ((https://doi.org/10.1007/s13225-020-00466-2)) |
| Otu04969 | 6 | k__Fungi(100);p__Ascomycota(100);c__Saccharomycetes(100);o__Saccharomycetales(100);f__Metschnikowiaceae(100);g__Metschnikowia(100);g__Metschnikowia_unclassified(100); | Metschnikowia | 13 | Saprotroph | \|Undefined Saprotroph\| | Yeast | NULL | Probable | As nectar-tap_saprotroph (Põlme et al. 2020); Decay sugar-rich substrates (Põlme et al. 2020) | Põlme S, et al. 2020. Fungal Diversity 105:1-16 ((https://doi.org/10.1007/s13225-020-00466-2)) |
| Otu04970 | 6 | k__Fungi(100);p__Ascomycota(100);c__Sordariomycetes(100);o__Xylariales(100);f__Diatrypaceae(100);g__Eutypa(100);s__Eutypa_spinosa(100); | Eutypa | 13 | Pathotroph | \|Plant Pathogen\| | Microfungus | NULL | Probable | NULL | Tedersoo L, et al. 2014. Science 346:e1256688 ((https://doi.org/10.1126/science.1256688)); Põlme S, et al. 2020. Fungal Diversity 105:1-16 ((https://doi.org/10.1007/s13225-020-00466-2)) |
| Otu04974 | 6 | k__Fungi(100);p__Basidiomycota(100);c__Agaricomycetes(100);o__Cantharellales(100);f__Ceratobasidiaceae(100);f__Ceratobasidiaceae_unclassified(100);f__Ceratobasidiaceae_unclassified(100); | Ceratobasidiaceae | 9 | Pathotroph-Saprotroph-Symbiotroph | Endomycorrhizal-Plant Pathogen-Undefined Saprotroph | Microfungus | NULL | Possible | Best known as plant pathogens, but also saprobic. Species are endomycorrhizal with a wide range of plant roots - especially of Orchidaceae but also even bryophytes (Cannon & Kirk 2007) | Cannon PF, Kirk PM. 2007. Fungal Families of the World. CAB International, Cambridge (ISBN: 978-0851998275) |
| Otu04976 | 6 | k__Fungi(100);p__Basidiomycota(100);c__Agaricomycetes(100);o__Cantharellales(100);f__Ceratobasidiaceae(100);g__Thanatephorus(100);s__Thanatephorus_cucumeris(100); | Thanatephorus | 13 | Pathotroph-Saprotroph | \|Plant Pathogen\|-Plant Saprotroph | Corticioid | NULL | Probable | As litter_saprotroph (Põlme et al. 2020); Decay leaf, fruit, seed (Põlme et al. 2020) | Tedersoo L, et al. 2014. Science 346:e1256688 ((https://doi.org/10.1126/science.1256688)); Põlme S, et al. 2020. Fungal Diversity 105:1-16 ((https://doi.org/10.1007/s13225-020-00466-2)) |
| Otu04977 | 6 | k__Fungi(100);p__Basidiomycota(100);c__Agaricomycetes(100);o__Agaricales(100);f__Strophariaceae(100);g__Psilocybe(100);g__Psilocybe_unclassified(100); | Psilocybe | 13 | Saprotroph | \|Plant Saprotroph\|-Undefined Saprotroph | Agaricoid | Poisonous | Probable | As litter_saprotroph (Põlme et al. 2020); bluing clade; Decay leaf, fruit, seed (Põlme et al. 2020); Typified by P. semilanceata (Norvell 2007); Psilocybe = blue staining hallucinogenic clade and Deconica = non | Norvell L. 2007. Taxon. 59: 291-293 ((https://doi.org/10.1002/tax.591029)); Tedersoo L, et al. 2014. Science 346:e1256688 ((https://doi.org/10.1126/science.1256688)); Põlme S, et al. 2020. Fungal Diversity 105:1-16 ((https://doi.org/10.1007/s13225-020-00466-2)) |
| Otu04978 | 6 | k__Fungi(100);p__Ascomycota(100);c__Saccharomycetes(100);o__Saccharomycetales(100);f__Metschnikowiaceae(100);g__Metschnikowia(100);g__Metschnikowia_unclassified(84); | Metschnikowia | 13 | Saprotroph | \|Undefined Saprotroph\| | Yeast | NULL | Probable | As nectar-tap_saprotroph (Põlme et al. 2020); Decay sugar-rich substrates (Põlme et al. 2020) | Põlme S, et al. 2020. Fungal Diversity 105:1-16 ((https://doi.org/10.1007/s13225-020-00466-2)) |
| Otu04979 | 6 | k__Fungi(100);p__Basidiomycota(100);c__Agaricomycetes(100);o__Agaricales(100);f__Agaricales_family_Incertae_sedis(100);g__unclassified_Agaricales(100);s__Agaricales_sp(100); | Agaricales | 7 | Pathotroph-Saprotroph-Symbiotroph | Bryophyte Parasite-Dung Saprotroph-Ectomycorrhizal-Fungal Parasite-Leaf Saprotroph-Plant Parasite-Undefined Saprotroph-Wood Saprotroph | Agaricoid-Gasteroid-Microfungus-Secotioid-Yeast | NULL | Possible | Mushrooms and toadstools, Gill fungi, Agarics... lignicolous, sometimes muscicolous or fungicolous, saprobic, mycorrhizal, rarely parasitic on plants or fungi (Kirk et al. 2008) | Kirk PM et al. 2008. Dictionary of the Fungi. Tenth Edition. CAB International, Wallingford (ISBN: 978-0851998268) |
| Otu04981 | 6 | k__Fungi(100);p__Basidiomycota(100);c__Agaricomycetes(100);o__Trechisporales(100);f__Hydnodontaceae(100);g__Trechispora(100);g__Trechispora_unclassified(100); | Trechispora | 13 | Saprotroph | \|Wood Saprotroph\| | Corticioid | White Rot | Highly Probable | As white_rot (Põlme et al. 2020); Decay wood (Põlme et al. 2020) | Gilbertson RL, Ryvarden L. 1987-1987. North American Polypores. Fungiflora, Oslo (ISBN: 978-0945345060); Rinaldi AC, et al. 2008. Fungal Diversity 33:1-45 ((http://www.fungaldiversity.org/fdp/sfdp/33-1.pdf)) (as Uncertain); Tedersoo L, et al. 2010. Mycorrhiza 20:217-263 ((https://doi.org/10.1007/s00572-009-0274-x)) (as not ECM).; Põlme S, et al. 2020. Fungal Diversity 105:1-16 ((https://doi.org/10.1007/s13225-020-00466-2)) |
| Otu04984 | 6 | k__Fungi(100);p__Basidiomycota(100);c__Agaricomycetes(100);o__Agaricales(100);f__Cortinariaceae(100);g__Cortinarius(100);g__Cortinarius_unclassified(67); | Cortinarius | 13 | Symbiotroph | \|Ectomycorrhizal\| | Agaricoid | NULL | Highly Probable | NULL | Rinaldi AC, et al. 2008. Fungal Diversity 33:1-45 ((http://www.fungaldiversity.org/fdp/sfdp/33-1.pdf)); Tedersoo L, et al. 2010. Mycorrhiza 20:217-263 ((https://doi.org/10.1007/s00572-009-0274-x)); Põlme S, et al. 2020. Fungal Diversity 105:1-16 ((https://doi.org/10.1007/s13225-020-00466-2)); http://mycorrhizas.info/ecmf.html ((http://mycorrhizas.info/ecmf.html)) |
| Otu04985 | 6 | k__Fungi(100);p__Ascomycota(100);c__Saccharomycetes(100);o__Saccharomycetales(100);f__Metschnikowiaceae(100);g__Metschnikowia(100);s__Metschnikowia_pulcherrima(100); | Metschnikowia | 13 | Saprotroph | \|Undefined Saprotroph\| | Yeast | NULL | Probable | As nectar-tap_saprotroph (Põlme et al. 2020); Decay sugar-rich substrates (Põlme et al. 2020) | Põlme S, et al. 2020. Fungal Diversity 105:1-16 ((https://doi.org/10.1007/s13225-020-00466-2)) |
| Otu04986 | 6 | k__Fungi(100);p__Ascomycota(100);c__Saccharomycetes(100);o__Saccharomycetales(100);f__Metschnikowiaceae(100);g__Metschnikowia(100);g__Metschnikowia_unclassified(100); | Metschnikowia | 13 | Saprotroph | \|Undefined Saprotroph\| | Yeast | NULL | Probable | As nectar-tap_saprotroph (Põlme et al. 2020); Decay sugar-rich substrates (Põlme et al. 2020) | Põlme S, et al. 2020. Fungal Diversity 105:1-16 ((https://doi.org/10.1007/s13225-020-00466-2)) |
| Otu04987 | 6 | k__Fungi(100);p__Ascomycota(100);c__Saccharomycetes(100);o__Saccharomycetales(100);f__Metschnikowiaceae(100);g__Metschnikowia(100);s__Metschnikowia_pulcherrima(100); | Metschnikowia | 13 | Saprotroph | \|Undefined Saprotroph\| | Yeast | NULL | Probable | As nectar-tap_saprotroph (Põlme et al. 2020); Decay sugar-rich substrates (Põlme et al. 2020) | Põlme S, et al. 2020. Fungal Diversity 105:1-16 ((https://doi.org/10.1007/s13225-020-00466-2)) |
| Otu04991 | 6 | k__Fungi(100);p__Ascomycota(100);c__Leotiomycetes(100);o__Helotiales(100);f__Sclerotiniaceae(100);g__Streptobotrys(100);s__Streptobotrys_streptothrix(100); | Streptobotrys | 13 | Pathotroph | \|Plant Pathogen\| | Microfungus | NULL | Probable | NULL | Tedersoo L, et al. 2014. Science 346:e1256688 ((https://doi.org/10.1126/science.1256688)); Põlme S, et al. 2020. Fungal Diversity 105:1-16 ((https://doi.org/10.1007/s13225-020-00466-2)) |
| Otu04992 | 6 | k__Fungi(100);p__Ascomycota(100);c__Saccharomycetes(100);o__Saccharomycetales(100);f__Metschnikowiaceae(100);g__Metschnikowia(100);s__Metschnikowia_pulcherrima(84); | Metschnikowia | 13 | Saprotroph | \|Undefined Saprotroph\| | Yeast | NULL | Probable | As nectar-tap_saprotroph (Põlme et al. 2020); Decay sugar-rich substrates (Põlme et al. 2020) | Põlme S, et al. 2020. Fungal Diversity 105:1-16 ((https://doi.org/10.1007/s13225-020-00466-2)) |
| Otu04993 | 6 | k__Fungi(100);p__Ascomycota(100);c__Saccharomycetes(100);o__Saccharomycetales(100);f__Metschnikowiaceae(100);g__Metschnikowia(100);s__Metschnikowia_pulcherrima(100); | Metschnikowia | 13 | Saprotroph | \|Undefined Saprotroph\| | Yeast | NULL | Probable | As nectar-tap_saprotroph (Põlme et al. 2020); Decay sugar-rich substrates (Põlme et al. 2020) | Põlme S, et al. 2020. Fungal Diversity 105:1-16 ((https://doi.org/10.1007/s13225-020-00466-2)) |
| Otu04994 | 6 | k__Fungi(100);p__Basidiomycota(100);c__Agaricomycetes(100);o__Agaricales(100);f__Cortinariaceae(100);g__Cortinarius(100);g__Cortinarius_unclassified(100); | Cortinarius | 13 | Symbiotroph | \|Ectomycorrhizal\| | Agaricoid | NULL | Highly Probable | NULL | Rinaldi AC, et al. 2008. Fungal Diversity 33:1-45 ((http://www.fungaldiversity.org/fdp/sfdp/33-1.pdf)); Tedersoo L, et al. 2010. Mycorrhiza 20:217-263 ((https://doi.org/10.1007/s00572-009-0274-x)); Põlme S, et al. 2020. Fungal Diversity 105:1-16 ((https://doi.org/10.1007/s13225-020-00466-2)); http://mycorrhizas.info/ecmf.html ((http://mycorrhizas.info/ecmf.html)) |
| Otu04995 | 6 | k__Fungi(100);p__Ascomycota(100);c__Saccharomycetes(100);o__Saccharomycetales(100);f__Metschnikowiaceae(100);g__Metschnikowia(100);s__Metschnikowia_pulcherrima(100); | Metschnikowia | 13 | Saprotroph | \|Undefined Saprotroph\| | Yeast | NULL | Probable | As nectar-tap_saprotroph (Põlme et al. 2020); Decay sugar-rich substrates (Põlme et al. 2020) | Põlme S, et al. 2020. Fungal Diversity 105:1-16 ((https://doi.org/10.1007/s13225-020-00466-2)) |
| Otu04997 | 6 | k__Fungi(100);p__Basidiomycota(100);c__Agaricomycetes(100);o__Agaricales(100);f__Cortinariaceae(100);g__Cortinarius(100);s__Cortinarius_bolaris(100); | Cortinarius | 13 | Symbiotroph | \|Ectomycorrhizal\| | Agaricoid | NULL | Highly Probable | NULL | Rinaldi AC, et al. 2008. Fungal Diversity 33:1-45 ((http://www.fungaldiversity.org/fdp/sfdp/33-1.pdf)); Tedersoo L, et al. 2010. Mycorrhiza 20:217-263 ((https://doi.org/10.1007/s00572-009-0274-x)); Põlme S, et al. 2020. Fungal Diversity 105:1-16 ((https://doi.org/10.1007/s13225-020-00466-2)); http://mycorrhizas.info/ecmf.html ((http://mycorrhizas.info/ecmf.html)) |
| Otu04998 | 6 | k__Fungi(100);p__Basidiomycota(100);c__Agaricomycetes(100);o__Agaricales(100);f__Agaricaceae(100);g__Agaricus(100);g__Agaricus_unclassified(67); | Agaricus | 13 | Saprotroph | \|Undefined Saprotroph\| | Agaricoid-Gasteroid-Secotioid | NULL | Probable | As soil_saprotroph (Põlme et al. 2020); Decay soil (Põlme et al. 2020) | Tedersoo L, et al. 2014. Science 346:e1256688 ((https://doi.org/10.1126/science.1256688)); Põlme S, et al. 2020. Fungal Diversity 105:1-16 ((https://doi.org/10.1007/s13225-020-00466-2)) |
| Otu05000 | 6 | k__Fungi(100);p__Ascomycota(100);c__Saccharomycetes(100);o__Saccharomycetales(100);f__Metschnikowiaceae(100);g__Metschnikowia(100);g__Metschnikowia_unclassified(67); | Metschnikowia | 13 | Saprotroph | \|Undefined Saprotroph\| | Yeast | NULL | Probable | As nectar-tap_saprotroph (Põlme et al. 2020); Decay sugar-rich substrates (Põlme et al. 2020) | Põlme S, et al. 2020. Fungal Diversity 105:1-16 ((https://doi.org/10.1007/s13225-020-00466-2)) |
| Otu05002 | 6 | k__Fungi(100);p__Ascomycota(100);c__Saccharomycetes(100);o__Saccharomycetales(100);f__Metschnikowiaceae(100);g__Metschnikowia(100);s__Metschnikowia_pulcherrima(100); | Metschnikowia | 13 | Saprotroph | \|Undefined Saprotroph\| | Yeast | NULL | Probable | As nectar-tap_saprotroph (Põlme et al. 2020); Decay sugar-rich substrates (Põlme et al. 2020) | Põlme S, et al. 2020. Fungal Diversity 105:1-16 ((https://doi.org/10.1007/s13225-020-00466-2)) |
| Otu05004 | 6 | k__Fungi(100);p__Basidiomycota(100);c__Agaricomycetes(100);o__Agaricales(100);f__Inocybaceae(100);g__Crepidotus(100);s__Crepidotus_sphaerosporus(100); | Crepidotus | 13 | Saprotroph | \|Wood Saprotroph\| | Agaricoid | NULL | Highly Probable | Decay wood (Põlme et al. 2020) | Grgurinovic CA, Simpson JA. 2001. Fungal Diversity 8:97-127 ((http://www.fungaldiversity.org/fdp/sfdp/FD_8_97-127.pdf)); Aime MC, et al. 2005. American Journal of Botany 92:74-82 ((https://doi.org/10.3732/ajb.92.1.74)); Tedersoo L, et al. 2014. Science 346:e1256688 ((https://doi.org/10.1126/science.1256688)); Põlme S, et al. 2020. Fungal Diversity 105:1-16 ((https://doi.org/10.1007/s13225-020-00466-2)) |
| Otu05007 | 6 | k__Fungi(100);p__Basidiomycota(100);c__Agaricomycetes(100);o__Trechisporales(100);f__Hydnodontaceae(100);g__Trechispora(100);s__Trechispora_sp(100); | Trechispora | 13 | Saprotroph | \|Wood Saprotroph\| | Corticioid | White Rot | Highly Probable | As white_rot (Põlme et al. 2020); Decay wood (Põlme et al. 2020) | Gilbertson RL, Ryvarden L. 1987-1987. North American Polypores. Fungiflora, Oslo (ISBN: 978-0945345060); Rinaldi AC, et al. 2008. Fungal Diversity 33:1-45 ((http://www.fungaldiversity.org/fdp/sfdp/33-1.pdf)) (as Uncertain); Tedersoo L, et al. 2010. Mycorrhiza 20:217-263 ((https://doi.org/10.1007/s00572-009-0274-x)) (as not ECM).; Põlme S, et al. 2020. Fungal Diversity 105:1-16 ((https://doi.org/10.1007/s13225-020-00466-2)) |
| Otu05008 | 6 | k__Fungi(100);p__Basidiomycota(100);c__Agaricomycetes(100);o__Trechisporales(100);f__Hydnodontaceae(100);g__Trechispora(100);g__Trechispora_unclassified(100); | Trechispora | 13 | Saprotroph | \|Wood Saprotroph\| | Corticioid | White Rot | Highly Probable | As white_rot (Põlme et al. 2020); Decay wood (Põlme et al. 2020) | Gilbertson RL, Ryvarden L. 1987-1987. North American Polypores. Fungiflora, Oslo (ISBN: 978-0945345060); Rinaldi AC, et al. 2008. Fungal Diversity 33:1-45 ((http://www.fungaldiversity.org/fdp/sfdp/33-1.pdf)) (as Uncertain); Tedersoo L, et al. 2010. Mycorrhiza 20:217-263 ((https://doi.org/10.1007/s00572-009-0274-x)) (as not ECM).; Põlme S, et al. 2020. Fungal Diversity 105:1-16 ((https://doi.org/10.1007/s13225-020-00466-2)) |
| Otu05009 | 6 | k__Fungi(100);p__Zygomycota(100);c__Zygomycota_class_Incertae_sedis(100);o__Mucorales(100);f__Mucoraceae(100);g__Mucor(100);s__Mucor_flavus(100); | Mucor | 13 | Saprotroph | \|Undefined Saprotroph\| | Microfungus | NULL | Probable | As soil_saprotroph (Põlme et al. 2020); Decay soil (Põlme et al. 2020) | Tedersoo L, et al. 2014. Science 346:e1256688 ((https://doi.org/10.1126/science.1256688)); Põlme S, et al. 2020. Fungal Diversity 105:1-16 ((https://doi.org/10.1007/s13225-020-00466-2)) |
| Otu05011 | 6 | k__Fungi(100);p__Ascomycota(100);c__Saccharomycetes(100);o__Saccharomycetales(100);f__Saccharomycetales_family_Incertae_sedis(100);g__Candida(84);g__Candida_unclassified(67); | Candida | 13 | Pathotroph-Saprotroph-Symbiotroph | Animal Pathogen-Endophyte-Endosymbiont-Epiphyte-\|Undefined Saprotroph\| | Dimorphic Yeast-Yeast | NULL | Probable | As nectar-tap_saprotroph (Põlme et al. 2020); Decay sugar-rich substrates (Põlme et al. 2020); This asexual genus is polyphyletic and contain species that function very differently depend on what they are. Caution should be taken when interpreting guilds for Candida species. | Manolakaki D, et al. 2010. Virulence. 1: 367-375 ((https://doi.org/10.4161/viru.1.5.12796)); Mekbib SB, et al. 2011. Tropical Plant Pathology 36:233-240 ((https://doi.org/10.1590/S1982-56762011000400004)); Põlme S, et al. 2020. Fungal Diversity 105:1-16 ((https://doi.org/10.1007/s13225-020-00466-2)) |
| Otu05013 | 6 | k__Fungi(100);p__Ascomycota(100);c__Eurotiomycetes(100);o__Ascosphaerales(100);f__Ascosphaeraceae(100);g__Ascosphaera(100);g__Ascosphaera_unclassified(100); | Ascosphaera | 13 | Pathotroph-Saprotroph | \|Animal Parasite\|-Animal Pathogen-Undefined Saprotroph | Microfungus | NULL | Probable | As animal_decomposer (Põlme et al. 2020); Decay animal material (Põlme et al. 2020); Host - Apis (Põlme et al. 2020) | Tedersoo L, et al. 2014. Science 346:e1256688 ((https://doi.org/10.1126/science.1256688)); Põlme S, et al. 2020. Fungal Diversity 105:1-16 ((https://doi.org/10.1007/s13225-020-00466-2)) |
| Otu05015 | 6 | k__Fungi(100);p__Ascomycota(100);c__Saccharomycetes(100);o__Saccharomycetales(100);f__Metschnikowiaceae(100);g__Metschnikowia(100);s__Metschnikowia_pulcherrima(100); | Metschnikowia | 13 | Saprotroph | \|Undefined Saprotroph\| | Yeast | NULL | Probable | As nectar-tap_saprotroph (Põlme et al. 2020); Decay sugar-rich substrates (Põlme et al. 2020) | Põlme S, et al. 2020. Fungal Diversity 105:1-16 ((https://doi.org/10.1007/s13225-020-00466-2)) |
| Otu05016 | 6 | k__Fungi(100);p__Ascomycota(100);c__Saccharomycetes(100);o__Saccharomycetales(100);f__Metschnikowiaceae(100);g__Metschnikowia(100);s__Metschnikowia_pulcherrima(100); | Metschnikowia | 13 | Saprotroph | \|Undefined Saprotroph\| | Yeast | NULL | Probable | As nectar-tap_saprotroph (Põlme et al. 2020); Decay sugar-rich substrates (Põlme et al. 2020) | Põlme S, et al. 2020. Fungal Diversity 105:1-16 ((https://doi.org/10.1007/s13225-020-00466-2)) |
| Otu05017 | 6 | k__Fungi(100);p__Ascomycota(100);c__Saccharomycetes(100);o__Saccharomycetales(100);f__Metschnikowiaceae(100);g__Metschnikowia(100);s__Metschnikowia_pulcherrima(100); | Metschnikowia | 13 | Saprotroph | \|Undefined Saprotroph\| | Yeast | NULL | Probable | As nectar-tap_saprotroph (Põlme et al. 2020); Decay sugar-rich substrates (Põlme et al. 2020) | Põlme S, et al. 2020. Fungal Diversity 105:1-16 ((https://doi.org/10.1007/s13225-020-00466-2)) |
| Otu05018 | 6 | k__Fungi(100);p__Ascomycota(100);c__Saccharomycetes(100);o__Saccharomycetales(100);f__Metschnikowiaceae(100);g__Metschnikowia(100);g__Metschnikowia_unclassified(100); | Metschnikowia | 13 | Saprotroph | \|Undefined Saprotroph\| | Yeast | NULL | Probable | As nectar-tap_saprotroph (Põlme et al. 2020); Decay sugar-rich substrates (Põlme et al. 2020) | Põlme S, et al. 2020. Fungal Diversity 105:1-16 ((https://doi.org/10.1007/s13225-020-00466-2)) |
| Otu05019 | 6 | k__Fungi(100);p__Ascomycota(100);c__Sordariomycetes(100);o__Xylariales(100);f__Xylariales_family_Incertae_sedis(100);g__Monographella(100);s__Monographella_cucumerina(100); | Monographella | 13 | Pathotroph | \|Plant Pathogen\| | Microfungus | NULL | Probable | NULL | Tedersoo L, et al. 2014. Science 346:e1256688 ((https://doi.org/10.1126/science.1256688)); Põlme S, et al. 2020. Fungal Diversity 105:1-16 ((https://doi.org/10.1007/s13225-020-00466-2)) |
| Otu05022 | 6 | k__Fungi(100);p__Ascomycota(100);c__Taphrinomycetes(100);o__Taphrinales(100);f__Taphrinaceae(100);f__Taphrinaceae_unclassified(100);f__Taphrinaceae_unclassified(100); | Taphrinaceae | 9 | Pathotroph | Plant Pathogen | Microfungus | NULL | Probable | Biotrophic on plants, usually causing hyperplasia (galls, witches broom) or lesions (Cannon & Kirk 2007) | Cannon PF, Kirk PM. 2007. Fungal Families of the World. CAB International, Cambridge (ISBN: 978-0851998275) |
| Otu05023 | 6 | k__Fungi(100);p__Ascomycota(100);c__Sordariomycetes(100);o__Hypocreales(100);f__Hypocreales_family_Incertae_sedis(84);g__Stachybotrys(84);g__Stachybotrys_unclassified(84); | Stachybotrys | 13 | Pathotroph-Saprotroph | Plant Pathogen-Plant Saprotroph-\|Wood Saprotroph\| | Microfungus | Soft Rot | Highly Probable | As litter_saprotroph (Põlme et al. 2020); Decay wood, leaf, fruit, seed (Põlme et al. 2020) | Seehann G, et al. 1975. List of Fungi in Soft-Rot Tissue Tests. Institute fur Holzbiologie und Holzschutz, Bundesforschungsanstalt fur Forst - und Holzwirtshaft, Hamburg - Lohbrugge, Germany; Costa IPMW, et al. 2012. Mycotaxon 119:e26 ((http://mycotaxon.com/resources/checklists/costa_v119_checklist.pdf)); Tedersoo L, et al. 2014. Science 346:e1256688 ((https://doi.org/10.1126/science.1256688)); Põlme S, et al. 2020. Fungal Diversity 105:1-16 ((https://doi.org/10.1007/s13225-020-00466-2)); The American Phytopathological Society Common Names of Plant Diseases ((http://www.apsnet.org/publications/commonnames/Pages/default.aspx)) |
| Otu05024 | 6 | k__Fungi(100);p__Ascomycota(100);c__Saccharomycetes(100);o__Saccharomycetales(100);f__Metschnikowiaceae(100);g__Metschnikowia(100);s__Metschnikowia_pulcherrima(84); | Metschnikowia | 13 | Saprotroph | \|Undefined Saprotroph\| | Yeast | NULL | Probable | As nectar-tap_saprotroph (Põlme et al. 2020); Decay sugar-rich substrates (Põlme et al. 2020) | Põlme S, et al. 2020. Fungal Diversity 105:1-16 ((https://doi.org/10.1007/s13225-020-00466-2)) |
| Otu05026 | 6 | k__Fungi(100);p__Ascomycota(100);c__Saccharomycetes(100);o__Saccharomycetales(100);f__Metschnikowiaceae(100);g__Metschnikowia(100);s__Metschnikowia_pulcherrima(84); | Metschnikowia | 13 | Saprotroph | \|Undefined Saprotroph\| | Yeast | NULL | Probable | As nectar-tap_saprotroph (Põlme et al. 2020); Decay sugar-rich substrates (Põlme et al. 2020) | Põlme S, et al. 2020. Fungal Diversity 105:1-16 ((https://doi.org/10.1007/s13225-020-00466-2)) |
| Otu05031 | 6 | k__Fungi(100);p__Ascomycota(100);c__Saccharomycetes(100);o__Saccharomycetales(100);f__Metschnikowiaceae(100);g__Metschnikowia(100);s__Metschnikowia_pulcherrima(100); | Metschnikowia | 13 | Saprotroph | \|Undefined Saprotroph\| | Yeast | NULL | Probable | As nectar-tap_saprotroph (Põlme et al. 2020); Decay sugar-rich substrates (Põlme et al. 2020) | Põlme S, et al. 2020. Fungal Diversity 105:1-16 ((https://doi.org/10.1007/s13225-020-00466-2)) |
| Otu05032 | 6 | k__Fungi(100);p__Basidiomycota(100);c__Agaricostilbomycetes(100);o__Agaricostilbales(100);f__Chionosphaeraceae(100);g__Kurtzmanomyces(84);g__Kurtzmanomyces_unclassified(84); | Kurtzmanomyces | 13 | Saprotroph | \|Undefined Saprotroph\| | Dimorphic | NULL | Probable | NULL | Põlme S, et al. 2020. Fungal Diversity 105:1-16 ((https://doi.org/10.1007/s13225-020-00466-2)) |
| Otu05037 | 6 | k__Fungi(100);p__Ascomycota(100);c__Sordariomycetes(100);o__Calosphaeriales(100);f__Calosphaeriaceae(100);g__Jattaea(100);g__Jattaea_unclassified(100); | Jattaea | 13 | Saprotroph | \|Undefined Saprotroph\| | Microfungus | NULL | Probable | NULL | Tedersoo L, et al. 2014. Science 346:e1256688 ((https://doi.org/10.1126/science.1256688)); Põlme S, et al. 2020. Fungal Diversity 105:1-16 ((https://doi.org/10.1007/s13225-020-00466-2)) |
| Otu05039 | 6 | k__Fungi(100);p__Ascomycota(100);c__Saccharomycetes(100);o__Saccharomycetales(100);f__Metschnikowiaceae(100);g__Metschnikowia(100);s__Metschnikowia_pulcherrima(100); | Metschnikowia | 13 | Saprotroph | \|Undefined Saprotroph\| | Yeast | NULL | Probable | As nectar-tap_saprotroph (Põlme et al. 2020); Decay sugar-rich substrates (Põlme et al. 2020) | Põlme S, et al. 2020. Fungal Diversity 105:1-16 ((https://doi.org/10.1007/s13225-020-00466-2)) |
| Otu05040 | 6 | k__Fungi(100);p__Ascomycota(100);c__Dothideomycetes(100);o__Pleosporales(100);f__Leptosphaeriaceae(100);g__Leptosphaeria(100);s__Leptosphaeria_doliolum(100); | Leptosphaeria | 13 | Pathotroph-Saprotroph | \|Plant Pathogen\|-Wood Saprotroph | Microfungus | NULL | Probable | As wood_pathogen (Põlme et al. 2020); Decay wood (Põlme et al. 2020) | Tedersoo L, et al. 2014. Science 346:e1256688 ((https://doi.org/10.1126/science.1256688)); Põlme S, et al. 2020. Fungal Diversity 105:1-16 ((https://doi.org/10.1007/s13225-020-00466-2)) |
| Otu05041 | 6 | k__Fungi(100);p__Ascomycota(100);c__Sordariomycetes(100);o__Xylariales(100);f__Diatrypaceae(100);g__Eutypella(100);s__Eutypella_parasitica(100); | Eutypella | 13 | Pathotroph | \|Plant Pathogen\| | Microfungus | NULL | Probable | NULL | Tedersoo L, et al. 2014. Science 346:e1256688 ((https://doi.org/10.1126/science.1256688)); Põlme S, et al. 2020. Fungal Diversity 105:1-16 ((https://doi.org/10.1007/s13225-020-00466-2)) |
| Otu05042 | 6 | k__Fungi(100);p__Ascomycota(100);c__Saccharomycetes(100);o__Saccharomycetales(100);f__Metschnikowiaceae(100);g__Metschnikowia(100);s__Metschnikowia_pulcherrima(100); | Metschnikowia | 13 | Saprotroph | \|Undefined Saprotroph\| | Yeast | NULL | Probable | As nectar-tap_saprotroph (Põlme et al. 2020); Decay sugar-rich substrates (Põlme et al. 2020) | Põlme S, et al. 2020. Fungal Diversity 105:1-16 ((https://doi.org/10.1007/s13225-020-00466-2)) |
| Otu05043 | 6 | k__Fungi(100);p__Ascomycota(100);c__Pezizomycetes(100);o__Pezizales(100);f__Tuberaceae(100);g__unclassified_Tuberaceae(100);s__Tuberaceae_sp(100); | Tuberaceae | 9 | Symbiotroph | Ectomycorrhizal | Gasteroid | Hypogeous | Probable | Common name - true truffle; Mycorrhizal with broadleaved trees, growing in the surface layer of soil and closely associated with roots. Highly specific pheromone-like chemicals are produced that play a role in dispersal by attracting mammals that uproot fruit bodies and ditribute spores (Cannon & Kirk 2007) | Cannon PF, Kirk PM. 2007. Fungal Families of the World. CAB International, Cambridge (ISBN: 978-0851998275) |
| Otu05045 | 6 | k__Fungi(100);p__Ascomycota(100);c__Sordariomycetes(100);o__Sordariomycetes_order_Incertae_sedis(100);f__Glomerellaceae(100);f__Glomerellaceae_unclassified(100);f__Glomerellaceae_unclassified(100); | Glomerellaceae | 9 | Pathotroph-Symbiotroph | Endophyte-Plant Pathogen | Microfungus | NULL | Probable | Species are hemibiotic, with an initial endophytic (symptomless) phase that leads to a necrotic phase were significant damage may be caused to the host (Cannon & Kirk 2007) | Cannon PF, Kirk PM. 2007. Fungal Families of the World. CAB International, Cambridge (ISBN: 978-0851998275) |
| Otu05050 | 6 | k__Fungi(100);p__Ascomycota(100);c__Sordariomycetes(100);o__Ophiostomatales(100);f__Ophiostomataceae(100);g__Ophiostoma(100);s__Ophiostoma_stenoceras(100); | Ophiostoma | 13 | Pathotroph-Saprotroph | \|Plant Pathogen\|-Wood Saprotroph | Microfungus | Blue-Staining | Probable | As wood_pathogen (Põlme et al. 2020); At least most species are transmitted via insect vectors, with mycelial colonies developing in bark; boring beetle galleries and spreading through the vascular system; they are highly specific to their beetle vectors. In some cases spores and mycelium are transmitted in specialized organs, termed mycangia (Cannon & Kirk 2007); Decay wood (Põlme et al. 2020) | Cannon PF, Kirk PM. 2007. Fungal Families of the World. CAB International, Cambridge (ISBN: 978-0851998275); Tedersoo L, et al. 2014. Science 346:e1256688 ((https://doi.org/10.1126/science.1256688)); Põlme S, et al. 2020. Fungal Diversity 105:1-16 ((https://doi.org/10.1007/s13225-020-00466-2)) |
| Otu05052 | 6 | k__Fungi(100);p__Ascomycota(100);c__Eurotiomycetes(100);o__Chaetothyriales(100);f__Chaetothyriaceae(100);g__Cyphellophora(100);s__Cyphellophora_europaea(100); | Cyphellophora | 13 | Pathotroph-Saprotroph | Animal Parasite-Animal Pathogen-\|Plant Saprotroph\|-Undefined Saprotroph | Dimorphic-Facultative Yeast-Microfungus | NULL | Probable | As litter_saprotroph (Põlme et al. 2020); Consult the current literature for updated names on this group that included the Black Yeasts; Some species are likely opportunistic human pathogen (Irinyi et al. 2015); Decay leaf, fruit, seed, animal material (Põlme et al. 2020) | Tedersoo L, et al. 2014. Science 346:e1256688 ((https://doi.org/10.1126/science.1256688)); Irinyi L, et al. 2015. Medical Mycology 53:313-337 ((https://doi.org/10.1093/mmy/myv008)); Põlme S, et al. 2020. Fungal Diversity 105:1-16 ((https://doi.org/10.1007/s13225-020-00466-2)) |
| Otu05053 | 6 | k__Fungi(100);p__Ascomycota(100);c__Saccharomycetes(100);o__Saccharomycetales(100);f__Metschnikowiaceae(100);g__Metschnikowia(100);s__Metschnikowia_pulcherrima(100); | Metschnikowia | 13 | Saprotroph | \|Undefined Saprotroph\| | Yeast | NULL | Probable | As nectar-tap_saprotroph (Põlme et al. 2020); Decay sugar-rich substrates (Põlme et al. 2020) | Põlme S, et al. 2020. Fungal Diversity 105:1-16 ((https://doi.org/10.1007/s13225-020-00466-2)) |
| Otu05055 | 6 | k__Fungi(100);p__Ascomycota(100);c__Leotiomycetes(100);o__Helotiales(100);f__Helotiaceae(100);g__Hymenoscyphus(100);s__Hymenoscyphus_caudatus(100); | Hymenoscyphus | 13 | Pathotroph-Saprotroph-Symbiotroph | Bryophyte Parasite-Ectomycorrhizal-Ericoid Mycorrhizal-Plant Pathogen-\|Plant Saprotroph\|-Undefined Saprotroph | Microfungus | NULL | Possible | As litter_saprotroph (Põlme et al. 2020); Dark Septate Microfungus; Syn. Pezoloma among other (see e.g., Pezoloma ericae); Decay leaf, fruit, seed (Põlme et al. 2020) | Baral HO, Kriegisteiner L. 2006. Acta Mycologica 41:11-20 ((https://doi.org/10.5586/am.2006.003)) (as a bryophyte parasite); Huhtinen S, et al. 2010. Nova Hedwigia 90:413-431 ((https://doi.org/10.1127/0029-5035/2010/0090-0413)); Bent E, et al. 2011. Applied and Environmental Microbiology 77:3351-3359 ((https://doi.org/10.1128/AEM.02575-10)) (as ectomycorrhizae); Tedersoo L, et al. 2014. Science 346:1256688 ((https://doi.org/10.1126/science.1256688)) (as saprotroph); Põlme S, et al. 2020. Fungal Diversity 105:1-16 ((https://doi.org/10.1007/s13225-020-00466-2)) |
| Otu05056 | 6 | k__Fungi(100);p__Ascomycota(100);c__Sordariomycetes(100);o__Hypocreales(100);o__Hypocreales_unclassified(100);o__Hypocreales_unclassified(100);o__Hypocreales_unclassified(100); | Hypocreales | 7 | Saprotroph | Undefined Saprotroph | Microfungus | NULL | Possible | NULL | Sterkenburg E, et al. 2015. New Phytologist 207:1145-1158 ((https://doi.org/10.1111/nph.13426)) |
| Otu05057 | 6 | k__Fungi(100);p__Basidiomycota(100);c__Agaricomycetes(100);o__Atheliales(100);f__Atheliaceae(100);g__unclassified_Atheliaceae(100);s__Atheliaceae_sp(100); | Atheliaceae | 9 | Pathotroph-Saprotroph-Symbiotroph | Ectomycorrhizal-Lichen Parasite-Lichenized-Plant Pathogen | Corticioid | NULL | Possible | Many species form ectomycorrhizal associations, often occupying a dominant position in mycorrhizal communities. Other species are pathogens of many different plants and even lichens, and one species has even been suggested to form a lichen association with cyanobacteria. Yet more species are sparse and active decomposers, and a few are aquatic (Cannon & Kirk 2007) | Cannon PF, Kirk PM. 2007. Fungal Families of the World. CAB International, Cambridge (ISBN: 978-0851998275) |
| Otu05060 | 6 | k__Fungi(100);p__Ascomycota(100);c__Saccharomycetes(100);o__Saccharomycetales(100);f__Metschnikowiaceae(100);g__Metschnikowia(100);s__Metschnikowia_pulcherrima(100); | Metschnikowia | 13 | Saprotroph | \|Undefined Saprotroph\| | Yeast | NULL | Probable | As nectar-tap_saprotroph (Põlme et al. 2020); Decay sugar-rich substrates (Põlme et al. 2020) | Põlme S, et al. 2020. Fungal Diversity 105:1-16 ((https://doi.org/10.1007/s13225-020-00466-2)) |
| Otu05062 | 6 | k__Fungi(100);p__Ascomycota(100);c__Saccharomycetes(100);o__Saccharomycetales(100);f__Metschnikowiaceae(100);g__Metschnikowia(100);s__Metschnikowia_pulcherrima(84); | Metschnikowia | 13 | Saprotroph | \|Undefined Saprotroph\| | Yeast | NULL | Probable | As nectar-tap_saprotroph (Põlme et al. 2020); Decay sugar-rich substrates (Põlme et al. 2020) | Põlme S, et al. 2020. Fungal Diversity 105:1-16 ((https://doi.org/10.1007/s13225-020-00466-2)) |
| Otu05063 | 6 | k__Fungi(100);p__Basidiomycota(100);c__Agaricomycetes(100);o__Thelephorales(100);f__Thelephoraceae(100);g__Tomentella(67);g__Tomentella_unclassified(67); | Tomentella | 13 | Symbiotroph | \|Ectomycorrhizal\| | Corticioid | NULL | Highly Probable | NULL | Rinaldi AC, et al. 2008. Fungal Diversity 33:1-45 ((http://www.fungaldiversity.org/fdp/sfdp/33-1.pdf)); Tedersoo L, et al. 2010. Mycorrhiza 20:217-263 ((https://doi.org/10.1007/s00572-009-0274-x)); Põlme S, et al. 2020. Fungal Diversity 105:1-16 ((https://doi.org/10.1007/s13225-020-00466-2)); http://mycorrhizas.info/ecmf.html ((http://mycorrhizas.info/ecmf.html)) |
| Otu05064 | 6 | k__Fungi(100);p__Ascomycota(100);c__Saccharomycetes(100);o__Saccharomycetales(100);f__Metschnikowiaceae(100);g__Metschnikowia(100);s__Metschnikowia_pulcherrima(100); | Metschnikowia | 13 | Saprotroph | \|Undefined Saprotroph\| | Yeast | NULL | Probable | As nectar-tap_saprotroph (Põlme et al. 2020); Decay sugar-rich substrates (Põlme et al. 2020) | Põlme S, et al. 2020. Fungal Diversity 105:1-16 ((https://doi.org/10.1007/s13225-020-00466-2)) |
| Otu05071 | 6 | k__Fungi(100);p__Ascomycota(100);c__Saccharomycetes(100);o__Saccharomycetales(100);f__Metschnikowiaceae(100);g__Metschnikowia(100);s__Metschnikowia_pulcherrima(100); | Metschnikowia | 13 | Saprotroph | \|Undefined Saprotroph\| | Yeast | NULL | Probable | As nectar-tap_saprotroph (Põlme et al. 2020); Decay sugar-rich substrates (Põlme et al. 2020) | Põlme S, et al. 2020. Fungal Diversity 105:1-16 ((https://doi.org/10.1007/s13225-020-00466-2)) |
| Otu05072 | 6 | k__Fungi(100);p__Ascomycota(100);c__Saccharomycetes(100);o__Saccharomycetales(100);f__Metschnikowiaceae(100);g__Metschnikowia(100);s__Metschnikowia_pulcherrima(100); | Metschnikowia | 13 | Saprotroph | \|Undefined Saprotroph\| | Yeast | NULL | Probable | As nectar-tap_saprotroph (Põlme et al. 2020); Decay sugar-rich substrates (Põlme et al. 2020) | Põlme S, et al. 2020. Fungal Diversity 105:1-16 ((https://doi.org/10.1007/s13225-020-00466-2)) |
| Otu05074 | 6 | k__Fungi(100);p__Basidiomycota(100);c__Agaricomycetes(100);o__Agaricales(100);f__Hydnangiaceae(100);g__Laccaria(100);g__Laccaria_unclassified(84); | Laccaria | 13 | Symbiotroph | \|Ectomycorrhizal\| | Agaricoid | NULL | Highly Probable | NULL | Rinaldi AC, et al. 2008. Fungal Diversity 33:1-45 ((http://www.fungaldiversity.org/fdp/sfdp/33-1.pdf)); Tedersoo L, et al. 2010. Mycorrhiza 20:217-263 ((https://doi.org/10.1007/s00572-009-0274-x)); Põlme S, et al. 2020. Fungal Diversity 105:1-16 ((https://doi.org/10.1007/s13225-020-00466-2)); http://mycorrhizas.info/ecmf.html ((http://mycorrhizas.info/ecmf.html)) |
| Otu05077 | 6 | k__Fungi(100);p__Basidiomycota(100);c__Tremellomycetes(100);o__Tremellales(100);f__Tremellales_family_Incertae_sedis(100);g__Cryptococcus(100);s__Cryptococcus_sp_QMW_2009a(100); | Cryptococcus | 13 | Pathotroph-Saprotroph-Symbiotroph | Animal Pathogen-Endophyte-Epiphyte-\|Undefined Saprotroph\| | Dimorphic Yeast-Yeast | NULL | Possible | As foliar_endophyte (Põlme et al. 2020); Decay leaf, fruit, seed, soil, dung, animal material (Põlme et al. 2020) | Andrews JH, et al. 1983. Phytopathology 73:228-234 ((https://doi.org/10.1094/Phyto-73-228)); Kurtzman CP, et al. (eds.) 2011. The Yeasts, a Taxonomic Study. Fifth Edition. Vols 1-3. Elsevier, San Diego (ISBN: 9780444521491); Põlme S, et al. 2020. Fungal Diversity 105:1-16 ((https://doi.org/10.1007/s13225-020-00466-2)) |
| Otu05078 | 6 | k__Fungi(100);p__Ascomycota(100);c__Sordariomycetes(100);o__Hypocreales(100);f__Cordycipitaceae(100);g__Simplicillium(100);s__Simplicillium_aogashimaense(100); | Simplicillium | 13 | Pathotroph-Saprotroph | \|Animal Parasite\|-Animal Pathogen-Fungal Parasite-Undefined Saprotroph | Microfungus | NULL | Probable | As animal_decomposer (Põlme et al. 2020); As mycoparasite (Põlme et al. 2020); Decay animal material (Põlme et al. 2020) | Tedersoo L, et al. 2014. Science 346:e1256688 ((https://doi.org/10.1126/science.1256688)); Põlme S, et al. 2020. Fungal Diversity 105:1-16 ((https://doi.org/10.1007/s13225-020-00466-2)) |
| Otu05079 | 6 | k__Fungi(100);p__Ascomycota(100);c__Saccharomycetes(100);o__Saccharomycetales(100);f__Metschnikowiaceae(100);g__Metschnikowia(100);g__Metschnikowia_unclassified(100); | Metschnikowia | 13 | Saprotroph | \|Undefined Saprotroph\| | Yeast | NULL | Probable | As nectar-tap_saprotroph (Põlme et al. 2020); Decay sugar-rich substrates (Põlme et al. 2020) | Põlme S, et al. 2020. Fungal Diversity 105:1-16 ((https://doi.org/10.1007/s13225-020-00466-2)) |
| Otu05083 | 6 | k__Fungi(100);p__Ascomycota(100);c__Saccharomycetes(100);o__Saccharomycetales(100);f__Metschnikowiaceae(100);g__Metschnikowia(100);s__Metschnikowia_pulcherrima(100); | Metschnikowia | 13 | Saprotroph | \|Undefined Saprotroph\| | Yeast | NULL | Probable | As nectar-tap_saprotroph (Põlme et al. 2020); Decay sugar-rich substrates (Põlme et al. 2020) | Põlme S, et al. 2020. Fungal Diversity 105:1-16 ((https://doi.org/10.1007/s13225-020-00466-2)) |
| Otu05084 | 6 | k__Fungi(100);p__Basidiomycota(100);c__Agaricomycetes(100);o__Cantharellales(100);f__Botryobasidiaceae(100);f__Botryobasidiaceae_unclassified(84);f__Botryobasidiaceae_unclassified(84); | Botryobasidiaceae | 9 | Saprotroph | Undefined Saprotroph-Wood Saprotroph | Corticioid | NULL | Probable | Saprobic, typically on rotten wood (Cannon & Kirk 2007) | Cannon PF, Kirk PM. 2007. Fungal Families of the World. CAB International, Cambridge (ISBN: 978-0851998275) |
| Otu05085 | 6 | k__Fungi(100);p__Ascomycota(100);c__Saccharomycetes(100);o__Saccharomycetales(100);f__Metschnikowiaceae(100);g__Metschnikowia(100);g__Metschnikowia_unclassified(67); | Metschnikowia | 13 | Saprotroph | \|Undefined Saprotroph\| | Yeast | NULL | Probable | As nectar-tap_saprotroph (Põlme et al. 2020); Decay sugar-rich substrates (Põlme et al. 2020) | Põlme S, et al. 2020. Fungal Diversity 105:1-16 ((https://doi.org/10.1007/s13225-020-00466-2)) |
| Otu05087 | 6 | k__Fungi(100);p__Ascomycota(100);c__Saccharomycetes(100);o__Saccharomycetales(100);f__Metschnikowiaceae(100);g__Metschnikowia(100);s__Metschnikowia_pulcherrima(100); | Metschnikowia | 13 | Saprotroph | \|Undefined Saprotroph\| | Yeast | NULL | Probable | As nectar-tap_saprotroph (Põlme et al. 2020); Decay sugar-rich substrates (Põlme et al. 2020) | Põlme S, et al. 2020. Fungal Diversity 105:1-16 ((https://doi.org/10.1007/s13225-020-00466-2)) |
| Otu05092 | 6 | k__Fungi(100);p__Basidiomycota(100);c__Cystobasidiomycetes(100);o__Cystobasidiales(100);f__Cystobasidiaceae(100);g__Occultifur(100);g__Occultifur_unclassified(100); | Occultifur | 13 | Pathotroph-Saprotroph | \|Fungal Parasite\|-Plant Saprotroph-Undefined Saprotroph | Dimorphic-Yeast | NULL | Probable | As litter_saprotroph (Põlme et al. 2020); As mycoparasite (Põlme et al. 2020); Decay leaf, fruit, seed, fungal material (Põlme et al. 2020) | Gomes FCO, et al. 2015. Antonie van Leeuwenhoek 107:597-611 ((https://doi.org/10.1007/s10482-014-0356-4)); Khunnamwong P, et al. 2017. International Journal of Systematic and Evolutionary Microbiology 67:2628-2633 ((https://doi.org/10.1099/ijsem.0.001988)); Põlme S, et al. 2020. Fungal Diversity 105:1-16 ((https://doi.org/10.1007/s13225-020-00466-2)) |
| Otu05093 | 6 | k__Fungi(100);p__Basidiomycota(100);c__Microbotryomycetes(100);o__Sporidiobolales(100);f__Sporidiobolales_family_Incertae_sedis(100);g__Rhodotorula(100);s__Rhodotorula_lamellibrachiae(67); | Rhodotorula | 13 | Pathotroph-Saprotroph-Symbiotroph | Animal Pathogen-Animal Symbiotroph-Endophyte-Plant Pathogen-\|Undefined Saprotroph\| | Yeast | NULL | Probable | As foliar_endophyte (Põlme et al. 2020); Opportunistic human pathogen | Põlme S, et al. 2020. Fungal Diversity 105:1-16 ((https://doi.org/10.1007/s13225-020-00466-2)); https://www.cdc.gov ((https://www.cdc.gov)) |
| Otu05096 | 6 | k__Fungi(100);p__Ascomycota(100);c__Lecanoromycetes(100);o__Lecanorales(100);f__Stereocaulaceae(100);g__Lepraria(100);s__Lepraria_lobificans(100); | Lepraria | 13 | Symbiotroph | \|Lichenized\| | Thallus | NULL | Highly Probable | NULL | Esslinger TL. 2014. North Dakota State University ((http://www.ndsu.edu/pubweb/~esslinge/chcklst/chcklst7.htm)); Põlme S, et al. 2020. Fungal Diversity 105:1-16 ((https://doi.org/10.1007/s13225-020-00466-2)) |
| Otu05102 | 6 | k__Fungi(100);p__Ascomycota(100);c__Saccharomycetes(100);o__Saccharomycetales(100);f__Metschnikowiaceae(100);g__Metschnikowia(100);g__Metschnikowia_unclassified(100); | Metschnikowia | 13 | Saprotroph | \|Undefined Saprotroph\| | Yeast | NULL | Probable | As nectar-tap_saprotroph (Põlme et al. 2020); Decay sugar-rich substrates (Põlme et al. 2020) | Põlme S, et al. 2020. Fungal Diversity 105:1-16 ((https://doi.org/10.1007/s13225-020-00466-2)) |
| Otu05103 | 6 | k__Fungi(100);p__Basidiomycota(100);c__Agaricomycetes(100);o__Agaricales(100);o__Agaricales_unclassified(100);o__Agaricales_unclassified(100);o__Agaricales_unclassified(100); | Agaricales | 7 | Pathotroph-Saprotroph-Symbiotroph | Bryophyte Parasite-Dung Saprotroph-Ectomycorrhizal-Fungal Parasite-Leaf Saprotroph-Plant Parasite-Undefined Saprotroph-Wood Saprotroph | Agaricoid-Gasteroid-Microfungus-Secotioid-Yeast | NULL | Possible | Mushrooms and toadstools, Gill fungi, Agarics... lignicolous, sometimes muscicolous or fungicolous, saprobic, mycorrhizal, rarely parasitic on plants or fungi (Kirk et al. 2008) | Kirk PM et al. 2008. Dictionary of the Fungi. Tenth Edition. CAB International, Wallingford (ISBN: 978-0851998268) |
| Otu05104 | 6 | k__Fungi(100);p__Ascomycota(100);c__Sordariomycetes(100);o__Xylariales(100);f__Amphisphaeriaceae(100);g__unclassified_Amphisphaeriaceae(100);s__Amphisphaeriaceae_sp(100); | Amphisphaeriaceae | 9 | Saprotroph | Undefined Saprotroph | Microfungus | NULL | Possible | Saprobes or weak parasites of leaves, stems, etc., often appearing as secondary colonizers following damage to plants by other agents (Cannon & Kirk 2007) | Cannon PF, Kirk PM. 2007. Fungal Families of the World. CAB International, Cambridge (ISBN: 978-0851998275) |
| Otu05105 | 6 | k__Fungi(100);p__Basidiomycota(100);c__Microbotryomycetes(100);o__Sporidiobolales(100);f__Sporidiobolales_family_Incertae_sedis(100);g__Sporobolomyces(100);s__Sporobolomyces_lactosus(100); | Sporobolomyces | 13 | Pathotroph-Saprotroph | \|Fungal Parasite\|-Plant Saprotroph | Yeast | NULL | Probable | As litter_saprotroph (Põlme et al. 2020); As mycoparasite (Põlme et al. 2020); Decay fungal material (Põlme et al. 2020) | Tedersoo L, et al. 2014. Science 346:e1256688 ((https://doi.org/10.1126/science.1256688)); Purahong W, et al. 2016. Molecular Ecology 25:4059-4074 ((https://doi.org/10.1111/mec.13739)); Põlme S, et al. 2020. Fungal Diversity 105:1-16 ((https://doi.org/10.1007/s13225-020-00466-2)) |
| Otu05106 | 6 | k__Fungi(100);p__Ascomycota(100);c__Saccharomycetes(100);o__Saccharomycetales(100);f__Metschnikowiaceae(100);g__Metschnikowia(100);g__Metschnikowia_unclassified(67); | Metschnikowia | 13 | Saprotroph | \|Undefined Saprotroph\| | Yeast | NULL | Probable | As nectar-tap_saprotroph (Põlme et al. 2020); Decay sugar-rich substrates (Põlme et al. 2020) | Põlme S, et al. 2020. Fungal Diversity 105:1-16 ((https://doi.org/10.1007/s13225-020-00466-2)) |
| Otu05110 | 6 | k__Fungi(100);p__Ascomycota(100);c__Saccharomycetes(100);o__Saccharomycetales(100);f__Metschnikowiaceae(100);g__Metschnikowia(100);s__Metschnikowia_pulcherrima(84); | Metschnikowia | 13 | Saprotroph | \|Undefined Saprotroph\| | Yeast | NULL | Probable | As nectar-tap_saprotroph (Põlme et al. 2020); Decay sugar-rich substrates (Põlme et al. 2020) | Põlme S, et al. 2020. Fungal Diversity 105:1-16 ((https://doi.org/10.1007/s13225-020-00466-2)) |
| Otu05114 | 6 | k__Fungi(100);p__Ascomycota(100);c__Saccharomycetes(100);o__Saccharomycetales(100);f__Metschnikowiaceae(100);g__Metschnikowia(100);g__Metschnikowia_unclassified(100); | Metschnikowia | 13 | Saprotroph | \|Undefined Saprotroph\| | Yeast | NULL | Probable | As nectar-tap_saprotroph (Põlme et al. 2020); Decay sugar-rich substrates (Põlme et al. 2020) | Põlme S, et al. 2020. Fungal Diversity 105:1-16 ((https://doi.org/10.1007/s13225-020-00466-2)) |
| Otu05119 | 6 | k__Fungi(100);p__Ascomycota(100);c__Dothideomycetes(100);o__Pleosporales(100);f__Phaeosphaeriaceae(100);f__Phaeosphaeriaceae_unclassified(67);f__Phaeosphaeriaceae_unclassified(67); | Phaeosphaeriaceae | 9 | Pathotroph-Saprotroph | Fungal Parasite-Plant Pathogen-Plant Saprotroph | Microfungus | NULL | Probable | Necrotrophic or saprobic on a wide range of plants, especially monocotyledons; a few species parasitic on other fungi (Cannon & Kirk 2007) | Cannon PF, Kirk PM. 2007. Fungal Families of the World. CAB International, Cambridge (ISBN: 978-0851998275) |
| Otu05120 | 6 | k__Fungi(100);p__Ascomycota(100);c__Sordariomycetes(100);o__Sordariomycetes_order_Incertae_sedis(100);f__Sordariomycetes_family_Incertae_sedis(100);g__Myrmecridium(100);s__Myrmecridium_schulzeri(100); | Myrmecridium | 13 | Saprotroph | \|Undefined Saprotroph\| | Microfungus | NULL | Probable | As soil_saprotroph (Põlme et al. 2020); Decay soil (Põlme et al. 2020) | Tedersoo L, et al. 2014. Science 346:e1256688 ((https://doi.org/10.1126/science.1256688)); Põlme S, et al. 2020. Fungal Diversity 105:1-16 ((https://doi.org/10.1007/s13225-020-00466-2)) |
| Otu05122 | 6 | k__Fungi(100);p__Ascomycota(100);c__Saccharomycetes(100);o__Saccharomycetales(100);f__Metschnikowiaceae(100);g__Metschnikowia(100);s__Metschnikowia_pulcherrima(84); | Metschnikowia | 13 | Saprotroph | \|Undefined Saprotroph\| | Yeast | NULL | Probable | As nectar-tap_saprotroph (Põlme et al. 2020); Decay sugar-rich substrates (Põlme et al. 2020) | Põlme S, et al. 2020. Fungal Diversity 105:1-16 ((https://doi.org/10.1007/s13225-020-00466-2)) |
| Otu05124 | 6 | k__Fungi(100);p__Basidiomycota(100);c__Microbotryomycetes(100);o__Sporidiobolales(100);f__Sporidiobolales_family_Incertae_sedis(100);g__Sporobolomyces(100);s__Sporobolomyces_aff_jilinensis_MCA_3774(100); | Sporobolomyces | 13 | Pathotroph-Saprotroph | \|Fungal Parasite\|-Plant Saprotroph | Yeast | NULL | Probable | As litter_saprotroph (Põlme et al. 2020); As mycoparasite (Põlme et al. 2020); Decay fungal material (Põlme et al. 2020) | Tedersoo L, et al. 2014. Science 346:e1256688 ((https://doi.org/10.1126/science.1256688)); Purahong W, et al. 2016. Molecular Ecology 25:4059-4074 ((https://doi.org/10.1111/mec.13739)); Põlme S, et al. 2020. Fungal Diversity 105:1-16 ((https://doi.org/10.1007/s13225-020-00466-2)) |
| Otu05125 | 6 | k__Fungi(100);p__Ascomycota(100);c__Saccharomycetes(100);o__Saccharomycetales(100);f__Metschnikowiaceae(100);g__Metschnikowia(100);s__Metschnikowia_pulcherrima(84); | Metschnikowia | 13 | Saprotroph | \|Undefined Saprotroph\| | Yeast | NULL | Probable | As nectar-tap_saprotroph (Põlme et al. 2020); Decay sugar-rich substrates (Põlme et al. 2020) | Põlme S, et al. 2020. Fungal Diversity 105:1-16 ((https://doi.org/10.1007/s13225-020-00466-2)) |
| Otu05126 | 6 | k__Fungi(100);p__Basidiomycota(100);c__Agaricostilbomycetes(100);o__Agaricostilbales(100);f__Agaricostilbaceae(100);g__Bensingtonia(100);g__Bensingtonia_unclassified(100); | Bensingtonia | 13 | Saprotroph | \|Undefined Saprotroph\| | Dimorphic-Facultative Yeast | NULL | Highly Probable | NULL | Kurtzman CP, et al. (eds.) 2011. The Yeasts, a Taxonomic Study. Fifth Edition. Vols 1-3. Elsevier, San Diego (ISBN: 9780444521491); Põlme S, et al. 2020. Fungal Diversity 105:1-16 ((https://doi.org/10.1007/s13225-020-00466-2)) |
| Otu05127 | 6 | k__Fungi(100);p__Ascomycota(100);c__Saccharomycetes(100);o__Saccharomycetales(100);f__Metschnikowiaceae(100);g__Metschnikowia(100);s__Metschnikowia_pulcherrima(84); | Metschnikowia | 13 | Saprotroph | \|Undefined Saprotroph\| | Yeast | NULL | Probable | As nectar-tap_saprotroph (Põlme et al. 2020); Decay sugar-rich substrates (Põlme et al. 2020) | Põlme S, et al. 2020. Fungal Diversity 105:1-16 ((https://doi.org/10.1007/s13225-020-00466-2)) |
| Otu05128 | 6 | k__Fungi(100);p__Basidiomycota(100);c__Agaricomycetes(100);o__Boletales(100);f__Coniophoraceae(100);g__Coniophora(100);s__Coniophora_olivacea(100); | Coniophora | 13 | Saprotroph | \|Wood Saprotroph\| | Corticioid | Brown Rot | Probable | As brown-rot (Põlme et al. 2020); Decay wood (Põlme et al. 2020) | Tedersoo L, et al. 2014. Science 346:e1256688 ((https://doi.org/10.1126/science.1256688)); Põlme S, et al. 2020. Fungal Diversity 105:1-16 ((https://doi.org/10.1007/s13225-020-00466-2)) |
| Otu05129 | 6 | k__Fungi(100);p__Ascomycota(100);c__Saccharomycetes(100);o__Saccharomycetales(100);f__Metschnikowiaceae(100);g__Metschnikowia(100);s__Metschnikowia_pulcherrima(84); | Metschnikowia | 13 | Saprotroph | \|Undefined Saprotroph\| | Yeast | NULL | Probable | As nectar-tap_saprotroph (Põlme et al. 2020); Decay sugar-rich substrates (Põlme et al. 2020) | Põlme S, et al. 2020. Fungal Diversity 105:1-16 ((https://doi.org/10.1007/s13225-020-00466-2)) |
| Otu05131 | 6 | k__Fungi(100);p__Ascomycota(100);c__Saccharomycetes(100);o__Saccharomycetales(100);f__Metschnikowiaceae(100);g__Metschnikowia(100);s__Metschnikowia_pulcherrima(67); | Metschnikowia | 13 | Saprotroph | \|Undefined Saprotroph\| | Yeast | NULL | Probable | As nectar-tap_saprotroph (Põlme et al. 2020); Decay sugar-rich substrates (Põlme et al. 2020) | Põlme S, et al. 2020. Fungal Diversity 105:1-16 ((https://doi.org/10.1007/s13225-020-00466-2)) |
| Otu05133 | 6 | k__Fungi(100);p__Ascomycota(100);c__Leotiomycetes(100);o__Helotiales(100);f__Helotiaceae(100);g__Hymenoscyphus(100);g__Hymenoscyphus_unclassified(100); | Hymenoscyphus | 13 | Pathotroph-Saprotroph-Symbiotroph | Bryophyte Parasite-Ectomycorrhizal-Ericoid Mycorrhizal-Plant Pathogen-\|Plant Saprotroph\|-Undefined Saprotroph | Microfungus | NULL | Possible | As litter_saprotroph (Põlme et al. 2020); Dark Septate Microfungus; Syn. Pezoloma among other (see e.g., Pezoloma ericae); Decay leaf, fruit, seed (Põlme et al. 2020) | Baral HO, Kriegisteiner L. 2006. Acta Mycologica 41:11-20 ((https://doi.org/10.5586/am.2006.003)) (as a bryophyte parasite); Huhtinen S, et al. 2010. Nova Hedwigia 90:413-431 ((https://doi.org/10.1127/0029-5035/2010/0090-0413)); Bent E, et al. 2011. Applied and Environmental Microbiology 77:3351-3359 ((https://doi.org/10.1128/AEM.02575-10)) (as ectomycorrhizae); Tedersoo L, et al. 2014. Science 346:1256688 ((https://doi.org/10.1126/science.1256688)) (as saprotroph); Põlme S, et al. 2020. Fungal Diversity 105:1-16 ((https://doi.org/10.1007/s13225-020-00466-2)) |
| Otu05136 | 6 | k__Fungi(100);p__Ascomycota(100);c__Dothideomycetes(100);o__Capnodiales(100);f__Teratosphaeriaceae(100);g__Teratosphaeria(84);g__Teratosphaeria_unclassified(84); | Teratosphaeria | 13 | Pathotroph-Saprotroph | \|Plant Pathogen\|-Plant Saprotroph-Wood Saprotroph | Microfungus | NULL | Probable | As litter_saprotroph (Põlme et al. 2020); As wood_pathogen (Põlme et al. 2020); Decay wood, leaf, fruit, seed (Põlme et al. 2020) | Yamazaki A, et al. 2010. Mycoscience 51:443-455 ((https://www.jstage.jst.go.jp/article/mycosci/51/6/51_MYC51443/_pdf)); Tedersoo L, et al. 2014. Science 346:e1256688 ((https://doi.org/10.1126/science.1256688)); Põlme S, et al. 2020. Fungal Diversity 105:1-16 ((https://doi.org/10.1007/s13225-020-00466-2)) |
| Otu05137 | 6 | k__Fungi(100);p__Ascomycota(100);c__Saccharomycetes(84);o__Saccharomycetales(84);f__Saccharomycetales_family_Incertae_sedis(84);g__Candida(67);g__Candida_unclassified(67); | Candida | 13 | Pathotroph-Saprotroph-Symbiotroph | Animal Pathogen-Endophyte-Endosymbiont-Epiphyte-\|Undefined Saprotroph\| | Dimorphic Yeast-Yeast | NULL | Probable | As nectar-tap_saprotroph (Põlme et al. 2020); Decay sugar-rich substrates (Põlme et al. 2020); This asexual genus is polyphyletic and contain species that function very differently depend on what they are. Caution should be taken when interpreting guilds for Candida species. | Manolakaki D, et al. 2010. Virulence. 1: 367-375 ((https://doi.org/10.4161/viru.1.5.12796)); Mekbib SB, et al. 2011. Tropical Plant Pathology 36:233-240 ((https://doi.org/10.1590/S1982-56762011000400004)); Põlme S, et al. 2020. Fungal Diversity 105:1-16 ((https://doi.org/10.1007/s13225-020-00466-2)) |
| Otu05138 | 6 | k__Fungi(100);p__Basidiomycota(100);c__Tremellomycetes(100);o__Tremellales(100);f__Tremellales_family_Incertae_sedis(100);g__Dioszegia(100);s__Dioszegia_hungarica(100); | Dioszegia | 13 | Saprotroph | \|Plant Saprotroph\|-Undefined Saprotroph | Yeast | NULL | Probable | As litter_saprotroph (Põlme et al. 2020); Decay leaf, fruit, seed (Põlme et al. 2020); The Dioszegia subclade sensu Inacio (Inacio et al. 20015) is characterized by orange; yellow or orange to deep orange colonies. Isolated from plant leaves and soil. | Põlme S, et al. 2020. Fungal Diversity 105:1-16 ((https://doi.org/10.1007/s13225-020-00466-2)); Takashima M, et al. 2011 International Journal of Systematic and Evolutionary Microbiology 61:1736-1739 ((https://doi.org/10.1099/ijs.0.025254-0)) |
| Otu05140 | 6 | k__Fungi(100);p__Ascomycota(100);c__Saccharomycetes(100);o__Saccharomycetales(100);f__Metschnikowiaceae(100);g__Metschnikowia(100);s__Metschnikowia_pulcherrima(84); | Metschnikowia | 13 | Saprotroph | \|Undefined Saprotroph\| | Yeast | NULL | Probable | As nectar-tap_saprotroph (Põlme et al. 2020); Decay sugar-rich substrates (Põlme et al. 2020) | Põlme S, et al. 2020. Fungal Diversity 105:1-16 ((https://doi.org/10.1007/s13225-020-00466-2)) |
| Otu05141 | 6 | k__Fungi(100);p__Basidiomycota(100);c__Agaricomycetes(100);o__Agaricales(100);f__Agaricaceae(100);g__Melanophyllum(84);s__Melanophyllum_haematospermum(84); | Melanophyllum | 13 | Saprotroph | \|Undefined Saprotroph\| | Agaricoid | NULL | Probable | As soil_saprotroph (Põlme et al. 2020); Decay soil (Põlme et al. 2020) | Tedersoo L, et al. 2014. Science 346:e1256688 ((https://doi.org/10.1126/science.1256688)); Põlme S, et al. 2020. Fungal Diversity 105:1-16 ((https://doi.org/10.1007/s13225-020-00466-2)) |
| Otu05143 | 6 | k__Fungi(100);p__Ascomycota(100);c__Saccharomycetes(100);o__Saccharomycetales(100);f__Metschnikowiaceae(100);g__Metschnikowia(100);s__Metschnikowia_pulcherrima(84); | Metschnikowia | 13 | Saprotroph | \|Undefined Saprotroph\| | Yeast | NULL | Probable | As nectar-tap_saprotroph (Põlme et al. 2020); Decay sugar-rich substrates (Põlme et al. 2020) | Põlme S, et al. 2020. Fungal Diversity 105:1-16 ((https://doi.org/10.1007/s13225-020-00466-2)) |
| Otu05144 | 6 | k__Fungi(100);p__Zygomycota(100);c__Zygomycota_class_Incertae_sedis(100);o__Mortierellales(100);f__Mortierellaceae(100);g__Mortierella(100);s__Mortierella_sp_1_RB_2011(100); | Mortierella | 13 | Saprotroph-Symbiotroph | Endophyte-Plant Saprotroph-\|Undefined Saprotroph\| | Microfungus | NULL | Probable | As chitinolytic (Põlme et al. 2020); As soil_saprotroph (Põlme et al. 2020); Decay roots, soil (Põlme et al. 2020); Saprobic in soil and thus frequently isolated from soil and material contaminated with soil such as hay or rotten silage and spoil tips (Cannon & Kirk 2007) | Cannon PF, Kirk PM. 2007. Fungal Families of the World. CAB International, Cambridge (ISBN: 978-0851998275); Tedersoo L, et al. 2014. Science 346:e1256688 ((https://doi.org/10.1126/science.1256688)); Purahong W, et al. 2016. Molecular Ecology 25:4059-4074 ((https://doi.org/10.1111/mec.13739)); Põlme S, et al. 2020. Fungal Diversity 105:1-16 ((https://doi.org/10.1007/s13225-020-00466-2)) |
| Otu05145 | 6 | k__Fungi(100);p__Ascomycota(100);c__Saccharomycetes(100);o__Saccharomycetales(100);f__Metschnikowiaceae(100);g__Metschnikowia(100);s__Metschnikowia_pulcherrima(100); | Metschnikowia | 13 | Saprotroph | \|Undefined Saprotroph\| | Yeast | NULL | Probable | As nectar-tap_saprotroph (Põlme et al. 2020); Decay sugar-rich substrates (Põlme et al. 2020) | Põlme S, et al. 2020. Fungal Diversity 105:1-16 ((https://doi.org/10.1007/s13225-020-00466-2)) |
| Otu05147 | 6 | k__Fungi(100);p__Ascomycota(100);c__Saccharomycetes(100);o__Saccharomycetales(100);f__Saccharomycetales_family_Incertae_sedis(84);g__Candida(84);g__Candida_unclassified(67); | Candida | 13 | Pathotroph-Saprotroph-Symbiotroph | Animal Pathogen-Endophyte-Endosymbiont-Epiphyte-\|Undefined Saprotroph\| | Dimorphic Yeast-Yeast | NULL | Probable | As nectar-tap_saprotroph (Põlme et al. 2020); Decay sugar-rich substrates (Põlme et al. 2020); This asexual genus is polyphyletic and contain species that function very differently depend on what they are. Caution should be taken when interpreting guilds for Candida species. | Manolakaki D, et al. 2010. Virulence. 1: 367-375 ((https://doi.org/10.4161/viru.1.5.12796)); Mekbib SB, et al. 2011. Tropical Plant Pathology 36:233-240 ((https://doi.org/10.1590/S1982-56762011000400004)); Põlme S, et al. 2020. Fungal Diversity 105:1-16 ((https://doi.org/10.1007/s13225-020-00466-2)) |
| Otu05148 | 6 | k__Fungi(100);p__Ascomycota(100);c__Dothideomycetes(100);o__Pleosporales(100);f__Venturiaceae(100);g__Venturia(100);g__Venturia_unclassified(100); | Venturia | 13 | Pathotroph-Pathotroph-Saprotroph | \|Plant Pathogen\|-Plant Saprotroph | Microfungus | NULL | Probable | As litter_saprotroph (Põlme et al. 2020); Decay leaf, fruit, seed (Põlme et al. 2020) | Tedersoo L, et al. 2014. Science 346:e1256688 ((https://doi.org/10.1126/science.1256688)); Põlme S, et al. 2020. Fungal Diversity 105:1-16 ((https://doi.org/10.1007/s13225-020-00466-2)) |
| Otu05149 | 6 | k__Fungi(100);p__Ascomycota(100);c__Saccharomycetes(100);o__Saccharomycetales(100);f__Metschnikowiaceae(100);g__Metschnikowia(100);g__Metschnikowia_unclassified(84); | Metschnikowia | 13 | Saprotroph | \|Undefined Saprotroph\| | Yeast | NULL | Probable | As nectar-tap_saprotroph (Põlme et al. 2020); Decay sugar-rich substrates (Põlme et al. 2020) | Põlme S, et al. 2020. Fungal Diversity 105:1-16 ((https://doi.org/10.1007/s13225-020-00466-2)) |
| Otu04614 | 6 | k__Fungi(100);p__Ascomycota(100);c__Dothideomycetes(100);o__Pleosporales(100);o__Pleosporales_unclassified(84);o__Pleosporales_unclassified(84);o__Pleosporales_unclassified(84); | - | - | - | - | - | - | - | Unassigned | - |
| Otu04615 | 6 | k__Fungi(100);p__Ascomycota(100);p__Ascomycota_unclassified(67);p__Ascomycota_unclassified(67);p__Ascomycota_unclassified(67);p__Ascomycota_unclassified(67);p__Ascomycota_unclassified(67); | - | - | - | - | - | - | - | Unassigned | - |
| Otu04617 | 6 | k__Fungi(100);k__Fungi_unclassified(67);k__Fungi_unclassified(67);k__Fungi_unclassified(67);k__Fungi_unclassified(67);k__Fungi_unclassified(67);k__Fungi_unclassified(67); | - | - | - | - | - | - | - | Unassigned | - |
| Otu04621 | 6 | k__Fungi(100);p__Basidiomycota(100);p__Basidiomycota_unclassified(100);p__Basidiomycota_unclassified(100);p__Basidiomycota_unclassified(100);p__Basidiomycota_unclassified(100);p__Basidiomycota_unclassified(100); | - | - | - | - | - | - | - | Unassigned | - |
| Otu04625 | 6 | k__Fungi(100);p__Ascomycota(100);c__Leotiomycetes(100);o__Helotiales(100);o__Helotiales_unclassified(100);o__Helotiales_unclassified(100);o__Helotiales_unclassified(100); | - | - | - | - | - | - | - | Unassigned | - |
| Otu04628 | 6 | k__Fungi(100);k__Fungi_unclassified(67);k__Fungi_unclassified(67);k__Fungi_unclassified(67);k__Fungi_unclassified(67);k__Fungi_unclassified(67);k__Fungi_unclassified(67); | - | - | - | - | - | - | - | Unassigned | - |
| Otu04629 | 6 | k__Fungi(100);p__Ascomycota(100);c__Sordariomycetes(100);c__Sordariomycetes_unclassified(100);c__Sordariomycetes_unclassified(100);c__Sordariomycetes_unclassified(100);c__Sordariomycetes_unclassified(100); | - | - | - | - | - | - | - | Unassigned | - |
| Otu04630 | 6 | k__Fungi(100);k__Fungi_unclassified(84);k__Fungi_unclassified(84);k__Fungi_unclassified(84);k__Fungi_unclassified(84);k__Fungi_unclassified(84);k__Fungi_unclassified(84); | - | - | - | - | - | - | - | Unassigned | - |
| Otu04632 | 6 | k__Fungi(100);k__Fungi_unclassified(67);k__Fungi_unclassified(67);k__Fungi_unclassified(67);k__Fungi_unclassified(67);k__Fungi_unclassified(67);k__Fungi_unclassified(67); | - | - | - | - | - | - | - | Unassigned | - |
| Otu04634 | 6 | k__Fungi(100);p__Ascomycota(67);p__Ascomycota_unclassified(67);p__Ascomycota_unclassified(67);p__Ascomycota_unclassified(67);p__Ascomycota_unclassified(67);p__Ascomycota_unclassified(67); | - | - | - | - | - | - | - | Unassigned | - |
| Otu04635 | 6 | k__Fungi(100);p__Ascomycota(67);p__Ascomycota_unclassified(67);p__Ascomycota_unclassified(67);p__Ascomycota_unclassified(67);p__Ascomycota_unclassified(67);p__Ascomycota_unclassified(67); | - | - | - | - | - | - | - | Unassigned | - |
| Otu04636 | 6 | k__Fungi(100);p__Ascomycota(100);p__Ascomycota_unclassified(100);p__Ascomycota_unclassified(100);p__Ascomycota_unclassified(100);p__Ascomycota_unclassified(100);p__Ascomycota_unclassified(100); | - | - | - | - | - | - | - | Unassigned | - |
| Otu04638 | 6 | k__Fungi(100);k__Fungi_unclassified(100);k__Fungi_unclassified(100);k__Fungi_unclassified(100);k__Fungi_unclassified(100);k__Fungi_unclassified(100);k__Fungi_unclassified(100); | - | - | - | - | - | - | - | Unassigned | - |
| Otu04639 | 6 | k__Fungi(100);p__Ascomycota(84);p__Ascomycota_unclassified(84);p__Ascomycota_unclassified(84);p__Ascomycota_unclassified(84);p__Ascomycota_unclassified(84);p__Ascomycota_unclassified(84); | - | - | - | - | - | - | - | Unassigned | - |

| | | | | | | | | | | | |
|---|---|---|---|---|---|---|---|---|---|---|---|
| Otu04642 | 6 | k__Fungi(100);p__Ascomycota(100);c__Sordariomycetes(100);c__Sordariomycetes_unclassified(100);c__Sordariomycetes_unclassified(100);c__Sordariomycetes_unclassified(100);c__Sordariomycetes_unclassified(100); | - | - | - | - | - | - | - | Unassigned | - |
| Otu04643 | 6 | k__Fungi(100);k__Fungi_unclassified(67);k__Fungi_unclassified(67);k__Fungi_unclassified(67);k__Fungi_unclassified(67);k__Fungi_unclassified(67);k__Fungi_unclassified(67); | - | - | - | - | - | - | - | Unassigned | - |
| Otu04644 | 6 | k__Fungi(100);p__Ascomycota(84);p__Ascomycota_unclassified(84);p__Ascomycota_unclassified(84);p__Ascomycota_unclassified(84);p__Ascomycota_unclassified(84);p__Ascomycota_unclassified(84); | - | - | - | - | - | - | - | Unassigned | - |
| Otu04648 | 6 | k__Fungi(100);k__Fungi_unclassified(100);k__Fungi_unclassified(100);k__Fungi_unclassified(100);k__Fungi_unclassified(100);k__Fungi_unclassified(100);k__Fungi_unclassified(100); | - | - | - | - | - | - | - | Unassigned | - |
| Otu04650 | 6 | k__Fungi(100);p__Basidiomycota(100);c__Agaricomycetes(100);o__Polyporales(100);o__Polyporales_unclassified(84);o__Polyporales_unclassified(84);o__Polyporales_unclassified(84); | - | - | - | - | - | - | - | Unassigned | - |
| Otu04652 | 6 | k__Fungi(100);p__Ascomycota(100);p__Ascomycota_unclassified(100);p__Ascomycota_unclassified(100);p__Ascomycota_unclassified(100);p__Ascomycota_unclassified(100);p__Ascomycota_unclassified(100); | - | - | - | - | - | - | - | Unassigned | - |
| Otu04657 | 6 | k__Fungi(100);p__Ascomycota(100);p__Ascomycota_unclassified(100);p__Ascomycota_unclassified(100);p__Ascomycota_unclassified(100);p__Ascomycota_unclassified(100);p__Ascomycota_unclassified(100); | - | - | - | - | - | - | - | Unassigned | - |
| Otu04661 | 6 | k__Fungi(100);p__Ascomycota(100);c__Leotiomycetes(100);o__Helotiales(100);f__Helotiales_family_Incertae_sedis(100);f__Helotiales_family_Incertae_sedis_unclassified(100);f__Helotiales_family_Incertae_sedis_unclassified(100); | - | - | - | - | - | - | - | Unassigned | - |
| Otu04662 | 6 | k__Fungi(100);p__Basidiomycota(100);c__Agaricomycetes(100);o__Agaricales(100);f__Typhulaceae(100);g__unclassified_Typhulaceae(100);s__Typhulaceae_sp(100); | - | - | - | - | - | - | - | Unassigned | - |
| Otu04664 | 6 | k__Fungi(100);k__Fungi_unclassified(67);k__Fungi_unclassified(67);k__Fungi_unclassified(67);k__Fungi_unclassified(67);k__Fungi_unclassified(67);k__Fungi_unclassified(67); | - | - | - | - | - | - | - | Unassigned | - |
| Otu04667 | 6 | k__Fungi(100);k__Fungi_unclassified(67);k__Fungi_unclassified(67);k__Fungi_unclassified(67);k__Fungi_unclassified(67);k__Fungi_unclassified(67);k__Fungi_unclassified(67); | - | - | - | - | - | - | - | Unassigned | - |
| Otu04668 | 6 | k__Fungi(100);p__Ascomycota(100);p__Ascomycota_unclassified(100);p__Ascomycota_unclassified(100);p__Ascomycota_unclassified(100);p__Ascomycota_unclassified(100);p__Ascomycota_unclassified(100); | - | - | - | - | - | - | - | Unassigned | - |
| Otu04669 | 6 | k__Fungi(100);p__Ascomycota(100);p__Ascomycota_unclassified(100);p__Ascomycota_unclassified(100);p__Ascomycota_unclassified(100);p__Ascomycota_unclassified(100);p__Ascomycota_unclassified(100); | - | - | - | - | - | - | - | Unassigned | - |
| Otu04670 | 6 | k__Fungi(100);p__Ascomycota(84);p__Ascomycota_unclassified(84);p__Ascomycota_unclassified(84);p__Ascomycota_unclassified(84);p__Ascomycota_unclassified(84);p__Ascomycota_unclassified(84); | - | - | - | - | - | - | - | Unassigned | - |
| Otu04672 | 6 | k__Fungi(100);k__Fungi_unclassified(100);k__Fungi_unclassified(100);k__Fungi_unclassified(100);k__Fungi_unclassified(100);k__Fungi_unclassified(100);k__Fungi_unclassified(100); | - | - | - | - | - | - | - | Unassigned | - |
| Otu04675 | 6 | k__Fungi(100);k__Fungi_unclassified(100);k__Fungi_unclassified(100);k__Fungi_unclassified(100);k__Fungi_unclassified(100);k__Fungi_unclassified(100);k__Fungi_unclassified(100); | - | - | - | - | - | - | - | Unassigned | - |
| Otu04676 | 6 | k__Fungi(100);p__Basidiomycota(100);c__Agaricomycetes(100);o__Cantharellales(100);f__unclassified_Cantharellales(100);g__unclassified_Cantharellales(100);s__Cantharellales_sp(100); | - | - | - | - | - | - | - | Unassigned | - |
| Otu04680 | 6 | k__Fungi(100);p__Ascomycota(100);p__Ascomycota_unclassified(100);p__Ascomycota_unclassified(100);p__Ascomycota_unclassified(100);p__Ascomycota_unclassified(100);p__Ascomycota_unclassified(100); | - | - | - | - | - | - | - | Unassigned | - |
| Otu04683 | 6 | k__Fungi(100);k__Fungi_unclassified(100);k__Fungi_unclassified(100);k__Fungi_unclassified(100);k__Fungi_unclassified(100);k__Fungi_unclassified(100);k__Fungi_unclassified(100); | - | - | - | - | - | - | - | Unassigned | - |
| Otu04685 | 6 | k__Plantae(100);p__unclassified_Plantae(100);c__unclassified_Plantae(100);o__unclassified_Plantae(100);f__unclassified_Plantae(100);g__unclassified_Plantae(100);s__Plantae_sp(100); | - | - | - | - | - | - | - | Unassigned | - |
| Otu04686 | 6 | k__Fungi(100);p__Ascomycota(100);c__Sordariomycetes(100);c__Sordariomycetes_unclassified(100);c__Sordariomycetes_unclassified(100);c__Sordariomycetes_unclassified(100);c__Sordariomycetes_unclassified(100); | - | - | - | - | - | - | - | Unassigned | - |
| Otu04688 | 6 | k__Fungi(100);k__Fungi_unclassified(100);k__Fungi_unclassified(100);k__Fungi_unclassified(100);k__Fungi_unclassified(100);k__Fungi_unclassified(100);k__Fungi_unclassified(100); | - | - | - | - | - | - | - | Unassigned | - |
| Otu04693 | 6 | k__Fungi(100);k__Fungi_unclassified(100);k__Fungi_unclassified(100);k__Fungi_unclassified(100);k__Fungi_unclassified(100);k__Fungi_unclassified(100);k__Fungi_unclassified(100); | - | - | - | - | - | - | - | Unassigned | - |
| Otu04700 | 6 | k__Fungi(100);p__Ascomycota(100);p__Ascomycota_unclassified(100);p__Ascomycota_unclassified(100);p__Ascomycota_unclassified(100);p__Ascomycota_unclassified(100);p__Ascomycota_unclassified(100); | - | - | - | - | - | - | - | Unassigned | - |
| Otu04703 | 6 | k__Fungi(100);p__Ascomycota(100);p__Ascomycota_unclassified(100);p__Ascomycota_unclassified(100);p__Ascomycota_unclassified(100);p__Ascomycota_unclassified(100);p__Ascomycota_unclassified(100); | - | - | - | - | - | - | - | Unassigned | - |
| Otu04704 | 6 | k__Fungi(100);k__Fungi_unclassified(100);k__Fungi_unclassified(100);k__Fungi_unclassified(100);k__Fungi_unclassified(100);k__Fungi_unclassified(100);k__Fungi_unclassified(100); | - | - | - | - | - | - | - | Unassigned | - |
| Otu04705 | 6 | k__Fungi(100);p__Ascomycota(67);p__Ascomycota_unclassified(67);p__Ascomycota_unclassified(67);p__Ascomycota_unclassified(67);p__Ascomycota_unclassified(67);p__Ascomycota_unclassified(67); | - | - | - | - | - | - | - | Unassigned | - |
| Otu04708 | 6 | k__Fungi(100);k__Fungi_unclassified(100);k__Fungi_unclassified(100);k__Fungi_unclassified(100);k__Fungi_unclassified(100);k__Fungi_unclassified(100);k__Fungi_unclassified(100); | - | - | - | - | - | - | - | Unassigned | - |
| Otu04710 | 6 | k__Fungi(100);k__Fungi_unclassified(84);k__Fungi_unclassified(84);k__Fungi_unclassified(84);k__Fungi_unclassified(84);k__Fungi_unclassified(84);k__Fungi_unclassified(84); | - | - | - | - | - | - | - | Unassigned | - |
| Otu04711 | 6 | k__Fungi(100);p__Basidiomycota(100);p__Basidiomycota_unclassified(100);p__Basidiomycota_unclassified(100);p__Basidiomycota_unclassified(100);p__Basidiomycota_unclassified(100);p__Basidiomycota_unclassified(100); | - | - | - | - | - | - | - | Unassigned | - |
| Otu04714 | 6 | k__Fungi(100);p__Basidiomycota(100);c__Agaricomycetes(100);o__Cantharellales(100);o__Cantharellales_unclassified(100);o__Cantharellales_unclassified(100);o__Cantharellales_unclassified(100); | - | - | - | - | - | - | - | Unassigned | - |
| Otu04715 | 6 | k__Fungi(100);k__Fungi_unclassified(100);k__Fungi_unclassified(100);k__Fungi_unclassified(100);k__Fungi_unclassified(100);k__Fungi_unclassified(100);k__Fungi_unclassified(100); | - | - | - | - | - | - | - | Unassigned | - |
| Otu04717 | 6 | k__Fungi(100);p__Ascomycota(84);p__Ascomycota_unclassified(84);p__Ascomycota_unclassified(84);p__Ascomycota_unclassified(84);p__Ascomycota_unclassified(84);p__Ascomycota_unclassified(84); | - | - | - | - | - | - | - | Unassigned | - |
| Otu04720 | 6 | k__Fungi(100);k__Fungi_unclassified(100);k__Fungi_unclassified(100);k__Fungi_unclassified(100);k__Fungi_unclassified(100);k__Fungi_unclassified(100);k__Fungi_unclassified(100); | - | - | - | - | - | - | - | Unassigned | - |
| Otu04723 | 6 | k__Fungi(100);k__Fungi_unclassified(84);k__Fungi_unclassified(84);k__Fungi_unclassified(84);k__Fungi_unclassified(84);k__Fungi_unclassified(84);k__Fungi_unclassified(84); | - | - | - | - | - | - | - | Unassigned | - |
| Otu04728 | 6 | k__Fungi(100);k__Fungi_unclassified(100);k__Fungi_unclassified(100);k__Fungi_unclassified(100);k__Fungi_unclassified(100);k__Fungi_unclassified(100);k__Fungi_unclassified(100); | - | - | - | - | - | - | - | Unassigned | - |
| Otu04730 | 6 | k__Fungi(100);k__Fungi_unclassified(67);k__Fungi_unclassified(67);k__Fungi_unclassified(67);k__Fungi_unclassified(67);k__Fungi_unclassified(67);k__Fungi_unclassified(67); | - | - | - | - | - | - | - | Unassigned | - |
| Otu04731 | 6 | k__Fungi(100);k__Fungi_unclassified(84);k__Fungi_unclassified(84);k__Fungi_unclassified(84);k__Fungi_unclassified(84);k__Fungi_unclassified(84);k__Fungi_unclassified(84); | - | - | - | - | - | - | - | Unassigned | - |
| Otu04732 | 6 | k__Fungi(100);k__Fungi_unclassified(67);k__Fungi_unclassified(67);k__Fungi_unclassified(67);k__Fungi_unclassified(67);k__Fungi_unclassified(67);k__Fungi_unclassified(67); | - | - | - | - | - | - | - | Unassigned | - |
| Otu04736 | 6 | k__Fungi(100);p__Basidiomycota(100);c__Agaricomycetes(100);c__Agaricomycetes_unclassified(100);c__Agaricomycetes_unclassified(100);c__Agaricomycetes_unclassified(100);c__Agaricomycetes_unclassified(100); | - | - | - | - | - | - | - | Unassigned | - |
| Otu04738 | 6 | k__Fungi(100);k__Fungi_unclassified(67);k__Fungi_unclassified(67);k__Fungi_unclassified(67);k__Fungi_unclassified(67);k__Fungi_unclassified(67);k__Fungi_unclassified(67); | - | - | - | - | - | - | - | Unassigned | - |
| Otu04740 | 6 | k__Fungi(100);k__Fungi_unclassified(67);k__Fungi_unclassified(67);k__Fungi_unclassified(67);k__Fungi_unclassified(67);k__Fungi_unclassified(67);k__Fungi_unclassified(67); | - | - | - | - | - | - | - | Unassigned | - |
| Otu04742 | 6 | k__Fungi(100);k__Fungi_unclassified(100);k__Fungi_unclassified(100);k__Fungi_unclassified(100);k__Fungi_unclassified(100);k__Fungi_unclassified(100);k__Fungi_unclassified(100); | - | - | - | - | - | - | - | Unassigned | - |
| Otu04745 | 6 | k__Fungi(100);p__Basidiomycota(100);c__Agaricomycetes(100);c__Agaricomycetes_unclassified(67);c__Agaricomycetes_unclassified(67);c__Agaricomycetes_unclassified(67);c__Agaricomycetes_unclassified(67); | - | - | - | - | - | - | - | Unassigned | - |
| Otu04750 | 6 | k__Fungi(100);p__Ascomycota(100);p__Ascomycota_unclassified(100);p__Ascomycota_unclassified(100);p__Ascomycota_unclassified(100);p__Ascomycota_unclassified(100);p__Ascomycota_unclassified(100); | - | - | - | - | - | - | - | Unassigned | - |
| Otu04752 | 6 | k__Fungi(100);p__Ascomycota(100);p__Ascomycota_unclassified(100);p__Ascomycota_unclassified(100);p__Ascomycota_unclassified(100);p__Ascomycota_unclassified(100);p__Ascomycota_unclassified(100); | - | - | - | - | - | - | - | Unassigned | - |
| Otu04754 | 6 | k__Fungi(100);p__Basidiomycota(100);c__Agaricomycetes(100);o__Sebacinales(100);o__Sebacinales_unclassified(100);o__Sebacinales_unclassified(100);o__Sebacinales_unclassified(100); | - | - | - | - | - | - | - | Unassigned | - |
| Otu04755 | 6 | k__Fungi(100);k__Fungi_unclassified(100);k__Fungi_unclassified(100);k__Fungi_unclassified(100);k__Fungi_unclassified(100);k__Fungi_unclassified(100);k__Fungi_unclassified(100); | - | - | - | - | - | - | - | Unassigned | - |
| Otu04756 | 6 | k__Fungi(100);p__Basidiomycota(100);c__Agaricomycetes(100);o__Agaricales(100);f__Niaceae(100);f__Niaceae_unclassified(100);f__Niaceae_unclassified(100); | - | - | - | - | - | - | - | Unassigned | - |
| Otu04758 | 6 | k__Fungi(100);k__Fungi_unclassified(67);k__Fungi_unclassified(67);k__Fungi_unclassified(67);k__Fungi_unclassified(67);k__Fungi_unclassified(67);k__Fungi_unclassified(67); | - | - | - | - | - | - | - | Unassigned | - |
| Otu04760 | 6 | k__Fungi(100);k__Fungi_unclassified(100);k__Fungi_unclassified(100);k__Fungi_unclassified(100);k__Fungi_unclassified(100);k__Fungi_unclassified(100);k__Fungi_unclassified(100); | - | - | - | - | - | - | - | Unassigned | - |
| Otu04765 | 6 | k__Fungi(100);p__Basidiomycota(100);c__Agaricomycetes(100);c__Agaricomycetes_unclassified(100);c__Agaricomycetes_unclassified(100);c__Agaricomycetes_unclassified(100);c__Agaricomycetes_unclassified(100); | - | - | - | - | - | - | - | Unassigned | - |
| Otu04767 | 6 | k__Fungi(100);p__Ascomycota(67);p__Ascomycota_unclassified(67);p__Ascomycota_unclassified(67);p__Ascomycota_unclassified(67);p__Ascomycota_unclassified(67);p__Ascomycota_unclassified(67); | - | - | - | - | - | - | - | Unassigned | - |
| Otu04769 | 6 | k__Fungi(100);k__Fungi_unclassified(100);k__Fungi_unclassified(100);k__Fungi_unclassified(100);k__Fungi_unclassified(100);k__Fungi_unclassified(100);k__Fungi_unclassified(100); | - | - | - | - | - | - | - | Unassigned | - |
| Otu04771 | 6 | k__Fungi(100);p__Ascomycota(100);p__Ascomycota_unclassified(84);p__Ascomycota_unclassified(84);p__Ascomycota_unclassified(84);p__Ascomycota_unclassified(84);p__Ascomycota_unclassified(84); | - | - | - | - | - | - | - | Unassigned | - |
| Otu04775 | 6 | k__Fungi(100);k__Fungi_unclassified(84);k__Fungi_unclassified(84);k__Fungi_unclassified(84);k__Fungi_unclassified(84);k__Fungi_unclassified(84);k__Fungi_unclassified(84); | - | - | - | - | - | - | - | Unassigned | - |
| Otu04776 | 6 | k__Fungi(100);k__Fungi_unclassified(67);k__Fungi_unclassified(67);k__Fungi_unclassified(67);k__Fungi_unclassified(67);k__Fungi_unclassified(67);k__Fungi_unclassified(67); | - | - | - | - | - | - | - | Unassigned | - |
| Otu04779 | 6 | k__Fungi(100);k__Fungi_unclassified(100);k__Fungi_unclassified(100);k__Fungi_unclassified(100);k__Fungi_unclassified(100);k__Fungi_unclassified(100);k__Fungi_unclassified(100); | - | - | - | - | - | - | - | Unassigned | - |
| Otu04780 | 6 | k__Fungi(100);k__Fungi_unclassified(100);k__Fungi_unclassified(100);k__Fungi_unclassified(100);k__Fungi_unclassified(100);k__Fungi_unclassified(100);k__Fungi_unclassified(100); | - | - | - | - | - | - | - | Unassigned | - |
| Otu04781 | 6 | k__Fungi(100);p__Basidiomycota(100);c__Agaricomycetes(100);o__Polyporales(100);o__Polyporales_unclassified(100);o__Polyporales_unclassified(100);o__Polyporales_unclassified(100); | - | - | - | - | - | - | - | Unassigned | - |
| Otu04782 | 6 | k__Fungi(100);k__Fungi_unclassified(100);k__Fungi_unclassified(100);k__Fungi_unclassified(100);k__Fungi_unclassified(100);k__Fungi_unclassified(100);k__Fungi_unclassified(100); | - | - | - | - | - | - | - | Unassigned | - |
| Otu04785 | 6 | k__Fungi(100);k__Fungi_unclassified(67);k__Fungi_unclassified(67);k__Fungi_unclassified(67);k__Fungi_unclassified(67);k__Fungi_unclassified(67);k__Fungi_unclassified(67); | - | - | - | - | - | - | - | Unassigned | - |
| Otu04787 | 6 | k__Fungi(100);p__Ascomycota(100);p__Ascomycota_unclassified(100);p__Ascomycota_unclassified(100);p__Ascomycota_unclassified(100);p__Ascomycota_unclassified(100);p__Ascomycota_unclassified(100); | - | - | - | - | - | - | - | Unassigned | - |
| Otu04789 | 6 | k__Fungi(100);k__Fungi_unclassified(100);k__Fungi_unclassified(100);k__Fungi_unclassified(100);k__Fungi_unclassified(100);k__Fungi_unclassified(100);k__Fungi_unclassified(100); | - | - | - | - | - | - | - | Unassigned | - |
| Otu04790 | 6 | k__Fungi(100);k__Fungi_unclassified(100);k__Fungi_unclassified(100);k__Fungi_unclassified(100);k__Fungi_unclassified(100);k__Fungi_unclassified(100);k__Fungi_unclassified(100); | - | - | - | - | - | - | - | Unassigned | - |
| Otu04793 | 6 | k__Fungi(100);p__Ascomycota(100);p__Ascomycota_unclassified(100);p__Ascomycota_unclassified(100);p__Ascomycota_unclassified(100);p__Ascomycota_unclassified(100);p__Ascomycota_unclassified(100); | - | - | - | - | - | - | - | Unassigned | - |
| Otu04795 | 6 | k__Fungi(100);p__Ascomycota(100);p__Ascomycota_unclassified(100);p__Ascomycota_unclassified(100);p__Ascomycota_unclassified(100);p__Ascomycota_unclassified(100);p__Ascomycota_unclassified(100); | - | - | - | - | - | - | - | Unassigned | - |
| Otu04796 | 6 | k__Fungi(100);k__Fungi_unclassified(100);k__Fungi_unclassified(100);k__Fungi_unclassified(100);k__Fungi_unclassified(100);k__Fungi_unclassified(100);k__Fungi_unclassified(100); | - | - | - | - | - | - | - | Unassigned | - |
| Otu04797 | 6 | k__Fungi(100);p__Ascomycota(67);p__Ascomycota_unclassified(67);p__Ascomycota_unclassified(67);p__Ascomycota_unclassified(67);p__Ascomycota_unclassified(67);p__Ascomycota_unclassified(67); | - | - | - | - | - | - | - | Unassigned | - |
| Otu04798 | 6 | k__Fungi(100);k__Fungi_unclassified(100);k__Fungi_unclassified(100);k__Fungi_unclassified(100);k__Fungi_unclassified(100);k__Fungi_unclassified(100);k__Fungi_unclassified(100); | - | - | - | - | - | - | - | Unassigned | - |
| Otu04801 | 6 | k__Fungi(100);p__Ascomycota(100);p__Ascomycota_unclassified(100);p__Ascomycota_unclassified(100);p__Ascomycota_unclassified(100);p__Ascomycota_unclassified(100);p__Ascomycota_unclassified(100); | - | - | - | - | - | - | - | Unassigned | - |
| Otu04803 | 6 | k__Fungi(100);p__Ascomycota(100);c__Sordariomycetes(100);c__Sordariomycetes_unclassified(100);c__Sordariomycetes_unclassified(100);c__Sordariomycetes_unclassified(100);c__Sordariomycetes_unclassified(100); | - | - | - | - | - | - | - | Unassigned | - |
| Otu04804 | 6 | k__Fungi(100);k__Fungi_unclassified(67);k__Fungi_unclassified(67);k__Fungi_unclassified(67);k__Fungi_unclassified(67);k__Fungi_unclassified(67);k__Fungi_unclassified(67); | - | - | - | - | - | - | - | Unassigned | - |
| Otu04808 | 6 | k__Fungi(100);k__Fungi_unclassified(67);k__Fungi_unclassified(67);k__Fungi_unclassified(67);k__Fungi_unclassified(67);k__Fungi_unclassified(67);k__Fungi_unclassified(67); | - | - | - | - | - | - | - | Unassigned | - |
| Otu04809 | 6 | k__Fungi(100);p__Ascomycota(100);p__Ascomycota_unclassified(100);p__Ascomycota_unclassified(100);p__Ascomycota_unclassified(100);p__Ascomycota_unclassified(100);p__Ascomycota_unclassified(100); | - | - | - | - | - | - | - | Unassigned | - |
| Otu04810 | 6 | k__Fungi(100);k__Fungi_unclassified(84);k__Fungi_unclassified(84);k__Fungi_unclassified(84);k__Fungi_unclassified(84);k__Fungi_unclassified(84);k__Fungi_unclassified(84); | - | - | - | - | - | - | - | Unassigned | - |
| Otu04812 | 6 | k__Fungi(100);k__Fungi_unclassified(67);k__Fungi_unclassified(67);k__Fungi_unclassified(67);k__Fungi_unclassified(67);k__Fungi_unclassified(67);k__Fungi_unclassified(67); | - | - | - | - | - | - | - | Unassigned | - |
| Otu04815 | 6 | k__Fungi(100);k__Fungi_unclassified(100);k__Fungi_unclassified(100);k__Fungi_unclassified(100);k__Fungi_unclassified(100);k__Fungi_unclassified(100);k__Fungi_unclassified(100); | - | - | - | - | - | - | - | Unassigned | - |
| Otu04818 | 6 | k__Fungi(100);k__Fungi_unclassified(100);k__Fungi_unclassified(100);k__Fungi_unclassified(100);k__Fungi_unclassified(100);k__Fungi_unclassified(100);k__Fungi_unclassified(100); | - | - | - | - | - | - | - | Unassigned | - |
| Otu04820 | 6 | k__Fungi(100);p__Basidiomycota(100);c__Agaricomycetes(100);o__Boletales(100);o__Boletales_unclassified(100);o__Boletales_unclassified(100);o__Boletales_unclassified(100); | - | - | - | - | - | - | - | Unassigned | - |
| Otu04823 | 6 | k__Fungi(100);k__Fungi_unclassified(100);k__Fungi_unclassified(100);k__Fungi_unclassified(100);k__Fungi_unclassified(100);k__Fungi_unclassified(100);k__Fungi_unclassified(100); | - | - | - | - | - | - | - | Unassigned | - |
| Otu04826 | 6 | k__Fungi(100);k__Fungi_unclassified(100);k__Fungi_unclassified(100);k__Fungi_unclassified(100);k__Fungi_unclassified(100);k__Fungi_unclassified(100);k__Fungi_unclassified(100); | - | - | - | - | - | - | - | Unassigned | - |
| Otu04828 | 6 | k__Fungi(100);p__Ascomycota(100);p__Ascomycota_unclassified(100);p__Ascomycota_unclassified(100);p__Ascomycota_unclassified(100);p__Ascomycota_unclassified(100);p__Ascomycota_unclassified(100); | - | - | - | - | - | - | - | Unassigned | - |
| Otu04831 | 6 | k__Fungi(100);p__Basidiomycota(100);c__Microbotryomycetes(100);o__Sporidiobolales(100);o__Sporidiobolales_unclassified(100);o__Sporidiobolales_unclassified(100);o__Sporidiobolales_unclassified(100); | - | - | - | - | - | - | - | Unassigned | - |
| Otu04835 | 6 | k__Fungi(100);k__Fungi_unclassified(100);k__Fungi_unclassified(100);k__Fungi_unclassified(100);k__Fungi_unclassified(100);k__Fungi_unclassified(100);k__Fungi_unclassified(100); | - | - | - | - | - | - | - | Unassigned | - |
| Otu04837 | 6 | k__Fungi(100);k__Fungi_unclassified(100);k__Fungi_unclassified(100);k__Fungi_unclassified(100);k__Fungi_unclassified(100);k__Fungi_unclassified(100);k__Fungi_unclassified(100); | - | - | - | - | - | - | - | Unassigned | - |
| Otu04840 | 6 | k__Fungi(100);p__Ascomycota(100);p__Ascomycota_unclassified(100);p__Ascomycota_unclassified(100);p__Ascomycota_unclassified(100);p__Ascomycota_unclassified(100);p__Ascomycota_unclassified(100); | - | - | - | - | - | - | - | Unassigned | - |
| Otu04841 | 6 | k__Fungi(100);k__Fungi_unclassified(84);k__Fungi_unclassified(84);k__Fungi_unclassified(84);k__Fungi_unclassified(84);k__Fungi_unclassified(84);k__Fungi_unclassified(84); | - | - | - | - | - | - | - | Unassigned | - |
| Otu04842 | 6 | k__Fungi(100);p__Basidiomycota(100);c__Agaricomycetes(100);c__Agaricomycetes_unclassified(100);c__Agaricomycetes_unclassified(100);c__Agaricomycetes_unclassified(100);c__Agaricomycetes_unclassified(100); | - | - | - | - | - | - | - | Unassigned | - |
| Otu04844 | 6 | k__Fungi(100);k__Fungi_unclassified(100);k__Fungi_unclassified(100);k__Fungi_unclassified(100);k__Fungi_unclassified(100);k__Fungi_unclassified(100);k__Fungi_unclassified(100); | - | - | - | - | - | - | - | Unassigned | - |
| Otu04848 | 6 | k__Fungi(100);p__Ascomycota(100);c__Eurotiomycetes(100);o__Chaetothyriales(100);o__Chaetothyriales_unclassified(100);o__Chaetothyriales_unclassified(100);o__Chaetothyriales_unclassified(100); | - | - | - | - | - | - | - | Unassigned | - |
| Otu04850 | 6 | k__Fungi(100);k__Fungi_unclassified(84);k__Fungi_unclassified(84);k__Fungi_unclassified(84);k__Fungi_unclassified(84);k__Fungi_unclassified(84);k__Fungi_unclassified(84); | - | - | - | - | - | - | - | Unassigned | - |
| Otu04851 | 6 | k__Fungi(100);p__Ascomycota(100);p__Ascomycota_unclassified(100);p__Ascomycota_unclassified(100);p__Ascomycota_unclassified(100);p__Ascomycota_unclassified(100);p__Ascomycota_unclassified(100); | - | - | - | - | - | - | - | Unassigned | - |
| Otu04852 | 6 | k__Fungi(100);k__Fungi_unclassified(67);k__Fungi_unclassified(67);k__Fungi_unclassified(67);k__Fungi_unclassified(67);k__Fungi_unclassified(67);k__Fungi_unclassified(67); | - | - | - | - | - | - | - | Unassigned | - |
| Otu04856 | 6 | k__Fungi(100);k__Fungi_unclassified(67);k__Fungi_unclassified(67);k__Fungi_unclassified(67);k__Fungi_unclassified(67);k__Fungi_unclassified(67);k__Fungi_unclassified(67); | - | - | - | - | - | - | - | Unassigned | - |
| Otu04857 | 6 | k__Fungi(100);p__Ascomycota(84);p__Ascomycota_unclassified(84);p__Ascomycota_unclassified(84);p__Ascomycota_unclassified(84);p__Ascomycota_unclassified(84);p__Ascomycota_unclassified(84); | - | - | - | - | - | - | - | Unassigned | - |
| Otu04863 | 6 | k__Fungi(100);k__Fungi_unclassified(67);k__Fungi_unclassified(67);k__Fungi_unclassified(67);k__Fungi_unclassified(67);k__Fungi_unclassified(67);k__Fungi_unclassified(67); | - | - | - | - | - | - | - | Unassigned | - |
| Otu04869 | 6 | k__Fungi(100);p__Ascomycota(84);p__Ascomycota_unclassified(84);p__Ascomycota_unclassified(84);p__Ascomycota_unclassified(84);p__Ascomycota_unclassified(84);p__Ascomycota_unclassified(84); | - | - | - | - | - | - | - | Unassigned | - |
| Otu04870 | 6 | k__Fungi(100);p__Basidiomycota(100);c__Agaricostilbomycetes(100);o__unclassified_Agaricostilbomycetes(100);f__unclassified_Agaricostilbomycetes(100);g__unclassified_Agaricostilbomycetes(100);s__Agaricostilbomycetes_sp(100); | - | - | - | - | - | - | - | Unassigned | - |
| Otu04872 | 6 | k__Fungi(100);p__Ascomycota(100);p__Ascomycota_unclassified(100);p__Ascomycota_unclassified(100);p__Ascomycota_unclassified(100);p__Ascomycota_unclassified(100);p__Ascomycota_unclassified(100); | - | - | - | - | - | - | - | Unassigned | - |
| Otu04873 | 6 | k__Fungi(100);p__Ascomycota(84);p__Ascomycota_unclassified(84);p__Ascomycota_unclassified(84);p__Ascomycota_unclassified(84);p__Ascomycota_unclassified(84);p__Ascomycota_unclassified(84); | - | - | - | - | - | - | - | Unassigned | - |
| Otu04874 | 6 | k__Fungi(100);p__Ascomycota(84);p__Ascomycota_unclassified(84);p__Ascomycota_unclassified(84);p__Ascomycota_unclassified(84);p__Ascomycota_unclassified(84);p__Ascomycota_unclassified(84); | - | - | - | - | - | - | - | Unassigned | - |
| Otu04881 | 6 | k__Fungi(100);k__Fungi_unclassified(100);k__Fungi_unclassified(100);k__Fungi_unclassified(100);k__Fungi_unclassified(100);k__Fungi_unclassified(100);k__Fungi_unclassified(100); | - | - | - | - | - | - | - | Unassigned | - |
| Otu04882 | 6 | k__Fungi(100);p__Ascomycota(100);c__Dothideomycetes(100);o__Pleosporales(100);o__Pleosporales_unclassified(100);o__Pleosporales_unclassified(100);o__Pleosporales_unclassified(100); | - | - | - | - | - | - | - | Unassigned | - |
| Otu04883 | 6 | k__Fungi(100);k__Fungi_unclassified(100);k__Fungi_unclassified(100);k__Fungi_unclassified(100);k__Fungi_unclassified(100);k__Fungi_unclassified(100);k__Fungi_unclassified(100); | - | - | - | - | - | - | - | Unassigned | - |
| Otu04884 | 6 | k__Fungi(100);k__Fungi_unclassified(67);k__Fungi_unclassified(67);k__Fungi_unclassified(67);k__Fungi_unclassified(67);k__Fungi_unclassified(67);k__Fungi_unclassified(67); | - | - | - | - | - | - | - | Unassigned | - |
| Otu04887 | 6 | k__Fungi(100);p__Ascomycota(100);c__Sordariomycetes(100);c__Sordariomycetes_unclassified(100);c__Sordariomycetes_unclassified(100);c__Sordariomycetes_unclassified(100);c__Sordariomycetes_unclassified(100); | - | - | - | - | - | - | - | Unassigned | - |
| Otu04891 | 6 | k__Fungi(100);p__Ascomycota(100);c__Sordariomycetes(100);c__Sordariomycetes_unclassified(100);c__Sordariomycetes_unclassified(100);c__Sordariomycetes_unclassified(100);c__Sordariomycetes_unclassified(100); | - | - | - | - | - | - | - | Unassigned | - |
| Otu04893 | 6 | k__Fungi(100);p__Ascomycota(100);p__Ascomycota_unclassified(100);p__Ascomycota_unclassified(100);p__Ascomycota_unclassified(100);p__Ascomycota_unclassified(100);p__Ascomycota_unclassified(100); | - | - | - | - | - | - | - | Unassigned | - |
| Otu04894 | 6 | k__Fungi(100);p__Ascomycota(67);p__Ascomycota_unclassified(67);p__Ascomycota_unclassified(67);p__Ascomycota_unclassified(67);p__Ascomycota_unclassified(67);p__Ascomycota_unclassified(67); | - | - | - | - | - | - | - | Unassigned | - |
| Otu04895 | 6 | k__Fungi(100);p__Ascomycota(67);p__Ascomycota_unclassified(67);p__Ascomycota_unclassified(67);p__Ascomycota_unclassified(67);p__Ascomycota_unclassified(67);p__Ascomycota_unclassified(67); | - | - | - | - | - | - | - | Unassigned | - |
| Otu04900 | 6 | k__Fungi(100);p__Basidiomycota(100);p__Basidiomycota_unclassified(100);p__Basidiomycota_unclassified(100);p__Basidiomycota_unclassified(100);p__Basidiomycota_unclassified(100);p__Basidiomycota_unclassified(100); | - | - | - | - | - | - | - | Unassigned | - |
| Otu04902 | 6 | k__Fungi(100);p__Basidiomycota(100);c__Agaricomycetes(100);c__Agaricomycetes_unclassified(100);c__Agaricomycetes_unclassified(100);c__Agaricomycetes_unclassified(100);c__Agaricomycetes_unclassified(100); | - | - | - | - | - | - | - | Unassigned | - |
| Otu04905 | 6 | k__Fungi(100);k__Fungi_unclassified(100);k__Fungi_unclassified(100);k__Fungi_unclassified(100);k__Fungi_unclassified(100);k__Fungi_unclassified(100);k__Fungi_unclassified(100); | - | - | - | - | - | - | - | Unassigned | - |
| Otu04906 | 6 | k__Fungi(100);k__Fungi_unclassified(100);k__Fungi_unclassified(100);k__Fungi_unclassified(100);k__Fungi_unclassified(100);k__Fungi_unclassified(100);k__Fungi_unclassified(100); | - | - | - | - | - | - | - | Unassigned | - |
| Otu04907 | 6 | k__Fungi(100);p__Ascomycota(84);p__Ascomycota_unclassified(84);p__Ascomycota_unclassified(84);p__Ascomycota_unclassified(84);p__Ascomycota_unclassified(84);p__Ascomycota_unclassified(84); | - | - | - | - | - | - | - | Unassigned | - |
| Otu04910 | 6 | k__Fungi(100);k__Fungi_unclassified(100);k__Fungi_unclassified(100);k__Fungi_unclassified(100);k__Fungi_unclassified(100);k__Fungi_unclassified(100);k__Fungi_unclassified(100); | - | - | - | - | - | - | - | Unassigned | - |
| Otu04911 | 6 | k__Fungi(100);p__Ascomycota(100);c__Dothideomycetes(100);o__Pleosporales(100);f__Pleosporales_family_Incertae_sedis(67);f__Pleosporales_family_Incertae_sedis_unclassified(67);f__Pleosporales_family_Incertae_sedis_unclassified(67); | - | - | - | - | - | - | - | Unassigned | - |
| Otu04915 | 6 | k__Fungi(100);k__Fungi_unclassified(100);k__Fungi_unclassified(100);k__Fungi_unclassified(100);k__Fungi_unclassified(100);k__Fungi_unclassified(100);k__Fungi_unclassified(100); | - | - | - | - | - | - | - | Unassigned | - |
| Otu04918 | 6 | k__Fungi(100);k__Fungi_unclassified(67);k__Fungi_unclassified(67);k__Fungi_unclassified(67);k__Fungi_unclassified(67);k__Fungi_unclassified(67);k__Fungi_unclassified(67); | - | - | - | - | - | - | - | Unassigned | - |
| Otu04920 | 6 | k__Fungi(100);p__Basidiomycota(100);c__Agaricomycetes(100);c__Agaricomycetes_unclassified(100);c__Agaricomycetes_unclassified(100);c__Agaricomycetes_unclassified(100);c__Agaricomycetes_unclassified(100); | - | - | - | - | - | - | - | Unassigned | - |
| Otu04923 | 6 | k__Fungi(100);k__Fungi_unclassified(67);k__Fungi_unclassified(67);k__Fungi_unclassified(67);k__Fungi_unclassified(67);k__Fungi_unclassified(67);k__Fungi_unclassified(67); | - | - | - | - | - | - | - | Unassigned | - |
| Otu04925 | 6 | k__Fungi(100);k__Fungi_unclassified(84);k__Fungi_unclassified(84);k__Fungi_unclassified(84);k__Fungi_unclassified(84);k__Fungi_unclassified(84);k__Fungi_unclassified(84); | - | - | - | - | - | - | - | Unassigned | - |
| Otu04927 | 6 | k__Fungi(100);k__Fungi_unclassified(100);k__Fungi_unclassified(100);k__Fungi_unclassified(100);k__Fungi_unclassified(100);k__Fungi_unclassified(100);k__Fungi_unclassified(100); | - | - | - | - | - | - | - | Unassigned | - |
| Otu04931 | 6 | k__Fungi(100);p__Ascomycota(100);c__Leotiomycetes(100);o__Helotiales(100);o__Helotiales_unclassified(84);o__Helotiales_unclassified(84);o__Helotiales_unclassified(84); | - | - | - | - | - | - | - | Unassigned | - |
| Otu04932 | 6 | k__Fungi(100);p__Basidiomycota(100);c__Agaricomycetes(100);c__Agaricomycetes_unclassified(100);c__Agaricomycetes_unclassified(100);c__Agaricomycetes_unclassified(100);c__Agaricomycetes_unclassified(100); | - | - | - | - | - | - | - | Unassigned | - |
| Otu04934 | 6 | k__Fungi(100);p__Basidiomycota(100);c__Agaricomycetes(100);o__Polyporales(100);o__Polyporales_unclassified(100);o__Polyporales_unclassified(100);o__Polyporales_unclassified(100); | - | - | - | - | - | - | - | Unassigned | - |
| Otu04935 | 6 | k__Fungi(100);p__Ascomycota(67);p__Ascomycota_unclassified(67);p__Ascomycota_unclassified(67);p__Ascomycota_unclassified(67);p__Ascomycota_unclassified(67);p__Ascomycota_unclassified(67); | - | - | - | - | - | - | - | Unassigned | - |
| Otu04936 | 6 | k__Fungi(100);k__Fungi_unclassified(67);k__Fungi_unclassified(67);k__Fungi_unclassified(67);k__Fungi_unclassified(67);k__Fungi_unclassified(67);k__Fungi_unclassified(67); | - | - | - | - | - | - | - | Unassigned | - |
| Otu04939 | 6 | k__Fungi(100);p__Ascomycota(100);c__Dothideomycetes(100);o__Capnodiales(100);o__Capnodiales_unclassified(84);o__Capnodiales_unclassified(84);o__Capnodiales_unclassified(84); | - | - | - | - | - | - | - | Unassigned | - |
| Otu04943 | 6 | k__Fungi(100);p__Ascomycota(67);p__Ascomycota_unclassified(67);p__Ascomycota_unclassified(67);p__Ascomycota_unclassified(67);p__Ascomycota_unclassified(67);p__Ascomycota_unclassified(67); | - | - | - | - | - | - | - | Unassigned | - |
| Otu04945 | 6 | k__Fungi(100);k__Fungi_unclassified(84);k__Fungi_unclassified(84);k__Fungi_unclassified(84);k__Fungi_unclassified(84);k__Fungi_unclassified(84);k__Fungi_unclassified(84); | - | - | - | - | - | - | - | Unassigned | - |
| Otu04946 | 6 | k__Fungi(100);k__Fungi_unclassified(84);k__Fungi_unclassified(84);k__Fungi_unclassified(84);k__Fungi_unclassified(84);k__Fungi_unclassified(84);k__Fungi_unclassified(84); | - | - | - | - | - | - | - | Unassigned | - |
| Otu04948 | 6 | k__Fungi(100);k__Fungi_unclassified(100);k__Fungi_unclassified(100);k__Fungi_unclassified(100);k__Fungi_unclassified(100);k__Fungi_unclassified(100);k__Fungi_unclassified(100); | - | - | - | - | - | - | - | Unassigned | - |
| Otu04950 | 6 | k__Fungi(100);p__Ascomycota(100);c__Dothideomycetes(100);o__Pleosporales(100);o__Pleosporales_unclassified(100);o__Pleosporales_unclassified(100);o__Pleosporales_unclassified(100); | - | - | - | - | - | - | - | Unassigned | - |
| Otu04953 | 6 | k__Fungi(100);p__Ascomycota(100);p__Ascomycota_unclassified(100);p__Ascomycota_unclassified(100);p__Ascomycota_unclassified(100);p__Ascomycota_unclassified(100);p__Ascomycota_unclassified(100); | - | - | - | - | - | - | - | Unassigned | - |
| Otu04954 | 6 | k__Fungi(100);p__Ascomycota(67);p__Ascomycota_unclassified(67);p__Ascomycota_unclassified(67);p__Ascomycota_unclassified(67);p__Ascomycota_unclassified(67);p__Ascomycota_unclassified(67); | - | - | - | - | - | - | - | Unassigned | - |
| Otu04957 | 6 | k__Fungi(100);k__Fungi_unclassified(100);k__Fungi_unclassified(100);k__Fungi_unclassified(100);k__Fungi_unclassified(100);k__Fungi_unclassified(100);k__Fungi_unclassified(100); | - | - | - | - | - | - | - | Unassigned | - |
| Otu04959 | 6 | k__Fungi(100);p__Ascomycota(67);p__Ascomycota_unclassified(67);p__Ascomycota_unclassified(67);p__Ascomycota_unclassified(67);p__Ascomycota_unclassified(67);p__Ascomycota_unclassified(67); | - | - | - | - | - | - | - | Unassigned | - |
| Otu04964 | 6 | k__Fungi(100);k__Fungi_unclassified(67);k__Fungi_unclassified(67);k__Fungi_unclassified(67);k__Fungi_unclassified(67);k__Fungi_unclassified(67);k__Fungi_unclassified(67); | - | - | - | - | - | - | - | Unassigned | - |
| Otu04965 | 6 | k__Fungi(100);k__Fungi_unclassified(100);k__Fungi_unclassified(100);k__Fungi_unclassified(100);k__Fungi_unclassified(100);k__Fungi_unclassified(100);k__Fungi_unclassified(100); | - | - | - | - | - | - | - | Unassigned | - |
| Otu04968 | 6 | k__Fungi(100);k__Fungi_unclassified(100);k__Fungi_unclassified(100);k__Fungi_unclassified(100);k__Fungi_unclassified(100);k__Fungi_unclassified(100);k__Fungi_unclassified(100); | - | - | - | - | - | - | - | Unassigned | - |
| Otu04971 | 6 | k__Fungi(100);p__Ascomycota(100);p__Ascomycota_unclassified(100);p__Ascomycota_unclassified(100);p__Ascomycota_unclassified(100);p__Ascomycota_unclassified(100);p__Ascomycota_unclassified(100); | - | - | - | - | - | - | - | Unassigned | - |
| Otu04972 | 6 | k__Fungi(100);k__Fungi_unclassified(100);k__Fungi_unclassified(100);k__Fungi_unclassified(100);k__Fungi_unclassified(100);k__Fungi_unclassified(100);k__Fungi_unclassified(100); | - | - | - | - | - | - | - | Unassigned | - |
| Otu04973 | 6 | k__Fungi(100);k__Fungi_unclassified(100);k__Fungi_unclassified(100);k__Fungi_unclassified(100);k__Fungi_unclassified(100);k__Fungi_unclassified(100);k__Fungi_unclassified(100); | - | - | - | - | - | - | - | Unassigned | - |
| Otu04975 | 6 | k__Fungi(100);p__Ascomycota(84);p__Ascomycota_unclassified(84);p__Ascomycota_unclassified(84);p__Ascomycota_unclassified(84);p__Ascomycota_unclassified(84);p__Ascomycota_unclassified(84); | - | - | - | - | - | - | - | Unassigned | - |
| Otu04980 | 6 | k__Fungi(100);p__Ascomycota(67);p__Ascomycota_unclassified(67);p__Ascomycota_unclassified(67);p__Ascomycota_unclassified(67);p__Ascomycota_unclassified(67);p__Ascomycota_unclassified(67); | - | - | - | - | - | - | - | Unassigned | - |
| Otu04982 | 6 | k__Fungi(100);k__Fungi_unclassified(100);k__Fungi_unclassified(100);k__Fungi_unclassified(100);k__Fungi_unclassified(100);k__Fungi_unclassified(100);k__Fungi_unclassified(100); | - | - | - | - | - | - | - | Unassigned | - |
| Otu04983 | 6 | k__Fungi(100);p__Basidiomycota(100);p__Basidiomycota_unclassified(100);p__Basidiomycota_unclassified(100);p__Basidiomycota_unclassified(100);p__Basidiomycota_unclassified(100);p__Basidiomycota_unclassified(100); | - | - | - | - | - | - | - | Unassigned | - |
| Otu04988 | 6 | k__Fungi(100);k__Fungi_unclassified(67);k__Fungi_unclassified(67);k__Fungi_unclassified(67);k__Fungi_unclassified(67);k__Fungi_unclassified(67);k__Fungi_unclassified(67); | - | - | - | - | - | - | - | Unassigned | - |
| Otu04989 | 6 | k__Fungi(100);k__Fungi_unclassified(84);k__Fungi_unclassified(84);k__Fungi_unclassified(84);k__Fungi_unclassified(84);k__Fungi_unclassified(84);k__Fungi_unclassified(84); | - | - | - | - | - | - | - | Unassigned | - |
| Otu04990 | 6 | k__Fungi(100);k__Fungi_unclassified(67);k__Fungi_unclassified(67);k__Fungi_unclassified(67);k__Fungi_unclassified(67);k__Fungi_unclassified(67);k__Fungi_unclassified(67); | - | - | - | - | - | - | - | Unassigned | - |
| Otu04996 | 6 | k__Fungi(100);k__Fungi_unclassified(67);k__Fungi_unclassified(67);k__Fungi_unclassified(67);k__Fungi_unclassified(67);k__Fungi_unclassified(67);k__Fungi_unclassified(67); | - | - | - | - | - | - | - | Unassigned | - |
| Otu04999 | 6 | k__Fungi(100);k__Fungi_unclassified(100);k__Fungi_unclassified(100);k__Fungi_unclassified(100);k__Fungi_unclassified(100);k__Fungi_unclassified(100);k__Fungi_unclassified(100); | - | - | - | - | - | - | - | Unassigned | - |
| Otu05001 | 6 | k__Fungi(100);p__Basidiomycota(100);c__Agaricomycetes(100);o__Trechisporales(100);f__unclassified_Trechisporales(100);g__unclassified_Trechisporales(100);s__Trechisporales_sp(100); | - | - | - | - | - | - | - | Unassigned | - |
| Otu05003 | 6 | k__Fungi(100);p__Ascomycota(67);p__Ascomycota_unclassified(67);p__Ascomycota_unclassified(67);p__Ascomycota_unclassified(67);p__Ascomycota_unclassified(67);p__Ascomycota_unclassified(67); | - | - | - | - | - | - | - | Unassigned | - |
| Otu05005 | 6 | k__Fungi(100);k__Fungi_unclassified(67);k__Fungi_unclassified(67);k__Fungi_unclassified(67);k__Fungi_unclassified(67);k__Fungi_unclassified(67);k__Fungi_unclassified(67); | - | - | - | - | - | - | - | Unassigned | - |
| Otu05006 | 6 | k__Fungi(100);p__Ascomycota(67);p__Ascomycota_unclassified(67);p__Ascomycota_unclassified(67);p__Ascomycota_unclassified(67);p__Ascomycota_unclassified(67);p__Ascomycota_unclassified(67); | - | - | - | - | - | - | - | Unassigned | - |
| Otu05010 | 6 | k__Fungi(100);p__Basidiomycota(100);c__Agaricomycetes(100);o__Trechisporales(100);f__Hydnodontaceae(100);g__unclassified_Hydnodontaceae(67);s__Hydnodontaceae_sp(67); | - | - | - | - | - | - | - | Unassigned | - |
| Otu05012 | 6 | k__Fungi(100);p__Ascomycota(100);p__Ascomycota_unclassified(100);p__Ascomycota_unclassified(100);p__Ascomycota_unclassified(100);p__Ascomycota_unclassified(100);p__Ascomycota_unclassified(100); | - | - | - | - | - | - | - | Unassigned | - |
| Otu05014 | 6 | k__Fungi(100);k__Fungi_unclassified(67);k__Fungi_unclassified(67);k__Fungi_unclassified(67);k__Fungi_unclassified(67);k__Fungi_unclassified(67);k__Fungi_unclassified(67); | - | - | - | - | - | - | - | Unassigned | - |
| Otu05020 | 6 | k__Fungi(100);k__Fungi_unclassified(100);k__Fungi_unclassified(100);k__Fungi_unclassified(100);k__Fungi_unclassified(100);k__Fungi_unclassified(100);k__Fungi_unclassified(100); | - | - | - | - | - | - | - | Unassigned | - |
| Otu05021 | 6 | k__Fungi(100);k__Fungi_unclassified(67);k__Fungi_unclassified(67);k__Fungi_unclassified(67);k__Fungi_unclassified(67);k__Fungi_unclassified(67);k__Fungi_unclassified(67); | - | - | - | - | - | - | - | Unassigned | - |
| Otu05025 | 6 | k__Fungi(100);p__Ascomycota(100);c__Sordariomycetes(100);c__Sordariomycetes_unclassified(100);c__Sordariomycetes_unclassified(100);c__Sordariomycetes_unclassified(100);c__Sordariomycetes_unclassified(100); | - | - | - | - | - | - | - | Unassigned | - |
| Otu05027 | 6 | k__Fungi(100);k__Fungi_unclassified(84);k__Fungi_unclassified(84);k__Fungi_unclassified(84);k__Fungi_unclassified(84);k__Fungi_unclassified(84);k__Fungi_unclassified(84); | - | - | - | - | - | - | - | Unassigned | - |
| Otu05028 | 6 | k__Fungi(100);p__Ascomycota(67);p__Ascomycota_unclassified(67);p__Ascomycota_unclassified(67);p__Ascomycota_unclassified(67);p__Ascomycota_unclassified(67);p__Ascomycota_unclassified(67); | - | - | - | - | - | - | - | Unassigned | - |
| Otu05029 | 6 | k__Fungi(100);k__Fungi_unclassified(100);k__Fungi_unclassified(100);k__Fungi_unclassified(100);k__Fungi_unclassified(100);k__Fungi_unclassified(100);k__Fungi_unclassified(100); | - | - | - | - | - | - | - | Unassigned | - |
| Otu05030 | 6 | k__Fungi(100);p__Ascomycota(100);c__Eurotiomycetes(67);c__Eurotiomycetes_unclassified(67);c__Eurotiomycetes_unclassified(67);c__Eurotiomycetes_unclassified(67);c__Eurotiomycetes_unclassified(67); | - | - | - | - | - | - | - | Unassigned | - |
| Otu05033 | 6 | k__Fungi(100);p__Ascomycota(100);p__Ascomycota_unclassified(100);p__Ascomycota_unclassified(100);p__Ascomycota_unclassified(100);p__Ascomycota_unclassified(100);p__Ascomycota_unclassified(100); | - | - | - | - | - | - | - | Unassigned | - |
| Otu05034 | 6 | k__Fungi(100);p__Ascomycota(100);c__Eurotiomycetes(100);c__Eurotiomycetes_unclassified(100);c__Eurotiomycetes_unclassified(100);c__Eurotiomycetes_unclassified(100);c__Eurotiomycetes_unclassified(100); | - | - | - | - | - | - | - | Unassigned | - |
| Otu05035 | 6 | k__Fungi(100);p__Ascomycota(100);p__Ascomycota_unclassified(100);p__Ascomycota_unclassified(100);p__Ascomycota_unclassified(100);p__Ascomycota_unclassified(100);p__Ascomycota_unclassified(100); | - | - | - | - | - | - | - | Unassigned | - |
| Otu05036 | 6 | k__Fungi(100);p__Ascomycota(100);c__Eurotiomycetes(100);o__Eurotiales(100);f__Trichocomaceae(100);f__Trichocomaceae_unclassified(100);f__Trichocomaceae_unclassified(100); | - | - | - | - | - | - | - | Unassigned | - |
| Otu05038 | 6 | k__Fungi(100);p__Ascomycota(100);p__Ascomycota_unclassified(100);p__Ascomycota_unclassified(100);p__Ascomycota_unclassified(100);p__Ascomycota_unclassified(100);p__Ascomycota_unclassified(100); | - | - | - | - | - | - | - | Unassigned | - |
| Otu05044 | 6 | k__Fungi(100);k__Fungi_unclassified(67);k__Fungi_unclassified(67);k__Fungi_unclassified(67);k__Fungi_unclassified(67);k__Fungi_unclassified(67);k__Fungi_unclassified(67); | - | - | - | - | - | - | - | Unassigned | - |
| Otu05046 | 6 | k__Fungi(100);p__Ascomycota(67);p__Ascomycota_unclassified(67);p__Ascomycota_unclassified(67);p__Ascomycota_unclassified(67);p__Ascomycota_unclassified(67);p__Ascomycota_unclassified(67); | - | - | - | - | - | - | - | Unassigned | - |
| Otu05047 | 6 | k__Fungi(100);p__Ascomycota(67);p__Ascomycota_unclassified(67);p__Ascomycota_unclassified(67);p__Ascomycota_unclassified(67);p__Ascomycota_unclassified(67);p__Ascomycota_unclassified(67); | - | - | - | - | - | - | - | Unassigned | - |
| Otu05048 | 6 | k__Fungi(100);p__Ascomycota(84);p__Ascomycota_unclassified(84);p__Ascomycota_unclassified(84);p__Ascomycota_unclassified(84);p__Ascomycota_unclassified(84);p__Ascomycota_unclassified(84); | - | - | - | - | - | - | - | Unassigned | - |
| Otu05049 | 6 | k__Fungi(100);p__Ascomycota(100);p__Ascomycota_unclassified(100);p__Ascomycota_unclassified(100);p__Ascomycota_unclassified(100);p__Ascomycota_unclassified(100);p__Ascomycota_unclassified(100); | - | - | - | - | - | - | - | Unassigned | - |
| Otu05051 | 6 | k__Fungi(100);k__Fungi_unclassified(67);k__Fungi_unclassified(67);k__Fungi_unclassified(67);k__Fungi_unclassified(67);k__Fungi_unclassified(67);k__Fungi_unclassified(67); | - | - | - | - | - | - | - | Unassigned | - |
| Otu05054 | 6 | k__Fungi(100);p__Basidiomycota(100);c__Agaricomycetes(100);c__Agaricomycetes_unclassified(100);c__Agaricomycetes_unclassified(100);c__Agaricomycetes_unclassified(100);c__Agaricomycetes_unclassified(100); | - | - | - | - | - | - | - | Unassigned | - |
| Otu05058 | 6 | k__Fungi(100);k__Fungi_unclassified(100);k__Fungi_unclassified(100);k__Fungi_unclassified(100);k__Fungi_unclassified(100);k__Fungi_unclassified(100);k__Fungi_unclassified(100); | - | - | - | - | - | - | - | Unassigned | - |
| Otu05059 | 6 | k__Fungi(100);k__Fungi_unclassified(84);k__Fungi_unclassified(84);k__Fungi_unclassified(84);k__Fungi_unclassified(84);k__Fungi_unclassified(84);k__Fungi_unclassified(84); | - | - | - | - | - | - | - | Unassigned | - |
| Otu05061 | 6 | k__Fungi(100);p__Basidiomycota(100);p__Basidiomycota_unclassified(100);p__Basidiomycota_unclassified(100);p__Basidiomycota_unclassified(100);p__Basidiomycota_unclassified(100);p__Basidiomycota_unclassified(100); | - | - | - | - | - | - | - | Unassigned | - |
| Otu05065 | 6 | k__Fungi(100);k__Fungi_unclassified(84);k__Fungi_unclassified(84);k__Fungi_unclassified(84);k__Fungi_unclassified(84);k__Fungi_unclassified(84);k__Fungi_unclassified(84); | - | - | - | - | - | - | - | Unassigned | - |
| Otu05066 | 6 | k__Fungi(100);k__Fungi_unclassified(100);k__Fungi_unclassified(100);k__Fungi_unclassified(100);k__Fungi_unclassified(100);k__Fungi_unclassified(100);k__Fungi_unclassified(100); | - | - | - | - | - | - | - | Unassigned | - |
| Otu05067 | 6 | k__Fungi(100);k__Fungi_unclassified(100);k__Fungi_unclassified(100);k__Fungi_unclassified(100);k__Fungi_unclassified(100);k__Fungi_unclassified(100);k__Fungi_unclassified(100); | - | - | - | - | - | - | - | Unassigned | - |
| Otu05068 | 6 | k__Fungi(100);k__Fungi_unclassified(100);k__Fungi_unclassified(100);k__Fungi_unclassified(100);k__Fungi_unclassified(100);k__Fungi_unclassified(100);k__Fungi_unclassified(100); | - | - | - | - | - | - | - | Unassigned | - |
| Otu05069 | 6 | k__Fungi(100);p__Ascomycota(67);p__Ascomycota_unclassified(67);p__Ascomycota_unclassified(67);p__Ascomycota_unclassified(67);p__Ascomycota_unclassified(67);p__Ascomycota_unclassified(67); | - | - | - | - | - | - | - | Unassigned | - |
| Otu05070 | 6 | k__Fungi(100);k__Fungi_unclassified(67);k__Fungi_unclassified(67);k__Fungi_unclassified(67);k__Fungi_unclassified(67);k__Fungi_unclassified(67);k__Fungi_unclassified(67); | - | - | - | - | - | - | - | Unassigned | - |
| Otu05073 | 6 | k__Fungi(100);p__Ascomycota(100);c__Dothideomycetes(100);o__Venturiales(100);o__Venturiales_unclassified(67);o__Venturiales_unclassified(67);o__Venturiales_unclassified(67); | - | - | - | - | - | - | - | Unassigned | - |
| Otu05075 | 6 | k__Fungi(100);p__Ascomycota(100);c__Dothideomycetes(100);o__Pleosporales(100);o__Pleosporales_unclassified(100);o__Pleosporales_unclassified(100);o__Pleosporales_unclassified(100); | - | - | - | - | - | - | - | Unassigned | - |
| Otu05076 | 6 | k__Fungi(100);p__Ascomycota(100);c__Dothideomycetes(100);o__Pleosporales(100);o__Pleosporales_unclassified(100);o__Pleosporales_unclassified(100);o__Pleosporales_unclassified(100); | - | - | - | - | - | - | - | Unassigned | - |
| Otu05080 | 6 | k__Fungi(100);k__Fungi_unclassified(100);k__Fungi_unclassified(100);k__Fungi_unclassified(100);k__Fungi_unclassified(100);k__Fungi_unclassified(100);k__Fungi_unclassified(100); | - | - | - | - | - | - | - | Unassigned | - |
| Otu05081 | 6 | k__Fungi(100);k__Fungi_unclassified(84);k__Fungi_unclassified(84);k__Fungi_unclassified(84);k__Fungi_unclassified(84);k__Fungi_unclassified(84);k__Fungi_unclassified(84); | - | - | - | - | - | - | - | Unassigned | - |
| Otu05082 | 6 | k__Fungi(100);p__Ascomycota(100);p__Ascomycota_unclassified(100);p__Ascomycota_unclassified(100);p__Ascomycota_unclassified(100);p__Ascomycota_unclassified(100);p__Ascomycota_unclassified(100); | - | - | - | - | - | - | - | Unassigned | - |
| Otu05086 | 6 | k__Fungi(100);k__Fungi_unclassified(67);k__Fungi_unclassified(67);k__Fungi_unclassified(67);k__Fungi_unclassified(67);k__Fungi_unclassified(67);k__Fungi_unclassified(67); | - | - | - | - | - | - | - | Unassigned | - |
| Otu05088 | 6 | k__Fungi(100);k__Fungi_unclassified(100);k__Fungi_unclassified(100);k__Fungi_unclassified(100);k__Fungi_unclassified(100);k__Fungi_unclassified(100);k__Fungi_unclassified(100); | - | - | - | - | - | - | - | Unassigned | - |
| Otu05089 | 6 | k__Fungi(100);k__Fungi_unclassified(100);k__Fungi_unclassified(100);k__Fungi_unclassified(100);k__Fungi_unclassified(100);k__Fungi_unclassified(100);k__Fungi_unclassified(100); | - | - | - | - | - | - | - | Unassigned | - |
| Otu05090 | 6 | k__Fungi(100);p__Basidiomycota(100);p__Basidiomycota_unclassified(100);p__Basidiomycota_unclassified(100);p__Basidiomycota_unclassified(100);p__Basidiomycota_unclassified(100);p__Basidiomycota_unclassified(100); | - | - | - | - | - | - | - | Unassigned | - |
| Otu05091 | 6 | k__Fungi(100);k__Fungi_unclassified(84);k__Fungi_unclassified(84);k__Fungi_unclassified(84);k__Fungi_unclassified(84);k__Fungi_unclassified(84);k__Fungi_unclassified(84); | - | - | - | - | - | - | - | Unassigned | - |
| Otu05094 | 6 | k__Fungi(100);p__Ascomycota(100);p__Ascomycota_unclassified(84);p__Ascomycota_unclassified(84);p__Ascomycota_unclassified(84);p__Ascomycota_unclassified(84);p__Ascomycota_unclassified(84); | - | - | - | - | - | - | - | Unassigned | - |
| Otu05095 | 6 | k__Fungi(100);k__Fungi_unclassified(67);k__Fungi_unclassified(67);k__Fungi_unclassified(67);k__Fungi_unclassified(67);k__Fungi_unclassified(67);k__Fungi_unclassified(67); | - | - | - | - | - | - | - | Unassigned | - |
| Otu05097 | 6 | k__Fungi(100);k__Fungi_unclassified(67);k__Fungi_unclassified(67);k__Fungi_unclassified(67);k__Fungi_unclassified(67);k__Fungi_unclassified(67);k__Fungi_unclassified(67); | - | - | - | - | - | - | - | Unassigned | - |
| Otu05098 | 6 | k__Fungi(100);p__Ascomycota(84);p__Ascomycota_unclassified(84);p__Ascomycota_unclassified(84);p__Ascomycota_unclassified(84);p__Ascomycota_unclassified(84);p__Ascomycota_unclassified(84); | - | - | - | - | - | - | - | Unassigned | - |
| Otu05099 | 6 | k__Fungi(100);p__Basidiomycota(100);c__Agaricomycetes(100);c__Agaricomycetes_unclassified(100);c__Agaricomycetes_unclassified(100);c__Agaricomycetes_unclassified(100);c__Agaricomycetes_unclassified(100); | - | - | - | - | - | - | - | Unassigned | - |
| Otu05100 | 6 | k__Fungi(100);p__Ascomycota(100);c__Dothideomycetes(100);o__Capnodiales(100);o__Capnodiales_unclassified(100);o__Capnodiales_unclassified(100);o__Capnodiales_unclassified(100); | - | - | - | - | - | - | - | Unassigned | - |
| Otu05101 | 6 | k__Fungi(100);k__Fungi_unclassified(100);k__Fungi_unclassified(100);k__Fungi_unclassified(100);k__Fungi_unclassified(100);k__Fungi_unclassified(100);k__Fungi_unclassified(100); | - | - | - | - | - | - | - | Unassigned | - |
| Otu05107 | 6 | k__Fungi(100);p__Basidiomycota(100);c__Agaricomycetes(100);o__unclassified_Agaricomycetes(67);f__unclassified_Agaricomycetes(67);g__unclassified_Agaricomycetes(67);s__Agaricomycetes_sp(67); | - | - | - | - | - | - | - | Unassigned | - |
| Otu05108 | 6 | k__Fungi(100);k__Fungi_unclassified(100);k__Fungi_unclassified(100);k__Fungi_unclassified(100);k__Fungi_unclassified(100);k__Fungi_unclassified(100);k__Fungi_unclassified(100); | - | - | - | - | - | - | - | Unassigned | - |
| Otu05109 | 6 | k__Fungi(100);p__Ascomycota(84);p__Ascomycota_unclassified(84);p__Ascomycota_unclassified(84);p__Ascomycota_unclassified(84);p__Ascomycota_unclassified(84);p__Ascomycota_unclassified(84); | - | - | - | - | - | - | - | Unassigned | - |
| Otu05111 | 6 | k__Fungi(100);p__Ascomycota(67);p__Ascomycota_unclassified(67);p__Ascomycota_unclassified(67);p__Ascomycota_unclassified(67);p__Ascomycota_unclassified(67);p__Ascomycota_unclassified(67); | - | - | - | - | - | - | - | Unassigned | - |
| Otu05112 | 6 | k__Fungi(100);p__Ascomycota(100);p__Ascomycota_unclassified(100);p__Ascomycota_unclassified(100);p__Ascomycota_unclassified(100);p__Ascomycota_unclassified(100);p__Ascomycota_unclassified(100); | - | - | - | - | - | - | - | Unassigned | - |
| Otu05113 | 6 | k__Fungi(100);p__Ascomycota(100);p__Ascomycota_unclassified(100);p__Ascomycota_unclassified(100);p__Ascomycota_unclassified(100);p__Ascomycota_unclassified(100);p__Ascomycota_unclassified(100); | - | - | - | - | - | - | - | Unassigned | - |
| Otu05115 | 6 | k__Fungi(100);p__Ascomycota(84);p__Ascomycota_unclassified(84);p__Ascomycota_unclassified(84);p__Ascomycota_unclassified(84);p__Ascomycota_unclassified(84);p__Ascomycota_unclassified(84); | - | - | - | - | - | - | - | Unassigned | - |
| Otu05116 | 6 | k__Fungi(100);p__Ascomycota(67);p__Ascomycota_unclassified(67);p__Ascomycota_unclassified(67);p__Ascomycota_unclassified(67);p__Ascomycota_unclassified(67);p__Ascomycota_unclassified(67); | - | - | - | - | - | - | - | Unassigned | - |
| Otu05117 | 6 | k__Fungi(100);k__Fungi_unclassified(100);k__Fungi_unclassified(100);k__Fungi_unclassified(100);k__Fungi_unclassified(100);k__Fungi_unclassified(100);k__Fungi_unclassified(100); | - | - | - | - | - | - | - | Unassigned | - |

| | | | | | | | | | | | |
|---|---|---|---|---|---|---|---|---|---|---|---|
| Otu05118 | 6 | k__Fungi(100);p__Ascomycota(100);p__Ascomycota_unclassified(100);p__Ascomycota_unclassified(100);p__Ascomycota_unclassified(100);p__Ascomycota_unclassified(100);p__Ascomycota_unclassified(100); | - | - | - | - | - | - | - | Unassigned | - |
| Otu05121 | 6 | k__Fungi(100);p__Ascomycota(100);p__Ascomycota_unclassified(100);p__Ascomycota_unclassified(100);p__Ascomycota_unclassified(100);p__Ascomycota_unclassified(100);p__Ascomycota_unclassified(100); | - | - | - | - | - | - | - | Unassigned | - |
| Otu05123 | 6 | k__Fungi(100);k__Fungi_unclassified(100);k__Fungi_unclassified(100);k__Fungi_unclassified(100);k__Fungi_unclassified(100);k__Fungi_unclassified(100);k__Fungi_unclassified(100); | - | - | - | - | - | - | - | Unassigned | - |
| Otu05130 | 6 | k__Fungi(100);p__Ascomycota(84);p__Ascomycota_unclassified(84);p__Ascomycota_unclassified(84);p__Ascomycota_unclassified(84);p__Ascomycota_unclassified(84);p__Ascomycota_unclassified(84); | - | - | - | - | - | - | - | Unassigned | - |
| Otu05132 | 6 | k__Fungi(100);p__Ascomycota(84);p__Ascomycota_unclassified(84);p__Ascomycota_unclassified(84);p__Ascomycota_unclassified(84);p__Ascomycota_unclassified(84);p__Ascomycota_unclassified(84); | - | - | - | - | - | - | - | Unassigned | - |
| Otu05134 | 6 | k__Fungi(100);p__Ascomycota(100);c__Dothideomycetes(67);c__Dothideomycetes_unclassified(67);c__Dothideomycetes_unclassified(67);c__Dothideomycetes_unclassified(67);c__Dothideomycetes_unclassified(67); | - | - | - | - | - | - | - | Unassigned | - |
| Otu05135 | 6 | k__Fungi(100);p__Ascomycota(100);c__Leotiomycetes(100);o__Helotiales(100);o__Helotiales_unclassified(100);o__Helotiales_unclassified(100);o__Helotiales_unclassified(100); | - | - | - | - | - | - | - | Unassigned | - |
| Otu05139 | 6 | k__Fungi(100);p__Ascomycota(100);c__Dothideomycetes(100);o__Pleosporales(100);o__Pleosporales_unclassified(100);o__Pleosporales_unclassified(100);o__Pleosporales_unclassified(100); | - | - | - | - | - | - | - | Unassigned | - |
| Otu05142 | 6 | k__Fungi(100);k__Fungi_unclassified(84);k__Fungi_unclassified(84);k__Fungi_unclassified(84);k__Fungi_unclassified(84);k__Fungi_unclassified(84);k__Fungi_unclassified(84); | - | - | - | - | - | - | - | Unassigned | - |
| Otu05146 | 6 | k__Fungi(100);k__Fungi_unclassified(67);k__Fungi_unclassified(67);k__Fungi_unclassified(67);k__Fungi_unclassified(67);k__Fungi_unclassified(67);k__Fungi_unclassified(67); | - | - | - | - | - | - | - | Unassigned | - |
| Otu05150 | 6 | k__Fungi(100);p__Ascomycota(100);p__Ascomycota_unclassified(100);p__Ascomycota_unclassified(100);p__Ascomycota_unclassified(100);p__Ascomycota_unclassified(100);p__Ascomycota_unclassified(100); | - | - | - | - | - | - | - | Unassigned | - |
| Otu05151 | 6 | k__Fungi(100);k__Fungi_unclassified(100);k__Fungi_unclassified(100);k__Fungi_unclassified(100);k__Fungi_unclassified(100);k__Fungi_unclassified(100);k__Fungi_unclassified(100); | - | - | - | - | - | - | - | Unassigned | - |
| Otu05156 | 5 | k__Fungi(100);p__Ascomycota(100);c__Saccharomycetes(100);o__Saccharomycetales(100);f__Metschnikowiaceae(100);g__Metschnikowia(100);s__Metschnikowia_pulcherrima(100); | Metschnikowia | 13 | Saprotroph | \|Undefined Saprotroph\| | Yeast | NULL | Probable | As nectar-tap_saprotroph (Põlme et al. 2020); Decay sugar-rich substrates (Põlme et al. 2020) | Põlme S, et al. 2020. Fungal Diversity 105:1-16 (https://doi.org/10.1007/s13225-020-00466-2) |
| Otu05158 | 5 | k__Fungi(100);p__Ascomycota(100);c__Saccharomycetes(100);o__Saccharomycetales(100);f__Metschnikowiaceae(100);g__Metschnikowia(100);s__Metschnikowia_pulcherrima(80); | Metschnikowia | 13 | Saprotroph | \|Undefined Saprotroph\| | Yeast | NULL | Probable | As nectar-tap_saprotroph (Põlme et al. 2020); Decay sugar-rich substrates (Põlme et al. 2020) | Põlme S, et al. 2020. Fungal Diversity 105:1-16 (https://doi.org/10.1007/s13225-020-00466-2) |
| Otu05159 | 5 | k__Fungi(100);p__Ascomycota(100);c__Sordariomycetes(100);o__Coniochaetales(100);f__Coniochaetaceae(100);g__Coniochaeta(80);s__Coniochaeta_lignicola(61); | Coniochaeta | 13 | Pathotroph-Saprotroph-Symbiotroph | Animal Pathogen-Dung Saprotroph-Endophyte-Lichen Parasite-Plant Pathogen-\|Undefined Saprotroph\| | Microfungus | NULL | Possible | As foliar_endophyte (Põlme et al. 2020); Decay leaf, fruit, seed, soil, dung, animal material (Põlme et al. 2020) | [illegible] |
| Otu05164 | 5 | k__Fungi(100);p__Ascomycota(100);c__Saccharomycetes(100);o__Saccharomycetales(100);f__Metschnikowiaceae(100);g__Metschnikowia(100);s__Metschnikowia_pulcherrima(100); | Metschnikowia | 13 | Saprotroph | \|Undefined Saprotroph\| | Yeast | NULL | Probable | As nectar-tap_saprotroph (Põlme et al. 2020); Decay sugar-rich substrates (Põlme et al. 2020) | Põlme S, et al. 2020. Fungal Diversity 105:1-16 (https://doi.org/10.1007/s13225-020-00466-2) |
| Otu05170 | 5 | k__Fungi(100);p__Ascomycota(100);c__Saccharomycetes(100);o__Saccharomycetales(100);f__Metschnikowiaceae(100);g__Metschnikowia(100);s__Metschnikowia_pulcherrima(80); | Metschnikowia | 13 | Saprotroph | \|Undefined Saprotroph\| | Yeast | NULL | Probable | As nectar-tap_saprotroph (Põlme et al. 2020); Decay sugar-rich substrates (Põlme et al. 2020) | Põlme S, et al. 2020. Fungal Diversity 105:1-16 (https://doi.org/10.1007/s13225-020-00466-2) |
| Otu05171 | 5 | k__Fungi(100);p__Basidiomycota(100);c__Agaricomycetes(100);o__Gomphales(100);f__Gomphaceae(100);g__Ramaria(100);s__Ramaria_rubella(100); | Ramaria | 13 | Saprotroph-Symbiotroph | \|Ectomycorrhizal\|-Plant Saprotroph | Clavarioid | NULL | Possible | As litter_saprotroph (Põlme et al. 2020) | [illegible] |
| Otu05172 | 5 | k__Fungi(100);p__Ascomycota(100);c__Saccharomycetes(100);o__Saccharomycetales(100);f__Metschnikowiaceae(100);g__Metschnikowia(100);s__Metschnikowia_pulcherrima(80); | Metschnikowia | 13 | Saprotroph | \|Undefined Saprotroph\| | Yeast | NULL | Probable | As nectar-tap_saprotroph (Põlme et al. 2020); Decay sugar-rich substrates (Põlme et al. 2020) | Põlme S, et al. 2020. Fungal Diversity 105:1-16 (https://doi.org/10.1007/s13225-020-00466-2) |
| Otu05173 | 5 | k__Fungi(100);p__Basidiomycota(100);c__Agaricostilbomycetes(100);o__Agaricostilbales(100);f__Agaricostilbaceae(100);g__Bensingtonia(100);g__Bensingtonia_unclassified(100); | Bensingtonia | 13 | Saprotroph | \|Undefined Saprotroph\| | Dimorphic-Facultative Yeast | NULL | Highly Probable | NULL | [illegible] |
| Otu05174 | 5 | k__Fungi(100);p__Ascomycota(100);c__Saccharomycetes(100);o__Saccharomycetales(100);f__Metschnikowiaceae(100);g__Metschnikowia(100);g__Metschnikowia_unclassified(61); | Metschnikowia | 13 | Saprotroph | \|Undefined Saprotroph\| | Yeast | NULL | Probable | As nectar-tap_saprotroph (Põlme et al. 2020); Decay sugar-rich substrates (Põlme et al. 2020) | Põlme S, et al. 2020. Fungal Diversity 105:1-16 (https://doi.org/10.1007/s13225-020-00466-2) |
| Otu05175 | 5 | k__Fungi(100);p__Ascomycota(100);c__Saccharomycetes(100);o__Saccharomycetales(100);f__Metschnikowiaceae(100);g__Metschnikowia(100);s__Metschnikowia_pulcherrima(61); | Metschnikowia | 13 | Saprotroph | \|Undefined Saprotroph\| | Yeast | NULL | Probable | As nectar-tap_saprotroph (Põlme et al. 2020); Decay sugar-rich substrates (Põlme et al. 2020) | Põlme S, et al. 2020. Fungal Diversity 105:1-16 (https://doi.org/10.1007/s13225-020-00466-2) |
| Otu05177 | 5 | k__Fungi(100);p__Ascomycota(100);c__Saccharomycetes(100);o__Saccharomycetales(100);f__Metschnikowiaceae(100);g__Metschnikowia(100);g__Metschnikowia_unclassified(61); | Metschnikowia | 13 | Saprotroph | \|Undefined Saprotroph\| | Yeast | NULL | Probable | As nectar-tap_saprotroph (Põlme et al. 2020); Decay sugar-rich substrates (Põlme et al. 2020) | Põlme S, et al. 2020. Fungal Diversity 105:1-16 (https://doi.org/10.1007/s13225-020-00466-2) |
| Otu05180 | 5 | k__Fungi(100);p__Ascomycota(100);c__Saccharomycetes(100);o__Saccharomycetales(100);f__Metschnikowiaceae(100);g__Metschnikowia(100);s__Metschnikowia_pulcherrima(80); | Metschnikowia | 13 | Saprotroph | \|Undefined Saprotroph\| | Yeast | NULL | Probable | As nectar-tap_saprotroph (Põlme et al. 2020); Decay sugar-rich substrates (Põlme et al. 2020) | Põlme S, et al. 2020. Fungal Diversity 105:1-16 (https://doi.org/10.1007/s13225-020-00466-2) |
| Otu05185 | 5 | k__Fungi(100);p__Ascomycota(100);c__Dothideomycetes(100);o__Capnodiales(100);f__Mycosphaerellaceae(100);g__Phaeoramularia(100);s__Phaeoramularia_weigelicola(100); | Phaeoramularia | 13 | Pathotroph | \|Plant Pathogen\| | Microfungus | NULL | Probable | NULL | [illegible] |
| Otu05189 | 5 | k__Fungi(100);p__Ascomycota(100);c__Saccharomycetes(100);o__Saccharomycetales(100);f__Metschnikowiaceae(100);g__Metschnikowia(100);s__Metschnikowia_pulcherrima(80); | Metschnikowia | 13 | Saprotroph | \|Undefined Saprotroph\| | Yeast | NULL | Probable | As nectar-tap_saprotroph (Põlme et al. 2020); Decay sugar-rich substrates (Põlme et al. 2020) | Põlme S, et al. 2020. Fungal Diversity 105:1-16 (https://doi.org/10.1007/s13225-020-00466-2) |
| Otu05191 | 5 | k__Fungi(100);p__Zygomycota(100);c__Zygomycota_class_Incertae_sedis(100);o__Mucorales(100);f__Mucoraceae(100);g__Mucor(100);s__Mucor_genevensis(100); | Mucor | 13 | Saprotroph | \|Undefined Saprotroph\| | Microfungus | NULL | Probable | As soil_saprotroph (Põlme et al. 2020); Decay soil (Põlme et al. 2020) | [illegible] |
| Otu05194 | 5 | k__Fungi(100);p__Ascomycota(100);c__Ascomycota_class_Incertae_sedis(100);o__Ascomycota_order_Incertae_sedis(100);f__Ascomycota_family_Incertae_sedis(100);g__Phialemoniopsis(100);s__Phialemoniopsis_cornearis(100); | Phialemoniopsis | 13 | Pathotroph-Saprotroph | \|Animal Parasite\|-Undefined Saprotroph | Microfungus | NULL | Probable | As animal_decomposer (Põlme et al. 2020); Decay animal material (Põlme et al. 2020) | [illegible] |
| Otu05195 | 5 | k__Fungi(100);p__Ascomycota(100);c__Saccharomycetes(100);o__Saccharomycetales(100);f__Metschnikowiaceae(100);g__Metschnikowia(100);s__Metschnikowia_pulcherrima(100); | Metschnikowia | 13 | Saprotroph | \|Undefined Saprotroph\| | Yeast | NULL | Probable | As nectar-tap_saprotroph (Põlme et al. 2020); Decay sugar-rich substrates (Põlme et al. 2020) | Põlme S, et al. 2020. Fungal Diversity 105:1-16 (https://doi.org/10.1007/s13225-020-00466-2) |
| Otu05197 | 5 | k__Fungi(100);p__Ascomycota(100);c__Saccharomycetes(100);o__Saccharomycetales(100);f__Metschnikowiaceae(100);g__Metschnikowia(100);s__Metschnikowia_unclassified(61); | Metschnikowia | 13 | Saprotroph | \|Undefined Saprotroph\| | Yeast | NULL | Probable | As nectar-tap_saprotroph (Põlme et al. 2020); Decay sugar-rich substrates (Põlme et al. 2020) | Põlme S, et al. 2020. Fungal Diversity 105:1-16 (https://doi.org/10.1007/s13225-020-00466-2) |
| Otu05199 | 5 | k__Fungi(100);p__Ascomycota(100);c__Saccharomycetes(100);o__Saccharomycetales(100);f__Metschnikowiaceae(100);g__Metschnikowia(100);s__Metschnikowia_pulcherrima(80); | Metschnikowia | 13 | Saprotroph | \|Undefined Saprotroph\| | Yeast | NULL | Probable | As nectar-tap_saprotroph (Põlme et al. 2020); Decay sugar-rich substrates (Põlme et al. 2020) | Põlme S, et al. 2020. Fungal Diversity 105:1-16 (https://doi.org/10.1007/s13225-020-00466-2) |
| Otu05204 | 5 | k__Fungi(100);p__Ascomycota(100);c__Saccharomycetes(100);o__Saccharomycetales(100);f__Metschnikowiaceae(100);g__Metschnikowia(100);s__Metschnikowia_pulcherrima(80); | Metschnikowia | 13 | Saprotroph | \|Undefined Saprotroph\| | Yeast | NULL | Probable | As nectar-tap_saprotroph (Põlme et al. 2020); Decay sugar-rich substrates (Põlme et al. 2020) | Põlme S, et al. 2020. Fungal Diversity 105:1-16 (https://doi.org/10.1007/s13225-020-00466-2) |
| Otu05206 | 5 | k__Fungi(100);p__Ascomycota(100);c__Saccharomycetes(100);o__Saccharomycetales(100);f__Metschnikowiaceae(100);g__Metschnikowia(100);s__Metschnikowia_pulcherrima(100); | Metschnikowia | 13 | Saprotroph | \|Undefined Saprotroph\| | Yeast | NULL | Probable | As nectar-tap_saprotroph (Põlme et al. 2020); Decay sugar-rich substrates (Põlme et al. 2020) | Põlme S, et al. 2020. Fungal Diversity 105:1-16 (https://doi.org/10.1007/s13225-020-00466-2) |
| Otu05208 | 5 | k__Fungi(100);p__Ascomycota(100);c__Saccharomycetes(100);o__Saccharomycetales(100);f__Saccharomycetales_family_Incertae_sedis(100);g__Candida(100);g__Candida_unclassified(61); | Candida | 13 | Pathotroph-Saprotroph-Symbiotroph | Animal Pathogen-Endophyte-Endosymbiont-Epiphyte-\|Undefined Saprotroph\| | Dimorphic Yeast-Yeast | NULL | Probable | As nectar-tap_saprotroph (Põlme et al. 2020); Decay sugar-rich substrates (Põlme et al. 2020). This asexual genus is polyphyletic and contain species that function very differently depend on what they are. Caution should be taken when interpreting guilds for Candida species. | [illegible] |
| Otu05210 | 5 | k__Fungi(100);p__Ascomycota(100);c__Saccharomycetes(100);o__Saccharomycetales(100);f__Metschnikowiaceae(100);g__Metschnikowia(100);s__Metschnikowia_pulcherrima(100); | Metschnikowia | 13 | Saprotroph | \|Undefined Saprotroph\| | Yeast | NULL | Probable | As nectar-tap_saprotroph (Põlme et al. 2020); Decay sugar-rich substrates (Põlme et al. 2020) | Põlme S, et al. 2020. Fungal Diversity 105:1-16 (https://doi.org/10.1007/s13225-020-00466-2) |
| Otu05211 | 5 | k__Fungi(100);p__Ascomycota(100);c__Saccharomycetes(100);o__Saccharomycetales(100);f__Metschnikowiaceae(100);g__Metschnikowia(100);s__Metschnikowia_pulcherrima(100); | Metschnikowia | 13 | Saprotroph | \|Undefined Saprotroph\| | Yeast | NULL | Probable | As nectar-tap_saprotroph (Põlme et al. 2020); Decay sugar-rich substrates (Põlme et al. 2020) | Põlme S, et al. 2020. Fungal Diversity 105:1-16 (https://doi.org/10.1007/s13225-020-00466-2) |
| Otu05212 | 5 | k__Fungi(100);p__Ascomycota(100);c__Sordariomycetes(100);o__Xylariales(100);f__Amphisphaeriaceae(100);g__Monochaetia(100);g__Monochaetia_unclassified(100); | Monochaetia | 13 | Pathotroph | \|Plant Pathogen\| | Microfungus | NULL | Probable | NULL | [illegible] |
| Otu05213 | 5 | k__Fungi(100);p__Basidiomycota(100);c__Tremellomycetes(100);o__Tremellales(100);f__Tremellales_family_Incertae_sedis(100);f__Tremellales_family_Incertae_sedis_unclassified(100);f__Tremellales_family_Incertae_sedis_unclassified(100); | Tremellales | 7 | Pathotroph-Saprotroph-Symbiotroph | Fungal Parasite-Undefined Saprotroph | Tremelloid-Yeast | NULL | Possible | The order contains both teleomorphic and anamorphic species, most of the latter being yeasts | [illegible] |
| Otu05214 | 5 | k__Fungi(100);p__Basidiomycota(100);c__Agaricomycetes(100);o__Polyporales(100);f__Fomitopsidaceae(100);g__Postia(100);s__Postia_caesia(100); | Postia | 13 | Saprotroph | \|Wood Saprotroph\| | Polyporoid | Brown Rot | Highly Probable | As brown-rot (Põlme et al. 2020); Decay wood (Põlme et al. 2020) | [illegible] |
| Otu05215 | 5 | k__Fungi(100);p__Ascomycota(100);c__Saccharomycetes(100);o__Saccharomycetales(100);f__Metschnikowiaceae(100);g__Metschnikowia(100);s__Metschnikowia_pulcherrima(61); | Metschnikowia | 13 | Saprotroph | \|Undefined Saprotroph\| | Yeast | NULL | Probable | As nectar-tap_saprotroph (Põlme et al. 2020); Decay sugar-rich substrates (Põlme et al. 2020) | Põlme S, et al. 2020. Fungal Diversity 105:1-16 (https://doi.org/10.1007/s13225-020-00466-2) |
| Otu05217 | 5 | k__Fungi(100);p__Ascomycota(100);c__Saccharomycetes(100);o__Saccharomycetales(100);f__Saccharomycetales_family_Incertae_sedis(80);g__Candida(80);g__Candida_unclassified(80); | Candida | 13 | Pathotroph-Saprotroph-Symbiotroph | Animal Pathogen-Endophyte-Endosymbiont-Epiphyte-\|Undefined Saprotroph\| | Dimorphic Yeast-Yeast | NULL | Probable | As nectar-tap_saprotroph (Põlme et al. 2020); Decay sugar-rich substrates (Põlme et al. 2020). This asexual genus is polyphyletic and contain species that function very differently depend on what they are. Caution should be taken when interpreting guilds for Candida species. | [illegible] |
| Otu05218 | 5 | k__Fungi(100);p__Ascomycota(100);c__Saccharomycetes(100);o__Saccharomycetales(100);f__Metschnikowiaceae(100);g__Metschnikowia(100);s__Metschnikowia_pulcherrima(100); | Metschnikowia | 13 | Saprotroph | \|Undefined Saprotroph\| | Yeast | NULL | Probable | As nectar-tap_saprotroph (Põlme et al. 2020); Decay sugar-rich substrates (Põlme et al. 2020) | Põlme S, et al. 2020. Fungal Diversity 105:1-16 (https://doi.org/10.1007/s13225-020-00466-2) |
| Otu05224 | 5 | k__Fungi(100);p__Ascomycota(100);c__Saccharomycetes(100);o__Saccharomycetales(100);f__Saccharomycetales_family_Incertae_sedis(100);g__Candida(100);g__Candida_unclassified(100); | Candida | 13 | Pathotroph-Saprotroph-Symbiotroph | Animal Pathogen-Endophyte-Endosymbiont-Epiphyte-\|Undefined Saprotroph\| | Dimorphic Yeast-Yeast | NULL | Probable | As nectar-tap_saprotroph (Põlme et al. 2020); Decay sugar-rich substrates (Põlme et al. 2020). This asexual genus is polyphyletic and contain species that function very differently depend on what they are. Caution should be taken when interpreting guilds for Candida species. | [illegible] |
| Otu05226 | 5 | k__Fungi(100);p__Ascomycota(100);c__Saccharomycetes(100);o__Saccharomycetales(100);f__Metschnikowiaceae(100);g__Metschnikowia(100);s__Metschnikowia_pulcherrima(61); | Metschnikowia | 13 | Saprotroph | \|Undefined Saprotroph\| | Yeast | NULL | Probable | As nectar-tap_saprotroph (Põlme et al. 2020); Decay sugar-rich substrates (Põlme et al. 2020) | Põlme S, et al. 2020. Fungal Diversity 105:1-16 (https://doi.org/10.1007/s13225-020-00466-2) |
| Otu05228 | 5 | k__Fungi(100);p__Ascomycota(100);c__Saccharomycetes(100);o__Saccharomycetales(100);f__Metschnikowiaceae(100);g__Metschnikowia(100);s__Metschnikowia_unclassified(80); | Metschnikowia | 13 | Saprotroph | \|Undefined Saprotroph\| | Yeast | NULL | Probable | As nectar-tap_saprotroph (Põlme et al. 2020); Decay sugar-rich substrates (Põlme et al. 2020) | Põlme S, et al. 2020. Fungal Diversity 105:1-16 (https://doi.org/10.1007/s13225-020-00466-2) |
| Otu05229 | 5 | k__Fungi(100);p__Ascomycota(100);c__Sordariomycetes(100);o__Boliniales(100);f__Boliniaceae(100);f__Boliniaceae_unclassified(80);f__Boliniaceae_unclassified(80); | Boliniaceae | 9 | Saprotroph | Wood Saprotroph | Microfungus | NULL | Probable | Saprobic on and in wood and bark (Cannon & Kirk 2007) | Cannon PF, Kirk PM. 2007. Fungal Families of the World. CAB International, Cambridge (ISBN: 9780851998275). |
| Otu05230 | 5 | k__Fungi(100);p__Ascomycota(100);c__Saccharomycetes(100);o__Saccharomycetales(100);f__Metschnikowiaceae(100);g__Metschnikowia(100);s__Metschnikowia_pulcherrima(61); | Metschnikowia | 13 | Saprotroph | \|Undefined Saprotroph\| | Yeast | NULL | Probable | As nectar-tap_saprotroph (Põlme et al. 2020); Decay sugar-rich substrates (Põlme et al. 2020) | Põlme S, et al. 2020. Fungal Diversity 105:1-16 (https://doi.org/10.1007/s13225-020-00466-2) |
| Otu05231 | 5 | k__Fungi(100);p__Ascomycota(100);c__Saccharomycetes(100);o__Saccharomycetales(100);f__Metschnikowiaceae(100);g__Metschnikowia(100);s__Metschnikowia_pulcherrima(100); | Metschnikowia | 13 | Saprotroph | \|Undefined Saprotroph\| | Yeast | NULL | Probable | As nectar-tap_saprotroph (Põlme et al. 2020); Decay sugar-rich substrates (Põlme et al. 2020) | Põlme S, et al. 2020. Fungal Diversity 105:1-16 (https://doi.org/10.1007/s13225-020-00466-2) |
| Otu05232 | 5 | k__Fungi(100);p__Ascomycota(100);c__Saccharomycetes(100);o__Saccharomycetales(100);f__Metschnikowiaceae(100);g__Metschnikowia(100);s__Metschnikowia_pulcherrima(80); | Metschnikowia | 13 | Saprotroph | \|Undefined Saprotroph\| | Yeast | NULL | Probable | As nectar-tap_saprotroph (Põlme et al. 2020); Decay sugar-rich substrates (Põlme et al. 2020) | Põlme S, et al. 2020. Fungal Diversity 105:1-16 (https://doi.org/10.1007/s13225-020-00466-2) |
| Otu05233 | 5 | k__Fungi(100);p__Ascomycota(100);c__Saccharomycetes(100);o__Saccharomycetales(100);f__Metschnikowiaceae(100);g__Metschnikowia(100);s__Metschnikowia_pulcherrima(61); | Metschnikowia | 13 | Saprotroph | \|Undefined Saprotroph\| | Yeast | NULL | Probable | As nectar-tap_saprotroph (Põlme et al. 2020); Decay sugar-rich substrates (Põlme et al. 2020) | Põlme S, et al. 2020. Fungal Diversity 105:1-16 (https://doi.org/10.1007/s13225-020-00466-2) |
| Otu05234 | 5 | k__Fungi(100);p__Ascomycota(100);c__Eurotiomycetes(100);o__Chaetothyriales(100);f__Herpotrichiellaceae(100);g__Cladophialophora(100);g__Cladophialophora_unclassified(100); | Cladophialophora | 13 | Pathotroph-Saprotroph-Symbiotroph | Animal Pathogen-Endophyte-Fungal Parasite-\|Undefined Saprotroph\| | Dimorphic-Microfungus | NULL | Probable | As mycoparasite (Põlme et al. 2020); As root_endophyte_dark_septate (Põlme et al. 2020); As soil_saprotroph (Põlme et al. 2020); Decay soil (Põlme et al. 2020) | [illegible] |
| Otu05236 | 5 | k__Fungi(100);p__Ascomycota(100);c__Saccharomycetes(100);o__Saccharomycetales(100);f__Metschnikowiaceae(100);g__Metschnikowia(100);s__Metschnikowia_pulcherrima(80); | Metschnikowia | 13 | Saprotroph | \|Undefined Saprotroph\| | Yeast | NULL | Probable | As nectar-tap_saprotroph (Põlme et al. 2020); Decay sugar-rich substrates (Põlme et al. 2020) | Põlme S, et al. 2020. Fungal Diversity 105:1-16 (https://doi.org/10.1007/s13225-020-00466-2) |
| Otu05238 | 5 | k__Fungi(100);p__Ascomycota(100);c__Saccharomycetes(100);o__Saccharomycetales(100);f__Saccharomycetaceae(100);g__Kazachstania(100);s__Kazachstania_intestinalis(100); | Kazachstania | 13 | Saprotroph | \|Undefined Saprotroph\| | Yeast | NULL | Probable | As nectar-tap_saprotroph (Põlme et al. 2020); Decay sugar-rich substrates (Põlme et al. 2020) | Põlme S, et al. 2020. Fungal Diversity 105:1-16 (https://doi.org/10.1007/s13225-020-00466-2) |
| Otu05241 | 5 | k__Fungi(100);p__Ascomycota(100);c__Saccharomycetes(100);o__Saccharomycetales(100);f__Metschnikowiaceae(100);g__Metschnikowia(100);s__Metschnikowia_unclassified(61); | Metschnikowia | 13 | Saprotroph | \|Undefined Saprotroph\| | Yeast | NULL | Probable | As nectar-tap_saprotroph (Põlme et al. 2020); Decay sugar-rich substrates (Põlme et al. 2020) | Põlme S, et al. 2020. Fungal Diversity 105:1-16 (https://doi.org/10.1007/s13225-020-00466-2) |
| Otu05242 | 5 | k__Fungi(100);p__unclassified_Fungi(100);c__unclassified_Fungi(100);o__unclassified_Fungi(100);f__unclassified_Fungi(100);g__unclassified_Fungi(100);s__Slopeiomyces_cylindrosporus(100); | Slopeiomyces | 13 | Symbiotroph | \|Endophyte\| | Microfungus | NULL | Probable | As root_endophyte (Põlme et al. 2020) | Põlme S, et al. 2020. Fungal Diversity 105:1-16 (https://doi.org/10.1007/s13225-020-00466-2) |
| Otu05243 | 5 | k__Fungi(100);p__Ascomycota(100);c__Saccharomycetes(100);o__Saccharomycetales(100);f__Metschnikowiaceae(100);g__Metschnikowia(100);s__Metschnikowia_pulcherrima(100); | Metschnikowia | 13 | Saprotroph | \|Undefined Saprotroph\| | Yeast | NULL | Probable | As nectar-tap_saprotroph (Põlme et al. 2020); Decay sugar-rich substrates (Põlme et al. 2020) | Põlme S, et al. 2020. Fungal Diversity 105:1-16 (https://doi.org/10.1007/s13225-020-00466-2) |
| Otu05245 | 5 | k__Fungi(100);p__Ascomycota(100);c__Saccharomycetes(100);o__Saccharomycetales(100);f__Metschnikowiaceae(100);g__Metschnikowia(100);s__Metschnikowia_pulcherrima(80); | Metschnikowia | 13 | Saprotroph | \|Undefined Saprotroph\| | Yeast | NULL | Probable | As nectar-tap_saprotroph (Põlme et al. 2020); Decay sugar-rich substrates (Põlme et al. 2020) | Põlme S, et al. 2020. Fungal Diversity 105:1-16 (https://doi.org/10.1007/s13225-020-00466-2) |
| Otu05249 | 5 | k__Fungi(100);p__Basidiomycota(100);c__Microbotryomycetes(100);o__Sporidiobolales(100);f__Sporidiobolales_family_Incertae_sedis(100);g__Sporobolomyces(100);s__Sporobolomyces_griseoflavus(100); | Sporobolomyces | 13 | Pathotroph-Saprotroph | \|Fungal Parasite\|-Plant Saprotroph | Yeast | NULL | Probable | As litter_saprotroph (Põlme et al. 2020); As mycoparasite (Põlme et al. 2020); Decay fungal material (Põlme et al. 2020) | [illegible] |
| Otu05256 | 5 | k__Fungi(100);p__Ascomycota(100);c__Saccharomycetes(100);o__Saccharomycetales(100);f__Metschnikowiaceae(100);g__Metschnikowia(100);s__Metschnikowia_pulcherrima(80); | Metschnikowia | 13 | Saprotroph | \|Undefined Saprotroph\| | Yeast | NULL | Probable | As nectar-tap_saprotroph (Põlme et al. 2020); Decay sugar-rich substrates (Põlme et al. 2020) | Põlme S, et al. 2020. Fungal Diversity 105:1-16 (https://doi.org/10.1007/s13225-020-00466-2) |
| Otu05259 | 5 | k__Fungi(100);p__Ascomycota(100);c__Dothideomycetes(100);o__Capnodiales(100);f__Teratosphaeriaceae(100);g__Devriesia(100);s__Devriesia_fraseriae(100); | Devriesia | 13 | Pathotroph-Saprotroph | \|Plant Pathogen\|-Plant Saprotroph | Microfungus | NULL | Probable | As litter_saprotroph (Põlme et al. 2020); Decay leaf, fruit, seed (Põlme et al. 2020) | [illegible] |
| Otu05260 | 5 | k__Fungi(100);p__Ascomycota(100);c__Saccharomycetes(100);o__Saccharomycetales(100);f__Metschnikowiaceae(100);g__Metschnikowia(100);s__Metschnikowia_pulcherrima(100); | Metschnikowia | 13 | Saprotroph | \|Undefined Saprotroph\| | Yeast | NULL | Probable | As nectar-tap_saprotroph (Põlme et al. 2020); Decay sugar-rich substrates (Põlme et al. 2020) | Põlme S, et al. 2020. Fungal Diversity 105:1-16 (https://doi.org/10.1007/s13225-020-00466-2) |
| Otu05265 | 5 | k__Fungi(100);p__Ascomycota(100);c__Saccharomycetes(100);o__Saccharomycetales(100);f__Metschnikowiaceae(100);g__Metschnikowia(100);s__Metschnikowia_pulcherrima(100); | Metschnikowia | 13 | Saprotroph | \|Undefined Saprotroph\| | Yeast | NULL | Probable | As nectar-tap_saprotroph (Põlme et al. 2020); Decay sugar-rich substrates (Põlme et al. 2020) | Põlme S, et al. 2020. Fungal Diversity 105:1-16 (https://doi.org/10.1007/s13225-020-00466-2) |
| Otu05272 | 5 | k__Fungi(100);p__Ascomycota(100);c__Saccharomycetes(100);o__Saccharomycetales(100);f__Saccharomycetales_family_Incertae_sedis(100);g__Candida(100);g__Candida_unclassified(80); | Candida | 13 | Pathotroph-Saprotroph-Symbiotroph | Animal Pathogen-Endophyte-Endosymbiont-Epiphyte-\|Undefined Saprotroph\| | Dimorphic Yeast-Yeast | NULL | Probable | As nectar-tap_saprotroph (Põlme et al. 2020); Decay sugar-rich substrates (Põlme et al. 2020). This asexual genus is polyphyletic and contain species that function very differently depend on what they are. Caution should be taken when interpreting guilds for Candida species. | [illegible] |
| Otu05274 | 5 | k__Fungi(100);p__Ascomycota(100);c__Saccharomycetes(100);o__Saccharomycetales(100);f__Metschnikowiaceae(100);g__Metschnikowia(100);s__Metschnikowia_pulcherrima(61); | Metschnikowia | 13 | Saprotroph | \|Undefined Saprotroph\| | Yeast | NULL | Probable | As nectar-tap_saprotroph (Põlme et al. 2020); Decay sugar-rich substrates (Põlme et al. 2020) | Põlme S, et al. 2020. Fungal Diversity 105:1-16 (https://doi.org/10.1007/s13225-020-00466-2) |
| Otu05275 | 5 | k__Fungi(100);p__Ascomycota(100);c__Saccharomycetes(100);o__Saccharomycetales(100);f__Metschnikowiaceae(100);g__Metschnikowia(100);s__Metschnikowia_pulcherrima(80); | Metschnikowia | 13 | Saprotroph | \|Undefined Saprotroph\| | Yeast | NULL | Probable | As nectar-tap_saprotroph (Põlme et al. 2020); Decay sugar-rich substrates (Põlme et al. 2020) | Põlme S, et al. 2020. Fungal Diversity 105:1-16 (https://doi.org/10.1007/s13225-020-00466-2) |
| Otu05278 | 5 | k__Fungi(100);p__Ascomycota(100);c__Saccharomycetes(100);o__Saccharomycetales(100);f__Metschnikowiaceae(100);g__Metschnikowia(100);s__Metschnikowia_pulcherrima(100); | Metschnikowia | 13 | Saprotroph | \|Undefined Saprotroph\| | Yeast | NULL | Probable | As nectar-tap_saprotroph (Põlme et al. 2020); Decay sugar-rich substrates (Põlme et al. 2020) | Põlme S, et al. 2020. Fungal Diversity 105:1-16 (https://doi.org/10.1007/s13225-020-00466-2) |
| Otu05279 | 5 | k__Fungi(100);p__Ascomycota(100);c__Eurotiomycetes(100);o__Chaetothyriales(100);f__Chaetothyriaceae(61);g__Cyphellophora(61);g__Cyphellophora_unclassified(61); | Cyphellophora | 13 | Pathotroph-Saprotroph | Animal Parasite-Animal Pathogen-\|Plant Saprotroph\|-Undefined Saprotroph | Dimorphic-Facultative Yeast-Microfungus | NULL | Probable | As litter_saprotroph (Põlme et al. 2020). Consult the current literature for updated names on this group that included the Black Yeasts; Some species are likely opportunistic human pathogen (Irinyi et al. 2015). Decay leaf, fruit, seed, animal material (Põlme et al. 2020) | [illegible] |
| Otu05283 | 5 | k__Fungi(100);p__Ascomycota(100);c__Saccharomycetes(100);o__Saccharomycetales(100);f__Metschnikowiaceae(100);g__Metschnikowia(100);s__Metschnikowia_pulcherrima(100); | Metschnikowia | 13 | Saprotroph | \|Undefined Saprotroph\| | Yeast | NULL | Probable | As nectar-tap_saprotroph (Põlme et al. 2020); Decay sugar-rich substrates (Põlme et al. 2020) | Põlme S, et al. 2020. Fungal Diversity 105:1-16 (https://doi.org/10.1007/s13225-020-00466-2) |
| Otu05284 | 5 | k__Fungi(100);p__Ascomycota(100);c__Saccharomycetes(100);o__Saccharomycetales(100);f__Metschnikowiaceae(100);g__Metschnikowia(100);s__Metschnikowia_pulcherrima(100); | Metschnikowia | 13 | Saprotroph | \|Undefined Saprotroph\| | Yeast | NULL | Probable | As nectar-tap_saprotroph (Põlme et al. 2020); Decay sugar-rich substrates (Põlme et al. 2020) | Põlme S, et al. 2020. Fungal Diversity 105:1-16 (https://doi.org/10.1007/s13225-020-00466-2) |
| Otu05286 | 5 | k__Fungi(100);p__Ascomycota(100);c__Saccharomycetes(100);o__Saccharomycetales(100);f__Metschnikowiaceae(100);g__Metschnikowia(100);s__Metschnikowia_pulcherrima(80); | Metschnikowia | 13 | Saprotroph | \|Undefined Saprotroph\| | Yeast | NULL | Probable | As nectar-tap_saprotroph (Põlme et al. 2020); Decay sugar-rich substrates (Põlme et al. 2020) | Põlme S, et al. 2020. Fungal Diversity 105:1-16 (https://doi.org/10.1007/s13225-020-00466-2) |
| Otu05287 | 5 | k__Fungi(100);p__Ascomycota(100);c__Saccharomycetes(100);o__Saccharomycetales(100);f__Metschnikowiaceae(100);g__Metschnikowia(100);s__Metschnikowia_pulcherrima(80); | Metschnikowia | 13 | Saprotroph | \|Undefined Saprotroph\| | Yeast | NULL | Probable | As nectar-tap_saprotroph (Põlme et al. 2020); Decay sugar-rich substrates (Põlme et al. 2020) | Põlme S, et al. 2020. Fungal Diversity 105:1-16 (https://doi.org/10.1007/s13225-020-00466-2) |
| Otu05289 | 5 | k__Fungi(100);p__Ascomycota(100);c__Sordariomycetes(100);o__Hypocreales(100);o__Hypocreales_unclassified(100);o__Hypocreales_unclassified(100);o__Hypocreales_unclassified(100); | Hypocreales | 7 | Saprotroph | Undefined Saprotroph | Microfungus | NULL | Possible | NULL | [illegible] |
| Otu05291 | 5 | k__Fungi(100);p__Ascomycota(100);c__Saccharomycetes(100);o__Saccharomycetales(100);f__Metschnikowiaceae(100);g__Metschnikowia(100);s__Metschnikowia_pulcherrima(61); | Metschnikowia | 13 | Saprotroph | \|Undefined Saprotroph\| | Yeast | NULL | Probable | As nectar-tap_saprotroph (Põlme et al. 2020); Decay sugar-rich substrates (Põlme et al. 2020) | Põlme S, et al. 2020. Fungal Diversity 105:1-16 (https://doi.org/10.1007/s13225-020-00466-2) |
| Otu05292 | 5 | k__Fungi(100);p__Ascomycota(100);c__Eurotiomycetes(100);o__Chaetothyriales(100);f__Herpotrichiellaceae(100);f__Herpotrichiellaceae_unclassified(100);f__Herpotrichiellaceae_unclassified(100); | Herpotrichiellaceae | 9 | Pathotroph-Saprotroph | Animal Pathogen-Fungal Parasite-Undefined Saprotroph | Facultative Yeast-Microfungus | NULL | Probable | Mostly saprobic on plants or other fungi; isolated from soil etc., and from medical sources (Cannon & Kirk 2007) | Cannon PF, Kirk PM. 2007. Fungal Families of the World. CAB International, Cambridge (ISBN: 9780851998275). |
| Otu05293 | 5 | k__Fungi(100);p__Ascomycota(100);c__Saccharomycetes(100);o__Saccharomycetales(100);f__Metschnikowiaceae(100);g__Metschnikowia(100);s__Metschnikowia_pulcherrima(100); | Metschnikowia | 13 | Saprotroph | \|Undefined Saprotroph\| | Yeast | NULL | Probable | As nectar-tap_saprotroph (Põlme et al. 2020); Decay sugar-rich substrates (Põlme et al. 2020) | Põlme S, et al. 2020. Fungal Diversity 105:1-16 (https://doi.org/10.1007/s13225-020-00466-2) |
| Otu05294 | 5 | k__Fungi(100);p__Ascomycota(100);c__Saccharomycetes(100);o__Saccharomycetales(100);f__Metschnikowiaceae(100);g__Metschnikowia(100);s__Metschnikowia_pulcherrima(61); | Metschnikowia | 13 | Saprotroph | \|Undefined Saprotroph\| | Yeast | NULL | Probable | As nectar-tap_saprotroph (Põlme et al. 2020); Decay sugar-rich substrates (Põlme et al. 2020) | Põlme S, et al. 2020. Fungal Diversity 105:1-16 (https://doi.org/10.1007/s13225-020-00466-2) |
| Otu05296 | 5 | k__Fungi(100);p__Ascomycota(100);c__Eurotiomycetes(100);o__Chaetothyriales(100);f__Herpotrichiellaceae(100);g__Cladophialophora(100);g__Cladophialophora_unclassified(100); | Cladophialophora | 13 | Pathotroph-Saprotroph-Symbiotroph | Animal Pathogen-Endophyte-Fungal Parasite-\|Undefined Saprotroph\| | Dimorphic-Microfungus | NULL | Probable | As mycoparasite (Põlme et al. 2020); As root_endophyte_dark_septate (Põlme et al. 2020); As soil_saprotroph (Põlme et al. 2020); Decay soil (Põlme et al. 2020) | [illegible] |
| Otu05300 | 5 | k__Fungi(100);p__Ascomycota(100);c__Eurotiomycetes(100);o__Chaetothyriales(100);f__Herpotrichiellaceae(100);f__Herpotrichiellaceae_unclassified(100);f__Herpotrichiellaceae_unclassified(100); | Herpotrichiellaceae | 9 | Pathotroph-Saprotroph | Animal Pathogen-Fungal Parasite-Undefined Saprotroph | Facultative Yeast-Microfungus | NULL | Probable | Mostly saprobic on plants or other fungi; isolated from soil etc., and from medical sources (Cannon & Kirk 2007) | Cannon PF, Kirk PM. 2007. Fungal Families of the World. CAB International, Cambridge (ISBN: 9780851998275). |
| Otu05302 | 5 | k__Fungi(100);p__Ascomycota(100);c__Saccharomycetes(61);o__Saccharomycetales(61);o__Saccharomycetales_unclassified(61);o__Saccharomycetales_unclassified(61);o__Saccharomycetales_unclassified(61); | Saccharomycetales | 7 | Saprotroph | Undefined Saprotroph | Yeast | NULL | Possible | NULL | [illegible] |
| Otu05304 | 5 | k__Fungi(100);p__Ascomycota(100);c__Saccharomycetes(100);o__Saccharomycetales(100);f__Metschnikowiaceae(100);g__Metschnikowia(100);s__Metschnikowia_pulcherrima(61); | Metschnikowia | 13 | Saprotroph | \|Undefined Saprotroph\| | Yeast | NULL | Probable | As nectar-tap_saprotroph (Põlme et al. 2020); Decay sugar-rich substrates (Põlme et al. 2020) | Põlme S, et al. 2020. Fungal Diversity 105:1-16 (https://doi.org/10.1007/s13225-020-00466-2) |
| Otu05308 | 5 | k__Fungi(100);p__Ascomycota(100);c__Dothideomycetes(100);o__Pleosporales(100);f__Pleosporales_family_Incertae_sedis(100);g__Periconia(100);s__Periconia_sp(100); | Periconia | 13 | Pathotroph-Saprotroph-Symbiotroph | Endophyte-\|Plant Pathogen\|-Wood Saprotroph | Microfungus | Soft Rot | Possible | Asteraceae, Musaceae (Costa et al. 2012); As foliar_endophyte (Põlme et al. 2020); Dark Septate Endophyte, known from plant roots (Mandyam et al. 2010); Endophyte Host : Decay leaf, fruit, seed (Põlme et al. 2020) | [illegible] |
| Otu05309 | 5 | k__Fungi(100);p__Basidiomycota(100);c__Microbotryomycetes(100);o__Microbotryomycetes_order_Incertae_sedis(80);f__Microbotryomycetes_family_Incertae_sedis(80);g__Curvibasidium(80);s__Curvibasidium_cygneicollum(80); | Curvibasidium | 13 | Saprotroph-Symbiotroph | Epiphyte-Plant Saprotroph-Undefined Saprotroph-\|Wood Saprotroph\| | Dimorphic-Microfungus | NULL | Possible | As litter_saprotroph (Põlme et al. 2020); Decay wood, leaf, fruit, seed, soil (Põlme et al. 2020) | [illegible] |
| Otu05311 | 5 | k__Fungi(100);p__Ascomycota(100);c__Saccharomycetes(100);o__Saccharomycetales(100);f__Metschnikowiaceae(100);g__Metschnikowia(100);s__Metschnikowia_pulcherrima(100); | Metschnikowia | 13 | Saprotroph | \|Undefined Saprotroph\| | Yeast | NULL | Probable | As nectar-tap_saprotroph (Põlme et al. 2020); Decay sugar-rich substrates (Põlme et al. 2020) | Põlme S, et al. 2020. Fungal Diversity 105:1-16 (https://doi.org/10.1007/s13225-020-00466-2) |
| Otu05312 | 5 | k__Fungi(100);p__Ascomycota(100);c__Saccharomycetes(100);o__Saccharomycetales(100);f__Metschnikowiaceae(100);g__Metschnikowia(100);s__Metschnikowia_pulcherrima(100); | Metschnikowia | 13 | Saprotroph | \|Undefined Saprotroph\| | Yeast | NULL | Probable | As nectar-tap_saprotroph (Põlme et al. 2020); Decay sugar-rich substrates (Põlme et al. 2020) | Põlme S, et al. 2020. Fungal Diversity 105:1-16 (https://doi.org/10.1007/s13225-020-00466-2) |
| Otu05313 | 5 | k__Fungi(100);p__Ascomycota(100);c__Sordariomycetes(100);o__Hypocreales(100);f__Cordycipitaceae(100);g__Beauveria(100);s__Beauveria_sp(100); | Beauveria | 13 | Pathotroph-Saprotroph-Symbiotroph | \|Animal Parasite\|-Animal Pathogen-Endophyte-Undefined Saprotroph | Microfungus | NULL | Probable | As animal_decomposer (Põlme et al. 2020); As foliar_endophyte (Põlme et al. 2020); Decay animal material (Põlme et al. 2020) | [illegible] |
| Otu05319 | 5 | k__Fungi(100);p__Basidiomycota(100);c__Microbotryomycetes(100);o__Sporidiobolales(100);f__Sporidiobolales_family_Incertae_sedis(100);g__Rhodotorula(100);s__Rhodotorula_cresolica(100); | Rhodotorula | 13 | Pathotroph-Saprotroph-Symbiotroph | Animal Pathogen-Animal Symbiotroph-Endophyte-Plant Pathogen-\|Undefined Saprotroph\| | Yeast | NULL | Probable | As foliar_endophyte (Põlme et al. 2020); Opportunistic human pathogen | [illegible] |
| Otu05321 | 5 | k__Fungi(100);p__Ascomycota(100);c__Sordariomycetes(100);o__Hypocreales(100);o__Hypocreales_unclassified(80);o__Hypocreales_unclassified(80);o__Hypocreales_unclassified(80); | Hypocreales | 7 | Saprotroph | Undefined Saprotroph | Microfungus | NULL | Possible | NULL | [illegible] |
| Otu05323 | 5 | k__Fungi(100);p__Ascomycota(100);c__Saccharomycetes(100);o__Saccharomycetales(100);f__Metschnikowiaceae(100);g__Metschnikowia(100);s__Metschnikowia_pulcherrima(100); | Metschnikowia | 13 | Saprotroph | \|Undefined Saprotroph\| | Yeast | NULL | Probable | As nectar-tap_saprotroph (Põlme et al. 2020); Decay sugar-rich substrates (Põlme et al. 2020) | Põlme S, et al. 2020. Fungal Diversity 105:1-16 (https://doi.org/10.1007/s13225-020-00466-2) |
| Otu05324 | 5 | k__Fungi(100);p__Basidiomycota(100);c__Agaricomycetes(100);o__Auriculariales(100);f__Auriculariales_family_Incertae_sedis(100);g__Auricularia(100);s__Auricularia_americana(80); | Auricularia | 13 | Saprotroph | Undefined Saprotroph-\|Wood Saprotroph\| | Tremelloid | NULL | Probable | Decay wood (Põlme et al. 2020) | [illegible] |
| Otu05325 | 5 | k__Fungi(100);p__Ascomycota(100);c__Saccharomycetes(100);o__Saccharomycetales(100);f__Metschnikowiaceae(100);g__Metschnikowia(100);s__Metschnikowia_pulcherrima(100); | Metschnikowia | 13 | Saprotroph | \|Undefined Saprotroph\| | Yeast | NULL | Probable | As nectar-tap_saprotroph (Põlme et al. 2020); Decay sugar-rich substrates (Põlme et al. 2020) | Põlme S, et al. 2020. Fungal Diversity 105:1-16 (https://doi.org/10.1007/s13225-020-00466-2) |
| Otu05326 | 5 | k__Fungi(100);p__Ascomycota(100);c__Saccharomycetes(100);o__Saccharomycetales(100);f__Saccharomycetales_family_Incertae_sedis(80);g__Candida(61);g__Candida_unclassified(61); | Candida | 13 | Pathotroph-Saprotroph-Symbiotroph | Animal Pathogen-Endophyte-Endosymbiont-Epiphyte-\|Undefined Saprotroph\| | Dimorphic Yeast-Yeast | NULL | Probable | As nectar-tap_saprotroph (Põlme et al. 2020); Decay sugar-rich substrates (Põlme et al. 2020). This asexual genus is polyphyletic and contain species that function very differently depend on what they are. Caution should be taken when interpreting guilds for Candida species. | [illegible] |
| Otu05327 | 5 | k__Fungi(100);p__Ascomycota(100);c__Dothideomycetes(100);o__Dothideomycetes_order_Incertae_sedis(100);f__Myxotrichaceae(100);g__Oidiodendron(100);s__Oidiodendron_sp(80); | Oidiodendron | 13 | Pathotroph-Saprotroph-Symbiotroph | Endophyte-Ericoid Mycorrhizal-\|Undefined Saprotroph\| | Microfungus | NULL | Probable | As root_endophyte (Põlme et al. 2020); As soil_saprotroph (Põlme et al. 2020); Dark Septate Endophyte; Decay soil (Põlme et al. 2020) | [illegible] |
| Otu05328 | 5 | k__Fungi(100);p__Ascomycota(100);c__Saccharomycetes(100);o__Saccharomycetales(100);f__Metschnikowiaceae(100);g__Metschnikowia(100);g__Metschnikowia_unclassified(61); | Metschnikowia | 13 | Saprotroph | \|Undefined Saprotroph\| | Yeast | NULL | Probable | As nectar-tap_saprotroph (Põlme et al. 2020); Decay sugar-rich substrates (Põlme et al. 2020) | Põlme S, et al. 2020. Fungal Diversity 105:1-16 (https://doi.org/10.1007/s13225-020-00466-2) |
| Otu05329 | 5 | k__Fungi(100);p__Zygomycota(100);c__Zygomycota_class_Incertae_sedis(100);o__Mucorales(100);f__Lentamycetaceae(80);g__Lentamyces(80);s__Lentamyces_parricida(80); | Lentamyces | 13 | Pathotroph-Symbiotroph | Animal Symbiotroph-\|Fungal Parasite\| | Microfungus | NULL | Probable | Animal gut fungus; As mycoparasite (Põlme et al. 2020); Decay fungal material (Põlme et al. 2020) | [illegible] |
| Otu05331 | 5 | k__Fungi(100);p__Ascomycota(100);c__Saccharomycetes(100);o__Saccharomycetales(100);f__Metschnikowiaceae(100);g__Metschnikowia(100);g__Metschnikowia_unclassified(80); | Metschnikowia | 13 | Saprotroph | \|Undefined Saprotroph\| | Yeast | NULL | Probable | As nectar-tap_saprotroph (Põlme et al. 2020); Decay sugar-rich substrates (Põlme et al. 2020) | Põlme S, et al. 2020. Fungal Diversity 105:1-16 (https://doi.org/10.1007/s13225-020-00466-2) |
| Otu05336 | 5 | k__Fungi(100);p__Ascomycota(100);c__Leotiomycetes(100);o__Helotiales(100);f__Vibrisseaceae(100);g__Phialocephala(100);s__Phialocephala_fusca(100); | Phialocephala | 13 | Saprotroph-Symbiotroph | Endophyte-\|Undefined Saprotroph\| | Microfungus | NULL | Highly Probable | As root_endophyte (Põlme et al. 2020); As root_endophyte_dark_septate (Põlme et al. 2020); As soil_saprotroph (Põlme et al. 2020); Dark Septate Endophyte, known from plant roots; Decay roots, soil (Põlme et al. 2020) | [illegible] |
| Otu05337 | 5 | k__Fungi(100);p__Ascomycota(100);c__Saccharomycetes(100);o__Saccharomycetales(100);f__Metschnikowiaceae(100);g__Metschnikowia(100);g__Metschnikowia_unclassified(61); | Metschnikowia | 13 | Saprotroph | \|Undefined Saprotroph\| | Yeast | NULL | Probable | As nectar-tap_saprotroph (Põlme et al. 2020); Decay sugar-rich substrates (Põlme et al. 2020) | Põlme S, et al. 2020. Fungal Diversity 105:1-16 (https://doi.org/10.1007/s13225-020-00466-2) |
| Otu05339 | 5 | k__Fungi(100);p__Ascomycota(100);c__Saccharomycetes(100);o__Saccharomycetales(100);f__Metschnikowiaceae(100);g__Metschnikowia(100);s__Metschnikowia_pulcherrima(100); | Metschnikowia | 13 | Saprotroph | \|Undefined Saprotroph\| | Yeast | NULL | Probable | As nectar-tap_saprotroph (Põlme et al. 2020); Decay sugar-rich substrates (Põlme et al. 2020) | Põlme S, et al. 2020. Fungal Diversity 105:1-16 (https://doi.org/10.1007/s13225-020-00466-2) |
| Otu05341 | 5 | k__Fungi(100);p__Ascomycota(100);c__Saccharomycetes(100);o__Saccharomycetales(100);f__Saccharomycetales_family_Incertae_sedis(100);g__Candida(100);g__Candida_unclassified(80); | Candida | 13 | Pathotroph-Saprotroph-Symbiotroph | Animal Pathogen-Endophyte-Endosymbiont-Epiphyte-\|Undefined Saprotroph\| | Dimorphic Yeast-Yeast | NULL | Probable | As nectar-tap_saprotroph (Põlme et al. 2020); Decay sugar-rich substrates (Põlme et al. 2020). This asexual genus is polyphyletic and contain species that function very differently depend on what they are. Caution should be taken when interpreting guilds for Candida species. | [illegible] |
| Otu05343 | 5 | k__Fungi(100);p__Ascomycota(100);c__Saccharomycetes(100);o__Saccharomycetales(100);f__Metschnikowiaceae(100);g__Metschnikowia(100);s__Metschnikowia_pulcherrima(61); | Metschnikowia | 13 | Saprotroph | \|Undefined Saprotroph\| | Yeast | NULL | Probable | As nectar-tap_saprotroph (Põlme et al. 2020); Decay sugar-rich substrates (Põlme et al. 2020) | Põlme S, et al. 2020. Fungal Diversity 105:1-16 (https://doi.org/10.1007/s13225-020-00466-2) |
| Otu05344 | 5 | k__Fungi(100);p__Ascomycota(100);c__Saccharomycetes(100);o__Saccharomycetales(100);f__Metschnikowiaceae(100);g__Metschnikowia(100);s__Metschnikowia_pulcherrima(80); | Metschnikowia | 13 | Saprotroph | \|Undefined Saprotroph\| | Yeast | NULL | Probable | As nectar-tap_saprotroph (Põlme et al. 2020); Decay sugar-rich substrates (Põlme et al. 2020) | Põlme S, et al. 2020. Fungal Diversity 105:1-16 (https://doi.org/10.1007/s13225-020-00466-2) |
| Otu05346 | 5 | k__Fungi(100);p__Ascomycota(100);c__Saccharomycetes(100);o__Saccharomycetales(100);f__Saccharomycetaceae(100);g__Kazachstania(100);s__Kazachstania_intestinalis(100); | Kazachstania | 13 | Saprotroph | \|Undefined Saprotroph\| | Yeast | NULL | Probable | As nectar-tap_saprotroph (Põlme et al. 2020); Decay sugar-rich substrates (Põlme et al. 2020) | Põlme S, et al. 2020. Fungal Diversity 105:1-16 (https://doi.org/10.1007/s13225-020-00466-2) |
| Otu05351 | 5 | k__Fungi(100);p__Ascomycota(100);c__Saccharomycetes(100);o__Saccharomycetales(100);f__Saccharomycetales_family_Incertae_sedis(100);g__Candida(100);g__Candida_unclassified(80); | Candida | 13 | Pathotroph-Saprotroph-Symbiotroph | Animal Pathogen-Endophyte-Endosymbiont-Epiphyte-\|Undefined Saprotroph\| | Dimorphic Yeast-Yeast | NULL | Probable | As nectar-tap_saprotroph (Põlme et al. 2020); Decay sugar-rich substrates (Põlme et al. 2020). This asexual genus is polyphyletic and contain species that function very differently depend on what they are. Caution should be taken when interpreting guilds for Candida species. | [illegible] |
| Otu05354 | 5 | k__Fungi(100);p__Ascomycota(100);c__Sordariomycetes(100);o__Boliniales(80);f__Boliniaceae(80);f__Boliniaceae_unclassified(80);f__Boliniaceae_unclassified(80); | Boliniaceae | 9 | Saprotroph | Wood Saprotroph | Microfungus | NULL | Probable | Saprobic on and in wood and bark (Cannon & Kirk 2007) | Cannon PF, Kirk PM. 2007. Fungal Families of the World. CAB International, Cambridge (ISBN: 9780851998275). |
| Otu05355 | 5 | k__Fungi(100);p__Ascomycota(100);c__Saccharomycetes(100);o__Saccharomycetales(100);f__Metschnikowiaceae(100);g__Metschnikowia(100);s__Metschnikowia_pulcherrima(80); | Metschnikowia | 13 | Saprotroph | \|Undefined Saprotroph\| | Yeast | NULL | Probable | As nectar-tap_saprotroph (Põlme et al. 2020); Decay sugar-rich substrates (Põlme et al. 2020) | Põlme S, et al. 2020. Fungal Diversity 105:1-16 (https://doi.org/10.1007/s13225-020-00466-2) |
| Otu05356 | 5 | k__Fungi(100);p__Ascomycota(100);c__Saccharomycetes(100);o__Saccharomycetales(100);f__Metschnikowiaceae(100);g__Metschnikowia(100);s__Metschnikowia_pulcherrima(100); | Metschnikowia | 13 | Saprotroph | \|Undefined Saprotroph\| | Yeast | NULL | Probable | As nectar-tap_saprotroph (Põlme et al. 2020); Decay sugar-rich substrates (Põlme et al. 2020) | Põlme S, et al. 2020. Fungal Diversity 105:1-16 (https://doi.org/10.1007/s13225-020-00466-2) |
| Otu05360 | 5 | k__Fungi(100);p__Ascomycota(100);c__Saccharomycetes(100);o__Saccharomycetales(100);f__Metschnikowiaceae(100);g__Metschnikowia(100);s__Metschnikowia_pulcherrima(100); | Metschnikowia | 13 | Saprotroph | \|Undefined Saprotroph\| | Yeast | NULL | Probable | As nectar-tap_saprotroph (Põlme et al. 2020); Decay sugar-rich substrates (Põlme et al. 2020) | Põlme S, et al. 2020. Fungal Diversity 105:1-16 (https://doi.org/10.1007/s13225-020-00466-2) |
| Otu05366 | 5 | k__Fungi(100);p__Basidiomycota(100);c__Exobasidiomycetes(100);o__Microstromatales(100);f__Microstromataceae(100);g__Microstroma(100);s__Microstroma_juglandis(100); | Microstroma | 13 | Pathotroph | \|Plant Pathogen\| | Dimorphic | NULL | Probable | NULL | [illegible] |
| Otu05367 | 5 | k__Fungi(100);p__Ascomycota(100);c__Saccharomycetes(100);o__Saccharomycetales(100);f__Saccharomycetaceae(100);g__Kazachstania(100);s__Kazachstania_intestinalis(100); | Kazachstania | 13 | Saprotroph | \|Undefined Saprotroph\| | Yeast | NULL | Probable | As nectar-tap_saprotroph (Põlme et al. 2020); Decay sugar-rich substrates (Põlme et al. 2020) | Põlme S, et al. 2020. Fungal Diversity 105:1-16 (https://doi.org/10.1007/s13225-020-00466-2) |
| Otu05368 | 5 | k__Fungi(100);p__Basidiomycota(100);c__Tremellomycetes(100);o__Cystofilobasidiales(100);f__Cystofilobasidiales_family_Incertae_sedis(100);g__Mrakiella(100);s__Mrakiella_aquatica(100); | Mrakiella | 13 | Saprotroph | \|Undefined Saprotroph\| | Dimorphic | NULL | Probable | As soil_saprotroph (Põlme et al. 2020); Decay soil (Põlme et al. 2020) | [illegible] |
| Otu05369 | 5 | k__Fungi(100);p__Ascomycota(100);c__Saccharomycetes(100);o__Saccharomycetales(100);f__Metschnikowiaceae(100);g__Metschnikowia(100);s__Metschnikowia_pulcherrima(80); | Metschnikowia | 13 | Saprotroph | \|Undefined Saprotroph\| | Yeast | NULL | Probable | As nectar-tap_saprotroph (Põlme et al. 2020); Decay sugar-rich substrates (Põlme et al. 2020) | Põlme S, et al. 2020. Fungal Diversity 105:1-16 (https://doi.org/10.1007/s13225-020-00466-2) |
| Otu05374 | 5 | k__Fungi(100);p__Basidiomycota(100);c__Microbotryomycetes(100);o__Sporidiobolales(100);f__Sporidiobolales_family_Incertae_sedis(100);g__Rhodotorula(61);s__Rhodotorula_taiwanensis(61); | Rhodotorula | 13 | Pathotroph-Saprotroph-Symbiotroph | Animal Pathogen-Animal Symbiotroph-Endophyte-Plant Pathogen-\|Undefined Saprotroph\| | Yeast | NULL | Probable | As foliar_endophyte (Põlme et al. 2020); Opportunistic human pathogen | [illegible] |
| Otu05377 | 5 | k__Fungi(100);p__Ascomycota(100);c__Saccharomycetes(80);o__Saccharomycetales(80);f__Saccharomycetales_family_Incertae_sedis(80);g__Candida(80);g__Candida_unclassified(80); | Candida | 13 | Pathotroph-Saprotroph-Symbiotroph | Animal Pathogen-Endophyte-Endosymbiont-Epiphyte-\|Undefined Saprotroph\| | Dimorphic Yeast-Yeast | NULL | Probable | As nectar-tap_saprotroph (Põlme et al. 2020); Decay sugar-rich substrates (Põlme et al. 2020). This asexual genus is polyphyletic and contain species that function very differently depend on what they are. Caution should be taken when interpreting guilds for Candida species. | [illegible] |
| Otu05378 | 5 | k__Fungi(100);p__Ascomycota(100);c__Saccharomycetes(100);o__Saccharomycetales(100);f__Metschnikowiaceae(100);g__Metschnikowia(100);s__Metschnikowia_pulcherrima(100); | Metschnikowia | 13 | Saprotroph | \|Undefined Saprotroph\| | Yeast | NULL | Probable | As nectar-tap_saprotroph (Põlme et al. 2020); Decay sugar-rich substrates (Põlme et al. 2020) | Põlme S, et al. 2020. Fungal Diversity 105:1-16 (https://doi.org/10.1007/s13225-020-00466-2) |
| Otu05380 | 5 | k__Fungi(100);p__Ascomycota(100);c__Saccharomycetes(100);o__Saccharomycetales(100);f__Metschnikowiaceae(100);g__Metschnikowia(100);s__Metschnikowia_pulcherrima(100); | Metschnikowia | 13 | Saprotroph | \|Undefined Saprotroph\| | Yeast | NULL | Probable | As nectar-tap_saprotroph (Põlme et al. 2020); Decay sugar-rich substrates (Põlme et al. 2020) | Põlme S, et al. 2020. Fungal Diversity 105:1-16 (https://doi.org/10.1007/s13225-020-00466-2) |
| Otu05381 | 5 | k__Fungi(100);p__Ascomycota(100);c__Saccharomycetes(100);o__Saccharomycetales(100);f__Metschnikowiaceae(100);g__Metschnikowia(100);s__Metschnikowia_pulcherrima(100); | Metschnikowia | 13 | Saprotroph | \|Undefined Saprotroph\| | Yeast | NULL | Probable | As nectar-tap_saprotroph (Põlme et al. 2020); Decay sugar-rich substrates (Põlme et al. 2020) | Põlme S, et al. 2020. Fungal Diversity 105:1-16 (https://doi.org/10.1007/s13225-020-00466-2) |
| Otu05384 | 5 | k__Fungi(100);p__Ascomycota(100);c__Saccharomycetes(100);o__Saccharomycetales(100);f__Metschnikowiaceae(100);g__Metschnikowia(100);s__Metschnikowia_pulcherrima(100); | Metschnikowia | 13 | Saprotroph | \|Undefined Saprotroph\| | Yeast | NULL | Probable | As nectar-tap_saprotroph (Põlme et al. 2020); Decay sugar-rich substrates (Põlme et al. 2020) | Põlme S, et al. 2020. Fungal Diversity 105:1-16 (https://doi.org/10.1007/s13225-020-00466-2) |
| Otu05386 | 5 | k__Fungi(100);p__Ascomycota(100);c__Saccharomycetes(100);o__Saccharomycetales(100);f__Metschnikowiaceae(100);g__Metschnikowia(100);s__Metschnikowia_pulcherrima(80); | Metschnikowia | 13 | Saprotroph | \|Undefined Saprotroph\| | Yeast | NULL | Probable | As nectar-tap_saprotroph (Põlme et al. 2020); Decay sugar-rich substrates (Põlme et al. 2020) | Põlme S, et al. 2020. Fungal Diversity 105:1-16 (https://doi.org/10.1007/s13225-020-00466-2) |
| Otu05387 | 5 | k__Fungi(100);p__Ascomycota(100);c__Saccharomycetes(100);o__Saccharomycetales(100);f__Metschnikowiaceae(100);g__Metschnikowia(100);s__Metschnikowia_pulcherrima(80); | Metschnikowia | 13 | Saprotroph | \|Undefined Saprotroph\| | Yeast | NULL | Probable | As nectar-tap_saprotroph (Põlme et al. 2020); Decay sugar-rich substrates (Põlme et al. 2020) | Põlme S, et al. 2020. Fungal Diversity 105:1-16 (https://doi.org/10.1007/s13225-020-00466-2) |
| Otu05388 | 5 | k__Fungi(100);p__Ascomycota(100);c__Saccharomycetes(100);o__Saccharomycetales(100);f__Metschnikowiaceae(100);g__Metschnikowia(100);s__Metschnikowia_pulcherrima(100); | Metschnikowia | 13 | Saprotroph | \|Undefined Saprotroph\| | Yeast | NULL | Probable | As nectar-tap_saprotroph (Põlme et al. 2020); Decay sugar-rich substrates (Põlme et al. 2020) | Põlme S, et al. 2020. Fungal Diversity 105:1-16 (https://doi.org/10.1007/s13225-020-00466-2) |
| Otu05390 | 5 | k__Fungi(100);p__Zygomycota(100);c__Zygomycota_class_Incertae_sedis(100);o__Mucorales(100);f__Mucoraceae(100);g__Mucor(100);s__Mucor_genevensis(100); | Mucor | 13 | Saprotroph | \|Undefined Saprotroph\| | Microfungus | NULL | Probable | As soil_saprotroph (Põlme et al. 2020); Decay soil (Põlme et al. 2020) | [illegible] |
| Otu05393 | 5 | k__Fungi(100);p__Ascomycota(100);c__Saccharomycetes(100);o__Saccharomycetales(100);f__Metschnikowiaceae(100);g__Metschnikowia(100);s__Metschnikowia_pulcherrima(61); | Metschnikowia | 13 | Saprotroph | \|Undefined Saprotroph\| | Yeast | NULL | Probable | As nectar-tap_saprotroph (Põlme et al. 2020); Decay sugar-rich substrates (Põlme et al. 2020) | Põlme S, et al. 2020. Fungal Diversity 105:1-16 (https://doi.org/10.1007/s13225-020-00466-2) |
| Otu05396 | 5 | k__Fungi(100);p__Basidiomycota(100);c__Agaricomycetes(100);o__Agaricales(100);f__Tricholomataceae(100);g__Tricholomopsis(100);s__Tricholomopsis_sp(100); | Tricholomopsis | 13 | Saprotroph | Undefined Saprotroph-\|Wood Saprotroph\| | Agaricoid | White Rot | Probable | Decay wood (Põlme et al. 2020) | [illegible] |
| Otu05397 | 5 | k__Fungi(100);p__Ascomycota(100);c__Saccharomycetes(100);o__Saccharomycetales(100);f__Metschnikowiaceae(100);g__Metschnikowia(100);s__Metschnikowia_pulcherrima(100); | Metschnikowia | 13 | Saprotroph | \|Undefined Saprotroph\| | Yeast | NULL | Probable | As nectar-tap_saprotroph (Põlme et al. 2020); Decay sugar-rich substrates (Põlme et al. 2020) | Põlme S, et al. 2020. Fungal Diversity 105:1-16 (https://doi.org/10.1007/s13225-020-00466-2) |
| Otu05399 | 5 | k__Fungi(100);p__Ascomycota(100);c__Saccharomycetes(100);o__Saccharomycetales(100);f__Metschnikowiaceae(100);g__Metschnikowia(100);s__Metschnikowia_pulcherrima(80); | Metschnikowia | 13 | Saprotroph | \|Undefined Saprotroph\| | Yeast | NULL | Probable | As nectar-tap_saprotroph (Põlme et al. 2020); Decay sugar-rich substrates (Põlme et al. 2020) | Põlme S, et al. 2020. Fungal Diversity 105:1-16 (https://doi.org/10.1007/s13225-020-00466-2) |
| Otu05401 | 5 | k__Fungi(100);p__Ascomycota(100);c__Saccharomycetes(100);o__Saccharomycetales(100);f__Metschnikowiaceae(100);g__Metschnikowia(100);s__Metschnikowia_pulcherrima(61); | Metschnikowia | 13 | Saprotroph | \|Undefined Saprotroph\| | Yeast | NULL | Probable | As nectar-tap_saprotroph (Põlme et al. 2020); Decay sugar-rich substrates (Põlme et al. 2020) | Põlme S, et al. 2020. Fungal Diversity 105:1-16 (https://doi.org/10.1007/s13225-020-00466-2) |
| Otu05402 | 5 | k__Fungi(100);p__Ascomycota(100);c__Saccharomycetes(100);o__Saccharomycetales(100);f__Metschnikowiaceae(100);g__Metschnikowia(100);s__Metschnikowia_pulcherrima(100); | Metschnikowia | 13 | Saprotroph | \|Undefined Saprotroph\| | Yeast | NULL | Probable | As nectar-tap_saprotroph (Põlme et al. 2020); Decay sugar-rich substrates (Põlme et al. 2020) | Põlme S, et al. 2020. Fungal Diversity 105:1-16 (https://doi.org/10.1007/s13225-020-00466-2) |
| Otu05405 | 5 | k__Fungi(100);p__Ascomycota(100);c__Saccharomycetes(100);o__Saccharomycetales(100);f__Metschnikowiaceae(100);g__Metschnikowia(100);g__Metschnikowia_unclassified(61); | Metschnikowia | 13 | Saprotroph | \|Undefined Saprotroph\| | Yeast | NULL | Probable | As nectar-tap_saprotroph (Põlme et al. 2020); Decay sugar-rich substrates (Põlme et al. 2020) | Põlme S, et al. 2020. Fungal Diversity 105:1-16 (https://doi.org/10.1007/s13225-020-00466-2) |
| Otu05406 | 5 | k__Fungi(100);p__Ascomycota(100);c__Saccharomycetes(100);o__Saccharomycetales(100);f__Metschnikowiaceae(100);g__Metschnikowia(100);s__Metschnikowia_pulcherrima(100); | Metschnikowia | 13 | Saprotroph | \|Undefined Saprotroph\| | Yeast | NULL | Probable | As nectar-tap_saprotroph (Põlme et al. 2020); Decay sugar-rich substrates (Põlme et al. 2020) | Põlme S, et al. 2020. Fungal Diversity 105:1-16 (https://doi.org/10.1007/s13225-020-00466-2) |
| Otu05407 | 5 | k__Fungi(100);p__Ascomycota(100);c__Sordariomycetes(80);o__Chaetosphaeriales(80);f__Chaetosphaeriaceae(80);g__unclassified_Chaetosphaeriaceae(80);s__Chaetosphaeriaceae_sp(80); | Chaetosphaeriaceae | 9 | Saprotroph | Plant Saprotroph-Wood Saprotroph | Microfungus | NULL | Probable | Saprobes associated with herbaceous plant material, including timber and timber products in damp buildings (Cannon & Kirk 2007) | Cannon PF, Kirk PM. 2007. Fungal Families of the World. CAB International, Cambridge (ISBN: 9780851998275). |
| Otu05412 | 5 | k__Fungi(100);p__Ascomycota(100);c__Saccharomycetes(100);o__Saccharomycetales(100);f__Metschnikowiaceae(100);g__Metschnikowia(100);s__Metschnikowia_pulcherrima(100); | Metschnikowia | 13 | Saprotroph | \|Undefined Saprotroph\| | Yeast | NULL | Probable | As nectar-tap_saprotroph (Põlme et al. 2020); Decay sugar-rich substrates (Põlme et al. 2020) | Põlme S, et al. 2020. Fungal Diversity 105:1-16 (https://doi.org/10.1007/s13225-020-00466-2) |
| Otu05415 | 5 | k__Fungi(100);p__Ascomycota(100);c__Saccharomycetes(100);o__Saccharomycetales(100);f__Saccharomycetaceae(100);g__Kazachstania(100);s__Kazachstania_intestinalis(100); | Kazachstania | 13 | Saprotroph | \|Undefined Saprotroph\| | Yeast | NULL | Probable | As nectar-tap_saprotroph (Põlme et al. 2020); Decay sugar-rich substrates (Põlme et al. 2020) | Põlme S, et al. 2020. Fungal Diversity 105:1-16 (https://doi.org/10.1007/s13225-020-00466-2) |
| Otu05416 | 5 | k__Fungi(100);p__Basidiomycota(100);c__Agaricomycetes(100);o__Agaricales(100);f__Psathyrellaceae(100);g__Parasola(100);s__Parasola_schroeterii(100); | Parasola | 13 | Saprotroph | \|Undefined Saprotroph\| | Agaricoid | NULL | Probable | As soil_saprotroph (Põlme et al. 2020); Decay soil (Põlme et al. 2020) | [illegible] |
| Otu05419 | 5 | k__Fungi(100);p__Ascomycota(100);c__Saccharomycetes(100);o__Saccharomycetales(100);f__Metschnikowiaceae(100);g__Metschnikowia(100);s__Metschnikowia_pulcherrima(100); | Metschnikowia | 13 | Saprotroph | \|Undefined Saprotroph\| | Yeast | NULL | Probable | As nectar-tap_saprotroph (Põlme et al. 2020); Decay sugar-rich substrates (Põlme et al. 2020) | Põlme S, et al. 2020. Fungal Diversity 105:1-16 (https://doi.org/10.1007/s13225-020-00466-2) |
| Otu05423 | 5 | k__Fungi(100);p__Basidiomycota(100);c__Agaricomycetes(100);o__Cantharellales(100);f__Clavulinaceae(100);g__Clavulina(100);g__Clavulina_unclassified(100); | Clavulina | 13 | Symbiotroph | \|Ectomycorrhizal\| | Clavarioid | NULL | Highly Probable | NULL | [illegible] |
| Otu05424 | 5 | k__Fungi(100);p__Ascomycota(100);c__Saccharomycetes(100);o__Saccharomycetales(100);f__Saccharomycetales_family_Incertae_sedis(100);g__Candida(80);g__Candida_unclassified(80); | Candida | 13 | Pathotroph-Saprotroph-Symbiotroph | Animal Pathogen-Endophyte-Endosymbiont-Epiphyte-\|Undefined Saprotroph\| | Dimorphic Yeast-Yeast | NULL | Probable | As nectar-tap_saprotroph (Põlme et al. 2020); Decay sugar-rich substrates (Põlme et al. 2020). This asexual genus is polyphyletic and contain species that function very differently depend on what they are. Caution should be taken when interpreting guilds for Candida species. | [illegible] |
| Otu05425 | 5 | k__Fungi(100);p__Ascomycota(100);c__Saccharomycetes(100);o__Saccharomycetales(100);f__Saccharomycetaceae(100);g__Kazachstania(100);s__Kazachstania_intestinalis(100); | Kazachstania | 13 | Saprotroph | \|Undefined Saprotroph\| | Yeast | NULL | Probable | As nectar-tap_saprotroph (Põlme et al. 2020); Decay sugar-rich substrates (Põlme et al. 2020) | Põlme S, et al. 2020. Fungal Diversity 105:1-16 (https://doi.org/10.1007/s13225-020-00466-2) |
| Otu05427 | 5 | k__Fungi(100);p__Ascomycota(100);c__Eurotiomycetes(100);o__Eurotiales(100);f__Elaphomycetaceae(100);g__Elaphomyces(100);s__Elaphomyces_sp_1_RB_2011(100); | Elaphomyces | 13 | Symbiotroph | \|Ectomycorrhizal\| | Gasteroid | Hypogeous | Highly Probable | false truffle; Common name | [illegible] |
| Otu05428 | 5 | k__Fungi(100);p__Ascomycota(100);c__Saccharomycetes(100);o__Saccharomycetales(100);f__Saccharomycetaceae(100);g__Kazachstania(100);s__Kazachstania_intestinalis(100); | Kazachstania | 13 | Saprotroph | \|Undefined Saprotroph\| | Yeast | NULL | Probable | As nectar-tap_saprotroph (Põlme et al. 2020); Decay sugar-rich substrates (Põlme et al. 2020) | Põlme S, et al. 2020. Fungal Diversity 105:1-16 (https://doi.org/10.1007/s13225-020-00466-2) |
| Otu05431 | 5 | k__Fungi(100);p__Ascomycota(100);c__Saccharomycetes(100);o__Saccharomycetales(100);f__Metschnikowiaceae(100);g__Metschnikowia(100);s__Metschnikowia_pulcherrima(61); | Metschnikowia | 13 | Saprotroph | \|Undefined Saprotroph\| | Yeast | NULL | Probable | As nectar-tap_saprotroph (Põlme et al. 2020); Decay sugar-rich substrates (Põlme et al. 2020) | Põlme S, et al. 2020. Fungal Diversity 105:1-16 (https://doi.org/10.1007/s13225-020-00466-2) |
| Otu05435 | 5 | k__Fungi(100);p__Ascomycota(100);c__Saccharomycetes(100);o__Saccharomycetales(100);f__Saccharomycetales_family_Incertae_sedis(80);g__Candida(80);g__Candida_unclassified(80); | Candida | 13 | Pathotroph-Saprotroph-Symbiotroph | Animal Pathogen-Endophyte-Endosymbiont-Epiphyte-\|Undefined Saprotroph\| | Dimorphic Yeast-Yeast | NULL | Probable | As nectar-tap_saprotroph (Põlme et al. 2020); Decay sugar-rich substrates (Põlme et al. 2020). This asexual genus is polyphyletic and contain species that function very differently depend on what they are. Caution should be taken when interpreting guilds for Candida species. | [illegible] |
| Otu05439 | 5 | k__Fungi(100);p__Ascomycota(100);c__Saccharomycetes(100);o__Saccharomycetales(100);f__Metschnikowiaceae(100);g__Metschnikowia(100);s__Metschnikowia_pulcherrima(100); | Metschnikowia | 13 | Saprotroph | \|Undefined Saprotroph\| | Yeast | NULL | Probable | As nectar-tap_saprotroph (Põlme et al. 2020); Decay sugar-rich substrates (Põlme et al. 2020) | Põlme S, et al. 2020. Fungal Diversity 105:1-16 (https://doi.org/10.1007/s13225-020-00466-2) |
| Otu05443 | 5 | k__Fungi(100);p__Ascomycota(100);c__Saccharomycetes(100);o__Saccharomycetales(100);f__Metschnikowiaceae(100);g__Metschnikowia(100);s__Metschnikowia_pulcherrima(100); | Metschnikowia | 13 | Saprotroph | \|Undefined Saprotroph\| | Yeast | NULL | Probable | As nectar-tap_saprotroph (Põlme et al. 2020); Decay sugar-rich substrates (Põlme et al. 2020) | Põlme S, et al. 2020. Fungal Diversity 105:1-16 (https://doi.org/10.1007/s13225-020-00466-2) |
| Otu05445 | 5 | k__Fungi(100);p__Ascomycota(100);c__Saccharomycetes(100);o__Saccharomycetales(100);f__Metschnikowiaceae(100);g__Metschnikowia(100);s__Metschnikowia_pulcherrima(100); | Metschnikowia | 13 | Saprotroph | \|Undefined Saprotroph\| | Yeast | NULL | Probable | As nectar-tap_saprotroph (Põlme et al. 2020); Decay sugar-rich substrates (Põlme et al. 2020) | Põlme S, et al. 2020. Fungal Diversity 105:1-16 (https://doi.org/10.1007/s13225-020-00466-2) |
| Otu05446 | 5 | k__Fungi(100);p__Ascomycota(100);c__Saccharomycetes(100);o__Saccharomycetales(100);f__Metschnikowiaceae(100);g__Metschnikowia(100);s__Metschnikowia_pulcherrima(100); | Metschnikowia | 13 | Saprotroph | \|Undefined Saprotroph\| | Yeast | NULL | Probable | As nectar-tap_saprotroph (Põlme et al. 2020); Decay sugar-rich substrates (Põlme et al. 2020) | Põlme S, et al. 2020. Fungal Diversity 105:1-16 (https://doi.org/10.1007/s13225-020-00466-2) |
| Otu05449 | 5 | k__Fungi(100);p__Ascomycota(100);c__Eurotiomycetes(100);o__Eurotiales(100);f__Trichocomaceae(100);g__Eupenicillium(80);s__Eupenicillium_sp_KB9(80); | Eupenicillium | 13 | Saprotroph | Undefined Saprotroph | NULL | NULL | Probable | NULL | [illegible] |
| Otu05455 | 5 | k__Fungi(100);p__Ascomycota(100);c__Saccharomycetes(100);o__Saccharomycetales(100);f__Metschnikowiaceae(100);g__Metschnikowia(100);s__Metschnikowia_pulcherrima(80); | Metschnikowia | 13 | Saprotroph | \|Undefined Saprotroph\| | Yeast | NULL | Probable | As nectar-tap_saprotroph (Põlme et al. 2020); Decay sugar-rich substrates (Põlme et al. 2020) | Põlme S, et al. 2020. Fungal Diversity 105:1-16 (https://doi.org/10.1007/s13225-020-00466-2) |
| Otu05462 | 5 | k__Fungi(100);p__Ascomycota(100);c__Saccharomycetes(100);o__Saccharomycetales(100);f__Metschnikowiaceae(100);g__Metschnikowia(100);s__Metschnikowia_pulcherrima(80); | Metschnikowia | 13 | Saprotroph | \|Undefined Saprotroph\| | Yeast | NULL | Probable | As nectar-tap_saprotroph (Põlme et al. 2020); Decay sugar-rich substrates (Põlme et al. 2020) | Põlme S, et al. 2020. Fungal Diversity 105:1-16 (https://doi.org/10.1007/s13225-020-00466-2) |
| Otu05464 | 5 | k__Fungi(100);p__Ascomycota(100);c__Saccharomycetes(100);o__Saccharomycetales(100);f__Metschnikowiaceae(100);g__Metschnikowia(100);s__Metschnikowia_pulcherrima(100); | Metschnikowia | 13 | Saprotroph | \|Undefined Saprotroph\| | Yeast | NULL | Probable | As nectar-tap_saprotroph (Põlme et al. 2020); Decay sugar-rich substrates (Põlme et al. 2020) | Põlme S, et al. 2020. Fungal Diversity 105:1-16 (https://doi.org/10.1007/s13225-020-00466-2) |
| Otu05467 | 5 | k__Fungi(100);p__Basidiomycota(100);c__Agaricomycetes(100);o__Agaricales(100);f__Strophariaceae(100);g__Kuehneromyces(100);s__Kuehneromyces_rostratus(100); | Kuehneromyces | 13 | Saprotroph | Undefined Saprotroph-\|Wood Saprotroph\| | Agaricoid | White Rot | Probable | As white_rot (Põlme et al. 2020); Decay wood (Põlme et al. 2020) | [illegible] |
| Otu05468 | 5 | k__Fungi(100);p__Ascomycota(100);c__Saccharomycetes(100);o__Saccharomycetales(100);f__Metschnikowiaceae(100);g__Metschnikowia(100);s__Metschnikowia_pulcherrima(80); | Metschnikowia | 13 | Saprotroph | \|Undefined Saprotroph\| | Yeast | NULL | Probable | As nectar-tap_saprotroph (Põlme et al. 2020); Decay sugar-rich substrates (Põlme et al. 2020) | Põlme S, et al. 2020. Fungal Diversity 105:1-16 (https://doi.org/10.1007/s13225-020-00466-2) |
| Otu05469 | 5 | k__Fungi(100);p__Ascomycota(100);c__Sordariomycetes(100);o__Hypocreales(100);f__Nectriaceae(100);f__Nectriaceae_unclassified(100);f__Nectriaceae_unclassified(100); | Nectriaceae | 9 | Pathotroph-Saprotroph-Symbiotroph | Animal Pathogen-Endophyte-Fungal Parasite-Lichen Parasite-Plant Pathogen-Wood Saprotroph | Microfungus | NULL | Possible | Fusarium species are amongst the economically significant plant pathogens (Cannon & Kirk 2007) | [illegible] |
| Otu05472 | 5 | k__Fungi(100);p__Ascomycota(100);c__Saccharomycetes(100);o__Saccharomycetales(100);f__Metschnikowiaceae(100);g__Metschnikowia(100);s__Metschnikowia_pulcherrima(80); | Metschnikowia | 13 | Saprotroph | \|Undefined Saprotroph\| | Yeast | NULL | Probable | As nectar-tap_saprotroph (Põlme et al. 2020); Decay sugar-rich substrates (Põlme et al. 2020) | Põlme S, et al. 2020. Fungal Diversity 105:1-16 (https://doi.org/10.1007/s13225-020-00466-2) |
| Otu05473 | 5 | k__Fungi(100);p__Ascomycota(100);c__Saccharomycetes(100);o__Saccharomycetales(100);f__Metschnikowiaceae(100);g__Metschnikowia(100);s__Metschnikowia_pulcherrima(80); | Metschnikowia | 13 | Saprotroph | \|Undefined Saprotroph\| | Yeast | NULL | Probable | As nectar-tap_saprotroph (Põlme et al. 2020); Decay sugar-rich substrates (Põlme et al. 2020) | Põlme S, et al. 2020. Fungal Diversity 105:1-16 (https://doi.org/10.1007/s13225-020-00466-2) |
| Otu05474 | 5 | k__Fungi(100);p__Ascomycota(100);c__Saccharomycetes(100);o__Saccharomycetales(100);f__Metschnikowiaceae(100);g__Metschnikowia(100);s__Metschnikowia_pulcherrima(100); | Metschnikowia | 13 | Saprotroph | \|Undefined Saprotroph\| | Yeast | NULL | Probable | As nectar-tap_saprotroph (Põlme et al. 2020); Decay sugar-rich substrates (Põlme et al. 2020) | Põlme S, et al. 2020. Fungal Diversity 105:1-16 (https://doi.org/10.1007/s13225-020-00466-2) |
| Otu05475 | 5 | k__Fungi(100);p__Ascomycota(100);c__Saccharomycetes(100);o__Saccharomycetales(100);f__Metschnikowiaceae(100);g__Metschnikowia(100);g__Metschnikowia_unclassified(61); | Metschnikowia | 13 | Saprotroph | \|Undefined Saprotroph\| | Yeast | NULL | Probable | As nectar-tap_saprotroph (Põlme et al. 2020); Decay sugar-rich substrates (Põlme et al. 2020) | Põlme S, et al. 2020. Fungal Diversity 105:1-16 (https://doi.org/10.1007/s13225-020-00466-2) |
| Otu05480 | 5 | k__Fungi(100);p__Ascomycota(100);c__Saccharomycetes(100);o__Saccharomycetales(100);f__Metschnikowiaceae(100);g__Metschnikowia(100);s__Metschnikowia_pulcherrima(100); | Metschnikowia | 13 | Saprotroph | \|Undefined Saprotroph\| | Yeast | NULL | Probable | As nectar-tap_saprotroph (Põlme et al. 2020); Decay sugar-rich substrates (Põlme et al. 2020) | Põlme S, et al. 2020. Fungal Diversity 105:1-16 (https://doi.org/10.1007/s13225-020-00466-2) |
| Otu05482 | 5 | k__Fungi(100);p__Ascomycota(100);c__Saccharomycetes(100);o__Saccharomycetales(100);f__Metschnikowiaceae(100);g__Metschnikowia(100);s__Metschnikowia_pulcherrima(100); | Metschnikowia | 13 | Saprotroph | \|Undefined Saprotroph\| | Yeast | NULL | Probable | As nectar-tap_saprotroph (Põlme et al. 2020); Decay sugar-rich substrates (Põlme et al. 2020) | Põlme S, et al. 2020. Fungal Diversity 105:1-16 (https://doi.org/10.1007/s13225-020-00466-2) |
| Otu05483 | 5 | k__Fungi(100);p__Ascomycota(100);c__Sordariomycetes(100);o__Hypocreales(100);f__Nectriaceae(100);g__Nectria(100);s__Nectria_sp(100); | Nectria | 13 | Pathotroph-Saprotroph-Symbiotroph | Animal Pathogen-Endophyte-Fungal Parasite-Lichen Parasite-\|Plant Pathogen\|-Plant Saprotroph-Wood Saprotroph | Microfungus | NULL | Possible | As foliar_endophyte (Põlme et al. 2020); As litter_saprotroph (Põlme et al. 2020); Decay wood, leaf, fruit, seed, roots, soil (Põlme et al. 2020) | [illegible] |
| Otu05487 | 5 | k__Fungi(100);p__Ascomycota(100);c__Saccharomycetes(100);o__Saccharomycetales(100);f__Saccharomycetales_family_Incertae_sedis(100);g__Candida(80);g__Candida_unclassified(80); | Candida | 13 | Pathotroph-Saprotroph-Symbiotroph | Animal Pathogen-Endophyte-Endosymbiont-Epiphyte-\|Undefined Saprotroph\| | Dimorphic Yeast-Yeast | NULL | Probable | As nectar-tap_saprotroph (Põlme et al. 2020); Decay sugar-rich substrates (Põlme et al. 2020). This asexual genus is polyphyletic and contain species that function very differently depend on what they are. Caution should be taken when interpreting guilds for Candida species. | [illegible] |
| Otu05488 | 5 | k__Fungi(100);p__Ascomycota(100);c__Lecanoromycetes(100);o__Lecanorales(100);f__Ramalinaceae(100);f__Ramalinaceae_unclassified(100);f__Ramalinaceae_unclassified(100); | Ramalinaceae | 9 | Symbiotroph | Lichenized | Thallus | NULL | Probable | Lichenized with green algae, on rocks, branches etc., often forming extensive carpets of thalli; often prominent in maritime communities (Cannon & Kirk 2007) | Cannon PF, Kirk PM. 2007. Fungal Families of the World. CAB International, Cambridge (ISBN: 9780851998275). |
| Otu05489 | 5 | k__Fungi(100);p__Ascomycota(100);c__Saccharomycetes(100);o__Saccharomycetales(100);f__Metschnikowiaceae(100);g__Metschnikowia(100);s__Metschnikowia_pulcherrima(100); | Metschnikowia | 13 | Saprotroph | \|Undefined Saprotroph\| | Yeast | NULL | Probable | As nectar-tap_saprotroph (Põlme et al. 2020); Decay sugar-rich substrates (Põlme et al. 2020) | Põlme S, et al. 2020. Fungal Diversity 105:1-16 (https://doi.org/10.1007/s13225-020-00466-2) |
| Otu05491 | 5 | k__Fungi(100);p__Ascomycota(100);c__Dothideomycetes(100);o__Pleosporales(100);f__Pleosporaceae(100);g__Stemphylium(100);s__Stemphylium_sp_1662(100); | Stemphylium | 13 | Pathotroph-Saprotroph | \|Plant Pathogen\|-Plant Saprotroph-Wood Saprotroph | Microfungus | Soft Rot | Highly Probable | As litter_saprotroph (Põlme et al. 2020); Decay leaf, fruit, seed (Põlme et al. 2020) | [illegible] |
| Otu05493 | 5 | k__Fungi(100);p__Ascomycota(100);c__Saccharomycetes(100);o__Saccharomycetales(100);f__Metschnikowiaceae(100);g__Metschnikowia(100);s__Metschnikowia_pulcherrima(61); | Metschnikowia | 13 | Saprotroph | \|Undefined Saprotroph\| | Yeast | NULL | Probable | As nectar-tap_saprotroph (Põlme et al. 2020); Decay sugar-rich substrates (Põlme et al. 2020) | Põlme S, et al. 2020. Fungal Diversity 105:1-16 (https://doi.org/10.1007/s13225-020-00466-2) |
| Otu05494 | 5 | k__Fungi(100);p__Ascomycota(100);c__Saccharomycetes(100);o__Saccharomycetales(100);f__Metschnikowiaceae(100);g__Metschnikowia(100);s__Metschnikowia_pulcherrima(61); | Metschnikowia | 13 | Saprotroph | \|Undefined Saprotroph\| | Yeast | NULL | Probable | As nectar-tap_saprotroph (Põlme et al. 2020); Decay sugar-rich substrates (Põlme et al. 2020) | Põlme S, et al. 2020. Fungal Diversity 105:1-16 (https://doi.org/10.1007/s13225-020-00466-2) |
| Otu05496 | 5 | k__Fungi(100);p__Ascomycota(100);c__Saccharomycetes(61);o__Saccharomycetales(61);o__Saccharomycetales_unclassified(61);o__Saccharomycetales_unclassified(61);o__Saccharomycetales_unclassified(61); | Saccharomycetales | 7 | Saprotroph | Undefined Saprotroph | Yeast | NULL | Possible | NULL | [illegible] |

| | | | | | | | | | | | |
|---|---|---|---|---|---|---|---|---|---|---|---|
| Otu05498 | 5 | k__Fungi(100);p__Ascomycota(100);c__Dothideomycetes(100);o__Pleosporales(100);f__Venturiaceae(100);g__Venturia(100);s__Venturia_macularis(100); | Venturia | 13 | Pathotroph-Pathotroph-Saprotroph | \|Plant Pathogen\|-Plant Saprotroph | Microfungus | NULL | Probable | As litter_saprotroph (Põlme et al. 2020); Decay leaf, fruit, seed (Põlme et al. 2020) | Tedersoo L, et al. 2014. Science 346:e1256688 ((https://doi.org/10.1126/science.1256688)); Põlme S, et al. 2020. Fungal Diversity 105:1-16 ((https://doi.org/10.1007/s13225-020-00466-2)) |
| Otu05501 | 5 | k__Fungi(100);p__Ascomycota(100);c__Saccharomycetes(100);o__Saccharomycetales(100);f__Metschnikowiaceae(100);g__Metschnikowia(100);g__Metschnikowia_unclassified(100); | Metschnikowia | 13 | Saprotroph | \|Undefined Saprotroph\| | Yeast | NULL | Probable | As nectar-tap_saprotroph (Põlme et al. 2020); Decay sugar-rich substrates (Põlme et al. 2020) | Põlme S, et al. 2020. Fungal Diversity 105:1-16 ((https://doi.org/10.1007/s13225-020-00466-2)) |
| Otu05504 | 5 | k__Fungi(100);p__Ascomycota(100);c__Saccharomycetes(100);o__Saccharomycetales(100);f__Metschnikowiaceae(100);g__Metschnikowia(100);g__Metschnikowia_unclassified(100); | Metschnikowia | 13 | Saprotroph | \|Undefined Saprotroph\| | Yeast | NULL | Probable | As nectar-tap_saprotroph (Põlme et al. 2020); Decay sugar-rich substrates (Põlme et al. 2020) | Põlme S, et al. 2020. Fungal Diversity 105:1-16 ((https://doi.org/10.1007/s13225-020-00466-2)) |
| Otu05505 | 5 | k__Fungi(100);p__Ascomycota(100);c__Saccharomycetes(100);o__Saccharomycetales(100);f__Metschnikowiaceae(100);g__Metschnikowia(100);s__Metschnikowia_pulcherrima(100); | Metschnikowia | 13 | Saprotroph | \|Undefined Saprotroph\| | Yeast | NULL | Probable | As nectar-tap_saprotroph (Põlme et al. 2020); Decay sugar-rich substrates (Põlme et al. 2020) | Põlme S, et al. 2020. Fungal Diversity 105:1-16 ((https://doi.org/10.1007/s13225-020-00466-2)) |
| Otu05508 | 5 | k__Fungi(100);p__Ascomycota(100);c__Saccharomycetes(100);o__Saccharomycetales(100);f__Saccharomycetales_family_Incertae_sedis(100);g__Candida(100);s__Candida_cretensis(61); | Candida | 13 | Pathotroph-Saprotroph-Symbiotroph | Animal Pathogen-Endophyte-Endosymbiont-Epiphyte-\|Undefined Saprotroph\| | Dimorphic Yeast-Yeast | NULL | Probable | As nectar-tap_saprotroph (Põlme et al. 2020); Decay sugar-rich substrates (Põlme et al. 2020); This asexual genus is polyphyletic and contain species that function very differently depend on what they are. Caution should be taken when interpreting guilds for Candida species. | Manolakaki D, et al. 2010. Virulence. 1: 367-375 (https://doi.org/10.4161/viru.1.5.12796)); Mekbib SB, et al. 2011. Tropical Plant Pathology 36:233-240 ((https://doi.org/10.1590/S1982-56762011000400004)); Põlme S, et al. 2020. Fungal Diversity 105:1-16 ((https://doi.org/10.1007/s13225-020-00466-2)) |
| Otu05509 | 5 | k__Fungi(100);p__Ascomycota(100);c__Dothideomycetes(100);o__Capnodiales(100);f__Mycosphaerellaceae(100);f__Mycosphaerellaceae_unclassified(100);f__Mycosphaerellaceae_unclassified(100); | Mycosphaerellaceae | 9 | Pathotroph-Saprotroph | Plant Pathogen-Undefined Saprotroph | Microfungus | NULL | Probable | Biotrophic, necrotrophic or saprobic on various plant tissues (Cannon & Kirk 2007) | Cannon PF, Kirk PM. 2007. Fungal Families of the World. CAB International, Cambridge (ISBN: 978-0851998275) |
| Otu05510 | 5 | k__Fungi(100);p__Zygomycota(100);c__Zygomycota_class_Incertae_sedis(100);o__Mucorales(100);f__Mucoraceae(100);g__Mucor(100);s__Mucor_genevensis(100); | Mucor | 13 | Saprotroph | \|Undefined Saprotroph\| | Microfungus | NULL | Probable | As soil_saprotroph (Põlme et al. 2020); Decay soil (Põlme et al. 2020) | Tedersoo L, et al. 2014. Science 346:e1256688 ((https://doi.org/10.1126/science.1256688)); Põlme S, et al. 2020. Fungal Diversity 105:1-16 ((https://doi.org/10.1007/s13225-020-00466-2)) |
| Otu05512 | 5 | k__Fungi(100);p__Ascomycota(100);c__Saccharomycetes(100);o__Saccharomycetales(100);f__Metschnikowiaceae(100);g__Metschnikowia(100);s__Metschnikowia_pulcherrima(100); | Metschnikowia | 13 | Saprotroph | \|Undefined Saprotroph\| | Yeast | NULL | Probable | As nectar-tap_saprotroph (Põlme et al. 2020); Decay sugar-rich substrates (Põlme et al. 2020) | Põlme S, et al. 2020. Fungal Diversity 105:1-16 ((https://doi.org/10.1007/s13225-020-00466-2)) |
| Otu05514 | 5 | k__Fungi(100);p__Ascomycota(100);c__Saccharomycetes(100);o__Saccharomycetales(100);f__Metschnikowiaceae(100);g__Metschnikowia(100);s__Metschnikowia_pulcherrima(61); | Metschnikowia | 13 | Saprotroph | \|Undefined Saprotroph\| | Yeast | NULL | Probable | As nectar-tap_saprotroph (Põlme et al. 2020); Decay sugar-rich substrates (Põlme et al. 2020) | Põlme S, et al. 2020. Fungal Diversity 105:1-16 ((https://doi.org/10.1007/s13225-020-00466-2)) |
| Otu05517 | 5 | k__Fungi(100);p__Ascomycota(100);c__Saccharomycetes(100);o__Saccharomycetales(100);f__Metschnikowiaceae(100);g__Metschnikowia(100);s__Metschnikowia_pulcherrima(80); | Metschnikowia | 13 | Saprotroph | \|Undefined Saprotroph\| | Yeast | NULL | Probable | As nectar-tap_saprotroph (Põlme et al. 2020); Decay sugar-rich substrates (Põlme et al. 2020) | Põlme S, et al. 2020. Fungal Diversity 105:1-16 ((https://doi.org/10.1007/s13225-020-00466-2)) |
| Otu05519 | 5 | k__Fungi(100);p__Ascomycota(100);c__Saccharomycetes(100);o__Saccharomycetales(100);f__Metschnikowiaceae(100);g__Metschnikowia(100);s__Metschnikowia_pulcherrima(80); | Metschnikowia | 13 | Saprotroph | \|Undefined Saprotroph\| | Yeast | NULL | Probable | As nectar-tap_saprotroph (Põlme et al. 2020); Decay sugar-rich substrates (Põlme et al. 2020) | Põlme S, et al. 2020. Fungal Diversity 105:1-16 ((https://doi.org/10.1007/s13225-020-00466-2)) |
| Otu05520 | 5 | k__Fungi(100);p__Ascomycota(100);c__Dothideomycetes(100);o__Dothideomycetes_order_Incertae_sedis(100);f__Myxotrichaceae(100);g__Oidiodendron(100);g__Oidiodendron_unclassified(100); | Oidiodendron | 13 | Pathotroph-Saprotroph-Symbiotroph | Endophyte-Ericoid Mycorrhizal-\|Undefined Saprotroph\| | Microfungus | NULL | Probable | As root_endophyte (Põlme et al. 2020); As soil_saprotroph (Põlme et al. 2020); Dark Septate Endophyte; Decay soil (Põlme et al. 2020) | Newsham KK. 2011. New Phytologist 190:783-793 ((https://doi.org/10.1111/j.1469-8137.2010.03611.x)) (as dark septate root endophytes); Põlme S, et al. 2020. Fungal Diversity 105:1-16 ((https://doi.org/10.1007/s13225-020-00466-2)) |
| Otu05521 | 5 | k__Fungi(100);p__Ascomycota(100);c__Saccharomycetes(100);o__Saccharomycetales(100);f__Metschnikowiaceae(100);g__Metschnikowia(100);s__Metschnikowia_pulcherrima(61); | Metschnikowia | 13 | Saprotroph | \|Undefined Saprotroph\| | Yeast | NULL | Probable | As nectar-tap_saprotroph (Põlme et al. 2020); Decay sugar-rich substrates (Põlme et al. 2020) | Põlme S, et al. 2020. Fungal Diversity 105:1-16 ((https://doi.org/10.1007/s13225-020-00466-2)) |
| Otu05524 | 5 | k__Fungi(100);p__Ascomycota(100);c__Leotiomycetes(100);o__Leotiomycetes_order_Incertae_sedis(100);f__Leotiomycetes_family_Incertae_sedis(80);g__Meliniomyces(80);g__Meliniomyces_unclassified(80); | Meliniomyces | 13 | Saprotroph-Symbiotroph | Ectomycorrhizal-\|Endophyte\|-Ericoid Mycorrhizal-Orchid Mycorrhizal-Plant Saprotroph-Undefined Saprotroph | Microfungus | NULL | Probable | As root_endophyte (Põlme et al. 2020); As soil_saprotroph (Põlme et al. 2020); Dark Septate Endophyte; Decay soil (Põlme et al. 2020) | Tedersoo L, et al. 2014. Science 346:e1256688 ((https://doi.org/10.1126/science.1256688)); Põlme S, et al. 2020. Fungal Diversity 105:1-16 ((https://doi.org/10.1007/s13225-020-00466-2)) |
| Otu05525 | 5 | k__Fungi(100);p__Ascomycota(100);c__Saccharomycetes(100);o__Saccharomycetales(100);f__Metschnikowiaceae(100);g__Metschnikowia(100);s__Metschnikowia_pulcherrima(80); | Metschnikowia | 13 | Saprotroph | \|Undefined Saprotroph\| | Yeast | NULL | Probable | As nectar-tap_saprotroph (Põlme et al. 2020); Decay sugar-rich substrates (Põlme et al. 2020) | Põlme S, et al. 2020. Fungal Diversity 105:1-16 ((https://doi.org/10.1007/s13225-020-00466-2)) |
| Otu05528 | 5 | k__Fungi(100);p__Ascomycota(100);c__Saccharomycetes(100);o__Saccharomycetales(100);f__Metschnikowiaceae(100);g__Metschnikowia(100);s__Metschnikowia_pulcherrima(80); | Metschnikowia | 13 | Saprotroph | \|Undefined Saprotroph\| | Yeast | NULL | Probable | As nectar-tap_saprotroph (Põlme et al. 2020); Decay sugar-rich substrates (Põlme et al. 2020) | Põlme S, et al. 2020. Fungal Diversity 105:1-16 ((https://doi.org/10.1007/s13225-020-00466-2)) |
| Otu05532 | 5 | k__Fungi(100);p__Ascomycota(100);c__Saccharomycetes(100);o__Saccharomycetales(100);f__Metschnikowiaceae(100);g__Metschnikowia(100);g__Metschnikowia_unclassified(80); | Metschnikowia | 13 | Saprotroph | \|Undefined Saprotroph\| | Yeast | NULL | Probable | As nectar-tap_saprotroph (Põlme et al. 2020); Decay sugar-rich substrates (Põlme et al. 2020) | Põlme S, et al. 2020. Fungal Diversity 105:1-16 ((https://doi.org/10.1007/s13225-020-00466-2)) |
| Otu05533 | 5 | k__Fungi(100);p__Ascomycota(100);c__Saccharomycetes(100);o__Saccharomycetales(100);f__Metschnikowiaceae(100);g__Metschnikowia(100);s__Metschnikowia_pulcherrima(100); | Metschnikowia | 13 | Saprotroph | \|Undefined Saprotroph\| | Yeast | NULL | Probable | As nectar-tap_saprotroph (Põlme et al. 2020); Decay sugar-rich substrates (Põlme et al. 2020) | Põlme S, et al. 2020. Fungal Diversity 105:1-16 ((https://doi.org/10.1007/s13225-020-00466-2)) |
| Otu05534 | 5 | k__Fungi(100);p__Ascomycota(100);c__Saccharomycetes(100);o__Saccharomycetales(100);f__Metschnikowiaceae(100);g__Metschnikowia(100);g__Metschnikowia_unclassified(61); | Metschnikowia | 13 | Saprotroph | \|Undefined Saprotroph\| | Yeast | NULL | Probable | As nectar-tap_saprotroph (Põlme et al. 2020); Decay sugar-rich substrates (Põlme et al. 2020) | Põlme S, et al. 2020. Fungal Diversity 105:1-16 ((https://doi.org/10.1007/s13225-020-00466-2)) |
| [illegible] | [illegible] | [illegible] | [illegible] | [illegible] | [illegible] | [illegible] | [illegible] | [illegible] | [illegible] | [illegible] | [illegible] |

| | | | | | | | | | | | |
|---|---|---|---|---|---|---|---|---|---|---|---|
| Otu05752 | 5 | k__Fungi(100);p__Ascomycota(100);c__Eurotiomycetes(100);o__Eurotiales(100);f__Trichocomaceae(100);g__Penicillium(100);g__Penicillium_unclassified(100); | Penicillium | 13 | Pathotroph-Saprotroph-Symbiotroph | Animal Parasite-Dung Saprotroph-Endophyte-Fungal Parasite-Plant Pathogen-\|Undefined Saprotroph\|-Wood Saprotroph | Microfungus | Soft Rot | Highly Probable | As foliar_endophyte (Põlme et al. 2020); As mycoparasite (Põlme et al. 2020); Decay soil (Põlme et al. 2020) | Duncan CG, Eslyn WE. 1966. Mycologia 58:642-664 ((https://doi.org/10.2307/3757045)); Seehann G, et al. 1975. List of Fungi in Soft-Rot Tissue Tests. Institute fur Holzbiologie und Holzschutz, Bundesforschungsanstalt fur Forst - und Holzwirtshaft, Hamburg - Lohbrugge, Germany ((https://www.irg-wp.com/irgdocs/details.php?d6493033-c9ea-40f3-8e3e-5697f32d640d)); Costa IPMW, et al. 2012. Mycotaxon 119:e26 ((http://mycotaxon.com/resources/checklists/costa_v119_checklist.pdf)); Bills GF, et al. 2013. Current Opinion in Microbiology 16: 549-565 ((https://doi.org/10.1016/j.mib.2013.08.001)); Tedersoo L, et al. 2014. Science 346:e1256688 ((https://doi.org/10.1126/science.1256688)); Põlme S, et al. 2020. Fungal Diversity 105:1-16 ((https://doi.org/10.1007/s13225-020-00466-2)) |
| Otu05753 | 5 | k__Fungi(100);p__Ascomycota(100);c__Eurotiomycetes(100);o__Onygenales(100);f__Onygenaceae(100);g__Aphanoascus(100);s__Aphanoascus_cinnabarinus(100); | Aphanoascus | 13 | Saprotroph | \|Undefined Saprotroph\| | Microfungus | NULL | Probable | As animal_decomposer (Põlme et al. 2020); As keratinolytic (Põlme et al. 2020); As soil_saprotroph (Põlme et al. 2020); Decay soil, animal material (Põlme et al. 2020) | Tedersoo L, et al. 2014. Science 346:e1256688 ((https://doi.org/10.1126/science.1256688)); Põlme S, et al. 2020. Fungal Diversity 105:1-16 ((https://doi.org/10.1007/s13225-020-00466-2)) |
| Otu05755 | 5 | k__Fungi(100);p__Basidiomycota(100);c__Agaricomycetes(100);o__Polyporales(100);f__Xenasmataceae(100);g__Xenasma(100);s__Xenasma_rimicola(100); | Xenasma | 13 | Saprotroph | Undefined Saprotroph-\|Wood Saprotroph\| | Corticioid | White Rot | Probable | As white_rot (Põlme et al. 2020); Decay wood (Põlme et al. 2020) | Tedersoo L, et al. 2014. Science 346:e1256688 ((https://doi.org/10.1126/science.1256688)); Põlme S, et al. 2020. Fungal Diversity 105:1-16 ((https://doi.org/10.1007/s13225-020-00466-2)) |
| Otu05757 | 5 | k__Fungi(100);p__Basidiomycota(100);c__Agaricomycetes(100);o__Polyporales(100);f__Phanerochaetaceae(100);g__Ceriporia(100);s__Ceriporia_aurantiocarnescens(80); | Ceriporia | 13 | Saprotroph | \|Wood Saprotroph\| | Corticioid | White Rot | Highly Probable | As white_rot (Põlme et al. 2020); Decay wood (Põlme et al. 2020) | Tedersoo L, et al. 2014. Science 346:e1256688 ((https://doi.org/10.1126/science.1256688)); Põlme S, et al. 2020. Fungal Diversity 105:1-16 ((https://doi.org/10.1007/s13225-020-00466-2)) |
| Otu05758 | 5 | k__Fungi(100);p__Ascomycota(100);c__Dothideomycetes(100);o__Capnodiales(100);f__Davidiellaceae(100);g__Cladosporium(100);s__Cladosporium_sphaerospermum(100); | Cladosporium | 13 | Pathotroph-Saprotroph-Symbiotroph | Animal Pathogen-Endophyte-Lichen Parasite-Plant Pathogen-\|Plant Saprotroph\|-Wood Saprotroph | Microfungus | NULL | Possible | As foliar_endophyte (Põlme et al. 2020); As litter_saprotroph (Põlme et al. 2020); Decay wood, leaf, fruit, seed, soil, dung, animal material (Põlme et al. 2020) | Seehann G, et al. 1975. List of Fungi in Soft-Rot Tissue Tests. Institute fur Holzbiologie und Holzschutz, Bundesforschungsanstalt fur Forst - und Holzwirtshaft, Hamburg - Lohbrugge, Germany; Lawrey JD, Diederich P. 2016. http://www.lichenicolous.net (accessed 24 May 2017) ((http://www.lichenicolous.net)); Põlme S, et al. 2020. Fungal Diversity 105:1-16 ((https://doi.org/10.1007/s13225-020-00466-2)); The American Phytopathological Society Common Names of Plant Diseases ((http://www.apsnet.org/publications/commonnames/Pages/default.aspx)) |
| Otu05759 | 5 | k__Fungi(100);p__Ascomycota(100);c__Saccharomycetes(100);o__Saccharomycetales(100);f__Saccharomycetaceae(100);g__Kazachstania(100);s__Kazachstania_intestinalis(100); | Kazachstania | 13 | Saprotroph | \|Undefined Saprotroph\| | Yeast | NULL | Probable | As nectar-tap_saprotroph (Põlme et al. 2020); Decay sugar-rich substrates (Põlme et al. 2020) | Põlme S, et al. 2020. Fungal Diversity 105:1-16 ((https://doi.org/10.1007/s13225-020-00466-2)) |
| Otu05761 | 5 | k__Fungi(100);p__Ascomycota(100);c__Saccharomycetes(100);o__Saccharomycetales(100);f__Metschnikowiaceae(100);g__Metschnikowia(100);s__Metschnikowia_pulcherrima(61); | Metschnikowia | 13 | Saprotroph | \|Undefined Saprotroph\| | Yeast | NULL | Probable | As nectar-tap_saprotroph (Põlme et al. 2020); Decay sugar-rich substrates (Põlme et al. 2020) | Põlme S, et al. 2020. Fungal Diversity 105:1-16 ((https://doi.org/10.1007/s13225-020-00466-2)) |
| Otu05762 | 5 | k__Fungi(100);p__Basidiomycota(100);c__Agaricomycetes(100);o__Agaricales(100);o__Agaricales_unclassified(100);o__Agaricales_unclassified(100);o__Agaricales_unclassified(100); | Agaricales | 7 | Pathotroph-Saprotroph-Symbiotroph | Bryophyte Parasite-Dung Saprotroph-Ectomycorrhizal-Fungal Parasite-Leaf Saprotroph-Plant Parasite-Undefined Saprotroph-Wood Saprotroph | Agaricoid-Gasteroid-Microfungus-Secotioid-Yeast | NULL | Possible | Mushrooms and toadstools, Gill fungi, Agarics... lignicolous, sometimes muscicolous or fungicolous, saprobic, mycorrhizal, rarely parasitic on plants or fungi (Kirk et al. 2008) | Kirk PM et al. 2008. Dictionary of the Fungi. Tenth Edition. CAB International, Wallingford (ISBN: 978-0851998268) |
| Otu05763 | 5 | k__Fungi(100);p__Ascomycota(100);c__Saccharomycetes(100);o__Saccharomycetales(100);f__Metschnikowiaceae(100);g__Metschnikowia(100);s__Metschnikowia_pulcherrima(80); | Metschnikowia | 13 | Saprotroph | \|Undefined Saprotroph\| | Yeast | NULL | Probable | As nectar-tap_saprotroph (Põlme et al. 2020); Decay sugar-rich substrates (Põlme et al. 2020) | Põlme S, et al. 2020. Fungal Diversity 105:1-16 ((https://doi.org/10.1007/s13225-020-00466-2)) |
| Otu05765 | 5 | k__Fungi(100);p__Ascomycota(100);c__Saccharomycetes(100);o__Saccharomycetales(100);f__Metschnikowiaceae(100);g__Metschnikowia(100);s__Metschnikowia_pulcherrima(80); | Metschnikowia | 13 | Saprotroph | \|Undefined Saprotroph\| | Yeast | NULL | Probable | As nectar-tap_saprotroph (Põlme et al. 2020); Decay sugar-rich substrates (Põlme et al. 2020) | Põlme S, et al. 2020. Fungal Diversity 105:1-16 ((https://doi.org/10.1007/s13225-020-00466-2)) |
| Otu05769 | 5 | k__Fungi(100);p__Basidiomycota(100);c__Agaricomycetes(100);o__Russulales(100);f__Russulaceae(100);g__unclassified_Russulaceae(80);s__Russulaceae_sp(80); | Russulaceae | 9 | Symbiotroph | Ectomycorrhizal | Agaricoid-Corticioid-Gasteroid-Secotioid | NULL | Probable | The genera Lactarius, Lactifluus, Multifurca and Russula form a mutualistic ectomycorrhizal root symbiosis with trees and shrubs, while the corticioid species in Boidinia, Gloeopeniophorella, and Pseudoxenasma are saprotrophic, wood-degrading fungi. | Larsson E, K-H Larsson. 2003. Mycologia 95:1037-1065 ((https://doi.org/10.2307/3761912)) |
| Otu05770 | 5 | k__Fungi(100);p__Basidiomycota(100);c__Agaricomycetes(100);o__Agaricales(100);f__Inocybaceae(100);g__Inocybe(100);g__Inocybe_unclassified(100); | Inocybe | 13 | Symbiotroph | \|Ectomycorrhizal\| | Agaricoid | NULL | Highly Probable | Sensu stricto (Matheny 2009). | Rinaldi AC, et al. 2008. Fungal Diversity 33:1-45 ((http://www.fungaldiversity.org/fdp/sfdp/33-1.pdf)); Matheny PB. 2009. McIlvainea 18:11-21 ((https://namyco.org/docs/3_Matheny.pdf)); Tedersoo L, et al. 2010. Mycorrhiza 20:217-263 ((https://doi.org/10.1007/s00572-009-0274-x)); Põlme S, et al. 2020. Fungal Diversity 105:1-16 ((https://doi.org/10.1007/s13225-020-00466-2)); http://mycorrhizas.info/ecmf.html ((http://mycorrhizas.info/ecmf.html)) |
| Otu05772 | 5 | k__Fungi(100);p__Basidiomycota(100);c__Agaricomycetes(100);o__Agaricales(100);o__Agaricales_unclassified(100);o__Agaricales_unclassified(100);o__Agaricales_unclassified(100); | Agaricales | 7 | Pathotroph-Saprotroph-Symbiotroph | Bryophyte Parasite-Dung Saprotroph-Ectomycorrhizal-Fungal Parasite-Leaf Saprotroph-Plant Parasite-Undefined Saprotroph-Wood Saprotroph | Agaricoid-Gasteroid-Microfungus-Secotioid-Yeast | NULL | Possible | Mushrooms and toadstools, Gill fungi, Agarics... lignicolous, sometimes muscicolous or fungicolous, saprobic, mycorrhizal, rarely parasitic on plants or fungi (Kirk et al. 2008) | Kirk PM et al. 2008. Dictionary of the Fungi. Tenth Edition. CAB International, Wallingford (ISBN: 978-0851998268) |
| Otu05773 | 5 | k__Fungi(100);p__Basidiomycota(100);c__Microbotryomycetes(100);o__Sporidiobolales(100);f__Sporidiobolales_family_Incertae_sedis(100);g__Rhodotorula(100);s__Rhodotorula_philyla(100); | Rhodotorula | 13 | Pathotroph-Saprotroph-Symbiotroph | Animal Pathogen-Animal Symbiotroph-Endophyte-Plant Pathogen-\|Undefined Saprotroph\| | Yeast | NULL | Probable | As foliar_endophyte (Põlme et al. 2020); Opportunistic human pathogen | Põlme S, et al. 2020. Fungal Diversity 105:1-16 ((https://doi.org/10.1007/s13225-020-00466-2)); https://www.cdc.gov ((https://www.cdc.gov)) |
| Otu05774 | 5 | k__Fungi(100);p__Ascomycota(100);c__Saccharomycetes(100);o__Saccharomycetales(100);f__Metschnikowiaceae(100);g__Metschnikowia(100);s__Metschnikowia_pulcherrima(100); | Metschnikowia | 13 | Saprotroph | \|Undefined Saprotroph\| | Yeast | NULL | Probable | As nectar-tap_saprotroph (Põlme et al. 2020); Decay sugar-rich substrates (Põlme et al. 2020) | Põlme S, et al. 2020. Fungal Diversity 105:1-16 ((https://doi.org/10.1007/s13225-020-00466-2)) |
| Otu05775 | 5 | k__Fungi(100);p__Ascomycota(100);c__Saccharomycetes(100);o__Saccharomycetales(100);f__Saccharomycetales_family_Incertae_sedis(100);g__Candida(100);g__Candida_unclassified(100); | Candida | 13 | Pathotroph-Saprotroph-Symbiotroph | Animal Pathogen-Endophyte-Endosymbiont-Epiphyte-\|Undefined Saprotroph\| | Dimorphic Yeast-Yeast | NULL | Probable | As nectar-tap_saprotroph (Põlme et al. 2020); Decay sugar-rich substrates (Põlme et al. 2020); This asexual genus is polyphyletic and contain species that function very differently depend on what they are. Caution should be taken when interpreting guilds for Candida species. | Manolakaki D, et al. 2010. Virulence. 1: 367-375 ((https://doi.org/10.4161/viru.1.5.12796)); Mekbib SB, et al. 2011. Tropical Plant Pathology 36:233-240 ((https://doi.org/10.1590/S1982-56762011000400004)); Põlme S, et al. 2020. Fungal Diversity 105:1-16 ((https://doi.org/10.1007/s13225-020-00466-2)) |
| Otu05776 | 5 | k__Fungi(100);p__Ascomycota(100);c__Eurotiomycetes(100);o__Eurotiales(100);f__Trichocomaceae(100);g__Paecilomyces(100);s__Paecilomyces_formosus(100); | Paecilomyces | 13 | Pathotroph-Saprotroph-Symbiotroph | Clavicipitaceous \|Endophyte\|-Plant Pathogen-Plant Saprotroph-Undefined Saprotroph-Wood Saprotroph | Microfungus | Soft Rot | Probable | As foliar_endophyte (Põlme et al. 2020); As litter_saprotroph (Põlme et al. 2020); culture (Lopez et al. 2014); Decay leaf, fruit, seed, animal material (Põlme et al. 2020); Endophyte detection method | Seehann G, et al. 1975. List of Fungi in Soft-Rot Tissue Tests. Institute fur Holzbiologie und Holzschutz, Bundesforschungsanstalt fur Forst - und Holzwirtshaft, Hamburg - Lohbrugge, Germany; Costa IPMW, et al. 2012. Mycotaxon 119:e26 ((http://mycotaxon.com/resources/checklists/costa_v119_checklist.pdf)); Tedersoo L, et al. 2014. Science 346:e1256688 ((https://doi.org/10.1126/science.1256688)); Põlme S, et al. 2020. Fungal Diversity 105:1-16 ((https://doi.org/10.1007/s13225-020-00466-2)); The American Phytopathological Society Common Names of Plant Diseases ((http://www.apsnet.org/publications/commonnames/Pages/default.aspx)); Lopez DC, et al. 2014. Plos One 9:e103891 ((https://doi.org/10.1371/journal.pone.0103891)) |
| Otu05778 | 5 | k__Fungi(100);p__Ascomycota(100);c__Saccharomycetes(100);o__Saccharomycetales(100);f__Metschnikowiaceae(100);g__Metschnikowia(100);s__Metschnikowia_pulcherrima(100); | Metschnikowia | 13 | Saprotroph | \|Undefined Saprotroph\| | Yeast | NULL | Probable | As nectar-tap_saprotroph (Põlme et al. 2020); Decay sugar-rich substrates (Põlme et al. 2020) | Põlme S, et al. 2020. Fungal Diversity 105:1-16 ((https://doi.org/10.1007/s13225-020-00466-2)) |
| Otu05779 | 5 | k__Fungi(100);p__Ascomycota(100);c__Eurotiomycetes(100);o__Eurotiales(100);f__Trichocomaceae(100);g__Penicillium(100);s__Penicillium_sp_SW_2014e(61); | Penicillium | 13 | Pathotroph-Saprotroph-Symbiotroph | Animal Parasite-Dung Saprotroph-Endophyte-Fungal Parasite-Plant Pathogen-\|Undefined Saprotroph\|-Wood Saprotroph | Microfungus | Soft Rot | Highly Probable | As foliar_endophyte (Põlme et al. 2020); As mycoparasite (Põlme et al. 2020); Decay soil (Põlme et al. 2020) | Duncan CG, Eslyn WE. 1966. Mycologia 58:642-664 ((https://doi.org/10.2307/3757045)); Seehann G, et al. 1975. List of Fungi in Soft-Rot Tissue Tests. Institute fur Holzbiologie und Holzschutz, Bundesforschungsanstalt fur Forst - und Holzwirtshaft, Hamburg - Lohbrugge, Germany ((https://www.irg-wp.com/irgdocs/details.php?d6493033-c9ea-40f3-8e3e-5697f32d640d)); Costa IPMW, et al. 2012. Mycotaxon 119:e26 ((http://mycotaxon.com/resources/checklists/costa_v119_checklist.pdf)); Bills GF, et al. 2013. Current Opinion in Microbiology 16: 549-565 ((https://doi.org/10.1016/j.mib.2013.08.001)); Tedersoo L, et al. 2014. Science 346:e1256688 ((https://doi.org/10.1126/science.1256688)); Põlme S, et al. 2020. Fungal Diversity 105:1-16 ((https://doi.org/10.1007/s13225-020-00466-2)) |
| Otu05780 | 5 | k__Fungi(100);p__Basidiomycota(100);c__Exobasidiomycetes(100);o__Exobasidiales(100);f__Exobasidiaceae(100);g__Exobasidium(100);g__Exobasidium_unclassified(61); | Exobasidium | 13 | Pathotroph | \|Plant Pathogen\| | Leaf Blister-Leaf Gall-Leaf Spot | NULL | Probable | NULL | Tedersoo L, et al. 2014. Science 346:e1256688 ((https://doi.org/10.1126/science.1256688)); Põlme S, et al. 2020. Fungal Diversity 105:1-16 ((https://doi.org/10.1007/s13225-020-00466-2)) |
| Otu05783 | 5 | k__Fungi(100);p__Basidiomycota(100);c__Agaricomycetes(100);o__Cantharellales(100);f__Hydnaceae(100);g__Sistotrema(100);s__Sistotrema_muscicola(100); | Sistotrema | 13 | Saprotroph-Symbiotroph | Ectomycorrhizal-\|Plant Saprotroph\|-Wood Saprotroph | Corticioid | White Rot | Possible | As litter_saprotroph (Põlme et al. 2020); Decay leaf, fruit, seed (Põlme et al. 2020) | Gilbertson RL, Ryvarden L. 1987-1987. North American Polypores. Fungiflora, Oslo (ISBN: 978-0945345060); Rinaldi AC, et al. 2008. Fungal Diversity 33:1-45 ((http://www.fungaldiversity.org/fdp/sfdp/33-1.pdf)) (as ECM/not ECM); Tedersoo L, et al. 2010. Mycorrhiza 20: 217-263 ((https://doi.org/10.1007/s00572-009-0274-x)) (pro parte); Põlme S, et al. 2020. Fungal Diversity 105:1-16 ((https://doi.org/10.1007/s13225-020-00466-2)); http://mycorrhizas.info/ecmf.html ((http://mycorrhizas.info/ecmf.html)) (as ECM/not ECM). |
| Otu05784 | 5 | k__Fungi(100);p__Basidiomycota(100);c__Microbotryomycetes(100);o__Sporidiobolales(100);f__Sporidiobolales_family_Incertae_sedis(100);g__Rhodotorula(100);g__Rhodotorula_unclassified(100); | Rhodotorula | 13 | Pathotroph-Saprotroph-Symbiotroph | Animal Pathogen-Animal Symbiotroph-Endophyte-Plant Pathogen-\|Undefined Saprotroph\| | Yeast | NULL | Probable | As foliar_endophyte (Põlme et al. 2020); Opportunistic human pathogen | Põlme S, et al. 2020. Fungal Diversity 105:1-16 ((https://doi.org/10.1007/s13225-020-00466-2)); https://www.cdc.gov ((https://www.cdc.gov)) |
| Otu05785 | 5 | k__Fungi(100);p__Basidiomycota(100);c__Agaricomycetes(100);o__Russulales(100);f__Russulaceae(100);g__Russula(100);s__Russula_mariae(100); | Russula | 13 | Symbiotroph | \|Ectomycorrhizal\| | Agaricoid | NULL | Highly Probable | NULL | Rinaldi AC, et al. 2008. Fungal Diversity 33:1-45 ((http://www.fungaldiversity.org/fdp/sfdp/33-1.pdf)); Tedersoo L, et al. 2010. Mycorrhiza 20:217-263 ((https://doi.org/10.1007/s00572-009-0274-x)); Põlme S, et al. 2020. Fungal Diversity 105:1-16 ((https://doi.org/10.1007/s13225-020-00466-2)); http://mycorrhizas.info/ecmf.html ((http://mycorrhizas.info/ecmf.html)) |
| Otu05786 | 5 | k__Fungi(100);p__Basidiomycota(100);c__Agaricomycetes(100);o__Cantharellales(100);f__Hydnaceae(100);g__Sistotrema(100);s__Sistotrema_sp_PC14(100); | Sistotrema | 13 | Saprotroph-Symbiotroph | Ectomycorrhizal-\|Plant Saprotroph\|-Wood Saprotroph | Corticioid | White Rot | Possible | As litter_saprotroph (Põlme et al. 2020); Decay leaf, fruit, seed (Põlme et al. 2020) | Gilbertson RL, Ryvarden L. 1987-1987. North American Polypores. Fungiflora, Oslo (ISBN: 978-0945345060); Rinaldi AC, et al. 2008. Fungal Diversity 33:1-45 ((http://www.fungaldiversity.org/fdp/sfdp/33-1.pdf)) (as ECM/not ECM); Tedersoo L, et al. 2010. Mycorrhiza 20: 217-263 ((https://doi.org/10.1007/s00572-009-0274-x)) (pro parte); Põlme S, et al. 2020. Fungal Diversity 105:1-16 ((https://doi.org/10.1007/s13225-020-00466-2)); http://mycorrhizas.info/ecmf.html ((http://mycorrhizas.info/ecmf.html)) (as ECM/not ECM). |
| Otu05790 | 5 | k__Fungi(100);p__Basidiomycota(100);c__Agaricomycetes(100);o__Polyporales(100);f__Ganodermataceae(100);f__Ganodermataceae_unclassified(80);f__Ganodermataceae_unclassified(80); | Ganodermataceae | 9 | Pathotroph-Saprotroph | Plant Pathogen-Wood Saprotroph | Polyporoid | White Rot | Probable | Lignicolous, causing sometimes agressive white rots of timber (Cannon & Kirk 2007) | Cannon PF, Kirk PM. 2007. Fungal Families of the World. CAB International, Cambridge (ISBN: 978-0851998275) |
| Otu05791 | 5 | k__Fungi(100);p__Basidiomycota(100);c__Agaricomycetes(100);o__Agaricales(100);f__Entolomataceae(100);f__Entolomataceae_unclassified(100);f__Entolomataceae_unclassified(100); | Entolomataceae | 9 | Pathotroph-Saprotroph-Symbiotroph | Ectomycorrhizal-Fungal Parasite-Soil Saprotroph-Undefined Saprotroph | Agaricoid | NULL | Possible | Saprobic or ectomycorrhizal, in both broadleaved and coniferous forests; a few species associated with other substrata such as dung and mosses (Cannon & Kirk 2007) | Cannon PF, Kirk PM. 2007. Fungal Families of the World. CAB International, Cambridge (ISBN: 978-0851998275); List compiled by John Plischke III ((https://goodmushroom.weebly.com/uploads/5/7/1/7/57176535/fungi_on_fungi_pdf_lq.pdf)) |
| Otu05792 | 5 | k__Fungi(100);p__Ascomycota(100);c__Eurotiomycetes(100);o__Chaetothyriales(100);f__Herpotrichiellaceae(100);g__Exophiala(100);s__Exophiala_heteromorpha(100); | Exophiala | 13 | Pathotroph-Saprotroph-Symbiotroph | \|Animal Parasite\|-Animal Pathogen-Endophyte-Plant Saprotroph-Undefined Saprotroph | Dimorphic-Facultative Yeast | NULL | Probable | As litter_saprotroph (Põlme et al. 2020); As root_endophyte_dark_septate (Põlme et al. 2020); Decay leaf, fruit, seed, animal material (Põlme et al. 2020) | Tedersoo L, et al. 2014. Science 346:e1256688 ((https://doi.org/10.1126/science.1256688)), Nagy LG, et al. 2014. Nature Communications 5:e4471 ((https://doi.org/10.1038/ncomms5471)); Põlme S, et al. 2020. Fungal Diversity 105:1-16 ((https://doi.org/10.1007/s13225-020-00466-2)) |
| Otu05794 | 5 | k__Fungi(100);p__Basidiomycota(100);c__Agaricomycetes(100);o__Boletales(100);f__Boletaceae(100);f__Boletaceae_unclassified(100);f__Boletaceae_unclassified(100); | Boletaceae | 9 | Pathotroph-Saprotroph-Symbiotroph | Ectomycorrhizal-Fungal Parasite-Plant Saprotroph-Wood Saprotroph | Boletoid | NULL | Probable | Most species form ectomycorrhizal associations with gymnosperm or angiosperm trees. Some are saprobes on decaying wood and leaf litter, a few are parasitic on other fungi (Cannon & Kirk 2007) | Cannon PF, Kirk PM. 2007. Fungal Families of the World. CAB International, Cambridge (ISBN: 978-0851998275) |
| Otu05795 | 5 | k__Fungi(100);p__Basidiomycota(100);c__Agaricomycetes(100);o__Agaricales(100);f__Cortinariaceae(100);g__Cortinarius(100);g__Cortinarius_unclassified(100); | Cortinarius | 13 | Symbiotroph | \|Ectomycorrhizal\| | Agaricoid | NULL | Highly Probable | NULL | Rinaldi AC, et al. 2008. Fungal Diversity 33:1-45 ((http://www.fungaldiversity.org/fdp/sfdp/33-1.pdf)); Tedersoo L, et al. 2010. Mycorrhiza 20:217-263 ((https://doi.org/10.1007/s00572-009-0274-x)); Põlme S, et al. 2020. Fungal Diversity 105:1-16 ((https://doi.org/10.1007/s13225-020-00466-2)); http://mycorrhizas.info/ecmf.html ((http://mycorrhizas.info/ecmf.html)) |
| Otu05796 | 5 | k__Fungi(100);p__Basidiomycota(100);c__Agaricomycetes(100);o__Polyporales(100);f__Fomitopsidaceae(100);g__Fibroporia(100);s__Fibroporia_norrlandica(100); | Fibroporia | 13 | Saprotroph | \|Wood Saprotroph\| | Corticioid | Brown Rot | Probable | As brown-rot (Põlme et al. 2020); Decay wood (Põlme et al. 2020) | Tedersoo L, et al. 2014. Science 346:e1256688 ((https://doi.org/10.1126/science.1256688)); Põlme S, et al. 2020. Fungal Diversity 105:1-16 ((https://doi.org/10.1007/s13225-020-00466-2)) |
| Otu05797 | 5 | k__Fungi(100);p__Ascomycota(100);c__Saccharomycetes(100);o__Saccharomycetales(100);f__Metschnikowiaceae(100);g__Metschnikowia(100);s__Metschnikowia_pulcherrima(100); | Metschnikowia | 13 | Saprotroph | \|Undefined Saprotroph\| | Yeast | NULL | Probable | As nectar-tap_saprotroph (Põlme et al. 2020); Decay sugar-rich substrates (Põlme et al. 2020) | Põlme S, et al. 2020. Fungal Diversity 105:1-16 ((https://doi.org/10.1007/s13225-020-00466-2)) |
| Otu05798 | 5 | k__Fungi(100);p__Basidiomycota(100);c__Agaricomycetes(100);o__Agaricales(100);o__Agaricales_unclassified(100);o__Agaricales_unclassified(100);o__Agaricales_unclassified(100); | Agaricales | 7 | Pathotroph-Saprotroph-Symbiotroph | Bryophyte Parasite-Dung Saprotroph-Ectomycorrhizal-Fungal Parasite-Leaf Saprotroph-Plant Parasite-Undefined Saprotroph-Wood Saprotroph | Agaricoid-Gasteroid-Microfungus-Secotioid-Yeast | NULL | Possible | Mushrooms and toadstools, Gill fungi, Agarics... lignicolous, sometimes muscicolous or fungicolous, saprobic, mycorrhizal, rarely parasitic on plants or fungi (Kirk et al. 2008) | Kirk PM et al. 2008. Dictionary of the Fungi. Tenth Edition. CAB International, Wallingford (ISBN: 978-0851998268) |
| Otu05799 | 5 | k__Fungi(100);p__Basidiomycota(100);c__Agaricomycetes(100);o__Agaricales(100);f__Cortinariaceae(100);g__Cortinarius(100);g__Cortinarius_unclassified(100); | Cortinarius | 13 | Symbiotroph | \|Ectomycorrhizal\| | Agaricoid | NULL | Highly Probable | NULL | Rinaldi AC, et al. 2008. Fungal Diversity 33:1-45 ((http://www.fungaldiversity.org/fdp/sfdp/33-1.pdf)); Tedersoo L, et al. 2010. Mycorrhiza 20:217-263 ((https://doi.org/10.1007/s00572-009-0274-x)); Põlme S, et al. 2020. Fungal Diversity 105:1-16 ((https://doi.org/10.1007/s13225-020-00466-2)); http://mycorrhizas.info/ecmf.html ((http://mycorrhizas.info/ecmf.html)) |
| Otu05800 | 5 | k__Fungi(100);p__Basidiomycota(100);c__Agaricomycetes(100);o__Agaricales(100);f__Strophariaceae(100);g__unclassified_Strophariaceae(100);s__Strophariaceae_sp(100); | Strophariaceae | 9 | Pathotroph-Saprotroph | Dung Saprotroph-Plant Parasite-Soil Saprotroph-Undefined Saprotroph-Wood Saprotroph | Agaricoid | NULL | Probable | Very variable ecology; Pholiota species are lignicolous and can act as weak parasites while most other taxa are either sparobes in grassy habitats or are found in dung. Species are not ectomycorrhizal (Cannon & Kirk 2007) | Cannon PF, Kirk PM. 2007. Fungal Families of the World. CAB International, Cambridge (ISBN: 978-0851998275) |
| Otu05801 | 5 | k__Fungi(100);p__Ascomycota(100);c__Saccharomycetes(100);o__Saccharomycetales(100);f__Saccharomycetales_family_Incertae_sedis(100);g__Candida(80);g__Candida_unclassified(80); | Candida | 13 | Pathotroph-Saprotroph-Symbiotroph | Animal Pathogen-Endophyte-Endosymbiont-Epiphyte-\|Undefined Saprotroph\| | Dimorphic Yeast-Yeast | NULL | Probable | As nectar-tap_saprotroph (Põlme et al. 2020); Decay sugar-rich substrates (Põlme et al. 2020); This asexual genus is polyphyletic and contain species that function very differently depend on what they are. Caution should be taken when interpreting guilds for Candida species. | Manolakaki D, et al. 2010. Virulence. 1: 367-375 ((https://doi.org/10.4161/viru.1.5.12796)); Mekbib SB, et al. 2011. Tropical Plant Pathology 36:233-240 ((https://doi.org/10.1590/S1982-56762011000400004)); Põlme S, et al. 2020. Fungal Diversity 105:1-16 ((https://doi.org/10.1007/s13225-020-00466-2)) |
| Otu05802 | 5 | k__Fungi(100);p__Ascomycota(100);c__Eurotiomycetes(100);o__Verrucariales(100);f__Verrucariaceae(100);g__Thelidium(100);s__Thelidium_rehmii(100); | Thelidium | 13 | Symbiotroph | \|Lichenized\| | Thallus | NULL | Highly Probable | NULL | Esslinger TL. 2014. North Dakota State University ((http://www.ndsu.edu/pubweb/~esslinge/chcklst/chcklst7.htm)); Põlme S, et al. 2020. Fungal Diversity 105:1-16 ((https://doi.org/10.1007/s13225-020-00466-2)) |
| Otu05803 | 5 | k__Fungi(100);p__Ascomycota(100);c__Saccharomycetes(100);o__Saccharomycetales(100);f__Metschnikowiaceae(100);g__Metschnikowia(100);s__Metschnikowia_pulcherrima(100); | Metschnikowia | 13 | Saprotroph | \|Undefined Saprotroph\| | Yeast | NULL | Probable | As nectar-tap_saprotroph (Põlme et al. 2020); Decay sugar-rich substrates (Põlme et al. 2020) | Põlme S, et al. 2020. Fungal Diversity 105:1-16 ((https://doi.org/10.1007/s13225-020-00466-2)) |
| Otu05804 | 5 | k__Fungi(100);p__Basidiomycota(100);c__Agaricomycetes(100);o__Russulales(100);f__Russulaceae(100);g__Lactarius(100);s__Lactarius_necator(100); | Lactarius | 13 | Symbiotroph | \|Ectomycorrhizal\| | Agaricoid | NULL | Highly Probable | milk; cap; Common name | Rinaldi AC, et al. 2008. Fungal Diversity 33:1-45 ((http://www.fungaldiversity.org/fdp/sfdp/33-1.pdf)); Tedersoo L, et al. 2010. Mycorrhiza 20:217-263 ((https://doi.org/10.1007/s00572-009-0274-x)); Põlme S, et al. 2020. Fungal Diversity 105:1-16 ((https://doi.org/10.1007/s13225-020-00466-2)); http://mycorrhizas.info/ecmf.html ((http://mycorrhizas.info/ecmf.html)) |
| Otu05806 | 5 | k__Fungi(100);p__Ascomycota(100);c__Sordariomycetes(100);o__Xylariales(100);f__Xylariaceae(100);g__unclassified_Xylariaceae(100);s__Xylariaceae_sp(100); | Xylariaceae | 9 | Pathotroph-Saprotroph-Symbiotroph | Dung Saprotroph-Endophyte-Plant Pathogen-Undefined Saprotroph | Xylarioid | NULL | Probable | Saprobes or weak pathogens, many endophytes, mainly in wood and bark, occasionally in soil and dung (Cannon & Kirk 2007) | Cannon PF, Kirk PM. 2007. Fungal Families of the World. CAB International, Cambridge (ISBN: 978-0851998275) |
| Otu05807 | 5 | k__Fungi(100);p__Basidiomycota(100);c__Microbotryomycetes(100);o__Sporidiobolales(100);f__Sporidiobolales_family_Incertae_sedis(100);g__Sporobolomyces(100);s__Sporobolomyces_lactosus(100); | Sporobolomyces | 13 | Pathotroph-Saprotroph | \|Fungal Parasite\|-Plant Saprotroph | Yeast | NULL | Probable | As litter_saprotroph (Põlme et al. 2020); As mycoparasite (Põlme et al. 2020); Decay fungal material (Põlme et al. 2020) | Tedersoo L, et al. 2014. Science 346:e1256688 ((https://doi.org/10.1126/science.1256688)); Purahong W, et al. 2016. Molecular Ecology 25:4059-4074 ((https://doi.org/10.1111/mec.13739)); Põlme S, et al. 2020. Fungal Diversity 105:1-16 ((https://doi.org/10.1007/s13225-020-00466-2)) |
| Otu05809 | 5 | k__Fungi(100);p__Ascomycota(100);c__Eurotiomycetes(100);o__Eurotiales(100);f__Trichocomaceae(100);g__Penicillium(100);g__Penicillium_unclassified(80); | Penicillium | 13 | Pathotroph-Saprotroph-Symbiotroph | Animal Parasite-Dung Saprotroph-Endophyte-Fungal Parasite-Plant Pathogen-\|Undefined Saprotroph\|-Wood Saprotroph | Microfungus | Soft Rot | Highly Probable | As foliar_endophyte (Põlme et al. 2020); As mycoparasite (Põlme et al. 2020); Decay soil (Põlme et al. 2020) | Duncan CG, Eslyn WE. 1966. Mycologia 58:642-664 ((https://doi.org/10.2307/3757045)); Seehann G, et al. 1975. List of Fungi in Soft-Rot Tissue Tests. Institute fur Holzbiologie und Holzschutz, Bundesforschungsanstalt fur Forst - und Holzwirtshaft, Hamburg - Lohbrugge, Germany ((https://www.irg-wp.com/irgdocs/details.php?d6493033-c9ea-40f3-8e3e-5697f32d640d)); Costa IPMW, et al. 2012. Mycotaxon 119:e26 ((http://mycotaxon.com/resources/checklists/costa_v119_checklist.pdf)); Bills GF, et al. 2013. Current Opinion in Microbiology 16: 549-565 ((https://doi.org/10.1016/j.mib.2013.08.001)); Tedersoo L, et al. 2014. Science 346:e1256688 ((https://doi.org/10.1126/science.1256688)); Põlme S, et al. 2020. Fungal Diversity 105:1-16 ((https://doi.org/10.1007/s13225-020-00466-2)) |
| Otu05812 | 5 | k__Fungi(100);p__Ascomycota(100);c__Lecanoromycetes(100);o__Teloschistales(100);f__Physciaceae(100);f__Physciaceae_unclassified(100);f__Physciaceae_unclassified(100); | Physciaceae | 9 | Symbiotroph | Lichenized | Thallus | NULL | Highly Probable | Lichenized with green algae, often on nitrogen-rich substrata (Cannon & Kirk 2007) | Cannon PF, Kirk PM. 2007. Fungal Families of the World. CAB International, Cambridge (ISBN: 978-0851998275) |
| Otu05813 | 5 | k__Fungi(100);p__Ascomycota(100);c__Lecanoromycetes(100);o__Acarosporales(100);f__Acarosporaceae(100);g__Acarospora(100);g__Acarospora_unclassified(100); | Acarospora | 13 | Symbiotroph | \|Lichenized\| | Thallus | NULL | Highly Probable | NULL | Esslinger TL. 2014. North Dakota State University ((http://www.ndsu.edu/pubweb/~esslinge/chcklst/chcklst7.htm)); Põlme S, et al. 2020. Fungal Diversity 105:1-16 ((https://doi.org/10.1007/s13225-020-00466-2)) |
| Otu05814 | 5 | k__Fungi(100);p__Basidiomycota(100);c__Agaricomycetes(100);o__Agaricales(100);f__Mycenaceae(100);g__Cruentomycena(100);g__Cruentomycena_unclassified(100); | Cruentomycena | 13 | Saprotroph | \|Plant Saprotroph\|-Undefined Saprotroph | Agaricoid | NULL | Probable | As litter_saprotroph (Põlme et al. 2020); Decay leaf, fruit, seed (Põlme et al. 2020) | Tedersoo L, et al. 2014. Science 346:e1256688 ((https://doi.org/10.1126/science.1256688)); Põlme S, et al. 2020. Fungal Diversity 105:1-16 ((https://doi.org/10.1007/s13225-020-00466-2)) |
| Otu05818 | 5 | k__Fungi(100);p__Ascomycota(100);c__Eurotiomycetes(100);o__Chaetothyriales(100);f__Herpotrichiellaceae(100);g__Exophiala(100);g__Exophiala_unclassified(100); | Exophiala | 13 | Pathotroph-Saprotroph-Symbiotroph | \|Animal Parasite\|-Animal Pathogen-Endophyte-Plant Saprotroph-Undefined Saprotroph | Dimorphic-Facultative Yeast | NULL | Probable | As litter_saprotroph (Põlme et al. 2020); As root_endophyte_dark_septate (Põlme et al. 2020); Decay leaf, fruit, seed, animal material (Põlme et al. 2020) | Tedersoo L, et al. 2014. Science 346:e1256688 ((https://doi.org/10.1126/science.1256688)), Nagy LG, et al. 2014. Nature Communications 5:e4471 ((https://doi.org/10.1038/ncomms5471)); Põlme S, et al. 2020. Fungal Diversity 105:1-16 ((https://doi.org/10.1007/s13225-020-00466-2)) |
| Otu05819 | 5 | k__Fungi(100);p__Basidiomycota(100);c__Agaricomycetes(100);o__Agaricales(100);f__Psathyrellaceae(100);g__Coprinopsis(100);s__Coprinopsis_villosa(100); | Coprinopsis | 13 | Saprotroph | \|Undefined Saprotroph\| | Agaricoid | NULL | Probable | As soil_saprotroph (Põlme et al. 2020); Decay soil (Põlme et al. 2020) | Tedersoo L, et al. 2014. Science 346:e1256688 ((https://doi.org/10.1126/science.1256688)); Põlme S, et al. 2020. Fungal Diversity 105:1-16 ((https://doi.org/10.1007/s13225-020-00466-2)) |
| Otu05821 | 5 | k__Fungi(100);p__Basidiomycota(100);c__Tremellomycetes(100);o__Tremellales(100);o__Tremellales_unclassified(100);o__Tremellales_unclassified(100);o__Tremellales_unclassified(100); | Tremellales | 7 | Pathotroph-Saprotroph-Symbiotroph | Fungal Parasite-Undefined Saprotroph | Tremelloid-Yeast | NULL | Possible | The order contains both teleomorphic and anamorphic species, most of the latter being yeasts | Sterkenburg E, et al. 2015. New Phytologist 207:1145-1158 ((https://doi.org/10.1111/nph.13426)) |
| Otu05822 | 5 | k__Fungi(100);p__Ascomycota(100);c__Saccharomycetes(100);o__Saccharomycetales(100);f__Metschnikowiaceae(100);g__Metschnikowia(100);s__Metschnikowia_pulcherrima(100); | Metschnikowia | 13 | Saprotroph | \|Undefined Saprotroph\| | Yeast | NULL | Probable | As nectar-tap_saprotroph (Põlme et al. 2020); Decay sugar-rich substrates (Põlme et al. 2020) | Põlme S, et al. 2020. Fungal Diversity 105:1-16 ((https://doi.org/10.1007/s13225-020-00466-2)) |
| Otu05824 | 5 | k__Fungi(100);p__Basidiomycota(100);c__Agaricomycetes(100);o__Cantharellales(100);f__Ceratobasidiaceae(100);g__Rhizoctonia(100);s__Rhizoctonia_sp_TC3(100); | Rhizoctonia | 13 | Pathotroph-Saprotroph-Symbiotroph | Ectomycorrhizal-Orchid Mycorrhizal-\|Plant Pathogen\|-Plant Saprotroph-Wood Saprotroph | Corticioid-Microfungus | Soft Rot | Probable | As litter_saprotroph (Põlme et al. 2020); Decay leaf, fruit, seed, roots, soil (Põlme et al. 2020); Only symbiotic (perhaps leaning parasitic) in special cases with orchids, otherwise typically a plant pathogen. | Seehann G, et al. 1975. List of Fungi in Soft-Rot Tissue Tests. Institute fur Holzbiologie und Holzschutz, Bundesforschungsanstalt fur Forst - und Holzwirtshaft, Hamburg - Lohbrugge, Germany; Costa IPMW, et al. 2012. Mycotaxon 119:e26 ((http://mycotaxon.com/resources/checklists/costa_v119_checklist.pdf)); Tedersoo L, et al. 2014. Science 346:e1256688 ((https://doi.org/10.1126/science.1256688)); Põlme S, et al. 2020. Fungal Diversity 105:1-16 ((https://doi.org/10.1007/s13225-020-00466-2)); List compiled by Sara Branco with input from Nhu Nguyen and Else Vellinga; The American Phytopathological Society Common Names of Plant Diseases ((http://www.apsnet.org/publications/commonnames/Pages/default.aspx)) |
| Otu05827 | 5 | k__Fungi(100);p__Basidiomycota(100);c__Agaricomycetes(100);o__Polyporales(100);f__Meruliaceae(100);g__Hypochnicium(100);s__Hypochnicium_albostramineum(100); | Hypochnicium | 13 | Saprotroph | Undefined Saprotroph-\|Wood Saprotroph\| | Corticioid | White Rot | Probable | As white_rot (Põlme et al. 2020); Decay wood (Põlme et al. 2020) | Tedersoo L, et al. 2014. Science 346:e1256688 ((https://doi.org/10.1126/science.1256688)); Põlme S, et al. 2020. Fungal Diversity 105:1-16 ((https://doi.org/10.1007/s13225-020-00466-2)) |
| Otu05829 | 5 | k__Fungi(100);p__Ascomycota(100);c__Saccharomycetes(100);o__Saccharomycetales(100);f__Metschnikowiaceae(100);g__Metschnikowia(100);s__Metschnikowia_pulcherrima(100); | Metschnikowia | 13 | Saprotroph | \|Undefined Saprotroph\| | Yeast | NULL | Probable | As nectar-tap_saprotroph (Põlme et al. 2020); Decay sugar-rich substrates (Põlme et al. 2020) | Põlme S, et al. 2020. Fungal Diversity 105:1-16 ((https://doi.org/10.1007/s13225-020-00466-2)) |
| Otu05831 | 5 | k__Fungi(100);p__Zygomycota(100);c__Zygomycota_class_Incertae_sedis(100);o__Mortierellales(100);f__Mortierellaceae(100);g__Mortierella(100);s__Mortierella_cystojenkinii(100); | Mortierella | 13 | Saprotroph-Symbiotroph | Endophyte-Plant Saprotroph-\|Undefined Saprotroph\| | Microfungus | NULL | Probable | As chitinolytic (Põlme et al. 2020); As soil_saprotroph (Põlme et al. 2020); Decay roots, soil (Põlme et al. 2020); Saprobic in soil and thus frequenctly isolated from soil and material contaminated with soil such as hay or rotten silage and spoil tips (Cannon & Kirk 2007) | Cannon PF, Kirk PM. 2007. Fungal Families of the World. CAB International, Cambridge (ISBN: 978-0851998275); Tedersoo L, et al. 2014. Science 346:e1256688 ((https://doi.org/10.1126/science.1256688)); Purahong W, et al. 2016. Molecular Ecology 25:4059-4074 ((https://doi.org/10.1111/mec.13739)); Põlme S, et al. 2020. Fungal Diversity 105:1-16 ((https://doi.org/10.1007/s13225-020-00466-2)) |
| Otu05832 | 5 | k__Fungi(100);p__Basidiomycota(100);c__Microbotryomycetes(100);o__Sporidiobolales(100);f__Sporidiobolales_family_Incertae_sedis(100);g__Sporobolomyces(100);s__Sporobolomyces_lactosus(100); | Sporobolomyces | 13 | Pathotroph-Saprotroph | \|Fungal Parasite\|-Plant Saprotroph | Yeast | NULL | Probable | As litter_saprotroph (Põlme et al. 2020); As mycoparasite (Põlme et al. 2020); Decay fungal material (Põlme et al. 2020) | Tedersoo L, et al. 2014. Science 346:e1256688 ((https://doi.org/10.1126/science.1256688)); Purahong W, et al. 2016. Molecular Ecology 25:4059-4074 ((https://doi.org/10.1111/mec.13739)); Põlme S, et al. 2020. Fungal Diversity 105:1-16 ((https://doi.org/10.1007/s13225-020-00466-2)) |
| Otu05833 | 5 | k__Fungi(100);p__Basidiomycota(100);c__Agaricomycetes(100);o__Agaricales(100);f__Clavariaceae(100);g__Clavaria(100);s__Clavaria_acuta(100); | Clavaria | 13 | Saprotroph-Symbiotroph | \|Undefined Saprotroph\|-Undefined Symbiotroph | Clavarioid | NULL | Probable | As soil_saprotroph (Põlme et al. 2020); Decay roots, soil (Põlme et al. 2020) | Tedersoo L, et al. 2014. Science 346:e1256688 ((https://doi.org/10.1126/science.1256688)); Põlme S, et al. 2020. Fungal Diversity 105:1-16 ((https://doi.org/10.1007/s13225-020-00466-2)) |
| Otu05837 | 5 | k__Fungi(100);p__Basidiomycota(100);c__Agaricomycetes(100);o__Agaricales(100);o__Agaricales_unclassified(100);o__Agaricales_unclassified(100);o__Agaricales_unclassified(100); | Agaricales | 7 | Pathotroph-Saprotroph-Symbiotroph | Bryophyte Parasite-Dung Saprotroph-Ectomycorrhizal-Fungal Parasite-Leaf Saprotroph-Plant Parasite-Undefined Saprotroph-Wood Saprotroph | Agaricoid-Gasteroid-Microfungus-Secotioid-Yeast | NULL | Possible | Mushrooms and toadstools, Gill fungi, Agarics... lignicolous, sometimes muscicolous or fungicolous, saprobic, mycorrhizal, rarely parasitic on plants or fungi (Kirk et al. 2008) | Kirk PM et al. 2008. Dictionary of the Fungi. Tenth Edition. CAB International, Wallingford (ISBN: 978-0851998268) |
| Otu05839 | 5 | k__Fungi(100);p__Ascomycota(100);c__Dothideomycetes(100);o__Pleosporales(100);f__Phaeosphaeriaceae(100);g__Phaeosphaeria(100);s__Phaeosphaeria_triglochinicola(100); | Phaeosphaeria | 13 | Pathotroph-Saprotroph-Symbiotroph | Endophyte-Plant Pathogen-\|Plant Saprotroph\| | Microfungus | NULL | Probable | As litter_saprotroph (Põlme et al. 2020); Decay leaf, fruit, seed (Põlme et al. 2020) | Tedersoo L, et al. 2014. Science 346:e1256688 ((https://doi.org/10.1126/science.1256688));David AS, et al. 2016. Microbial Ecology 71: 912-926 ((https://doi.org/10.1007/s00248-015-0712-6)); Põlme S, et al. 2020. Fungal Diversity 105:1-16 ((https://doi.org/10.1007/s13225-020-00466-2)) |
| Otu05840 | 5 | k__Fungi(100);p__Ascomycota(100);c__Eurotiomycetes(100);o__Onygenales(100);f__Gymnoascaceae(100);g__Gymnoascus(100);s__Gymnoascus_sp_02NH07(100); | Gymnoascus | 13 | Saprotroph | \|Undefined Saprotroph\| | Microfungus | NULL | Probable | As soil_saprotroph (Põlme et al. 2020); Decay soil (Põlme et al. 2020) | Tedersoo L, et al. 2014. Science 346:e1256688 ((https://doi.org/10.1126/science.1256688)); Põlme S, et al. 2020. Fungal Diversity 105:1-16 ((https://doi.org/10.1007/s13225-020-00466-2)) |
| Otu05841 | 5 | k__Fungi(100);p__Basidiomycota(100);c__Agaricomycetes(100);o__Thelephorales(100);f__Thelephoraceae(100);g__Tomentella(100);g__Tomentella_unclassified(100); | Tomentella | 13 | Symbiotroph | \|Ectomycorrhizal\| | Corticioid | NULL | Highly Probable | NULL | Rinaldi AC, et al. 2008. Fungal Diversity 33:1-45 ((http://www.fungaldiversity.org/fdp/sfdp/33-1.pdf)); Tedersoo L, et al. 2010. Mycorrhiza 20:217-263 ((https://doi.org/10.1007/s00572-009-0274-x)); Põlme S, et al. 2020. Fungal Diversity 105:1-16 ((https://doi.org/10.1007/s13225-020-00466-2)); http://mycorrhizas.info/ecmf.html ((http://mycorrhizas.info/ecmf.html)) |
| Otu05842 | 5 | k__Fungi(100);p__Ascomycota(100);c__Sordariomycetes(100);o__Hypocreales(100);o__Hypocreales_unclassified(100);o__Hypocreales_unclassified(100);o__Hypocreales_unclassified(100); | Hypocreales | 7 | Saprotroph | Undefined Saprotroph | Microfungus | NULL | Possible | NULL | Sterkenburg E, et al. 2015. New Phytologist 207:1145-1158 ((https://doi.org/10.1111/nph.13426)) |
| Otu05844 | 5 | k__Fungi(100);p__Ascomycota(100);c__Saccharomycetes(100);o__Saccharomycetales(100);f__Metschnikowiaceae(100);g__Metschnikowia(100);s__Metschnikowia_pulcherrima(80); | Metschnikowia | 13 | Saprotroph | \|Undefined Saprotroph\| | Yeast | NULL | Probable | As nectar-tap_saprotroph (Põlme et al. 2020); Decay sugar-rich substrates (Põlme et al. 2020) | Põlme S, et al. 2020. Fungal Diversity 105:1-16 ((https://doi.org/10.1007/s13225-020-00466-2)) |
| Otu05847 | 5 | k__Fungi(100);p__Ascomycota(100);c__Saccharomycetes(100);o__Saccharomycetales(100);f__Metschnikowiaceae(100);g__Metschnikowia(100);s__Metschnikowia_pulcherrima(100); | Metschnikowia | 13 | Saprotroph | \|Undefined Saprotroph\| | Yeast | NULL | Probable | As nectar-tap_saprotroph (Põlme et al. 2020); Decay sugar-rich substrates (Põlme et al. 2020) | Põlme S, et al. 2020. Fungal Diversity 105:1-16 ((https://doi.org/10.1007/s13225-020-00466-2)) |
| Otu05849 | 5 | k__Fungi(100);p__Chytridiomycota(100);c__Chytridiomycetes(100);o__Rhizophlyctidales(100);f__Rhizophlyctidaceae(100);g__Rhizophlyctis(100);s__Rhizophlyctis_rosea(100); | Rhizophlyctis | 13 | Pathotroph-Saprotroph | Algal Parasite-\|Plant Saprotroph\|-Undefined Saprotroph | Chytrid-Microfungus | NULL | Probable | As litter_saprotroph (Põlme et al. 2020); Decay leaf, fruit, seed, algal material (Põlme et al. 2020) | James TY, et al. 2006. Nature 443:818-822 ((https://doi.org/10.1038/nature05110)); Põlme S, et al. 2020. Fungal Diversity 105:1-16 ((https://doi.org/10.1007/s13225-020-00466-2)) |
| Otu05850 | 5 | k__Fungi(100);p__Ascomycota(100);c__Dothideomycetes(100);o__Pleosporales(100);f__Phaeosphaeriaceae(100);f__Phaeosphaeriaceae_unclassified(100);f__Phaeosphaeriaceae_unclassified(100); | Phaeosphaeriaceae | 9 | Pathotroph-Saprotroph | Fungal Parasite-Plant Pathogen-Plant Saprotroph | Microfungus | NULL | Probable | Necrotrophic or saprobic on a wide range of plants, especially monocotyledons; a few species parasitic on other fungi (Cannon & Kirk 2007) | Cannon PF, Kirk PM. 2007. Fungal Families of the World. CAB International, Cambridge (ISBN: 978-0851998275) |
| Otu05855 | 5 | k__Fungi(100);p__Ascomycota(100);c__Eurotiomycetes(100);o__Chaetothyriales(100);f__Herpotrichiellaceae(100);g__Exophiala(100);g__Exophiala_unclassified(100); | Exophiala | 13 | Pathotroph-Saprotroph-Symbiotroph | \|Animal Parasite\|-Animal Pathogen-Endophyte-Plant Saprotroph-Undefined Saprotroph | Dimorphic-Facultative Yeast | NULL | Probable | As litter_saprotroph (Põlme et al. 2020); As root_endophyte_dark_septate (Põlme et al. 2020); Decay leaf, fruit, seed, animal material (Põlme et al. 2020) | Tedersoo L, et al. 2014. Science 346:e1256688 ((https://doi.org/10.1126/science.1256688)), Nagy LG, et al. 2014. Nature Communications 5:e4471 ((https://doi.org/10.1038/ncomms5471)); Põlme S, et al. 2020. Fungal Diversity 105:1-16 ((https://doi.org/10.1007/s13225-020-00466-2)) |
| Otu05859 | 5 | k__Fungi(100);p__Ascomycota(100);c__Saccharomycetes(100);o__Saccharomycetales(100);f__Metschnikowiaceae(100);g__Metschnikowia(100);s__Metschnikowia_pulcherrima(100); | Metschnikowia | 13 | Saprotroph | \|Undefined Saprotroph\| | Yeast | NULL | Probable | As nectar-tap_saprotroph (Põlme et al. 2020); Decay sugar-rich substrates (Põlme et al. 2020) | Põlme S, et al. 2020. Fungal Diversity 105:1-16 ((https://doi.org/10.1007/s13225-020-00466-2)) |
| Otu05860 | 5 | k__Fungi(100);p__Ascomycota(100);c__Saccharomycetes(100);o__Saccharomycetales(100);f__Metschnikowiaceae(100);g__Metschnikowia(100);s__Metschnikowia_pulcherrima(100); | Metschnikowia | 13 | Saprotroph | \|Undefined Saprotroph\| | Yeast | NULL | Probable | As nectar-tap_saprotroph (Põlme et al. 2020); Decay sugar-rich substrates (Põlme et al. 2020) | Põlme S, et al. 2020. Fungal Diversity 105:1-16 ((https://doi.org/10.1007/s13225-020-00466-2)) |
| Otu05862 | 5 | k__Fungi(100);p__Ascomycota(100);c__Sordariomycetes(100);o__Xylariales(100);f__Xylariaceae(100);g__Xylaria(100);s__Xylaria_oxyacanthae(100); | Xylaria | 13 | Saprotroph-Symbiotroph | Endophyte-Plant Saprotroph-Undefined Saprotroph-\|Wood Saprotroph\| | Xylarioid | Soft Rot | Probable | As foliar_endophyte (Põlme et al. 2020); As litter_saprotroph (Põlme et al. 2020); Decay wood, leaf, fruit, seed (Põlme et al. 2020); Host - angiosperms (Põlme et al. 2020) | Duncan CG, Eslyn WE. 1966. Mycologia 58:642-645 ((https://doi.org/10.2307/3757045)); Costa IPMW, et al. 2012. Mycotaxon 119:e26 ((http://mycotaxon.com/resources/checklists/costa_v119_checklist.pdf)); Põlme S, et al. 2020. Fungal Diversity 105:1-16 ((https://doi.org/10.1007/s13225-020-00466-2)); The American Phytopathological Society Common Names of Plant Diseases ((http://www.apsnet.org/publications/commonnames/Pages/default.aspx)) |
| Otu05863 | 5 | k__Fungi(100);p__Basidiomycota(100);c__Agaricomycetes(100);o__Cantharellales(100);f__Tulasnellaceae(100);g__Tulasnella(100);s__Tulasnella_irregularis(100); | Tulasnella | 13 | Saprotroph-Symbiotroph | Ectomycorrhizal-\|Plant Saprotroph\|-Undefined Saprotroph | Corticioid | NULL | Possible | As litter_saprotroph (Põlme et al. 2020); Decay leaf, fruit, seed (Põlme et al. 2020) | Rinaldi AC, et al. 2008. Fungal Diversity 33:1-45 ((http://www.fungaldiversity.org/fdp/sfdp/33-1.pdf)); Tedersoo L, et al. 2010. Mycorrhiza 20:217-263 ((https://doi.org/10.1007/s00572-009-0274-x)) (pro parte); Põlme S, et al. 2020. Fungal Diversity 105:1-16 ((https://doi.org/10.1007/s13225-020-00466-2)) |
| Otu05865 | 5 | k__Fungi(100);p__Ascomycota(100);c__Saccharomycetes(100);o__Saccharomycetales(100);f__Metschnikowiaceae(100);g__Metschnikowia(100);s__Metschnikowia_pulcherrima(100); | Metschnikowia | 13 | Saprotroph | \|Undefined Saprotroph\| | Yeast | NULL | Probable | As nectar-tap_saprotroph (Põlme et al. 2020); Decay sugar-rich substrates (Põlme et al. 2020) | Põlme S, et al. 2020. Fungal Diversity 105:1-16 ((https://doi.org/10.1007/s13225-020-00466-2)) |
| Otu05867 | 5 | k__Fungi(100);p__Ascomycota(100);c__Dothideomycetes(100);o__Botryosphaeriales(100);f__Botryosphaeriaceae(100);g__Botryosphaeria(100);s__Botryosphaeria_agaves(100); | Botryosphaeria | 13 | Pathotroph-Saprotroph-Symbiotroph | Endophyte-\|Plant Pathogen\|-Plant Saprotroph | Microfungus | NULL | Probable | As foliar_endophyte (Põlme et al. 2020); As litter_saprotroph (Põlme et al. 2020); Decay leaf, fruit, seed (Põlme et al. 2020) | Tedersoo L, et al. 2014. Science 346:e1256688 ((https://doi.org/10.1126/science.1256688)); Christian N, et al. 2016. Microbial Ecology 72: 621-632 ((https://doi.org/10.1007/s00248-016-0804-y)); Põlme S, et al. 2020. Fungal Diversity 105:1-16 ((https://doi.org/10.1007/s13225-020-00466-2)); The American Phytopathological Society Common Names of Plant Diseases ((http://www.apsnet.org/publications/commonnames/Pages/default.aspx)) |
| Otu05868 | 5 | k__Fungi(100);p__Ascomycota(100);c__Saccharomycetes(100);o__Saccharomycetales(100);f__Metschnikowiaceae(100);g__Metschnikowia(100);s__Metschnikowia_pulcherrima(80); | Metschnikowia | 13 | Saprotroph | \|Undefined Saprotroph\| | Yeast | NULL | Probable | As nectar-tap_saprotroph (Põlme et al. 2020); Decay sugar-rich substrates (Põlme et al. 2020) | Põlme S, et al. 2020. Fungal Diversity 105:1-16 ((https://doi.org/10.1007/s13225-020-00466-2)) |
| Otu05869 | 5 | k__Fungi(100);p__Ascomycota(100);c__Leotiomycetes(100);o__Rhytismatales(100);f__Rhytismataceae(100);g__unclassified_Rhytismataceae(100);s__Rhytismataceae_sp(100); | Rhytismataceae | 9 | Pathotroph-Saprotroph-Symbiotroph | Endophyte-Leaf Saprotroph-Plant Pathogen | Microfungus | NULL | Probable | Biotrophic, necrotrophic or saprobic on leaves and bark, often initially as endophytes, causing tar spots, premature leaf fall and occasionally witches broom symptoms (Cannon & Kirk 2007) | Cannon PF, Kirk PM. 2007. Fungal Families of the World. CAB International, Cambridge (ISBN: 978-0851998275) |
| Otu05870 | 5 | k__Fungi(100);p__Ascomycota(100);c__Taphrinomycetes(100);o__Taphrinales(100);f__Taphrinaceae(100);g__unclassified_Taphrinaceae(100);s__Taphrinaceae_sp(100); | Taphrinaceae | 9 | Pathotroph | Plant Pathogen | Microfungus | NULL | Probable | Biotrophic on plants, usually causing hyperplasia (galls, witches broom) or lesions (Cannon & Kirk 2007) | Cannon PF, Kirk PM. 2007. Fungal Families of the World. CAB International, Cambridge (ISBN: 978-0851998275) |
| Otu05871 | 5 | k__Fungi(100);p__Ascomycota(100);c__Saccharomycetes(100);o__Saccharomycetales(100);f__Metschnikowiaceae(100);g__Metschnikowia(100);s__Metschnikowia_pulcherrima(80); | Metschnikowia | 13 | Saprotroph | \|Undefined Saprotroph\| | Yeast | NULL | Probable | As nectar-tap_saprotroph (Põlme et al. 2020); Decay sugar-rich substrates (Põlme et al. 2020) | Põlme S, et al. 2020. Fungal Diversity 105:1-16 ((https://doi.org/10.1007/s13225-020-00466-2)) |
| Otu05873 | 5 | k__Fungi(100);p__Basidiomycota(100);c__Tremellomycetes(100);o__Tremellales(100);f__Tremellales_family_Incertae_sedis(100);g__Tremella(100);s__Tremella_candelariellae(100); | Tremella | 13 | Pathotroph | \|Fungal Parasite\|-Lichen Parasite | Dimorphic-Facultative Yeast-Microfungus-Tremelloid | NULL | Probable | As mycoparasite (Põlme et al. 2020); Decay fungal material (Põlme et al. 2020) | Tedersoo L, et al. 2014. Science 346:e1256688 ((https://doi.org/10.1126/science.1256688)); Põlme S, et al. 2020. Fungal Diversity 105:1-16 ((https://doi.org/10.1007/s13225-020-00466-2)) |
| Otu05874 | 5 | k__Fungi(100);p__Ascomycota(100);c__Saccharomycetes(100);o__Saccharomycetales(100);f__Metschnikowiaceae(100);g__Metschnikowia(100);s__Metschnikowia_pulcherrima(100); | Metschnikowia | 13 | Saprotroph | \|Undefined Saprotroph\| | Yeast | NULL | Probable | As nectar-tap_saprotroph (Põlme et al. 2020); Decay sugar-rich substrates (Põlme et al. 2020) | Põlme S, et al. 2020. Fungal Diversity 105:1-16 ((https://doi.org/10.1007/s13225-020-00466-2)) |
| Otu05875 | 5 | k__Fungi(100);p__Ascomycota(100);c__Sordariomycetes(100);o__Diaporthales(100);f__Melanconidaceae(100);g__Melanconis(100);s__Melanconis_desmazieri(100); | Melanconis | 13 | Pathotroph | \|Plant Pathogen\| | Microfungus | NULL | Probable | NULL | Tedersoo L, et al. 2014. Science 346:e1256688 ((https://doi.org/10.1126/science.1256688)); Põlme S, et al. 2020. Fungal Diversity 105:1-16 ((https://doi.org/10.1007/s13225-020-00466-2)) |
| Otu05876 | 5 | k__Fungi(100);p__Basidiomycota(80);c__Microbotryomycetes(80);o__Sporidiobolales(80);f__Sporidiobolales_family_Incertae_sedis(80);g__Sporobolomyces(80);s__Sporobolomyces_lactosus(80); | Sporobolomyces | 13 | Pathotroph-Saprotroph | \|Fungal Parasite\|-Plant Saprotroph | Yeast | NULL | Probable | As litter_saprotroph (Põlme et al. 2020); As mycoparasite (Põlme et al. 2020); Decay fungal material (Põlme et al. 2020) | Tedersoo L, et al. 2014. Science 346:e1256688 ((https://doi.org/10.1126/science.1256688)); Purahong W, et al. 2016. Molecular Ecology 25:4059-4074 ((https://doi.org/10.1111/mec.13739)); Põlme S, et al. 2020. Fungal Diversity 105:1-16 ((https://doi.org/10.1007/s13225-020-00466-2)) |
| Otu05877 | 5 | k__Fungi(100);p__Ascomycota(100);c__Saccharomycetes(100);o__Saccharomycetales(100);f__Metschnikowiaceae(100);g__Metschnikowia(100);g__Metschnikowia_unclassified(80); | Metschnikowia | 13 | Saprotroph | \|Undefined Saprotroph\| | Yeast | NULL | Probable | As nectar-tap_saprotroph (Põlme et al. 2020); Decay sugar-rich substrates (Põlme et al. 2020) | Põlme S, et al. 2020. Fungal Diversity 105:1-16 ((https://doi.org/10.1007/s13225-020-00466-2)) |
| Otu05878 | 5 | k__Fungi(100);p__Ascomycota(100);c__Saccharomycetes(100);o__Saccharomycetales(100);f__Metschnikowiaceae(100);g__Metschnikowia(100);g__Metschnikowia_unclassified(61); | Metschnikowia | 13 | Saprotroph | \|Undefined Saprotroph\| | Yeast | NULL | Probable | As nectar-tap_saprotroph (Põlme et al. 2020); Decay sugar-rich substrates (Põlme et al. 2020) | Põlme S, et al. 2020. Fungal Diversity 105:1-16 ((https://doi.org/10.1007/s13225-020-00466-2)) |
| Otu05152 | 5 | k__Fungi(100);p__Basidiomycota(100);p__Basidiomycota_unclassified(100);p__Basidiomycota_unclassified(100);p__Basidiomycota_unclassified(100);p__Basidiomycota_unclassified(100);p__Basidiomycota_unclassified(100); | - | - | - | - | - | - | - | Unassigned | - |
| Otu05153 | 5 | k__Plantae(100);p__unclassified_Plantae(100);c__unclassified_Plantae(100);o__unclassified_Plantae(100);f__unclassified_Plantae(100);g__unclassified_Plantae(100);s__Plantae_sp(100); | - | - | - | - | - | - | - | Unassigned | - |
| Otu05154 | 5 | k__Fungi(100);p__Ascomycota(61);p__Ascomycota_unclassified(61);p__Ascomycota_unclassified(61);p__Ascomycota_unclassified(61);p__Ascomycota_unclassified(61);p__Ascomycota_unclassified(61); | - | - | - | - | - | - | - | Unassigned | - |
| Otu05155 | 5 | k__Fungi(100);k__Fungi_unclassified(61);k__Fungi_unclassified(61);k__Fungi_unclassified(61);k__Fungi_unclassified(61);k__Fungi_unclassified(61);k__Fungi_unclassified(61); | - | - | - | - | - | - | - | Unassigned | - |
| Otu05157 | 5 | k__Fungi(100);k__Fungi_unclassified(61);k__Fungi_unclassified(61);k__Fungi_unclassified(61);k__Fungi_unclassified(61);k__Fungi_unclassified(61);k__Fungi_unclassified(61); | - | - | - | - | - | - | - | Unassigned | - |
| Otu05160 | 5 | k__Fungi(100);k__Fungi_unclassified(61);k__Fungi_unclassified(61);k__Fungi_unclassified(61);k__Fungi_unclassified(61);k__Fungi_unclassified(61);k__Fungi_unclassified(61); | - | - | - | - | - | - | - | Unassigned | - |
| Otu05161 | 5 | k__Fungi(100);k__Fungi_unclassified(61);k__Fungi_unclassified(61);k__Fungi_unclassified(61);k__Fungi_unclassified(61);k__Fungi_unclassified(61);k__Fungi_unclassified(61); | - | - | - | - | - | - | - | Unassigned | - |
| Otu05162 | 5 | k__Fungi(100);k__Fungi_unclassified(61);k__Fungi_unclassified(61);k__Fungi_unclassified(61);k__Fungi_unclassified(61);k__Fungi_unclassified(61);k__Fungi_unclassified(61); | - | - | - | - | - | - | - | Unassigned | - |
| Otu05163 | 5 | k__Fungi(100);k__Fungi_unclassified(100);k__Fungi_unclassified(100);k__Fungi_unclassified(100);k__Fungi_unclassified(100);k__Fungi_unclassified(100);k__Fungi_unclassified(100); | - | - | - | - | - | - | - | Unassigned | - |
| Otu05165 | 5 | k__Fungi(100);p__Ascomycota(80);p__Ascomycota_unclassified(80);p__Ascomycota_unclassified(80);p__Ascomycota_unclassified(80);p__Ascomycota_unclassified(80);p__Ascomycota_unclassified(80); | - | - | - | - | - | - | - | Unassigned | - |
| Otu05166 | 5 | k__Fungi(100);p__Ascomycota(100);c__Dothideomycetes(100);o__Pleosporales(100);o__Pleosporales_unclassified(100);o__Pleosporales_unclassified(100);o__Pleosporales_unclassified(100); | - | - | - | - | - | - | - | Unassigned | - |
| Otu05167 | 5 | k__Fungi(100);k__Fungi_unclassified(100);k__Fungi_unclassified(100);k__Fungi_unclassified(100);k__Fungi_unclassified(100);k__Fungi_unclassified(100);k__Fungi_unclassified(100); | - | - | - | - | - | - | - | Unassigned | - |
| Otu05168 | 5 | k__Fungi(100);p__Ascomycota(100);c__Dothideomycetes(100);o__Pleosporales(100);o__Pleosporales_unclassified(100);o__Pleosporales_unclassified(100);o__Pleosporales_unclassified(100); | - | - | - | - | - | - | - | Unassigned | - |
| Otu05169 | 5 | k__Fungi(100);p__Ascomycota(80);p__Ascomycota_unclassified(80);p__Ascomycota_unclassified(80);p__Ascomycota_unclassified(80);p__Ascomycota_unclassified(80);p__Ascomycota_unclassified(80); | - | - | - | - | - | - | - | Unassigned | - |
| Otu05176 | 5 | k__Fungi(100);k__Fungi_unclassified(80);k__Fungi_unclassified(80);k__Fungi_unclassified(80);k__Fungi_unclassified(80);k__Fungi_unclassified(80);k__Fungi_unclassified(80); | - | - | - | - | - | - | - | Unassigned | - |
| Otu05178 | 5 | k__Fungi(100);p__Ascomycota(100);p__Ascomycota_unclassified(100);p__Ascomycota_unclassified(100);p__Ascomycota_unclassified(100);p__Ascomycota_unclassified(100);p__Ascomycota_unclassified(100); | - | - | - | - | - | - | - | Unassigned | - |
| Otu05179 | 5 | k__Fungi(100);p__Ascomycota(61);p__Ascomycota_unclassified(61);p__Ascomycota_unclassified(61);p__Ascomycota_unclassified(61);p__Ascomycota_unclassified(61);p__Ascomycota_unclassified(61); | - | - | - | - | - | - | - | Unassigned | - |
| Otu05181 | 5 | k__Fungi(100);k__Fungi_unclassified(100);k__Fungi_unclassified(100);k__Fungi_unclassified(100);k__Fungi_unclassified(100);k__Fungi_unclassified(100);k__Fungi_unclassified(100); | - | - | - | - | - | - | - | Unassigned | - |
| Otu05182 | 5 | k__Fungi(100);k__Fungi_unclassified(61);k__Fungi_unclassified(61);k__Fungi_unclassified(61);k__Fungi_unclassified(61);k__Fungi_unclassified(61);k__Fungi_unclassified(61); | - | - | - | - | - | - | - | Unassigned | - |
| Otu05183 | 5 | k__Fungi(100);p__Ascomycota(100);c__Sordariomycetes(100);o__Diaporthales(80);o__Diaporthales_unclassified(80);o__Diaporthales_unclassified(80);o__Diaporthales_unclassified(80); | - | - | - | - | - | - | - | Unassigned | - |
| Otu05184 | 5 | k__Fungi(100);p__Ascomycota(80);p__Ascomycota_unclassified(80);p__Ascomycota_unclassified(80);p__Ascomycota_unclassified(80);p__Ascomycota_unclassified(80);p__Ascomycota_unclassified(80); | - | - | - | - | - | - | - | Unassigned | - |
| Otu05186 | 5 | k__Fungi(100);k__Fungi_unclassified(80);k__Fungi_unclassified(80);k__Fungi_unclassified(80);k__Fungi_unclassified(80);k__Fungi_unclassified(80);k__Fungi_unclassified(80); | - | - | - | - | - | - | - | Unassigned | - |
| Otu05187 | 5 | k__Fungi(100);p__Ascomycota(61);p__Ascomycota_unclassified(61);p__Ascomycota_unclassified(61);p__Ascomycota_unclassified(61);p__Ascomycota_unclassified(61);p__Ascomycota_unclassified(61); | - | - | - | - | - | - | - | Unassigned | - |
| Otu05188 | 5 | k__Fungi(100);k__Fungi_unclassified(61);k__Fungi_unclassified(61);k__Fungi_unclassified(61);k__Fungi_unclassified(61);k__Fungi_unclassified(61);k__Fungi_unclassified(61); | - | - | - | - | - | - | - | Unassigned | - |
| Otu05190 | 5 | k__Fungi(100);p__Ascomycota(61);p__Ascomycota_unclassified(61);p__Ascomycota_unclassified(61);p__Ascomycota_unclassified(61);p__Ascomycota_unclassified(61);p__Ascomycota_unclassified(61); | - | - | - | - | - | - | - | Unassigned | - |
| Otu05192 | 5 | k__Fungi(100);k__Fungi_unclassified(100);k__Fungi_unclassified(100);k__Fungi_unclassified(100);k__Fungi_unclassified(100);k__Fungi_unclassified(100);k__Fungi_unclassified(100); | - | - | - | - | - | - | - | Unassigned | - |
| Otu05193 | 5 | k__Fungi(100);k__Fungi_unclassified(80);k__Fungi_unclassified(80);k__Fungi_unclassified(80);k__Fungi_unclassified(80);k__Fungi_unclassified(80);k__Fungi_unclassified(80); | - | - | - | - | - | - | - | Unassigned | - |
| Otu05196 | 5 | k__Fungi(100);k__Fungi_unclassified(80);k__Fungi_unclassified(80);k__Fungi_unclassified(80);k__Fungi_unclassified(80);k__Fungi_unclassified(80);k__Fungi_unclassified(80); | - | - | - | - | - | - | - | Unassigned | - |
| Otu05198 | 5 | k__Fungi(100);p__Ascomycota(80);p__Ascomycota_unclassified(80);p__Ascomycota_unclassified(80);p__Ascomycota_unclassified(80);p__Ascomycota_unclassified(80);p__Ascomycota_unclassified(80); | - | - | - | - | - | - | - | Unassigned | - |
| Otu05200 | 5 | k__Fungi(100);p__Ascomycota(100);p__Ascomycota_unclassified(100);p__Ascomycota_unclassified(100);p__Ascomycota_unclassified(100);p__Ascomycota_unclassified(100);p__Ascomycota_unclassified(100); | - | - | - | - | - | - | - | Unassigned | - |
| Otu05201 | 5 | k__Fungi(100);p__Ascomycota(100);p__Ascomycota_unclassified(100);p__Ascomycota_unclassified(100);p__Ascomycota_unclassified(100);p__Ascomycota_unclassified(100);p__Ascomycota_unclassified(100); | - | - | - | - | - | - | - | Unassigned | - |
| Otu05202 | 5 | k__Fungi(100);p__Ascomycota(61);p__Ascomycota_unclassified(61);p__Ascomycota_unclassified(61);p__Ascomycota_unclassified(61);p__Ascomycota_unclassified(61);p__Ascomycota_unclassified(61); | - | - | - | - | - | - | - | Unassigned | - |
| Otu05203 | 5 | k__Fungi(100);k__Fungi_unclassified(100);k__Fungi_unclassified(100);k__Fungi_unclassified(100);k__Fungi_unclassified(100);k__Fungi_unclassified(100);k__Fungi_unclassified(100); | - | - | - | - | - | - | - | Unassigned | - |
| Otu05205 | 5 | k__Fungi(100);p__Ascomycota(100);p__Ascomycota_unclassified(100);p__Ascomycota_unclassified(100);p__Ascomycota_unclassified(100);p__Ascomycota_unclassified(100);p__Ascomycota_unclassified(100); | - | - | - | - | - | - | - | Unassigned | - |
| Otu05207 | 5 | k__Fungi(100);p__Ascomycota(100);p__Ascomycota_unclassified(100);p__Ascomycota_unclassified(100);p__Ascomycota_unclassified(100);p__Ascomycota_unclassified(100);p__Ascomycota_unclassified(100); | - | - | - | - | - | - | - | Unassigned | - |
| Otu05209 | 5 | k__Fungi(100);p__Ascomycota(80);p__Ascomycota_unclassified(80);p__Ascomycota_unclassified(80);p__Ascomycota_unclassified(80);p__Ascomycota_unclassified(80);p__Ascomycota_unclassified(80); | - | - | - | - | - | - | - | Unassigned | - |
| Otu05216 | 5 | k__Fungi(100);k__Fungi_unclassified(100);k__Fungi_unclassified(100);k__Fungi_unclassified(100);k__Fungi_unclassified(100);k__Fungi_unclassified(100);k__Fungi_unclassified(100); | - | - | - | - | - | - | - | Unassigned | - |
| Otu05219 | 5 | k__Fungi(100);k__Fungi_unclassified(80);k__Fungi_unclassified(80);k__Fungi_unclassified(80);k__Fungi_unclassified(80);k__Fungi_unclassified(80);k__Fungi_unclassified(80); | - | - | - | - | - | - | - | Unassigned | - |
| Otu05220 | 5 | k__Fungi(100);p__Ascomycota(100);c__Sordariomycetes(100);c__Sordariomycetes_unclassified(80);c__Sordariomycetes_unclassified(80);c__Sordariomycetes_unclassified(80);c__Sordariomycetes_unclassified(80); | - | - | - | - | - | - | - | Unassigned | - |
| Otu05221 | 5 | k__Fungi(100);p__Ascomycota(80);p__Ascomycota_unclassified(80);p__Ascomycota_unclassified(80);p__Ascomycota_unclassified(80);p__Ascomycota_unclassified(80);p__Ascomycota_unclassified(80); | - | - | - | - | - | - | - | Unassigned | - |
| Otu05222 | 5 | k__Fungi(100);p__Ascomycota(80);p__Ascomycota_unclassified(80);p__Ascomycota_unclassified(80);p__Ascomycota_unclassified(80);p__Ascomycota_unclassified(80);p__Ascomycota_unclassified(80); | - | - | - | - | - | - | - | Unassigned | - |
| Otu05223 | 5 | k__Fungi(100);k__Fungi_unclassified(61);k__Fungi_unclassified(61);k__Fungi_unclassified(61);k__Fungi_unclassified(61);k__Fungi_unclassified(61);k__Fungi_unclassified(61); | - | - | - | - | - | - | - | Unassigned | - |
| Otu05225 | 5 | k__Fungi(100);p__Ascomycota(100);p__Ascomycota_unclassified(100);p__Ascomycota_unclassified(100);p__Ascomycota_unclassified(100);p__Ascomycota_unclassified(100);p__Ascomycota_unclassified(100); | - | - | - | - | - | - | - | Unassigned | - |
| Otu05227 | 5 | k__Fungi(100);p__Ascomycota(100);p__Ascomycota_unclassified(100);p__Ascomycota_unclassified(100);p__Ascomycota_unclassified(100);p__Ascomycota_unclassified(100);p__Ascomycota_unclassified(100); | - | - | - | - | - | - | - | Unassigned | - |
| Otu05235 | 5 | k__Fungi(100);p__Ascomycota(100);p__Ascomycota_unclassified(100);p__Ascomycota_unclassified(100);p__Ascomycota_unclassified(100);p__Ascomycota_unclassified(100);p__Ascomycota_unclassified(100); | - | - | - | - | - | - | - | Unassigned | - |
| Otu05237 | 5 | k__Fungi(100);k__Fungi_unclassified(61);k__Fungi_unclassified(61);k__Fungi_unclassified(61);k__Fungi_unclassified(61);k__Fungi_unclassified(61);k__Fungi_unclassified(61); | - | - | - | - | - | - | - | Unassigned | - |
| Otu05239 | 5 | k__Fungi(100);p__Ascomycota(61);p__Ascomycota_unclassified(61);p__Ascomycota_unclassified(61);p__Ascomycota_unclassified(61);p__Ascomycota_unclassified(61);p__Ascomycota_unclassified(61); | - | - | - | - | - | - | - | Unassigned | - |
| Otu05240 | 5 | k__Fungi(100);k__Fungi_unclassified(80);k__Fungi_unclassified(80);k__Fungi_unclassified(80);k__Fungi_unclassified(80);k__Fungi_unclassified(80);k__Fungi_unclassified(80); | - | - | - | - | - | - | - | Unassigned | - |
| Otu05244 | 5 | k__Fungi(100);p__Ascomycota(61);p__Ascomycota_unclassified(61);p__Ascomycota_unclassified(61);p__Ascomycota_unclassified(61);p__Ascomycota_unclassified(61);p__Ascomycota_unclassified(61); | - | - | - | - | - | - | - | Unassigned | - |
| Otu05246 | 5 | k__Fungi(100);p__Ascomycota(61);p__Ascomycota_unclassified(61);p__Ascomycota_unclassified(61);p__Ascomycota_unclassified(61);p__Ascomycota_unclassified(61);p__Ascomycota_unclassified(61); | - | - | - | - | - | - | - | Unassigned | - |
| Otu05247 | 5 | k__Fungi(100);k__Fungi_unclassified(61);k__Fungi_unclassified(61);k__Fungi_unclassified(61);k__Fungi_unclassified(61);k__Fungi_unclassified(61);k__Fungi_unclassified(61); | - | - | - | - | - | - | - | Unassigned | - |
| Otu05248 | 5 | k__Fungi(100);k__Fungi_unclassified(100);k__Fungi_unclassified(100);k__Fungi_unclassified(100);k__Fungi_unclassified(100);k__Fungi_unclassified(100);k__Fungi_unclassified(100); | - | - | - | - | - | - | - | Unassigned | - |
| Otu05250 | 5 | k__Fungi(100);p__Ascomycota(80);p__Ascomycota_unclassified(80);p__Ascomycota_unclassified(80);p__Ascomycota_unclassified(80);p__Ascomycota_unclassified(80);p__Ascomycota_unclassified(80); | - | - | - | - | - | - | - | Unassigned | - |
| Otu05251 | 5 | k__Fungi(100);p__Ascomycota(61);p__Ascomycota_unclassified(61);p__Ascomycota_unclassified(61);p__Ascomycota_unclassified(61);p__Ascomycota_unclassified(61);p__Ascomycota_unclassified(61); | - | - | - | - | - | - | - | Unassigned | - |
| Otu05252 | 5 | k__Fungi(100);k__Fungi_unclassified(100);k__Fungi_unclassified(100);k__Fungi_unclassified(100);k__Fungi_unclassified(100);k__Fungi_unclassified(100);k__Fungi_unclassified(100); | - | - | - | - | - | - | - | Unassigned | - |
| Otu05253 | 5 | k__Fungi(100);p__Ascomycota(61);p__Ascomycota_unclassified(61);p__Ascomycota_unclassified(61);p__Ascomycota_unclassified(61);p__Ascomycota_unclassified(61);p__Ascomycota_unclassified(61); | - | - | - | - | - | - | - | Unassigned | - |
| Otu05254 | 5 | k__Fungi(100);k__Fungi_unclassified(80);k__Fungi_unclassified(80);k__Fungi_unclassified(80);k__Fungi_unclassified(80);k__Fungi_unclassified(80);k__Fungi_unclassified(80); | - | - | - | - | - | - | - | Unassigned | - |
| Otu05255 | 5 | k__Fungi(100);k__Fungi_unclassified(61);k__Fungi_unclassified(61);k__Fungi_unclassified(61);k__Fungi_unclassified(61);k__Fungi_unclassified(61);k__Fungi_unclassified(61); | - | - | - | - | - | - | - | Unassigned | - |
| Otu05257 | 5 | k__Fungi(100);p__Ascomycota(61);p__Ascomycota_unclassified(61);p__Ascomycota_unclassified(61);p__Ascomycota_unclassified(61);p__Ascomycota_unclassified(61);p__Ascomycota_unclassified(61); | - | - | - | - | - | - | - | Unassigned | - |
| Otu05258 | 5 | k__Fungi(100);p__Ascomycota(61);p__Ascomycota_unclassified(61);p__Ascomycota_unclassified(61);p__Ascomycota_unclassified(61);p__Ascomycota_unclassified(61);p__Ascomycota_unclassified(61); | - | - | - | - | - | - | - | Unassigned | - |
| Otu05261 | 5 | k__Fungi(100);p__Ascomycota(61);p__Ascomycota_unclassified(61);p__Ascomycota_unclassified(61);p__Ascomycota_unclassified(61);p__Ascomycota_unclassified(61);p__Ascomycota_unclassified(61); | - | - | - | - | - | - | - | Unassigned | - |
| Otu05262 | 5 | k__Fungi(100);k__Fungi_unclassified(80);k__Fungi_unclassified(80);k__Fungi_unclassified(80);k__Fungi_unclassified(80);k__Fungi_unclassified(80);k__Fungi_unclassified(80); | - | - | - | - | - | - | - | Unassigned | - |
| Otu05263 | 5 | k__Plantae(100);p__unclassified_Plantae(100);c__unclassified_Plantae(100);o__unclassified_Plantae(100);f__unclassified_Plantae(100);g__unclassified_Plantae(100);s__Plantae_sp(100); | - | - | - | - | - | - | - | Unassigned | - |
| Otu05264 | 5 | k__Fungi(100);p__Ascomycota(80);p__Ascomycota_unclassified(80);p__Ascomycota_unclassified(80);p__Ascomycota_unclassified(80);p__Ascomycota_unclassified(80);p__Ascomycota_unclassified(80); | - | - | - | - | - | - | - | Unassigned | - |
| Otu05266 | 5 | k__Fungi(100);k__Fungi_unclassified(80);k__Fungi_unclassified(80);k__Fungi_unclassified(80);k__Fungi_unclassified(80);k__Fungi_unclassified(80);k__Fungi_unclassified(80); | - | - | - | - | - | - | - | Unassigned | - |
| Otu05267 | 5 | k__Fungi(100);p__Ascomycota(61);p__Ascomycota_unclassified(61);p__Ascomycota_unclassified(61);p__Ascomycota_unclassified(61);p__Ascomycota_unclassified(61);p__Ascomycota_unclassified(61); | - | - | - | - | - | - | - | Unassigned | - |
| Otu05268 | 5 | k__Fungi(100);k__Fungi_unclassified(61);k__Fungi_unclassified(61);k__Fungi_unclassified(61);k__Fungi_unclassified(61);k__Fungi_unclassified(61);k__Fungi_unclassified(61); | - | - | - | - | - | - | - | Unassigned | - |
| Otu05269 | 5 | k__Fungi(100);k__Fungi_unclassified(80);k__Fungi_unclassified(80);k__Fungi_unclassified(80);k__Fungi_unclassified(80);k__Fungi_unclassified(80);k__Fungi_unclassified(80); | - | - | - | - | - | - | - | Unassigned | - |
| Otu05270 | 5 | k__Fungi(100);k__Fungi_unclassified(100);k__Fungi_unclassified(100);k__Fungi_unclassified(100);k__Fungi_unclassified(100);k__Fungi_unclassified(100);k__Fungi_unclassified(100); | - | - | - | - | - | - | - | Unassigned | - |
| Otu05271 | 5 | k__Fungi(100);k__Fungi_unclassified(100);k__Fungi_unclassified(100);k__Fungi_unclassified(100);k__Fungi_unclassified(100);k__Fungi_unclassified(100);k__Fungi_unclassified(100); | - | - | - | - | - | - | - | Unassigned | - |
| Otu05273 | 5 | k__Fungi(100);k__Fungi_unclassified(80);k__Fungi_unclassified(80);k__Fungi_unclassified(80);k__Fungi_unclassified(80);k__Fungi_unclassified(80);k__Fungi_unclassified(80); | - | - | - | - | - | - | - | Unassigned | - |

| | | | | | | | | | | | |
|---|---|---|---|---|---|---|---|---|---|---|---|
| Otu05276 | 5 | k__Fungi(100);p__Ascomycota(100);c__Sordariomycetes(100);c__Sordariomycetes_unclassified(100);c__Sordariomycetes_unclassified(100);c__Sordariomycetes_unclassified(100);c__Sordariomycetes_unclassified(100); | - | - | - | - | - | - | - | Unassigned | - |
| Otu05277 | 5 | k__Fungi(100);k__Fungi_unclassified(61);k__Fungi_unclassified(61);k__Fungi_unclassified(61);k__Fungi_unclassified(61);k__Fungi_unclassified(61);k__Fungi_unclassified(61); | - | - | - | - | - | - | - | Unassigned | - |
| Otu05280 | 5 | k__Fungi(100);p__Ascomycota(100);p__Ascomycota_unclassified(61);p__Ascomycota_unclassified(61);p__Ascomycota_unclassified(61);p__Ascomycota_unclassified(61);p__Ascomycota_unclassified(61); | - | - | - | - | - | - | - | Unassigned | - |
| Otu05281 | 5 | k__Fungi(100);k__Fungi_unclassified(100);k__Fungi_unclassified(100);k__Fungi_unclassified(100);k__Fungi_unclassified(100);k__Fungi_unclassified(100);k__Fungi_unclassified(100); | - | - | - | - | - | - | - | Unassigned | - |
| Otu05282 | 5 | k__Fungi(100);p__Ascomycota(80);p__Ascomycota_unclassified(80);p__Ascomycota_unclassified(80);p__Ascomycota_unclassified(80);p__Ascomycota_unclassified(80);p__Ascomycota_unclassified(80); | - | - | - | - | - | - | - | Unassigned | - |
| Otu05285 | 5 | k__Fungi(100);p__Ascomycota(61);p__Ascomycota_unclassified(61);p__Ascomycota_unclassified(61);p__Ascomycota_unclassified(61);p__Ascomycota_unclassified(61);p__Ascomycota_unclassified(61); | - | - | - | - | - | - | - | Unassigned | - |
| Otu05288 | 5 | k__Fungi(100);p__Ascomycota(80);p__Ascomycota_unclassified(80);p__Ascomycota_unclassified(80);p__Ascomycota_unclassified(80);p__Ascomycota_unclassified(80);p__Ascomycota_unclassified(80); | - | - | - | - | - | - | - | Unassigned | - |
| Otu05290 | 5 | k__Fungi(100);p__Ascomycota(100);p__Ascomycota_unclassified(100);p__Ascomycota_unclassified(100);p__Ascomycota_unclassified(100);p__Ascomycota_unclassified(100);p__Ascomycota_unclassified(100); | - | - | - | - | - | - | - | Unassigned | - |
| Otu05295 | 5 | k__Fungi(100);k__Fungi_unclassified(80);k__Fungi_unclassified(80);k__Fungi_unclassified(80);k__Fungi_unclassified(80);k__Fungi_unclassified(80);k__Fungi_unclassified(80); | - | - | - | - | - | - | - | Unassigned | - |
| Otu05297 | 5 | k__Fungi(100);k__Fungi_unclassified(100);k__Fungi_unclassified(100);k__Fungi_unclassified(100);k__Fungi_unclassified(100);k__Fungi_unclassified(100);k__Fungi_unclassified(100); | - | - | - | - | - | - | - | Unassigned | - |
| Otu05298 | 5 | k__Fungi(100);p__Ascomycota(61);p__Ascomycota_unclassified(61);p__Ascomycota_unclassified(61);p__Ascomycota_unclassified(61);p__Ascomycota_unclassified(61);p__Ascomycota_unclassified(61); | - | - | - | - | - | - | - | Unassigned | - |
| Otu05299 | 5 | k__Fungi(100);k__Fungi_unclassified(61);k__Fungi_unclassified(61);k__Fungi_unclassified(61);k__Fungi_unclassified(61);k__Fungi_unclassified(61);k__Fungi_unclassified(61); | - | - | - | - | - | - | - | Unassigned | - |
| Otu05301 | 5 | k__Fungi(100);p__Ascomycota(100);p__Ascomycota_unclassified(100);p__Ascomycota_unclassified(100);p__Ascomycota_unclassified(100);p__Ascomycota_unclassified(100);p__Ascomycota_unclassified(100); | - | - | - | - | - | - | - | Unassigned | - |
| Otu05303 | 5 | k__Fungi(100);k__Fungi_unclassified(100);k__Fungi_unclassified(100);k__Fungi_unclassified(100);k__Fungi_unclassified(100);k__Fungi_unclassified(100);k__Fungi_unclassified(100); | - | - | - | - | - | - | - | Unassigned | - |
| Otu05305 | 5 | k__Fungi(100);p__Ascomycota(61);p__Ascomycota_unclassified(61);p__Ascomycota_unclassified(61);p__Ascomycota_unclassified(61);p__Ascomycota_unclassified(61);p__Ascomycota_unclassified(61); | - | - | - | - | - | - | - | Unassigned | - |
| Otu05306 | 5 | k__Fungi(100);p__Ascomycota(100);p__Ascomycota_unclassified(100);p__Ascomycota_unclassified(100);p__Ascomycota_unclassified(100);p__Ascomycota_unclassified(100);p__Ascomycota_unclassified(100); | - | - | - | - | - | - | - | Unassigned | - |
| Otu05307 | 5 | k__Fungi(100);p__Ascomycota(100);p__Ascomycota_unclassified(100);p__Ascomycota_unclassified(100);p__Ascomycota_unclassified(100);p__Ascomycota_unclassified(100);p__Ascomycota_unclassified(100); | - | - | - | - | - | - | - | Unassigned | - |
| Otu05310 | 5 | k__Fungi(100);k__Fungi_unclassified(61);k__Fungi_unclassified(61);k__Fungi_unclassified(61);k__Fungi_unclassified(61);k__Fungi_unclassified(61);k__Fungi_unclassified(61); | - | - | - | - | - | - | - | Unassigned | - |
| Otu05314 | 5 | k__Fungi(100);p__Ascomycota(61);p__Ascomycota_unclassified(61);p__Ascomycota_unclassified(61);p__Ascomycota_unclassified(61);p__Ascomycota_unclassified(61);p__Ascomycota_unclassified(61); | - | - | - | - | - | - | - | Unassigned | - |
| Otu05315 | 5 | k__Fungi(100);k__Fungi_unclassified(80);k__Fungi_unclassified(80);k__Fungi_unclassified(80);k__Fungi_unclassified(80);k__Fungi_unclassified(80);k__Fungi_unclassified(80); | - | - | - | - | - | - | - | Unassigned | - |
| Otu05316 | 5 | k__Fungi(100);p__Ascomycota(100);p__Ascomycota_unclassified(80);p__Ascomycota_unclassified(80);p__Ascomycota_unclassified(80);p__Ascomycota_unclassified(80);p__Ascomycota_unclassified(80); | - | - | - | - | - | - | - | Unassigned | - |
| Otu05317 | 5 | k__Fungi(100);p__Basidiomycota(100);c__Microbotryomycetes(100);o__Sporidiobolales(100);f__Sporidiobolales_family_Incertae_sedis(100);f__Sporidiobolales_family_Incertae_sedis_unclassified(100);f__Sporidiobolales_family_Incertae_sedis_unclassified(100); | - | - | - | - | - | - | - | Unassigned | - |
| Otu05318 | 5 | k__Fungi(100);p__Ascomycota(61);p__Ascomycota_unclassified(61);p__Ascomycota_unclassified(61);p__Ascomycota_unclassified(61);p__Ascomycota_unclassified(61);p__Ascomycota_unclassified(61); | - | - | - | - | - | - | - | Unassigned | - |
| Otu05320 | 5 | k__Fungi(100);k__Fungi_unclassified(80);k__Fungi_unclassified(80);k__Fungi_unclassified(80);k__Fungi_unclassified(80);k__Fungi_unclassified(80);k__Fungi_unclassified(80); | - | - | - | - | - | - | - | Unassigned | - |
| Otu05322 | 5 | k__Fungi(100);p__Ascomycota(100);c__Dothideomycetes(100);o__Capnodiales(100);o__Capnodiales_unclassified(61);o__Capnodiales_unclassified(61);o__Capnodiales_unclassified(61); | - | - | - | - | - | - | - | Unassigned | - |
| Otu05330 | 5 | k__Fungi(100);p__Ascomycota(100);p__Ascomycota_unclassified(100);p__Ascomycota_unclassified(100);p__Ascomycota_unclassified(100);p__Ascomycota_unclassified(100);p__Ascomycota_unclassified(100); | - | - | - | - | - | - | - | Unassigned | - |
| Otu05332 | 5 | k__Plantae(100);p__unclassified_Plantae(100);c__unclassified_Plantae(100);o__unclassified_Plantae(100);f__unclassified_Plantae(100);g__unclassified_Plantae(100);s__Plantae_sp(100); | - | - | - | - | - | - | - | Unassigned | - |
| Otu05333 | 5 | k__Fungi(100);p__Ascomycota(80);p__Ascomycota_unclassified(80);p__Ascomycota_unclassified(80);p__Ascomycota_unclassified(80);p__Ascomycota_unclassified(80);p__Ascomycota_unclassified(80); | - | - | - | - | - | - | - | Unassigned | - |
| Otu05334 | 5 | k__Fungi(100);p__Ascomycota(100);p__Ascomycota_unclassified(100);p__Ascomycota_unclassified(100);p__Ascomycota_unclassified(100);p__Ascomycota_unclassified(100);p__Ascomycota_unclassified(100); | - | - | - | - | - | - | - | Unassigned | - |
| Otu05335 | 5 | k__Fungi(100);k__Fungi_unclassified(61);k__Fungi_unclassified(61);k__Fungi_unclassified(61);k__Fungi_unclassified(61);k__Fungi_unclassified(61);k__Fungi_unclassified(61); | - | - | - | - | - | - | - | Unassigned | - |
| Otu05338 | 5 | k__Fungi(100);k__Fungi_unclassified(61);k__Fungi_unclassified(61);k__Fungi_unclassified(61);k__Fungi_unclassified(61);k__Fungi_unclassified(61);k__Fungi_unclassified(61); | - | - | - | - | - | - | - | Unassigned | - |
| Otu05340 | 5 | k__Fungi(100);p__Ascomycota(100);p__Ascomycota_unclassified(61);p__Ascomycota_unclassified(61);p__Ascomycota_unclassified(61);p__Ascomycota_unclassified(61);p__Ascomycota_unclassified(61); | - | - | - | - | - | - | - | Unassigned | - |
| Otu05342 | 5 | k__Fungi(100);p__Ascomycota(100);p__Ascomycota_unclassified(100);p__Ascomycota_unclassified(100);p__Ascomycota_unclassified(100);p__Ascomycota_unclassified(100);p__Ascomycota_unclassified(100); | - | - | - | - | - | - | - | Unassigned | - |
| Otu05345 | 5 | k__Fungi(100);p__Ascomycota(80);p__Ascomycota_unclassified(80);p__Ascomycota_unclassified(80);p__Ascomycota_unclassified(80);p__Ascomycota_unclassified(80);p__Ascomycota_unclassified(80); | - | - | - | - | - | - | - | Unassigned | - |
| Otu05347 | 5 | k__Fungi(100);k__Fungi_unclassified(80);k__Fungi_unclassified(80);k__Fungi_unclassified(80);k__Fungi_unclassified(80);k__Fungi_unclassified(80);k__Fungi_unclassified(80); | - | - | - | - | - | - | - | Unassigned | - |
| Otu05348 | 5 | k__Fungi(100);k__Fungi_unclassified(61);k__Fungi_unclassified(61);k__Fungi_unclassified(61);k__Fungi_unclassified(61);k__Fungi_unclassified(61);k__Fungi_unclassified(61); | - | - | - | - | - | - | - | Unassigned | - |
| Otu05349 | 5 | k__Fungi(100);p__Ascomycota(100);p__Ascomycota_unclassified(100);p__Ascomycota_unclassified(100);p__Ascomycota_unclassified(100);p__Ascomycota_unclassified(100);p__Ascomycota_unclassified(100); | - | - | - | - | - | - | - | Unassigned | - |
| Otu05350 | 5 | k__Fungi(100);k__Fungi_unclassified(100);k__Fungi_unclassified(100);k__Fungi_unclassified(100);k__Fungi_unclassified(100);k__Fungi_unclassified(100);k__Fungi_unclassified(100); | - | - | - | - | - | - | - | Unassigned | - |
| Otu05352 | 5 | k__Fungi(100);k__Fungi_unclassified(80);k__Fungi_unclassified(80);k__Fungi_unclassified(80);k__Fungi_unclassified(80);k__Fungi_unclassified(80);k__Fungi_unclassified(80); | - | - | - | - | - | - | - | Unassigned | - |
| Otu05353 | 5 | k__Fungi(100);p__Ascomycota(100);c__Dothideomycetes(100);o__Pleosporales(100);o__Pleosporales_unclassified(100);o__Pleosporales_unclassified(100);o__Pleosporales_unclassified(100); | - | - | - | - | - | - | - | Unassigned | - |
| Otu05357 | 5 | k__Fungi(100);p__Ascomycota(100);p__Ascomycota_unclassified(100);p__Ascomycota_unclassified(100);p__Ascomycota_unclassified(100);p__Ascomycota_unclassified(100);p__Ascomycota_unclassified(100); | - | - | - | - | - | - | - | Unassigned | - |
| Otu05358 | 5 | k__Fungi(100);p__Ascomycota(100);p__Ascomycota_unclassified(100);p__Ascomycota_unclassified(100);p__Ascomycota_unclassified(100);p__Ascomycota_unclassified(100);p__Ascomycota_unclassified(100); | - | - | - | - | - | - | - | Unassigned | - |
| Otu05359 | 5 | k__Fungi(100);p__Ascomycota(100);p__Ascomycota_unclassified(100);p__Ascomycota_unclassified(100);p__Ascomycota_unclassified(100);p__Ascomycota_unclassified(100);p__Ascomycota_unclassified(100); | - | - | - | - | - | - | - | Unassigned | - |
| Otu05361 | 5 | k__Fungi(100);p__Ascomycota(100);p__Ascomycota_unclassified(100);p__Ascomycota_unclassified(100);p__Ascomycota_unclassified(100);p__Ascomycota_unclassified(100);p__Ascomycota_unclassified(100); | - | - | - | - | - | - | - | Unassigned | - |
| Otu05362 | 5 | k__Fungi(100);k__Fungi_unclassified(61);k__Fungi_unclassified(61);k__Fungi_unclassified(61);k__Fungi_unclassified(61);k__Fungi_unclassified(61);k__Fungi_unclassified(61); | - | - | - | - | - | - | - | Unassigned | - |
| Otu05363 | 5 | k__Fungi(100);p__Ascomycota(100);p__Ascomycota_unclassified(100);p__Ascomycota_unclassified(100);p__Ascomycota_unclassified(100);p__Ascomycota_unclassified(100);p__Ascomycota_unclassified(100); | - | - | - | - | - | - | - | Unassigned | - |
| Otu05364 | 5 | k__Fungi(100);k__Fungi_unclassified(80);k__Fungi_unclassified(80);k__Fungi_unclassified(80);k__Fungi_unclassified(80);k__Fungi_unclassified(80);k__Fungi_unclassified(80); | - | - | - | - | - | - | - | Unassigned | - |
| Otu05365 | 5 | k__Fungi(100);p__Ascomycota(100);p__Ascomycota_unclassified(100);p__Ascomycota_unclassified(100);p__Ascomycota_unclassified(100);p__Ascomycota_unclassified(100);p__Ascomycota_unclassified(100); | - | - | - | - | - | - | - | Unassigned | - |
| Otu05370 | 5 | k__Fungi(100);p__Ascomycota(100);p__Ascomycota_unclassified(100);p__Ascomycota_unclassified(100);p__Ascomycota_unclassified(100);p__Ascomycota_unclassified(100);p__Ascomycota_unclassified(100); | - | - | - | - | - | - | - | Unassigned | - |
| Otu05371 | 5 | k__Fungi(100);p__Ascomycota(61);p__Ascomycota_unclassified(61);p__Ascomycota_unclassified(61);p__Ascomycota_unclassified(61);p__Ascomycota_unclassified(61);p__Ascomycota_unclassified(61); | - | - | - | - | - | - | - | Unassigned | - |
| Otu05372 | 5 | k__Fungi(100);p__Ascomycota(80);p__Ascomycota_unclassified(80);p__Ascomycota_unclassified(80);p__Ascomycota_unclassified(80);p__Ascomycota_unclassified(80);p__Ascomycota_unclassified(80); | - | - | - | - | - | - | - | Unassigned | - |
| Otu05373 | 5 | k__Fungi(100);k__Fungi_unclassified(80);k__Fungi_unclassified(80);k__Fungi_unclassified(80);k__Fungi_unclassified(80);k__Fungi_unclassified(80);k__Fungi_unclassified(80); | - | - | - | - | - | - | - | Unassigned | - |
| Otu05375 | 5 | k__Fungi(100);k__Fungi_unclassified(100);k__Fungi_unclassified(100);k__Fungi_unclassified(100);k__Fungi_unclassified(100);k__Fungi_unclassified(100);k__Fungi_unclassified(100); | - | - | - | - | - | - | - | Unassigned | - |
| Otu05376 | 5 | k__Fungi(100);p__Ascomycota(100);c__Dothideomycetes(100);o__Capnodiales(100);o__Capnodiales_unclassified(100);o__Capnodiales_unclassified(100);o__Capnodiales_unclassified(100); | - | - | - | - | - | - | - | Unassigned | - |
| Otu05379 | 5 | k__Fungi(100);k__Fungi_unclassified(100);k__Fungi_unclassified(100);k__Fungi_unclassified(100);k__Fungi_unclassified(100);k__Fungi_unclassified(100);k__Fungi_unclassified(100); | - | - | - | - | - | - | - | Unassigned | - |
| Otu05382 | 5 | k__Fungi(100);p__Ascomycota(100);p__Ascomycota_unclassified(100);p__Ascomycota_unclassified(100);p__Ascomycota_unclassified(100);p__Ascomycota_unclassified(100);p__Ascomycota_unclassified(100); | - | - | - | - | - | - | - | Unassigned | - |
| Otu05383 | 5 | k__Fungi(100);k__Fungi_unclassified(61);k__Fungi_unclassified(61);k__Fungi_unclassified(61);k__Fungi_unclassified(61);k__Fungi_unclassified(61);k__Fungi_unclassified(61); | - | - | - | - | - | - | - | Unassigned | - |
| Otu05385 | 5 | k__Fungi(100);p__Ascomycota(61);p__Ascomycota_unclassified(61);p__Ascomycota_unclassified(61);p__Ascomycota_unclassified(61);p__Ascomycota_unclassified(61);p__Ascomycota_unclassified(61); | - | - | - | - | - | - | - | Unassigned | - |
| Otu05389 | 5 | k__Fungi(100);k__Fungi_unclassified(100);k__Fungi_unclassified(100);k__Fungi_unclassified(100);k__Fungi_unclassified(100);k__Fungi_unclassified(100);k__Fungi_unclassified(100); | - | - | - | - | - | - | - | Unassigned | - |
| Otu05391 | 5 | k__Fungi(100);k__Fungi_unclassified(80);k__Fungi_unclassified(80);k__Fungi_unclassified(80);k__Fungi_unclassified(80);k__Fungi_unclassified(80);k__Fungi_unclassified(80); | - | - | - | - | - | - | - | Unassigned | - |
| Otu05392 | 5 | k__Fungi(100);k__Fungi_unclassified(100);k__Fungi_unclassified(100);k__Fungi_unclassified(100);k__Fungi_unclassified(100);k__Fungi_unclassified(100);k__Fungi_unclassified(100); | - | - | - | - | - | - | - | Unassigned | - |
| Otu05394 | 5 | k__Fungi(100);p__Ascomycota(80);p__Ascomycota_unclassified(80);p__Ascomycota_unclassified(80);p__Ascomycota_unclassified(80);p__Ascomycota_unclassified(80);p__Ascomycota_unclassified(80); | - | - | - | - | - | - | - | Unassigned | - |
| Otu05395 | 5 | k__Fungi(100);p__Ascomycota(80);p__Ascomycota_unclassified(80);p__Ascomycota_unclassified(80);p__Ascomycota_unclassified(80);p__Ascomycota_unclassified(80);p__Ascomycota_unclassified(80); | - | - | - | - | - | - | - | Unassigned | - |
| Otu05398 | 5 | k__Fungi(100);p__Ascomycota(61);p__Ascomycota_unclassified(61);p__Ascomycota_unclassified(61);p__Ascomycota_unclassified(61);p__Ascomycota_unclassified(61);p__Ascomycota_unclassified(61); | - | - | - | - | - | - | - | Unassigned | - |
| Otu05400 | 5 | k__Fungi(100);k__Fungi_unclassified(100);k__Fungi_unclassified(100);k__Fungi_unclassified(100);k__Fungi_unclassified(100);k__Fungi_unclassified(100);k__Fungi_unclassified(100); | - | - | - | - | - | - | - | Unassigned | - |
| Otu05403 | 5 | k__Fungi(100);p__Ascomycota(80);p__Ascomycota_unclassified(80);p__Ascomycota_unclassified(80);p__Ascomycota_unclassified(80);p__Ascomycota_unclassified(80);p__Ascomycota_unclassified(80); | - | - | - | - | - | - | - | Unassigned | - |
| Otu05404 | 5 | k__Fungi(100);k__Fungi_unclassified(61);k__Fungi_unclassified(61);k__Fungi_unclassified(61);k__Fungi_unclassified(61);k__Fungi_unclassified(61);k__Fungi_unclassified(61); | - | - | - | - | - | - | - | Unassigned | - |
| Otu05408 | 5 | k__Fungi(100);p__Ascomycota(80);p__Ascomycota_unclassified(80);p__Ascomycota_unclassified(80);p__Ascomycota_unclassified(80);p__Ascomycota_unclassified(80);p__Ascomycota_unclassified(80); | - | - | - | - | - | - | - | Unassigned | - |
| Otu05409 | 5 | k__Fungi(100);p__Ascomycota(80);p__Ascomycota_unclassified(80);p__Ascomycota_unclassified(80);p__Ascomycota_unclassified(80);p__Ascomycota_unclassified(80);p__Ascomycota_unclassified(80); | - | - | - | - | - | - | - | Unassigned | - |
| Otu05410 | 5 | k__Fungi(100);k__Fungi_unclassified(100);k__Fungi_unclassified(100);k__Fungi_unclassified(100);k__Fungi_unclassified(100);k__Fungi_unclassified(100);k__Fungi_unclassified(100); | - | - | - | - | - | - | - | Unassigned | - |
| Otu05411 | 5 | k__Fungi(100);k__Fungi_unclassified(100);k__Fungi_unclassified(100);k__Fungi_unclassified(100);k__Fungi_unclassified(100);k__Fungi_unclassified(100);k__Fungi_unclassified(100); | - | - | - | - | - | - | - | Unassigned | - |
| Otu05413 | 5 | k__Fungi(100);p__Ascomycota(80);p__Ascomycota_unclassified(80);p__Ascomycota_unclassified(80);p__Ascomycota_unclassified(80);p__Ascomycota_unclassified(80);p__Ascomycota_unclassified(80); | - | - | - | - | - | - | - | Unassigned | - |
| Otu05414 | 5 | k__Fungi(100);p__Ascomycota(100);c__Lecanoromycetes(100);o__Teloschistales(100);o__Teloschistales_unclassified(100);o__Teloschistales_unclassified(100);o__Teloschistales_unclassified(100); | - | - | - | - | - | - | - | Unassigned | - |
| Otu05417 | 5 | k__Fungi(100);k__Fungi_unclassified(100);k__Fungi_unclassified(100);k__Fungi_unclassified(100);k__Fungi_unclassified(100);k__Fungi_unclassified(100);k__Fungi_unclassified(100); | - | - | - | - | - | - | - | Unassigned | - |
| Otu05418 | 5 | k__Fungi(100);p__Ascomycota(100);p__Ascomycota_unclassified(100);p__Ascomycota_unclassified(100);p__Ascomycota_unclassified(100);p__Ascomycota_unclassified(100);p__Ascomycota_unclassified(100); | - | - | - | - | - | - | - | Unassigned | - |
| Otu05420 | 5 | k__Fungi(100);k__Fungi_unclassified(80);k__Fungi_unclassified(80);k__Fungi_unclassified(80);k__Fungi_unclassified(80);k__Fungi_unclassified(80);k__Fungi_unclassified(80); | - | - | - | - | - | - | - | Unassigned | - |
| Otu05421 | 5 | k__Fungi(100);p__Ascomycota(80);p__Ascomycota_unclassified(80);p__Ascomycota_unclassified(80);p__Ascomycota_unclassified(80);p__Ascomycota_unclassified(80);p__Ascomycota_unclassified(80); | - | - | - | - | - | - | - | Unassigned | - |
| Otu05422 | 5 | k__Fungi(100);p__Ascomycota(61);p__Ascomycota_unclassified(61);p__Ascomycota_unclassified(61);p__Ascomycota_unclassified(61);p__Ascomycota_unclassified(61);p__Ascomycota_unclassified(61); | - | - | - | - | - | - | - | Unassigned | - |
| Otu05426 | 5 | k__Fungi(100);k__Fungi_unclassified(100);k__Fungi_unclassified(100);k__Fungi_unclassified(100);k__Fungi_unclassified(100);k__Fungi_unclassified(100);k__Fungi_unclassified(100); | - | - | - | - | - | - | - | Unassigned | - |
| Otu05429 | 5 | k__Fungi(100);k__Fungi_unclassified(100);k__Fungi_unclassified(100);k__Fungi_unclassified(100);k__Fungi_unclassified(100);k__Fungi_unclassified(100);k__Fungi_unclassified(100); | - | - | - | - | - | - | - | Unassigned | - |
| Otu05430 | 5 | k__Fungi(100);p__Ascomycota(61);p__Ascomycota_unclassified(61);p__Ascomycota_unclassified(61);p__Ascomycota_unclassified(61);p__Ascomycota_unclassified(61);p__Ascomycota_unclassified(61); | - | - | - | - | - | - | - | Unassigned | - |
| Otu05432 | 5 | k__Fungi(100);k__Fungi_unclassified(100);k__Fungi_unclassified(100);k__Fungi_unclassified(100);k__Fungi_unclassified(100);k__Fungi_unclassified(100);k__Fungi_unclassified(100); | - | - | - | - | - | - | - | Unassigned | - |
| Otu05433 | 5 | k__Fungi(100);p__Ascomycota(61);p__Ascomycota_unclassified(61);p__Ascomycota_unclassified(61);p__Ascomycota_unclassified(61);p__Ascomycota_unclassified(61);p__Ascomycota_unclassified(61); | - | - | - | - | - | - | - | Unassigned | - |
| Otu05434 | 5 | k__Fungi(100);p__Ascomycota(100);p__Ascomycota_unclassified(100);p__Ascomycota_unclassified(100);p__Ascomycota_unclassified(100);p__Ascomycota_unclassified(100);p__Ascomycota_unclassified(100); | - | - | - | - | - | - | - | Unassigned | - |
| Otu05436 | 5 | k__Fungi(100);k__Fungi_unclassified(61);k__Fungi_unclassified(61);k__Fungi_unclassified(61);k__Fungi_unclassified(61);k__Fungi_unclassified(61);k__Fungi_unclassified(61); | - | - | - | - | - | - | - | Unassigned | - |
| Otu05437 | 5 | k__Fungi(100);k__Fungi_unclassified(80);k__Fungi_unclassified(80);k__Fungi_unclassified(80);k__Fungi_unclassified(80);k__Fungi_unclassified(80);k__Fungi_unclassified(80); | - | - | - | - | - | - | - | Unassigned | - |
| Otu05438 | 5 | k__Fungi(100);k__Fungi_unclassified(80);k__Fungi_unclassified(80);k__Fungi_unclassified(80);k__Fungi_unclassified(80);k__Fungi_unclassified(80);k__Fungi_unclassified(80); | - | - | - | - | - | - | - | Unassigned | - |
| Otu05440 | 5 | k__Fungi(100);k__Fungi_unclassified(61);k__Fungi_unclassified(61);k__Fungi_unclassified(61);k__Fungi_unclassified(61);k__Fungi_unclassified(61);k__Fungi_unclassified(61); | - | - | - | - | - | - | - | Unassigned | - |
| Otu05441 | 5 | k__Fungi(100);k__Fungi_unclassified(80);k__Fungi_unclassified(80);k__Fungi_unclassified(80);k__Fungi_unclassified(80);k__Fungi_unclassified(80);k__Fungi_unclassified(80); | - | - | - | - | - | - | - | Unassigned | - |
| Otu05442 | 5 | k__Fungi(100);p__Ascomycota(100);c__Dothideomycetes(61);c__Dothideomycetes_unclassified(61);c__Dothideomycetes_unclassified(61);c__Dothideomycetes_unclassified(61);c__Dothideomycetes_unclassified(61); | - | - | - | - | - | - | - | Unassigned | - |
| Otu05444 | 5 | k__Fungi(100);k__Fungi_unclassified(100);k__Fungi_unclassified(100);k__Fungi_unclassified(100);k__Fungi_unclassified(100);k__Fungi_unclassified(100);k__Fungi_unclassified(100); | - | - | - | - | - | - | - | Unassigned | - |
| Otu05447 | 5 | k__Fungi(100);p__Ascomycota(100);p__Ascomycota_unclassified(100);p__Ascomycota_unclassified(100);p__Ascomycota_unclassified(100);p__Ascomycota_unclassified(100);p__Ascomycota_unclassified(100); | - | - | - | - | - | - | - | Unassigned | - |
| Otu05448 | 5 | k__Fungi(100);k__Fungi_unclassified(61);k__Fungi_unclassified(61);k__Fungi_unclassified(61);k__Fungi_unclassified(61);k__Fungi_unclassified(61);k__Fungi_unclassified(61); | - | - | - | - | - | - | - | Unassigned | - |
| Otu05450 | 5 | k__Fungi(100);k__Fungi_unclassified(61);k__Fungi_unclassified(61);k__Fungi_unclassified(61);k__Fungi_unclassified(61);k__Fungi_unclassified(61);k__Fungi_unclassified(61); | - | - | - | - | - | - | - | Unassigned | - |
| Otu05451 | 5 | k__Plantae(100);p__unclassified_Plantae(100);c__unclassified_Plantae(100);o__unclassified_Plantae(100);f__unclassified_Plantae(100);g__unclassified_Plantae(100);s__Plantae_sp(100); | - | - | - | - | - | - | - | Unassigned | - |
| Otu05452 | 5 | k__Fungi(100);p__Ascomycota(100);p__Ascomycota_unclassified(100);p__Ascomycota_unclassified(100);p__Ascomycota_unclassified(100);p__Ascomycota_unclassified(100);p__Ascomycota_unclassified(100); | - | - | - | - | - | - | - | Unassigned | - |
| Otu05453 | 5 | k__Fungi(100);p__Ascomycota(100);p__Ascomycota_unclassified(100);p__Ascomycota_unclassified(100);p__Ascomycota_unclassified(100);p__Ascomycota_unclassified(100);p__Ascomycota_unclassified(100); | - | - | - | - | - | - | - | Unassigned | - |
| Otu05454 | 5 | k__Fungi(100);k__Fungi_unclassified(61);k__Fungi_unclassified(61);k__Fungi_unclassified(61);k__Fungi_unclassified(61);k__Fungi_unclassified(61);k__Fungi_unclassified(61); | - | - | - | - | - | - | - | Unassigned | - |
| Otu05456 | 5 | k__Fungi(100);k__Fungi_unclassified(80);k__Fungi_unclassified(80);k__Fungi_unclassified(80);k__Fungi_unclassified(80);k__Fungi_unclassified(80);k__Fungi_unclassified(80); | - | - | - | - | - | - | - | Unassigned | - |
| Otu05457 | 5 | k__Fungi(100);p__Ascomycota(61);p__Ascomycota_unclassified(61);p__Ascomycota_unclassified(61);p__Ascomycota_unclassified(61);p__Ascomycota_unclassified(61);p__Ascomycota_unclassified(61); | - | - | - | - | - | - | - | Unassigned | - |
| Otu05458 | 5 | k__Fungi(100);p__Basidiomycota(100);c__Tremellomycetes(100);c__Tremellomycetes_unclassified(100);c__Tremellomycetes_unclassified(100);c__Tremellomycetes_unclassified(100);c__Tremellomycetes_unclassified(100); | - | - | - | - | - | - | - | Unassigned | - |
| Otu05459 | 5 | k__Fungi(100);k__Fungi_unclassified(61);k__Fungi_unclassified(61);k__Fungi_unclassified(61);k__Fungi_unclassified(61);k__Fungi_unclassified(61);k__Fungi_unclassified(61); | - | - | - | - | - | - | - | Unassigned | - |
| Otu05460 | 5 | k__Fungi(100);p__Ascomycota(100);p__Ascomycota_unclassified(100);p__Ascomycota_unclassified(100);p__Ascomycota_unclassified(100);p__Ascomycota_unclassified(100);p__Ascomycota_unclassified(100); | - | - | - | - | - | - | - | Unassigned | - |
| Otu05461 | 5 | k__Fungi(100);p__Ascomycota(100);p__Ascomycota_unclassified(100);p__Ascomycota_unclassified(100);p__Ascomycota_unclassified(100);p__Ascomycota_unclassified(100);p__Ascomycota_unclassified(100); | - | - | - | - | - | - | - | Unassigned | - |
| Otu05463 | 5 | k__Fungi(100);p__Ascomycota(100);p__Ascomycota_unclassified(100);p__Ascomycota_unclassified(100);p__Ascomycota_unclassified(100);p__Ascomycota_unclassified(100);p__Ascomycota_unclassified(100); | - | - | - | - | - | - | - | Unassigned | - |
| Otu05465 | 5 | k__Fungi(100);k__Fungi_unclassified(80);k__Fungi_unclassified(80);k__Fungi_unclassified(80);k__Fungi_unclassified(80);k__Fungi_unclassified(80);k__Fungi_unclassified(80); | - | - | - | - | - | - | - | Unassigned | - |
| Otu05466 | 5 | k__Fungi(100);p__Ascomycota(100);p__Ascomycota_unclassified(61);p__Ascomycota_unclassified(61);p__Ascomycota_unclassified(61);p__Ascomycota_unclassified(61);p__Ascomycota_unclassified(61); | - | - | - | - | - | - | - | Unassigned | - |
| Otu05470 | 5 | k__Fungi(100);k__Fungi_unclassified(80);k__Fungi_unclassified(80);k__Fungi_unclassified(80);k__Fungi_unclassified(80);k__Fungi_unclassified(80);k__Fungi_unclassified(80); | - | - | - | - | - | - | - | Unassigned | - |
| Otu05471 | 5 | k__Fungi(100);p__Ascomycota(80);p__Ascomycota_unclassified(80);p__Ascomycota_unclassified(80);p__Ascomycota_unclassified(80);p__Ascomycota_unclassified(80);p__Ascomycota_unclassified(80); | - | - | - | - | - | - | - | Unassigned | - |
| Otu05476 | 5 | k__Fungi(100);p__Ascomycota(61);p__Ascomycota_unclassified(61);p__Ascomycota_unclassified(61);p__Ascomycota_unclassified(61);p__Ascomycota_unclassified(61);p__Ascomycota_unclassified(61); | - | - | - | - | - | - | - | Unassigned | - |
| Otu05477 | 5 | k__Fungi(100);k__Fungi_unclassified(100);k__Fungi_unclassified(100);k__Fungi_unclassified(100);k__Fungi_unclassified(100);k__Fungi_unclassified(100);k__Fungi_unclassified(100); | - | - | - | - | - | - | - | Unassigned | - |
| Otu05478 | 5 | k__Fungi(100);k__Fungi_unclassified(61);k__Fungi_unclassified(61);k__Fungi_unclassified(61);k__Fungi_unclassified(61);k__Fungi_unclassified(61);k__Fungi_unclassified(61); | - | - | - | - | - | - | - | Unassigned | - |
| Otu05479 | 5 | k__Fungi(100);k__Fungi_unclassified(100);k__Fungi_unclassified(100);k__Fungi_unclassified(100);k__Fungi_unclassified(100);k__Fungi_unclassified(100);k__Fungi_unclassified(100); | - | - | - | - | - | - | - | Unassigned | - |
| Otu05481 | 5 | k__Fungi(100);p__Ascomycota(100);p__Ascomycota_unclassified(100);p__Ascomycota_unclassified(100);p__Ascomycota_unclassified(100);p__Ascomycota_unclassified(100);p__Ascomycota_unclassified(100); | - | - | - | - | - | - | - | Unassigned | - |
| Otu05484 | 5 | k__Fungi(100);p__Ascomycota(100);c__Eurotiomycetes(100);c__Eurotiomycetes_unclassified(80);c__Eurotiomycetes_unclassified(80);c__Eurotiomycetes_unclassified(80);c__Eurotiomycetes_unclassified(80); | - | - | - | - | - | - | - | Unassigned | - |
| Otu05485 | 5 | k__Fungi(100);p__Ascomycota(80);p__Ascomycota_unclassified(80);p__Ascomycota_unclassified(80);p__Ascomycota_unclassified(80);p__Ascomycota_unclassified(80);p__Ascomycota_unclassified(80); | - | - | - | - | - | - | - | Unassigned | - |
| Otu05486 | 5 | k__Fungi(100);k__Fungi_unclassified(61);k__Fungi_unclassified(61);k__Fungi_unclassified(61);k__Fungi_unclassified(61);k__Fungi_unclassified(61);k__Fungi_unclassified(61); | - | - | - | - | - | - | - | Unassigned | - |
| Otu05490 | 5 | k__Fungi(100);k__Fungi_unclassified(100);k__Fungi_unclassified(100);k__Fungi_unclassified(100);k__Fungi_unclassified(100);k__Fungi_unclassified(100);k__Fungi_unclassified(100); | - | - | - | - | - | - | - | Unassigned | - |
| Otu05492 | 5 | k__Fungi(100);k__Fungi_unclassified(61);k__Fungi_unclassified(61);k__Fungi_unclassified(61);k__Fungi_unclassified(61);k__Fungi_unclassified(61);k__Fungi_unclassified(61); | - | - | - | - | - | - | - | Unassigned | - |
| Otu05495 | 5 | k__Fungi(100);k__Fungi_unclassified(100);k__Fungi_unclassified(100);k__Fungi_unclassified(100);k__Fungi_unclassified(100);k__Fungi_unclassified(100);k__Fungi_unclassified(100); | - | - | - | - | - | - | - | Unassigned | - |
| Otu05497 | 5 | k__Fungi(100);k__Fungi_unclassified(100);k__Fungi_unclassified(100);k__Fungi_unclassified(100);k__Fungi_unclassified(100);k__Fungi_unclassified(100);k__Fungi_unclassified(100); | - | - | - | - | - | - | - | Unassigned | - |
| Otu05499 | 5 | k__Fungi(100);k__Fungi_unclassified(100);k__Fungi_unclassified(100);k__Fungi_unclassified(100);k__Fungi_unclassified(100);k__Fungi_unclassified(100);k__Fungi_unclassified(100); | - | - | - | - | - | - | - | Unassigned | - |
| Otu05500 | 5 | k__Fungi(100);p__Ascomycota(61);p__Ascomycota_unclassified(61);p__Ascomycota_unclassified(61);p__Ascomycota_unclassified(61);p__Ascomycota_unclassified(61);p__Ascomycota_unclassified(61); | - | - | - | - | - | - | - | Unassigned | - |
| Otu05502 | 5 | k__Fungi(100);p__Ascomycota(100);c__Dothideomycetes(100);o__Capnodiales(100);o__Capnodiales_unclassified(80);o__Capnodiales_unclassified(80);o__Capnodiales_unclassified(80); | - | - | - | - | - | - | - | Unassigned | - |
| Otu05503 | 5 | k__Fungi(100);p__Ascomycota(61);p__Ascomycota_unclassified(61);p__Ascomycota_unclassified(61);p__Ascomycota_unclassified(61);p__Ascomycota_unclassified(61);p__Ascomycota_unclassified(61); | - | - | - | - | - | - | - | Unassigned | - |
| Otu05506 | 5 | k__Fungi(100);p__Ascomycota(100);p__Ascomycota_unclassified(100);p__Ascomycota_unclassified(100);p__Ascomycota_unclassified(100);p__Ascomycota_unclassified(100);p__Ascomycota_unclassified(100); | - | - | - | - | - | - | - | Unassigned | - |
| Otu05507 | 5 | k__Fungi(100);k__Fungi_unclassified(61);k__Fungi_unclassified(61);k__Fungi_unclassified(61);k__Fungi_unclassified(61);k__Fungi_unclassified(61);k__Fungi_unclassified(61); | - | - | - | - | - | - | - | Unassigned | - |
| Otu05511 | 5 | k__Fungi(100);k__Fungi_unclassified(61);k__Fungi_unclassified(61);k__Fungi_unclassified(61);k__Fungi_unclassified(61);k__Fungi_unclassified(61);k__Fungi_unclassified(61); | - | - | - | - | - | - | - | Unassigned | - |
| Otu05513 | 5 | k__Fungi(100);p__Ascomycota(61);p__Ascomycota_unclassified(61);p__Ascomycota_unclassified(61);p__Ascomycota_unclassified(61);p__Ascomycota_unclassified(61);p__Ascomycota_unclassified(61); | - | - | - | - | - | - | - | Unassigned | - |
| Otu05515 | 5 | k__Fungi(100);p__Ascomycota(80);p__Ascomycota_unclassified(80);p__Ascomycota_unclassified(80);p__Ascomycota_unclassified(80);p__Ascomycota_unclassified(80);p__Ascomycota_unclassified(80); | - | - | - | - | - | - | - | Unassigned | - |
| Otu05516 | 5 | k__Fungi(100);k__Fungi_unclassified(61);k__Fungi_unclassified(61);k__Fungi_unclassified(61);k__Fungi_unclassified(61);k__Fungi_unclassified(61);k__Fungi_unclassified(61); | - | - | - | - | - | - | - | Unassigned | - |
| Otu05518 | 5 | k__Fungi(100);k__Fungi_unclassified(61);k__Fungi_unclassified(61);k__Fungi_unclassified(61);k__Fungi_unclassified(61);k__Fungi_unclassified(61);k__Fungi_unclassified(61); | - | - | - | - | - | - | - | Unassigned | - |
| Otu05522 | 5 | k__Fungi(100);k__Fungi_unclassified(61);k__Fungi_unclassified(61);k__Fungi_unclassified(61);k__Fungi_unclassified(61);k__Fungi_unclassified(61);k__Fungi_unclassified(61); | - | - | - | - | - | - | - | Unassigned | - |
| Otu05523 | 5 | k__Fungi(100);p__Ascomycota(100);c__Dothideomycetes(100);c__Dothideomycetes_unclassified(80);c__Dothideomycetes_unclassified(80);c__Dothideomycetes_unclassified(80);c__Dothideomycetes_unclassified(80); | - | - | - | - | - | - | - | Unassigned | - |
| Otu05526 | 5 | k__Fungi(100);p__Ascomycota(80);p__Ascomycota_unclassified(80);p__Ascomycota_unclassified(80);p__Ascomycota_unclassified(80);p__Ascomycota_unclassified(80);p__Ascomycota_unclassified(80); | - | - | - | - | - | - | - | Unassigned | - |
| Otu05527 | 5 | k__Fungi(100);p__Ascomycota(80);p__Ascomycota_unclassified(80);p__Ascomycota_unclassified(80);p__Ascomycota_unclassified(80);p__Ascomycota_unclassified(80);p__Ascomycota_unclassified(80); | - | - | - | - | - | - | - | Unassigned | - |
| Otu05529 | 5 | k__Fungi(100);p__Ascomycota(80);p__Ascomycota_unclassified(80);p__Ascomycota_unclassified(80);p__Ascomycota_unclassified(80);p__Ascomycota_unclassified(80);p__Ascomycota_unclassified(80); | - | - | - | - | - | - | - | Unassigned | - |
| Otu05530 | 5 | k__Fungi(100);p__Ascomycota(100);p__Ascomycota_unclassified(100);p__Ascomycota_unclassified(100);p__Ascomycota_unclassified(100);p__Ascomycota_unclassified(100);p__Ascomycota_unclassified(100); | - | - | - | - | - | - | - | Unassigned | - |
| Otu05531 | 5 | k__Fungi(100);p__Ascomycota(100);c__Dothideomycetes(100);o__Pleosporales(100);o__Pleosporales_unclassified(61);o__Pleosporales_unclassified(61);o__Pleosporales_unclassified(61); | - | - | - | - | - | - | - | Unassigned | - |
| Otu05535 | 5 | k__Fungi(100);k__Fungi_unclassified(100);k__Fungi_unclassified(100);k__Fungi_unclassified(100);k__Fungi_unclassified(100);k__Fungi_unclassified(100);k__Fungi_unclassified(100); | - | - | - | - | - | - | - | Unassigned | - |
| Otu05544 | 5 | k__Fungi(100);p__Basidiomycota(100);c__Agaricomycetes(100);o__Cantharellales(100);f__unclassified_Cantharellales(100);g__unclassified_Cantharellales(100);s__Cantharellales_sp(100); | - | - | - | - | - | - | - | Unassigned | - |
| Otu05545 | 5 | k__Fungi(100);p__Ascomycota(100);c__Dothideomycetes(100);o__Capnodiales(100);o__Capnodiales_unclassified(100);o__Capnodiales_unclassified(100);o__Capnodiales_unclassified(100); | - | - | - | - | - | - | - | Unassigned | - |
| Otu05546 | 5 | k__Fungi(100);p__Ascomycota(100);p__Ascomycota_unclassified(100);p__Ascomycota_unclassified(100);p__Ascomycota_unclassified(100);p__Ascomycota_unclassified(100);p__Ascomycota_unclassified(100); | - | - | - | - | - | - | - | Unassigned | - |
| Otu05549 | 5 | k__Fungi(100);k__Fungi_unclassified(61);k__Fungi_unclassified(61);k__Fungi_unclassified(61);k__Fungi_unclassified(61);k__Fungi_unclassified(61);k__Fungi_unclassified(61); | - | - | - | - | - | - | - | Unassigned | - |
| Otu05553 | 5 | k__Fungi(100);p__Ascomycota(100);p__Ascomycota_unclassified(100);p__Ascomycota_unclassified(100);p__Ascomycota_unclassified(100);p__Ascomycota_unclassified(100);p__Ascomycota_unclassified(100); | - | - | - | - | - | - | - | Unassigned | - |
| Otu05555 | 5 | k__Fungi(100);k__Fungi_unclassified(80);k__Fungi_unclassified(80);k__Fungi_unclassified(80);k__Fungi_unclassified(80);k__Fungi_unclassified(80);k__Fungi_unclassified(80); | - | - | - | - | - | - | - | Unassigned | - |
| Otu05556 | 5 | k__Fungi(100);p__Ascomycota(100);p__Ascomycota_unclassified(100);p__Ascomycota_unclassified(100);p__Ascomycota_unclassified(100);p__Ascomycota_unclassified(100);p__Ascomycota_unclassified(100); | - | - | - | - | - | - | - | Unassigned | - |
| Otu05558 | 5 | k__Fungi(100);p__Basidiomycota(100);p__Basidiomycota_unclassified(100);p__Basidiomycota_unclassified(100);p__Basidiomycota_unclassified(100);p__Basidiomycota_unclassified(100);p__Basidiomycota_unclassified(100); | - | - | - | - | - | - | - | Unassigned | - |
| Otu05560 | 5 | k__Fungi(100);p__Ascomycota(61);p__Ascomycota_unclassified(61);p__Ascomycota_unclassified(61);p__Ascomycota_unclassified(61);p__Ascomycota_unclassified(61);p__Ascomycota_unclassified(61); | - | - | - | - | - | - | - | Unassigned | - |
| Otu05562 | 5 | k__Fungi(100);p__Ascomycota(100);p__Ascomycota_unclassified(100);p__Ascomycota_unclassified(100);p__Ascomycota_unclassified(100);p__Ascomycota_unclassified(100);p__Ascomycota_unclassified(100); | - | - | - | - | - | - | - | Unassigned | - |
| Otu05566 | 5 | k__Fungi(100);p__Ascomycota(100);p__Ascomycota_unclassified(100);p__Ascomycota_unclassified(100);p__Ascomycota_unclassified(100);p__Ascomycota_unclassified(100);p__Ascomycota_unclassified(100); | - | - | - | - | - | - | - | Unassigned | - |
| Otu05569 | 5 | k__Fungi(100);p__Basidiomycota(100);c__Agaricomycetes(100);c__Agaricomycetes_unclassified(100);c__Agaricomycetes_unclassified(100);c__Agaricomycetes_unclassified(100);c__Agaricomycetes_unclassified(100); | - | - | - | - | - | - | - | Unassigned | - |
| Otu05572 | 5 | k__Fungi(100);p__Ascomycota(100);p__Ascomycota_unclassified(61);p__Ascomycota_unclassified(61);p__Ascomycota_unclassified(61);p__Ascomycota_unclassified(61);p__Ascomycota_unclassified(61); | - | - | - | - | - | - | - | Unassigned | - |
| Otu05573 | 5 | k__Fungi(100);k__Fungi_unclassified(61);k__Fungi_unclassified(61);k__Fungi_unclassified(61);k__Fungi_unclassified(61);k__Fungi_unclassified(61);k__Fungi_unclassified(61); | - | - | - | - | - | - | - | Unassigned | - |
| Otu05576 | 5 | k__Fungi(100);p__Basidiomycota(80);p__Basidiomycota_unclassified(80);p__Basidiomycota_unclassified(80);p__Basidiomycota_unclassified(80);p__Basidiomycota_unclassified(80);p__Basidiomycota_unclassified(80); | - | - | - | - | - | - | - | Unassigned | - |
| Otu05577 | 5 | k__Fungi(100);k__Fungi_unclassified(61);k__Fungi_unclassified(61);k__Fungi_unclassified(61);k__Fungi_unclassified(61);k__Fungi_unclassified(61);k__Fungi_unclassified(61); | - | - | - | - | - | - | - | Unassigned | - |
| Otu05578 | 5 | k__Fungi(100);p__Ascomycota(100);c__Dothideomycetes(100);o__Pleosporales(100);o__Pleosporales_unclassified(100);o__Pleosporales_unclassified(100);o__Pleosporales_unclassified(100); | - | - | - | - | - | - | - | Unassigned | - |
| Otu05580 | 5 | k__Fungi(100);p__Basidiomycota(100);c__Tremellomycetes(100);c__Tremellomycetes_unclassified(80);c__Tremellomycetes_unclassified(80);c__Tremellomycetes_unclassified(80);c__Tremellomycetes_unclassified(80); | - | - | - | - | - | - | - | Unassigned | - |
| Otu05584 | 5 | k__Fungi(100);p__Ascomycota(100);c__Sordariomycetes(100);c__Sordariomycetes_unclassified(100);c__Sordariomycetes_unclassified(100);c__Sordariomycetes_unclassified(100);c__Sordariomycetes_unclassified(100); | - | - | - | - | - | - | - | Unassigned | - |
| Otu05585 | 5 | k__Fungi(100);p__Basidiomycota(100);p__Basidiomycota_unclassified(100);p__Basidiomycota_unclassified(100);p__Basidiomycota_unclassified(100);p__Basidiomycota_unclassified(100);p__Basidiomycota_unclassified(100); | - | - | - | - | - | - | - | Unassigned | - |
| Otu05591 | 5 | k__Fungi(100);k__Fungi_unclassified(80);k__Fungi_unclassified(80);k__Fungi_unclassified(80);k__Fungi_unclassified(80);k__Fungi_unclassified(80);k__Fungi_unclassified(80); | - | - | - | - | - | - | - | Unassigned | - |
| Otu05597 | 5 | k__Plantae(100);p__unclassified_Plantae(100);c__unclassified_Plantae(100);o__unclassified_Plantae(100);f__unclassified_Plantae(100);g__unclassified_Plantae(100);s__Plantae_sp(100); | - | - | - | - | - | - | - | Unassigned | - |
| Otu05598 | 5 | k__Fungi(100);p__Ascomycota(100);p__Ascomycota_unclassified(100);p__Ascomycota_unclassified(100);p__Ascomycota_unclassified(100);p__Ascomycota_unclassified(100);p__Ascomycota_unclassified(100); | - | - | - | - | - | - | - | Unassigned | - |
| Otu05601 | 5 | k__Fungi(100);p__Basidiomycota(100);p__Basidiomycota_unclassified(100);p__Basidiomycota_unclassified(100);p__Basidiomycota_unclassified(100);p__Basidiomycota_unclassified(100);p__Basidiomycota_unclassified(100); | - | - | - | - | - | - | - | Unassigned | - |
| Otu05602 | 5 | k__Fungi(100);p__Ascomycota(100);p__Ascomycota_unclassified(100);p__Ascomycota_unclassified(100);p__Ascomycota_unclassified(100);p__Ascomycota_unclassified(100);p__Ascomycota_unclassified(100); | - | - | - | - | - | - | - | Unassigned | - |
| Otu05604 | 5 | k__Fungi(100);p__Ascomycota(100);c__Dothideomycetes(100);o__Pleosporales(100);o__Pleosporales_unclassified(100);o__Pleosporales_unclassified(100);o__Pleosporales_unclassified(100); | - | - | - | - | - | - | - | Unassigned | - |
| Otu05605 | 5 | k__Fungi(100);k__Fungi_unclassified(61);k__Fungi_unclassified(61);k__Fungi_unclassified(61);k__Fungi_unclassified(61);k__Fungi_unclassified(61);k__Fungi_unclassified(61); | - | - | - | - | - | - | - | Unassigned | - |
| Otu05611 | 5 | k__Fungi(100);p__Ascomycota(100);p__Ascomycota_unclassified(100);p__Ascomycota_unclassified(100);p__Ascomycota_unclassified(100);p__Ascomycota_unclassified(100);p__Ascomycota_unclassified(100); | - | - | - | - | - | - | - | Unassigned | - |
| Otu05618 | 5 | k__Fungi(100);p__Ascomycota(61);p__Ascomycota_unclassified(61);p__Ascomycota_unclassified(61);p__Ascomycota_unclassified(61);p__Ascomycota_unclassified(61);p__Ascomycota_unclassified(61); | - | - | - | - | - | - | - | Unassigned | - |
| Otu05620 | 5 | k__Fungi(100);k__Fungi_unclassified(100);k__Fungi_unclassified(100);k__Fungi_unclassified(100);k__Fungi_unclassified(100);k__Fungi_unclassified(100);k__Fungi_unclassified(100); | - | - | - | - | - | - | - | Unassigned | - |
| Otu05621 | 5 | k__Fungi(100);p__Ascomycota(61);p__Ascomycota_unclassified(61);p__Ascomycota_unclassified(61);p__Ascomycota_unclassified(61);p__Ascomycota_unclassified(61);p__Ascomycota_unclassified(61); | - | - | - | - | - | - | - | Unassigned | - |
| Otu05622 | 5 | k__Fungi(100);p__Basidiomycota(100);c__Agaricomycetes(100);c__Agaricomycetes_unclassified(100);c__Agaricomycetes_unclassified(100);c__Agaricomycetes_unclassified(100);c__Agaricomycetes_unclassified(100); | - | - | - | - | - | - | - | Unassigned | - |
| Otu05623 | 5 | k__Fungi(100);k__Fungi_unclassified(100);k__Fungi_unclassified(100);k__Fungi_unclassified(100);k__Fungi_unclassified(100);k__Fungi_unclassified(100);k__Fungi_unclassified(100); | - | - | - | - | - | - | - | Unassigned | - |
| Otu05627 | 5 | k__Fungi(100);k__Fungi_unclassified(80);k__Fungi_unclassified(80);k__Fungi_unclassified(80);k__Fungi_unclassified(80);k__Fungi_unclassified(80);k__Fungi_unclassified(80); | - | - | - | - | - | - | - | Unassigned | - |
| Otu05628 | 5 | k__Fungi(100);k__Fungi_unclassified(100);k__Fungi_unclassified(100);k__Fungi_unclassified(100);k__Fungi_unclassified(100);k__Fungi_unclassified(100);k__Fungi_unclassified(100); | - | - | - | - | - | - | - | Unassigned | - |
| Otu05629 | 5 | k__Fungi(100);p__Basidiomycota(100);c__Agaricomycetes(100);o__Auriculariales(100);f__Auriculariales_family_Incertae_sedis(100);f__Auriculariales_family_Incertae_sedis_unclassified(100);f__Auriculariales_family_Incertae_sedis_unclassified(100); | - | - | - | - | - | - | - | Unassigned | - |
| Otu05631 | 5 | k__Fungi(100);p__Ascomycota(100);c__Sordariomycetes(100);c__Sordariomycetes_unclassified(100);c__Sordariomycetes_unclassified(100);c__Sordariomycetes_unclassified(100);c__Sordariomycetes_unclassified(100); | - | - | - | - | - | - | - | Unassigned | - |
| Otu05633 | 5 | k__Fungi(100);k__Fungi_unclassified(100);k__Fungi_unclassified(100);k__Fungi_unclassified(100);k__Fungi_unclassified(100);k__Fungi_unclassified(100);k__Fungi_unclassified(100); | - | - | - | - | - | - | - | Unassigned | - |
| Otu05635 | 5 | k__Fungi(100);k__Fungi_unclassified(100);k__Fungi_unclassified(100);k__Fungi_unclassified(100);k__Fungi_unclassified(100);k__Fungi_unclassified(100);k__Fungi_unclassified(100); | - | - | - | - | - | - | - | Unassigned | - |
| Otu05636 | 5 | k__Fungi(100);k__Fungi_unclassified(100);k__Fungi_unclassified(100);k__Fungi_unclassified(100);k__Fungi_unclassified(100);k__Fungi_unclassified(100);k__Fungi_unclassified(100); | - | - | - | - | - | - | - | Unassigned | - |
| Otu05638 | 5 | k__Fungi(100);p__Ascomycota(80);p__Ascomycota_unclassified(80);p__Ascomycota_unclassified(80);p__Ascomycota_unclassified(80);p__Ascomycota_unclassified(80);p__Ascomycota_unclassified(80); | - | - | - | - | - | - | - | Unassigned | - |
| Otu05639 | 5 | k__Fungi(100);p__Ascomycota(100);p__Ascomycota_unclassified(100);p__Ascomycota_unclassified(100);p__Ascomycota_unclassified(100);p__Ascomycota_unclassified(100);p__Ascomycota_unclassified(100); | - | - | - | - | - | - | - | Unassigned | - |
| Otu05641 | 5 | k__Fungi(100);p__Ascomycota(100);c__Eurotiomycetes(100);o__Chaetothyriales(100);o__Chaetothyriales_unclassified(100);o__Chaetothyriales_unclassified(100);o__Chaetothyriales_unclassified(100); | - | - | - | - | - | - | - | Unassigned | - |
| Otu05642 | 5 | k__Fungi(100);p__Ascomycota(100);c__Leotiomycetes(100);o__Helotiales(100);f__Helotiales_family_Incertae_sedis(100);f__Helotiales_family_Incertae_sedis_unclassified(100);f__Helotiales_family_Incertae_sedis_unclassified(100); | - | - | - | - | - | - | - | Unassigned | - |
| Otu05649 | 5 | k__Fungi(100);p__Ascomycota(100);p__Ascomycota_unclassified(100);p__Ascomycota_unclassified(100);p__Ascomycota_unclassified(100);p__Ascomycota_unclassified(100);p__Ascomycota_unclassified(100); | - | - | - | - | - | - | - | Unassigned | - |
| Otu05650 | 5 | k__Fungi(100);p__Basidiomycota(100);p__Basidiomycota_unclassified(100);p__Basidiomycota_unclassified(100);p__Basidiomycota_unclassified(100);p__Basidiomycota_unclassified(100);p__Basidiomycota_unclassified(100); | - | - | - | - | - | - | - | Unassigned | - |
| Otu05651 | 5 | k__Fungi(100);k__Fungi_unclassified(100);k__Fungi_unclassified(100);k__Fungi_unclassified(100);k__Fungi_unclassified(100);k__Fungi_unclassified(100);k__Fungi_unclassified(100); | - | - | - | - | - | - | - | Unassigned | - |
| Otu05656 | 5 | k__Fungi(100);p__Basidiomycota(100);c__Agaricomycetes(100);c__Agaricomycetes_unclassified(100);c__Agaricomycetes_unclassified(100);c__Agaricomycetes_unclassified(100);c__Agaricomycetes_unclassified(100); | - | - | - | - | - | - | - | Unassigned | - |
| Otu05657 | 5 | k__Fungi(100);p__Basidiomycota(100);c__Agaricomycetes(100);o__Corticiales(100);o__Corticiales_unclassified(80);o__Corticiales_unclassified(80);o__Corticiales_unclassified(80); | - | - | - | - | - | - | - | Unassigned | - |
| Otu05659 | 5 | k__Fungi(100);p__Ascomycota(100);p__Ascomycota_unclassified(100);p__Ascomycota_unclassified(100);p__Ascomycota_unclassified(100);p__Ascomycota_unclassified(100);p__Ascomycota_unclassified(100); | - | - | - | - | - | - | - | Unassigned | - |
| Otu05660 | 5 | k__Fungi(100);p__Ascomycota(100);c__Leotiomycetes(80);c__Leotiomycetes_unclassified(80);c__Leotiomycetes_unclassified(80);c__Leotiomycetes_unclassified(80);c__Leotiomycetes_unclassified(80); | - | - | - | - | - | - | - | Unassigned | - |
| Otu05661 | 5 | k__Fungi(100);p__Basidiomycota(100);c__Agaricomycetes(100);c__Agaricomycetes_unclassified(100);c__Agaricomycetes_unclassified(100);c__Agaricomycetes_unclassified(100);c__Agaricomycetes_unclassified(100); | - | - | - | - | - | - | - | Unassigned | - |
| Otu05666 | 5 | k__Fungi(100);p__Ascomycota(80);p__Ascomycota_unclassified(80);p__Ascomycota_unclassified(80);p__Ascomycota_unclassified(80);p__Ascomycota_unclassified(80);p__Ascomycota_unclassified(80); | - | - | - | - | - | - | - | Unassigned | - |
| Otu05667 | 5 | k__Fungi(100);p__Ascomycota(100);p__Ascomycota_unclassified(100);p__Ascomycota_unclassified(100);p__Ascomycota_unclassified(100);p__Ascomycota_unclassified(100);p__Ascomycota_unclassified(100); | - | - | - | - | - | - | - | Unassigned | - |
| Otu05669 | 5 | k__Fungi(100);p__Basidiomycota(100);p__Basidiomycota_unclassified(100);p__Basidiomycota_unclassified(100);p__Basidiomycota_unclassified(100);p__Basidiomycota_unclassified(100);p__Basidiomycota_unclassified(100); | - | - | - | - | - | - | - | Unassigned | - |
| Otu05673 | 5 | k__Fungi(100);p__Ascomycota(100);c__Leotiomycetes(80);c__Leotiomycetes_unclassified(80);c__Leotiomycetes_unclassified(80);c__Leotiomycetes_unclassified(80);c__Leotiomycetes_unclassified(80); | - | - | - | - | - | - | - | Unassigned | - |
| Otu05677 | 5 | k__Fungi(100);p__Ascomycota(80);p__Ascomycota_unclassified(80);p__Ascomycota_unclassified(80);p__Ascomycota_unclassified(80);p__Ascomycota_unclassified(80);p__Ascomycota_unclassified(80); | - | - | - | - | - | - | - | Unassigned | - |
| Otu05678 | 5 | k__Fungi(100);k__Fungi_unclassified(61);k__Fungi_unclassified(61);k__Fungi_unclassified(61);k__Fungi_unclassified(61);k__Fungi_unclassified(61);k__Fungi_unclassified(61); | - | - | - | - | - | - | - | Unassigned | - |
| Otu05679 | 5 | k__Fungi(100);p__Basidiomycota(100);c__Agaricomycetes(100);o__Corticiales(100);f__unclassified_Corticiales(100);g__unclassified_Corticiales(100);s__Corticiales_sp(100); | - | - | - | - | - | - | - | Unassigned | - |
| Otu05681 | 5 | k__Fungi(100);p__Basidiomycota(100);c__Agaricomycetes(100);c__Agaricomycetes_unclassified(100);c__Agaricomycetes_unclassified(100);c__Agaricomycetes_unclassified(100);c__Agaricomycetes_unclassified(100); | - | - | - | - | - | - | - | Unassigned | - |
| Otu05689 | 5 | k__Fungi(100);k__Fungi_unclassified(100);k__Fungi_unclassified(100);k__Fungi_unclassified(100);k__Fungi_unclassified(100);k__Fungi_unclassified(100);k__Fungi_unclassified(100); | - | - | - | - | - | - | - | Unassigned | - |
| Otu05691 | 5 | k__Fungi(100);p__Ascomycota(61);p__Ascomycota_unclassified(61);p__Ascomycota_unclassified(61);p__Ascomycota_unclassified(61);p__Ascomycota_unclassified(61);p__Ascomycota_unclassified(61); | - | - | - | - | - | - | - | Unassigned | - |
| Otu05694 | 5 | k__Fungi(100);p__Basidiomycota(100);p__Basidiomycota_unclassified(100);p__Basidiomycota_unclassified(100);p__Basidiomycota_unclassified(100);p__Basidiomycota_unclassified(100);p__Basidiomycota_unclassified(100); | - | - | - | - | - | - | - | Unassigned | - |
| Otu05695 | 5 | k__Fungi(100);p__Ascomycota(100);c__Leotiomycetes(61);o__Helotiales(61);f__Helotiales_family_Incertae_sedis(61);f__Helotiales_family_Incertae_sedis_unclassified(61);f__Helotiales_family_Incertae_sedis_unclassified(61); | - | - | - | - | - | - | - | Unassigned | - |
| Otu05698 | 5 | k__Fungi(100);p__Basidiomycota(100);p__Basidiomycota_unclassified(100);p__Basidiomycota_unclassified(100);p__Basidiomycota_unclassified(100);p__Basidiomycota_unclassified(100);p__Basidiomycota_unclassified(100); | - | - | - | - | - | - | - | Unassigned | - |
| Otu05701 | 5 | k__Fungi(100);p__Ascomycota(61);p__Ascomycota_unclassified(61);p__Ascomycota_unclassified(61);p__Ascomycota_unclassified(61);p__Ascomycota_unclassified(61);p__Ascomycota_unclassified(61); | - | - | - | - | - | - | - | Unassigned | - |
| Otu05702 | 5 | k__Fungi(100);k__Fungi_unclassified(80);k__Fungi_unclassified(80);k__Fungi_unclassified(80);k__Fungi_unclassified(80);k__Fungi_unclassified(80);k__Fungi_unclassified(80); | - | - | - | - | - | - | - | Unassigned | - |
| Otu05705 | 5 | k__Fungi(100);k__Fungi_unclassified(100);k__Fungi_unclassified(100);k__Fungi_unclassified(100);k__Fungi_unclassified(100);k__Fungi_unclassified(100);k__Fungi_unclassified(100); | - | - | - | - | - | - | - | Unassigned | - |
| Otu05706 | 5 | k__Fungi(100);p__Ascomycota(61);p__Ascomycota_unclassified(61);p__Ascomycota_unclassified(61);p__Ascomycota_unclassified(61);p__Ascomycota_unclassified(61);p__Ascomycota_unclassified(61); | - | - | - | - | - | - | - | Unassigned | - |
| Otu05708 | 5 | k__Fungi(100);p__Ascomycota(100);p__Ascomycota_unclassified(80);p__Ascomycota_unclassified(80);p__Ascomycota_unclassified(80);p__Ascomycota_unclassified(80);p__Ascomycota_unclassified(80); | - | - | - | - | - | - | - | Unassigned | - |
| Otu05710 | 5 | k__Fungi(100);p__Ascomycota(100);c__Leotiomycetes(80);o__Helotiales(80);o__Helotiales_unclassified(80);o__Helotiales_unclassified(80);o__Helotiales_unclassified(80); | - | - | - | - | - | - | - | Unassigned | - |
| Otu05712 | 5 | k__Fungi(100);p__Basidiomycota(100);c__Agaricomycetes(100);o__Russulales(80);o__Russulales_unclassified(80);o__Russulales_unclassified(80);o__Russulales_unclassified(80); | - | - | - | - | - | - | - | Unassigned | - |
| Otu05715 | 5 | k__Fungi(100);p__Ascomycota(100);p__Ascomycota_unclassified(100);p__Ascomycota_unclassified(100);p__Ascomycota_unclassified(100);p__Ascomycota_unclassified(100);p__Ascomycota_unclassified(100); | - | - | - | - | - | - | - | Unassigned | - |
| Otu05716 | 5 | k__Fungi(100);p__Ascomycota(61);p__Ascomycota_unclassified(61);p__Ascomycota_unclassified(61);p__Ascomycota_unclassified(61);p__Ascomycota_unclassified(61);p__Ascomycota_unclassified(61); | - | - | - | - | - | - | - | Unassigned | - |
| Otu05720 | 5 | k__Fungi(100);p__Ascomycota(100);p__Ascomycota_unclassified(100);p__Ascomycota_unclassified(100);p__Ascomycota_unclassified(100);p__Ascomycota_unclassified(100);p__Ascomycota_unclassified(100); | - | - | - | - | - | - | - | Unassigned | - |
| Otu05722 | 5 | k__Fungi(100);p__Basidiomycota(100);c__Agaricomycetes(100);o__Trechisporales(100);f__unclassified_Trechisporales(100);g__unclassified_Trechisporales(100);s__Trechisporales_sp(100); | - | - | - | - | - | - | - | Unassigned | - |

| | | | | | | | | | | | |
|---|---|---|---|---|---|---|---|---|---|---|---|
| Otu05724 | 5 | k__Fungi(100);p__Ascomycota(80);p__Ascomycota_unclassified(80);p__Ascomycota_unclassified(80);p__Ascomycota_unclassified(80);p__Ascomycota_unclassified(80);p__Ascomycota_unclassified(80); | - | - | - | - | - | - | - | Unassigned | - |
| Otu05729 | 5 | k__Fungi(100);k__Fungi_unclassified(80);k__Fungi_unclassified(80);k__Fungi_unclassified(80);k__Fungi_unclassified(80);k__Fungi_unclassified(80);k__Fungi_unclassified(80); | - | - | - | - | - | - | - | Unassigned | - |
| Otu05730 | 5 | k__Fungi(100);k__Fungi_unclassified(61);k__Fungi_unclassified(61);k__Fungi_unclassified(61);k__Fungi_unclassified(61);k__Fungi_unclassified(61);k__Fungi_unclassified(61); | - | - | - | - | - | - | - | Unassigned | - |
| Otu05732 | 5 | k__Fungi(100);p__Basidiomycota(100);c__Tremellomycetes(100);c__Tremellomycetes_unclassified(100);c__Tremellomycetes_unclassified(100);c__Tremellomycetes_unclassified(100);c__Tremellomycetes_unclassified(100); | - | - | - | - | - | - | - | Unassigned | - |
| Otu05734 | 5 | k__Fungi(100);k__Fungi_unclassified(61);k__Fungi_unclassified(61);k__Fungi_unclassified(61);k__Fungi_unclassified(61);k__Fungi_unclassified(61);k__Fungi_unclassified(61); | - | - | - | - | - | - | - | Unassigned | - |
| Otu05735 | 5 | k__Fungi(100);p__Ascomycota(80);p__Ascomycota_unclassified(80);p__Ascomycota_unclassified(80);p__Ascomycota_unclassified(80);p__Ascomycota_unclassified(80);p__Ascomycota_unclassified(80); | - | - | - | - | - | - | - | Unassigned | - |
| Otu05737 | 5 | k__Fungi(100);p__Ascomycota(100);c__Dothideomycetes(100);o__Pleosporales(100);o__Pleosporales_unclassified(100);o__Pleosporales_unclassified(100);o__Pleosporales_unclassified(100); | - | - | - | - | - | - | - | Unassigned | - |
| Otu05745 | 5 | k__Fungi(100);p__Ascomycota(80);p__Ascomycota_unclassified(80);p__Ascomycota_unclassified(80);p__Ascomycota_unclassified(80);p__Ascomycota_unclassified(80);p__Ascomycota_unclassified(80); | - | - | - | - | - | - | - | Unassigned | - |
| Otu05748 | 5 | k__Fungi(100);p__Ascomycota(100);c__Dothideomycetes(80);c__Dothideomycetes_unclassified(80);c__Dothideomycetes_unclassified(80);c__Dothideomycetes_unclassified(80);c__Dothideomycetes_unclassified(80); | - | - | - | - | - | - | - | Unassigned | - |
| Otu05754 | 5 | k__Fungi(100);p__Basidiomycota(100);c__Agaricomycetes(100);c__Agaricomycetes_unclassified(100);c__Agaricomycetes_unclassified(100);c__Agaricomycetes_unclassified(100);c__Agaricomycetes_unclassified(100); | - | - | - | - | - | - | - | Unassigned | - |
| Otu05756 | 5 | k__Fungi(100);p__Ascomycota(100);c__Dothideomycetes(100);c__Dothideomycetes_unclassified(80);c__Dothideomycetes_unclassified(80);c__Dothideomycetes_unclassified(80);c__Dothideomycetes_unclassified(80); | - | - | - | - | - | - | - | Unassigned | - |
| Otu05760 | 5 | k__Fungi(100);k__Fungi_unclassified(100);k__Fungi_unclassified(100);k__Fungi_unclassified(100);k__Fungi_unclassified(100);k__Fungi_unclassified(100);k__Fungi_unclassified(100); | - | - | - | - | - | - | - | Unassigned | - |
| Otu05764 | 5 | k__Fungi(100);p__Basidiomycota(100);c__Agaricomycetes(100);o__Cantharellales(100);o__Cantharellales_unclassified(100);o__Cantharellales_unclassified(100);o__Cantharellales_unclassified(100); | - | - | - | - | - | - | - | Unassigned | - |
| Otu05766 | 5 | k__Fungi(100);p__Ascomycota(100);c__Leotiomycetes(100);o__Helotiales(100);o__Helotiales_unclassified(100);o__Helotiales_unclassified(100);o__Helotiales_unclassified(100); | - | - | - | - | - | - | - | Unassigned | - |
| Otu05767 | 5 | k__Fungi(100);p__Ascomycota(100);p__Ascomycota_unclassified(100);p__Ascomycota_unclassified(100);p__Ascomycota_unclassified(100);p__Ascomycota_unclassified(100);p__Ascomycota_unclassified(100); | - | - | - | - | - | - | - | Unassigned | - |
| Otu05768 | 5 | k__Fungi(100);k__Fungi_unclassified(61);k__Fungi_unclassified(61);k__Fungi_unclassified(61);k__Fungi_unclassified(61);k__Fungi_unclassified(61);k__Fungi_unclassified(61); | - | - | - | - | - | - | - | Unassigned | - |
| Otu05771 | 5 | k__Fungi(100);p__Ascomycota(100);c__Leotiomycetes(100);o__Helotiales(100);f__Helotiales_family_Incertae_sedis(100);f__Helotiales_family_Incertae_sedis_unclassified(100);f__Helotiales_family_Incertae_sedis_unclassified(100); | - | - | - | - | - | - | - | Unassigned | - |
| Otu05777 | 5 | k__Fungi(100);p__Basidiomycota(100);c__Agaricomycetes(100);c__Agaricomycetes_unclassified(100);c__Agaricomycetes_unclassified(100);c__Agaricomycetes_unclassified(100);c__Agaricomycetes_unclassified(100); | - | - | - | - | - | - | - | Unassigned | - |
| Otu05781 | 5 | k__Fungi(100);p__Basidiomycota(100);p__Basidiomycota_unclassified(100);p__Basidiomycota_unclassified(100);p__Basidiomycota_unclassified(100);p__Basidiomycota_unclassified(100);p__Basidiomycota_unclassified(100); | - | - | - | - | - | - | - | Unassigned | - |
| Otu05782 | 5 | k__Fungi(100);p__Ascomycota(80);p__Ascomycota_unclassified(80);p__Ascomycota_unclassified(80);p__Ascomycota_unclassified(80);p__Ascomycota_unclassified(80);p__Ascomycota_unclassified(80); | - | - | - | - | - | - | - | Unassigned | - |
| Otu05787 | 5 | k__Fungi(100);k__Fungi_unclassified(61);k__Fungi_unclassified(61);k__Fungi_unclassified(61);k__Fungi_unclassified(61);k__Fungi_unclassified(61);k__Fungi_unclassified(61); | - | - | - | - | - | - | - | Unassigned | - |
| Otu05788 | 5 | k__Fungi(100);k__Fungi_unclassified(61);k__Fungi_unclassified(61);k__Fungi_unclassified(61);k__Fungi_unclassified(61);k__Fungi_unclassified(61);k__Fungi_unclassified(61); | - | - | - | - | - | - | - | Unassigned | - |
| Otu05789 | 5 | k__Fungi(100);p__Basidiomycota(100);c__Agaricomycetes(100);o__Boletales(100);o__Boletales_unclassified(100);o__Boletales_unclassified(100);o__Boletales_unclassified(100); | - | - | - | - | - | - | - | Unassigned | - |
| Otu05793 | 5 | k__Fungi(100);p__Basidiomycota(100);c__Agaricomycetes(100);o__Cantharellales(100);f__Cantharellaceae(80);g__unclassified_Cantharellaceae(80);s__Cantharellaceae_sp(80); | - | - | - | - | - | - | - | Unassigned | - |
| Otu05805 | 5 | k__Fungi(100);p__Basidiomycota(100);c__Agaricomycetes(100);c__Agaricomycetes_unclassified(100);c__Agaricomycetes_unclassified(100);c__Agaricomycetes_unclassified(100);c__Agaricomycetes_unclassified(100); | - | - | - | - | - | - | - | Unassigned | - |
| Otu05808 | 5 | k__Fungi(100);k__Fungi_unclassified(100);k__Fungi_unclassified(100);k__Fungi_unclassified(100);k__Fungi_unclassified(100);k__Fungi_unclassified(100);k__Fungi_unclassified(100); | - | - | - | - | - | - | - | Unassigned | - |
| Otu05810 | 5 | k__Fungi(100);k__Fungi_unclassified(61);k__Fungi_unclassified(61);k__Fungi_unclassified(61);k__Fungi_unclassified(61);k__Fungi_unclassified(61);k__Fungi_unclassified(61); | - | - | - | - | - | - | - | Unassigned | - |
| Otu05811 | 5 | k__Fungi(100);p__Basidiomycota(100);c__Agaricomycetes(100);c__Agaricomycetes_unclassified(100);c__Agaricomycetes_unclassified(100);c__Agaricomycetes_unclassified(100);c__Agaricomycetes_unclassified(100); | - | - | - | - | - | - | - | Unassigned | - |
| Otu05815 | 5 | k__Fungi(100);p__Ascomycota(100);p__Ascomycota_unclassified(100);p__Ascomycota_unclassified(100);p__Ascomycota_unclassified(100);p__Ascomycota_unclassified(100);p__Ascomycota_unclassified(100); | - | - | - | - | - | - | - | Unassigned | - |
| Otu05816 | 5 | k__Fungi(100);p__Basidiomycota(100);p__Basidiomycota_unclassified(100);p__Basidiomycota_unclassified(100);p__Basidiomycota_unclassified(100);p__Basidiomycota_unclassified(100);p__Basidiomycota_unclassified(100); | - | - | - | - | - | - | - | Unassigned | - |
| Otu05817 | 5 | k__Fungi(100);p__Basidiomycota(100);p__Basidiomycota_unclassified(100);p__Basidiomycota_unclassified(100);p__Basidiomycota_unclassified(100);p__Basidiomycota_unclassified(100);p__Basidiomycota_unclassified(100); | - | - | - | - | - | - | - | Unassigned | - |
| Otu05820 | 5 | k__Fungi(100);p__Ascomycota(61);p__Ascomycota_unclassified(61);p__Ascomycota_unclassified(61);p__Ascomycota_unclassified(61);p__Ascomycota_unclassified(61);p__Ascomycota_unclassified(61); | - | - | - | - | - | - | - | Unassigned | - |
| Otu05823 | 5 | k__Fungi(100);p__Basidiomycota(100);c__Agaricomycetes(100);c__Agaricomycetes_unclassified(100);c__Agaricomycetes_unclassified(100);c__Agaricomycetes_unclassified(100);c__Agaricomycetes_unclassified(100); | - | - | - | - | - | - | - | Unassigned | - |
| Otu05825 | 5 | k__Fungi(100);p__Basidiomycota(100);p__Basidiomycota_unclassified(100);p__Basidiomycota_unclassified(100);p__Basidiomycota_unclassified(100);p__Basidiomycota_unclassified(100);p__Basidiomycota_unclassified(100); | - | - | - | - | - | - | - | Unassigned | - |
| Otu05826 | 5 | k__Fungi(100);k__Fungi_unclassified(61);k__Fungi_unclassified(61);k__Fungi_unclassified(61);k__Fungi_unclassified(61);k__Fungi_unclassified(61);k__Fungi_unclassified(61); | - | - | - | - | - | - | - | Unassigned | - |
| Otu05828 | 5 | k__Fungi(100);k__Fungi_unclassified(100);k__Fungi_unclassified(100);k__Fungi_unclassified(100);k__Fungi_unclassified(100);k__Fungi_unclassified(100);k__Fungi_unclassified(100); | - | - | - | - | - | - | - | Unassigned | - |
| Otu05830 | 5 | k__Fungi(100);k__Fungi_unclassified(61);k__Fungi_unclassified(61);k__Fungi_unclassified(61);k__Fungi_unclassified(61);k__Fungi_unclassified(61);k__Fungi_unclassified(61); | - | - | - | - | - | - | - | Unassigned | - |
| Otu05834 | 5 | k__Fungi(100);p__Ascomycota(100);c__Eurotiomycetes(100);c__Eurotiomycetes_unclassified(100);c__Eurotiomycetes_unclassified(100);c__Eurotiomycetes_unclassified(100);c__Eurotiomycetes_unclassified(100); | - | - | - | - | - | - | - | Unassigned | - |
| Otu05835 | 5 | k__Fungi(100);p__Ascomycota(100);c__Lecanoromycetes(100);c__Lecanoromycetes_unclassified(100);c__Lecanoromycetes_unclassified(100);c__Lecanoromycetes_unclassified(100);c__Lecanoromycetes_unclassified(100); | - | - | - | - | - | - | - | Unassigned | - |
| Otu05836 | 5 | k__Fungi(100);p__Basidiomycota(100);p__Basidiomycota_unclassified(100);p__Basidiomycota_unclassified(100);p__Basidiomycota_unclassified(100);p__Basidiomycota_unclassified(100);p__Basidiomycota_unclassified(100); | - | - | - | - | - | - | - | Unassigned | - |
| Otu05838 | 5 | k__Fungi(100);k__Fungi_unclassified(100);k__Fungi_unclassified(100);k__Fungi_unclassified(100);k__Fungi_unclassified(100);k__Fungi_unclassified(100);k__Fungi_unclassified(100); | - | - | - | - | - | - | - | Unassigned | - |
| Otu05843 | 5 | k__Fungi(100);p__Basidiomycota(100);p__Basidiomycota_unclassified(100);p__Basidiomycota_unclassified(100);p__Basidiomycota_unclassified(100);p__Basidiomycota_unclassified(100);p__Basidiomycota_unclassified(100); | - | - | - | - | - | - | - | Unassigned | - |
| Otu05845 | 5 | k__Fungi(100);k__Fungi_unclassified(80);k__Fungi_unclassified(80);k__Fungi_unclassified(80);k__Fungi_unclassified(80);k__Fungi_unclassified(80);k__Fungi_unclassified(80); | - | - | - | - | - | - | - | Unassigned | - |
| Otu05846 | 5 | k__Fungi(100);p__Ascomycota(100);c__Dothideomycetes(100);o__Pleosporales(100);o__Pleosporales_unclassified(100);o__Pleosporales_unclassified(100);o__Pleosporales_unclassified(100); | - | - | - | - | - | - | - | Unassigned | - |
| Otu05848 | 5 | k__Fungi(100);k__Fungi_unclassified(61);k__Fungi_unclassified(61);k__Fungi_unclassified(61);k__Fungi_unclassified(61);k__Fungi_unclassified(61);k__Fungi_unclassified(61); | - | - | - | - | - | - | - | Unassigned | - |
| Otu05851 | 5 | k__Fungi(100);p__Ascomycota(61);p__Ascomycota_unclassified(61);p__Ascomycota_unclassified(61);p__Ascomycota_unclassified(61);p__Ascomycota_unclassified(61);p__Ascomycota_unclassified(61); | - | - | - | - | - | - | - | Unassigned | - |
| Otu05852 | 5 | k__Fungi(100);p__Ascomycota(61);p__Ascomycota_unclassified(61);p__Ascomycota_unclassified(61);p__Ascomycota_unclassified(61);p__Ascomycota_unclassified(61);p__Ascomycota_unclassified(61); | - | - | - | - | - | - | - | Unassigned | - |
| Otu05853 | 5 | k__Fungi(100);p__Ascomycota(100);c__Dothideomycetes(100);c__Dothideomycetes_unclassified(100);c__Dothideomycetes_unclassified(100);c__Dothideomycetes_unclassified(100);c__Dothideomycetes_unclassified(100); | - | - | - | - | - | - | - | Unassigned | - |
| Otu05854 | 5 | k__Fungi(100);p__Basidiomycota(100);p__Basidiomycota_unclassified(80);p__Basidiomycota_unclassified(80);p__Basidiomycota_unclassified(80);p__Basidiomycota_unclassified(80);p__Basidiomycota_unclassified(80); | - | - | - | - | - | - | - | Unassigned | - |
| Otu05856 | 5 | k__Fungi(100);k__Fungi_unclassified(80);k__Fungi_unclassified(80);k__Fungi_unclassified(80);k__Fungi_unclassified(80);k__Fungi_unclassified(80);k__Fungi_unclassified(80); | - | - | - | - | - | - | - | Unassigned | - |
| Otu05857 | 5 | k__Fungi(100);k__Fungi_unclassified(61);k__Fungi_unclassified(61);k__Fungi_unclassified(61);k__Fungi_unclassified(61);k__Fungi_unclassified(61);k__Fungi_unclassified(61); | - | - | - | - | - | - | - | Unassigned | - |
| Otu05858 | 5 | k__Fungi(100);k__Fungi_unclassified(61);k__Fungi_unclassified(61);k__Fungi_unclassified(61);k__Fungi_unclassified(61);k__Fungi_unclassified(61);k__Fungi_unclassified(61); | - | - | - | - | - | - | - | Unassigned | - |
| Otu05861 | 5 | k__Fungi(100);p__Ascomycota(100);c__Lecanoromycetes(100);c__Lecanoromycetes_unclassified(61);c__Lecanoromycetes_unclassified(61);c__Lecanoromycetes_unclassified(61);c__Lecanoromycetes_unclassified(61); | - | - | - | - | - | - | - | Unassigned | - |
| Otu05864 | 5 | k__Fungi(100);p__Ascomycota(100);p__Ascomycota_unclassified(100);p__Ascomycota_unclassified(100);p__Ascomycota_unclassified(100);p__Ascomycota_unclassified(100);p__Ascomycota_unclassified(100); | - | - | - | - | - | - | - | Unassigned | - |
| Otu05866 | 5 | k__Fungi(100);p__Ascomycota(100);c__Leotiomycetes(100);o__Helotiales(100);o__Helotiales_unclassified(100);o__Helotiales_unclassified(100);o__Helotiales_unclassified(100); | - | - | - | - | - | - | - | Unassigned | - |
| Otu05872 | 5 | k__Fungi(100);p__Ascomycota(80);p__Ascomycota_unclassified(80);p__Ascomycota_unclassified(80);p__Ascomycota_unclassified(80);p__Ascomycota_unclassified(80);p__Ascomycota_unclassified(80); | - | - | - | - | - | - | - | Unassigned | - |
| Otu05879 | 5 | k__Fungi(100);k__Fungi_unclassified(100);k__Fungi_unclassified(100);k__Fungi_unclassified(100);k__Fungi_unclassified(100);k__Fungi_unclassified(100);k__Fungi_unclassified(100); | - | - | - | - | - | - | - | Unassigned | - |
| Otu05880 | 4 | k__Fungi(100);p__Ascomycota(100);c__Saccharomycetes(100);o__Saccharomycetales(100);f__Metschnikowiaceae(100);g__Metschnikowia(100);s__Metschnikowia_pulcherrima(100); | Metschnikowia | 13 | Saprotroph | \|Undefined Saprotroph\| | Yeast | NULL | Probable | As nectar-tap_saprotroph (Põlme et al. 2020); Decay sugar-rich substrates (Põlme et al. 2020) | Põlme S, et al. 2020. Fungal Diversity 105:1-16 ((https://doi.org/10.1007/s13225-020-00466-2)) |
| Otu05881 | 4 | k__Fungi(100);p__Ascomycota(100);c__Dothideomycetes(100);o__Pleosporales(100);f__Venturiaceae(100);f__Venturiaceae_unclassified(100);f__Venturiaceae_unclassified(100); | Venturiaceae | 9 | Pathotroph-Saprotroph-Symbiotroph | Endophyte-Plant Pathogen-Undefined Saprotroph | Microfungus | NULL | Probable | Mostly endophytes and weak parasites of aerial plant parts, including bark and wood, leaf and fruit tissue; rarely biotrophic. Little is known except for the important pathogens (Cannon & Kirk 2007) | Cannon PF, Kirk PM. 2007. Fungal Families of the World. CAB International, Cambridge (ISBN: 978-0851998275) |
| Otu05882 | 4 | k__Fungi(100);p__Ascomycota(100);c__Saccharomycetes(100);o__Saccharomycetales(100);f__Metschnikowiaceae(100);g__Metschnikowia(100);s__Metschnikowia_pulcherrima(75); | Metschnikowia | 13 | Saprotroph | \|Undefined Saprotroph\| | Yeast | NULL | Probable | As nectar-tap_saprotroph (Põlme et al. 2020); Decay sugar-rich substrates (Põlme et al. 2020) | Põlme S, et al. 2020. Fungal Diversity 105:1-16 ((https://doi.org/10.1007/s13225-020-00466-2)) |
| Otu05883 | 4 | k__Fungi(100);p__Ascomycota(100);c__Saccharomycetes(100);o__Saccharomycetales(100);f__Metschnikowiaceae(100);g__Metschnikowia(100);g__Metschnikowia_unclassified(100); | Metschnikowia | 13 | Saprotroph | \|Undefined Saprotroph\| | Yeast | NULL | Probable | As nectar-tap_saprotroph (Põlme et al. 2020); Decay sugar-rich substrates (Põlme et al. 2020) | Põlme S, et al. 2020. Fungal Diversity 105:1-16 ((https://doi.org/10.1007/s13225-020-00466-2)) |
| Otu05884 | 4 | k__Fungi(100);p__Ascomycota(100);c__Saccharomycetes(100);o__Saccharomycetales(100);f__Metschnikowiaceae(100);g__Metschnikowia(100);g__Metschnikowia_unclassified(100); | Metschnikowia | 13 | Saprotroph | \|Undefined Saprotroph\| | Yeast | NULL | Probable | As nectar-tap_saprotroph (Põlme et al. 2020); Decay sugar-rich substrates (Põlme et al. 2020) | Põlme S, et al. 2020. Fungal Diversity 105:1-16 ((https://doi.org/10.1007/s13225-020-00466-2)) |
| Otu05885 | 4 | k__Fungi(100);p__Basidiomycota(100);c__Microbotryomycetes(100);o__Sporidiobolales(100);f__Sporidiobolales_family_Incertae_sedis(100);g__Sporobolomyces(100);s__Sporobolomyces_lactosus(100); | Sporobolomyces | 13 | Pathotroph-Saprotroph | \|Fungal Parasite\|-Plant Saprotroph | Yeast | NULL | Probable | As litter_saprotroph (Põlme et al. 2020); As mycoparasite (Põlme et al. 2020); Decay fungal material (Põlme et al. 2020) | Tedersoo L, et al. 2014. Science 346:e1256688 ((https://doi.org/10.1126/science.1256688)); Purahong W, et al. 2016. Molecular Ecology 25:4059-4074 ((https://doi.org/10.1111/mec.13739)); Põlme S, et al. 2020. Fungal Diversity 105:1-16 ((https://doi.org/10.1007/s13225-020-00466-2)) |
| Otu05886 | 4 | k__Fungi(100);p__Ascomycota(100);c__Saccharomycetes(100);o__Saccharomycetales(100);f__Metschnikowiaceae(100);g__Metschnikowia(100);s__Metschnikowia_pulcherrima(100); | Metschnikowia | 13 | Saprotroph | \|Undefined Saprotroph\| | Yeast | NULL | Probable | As nectar-tap_saprotroph (Põlme et al. 2020); Decay sugar-rich substrates (Põlme et al. 2020) | Põlme S, et al. 2020. Fungal Diversity 105:1-16 ((https://doi.org/10.1007/s13225-020-00466-2)) |
| Otu05887 | 4 | k__Fungi(100);p__Basidiomycota(100);c__Tremellomycetes(100);o__Filobasidiales(100);f__Filobasidiaceae(100);g__Cryptococcus(100);s__Cryptococcus_arrabidensis(100); | Cryptococcus | 13 | Pathotroph-Saprotroph-Symbiotroph | Animal Pathogen-Endophyte-Epiphyte-\|Undefined Saprotroph\| | Dimorphic Yeast-Yeast | NULL | Possible | As foliar_endophyte (Põlme et al. 2020); Decay leaf, fruit, seed, soil, dung, animal material (Põlme et al. 2020) | Andrews JH, et al. 1983. Phytopathology 73:228-234 ((https://doi.org/10.1094/Phyto-73-228)); Kurtzman CP, et al. (eds.) 2011. The Yeasts, a Taxonomic Study. Fifth Edition. Vols 1-3. Elsevier, San Diego (ISBN: 9780444521491); Põlme S, et al. 2020. Fungal Diversity 105:1-16 ((https://doi.org/10.1007/s13225-020-00466-2)) |
| Otu05888 | 4 | k__Fungi(100);p__Ascomycota(100);c__Saccharomycetes(100);o__Saccharomycetales(100);f__Saccharomycetales_family_Incertae_sedis(100);g__Candida(100);g__Candida_unclassified(100); | Candida | 13 | Pathotroph-Saprotroph-Symbiotroph | Animal Pathogen-Endophyte-Endosymbiont-Epiphyte-\|Undefined Saprotroph\| | Dimorphic Yeast-Yeast | NULL | Probable | As nectar-tap_saprotroph (Põlme et al. 2020); Decay sugar-rich substrates (Põlme et al. 2020); This asexual genus is polyphyletic and contain species that function very differently depend on what they are. Caution should be taken when interpreting guilds for Candida species. | Manolakaki D, et al. 2010. Virulence. 1: 367-375 ((https://doi.org/10.4161/viru.1.5.12796)); Mekbib SB, et al. 2011. Tropical Plant Pathology 36:233-240 ((https://doi.org/10.1590/S1982-56762011000400004)); Põlme S, et al. 2020. Fungal Diversity 105:1-16 ((https://doi.org/10.1007/s13225-020-00466-2)) |
| Otu05891 | 4 | k__Fungi(100);p__Ascomycota(100);c__Saccharomycetes(100);o__Saccharomycetales(100);f__Metschnikowiaceae(100);g__Metschnikowia(100);g__Metschnikowia_unclassified(100); | Metschnikowia | 13 | Saprotroph | \|Undefined Saprotroph\| | Yeast | NULL | Probable | As nectar-tap_saprotroph (Põlme et al. 2020); Decay sugar-rich substrates (Põlme et al. 2020) | Põlme S, et al. 2020. Fungal Diversity 105:1-16 ((https://doi.org/10.1007/s13225-020-00466-2)) |
| Otu05892 | 4 | k__Fungi(100);p__Ascomycota(100);c__Sordariomycetes(100);o__Ophiostomatales(100);f__Ophiostomataceae(100);f__Ophiostomataceae_unclassified(100);f__Ophiostomataceae_unclassified(100); | Ophiostomataceae | 9 | Pathotroph | Plant Pathogen | Microfungus | Blue-Staining | Probable | At least most species are transmitted via insect vectors, with mycelial colonies developing in bark-boring beetle galleries and spreading through the vascular system; they are highly specific to their beetle vectors. In some cases spores and mycelium are transmitted in specialized organs, termed mycangia (Cannon & Kirk 2007) | Cannon PF, Kirk PM. 2007. Fungal Families of the World. CAB International, Cambridge (ISBN: 978-0851998275); Tedersoo L, et al. 2014. Science 346:e1256688 ((https://doi.org/10.1126/science.1256688)) |
| Otu05893 | 4 | k__Fungi(100);p__Ascomycota(100);c__Pezizomycetes(100);o__Pezizales(100);f__Sarcosomataceae(100);g__Pseudoplectania(100);s__Pseudoplectania_sp(100); | Pseudoplectania | 13 | Saprotroph | \|Plant Saprotroph\|-Undefined Saprotroph | Microfungus-Pezizoid | NULL | Probable | As litter_saprotroph (Põlme et al. 2020); Decay leaf, fruit, seed (Põlme et al. 2020) | Tedersoo L, et al. 2014. Science 346:e1256688 ((https://doi.org/10.1126/science.1256688)); Põlme S, et al. 2020. Fungal Diversity 105:1-16 ((https://doi.org/10.1007/s13225-020-00466-2)) |
| Otu05895 | 4 | k__Fungi(100);p__Ascomycota(100);c__Saccharomycetes(100);o__Saccharomycetales(100);f__Metschnikowiaceae(100);g__Metschnikowia(100);g__Metschnikowia_unclassified(100); | Metschnikowia | 13 | Saprotroph | \|Undefined Saprotroph\| | Yeast | NULL | Probable | As nectar-tap_saprotroph (Põlme et al. 2020); Decay sugar-rich substrates (Põlme et al. 2020) | Põlme S, et al. 2020. Fungal Diversity 105:1-16 ((https://doi.org/10.1007/s13225-020-00466-2)) |
| Otu05896 | 4 | k__Fungi(100);p__Ascomycota(100);c__Saccharomycetes(100);o__Saccharomycetales(100);f__Metschnikowiaceae(100);g__Metschnikowia(100);g__Metschnikowia_unclassified(100); | Metschnikowia | 13 | Saprotroph | \|Undefined Saprotroph\| | Yeast | NULL | Probable | As nectar-tap_saprotroph (Põlme et al. 2020); Decay sugar-rich substrates (Põlme et al. 2020) | Põlme S, et al. 2020. Fungal Diversity 105:1-16 ((https://doi.org/10.1007/s13225-020-00466-2)) |
| Otu05900 | 4 | k__Fungi(100);p__Ascomycota(75);c__Saccharomycetes(75);o__Saccharomycetales(75);f__Metschnikowiaceae(75);g__Metschnikowia(75);g__Metschnikowia_unclassified(75); | Metschnikowia | 13 | Saprotroph | \|Undefined Saprotroph\| | Yeast | NULL | Probable | As nectar-tap_saprotroph (Põlme et al. 2020); Decay sugar-rich substrates (Põlme et al. 2020) | Põlme S, et al. 2020. Fungal Diversity 105:1-16 ((https://doi.org/10.1007/s13225-020-00466-2)) |
| Otu05901 | 4 | k__Fungi(100);p__Ascomycota(100);c__Leotiomycetes(100);o__Helotiales(100);f__Hyaloscyphaceae(100);g__Lachnum(100);g__Lachnum_unclassified(75); | Lachnum | 13 | Saprotroph-Symbiotroph | Endophyte-\|Plant Saprotroph\|-Undefined Saprotroph-Wood Saprotroph | Microfungus | NULL | Probable | As litter_saprotroph (Põlme et al. 2020); As root_endophyte (Põlme et al. 2020); Decay wood, leaf, fruit, seed (Põlme et al. 2020) | Tedersoo L, et al. 2014. Science 346:e1256688 ((https://doi.org/10.1126/science.1256688)); Põlme S, et al. 2020. Fungal Diversity 105:1-16 ((https://doi.org/10.1007/s13225-020-00466-2)) |
| Otu05905 | 4 | k__Fungi(100);p__Ascomycota(100);c__Saccharomycetes(100);o__Saccharomycetales(100);f__Saccharomycetaceae(100);g__Kazachstania(100);s__Kazachstania_intestinalis(100); | Kazachstania | 13 | Saprotroph | \|Undefined Saprotroph\| | Yeast | NULL | Probable | As nectar-tap_saprotroph (Põlme et al. 2020); Decay sugar-rich substrates (Põlme et al. 2020) | Põlme S, et al. 2020. Fungal Diversity 105:1-16 ((https://doi.org/10.1007/s13225-020-00466-2)) |
| Otu05909 | 4 | k__Fungi(100);p__Ascomycota(100);c__Saccharomycetes(100);o__Saccharomycetales(100);f__Metschnikowiaceae(100);g__Metschnikowia(100);s__Metschnikowia_pulcherrima(100); | Metschnikowia | 13 | Saprotroph | \|Undefined Saprotroph\| | Yeast | NULL | Probable | As nectar-tap_saprotroph (Põlme et al. 2020); Decay sugar-rich substrates (Põlme et al. 2020) | Põlme S, et al. 2020. Fungal Diversity 105:1-16 ((https://doi.org/10.1007/s13225-020-00466-2)) |
| Otu05912 | 4 | k__Fungi(100);p__Ascomycota(100);c__Saccharomycetes(100);o__Saccharomycetales(100);f__Metschnikowiaceae(100);g__Metschnikowia(100);s__Metschnikowia_pulcherrima(100); | Metschnikowia | 13 | Saprotroph | \|Undefined Saprotroph\| | Yeast | NULL | Probable | As nectar-tap_saprotroph (Põlme et al. 2020); Decay sugar-rich substrates (Põlme et al. 2020) | Põlme S, et al. 2020. Fungal Diversity 105:1-16 ((https://doi.org/10.1007/s13225-020-00466-2)) |
| Otu05919 | 4 | k__Fungi(100);p__Basidiomycota(100);c__Agaricomycetes(100);o__Agaricales(100);f__Cortinariaceae(100);g__Cortinarius(100);g__Cortinarius_unclassified(100); | Cortinarius | 13 | Symbiotroph | \|Ectomycorrhizal\| | Agaricoid | NULL | Highly Probable | NULL | Rinaldi AC, et al. 2008. Fungal Diversity 33:1-45 ((http://www.fungaldiversity.org/fdp/sfdp/33-1.pdf)); Tedersoo L, et al. 2010. Mycorrhiza 20:217-263 ((https://doi.org/10.1007/s00572-009-0274-x)); Põlme S, et al. 2020. Fungal Diversity 105:1-16 ((https://doi.org/10.1007/s13225-020-00466-2)); http://mycorrhizas.info/ecmf.html ((http://mycorrhizas.info/ecmf.html)) |
| Otu05922 | 4 | k__Fungi(100);p__Basidiomycota(100);c__Pucciniomycetes(100);o__Platygloeales(100);f__Eocronartiaceae(100);g__Eocronartium(100);s__Eocronartium_sp(100); | Eocronartium | 13 | Pathotroph | \|Plant Pathogen\| | Clavarioid | NULL | Probable | Host - mosses (Põlme et al. 2020) | Tedersoo L, et al. 2014. Science 346:e1256688 ((https://doi.org/10.1126/science.1256688)); Põlme S, et al. 2020. Fungal Diversity 105:1-16 ((https://doi.org/10.1007/s13225-020-00466-2)) |
| Otu05923 | 4 | k__Fungi(100);p__Ascomycota(100);c__Dothideomycetes(100);o__Pleosporales(100);f__Venturiaceae(100);g__Venturia(100);g__Venturia_unclassified(100); | Venturia | 13 | Pathotroph-Pathotroph-Saprotroph | \|Plant Pathogen\|-Plant Saprotroph | Microfungus | NULL | Probable | As litter_saprotroph (Põlme et al. 2020); Decay leaf, fruit, seed (Põlme et al. 2020) | Tedersoo L, et al. 2014. Science 346:e1256688 ((https://doi.org/10.1126/science.1256688)); Põlme S, et al. 2020. Fungal Diversity 105:1-16 ((https://doi.org/10.1007/s13225-020-00466-2)) |
| Otu05927 | 4 | k__Fungi(100);p__Ascomycota(100);c__Saccharomycetes(100);o__Saccharomycetales(100);f__Saccharomycetales_family_Incertae_sedis(100);g__Candida(100);g__Candida_unclassified(100); | Candida | 13 | Pathotroph-Saprotroph-Symbiotroph | Animal Pathogen-Endophyte-Endosymbiont-Epiphyte-\|Undefined Saprotroph\| | Dimorphic Yeast-Yeast | NULL | Probable | As nectar-tap_saprotroph (Põlme et al. 2020); Decay sugar-rich substrates (Põlme et al. 2020); This asexual genus is polyphyletic and contain species that function very differently depend on what they are. Caution should be taken when interpreting guilds for Candida species. | Manolakaki D, et al. 2010. Virulence. 1: 367-375 ((https://doi.org/10.4161/viru.1.5.12796)); Mekbib SB, et al. 2011. Tropical Plant Pathology 36:233-240 ((https://doi.org/10.1590/S1982-56762011000400004)); Põlme S, et al. 2020. Fungal Diversity 105:1-16 ((https://doi.org/10.1007/s13225-020-00466-2)) |
| Otu05928 | 4 | k__Fungi(100);p__Ascomycota(100);c__Saccharomycetes(100);o__Saccharomycetales(100);f__Metschnikowiaceae(100);g__Metschnikowia(100);s__Metschnikowia_pulcherrima(75); | Metschnikowia | 13 | Saprotroph | \|Undefined Saprotroph\| | Yeast | NULL | Probable | As nectar-tap_saprotroph (Põlme et al. 2020); Decay sugar-rich substrates (Põlme et al. 2020) | Põlme S, et al. 2020. Fungal Diversity 105:1-16 ((https://doi.org/10.1007/s13225-020-00466-2)) |
| Otu05930 | 4 | k__Fungi(100);p__Ascomycota(100);c__Saccharomycetes(100);o__Saccharomycetales(100);f__Metschnikowiaceae(100);g__Metschnikowia(100);s__Metschnikowia_pulcherrima(75); | Metschnikowia | 13 | Saprotroph | \|Undefined Saprotroph\| | Yeast | NULL | Probable | As nectar-tap_saprotroph (Põlme et al. 2020); Decay sugar-rich substrates (Põlme et al. 2020) | Põlme S, et al. 2020. Fungal Diversity 105:1-16 ((https://doi.org/10.1007/s13225-020-00466-2)) |
| Otu05932 | 4 | k__Fungi(100);p__Ascomycota(100);c__Saccharomycetes(100);o__Saccharomycetales(100);f__Metschnikowiaceae(100);g__Metschnikowia(100);s__Metschnikowia_pulcherrima(100); | Metschnikowia | 13 | Saprotroph | \|Undefined Saprotroph\| | Yeast | NULL | Probable | As nectar-tap_saprotroph (Põlme et al. 2020); Decay sugar-rich substrates (Põlme et al. 2020) | Põlme S, et al. 2020. Fungal Diversity 105:1-16 ((https://doi.org/10.1007/s13225-020-00466-2)) |
| Otu05933 | 4 | k__Fungi(100);p__Basidiomycota(100);c__Wallemiomycetes(100);o__Wallemiales(100);f__Wallemiaceae(100);g__Wallemia(100);s__Wallemia_sebi(100); | Wallemia | 13 | Saprotroph | \|Undefined Saprotroph\| | Microfungus | NULL | Probable | Decay sugar-rich substrates (Põlme et al. 2020) | Tedersoo L, et al. 2014. Science 346:e1256688 ((https://doi.org/10.1126/science.1256688)); Põlme S, et al. 2020. Fungal Diversity 105:1-16 ((https://doi.org/10.1007/s13225-020-00466-2)) |
| Otu05934 | 4 | k__Fungi(100);p__Ascomycota(100);c__Saccharomycetes(100);o__Saccharomycetales(100);f__Metschnikowiaceae(100);g__Metschnikowia(100);s__Metschnikowia_pulcherrima(75); | Metschnikowia | 13 | Saprotroph | \|Undefined Saprotroph\| | Yeast | NULL | Probable | As nectar-tap_saprotroph (Põlme et al. 2020); Decay sugar-rich substrates (Põlme et al. 2020) | Põlme S, et al. 2020. Fungal Diversity 105:1-16 ((https://doi.org/10.1007/s13225-020-00466-2)) |
| Otu05936 | 4 | k__Fungi(100);p__Ascomycota(100);c__Saccharomycetes(100);o__Saccharomycetales(100);f__Metschnikowiaceae(100);g__Metschnikowia(100);s__Metschnikowia_pulcherrima(100); | Metschnikowia | 13 | Saprotroph | \|Undefined Saprotroph\| | Yeast | NULL | Probable | As nectar-tap_saprotroph (Põlme et al. 2020); Decay sugar-rich substrates (Põlme et al. 2020) | Põlme S, et al. 2020. Fungal Diversity 105:1-16 ((https://doi.org/10.1007/s13225-020-00466-2)) |
| Otu05937 | 4 | k__Fungi(100);p__Ascomycota(100);c__Sordariomycetes(100);o__Hypocreales(100);o__Hypocreales_unclassified(100);o__Hypocreales_unclassified(100);o__Hypocreales_unclassified(100); | Hypocreales | 7 | Saprotroph | Undefined Saprotroph | Microfungus | NULL | Possible | NULL | Sterkenburg E, et al. 2015. New Phytologist 207:1145-1158 ((https://doi.org/10.1111/nph.13426)) |
| Otu05941 | 4 | k__Fungi(100);p__Ascomycota(100);c__Saccharomycetes(100);o__Saccharomycetales(100);f__Metschnikowiaceae(100);g__Metschnikowia(100);s__Metschnikowia_pulcherrima(100); | Metschnikowia | 13 | Saprotroph | \|Undefined Saprotroph\| | Yeast | NULL | Probable | As nectar-tap_saprotroph (Põlme et al. 2020); Decay sugar-rich substrates (Põlme et al. 2020) | Põlme S, et al. 2020. Fungal Diversity 105:1-16 ((https://doi.org/10.1007/s13225-020-00466-2)) |
| Otu05942 | 4 | k__Fungi(100);p__Ascomycota(100);c__Pezizomycetes(100);o__Pezizales(100);f__Pezizaceae(100);g__Peziza(100);s__Peziza_sp_HK_87_273(100); | Peziza | 13 | Saprotroph-Symbiotroph | Dung Saprotroph-Ectomycorrhizal-Endophyte-Plant Saprotroph-\|Undefined Saprotroph\| | Pezizoid | NULL | Possible | As foliar_endophyte (Põlme et al. 2020); As soil_saprotroph (Põlme et al. 2020); Decay soil (Põlme et al. 2020); Saprobic or ectomycorrhizal, on or in soil, leafmould etc., some species associated with dung or fire sites (Cannon & Kirk 2007) | Cannon PF, Kirk PM. 2007. Fungal Families of the World. CAB International, Cambridge (ISBN: 978-0851998275); Rinaldi AC, et al. 2008. Fungal Diversity 33:1-45 ((http://www.fungaldiversity.org/fdp/sfdp/33-1.pdf)); Tedersoo L, et al. 2010. Mycorrhiza 20:217-263 ((https://doi.org/10.1007/s00572-009-0274-x)) (pro parte); Põlme S, et al. 2020. Fungal Diversity 105:1-16 ((https://doi.org/10.1007/s13225-020-00466-2)) |
| Otu05943 | 4 | k__Fungi(100);p__Ascomycota(100);c__Saccharomycetes(100);o__Saccharomycetales(100);f__Metschnikowiaceae(100);g__Metschnikowia(100);s__Metschnikowia_pulcherrima(75); | Metschnikowia | 13 | Saprotroph | \|Undefined Saprotroph\| | Yeast | NULL | Probable | As nectar-tap_saprotroph (Põlme et al. 2020); Decay sugar-rich substrates (Põlme et al. 2020) | Põlme S, et al. 2020. Fungal Diversity 105:1-16 ((https://doi.org/10.1007/s13225-020-00466-2)) |
| Otu05944 | 4 | k__Fungi(100);p__Ascomycota(100);c__Saccharomycetes(100);o__Saccharomycetales(100);f__Saccharomycetaceae(100);g__Kazachstania(100);s__Kazachstania_intestinalis(100); | Kazachstania | 13 | Saprotroph | \|Undefined Saprotroph\| | Yeast | NULL | Probable | As nectar-tap_saprotroph (Põlme et al. 2020); Decay sugar-rich substrates (Põlme et al. 2020) | Põlme S, et al. 2020. Fungal Diversity 105:1-16 ((https://doi.org/10.1007/s13225-020-00466-2)) |
| Otu05945 | 4 | k__Fungi(100);p__Basidiomycota(100);c__Wallemiomycetes(100);o__Wallemiales(100);f__Wallemiaceae(100);g__Wallemia(100);s__Wallemia_sebi(100); | Wallemia | 13 | Saprotroph | \|Undefined Saprotroph\| | Microfungus | NULL | Probable | Decay sugar-rich substrates (Põlme et al. 2020) | Tedersoo L, et al. 2014. Science 346:e1256688 ((https://doi.org/10.1126/science.1256688)); Põlme S, et al. 2020. Fungal Diversity 105:1-16 ((https://doi.org/10.1007/s13225-020-00466-2)) |
| Otu05946 | 4 | k__Fungi(100);p__Basidiomycota(100);c__Agaricomycetes(100);o__Cantharellales(100);f__Ceratobasidiaceae(100);g__Rhizoctonia(75);g__Rhizoctonia_unclassified(75); | Rhizoctonia | 13 | Pathotroph-Saprotroph-Symbiotroph | Ectomycorrhizal-Orchid Mycorrhizal-\|Plant Pathogen\|-Plant Saprotroph-Wood Saprotroph | Corticioid-Microfungus | Soft Rot | Probable | As litter_saprotroph (Põlme et al. 2020); Decay leaf, fruit, seed, roots, soil (Põlme et al. 2020); Only symbiotic (perhaps leaning parasitic) in special cases with orchids, otherwise typically a plant pathogen. | Seehann G, et al. 1975. List of Fungi in Soft-Rot Tissue Tests. Institute fur Holzbiologie und Holzschutz, Bundesforschungsanstalt fur Forst - und Holzwirtshaft, Hamburg - Lohbrugge, Germany; Costa IPMW, et al. 2012. Mycotaxon 119:e26 ((http://mycotaxon.com/resources/checklists/costa_v119_checklist.pdf)); Tedersoo L, et al. 2014. Science 346:e1256688 ((https://doi.org/10.1126/science.1256688)); Põlme S, et al. 2020. Fungal Diversity 105:1-16 ((https://doi.org/10.1007/s13225-020-00466-2)); List compiled by Sara Branco with input from Nhu Nguyen and Else Vellinga; The American Phytopathological Society Common Names of Plant Diseases ((http://www.apsnet.org/publications/commonnames/Pages/default.aspx)) |
| Otu05947 | 4 | k__Fungi(100);p__Ascomycota(100);c__Saccharomycetes(100);o__Saccharomycetales(100);f__Metschnikowiaceae(100);g__Metschnikowia(100);s__Metschnikowia_pulcherrima(75); | Metschnikowia | 13 | Saprotroph | \|Undefined Saprotroph\| | Yeast | NULL | Probable | As nectar-tap_saprotroph (Põlme et al. 2020); Decay sugar-rich substrates (Põlme et al. 2020) | Põlme S, et al. 2020. Fungal Diversity 105:1-16 ((https://doi.org/10.1007/s13225-020-00466-2)) |
| Otu05951 | 4 | k__Fungi(100);p__Ascomycota(100);c__Saccharomycetes(100);o__Saccharomycetales(100);f__Metschnikowiaceae(100);g__Metschnikowia(100);g__Metschnikowia_unclassified(100); | Metschnikowia | 13 | Saprotroph | \|Undefined Saprotroph\| | Yeast | NULL | Probable | As nectar-tap_saprotroph (Põlme et al. 2020); Decay sugar-rich substrates (Põlme et al. 2020) | Põlme S, et al. 2020. Fungal Diversity 105:1-16 ((https://doi.org/10.1007/s13225-020-00466-2)) |
| Otu05953 | 4 | k__Fungi(100);p__Ascomycota(100);c__Saccharomycetes(100);o__Saccharomycetales(100);f__Metschnikowiaceae(100);g__Metschnikowia(100);s__Metschnikowia_pulcherrima(100); | Metschnikowia | 13 | Saprotroph | \|Undefined Saprotroph\| | Yeast | NULL | Probable | As nectar-tap_saprotroph (Põlme et al. 2020); Decay sugar-rich substrates (Põlme et al. 2020) | Põlme S, et al. 2020. Fungal Diversity 105:1-16 ((https://doi.org/10.1007/s13225-020-00466-2)) |
| Otu05954 | 4 | k__Fungi(100);p__Ascomycota(100);c__Saccharomycetes(100);o__Saccharomycetales(100);f__Metschnikowiaceae(100);g__Metschnikowia(100);s__Metschnikowia_pulcherrima(100); | Metschnikowia | 13 | Saprotroph | \|Undefined Saprotroph\| | Yeast | NULL | Probable | As nectar-tap_saprotroph (Põlme et al. 2020); Decay sugar-rich substrates (Põlme et al. 2020) | Põlme S, et al. 2020. Fungal Diversity 105:1-16 ((https://doi.org/10.1007/s13225-020-00466-2)) |
| Otu05958 | 4 | k__Fungi(100);p__Ascomycota(100);c__Saccharomycetes(100);o__Saccharomycetales(100);f__Metschnikowiaceae(100);g__Metschnikowia(100);g__Metschnikowia_unclassified(100); | Metschnikowia | 13 | Saprotroph | \|Undefined Saprotroph\| | Yeast | NULL | Probable | As nectar-tap_saprotroph (Põlme et al. 2020); Decay sugar-rich substrates (Põlme et al. 2020) | Põlme S, et al. 2020. Fungal Diversity 105:1-16 ((https://doi.org/10.1007/s13225-020-00466-2)) |
| Otu05959 | 4 | k__Fungi(100);p__Ascomycota(100);c__Saccharomycetes(100);o__Saccharomycetales(100);f__Metschnikowiaceae(100);g__Metschnikowia(100);s__Metschnikowia_pulcherrima(75); | Metschnikowia | 13 | Saprotroph | \|Undefined Saprotroph\| | Yeast | NULL | Probable | As nectar-tap_saprotroph (Põlme et al. 2020); Decay sugar-rich substrates (Põlme et al. 2020) | Põlme S, et al. 2020. Fungal Diversity 105:1-16 ((https://doi.org/10.1007/s13225-020-00466-2)) |
| Otu05962 | 4 | k__Fungi(100);p__Ascomycota(100);c__Saccharomycetes(100);o__Saccharomycetales(100);f__Metschnikowiaceae(100);g__Metschnikowia(100);s__Metschnikowia_pulcherrima(100); | Metschnikowia | 13 | Saprotroph | \|Undefined Saprotroph\| | Yeast | NULL | Probable | As nectar-tap_saprotroph (Põlme et al. 2020); Decay sugar-rich substrates (Põlme et al. 2020) | Põlme S, et al. 2020. Fungal Diversity 105:1-16 ((https://doi.org/10.1007/s13225-020-00466-2)) |
| Otu05963 | 4 | k__Fungi(100);p__Zygomycota(100);c__Zygomycota_class_Incertae_sedis(100);o__Mucorales(100);f__Mucoraceae(100);g__Mucor(100);s__Mucor_circinelloides_f._circinelloides(75); | Mucor | 13 | Saprotroph | \|Undefined Saprotroph\| | Microfungus | NULL | Probable | As soil_saprotroph (Põlme et al. 2020); Decay soil (Põlme et al. 2020) | Tedersoo L, et al. 2014. Science 346:e1256688 ((https://doi.org/10.1126/science.1256688)); Põlme S, et al. 2020. Fungal Diversity 105:1-16 ((https://doi.org/10.1007/s13225-020-00466-2)) |
| Otu05964 | 4 | k__Fungi(100);p__Zygomycota(100);c__Zygomycota_class_Incertae_sedis(100);o__Mucorales(100);f__Mucoraceae(100);g__Mucor(100);g__Mucor_unclassified(100); | Mucor | 13 | Saprotroph | \|Undefined Saprotroph\| | Microfungus | NULL | Probable | As soil_saprotroph (Põlme et al. 2020); Decay soil (Põlme et al. 2020) | Tedersoo L, et al. 2014. Science 346:e1256688 ((https://doi.org/10.1126/science.1256688)); Põlme S, et al. 2020. Fungal Diversity 105:1-16 ((https://doi.org/10.1007/s13225-020-00466-2)) |
| Otu05965 | 4 | k__Fungi(100);p__Ascomycota(100);c__Eurotiomycetes(100);o__Chaetothyriales(100);f__Herpotrichiellaceae(100);g__Cladophialophora(100);g__Cladophialophora_unclassified(75); | Cladophialophora | 13 | Pathotroph-Saprotroph-Symbiotroph | Animal Pathogen-Endophyte-Fungal Parasite-\|Undefined Saprotroph\| | Dimorphic-Microfungus | NULL | Probable | As mycoparasite (Põlme et al. 2020); As root_endophyte_dark_septate (Põlme et al. 2020); As soil_saprotroph (Põlme et al. 2020); Decay soil (Põlme et al. 2020) | James TY, et al. 2006. Nature 443:818-822 ((https://doi.org/10.1038/nature05110)); Põlme S, et al. 2020. Fungal Diversity 105:1-16 ((https://doi.org/10.1007/s13225-020-00466-2)) |
| Otu05966 | 4 | k__Fungi(100);p__Ascomycota(100);c__Pezizomycetes(100);o__Pezizales(100);f__Pezizaceae(100);g__Peziza(100);s__Peziza_phyllogena(100); | Peziza | 13 | Saprotroph-Symbiotroph | Dung Saprotroph-Ectomycorrhizal-Endophyte-Plant Saprotroph-\|Undefined Saprotroph\| | Pezizoid | NULL | Possible | As foliar_endophyte (Põlme et al. 2020); As soil_saprotroph (Põlme et al. 2020); Decay soil (Põlme et al. 2020); Saprobic or ectomycorrhizal, on or in soil, leafmould etc., some species associated with dung or fire sites (Cannon & Kirk 2007) | Cannon PF, Kirk PM. 2007. Fungal Families of the World. CAB International, Cambridge (ISBN: 978-0851998275); Rinaldi AC, et al. 2008. Fungal Diversity 33:1-45 ((http://www.fungaldiversity.org/fdp/sfdp/33-1.pdf)); Tedersoo L, et al. 2010. Mycorrhiza 20:217-263 ((https://doi.org/10.1007/s00572-009-0274-x)) (pro parte); Põlme S, et al. 2020. Fungal Diversity 105:1-16 ((https://doi.org/10.1007/s13225-020-00466-2)) |
| Otu05967 | 4 | k__Fungi(100);p__Ascomycota(100);c__Saccharomycetes(100);o__Saccharomycetales(100);f__Metschnikowiaceae(100);g__Metschnikowia(100);g__Metschnikowia_unclassified(100); | Metschnikowia | 13 | Saprotroph | \|Undefined Saprotroph\| | Yeast | NULL | Probable | As nectar-tap_saprotroph (Põlme et al. 2020); Decay sugar-rich substrates (Põlme et al. 2020) | Põlme S, et al. 2020. Fungal Diversity 105:1-16 ((https://doi.org/10.1007/s13225-020-00466-2)) |
| Otu05968 | 4 | k__Fungi(100);p__Ascomycota(100);c__Lecanoromycetes(100);o__Lecanorales(100);f__Byssolomataceae(100);g__Micarea(100);s__Micarea_sp(100); | Micarea | 13 | Pathotroph-Symbiotroph | Lichen Parasite-\|Lichenized\| | Thallus | NULL | Highly Probable | NULL | Esslinger TL. 2014. North Dakota State University ((http://www.ndsu.edu/pubweb/~esslinge/chcklst/chcklst7.htm)); Põlme S, et al. 2020. Fungal Diversity 105:1-16 ((https://doi.org/10.1007/s13225-020-00466-2)) |
| Otu05971 | 4 | k__Fungi(100);p__Basidiomycota(100);c__Agaricomycetes(100);o__Hymenochaetales(100);f__Hymenochaetaceae(100);g__Phellinus(100);s__Phellinus_hartigii(100); | Phellinus | 13 | Pathotroph-Saprotroph | \|Plant Pathogen\|-Wood Saprotroph | Polyporoid | White Rot | Highly Probable | As white_rot (Põlme et al. 2020); As wood_pathogen (Põlme et al. 2020); Decay wood (Põlme et al. 2020) | Gilbertson RL, Ryvarden L. 1987-1987. North American Polypores. Fungiflora, Oslo (ISBN: 978-0945345060); Põlme S, et al. 2020. Fungal Diversity 105:1-16 ((https://doi.org/10.1007/s13225-020-00466-2)) |
| Otu05977 | 4 | k__Fungi(100);p__Ascomycota(100);c__Saccharomycetes(100);o__Saccharomycetales(100);f__Metschnikowiaceae(100);g__Metschnikowia(100);s__Metschnikowia_pulcherrima(100); | Metschnikowia | 13 | Saprotroph | \|Undefined Saprotroph\| | Yeast | NULL | Probable | As nectar-tap_saprotroph (Põlme et al. 2020); Decay sugar-rich substrates (Põlme et al. 2020) | Põlme S, et al. 2020. Fungal Diversity 105:1-16 ((https://doi.org/10.1007/s13225-020-00466-2)) |
| Otu05979 | 4 | k__Fungi(100);p__Ascomycota(100);c__Eurotiomycetes(100);o__Chaetothyriales(100);f__Chaetothyriaceae(100);g__unclassified_Chaetothyriaceae(100);s__Chaetothyriaceae_sp(100); | Chaetothyriaceae | 9 | Pathotroph-Symbiotroph | Epiphyte-Plant Pathogen | Microfungus | NULL | Possible | Epiphytic or biotrophic on living leaves, very little is known (Cannon & Kirk 2007) | Cannon PF, Kirk PM. 2007. Fungal Families of the World. CAB International, Cambridge (ISBN: 978-0851998275) |
| Otu05980 | 4 | k__Fungi(100);p__Ascomycota(100);c__Dothideomycetes(100);o__Capnodiales(100);f__Mycosphaerellaceae(100);f__Mycosphaerellaceae_unclassified(100);f__Mycosphaerellaceae_unclassified(100); | Mycosphaerellaceae | 9 | Pathotroph-Saprotroph | Plant Pathogen-Undefined Saprotroph | Microfungus | NULL | Probable | Biotrophic, necrotrophic or saprobic on various plant tissues (Cannon & Kirk 2007) | Cannon PF, Kirk PM. 2007. Fungal Families of the World. CAB International, Cambridge (ISBN: 978-0851998275) |
| Otu05981 | 4 | k__Fungi(100);p__Ascomycota(100);c__Saccharomycetes(100);o__Saccharomycetales(100);f__Metschnikowiaceae(100);g__Metschnikowia(100);g__Metschnikowia_unclassified(100); | Metschnikowia | 13 | Saprotroph | \|Undefined Saprotroph\| | Yeast | NULL | Probable | As nectar-tap_saprotroph (Põlme et al. 2020); Decay sugar-rich substrates (Põlme et al. 2020) | Põlme S, et al. 2020. Fungal Diversity 105:1-16 ((https://doi.org/10.1007/s13225-020-00466-2)) |
| Otu05986 | 4 | k__Fungi(100);p__Ascomycota(100);c__Saccharomycetes(100);o__Saccharomycetales(100);f__Saccharomycetaceae(100);g__Kazachstania(100);s__Kazachstania_intestinalis(100); | Kazachstania | 13 | Saprotroph | \|Undefined Saprotroph\| | Yeast | NULL | Probable | As nectar-tap_saprotroph (Põlme et al. 2020); Decay sugar-rich substrates (Põlme et al. 2020) | Põlme S, et al. 2020. Fungal Diversity 105:1-16 ((https://doi.org/10.1007/s13225-020-00466-2)) |
| Otu05987 | 4 | k__Fungi(100);p__Ascomycota(100);c__Sordariomycetes(100);o__Hypocreales(100);f__Ophiocordycipitaceae(100);g__Tolypocladium(100);s__Tolypocladium_RG_2013a(100); | Tolypocladium | 13 | Pathotroph-Symbiotroph | \|Animal Parasite\|-Animal Pathogen-Clavicipitaceous Endophyte-Fungal Parasite | Microfungus | NULL | Possible | As foliar_endophyte (Põlme et al. 2020); As mycoparasite (Põlme et al. 2020); culture (Gazis et al. 2014); Decay animal material (Põlme et al. 2020); Endophyte detection method | Gazis R, et al. 2014. Mycologia 106:1090-1105 ((https://doi.org/10.3852/13-346)); Põlme S, et al. 2020. Fungal Diversity 105:1-16 ((https://doi.org/10.1007/s13225-020-00466-2)) |
| Otu05991 | 4 | k__Fungi(100);p__Ascomycota(100);c__Sordariomycetes(100);o__Hypocreales(100);f__Clavicipitaceae(100);g__Pochonia(100);s__Pochonia_sp(100); | Pochonia | 13 | Pathotroph-Saprotroph | \|Animal Parasite\|-Animal Pathogen-Undefined Saprotroph | Microfungus | NULL | Probable | As animal_decomposer (Põlme et al. 2020); Decay animal material (Põlme et al. 2020) | Tedersoo L, et al. 2014. Science 346:e1256688 ((https://doi.org/10.1126/science.1256688)); Põlme S, et al. 2020. Fungal Diversity 105:1-16 ((https://doi.org/10.1007/s13225-020-00466-2)) |
| Otu05993 | 4 | k__Fungi(100);p__Ascomycota(100);c__Sordariomycetes(100);o__Hypocreales(100);f__Cordycipitaceae(100);g__Cordyceps(100);g__Cordyceps_unclassified(100); | Cordyceps | 13 | Pathotroph-Symbiotroph | Clavicipitaceous Endophyte-Fungal Parasite-\|Insect Pathogen\| | Cordyceptoid | NULL | Possible | As foliar_endophyte (Põlme et al. 2020); Decay animal material (Põlme et al. 2020) | Samuels GJ. 2007. Self-published; Costa IPMW, et al. 2012. Mycotaxon 119:e26 ((http://mycotaxon.com/resources/checklists/costa_v119_checklist.pdf)); Tedersoo L, et al. 2014. Science 346:e1256688 ((https://doi.org/10.1126/science.1256688)); Põlme S, et al. 2020. Fungal Diversity 105:1-16 ((https://doi.org/10.1007/s13225-020-00466-2)) |
| Otu05994 | 4 | k__Fungi(100);p__Ascomycota(100);c__Saccharomycetes(100);o__Saccharomycetales(100);f__Metschnikowiaceae(100);g__Metschnikowia(100);g__Metschnikowia_unclassified(75); | Metschnikowia | 13 | Saprotroph | \|Undefined Saprotroph\| | Yeast | NULL | Probable | As nectar-tap_saprotroph (Põlme et al. 2020); Decay sugar-rich substrates (Põlme et al. 2020) | Põlme S, et al. 2020. Fungal Diversity 105:1-16 ((https://doi.org/10.1007/s13225-020-00466-2)) |
| Otu05996 | 4 | k__Fungi(100);p__Ascomycota(100);c__Leotiomycetes(100);o__Leotiomycetes_order_Incertae_sedis(100);f__Leotiomycetes_family_Incertae_sedis(100);g__Leohumicola(100);g__Leohumicola_unclassified(100); | Leohumicola | 13 | Saprotroph-Symbiotroph | Endophyte-Ericoid Mycorrhizal-\|Undefined Saprotroph\| | Microfungus | NULL | Probable | As root_endophyte_dark_septate (Põlme et al. 2020); As soil_saprotroph (Põlme et al. 2020); Decay soil, roots (Põlme et al. 2020) | Tedersoo L, et al. 2014. Science 346:e1256688 ((https://doi.org/10.1126/science.1256688)); Põlme S, et al. 2020. Fungal Diversity 105:1-16 ((https://doi.org/10.1007/s13225-020-00466-2)) |
| Otu05997 | 4 | k__Fungi(100);p__Ascomycota(100);c__Saccharomycetes(100);o__Saccharomycetales(100);f__Metschnikowiaceae(100);g__Metschnikowia(100);s__Metschnikowia_pulcherrima(100); | Metschnikowia | 13 | Saprotroph | \|Undefined Saprotroph\| | Yeast | NULL | Probable | As nectar-tap_saprotroph (Põlme et al. 2020); Decay sugar-rich substrates (Põlme et al. 2020) | Põlme S, et al. 2020. Fungal Diversity 105:1-16 ((https://doi.org/10.1007/s13225-020-00466-2)) |
| Otu05998 | 4 | k__Fungi(100);p__Ascomycota(100);c__Saccharomycetes(100);o__Saccharomycetales(100);f__Metschnikowiaceae(100);g__Metschnikowia(100);s__Metschnikowia_pulcherrima(100); | Metschnikowia | 13 | Saprotroph | \|Undefined Saprotroph\| | Yeast | NULL | Probable | As nectar-tap_saprotroph (Põlme et al. 2020); Decay sugar-rich substrates (Põlme et al. 2020) | Põlme S, et al. 2020. Fungal Diversity 105:1-16 ((https://doi.org/10.1007/s13225-020-00466-2)) |
| Otu05999 | 4 | k__Fungi(100);p__Ascomycota(100);c__Saccharomycetes(100);o__Saccharomycetales(100);f__Metschnikowiaceae(100);g__Metschnikowia(100);s__Metschnikowia_pulcherrima(100); | Metschnikowia | 13 | Saprotroph | \|Undefined Saprotroph\| | Yeast | NULL | Probable | As nectar-tap_saprotroph (Põlme et al. 2020); Decay sugar-rich substrates (Põlme et al. 2020) | Põlme S, et al. 2020. Fungal Diversity 105:1-16 ((https://doi.org/10.1007/s13225-020-00466-2)) |
| Otu06000 | 4 | k__Fungi(100);p__Zygomycota(100);c__Zygomycota_class_Incertae_sedis(100);o__Mortierellales(100);f__Mortierellaceae(100);f__Mortierellaceae_unclassified(75);f__Mortierellaceae_unclassified(75); | Mortierellaceae | 9 | Saprotroph-Symbiotroph | Endophyte-Litter Saprotroph-Soil Saprotroph-Undefined Saprotroph | Microfungus | NULL | Possible | Saprobic in soil and thus frequently isolated from soil and material contaminated with soil such as hay or rotten silage and spoil tips (Cannon & Kirk 2007) | Cannon PF, Kirk PM. 2007. Fungal Families of the World. CAB International, Cambridge (ISBN: 978-0851998275); Purahong W, et al. 2016. Molecular Ecology 25:4059-4074 ((https://doi.org/10.1111/mec.13739)) |
| Otu06002 | 4 | k__Fungi(100);p__Ascomycota(100);c__Eurotiomycetes(100);o__Chaetothyriales(100);f__Chaetothyriaceae(100);g__Cyphellophora(100);s__Cyphellophora_europaea(100); | Cyphellophora | 13 | Pathotroph-Saprotroph | Animal Parasite-Animal Pathogen-\|Plant Saprotroph\|-Undefined Saprotroph | Dimorphic-Facultative Yeast-Microfungus | NULL | Probable | As litter_saprotroph (Põlme et al. 2020); Consult the current literature for updated names on this group that included the Black Yeasts; Some species are likely opportunistic human pathogen (Irinyi et al. 2015); Decay leaf, fruit, seed, animal material (Põlme et al. 2020) | Tedersoo L, et al. 2014. Science 346:e1256688 ((https://doi.org/10.1126/science.1256688)); Irinyi L, et al. 2015. Medical Mycology 53:313-337 ((https://doi.org/10.1093/mmy/myv008)); Põlme S, et al. 2020. Fungal Diversity 105:1-16 ((https://doi.org/10.1007/s13225-020-00466-2)) |
| Otu06003 | 4 | k__Fungi(100);p__Ascomycota(100);c__Eurotiomycetes(100);o__Eurotiales(100);f__Trichocomaceae(100);g__Paecilomyces(100);s__Paecilomyces_lilacinus(100); | Paecilomyces | 13 | Pathotroph-Saprotroph-Symbiotroph | Clavicipitaceous \|Endophyte\|-Plant Pathogen-Plant Saprotroph-Undefined Saprotroph-Wood Saprotroph | Microfungus | Soft Rot | Probable | As foliar_endophyte (Põlme et al. 2020); As litter_saprotroph (Põlme et al. 2020); culture (Lopez et al. 2014); Decay leaf, fruit, seed, animal material (Põlme et al. 2020); Endophyte detection method | Seehann G, et al. 1975. List of Fungi in Soft-Rot Tissue Tests. Institute fur Holzbiologie und Holzschutz, Bundesforschungsanstalt fur Forst - und Holzwirtshaft, Hamburg - Lohbrugge, Germany; Costa IPMW, et al. 2012. Mycotaxon 119:e26 ((http://mycotaxon.com/resources/checklists/costa_v119_checklist.pdf)); Tedersoo L, et al. 2014. Science 346:e1256688 ((https://doi.org/10.1126/science.1256688)); Põlme S, et al. 2020. Fungal Diversity 105:1-16 ((https://doi.org/10.1007/s13225-020-00466-2)); The American Phytopathological Society Common Names of Plant Diseases ((http://www.apsnet.org/publications/commonnames/Pages/default.aspx)); Lopez DC, et al. 2014. Plos One 9:e103891 ((https://doi.org/10.1371/journal.pone.0103891)) |
| Otu06006 | 4 | k__Fungi(100);p__Ascomycota(100);c__Sordariomycetes(100);o__Diaporthales(100);f__Melanconidaceae(100);g__Melanconiella(100);g__Melanconiella_unclassified(100); | Melanconiella | 13 | Saprotroph | Undefined Saprotroph-\|Wood Saprotroph\| | Microfungus | NULL | Probable | Decay wood (Põlme et al. 2020) | Tedersoo L, et al. 2014. Science 346:e1256688 ((https://doi.org/10.1126/science.1256688)); Põlme S, et al. 2020. Fungal Diversity 105:1-16 ((https://doi.org/10.1007/s13225-020-00466-2)) |
| Otu06007 | 4 | k__Fungi(100);p__Ascomycota(100);c__Sordariomycetes(100);o__Hypocreales(100);o__Hypocreales_unclassified(100);o__Hypocreales_unclassified(100);o__Hypocreales_unclassified(100); | Hypocreales | 7 | Saprotroph | Undefined Saprotroph | Microfungus | NULL | Possible | NULL | Sterkenburg E, et al. 2015. New Phytologist 207:1145-1158 ((https://doi.org/10.1111/nph.13426)) |
| Otu06009 | 4 | k__Fungi(100);p__Ascomycota(100);c__Saccharomycetes(100);o__Saccharomycetales(100);f__Metschnikowiaceae(100);g__Metschnikowia(100);s__Metschnikowia_pulcherrima(100); | Metschnikowia | 13 | Saprotroph | \|Undefined Saprotroph\| | Yeast | NULL | Probable | As nectar-tap_saprotroph (Põlme et al. 2020); Decay sugar-rich substrates (Põlme et al. 2020) | Põlme S, et al. 2020. Fungal Diversity 105:1-16 ((https://doi.org/10.1007/s13225-020-00466-2)) |
| Otu06011 | 4 | k__Fungi(100);p__Ascomycota(100);c__Sordariomycetes(100);o__Microascales(100);f__Ceratocystidaceae(100);g__Sporendocladia(100);s__Sporendocladia_bactrospora(100); | Sporendocladia | 13 | Pathotroph | \|Plant Pathogen\| | Microfungus | NULL | Probable | NULL | Tedersoo L, et al. 2014. Science 346:e1256688 ((https://doi.org/10.1126/science.1256688)); Põlme S, et al. 2020. Fungal Diversity 105:1-16 ((https://doi.org/10.1007/s13225-020-00466-2)) |
| Otu06012 | 4 | k__Fungi(100);p__Basidiomycota(100);c__Agaricomycetes(100);o__Trechisporales(100);f__Hydnodontaceae(100);g__Trechispora(100);g__Trechispora_unclassified(75); | Trechispora | 13 | Saprotroph | \|Wood Saprotroph\| | Corticioid | White Rot | Highly Probable | As white_rot (Põlme et al. 2020); Decay wood (Põlme et al. 2020) | Gilbertson RL, Ryvarden L. 1987-1987. North American Polypores. Fungiflora, Oslo (ISBN: 978-0945345060); Rinaldi AC, et al. 2008. Fungal Diversity 33:1-45 ((http://www.fungaldiversity.org/fdp/sfdp/33-1.pdf)) (as Uncertain); Tedersoo L, et al. 2010. Mycorrhiza 20:217-263 ((https://doi.org/10.1007/s00572-009-0274-x)) (as not ECM).; Põlme S, et al. 2020. Fungal Diversity 105:1-16 ((https://doi.org/10.1007/s13225-020-00466-2)) |
| Otu06014 | 4 | k__Fungi(100);p__Basidiomycota(100);c__Agaricomycetes(100);o__Russulales(100);f__Stereaceae(100);g__Gloeocystidiellum(100);s__Gloeocystidiellum_kenyense(100); | Gloeocystidiellum | 13 | Saprotroph | Undefined Saprotroph-\|Wood Saprotroph\| | Corticioid | White Rot | Probable | As white_rot (Põlme et al. 2020); Decay wood (Põlme et al. 2020) | Tedersoo L, et al. 2014. Science 346:e1256688 ((https://doi.org/10.1126/science.1256688)); Põlme S, et al. 2020. Fungal Diversity 105:1-16 ((https://doi.org/10.1007/s13225-020-00466-2)) |
| Otu06016 | 4 | k__Fungi(100);p__Ascomycota(100);c__Saccharomycetes(100);o__Saccharomycetales(100);f__Saccharomycetales_family_Incertae_sedis(75);f__Saccharomycetales_family_Incertae_sedis_unclassified(75);f__Saccharomycetales_family_Incertae_sedis_unclassified(75); | Saccharomycetales | 7 | Saprotroph | Undefined Saprotroph | Yeast | NULL | Possible | NULL | Sterkenburg E, et al. 2015. New Phytologist 207:1145-1158 ((https://doi.org/10.1111/nph.13426)) |
| Otu06017 | 4 | k__Fungi(100);p__Ascomycota(100);c__Sordariomycetes(100);o__Ophiostomatales(100);f__Ophiostomataceae(100);g__Ophiostoma(100);s__Ophiostoma_piliferum(100); | Ophiostoma | 13 | Pathotroph-Saprotroph | \|Plant Pathogen\|-Wood Saprotroph | Microfungus | Blue-Staining | Probable | As wood_pathogen (Põlme et al. 2020); At least most species are transmitted via insect vectors, with mycelial colonies developing in bark; boring beetle galleries and spreading through the vascular system; they are highly specific to their beetle vectors. In some cases spores and mycelium are transmitted in specialized organs, termed mycangia (Cannon & Kirk 2007); Decay wood (Põlme et al. 2020) | Cannon PF, Kirk PM. 2007. Fungal Families of the World. CAB International, Cambridge (ISBN: 978-0851998275); Tedersoo L, et al. 2014. Science 346:e1256688 ((https://doi.org/10.1126/science.1256688)); Põlme S, et al. 2020. Fungal Diversity 105:1-16 ((https://doi.org/10.1007/s13225-020-00466-2)) |
| Otu06021 | 4 | k__Fungi(100);p__Basidiomycota(100);c__Tremellomycetes(100);o__Tremellales(100);f__Tremellales_family_Incertae_sedis(100);g__Fellomyces(100);s__Fellomyces_horovitziae(100); | Fellomyces | 13 | Pathotroph-Saprotroph | Fungal Parasite-\|Undefined Saprotroph\| | Yeast | NULL | Probable | As mycoparasite (Põlme et al. 2020) | Põlme S, et al. 2020. Fungal Diversity 105:1-16 ((https://doi.org/10.1007/s13225-020-00466-2)) |
| Otu06022 | 4 | k__Fungi(100);p__Ascomycota(100);c__Saccharomycetes(100);o__Saccharomycetales(100);f__Metschnikowiaceae(100);g__Metschnikowia(100);s__Metschnikowia_pulcherrima(100); | Metschnikowia | 13 | Saprotroph | \|Undefined Saprotroph\| | Yeast | NULL | Probable | As nectar-tap_saprotroph (Põlme et al. 2020); Decay sugar-rich substrates (Põlme et al. 2020) | Põlme S, et al. 2020. Fungal Diversity 105:1-16 ((https://doi.org/10.1007/s13225-020-00466-2)) |
| Otu06025 | 4 | k__Fungi(100);p__Ascomycota(75);c__Saccharomycetes(75);o__Saccharomycetales(75);f__Metschnikowiaceae(75);g__Metschnikowia(75);s__Metschnikowia_pulcherrima(75); | Metschnikowia | 13 | Saprotroph | \|Undefined Saprotroph\| | Yeast | NULL | Probable | As nectar-tap_saprotroph (Põlme et al. 2020); Decay sugar-rich substrates (Põlme et al. 2020) | Põlme S, et al. 2020. Fungal Diversity 105:1-16 ((https://doi.org/10.1007/s13225-020-00466-2)) |
| Otu06027 | 4 | k__Fungi(100);p__Ascomycota(100);c__Saccharomycetes(100);o__Saccharomycetales(100);f__Metschnikowiaceae(100);g__Metschnikowia(100);g__Metschnikowia_unclassified(100); | Metschnikowia | 13 | Saprotroph | \|Undefined Saprotroph\| | Yeast | NULL | Probable | As nectar-tap_saprotroph (Põlme et al. 2020); Decay sugar-rich substrates (Põlme et al. 2020) | Põlme S, et al. 2020. Fungal Diversity 105:1-16 ((https://doi.org/10.1007/s13225-020-00466-2)) |
| Otu06028 | 4 | k__Fungi(100);p__Ascomycota(100);c__Sordariomycetes(100);o__Coniochaetales(75);f__Coniochaetaceae(75);f__Coniochaetaceae_unclassified(75);f__Coniochaetaceae_unclassified(75); | Coniochaetaceae | 9 | Pathotroph-Saprotroph-Symbiotroph | Animal Pathogen-Dung Saprotroph-Endophyte-Lichen Parasite-Plant Pathogen-Undefined Saprotroph | Microfungus | NULL | Possible | Saprobic on wood and bark (especially in dry habitats), coprophilous or isolated from soil and similar substrata (Cannon & Kirk 2007) | Cannon PF, Kirk PM. 2007. Fungal Families of the World. CAB International, Cambridge (ISBN: 978-0851998275); Damm U, et al. 2010. Persoonia 24:60-80 ((http://doi.org/10.3767/003158510X500705)); Irinyi L, et al. 2015. Medical Mycology 53:313-337 ((https://doi.org/10.1093/mmy/myv008)); Lawrey JD, Diederich P. 2016. http://www.lichenicolous.net (accessed 24 May 2017) ((http://www.lichenicolous.net)) |
| Otu06030 | 4 | k__Fungi(100);p__Ascomycota(100);c__Dothideomycetes(100);o__Pleosporales(100);f__Sporormiaceae(100);g__Preussia(100);g__Preussia_unclassified(100); | Preussia | 13 | Saprotroph | \|Dung Saprotroph\|-Undefined Saprotroph | Microfungus | NULL | Probable | Decay dung (Põlme et al. 2020) | Tedersoo L, et al. 2014. Science 346:e1256688 ((https://doi.org/10.1126/science.1256688)); Põlme S, et al. 2020. Fungal Diversity 105:1-16 ((https://doi.org/10.1007/s13225-020-00466-2)) |
| Otu06031 | 4 | k__Fungi(100);p__Basidiomycota(100);c__Tremellomycetes(100);o__Cystofilobasidiales(100);f__Cystofilobasidiaceae(100);g__Cystofilobasidium(100);s__Cystofilobasidium_capitatum(100); | Cystofilobasidium | 13 | Saprotroph | \|Plant Saprotroph\| | Yeast | NULL | Possible | As litter_saprotroph (Põlme et al. 2020); Decay leaf, fruit, seed (Põlme et al. 2020) | Oberwinkler F, et al. 1983. Systematic and Applied Microbiology 41:14-122 ((https://doi.org/10.1016/S0723-2020((83))80039-3)) |
| Otu06036 | 4 | k__Fungi(100);p__Ascomycota(100);c__Eurotiomycetes(100);o__Chaetothyriales(100);f__Herpotrichiellaceae(100);f__Herpotrichiellaceae_unclassified(100);f__Herpotrichiellaceae_unclassified(100); | Herpotrichiellaceae | 9 | Pathotroph-Saprotroph | Animal Pathogen-Fungal Parasite-Undefined Saprotroph | Facultative Yeast-Microfungus | NULL | Probable | Mostly saprobic on plants or other fungi, isolated from soil etc., and from medical sources (Cannon & Kirk 2007) | Cannon PF, Kirk PM. 2007. Fungal Families of the World. CAB International, Cambridge (ISBN: 978-0851998275) |
| Otu06037 | 4 | k__Fungi(100);p__Ascomycota(100);c__Saccharomycetes(100);o__Saccharomycetales(100);f__Metschnikowiaceae(100);g__Metschnikowia(100);s__Metschnikowia_pulcherrima(75); | Metschnikowia | 13 | Saprotroph | \|Undefined Saprotroph\| | Yeast | NULL | Probable | As nectar-tap_saprotroph (Põlme et al. 2020); Decay sugar-rich substrates (Põlme et al. 2020) | Põlme S, et al. 2020. Fungal Diversity 105:1-16 ((https://doi.org/10.1007/s13225-020-00466-2)) |
| Otu06038 | 4 | k__Fungi(100);p__Ascomycota(100);c__Saccharomycetes(100);o__Saccharomycetales(100);f__Metschnikowiaceae(100);g__Metschnikowia(100);s__Metschnikowia_pulcherrima(100); | Metschnikowia | 13 | Saprotroph | \|Undefined Saprotroph\| | Yeast | NULL | Probable | As nectar-tap_saprotroph (Põlme et al. 2020); Decay sugar-rich substrates (Põlme et al. 2020) | Põlme S, et al. 2020. Fungal Diversity 105:1-16 ((https://doi.org/10.1007/s13225-020-00466-2)) |
| Otu06039 | 4 | k__Fungi(100);p__Ascomycota(100);c__Saccharomycetes(100);o__Saccharomycetales(100);f__Metschnikowiaceae(100);g__Metschnikowia(100);g__Metschnikowia_unclassified(100); | Metschnikowia | 13 | Saprotroph | \|Undefined Saprotroph\| | Yeast | NULL | Probable | As nectar-tap_saprotroph (Põlme et al. 2020); Decay sugar-rich substrates (Põlme et al. 2020) | Põlme S, et al. 2020. Fungal Diversity 105:1-16 ((https://doi.org/10.1007/s13225-020-00466-2)) |
| Otu06040 | 4 | k__Fungi(100);p__Ascomycota(100);c__Eurotiomycetes(100);o__Chaetothyriales(100);f__Herpotrichiellaceae(100);g__Exophiala(100);s__Exophiala_sp_NO170(100); | Exophiala | 13 | Pathotroph-Saprotroph-Symbiotroph | \|Animal Parasite\|-Animal Pathogen-Endophyte-Plant Saprotroph-Undefined Saprotroph | Dimorphic-Facultative Yeast | NULL | Probable | As litter_saprotroph (Põlme et al. 2020); As root_endophyte_dark_septate (Põlme et al. 2020); Decay leaf, fruit, seed, animal material (Põlme et al. 2020) | Tedersoo L, et al. 2014. Science 346:e1256688 ((https://doi.org/10.1126/science.1256688)), Nagy LG, et al. 2014. Nature Communications 5:e4471 ((https://doi.org/10.1038/ncomms5471)); Põlme S, et al. 2020. Fungal Diversity 105:1-16 ((https://doi.org/10.1007/s13225-020-00466-2)) |
| Otu06041 | 4 | k__Fungi(100);p__Ascomycota(100);c__Saccharomycetes(100);o__Saccharomycetales(100);f__Metschnikowiaceae(100);g__Metschnikowia(100);g__Metschnikowia_unclassified(75); | Metschnikowia | 13 | Saprotroph | \|Undefined Saprotroph\| | Yeast | NULL | Probable | As nectar-tap_saprotroph (Põlme et al. 2020); Decay sugar-rich substrates (Põlme et al. 2020) | Põlme S, et al. 2020. Fungal Diversity 105:1-16 ((https://doi.org/10.1007/s13225-020-00466-2)) |
| Otu06044 | 4 | k__Fungi(100);p__Ascomycota(100);c__Eurotiomycetes(100);o__Chaetothyriales(100);f__Herpotrichiellaceae(100);g__Cladophialophora(100);g__Cladophialophora_unclassified(100); | Cladophialophora | 13 | Pathotroph-Saprotroph-Symbiotroph | Animal Pathogen-Endophyte-Fungal Parasite-\|Undefined Saprotroph\| | Dimorphic-Microfungus | NULL | Probable | As mycoparasite (Põlme et al. 2020); As root_endophyte_dark_septate (Põlme et al. 2020); As soil_saprotroph (Põlme et al. 2020); Decay soil (Põlme et al. 2020) | James TY, et al. 2006. Nature 443:818-822 ((https://doi.org/10.1038/nature05110)); Põlme S, et al. 2020. Fungal Diversity 105:1-16 ((https://doi.org/10.1007/s13225-020-00466-2)) |
| Otu06045 | 4 | k__Fungi(100);p__Ascomycota(100);c__Saccharomycetes(100);o__Saccharomycetales(100);f__Saccharomycetales_family_Incertae_sedis(100);g__Candida(100);g__Candida_unclassified(75); | Candida | 13 | Pathotroph-Saprotroph-Symbiotroph | Animal Pathogen-Endophyte-Endosymbiont-Epiphyte-\|Undefined Saprotroph\| | Dimorphic Yeast-Yeast | NULL | Probable | As nectar-tap_saprotroph (Põlme et al. 2020); Decay sugar-rich substrates (Põlme et al. 2020); This asexual genus is polyphyletic and contain species that function very differently depend on what they are. Caution should be taken when interpreting guilds for Candida species. | Manolakaki D, et al. 2010. Virulence. 1: 367-375 ((https://doi.org/10.4161/viru.1.5.12796)); Mekbib SB, et al. 2011. Tropical Plant Pathology 36:233-240 ((https://doi.org/10.1590/S1982-56762011000400004)); Põlme S, et al. 2020. Fungal Diversity 105:1-16 ((https://doi.org/10.1007/s13225-020-00466-2)) |
| Otu06048 | 4 | k__Fungi(100);p__Ascomycota(100);c__Saccharomycetes(100);o__Saccharomycetales(100);f__Metschnikowiaceae(100);g__Metschnikowia(100);s__Metschnikowia_pulcherrima(100); | Metschnikowia | 13 | Saprotroph | \|Undefined Saprotroph\| | Yeast | NULL | Probable | As nectar-tap_saprotroph (Põlme et al. 2020); Decay sugar-rich substrates (Põlme et al. 2020) | Põlme S, et al. 2020. Fungal Diversity 105:1-16 ((https://doi.org/10.1007/s13225-020-00466-2)) |
| Otu06050 | 4 | k__Fungi(100);p__Ascomycota(100);c__Saccharomycetes(100);o__Saccharomycetales(100);f__Metschnikowiaceae(100);g__Metschnikowia(100);s__Metschnikowia_pulcherrima(100); | Metschnikowia | 13 | Saprotroph | \|Undefined Saprotroph\| | Yeast | NULL | Probable | As nectar-tap_saprotroph (Põlme et al. 2020); Decay sugar-rich substrates (Põlme et al. 2020) | Põlme S, et al. 2020. Fungal Diversity 105:1-16 ((https://doi.org/10.1007/s13225-020-00466-2)) |
| Otu06051 | 4 | k__Fungi(100);p__Ascomycota(100);c__Saccharomycetes(100);o__Saccharomycetales(100);f__Metschnikowiaceae(100);g__Metschnikowia(100);s__Metschnikowia_pulcherrima(100); | Metschnikowia | 13 | Saprotroph | \|Undefined Saprotroph\| | Yeast | NULL | Probable | As nectar-tap_saprotroph (Põlme et al. 2020); Decay sugar-rich substrates (Põlme et al. 2020) | Põlme S, et al. 2020. Fungal Diversity 105:1-16 ((https://doi.org/10.1007/s13225-020-00466-2)) |
| Otu06052 | 4 | k__Fungi(100);p__Ascomycota(100);c__Dothideomycetes(100);o__Capnodiales(100);f__Mycosphaerellaceae(100);f__Mycosphaerellaceae_unclassified(100);f__Mycosphaerellaceae_unclassified(100); | Mycosphaerellaceae | 9 | Pathotroph-Saprotroph | Plant Pathogen-Undefined Saprotroph | Microfungus | NULL | Probable | Biotrophic, necrotrophic or saprobic on various plant tissues (Cannon & Kirk 2007) | Cannon PF, Kirk PM. 2007. Fungal Families of the World. CAB International, Cambridge (ISBN: 978-0851998275) |
| Otu06054 | 4 | k__Fungi(100);p__Ascomycota(100);c__Saccharomycetes(100);o__Saccharomycetales(100);f__Metschnikowiaceae(100);g__Metschnikowia(100);g__Metschnikowia_unclassified(75); | Metschnikowia | 13 | Saprotroph | \|Undefined Saprotroph\| | Yeast | NULL | Probable | As nectar-tap_saprotroph (Põlme et al. 2020); Decay sugar-rich substrates (Põlme et al. 2020) | Põlme S, et al. 2020. Fungal Diversity 105:1-16 ((https://doi.org/10.1007/s13225-020-00466-2)) |
| Otu06056 | 4 | k__Fungi(100);p__Ascomycota(100);c__Saccharomycetes(100);o__Saccharomycetales(100);f__Saccharomycetaceae(100);g__Kazachstania(100);s__Kazachstania_intestinalis(100); | Kazachstania | 13 | Saprotroph | \|Undefined Saprotroph\| | Yeast | NULL | Probable | As nectar-tap_saprotroph (Põlme et al. 2020); Decay sugar-rich substrates (Põlme et al. 2020) | Põlme S, et al. 2020. Fungal Diversity 105:1-16 ((https://doi.org/10.1007/s13225-020-00466-2)) |
| Otu06057 | 4 | k__Fungi(100);p__Ascomycota(100);c__Saccharomycetes(100);o__Saccharomycetales(100);f__Metschnikowiaceae(100);g__Metschnikowia(100);s__Metschnikowia_pulcherrima(100); | Metschnikowia | 13 | Saprotroph | \|Undefined Saprotroph\| | Yeast | NULL | Probable | As nectar-tap_saprotroph (Põlme et al. 2020); Decay sugar-rich substrates (Põlme et al. 2020) | Põlme S, et al. 2020. Fungal Diversity 105:1-16 ((https://doi.org/10.1007/s13225-020-00466-2)) |
| Otu06058 | 4 | k__Fungi(100);p__Basidiomycota(100);c__Agaricomycetes(100);o__Russulales(100);f__Hericiaceae(100);g__Hericium(100);s__Hericium_coralloides(100); | Hericium | 13 | Saprotroph-Pathotroph-Saprotroph | Plant Pathogen-Undefined Saprotroph-\|Wood Saprotroph\| | Hydnoid-Polyporoid | White Rot | Probable | As root_pathogen (Põlme et al. 2020); As white_rot (Põlme et al. 2020); Decay wood, roots (Põlme et al. 2020) | Tedersoo L, et al. 2014. Science 346:e1256688 ((https://doi.org/10.1126/science.1256688)); Põlme S, et al. 2020. Fungal Diversity 105:1-16 ((https://doi.org/10.1007/s13225-020-00466-2)) |
| Otu06061 | 4 | k__Fungi(100);p__Ascomycota(100);c__Saccharomycetes(100);o__Saccharomycetales(100);f__Metschnikowiaceae(100);g__Metschnikowia(100);s__Metschnikowia_pulcherrima(100); | Metschnikowia | 13 | Saprotroph | \|Undefined Saprotroph\| | Yeast | NULL | Probable | As nectar-tap_saprotroph (Põlme et al. 2020); Decay sugar-rich substrates (Põlme et al. 2020) | Põlme S, et al. 2020. Fungal Diversity 105:1-16 ((https://doi.org/10.1007/s13225-020-00466-2)) |
| Otu06062 | 4 | k__Fungi(100);p__Ascomycota(100);c__Saccharomycetes(100);o__Saccharomycetales(100);f__Metschnikowiaceae(100);g__Metschnikowia(100);s__Metschnikowia_pulcherrima(100); | Metschnikowia | 13 | Saprotroph | \|Undefined Saprotroph\| | Yeast | NULL | Probable | As nectar-tap_saprotroph (Põlme et al. 2020); Decay sugar-rich substrates (Põlme et al. 2020) | Põlme S, et al. 2020. Fungal Diversity 105:1-16 ((https://doi.org/10.1007/s13225-020-00466-2)) |
| Otu06063 | 4 | k__Fungi(100);p__Ascomycota(100);c__Saccharomycetes(100);o__Saccharomycetales(100);f__Metschnikowiaceae(100);g__Metschnikowia(100);s__Metschnikowia_pulcherrima(75); | Metschnikowia | 13 | Saprotroph | \|Undefined Saprotroph\| | Yeast | NULL | Probable | As nectar-tap_saprotroph (Põlme et al. 2020); Decay sugar-rich substrates (Põlme et al. 2020) | Põlme S, et al. 2020. Fungal Diversity 105:1-16 ((https://doi.org/10.1007/s13225-020-00466-2)) |
| Otu06064 | 4 | k__Fungi(100);p__Ascomycota(100);c__Leotiomycetes(100);o__Helotiales(100);f__Hyaloscyphaceae(100);f__Hyaloscyphaceae_unclassified(100);f__Hyaloscyphaceae_unclassified(100); | Hyaloscyphaceae | 9 | Saprotroph | Plant Saprotroph-Wood Saprotroph | Microfungus | NULL | Probable | Saprobic on woody and herbaceous material (Cannon & Kirk 2007) | Cannon PF, Kirk PM. 2007. Fungal Families of the World. CAB International, Cambridge (ISBN: 978-0851998275) |
| Otu06065 | 4 | k__Fungi(100);p__Ascomycota(100);c__Leotiomycetes(100);o__Helotiales(100);f__Vibrisseaceae(100);g__Phialocephala(100);s__Phialocephala_sp_CM16s1(100); | Phialocephala | 13 | Saprotroph-Symbiotroph | Endophyte-\|Undefined Saprotroph\| | Microfungus | NULL | Highly Probable | As root_endophyte (Põlme et al. 2020); As root_endophyte_dark_septate (Põlme et al. 2020); As soil_saprotroph (Põlme et al. 2020); Dark Septate Endophyte, known from plant roots.; Decay roots, soil (Põlme et al. 2020) | Newsham KK. 2011. New Phytologist 190:783-793 ((https://doi.org/10.1111/j.1469-8137.2010.03611.x)) (as dark septate root endophytes); Põlme S, et al. 2020. Fungal Diversity 105:1-16 ((https://doi.org/10.1007/s13225-020-00466-2)) |
| Otu06066 | 4 | k__Fungi(100);p__Ascomycota(100);c__Saccharomycetes(100);o__Saccharomycetales(100);f__Metschnikowiaceae(100);g__Metschnikowia(100);s__Metschnikowia_pulcherrima(100); | Metschnikowia | 13 | Saprotroph | \|Undefined Saprotroph\| | Yeast | NULL | Probable | As nectar-tap_saprotroph (Põlme et al. 2020); Decay sugar-rich substrates (Põlme et al. 2020) | Põlme S, et al. 2020. Fungal Diversity 105:1-16 ((https://doi.org/10.1007/s13225-020-00466-2)) |
| Otu06068 | 4 | k__Fungi(100);p__Zygomycota(100);c__Zygomycota_class_Incertae_sedis(100);o__Mucorales(100);f__Mucoraceae(100);g__Mucor(100);s__Mucor_genevensis(100); | Mucor | 13 | Saprotroph | \|Undefined Saprotroph\| | Microfungus | NULL | Probable | As soil_saprotroph (Põlme et al. 2020); Decay soil (Põlme et al. 2020) | Tedersoo L, et al. 2014. Science 346:e1256688 ((https://doi.org/10.1126/science.1256688)); Põlme S, et al. 2020. Fungal Diversity 105:1-16 ((https://doi.org/10.1007/s13225-020-00466-2)) |
| Otu06070 | 4 | k__Fungi(100);p__Basidiomycota(100);c__Agaricomycetes(100);o__Boletales(100);f__Hygrophoropsidaceae(100);g__Hygrophoropsis(100);s__Hygrophoropsis_aurantiaca(100); | Hygrophoropsis | 13 | Saprotroph | \|Plant Saprotroph\|-Wood Saprotroph | Agaricoid | Brown Rot | Probable | As litter_saprotroph (Põlme et al. 2020); Decay wood, leaf, fruit, seed (Põlme et al. 2020) | Tedersoo L, et al. 2014. Science 346:e1256688 ((https://doi.org/10.1126/science.1256688)); Põlme S, et al. 2020. Fungal Diversity 105:1-16 ((https://doi.org/10.1007/s13225-020-00466-2)) |
| Otu06071 | 4 | k__Fungi(100);p__Ascomycota(100);c__Saccharomycetes(100);o__Saccharomycetales(100);f__Metschnikowiaceae(100);g__Metschnikowia(100);s__Metschnikowia_pulcherrima(100); | Metschnikowia | 13 | Saprotroph | \|Undefined Saprotroph\| | Yeast | NULL | Probable | As nectar-tap_saprotroph (Põlme et al. 2020); Decay sugar-rich substrates (Põlme et al. 2020) | Põlme S, et al. 2020. Fungal Diversity 105:1-16 ((https://doi.org/10.1007/s13225-020-00466-2)) |
| Otu06074 | 4 | k__Fungi(100);p__Basidiomycota(100);c__Agaricomycetes(100);o__Auriculariales(100);f__Auriculariales_family_Incertae_sedis(100);g__Exidia(100);g__Exidia_unclassified(100); | Exidia | 13 | Saprotroph-Symbiotroph | Endophyte-Plant Saprotroph-Undefined Saprotroph-\|Wood Saprotroph\| | Tremelloid | NULL | Probable | As foliar_endophyte (Põlme et al. 2020); As litter_saprotroph (Põlme et al. 2020); Decay wood, leaf, fruit, seed (Põlme et al. 2020) | Tedersoo L, et al. 2014. Science 346:e1256688 ((https://doi.org/10.1126/science.1256688)); David AS, et al. 2016. Microbial Ecology 71:912-926 ((https://doi.org/10.1007/s00248-015-0712-6)); Põlme S, et al. 2020. Fungal Diversity 105:1-16 ((https://doi.org/10.1007/s13225-020-00466-2)) |
| Otu06075 | 4 | k__Fungi(100);p__Ascomycota(100);c__Saccharomycetes(100);o__Saccharomycetales(100);f__Metschnikowiaceae(100);g__Metschnikowia(100);s__Metschnikowia_pulcherrima(100); | Metschnikowia | 13 | Saprotroph | \|Undefined Saprotroph\| | Yeast | NULL | Probable | As nectar-tap_saprotroph (Põlme et al. 2020); Decay sugar-rich substrates (Põlme et al. 2020) | Põlme S, et al. 2020. Fungal Diversity 105:1-16 ((https://doi.org/10.1007/s13225-020-00466-2)) |
| Otu06076 | 4 | k__Fungi(100);p__Ascomycota(100);c__Saccharomycetes(100);o__Saccharomycetales(100);f__Metschnikowiaceae(100);g__Metschnikowia(100);s__Metschnikowia_pulcherrima(100); | Metschnikowia | 13 | Saprotroph | \|Undefined Saprotroph\| | Yeast | NULL | Probable | As nectar-tap_saprotroph (Põlme et al. 2020); Decay sugar-rich substrates (Põlme et al. 2020) | Põlme S, et al. 2020. Fungal Diversity 105:1-16 ((https://doi.org/10.1007/s13225-020-00466-2)) |
| Otu06079 | 4 | k__Fungi(100);p__Ascomycota(100);c__Saccharomycetes(100);o__Saccharomycetales(100);f__Metschnikowiaceae(100);g__Metschnikowia(100);g__Metschnikowia_unclassified(75); | Metschnikowia | 13 | Saprotroph | \|Undefined Saprotroph\| | Yeast | NULL | Probable | As nectar-tap_saprotroph (Põlme et al. 2020); Decay sugar-rich substrates (Põlme et al. 2020) | Põlme S, et al. 2020. Fungal Diversity 105:1-16 ((https://doi.org/10.1007/s13225-020-00466-2)) |
| Otu06082 | 4 | k__Fungi(100);p__Ascomycota(75);c__Saccharomycetes(75);o__Saccharomycetales(75);f__Metschnikowiaceae(75);g__Metschnikowia(75);s__Metschnikowia_pulcherrima(75); | Metschnikowia | 13 | Saprotroph | \|Undefined Saprotroph\| | Yeast | NULL | Probable | As nectar-tap_saprotroph (Põlme et al. 2020); Decay sugar-rich substrates (Põlme et al. 2020) | Põlme S, et al. 2020. Fungal Diversity 105:1-16 ((https://doi.org/10.1007/s13225-020-00466-2)) |
| Otu06084 | 4 | k__Fungi(100);p__Basidiomycota(100);c__Microbotryomycetes(100);o__Sporidiobolales(100);f__Sporidiobolales_family_Incertae_sedis(100);g__Rhodotorula(100);s__Rhodotorula_minuta(100); | Rhodotorula | 13 | Pathotroph-Saprotroph-Symbiotroph | Animal Pathogen-Animal Symbiotroph-Endophyte-Plant Pathogen-\|Undefined Saprotroph\| | Yeast | NULL | Probable | As foliar_endophyte (Põlme et al. 2020); Opportunistic human pathogen | Põlme S, et al. 2020. Fungal Diversity 105:1-16 ((https://doi.org/10.1007/s13225-020-00466-2)); https://www.cdc.gov ((https://www.cdc.gov)) |

| | | | | | | | | | | | |
|---|---|---|---|---|---|---|---|---|---|---|---|
| Otu06085 | 4 | k__Fungi(100);p__Ascomycota(100);c__Dothideomycetes(100);o__Pleosporales(100);f__Pleosporales_family_Incertae_sedis(100);g__Phoma(100);s__Phoma_sp(100); | Phoma | 13 | Pathotroph-Saprotroph-Symbiotroph | Dung Saprotroph-Endophyte-Lichen Parasite-\|Plant Pathogen\|-Plant Saprotroph-Undefined Saprotroph-Wood Saprotroph | Microfungus | Soft Rot | Possible | As foliar_endophyte (Põlme et al. 2020); As litter_saprotroph (Põlme et al. 2020); Decay leaf, fruit, seed, roots, soil (Põlme et al. 2020) | Richardson MJ. 1972. Transactions of the British Mycological Society 58:37-48 ((https://doi.org/10.1016/S0007-1536(72)80069-X)); Seehann G, et al. 1975. List of Fungi in Soft-Rot Tissue Tests. Institute fur Holzbiologie und Holzschutz, Bundesforschungsanstalt fur Forst - und Holzwirtshaft, Hamburg - Lohbrugge, Germany; de Gruyter J, et al. 2010. Mycologia 102:1066-1081 ((https://doi.org/10.3852/09-240)); Costa IPMW, et al. 2012. Mycotaxon 119:e26 ((http://mycotaxon.com/resources/checklists/costa_v119_checklist.pdf)); Tedersoo L, et al. 2014. Science 346:e1256688 ((https://doi.org/10.1126/science.1256688)); Lawrey JD, Diederich P. 2016. http://www.lichenicolous.net (accessed 24 May 2017) ((http://www.lichenicolous.net)); Põlme S, et al. 2020. Fungal Diversity 105:1-16 ((https://doi.org/10.1007/s13225-020-00466-2)) |
| Otu06086 | 4 | k__Fungi(100);p__Ascomycota(100);c__Saccharomycetes(100);o__Saccharomycetales(100);f__Metschnikowiaceae(100);g__Metschnikowia(100);s__Metschnikowia_pulcherrima(100); | Metschnikowia | 13 | Saprotroph | \|Undefined Saprotroph\| | Yeast | NULL | Probable | As nectar-tap_saprotroph (Põlme et al. 2020); Decay sugar-rich substrates (Põlme et al. 2020) | Põlme S, et al. 2020. Fungal Diversity 105:1-16 ((https://doi.org/10.1007/s13225-020-00466-2)) |
| Otu06087 | 4 | k__Fungi(100);p__Ascomycota(100);c__Saccharomycetes(100);o__Saccharomycetales(100);f__Metschnikowiaceae(100);g__Metschnikowia(100);s__Metschnikowia_pulcherrima(100); | Metschnikowia | 13 | Saprotroph | \|Undefined Saprotroph\| | Yeast | NULL | Probable | As nectar-tap_saprotroph (Põlme et al. 2020); Decay sugar-rich substrates (Põlme et al. 2020) | Põlme S, et al. 2020. Fungal Diversity 105:1-16 ((https://doi.org/10.1007/s13225-020-00466-2)) |
| Otu06088 | 4 | k__Fungi(100);p__Ascomycota(100);c__Eurotiomycetes(100);o__Eurotiales(100);f__Trichocomaceae(100);g__Penicillium(100);s__Penicillium_maximae(100); | Penicillium | 13 | Pathotroph-Saprotroph-Symbiotroph | Animal Parasite-Dung Saprotroph-Endophyte-Fungal Parasite-Plant Pathogen-\|Undefined Saprotroph\|-Wood Saprotroph | Microfungus | Soft Rot | Highly Probable | As foliar_endophyte (Põlme et al. 2020); As mycoparasite (Põlme et al. 2020); Decay soil (Põlme et al. 2020) | Duncan CG, Eslyn WE. 1966. Mycologia 58:642-664 ((https://doi.org/10.2307/3757045)); Seehann G, et al. 1975. List of Fungi in Soft-Rot Tissue Tests. Institute fur Holzbiologie und Holzschutz, Bundesforschungsanstalt fur Forst - und Holzwirtshaft, Hamburg - Lohbrugge, Germany ((https://www.irg-wp.com/irgdocs/details.php?d6493033-c9ea-40f3-8e3e-5697f32d640d)); Costa IPMW, et al. 2012. Mycotaxon 119:e26 ((http://mycotaxon.com/resources/checklists/costa_v119_checklist.pdf)); Bills GF, et al. 2013. Current Opinion in Microbiology 16:549-565 ((https://doi.org/10.1016/j.mib.2013.08.001)); Tedersoo L, et al. 2014. Science 346:e1256688 ((https://doi.org/10.1126/science.1256688)); Põlme S, et al. 2020. Fungal Diversity 105:1-16 ((https://doi.org/10.1007/s13225-020-00466-2)) |
| Otu06090 | 4 | k__Fungi(100);p__Ascomycota(100);c__Saccharomycetes(100);o__Saccharomycetales(100);f__Metschnikowiaceae(100);g__Metschnikowia(100);s__Metschnikowia_pulcherrima(100); | Metschnikowia | 13 | Saprotroph | \|Undefined Saprotroph\| | Yeast | NULL | Probable | As nectar-tap_saprotroph (Põlme et al. 2020); Decay sugar-rich substrates (Põlme et al. 2020) | Põlme S, et al. 2020. Fungal Diversity 105:1-16 ((https://doi.org/10.1007/s13225-020-00466-2)) |
| Otu06094 | 4 | k__Fungi(100);p__Ascomycota(100);c__Saccharomycetes(100);o__Saccharomycetales(100);f__Metschnikowiaceae(100);g__Metschnikowia(100);s__Metschnikowia_pulcherrima(75); | Metschnikowia | 13 | Saprotroph | \|Undefined Saprotroph\| | Yeast | NULL | Probable | As nectar-tap_saprotroph (Põlme et al. 2020); Decay sugar-rich substrates (Põlme et al. 2020) | Põlme S, et al. 2020. Fungal Diversity 105:1-16 ((https://doi.org/10.1007/s13225-020-00466-2)) |
| Otu06095 | 4 | k__Fungi(100);p__Ascomycota(100);c__Saccharomycetes(100);o__Saccharomycetales(100);f__Metschnikowiaceae(100);g__Metschnikowia(100);s__Metschnikowia_pulcherrima(75); | Metschnikowia | 13 | Saprotroph | \|Undefined Saprotroph\| | Yeast | NULL | Probable | As nectar-tap_saprotroph (Põlme et al. 2020); Decay sugar-rich substrates (Põlme et al. 2020) | Põlme S, et al. 2020. Fungal Diversity 105:1-16 ((https://doi.org/10.1007/s13225-020-00466-2)) |
| Otu06099 | 4 | k__Fungi(100);p__Ascomycota(100);c__Leotiomycetes(100);o__Helotiales(100);f__Hyaloscyphaceae(100);g__Lachnellula(100);s__Lachnellula_sp(100); | Lachnellula | 13 | Saprotroph | Undefined Saprotroph-\|Wood Saprotroph\| | Microfungus-Pezizoid | NULL | Probable | Decay: wood (Põlme et al. 2020) | Tedersoo L, et al. 2014. Science 346:e1256688 ((https://doi.org/10.1126/science.1256688)); Põlme S, et al. 2020. Fungal Diversity 105:1-16 ((https://doi.org/10.1007/s13225-020-00466-2)) |
| Otu06100 | 4 | k__Fungi(100);p__Ascomycota(100);c__Saccharomycetes(100);o__Saccharomycetales(100);f__Metschnikowiaceae(100);g__Metschnikowia(100);s__Metschnikowia_pulcherrima(100); | Metschnikowia | 13 | Saprotroph | \|Undefined Saprotroph\| | Yeast | NULL | Probable | As nectar-tap_saprotroph (Põlme et al. 2020); Decay sugar-rich substrates (Põlme et al. 2020) | Põlme S, et al. 2020. Fungal Diversity 105:1-16 ((https://doi.org/10.1007/s13225-020-00466-2)) |
| Otu06101 | 4 | k__Fungi(100);p__Ascomycota(100);c__Saccharomycetes(100);o__Saccharomycetales(100);f__Metschnikowiaceae(100);g__Metschnikowia(100);s__Metschnikowia_pulcherrima(75); | Metschnikowia | 13 | Saprotroph | \|Undefined Saprotroph\| | Yeast | NULL | Probable | As nectar-tap_saprotroph (Põlme et al. 2020); Decay sugar-rich substrates (Põlme et al. 2020) | Põlme S, et al. 2020. Fungal Diversity 105:1-16 ((https://doi.org/10.1007/s13225-020-00466-2)) |
| Otu06102 | 4 | k__Fungi(100);p__Ascomycota(100);c__Saccharomycetes(100);o__Saccharomycetales(100);f__Metschnikowiaceae(100);g__Metschnikowia(100);s__Metschnikowia_pulcherrima(100); | Metschnikowia | 13 | Saprotroph | \|Undefined Saprotroph\| | Yeast | NULL | Probable | As nectar-tap_saprotroph (Põlme et al. 2020); Decay sugar-rich substrates (Põlme et al. 2020) | Põlme S, et al. 2020. Fungal Diversity 105:1-16 ((https://doi.org/10.1007/s13225-020-00466-2)) |
| Otu06106 | 4 | k__Fungi(100);p__Basidiomycota(100);c__Agaricomycetes(100);o__Cantharellales(100);f__Tulasnellaceae(100);g__Epulorhiza(100);s__Epulorhiza_sp_Ec1B(75); | Epulorhiza | 13 | Saprotroph-Symbiotroph | Ectomycorrhizal-\|Plant Saprotroph\| | Corticioid-Microfungus | NULL | Possible | As litter_saprotroph (Põlme et al. 2020); Decay leaf, fruit, seed (Põlme et al. 2020) | Pereira OL, et al. 2003. Mycoscience 44:153-155 ((https://doi.org/10.1007/S10267-002-0087-7)); Põlme S, et al. 2020. Fungal Diversity 105:1-16 ((https://doi.org/10.1007/s13225-020-00466-2)) |
| Otu06107 | 4 | k__Fungi(100);p__Ascomycota(100);c__Saccharomycetes(100);o__Saccharomycetales(100);f__Metschnikowiaceae(100);g__Metschnikowia(100);g__Metschnikowia_unclassified(75); | Metschnikowia | 13 | Saprotroph | \|Undefined Saprotroph\| | Yeast | NULL | Probable | As nectar-tap_saprotroph (Põlme et al. 2020); Decay sugar-rich substrates (Põlme et al. 2020) | Põlme S, et al. 2020. Fungal Diversity 105:1-16 ((https://doi.org/10.1007/s13225-020-00466-2)) |
| Otu06109 | 4 | k__Fungi(100);p__Ascomycota(100);c__Saccharomycetes(100);o__Saccharomycetales(100);f__Metschnikowiaceae(100);g__Metschnikowia(100);s__Metschnikowia_pulcherrima(75); | Metschnikowia | 13 | Saprotroph | \|Undefined Saprotroph\| | Yeast | NULL | Probable | As nectar-tap_saprotroph (Põlme et al. 2020); Decay sugar-rich substrates (Põlme et al. 2020) | Põlme S, et al. 2020. Fungal Diversity 105:1-16 ((https://doi.org/10.1007/s13225-020-00466-2)) |
| Otu06110 | 4 | k__Fungi(100);p__Ascomycota(100);c__Saccharomycetes(100);o__Saccharomycetales(100);f__Metschnikowiaceae(100);g__Metschnikowia(100);s__Metschnikowia_pulcherrima(100); | Metschnikowia | 13 | Saprotroph | \|Undefined Saprotroph\| | Yeast | NULL | Probable | As nectar-tap_saprotroph (Põlme et al. 2020); Decay sugar-rich substrates (Põlme et al. 2020) | Põlme S, et al. 2020. Fungal Diversity 105:1-16 ((https://doi.org/10.1007/s13225-020-00466-2)) |
| Otu06115 | 4 | k__Fungi(100);p__Basidiomycota(100);c__Exobasidiomycetes(100);o__Exobasidiales(100);f__Exobasidiaceae(100);g__Exobasidium(100);s__Exobasidium_arescens(100); | Exobasidium | 13 | Pathotroph | \|Plant Pathogen\| | Leaf Blister-Leaf Gall-Leaf Spot | NULL | Probable | NULL | Tedersoo L, et al. 2014. Science 346:e1256688 ((https://doi.org/10.1126/science.1256688)); Põlme S, et al. 2020. Fungal Diversity 105:1-16 ((https://doi.org/10.1007/s13225-020-00466-2)) |
| Otu06117 | 4 | k__Fungi(100);p__Ascomycota(100);c__Saccharomycetes(100);o__Saccharomycetales(100);f__Saccharomycetales_family_Incertae_sedis(100);f__Saccharomycetales_family_Incertae_sedis_unclassified(100);f__Saccharomycetales_family_Incertae_sedis_unclassified(100); | Saccharomycetales | 7 | Saprotroph | Undefined Saprotroph | Yeast | NULL | Possible | NULL | Sterkenburg E, et al. 2015. New Phytologist 207:1145-1158 ((https://doi.org/10.1111/nph.13426)) |
| Otu06120 | 4 | k__Fungi(100);p__Ascomycota(100);c__Saccharomycetes(100);o__Saccharomycetales(100);f__Metschnikowiaceae(100);g__Metschnikowia(100);s__Metschnikowia_pulcherrima(75); | Metschnikowia | 13 | Saprotroph | \|Undefined Saprotroph\| | Yeast | NULL | Probable | As nectar-tap_saprotroph (Põlme et al. 2020); Decay sugar-rich substrates (Põlme et al. 2020) | Põlme S, et al. 2020. Fungal Diversity 105:1-16 ((https://doi.org/10.1007/s13225-020-00466-2)) |
| Otu06122 | 4 | k__Fungi(100);p__Ascomycota(100);c__Saccharomycetes(100);o__Saccharomycetales(100);f__Metschnikowiaceae(100);g__Metschnikowia(100);s__Metschnikowia_pulcherrima(100); | Metschnikowia | 13 | Saprotroph | \|Undefined Saprotroph\| | Yeast | NULL | Probable | As nectar-tap_saprotroph (Põlme et al. 2020); Decay sugar-rich substrates (Põlme et al. 2020) | Põlme S, et al. 2020. Fungal Diversity 105:1-16 ((https://doi.org/10.1007/s13225-020-00466-2)) |
| Otu06123 | 4 | k__Fungi(100);p__Ascomycota(100);c__Saccharomycetes(100);o__Saccharomycetales(100);f__Metschnikowiaceae(100);g__Metschnikowia(100);s__Metschnikowia_pulcherrima(100); | Metschnikowia | 13 | Saprotroph | \|Undefined Saprotroph\| | Yeast | NULL | Probable | As nectar-tap_saprotroph (Põlme et al. 2020); Decay sugar-rich substrates (Põlme et al. 2020) | Põlme S, et al. 2020. Fungal Diversity 105:1-16 ((https://doi.org/10.1007/s13225-020-00466-2)) |
| Otu06124 | 4 | k__Fungi(100);p__Basidiomycota(100);c__Microbotryomycetes(100);o__Sporidiobolales(100);f__Sporidiobolales_family_Incertae_sedis(100);g__Sporobolomyces(100);s__Sporobolomyces_lactosus(100); | Sporobolomyces | 13 | Pathotroph-Saprotroph | \|Fungal Parasite\|-Plant Saprotroph | Yeast | NULL | Probable | As litter_saprotroph (Põlme et al. 2020); As mycoparasite (Põlme et al. 2020); Decay fungal material (Põlme et al. 2020) | Tedersoo L, et al. 2014. Science 346:e1256688 ((https://doi.org/10.1126/science.1256688)); Purahong W, et al. 2016. Molecular Ecology 25:4059-4074 ((https://doi.org/10.1111/mec.13739)); Põlme S, et al. 2020. Fungal Diversity 105:1-16 ((https://doi.org/10.1007/s13225-020-00466-2)) |
| Otu06125 | 4 | k__Fungi(100);p__Ascomycota(100);c__Saccharomycetes(100);o__Saccharomycetales(100);f__Metschnikowiaceae(100);g__Metschnikowia(100);s__Metschnikowia_pulcherrima(100); | Metschnikowia | 13 | Saprotroph | \|Undefined Saprotroph\| | Yeast | NULL | Probable | As nectar-tap_saprotroph (Põlme et al. 2020); Decay sugar-rich substrates (Põlme et al. 2020) | Põlme S, et al. 2020. Fungal Diversity 105:1-16 ((https://doi.org/10.1007/s13225-020-00466-2)) |
| Otu06128 | 4 | k__Fungi(100);p__Ascomycota(100);c__Saccharomycetes(100);o__Saccharomycetales(100);f__Metschnikowiaceae(100);g__Metschnikowia(100);s__Metschnikowia_pulcherrima(75); | Metschnikowia | 13 | Saprotroph | \|Undefined Saprotroph\| | Yeast | NULL | Probable | As nectar-tap_saprotroph (Põlme et al. 2020); Decay sugar-rich substrates (Põlme et al. 2020) | Põlme S, et al. 2020. Fungal Diversity 105:1-16 ((https://doi.org/10.1007/s13225-020-00466-2)) |
| Otu06130 | 4 | k__Fungi(100);p__Ascomycota(100);c__Sordariomycetes(100);o__Xylariales(100);f__Amphisphaeriaceae(100);f__Amphisphaeriaceae_unclassified(100);f__Amphisphaeriaceae_unclassified(100); | Amphisphaeriaceae | 9 | Saprotroph | Undefined Saprotroph | Microfungus | NULL | Possible | Saprobes or weak parasites of leaves, stems, etc., often appearing as secondary colonizers following damage to plants by other agents (Cannon & Kirk 2007) | Cannon PF, Kirk PM. 2007. Fungal Families of the World. CAB International, Cambridge (ISBN: 978-0851998275) |
| Otu06131 | 4 | k__Fungi(100);p__Ascomycota(100);c__Saccharomycetes(100);o__Saccharomycetales(100);f__Metschnikowiaceae(100);g__Metschnikowia(100);s__Metschnikowia_pulcherrima(100); | Metschnikowia | 13 | Saprotroph | \|Undefined Saprotroph\| | Yeast | NULL | Probable | As nectar-tap_saprotroph (Põlme et al. 2020); Decay sugar-rich substrates (Põlme et al. 2020) | Põlme S, et al. 2020. Fungal Diversity 105:1-16 ((https://doi.org/10.1007/s13225-020-00466-2)) |
| Otu06132 | 4 | k__Fungi(100);p__Ascomycota(100);c__Saccharomycetes(100);o__Saccharomycetales(100);f__Metschnikowiaceae(100);g__Metschnikowia(100);s__Metschnikowia_pulcherrima(100); | Metschnikowia | 13 | Saprotroph | \|Undefined Saprotroph\| | Yeast | NULL | Probable | As nectar-tap_saprotroph (Põlme et al. 2020); Decay sugar-rich substrates (Põlme et al. 2020) | Põlme S, et al. 2020. Fungal Diversity 105:1-16 ((https://doi.org/10.1007/s13225-020-00466-2)) |
| Otu06133 | 4 | k__Fungi(100);p__Ascomycota(100);c__Saccharomycetes(100);o__Saccharomycetales(100);f__Metschnikowiaceae(100);g__Metschnikowia(100);s__Metschnikowia_pulcherrima(100); | Metschnikowia | 13 | Saprotroph | \|Undefined Saprotroph\| | Yeast | NULL | Probable | As nectar-tap_saprotroph (Põlme et al. 2020); Decay sugar-rich substrates (Põlme et al. 2020) | Põlme S, et al. 2020. Fungal Diversity 105:1-16 ((https://doi.org/10.1007/s13225-020-00466-2)) |
| Otu06134 | 4 | k__Fungi(100);p__Ascomycota(100);c__Saccharomycetes(100);o__Saccharomycetales(100);f__Metschnikowiaceae(100);g__Metschnikowia(100);s__Metschnikowia_pulcherrima(100); | Metschnikowia | 13 | Saprotroph | \|Undefined Saprotroph\| | Yeast | NULL | Probable | As nectar-tap_saprotroph (Põlme et al. 2020); Decay sugar-rich substrates (Põlme et al. 2020) | Põlme S, et al. 2020. Fungal Diversity 105:1-16 ((https://doi.org/10.1007/s13225-020-00466-2)) |
| Otu06135 | 4 | k__Fungi(100);p__Ascomycota(100);c__Sordariomycetes(100);o__Ophiostomatales(100);f__Ophiostomataceae(100);f__Ophiostomataceae_unclassified(100);f__Ophiostomataceae_unclassified(100); | Ophiostomataceae | 9 | Pathotroph | Plant Pathogen | Microfungus | Blue-Staining | Probable | At least most species are transmitted via insect vectors, with mycelial colonies developing in bark-boring beetle galleries and spreading through the vascular system; they are highly specific to their beetle vectors. In some cases spores and mycelium are transmitted in specialized organs, termed mycangia (Cannon & Kirk 2007) | Cannon PF, Kirk PM. 2007. Fungal Families of the World. CAB International, Cambridge (ISBN: 978-0851998275); Tedersoo L, et al. 2014. Science 346:e1256688 ((https://doi.org/10.1126/science.1256688)) |
| Otu06136 | 4 | k__Fungi(100);p__Basidiomycota(100);c__Agaricomycetes(100);o__Agaricales(100);f__Marasmiaceae(100);g__Rhodocollybia(100);g__Rhodocollybia_unclassified(100); | Rhodocollybia | 13 | Saprotroph | \|Plant Saprotroph\|-Undefined Saprotroph | Agaricoid | NULL | Probable | As litter_saprotroph (Põlme et al. 2020); Decay leaf, fruit, seed (Põlme et al. 2020) | Tedersoo L, et al. 2014. Science 346:e1256688 ((https://doi.org/10.1126/science.1256688)); Põlme S, et al. 2020. Fungal Diversity 105:1-16 ((https://doi.org/10.1007/s13225-020-00466-2)) |
| Otu06137 | 4 | k__Fungi(100);p__Ascomycota(100);c__Saccharomycetes(100);o__Saccharomycetales(100);f__Metschnikowiaceae(100);g__Metschnikowia(100);s__Metschnikowia_pulcherrima(75); | Metschnikowia | 13 | Saprotroph | \|Undefined Saprotroph\| | Yeast | NULL | Probable | As nectar-tap_saprotroph (Põlme et al. 2020); Decay sugar-rich substrates (Põlme et al. 2020) | Põlme S, et al. 2020. Fungal Diversity 105:1-16 ((https://doi.org/10.1007/s13225-020-00466-2)) |
| Otu06139 | 4 | k__Fungi(100);p__Ascomycota(100);c__Sordariomycetes(100);o__Hypocreales(100);f__Nectriaceae(100);g__Fusarium(100);s__Fusarium_acetilerea(100); | Fusarium | 13 | Pathotroph-Saprotroph-Symbiotroph | Animal Pathogen-Endophyte-Lichen Parasite-\|Plant Pathogen\|-Plant Saprotroph-Undefined Saprotroph-Wood Saprotroph | Microfungus | Soft Rot | Possible | As foliar_endophyte (Põlme et al. 2020); As litter_saprotroph (Põlme et al. 2020); Causing damping-off disease (Põlme et al. 2020); Decay leaf, fruit, seed, roots, soil, animal material (Põlme et al. 2020); Likely opportunistic human pathogen (Irinyi et al. 2015) | Duncan CG, Eslyn WE. 1966. Mycologia 58:642-664 ((https://doi.org/10.2307/3757045)); Seehann G, et al. 1975. List of Fungi in Soft-Rot Tissue Tests. Institute fur Holzbiologie und Holzschutz, Bundesforschungsanstalt fur Forst - und Holzwirtshaft, Hamburg - Lohbrugge, Germany; Costa IPMW, et al. 2012. Mycotaxon 119:e26 ((http://mycotaxon.com/resources/checklists/costa_v119_checklist.pdf)); Tedersoo L, et al. 2014. Science 346:e1256688 ((https://doi.org/10.1126/science.1256688)); Irinyi L, et al. 2015. Medical Mycology 53:313-337 ((https://doi.org/10.1093/mmy/myv008)); Schroers, et al. 2016. Mycologia 108:806-819 ((https://doi.org/10.3852/15-255)); Põlme S, et al. 2020. Fungal Diversity 105:1-16 ((https://doi.org/10.1007/s13225-020-00466-2)); The American Phytopathological Society Common Names of Plant Diseases ((http://www.apsnet.org/publications/commonnames/Pages/default.aspx)); Lawrey JD, Diederich P. 2016. http://www.lichenicolous.net (accessed 24 May 2017) ((http://www.lichenicolous.net)) |
| Otu06140 | 4 | k__Fungi(100);p__Ascomycota(100);c__Saccharomycetes(100);o__Saccharomycetales(100);f__Metschnikowiaceae(100);g__Metschnikowia(100);g__Metschnikowia_unclassified(100); | Metschnikowia | 13 | Saprotroph | \|Undefined Saprotroph\| | Yeast | NULL | Probable | As nectar-tap_saprotroph (Põlme et al. 2020); Decay sugar-rich substrates (Põlme et al. 2020) | Põlme S, et al. 2020. Fungal Diversity 105:1-16 ((https://doi.org/10.1007/s13225-020-00466-2)) |
| Otu06141 | 4 | k__Fungi(100);p__Ascomycota(100);c__Saccharomycetes(100);o__Saccharomycetales(100);f__Metschnikowiaceae(100);g__Metschnikowia(100);g__Metschnikowia_unclassified(75); | Metschnikowia | 13 | Saprotroph | \|Undefined Saprotroph\| | Yeast | NULL | Probable | As nectar-tap_saprotroph (Põlme et al. 2020); Decay sugar-rich substrates (Põlme et al. 2020) | Põlme S, et al. 2020. Fungal Diversity 105:1-16 ((https://doi.org/10.1007/s13225-020-00466-2)) |
| Otu06142 | 4 | k__Fungi(100);p__Basidiomycota(100);c__Agaricomycetes(100);o__Cantharellales(100);f__Ceratobasidiaceae(100);g__Ceratobasidium(100);s__Ceratobasidium_sp_AG_I(100); | Ceratobasidium | 13 | Pathotroph-Saprotroph-Symbiotroph | Ectomycorrhizal-Endomycorrhizal-Orchid Mycorrhizal-\|Plant Pathogen\|-Plant Saprotroph-Undefined Saprotroph | Corticioid-Microfungus | NULL | Possible | especially of Orchidaceae but also even bryophytes (Cannon & Kirk 2007); As litter_saprotroph (Põlme et al. 2020); Best known as plant pathogens, but also saprobic. Species are endomycorrhizal with a wide range of plant roots ; Decay leaf, fruit, seed, roots, soil (Põlme et al. 2020) | Cannon PF, Kirk PM. 2007. Fungal Families of the World. CAB International, Cambridge (ISBN: 978-0851998275); Tedersoo L, et al. 2014. Science 346:e1256688 ((https://doi.org/10.1126/science.1256688)); Põlme S, et al. 2020. Fungal Diversity 105:1-16 ((https://doi.org/10.1007/s13225-020-00466-2)) |
| Otu06143 | 4 | k__Fungi(100);p__Ascomycota(100);c__Saccharomycetes(100);o__Saccharomycetales(100);f__Metschnikowiaceae(100);g__Metschnikowia(100);s__Metschnikowia_pulcherrima(75); | Metschnikowia | 13 | Saprotroph | \|Undefined Saprotroph\| | Yeast | NULL | Probable | As nectar-tap_saprotroph (Põlme et al. 2020); Decay sugar-rich substrates (Põlme et al. 2020) | Põlme S, et al. 2020. Fungal Diversity 105:1-16 ((https://doi.org/10.1007/s13225-020-00466-2)) |
| Otu06145 | 4 | k__Fungi(100);p__Ascomycota(100);c__Saccharomycetes(100);o__Saccharomycetales(100);f__Metschnikowiaceae(100);g__Metschnikowia(100);g__Metschnikowia_unclassified(100); | Metschnikowia | 13 | Saprotroph | \|Undefined Saprotroph\| | Yeast | NULL | Probable | As nectar-tap_saprotroph (Põlme et al. 2020); Decay sugar-rich substrates (Põlme et al. 2020) | Põlme S, et al. 2020. Fungal Diversity 105:1-16 ((https://doi.org/10.1007/s13225-020-00466-2)) |
| Otu06148 | 4 | k__Fungi(100);p__Ascomycota(100);c__Saccharomycetes(100);o__Saccharomycetales(100);f__Metschnikowiaceae(100);g__Metschnikowia(100);s__Metschnikowia_pulcherrima(100); | Metschnikowia | 13 | Saprotroph | \|Undefined Saprotroph\| | Yeast | NULL | Probable | As nectar-tap_saprotroph (Põlme et al. 2020); Decay sugar-rich substrates (Põlme et al. 2020) | Põlme S, et al. 2020. Fungal Diversity 105:1-16 ((https://doi.org/10.1007/s13225-020-00466-2)) |
| Otu06150 | 4 | k__Fungi(100);p__Ascomycota(100);c__Lecanoromycetes(100);o__Lecanorales(100);o__Lecanorales_unclassified(100);o__Lecanorales_unclassified(100);o__Lecanorales_unclassified(100); | Lecanorales | 7 | Pathotroph-Symbiotroph | Lichen Parasite-Lichenized | Microfungus-Thallus | NULL | Probable | Lichenized with green algae, occasionally lichenicolous, on rocks, mosses and wood (Cannon & Kirk 2007); ...an order of mostly lichen-forming fungi (Kirk et al. 2008) | Cannon PF, Kirk PM. 2007. Fungal Families of the World. CAB International, Cambridge (ISBN: 978-0851998275); Kirk PM et al. 2008. Dictionary of the Fungi. Tenth Edition. CAB International, Wallingford (ISBN: 978-0851998268); Esslinger TL. 2014. North Dakota State University ((http://www.ndsu.edu/pubweb/~esslinge/chcklst/chcklst7.htm)) |
| Otu06151 | 4 | k__Fungi(100);p__Ascomycota(100);c__Eurotiomycetes(100);o__Arachnomycetales(100);f__Arachnomycetaceae(100);g__Arachnomyces(100);s__Arachnomyces_nodosetosus(100); | Arachnomyces | 13 | Pathotroph-Saprotroph | \|Animal Parasite\|-Dung Saprotroph-Undefined Saprotroph | Microfungus | NULL | Highly Probable | As keratinolytic (Põlme et al. 2020); Decay animal material, dung (Põlme et al. 2020) | Bills GF, et al. 2013. Current Opinion in Microbiology 16:549-565 ((https://doi.org/10.1016/j.mib.2013.08.001)); Põlme S, et al. 2020. Fungal Diversity 105:1-16 ((https://doi.org/10.1007/s13225-020-00466-2)) |
| Otu06154 | 4 | k__Fungi(100);p__Ascomycota(100);c__Saccharomycetes(100);o__Saccharomycetales(100);f__Metschnikowiaceae(100);g__Metschnikowia(100);s__Metschnikowia_pulcherrima(100); | Metschnikowia | 13 | Saprotroph | \|Undefined Saprotroph\| | Yeast | NULL | Probable | As nectar-tap_saprotroph (Põlme et al. 2020); Decay sugar-rich substrates (Põlme et al. 2020) | Põlme S, et al. 2020. Fungal Diversity 105:1-16 ((https://doi.org/10.1007/s13225-020-00466-2)) |
| Otu06158 | 4 | k__Fungi(100);p__Basidiomycota(100);c__Agaricomycetes(100);o__Russulales(100);f__Russulaceae(100);g__Russula(100);s__Russula_sp_GAL17323(100); | Russula | 13 | Symbiotroph | \|Ectomycorrhizal\| | Agaricoid | NULL | Highly Probable | NULL | Rinaldi AC, et al. 2008. Fungal Diversity 33:1-45 ((http://www.fungaldiversity.org/fdp/sfdp/33-1.pdf)); Tedersoo L, et al. 2010. Mycorrhiza 20:217-263 ((https://doi.org/10.1007/s00572-009-0274-x)); Põlme S, et al. 2020. Fungal Diversity 105:1-16 ((https://doi.org/10.1007/s13225-020-00466-2)); http://mycorrhizas.info/ecmf.html ((http://mycorrhizas.info/ecmf.html)) |
| Otu06159 | 4 | k__Fungi(100);p__Ascomycota(100);c__Saccharomycetes(100);o__Saccharomycetales(100);f__Metschnikowiaceae(100);g__Metschnikowia(100);s__Metschnikowia_pulcherrima(100); | Metschnikowia | 13 | Saprotroph | \|Undefined Saprotroph\| | Yeast | NULL | Probable | As nectar-tap_saprotroph (Põlme et al. 2020); Decay sugar-rich substrates (Põlme et al. 2020) | Põlme S, et al. 2020. Fungal Diversity 105:1-16 ((https://doi.org/10.1007/s13225-020-00466-2)) |
| Otu06160 | 4 | k__Fungi(100);p__Ascomycota(100);c__Saccharomycetes(100);o__Saccharomycetales(100);f__Saccharomycetales_family_Incertae_sedis(100);g__Candida(75);g__Candida_unclassified(75); | Candida | 13 | Pathotroph-Saprotroph-Symbiotroph | Animal Pathogen-Endophyte-Endosymbiont-Epiphyte-\|Undefined Saprotroph\| | Dimorphic Yeast-Yeast | NULL | Probable | As nectar-tap_saprotroph (Põlme et al. 2020); Decay sugar-rich substrates (Põlme et al. 2020); This asexual genus is polyphyletic and contain species that function very differently depend on what they are. Caution should be taken when interpreting guilds for Candida species. | Manolakaki D, et al. 2010. Virulence. 1: 367-375 ((https://doi.org/10.4161/viru.1.5.12796)); Mekbib SB, et al. 2011. Tropical Plant Pathology 36:233-240 ((https://doi.org/10.1590/S1982-56762011000400004)); Põlme S, et al. 2020. Fungal Diversity 105:1-16 ((https://doi.org/10.1007/s13225-020-00466-2)) |
| Otu06161 | 4 | k__Fungi(100);p__Ascomycota(100);c__Sordariomycetes(100);o__Xylariales(100);f__Xylariaceae(100);g__Xylaria(100);s__Xylaria_polymorpha(100); | Xylaria | 13 | Saprotroph-Symbiotroph | Endophyte-Plant Saprotroph-Undefined Saprotroph-\|Wood Saprotroph\| | Xylarioid | Soft Rot | Probable | As foliar_endophyte (Põlme et al. 2020); As litter_saprotroph (Põlme et al. 2020); Decay wood, leaf, fruit, seed (Põlme et al. 2020); Host - angiosperms (Põlme et al. 2020) | Duncan CG, Eslyn WE. 1966. Mycologia 58:642-645 ((https://doi.org/10.2307/3757045)); Costa IPMW, et al. 2012. Mycotaxon 119:e26 ((http://mycotaxon.com/resources/checklists/costa_v119_checklist.pdf)); Põlme S, et al. 2020. Fungal Diversity 105:1-16 ((https://doi.org/10.1007/s13225-020-00466-2)); The American Phytopathological Society Common Names of Plant Diseases ((http://www.apsnet.org/publications/commonnames/Pages/default.aspx)) |
| Otu06162 | 4 | k__Fungi(100);p__Basidiomycota(100);c__Agaricomycetes(100);o__Trechisporales(100);f__Hydnodontaceae(100);g__Trechispora(100);g__Trechispora_unclassified(100); | Trechispora | 13 | Saprotroph | \|Wood Saprotroph\| | Corticioid | White Rot | Highly Probable | As white_rot (Põlme et al. 2020); Decay wood (Põlme et al. 2020) | Gilbertson RL, Ryvarden L. 1987-1987. North American Polypores. Fungiflora, Oslo (ISBN: 978-0945345060); Rinaldi AC, et al. 2008. Fungal Diversity 33:1-45 ((http://www.fungaldiversity.org/fdp/sfdp/33-1.pdf)) (as Uncertain); Tedersoo L, et al. 2010. Mycorrhiza 20:217-263 ((https://doi.org/10.1007/s00572-009-0274-x)) (as not ECM).; Põlme S, et al. 2020. Fungal Diversity 105:1-16 ((https://doi.org/10.1007/s13225-020-00466-2)) |
| Otu06164 | 4 | k__Fungi(100);p__Ascomycota(100);c__Saccharomycetes(100);o__Saccharomycetales(100);f__Metschnikowiaceae(100);g__Metschnikowia(100);s__Metschnikowia_pulcherrima(100); | Metschnikowia | 13 | Saprotroph | \|Undefined Saprotroph\| | Yeast | NULL | Probable | As nectar-tap_saprotroph (Põlme et al. 2020); Decay sugar-rich substrates (Põlme et al. 2020) | Põlme S, et al. 2020. Fungal Diversity 105:1-16 ((https://doi.org/10.1007/s13225-020-00466-2)) |
| Otu06165 | 4 | k__Fungi(100);p__Ascomycota(100);c__Saccharomycetes(100);o__Saccharomycetales(100);f__Metschnikowiaceae(100);g__Metschnikowia(100);s__Metschnikowia_pulcherrima(100); | Metschnikowia | 13 | Saprotroph | \|Undefined Saprotroph\| | Yeast | NULL | Probable | As nectar-tap_saprotroph (Põlme et al. 2020); Decay sugar-rich substrates (Põlme et al. 2020) | Põlme S, et al. 2020. Fungal Diversity 105:1-16 ((https://doi.org/10.1007/s13225-020-00466-2)) |
| Otu06168 | 4 | k__Fungi(100);p__Ascomycota(100);c__Leotiomycetes(100);o__Helotiales(100);f__Helotiales_family_Incertae_sedis(100);g__Leptodontidium(100);s__Leptodontidium_sp_Di124_4(100); | Leptodontidium | 13 | Saprotroph-Symbiotroph | Endophyte-Ericoid Mycorrhizal-\|Plant Saprotroph\| | Microfungus | NULL | Highly Probable | As litter_saprotroph (Põlme et al. 2020); As root_endophyte_dark_septate (Põlme et al. 2020); Dark Septate Endophyte, known from plant roots.; Decay leaf, fruit, seed (Põlme et al. 2020) | Jumpponen A, Trappe J. 1998. New Phytologist 140:295-310 ((https://doi.org/10.1046/j.1469-8137.1998.00265.x)) (as DSE); Taylor DL, et al. 2014. Ecological Monographs 8:43-20 ((https://esajournals.onlinelibrary.wiley.com/doi/10.1890/12-1693.1)); Põlme S, et al. 2020. Fungal Diversity 105:1-16 ((https://doi.org/10.1007/s13225-020-00466-2)) |
| Otu06173 | 4 | k__Fungi(100);p__Ascomycota(100);c__Saccharomycetes(100);o__Saccharomycetales(100);f__Metschnikowiaceae(100);g__Metschnikowia(100);s__Metschnikowia_pulcherrima(100); | Metschnikowia | 13 | Saprotroph | \|Undefined Saprotroph\| | Yeast | NULL | Probable | As nectar-tap_saprotroph (Põlme et al. 2020); Decay sugar-rich substrates (Põlme et al. 2020) | Põlme S, et al. 2020. Fungal Diversity 105:1-16 ((https://doi.org/10.1007/s13225-020-00466-2)) |
| Otu06174 | 4 | k__Fungi(100);p__Basidiomycota(100);c__Agaricomycetes(100);o__Cantharellales(100);f__Ceratobasidiaceae(100);f__Ceratobasidiaceae_unclassified(100);f__Ceratobasidiaceae_unclassified(100); | Ceratobasidiaceae | 9 | Pathotroph-Saprotroph-Symbiotroph | Endomycorrhizal-Plant Pathogen-Undefined Saprotroph | Microfungus | NULL | Possible | Best known as plant pathogens, but also saprobic. Species are endomycorrhizal with a wide range of plant roots - especially of Orchidaceae but also even bryophytes (Cannon & Kirk 2007) | Cannon PF, Kirk PM. 2007. Fungal Families of the World. CAB International, Cambridge (ISBN: 978-0851998275) |
| Otu06175 | 4 | k__Fungi(100);p__Basidiomycota(100);c__Agaricomycetes(100);o__Boletales(100);f__Boletaceae(100);g__Leccinum(100);s__Leccinum_rotundifoliae(100); | Leccinum | 13 | Symbiotroph | \|Ectomycorrhizal\| | Agaricoid-Boletoid | NULL | Highly Probable | NULL | Rinaldi AC, et al. 2008. Fungal Diversity 33:1-45 ((http://www.fungaldiversity.org/fdp/sfdp/33-1.pdf)); Tedersoo L, et al. 2010. Mycorrhiza 20:217-263 ((https://doi.org/10.1007/s00572-009-0274-x)); Põlme S, et al. 2020. Fungal Diversity 105:1-16 ((https://doi.org/10.1007/s13225-020-00466-2)); http://mycorrhizas.info/ecmf.html ((http://mycorrhizas.info/ecmf.html)) |
| Otu06176 | 4 | k__Fungi(100);p__Ascomycota(100);c__Saccharomycetes(100);o__Saccharomycetales(100);f__Metschnikowiaceae(100);g__Metschnikowia(100);s__Metschnikowia_pulcherrima(100); | Metschnikowia | 13 | Saprotroph | \|Undefined Saprotroph\| | Yeast | NULL | Probable | As nectar-tap_saprotroph (Põlme et al. 2020); Decay sugar-rich substrates (Põlme et al. 2020) | Põlme S, et al. 2020. Fungal Diversity 105:1-16 ((https://doi.org/10.1007/s13225-020-00466-2)) |
| Otu06178 | 4 | k__Fungi(100);p__Ascomycota(100);c__Saccharomycetes(100);o__Saccharomycetales(100);f__Metschnikowiaceae(100);g__Metschnikowia(100);s__Metschnikowia_pulcherrima(100); | Metschnikowia | 13 | Saprotroph | \|Undefined Saprotroph\| | Yeast | NULL | Probable | As nectar-tap_saprotroph (Põlme et al. 2020); Decay sugar-rich substrates (Põlme et al. 2020) | Põlme S, et al. 2020. Fungal Diversity 105:1-16 ((https://doi.org/10.1007/s13225-020-00466-2)) |
| Otu06180 | 4 | k__Fungi(100);p__Ascomycota(100);c__Dothideomycetes(100);o__Dothideales(100);f__Dothideaceae(100);g__Endoconidioma(75);s__Endoconidioma_populi(75); | Endoconidioma | 13 | Saprotroph | Undefined Saprotroph-\|Wood Saprotroph\| | Microfungus | NULL | Probable | Decay: wood (Põlme et al. 2020) | Tedersoo L, et al. 2014. Science 346:e1256688 ((https://doi.org/10.1126/science.1256688)); Põlme S, et al. 2020. Fungal Diversity 105:1-16 ((https://doi.org/10.1007/s13225-020-00466-2)) |
| Otu06181 | 4 | k__Fungi(100);p__Ascomycota(100);c__Saccharomycetes(100);o__Saccharomycetales(100);f__Metschnikowiaceae(100);g__Metschnikowia(100);s__Metschnikowia_pulcherrima(75); | Metschnikowia | 13 | Saprotroph | \|Undefined Saprotroph\| | Yeast | NULL | Probable | As nectar-tap_saprotroph (Põlme et al. 2020); Decay sugar-rich substrates (Põlme et al. 2020) | Põlme S, et al. 2020. Fungal Diversity 105:1-16 ((https://doi.org/10.1007/s13225-020-00466-2)) |
| Otu06182 | 4 | k__Fungi(100);p__Basidiomycota(100);c__Agaricomycetes(100);o__Agaricales(100);f__Pleurotaceae(100);g__Pleurotus(100);s__Pleurotus_dryinus(100); | Pleurotus | 13 | Pathotroph-Saprotroph | Endophyte-Nematophagous-Plant Pathogen-\|Wood Saprotroph\| | Agaricoid | White Rot | Probable | As white_rot (Põlme et al. 2020); As wood_pathogen (Põlme et al. 2020); Decay wood, animal material (Põlme et al. 2020); This genus is known to be nematophagous, capturing and consuming nematodes. | James TY, et al. 2006. Nature 443:818-822 ((https://doi.org/10.1038/nature05110)); Costa IPMW, et al. 2012. Mycotaxon 119:e26 ((http://mycotaxon.com/resources/checklists/costa_v119_checklist.pdf)); Tedersoo L, et al. 2014. Science 346:e1256688 ((https://doi.org/10.1126/science.1256688)); Põlme S, et al. 2020. Fungal Diversity 105:1-16 ((https://doi.org/10.1007/s13225-020-00466-2)); The American Phytopathological Society Common Names of Plant Diseases ((http://www.apsnet.org/publications/commonnames/Pages/default.aspx)) |
| Otu06186 | 4 | k__Fungi(100);p__Basidiomycota(100);c__Tremellomycetes(100);o__Tremellales(100);f__Tremellales_family_Incertae_sedis(100);g__Tremella(100);s__Tremella_moriformis(100); | Tremella | 13 | Pathotroph | \|Fungal Parasite\|-Lichen Parasite | Dimorphic-Facultative Yeast-Microfungus-Tremelloid | NULL | Probable | As mycoparasite (Põlme et al. 2020); Decay fungal material (Põlme et al. 2020) | Tedersoo L, et al. 2014. Science 346:e1256688 ((https://doi.org/10.1126/science.1256688)); Põlme S, et al. 2020. Fungal Diversity 105:1-16 ((https://doi.org/10.1007/s13225-020-00466-2)) |
| Otu06187 | 4 | k__Fungi(100);p__Ascomycota(100);c__Saccharomycetes(100);o__Saccharomycetales(100);f__Metschnikowiaceae(100);g__Metschnikowia(100);g__Metschnikowia_unclassified(75); | Metschnikowia | 13 | Saprotroph | \|Undefined Saprotroph\| | Yeast | NULL | Probable | As nectar-tap_saprotroph (Põlme et al. 2020); Decay sugar-rich substrates (Põlme et al. 2020) | Põlme S, et al. 2020. Fungal Diversity 105:1-16 ((https://doi.org/10.1007/s13225-020-00466-2)) |
| Otu06188 | 4 | k__Fungi(100);p__Basidiomycota(100);c__Microbotryomycetes(100);o__Sporidiobolales(100);f__Sporidiobolales_family_Incertae_sedis(100);g__Rhodotorula(100);g__Rhodotorula_unclassified(100); | Rhodotorula | 13 | Pathotroph-Saprotroph-Symbiotroph | Animal Pathogen-Animal Symbiotroph-Endophyte-Plant Pathogen-\|Undefined Saprotroph\| | Yeast | NULL | Probable | As foliar_endophyte (Põlme et al. 2020); Opportunistic human pathogen | Põlme S, et al. 2020. Fungal Diversity 105:1-16 ((https://doi.org/10.1007/s13225-020-00466-2)); https://www.cdc.gov ((https://www.cdc.gov)) |
| Otu06195 | 4 | k__Fungi(100);p__Ascomycota(100);c__Saccharomycetes(100);o__Saccharomycetales(100);f__Metschnikowiaceae(100);g__Metschnikowia(100);g__Metschnikowia_unclassified(100); | Metschnikowia | 13 | Saprotroph | \|Undefined Saprotroph\| | Yeast | NULL | Probable | As nectar-tap_saprotroph (Põlme et al. 2020); Decay sugar-rich substrates (Põlme et al. 2020) | Põlme S, et al. 2020. Fungal Diversity 105:1-16 ((https://doi.org/10.1007/s13225-020-00466-2)) |
| Otu06196 | 4 | k__Fungi(100);p__Ascomycota(100);c__Saccharomycetes(100);o__Saccharomycetales(100);f__Metschnikowiaceae(100);g__Metschnikowia(100);s__Metschnikowia_pulcherrima(75); | Metschnikowia | 13 | Saprotroph | \|Undefined Saprotroph\| | Yeast | NULL | Probable | As nectar-tap_saprotroph (Põlme et al. 2020); Decay sugar-rich substrates (Põlme et al. 2020) | Põlme S, et al. 2020. Fungal Diversity 105:1-16 ((https://doi.org/10.1007/s13225-020-00466-2)) |
| Otu06197 | 4 | k__Fungi(100);p__Basidiomycota(100);c__Agaricomycetes(100);o__Agaricales(100);f__Strophariaceae(100);g__Agrocybe(100);s__Agrocybe_praecox(100); | Agrocybe | 13 | Saprotroph | Dung Saprotroph-\|Undefined Saprotroph\| | Agaricoid | NULL | Probable | As soil_saprotroph (Põlme et al. 2020); Decay soil (Põlme et al. 2020) | Tedersoo L, et al. 2014. Science 346:e1256688 ((https://doi.org/10.1126/science.1256688)); Põlme S, et al. 2020. Fungal Diversity 105:1-16 ((https://doi.org/10.1007/s13225-020-00466-2)) |
| Otu06199 | 4 | k__Fungi(100);p__Basidiomycota(100);c__Microbotryomycetes(100);o__Sporidiobolales(100);f__Sporidiobolales_family_Incertae_sedis(100);g__Rhodotorula(100);s__Rhodotorula_nothofagi(100); | Rhodotorula | 13 | Pathotroph-Saprotroph-Symbiotroph | Animal Pathogen-Animal Symbiotroph-Endophyte-Plant Pathogen-\|Undefined Saprotroph\| | Yeast | NULL | Probable | As foliar_endophyte (Põlme et al. 2020); Opportunistic human pathogen | Põlme S, et al. 2020. Fungal Diversity 105:1-16 ((https://doi.org/10.1007/s13225-020-00466-2)); https://www.cdc.gov ((https://www.cdc.gov)) |
| Otu06200 | 4 | k__Fungi(100);p__Basidiomycota(100);c__Agaricomycetes(100);o__Auriculariales(100);f__Auriculariales_family_Incertae_sedis(100);g__Basidiodendron(100);s__Basidiodendron_caesiocinereum(100); | Basidiodendron | 13 | Saprotroph | Undefined Saprotroph-\|Wood Saprotroph\| | Corticioid | NULL | Probable | Decay wood (Põlme et al. 2020) | Tedersoo L, et al. 2014. Science 346:e1256688 ((https://doi.org/10.1126/science.1256688)); Põlme S, et al. 2020. Fungal Diversity 105:1-16 ((https://doi.org/10.1007/s13225-020-00466-2)) |
| Otu06201 | 4 | k__Fungi(100);p__Ascomycota(100);c__Saccharomycetes(75);o__Saccharomycetales(75);f__Saccharomycetales_family_Incertae_sedis(75);g__Candida(75);g__Candida_unclassified(75); | Candida | 13 | Pathotroph-Saprotroph-Symbiotroph | Animal Pathogen-Endophyte-Endosymbiont-Epiphyte-\|Undefined Saprotroph\| | Dimorphic Yeast-Yeast | NULL | Probable | As nectar-tap_saprotroph (Põlme et al. 2020); Decay sugar-rich substrates (Põlme et al. 2020); This asexual genus is polyphyletic and contain species that function very differently depend on what they are. Caution should be taken when interpreting guilds for Candida species. | Manolakaki D, et al. 2010. Virulence. 1: 367-375 ((https://doi.org/10.4161/viru.1.5.12796)); Mekbib SB, et al. 2011. Tropical Plant Pathology 36:233-240 ((https://doi.org/10.1590/S1982-56762011000400004)); Põlme S, et al. 2020. Fungal Diversity 105:1-16 ((https://doi.org/10.1007/s13225-020-00466-2)) |
| Otu06203 | 4 | k__Fungi(100);p__Ascomycota(100);c__Saccharomycetes(100);o__Saccharomycetales(100);f__Metschnikowiaceae(100);g__Metschnikowia(100);s__Metschnikowia_pulcherrima(100); | Metschnikowia | 13 | Saprotroph | \|Undefined Saprotroph\| | Yeast | NULL | Probable | As nectar-tap_saprotroph (Põlme et al. 2020); Decay sugar-rich substrates (Põlme et al. 2020) | Põlme S, et al. 2020. Fungal Diversity 105:1-16 ((https://doi.org/10.1007/s13225-020-00466-2)) |
| Otu06206 | 4 | k__Fungi(100);p__Ascomycota(100);c__Saccharomycetes(100);o__Saccharomycetales(100);f__Metschnikowiaceae(100);g__Metschnikowia(100);s__Metschnikowia_pulcherrima(75); | Metschnikowia | 13 | Saprotroph | \|Undefined Saprotroph\| | Yeast | NULL | Probable | As nectar-tap_saprotroph (Põlme et al. 2020); Decay sugar-rich substrates (Põlme et al. 2020) | Põlme S, et al. 2020. Fungal Diversity 105:1-16 ((https://doi.org/10.1007/s13225-020-00466-2)) |
| Otu06208 | 4 | k__Fungi(100);p__Ascomycota(100);c__Saccharomycetes(100);o__Saccharomycetales(100);f__Metschnikowiaceae(100);g__Metschnikowia(100);s__Metschnikowia_pulcherrima(100); | Metschnikowia | 13 | Saprotroph | \|Undefined Saprotroph\| | Yeast | NULL | Probable | As nectar-tap_saprotroph (Põlme et al. 2020); Decay sugar-rich substrates (Põlme et al. 2020) | Põlme S, et al. 2020. Fungal Diversity 105:1-16 ((https://doi.org/10.1007/s13225-020-00466-2)) |
| Otu06209 | 4 | k__Fungi(100);p__Basidiomycota(100);c__Agaricomycetes(100);o__Polyporales(100);f__Phanerochaetaceae(100);g__Ceriporia(100);g__Ceriporia_unclassified(100); | Ceriporia | 13 | Saprotroph | \|Wood Saprotroph\| | Corticioid | White Rot | Highly Probable | As white_rot (Põlme et al. 2020); Decay wood (Põlme et al. 2020) | Tedersoo L, et al. 2014. Science 346:e1256688 ((https://doi.org/10.1126/science.1256688)); Põlme S, et al. 2020. Fungal Diversity 105:1-16 ((https://doi.org/10.1007/s13225-020-00466-2)) |
| Otu06211 | 4 | k__Fungi(100);p__Ascomycota(100);c__Saccharomycetes(100);o__Saccharomycetales(100);f__Metschnikowiaceae(100);g__Metschnikowia(100);s__Metschnikowia_pulcherrima(100); | Metschnikowia | 13 | Saprotroph | \|Undefined Saprotroph\| | Yeast | NULL | Probable | As nectar-tap_saprotroph (Põlme et al. 2020); Decay sugar-rich substrates (Põlme et al. 2020) | Põlme S, et al. 2020. Fungal Diversity 105:1-16 ((https://doi.org/10.1007/s13225-020-00466-2)) |
| Otu06213 | 4 | k__Fungi(100);p__Ascomycota(100);c__Eurotiomycetes(100);o__Chaetothyriales(100);f__Herpotrichiellaceae(100);g__Exophiala(100);s__Exophiala_sp(100); | Exophiala | 13 | Pathotroph-Saprotroph-Symbiotroph | \|Animal Parasite\|-Animal Pathogen-Endophyte-Plant Saprotroph-Undefined Saprotroph | Dimorphic-Facultative Yeast | NULL | Probable | As litter_saprotroph (Põlme et al. 2020); As root_endophyte_dark_septate (Põlme et al. 2020); Decay leaf, fruit, seed, animal material (Põlme et al. 2020) | Tedersoo L, et al. 2014. Science 346:e1256688 ((https://doi.org/10.1126/science.1256688)); Nagy LG, et al. 2014. Nature Communications 5:e4471 ((https://doi.org/10.1038/ncomms5471)); Põlme S, et al. 2020. Fungal Diversity 105:1-16 ((https://doi.org/10.1007/s13225-020-00466-2)) |
| Otu06214 | 4 | k__Fungi(100);p__Ascomycota(100);c__Saccharomycetes(100);o__Saccharomycetales(100);f__Metschnikowiaceae(100);g__Metschnikowia(100);s__Metschnikowia_pulcherrima(75); | Metschnikowia | 13 | Saprotroph | \|Undefined Saprotroph\| | Yeast | NULL | Probable | As nectar-tap_saprotroph (Põlme et al. 2020); Decay sugar-rich substrates (Põlme et al. 2020) | Põlme S, et al. 2020. Fungal Diversity 105:1-16 ((https://doi.org/10.1007/s13225-020-00466-2)) |
| Otu06216 | 4 | k__Fungi(100);p__Ascomycota(100);c__Saccharomycetes(100);o__Saccharomycetales(100);f__Metschnikowiaceae(100);g__Metschnikowia(100);s__Metschnikowia_pulcherrima(75); | Metschnikowia | 13 | Saprotroph | \|Undefined Saprotroph\| | Yeast | NULL | Probable | As nectar-tap_saprotroph (Põlme et al. 2020); Decay sugar-rich substrates (Põlme et al. 2020) | Põlme S, et al. 2020. Fungal Diversity 105:1-16 ((https://doi.org/10.1007/s13225-020-00466-2)) |
| Otu06217 | 4 | k__Fungi(100);p__Ascomycota(100);c__Saccharomycetes(100);o__Saccharomycetales(100);f__Metschnikowiaceae(100);g__Metschnikowia(100);s__Metschnikowia_pulcherrima(75); | Metschnikowia | 13 | Saprotroph | \|Undefined Saprotroph\| | Yeast | NULL | Probable | As nectar-tap_saprotroph (Põlme et al. 2020); Decay sugar-rich substrates (Põlme et al. 2020) | Põlme S, et al. 2020. Fungal Diversity 105:1-16 ((https://doi.org/10.1007/s13225-020-00466-2)) |
| Otu06218 | 4 | k__Fungi(100);p__Ascomycota(100);c__Saccharomycetes(100);o__Saccharomycetales(100);f__Metschnikowiaceae(100);g__Metschnikowia(100);s__Metschnikowia_pulcherrima(100); | Metschnikowia | 13 | Saprotroph | \|Undefined Saprotroph\| | Yeast | NULL | Probable | As nectar-tap_saprotroph (Põlme et al. 2020); Decay sugar-rich substrates (Põlme et al. 2020) | Põlme S, et al. 2020. Fungal Diversity 105:1-16 ((https://doi.org/10.1007/s13225-020-00466-2)) |
| Otu06219 | 4 | k__Fungi(100);p__Ascomycota(100);c__Saccharomycetes(100);o__Saccharomycetales(100);f__Saccharomycetales_family_Incertae_sedis(100);g__Candida(100);g__Candida_unclassified(100); | Candida | 13 | Pathotroph-Saprotroph-Symbiotroph | Animal Pathogen-Endophyte-Endosymbiont-Epiphyte-\|Undefined Saprotroph\| | Dimorphic Yeast-Yeast | NULL | Probable | As nectar-tap_saprotroph (Põlme et al. 2020); Decay sugar-rich substrates (Põlme et al. 2020); This asexual genus is polyphyletic and contain species that function very differently depend on what they are. Caution should be taken when interpreting guilds for Candida species. | Manolakaki D, et al. 2010. Virulence. 1: 367-375 ((https://doi.org/10.4161/viru.1.5.12796)); Mekbib SB, et al. 2011. Tropical Plant Pathology 36:233-240 ((https://doi.org/10.1590/S1982-56762011000400004)); Põlme S, et al. 2020. Fungal Diversity 105:1-16 ((https://doi.org/10.1007/s13225-020-00466-2)) |
| Otu06220 | 4 | k__Fungi(100);p__Ascomycota(100);c__Saccharomycetes(100);o__Saccharomycetales(100);f__Metschnikowiaceae(100);g__Metschnikowia(100);s__Metschnikowia_pulcherrima(100); | Metschnikowia | 13 | Saprotroph | \|Undefined Saprotroph\| | Yeast | NULL | Probable | As nectar-tap_saprotroph (Põlme et al. 2020); Decay sugar-rich substrates (Põlme et al. 2020) | Põlme S, et al. 2020. Fungal Diversity 105:1-16 ((https://doi.org/10.1007/s13225-020-00466-2)) |
| Otu06221 | 4 | k__Fungi(100);p__Ascomycota(100);c__Saccharomycetes(100);o__Saccharomycetales(100);f__Metschnikowiaceae(100);g__Metschnikowia(100);g__Metschnikowia_unclassified(100); | Metschnikowia | 13 | Saprotroph | \|Undefined Saprotroph\| | Yeast | NULL | Probable | As nectar-tap_saprotroph (Põlme et al. 2020); Decay sugar-rich substrates (Põlme et al. 2020) | Põlme S, et al. 2020. Fungal Diversity 105:1-16 ((https://doi.org/10.1007/s13225-020-00466-2)) |
| Otu06224 | 4 | k__Fungi(100);p__Ascomycota(100);c__Saccharomycetes(100);o__Saccharomycetales(100);f__Saccharomycetales_family_Incertae_sedis(100);g__Candida(75);g__Candida_unclassified(75); | Candida | 13 | Pathotroph-Saprotroph-Symbiotroph | Animal Pathogen-Endophyte-Endosymbiont-Epiphyte-\|Undefined Saprotroph\| | Dimorphic Yeast-Yeast | NULL | Probable | As nectar-tap_saprotroph (Põlme et al. 2020); Decay sugar-rich substrates (Põlme et al. 2020); This asexual genus is polyphyletic and contain species that function very differently depend on what they are. Caution should be taken when interpreting guilds for Candida species. | Manolakaki D, et al. 2010. Virulence. 1: 367-375 ((https://doi.org/10.4161/viru.1.5.12796)); Mekbib SB, et al. 2011. Tropical Plant Pathology 36:233-240 ((https://doi.org/10.1590/S1982-56762011000400004)); Põlme S, et al. 2020. Fungal Diversity 105:1-16 ((https://doi.org/10.1007/s13225-020-00466-2)) |
| Otu06227 | 4 | k__Fungi(100);p__Basidiomycota(100);c__Agaricomycetes(100);o__Boletales(100);f__Coniophoraceae(100);g__Coniophora(100);s__Coniophora_puteana(100); | Coniophora | 13 | Saprotroph | \|Wood Saprotroph\| | Corticioid | Brown Rot | Probable | As brown-rot (Põlme et al. 2020); Decay wood (Põlme et al. 2020) | Tedersoo L, et al. 2014. Science 346:e1256688 ((https://doi.org/10.1126/science.1256688)); Põlme S, et al. 2020. Fungal Diversity 105:1-16 ((https://doi.org/10.1007/s13225-020-00466-2)) |
| Otu06228 | 4 | k__Fungi(100);p__Ascomycota(100);c__Saccharomycetes(100);o__Saccharomycetales(100);f__Metschnikowiaceae(100);g__Metschnikowia(100);s__Metschnikowia_pulcherrima(75); | Metschnikowia | 13 | Saprotroph | \|Undefined Saprotroph\| | Yeast | NULL | Probable | As nectar-tap_saprotroph (Põlme et al. 2020); Decay sugar-rich substrates (Põlme et al. 2020) | Põlme S, et al. 2020. Fungal Diversity 105:1-16 ((https://doi.org/10.1007/s13225-020-00466-2)) |
| Otu06230 | 4 | k__Fungi(100);p__Ascomycota(100);c__Saccharomycetes(100);o__Saccharomycetales(100);f__Metschnikowiaceae(100);g__Metschnikowia(100);g__Metschnikowia_unclassified(100); | Metschnikowia | 13 | Saprotroph | \|Undefined Saprotroph\| | Yeast | NULL | Probable | As nectar-tap_saprotroph (Põlme et al. 2020); Decay sugar-rich substrates (Põlme et al. 2020) | Põlme S, et al. 2020. Fungal Diversity 105:1-16 ((https://doi.org/10.1007/s13225-020-00466-2)) |
| Otu06234 | 4 | k__Fungi(100);p__Basidiomycota(100);c__Agaricostilbomycetes(100);o__Agaricostilbales(100);f__Agaricostilbaceae(100);g__Bensingtonia(100);g__Bensingtonia_unclassified(75); | Bensingtonia | 13 | Saprotroph | \|Undefined Saprotroph\| | Dimorphic-Facultative Yeast | NULL | Highly Probable | NULL | Kurtzman CP, et al. (eds.) 2011. The Yeasts, a Taxonomic Study. Fifth Edition. Vols 1-3. Elsevier, San Diego (ISBN: 9780444521491); Põlme S, et al. 2020. Fungal Diversity 105:1-16 ((https://doi.org/10.1007/s13225-020-00466-2)) |
| Otu06235 | 4 | k__Fungi(100);p__Ascomycota(100);c__Saccharomycetes(100);o__Saccharomycetales(100);f__Metschnikowiaceae(100);g__Metschnikowia(100);s__Metschnikowia_pulcherrima(100); | Metschnikowia | 13 | Saprotroph | \|Undefined Saprotroph\| | Yeast | NULL | Probable | As nectar-tap_saprotroph (Põlme et al. 2020); Decay sugar-rich substrates (Põlme et al. 2020) | Põlme S, et al. 2020. Fungal Diversity 105:1-16 ((https://doi.org/10.1007/s13225-020-00466-2)) |
| Otu06237 | 4 | k__Fungi(100);p__Basidiomycota(100);c__Agaricomycetes(100);o__Agaricales(100);f__Inocybaceae(100);g__Inocybe(100);s__Inocybe_assimilata(100); | Inocybe | 13 | Symbiotroph | \|Ectomycorrhizal\| | Agaricoid | NULL | Highly Probable | Sensu stricto (Matheny 2009). | Rinaldi AC, et al. 2008. Fungal Diversity 33:1-45 ((http://www.fungaldiversity.org/fdp/sfdp/33-1.pdf)); Matheny PB. 2009. McIlvainea 18:11-21 ((https://namyco.org/docs/3_Matheny.pdf)); Tedersoo L, et al. 2010. Mycorrhiza 20:217-263 ((https://doi.org/10.1007/s00572-009-0274-x)); Põlme S, et al. 2020. Fungal Diversity 105:1-16 ((https://doi.org/10.1007/s13225-020-00466-2)); http://mycorrhizas.info/ecmf.html ((http://mycorrhizas.info/ecmf.html)) |
| Otu06238 | 4 | k__Fungi(100);p__Ascomycota(100);c__Saccharomycetes(100);o__Saccharomycetales(100);f__Metschnikowiaceae(100);g__Metschnikowia(100);s__Metschnikowia_pulcherrima(100); | Metschnikowia | 13 | Saprotroph | \|Undefined Saprotroph\| | Yeast | NULL | Probable | As nectar-tap_saprotroph (Põlme et al. 2020); Decay sugar-rich substrates (Põlme et al. 2020) | Põlme S, et al. 2020. Fungal Diversity 105:1-16 ((https://doi.org/10.1007/s13225-020-00466-2)) |
| Otu06239 | 4 | k__Fungi(100);p__Ascomycota(100);c__Saccharomycetes(100);o__Saccharomycetales(100);f__Metschnikowiaceae(100);g__Metschnikowia(100);g__Metschnikowia_unclassified(75); | Metschnikowia | 13 | Saprotroph | \|Undefined Saprotroph\| | Yeast | NULL | Probable | As nectar-tap_saprotroph (Põlme et al. 2020); Decay sugar-rich substrates (Põlme et al. 2020) | Põlme S, et al. 2020. Fungal Diversity 105:1-16 ((https://doi.org/10.1007/s13225-020-00466-2)) |
| Otu06240 | 4 | k__Fungi(100);p__Ascomycota(100);c__Saccharomycetes(100);o__Saccharomycetales(100);f__Metschnikowiaceae(100);g__Metschnikowia(100);s__Metschnikowia_pulcherrima(100); | Metschnikowia | 13 | Saprotroph | \|Undefined Saprotroph\| | Yeast | NULL | Probable | As nectar-tap_saprotroph (Põlme et al. 2020); Decay sugar-rich substrates (Põlme et al. 2020) | Põlme S, et al. 2020. Fungal Diversity 105:1-16 ((https://doi.org/10.1007/s13225-020-00466-2)) |
| Otu06241 | 4 | k__Fungi(100);p__Ascomycota(100);c__Saccharomycetes(100);o__Saccharomycetales(100);f__Metschnikowiaceae(100);g__Metschnikowia(100);s__Metschnikowia_pulcherrima(100); | Metschnikowia | 13 | Saprotroph | \|Undefined Saprotroph\| | Yeast | NULL | Probable | As nectar-tap_saprotroph (Põlme et al. 2020); Decay sugar-rich substrates (Põlme et al. 2020) | Põlme S, et al. 2020. Fungal Diversity 105:1-16 ((https://doi.org/10.1007/s13225-020-00466-2)) |
| Otu06242 | 4 | k__Fungi(100);p__Ascomycota(100);c__Saccharomycetes(100);o__Saccharomycetales(100);f__Saccharomycetales_family_Incertae_sedis(100);g__Candida(100);g__Candida_unclassified(100); | Candida | 13 | Pathotroph-Saprotroph-Symbiotroph | Animal Pathogen-Endophyte-Endosymbiont-Epiphyte-\|Undefined Saprotroph\| | Dimorphic Yeast-Yeast | NULL | Probable | As nectar-tap_saprotroph (Põlme et al. 2020); Decay sugar-rich substrates (Põlme et al. 2020); This asexual genus is polyphyletic and contain species that function very differently depend on what they are. Caution should be taken when interpreting guilds for Candida species. | Manolakaki D, et al. 2010. Virulence. 1: 367-375 ((https://doi.org/10.4161/viru.1.5.12796)); Mekbib SB, et al. 2011. Tropical Plant Pathology 36:233-240 ((https://doi.org/10.1590/S1982-56762011000400004)); Põlme S, et al. 2020. Fungal Diversity 105:1-16 ((https://doi.org/10.1007/s13225-020-00466-2)) |
| Otu06246 | 4 | k__Fungi(100);p__Zygomycota(100);c__Zygomycota_class_Incertae_sedis(100);o__Mortierellales(100);f__Mortierellaceae(100);g__Mortierella(100);s__Mortierella_globulifera(100); | Mortierella | 13 | Saprotroph-Symbiotroph | Endophyte-Plant Saprotroph-\|Undefined Saprotroph\| | Microfungus | NULL | Probable | As chitinolytic (Põlme et al. 2020); As soil_saprotroph (Põlme et al. 2020); Decay roots, soil (Põlme et al. 2020); Saprobic in soil and thus frequently isolated from soil and material contaminated with soil such as hay or rotten silage and spoil tips (Cannon & Kirk 2007) | Cannon PF, Kirk PM. 2007. Fungal Families of the World. CAB International, Cambridge (ISBN: 978-0851998275); Tedersoo L, et al. 2014. Science 346:e1256688 ((https://doi.org/10.1126/science.1256688)); Purahong W, et al. 2016. Molecular Ecology 25:4059-4074 ((https://doi.org/10.1111/mec.13739)); Põlme S, et al. 2020. Fungal Diversity 105:1-16 ((https://doi.org/10.1007/s13225-020-00466-2)) |
| Otu06247 | 4 | k__Fungi(100);p__Ascomycota(100);c__Saccharomycetes(100);o__Saccharomycetales(100);f__Saccharomycetales_family_Incertae_sedis(100);g__Candida(100);g__Candida_unclassified(75); | Candida | 13 | Pathotroph-Saprotroph-Symbiotroph | Animal Pathogen-Endophyte-Endosymbiont-Epiphyte-\|Undefined Saprotroph\| | Dimorphic Yeast-Yeast | NULL | Probable | As nectar-tap_saprotroph (Põlme et al. 2020); Decay sugar-rich substrates (Põlme et al. 2020); This asexual genus is polyphyletic and contain species that function very differently depend on what they are. Caution should be taken when interpreting guilds for Candida species. | Manolakaki D, et al. 2010. Virulence. 1: 367-375 ((https://doi.org/10.4161/viru.1.5.12796)); Mekbib SB, et al. 2011. Tropical Plant Pathology 36:233-240 ((https://doi.org/10.1590/S1982-56762011000400004)); Põlme S, et al. 2020. Fungal Diversity 105:1-16 ((https://doi.org/10.1007/s13225-020-00466-2)) |
| Otu06252 | 4 | k__Fungi(100);p__Ascomycota(100);c__Sordariomycetes(100);o__Sordariomycetes_order_Incertae_sedis(100);f__Sordariomycetes_family_Incertae_sedis(100);g__Myrmecridium(100);s__Myrmecridium_schulzeri(100); | Myrmecridium | 13 | Saprotroph | \|Undefined Saprotroph\| | Microfungus | NULL | Probable | As soil_saprotroph (Põlme et al. 2020); Decay soil (Põlme et al. 2020) | Tedersoo L, et al. 2014. Science 346:e1256688 ((https://doi.org/10.1126/science.1256688)); Põlme S, et al. 2020. Fungal Diversity 105:1-16 ((https://doi.org/10.1007/s13225-020-00466-2)) |
| Otu06254 | 4 | k__Fungi(100);p__Ascomycota(100);c__Saccharomycetes(100);o__Saccharomycetales(100);f__Metschnikowiaceae(100);g__Metschnikowia(100);s__Metschnikowia_pulcherrima(75); | Metschnikowia | 13 | Saprotroph | \|Undefined Saprotroph\| | Yeast | NULL | Probable | As nectar-tap_saprotroph (Põlme et al. 2020); Decay sugar-rich substrates (Põlme et al. 2020) | Põlme S, et al. 2020. Fungal Diversity 105:1-16 ((https://doi.org/10.1007/s13225-020-00466-2)) |
| Otu06256 | 4 | k__Fungi(100);p__Ascomycota(100);c__Saccharomycetes(100);o__Saccharomycetales(100);f__Metschnikowiaceae(100);g__Metschnikowia(100);s__Metschnikowia_pulcherrima(100); | Metschnikowia | 13 | Saprotroph | \|Undefined Saprotroph\| | Yeast | NULL | Probable | As nectar-tap_saprotroph (Põlme et al. 2020); Decay sugar-rich substrates (Põlme et al. 2020) | Põlme S, et al. 2020. Fungal Diversity 105:1-16 ((https://doi.org/10.1007/s13225-020-00466-2)) |
| Otu06257 | 4 | k__Fungi(100);p__Ascomycota(100);c__Saccharomycetes(100);o__Saccharomycetales(100);f__Metschnikowiaceae(100);g__Metschnikowia(100);s__Metschnikowia_pulcherrima(100); | Metschnikowia | 13 | Saprotroph | \|Undefined Saprotroph\| | Yeast | NULL | Probable | As nectar-tap_saprotroph (Põlme et al. 2020); Decay sugar-rich substrates (Põlme et al. 2020) | Põlme S, et al. 2020. Fungal Diversity 105:1-16 ((https://doi.org/10.1007/s13225-020-00466-2)) |
| Otu06261 | 4 | k__Fungi(100);p__Ascomycota(100);c__Saccharomycetes(100);o__Saccharomycetales(100);f__Metschnikowiaceae(100);g__Metschnikowia(100);s__Metschnikowia_pulcherrima(100); | Metschnikowia | 13 | Saprotroph | \|Undefined Saprotroph\| | Yeast | NULL | Probable | As nectar-tap_saprotroph (Põlme et al. 2020); Decay sugar-rich substrates (Põlme et al. 2020) | Põlme S, et al. 2020. Fungal Diversity 105:1-16 ((https://doi.org/10.1007/s13225-020-00466-2)) |
| Otu06262 | 4 | k__Fungi(100);p__Ascomycota(100);c__Saccharomycetes(100);o__Saccharomycetales(100);f__Metschnikowiaceae(100);g__Metschnikowia(100);s__Metschnikowia_pulcherrima(100); | Metschnikowia | 13 | Saprotroph | \|Undefined Saprotroph\| | Yeast | NULL | Probable | As nectar-tap_saprotroph (Põlme et al. 2020); Decay sugar-rich substrates (Põlme et al. 2020) | Põlme S, et al. 2020. Fungal Diversity 105:1-16 ((https://doi.org/10.1007/s13225-020-00466-2)) |
| Otu06263 | 4 | k__Fungi(100);p__Ascomycota(100);c__Saccharomycetes(100);o__Saccharomycetales(100);f__Metschnikowiaceae(100);g__Metschnikowia(100);s__Metschnikowia_pulcherrima(100); | Metschnikowia | 13 | Saprotroph | \|Undefined Saprotroph\| | Yeast | NULL | Probable | As nectar-tap_saprotroph (Põlme et al. 2020); Decay sugar-rich substrates (Põlme et al. 2020) | Põlme S, et al. 2020. Fungal Diversity 105:1-16 ((https://doi.org/10.1007/s13225-020-00466-2)) |
| Otu06264 | 4 | k__Fungi(100);p__Ascomycota(100);c__Saccharomycetes(100);o__Saccharomycetales(100);f__Metschnikowiaceae(100);g__Metschnikowia(100);s__Metschnikowia_pulcherrima(100); | Metschnikowia | 13 | Saprotroph | \|Undefined Saprotroph\| | Yeast | NULL | Probable | As nectar-tap_saprotroph (Põlme et al. 2020); Decay sugar-rich substrates (Põlme et al. 2020) | Põlme S, et al. 2020. Fungal Diversity 105:1-16 ((https://doi.org/10.1007/s13225-020-00466-2)) |
| Otu06265 | 4 | k__Fungi(100);p__Ascomycota(100);c__Saccharomycetes(100);o__Saccharomycetales(100);f__Metschnikowiaceae(100);g__Metschnikowia(100);s__Metschnikowia_pulcherrima(75); | Metschnikowia | 13 | Saprotroph | \|Undefined Saprotroph\| | Yeast | NULL | Probable | As nectar-tap_saprotroph (Põlme et al. 2020); Decay sugar-rich substrates (Põlme et al. 2020) | Põlme S, et al. 2020. Fungal Diversity 105:1-16 ((https://doi.org/10.1007/s13225-020-00466-2)) |
| Otu06269 | 4 | k__Fungi(100);p__Ascomycota(100);c__Dothideomycetes(100);o__Pleosporales(100);f__Pleosporales_family_Incertae_sedis(75);g__Pyrenochaeta(75);s__Pyrenochaeta_sp_MUT_4382(75); | Pyrenochaeta | 13 | Pathotroph-Saprotroph | Plant Pathogen-Undefined Saprotroph-\|Wood Saprotroph\| | Microfungus | Soft Rot | Highly Probable | Decay wood (Põlme et al. 2020) | Seehann G, et al. 1975. List of Fungi in Soft-Rot Tissue Tests. Institute fur Holzbiologie und Holzschutz, Bundesforschungsanstalt fur Forst - und Holzwirtshaft, Hamburg - Lohbrugge, Germany; Costa IPMW, et al. 2012. Mycotaxon 119:e26 ((http://mycotaxon.com/resources/checklists/costa_v119_checklist.pdf)); Tedersoo L, et al. 2014. Science 346:e1256688 ((https://doi.org/10.1126/science.1256688)); Põlme S, et al. 2020. Fungal Diversity 105:1-16 ((https://doi.org/10.1007/s13225-020-00466-2)) |
| Otu06272 | 4 | k__Fungi(100);p__Ascomycota(100);c__Saccharomycetes(100);o__Saccharomycetales(100);f__Metschnikowiaceae(100);g__Metschnikowia(100);g__Metschnikowia_unclassified(100); | Metschnikowia | 13 | Saprotroph | \|Undefined Saprotroph\| | Yeast | NULL | Probable | As nectar-tap_saprotroph (Põlme et al. 2020); Decay sugar-rich substrates (Põlme et al. 2020) | Põlme S, et al. 2020. Fungal Diversity 105:1-16 ((https://doi.org/10.1007/s13225-020-00466-2)) |
| Otu06273 | 4 | k__Fungi(100);p__Ascomycota(100);c__Saccharomycetes(100);o__Saccharomycetales(100);f__Metschnikowiaceae(100);g__Metschnikowia(100);s__Metschnikowia_pulcherrima(100); | Metschnikowia | 13 | Saprotroph | \|Undefined Saprotroph\| | Yeast | NULL | Probable | As nectar-tap_saprotroph (Põlme et al. 2020); Decay sugar-rich substrates (Põlme et al. 2020) | Põlme S, et al. 2020. Fungal Diversity 105:1-16 ((https://doi.org/10.1007/s13225-020-00466-2)) |
| Otu06274 | 4 | k__Fungi(100);p__Ascomycota(100);c__Saccharomycetes(100);o__Saccharomycetales(100);f__Metschnikowiaceae(100);g__Metschnikowia(100);s__Metschnikowia_pulcherrima(100); | Metschnikowia | 13 | Saprotroph | \|Undefined Saprotroph\| | Yeast | NULL | Probable | As nectar-tap_saprotroph (Põlme et al. 2020); Decay sugar-rich substrates (Põlme et al. 2020) | Põlme S, et al. 2020. Fungal Diversity 105:1-16 ((https://doi.org/10.1007/s13225-020-00466-2)) |
| Otu06275 | 4 | k__Fungi(100);p__Ascomycota(100);c__Leotiomycetes(100);o__Helotiales(100);f__Hyaloscyphaceae(100);g__Hyaloscypha(100);s__Hyaloscypha_aureliella(100); | Hyaloscypha | 13 | Pathotroph-Saprotroph-Symbiotroph | Ectomycorrhizal-Ericoid Mycorrhizal-Plant Pathogen-\|Plant Saprotroph\|-Undefined Saprotroph-Wood Saprotroph | Microfungus | NULL | Probable | As litter_saprotroph (Põlme et al. 2020); Decay wood, leaf, fruit, seed (Põlme et al. 2020); Host - mosses (Põlme et al. 2020) | Tedersoo L, et al. 2014. Science 346:e1256688 ((https://doi.org/10.1126/science.1256688)); Põlme S, et al. 2020. Fungal Diversity 105:1-16 ((https://doi.org/10.1007/s13225-020-00466-2)) |
| Otu06278 | 4 | k__Fungi(100);p__Basidiomycota(100);c__Agaricomycetes(100);o__Boletales(100);f__Hygrophoropsidaceae(100);g__Hygrophoropsis(100);s__Hygrophoropsis_rufa(100); | Hygrophoropsis | 13 | Saprotroph | \|Plant Saprotroph\|-Wood Saprotroph | Agaricoid | Brown Rot | Probable | As litter_saprotroph (Põlme et al. 2020); Decay wood, leaf, fruit, seed (Põlme et al. 2020) | Tedersoo L, et al. 2014. Science 346:e1256688 ((https://doi.org/10.1126/science.1256688)); Põlme S, et al. 2020. Fungal Diversity 105:1-16 ((https://doi.org/10.1007/s13225-020-00466-2)) |
| Otu06279 | 4 | k__Fungi(100);p__Ascomycota(100);c__Saccharomycetes(100);o__Saccharomycetales(100);f__Metschnikowiaceae(100);g__Metschnikowia(100);s__Metschnikowia_pulcherrima(100); | Metschnikowia | 13 | Saprotroph | \|Undefined Saprotroph\| | Yeast | NULL | Probable | As nectar-tap_saprotroph (Põlme et al. 2020); Decay sugar-rich substrates (Põlme et al. 2020) | Põlme S, et al. 2020. Fungal Diversity 105:1-16 ((https://doi.org/10.1007/s13225-020-00466-2)) |
| Otu06281 | 4 | k__Fungi(100);p__Ascomycota(100);c__Saccharomycetes(100);o__Saccharomycetales(100);f__Metschnikowiaceae(100);g__Metschnikowia(100);s__Metschnikowia_pulcherrima(100); | Metschnikowia | 13 | Saprotroph | \|Undefined Saprotroph\| | Yeast | NULL | Probable | As nectar-tap_saprotroph (Põlme et al. 2020); Decay sugar-rich substrates (Põlme et al. 2020) | Põlme S, et al. 2020. Fungal Diversity 105:1-16 ((https://doi.org/10.1007/s13225-020-00466-2)) |
| Otu06282 | 4 | k__Fungi(100);p__Basidiomycota(100);c__Microbotryomycetes(100);o__Sporidiobolales(100);f__Sporidiobolales_family_Incertae_sedis(100);g__Sporobolomyces(100);s__Sporobolomyces_lactosus(100); | Sporobolomyces | 13 | Pathotroph-Saprotroph | \|Fungal Parasite\|-Plant Saprotroph | Yeast | NULL | Probable | As litter_saprotroph (Põlme et al. 2020); As mycoparasite (Põlme et al. 2020); Decay fungal material (Põlme et al. 2020) | Tedersoo L, et al. 2014. Science 346:e1256688 ((https://doi.org/10.1126/science.1256688)); Purahong W, et al. 2016. Molecular Ecology 25:4059-4074 ((https://doi.org/10.1111/mec.13739)); Põlme S, et al. 2020. Fungal Diversity 105:1-16 ((https://doi.org/10.1007/s13225-020-00466-2)) |
| Otu06283 | 4 | k__Fungi(100);p__Ascomycota(100);c__Saccharomycetes(100);o__Saccharomycetales(100);f__Metschnikowiaceae(100);g__Metschnikowia(100);s__Metschnikowia_pulcherrima(100); | Metschnikowia | 13 | Saprotroph | \|Undefined Saprotroph\| | Yeast | NULL | Probable | As nectar-tap_saprotroph (Põlme et al. 2020); Decay sugar-rich substrates (Põlme et al. 2020) | Põlme S, et al. 2020. Fungal Diversity 105:1-16 ((https://doi.org/10.1007/s13225-020-00466-2)) |
| Otu06284 | 4 | k__Fungi(100);p__Ascomycota(100);c__Orbiliomycetes(100);o__Orbiliales(100);f__Orbiliaceae(100);g__unclassified_Orbiliaceae(100);s__Orbiliaceae_sp(100); | Orbiliaceae | 9 | Saprotroph | Wood Saprotroph | Microfungus | NULL | Probable | Saprobic, found especially on wood in both wet and dry habitats. Many species are nematophagous, forming adhesive knobs, nets or ring-like traps in the presence of a food source; perhaps a strategy for life in low-nitrogen environments. There have been reports of associations with algae for a few species, but an established lichen association appears not to exist (Cannon & Kirk 2007) | Cannon PF, Kirk PM. 2007. Fungal Families of the World. CAB International, Cambridge (ISBN: 978-0851998275) |
| Otu06286 | 4 | k__Fungi(100);p__Ascomycota(100);c__Dothideomycetes(100);o__Capnodiales(100);f__Schizothyriaceae(100);g__Zygophiala(100);s__Zygophiala_wisconsinensis(100); | Zygophiala | 13 | Pathotroph-Saprotroph | \|Plant Pathogen\|-Plant Saprotroph | Microfungus | NULL | Probable | As Biotroph (Tedersoo et al. 2014); As litter_saprotroph (Põlme et al. 2020); Decay leaf, fruit, seed (Põlme et al. 2020); Z. jamaicensis (Syn. Schizothyrium pomi - teleomorph) causes flyspeck and greasy blotch disease on apples (Baker, et al. 1977) | Baker, et al. 1977. Phytopathology 67:580-588 ((https://www.apsnet.org/publications/phytopathology/backissues/Documents/1977Articles/Phyto67n05_580.PDF)); Tedersoo L, et al. 2014. Science 346:e1256688 ((https://doi.org/10.1126/science.1256688)); Põlme S, et al. 2020. Fungal Diversity 105:1-16 ((https://doi.org/10.1007/s13225-020-00466-2)); The American Phytopathological Society Common Names of Plant Diseases ((https://www.apsnet.org/edcenter/resources/commonnames/Pages/default.aspx)) |
| Otu06288 | 4 | k__Fungi(100);p__Ascomycota(100);c__Saccharomycetes(100);o__Saccharomycetales(100);f__Metschnikowiaceae(100);g__Metschnikowia(100);s__Metschnikowia_pulcherrima(100); | Metschnikowia | 13 | Saprotroph | \|Undefined Saprotroph\| | Yeast | NULL | Probable | As nectar-tap_saprotroph (Põlme et al. 2020); Decay sugar-rich substrates (Põlme et al. 2020) | Põlme S, et al. 2020. Fungal Diversity 105:1-16 ((https://doi.org/10.1007/s13225-020-00466-2)) |
| Otu06291 | 4 | k__Fungi(100);p__Ascomycota(100);c__Saccharomycetes(100);o__Saccharomycetales(100);f__Metschnikowiaceae(100);g__Metschnikowia(100);g__Metschnikowia_unclassified(100); | Metschnikowia | 13 | Saprotroph | \|Undefined Saprotroph\| | Yeast | NULL | Probable | As nectar-tap_saprotroph (Põlme et al. 2020); Decay sugar-rich substrates (Põlme et al. 2020) | Põlme S, et al. 2020. Fungal Diversity 105:1-16 ((https://doi.org/10.1007/s13225-020-00466-2)) |
| Otu06292 | 4 | k__Fungi(100);p__Ascomycota(100);c__Saccharomycetes(100);o__Saccharomycetales(100);f__Metschnikowiaceae(100);g__Metschnikowia(100);g__Metschnikowia_unclassified(100); | Metschnikowia | 13 | Saprotroph | \|Undefined Saprotroph\| | Yeast | NULL | Probable | As nectar-tap_saprotroph (Põlme et al. 2020); Decay sugar-rich substrates (Põlme et al. 2020) | Põlme S, et al. 2020. Fungal Diversity 105:1-16 ((https://doi.org/10.1007/s13225-020-00466-2)) |
| Otu06294 | 4 | k__Fungi(100);p__Ascomycota(100);c__Saccharomycetes(100);o__Saccharomycetales(100);f__Metschnikowiaceae(100);g__Metschnikowia(100);s__Metschnikowia_pulcherrima(100); | Metschnikowia | 13 | Saprotroph | \|Undefined Saprotroph\| | Yeast | NULL | Probable | As nectar-tap_saprotroph (Põlme et al. 2020); Decay sugar-rich substrates (Põlme et al. 2020) | Põlme S, et al. 2020. Fungal Diversity 105:1-16 ((https://doi.org/10.1007/s13225-020-00466-2)) |
| Otu06296 | 4 | k__Fungi(100);p__Basidiomycota(100);c__Agaricomycetes(100);o__Agaricales(100);f__Psathyrellaceae(100);g__Coprinopsis(100);s__Coprinopsis_phlyctidospora(100); | Coprinopsis | 13 | Saprotroph | \|Undefined Saprotroph\| | Agaricoid | NULL | Probable | As soil_saprotroph (Põlme et al. 2020); Decay soil (Põlme et al. 2020) | Tedersoo L, et al. 2014. Science 346:e1256688 ((https://doi.org/10.1126/science.1256688)); Põlme S, et al. 2020. Fungal Diversity 105:1-16 ((https://doi.org/10.1007/s13225-020-00466-2)) |
| Otu06298 | 4 | k__Fungi(100);p__Ascomycota(100);c__Saccharomycetes(100);o__Saccharomycetales(100);f__Metschnikowiaceae(100);g__Metschnikowia(100);s__Metschnikowia_pulcherrima(100); | Metschnikowia | 13 | Saprotroph | \|Undefined Saprotroph\| | Yeast | NULL | Probable | As nectar-tap_saprotroph (Põlme et al. 2020); Decay sugar-rich substrates (Põlme et al. 2020) | Põlme S, et al. 2020. Fungal Diversity 105:1-16 ((https://doi.org/10.1007/s13225-020-00466-2)) |
| Otu06299 | 4 | k__Fungi(100);p__Ascomycota(100);c__Saccharomycetes(100);o__Saccharomycetales(100);f__Metschnikowiaceae(100);g__Metschnikowia(100);s__Metschnikowia_pulcherrima(100); | Metschnikowia | 13 | Saprotroph | \|Undefined Saprotroph\| | Yeast | NULL | Probable | As nectar-tap_saprotroph (Põlme et al. 2020); Decay sugar-rich substrates (Põlme et al. 2020) | Põlme S, et al. 2020. Fungal Diversity 105:1-16 ((https://doi.org/10.1007/s13225-020-00466-2)) |
| Otu06300 | 4 | k__Fungi(100);p__Basidiomycota(100);c__Microbotryomycetes(100);o__Sporidiobolales(100);f__Sporidiobolales_family_Incertae_sedis(100);g__Sporobolomyces(100);s__Sporobolomyces_inositophilus(100); | Sporobolomyces | 13 | Pathotroph-Saprotroph | \|Fungal Parasite\|-Plant Saprotroph | Yeast | NULL | Probable | As litter_saprotroph (Põlme et al. 2020); As mycoparasite (Põlme et al. 2020); Decay fungal material (Põlme et al. 2020) | Tedersoo L, et al. 2014. Science 346:e1256688 ((https://doi.org/10.1126/science.1256688)); Purahong W, et al. 2016. Molecular Ecology 25:4059-4074 ((https://doi.org/10.1111/mec.13739)); Põlme S, et al. 2020. Fungal Diversity 105:1-16 ((https://doi.org/10.1007/s13225-020-00466-2)) |
| Otu06301 | 4 | k__Fungi(100);p__Ascomycota(100);c__Saccharomycetes(100);o__Saccharomycetales(100);f__Metschnikowiaceae(100);g__Metschnikowia(100);s__Metschnikowia_pulcherrima(100); | Metschnikowia | 13 | Saprotroph | \|Undefined Saprotroph\| | Yeast | NULL | Probable | As nectar-tap_saprotroph (Põlme et al. 2020); Decay sugar-rich substrates (Põlme et al. 2020) | Põlme S, et al. 2020. Fungal Diversity 105:1-16 ((https://doi.org/10.1007/s13225-020-00466-2)) |
| Otu06304 | 4 | k__Fungi(100);p__Basidiomycota(100);c__Agaricomycetes(100);o__Hymenochaetales(100);f__Hymenochaetaceae(100);g__Hymenochaete(100);s__Hymenochaete_tenuis(100); | Hymenochaete | 13 | Pathotroph-Saprotroph | Plant Pathogen-Undefined Saprotroph-\|Wood Saprotroph\| | Corticioid-Polyporoid | White Rot | Probable | As white_rot (Põlme et al. 2020); As wood_pathogen (Põlme et al. 2020); Decay wood (Põlme et al. 2020) | Tedersoo L, et al. 2014. Science 346:e1256688 ((https://doi.org/10.1126/science.1256688)); Põlme S, et al. 2020. Fungal Diversity 105:1-16 ((https://doi.org/10.1007/s13225-020-00466-2)) |
| Otu06307 | 4 | k__Fungi(100);p__Ascomycota(100);c__Saccharomycetes(100);o__Saccharomycetales(100);f__Metschnikowiaceae(100);g__Metschnikowia(100);g__Metschnikowia_unclassified(100); | Metschnikowia | 13 | Saprotroph | \|Undefined Saprotroph\| | Yeast | NULL | Probable | As nectar-tap_saprotroph (Põlme et al. 2020); Decay sugar-rich substrates (Põlme et al. 2020) | Põlme S, et al. 2020. Fungal Diversity 105:1-16 ((https://doi.org/10.1007/s13225-020-00466-2)) |
| Otu06308 | 4 | k__Fungi(100);p__Ascomycota(100);c__Leotiomycetes(100);o__Helotiales(100);f__Hyaloscyphaceae(100);g__Lachnum(100);g__Lachnum_unclassified(100); | Lachnum | 13 | Saprotroph-Symbiotroph | Endophyte-\|Plant Saprotroph\|-Undefined Saprotroph-Wood Saprotroph | Microfungus | NULL | Probable | As litter_saprotroph (Põlme et al. 2020); As root_endophyte (Põlme et al. 2020); Decay wood, leaf, fruit, seed (Põlme et al. 2020) | Tedersoo L, et al. 2014. Science 346:e1256688 ((https://doi.org/10.1126/science.1256688)); Põlme S, et al. 2020. Fungal Diversity 105:1-16 ((https://doi.org/10.1007/s13225-020-00466-2)) |
| Otu06309 | 4 | k__Fungi(100);p__Zygomycota(100);c__Zygomycota_class_Incertae_sedis(100);o__Mucorales(100);f__Cunninghamellaceae(100);g__Absidia(100);s__Absidia_cylindrospora_var._cylindrospora(100); | Absidia | 13 | Saprotroph | \|Undefined Saprotroph\| | Microfungus | NULL | Probable | As soil_saprotroph (Põlme et al. 2020); Decay soil (Põlme et al. 2020) | Tedersoo L, et al. 2014. Science 346:e1256688 ((https://doi.org/10.1126/science.1256688)); Põlme S, et al. 2020. Fungal Diversity 105:1-16 ((https://doi.org/10.1007/s13225-020-00466-2)) |
| Otu06310 | 4 | k__Fungi(100);p__Ascomycota(100);c__Saccharomycetes(100);o__Saccharomycetales(100);f__Metschnikowiaceae(100);g__Metschnikowia(100);g__Metschnikowia_unclassified(100); | Metschnikowia | 13 | Saprotroph | \|Undefined Saprotroph\| | Yeast | NULL | Probable | As nectar-tap_saprotroph (Põlme et al. 2020); Decay sugar-rich substrates (Põlme et al. 2020) | Põlme S, et al. 2020. Fungal Diversity 105:1-16 ((https://doi.org/10.1007/s13225-020-00466-2)) |
| Otu06312 | 4 | k__Fungi(100);p__Chytridiomycota(75);c__Chytridiomycetes(75);o__Spizellomycetales(75);f__Spizellomycetaceae(75);f__Spizellomycetaceae_unclassified(75);f__Spizellomycetaceae_unclassified(75); | Spizellomycetaceae | 9 | Pathotroph | Plant Pathogen-Undefined Parasite-Undefined Saprotroph | Microfungus | NULL | Probable | Common in the soil and also as parasites of a range of soil organisms and plants (Cannon & Kirk 2007) | Cannon PF, Kirk PM. 2007. Fungal Families of the World. CAB International, Cambridge (ISBN: 978-0851998275) |
| Otu06316 | 4 | k__Fungi(100);p__Ascomycota(100);c__Saccharomycetes(100);o__Saccharomycetales(100);f__Metschnikowiaceae(100);g__Metschnikowia(100);s__Metschnikowia_pulcherrima(100); | Metschnikowia | 13 | Saprotroph | \|Undefined Saprotroph\| | Yeast | NULL | Probable | As nectar-tap_saprotroph (Põlme et al. 2020); Decay sugar-rich substrates (Põlme et al. 2020) | Põlme S, et al. 2020. Fungal Diversity 105:1-16 ((https://doi.org/10.1007/s13225-020-00466-2)) |
| Otu06317 | 4 | k__Fungi(100);p__Basidiomycota(100);c__Agaricomycetes(100);o__Trechisporales(100);f__Hydnodontaceae(100);g__Trechispora(100);g__Trechispora_unclassified(100); | Trechispora | 13 | Saprotroph | \|Wood Saprotroph\| | Corticioid | White Rot | Highly Probable | As white_rot (Põlme et al. 2020); Decay wood (Põlme et al. 2020) | Gilbertson RL, Ryvarden L. 1987-1987. North American Polypores. Fungiflora, Oslo (ISBN: 978-0945345060); Rinaldi AC, et al. 2008. Fungal Diversity 33:1-45 ((http://www.fungaldiversity.org/fdp/sfdp/33-1.pdf)) (as Uncertain); Tedersoo L, et al. 2010. Mycorrhiza 20:217-263 ((https://doi.org/10.1007/s00572-009-0274-x)) (as not ECM).; Põlme S, et al. 2020. Fungal Diversity 105:1-16 ((https://doi.org/10.1007/s13225-020-00466-2)) |
| Otu06323 | 4 | k__Fungi(100);p__Ascomycota(100);c__Saccharomycetes(100);o__Saccharomycetales(100);f__Saccharomycetales_family_Incertae_sedis(100);g__Candida(75);g__Candida_unclassified(75); | Candida | 13 | Pathotroph-Saprotroph-Symbiotroph | Animal Pathogen-Endophyte-Endosymbiont-Epiphyte-\|Undefined Saprotroph\| | Dimorphic Yeast-Yeast | NULL | Probable | As nectar-tap_saprotroph (Põlme et al. 2020); Decay sugar-rich substrates (Põlme et al. 2020); This asexual genus is polyphyletic and contain species that function very differently depend on what they are. Caution should be taken when interpreting guilds for Candida species. | Manolakaki D, et al. 2010. Virulence. 1: 367-375 ((https://doi.org/10.4161/viru.1.5.12796)); Mekbib SB, et al. 2011. Tropical Plant Pathology 36:233-240 ((https://doi.org/10.1590/S1982-56762011000400004)); Põlme S, et al. 2020. Fungal Diversity 105:1-16 ((https://doi.org/10.1007/s13225-020-00466-2)) |
| Otu06324 | 4 | k__Fungi(100);p__Basidiomycota(100);c__Basidiomycota_class_Incertae_sedis(100);o__Malasseziales(100);f__Malasseziales_family_Incertae_sedis(100);g__Malassezia(100);s__Malassezia_globosa(100); | Malassezia | 13 | Pathotroph-Saprotroph | Animal Pathogen-\|Undefined Saprotroph\| | Facultative Yeast-Yeast | NULL | Probable | A lipid; As soil_saprotroph (Põlme et al. 2020); Decay roots, soil (Põlme et al. 2020); dependent fungus growing on the sebaceous areas of human skin that may cause or exacerbate several skin diseases (Pityrosporum is a synonym that has been used for the yeast forms) | Kurtzman CP, et al. (eds.) 2011. The Yeasts, a Taxonomic Study. Fifth Edition. Vols 1-3. Elsevier, San Diego (ISBN: 9780444521491); Findley K, et al. 2013. Nature 498:367-370 ((https://doi.org/10.1038/nature12171)); Põlme S, et al. 2020. Fungal Diversity 105:1-16 ((https://doi.org/10.1007/s13225-020-00466-2)) |
| Otu06325 | 4 | k__Fungi(100);p__Ascomycota(100);c__Saccharomycetes(100);o__Saccharomycetales(100);f__Metschnikowiaceae(100);g__Metschnikowia(100);s__Metschnikowia_pulcherrima(100); | Metschnikowia | 13 | Saprotroph | \|Undefined Saprotroph\| | Yeast | NULL | Probable | As nectar-tap_saprotroph (Põlme et al. 2020); Decay sugar-rich substrates (Põlme et al. 2020) | Põlme S, et al. 2020. Fungal Diversity 105:1-16 ((https://doi.org/10.1007/s13225-020-00466-2)) |
| Otu06329 | 4 | k__Fungi(100);p__Ascomycota(100);c__Sordariomycetes(100);o__Xylariales(100);f__Xylariaceae(100);g__Nemania(100);s__Nemania_beaumontii(100); | Nemania | 13 | Saprotroph | Undefined Saprotroph-\|Wood Saprotroph\| | Microfungus | NULL | Probable | Decay wood (Põlme et al. 2020) | Tedersoo L, et al. 2014. Science 346:e1256688 ((https://doi.org/10.1126/science.1256688)); Põlme S, et al. 2020. Fungal Diversity 105:1-16 ((https://doi.org/10.1007/s13225-020-00466-2)) |
| Otu06330 | 4 | k__Fungi(100);p__Ascomycota(100);c__Saccharomycetes(100);o__Saccharomycetales(100);f__Metschnikowiaceae(100);g__Metschnikowia(100);g__Metschnikowia_unclassified(75); | Metschnikowia | 13 | Saprotroph | \|Undefined Saprotroph\| | Yeast | NULL | Probable | As nectar-tap_saprotroph (Põlme et al. 2020); Decay sugar-rich substrates (Põlme et al. 2020) | Põlme S, et al. 2020. Fungal Diversity 105:1-16 ((https://doi.org/10.1007/s13225-020-00466-2)) |
| Otu06331 | 4 | k__Fungi(100);p__Ascomycota(100);c__Saccharomycetes(100);o__Saccharomycetales(100);f__Saccharomycetales_family_Incertae_sedis(100);g__Candida(75);g__Candida_unclassified(75); | Candida | 13 | Pathotroph-Saprotroph-Symbiotroph | Animal Pathogen-Endophyte-Endosymbiont-Epiphyte-\|Undefined Saprotroph\| | Dimorphic Yeast-Yeast | NULL | Probable | As nectar-tap_saprotroph (Põlme et al. 2020); Decay sugar-rich substrates (Põlme et al. 2020); This asexual genus is polyphyletic and contain species that function very differently depend on what they are. Caution should be taken when interpreting guilds for Candida species. | Manolakaki D, et al. 2010. Virulence. 1: 367-375 ((https://doi.org/10.4161/viru.1.5.12796)); Mekbib SB, et al. 2011. Tropical Plant Pathology 36:233-240 ((https://doi.org/10.1590/S1982-56762011000400004)); Põlme S, et al. 2020. Fungal Diversity 105:1-16 ((https://doi.org/10.1007/s13225-020-00466-2)) |
| Otu06332 | 4 | k__Fungi(100);p__Ascomycota(100);c__Saccharomycetes(100);o__Saccharomycetales(100);f__Saccharomycetales_family_Incertae_sedis(100);f__Saccharomycetales_family_Incertae_sedis_unclassified(100);f__Saccharomycetales_family_Incertae_sedis_unclassified(100); | Saccharomycetales | 7 | Saprotroph | Undefined Saprotroph | Yeast | NULL | Possible | NULL | Sterkenburg E, et al. 2015. New Phytologist 207:1145-1158 ((https://doi.org/10.1111/nph.13426)) |
| Otu06334 | 4 | k__Fungi(100);p__Ascomycota(100);c__Saccharomycetes(100);o__Saccharomycetales(100);f__Metschnikowiaceae(100);g__Metschnikowia(100);g__Metschnikowia_unclassified(75); | Metschnikowia | 13 | Saprotroph | \|Undefined Saprotroph\| | Yeast | NULL | Probable | As nectar-tap_saprotroph (Põlme et al. 2020); Decay sugar-rich substrates (Põlme et al. 2020) | Põlme S, et al. 2020. Fungal Diversity 105:1-16 ((https://doi.org/10.1007/s13225-020-00466-2)) |
| Otu06339 | 4 | k__Fungi(100);p__Ascomycota(100);c__Saccharomycetes(100);o__Saccharomycetales(100);f__Metschnikowiaceae(100);g__Metschnikowia(100);s__Metschnikowia_pulcherrima(75); | Metschnikowia | 13 | Saprotroph | \|Undefined Saprotroph\| | Yeast | NULL | Probable | As nectar-tap_saprotroph (Põlme et al. 2020); Decay sugar-rich substrates (Põlme et al. 2020) | Põlme S, et al. 2020. Fungal Diversity 105:1-16 ((https://doi.org/10.1007/s13225-020-00466-2)) |
| Otu06340 | 4 | k__Fungi(100);p__Zygomycota(100);c__Zygomycota_class_Incertae_sedis(100);o__Mortierellales(100);f__Mortierellaceae(100);g__Mortierella(100);s__Mortierella_sp(100); | Mortierella | 13 | Saprotroph-Symbiotroph | Endophyte-Plant Saprotroph-\|Undefined Saprotroph\| | Microfungus | NULL | Probable | As chitinolytic (Põlme et al. 2020); As soil_saprotroph (Põlme et al. 2020); Decay roots, soil (Põlme et al. 2020); Saprobic in soil and thus frequently isolated from soil and material contaminated with soil such as hay or rotten silage and spoil tips (Cannon & Kirk 2007) | Cannon PF, Kirk PM. 2007. Fungal Families of the World. CAB International, Cambridge (ISBN: 978-0851998275); Tedersoo L, et al. 2014. Science 346:e1256688 ((https://doi.org/10.1126/science.1256688)); Purahong W, et al. 2016. Molecular Ecology 25:4059-4074 ((https://doi.org/10.1111/mec.13739)); Põlme S, et al. 2020. Fungal Diversity 105:1-16 ((https://doi.org/10.1007/s13225-020-00466-2)) |
| Otu06342 | 4 | k__Fungi(100);p__Ascomycota(100);c__Saccharomycetes(100);o__Saccharomycetales(100);f__Metschnikowiaceae(100);g__Metschnikowia(100);s__Metschnikowia_pulcherrima(100); | Metschnikowia | 13 | Saprotroph | \|Undefined Saprotroph\| | Yeast | NULL | Probable | As nectar-tap_saprotroph (Põlme et al. 2020); Decay sugar-rich substrates (Põlme et al. 2020) | Põlme S, et al. 2020. Fungal Diversity 105:1-16 ((https://doi.org/10.1007/s13225-020-00466-2)) |
| Otu06343 | 4 | k__Fungi(100);p__Basidiomycota(100);c__Microbotryomycetes(100);o__Sporidiobolales(100);f__Sporidiobolales_family_Incertae_sedis(100);g__Sporobolomyces(100);s__Sporobolomyces_lactosus(100); | Sporobolomyces | 13 | Pathotroph-Saprotroph | \|Fungal Parasite\|-Plant Saprotroph | Yeast | NULL | Probable | As litter_saprotroph (Põlme et al. 2020); As mycoparasite (Põlme et al. 2020); Decay fungal material (Põlme et al. 2020) | Tedersoo L, et al. 2014. Science 346:e1256688 ((https://doi.org/10.1126/science.1256688)); Purahong W, et al. 2016. Molecular Ecology 25:4059-4074 ((https://doi.org/10.1111/mec.13739)); Põlme S, et al. 2020. Fungal Diversity 105:1-16 ((https://doi.org/10.1007/s13225-020-00466-2)) |

| | | | | | | | | | | | |
|---|---|---|---|---|---|---|---|---|---|---|---|
| Otu06345 | 4 | k__Fungi(100);p__Ascomycota(100);c__Saccharomycetes(100);o__Saccharomycetales(100);f__Metschnikowiaceae(100);g__Metschnikowia(100);s__Metschnikowia_pulcherrima(100); | Metschnikowia | 13 | Saprotroph | \|Undefined Saprotroph\| | Yeast | NULL | Probable | As nectar-tap_saprotroph (Põlme et al. 2020); Decay sugar-rich substrates (Põlme et al. 2020) | Põlme S, et al. 2020. Fungal Diversity 105:1-16 ((https://doi.org/10.1007/s13225-020-00466-2)) |
| Otu06347 | 4 | k__Fungi(100);p__Basidiomycota(100);c__Agaricomycetes(100);o__Agaricales(100);f__Inocybaceae(100);g__Inocybe(100);s__Inocybe_semifulva(100); | Inocybe | 13 | Symbiotroph | \|Ectomycorrhizal\| | Agaricoid | NULL | Highly Probable | Sensu stricto (Matheny 2009). | Rinaldi AC, et al. 2008. Fungal Diversity 33:1-45 ((http://www.fungaldiversity.org/fdp/sfdp/33-1.pdf)); Matheny PB. 2009. McIlvainea 18:11-21 ((https://namyco.org/docs/3_Matheny.pdf)); Tedersoo L, et al. 2010. Mycorrhiza 20:217-263 ((https://doi.org/10.1007/s00572-009-0274-x)); Põlme S, et al. 2020. Fungal Diversity 105:1-16 ((https://doi.org/10.1007/s13225-020-00466-2)); http://mycorrhizas.info/ecmf.html ((http://mycorrhizas.info/ecmf.html)) |
| Otu06348 | 4 | k__Fungi(100);p__Ascomycota(100);c__Saccharomycetes(100);o__Saccharomycetales(100);f__Metschnikowiaceae(100);g__Metschnikowia(100);s__Metschnikowia_pulcherrima(75); | Metschnikowia | 13 | Saprotroph | \|Undefined Saprotroph\| | Yeast | NULL | Probable | As nectar-tap_saprotroph (Põlme et al. 2020); Decay sugar-rich substrates (Põlme et al. 2020) | Põlme S, et al. 2020. Fungal Diversity 105:1-16 ((https://doi.org/10.1007/s13225-020-00466-2)) |
| Otu06350 | 4 | k__Fungi(100);p__Ascomycota(100);c__Saccharomycetes(100);o__Saccharomycetales(100);f__Metschnikowiaceae(100);g__Metschnikowia(100);s__Metschnikowia_pulcherrima(100); | Metschnikowia | 13 | Saprotroph | \|Undefined Saprotroph\| | Yeast | NULL | Probable | As nectar-tap_saprotroph (Põlme et al. 2020); Decay sugar-rich substrates (Põlme et al. 2020) | Põlme S, et al. 2020. Fungal Diversity 105:1-16 ((https://doi.org/10.1007/s13225-020-00466-2)) |
| Otu06351 | 4 | k__Fungi(100);p__Ascomycota(100);c__Saccharomycetes(100);o__Saccharomycetales(100);f__Metschnikowiaceae(100);g__Metschnikowia(100);s__Metschnikowia_pulcherrima(75); | Metschnikowia | 13 | Saprotroph | \|Undefined Saprotroph\| | Yeast | NULL | Probable | As nectar-tap_saprotroph (Põlme et al. 2020); Decay sugar-rich substrates (Põlme et al. 2020) | Põlme S, et al. 2020. Fungal Diversity 105:1-16 ((https://doi.org/10.1007/s13225-020-00466-2)) |
| Otu06355 | 4 | k__Fungi(100);p__Ascomycota(100);c__Saccharomycetes(100);o__Saccharomycetales(100);f__Metschnikowiaceae(100);g__Metschnikowia(100);s__Metschnikowia_pulcherrima(100); | Metschnikowia | 13 | Saprotroph | \|Undefined Saprotroph\| | Yeast | NULL | Probable | As nectar-tap_saprotroph (Põlme et al. 2020); Decay sugar-rich substrates (Põlme et al. 2020) | Põlme S, et al. 2020. Fungal Diversity 105:1-16 ((https://doi.org/10.1007/s13225-020-00466-2)) |
| Otu06357 | 4 | k__Fungi(100);p__Basidiomycota(100);c__Microbotryomycetes(100);o__Sporidiobolales(100);f__Sporidiobolales_family_Incertae_sedis(100);g__Rhodotorula(100);s__Rhodotorula_hordea(100); | Rhodotorula | 13 | Pathotroph-Saprotroph-Symbiotroph | Animal Pathogen-Animal Symbiotroph-Endophyte-Plant Pathogen-\|Undefined Saprotroph\| | Yeast | NULL | Probable | As foliar_endophyte (Põlme et al. 2020); Opportunistic human pathogen | Põlme S, et al. 2020. Fungal Diversity 105:1-16 ((https://doi.org/10.1007/s13225-020-00466-2)); https://www.cdc.gov ((https://www.cdc.gov)) |
| Otu06359 | 4 | k__Fungi(100);p__Ascomycota(100);c__Ascomycota_class_Incertae_sedis(100);o__Ascomycota_order_Incertae_sedis(100);f__Ascomycota_family_Incertae_sedis(100);g__Pyrigemmula(100);s__Pyrigemmula_aurantiaca(100); | Pyrigemmula | 13 | Saprotroph | \|Undefined Saprotroph\|-Wood Saprotroph | Microfungus | NULL | Probable | As nectar-tap_saprotroph (Põlme et al. 2020); Decay wood, sugar-rich substrates (Põlme et al. 2020) | Põlme S, et al. 2020. Fungal Diversity 105:1-16 ((https://doi.org/10.1007/s13225-020-00466-2)) |
| Otu06361 | 4 | k__Fungi(100);p__Ascomycota(100);c__Saccharomycetes(100);o__Saccharomycetales(100);f__Metschnikowiaceae(100);g__Metschnikowia(100);s__Metschnikowia_viticola(100); | Metschnikowia | 13 | Saprotroph | \|Undefined Saprotroph\| | Yeast | NULL | Probable | As nectar-tap_saprotroph (Põlme et al. 2020); Decay sugar-rich substrates (Põlme et al. 2020) | Põlme S, et al. 2020. Fungal Diversity 105:1-16 ((https://doi.org/10.1007/s13225-020-00466-2)) |
| Otu06363 | 4 | k__Fungi(100);p__Basidiomycota(100);c__Agaricomycetes(100);o__Agaricales(100);f__Inocybaceae(100);g__Inocybe(100);g__Inocybe_unclassified(100); | Inocybe | 13 | Symbiotroph | \|Ectomycorrhizal\| | Agaricoid | NULL | Highly Probable | Sensu stricto (Matheny 2009). | Rinaldi AC, et al. 2008. Fungal Diversity 33:1-45 ((http://www.fungaldiversity.org/fdp/sfdp/33-1.pdf)); Matheny PB. 2009. McIlvainea 18:11-21 ((https://namyco.org/docs/3_Matheny.pdf)); Tedersoo L, et al. 2010. Mycorrhiza 20:217-263 ((https://doi.org/10.1007/s00572-009-0274-x)); Põlme S, et al. 2020. Fungal Diversity 105:1-16 ((https://doi.org/10.1007/s13225-020-00466-2)); http://mycorrhizas.info/ecmf.html ((http://mycorrhizas.info/ecmf.html)) |
| Otu06366 | 4 | k__Fungi(100);p__Ascomycota(100);c__Saccharomycetes(100);o__Saccharomycetales(100);f__Metschnikowiaceae(100);g__Metschnikowia(100);s__Metschnikowia_pulcherrima(75); | Metschnikowia | 13 | Saprotroph | \|Undefined Saprotroph\| | Yeast | NULL | Probable | As nectar-tap_saprotroph (Põlme et al. 2020); Decay sugar-rich substrates (Põlme et al. 2020) | Põlme S, et al. 2020. Fungal Diversity 105:1-16 ((https://doi.org/10.1007/s13225-020-00466-2)) |
| Otu06369 | 4 | k__Fungi(100);p__Ascomycota(100);c__Saccharomycetes(100);o__Saccharomycetales(100);f__Metschnikowiaceae(100);g__Metschnikowia(100);s__Metschnikowia_pulcherrima(75); | Metschnikowia | 13 | Saprotroph | \|Undefined Saprotroph\| | Yeast | NULL | Probable | As nectar-tap_saprotroph (Põlme et al. 2020); Decay sugar-rich substrates (Põlme et al. 2020) | Põlme S, et al. 2020. Fungal Diversity 105:1-16 ((https://doi.org/10.1007/s13225-020-00466-2)) |
| Otu06371 | 4 | k__Fungi(100);p__Ascomycota(100);c__Saccharomycetes(100);o__Saccharomycetales(100);f__Metschnikowiaceae(100);g__Metschnikowia(100);s__Metschnikowia_pulcherrima(100); | Metschnikowia | 13 | Saprotroph | \|Undefined Saprotroph\| | Yeast | NULL | Probable | As nectar-tap_saprotroph (Põlme et al. 2020); Decay sugar-rich substrates (Põlme et al. 2020) | Põlme S, et al. 2020. Fungal Diversity 105:1-16 ((https://doi.org/10.1007/s13225-020-00466-2)) |
| Otu06372 | 4 | k__Fungi(100);p__Ascomycota(100);c__Saccharomycetes(100);o__Saccharomycetales(100);f__Metschnikowiaceae(100);g__Metschnikowia(100);s__Metschnikowia_pulcherrima(75); | Metschnikowia | 13 | Saprotroph | \|Undefined Saprotroph\| | Yeast | NULL | Probable | As nectar-tap_saprotroph (Põlme et al. 2020); Decay sugar-rich substrates (Põlme et al. 2020) | Põlme S, et al. 2020. Fungal Diversity 105:1-16 ((https://doi.org/10.1007/s13225-020-00466-2)) |
| Otu06373 | 4 | k__Fungi(100);p__Ascomycota(100);c__Saccharomycetes(100);o__Saccharomycetales(100);f__Metschnikowiaceae(100);g__Metschnikowia(100);s__Metschnikowia_pulcherrima(75); | Metschnikowia | 13 | Saprotroph | \|Undefined Saprotroph\| | Yeast | NULL | Probable | As nectar-tap_saprotroph (Põlme et al. 2020); Decay sugar-rich substrates (Põlme et al. 2020) | Põlme S, et al. 2020. Fungal Diversity 105:1-16 ((https://doi.org/10.1007/s13225-020-00466-2)) |
| Otu06376 | 4 | k__Fungi(100);p__Ascomycota(100);c__Dothideomycetes(100);o__Pleosporales(100);f__Montagnulaceae(100);g__unclassified_Montagnulaceae(100);s__Montagnulaceae_sp(100); | Montagnulaceae | 9 | Pathotroph-Saprotroph | Fungal Parasite-Plant Pathogen-Undefined Saprotroph | Microfungus | NULL | Probable | Species in this clade can be saprobic, plant pathogenic, or mycoparasitic (Zhang et al. 2009) | Zhang Y, et al. 2009. Studies in Mycology 64:85-102 ((https://doi.org/10.3114/sim.2009.64.04)) |
| Otu06377 | 4 | k__Fungi(100);p__Ascomycota(100);c__Sordariomycetes(100);o__Hypocreales(100);f__Hypocreales_family_Incertae_sedis(100);g__Acremonium(100);s__Acremonium_furcatum(100); | Acremonium | 13 | Pathotroph-Saprotroph-Symbiotroph | Animal Pathogen-Endophyte-Fungal Parasite-Plant Pathogen-\|Undefined Saprotroph\|-Wood Saprotroph | Microfungus | Soft Rot | Possible | As foliar_endophyte (Põlme et al. 2020); culture (Hanada et al. 2010, Del Olmo; Decay leaf, fruit, seed, soil, dung, animal material (Põlme et al. 2020); Endophyte detection method; Ruiz & Arnold 2014); Likely opportunistic human pathogen (Irinyi et al. 2015) | Duncan CG, Eslyn WE. 1966. Mycologia 58:642-645 ((https://doi.org/10.2307/3757045)); Nilsson T. 1973. Studia Forestalia Suecica 104: 1-40 ((https://pub.epsilon.slu.se/5711/)); Worrall JJ, et al. 1997. Mycologia 89:199-219 ((https://doi.org/10.2307/3761073)); Hanada RE, et al. 2010. Fungal Biology 114:901-910 ((https://doi.org/10.1016/j.funbio.2010.08.006)); Del Olmo-Ruiz M, Arnold AE. 2014. Mycologia 106: 8-21 ((https://doi.org/10.3852/13-098)); Irinyi L, et al. 2015. Medical Mycology 53313-337 ((https://doi.org/10.1093/mmy/myv008)); Põlme S, et al. 2020. Fungal Diversity 105:1-16 ((https://doi.org/10.1007/s13225-020-00466-2)) |
| Otu06378 | 4 | k__Fungi(100);p__Ascomycota(100);c__Saccharomycetes(100);o__Saccharomycetales(100);f__Metschnikowiaceae(100);g__Metschnikowia(100);s__Metschnikowia_pulcherrima(100); | Metschnikowia | 13 | Saprotroph | \|Undefined Saprotroph\| | Yeast | NULL | Probable | As nectar-tap_saprotroph (Põlme et al. 2020); Decay sugar-rich substrates (Põlme et al. 2020) | Põlme S, et al. 2020. Fungal Diversity 105:1-16 ((https://doi.org/10.1007/s13225-020-00466-2)) |
| Otu06379 | 4 | k__Fungi(100);p__Ascomycota(100);c__Sordariomycetes(100);o__Sordariales(100);f__Chaetomiaceae(100);g__Chaetomium(100);s__Chaetomium_sp_8444(100); | Chaetomium | 13 | Pathotroph-Saprotroph-Symbiotroph | Animal Pathogen-Dung Saprotroph-Endophyte-Epiphyte-Plant Pathogen-\|Plant Saprotroph\|-Wood Saprotroph | Microfungus | Soft Rot | Possible | As foliar_endophyte (Põlme et al. 2020); As litter_saprotroph (Põlme et al. 2020); culture (Busby et al. 2016, David et al. 2016); Likely opportunistic human pathogen (Irinyi et al. 2015); culture (Massimo et al. 2015); Endophyte detection method; Decay wood, leaf, fruit, seed, soil, dung (Põlme et al. 2020); derived commodities (Cannon & Kirk 2007); Epiphyte detection method; Saprobic, acting as key decay organisms of plant materials and plant | Seehann G, et al. 1975. List of Fungi in Soft-Rot Tissue Tests. Institute fur Holzbiologie und Holzschutz, Bundesforschungsanstalt fur Forst - und Holzwirtshaft, Hamburg - Lohbrugge, Germany; Cannon PF, Kirk PM. 2007. Fungal Families of the World. CAB International, Cambridge ((https://isbnsearch.org/isbn/9780851998275)); Bills GF, et al. 2013. Current Opinion in Microbiology 16:549-565 ((https://doi.org/10.1016/j.mib.2013.08.001)); Tedersoo L, et al. 2014. Science 346:e1256688 ((https://doi.org/10.1126/science.1256688)); Irinyi L, et al. 2015. Medical Mycology 53:313-337 ((https://doi.org/10.1093/mmy/myv008)); Busby PE, et al. 2016. New Phytologist 209:1681-1692 ((https://doi.org/10.1111/nph.13742)); Põlme S, et al. 2020. Fungal Diversity 105:1-16 ((https://doi.org/10.1007/s13225-020-00466-2)) |
| Otu06381 | 4 | k__Fungi(100);p__Ascomycota(100);c__Eurotiomycetes(100);o__Eurotiales(100);f__Trichocomaceae(100);g__Penicillium(100);s__Penicillium_sclerotiorum(100); | Penicillium | 13 | Pathotroph-Saprotroph-Symbiotroph | Animal Parasite-Dung Saprotroph-Endophyte-Fungal Parasite-Plant Pathogen-\|Undefined Saprotroph\|-Wood Saprotroph | Microfungus | Soft Rot | Highly Probable | As foliar_endophyte (Põlme et al. 2020); As mycoparasite (Põlme et al. 2020); Decay soil (Põlme et al. 2020) | Duncan CG, Eslyn WE. 1966. Mycologia 58:642-664 ((https://doi.org/10.2307/3757045)); Seehann G, et al. 1975. List of Fungi in Soft-Rot Tissue Tests. Institute fur Holzbiologie und Holzschutz, Bundesforschungsanstalt fur Forst - und Holzwirtshaft, Hamburg - Lohbrugge, Germany ((https://www.irg-wp.com/irgdocs/details.php?d6493033-c9ea-40f3-8e3e-5697f32d640d)); Costa IPMW, et al. 2012. Mycotaxon 119:e26 ((http://mycotaxon.com/resources/checklists/costa_v119_checklist.pdf)); Bills GF, et al. 2013. Current Opinion in Microbiology 16: 549-565 ((https://doi.org/10.1016/j.mib.2013.08.001)); Tedersoo L, et al. 2014. Science 346:e1256688 ((https://doi.org/10.1126/science.1256688)); Põlme S, et al. 2020. Fungal Diversity 105:1-16 ((https://doi.org/10.1007/s13225-020-00466-2)) |
| Otu06382 | 4 | k__Fungi(100);p__Ascomycota(100);c__Sordariomycetes(100);o__Sordariales(100);f__Cephalothecaceae(100);g__Phialemonium(100);s__Phialemonium_atrogriseum(100); | Phialemonium | 13 | Pathotroph-Saprotroph | \|Animal Parasite\|-Undefined Saprotroph | Microfungus | NULL | Probable | As animal_decomposer (Põlme et al. 2020); Decay animal material (Põlme et al. 2020) | Tedersoo L, et al. 2014. Science 346:e1256688 ((https://doi.org/10.1126/science.1256688)); Põlme S, et al. 2020. Fungal Diversity 105:1-16 ((https://doi.org/10.1007/s13225-020-00466-2)) |
| Otu06384 | 4 | k__Fungi(100);p__Basidiomycota(100);c__Agaricomycetes(100);o__Cantharellales(100);f__Hydnaceae(100);g__Sistotrema(75);g__Sistotrema_unclassified(75); | Sistotrema | 13 | Saprotroph-Symbiotroph | Ectomycorrhizal-\|Plant Saprotroph\|-Wood Saprotroph | Corticioid | White Rot | Possible | As litter_saprotroph (Põlme et al. 2020); Decay leaf, fruit, seed (Põlme et al. 2020) | Gilbertson RL, Ryvarden L. 1987-1987. North American Polypores. Fungiflora, Oslo (ISBN: 978-0945345060); Rinaldi AC, et al. 2008. Fungal Diversity 33:1-45 ((http://www.fungaldiversity.org/fdp/sfdp/33-1.pdf)) (as ECM/not ECM); Tedersoo L, et al. 2010. Mycorrhiza 20: 217-263 ((https://doi.org/10.1007/s00572-009-0274-x)) (pro parte); Põlme S, et al. 2020. Fungal Diversity 105:1-16 ((https://doi.org/10.1007/s13225-020-00466-2)); http://mycorrhizas.info/ecmf.html ((http://mycorrhizas.info/ecmf.html)) (as ECM/not ECM). |
| Otu06385 | 4 | k__Fungi(100);p__Ascomycota(100);c__Saccharomycetes(100);o__Saccharomycetales(100);f__Metschnikowiaceae(100);g__Metschnikowia(100);s__Metschnikowia_pulcherrima(100); | Metschnikowia | 13 | Saprotroph | \|Undefined Saprotroph\| | Yeast | NULL | Probable | As nectar-tap_saprotroph (Põlme et al. 2020); Decay sugar-rich substrates (Põlme et al. 2020) | Põlme S, et al. 2020. Fungal Diversity 105:1-16 ((https://doi.org/10.1007/s13225-020-00466-2)) |
| Otu06388 | 4 | k__Fungi(100);p__Ascomycota(100);c__Saccharomycetes(100);o__Saccharomycetales(100);f__Metschnikowiaceae(100);g__Metschnikowia(100);g__Metschnikowia_unclassified(100); | Metschnikowia | 13 | Saprotroph | \|Undefined Saprotroph\| | Yeast | NULL | Probable | As nectar-tap_saprotroph (Põlme et al. 2020); Decay sugar-rich substrates (Põlme et al. 2020) | Põlme S, et al. 2020. Fungal Diversity 105:1-16 ((https://doi.org/10.1007/s13225-020-00466-2)) |
| Otu06393 | 4 | k__Fungi(100);p__Ascomycota(100);c__Saccharomycetes(100);o__Saccharomycetales(100);f__Saccharomycetales_family_Incertae_sedis(75);g__Candida(75);g__Candida_unclassified(75); | Candida | 13 | Pathotroph-Saprotroph-Symbiotroph | Animal Pathogen-Endophyte-Endosymbiont-Epiphyte-\|Undefined Saprotroph\| | Dimorphic Yeast-Yeast | NULL | Probable | As nectar-tap_saprotroph (Põlme et al. 2020); Decay sugar-rich substrates (Põlme et al. 2020); This asexual genus is polyphyletic and contain species that function very differently depend on what they are. Caution should be taken when interpreting guilds for Candida species. | Manolakaki D, et al. 2010. Virulence. 1: 367-375 ((https://doi.org/10.4161/viru.1.5.12796)); Mekbib SB, et al. 2011. Tropical Plant Pathology 36:233-240 ((https://doi.org/10.1590/S1982-56762011000400004)); Põlme S, et al. 2020. Fungal Diversity 105:1-16 ((https://doi.org/10.1007/s13225-020-00466-2)) |
| Otu06397 | 4 | k__Fungi(100);p__Ascomycota(100);c__Ascomycota_class_Incertae_sedis(100);o__Ascomycota_order_Incertae_sedis(100);f__Ascomycota_family_Incertae_sedis(100);g__Scolecobasidium(100);g__Scolecobasidium_unclassified(100); | Scolecobasidium | 13 | Pathotroph-Saprotroph-Symbiotroph | \|Animal Parasite\|-Endophyte-Plant Pathogen-Undefined Saprotroph | Microfungus | NULL | Probable | As foliar_endophyte (Põlme et al. 2020); Decay leaf, fruit, seed, animal material (Põlme et al. 2020) | Tedersoo L, et al. 2014. Science 346:e1256688 ((https://doi.org/10.1126/science.1256688)); Põlme S, et al. 2020. Fungal Diversity 105:1-16 ((https://doi.org/10.1007/s13225-020-00466-2)) |
| Otu06398 | 4 | k__Fungi(100);p__Ascomycota(100);c__Sordariomycetes(100);o__Sordariales(100);f__Chaetomiaceae(100);f__Chaetomiaceae_unclassified(75);f__Chaetomiaceae_unclassified(75); | Chaetomiaceae | 9 | Pathotroph-Saprotroph-Symbiotroph | Animal Pathogen-Dung Saprotroph-Endophyte-Epiphyte-Plant Saprotroph-Wood Saprotroph | Microfungus | Soft Rot | Possible | Saprobic, acting as key decay organisms of plant materials and plant-derived commodities (Cannon & Kirk 2007); Epiphyte detection method-culture (Massimo et al. 2015); Endophyte detection method-culture (Busby et al. 2016, David et al. 2016); Likely opportunistic human pathogen (Irinyi et al. 2015) | Seehann G, et al. 1975. List of Fungi in Soft-Rot Tissue Tests. Institute fur Holzbiologie und Holzschutz, Bundesforschungsanstalt fur Forst - und Holzwirtshaft, Hamburg - Lohbrugge, Germany; Cannon PF, Kirk PM. 2007. Fungal Families of the World. CAB International, Cambridge ((https://isbnsearch.org/isbn/9780851998275)); Bills GF, et al. 2013. Current Opinion in Microbiology 16:549-565 ((https://doi.org/10.1016/j.mib.2013.08.001)); Tedersoo L, et al. 2014. Science 346:e1256688 ((https://doi.org/10.1126/science.1256688)); Irinyi L, et al. 2015. Medical Mycology 53:313-337 ((https://doi.org/10.1093/mmy/myv008)); Massimo NC, et al. 2015. Microbial Ecology 70:61-76 ((https://doi.org/10.1007/s00248-014-0563-6)); Busby PE, et al. 2016. New Phytologist 209:1681-1692 ((https://doi.org/10.1111/nph.13742)); David AS, et al. 2016. Microbial Ecology 71:912-926 ((https://doi.org/10.1007/s00248-015-0712-6)) |
| Otu06400 | 4 | k__Fungi(100);p__Ascomycota(100);c__Saccharomycetes(100);o__Saccharomycetales(100);f__Metschnikowiaceae(100);g__Metschnikowia(100);s__Metschnikowia_pulcherrima(75); | Metschnikowia | 13 | Saprotroph | \|Undefined Saprotroph\| | Yeast | NULL | Probable | As nectar-tap_saprotroph (Põlme et al. 2020); Decay sugar-rich substrates (Põlme et al. 2020) | Põlme S, et al. 2020. Fungal Diversity 105:1-16 ((https://doi.org/10.1007/s13225-020-00466-2)) |
| Otu06402 | 4 | k__Fungi(100);p__Ascomycota(100);c__Saccharomycetes(100);o__Saccharomycetales(100);f__Metschnikowiaceae(100);g__Metschnikowia(100);s__Metschnikowia_pulcherrima(75); | Metschnikowia | 13 | Saprotroph | \|Undefined Saprotroph\| | Yeast | NULL | Probable | As nectar-tap_saprotroph (Põlme et al. 2020); Decay sugar-rich substrates (Põlme et al. 2020) | Põlme S, et al. 2020. Fungal Diversity 105:1-16 ((https://doi.org/10.1007/s13225-020-00466-2)) |
| Otu06406 | 4 | k__Fungi(100);p__Basidiomycota(100);c__Tremellomycetes(100);o__Tremellales(100);f__Tremellales_family_Incertae_sedis(100);f__Tremellales_family_Incertae_sedis_unclassified(100);f__Tremellales_family_Incertae_sedis_unclassified(100); | Tremellales | 7 | Pathotroph-Saprotroph-Symbiotroph | Fungal Parasite-Undefined Saprotroph | Tremelloid-Yeast | NULL | Possible | The order contains both teleomorphic and anamorphic species, most of the latter being yeasts | Sterkenburg E, et al. 2015. New Phytologist 207:1145-1158 ((https://doi.org/10.1111/nph.13426)) |
| Otu06407 | 4 | k__Fungi(100);p__Ascomycota(100);c__Saccharomycetes(100);o__Saccharomycetales(100);f__Saccharomycetales_family_Incertae_sedis(75);g__Candida(75);g__Candida_unclassified(75); | Candida | 13 | Pathotroph-Saprotroph-Symbiotroph | Animal Pathogen-Endophyte-Endosymbiont-Epiphyte-\|Undefined Saprotroph\| | Dimorphic Yeast-Yeast | NULL | Probable | As nectar-tap_saprotroph (Põlme et al. 2020); Decay sugar-rich substrates (Põlme et al. 2020); This asexual genus is polyphyletic and contain species that function very differently depend on what they are. Caution should be taken when interpreting guilds for Candida species. | Manolakaki D, et al. 2010. Virulence. 1: 367-375 ((https://doi.org/10.4161/viru.1.5.12796)); Mekbib SB, et al. 2011. Tropical Plant Pathology 36:233-240 ((https://doi.org/10.1590/S1982-56762011000400004)); Põlme S, et al. 2020. Fungal Diversity 105:1-16 ((https://doi.org/10.1007/s13225-020-00466-2)) |
| Otu06409 | 4 | k__Fungi(100);p__Ascomycota(100);c__Lecanoromycetes(100);o__Ostropales(100);f__Stictidaceae(100);g__Cryptodiscus(100);s__Cryptodiscus_pini(75); | Cryptodiscus | 13 | Pathotroph-Saprotroph | \|Lichen Parasite\|-Undefined Saprotroph | Microfungus | NULL | Probable | Decay fungal material, algal material (Põlme et al. 2020) | Tedersoo L, et al. 2014. Science 346:e1256688 ((https://doi.org/10.1126/science.1256688)); Põlme S, et al. 2020. Fungal Diversity 105:1-16 ((https://doi.org/10.1007/s13225-020-00466-2)) |
| Otu06411 | 4 | k__Fungi(100);p__Ascomycota(100);c__Saccharomycetes(100);o__Saccharomycetales(100);f__Saccharomycetales_family_Incertae_sedis(100);g__Candida(100);g__Candida_unclassified(100); | Candida | 13 | Pathotroph-Saprotroph-Symbiotroph | Animal Pathogen-Endophyte-Endosymbiont-Epiphyte-\|Undefined Saprotroph\| | Dimorphic Yeast-Yeast | NULL | Probable | As nectar-tap_saprotroph (Põlme et al. 2020); Decay sugar-rich substrates (Põlme et al. 2020); This asexual genus is polyphyletic and contain species that function very differently depend on what they are. Caution should be taken when interpreting guilds for Candida species. | Manolakaki D, et al. 2010. Virulence. 1: 367-375 ((https://doi.org/10.4161/viru.1.5.12796)); Mekbib SB, et al. 2011. Tropical Plant Pathology 36:233-240 ((https://doi.org/10.1590/S1982-56762011000400004)); Põlme S, et al. 2020. Fungal Diversity 105:1-16 ((https://doi.org/10.1007/s13225-020-00466-2)) |
| Otu06413 | 4 | k__Fungi(100);p__Basidiomycota(100);c__Agaricomycetes(100);o__Trechisporales(100);f__Hydnodontaceae(100);g__Trechispora(100);g__Trechispora_unclassified(75); | Trechispora | 13 | Saprotroph | \|Wood Saprotroph\| | Corticioid | White Rot | Highly Probable | As white_rot (Põlme et al. 2020); Decay wood (Põlme et al. 2020) | Gilbertson RL, Ryvarden L. [illegible] |
| Otu06414 | 4 | k__Fungi(100);p__Zygomycota(100);c__Zygomycota_class_Incertae_sedis(100);o__Mortierellales(100);f__Mortierellaceae(100);g__Mortierella(100);s__Mortierella_sp(100); | Mortierella | 13 | Saprotroph-Symbiotroph | Endophyte-Plant Saprotroph-\|Undefined Saprotroph\| | Microfungus | NULL | Probable | As chitinolytic (Põlme et al. 2020); As soil_saprotroph (Põlme et al. 2020); Decay roots, soil (Põlme et al. 2020); Saprobic in soil and thus frequently isolated from soil and material contaminated with soil such as hay or rotten silage and spoil tips (Cannon & Kirk 2007) | Cannon PF, Kirk PM. [illegible] |
| Otu06415 | 4 | k__Fungi(100);p__Ascomycota(100);c__Eurotiomycetes(100);o__Chaetothyriales(100);f__Herpotrichiellaceae(100);g__Phialophora(100);s__Phialophora_hyalina(100); | Phialophora | 13 | Pathotroph-Saprotroph-Symbiotroph | Animal Pathogen-Endophyte-Ericoid Mycorrhizal-Plant Pathogen-\|Plant Saprotroph\|-Wood Saprotroph | Dimorphic-Facultative Yeast-Microfungus | NULL | Possible | As foliar_endophyte (Põlme et al. 2020); As litter_saprotroph (Põlme et al. 2020); Consult the current literature for updated names on this paraphyletic grouping that included the Black Yeasts, Dark Septate Endophytes, etc.; Decay wood, leaf, fruit, seed, soil, dung, animal material (Põlme et al. 2020) | Nilsson T. 1973. Studia [illegible] |
| Otu06416 | 4 | k__Fungi(100);p__Ascomycota(100);c__Sordariomycetes(100);o__Hypocreales(100);f__Nectriaceae(100);g__Gibberella(100);s__Gibberella_intricans(100); | Gibberella | 13 | Pathotroph-Saprotroph-Symbiotroph | Endophyte-\|Plant Pathogen\|-Plant Saprotroph-Undefined Saprotroph | Microfungus | Soft Rot | Probable | As foliar_endophyte (Põlme et al. 2020); As litter_saprotroph (Põlme et al. 2020); Causing damping-off disease (Põlme et al. 2020); Decay leaf, fruit, seed, roots, soil, animal material (Põlme et al. 2020) | Tedersoo L, et al. 2014. Science 346:e1256688 ((https://doi.org/10.1126/science.1256688)); Põlme S, et al. 2020. Fungal Diversity 105:1-16 ((https://doi.org/10.1007/s13225-020-00466-2)) |
| Otu06418 | 4 | k__Fungi(100);p__Ascomycota(100);c__Saccharomycetes(100);o__Saccharomycetales(100);f__Metschnikowiaceae(100);g__Metschnikowia(100);s__Metschnikowia_pulcherrima(75); | Metschnikowia | 13 | Saprotroph | \|Undefined Saprotroph\| | Yeast | NULL | Probable | As nectar-tap_saprotroph (Põlme et al. 2020); Decay sugar-rich substrates (Põlme et al. 2020) | Põlme S, et al. 2020. Fungal Diversity 105:1-16 ((https://doi.org/10.1007/s13225-020-00466-2)) |
| Otu06419 | 4 | k__Fungi(100);p__Ascomycota(100);c__Saccharomycetes(100);o__Saccharomycetales(100);f__Metschnikowiaceae(100);g__Metschnikowia(100);s__Metschnikowia_pulcherrima(100); | Metschnikowia | 13 | Saprotroph | \|Undefined Saprotroph\| | Yeast | NULL | Probable | As nectar-tap_saprotroph (Põlme et al. 2020); Decay sugar-rich substrates (Põlme et al. 2020) | Põlme S, et al. 2020. Fungal Diversity 105:1-16 ((https://doi.org/10.1007/s13225-020-00466-2)) |
| Otu06420 | 4 | k__Fungi(100);p__Ascomycota(100);c__Saccharomycetes(100);o__Saccharomycetales(100);f__Metschnikowiaceae(100);g__Metschnikowia(100);s__Metschnikowia_pulcherrima(100); | Metschnikowia | 13 | Saprotroph | \|Undefined Saprotroph\| | Yeast | NULL | Probable | As nectar-tap_saprotroph (Põlme et al. 2020); Decay sugar-rich substrates (Põlme et al. 2020) | Põlme S, et al. 2020. Fungal Diversity 105:1-16 ((https://doi.org/10.1007/s13225-020-00466-2)) |
| Otu06421 | 4 | k__Fungi(100);p__Ascomycota(100);c__Saccharomycetes(100);o__Saccharomycetales(100);f__Metschnikowiaceae(100);g__Metschnikowia(100);g__Metschnikowia_unclassified(100); | Metschnikowia | 13 | Saprotroph | \|Undefined Saprotroph\| | Yeast | NULL | Probable | As nectar-tap_saprotroph (Põlme et al. 2020); Decay sugar-rich substrates (Põlme et al. 2020) | Põlme S, et al. 2020. Fungal Diversity 105:1-16 ((https://doi.org/10.1007/s13225-020-00466-2)) |
| Otu06424 | 4 | k__Fungi(100);p__Ascomycota(100);c__Saccharomycetes(100);o__Saccharomycetales(100);f__Saccharomycetales_family_Incertae_sedis(100);g__Candida(100);g__Candida_unclassified(75); | Candida | 13 | Pathotroph-Saprotroph-Symbiotroph | Animal Pathogen-Endophyte-Endosymbiont-Epiphyte-\|Undefined Saprotroph\| | Dimorphic Yeast-Yeast | NULL | Probable | As nectar-tap_saprotroph (Põlme et al. 2020); Decay sugar-rich substrates (Põlme et al. 2020); This asexual genus is polyphyletic and contain species that function very differently depend on what they are. Caution should be taken when interpreting guilds for Candida species. | Manolakaki D, et al. 2010. Virulence. 1: 367-375 ((https://doi.org/10.4161/viru.1.5.12796)); Mekbib SB, et al. 2011. Tropical Plant Pathology 36:233-240 ((https://doi.org/10.1590/S1982-56762011000400004)); Põlme S, et al. 2020. Fungal Diversity 105:1-16 ((https://doi.org/10.1007/s13225-020-00466-2)) |
| Otu06425 | 4 | k__Fungi(100);p__Ascomycota(100);c__Saccharomycetes(100);o__Saccharomycetales(100);f__Metschnikowiaceae(100);g__Metschnikowia(100);s__Metschnikowia_pulcherrima(100); | Metschnikowia | 13 | Saprotroph | \|Undefined Saprotroph\| | Yeast | NULL | Probable | As nectar-tap_saprotroph (Põlme et al. 2020); Decay sugar-rich substrates (Põlme et al. 2020) | Põlme S, et al. 2020. Fungal Diversity 105:1-16 ((https://doi.org/10.1007/s13225-020-00466-2)) |
| Otu06426 | 4 | k__Fungi(100);p__Ascomycota(100);c__Saccharomycetes(100);o__Saccharomycetales(100);f__Metschnikowiaceae(100);g__Metschnikowia(100);s__Metschnikowia_pulcherrima(100); | Metschnikowia | 13 | Saprotroph | \|Undefined Saprotroph\| | Yeast | NULL | Probable | As nectar-tap_saprotroph (Põlme et al. 2020); Decay sugar-rich substrates (Põlme et al. 2020) | Põlme S, et al. 2020. Fungal Diversity 105:1-16 ((https://doi.org/10.1007/s13225-020-00466-2)) |
| Otu06427 | 4 | k__Fungi(100);p__Basidiomycota(100);c__Agaricomycetes(100);o__Geastrales(100);f__Geastraceae(100);g__Geastrum(100);s__Geastrum_triplex(100); | Geastrum | 13 | Saprotroph | \|Plant Saprotroph\|-Undefined Saprotroph | Gasteroid | NULL | Probable | earthstar; Terrestrial, rarely on wood or coprophilous. Most species are presumed saprobes, but at least some may be ectomycorrhizal (Cannon & Kirk 2007); As litter_saprotroph (Põlme et al. 2020); Common name ; Decay leaf, fruit, seed (Põlme et al. 2020) | Grgurinovic CA, Simpson [illegible] |
| Otu06430 | 4 | k__Fungi(100);p__Ascomycota(100);c__Saccharomycetes(100);o__Saccharomycetales(100);f__Metschnikowiaceae(100);g__Metschnikowia(100);s__Metschnikowia_pulcherrima(100); | Metschnikowia | 13 | Saprotroph | \|Undefined Saprotroph\| | Yeast | NULL | Probable | As nectar-tap_saprotroph (Põlme et al. 2020); Decay sugar-rich substrates (Põlme et al. 2020) | Põlme S, et al. 2020. Fungal Diversity 105:1-16 ((https://doi.org/10.1007/s13225-020-00466-2)) |
| Otu06432 | 4 | k__Fungi(100);p__Ascomycota(100);c__Dothideomycetes(100);o__Pleosporales(100);f__Phaeosphaeriaceae(100);f__Phaeosphaeriaceae_unclassified(100);f__Phaeosphaeriaceae_unclassified(100); | Phaeosphaeriaceae | 9 | Pathotroph-Saprotroph | Fungal Parasite-Plant Pathogen-Plant Saprotroph | Microfungus | NULL | Probable | Necrotrophic or saprobic on a wide range of plants, especially monocotyledons; a few species parasitic on other fungi (Cannon & Kirk 2007) | Cannon PF, Kirk PM. [illegible] |
| Otu06436 | 4 | k__Fungi(100);p__Ascomycota(100);c__Saccharomycetes(100);o__Saccharomycetales(100);f__Saccharomycetales_family_Incertae_sedis(100);g__Candida(100);g__Candida_unclassified(100); | Candida | 13 | Pathotroph-Saprotroph-Symbiotroph | Animal Pathogen-Endophyte-Endosymbiont-Epiphyte-\|Undefined Saprotroph\| | Dimorphic Yeast-Yeast | NULL | Probable | As nectar-tap_saprotroph (Põlme et al. 2020); Decay sugar-rich substrates (Põlme et al. 2020); This asexual genus is polyphyletic and contain species that function very differently depend on what they are. Caution should be taken when interpreting guilds for Candida species. | Manolakaki D, et al. 2010. Virulence. 1: 367-375 ((https://doi.org/10.4161/viru.1.5.12796)); Mekbib SB, et al. 2011. Tropical Plant Pathology 36:233-240 ((https://doi.org/10.1590/S1982-56762011000400004)); Põlme S, et al. 2020. Fungal Diversity 105:1-16 ((https://doi.org/10.1007/s13225-020-00466-2)) |
| Otu06437 | 4 | k__Fungi(100);p__Zygomycota(100);c__Zygomycota_class_Incertae_sedis(100);o__Mortierellales(100);f__Mortierellaceae(100);g__Mortierella(100);s__Mortierella_sp(100); | Mortierella | 13 | Saprotroph-Symbiotroph | Endophyte-Plant Saprotroph-\|Undefined Saprotroph\| | Microfungus | NULL | Probable | As chitinolytic (Põlme et al. 2020); As soil_saprotroph (Põlme et al. 2020); Decay roots, soil (Põlme et al. 2020); Saprobic in soil and thus frequently isolated from soil and material contaminated with soil such as hay or rotten silage and spoil tips (Cannon & Kirk 2007) | Cannon PF, Kirk PM. [illegible] |
| Otu06441 | 4 | k__Fungi(100);p__Ascomycota(100);c__Dothideomycetes(100);o__Dothideomycetes_order_Incertae_sedis(100);f__Myxotrichaceae(100);g__Oidiodendron(100);s__Oidiodendron_myxotrichoides(100); | Oidiodendron | 13 | Pathotroph-Saprotroph-Symbiotroph | Endophyte-Ericoid Mycorrhizal-\|Undefined Saprotroph\| | Microfungus | NULL | Probable | As root_endophyte (Põlme et al. 2020); As soil_saprotroph (Põlme et al. 2020); Dark Septate Endophyte; Decay soil (Põlme et al. 2020) | Newsham KK. 2011. [illegible] |
| Otu06442 | 4 | k__Fungi(100);p__Basidiomycota(100);c__Agaricomycetes(100);o__Agaricales(100);f__Agaricales_family_Incertae_sedis(100);g__unclassified_Agaricales(100);s__Agaricales_sp(100); | Agaricales | 7 | Pathotroph-Saprotroph-Symbiotroph | Bryophyte Parasite-Dung Saprotroph-Ectomycorrhizal-Fungal Parasite-Leaf Saprotroph-Plant Parasite-Undefined Saprotroph-Wood Saprotroph | Agaricoid-Gasteroid-Microfungus-Secotioid-Yeast | NULL | Possible | Mushrooms and toadstools, Gill fungi, Agarics... lignicolous, sometimes muscicolous or fungicolous, saprobic, mycorrhizal, rarely parasitic on plants or fungi (Kirk et al. 2008) | Kirk PM et al. 2008. [illegible] |
| Otu06443 | 4 | k__Fungi(100);p__Basidiomycota(100);c__Agaricomycetes(100);o__Agaricales(100);f__Inocybaceae(100);g__Inocybe(100);g__Inocybe_unclassified(100); | Inocybe | 13 | Symbiotroph | \|Ectomycorrhizal\| | Agaricoid | NULL | Highly Probable | Sensu stricto (Matheny 2009). | Rinaldi AC, et al. 2008. [illegible] |
| Otu06444 | 4 | k__Fungi(100);p__Ascomycota(100);c__Leotiomycetes(100);o__Helotiales(100);f__Hyaloscyphaceae(75);f__Hyaloscyphaceae_unclassified(75);f__Hyaloscyphaceae_unclassified(75); | Hyaloscyphaceae | 9 | Saprotroph | Plant Saprotroph-Wood Saprotroph | Microfungus | NULL | Probable | Saprobic on woody and herbaceous material (Cannon & Kirk 2007) | Cannon PF, Kirk PM. [illegible] |
| Otu06447 | 4 | k__Fungi(100);p__Ascomycota(100);c__Saccharomycetes(100);o__Saccharomycetales(100);f__Metschnikowiaceae(100);g__Metschnikowia(100);s__Metschnikowia_pulcherrima(75); | Metschnikowia | 13 | Saprotroph | \|Undefined Saprotroph\| | Yeast | NULL | Probable | As nectar-tap_saprotroph (Põlme et al. 2020); Decay sugar-rich substrates (Põlme et al. 2020) | Põlme S, et al. 2020. Fungal Diversity 105:1-16 ((https://doi.org/10.1007/s13225-020-00466-2)) |
| Otu06449 | 4 | k__Fungi(100);p__Ascomycota(100);c__Saccharomycetes(100);o__Saccharomycetales(100);f__Metschnikowiaceae(100);g__Metschnikowia(100);s__Metschnikowia_pulcherrima(75); | Metschnikowia | 13 | Saprotroph | \|Undefined Saprotroph\| | Yeast | NULL | Probable | As nectar-tap_saprotroph (Põlme et al. 2020); Decay sugar-rich substrates (Põlme et al. 2020) | Põlme S, et al. 2020. Fungal Diversity 105:1-16 ((https://doi.org/10.1007/s13225-020-00466-2)) |
| Otu06450 | 4 | k__Fungi(100);p__Ascomycota(100);c__Saccharomycetes(100);o__Saccharomycetales(100);f__Metschnikowiaceae(100);g__Metschnikowia(100);g__Metschnikowia_unclassified(100); | Metschnikowia | 13 | Saprotroph | \|Undefined Saprotroph\| | Yeast | NULL | Probable | As nectar-tap_saprotroph (Põlme et al. 2020); Decay sugar-rich substrates (Põlme et al. 2020) | Põlme S, et al. 2020. Fungal Diversity 105:1-16 ((https://doi.org/10.1007/s13225-020-00466-2)) |
| Otu06451 | 4 | k__Fungi(100);p__Ascomycota(100);c__Leotiomycetes(100);o__Helotiales(100);f__Helotiaceae(75);g__Hymenoscyphus(75);g__Hymenoscyphus_unclassified(75); | Hymenoscyphus | 13 | Pathotroph-Saprotroph-Symbiotroph | Bryophyte Parasite-Ectomycorrhizal-Ericoid Mycorrhizal-Plant Pathogen-\|Plant Saprotroph\|-Undefined Saprotroph | Microfungus | NULL | Possible | As litter_saprotroph (Põlme et al. 2020); Dark Septate Microfungus; Syn. Pezoloma among other (see e.g., Pezoloma ericae); Decay leaf, fruit, seed (Põlme et al. 2020) | Baral HO, Krieglsteiner L [illegible] |
| Otu06453 | 4 | k__Fungi(100);p__Basidiomycota(100);c__Agaricomycetes(100);o__Agaricales(100);f__Cortinariaceae(100);g__Cortinarius(100);g__Cortinarius_unclassified(100); | Cortinarius | 13 | Symbiotroph | \|Ectomycorrhizal\| | Agaricoid | NULL | Highly Probable | NULL | Rinaldi AC, et al. 2008. [illegible] |
| Otu06454 | 4 | k__Fungi(100);p__Ascomycota(100);c__Sordariomycetes(100);o__Xylariales(100);f__Xylariaceae(100);g__Nemania(100);s__Nemania_plumbea(100); | Nemania | 13 | Saprotroph | Undefined Saprotroph-\|Wood Saprotroph\| | Microfungus | NULL | Probable | Decay wood (Põlme et al. 2020) | Tedersoo L, et al. 2014. Science 346:e1256688 ((https://doi.org/10.1126/science.1256688)); Põlme S, et al. 2020. Fungal Diversity 105:1-16 ((https://doi.org/10.1007/s13225-020-00466-2)) |
| Otu06455 | 4 | k__Fungi(100);p__Ascomycota(100);c__Leotiomycetes(100);o__Helotiales(100);f__Vibrisseaceae(100);g__Phialocephala(100);g__Phialocephala_unclassified(75); | Phialocephala | 13 | Saprotroph-Symbiotroph | Endophyte-\|Undefined Saprotroph\| | Microfungus | NULL | Highly Probable | As root_endophyte (Põlme et al. 2020); As root_endophyte_dark_septate (Põlme et al. 2020); As soil_saprotroph (Põlme et al. 2020); Dark Septate Endophyte, known from plant roots.; Decay roots, soil (Põlme et al. 2020) | Newsham KK. 2011. [illegible] |
| Otu06456 | 4 | k__Fungi(100);p__Basidiomycota(100);c__Microbotryomycetes(100);o__Sporidiobolales(100);f__Sporidiobolales_family_Incertae_sedis(100);g__Rhodotorula(100);s__Rhodotorula_mucilaginosa(100); | Rhodotorula | 13 | Pathotroph-Saprotroph-Symbiotroph | Animal Pathogen-Animal Symbiotroph-Endophyte-Plant Pathogen-\|Undefined Saprotroph\| | Yeast | NULL | Probable | As foliar_endophyte (Põlme et al. 2020); Opportunistic human pathogen | Põlme S, et al. 2020. Fungal Diversity 105:1-16 ((https://doi.org/10.1007/s13225-020-00466-2)); https://www.cdc.gov ((https://www.cdc.gov)) |
| Otu06457 | 4 | k__Fungi(100);p__Basidiomycota(100);c__Agaricomycetes(100);o__Agaricales(100);o__Agaricales_unclassified(100);o__Agaricales_unclassified(100);o__Agaricales_unclassified(100); | Agaricales | 7 | Pathotroph-Saprotroph-Symbiotroph | Bryophyte Parasite-Dung Saprotroph-Ectomycorrhizal-Fungal Parasite-Leaf Saprotroph-Plant Parasite-Undefined Saprotroph-Wood Saprotroph | Agaricoid-Gasteroid-Microfungus-Secotioid-Yeast | NULL | Possible | Mushrooms and toadstools, Gill fungi, Agarics... lignicolous, sometimes muscicolous or fungicolous, saprobic, mycorrhizal, rarely parasitic on plants or fungi (Kirk et al. 2008) | Kirk PM et al. 2008. [illegible] |
| Otu06458 | 4 | k__Fungi(100);p__Basidiomycota(100);c__Agaricomycetes(100);o__Agaricales(100);f__Psathyrellaceae(100);g__Coprinopsis(100);s__Coprinopsis_insignis(100); | Coprinopsis | 13 | Saprotroph | \|Undefined Saprotroph\| | Agaricoid | NULL | Probable | As soil_saprotroph (Põlme et al. 2020); Decay soil (Põlme et al. 2020) | Tedersoo L, et al. 2014. Science 346:e1256688 ((https://doi.org/10.1126/science.1256688)); Põlme S, et al. 2020. Fungal Diversity 105:1-16 ((https://doi.org/10.1007/s13225-020-00466-2)) |
| Otu06459 | 4 | k__Fungi(100);p__Basidiomycota(100);c__Agaricomycetes(100);o__Agaricales(100);f__Strophariaceae(100);f__Strophariaceae_unclassified(100);f__Strophariaceae_unclassified(100); | Strophariaceae | 9 | Pathotroph-Saprotroph | Dung Saprotroph-Plant Parasite-Soil Saprotroph-Undefined Saprotroph-Wood Saprotroph | Agaricoid | NULL | Probable | Very variable ecology; Pholiota species are lignicolous and can act as weak parasites while most other taxa are either sparobes in grassy habitats or are found in dung. Species are not ectomycorrhizal (Cannon & Kirk 2007) | Cannon PF, Kirk PM. [illegible] |
| Otu06460 | 4 | k__Fungi(100);p__Basidiomycota(100);c__Agaricomycetes(100);o__Agaricales(100);f__Cortinariaceae(100);g__Cortinarius(100);g__Cortinarius_unclassified(100); | Cortinarius | 13 | Symbiotroph | \|Ectomycorrhizal\| | Agaricoid | NULL | Highly Probable | NULL | Rinaldi AC, et al. 2008. [illegible] |
| Otu06465 | 4 | k__Fungi(100);p__Ascomycota(100);c__Sordariomycetes(100);o__Hypocreales(100);f__Nectriaceae(100);g__Microcera(100);s__Microcera_rubra(100); | Microcera | 13 | Pathotroph-Saprotroph | \|Animal Parasite\|-Undefined Saprotroph | Microfungus | NULL | Probable | As animal_decomposer (Põlme et al. 2020); Decay animal material (Põlme et al. 2020) | Põlme S, et al. 2020. Fungal Diversity 105:1-16 ((https://doi.org/10.1007/s13225-020-00466-2)) |
| Otu06467 | 4 | k__Fungi(100);p__Ascomycota(100);c__Saccharomycetes(100);o__Saccharomycetales(100);f__Metschnikowiaceae(100);g__Metschnikowia(100);g__Metschnikowia_unclassified(100); | Metschnikowia | 13 | Saprotroph | \|Undefined Saprotroph\| | Yeast | NULL | Probable | As nectar-tap_saprotroph (Põlme et al. 2020); Decay sugar-rich substrates (Põlme et al. 2020) | Põlme S, et al. 2020. Fungal Diversity 105:1-16 ((https://doi.org/10.1007/s13225-020-00466-2)) |
| Otu06468 | 4 | k__Fungi(100);p__Basidiomycota(100);c__Agaricomycetes(100);o__Hymenochaetales(100);f__Hymenochaetaceae(100);g__Coltriciella(100);s__Coltriciella_sp(100); | Coltriciella | 13 | Symbiotroph | \|Ectomycorrhizal\| | Polyporoid | NULL | Probable | NULL | Gilbertson RL, Ryvarden L. [illegible] |
| Otu06473 | 4 | k__Fungi(100);p__Basidiomycota(100);c__Microbotryomycetes(100);o__Sporidiobolales(100);f__Sporidiobolales_family_Incertae_sedis(100);g__Sporobolomyces(100);s__Sporobolomyces_lactosus(100); | Sporobolomyces | 13 | Pathotroph-Saprotroph | \|Fungal Parasite\|-Plant Saprotroph | Yeast | NULL | Probable | As litter_saprotroph (Põlme et al. 2020); As mycoparasite (Põlme et al. 2020); Decay fungal material (Põlme et al. 2020) | Tedersoo L, et al. 2014. [illegible] |
| Otu06474 | 4 | k__Fungi(100);p__Basidiomycota(100);c__Agaricomycetes(100);o__Agaricales(100);f__Agaricaceae(100);f__Agaricaceae_unclassified(100);f__Agaricaceae_unclassified(100); | Agaricaceae | 9 | Saprotroph | Undefined Saprotroph | Agaricoid-Gasteroid-Secotioid | NULL | Probable | Primarily saprobes in grassland and woodland situations (Cannon & Kirk 2007) | Cannon PF, Kirk PM. [illegible] |
| Otu06484 | 4 | k__Fungi(100);p__Ascomycota(100);c__Saccharomycetes(100);o__Saccharomycetales(100);f__Metschnikowiaceae(100);g__Metschnikowia(100);g__Metschnikowia_unclassified(100); | Metschnikowia | 13 | Saprotroph | \|Undefined Saprotroph\| | Yeast | NULL | Probable | As nectar-tap_saprotroph (Põlme et al. 2020); Decay sugar-rich substrates (Põlme et al. 2020) | Põlme S, et al. 2020. Fungal Diversity 105:1-16 ((https://doi.org/10.1007/s13225-020-00466-2)) |
| Otu06485 | 4 | k__Fungi(100);p__Ascomycota(100);c__Saccharomycetes(100);o__Saccharomycetales(100);f__Metschnikowiaceae(100);g__Metschnikowia(100);s__Metschnikowia_pulcherrima(100); | Metschnikowia | 13 | Saprotroph | \|Undefined Saprotroph\| | Yeast | NULL | Probable | As nectar-tap_saprotroph (Põlme et al. 2020); Decay sugar-rich substrates (Põlme et al. 2020) | Põlme S, et al. 2020. Fungal Diversity 105:1-16 ((https://doi.org/10.1007/s13225-020-00466-2)) |
| Otu06488 | 4 | k__Fungi(100);p__Ascomycota(100);c__Dothideomycetes(100);o__Pleosporales(100);f__Pleosporales_family_Incertae_sedis(100);g__Phoma(75);g__Phoma_unclassified(75); | Phoma | 13 | Pathotroph-Saprotroph-Symbiotroph | Dung Saprotroph-Endophyte-Lichen Parasite-\|Plant Pathogen\|-Plant Saprotroph-Undefined Saprotroph-Wood Saprotroph | Microfungus | Soft Rot | Possible | As foliar_endophyte (Põlme et al. 2020); As litter_saprotroph (Põlme et al. 2020); Decay leaf, fruit, seed, roots, soil (Põlme et al. 2020) | Richardson MJ. 1972. [illegible] |
| Otu06494 | 4 | k__Fungi(100);p__Basidiomycota(100);c__Microbotryomycetes(100);o__Sporidiobolales(100);f__Sporidiobolales_family_Incertae_sedis(100);g__Rhodotorula(100);g__Rhodotorula_unclassified(100); | Rhodotorula | 13 | Pathotroph-Saprotroph-Symbiotroph | Animal Pathogen-Animal Symbiotroph-Endophyte-Plant Pathogen-\|Undefined Saprotroph\| | Yeast | NULL | Probable | As foliar_endophyte (Põlme et al. 2020); Opportunistic human pathogen | Põlme S, et al. 2020. Fungal Diversity 105:1-16 ((https://doi.org/10.1007/s13225-020-00466-2)); https://www.cdc.gov ((https://www.cdc.gov)) |
| Otu06496 | 4 | k__Fungi(100);p__Basidiomycota(100);c__Agaricomycetes(100);o__Cantharellales(100);f__Ceratobasidiaceae(100);g__Ceratobasidium(100);g__Ceratobasidium_unclassified(100); | Ceratobasidium | 13 | Pathotroph-Saprotroph-Symbiotroph | Ectomycorrhizal-Endomycorrhizal-Orchid Mycorrhizal-\|Plant Pathogen\|-Plant Saprotroph-Undefined Saprotroph | Corticioid-Microfungus | NULL | Possible | especially of Orchidaceae but also even bryophytes (Cannon & Kirk 2007); As litter_saprotroph (Põlme et al. 2020); Best known as plant pathogens, but also saprobic. Species are endomycorrhizal with a wide range of plant roots ; Decay leaf, fruit, seed, roots, soil (Põlme et al. 2020) | Cannon PF, Kirk PM. [illegible] |
| Otu06501 | 4 | k__Fungi(100);p__Basidiomycota(100);c__Agaricomycetes(100);o__Polyporales(100);f__Meruliaceae(100);g__Hyphoderma(100);s__Hyphoderma_mutatum(100); | Hyphoderma | 13 | Saprotroph | Undefined Saprotroph-\|Wood Saprotroph\| | Corticioid | White Rot | Probable | As white_rot (Põlme et al. 2020); Decay wood (Põlme et al. 2020) | Tedersoo L, et al. 2014. Science 346:e1256688 ((https://doi.org/10.1126/science.1256688)); Põlme S, et al. 2020. Fungal Diversity 105:1-16 ((https://doi.org/10.1007/s13225-020-00466-2)) |
| Otu06505 | 4 | k__Fungi(100);p__Ascomycota(100);c__Saccharomycetes(100);o__Saccharomycetales(100);f__Metschnikowiaceae(100);g__Metschnikowia(100);g__Metschnikowia_unclassified(100); | Metschnikowia | 13 | Saprotroph | \|Undefined Saprotroph\| | Yeast | NULL | Probable | As nectar-tap_saprotroph (Põlme et al. 2020); Decay sugar-rich substrates (Põlme et al. 2020) | Põlme S, et al. 2020. Fungal Diversity 105:1-16 ((https://doi.org/10.1007/s13225-020-00466-2)) |
| Otu06508 | 4 | k__Fungi(100);p__Basidiomycota(100);c__Microbotryomycetes(100);o__Sporidiobolales(100);f__Sporidiobolales_family_Incertae_sedis(100);g__Rhodotorula(100);s__Rhodotorula_taiwanensis(100); | Rhodotorula | 13 | Pathotroph-Saprotroph-Symbiotroph | Animal Pathogen-Animal Symbiotroph-Endophyte-Plant Pathogen-\|Undefined Saprotroph\| | Yeast | NULL | Probable | As foliar_endophyte (Põlme et al. 2020); Opportunistic human pathogen | Põlme S, et al. 2020. Fungal Diversity 105:1-16 ((https://doi.org/10.1007/s13225-020-00466-2)); https://www.cdc.gov ((https://www.cdc.gov)) |
| Otu06510 | 4 | k__Fungi(100);p__Basidiomycota(100);c__Exobasidiomycetes(100);o__Microstromatales(100);f__Microstromataceae(100);g__Microstroma(100);s__Microstroma_juglandis(100); | Microstroma | 13 | Pathotroph | \|Plant Pathogen\| | Dimorphic | NULL | Probable | NULL | Tedersoo L, et al. 2014. Science 346:e1256688 ((https://doi.org/10.1126/science.1256688)); Põlme S, et al. 2020. Fungal Diversity 105:1-16 ((https://doi.org/10.1007/s13225-020-00466-2)) |
| Otu06511 | 4 | k__Fungi(100);p__Ascomycota(100);c__Saccharomycetes(100);o__Saccharomycetales(100);f__Saccharomycetales_family_Incertae_sedis(75);g__Candida(75);g__Candida_unclassified(75); | Candida | 13 | Pathotroph-Saprotroph-Symbiotroph | Animal Pathogen-Endophyte-Endosymbiont-Epiphyte-\|Undefined Saprotroph\| | Dimorphic Yeast-Yeast | NULL | Probable | As nectar-tap_saprotroph (Põlme et al. 2020); Decay sugar-rich substrates (Põlme et al. 2020); This asexual genus is polyphyletic and contain species that function very differently depend on what they are. Caution should be taken when interpreting guilds for Candida species. | Manolakaki D, et al. 2010. Virulence. 1: 367-375 ((https://doi.org/10.4161/viru.1.5.12796)); Mekbib SB, et al. 2011. Tropical Plant Pathology 36:233-240 ((https://doi.org/10.1590/S1982-56762011000400004)); Põlme S, et al. 2020. Fungal Diversity 105:1-16 ((https://doi.org/10.1007/s13225-020-00466-2)) |
| Otu06523 | 4 | k__Fungi(100);p__Ascomycota(100);c__Saccharomycetes(100);o__Saccharomycetales(100);f__Metschnikowiaceae(100);g__Metschnikowia(100);s__Metschnikowia_pulcherrima(100); | Metschnikowia | 13 | Saprotroph | \|Undefined Saprotroph\| | Yeast | NULL | Probable | As nectar-tap_saprotroph (Põlme et al. 2020); Decay sugar-rich substrates (Põlme et al. 2020) | Põlme S, et al. 2020. Fungal Diversity 105:1-16 ((https://doi.org/10.1007/s13225-020-00466-2)) |
| Otu06527 | 4 | k__Fungi(100);p__Ascomycota(100);c__Dothideomycetes(100);o__Pleosporales(100);f__Venturiaceae(100);f__Venturiaceae_unclassified(75);f__Venturiaceae_unclassified(75); | Venturiaceae | 9 | Pathotroph-Saprotroph-Symbiotroph | Endophyte-Plant Pathogen-Undefined Saprotroph | Microfungus | NULL | Probable | Mostly endophytes and weak parasites of aerial plant parts, including bark and wood, leaf and fruit tissue; rarely biotrophic. Little is known except for the important pathogens (Cannon & Kirk 2007) | Cannon PF, Kirk PM. [illegible] |
| Otu06528 | 4 | k__Fungi(100);p__Ascomycota(100);c__Sordariomycetes(100);o__Diaporthales(100);f__Valsaceae(100);f__Valsaceae_unclassified(100);f__Valsaceae_unclassified(100); | Valsaceae | 9 | Pathotroph-Saprotroph-Symbiotroph | Endophyte-Plant Pathogen-Undefined Saprotroph | Microfungus | NULL | Probable | Saprobes, endophytes and weak to virulent parasites of woody plants (Cannon & Kirk 2007) | Cannon PF, Kirk PM. [illegible] |
| Otu06529 | 4 | k__Fungi(100);p__Ascomycota(100);c__Saccharomycetes(100);o__Saccharomycetales(100);f__Metschnikowiaceae(100);g__Metschnikowia(100);g__Metschnikowia_unclassified(100); | Metschnikowia | 13 | Saprotroph | \|Undefined Saprotroph\| | Yeast | NULL | Probable | As nectar-tap_saprotroph (Põlme et al. 2020); Decay sugar-rich substrates (Põlme et al. 2020) | Põlme S, et al. 2020. Fungal Diversity 105:1-16 ((https://doi.org/10.1007/s13225-020-00466-2)) |
| Otu06531 | 4 | k__Fungi(100);p__Basidiomycota(100);c__Microbotryomycetes(100);o__Sporidiobolales(100);f__Sporidiobolales_family_Incertae_sedis(100);g__Rhodotorula(75);g__Rhodotorula_unclassified(75); | Rhodotorula | 13 | Pathotroph-Saprotroph-Symbiotroph | Animal Pathogen-Animal Symbiotroph-Endophyte-Plant Pathogen-\|Undefined Saprotroph\| | Yeast | NULL | Probable | As foliar_endophyte (Põlme et al. 2020); Opportunistic human pathogen | Põlme S, et al. 2020. Fungal Diversity 105:1-16 ((https://doi.org/10.1007/s13225-020-00466-2)); https://www.cdc.gov ((https://www.cdc.gov)) |
| Otu06533 | 4 | k__Fungi(100);p__Basidiomycota(100);c__Agaricomycetes(100);o__Agaricales(100);o__Agaricales_unclassified(100);o__Agaricales_unclassified(100);o__Agaricales_unclassified(100); | Agaricales | 7 | Pathotroph-Saprotroph-Symbiotroph | Bryophyte Parasite-Dung Saprotroph-Ectomycorrhizal-Fungal Parasite-Leaf Saprotroph-Plant Parasite-Undefined Saprotroph-Wood Saprotroph | Agaricoid-Gasteroid-Microfungus-Secotioid-Yeast | NULL | Possible | Mushrooms and toadstools, Gill fungi, Agarics... lignicolous, sometimes muscicolous or fungicolous, saprobic, mycorrhizal, rarely parasitic on plants or fungi (Kirk et al. 2008) | Kirk PM et al. 2008. [illegible] |
| Otu06534 | 4 | k__Fungi(100);p__Basidiomycota(100);c__Tremellomycetes(100);o__Tremellales(100);f__Tremellales_family_Incertae_sedis(100);g__Dioszegia(100);s__Dioszegia_zsoltii_var._zsoltii(100); | Dioszegia | 13 | Saprotroph | \|Plant Saprotroph\|-Undefined Saprotroph | Yeast | NULL | Probable | As litter_saprotroph (Põlme et al. 2020); Decay leaf, fruit, seed (Põlme et al. 2020); The Dioszegia subclade sensu Inacio (Inacio et al. 20015) is characterized by orange; yellow or orange to deep orange colonies. Isolated from plant leaves and soil. | Põlme S, et al. 2020. [illegible] |
| Otu06535 | 4 | k__Fungi(100);p__Ascomycota(100);c__Saccharomycetes(100);o__Saccharomycetales(100);f__Metschnikowiaceae(100);g__Metschnikowia(100);g__Metschnikowia_unclassified(75); | Metschnikowia | 13 | Saprotroph | \|Undefined Saprotroph\| | Yeast | NULL | Probable | As nectar-tap_saprotroph (Põlme et al. 2020); Decay sugar-rich substrates (Põlme et al. 2020) | Põlme S, et al. 2020. Fungal Diversity 105:1-16 ((https://doi.org/10.1007/s13225-020-00466-2)) |
| Otu06536 | 4 | k__Fungi(100);p__Ascomycota(100);c__Saccharomycetes(100);o__Saccharomycetales(100);f__Saccharomycetales_family_Incertae_sedis(100);f__Saccharomycetales_family_Incertae_sedis_unclassified(75);f__Saccharomycetales_family_Incertae_sedis_unclassified(75); | Saccharomycetales | 7 | Saprotroph | Undefined Saprotroph | Yeast | NULL | Possible | NULL | Sterkenburg E, et al. 201[illegible] |
| Otu06537 | 4 | k__Fungi(100);p__Ascomycota(100);c__Sordariomycetes(100);o__Sordariales(100);f__Cephalothecaceae(100);g__Phialemonium(100);s__Phialemonium_atrogriseum(100); | Phialemonium | 13 | Pathotroph-Saprotroph | \|Animal Parasite\|-Undefined Saprotroph | Microfungus | NULL | Probable | As animal_decomposer (Põlme et al. 2020); Decay animal material (Põlme et al. 2020) | Tedersoo L, et al. 2014. Science 346:e1256688 ((https://doi.org/10.1126/science.1256688)); Põlme S, et al. 2020. Fungal Diversity 105:1-16 ((https://doi.org/10.1007/s13225-020-00466-2)) |
| Otu06538 | 4 | k__Fungi(100);p__Ascomycota(100);c__Saccharomycetes(100);o__Saccharomycetales(100);f__Metschnikowiaceae(100);g__Metschnikowia(100);g__Metschnikowia_unclassified(100); | Metschnikowia | 13 | Saprotroph | \|Undefined Saprotroph\| | Yeast | NULL | Probable | As nectar-tap_saprotroph (Põlme et al. 2020); Decay sugar-rich substrates (Põlme et al. 2020) | Põlme S, et al. 2020. Fungal Diversity 105:1-16 ((https://doi.org/10.1007/s13225-020-00466-2)) |
| Otu06539 | 4 | k__Fungi(100);p__Ascomycota(100);c__Saccharomycetes(100);o__Saccharomycetales(100);f__Metschnikowiaceae(100);g__Metschnikowia(100);g__Metschnikowia_unclassified(75); | Metschnikowia | 13 | Saprotroph | \|Undefined Saprotroph\| | Yeast | NULL | Probable | As nectar-tap_saprotroph (Põlme et al. 2020); Decay sugar-rich substrates (Põlme et al. 2020) | Põlme S, et al. 2020. Fungal Diversity 105:1-16 ((https://doi.org/10.1007/s13225-020-00466-2)) |
| Otu06540 | 4 | k__Fungi(100);p__Ascomycota(100);c__Saccharomycetes(100);o__Saccharomycetales(100);f__Metschnikowiaceae(100);g__Metschnikowia(100);s__Metschnikowia_pulcherrima(100); | Metschnikowia | 13 | Saprotroph | \|Undefined Saprotroph\| | Yeast | NULL | Probable | As nectar-tap_saprotroph (Põlme et al. 2020); Decay sugar-rich substrates (Põlme et al. 2020) | Põlme S, et al. 2020. Fungal Diversity 105:1-16 ((https://doi.org/10.1007/s13225-020-00466-2)) |
| Otu06541 | 4 | k__Fungi(100);p__Ascomycota(100);c__Saccharomycetes(100);o__Saccharomycetales(100);f__Saccharomycetales_family_Incertae_sedis(100);g__Candida(100);g__Candida_unclassified(100); | Candida | 13 | Pathotroph-Saprotroph-Symbiotroph | Animal Pathogen-Endophyte-Endosymbiont-Epiphyte-\|Undefined Saprotroph\| | Dimorphic Yeast-Yeast | NULL | Probable | As nectar-tap_saprotroph (Põlme et al. 2020); Decay sugar-rich substrates (Põlme et al. 2020); This asexual genus is polyphyletic and contain species that function very differently depend on what they are. Caution should be taken when interpreting guilds for Candida species. | Manolakaki D, et al. 2010. Virulence. 1: 367-375 ((https://doi.org/10.4161/viru.1.5.12796)); Mekbib SB, et al. 2011. Tropical Plant Pathology 36:233-240 ((https://doi.org/10.1590/S1982-56762011000400004)); Põlme S, et al. 2020. Fungal Diversity 105:1-16 ((https://doi.org/10.1007/s13225-020-00466-2)) |
| Otu06543 | 4 | k__Fungi(100);p__Ascomycota(100);c__Dothideomycetes(100);o__Dothideomycetes_order_Incertae_sedis(100);f__Dothideomycetes_family_Incertae_sedis(100);g__Leptospora(100);s__Leptospora_rubella(100); | Leptospora | 13 | Saprotroph | \|Plant Saprotroph\| | Microfungus | NULL | Probable | As litter_saprotroph (Põlme et al. 2020); Decay leaf, fruit, seed (Põlme et al. 2020) | Põlme S, et al. 2020. Fungal Diversity 105:1-16 ((https://doi.org/10.1007/s13225-020-00466-2)) |
| Otu06544 | 4 | k__Fungi(100);p__Ascomycota(100);c__Saccharomycetes(100);o__Saccharomycetales(100);f__Metschnikowiaceae(100);g__Metschnikowia(100);g__Metschnikowia_unclassified(100); | Metschnikowia | 13 | Saprotroph | \|Undefined Saprotroph\| | Yeast | NULL | Probable | As nectar-tap_saprotroph (Põlme et al. 2020); Decay sugar-rich substrates (Põlme et al. 2020) | Põlme S, et al. 2020. Fungal Diversity 105:1-16 ((https://doi.org/10.1007/s13225-020-00466-2)) |
| Otu06545 | 4 | k__Fungi(100);p__Ascomycota(100);c__Saccharomycetes(100);o__Saccharomycetales(100);f__Metschnikowiaceae(100);g__Metschnikowia(100);s__Metschnikowia_pulcherrima(75); | Metschnikowia | 13 | Saprotroph | \|Undefined Saprotroph\| | Yeast | NULL | Probable | As nectar-tap_saprotroph (Põlme et al. 2020); Decay sugar-rich substrates (Põlme et al. 2020) | Põlme S, et al. 2020. Fungal Diversity 105:1-16 ((https://doi.org/10.1007/s13225-020-00466-2)) |
| Otu06548 | 4 | k__Fungi(100);p__Ascomycota(100);c__Saccharomycetes(100);o__Saccharomycetales(100);f__Metschnikowiaceae(100);g__Metschnikowia(100);s__Metschnikowia_pulcherrima(75); | Metschnikowia | 13 | Saprotroph | \|Undefined Saprotroph\| | Yeast | NULL | Probable | As nectar-tap_saprotroph (Põlme et al. 2020); Decay sugar-rich substrates (Põlme et al. 2020) | Põlme S, et al. 2020. Fungal Diversity 105:1-16 ((https://doi.org/10.1007/s13225-020-00466-2)) |
| Otu06549 | 4 | k__Fungi(100);p__Basidiomycota(100);c__Agaricomycetes(100);o__Agaricales(100);o__Agaricales_unclassified(100);o__Agaricales_unclassified(100);o__Agaricales_unclassified(100); | Agaricales | 7 | Pathotroph-Saprotroph-Symbiotroph | Bryophyte Parasite-Dung Saprotroph-Ectomycorrhizal-Fungal Parasite-Leaf Saprotroph-Plant Parasite-Undefined Saprotroph-Wood Saprotroph | Agaricoid-Gasteroid-Microfungus-Secotioid-Yeast | NULL | Possible | Mushrooms and toadstools, Gill fungi, Agarics... lignicolous, sometimes muscicolous or fungicolous, saprobic, mycorrhizal, rarely parasitic on plants or fungi (Kirk et al. 2008) | Kirk PM et al. 2008. [illegible] |
| Otu06550 | 4 | k__Fungi(100);p__Ascomycota(100);c__Eurotiomycetes(100);o__Chaetothyriales(100);f__Herpotrichiellaceae(100);g__Cladophialophora(100);g__Cladophialophora_unclassified(100); | Cladophialophora | 13 | Pathotroph-Saprotroph-Symbiotroph | Animal Pathogen-Endophyte-Fungal Parasite-\|Undefined Saprotroph\| | Dimorphic-Microfungus | NULL | Probable | As mycoparasite (Põlme et al. 2020); As root_endophyte_dark_septate (Põlme et al. 2020); As soil_saprotroph (Põlme et al. 2020); Decay soil (Põlme et al. 2020) | James TY, et al. 2006. [illegible] |
| Otu06551 | 4 | k__Fungi(100);p__Ascomycota(100);c__Saccharomycetes(100);o__Saccharomycetales(100);f__Metschnikowiaceae(100);g__Metschnikowia(100);g__Metschnikowia_unclassified(100); | Metschnikowia | 13 | Saprotroph | \|Undefined Saprotroph\| | Yeast | NULL | Probable | As nectar-tap_saprotroph (Põlme et al. 2020); Decay sugar-rich substrates (Põlme et al. 2020) | Põlme S, et al. 2020. Fungal Diversity 105:1-16 ((https://doi.org/10.1007/s13225-020-00466-2)) |
| Otu06553 | 4 | k__Fungi(100);p__Ascomycota(100);c__Sordariomycetes(100);o__Xylariales(100);f__Xylariaceae(100);g__Nodulisporium(100);s__Nodulisporium_sp(100); | Nodulisporium | 13 | Saprotroph | \|Undefined Saprotroph\| | Microfungus | NULL | Probable | NULL | Tedersoo L, et al. 2014. Science 346:e1256688 ((https://doi.org/10.1126/science.1256688)); Põlme S, et al. 2020. Fungal Diversity 105:1-16 ((https://doi.org/10.1007/s13225-020-00466-2)) |
| Otu06554 | 4 | k__Fungi(100);p__Basidiomycota(100);c__Agaricomycetes(100);o__Agaricales(100);f__Entolomataceae(100);g__Nolanea(100);s__Nolanea_strictior(100); | Nolanea | 13 | Saprotroph-Symbiotroph | \|Undefined Saprotroph\|-Undefined Symbiotroph | Agaricoid | NULL | Probable | As soil_saprotroph (Põlme et al. 2020); Decay roots, soil (Põlme et al. 2020) | Tedersoo L, et al. 2014. Science 346:e1256688 ((https://doi.org/10.1126/science.1256688)); Põlme S, et al. 2020. Fungal Diversity 105:1-16 ((https://doi.org/10.1007/s13225-020-00466-2)) |
| Otu06555 | 4 | k__Fungi(100);p__Ascomycota(100);c__Sordariomycetes(100);o__Sordariales(100);f__Lasiosphaeriaceae(100);g__Cladorrhinum(100);s__Cladorrhinum_sp(100); | Cladorrhinum | 13 | Pathotroph-Saprotroph-Symbiotroph | Animal Pathogen-Dung Saprotroph-Endophyte-Plant Pathogen-Plant Saprotroph-\|Undefined Saprotroph\|-Wood Saprotroph | Microfungus | Soft Rot | Probable | As foliar_endophyte (Põlme et al. 2020); Decay leaf, fruit, seed (Põlme et al. 2020); Reported in soils (including agricultural) as saprotrophs on dung or plant material, or in roots as endophytes, while some species are associated with human and animal opportunistic diseases (Carmaran et al.2015) | Seehann G, et al. 1975. [illegible] |
| Otu06557 | 4 | k__Fungi(100);p__Basidiomycota(100);c__Microbotryomycetes(100);o__Sporidiobolales(100);f__Sporidiobolales_family_Incertae_sedis(100);g__Rhodotorula(100);g__Rhodotorula_unclassified(100); | Rhodotorula | 13 | Pathotroph-Saprotroph-Symbiotroph | Animal Pathogen-Animal Symbiotroph-Endophyte-Plant Pathogen-\|Undefined Saprotroph\| | Yeast | NULL | Probable | As foliar_endophyte (Põlme et al. 2020); Opportunistic human pathogen | Põlme S, et al. 2020. Fungal Diversity 105:1-16 ((https://doi.org/10.1007/s13225-020-00466-2)); https://www.cdc.gov ((https://www.cdc.gov)) |
| Otu06559 | 4 | k__Fungi(100);p__Basidiomycota(100);c__Agaricomycetes(100);o__Hymenochaetales(100);f__Hymenochaetaceae(100);g__Mensularia(100);s__Mensularia_radiata(100); | Mensularia | 13 | Saprotroph | \|Wood Saprotroph\| | Polyporoid | White Rot | Probable | As white_rot (Põlme et al. 2020); Decay wood (Põlme et al. 2020) | Tedersoo L, et al. 2014. [illegible] |
| Otu06560 | 4 | k__Fungi(100);p__Basidiomycota(100);c__Agaricomycetes(100);o__Agaricales(100);f__Marasmiaceae(100);g__Gymnopus(100);g__Gymnopus_unclassified(100); | Gymnopus | 13 | Saprotroph | \|Plant Saprotroph\| | Agaricoid | NULL | Probable | As litter_saprotroph (Põlme et al. 2020); Decay leaf, fruit, seed (Põlme et al. 2020) | Põlme S, et al. 2020. Fungal Diversity 105:1-16 ((https://doi.org/10.1007/s13225-020-00466-2)) |
| Otu06561 | 4 | k__Fungi(100);p__Ascomycota(100);c__Dothideomycetes(100);o__Pleosporales(100);f__Pleosporales_family_Incertae_sedis(75);g__Boeremia(75);s__Boeremia_exigua_var._populi(75); | Boeremia | 13 | Pathotroph-Symbiotroph | Endophyte-\|Plant Pathogen\| | Microfungus | NULL | Possible | Asteraceae; As foliar_endophyte (Põlme et al. 2020); Decay leaf, fruit, seed (Põlme et al. 2020); Host | Costa IPMW, et al. 2012. [illegible] |
| Otu06563 | 4 | k__Fungi(100);p__Ascomycota(100);c__Saccharomycetes(100);o__Saccharomycetales(100);f__Saccharomycetales_family_Incertae_sedis(100);g__Candida(100);g__Candida_unclassified(100); | Candida | 13 | Pathotroph-Saprotroph-Symbiotroph | Animal Pathogen-Endophyte-Endosymbiont-Epiphyte-\|Undefined Saprotroph\| | Dimorphic Yeast-Yeast | NULL | Probable | As nectar-tap_saprotroph (Põlme et al. 2020); Decay sugar-rich substrates (Põlme et al. 2020); This asexual genus is polyphyletic and contain species that function very differently depend on what they are. Caution should be taken when interpreting guilds for Candida species. | Manolakaki D, et al. 2010. Virulence. 1: 367-375 ((https://doi.org/10.4161/viru.1.5.12796)); Mekbib SB, et al. 2011. Tropical Plant Pathology 36:233-240 ((https://doi.org/10.1590/S1982-56762011000400004)); Põlme S, et al. 2020. Fungal Diversity 105:1-16 ((https://doi.org/10.1007/s13225-020-00466-2)) |
| Otu06564 | 4 | k__Fungi(100);p__Ascomycota(100);c__Saccharomycetes(100);o__Saccharomycetales(100);f__Metschnikowiaceae(100);g__Metschnikowia(100);g__Metschnikowia_unclassified(75); | Metschnikowia | 13 | Saprotroph | \|Undefined Saprotroph\| | Yeast | NULL | Probable | As nectar-tap_saprotroph (Põlme et al. 2020); Decay sugar-rich substrates (Põlme et al. 2020) | Põlme S, et al. 2020. Fungal Diversity 105:1-16 ((https://doi.org/10.1007/s13225-020-00466-2)) |
| Otu06567 | 4 | k__Fungi(100);p__Basidiomycota(100);c__Exobasidiomycetes(100);o__Microstromatales(100);f__Microstromataceae(100);g__Microstroma(100);s__Microstroma_juglandis(100); | Microstroma | 13 | Pathotroph | \|Plant Pathogen\| | Dimorphic | NULL | Probable | NULL | Tedersoo L, et al. 2014. Science 346:e1256688 ((https://doi.org/10.1126/science.1256688)); Põlme S, et al. 2020. Fungal Diversity 105:1-16 ((https://doi.org/10.1007/s13225-020-00466-2)) |
| Otu06568 | 4 | k__Fungi(100);p__Ascomycota(100);c__Saccharomycetes(100);o__Saccharomycetales(100);f__Metschnikowiaceae(100);g__Metschnikowia(100);s__Metschnikowia_pulcherrima(75); | Metschnikowia | 13 | Saprotroph | \|Undefined Saprotroph\| | Yeast | NULL | Probable | As nectar-tap_saprotroph (Põlme et al. 2020); Decay sugar-rich substrates (Põlme et al. 2020) | Põlme S, et al. 2020. Fungal Diversity 105:1-16 ((https://doi.org/10.1007/s13225-020-00466-2)) |
| Otu06570 | 4 | k__Fungi(100);p__Ascomycota(100);c__Saccharomycetes(100);o__Saccharomycetales(100);f__Metschnikowiaceae(100);g__Metschnikowia(100);g__Metschnikowia_unclassified(100); | Metschnikowia | 13 | Saprotroph | \|Undefined Saprotroph\| | Yeast | NULL | Probable | As nectar-tap_saprotroph (Põlme et al. 2020); Decay sugar-rich substrates (Põlme et al. 2020) | Põlme S, et al. 2020. Fungal Diversity 105:1-16 ((https://doi.org/10.1007/s13225-020-00466-2)) |
| Otu06571 | 4 | k__Fungi(100);p__Ascomycota(100);c__Saccharomycetes(100);o__Saccharomycetales(100);f__Metschnikowiaceae(100);g__Metschnikowia(100);g__Metschnikowia_unclassified(100); | Metschnikowia | 13 | Saprotroph | \|Undefined Saprotroph\| | Yeast | NULL | Probable | As nectar-tap_saprotroph (Põlme et al. 2020); Decay sugar-rich substrates (Põlme et al. 2020) | Põlme S, et al. 2020. Fungal Diversity 105:1-16 ((https://doi.org/10.1007/s13225-020-00466-2)) |
| Otu06574 | 4 | k__Fungi(100);p__Ascomycota(100);c__Saccharomycetes(100);o__Saccharomycetales(100);f__Metschnikowiaceae(100);g__Metschnikowia(100);s__Metschnikowia_pulcherrima(75); | Metschnikowia | 13 | Saprotroph | \|Undefined Saprotroph\| | Yeast | NULL | Probable | As nectar-tap_saprotroph (Põlme et al. 2020); Decay sugar-rich substrates (Põlme et al. 2020) | Põlme S, et al. 2020. Fungal Diversity 105:1-16 ((https://doi.org/10.1007/s13225-020-00466-2)) |
| Otu06577 | 4 | k__Fungi(100);p__Ascomycota(100);c__Saccharomycetes(100);o__Saccharomycetales(100);f__Metschnikowiaceae(100);g__Metschnikowia(100);s__Metschnikowia_pulcherrima(75); | Metschnikowia | 13 | Saprotroph | \|Undefined Saprotroph\| | Yeast | NULL | Probable | As nectar-tap_saprotroph (Põlme et al. 2020); Decay sugar-rich substrates (Põlme et al. 2020) | Põlme S, et al. 2020. Fungal Diversity 105:1-16 ((https://doi.org/10.1007/s13225-020-00466-2)) |
| Otu06578 | 4 | k__Fungi(100);p__Basidiomycota(100);c__Agaricomycetes(100);o__Agaricales(100);f__Tricholomataceae(100);g__Callistosporium(100);s__Callistosporium_xanthophyllum(100); | Callistosporium | 13 | Saprotroph | \|Undefined Saprotroph\| | Agaricoid | NULL | Probable | As soil_saprotroph (Põlme et al. 2020); Decay soil (Põlme et al. 2020) | Tedersoo L, et al. 2014. Science 346:e1256688 ((https://doi.org/10.1126/science.1256688)); Põlme S, et al. 2020. Fungal Diversity 105:1-16 ((https://doi.org/10.1007/s13225-020-00466-2)) |
| Otu06579 | 4 | k__Fungi(100);p__Ascomycota(100);c__Saccharomycetes(100);o__Saccharomycetales(100);f__Metschnikowiaceae(100);g__Metschnikowia(100);g__Metschnikowia_unclassified(75); | Metschnikowia | 13 | Saprotroph | \|Undefined Saprotroph\| | Yeast | NULL | Probable | As nectar-tap_saprotroph (Põlme et al. 2020); Decay sugar-rich substrates (Põlme et al. 2020) | Põlme S, et al. 2020. Fungal Diversity 105:1-16 ((https://doi.org/10.1007/s13225-020-00466-2)) |
| Otu06581 | 4 | k__Fungi(100);p__Basidiomycota(100);c__Agaricomycetes(100);o__Auriculariales(100);f__Auriculariales_family_Incertae_sedis(100);g__Basidiodendron(100);s__Basidiodendron_caesiocinereum(100); | Basidiodendron | 13 | Saprotroph | Undefined Saprotroph-\|Wood Saprotroph\| | Corticioid | NULL | Probable | Decay wood (Põlme et al. 2020) | Tedersoo L, et al. 2014. Science 346:e1256688 ((https://doi.org/10.1126/science.1256688)); Põlme S, et al. 2020. Fungal Diversity 105:1-16 ((https://doi.org/10.1007/s13225-020-00466-2)) |
| Otu06582 | 4 | k__Fungi(100);p__Ascomycota(100);c__Saccharomycetes(100);o__Saccharomycetales(100);f__Metschnikowiaceae(100);g__Metschnikowia(100);s__Metschnikowia_pulcherrima(100); | Metschnikowia | 13 | Saprotroph | \|Undefined Saprotroph\| | Yeast | NULL | Probable | As nectar-tap_saprotroph (Põlme et al. 2020); Decay sugar-rich substrates (Põlme et al. 2020) | Põlme S, et al. 2020. Fungal Diversity 105:1-16 ((https://doi.org/10.1007/s13225-020-00466-2)) |
| Otu06583 | 4 | k__Fungi(100);p__Ascomycota(100);c__Saccharomycetes(100);o__Saccharomycetales(100);f__Metschnikowiaceae(100);g__Metschnikowia(100);g__Metschnikowia_unclassified(75); | Metschnikowia | 13 | Saprotroph | \|Undefined Saprotroph\| | Yeast | NULL | Probable | As nectar-tap_saprotroph (Põlme et al. 2020); Decay sugar-rich substrates (Põlme et al. 2020) | Põlme S, et al. 2020. Fungal Diversity 105:1-16 ((https://doi.org/10.1007/s13225-020-00466-2)) |
| Otu06586 | 4 | k__Fungi(100);p__Ascomycota(100);c__Saccharomycetes(100);o__Saccharomycetales(100);f__Metschnikowiaceae(100);g__Metschnikowia(100);g__Metschnikowia_unclassified(100); | Metschnikowia | 13 | Saprotroph | \|Undefined Saprotroph\| | Yeast | NULL | Probable | As nectar-tap_saprotroph (Põlme et al. 2020); Decay sugar-rich substrates (Põlme et al. 2020) | Põlme S, et al. 2020. Fungal Diversity 105:1-16 ((https://doi.org/10.1007/s13225-020-00466-2)) |
| Otu06587 | 4 | k__Fungi(100);p__Ascomycota(100);c__Dothideomycetes(100);o__Capnodiales(100);f__Mycosphaerellaceae(100);f__Mycosphaerellaceae_unclassified(100);f__Mycosphaerellaceae_unclassified(100); | Mycosphaerellaceae | 9 | Pathotroph-Saprotroph | Plant Pathogen-Undefined Saprotroph | Microfungus | NULL | Probable | Biotrophic, necrotrophic or saprobic on various plant tissues (Cannon & Kirk 2007) | Cannon PF, Kirk PM. [illegible] |
| Otu06588 | 4 | k__Fungi(100);p__Ascomycota(100);c__Saccharomycetes(100);o__Saccharomycetales(100);f__Metschnikowiaceae(100);g__Metschnikowia(100);s__Metschnikowia_pulcherrima(100); | Metschnikowia | 13 | Saprotroph | \|Undefined Saprotroph\| | Yeast | NULL | Probable | As nectar-tap_saprotroph (Põlme et al. 2020); Decay sugar-rich substrates (Põlme et al. 2020) | Põlme S, et al. 2020. Fungal Diversity 105:1-16 ((https://doi.org/10.1007/s13225-020-00466-2)) |
| Otu06589 | 4 | k__Fungi(100);p__Ascomycota(100);c__Saccharomycetes(100);o__Saccharomycetales(100);f__Metschnikowiaceae(100);g__Metschnikowia(100);s__Metschnikowia_pulcherrima(100); | Metschnikowia | 13 | Saprotroph | \|Undefined Saprotroph\| | Yeast | NULL | Probable | As nectar-tap_saprotroph (Põlme et al. 2020); Decay sugar-rich substrates (Põlme et al. 2020) | Põlme S, et al. 2020. Fungal Diversity 105:1-16 ((https://doi.org/10.1007/s13225-020-00466-2)) |
| Otu06592 | 4 | k__Fungi(100);p__Basidiomycota(100);c__Agaricomycetes(100);o__Agaricales(100);f__Fistulinaceae(100);g__Fistulina(100);s__Fistulina_sp_FP_103444_T(100); | Fistulina | 13 | Pathotroph-Saprotroph | \|Plant Pathogen\|-Wood Saprotroph | Polyporoid | Brown Rot | Highly Probable | As brown-rot (Põlme et al. 2020); As wood_pathogen (Põlme et al. 2020); Decay wood (Põlme et al. 2020) | Gilbertson RL, Ryvarden [illegible] |
| Otu06594 | 4 | k__Fungi(100);p__Ascomycota(100);c__Saccharomycetes(100);o__Saccharomycetales(100);f__Metschnikowiaceae(100);g__Metschnikowia(100);s__Metschnikowia_pulcherrima(100); | Metschnikowia | 13 | Saprotroph | \|Undefined Saprotroph\| | Yeast | NULL | Probable | As nectar-tap_saprotroph (Põlme et al. 2020); Decay sugar-rich substrates (Põlme et al. 2020) | Põlme S, et al. 2020. Fungal Diversity 105:1-16 ((https://doi.org/10.1007/s13225-020-00466-2)) |
| Otu06597 | 4 | k__Fungi(100);p__Ascomycota(100);c__Saccharomycetes(100);o__Saccharomycetales(100);f__Metschnikowiaceae(100);g__Metschnikowia(100);s__Metschnikowia_pulcherrima(100); | Metschnikowia | 13 | Saprotroph | \|Undefined Saprotroph\| | Yeast | NULL | Probable | As nectar-tap_saprotroph (Põlme et al. 2020); Decay sugar-rich substrates (Põlme et al. 2020) | Põlme S, et al. 2020. Fungal Diversity 105:1-16 ((https://doi.org/10.1007/s13225-020-00466-2)) |
| Otu06598 | 4 | k__Fungi(100);p__Ascomycota(100);c__Saccharomycetes(100);o__Saccharomycetales(100);f__Metschnikowiaceae(100);g__Metschnikowia(100);s__Metschnikowia_pulcherrima(75); | Metschnikowia | 13 | Saprotroph | \|Undefined Saprotroph\| | Yeast | NULL | Probable | As nectar-tap_saprotroph (Põlme et al. 2020); Decay sugar-rich substrates (Põlme et al. 2020) | Põlme S, et al. 2020. Fungal Diversity 105:1-16 ((https://doi.org/10.1007/s13225-020-00466-2)) |
| Otu06599 | 4 | k__Fungi(100);p__Ascomycota(100);c__Saccharomycetes(100);o__Saccharomycetales(100);f__Metschnikowiaceae(100);g__Metschnikowia(100);s__Metschnikowia_pulcherrima(100); | Metschnikowia | 13 | Saprotroph | \|Undefined Saprotroph\| | Yeast | NULL | Probable | As nectar-tap_saprotroph (Põlme et al. 2020); Decay sugar-rich substrates (Põlme et al. 2020) | Põlme S, et al. 2020. Fungal Diversity 105:1-16 ((https://doi.org/10.1007/s13225-020-00466-2)) |
| Otu06600 | 4 | k__Fungi(100);p__Ascomycota(100);c__Saccharomycetes(100);o__Saccharomycetales(100);f__Metschnikowiaceae(100);g__Metschnikowia(100);s__Metschnikowia_pulcherrima(100); | Metschnikowia | 13 | Saprotroph | \|Undefined Saprotroph\| | Yeast | NULL | Probable | As nectar-tap_saprotroph (Põlme et al. 2020); Decay sugar-rich substrates (Põlme et al. 2020) | Põlme S, et al. 2020. Fungal Diversity 105:1-16 ((https://doi.org/10.1007/s13225-020-00466-2)) |
| Otu06601 | 4 | k__Fungi(100);p__Ascomycota(100);c__Saccharomycetes(100);o__Saccharomycetales(100);f__Metschnikowiaceae(100);g__Metschnikowia(100);s__Metschnikowia_pulcherrima(100); | Metschnikowia | 13 | Saprotroph | \|Undefined Saprotroph\| | Yeast | NULL | Probable | As nectar-tap_saprotroph (Põlme et al. 2020); Decay sugar-rich substrates (Põlme et al. 2020) | Põlme S, et al. 2020. Fungal Diversity 105:1-16 ((https://doi.org/10.1007/s13225-020-00466-2)) |
| Otu06602 | 4 | k__Fungi(100);p__Ascomycota(100);c__Dothideomycetes(100);o__Pleosporales(100);f__Pleosporales_family_Incertae_sedis(100);g__Periconia(100);s__Periconia_sp_CY137(100); | Periconia | 13 | Pathotroph-Saprotroph-Symbiotroph | Endophyte-\|Plant Pathogen\|-Wood Saprotroph | Microfungus | Soft Rot | Possible | Asteraceae, Musaceae (Costa et al. 2012); As foliar_endophyte (Põlme et al. 2020); Dark Septate Endophyte, known from plant roots (Mandyam et al. 2010); Endophyte Host ; Decay leaf, fruit, seed (Põlme et al. 2020) | Duncan CG, Eslyn WE. [illegible] |
| Otu06603 | 4 | k__Fungi(100);p__Ascomycota(100);c__Saccharomycetes(100);o__Saccharomycetales(100);f__Metschnikowiaceae(100);g__Metschnikowia(100);s__Metschnikowia_pulcherrima(75); | Metschnikowia | 13 | Saprotroph | \|Undefined Saprotroph\| | Yeast | NULL | Probable | As nectar-tap_saprotroph (Põlme et al. 2020); Decay sugar-rich substrates (Põlme et al. 2020) | Põlme S, et al. 2020. Fungal Diversity 105:1-16 ((https://doi.org/10.1007/s13225-020-00466-2)) |
| Otu06605 | 4 | k__Fungi(100);p__Ascomycota(100);c__Saccharomycetes(100);o__Saccharomycetales(100);f__Metschnikowiaceae(100);g__Metschnikowia(100);s__Metschnikowia_pulcherrima(100); | Metschnikowia | 13 | Saprotroph | \|Undefined Saprotroph\| | Yeast | NULL | Probable | As nectar-tap_saprotroph (Põlme et al. 2020); Decay sugar-rich substrates (Põlme et al. 2020) | Põlme S, et al. 2020. Fungal Diversity 105:1-16 ((https://doi.org/10.1007/s13225-020-00466-2)) |
| Otu06606 | 4 | k__Fungi(100);p__Basidiomycota(100);c__Agaricomycetes(100);o__Russulales(100);f__Lachnocladiaceae(100);g__Scytinostroma(100);s__Scytinostroma_sp_DLL2011_1(100); | Scytinostroma | 13 | Saprotroph | Undefined Saprotroph-\|Wood Saprotroph\| | Corticioid | White Rot | Probable | As white_rot (Põlme et al. 2020); Decay wood (Põlme et al. 2020) | Tedersoo L, et al. 2014. Science 346:e1256688 ((https://doi.org/10.1126/science.1256688)); Põlme S, et al. 2020. Fungal Diversity 105:1-16 ((https://doi.org/10.1007/s13225-020-00466-2)) |

| | | | | | | | | | | | |
|---|---|---|---|---|---|---|---|---|---|---|---|
| Otu06607 | 4 | k__Fungi(100);p__Basidiomycota(100);c__Agaricomycetes(100);o__Agaricales(100);o__Agaricales_unclassified(100);o__Agaricales_unclassified(100);o__Agaricales_unclassified(100); | Agaricales | 7 | Pathotroph-Saprotroph-Symbiotroph | Bryophyte Parasite-Dung Saprotroph-Ectomycorrhizal-Fungal Parasite-Leaf Saprotroph-Plant Parasite-Undefined Saprotroph-Wood Saprotroph | Agaricoid-Gasteroid-Microfungus-Secotioid-Yeast | NULL | Possible | Mushrooms and toadstools, Gill fungi, Agarics... lignicolous, sometimes muscicolous or fungicolous, saprobic, mycorrhizal, rarely parasitic on plants or fungi (Kirk et al. 2008) | [illegible] |
| Otu06608 | 4 | k__Fungi(100);p__Ascomycota(100);c__Saccharomycetes(100);o__Saccharomycetales(100);f__Metschnikowiaceae(100);g__Metschnikowia(100);s__Metschnikowia_pulcherrima(75); | Metschnikowia | 13 | Saprotroph | \|Undefined Saprotroph\| | Yeast | NULL | Probable | As nectar-tap_saprotroph (Põlme et al. 2020); Decay sugar-rich substrates (Põlme et al. 2020) | Põlme S, et al. 2020. Fungal Diversity 105:1-16 ((https://doi.org/10.1007/s13225-020-00466-2)) |
| Otu06610 | 4 | k__Fungi(100);p__Ascomycota(100);c__Saccharomycetes(100);o__Saccharomycetales(100);f__Metschnikowiaceae(100);g__Metschnikowia(100);g__Metschnikowia_unclassified(100); | Metschnikowia | 13 | Saprotroph | \|Undefined Saprotroph\| | Yeast | NULL | Probable | As nectar-tap_saprotroph (Põlme et al. 2020); Decay sugar-rich substrates (Põlme et al. 2020) | Põlme S, et al. 2020. Fungal Diversity 105:1-16 ((https://doi.org/10.1007/s13225-020-00466-2)) |
| Otu06614 | 4 | k__Fungi(100);p__Ascomycota(100);c__Saccharomycetes(100);o__Saccharomycetales(100);f__Metschnikowiaceae(100);g__Metschnikowia(100);s__Metschnikowia_pulcherrima(100); | Metschnikowia | 13 | Saprotroph | \|Undefined Saprotroph\| | Yeast | NULL | Probable | As nectar-tap_saprotroph (Põlme et al. 2020); Decay sugar-rich substrates (Põlme et al. 2020) | Põlme S, et al. 2020. Fungal Diversity 105:1-16 ((https://doi.org/10.1007/s13225-020-00466-2)) |
| Otu06615 | 4 | k__Fungi(100);p__Ascomycota(100);c__Saccharomycetes(100);o__Saccharomycetales(100);f__Metschnikowiaceae(100);g__Metschnikowia(100);s__Metschnikowia_pulcherrima(100); | Metschnikowia | 13 | Saprotroph | \|Undefined Saprotroph\| | Yeast | NULL | Probable | As nectar-tap_saprotroph (Põlme et al. 2020); Decay sugar-rich substrates (Põlme et al. 2020) | Põlme S, et al. 2020. Fungal Diversity 105:1-16 ((https://doi.org/10.1007/s13225-020-00466-2)) |
| Otu06617 | 4 | k__Fungi(100);p__Ascomycota(100);c__Saccharomycetes(100);o__Saccharomycetales(100);f__Metschnikowiaceae(100);g__Metschnikowia(100);s__Metschnikowia_pulcherrima(75); | Metschnikowia | 13 | Saprotroph | \|Undefined Saprotroph\| | Yeast | NULL | Probable | As nectar-tap_saprotroph (Põlme et al. 2020); Decay sugar-rich substrates (Põlme et al. 2020) | Põlme S, et al. 2020. Fungal Diversity 105:1-16 ((https://doi.org/10.1007/s13225-020-00466-2)) |
| Otu06620 | 4 | k__Fungi(100);p__Ascomycota(100);c__Dothideomycetes(100);o__Dothideomycetes_order_Incertae_sedis(100);f__Myxotrichaceae(100);g__Oidiodendron(100);s__Oidiodendron_sp_18VA18(100); | Oidiodendron | 13 | Pathotroph-Saprotroph-Symbiotroph | Endophyte-Ericoid Mycorrhizal-\|Undefined Saprotroph\| | Microfungus | NULL | Probable | As root_endophyte (Põlme et al. 2020); As soil_saprotroph (Põlme et al. 2020); Dark Septate Endophyte; Decay soil (Põlme et al. 2020) | [illegible] |
| Otu06621 | 4 | k__Fungi(100);p__Ascomycota(100);c__Saccharomycetes(100);o__Saccharomycetales(100);f__Metschnikowiaceae(100);g__Metschnikowia(100);s__Metschnikowia_pulcherrima(100); | Metschnikowia | 13 | Saprotroph | \|Undefined Saprotroph\| | Yeast | NULL | Probable | As nectar-tap_saprotroph (Põlme et al. 2020); Decay sugar-rich substrates (Põlme et al. 2020) | Põlme S, et al. 2020. Fungal Diversity 105:1-16 ((https://doi.org/10.1007/s13225-020-00466-2)) |
| Otu06622 | 4 | k__Fungi(100);p__Ascomycota(100);c__Saccharomycetes(100);o__Saccharomycetales(100);f__Metschnikowiaceae(100);g__Metschnikowia(100);s__Metschnikowia_pulcherrima(100); | Metschnikowia | 13 | Saprotroph | \|Undefined Saprotroph\| | Yeast | NULL | Probable | As nectar-tap_saprotroph (Põlme et al. 2020); Decay sugar-rich substrates (Põlme et al. 2020) | Põlme S, et al. 2020. Fungal Diversity 105:1-16 ((https://doi.org/10.1007/s13225-020-00466-2)) |
| Otu06624 | 4 | k__Fungi(100);p__Ascomycota(100);c__Saccharomycetes(100);o__Saccharomycetales(100);f__Metschnikowiaceae(100);g__Metschnikowia(100);s__Metschnikowia_pulcherrima(75); | Metschnikowia | 13 | Saprotroph | \|Undefined Saprotroph\| | Yeast | NULL | Probable | As nectar-tap_saprotroph (Põlme et al. 2020); Decay sugar-rich substrates (Põlme et al. 2020) | Põlme S, et al. 2020. Fungal Diversity 105:1-16 ((https://doi.org/10.1007/s13225-020-00466-2)) |
| Otu06625 | 4 | k__Fungi(100);p__Ascomycota(100);c__Saccharomycetes(100);o__Saccharomycetales(100);f__Metschnikowiaceae(100);g__Metschnikowia(100);g__Metschnikowia_unclassified(100); | Metschnikowia | 13 | Saprotroph | \|Undefined Saprotroph\| | Yeast | NULL | Probable | As nectar-tap_saprotroph (Põlme et al. 2020); Decay sugar-rich substrates (Põlme et al. 2020) | Põlme S, et al. 2020. Fungal Diversity 105:1-16 ((https://doi.org/10.1007/s13225-020-00466-2)) |
| Otu06628 | 4 | k__Fungi(100);p__Ascomycota(100);c__Saccharomycetes(100);o__Saccharomycetales(100);f__Metschnikowiaceae(100);g__Metschnikowia(100);s__Metschnikowia_pulcherrima(100); | Metschnikowia | 13 | Saprotroph | \|Undefined Saprotroph\| | Yeast | NULL | Probable | As nectar-tap_saprotroph (Põlme et al. 2020); Decay sugar-rich substrates (Põlme et al. 2020) | Põlme S, et al. 2020. Fungal Diversity 105:1-16 ((https://doi.org/10.1007/s13225-020-00466-2)) |
| Otu06631 | 4 | k__Fungi(100);p__Ascomycota(100);c__Saccharomycetes(100);o__Saccharomycetales(100);f__Metschnikowiaceae(100);g__Metschnikowia(100);s__Metschnikowia_pulcherrima(100); | Metschnikowia | 13 | Saprotroph | \|Undefined Saprotroph\| | Yeast | NULL | Probable | As nectar-tap_saprotroph (Põlme et al. 2020); Decay sugar-rich substrates (Põlme et al. 2020) | Põlme S, et al. 2020. Fungal Diversity 105:1-16 ((https://doi.org/10.1007/s13225-020-00466-2)) |
| Otu06632 | 4 | k__Fungi(100);p__Basidiomycota(100);c__Microbotryomycetes(100);o__Sporidiobolales(100);f__Sporidiobolales_family_Incertae_sedis(100);g__Rhodotorula(100);g__Rhodotorula_unclassified(100); | Rhodotorula | 13 | Pathotroph-Saprotroph-Symbiotroph | Animal Pathogen-Animal Symbiotroph-Endophyte-Plant Pathogen-\|Undefined Saprotroph\| | Yeast | NULL | Probable | As foliar_endophyte (Põlme et al. 2020); Opportunistic human pathogen | [illegible] |
| Otu06634 | 4 | k__Fungi(100);p__Basidiomycota(100);c__Agaricomycetes(100);o__Agaricales(100);o__Agaricales_unclassified(100);o__Agaricales_unclassified(100);o__Agaricales_unclassified(100); | Agaricales | 7 | Pathotroph-Saprotroph-Symbiotroph | Bryophyte Parasite-Dung Saprotroph-Ectomycorrhizal-Fungal Parasite-Leaf Saprotroph-Plant Parasite-Undefined Saprotroph-Wood Saprotroph | Agaricoid-Gasteroid-Microfungus-Secotioid-Yeast | NULL | Possible | Mushrooms and toadstools, Gill fungi, Agarics... lignicolous, sometimes muscicolous or fungicolous, saprobic, mycorrhizal, rarely parasitic on plants or fungi (Kirk et al. 2008) | [illegible] |
| Otu06636 | 4 | k__Fungi(100);p__Ascomycota(100);c__Sordariomycetes(100);o__Melanosporales(100);f__Ceratostomataceae(100);g__Harzia(100);s__Harzia_acremonioides(100); | Harzia | 13 | Saprotroph | Plant Saprotroph-Undefined Saprotroph-\|Wood Saprotroph\| | Microfungus | NULL | Probable | As litter_saprotroph (Põlme et al. 2020); Decay wood, leaf, fruit, seed (Põlme et al. 2020) | [illegible] |
| Otu06637 | 4 | k__Fungi(100);p__Ascomycota(100);c__Dothideomycetes(100);o__Capnodiales(100);f__Mycosphaerellaceae(100);f__Mycosphaerellaceae_unclassified(100);f__Mycosphaerellaceae_unclassified(100); | Mycosphaerellaceae | 9 | Pathotroph-Saprotroph | Plant Pathogen-Undefined Saprotroph | Microfungus | NULL | Probable | Biotrophic, necrotrophic or saprobic on various plant tissues (Cannon & Kirk 2007) | [illegible] |
| Otu06642 | 4 | k__Fungi(100);p__Ascomycota(100);c__Saccharomycetes(100);o__Saccharomycetales(100);f__Metschnikowiaceae(100);g__Metschnikowia(100);g__Metschnikowia_unclassified(75); | Metschnikowia | 13 | Saprotroph | \|Undefined Saprotroph\| | Yeast | NULL | Probable | As nectar-tap_saprotroph (Põlme et al. 2020); Decay sugar-rich substrates (Põlme et al. 2020) | Põlme S, et al. 2020. Fungal Diversity 105:1-16 ((https://doi.org/10.1007/s13225-020-00466-2)) |
| Otu06643 | 4 | k__Fungi(100);p__Basidiomycota(100);c__Agaricomycetes(100);o__Trechisporales(100);f__Hydnodontaceae(100);g__Trechispora(100);g__Trechispora_unclassified(100); | Trechispora | 13 | Saprotroph | \|Wood Saprotroph\| | Corticioid | White Rot | Highly Probable | As white_rot (Põlme et al. 2020); Decay wood (Põlme et al. 2020) | [illegible] |
| Otu06645 | 4 | k__Fungi(100);p__Ascomycota(100);c__Dothideomycetes(100);o__Capnodiales(100);f__Teratosphaeriaceae(100);g__Teratosphaeria(100);s__Teratosphaeria_knoxdavesii(100); | Teratosphaeria | 13 | Pathotroph-Saprotroph | \|Plant Pathogen\|-Plant Saprotroph-Wood Saprotroph | Microfungus | NULL | Probable | As litter_saprotroph (Põlme et al. 2020); As wood_pathogen (Põlme et al. 2020); Decay wood, leaf, fruit, seed (Põlme et al. 2020) | [illegible] |
| Otu06648 | 4 | k__Fungi(100);p__Ascomycota(100);c__Lecanoromycetes(100);o__Lecanorales(100);f__Parmeliaceae(100);g__Punctelia(100);s__Punctelia_jeckeri(100); | Punctelia | 13 | Symbiotroph | \|Lichenized\| | Thallus | NULL | Highly Probable | NULL | [illegible] |
| Otu06649 | 4 | k__Fungi(100);p__Ascomycota(100);c__Saccharomycetes(100);o__Saccharomycetales(100);f__Metschnikowiaceae(100);g__Metschnikowia(100);s__Metschnikowia_pulcherrima(75); | Metschnikowia | 13 | Saprotroph | \|Undefined Saprotroph\| | Yeast | NULL | Probable | As nectar-tap_saprotroph (Põlme et al. 2020); Decay sugar-rich substrates (Põlme et al. 2020) | Põlme S, et al. 2020. Fungal Diversity 105:1-16 ((https://doi.org/10.1007/s13225-020-00466-2)) |
| Otu06652 | 4 | k__Fungi(100);p__Ascomycota(100);c__Saccharomycetes(100);o__Saccharomycetales(100);f__Metschnikowiaceae(100);g__Metschnikowia(100);s__Metschnikowia_pulcherrima(100); | Metschnikowia | 13 | Saprotroph | \|Undefined Saprotroph\| | Yeast | NULL | Probable | As nectar-tap_saprotroph (Põlme et al. 2020); Decay sugar-rich substrates (Põlme et al. 2020) | Põlme S, et al. 2020. Fungal Diversity 105:1-16 ((https://doi.org/10.1007/s13225-020-00466-2)) |
| Otu06653 | 4 | k__Fungi(100);p__Ascomycota(100);c__Sordariomycetes(100);o__Hypocreales(100);f__Hypocreales_family_Incertae_sedis(100);g__Ilyonectria(100);s__Ilyonectria_macrodidyma(100); | Ilyonectria | 13 | Pathotroph | \|Plant Pathogen\| | Microfungus | NULL | Probable | NULL | [illegible] |
| Otu06654 | 4 | k__Fungi(100);p__Ascomycota(100);c__Saccharomycetes(100);o__Saccharomycetales(100);f__Metschnikowiaceae(100);g__Metschnikowia(100);s__Metschnikowia_pulcherrima(100); | Metschnikowia | 13 | Saprotroph | \|Undefined Saprotroph\| | Yeast | NULL | Probable | As nectar-tap_saprotroph (Põlme et al. 2020); Decay sugar-rich substrates (Põlme et al. 2020) | Põlme S, et al. 2020. Fungal Diversity 105:1-16 ((https://doi.org/10.1007/s13225-020-00466-2)) |
| Otu06657 | 4 | k__Fungi(100);p__Ascomycota(100);c__Sordariomycetes(100);o__Hypocreales(100);f__Nectriaceae(100);f__Nectriaceae_unclassified(75);f__Nectriaceae_unclassified(75); | Nectriaceae | 9 | Pathotroph-Saprotroph-Symbiotroph | Animal Pathogen-Endophyte-Fungal Parasite-Lichen Parasite-Plant Pathogen-Wood Saprotroph | Microfungus | NULL | Possible | Fusarium species are amongst the economically significant plant pathogens (Cannon & Kirk 2007) | [illegible] |
| Otu06661 | 4 | k__Fungi(100);p__Ascomycota(100);c__Dothideomycetes(100);o__Dothideomycetes_order_Incertae_sedis(100);f__Myxotrichaceae(100);g__Oidiodendron(100);g__Oidiodendron_unclassified(100); | Oidiodendron | 13 | Pathotroph-Saprotroph-Symbiotroph | Endophyte-Ericoid Mycorrhizal-\|Undefined Saprotroph\| | Microfungus | NULL | Probable | As root_endophyte (Põlme et al. 2020); As soil_saprotroph (Põlme et al. 2020); Dark Septate Endophyte; Decay soil (Põlme et al. 2020) | [illegible] |
| Otu06662 | 4 | k__Fungi(100);p__Basidiomycota(100);c__Agaricomycetes(100);o__Agaricales(100);f__Mycenaceae(100);g__Mycena(100);s__Mycena_sp(75); | Mycena | 13 | Pathotroph-Saprotroph | Plant Pathogen-\|Plant Saprotroph\|-Undefined Saprotroph-Wood Saprotroph | Agaricoid | NULL | Probable | As litter_saprotroph (Põlme et al. 2020); Decay leaf, fruit, seed (Põlme et al. 2020) | [illegible] |
| Otu06665 | 4 | k__Fungi(100);p__Ascomycota(100);c__Saccharomycetes(100);o__Saccharomycetales(100);f__Metschnikowiaceae(100);g__Metschnikowia(100);s__Metschnikowia_pulcherrima(100); | Metschnikowia | 13 | Saprotroph | \|Undefined Saprotroph\| | Yeast | NULL | Probable | As nectar-tap_saprotroph (Põlme et al. 2020); Decay sugar-rich substrates (Põlme et al. 2020) | Põlme S, et al. 2020. Fungal Diversity 105:1-16 ((https://doi.org/10.1007/s13225-020-00466-2)) |
| Otu06668 | 4 | k__Fungi(100);p__Basidiomycota(100);c__Agaricomycetes(100);o__Agaricales(100);o__Agaricales_unclassified(100);o__Agaricales_unclassified(100);o__Agaricales_unclassified(100); | Agaricales | 7 | Pathotroph-Saprotroph-Symbiotroph | Bryophyte Parasite-Dung Saprotroph-Ectomycorrhizal-Fungal Parasite-Leaf Saprotroph-Plant Parasite-Undefined Saprotroph-Wood Saprotroph | Agaricoid-Gasteroid-Microfungus-Secotioid-Yeast | NULL | Possible | Mushrooms and toadstools, Gill fungi, Agarics... lignicolous, sometimes muscicolous or fungicolous, saprobic, mycorrhizal, rarely parasitic on plants or fungi (Kirk et al. 2008) | [illegible] |
| Otu06669 | 4 | k__Fungi(100);p__Basidiomycota(100);c__Agaricomycetes(100);o__Russulales(100);f__Stereaceae(100);f__Stereaceae_unclassified(100);f__Stereaceae_unclassified(100); | Stereaceae | 9 | Saprotroph | Undefined Saprotroph-Wood Saprotroph | Corticioid-Polyporoid | NULL | Probable | Lignicolous or terrestrial (in leaf litter); typically saprobic (Cannon & Kirk 2007) | [illegible] |
| Otu06670 | 4 | k__Fungi(100);p__Basidiomycota(100);c__Microbotryomycetes(100);o__Sporidiobolales(100);f__Sporidiobolales_family_Incertae_sedis(100);g__Sporobolomyces(100);s__Sporobolomyces_lactosus(100); | Sporobolomyces | 13 | Pathotroph-Saprotroph | \|Fungal Parasite\|-Plant Saprotroph | Yeast | NULL | Probable | As litter_saprotroph (Põlme et al. 2020); As mycoparasite (Põlme et al. 2020); Decay fungal material (Põlme et al. 2020) | [illegible] |
| Otu06671 | 4 | k__Fungi(100);p__Ascomycota(100);c__Saccharomycetes(100);o__Saccharomycetales(100);f__Metschnikowiaceae(100);g__Metschnikowia(100);s__Metschnikowia_pulcherrima(100); | Metschnikowia | 13 | Saprotroph | \|Undefined Saprotroph\| | Yeast | NULL | Probable | As nectar-tap_saprotroph (Põlme et al. 2020); Decay sugar-rich substrates (Põlme et al. 2020) | Põlme S, et al. 2020. Fungal Diversity 105:1-16 ((https://doi.org/10.1007/s13225-020-00466-2)) |
| Otu06673 | 4 | k__Fungi(100);p__Ascomycota(100);c__Saccharomycetes(100);o__Saccharomycetales(100);f__Metschnikowiaceae(100);g__Metschnikowia(100);s__Metschnikowia_pulcherrima(100); | Metschnikowia | 13 | Saprotroph | \|Undefined Saprotroph\| | Yeast | NULL | Probable | As nectar-tap_saprotroph (Põlme et al. 2020); Decay sugar-rich substrates (Põlme et al. 2020) | Põlme S, et al. 2020. Fungal Diversity 105:1-16 ((https://doi.org/10.1007/s13225-020-00466-2)) |
| Otu06675 | 4 | k__Fungi(100);p__Ascomycota(100);c__Saccharomycetes(100);o__Saccharomycetales(100);f__Metschnikowiaceae(100);g__Metschnikowia(100);s__Metschnikowia_pulcherrima(75); | Metschnikowia | 13 | Saprotroph | \|Undefined Saprotroph\| | Yeast | NULL | Probable | As nectar-tap_saprotroph (Põlme et al. 2020); Decay sugar-rich substrates (Põlme et al. 2020) | Põlme S, et al. 2020. Fungal Diversity 105:1-16 ((https://doi.org/10.1007/s13225-020-00466-2)) |
| Otu06678 | 4 | k__Fungi(100);p__Ascomycota(100);c__Sordariomycetes(100);o__Hypocreales(100);o__Hypocreales_unclassified(100);o__Hypocreales_unclassified(100);o__Hypocreales_unclassified(100); | Hypocreales | 7 | Saprotroph | Undefined Saprotroph | Microfungus | NULL | Possible | NULL | [illegible] |
| Otu06679 | 4 | k__Fungi(100);p__Basidiomycota(100);c__Tremellomycetes(100);o__Tremellales(100);f__Tremellales_family_Incertae_sedis(100);g__Dioszegia(100);g__Dioszegia_unclassified(100); | Dioszegia | 13 | Saprotroph | \|Plant Saprotroph\|-Undefined Saprotroph | Yeast | NULL | Probable | As litter_saprotroph (Põlme et al. 2020); Decay leaf, fruit, seed (Põlme et al. 2020); The Dioszegia subclade sensu Inacio (Inacio et al. 20015) is characterized by orange; yellow or orange to deep orange colonies. Isolated from plant leaves and soil. | [illegible] |
| Otu06680 | 4 | k__Fungi(100);p__Basidiomycota(100);c__Tremellomycetes(100);o__Tremellales(100);f__Tremellales_family_Incertae_sedis(100);g__Fellomyces(100);s__Fellomyces_fuzhouensis(100); | Fellomyces | 13 | Pathotroph-Saprotroph | Fungal Parasite-\|Undefined Saprotroph\| | Yeast | NULL | Probable | As mycoparasite (Põlme et al. 2020) | [illegible] |
| Otu06681 | 4 | k__Fungi(100);p__Ascomycota(100);c__Saccharomycetes(100);o__Saccharomycetales(100);f__Metschnikowiaceae(100);g__Metschnikowia(100);s__Metschnikowia_pulcherrima(75); | Metschnikowia | 13 | Saprotroph | \|Undefined Saprotroph\| | Yeast | NULL | Probable | As nectar-tap_saprotroph (Põlme et al. 2020); Decay sugar-rich substrates (Põlme et al. 2020) | Põlme S, et al. 2020. Fungal Diversity 105:1-16 ((https://doi.org/10.1007/s13225-020-00466-2)) |
| Otu06682 | 4 | k__Fungi(100);p__Ascomycota(100);c__Lecanoromycetes(100);o__Lecanorales(100);o__Lecanorales_unclassified(100);o__Lecanorales_unclassified(100);o__Lecanorales_unclassified(100); | Lecanorales | 7 | Pathotroph-Symbiotroph | Lichen Parasite-Lichenized | Microfungus-Thallus | NULL | Probable | Lichenized with green algae, occasionally lichenicolous, on rocks, mosses and wood (Cannon & Kirk 2007); ...an order of mostly lichen-forming fungi (Kirk et al. 2008) | [illegible] |
| Otu06683 | 4 | k__Fungi(100);p__Basidiomycota(100);c__Microbotryomycetes(100);o__Sporidiobolales(100);f__Sporidiobolales_family_Incertae_sedis(100);g__Sporobolomyces(100);s__Sporobolomyces_lactosus(100); | Sporobolomyces | 13 | Pathotroph-Saprotroph | \|Fungal Parasite\|-Plant Saprotroph | Yeast | NULL | Probable | As litter_saprotroph (Põlme et al. 2020); As mycoparasite (Põlme et al. 2020); Decay fungal material (Põlme et al. 2020) | [illegible] |
| Otu06685 | 4 | k__Fungi(100);p__Ascomycota(100);c__Lecanoromycetes(100);o__Lecanorales(100);f__Ramalinaceae(100);g__Bacidina(100);s__Bacidina_sulphurella(100); | Bacidina | 13 | Symbiotroph | \|Lichenized\| | Thallus | NULL | Highly Probable | NULL | [illegible] |
| Otu06686 | 4 | k__Fungi(100);p__Ascomycota(100);c__Saccharomycetes(100);o__Saccharomycetales(100);f__Saccharomycetales_family_Incertae_sedis(100);g__Candida(100);g__Candida_unclassified(100); | Candida | 13 | Pathotroph-Saprotroph-Symbiotroph | Animal Pathogen-Endophyte-Endosymbiont-Epiphyte-\|Undefined Saprotroph\| | Dimorphic Yeast-Yeast | NULL | Probable | As nectar-tap_saprotroph (Põlme et al. 2020); Decay sugar-rich substrates (Põlme et al. 2020); This asexual genus is polyphyletic and contain species that function very differently depend on what they are. Caution should be taken when interpreting guilds for Candida species. | [illegible] |
| Otu06689 | 4 | k__Fungi(100);p__Ascomycota(100);c__Saccharomycetes(100);o__Saccharomycetales(100);f__Metschnikowiaceae(100);g__Metschnikowia(100);g__Metschnikowia_unclassified(75); | Metschnikowia | 13 | Saprotroph | \|Undefined Saprotroph\| | Yeast | NULL | Probable | As nectar-tap_saprotroph (Põlme et al. 2020); Decay sugar-rich substrates (Põlme et al. 2020) | Põlme S, et al. 2020. Fungal Diversity 105:1-16 ((https://doi.org/10.1007/s13225-020-00466-2)) |
| Otu06690 | 4 | k__Fungi(100);p__Basidiomycota(100);c__Agaricomycetes(100);o__Trechisporales(100);f__Hydnodontaceae(100);g__Trechispora(100);s__Trechispora_sp(100); | Trechispora | 13 | Saprotroph | \|Wood Saprotroph\| | Corticioid | White Rot | Highly Probable | As white_rot (Põlme et al. 2020); Decay wood (Põlme et al. 2020) | [illegible] |
| Otu06691 | 4 | k__Fungi(100);p__Ascomycota(100);c__Saccharomycetes(100);o__Saccharomycetales(100);f__Metschnikowiaceae(100);g__Metschnikowia(100);s__Metschnikowia_pulcherrima(75); | Metschnikowia | 13 | Saprotroph | \|Undefined Saprotroph\| | Yeast | NULL | Probable | As nectar-tap_saprotroph (Põlme et al. 2020); Decay sugar-rich substrates (Põlme et al. 2020) | Põlme S, et al. 2020. Fungal Diversity 105:1-16 ((https://doi.org/10.1007/s13225-020-00466-2)) |
| Otu06692 | 4 | k__Fungi(100);p__Ascomycota(100);c__Dothideomycetes(100);o__Pleosporales(100);f__Pleosporales_family_Incertae_sedis(100);g__Phoma(100);s__Phoma_paspali(100); | Phoma | 13 | Pathotroph-Saprotroph-Symbiotroph | Dung Saprotroph-Endophyte-Lichen Parasite-\|Plant Pathogen\|-Plant Saprotroph-Undefined Saprotroph-Wood Saprotroph | Microfungus | Soft Rot | Possible | As foliar_endophyte (Põlme et al. 2020); As litter_saprotroph (Põlme et al. 2020); Decay leaf, fruit, seed, roots, soil (Põlme et al. 2020) | [illegible] |
| Otu06693 | 4 | k__Fungi(100);p__Ascomycota(100);c__Saccharomycetes(100);o__Saccharomycetales(100);f__Metschnikowiaceae(100);g__Metschnikowia(100);s__Metschnikowia_pulcherrima(100); | Metschnikowia | 13 | Saprotroph | \|Undefined Saprotroph\| | Yeast | NULL | Probable | As nectar-tap_saprotroph (Põlme et al. 2020); Decay sugar-rich substrates (Põlme et al. 2020) | Põlme S, et al. 2020. Fungal Diversity 105:1-16 ((https://doi.org/10.1007/s13225-020-00466-2)) |
| Otu06694 | 4 | k__Fungi(100);p__Ascomycota(100);c__Saccharomycetes(100);o__Saccharomycetales(100);f__Metschnikowiaceae(100);g__Metschnikowia(100);g__Metschnikowia_unclassified(75); | Metschnikowia | 13 | Saprotroph | \|Undefined Saprotroph\| | Yeast | NULL | Probable | As nectar-tap_saprotroph (Põlme et al. 2020); Decay sugar-rich substrates (Põlme et al. 2020) | Põlme S, et al. 2020. Fungal Diversity 105:1-16 ((https://doi.org/10.1007/s13225-020-00466-2)) |
| Otu06695 | 4 | k__Fungi(100);p__Basidiomycota(100);c__Agaricomycetes(100);o__Cantharellales(100);f__Botryobasidiaceae(100);g__Haplotrichum(100);s__Haplotrichum_conspersum(100); | Haplotrichum | 13 | Saprotroph | \|Wood Saprotroph\| | Corticioid | White Rot | Probable | As white_rot (Põlme et al. 2020); Decay wood (Põlme et al. 2020) | [illegible] |
| Otu06697 | 4 | k__Fungi(100);p__Ascomycota(100);c__Saccharomycetes(100);o__Saccharomycetales(100);f__Metschnikowiaceae(100);g__Metschnikowia(100);s__Metschnikowia_pulcherrima(75); | Metschnikowia | 13 | Saprotroph | \|Undefined Saprotroph\| | Yeast | NULL | Probable | As nectar-tap_saprotroph (Põlme et al. 2020); Decay sugar-rich substrates (Põlme et al. 2020) | Põlme S, et al. 2020. Fungal Diversity 105:1-16 ((https://doi.org/10.1007/s13225-020-00466-2)) |
| Otu06699 | 4 | k__Fungi(100);p__Basidiomycota(100);c__Microbotryomycetes(100);o__Sporidiobolales(100);f__Sporidiobolales_family_Incertae_sedis(100);g__Sporobolomyces(100);s__Sporobolomyces_lactosus(100); | Sporobolomyces | 13 | Pathotroph-Saprotroph | \|Fungal Parasite\|-Plant Saprotroph | Yeast | NULL | Probable | As litter_saprotroph (Põlme et al. 2020); As mycoparasite (Põlme et al. 2020); Decay fungal material (Põlme et al. 2020) | [illegible] |
| Otu06700 | 4 | k__Fungi(100);p__Basidiomycota(100);c__Agaricomycetes(100);o__Thelephorales(100);f__Thelephoraceae(100);f__Thelephoraceae_unclassified(100);f__Thelephoraceae_unclassified(100); | Thelephoraceae | 9 | Saprotroph-Symbiotroph | Ectomycorrhizal-Undefined Saprotroph | Clavarioid | NULL | Probable | On the ground, on soil, a few on wood. At least some speices ectomycorrhizal (Cannon & Kirk 2007) | [illegible] |
| Otu06702 | 4 | k__Fungi(100);p__Ascomycota(100);c__Eurotiomycetes(100);o__Chaetothyriales(100);f__Chaetothyriaceae(100);g__Ceramothyrium(100);s__Ceramothyrium_carniolicum(100); | Ceramothyrium | 13 | Pathotroph-Saprotroph-Symbiotroph | Endophyte-Plant Pathogen-\|Plant Saprotroph\|-Undefined Saprotroph | Microfungus | NULL | Probable | As foliar_endophyte (Põlme et al. 2020); As litter_saprotroph (Põlme et al. 2020); Causes sooty mold (Põlme et al. 2020); Decay leaf, fruit, seed, sugar-rich substrates (Põlme et al. 2020) | [illegible] |
| Otu06703 | 4 | k__Fungi(100);p__Ascomycota(100);c__Dothideomycetes(100);o__Capnodiales(100);f__Davidiellaceae(100);g__Cladosporium(100);g__Cladosporium_unclassified(100); | Cladosporium | 13 | Pathotroph-Saprotroph-Symbiotroph | Animal Pathogen-Endophyte-Lichen Parasite-Plant Pathogen-\|Plant Saprotroph\|-Wood Saprotroph | Microfungus | NULL | Possible | As foliar_endophyte (Põlme et al. 2020); As litter_saprotroph (Põlme et al. 2020); Decay wood, leaf, fruit, seed, soil, dung, animal material (Põlme et al. 2020) | [illegible] |
| Otu06704 | 4 | k__Fungi(100);p__Ascomycota(100);c__Dothideomycetes(100);o__Pleosporales(100);f__Phaeosphaeriaceae(100);g__Eudarluca(100);s__Eudarluca_caricis(100); | Eudarluca | 13 | Pathotroph-Saprotroph | \|Fungal Parasite\|-Undefined Saprotroph | Microfungus | NULL | Probable | As mycoparasite (Põlme et al. 2020); Decay fungal material (Põlme et al. 2020); Host - rusts (Põlme et al. 2020) | [illegible] |
| Otu06705 | 4 | k__Fungi(100);p__Ascomycota(100);c__Leotiomycetes(100);o__Helotiales(100);f__Vibrisseaceae(100);g__Phialocephala(100);s__Phialocephala_fusca(100); | Phialocephala | 13 | Saprotroph-Symbiotroph | Endophyte-\|Undefined Saprotroph\| | Microfungus | NULL | Highly Probable | As root_endophyte (Põlme et al. 2020); As root_endophyte_dark_septate (Põlme et al. 2020); As soil_saprotroph (Põlme et al. 2020); Dark Septate Endophyte, known from plant roots.; Decay roots, soil (Põlme et al. 2020) | [illegible] |
| Otu06708 | 4 | k__Fungi(100);p__Ascomycota(100);c__Saccharomycetes(100);o__Saccharomycetales(100);f__Metschnikowiaceae(100);g__Metschnikowia(100);g__Metschnikowia_unclassified(75); | Metschnikowia | 13 | Saprotroph | \|Undefined Saprotroph\| | Yeast | NULL | Probable | As nectar-tap_saprotroph (Põlme et al. 2020); Decay sugar-rich substrates (Põlme et al. 2020) | Põlme S, et al. 2020. Fungal Diversity 105:1-16 ((https://doi.org/10.1007/s13225-020-00466-2)) |
| Otu06712 | 4 | k__Fungi(100);p__Ascomycota(100);c__Sordariomycetes(100);o__Hypocreales(100);f__Cordycipitaceae(100);g__Gibellula(100);s__Gibellula_sp_WK_2013(100); | Gibellula | 13 | Pathotroph | \|Animal Parasite\|-Animal Pathogen-Arthropod Parasite | Microfungus | NULL | Probable | Decay animal material (Põlme et al. 2020); Host - spiders (Põlme et al. 2020) | [illegible] |
| Otu06713 | 4 | k__Fungi(100);p__Ascomycota(100);c__Sordariomycetes(100);o__Sordariomycetes_order_Incertae_sedis(100);f__Apiosporaceae(100);g__Arthrinium(100);g__Arthrinium_unclassified(100); | Arthrinium | 13 | Pathotroph-Saprotroph-Symbiotroph | Dung Saprotroph-Endophyte-\|Plant Pathogen\|-Undefined Saprotroph | Microfungus | NULL | Probable | As foliar_endophyte (Põlme et al. 2020); Decay leaf, fruit, seed (Põlme et al. 2020) | [illegible] |
| Otu06715 | 4 | k__Fungi(100);p__Ascomycota(100);c__Ascomycota_class_Incertae_sedis(100);o__Ascomycota_order_Incertae_sedis(100);f__Ascomycota_family_Incertae_sedis(100);g__Phialemoniopsis(100);s__Phialemoniopsis_cornearis(100); | Phialemoniopsis | 13 | Pathotroph-Saprotroph | \|Animal Parasite\|-Undefined Saprotroph | Microfungus | NULL | Probable | As animal_decomposer (Põlme et al. 2020); Decay animal material (Põlme et al. 2020) | [illegible] |
| Otu06717 | 4 | k__Fungi(100);p__Basidiomycota(100);c__Microbotryomycetes(100);o__Sporidiobolales(100);f__Sporidiobolales_family_Incertae_sedis(100);g__Rhodotorula(100);g__Rhodotorula_unclassified(100); | Rhodotorula | 13 | Pathotroph-Saprotroph-Symbiotroph | Animal Pathogen-Animal Symbiotroph-Endophyte-Plant Pathogen-\|Undefined Saprotroph\| | Yeast | NULL | Probable | As foliar_endophyte (Põlme et al. 2020); Opportunistic human pathogen | [illegible] |
| Otu06718 | 4 | k__Fungi(100);p__Ascomycota(100);c__Saccharomycetes(100);o__Saccharomycetales(100);f__Metschnikowiaceae(100);g__Metschnikowia(100);s__Metschnikowia_pulcherrima(75); | Metschnikowia | 13 | Saprotroph | \|Undefined Saprotroph\| | Yeast | NULL | Probable | As nectar-tap_saprotroph (Põlme et al. 2020); Decay sugar-rich substrates (Põlme et al. 2020) | Põlme S, et al. 2020. Fungal Diversity 105:1-16 ((https://doi.org/10.1007/s13225-020-00466-2)) |
| Otu06720 | 4 | k__Fungi(100);p__Ascomycota(100);c__Dothideomycetes(100);o__Pleosporales(100);f__Leptosphaeriaceae(100);g__Leptosphaeria(100);g__Leptosphaeria_unclassified(100); | Leptosphaeria | 13 | Pathotroph-Saprotroph | \|Plant Pathogen\|-Wood Saprotroph | Microfungus | NULL | Probable | As wood_pathogen (Põlme et al. 2020); Decay wood (Põlme et al. 2020) | [illegible] |
| Otu06723 | 4 | k__Fungi(100);p__Ascomycota(100);c__Saccharomycetes(100);o__Saccharomycetales(100);o__Saccharomycetales_unclassified(100);o__Saccharomycetales_unclassified(100);o__Saccharomycetales_unclassified(100); | Saccharomycetales | 7 | Saprotroph | Undefined Saprotroph | Yeast | NULL | Possible | NULL | [illegible] |
| Otu06724 | 4 | k__Fungi(100);p__Ascomycota(100);c__Saccharomycetes(100);o__Saccharomycetales(100);f__Metschnikowiaceae(100);g__Metschnikowia(100);s__Metschnikowia_pulcherrima(100); | Metschnikowia | 13 | Saprotroph | \|Undefined Saprotroph\| | Yeast | NULL | Probable | As nectar-tap_saprotroph (Põlme et al. 2020); Decay sugar-rich substrates (Põlme et al. 2020) | Põlme S, et al. 2020. Fungal Diversity 105:1-16 ((https://doi.org/10.1007/s13225-020-00466-2)) |
| Otu06725 | 4 | k__Fungi(100);p__Basidiomycota(100);c__Tremellomycetes(100);o__Tremellales(100);f__Tremellales_family_Incertae_sedis(100);g__Kockovaella(100);s__Kockovaella_cucphuongensis(100); | Kockovaella | 13 | Saprotroph-Symbiotroph | Endophyte-\|Epiphyte\|-Undefined Saprotroph | Yeast | NULL | Probable | As foliar_endophyte (Põlme et al. 2020); Decay leaf, fruit, seed (Põlme et al. 2020); Foliar, isolated from a plant sample collected in a tropical rain forest in Thailand. | [illegible] |
| Otu06726 | 4 | k__Fungi(100);p__Ascomycota(100);c__Saccharomycetes(100);o__Saccharomycetales(100);f__Metschnikowiaceae(100);g__Metschnikowia(100);s__Metschnikowia_pulcherrima(75); | Metschnikowia | 13 | Saprotroph | \|Undefined Saprotroph\| | Yeast | NULL | Probable | As nectar-tap_saprotroph (Põlme et al. 2020); Decay sugar-rich substrates (Põlme et al. 2020) | Põlme S, et al. 2020. Fungal Diversity 105:1-16 ((https://doi.org/10.1007/s13225-020-00466-2)) |
| Otu06727 | 4 | k__Fungi(100);p__Ascomycota(100);c__Saccharomycetes(100);o__Saccharomycetales(100);f__Metschnikowiaceae(100);g__Metschnikowia(100);s__Metschnikowia_pulcherrima(100); | Metschnikowia | 13 | Saprotroph | \|Undefined Saprotroph\| | Yeast | NULL | Probable | As nectar-tap_saprotroph (Põlme et al. 2020); Decay sugar-rich substrates (Põlme et al. 2020) | Põlme S, et al. 2020. Fungal Diversity 105:1-16 ((https://doi.org/10.1007/s13225-020-00466-2)) |
| Otu06728 | 4 | k__Fungi(100);p__Basidiomycota(100);c__Tremellomycetes(100);o__Tremellales(100);f__Tremellales_family_Incertae_sedis(100);f__Tremellales_family_Incertae_sedis_unclassified(100);f__Tremellales_family_Incertae_sedis_unclassified(100); | Tremellales | 7 | Pathotroph-Saprotroph-Symbiotroph | Fungal Parasite-Undefined Saprotroph | Tremelloid-Yeast | NULL | Possible | The order contains both teleomorphic and anamorphic species, most of the latter being yeasts | [illegible] |
| Otu06730 | 4 | k__Fungi(100);p__Ascomycota(100);c__Saccharomycetes(100);o__Saccharomycetales(100);f__Saccharomycetaceae(100);g__Kazachstania(100);s__Kazachstania_intestinalis(100); | Kazachstania | 13 | Saprotroph | \|Undefined Saprotroph\| | Yeast | NULL | Probable | As nectar-tap_saprotroph (Põlme et al. 2020); Decay sugar-rich substrates (Põlme et al. 2020) | Põlme S, et al. 2020. Fungal Diversity 105:1-16 ((https://doi.org/10.1007/s13225-020-00466-2)) |
| Otu06731 | 4 | k__Fungi(100);p__Ascomycota(100);c__Saccharomycetes(100);o__Saccharomycetales(100);f__Metschnikowiaceae(100);g__Metschnikowia(100);s__Metschnikowia_pulcherrima(100); | Metschnikowia | 13 | Saprotroph | \|Undefined Saprotroph\| | Yeast | NULL | Probable | As nectar-tap_saprotroph (Põlme et al. 2020); Decay sugar-rich substrates (Põlme et al. 2020) | Põlme S, et al. 2020. Fungal Diversity 105:1-16 ((https://doi.org/10.1007/s13225-020-00466-2)) |
| Otu06732 | 4 | k__Fungi(100);p__Basidiomycota(100);c__Agaricomycetes(100);o__Agaricales(100);f__Marasmiaceae(100);g__Marasmius(100);s__Marasmius_oreades(100); | Marasmius | 13 | Pathotroph-Saprotroph-Symbiotroph | Endophyte-Plant Pathogen-\|Plant Saprotroph\|-Undefined Saprotroph | Agaricoid | NULL | Probable | As litter_saprotroph (Põlme et al. 2020); Decay leaf, fruit, seed (Põlme et al. 2020); Often one of the most prominent fungal components of tropical forests, playing a major role in nutrient cycling (Cannon & Kirk 2007) | [illegible] |
| Otu06733 | 4 | k__Fungi(100);p__Ascomycota(100);c__Sordariomycetes(100);o__Hypocreales(100);o__Hypocreales_unclassified(100);o__Hypocreales_unclassified(100);o__Hypocreales_unclassified(100); | Hypocreales | 7 | Saprotroph | Undefined Saprotroph | Microfungus | NULL | Possible | NULL | [illegible] |
| Otu06734 | 4 | k__Fungi(100);p__Ascomycota(100);c__Dothideomycetes(100);o__Capnodiales(100);f__Teratosphaeriaceae(100);f__Teratosphaeriaceae_unclassified(100);f__Teratosphaeriaceae_unclassified(100); | Teratosphaeriaceae | 9 | Pathotroph-Saprotroph | Animal Pathogen-Plant Pathogen-Undefined Saprotroph | NULL | NULL | Probable | Includes numerous saprobic, extremophilic, human opportunistic, and plant pathogenic fungi (Quaedvlieg et al. 2014) | [illegible] |
| Otu06735 | 4 | k__Fungi(100);p__Basidiomycota(100);c__Tremellomycetes(100);o__Tremellales(100);f__Tremellales_family_Incertae_sedis(100);g__Hannaella(100);s__Hannaella_luteola(100); | Hannaella | 13 | Pathotroph-Saprotroph | \|Animal Parasite\|-Undefined Saprotroph | Yeast | NULL | Probable | As animal_decomposer (Põlme et al. 2020); Decay animal material (Põlme et al. 2020) | [illegible] |
| Otu06739 | 4 | k__Fungi(100);p__Ascomycota(100);c__Eurotiomycetes(100);o__Eurotiales(100);o__Eurotiales_unclassified(100);o__Eurotiales_unclassified(100);o__Eurotiales_unclassified(100); | Eurotiales | 7 | Saprotroph | Undefined Saprotroph | Microfungus | NULL | Possible | NULL | [illegible] |
| Otu06740 | 4 | k__Fungi(100);p__Ascomycota(100);c__Saccharomycetes(100);o__Saccharomycetales(100);f__Metschnikowiaceae(100);g__Metschnikowia(100);s__Metschnikowia_pulcherrima(100); | Metschnikowia | 13 | Saprotroph | \|Undefined Saprotroph\| | Yeast | NULL | Probable | As nectar-tap_saprotroph (Põlme et al. 2020); Decay sugar-rich substrates (Põlme et al. 2020) | Põlme S, et al. 2020. Fungal Diversity 105:1-16 ((https://doi.org/10.1007/s13225-020-00466-2)) |
| Otu06744 | 4 | k__Fungi(100);p__Ascomycota(100);c__Sordariomycetes(100);o__Diaporthales(100);f__Diaporthaceae(100);g__Diaporthe(100);s__Diaporthe_kochmanii(75); | Diaporthe | 13 | Pathotroph-Saprotroph-Symbiotroph | Endophyte-\|Plant Pathogen\|-Plant Saprotroph | Microfungus | NULL | Possible | As foliar_endophyte (Põlme et al. 2020); As litter_saprotroph (Põlme et al. 2020); Decay leaf, fruit, seed (Põlme et al. 2020) | [illegible] |
| Otu06746 | 4 | k__Fungi(100);p__Basidiomycota(100);c__Tremellomycetes(100);o__Tremellales(100);f__Tremellales_family_Incertae_sedis(100);f__Tremellales_family_Incertae_sedis_unclassified(100);f__Tremellales_family_Incertae_sedis_unclassified(100); | Tremellales | 7 | Pathotroph-Saprotroph-Symbiotroph | Fungal Parasite-Undefined Saprotroph | Tremelloid-Yeast | NULL | Possible | The order contains both teleomorphic and anamorphic species, most of the latter being yeasts | [illegible] |
| Otu06748 | 4 | k__Fungi(100);p__Ascomycota(100);c__Saccharomycetes(100);o__Saccharomycetales(100);f__Metschnikowiaceae(100);g__Metschnikowia(100);s__Metschnikowia_pulcherrima(100); | Metschnikowia | 13 | Saprotroph | \|Undefined Saprotroph\| | Yeast | NULL | Probable | As nectar-tap_saprotroph (Põlme et al. 2020); Decay sugar-rich substrates (Põlme et al. 2020) | Põlme S, et al. 2020. Fungal Diversity 105:1-16 ((https://doi.org/10.1007/s13225-020-00466-2)) |
| Otu06749 | 4 | k__Fungi(100);p__Ascomycota(100);c__Saccharomycetes(100);o__Saccharomycetales(100);f__Metschnikowiaceae(100);g__Metschnikowia(100);s__Metschnikowia_pulcherrima(100); | Metschnikowia | 13 | Saprotroph | \|Undefined Saprotroph\| | Yeast | NULL | Probable | As nectar-tap_saprotroph (Põlme et al. 2020); Decay sugar-rich substrates (Põlme et al. 2020) | Põlme S, et al. 2020. Fungal Diversity 105:1-16 ((https://doi.org/10.1007/s13225-020-00466-2)) |
| Otu06750 | 4 | k__Fungi(100);p__Ascomycota(100);c__Saccharomycetes(100);o__Saccharomycetales(100);f__Metschnikowiaceae(100);g__Metschnikowia(100);s__Metschnikowia_pulcherrima(100); | Metschnikowia | 13 | Saprotroph | \|Undefined Saprotroph\| | Yeast | NULL | Probable | As nectar-tap_saprotroph (Põlme et al. 2020); Decay sugar-rich substrates (Põlme et al. 2020) | Põlme S, et al. 2020. Fungal Diversity 105:1-16 ((https://doi.org/10.1007/s13225-020-00466-2)) |
| Otu06751 | 4 | k__Fungi(100);p__Ascomycota(100);c__Saccharomycetes(100);o__Saccharomycetales(100);f__Metschnikowiaceae(100);g__Metschnikowia(100);g__Metschnikowia_unclassified(100); | Metschnikowia | 13 | Saprotroph | \|Undefined Saprotroph\| | Yeast | NULL | Probable | As nectar-tap_saprotroph (Põlme et al. 2020); Decay sugar-rich substrates (Põlme et al. 2020) | Põlme S, et al. 2020. Fungal Diversity 105:1-16 ((https://doi.org/10.1007/s13225-020-00466-2)) |
| Otu06752 | 4 | k__Fungi(100);p__Ascomycota(100);c__Saccharomycetes(100);o__Saccharomycetales(100);f__Metschnikowiaceae(100);g__Metschnikowia(100);s__Metschnikowia_pulcherrima(75); | Metschnikowia | 13 | Saprotroph | \|Undefined Saprotroph\| | Yeast | NULL | Probable | As nectar-tap_saprotroph (Põlme et al. 2020); Decay sugar-rich substrates (Põlme et al. 2020) | Põlme S, et al. 2020. Fungal Diversity 105:1-16 ((https://doi.org/10.1007/s13225-020-00466-2)) |
| Otu06753 | 4 | k__Fungi(100);p__Ascomycota(100);c__Saccharomycetes(100);o__Saccharomycetales(100);f__Metschnikowiaceae(100);g__Metschnikowia(100);s__Metschnikowia_pulcherrima(100); | Metschnikowia | 13 | Saprotroph | \|Undefined Saprotroph\| | Yeast | NULL | Probable | As nectar-tap_saprotroph (Põlme et al. 2020); Decay sugar-rich substrates (Põlme et al. 2020) | Põlme S, et al. 2020. Fungal Diversity 105:1-16 ((https://doi.org/10.1007/s13225-020-00466-2)) |
| Otu06755 | 4 | k__Fungi(100);p__Ascomycota(100);c__Saccharomycetes(100);o__Saccharomycetales(100);f__Metschnikowiaceae(100);g__Metschnikowia(100);s__Metschnikowia_pulcherrima(100); | Metschnikowia | 13 | Saprotroph | \|Undefined Saprotroph\| | Yeast | NULL | Probable | As nectar-tap_saprotroph (Põlme et al. 2020); Decay sugar-rich substrates (Põlme et al. 2020) | Põlme S, et al. 2020. Fungal Diversity 105:1-16 ((https://doi.org/10.1007/s13225-020-00466-2)) |
| Otu06760 | 4 | k__Fungi(100);p__Ascomycota(100);c__Sordariomycetes(100);o__Melanosporales(100);f__Ceratostomataceae(100);g__Harzia(100);s__Harzia_acremonioides(100); | Harzia | 13 | Saprotroph | Plant Saprotroph-Undefined Saprotroph-\|Wood Saprotroph\| | Microfungus | NULL | Probable | As litter_saprotroph (Põlme et al. 2020); Decay wood, leaf, fruit, seed (Põlme et al. 2020) | [illegible] |
| Otu06763 | 4 | k__Fungi(100);p__Basidiomycota(100);c__Agaricomycetes(100);o__Agaricales(100);o__Agaricales_unclassified(100);o__Agaricales_unclassified(100);o__Agaricales_unclassified(100); | Agaricales | 7 | Pathotroph-Saprotroph-Symbiotroph | Bryophyte Parasite-Dung Saprotroph-Ectomycorrhizal-Fungal Parasite-Leaf Saprotroph-Plant Parasite-Undefined Saprotroph-Wood Saprotroph | Agaricoid-Gasteroid-Microfungus-Secotioid-Yeast | NULL | Possible | Mushrooms and toadstools, Gill fungi, Agarics... lignicolous, sometimes muscicolous or fungicolous, saprobic, mycorrhizal, rarely parasitic on plants or fungi (Kirk et al. 2008) | [illegible] |
| Otu06764 | 4 | k__Fungi(100);p__Ascomycota(100);c__Dothideomycetes(100);o__Pleosporales(100);f__Lophiostomataceae(100);g__Lophiostoma(100);s__Lophiostoma_sp_CBS_109932(100); | Lophiostoma | 13 | Saprotroph | Plant Saprotroph-Undefined Saprotroph-\|Wood Saprotroph\| | Microfungus | NULL | Probable | As litter_saprotroph (Põlme et al. 2020); Decay wood, leaf, fruit, seed (Põlme et al. 2020) | [illegible] |
| Otu06765 | 4 | k__Fungi(100);p__Ascomycota(100);c__Eurotiomycetes(100);o__Chaetothyriales(100);f__Herpotrichiellaceae(100);g__Phaeomoniella(100);g__Phaeomoniella_unclassified(100); | Phaeomoniella | 13 | Pathotroph | \|Plant Pathogen\| | Dimorphic-Microfungus; Facultative Yeast (Tedersoo et | NULL | Highly Probable | Pathotroph isolated from necrosis in Prunus wood (Damm et al. 2010) | [illegible] |
| Otu06767 | 4 | k__Fungi(100);p__Ascomycota(100);c__Saccharomycetes(100);o__Saccharomycetales(100);f__Metschnikowiaceae(100);g__Metschnikowia(100);s__Metschnikowia_pulcherrima(100); | Metschnikowia | 13 | Saprotroph | \|Undefined Saprotroph\| | Yeast | NULL | Probable | As nectar-tap_saprotroph (Põlme et al. 2020); Decay sugar-rich substrates (Põlme et al. 2020) | Põlme S, et al. 2020. Fungal Diversity 105:1-16 ((https://doi.org/10.1007/s13225-020-00466-2)) |
| Otu06768 | 4 | k__Fungi(100);p__Basidiomycota(100);c__Agaricomycetes(100);o__Polyporales(100);f__Meruliaceae(100);g__Phlebia(100);s__Phlebia_albomellea(100); | Phlebia | 13 | Saprotroph | \|Wood Saprotroph\| | Corticioid | White Rot | Highly Probable | As white_rot (Põlme et al. 2020); Decay wood (Põlme et al. 2020) | [illegible] |
| Otu06771 | 4 | k__Fungi(100);p__Basidiomycota(100);c__Tremellomycetes(100);o__Tremellales(100);f__Tremellales_family_Incertae_sedis(100);f__Tremellales_family_Incertae_sedis_unclassified(75);f__Tremellales_family_Incertae_sedis_unclassified(75); | Tremellales | 7 | Pathotroph-Saprotroph-Symbiotroph | Fungal Parasite-Undefined Saprotroph | Tremelloid-Yeast | NULL | Possible | The order contains both teleomorphic and anamorphic species, most of the latter being yeasts | [illegible] |
| Otu06772 | 4 | k__Fungi(100);p__Ascomycota(100);c__Dothideomycetes(100);o__Pleosporales(100);f__Pleosporaceae(100);g__Cochliobolus(100);s__Cochliobolus_kusanoi(100); | Cochliobolus | 13 | Pathotroph | \|Plant Pathogen\| | Microfungus | NULL | Probable | NULL | [illegible] |
| Otu06773 | 4 | k__Fungi(100);p__Basidiomycota(100);c__Agaricomycetes(100);o__Agaricales(100);f__Strophariaceae(100);g__Gymnopilus(100);g__Gymnopilus_unclassified(100); | Gymnopilus | 13 | Saprotroph | Plant Saprotroph-\|Wood Saprotroph\| | Agaricoid | White Rot | Probable | As litter_saprotroph (Põlme et al. 2020); As white_rot (Põlme et al. 2020); Decay wood (Põlme et al. 2020) | [illegible] |
| Otu06774 | 4 | k__Fungi(100);p__Ascomycota(100);c__Saccharomycetes(100);o__Saccharomycetales(100);f__Saccharomycetales_family_Incertae_sedis(100);g__Candida(75);g__Candida_unclassified(75); | Candida | 13 | Pathotroph-Saprotroph-Symbiotroph | Animal Pathogen-Endophyte-Endosymbiont-Epiphyte-\|Undefined Saprotroph\| | Dimorphic Yeast-Yeast | NULL | Probable | As nectar-tap_saprotroph (Põlme et al. 2020); Decay sugar-rich substrates (Põlme et al. 2020); This asexual genus is polyphyletic and contain species that function very differently depend on what they are. Caution should be taken when interpreting guilds for Candida species. | [illegible] |
| Otu06775 | 4 | k__Fungi(100);p__Ascomycota(100);c__Pezizomycetes(100);o__Pezizales(100);f__Discinaceae(100);g__Hydnotrya(100);s__Hydnotrya_michaelis(100); | Hydnotrya | 13 | Symbiotroph | \|Ectomycorrhizal\| | Gasteroid | Hypogeous | Highly Probable | Common name - false truffle; Syn. Geoporella | [illegible] |
| Otu06776 | 4 | k__Fungi(100);p__Basidiomycota(100);c__Agaricomycetes(100);o__Russulales(100);f__Peniophoraceae(100);g__Peniophora(100);s__Peniophora_sp_3_RG_2014(100); | Peniophora | 13 | Pathotroph-Saprotroph | Plant Pathogen-\|Wood Saprotroph\| | Corticioid | White Rot | Probable | As white_rot (Põlme et al. 2020); As wood_pathogen (Põlme et al. 2020); Decay wood (Põlme et al. 2020) | [illegible] |
| Otu06777 | 4 | k__Fungi(100);p__Ascomycota(100);c__Dothideomycetes(100);o__Dothideomycetes_order_Incertae_sedis(100);f__Pseudeurotiaceae(100);g__Pseudeurotium(100);g__Pseudeurotium_unclassified(75); | Pseudeurotium | 13 | Saprotroph | \|Dung Saprotroph\|-Undefined Saprotroph | Microfungus | NULL | Probable | As soil_saprotroph (Põlme et al. 2020); Decay soil, dung (Põlme et al. 2020) | [illegible] |
| Otu06778 | 4 | k__Fungi(100);p__Ascomycota(100);c__Saccharomycetes(100);o__Saccharomycetales(100);f__Metschnikowiaceae(100);g__Metschnikowia(100);s__Metschnikowia_pulcherrima(100); | Metschnikowia | 13 | Saprotroph | \|Undefined Saprotroph\| | Yeast | NULL | Probable | As nectar-tap_saprotroph (Põlme et al. 2020); Decay sugar-rich substrates (Põlme et al. 2020) | Põlme S, et al. 2020. Fungal Diversity 105:1-16 ((https://doi.org/10.1007/s13225-020-00466-2)) |
| Otu06781 | 4 | k__Fungi(100);p__Basidiomycota(100);c__Agaricomycetes(100);o__Agaricales(100);f__Cortinariaceae(100);g__Cortinarius(100);g__Cortinarius_unclassified(100); | Cortinarius | 13 | Symbiotroph | \|Ectomycorrhizal\| | Agaricoid | NULL | Highly Probable | NULL | [illegible] |
| Otu06784 | 4 | k__Fungi(100);p__Ascomycota(100);c__Saccharomycetes(100);o__Saccharomycetales(100);f__Metschnikowiaceae(100);g__Metschnikowia(100);s__Metschnikowia_pulcherrima(100); | Metschnikowia | 13 | Saprotroph | \|Undefined Saprotroph\| | Yeast | NULL | Probable | As nectar-tap_saprotroph (Põlme et al. 2020); Decay sugar-rich substrates (Põlme et al. 2020) | Põlme S, et al. 2020. Fungal Diversity 105:1-16 ((https://doi.org/10.1007/s13225-020-00466-2)) |
| Otu06787 | 4 | k__Fungi(100);p__Ascomycota(100);c__Saccharomycetes(100);o__Saccharomycetales(100);f__Metschnikowiaceae(100);g__Metschnikowia(100);s__Metschnikowia_pulcherrima(100); | Metschnikowia | 13 | Saprotroph | \|Undefined Saprotroph\| | Yeast | NULL | Probable | As nectar-tap_saprotroph (Põlme et al. 2020); Decay sugar-rich substrates (Põlme et al. 2020) | Põlme S, et al. 2020. Fungal Diversity 105:1-16 ((https://doi.org/10.1007/s13225-020-00466-2)) |
| Otu06788 | 4 | k__Fungi(100);p__Basidiomycota(100);c__Agaricomycetes(100);o__Agaricales(100);f__Pleurotaceae(100);g__Nematoctonus(100);s__Nematoctonus_subreniformis(100); | Nematoctonus | 13 | Pathotroph-Saprotroph | Nematophagous-Undefined Saprotroph-\|Wood Saprotroph\| | Microfungus | White Rot | Probable | Decay wood, animal material (Põlme et al. 2020) | [illegible] |
| Otu06792 | 4 | k__Fungi(100);p__Ascomycota(100);c__Sordariomycetes(100);o__Hypocreales(100);f__Nectriaceae(100);g__Gibberella(100);s__Gibberella_intricans(100); | Gibberella | 13 | Pathotroph-Saprotroph-Symbiotroph | Endophyte-\|Plant Pathogen\|-Plant Saprotroph-Undefined Saprotroph | Microfungus | Soft Rot | Probable | As foliar_endophyte (Põlme et al. 2020); As litter_saprotroph (Põlme et al. 2020); Causing damping-off disease (Põlme et al. 2020); Decay leaf, fruit, seed, roots, soil, animal material (Põlme et al. 2020) | [illegible] |
| Otu06795 | 4 | k__Fungi(100);p__Basidiomycota(100);c__Microbotryomycetes(100);o__Sporidiobolales(100);f__Sporidiobolales_family_Incertae_sedis(100);g__Sporobolomyces(100);s__Sporobolomyces_lactosus(100); | Sporobolomyces | 13 | Pathotroph-Saprotroph | \|Fungal Parasite\|-Plant Saprotroph | Yeast | NULL | Probable | As litter_saprotroph (Põlme et al. 2020); As mycoparasite (Põlme et al. 2020); Decay fungal material (Põlme et al. 2020) | [illegible] |
| Otu06796 | 4 | k__Fungi(100);p__Basidiomycota(100);c__Pucciniomycetes(100);o__Pucciniales(100);f__Pucciniaceae(100);f__Pucciniaceae_unclassified(100);f__Pucciniaceae_unclassified(100); | Pucciniaceae | 9 | Pathotroph | Plant Pathogen | Rust | NULL | Probable | Biotrophic, heteroecious or autoecious, associated with leaf and stem tissue of a very wide range of plants (Cannon & Kirk 2007) | [illegible] |
| Otu06798 | 4 | k__Fungi(100);p__Basidiomycota(100);c__Agaricomycetes(100);o__Agaricales(100);f__Mycenaceae(100);g__Mycena(100);s__Mycena_speirea(100); | Mycena | 13 | Pathotroph-Saprotroph | Plant Pathogen-\|Plant Saprotroph\|-Undefined Saprotroph-Wood Saprotroph | Agaricoid | NULL | Probable | As litter_saprotroph (Põlme et al. 2020); Decay leaf, fruit, seed (Põlme et al. 2020) | [illegible] |
| Otu06799 | 4 | k__Fungi(100);p__Basidiomycota(100);c__Agaricomycetes(100);o__Cantharellales(100);f__Botryobasidiaceae(100);g__Botryobasidium(100);s__Botryobasidium_subcoronatum(100); | Botryobasidium | 13 | Saprotroph | Undefined Saprotroph-\|Wood Saprotroph\| | Corticioid | White Rot | Probable | As white_rot (Põlme et al. 2020); Decay wood (Põlme et al. 2020) | [illegible] |
| Otu06800 | 4 | k__Fungi(100);p__Ascomycota(100);c__Saccharomycetes(100);o__Saccharomycetales(100);f__Metschnikowiaceae(100);g__Metschnikowia(100);s__Metschnikowia_pulcherrima(75); | Metschnikowia | 13 | Saprotroph | \|Undefined Saprotroph\| | Yeast | NULL | Probable | As nectar-tap_saprotroph (Põlme et al. 2020); Decay sugar-rich substrates (Põlme et al. 2020) | Põlme S, et al. 2020. Fungal Diversity 105:1-16 ((https://doi.org/10.1007/s13225-020-00466-2)) |
| Otu06801 | 4 | k__Fungi(100);p__Ascomycota(100);c__Leotiomycetes(100);o__Helotiales(100);f__Helotiales_family_Incertae_sedis(100);g__Rhexocercosporidium(100);s__Rhexocercosporidium_carotae(100); | Rhexocercosporidium | 13 | Pathotroph-Saprotroph | \|Plant Pathogen\|-Plant Saprotroph | Microfungus | NULL | Probable | As litter_saprotroph (Põlme et al. 2020); As root_pathogen (Põlme et al. 2020); Decay leaf, fruit, seed, roots (Põlme et al. 2020) | [illegible] |
| Otu06802 | 4 | k__Fungi(100);p__Ascomycota(100);c__Leotiomycetes(100);o__Helotiales(100);f__Vibrisseaceae(100);g__Phialocephala(100);g__Phialocephala_unclassified(100); | Phialocephala | 13 | Saprotroph-Symbiotroph | Endophyte-\|Undefined Saprotroph\| | Microfungus | NULL | Highly Probable | As root_endophyte (Põlme et al. 2020); As root_endophyte_dark_septate (Põlme et al. 2020); As soil_saprotroph (Põlme et al. 2020); Dark Septate Endophyte, known from plant roots.; Decay roots, soil (Põlme et al. 2020) | [illegible] |
| Otu06805 | 4 | k__Fungi(100);p__Basidiomycota(100);c__Microbotryomycetes(100);o__Sporidiobolales(100);f__Sporidiobolales_family_Incertae_sedis(100);g__Rhodotorula(100);g__Rhodotorula_unclassified(100); | Rhodotorula | 13 | Pathotroph-Saprotroph-Symbiotroph | Animal Pathogen-Animal Symbiotroph-Endophyte-Plant Pathogen-\|Undefined Saprotroph\| | Yeast | NULL | Probable | As foliar_endophyte (Põlme et al. 2020); Opportunistic human pathogen | [illegible] |
| Otu06807 | 4 | k__Fungi(100);p__Ascomycota(100);c__Saccharomycetes(100);o__Saccharomycetales(100);f__Metschnikowiaceae(100);g__Metschnikowia(100);s__Metschnikowia_pulcherrima(100); | Metschnikowia | 13 | Saprotroph | \|Undefined Saprotroph\| | Yeast | NULL | Probable | As nectar-tap_saprotroph (Põlme et al. 2020); Decay sugar-rich substrates (Põlme et al. 2020) | Põlme S, et al. 2020. Fungal Diversity 105:1-16 ((https://doi.org/10.1007/s13225-020-00466-2)) |
| Otu06810 | 4 | k__Fungi(100);p__Ascomycota(100);c__Saccharomycetes(100);o__Saccharomycetales(100);f__Metschnikowiaceae(100);g__Metschnikowia(100);s__Metschnikowia_pulcherrima(100); | Metschnikowia | 13 | Saprotroph | \|Undefined Saprotroph\| | Yeast | NULL | Probable | As nectar-tap_saprotroph (Põlme et al. 2020); Decay sugar-rich substrates (Põlme et al. 2020) | Põlme S, et al. 2020. Fungal Diversity 105:1-16 ((https://doi.org/10.1007/s13225-020-00466-2)) |
| Otu06811 | 4 | k__Fungi(100);p__Ascomycota(100);c__Sordariomycetes(100);o__Diaporthales(100);f__Diaporthales_family_Incertae_sedis(100);g__Valsaria(100);s__Valsaria_ceratoniae(100); | Valsaria | 13 | Saprotroph | Plant Saprotroph-\|Wood Saprotroph\| | Microfungus | NULL | Highly Probable | As litter_saprotroph (Põlme et al. 2020); Decay wood, leaf, fruit, seed (Põlme et al. 2020); Generally occur in little to moderately decayed bark (Jaklitsch et al. 2015) | [illegible] |
| Otu06813 | 4 | k__Fungi(100);p__Basidiomycota(100);c__Agaricomycetes(100);o__Cantharellales(100);f__Clavulinaceae(100);f__Clavulinaceae_unclassified(100);f__Clavulinaceae_unclassified(100); | Clavulinaceae | 9 | Saprotroph-Symbiotroph | Ectomycorrhizal-Lichenized-Wood Saprotroph | Clavarioid | NULL | Possible | Some species form ectomycorrhizas, with fruit bodies in leaf litter and on soil. Others are probably saprobes associated with rotten wood (Cannon & Kirk 2007) | [illegible] |
| Otu06815 | 4 | k__Fungi(100);p__Ascomycota(100);c__Saccharomycetes(100);o__Saccharomycetales(100);f__Metschnikowiaceae(100);g__Metschnikowia(100);g__Metschnikowia_unclassified(75); | Metschnikowia | 13 | Saprotroph | \|Undefined Saprotroph\| | Yeast | NULL | Probable | As nectar-tap_saprotroph (Põlme et al. 2020); Decay sugar-rich substrates (Põlme et al. 2020) | Põlme S, et al. 2020. Fungal Diversity 105:1-16 ((https://doi.org/10.1007/s13225-020-00466-2)) |
| Otu06817 | 4 | k__Fungi(100);p__Ascomycota(100);c__Saccharomycetes(100);o__Saccharomycetales(100);f__Metschnikowiaceae(100);g__Metschnikowia(100);s__Metschnikowia_pulcherrima(75); | Metschnikowia | 13 | Saprotroph | \|Undefined Saprotroph\| | Yeast | NULL | Probable | As nectar-tap_saprotroph (Põlme et al. 2020); Decay sugar-rich substrates (Põlme et al. 2020) | Põlme S, et al. 2020. Fungal Diversity 105:1-16 ((https://doi.org/10.1007/s13225-020-00466-2)) |
| Otu06819 | 4 | k__Fungi(100);p__Ascomycota(100);c__Saccharomycetes(100);o__Saccharomycetales(100);f__Metschnikowiaceae(100);g__Metschnikowia(100);s__Metschnikowia_pulcherrima(100); | Metschnikowia | 13 | Saprotroph | \|Undefined Saprotroph\| | Yeast | NULL | Probable | As nectar-tap_saprotroph (Põlme et al. 2020); Decay sugar-rich substrates (Põlme et al. 2020) | Põlme S, et al. 2020. Fungal Diversity 105:1-16 ((https://doi.org/10.1007/s13225-020-00466-2)) |
| Otu06820 | 4 | k__Fungi(100);p__Ascomycota(100);c__Saccharomycetes(100);o__Saccharomycetales(100);f__Metschnikowiaceae(100);g__Metschnikowia(100);s__Metschnikowia_pulcherrima(75); | Metschnikowia | 13 | Saprotroph | \|Undefined Saprotroph\| | Yeast | NULL | Probable | As nectar-tap_saprotroph (Põlme et al. 2020); Decay sugar-rich substrates (Põlme et al. 2020) | Põlme S, et al. 2020. Fungal Diversity 105:1-16 ((https://doi.org/10.1007/s13225-020-00466-2)) |
| Otu06826 | 4 | k__Fungi(100);p__Ascomycota(100);c__Saccharomycetes(100);o__Saccharomycetales(100);f__Metschnikowiaceae(100);g__Metschnikowia(100);s__Metschnikowia_pulcherrima(100); | Metschnikowia | 13 | Saprotroph | \|Undefined Saprotroph\| | Yeast | NULL | Probable | As nectar-tap_saprotroph (Põlme et al. 2020); Decay sugar-rich substrates (Põlme et al. 2020) | Põlme S, et al. 2020. Fungal Diversity 105:1-16 ((https://doi.org/10.1007/s13225-020-00466-2)) |
| Otu06828 | 4 | k__Fungi(100);p__Ascomycota(100);c__Saccharomycetes(100);o__Saccharomycetales(100);f__Metschnikowiaceae(100);g__Metschnikowia(100);g__Metschnikowia_unclassified(100); | Metschnikowia | 13 | Saprotroph | \|Undefined Saprotroph\| | Yeast | NULL | Probable | As nectar-tap_saprotroph (Põlme et al. 2020); Decay sugar-rich substrates (Põlme et al. 2020) | Põlme S, et al. 2020. Fungal Diversity 105:1-16 ((https://doi.org/10.1007/s13225-020-00466-2)) |
| Otu06831 | 4 | k__Fungi(100);p__Ascomycota(100);c__Sordariomycetes(100);o__Hypocreales(100);f__Hypocreales_family_Incertae_sedis(100);g__Acremonium(100);g__Acremonium_unclassified(100); | Acremonium | 13 | Pathotroph-Saprotroph-Symbiotroph | Animal Pathogen-Endophyte-Fungal Parasite-Plant Pathogen-\|Undefined Saprotroph\|-Wood Saprotroph | Microfungus | Soft Rot | Possible | As foliar_endophyte (Põlme et al. 2020); culture (Hanada et al. 2010, Del Olmo; Decay leaf, fruit, seed, soil, dung, animal material (Põlme et al. 2020); Endophyte detection method; Ruiz & Arnold 2014); Likely opportunistic human pathogen (Irinyi et al. 2015) | [illegible] |
| Otu06832 | 4 | k__Fungi(100);p__Ascomycota(100);c__Sordariomycetes(100);o__Hypocreales(100);o__Hypocreales_unclassified(75);o__Hypocreales_unclassified(75);o__Hypocreales_unclassified(75); | Hypocreales | 7 | Saprotroph | Undefined Saprotroph | Microfungus | NULL | Possible | NULL | [illegible] |
| Otu06835 | 4 | k__Fungi(100);p__Basidiomycota(100);c__Agaricomycetes(100);o__Polyporales(100);f__Fomitopsidaceae(100);g__Antrodia(100);s__Antrodia_macra(100); | Antrodia | 13 | Saprotroph | \|Wood Saprotroph\| | Corticioid | Brown Rot | Highly Probable | As brown-rot (Põlme et al. 2020); Decay wood (Põlme et al. 2020) | [illegible] |
| Otu06837 | 4 | k__Fungi(100);p__Basidiomycota(100);c__Agaricomycetes(100);o__Polyporales(100);f__Fomitopsidaceae(100);g__Postia(100);s__Postia_sp(100); | Postia | 13 | Saprotroph | \|Wood Saprotroph\| | Polyporoid | Brown Rot | Highly Probable | As brown-rot (Põlme et al. 2020); Decay wood (Põlme et al. 2020) | [illegible] |
| Otu06838 | 4 | k__Fungi(100);p__Basidiomycota(100);c__Agaricomycetes(100);o__Agaricales(100);f__Lyophyllaceae(100);g__Lyophyllum(100);s__Lyophyllum_semitale(100); | Lyophyllum | 13 | Saprotroph-Symbiotroph | \|Ectomycorrhizal\|-Undefined Saprotroph | Agaricoid | NULL | Possible | As soil_saprotroph (Põlme et al. 2020) | [illegible] |
| Otu06841 | 4 | k__Fungi(100);p__Basidiomycota(100);c__Agaricomycetes(100);o__Cantharellales(100);f__Clavulinaceae(100);g__unclassified_Clavulinaceae(100);s__Clavulinaceae_sp(100); | Clavulinaceae | 9 | Saprotroph-Symbiotroph | Ectomycorrhizal-Lichenized-Wood Saprotroph | Clavarioid | NULL | Possible | Some species form ectomycorrhizas, with fruit bodies in leaf litter and on soil. Others are probably saprobes associated with rotten wood (Cannon & Kirk 2007) | [illegible] |
| Otu06846 | 4 | k__Fungi(100);p__Ascomycota(100);c__Saccharomycetes(100);o__Saccharomycetales(100);f__Metschnikowiaceae(100);g__Metschnikowia(100);g__Metschnikowia_unclassified(100); | Metschnikowia | 13 | Saprotroph | \|Undefined Saprotroph\| | Yeast | NULL | Probable | As nectar-tap_saprotroph (Põlme et al. 2020); Decay sugar-rich substrates (Põlme et al. 2020) | Põlme S, et al. 2020. Fungal Diversity 105:1-16 ((https://doi.org/10.1007/s13225-020-00466-2)) |
| Otu06847 | 4 | k__Fungi(100);p__Basidiomycota(100);c__Exobasidiomycetes(100);o__Microstromatales(100);f__Microstromataceae(100);g__Microstroma(100);s__Microstroma_juglandis(100); | Microstroma | 13 | Pathotroph | \|Plant Pathogen\| | Dimorphic | NULL | Probable | NULL | [illegible] |
| Otu06848 | 4 | k__Fungi(100);p__Ascomycota(100);c__Saccharomycetes(100);o__Saccharomycetales(100);f__Metschnikowiaceae(100);g__Metschnikowia(100);g__Metschnikowia_unclassified(100); | Metschnikowia | 13 | Saprotroph | \|Undefined Saprotroph\| | Yeast | NULL | Probable | As nectar-tap_saprotroph (Põlme et al. 2020); Decay sugar-rich substrates (Põlme et al. 2020) | Põlme S, et al. 2020. Fungal Diversity 105:1-16 ((https://doi.org/10.1007/s13225-020-00466-2)) |
| Otu06849 | 4 | k__Fungi(100);p__Ascomycota(100);c__Saccharomycetes(100);o__Saccharomycetales(100);f__Metschnikowiaceae(100);g__Metschnikowia(100);s__Metschnikowia_pulcherrima(100); | Metschnikowia | 13 | Saprotroph | \|Undefined Saprotroph\| | Yeast | NULL | Probable | As nectar-tap_saprotroph (Põlme et al. 2020); Decay sugar-rich substrates (Põlme et al. 2020) | Põlme S, et al. 2020. Fungal Diversity 105:1-16 ((https://doi.org/10.1007/s13225-020-00466-2)) |
| Otu06850 | 4 | k__Fungi(100);p__Basidiomycota(100);c__Ustilaginomycetes(100);o__Ustilaginales(100);f__Ustilaginaceae(100);g__Pseudozyma(100);s__Pseudozyma_sp(100); | Pseudozyma | 13 | Pathotroph-Symbiotroph | Endophyte-\|Epiphyte\|-Fungal Parasite | Yeast | NULL | Probable | As foliar_endophyte (Põlme et al. 2020); As mycoparasite (Põlme et al. 2020); Decay leaf, fruit, seed, fungal material (Põlme et al. 2020) | [illegible] |
| Otu06851 | 4 | k__Fungi(100);p__Ascomycota(100);c__Saccharomycetes(100);o__Saccharomycetales(100);f__Metschnikowiaceae(100);g__Metschnikowia(100);s__Metschnikowia_pulcherrima(75); | Metschnikowia | 13 | Saprotroph | \|Undefined Saprotroph\| | Yeast | NULL | Probable | As nectar-tap_saprotroph (Põlme et al. 2020); Decay sugar-rich substrates (Põlme et al. 2020) | Põlme S, et al. 2020. Fungal Diversity 105:1-16 ((https://doi.org/10.1007/s13225-020-00466-2)) |
| Otu06853 | 4 | k__Fungi(100);p__Basidiomycota(100);c__Agaricomycetes(100);o__Agaricales(100);f__Agaricaceae(100);g__Lycoperdon(100);g__Lycoperdon_unclassified(100); | Lycoperdon | 13 | Saprotroph | \|Plant Saprotroph\|-Undefined Saprotroph | Gasteroid | NULL | Probable | puffball; As litter_saprotroph (Põlme et al. 2020); Common name ; Decay leaf, fruit, seed (Põlme et al. 2020) | [illegible] |
| Otu06854 | 4 | k__Fungi(100);p__Ascomycota(100);c__Saccharomycetes(100);o__Saccharomycetales(100);f__Metschnikowiaceae(100);g__Metschnikowia(100);s__Metschnikowia_pulcherrima(75); | Metschnikowia | 13 | Saprotroph | \|Undefined Saprotroph\| | Yeast | NULL | Probable | As nectar-tap_saprotroph (Põlme et al. 2020); Decay sugar-rich substrates (Põlme et al. 2020) | Põlme S, et al. 2020. Fungal Diversity 105:1-16 ((https://doi.org/10.1007/s13225-020-00466-2)) |
| Otu06856 | 4 | k__Fungi(100);p__Ascomycota(100);c__Sordariomycetes(100);o__Hypocreales(100);f__Hypocreales_family_Incertae_sedis(100);f__Hypocreales_family_Incertae_sedis_unclassified(75);f__Hypocreales_family_Incertae_sedis_unclassified(75); | Hypocreales | 7 | Saprotroph | Undefined Saprotroph | Microfungus | NULL | Possible | NULL | [illegible] |
| Otu06857 | 4 | k__Fungi(100);p__Basidiomycota(100);c__Agaricomycetes(100);o__Agaricales(100);f__Psathyrellaceae(100);g__Coprinopsis(100);g__Coprinopsis_unclassified(100); | Coprinopsis | 13 | Saprotroph | \|Undefined Saprotroph\| | Agaricoid | NULL | Probable | As soil_saprotroph (Põlme et al. 2020); Decay soil (Põlme et al. 2020) | [illegible] |
| Otu06858 | 4 | k__Fungi(100);p__Ascomycota(100);c__Saccharomycetes(100);o__Saccharomycetales(100);f__Metschnikowiaceae(100);g__Metschnikowia(100);s__Metschnikowia_pulcherrima(100); | Metschnikowia | 13 | Saprotroph | \|Undefined Saprotroph\| | Yeast | NULL | Probable | As nectar-tap_saprotroph (Põlme et al. 2020); Decay sugar-rich substrates (Põlme et al. 2020) | Põlme S, et al. 2020. Fungal Diversity 105:1-16 ((https://doi.org/10.1007/s13225-020-00466-2)) |
| Otu06860 | 4 | k__Fungi(100);p__Ascomycota(100);c__Sordariomycetes(100);o__Sordariales(100);f__Cephalothecaceae(100);g__Phialemonium(100);s__Phialemonium_atrogriseum(100); | Phialemonium | 13 | Pathotroph-Saprotroph | \|Animal Parasite\|-Undefined Saprotroph | Microfungus | NULL | Probable | As animal_decomposer (Põlme et al. 2020); Decay animal material (Põlme et al. 2020) | [illegible] |
| Otu06861 | 4 | k__Fungi(100);p__Basidiomycota(100);c__Agaricomycetes(100);o__Agaricales(100);f__Psathyrellaceae(100);g__Coprinopsis(100);g__Coprinopsis_unclassified(100); | Coprinopsis | 13 | Saprotroph | \|Undefined Saprotroph\| | Agaricoid | NULL | Probable | As soil_saprotroph (Põlme et al. 2020); Decay soil (Põlme et al. 2020) | [illegible] |
| Otu06862 | 4 | k__Fungi(100);p__Ascomycota(100);c__Dothideomycetes(100);o__Pleosporales(100);f__Phaeosphaeriaceae(75);f__Phaeosphaeriaceae_unclassified(75);f__Phaeosphaeriaceae_unclassified(75); | Phaeosphaeriaceae | 9 | Pathotroph-Saprotroph | Fungal Parasite-Plant Pathogen-Plant Saprotroph | Microfungus | NULL | Probable | Necrotrophic or saprobic on a wide range of plants, especially monocotyledons; a few species parasitic on other fungi (Cannon & Kirk 2007) | [illegible] |
| Otu06865 | 4 | k__Fungi(100);p__Basidiomycota(100);c__Agaricomycetes(100);o__Polyporales(100);f__Meruliaceae(100);g__Hypochnicium(100);s__Hypochnicium_geogenium(100); | Hypochnicium | 13 | Saprotroph | Undefined Saprotroph-\|Wood Saprotroph\| | Corticioid | White Rot | Probable | As white_rot (Põlme et al. 2020); Decay wood (Põlme et al. 2020) | [illegible] |
| Otu06867 | 4 | k__Fungi(100);p__Ascomycota(100);c__Saccharomycetes(100);o__Saccharomycetales(100);f__Metschnikowiaceae(100);g__Metschnikowia(100);s__Metschnikowia_pulcherrima(100); | Metschnikowia | 13 | Saprotroph | \|Undefined Saprotroph\| | Yeast | NULL | Probable | As nectar-tap_saprotroph (Põlme et al. 2020); Decay sugar-rich substrates (Põlme et al. 2020) | Põlme S, et al. 2020. Fungal Diversity 105:1-16 ((https://doi.org/10.1007/s13225-020-00466-2)) |
| Otu06871 | 4 | k__Fungi(100);p__Ascomycota(100);c__Sordariomycetes(100);o__Diaporthales(100);f__Gnomoniaceae(100);g__Gnomonia(100);s__Gnomonia_mirabilis(100); | Gnomonia | 13 | Pathotroph-Saprotroph-Symbiotroph | \|Endophyte\|-Plant Pathogen-Plant Saprotroph-Undefined Saprotroph | Microfungus | NULL | Probable | As foliar_endophyte (Põlme et al. 2020); As litter_saprotroph (Põlme et al. 2020); Decay leaf, fruit, seed (Põlme et al. 2020); Most species are saprobes or endophytes in leaves and bark, but a few are neurotrophic pathogens (Cannon & Kirk 2007) | [illegible] |

| | | | | | | | | | | | |
|---|---|---|---|---|---|---|---|---|---|---|---|
| Otu06872 | 4 | k__Fungi(100);p__Basidiomycota(100);c__Agaricomycetes(100);o__Polyporales(100);f__Meruliaceae(100);g__Phlebia(100);s__Phlebia_centrifuga(100); | Phlebia | 13 | Saprotroph | \|Wood Saprotroph\| | Corticioid | White Rot | Highly Probable | As white_rot (Põlme et al. 2020); Decay wood (Põlme et al. 2020) | Kirk PM et al. 2008. Dictionary of the Fungi. Tenth Edition. CAB International, Wallingford (ISBN: 978-0851998268); Põlme S, et al. 2020. Fungal Diversity 105:1-16 ((https://doi.org/10.1007/s13225-020-00466-2)) |
| Otu06873 | 4 | k__Fungi(100);p__Ascomycota(100);c__Saccharomycetes(100);o__Saccharomycetales(100);f__Metschnikowiaceae(100);g__Metschnikowia(100);s__Metschnikowia_pulcherrima(100); | Metschnikowia | 13 | Saprotroph | \|Undefined Saprotroph\| | Yeast | NULL | Probable | As nectar-tap_saprotroph (Põlme et al. 2020); Decay sugar-rich substrates (Põlme et al. 2020) | Põlme S, et al. 2020. Fungal Diversity 105:1-16 ((https://doi.org/10.1007/s13225-020-00466-2)) |
| Otu06874 | 4 | k__Fungi(100);p__Basidiomycota(100);c__Agaricomycetes(100);o__Agaricales(100);o__Agaricales_unclassified(100);o__Agaricales_unclassified(100);o__Agaricales_unclassified(100); | Agaricales | 7 | Pathotroph-Saprotroph-Symbiotroph | Bryophyte Parasite-Dung Saprotroph-Ectomycorrhizal-Fungal Parasite-Leaf Saprotroph-Plant Parasite-Undefined Saprotroph-Wood Saprotroph | Agaricoid-Gasteroid-Microfungus-Secotioid-Yeast | NULL | Possible | Mushrooms and toadstools, Gill fungi, Agarics... lignicolous, sometimes muscicolous or fungicolous, saprobic, mycorrhizal, rarely parasitic on plants or fungi (Kirk et al. 2008) | Kirk PM et al. 2008. Dictionary of the Fungi. Tenth Edition. CAB International, Wallingford (ISBN: 978-0851998268) |
| Otu06875 | 4 | k__Fungi(100);p__Ascomycota(100);c__Saccharomycetes(100);o__Saccharomycetales(100);f__Saccharomycetales_family_Incertae_sedis(75);f__Saccharomycetales_family_Incertae_sedis_unclassified(75);f__Saccharomycetales_family_Incertae_sedis_unclassified(75); | Saccharomycetales | 7 | Saprotroph | Undefined Saprotroph | Yeast | NULL | Possible | NULL | Sterkenburg E, et al. 2015. New Phytologist 207:1145-1158 ((https://doi.org/10.1111/nph.13426)) |
| Otu06876 | 4 | k__Fungi(100);p__Basidiomycota(100);c__Agaricomycetes(100);o__Agaricales(100);o__Agaricales_unclassified(100);o__Agaricales_unclassified(100);o__Agaricales_unclassified(100); | Agaricales | 7 | Pathotroph-Saprotroph-Symbiotroph | Bryophyte Parasite-Dung Saprotroph-Ectomycorrhizal-Fungal Parasite-Leaf Saprotroph-Plant Parasite-Undefined Saprotroph-Wood Saprotroph | Agaricoid-Gasteroid-Microfungus-Secotioid-Yeast | NULL | Possible | Mushrooms and toadstools, Gill fungi, Agarics... lignicolous, sometimes muscicolous or fungicolous, saprobic, mycorrhizal, rarely parasitic on plants or fungi (Kirk et al. 2008) | Kirk PM et al. 2008. Dictionary of the Fungi. Tenth Edition. CAB International, Wallingford (ISBN: 978-0851998268) |
| Otu06878 | 4 | k__Fungi(100);p__Ascomycota(100);c__Dothideomycetes(100);o__Capnodiales(100);f__Capnodiales_family_Incertae_sedis(100);g__Rachicladosporium(75);g__Rachicladosporium_unclassified(75); | Rachicladosporium | 13 | Saprotroph | \|Plant Saprotroph\| | Microfungus | NULL | Probable | As litter_saprotroph (Põlme et al. 2020); Decay leaf, fruit, seed (Põlme et al. 2020) | Põlme S, et al. 2020. Fungal Diversity 105:1-16 ((https://doi.org/10.1007/s13225-020-00466-2)) |
| Otu06879 | 4 | k__Fungi(100);p__Ascomycota(100);c__Saccharomycetes(100);o__Saccharomycetales(100);f__Metschnikowiaceae(100);g__Metschnikowia(100);g__Metschnikowia_unclassified(100); | Metschnikowia | 13 | Saprotroph | \|Undefined Saprotroph\| | Yeast | NULL | Probable | As nectar-tap_saprotroph (Põlme et al. 2020); Decay sugar-rich substrates (Põlme et al. 2020) | Põlme S, et al. 2020. Fungal Diversity 105:1-16 ((https://doi.org/10.1007/s13225-020-00466-2)) |
| Otu06883 | 4 | k__Fungi(100);p__Ascomycota(100);c__Saccharomycetes(100);o__Saccharomycetales(100);f__Metschnikowiaceae(100);g__Metschnikowia(100);s__Metschnikowia_pulcherrima(100); | Metschnikowia | 13 | Saprotroph | \|Undefined Saprotroph\| | Yeast | NULL | Probable | As nectar-tap_saprotroph (Põlme et al. 2020); Decay sugar-rich substrates (Põlme et al. 2020) | Põlme S, et al. 2020. Fungal Diversity 105:1-16 ((https://doi.org/10.1007/s13225-020-00466-2)) |
| Otu06884 | 4 | k__Fungi(100);p__Ascomycota(100);c__Saccharomycetes(100);o__Saccharomycetales(100);f__Metschnikowiaceae(100);g__Metschnikowia(100);g__Metschnikowia_unclassified(100); | Metschnikowia | 13 | Saprotroph | \|Undefined Saprotroph\| | Yeast | NULL | Probable | As nectar-tap_saprotroph (Põlme et al. 2020); Decay sugar-rich substrates (Põlme et al. 2020) | Põlme S, et al. 2020. Fungal Diversity 105:1-16 ((https://doi.org/10.1007/s13225-020-00466-2)) |
| Otu06885 | 4 | k__Fungi(100);p__Ascomycota(100);c__Sordariomycetes(100);o__Hypocreales(100);f__Hypocreales_family_Incertae_sedis(100);g__Trichothecium(100);s__Trichothecium_roseum(100); | Trichothecium | 13 | Pathotroph-Saprotroph | Fungal Parasite-\|Plant Pathogen\|-Plant Saprotroph | Microfungus | NULL | Probable | As litter_saprotroph (Põlme et al. 2020); As mycoparasite (Põlme et al. 2020); Decay leaf, fruit, seed (Põlme et al. 2020) | Põlme S, et al. 2020. Fungal Diversity 105:1-16 ((https://doi.org/10.1007/s13225-020-00466-2)); The American Phytopathological Society Common Names of Plant Diseases ((http://www.apsnet.org/publications/commonnames/Pages/default.aspx)) |
| Otu06886 | 4 | k__Fungi(100);p__Basidiomycota(100);c__Agaricomycetes(100);o__Polyporales(100);f__Phanerochaetaceae(100);g__Ceriporia(100);s__Ceriporia_alachuana(100); | Ceriporia | 13 | Saprotroph | \|Wood Saprotroph\| | Corticioid | White Rot | Highly Probable | As white_rot (Põlme et al. 2020); Decay wood (Põlme et al. 2020) | Tedersoo L, et al. 2014. Science 346:e1256688 ((https://doi.org/10.1126/science.1256688)); Põlme S, et al. 2020. Fungal Diversity 105:1-16 ((https://doi.org/10.1007/s13225-020-00466-2)) |
| Otu06888 | 4 | k__Fungi(100);p__Ascomycota(100);c__Saccharomycetes(100);o__Saccharomycetales(100);f__Metschnikowiaceae(100);g__Metschnikowia(100);g__Metschnikowia_unclassified(100); | Metschnikowia | 13 | Saprotroph | \|Undefined Saprotroph\| | Yeast | NULL | Probable | As nectar-tap_saprotroph (Põlme et al. 2020); Decay sugar-rich substrates (Põlme et al. 2020) | Põlme S, et al. 2020. Fungal Diversity 105:1-16 ((https://doi.org/10.1007/s13225-020-00466-2)) |
| Otu06889 | 4 | k__Fungi(100);p__Basidiomycota(100);c__Agaricomycetes(100);o__Polyporales(100);f__Polyporaceae(100);f__Polyporaceae_unclassified(75);f__Polyporaceae_unclassified(75); | Polyporaceae | 9 | Saprotroph | Wood Saprotroph | Polyporoid | Brown Rot-White | Probable | Lignicolous, causing brown or white rot, mostly of broadleaved trees (Cannon & Kirk 2007) | Cannon PF, Kirk PM. 2007. Fungal Families of the World. CAB International, Cambridge (ISBN: 978-0851998275) |
| Otu06890 | 4 | k__Fungi(100);p__Basidiomycota(100);c__Agaricomycetes(100);o__Agaricales(100);f__Bolbitiaceae(100);g__Conocybe(100);s__Conocybe_hausknechtii(100); | Conocybe | 13 | Saprotroph | Dung Saprotroph-Plant Saprotroph-\|Undefined Saprotroph\| | Agaricoid | NULL | Probable | As litter_saprotroph (Põlme et al. 2020); As soil_saprotroph (Põlme et al. 2020); Decay soil (Põlme et al. 2020) | Tedersoo L, et al. 2014. Science 346:e1256688 ((https://doi.org/10.1126/science.1256688)); Põlme S, et al. 2020. Fungal Diversity 105:1-16 ((https://doi.org/10.1007/s13225-020-00466-2)) |
| Otu06893 | 4 | k__Fungi(100);p__Basidiomycota(100);c__Microbotryomycetes(100);o__Sporidiobolales(100);f__Sporidiobolales_family_Incertae_sedis(100);g__Sporobolomyces(100);s__Sporobolomyces_lactosus(100); | Sporobolomyces | 13 | Pathotroph-Saprotroph | \|Fungal Parasite\|-Plant Saprotroph | Yeast | NULL | Probable | As litter_saprotroph (Põlme et al. 2020); As mycoparasite (Põlme et al. 2020); Decay fungal material (Põlme et al. 2020) | Tedersoo L, et al. 2014. Science 346:e1256688 ((https://doi.org/10.1126/science.1256688)); Purahong W, et al. 2016. Molecular Ecology 25:4059-4074 ((https://doi.org/10.1111/mec.13739)); Põlme S, et al. 2020. Fungal Diversity 105:1-16 ((https://doi.org/10.1007/s13225-020-00466-2)) |
| Otu06894 | 4 | k__Fungi(100);p__Ascomycota(100);c__Sordariomycetes(100);o__Sordariales(100);f__Lasiosphaeriaceae(100);g__Podospora(75);s__Podospora_inflatula(75); | Podospora | 13 | Saprotroph-Symbiotroph | \|Dung Saprotroph\|-Endophyte-Plant Saprotroph-Undefined Saprotroph | Microfungus | NULL | Possible | As foliar_endophyte (Põlme et al. 2020); As litter_saprotroph (Põlme et al. 2020); culture (David et al. 2016); Decay leaf, fruit, seed, dung (Põlme et al. 2020); Endophyte detection method | Bell A. 1983. Dung Fungi: An Illustrated Guide to Coprophilous Fungi in New Zealand. Victoria University Press, Wellington (ISBN: 978-0864730015); Tedersoo L., et al. 2014. Science 346:e1256688 ((https://doi.org/10.1126/science.1256688)); David AS, et al. 2016. Microbial Ecology 71:912-926 ((https://doi.org/10.1007/s00248-015-0712-6)); Põlme S, et al. 2020. Fungal Diversity 105:1-16 ((https://doi.org/10.1007/s13225-020-00466-2)) |
| Otu06895 | 4 | k__Fungi(100);p__Ascomycota(100);c__Saccharomycetes(100);o__Saccharomycetales(100);f__Metschnikowiaceae(100);g__Metschnikowia(100);g__Metschnikowia_unclassified(100); | Metschnikowia | 13 | Saprotroph | \|Undefined Saprotroph\| | Yeast | NULL | Probable | As nectar-tap_saprotroph (Põlme et al. 2020); Decay sugar-rich substrates (Põlme et al. 2020) | Põlme S, et al. 2020. Fungal Diversity 105:1-16 ((https://doi.org/10.1007/s13225-020-00466-2)) |
| Otu06896 | 4 | k__Fungi(100);p__Ascomycota(100);c__Sordariomycetes(100);o__Xylariales(100);f__Xylariaceae(100);g__Nodulisporium(100);s__Nodulisporium_sp(100); | Nodulisporium | 13 | Saprotroph | \|Undefined Saprotroph\| | Microfungus | NULL | Probable | NULL | Tedersoo L, et al. 2014. Science 346:e1256688 ((https://doi.org/10.1126/science.1256688)); Põlme S, et al. 2020. Fungal Diversity 105:1-16 ((https://doi.org/10.1007/s13225-020-00466-2)) |
| Otu06899 | 4 | k__Fungi(100);p__Basidiomycota(100);c__Microbotryomycetes(100);o__Sporidiobolales(100);f__Sporidiobolales_family_Incertae_sedis(100);g__Rhodotorula(100);s__Rhodotorula_mucilaginosa(100); | Rhodotorula | 13 | Pathotroph-Saprotroph-Symbiotroph | Animal Pathogen-Animal Symbiotroph-Endophyte-Plant Pathogen-\|Undefined Saprotroph\| | Yeast | NULL | Probable | As foliar_endophyte (Põlme et al. 2020); Opportunistic human pathogen | Põlme S, et al. 2020. Fungal Diversity 105:1-16 ((https://doi.org/10.1007/s13225-020-00466-2)); https://www.cdc.gov ((https://www.cdc.gov)) |
| Otu06900 | 4 | k__Fungi(100);p__Ascomycota(100);c__Saccharomycetes(100);o__Saccharomycetales(100);f__Metschnikowiaceae(100);g__Metschnikowia(100);s__Metschnikowia_pulcherrima(100); | Metschnikowia | 13 | Saprotroph | \|Undefined Saprotroph\| | Yeast | NULL | Probable | As nectar-tap_saprotroph (Põlme et al. 2020); Decay sugar-rich substrates (Põlme et al. 2020) | Põlme S, et al. 2020. Fungal Diversity 105:1-16 ((https://doi.org/10.1007/s13225-020-00466-2)) |
| Otu06901 | 4 | k__Fungi(100);p__Ascomycota(100);c__Saccharomycetes(100);o__Saccharomycetales(100);f__Metschnikowiaceae(100);g__Metschnikowia(100);s__Metschnikowia_pulcherrima(100); | Metschnikowia | 13 | Saprotroph | \|Undefined Saprotroph\| | Yeast | NULL | Probable | As nectar-tap_saprotroph (Põlme et al. 2020); Decay sugar-rich substrates (Põlme et al. 2020) | Põlme S, et al. 2020. Fungal Diversity 105:1-16 ((https://doi.org/10.1007/s13225-020-00466-2)) |
| Otu06903 | 4 | k__Fungi(100);p__Ascomycota(100);c__Saccharomycetes(100);o__Saccharomycetales(100);f__Metschnikowiaceae(100);g__Metschnikowia(100);s__Metschnikowia_pulcherrima(100); | Metschnikowia | 13 | Saprotroph | \|Undefined Saprotroph\| | Yeast | NULL | Probable | As nectar-tap_saprotroph (Põlme et al. 2020); Decay sugar-rich substrates (Põlme et al. 2020) | Põlme S, et al. 2020. Fungal Diversity 105:1-16 ((https://doi.org/10.1007/s13225-020-00466-2)) |
| Otu06905 | 4 | k__Fungi(100);p__Ascomycota(100);c__Saccharomycetes(100);o__Saccharomycetales(100);f__Metschnikowiaceae(100);g__Metschnikowia(100);s__Metschnikowia_pulcherrima(100); | Metschnikowia | 13 | Saprotroph | \|Undefined Saprotroph\| | Yeast | NULL | Probable | As nectar-tap_saprotroph (Põlme et al. 2020); Decay sugar-rich substrates (Põlme et al. 2020) | Põlme S, et al. 2020. Fungal Diversity 105:1-16 ((https://doi.org/10.1007/s13225-020-00466-2)) |
| Otu06907 | 4 | k__Fungi(100);p__Ascomycota(100);c__Sordariomycetes(100);o__Hypocreales(75);o__Hypocreales_unclassified(75);o__Hypocreales_unclassified(75);o__Hypocreales_unclassified(75); | Hypocreales | 7 | Saprotroph | Undefined Saprotroph | Microfungus | NULL | Possible | NULL | Sterkenburg E, et al. 2015. New Phytologist 207:1145-1158 ((https://doi.org/10.1111/nph.13426)) |
| Otu06908 | 4 | k__Fungi(100);p__Basidiomycota(100);c__Agaricomycetes(100);o__Auriculariales(100);f__Auriculariales_family_Incertae_sedis(100);g__Exidia(100);s__Exidia_saccharina(100); | Exidia | 13 | Saprotroph-Symbiotroph | Endophyte-Plant Saprotroph-Undefined Saprotroph-\|Wood Saprotroph\| | Tremelloid | NULL | Probable | As foliar_endophyte (Põlme et al. 2020); As litter_saprotroph (Põlme et al. 2020); Decay wood, leaf, fruit, seed (Põlme et al. 2020) | Tedersoo L, et al. 2014. Science 346:e1256688 ((https://doi.org/10.1126/science.1256688)); David AS, et al. 2016. Microbial Ecology 71:912-926 ((https://doi.org/10.1007/s00248-015-0712-6)); Põlme S, et al. 2020. Fungal Diversity 105:1-16 ((https://doi.org/10.1007/s13225-020-00466-2)) |
| Otu06909 | 4 | k__Fungi(100);p__Basidiomycota(100);c__Agaricomycetes(100);o__Thelephorales(100);f__Thelephoraceae(100);g__Tomentellopsis(100);g__Tomentellopsis_unclassified(100); | Tomentellopsis | 13 | Symbiotroph | \|Ectomycorrhizal\| | Corticioid | NULL | Highly Probable | NULL | Rinaldi AC, et al. 2008. Fungal Diversity 33:1-45 ((http://www.fungaldiversity.org/fdp/sfdp/33-1.pdf)); Tedersoo L, et al. 2010. Mycorrhiza 20:217-263 ((https://doi.org/10.1007/s00572-009-0274-x)); Põlme S, et al. 2020. Fungal Diversity 105:1-16 ((https://doi.org/10.1007/s13225-020-00466-2)); http://mycorrhizas.info/ecmf.html ((http://mycorrhizas.info/ecmf.html)) |
| Otu06912 | 4 | k__Fungi(100);p__Basidiomycota(100);c__Agaricomycetes(100);o__Corticiales(100);f__Corticiaceae(100);g__Corticium(100);s__Corticium_roseum(100); | Corticium | 13 | Saprotroph | Undefined Saprotroph-\|Wood Saprotroph\| | Corticioid | White Rot | Probable | As white_rot (Põlme et al. 2020); Decay wood (Põlme et al. 2020) | Tedersoo L, et al. 2014. Science 346:e1256688 ((https://doi.org/10.1126/science.1256688)); Põlme S, et al. 2020. Fungal Diversity 105:1-16 ((https://doi.org/10.1007/s13225-020-00466-2)) |
| Otu06913 | 4 | k__Fungi(100);p__Basidiomycota(100);c__Agaricomycetes(100);o__Polyporales(100);f__Meruliaceae(100);g__Steccherinum(100);g__Steccherinum_unclassified(100); | Steccherinum | 13 | Saprotroph | Undefined Saprotroph-\|Wood Saprotroph\| | Corticioid | White Rot | Probable | As white_rot (Põlme et al. 2020); Decay wood (Põlme et al. 2020) | Tedersoo L, et al. 2014. Science 346:e1256688 ((https://doi.org/10.1126/science.1256688)); Põlme S, et al. 2020. Fungal Diversity 105:1-16 ((https://doi.org/10.1007/s13225-020-00466-2)) |
| Otu06914 | 4 | k__Fungi(100);p__Ascomycota(100);c__Saccharomycetes(100);o__Saccharomycetales(100);f__Metschnikowiaceae(100);g__Metschnikowia(100);s__Metschnikowia_pulcherrima(100); | Metschnikowia | 13 | Saprotroph | \|Undefined Saprotroph\| | Yeast | NULL | Probable | As nectar-tap_saprotroph (Põlme et al. 2020); Decay sugar-rich substrates (Põlme et al. 2020) | Põlme S, et al. 2020. Fungal Diversity 105:1-16 ((https://doi.org/10.1007/s13225-020-00466-2)) |
| Otu06916 | 4 | k__Fungi(100);p__Basidiomycota(100);c__Agaricomycetes(100);o__Polyporales(100);f__Meruliaceae(100);g__Phlebia(100);s__Phlebia_acanthocystis(100); | Phlebia | 13 | Saprotroph | \|Wood Saprotroph\| | Corticioid | White Rot | Highly Probable | As white_rot (Põlme et al. 2020); Decay wood (Põlme et al. 2020) | Kirk PM et al. 2008. Dictionary of the Fungi. Tenth Edition. CAB International, Wallingford (ISBN: 978-0851998268); Põlme S, et al. 2020. Fungal Diversity 105:1-16 ((https://doi.org/10.1007/s13225-020-00466-2)) |
| Otu06917 | 4 | k__Fungi(100);p__Ascomycota(100);c__Dothideomycetes(100);o__Pleosporales(100);f__Phaeosphaeriaceae(100);f__Phaeosphaeriaceae_unclassified(100);f__Phaeosphaeriaceae_unclassified(100); | Phaeosphaeriaceae | 9 | Pathotroph-Saprotroph | Fungal Parasite-Plant Pathogen-Plant Saprotroph | Microfungus | NULL | Probable | Necrotrophic or saprobic on a wide range of plants, especially monocotyledons; a few species parasitic on other fungi (Cannon & Kirk 2007) | Cannon PF, Kirk PM. 2007. Fungal Families of the World. CAB International, Cambridge (ISBN: 978-0851998275) |
| Otu06920 | 4 | k__Fungi(100);p__Ascomycota(100);c__Saccharomycetes(100);o__Saccharomycetales(100);f__Metschnikowiaceae(100);g__Metschnikowia(100);g__Metschnikowia_unclassified(75); | Metschnikowia | 13 | Saprotroph | \|Undefined Saprotroph\| | Yeast | NULL | Probable | As nectar-tap_saprotroph (Põlme et al. 2020); Decay sugar-rich substrates (Põlme et al. 2020) | Põlme S, et al. 2020. Fungal Diversity 105:1-16 ((https://doi.org/10.1007/s13225-020-00466-2)) |
| Otu06921 | 4 | k__Fungi(100);p__Basidiomycota(100);c__Agaricomycetes(100);o__Agaricales(100);f__Entolomataceae(100);g__Entoloma(100);s__Entoloma_minutisporum(100); | Entoloma | 13 | Pathotroph-Saprotroph-Symbiotroph | Ectomycorrhizal-Fungal Parasite-\|Undefined Saprotroph\|-Undefined Symbiotroph | Agaricoid | NULL | Possible | As soil_saprotroph (Põlme et al. 2020); Decay roots, soil (Põlme et al. 2020) | Rinaldi AC, et al. 2008. Fungal Diversity 33:1-45 ((http://www.fungaldiversity.org/fdp/sfdp/33-1.pdf)) (as ECM/not ECM); Tedersoo L, et al. 2010. Mycorrhiza 20:217-263 ((https://doi.org/10.1007/s00572-009-0274-x; Põlme S, et al. 2020. Fungal Diversity 105:1-16 ((https://doi.org/10.1007/s13225-020-00466-2)); sensu stricto)); http://mycorrhizas.info/ecmf.html ((http://mycorrhizas.info/ecmf.html)) ((as ECM/not ECM) |
| Otu06923 | 4 | k__Fungi(100);p__Ascomycota(100);c__Sordariomycetes(100);o__Sordariales(100);f__Chaetomiaceae(100);g__Chaetomium(100);s__Chaetomium_sp_CGMCC_314188(100); | Chaetomium | 13 | Pathotroph-Saprotroph-Symbiotroph | Animal Pathogen-Dung Saprotroph-Endophyte-Epiphyte-Plant Pathogen-\|Plant Saprotroph\|-Wood Saprotroph | Microfungus | Soft Rot | Possible | As foliar_endophyte (Põlme et al. 2020); As litter_saprotroph (Põlme et al. 2020); culture (Busby et al. 2016, David et al. 2016); Likely opportunistic human pathogen (Irinyi et al. 2015); culture (Massimo et al. 2015); Endophyte detection method; Decay wood, leaf, fruit, seed, soil, dung (Põlme et al. 2020); derived commodities (Cannon & Kirk 2007); Epiphyte detection method; Saprobic, acting as key decay organisms of plant materials and plant | Seehann G, et al. 1975. List of Fungi in Soft-Rot Tissue Tests. Institute fur Holzbiologie und Holzschutz, Bundesforschungsanstalt fur Forst - und Holzwirtshaft, Hamburg - Lohbrugge, Germany; Cannon PF, Kirk PM. 2007. Fungal Families of the World. CAB International, Cambridge ((https://isbnsearch.org/isbn/9780851998275)); Bills GF, et al. 2013. Current Opinion in Microbiology 16:549-565 ((https://doi.org/10.1016/j.mib.2013.08.001)); Tedersoo L, et al. 2014. Science 346:e1256688 ((https://doi.org/10.1126/science.1256688)); Irinyi L, et al. 2015. Medical Mycology 53:313-337 ((https://doi.org/10.1093/mmy/myv008)); Busby PE, et al. 2016. New Phytologist 209:1681-1692 ((https://doi.org/10.1111/nph.13742)); Põlme S, et al. 2020. Fungal Diversity 105:1-16 ((https://doi.org/10.1007/s13225-020-00466-2)) |
| Otu06924 | 4 | k__Fungi(100);p__Ascomycota(100);c__Saccharomycetes(100);o__Saccharomycetales(100);f__Metschnikowiaceae(100);g__Metschnikowia(100);s__Metschnikowia_pulcherrima(100); | Metschnikowia | 13 | Saprotroph | \|Undefined Saprotroph\| | Yeast | NULL | Probable | As nectar-tap_saprotroph (Põlme et al. 2020); Decay sugar-rich substrates (Põlme et al. 2020) | Põlme S, et al. 2020. Fungal Diversity 105:1-16 ((https://doi.org/10.1007/s13225-020-00466-2)) |
| Otu06925 | 4 | k__Fungi(100);p__Ascomycota(100);c__Sordariomycetes(100);o__Ophiostomatales(100);f__Ophiostomataceae(100);g__Sporothrix(100);s__Sporothrix_inflata(100); | Sporothrix | 13 | Pathotroph-Saprotroph-Symbiotroph | Animal Pathogen-Endophyte-\|Plant Pathogen\|-Plant Saprotroph-Undefined Saprotroph-Wood Saprotroph | Microfungus | NULL | Probable | As foliar_endophyte (Põlme et al. 2020); As wood_pathogen (Põlme et al. 2020); culture (Dik et al. 1998); The fungal genus Sporothrix comprises a group of thermodimorphic pathogens that cause skin infections in humans and other mammals. Rose gardenerâ€™s disease (Sporotrichosis) occurs worldwide, with hyperendemic areas in Brazil, China, and South Africa (Rodrigues et al. 2016); The fungus lives throughout the world in soil and on plant matter such as sphagnum moss, rose bushes, and hay (www.cdc.gov); Decay wood (Põlme et al. 2020); Endophyte detection method | Dik AJ, et al. 1998. European Journal of Plant Pathology 104:413-423 ((https://doi.org/10.1023/A:1008025416672)); Rodrigues AM, et al. 2016. PLoS Pathogens 12:e1005638 ((https://doi.org/10.1371/journal.ppat.1005638)); Põlme S, et al. 2020. Fungal Diversity 105:1-16 ((https://doi.org/10.1007/s13225-020-00466-2)); https://www.cdc.gov ((https://www.cdc.gov)) |
| Otu06926 | 4 | k__Fungi(100);p__Ascomycota(100);c__Sordariomycetes(100);o__Hypocreales(100);f__Nectriaceae(100);g__Cylindrocarpon(100);s__Cylindrocarpon_ianthothele_var_majus(75); | Cylindrocarpon | 13 | Pathotroph | \|Plant Pathogen\| | Microfungus | NULL | Probable | NULL | Tedersoo L, et al. 2014. Science 346:e1256688 ((https://doi.org/10.1126/science.1256688)); Põlme S, et al. 2020. Fungal Diversity 105:1-16 ((https://doi.org/10.1007/s13225-020-00466-2)) |
| Otu06927 | 4 | k__Fungi(100);p__Ascomycota(100);c__Saccharomycetes(100);o__Saccharomycetales(100);f__Metschnikowiaceae(100);g__Metschnikowia(100);s__Metschnikowia_pulcherrima(100); | Metschnikowia | 13 | Saprotroph | \|Undefined Saprotroph\| | Yeast | NULL | Probable | As nectar-tap_saprotroph (Põlme et al. 2020); Decay sugar-rich substrates (Põlme et al. 2020) | Põlme S, et al. 2020. Fungal Diversity 105:1-16 ((https://doi.org/10.1007/s13225-020-00466-2)) |
| Otu06928 | 4 | k__Fungi(100);p__Ascomycota(100);c__Saccharomycetes(100);o__Saccharomycetales(100);f__Metschnikowiaceae(100);g__Metschnikowia(100);s__Metschnikowia_pulcherrima(100); | Metschnikowia | 13 | Saprotroph | \|Undefined Saprotroph\| | Yeast | NULL | Probable | As nectar-tap_saprotroph (Põlme et al. 2020); Decay sugar-rich substrates (Põlme et al. 2020) | Põlme S, et al. 2020. Fungal Diversity 105:1-16 ((https://doi.org/10.1007/s13225-020-00466-2)) |
| Otu06929 | 4 | k__Fungi(100);p__Basidiomycota(100);c__Agaricomycetes(100);o__Boletales(100);f__Boletaceae(100);g__Buchwaldoboletus(100);s__Buchwaldoboletus_lignicola(100); | Buchwaldoboletus | 13 | Pathotroph-Saprotroph | \|Fungal Parasite\|-Wood Saprotroph | Agaricoid-Boletoid | Brown Rot | Probable | As mycoparasite (Põlme et al. 2020); Decay fungal material (Põlme et al. 2020); Putatively mycoparasitic (Nuhn et al. 2013) | Nuhn ME et al. 2013. Fungal Biology 117:7-8 ((https://doi.org/10.1016/j.funbio.2013.04.008)); Tedersoo L, et al. 2014. Science 346:e1256688 ((https://doi.org/10.1126/science.1256688)); Põlme S, et al. 2020. Fungal Diversity 105:1-16 ((https://doi.org/10.1007/s13225-020-00466-2)) |
| Otu06930 | 4 | k__Fungi(100);p__Ascomycota(100);c__Sordariomycetes(100);o__Diaporthales(100);f__Gnomoniaceae(100);g__Greeneria(100);s__Greeneria_uvicola(100); | Greeneria | 13 | Pathotroph-Saprotroph | \|Plant Pathogen\|-Undefined Saprotroph | Microfungus | NULL | Probable | NULL | Tedersoo L, et al. 2014. Science 346:e1256688 ((https://doi.org/10.1126/science.1256688)); Põlme S, et al. 2020. Fungal Diversity 105:1-16 ((https://doi.org/10.1007/s13225-020-00466-2)) |
| Otu06931 | 4 | k__Fungi(100);p__Ascomycota(100);c__Saccharomycetes(100);o__Saccharomycetales(100);f__Metschnikowiaceae(100);g__Metschnikowia(100);g__Metschnikowia_unclassified(100); | Metschnikowia | 13 | Saprotroph | \|Undefined Saprotroph\| | Yeast | NULL | Probable | As nectar-tap_saprotroph (Põlme et al. 2020); Decay sugar-rich substrates (Põlme et al. 2020) | Põlme S, et al. 2020. Fungal Diversity 105:1-16 ((https://doi.org/10.1007/s13225-020-00466-2)) |
| Otu05889 | 4 | k__Fungi(100);k__Fungi_unclassified(100);k__Fungi_unclassified(100);k__Fungi_unclassified(100);k__Fungi_unclassified(100);k__Fungi_unclassified(100);k__Fungi_unclassified(100); | - | - | - | - | - | - | - | Unassigned | - |
| Otu05890 | 4 | k__Fungi(100);p__Ascomycota(100);p__Ascomycota_unclassified(100);p__Ascomycota_unclassified(100);p__Ascomycota_unclassified(100);p__Ascomycota_unclassified(100);p__Ascomycota_unclassified(100); | - | - | - | - | - | - | - | Unassigned | - |
| Otu05894 | 4 | k__Fungi(100);p__Ascomycota(100);c__Leotiomycetes(100);c__Leotiomycetes_unclassified(100);c__Leotiomycetes_unclassified(100);c__Leotiomycetes_unclassified(100);c__Leotiomycetes_unclassified(100); | - | - | - | - | - | - | - | Unassigned | - |
| Otu05897 | 4 | k__Fungi(100);k__Fungi_unclassified(100);k__Fungi_unclassified(100);k__Fungi_unclassified(100);k__Fungi_unclassified(100);k__Fungi_unclassified(100);k__Fungi_unclassified(100); | - | - | - | - | - | - | - | Unassigned | - |
| Otu05898 | 4 | k__Fungi(100);p__Ascomycota(75);p__Ascomycota_unclassified(75);p__Ascomycota_unclassified(75);p__Ascomycota_unclassified(75);p__Ascomycota_unclassified(75);p__Ascomycota_unclassified(75); | - | - | - | - | - | - | - | Unassigned | - |
| Otu05899 | 4 | k__Fungi(100);k__Fungi_unclassified(75);k__Fungi_unclassified(75);k__Fungi_unclassified(75);k__Fungi_unclassified(75);k__Fungi_unclassified(75);k__Fungi_unclassified(75); | - | - | - | - | - | - | - | Unassigned | - |
| Otu05902 | 4 | k__Plantae(100);p__unclassified_Plantae(100);c__unclassified_Plantae(100);o__unclassified_Plantae(100);f__unclassified_Plantae(100);g__unclassified_Plantae(100);s__Plantae_sp(100); | - | - | - | - | - | - | - | Unassigned | - |
| Otu05903 | 4 | k__Fungi(100);p__Basidiomycota(100);p__Basidiomycota_unclassified(100);p__Basidiomycota_unclassified(100);p__Basidiomycota_unclassified(100);p__Basidiomycota_unclassified(100);p__Basidiomycota_unclassified(100); | - | - | - | - | - | - | - | Unassigned | - |
| Otu05904 | 4 | k__Plantae(100);p__unclassified_Plantae(100);c__unclassified_Plantae(100);o__unclassified_Plantae(100);f__unclassified_Plantae(100);g__unclassified_Plantae(100);s__Plantae_sp(100); | - | - | - | - | - | - | - | Unassigned | - |
| Otu05906 | 4 | k__Fungi(100);p__Ascomycota(100);c__Dothideomycetes(100);o__Capnodiales(100);o__Capnodiales_unclassified(100);o__Capnodiales_unclassified(100);o__Capnodiales_unclassified(100); | - | - | - | - | - | - | - | Unassigned | - |
| Otu05907 | 4 | k__Fungi(100);p__Ascomycota(100);c__Eurotiomycetes(100);o__Eurotiales(100);f__Trichocomaceae(100);f__Trichocomaceae_unclassified(100);f__Trichocomaceae_unclassified(100); | - | - | - | - | - | - | - | Unassigned | - |
| Otu05908 | 4 | k__Fungi(100);p__Ascomycota(100);p__Ascomycota_unclassified(100);p__Ascomycota_unclassified(100);p__Ascomycota_unclassified(100);p__Ascomycota_unclassified(100);p__Ascomycota_unclassified(100); | - | - | - | - | - | - | - | Unassigned | - |
| Otu05910 | 4 | k__Fungi(100);k__Fungi_unclassified(100);k__Fungi_unclassified(100);k__Fungi_unclassified(100);k__Fungi_unclassified(100);k__Fungi_unclassified(100);k__Fungi_unclassified(100); | - | - | - | - | - | - | - | Unassigned | - |
| Otu05911 | 4 | k__Fungi(100);p__Ascomycota(75);p__Ascomycota_unclassified(75);p__Ascomycota_unclassified(75);p__Ascomycota_unclassified(75);p__Ascomycota_unclassified(75);p__Ascomycota_unclassified(75); | - | - | - | - | - | - | - | Unassigned | - |
| Otu05913 | 4 | k__Fungi(100);k__Fungi_unclassified(100);k__Fungi_unclassified(100);k__Fungi_unclassified(100);k__Fungi_unclassified(100);k__Fungi_unclassified(100);k__Fungi_unclassified(100); | - | - | - | - | - | - | - | Unassigned | - |
| Otu05914 | 4 | k__Fungi(100);p__Ascomycota(100);p__Ascomycota_unclassified(100);p__Ascomycota_unclassified(100);p__Ascomycota_unclassified(100);p__Ascomycota_unclassified(100);p__Ascomycota_unclassified(100); | - | - | - | - | - | - | - | Unassigned | - |
| Otu05915 | 4 | k__Fungi(100);p__Ascomycota(75);p__Ascomycota_unclassified(75);p__Ascomycota_unclassified(75);p__Ascomycota_unclassified(75);p__Ascomycota_unclassified(75);p__Ascomycota_unclassified(75); | - | - | - | - | - | - | - | Unassigned | - |
| Otu05916 | 4 | k__Fungi(100);p__Ascomycota(100);c__Leotiomycetes(100);o__Helotiales(100);o__Helotiales_unclassified(100);o__Helotiales_unclassified(100);o__Helotiales_unclassified(100); | - | - | - | - | - | - | - | Unassigned | - |
| Otu05917 | 4 | k__Fungi(100);k__Fungi_unclassified(100);k__Fungi_unclassified(100);k__Fungi_unclassified(100);k__Fungi_unclassified(100);k__Fungi_unclassified(100);k__Fungi_unclassified(100); | - | - | - | - | - | - | - | Unassigned | - |
| Otu05918 | 4 | k__Fungi(100);p__Ascomycota(75);p__Ascomycota_unclassified(75);p__Ascomycota_unclassified(75);p__Ascomycota_unclassified(75);p__Ascomycota_unclassified(75);p__Ascomycota_unclassified(75); | - | - | - | - | - | - | - | Unassigned | - |
| Otu05920 | 4 | k__Fungi(100);p__Ascomycota(100);p__Ascomycota_unclassified(75);p__Ascomycota_unclassified(75);p__Ascomycota_unclassified(75);p__Ascomycota_unclassified(75);p__Ascomycota_unclassified(75); | - | - | - | - | - | - | - | Unassigned | - |
| Otu05921 | 4 | k__Fungi(100);p__Ascomycota(100);p__Ascomycota_unclassified(100);p__Ascomycota_unclassified(100);p__Ascomycota_unclassified(100);p__Ascomycota_unclassified(100);p__Ascomycota_unclassified(100); | - | - | - | - | - | - | - | Unassigned | - |
| Otu05924 | 4 | k__Fungi(100);p__Ascomycota(75);p__Ascomycota_unclassified(75);p__Ascomycota_unclassified(75);p__Ascomycota_unclassified(75);p__Ascomycota_unclassified(75);p__Ascomycota_unclassified(75); | - | - | - | - | - | - | - | Unassigned | - |
| Otu05925 | 4 | k__Fungi(100);k__Fungi_unclassified(75);k__Fungi_unclassified(75);k__Fungi_unclassified(75);k__Fungi_unclassified(75);k__Fungi_unclassified(75);k__Fungi_unclassified(75); | - | - | - | - | - | - | - | Unassigned | - |
| Otu05926 | 4 | k__Fungi(100);p__Ascomycota(100);c__Sordariomycetes(100);c__Sordariomycetes_unclassified(100);c__Sordariomycetes_unclassified(100);c__Sordariomycetes_unclassified(100);c__Sordariomycetes_unclassified(100); | - | - | - | - | - | - | - | Unassigned | - |
| Otu05929 | 4 | k__Fungi(100);p__Basidiomycota(100);c__Agaricomycetes(100);o__Russulales(100);o__Russulales_unclassified(75);o__Russulales_unclassified(75);o__Russulales_unclassified(75); | - | - | - | - | - | - | - | Unassigned | - |
| Otu05931 | 4 | k__Fungi(100);k__Fungi_unclassified(100);k__Fungi_unclassified(100);k__Fungi_unclassified(100);k__Fungi_unclassified(100);k__Fungi_unclassified(100);k__Fungi_unclassified(100); | - | - | - | - | - | - | - | Unassigned | - |
| Otu05935 | 4 | k__Fungi(100);p__Ascomycota(100);p__Ascomycota_unclassified(100);p__Ascomycota_unclassified(100);p__Ascomycota_unclassified(100);p__Ascomycota_unclassified(100);p__Ascomycota_unclassified(100); | - | - | - | - | - | - | - | Unassigned | - |
| Otu05938 | 4 | k__Fungi(100);p__Ascomycota(100);c__Dothideomycetes(100);o__Pleosporales(100);o__Pleosporales_unclassified(100);o__Pleosporales_unclassified(100);o__Pleosporales_unclassified(100); | - | - | - | - | - | - | - | Unassigned | - |
| Otu05939 | 4 | k__Fungi(100);k__Fungi_unclassified(100);k__Fungi_unclassified(100);k__Fungi_unclassified(100);k__Fungi_unclassified(100);k__Fungi_unclassified(100);k__Fungi_unclassified(100); | - | - | - | - | - | - | - | Unassigned | - |
| Otu05940 | 4 | k__Fungi(100);k__Fungi_unclassified(100);k__Fungi_unclassified(100);k__Fungi_unclassified(100);k__Fungi_unclassified(100);k__Fungi_unclassified(100);k__Fungi_unclassified(100); | - | - | - | - | - | - | - | Unassigned | - |
| Otu05948 | 4 | k__Fungi(100);k__Fungi_unclassified(100);k__Fungi_unclassified(100);k__Fungi_unclassified(100);k__Fungi_unclassified(100);k__Fungi_unclassified(100);k__Fungi_unclassified(100); | - | - | - | - | - | - | - | Unassigned | - |
| Otu05949 | 4 | k__Fungi(100);k__Fungi_unclassified(75);k__Fungi_unclassified(75);k__Fungi_unclassified(75);k__Fungi_unclassified(75);k__Fungi_unclassified(75);k__Fungi_unclassified(75); | - | - | - | - | - | - | - | Unassigned | - |
| Otu05950 | 4 | k__Fungi(100);k__Fungi_unclassified(75);k__Fungi_unclassified(75);k__Fungi_unclassified(75);k__Fungi_unclassified(75);k__Fungi_unclassified(75);k__Fungi_unclassified(75); | - | - | - | - | - | - | - | Unassigned | - |
| Otu05952 | 4 | k__Fungi(100);k__Fungi_unclassified(75);k__Fungi_unclassified(75);k__Fungi_unclassified(75);k__Fungi_unclassified(75);k__Fungi_unclassified(75);k__Fungi_unclassified(75); | - | - | - | - | - | - | - | Unassigned | - |
| Otu05955 | 4 | k__Fungi(100);p__Ascomycota(100);p__Ascomycota_unclassified(100);p__Ascomycota_unclassified(100);p__Ascomycota_unclassified(100);p__Ascomycota_unclassified(100);p__Ascomycota_unclassified(100); | - | - | - | - | - | - | - | Unassigned | - |
| Otu05956 | 4 | k__Fungi(100);p__Ascomycota(100);p__Ascomycota_unclassified(100);p__Ascomycota_unclassified(100);p__Ascomycota_unclassified(100);p__Ascomycota_unclassified(100);p__Ascomycota_unclassified(100); | - | - | - | - | - | - | - | Unassigned | - |
| Otu05957 | 4 | k__Fungi(100);p__Ascomycota(100);c__Dothideomycetes(100);c__Dothideomycetes_unclassified(75);c__Dothideomycetes_unclassified(75);c__Dothideomycetes_unclassified(75);c__Dothideomycetes_unclassified(75); | - | - | - | - | - | - | - | Unassigned | - |
| Otu05960 | 4 | k__Fungi(100);k__Fungi_unclassified(100);k__Fungi_unclassified(100);k__Fungi_unclassified(100);k__Fungi_unclassified(100);k__Fungi_unclassified(100);k__Fungi_unclassified(100); | - | - | - | - | - | - | - | Unassigned | - |
| Otu05961 | 4 | k__Fungi(100);p__Ascomycota(100);c__Dothideomycetes(100);c__Dothideomycetes_unclassified(100);c__Dothideomycetes_unclassified(100);c__Dothideomycetes_unclassified(100);c__Dothideomycetes_unclassified(100); | - | - | - | - | - | - | - | Unassigned | - |
| Otu05969 | 4 | k__Fungi(100);p__Ascomycota(75);p__Ascomycota_unclassified(75);p__Ascomycota_unclassified(75);p__Ascomycota_unclassified(75);p__Ascomycota_unclassified(75);p__Ascomycota_unclassified(75); | - | - | - | - | - | - | - | Unassigned | - |
| Otu05970 | 4 | k__Fungi(100);p__Ascomycota(75);p__Ascomycota_unclassified(75);p__Ascomycota_unclassified(75);p__Ascomycota_unclassified(75);p__Ascomycota_unclassified(75);p__Ascomycota_unclassified(75); | - | - | - | - | - | - | - | Unassigned | - |
| Otu05972 | 4 | k__Fungi(100);k__Fungi_unclassified(75);k__Fungi_unclassified(75);k__Fungi_unclassified(75);k__Fungi_unclassified(75);k__Fungi_unclassified(75);k__Fungi_unclassified(75); | - | - | - | - | - | - | - | Unassigned | - |
| Otu05973 | 4 | k__Fungi(100);k__Fungi_unclassified(100);k__Fungi_unclassified(100);k__Fungi_unclassified(100);k__Fungi_unclassified(100);k__Fungi_unclassified(100);k__Fungi_unclassified(100); | - | - | - | - | - | - | - | Unassigned | - |
| Otu05974 | 4 | k__Fungi(100);p__Ascomycota(100);p__Ascomycota_unclassified(100);p__Ascomycota_unclassified(100);p__Ascomycota_unclassified(100);p__Ascomycota_unclassified(100);p__Ascomycota_unclassified(100); | - | - | - | - | - | - | - | Unassigned | - |
| Otu05975 | 4 | k__Fungi(100);p__Ascomycota(100);c__Dothideomycetes(100);o__Botryosphaeriales(100);o__Botryosphaeriales_unclassified(100);o__Botryosphaeriales_unclassified(100);o__Botryosphaeriales_unclassified(100); | - | - | - | - | - | - | - | Unassigned | - |
| Otu05976 | 4 | k__Fungi(100);p__Ascomycota(100);p__Ascomycota_unclassified(100);p__Ascomycota_unclassified(100);p__Ascomycota_unclassified(100);p__Ascomycota_unclassified(100);p__Ascomycota_unclassified(100); | - | - | - | - | - | - | - | Unassigned | - |
| Otu05978 | 4 | k__Fungi(100);p__Ascomycota(100);p__Ascomycota_unclassified(100);p__Ascomycota_unclassified(100);p__Ascomycota_unclassified(100);p__Ascomycota_unclassified(100);p__Ascomycota_unclassified(100); | - | - | - | - | - | - | - | Unassigned | - |
| Otu05982 | 4 | k__Fungi(100);p__Ascomycota(100);p__Ascomycota_unclassified(100);p__Ascomycota_unclassified(100);p__Ascomycota_unclassified(100);p__Ascomycota_unclassified(100);p__Ascomycota_unclassified(100); | - | - | - | - | - | - | - | Unassigned | - |
| Otu05983 | 4 | k__Fungi(100);k__Fungi_unclassified(75);k__Fungi_unclassified(75);k__Fungi_unclassified(75);k__Fungi_unclassified(75);k__Fungi_unclassified(75);k__Fungi_unclassified(75); | - | - | - | - | - | - | - | Unassigned | - |
| Otu05984 | 4 | k__Fungi(100);p__Ascomycota(100);p__Ascomycota_unclassified(100);p__Ascomycota_unclassified(100);p__Ascomycota_unclassified(100);p__Ascomycota_unclassified(100);p__Ascomycota_unclassified(100); | - | - | - | - | - | - | - | Unassigned | - |
| Otu05985 | 4 | k__Fungi(100);p__Ascomycota(100);p__Ascomycota_unclassified(75);p__Ascomycota_unclassified(75);p__Ascomycota_unclassified(75);p__Ascomycota_unclassified(75);p__Ascomycota_unclassified(75); | - | - | - | - | - | - | - | Unassigned | - |
| Otu05988 | 4 | k__Fungi(100);p__Ascomycota(75);p__Ascomycota_unclassified(75);p__Ascomycota_unclassified(75);p__Ascomycota_unclassified(75);p__Ascomycota_unclassified(75);p__Ascomycota_unclassified(75); | - | - | - | - | - | - | - | Unassigned | - |
| Otu05989 | 4 | k__Fungi(100);p__Ascomycota(100);p__Ascomycota_unclassified(100);p__Ascomycota_unclassified(100);p__Ascomycota_unclassified(100);p__Ascomycota_unclassified(100);p__Ascomycota_unclassified(100); | - | - | - | - | - | - | - | Unassigned | - |
| Otu05990 | 4 | k__Fungi(100);k__Fungi_unclassified(100);k__Fungi_unclassified(100);k__Fungi_unclassified(100);k__Fungi_unclassified(100);k__Fungi_unclassified(100);k__Fungi_unclassified(100); | - | - | - | - | - | - | - | Unassigned | - |
| Otu05992 | 4 | k__Fungi(100);k__Fungi_unclassified(75);k__Fungi_unclassified(75);k__Fungi_unclassified(75);k__Fungi_unclassified(75);k__Fungi_unclassified(75);k__Fungi_unclassified(75); | - | - | - | - | - | - | - | Unassigned | - |
| Otu05995 | 4 | k__Fungi(100);k__Fungi_unclassified(75);k__Fungi_unclassified(75);k__Fungi_unclassified(75);k__Fungi_unclassified(75);k__Fungi_unclassified(75);k__Fungi_unclassified(75); | - | - | - | - | - | - | - | Unassigned | - |
| Otu06001 | 4 | k__Fungi(100);p__Ascomycota(100);c__Sordariomycetes(100);c__Sordariomycetes_unclassified(100);c__Sordariomycetes_unclassified(100);c__Sordariomycetes_unclassified(100);c__Sordariomycetes_unclassified(100); | - | - | - | - | - | - | - | Unassigned | - |
| Otu06004 | 4 | k__Fungi(100);p__Ascomycota(75);p__Ascomycota_unclassified(75);p__Ascomycota_unclassified(75);p__Ascomycota_unclassified(75);p__Ascomycota_unclassified(75);p__Ascomycota_unclassified(75); | - | - | - | - | - | - | - | Unassigned | - |
| Otu06005 | 4 | k__Fungi(100);p__Ascomycota(100);c__Sordariomycetes(100);c__Sordariomycetes_unclassified(100);c__Sordariomycetes_unclassified(100);c__Sordariomycetes_unclassified(100);c__Sordariomycetes_unclassified(100); | - | - | - | - | - | - | - | Unassigned | - |
| Otu06008 | 4 | k__Fungi(100);k__Fungi_unclassified(100);k__Fungi_unclassified(100);k__Fungi_unclassified(100);k__Fungi_unclassified(100);k__Fungi_unclassified(100);k__Fungi_unclassified(100); | - | - | - | - | - | - | - | Unassigned | - |
| Otu06010 | 4 | k__Fungi(100);k__Fungi_unclassified(100);k__Fungi_unclassified(100);k__Fungi_unclassified(100);k__Fungi_unclassified(100);k__Fungi_unclassified(100);k__Fungi_unclassified(100); | - | - | - | - | - | - | - | Unassigned | - |
| Otu06013 | 4 | k__Plantae(100);p__unclassified_Plantae(100);c__unclassified_Plantae(100);o__unclassified_Plantae(100);f__unclassified_Plantae(100);g__unclassified_Plantae(100);s__Plantae_sp(100); | - | - | - | - | - | - | - | Unassigned | - |
| Otu06015 | 4 | k__Fungi(100);p__Ascomycota(100);c__Dothideomycetes(100);o__Capnodiales(100);o__Capnodiales_unclassified(75);o__Capnodiales_unclassified(75);o__Capnodiales_unclassified(75); | - | - | - | - | - | - | - | Unassigned | - |
| Otu06018 | 4 | k__Fungi(100);p__Ascomycota(100);c__Eurotiomycetes(100);o__Chaetothyriales(100);o__Chaetothyriales_unclassified(100);o__Chaetothyriales_unclassified(100);o__Chaetothyriales_unclassified(100); | - | - | - | - | - | - | - | Unassigned | - |
| Otu06019 | 4 | k__Fungi(100);p__Basidiomycota(100);c__Agaricomycetes(100);c__Agaricomycetes_unclassified(100);c__Agaricomycetes_unclassified(100);c__Agaricomycetes_unclassified(100);c__Agaricomycetes_unclassified(100); | - | - | - | - | - | - | - | Unassigned | - |
| Otu06020 | 4 | k__Fungi(100);k__Fungi_unclassified(100);k__Fungi_unclassified(100);k__Fungi_unclassified(100);k__Fungi_unclassified(100);k__Fungi_unclassified(100);k__Fungi_unclassified(100); | - | - | - | - | - | - | - | Unassigned | - |
| Otu06023 | 4 | k__Fungi(100);p__Basidiomycota(100);p__Basidiomycota_unclassified(100);p__Basidiomycota_unclassified(100);p__Basidiomycota_unclassified(100);p__Basidiomycota_unclassified(100);p__Basidiomycota_unclassified(100); | - | - | - | - | - | - | - | Unassigned | - |
| Otu06024 | 4 | k__Fungi(100);k__Fungi_unclassified(100);k__Fungi_unclassified(100);k__Fungi_unclassified(100);k__Fungi_unclassified(100);k__Fungi_unclassified(100);k__Fungi_unclassified(100); | - | - | - | - | - | - | - | Unassigned | - |
| Otu06026 | 4 | k__Fungi(100);p__Ascomycota(100);c__Leotiomycetes(100);o__Helotiales(100);f__Helotiales_family_Incertae_sedis(100);f__Helotiales_family_Incertae_sedis_unclassified(100);f__Helotiales_family_Incertae_sedis_unclassified(100); | - | - | - | - | - | - | - | Unassigned | - |
| Otu06029 | 4 | k__Fungi(100);k__Fungi_unclassified(75);k__Fungi_unclassified(75);k__Fungi_unclassified(75);k__Fungi_unclassified(75);k__Fungi_unclassified(75);k__Fungi_unclassified(75); | - | - | - | - | - | - | - | Unassigned | - |
| Otu06032 | 4 | k__Fungi(100);k__Fungi_unclassified(100);k__Fungi_unclassified(100);k__Fungi_unclassified(100);k__Fungi_unclassified(100);k__Fungi_unclassified(100);k__Fungi_unclassified(100); | - | - | - | - | - | - | - | Unassigned | - |
| Otu06033 | 4 | k__Fungi(100);k__Fungi_unclassified(100);k__Fungi_unclassified(100);k__Fungi_unclassified(100);k__Fungi_unclassified(100);k__Fungi_unclassified(100);k__Fungi_unclassified(100); | - | - | - | - | - | - | - | Unassigned | - |
| Otu06034 | 4 | k__Fungi(100);p__Ascomycota(75);p__Ascomycota_unclassified(75);p__Ascomycota_unclassified(75);p__Ascomycota_unclassified(75);p__Ascomycota_unclassified(75);p__Ascomycota_unclassified(75); | - | - | - | - | - | - | - | Unassigned | - |
| Otu06035 | 4 | k__Fungi(100);k__Fungi_unclassified(100);k__Fungi_unclassified(100);k__Fungi_unclassified(100);k__Fungi_unclassified(100);k__Fungi_unclassified(100);k__Fungi_unclassified(100); | - | - | - | - | - | - | - | Unassigned | - |
| Otu06042 | 4 | k__Fungi(100);k__Fungi_unclassified(100);k__Fungi_unclassified(100);k__Fungi_unclassified(100);k__Fungi_unclassified(100);k__Fungi_unclassified(100);k__Fungi_unclassified(100); | - | - | - | - | - | - | - | Unassigned | - |
| Otu06043 | 4 | k__Fungi(100);k__Fungi_unclassified(100);k__Fungi_unclassified(100);k__Fungi_unclassified(100);k__Fungi_unclassified(100);k__Fungi_unclassified(100);k__Fungi_unclassified(100); | - | - | - | - | - | - | - | Unassigned | - |
| Otu06046 | 4 | k__Fungi(100);k__Fungi_unclassified(100);k__Fungi_unclassified(100);k__Fungi_unclassified(100);k__Fungi_unclassified(100);k__Fungi_unclassified(100);k__Fungi_unclassified(100); | - | - | - | - | - | - | - | Unassigned | - |
| Otu06047 | 4 | k__Fungi(100);p__Ascomycota(75);p__Ascomycota_unclassified(75);p__Ascomycota_unclassified(75);p__Ascomycota_unclassified(75);p__Ascomycota_unclassified(75);p__Ascomycota_unclassified(75); | - | - | - | - | - | - | - | Unassigned | - |
| Otu06049 | 4 | k__Fungi(100);k__Fungi_unclassified(100);k__Fungi_unclassified(100);k__Fungi_unclassified(100);k__Fungi_unclassified(100);k__Fungi_unclassified(100);k__Fungi_unclassified(100); | - | - | - | - | - | - | - | Unassigned | - |
| Otu06053 | 4 | k__Fungi(100);k__Fungi_unclassified(100);k__Fungi_unclassified(100);k__Fungi_unclassified(100);k__Fungi_unclassified(100);k__Fungi_unclassified(100);k__Fungi_unclassified(100); | - | - | - | - | - | - | - | Unassigned | - |
| Otu06055 | 4 | k__Fungi(100);k__Fungi_unclassified(100);k__Fungi_unclassified(100);k__Fungi_unclassified(100);k__Fungi_unclassified(100);k__Fungi_unclassified(100);k__Fungi_unclassified(100); | - | - | - | - | - | - | - | Unassigned | - |
| Otu06059 | 4 | k__Fungi(100);k__Fungi_unclassified(100);k__Fungi_unclassified(100);k__Fungi_unclassified(100);k__Fungi_unclassified(100);k__Fungi_unclassified(100);k__Fungi_unclassified(100); | - | - | - | - | - | - | - | Unassigned | - |
| Otu06060 | 4 | k__Fungi(100);k__Fungi_unclassified(100);k__Fungi_unclassified(100);k__Fungi_unclassified(100);k__Fungi_unclassified(100);k__Fungi_unclassified(100);k__Fungi_unclassified(100); | - | - | - | - | - | - | - | Unassigned | - |
| Otu06067 | 4 | k__Fungi(100);k__Fungi_unclassified(100);k__Fungi_unclassified(100);k__Fungi_unclassified(100);k__Fungi_unclassified(100);k__Fungi_unclassified(100);k__Fungi_unclassified(100); | - | - | - | - | - | - | - | Unassigned | - |
| Otu06069 | 4 | k__Fungi(100);k__Fungi_unclassified(100);k__Fungi_unclassified(100);k__Fungi_unclassified(100);k__Fungi_unclassified(100);k__Fungi_unclassified(100);k__Fungi_unclassified(100); | - | - | - | - | - | - | - | Unassigned | - |
| Otu06072 | 4 | k__Fungi(100);p__Ascomycota(75);p__Ascomycota_unclassified(75);p__Ascomycota_unclassified(75);p__Ascomycota_unclassified(75);p__Ascomycota_unclassified(75);p__Ascomycota_unclassified(75); | - | - | - | - | - | - | - | Unassigned | - |
| Otu06073 | 4 | k__Fungi(100);k__Fungi_unclassified(75);k__Fungi_unclassified(75);k__Fungi_unclassified(75);k__Fungi_unclassified(75);k__Fungi_unclassified(75);k__Fungi_unclassified(75); | - | - | - | - | - | - | - | Unassigned | - |
| Otu06077 | 4 | k__Fungi(100);k__Fungi_unclassified(100);k__Fungi_unclassified(100);k__Fungi_unclassified(100);k__Fungi_unclassified(100);k__Fungi_unclassified(100);k__Fungi_unclassified(100); | - | - | - | - | - | - | - | Unassigned | - |
| Otu06078 | 4 | k__Fungi(100);p__Basidiomycota(100);c__Tremellomycetes(100);c__Tremellomycetes_unclassified(100);c__Tremellomycetes_unclassified(100);c__Tremellomycetes_unclassified(100);c__Tremellomycetes_unclassified(100); | - | - | - | - | - | - | - | Unassigned | - |
| Otu06080 | 4 | k__Fungi(100);p__Ascomycota(75);p__Ascomycota_unclassified(75);p__Ascomycota_unclassified(75);p__Ascomycota_unclassified(75);p__Ascomycota_unclassified(75);p__Ascomycota_unclassified(75); | - | - | - | - | - | - | - | Unassigned | - |
| Otu06081 | 4 | k__Fungi(100);k__Fungi_unclassified(100);k__Fungi_unclassified(100);k__Fungi_unclassified(100);k__Fungi_unclassified(100);k__Fungi_unclassified(100);k__Fungi_unclassified(100); | - | - | - | - | - | - | - | Unassigned | - |
| Otu06083 | 4 | k__Fungi(100);p__Basidiomycota(100);c__Microbotryomycetes(100);o__Sporidiobolales(100);f__Sporidiobolales_family_Incertae_sedis(100);f__Sporidiobolales_family_Incertae_sedis_unclassified(100);f__Sporidiobolales_family_Incertae_sedis_unclassified(100); | - | - | - | - | - | - | - | Unassigned | - |
| Otu06089 | 4 | k__Fungi(100);k__Fungi_unclassified(100);k__Fungi_unclassified(100);k__Fungi_unclassified(100);k__Fungi_unclassified(100);k__Fungi_unclassified(100);k__Fungi_unclassified(100); | - | - | - | - | - | - | - | Unassigned | - |
| Otu06091 | 4 | k__Fungi(100);p__Ascomycota(100);c__Dothideomycetes(100);o__Pleosporales(100);f__Pleosporales_family_Incertae_sedis(100);f__Pleosporales_family_Incertae_sedis_unclassified(75);f__Pleosporales_family_Incertae_sedis_unclassified(75); | - | - | - | - | - | - | - | Unassigned | - |
| Otu06092 | 4 | k__Fungi(100);p__Ascomycota(100);p__Ascomycota_unclassified(100);p__Ascomycota_unclassified(100);p__Ascomycota_unclassified(100);p__Ascomycota_unclassified(100);p__Ascomycota_unclassified(100); | - | - | - | - | - | - | - | Unassigned | - |
| Otu06093 | 4 | k__Fungi(100);k__Fungi_unclassified(100);k__Fungi_unclassified(100);k__Fungi_unclassified(100);k__Fungi_unclassified(100);k__Fungi_unclassified(100);k__Fungi_unclassified(100); | - | - | - | - | - | - | - | Unassigned | - |
| Otu06096 | 4 | k__Fungi(100);k__Fungi_unclassified(100);k__Fungi_unclassified(100);k__Fungi_unclassified(100);k__Fungi_unclassified(100);k__Fungi_unclassified(100);k__Fungi_unclassified(100); | - | - | - | - | - | - | - | Unassigned | - |
| Otu06097 | 4 | k__Fungi(100);k__Fungi_unclassified(100);k__Fungi_unclassified(100);k__Fungi_unclassified(100);k__Fungi_unclassified(100);k__Fungi_unclassified(100);k__Fungi_unclassified(100); | - | - | - | - | - | - | - | Unassigned | - |
| Otu06098 | 4 | k__Fungi(100);p__Ascomycota(100);p__Ascomycota_unclassified(75);p__Ascomycota_unclassified(75);p__Ascomycota_unclassified(75);p__Ascomycota_unclassified(75);p__Ascomycota_unclassified(75); | - | - | - | - | - | - | - | Unassigned | - |
| Otu06103 | 4 | k__Fungi(100);k__Fungi_unclassified(100);k__Fungi_unclassified(100);k__Fungi_unclassified(100);k__Fungi_unclassified(100);k__Fungi_unclassified(100);k__Fungi_unclassified(100); | - | - | - | - | - | - | - | Unassigned | - |
| Otu06104 | 4 | k__Fungi(100);k__Fungi_unclassified(100);k__Fungi_unclassified(100);k__Fungi_unclassified(100);k__Fungi_unclassified(100);k__Fungi_unclassified(100);k__Fungi_unclassified(100); | - | - | - | - | - | - | - | Unassigned | - |
| Otu06105 | 4 | k__Fungi(100);k__Fungi_unclassified(100);k__Fungi_unclassified(100);k__Fungi_unclassified(100);k__Fungi_unclassified(100);k__Fungi_unclassified(100);k__Fungi_unclassified(100); | - | - | - | - | - | - | - | Unassigned | - |
| Otu06108 | 4 | k__Fungi(100);k__Fungi_unclassified(100);k__Fungi_unclassified(100);k__Fungi_unclassified(100);k__Fungi_unclassified(100);k__Fungi_unclassified(100);k__Fungi_unclassified(100); | - | - | - | - | - | - | - | Unassigned | - |
| Otu06111 | 4 | k__Fungi(100);k__Fungi_unclassified(75);k__Fungi_unclassified(75);k__Fungi_unclassified(75);k__Fungi_unclassified(75);k__Fungi_unclassified(75);k__Fungi_unclassified(75); | - | - | - | - | - | - | - | Unassigned | - |
| Otu06112 | 4 | k__Fungi(100);k__Fungi_unclassified(100);k__Fungi_unclassified(100);k__Fungi_unclassified(100);k__Fungi_unclassified(100);k__Fungi_unclassified(100);k__Fungi_unclassified(100); | - | - | - | - | - | - | - | Unassigned | - |
| Otu06113 | 4 | k__Fungi(100);k__Fungi_unclassified(100);k__Fungi_unclassified(100);k__Fungi_unclassified(100);k__Fungi_unclassified(100);k__Fungi_unclassified(100);k__Fungi_unclassified(100); | - | - | - | - | - | - | - | Unassigned | - |
| Otu06114 | 4 | k__Fungi(100);k__Fungi_unclassified(75);k__Fungi_unclassified(75);k__Fungi_unclassified(75);k__Fungi_unclassified(75);k__Fungi_unclassified(75);k__Fungi_unclassified(75); | - | - | - | - | - | - | - | Unassigned | - |
| Otu06116 | 4 | k__Fungi(100);k__Fungi_unclassified(100);k__Fungi_unclassified(100);k__Fungi_unclassified(100);k__Fungi_unclassified(100);k__Fungi_unclassified(100);k__Fungi_unclassified(100); | - | - | - | - | - | - | - | Unassigned | - |
| Otu06118 | 4 | k__Fungi(100);k__Fungi_unclassified(75);k__Fungi_unclassified(75);k__Fungi_unclassified(75);k__Fungi_unclassified(75);k__Fungi_unclassified(75);k__Fungi_unclassified(75); | - | - | - | - | - | - | - | Unassigned | - |
| Otu06119 | 4 | k__Fungi(100);k__Fungi_unclassified(100);k__Fungi_unclassified(100);k__Fungi_unclassified(100);k__Fungi_unclassified(100);k__Fungi_unclassified(100);k__Fungi_unclassified(100); | - | - | - | - | - | - | - | Unassigned | - |
| Otu06121 | 4 | k__Fungi(100);p__Ascomycota(100);c__Dothideomycetes(100);c__Dothideomycetes_unclassified(75);c__Dothideomycetes_unclassified(75);c__Dothideomycetes_unclassified(75);c__Dothideomycetes_unclassified(75); | - | - | - | - | - | - | - | Unassigned | - |
| Otu06126 | 4 | k__Fungi(100);p__Basidiomycota(100);c__Agaricomycetes(100);o__Trechisporales(75);f__Hydnodontaceae(75);f__Hydnodontaceae_unclassified(75);f__Hydnodontaceae_unclassified(75); | - | - | - | - | - | - | - | Unassigned | - |
| Otu06127 | 4 | k__Fungi(100);k__Fungi_unclassified(100);k__Fungi_unclassified(100);k__Fungi_unclassified(100);k__Fungi_unclassified(100);k__Fungi_unclassified(100);k__Fungi_unclassified(100); | - | - | - | - | - | - | - | Unassigned | - |
| Otu06129 | 4 | k__Fungi(100);p__Ascomycota(100);p__Ascomycota_unclassified(100);p__Ascomycota_unclassified(100);p__Ascomycota_unclassified(100);p__Ascomycota_unclassified(100);p__Ascomycota_unclassified(100); | - | - | - | - | - | - | - | Unassigned | - |
| Otu06138 | 4 | k__Fungi(100);k__Fungi_unclassified(75);k__Fungi_unclassified(75);k__Fungi_unclassified(75);k__Fungi_unclassified(75);k__Fungi_unclassified(75);k__Fungi_unclassified(75); | - | - | - | - | - | - | - | Unassigned | - |
| Otu06144 | 4 | k__Fungi(100);p__Ascomycota(75);p__Ascomycota_unclassified(75);p__Ascomycota_unclassified(75);p__Ascomycota_unclassified(75);p__Ascomycota_unclassified(75);p__Ascomycota_unclassified(75); | - | - | - | - | - | - | - | Unassigned | - |
| Otu06146 | 4 | k__Fungi(100);k__Fungi_unclassified(100);k__Fungi_unclassified(100);k__Fungi_unclassified(100);k__Fungi_unclassified(100);k__Fungi_unclassified(100);k__Fungi_unclassified(100); | - | - | - | - | - | - | - | Unassigned | - |
| Otu06147 | 4 | k__Fungi(100);k__Fungi_unclassified(100);k__Fungi_unclassified(100);k__Fungi_unclassified(100);k__Fungi_unclassified(100);k__Fungi_unclassified(100);k__Fungi_unclassified(100); | - | - | - | - | - | - | - | Unassigned | - |
| Otu06149 | 4 | k__Plantae(100);p__unclassified_Plantae(100);c__unclassified_Plantae(100);o__unclassified_Plantae(100);f__unclassified_Plantae(100);g__unclassified_Plantae(100);s__Plantae_sp(100); | - | - | - | - | - | - | - | Unassigned | - |
| Otu06152 | 4 | k__Fungi(100);p__Ascomycota(100);p__Ascomycota_unclassified(75);p__Ascomycota_unclassified(75);p__Ascomycota_unclassified(75);p__Ascomycota_unclassified(75);p__Ascomycota_unclassified(75); | - | - | - | - | - | - | - | Unassigned | - |
| Otu06153 | 4 | k__Fungi(100);k__Fungi_unclassified(100);k__Fungi_unclassified(100);k__Fungi_unclassified(100);k__Fungi_unclassified(100);k__Fungi_unclassified(100);k__Fungi_unclassified(100); | - | - | - | - | - | - | - | Unassigned | - |
| Otu06155 | 4 | k__Fungi(100);p__Ascomycota(100);p__Ascomycota_unclassified(100);p__Ascomycota_unclassified(100);p__Ascomycota_unclassified(100);p__Ascomycota_unclassified(100);p__Ascomycota_unclassified(100); | - | - | - | - | - | - | - | Unassigned | - |
| Otu06156 | 4 | k__Fungi(100);k__Fungi_unclassified(100);k__Fungi_unclassified(100);k__Fungi_unclassified(100);k__Fungi_unclassified(100);k__Fungi_unclassified(100);k__Fungi_unclassified(100); | - | - | - | - | - | - | - | Unassigned | - |
| Otu06157 | 4 | k__Fungi(100);k__Fungi_unclassified(100);k__Fungi_unclassified(100);k__Fungi_unclassified(100);k__Fungi_unclassified(100);k__Fungi_unclassified(100);k__Fungi_unclassified(100); | - | - | - | - | - | - | - | Unassigned | - |
| Otu06163 | 4 | k__Fungi(100);k__Fungi_unclassified(75);k__Fungi_unclassified(75);k__Fungi_unclassified(75);k__Fungi_unclassified(75);k__Fungi_unclassified(75);k__Fungi_unclassified(75); | - | - | - | - | - | - | - | Unassigned | - |
| Otu06166 | 4 | k__Fungi(100);p__Ascomycota(100);c__Leotiomycetes(100);o__Helotiales(100);o__Helotiales_unclassified(100);o__Helotiales_unclassified(100);o__Helotiales_unclassified(100); | - | - | - | - | - | - | - | Unassigned | - |
| Otu06167 | 4 | k__Fungi(100);p__Basidiomycota(100);c__Agaricomycetes(100);o__Trechisporales(100);f__unclassified_Trechisporales(100);g__unclassified_Trechisporales(100);s__Trechisporales_sp(100); | - | - | - | - | - | - | - | Unassigned | - |
| Otu06169 | 4 | k__Fungi(100);k__Fungi_unclassified(100);k__Fungi_unclassified(100);k__Fungi_unclassified(100);k__Fungi_unclassified(100);k__Fungi_unclassified(100);k__Fungi_unclassified(100); | - | - | - | - | - | - | - | Unassigned | - |
| Otu06170 | 4 | k__Fungi(100);p__Ascomycota(100);c__Sordariomycetes(100);c__Sordariomycetes_unclassified(100);c__Sordariomycetes_unclassified(100);c__Sordariomycetes_unclassified(100);c__Sordariomycetes_unclassified(100); | - | - | - | - | - | - | - | Unassigned | - |
| Otu06171 | 4 | k__Fungi(100);k__Fungi_unclassified(75);k__Fungi_unclassified(75);k__Fungi_unclassified(75);k__Fungi_unclassified(75);k__Fungi_unclassified(75);k__Fungi_unclassified(75); | - | - | - | - | - | - | - | Unassigned | - |
| Otu06172 | 4 | k__Fungi(100);k__Fungi_unclassified(100);k__Fungi_unclassified(100);k__Fungi_unclassified(100);k__Fungi_unclassified(100);k__Fungi_unclassified(100);k__Fungi_unclassified(100); | - | - | - | - | - | - | - | Unassigned | - |
| Otu06177 | 4 | k__Fungi(100);k__Fungi_unclassified(100);k__Fungi_unclassified(100);k__Fungi_unclassified(100);k__Fungi_unclassified(100);k__Fungi_unclassified(100);k__Fungi_unclassified(100); | - | - | - | - | - | - | - | Unassigned | - |
| Otu06179 | 4 | k__Fungi(100);k__Fungi_unclassified(100);k__Fungi_unclassified(100);k__Fungi_unclassified(100);k__Fungi_unclassified(100);k__Fungi_unclassified(100);k__Fungi_unclassified(100); | - | - | - | - | - | - | - | Unassigned | - |
| Otu06183 | 4 | k__Fungi(100);p__Ascomycota(100);c__Sordariomycetes(100);c__Sordariomycetes_unclassified(100);c__Sordariomycetes_unclassified(100);c__Sordariomycetes_unclassified(100);c__Sordariomycetes_unclassified(100); | - | - | - | - | - | - | - | Unassigned | - |
| Otu06184 | 4 | k__Fungi(100);k__Fungi_unclassified(100);k__Fungi_unclassified(100);k__Fungi_unclassified(100);k__Fungi_unclassified(100);k__Fungi_unclassified(100);k__Fungi_unclassified(100); | - | - | - | - | - | - | - | Unassigned | - |
| Otu06185 | 4 | k__Fungi(100);p__Basidiomycota(100);p__Basidiomycota_unclassified(100);p__Basidiomycota_unclassified(100);p__Basidiomycota_unclassified(100);p__Basidiomycota_unclassified(100);p__Basidiomycota_unclassified(100); | - | - | - | - | - | - | - | Unassigned | - |
| Otu06189 | 4 | k__Fungi(100);p__Ascomycota(75);p__Ascomycota_unclassified(75);p__Ascomycota_unclassified(75);p__Ascomycota_unclassified(75);p__Ascomycota_unclassified(75);p__Ascomycota_unclassified(75); | - | - | - | - | - | - | - | Unassigned | - |
| Otu06190 | 4 | k__Fungi(100);p__Ascomycota(100);p__Ascomycota_unclassified(100);p__Ascomycota_unclassified(100);p__Ascomycota_unclassified(100);p__Ascomycota_unclassified(100);p__Ascomycota_unclassified(100); | - | - | - | - | - | - | - | Unassigned | - |
| Otu06191 | 4 | k__Fungi(100);p__Ascomycota(75);p__Ascomycota_unclassified(75);p__Ascomycota_unclassified(75);p__Ascomycota_unclassified(75);p__Ascomycota_unclassified(75);p__Ascomycota_unclassified(75); | - | - | - | - | - | - | - | Unassigned | - |
| Otu06192 | 4 | k__Fungi(100);k__Fungi_unclassified(75);k__Fungi_unclassified(75);k__Fungi_unclassified(75);k__Fungi_unclassified(75);k__Fungi_unclassified(75);k__Fungi_unclassified(75); | - | - | - | - | - | - | - | Unassigned | - |
| Otu06193 | 4 | k__Fungi(100);p__Ascomycota(100);p__Ascomycota_unclassified(100);p__Ascomycota_unclassified(100);p__Ascomycota_unclassified(100);p__Ascomycota_unclassified(100);p__Ascomycota_unclassified(100); | - | - | - | - | - | - | - | Unassigned | - |
| Otu06194 | 4 | k__Fungi(100);p__Ascomycota(75);p__Ascomycota_unclassified(75);p__Ascomycota_unclassified(75);p__Ascomycota_unclassified(75);p__Ascomycota_unclassified(75);p__Ascomycota_unclassified(75); | - | - | - | - | - | - | - | Unassigned | - |
| Otu06198 | 4 | k__Fungi(100);k__Fungi_unclassified(100);k__Fungi_unclassified(100);k__Fungi_unclassified(100);k__Fungi_unclassified(100);k__Fungi_unclassified(100);k__Fungi_unclassified(100); | - | - | - | - | - | - | - | Unassigned | - |
| Otu06202 | 4 | k__Fungi(100);k__Fungi_unclassified(100);k__Fungi_unclassified(100);k__Fungi_unclassified(100);k__Fungi_unclassified(100);k__Fungi_unclassified(100);k__Fungi_unclassified(100); | - | - | - | - | - | - | - | Unassigned | - |
| Otu06204 | 4 | k__Fungi(100);k__Fungi_unclassified(100);k__Fungi_unclassified(100);k__Fungi_unclassified(100);k__Fungi_unclassified(100);k__Fungi_unclassified(100);k__Fungi_unclassified(100); | - | - | - | - | - | - | - | Unassigned | - |
| Otu06205 | 4 | k__Fungi(100);p__Ascomycota(100);c__Eurotiomycetes(75);o__Chaetothyriales(75);o__Chaetothyriales_unclassified(75);o__Chaetothyriales_unclassified(75);o__Chaetothyriales_unclassified(75); | - | - | - | - | - | - | - | Unassigned | - |
| Otu06207 | 4 | k__Fungi(100);k__Fungi_unclassified(100);k__Fungi_unclassified(100);k__Fungi_unclassified(100);k__Fungi_unclassified(100);k__Fungi_unclassified(100);k__Fungi_unclassified(100); | - | - | - | - | - | - | - | Unassigned | - |
| Otu06210 | 4 | k__Fungi(100);p__Ascomycota(75);p__Ascomycota_unclassified(75);p__Ascomycota_unclassified(75);p__Ascomycota_unclassified(75);p__Ascomycota_unclassified(75);p__Ascomycota_unclassified(75); | - | - | - | - | - | - | - | Unassigned | - |
| Otu06212 | 4 | k__Fungi(100);k__Fungi_unclassified(75);k__Fungi_unclassified(75);k__Fungi_unclassified(75);k__Fungi_unclassified(75);k__Fungi_unclassified(75);k__Fungi_unclassified(75); | - | - | - | - | - | - | - | Unassigned | - |
| Otu06215 | 4 | k__Fungi(100);k__Fungi_unclassified(75);k__Fungi_unclassified(75);k__Fungi_unclassified(75);k__Fungi_unclassified(75);k__Fungi_unclassified(75);k__Fungi_unclassified(75); | - | - | - | - | - | - | - | Unassigned | - |
| Otu06222 | 4 | k__Fungi(100);p__Ascomycota(100);p__Ascomycota_unclassified(100);p__Ascomycota_unclassified(100);p__Ascomycota_unclassified(100);p__Ascomycota_unclassified(100);p__Ascomycota_unclassified(100); | - | - | - | - | - | - | - | Unassigned | - |

| | | | | | | | | | | | |
|---|---|---|---|---|---|---|---|---|---|---|---|
| Otu06223 | 4 | k__Fungi(100);k__Fungi_unclassified(75);k__Fungi_unclassified(75);k__Fungi_unclassified(75);k__Fungi_unclassified(75);k__Fungi_unclassified(75);k__Fungi_unclassified(75); | - | - | - | - | - | - | - | Unassigned | - |
| Otu06225 | 4 | k__Fungi(100);k__Fungi_unclassified(100);k__Fungi_unclassified(100);k__Fungi_unclassified(100);k__Fungi_unclassified(100);k__Fungi_unclassified(100);k__Fungi_unclassified(100); | - | - | - | - | - | - | - | Unassigned | - |
| Otu06226 | 4 | k__Fungi(100);p__Basidiomycota(100);c__Microbotryomycetes(100);o__Sporidiobolales(100);f__Sporidiobolales_family_Incertae_sedis(100);f__Sporidiobolales_family_Incertae_sedis_unclassified(75);f__Sporidiobolales_family_Incertae_sedis_unclassified(75); | - | - | - | - | - | - | - | Unassigned | - |
| Otu06229 | 4 | k__Fungi(100);k__Fungi_unclassified(100);k__Fungi_unclassified(100);k__Fungi_unclassified(100);k__Fungi_unclassified(100);k__Fungi_unclassified(100);k__Fungi_unclassified(100); | - | - | - | - | - | - | - | Unassigned | - |
| Otu06231 | 4 | k__Fungi(100);k__Fungi_unclassified(100);k__Fungi_unclassified(100);k__Fungi_unclassified(100);k__Fungi_unclassified(100);k__Fungi_unclassified(100);k__Fungi_unclassified(100); | - | - | - | - | - | - | - | Unassigned | - |
| Otu06232 | 4 | k__Fungi(100);k__Fungi_unclassified(100);k__Fungi_unclassified(100);k__Fungi_unclassified(100);k__Fungi_unclassified(100);k__Fungi_unclassified(100);k__Fungi_unclassified(100); | - | - | - | - | - | - | - | Unassigned | - |
| Otu06233 | 4 | k__Fungi(100);k__Fungi_unclassified(100);k__Fungi_unclassified(100);k__Fungi_unclassified(100);k__Fungi_unclassified(100);k__Fungi_unclassified(100);k__Fungi_unclassified(100); | - | - | - | - | - | - | - | Unassigned | - |
| Otu06236 | 4 | k__Fungi(100);k__Fungi_unclassified(100);k__Fungi_unclassified(100);k__Fungi_unclassified(100);k__Fungi_unclassified(100);k__Fungi_unclassified(100);k__Fungi_unclassified(100); | - | - | - | - | - | - | - | Unassigned | - |
| Otu06243 | 4 | k__Fungi(100);p__Basidiomycota(100);c__Agaricomycetes(100);c__Agaricomycetes_unclassified(100);c__Agaricomycetes_unclassified(100);c__Agaricomycetes_unclassified(100);c__Agaricomycetes_unclassified(100); | - | - | - | - | - | - | - | Unassigned | - |
| Otu06244 | 4 | k__Fungi(100);k__Fungi_unclassified(100);k__Fungi_unclassified(100);k__Fungi_unclassified(100);k__Fungi_unclassified(100);k__Fungi_unclassified(100);k__Fungi_unclassified(100); | - | - | - | - | - | - | - | Unassigned | - |
| Otu06245 | 4 | k__Fungi(100);k__Fungi_unclassified(100);k__Fungi_unclassified(100);k__Fungi_unclassified(100);k__Fungi_unclassified(100);k__Fungi_unclassified(100);k__Fungi_unclassified(100); | - | - | - | - | - | - | - | Unassigned | - |
| Otu06248 | 4 | k__Fungi(100);k__Fungi_unclassified(75);k__Fungi_unclassified(75);k__Fungi_unclassified(75);k__Fungi_unclassified(75);k__Fungi_unclassified(75);k__Fungi_unclassified(75); | - | - | - | - | - | - | - | Unassigned | - |
| Otu06249 | 4 | k__Fungi(100);p__Ascomycota(75);p__Ascomycota_unclassified(75);p__Ascomycota_unclassified(75);p__Ascomycota_unclassified(75);p__Ascomycota_unclassified(75);p__Ascomycota_unclassified(75); | - | - | - | - | - | - | - | Unassigned | - |
| Otu06250 | 4 | k__Fungi(100);k__Fungi_unclassified(100);k__Fungi_unclassified(100);k__Fungi_unclassified(100);k__Fungi_unclassified(100);k__Fungi_unclassified(100);k__Fungi_unclassified(100); | - | - | - | - | - | - | - | Unassigned | - |
| Otu06251 | 4 | k__Fungi(100);k__Fungi_unclassified(75);k__Fungi_unclassified(75);k__Fungi_unclassified(75);k__Fungi_unclassified(75);k__Fungi_unclassified(75);k__Fungi_unclassified(75); | - | - | - | - | - | - | - | Unassigned | - |
| Otu06253 | 4 | k__Plantae(100);p__unclassified_Plantae(100);c__unclassified_Plantae(100);o__unclassified_Plantae(100);f__unclassified_Plantae(100);g__unclassified_Plantae(100);s__Plantae_sp(100); | - | - | - | - | - | - | - | Unassigned | - |
| Otu06255 | 4 | k__Fungi(100);k__Fungi_unclassified(100);k__Fungi_unclassified(100);k__Fungi_unclassified(100);k__Fungi_unclassified(100);k__Fungi_unclassified(100);k__Fungi_unclassified(100); | - | - | - | - | - | - | - | Unassigned | - |
| Otu06258 | 4 | k__Fungi(100);k__Fungi_unclassified(100);k__Fungi_unclassified(100);k__Fungi_unclassified(100);k__Fungi_unclassified(100);k__Fungi_unclassified(100);k__Fungi_unclassified(100); | - | - | - | - | - | - | - | Unassigned | - |
| Otu06259 | 4 | k__Fungi(100);k__Fungi_unclassified(75);k__Fungi_unclassified(75);k__Fungi_unclassified(75);k__Fungi_unclassified(75);k__Fungi_unclassified(75);k__Fungi_unclassified(75); | - | - | - | - | - | - | - | Unassigned | - |
| Otu06260 | 4 | k__Fungi(100);p__Ascomycota(100);c__Dothideomycetes(100);o__Capnodiales(75);o__Capnodiales_unclassified(75);o__Capnodiales_unclassified(75);o__Capnodiales_unclassified(75); | - | - | - | - | - | - | - | Unassigned | - |
| Otu06266 | 4 | k__Fungi(100);k__Fungi_unclassified(100);k__Fungi_unclassified(100);k__Fungi_unclassified(100);k__Fungi_unclassified(100);k__Fungi_unclassified(100);k__Fungi_unclassified(100); | - | - | - | - | - | - | - | Unassigned | - |
| Otu06267 | 4 | k__Fungi(100);p__Ascomycota(100);p__Ascomycota_unclassified(100);p__Ascomycota_unclassified(100);p__Ascomycota_unclassified(100);p__Ascomycota_unclassified(100);p__Ascomycota_unclassified(100); | - | - | - | - | - | - | - | Unassigned | - |
| Otu06268 | 4 | k__Fungi(100);k__Fungi_unclassified(100);k__Fungi_unclassified(100);k__Fungi_unclassified(100);k__Fungi_unclassified(100);k__Fungi_unclassified(100);k__Fungi_unclassified(100); | - | - | - | - | - | - | - | Unassigned | - |
| Otu06270 | 4 | k__Plantae(100);p__unclassified_Plantae(100);c__unclassified_Plantae(100);o__unclassified_Plantae(100);f__unclassified_Plantae(100);g__unclassified_Plantae(100);s__Plantae_sp(100); | - | - | - | - | - | - | - | Unassigned | - |
| Otu06271 | 4 | k__Fungi(100);k__Fungi_unclassified(75);k__Fungi_unclassified(75);k__Fungi_unclassified(75);k__Fungi_unclassified(75);k__Fungi_unclassified(75);k__Fungi_unclassified(75); | - | - | - | - | - | - | - | Unassigned | - |
| Otu06276 | 4 | k__Fungi(100);k__Fungi_unclassified(100);k__Fungi_unclassified(100);k__Fungi_unclassified(100);k__Fungi_unclassified(100);k__Fungi_unclassified(100);k__Fungi_unclassified(100); | - | - | - | - | - | - | - | Unassigned | - |
| Otu06277 | 4 | k__Fungi(100);k__Fungi_unclassified(75);k__Fungi_unclassified(75);k__Fungi_unclassified(75);k__Fungi_unclassified(75);k__Fungi_unclassified(75);k__Fungi_unclassified(75); | - | - | - | - | - | - | - | Unassigned | - |
| Otu06280 | 4 | k__Fungi(100);k__Fungi_unclassified(100);k__Fungi_unclassified(100);k__Fungi_unclassified(100);k__Fungi_unclassified(100);k__Fungi_unclassified(100);k__Fungi_unclassified(100); | - | - | - | - | - | - | - | Unassigned | - |
| Otu06285 | 4 | k__Fungi(100);k__Fungi_unclassified(75);k__Fungi_unclassified(75);k__Fungi_unclassified(75);k__Fungi_unclassified(75);k__Fungi_unclassified(75);k__Fungi_unclassified(75); | - | - | - | - | - | - | - | Unassigned | - |
| Otu06287 | 4 | k__Fungi(100);k__Fungi_unclassified(75);k__Fungi_unclassified(75);k__Fungi_unclassified(75);k__Fungi_unclassified(75);k__Fungi_unclassified(75);k__Fungi_unclassified(75); | - | - | - | - | - | - | - | Unassigned | - |
| Otu06289 | 4 | k__Fungi(100);p__Ascomycota(75);p__Ascomycota_unclassified(75);p__Ascomycota_unclassified(75);p__Ascomycota_unclassified(75);p__Ascomycota_unclassified(75);p__Ascomycota_unclassified(75); | - | - | - | - | - | - | - | Unassigned | - |
| Otu06290 | 4 | k__Fungi(100);p__Ascomycota(100);p__Ascomycota_unclassified(100);p__Ascomycota_unclassified(100);p__Ascomycota_unclassified(100);p__Ascomycota_unclassified(100);p__Ascomycota_unclassified(100); | - | - | - | - | - | - | - | Unassigned | - |
| Otu06293 | 4 | k__Fungi(100);p__Basidiomycota(100);c__Agaricomycetes(100);o__Polyporales(100);o__Polyporales_unclassified(100);o__Polyporales_unclassified(100);o__Polyporales_unclassified(100); | - | - | - | - | - | - | - | Unassigned | - |
| Otu06295 | 4 | k__Fungi(100);p__Basidiomycota(100);c__Agaricomycetes(100);c__Agaricomycetes_unclassified(100);c__Agaricomycetes_unclassified(100);c__Agaricomycetes_unclassified(100);c__Agaricomycetes_unclassified(100); | - | - | - | - | - | - | - | Unassigned | - |
| Otu06297 | 4 | k__Fungi(100);k__Fungi_unclassified(100);k__Fungi_unclassified(100);k__Fungi_unclassified(100);k__Fungi_unclassified(100);k__Fungi_unclassified(100);k__Fungi_unclassified(100); | - | - | - | - | - | - | - | Unassigned | - |
| Otu06302 | 4 | k__Fungi(100);p__Ascomycota(100);c__Leotiomycetes(100);o__Helotiales(100);o__Helotiales_unclassified(75);o__Helotiales_unclassified(75);o__Helotiales_unclassified(75); | - | - | - | - | - | - | - | Unassigned | - |
| Otu06303 | 4 | k__Fungi(100);k__Fungi_unclassified(75);k__Fungi_unclassified(75);k__Fungi_unclassified(75);k__Fungi_unclassified(75);k__Fungi_unclassified(75);k__Fungi_unclassified(75); | - | - | - | - | - | - | - | Unassigned | - |
| Otu06305 | 4 | k__Fungi(100);p__Ascomycota(100);p__Ascomycota_unclassified(100);p__Ascomycota_unclassified(100);p__Ascomycota_unclassified(100);p__Ascomycota_unclassified(100);p__Ascomycota_unclassified(100); | - | - | - | - | - | - | - | Unassigned | - |
| Otu06306 | 4 | k__Fungi(100);p__Basidiomycota(100);c__Pucciniomycetes(100);o__Pucciniales(100);o__Pucciniales_unclassified(100);o__Pucciniales_unclassified(100);o__Pucciniales_unclassified(100); | - | - | - | - | - | - | - | Unassigned | - |
| Otu06311 | 4 | k__Fungi(100);k__Fungi_unclassified(100);k__Fungi_unclassified(100);k__Fungi_unclassified(100);k__Fungi_unclassified(100);k__Fungi_unclassified(100);k__Fungi_unclassified(100); | - | - | - | - | - | - | - | Unassigned | - |
| Otu06313 | 4 | k__Fungi(100);k__Fungi_unclassified(75);k__Fungi_unclassified(75);k__Fungi_unclassified(75);k__Fungi_unclassified(75);k__Fungi_unclassified(75);k__Fungi_unclassified(75); | - | - | - | - | - | - | - | Unassigned | - |
| Otu06314 | 4 | k__Fungi(100);p__Basidiomycota(100);c__Agaricomycetes(100);o__Trechisporales(100);f__Hydnodontaceae(100);f__Hydnodontaceae_unclassified(100);f__Hydnodontaceae_unclassified(100); | - | - | - | - | - | - | - | Unassigned | - |
| Otu06315 | 4 | k__Fungi(100);p__Ascomycota(75);p__Ascomycota_unclassified(75);p__Ascomycota_unclassified(75);p__Ascomycota_unclassified(75);p__Ascomycota_unclassified(75);p__Ascomycota_unclassified(75); | - | - | - | - | - | - | - | Unassigned | - |
| Otu06318 | 4 | k__Fungi(100);p__Basidiomycota(100);c__Agaricomycetes(100);c__Agaricomycetes_unclassified(100);c__Agaricomycetes_unclassified(100);c__Agaricomycetes_unclassified(100);c__Agaricomycetes_unclassified(100); | - | - | - | - | - | - | - | Unassigned | - |
| Otu06319 | 4 | k__Fungi(100);p__Ascomycota(100);c__Sordariomycetes(100);o__Xylariales(75);o__Xylariales_unclassified(75);o__Xylariales_unclassified(75);o__Xylariales_unclassified(75); | - | - | - | - | - | - | - | Unassigned | - |
| Otu06320 | 4 | k__Fungi(100);p__Ascomycota(100);p__Ascomycota_unclassified(100);p__Ascomycota_unclassified(100);p__Ascomycota_unclassified(100);p__Ascomycota_unclassified(100);p__Ascomycota_unclassified(100); | - | - | - | - | - | - | - | Unassigned | - |
| Otu06321 | 4 | k__Fungi(100);p__Ascomycota(75);p__Ascomycota_unclassified(75);p__Ascomycota_unclassified(75);p__Ascomycota_unclassified(75);p__Ascomycota_unclassified(75);p__Ascomycota_unclassified(75); | - | - | - | - | - | - | - | Unassigned | - |
| Otu06322 | 4 | k__Fungi(100);k__Fungi_unclassified(75);k__Fungi_unclassified(75);k__Fungi_unclassified(75);k__Fungi_unclassified(75);k__Fungi_unclassified(75);k__Fungi_unclassified(75); | - | - | - | - | - | - | - | Unassigned | - |
| Otu06326 | 4 | k__Fungi(100);k__Fungi_unclassified(100);k__Fungi_unclassified(100);k__Fungi_unclassified(100);k__Fungi_unclassified(100);k__Fungi_unclassified(100);k__Fungi_unclassified(100); | - | - | - | - | - | - | - | Unassigned | - |
| Otu06327 | 4 | k__Fungi(100);k__Fungi_unclassified(100);k__Fungi_unclassified(100);k__Fungi_unclassified(100);k__Fungi_unclassified(100);k__Fungi_unclassified(100);k__Fungi_unclassified(100); | - | - | - | - | - | - | - | Unassigned | - |
| Otu06328 | 4 | k__Fungi(100);k__Fungi_unclassified(100);k__Fungi_unclassified(100);k__Fungi_unclassified(100);k__Fungi_unclassified(100);k__Fungi_unclassified(100);k__Fungi_unclassified(100); | - | - | - | - | - | - | - | Unassigned | - |
| Otu06333 | 4 | k__Fungi(100);k__Fungi_unclassified(75);k__Fungi_unclassified(75);k__Fungi_unclassified(75);k__Fungi_unclassified(75);k__Fungi_unclassified(75);k__Fungi_unclassified(75); | - | - | - | - | - | - | - | Unassigned | - |
| Otu06335 | 4 | k__Fungi(100);k__Fungi_unclassified(100);k__Fungi_unclassified(100);k__Fungi_unclassified(100);k__Fungi_unclassified(100);k__Fungi_unclassified(100);k__Fungi_unclassified(100); | - | - | - | - | - | - | - | Unassigned | - |
| Otu06336 | 4 | k__Fungi(100);k__Fungi_unclassified(100);k__Fungi_unclassified(100);k__Fungi_unclassified(100);k__Fungi_unclassified(100);k__Fungi_unclassified(100);k__Fungi_unclassified(100); | - | - | - | - | - | - | - | Unassigned | - |
| Otu06337 | 4 | k__Fungi(100);p__Basidiomycota(100);c__Agaricomycetes(100);c__Agaricomycetes_unclassified(100);c__Agaricomycetes_unclassified(100);c__Agaricomycetes_unclassified(100);c__Agaricomycetes_unclassified(100); | - | - | - | - | - | - | - | Unassigned | - |
| Otu06338 | 4 | k__Fungi(100);p__Ascomycota(100);p__Ascomycota_unclassified(100);p__Ascomycota_unclassified(100);p__Ascomycota_unclassified(100);p__Ascomycota_unclassified(100);p__Ascomycota_unclassified(100); | - | - | - | - | - | - | - | Unassigned | - |
| Otu06341 | 4 | k__Fungi(100);k__Fungi_unclassified(100);k__Fungi_unclassified(100);k__Fungi_unclassified(100);k__Fungi_unclassified(100);k__Fungi_unclassified(100);k__Fungi_unclassified(100); | - | - | - | - | - | - | - | Unassigned | - |
| Otu06344 | 4 | k__Fungi(100);k__Fungi_unclassified(75);k__Fungi_unclassified(75);k__Fungi_unclassified(75);k__Fungi_unclassified(75);k__Fungi_unclassified(75);k__Fungi_unclassified(75); | - | - | - | - | - | - | - | Unassigned | - |
| Otu06346 | 4 | k__Fungi(100);p__Ascomycota(100);p__Ascomycota_unclassified(100);p__Ascomycota_unclassified(100);p__Ascomycota_unclassified(100);p__Ascomycota_unclassified(100);p__Ascomycota_unclassified(100); | - | - | - | - | - | - | - | Unassigned | - |
| Otu06349 | 4 | k__Fungi(100);p__Ascomycota(75);p__Ascomycota_unclassified(75);p__Ascomycota_unclassified(75);p__Ascomycota_unclassified(75);p__Ascomycota_unclassified(75);p__Ascomycota_unclassified(75); | - | - | - | - | - | - | - | Unassigned | - |
| Otu06352 | 4 | k__Fungi(100);k__Fungi_unclassified(100);k__Fungi_unclassified(100);k__Fungi_unclassified(100);k__Fungi_unclassified(100);k__Fungi_unclassified(100);k__Fungi_unclassified(100); | - | - | - | - | - | - | - | Unassigned | - |
| Otu06353 | 4 | k__Fungi(100);p__Basidiomycota(100);c__Agaricomycetes(100);c__Agaricomycetes_unclassified(100);c__Agaricomycetes_unclassified(100);c__Agaricomycetes_unclassified(100);c__Agaricomycetes_unclassified(100); | - | - | - | - | - | - | - | Unassigned | - |
| Otu06354 | 4 | k__Fungi(100);k__Fungi_unclassified(75);k__Fungi_unclassified(75);k__Fungi_unclassified(75);k__Fungi_unclassified(75);k__Fungi_unclassified(75);k__Fungi_unclassified(75); | - | - | - | - | - | - | - | Unassigned | - |
| Otu06356 | 4 | k__Fungi(100);p__Ascomycota(100);p__Ascomycota_unclassified(100);p__Ascomycota_unclassified(100);p__Ascomycota_unclassified(100);p__Ascomycota_unclassified(100);p__Ascomycota_unclassified(100); | - | - | - | - | - | - | - | Unassigned | - |
| Otu06358 | 4 | k__Fungi(100);p__Basidiomycota(100);p__Basidiomycota_unclassified(100);p__Basidiomycota_unclassified(100);p__Basidiomycota_unclassified(100);p__Basidiomycota_unclassified(100);p__Basidiomycota_unclassified(100); | - | - | - | - | - | - | - | Unassigned | - |
| Otu06360 | 4 | k__Fungi(100);p__Ascomycota(75);p__Ascomycota_unclassified(75);p__Ascomycota_unclassified(75);p__Ascomycota_unclassified(75);p__Ascomycota_unclassified(75);p__Ascomycota_unclassified(75); | - | - | - | - | - | - | - | Unassigned | - |
| Otu06362 | 4 | k__Fungi(100);k__Fungi_unclassified(75);k__Fungi_unclassified(75);k__Fungi_unclassified(75);k__Fungi_unclassified(75);k__Fungi_unclassified(75);k__Fungi_unclassified(75); | - | - | - | - | - | - | - | Unassigned | - |
| Otu06364 | 4 | k__Fungi(100);p__Ascomycota(75);p__Ascomycota_unclassified(75);p__Ascomycota_unclassified(75);p__Ascomycota_unclassified(75);p__Ascomycota_unclassified(75);p__Ascomycota_unclassified(75); | - | - | - | - | - | - | - | Unassigned | - |
| Otu06365 | 4 | k__Fungi(100);k__Fungi_unclassified(100);k__Fungi_unclassified(100);k__Fungi_unclassified(100);k__Fungi_unclassified(100);k__Fungi_unclassified(100);k__Fungi_unclassified(100); | - | - | - | - | - | - | - | Unassigned | - |
| Otu06367 | 4 | k__Fungi(100);k__Fungi_unclassified(100);k__Fungi_unclassified(100);k__Fungi_unclassified(100);k__Fungi_unclassified(100);k__Fungi_unclassified(100);k__Fungi_unclassified(100); | - | - | - | - | - | - | - | Unassigned | - |
| Otu06368 | 4 | k__Fungi(100);p__Basidiomycota(100);c__Agaricomycetes(100);c__Agaricomycetes_unclassified(100);c__Agaricomycetes_unclassified(100);c__Agaricomycetes_unclassified(100);c__Agaricomycetes_unclassified(100); | - | - | - | - | - | - | - | Unassigned | - |
| Otu06370 | 4 | k__Fungi(100);p__Ascomycota(75);p__Ascomycota_unclassified(75);p__Ascomycota_unclassified(75);p__Ascomycota_unclassified(75);p__Ascomycota_unclassified(75);p__Ascomycota_unclassified(75); | - | - | - | - | - | - | - | Unassigned | - |
| Otu06374 | 4 | k__Fungi(100);k__Fungi_unclassified(75);k__Fungi_unclassified(75);k__Fungi_unclassified(75);k__Fungi_unclassified(75);k__Fungi_unclassified(75);k__Fungi_unclassified(75); | - | - | - | - | - | - | - | Unassigned | - |
| Otu06375 | 4 | k__Fungi(100);p__Ascomycota(75);p__Ascomycota_unclassified(75);p__Ascomycota_unclassified(75);p__Ascomycota_unclassified(75);p__Ascomycota_unclassified(75);p__Ascomycota_unclassified(75); | - | - | - | - | - | - | - | Unassigned | - |
| Otu06380 | 4 | k__Fungi(100);p__Ascomycota(100);p__Ascomycota_unclassified(100);p__Ascomycota_unclassified(100);p__Ascomycota_unclassified(100);p__Ascomycota_unclassified(100);p__Ascomycota_unclassified(100); | - | - | - | - | - | - | - | Unassigned | - |
| Otu06383 | 4 | k__Fungi(100);k__Fungi_unclassified(100);k__Fungi_unclassified(100);k__Fungi_unclassified(100);k__Fungi_unclassified(100);k__Fungi_unclassified(100);k__Fungi_unclassified(100); | - | - | - | - | - | - | - | Unassigned | - |
| Otu06386 | 4 | k__Fungi(100);p__Ascomycota(100);c__Orbiliomycetes(100);o__unclassified_Orbiliomycetes(100);f__unclassified_Orbiliomycetes(100);g__unclassified_Orbiliomycetes(100);s__Orbiliomycetes_sp(100); | - | - | - | - | - | - | - | Unassigned | - |
| Otu06387 | 4 | k__Fungi(100);k__Fungi_unclassified(100);k__Fungi_unclassified(100);k__Fungi_unclassified(100);k__Fungi_unclassified(100);k__Fungi_unclassified(100);k__Fungi_unclassified(100); | - | - | - | - | - | - | - | Unassigned | - |
| Otu06389 | 4 | k__Fungi(100);p__Ascomycota(100);c__Sordariomycetes(100);c__Sordariomycetes_unclassified(100);c__Sordariomycetes_unclassified(100);c__Sordariomycetes_unclassified(100);c__Sordariomycetes_unclassified(100); | - | - | - | - | - | - | - | Unassigned | - |
| Otu06390 | 4 | k__Fungi(100);p__Ascomycota(100);p__Ascomycota_unclassified(75);p__Ascomycota_unclassified(75);p__Ascomycota_unclassified(75);p__Ascomycota_unclassified(75);p__Ascomycota_unclassified(75); | - | - | - | - | - | - | - | Unassigned | - |
| Otu06391 | 4 | k__Fungi(100);k__Fungi_unclassified(75);k__Fungi_unclassified(75);k__Fungi_unclassified(75);k__Fungi_unclassified(75);k__Fungi_unclassified(75);k__Fungi_unclassified(75); | - | - | - | - | - | - | - | Unassigned | - |
| Otu06392 | 4 | k__Fungi(100);k__Fungi_unclassified(100);k__Fungi_unclassified(100);k__Fungi_unclassified(100);k__Fungi_unclassified(100);k__Fungi_unclassified(100);k__Fungi_unclassified(100); | - | - | - | - | - | - | - | Unassigned | - |
| Otu06394 | 4 | k__Fungi(100);p__Ascomycota(100);c__Dothideomycetes(100);o__Pleosporales(100);o__Pleosporales_unclassified(100);o__Pleosporales_unclassified(100);o__Pleosporales_unclassified(100); | - | - | - | - | - | - | - | Unassigned | - |
| Otu06395 | 4 | k__Fungi(100);p__Ascomycota(75);p__Ascomycota_unclassified(75);p__Ascomycota_unclassified(75);p__Ascomycota_unclassified(75);p__Ascomycota_unclassified(75);p__Ascomycota_unclassified(75); | - | - | - | - | - | - | - | Unassigned | - |
| Otu06396 | 4 | k__Fungi(100);p__Ascomycota(100);c__Ascomycota_class_Incertae_sedis(100);o__Ascomycota_order_Incertae_sedis(100);f__Ascomycota_family_Incertae_sedis(100);g__unclassified_Ascomycota(100);s__Ascomycota_sp(100); | - | - | - | - | - | - | - | Unassigned | - |
| Otu06399 | 4 | k__Fungi(100);k__Fungi_unclassified(100);k__Fungi_unclassified(100);k__Fungi_unclassified(100);k__Fungi_unclassified(100);k__Fungi_unclassified(100);k__Fungi_unclassified(100); | - | - | - | - | - | - | - | Unassigned | - |
| Otu06401 | 4 | k__Fungi(100);p__Basidiomycota(100);p__Basidiomycota_unclassified(100);p__Basidiomycota_unclassified(100);p__Basidiomycota_unclassified(100);p__Basidiomycota_unclassified(100);p__Basidiomycota_unclassified(100); | - | - | - | - | - | - | - | Unassigned | - |
| Otu06403 | 4 | k__Fungi(100);k__Fungi_unclassified(100);k__Fungi_unclassified(100);k__Fungi_unclassified(100);k__Fungi_unclassified(100);k__Fungi_unclassified(100);k__Fungi_unclassified(100); | - | - | - | - | - | - | - | Unassigned | - |
| Otu06404 | 4 | k__Fungi(100);p__Basidiomycota(100);p__Basidiomycota_unclassified(100);p__Basidiomycota_unclassified(100);p__Basidiomycota_unclassified(100);p__Basidiomycota_unclassified(100);p__Basidiomycota_unclassified(100); | - | - | - | - | - | - | - | Unassigned | - |
| Otu06405 | 4 | k__Fungi(100);p__Ascomycota(75);p__Ascomycota_unclassified(75);p__Ascomycota_unclassified(75);p__Ascomycota_unclassified(75);p__Ascomycota_unclassified(75);p__Ascomycota_unclassified(75); | - | - | - | - | - | - | - | Unassigned | - |
| Otu06408 | 4 | k__Fungi(100);p__Ascomycota(100);p__Ascomycota_unclassified(100);p__Ascomycota_unclassified(100);p__Ascomycota_unclassified(100);p__Ascomycota_unclassified(100);p__Ascomycota_unclassified(100); | - | - | - | - | - | - | - | Unassigned | - |
| Otu06410 | 4 | k__Fungi(100);p__Ascomycota(75);p__Ascomycota_unclassified(75);p__Ascomycota_unclassified(75);p__Ascomycota_unclassified(75);p__Ascomycota_unclassified(75);p__Ascomycota_unclassified(75); | - | - | - | - | - | - | - | Unassigned | - |
| Otu06412 | 4 | k__Fungi(100);p__Ascomycota(100);c__Dothideomycetes(100);o__Pleosporales(100);o__Pleosporales_unclassified(100);o__Pleosporales_unclassified(100);o__Pleosporales_unclassified(100); | - | - | - | - | - | - | - | Unassigned | - |
| Otu06417 | 4 | k__Fungi(100);k__Fungi_unclassified(100);k__Fungi_unclassified(100);k__Fungi_unclassified(100);k__Fungi_unclassified(100);k__Fungi_unclassified(100);k__Fungi_unclassified(100); | - | - | - | - | - | - | - | Unassigned | - |
| Otu06422 | 4 | k__Fungi(100);p__Ascomycota(75);p__Ascomycota_unclassified(75);p__Ascomycota_unclassified(75);p__Ascomycota_unclassified(75);p__Ascomycota_unclassified(75);p__Ascomycota_unclassified(75); | - | - | - | - | - | - | - | Unassigned | - |
| Otu06423 | 4 | k__Fungi(100);p__Ascomycota(100);p__Ascomycota_unclassified(100);p__Ascomycota_unclassified(100);p__Ascomycota_unclassified(100);p__Ascomycota_unclassified(100);p__Ascomycota_unclassified(100); | - | - | - | - | - | - | - | Unassigned | - |
| Otu06428 | 4 | k__Fungi(100);p__Basidiomycota(100);p__Basidiomycota_unclassified(100);p__Basidiomycota_unclassified(100);p__Basidiomycota_unclassified(100);p__Basidiomycota_unclassified(100);p__Basidiomycota_unclassified(100); | - | - | - | - | - | - | - | Unassigned | - |
| Otu06429 | 4 | k__Fungi(100);p__Ascomycota(100);c__Sordariomycetes(100);o__Hypocreales(100);f__Hypocreaceae(100);f__Hypocreaceae_unclassified(100);f__Hypocreaceae_unclassified(100); | - | - | - | - | - | - | - | Unassigned | - |
| Otu06431 | 4 | k__Fungi(100);p__Ascomycota(100);p__Ascomycota_unclassified(100);p__Ascomycota_unclassified(100);p__Ascomycota_unclassified(100);p__Ascomycota_unclassified(100);p__Ascomycota_unclassified(100); | - | - | - | - | - | - | - | Unassigned | - |
| Otu06433 | 4 | k__Fungi(100);k__Fungi_unclassified(100);k__Fungi_unclassified(100);k__Fungi_unclassified(100);k__Fungi_unclassified(100);k__Fungi_unclassified(100);k__Fungi_unclassified(100); | - | - | - | - | - | - | - | Unassigned | - |
| Otu06434 | 4 | k__Fungi(100);k__Fungi_unclassified(100);k__Fungi_unclassified(100);k__Fungi_unclassified(100);k__Fungi_unclassified(100);k__Fungi_unclassified(100);k__Fungi_unclassified(100); | - | - | - | - | - | - | - | Unassigned | - |
| Otu06435 | 4 | k__Fungi(100);p__Ascomycota(100);c__Sordariomycetes(100);c__Sordariomycetes_unclassified(100);c__Sordariomycetes_unclassified(100);c__Sordariomycetes_unclassified(100);c__Sordariomycetes_unclassified(100); | - | - | - | - | - | - | - | Unassigned | - |
| Otu06438 | 4 | k__Fungi(100);p__Ascomycota(75);p__Ascomycota_unclassified(75);p__Ascomycota_unclassified(75);p__Ascomycota_unclassified(75);p__Ascomycota_unclassified(75);p__Ascomycota_unclassified(75); | - | - | - | - | - | - | - | Unassigned | - |
| Otu06439 | 4 | k__Fungi(100);k__Fungi_unclassified(100);k__Fungi_unclassified(100);k__Fungi_unclassified(100);k__Fungi_unclassified(100);k__Fungi_unclassified(100);k__Fungi_unclassified(100); | - | - | - | - | - | - | - | Unassigned | - |
| Otu06440 | 4 | k__Fungi(100);p__Ascomycota(100);c__Dothideomycetes(100);o__Capnodiales(100);o__Capnodiales_unclassified(100);o__Capnodiales_unclassified(100);o__Capnodiales_unclassified(100); | - | - | - | - | - | - | - | Unassigned | - |
| Otu06445 | 4 | k__Fungi(100);k__Fungi_unclassified(75);k__Fungi_unclassified(75);k__Fungi_unclassified(75);k__Fungi_unclassified(75);k__Fungi_unclassified(75);k__Fungi_unclassified(75); | - | - | - | - | - | - | - | Unassigned | - |
| Otu06446 | 4 | k__Fungi(100);k__Fungi_unclassified(75);k__Fungi_unclassified(75);k__Fungi_unclassified(75);k__Fungi_unclassified(75);k__Fungi_unclassified(75);k__Fungi_unclassified(75); | - | - | - | - | - | - | - | Unassigned | - |
| Otu06448 | 4 | k__Fungi(100);p__Ascomycota(100);p__Ascomycota_unclassified(100);p__Ascomycota_unclassified(100);p__Ascomycota_unclassified(100);p__Ascomycota_unclassified(100);p__Ascomycota_unclassified(100); | - | - | - | - | - | - | - | Unassigned | - |
| Otu06452 | 4 | k__Fungi(100);k__Fungi_unclassified(100);k__Fungi_unclassified(100);k__Fungi_unclassified(100);k__Fungi_unclassified(100);k__Fungi_unclassified(100);k__Fungi_unclassified(100); | - | - | - | - | - | - | - | Unassigned | - |
| Otu06461 | 4 | k__Fungi(100);k__Fungi_unclassified(100);k__Fungi_unclassified(100);k__Fungi_unclassified(100);k__Fungi_unclassified(100);k__Fungi_unclassified(100);k__Fungi_unclassified(100); | - | - | - | - | - | - | - | Unassigned | - |
| Otu06462 | 4 | k__Fungi(100);p__Ascomycota(75);p__Ascomycota_unclassified(75);p__Ascomycota_unclassified(75);p__Ascomycota_unclassified(75);p__Ascomycota_unclassified(75);p__Ascomycota_unclassified(75); | - | - | - | - | - | - | - | Unassigned | - |
| Otu06463 | 4 | k__Fungi(100);k__Fungi_unclassified(100);k__Fungi_unclassified(100);k__Fungi_unclassified(100);k__Fungi_unclassified(100);k__Fungi_unclassified(100);k__Fungi_unclassified(100); | - | - | - | - | - | - | - | Unassigned | - |
| Otu06464 | 4 | k__Fungi(100);p__Basidiomycota(100);c__Microbotryomycetes(100);o__Sporidiobolales(100);f__Sporidiobolales_family_Incertae_sedis(100);g__unclassified_Sporidiobolales(75);s__Sporidiobolales_sp(75); | - | - | - | - | - | - | - | Unassigned | - |
| Otu06466 | 4 | k__Fungi(100);k__Fungi_unclassified(100);k__Fungi_unclassified(100);k__Fungi_unclassified(100);k__Fungi_unclassified(100);k__Fungi_unclassified(100);k__Fungi_unclassified(100); | - | - | - | - | - | - | - | Unassigned | - |
| Otu06469 | 4 | k__Fungi(100);p__Basidiomycota(100);c__Agaricostilbomycetes(100);o__Agaricostilbales(100);o__Agaricostilbales_unclassified(75);o__Agaricostilbales_unclassified(75);o__Agaricostilbales_unclassified(75); | - | - | - | - | - | - | - | Unassigned | - |
| Otu06470 | 4 | k__Fungi(100);p__Basidiomycota(100);c__Agaricomycetes(100);o__Cantharellales(100);o__Cantharellales_unclassified(100);o__Cantharellales_unclassified(100);o__Cantharellales_unclassified(100); | - | - | - | - | - | - | - | Unassigned | - |
| Otu06471 | 4 | k__Fungi(100);k__Fungi_unclassified(100);k__Fungi_unclassified(100);k__Fungi_unclassified(100);k__Fungi_unclassified(100);k__Fungi_unclassified(100);k__Fungi_unclassified(100); | - | - | - | - | - | - | - | Unassigned | - |
| Otu06472 | 4 | k__Fungi(100);p__Ascomycota(100);c__Dothideomycetes(100);o__Capnodiales(100);f__Capnodiales_family_Incertae_sedis(100);f__Capnodiales_family_Incertae_sedis_unclassified(75);f__Capnodiales_family_Incertae_sedis_unclassified(75); | - | - | - | - | - | - | - | Unassigned | - |
| Otu06475 | 4 | k__Fungi(100);k__Fungi_unclassified(100);k__Fungi_unclassified(100);k__Fungi_unclassified(100);k__Fungi_unclassified(100);k__Fungi_unclassified(100);k__Fungi_unclassified(100); | - | - | - | - | - | - | - | Unassigned | - |
| Otu06476 | 4 | k__Fungi(100);p__Ascomycota(75);p__Ascomycota_unclassified(75);p__Ascomycota_unclassified(75);p__Ascomycota_unclassified(75);p__Ascomycota_unclassified(75);p__Ascomycota_unclassified(75); | - | - | - | - | - | - | - | Unassigned | - |
| Otu06477 | 4 | k__Fungi(100);k__Fungi_unclassified(75);k__Fungi_unclassified(75);k__Fungi_unclassified(75);k__Fungi_unclassified(75);k__Fungi_unclassified(75);k__Fungi_unclassified(75); | - | - | - | - | - | - | - | Unassigned | - |
| Otu06478 | 4 | k__Fungi(100);p__Ascomycota(75);p__Ascomycota_unclassified(75);p__Ascomycota_unclassified(75);p__Ascomycota_unclassified(75);p__Ascomycota_unclassified(75);p__Ascomycota_unclassified(75); | - | - | - | - | - | - | - | Unassigned | - |
| Otu06479 | 4 | k__Fungi(100);k__Fungi_unclassified(75);k__Fungi_unclassified(75);k__Fungi_unclassified(75);k__Fungi_unclassified(75);k__Fungi_unclassified(75);k__Fungi_unclassified(75); | - | - | - | - | - | - | - | Unassigned | - |
| Otu06480 | 4 | k__Fungi(100);k__Fungi_unclassified(100);k__Fungi_unclassified(100);k__Fungi_unclassified(100);k__Fungi_unclassified(100);k__Fungi_unclassified(100);k__Fungi_unclassified(100); | - | - | - | - | - | - | - | Unassigned | - |
| Otu06481 | 4 | k__Fungi(100);p__Ascomycota(75);p__Ascomycota_unclassified(75);p__Ascomycota_unclassified(75);p__Ascomycota_unclassified(75);p__Ascomycota_unclassified(75);p__Ascomycota_unclassified(75); | - | - | - | - | - | - | - | Unassigned | - |
| Otu06482 | 4 | k__Fungi(100);k__Fungi_unclassified(75);k__Fungi_unclassified(75);k__Fungi_unclassified(75);k__Fungi_unclassified(75);k__Fungi_unclassified(75);k__Fungi_unclassified(75); | - | - | - | - | - | - | - | Unassigned | - |
| Otu06483 | 4 | k__Fungi(100);k__Fungi_unclassified(100);k__Fungi_unclassified(100);k__Fungi_unclassified(100);k__Fungi_unclassified(100);k__Fungi_unclassified(100);k__Fungi_unclassified(100); | - | - | - | - | - | - | - | Unassigned | - |
| Otu06486 | 4 | k__Fungi(100);p__Ascomycota(100);p__Ascomycota_unclassified(100);p__Ascomycota_unclassified(100);p__Ascomycota_unclassified(100);p__Ascomycota_unclassified(100);p__Ascomycota_unclassified(100); | - | - | - | - | - | - | - | Unassigned | - |
| Otu06487 | 4 | k__Fungi(100);k__Fungi_unclassified(100);k__Fungi_unclassified(100);k__Fungi_unclassified(100);k__Fungi_unclassified(100);k__Fungi_unclassified(100);k__Fungi_unclassified(100); | - | - | - | - | - | - | - | Unassigned | - |
| Otu06489 | 4 | k__Fungi(100);p__Basidiomycota(100);c__Agaricomycetes(100);o__Polyporales(100);o__Polyporales_unclassified(100);o__Polyporales_unclassified(100);o__Polyporales_unclassified(100); | - | - | - | - | - | - | - | Unassigned | - |
| Otu06490 | 4 | k__Fungi(100);k__Fungi_unclassified(75);k__Fungi_unclassified(75);k__Fungi_unclassified(75);k__Fungi_unclassified(75);k__Fungi_unclassified(75);k__Fungi_unclassified(75); | - | - | - | - | - | - | - | Unassigned | - |
| Otu06491 | 4 | k__Fungi(100);p__Ascomycota(75);p__Ascomycota_unclassified(75);p__Ascomycota_unclassified(75);p__Ascomycota_unclassified(75);p__Ascomycota_unclassified(75);p__Ascomycota_unclassified(75); | - | - | - | - | - | - | - | Unassigned | - |
| Otu06492 | 4 | k__Fungi(100);k__Fungi_unclassified(75);k__Fungi_unclassified(75);k__Fungi_unclassified(75);k__Fungi_unclassified(75);k__Fungi_unclassified(75);k__Fungi_unclassified(75); | - | - | - | - | - | - | - | Unassigned | - |
| Otu06493 | 4 | k__Fungi(100);p__Ascomycota(100);c__Dothideomycetes(100);o__Capnodiales(100);f__Davidiellaceae(100);f__Davidiellaceae_unclassified(100);f__Davidiellaceae_unclassified(100); | - | - | - | - | - | - | - | Unassigned | - |
| Otu06495 | 4 | k__Fungi(100);k__Fungi_unclassified(75);k__Fungi_unclassified(75);k__Fungi_unclassified(75);k__Fungi_unclassified(75);k__Fungi_unclassified(75);k__Fungi_unclassified(75); | - | - | - | - | - | - | - | Unassigned | - |
| Otu06497 | 4 | k__Fungi(100);p__Ascomycota(100);c__Sordariomycetes(100);o__Sordariales(100);o__Sordariales_unclassified(100);o__Sordariales_unclassified(100);o__Sordariales_unclassified(100); | - | - | - | - | - | - | - | Unassigned | - |
| Otu06498 | 4 | k__Fungi(100);p__Basidiomycota(100);p__Basidiomycota_unclassified(100);p__Basidiomycota_unclassified(100);p__Basidiomycota_unclassified(100);p__Basidiomycota_unclassified(100);p__Basidiomycota_unclassified(100); | - | - | - | - | - | - | - | Unassigned | - |
| Otu06499 | 4 | k__Fungi(100);p__Ascomycota(100);p__Ascomycota_unclassified(100);p__Ascomycota_unclassified(100);p__Ascomycota_unclassified(100);p__Ascomycota_unclassified(100);p__Ascomycota_unclassified(100); | - | - | - | - | - | - | - | Unassigned | - |
| Otu06500 | 4 | k__Fungi(100);p__Basidiomycota(100);c__Agaricomycetes(100);c__Agaricomycetes_unclassified(100);c__Agaricomycetes_unclassified(100);c__Agaricomycetes_unclassified(100);c__Agaricomycetes_unclassified(100); | - | - | - | - | - | - | - | Unassigned | - |
| Otu06502 | 4 | k__Fungi(100);k__Fungi_unclassified(100);k__Fungi_unclassified(100);k__Fungi_unclassified(100);k__Fungi_unclassified(100);k__Fungi_unclassified(100);k__Fungi_unclassified(100); | - | - | - | - | - | - | - | Unassigned | - |
| Otu06503 | 4 | k__Fungi(100);k__Fungi_unclassified(100);k__Fungi_unclassified(100);k__Fungi_unclassified(100);k__Fungi_unclassified(100);k__Fungi_unclassified(100);k__Fungi_unclassified(100); | - | - | - | - | - | - | - | Unassigned | - |
| Otu06504 | 4 | k__Fungi(100);k__Fungi_unclassified(100);k__Fungi_unclassified(100);k__Fungi_unclassified(100);k__Fungi_unclassified(100);k__Fungi_unclassified(100);k__Fungi_unclassified(100); | - | - | - | - | - | - | - | Unassigned | - |
| Otu06506 | 4 | k__Fungi(100);k__Fungi_unclassified(75);k__Fungi_unclassified(75);k__Fungi_unclassified(75);k__Fungi_unclassified(75);k__Fungi_unclassified(75);k__Fungi_unclassified(75); | - | - | - | - | - | - | - | Unassigned | - |
| Otu06507 | 4 | k__Fungi(100);p__Ascomycota(100);c__Lecanoromycetes(100);c__Lecanoromycetes_unclassified(100);c__Lecanoromycetes_unclassified(100);c__Lecanoromycetes_unclassified(100);c__Lecanoromycetes_unclassified(100); | - | - | - | - | - | - | - | Unassigned | - |
| Otu06509 | 4 | k__Fungi(100);p__Ascomycota(75);p__Ascomycota_unclassified(75);p__Ascomycota_unclassified(75);p__Ascomycota_unclassified(75);p__Ascomycota_unclassified(75);p__Ascomycota_unclassified(75); | - | - | - | - | - | - | - | Unassigned | - |
| Otu06512 | 4 | k__Fungi(100);p__Ascomycota(100);p__Ascomycota_unclassified(100);p__Ascomycota_unclassified(100);p__Ascomycota_unclassified(100);p__Ascomycota_unclassified(100);p__Ascomycota_unclassified(100); | - | - | - | - | - | - | - | Unassigned | - |
| Otu06513 | 4 | k__Fungi(100);k__Fungi_unclassified(75);k__Fungi_unclassified(75);k__Fungi_unclassified(75);k__Fungi_unclassified(75);k__Fungi_unclassified(75);k__Fungi_unclassified(75); | - | - | - | - | - | - | - | Unassigned | - |
| Otu06514 | 4 | k__Fungi(100);p__Ascomycota(100);c__Sordariomycetes(100);c__Sordariomycetes_unclassified(100);c__Sordariomycetes_unclassified(100);c__Sordariomycetes_unclassified(100);c__Sordariomycetes_unclassified(100); | - | - | - | - | - | - | - | Unassigned | - |
| Otu06515 | 4 | k__Fungi(100);p__Ascomycota(100);p__Ascomycota_unclassified(100);p__Ascomycota_unclassified(100);p__Ascomycota_unclassified(100);p__Ascomycota_unclassified(100);p__Ascomycota_unclassified(100); | - | - | - | - | - | - | - | Unassigned | - |
| Otu06516 | 4 | k__Fungi(100);k__Fungi_unclassified(100);k__Fungi_unclassified(100);k__Fungi_unclassified(100);k__Fungi_unclassified(100);k__Fungi_unclassified(100);k__Fungi_unclassified(100); | - | - | - | - | - | - | - | Unassigned | - |
| Otu06517 | 4 | k__Fungi(100);p__Ascomycota(100);p__Ascomycota_unclassified(100);p__Ascomycota_unclassified(100);p__Ascomycota_unclassified(100);p__Ascomycota_unclassified(100);p__Ascomycota_unclassified(100); | - | - | - | - | - | - | - | Unassigned | - |
| Otu06518 | 4 | k__Fungi(100);k__Fungi_unclassified(75);k__Fungi_unclassified(75);k__Fungi_unclassified(75);k__Fungi_unclassified(75);k__Fungi_unclassified(75);k__Fungi_unclassified(75); | - | - | - | - | - | - | - | Unassigned | - |
| Otu06519 | 4 | k__Fungi(100);p__Basidiomycota(100);c__Agaricomycetes(100);c__Agaricomycetes_unclassified(100);c__Agaricomycetes_unclassified(100);c__Agaricomycetes_unclassified(100);c__Agaricomycetes_unclassified(100); | - | - | - | - | - | - | - | Unassigned | - |
| Otu06520 | 4 | k__Fungi(100);k__Fungi_unclassified(75);k__Fungi_unclassified(75);k__Fungi_unclassified(75);k__Fungi_unclassified(75);k__Fungi_unclassified(75);k__Fungi_unclassified(75); | - | - | - | - | - | - | - | Unassigned | - |
| Otu06521 | 4 | k__Fungi(100);p__Basidiomycota(100);c__Agaricomycetes(100);c__Agaricomycetes_unclassified(100);c__Agaricomycetes_unclassified(100);c__Agaricomycetes_unclassified(100);c__Agaricomycetes_unclassified(100); | - | - | - | - | - | - | - | Unassigned | - |
| Otu06522 | 4 | k__Fungi(100);k__Fungi_unclassified(100);k__Fungi_unclassified(100);k__Fungi_unclassified(100);k__Fungi_unclassified(100);k__Fungi_unclassified(100);k__Fungi_unclassified(100); | - | - | - | - | - | - | - | Unassigned | - |
| Otu06524 | 4 | k__Fungi(100);p__Ascomycota(100);p__Ascomycota_unclassified(100);p__Ascomycota_unclassified(100);p__Ascomycota_unclassified(100);p__Ascomycota_unclassified(100);p__Ascomycota_unclassified(100); | - | - | - | - | - | - | - | Unassigned | - |
| Otu06525 | 4 | k__Fungi(100);p__Basidiomycota(100);c__Microbotryomycetes(100);o__Sporidiobolales(100);f__Sporidiobolales_family_Incertae_sedis(100);f__Sporidiobolales_family_Incertae_sedis_unclassified(100);f__Sporidiobolales_family_Incertae_sedis_unclassified(100); | - | - | - | - | - | - | - | Unassigned | - |
| Otu06526 | 4 | k__Fungi(100);k__Fungi_unclassified(75);k__Fungi_unclassified(75);k__Fungi_unclassified(75);k__Fungi_unclassified(75);k__Fungi_unclassified(75);k__Fungi_unclassified(75); | - | - | - | - | - | - | - | Unassigned | - |
| Otu06530 | 4 | k__Fungi(100);p__Ascomycota(100);c__Lecanoromycetes(100);c__Lecanoromycetes_unclassified(100);c__Lecanoromycetes_unclassified(100);c__Lecanoromycetes_unclassified(100);c__Lecanoromycetes_unclassified(100); | - | - | - | - | - | - | - | Unassigned | - |
| Otu06532 | 4 | k__Fungi(100);k__Fungi_unclassified(75);k__Fungi_unclassified(75);k__Fungi_unclassified(75);k__Fungi_unclassified(75);k__Fungi_unclassified(75);k__Fungi_unclassified(75); | - | - | - | - | - | - | - | Unassigned | - |
| Otu06542 | 4 | k__Fungi(100);p__Ascomycota(100);p__Ascomycota_unclassified(100);p__Ascomycota_unclassified(100);p__Ascomycota_unclassified(100);p__Ascomycota_unclassified(100);p__Ascomycota_unclassified(100); | - | - | - | - | - | - | - | Unassigned | - |
| Otu06546 | 4 | k__Fungi(100);k__Fungi_unclassified(75);k__Fungi_unclassified(75);k__Fungi_unclassified(75);k__Fungi_unclassified(75);k__Fungi_unclassified(75);k__Fungi_unclassified(75); | - | - | - | - | - | - | - | Unassigned | - |
| Otu06547 | 4 | k__Fungi(100);k__Fungi_unclassified(100);k__Fungi_unclassified(100);k__Fungi_unclassified(100);k__Fungi_unclassified(100);k__Fungi_unclassified(100);k__Fungi_unclassified(100); | - | - | - | - | - | - | - | Unassigned | - |
| Otu06552 | 4 | k__Fungi(100);p__Ascomycota(100);c__Leotiomycetes(100);c__Leotiomycetes_unclassified(100);c__Leotiomycetes_unclassified(100);c__Leotiomycetes_unclassified(100);c__Leotiomycetes_unclassified(100); | - | - | - | - | - | - | - | Unassigned | - |
| Otu06556 | 4 | k__Fungi(100);k__Fungi_unclassified(100);k__Fungi_unclassified(100);k__Fungi_unclassified(100);k__Fungi_unclassified(100);k__Fungi_unclassified(100);k__Fungi_unclassified(100); | - | - | - | - | - | - | - | Unassigned | - |
| Otu06558 | 4 | k__Fungi(100);k__Fungi_unclassified(100);k__Fungi_unclassified(100);k__Fungi_unclassified(100);k__Fungi_unclassified(100);k__Fungi_unclassified(100);k__Fungi_unclassified(100); | - | - | - | - | - | - | - | Unassigned | - |
| Otu06562 | 4 | k__Fungi(100);k__Fungi_unclassified(100);k__Fungi_unclassified(100);k__Fungi_unclassified(100);k__Fungi_unclassified(100);k__Fungi_unclassified(100);k__Fungi_unclassified(100); | - | - | - | - | - | - | - | Unassigned | - |
| Otu06565 | 4 | k__Fungi(100);p__Ascomycota(75);p__Ascomycota_unclassified(75);p__Ascomycota_unclassified(75);p__Ascomycota_unclassified(75);p__Ascomycota_unclassified(75);p__Ascomycota_unclassified(75); | - | - | - | - | - | - | - | Unassigned | - |
| Otu06566 | 4 | k__Fungi(100);k__Fungi_unclassified(75);k__Fungi_unclassified(75);k__Fungi_unclassified(75);k__Fungi_unclassified(75);k__Fungi_unclassified(75);k__Fungi_unclassified(75); | - | - | - | - | - | - | - | Unassigned | - |
| Otu06569 | 4 | k__Fungi(100);p__Ascomycota(75);p__Ascomycota_unclassified(75);p__Ascomycota_unclassified(75);p__Ascomycota_unclassified(75);p__Ascomycota_unclassified(75);p__Ascomycota_unclassified(75); | - | - | - | - | - | - | - | Unassigned | - |
| Otu06572 | 4 | k__Fungi(100);k__Fungi_unclassified(75);k__Fungi_unclassified(75);k__Fungi_unclassified(75);k__Fungi_unclassified(75);k__Fungi_unclassified(75);k__Fungi_unclassified(75); | - | - | - | - | - | - | - | Unassigned | - |
| Otu06573 | 4 | k__Fungi(100);k__Fungi_unclassified(75);k__Fungi_unclassified(75);k__Fungi_unclassified(75);k__Fungi_unclassified(75);k__Fungi_unclassified(75);k__Fungi_unclassified(75); | - | - | - | - | - | - | - | Unassigned | - |
| Otu06575 | 4 | k__Fungi(100);k__Fungi_unclassified(100);k__Fungi_unclassified(100);k__Fungi_unclassified(100);k__Fungi_unclassified(100);k__Fungi_unclassified(100);k__Fungi_unclassified(100); | - | - | - | - | - | - | - | Unassigned | - |
| Otu06576 | 4 | k__Fungi(100);p__Ascomycota(100);c__Eurotiomycetes(100);o__Chaetothyriales(100);o__Chaetothyriales_unclassified(100);o__Chaetothyriales_unclassified(100);o__Chaetothyriales_unclassified(100); | - | - | - | - | - | - | - | Unassigned | - |
| Otu06580 | 4 | k__Fungi(100);p__Ascomycota(100);p__Ascomycota_unclassified(100);p__Ascomycota_unclassified(100);p__Ascomycota_unclassified(100);p__Ascomycota_unclassified(100);p__Ascomycota_unclassified(100); | - | - | - | - | - | - | - | Unassigned | - |
| Otu06584 | 4 | k__Fungi(100);p__Ascomycota(100);c__Dothideomycetes(100);o__Capnodiales(100);o__Capnodiales_unclassified(100);o__Capnodiales_unclassified(100);o__Capnodiales_unclassified(100); | - | - | - | - | - | - | - | Unassigned | - |
| Otu06585 | 4 | k__Fungi(100);p__Ascomycota(100);p__Ascomycota_unclassified(100);p__Ascomycota_unclassified(100);p__Ascomycota_unclassified(100);p__Ascomycota_unclassified(100);p__Ascomycota_unclassified(100); | - | - | - | - | - | - | - | Unassigned | - |
| Otu06590 | 4 | k__Fungi(100);k__Fungi_unclassified(100);k__Fungi_unclassified(100);k__Fungi_unclassified(100);k__Fungi_unclassified(100);k__Fungi_unclassified(100);k__Fungi_unclassified(100); | - | - | - | - | - | - | - | Unassigned | - |
| Otu06591 | 4 | k__Fungi(100);k__Fungi_unclassified(100);k__Fungi_unclassified(100);k__Fungi_unclassified(100);k__Fungi_unclassified(100);k__Fungi_unclassified(100);k__Fungi_unclassified(100); | - | - | - | - | - | - | - | Unassigned | - |
| Otu06593 | 4 | k__Fungi(100);k__Fungi_unclassified(100);k__Fungi_unclassified(100);k__Fungi_unclassified(100);k__Fungi_unclassified(100);k__Fungi_unclassified(100);k__Fungi_unclassified(100); | - | - | - | - | - | - | - | Unassigned | - |
| Otu06595 | 4 | k__Fungi(100);p__Ascomycota(100);p__Ascomycota_unclassified(100);p__Ascomycota_unclassified(100);p__Ascomycota_unclassified(100);p__Ascomycota_unclassified(100);p__Ascomycota_unclassified(100); | - | - | - | - | - | - | - | Unassigned | - |
| Otu06596 | 4 | k__Fungi(100);p__Ascomycota(100);p__Ascomycota_unclassified(100);p__Ascomycota_unclassified(100);p__Ascomycota_unclassified(100);p__Ascomycota_unclassified(100);p__Ascomycota_unclassified(100); | - | - | - | - | - | - | - | Unassigned | - |
| Otu06604 | 4 | k__Fungi(100);k__Fungi_unclassified(75);k__Fungi_unclassified(75);k__Fungi_unclassified(75);k__Fungi_unclassified(75);k__Fungi_unclassified(75);k__Fungi_unclassified(75); | - | - | - | - | - | - | - | Unassigned | - |
| Otu06609 | 4 | k__Fungi(100);k__Fungi_unclassified(100);k__Fungi_unclassified(100);k__Fungi_unclassified(100);k__Fungi_unclassified(100);k__Fungi_unclassified(100);k__Fungi_unclassified(100); | - | - | - | - | - | - | - | Unassigned | - |
| Otu06611 | 4 | k__Fungi(100);p__Ascomycota(75);p__Ascomycota_unclassified(75);p__Ascomycota_unclassified(75);p__Ascomycota_unclassified(75);p__Ascomycota_unclassified(75);p__Ascomycota_unclassified(75); | - | - | - | - | - | - | - | Unassigned | - |
| Otu06612 | 4 | k__Fungi(100);p__Ascomycota(100);p__Ascomycota_unclassified(100);p__Ascomycota_unclassified(100);p__Ascomycota_unclassified(100);p__Ascomycota_unclassified(100);p__Ascomycota_unclassified(100); | - | - | - | - | - | - | - | Unassigned | - |
| Otu06613 | 4 | k__Fungi(100);k__Fungi_unclassified(100);k__Fungi_unclassified(100);k__Fungi_unclassified(100);k__Fungi_unclassified(100);k__Fungi_unclassified(100);k__Fungi_unclassified(100); | - | - | - | - | - | - | - | Unassigned | - |
| Otu06616 | 4 | k__Fungi(100);p__Ascomycota(100);c__Dothideomycetes(100);o__Capnodiales(100);o__Capnodiales_unclassified(100);o__Capnodiales_unclassified(100);o__Capnodiales_unclassified(100); | - | - | - | - | - | - | - | Unassigned | - |
| Otu06618 | 4 | k__Fungi(100);p__Ascomycota(100);p__Ascomycota_unclassified(100);p__Ascomycota_unclassified(100);p__Ascomycota_unclassified(100);p__Ascomycota_unclassified(100);p__Ascomycota_unclassified(100); | - | - | - | - | - | - | - | Unassigned | - |
| Otu06619 | 4 | k__Fungi(100);p__Ascomycota(100);p__Ascomycota_unclassified(100);p__Ascomycota_unclassified(100);p__Ascomycota_unclassified(100);p__Ascomycota_unclassified(100);p__Ascomycota_unclassified(100); | - | - | - | - | - | - | - | Unassigned | - |
| Otu06623 | 4 | k__Fungi(100);p__Ascomycota(100);p__Ascomycota_unclassified(100);p__Ascomycota_unclassified(100);p__Ascomycota_unclassified(100);p__Ascomycota_unclassified(100);p__Ascomycota_unclassified(100); | - | - | - | - | - | - | - | Unassigned | - |
| Otu06626 | 4 | k__Fungi(100);p__Ascomycota(75);p__Ascomycota_unclassified(75);p__Ascomycota_unclassified(75);p__Ascomycota_unclassified(75);p__Ascomycota_unclassified(75);p__Ascomycota_unclassified(75); | - | - | - | - | - | - | - | Unassigned | - |
| Otu06627 | 4 | k__Fungi(100);p__Basidiomycota(100);c__Agaricomycetes(100);o__Polyporales(100);f__Phanerochaetaceae(100);f__Phanerochaetaceae_unclassified(100);f__Phanerochaetaceae_unclassified(100); | - | - | - | - | - | - | - | Unassigned | - |
| Otu06629 | 4 | k__Fungi(100);k__Fungi_unclassified(75);k__Fungi_unclassified(75);k__Fungi_unclassified(75);k__Fungi_unclassified(75);k__Fungi_unclassified(75);k__Fungi_unclassified(75); | - | - | - | - | - | - | - | Unassigned | - |
| Otu06630 | 4 | k__Fungi(100);k__Fungi_unclassified(75);k__Fungi_unclassified(75);k__Fungi_unclassified(75);k__Fungi_unclassified(75);k__Fungi_unclassified(75);k__Fungi_unclassified(75); | - | - | - | - | - | - | - | Unassigned | - |
| Otu06633 | 4 | k__Fungi(100);p__Ascomycota(100);c__Dothideomycetes(100);o__Pleosporales(100);o__Pleosporales_unclassified(100);o__Pleosporales_unclassified(100);o__Pleosporales_unclassified(100); | - | - | - | - | - | - | - | Unassigned | - |
| Otu06635 | 4 | k__Fungi(100);k__Fungi_unclassified(100);k__Fungi_unclassified(100);k__Fungi_unclassified(100);k__Fungi_unclassified(100);k__Fungi_unclassified(100);k__Fungi_unclassified(100); | - | - | - | - | - | - | - | Unassigned | - |
| Otu06638 | 4 | k__Fungi(100);p__Ascomycota(100);c__Dothideomycetes(100);o__Myriangiales(100);f__unclassified_Myriangiales(100);g__unclassified_Myriangiales(100);s__Myriangiales_sp(100); | - | - | - | - | - | - | - | Unassigned | - |
| Otu06639 | 4 | k__Fungi(100);k__Fungi_unclassified(75);k__Fungi_unclassified(75);k__Fungi_unclassified(75);k__Fungi_unclassified(75);k__Fungi_unclassified(75);k__Fungi_unclassified(75); | - | - | - | - | - | - | - | Unassigned | - |
| Otu06640 | 4 | k__Fungi(100);k__Fungi_unclassified(100);k__Fungi_unclassified(100);k__Fungi_unclassified(100);k__Fungi_unclassified(100);k__Fungi_unclassified(100);k__Fungi_unclassified(100); | - | - | - | - | - | - | - | Unassigned | - |
| Otu06641 | 4 | k__Plantae(100);p__unclassified_Plantae(100);c__unclassified_Plantae(100);o__unclassified_Plantae(100);f__unclassified_Plantae(100);g__unclassified_Plantae(100);s__Plantae_sp(100); | - | - | - | - | - | - | - | Unassigned | - |
| Otu06644 | 4 | k__Fungi(100);p__Ascomycota(100);c__Eurotiomycetes(100);c__Eurotiomycetes_unclassified(100);c__Eurotiomycetes_unclassified(100);c__Eurotiomycetes_unclassified(100);c__Eurotiomycetes_unclassified(100); | - | - | - | - | - | - | - | Unassigned | - |
| Otu06646 | 4 | k__Fungi(100);p__Ascomycota(100);p__Ascomycota_unclassified(100);p__Ascomycota_unclassified(100);p__Ascomycota_unclassified(100);p__Ascomycota_unclassified(100);p__Ascomycota_unclassified(100); | - | - | - | - | - | - | - | Unassigned | - |
| Otu06647 | 4 | k__Fungi(100);p__Basidiomycota(100);c__Pucciniomycetes(100);c__Pucciniomycetes_unclassified(100);c__Pucciniomycetes_unclassified(100);c__Pucciniomycetes_unclassified(100);c__Pucciniomycetes_unclassified(100); | - | - | - | - | - | - | - | Unassigned | - |
| Otu06650 | 4 | k__Fungi(100);p__Ascomycota(100);p__Ascomycota_unclassified(100);p__Ascomycota_unclassified(100);p__Ascomycota_unclassified(100);p__Ascomycota_unclassified(100);p__Ascomycota_unclassified(100); | - | - | - | - | - | - | - | Unassigned | - |
| Otu06651 | 4 | k__Fungi(100);k__Fungi_unclassified(75);k__Fungi_unclassified(75);k__Fungi_unclassified(75);k__Fungi_unclassified(75);k__Fungi_unclassified(75);k__Fungi_unclassified(75); | - | - | - | - | - | - | - | Unassigned | - |
| Otu06655 | 4 | k__Fungi(100);p__Basidiomycota(100);c__Agaricomycetes(100);c__Agaricomycetes_unclassified(100);c__Agaricomycetes_unclassified(100);c__Agaricomycetes_unclassified(100);c__Agaricomycetes_unclassified(100); | - | - | - | - | - | - | - | Unassigned | - |
| Otu06656 | 4 | k__Fungi(100);p__Ascomycota(75);p__Ascomycota_unclassified(75);p__Ascomycota_unclassified(75);p__Ascomycota_unclassified(75);p__Ascomycota_unclassified(75);p__Ascomycota_unclassified(75); | - | - | - | - | - | - | - | Unassigned | - |
| Otu06658 | 4 | k__Fungi(100);p__Ascomycota(100);c__Dothideomycetes(100);c__Dothideomycetes_unclassified(100);c__Dothideomycetes_unclassified(100);c__Dothideomycetes_unclassified(100);c__Dothideomycetes_unclassified(100); | - | - | - | - | - | - | - | Unassigned | - |
| Otu06659 | 4 | k__Fungi(100);p__Ascomycota(100);p__Ascomycota_unclassified(100);p__Ascomycota_unclassified(100);p__Ascomycota_unclassified(100);p__Ascomycota_unclassified(100);p__Ascomycota_unclassified(100); | - | - | - | - | - | - | - | Unassigned | - |
| Otu06660 | 4 | k__Fungi(100);p__Ascomycota(100);p__Ascomycota_unclassified(100);p__Ascomycota_unclassified(100);p__Ascomycota_unclassified(100);p__Ascomycota_unclassified(100);p__Ascomycota_unclassified(100); | - | - | - | - | - | - | - | Unassigned | - |
| Otu06663 | 4 | k__Fungi(100);k__Fungi_unclassified(100);k__Fungi_unclassified(100);k__Fungi_unclassified(100);k__Fungi_unclassified(100);k__Fungi_unclassified(100);k__Fungi_unclassified(100); | - | - | - | - | - | - | - | Unassigned | - |
| Otu06664 | 4 | k__Fungi(100);p__Ascomycota(100);p__Ascomycota_unclassified(100);p__Ascomycota_unclassified(100);p__Ascomycota_unclassified(100);p__Ascomycota_unclassified(100);p__Ascomycota_unclassified(100); | - | - | - | - | - | - | - | Unassigned | - |
| Otu06666 | 4 | k__Fungi(100);p__Ascomycota(100);p__Ascomycota_unclassified(100);p__Ascomycota_unclassified(100);p__Ascomycota_unclassified(100);p__Ascomycota_unclassified(100);p__Ascomycota_unclassified(100); | - | - | - | - | - | - | - | Unassigned | - |
| Otu06667 | 4 | k__Fungi(100);k__Fungi_unclassified(100);k__Fungi_unclassified(100);k__Fungi_unclassified(100);k__Fungi_unclassified(100);k__Fungi_unclassified(100);k__Fungi_unclassified(100); | - | - | - | - | - | - | - | Unassigned | - |

| | | | | | | | | | | | |
|---|---|---|---|---|---|---|---|---|---|---|---|
| Otu06672 | 4 | k__Fungi(100);k__Fungi_unclassified(100);k__Fungi_unclassified(100);k__Fungi_unclassified(100);k__Fungi_unclassified(100);k__Fungi_unclassified(100);k__Fungi_unclassified(100); | - | - | - | - | - | - | - | Unassigned | - |
| Otu06674 | 4 | k__Fungi(100);p__Basidiomycota(100);p__Basidiomycota_unclassified(100);p__Basidiomycota_unclassified(100);p__Basidiomycota_unclassified(100);p__Basidiomycota_unclassified(100);p__Basidiomycota_unclassified(100); | - | - | - | - | - | - | - | Unassigned | - |
| Otu06676 | 4 | k__Fungi(100);p__Basidiomycota(100);c__Agaricomycetes(100);c__Agaricomycetes_unclassified(100);c__Agaricomycetes_unclassified(100);c__Agaricomycetes_unclassified(100);c__Agaricomycetes_unclassified(100); | - | - | - | - | - | - | - | Unassigned | - |
| Otu06677 | 4 | k__Fungi(100);p__Ascomycota(100);p__Ascomycota_unclassified(100);p__Ascomycota_unclassified(100);p__Ascomycota_unclassified(100);p__Ascomycota_unclassified(100);p__Ascomycota_unclassified(100); | - | - | - | - | - | - | - | Unassigned | - |
| Otu06684 | 4 | k__Fungi(100);k__Fungi_unclassified(75);k__Fungi_unclassified(75);k__Fungi_unclassified(75);k__Fungi_unclassified(75);k__Fungi_unclassified(75);k__Fungi_unclassified(75); | - | - | - | - | - | - | - | Unassigned | - |
| Otu06687 | 4 | k__Fungi(100);p__Basidiomycota(100);c__Agaricomycetes(100);o__Auriculariales(100);f__Auriculariales_family_Incertae_sedis(100);f__Auriculariales_family_Incertae_sedis_unclassified(100);f__Auriculariales_family_Incertae_sedis_unclassified(100); | - | - | - | - | - | - | - | Unassigned | - |
| Otu06688 | 4 | k__Fungi(100);k__Fungi_unclassified(100);k__Fungi_unclassified(100);k__Fungi_unclassified(100);k__Fungi_unclassified(100);k__Fungi_unclassified(100);k__Fungi_unclassified(100); | - | - | - | - | - | - | - | Unassigned | - |
| Otu06696 | 4 | k__Fungi(100);k__Fungi_unclassified(100);k__Fungi_unclassified(100);k__Fungi_unclassified(100);k__Fungi_unclassified(100);k__Fungi_unclassified(100);k__Fungi_unclassified(100); | - | - | - | - | - | - | - | Unassigned | - |
| Otu06698 | 4 | k__Fungi(100);k__Fungi_unclassified(75);k__Fungi_unclassified(75);k__Fungi_unclassified(75);k__Fungi_unclassified(75);k__Fungi_unclassified(75);k__Fungi_unclassified(75); | - | - | - | - | - | - | - | Unassigned | - |
| Otu06701 | 4 | k__Fungi(100);k__Fungi_unclassified(100);k__Fungi_unclassified(100);k__Fungi_unclassified(100);k__Fungi_unclassified(100);k__Fungi_unclassified(100);k__Fungi_unclassified(100); | - | - | - | - | - | - | - | Unassigned | - |
| Otu06706 | 4 | k__Fungi(100);p__Ascomycota(100);p__Ascomycota_unclassified(100);p__Ascomycota_unclassified(100);p__Ascomycota_unclassified(100);p__Ascomycota_unclassified(100);p__Ascomycota_unclassified(100); | - | - | - | - | - | - | - | Unassigned | - |
| Otu06707 | 4 | k__Fungi(100);p__Ascomycota(75);p__Ascomycota_unclassified(75);p__Ascomycota_unclassified(75);p__Ascomycota_unclassified(75);p__Ascomycota_unclassified(75);p__Ascomycota_unclassified(75); | - | - | - | - | - | - | - | Unassigned | - |
| Otu06709 | 4 | k__Fungi(100);p__Ascomycota(100);c__Sordariomycetes(100);o__Hypocreales(100);f__Cordycipitaceae(100);f__Cordycipitaceae_unclassified(100);f__Cordycipitaceae_unclassified(100); | - | - | - | - | - | - | - | Unassigned | - |
| Otu06710 | 4 | k__Fungi(100);k__Fungi_unclassified(75);k__Fungi_unclassified(75);k__Fungi_unclassified(75);k__Fungi_unclassified(75);k__Fungi_unclassified(75);k__Fungi_unclassified(75); | - | - | - | - | - | - | - | Unassigned | - |
| Otu06711 | 4 | k__Fungi(100);p__Basidiomycota(100);c__Agaricomycetes(100);o__Polyporales(100);o__Polyporales_unclassified(100);o__Polyporales_unclassified(100);o__Polyporales_unclassified(100); | - | - | - | - | - | - | - | Unassigned | - |
| Otu06714 | 4 | k__Fungi(100);p__Ascomycota(100);c__Leotiomycetes(75);c__Leotiomycetes_unclassified(75);c__Leotiomycetes_unclassified(75);c__Leotiomycetes_unclassified(75);c__Leotiomycetes_unclassified(75); | - | - | - | - | - | - | - | Unassigned | - |
| Otu06716 | 4 | k__Fungi(100);k__Fungi_unclassified(100);k__Fungi_unclassified(100);k__Fungi_unclassified(100);k__Fungi_unclassified(100);k__Fungi_unclassified(100);k__Fungi_unclassified(100); | - | - | - | - | - | - | - | Unassigned | - |
| Otu06719 | 4 | k__Fungi(100);k__Fungi_unclassified(100);k__Fungi_unclassified(100);k__Fungi_unclassified(100);k__Fungi_unclassified(100);k__Fungi_unclassified(100);k__Fungi_unclassified(100); | - | - | - | - | - | - | - | Unassigned | - |
| Otu06721 | 4 | k__Fungi(100);k__Fungi_unclassified(100);k__Fungi_unclassified(100);k__Fungi_unclassified(100);k__Fungi_unclassified(100);k__Fungi_unclassified(100);k__Fungi_unclassified(100); | - | - | - | - | - | - | - | Unassigned | - |
| Otu06722 | 4 | k__Fungi(100);p__Ascomycota(100);c__Dothideomycetes(100);o__Pleosporales(100);o__Pleosporales_unclassified(100);o__Pleosporales_unclassified(100);o__Pleosporales_unclassified(100); | - | - | - | - | - | - | - | Unassigned | - |
| Otu06729 | 4 | k__Fungi(100);k__Fungi_unclassified(100);k__Fungi_unclassified(100);k__Fungi_unclassified(100);k__Fungi_unclassified(100);k__Fungi_unclassified(100);k__Fungi_unclassified(100); | - | - | - | - | - | - | - | Unassigned | - |
| Otu06736 | 4 | k__Fungi(100);p__Ascomycota(100);c__Leotiomycetes(100);o__Helotiales(75);o__Helotiales_unclassified(75);o__Helotiales_unclassified(75);o__Helotiales_unclassified(75); | - | - | - | - | - | - | - | Unassigned | - |
| Otu06737 | 4 | k__Fungi(100);p__Ascomycota(100);p__Ascomycota_unclassified(100);p__Ascomycota_unclassified(100);p__Ascomycota_unclassified(100);p__Ascomycota_unclassified(100);p__Ascomycota_unclassified(100); | - | - | - | - | - | - | - | Unassigned | - |
| Otu06738 | 4 | k__Fungi(100);k__Fungi_unclassified(100);k__Fungi_unclassified(100);k__Fungi_unclassified(100);k__Fungi_unclassified(100);k__Fungi_unclassified(100);k__Fungi_unclassified(100); | - | - | - | - | - | - | - | Unassigned | - |
| Otu06741 | 4 | k__Fungi(100);p__Basidiomycota(100);c__Agaricomycetes(100);c__Agaricomycetes_unclassified(100);c__Agaricomycetes_unclassified(100);c__Agaricomycetes_unclassified(100);c__Agaricomycetes_unclassified(100); | - | - | - | - | - | - | - | Unassigned | - |
| Otu06742 | 4 | k__Fungi(100);p__Basidiomycota(100);c__Agaricomycetes(100);c__Agaricomycetes_unclassified(100);c__Agaricomycetes_unclassified(100);c__Agaricomycetes_unclassified(100);c__Agaricomycetes_unclassified(100); | - | - | - | - | - | - | - | Unassigned | - |
| Otu06743 | 4 | k__Fungi(100);k__Fungi_unclassified(100);k__Fungi_unclassified(100);k__Fungi_unclassified(100);k__Fungi_unclassified(100);k__Fungi_unclassified(100);k__Fungi_unclassified(100); | - | - | - | - | - | - | - | Unassigned | - |
| Otu06745 | 4 | k__Fungi(100);k__Fungi_unclassified(75);k__Fungi_unclassified(75);k__Fungi_unclassified(75);k__Fungi_unclassified(75);k__Fungi_unclassified(75);k__Fungi_unclassified(75); | - | - | - | - | - | - | - | Unassigned | - |
| Otu06747 | 4 | k__Fungi(100);k__Fungi_unclassified(100);k__Fungi_unclassified(100);k__Fungi_unclassified(100);k__Fungi_unclassified(100);k__Fungi_unclassified(100);k__Fungi_unclassified(100); | - | - | - | - | - | - | - | Unassigned | - |
| Otu06754 | 4 | k__Fungi(100);p__Ascomycota(100);c__Leotiomycetes(100);c__Leotiomycetes_unclassified(75);c__Leotiomycetes_unclassified(75);c__Leotiomycetes_unclassified(75);c__Leotiomycetes_unclassified(75); | - | - | - | - | - | - | - | Unassigned | - |
| Otu06756 | 4 | k__Fungi(100);p__Ascomycota(100);c__Leotiomycetes(100);o__Helotiales(75);o__Helotiales_unclassified(75);o__Helotiales_unclassified(75);o__Helotiales_unclassified(75); | - | - | - | - | - | - | - | Unassigned | - |
| Otu06757 | 4 | k__Fungi(100);k__Fungi_unclassified(100);k__Fungi_unclassified(100);k__Fungi_unclassified(100);k__Fungi_unclassified(100);k__Fungi_unclassified(100);k__Fungi_unclassified(100); | - | - | - | - | - | - | - | Unassigned | - |
| Otu06758 | 4 | k__Fungi(100);k__Fungi_unclassified(75);k__Fungi_unclassified(75);k__Fungi_unclassified(75);k__Fungi_unclassified(75);k__Fungi_unclassified(75);k__Fungi_unclassified(75); | - | - | - | - | - | - | - | Unassigned | - |
| Otu06759 | 4 | k__Fungi(100);k__Fungi_unclassified(100);k__Fungi_unclassified(100);k__Fungi_unclassified(100);k__Fungi_unclassified(100);k__Fungi_unclassified(100);k__Fungi_unclassified(100); | - | - | - | - | - | - | - | Unassigned | - |
| Otu06761 | 4 | k__Fungi(100);k__Fungi_unclassified(100);k__Fungi_unclassified(100);k__Fungi_unclassified(100);k__Fungi_unclassified(100);k__Fungi_unclassified(100);k__Fungi_unclassified(100); | - | - | - | - | - | - | - | Unassigned | - |
| Otu06762 | 4 | k__Fungi(100);k__Fungi_unclassified(75);k__Fungi_unclassified(75);k__Fungi_unclassified(75);k__Fungi_unclassified(75);k__Fungi_unclassified(75);k__Fungi_unclassified(75); | - | - | - | - | - | - | - | Unassigned | - |
| Otu06766 | 4 | k__Fungi(100);p__Ascomycota(100);p__Ascomycota_unclassified(100);p__Ascomycota_unclassified(100);p__Ascomycota_unclassified(100);p__Ascomycota_unclassified(100);p__Ascomycota_unclassified(100); | - | - | - | - | - | - | - | Unassigned | - |
| Otu06769 | 4 | k__Fungi(100);p__Ascomycota(100);p__Ascomycota_unclassified(100);p__Ascomycota_unclassified(100);p__Ascomycota_unclassified(100);p__Ascomycota_unclassified(100);p__Ascomycota_unclassified(100); | - | - | - | - | - | - | - | Unassigned | - |
| Otu06770 | 4 | k__Fungi(100);k__Fungi_unclassified(75);k__Fungi_unclassified(75);k__Fungi_unclassified(75);k__Fungi_unclassified(75);k__Fungi_unclassified(75);k__Fungi_unclassified(75); | - | - | - | - | - | - | - | Unassigned | - |
| Otu06779 | 4 | k__Fungi(100);p__Ascomycota(100);p__Ascomycota_unclassified(75);p__Ascomycota_unclassified(75);p__Ascomycota_unclassified(75);p__Ascomycota_unclassified(75);p__Ascomycota_unclassified(75); | - | - | - | - | - | - | - | Unassigned | - |
| Otu06780 | 4 | k__Fungi(100);p__Basidiomycota(100);c__Agaricomycetes(100);o__Polyporales(100);o__Polyporales_unclassified(100);o__Polyporales_unclassified(100);o__Polyporales_unclassified(100); | - | - | - | - | - | - | - | Unassigned | - |
| Otu06782 | 4 | k__Fungi(100);k__Fungi_unclassified(100);k__Fungi_unclassified(100);k__Fungi_unclassified(100);k__Fungi_unclassified(100);k__Fungi_unclassified(100);k__Fungi_unclassified(100); | - | - | - | - | - | - | - | Unassigned | - |
| Otu06783 | 4 | k__Fungi(100);p__Ascomycota(100);c__Leotiomycetes(100);o__Helotiales(100);f__Helotiales_family_Incertae_sedis(100);f__Helotiales_family_Incertae_sedis_unclassified(100);f__Helotiales_family_Incertae_sedis_unclassified(100); | - | - | - | - | - | - | - | Unassigned | - |
| Otu06785 | 4 | k__Fungi(100);k__Fungi_unclassified(100);k__Fungi_unclassified(100);k__Fungi_unclassified(100);k__Fungi_unclassified(100);k__Fungi_unclassified(100);k__Fungi_unclassified(100); | - | - | - | - | - | - | - | Unassigned | - |
| Otu06786 | 4 | k__Fungi(100);k__Fungi_unclassified(100);k__Fungi_unclassified(100);k__Fungi_unclassified(100);k__Fungi_unclassified(100);k__Fungi_unclassified(100);k__Fungi_unclassified(100); | - | - | - | - | - | - | - | Unassigned | - |
| Otu06789 | 4 | k__Fungi(100);p__Basidiomycota(100);p__Basidiomycota_unclassified(100);p__Basidiomycota_unclassified(100);p__Basidiomycota_unclassified(100);p__Basidiomycota_unclassified(100);p__Basidiomycota_unclassified(100); | - | - | - | - | - | - | - | Unassigned | - |
| Otu06790 | 4 | k__Fungi(100);p__Ascomycota(100);p__Ascomycota_unclassified(100);p__Ascomycota_unclassified(100);p__Ascomycota_unclassified(100);p__Ascomycota_unclassified(100);p__Ascomycota_unclassified(100); | - | - | - | - | - | - | - | Unassigned | - |
| Otu06791 | 4 | k__Fungi(100);k__Fungi_unclassified(100);k__Fungi_unclassified(100);k__Fungi_unclassified(100);k__Fungi_unclassified(100);k__Fungi_unclassified(100);k__Fungi_unclassified(100); | - | - | - | - | - | - | - | Unassigned | - |
| Otu06793 | 4 | k__Fungi(100);p__Basidiomycota(100);c__Agaricomycetes(100);c__Agaricomycetes_unclassified(100);c__Agaricomycetes_unclassified(100);c__Agaricomycetes_unclassified(100);c__Agaricomycetes_unclassified(100); | - | - | - | - | - | - | - | Unassigned | - |
| Otu06794 | 4 | k__Fungi(100);p__Basidiomycota(100);c__Agaricomycetes(100);o__Sebacinales(100);f__Sebacinales_Group_B(100);g__unclassified_Sebacinales_Group_B(100);s__Sebacinales_Group_B_sp(100); | - | - | - | - | - | - | - | Unassigned | - |
| Otu06797 | 4 | k__Fungi(100);p__Ascomycota(100);p__Ascomycota_unclassified(100);p__Ascomycota_unclassified(100);p__Ascomycota_unclassified(100);p__Ascomycota_unclassified(100);p__Ascomycota_unclassified(100); | - | - | - | - | - | - | - | Unassigned | - |
| Otu06803 | 4 | k__Fungi(100);p__Ascomycota(75);p__Ascomycota_unclassified(75);p__Ascomycota_unclassified(75);p__Ascomycota_unclassified(75);p__Ascomycota_unclassified(75);p__Ascomycota_unclassified(75); | - | - | - | - | - | - | - | Unassigned | - |
| Otu06804 | 4 | k__Fungi(100);k__Fungi_unclassified(100);k__Fungi_unclassified(100);k__Fungi_unclassified(100);k__Fungi_unclassified(100);k__Fungi_unclassified(100);k__Fungi_unclassified(100); | - | - | - | - | - | - | - | Unassigned | - |
| Otu06806 | 4 | k__Fungi(100);p__Ascomycota(100);p__Ascomycota_unclassified(100);p__Ascomycota_unclassified(100);p__Ascomycota_unclassified(100);p__Ascomycota_unclassified(100);p__Ascomycota_unclassified(100); | - | - | - | - | - | - | - | Unassigned | - |
| Otu06808 | 4 | k__Fungi(100);k__Fungi_unclassified(75);k__Fungi_unclassified(75);k__Fungi_unclassified(75);k__Fungi_unclassified(75);k__Fungi_unclassified(75);k__Fungi_unclassified(75); | - | - | - | - | - | - | - | Unassigned | - |
| Otu06809 | 4 | k__Fungi(100);k__Fungi_unclassified(100);k__Fungi_unclassified(100);k__Fungi_unclassified(100);k__Fungi_unclassified(100);k__Fungi_unclassified(100);k__Fungi_unclassified(100); | - | - | - | - | - | - | - | Unassigned | - |
| Otu06812 | 4 | k__Fungi(100);k__Fungi_unclassified(100);k__Fungi_unclassified(100);k__Fungi_unclassified(100);k__Fungi_unclassified(100);k__Fungi_unclassified(100);k__Fungi_unclassified(100); | - | - | - | - | - | - | - | Unassigned | - |
| Otu06814 | 4 | k__Fungi(100);k__Fungi_unclassified(75);k__Fungi_unclassified(75);k__Fungi_unclassified(75);k__Fungi_unclassified(75);k__Fungi_unclassified(75);k__Fungi_unclassified(75); | - | - | - | - | - | - | - | Unassigned | - |
| Otu06816 | 4 | k__Fungi(100);p__Ascomycota(75);p__Ascomycota_unclassified(75);p__Ascomycota_unclassified(75);p__Ascomycota_unclassified(75);p__Ascomycota_unclassified(75);p__Ascomycota_unclassified(75); | - | - | - | - | - | - | - | Unassigned | - |
| Otu06818 | 4 | k__Fungi(100);k__Fungi_unclassified(100);k__Fungi_unclassified(100);k__Fungi_unclassified(100);k__Fungi_unclassified(100);k__Fungi_unclassified(100);k__Fungi_unclassified(100); | - | - | - | - | - | - | - | Unassigned | - |
| Otu06821 | 4 | k__Fungi(100);p__Basidiomycota(100);c__Agaricomycetes(100);c__Agaricomycetes_unclassified(100);c__Agaricomycetes_unclassified(100);c__Agaricomycetes_unclassified(100);c__Agaricomycetes_unclassified(100); | - | - | - | - | - | - | - | Unassigned | - |
| Otu06822 | 4 | k__Fungi(100);k__Fungi_unclassified(100);k__Fungi_unclassified(100);k__Fungi_unclassified(100);k__Fungi_unclassified(100);k__Fungi_unclassified(100);k__Fungi_unclassified(100); | - | - | - | - | - | - | - | Unassigned | - |
| Otu06823 | 4 | k__Fungi(100);p__Basidiomycota(100);c__Agaricomycetes(100);o__Polyporales(100);o__Polyporales_unclassified(100);o__Polyporales_unclassified(100);o__Polyporales_unclassified(100); | - | - | - | - | - | - | - | Unassigned | - |
| Otu06824 | 4 | k__Fungi(100);p__Ascomycota(75);p__Ascomycota_unclassified(75);p__Ascomycota_unclassified(75);p__Ascomycota_unclassified(75);p__Ascomycota_unclassified(75);p__Ascomycota_unclassified(75); | - | - | - | - | - | - | - | Unassigned | - |
| Otu06825 | 4 | k__Fungi(100);k__Fungi_unclassified(100);k__Fungi_unclassified(100);k__Fungi_unclassified(100);k__Fungi_unclassified(100);k__Fungi_unclassified(100);k__Fungi_unclassified(100); | - | - | - | - | - | - | - | Unassigned | - |
| Otu06827 | 4 | k__Fungi(100);p__Ascomycota(100);p__Ascomycota_unclassified(100);p__Ascomycota_unclassified(100);p__Ascomycota_unclassified(100);p__Ascomycota_unclassified(100);p__Ascomycota_unclassified(100); | - | - | - | - | - | - | - | Unassigned | - |
| Otu06829 | 4 | k__Fungi(100);k__Fungi_unclassified(100);k__Fungi_unclassified(100);k__Fungi_unclassified(100);k__Fungi_unclassified(100);k__Fungi_unclassified(100);k__Fungi_unclassified(100); | - | - | - | - | - | - | - | Unassigned | - |
| Otu06830 | 4 | k__Fungi(100);p__Ascomycota(100);c__Dothideomycetes(75);o__Capnodiales(75);o__Capnodiales_unclassified(75);o__Capnodiales_unclassified(75);o__Capnodiales_unclassified(75); | - | - | - | - | - | - | - | Unassigned | - |
| Otu06833 | 4 | k__Fungi(100);p__Ascomycota(100);c__Dothideomycetes(100);o__Pleosporales(100);o__Pleosporales_unclassified(100);o__Pleosporales_unclassified(100);o__Pleosporales_unclassified(100); | - | - | - | - | - | - | - | Unassigned | - |
| Otu06834 | 4 | k__Fungi(100);k__Fungi_unclassified(75);k__Fungi_unclassified(75);k__Fungi_unclassified(75);k__Fungi_unclassified(75);k__Fungi_unclassified(75);k__Fungi_unclassified(75); | - | - | - | - | - | - | - | Unassigned | - |
| Otu06836 | 4 | k__Fungi(100);k__Fungi_unclassified(100);k__Fungi_unclassified(100);k__Fungi_unclassified(100);k__Fungi_unclassified(100);k__Fungi_unclassified(100);k__Fungi_unclassified(100); | - | - | - | - | - | - | - | Unassigned | - |
| Otu06839 | 4 | k__Fungi(100);k__Fungi_unclassified(100);k__Fungi_unclassified(100);k__Fungi_unclassified(100);k__Fungi_unclassified(100);k__Fungi_unclassified(100);k__Fungi_unclassified(100); | - | - | - | - | - | - | - | Unassigned | - |
| Otu06840 | 4 | k__Fungi(100);k__Fungi_unclassified(100);k__Fungi_unclassified(100);k__Fungi_unclassified(100);k__Fungi_unclassified(100);k__Fungi_unclassified(100);k__Fungi_unclassified(100); | - | - | - | - | - | - | - | Unassigned | - |
| Otu06842 | 4 | k__Fungi(100);p__Ascomycota(75);p__Ascomycota_unclassified(75);p__Ascomycota_unclassified(75);p__Ascomycota_unclassified(75);p__Ascomycota_unclassified(75);p__Ascomycota_unclassified(75); | - | - | - | - | - | - | - | Unassigned | - |
| Otu06843 | 4 | k__Fungi(100);k__Fungi_unclassified(100);k__Fungi_unclassified(100);k__Fungi_unclassified(100);k__Fungi_unclassified(100);k__Fungi_unclassified(100);k__Fungi_unclassified(100); | - | - | - | - | - | - | - | Unassigned | - |
| Otu06844 | 4 | k__Fungi(100);p__Ascomycota(75);p__Ascomycota_unclassified(75);p__Ascomycota_unclassified(75);p__Ascomycota_unclassified(75);p__Ascomycota_unclassified(75);p__Ascomycota_unclassified(75); | - | - | - | - | - | - | - | Unassigned | - |
| Otu06845 | 4 | k__Fungi(100);k__Fungi_unclassified(100);k__Fungi_unclassified(100);k__Fungi_unclassified(100);k__Fungi_unclassified(100);k__Fungi_unclassified(100);k__Fungi_unclassified(100); | - | - | - | - | - | - | - | Unassigned | - |
| Otu06852 | 4 | k__Fungi(100);p__Ascomycota(75);p__Ascomycota_unclassified(75);p__Ascomycota_unclassified(75);p__Ascomycota_unclassified(75);p__Ascomycota_unclassified(75);p__Ascomycota_unclassified(75); | - | - | - | - | - | - | - | Unassigned | - |
| Otu06855 | 4 | k__Fungi(100);p__Ascomycota(100);p__Ascomycota_unclassified(100);p__Ascomycota_unclassified(100);p__Ascomycota_unclassified(100);p__Ascomycota_unclassified(100);p__Ascomycota_unclassified(100); | - | - | - | - | - | - | - | Unassigned | - |
| Otu06859 | 4 | k__Fungi(100);k__Fungi_unclassified(75);k__Fungi_unclassified(75);k__Fungi_unclassified(75);k__Fungi_unclassified(75);k__Fungi_unclassified(75);k__Fungi_unclassified(75); | - | - | - | - | - | - | - | Unassigned | - |
| Otu06863 | 4 | k__Fungi(100);p__Basidiomycota(100);c__Agaricomycetes(100);c__Agaricomycetes_unclassified(100);c__Agaricomycetes_unclassified(100);c__Agaricomycetes_unclassified(100);c__Agaricomycetes_unclassified(100); | - | - | - | - | - | - | - | Unassigned | - |
| Otu06864 | 4 | k__Fungi(100);k__Fungi_unclassified(100);k__Fungi_unclassified(100);k__Fungi_unclassified(100);k__Fungi_unclassified(100);k__Fungi_unclassified(100);k__Fungi_unclassified(100); | - | - | - | - | - | - | - | Unassigned | - |
| Otu06866 | 4 | k__Fungi(100);p__Ascomycota(100);p__Ascomycota_unclassified(100);p__Ascomycota_unclassified(100);p__Ascomycota_unclassified(100);p__Ascomycota_unclassified(100);p__Ascomycota_unclassified(100); | - | - | - | - | - | - | - | Unassigned | - |
| Otu06868 | 4 | k__Fungi(100);p__Ascomycota(75);p__Ascomycota_unclassified(75);p__Ascomycota_unclassified(75);p__Ascomycota_unclassified(75);p__Ascomycota_unclassified(75);p__Ascomycota_unclassified(75); | - | - | - | - | - | - | - | Unassigned | - |
| Otu06869 | 4 | k__Fungi(100);p__Ascomycota(75);p__Ascomycota_unclassified(75);p__Ascomycota_unclassified(75);p__Ascomycota_unclassified(75);p__Ascomycota_unclassified(75);p__Ascomycota_unclassified(75); | - | - | - | - | - | - | - | Unassigned | - |
| Otu06870 | 4 | k__Fungi(100);k__Fungi_unclassified(100);k__Fungi_unclassified(100);k__Fungi_unclassified(100);k__Fungi_unclassified(100);k__Fungi_unclassified(100);k__Fungi_unclassified(100); | - | - | - | - | - | - | - | Unassigned | - |
| Otu06877 | 4 | k__Fungi(100);k__Fungi_unclassified(75);k__Fungi_unclassified(75);k__Fungi_unclassified(75);k__Fungi_unclassified(75);k__Fungi_unclassified(75);k__Fungi_unclassified(75); | - | - | - | - | - | - | - | Unassigned | - |
| Otu06880 | 4 | k__Fungi(100);p__Ascomycota(100);p__Ascomycota_unclassified(100);p__Ascomycota_unclassified(100);p__Ascomycota_unclassified(100);p__Ascomycota_unclassified(100);p__Ascomycota_unclassified(100); | - | - | - | - | - | - | - | Unassigned | - |
| Otu06881 | 4 | k__Fungi(100);k__Fungi_unclassified(100);k__Fungi_unclassified(100);k__Fungi_unclassified(100);k__Fungi_unclassified(100);k__Fungi_unclassified(100);k__Fungi_unclassified(100); | - | - | - | - | - | - | - | Unassigned | - |
| Otu06882 | 4 | k__Fungi(100);k__Fungi_unclassified(100);k__Fungi_unclassified(100);k__Fungi_unclassified(100);k__Fungi_unclassified(100);k__Fungi_unclassified(100);k__Fungi_unclassified(100); | - | - | - | - | - | - | - | Unassigned | - |
| Otu06887 | 4 | k__Fungi(100);p__Ascomycota(100);c__Eurotiomycetes(100);c__Eurotiomycetes_unclassified(100);c__Eurotiomycetes_unclassified(100);c__Eurotiomycetes_unclassified(100);c__Eurotiomycetes_unclassified(100); | - | - | - | - | - | - | - | Unassigned | - |
| Otu06891 | 4 | k__Fungi(100);p__Ascomycota(100);p__Ascomycota_unclassified(100);p__Ascomycota_unclassified(100);p__Ascomycota_unclassified(100);p__Ascomycota_unclassified(100);p__Ascomycota_unclassified(100); | - | - | - | - | - | - | - | Unassigned | - |
| Otu06892 | 4 | k__Fungi(100);k__Fungi_unclassified(100);k__Fungi_unclassified(100);k__Fungi_unclassified(100);k__Fungi_unclassified(100);k__Fungi_unclassified(100);k__Fungi_unclassified(100); | - | - | - | - | - | - | - | Unassigned | - |
| Otu06897 | 4 | k__Fungi(100);p__Ascomycota(75);p__Ascomycota_unclassified(75);p__Ascomycota_unclassified(75);p__Ascomycota_unclassified(75);p__Ascomycota_unclassified(75);p__Ascomycota_unclassified(75); | - | - | - | - | - | - | - | Unassigned | - |
| Otu06898 | 4 | k__Fungi(100);k__Fungi_unclassified(75);k__Fungi_unclassified(75);k__Fungi_unclassified(75);k__Fungi_unclassified(75);k__Fungi_unclassified(75);k__Fungi_unclassified(75); | - | - | - | - | - | - | - | Unassigned | - |
| Otu06902 | 4 | k__Fungi(100);k__Fungi_unclassified(75);k__Fungi_unclassified(75);k__Fungi_unclassified(75);k__Fungi_unclassified(75);k__Fungi_unclassified(75);k__Fungi_unclassified(75); | - | - | - | - | - | - | - | Unassigned | - |
| Otu06904 | 4 | k__Fungi(100);p__Ascomycota(100);p__Ascomycota_unclassified(100);p__Ascomycota_unclassified(100);p__Ascomycota_unclassified(100);p__Ascomycota_unclassified(100);p__Ascomycota_unclassified(100); | - | - | - | - | - | - | - | Unassigned | - |
| Otu06906 | 4 | k__Fungi(100);p__Ascomycota(75);p__Ascomycota_unclassified(75);p__Ascomycota_unclassified(75);p__Ascomycota_unclassified(75);p__Ascomycota_unclassified(75);p__Ascomycota_unclassified(75); | - | - | - | - | - | - | - | Unassigned | - |
| Otu06910 | 4 | k__Fungi(100);p__Basidiomycota(100);c__Exobasidiomycetes(100);o__Entylomatales(100);f__unclassified_Entylomatales(100);g__unclassified_Entylomatales(100);s__Entylomatales_sp(100); | - | - | - | - | - | - | - | Unassigned | - |
| Otu06911 | 4 | k__Fungi(100);k__Fungi_unclassified(75);k__Fungi_unclassified(75);k__Fungi_unclassified(75);k__Fungi_unclassified(75);k__Fungi_unclassified(75);k__Fungi_unclassified(75); | - | - | - | - | - | - | - | Unassigned | - |
| Otu06915 | 4 | k__Fungi(100);k__Fungi_unclassified(100);k__Fungi_unclassified(100);k__Fungi_unclassified(100);k__Fungi_unclassified(100);k__Fungi_unclassified(100);k__Fungi_unclassified(100); | - | - | - | - | - | - | - | Unassigned | - |
| Otu06918 | 4 | k__Fungi(100);p__Ascomycota(75);p__Ascomycota_unclassified(75);p__Ascomycota_unclassified(75);p__Ascomycota_unclassified(75);p__Ascomycota_unclassified(75);p__Ascomycota_unclassified(75); | - | - | - | - | - | - | - | Unassigned | - |
| Otu06919 | 4 | k__Fungi(100);k__Fungi_unclassified(100);k__Fungi_unclassified(100);k__Fungi_unclassified(100);k__Fungi_unclassified(100);k__Fungi_unclassified(100);k__Fungi_unclassified(100); | - | - | - | - | - | - | - | Unassigned | - |
| Otu06922 | 4 | k__Fungi(100);k__Fungi_unclassified(100);k__Fungi_unclassified(100);k__Fungi_unclassified(100);k__Fungi_unclassified(100);k__Fungi_unclassified(100);k__Fungi_unclassified(100); | - | - | - | - | - | - | - | Unassigned | - |
| Otu06932 | 3 | k__Fungi(100);p__Ascomycota(100);c__Saccharomycetes(100);o__Saccharomycetales(100);f__Metschnikowiaceae(100);g__Metschnikowia(100);s__Metschnikowia_pulcherrima(100); | Metschnikowia | 13 | Saprotroph | \|Undefined Saprotroph\| | Yeast | NULL | Probable | As nectar-tap_saprotroph (Põlme et al. 2020); Decay sugar-rich substrates (Põlme et al. 2020) | Põlme S, et al. 2020. Fungal Diversity 105:1-16 ((https://doi.org/10.1007/s13225-020-00466-2)) |
| Otu06935 | 3 | k__Fungi(100);p__Ascomycota(100);c__Saccharomycetes(100);o__Saccharomycetales(100);f__Metschnikowiaceae(100);g__Metschnikowia(100);s__Metschnikowia_pulcherrima(100); | Metschnikowia | 13 | Saprotroph | \|Undefined Saprotroph\| | Yeast | NULL | Probable | As nectar-tap_saprotroph (Põlme et al. 2020); Decay sugar-rich substrates (Põlme et al. 2020) | Põlme S, et al. 2020. Fungal Diversity 105:1-16 ((https://doi.org/10.1007/s13225-020-00466-2)) |
| Otu06936 | 3 | k__Fungi(100);p__Ascomycota(100);c__Saccharomycetes(100);o__Saccharomycetales(100);f__Metschnikowiaceae(100);g__Metschnikowia(100);s__Metschnikowia_pulcherrima(100); | Metschnikowia | 13 | Saprotroph | \|Undefined Saprotroph\| | Yeast | NULL | Probable | As nectar-tap_saprotroph (Põlme et al. 2020); Decay sugar-rich substrates (Põlme et al. 2020) | Põlme S, et al. 2020. Fungal Diversity 105:1-16 ((https://doi.org/10.1007/s13225-020-00466-2)) |
| Otu06940 | 3 | k__Fungi(100);p__Basidiomycota(100);c__Agaricomycetes(100);o__Cantharellales(100);f__Ceratobasidiaceae(100);g__Thanatephorus(100);s__Thanatephorus_fusisporus(100); | Thanatephorus | 13 | Pathotroph-Saprotroph | \|Plant Pathogen\|-Plant Saprotroph | Corticioid | NULL | Probable | As litter_saprotroph (Põlme et al. 2020); Decay leaf, fruit, seed (Põlme et al. 2020) | Tedersoo L, et al. 2014. Science 346:e1256688 ((https://doi.org/10.1126/science.1256688)); Põlme S, et al. 2020. Fungal Diversity 105:1-16 ((https://doi.org/10.1007/s13225-020-00466-2)) |
| Otu06943 | 3 | k__Fungi(100);p__Ascomycota(100);c__Dothideomycetes(100);o__Pleosporales(100);f__Phaeosphaeriaceae(100);g__unclassified_Phaeosphaeriaceae(67);s__Phaeosphaeriaceae_sp(67); | Phaeosphaeriaceae | 9 | Pathotroph-Saprotroph | Fungal Parasite-Plant Pathogen-Plant Saprotroph | Microfungus | NULL | Probable | Necrotrophic or saprobic on a wide range of plants, especially monocotyledons; a few species parasitic on other fungi (Cannon & Kirk 2007) | Cannon PF, Kirk PM. 2007. Fungal Families of the World. CAB International, Cambridge (ISBN: 978-0851998275) |
| Otu06946 | 3 | k__Fungi(100);p__Ascomycota(100);c__Leotiomycetes(100);o__Helotiales(100);f__Rutstroemiaceae(100);g__Scleromitrula(100);g__Scleromitrula_unclassified(100); | Scleromitrula | 13 | Pathotroph-Saprotroph | \|Plant Pathogen\|-Undefined Saprotroph | Microfungus | NULL | Probable | NULL | Tedersoo L, et al. 2014. Science 346:e1256688 ((https://doi.org/10.1126/science.1256688)); Põlme S, et al. 2020. Fungal Diversity 105:1-16 ((https://doi.org/10.1007/s13225-020-00466-2)) |
| Otu06950 | 3 | k__Fungi(100);p__Ascomycota(100);c__Dothideomycetes(100);o__Pleosporales(100);f__Phaeosphaeriaceae(100);g__Ophiosphaerella(100);g__Ophiosphaerella_unclassified(67); | Ophiosphaerella | 13 | Pathotroph-Saprotroph | \|Plant Pathogen\|-Plant Saprotroph-Undefined Saprotroph | Microfungus | NULL | Probable | As litter_saprotroph (Põlme et al. 2020); Decay leaf, fruit, seed (Põlme et al. 2020) | Tedersoo L, et al. 2014. Science 346:e1256688 ((https://doi.org/10.1126/science.1256688)); Põlme S, et al. 2020. Fungal Diversity 105:1-16 ((https://doi.org/10.1007/s13225-020-00466-2)) |
| Otu06951 | 3 | k__Fungi(100);p__Ascomycota(100);c__Sordariomycetes(100);o__Hypocreales(100);f__Clavicipitaceae(100);g__Metarhizium(100);s__Metarhizium_lepidiotae(100); | Metarhizium | 13 | Pathotroph-Saprotroph | \|Animal Parasite\|-Animal Pathogen-Undefined Saprotroph | Microfungus | NULL | Probable | As animal_decomposer (Põlme et al. 2020); Decay animal material (Põlme et al. 2020) | Tedersoo L, et al. 2014. Science 346:e1256688 ((https://doi.org/10.1126/science.1256688)); Põlme S, et al. 2020. Fungal Diversity 105:1-16 ((https://doi.org/10.1007/s13225-020-00466-2)) |
| Otu06953 | 3 | k__Fungi(100);p__Ascomycota(100);c__Saccharomycetes(100);o__Saccharomycetales(100);f__Metschnikowiaceae(100);g__Metschnikowia(100);s__Metschnikowia_pulcherrima(67); | Metschnikowia | 13 | Saprotroph | \|Undefined Saprotroph\| | Yeast | NULL | Probable | As nectar-tap_saprotroph (Põlme et al. 2020); Decay sugar-rich substrates (Põlme et al. 2020) | Põlme S, et al. 2020. Fungal Diversity 105:1-16 ((https://doi.org/10.1007/s13225-020-00466-2)) |
| Otu06954 | 3 | k__Fungi(100);p__Ascomycota(100);c__Saccharomycetes(100);o__Saccharomycetales(100);f__Metschnikowiaceae(100);g__Metschnikowia(100);s__Metschnikowia_pulcherrima(100); | Metschnikowia | 13 | Saprotroph | \|Undefined Saprotroph\| | Yeast | NULL | Probable | As nectar-tap_saprotroph (Põlme et al. 2020); Decay sugar-rich substrates (Põlme et al. 2020) | Põlme S, et al. 2020. Fungal Diversity 105:1-16 ((https://doi.org/10.1007/s13225-020-00466-2)) |
| Otu06955 | 3 | k__Fungi(100);p__Ascomycota(100);c__Saccharomycetes(100);o__Saccharomycetales(100);f__Saccharomycetales_family_Incertae_sedis(100);g__Candida(100);s__Candida_sp_AS23486(100); | Candida | 13 | Pathotroph-Saprotroph-Symbiotroph | Animal Pathogen-Endophyte-Endosymbiont-Epiphyte-\|Undefined Saprotroph\| | Dimorphic Yeast-Yeast | NULL | Probable | As nectar-tap_saprotroph (Põlme et al. 2020); Decay sugar-rich substrates (Põlme et al. 2020); This asexual genus is polyphyletic and contain species that function very differently depend on what they are. Caution should be taken when interpreting guilds for Candida species. | Manolakaki D, et al. 2010. Virulence. 1: 367-375 ((https://doi.org/10.4161/viru.1.5.12796)); Mekbib SB, et al. 2011. Tropical Plant Pathology 36:233-240 ((https://doi.org/10.1590/S1982-56762011000400004)); Põlme S, et al. 2020. Fungal Diversity 105:1-16 ((https://doi.org/10.1007/s13225-020-00466-2)) |
| Otu06956 | 3 | k__Fungi(100);p__Ascomycota(100);c__Saccharomycetes(100);o__Saccharomycetales(100);f__Metschnikowiaceae(100);g__Metschnikowia(100);s__Metschnikowia_pulcherrima(100); | Metschnikowia | 13 | Saprotroph | \|Undefined Saprotroph\| | Yeast | NULL | Probable | As nectar-tap_saprotroph (Põlme et al. 2020); Decay sugar-rich substrates (Põlme et al. 2020) | Põlme S, et al. 2020. Fungal Diversity 105:1-16 ((https://doi.org/10.1007/s13225-020-00466-2)) |
| Otu06958 | 3 | k__Fungi(100);p__Ascomycota(100);c__Sordariomycetes(100);o__Chaetosphaeriales(100);f__Chaetosphaeriaceae(100);g__Chloridium(67);g__Chloridium_unclassified(67); | Chloridium | 13 | Pathotroph-Saprotroph-Symbiotroph | Ectomycorrhizal-Endophyte-Plant Pathogen-\|Plant Saprotroph\|-Wood Saprotroph | Microfungus | NULL | Possible | As litter_saprotroph (Põlme et al. 2020); As root_pathogen (Põlme et al. 2020); Decay wood, leaf, fruit, seed, roots, animal material (Põlme et al. 2020) | Duncan CG, Eslyn WE. 1966. Mycologia 58:642-645 ((https://doi.org/10.2307/3757045)); Jumpponen A, Trappe J. 1998. New Phytologist 140:295-310 ((https://doi.org/10.1046/j.1469-8137.1998.00265.x)); Rinaldi AC, et al. 2008. Fungal Diversity 33:1-45 ((http://www.fungaldiversity.org/fdp/sfdp/33-1.pdf)); Tedersoo L, et al. 2014. Science 346:e1256688 ((https://doi.org/10.1126/science.1256688)); Põlme S, et al. 2020. Fungal Diversity 105:1-16 ((https://doi.org/10.1007/s13225-020-00466-2)) |
| Otu06959 | 3 | k__Fungi(100);p__Ascomycota(100);c__Saccharomycetes(100);o__Saccharomycetales(100);f__Metschnikowiaceae(100);g__Metschnikowia(100);s__Metschnikowia_pulcherrima(100); | Metschnikowia | 13 | Saprotroph | \|Undefined Saprotroph\| | Yeast | NULL | Probable | As nectar-tap_saprotroph (Põlme et al. 2020); Decay sugar-rich substrates (Põlme et al. 2020) | Põlme S, et al. 2020. Fungal Diversity 105:1-16 ((https://doi.org/10.1007/s13225-020-00466-2)) |
| Otu06960 | 3 | k__Fungi(100);p__Ascomycota(100);c__Saccharomycetes(100);o__Saccharomycetales(100);f__Metschnikowiaceae(100);g__Metschnikowia(100);s__Metschnikowia_pulcherrima(67); | Metschnikowia | 13 | Saprotroph | \|Undefined Saprotroph\| | Yeast | NULL | Probable | As nectar-tap_saprotroph (Põlme et al. 2020); Decay sugar-rich substrates (Põlme et al. 2020) | Põlme S, et al. 2020. Fungal Diversity 105:1-16 ((https://doi.org/10.1007/s13225-020-00466-2)) |
| Otu06962 | 3 | k__Fungi(100);p__Ascomycota(100);c__Saccharomycetes(100);o__Saccharomycetales(100);f__Saccharomycetales_family_Incertae_sedis(67);g__Candida(67);g__Candida_unclassified(67); | Candida | 13 | Pathotroph-Saprotroph-Symbiotroph | Animal Pathogen-Endophyte-Endosymbiont-Epiphyte-\|Undefined Saprotroph\| | Dimorphic Yeast-Yeast | NULL | Probable | As nectar-tap_saprotroph (Põlme et al. 2020); Decay sugar-rich substrates (Põlme et al. 2020); This asexual genus is polyphyletic and contain species that function very differently depend on what they are. Caution should be taken when interpreting guilds for Candida species. | Manolakaki D, et al. 2010. Virulence. 1: 367-375 ((https://doi.org/10.4161/viru.1.5.12796)); Mekbib SB, et al. 2011. Tropical Plant Pathology 36:233-240 ((https://doi.org/10.1590/S1982-56762011000400004)); Põlme S, et al. 2020. Fungal Diversity 105:1-16 ((https://doi.org/10.1007/s13225-020-00466-2)) |
| Otu06964 | 3 | k__Fungi(100);p__Ascomycota(100);c__Saccharomycetes(100);o__Saccharomycetales(100);f__Metschnikowiaceae(100);g__Metschnikowia(100);s__Metschnikowia_pulcherrima(100); | Metschnikowia | 13 | Saprotroph | \|Undefined Saprotroph\| | Yeast | NULL | Probable | As nectar-tap_saprotroph (Põlme et al. 2020); Decay sugar-rich substrates (Põlme et al. 2020) | Põlme S, et al. 2020. Fungal Diversity 105:1-16 ((https://doi.org/10.1007/s13225-020-00466-2)) |
| Otu06966 | 3 | k__Fungi(100);p__Ascomycota(100);c__Saccharomycetes(100);o__Saccharomycetales(100);f__Metschnikowiaceae(100);g__Metschnikowia(100);s__Metschnikowia_pulcherrima(100); | Metschnikowia | 13 | Saprotroph | \|Undefined Saprotroph\| | Yeast | NULL | Probable | As nectar-tap_saprotroph (Põlme et al. 2020); Decay sugar-rich substrates (Põlme et al. 2020) | Põlme S, et al. 2020. Fungal Diversity 105:1-16 ((https://doi.org/10.1007/s13225-020-00466-2)) |
| Otu06967 | 3 | k__Fungi(100);p__Ascomycota(100);c__Saccharomycetes(100);o__Saccharomycetales(100);f__Metschnikowiaceae(100);g__Metschnikowia(100);s__Metschnikowia_pulcherrima(67); | Metschnikowia | 13 | Saprotroph | \|Undefined Saprotroph\| | Yeast | NULL | Probable | As nectar-tap_saprotroph (Põlme et al. 2020); Decay sugar-rich substrates (Põlme et al. 2020) | Põlme S, et al. 2020. Fungal Diversity 105:1-16 ((https://doi.org/10.1007/s13225-020-00466-2)) |
| Otu06968 | 3 | k__Fungi(100);p__Ascomycota(100);c__Dothideomycetes(100);o__Pleosporales(100);f__Pleosporales_family_Incertae_sedis(100);g__Periconia(100);s__Periconia_sp(100); | Periconia | 13 | Pathotroph-Saprotroph-Symbiotroph | Endophyte-\|Plant Pathogen\|-Wood Saprotroph | Microfungus | Soft Rot | Possible | Asteraceae, Musaceae (Costa et al. 2012); As foliar_endophyte (Põlme et al. 2020); Dark Septate Endophyte, known from plant roots (Mandyam et al. 2010); Endophyte Host ; Decay leaf, fruit, seed (Põlme et al. 2020) | Duncan CG, Eslyn WE. 1966. Mycologia 58:642-645 ((https://doi.org/10.2307/3757045)); Costa IPMW, et al. 2012. Mycotaxon 119:e26 ((http://mycotaxon.com/resources/checklists/costa_v119_checklist.pdf)); Põlme S, et al. 2020. Fungal Diversity 105:1-16 ((https://doi.org/10.1007/s13225-020-00466-2)); Mandyam K, et al. 2012. Mycorrhiza 22:109-119 ((https://doi.org/10.1007/s00572-011-0386-y)); Knapp DG, et al. 2012. PLOS ONE 7:e32570 (( https://doi.org/10.1371/journal.pone.0032570)); The American Phytopathological Society Common Names of Plant Diseases ((http://www.apsnet.org/publications/commonnames/Pages/default.aspx)) |
| Otu06969 | 3 | k__Fungi(100);p__Ascomycota(100);c__Saccharomycetes(100);o__Saccharomycetales(100);f__Metschnikowiaceae(100);g__Metschnikowia(100);s__Metschnikowia_pulcherrima(67); | Metschnikowia | 13 | Saprotroph | \|Undefined Saprotroph\| | Yeast | NULL | Probable | As nectar-tap_saprotroph (Põlme et al. 2020); Decay sugar-rich substrates (Põlme et al. 2020) | Põlme S, et al. 2020. Fungal Diversity 105:1-16 ((https://doi.org/10.1007/s13225-020-00466-2)) |
| Otu06970 | 3 | k__Fungi(100);p__Ascomycota(100);c__Saccharomycetes(100);o__Saccharomycetales(100);f__Metschnikowiaceae(100);g__Metschnikowia(100);s__Metschnikowia_pulcherrima(67); | Metschnikowia | 13 | Saprotroph | \|Undefined Saprotroph\| | Yeast | NULL | Probable | As nectar-tap_saprotroph (Põlme et al. 2020); Decay sugar-rich substrates (Põlme et al. 2020) | Põlme S, et al. 2020. Fungal Diversity 105:1-16 ((https://doi.org/10.1007/s13225-020-00466-2)) |
| Otu06971 | 3 | k__Fungi(100);p__Ascomycota(100);c__Dothideomycetes(100);o__Pleosporales(100);f__Cucurbitariaceae(100);g__Pyrenochaetopsis(100);s__Pyrenochaetopsis_sp(100); | Pyrenochaetopsis | 13 | Pathotroph-Saprotroph-Symbiotroph | \|Endophyte\|-Lichen Parasite-Plant Saprotroph-Undefined Saprotroph | Microfungus | NULL | Possible | As foliar_endophyte (Põlme et al. 2020); As litter_saprotroph (Põlme et al. 2020); Decay leaf, fruit, seed (Põlme et al. 2020) | Tedersoo L, et al. 2014. Science 346:e1256688 ((https://doi.org/10.1126/science.1256688)); Massimo NC, et al. 2015. Microbial Ecology 70:61-76 ((https://doi.org/10.1007/s00248-014-0563-6)); Lawrey JD, Diederich P. 2016. http://www.lichenicolous.net (accessed 24 May 2017) ((http://www.lichenicolous.net)); Põlme S, et al. 2020. Fungal Diversity 105:1-16 ((https://doi.org/10.1007/s13225-020-00466-2)) |
| Otu06974 | 3 | k__Fungi(100);p__Ascomycota(100);c__Saccharomycetes(100);o__Saccharomycetales(100);f__Saccharomycetaceae(100);g__Kazachstania(100);s__Kazachstania_intestinalis(100); | Kazachstania | 13 | Saprotroph | \|Undefined Saprotroph\| | Yeast | NULL | Probable | As nectar-tap_saprotroph (Põlme et al. 2020); Decay sugar-rich substrates (Põlme et al. 2020) | Põlme S, et al. 2020. Fungal Diversity 105:1-16 ((https://doi.org/10.1007/s13225-020-00466-2)) |
| Otu06975 | 3 | k__Fungi(100);p__Basidiomycota(100);c__Agaricomycetes(100);o__Hymenochaetales(100);f__Hymenochaetaceae(100);g__Phellinus(100);s__Phellinus_ribis_f_ulicis(100); | Phellinus | 13 | Pathotroph-Saprotroph | \|Plant Pathogen\|-Wood Saprotroph | Polyporoid | White Rot | Highly Probable | As white_rot (Põlme et al. 2020); As wood_pathogen (Põlme et al. 2020); Decay wood (Põlme et al. 2020) | Gilbertson RL, Ryvarden L. 1987-1987. North American Polypores. Fungiflora, Oslo (ISBN: 978-0945345060); Põlme S, et al. 2020. Fungal Diversity 105:1-16 ((https://doi.org/10.1007/s13225-020-00466-2)) |
| Otu06980 | 3 | k__Fungi(100);p__Ascomycota(100);c__Sordariomycetes(100);o__Diaporthales(100);f__Diaporthaceae(100);f__Diaporthaceae_unclassified(67);f__Diaporthaceae_unclassified(67); | Diaporthaceae | 9 | Pathotroph-Saprotroph-Symbiotroph | Endophyte-Plant Pathogen-Plant Saprotroph | Microfungus | NULL | Probable | Most species are weak pathogens or endophytes in leaf or stem tissue, growing saprobically after the plant dies (Cannon & Kirk 2007) | Cannon PF, Kirk PM. 2007. Fungal Families of the World. CAB International, Cambridge (ISBN: 978-0851998275) |
| Otu06981 | 3 | k__Fungi(100);p__Basidiomycota(100);c__Agaricomycetes(100);o__Agaricales(100);f__Mycenaceae(100);g__Mycena(100);s__Mycena_zephirus(100); | Mycena | 13 | Pathotroph-Saprotroph | Plant Pathogen-\|Plant Saprotroph\|-Undefined Saprotroph-Wood Saprotroph | Agaricoid | NULL | Probable | As litter_saprotroph (Põlme et al. 2020); Decay leaf, fruit, seed (Põlme et al. 2020) | Grgurinovic CA, Simpson JA. 2001. Fungal Diversity 8:97-127 ((http://www.fungaldiversity.org/fdp/sfdp/FD_8_97-127.pdf)); Tedersoo L, et al. 2014. Science 346:e1256688 ((https://doi.org/10.1126/science.1256688)); Põlme S, et al. 2020. Fungal Diversity 105:1-16 ((https://doi.org/10.1007/s13225-020-00466-2)); The American Phytopathological Society Common Names of Plant Diseases ((http://www.apsnet.org/publications/commonnames/Pages/default.aspx)) |
| Otu06983 | 3 | k__Fungi(100);p__Ascomycota(100);c__Saccharomycetes(100);o__Saccharomycetales(100);f__Metschnikowiaceae(100);g__Metschnikowia(100);s__Metschnikowia_pulcherrima(100); | Metschnikowia | 13 | Saprotroph | \|Undefined Saprotroph\| | Yeast | NULL | Probable | As nectar-tap_saprotroph (Põlme et al. 2020); Decay sugar-rich substrates (Põlme et al. 2020) | Põlme S, et al. 2020. Fungal Diversity 105:1-16 ((https://doi.org/10.1007/s13225-020-00466-2)) |
| Otu06985 | 3 | k__Fungi(100);p__Ascomycota(100);c__Saccharomycetes(100);o__Saccharomycetales(100);f__Metschnikowiaceae(100);g__Metschnikowia(100);s__Metschnikowia_pulcherrima(67); | Metschnikowia | 13 | Saprotroph | \|Undefined Saprotroph\| | Yeast | NULL | Probable | As nectar-tap_saprotroph (Põlme et al. 2020); Decay sugar-rich substrates (Põlme et al. 2020) | Põlme S, et al. 2020. Fungal Diversity 105:1-16 ((https://doi.org/10.1007/s13225-020-00466-2)) |
| Otu06988 | 3 | k__Fungi(100);p__Ascomycota(100);c__Saccharomycetes(100);o__Saccharomycetales(100);f__Saccharomycetales_family_Incertae_sedis(100);g__Candida(100);g__Candida_unclassified(67); | Candida | 13 | Pathotroph-Saprotroph-Symbiotroph | Animal Pathogen-Endophyte-Endosymbiont-Epiphyte-\|Undefined Saprotroph\| | Dimorphic Yeast-Yeast | NULL | Probable | As nectar-tap_saprotroph (Põlme et al. 2020); Decay sugar-rich substrates (Põlme et al. 2020); This asexual genus is polyphyletic and contain species that function very differently depend on what they are. Caution should be taken when interpreting guilds for Candida species. | Manolakaki D, et al. 2010. Virulence. 1: 367-375 ((https://doi.org/10.4161/viru.1.5.12796)); Mekbib SB, et al. 2011. Tropical Plant Pathology 36:233-240 ((https://doi.org/10.1590/S1982-56762011000400004)); Põlme S, et al. 2020. Fungal Diversity 105:1-16 ((https://doi.org/10.1007/s13225-020-00466-2)) |
| Otu06989 | 3 | k__Fungi(100);p__Ascomycota(100);c__Saccharomycetes(100);o__Saccharomycetales(100);f__Metschnikowiaceae(100);g__Metschnikowia(100);g__Metschnikowia_unclassified(100); | Metschnikowia | 13 | Saprotroph | \|Undefined Saprotroph\| | Yeast | NULL | Probable | As nectar-tap_saprotroph (Põlme et al. 2020); Decay sugar-rich substrates (Põlme et al. 2020) | Põlme S, et al. 2020. Fungal Diversity 105:1-16 ((https://doi.org/10.1007/s13225-020-00466-2)) |
| Otu06990 | 3 | k__Fungi(100);p__Ascomycota(100);c__Leotiomycetes(100);o__Helotiales(100);f__Helotiaceae(100);g__Articulospora(100);s__Articulospora_sp(100); | Articulospora | 13 | Saprotroph | \|Plant Saprotroph\|-Undefined Saprotroph | Microfungus | NULL | Probable | As litter_saprotroph (Põlme et al. 2020); Decay leaf, fruit, seed (Põlme et al. 2020) | Tedersoo L, et al. 2014. Science 346:e1256688 ((https://doi.org/10.1126/science.1256688)); Põlme S, et al. 2020. Fungal Diversity 105:1-16 ((https://doi.org/10.1007/s13225-020-00466-2)) |
| Otu06995 | 3 | k__Fungi(100);p__Ascomycota(100);c__Saccharomycetes(100);o__Saccharomycetales(100);f__Metschnikowiaceae(100);g__Metschnikowia(100);s__Metschnikowia_pulcherrima(100); | Metschnikowia | 13 | Saprotroph | \|Undefined Saprotroph\| | Yeast | NULL | Probable | As nectar-tap_saprotroph (Põlme et al. 2020); Decay sugar-rich substrates (Põlme et al. 2020) | Põlme S, et al. 2020. Fungal Diversity 105:1-16 ((https://doi.org/10.1007/s13225-020-00466-2)) |
| Otu06996 | 3 | k__Fungi(100);p__Ascomycota(100);c__Sordariomycetes(100);o__Hypocreales(100);f__Nectriaceae(100);g__Gibberella(100);s__Gibberella_intricans(100); | Gibberella | 13 | Pathotroph-Saprotroph-Symbiotroph | Endophyte-\|Plant Pathogen\|-Plant Saprotroph-Undefined Saprotroph | Microfungus | Soft Rot | Probable | As foliar_endophyte (Põlme et al. 2020); As litter_saprotroph (Põlme et al. 2020); Causing damping-off disease (Põlme et al. 2020); Decay leaf, fruit, seed, roots, soil, animal material (Põlme et al. 2020) | Tedersoo L, et al. 2014. Science 346:e1256688 ((https://doi.org/10.1126/science.1256688)); Põlme S, et al. 2020. Fungal Diversity 105:1-16 ((https://doi.org/10.1007/s13225-020-00466-2)) |
| Otu06997 | 3 | k__Fungi(100);p__Ascomycota(100);c__Saccharomycetes(100);o__Saccharomycetales(100);f__Metschnikowiaceae(100);g__Metschnikowia(100);s__Metschnikowia_pulcherrima(100); | Metschnikowia | 13 | Saprotroph | \|Undefined Saprotroph\| | Yeast | NULL | Probable | As nectar-tap_saprotroph (Põlme et al. 2020); Decay sugar-rich substrates (Põlme et al. 2020) | Põlme S, et al. 2020. Fungal Diversity 105:1-16 ((https://doi.org/10.1007/s13225-020-00466-2)) |
| Otu06998 | 3 | k__Fungi(100);p__Basidiomycota(100);c__Agaricomycetes(100);o__Polyporales(100);f__Phanerochaetaceae(100);g__Rhizochaete(100);s__Rhizochaete_filamentosa(100); | Rhizochaete | 13 | Saprotroph | Undefined Saprotroph-\|Wood Saprotroph\| | Corticioid | White Rot | Probable | As white_rot (Põlme et al. 2020); Decay wood (Põlme et al. 2020) | Tedersoo L, et al. 2014. Science 346:e1256688 ((https://doi.org/10.1126/science.1256688)); Põlme S, et al. 2020. Fungal Diversity 105:1-16 ((https://doi.org/10.1007/s13225-020-00466-2)) |
| Otu06999 | 3 | k__Fungi(100);p__Basidiomycota(100);c__Agaricostilbomycetes(100);o__Agaricostilbales(100);f__Chionosphaeraceae(100);f__Chionosphaeraceae_unclassified(100);f__Chionosphaeraceae_unclassified(100); | Chionosphaeraceae | 9 | Pathotroph-Saprotroph | Fungal Parasite-Undefined Saprotroph | Microfungus | NULL | Possible | Species appear to be mycoparasitic, associated with fungi of bark beetle galleries, wood-inhabiting Xylariaceae or lichens. Others are found erumpent from bark or decaying wood and may be sabrobic (Cannon & Kirk 2007) | Cannon PF, Kirk PM. 2007. Fungal Families of the World. CAB International, Cambridge (ISBN: 978-0851998275) |
| Otu07000 | 3 | k__Fungi(100);p__Ascomycota(100);c__Dothideomycetes(100);o__Dothideomycetes_order_Incertae_sedis(100);f__Myxotrichaceae(100);g__unclassified_Myxotrichaceae(100);s__Myxotrichaceae_sp(100); | Myxotrichaceae | 9 | Saprotroph-Symbiotroph | Ectomycorrhizal-Undefined Saprotroph | Microfungus | NULL | Probable | Many species are cellulolytic, occurring on paper, straw etc. Others are mycorrhizal associated with roots of Ericaceae and Epacridaceae (Cannon & Kirk 2007) | Cannon PF, Kirk PM. 2007. Fungal Families of the World. CAB International, Cambridge (ISBN: 978-0851998275) |
| Otu07005 | 3 | k__Fungi(100);p__Ascomycota(100);c__Saccharomycetes(100);o__Saccharomycetales(100);f__Metschnikowiaceae(100);g__Metschnikowia(100);s__Metschnikowia_pulcherrima(100); | Metschnikowia | 13 | Saprotroph | \|Undefined Saprotroph\| | Yeast | NULL | Probable | As nectar-tap_saprotroph (Põlme et al. 2020); Decay sugar-rich substrates (Põlme et al. 2020) | Põlme S, et al. 2020. Fungal Diversity 105:1-16 ((https://doi.org/10.1007/s13225-020-00466-2)) |
| Otu07006 | 3 | k__Fungi(100);p__Ascomycota(100);c__Saccharomycetes(100);o__Saccharomycetales(100);f__Metschnikowiaceae(100);g__Metschnikowia(100);g__Metschnikowia_unclassified(67); | Metschnikowia | 13 | Saprotroph | \|Undefined Saprotroph\| | Yeast | NULL | Probable | As nectar-tap_saprotroph (Põlme et al. 2020); Decay sugar-rich substrates (Põlme et al. 2020) | Põlme S, et al. 2020. Fungal Diversity 105:1-16 ((https://doi.org/10.1007/s13225-020-00466-2)) |
| Otu07007 | 3 | k__Fungi(100);p__Ascomycota(100);c__Saccharomycetes(100);o__Saccharomycetales(100);f__Saccharomycetales_family_Incertae_sedis(100);g__Candida(100);s__Candida_germanica(100); | Candida | 13 | Pathotroph-Saprotroph-Symbiotroph | Animal Pathogen-Endophyte-Endosymbiont-Epiphyte-\|Undefined Saprotroph\| | Dimorphic Yeast-Yeast | NULL | Probable | As nectar-tap_saprotroph (Põlme et al. 2020); Decay sugar-rich substrates (Põlme et al. 2020); This asexual genus is polyphyletic and contain species that function very differently depend on what they are. Caution should be taken when interpreting guilds for Candida species. | Manolakaki D, et al. 2010. Virulence. 1: 367-375 ((https://doi.org/10.4161/viru.1.5.12796)); Mekbib SB, et al. 2011. Tropical Plant Pathology 36:233-240 ((https://doi.org/10.1590/S1982-56762011000400004)); Põlme S, et al. 2020. Fungal Diversity 105:1-16 ((https://doi.org/10.1007/s13225-020-00466-2)) |
| Otu07008 | 3 | k__Fungi(100);p__Ascomycota(100);c__Saccharomycetes(100);o__Saccharomycetales(100);f__Metschnikowiaceae(100);g__Metschnikowia(100);s__Metschnikowia_pulcherrima(100); | Metschnikowia | 13 | Saprotroph | \|Undefined Saprotroph\| | Yeast | NULL | Probable | As nectar-tap_saprotroph (Põlme et al. 2020); Decay sugar-rich substrates (Põlme et al. 2020) | Põlme S, et al. 2020. Fungal Diversity 105:1-16 ((https://doi.org/10.1007/s13225-020-00466-2)) |
| Otu07009 | 3 | k__Fungi(100);p__Ascomycota(100);c__Saccharomycetes(100);o__Saccharomycetales(100);f__Metschnikowiaceae(100);g__Metschnikowia(100);s__Metschnikowia_pulcherrima(100); | Metschnikowia | 13 | Saprotroph | \|Undefined Saprotroph\| | Yeast | NULL | Probable | As nectar-tap_saprotroph (Põlme et al. 2020); Decay sugar-rich substrates (Põlme et al. 2020) | Põlme S, et al. 2020. Fungal Diversity 105:1-16 ((https://doi.org/10.1007/s13225-020-00466-2)) |
| Otu07010 | 3 | k__Fungi(100);p__Ascomycota(100);c__Saccharomycetes(100);o__Saccharomycetales(100);f__Saccharomycetaceae(100);g__Kazachstania(100);s__Kazachstania_intestinalis(100); | Kazachstania | 13 | Saprotroph | \|Undefined Saprotroph\| | Yeast | NULL | Probable | As nectar-tap_saprotroph (Põlme et al. 2020); Decay sugar-rich substrates (Põlme et al. 2020) | Põlme S, et al. 2020. Fungal Diversity 105:1-16 ((https://doi.org/10.1007/s13225-020-00466-2)) |
| Otu07012 | 3 | k__Fungi(100);p__Basidiomycota(100);c__Agaricomycetes(100);o__Agaricales(100);f__Marasmiaceae(100);g__Marasmius(100);s__Marasmius_sp_SM2425(100); | Marasmius | 13 | Pathotroph-Saprotroph-Symbiotroph | Endophyte-Plant Pathogen-\|Plant Saprotroph\|-Undefined Saprotroph | Agaricoid | NULL | Probable | As litter_saprotroph (Põlme et al. 2020); Decay leaf, fruit, seed (Põlme et al. 2020); Often one of the most prominent fungal components of tropical forests, playing a major role in nutrient cycling (Cannon & Kirk 2007) | Cannon PF, Kirk PM. 2007. Fungal Families of the World. CAB International, Cambridge (ISBN: 978-0851998275); Tedersoo L, et al. 2014. Science 346:e1256688 ((https://doi.org/10.1126/science.1256688)); David AS, et al. 2016. Microbial Ecology 71:912-926 ((https://doi.org/10.1007/s00248-015-0712-6)); Põlme S, et al. 2020. Fungal Diversity 105:1-16 ((https://doi.org/10.1007/s13225-020-00466-2)) |
| Otu07014 | 3 | k__Fungi(100);p__Ascomycota(100);c__Dothideomycetes(100);o__Pleosporales(100);f__Tubeufiaceae(100);f__Tubeufiaceae_unclassified(100);f__Tubeufiaceae_unclassified(100); | Tubeufiaceae | 9 | Pathotroph-Saprotroph | Animal Pathogen-Undefined Saprotroph | NULL | NULL | Probable | Biotrophic or saprobic, especially on scale insects or other fungi. The anamorphic forms are fequently aero-aquatic (Cannon & Kirk 2007) | Cannon PF, Kirk PM. 2007. Fungal Families of the World. CAB International, Cambridge (ISBN: 978-0851998275) |
| Otu07016 | 3 | k__Fungi(100);p__Ascomycota(100);c__Saccharomycetes(100);o__Saccharomycetales(100);f__Metschnikowiaceae(100);g__Metschnikowia(100);s__Metschnikowia_pulcherrima(67); | Metschnikowia | 13 | Saprotroph | \|Undefined Saprotroph\| | Yeast | NULL | Probable | As nectar-tap_saprotroph (Põlme et al. 2020); Decay sugar-rich substrates (Põlme et al. 2020) | Põlme S, et al. 2020. Fungal Diversity 105:1-16 ((https://doi.org/10.1007/s13225-020-00466-2)) |
| Otu07017 | 3 | k__Fungi(100);p__Basidiomycota(100);c__Agaricomycetes(100);o__Agaricales(100);f__Strophariaceae(100);g__Agrocybe(100);s__Agrocybe_erebia(100); | Agrocybe | 13 | Saprotroph | Dung Saprotroph-\|Undefined Saprotroph\| | Agaricoid | NULL | Probable | As soil_saprotroph (Põlme et al. 2020); Decay soil (Põlme et al. 2020) | Tedersoo L, et al. 2014. Science 346:e1256688 ((https://doi.org/10.1126/science.1256688)); Põlme S, et al. 2020. Fungal Diversity 105:1-16 ((https://doi.org/10.1007/s13225-020-00466-2)) |
| Otu07018 | 3 | k__Fungi(100);p__Ascomycota(100);c__Saccharomycetes(100);o__Saccharomycetales(100);f__Metschnikowiaceae(100);g__Metschnikowia(100);g__Metschnikowia_unclassified(67); | Metschnikowia | 13 | Saprotroph | \|Undefined Saprotroph\| | Yeast | NULL | Probable | As nectar-tap_saprotroph (Põlme et al. 2020); Decay sugar-rich substrates (Põlme et al. 2020) | Põlme S, et al. 2020. Fungal Diversity 105:1-16 ((https://doi.org/10.1007/s13225-020-00466-2)) |
| Otu07019 | 3 | k__Fungi(100);p__Ascomycota(100);c__Saccharomycetes(100);o__Saccharomycetales(100);f__Metschnikowiaceae(100);g__Metschnikowia(100);g__Metschnikowia_unclassified(67); | Metschnikowia | 13 | Saprotroph | \|Undefined Saprotroph\| | Yeast | NULL | Probable | As nectar-tap_saprotroph (Põlme et al. 2020); Decay sugar-rich substrates (Põlme et al. 2020) | Põlme S, et al. 2020. Fungal Diversity 105:1-16 ((https://doi.org/10.1007/s13225-020-00466-2)) |
| Otu07020 | 3 | k__Fungi(100);p__Ascomycota(100);c__Saccharomycetes(100);o__Saccharomycetales(100);f__Metschnikowiaceae(100);g__Metschnikowia(100);g__Metschnikowia_unclassified(100); | Metschnikowia | 13 | Saprotroph | \|Undefined Saprotroph\| | Yeast | NULL | Probable | As nectar-tap_saprotroph (Põlme et al. 2020); Decay sugar-rich substrates (Põlme et al. 2020) | Põlme S, et al. 2020. Fungal Diversity 105:1-16 ((https://doi.org/10.1007/s13225-020-00466-2)) |
| Otu07022 | 3 | k__Fungi(100);p__Ascomycota(100);c__Saccharomycetes(100);o__Saccharomycetales(100);f__Metschnikowiaceae(100);g__Metschnikowia(100);s__Metschnikowia_pulcherrima(67); | Metschnikowia | 13 | Saprotroph | \|Undefined Saprotroph\| | Yeast | NULL | Probable | As nectar-tap_saprotroph (Põlme et al. 2020); Decay sugar-rich substrates (Põlme et al. 2020) | Põlme S, et al. 2020. Fungal Diversity 105:1-16 ((https://doi.org/10.1007/s13225-020-00466-2)) |
| Otu07024 | 3 | k__Fungi(100);p__Basidiomycota(100);c__Agaricomycetes(100);o__Agaricales(100);f__Marasmiaceae(100);g__Marasmius(100);s__Marasmius_sp_NW_2008g(100); | Marasmius | 13 | Pathotroph-Saprotroph-Symbiotroph | Endophyte-Plant Pathogen-\|Plant Saprotroph\|-Undefined Saprotroph | Agaricoid | NULL | Probable | As litter_saprotroph (Põlme et al. 2020); Decay leaf, fruit, seed (Põlme et al. 2020); Often one of the most prominent fungal components of tropical forests, playing a major role in nutrient cycling (Cannon & Kirk 2007) | Cannon PF, Kirk PM. 2007. Fungal Families of the World. CAB International, Cambridge (ISBN: 978-0851998275); Tedersoo L, et al. 2014. Science 346:e1256688 ((https://doi.org/10.1126/science.1256688)); David AS, et al. 2016. Microbial Ecology 71:912-926 ((https://doi.org/10.1007/s00248-015-0712-6)); Põlme S, et al. 2020. Fungal Diversity 105:1-16 ((https://doi.org/10.1007/s13225-020-00466-2)) |
| Otu07025 | 3 | k__Fungi(100);p__Ascomycota(100);c__Saccharomycetes(100);o__Saccharomycetales(100);f__Metschnikowiaceae(100);g__Metschnikowia(100);s__Metschnikowia_pulcherrima(100); | Metschnikowia | 13 | Saprotroph | \|Undefined Saprotroph\| | Yeast | NULL | Probable | As nectar-tap_saprotroph (Põlme et al. 2020); Decay sugar-rich substrates (Põlme et al. 2020) | Põlme S, et al. 2020. Fungal Diversity 105:1-16 ((https://doi.org/10.1007/s13225-020-00466-2)) |
| Otu07027 | 3 | k__Fungi(100);p__Ascomycota(100);c__Saccharomycetes(100);o__Saccharomycetales(100);f__Metschnikowiaceae(100);g__Metschnikowia(100);s__Metschnikowia_pulcherrima(100); | Metschnikowia | 13 | Saprotroph | \|Undefined Saprotroph\| | Yeast | NULL | Probable | As nectar-tap_saprotroph (Põlme et al. 2020); Decay sugar-rich substrates (Põlme et al. 2020) | Põlme S, et al. 2020. Fungal Diversity 105:1-16 ((https://doi.org/10.1007/s13225-020-00466-2)) |
| Otu07029 | 3 | k__Fungi(100);p__Ascomycota(100);c__Saccharomycetes(100);o__Saccharomycetales(100);f__Metschnikowiaceae(100);g__Metschnikowia(100);s__Metschnikowia_pulcherrima(100); | Metschnikowia | 13 | Saprotroph | \|Undefined Saprotroph\| | Yeast | NULL | Probable | As nectar-tap_saprotroph (Põlme et al. 2020); Decay sugar-rich substrates (Põlme et al. 2020) | Põlme S, et al. 2020. Fungal Diversity 105:1-16 ((https://doi.org/10.1007/s13225-020-00466-2)) |
| Otu07032 | 3 | k__Fungi(100);p__Basidiomycota(100);c__Microbotryomycetes(100);o__Sporidiobolales(100);f__Sporidiobolales_family_Incertae_sedis(100);g__Rhodotorula(100);s__Rhodotorula_taiwanensis(100); | Rhodotorula | 13 | Pathotroph-Saprotroph-Symbiotroph | Animal Pathogen-Animal Symbiotroph-Endophyte-Plant Pathogen-\|Undefined Saprotroph\| | Yeast | NULL | Probable | As foliar_endophyte (Põlme et al. 2020); Opportunistic human pathogen | Põlme S, et al. 2020. Fungal Diversity 105:1-16 ((https://doi.org/10.1007/s13225-020-00466-2)); https://www.cdc.gov ((https://www.cdc.gov)) |
| Otu07035 | 3 | k__Fungi(100);p__Basidiomycota(100);c__Tremellomycetes(100);o__Tremellales(100);f__Tremellales_family_Incertae_sedis(100);f__Tremellales_family_Incertae_sedis_unclassified(100);f__Tremellales_family_Incertae_sedis_unclassified(100); | Tremellales | 7 | Pathotroph-Saprotroph-Symbiotroph | Fungal Parasite-Undefined Saprotroph | Tremelloid-Yeast | NULL | Possible | The order contains both teleomorphic and anamorphic species, most of the latter being yeasts | Sterkenburg E, et al. 2015. New Phytologist 207:1145-1158 ((https://doi.org/10.1111/nph.13426)) |
| Otu07036 | 3 | k__Fungi(100);p__Ascomycota(100);c__Saccharomycetes(100);o__Saccharomycetales(100);f__Metschnikowiaceae(100);g__Metschnikowia(100);s__Metschnikowia_pulcherrima(100); | Metschnikowia | 13 | Saprotroph | \|Undefined Saprotroph\| | Yeast | NULL | Probable | As nectar-tap_saprotroph (Põlme et al. 2020); Decay sugar-rich substrates (Põlme et al. 2020) | Põlme S, et al. 2020. Fungal Diversity 105:1-16 ((https://doi.org/10.1007/s13225-020-00466-2)) |
| Otu07037 | 3 | k__Fungi(100);p__Basidiomycota(100);c__Tremellomycetes(100);o__Tremellales(100);f__Tremellales_family_Incertae_sedis(100);g__Cryptococcus(100);g__Cryptococcus_unclassified(100); | Cryptococcus | 13 | Pathotroph-Saprotroph-Symbiotroph | Animal Pathogen-Endophyte-Epiphyte-\|Undefined Saprotroph\| | Dimorphic Yeast-Yeast | NULL | Possible | As foliar_endophyte (Põlme et al. 2020); Decay leaf, fruit, seed, soil, dung, animal material (Põlme et al. 2020) | Andrews JH, et al. 1983. Phytopathology 73:228-234 ((https://doi.org/10.1094/Phyto-73-228)); Kurtzman CP, et al. (eds.) 2011. The Yeasts, a Taxonomic Study. Fifth Edition. Vols 1-3. Elsevier, San Diego (ISBN: 9780444521491); Põlme S, et al. 2020. Fungal Diversity 105:1-16 ((https://doi.org/10.1007/s13225-020-00466-2)) |
| Otu07039 | 3 | k__Fungi(100);p__Basidiomycota(100);c__Tremellomycetes(100);o__Tremellales(100);f__Tremellales_family_Incertae_sedis(100);f__Tremellales_family_Incertae_sedis_unclassified(100);f__Tremellales_family_Incertae_sedis_unclassified(100); | Tremellales | 7 | Pathotroph-Saprotroph-Symbiotroph | Fungal Parasite-Undefined Saprotroph | Tremelloid-Yeast | NULL | Possible | The order contains both teleomorphic and anamorphic species, most of the latter being yeasts | Sterkenburg E, et al. 2015. New Phytologist 207:1145-1158 ((https://doi.org/10.1111/nph.13426)) |
| Otu07040 | 3 | k__Fungi(100);p__Ascomycota(100);c__Saccharomycetes(100);o__Saccharomycetales(100);f__Metschnikowiaceae(100);g__Metschnikowia(100);g__Metschnikowia_unclassified(100); | Metschnikowia | 13 | Saprotroph | \|Undefined Saprotroph\| | Yeast | NULL | Probable | As nectar-tap_saprotroph (Põlme et al. 2020); Decay sugar-rich substrates (Põlme et al. 2020) | Põlme S, et al. 2020. Fungal Diversity 105:1-16 ((https://doi.org/10.1007/s13225-020-00466-2)) |
| Otu07041 | 3 | k__Fungi(100);p__Ascomycota(100);c__Saccharomycetes(100);o__Saccharomycetales(100);f__Metschnikowiaceae(100);g__Metschnikowia(100);g__Metschnikowia_unclassified(100); | Metschnikowia | 13 | Saprotroph | \|Undefined Saprotroph\| | Yeast | NULL | Probable | As nectar-tap_saprotroph (Põlme et al. 2020); Decay sugar-rich substrates (Põlme et al. 2020) | Põlme S, et al. 2020. Fungal Diversity 105:1-16 ((https://doi.org/10.1007/s13225-020-00466-2)) |
| Otu07042 | 3 | k__Fungi(100);p__Ascomycota(100);c__Saccharomycetes(100);o__Saccharomycetales(100);f__Metschnikowiaceae(100);g__Metschnikowia(100);g__Metschnikowia_unclassified(67); | Metschnikowia | 13 | Saprotroph | \|Undefined Saprotroph\| | Yeast | NULL | Probable | As nectar-tap_saprotroph (Põlme et al. 2020); Decay sugar-rich substrates (Põlme et al. 2020) | Põlme S, et al. 2020. Fungal Diversity 105:1-16 ((https://doi.org/10.1007/s13225-020-00466-2)) |
| Otu07043 | 3 | k__Fungi(100);p__Ascomycota(100);c__Saccharomycetes(100);o__Saccharomycetales(100);f__Saccharomycetaceae(100);g__Kazachstania(100);s__Kazachstania_intestinalis(100); | Kazachstania | 13 | Saprotroph | \|Undefined Saprotroph\| | Yeast | NULL | Probable | As nectar-tap_saprotroph (Põlme et al. 2020); Decay sugar-rich substrates (Põlme et al. 2020) | Põlme S, et al. 2020. Fungal Diversity 105:1-16 ((https://doi.org/10.1007/s13225-020-00466-2)) |
| Otu07044 | 3 | k__Fungi(100);p__Ascomycota(100);c__Saccharomycetes(100);o__Saccharomycetales(100);f__Metschnikowiaceae(100);g__Metschnikowia(100);g__Metschnikowia_unclassified(67); | Metschnikowia | 13 | Saprotroph | \|Undefined Saprotroph\| | Yeast | NULL | Probable | As nectar-tap_saprotroph (Põlme et al. 2020); Decay sugar-rich substrates (Põlme et al. 2020) | Põlme S, et al. 2020. Fungal Diversity 105:1-16 ((https://doi.org/10.1007/s13225-020-00466-2)) |
| Otu07045 | 3 | k__Fungi(100);p__Ascomycota(100);c__Saccharomycetes(100);o__Saccharomycetales(100);f__Metschnikowiaceae(100);g__Metschnikowia(100);s__Metschnikowia_pulcherrima(100); | Metschnikowia | 13 | Saprotroph | \|Undefined Saprotroph\| | Yeast | NULL | Probable | As nectar-tap_saprotroph (Põlme et al. 2020); Decay sugar-rich substrates (Põlme et al. 2020) | Põlme S, et al. 2020. Fungal Diversity 105:1-16 ((https://doi.org/10.1007/s13225-020-00466-2)) |
| Otu07047 | 3 | k__Fungi(100);p__Ascomycota(100);c__Saccharomycetes(100);o__Saccharomycetales(100);f__Metschnikowiaceae(100);g__Metschnikowia(100);s__Metschnikowia_pulcherrima(67); | Metschnikowia | 13 | Saprotroph | \|Undefined Saprotroph\| | Yeast | NULL | Probable | As nectar-tap_saprotroph (Põlme et al. 2020); Decay sugar-rich substrates (Põlme et al. 2020) | Põlme S, et al. 2020. Fungal Diversity 105:1-16 ((https://doi.org/10.1007/s13225-020-00466-2)) |
| Otu07048 | 3 | k__Fungi(100);p__Basidiomycota(100);c__Agaricomycetes(100);o__Thelephorales(100);f__Thelephoraceae(100);g__Tomentella(100);s__Tomentella_ellisii(100); | Tomentella | 13 | Symbiotroph | \|Ectomycorrhizal\| | Corticioid | NULL | Highly Probable | NULL | Rinaldi AC, et al. 2008. Fungal Diversity 33:1-45 ((http://www.fungaldiversity.org/fdp/sfdp/33-1.pdf)); Tedersoo L, et al. 2010. Mycorrhiza 20:217-263 ((https://doi.org/10.1007/s00572-009-0274-x)); Põlme S, et al. 2020. Fungal Diversity 105:1-16 ((https://doi.org/10.1007/s13225-020-00466-2)); http://mycorrhizas.info/ecmf.html ((http://mycorrhizas.info/ecmf.html)) |
| Otu07049 | 3 | k__Fungi(100);p__Ascomycota(100);c__Dothideomycetes(100);o__Dothideales(100);f__Dothioraceae(100);g__Aureobasidium(100);s__Aureobasidium_pullulans(100); | Aureobasidium | 13 | Pathotroph-Saprotroph-Symbiotroph | Animal Pathogen-Endophyte-Epiphyte-\|Plant Pathogen\|-Plant Saprotroph-Undefined Saprotroph | Dimorphic-Facultative Yeast | NULL | Possible | As foliar_endophyte (Põlme et al. 2020); As litter_saprotroph (Põlme et al. 2020); Causes sooty mold (Põlme et al. 2020); Decay leaf, fruit, seed, roots, animal material, sugar-rich substrates (Põlme et al. 2020) | Costa IPMW, et al. 2012. Mycotaxon 119:e26 ((http://mycotaxon.com/resources/checklists/costa_v119_checklist.pdf)); Tedersoo L, et al. 2014. Science 346:e1256688 ((https://doi.org/10.1126/science.1256688)); Irinyia L, et al. 2016. Fungal Biology 120:125-136 ((https://doi.org/10.1016/j.funbio.2015.04.007)); Põlme S, et al. 2020. Fungal Diversity 105:1-16 ((https://doi.org/10.1007/s13225-020-00466-2)) |
| Otu07051 | 3 | k__Fungi(100);p__Ascomycota(100);c__Saccharomycetes(100);o__Saccharomycetales(100);f__Metschnikowiaceae(100);g__Metschnikowia(100);s__Metschnikowia_pulcherrima(100); | Metschnikowia | 13 | Saprotroph | \|Undefined Saprotroph\| | Yeast | NULL | Probable | As nectar-tap_saprotroph (Põlme et al. 2020); Decay sugar-rich substrates (Põlme et al. 2020) | Põlme S, et al. 2020. Fungal Diversity 105:1-16 ((https://doi.org/10.1007/s13225-020-00466-2)) |
| Otu07053 | 3 | k__Fungi(100);p__Ascomycota(100);c__Sordariomycetes(100);o__Diaporthales(100);f__Diaporthaceae(100);g__Diaporthe(100);g__Diaporthe_unclassified(100); | Diaporthe | 13 | Pathotroph-Saprotroph-Symbiotroph | Endophyte-\|Plant Pathogen\|-Plant Saprotroph | Microfungus | NULL | Possible | As foliar_endophyte (Põlme et al. 2020); As litter_saprotroph (Põlme et al. 2020); Decay leaf, fruit, seed (Põlme et al. 2020) | Tedersoo L, et al. 2014. Science 346:e1256688 ((https://doi.org/10.1126/science.1256688)); Põlme S, et al. 2020. Fungal Diversity 105:1-16 ((https://doi.org/10.1007/s13225-020-00466-2)) |
| Otu07054 | 3 | k__Fungi(100);p__Basidiomycota(100);c__Agaricomycetes(100);o__Thelephorales(100);f__Thelephoraceae(100);g__Tomentella(100);s__Tomentella_badia(100); | Tomentella | 13 | Symbiotroph | \|Ectomycorrhizal\| | Corticioid | NULL | Highly Probable | NULL | Rinaldi AC, et al. 2008. Fungal Diversity 33:1-45 ((http://www.fungaldiversity.org/fdp/sfdp/33-1.pdf)); Tedersoo L, et al. 2010. Mycorrhiza 20:217-263 ((https://doi.org/10.1007/s00572-009-0274-x)); Põlme S, et al. 2020. Fungal Diversity 105:1-16 ((https://doi.org/10.1007/s13225-020-00466-2)); http://mycorrhizas.info/ecmf.html ((http://mycorrhizas.info/ecmf.html)) |
| Otu07055 | 3 | k__Fungi(100);p__Basidiomycota(100);c__Agaricomycetes(100);o__Thelephorales(100);f__Thelephoraceae(100);f__Thelephoraceae_unclassified(100);f__Thelephoraceae_unclassified(100); | Thelephoraceae | 9 | Saprotroph-Symbiotroph | Ectomycorrhizal-Undefined Saprotroph | Clavarioid | NULL | Probable | On the ground, on soil, a few on wood. At least some speices ectomycorrhizal (Cannon & Kirk 2007) | Cannon PF, Kirk PM. 2007. Fungal Families of the World. CAB International, Cambridge (ISBN: 978-0851998275) |
| Otu07057 | 3 | k__Fungi(100);p__Ascomycota(100);c__Saccharomycetes(100);o__Saccharomycetales(100);f__Metschnikowiaceae(100);g__Metschnikowia(100);s__Metschnikowia_pulcherrima(100); | Metschnikowia | 13 | Saprotroph | \|Undefined Saprotroph\| | Yeast | NULL | Probable | As nectar-tap_saprotroph (Põlme et al. 2020); Decay sugar-rich substrates (Põlme et al. 2020) | Põlme S, et al. 2020. Fungal Diversity 105:1-16 ((https://doi.org/10.1007/s13225-020-00466-2)) |
| Otu07058 | 3 | k__Fungi(100);p__Ascomycota(100);c__Saccharomycetes(100);o__Saccharomycetales(100);f__Metschnikowiaceae(100);g__Metschnikowia(100);s__Metschnikowia_pulcherrima(100); | Metschnikowia | 13 | Saprotroph | \|Undefined Saprotroph\| | Yeast | NULL | Probable | As nectar-tap_saprotroph (Põlme et al. 2020); Decay sugar-rich substrates (Põlme et al. 2020) | Põlme S, et al. 2020. Fungal Diversity 105:1-16 ((https://doi.org/10.1007/s13225-020-00466-2)) |
| Otu07059 | 3 | k__Fungi(100);p__Ascomycota(100);c__Saccharomycetes(100);o__Saccharomycetales(100);f__Metschnikowiaceae(100);g__Metschnikowia(100);s__Metschnikowia_pulcherrima(100); | Metschnikowia | 13 | Saprotroph | \|Undefined Saprotroph\| | Yeast | NULL | Probable | As nectar-tap_saprotroph (Põlme et al. 2020); Decay sugar-rich substrates (Põlme et al. 2020) | Põlme S, et al. 2020. Fungal Diversity 105:1-16 ((https://doi.org/10.1007/s13225-020-00466-2)) |
| Otu07064 | 3 | k__Fungi(100);p__Ascomycota(100);c__Saccharomycetes(100);o__Saccharomycetales(100);f__Metschnikowiaceae(100);g__Metschnikowia(100);s__Metschnikowia_pulcherrima(100); | Metschnikowia | 13 | Saprotroph | \|Undefined Saprotroph\| | Yeast | NULL | Probable | As nectar-tap_saprotroph (Põlme et al. 2020); Decay sugar-rich substrates (Põlme et al. 2020) | Põlme S, et al. 2020. Fungal Diversity 105:1-16 ((https://doi.org/10.1007/s13225-020-00466-2)) |
| Otu07065 | 3 | k__Fungi(100);p__Ascomycota(100);c__Saccharomycetes(100);o__Saccharomycetales(100);f__Metschnikowiaceae(100);g__Metschnikowia(100);s__Metschnikowia_pulcherrima(100); | Metschnikowia | 13 | Saprotroph | \|Undefined Saprotroph\| | Yeast | NULL | Probable | As nectar-tap_saprotroph (Põlme et al. 2020); Decay sugar-rich substrates (Põlme et al. 2020) | Põlme S, et al. 2020. Fungal Diversity 105:1-16 ((https://doi.org/10.1007/s13225-020-00466-2)) |
| Otu07068 | 3 | k__Fungi(100);p__Ascomycota(100);c__Saccharomycetes(100);o__Saccharomycetales(100);f__Metschnikowiaceae(100);g__Metschnikowia(100);s__Metschnikowia_pulcherrima(100); | Metschnikowia | 13 | Saprotroph | \|Undefined Saprotroph\| | Yeast | NULL | Probable | As nectar-tap_saprotroph (Põlme et al. 2020); Decay sugar-rich substrates (Põlme et al. 2020) | Põlme S, et al. 2020. Fungal Diversity 105:1-16 ((https://doi.org/10.1007/s13225-020-00466-2)) |
| Otu07069 | 3 | k__Fungi(100);p__Basidiomycota(100);c__Agaricomycetes(100);o__Agaricales(100);f__Marasmiaceae(100);g__Gymnopus(100);s__Gymnopus_iocephalus(100); | Gymnopus | 13 | Saprotroph | \|Plant Saprotroph\| | Agaricoid | NULL | Probable | As litter_saprotroph (Põlme et al. 2020); Decay leaf, fruit, seed (Põlme et al. 2020) | Põlme S, et al. 2020. Fungal Diversity 105:1-16 ((https://doi.org/10.1007/s13225-020-00466-2)) |
| Otu07070 | 3 | k__Fungi(100);p__Ascomycota(100);c__Saccharomycetes(100);o__Saccharomycetales(100);f__Metschnikowiaceae(100);g__Metschnikowia(100);s__Metschnikowia_pulcherrima(100); | Metschnikowia | 13 | Saprotroph | \|Undefined Saprotroph\| | Yeast | NULL | Probable | As nectar-tap_saprotroph (Põlme et al. 2020); Decay sugar-rich substrates (Põlme et al. 2020) | Põlme S, et al. 2020. Fungal Diversity 105:1-16 ((https://doi.org/10.1007/s13225-020-00466-2)) |
| Otu07072 | 3 | k__Fungi(100);p__Ascomycota(100);c__Saccharomycetes(100);o__Saccharomycetales(100);f__Metschnikowiaceae(100);g__Metschnikowia(100);g__Metschnikowia_unclassified(100); | Metschnikowia | 13 | Saprotroph | \|Undefined Saprotroph\| | Yeast | NULL | Probable | As nectar-tap_saprotroph (Põlme et al. 2020); Decay sugar-rich substrates (Põlme et al. 2020) | Põlme S, et al. 2020. Fungal Diversity 105:1-16 ((https://doi.org/10.1007/s13225-020-00466-2)) |

| | | | | | | | | | | | |
|---|---|---|---|---|---|---|---|---|---|---|---|
| Otu07074 | 3 | k__Fungi(100);p__Ascomycota(100);c__Saccharomycetes(100);o__Saccharomycetales(100);f__Metschnikowiaceae(100);g__Metschnikowia(100);s__Metschnikowia_pulcherrima(100); | Metschnikowia | 13 | Saprotroph | \|Undefined Saprotroph\| | Yeast | NULL | Probable | [illegible] | [illegible] |
| Otu07075 | 3 | k__Fungi(100);p__Ascomycota(100);c__Sordariomycetes(100);o__Diaporthales(100);f__Valsaceae(100);f__Valsaceae_unclassified(100);f__Valsaceae_unclassified(100); | Valsaceae | 9 | Pathotroph-Saprotroph-Symbiotroph | Endophyte-Plant Pathogen-Undefined Saprotroph | Microfungus | NULL | Probable | [illegible] | [illegible] |
| Otu07076 | 3 | k__Fungi(100);p__Ascomycota(100);c__Saccharomycetes(100);o__Saccharomycetales(100);f__Metschnikowiaceae(100);g__Metschnikowia(100);s__Metschnikowia_pulcherrima(100); | Metschnikowia | 13 | Saprotroph | \|Undefined Saprotroph\| | Yeast | NULL | Probable | [illegible] | [illegible] |
| Otu07077 | 3 | k__Fungi(100);p__Ascomycota(100);c__Saccharomycetes(100);o__Saccharomycetales(100);f__Saccharomycetales_family_Incertae_sedis(100);g__Candida(100);s__Candida_sp_AS23486(100); | Candida | 13 | Pathotroph-Saprotroph-Symbiotroph | Animal Pathogen-Endophyte-Endosymbiont-Epiphyte-\|Undefined Saprotroph\| | Dimorphic Yeast-Yeast | NULL | Probable | [illegible] | [illegible] |
| Otu07079 | 3 | k__Fungi(100);p__Ascomycota(100);c__Saccharomycetes(100);o__Saccharomycetales(100);f__Metschnikowiaceae(100);g__Metschnikowia(100);s__Metschnikowia_pulcherrima(100); | Metschnikowia | 13 | Saprotroph | \|Undefined Saprotroph\| | Yeast | NULL | Probable | [illegible] | [illegible] |
| Otu07080 | 3 | k__Fungi(100);p__Ascomycota(100);c__Saccharomycetes(100);o__Saccharomycetales(100);f__Metschnikowiaceae(100);g__Metschnikowia(100);g__Metschnikowia_unclassified(67); | Metschnikowia | 13 | Saprotroph | \|Undefined Saprotroph\| | Yeast | NULL | Probable | [illegible] | [illegible] |
| Otu07083 | 3 | k__Fungi(100);p__Ascomycota(100);c__Saccharomycetes(100);o__Saccharomycetales(100);f__Metschnikowiaceae(100);g__Metschnikowia(100);s__Metschnikowia_pulcherrima(100); | Metschnikowia | 13 | Saprotroph | \|Undefined Saprotroph\| | Yeast | NULL | Probable | [illegible] | [illegible] |
| Otu07084 | 3 | k__Fungi(100);p__Basidiomycota(100);c__Agaricomycetes(100);o__Polyporales(100);f__Meruliaceae(100);g__Phlebia(100);s__Phlebia_fuscoatra(100); | Phlebia | 13 | Saprotroph | \|Wood Saprotroph\| | Corticioid | White Rot | Highly Probable | [illegible] | [illegible] |
| Otu07085 | 3 | k__Fungi(100);p__Basidiomycota(100);c__Agaricomycetes(100);o__Agaricales(100);f__Marasmiaceae(100);g__Gymnopus(100);s__Gymnopus_androsaceus(100); | Gymnopus | 13 | Saprotroph | \|Plant Saprotroph\| | Agaricoid | NULL | Probable | [illegible] | [illegible] |
| Otu07086 | 3 | k__Fungi(100);p__Ascomycota(100);c__Saccharomycetes(100);o__Saccharomycetales(100);f__Metschnikowiaceae(100);g__Metschnikowia(100);g__Metschnikowia_unclassified(100); | Metschnikowia | 13 | Saprotroph | \|Undefined Saprotroph\| | Yeast | NULL | Probable | [illegible] | [illegible] |
| Otu07087 | 3 | k__Fungi(100);p__Ascomycota(100);c__Dothideomycetes(100);o__Pleosporales(100);f__Massarinaceae(100);g__Helminthosporium(100);s__Helminthosporium_sp(100); | Helminthosporium | 13 | Pathotroph | Fungal Parasite-\|Plant Pathogen\| | Microfungus | NULL | Probable | NULL | [illegible] |
| Otu07088 | 3 | k__Fungi(100);p__Basidiomycota(100);c__Agaricomycetes(100);o__Polyporales(100);f__Phanerochaetaceae(100);g__Phanerochaete(100);s__Phanerochaete_sordida(100); | Phanerochaete | 13 | Saprotroph | \|Wood Saprotroph\| | Corticioid | White Rot | Highly Probable | [illegible] | [illegible] |
| Otu07089 | 3 | k__Fungi(100);p__Ascomycota(100);c__Taphrinomycetes(100);o__Taphrinales(100);f__Taphrinaceae(100);f__Taphrinaceae_unclassified(67);f__Taphrinaceae_unclassified(67); | Taphrinaceae | 9 | Pathotroph | Plant Pathogen | Microfungus | NULL | Probable | [illegible] | [illegible] |
| Otu07091 | 3 | k__Fungi(100);p__Ascomycota(100);c__Saccharomycetes(100);o__Saccharomycetales(100);f__Metschnikowiaceae(100);g__Metschnikowia(100);s__Metschnikowia_pulcherrima(100); | Metschnikowia | 13 | Saprotroph | \|Undefined Saprotroph\| | Yeast | NULL | Probable | [illegible] | [illegible] |
| Otu07093 | 3 | k__Fungi(100);p__Ascomycota(100);c__Dothideomycetes(100);o__Capnodiales(100);f__Mycosphaerellaceae(100);f__Mycosphaerellaceae_unclassified(67);f__Mycosphaerellaceae_unclassified(67); | Mycosphaerellaceae | 9 | Pathotroph-Saprotroph | Plant Pathogen-Undefined Saprotroph | Microfungus | NULL | Probable | [illegible] | [illegible] |
| Otu07094 | 3 | k__Fungi(100);p__Ascomycota(100);c__Saccharomycetes(100);o__Saccharomycetales(100);f__Metschnikowiaceae(100);g__Metschnikowia(100);g__Metschnikowia_unclassified(67); | Metschnikowia | 13 | Saprotroph | \|Undefined Saprotroph\| | Yeast | NULL | Probable | [illegible] | [illegible] |
| Otu07095 | 3 | k__Fungi(100);p__Ascomycota(100);c__Saccharomycetes(100);o__Saccharomycetales(100);f__Saccharomycetales_family_Incertae_sedis(100);g__Candida(67);g__Candida_unclassified(67); | Candida | 13 | Pathotroph-Saprotroph-Symbiotroph | Animal Pathogen-Endophyte-Endosymbiont-Epiphyte-\|Undefined Saprotroph\| | Dimorphic Yeast-Yeast | NULL | Probable | [illegible] | [illegible] |
| Otu07096 | 3 | k__Fungi(100);p__Basidiomycota(100);c__Tremellomycetes(100);o__Tremellales(100);f__Tremellales_family_Incertae_sedis(100);f__Tremellales_family_Incertae_sedis_unclassified(100);f__Tremellales_family_Incertae_sedis_unclassified(100); | Tremellales | 7 | Pathotroph-Saprotroph-Symbiotroph | Fungal Parasite-Undefined Saprotroph | Tremelloid-Yeast | NULL | Possible | [illegible] | [illegible] |
| Otu07097 | 3 | k__Fungi(100);p__Basidiomycota(100);c__Microbotryomycetes(100);o__Sporidiobolales(100);f__Sporidiobolales_family_Incertae_sedis(100);g__Rhodotorula(100);s__Rhodotorula_cresolica(100); | Rhodotorula | 13 | Pathotroph-Saprotroph-Symbiotroph | Animal Pathogen-Animal Symbiotroph-Endophyte-Plant Pathogen-\|Undefined Saprotroph\| | Yeast | NULL | Probable | [illegible] | [illegible] |
| Otu07098 | 3 | k__Fungi(100);p__Ascomycota(100);c__Saccharomycetes(100);o__Saccharomycetales(100);f__Saccharomycetaceae(100);g__Kazachstania(100);s__Kazachstania_intestinalis(100); | Kazachstania | 13 | Saprotroph | \|Undefined Saprotroph\| | Yeast | NULL | Probable | [illegible] | [illegible] |
| Otu07099 | 3 | k__Fungi(100);p__Ascomycota(100);c__Sordariomycetes(100);o__Sordariomycetes_order_Incertae_sedis(100);f__Sordariomycetes_family_Incertae_sedis(100);g__Myrmecridium(100);s__Myrmecridium_schulzeri(100); | Myrmecridium | 13 | Saprotroph | \|Undefined Saprotroph\| | Microfungus | NULL | Probable | [illegible] | [illegible] |
| Otu07105 | 3 | k__Fungi(100);p__Basidiomycota(100);c__Tremellomycetes(100);o__Tremellales(100);f__Tremellales_family_Incertae_sedis(100);g__Cryptococcus(100);s__Cryptococcus_dejecticola(100); | Cryptococcus | 13 | Pathotroph-Saprotroph-Symbiotroph | Animal Pathogen-Endophyte-Epiphyte-\|Undefined Saprotroph\| | Dimorphic Yeast-Yeast | NULL | Possible | [illegible] | [illegible] |
| Otu07106 | 3 | k__Fungi(100);p__Ascomycota(100);c__Sordariomycetes(100);o__Sordariales(100);f__Chaetomiaceae(100);f__Chaetomiaceae_unclassified(67);f__Chaetomiaceae_unclassified(67); | Chaetomiaceae | 9 | Pathotroph-Saprotroph-Symbiotroph | Animal Pathogen-Dung Saprotroph-Endophyte-Epiphyte-Plant Saprotroph-Wood Saprotroph | Microfungus | Soft Rot | Possible | [illegible] | [illegible] |
| Otu07107 | 3 | k__Fungi(100);p__Basidiomycota(100);c__Agaricomycetes(100);o__Russulales(100);f__Russulaceae(100);g__Russula(100);g__Russula_unclassified(100); | Russula | 13 | Symbiotroph | \|Ectomycorrhizal\| | Agaricoid | NULL | Highly Probable | NULL | [illegible] |
| Otu07108 | 3 | k__Fungi(100);p__Ascomycota(100);c__Saccharomycetes(100);o__Saccharomycetales(100);f__Metschnikowiaceae(100);g__Metschnikowia(100);s__Metschnikowia_pulcherrima(67); | Metschnikowia | 13 | Saprotroph | \|Undefined Saprotroph\| | Yeast | NULL | Probable | [illegible] | [illegible] |
| Otu07109 | 3 | k__Fungi(100);p__Basidiomycota(100);c__Agaricomycetes(100);o__Agaricales(100);f__Tricholomataceae(100);g__Melanoleuca(100);s__Melanoleuca_sp_MSG_2013a(100); | Melanoleuca | 13 | Saprotroph | Plant Saprotroph-\|Undefined Saprotroph\| | Agaricoid | NULL | Probable | [illegible] | [illegible] |
| Otu07111 | 3 | k__Fungi(100);p__Ascomycota(100);c__Saccharomycetes(100);o__Saccharomycetales(100);f__Metschnikowiaceae(100);g__Metschnikowia(100);s__Metschnikowia_pulcherrima(100); | Metschnikowia | 13 | Saprotroph | \|Undefined Saprotroph\| | Yeast | NULL | Probable | [illegible] | [illegible] |
| Otu07112 | 3 | k__Fungi(100);p__Ascomycota(100);c__Saccharomycetes(100);o__Saccharomycetales(100);f__Metschnikowiaceae(100);g__Metschnikowia(100);s__Metschnikowia_pulcherrima(67); | Metschnikowia | 13 | Saprotroph | \|Undefined Saprotroph\| | Yeast | NULL | Probable | [illegible] | [illegible] |
| Otu07113 | 3 | k__Fungi(100);p__Ascomycota(100);c__Saccharomycetes(100);o__Saccharomycetales(100);f__Metschnikowiaceae(100);g__Metschnikowia(100);g__Metschnikowia_unclassified(67); | Metschnikowia | 13 | Saprotroph | \|Undefined Saprotroph\| | Yeast | NULL | Probable | [illegible] | [illegible] |
| Otu07115 | 3 | k__Fungi(100);p__Basidiomycota(100);c__Agaricomycetes(100);o__Russulales(100);f__Stereaceae(100);g__Gloeocystidiellum(100);g__Gloeocystidiellum_unclassified(100); | Gloeocystidiellum | 13 | Saprotroph | Undefined Saprotroph-\|Wood Saprotroph\| | Corticioid | White Rot | Probable | [illegible] | [illegible] |
| Otu07116 | 3 | k__Fungi(100);p__Ascomycota(100);c__Saccharomycetes(100);o__Saccharomycetales(100);f__Metschnikowiaceae(100);g__Metschnikowia(100);s__Metschnikowia_pulcherrima(67); | Metschnikowia | 13 | Saprotroph | \|Undefined Saprotroph\| | Yeast | NULL | Probable | [illegible] | [illegible] |
| Otu07117 | 3 | k__Fungi(100);p__Basidiomycota(100);c__Agaricomycetes(100);o__Hymenochaetales(100);f__Hymenochaetaceae(100);g__Hymenochaete(100);g__Hymenochaete_unclassified(100); | Hymenochaete | 13 | Pathotroph-Saprotroph | Plant Pathogen-Undefined Saprotroph-\|Wood Saprotroph\| | Corticioid-Polyporoid | White Rot | Probable | [illegible] | [illegible] |
| Otu07118 | 3 | k__Fungi(100);p__Ascomycota(100);c__Saccharomycetes(100);o__Saccharomycetales(100);f__Metschnikowiaceae(100);g__Metschnikowia(100);s__Metschnikowia_pulcherrima(100); | Metschnikowia | 13 | Saprotroph | \|Undefined Saprotroph\| | Yeast | NULL | Probable | [illegible] | [illegible] |
| Otu07120 | 3 | k__Fungi(100);p__Ascomycota(100);c__Saccharomycetes(100);o__Saccharomycetales(100);f__Saccharomycetales_family_Incertae_sedis(100);g__Candida(100);g__Candida_unclassified(67); | Candida | 13 | Pathotroph-Saprotroph-Symbiotroph | Animal Pathogen-Endophyte-Endosymbiont-Epiphyte-\|Undefined Saprotroph\| | Dimorphic Yeast-Yeast | NULL | Probable | [illegible] | [illegible] |
| Otu07121 | 3 | k__Fungi(100);p__Ascomycota(100);c__Saccharomycetes(100);o__Saccharomycetales(100);f__Metschnikowiaceae(100);g__Metschnikowia(100);s__Metschnikowia_pulcherrima(100); | Metschnikowia | 13 | Saprotroph | \|Undefined Saprotroph\| | Yeast | NULL | Probable | [illegible] | [illegible] |
| Otu07125 | 3 | k__Fungi(100);p__Ascomycota(100);c__Sordariomycetes(100);o__Hypocreales(100);f__Clavicipitaceae(100);g__Conoideocrella(100);s__Conoideocrella_tenuis(100); | Conoideocrella | 13 | Pathotroph-Saprotroph | \|Animal Parasite\|-Undefined Saprotroph | Microfungus | NULL | Probable | [illegible] | [illegible] |
| Otu07126 | 3 | k__Fungi(100);p__Ascomycota(100);c__Saccharomycetes(100);o__Saccharomycetales(100);f__Metschnikowiaceae(100);g__Metschnikowia(100);g__Metschnikowia_unclassified(100); | Metschnikowia | 13 | Saprotroph | \|Undefined Saprotroph\| | Yeast | NULL | Probable | [illegible] | [illegible] |
| Otu07127 | 3 | k__Fungi(100);p__Ascomycota(100);c__Sordariomycetes(100);o__Xylariales(100);f__Diatrypaceae(100);f__Diatrypaceae_unclassified(100);f__Diatrypaceae_unclassified(100); | Diatrypaceae | 9 | Pathotroph-Saprotroph | Plant Pathogen-Plant Saprotroph | Microfungus | NULL | Probable | [illegible] | [illegible] |
| Otu07129 | 3 | k__Fungi(100);p__Ascomycota(100);c__Saccharomycetes(100);o__Saccharomycetales(100);f__Saccharomycetales_family_Incertae_sedis(100);g__Candida(100);s__Candida_cretensis(67); | Candida | 13 | Pathotroph-Saprotroph-Symbiotroph | Animal Pathogen-Endophyte-Endosymbiont-Epiphyte-\|Undefined Saprotroph\| | Dimorphic Yeast-Yeast | NULL | Probable | [illegible] | [illegible] |
| Otu07132 | 3 | k__Fungi(100);p__Ascomycota(100);c__Eurotiomycetes(100);o__Chaetothyriales(100);f__Herpotrichiellaceae(100);f__Herpotrichiellaceae_unclassified(67);f__Herpotrichiellaceae_unclassified(67); | Herpotrichiellaceae | 9 | Pathotroph-Saprotroph | Animal Pathogen-Fungal Parasite-Undefined Saprotroph | Facultative Yeast-Microfungus | NULL | Probable | [illegible] | [illegible] |
| Otu07133 | 3 | k__Fungi(100);p__Ascomycota(100);c__Saccharomycetes(100);o__Saccharomycetales(100);f__Saccharomycetales_family_Incertae_sedis(100);g__Candida(100);g__Candida_unclassified(100); | Candida | 13 | Pathotroph-Saprotroph-Symbiotroph | Animal Pathogen-Endophyte-Endosymbiont-Epiphyte-\|Undefined Saprotroph\| | Dimorphic Yeast-Yeast | NULL | Probable | [illegible] | [illegible] |
| Otu07134 | 3 | k__Fungi(100);p__Ascomycota(100);c__Eurotiomycetes(100);o__Eurotiales(100);f__Trichocomaceae(100);g__Talaromyces(100);s__Talaromyces_proteolyticus(100); | Talaromyces | 13 | Pathotroph-Saprotroph | Animal Parasite-\|Undefined Saprotroph\| | Microfungus | NULL | Probable | [illegible] | [illegible] |
| Otu07138 | 3 | k__Fungi(100);p__Ascomycota(100);c__Leotiomycetes(100);o__Rhytismatales(100);f__Rhytismataceae(100);f__Rhytismataceae_unclassified(100);f__Rhytismataceae_unclassified(100); | Rhytismataceae | 9 | Pathotroph-Saprotroph-Symbiotroph | Endophyte-Leaf Saprotroph-Plant Pathogen | Microfungus | NULL | Probable | [illegible] | [illegible] |
| Otu07139 | 3 | k__Fungi(100);p__Ascomycota(100);c__Eurotiomycetes(100);o__Onygenales(100);f__Onygenaceae(100);g__Chrysosporium(100);s__Chrysosporium_speluncarum(100); | Chrysosporium | 13 | Pathotroph-Saprotroph | Plant Pathogen-Undefined Saprotroph-\|Wood Saprotroph\| | Microfungus | Soft Rot | Highly Probable | [illegible] | [illegible] |
| Otu07140 | 3 | k__Fungi(100);p__Basidiomycota(100);c__Tremellomycetes(100);o__Tremellales(100);f__Tremellales_family_Incertae_sedis(100);g__Hannaella(100);s__Hannaella_surugaensis(100); | Hannaella | 13 | Pathotroph-Saprotroph | \|Animal Parasite\|-Undefined Saprotroph | Yeast | NULL | Probable | [illegible] | [illegible] |
| Otu07141 | 3 | k__Fungi(100);p__Ascomycota(100);c__Taphrinomycetes(100);o__Taphrinales(100);f__Taphrinaceae(100);g__Taphrina(100);s__Taphrina_betulina(100); | Taphrina | 13 | Pathotroph | \|Plant Pathogen\| | Dimorphic-Microfungus | NULL | Probable | NULL | [illegible] |
| Otu07142 | 3 | k__Fungi(100);p__Ascomycota(100);c__Saccharomycetes(100);o__Saccharomycetales(100);f__Metschnikowiaceae(100);g__Metschnikowia(100);s__Metschnikowia_pulcherrima(67); | Metschnikowia | 13 | Saprotroph | \|Undefined Saprotroph\| | Yeast | NULL | Probable | [illegible] | [illegible] |
| Otu07146 | 3 | k__Fungi(100);p__Ascomycota(100);c__Saccharomycetes(100);o__Saccharomycetales(100);f__Metschnikowiaceae(100);g__Metschnikowia(100);s__Metschnikowia_pulcherrima(100); | Metschnikowia | 13 | Saprotroph | \|Undefined Saprotroph\| | Yeast | NULL | Probable | [illegible] | [illegible] |
| Otu07148 | 3 | k__Fungi(100);p__Basidiomycota(100);c__Agaricomycetes(100);o__Agaricales(100);f__Strophariaceae(100);g__Pholiota(100);s__Pholiota_abieticola(100); | Pholiota | 13 | Pathotroph-Saprotroph | Plant Pathogen-Plant Saprotroph-Undefined Saprotroph-\|Wood Saprotroph\| | Agaricoid | White Rot | Probable | [illegible] | [illegible] |
| Otu07149 | 3 | k__Fungi(100);p__Ascomycota(100);c__Dothideomycetes(100);o__Capnodiales(100);f__Teratosphaeriaceae(100);g__Devriesia(100);s__Devriesia_sp(100); | Devriesia | 13 | Pathotroph-Saprotroph | \|Plant Pathogen\|-Plant Saprotroph | Microfungus | NULL | Probable | [illegible] | [illegible] |
| Otu07152 | 3 | k__Fungi(100);p__Ascomycota(100);c__Eurotiomycetes(100);o__Chaetothyriales(100);f__Herpotrichiellaceae(100);g__Exophiala(100);g__Exophiala_unclassified(100); | Exophiala | 13 | Pathotroph-Saprotroph-Symbiotroph | \|Animal Parasite\|-Animal Pathogen-Endophyte-Plant Saprotroph-Undefined Saprotroph | Dimorphic-Facultative Yeast | NULL | Probable | [illegible] | [illegible] |
| Otu07155 | 3 | k__Fungi(100);p__Ascomycota(100);c__Saccharomycetes(100);o__Saccharomycetales(100);f__Metschnikowiaceae(100);g__Metschnikowia(100);s__Metschnikowia_pulcherrima(100); | Metschnikowia | 13 | Saprotroph | \|Undefined Saprotroph\| | Yeast | NULL | Probable | [illegible] | [illegible] |
| Otu07157 | 3 | k__Fungi(100);p__Ascomycota(100);c__Saccharomycetes(100);o__Saccharomycetales(100);f__Metschnikowiaceae(100);g__Metschnikowia(100);s__Metschnikowia_pulcherrima(67); | Metschnikowia | 13 | Saprotroph | \|Undefined Saprotroph\| | Yeast | NULL | Probable | [illegible] | [illegible] |
| Otu07158 | 3 | k__Fungi(100);p__Ascomycota(100);c__Sordariomycetes(100);o__Microascales(100);f__Microascaceae(100);g__Microascus(100);s__Microascus_sp_r249(100); | Microascus | 13 | Pathotroph-Saprotroph-Symbiotroph | Animal Pathogen-Dung Saprotroph-Endophyte-Plant Pathogen-Plant Saprotroph-\|Undefined Saprotroph\| | Microfungus | NULL | Possible | [illegible] | [illegible] |
| Otu07159 | 3 | k__Fungi(100);p__Basidiomycota(100);c__Microbotryomycetes(100);o__Sporidiobolales(100);f__Sporidiobolales_family_Incertae_sedis(100);g__Rhodotorula(100);s__Rhodotorula_mucilaginosa(100); | Rhodotorula | 13 | Pathotroph-Saprotroph-Symbiotroph | Animal Pathogen-Animal Symbiotroph-Endophyte-Plant Pathogen-\|Undefined Saprotroph\| | Yeast | NULL | Probable | [illegible] | [illegible] |
| Otu07161 | 3 | k__Fungi(100);p__Ascomycota(100);c__Saccharomycetes(100);o__Saccharomycetales(100);f__Metschnikowiaceae(100);g__Metschnikowia(100);s__Metschnikowia_pulcherrima(100); | Metschnikowia | 13 | Saprotroph | \|Undefined Saprotroph\| | Yeast | NULL | Probable | [illegible] | [illegible] |
| Otu07164 | 3 | k__Fungi(100);p__Basidiomycota(100);c__Microbotryomycetes(100);o__Sporidiobolales(100);f__Sporidiobolales_family_Incertae_sedis(100);g__Rhodotorula(67);s__Rhodotorula_graminis(67); | Rhodotorula | 13 | Pathotroph-Saprotroph-Symbiotroph | Animal Pathogen-Animal Symbiotroph-Endophyte-Plant Pathogen-\|Undefined Saprotroph\| | Yeast | NULL | Probable | [illegible] | [illegible] |
| Otu07168 | 3 | k__Fungi(100);p__Ascomycota(100);c__Saccharomycetes(100);o__Saccharomycetales(100);f__Metschnikowiaceae(100);g__Metschnikowia(100);s__Metschnikowia_pulcherrima(67); | Metschnikowia | 13 | Saprotroph | \|Undefined Saprotroph\| | Yeast | NULL | Probable | [illegible] | [illegible] |
| Otu07170 | 3 | k__Fungi(100);p__Ascomycota(100);c__Eurotiomycetes(100);o__Chaetothyriales(100);f__Herpotrichiellaceae(67);f__Herpotrichiellaceae_unclassified(67);f__Herpotrichiellaceae_unclassified(67); | Herpotrichiellaceae | 9 | Pathotroph-Saprotroph | Animal Pathogen-Fungal Parasite-Undefined Saprotroph | Facultative Yeast-Microfungus | NULL | Probable | [illegible] | [illegible] |
| Otu07171 | 3 | k__Fungi(100);p__Ascomycota(100);c__Saccharomycetes(100);o__Saccharomycetales(100);f__Metschnikowiaceae(100);g__Metschnikowia(100);s__Metschnikowia_pulcherrima(100); | Metschnikowia | 13 | Saprotroph | \|Undefined Saprotroph\| | Yeast | NULL | Probable | [illegible] | [illegible] |
| Otu07176 | 3 | k__Fungi(100);p__Ascomycota(100);c__Eurotiomycetes(100);o__Chaetothyriales(100);f__Herpotrichiellaceae(100);g__Phaeomoniella(100);s__Phaeomoniella_prunicola(100); | Phaeomoniella | 13 | Pathotroph | \|Plant Pathogen\| | Dimorphic-Microfungus; Facultative Yeast (Tedersoo et | NULL | Highly Probable | [illegible] | [illegible] |
| Otu07178 | 3 | k__Fungi(100);p__Basidiomycota(100);c__Microbotryomycetes(100);o__Sporidiobolales(100);f__Sporidiobolales_family_Incertae_sedis(100);g__Rhodotorula(100);s__Rhodotorula_taiwanensis(100); | Rhodotorula | 13 | Pathotroph-Saprotroph-Symbiotroph | Animal Pathogen-Animal Symbiotroph-Endophyte-Plant Pathogen-\|Undefined Saprotroph\| | Yeast | NULL | Probable | [illegible] | [illegible] |
| Otu07179 | 3 | k__Fungi(100);p__Ascomycota(100);c__Saccharomycetes(100);o__Saccharomycetales(100);f__Metschnikowiaceae(100);g__Metschnikowia(100);s__Metschnikowia_pulcherrima(100); | Metschnikowia | 13 | Saprotroph | \|Undefined Saprotroph\| | Yeast | NULL | Probable | [illegible] | [illegible] |
| Otu07180 | 3 | k__Fungi(100);p__Ascomycota(100);c__Ascomycota_class_Incertae_sedis(100);o__Ascomycota_order_Incertae_sedis(100);f__Ascomycota_family_Incertae_sedis(100);g__Veronaea(100);s__Veronaea_compacta(100); | Veronaea | 13 | Pathotroph-Saprotroph | Dung Saprotroph-\|Plant Pathogen\|-Plant Saprotroph | Dimorphic | NULL | Probable | [illegible] | [illegible] |
| Otu07182 | 3 | k__Fungi(100);p__Ascomycota(100);c__Saccharomycetes(100);o__Saccharomycetales(100);f__Metschnikowiaceae(100);g__Metschnikowia(100);g__Metschnikowia_unclassified(67); | Metschnikowia | 13 | Saprotroph | \|Undefined Saprotroph\| | Yeast | NULL | Probable | [illegible] | [illegible] |
| Otu07184 | 3 | k__Fungi(100);p__Ascomycota(100);c__Saccharomycetes(100);o__Saccharomycetales(100);f__Metschnikowiaceae(100);g__Metschnikowia(100);s__Metschnikowia_pulcherrima(100); | Metschnikowia | 13 | Saprotroph | \|Undefined Saprotroph\| | Yeast | NULL | Probable | [illegible] | [illegible] |
| Otu07186 | 3 | k__Fungi(100);p__Ascomycota(100);c__Saccharomycetes(100);o__Saccharomycetales(100);f__Metschnikowiaceae(100);g__Metschnikowia(100);g__Metschnikowia_unclassified(67); | Metschnikowia | 13 | Saprotroph | \|Undefined Saprotroph\| | Yeast | NULL | Probable | [illegible] | [illegible] |
| Otu07187 | 3 | k__Fungi(100);p__Ascomycota(100);c__Saccharomycetes(100);o__Saccharomycetales(100);f__Metschnikowiaceae(100);g__Metschnikowia(100);s__Metschnikowia_pulcherrima(67); | Metschnikowia | 13 | Saprotroph | \|Undefined Saprotroph\| | Yeast | NULL | Probable | [illegible] | [illegible] |
| Otu07188 | 3 | k__Fungi(100);p__Ascomycota(100);c__Saccharomycetes(100);o__Saccharomycetales(100);f__Metschnikowiaceae(100);g__Metschnikowia(100);s__Metschnikowia_pulcherrima(67); | Metschnikowia | 13 | Saprotroph | \|Undefined Saprotroph\| | Yeast | NULL | Probable | [illegible] | [illegible] |
| Otu07192 | 3 | k__Fungi(100);p__Ascomycota(100);c__Sordariomycetes(100);o__Xylariales(100);f__Xylariaceae(100);g__Xylaria(100);g__Xylaria_unclassified(100); | Xylaria | 13 | Saprotroph-Symbiotroph | Endophyte-Plant Saprotroph-Undefined Saprotroph-\|Wood Saprotroph\| | Xylarioid | Soft Rot | Probable | [illegible] | [illegible] |
| Otu07194 | 3 | k__Fungi(100);p__Ascomycota(100);c__Saccharomycetes(100);o__Saccharomycetales(100);f__Saccharomycetales_family_Incertae_sedis(100);g__Candida(100);g__Candida_unclassified(100); | Candida | 13 | Pathotroph-Saprotroph-Symbiotroph | Animal Pathogen-Endophyte-Endosymbiont-Epiphyte-\|Undefined Saprotroph\| | Dimorphic Yeast-Yeast | NULL | Probable | [illegible] | [illegible] |
| Otu07197 | 3 | k__Fungi(100);p__Ascomycota(100);c__Saccharomycetes(100);o__Saccharomycetales(100);f__Metschnikowiaceae(100);g__Metschnikowia(100);s__Metschnikowia_pulcherrima(67); | Metschnikowia | 13 | Saprotroph | \|Undefined Saprotroph\| | Yeast | NULL | Probable | [illegible] | [illegible] |
| Otu07200 | 3 | k__Fungi(100);p__Ascomycota(100);c__Eurotiomycetes(100);o__Chaetothyriales(100);f__Herpotrichiellaceae(100);g__Exophiala(100);s__Exophiala_moniliae(100); | Exophiala | 13 | Pathotroph-Saprotroph-Symbiotroph | \|Animal Parasite\|-Animal Pathogen-Endophyte-Plant Saprotroph-Undefined Saprotroph | Dimorphic-Facultative Yeast | NULL | Probable | [illegible] | [illegible] |
| Otu07203 | 3 | k__Fungi(100);p__Ascomycota(100);c__Saccharomycetes(100);o__Saccharomycetales(100);f__Metschnikowiaceae(100);g__Metschnikowia(100);g__Metschnikowia_unclassified(67); | Metschnikowia | 13 | Saprotroph | \|Undefined Saprotroph\| | Yeast | NULL | Probable | [illegible] | [illegible] |
| Otu07206 | 3 | k__Fungi(100);p__Basidiomycota(100);c__Agaricomycetes(100);o__Agaricales(100);f__Mycenaceae(100);g__Mycena(100);g__Mycena_unclassified(100); | Mycena | 13 | Pathotroph-Saprotroph | Plant Pathogen-\|Plant Saprotroph\|-Undefined Saprotroph-Wood Saprotroph | Agaricoid | NULL | Probable | [illegible] | [illegible] |
| Otu07207 | 3 | k__Fungi(100);p__Ascomycota(100);c__Eurotiomycetes(100);o__Chaetothyriales(100);f__Herpotrichiellaceae(100);g__Exophiala(100);g__Exophiala_unclassified(100); | Exophiala | 13 | Pathotroph-Saprotroph-Symbiotroph | \|Animal Parasite\|-Animal Pathogen-Endophyte-Plant Saprotroph-Undefined Saprotroph | Dimorphic-Facultative Yeast | NULL | Probable | [illegible] | [illegible] |
| Otu07209 | 3 | k__Fungi(100);p__Basidiomycota(100);c__Agaricomycetes(100);o__Agaricales(100);f__Bolbitiaceae(100);g__Conocybe(100);s__Conocybe_aff_ochrostriata_NL_0830(100); | Conocybe | 13 | Saprotroph | Dung Saprotroph-Plant Saprotroph-\|Undefined Saprotroph\| | Agaricoid | NULL | Probable | [illegible] | [illegible] |
| Otu07210 | 3 | k__Fungi(100);p__Ascomycota(100);c__Saccharomycetes(100);o__Saccharomycetales(100);f__Metschnikowiaceae(100);g__Metschnikowia(100);s__Metschnikowia_pulcherrima(100); | Metschnikowia | 13 | Saprotroph | \|Undefined Saprotroph\| | Yeast | NULL | Probable | [illegible] | [illegible] |
| Otu07211 | 3 | k__Fungi(100);p__Zygomycota(100);c__Zygomycota_class_Incertae_sedis(100);o__Mortierellales(100);f__Mortierellaceae(100);g__Mortierella(100);g__Mortierella_unclassified(100); | Mortierella | 13 | Saprotroph-Symbiotroph | Endophyte-Plant Saprotroph-\|Undefined Saprotroph\| | Microfungus | NULL | Probable | [illegible] | [illegible] |
| Otu07212 | 3 | k__Fungi(100);p__Zygomycota(100);c__Zygomycota_class_Incertae_sedis(100);o__Mortierellales(100);f__Mortierellaceae(100);g__Mortierella(100);s__Mortierella_parvispora(100); | Mortierella | 13 | Saprotroph-Symbiotroph | Endophyte-Plant Saprotroph-\|Undefined Saprotroph\| | Microfungus | NULL | Probable | [illegible] | [illegible] |
| Otu07213 | 3 | k__Fungi(100);p__Ascomycota(100);c__Saccharomycetes(100);o__Saccharomycetales(100);f__Metschnikowiaceae(100);g__Metschnikowia(100);s__Metschnikowia_pulcherrima(67); | Metschnikowia | 13 | Saprotroph | \|Undefined Saprotroph\| | Yeast | NULL | Probable | [illegible] | [illegible] |
| Otu07217 | 3 | k__Fungi(100);p__Ascomycota(100);c__Saccharomycetes(100);o__Saccharomycetales(100);f__Saccharomycetales_family_Incertae_sedis(100);f__Saccharomycetales_family_Incertae_sedis_unclassified(67);f__Saccharomycetales_family_Incertae_sedis_unclassified(67); | Saccharomycetales | 7 | Saprotroph | Undefined Saprotroph | Yeast | NULL | Possible | NULL | [illegible] |
| Otu07219 | 3 | k__Fungi(100);p__Ascomycota(100);c__Eurotiomycetes(100);o__Chaetothyriales(100);f__Herpotrichiellaceae(100);f__Herpotrichiellaceae_unclassified(100);f__Herpotrichiellaceae_unclassified(100); | Herpotrichiellaceae | 9 | Pathotroph-Saprotroph | Animal Pathogen-Fungal Parasite-Undefined Saprotroph | Facultative Yeast-Microfungus | NULL | Probable | [illegible] | [illegible] |
| Otu07220 | 3 | k__Fungi(100);p__Ascomycota(100);c__Saccharomycetes(100);o__Saccharomycetales(100);f__Metschnikowiaceae(100);g__Metschnikowia(100);s__Metschnikowia_pulcherrima(100); | Metschnikowia | 13 | Saprotroph | \|Undefined Saprotroph\| | Yeast | NULL | Probable | [illegible] | [illegible] |
| Otu07224 | 3 | k__Fungi(100);p__Ascomycota(100);c__Saccharomycetes(100);o__Saccharomycetales(100);f__Metschnikowiaceae(100);g__Metschnikowia(100);s__Metschnikowia_pulcherrima(100); | Metschnikowia | 13 | Saprotroph | \|Undefined Saprotroph\| | Yeast | NULL | Probable | [illegible] | [illegible] |
| Otu07226 | 3 | k__Fungi(100);p__Ascomycota(100);c__Saccharomycetes(100);o__Saccharomycetales(100);f__Metschnikowiaceae(100);g__Metschnikowia(100);s__Metschnikowia_pulcherrima(100); | Metschnikowia | 13 | Saprotroph | \|Undefined Saprotroph\| | Yeast | NULL | Probable | [illegible] | [illegible] |
| Otu07228 | 3 | k__Fungi(100);p__Ascomycota(100);c__Lecanoromycetes(100);o__Teloschistales(100);f__Caliciaceae(100);g__Amandinea(100);s__Amandinea_punctata(100); | Amandinea | 13 | Symbiotroph | \|Lichenized\| | Thallus | NULL | Highly Probable | NULL | [illegible] |
| Otu07230 | 3 | k__Fungi(100);p__Ascomycota(100);c__Saccharomycetes(100);o__Saccharomycetales(100);f__Metschnikowiaceae(100);g__Metschnikowia(100);s__Metschnikowia_pulcherrima(67); | Metschnikowia | 13 | Saprotroph | \|Undefined Saprotroph\| | Yeast | NULL | Probable | [illegible] | [illegible] |
| Otu07231 | 3 | k__Fungi(100);p__Ascomycota(100);c__Saccharomycetes(100);o__Saccharomycetales(100);f__Metschnikowiaceae(100);g__Metschnikowia(100);s__Metschnikowia_pulcherrima(100); | Metschnikowia | 13 | Saprotroph | \|Undefined Saprotroph\| | Yeast | NULL | Probable | [illegible] | [illegible] |
| Otu07232 | 3 | k__Fungi(100);p__Basidiomycota(100);c__Microbotryomycetes(100);o__Sporidiobolales(100);f__Sporidiobolales_family_Incertae_sedis(100);g__Rhodotorula(100);s__Rhodotorula_mucilaginosa(100); | Rhodotorula | 13 | Pathotroph-Saprotroph-Symbiotroph | Animal Pathogen-Animal Symbiotroph-Endophyte-Plant Pathogen-\|Undefined Saprotroph\| | Yeast | NULL | Probable | [illegible] | [illegible] |
| Otu07234 | 3 | k__Fungi(100);p__Ascomycota(100);c__Saccharomycetes(100);o__Saccharomycetales(100);f__Metschnikowiaceae(100);g__Metschnikowia(100);s__Metschnikowia_pulcherrima(100); | Metschnikowia | 13 | Saprotroph | \|Undefined Saprotroph\| | Yeast | NULL | Probable | [illegible] | [illegible] |
| Otu07236 | 3 | k__Fungi(100);p__Ascomycota(100);c__Dothideomycetes(100);o__Pleosporales(100);f__Pleosporaceae(100);f__Pleosporaceae_unclassified(100);f__Pleosporaceae_unclassified(100); | Pleosporaceae | 9 | Pathotroph-Saprotroph | Endophyte-Lichen Parasite-Plant Pathogen-Undefined Saprotroph | Microfungus | NULL | Probable | [illegible] | [illegible] |
| Otu07237 | 3 | k__Fungi(100);p__Ascomycota(100);c__Saccharomycetes(67);o__Saccharomycetales(67);f__Saccharomycetales_family_Incertae_sedis(67);g__Candida(67);g__Candida_unclassified(67); | Candida | 13 | Pathotroph-Saprotroph-Symbiotroph | Animal Pathogen-Endophyte-Endosymbiont-Epiphyte-\|Undefined Saprotroph\| | Dimorphic Yeast-Yeast | NULL | Probable | [illegible] | [illegible] |
| Otu07238 | 3 | k__Fungi(100);p__Ascomycota(100);c__Saccharomycetes(100);o__Saccharomycetales(100);f__Metschnikowiaceae(100);g__Metschnikowia(100);s__Metschnikowia_pulcherrima(100); | Metschnikowia | 13 | Saprotroph | \|Undefined Saprotroph\| | Yeast | NULL | Probable | [illegible] | [illegible] |
| Otu07239 | 3 | k__Fungi(100);p__Ascomycota(100);c__Dothideomycetes(100);o__Pleosporales(100);f__Pleosporaceae(100);g__Stemphylium(100);s__Stemphylium_loti(100); | Stemphylium | 13 | Pathotroph-Saprotroph | \|Plant Pathogen\|-Plant Saprotroph-Wood Saprotroph | Microfungus | Soft Rot | Highly Probable | [illegible] | [illegible] |
| Otu07240 | 3 | k__Fungi(100);p__Ascomycota(100);c__Dothideomycetes(100);o__Pleosporales(100);f__Pleosporaceae(100);g__Pleospora(100);s__Pleospora_herbarum(100); | Pleospora | 13 | Pathotroph-Saprotroph-Symbiotroph | Endophyte-Lichen Parasite-\|Plant Pathogen\|-Undefined Saprotroph | Microfungus | NULL | Probable | [illegible] | [illegible] |
| Otu07241 | 3 | k__Fungi(100);p__Ascomycota(100);c__Saccharomycetes(100);o__Saccharomycetales(100);f__Metschnikowiaceae(100);g__Metschnikowia(100);s__Metschnikowia_pulcherrima(100); | Metschnikowia | 13 | Saprotroph | \|Undefined Saprotroph\| | Yeast | NULL | Probable | [illegible] | [illegible] |
| Otu07243 | 3 | k__Fungi(100);p__Ascomycota(100);c__Saccharomycetes(100);o__Saccharomycetales(100);f__Metschnikowiaceae(100);g__Metschnikowia(100);s__Metschnikowia_pulcherrima(100); | Metschnikowia | 13 | Saprotroph | \|Undefined Saprotroph\| | Yeast | NULL | Probable | [illegible] | [illegible] |
| Otu07246 | 3 | k__Fungi(100);p__Ascomycota(100);c__Saccharomycetes(100);o__Saccharomycetales(100);f__Metschnikowiaceae(100);g__Metschnikowia(100);s__Metschnikowia_pulcherrima(100); | Metschnikowia | 13 | Saprotroph | \|Undefined Saprotroph\| | Yeast | NULL | Probable | [illegible] | [illegible] |
| Otu07247 | 3 | k__Fungi(100);p__Basidiomycota(100);c__Agaricomycetes(100);o__Russulales(100);f__Stephanosporaceae(100);g__unclassified_Stephanosporaceae(100);s__Stephanosporaceae_sp(100); | Stephanosporaceae | 9 | Saprotroph | Plant Saprotroph-Wood Saprotroph | Corticioid-Gasteroid | NULL | Probable | [illegible] | [illegible] |
| Otu07251 | 3 | k__Fungi(100);p__Basidiomycota(100);c__Agaricomycetes(100);o__Agaricales(100);f__Marasmiaceae(100);g__Marasmius(100);s__Marasmius_sp_SM2425(100); | Marasmius | 13 | Pathotroph-Saprotroph-Symbiotroph | Endophyte-Plant Pathogen-\|Plant Saprotroph\|-Undefined Saprotroph | Agaricoid | NULL | Probable | [illegible] | [illegible] |
| Otu07253 | 3 | k__Fungi(100);p__Ascomycota(100);c__Saccharomycetes(100);o__Saccharomycetales(100);f__Metschnikowiaceae(100);g__Metschnikowia(100);s__Metschnikowia_pulcherrima(67); | Metschnikowia | 13 | Saprotroph | \|Undefined Saprotroph\| | Yeast | NULL | Probable | [illegible] | [illegible] |
| Otu07256 | 3 | k__Fungi(100);p__Ascomycota(100);c__Saccharomycetes(100);o__Saccharomycetales(100);f__Metschnikowiaceae(100);g__Metschnikowia(100);s__Metschnikowia_pulcherrima(100); | Metschnikowia | 13 | Saprotroph | \|Undefined Saprotroph\| | Yeast | NULL | Probable | [illegible] | [illegible] |
| Otu07257 | 3 | k__Fungi(100);p__Ascomycota(100);c__Saccharomycetes(100);o__Saccharomycetales(100);f__Metschnikowiaceae(100);g__Metschnikowia(100);s__Metschnikowia_pulcherrima(67); | Metschnikowia | 13 | Saprotroph | \|Undefined Saprotroph\| | Yeast | NULL | Probable | [illegible] | [illegible] |
| Otu07258 | 3 | k__Fungi(100);p__Ascomycota(100);c__Saccharomycetes(100);o__Saccharomycetales(100);f__Metschnikowiaceae(100);g__Metschnikowia(100);g__Metschnikowia_unclassified(67); | Metschnikowia | 13 | Saprotroph | \|Undefined Saprotroph\| | Yeast | NULL | Probable | [illegible] | [illegible] |
| Otu07261 | 3 | k__Fungi(100);p__Ascomycota(100);c__Ascomycota_class_Incertae_sedis(100);o__Ascomycota_order_Incertae_sedis(100);f__Ascomycota_family_Incertae_sedis(100);g__Slimacomyces(100);s__Slimacomyces_isiola(100); | Slimacomyces | 13 | Saprotroph | \|Plant Saprotroph\| | Microfungus | NULL | Probable | [illegible] | [illegible] |
| Otu07262 | 3 | k__Fungi(100);p__Basidiomycota(100);c__Tremellomycetes(100);o__Tremellales(100);f__Tremellales_family_Incertae_sedis(100);g__Fellomyces(100);s__Fellomyces_penicillatus(100); | Fellomyces | 13 | Pathotroph-Saprotroph | Fungal Parasite-\|Undefined Saprotroph\| | Yeast | NULL | Probable | [illegible] | [illegible] |
| Otu07263 | 3 | k__Fungi(100);p__Basidiomycota(100);c__Exobasidiomycetes(100);o__Microstromatales(100);f__Microstromataceae(100);g__Microstroma(100);s__Microstroma_juglandis(100); | Microstroma | 13 | Pathotroph | \|Plant Pathogen\| | Dimorphic | NULL | Probable | NULL | [illegible] |
| Otu07264 | 3 | k__Fungi(100);p__Ascomycota(100);c__Saccharomycetes(100);o__Saccharomycetales(100);f__Saccharomycetales_family_Incertae_sedis(100);g__Candida(100);g__Candida_unclassified(67); | Candida | 13 | Pathotroph-Saprotroph-Symbiotroph | Animal Pathogen-Endophyte-Endosymbiont-Epiphyte-\|Undefined Saprotroph\| | Dimorphic Yeast-Yeast | NULL | Probable | [illegible] | [illegible] |
| Otu07266 | 3 | k__Fungi(100);p__Basidiomycota(100);c__Agaricomycetes(100);o__Agaricales(100);f__Pleurotaceae(100);g__unclassified_Pleurotaceae(100);s__Pleurotaceae_sp(100); | Pleurotaceae | 9 | Pathotroph-Saprotroph | Endophyte-Plant Pathogen-Wood Saprotroph | Agaricoid | White Rot | Probable | [illegible] | [illegible] |
| Otu07267 | 3 | k__Fungi(100);p__Ascomycota(100);c__Saccharomycetes(100);o__Saccharomycetales(100);f__Saccharomycetales_family_Incertae_sedis(100);g__Candida(100);g__Candida_unclassified(100); | Candida | 13 | Pathotroph-Saprotroph-Symbiotroph | Animal Pathogen-Endophyte-Endosymbiont-Epiphyte-\|Undefined Saprotroph\| | Dimorphic Yeast-Yeast | NULL | Probable | [illegible] | [illegible] |
| Otu07268 | 3 | k__Fungi(100);p__Basidiomycota(100);c__Agaricomycetes(100);o__Boletales(100);f__Suillaceae(100);g__Suillus(100);s__Suillus_flavidus(100); | Suillus | 13 | Symbiotroph | \|Ectomycorrhizal\| | Agaricoid-Boletoid | NULL | Highly Probable | [illegible] | [illegible] |
| Otu07272 | 3 | k__Fungi(100);p__Basidiomycota(100);c__Agaricomycetes(100);o__Agaricales(100);f__Mycenaceae(100);g__Mycena(100);s__Mycena_stipata(100); | Mycena | 13 | Pathotroph-Saprotroph | Plant Pathogen-\|Plant Saprotroph\|-Undefined Saprotroph-Wood Saprotroph | Agaricoid | NULL | Probable | [illegible] | [illegible] |
| Otu07273 | 3 | k__Fungi(100);p__Ascomycota(100);c__Saccharomycetes(100);o__Saccharomycetales(100);f__Metschnikowiaceae(100);g__Metschnikowia(100);s__Metschnikowia_pulcherrima(67); | Metschnikowia | 13 | Saprotroph | \|Undefined Saprotroph\| | Yeast | NULL | Probable | [illegible] | [illegible] |
| Otu07274 | 3 | k__Fungi(100);p__Basidiomycota(100);c__Pucciniomycetes(100);o__Platygloeales(100);f__Eocronartiaceae(100);g__Eocronartium(100);s__Eocronartium_sp(100); | Eocronartium | 13 | Pathotroph | \|Plant Pathogen\| | Clavarioid | NULL | Probable | [illegible] | [illegible] |
| Otu07277 | 3 | k__Fungi(100);p__Ascomycota(100);c__Saccharomycetes(100);o__Saccharomycetales(100);f__Metschnikowiaceae(100);g__Metschnikowia(100);s__Metschnikowia_pulcherrima(67); | Metschnikowia | 13 | Saprotroph | \|Undefined Saprotroph\| | Yeast | NULL | Probable | [illegible] | [illegible] |
| Otu07279 | 3 | k__Fungi(100);p__Ascomycota(100);c__Saccharomycetes(100);o__Saccharomycetales(100);f__Saccharomycetales_family_Incertae_sedis(100);g__Candida(100);g__Candida_unclassified(67); | Candida | 13 | Pathotroph-Saprotroph-Symbiotroph | Animal Pathogen-Endophyte-Endosymbiont-Epiphyte-\|Undefined Saprotroph\| | Dimorphic Yeast-Yeast | NULL | Probable | [illegible] | [illegible] |
| Otu07283 | 3 | k__Fungi(100);p__Basidiomycota(100);c__Exobasidiomycetes(100);o__Microstromatales(100);f__Microstromataceae(100);g__Microstroma(100);s__Microstroma_juglandis(100); | Microstroma | 13 | Pathotroph | \|Plant Pathogen\| | Dimorphic | NULL | Probable | NULL | [illegible] |
| Otu07284 | 3 | k__Fungi(100);p__Basidiomycota(100);c__Microbotryomycetes(100);o__Sporidiobolales(100);f__Sporidiobolales_family_Incertae_sedis(100);g__Sporobolomyces(100);s__Sporobolomyces_lactosus(100); | Sporobolomyces | 13 | Pathotroph-Saprotroph | \|Fungal Parasite\|-Plant Saprotroph | Yeast | NULL | Probable | [illegible] | [illegible] |
| Otu07285 | 3 | k__Fungi(100);p__Ascomycota(100);c__Saccharomycetes(100);o__Saccharomycetales(100);f__Metschnikowiaceae(100);g__Metschnikowia(100);g__Metschnikowia_unclassified(67); | Metschnikowia | 13 | Saprotroph | \|Undefined Saprotroph\| | Yeast | NULL | Probable | [illegible] | [illegible] |
| Otu07289 | 3 | k__Fungi(100);p__Ascomycota(100);c__Saccharomycetes(100);o__Saccharomycetales(100);f__Metschnikowiaceae(100);g__Metschnikowia(100);s__Metschnikowia_pulcherrima(100); | Metschnikowia | 13 | Saprotroph | \|Undefined Saprotroph\| | Yeast | NULL | Probable | [illegible] | [illegible] |
| Otu07291 | 3 | k__Fungi(100);p__Ascomycota(100);c__Dothideomycetes(100);o__Dothideomycetes_order_Incertae_sedis(100);f__Myxotrichaceae(100);g__Oidiodendron(100);s__Oidiodendron_cereale(100); | Oidiodendron | 13 | Pathotroph-Saprotroph-Symbiotroph | Endophyte-Ericoid Mycorrhizal-\|Undefined Saprotroph\| | Microfungus | NULL | Probable | [illegible] | [illegible] |
| Otu07294 | 3 | k__Fungi(100);p__Basidiomycota(100);c__Agaricomycetes(100);o__Polyporales(100);f__Fomitopsidaceae(100);g__Postia(100);s__Postia_sp_DLL2009_106(100); | Postia | 13 | Saprotroph | \|Wood Saprotroph\| | Polyporoid | Brown Rot | Highly Probable | [illegible] | [illegible] |
| Otu07296 | 3 | k__Fungi(100);p__Ascomycota(100);c__Saccharomycetes(100);o__Saccharomycetales(100);f__Metschnikowiaceae(100);g__Metschnikowia(100);s__Metschnikowia_pulcherrima(100); | Metschnikowia | 13 | Saprotroph | \|Undefined Saprotroph\| | Yeast | NULL | Probable | [illegible] | [illegible] |
| Otu07303 | 3 | k__Fungi(100);p__Ascomycota(100);c__Saccharomycetes(100);o__Saccharomycetales(100);f__Metschnikowiaceae(100);g__Metschnikowia(100);s__Metschnikowia_pulcherrima(100); | Metschnikowia | 13 | Saprotroph | \|Undefined Saprotroph\| | Yeast | NULL | Probable | [illegible] | [illegible] |
| Otu07304 | 3 | k__Fungi(100);p__Basidiomycota(100);c__Agaricomycetes(100);o__Cantharellales(100);f__Hydnaceae(100);g__Sistotrema(100);s__Sistotrema_sp(100); | Sistotrema | 13 | Saprotroph-Symbiotroph | Ectomycorrhizal-\|Plant Saprotroph\|-Wood Saprotroph | Corticioid | White Rot | Possible | [illegible] | [illegible] |
| Otu07305 | 3 | k__Fungi(100);p__Ascomycota(100);c__Saccharomycetes(100);o__Saccharomycetales(100);f__Saccharomycetales_family_Incertae_sedis(100);g__Candida(100);s__Candida_sp_AS23486(100); | Candida | 13 | Pathotroph-Saprotroph-Symbiotroph | Animal Pathogen-Endophyte-Endosymbiont-Epiphyte-\|Undefined Saprotroph\| | Dimorphic Yeast-Yeast | NULL | Probable | [illegible] | [illegible] |
| Otu07306 | 3 | k__Fungi(100);p__Ascomycota(100);c__Saccharomycetes(100);o__Saccharomycetales(100);f__Metschnikowiaceae(100);g__Metschnikowia(100);s__Metschnikowia_pulcherrima(100); | Metschnikowia | 13 | Saprotroph | \|Undefined Saprotroph\| | Yeast | NULL | Probable | [illegible] | [illegible] |
| Otu07308 | 3 | k__Fungi(100);p__Basidiomycota(100);c__Exobasidiomycetes(100);o__Exobasidiales(100);f__Exobasidiaceae(100);g__Exobasidium(100);s__Exobasidium_kishianum(100); | Exobasidium | 13 | Pathotroph | \|Plant Pathogen\| | Leaf Blister-Leaf Gall-Leaf Spot | NULL | Probable | NULL | [illegible] |
| Otu07309 | 3 | k__Fungi(100);p__Basidiomycota(100);c__Agaricomycetes(100);o__Agaricales(100);f__Pluteaceae(100);g__Pluteus(100);g__Pluteus_unclassified(100); | Pluteus | 13 | Pathotroph-Saprotroph | Bryophyte Parasite-\|Plant Saprotroph\|-Wood Saprotroph | Agaricoid | White Rot | Probable | [illegible] | [illegible] |
| Otu07311 | 3 | k__Fungi(100);p__Basidiomycota(100);c__Agaricomycetes(100);o__Atheliales(100);f__Atheliaceae(100);g__Athelia(100);s__Athelia_decipiens(100); | Athelia | 13 | Pathotroph-Saprotroph-Symbiotroph | Algal Parasite-Epiphyte-\|Lichen Parasite\|-Lichenized-Plant Pathogen-Plant Saprotroph-Wood Saprotroph | Corticioid-Thallus | NULL | Possible | [illegible] | [illegible] |
| Otu07312 | 3 | k__Fungi(100);p__Ascomycota(100);c__Eurotiomycetes(100);o__Eurotiales(100);f__Trichocomaceae(100);g__Sagenomella(100);s__Sagenomella_diversispora(100); | Sagenomella | 13 | Pathotroph-Saprotroph | Animal Parasite-\|Undefined Saprotroph\| | Microfungus | NULL | Probable | NULL | [illegible] |
| Otu07313 | 3 | k__Fungi(100);p__Basidiomycota(100);c__Agaricomycetes(100);o__Hymenochaetales(100);f__Schizoporaceae(100);g__Hyphodontia(100);s__Hyphodontia_sp_DLL2011_1(100); | Hyphodontia | 13 | Saprotroph | Undefined Saprotroph-\|Wood Saprotroph\| | Corticioid | White Rot | Probable | [illegible] | [illegible] |
| Otu07314 | 3 | k__Fungi(100);p__Ascomycota(100);c__Saccharomycetes(100);o__Saccharomycetales(100);f__Metschnikowiaceae(100);g__Metschnikowia(100);s__Metschnikowia_pulcherrima(100); | Metschnikowia | 13 | Saprotroph | \|Undefined Saprotroph\| | Yeast | NULL | Probable | [illegible] | [illegible] |
| Otu07316 | 3 | k__Fungi(100);p__Ascomycota(100);c__Saccharomycetes(100);o__Saccharomycetales(100);f__Metschnikowiaceae(100);g__Metschnikowia(100);s__Metschnikowia_pulcherrima(100); | Metschnikowia | 13 | Saprotroph | \|Undefined Saprotroph\| | Yeast | NULL | Probable | [illegible] | [illegible] |
| Otu07317 | 3 | k__Fungi(100);p__Ascomycota(100);c__Saccharomycetes(100);o__Saccharomycetales(100);f__Saccharomycetales_family_Incertae_sedis(100);g__Candida(100);g__Candida_unclassified(67); | Candida | 13 | Pathotroph-Saprotroph-Symbiotroph | Animal Pathogen-Endophyte-Endosymbiont-Epiphyte-\|Undefined Saprotroph\| | Dimorphic Yeast-Yeast | NULL | Probable | [illegible] | [illegible] |
| Otu07319 | 3 | k__Fungi(100);p__Ascomycota(100);c__Sordariomycetes(100);o__Hypocreales(100);f__Hypocreales_family_Incertae_sedis(100);g__Myrothecium(100);g__Myrothecium_unclassified(100); | Myrothecium | 13 | Pathotroph-Saprotroph | Fungal Parasite-\|Plant Saprotroph\|-Undefined Saprotroph | Microfungus | NULL | Probable | [illegible] | [illegible] |

| | | | | | | | | | | | |
|---|---|---|---|---|---|---|---|---|---|---|---|
| Otu07320 | 3 | k__Fungi(100);p__Ascomycota(100);c__Saccharomycetes(100);o__Saccharomycetales(100);f__Metschnikowiaceae(100);g__Metschnikowia(100);s__Metschnikowia_pulcherrima(100); | Metschnikowia | 13 | Saprotroph | \|Undefined Saprotroph\| | Yeast | NULL | Probable | As nectar-tap_saprotroph (Põlme et al. 2020); Decay sugar-rich substrates (Põlme et al. 2020) | Põlme S, et al. 2020. Fungal Diversity 105:1-16 ((https://doi.org/10.1007/s13225-020-00466-2)) |
| Otu07322 | 3 | k__Fungi(100);p__Ascomycota(100);c__Saccharomycetes(100);o__Saccharomycetales(100);f__Metschnikowiaceae(100);g__Metschnikowia(100);s__Metschnikowia_pulcherrima(100); | Metschnikowia | 13 | Saprotroph | \|Undefined Saprotroph\| | Yeast | NULL | Probable | As nectar-tap_saprotroph (Põlme et al. 2020); Decay sugar-rich substrates (Põlme et al. 2020) | Põlme S, et al. 2020. Fungal Diversity 105:1-16 ((https://doi.org/10.1007/s13225-020-00466-2)) |
| Otu07323 | 3 | k__Fungi(100);p__Ascomycota(100);c__Sordariomycetes(100);o__Ophiostomatales(100);f__Ophiostomataceae(100);f__Ophiostomataceae_unclassified(100);f__Ophiostomataceae_unclassified(100); | Ophiostomataceae | 9 | Pathotroph | Plant Pathogen | Microfungus | Blue-Staining | Probable | At least most species are transmitted via insect vectors, with mycelial colonies developing in bark-boring beetle galleries and spreading through the vascular system; they are highly specific to their beetle vectors. In some cases spores and mycelium are transmitted in specialized organs, termed mycangia (Cannon & Kirk 2007) | Cannon PF, Kirk PM. 2007. Fungal Families of the World. CAB International, Cambridge (ISBN: 978-0851998275); Tedersoo L, et al. 2014. Science 346:e1256688 ((https://doi.org/10.1126/science.1256688)) |
| Otu07326 | 3 | k__Fungi(100);p__Ascomycota(100);c__Saccharomycetes(100);o__Saccharomycetales(100);f__Metschnikowiaceae(100);g__Metschnikowia(100);s__Metschnikowia_pulcherrima(100); | Metschnikowia | 13 | Saprotroph | \|Undefined Saprotroph\| | Yeast | NULL | Probable | As nectar-tap_saprotroph (Põlme et al. 2020); Decay sugar-rich substrates (Põlme et al. 2020) | Põlme S, et al. 2020. Fungal Diversity 105:1-16 ((https://doi.org/10.1007/s13225-020-00466-2)) |
| Otu07329 | 3 | k__Fungi(100);p__Ascomycota(100);c__Saccharomycetes(100);o__Saccharomycetales(100);f__Metschnikowiaceae(100);g__Metschnikowia(100);s__Metschnikowia_pulcherrima(100); | Metschnikowia | 13 | Saprotroph | \|Undefined Saprotroph\| | Yeast | NULL | Probable | As nectar-tap_saprotroph (Põlme et al. 2020); Decay sugar-rich substrates (Põlme et al. 2020) | Põlme S, et al. 2020. Fungal Diversity 105:1-16 ((https://doi.org/10.1007/s13225-020-00466-2)) |
| Otu07330 | 3 | k__Fungi(100);p__Ascomycota(100);c__Saccharomycetes(100);o__Saccharomycetales(100);f__Metschnikowiaceae(100);g__Metschnikowia(100);s__Metschnikowia_pulcherrima(67); | Metschnikowia | 13 | Saprotroph | \|Undefined Saprotroph\| | Yeast | NULL | Probable | As nectar-tap_saprotroph (Põlme et al. 2020); Decay sugar-rich substrates (Põlme et al. 2020) | Põlme S, et al. 2020. Fungal Diversity 105:1-16 ((https://doi.org/10.1007/s13225-020-00466-2)) |
| Otu07331 | 3 | k__Fungi(100);p__Basidiomycota(100);c__Ustilaginomycetes(100);o__Ustilaginales(100);f__Ustilaginaceae(100);g__unclassified_Ustilaginaceae(100);s__Ustilaginaceae_sp(100); | Ustilaginaceae | 9 | Pathotroph | Plant Pathogen | Smut | NULL | Probable | Biotrophic in living tissue of Gramineae and Cyperaceae (Cannon & Kirk 2007) | Cannon PF, Kirk PM. 2007. Fungal Families of the World. CAB International, Cambridge (ISBN: 978-0851998275) |
| Otu07333 | 3 | k__Fungi(100);p__Ascomycota(100);c__Saccharomycetes(100);o__Saccharomycetales(100);f__Metschnikowiaceae(100);g__Metschnikowia(100);s__Metschnikowia_pulcherrima(100); | Metschnikowia | 13 | Saprotroph | \|Undefined Saprotroph\| | Yeast | NULL | Probable | As nectar-tap_saprotroph (Põlme et al. 2020); Decay sugar-rich substrates (Põlme et al. 2020) | Põlme S, et al. 2020. Fungal Diversity 105:1-16 ((https://doi.org/10.1007/s13225-020-00466-2)) |
| Otu07334 | 3 | k__Fungi(100);p__Ascomycota(100);c__Saccharomycetes(100);o__Saccharomycetales(100);f__Metschnikowiaceae(100);g__Metschnikowia(100);g__Metschnikowia_unclassified(67); | Metschnikowia | 13 | Saprotroph | \|Undefined Saprotroph\| | Yeast | NULL | Probable | As nectar-tap_saprotroph (Põlme et al. 2020); Decay sugar-rich substrates (Põlme et al. 2020) | Põlme S, et al. 2020. Fungal Diversity 105:1-16 ((https://doi.org/10.1007/s13225-020-00466-2)) |
| Otu07335 | 3 | k__Fungi(100);p__Ascomycota(100);c__Eurotiomycetes(100);o__Chaetothyriales(100);f__Herpotrichiellaceae(100);g__Phaeomoniella(100);s__Phaeomoniella_prunicola(100); | Phaeomoniella | 13 | Pathotroph | \|Plant Pathogen\| | Dimorphic-Microfungus; Facultative Yeast (Tedersoo et | NULL | Highly Probable | Pathotroph isolated from necrosis in Prunus wood (Damm et al. 2010) | Damm U, et al. 2010. Persoonia 24:60-80 ((http://doi.org/10.3767/003158510X500705)); Tedersoo L, et al. 2014. Science 346:e1256688 ((https://doi.org/10.1126/science.1256688)); Põlme S, et al. 2020. Fungal Diversity 105:1-16 ((https://doi.org/10.1007/s13225-020-00466-2)) |
| Otu07336 | 3 | k__Fungi(100);p__Ascomycota(100);c__Sordariomycetes(100);o__Hypocreales(100);f__Clavicipitaceae(100);g__Pochonia(100);s__Pochonia_bulbillosa(100); | Pochonia | 13 | Pathotroph-Saprotroph | \|Animal Parasite\|-Animal Pathogen-Undefined Saprotroph | Microfungus | NULL | Probable | As animal_decomposer (Põlme et al. 2020); Decay animal material (Põlme et al. 2020) | Tedersoo L, et al. 2014. Science 346:e1256688 ((https://doi.org/10.1126/science.1256688)); Põlme S, et al. 2020. Fungal Diversity 105:1-16 ((https://doi.org/10.1007/s13225-020-00466-2)) |
| Otu07337 | 3 | k__Fungi(100);p__Ascomycota(100);c__Sordariomycetes(100);o__Hypocreales(100);f__Hypocreales_family_Incertae_sedis(100);g__Myrothecium(100);s__Myrothecium_verrucaria(100); | Myrothecium | 13 | Pathotroph-Saprotroph | Fungal Parasite-\|Plant Saprotroph\|-Undefined Saprotroph | Microfungus | NULL | Probable | As litter_saprotroph (Põlme et al. 2020); As mycoparasite (Põlme et al. 2020); Decay leaf, fruit, seed (Põlme et al. 2020) | Tedersoo L, et al. 2014. Science 346:e1256688 ((https://doi.org/10.1126/science.1256688)); Põlme S, et al. 2020. Fungal Diversity 105:1-16 ((https://doi.org/10.1007/s13225-020-00466-2)) |
| Otu07338 | 3 | k__Fungi(100);p__Basidiomycota(100);c__Microbotryomycetes(100);o__Sporidiobolales(100);f__Sporidiobolales_family_Incertae_sedis(100);g__Rhodotorula(100);s__Rhodotorula_taiwanensis(100); | Rhodotorula | 13 | Pathotroph-Saprotroph-Symbiotroph | Animal Pathogen-Animal Symbiotroph-Endophyte-Plant Pathogen-\|Undefined Saprotroph\| | Yeast | NULL | Probable | As foliar_endophyte (Põlme et al. 2020); Opportunistic human pathogen | Põlme S, et al. 2020. Fungal Diversity 105:1-16 ((https://doi.org/10.1007/s13225-020-00466-2)); https://www.cdc.gov ((https://www.cdc.gov)) |
| Otu07341 | 3 | k__Fungi(100);p__Basidiomycota(100);c__Microbotryomycetes(100);o__Sporidiobolales(100);f__Sporidiobolales_family_Incertae_sedis(100);g__Rhodotorula(100);s__Rhodotorula_taiwanensis(100); | Rhodotorula | 13 | Pathotroph-Saprotroph-Symbiotroph | Animal Pathogen-Animal Symbiotroph-Endophyte-Plant Pathogen-\|Undefined Saprotroph\| | Yeast | NULL | Probable | As foliar_endophyte (Põlme et al. 2020); Opportunistic human pathogen | Põlme S, et al. 2020. Fungal Diversity 105:1-16 ((https://doi.org/10.1007/s13225-020-00466-2)); https://www.cdc.gov ((https://www.cdc.gov)) |
| Otu07343 | 3 | k__Fungi(100);p__Ascomycota(100);c__Saccharomycetes(100);o__Saccharomycetales(100);f__Metschnikowiaceae(100);g__Metschnikowia(100);s__Metschnikowia_pulcherrima(67); | Metschnikowia | 13 | Saprotroph | \|Undefined Saprotroph\| | Yeast | NULL | Probable | As nectar-tap_saprotroph (Põlme et al. 2020); Decay sugar-rich substrates (Põlme et al. 2020) | Põlme S, et al. 2020. Fungal Diversity 105:1-16 ((https://doi.org/10.1007/s13225-020-00466-2)) |
| Otu07345 | 3 | k__Fungi(100);p__Ascomycota(100);c__Sordariomycetes(100);o__Hypocreales(100);o__Hypocreales_unclassified(100);o__Hypocreales_unclassified(100);o__Hypocreales_unclassified(100); | Hypocreales | 7 | Saprotroph | Undefined Saprotroph | Microfungus | NULL | Possible | NULL | Sterkenburg E, et al. 2015. New Phytologist 207:1145-1158 ((https://doi.org/10.1111/nph.13426)) |
| Otu07347 | 3 | k__Fungi(100);p__Ascomycota(100);c__Ascomycota_class_Incertae_sedis(67);o__Ascomycota_order_Incertae_sedis(67);f__Ascomycota_family_Incertae_sedis(67);g__Veronaea(67);s__Veronaea_compacta(67); | Veronaea | 13 | Pathotroph-Saprotroph | Dung Saprotroph-\|Plant Pathogen\|-Plant Saprotroph | Dimorphic | NULL | Probable | As litter_saprotroph (Põlme et al. 2020); Decay leaf, fruit, seed (Põlme et al. 2020) | Põlme S, et al. 2020. Fungal Diversity 105:1-16 ((https://doi.org/10.1007/s13225-020-00466-2)); The American Phytopathological Society Common Names of Plant Diseases ((http://www.apsnet.org/publications/commonnames/Pages/default.aspx)) |
| Otu07349 | 3 | k__Fungi(100);p__Basidiomycota(100);c__Agaricomycetes(100);o__Boletales(100);f__Boletaceae(100);g__Boletus(100);s__Boletus_edulis(100); | Boletus | 13 | Pathotroph-Symbiotroph | \|Ectomycorrhizal\|-Fungal Parasite | Agaricoid-Boletoid | NULL | Highly Probable | NULL | Rinaldi AC, et al. 2008. Fungal Diversity 33:1-45 ((http://www.fungaldiversity.org/fdp/sfdp/33-1.pdf)); Tedersoo L, et al. 2010. Mycorrhiza 20:217-263 ((https://doi.org/10.1007/s00572-009-0274-x)); Põlme S, et al. 2020. Fungal Diversity 105:1-16 ((https://doi.org/10.1007/s13225-020-00466-2)); http://mycorrhizas.info/ecmf.html ((http://mycorrhizas.info/ecmf.html)); List compiled by John Plischke III ((https://goodmushroom.weebly.com/uploads/5/7/1/7/57176535/fungi_on_fungi_pdf_lq.pdf)) |
| Otu07351 | 3 | k__Fungi(100);p__Basidiomycota(100);c__Microbotryomycetes(67);o__Sporidiobolales(67);f__Sporidiobolales_family_Incertae_sedis(67);g__Rhodotorula(67);s__Rhodotorula_sp(67); | Rhodotorula | 13 | Pathotroph-Saprotroph-Symbiotroph | Animal Pathogen-Animal Symbiotroph-Endophyte-Plant Pathogen-\|Undefined Saprotroph\| | Yeast | NULL | Probable | As foliar_endophyte (Põlme et al. 2020); Opportunistic human pathogen | Põlme S, et al. 2020. Fungal Diversity 105:1-16 ((https://doi.org/10.1007/s13225-020-00466-2)); https://www.cdc.gov ((https://www.cdc.gov)) |
| Otu07352 | 3 | k__Fungi(100);p__Ascomycota(100);c__Ascomycota_class_Incertae_sedis(100);o__Ascomycota_order_Incertae_sedis(100);f__Ascomycota_family_Incertae_sedis(100);g__Zymoseptoria(100);s__Zymoseptoria_verkleyi(100); | Zymoseptoria | 13 | Pathotroph-Saprotroph | \|Plant Pathogen\|-Plant Saprotroph | Microfungus | NULL | Probable | As litter_saprotroph (Põlme et al. 2020); Decay leaf, fruit, seed (Põlme et al. 2020) | Põlme S, et al. 2020. Fungal Diversity 105:1-16 ((https://doi.org/10.1007/s13225-020-00466-2)) |
| Otu07353 | 3 | k__Fungi(100);p__Basidiomycota(100);c__Agaricomycetes(100);o__Boletales(100);f__Boletaceae(100);g__Boletus(100);s__Boletus_eastwoodiae(100); | Boletus | 13 | Pathotroph-Symbiotroph | \|Ectomycorrhizal\|-Fungal Parasite | Agaricoid-Boletoid | NULL | Highly Probable | NULL | Rinaldi AC, et al. 2008. Fungal Diversity 33:1-45 ((http://www.fungaldiversity.org/fdp/sfdp/33-1.pdf)); Tedersoo L, et al. 2010. Mycorrhiza 20:217-263 ((https://doi.org/10.1007/s00572-009-0274-x)); Põlme S, et al. 2020. Fungal Diversity 105:1-16 ((https://doi.org/10.1007/s13225-020-00466-2)); http://mycorrhizas.info/ecmf.html ((http://mycorrhizas.info/ecmf.html)); List compiled by John Plischke III ((https://goodmushroom.weebly.com/uploads/5/7/1/7/57176535/fungi_on_fungi_pdf_lq.pdf)) |
| Otu07355 | 3 | k__Fungi(100);p__Ascomycota(100);c__Saccharomycetes(100);o__Saccharomycetales(100);f__Metschnikowiaceae(100);g__Metschnikowia(100);s__Metschnikowia_pulcherrima(67); | Metschnikowia | 13 | Saprotroph | \|Undefined Saprotroph\| | Yeast | NULL | Probable | As nectar-tap_saprotroph (Põlme et al. 2020); Decay sugar-rich substrates (Põlme et al. 2020) | Põlme S, et al. 2020. Fungal Diversity 105:1-16 ((https://doi.org/10.1007/s13225-020-00466-2)) |
| Otu07356 | 3 | k__Fungi(100);p__Ascomycota(100);c__Dothideomycetes(100);o__Pleosporales(100);f__Montagnulaceae(100);g__Paraconiothyrium(100);s__Paraconiothyrium_variabile(100); | Paraconiothyrium | 13 | Saprotroph | Undefined Saprotroph-\|Wood Saprotroph\| | Microfungus | NULL | Probable | Decay wood (Põlme et al. 2020) | Tedersoo L, et al. 2014. Science 346:e1256688 ((https://doi.org/10.1126/science.1256688)); Põlme S, et al. 2020. Fungal Diversity 105:1-16 ((https://doi.org/10.1007/s13225-020-00466-2)) |
| Otu07357 | 3 | k__Fungi(100);p__Ascomycota(100);c__Saccharomycetes(100);o__Saccharomycetales(100);f__Metschnikowiaceae(100);g__Metschnikowia(100);s__Metschnikowia_pulcherrima(67); | Metschnikowia | 13 | Saprotroph | \|Undefined Saprotroph\| | Yeast | NULL | Probable | As nectar-tap_saprotroph (Põlme et al. 2020); Decay sugar-rich substrates (Põlme et al. 2020) | Põlme S, et al. 2020. Fungal Diversity 105:1-16 ((https://doi.org/10.1007/s13225-020-00466-2)) |
| Otu07358 | 3 | k__Fungi(100);p__Ascomycota(100);c__Saccharomycetes(100);o__Saccharomycetales(100);f__Metschnikowiaceae(100);g__Metschnikowia(100);s__Metschnikowia_pulcherrima(67); | Metschnikowia | 13 | Saprotroph | \|Undefined Saprotroph\| | Yeast | NULL | Probable | As nectar-tap_saprotroph (Põlme et al. 2020); Decay sugar-rich substrates (Põlme et al. 2020) | Põlme S, et al. 2020. Fungal Diversity 105:1-16 ((https://doi.org/10.1007/s13225-020-00466-2)) |
| Otu07360 | 3 | k__Fungi(100);p__Basidiomycota(100);c__Agaricomycetes(100);o__Cantharellales(100);f__Ceratobasidiaceae(100);g__unclassified_Ceratobasidiaceae(67);g__unclassified_Ceratobasidiaceae_unclassified(67); | Ceratobasidiaceae | 9 | Pathotroph-Saprotroph-Symbiotroph | Endomycorrhizal-Plant Pathogen-Undefined Saprotroph | Microfungus | NULL | Possible | Best known as plant pathogens, but also saprobic. Species are endomycorrhizal with a wide range of plant roots - especially of Orchidaceae but also even bryophytes (Cannon & Kirk 2007) | Cannon PF, Kirk PM. 2007. Fungal Families of the World. CAB International, Cambridge (ISBN: 978-0851998275) |
| Otu07361 | 3 | k__Fungi(100);p__Ascomycota(100);c__Dothideomycetes(100);o__Pleosporales(100);f__Phaeosphaeriaceae(100);f__Phaeosphaeriaceae_unclassified(100);f__Phaeosphaeriaceae_unclassified(100); | Phaeosphaeriaceae | 9 | Pathotroph-Saprotroph | Fungal Parasite-Plant Pathogen-Plant Saprotroph | Microfungus | NULL | Probable | Necrotrophic or saprobic on a wide range of plants, especially monocotyledons; a few species parasitic on other fungi (Cannon & Kirk 2007) | Cannon PF, Kirk PM. 2007. Fungal Families of the World. CAB International, Cambridge (ISBN: 978-0851998275) |
| Otu07364 | 3 | k__Fungi(100);p__Ascomycota(100);c__Saccharomycetes(100);o__Saccharomycetales(100);f__Metschnikowiaceae(100);g__Metschnikowia(100);g__Metschnikowia_unclassified(100); | Metschnikowia | 13 | Saprotroph | \|Undefined Saprotroph\| | Yeast | NULL | Probable | As nectar-tap_saprotroph (Põlme et al. 2020); Decay sugar-rich substrates (Põlme et al. 2020) | Põlme S, et al. 2020. Fungal Diversity 105:1-16 ((https://doi.org/10.1007/s13225-020-00466-2)) |
| Otu07366 | 3 | k__Fungi(100);p__Ascomycota(100);c__Saccharomycetes(100);o__Saccharomycetales(100);f__Metschnikowiaceae(100);g__Metschnikowia(100);g__Metschnikowia_unclassified(67); | Metschnikowia | 13 | Saprotroph | \|Undefined Saprotroph\| | Yeast | NULL | Probable | As nectar-tap_saprotroph (Põlme et al. 2020); Decay sugar-rich substrates (Põlme et al. 2020) | Põlme S, et al. 2020. Fungal Diversity 105:1-16 ((https://doi.org/10.1007/s13225-020-00466-2)) |
| Otu07369 | 3 | k__Fungi(100);p__Ascomycota(100);c__Saccharomycetes(100);o__Saccharomycetales(100);f__Metschnikowiaceae(100);g__Metschnikowia(100);s__Metschnikowia_pulcherrima(100); | Metschnikowia | 13 | Saprotroph | \|Undefined Saprotroph\| | Yeast | NULL | Probable | As nectar-tap_saprotroph (Põlme et al. 2020); Decay sugar-rich substrates (Põlme et al. 2020) | Põlme S, et al. 2020. Fungal Diversity 105:1-16 ((https://doi.org/10.1007/s13225-020-00466-2)) |
| Otu07372 | 3 | k__Fungi(100);p__Ascomycota(100);c__Saccharomycetes(100);o__Saccharomycetales(100);f__Metschnikowiaceae(100);g__Metschnikowia(100);g__Metschnikowia_unclassified(67); | Metschnikowia | 13 | Saprotroph | \|Undefined Saprotroph\| | Yeast | NULL | Probable | As nectar-tap_saprotroph (Põlme et al. 2020); Decay sugar-rich substrates (Põlme et al. 2020) | Põlme S, et al. 2020. Fungal Diversity 105:1-16 ((https://doi.org/10.1007/s13225-020-00466-2)) |
| Otu07373 | 3 | k__Fungi(100);p__Ascomycota(100);c__Saccharomycetes(100);o__Saccharomycetales(100);f__Metschnikowiaceae(100);g__Metschnikowia(100);s__Metschnikowia_pulcherrima(67); | Metschnikowia | 13 | Saprotroph | \|Undefined Saprotroph\| | Yeast | NULL | Probable | As nectar-tap_saprotroph (Põlme et al. 2020); Decay sugar-rich substrates (Põlme et al. 2020) | Põlme S, et al. 2020. Fungal Diversity 105:1-16 ((https://doi.org/10.1007/s13225-020-00466-2)) |
| Otu07374 | 3 | k__Fungi(100);p__Chytridiomycota(100);c__Chytridiomycetes(100);o__Spizellomycetales(100);f__Spizellomycetaceae(100);g__Spizellomyces(100);s__Spizellomyces_sp(100); | Spizellomyces | 13 | Pathotroph-Saprotroph | Plant Pathogen-\|Pollen Saprotroph\|-Undefined Parasite-Undefined Saprotroph | Chytrid-Microfungus | NULL | Probable | Decay pollen (Põlme et al. 2020); Pollen degrader; Common in the soil and also as parasites of a range of soil organisms and plants (Cannon & Kirk 2007) | Cannon PF, Kirk PM. 2007. Fungal Families of the World. CAB International, Cambridge (ISBN: 978-0851998275); Tedersoo L, et al. 2014. Science 346:e1256688 ((https://doi.org/10.1126/science.1256688)); Põlme S, et al. 2020. Fungal Diversity 105:1-16 ((https://doi.org/10.1007/s13225-020-00466-2)) |
| Otu07377 | 3 | k__Fungi(100);p__Ascomycota(100);c__Saccharomycetes(100);o__Saccharomycetales(100);f__Metschnikowiaceae(100);g__Metschnikowia(100);s__Metschnikowia_pulcherrima(100); | Metschnikowia | 13 | Saprotroph | \|Undefined Saprotroph\| | Yeast | NULL | Probable | As nectar-tap_saprotroph (Põlme et al. 2020); Decay sugar-rich substrates (Põlme et al. 2020) | Põlme S, et al. 2020. Fungal Diversity 105:1-16 ((https://doi.org/10.1007/s13225-020-00466-2)) |
| Otu07378 | 3 | k__Fungi(100);p__Zygomycota(100);c__Zygomycota_class_Incertae_sedis(100);o__Mucorales(100);f__Mucoraceae(100);g__Mucor(100);s__Mucor_circinelloides_f._lusitanicus(67); | Mucor | 13 | Saprotroph | \|Undefined Saprotroph\| | Microfungus | NULL | Probable | As soil_saprotroph (Põlme et al. 2020); Decay soil (Põlme et al. 2020) | Tedersoo L, et al. 2014. Science 346:e1256688 ((https://doi.org/10.1126/science.1256688)); Põlme S, et al. 2020. Fungal Diversity 105:1-16 ((https://doi.org/10.1007/s13225-020-00466-2)) |
| Otu07380 | 3 | k__Fungi(100);p__Ascomycota(100);c__Saccharomycetes(100);o__Saccharomycetales(100);f__Metschnikowiaceae(100);g__Metschnikowia(100);s__Metschnikowia_pulcherrima(67); | Metschnikowia | 13 | Saprotroph | \|Undefined Saprotroph\| | Yeast | NULL | Probable | As nectar-tap_saprotroph (Põlme et al. 2020); Decay sugar-rich substrates (Põlme et al. 2020) | Põlme S, et al. 2020. Fungal Diversity 105:1-16 ((https://doi.org/10.1007/s13225-020-00466-2)) |
| Otu07381 | 3 | k__Fungi(100);p__Ascomycota(100);c__Sordariomycetes(100);o__Sordariomycetes_order_Incertae_sedis(100);f__Annulatascaceae(100);g__Rhodoveronaea(100);s__Rhodoveronaea_varioseptata(100); | Rhodoveronaea | 13 | Saprotroph | \|Undefined Saprotroph\| | Microfungus | NULL | Probable | NULL | Tedersoo L, et al. 2014. Science 346:e1256688 ((https://doi.org/10.1126/science.1256688)); Põlme S, et al. 2020. Fungal Diversity 105:1-16 ((https://doi.org/10.1007/s13225-020-00466-2)) |
| Otu07382 | 3 | k__Fungi(100);p__Ascomycota(100);c__Saccharomycetes(100);o__Saccharomycetales(100);f__Metschnikowiaceae(100);g__Metschnikowia(100);g__Metschnikowia_unclassified(100); | Metschnikowia | 13 | Saprotroph | \|Undefined Saprotroph\| | Yeast | NULL | Probable | As nectar-tap_saprotroph (Põlme et al. 2020); Decay sugar-rich substrates (Põlme et al. 2020) | Põlme S, et al. 2020. Fungal Diversity 105:1-16 ((https://doi.org/10.1007/s13225-020-00466-2)) |
| Otu07385 | 3 | k__Fungi(100);p__Basidiomycota(100);c__Tremellomycetes(100);o__Cystofilobasidiales(100);f__Cystofilobasidiales_family_Incertae_sedis(100);g__Syzygospora(100);s__Syzygospora_sp(100); | Syzygospora | 13 | Pathotroph | \|Fungal Parasite\|-Lichen Parasite | Dimorphic-Tremelloid | NULL | Highly Probable | As mycoparasite (Põlme et al. 2020); Decay fungal material (Põlme et al. 2020) | Põlme S, et al. 2020. Fungal Diversity 105:1-16 ((https://doi.org/10.1007/s13225-020-00466-2)); List compiled by John Plischke III ((https://goodmushroom.weebly.com/uploads/5/7/1/7/57176535/fungi_on_fungi_pdf_lq.pdf)) |
| Otu07387 | 3 | k__Fungi(100);p__Basidiomycota(100);c__Agaricomycetes(100);o__Agaricales(100);f__Psathyrellaceae(100);g__Coprinellus(100);s__Coprinellus_micaceus(100); | Coprinellus | 13 | Saprotroph | \|Undefined Saprotroph\| | Agaricoid | NULL | Probable | As soil_saprotroph (Põlme et al. 2020); Decay soil (Põlme et al. 2020) | Tedersoo L, et al. 2014. Science 346:e1256688 ((https://doi.org/10.1126/science.1256688)); Põlme S, et al. 2020. Fungal Diversity 105:1-16 ((https://doi.org/10.1007/s13225-020-00466-2)) |
| Otu07392 | 3 | k__Fungi(100);p__Basidiomycota(100);c__Agaricomycetes(100);o__Polyporales(100);f__Meruliaceae(100);g__Crustoderma(100);s__Crustoderma_corneum(100); | Crustoderma | 13 | Saprotroph | \|Wood Saprotroph\| | Corticioid | Brown Rot | Probable | As brown-rot (Põlme et al. 2020); Decay wood (Põlme et al. 2020) | Tedersoo L, et al. 2014. Science 346:e1256688 ((https://doi.org/10.1126/science.1256688)); Põlme S, et al. 2020. Fungal Diversity 105:1-16 ((https://doi.org/10.1007/s13225-020-00466-2)) |
| Otu07393 | 3 | k__Fungi(100);p__Ascomycota(100);c__Sordariomycetes(100);o__Coniochaetales(100);f__Coniochaetaceae(100);f__Coniochaetaceae_unclassified(100);f__Coniochaetaceae_unclassified(100); | Coniochaetaceae | 9 | Pathotroph-Saprotroph-Symbiotroph | Animal Pathogen-Dung Saprotroph-Endophyte-Lichen Parasite-Plant Pathogen-Undefined Saprotroph | Microfungus | NULL | Possible | Saprobic on wood and bark (especially in dry habitats), coprophilous or isolated from soil and similar substrata (Cannon & Kirk 2007) | Cannon PF, Kirk PM. 2007. Fungal Families of the World. CAB International, Cambridge (ISBN: 978-0851998275); Damm U, et al. 2010. Persoonia 24:60-80 ((http://doi.org/10.3767/003158510X500705)); Irinyi L, et al. 2015. Medical Mycology 53:313-337 ((https://doi.org/10.1093/mmy/myv008)); Lawrey JD, Diederich P. 2016. http://www.lichenicolous.net (accessed 24 May 2017) ((http://www.lichenicolous.net)) |
| Otu07395 | 3 | k__Fungi(100);p__Basidiomycota(100);c__Tremellomycetes(100);o__Tremellales(100);f__Sirobasidiaceae(100);g__Sirobasidium(100);s__Sirobasidium_intermedium(100); | Sirobasidium | 13 | Pathotroph | \|Fungal Parasite\| | Dimorphic | NULL | Probable | As mycoparasite (Põlme et al. 2020); Decay fungal material (Põlme et al. 2020) | Põlme S, et al. 2020. Fungal Diversity 105:1-16 ((https://doi.org/10.1007/s13225-020-00466-2)) |
| Otu07397 | 3 | k__Fungi(100);p__Ascomycota(100);c__Saccharomycetes(100);o__Saccharomycetales(100);f__Metschnikowiaceae(100);g__Metschnikowia(100);s__Metschnikowia_pulcherrima(100); | Metschnikowia | 13 | Saprotroph | \|Undefined Saprotroph\| | Yeast | NULL | Probable | As nectar-tap_saprotroph (Põlme et al. 2020); Decay sugar-rich substrates (Põlme et al. 2020) | Põlme S, et al. 2020. Fungal Diversity 105:1-16 ((https://doi.org/10.1007/s13225-020-00466-2)) |
| Otu07398 | 3 | k__Fungi(100);p__Ascomycota(100);c__Saccharomycetes(100);o__Saccharomycetales(100);f__Metschnikowiaceae(100);g__Metschnikowia(100);s__Metschnikowia_pulcherrima(100); | Metschnikowia | 13 | Saprotroph | \|Undefined Saprotroph\| | Yeast | NULL | Probable | As nectar-tap_saprotroph (Põlme et al. 2020); Decay sugar-rich substrates (Põlme et al. 2020) | Põlme S, et al. 2020. Fungal Diversity 105:1-16 ((https://doi.org/10.1007/s13225-020-00466-2)) |
| Otu07399 | 3 | k__Fungi(100);p__Basidiomycota(100);c__Agaricomycetes(100);o__Polyporales(100);f__Phanerochaetaceae(100);g__Ceriporia(100);s__Ceriporia_bubalinomarginata(100); | Ceriporia | 13 | Saprotroph | \|Wood Saprotroph\| | Corticioid | White Rot | Highly Probable | As white_rot (Põlme et al. 2020); Decay wood (Põlme et al. 2020) | Tedersoo L, et al. 2014. Science 346:e1256688 ((https://doi.org/10.1126/science.1256688)); Põlme S, et al. 2020. Fungal Diversity 105:1-16 ((https://doi.org/10.1007/s13225-020-00466-2)) |
| Otu07403 | 3 | k__Fungi(100);p__Ascomycota(100);c__Saccharomycetes(100);o__Saccharomycetales(100);f__Metschnikowiaceae(100);g__Metschnikowia(100);s__Metschnikowia_pulcherrima(100); | Metschnikowia | 13 | Saprotroph | \|Undefined Saprotroph\| | Yeast | NULL | Probable | As nectar-tap_saprotroph (Põlme et al. 2020); Decay sugar-rich substrates (Põlme et al. 2020) | Põlme S, et al. 2020. Fungal Diversity 105:1-16 ((https://doi.org/10.1007/s13225-020-00466-2)) |
| Otu07404 | 3 | k__Fungi(100);p__Ascomycota(100);c__Saccharomycetes(100);o__Saccharomycetales(100);f__Metschnikowiaceae(100);g__Metschnikowia(100);s__Metschnikowia_pulcherrima(100); | Metschnikowia | 13 | Saprotroph | \|Undefined Saprotroph\| | Yeast | NULL | Probable | As nectar-tap_saprotroph (Põlme et al. 2020); Decay sugar-rich substrates (Põlme et al. 2020) | Põlme S, et al. 2020. Fungal Diversity 105:1-16 ((https://doi.org/10.1007/s13225-020-00466-2)) |
| Otu07406 | 3 | k__Fungi(100);p__Ascomycota(100);c__Sordariomycetes(100);o__Hypocreales(100);f__Nectriaceae(100);g__Fusarium(100);g__Fusarium_unclassified(100); | Fusarium | 13 | Pathotroph-Saprotroph-Symbiotroph | Animal Pathogen-Endophyte-Lichen Parasite-\|Plant Pathogen\|-Plant Saprotroph-Undefined Saprotroph-Wood Saprotroph | Microfungus | Soft Rot | Possible | As foliar_endophyte (Põlme et al. 2020); As litter_saprotroph (Põlme et al. 2020); Causing damping-off disease (Põlme et al. 2020); Decay leaf, fruit, seed, roots, soil, animal material (Põlme et al. 2020); Likely opportunistic human pathogen (Irinyi et al. 2015) | Duncan CG, Eslyn WE. 1966. Mycologia 58:642-664 ((https://doi.org/10.2307/3757045)); Seehann G, et al. 1975. List of Fungi in Soft-Rot Tissue Tests. Institute fur Holzbiologie und Holzschutz, Bundesforschungsanstalt fur Forst - und Holzwirtshaft, Hamburg - Lohbrugge, Germany; Costa IPMW, et al. 2012. Mycotaxon 119:e26 ((http://mycotaxon.com/resources/checklists/costa_v119_checklist.pdf)); Tedersoo L, et al. 2014. Science 346:e1256688 ((https://doi.org/10.1126/science.1256688)); Irinyi L, et al. 2015. Medical Mycology 53:313-337 ((https://doi.org/10.1093/mmy/myv008)); Schroers, et al. 2016. Mycologia 108:806-819 ((https://doi.org/10.3852/15-255)); Põlme S, et al. 2020. Fungal Diversity 105:1-16 ((https://doi.org/10.1007/s13225-020-00466-2)); The American Phytopathological Society Common Names of Plant Diseases ((http://www.apsnet.org/publications/commonnames/Pages/default.aspx)); Lawrey JD, Diederich P. 2016. http://www.lichenicolous.net (accessed 24 May 2017) ((http://www.lichenicolous.net)) |
| Otu07408 | 3 | k__Fungi(100);p__Ascomycota(100);c__Saccharomycetes(100);o__Saccharomycetales(100);f__Metschnikowiaceae(100);g__Metschnikowia(100);g__Metschnikowia_unclassified(67); | Metschnikowia | 13 | Saprotroph | \|Undefined Saprotroph\| | Yeast | NULL | Probable | As nectar-tap_saprotroph (Põlme et al. 2020); Decay sugar-rich substrates (Põlme et al. 2020) | Põlme S, et al. 2020. Fungal Diversity 105:1-16 ((https://doi.org/10.1007/s13225-020-00466-2)) |
| Otu07409 | 3 | k__Fungi(100);p__Basidiomycota(100);c__Microbotryomycetes(100);o__Sporidiobolales(100);f__Sporidiobolales_family_Incertae_sedis(100);g__Rhodotorula(100);s__Rhodotorula_minuta(100); | Rhodotorula | 13 | Pathotroph-Saprotroph-Symbiotroph | Animal Pathogen-Animal Symbiotroph-Endophyte-Plant Pathogen-\|Undefined Saprotroph\| | Yeast | NULL | Probable | As foliar_endophyte (Põlme et al. 2020); Opportunistic human pathogen | Põlme S, et al. 2020. Fungal Diversity 105:1-16 ((https://doi.org/10.1007/s13225-020-00466-2)); https://www.cdc.gov ((https://www.cdc.gov)) |
| Otu07410 | 3 | k__Fungi(100);p__Ascomycota(100);c__Saccharomycetes(100);o__Saccharomycetales(100);f__Metschnikowiaceae(100);g__Metschnikowia(100);g__Metschnikowia_unclassified(100); | Metschnikowia | 13 | Saprotroph | \|Undefined Saprotroph\| | Yeast | NULL | Probable | As nectar-tap_saprotroph (Põlme et al. 2020); Decay sugar-rich substrates (Põlme et al. 2020) | Põlme S, et al. 2020. Fungal Diversity 105:1-16 ((https://doi.org/10.1007/s13225-020-00466-2)) |
| Otu07411 | 3 | k__Fungi(100);p__Ascomycota(100);c__Saccharomycetes(100);o__Saccharomycetales(100);f__Metschnikowiaceae(100);g__Metschnikowia(100);s__Metschnikowia_pulcherrima(100); | Metschnikowia | 13 | Saprotroph | \|Undefined Saprotroph\| | Yeast | NULL | Probable | As nectar-tap_saprotroph (Põlme et al. 2020); Decay sugar-rich substrates (Põlme et al. 2020) | Põlme S, et al. 2020. Fungal Diversity 105:1-16 ((https://doi.org/10.1007/s13225-020-00466-2)) |
| Otu07413 | 3 | k__Fungi(100);p__Ascomycota(100);c__Sordariomycetes(100);o__Coniochaetales(100);f__Coniochaetaceae(100);g__Coniochaeta(67);s__Coniochaeta_sp(67); | Coniochaeta | 13 | Pathotroph-Saprotroph-Symbiotroph | Animal Pathogen-Dung Saprotroph-Endophyte-Lichen Parasite-Plant Pathogen-\|Undefined Saprotroph\| | Microfungus | NULL | Possible | As foliar_endophyte (Põlme et al. 2020); Decay leaf, fruit, seed, soil, dung, animal material (Põlme et al. 2020) | Bell A. 1983. Dung Fungi: An Illustrated Guide to Coprophilous Fungi in New Zealand. Victoria University Press, Wellington (ISBN: 978-0864730015); Damm U, et al. 2010. Persoonia 24:60-80 ((http://doi.org/10.3767/003158510X500705)); Costa IPMW, et al. 2012. Mycotaxon 119:e26 ((http://mycotaxon.com/resources/checklists/costa_v119_checklist.pdf)); Tedersoo L, et al. 2014. Science 346: 1256688 ((https://doi.org/:10.1126/science.1256688)); Irinyi L, et al. 2015. Medical Mycology 53:313-337 ((https://doi.org/10.1093/mmy/myv008)); Lawrey JD, Diederich P. 2016. http://www.lichenicolous.net (accessed 24 May 2017) ((http://www.lichenicolous.net)); Põlme S, et al. 2020. Fungal Diversity 105:1-16 ((https://doi.org/10.1007/s13225-020-00466-2)) |
| Otu07418 | 3 | k__Fungi(100);p__Ascomycota(100);c__Saccharomycetes(100);o__Saccharomycetales(100);f__Metschnikowiaceae(100);g__Metschnikowia(100);g__Metschnikowia_unclassified(67); | Metschnikowia | 13 | Saprotroph | \|Undefined Saprotroph\| | Yeast | NULL | Probable | As nectar-tap_saprotroph (Põlme et al. 2020); Decay sugar-rich substrates (Põlme et al. 2020) | Põlme S, et al. 2020. Fungal Diversity 105:1-16 ((https://doi.org/10.1007/s13225-020-00466-2)) |
| Otu07420 | 3 | k__Fungi(100);p__Basidiomycota(100);c__Tremellomycetes(100);o__Tremellales(100);f__Tremellales_family_Incertae_sedis(100);g__Cryptococcus(100);s__Cryptococcus_sp_D162_2(100); | Cryptococcus | 13 | Pathotroph-Saprotroph-Symbiotroph | Animal Pathogen-Endophyte-Epiphyte-\|Undefined Saprotroph\| | Dimorphic Yeast-Yeast | NULL | Possible | As foliar_endophyte (Põlme et al. 2020); Decay leaf, fruit, seed, soil, dung, animal material (Põlme et al. 2020) | Andrews JH, et al. 1983. Phytopathology 73:228-234 ((https://doi.org/10.1094/Phyto-73-228)); Kurtzman CP, et al. (eds.) 2011. The Yeasts, a Taxonomic Study. Fifth Edition. Vols 1-3. Elsevier, San Diego (ISBN: 9780444521491); Põlme S, et al. 2020. Fungal Diversity 105:1-16 ((https://doi.org/10.1007/s13225-020-00466-2)) |
| Otu07421 | 3 | k__Fungi(100);p__Basidiomycota(100);c__Agaricomycetes(100);o__Thelephorales(100);f__Thelephoraceae(100);g__Odontia(100);s__Odontia_fibrosa(100); | Odontia | 13 | Saprotroph | \|Plant Saprotroph\| | Corticioid | NULL | Probable | As litter_saprotroph (Põlme et al. 2020); Decay leaf, fruit, seed (Põlme et al. 2020) | Põlme S, et al. 2020. Fungal Diversity 105:1-16 ((https://doi.org/10.1007/s13225-020-00466-2)) |
| Otu07425 | 3 | k__Fungi(100);p__Ascomycota(100);c__Saccharomycetes(100);o__Saccharomycetales(100);f__Metschnikowiaceae(100);g__Metschnikowia(100);s__Metschnikowia_pulcherrima(67); | Metschnikowia | 13 | Saprotroph | \|Undefined Saprotroph\| | Yeast | NULL | Probable | As nectar-tap_saprotroph (Põlme et al. 2020); Decay sugar-rich substrates (Põlme et al. 2020) | Põlme S, et al. 2020. Fungal Diversity 105:1-16 ((https://doi.org/10.1007/s13225-020-00466-2)) |
| Otu07427 | 3 | k__Fungi(100);p__Ascomycota(100);c__Dothideomycetes(100);o__Capnodiales(100);f__Mycosphaerellaceae(100);g__Pseudocercosporella(100);s__Pseudocercosporella_sp(100); | Pseudocercosporella | 13 | Pathotroph | \|Plant Pathogen\| | Microfungus | NULL | Probable | NULL | Tedersoo L, et al. 2014. Science 346:e1256688 ((https://doi.org/10.1126/science.1256688)); Põlme S, et al. 2020. Fungal Diversity 105:1-16 ((https://doi.org/10.1007/s13225-020-00466-2)) |
| Otu07428 | 3 | k__Fungi(100);p__Ascomycota(100);c__Sordariomycetes(100);o__Hypocreales(100);o__Hypocreales_unclassified(100);o__Hypocreales_unclassified(100);o__Hypocreales_unclassified(100); | Hypocreales | 7 | Saprotroph | Undefined Saprotroph | Microfungus | NULL | Possible | NULL | Sterkenburg E, et al. 2015. New Phytologist 207:1145-1158 ((https://doi.org/10.1111/nph.13426)) |
| Otu07429 | 3 | k__Fungi(100);p__Ascomycota(100);c__Saccharomycetes(100);o__Saccharomycetales(100);f__Metschnikowiaceae(100);g__Metschnikowia(100);s__Metschnikowia_pulcherrima(100); | Metschnikowia | 13 | Saprotroph | \|Undefined Saprotroph\| | Yeast | NULL | Probable | As nectar-tap_saprotroph (Põlme et al. 2020); Decay sugar-rich substrates (Põlme et al. 2020) | Põlme S, et al. 2020. Fungal Diversity 105:1-16 ((https://doi.org/10.1007/s13225-020-00466-2)) |
| Otu07431 | 3 | k__Fungi(100);p__Ascomycota(100);c__Saccharomycetes(100);o__Saccharomycetales(100);f__Metschnikowiaceae(100);g__Metschnikowia(100);g__Metschnikowia_unclassified(67); | Metschnikowia | 13 | Saprotroph | \|Undefined Saprotroph\| | Yeast | NULL | Probable | As nectar-tap_saprotroph (Põlme et al. 2020); Decay sugar-rich substrates (Põlme et al. 2020) | Põlme S, et al. 2020. Fungal Diversity 105:1-16 ((https://doi.org/10.1007/s13225-020-00466-2)) |
| Otu07432 | 3 | k__Fungi(100);p__Ascomycota(100);c__Saccharomycetes(100);o__Saccharomycetales(100);f__Metschnikowiaceae(100);g__Metschnikowia(100);s__Metschnikowia_pulcherrima(100); | Metschnikowia | 13 | Saprotroph | \|Undefined Saprotroph\| | Yeast | NULL | Probable | As nectar-tap_saprotroph (Põlme et al. 2020); Decay sugar-rich substrates (Põlme et al. 2020) | Põlme S, et al. 2020. Fungal Diversity 105:1-16 ((https://doi.org/10.1007/s13225-020-00466-2)) |
| Otu07433 | 3 | k__Fungi(100);p__Basidiomycota(100);c__Agaricomycetes(100);o__Agaricales(100);f__Marasmiaceae(100);f__Marasmiaceae_unclassified(100);f__Marasmiaceae_unclassified(100); | Marasmiaceae | 9 | Pathotroph-Saprotroph-Symbiotroph | Endophyte-Plant Pathogen-Undefined Saprotroph | Agaricoid | NULL | Probable | Species in Moniliophthora are devastating pathogens of tree crops, causing witches broom disease; Saprobic on litter and dead wood, or parasitic on woody plants - often one of the most prominent fungal components of tropical forests, playing a major role in nutrient cycling (Cannon & Kirk 2007) | Cannon PF, Kirk PM. 2007. Fungal Families of the World. CAB International, Cambridge (ISBN: 978-0851998275); David AS, et al. 2016. Microbial Ecology 71:912-926 ((https://doi.org/10.1007/s00248-015-0712-6)) |
| Otu07434 | 3 | k__Fungi(100);p__Basidiomycota(100);c__Agaricomycetes(100);o__Agaricales(100);f__Agaricaceae(100);g__Langermannia(100);s__Langermannia_gigantea(100); | Langermannia | 13 | Saprotroph | \|Undefined Saprotroph\| | Gasteroid | NULL | Probable | As soil_saprotroph (Põlme et al. 2020); Decay soil (Põlme et al. 2020) | Tedersoo L, et al. 2014. Science 346:e1256688 ((https://doi.org/10.1126/science.1256688)); Põlme S, et al. 2020. Fungal Diversity 105:1-16 ((https://doi.org/10.1007/s13225-020-00466-2)) |
| Otu07435 | 3 | k__Fungi(100);p__Chytridiomycota(67);c__Chytridiomycetes(67);o__Rhizophlyctidales(67);f__Rhizophlyctidaceae(67);g__Rhizophlyctis(67);s__Rhizophlyctis_rosea(67); | Rhizophlyctis | 13 | Pathotroph-Saprotroph | Algal Parasite-\|Plant Saprotroph\|-Undefined Saprotroph | Chytrid-Microfungus | NULL | Probable | As litter_saprotroph (Põlme et al. 2020); Decay leaf, fruit, seed, algal material (Põlme et al. 2020) | James TY, et al. 2006. Nature 443:818-822 ((https://doi.org/10.1038/nature05110)); Põlme S, et al. 2020. Fungal Diversity 105:1-16 ((https://doi.org/10.1007/s13225-020-00466-2)) |
| Otu07436 | 3 | k__Fungi(100);p__Ascomycota(100);c__Saccharomycetes(100);o__Saccharomycetales(100);f__Metschnikowiaceae(100);g__Metschnikowia(100);g__Metschnikowia_unclassified(100); | Metschnikowia | 13 | Saprotroph | \|Undefined Saprotroph\| | Yeast | NULL | Probable | As nectar-tap_saprotroph (Põlme et al. 2020); Decay sugar-rich substrates (Põlme et al. 2020) | Põlme S, et al. 2020. Fungal Diversity 105:1-16 ((https://doi.org/10.1007/s13225-020-00466-2)) |
| Otu07437 | 3 | k__Fungi(100);p__Ascomycota(100);c__Saccharomycetes(100);o__Saccharomycetales(100);f__Metschnikowiaceae(100);g__Metschnikowia(100);g__Metschnikowia_unclassified(100); | Metschnikowia | 13 | Saprotroph | \|Undefined Saprotroph\| | Yeast | NULL | Probable | As nectar-tap_saprotroph (Põlme et al. 2020); Decay sugar-rich substrates (Põlme et al. 2020) | Põlme S, et al. 2020. Fungal Diversity 105:1-16 ((https://doi.org/10.1007/s13225-020-00466-2)) |
| Otu07439 | 3 | k__Fungi(100);p__Ascomycota(100);c__Saccharomycetes(100);o__Saccharomycetales(100);f__Metschnikowiaceae(100);g__Metschnikowia(100);g__Metschnikowia_unclassified(67); | Metschnikowia | 13 | Saprotroph | \|Undefined Saprotroph\| | Yeast | NULL | Probable | As nectar-tap_saprotroph (Põlme et al. 2020); Decay sugar-rich substrates (Põlme et al. 2020) | Põlme S, et al. 2020. Fungal Diversity 105:1-16 ((https://doi.org/10.1007/s13225-020-00466-2)) |
| Otu07440 | 3 | k__Fungi(100);p__Zygomycota(100);c__Zygomycota_class_Incertae_sedis(100);o__Basidiobolales(100);f__Basidiobolaceae(100);g__Basidiobolus(100);s__Basidiobolus_ranarum(100); | Basidiobolus | 13 | Pathotroph-Saprotroph-Symbiotroph | Animal Pathogen-Animal Symbiotroph-\|Undefined Saprotroph\| | Microfungus | NULL | Probable | Cause of subcutaneous zygomycosis in humans (Cannon & Kirk 2007); Basidiobolus ranarum is commonly present in decaying fruit and vegetable matter, and as a commensal in the intestinal tract of frogs, toads, and lizards ; Decay leaf, fruit, seed, soil, dung, animal material (Põlme et al. 2020) | Cannon PF, Kirk PM. 2007. Fungal Families of the World. CAB International, Cambridge (ISBN: 978-0851998275); Tedersoo L, et al. 2014. Science 346:e1256688 ((https://doi.org/10.1126/science.1256688)); Põlme S, et al. 2020. Fungal Diversity 105:1-16 ((https://doi.org/10.1007/s13225-020-00466-2)) |
| Otu07444 | 3 | k__Fungi(100);p__Ascomycota(100);c__Saccharomycetes(100);o__Saccharomycetales(100);f__Metschnikowiaceae(100);g__Metschnikowia(100);s__Metschnikowia_pulcherrima(100); | Metschnikowia | 13 | Saprotroph | \|Undefined Saprotroph\| | Yeast | NULL | Probable | As nectar-tap_saprotroph (Põlme et al. 2020); Decay sugar-rich substrates (Põlme et al. 2020) | Põlme S, et al. 2020. Fungal Diversity 105:1-16 ((https://doi.org/10.1007/s13225-020-00466-2)) |
| Otu07446 | 3 | k__Fungi(100);p__Ascomycota(100);c__Saccharomycetes(100);o__Saccharomycetales(100);f__Metschnikowiaceae(100);g__Metschnikowia(100);s__Metschnikowia_pulcherrima(100); | Metschnikowia | 13 | Saprotroph | \|Undefined Saprotroph\| | Yeast | NULL | Probable | As nectar-tap_saprotroph (Põlme et al. 2020); Decay sugar-rich substrates (Põlme et al. 2020) | Põlme S, et al. 2020. Fungal Diversity 105:1-16 ((https://doi.org/10.1007/s13225-020-00466-2)) |
| Otu07448 | 3 | k__Fungi(100);p__Ascomycota(100);c__Saccharomycetes(100);o__Saccharomycetales(100);f__Metschnikowiaceae(100);g__Metschnikowia(100);s__Metschnikowia_pulcherrima(67); | Metschnikowia | 13 | Saprotroph | \|Undefined Saprotroph\| | Yeast | NULL | Probable | As nectar-tap_saprotroph (Põlme et al. 2020); Decay sugar-rich substrates (Põlme et al. 2020) | Põlme S, et al. 2020. Fungal Diversity 105:1-16 ((https://doi.org/10.1007/s13225-020-00466-2)) |
| Otu07451 | 3 | k__Fungi(100);p__Ascomycota(100);c__Saccharomycetes(100);o__Saccharomycetales(100);f__Metschnikowiaceae(100);g__Metschnikowia(100);s__Metschnikowia_agaves(100); | Metschnikowia | 13 | Saprotroph | \|Undefined Saprotroph\| | Yeast | NULL | Probable | As nectar-tap_saprotroph (Põlme et al. 2020); Decay sugar-rich substrates (Põlme et al. 2020) | Põlme S, et al. 2020. Fungal Diversity 105:1-16 ((https://doi.org/10.1007/s13225-020-00466-2)) |
| Otu07452 | 3 | k__Fungi(100);p__Ascomycota(100);c__Saccharomycetes(100);o__Saccharomycetales(100);f__Metschnikowiaceae(100);g__Metschnikowia(100);s__Metschnikowia_pulcherrima(67); | Metschnikowia | 13 | Saprotroph | \|Undefined Saprotroph\| | Yeast | NULL | Probable | As nectar-tap_saprotroph (Põlme et al. 2020); Decay sugar-rich substrates (Põlme et al. 2020) | Põlme S, et al. 2020. Fungal Diversity 105:1-16 ((https://doi.org/10.1007/s13225-020-00466-2)) |
| Otu07453 | 3 | k__Fungi(100);p__Ascomycota(100);c__Saccharomycetes(100);o__Saccharomycetales(100);f__Metschnikowiaceae(100);g__Metschnikowia(100);g__Metschnikowia_unclassified(67); | Metschnikowia | 13 | Saprotroph | \|Undefined Saprotroph\| | Yeast | NULL | Probable | As nectar-tap_saprotroph (Põlme et al. 2020); Decay sugar-rich substrates (Põlme et al. 2020) | Põlme S, et al. 2020. Fungal Diversity 105:1-16 ((https://doi.org/10.1007/s13225-020-00466-2)) |
| Otu07456 | 3 | k__Fungi(100);p__Basidiomycota(100);c__Agaricomycetes(100);o__Boletales(100);f__Suillaceae(100);g__Suillus(100);s__Suillus_flavidus(100); | Suillus | 13 | Symbiotroph | \|Ectomycorrhizal\| | Agaricoid-Boletoid | NULL | Highly Probable | Host - Pinaceae (Põlme et al. 2020) | Rinaldi AC, et al. 2008. Fungal Diversity 33:1-45 ((http://www.fungaldiversity.org/fdp/sfdp/33-1.pdf)); Tedersoo L, et al. 2010. Mycorrhiza 20:217-263 ((https://doi.org/10.1007/s00572-009-0274-x)); Põlme S, et al. 2020. Fungal Diversity 105:1-16 ((https://doi.org/10.1007/s13225-020-00466-2)); http://mycorrhizas.info/ecmf.html ((http://mycorrhizas.info/ecmf.html)) |
| Otu07458 | 3 | k__Fungi(100);p__Ascomycota(100);c__Pezizomycetes(100);o__Pezizales(100);f__Tuberaceae(100);g__Tuber(100);g__Tuber_unclassified(100); | Tuber | 13 | Symbiotroph | \|Ectomycorrhizal\| | Gasteroid | Hypogeous | Highly Probable | true truffle; Common name | Rinaldi AC, et al. 2008. Fungal Diversity 33:1-45 ((http://www.fungaldiversity.org/fdp/sfdp/33-1.pdf)); Tedersoo L, et al. 2010. Mycorrhiza 20:217-263 ((https://doi.org/10.1007/s00572-009-0274-x)); Põlme S, et al. 2020. Fungal Diversity 105:1-16 ((https://doi.org/10.1007/s13225-020-00466-2)); http://mycorrhizas.info/ecmf.html ((http://mycorrhizas.info/ecmf.html)) |
| Otu07460 | 3 | k__Fungi(100);p__Ascomycota(100);c__Saccharomycetes(100);o__Saccharomycetales(100);f__Metschnikowiaceae(100);g__Metschnikowia(100);s__Metschnikowia_pulcherrima(100); | Metschnikowia | 13 | Saprotroph | \|Undefined Saprotroph\| | Yeast | NULL | Probable | As nectar-tap_saprotroph (Põlme et al. 2020); Decay sugar-rich substrates (Põlme et al. 2020) | Põlme S, et al. 2020. Fungal Diversity 105:1-16 ((https://doi.org/10.1007/s13225-020-00466-2)) |
| Otu07462 | 3 | k__Fungi(100);p__Basidiomycota(100);c__Agaricomycetes(100);o__Russulales(100);f__Russulaceae(100);g__Russula(100);g__Russula_unclassified(100); | Russula | 13 | Symbiotroph | \|Ectomycorrhizal\| | Agaricoid | NULL | Highly Probable | NULL | Rinaldi AC, et al. 2008. Fungal Diversity 33:1-45 ((http://www.fungaldiversity.org/fdp/sfdp/33-1.pdf)); Tedersoo L, et al. 2010. Mycorrhiza 20:217-263 ((https://doi.org/10.1007/s00572-009-0274-x)); Põlme S, et al. 2020. Fungal Diversity 105:1-16 ((https://doi.org/10.1007/s13225-020-00466-2)); http://mycorrhizas.info/ecmf.html ((http://mycorrhizas.info/ecmf.html)) |
| Otu07463 | 3 | k__Fungi(100);p__Ascomycota(100);c__Eurotiomycetes(100);o__Pyrenulales(100);f__Massariaceae(100);g__Massaria(100);s__Massaria_pyri(100); | Massaria | 13 | Saprotroph | Undefined Saprotroph-\|Wood Saprotroph\| | Microfungus | NULL | Probable | Decay wood (Põlme et al. 2020) | Tedersoo L, et al. 2014. Science 346:e1256688 ((https://doi.org/10.1126/science.1256688)); Põlme S, et al. 2020. Fungal Diversity 105:1-16 ((https://doi.org/10.1007/s13225-020-00466-2)) |
| Otu07464 | 3 | k__Fungi(100);p__Ascomycota(100);c__Saccharomycetes(100);o__Saccharomycetales(100);f__Metschnikowiaceae(100);g__Metschnikowia(100);s__Metschnikowia_pulcherrima(67); | Metschnikowia | 13 | Saprotroph | \|Undefined Saprotroph\| | Yeast | NULL | Probable | As nectar-tap_saprotroph (Põlme et al. 2020); Decay sugar-rich substrates (Põlme et al. 2020) | Põlme S, et al. 2020. Fungal Diversity 105:1-16 ((https://doi.org/10.1007/s13225-020-00466-2)) |
| Otu07467 | 3 | k__Fungi(100);p__Basidiomycota(100);c__Agaricomycetes(100);o__Russulales(100);f__Amylostereaceae(100);g__Amylostereum(100);s__Amylostereum_chailletii(100); | Amylostereum | 13 | Saprotroph | Undefined Saprotroph-\|Wood Saprotroph\| | Corticioid | White Rot | Probable | As white_rot (Põlme et al. 2020); Decay wood (Põlme et al. 2020) | Tedersoo L, et al. 2014. Science 346:e1256688 ((https://doi.org/10.1126/science.1256688)); Põlme S, et al. 2020. Fungal Diversity 105:1-16 ((https://doi.org/10.1007/s13225-020-00466-2)) |
| Otu07468 | 3 | k__Fungi(100);p__Ascomycota(100);c__Saccharomycetes(100);o__Saccharomycetales(100);f__Metschnikowiaceae(100);g__Metschnikowia(100);s__Metschnikowia_pulcherrima(100); | Metschnikowia | 13 | Saprotroph | \|Undefined Saprotroph\| | Yeast | NULL | Probable | As nectar-tap_saprotroph (Põlme et al. 2020); Decay sugar-rich substrates (Põlme et al. 2020) | Põlme S, et al. 2020. Fungal Diversity 105:1-16 ((https://doi.org/10.1007/s13225-020-00466-2)) |
| Otu07469 | 3 | k__Fungi(100);p__Ascomycota(100);c__Saccharomycetes(100);o__Saccharomycetales(100);f__Metschnikowiaceae(100);g__Metschnikowia(100);s__Metschnikowia_pulcherrima(100); | Metschnikowia | 13 | Saprotroph | \|Undefined Saprotroph\| | Yeast | NULL | Probable | As nectar-tap_saprotroph (Põlme et al. 2020); Decay sugar-rich substrates (Põlme et al. 2020) | Põlme S, et al. 2020. Fungal Diversity 105:1-16 ((https://doi.org/10.1007/s13225-020-00466-2)) |
| Otu07470 | 3 | k__Fungi(100);p__Ascomycota(100);c__Saccharomycetes(100);o__Saccharomycetales(100);f__Metschnikowiaceae(100);g__Metschnikowia(100);s__Metschnikowia_pulcherrima(100); | Metschnikowia | 13 | Saprotroph | \|Undefined Saprotroph\| | Yeast | NULL | Probable | As nectar-tap_saprotroph (Põlme et al. 2020); Decay sugar-rich substrates (Põlme et al. 2020) | Põlme S, et al. 2020. Fungal Diversity 105:1-16 ((https://doi.org/10.1007/s13225-020-00466-2)) |
| Otu07471 | 3 | k__Fungi(100);p__Basidiomycota(100);c__Agaricomycetes(100);o__Atheliales(100);f__Atheliaceae(100);g__Piloderma(100);s__Piloderma_lanatum(100); | Piloderma | 13 | Symbiotroph | \|Ectomycorrhizal\| | Corticioid | NULL | Highly Probable | NULL | Rinaldi AC, et al. 2008. Fungal Diversity 33:1-45 ((http://www.fungaldiversity.org/fdp/sfdp/33-1.pdf)); Tedersoo L, et al. 2010. Mycorrhiza 20:217-263 ((https://doi.org/10.1007/s00572-009-0274-x)); Põlme S, et al. 2020. Fungal Diversity 105:1-16 ((https://doi.org/10.1007/s13225-020-00466-2)); http://mycorrhizas.info/ecmf.html ((http://mycorrhizas.info/ecmf.html)) |
| Otu07474 | 3 | k__Fungi(100);p__Ascomycota(100);c__Saccharomycetes(100);o__Saccharomycetales(100);f__Metschnikowiaceae(100);g__Metschnikowia(100);s__Metschnikowia_pulcherrima(67); | Metschnikowia | 13 | Saprotroph | \|Undefined Saprotroph\| | Yeast | NULL | Probable | As nectar-tap_saprotroph (Põlme et al. 2020); Decay sugar-rich substrates (Põlme et al. 2020) | Põlme S, et al. 2020. Fungal Diversity 105:1-16 ((https://doi.org/10.1007/s13225-020-00466-2)) |
| Otu07477 | 3 | k__Fungi(100);p__Ascomycota(100);c__Saccharomycetes(100);o__Saccharomycetales(100);f__Metschnikowiaceae(100);g__Metschnikowia(100);g__Metschnikowia_unclassified(67); | Metschnikowia | 13 | Saprotroph | \|Undefined Saprotroph\| | Yeast | NULL | Probable | As nectar-tap_saprotroph (Põlme et al. 2020); Decay sugar-rich substrates (Põlme et al. 2020) | Põlme S, et al. 2020. Fungal Diversity 105:1-16 ((https://doi.org/10.1007/s13225-020-00466-2)) |
| Otu07480 | 3 | k__Fungi(100);p__Ascomycota(100);c__Saccharomycetes(100);o__Saccharomycetales(100);f__Metschnikowiaceae(100);g__Metschnikowia(100);g__Metschnikowia_unclassified(100); | Metschnikowia | 13 | Saprotroph | \|Undefined Saprotroph\| | Yeast | NULL | Probable | As nectar-tap_saprotroph (Põlme et al. 2020); Decay sugar-rich substrates (Põlme et al. 2020) | Põlme S, et al. 2020. Fungal Diversity 105:1-16 ((https://doi.org/10.1007/s13225-020-00466-2)) |
| Otu07481 | 3 | k__Fungi(100);p__Basidiomycota(100);c__Microbotryomycetes(100);o__Sporidiobolales(100);f__Sporidiobolales_family_Incertae_sedis(100);g__Rhodotorula(100);g__Rhodotorula_unclassified(100); | Rhodotorula | 13 | Pathotroph-Saprotroph-Symbiotroph | Animal Pathogen-Animal Symbiotroph-Endophyte-Plant Pathogen-\|Undefined Saprotroph\| | Yeast | NULL | Probable | As foliar_endophyte (Põlme et al. 2020); Opportunistic human pathogen | Põlme S, et al. 2020. Fungal Diversity 105:1-16 ((https://doi.org/10.1007/s13225-020-00466-2)); https://www.cdc.gov ((https://www.cdc.gov)) |
| Otu07482 | 3 | k__Fungi(100);p__Ascomycota(100);c__Saccharomycetes(100);o__Saccharomycetales(100);f__Metschnikowiaceae(100);g__Metschnikowia(100);g__Metschnikowia_unclassified(67); | Metschnikowia | 13 | Saprotroph | \|Undefined Saprotroph\| | Yeast | NULL | Probable | As nectar-tap_saprotroph (Põlme et al. 2020); Decay sugar-rich substrates (Põlme et al. 2020) | Põlme S, et al. 2020. Fungal Diversity 105:1-16 ((https://doi.org/10.1007/s13225-020-00466-2)) |
| Otu07483 | 3 | k__Fungi(100);p__Ascomycota(100);c__Saccharomycetes(100);o__Saccharomycetales(100);f__Metschnikowiaceae(100);g__Metschnikowia(100);s__Metschnikowia_pulcherrima(100); | Metschnikowia | 13 | Saprotroph | \|Undefined Saprotroph\| | Yeast | NULL | Probable | As nectar-tap_saprotroph (Põlme et al. 2020); Decay sugar-rich substrates (Põlme et al. 2020) | Põlme S, et al. 2020. Fungal Diversity 105:1-16 ((https://doi.org/10.1007/s13225-020-00466-2)) |
| Otu07486 | 3 | k__Fungi(100);p__Ascomycota(100);c__Saccharomycetes(100);o__Saccharomycetales(100);f__Metschnikowiaceae(100);g__Metschnikowia(100);s__Metschnikowia_pulcherrima(67); | Metschnikowia | 13 | Saprotroph | \|Undefined Saprotroph\| | Yeast | NULL | Probable | As nectar-tap_saprotroph (Põlme et al. 2020); Decay sugar-rich substrates (Põlme et al. 2020) | Põlme S, et al. 2020. Fungal Diversity 105:1-16 ((https://doi.org/10.1007/s13225-020-00466-2)) |
| Otu07487 | 3 | k__Fungi(100);p__Ascomycota(100);c__Saccharomycetes(100);o__Saccharomycetales(100);f__Saccharomycetales_family_Incertae_sedis(100);g__Candida(100);g__Candida_unclassified(100); | Candida | 13 | Pathotroph-Saprotroph-Symbiotroph | Animal Pathogen-Endophyte-Endosymbiont-Epiphyte-\|Undefined Saprotroph\| | Dimorphic Yeast-Yeast | NULL | Probable | As nectar-tap_saprotroph (Põlme et al. 2020); Decay sugar-rich substrates (Põlme et al. 2020); This asexual genus is polyphyletic and contain species that function very differently depend on what they are. Caution should be taken when interpreting guilds for Candida species. | Manolakaki D, et al. 2010. Virulence. 1: 367-375 ((https://doi.org/10.4161/viru.1.5.12796)); Mekbib SB, et al. 2011. Tropical Plant Pathology 36:233-240 ((https://doi.org/10.1590/S1982-56762011000400004)); Põlme S, et al. 2020. Fungal Diversity 105:1-16 ((https://doi.org/10.1007/s13225-020-00466-2)) |
| Otu07489 | 3 | k__Fungi(100);p__Basidiomycota(100);c__Agaricomycetes(100);o__Cantharellales(100);f__Ceratobasidiaceae(100);g__Ceratobasidium(100);s__Ceratobasidium_sp_B545cShBmS2(67); | Ceratobasidium | 13 | Pathotroph-Saprotroph-Symbiotroph | Ectomycorrhizal-Endomycorrhizal-Orchid Mycorrhizal-\|Plant Pathogen\|-Plant Saprotroph-Undefined Saprotroph | Corticioid-Microfungus | NULL | Possible | especially of Orchidaceae but also even bryophytes (Cannon & Kirk 2007); As litter_saprotroph (Põlme et al. 2020); Best known as plant pathogens, but also saprobic. Species are endomycorrhizal with a wide range of plant roots ; Decay leaf, fruit, seed, roots, soil (Põlme et al. 2020) | Cannon PF, Kirk PM. 2007. Fungal Families of the World. CAB International, Cambridge (ISBN: 978-0851998275); Tedersoo L, et al. 2014. Science 346:e1256688 ((https://doi.org/10.1126/science.1256688)); Põlme S, et al. 2020. Fungal Diversity 105:1-16 ((https://doi.org/10.1007/s13225-020-00466-2)) |
| Otu07493 | 3 | k__Fungi(100);p__Ascomycota(100);c__Saccharomycetes(100);o__Saccharomycetales(100);f__Metschnikowiaceae(100);g__Metschnikowia(100);s__Metschnikowia_pulcherrima(100); | Metschnikowia | 13 | Saprotroph | \|Undefined Saprotroph\| | Yeast | NULL | Probable | As nectar-tap_saprotroph (Põlme et al. 2020); Decay sugar-rich substrates (Põlme et al. 2020) | Põlme S, et al. 2020. Fungal Diversity 105:1-16 ((https://doi.org/10.1007/s13225-020-00466-2)) |
| Otu07494 | 3 | k__Fungi(100);p__Ascomycota(100);c__Saccharomycetes(100);o__Saccharomycetales(100);f__Metschnikowiaceae(100);g__Metschnikowia(100);s__Metschnikowia_pulcherrima(100); | Metschnikowia | 13 | Saprotroph | \|Undefined Saprotroph\| | Yeast | NULL | Probable | As nectar-tap_saprotroph (Põlme et al. 2020); Decay sugar-rich substrates (Põlme et al. 2020) | Põlme S, et al. 2020. Fungal Diversity 105:1-16 ((https://doi.org/10.1007/s13225-020-00466-2)) |
| Otu07495 | 3 | k__Fungi(100);p__Ascomycota(100);c__Saccharomycetes(100);o__Saccharomycetales(100);f__Metschnikowiaceae(100);g__Metschnikowia(100);s__Metschnikowia_pulcherrima(100); | Metschnikowia | 13 | Saprotroph | \|Undefined Saprotroph\| | Yeast | NULL | Probable | As nectar-tap_saprotroph (Põlme et al. 2020); Decay sugar-rich substrates (Põlme et al. 2020) | Põlme S, et al. 2020. Fungal Diversity 105:1-16 ((https://doi.org/10.1007/s13225-020-00466-2)) |
| Otu07498 | 3 | k__Fungi(100);p__Ascomycota(100);c__Dothideomycetes(100);o__Pleosporales(100);f__Venturiaceae(100);g__Fusicladium(100);s__Fusicladium_caryigenum(100); | Fusicladium | 13 | Pathotroph-Symbiotroph | Endophyte-Lichen Parasite-\|Plant Pathogen\| | Microfungus | NULL | Probable | NULL | Tedersoo L, et al. 2014. Science 346:e1256688 ((https://doi.org/10.1126/science.1256688)); Põlme S, et al. 2020. Fungal Diversity 105:1-16 ((https://doi.org/10.1007/s13225-020-00466-2)) |
| Otu07499 | 3 | k__Fungi(100);p__Ascomycota(100);c__Dothideomycetes(100);o__Pleosporales(100);f__Venturiaceae(100);g__Fusicladium(100);s__Fusicladium_caryigenum(100); | Fusicladium | 13 | Pathotroph-Symbiotroph | Endophyte-Lichen Parasite-\|Plant Pathogen\| | Microfungus | NULL | Probable | NULL | Tedersoo L, et al. 2014. Science 346:e1256688 ((https://doi.org/10.1126/science.1256688)); Põlme S, et al. 2020. Fungal Diversity 105:1-16 ((https://doi.org/10.1007/s13225-020-00466-2)) |
| Otu07502 | 3 | k__Fungi(100);p__Ascomycota(100);c__Saccharomycetes(100);o__Saccharomycetales(100);f__Metschnikowiaceae(100);g__Metschnikowia(100);s__Metschnikowia_pulcherrima(67); | Metschnikowia | 13 | Saprotroph | \|Undefined Saprotroph\| | Yeast | NULL | Probable | As nectar-tap_saprotroph (Põlme et al. 2020); Decay sugar-rich substrates (Põlme et al. 2020) | Põlme S, et al. 2020. Fungal Diversity 105:1-16 ((https://doi.org/10.1007/s13225-020-00466-2)) |
| Otu07503 | 3 | k__Fungi(100);p__Ascomycota(100);c__Saccharomycetes(100);o__Saccharomycetales(100);f__Metschnikowiaceae(100);g__Metschnikowia(100);s__Metschnikowia_pulcherrima(67); | Metschnikowia | 13 | Saprotroph | \|Undefined Saprotroph\| | Yeast | NULL | Probable | As nectar-tap_saprotroph (Põlme et al. 2020); Decay sugar-rich substrates (Põlme et al. 2020) | Põlme S, et al. 2020. Fungal Diversity 105:1-16 ((https://doi.org/10.1007/s13225-020-00466-2)) |
| Otu07504 | 3 | k__Fungi(100);p__Ascomycota(100);c__Saccharomycetes(100);o__Saccharomycetales(100);f__Metschnikowiaceae(100);g__Metschnikowia(100);s__Metschnikowia_pulcherrima(67); | Metschnikowia | 13 | Saprotroph | \|Undefined Saprotroph\| | Yeast | NULL | Probable | As nectar-tap_saprotroph (Põlme et al. 2020); Decay sugar-rich substrates (Põlme et al. 2020) | Põlme S, et al. 2020. Fungal Diversity 105:1-16 ((https://doi.org/10.1007/s13225-020-00466-2)) |
| Otu07505 | 3 | k__Fungi(100);p__Basidiomycota(100);c__Agaricomycetes(100);o__Russulales(100);f__Russulaceae(100);g__Russula(100);s__Russula_xerampelina(100); | Russula | 13 | Symbiotroph | \|Ectomycorrhizal\| | Agaricoid | NULL | Highly Probable | NULL | Rinaldi AC, et al. 2008. Fungal Diversity 33:1-45 ((http://www.fungaldiversity.org/fdp/sfdp/33-1.pdf)); Tedersoo L, et al. 2010. Mycorrhiza 20:217-263 ((https://doi.org/10.1007/s00572-009-0274-x)); Põlme S, et al. 2020. Fungal Diversity 105:1-16 ((https://doi.org/10.1007/s13225-020-00466-2)); http://mycorrhizas.info/ecmf.html ((http://mycorrhizas.info/ecmf.html)) |
| Otu07506 | 3 | k__Fungi(100);p__Ascomycota(100);c__Saccharomycetes(100);o__Saccharomycetales(100);f__Metschnikowiaceae(100);g__Metschnikowia(100);g__Metschnikowia_unclassified(100); | Metschnikowia | 13 | Saprotroph | \|Undefined Saprotroph\| | Yeast | NULL | Probable | As nectar-tap_saprotroph (Põlme et al. 2020); Decay sugar-rich substrates (Põlme et al. 2020) | Põlme S, et al. 2020. Fungal Diversity 105:1-16 ((https://doi.org/10.1007/s13225-020-00466-2)) |
| Otu07507 | 3 | k__Fungi(100);p__Ascomycota(100);c__Saccharomycetes(100);o__Saccharomycetales(100);f__Metschnikowiaceae(100);g__Metschnikowia(100);g__Metschnikowia_unclassified(67); | Metschnikowia | 13 | Saprotroph | \|Undefined Saprotroph\| | Yeast | NULL | Probable | As nectar-tap_saprotroph (Põlme et al. 2020); Decay sugar-rich substrates (Põlme et al. 2020) | Põlme S, et al. 2020. Fungal Diversity 105:1-16 ((https://doi.org/10.1007/s13225-020-00466-2)) |
| Otu07509 | 3 | k__Fungi(100);p__Basidiomycota(100);c__Agaricomycetes(100);o__Russulales(100);f__Peniophoraceae(100);g__Peniophora(67);g__Peniophora_unclassified(67); | Peniophora | 13 | Pathotroph-Saprotroph | Plant Pathogen-\|Wood Saprotroph\| | Corticioid | White Rot | Probable | As white_rot (Põlme et al. 2020); As wood_pathogen (Põlme et al. 2020); Decay wood (Põlme et al. 2020) | Tedersoo L, et al. 2014. Science 346:e1256688 ((https://doi.org/10.1126/science.1256688)); Põlme S, et al. 2020. Fungal Diversity 105:1-16 ((https://doi.org/10.1007/s13225-020-00466-2)) |
| Otu07510 | 3 | k__Fungi(100);p__Ascomycota(100);c__Pezizomycetes(100);o__Pezizales(100);f__Tuberaceae(100);f__Tuberaceae_unclassified(100);f__Tuberaceae_unclassified(100); | Tuberaceae | 9 | Symbiotroph | Ectomycorrhizal | Gasteroid | Hypogeous | Probable | Common name - true truffle; Mycorrhizal with broadleaved trees, growing in the surface layer of soil and closely associated with roots. Highly specific pheromone-like chemicals are produced that play a role in dispersal by attracting mammals that uproot fruit bodies and ditribute spores (Cannon & Kirk 2007) | Cannon PF, Kirk PM. 2007. Fungal Families of the World. CAB International, Cambridge (ISBN: 978-0851998275) |
| Otu07512 | 3 | k__Fungi(100);p__Ascomycota(100);c__Saccharomycetes(100);o__Saccharomycetales(100);f__Metschnikowiaceae(100);g__Metschnikowia(100);g__Metschnikowia_unclassified(100); | Metschnikowia | 13 | Saprotroph | \|Undefined Saprotroph\| | Yeast | NULL | Probable | As nectar-tap_saprotroph (Põlme et al. 2020); Decay sugar-rich substrates (Põlme et al. 2020) | Põlme S, et al. 2020. Fungal Diversity 105:1-16 ((https://doi.org/10.1007/s13225-020-00466-2)) |
| Otu07513 | 3 | k__Fungi(100);p__Ascomycota(100);c__Saccharomycetes(100);o__Saccharomycetales(100);f__Metschnikowiaceae(100);g__Metschnikowia(100);s__Metschnikowia_pulcherrima(100); | Metschnikowia | 13 | Saprotroph | \|Undefined Saprotroph\| | Yeast | NULL | Probable | As nectar-tap_saprotroph (Põlme et al. 2020); Decay sugar-rich substrates (Põlme et al. 2020) | Põlme S, et al. 2020. Fungal Diversity 105:1-16 ((https://doi.org/10.1007/s13225-020-00466-2)) |
| Otu07517 | 3 | k__Fungi(100);p__Ascomycota(100);c__Saccharomycetes(100);o__Saccharomycetales(100);f__Metschnikowiaceae(100);g__Metschnikowia(100);g__Metschnikowia_unclassified(100); | Metschnikowia | 13 | Saprotroph | \|Undefined Saprotroph\| | Yeast | NULL | Probable | As nectar-tap_saprotroph (Põlme et al. 2020); Decay sugar-rich substrates (Põlme et al. 2020) | Põlme S, et al. 2020. Fungal Diversity 105:1-16 ((https://doi.org/10.1007/s13225-020-00466-2)) |
| Otu07521 | 3 | k__Fungi(100);p__Ascomycota(100);c__Saccharomycetes(100);o__Saccharomycetales(100);f__Metschnikowiaceae(100);g__Metschnikowia(100);s__Metschnikowia_pulcherrima(67); | Metschnikowia | 13 | Saprotroph | \|Undefined Saprotroph\| | Yeast | NULL | Probable | As nectar-tap_saprotroph (Põlme et al. 2020); Decay sugar-rich substrates (Põlme et al. 2020) | Põlme S, et al. 2020. Fungal Diversity 105:1-16 ((https://doi.org/10.1007/s13225-020-00466-2)) |
| Otu07523 | 3 | k__Fungi(100);p__Basidiomycota(100);c__Agaricomycetes(100);o__Russulales(100);f__Russulaceae(100);g__Lactifluus(100);s__Lactifluus_hygrophoroides(100); | Lactifluus | 13 | Symbiotroph | \|Ectomycorrhizal\| | Agaricoid | NULL | Highly Probable | NULL | Põlme S, et al. 2020. Fungal Diversity 105:1-16 ((https://doi.org/10.1007/s13225-020-00466-2)); List compiled by Sara Branco with input from Nhu Nguyen and Else Vellinga |
| Otu07525 | 3 | k__Fungi(100);p__Ascomycota(100);c__Pezizomycetes(100);o__Pezizales(100);f__Pezizaceae(100);g__Peziza(100);s__Peziza_ammophila(100); | Peziza | 13 | Saprotroph-Symbiotroph | Dung Saprotroph-Ectomycorrhizal-Endophyte-Plant Saprotroph-\|Undefined Saprotroph\| | Pezizoid | NULL | Possible | As foliar_endophyte (Põlme et al. 2020); As soil_saprotroph (Põlme et al. 2020); Decay soil (Põlme et al. 2020); Saprobic or ectomycorrhizal, on or in soil, leafmould etc., some species associated with dung or fire sites (Cannon & Kirk 2007) | Cannon PF, Kirk PM. 2007. Fungal Families of the World. CAB International, Cambridge (ISBN: 978-0851998275); Rinaldi AC, et al. 2008. Fungal Diversity 33:1-45 ((http://www.fungaldiversity.org/fdp/sfdp/33-1.pdf)); Tedersoo L, et al. 2010. Mycorrhiza 20:217-263 ((https://doi.org/10.1007/s00572-009-0274-x)) (pro parte); Põlme S, et al. 2020. Fungal Diversity 105:1-16 ((https://doi.org/10.1007/s13225-020-00466-2)) |
| Otu07526 | 3 | k__Fungi(100);p__Ascomycota(100);c__Saccharomycetes(100);o__Saccharomycetales(100);f__Metschnikowiaceae(100);g__Metschnikowia(100);g__Metschnikowia_unclassified(67); | Metschnikowia | 13 | Saprotroph | \|Undefined Saprotroph\| | Yeast | NULL | Probable | As nectar-tap_saprotroph (Põlme et al. 2020); Decay sugar-rich substrates (Põlme et al. 2020) | Põlme S, et al. 2020. Fungal Diversity 105:1-16 ((https://doi.org/10.1007/s13225-020-00466-2)) |
| Otu07531 | 3 | k__Fungi(100);p__Ascomycota(100);c__Saccharomycetes(100);o__Saccharomycetales(100);f__Metschnikowiaceae(100);g__Metschnikowia(100);g__Metschnikowia_unclassified(67); | Metschnikowia | 13 | Saprotroph | \|Undefined Saprotroph\| | Yeast | NULL | Probable | As nectar-tap_saprotroph (Põlme et al. 2020); Decay sugar-rich substrates (Põlme et al. 2020) | Põlme S, et al. 2020. Fungal Diversity 105:1-16 ((https://doi.org/10.1007/s13225-020-00466-2)) |
| Otu07532 | 3 | k__Fungi(100);p__Ascomycota(100);c__Saccharomycetes(100);o__Saccharomycetales(100);f__Metschnikowiaceae(100);g__Metschnikowia(100);g__Metschnikowia_unclassified(100); | Metschnikowia | 13 | Saprotroph | \|Undefined Saprotroph\| | Yeast | NULL | Probable | As nectar-tap_saprotroph (Põlme et al. 2020); Decay sugar-rich substrates (Põlme et al. 2020) | Põlme S, et al. 2020. Fungal Diversity 105:1-16 ((https://doi.org/10.1007/s13225-020-00466-2)) |
| Otu07534 | 3 | k__Fungi(100);p__Basidiomycota(100);c__Agaricomycetes(100);o__Agaricales(100);f__Entolomataceae(100);g__Clitopilus(100);g__Clitopilus_unclassified(100); | Clitopilus | 13 | Pathotroph-Saprotroph | Fungal Parasite-\|Plant Saprotroph\|-Undefined Saprotroph | Agaricoid | NULL | Probable | As litter_saprotroph (Põlme et al. 2020); As mycoparasite (Põlme et al. 2020); Decay leaf, fruit, seed (Põlme et al. 2020) | Tedersoo L, et al. 2014. Science 346:e1256688 ((https://doi.org/10.1126/science.1256688)); Põlme S, et al. 2020. Fungal Diversity 105:1-16 ((https://doi.org/10.1007/s13225-020-00466-2)) |
| Otu07538 | 3 | k__Fungi(100);p__Basidiomycota(100);c__Agaricomycetes(100);o__Russulales(100);f__Hericiaceae(100);g__Dentipellis(100);s__Dentipellis_fragilis(100); | Dentipellis | 13 | Saprotroph | Undefined Saprotroph-\|Wood Saprotroph\| | Corticioid-Hydnoid | White Rot | Probable | As white_rot (Põlme et al. 2020); Decay wood (Põlme et al. 2020) | Tedersoo L, et al. 2014. Science 346:e1256688 ((https://doi.org/10.1126/science.1256688)); Põlme S, et al. 2020. Fungal Diversity 105:1-16 ((https://doi.org/10.1007/s13225-020-00466-2)) |
| Otu07539 | 3 | k__Fungi(100);p__Ascomycota(100);c__Sordariomycetes(100);o__Hypocreales(100);f__Hypocreales_family_Incertae_sedis(100);g__Acremonium(100);s__Acremonium_nepalense(100); | Acremonium | 13 | Pathotroph-Saprotroph-Symbiotroph | Animal Pathogen-Endophyte-Fungal Parasite-Plant Pathogen-\|Undefined Saprotroph\|-Wood Saprotroph | Microfungus | Soft Rot | Possible | As foliar_endophyte (Põlme et al. 2020); culture (Hanada et al. 2010, Del Olmo; Decay leaf, fruit, seed, soil, dung, animal material (Põlme et al. 2020); Endophyte detection method; Ruiz & Arnold 2014); Likely opportunistic human pathogen (Irinyi et al. 2015) | Duncan CG, Eslyn WE. 1966. Mycologia 58:642-645 ((https://doi.org/10.2307/3757045)); Nilsson T. 1973. Studia Forestalia Suecica 104: 1-40 ((https://pub.epsilon.slu.se/5711/)); Worrall JJ, et al. 1997. Mycologia 89:199-219 ((https://doi.org/10.2307/3761073)); Hanada RE, et al. 2010. Fungal Biology 114:901-910 ((https://doi.org/10.1016/j.funbio.2010.08.006)); Del Olmo-Ruiz M, Arnold AE. 2014. Mycologia 106: 8-21 ((https://doi.org/10.3852/13-098)); Irinyi L, et al. 2015. Medical Mycology 53313-337 ((https://doi.org/10.1093/mmy/myv008)); Põlme S, et al. 2020. Fungal Diversity 105:1-16 ((https://doi.org/10.1007/s13225-020-00466-2)) |
| Otu07540 | 3 | k__Fungi(100);p__Ascomycota(100);c__Eurotiomycetes(100);o__Chaetothyriales(100);f__Herpotrichiellaceae(100);f__Herpotrichiellaceae_unclassified(100);f__Herpotrichiellaceae_unclassified(100); | Herpotrichiellaceae | 9 | Pathotroph-Saprotroph | Animal Pathogen-Fungal Parasite-Undefined Saprotroph | Facultative Yeast-Microfungus | NULL | Probable | Mostly saprobic on plants or other fungi, isolated from soil etc., and from medical sources (Cannon & Kirk 2007) | Cannon PF, Kirk PM. 2007. Fungal Families of the World. CAB International, Cambridge (ISBN: 978-0851998275) |
| Otu07542 | 3 | k__Fungi(100);p__Basidiomycota(100);c__Agaricomycetes(100);o__Agaricales(100);f__Cortinariaceae(100);g__Cortinarius(100);g__Cortinarius_unclassified(67); | Cortinarius | 13 | Symbiotroph | \|Ectomycorrhizal\| | Agaricoid | NULL | Highly Probable | NULL | Rinaldi AC, et al. 2008. Fungal Diversity 33:1-45 ((http://www.fungaldiversity.org/fdp/sfdp/33-1.pdf)); Tedersoo L, et al. 2010. Mycorrhiza 20:217-263 ((https://doi.org/10.1007/s00572-009-0274-x)); Põlme S, et al. 2020. Fungal Diversity 105:1-16 ((https://doi.org/10.1007/s13225-020-00466-2)); http://mycorrhizas.info/ecmf.html ((http://mycorrhizas.info/ecmf.html)) |
| Otu07543 | 3 | k__Fungi(100);p__Ascomycota(100);c__Eurotiomycetes(100);o__Chaetothyriales(100);f__Herpotrichiellaceae(100);f__Herpotrichiellaceae_unclassified(100);f__Herpotrichiellaceae_unclassified(100); | Herpotrichiellaceae | 9 | Pathotroph-Saprotroph | Animal Pathogen-Fungal Parasite-Undefined Saprotroph | Facultative Yeast-Microfungus | NULL | Probable | Mostly saprobic on plants or other fungi, isolated from soil etc., and from medical sources (Cannon & Kirk 2007) | Cannon PF, Kirk PM. 2007. Fungal Families of the World. CAB International, Cambridge (ISBN: 978-0851998275) |
| Otu07546 | 3 | k__Fungi(100);p__Ascomycota(100);c__Saccharomycetes(100);o__Saccharomycetales(100);f__Metschnikowiaceae(100);g__Metschnikowia(100);s__Metschnikowia_pulcherrima(100); | Metschnikowia | 13 | Saprotroph | \|Undefined Saprotroph\| | Yeast | NULL | Probable | As nectar-tap_saprotroph (Põlme et al. 2020); Decay sugar-rich substrates (Põlme et al. 2020) | Põlme S, et al. 2020. Fungal Diversity 105:1-16 ((https://doi.org/10.1007/s13225-020-00466-2)) |
| Otu07549 | 3 | k__Fungi(100);p__Ascomycota(100);c__Saccharomycetes(100);o__Saccharomycetales(100);f__Metschnikowiaceae(100);g__Metschnikowia(100);s__Metschnikowia_pulcherrima(67); | Metschnikowia | 13 | Saprotroph | \|Undefined Saprotroph\| | Yeast | NULL | Probable | As nectar-tap_saprotroph (Põlme et al. 2020); Decay sugar-rich substrates (Põlme et al. 2020) | Põlme S, et al. 2020. Fungal Diversity 105:1-16 ((https://doi.org/10.1007/s13225-020-00466-2)) |
| Otu07550 | 3 | k__Fungi(100);p__Ascomycota(100);c__Saccharomycetes(100);o__Saccharomycetales(100);f__Metschnikowiaceae(100);g__Metschnikowia(100);g__Metschnikowia_unclassified(67); | Metschnikowia | 13 | Saprotroph | \|Undefined Saprotroph\| | Yeast | NULL | Probable | As nectar-tap_saprotroph (Põlme et al. 2020); Decay sugar-rich substrates (Põlme et al. 2020) | Põlme S, et al. 2020. Fungal Diversity 105:1-16 ((https://doi.org/10.1007/s13225-020-00466-2)) |
| Otu07554 | 3 | k__Fungi(100);p__Basidiomycota(100);c__Microbotryomycetes(100);o__Sporidiobolales(100);f__Sporidiobolales_family_Incertae_sedis(100);g__Rhodotorula(100);g__Rhodotorula_unclassified(100); | Rhodotorula | 13 | Pathotroph-Saprotroph-Symbiotroph | Animal Pathogen-Animal Symbiotroph-Endophyte-Plant Pathogen-\|Undefined Saprotroph\| | Yeast | NULL | Probable | As foliar_endophyte (Põlme et al. 2020); Opportunistic human pathogen | Põlme S, et al. 2020. Fungal Diversity 105:1-16 ((https://doi.org/10.1007/s13225-020-00466-2)); https://www.cdc.gov ((https://www.cdc.gov)) |
| Otu07555 | 3 | k__Fungi(100);p__Ascomycota(100);c__Saccharomycetes(100);o__Saccharomycetales(100);f__Metschnikowiaceae(100);g__Metschnikowia(100);s__Metschnikowia_pulcherrima(67); | Metschnikowia | 13 | Saprotroph | \|Undefined Saprotroph\| | Yeast | NULL | Probable | As nectar-tap_saprotroph (Põlme et al. 2020); Decay sugar-rich substrates (Põlme et al. 2020) | Põlme S, et al. 2020. Fungal Diversity 105:1-16 ((https://doi.org/10.1007/s13225-020-00466-2)) |
| Otu07558 | 3 | k__Fungi(100);p__Ascomycota(100);c__Saccharomycetes(100);o__Saccharomycetales(100);f__Metschnikowiaceae(100);g__Metschnikowia(100);s__Metschnikowia_pulcherrima(67); | Metschnikowia | 13 | Saprotroph | \|Undefined Saprotroph\| | Yeast | NULL | Probable | As nectar-tap_saprotroph (Põlme et al. 2020); Decay sugar-rich substrates (Põlme et al. 2020) | Põlme S, et al. 2020. Fungal Diversity 105:1-16 ((https://doi.org/10.1007/s13225-020-00466-2)) |
| Otu07559 | 3 | k__Fungi(100);p__Ascomycota(100);c__Saccharomycetes(100);o__Saccharomycetales(100);f__Metschnikowiaceae(100);g__Metschnikowia(100);g__Metschnikowia_unclassified(100); | Metschnikowia | 13 | Saprotroph | \|Undefined Saprotroph\| | Yeast | NULL | Probable | As nectar-tap_saprotroph (Põlme et al. 2020); Decay sugar-rich substrates (Põlme et al. 2020) | Põlme S, et al. 2020. Fungal Diversity 105:1-16 ((https://doi.org/10.1007/s13225-020-00466-2)) |
| Otu07561 | 3 | k__Fungi(100);p__Ascomycota(100);c__Dothideomycetes(100);o__Pleosporales(100);f__Pleosporaceae(100);f__Pleosporaceae_unclassified(100);f__Pleosporaceae_unclassified(100); | Pleosporaceae | 9 | Pathotroph-Saprotroph | Endophyte-Lichen Parasite-Plant Pathogen-Undefined Saprotroph | Microfungus | NULL | Probable | Necrotrophic pathogens and saprobes, especially associated with grasses. Many species are seed-borne (Cannon & Kirk 2007) | Cannon PF, Kirk PM. 2007. Fungal Families of the World. CAB International, Cambridge (ISBN: 978-0851998275); David AS, et al. 2016. Microbial Ecology 71:912-926 ((https://doi.org/10.1007/s00248-015-0712-6)); Lawrey JD, Diederich P. 2016. http://www.lichenicolous.net (accessed 24 May 2017) ((http://www.lichenicolous.net)) |
| Otu07562 | 3 | k__Fungi(100);p__Ascomycota(100);c__Saccharomycetes(100);o__Saccharomycetales(100);f__Metschnikowiaceae(100);g__Metschnikowia(100);s__Metschnikowia_pulcherrima(67); | Metschnikowia | 13 | Saprotroph | \|Undefined Saprotroph\| | Yeast | NULL | Probable | As nectar-tap_saprotroph (Põlme et al. 2020); Decay sugar-rich substrates (Põlme et al. 2020) | Põlme S, et al. 2020. Fungal Diversity 105:1-16 ((https://doi.org/10.1007/s13225-020-00466-2)) |
| Otu07565 | 3 | k__Fungi(100);p__Ascomycota(100);c__Saccharomycetes(100);o__Saccharomycetales(100);f__Metschnikowiaceae(100);g__Metschnikowia(100);s__Metschnikowia_pulcherrima(100); | Metschnikowia | 13 | Saprotroph | \|Undefined Saprotroph\| | Yeast | NULL | Probable | As nectar-tap_saprotroph (Põlme et al. 2020); Decay sugar-rich substrates (Põlme et al. 2020) | Põlme S, et al. 2020. Fungal Diversity 105:1-16 ((https://doi.org/10.1007/s13225-020-00466-2)) |
| Otu07573 | 3 | k__Fungi(100);p__Basidiomycota(100);c__Microbotryomycetes(100);o__Sporidiobolales(100);f__Sporidiobolales_family_Incertae_sedis(100);g__Rhodotorula(100);s__Rhodotorula_hordea(100); | Rhodotorula | 13 | Pathotroph-Saprotroph-Symbiotroph | Animal Pathogen-Animal Symbiotroph-Endophyte-Plant Pathogen-\|Undefined Saprotroph\| | Yeast | NULL | Probable | As foliar_endophyte (Põlme et al. 2020); Opportunistic human pathogen | Põlme S, et al. 2020. Fungal Diversity 105:1-16 ((https://doi.org/10.1007/s13225-020-00466-2)); https://www.cdc.gov ((https://www.cdc.gov)) |
| Otu07574 | 3 | k__Fungi(100);p__Ascomycota(100);c__Saccharomycetes(100);o__Saccharomycetales(100);f__Metschnikowiaceae(100);g__Metschnikowia(100);g__Metschnikowia_unclassified(100); | Metschnikowia | 13 | Saprotroph | \|Undefined Saprotroph\| | Yeast | NULL | Probable | As nectar-tap_saprotroph (Põlme et al. 2020); Decay sugar-rich substrates (Põlme et al. 2020) | Põlme S, et al. 2020. Fungal Diversity 105:1-16 ((https://doi.org/10.1007/s13225-020-00466-2)) |
| Otu07575 | 3 | k__Fungi(100);p__Ascomycota(100);c__Saccharomycetes(100);o__Saccharomycetales(100);f__Metschnikowiaceae(100);g__Metschnikowia(100);g__Metschnikowia_unclassified(100); | Metschnikowia | 13 | Saprotroph | \|Undefined Saprotroph\| | Yeast | NULL | Probable | As nectar-tap_saprotroph (Põlme et al. 2020); Decay sugar-rich substrates (Põlme et al. 2020) | Põlme S, et al. 2020. Fungal Diversity 105:1-16 ((https://doi.org/10.1007/s13225-020-00466-2)) |
| Otu07578 | 3 | k__Fungi(100);p__Basidiomycota(100);c__Agaricomycetes(100);o__Agaricales(100);f__Clavariaceae(100);f__Clavariaceae_unclassified(100);f__Clavariaceae_unclassified(100); | Clavariaceae | 9 | Saprotroph-Symbiotroph | Lichenized-Undefined Saprotroph | Clavarioid | NULL | Probable | Most species are presumed saprobic on the ground or rotten wood, but many Multiclavula species are lichenized with green algae (Cannon & Kirk 2007) | Cannon PF, Kirk PM. 2007. Fungal Families of the World. CAB International, Cambridge (ISBN: 978-0851998275) |
| Otu07579 | 3 | k__Fungi(100);p__Ascomycota(100);c__Saccharomycetes(100);o__Saccharomycetales(100);f__Metschnikowiaceae(100);g__Metschnikowia(100);s__Metschnikowia_pulcherrima(100); | Metschnikowia | 13 | Saprotroph | \|Undefined Saprotroph\| | Yeast | NULL | Probable | As nectar-tap_saprotroph (Põlme et al. 2020); Decay sugar-rich substrates (Põlme et al. 2020) | Põlme S, et al. 2020. Fungal Diversity 105:1-16 ((https://doi.org/10.1007/s13225-020-00466-2)) |
| Otu07580 | 3 | k__Fungi(100);p__Ascomycota(100);c__Saccharomycetes(100);o__Saccharomycetales(100);f__Saccharomycetales_family_Incertae_sedis(100);g__Candida(100);g__Candida_unclassified(67); | Candida | 13 | Pathotroph-Saprotroph-Symbiotroph | Animal Pathogen-Endophyte-Endosymbiont-Epiphyte-\|Undefined Saprotroph\| | Dimorphic Yeast-Yeast | NULL | Probable | As nectar-tap_saprotroph (Põlme et al. 2020); Decay sugar-rich substrates (Põlme et al. 2020); This asexual genus is polyphyletic and contain species that function very differently depend on what they are. Caution should be taken when interpreting guilds for Candida species. | Manolakaki D, et al. 2010. Virulence. 1: 367-375 ((https://doi.org/10.4161/viru.1.5.12796)); Mekbib SB, et al. 2011. Tropical Plant Pathology 36:233-240 ((https://doi.org/10.1590/S1982-56762011000400004)); Põlme S, et al. 2020. Fungal Diversity 105:1-16 ((https://doi.org/10.1007/s13225-020-00466-2)) |
| Otu07581 | 3 | k__Fungi(100);p__Chytridiomycota(100);c__Chytridiomycetes(100);o__Spizellomycetales(100);f__Spizellomycetaceae(100);f__Spizellomycetaceae_unclassified(100);f__Spizellomycetaceae_unclassified(100); | Spizellomycetaceae | 9 | Pathotroph | Plant Pathogen-Undefined Parasite-Undefined Saprotroph | Microfungus | NULL | Probable | Common in the soil and also as parasites of a range of soil organisms and plants (Cannon & Kirk 2007) | Cannon PF, Kirk PM. 2007. Fungal Families of the World. CAB International, Cambridge (ISBN: 978-0851998275) |
| Otu07583 | 3 | k__Fungi(100);p__Ascomycota(100);c__Saccharomycetes(100);o__Saccharomycetales(100);f__Metschnikowiaceae(100);g__Metschnikowia(100);s__Metschnikowia_pulcherrima(100); | Metschnikowia | 13 | Saprotroph | \|Undefined Saprotroph\| | Yeast | NULL | Probable | As nectar-tap_saprotroph (Põlme et al. 2020); Decay sugar-rich substrates (Põlme et al. 2020) | Põlme S, et al. 2020. Fungal Diversity 105:1-16 ((https://doi.org/10.1007/s13225-020-00466-2)) |
| Otu07584 | 3 | k__Fungi(100);p__Ascomycota(100);c__Saccharomycetes(100);o__Saccharomycetales(100);f__Metschnikowiaceae(100);g__Metschnikowia(100);s__Metschnikowia_pulcherrima(100); | Metschnikowia | 13 | Saprotroph | \|Undefined Saprotroph\| | Yeast | NULL | Probable | As nectar-tap_saprotroph (Põlme et al. 2020); Decay sugar-rich substrates (Põlme et al. 2020) | Põlme S, et al. 2020. Fungal Diversity 105:1-16 ((https://doi.org/10.1007/s13225-020-00466-2)) |
| Otu07587 | 3 | k__Fungi(100);p__Basidiomycota(100);c__Agaricomycetes(100);o__Agaricales(100);f__Inocybaceae(100);g__Tubaria(100);s__Tubaria_minutalis(100); | Tubaria | 13 | Saprotroph | \|Plant Saprotroph\|-Wood Saprotroph | Agaricoid | NULL | Probable | As litter_saprotroph (Põlme et al. 2020); Decay leaf, fruit, seed (Põlme et al. 2020) | Tedersoo L, et al. 2014. Science 346:e1256688 ((https://doi.org/10.1126/science.1256688)); Põlme S, et al. 2020. Fungal Diversity 105:1-16 ((https://doi.org/10.1007/s13225-020-00466-2)) |
| Otu07592 | 3 | k__Fungi(100);p__Ascomycota(100);c__Saccharomycetes(100);o__Saccharomycetales(100);f__Metschnikowiaceae(100);g__Metschnikowia(100);g__Metschnikowia_unclassified(67); | Metschnikowia | 13 | Saprotroph | \|Undefined Saprotroph\| | Yeast | NULL | Probable | As nectar-tap_saprotroph (Põlme et al. 2020); Decay sugar-rich substrates (Põlme et al. 2020) | Põlme S, et al. 2020. Fungal Diversity 105:1-16 ((https://doi.org/10.1007/s13225-020-00466-2)) |
| Otu07594 | 3 | k__Fungi(100);p__Ascomycota(100);c__Saccharomycetes(100);o__Saccharomycetales(100);f__Metschnikowiaceae(100);g__Metschnikowia(100);g__Metschnikowia_unclassified(100); | Metschnikowia | 13 | Saprotroph | \|Undefined Saprotroph\| | Yeast | NULL | Probable | As nectar-tap_saprotroph (Põlme et al. 2020); Decay sugar-rich substrates (Põlme et al. 2020) | Põlme S, et al. 2020. Fungal Diversity 105:1-16 ((https://doi.org/10.1007/s13225-020-00466-2)) |
| Otu07598 | 3 | k__Fungi(100);p__Basidiomycota(100);c__Microbotryomycetes(100);o__Sporidiobolales(100);f__Sporidiobolales_family_Incertae_sedis(67);g__Sporobolomyces(67);g__Sporobolomyces_unclassified(67); | Sporobolomyces | 13 | Pathotroph-Saprotroph | \|Fungal Parasite\|-Plant Saprotroph | Yeast | NULL | Probable | As litter_saprotroph (Põlme et al. 2020); As mycoparasite (Põlme et al. 2020); Decay fungal material (Põlme et al. 2020) | Tedersoo L, et al. 2014. Science 346:e1256688 ((https://doi.org/10.1126/science.1256688)); Purahong W, et al. 2016. Molecular Ecology 25:4059-4074 ((https://doi.org/10.1111/mec.13739)); Põlme S, et al. 2020. Fungal Diversity 105:1-16 ((https://doi.org/10.1007/s13225-020-00466-2)) |
| Otu07600 | 3 | k__Fungi(100);p__Ascomycota(100);c__Sordariomycetes(100);o__Xylariales(100);f__Xylariaceae(100);g__Hypoxylon(100);s__Hypoxylon_sp(100); | Hypoxylon | 13 | Saprotroph-Saprotroph-Symbiotroph | Endophyte-Undefined Saprotroph-\|Wood Saprotroph\| | Corticioid-Microfungus-Xylarioid | NULL | Probable | As foliar_endophyte (Põlme et al. 2020); Decay wood, leaf, fruit, seed (Põlme et al. 2020); Host - angiosperms (Põlme et al. 2020) | Tedersoo L, et al. 2014. Science 346:e1256688 ((https://doi.org/10.1126/science.1256688)); Põlme S, et al. 2020. Fungal Diversity 105:1-16 ((https://doi.org/10.1007/s13225-020-00466-2)) |
| Otu07602 | 3 | k__Fungi(100);p__Ascomycota(100);c__Saccharomycetes(100);o__Saccharomycetales(100);f__Metschnikowiaceae(100);g__Metschnikowia(100);s__Metschnikowia_pulcherrima(67); | Metschnikowia | 13 | Saprotroph | \|Undefined Saprotroph\| | Yeast | NULL | Probable | As nectar-tap_saprotroph (Põlme et al. 2020); Decay sugar-rich substrates (Põlme et al. 2020) | Põlme S, et al. 2020. Fungal Diversity 105:1-16 ((https://doi.org/10.1007/s13225-020-00466-2)) |
| Otu07604 | 3 | k__Fungi(100);p__Basidiomycota(100);c__Agaricomycetes(100);o__Gomphales(100);f__Gomphaceae(100);f__Gomphaceae_unclassified(67);f__Gomphaceae_unclassified(67); | Gomphaceae | 9 | Saprotroph-Symbiotroph | Ectomycorrhizal-Undefined Saprotroph-Wood Saprotroph | Clavarioid-Gomphoid | NULL | Probable | Terrestrial or lignicolous, either saprobic or forming ectomycorrhizal relationships with woody plants (Cannon & Kirk 2007) | Cannon PF, Kirk PM. 2007. Fungal Families of the World. CAB International, Cambridge (ISBN: 978-0851998275) |
| Otu07608 | 3 | k__Fungi(100);p__Ascomycota(100);c__Dothideomycetes(100);o__Capnodiales(100);f__Teratosphaeriaceae(100);g__Teratosphaeria(100);s__Teratosphaeria_knoxdavesii(100); | Teratosphaeria | 13 | Pathotroph-Saprotroph | \|Plant Pathogen\|-Plant Saprotroph-Wood Saprotroph | Microfungus | NULL | Probable | As litter_saprotroph (Põlme et al. 2020); As wood_pathogen (Põlme et al. 2020); Decay wood, leaf, fruit, seed (Põlme et al. 2020) | Yamazaki A, et al. 2010. Mycoscience 51:443-455 ((https://www.jstage.jst.go.jp/article/mycosci/51/6/51_MYC51443/_pdf)); Tedersoo L, et al. 2014. Science 346:e1256688 ((https://doi.org/10.1126/science.1256688)); Põlme S, et al. 2020. Fungal Diversity 105:1-16 ((https://doi.org/10.1007/s13225-020-00466-2)) |
| Otu07609 | 3 | k__Fungi(100);p__Ascomycota(100);c__Saccharomycetes(100);o__Saccharomycetales(100);f__Metschnikowiaceae(100);g__Metschnikowia(100);s__Metschnikowia_pulcherrima(100); | Metschnikowia | 13 | Saprotroph | \|Undefined Saprotroph\| | Yeast | NULL | Probable | As nectar-tap_saprotroph (Põlme et al. 2020); Decay sugar-rich substrates (Põlme et al. 2020) | Põlme S, et al. 2020. Fungal Diversity 105:1-16 ((https://doi.org/10.1007/s13225-020-00466-2)) |
| Otu07612 | 3 | k__Fungi(100);p__Ascomycota(100);c__Saccharomycetes(100);o__Saccharomycetales(100);f__Metschnikowiaceae(100);g__Metschnikowia(100);g__Metschnikowia_unclassified(67); | Metschnikowia | 13 | Saprotroph | \|Undefined Saprotroph\| | Yeast | NULL | Probable | As nectar-tap_saprotroph (Põlme et al. 2020); Decay sugar-rich substrates (Põlme et al. 2020) | Põlme S, et al. 2020. Fungal Diversity 105:1-16 ((https://doi.org/10.1007/s13225-020-00466-2)) |

| | | | | | | | | | | | |
|---|---|---|---|---|---|---|---|---|---|---|---|
| Otu07613 | 3 | k__Fungi(100);p__Ascomycota(100);c__Saccharomycetes(100);o__Saccharomycetales(100);f__Metschnikowiaceae(100);g__Metschnikowia(100);s__Metschnikowia_pulcherrima(100); | Metschnikowia | 13 | Saprotroph | \|Undefined Saprotroph\| | Yeast | NULL | Probable | As nectar-tap_saprotroph (Põlme et al. 2020); Decay sugar-rich substrates (Põlme et al. 2020) | Põlme S, et al. 2020. Fungal Diversity 105:1-16 ((https://doi.org/10.1007/s13225-020-00466-2)) |
| Otu07614 | 3 | k__Fungi(100);p__Basidiomycota(100);c__Agaricomycetes(100);o__Gomphales(100);f__Gomphaceae(100);g__Ramaria(100);s__Ramaria_sp_5_RG_2012(100); | Ramaria | 13 | Saprotroph-Symbiotroph | \|Ectomycorrhizal\|-Plant Saprotroph | Clavarioid | NULL | Possible | As litter_saprotroph (Põlme et al. 2020) | Rinaldi AC, et al. 2008. Fungal Diversity 33:1-45 ((http://www.fungaldiversity.org/fdp/sfdp/33-1.pdf)) (as ECM/not ECM); Tedersoo L, et al. 2010. Mycorrhiza 20:217-263 ((10.1007/s00572-009-0274-x)) (pro parte); Põlme S, et al. 2020. Fungal Diversity 105:1-16 ((https://doi.org/10.1007/s13225-020-00466-2)); http://mycorrhizas.info/ecmf.html ((http://mycorrhizas.info/ecmf.html)) (as ECM) |
| Otu07615 | 3 | k__Fungi(100);p__Ascomycota(100);c__Lecanoromycetes(100);o__Lecanorales(100);f__Parmeliaceae(100);g__Melanelixia(100);s__Melanelixia_subaurifera(100); | Melanelixia | 13 | Symbiotroph | \|Lichenized\| | Thallus | NULL | Highly Probable | NULL | Esslinger TL. 2014. North Dakota State University ((http://www.ndsu.edu/pubweb/~esslinge/chcklst/chcklst7.htm)); Põlme S, et al. 2020. Fungal Diversity 105:1-16 ((https://doi.org/10.1007/s13225-020-00466-2)) |
| Otu07617 | 3 | k__Fungi(100);p__Ascomycota(100);c__Ascomycota_class_Incertae_sedis(100);o__Ascomycota_order_Incertae_sedis(100);f__Ascomycota_family_Incertae_sedis(100);g__Veronaea(100);s__Veronaea_compacta(100); | Veronaea | 13 | Pathotroph-Saprotroph | Dung Saprotroph-\|Plant Pathogen\|-Plant Saprotroph | Dimorphic | NULL | Probable | As litter_saprotroph (Põlme et al. 2020); Decay leaf, fruit, seed (Põlme et al. 2020) | Põlme S, et al. 2020. Fungal Diversity 105:1-16 ((https://doi.org/10.1007/s13225-020-00466-2)); The American Phytopathological Society Common Names of Plant Diseases ((http://www.apsnet.org/publications/commonnames/Pages/default.aspx)) |
| Otu07619 | 3 | k__Fungi(100);p__Basidiomycota(100);c__Agaricomycetes(100);o__Polyporales(100);f__Meripilaceae(100);g__Hydnopolyporus(100);s__Hydnopolyporus_fimbriatus(100); | Hydnopolyporus | 13 | Pathotroph-Saprotroph | \|Plant Pathogen\|-Wood Saprotroph | Clavarioid | White Rot | Highly Probable | As white_rot (Põlme et al. 2020); As wood_pathogen (Põlme et al. 2020); Decay wood (Põlme et al. 2020) | Gilbertson RL, Ryvarden L. 1987-1987. North American Polypores. Fungiflora, Oslo (ISBN: 978-0945345060); Põlme S, et al. 2020. Fungal Diversity 105:1-16 ((https://doi.org/10.1007/s13225-020-00466-2)) |
| Otu07620 | 3 | k__Fungi(100);p__Basidiomycota(100);c__Agaricomycetes(100);o__Cantharellales(100);f__Cantharellales_family_Incertae_sedis(100);g__Minimedusa(100);s__Minimedusa_sp(100); | Minimedusa | 13 | Pathotroph | \|Lichen Parasite\| | Microfungus | NULL | Probable | Decay fungal material, algal material (Põlme et al. 2020) | Põlme S, et al. 2020. Fungal Diversity 105:1-16 ((https://doi.org/10.1007/s13225-020-00466-2)) |
| Otu07625 | 3 | k__Fungi(100);p__Basidiomycota(100);c__Tremellomycetes(100);o__Cystofilobasidiales(100);f__Cystofilobasidiales_family_Incertae_sedis(100);g__Syzygospora(100);s__Syzygospora_sp(100); | Syzygospora | 13 | Pathotroph | \|Fungal Parasite\|-Lichen Parasite | Dimorphic-Tremelloid | NULL | Highly Probable | As mycoparasite (Põlme et al. 2020); Decay fungal material (Põlme et al. 2020) | Põlme S, et al. 2020. Fungal Diversity 105:1-16 ((https://doi.org/10.1007/s13225-020-00466-2)); List compiled by John Plischke III ((https://goodmushroom.weebly.com/uploads/5/7/1/7/57176535/fungi_on_fungi_pdf_lq.pdf)) |
| Otu07626 | 3 | k__Fungi(100);p__Ascomycota(100);c__Saccharomycetes(100);o__Saccharomycetales(100);f__Metschnikowiaceae(100);g__Metschnikowia(100);s__Metschnikowia_pulcherrima(100); | Metschnikowia | 13 | Saprotroph | \|Undefined Saprotroph\| | Yeast | NULL | Probable | As nectar-tap_saprotroph (Põlme et al. 2020); Decay sugar-rich substrates (Põlme et al. 2020) | Põlme S, et al. 2020. Fungal Diversity 105:1-16 ((https://doi.org/10.1007/s13225-020-00466-2)) |
| Otu07628 | 3 | k__Fungi(100);p__Basidiomycota(100);c__Pucciniomycetes(100);o__Pucciniales(100);f__Pucciniaceae(100);g__Puccinia(100);s__Puccinia_bartholomaei(100); | Puccinia | 13 | Pathotroph | \|Plant Pathogen\| | Rust | NULL | Highly Probable | NULL | Tedersoo L, et al. 2014. Science 346:e1256688 ((https://doi.org/10.1126/science.1256688)); Põlme S, et al. 2020. Fungal Diversity 105:1-16 ((https://doi.org/10.1007/s13225-020-00466-2)) |
| Otu07630 | 3 | k__Fungi(100);p__Ascomycota(100);c__Saccharomycetes(100);o__Saccharomycetales(100);f__Metschnikowiaceae(100);g__Metschnikowia(100);g__Metschnikowia_unclassified(67); | Metschnikowia | 13 | Saprotroph | \|Undefined Saprotroph\| | Yeast | NULL | Probable | As nectar-tap_saprotroph (Põlme et al. 2020); Decay sugar-rich substrates (Põlme et al. 2020) | Põlme S, et al. 2020. Fungal Diversity 105:1-16 ((https://doi.org/10.1007/s13225-020-00466-2)) |
| Otu07631 | 3 | k__Fungi(100);p__Ascomycota(100);c__Saccharomycetes(100);o__Saccharomycetales(100);f__Metschnikowiaceae(100);g__Metschnikowia(100);s__Metschnikowia_pulcherrima(100); | Metschnikowia | 13 | Saprotroph | \|Undefined Saprotroph\| | Yeast | NULL | Probable | As nectar-tap_saprotroph (Põlme et al. 2020); Decay sugar-rich substrates (Põlme et al. 2020) | Põlme S, et al. 2020. Fungal Diversity 105:1-16 ((https://doi.org/10.1007/s13225-020-00466-2)) |
| Otu07632 | 3 | k__Fungi(100);p__Ascomycota(100);c__Saccharomycetes(100);o__Saccharomycetales(100);f__Metschnikowiaceae(100);g__Metschnikowia(100);s__Metschnikowia_pulcherrima(100); | Metschnikowia | 13 | Saprotroph | \|Undefined Saprotroph\| | Yeast | NULL | Probable | As nectar-tap_saprotroph (Põlme et al. 2020); Decay sugar-rich substrates (Põlme et al. 2020) | Põlme S, et al. 2020. Fungal Diversity 105:1-16 ((https://doi.org/10.1007/s13225-020-00466-2)) |
| Otu07638 | 3 | k__Fungi(100);p__Ascomycota(100);c__Saccharomycetes(100);o__Saccharomycetales(100);f__Metschnikowiaceae(100);g__Metschnikowia(100);s__Metschnikowia_pulcherrima(67); | Metschnikowia | 13 | Saprotroph | \|Undefined Saprotroph\| | Yeast | NULL | Probable | As nectar-tap_saprotroph (Põlme et al. 2020); Decay sugar-rich substrates (Põlme et al. 2020) | Põlme S, et al. 2020. Fungal Diversity 105:1-16 ((https://doi.org/10.1007/s13225-020-00466-2)) |
| Otu07640 | 3 | k__Fungi(100);p__Ascomycota(100);c__Saccharomycetes(100);o__Saccharomycetales(100);f__Metschnikowiaceae(100);g__Metschnikowia(100);s__Metschnikowia_pulcherrima(100); | Metschnikowia | 13 | Saprotroph | \|Undefined Saprotroph\| | Yeast | NULL | Probable | As nectar-tap_saprotroph (Põlme et al. 2020); Decay sugar-rich substrates (Põlme et al. 2020) | Põlme S, et al. 2020. Fungal Diversity 105:1-16 ((https://doi.org/10.1007/s13225-020-00466-2)) |
| Otu07642 | 3 | k__Fungi(100);p__Ascomycota(100);c__Dothideomycetes(100);o__Capnodiales(100);f__Capnodiales_family_Incertae_sedis(67);g__Hispidoconidioma(67);s__Hispidoconidioma_sp(67); | Hispidoconidioma | 13 | Saprotroph | \|Wood Saprotroph\| | Microfungus | NULL | Probable | Decay wood (Põlme et al. 2020) | Põlme S, et al. 2020. Fungal Diversity 105:1-16 ((https://doi.org/10.1007/s13225-020-00466-2)) |
| Otu07643 | 3 | k__Fungi(100);p__Ascomycota(100);c__Saccharomycetes(100);o__Saccharomycetales(100);f__Saccharomycetales_family_Incertae_sedis(100);g__Candida(100);g__Candida_unclassified(100); | Candida | 13 | Pathotroph-Saprotroph-Symbiotroph | Animal Pathogen-Endophyte-Endosymbiont-Epiphyte-\|Undefined Saprotroph\| | Dimorphic Yeast-Yeast | NULL | Probable | As nectar-tap_saprotroph (Põlme et al. 2020); Decay sugar-rich substrates (Põlme et al. 2020); This asexual genus is polyphyletic and contain species that function very differently depend on what they are. Caution should be taken when interpreting guilds for Candida species. | Manolakaki D, et al. 2010. Virulence. 1: 367-375 ((https://doi.org/10.4161/viru.1.5.12796)); Mekbib SB, et al. 2011. Tropical Plant Pathology 36:233-240 ((https://doi.org/10.1590/S1982-56762011000400004)); Põlme S, et al. 2020. Fungal Diversity 105:1-16 ((https://doi.org/10.1007/s13225-020-00466-2)) |
| Otu07645 | 3 | k__Fungi(100);p__Ascomycota(100);c__Dothideomycetes(100);o__Capnodiales(100);f__Mycosphaerellaceae(100);g__Mycosphaerella(100);s__Mycosphaerella_ulmi(100); | Mycosphaerella | 13 | Pathotroph-Saprotroph | \|Plant Pathogen\|-Plant Saprotroph | Microfungus | NULL | Probable | As litter_saprotroph (Põlme et al. 2020); Decay leaf, fruit, seed (Põlme et al. 2020) | Tedersoo L, et al. 2014. Science 346:e1256688 ((https://doi.org/10.1126/science.1256688)); Põlme S, et al. 2020. Fungal Diversity 105:1-16 ((https://doi.org/10.1007/s13225-020-00466-2)) |
| Otu07646 | 3 | k__Fungi(100);p__Ascomycota(100);c__Leotiomycetes(100);o__Helotiales(100);f__Helotiales_family_Incertae_sedis(100);g__Clathrosporium(100);s__Clathrosporium_intricatum(100); | Clathrosporium | 13 | Saprotroph | \|Plant Saprotroph\|-Undefined Saprotroph | Microfungus | NULL | Probable | As litter_saprotroph (Põlme et al. 2020); Decay leaf, fruit, seed (Põlme et al. 2020) | Põlme S, et al. 2020. Fungal Diversity 105:1-16 ((https://doi.org/10.1007/s13225-020-00466-2)); The American Phytopathological Society Common Names of Plant Diseases ((http://www.apsnet.org/publications/commonnames/Pages/default.aspx)) |
| Otu07648 | 3 | k__Fungi(100);p__Ascomycota(100);c__Saccharomycetes(100);o__Saccharomycetales(100);f__Metschnikowiaceae(100);g__Metschnikowia(100);s__Metschnikowia_pulcherrima(100); | Metschnikowia | 13 | Saprotroph | \|Undefined Saprotroph\| | Yeast | NULL | Probable | As nectar-tap_saprotroph (Põlme et al. 2020); Decay sugar-rich substrates (Põlme et al. 2020) | Põlme S, et al. 2020. Fungal Diversity 105:1-16 ((https://doi.org/10.1007/s13225-020-00466-2)) |
| Otu07650 | 3 | k__Fungi(100);p__Ascomycota(100);c__Dothideomycetes(100);o__Pleosporales(100);f__Pleosporaceae(100);f__Pleosporaceae_unclassified(100);f__Pleosporaceae_unclassified(100); | Pleosporaceae | 9 | Pathotroph-Saprotroph | Endophyte-Lichen Parasite-Plant Pathogen-Undefined Saprotroph | Microfungus | NULL | Probable | Necrotrophic pathogens and saprobes, especially associated with grasses. Many species are seed-borne (Cannon & Kirk 2007) | Cannon PF, Kirk PM. 2007. Fungal Families of the World. CAB International, Cambridge (ISBN: 978-0851998275); David AS, et al. 2016. Microbial Ecology 71:912-926 ((https://doi.org/10.1007/s00248-015-0712-6)); Lawrey JD, Diederich P. 2016. http://www.lichenicolous.net (accessed 24 May 2017) ((http://www.lichenicolous.net)) |
| Otu07651 | 3 | k__Fungi(100);p__Ascomycota(100);c__Leotiomycetes(100);o__Helotiales(100);f__Helotiales_family_Incertae_sedis(100);g__Rhexocercosporidium(100);g__Rhexocercosporidium_unclassified(67); | Rhexocercosporidium | 13 | Pathotroph-Saprotroph | \|Plant Pathogen\|-Plant Saprotroph | Microfungus | NULL | Probable | As litter_saprotroph (Põlme et al. 2020); As root_pathogen (Põlme et al. 2020); Decay leaf, fruit, seed, roots (Põlme et al. 2020) | Tedersoo L, et al. 2014. Science 346:e1256688 ((https://doi.org/10.1126/science.1256688)); Põlme S, et al. 2020. Fungal Diversity 105:1-16 ((https://doi.org/10.1007/s13225-020-00466-2)) |
| Otu07654 | 3 | k__Fungi(100);p__Ascomycota(100);c__Saccharomycetes(100);o__Saccharomycetales(100);f__Metschnikowiaceae(100);g__Metschnikowia(100);s__Metschnikowia_pulcherrima(100); | Metschnikowia | 13 | Saprotroph | \|Undefined Saprotroph\| | Yeast | NULL | Probable | As nectar-tap_saprotroph (Põlme et al. 2020); Decay sugar-rich substrates (Põlme et al. 2020) | Põlme S, et al. 2020. Fungal Diversity 105:1-16 ((https://doi.org/10.1007/s13225-020-00466-2)) |
| Otu07658 | 3 | k__Fungi(100);p__Ascomycota(100);c__Saccharomycetes(100);o__Saccharomycetales(100);f__Metschnikowiaceae(100);g__Metschnikowia(100);s__Metschnikowia_pulcherrima(67); | Metschnikowia | 13 | Saprotroph | \|Undefined Saprotroph\| | Yeast | NULL | Probable | As nectar-tap_saprotroph (Põlme et al. 2020); Decay sugar-rich substrates (Põlme et al. 2020) | Põlme S, et al. 2020. Fungal Diversity 105:1-16 ((https://doi.org/10.1007/s13225-020-00466-2)) |
| Otu07659 | 3 | k__Fungi(100);p__Ascomycota(100);c__Saccharomycetes(67);o__Saccharomycetales(67);f__Saccharomycetales_family_Incertae_sedis(67);g__Candida(67);g__Candida_unclassified(67); | Candida | 13 | Pathotroph-Saprotroph-Symbiotroph | Animal Pathogen-Endophyte-Endosymbiont-Epiphyte-\|Undefined Saprotroph\| | Dimorphic Yeast-Yeast | NULL | Probable | As nectar-tap_saprotroph (Põlme et al. 2020); Decay sugar-rich substrates (Põlme et al. 2020); This asexual genus is polyphyletic and contain species that function very differently depend on what they are. Caution should be taken when interpreting guilds for Candida species. | Manolakaki D, et al. 2010. Virulence. 1: 367-375 ((https://doi.org/10.4161/viru.1.5.12796)); Mekbib SB, et al. 2011. Tropical Plant Pathology 36:233-240 ((https://doi.org/10.1590/S1982-56762011000400004)); Põlme S, et al. 2020. Fungal Diversity 105:1-16 ((https://doi.org/10.1007/s13225-020-00466-2)) |
| Otu07660 | 3 | k__Fungi(100);p__Ascomycota(100);c__Saccharomycetes(100);o__Saccharomycetales(100);f__Metschnikowiaceae(100);g__Metschnikowia(100);s__Metschnikowia_pulcherrima(67); | Metschnikowia | 13 | Saprotroph | \|Undefined Saprotroph\| | Yeast | NULL | Probable | As nectar-tap_saprotroph (Põlme et al. 2020); Decay sugar-rich substrates (Põlme et al. 2020) | Põlme S, et al. 2020. Fungal Diversity 105:1-16 ((https://doi.org/10.1007/s13225-020-00466-2)) |
| Otu07662 | 3 | k__Fungi(100);p__Ascomycota(100);c__Saccharomycetes(100);o__Saccharomycetales(100);f__Metschnikowiaceae(100);g__Metschnikowia(100);g__Metschnikowia_unclassified(100); | Metschnikowia | 13 | Saprotroph | \|Undefined Saprotroph\| | Yeast | NULL | Probable | As nectar-tap_saprotroph (Põlme et al. 2020); Decay sugar-rich substrates (Põlme et al. 2020) | Põlme S, et al. 2020. Fungal Diversity 105:1-16 ((https://doi.org/10.1007/s13225-020-00466-2)) |
| Otu07664 | 3 | k__Fungi(100);p__Ascomycota(100);c__Saccharomycetes(100);o__Saccharomycetales(100);f__Metschnikowiaceae(100);g__Metschnikowia(100);s__Metschnikowia_pulcherrima(67); | Metschnikowia | 13 | Saprotroph | \|Undefined Saprotroph\| | Yeast | NULL | Probable | As nectar-tap_saprotroph (Põlme et al. 2020); Decay sugar-rich substrates (Põlme et al. 2020) | Põlme S, et al. 2020. Fungal Diversity 105:1-16 ((https://doi.org/10.1007/s13225-020-00466-2)) |
| Otu07666 | 3 | k__Fungi(100);p__Ascomycota(100);c__Saccharomycetes(100);o__Saccharomycetales(100);f__Metschnikowiaceae(100);g__Metschnikowia(100);s__Metschnikowia_pulcherrima(67); | Metschnikowia | 13 | Saprotroph | \|Undefined Saprotroph\| | Yeast | NULL | Probable | As nectar-tap_saprotroph (Põlme et al. 2020); Decay sugar-rich substrates (Põlme et al. 2020) | Põlme S, et al. 2020. Fungal Diversity 105:1-16 ((https://doi.org/10.1007/s13225-020-00466-2)) |
| Otu07667 | 3 | k__Fungi(100);p__Ascomycota(100);c__Saccharomycetes(100);o__Saccharomycetales(100);f__Metschnikowiaceae(100);g__Metschnikowia(100);s__Metschnikowia_pulcherrima(67); | Metschnikowia | 13 | Saprotroph | \|Undefined Saprotroph\| | Yeast | NULL | Probable | As nectar-tap_saprotroph (Põlme et al. 2020); Decay sugar-rich substrates (Põlme et al. 2020) | Põlme S, et al. 2020. Fungal Diversity 105:1-16 ((https://doi.org/10.1007/s13225-020-00466-2)) |
| Otu07669 | 3 | k__Fungi(100);p__Ascomycota(100);c__Saccharomycetes(100);o__Saccharomycetales(100);f__Metschnikowiaceae(100);g__Metschnikowia(100);s__Metschnikowia_pulcherrima(100); | Metschnikowia | 13 | Saprotroph | \|Undefined Saprotroph\| | Yeast | NULL | Probable | As nectar-tap_saprotroph (Põlme et al. 2020); Decay sugar-rich substrates (Põlme et al. 2020) | Põlme S, et al. 2020. Fungal Diversity 105:1-16 ((https://doi.org/10.1007/s13225-020-00466-2)) |
| Otu07670 | 3 | k__Fungi(100);p__Ascomycota(100);c__Saccharomycetes(100);o__Saccharomycetales(100);f__Metschnikowiaceae(100);g__Metschnikowia(100);s__Metschnikowia_pulcherrima(67); | Metschnikowia | 13 | Saprotroph | \|Undefined Saprotroph\| | Yeast | NULL | Probable | As nectar-tap_saprotroph (Põlme et al. 2020); Decay sugar-rich substrates (Põlme et al. 2020) | Põlme S, et al. 2020. Fungal Diversity 105:1-16 ((https://doi.org/10.1007/s13225-020-00466-2)) |
| Otu07672 | 3 | k__Fungi(100);p__Ascomycota(100);c__Saccharomycetes(100);o__Saccharomycetales(100);f__Metschnikowiaceae(100);g__Metschnikowia(100);s__Metschnikowia_pulcherrima(100); | Metschnikowia | 13 | Saprotroph | \|Undefined Saprotroph\| | Yeast | NULL | Probable | As nectar-tap_saprotroph (Põlme et al. 2020); Decay sugar-rich substrates (Põlme et al. 2020) | Põlme S, et al. 2020. Fungal Diversity 105:1-16 ((https://doi.org/10.1007/s13225-020-00466-2)) |
| Otu07673 | 3 | k__Fungi(100);p__Ascomycota(100);c__Leotiomycetes(100);o__Helotiales(100);f__Hyaloscyphaceae(100);f__Hyaloscyphaceae_unclassified(100);f__Hyaloscyphaceae_unclassified(100); | Hyaloscyphaceae | 9 | Saprotroph | Plant Saprotroph-Wood Saprotroph | Microfungus | NULL | Probable | Saprobic on woody and herbaceous material (Cannon & Kirk 2007) | Cannon PF, Kirk PM. 2007. Fungal Families of the World. CAB International, Cambridge (ISBN: 978-0851998275) |
| Otu07674 | 3 | k__Fungi(100);p__Ascomycota(100);c__Saccharomycetes(100);o__Saccharomycetales(100);f__Metschnikowiaceae(100);g__Metschnikowia(100);s__Metschnikowia_pulcherrima(67); | Metschnikowia | 13 | Saprotroph | \|Undefined Saprotroph\| | Yeast | NULL | Probable | As nectar-tap_saprotroph (Põlme et al. 2020); Decay sugar-rich substrates (Põlme et al. 2020) | Põlme S, et al. 2020. Fungal Diversity 105:1-16 ((https://doi.org/10.1007/s13225-020-00466-2)) |
| Otu07675 | 3 | k__Fungi(100);p__Ascomycota(100);c__Saccharomycetes(100);o__Saccharomycetales(100);f__Metschnikowiaceae(100);g__Metschnikowia(100);s__Metschnikowia_pulcherrima(100); | Metschnikowia | 13 | Saprotroph | \|Undefined Saprotroph\| | Yeast | NULL | Probable | As nectar-tap_saprotroph (Põlme et al. 2020); Decay sugar-rich substrates (Põlme et al. 2020) | Põlme S, et al. 2020. Fungal Diversity 105:1-16 ((https://doi.org/10.1007/s13225-020-00466-2)) |
| Otu07676 | 3 | k__Fungi(100);p__Ascomycota(100);c__Saccharomycetes(100);o__Saccharomycetales(100);f__Metschnikowiaceae(100);g__Metschnikowia(100);g__Metschnikowia_unclassified(100); | Metschnikowia | 13 | Saprotroph | \|Undefined Saprotroph\| | Yeast | NULL | Probable | As nectar-tap_saprotroph (Põlme et al. 2020); Decay sugar-rich substrates (Põlme et al. 2020) | Põlme S, et al. 2020. Fungal Diversity 105:1-16 ((https://doi.org/10.1007/s13225-020-00466-2)) |
| Otu07678 | 3 | k__Fungi(100);p__Ascomycota(100);c__Saccharomycetes(100);o__Saccharomycetales(100);f__Metschnikowiaceae(100);g__Metschnikowia(100);s__Metschnikowia_pulcherrima(67); | Metschnikowia | 13 | Saprotroph | \|Undefined Saprotroph\| | Yeast | NULL | Probable | As nectar-tap_saprotroph (Põlme et al. 2020); Decay sugar-rich substrates (Põlme et al. 2020) | Põlme S, et al. 2020. Fungal Diversity 105:1-16 ((https://doi.org/10.1007/s13225-020-00466-2)) |
| Otu07679 | 3 | k__Fungi(100);p__Ascomycota(100);c__Leotiomycetes(100);o__Helotiales(100);f__Helotiaceae(100);g__Hymenoscyphus(100);s__Hymenoscyphus_fructigenus(100); | Hymenoscyphus | 13 | Pathotroph-Saprotroph-Symbiotroph | Bryophyte Parasite-Ectomycorrhizal-Ericoid Mycorrhizal-Plant Pathogen-\|Plant Saprotroph\|-Undefined Saprotroph | Microfungus | NULL | Possible | As litter_saprotroph (Põlme et al. 2020); Dark Septate Microfungus; Syn. Pezoloma among other (see e.g., Pezoloma ericae); Decay leaf, fruit, seed (Põlme et al. 2020) | Baral HO, Krieglsteiner L. 2006. Acta Mycologica 41:11-20 ((https://doi.org/10.5586/am.2006.003)) (as a bryophyte parasite); Huhtinen S, et al. 2010. Nova Hedwigia 90:413-431 ((https://doi.org/10.1127/0029-5035/2010/0090-0413)); Bent E, et al. 2011. Applied and Environmental Microbiology 77:3351-3359 ((https://doi.org/10.1128/AEM.02575-10)) (as ectomycorrhizae); Tedersoo L, et al. 2014. Science 346:1256688 ((https://doi.org/10.1126/science.1256688)) (as saprotroph); Põlme S, et al. 2020. Fungal Diversity 105:1-16 ((https://doi.org/10.1007/s13225-020-00466-2)) |
| Otu07683 | 3 | k__Fungi(100);p__Basidiomycota(100);c__Agaricomycetes(100);o__Agaricales(100);f__Psathyrellaceae(100);g__Coprinellus(100);s__Coprinellus_sp(100); | Coprinellus | 13 | Saprotroph | \|Undefined Saprotroph\| | Agaricoid | NULL | Probable | As soil_saprotroph (Põlme et al. 2020); Decay soil (Põlme et al. 2020) | Tedersoo L, et al. 2014. Science 346:e1256688 ((https://doi.org/10.1126/science.1256688)); Põlme S, et al. 2020. Fungal Diversity 105:1-16 ((https://doi.org/10.1007/s13225-020-00466-2)) |
| Otu07686 | 3 | k__Fungi(100);p__Basidiomycota(100);c__Microbotryomycetes(100);o__Sporidiobolales(100);f__Sporidiobolales_family_Incertae_sedis(100);g__Sporobolomyces(100);s__Sporobolomyces_lactosus(100); | Sporobolomyces | 13 | Pathotroph-Saprotroph | \|Fungal Parasite\|-Plant Saprotroph | Yeast | NULL | Probable | As litter_saprotroph (Põlme et al. 2020); As mycoparasite (Põlme et al. 2020); Decay fungal material (Põlme et al. 2020) | Tedersoo L, et al. 2014. Science 346:e1256688 ((https://doi.org/10.1126/science.1256688)); Purahong W, et al. 2016. Molecular Ecology 25:4059-4074 ((https://doi.org/10.1111/mec.13739)); Põlme S, et al. 2020. Fungal Diversity 105:1-16 ((https://doi.org/10.1007/s13225-020-00466-2)) |
| Otu07687 | 3 | k__Fungi(100);p__Ascomycota(100);c__Saccharomycetes(100);o__Saccharomycetales(100);f__Metschnikowiaceae(100);g__Metschnikowia(100);g__Metschnikowia_unclassified(100); | Metschnikowia | 13 | Saprotroph | \|Undefined Saprotroph\| | Yeast | NULL | Probable | As nectar-tap_saprotroph (Põlme et al. 2020); Decay sugar-rich substrates (Põlme et al. 2020) | Põlme S, et al. 2020. Fungal Diversity 105:1-16 ((https://doi.org/10.1007/s13225-020-00466-2)) |
| Otu07688 | 3 | k__Fungi(100);p__Ascomycota(100);c__Leotiomycetes(100);o__Helotiales(100);f__Hyaloscyphaceae(100);g__Lachnum(100);s__Lachnum_nudipes(100); | Lachnum | 13 | Saprotroph-Symbiotroph | Endophyte-\|Plant Saprotroph\|-Undefined Saprotroph-Wood Saprotroph | Microfungus | NULL | Probable | As litter_saprotroph (Põlme et al. 2020); As root_endophyte (Põlme et al. 2020); Decay wood, leaf, fruit, seed (Põlme et al. 2020) | Tedersoo L, et al. 2014. Science 346:e1256688 ((https://doi.org/10.1126/science.1256688)); Põlme S, et al. 2020. Fungal Diversity 105:1-16 ((https://doi.org/10.1007/s13225-020-00466-2)) |
| Otu07689 | 3 | k__Fungi(100);p__Ascomycota(100);c__Saccharomycetes(100);o__Saccharomycetales(100);f__Metschnikowiaceae(100);g__Metschnikowia(100);s__Metschnikowia_pulcherrima(67); | Metschnikowia | 13 | Saprotroph | \|Undefined Saprotroph\| | Yeast | NULL | Probable | As nectar-tap_saprotroph (Põlme et al. 2020); Decay sugar-rich substrates (Põlme et al. 2020) | Põlme S, et al. 2020. Fungal Diversity 105:1-16 ((https://doi.org/10.1007/s13225-020-00466-2)) |
| Otu07690 | 3 | k__Fungi(100);p__Ascomycota(100);c__Saccharomycetes(100);o__Saccharomycetales(100);f__Metschnikowiaceae(100);g__Metschnikowia(100);s__Metschnikowia_pulcherrima(100); | Metschnikowia | 13 | Saprotroph | \|Undefined Saprotroph\| | Yeast | NULL | Probable | As nectar-tap_saprotroph (Põlme et al. 2020); Decay sugar-rich substrates (Põlme et al. 2020) | Põlme S, et al. 2020. Fungal Diversity 105:1-16 ((https://doi.org/10.1007/s13225-020-00466-2)) |
| Otu07691 | 3 | k__Fungi(100);p__Ascomycota(100);c__Saccharomycetes(100);o__Saccharomycetales(100);f__Metschnikowiaceae(100);g__Metschnikowia(100);g__Metschnikowia_unclassified(67); | Metschnikowia | 13 | Saprotroph | \|Undefined Saprotroph\| | Yeast | NULL | Probable | As nectar-tap_saprotroph (Põlme et al. 2020); Decay sugar-rich substrates (Põlme et al. 2020) | Põlme S, et al. 2020. Fungal Diversity 105:1-16 ((https://doi.org/10.1007/s13225-020-00466-2)) |
| Otu07693 | 3 | k__Fungi(100);p__Basidiomycota(100);c__Wallemiomycetes(100);o__Wallemiales(100);f__Wallemiaceae(100);g__Wallemia(100);s__Wallemia_sebi(100); | Wallemia | 13 | Saprotroph | \|Undefined Saprotroph\| | Microfungus | NULL | Probable | Decay sugar-rich substrates (Põlme et al. 2020) | Tedersoo L, et al. 2014. Science 346:e1256688 ((https://doi.org/10.1126/science.1256688)); Põlme S, et al. 2020. Fungal Diversity 105:1-16 ((https://doi.org/10.1007/s13225-020-00466-2)) |
| Otu07695 | 3 | k__Fungi(100);p__Ascomycota(100);c__Sordariomycetes(100);o__Sordariales(100);f__Lasiosphaeriaceae(100);g__Podospora(100);s__Podospora_longicollis(100); | Podospora | 13 | Saprotroph-Symbiotroph | \|Dung Saprotroph\|-Endophyte-Plant Saprotroph-Undefined Saprotroph | Microfungus | NULL | Possible | As foliar_endophyte (Põlme et al. 2020); As litter_saprotroph (Põlme et al. 2020); culture (David et al. 2016); Decay leaf, fruit, seed, dung (Põlme et al. 2020); Endophyte detection method | Bell A. 1983. Dung Fungi: An Illustrated Guide to Coprophilous Fungi in New Zealand. Victoria University Press, Wellington (ISBN: 978-0864730015); Tedersoo L, et al. 2014. Science 346:e1256688 ((https://doi.org/10.1126/science.1256688)); David AS, et al. 2016. Microbial Ecology 71:912-926 ((https://doi.org/10.1007/s00248-015-0712-6)); Põlme S, et al. 2020. Fungal Diversity 105:1-16 ((https://doi.org/10.1007/s13225-020-00466-2)) |
| Otu07696 | 3 | k__Fungi(100);p__Ascomycota(100);c__Saccharomycetes(100);o__Saccharomycetales(100);f__Metschnikowiaceae(100);g__Metschnikowia(100);g__Metschnikowia_unclassified(67); | Metschnikowia | 13 | Saprotroph | \|Undefined Saprotroph\| | Yeast | NULL | Probable | As nectar-tap_saprotroph (Põlme et al. 2020); Decay sugar-rich substrates (Põlme et al. 2020) | Põlme S, et al. 2020. Fungal Diversity 105:1-16 ((https://doi.org/10.1007/s13225-020-00466-2)) |
| Otu07698 | 3 | k__Fungi(100);p__Ascomycota(100);c__Saccharomycetes(100);o__Saccharomycetales(100);f__Metschnikowiaceae(100);g__Metschnikowia(100);s__Metschnikowia_pulcherrima(67); | Metschnikowia | 13 | Saprotroph | \|Undefined Saprotroph\| | Yeast | NULL | Probable | As nectar-tap_saprotroph (Põlme et al. 2020); Decay sugar-rich substrates (Põlme et al. 2020) | Põlme S, et al. 2020. Fungal Diversity 105:1-16 ((https://doi.org/10.1007/s13225-020-00466-2)) |
| Otu07701 | 3 | k__Fungi(100);p__Basidiomycota(100);c__Wallemiomycetes(100);o__Wallemiales(100);f__Wallemiaceae(100);g__Wallemia(100);s__Wallemia_sebi(100); | Wallemia | 13 | Saprotroph | \|Undefined Saprotroph\| | Microfungus | NULL | Probable | Decay sugar-rich substrates (Põlme et al. 2020) | Tedersoo L, et al. 2014. Science 346:e1256688 ((https://doi.org/10.1126/science.1256688)); Põlme S, et al. 2020. Fungal Diversity 105:1-16 ((https://doi.org/10.1007/s13225-020-00466-2)) |
| Otu07704 | 3 | k__Fungi(100);p__Ascomycota(100);c__Saccharomycetes(100);o__Saccharomycetales(100);f__Metschnikowiaceae(100);g__Metschnikowia(100);s__Metschnikowia_pulcherrima(100); | Metschnikowia | 13 | Saprotroph | \|Undefined Saprotroph\| | Yeast | NULL | Probable | As nectar-tap_saprotroph (Põlme et al. 2020); Decay sugar-rich substrates (Põlme et al. 2020) | Põlme S, et al. 2020. Fungal Diversity 105:1-16 ((https://doi.org/10.1007/s13225-020-00466-2)) |
| Otu07706 | 3 | k__Fungi(100);p__Ascomycota(100);c__Saccharomycetes(100);o__Saccharomycetales(100);f__Metschnikowiaceae(100);g__Metschnikowia(100);s__Metschnikowia_pulcherrima(100); | Metschnikowia | 13 | Saprotroph | \|Undefined Saprotroph\| | Yeast | NULL | Probable | As nectar-tap_saprotroph (Põlme et al. 2020); Decay sugar-rich substrates (Põlme et al. 2020) | Põlme S, et al. 2020. Fungal Diversity 105:1-16 ((https://doi.org/10.1007/s13225-020-00466-2)) |
| Otu07707 | 3 | k__Fungi(100);p__Basidiomycota(100);c__Microbotryomycetes(100);o__Sporidiobolales(100);f__Sporidiobolales_family_Incertae_sedis(100);g__Sporobolomyces(100);s__Sporobolomyces_lactosus(100); | Sporobolomyces | 13 | Pathotroph-Saprotroph | \|Fungal Parasite\|-Plant Saprotroph | Yeast | NULL | Probable | As litter_saprotroph (Põlme et al. 2020); As mycoparasite (Põlme et al. 2020); Decay fungal material (Põlme et al. 2020) | Tedersoo L, et al. 2014. Science 346:e1256688 ((https://doi.org/10.1126/science.1256688)); Purahong W, et al. 2016. Molecular Ecology 25:4059-4074 ((https://doi.org/10.1111/mec.13739)); Põlme S, et al. 2020. Fungal Diversity 105:1-16 ((https://doi.org/10.1007/s13225-020-00466-2)) |
| Otu07709 | 3 | k__Fungi(100);p__Ascomycota(100);c__Sordariomycetes(100);o__Sordariales(100);f__Lasiosphaeriaceae(67);g__Cladorrhinum(67);s__Cladorrhinum_sp(67); | Cladorrhinum | 13 | Pathotroph-Saprotroph-Symbiotroph | Animal Pathogen-Dung Saprotroph-Endophyte-Plant Pathogen-Plant Saprotroph-\|Undefined Saprotroph\|-Wood Saprotroph | Microfungus | Soft Rot | Probable | As foliar_endophyte (Põlme et al. 2020); Decay leaf, fruit, seed (Põlme et al. 2020); Reported in soils (including agricultural) as saprotrophs on dung or plant material, or in roots as endophytes, while some species are associated with human and animal opportunistic diseases (Carmaran et al.2015) | Seehann G, et al. 1975. List of Fungi in Soft-Rot Tissue Tests. Institute fur Holzbiologie und Holzschutz, Bundesforschungsanstalt fur Forst - und Holzwirtshaft, Hamburg - Lohbrugge, Germany; Carmaran CC, et al. 2015. Mycological Progress 14:e94 ((https://doi.org/10.1007/s11557-015-1106-3)); Põlme S, et al. 2020. Fungal Diversity 105:1-16 ((https://doi.org/10.1007/s13225-020-00466-2)) |
| Otu07711 | 3 | k__Fungi(100);p__Ascomycota(100);c__Saccharomycetes(100);o__Saccharomycetales(100);f__Metschnikowiaceae(100);g__Metschnikowia(100);s__Metschnikowia_pulcherrima(100); | Metschnikowia | 13 | Saprotroph | \|Undefined Saprotroph\| | Yeast | NULL | Probable | As nectar-tap_saprotroph (Põlme et al. 2020); Decay sugar-rich substrates (Põlme et al. 2020) | Põlme S, et al. 2020. Fungal Diversity 105:1-16 ((https://doi.org/10.1007/s13225-020-00466-2)) |
| Otu07713 | 3 | k__Fungi(100);p__Ascomycota(100);c__Saccharomycetes(100);o__Saccharomycetales(100);f__Metschnikowiaceae(100);g__Metschnikowia(100);s__Metschnikowia_pulcherrima(67); | Metschnikowia | 13 | Saprotroph | \|Undefined Saprotroph\| | Yeast | NULL | Probable | As nectar-tap_saprotroph (Põlme et al. 2020); Decay sugar-rich substrates (Põlme et al. 2020) | Põlme S, et al. 2020. Fungal Diversity 105:1-16 ((https://doi.org/10.1007/s13225-020-00466-2)) |
| Otu07714 | 3 | k__Fungi(100);p__Ascomycota(100);c__Saccharomycetes(100);o__Saccharomycetales(100);f__Metschnikowiaceae(100);g__Metschnikowia(100);s__Metschnikowia_pulcherrima(100); | Metschnikowia | 13 | Saprotroph | \|Undefined Saprotroph\| | Yeast | NULL | Probable | As nectar-tap_saprotroph (Põlme et al. 2020); Decay sugar-rich substrates (Põlme et al. 2020) | Põlme S, et al. 2020. Fungal Diversity 105:1-16 ((https://doi.org/10.1007/s13225-020-00466-2)) |
| Otu07716 | 3 | k__Fungi(100);p__Ascomycota(100);c__Saccharomycetes(100);o__Saccharomycetales(100);f__Metschnikowiaceae(100);g__Metschnikowia(100);s__Metschnikowia_pulcherrima(67); | Metschnikowia | 13 | Saprotroph | \|Undefined Saprotroph\| | Yeast | NULL | Probable | As nectar-tap_saprotroph (Põlme et al. 2020); Decay sugar-rich substrates (Põlme et al. 2020) | Põlme S, et al. 2020. Fungal Diversity 105:1-16 ((https://doi.org/10.1007/s13225-020-00466-2)) |
| Otu07718 | 3 | k__Fungi(100);p__Ascomycota(100);c__Saccharomycetes(100);o__Saccharomycetales(100);f__Metschnikowiaceae(100);g__Metschnikowia(100);s__Metschnikowia_pulcherrima(100); | Metschnikowia | 13 | Saprotroph | \|Undefined Saprotroph\| | Yeast | NULL | Probable | As nectar-tap_saprotroph (Põlme et al. 2020); Decay sugar-rich substrates (Põlme et al. 2020) | Põlme S, et al. 2020. Fungal Diversity 105:1-16 ((https://doi.org/10.1007/s13225-020-00466-2)) |
| Otu07719 | 3 | k__Fungi(100);p__Ascomycota(100);c__Saccharomycetes(100);o__Saccharomycetales(100);f__Metschnikowiaceae(100);g__Metschnikowia(100);s__Metschnikowia_pulcherrima(67); | Metschnikowia | 13 | Saprotroph | \|Undefined Saprotroph\| | Yeast | NULL | Probable | As nectar-tap_saprotroph (Põlme et al. 2020); Decay sugar-rich substrates (Põlme et al. 2020) | Põlme S, et al. 2020. Fungal Diversity 105:1-16 ((https://doi.org/10.1007/s13225-020-00466-2)) |
| Otu07720 | 3 | k__Fungi(100);p__Ascomycota(100);c__Saccharomycetes(100);o__Saccharomycetales(100);f__Metschnikowiaceae(100);g__Metschnikowia(100);g__Metschnikowia_unclassified(67); | Metschnikowia | 13 | Saprotroph | \|Undefined Saprotroph\| | Yeast | NULL | Probable | As nectar-tap_saprotroph (Põlme et al. 2020); Decay sugar-rich substrates (Põlme et al. 2020) | Põlme S, et al. 2020. Fungal Diversity 105:1-16 ((https://doi.org/10.1007/s13225-020-00466-2)) |
| Otu07721 | 3 | k__Fungi(100);p__Ascomycota(100);c__Saccharomycetes(100);o__Saccharomycetales(100);f__Metschnikowiaceae(100);g__Metschnikowia(100);s__Metschnikowia_pulcherrima(100); | Metschnikowia | 13 | Saprotroph | \|Undefined Saprotroph\| | Yeast | NULL | Probable | As nectar-tap_saprotroph (Põlme et al. 2020); Decay sugar-rich substrates (Põlme et al. 2020) | Põlme S, et al. 2020. Fungal Diversity 105:1-16 ((https://doi.org/10.1007/s13225-020-00466-2)) |
| Otu07722 | 3 | k__Fungi(100);p__Ascomycota(100);c__Saccharomycetes(100);o__Saccharomycetales(100);f__Metschnikowiaceae(100);g__Metschnikowia(100);s__Metschnikowia_pulcherrima(67); | Metschnikowia | 13 | Saprotroph | \|Undefined Saprotroph\| | Yeast | NULL | Probable | As nectar-tap_saprotroph (Põlme et al. 2020); Decay sugar-rich substrates (Põlme et al. 2020) | Põlme S, et al. 2020. Fungal Diversity 105:1-16 ((https://doi.org/10.1007/s13225-020-00466-2)) |
| Otu07726 | 3 | k__Fungi(100);p__Ascomycota(100);c__Saccharomycetes(100);o__Saccharomycetales(100);f__Metschnikowiaceae(100);g__Metschnikowia(100);s__Metschnikowia_pulcherrima(100); | Metschnikowia | 13 | Saprotroph | \|Undefined Saprotroph\| | Yeast | NULL | Probable | As nectar-tap_saprotroph (Põlme et al. 2020); Decay sugar-rich substrates (Põlme et al. 2020) | Põlme S, et al. 2020. Fungal Diversity 105:1-16 ((https://doi.org/10.1007/s13225-020-00466-2)) |
| Otu07727 | 3 | k__Fungi(100);p__Ascomycota(100);c__Saccharomycetes(100);o__Saccharomycetales(100);f__Metschnikowiaceae(100);g__Metschnikowia(100);s__Metschnikowia_pulcherrima(100); | Metschnikowia | 13 | Saprotroph | \|Undefined Saprotroph\| | Yeast | NULL | Probable | As nectar-tap_saprotroph (Põlme et al. 2020); Decay sugar-rich substrates (Põlme et al. 2020) | Põlme S, et al. 2020. Fungal Diversity 105:1-16 ((https://doi.org/10.1007/s13225-020-00466-2)) |
| Otu07729 | 3 | k__Fungi(100);p__Ascomycota(100);c__Saccharomycetes(100);o__Saccharomycetales(100);f__Metschnikowiaceae(100);g__Metschnikowia(100);g__Metschnikowia_unclassified(67); | Metschnikowia | 13 | Saprotroph | \|Undefined Saprotroph\| | Yeast | NULL | Probable | As nectar-tap_saprotroph (Põlme et al. 2020); Decay sugar-rich substrates (Põlme et al. 2020) | Põlme S, et al. 2020. Fungal Diversity 105:1-16 ((https://doi.org/10.1007/s13225-020-00466-2)) |
| Otu07732 | 3 | k__Fungi(100);p__Ascomycota(100);c__Saccharomycetes(100);o__Saccharomycetales(100);f__Metschnikowiaceae(100);g__Metschnikowia(100);g__Metschnikowia_unclassified(67); | Metschnikowia | 13 | Saprotroph | \|Undefined Saprotroph\| | Yeast | NULL | Probable | As nectar-tap_saprotroph (Põlme et al. 2020); Decay sugar-rich substrates (Põlme et al. 2020) | Põlme S, et al. 2020. Fungal Diversity 105:1-16 ((https://doi.org/10.1007/s13225-020-00466-2)) |
| Otu07733 | 3 | k__Fungi(100);p__Ascomycota(100);c__Saccharomycetes(100);o__Saccharomycetales(100);f__Saccharomycetales_family_Incertae_sedis(100);f__Saccharomycetales_family_Incertae_sedis_unclassified(100);f__Saccharomycetales_family_Incertae_sedis_unclassified(100); | Saccharomycetales | 7 | Saprotroph | Undefined Saprotroph | Yeast | NULL | Possible | NULL | Sterkenburg E, et al. 2015. New Phytologist 207:1145-1158 ((https://doi.org/10.1111/nph.13426)) |
| Otu07734 | 3 | k__Fungi(100);p__Ascomycota(100);c__Saccharomycetes(100);o__Saccharomycetales(100);f__Metschnikowiaceae(100);g__Metschnikowia(100);s__Metschnikowia_pulcherrima(100); | Metschnikowia | 13 | Saprotroph | \|Undefined Saprotroph\| | Yeast | NULL | Probable | As nectar-tap_saprotroph (Põlme et al. 2020); Decay sugar-rich substrates (Põlme et al. 2020) | Põlme S, et al. 2020. Fungal Diversity 105:1-16 ((https://doi.org/10.1007/s13225-020-00466-2)) |
| Otu07736 | 3 | k__Fungi(100);p__Ascomycota(100);c__Saccharomycetes(100);o__Saccharomycetales(100);f__Metschnikowiaceae(100);g__Metschnikowia(100);s__Metschnikowia_pulcherrima(67); | Metschnikowia | 13 | Saprotroph | \|Undefined Saprotroph\| | Yeast | NULL | Probable | As nectar-tap_saprotroph (Põlme et al. 2020); Decay sugar-rich substrates (Põlme et al. 2020) | Põlme S, et al. 2020. Fungal Diversity 105:1-16 ((https://doi.org/10.1007/s13225-020-00466-2)) |
| Otu07738 | 3 | k__Fungi(100);p__Ascomycota(100);c__Saccharomycetes(100);o__Saccharomycetales(100);f__Metschnikowiaceae(100);g__Metschnikowia(100);s__Metschnikowia_pulcherrima(67); | Metschnikowia | 13 | Saprotroph | \|Undefined Saprotroph\| | Yeast | NULL | Probable | As nectar-tap_saprotroph (Põlme et al. 2020); Decay sugar-rich substrates (Põlme et al. 2020) | Põlme S, et al. 2020. Fungal Diversity 105:1-16 ((https://doi.org/10.1007/s13225-020-00466-2)) |
| Otu07739 | 3 | k__Fungi(100);p__Ascomycota(100);c__Saccharomycetes(100);o__Saccharomycetales(100);f__Metschnikowiaceae(100);g__Metschnikowia(100);s__Metschnikowia_pulcherrima(100); | Metschnikowia | 13 | Saprotroph | \|Undefined Saprotroph\| | Yeast | NULL | Probable | As nectar-tap_saprotroph (Põlme et al. 2020); Decay sugar-rich substrates (Põlme et al. 2020) | Põlme S, et al. 2020. Fungal Diversity 105:1-16 ((https://doi.org/10.1007/s13225-020-00466-2)) |
| Otu07740 | 3 | k__Fungi(100);p__Ascomycota(100);c__Saccharomycetes(100);o__Saccharomycetales(100);f__Metschnikowiaceae(100);g__Metschnikowia(100);s__Metschnikowia_pulcherrima(67); | Metschnikowia | 13 | Saprotroph | \|Undefined Saprotroph\| | Yeast | NULL | Probable | As nectar-tap_saprotroph (Põlme et al. 2020); Decay sugar-rich substrates (Põlme et al. 2020) | Põlme S, et al. 2020. Fungal Diversity 105:1-16 ((https://doi.org/10.1007/s13225-020-00466-2)) |
| Otu07741 | 3 | k__Fungi(100);p__Ascomycota(100);c__Saccharomycetes(100);o__Saccharomycetales(100);f__Saccharomycetales_family_Incertae_sedis(67);g__Candida(67);g__Candida_unclassified(67); | Candida | 13 | Pathotroph-Saprotroph-Symbiotroph | Animal Pathogen-Endophyte-Endosymbiont-Epiphyte-\|Undefined Saprotroph\| | Dimorphic Yeast-Yeast | NULL | Probable | As nectar-tap_saprotroph (Põlme et al. 2020); Decay sugar-rich substrates (Põlme et al. 2020); This asexual genus is polyphyletic and contain species that function very differently depend on what they are. Caution should be taken when interpreting guilds for Candida species. | Manolakaki D, et al. 2010. Virulence. 1: 367-375 ((https://doi.org/10.4161/viru.1.5.12796)); Mekbib SB, et al. 2011. Tropical Plant Pathology 36:233-240 ((https://doi.org/10.1590/S1982-56762011000400004)); Põlme S, et al. 2020. Fungal Diversity 105:1-16 ((https://doi.org/10.1007/s13225-020-00466-2)) |
| Otu07743 | 3 | k__Fungi(100);p__Ascomycota(100);c__Saccharomycetes(100);o__Saccharomycetales(100);f__Metschnikowiaceae(100);g__Metschnikowia(100);s__Metschnikowia_pulcherrima(100); | Metschnikowia | 13 | Saprotroph | \|Undefined Saprotroph\| | Yeast | NULL | Probable | As nectar-tap_saprotroph (Põlme et al. 2020); Decay sugar-rich substrates (Põlme et al. 2020) | Põlme S, et al. 2020. Fungal Diversity 105:1-16 ((https://doi.org/10.1007/s13225-020-00466-2)) |
| Otu07746 | 3 | k__Fungi(100);p__Ascomycota(100);c__Saccharomycetes(100);o__Saccharomycetales(100);f__Metschnikowiaceae(100);g__Metschnikowia(100);g__Metschnikowia_unclassified(67); | Metschnikowia | 13 | Saprotroph | \|Undefined Saprotroph\| | Yeast | NULL | Probable | As nectar-tap_saprotroph (Põlme et al. 2020); Decay sugar-rich substrates (Põlme et al. 2020) | Põlme S, et al. 2020. Fungal Diversity 105:1-16 ((https://doi.org/10.1007/s13225-020-00466-2)) |
| Otu07748 | 3 | k__Fungi(100);p__Ascomycota(100);c__Saccharomycetes(100);o__Saccharomycetales(100);f__Metschnikowiaceae(100);g__Metschnikowia(100);s__Metschnikowia_pulcherrima(100); | Metschnikowia | 13 | Saprotroph | \|Undefined Saprotroph\| | Yeast | NULL | Probable | As nectar-tap_saprotroph (Põlme et al. 2020); Decay sugar-rich substrates (Põlme et al. 2020) | Põlme S, et al. 2020. Fungal Diversity 105:1-16 ((https://doi.org/10.1007/s13225-020-00466-2)) |
| Otu07749 | 3 | k__Fungi(100);p__Ascomycota(100);c__Saccharomycetes(100);o__Saccharomycetales(100);f__Metschnikowiaceae(100);g__Metschnikowia(100);g__Metschnikowia_unclassified(67); | Metschnikowia | 13 | Saprotroph | \|Undefined Saprotroph\| | Yeast | NULL | Probable | As nectar-tap_saprotroph (Põlme et al. 2020); Decay sugar-rich substrates (Põlme et al. 2020) | Põlme S, et al. 2020. Fungal Diversity 105:1-16 ((https://doi.org/10.1007/s13225-020-00466-2)) |
| Otu07750 | 3 | k__Fungi(100);p__Ascomycota(100);c__Saccharomycetes(100);o__Saccharomycetales(100);f__Metschnikowiaceae(100);g__Metschnikowia(100);s__Metschnikowia_pulcherrima(100); | Metschnikowia | 13 | Saprotroph | \|Undefined Saprotroph\| | Yeast | NULL | Probable | As nectar-tap_saprotroph (Põlme et al. 2020); Decay sugar-rich substrates (Põlme et al. 2020) | Põlme S, et al. 2020. Fungal Diversity 105:1-16 ((https://doi.org/10.1007/s13225-020-00466-2)) |
| Otu07751 | 3 | k__Fungi(100);p__Ascomycota(100);c__Saccharomycetes(100);o__Saccharomycetales(100);f__Metschnikowiaceae(100);g__Metschnikowia(100);s__Metschnikowia_pulcherrima(100); | Metschnikowia | 13 | Saprotroph | \|Undefined Saprotroph\| | Yeast | NULL | Probable | As nectar-tap_saprotroph (Põlme et al. 2020); Decay sugar-rich substrates (Põlme et al. 2020) | Põlme S, et al. 2020. Fungal Diversity 105:1-16 ((https://doi.org/10.1007/s13225-020-00466-2)) |
| Otu07752 | 3 | k__Fungi(100);p__Ascomycota(100);c__Saccharomycetes(100);o__Saccharomycetales(100);f__Metschnikowiaceae(100);g__Metschnikowia(100);s__Metschnikowia_pulcherrima(67); | Metschnikowia | 13 | Saprotroph | \|Undefined Saprotroph\| | Yeast | NULL | Probable | As nectar-tap_saprotroph (Põlme et al. 2020); Decay sugar-rich substrates (Põlme et al. 2020) | Põlme S, et al. 2020. Fungal Diversity 105:1-16 ((https://doi.org/10.1007/s13225-020-00466-2)) |
| Otu07753 | 3 | k__Fungi(100);p__Ascomycota(100);c__Saccharomycetes(100);o__Saccharomycetales(100);f__Metschnikowiaceae(100);g__Metschnikowia(100);g__Metschnikowia_unclassified(67); | Metschnikowia | 13 | Saprotroph | \|Undefined Saprotroph\| | Yeast | NULL | Probable | As nectar-tap_saprotroph (Põlme et al. 2020); Decay sugar-rich substrates (Põlme et al. 2020) | Põlme S, et al. 2020. Fungal Diversity 105:1-16 ((https://doi.org/10.1007/s13225-020-00466-2)) |
| Otu07754 | 3 | k__Fungi(100);p__Ascomycota(100);c__Saccharomycetes(100);o__Saccharomycetales(100);f__Metschnikowiaceae(100);g__Metschnikowia(100);g__Metschnikowia_unclassified(67); | Metschnikowia | 13 | Saprotroph | \|Undefined Saprotroph\| | Yeast | NULL | Probable | As nectar-tap_saprotroph (Põlme et al. 2020); Decay sugar-rich substrates (Põlme et al. 2020) | Põlme S, et al. 2020. Fungal Diversity 105:1-16 ((https://doi.org/10.1007/s13225-020-00466-2)) |
| Otu07755 | 3 | k__Fungi(100);p__Ascomycota(100);c__Dothideomycetes(100);o__Pleosporales(100);f__Massarinaceae(100);g__Massarina(100);s__Massarina_sp_HF12700(100); | Massarina | 13 | Pathotroph-Saprotroph | \|Plant Pathogen\|-Undefined Saprotroph | Microfungus | NULL | Probable | NULL | Tedersoo L, et al. 2014. Science 346:e1256688 ((https://doi.org/10.1126/science.1256688)); Põlme S, et al. 2020. Fungal Diversity 105:1-16 ((https://doi.org/10.1007/s13225-020-00466-2)) |
| Otu07757 | 3 | k__Fungi(100);p__Ascomycota(100);c__Saccharomycetes(100);o__Saccharomycetales(100);f__Metschnikowiaceae(100);g__Metschnikowia(100);s__Metschnikowia_pulcherrima(100); | Metschnikowia | 13 | Saprotroph | \|Undefined Saprotroph\| | Yeast | NULL | Probable | As nectar-tap_saprotroph (Põlme et al. 2020); Decay sugar-rich substrates (Põlme et al. 2020) | Põlme S, et al. 2020. Fungal Diversity 105:1-16 ((https://doi.org/10.1007/s13225-020-00466-2)) |
| Otu07758 | 3 | k__Fungi(100);p__Basidiomycota(100);c__Microbotryomycetes(100);o__Sporidiobolales(100);f__Sporidiobolales_family_Incertae_sedis(100);g__Sporobolomyces(100);s__Sporobolomyces_lactosus(100); | Sporobolomyces | 13 | Pathotroph-Saprotroph | \|Fungal Parasite\|-Plant Saprotroph | Yeast | NULL | Probable | As litter_saprotroph (Põlme et al. 2020); As mycoparasite (Põlme et al. 2020); Decay fungal material (Põlme et al. 2020) | Tedersoo L, et al. 2014. Science 346:e1256688 ((https://doi.org/10.1126/science.1256688)); Purahong W, et al. 2016. Molecular Ecology 25:4059-4074 ((https://doi.org/10.1111/mec.13739)); Põlme S, et al. 2020. Fungal Diversity 105:1-16 ((https://doi.org/10.1007/s13225-020-00466-2)) |
| Otu07759 | 3 | k__Fungi(100);p__Basidiomycota(100);c__Microbotryomycetes(100);o__Sporidiobolales(100);f__Sporidiobolales_family_Incertae_sedis(100);g__Sporobolomyces(100);s__Sporobolomyces_lactosus(100); | Sporobolomyces | 13 | Pathotroph-Saprotroph | \|Fungal Parasite\|-Plant Saprotroph | Yeast | NULL | Probable | As litter_saprotroph (Põlme et al. 2020); As mycoparasite (Põlme et al. 2020); Decay fungal material (Põlme et al. 2020) | Tedersoo L, et al. 2014. Science 346:e1256688 ((https://doi.org/10.1126/science.1256688)); Purahong W, et al. 2016. Molecular Ecology 25:4059-4074 ((https://doi.org/10.1111/mec.13739)); Põlme S, et al. 2020. Fungal Diversity 105:1-16 ((https://doi.org/10.1007/s13225-020-00466-2)) |
| Otu07762 | 3 | k__Fungi(100);p__Ascomycota(100);c__Dothideomycetes(100);o__Pleosporales(100);f__Montagnulaceae(67);f__Montagnulaceae_unclassified(67);f__Montagnulaceae_unclassified(67); | Montagnulaceae | 9 | Pathotroph-Saprotroph | Fungal Parasite-Plant Pathogen-Undefined Saprotroph | Microfungus | NULL | Probable | Species in this clade can be saprobic, plant pathogenic, or mycoparasitic (Zhang et al. 2009) | Zhang Y, et al. 2009. Studies in Mycology 64:85-102 ((https://doi.org/10.3114/sim.2009.64.04)) |
| Otu07767 | 3 | k__Fungi(100);p__Ascomycota(100);c__Saccharomycetes(100);o__Saccharomycetales(100);f__Metschnikowiaceae(100);g__Metschnikowia(100);g__Metschnikowia_unclassified(67); | Metschnikowia | 13 | Saprotroph | \|Undefined Saprotroph\| | Yeast | NULL | Probable | As nectar-tap_saprotroph (Põlme et al. 2020); Decay sugar-rich substrates (Põlme et al. 2020) | Põlme S, et al. 2020. Fungal Diversity 105:1-16 ((https://doi.org/10.1007/s13225-020-00466-2)) |
| Otu07768 | 3 | k__Fungi(100);p__Ascomycota(100);c__Saccharomycetes(100);o__Saccharomycetales(100);f__Metschnikowiaceae(100);g__Metschnikowia(100);s__Metschnikowia_pulcherrima(67); | Metschnikowia | 13 | Saprotroph | \|Undefined Saprotroph\| | Yeast | NULL | Probable | As nectar-tap_saprotroph (Põlme et al. 2020); Decay sugar-rich substrates (Põlme et al. 2020) | Põlme S, et al. 2020. Fungal Diversity 105:1-16 ((https://doi.org/10.1007/s13225-020-00466-2)) |
| Otu07769 | 3 | k__Fungi(100);p__Ascomycota(100);c__Saccharomycetes(100);o__Saccharomycetales(100);f__Metschnikowiaceae(100);g__Metschnikowia(100);s__Metschnikowia_pulcherrima(100); | Metschnikowia | 13 | Saprotroph | \|Undefined Saprotroph\| | Yeast | NULL | Probable | As nectar-tap_saprotroph (Põlme et al. 2020); Decay sugar-rich substrates (Põlme et al. 2020) | Põlme S, et al. 2020. Fungal Diversity 105:1-16 ((https://doi.org/10.1007/s13225-020-00466-2)) |
| Otu07770 | 3 | k__Fungi(100);p__Basidiomycota(100);c__Tremellomycetes(100);o__Cystofilobasidiales(100);f__Cystofilobasidiales_family_Incertae_sedis(100);g__Syzygospora(100);s__Syzygospora_bachmannii(100); | Syzygospora | 13 | Pathotroph | \|Fungal Parasite\|-Lichen Parasite | Dimorphic-Tremelloid | NULL | Highly Probable | As mycoparasite (Põlme et al. 2020); Decay fungal material (Põlme et al. 2020) | Põlme S, et al. 2020. Fungal Diversity 105:1-16 ((https://doi.org/10.1007/s13225-020-00466-2)); List compiled by John Plischke III ((https://goodmushroom.weebly.com/uploads/5/7/1/7/57176535/fungi_on_fungi_pdf_lq.pdf)) |
| Otu07772 | 3 | k__Fungi(100);p__Ascomycota(100);c__Saccharomycetes(100);o__Saccharomycetales(100);f__Metschnikowiaceae(100);g__Metschnikowia(100);s__Metschnikowia_pulcherrima(100); | Metschnikowia | 13 | Saprotroph | \|Undefined Saprotroph\| | Yeast | NULL | Probable | As nectar-tap_saprotroph (Põlme et al. 2020); Decay sugar-rich substrates (Põlme et al. 2020) | Põlme S, et al. 2020. Fungal Diversity 105:1-16 ((https://doi.org/10.1007/s13225-020-00466-2)) |
| Otu07773 | 3 | k__Fungi(100);p__Ascomycota(100);c__Sordariomycetes(100);o__Sordariales(100);f__Cephalothecaceae(100);g__Phialemonium(100);s__Phialemonium_globosum(100); | Phialemonium | 13 | Pathotroph-Saprotroph | \|Animal Parasite\|-Undefined Saprotroph | Microfungus | NULL | Probable | As animal_decomposer (Põlme et al. 2020); Decay animal material (Põlme et al. 2020) | Tedersoo L, et al. 2014. Science 346:e1256688 ((https://doi.org/10.1126/science.1256688)); Põlme S, et al. 2020. Fungal Diversity 105:1-16 ((https://doi.org/10.1007/s13225-020-00466-2)) |
| Otu07776 | 3 | k__Fungi(100);p__Ascomycota(100);c__Dothideomycetes(100);o__Pleosporales(100);f__Tubeufiaceae(100);g__Chlamydotubeufia(100);s__Chlamydotubeufia_khunkornensis(100); | Chlamydotubeufia | 13 | Saprotroph | Undefined Saprotroph-\|Wood Saprotroph\| | Microfungus | NULL | Probable | Decay wood (Põlme et al. 2020) | Tedersoo L, et al. 2014. Science 346:e1256688 ((https://doi.org/10.1126/science.1256688)); Põlme S, et al. 2020. Fungal Diversity 105:1-16 ((https://doi.org/10.1007/s13225-020-00466-2)) |
| Otu07778 | 3 | k__Fungi(100);p__Ascomycota(100);c__Saccharomycetes(100);o__Saccharomycetales(100);f__Metschnikowiaceae(100);g__Metschnikowia(100);s__Metschnikowia_pulcherrima(100); | Metschnikowia | 13 | Saprotroph | \|Undefined Saprotroph\| | Yeast | NULL | Probable | As nectar-tap_saprotroph (Põlme et al. 2020); Decay sugar-rich substrates (Põlme et al. 2020) | Põlme S, et al. 2020. Fungal Diversity 105:1-16 ((https://doi.org/10.1007/s13225-020-00466-2)) |
| Otu07780 | 3 | k__Fungi(100);p__Ascomycota(100);c__Saccharomycetes(100);o__Saccharomycetales(100);f__Metschnikowiaceae(100);g__Metschnikowia(100);s__Metschnikowia_pulcherrima(67); | Metschnikowia | 13 | Saprotroph | \|Undefined Saprotroph\| | Yeast | NULL | Probable | As nectar-tap_saprotroph (Põlme et al. 2020); Decay sugar-rich substrates (Põlme et al. 2020) | Põlme S, et al. 2020. Fungal Diversity 105:1-16 ((https://doi.org/10.1007/s13225-020-00466-2)) |
| Otu07786 | 3 | k__Fungi(100);p__Ascomycota(100);c__Saccharomycetes(100);o__Saccharomycetales(100);f__Metschnikowiaceae(100);g__Metschnikowia(100);s__Metschnikowia_pulcherrima(100); | Metschnikowia | 13 | Saprotroph | \|Undefined Saprotroph\| | Yeast | NULL | Probable | As nectar-tap_saprotroph (Põlme et al. 2020); Decay sugar-rich substrates (Põlme et al. 2020) | Põlme S, et al. 2020. Fungal Diversity 105:1-16 ((https://doi.org/10.1007/s13225-020-00466-2)) |
| Otu07789 | 3 | k__Fungi(100);p__Ascomycota(100);c__Saccharomycetes(100);o__Saccharomycetales(100);f__Metschnikowiaceae(100);g__Metschnikowia(100);s__Metschnikowia_pulcherrima(100); | Metschnikowia | 13 | Saprotroph | \|Undefined Saprotroph\| | Yeast | NULL | Probable | As nectar-tap_saprotroph (Põlme et al. 2020); Decay sugar-rich substrates (Põlme et al. 2020) | Põlme S, et al. 2020. Fungal Diversity 105:1-16 ((https://doi.org/10.1007/s13225-020-00466-2)) |
| Otu07790 | 3 | k__Fungi(100);p__Ascomycota(100);c__Dothideomycetes(100);o__Capnodiales(100);f__Teratosphaeriaceae(100);g__Teratosphaeria(100);g__Teratosphaeria_unclassified(100); | Teratosphaeria | 13 | Pathotroph-Saprotroph | \|Plant Pathogen\|-Plant Saprotroph-Wood Saprotroph | Microfungus | NULL | Probable | As litter_saprotroph (Põlme et al. 2020); As wood_pathogen (Põlme et al. 2020); Decay wood, leaf, fruit, seed (Põlme et al. 2020) | Yamazaki A, et al. 2010. Mycoscience 51:443-455 ((https://www.jstage.jst.go.jp/article/mycosci/51/6/51_MYC51443/_pdf)); Tedersoo L, et al. 2014. Science 346:e1256688 ((https://doi.org/10.1126/science.1256688)); Põlme S, et al. 2020. Fungal Diversity 105:1-16 ((https://doi.org/10.1007/s13225-020-00466-2)) |
| Otu07791 | 3 | k__Fungi(100);p__Basidiomycota(100);c__Agaricostilbomycetes(100);o__Agaricostilbales(100);f__Agaricostilbaceae(100);g__Agaricostilbum(100);s__Agaricostilbum_hyphaenes(100); | Agaricostilbum | 13 | Saprotroph | \|Plant Saprotroph\|-Undefined Saprotroph | Dimorphic-Facultative Yeast | NULL | Highly Probable | As litter_saprotroph (Põlme et al. 2020); Decay leaf, fruit, seed (Põlme et al. 2020); Host - Arecaceae (Põlme et al. 2020) | Kurtzman CP, et al. (eds.) 2011. The Yeasts, a Taxonomic Study. Fifth Edition. Vols 1-3. Elsevier, San Diego (ISBN: 9780444521491); Põlme S, et al. 2020. Fungal Diversity 105:1-16 ((https://doi.org/10.1007/s13225-020-00466-2)) |
| Otu07794 | 3 | k__Fungi(100);p__Ascomycota(100);c__Saccharomycetes(100);o__Saccharomycetales(100);f__Metschnikowiaceae(100);g__Metschnikowia(100);g__Metschnikowia_unclassified(67); | Metschnikowia | 13 | Saprotroph | \|Undefined Saprotroph\| | Yeast | NULL | Probable | As nectar-tap_saprotroph (Põlme et al. 2020); Decay sugar-rich substrates (Põlme et al. 2020) | Põlme S, et al. 2020. Fungal Diversity 105:1-16 ((https://doi.org/10.1007/s13225-020-00466-2)) |
| Otu07801 | 3 | k__Fungi(100);p__Basidiomycota(100);c__Tremellomycetes(100);o__Tremellales(100);f__Tremellales_family_Incertae_sedis(100);g__Tremella(100);s__Tremella_aff_globispora_IMUFRJ_52033(100); | Tremella | 13 | Pathotroph | \|Fungal Parasite\|-Lichen Parasite | Dimorphic-Facultative Yeast-Microfungus-Tremelloid | NULL | Probable | As mycoparasite (Põlme et al. 2020); Decay fungal material (Põlme et al. 2020) | Tedersoo L, et al. 2014. Science 346:e1256688 ((https://doi.org/10.1126/science.1256688)); Põlme S, et al. 2020. Fungal Diversity 105:1-16 ((https://doi.org/10.1007/s13225-020-00466-2)) |
| Otu07804 | 3 | k__Fungi(100);p__Basidiomycota(100);c__Agaricomycetes(100);o__Polyporales(100);f__Meruliaceae(100);g__Junghuhnia(100);g__Junghuhnia_unclassified(100); | Junghuhnia | 13 | Saprotroph | \|Wood Saprotroph\| | Corticioid | White Rot | Highly Probable | As white_rot (Põlme et al. 2020); Decay wood (Põlme et al. 2020) | Gilbertson RL, Ryvarden L. 1987-1987. North American Polypores. Fungiflora, Oslo (ISBN: 978-0945345060); Põlme S, et al. 2020. Fungal Diversity 105:1-16 ((https://doi.org/10.1007/s13225-020-00466-2)) |
| Otu07805 | 3 | k__Fungi(100);p__Ascomycota(100);c__Saccharomycetes(100);o__Saccharomycetales(100);f__Metschnikowiaceae(100);g__Metschnikowia(100);s__Metschnikowia_pulcherrima(100); | Metschnikowia | 13 | Saprotroph | \|Undefined Saprotroph\| | Yeast | NULL | Probable | As nectar-tap_saprotroph (Põlme et al. 2020); Decay sugar-rich substrates (Põlme et al. 2020) | Põlme S, et al. 2020. Fungal Diversity 105:1-16 ((https://doi.org/10.1007/s13225-020-00466-2)) |
| Otu07806 | 3 | k__Fungi(100);p__Ascomycota(100);c__Saccharomycetes(100);o__Saccharomycetales(100);f__Metschnikowiaceae(100);g__Metschnikowia(100);s__Metschnikowia_pulcherrima(67); | Metschnikowia | 13 | Saprotroph | \|Undefined Saprotroph\| | Yeast | NULL | Probable | As nectar-tap_saprotroph (Põlme et al. 2020); Decay sugar-rich substrates (Põlme et al. 2020) | Põlme S, et al. 2020. Fungal Diversity 105:1-16 ((https://doi.org/10.1007/s13225-020-00466-2)) |
| Otu07807 | 3 | k__Fungi(100);p__Basidiomycota(100);c__Tremellomycetes(100);o__Tremellales(100);f__Tremellales_family_Incertae_sedis(100);g__Cryptococcus(100);s__Cryptococcus_sp_JCM_11356(100); | Cryptococcus | 13 | Pathotroph-Saprotroph-Symbiotroph | Animal Pathogen-Endophyte-Epiphyte-\|Undefined Saprotroph\| | Dimorphic Yeast-Yeast | NULL | Possible | As foliar_endophyte (Põlme et al. 2020); Decay leaf, fruit, seed, soil, dung, animal material (Põlme et al. 2020) | Andrews JH, et al. 1983. Phytopathology 73:228-234 ((https://doi.org/10.1094/Phyto-73-228)); Kurtzman CP, et al. (eds.) 2011. The Yeasts, a Taxonomic Study. Fifth Edition. Vols 1-3. Elsevier, San Diego (ISBN: 9780444521491); Põlme S, et al. 2020. Fungal Diversity 105:1-16 ((https://doi.org/10.1007/s13225-020-00466-2)) |
| Otu07808 | 3 | k__Fungi(100);p__Basidiomycota(100);c__Agaricomycetes(100);o__Polyporales(100);f__Fomitopsidaceae(100);g__Fomitopsis(100);g__Fomitopsis_unclassified(100); | Fomitopsis | 13 | Pathotroph-Saprotroph | Plant Pathogen-\|Wood Saprotroph\| | Polyporoid | Brown Rot | Highly Probable | As brown-rot (Põlme et al. 2020); As wood_pathogen (Põlme et al. 2020); Decay wood (Põlme et al. 2020); Most species are parasitic on woody plants, causing brown rots (Cannon & Kirk 2007) | Gilbertson RL, Ryvarden L. 1987-1987. North American Polypores. Fungiflora, Oslo (ISBN: 978-0945345060); Cannon PF, Kirk PM. 2007. Fungal Families of the World. CAB International, Cambridge (ISBN: 978-0851998275); Põlme S, et al. 2020. Fungal Diversity 105:1-16 ((https://doi.org/10.1007/s13225-020-00466-2)) |
| Otu07809 | 3 | k__Fungi(100);p__Ascomycota(100);c__Saccharomycetes(100);o__Saccharomycetales(100);f__Metschnikowiaceae(100);g__Metschnikowia(100);g__Metschnikowia_unclassified(67); | Metschnikowia | 13 | Saprotroph | \|Undefined Saprotroph\| | Yeast | NULL | Probable | As nectar-tap_saprotroph (Põlme et al. 2020); Decay sugar-rich substrates (Põlme et al. 2020) | Põlme S, et al. 2020. Fungal Diversity 105:1-16 ((https://doi.org/10.1007/s13225-020-00466-2)) |
| Otu07810 | 3 | k__Fungi(100);p__Ascomycota(100);c__Saccharomycetes(100);o__Saccharomycetales(100);f__Metschnikowiaceae(100);g__Metschnikowia(100);s__Metschnikowia_pulcherrima(100); | Metschnikowia | 13 | Saprotroph | \|Undefined Saprotroph\| | Yeast | NULL | Probable | As nectar-tap_saprotroph (Põlme et al. 2020); Decay sugar-rich substrates (Põlme et al. 2020) | Põlme S, et al. 2020. Fungal Diversity 105:1-16 ((https://doi.org/10.1007/s13225-020-00466-2)) |
| Otu07816 | 3 | k__Fungi(100);p__Ascomycota(100);c__Lecanoromycetes(100);o__Lecanorales(100);o__Lecanorales_unclassified(100);o__Lecanorales_unclassified(100);o__Lecanorales_unclassified(100); | Lecanorales | 7 | Pathotroph-Symbiotroph | Lichen Parasite-Lichenized | Microfungus-Thallus | NULL | Probable | Lichenized with green algae, occasionally lichenicolous, on rocks, mosses and wood (Cannon & Kirk 2007); ...an order of mostly lichen-forming fungi (Kirk et al. 2008) | Cannon PF, Kirk PM. 2007. Fungal Families of the World. CAB International, Cambridge (ISBN: 978-0851998275); Kirk PM et al. 2008. Dictionary of the Fungi. Tenth Edition. CAB International, Wallingford (ISBN: 978-0851998268); Esslinger TL. 2014. North Dakota State University ((http://www.ndsu.edu/pubweb/~esslinge/chcklst/chcklst7.htm)) |
| Otu07824 | 3 | k__Fungi(100);p__Ascomycota(100);c__Dothideomycetes(100);o__Pleosporales(100);f__Leptosphaeriaceae(100);g__Leptosphaeria(100);s__Leptosphaeria_sp(100); | Leptosphaeria | 13 | Pathotroph-Saprotroph | \|Plant Pathogen\|-Wood Saprotroph | Microfungus | NULL | Probable | As wood_pathogen (Põlme et al. 2020); Decay wood (Põlme et al. 2020) | Tedersoo L, et al. 2014. Science 346:e1256688 ((https://doi.org/10.1126/science.1256688)); Põlme S, et al. 2020. Fungal Diversity 105:1-16 ((https://doi.org/10.1007/s13225-020-00466-2)) |
| Otu07829 | 3 | k__Fungi(100);p__Ascomycota(100);c__Saccharomycetes(100);o__Saccharomycetales(100);f__Saccharomycetales_family_Incertae_sedis(100);g__Candida(100);g__Candida_unclassified(67); | Candida | 13 | Pathotroph-Saprotroph-Symbiotroph | Animal Pathogen-Endophyte-Endosymbiont-Epiphyte-\|Undefined Saprotroph\| | Dimorphic Yeast-Yeast | NULL | Probable | As nectar-tap_saprotroph (Põlme et al. 2020); Decay sugar-rich substrates (Põlme et al. 2020); This asexual genus is polyphyletic and contain species that function very differently depend on what they are. Caution should be taken when interpreting guilds for Candida species. | Manolakaki D, et al. 2010. Virulence. 1: 367-375 ((https://doi.org/10.4161/viru.1.5.12796)); Mekbib SB, et al. 2011. Tropical Plant Pathology 36:233-240 ((https://doi.org/10.1590/S1982-56762011000400004)); Põlme S, et al. 2020. Fungal Diversity 105:1-16 ((https://doi.org/10.1007/s13225-020-00466-2)) |
| Otu07831 | 3 | k__Fungi(100);p__Ascomycota(100);c__Dothideomycetes(100);o__Pleosporales(100);f__Venturiaceae(100);g__Venturia(100);g__Venturia_unclassified(100); | Venturia | 13 | Pathotroph-Pathotroph-Saprotroph | \|Plant Pathogen\|-Plant Saprotroph | Microfungus | NULL | Probable | As litter_saprotroph (Põlme et al. 2020); Decay leaf, fruit, seed (Põlme et al. 2020) | Tedersoo L, et al. 2014. Science 346:e1256688 ((https://doi.org/10.1126/science.1256688)); Põlme S, et al. 2020. Fungal Diversity 105:1-16 ((https://doi.org/10.1007/s13225-020-00466-2)) |
| Otu07835 | 3 | k__Fungi(100);p__Ascomycota(100);c__Sordariomycetes(100);o__Hypocreales(100);f__Nectriaceae(100);f__Nectriaceae_unclassified(100);f__Nectriaceae_unclassified(100); | Nectriaceae | 9 | Pathotroph-Saprotroph-Symbiotroph | Animal Pathogen-Endophyte-Fungal Parasite-Lichen Parasite-Plant Pathogen-Wood Saprotroph | Microfungus | NULL | Possible | Fusarium species are amongst the economically significant plant pathogens (Cannon & Kirk 2007) | Cannon PF, Kirk PM. 2007. Fungal Families of the World. CAB International, Cambridge (ISBN: 978-0851998275); Hirooka Y, et al. 2012. Studies in Mycology 711-210 ((https://doi.org/10.3114/sim0001)); Tedersoo L, et al. 2014. Science 346:e1256688 ((https://doi.org/10.1126/science.1256688)); The American Phytopathological Society Common Names of Plant Diseases ((http://www.apsnet.org/publications/commonnames/Pages/default.aspx)); https://www.cdc.gov ((https://www.cdc.gov)) |
| Otu07836 | 3 | k__Fungi(100);p__Ascomycota(100);c__Dothideomycetes(100);o__Pleosporales(100);f__Tubeufiaceae(100);f__Tubeufiaceae_unclassified(100);f__Tubeufiaceae_unclassified(100); | Tubeufiaceae | 9 | Pathotroph-Saprotroph | Animal Pathogen-Undefined Saprotroph | NULL | NULL | Probable | Biotrophic or saprobic, especially on scale insects or other fungi. The anamorphic forms are fequently aero-aquatic (Cannon & Kirk 2007) | Cannon PF, Kirk PM. 2007. Fungal Families of the World. CAB International, Cambridge (ISBN: 978-0851998275) |
| Otu07838 | 3 | k__Fungi(100);p__Ascomycota(100);c__Saccharomycetes(100);o__Saccharomycetales(100);f__Metschnikowiaceae(100);g__Metschnikowia(100);g__Metschnikowia_unclassified(100); | Metschnikowia | 13 | Saprotroph | \|Undefined Saprotroph\| | Yeast | NULL | Probable | As nectar-tap_saprotroph (Põlme et al. 2020); Decay sugar-rich substrates (Põlme et al. 2020) | Põlme S, et al. 2020. Fungal Diversity 105:1-16 ((https://doi.org/10.1007/s13225-020-00466-2)) |
| Otu07841 | 3 | k__Fungi(100);p__Ascomycota(100);c__Saccharomycetes(100);o__Saccharomycetales(100);f__Metschnikowiaceae(100);g__Metschnikowia(100);g__Metschnikowia_unclassified(100); | Metschnikowia | 13 | Saprotroph | \|Undefined Saprotroph\| | Yeast | NULL | Probable | As nectar-tap_saprotroph (Põlme et al. 2020); Decay sugar-rich substrates (Põlme et al. 2020) | Põlme S, et al. 2020. Fungal Diversity 105:1-16 ((https://doi.org/10.1007/s13225-020-00466-2)) |
| Otu07844 | 3 | k__Fungi(100);p__Ascomycota(100);c__Sordariomycetes(100);o__Xylariales(100);f__Amphisphaeriaceae(100);f__Amphisphaeriaceae_unclassified(100);f__Amphisphaeriaceae_unclassified(100); | Amphisphaeriaceae | 9 | Saprotroph | Undefined Saprotroph | Microfungus | NULL | Possible | Saprobes or weak parasites of leaves, stems, etc., often appearing as secondary colonizers following damage to plants by other agents (Cannon & Kirk 2007) | Cannon PF, Kirk PM. 2007. Fungal Families of the World. CAB International, Cambridge (ISBN: 978-0851998275) |
| Otu07846 | 3 | k__Fungi(100);p__Ascomycota(100);c__Saccharomycetes(100);o__Saccharomycetales(100);f__Metschnikowiaceae(100);g__Metschnikowia(100);s__Metschnikowia_pulcherrima(100); | Metschnikowia | 13 | Saprotroph | \|Undefined Saprotroph\| | Yeast | NULL | Probable | As nectar-tap_saprotroph (Põlme et al. 2020); Decay sugar-rich substrates (Põlme et al. 2020) | Põlme S, et al. 2020. Fungal Diversity 105:1-16 ((https://doi.org/10.1007/s13225-020-00466-2)) |
| Otu07852 | 3 | k__Fungi(100);p__Ascomycota(100);c__Saccharomycetes(100);o__Saccharomycetales(100);f__Metschnikowiaceae(100);g__Metschnikowia(100);g__Metschnikowia_unclassified(100); | Metschnikowia | 13 | Saprotroph | \|Undefined Saprotroph\| | Yeast | NULL | Probable | As nectar-tap_saprotroph (Põlme et al. 2020); Decay sugar-rich substrates (Põlme et al. 2020) | Põlme S, et al. 2020. Fungal Diversity 105:1-16 ((https://doi.org/10.1007/s13225-020-00466-2)) |
| Otu07857 | 3 | k__Fungi(100);p__Basidiomycota(100);c__Tremellomycetes(100);o__Tremellales(100);f__Tremellales_family_Incertae_sedis(100);g__Fellomyces(100);s__Fellomyces_horovitziae(100); | Fellomyces | 13 | Pathotroph-Saprotroph | Fungal Parasite-\|Undefined Saprotroph\| | Yeast | NULL | Probable | As mycoparasite (Põlme et al. 2020) | Põlme S, et al. 2020. Fungal Diversity 105:1-16 ((https://doi.org/10.1007/s13225-020-00466-2)) |
| Otu07859 | 3 | k__Fungi(100);p__Zygomycota(100);c__Zygomycota_class_Incertae_sedis(100);o__Mucorales(100);f__Mucoraceae(100);g__Mucor(100);s__Mucor_circinelloides_f._janssenii(100); | Mucor | 13 | Saprotroph | \|Undefined Saprotroph\| | Microfungus | NULL | Probable | As soil_saprotroph (Põlme et al. 2020); Decay soil (Põlme et al. 2020) | Tedersoo L, et al. 2014. Science 346:e1256688 ((https://doi.org/10.1126/science.1256688)); Põlme S, et al. 2020. Fungal Diversity 105:1-16 ((https://doi.org/10.1007/s13225-020-00466-2)) |
| Otu07860 | 3 | k__Fungi(100);p__Ascomycota(100);c__Saccharomycetes(100);o__Saccharomycetales(100);f__Metschnikowiaceae(100);g__Metschnikowia(100);g__Metschnikowia_unclassified(67); | Metschnikowia | 13 | Saprotroph | \|Undefined Saprotroph\| | Yeast | NULL | Probable | As nectar-tap_saprotroph (Põlme et al. 2020); Decay sugar-rich substrates (Põlme et al. 2020) | Põlme S, et al. 2020. Fungal Diversity 105:1-16 ((https://doi.org/10.1007/s13225-020-00466-2)) |
| Otu07862 | 3 | k__Fungi(100);p__Ascomycota(100);c__Saccharomycetes(100);o__Saccharomycetales(100);f__Metschnikowiaceae(100);g__Metschnikowia(100);s__Metschnikowia_pulcherrima(100); | Metschnikowia | 13 | Saprotroph | \|Undefined Saprotroph\| | Yeast | NULL | Probable | As nectar-tap_saprotroph (Põlme et al. 2020); Decay sugar-rich substrates (Põlme et al. 2020) | Põlme S, et al. 2020. Fungal Diversity 105:1-16 ((https://doi.org/10.1007/s13225-020-00466-2)) |
| Otu07863 | 3 | k__Fungi(100);p__Ascomycota(100);c__Saccharomycetes(100);o__Saccharomycetales(100);f__Metschnikowiaceae(100);g__Metschnikowia(100);s__Metschnikowia_pulcherrima(100); | Metschnikowia | 13 | Saprotroph | \|Undefined Saprotroph\| | Yeast | NULL | Probable | As nectar-tap_saprotroph (Põlme et al. 2020); Decay sugar-rich substrates (Põlme et al. 2020) | Põlme S, et al. 2020. Fungal Diversity 105:1-16 ((https://doi.org/10.1007/s13225-020-00466-2)) |
| Otu07864 | 3 | k__Fungi(100);p__Basidiomycota(100);c__Agaricomycetes(100);o__Polyporales(100);f__Fomitopsidaceae(100);g__Auriporia(100);s__Auriporia_aurea(100); | Auriporia | 13 | Saprotroph | \|Wood Saprotroph\| | Corticioid | Brown Rot | Highly Probable | As brown-rot (Põlme et al. 2020); Decay wood (Põlme et al. 2020) | Gilbertson RL, Ryvarden L. 1987-1987. North American Polypores. Fungiflora, Oslo (ISBN: 978-0945345060); Põlme S, et al. 2020. Fungal Diversity 105:1-16 ((https://doi.org/10.1007/s13225-020-00466-2)) |
| Otu07867 | 3 | k__Fungi(100);p__Ascomycota(100);c__Saccharomycetes(100);o__Saccharomycetales(100);f__Metschnikowiaceae(100);g__Metschnikowia(100);g__Metschnikowia_unclassified(100); | Metschnikowia | 13 | Saprotroph | \|Undefined Saprotroph\| | Yeast | NULL | Probable | As nectar-tap_saprotroph (Põlme et al. 2020); Decay sugar-rich substrates (Põlme et al. 2020) | Põlme S, et al. 2020. Fungal Diversity 105:1-16 ((https://doi.org/10.1007/s13225-020-00466-2)) |
| Otu07869 | 3 | k__Fungi(100);p__Ascomycota(100);c__Saccharomycetes(100);o__Saccharomycetales(100);f__Metschnikowiaceae(100);g__Metschnikowia(100);s__Metschnikowia_pulcherrima(67); | Metschnikowia | 13 | Saprotroph | \|Undefined Saprotroph\| | Yeast | NULL | Probable | As nectar-tap_saprotroph (Põlme et al. 2020); Decay sugar-rich substrates (Põlme et al. 2020) | Põlme S, et al. 2020. Fungal Diversity 105:1-16 ((https://doi.org/10.1007/s13225-020-00466-2)) |
| Otu07870 | 3 | k__Fungi(100);p__Ascomycota(100);c__Saccharomycetes(100);o__Saccharomycetales(100);f__Metschnikowiaceae(100);g__Metschnikowia(100);s__Metschnikowia_pulcherrima(100); | Metschnikowia | 13 | Saprotroph | \|Undefined Saprotroph\| | Yeast | NULL | Probable | As nectar-tap_saprotroph (Põlme et al. 2020); Decay sugar-rich substrates (Põlme et al. 2020) | Põlme S, et al. 2020. Fungal Diversity 105:1-16 ((https://doi.org/10.1007/s13225-020-00466-2)) |
| Otu07871 | 3 | k__Fungi(100);p__Ascomycota(100);c__Saccharomycetes(100);o__Saccharomycetales(100);f__Metschnikowiaceae(100);g__Metschnikowia(100);g__Metschnikowia_unclassified(67); | Metschnikowia | 13 | Saprotroph | \|Undefined Saprotroph\| | Yeast | NULL | Probable | As nectar-tap_saprotroph (Põlme et al. 2020); Decay sugar-rich substrates (Põlme et al. 2020) | Põlme S, et al. 2020. Fungal Diversity 105:1-16 ((https://doi.org/10.1007/s13225-020-00466-2)) |
| Otu07872 | 3 | k__Fungi(100);p__Ascomycota(100);c__Saccharomycetes(100);o__Saccharomycetales(100);f__Metschnikowiaceae(100);g__Metschnikowia(100);s__Metschnikowia_pulcherrima(100); | Metschnikowia | 13 | Saprotroph | \|Undefined Saprotroph\| | Yeast | NULL | Probable | As nectar-tap_saprotroph (Põlme et al. 2020); Decay sugar-rich substrates (Põlme et al. 2020) | Põlme S, et al. 2020. Fungal Diversity 105:1-16 ((https://doi.org/10.1007/s13225-020-00466-2)) |
| Otu07876 | 3 | k__Fungi(100);p__Ascomycota(100);c__Saccharomycetes(100);o__Saccharomycetales(100);f__Metschnikowiaceae(100);g__Metschnikowia(100);s__Metschnikowia_pulcherrima(67); | Metschnikowia | 13 | Saprotroph | \|Undefined Saprotroph\| | Yeast | NULL | Probable | As nectar-tap_saprotroph (Põlme et al. 2020); Decay sugar-rich substrates (Põlme et al. 2020) | Põlme S, et al. 2020. Fungal Diversity 105:1-16 ((https://doi.org/10.1007/s13225-020-00466-2)) |
| Otu07877 | 3 | k__Fungi(100);p__Ascomycota(100);c__Saccharomycetes(100);o__Saccharomycetales(100);f__Metschnikowiaceae(100);g__Metschnikowia(100);s__Metschnikowia_pulcherrima(100); | Metschnikowia | 13 | Saprotroph | \|Undefined Saprotroph\| | Yeast | NULL | Probable | As nectar-tap_saprotroph (Põlme et al. 2020); Decay sugar-rich substrates (Põlme et al. 2020) | Põlme S, et al. 2020. Fungal Diversity 105:1-16 ((https://doi.org/10.1007/s13225-020-00466-2)) |
| Otu07880 | 3 | k__Fungi(100);p__Ascomycota(100);c__Saccharomycetes(100);o__Saccharomycetales(100);f__Metschnikowiaceae(100);g__Metschnikowia(100);s__Metschnikowia_pulcherrima(100); | Metschnikowia | 13 | Saprotroph | \|Undefined Saprotroph\| | Yeast | NULL | Probable | As nectar-tap_saprotroph (Põlme et al. 2020); Decay sugar-rich substrates (Põlme et al. 2020) | Põlme S, et al. 2020. Fungal Diversity 105:1-16 ((https://doi.org/10.1007/s13225-020-00466-2)) |
| Otu07882 | 3 | k__Fungi(100);p__Ascomycota(100);c__Saccharomycetes(100);o__Saccharomycetales(100);f__Metschnikowiaceae(100);g__Metschnikowia(100);s__Metschnikowia_pulcherrima(100); | Metschnikowia | 13 | Saprotroph | \|Undefined Saprotroph\| | Yeast | NULL | Probable | As nectar-tap_saprotroph (Põlme et al. 2020); Decay sugar-rich substrates (Põlme et al. 2020) | Põlme S, et al. 2020. Fungal Diversity 105:1-16 ((https://doi.org/10.1007/s13225-020-00466-2)) |
| Otu07884 | 3 | k__Fungi(100);p__Ascomycota(100);c__Saccharomycetes(100);o__Saccharomycetales(100);f__Metschnikowiaceae(100);g__Metschnikowia(100);s__Metschnikowia_pulcherrima(100); | Metschnikowia | 13 | Saprotroph | \|Undefined Saprotroph\| | Yeast | NULL | Probable | As nectar-tap_saprotroph (Põlme et al. 2020); Decay sugar-rich substrates (Põlme et al. 2020) | Põlme S, et al. 2020. Fungal Diversity 105:1-16 ((https://doi.org/10.1007/s13225-020-00466-2)) |
| Otu07886 | 3 | k__Fungi(100);p__Ascomycota(100);c__Saccharomycetes(100);o__Saccharomycetales(100);f__Metschnikowiaceae(100);g__Metschnikowia(100);s__Metschnikowia_pulcherrima(100); | Metschnikowia | 13 | Saprotroph | \|Undefined Saprotroph\| | Yeast | NULL | Probable | As nectar-tap_saprotroph (Põlme et al. 2020); Decay sugar-rich substrates (Põlme et al. 2020) | Põlme S, et al. 2020. Fungal Diversity 105:1-16 ((https://doi.org/10.1007/s13225-020-00466-2)) |
| Otu07887 | 3 | k__Fungi(100);p__Ascomycota(100);c__Saccharomycetes(100);o__Saccharomycetales(100);f__Metschnikowiaceae(100);g__Metschnikowia(100);s__Metschnikowia_pulcherrima(100); | Metschnikowia | 13 | Saprotroph | \|Undefined Saprotroph\| | Yeast | NULL | Probable | As nectar-tap_saprotroph (Põlme et al. 2020); Decay sugar-rich substrates (Põlme et al. 2020) | Põlme S, et al. 2020. Fungal Diversity 105:1-16 ((https://doi.org/10.1007/s13225-020-00466-2)) |
| Otu07889 | 3 | k__Fungi(100);p__Ascomycota(100);c__Saccharomycetes(100);o__Saccharomycetales(100);f__Metschnikowiaceae(100);g__Metschnikowia(100);s__Metschnikowia_pulcherrima(100); | Metschnikowia | 13 | Saprotroph | \|Undefined Saprotroph\| | Yeast | NULL | Probable | As nectar-tap_saprotroph (Põlme et al. 2020); Decay sugar-rich substrates (Põlme et al. 2020) | Põlme S, et al. 2020. Fungal Diversity 105:1-16 ((https://doi.org/10.1007/s13225-020-00466-2)) |
| Otu07891 | 3 | k__Fungi(100);p__Ascomycota(100);c__Saccharomycetes(100);o__Saccharomycetales(100);f__Metschnikowiaceae(100);g__Metschnikowia(100);s__Metschnikowia_pulcherrima(67); | Metschnikowia | 13 | Saprotroph | \|Undefined Saprotroph\| | Yeast | NULL | Probable | As nectar-tap_saprotroph (Põlme et al. 2020); Decay sugar-rich substrates (Põlme et al. 2020) | Põlme S, et al. 2020. Fungal Diversity 105:1-16 ((https://doi.org/10.1007/s13225-020-00466-2)) |
| Otu07896 | 3 | k__Fungi(100);p__Ascomycota(100);c__Saccharomycetes(100);o__Saccharomycetales(100);f__Metschnikowiaceae(100);g__Metschnikowia(100);s__Metschnikowia_pulcherrima(67); | Metschnikowia | 13 | Saprotroph | \|Undefined Saprotroph\| | Yeast | NULL | Probable | As nectar-tap_saprotroph (Põlme et al. 2020); Decay sugar-rich substrates (Põlme et al. 2020) | Põlme S, et al. 2020. Fungal Diversity 105:1-16 ((https://doi.org/10.1007/s13225-020-00466-2)) |
| Otu07897 | 3 | k__Fungi(100);p__Ascomycota(100);c__Saccharomycetes(100);o__Saccharomycetales(100);f__Metschnikowiaceae(100);g__Metschnikowia(100);s__Metschnikowia_pulcherrima(100); | Metschnikowia | 13 | Saprotroph | \|Undefined Saprotroph\| | Yeast | NULL | Probable | As nectar-tap_saprotroph (Põlme et al. 2020); Decay sugar-rich substrates (Põlme et al. 2020) | Põlme S, et al. 2020. Fungal Diversity 105:1-16 ((https://doi.org/10.1007/s13225-020-00466-2)) |
| Otu07899 | 3 | k__Fungi(100);p__Ascomycota(100);c__Saccharomycetes(100);o__Saccharomycetales(100);f__Metschnikowiaceae(100);g__Metschnikowia(100);s__Metschnikowia_pulcherrima(100); | Metschnikowia | 13 | Saprotroph | \|Undefined Saprotroph\| | Yeast | NULL | Probable | As nectar-tap_saprotroph (Põlme et al. 2020); Decay sugar-rich substrates (Põlme et al. 2020) | Põlme S, et al. 2020. Fungal Diversity 105:1-16 ((https://doi.org/10.1007/s13225-020-00466-2)) |
| Otu07900 | 3 | k__Fungi(100);p__Ascomycota(100);c__Saccharomycetes(100);o__Saccharomycetales(100);f__Saccharomycetales_family_Incertae_sedis(100);g__Candida(100);g__Candida_unclassified(67); | Candida | 13 | Pathotroph-Saprotroph-Symbiotroph | Animal Pathogen-Endophyte-Endosymbiont-Epiphyte-\|Undefined Saprotroph\| | Dimorphic Yeast-Yeast | NULL | Probable | As nectar-tap_saprotroph (Põlme et al. 2020); Decay sugar-rich substrates (Põlme et al. 2020); This asexual genus is polyphyletic and contain species that function very differently depend on what they are. Caution should be taken when interpreting guilds for Candida species. | Manolakaki D, et al. 2010. Virulence. 1: 367-375 ((https://doi.org/10.4161/viru.1.5.12796)); Mekbib SB, et al. 2011. Tropical Plant Pathology 36:233-240 ((https://doi.org/10.1590/S1982-56762011000400004)); Põlme S, et al. 2020. Fungal Diversity 105:1-16 ((https://doi.org/10.1007/s13225-020-00466-2)) |
| Otu07901 | 3 | k__Fungi(100);p__Ascomycota(100);c__Saccharomycetes(100);o__Saccharomycetales(100);f__Metschnikowiaceae(100);g__Metschnikowia(100);s__Metschnikowia_pulcherrima(67); | Metschnikowia | 13 | Saprotroph | \|Undefined Saprotroph\| | Yeast | NULL | Probable | As nectar-tap_saprotroph (Põlme et al. 2020); Decay sugar-rich substrates (Põlme et al. 2020) | Põlme S, et al. 2020. Fungal Diversity 105:1-16 ((https://doi.org/10.1007/s13225-020-00466-2)) |
| Otu07902 | 3 | k__Fungi(100);p__Ascomycota(100);c__Saccharomycetes(100);o__Saccharomycetales(100);f__Metschnikowiaceae(100);g__Metschnikowia(100);s__Metschnikowia_pulcherrima(67); | Metschnikowia | 13 | Saprotroph | \|Undefined Saprotroph\| | Yeast | NULL | Probable | As nectar-tap_saprotroph (Põlme et al. 2020); Decay sugar-rich substrates (Põlme et al. 2020) | Põlme S, et al. 2020. Fungal Diversity 105:1-16 ((https://doi.org/10.1007/s13225-020-00466-2)) |
| Otu07906 | 3 | k__Fungi(100);p__Ascomycota(100);c__Saccharomycetes(100);o__Saccharomycetales(100);f__Metschnikowiaceae(100);g__Metschnikowia(100);s__Metschnikowia_pulcherrima(100); | Metschnikowia | 13 | Saprotroph | \|Undefined Saprotroph\| | Yeast | NULL | Probable | As nectar-tap_saprotroph (Põlme et al. 2020); Decay sugar-rich substrates (Põlme et al. 2020) | Põlme S, et al. 2020. Fungal Diversity 105:1-16 ((https://doi.org/10.1007/s13225-020-00466-2)) |
| Otu07909 | 3 | k__Fungi(100);p__Zygomycota(100);c__Zygomycota_class_Incertae_sedis(100);o__Mucorales(100);f__Mucoraceae(100);g__Mucor(100);s__Mucor_genevensis(100); | Mucor | 13 | Saprotroph | \|Undefined Saprotroph\| | Microfungus | NULL | Probable | As soil_saprotroph (Põlme et al. 2020); Decay soil (Põlme et al. 2020) | Tedersoo L, et al. 2014. Science 346:e1256688 ((https://doi.org/10.1126/science.1256688)); Põlme S, et al. 2020. Fungal Diversity 105:1-16 ((https://doi.org/10.1007/s13225-020-00466-2)) |
| Otu07910 | 3 | k__Fungi(100);p__Ascomycota(100);c__Saccharomycetes(100);o__Saccharomycetales(100);f__Metschnikowiaceae(100);g__Metschnikowia(100);g__Metschnikowia_unclassified(67); | Metschnikowia | 13 | Saprotroph | \|Undefined Saprotroph\| | Yeast | NULL | Probable | As nectar-tap_saprotroph (Põlme et al. 2020); Decay sugar-rich substrates (Põlme et al. 2020) | Põlme S, et al. 2020. Fungal Diversity 105:1-16 ((https://doi.org/10.1007/s13225-020-00466-2)) |
| Otu07912 | 3 | k__Fungi(100);p__Ascomycota(100);c__Saccharomycetes(100);o__Saccharomycetales(100);f__Metschnikowiaceae(100);g__Metschnikowia(100);s__Metschnikowia_pulcherrima(100); | Metschnikowia | 13 | Saprotroph | \|Undefined Saprotroph\| | Yeast | NULL | Probable | As nectar-tap_saprotroph (Põlme et al. 2020); Decay sugar-rich substrates (Põlme et al. 2020) | Põlme S, et al. 2020. Fungal Diversity 105:1-16 ((https://doi.org/10.1007/s13225-020-00466-2)) |
| Otu07913 | 3 | k__Fungi(100);p__Ascomycota(100);c__Saccharomycetes(100);o__Saccharomycetales(100);f__Metschnikowiaceae(100);g__Metschnikowia(100);s__Metschnikowia_pulcherrima(100); | Metschnikowia | 13 | Saprotroph | \|Undefined Saprotroph\| | Yeast | NULL | Probable | As nectar-tap_saprotroph (Põlme et al. 2020); Decay sugar-rich substrates (Põlme et al. 2020) | Põlme S, et al. 2020. Fungal Diversity 105:1-16 ((https://doi.org/10.1007/s13225-020-00466-2)) |
| Otu07915 | 3 | k__Fungi(100);p__Basidiomycota(100);c__Tremellomycetes(100);o__Filobasidiales(100);f__Filobasidiaceae(100);g__Cryptococcus(100);s__Cryptococcus_sp(100); | Cryptococcus | 13 | Pathotroph-Saprotroph-Symbiotroph | Animal Pathogen-Endophyte-Epiphyte-\|Undefined Saprotroph\| | Dimorphic Yeast-Yeast | NULL | Possible | As foliar_endophyte (Põlme et al. 2020); Decay leaf, fruit, seed, soil, dung, animal material (Põlme et al. 2020) | Andrews JH, et al. 1983. Phytopathology 73:228-234 ((https://doi.org/10.1094/Phyto-73-228)); Kurtzman CP, et al. (eds.) 2011. The Yeasts, a Taxonomic Study. Fifth Edition. Vols 1-3. Elsevier, San Diego (ISBN: 9780444521491); Põlme S, et al. 2020. Fungal Diversity 105:1-16 ((https://doi.org/10.1007/s13225-020-00466-2)) |
| Otu07917 | 3 | k__Fungi(100);p__Basidiomycota(100);c__Agaricomycetes(100);o__Agaricales(100);f__Inocybaceae(100);g__Inocybe(100);g__Inocybe_unclassified(100); | Inocybe | 13 | Symbiotroph | \|Ectomycorrhizal\| | Agaricoid | NULL | Highly Probable | Sensu stricto (Matheny 2009). | Rinaldi AC, et al. 2008. Fungal Diversity 33:1-45 ((http://www.fungaldiversity.org/fdp/sfdp/33-1.pdf)); Matheny PB. 2009. McIlvainea 18:11-21 ((https://namyco.org/docs/3_Matheny.pdf)); Tedersoo L, et al. 2010. Mycorrhiza 20:217-263 ((https://doi.org/10.1007/s00572-009-0274-x)); Põlme S, et al. 2020. Fungal Diversity 105:1-16 ((https://doi.org/10.1007/s13225-020-00466-2)); http://mycorrhizas.info/ecmf.html ((http://mycorrhizas.info/ecmf.html)) |
| Otu07927 | 3 | k__Fungi(100);p__Ascomycota(100);c__Saccharomycetes(100);o__Saccharomycetales(100);f__Metschnikowiaceae(100);g__Metschnikowia(100);s__Metschnikowia_pulcherrima(67); | Metschnikowia | 13 | Saprotroph | \|Undefined Saprotroph\| | Yeast | NULL | Probable | As nectar-tap_saprotroph (Põlme et al. 2020); Decay sugar-rich substrates (Põlme et al. 2020) | Põlme S, et al. 2020. Fungal Diversity 105:1-16 ((https://doi.org/10.1007/s13225-020-00466-2)) |
| Otu07930 | 3 | k__Fungi(100);p__Ascomycota(100);c__Saccharomycetes(100);o__Saccharomycetales(100);f__Metschnikowiaceae(100);g__Metschnikowia(100);s__Metschnikowia_pulcherrima(100); | Metschnikowia | 13 | Saprotroph | \|Undefined Saprotroph\| | Yeast | NULL | Probable | As nectar-tap_saprotroph (Põlme et al. 2020); Decay sugar-rich substrates (Põlme et al. 2020) | Põlme S, et al. 2020. Fungal Diversity 105:1-16 ((https://doi.org/10.1007/s13225-020-00466-2)) |
| Otu07932 | 3 | k__Fungi(100);p__Ascomycota(100);c__Saccharomycetes(100);o__Saccharomycetales(100);f__Metschnikowiaceae(100);g__Metschnikowia(100);s__Metschnikowia_pulcherrima(100); | Metschnikowia | 13 | Saprotroph | \|Undefined Saprotroph\| | Yeast | NULL | Probable | As nectar-tap_saprotroph (Põlme et al. 2020); Decay sugar-rich substrates (Põlme et al. 2020) | Põlme S, et al. 2020. Fungal Diversity 105:1-16 ((https://doi.org/10.1007/s13225-020-00466-2)) |
| Otu07933 | 3 | k__Fungi(100);p__Ascomycota(100);c__Saccharomycetes(100);o__Saccharomycetales(100);f__Metschnikowiaceae(100);g__Metschnikowia(100);g__Metschnikowia_unclassified(67); | Metschnikowia | 13 | Saprotroph | \|Undefined Saprotroph\| | Yeast | NULL | Probable | As nectar-tap_saprotroph (Põlme et al. 2020); Decay sugar-rich substrates (Põlme et al. 2020) | Põlme S, et al. 2020. Fungal Diversity 105:1-16 ((https://doi.org/10.1007/s13225-020-00466-2)) |
| Otu07934 | 3 | k__Fungi(100);p__Ascomycota(100);c__Saccharomycetes(100);o__Saccharomycetales(100);f__Metschnikowiaceae(100);g__Metschnikowia(100);s__Metschnikowia_pulcherrima(67); | Metschnikowia | 13 | Saprotroph | \|Undefined Saprotroph\| | Yeast | NULL | Probable | As nectar-tap_saprotroph (Põlme et al. 2020); Decay sugar-rich substrates (Põlme et al. 2020) | Põlme S, et al. 2020. Fungal Diversity 105:1-16 ((https://doi.org/10.1007/s13225-020-00466-2)) |
| Otu07936 | 3 | k__Fungi(100);p__Ascomycota(100);c__Saccharomycetes(100);o__Saccharomycetales(100);f__Metschnikowiaceae(100);g__Metschnikowia(100);g__Metschnikowia_unclassified(67); | Metschnikowia | 13 | Saprotroph | \|Undefined Saprotroph\| | Yeast | NULL | Probable | As nectar-tap_saprotroph (Põlme et al. 2020); Decay sugar-rich substrates (Põlme et al. 2020) | Põlme S, et al. 2020. Fungal Diversity 105:1-16 ((https://doi.org/10.1007/s13225-020-00466-2)) |
| Otu07938 | 3 | k__Fungi(100);p__Ascomycota(100);c__Saccharomycetes(100);o__Saccharomycetales(100);f__Metschnikowiaceae(100);g__Metschnikowia(100);s__Metschnikowia_pulcherrima(67); | Metschnikowia | 13 | Saprotroph | \|Undefined Saprotroph\| | Yeast | NULL | Probable | As nectar-tap_saprotroph (Põlme et al. 2020); Decay sugar-rich substrates (Põlme et al. 2020) | Põlme S, et al. 2020. Fungal Diversity 105:1-16 ((https://doi.org/10.1007/s13225-020-00466-2)) |
| Otu07940 | 3 | k__Fungi(100);p__Ascomycota(100);c__Saccharomycetes(100);o__Saccharomycetales(100);f__Metschnikowiaceae(100);g__Metschnikowia(100);s__Metschnikowia_pulcherrima(100); | Metschnikowia | 13 | Saprotroph | \|Undefined Saprotroph\| | Yeast | NULL | Probable | As nectar-tap_saprotroph (Põlme et al. 2020); Decay sugar-rich substrates (Põlme et al. 2020) | Põlme S, et al. 2020. Fungal Diversity 105:1-16 ((https://doi.org/10.1007/s13225-020-00466-2)) |
| Otu07943 | 3 | k__Fungi(100);p__Ascomycota(100);c__Saccharomycetes(100);o__Saccharomycetales(100);f__Metschnikowiaceae(100);g__Metschnikowia(100);g__Metschnikowia_unclassified(100); | Metschnikowia | 13 | Saprotroph | \|Undefined Saprotroph\| | Yeast | NULL | Probable | As nectar-tap_saprotroph (Põlme et al. 2020); Decay sugar-rich substrates (Põlme et al. 2020) | Põlme S, et al. 2020. Fungal Diversity 105:1-16 ((https://doi.org/10.1007/s13225-020-00466-2)) |
| Otu07944 | 3 | k__Fungi(100);p__Ascomycota(100);c__Saccharomycetes(100);o__Saccharomycetales(100);f__Metschnikowiaceae(100);g__Metschnikowia(100);s__Metschnikowia_pulcherrima(100); | Metschnikowia | 13 | Saprotroph | \|Undefined Saprotroph\| | Yeast | NULL | Probable | As nectar-tap_saprotroph (Põlme et al. 2020); Decay sugar-rich substrates (Põlme et al. 2020) | Põlme S, et al. 2020. Fungal Diversity 105:1-16 ((https://doi.org/10.1007/s13225-020-00466-2)) |
| Otu07947 | 3 | k__Fungi(100);p__Ascomycota(100);c__Saccharomycetes(100);o__Saccharomycetales(100);f__Metschnikowiaceae(100);g__Metschnikowia(100);g__Metschnikowia_unclassified(67); | Metschnikowia | 13 | Saprotroph | \|Undefined Saprotroph\| | Yeast | NULL | Probable | As nectar-tap_saprotroph (Põlme et al. 2020); Decay sugar-rich substrates (Põlme et al. 2020) | Põlme S, et al. 2020. Fungal Diversity 105:1-16 ((https://doi.org/10.1007/s13225-020-00466-2)) |
| Otu07950 | 3 | k__Fungi(100);p__Ascomycota(100);c__Saccharomycetes(100);o__Saccharomycetales(100);f__Saccharomycetales_family_Incertae_sedis(100);g__Candida(67);g__Candida_unclassified(67); | Candida | 13 | Pathotroph-Saprotroph-Symbiotroph | Animal Pathogen-Endophyte-Endosymbiont-Epiphyte-\|Undefined Saprotroph\| | Dimorphic Yeast-Yeast | NULL | Probable | As nectar-tap_saprotroph (Põlme et al. 2020); Decay sugar-rich substrates (Põlme et al. 2020); This asexual genus is polyphyletic and contain species that function very differently depend on what they are. Caution should be taken when interpreting guilds for Candida species. | Manolakaki D, et al. 2010. Virulence. 1: 367-375 ((https://doi.org/10.4161/viru.1.5.12796)); Mekbib SB, et al. 2011. Tropical Plant Pathology 36:233-240 ((https://doi.org/10.1590/S1982-56762011000400004)); Põlme S, et al. 2020. Fungal Diversity 105:1-16 ((https://doi.org/10.1007/s13225-020-00466-2)) |

| | | | | | | | | | | | |
|---|---|---|---|---|---|---|---|---|---|---|---|
| Otu07951 | 3 | k__Fungi(100);p__Ascomycota(100);c__Saccharomycetes(100);o__Saccharomycetales(100);f__Metschnikowiaceae(100);g__Metschnikowia(100);s__Metschnikowia_pulcherrima(67); | Metschnikowia | 13 | Saprotroph | \|Undefined Saprotroph\| | Yeast | NULL | Probable | As nectar-tap_saprotroph (Põlme et al. 2020); Decay sugar-rich substrates (Põlme et al. 2020) | Põlme S, et al. 2020. Fungal Diversity 105:1-16 ((https://doi.org/10.1007/s13225-020-00466-2)) |
| Otu07955 | 3 | k__Fungi(100);p__Ascomycota(100);c__Saccharomycetes(100);o__Saccharomycetales(100);f__Metschnikowiaceae(100);g__Metschnikowia(100);s__Metschnikowia_pulcherrima(100); | Metschnikowia | 13 | Saprotroph | \|Undefined Saprotroph\| | Yeast | NULL | Probable | As nectar-tap_saprotroph (Põlme et al. 2020); Decay sugar-rich substrates (Põlme et al. 2020) | Põlme S, et al. 2020. Fungal Diversity 105:1-16 ((https://doi.org/10.1007/s13225-020-00466-2)) |
| Otu07957 | 3 | k__Fungi(100);p__Ascomycota(100);c__Saccharomycetes(100);o__Saccharomycetales(100);f__Metschnikowiaceae(100);g__Metschnikowia(100);s__Metschnikowia_pulcherrima(67); | Metschnikowia | 13 | Saprotroph | \|Undefined Saprotroph\| | Yeast | NULL | Probable | As nectar-tap_saprotroph (Põlme et al. 2020); Decay sugar-rich substrates (Põlme et al. 2020) | Põlme S, et al. 2020. Fungal Diversity 105:1-16 ((https://doi.org/10.1007/s13225-020-00466-2)) |
| Otu07960 | 3 | k__Fungi(100);p__Ascomycota(100);c__Saccharomycetes(100);o__Saccharomycetales(100);f__Saccharomycetales_family_Incertae_sedis(100);g__Candida(100);g__Candida_unclassified(100); | Candida | 13 | Pathotroph-Saprotroph-Symbiotroph | Animal Pathogen-Endophyte-Endosymbiont-Epiphyte-\|Undefined Saprotroph\| | Dimorphic Yeast-Yeast | NULL | Probable | As nectar-tap_saprotroph (Põlme et al. 2020); Decay sugar-rich substrates (Põlme et al. 2020); This asexual genus is polyphyletic and contain species that function very differently depend on what they are. Caution should be taken when interpreting guilds for Candida species. | Manolakaki D, et al. 2010. Virulence. 1: 367-375 ((https://doi.org/10.4161/viru.1.5.12796)); Mekbib SB, et al. 2011. Tropical Plant Pathology 36:233-240 ((https://doi.org/10.1590/S1982-56762011000400004)); Põlme S, et al. 2020. Fungal Diversity 105:1-16 ((https://doi.org/10.1007/s13225-020-00466-2)) |
| Otu07961 | 3 | k__Fungi(100);p__Ascomycota(100);c__Saccharomycetes(100);o__Saccharomycetales(100);f__Metschnikowiaceae(100);g__Metschnikowia(100);g__Metschnikowia_unclassified(100); | Metschnikowia | 13 | Saprotroph | \|Undefined Saprotroph\| | Yeast | NULL | Probable | As nectar-tap_saprotroph (Põlme et al. 2020); Decay sugar-rich substrates (Põlme et al. 2020) | Põlme S, et al. 2020. Fungal Diversity 105:1-16 ((https://doi.org/10.1007/s13225-020-00466-2)) |
| Otu07963 | 3 | k__Fungi(100);p__Ascomycota(100);c__Saccharomycetes(100);o__Saccharomycetales(100);f__Metschnikowiaceae(100);g__Metschnikowia(100);s__Metschnikowia_pulcherrima(100); | Metschnikowia | 13 | Saprotroph | \|Undefined Saprotroph\| | Yeast | NULL | Probable | As nectar-tap_saprotroph (Põlme et al. 2020); Decay sugar-rich substrates (Põlme et al. 2020) | Põlme S, et al. 2020. Fungal Diversity 105:1-16 ((https://doi.org/10.1007/s13225-020-00466-2)) |
| Otu07966 | 3 | k__Fungi(100);p__Ascomycota(100);c__Saccharomycetes(100);o__Saccharomycetales(100);f__Metschnikowiaceae(100);g__Metschnikowia(100);s__Metschnikowia_pulcherrima(67); | Metschnikowia | 13 | Saprotroph | \|Undefined Saprotroph\| | Yeast | NULL | Probable | As nectar-tap_saprotroph (Põlme et al. 2020); Decay sugar-rich substrates (Põlme et al. 2020) | Põlme S, et al. 2020. Fungal Diversity 105:1-16 ((https://doi.org/10.1007/s13225-020-00466-2)) |
| Otu07968 | 3 | k__Fungi(100);p__Ascomycota(100);c__Saccharomycetes(100);o__Saccharomycetales(100);f__Metschnikowiaceae(100);g__Metschnikowia(100);s__Metschnikowia_pulcherrima(100); | Metschnikowia | 13 | Saprotroph | \|Undefined Saprotroph\| | Yeast | NULL | Probable | As nectar-tap_saprotroph (Põlme et al. 2020); Decay sugar-rich substrates (Põlme et al. 2020) | Põlme S, et al. 2020. Fungal Diversity 105:1-16 ((https://doi.org/10.1007/s13225-020-00466-2)) |
| Otu07969 | 3 | k__Fungi(100);p__Ascomycota(100);c__Saccharomycetes(100);o__Saccharomycetales(100);f__Metschnikowiaceae(100);g__Metschnikowia(100);g__Metschnikowia_unclassified(67); | Metschnikowia | 13 | Saprotroph | \|Undefined Saprotroph\| | Yeast | NULL | Probable | As nectar-tap_saprotroph (Põlme et al. 2020); Decay sugar-rich substrates (Põlme et al. 2020) | Põlme S, et al. 2020. Fungal Diversity 105:1-16 ((https://doi.org/10.1007/s13225-020-00466-2)) |
| Otu07970 | 3 | k__Fungi(100);p__Ascomycota(100);c__Saccharomycetes(100);o__Saccharomycetales(100);f__Metschnikowiaceae(100);g__Metschnikowia(100);s__Metschnikowia_pulcherrima(67); | Metschnikowia | 13 | Saprotroph | \|Undefined Saprotroph\| | Yeast | NULL | Probable | As nectar-tap_saprotroph (Põlme et al. 2020); Decay sugar-rich substrates (Põlme et al. 2020) | Põlme S, et al. 2020. Fungal Diversity 105:1-16 ((https://doi.org/10.1007/s13225-020-00466-2)) |
| Otu07971 | 3 | k__Fungi(100);p__Ascomycota(100);c__Saccharomycetes(100);o__Saccharomycetales(100);f__Metschnikowiaceae(100);g__Metschnikowia(100);s__Metschnikowia_pulcherrima(100); | Metschnikowia | 13 | Saprotroph | \|Undefined Saprotroph\| | Yeast | NULL | Probable | As nectar-tap_saprotroph (Põlme et al. 2020); Decay sugar-rich substrates (Põlme et al. 2020) | Põlme S, et al. 2020. Fungal Diversity 105:1-16 ((https://doi.org/10.1007/s13225-020-00466-2)) |
| Otu07972 | 3 | k__Fungi(100);p__Ascomycota(100);c__Saccharomycetes(100);o__Saccharomycetales(100);f__Metschnikowiaceae(100);g__Metschnikowia(100);g__Metschnikowia_unclassified(100); | Metschnikowia | 13 | Saprotroph | \|Undefined Saprotroph\| | Yeast | NULL | Probable | As nectar-tap_saprotroph (Põlme et al. 2020); Decay sugar-rich substrates (Põlme et al. 2020) | Põlme S, et al. 2020. Fungal Diversity 105:1-16 ((https://doi.org/10.1007/s13225-020-00466-2)) |
| Otu07973 | 3 | k__Fungi(100);p__Ascomycota(100);c__Saccharomycetes(100);o__Saccharomycetales(100);f__Metschnikowiaceae(100);g__Metschnikowia(100);s__Metschnikowia_pulcherrima(100); | Metschnikowia | 13 | Saprotroph | \|Undefined Saprotroph\| | Yeast | NULL | Probable | As nectar-tap_saprotroph (Põlme et al. 2020); Decay sugar-rich substrates (Põlme et al. 2020) | Põlme S, et al. 2020. Fungal Diversity 105:1-16 ((https://doi.org/10.1007/s13225-020-00466-2)) |
| Otu07978 | 3 | k__Fungi(100);p__Ascomycota(100);c__Saccharomycetes(100);o__Saccharomycetales(100);f__Metschnikowiaceae(100);g__Metschnikowia(100);s__Metschnikowia_pulcherrima(100); | Metschnikowia | 13 | Saprotroph | \|Undefined Saprotroph\| | Yeast | NULL | Probable | As nectar-tap_saprotroph (Põlme et al. 2020); Decay sugar-rich substrates (Põlme et al. 2020) | Põlme S, et al. 2020. Fungal Diversity 105:1-16 ((https://doi.org/10.1007/s13225-020-00466-2)) |
| Otu07979 | 3 | k__Fungi(100);p__Ascomycota(100);c__Saccharomycetes(100);o__Saccharomycetales(100);f__Metschnikowiaceae(100);g__Metschnikowia(100);s__Metschnikowia_pulcherrima(67); | Metschnikowia | 13 | Saprotroph | \|Undefined Saprotroph\| | Yeast | NULL | Probable | As nectar-tap_saprotroph (Põlme et al. 2020); Decay sugar-rich substrates (Põlme et al. 2020) | Põlme S, et al. 2020. Fungal Diversity 105:1-16 ((https://doi.org/10.1007/s13225-020-00466-2)) |
| Otu07980 | 3 | k__Fungi(100);p__Ascomycota(100);c__Saccharomycetes(100);o__Saccharomycetales(100);f__Metschnikowiaceae(100);g__Metschnikowia(100);s__Metschnikowia_pulcherrima(100); | Metschnikowia | 13 | Saprotroph | \|Undefined Saprotroph\| | Yeast | NULL | Probable | As nectar-tap_saprotroph (Põlme et al. 2020); Decay sugar-rich substrates (Põlme et al. 2020) | Põlme S, et al. 2020. Fungal Diversity 105:1-16 ((https://doi.org/10.1007/s13225-020-00466-2)) |
| Otu07981 | 3 | k__Fungi(100);p__Ascomycota(100);c__Dothideomycetes(100);o__Capnodiales(100);f__Davidiellaceae(100);g__Cladosporium(100);g__Cladosporium_unclassified(100); | Cladosporium | 13 | Pathotroph-Saprotroph-Symbiotroph | Animal Pathogen-Endophyte-Lichen Parasite-Plant Pathogen-\|Plant Saprotroph\|-Wood Saprotroph | Microfungus | NULL | Possible | As foliar_endophyte (Põlme et al. 2020); As litter_saprotroph (Põlme et al. 2020); Decay wood, leaf, fruit, seed, soil, dung, animal material (Põlme et al. 2020) | Seehann G, et al. 1975. List of Fungi in Soft-Rot Tissue Tests. Institute fur Holzbiologie und Holzschutz, Bundesforschungsanstalt fur Forst - und Holzwirtshaft, Hamburg - Lohbrugge, Germany; Lawrey JD, Diederich P. 2016. http://www.lichenicolous.net (accessed 24 May 2017) ((http://www.lichenicolous.net)); Põlme S, et al. 2020. Fungal Diversity 105:1-16 ((https://doi.org/10.1007/s13225-020-00466-2)); The American Phytopathological Society Common Names of Plant Diseases ((http://www.apsnet.org/publications/commonnames/Pages/default.aspx)) |
| Otu07982 | 3 | k__Fungi(100);p__Ascomycota(100);c__Saccharomycetes(100);o__Saccharomycetales(100);f__Metschnikowiaceae(100);g__Metschnikowia(100);s__Metschnikowia_pulcherrima(67); | Metschnikowia | 13 | Saprotroph | \|Undefined Saprotroph\| | Yeast | NULL | Probable | As nectar-tap_saprotroph (Põlme et al. 2020); Decay sugar-rich substrates (Põlme et al. 2020) | Põlme S, et al. 2020. Fungal Diversity 105:1-16 ((https://doi.org/10.1007/s13225-020-00466-2)) |
| Otu07984 | 3 | k__Fungi(100);p__Basidiomycota(100);c__Agaricomycetes(100);o__Cantharellales(100);f__Ceratobasidiaceae(100);g__Thanatephorus(100);s__Thanatephorus_cucumeris(100); | Thanatephorus | 13 | Pathotroph-Saprotroph | \|Plant Pathogen\|-Plant Saprotroph | Corticioid | NULL | Probable | As litter_saprotroph (Põlme et al. 2020); Decay leaf, fruit, seed (Põlme et al. 2020) | Tedersoo L, et al. 2014. Science 346:e1256688 ((https://doi.org/10.1126/science.1256688)); Põlme S, et al. 2020. Fungal Diversity 105:1-16 ((https://doi.org/10.1007/s13225-020-00466-2)) |
| Otu07986 | 3 | k__Fungi(100);p__Ascomycota(100);c__Saccharomycetes(100);o__Saccharomycetales(100);f__Metschnikowiaceae(100);g__Metschnikowia(100);s__Metschnikowia_pulcherrima(67); | Metschnikowia | 13 | Saprotroph | \|Undefined Saprotroph\| | Yeast | NULL | Probable | As nectar-tap_saprotroph (Põlme et al. 2020); Decay sugar-rich substrates (Põlme et al. 2020) | Põlme S, et al. 2020. Fungal Diversity 105:1-16 ((https://doi.org/10.1007/s13225-020-00466-2)) |
| Otu07988 | 3 | k__Fungi(100);p__Ascomycota(100);c__Saccharomycetes(100);o__Saccharomycetales(100);f__Metschnikowiaceae(100);g__Metschnikowia(100);g__Metschnikowia_unclassified(67); | Metschnikowia | 13 | Saprotroph | \|Undefined Saprotroph\| | Yeast | NULL | Probable | As nectar-tap_saprotroph (Põlme et al. 2020); Decay sugar-rich substrates (Põlme et al. 2020) | Põlme S, et al. 2020. Fungal Diversity 105:1-16 ((https://doi.org/10.1007/s13225-020-00466-2)) |
| Otu07990 | 3 | k__Fungi(100);p__Ascomycota(100);c__Saccharomycetes(100);o__Saccharomycetales(100);f__Metschnikowiaceae(100);g__Metschnikowia(100);s__Metschnikowia_pulcherrima(100); | Metschnikowia | 13 | Saprotroph | \|Undefined Saprotroph\| | Yeast | NULL | Probable | As nectar-tap_saprotroph (Põlme et al. 2020); Decay sugar-rich substrates (Põlme et al. 2020) | Põlme S, et al. 2020. Fungal Diversity 105:1-16 ((https://doi.org/10.1007/s13225-020-00466-2)) |
| Otu07991 | 3 | k__Fungi(100);p__Ascomycota(100);c__Saccharomycetes(100);o__Saccharomycetales(100);f__Metschnikowiaceae(100);g__Metschnikowia(100);s__Metschnikowia_pulcherrima(100); | Metschnikowia | 13 | Saprotroph | \|Undefined Saprotroph\| | Yeast | NULL | Probable | As nectar-tap_saprotroph (Põlme et al. 2020); Decay sugar-rich substrates (Põlme et al. 2020) | Põlme S, et al. 2020. Fungal Diversity 105:1-16 ((https://doi.org/10.1007/s13225-020-00466-2)) |
| Otu07993 | 3 | k__Fungi(100);p__Basidiomycota(100);c__Agaricomycetes(100);o__Cantharellales(100);f__Ceratobasidiaceae(100);f__Ceratobasidiaceae_unclassified(100);f__Ceratobasidiaceae_unclassified(100); | Ceratobasidiaceae | 9 | Pathotroph-Saprotroph-Symbiotroph | Endomycorrhizal-Plant Pathogen-Undefined Saprotroph | Microfungus | NULL | Possible | Best known as plant pathogens, but also saprobic. Species are endomycorrhizal with a wide range of plant roots - especially of Orchidaceae but also even bryophytes (Cannon & Kirk 2007) | Cannon PF, Kirk PM. 2007. Fungal Families of the World. CAB International, Cambridge (ISBN: 978-0851998275) |
| Otu07994 | 3 | k__Fungi(100);p__Ascomycota(100);c__Saccharomycetes(100);o__Saccharomycetales(100);f__Metschnikowiaceae(100);g__Metschnikowia(100);s__Metschnikowia_pulcherrima(100); | Metschnikowia | 13 | Saprotroph | \|Undefined Saprotroph\| | Yeast | NULL | Probable | As nectar-tap_saprotroph (Põlme et al. 2020); Decay sugar-rich substrates (Põlme et al. 2020) | Põlme S, et al. 2020. Fungal Diversity 105:1-16 ((https://doi.org/10.1007/s13225-020-00466-2)) |
| Otu07999 | 3 | k__Fungi(100);p__Ascomycota(100);c__Saccharomycetes(100);o__Saccharomycetales(100);f__Metschnikowiaceae(100);g__Metschnikowia(100);s__Metschnikowia_pulcherrima(67); | Metschnikowia | 13 | Saprotroph | \|Undefined Saprotroph\| | Yeast | NULL | Probable | As nectar-tap_saprotroph (Põlme et al. 2020); Decay sugar-rich substrates (Põlme et al. 2020) | Põlme S, et al. 2020. Fungal Diversity 105:1-16 ((https://doi.org/10.1007/s13225-020-00466-2)) |
| Otu08002 | 3 | k__Fungi(100);p__Ascomycota(100);c__Saccharomycetes(100);o__Saccharomycetales(100);f__Metschnikowiaceae(100);g__Metschnikowia(100);s__Metschnikowia_pulcherrima(67); | Metschnikowia | 13 | Saprotroph | \|Undefined Saprotroph\| | Yeast | NULL | Probable | As nectar-tap_saprotroph (Põlme et al. 2020); Decay sugar-rich substrates (Põlme et al. 2020) | Põlme S, et al. 2020. Fungal Diversity 105:1-16 ((https://doi.org/10.1007/s13225-020-00466-2)) |
| Otu08003 | 3 | k__Fungi(100);p__Ascomycota(100);c__Saccharomycetes(100);o__Saccharomycetales(100);f__Metschnikowiaceae(100);g__Metschnikowia(100);s__Metschnikowia_pulcherrima(100); | Metschnikowia | 13 | Saprotroph | \|Undefined Saprotroph\| | Yeast | NULL | Probable | As nectar-tap_saprotroph (Põlme et al. 2020); Decay sugar-rich substrates (Põlme et al. 2020) | Põlme S, et al. 2020. Fungal Diversity 105:1-16 ((https://doi.org/10.1007/s13225-020-00466-2)) |
| Otu08004 | 3 | k__Fungi(100);p__Ascomycota(100);c__Saccharomycetes(100);o__Saccharomycetales(100);f__Metschnikowiaceae(100);g__Metschnikowia(100);s__Metschnikowia_pulcherrima(100); | Metschnikowia | 13 | Saprotroph | \|Undefined Saprotroph\| | Yeast | NULL | Probable | As nectar-tap_saprotroph (Põlme et al. 2020); Decay sugar-rich substrates (Põlme et al. 2020) | Põlme S, et al. 2020. Fungal Diversity 105:1-16 ((https://doi.org/10.1007/s13225-020-00466-2)) |
| Otu08005 | 3 | k__Fungi(100);p__Ascomycota(100);c__Saccharomycetes(100);o__Saccharomycetales(100);f__Metschnikowiaceae(100);g__Metschnikowia(100);s__Metschnikowia_pulcherrima(100); | Metschnikowia | 13 | Saprotroph | \|Undefined Saprotroph\| | Yeast | NULL | Probable | As nectar-tap_saprotroph (Põlme et al. 2020); Decay sugar-rich substrates (Põlme et al. 2020) | Põlme S, et al. 2020. Fungal Diversity 105:1-16 ((https://doi.org/10.1007/s13225-020-00466-2)) |
| Otu08006 | 3 | k__Fungi(100);p__Ascomycota(100);c__Saccharomycetes(100);o__Saccharomycetales(100);f__Metschnikowiaceae(100);g__Metschnikowia(100);s__Metschnikowia_pulcherrima(100); | Metschnikowia | 13 | Saprotroph | \|Undefined Saprotroph\| | Yeast | NULL | Probable | As nectar-tap_saprotroph (Põlme et al. 2020); Decay sugar-rich substrates (Põlme et al. 2020) | Põlme S, et al. 2020. Fungal Diversity 105:1-16 ((https://doi.org/10.1007/s13225-020-00466-2)) |
| Otu08008 | 3 | k__Fungi(100);p__Ascomycota(100);c__Saccharomycetes(100);o__Saccharomycetales(100);f__Metschnikowiaceae(100);g__Metschnikowia(100);g__Metschnikowia_unclassified(67); | Metschnikowia | 13 | Saprotroph | \|Undefined Saprotroph\| | Yeast | NULL | Probable | As nectar-tap_saprotroph (Põlme et al. 2020); Decay sugar-rich substrates (Põlme et al. 2020) | Põlme S, et al. 2020. Fungal Diversity 105:1-16 ((https://doi.org/10.1007/s13225-020-00466-2)) |
| Otu08011 | 3 | k__Fungi(100);p__Ascomycota(100);c__Eurotiomycetes(100);o__Eurotiales(100);f__Trichocomaceae(100);g__Penicillium(100);g__Penicillium_unclassified(100); | Penicillium | 13 | Pathotroph-Saprotroph-Symbiotroph | Animal Parasite-Dung Saprotroph-Endophyte-Fungal Parasite-Plant Pathogen-\|Undefined Saprotroph\|-Wood Saprotroph | Microfungus | Soft Rot | Highly Probable | As foliar_endophyte (Põlme et al. 2020); As mycoparasite (Põlme et al. 2020); Decay soil (Põlme et al. 2020) | Duncan CG, Eslyn WE. 1966. Mycologia 58:642-664 ((https://doi.org/10.2307/3757045)); Seehann G, et al. 1975. List of Fungi in Soft-Rot Tissue Tests. Institute fur Holzbiologie und Holzschutz, Bundesforschungsanstalt fur Forst - und Holzwirtshaft, Hamburg - Lohbrugge, Germany ((https://www.irg-wp.com/irgdocs/details.php?d6493033-c9ea-40f3-8e3e-5697f32d640d)); Costa IPMW, et al. 2012. Mycotaxon 119:e26 ((http://mycotaxon.com/resources/checklists/costa_v119_checklist.pdf)); Bills GF, et al. 2013. Current Opinion in Microbiology 16: 549-565 ((https://doi.org/10.1016/j.mib.2013.08.001)); Tedersoo L, et al. 2014. Science 346:e1256688 ((https://doi.org/10.1126/science.1256688)); Põlme S, et al. 2020. Fungal Diversity 105:1-16 ((https://doi.org/10.1007/s13225-020-00466-2)) |
| Otu08012 | 3 | k__Fungi(100);p__Ascomycota(100);c__Saccharomycetes(100);o__Saccharomycetales(100);f__Metschnikowiaceae(100);g__Metschnikowia(100);s__Metschnikowia_pulcherrima(67); | Metschnikowia | 13 | Saprotroph | \|Undefined Saprotroph\| | Yeast | NULL | Probable | As nectar-tap_saprotroph (Põlme et al. 2020); Decay sugar-rich substrates (Põlme et al. 2020) | Põlme S, et al. 2020. Fungal Diversity 105:1-16 ((https://doi.org/10.1007/s13225-020-00466-2)) |
| Otu08013 | 3 | k__Fungi(100);p__Ascomycota(100);c__Saccharomycetes(100);o__Saccharomycetales(100);f__Metschnikowiaceae(100);g__Metschnikowia(100);s__Metschnikowia_pulcherrima(100); | Metschnikowia | 13 | Saprotroph | \|Undefined Saprotroph\| | Yeast | NULL | Probable | As nectar-tap_saprotroph (Põlme et al. 2020); Decay sugar-rich substrates (Põlme et al. 2020) | Põlme S, et al. 2020. Fungal Diversity 105:1-16 ((https://doi.org/10.1007/s13225-020-00466-2)) |
| Otu08014 | 3 | k__Fungi(100);p__Ascomycota(100);c__Saccharomycetes(100);o__Saccharomycetales(100);f__Metschnikowiaceae(100);g__Metschnikowia(100);s__Metschnikowia_pulcherrima(100); | Metschnikowia | 13 | Saprotroph | \|Undefined Saprotroph\| | Yeast | NULL | Probable | As nectar-tap_saprotroph (Põlme et al. 2020); Decay sugar-rich substrates (Põlme et al. 2020) | Põlme S, et al. 2020. Fungal Diversity 105:1-16 ((https://doi.org/10.1007/s13225-020-00466-2)) |
| Otu08015 | 3 | k__Fungi(100);p__Ascomycota(100);c__Saccharomycetes(100);o__Saccharomycetales(100);f__Metschnikowiaceae(100);g__Metschnikowia(100);s__Metschnikowia_pulcherrima(67); | Metschnikowia | 13 | Saprotroph | \|Undefined Saprotroph\| | Yeast | NULL | Probable | As nectar-tap_saprotroph (Põlme et al. 2020); Decay sugar-rich substrates (Põlme et al. 2020) | Põlme S, et al. 2020. Fungal Diversity 105:1-16 ((https://doi.org/10.1007/s13225-020-00466-2)) |
| Otu08016 | 3 | k__Fungi(100);p__Ascomycota(100);c__Saccharomycetes(100);o__Saccharomycetales(100);f__Metschnikowiaceae(100);g__Metschnikowia(100);s__Metschnikowia_pulcherrima(100); | Metschnikowia | 13 | Saprotroph | \|Undefined Saprotroph\| | Yeast | NULL | Probable | As nectar-tap_saprotroph (Põlme et al. 2020); Decay sugar-rich substrates (Põlme et al. 2020) | Põlme S, et al. 2020. Fungal Diversity 105:1-16 ((https://doi.org/10.1007/s13225-020-00466-2)) |
| Otu08017 | 3 | k__Fungi(100);p__Ascomycota(100);c__Saccharomycetes(100);o__Saccharomycetales(100);f__Metschnikowiaceae(100);g__Metschnikowia(100);g__Metschnikowia_unclassified(100); | Metschnikowia | 13 | Saprotroph | \|Undefined Saprotroph\| | Yeast | NULL | Probable | As nectar-tap_saprotroph (Põlme et al. 2020); Decay sugar-rich substrates (Põlme et al. 2020) | Põlme S, et al. 2020. Fungal Diversity 105:1-16 ((https://doi.org/10.1007/s13225-020-00466-2)) |
| Otu08020 | 3 | k__Fungi(100);p__Ascomycota(100);c__Saccharomycetes(100);o__Saccharomycetales(100);f__Metschnikowiaceae(100);g__Metschnikowia(100);s__Metschnikowia_pulcherrima(100); | Metschnikowia | 13 | Saprotroph | \|Undefined Saprotroph\| | Yeast | NULL | Probable | As nectar-tap_saprotroph (Põlme et al. 2020); Decay sugar-rich substrates (Põlme et al. 2020) | Põlme S, et al. 2020. Fungal Diversity 105:1-16 ((https://doi.org/10.1007/s13225-020-00466-2)) |
| Otu08021 | 3 | k__Fungi(100);p__Ascomycota(100);c__Dothideomycetes(100);o__Capnodiales(100);f__Teratosphaeriaceae(100);g__Devriesia(100);s__Devriesia_fraseriae(100); | Devriesia | 13 | Pathotroph-Saprotroph | \|Plant Pathogen\|-Plant Saprotroph | Microfungus | NULL | Probable | As litter_saprotroph (Põlme et al. 2020); Decay leaf, fruit, seed (Põlme et al. 2020) | Tedersoo L, et al. 2014. Science 346:e1256688 ((https://doi.org/10.1126/science.1256688)); Põlme S, et al. 2020. Fungal Diversity 105:1-16 ((https://doi.org/10.1007/s13225-020-00466-2)) |
| Otu08022 | 3 | k__Fungi(100);p__Ascomycota(100);c__Saccharomycetes(100);o__Saccharomycetales(100);f__Metschnikowiaceae(100);g__Metschnikowia(100);s__Metschnikowia_pulcherrima(100); | Metschnikowia | 13 | Saprotroph | \|Undefined Saprotroph\| | Yeast | NULL | Probable | As nectar-tap_saprotroph (Põlme et al. 2020); Decay sugar-rich substrates (Põlme et al. 2020) | Põlme S, et al. 2020. Fungal Diversity 105:1-16 ((https://doi.org/10.1007/s13225-020-00466-2)) |
| Otu08024 | 3 | k__Fungi(100);p__Ascomycota(100);c__Lecanoromycetes(100);o__Lecanorales(100);f__Parmeliaceae(100);g__Hypogymnia(100);s__Hypogymnia_physodes(100); | Hypogymnia | 13 | Symbiotroph | \|Lichenized\| | Thallus | NULL | Highly Probable | NULL | Esslinger TL. 2014. North Dakota State University ((http://www.ndsu.edu/pubweb/~esslinge/chcklst/chcklst7.htm)); Põlme S, et al. 2020. Fungal Diversity 105:1-16 ((https://doi.org/10.1007/s13225-020-00466-2)) |
| Otu08025 | 3 | k__Fungi(100);p__Basidiomycota(100);c__Tremellomycetes(100);o__Tremellales(100);f__Tremellales_family_Incertae_sedis(100);f__Tremellales_family_Incertae_sedis_unclassified(100);f__Tremellales_family_Incertae_sedis_unclassified(100); | Tremellales | 7 | Pathotroph-Saprotroph-Symbiotroph | Fungal Parasite-Undefined Saprotroph | Tremelloid-Yeast | NULL | Possible | The order contains both teleomorphic and anamorphic species, most of the latter being yeasts | Sterkenburg E, et al. 2015. New Phytologist 207:1145-1158 ((https://doi.org/10.1111/nph.13426)) |
| Otu08027 | 3 | k__Fungi(100);p__Ascomycota(100);c__Saccharomycetes(100);o__Saccharomycetales(100);f__Saccharomycetales_family_Incertae_sedis(100);g__Candida(100);s__Candida_sp_NG_31(100); | Candida | 13 | Pathotroph-Saprotroph-Symbiotroph | Animal Pathogen-Endophyte-Endosymbiont-Epiphyte-\|Undefined Saprotroph\| | Dimorphic Yeast-Yeast | NULL | Probable | As nectar-tap_saprotroph (Põlme et al. 2020); Decay sugar-rich substrates (Põlme et al. 2020); This asexual genus is polyphyletic and contain species that function very differently depend on what they are. Caution should be taken when interpreting guilds for Candida species. | Manolakaki D, et al. 2010. Virulence. 1: 367-375 ((https://doi.org/10.4161/viru.1.5.12796)); Mekbib SB, et al. 2011. Tropical Plant Pathology 36:233-240 ((https://doi.org/10.1590/S1982-56762011000400004)); Põlme S, et al. 2020. Fungal Diversity 105:1-16 ((https://doi.org/10.1007/s13225-020-00466-2)) |
| Otu08030 | 3 | k__Fungi(100);p__Ascomycota(100);c__Dothideomycetes(100);o__Capnodiales(100);f__Teratosphaeriaceae(100);g__Teratosphaeria(100);s__Teratosphaeria_knoxdavesii(100); | Teratosphaeria | 13 | Pathotroph-Saprotroph | \|Plant Pathogen\|-Plant Saprotroph-Wood Saprotroph | Microfungus | NULL | Probable | As litter_saprotroph (Põlme et al. 2020); As wood_pathogen (Põlme et al. 2020); Decay wood, leaf, fruit, seed (Põlme et al. 2020) | Yamazaki A, et al. 2010. Mycoscience 51:443-455 ((https://www.jstage.jst.go.jp/article/mycosci/51/6/51_MYC51443/_pdf)); Tedersoo L, et al. 2014. Science 346:e1256688 ((https://doi.org/10.1126/science.1256688)); Põlme S, et al. 2020. Fungal Diversity 105:1-16 ((https://doi.org/10.1007/s13225-020-00466-2)) |
| Otu08031 | 3 | k__Fungi(100);p__Ascomycota(100);c__Saccharomycetes(100);o__Saccharomycetales(100);f__Metschnikowiaceae(100);g__Metschnikowia(100);s__Metschnikowia_pulcherrima(67); | Metschnikowia | 13 | Saprotroph | \|Undefined Saprotroph\| | Yeast | NULL | Probable | As nectar-tap_saprotroph (Põlme et al. 2020); Decay sugar-rich substrates (Põlme et al. 2020) | Põlme S, et al. 2020. Fungal Diversity 105:1-16 ((https://doi.org/10.1007/s13225-020-00466-2)) |
| Otu08034 | 3 | k__Fungi(100);p__Ascomycota(100);c__Saccharomycetes(100);o__Saccharomycetales(100);f__Metschnikowiaceae(100);g__Metschnikowia(100);s__Metschnikowia_pulcherrima(100); | Metschnikowia | 13 | Saprotroph | \|Undefined Saprotroph\| | Yeast | NULL | Probable | As nectar-tap_saprotroph (Põlme et al. 2020); Decay sugar-rich substrates (Põlme et al. 2020) | Põlme S, et al. 2020. Fungal Diversity 105:1-16 ((https://doi.org/10.1007/s13225-020-00466-2)) |
| Otu08036 | 3 | k__Fungi(100);p__Ascomycota(100);c__Saccharomycetes(100);o__Saccharomycetales(100);f__Metschnikowiaceae(100);g__Metschnikowia(100);s__Metschnikowia_pulcherrima(67); | Metschnikowia | 13 | Saprotroph | \|Undefined Saprotroph\| | Yeast | NULL | Probable | As nectar-tap_saprotroph (Põlme et al. 2020); Decay sugar-rich substrates (Põlme et al. 2020) | Põlme S, et al. 2020. Fungal Diversity 105:1-16 ((https://doi.org/10.1007/s13225-020-00466-2)) |
| Otu08038 | 3 | k__Fungi(100);p__Ascomycota(100);c__Saccharomycetes(100);o__Saccharomycetales(100);f__Metschnikowiaceae(100);g__Metschnikowia(100);g__Metschnikowia_unclassified(67); | Metschnikowia | 13 | Saprotroph | \|Undefined Saprotroph\| | Yeast | NULL | Probable | As nectar-tap_saprotroph (Põlme et al. 2020); Decay sugar-rich substrates (Põlme et al. 2020) | Põlme S, et al. 2020. Fungal Diversity 105:1-16 ((https://doi.org/10.1007/s13225-020-00466-2)) |
| Otu08039 | 3 | k__Fungi(100);p__Ascomycota(100);c__Saccharomycetes(100);o__Saccharomycetales(100);f__Metschnikowiaceae(100);g__Metschnikowia(100);s__Metschnikowia_pulcherrima(67); | Metschnikowia | 13 | Saprotroph | \|Undefined Saprotroph\| | Yeast | NULL | Probable | As nectar-tap_saprotroph (Põlme et al. 2020); Decay sugar-rich substrates (Põlme et al. 2020) | Põlme S, et al. 2020. Fungal Diversity 105:1-16 ((https://doi.org/10.1007/s13225-020-00466-2)) |
| Otu08040 | 3 | k__Fungi(100);p__Ascomycota(100);c__Ascomycota_class_Incertae_sedis(100);o__Ascomycota_order_Incertae_sedis(100);f__Ascomycota_family_Incertae_sedis(100);g__Torula(100);s__Torula_sp(100); | Torula | 13 | Saprotroph-Symbiotroph | \|Endophyte\|-Plant Saprotroph-Wood Saprotroph | Facultative Yeast-Microfungus | Soft Rot | Probable | As foliar_endophyte (Põlme et al. 2020); Decay wood, leaf, fruit, seed (Põlme et al. 2020); Yeast can be propagated for long periods in wood sugar solutions (Fink & Lechner 1936) | Fink H, Lechner R. 1936. Biochemische Zeitschrift 286:83-90 ((https://archive.org/details/sim_biochemische-zeitschrift-beitraege-zur-chemischen_1936_286)); Crous PW, et al. 2015. IMA Fungus 6: 163-198 ((https://doi.org/10.5598/imafungus.2015.06.01.11)); Põlme S, et al. 2020. Fungal Diversity 105:1-16 ((https://doi.org/10.1007/s13225-020-00466-2)) |
| Otu08043 | 3 | k__Fungi(100);p__Ascomycota(100);c__Saccharomycetes(100);o__Saccharomycetales(100);f__Metschnikowiaceae(100);g__Metschnikowia(100);s__Metschnikowia_pulcherrima(67); | Metschnikowia | 13 | Saprotroph | \|Undefined Saprotroph\| | Yeast | NULL | Probable | As nectar-tap_saprotroph (Põlme et al. 2020); Decay sugar-rich substrates (Põlme et al. 2020) | Põlme S, et al. 2020. Fungal Diversity 105:1-16 ((https://doi.org/10.1007/s13225-020-00466-2)) |
| Otu08046 | 3 | k__Fungi(100);p__Ascomycota(100);c__Saccharomycetes(100);o__Saccharomycetales(100);f__Metschnikowiaceae(100);g__Metschnikowia(100);s__Metschnikowia_pulcherrima(100); | Metschnikowia | 13 | Saprotroph | \|Undefined Saprotroph\| | Yeast | NULL | Probable | As nectar-tap_saprotroph (Põlme et al. 2020); Decay sugar-rich substrates (Põlme et al. 2020) | Põlme S, et al. 2020. Fungal Diversity 105:1-16 ((https://doi.org/10.1007/s13225-020-00466-2)) |
| Otu08047 | 3 | k__Fungi(100);p__Ascomycota(100);c__Saccharomycetes(100);o__Saccharomycetales(100);f__Metschnikowiaceae(100);g__Metschnikowia(100);s__Metschnikowia_pulcherrima(67); | Metschnikowia | 13 | Saprotroph | \|Undefined Saprotroph\| | Yeast | NULL | Probable | As nectar-tap_saprotroph (Põlme et al. 2020); Decay sugar-rich substrates (Põlme et al. 2020) | Põlme S, et al. 2020. Fungal Diversity 105:1-16 ((https://doi.org/10.1007/s13225-020-00466-2)) |
| Otu08050 | 3 | k__Fungi(100);p__Ascomycota(100);c__Saccharomycetes(100);o__Saccharomycetales(100);f__Metschnikowiaceae(100);g__Metschnikowia(100);s__Metschnikowia_pulcherrima(67); | Metschnikowia | 13 | Saprotroph | \|Undefined Saprotroph\| | Yeast | NULL | Probable | As nectar-tap_saprotroph (Põlme et al. 2020); Decay sugar-rich substrates (Põlme et al. 2020) | Põlme S, et al. 2020. Fungal Diversity 105:1-16 ((https://doi.org/10.1007/s13225-020-00466-2)) |
| Otu08051 | 3 | k__Fungi(100);p__Ascomycota(100);c__Saccharomycetes(100);o__Saccharomycetales(100);f__Saccharomycetales_family_Incertae_sedis(100);g__Candida(100);g__Candida_unclassified(67); | Candida | 13 | Pathotroph-Saprotroph-Symbiotroph | Animal Pathogen-Endophyte-Endosymbiont-Epiphyte-\|Undefined Saprotroph\| | Dimorphic Yeast-Yeast | NULL | Probable | As nectar-tap_saprotroph (Põlme et al. 2020); Decay sugar-rich substrates (Põlme et al. 2020); This asexual genus is polyphyletic and contain species that function very differently depend on what they are. Caution should be taken when interpreting guilds for Candida species. | Manolakaki D, et al. 2010. Virulence. 1: 367-375 ((https://doi.org/10.4161/viru.1.5.12796)); Mekbib SB, et al. 2011. Tropical Plant Pathology 36:233-240 ((https://doi.org/10.1590/S1982-56762011000400004)); Põlme S, et al. 2020. Fungal Diversity 105:1-16 ((https://doi.org/10.1007/s13225-020-00466-2)) |
| Otu08052 | 3 | k__Fungi(100);p__Ascomycota(100);c__Saccharomycetes(100);o__Saccharomycetales(100);f__Metschnikowiaceae(100);g__Metschnikowia(100);s__Metschnikowia_pulcherrima(100); | Metschnikowia | 13 | Saprotroph | \|Undefined Saprotroph\| | Yeast | NULL | Probable | As nectar-tap_saprotroph (Põlme et al. 2020); Decay sugar-rich substrates (Põlme et al. 2020) | Põlme S, et al. 2020. Fungal Diversity 105:1-16 ((https://doi.org/10.1007/s13225-020-00466-2)) |
| Otu08054 | 3 | k__Fungi(100);p__Ascomycota(100);c__Saccharomycetes(100);o__Saccharomycetales(100);f__Metschnikowiaceae(100);g__Metschnikowia(100);s__Metschnikowia_pulcherrima(100); | Metschnikowia | 13 | Saprotroph | \|Undefined Saprotroph\| | Yeast | NULL | Probable | As nectar-tap_saprotroph (Põlme et al. 2020); Decay sugar-rich substrates (Põlme et al. 2020) | Põlme S, et al. 2020. Fungal Diversity 105:1-16 ((https://doi.org/10.1007/s13225-020-00466-2)) |
| Otu08055 | 3 | k__Fungi(100);p__Ascomycota(100);c__Saccharomycetes(100);o__Saccharomycetales(100);f__Metschnikowiaceae(100);g__Metschnikowia(100);g__Metschnikowia_unclassified(67); | Metschnikowia | 13 | Saprotroph | \|Undefined Saprotroph\| | Yeast | NULL | Probable | As nectar-tap_saprotroph (Põlme et al. 2020); Decay sugar-rich substrates (Põlme et al. 2020) | Põlme S, et al. 2020. Fungal Diversity 105:1-16 ((https://doi.org/10.1007/s13225-020-00466-2)) |
| Otu08056 | 3 | k__Fungi(100);p__Ascomycota(100);c__Saccharomycetes(100);o__Saccharomycetales(100);f__Metschnikowiaceae(100);g__Metschnikowia(100);s__Metschnikowia_pulcherrima(100); | Metschnikowia | 13 | Saprotroph | \|Undefined Saprotroph\| | Yeast | NULL | Probable | As nectar-tap_saprotroph (Põlme et al. 2020); Decay sugar-rich substrates (Põlme et al. 2020) | Põlme S, et al. 2020. Fungal Diversity 105:1-16 ((https://doi.org/10.1007/s13225-020-00466-2)) |
| Otu08057 | 3 | k__Fungi(100);p__Ascomycota(100);c__Saccharomycetes(100);o__Saccharomycetales(100);f__Metschnikowiaceae(100);g__Metschnikowia(100);g__Metschnikowia_unclassified(67); | Metschnikowia | 13 | Saprotroph | \|Undefined Saprotroph\| | Yeast | NULL | Probable | As nectar-tap_saprotroph (Põlme et al. 2020); Decay sugar-rich substrates (Põlme et al. 2020) | Põlme S, et al. 2020. Fungal Diversity 105:1-16 ((https://doi.org/10.1007/s13225-020-00466-2)) |
| Otu08058 | 3 | k__Fungi(100);p__Ascomycota(100);c__Saccharomycetes(100);o__Saccharomycetales(100);f__Metschnikowiaceae(100);g__Metschnikowia(100);g__Metschnikowia_unclassified(67); | Metschnikowia | 13 | Saprotroph | \|Undefined Saprotroph\| | Yeast | NULL | Probable | As nectar-tap_saprotroph (Põlme et al. 2020); Decay sugar-rich substrates (Põlme et al. 2020) | Põlme S, et al. 2020. Fungal Diversity 105:1-16 ((https://doi.org/10.1007/s13225-020-00466-2)) |
| Otu08059 | 3 | k__Fungi(100);p__Ascomycota(100);c__Saccharomycetes(100);o__Saccharomycetales(100);f__Metschnikowiaceae(100);g__Metschnikowia(100);g__Metschnikowia_unclassified(100); | Metschnikowia | 13 | Saprotroph | \|Undefined Saprotroph\| | Yeast | NULL | Probable | As nectar-tap_saprotroph (Põlme et al. 2020); Decay sugar-rich substrates (Põlme et al. 2020) | Põlme S, et al. 2020. Fungal Diversity 105:1-16 ((https://doi.org/10.1007/s13225-020-00466-2)) |
| Otu08061 | 3 | k__Fungi(100);p__Ascomycota(100);c__Saccharomycetes(100);o__Saccharomycetales(100);f__Metschnikowiaceae(100);g__Metschnikowia(100);s__Metschnikowia_pulcherrima(100); | Metschnikowia | 13 | Saprotroph | \|Undefined Saprotroph\| | Yeast | NULL | Probable | As nectar-tap_saprotroph (Põlme et al. 2020); Decay sugar-rich substrates (Põlme et al. 2020) | Põlme S, et al. 2020. Fungal Diversity 105:1-16 ((https://doi.org/10.1007/s13225-020-00466-2)) |
| Otu08062 | 3 | k__Fungi(100);p__Ascomycota(100);c__Saccharomycetes(100);o__Saccharomycetales(100);f__Metschnikowiaceae(100);g__Metschnikowia(100);s__Metschnikowia_pulcherrima(100); | Metschnikowia | 13 | Saprotroph | \|Undefined Saprotroph\| | Yeast | NULL | Probable | As nectar-tap_saprotroph (Põlme et al. 2020); Decay sugar-rich substrates (Põlme et al. 2020) | Põlme S, et al. 2020. Fungal Diversity 105:1-16 ((https://doi.org/10.1007/s13225-020-00466-2)) |
| Otu08063 | 3 | k__Fungi(100);p__Ascomycota(100);c__Saccharomycetes(100);o__Saccharomycetales(100);f__Metschnikowiaceae(100);g__Metschnikowia(100);s__Metschnikowia_pulcherrima(100); | Metschnikowia | 13 | Saprotroph | \|Undefined Saprotroph\| | Yeast | NULL | Probable | As nectar-tap_saprotroph (Põlme et al. 2020); Decay sugar-rich substrates (Põlme et al. 2020) | Põlme S, et al. 2020. Fungal Diversity 105:1-16 ((https://doi.org/10.1007/s13225-020-00466-2)) |
| Otu08064 | 3 | k__Fungi(100);p__Ascomycota(100);c__Saccharomycetes(100);o__Saccharomycetales(100);f__Metschnikowiaceae(100);g__Metschnikowia(100);g__Metschnikowia_unclassified(67); | Metschnikowia | 13 | Saprotroph | \|Undefined Saprotroph\| | Yeast | NULL | Probable | As nectar-tap_saprotroph (Põlme et al. 2020); Decay sugar-rich substrates (Põlme et al. 2020) | Põlme S, et al. 2020. Fungal Diversity 105:1-16 ((https://doi.org/10.1007/s13225-020-00466-2)) |
| Otu08067 | 3 | k__Fungi(100);p__Ascomycota(100);c__Saccharomycetes(100);o__Saccharomycetales(100);f__Metschnikowiaceae(100);g__Metschnikowia(100);s__Metschnikowia_pulcherrima(67); | Metschnikowia | 13 | Saprotroph | \|Undefined Saprotroph\| | Yeast | NULL | Probable | As nectar-tap_saprotroph (Põlme et al. 2020); Decay sugar-rich substrates (Põlme et al. 2020) | Põlme S, et al. 2020. Fungal Diversity 105:1-16 ((https://doi.org/10.1007/s13225-020-00466-2)) |
| Otu08068 | 3 | k__Fungi(100);p__Ascomycota(100);c__Saccharomycetes(100);o__Saccharomycetales(100);f__Saccharomycetales_family_Incertae_sedis(100);g__Candida(67);g__Candida_unclassified(67); | Candida | 13 | Pathotroph-Saprotroph-Symbiotroph | Animal Pathogen-Endophyte-Endosymbiont-Epiphyte-\|Undefined Saprotroph\| | Dimorphic Yeast-Yeast | NULL | Probable | As nectar-tap_saprotroph (Põlme et al. 2020); Decay sugar-rich substrates (Põlme et al. 2020); This asexual genus is polyphyletic and contain species that function very differently depend on what they are. Caution should be taken when interpreting guilds for Candida species. | Manolakaki D, et al. 2010. Virulence. 1: 367-375 ((https://doi.org/10.4161/viru.1.5.12796)); Mekbib SB, et al. 2011. Tropical Plant Pathology 36:233-240 ((https://doi.org/10.1590/S1982-56762011000400004)); Põlme S, et al. 2020. Fungal Diversity 105:1-16 ((https://doi.org/10.1007/s13225-020-00466-2)) |
| Otu08072 | 3 | k__Fungi(100);p__Ascomycota(100);c__Saccharomycetes(100);o__Saccharomycetales(100);f__Metschnikowiaceae(100);g__Metschnikowia(100);s__Metschnikowia_pulcherrima(100); | Metschnikowia | 13 | Saprotroph | \|Undefined Saprotroph\| | Yeast | NULL | Probable | As nectar-tap_saprotroph (Põlme et al. 2020); Decay sugar-rich substrates (Põlme et al. 2020) | Põlme S, et al. 2020. Fungal Diversity 105:1-16 ((https://doi.org/10.1007/s13225-020-00466-2)) |
| Otu08074 | 3 | k__Fungi(100);p__Ascomycota(100);c__Saccharomycetes(100);o__Saccharomycetales(100);f__Metschnikowiaceae(100);g__Metschnikowia(100);g__Metschnikowia_unclassified(67); | Metschnikowia | 13 | Saprotroph | \|Undefined Saprotroph\| | Yeast | NULL | Probable | As nectar-tap_saprotroph (Põlme et al. 2020); Decay sugar-rich substrates (Põlme et al. 2020) | Põlme S, et al. 2020. Fungal Diversity 105:1-16 ((https://doi.org/10.1007/s13225-020-00466-2)) |
| Otu08075 | 3 | k__Fungi(100);p__Ascomycota(100);c__Sordariomycetes(100);o__Coniochaetales(100);f__Coniochaetaceae(100);g__Coniochaeta(67);g__Coniochaeta_unclassified(67); | Coniochaeta | 13 | Pathotroph-Saprotroph-Symbiotroph | Animal Pathogen-Dung Saprotroph-Endophyte-Lichen Parasite-Plant Pathogen-\|Undefined Saprotroph\| | Microfungus | NULL | Possible | As foliar_endophyte (Põlme et al. 2020); Decay leaf, fruit, seed, soil, dung, animal material (Põlme et al. 2020) | Bell A. 1983. Dung Fungi: An Illustrated Guide to Coprophilous Fungi in New Zealand. Victoria University Press, Wellington (ISBN: 978-0864730015); Damm U, et al. 2010. Persoonia 24:60-80 ((http://doi.org/10.3767/003158510X500705)); Costa IPMW, et al. 2012. Mycotaxon 119:e26 ((http://mycotaxon.com/resources/checklists/costa_v119_checklist.pdf)); Tedersoo L, et al. 2014. Science 346: 1256688 ((https://doi.org/ 10.1126/science.1256688)); Irinyi L, et al. 2015. Medical Mycology 53:313-337 ((https://doi.org/10.1093/mmy/myv008)); Lawrey JD, Diederich P. 2016. http://www.lichenicolous.net (accessed 24 May 2017) ((http://www.lichenicolous.net)); Põlme S, et al. 2020. Fungal Diversity 105:1-16 ((https://doi.org/10.1007/s13225-020-00466-2)) |
| Otu08078 | 3 | k__Fungi(100);p__Ascomycota(100);c__Saccharomycetes(100);o__Saccharomycetales(100);f__Metschnikowiaceae(100);g__Metschnikowia(100);s__Metschnikowia_pulcherrima(100); | Metschnikowia | 13 | Saprotroph | \|Undefined Saprotroph\| | Yeast | NULL | Probable | As nectar-tap_saprotroph (Põlme et al. 2020); Decay sugar-rich substrates (Põlme et al. 2020) | Põlme S, et al. 2020. Fungal Diversity 105:1-16 ((https://doi.org/10.1007/s13225-020-00466-2)) |
| Otu08080 | 3 | k__Fungi(100);p__Ascomycota(100);c__Saccharomycetes(100);o__Saccharomycetales(100);f__Metschnikowiaceae(100);g__Metschnikowia(100);s__Metschnikowia_pulcherrima(100); | Metschnikowia | 13 | Saprotroph | \|Undefined Saprotroph\| | Yeast | NULL | Probable | As nectar-tap_saprotroph (Põlme et al. 2020); Decay sugar-rich substrates (Põlme et al. 2020) | Põlme S, et al. 2020. Fungal Diversity 105:1-16 ((https://doi.org/10.1007/s13225-020-00466-2)) |
| Otu08082 | 3 | k__Fungi(100);p__Ascomycota(100);c__Saccharomycetes(100);o__Saccharomycetales(100);f__Metschnikowiaceae(100);g__Metschnikowia(100);g__Metschnikowia_unclassified(67); | Metschnikowia | 13 | Saprotroph | \|Undefined Saprotroph\| | Yeast | NULL | Probable | As nectar-tap_saprotroph (Põlme et al. 2020); Decay sugar-rich substrates (Põlme et al. 2020) | Põlme S, et al. 2020. Fungal Diversity 105:1-16 ((https://doi.org/10.1007/s13225-020-00466-2)) |
| Otu08083 | 3 | k__Fungi(100);p__Ascomycota(100);c__Saccharomycetes(100);o__Saccharomycetales(100);f__Metschnikowiaceae(100);g__Metschnikowia(100);s__Metschnikowia_pulcherrima(67); | Metschnikowia | 13 | Saprotroph | \|Undefined Saprotroph\| | Yeast | NULL | Probable | As nectar-tap_saprotroph (Põlme et al. 2020); Decay sugar-rich substrates (Põlme et al. 2020) | Põlme S, et al. 2020. Fungal Diversity 105:1-16 ((https://doi.org/10.1007/s13225-020-00466-2)) |
| Otu08088 | 3 | k__Fungi(100);p__Ascomycota(100);c__Saccharomycetes(100);o__Saccharomycetales(100);f__Metschnikowiaceae(100);g__Metschnikowia(100);s__Metschnikowia_pulcherrima(100); | Metschnikowia | 13 | Saprotroph | \|Undefined Saprotroph\| | Yeast | NULL | Probable | As nectar-tap_saprotroph (Põlme et al. 2020); Decay sugar-rich substrates (Põlme et al. 2020) | Põlme S, et al. 2020. Fungal Diversity 105:1-16 ((https://doi.org/10.1007/s13225-020-00466-2)) |
| Otu08090 | 3 | k__Fungi(100);p__Ascomycota(100);c__Saccharomycetes(100);o__Saccharomycetales(100);f__Saccharomycetales_family_Incertae_sedis(100);g__Candida(67);g__Candida_unclassified(67); | Candida | 13 | Pathotroph-Saprotroph-Symbiotroph | Animal Pathogen-Endophyte-Endosymbiont-Epiphyte-\|Undefined Saprotroph\| | Dimorphic Yeast-Yeast | NULL | Probable | As nectar-tap_saprotroph (Põlme et al. 2020); Decay sugar-rich substrates (Põlme et al. 2020); This asexual genus is polyphyletic and contain species that function very differently depend on what they are. Caution should be taken when interpreting guilds for Candida species. | Manolakaki D, et al. 2010. Virulence. 1: 367-375 ((https://doi.org/10.4161/viru.1.5.12796)); Mekbib SB, et al. 2011. Tropical Plant Pathology 36:233-240 ((https://doi.org/10.1590/S1982-56762011000400004)); Põlme S, et al. 2020. Fungal Diversity 105:1-16 ((https://doi.org/10.1007/s13225-020-00466-2)) |
| Otu08092 | 3 | k__Fungi(100);p__Ascomycota(100);c__Saccharomycetes(100);o__Saccharomycetales(100);f__Metschnikowiaceae(100);g__Metschnikowia(100);s__Metschnikowia_pulcherrima(100); | Metschnikowia | 13 | Saprotroph | \|Undefined Saprotroph\| | Yeast | NULL | Probable | As nectar-tap_saprotroph (Põlme et al. 2020); Decay sugar-rich substrates (Põlme et al. 2020) | Põlme S, et al. 2020. Fungal Diversity 105:1-16 ((https://doi.org/10.1007/s13225-020-00466-2)) |
| Otu08094 | 3 | k__Fungi(100);p__Ascomycota(100);c__Saccharomycetes(100);o__Saccharomycetales(100);f__Metschnikowiaceae(100);g__Metschnikowia(100);s__Metschnikowia_pulcherrima(100); | Metschnikowia | 13 | Saprotroph | \|Undefined Saprotroph\| | Yeast | NULL | Probable | As nectar-tap_saprotroph (Põlme et al. 2020); Decay sugar-rich substrates (Põlme et al. 2020) | Põlme S, et al. 2020. Fungal Diversity 105:1-16 ((https://doi.org/10.1007/s13225-020-00466-2)) |
| Otu08095 | 3 | k__Fungi(100);p__Ascomycota(100);c__Saccharomycetes(100);o__Saccharomycetales(100);f__Metschnikowiaceae(100);g__Metschnikowia(100);s__Metschnikowia_pulcherrima(67); | Metschnikowia | 13 | Saprotroph | \|Undefined Saprotroph\| | Yeast | NULL | Probable | As nectar-tap_saprotroph (Põlme et al. 2020); Decay sugar-rich substrates (Põlme et al. 2020) | Põlme S, et al. 2020. Fungal Diversity 105:1-16 ((https://doi.org/10.1007/s13225-020-00466-2)) |
| Otu08096 | 3 | k__Fungi(100);p__Ascomycota(100);c__Saccharomycetes(100);o__Saccharomycetales(100);f__Metschnikowiaceae(100);g__Metschnikowia(100);s__Metschnikowia_pulcherrima(100); | Metschnikowia | 13 | Saprotroph | \|Undefined Saprotroph\| | Yeast | NULL | Probable | As nectar-tap_saprotroph (Põlme et al. 2020); Decay sugar-rich substrates (Põlme et al. 2020) | Põlme S, et al. 2020. Fungal Diversity 105:1-16 ((https://doi.org/10.1007/s13225-020-00466-2)) |
| Otu08098 | 3 | k__Fungi(100);p__Ascomycota(100);c__Saccharomycetes(100);o__Saccharomycetales(100);f__Metschnikowiaceae(100);g__Metschnikowia(100);s__Metschnikowia_pulcherrima(100); | Metschnikowia | 13 | Saprotroph | \|Undefined Saprotroph\| | Yeast | NULL | Probable | As nectar-tap_saprotroph (Põlme et al. 2020); Decay sugar-rich substrates (Põlme et al. 2020) | Põlme S, et al. 2020. Fungal Diversity 105:1-16 ((https://doi.org/10.1007/s13225-020-00466-2)) |
| Otu08099 | 3 | k__Fungi(100);p__Ascomycota(100);c__Saccharomycetes(100);o__Saccharomycetales(100);f__Saccharomycetales_family_Incertae_sedis(100);g__Candida(100);g__Candida_unclassified(67); | Candida | 13 | Pathotroph-Saprotroph-Symbiotroph | Animal Pathogen-Endophyte-Endosymbiont-Epiphyte-\|Undefined Saprotroph\| | Dimorphic Yeast-Yeast | NULL | Probable | As nectar-tap_saprotroph (Põlme et al. 2020); Decay sugar-rich substrates (Põlme et al. 2020); This asexual genus is polyphyletic and contain species that function very differently depend on what they are. Caution should be taken when interpreting guilds for Candida species. | Manolakaki D, et al. 2010. Virulence. 1: 367-375 ((https://doi.org/10.4161/viru.1.5.12796)); Mekbib SB, et al. 2011. Tropical Plant Pathology 36:233-240 ((https://doi.org/10.1590/S1982-56762011000400004)); Põlme S, et al. 2020. Fungal Diversity 105:1-16 ((https://doi.org/10.1007/s13225-020-00466-2)) |
| Otu08101 | 3 | k__Fungi(100);p__Ascomycota(100);c__Saccharomycetes(100);o__Saccharomycetales(100);f__Metschnikowiaceae(100);g__Metschnikowia(100);s__Metschnikowia_pulcherrima(100); | Metschnikowia | 13 | Saprotroph | \|Undefined Saprotroph\| | Yeast | NULL | Probable | As nectar-tap_saprotroph (Põlme et al. 2020); Decay sugar-rich substrates (Põlme et al. 2020) | Põlme S, et al. 2020. Fungal Diversity 105:1-16 ((https://doi.org/10.1007/s13225-020-00466-2)) |
| Otu08103 | 3 | k__Fungi(100);p__Ascomycota(100);c__Saccharomycetes(100);o__Saccharomycetales(100);f__Metschnikowiaceae(100);g__Metschnikowia(100);s__Metschnikowia_pulcherrima(100); | Metschnikowia | 13 | Saprotroph | \|Undefined Saprotroph\| | Yeast | NULL | Probable | As nectar-tap_saprotroph (Põlme et al. 2020); Decay sugar-rich substrates (Põlme et al. 2020) | Põlme S, et al. 2020. Fungal Diversity 105:1-16 ((https://doi.org/10.1007/s13225-020-00466-2)) |
| Otu08104 | 3 | k__Fungi(100);p__Ascomycota(100);c__Saccharomycetes(100);o__Saccharomycetales(100);f__Metschnikowiaceae(100);g__Metschnikowia(100);g__Metschnikowia_unclassified(100); | Metschnikowia | 13 | Saprotroph | \|Undefined Saprotroph\| | Yeast | NULL | Probable | As nectar-tap_saprotroph (Põlme et al. 2020); Decay sugar-rich substrates (Põlme et al. 2020) | Põlme S, et al. 2020. Fungal Diversity 105:1-16 ((https://doi.org/10.1007/s13225-020-00466-2)) |
| Otu08107 | 3 | k__Fungi(100);p__Ascomycota(100);c__Saccharomycetes(100);o__Saccharomycetales(100);f__Metschnikowiaceae(100);g__Metschnikowia(100);g__Metschnikowia_unclassified(67); | Metschnikowia | 13 | Saprotroph | \|Undefined Saprotroph\| | Yeast | NULL | Probable | As nectar-tap_saprotroph (Põlme et al. 2020); Decay sugar-rich substrates (Põlme et al. 2020) | Põlme S, et al. 2020. Fungal Diversity 105:1-16 ((https://doi.org/10.1007/s13225-020-00466-2)) |
| Otu08108 | 3 | k__Fungi(100);p__Ascomycota(100);c__Saccharomycetes(100);o__Saccharomycetales(100);f__Metschnikowiaceae(100);g__Metschnikowia(100);s__Metschnikowia_pulcherrima(100); | Metschnikowia | 13 | Saprotroph | \|Undefined Saprotroph\| | Yeast | NULL | Probable | As nectar-tap_saprotroph (Põlme et al. 2020); Decay sugar-rich substrates (Põlme et al. 2020) | Põlme S, et al. 2020. Fungal Diversity 105:1-16 ((https://doi.org/10.1007/s13225-020-00466-2)) |
| Otu08109 | 3 | k__Fungi(100);p__Ascomycota(100);c__Saccharomycetes(100);o__Saccharomycetales(100);f__Metschnikowiaceae(100);g__Metschnikowia(100);g__Metschnikowia_unclassified(67); | Metschnikowia | 13 | Saprotroph | \|Undefined Saprotroph\| | Yeast | NULL | Probable | As nectar-tap_saprotroph (Põlme et al. 2020); Decay sugar-rich substrates (Põlme et al. 2020) | Põlme S, et al. 2020. Fungal Diversity 105:1-16 ((https://doi.org/10.1007/s13225-020-00466-2)) |
| Otu08110 | 3 | k__Fungi(100);p__Basidiomycota(100);c__Exobasidiomycetes(100);o__Exobasidiales(100);f__Exobasidiaceae(100);g__Exobasidium(100);g__Exobasidium_unclassified(100); | Exobasidium | 13 | Pathotroph | \|Plant Pathogen\| | Leaf Blister-Leaf Gall-Leaf Spot | NULL | Probable | NULL | Tedersoo L, et al. 2014. Science 346:e1256688 ((https://doi.org/10.1126/science.1256688)); Põlme S, et al. 2020. Fungal Diversity 105:1-16 ((https://doi.org/10.1007/s13225-020-00466-2)) |
| Otu08111 | 3 | k__Fungi(100);p__Ascomycota(100);c__Saccharomycetes(100);o__Saccharomycetales(100);f__Metschnikowiaceae(100);g__Metschnikowia(100);s__Metschnikowia_pulcherrima(67); | Metschnikowia | 13 | Saprotroph | \|Undefined Saprotroph\| | Yeast | NULL | Probable | As nectar-tap_saprotroph (Põlme et al. 2020); Decay sugar-rich substrates (Põlme et al. 2020) | Põlme S, et al. 2020. Fungal Diversity 105:1-16 ((https://doi.org/10.1007/s13225-020-00466-2)) |
| Otu08113 | 3 | k__Fungi(100);p__Ascomycota(100);c__Saccharomycetes(100);o__Saccharomycetales(100);f__Metschnikowiaceae(100);g__Metschnikowia(100);g__Metschnikowia_unclassified(67); | Metschnikowia | 13 | Saprotroph | \|Undefined Saprotroph\| | Yeast | NULL | Probable | As nectar-tap_saprotroph (Põlme et al. 2020); Decay sugar-rich substrates (Põlme et al. 2020) | Põlme S, et al. 2020. Fungal Diversity 105:1-16 ((https://doi.org/10.1007/s13225-020-00466-2)) |
| Otu08114 | 3 | k__Fungi(100);p__Ascomycota(100);c__Saccharomycetes(100);o__Saccharomycetales(100);f__Metschnikowiaceae(100);g__Metschnikowia(100);s__Metschnikowia_pulcherrima(100); | Metschnikowia | 13 | Saprotroph | \|Undefined Saprotroph\| | Yeast | NULL | Probable | As nectar-tap_saprotroph (Põlme et al. 2020); Decay sugar-rich substrates (Põlme et al. 2020) | Põlme S, et al. 2020. Fungal Diversity 105:1-16 ((https://doi.org/10.1007/s13225-020-00466-2)) |
| Otu08116 | 3 | k__Fungi(100);p__Ascomycota(100);c__Saccharomycetes(100);o__Saccharomycetales(100);f__Metschnikowiaceae(100);g__Metschnikowia(100);s__Metschnikowia_pulcherrima(100); | Metschnikowia | 13 | Saprotroph | \|Undefined Saprotroph\| | Yeast | NULL | Probable | As nectar-tap_saprotroph (Põlme et al. 2020); Decay sugar-rich substrates (Põlme et al. 2020) | Põlme S, et al. 2020. Fungal Diversity 105:1-16 ((https://doi.org/10.1007/s13225-020-00466-2)) |
| Otu08118 | 3 | k__Fungi(100);p__Ascomycota(100);c__Saccharomycetes(100);o__Saccharomycetales(100);f__Metschnikowiaceae(100);g__Metschnikowia(100);s__Metschnikowia_pulcherrima(67); | Metschnikowia | 13 | Saprotroph | \|Undefined Saprotroph\| | Yeast | NULL | Probable | As nectar-tap_saprotroph (Põlme et al. 2020); Decay sugar-rich substrates (Põlme et al. 2020) | Põlme S, et al. 2020. Fungal Diversity 105:1-16 ((https://doi.org/10.1007/s13225-020-00466-2)) |
| Otu08121 | 3 | k__Fungi(100);p__Ascomycota(100);c__Saccharomycetes(100);o__Saccharomycetales(100);f__Metschnikowiaceae(100);g__Metschnikowia(100);s__Metschnikowia_pulcherrima(67); | Metschnikowia | 13 | Saprotroph | \|Undefined Saprotroph\| | Yeast | NULL | Probable | As nectar-tap_saprotroph (Põlme et al. 2020); Decay sugar-rich substrates (Põlme et al. 2020) | Põlme S, et al. 2020. Fungal Diversity 105:1-16 ((https://doi.org/10.1007/s13225-020-00466-2)) |
| Otu08122 | 3 | k__Fungi(100);p__Ascomycota(100);c__Saccharomycetes(100);o__Saccharomycetales(100);f__Metschnikowiaceae(100);g__Metschnikowia(100);s__Metschnikowia_pulcherrima(67); | Metschnikowia | 13 | Saprotroph | \|Undefined Saprotroph\| | Yeast | NULL | Probable | As nectar-tap_saprotroph (Põlme et al. 2020); Decay sugar-rich substrates (Põlme et al. 2020) | Põlme S, et al. 2020. Fungal Diversity 105:1-16 ((https://doi.org/10.1007/s13225-020-00466-2)) |
| Otu08127 | 3 | k__Fungi(100);p__Ascomycota(100);c__Saccharomycetes(100);o__Saccharomycetales(100);f__Metschnikowiaceae(100);g__Metschnikowia(100);s__Metschnikowia_pulcherrima(100); | Metschnikowia | 13 | Saprotroph | \|Undefined Saprotroph\| | Yeast | NULL | Probable | As nectar-tap_saprotroph (Põlme et al. 2020); Decay sugar-rich substrates (Põlme et al. 2020) | Põlme S, et al. 2020. Fungal Diversity 105:1-16 ((https://doi.org/10.1007/s13225-020-00466-2)) |
| Otu08128 | 3 | k__Fungi(100);p__Ascomycota(100);c__Saccharomycetes(100);o__Saccharomycetales(100);f__Metschnikowiaceae(100);g__Metschnikowia(100);s__Metschnikowia_pulcherrima(67); | Metschnikowia | 13 | Saprotroph | \|Undefined Saprotroph\| | Yeast | NULL | Probable | As nectar-tap_saprotroph (Põlme et al. 2020); Decay sugar-rich substrates (Põlme et al. 2020) | Põlme S, et al. 2020. Fungal Diversity 105:1-16 ((https://doi.org/10.1007/s13225-020-00466-2)) |
| Otu08129 | 3 | k__Fungi(100);p__Ascomycota(100);c__Saccharomycetes(100);o__Saccharomycetales(100);f__Metschnikowiaceae(100);g__Metschnikowia(100);g__Metschnikowia_unclassified(100); | Metschnikowia | 13 | Saprotroph | \|Undefined Saprotroph\| | Yeast | NULL | Probable | As nectar-tap_saprotroph (Põlme et al. 2020); Decay sugar-rich substrates (Põlme et al. 2020) | Põlme S, et al. 2020. Fungal Diversity 105:1-16 ((https://doi.org/10.1007/s13225-020-00466-2)) |
| Otu08133 | 3 | k__Fungi(100);p__Ascomycota(100);c__Saccharomycetes(100);o__Saccharomycetales(100);f__Metschnikowiaceae(100);g__Metschnikowia(100);s__Metschnikowia_pulcherrima(100); | Metschnikowia | 13 | Saprotroph | \|Undefined Saprotroph\| | Yeast | NULL | Probable | As nectar-tap_saprotroph (Põlme et al. 2020); Decay sugar-rich substrates (Põlme et al. 2020) | Põlme S, et al. 2020. Fungal Diversity 105:1-16 ((https://doi.org/10.1007/s13225-020-00466-2)) |
| Otu08134 | 3 | k__Fungi(100);p__Ascomycota(100);c__Saccharomycetes(100);o__Saccharomycetales(100);f__Metschnikowiaceae(100);g__Metschnikowia(100);s__Metschnikowia_pulcherrima(100); | Metschnikowia | 13 | Saprotroph | \|Undefined Saprotroph\| | Yeast | NULL | Probable | As nectar-tap_saprotroph (Põlme et al. 2020); Decay sugar-rich substrates (Põlme et al. 2020) | Põlme S, et al. 2020. Fungal Diversity 105:1-16 ((https://doi.org/10.1007/s13225-020-00466-2)) |
| Otu08137 | 3 | k__Fungi(100);p__Ascomycota(100);c__Saccharomycetes(100);o__Saccharomycetales(100);f__Metschnikowiaceae(100);g__Metschnikowia(100);g__Metschnikowia_unclassified(67); | Metschnikowia | 13 | Saprotroph | \|Undefined Saprotroph\| | Yeast | NULL | Probable | As nectar-tap_saprotroph (Põlme et al. 2020); Decay sugar-rich substrates (Põlme et al. 2020) | Põlme S, et al. 2020. Fungal Diversity 105:1-16 ((https://doi.org/10.1007/s13225-020-00466-2)) |
| Otu08139 | 3 | k__Fungi(100);p__Ascomycota(100);c__Dothideomycetes(100);o__Capnodiales(100);f__Mycosphaerellaceae(100);g__Mycosphaerella(67);s__Mycosphaerella_nyssicola(67); | Mycosphaerella | 13 | Pathotroph-Saprotroph | \|Plant Pathogen\|-Plant Saprotroph | Microfungus | NULL | Probable | As litter_saprotroph (Põlme et al. 2020); Decay leaf, fruit, seed (Põlme et al. 2020) | Tedersoo L, et al. 2014. Science 346:e1256688 ((https://doi.org/10.1126/science.1256688)); Põlme S, et al. 2020. Fungal Diversity 105:1-16 ((https://doi.org/10.1007/s13225-020-00466-2)) |
| Otu08140 | 3 | k__Fungi(100);p__Ascomycota(100);c__Saccharomycetes(100);o__Saccharomycetales(100);f__Metschnikowiaceae(100);g__Metschnikowia(100);s__Metschnikowia_pulcherrima(100); | Metschnikowia | 13 | Saprotroph | \|Undefined Saprotroph\| | Yeast | NULL | Probable | As nectar-tap_saprotroph (Põlme et al. 2020); Decay sugar-rich substrates (Põlme et al. 2020) | Põlme S, et al. 2020. Fungal Diversity 105:1-16 ((https://doi.org/10.1007/s13225-020-00466-2)) |
| Otu08144 | 3 | k__Fungi(100);p__Ascomycota(100);c__Saccharomycetes(100);o__Saccharomycetales(100);f__Metschnikowiaceae(100);g__Metschnikowia(100);s__Metschnikowia_pulcherrima(67); | Metschnikowia | 13 | Saprotroph | \|Undefined Saprotroph\| | Yeast | NULL | Probable | As nectar-tap_saprotroph (Põlme et al. 2020); Decay sugar-rich substrates (Põlme et al. 2020) | Põlme S, et al. 2020. Fungal Diversity 105:1-16 ((https://doi.org/10.1007/s13225-020-00466-2)) |
| Otu08147 | 3 | k__Fungi(100);p__Ascomycota(100);c__Saccharomycetes(100);o__Saccharomycetales(100);f__Metschnikowiaceae(100);g__Metschnikowia(100);s__Metschnikowia_pulcherrima(100); | Metschnikowia | 13 | Saprotroph | \|Undefined Saprotroph\| | Yeast | NULL | Probable | As nectar-tap_saprotroph (Põlme et al. 2020); Decay sugar-rich substrates (Põlme et al. 2020) | Põlme S, et al. 2020. Fungal Diversity 105:1-16 ((https://doi.org/10.1007/s13225-020-00466-2)) |
| Otu08148 | 3 | k__Fungi(100);p__Ascomycota(100);c__Saccharomycetes(100);o__Saccharomycetales(100);f__Metschnikowiaceae(100);g__Metschnikowia(100);s__Metschnikowia_pulcherrima(100); | Metschnikowia | 13 | Saprotroph | \|Undefined Saprotroph\| | Yeast | NULL | Probable | As nectar-tap_saprotroph (Põlme et al. 2020); Decay sugar-rich substrates (Põlme et al. 2020) | Põlme S, et al. 2020. Fungal Diversity 105:1-16 ((https://doi.org/10.1007/s13225-020-00466-2)) |
| Otu08150 | 3 | k__Fungi(100);p__Ascomycota(100);c__Saccharomycetes(100);o__Saccharomycetales(100);f__Metschnikowiaceae(100);g__Metschnikowia(100);s__Metschnikowia_pulcherrima(67); | Metschnikowia | 13 | Saprotroph | \|Undefined Saprotroph\| | Yeast | NULL | Probable | As nectar-tap_saprotroph (Põlme et al. 2020); Decay sugar-rich substrates (Põlme et al. 2020) | Põlme S, et al. 2020. Fungal Diversity 105:1-16 ((https://doi.org/10.1007/s13225-020-00466-2)) |
| Otu08152 | 3 | k__Fungi(100);p__Ascomycota(100);c__Saccharomycetes(100);o__Saccharomycetales(100);f__Metschnikowiaceae(100);g__Metschnikowia(100);s__Metschnikowia_pulcherrima(67); | Metschnikowia | 13 | Saprotroph | \|Undefined Saprotroph\| | Yeast | NULL | Probable | As nectar-tap_saprotroph (Põlme et al. 2020); Decay sugar-rich substrates (Põlme et al. 2020) | Põlme S, et al. 2020. Fungal Diversity 105:1-16 ((https://doi.org/10.1007/s13225-020-00466-2)) |
| Otu08153 | 3 | k__Fungi(100);p__Ascomycota(100);c__Saccharomycetes(100);o__Saccharomycetales(100);f__Metschnikowiaceae(100);g__Metschnikowia(100);s__Metschnikowia_pulcherrima(67); | Metschnikowia | 13 | Saprotroph | \|Undefined Saprotroph\| | Yeast | NULL | Probable | As nectar-tap_saprotroph (Põlme et al. 2020); Decay sugar-rich substrates (Põlme et al. 2020) | Põlme S, et al. 2020. Fungal Diversity 105:1-16 ((https://doi.org/10.1007/s13225-020-00466-2)) |
| Otu08156 | 3 | k__Fungi(100);p__Ascomycota(100);c__Saccharomycetes(100);o__Saccharomycetales(100);f__Metschnikowiaceae(100);g__Metschnikowia(100);s__Metschnikowia_pulcherrima(100); | Metschnikowia | 13 | Saprotroph | \|Undefined Saprotroph\| | Yeast | NULL | Probable | As nectar-tap_saprotroph (Põlme et al. 2020); Decay sugar-rich substrates (Põlme et al. 2020) | Põlme S, et al. 2020. Fungal Diversity 105:1-16 ((https://doi.org/10.1007/s13225-020-00466-2)) |
| Otu08158 | 3 | k__Fungi(100);p__Ascomycota(100);c__Saccharomycetes(100);o__Saccharomycetales(100);f__Metschnikowiaceae(100);g__Metschnikowia(100);s__Metschnikowia_pulcherrima(100); | Metschnikowia | 13 | Saprotroph | \|Undefined Saprotroph\| | Yeast | NULL | Probable | As nectar-tap_saprotroph (Põlme et al. 2020); Decay sugar-rich substrates (Põlme et al. 2020) | Põlme S, et al. 2020. Fungal Diversity 105:1-16 ((https://doi.org/10.1007/s13225-020-00466-2)) |
| Otu08160 | 3 | k__Fungi(100);p__Ascomycota(100);c__Dothideomycetes(100);o__Pleosporales(100);f__Montagnulaceae(100);f__Montagnulaceae_unclassified(100);f__Montagnulaceae_unclassified(100); | Montagnulaceae | 9 | Pathotroph-Saprotroph | Fungal Parasite-Plant Pathogen-Undefined Saprotroph | Microfungus | NULL | Probable | Species in this clade can be saprobic, plant pathogenic, or mycoparasitic (Zhang et al. 2009) | Zhang Y, et al. 2009. Studies in Mycology 64:85-102 ((https://doi.org/10.3114/sim.2009.64.04)) |
| Otu08161 | 3 | k__Fungi(100);p__Ascomycota(100);c__Saccharomycetes(100);o__Saccharomycetales(100);f__Metschnikowiaceae(100);g__Metschnikowia(100);s__Metschnikowia_pulcherrima(67); | Metschnikowia | 13 | Saprotroph | \|Undefined Saprotroph\| | Yeast | NULL | Probable | As nectar-tap_saprotroph (Põlme et al. 2020); Decay sugar-rich substrates (Põlme et al. 2020) | Põlme S, et al. 2020. Fungal Diversity 105:1-16 ((https://doi.org/10.1007/s13225-020-00466-2)) |
| Otu08162 | 3 | k__Fungi(100);p__Ascomycota(100);c__Dothideomycetes(100);o__Dothideales(100);f__Dothideales_family_Incertae_sedis(100);g__Phaeocryptopus(100);s__Phaeocryptopus_gaeumannii(100); | Phaeocryptopus | 13 | Pathotroph-Saprotroph | \|Plant Pathogen\|-Plant Saprotroph | Microfungus | NULL | Probable | As litter_saprotroph (Põlme et al. 2020); Decay leaf, fruit, seed (Põlme et al. 2020) | Põlme S, et al. 2020. Fungal Diversity 105:1-16 ((https://doi.org/10.1007/s13225-020-00466-2)); The American Phytopathological Society Common Names of Plant Diseases ((http://www.apsnet.org/publications/commonnames/Pages/default.aspx)) |
| Otu08163 | 3 | k__Fungi(100);p__Ascomycota(100);c__Saccharomycetes(100);o__Saccharomycetales(100);f__Metschnikowiaceae(100);g__Metschnikowia(100);s__Metschnikowia_pulcherrima(100); | Metschnikowia | 13 | Saprotroph | \|Undefined Saprotroph\| | Yeast | NULL | Probable | As nectar-tap_saprotroph (Põlme et al. 2020); Decay sugar-rich substrates (Põlme et al. 2020) | Põlme S, et al. 2020. Fungal Diversity 105:1-16 ((https://doi.org/10.1007/s13225-020-00466-2)) |
| Otu08167 | 3 | k__Fungi(100);p__Basidiomycota(100);c__Exobasidiomycetes(100);o__Exobasidiales(100);f__Exobasidiaceae(100);g__Exobasidium(100);g__Exobasidium_unclassified(100); | Exobasidium | 13 | Pathotroph | \|Plant Pathogen\| | Leaf Blister-Leaf Gall-Leaf Spot | NULL | Probable | NULL | Tedersoo L, et al. 2014. Science 346:e1256688 ((https://doi.org/10.1126/science.1256688)); Põlme S, et al. 2020. Fungal Diversity 105:1-16 ((https://doi.org/10.1007/s13225-020-00466-2)) |
| Otu08170 | 3 | k__Fungi(100);p__Ascomycota(100);c__Saccharomycetes(100);o__Saccharomycetales(100);f__Metschnikowiaceae(100);g__Metschnikowia(100);g__Metschnikowia_unclassified(100); | Metschnikowia | 13 | Saprotroph | \|Undefined Saprotroph\| | Yeast | NULL | Probable | As nectar-tap_saprotroph (Põlme et al. 2020); Decay sugar-rich substrates (Põlme et al. 2020) | Põlme S, et al. 2020. Fungal Diversity 105:1-16 ((https://doi.org/10.1007/s13225-020-00466-2)) |
| Otu08172 | 3 | k__Fungi(100);p__Ascomycota(100);c__Saccharomycetes(100);o__Saccharomycetales(100);f__Metschnikowiaceae(100);g__Metschnikowia(100);s__Metschnikowia_pulcherrima(67); | Metschnikowia | 13 | Saprotroph | \|Undefined Saprotroph\| | Yeast | NULL | Probable | As nectar-tap_saprotroph (Põlme et al. 2020); Decay sugar-rich substrates (Põlme et al. 2020) | Põlme S, et al. 2020. Fungal Diversity 105:1-16 ((https://doi.org/10.1007/s13225-020-00466-2)) |
| Otu08174 | 3 | k__Fungi(100);p__Ascomycota(100);c__Saccharomycetes(100);o__Saccharomycetales(100);f__Metschnikowiaceae(100);g__Metschnikowia(100);s__Metschnikowia_pulcherrima(67); | Metschnikowia | 13 | Saprotroph | \|Undefined Saprotroph\| | Yeast | NULL | Probable | As nectar-tap_saprotroph (Põlme et al. 2020); Decay sugar-rich substrates (Põlme et al. 2020) | Põlme S, et al. 2020. Fungal Diversity 105:1-16 ((https://doi.org/10.1007/s13225-020-00466-2)) |
| Otu08178 | 3 | k__Fungi(100);p__Ascomycota(100);c__Dothideomycetes(100);o__Pleosporales(100);f__Cucurbitariaceae(100);g__Curreya(100);s__Curreya_austroafricana(100); | Curreya | 13 | Saprotroph | \|Plant Saprotroph\|-Undefined Saprotroph | Microfungus | NULL | Probable | As litter_saprotroph (Põlme et al. 2020); Decay leaf, fruit, seed (Põlme et al. 2020) | Tedersoo L, et al. 2014. Science 346:e1256688 ((https://doi.org/10.1126/science.1256688)); Põlme S, et al. 2020. Fungal Diversity 105:1-16 ((https://doi.org/10.1007/s13225-020-00466-2)) |
| Otu08179 | 3 | k__Fungi(100);p__Ascomycota(100);c__Saccharomycetes(100);o__Saccharomycetales(100);f__Metschnikowiaceae(100);g__Metschnikowia(100);s__Metschnikowia_pulcherrima(100); | Metschnikowia | 13 | Saprotroph | \|Undefined Saprotroph\| | Yeast | NULL | Probable | As nectar-tap_saprotroph (Põlme et al. 2020); Decay sugar-rich substrates (Põlme et al. 2020) | Põlme S, et al. 2020. Fungal Diversity 105:1-16 ((https://doi.org/10.1007/s13225-020-00466-2)) |
| Otu08185 | 3 | k__Fungi(100);p__Ascomycota(100);c__Saccharomycetes(100);o__Saccharomycetales(100);f__Metschnikowiaceae(100);g__Metschnikowia(100);s__Metschnikowia_pulcherrima(100); | Metschnikowia | 13 | Saprotroph | \|Undefined Saprotroph\| | Yeast | NULL | Probable | As nectar-tap_saprotroph (Põlme et al. 2020); Decay sugar-rich substrates (Põlme et al. 2020) | Põlme S, et al. 2020. Fungal Diversity 105:1-16 ((https://doi.org/10.1007/s13225-020-00466-2)) |
| Otu08186 | 3 | k__Fungi(100);p__Ascomycota(100);c__Saccharomycetes(100);o__Saccharomycetales(100);f__Metschnikowiaceae(100);g__Metschnikowia(100);g__Metschnikowia_unclassified(100); | Metschnikowia | 13 | Saprotroph | \|Undefined Saprotroph\| | Yeast | NULL | Probable | As nectar-tap_saprotroph (Põlme et al. 2020); Decay sugar-rich substrates (Põlme et al. 2020) | Põlme S, et al. 2020. Fungal Diversity 105:1-16 ((https://doi.org/10.1007/s13225-020-00466-2)) |
| Otu08188 | 3 | k__Fungi(100);p__Ascomycota(100);c__Saccharomycetes(100);o__Saccharomycetales(100);f__Metschnikowiaceae(100);g__Metschnikowia(100);s__Metschnikowia_pulcherrima(100); | Metschnikowia | 13 | Saprotroph | \|Undefined Saprotroph\| | Yeast | NULL | Probable | As nectar-tap_saprotroph (Põlme et al. 2020); Decay sugar-rich substrates (Põlme et al. 2020) | Põlme S, et al. 2020. Fungal Diversity 105:1-16 ((https://doi.org/10.1007/s13225-020-00466-2)) |
| Otu08189 | 3 | k__Fungi(100);p__Ascomycota(100);c__Saccharomycetes(100);o__Saccharomycetales(100);f__Metschnikowiaceae(100);g__Metschnikowia(100);s__Metschnikowia_pulcherrima(67); | Metschnikowia | 13 | Saprotroph | \|Undefined Saprotroph\| | Yeast | NULL | Probable | As nectar-tap_saprotroph (Põlme et al. 2020); Decay sugar-rich substrates (Põlme et al. 2020) | Põlme S, et al. 2020. Fungal Diversity 105:1-16 ((https://doi.org/10.1007/s13225-020-00466-2)) |
| Otu08190 | 3 | k__Fungi(100);p__Ascomycota(100);c__Saccharomycetes(100);o__Saccharomycetales(100);f__Metschnikowiaceae(100);g__Metschnikowia(100);s__Metschnikowia_pulcherrima(100); | Metschnikowia | 13 | Saprotroph | \|Undefined Saprotroph\| | Yeast | NULL | Probable | As nectar-tap_saprotroph (Põlme et al. 2020); Decay sugar-rich substrates (Põlme et al. 2020) | Põlme S, et al. 2020. Fungal Diversity 105:1-16 ((https://doi.org/10.1007/s13225-020-00466-2)) |
| Otu08191 | 3 | k__Fungi(100);p__Ascomycota(100);c__Saccharomycetes(100);o__Saccharomycetales(100);f__Metschnikowiaceae(100);g__Metschnikowia(100);s__Metschnikowia_pulcherrima(100); | Metschnikowia | 13 | Saprotroph | \|Undefined Saprotroph\| | Yeast | NULL | Probable | As nectar-tap_saprotroph (Põlme et al. 2020); Decay sugar-rich substrates (Põlme et al. 2020) | Põlme S, et al. 2020. Fungal Diversity 105:1-16 ((https://doi.org/10.1007/s13225-020-00466-2)) |
| Otu08192 | 3 | k__Fungi(100);p__Ascomycota(100);c__Saccharomycetes(100);o__Saccharomycetales(100);f__Metschnikowiaceae(100);g__Metschnikowia(100);s__Metschnikowia_pulcherrima(100); | Metschnikowia | 13 | Saprotroph | \|Undefined Saprotroph\| | Yeast | NULL | Probable | As nectar-tap_saprotroph (Põlme et al. 2020); Decay sugar-rich substrates (Põlme et al. 2020) | Põlme S, et al. 2020. Fungal Diversity 105:1-16 ((https://doi.org/10.1007/s13225-020-00466-2)) |
| Otu08194 | 3 | k__Fungi(100);p__Ascomycota(100);c__Saccharomycetes(100);o__Saccharomycetales(100);f__Metschnikowiaceae(100);g__Metschnikowia(100);s__Metschnikowia_pulcherrima(67); | Metschnikowia | 13 | Saprotroph | \|Undefined Saprotroph\| | Yeast | NULL | Probable | As nectar-tap_saprotroph (Põlme et al. 2020); Decay sugar-rich substrates (Põlme et al. 2020) | Põlme S, et al. 2020. Fungal Diversity 105:1-16 ((https://doi.org/10.1007/s13225-020-00466-2)) |
| Otu08195 | 3 | k__Fungi(100);p__Ascomycota(100);c__Saccharomycetes(100);o__Saccharomycetales(100);f__Metschnikowiaceae(100);g__Metschnikowia(100);s__Metschnikowia_pulcherrima(67); | Metschnikowia | 13 | Saprotroph | \|Undefined Saprotroph\| | Yeast | NULL | Probable | As nectar-tap_saprotroph (Põlme et al. 2020); Decay sugar-rich substrates (Põlme et al. 2020) | Põlme S, et al. 2020. Fungal Diversity 105:1-16 ((https://doi.org/10.1007/s13225-020-00466-2)) |
| Otu08196 | 3 | k__Fungi(100);p__Ascomycota(100);c__Saccharomycetes(100);o__Saccharomycetales(100);f__Saccharomycetales_family_Incertae_sedis(100);g__Candida(100);s__Candida_cretensis(67); | Candida | 13 | Pathotroph-Saprotroph-Symbiotroph | Animal Pathogen-Endophyte-Endosymbiont-Epiphyte-\|Undefined Saprotroph\| | Dimorphic Yeast-Yeast | NULL | Probable | As nectar-tap_saprotroph (Põlme et al. 2020); Decay sugar-rich substrates (Põlme et al. 2020); This asexual genus is polyphyletic and contain species that function very differently depend on what they are. Caution should be taken when interpreting guilds for Candida species. | Manolakaki D, et al. 2010. Virulence. 1: 367-375 ((https://doi.org/10.4161/viru.1.5.12796)); Mekbib SB, et al. 2011. Tropical Plant Pathology 36:233-240 ((https://doi.org/10.1590/S1982-56762011000400004)); Põlme S, et al. 2020. Fungal Diversity 105:1-16 ((https://doi.org/10.1007/s13225-020-00466-2)) |
| Otu08198 | 3 | k__Fungi(100);p__Ascomycota(100);c__Saccharomycetes(100);o__Saccharomycetales(100);f__Metschnikowiaceae(100);g__Metschnikowia(100);s__Metschnikowia_pulcherrima(67); | Metschnikowia | 13 | Saprotroph | \|Undefined Saprotroph\| | Yeast | NULL | Probable | As nectar-tap_saprotroph (Põlme et al. 2020); Decay sugar-rich substrates (Põlme et al. 2020) | Põlme S, et al. 2020. Fungal Diversity 105:1-16 ((https://doi.org/10.1007/s13225-020-00466-2)) |
| Otu08199 | 3 | k__Fungi(100);p__Ascomycota(100);c__Saccharomycetes(100);o__Saccharomycetales(100);f__Metschnikowiaceae(100);g__Metschnikowia(100);s__Metschnikowia_pulcherrima(67); | Metschnikowia | 13 | Saprotroph | \|Undefined Saprotroph\| | Yeast | NULL | Probable | As nectar-tap_saprotroph (Põlme et al. 2020); Decay sugar-rich substrates (Põlme et al. 2020) | Põlme S, et al. 2020. Fungal Diversity 105:1-16 ((https://doi.org/10.1007/s13225-020-00466-2)) |
| Otu08207 | 3 | k__Fungi(100);p__Ascomycota(100);c__Saccharomycetes(100);o__Saccharomycetales(100);f__Metschnikowiaceae(100);g__Metschnikowia(100);s__Metschnikowia_pulcherrima(100); | Metschnikowia | 13 | Saprotroph | \|Undefined Saprotroph\| | Yeast | NULL | Probable | As nectar-tap_saprotroph (Põlme et al. 2020); Decay sugar-rich substrates (Põlme et al. 2020) | Põlme S, et al. 2020. Fungal Diversity 105:1-16 ((https://doi.org/10.1007/s13225-020-00466-2)) |
| Otu08208 | 3 | k__Fungi(100);p__Ascomycota(100);c__Saccharomycetes(100);o__Saccharomycetales(100);f__Metschnikowiaceae(100);g__Metschnikowia(100);s__Metschnikowia_pulcherrima(67); | Metschnikowia | 13 | Saprotroph | \|Undefined Saprotroph\| | Yeast | NULL | Probable | As nectar-tap_saprotroph (Põlme et al. 2020); Decay sugar-rich substrates (Põlme et al. 2020) | Põlme S, et al. 2020. Fungal Diversity 105:1-16 ((https://doi.org/10.1007/s13225-020-00466-2)) |
| Otu08209 | 3 | k__Fungi(100);p__Ascomycota(100);c__Saccharomycetes(100);o__Saccharomycetales(100);f__Metschnikowiaceae(100);g__Metschnikowia(100);s__Metschnikowia_pulcherrima(100); | Metschnikowia | 13 | Saprotroph | \|Undefined Saprotroph\| | Yeast | NULL | Probable | As nectar-tap_saprotroph (Põlme et al. 2020); Decay sugar-rich substrates (Põlme et al. 2020) | Põlme S, et al. 2020. Fungal Diversity 105:1-16 ((https://doi.org/10.1007/s13225-020-00466-2)) |
| Otu08212 | 3 | k__Fungi(100);p__Ascomycota(100);c__Saccharomycetes(100);o__Saccharomycetales(100);f__Metschnikowiaceae(100);g__Metschnikowia(100);s__Metschnikowia_pulcherrima(67); | Metschnikowia | 13 | Saprotroph | \|Undefined Saprotroph\| | Yeast | NULL | Probable | As nectar-tap_saprotroph (Põlme et al. 2020); Decay sugar-rich substrates (Põlme et al. 2020) | Põlme S, et al. 2020. Fungal Diversity 105:1-16 ((https://doi.org/10.1007/s13225-020-00466-2)) |
| Otu08213 | 3 | k__Fungi(100);p__Basidiomycota(100);c__Agaricomycetes(100);o__Polyporales(100);f__Polyporaceae(100);g__Trametes(100);g__Trametes_unclassified(100); | Trametes | 13 | Pathotroph-Saprotroph | Plant Pathogen-\|Wood Saprotroph\| | Polyporoid | White Rot | Highly Probable | As white_rot (Põlme et al. 2020); As wood_pathogen (Põlme et al. 2020); Decay wood (Põlme et al. 2020) | Gilbertson RL, Ryvarden L. 1987-1987. North American Polypores. Fungiflora, Oslo (ISBN: 978-0945345060); Põlme S, et al. 2020. Fungal Diversity 105:1-16 ((https://doi.org/10.1007/s13225-020-00466-2)) |
| Otu08214 | 3 | k__Fungi(100);p__Ascomycota(100);c__Saccharomycetes(100);o__Saccharomycetales(100);f__Metschnikowiaceae(100);g__Metschnikowia(100);g__Metschnikowia_unclassified(100); | Metschnikowia | 13 | Saprotroph | \|Undefined Saprotroph\| | Yeast | NULL | Probable | As nectar-tap_saprotroph (Põlme et al. 2020); Decay sugar-rich substrates (Põlme et al. 2020) | Põlme S, et al. 2020. Fungal Diversity 105:1-16 ((https://doi.org/10.1007/s13225-020-00466-2)) |
| Otu08215 | 3 | k__Fungi(100);p__Ascomycota(100);c__Saccharomycetes(100);o__Saccharomycetales(100);f__Saccharomycetales_family_Incertae_sedis(100);g__Candida(100);g__Candida_unclassified(100); | Candida | 13 | Pathotroph-Saprotroph-Symbiotroph | Animal Pathogen-Endophyte-Endosymbiont-Epiphyte-\|Undefined Saprotroph\| | Dimorphic Yeast-Yeast | NULL | Probable | As nectar-tap_saprotroph (Põlme et al. 2020); Decay sugar-rich substrates (Põlme et al. 2020); This asexual genus is polyphyletic and contain species that function very differently depend on what they are. Caution should be taken when interpreting guilds for Candida species. | Manolakaki D, et al. 2010. Virulence. 1: 367-375 ((https://doi.org/10.4161/viru.1.5.12796)); Mekbib SB, et al. 2011. Tropical Plant Pathology 36:233-240 ((https://doi.org/10.1590/S1982-56762011000400004)); Põlme S, et al. 2020. Fungal Diversity 105:1-16 ((https://doi.org/10.1007/s13225-020-00466-2)) |
| Otu08216 | 3 | k__Fungi(100);p__Ascomycota(100);c__Saccharomycetes(100);o__Saccharomycetales(100);f__Metschnikowiaceae(100);g__Metschnikowia(100);s__Metschnikowia_pulcherrima(67); | Metschnikowia | 13 | Saprotroph | \|Undefined Saprotroph\| | Yeast | NULL | Probable | As nectar-tap_saprotroph (Põlme et al. 2020); Decay sugar-rich substrates (Põlme et al. 2020) | Põlme S, et al. 2020. Fungal Diversity 105:1-16 ((https://doi.org/10.1007/s13225-020-00466-2)) |
| Otu08217 | 3 | k__Fungi(100);p__Ascomycota(100);c__Saccharomycetes(100);o__Saccharomycetales(100);f__Metschnikowiaceae(100);g__Metschnikowia(100);s__Metschnikowia_pulcherrima(67); | Metschnikowia | 13 | Saprotroph | \|Undefined Saprotroph\| | Yeast | NULL | Probable | As nectar-tap_saprotroph (Põlme et al. 2020); Decay sugar-rich substrates (Põlme et al. 2020) | Põlme S, et al. 2020. Fungal Diversity 105:1-16 ((https://doi.org/10.1007/s13225-020-00466-2)) |
| Otu08218 | 3 | k__Fungi(100);p__Ascomycota(100);c__Saccharomycetes(100);o__Saccharomycetales(100);f__Metschnikowiaceae(100);g__Metschnikowia(100);s__Metschnikowia_pulcherrima(67); | Metschnikowia | 13 | Saprotroph | \|Undefined Saprotroph\| | Yeast | NULL | Probable | As nectar-tap_saprotroph (Põlme et al. 2020); Decay sugar-rich substrates (Põlme et al. 2020) | Põlme S, et al. 2020. Fungal Diversity 105:1-16 ((https://doi.org/10.1007/s13225-020-00466-2)) |
| Otu08220 | 3 | k__Fungi(100);p__Ascomycota(100);c__Saccharomycetes(100);o__Saccharomycetales(100);f__Saccharomycetales_family_Incertae_sedis(100);g__Candida(100);s__Candida_cretensis(67); | Candida | 13 | Pathotroph-Saprotroph-Symbiotroph | Animal Pathogen-Endophyte-Endosymbiont-Epiphyte-\|Undefined Saprotroph\| | Dimorphic Yeast-Yeast | NULL | Probable | As nectar-tap_saprotroph (Põlme et al. 2020); Decay sugar-rich substrates (Põlme et al. 2020); This asexual genus is polyphyletic and contain species that function very differently depend on what they are. Caution should be taken when interpreting guilds for Candida species. | Manolakaki D, et al. 2010. Virulence. 1: 367-375 ((https://doi.org/10.4161/viru.1.5.12796)); Mekbib SB, et al. 2011. Tropical Plant Pathology 36:233-240 ((https://doi.org/10.1590/S1982-56762011000400004)); Põlme S, et al. 2020. Fungal Diversity 105:1-16 ((https://doi.org/10.1007/s13225-020-00466-2)) |
| Otu08221 | 3 | k__Fungi(100);p__Ascomycota(100);c__Saccharomycetes(100);o__Saccharomycetales(100);f__Metschnikowiaceae(100);g__Metschnikowia(100);g__Metschnikowia_unclassified(67); | Metschnikowia | 13 | Saprotroph | \|Undefined Saprotroph\| | Yeast | NULL | Probable | As nectar-tap_saprotroph (Põlme et al. 2020); Decay sugar-rich substrates (Põlme et al. 2020) | Põlme S, et al. 2020. Fungal Diversity 105:1-16 ((https://doi.org/10.1007/s13225-020-00466-2)) |
| Otu08222 | 3 | k__Fungi(100);p__Basidiomycota(100);c__Agaricomycetes(100);o__Agaricales(100);f__Cortinariaceae(100);g__Cortinarius(100);g__Cortinarius_unclassified(100); | Cortinarius | 13 | Symbiotroph | \|Ectomycorrhizal\| | Agaricoid | NULL | Highly Probable | NULL | Rinaldi AC, et al. 2008. Fungal Diversity 33:1-45 ((http://www.fungaldiversity.org/fdp/sfdp/33-1.pdf)); Tedersoo L, et al. 2010. Mycorrhiza 20:217-263 ((https://doi.org/10.1007/s00572-009-0274-x)); Põlme S, et al. 2020. Fungal Diversity 105:1-16 ((https://doi.org/10.1007/s13225-020-00466-2)); http://mycorrhizas.info/ecmf.html ((http://mycorrhizas.info/ecmf.html)) |
| Otu08223 | 3 | k__Fungi(100);p__Ascomycota(100);c__Saccharomycetes(100);o__Saccharomycetales(100);f__Metschnikowiaceae(100);g__Metschnikowia(100);s__Metschnikowia_pulcherrima(67); | Metschnikowia | 13 | Saprotroph | \|Undefined Saprotroph\| | Yeast | NULL | Probable | As nectar-tap_saprotroph (Põlme et al. 2020); Decay sugar-rich substrates (Põlme et al. 2020) | Põlme S, et al. 2020. Fungal Diversity 105:1-16 ((https://doi.org/10.1007/s13225-020-00466-2)) |
| Otu08226 | 3 | k__Fungi(100);p__Basidiomycota(100);c__Agaricomycetes(100);o__Agaricales(100);f__Cortinariaceae(100);g__Cortinarius(100);g__Cortinarius_unclassified(100); | Cortinarius | 13 | Symbiotroph | \|Ectomycorrhizal\| | Agaricoid | NULL | Highly Probable | NULL | Rinaldi AC, et al. 2008. Fungal Diversity 33:1-45 ((http://www.fungaldiversity.org/fdp/sfdp/33-1.pdf)); Tedersoo L, et al. 2010. Mycorrhiza 20:217-263 ((https://doi.org/10.1007/s00572-009-0274-x)); Põlme S, et al. 2020. Fungal Diversity 105:1-16 ((https://doi.org/10.1007/s13225-020-00466-2)); http://mycorrhizas.info/ecmf.html ((http://mycorrhizas.info/ecmf.html)) |
| Otu08227 | 3 | k__Fungi(100);p__Basidiomycota(100);c__Agaricomycetes(100);o__Agaricales(100);f__Cortinariaceae(100);g__Cortinarius(100);g__Cortinarius_unclassified(100); | Cortinarius | 13 | Symbiotroph | \|Ectomycorrhizal\| | Agaricoid | NULL | Highly Probable | NULL | Rinaldi AC, et al. 2008. Fungal Diversity 33:1-45 ((http://www.fungaldiversity.org/fdp/sfdp/33-1.pdf)); Tedersoo L, et al. 2010. Mycorrhiza 20:217-263 ((https://doi.org/10.1007/s00572-009-0274-x)); Põlme S, et al. 2020. Fungal Diversity 105:1-16 ((https://doi.org/10.1007/s13225-020-00466-2)); http://mycorrhizas.info/ecmf.html ((http://mycorrhizas.info/ecmf.html)) |
| Otu08229 | 3 | k__Fungi(100);p__Ascomycota(100);c__Saccharomycetes(100);o__Saccharomycetales(100);f__Metschnikowiaceae(100);g__Metschnikowia(100);s__Metschnikowia_pulcherrima(100); | Metschnikowia | 13 | Saprotroph | \|Undefined Saprotroph\| | Yeast | NULL | Probable | As nectar-tap_saprotroph (Põlme et al. 2020); Decay sugar-rich substrates (Põlme et al. 2020) | Põlme S, et al. 2020. Fungal Diversity 105:1-16 ((https://doi.org/10.1007/s13225-020-00466-2)) |
| Otu08232 | 3 | k__Fungi(100);p__Ascomycota(100);c__Saccharomycetes(100);o__Saccharomycetales(100);f__Metschnikowiaceae(100);g__Metschnikowia(100);s__Metschnikowia_pulcherrima(100); | Metschnikowia | 13 | Saprotroph | \|Undefined Saprotroph\| | Yeast | NULL | Probable | As nectar-tap_saprotroph (Põlme et al. 2020); Decay sugar-rich substrates (Põlme et al. 2020) | Põlme S, et al. 2020. Fungal Diversity 105:1-16 ((https://doi.org/10.1007/s13225-020-00466-2)) |
| Otu08233 | 3 | k__Fungi(100);p__Ascomycota(100);c__Saccharomycetes(100);o__Saccharomycetales(100);f__Metschnikowiaceae(100);g__Metschnikowia(100);s__Metschnikowia_pulcherrima(100); | Metschnikowia | 13 | Saprotroph | \|Undefined Saprotroph\| | Yeast | NULL | Probable | As nectar-tap_saprotroph (Põlme et al. 2020); Decay sugar-rich substrates (Põlme et al. 2020) | Põlme S, et al. 2020. Fungal Diversity 105:1-16 ((https://doi.org/10.1007/s13225-020-00466-2)) |
| Otu08234 | 3 | k__Fungi(100);p__Ascomycota(100);c__Saccharomycetes(100);o__Saccharomycetales(100);f__Metschnikowiaceae(100);g__Metschnikowia(100);g__Metschnikowia_unclassified(100); | Metschnikowia | 13 | Saprotroph | \|Undefined Saprotroph\| | Yeast | NULL | Probable | As nectar-tap_saprotroph (Põlme et al. 2020); Decay sugar-rich substrates (Põlme et al. 2020) | Põlme S, et al. 2020. Fungal Diversity 105:1-16 ((https://doi.org/10.1007/s13225-020-00466-2)) |
| Otu08235 | 3 | k__Fungi(100);p__Ascomycota(100);c__Saccharomycetes(100);o__Saccharomycetales(100);f__Metschnikowiaceae(100);g__Metschnikowia(100);s__Metschnikowia_pulcherrima(67); | Metschnikowia | 13 | Saprotroph | \|Undefined Saprotroph\| | Yeast | NULL | Probable | As nectar-tap_saprotroph (Põlme et al. 2020); Decay sugar-rich substrates (Põlme et al. 2020) | Põlme S, et al. 2020. Fungal Diversity 105:1-16 ((https://doi.org/10.1007/s13225-020-00466-2)) |
| Otu08239 | 3 | k__Fungi(100);p__Ascomycota(100);c__Sordariomycetes(100);o__Hypocreales(100);f__Ophiocordycipitaceae(100);f__Ophiocordycipitaceae_unclassified(100);f__Ophiocordycipitaceae_unclassified(100); | Ophiocordycipitaceae | 9 | Pathotroph-Symbiotroph | Endophyte-Insect Pathogen | Microfungus | NULL | Probable | A family of parasitic fungi (Sung et al. 2007) | Sung GH, et al. 2007. Studies in Mycology 57:5-59 ((https://doi.org/10.3114/sim.2007.57.01)); Costa IPMW, et al. 2012. Mycotaxon 119: e26 ((http://mycotaxon.com/resources/checklists/costa_v119_checklist.pdf)) |
| Otu08240 | 3 | k__Fungi(100);p__Ascomycota(100);c__Saccharomycetes(100);o__Saccharomycetales(100);f__Metschnikowiaceae(100);g__Metschnikowia(100);s__Metschnikowia_pulcherrima(100); | Metschnikowia | 13 | Saprotroph | \|Undefined Saprotroph\| | Yeast | NULL | Probable | As nectar-tap_saprotroph (Põlme et al. 2020); Decay sugar-rich substrates (Põlme et al. 2020) | Põlme S, et al. 2020. Fungal Diversity 105:1-16 ((https://doi.org/10.1007/s13225-020-00466-2)) |
| Otu08243 | 3 | k__Fungi(100);p__Basidiomycota(100);c__Agaricomycetes(100);o__Agaricales(100);f__Entolomataceae(100);g__Entoloma(100);g__Entoloma_unclassified(100); | Entoloma | 13 | Pathotroph-Saprotroph-Symbiotroph | Ectomycorrhizal-Fungal Parasite-\|Undefined Saprotroph\|-Undefined Symbiotroph | Agaricoid | NULL | Possible | As soil_saprotroph (Põlme et al. 2020); Decay roots, soil (Põlme et al. 2020) | Rinaldi AC, et al. 2008. Fungal Diversity 33:1-45 ((http://www.fungaldiversity.org/fdp/sfdp/33-1.pdf)) (as ECM/not ECM); Tedersoo L, et al. 2010. Mycorrhiza 20:217-263 ((https://doi.org/10.1007/s00572-009-0274-x; Põlme S, et al. 2020. Fungal Diversity 105:1-16 ((https://doi.org/10.1007/s13225-020-00466-2)); sensu stricto)); http://mycorrhizas.info/ecmf.html ((http://mycorrhizas.info/ecmf.html)) ((as ECM/not ECM)) |
| Otu08245 | 3 | k__Fungi(100);p__Ascomycota(100);c__Saccharomycetes(100);o__Saccharomycetales(100);f__Saccharomycetales_family_Incertae_sedis(100);g__Candida(100);g__Candida_unclassified(100); | Candida | 13 | Pathotroph-Saprotroph-Symbiotroph | Animal Pathogen-Endophyte-Endosymbiont-Epiphyte-\|Undefined Saprotroph\| | Dimorphic Yeast-Yeast | NULL | Probable | As nectar-tap_saprotroph (Põlme et al. 2020); Decay sugar-rich substrates (Põlme et al. 2020); This asexual genus is polyphyletic and contain species that function very differently depend on what they are. Caution should be taken when interpreting guilds for Candida species. | Manolakaki D, et al. 2010. Virulence. 1: 367-375 ((https://doi.org/10.4161/viru.1.5.12796)); Mekbib SB, et al. 2011. Tropical Plant Pathology 36:233-240 ((https://doi.org/10.1590/S1982-56762011000400004)); Põlme S, et al. 2020. Fungal Diversity 105:1-16 ((https://doi.org/10.1007/s13225-020-00466-2)) |
| Otu08246 | 3 | k__Fungi(100);p__Ascomycota(100);c__Saccharomycetes(100);o__Saccharomycetales(100);f__Metschnikowiaceae(100);g__Metschnikowia(100);g__Metschnikowia_unclassified(67); | Metschnikowia | 13 | Saprotroph | \|Undefined Saprotroph\| | Yeast | NULL | Probable | As nectar-tap_saprotroph (Põlme et al. 2020); Decay sugar-rich substrates (Põlme et al. 2020) | Põlme S, et al. 2020. Fungal Diversity 105:1-16 ((https://doi.org/10.1007/s13225-020-00466-2)) |
| Otu08247 | 3 | k__Fungi(100);p__Ascomycota(100);c__Saccharomycetes(100);o__Saccharomycetales(100);f__Metschnikowiaceae(100);g__Metschnikowia(100);s__Metschnikowia_pulcherrima(100); | Metschnikowia | 13 | Saprotroph | \|Undefined Saprotroph\| | Yeast | NULL | Probable | As nectar-tap_saprotroph (Põlme et al. 2020); Decay sugar-rich substrates (Põlme et al. 2020) | Põlme S, et al. 2020. Fungal Diversity 105:1-16 ((https://doi.org/10.1007/s13225-020-00466-2)) |
| Otu08250 | 3 | k__Fungi(100);p__Ascomycota(100);c__Saccharomycetes(100);o__Saccharomycetales(100);f__Metschnikowiaceae(100);g__Metschnikowia(100);s__Metschnikowia_pulcherrima(67); | Metschnikowia | 13 | Saprotroph | \|Undefined Saprotroph\| | Yeast | NULL | Probable | As nectar-tap_saprotroph (Põlme et al. 2020); Decay sugar-rich substrates (Põlme et al. 2020) | Põlme S, et al. 2020. Fungal Diversity 105:1-16 ((https://doi.org/10.1007/s13225-020-00466-2)) |
| Otu08252 | 3 | k__Fungi(100);p__Ascomycota(100);c__Saccharomycetes(100);o__Saccharomycetales(100);f__Metschnikowiaceae(100);g__Metschnikowia(100);s__Metschnikowia_pulcherrima(100); | Metschnikowia | 13 | Saprotroph | \|Undefined Saprotroph\| | Yeast | NULL | Probable | As nectar-tap_saprotroph (Põlme et al. 2020); Decay sugar-rich substrates (Põlme et al. 2020) | Põlme S, et al. 2020. Fungal Diversity 105:1-16 ((https://doi.org/10.1007/s13225-020-00466-2)) |
| Otu08254 | 3 | k__Fungi(100);p__Basidiomycota(100);c__Agaricomycetes(100);o__Boletales(100);f__Boletaceae(100);g__Aureoboletus(100);s__Aureoboletus_gentilis(100); | Aureoboletus | 13 | Symbiotroph | \|Ectomycorrhizal\| | Agaricoid-Boletoid | NULL | Highly Probable | NULL | Rinaldi AC, et al. 2008. Fungal Diversity 33:1-45 ((http://www.fungaldiversity.org/fdp/sfdp/33-1.pdf)); Tedersoo L, et al. 2010. Mycorrhiza 20:217-263 ((https://doi.org/10.1007/s00572-009-0274-x)); Põlme S, et al. 2020. Fungal Diversity 105:1-16 ((https://doi.org/10.1007/s13225-020-00466-2)); http://mycorrhizas.info/ecmf.html ((http://mycorrhizas.info/ecmf.html)) |
| Otu08255 | 3 | k__Fungi(100);p__Ascomycota(100);c__Saccharomycetes(100);o__Saccharomycetales(100);f__Metschnikowiaceae(100);g__Metschnikowia(100);s__Metschnikowia_pulcherrima(100); | Metschnikowia | 13 | Saprotroph | \|Undefined Saprotroph\| | Yeast | NULL | Probable | As nectar-tap_saprotroph (Põlme et al. 2020); Decay sugar-rich substrates (Põlme et al. 2020) | Põlme S, et al. 2020. Fungal Diversity 105:1-16 ((https://doi.org/10.1007/s13225-020-00466-2)) |
| Otu08257 | 3 | k__Fungi(100);p__Ascomycota(100);c__Saccharomycetes(100);o__Saccharomycetales(100);f__Metschnikowiaceae(100);g__Metschnikowia(100);s__Metschnikowia_pulcherrima(100); | Metschnikowia | 13 | Saprotroph | \|Undefined Saprotroph\| | Yeast | NULL | Probable | As nectar-tap_saprotroph (Põlme et al. 2020); Decay sugar-rich substrates (Põlme et al. 2020) | Põlme S, et al. 2020. Fungal Diversity 105:1-16 ((https://doi.org/10.1007/s13225-020-00466-2)) |
| Otu08259 | 3 | k__Fungi(100);p__Ascomycota(100);c__Saccharomycetes(100);o__Saccharomycetales(100);f__Metschnikowiaceae(100);g__Metschnikowia(100);s__Metschnikowia_pulcherrima(67); | Metschnikowia | 13 | Saprotroph | \|Undefined Saprotroph\| | Yeast | NULL | Probable | As nectar-tap_saprotroph (Põlme et al. 2020); Decay sugar-rich substrates (Põlme et al. 2020) | Põlme S, et al. 2020. Fungal Diversity 105:1-16 ((https://doi.org/10.1007/s13225-020-00466-2)) |
| Otu08260 | 3 | k__Fungi(100);p__Ascomycota(100);c__Saccharomycetes(100);o__Saccharomycetales(100);f__Metschnikowiaceae(100);g__Metschnikowia(100);g__Metschnikowia_unclassified(67); | Metschnikowia | 13 | Saprotroph | \|Undefined Saprotroph\| | Yeast | NULL | Probable | As nectar-tap_saprotroph (Põlme et al. 2020); Decay sugar-rich substrates (Põlme et al. 2020) | Põlme S, et al. 2020. Fungal Diversity 105:1-16 ((https://doi.org/10.1007/s13225-020-00466-2)) |
| Otu08262 | 3 | k__Fungi(100);p__Basidiomycota(100);c__Agaricomycetes(100);o__Agaricales(100);f__Bolbitiaceae(100);g__Bolbitius(100);s__Bolbitius_reticulatus(100); | Bolbitius | 13 | Saprotroph | \|Dung Saprotroph\|-Plant Saprotroph | Agaricoid | NULL | Probable | As litter_saprotroph (Põlme et al. 2020); Decay leaf, fruit, seed, dung (Põlme et al. 2020) | Tedersoo L, et al. 2014. Science 346:e1256688 ((https://doi.org/10.1126/science.1256688)); Põlme S, et al. 2020. Fungal Diversity 105:1-16 ((https://doi.org/10.1007/s13225-020-00466-2)) |
| Otu08264 | 3 | k__Fungi(100);p__Ascomycota(100);c__Saccharomycetes(100);o__Saccharomycetales(100);f__Metschnikowiaceae(100);g__Metschnikowia(100);s__Metschnikowia_pulcherrima(100); | Metschnikowia | 13 | Saprotroph | \|Undefined Saprotroph\| | Yeast | NULL | Probable | As nectar-tap_saprotroph (Põlme et al. 2020); Decay sugar-rich substrates (Põlme et al. 2020) | Põlme S, et al. 2020. Fungal Diversity 105:1-16 ((https://doi.org/10.1007/s13225-020-00466-2)) |
| Otu08266 | 3 | k__Fungi(100);p__Basidiomycota(100);c__Agaricomycetes(100);o__Agaricales(100);f__Entolomataceae(100);g__Clitopilus(100);s__Clitopilus_hobsonii(100); | Clitopilus | 13 | Pathotroph-Saprotroph | Fungal Parasite-\|Plant Saprotroph\|-Undefined Saprotroph | Agaricoid | NULL | Probable | As litter_saprotroph (Põlme et al. 2020); As mycoparasite (Põlme et al. 2020); Decay leaf, fruit, seed (Põlme et al. 2020) | Tedersoo L, et al. 2014. Science 346:e1256688 ((https://doi.org/10.1126/science.1256688)); Põlme S, et al. 2020. Fungal Diversity 105:1-16 ((https://doi.org/10.1007/s13225-020-00466-2)) |

| | | | | | | | | | | | |
|---|---|---|---|---|---|---|---|---|---|---|---|
| Otu08267 | 3 | k__Fungi(100);p__Ascomycota(100);c__Saccharomycetes(100);o__Saccharomycetales(100);f__Metschnikowiaceae(100);g__Metschnikowia(100);g__Metschnikowia_unclassified(67); | Metschnikowia | 13 | Saprotroph | \|Undefined Saprotroph\| | Yeast | NULL | Probable | As nectar-tap_saprotroph (Põlme et al. 2020); Decay sugar-rich substrates (Põlme et al. 2020) | Põlme S, et al. 2020. Fungal Diversity 105:1-16 ((https://doi.org/10.1007/s13225-020-00466-2)) |
| Otu08268 | 3 | k__Fungi(100);p__Basidiomycota(100);c__Agaricomycetes(100);o__Agaricales(100);f__Strophariaceae(100);g__Galerina(100);g__Galerina_unclassified(100); | Galerina | 13 | Pathotroph-Saprotroph | Bryophyte Parasite-Plant Pathogen-Plant Saprotroph-Undefined Saprotroph-\|Wood Saprotroph\| | Agaricoid | White Rot | Probable | As litter_saprotroph (Põlme et al. 2020); Decay wood, leaf, fruit, seed (Põlme et al. 2020) | Tedersoo L, et al. 2014. Science 346:e1256688 ((https://doi.org/10.1126/science.1256688)); Põlme S, et al. 2020. Fungal Diversity 105:1-16 ((https://doi.org/10.1007/s13225-020-00466-2)) |
| Otu08270 | 3 | k__Fungi(100);p__Ascomycota(100);c__Saccharomycetes(100);o__Saccharomycetales(100);f__Metschnikowiaceae(100);g__Metschnikowia(100);s__Metschnikowia_pulcherrima(100); | Metschnikowia | 13 | Saprotroph | \|Undefined Saprotroph\| | Yeast | NULL | Probable | As nectar-tap_saprotroph (Põlme et al. 2020); Decay sugar-rich substrates (Põlme et al. 2020) | Põlme S, et al. 2020. Fungal Diversity 105:1-16 ((https://doi.org/10.1007/s13225-020-00466-2)) |
| Otu08272 | 3 | k__Fungi(100);p__Ascomycota(100);c__Dothideomycetes(100);o__Dothideomycetes_order_Incertae_sedis(100);f__Myxotrichaceae(100);g__Oidiodendron(100);g__Oidiodendron_unclassified(100); | Oidiodendron | 13 | Pathotroph-Saprotroph-Symbiotroph | Endophyte-Ericoid Mycorrhizal-\|Undefined Saprotroph\| | Microfungus | NULL | Probable | As root_endophyte (Põlme et al. 2020); As soil_saprotroph (Põlme et al. 2020); Dark Septate Endophyte; Decay soil (Põlme et al. 2020) | Newsham KK. 2011. New Phytologist 190:783-793 ((https://doi.org/10.1111/j.1469-8137.2010.03611.x)) (as dark septate root endophytes); Põlme S, et al. 2020. Fungal Diversity 105:1-16 ((https://doi.org/10.1007/s13225-020-00466-2)) |
| Otu08273 | 3 | k__Fungi(100);p__Ascomycota(100);c__Dothideomycetes(100);o__Pleosporales(100);f__Venturiaceae(100);g__Venturia(100);s__Venturia_chlorospora(100); | Venturia | 13 | Pathotroph-Pathotroph-Saprotroph | \|Plant Pathogen\|-Plant Saprotroph | Microfungus | NULL | Probable | As litter_saprotroph (Põlme et al. 2020); Decay leaf, fruit, seed (Põlme et al. 2020) | Tedersoo L, et al. 2014. Science 346:e1256688 ((https://doi.org/10.1126/science.1256688)); Põlme S, et al. 2020. Fungal Diversity 105:1-16 ((https://doi.org/10.1007/s13225-020-00466-2)) |
| Otu08277 | 3 | k__Fungi(100);p__Ascomycota(100);c__Saccharomycetes(100);o__Saccharomycetales(100);f__Metschnikowiaceae(100);g__Metschnikowia(100);g__Metschnikowia_unclassified(67); | Metschnikowia | 13 | Saprotroph | \|Undefined Saprotroph\| | Yeast | NULL | Probable | As nectar-tap_saprotroph (Põlme et al. 2020); Decay sugar-rich substrates (Põlme et al. 2020) | Põlme S, et al. 2020. Fungal Diversity 105:1-16 ((https://doi.org/10.1007/s13225-020-00466-2)) |
| Otu08278 | 3 | k__Fungi(100);p__Ascomycota(100);c__Saccharomycetes(100);o__Saccharomycetales(100);f__Metschnikowiaceae(100);g__Metschnikowia(100);g__Metschnikowia_unclassified(67); | Metschnikowia | 13 | Saprotroph | \|Undefined Saprotroph\| | Yeast | NULL | Probable | As nectar-tap_saprotroph (Põlme et al. 2020); Decay sugar-rich substrates (Põlme et al. 2020) | Põlme S, et al. 2020. Fungal Diversity 105:1-16 ((https://doi.org/10.1007/s13225-020-00466-2)) |
| Otu08279 | 3 | k__Fungi(100);p__Ascomycota(100);c__Saccharomycetes(100);o__Saccharomycetales(100);f__Metschnikowiaceae(100);g__Metschnikowia(100);g__Metschnikowia_unclassified(100); | Metschnikowia | 13 | Saprotroph | \|Undefined Saprotroph\| | Yeast | NULL | Probable | As nectar-tap_saprotroph (Põlme et al. 2020); Decay sugar-rich substrates (Põlme et al. 2020) | Põlme S, et al. 2020. Fungal Diversity 105:1-16 ((https://doi.org/10.1007/s13225-020-00466-2)) |
| Otu08283 | 3 | k__Fungi(100);p__Ascomycota(100);c__Saccharomycetes(100);o__Saccharomycetales(100);f__Metschnikowiaceae(100);g__Metschnikowia(100);g__Metschnikowia_unclassified(100); | Metschnikowia | 13 | Saprotroph | \|Undefined Saprotroph\| | Yeast | NULL | Probable | As nectar-tap_saprotroph (Põlme et al. 2020); Decay sugar-rich substrates (Põlme et al. 2020) | Põlme S, et al. 2020. Fungal Diversity 105:1-16 ((https://doi.org/10.1007/s13225-020-00466-2)) |
| Otu08284 | 3 | k__Fungi(100);p__Ascomycota(100);c__Saccharomycetes(100);o__Saccharomycetales(100);f__Metschnikowiaceae(100);g__Metschnikowia(100);s__Metschnikowia_pulcherrima(100); | Metschnikowia | 13 | Saprotroph | \|Undefined Saprotroph\| | Yeast | NULL | Probable | As nectar-tap_saprotroph (Põlme et al. 2020); Decay sugar-rich substrates (Põlme et al. 2020) | Põlme S, et al. 2020. Fungal Diversity 105:1-16 ((https://doi.org/10.1007/s13225-020-00466-2)) |
| Otu08286 | 3 | k__Fungi(100);p__Basidiomycota(100);c__Agaricomycetes(100);o__Agaricales(100);f__Hygrophoraceae(100);g__Gloioxanthomyces(100);s__Gloioxanthomyces_nitidus(100); | Gloioxanthomyces | 13 | Saprotroph-Symbiotroph | \|Undefined Saprotroph\|-Undefined Symbiotroph | Agaricoid | NULL | Highly Probable | As soil_saprotroph (Põlme et al. 2020); Decay roots, soil (Põlme et al. 2020); Previously classified as Hygrocybe; carbon signature suggests biotrophy with bryophytes, grasses, or understory plants (Seitzman et al. 2011) | Seitzman BH, et al. 2011. Mycologia 103:280-290 ((https://doi.org/10.3852/10-195)); Lodge J., et al. 2014. Fungal Diversity 64:1-99 ((https://doi.org/10.1007/s13225-013-0259-0)); Põlme S, et al. 2020. Fungal Diversity 105:1-16 ((https://doi.org/10.1007/s13225-020-00466-2)) |
| Otu08287 | 3 | k__Fungi(100);p__Ascomycota(100);c__Saccharomycetes(100);o__Saccharomycetales(100);f__Metschnikowiaceae(100);g__Metschnikowia(100);g__Metschnikowia_unclassified(67); | Metschnikowia | 13 | Saprotroph | \|Undefined Saprotroph\| | Yeast | NULL | Probable | As nectar-tap_saprotroph (Põlme et al. 2020); Decay sugar-rich substrates (Põlme et al. 2020) | Põlme S, et al. 2020. Fungal Diversity 105:1-16 ((https://doi.org/10.1007/s13225-020-00466-2)) |
| Otu08288 | 3 | k__Fungi(100);p__Ascomycota(100);c__Saccharomycetes(100);o__Saccharomycetales(100);f__Metschnikowiaceae(100);g__Metschnikowia(100);s__Metschnikowia_pulcherrima(100); | Metschnikowia | 13 | Saprotroph | \|Undefined Saprotroph\| | Yeast | NULL | Probable | As nectar-tap_saprotroph (Põlme et al. 2020); Decay sugar-rich substrates (Põlme et al. 2020) | Põlme S, et al. 2020. Fungal Diversity 105:1-16 ((https://doi.org/10.1007/s13225-020-00466-2)) |
| Otu08289 | 3 | k__Fungi(100);p__Ascomycota(100);c__Saccharomycetes(100);o__Saccharomycetales(100);f__Metschnikowiaceae(100);g__Metschnikowia(100);s__Metschnikowia_pulcherrima(100); | Metschnikowia | 13 | Saprotroph | \|Undefined Saprotroph\| | Yeast | NULL | Probable | As nectar-tap_saprotroph (Põlme et al. 2020); Decay sugar-rich substrates (Põlme et al. 2020) | Põlme S, et al. 2020. Fungal Diversity 105:1-16 ((https://doi.org/10.1007/s13225-020-00466-2)) |
| Otu08290 | 3 | k__Fungi(100);p__Ascomycota(100);c__Sordariomycetes(100);o__Hypocreales(100);f__Cordycipitaceae(100);g__Cordyceps(100);g__Cordyceps_unclassified(100); | Cordyceps | 13 | Pathotroph-Symbiotroph | Clavicipitaceous Endophyte-Fungal Parasite-\|Insect Pathogen\| | Cordyceptoid | NULL | Possible | As foliar_endophyte (Põlme et al. 2020); Decay animal material (Põlme et al. 2020) | Samuels GJ. 2007. Self-published; Costa IPMW, et al. 2012. Mycotaxon 119:e26 ((http://mycotaxon.com/resources/checklists/costa_v119_checklist.pdf)); Tedersoo L, et al. 2014. Science 346:e1256688 ((https://doi.org/10.1126/science.1256688)); Põlme S, et al. 2020. Fungal Diversity 105:1-16 ((https://doi.org/10.1007/s13225-020-00466-2)) |
| Otu08291 | 3 | k__Fungi(100);p__Basidiomycota(100);c__Agaricomycetes(100);o__Agaricales(100);f__Psathyrellaceae(100);f__Psathyrellaceae_unclassified(100);f__Psathyrellaceae_unclassified(100); | Psathyrellaceae | 9 | Saprotroph | Dung Saprotroph-Plant Saprotroph-Wood Saprotroph | Agaricoid | NULL | Probable | Saprobic, in grass, on dung, rotten wood and similar substrata (Cannon & Kirk 2007) | Cannon PF, Kirk PM. 2007. Fungal Families of the World. CAB International, Cambridge (ISBN: 978-0851998275) |
| Otu08292 | 3 | k__Fungi(100);p__Ascomycota(100);c__Sordariomycetes(100);o__Sordariomycetes_order_Incertae_sedis(100);f__Plectosphaerellaceae(100);g__Verticillium(100);s__Verticillium_sp(100); | Verticillium | 13 | Pathotroph-Saprotroph-Symbiotroph | Endophyte-Fungal Parasite-\|Plant Pathogen\|-Plant Saprotroph | Microfungus | NULL | Probable | As foliar_endophyte (Põlme et al. 2020); As litter_saprotroph (Põlme et al. 2020); Decay leaf, fruit, seed, animal material (Põlme et al. 2020) | Costa IPMW, et al. 2012. Mycotaxon 119:e26 ((http://mycotaxon.com/resources/checklists/costa_v119_checklist.pdf)); Tedersoo L, et al. 2014. Science 346:e1256688 ((https://doi.org/10.1126/science.1256688)); Põlme S, et al. 2020. Fungal Diversity 105:1-16 ((https://doi.org/10.1007/s13225-020-00466-2)); List compiled by John Plischke III ((https://goodmushroom.weebly.com/uploads/5/7/1/7/57176535/fungi_on_fungi_pdf_lq.pdf)) |
| Otu08294 | 3 | k__Fungi(100);p__Ascomycota(100);c__Sordariomycetes(100);o__Hypocreales(100);f__Cordycipitaceae(100);g__Lecanicillium(100);s__Lecanicillium_fungicola_var._aleophilum(67); | Lecanicillium | 13 | Pathotroph-Saprotroph-Symbiotroph | \|Animal Parasite\|-Animal Pathogen-Endophyte-Undefined Saprotroph | Microfungus | NULL | Probable | As animal_decomposer (Põlme et al. 2020); As foliar_endophyte (Põlme et al. 2020); Decay animal material (Põlme et al. 2020) | Tedersoo L, et al. 2014. Science 346:e1256688 ((https://doi.org/10.1126/science.1256688)); Põlme S, et al. 2020. Fungal Diversity 105:1-16 ((https://doi.org/10.1007/s13225-020-00466-2)) |
| Otu08296 | 3 | k__Fungi(100);p__Ascomycota(100);c__Sordariomycetes(100);o__Xylariales(100);f__Xylariaceae(100);g__Hypoxylon(67);g__Hypoxylon_unclassified(67); | Hypoxylon | 13 | Saprotroph-Saprotroph-Symbiotroph | Endophyte-Undefined Saprotroph-\|Wood Saprotroph\| | Corticioid-Microfungus-Xylarioid | NULL | Probable | As foliar_endophyte (Põlme et al. 2020); Decay wood, leaf, fruit, seed (Põlme et al. 2020); Host - angiosperms (Põlme et al. 2020) | Tedersoo L, et al. 2014. Science 346:e1256688 ((https://doi.org/10.1126/science.1256688)); Põlme S, et al. 2020. Fungal Diversity 105:1-16 ((https://doi.org/10.1007/s13225-020-00466-2)) |
| Otu08297 | 3 | k__Fungi(100);p__Ascomycota(100);c__Saccharomycetes(100);o__Saccharomycetales(100);f__Saccharomycetales_family_Incertae_sedis(100);g__Candida(100);g__Candida_unclassified(67); | Candida | 13 | Pathotroph-Saprotroph-Symbiotroph | Animal Pathogen-Endophyte-Endosymbiont-Epiphyte-\|Undefined Saprotroph\| | Dimorphic Yeast-Yeast | NULL | Probable | As nectar-tap_saprotroph (Põlme et al. 2020); Decay sugar-rich substrates (Põlme et al. 2020); This asexual genus is polyphyletic and contain species that function very differently depend on what they are. Caution should be taken when interpreting guilds for Candida species. | Manolakaki D, et al. 2010. Virulence. 1: 367-375 ((https://doi.org/10.4161/viru.1.5.12796)); Mekbib SB, et al. 2011. Tropical Plant Pathology 36:233-240 ((https://doi.org/10.1590/S1982-56762011000400004)); Põlme S, et al. 2020. Fungal Diversity 105:1-16 ((https://doi.org/10.1007/s13225-020-00466-2)) |
| Otu08298 | 3 | k__Fungi(100);p__Basidiomycota(100);c__Agaricomycetes(100);o__Polyporales(100);f__Phanerochaetaceae(100);g__Ceriporia(100);s__Ceriporia_sp_DLL2011_1(100); | Ceriporia | 13 | Saprotroph | \|Wood Saprotroph\| | Corticioid | White Rot | Highly Probable | As white_rot (Põlme et al. 2020); Decay wood (Põlme et al. 2020) | Tedersoo L, et al. 2014. Science 346:e1256688 ((https://doi.org/10.1126/science.1256688)); Põlme S, et al. 2020. Fungal Diversity 105:1-16 ((https://doi.org/10.1007/s13225-020-00466-2)) |
| Otu08300 | 3 | k__Fungi(100);p__Ascomycota(100);c__Saccharomycetes(100);o__Saccharomycetales(100);f__Saccharomycetales_family_Incertae_sedis(100);g__Candida(100);g__Candida_unclassified(100); | Candida | 13 | Pathotroph-Saprotroph-Symbiotroph | Animal Pathogen-Endophyte-Endosymbiont-Epiphyte-\|Undefined Saprotroph\| | Dimorphic Yeast-Yeast | NULL | Probable | As nectar-tap_saprotroph (Põlme et al. 2020); Decay sugar-rich substrates (Põlme et al. 2020); This asexual genus is polyphyletic and contain species that function very differently depend on what they are. Caution should be taken when interpreting guilds for Candida species. | Manolakaki D, et al. 2010. Virulence. 1: 367-375 ((https://doi.org/10.4161/viru.1.5.12796)); Mekbib SB, et al. 2011. Tropical Plant Pathology 36:233-240 ((https://doi.org/10.1590/S1982-56762011000400004)); Põlme S, et al. 2020. Fungal Diversity 105:1-16 ((https://doi.org/10.1007/s13225-020-00466-2)) |
| Otu08304 | 3 | k__Fungi(100);p__Ascomycota(100);c__Saccharomycetes(100);o__Saccharomycetales(100);f__Metschnikowiaceae(100);g__Metschnikowia(100);s__Metschnikowia_pulcherrima(100); | Metschnikowia | 13 | Saprotroph | \|Undefined Saprotroph\| | Yeast | NULL | Probable | As nectar-tap_saprotroph (Põlme et al. 2020); Decay sugar-rich substrates (Põlme et al. 2020) | Põlme S, et al. 2020. Fungal Diversity 105:1-16 ((https://doi.org/10.1007/s13225-020-00466-2)) |
| Otu08305 | 3 | k__Fungi(100);p__Basidiomycota(100);c__Tremellomycetes(100);o__Tremellales(100);f__Tremellales_family_Incertae_sedis(100);g__Cryptococcus(100);s__Cryptococcus_sp_SJ4L02(100); | Cryptococcus | 13 | Pathotroph-Saprotroph-Symbiotroph | Animal Pathogen-Endophyte-Epiphyte-\|Undefined Saprotroph\| | Dimorphic Yeast-Yeast | NULL | Possible | As foliar_endophyte (Põlme et al. 2020); Decay leaf, fruit, seed, soil, dung, animal material (Põlme et al. 2020) | Andrews JH, et al. 1983. Phytopathology 73:228-234 ((https://doi.org/10.1094/Phyto-73-228)); Kurtzman CP, et al. (eds.) 2011. The Yeasts, a Taxonomic Study. Fifth Edition. Vols 1-3. Elsevier, San Diego (ISBN: 9780444521491); Põlme S, et al. 2020. Fungal Diversity 105:1-16 ((https://doi.org/10.1007/s13225-020-00466-2)) |
| Otu08306 | 3 | k__Fungi(100);p__Ascomycota(100);c__Saccharomycetes(100);o__Saccharomycetales(100);f__Saccharomycetales_family_Incertae_sedis(100);g__Candida(67);g__Candida_unclassified(67); | Candida | 13 | Pathotroph-Saprotroph-Symbiotroph | Animal Pathogen-Endophyte-Endosymbiont-Epiphyte-\|Undefined Saprotroph\| | Dimorphic Yeast-Yeast | NULL | Probable | As nectar-tap_saprotroph (Põlme et al. 2020); Decay sugar-rich substrates (Põlme et al. 2020); This asexual genus is polyphyletic and contain species that function very differently depend on what they are. Caution should be taken when interpreting guilds for Candida species. | Manolakaki D, et al. 2010. Virulence. 1: 367-375 ((https://doi.org/10.4161/viru.1.5.12796)); Mekbib SB, et al. 2011. Tropical Plant Pathology 36:233-240 ((https://doi.org/10.1590/S1982-56762011000400004)); Põlme S, et al. 2020. Fungal Diversity 105:1-16 ((https://doi.org/10.1007/s13225-020-00466-2)) |
| Otu08309 | 3 | k__Fungi(100);p__Basidiomycota(100);c__Agaricomycetes(100);o__Polyporales(100);f__Phanerochaetaceae(100);g__Antrodiella(100);s__Antrodiella_micra(100); | Antrodiella | 13 | Saprotroph | \|Wood Saprotroph\| | Corticioid-Polyporoid | White Rot | Highly Probable | As white_rot (Põlme et al. 2020); Decay wood (Põlme et al. 2020) | Gilbertson RL, Ryvarden L. 1987-1987. North American Polypores. Fungiflora, Oslo (ISBN: 978-0945345060); Põlme S, et al. 2020. Fungal Diversity 105:1-16 ((https://doi.org/10.1007/s13225-020-00466-2)) |
| Otu08310 | 3 | k__Fungi(100);p__Basidiomycota(100);c__Agaricomycetes(100);o__Boletales(100);f__Tapinellaceae(100);g__Tapinella(100);s__Tapinella_panuoides(100); | Tapinella | 13 | Saprotroph | \|Wood Saprotroph\| | Agaricoid | Brown Rot | Probable | As brown-rot (Põlme et al. 2020); Decay wood (Põlme et al. 2020) | Grgurinovic CA, Simpson JA. 2001. Fungal Diversity 8:97-127 ((http://www.fungaldiversity.org/fdp/sfdp/FD_8_97-127.pdf)); Tedersoo L, et al. 2014. Science 346:e1256688 ((https://doi.org/10.1126/science.1256688)); Põlme S, et al. 2020. Fungal Diversity 105:1-16 ((https://doi.org/10.1007/s13225-020-00466-2)) |
| Otu08312 | 3 | k__Fungi(100);p__Ascomycota(100);c__Saccharomycetes(100);o__Saccharomycetales(100);f__Metschnikowiaceae(100);g__Metschnikowia(100);s__Metschnikowia_pulcherrima(100); | Metschnikowia | 13 | Saprotroph | \|Undefined Saprotroph\| | Yeast | NULL | Probable | As nectar-tap_saprotroph (Põlme et al. 2020); Decay sugar-rich substrates (Põlme et al. 2020) | Põlme S, et al. 2020. Fungal Diversity 105:1-16 ((https://doi.org/10.1007/s13225-020-00466-2)) |
| Otu08315 | 3 | k__Fungi(100);p__Basidiomycota(100);c__Agaricomycetes(100);o__Hymenochaetales(100);f__Hymenochaetaceae(100);g__Fomitiporia(100);s__Fomitiporia_punctata(100); | Fomitiporia | 13 | Pathotroph-Saprotroph | \|Plant Pathogen\|-Wood Saprotroph | Polyporoid | White Rot | Probable | As white_rot (Põlme et al. 2020); As wood_pathogen (Põlme et al. 2020); Decay wood (Põlme et al. 2020) | Tedersoo L, et al. 2014. Science 346:e1256688 ((https://doi.org/10.1126/science.1256688)); Põlme S, et al. 2020. Fungal Diversity 105:1-16 ((https://doi.org/10.1007/s13225-020-00466-2)) |
| Otu08316 | 3 | k__Fungi(100);p__Ascomycota(100);c__Saccharomycetes(100);o__Saccharomycetales(100);f__Metschnikowiaceae(100);g__Metschnikowia(100);s__Metschnikowia_pulcherrima(67); | Metschnikowia | 13 | Saprotroph | \|Undefined Saprotroph\| | Yeast | NULL | Probable | As nectar-tap_saprotroph (Põlme et al. 2020); Decay sugar-rich substrates (Põlme et al. 2020) | Põlme S, et al. 2020. Fungal Diversity 105:1-16 ((https://doi.org/10.1007/s13225-020-00466-2)) |
| Otu08318 | 3 | k__Fungi(100);p__Ascomycota(100);c__Leotiomycetes(100);o__Helotiales(100);f__Dermateaceae(100);f__Dermateaceae_unclassified(67);f__Dermateaceae_unclassified(67); | Dermateaceae | 9 | Pathotroph-Saprotroph | Endophyte-\|Plant Pathogen\|-Plant Saprotroph-Wood Saprotroph | Microfungus-Pezizoid | NULL | Possible | As foliar_endophyte (Põlme et al. 2020); As litter_saprotroph (Põlme et al. 2020); Dark Septate Endophyte, known from plant roots.; Decay leaf, fruit, seed (Põlme et al. 2020); Saprobic or parasitic on herbaceous and woody material (Cannon & Kirk 2007) | Cannon PF, Kirk PM. 2007. Fungal Families of the World. CAB International, Cambridge (ISBN: 978-0851998275); Põlme S, et al. 2020. Fungal Diversity 105:1-16 ((https://doi.org/10.1007/s13225-020-00466-2)) |
| Otu08319 | 3 | k__Fungi(100);p__Ascomycota(100);c__Saccharomycetes(100);o__Saccharomycetales(100);f__Metschnikowiaceae(100);g__Metschnikowia(100);g__Metschnikowia_unclassified(67); | Metschnikowia | 13 | Saprotroph | \|Undefined Saprotroph\| | Yeast | NULL | Probable | As nectar-tap_saprotroph (Põlme et al. 2020); Decay sugar-rich substrates (Põlme et al. 2020) | Põlme S, et al. 2020. Fungal Diversity 105:1-16 ((https://doi.org/10.1007/s13225-020-00466-2)) |
| Otu08320 | 3 | k__Fungi(100);p__Basidiomycota(100);c__Agaricomycetes(100);o__Polyporales(100);f__Fomitopsidaceae(100);g__Fibroporia(100);s__Fibroporia_citrina(100); | Fibroporia | 13 | Saprotroph | \|Wood Saprotroph\| | Corticioid | Brown Rot | Probable | As brown-rot (Põlme et al. 2020); Decay wood (Põlme et al. 2020) | Tedersoo L, et al. 2014. Science 346:e1256688 ((https://doi.org/10.1126/science.1256688)); Põlme S, et al. 2020. Fungal Diversity 105:1-16 ((https://doi.org/10.1007/s13225-020-00466-2)) |
| Otu08321 | 3 | k__Fungi(100);p__Ascomycota(100);c__Saccharomycetes(100);o__Saccharomycetales(100);f__Metschnikowiaceae(100);g__Metschnikowia(100);s__Metschnikowia_pulcherrima(100); | Metschnikowia | 13 | Saprotroph | \|Undefined Saprotroph\| | Yeast | NULL | Probable | As nectar-tap_saprotroph (Põlme et al. 2020); Decay sugar-rich substrates (Põlme et al. 2020) | Põlme S, et al. 2020. Fungal Diversity 105:1-16 ((https://doi.org/10.1007/s13225-020-00466-2)) |
| Otu08324 | 3 | k__Fungi(100);p__Basidiomycota(100);c__Agaricomycetes(100);o__Agaricales(100);o__Agaricales_unclassified(100);o__Agaricales_unclassified(100);o__Agaricales_unclassified(100); | Agaricales | 7 | Pathotroph-Saprotroph-Symbiotroph | Bryophyte Parasite-Dung Saprotroph-Ectomycorrhizal-Fungal Parasite-Leaf Saprotroph-Plant Parasite-Undefined Saprotroph-Wood Saprotroph | Agaricoid-Gasteroid-Microfungus-Secotioid-Yeast | NULL | Possible | Mushrooms and toadstools, Gill fungi, Agarics... lignicolous, sometimes muscicolous or fungicolous, saprobic, mycorrhizal, rarely parasitic on plants or fungi (Kirk et al. 2008) | Kirk PM et al. 2008. Dictionary of the Fungi. Tenth Edition. CAB International, Wallingford (ISBN: 978-0851998268) |
| Otu08325 | 3 | k__Fungi(100);p__Basidiomycota(100);c__Tremellomycetes(100);o__Tremellales(100);f__Tremellales_family_Incertae_sedis(100);g__Tremella(100);s__Tremella_foliacea(100); | Tremella | 13 | Pathotroph | \|Fungal Parasite\|-Lichen Parasite | Dimorphic-Facultative Yeast-Microfungus-Tremelloid | NULL | Probable | As mycoparasite (Põlme et al. 2020); Decay fungal material (Põlme et al. 2020) | Tedersoo L, et al. 2014. Science 346:e1256688 ((https://doi.org/10.1126/science.1256688)); Põlme S, et al. 2020. Fungal Diversity 105:1-16 ((https://doi.org/10.1007/s13225-020-00466-2)) |
| Otu08329 | 3 | k__Fungi(100);p__Ascomycota(100);c__Saccharomycetes(100);o__Saccharomycetales(100);f__Saccharomycetales_family_Incertae_sedis(100);g__Candida(100);s__Candida_sp_AS23486(67); | Candida | 13 | Pathotroph-Saprotroph-Symbiotroph | Animal Pathogen-Endophyte-Endosymbiont-Epiphyte-\|Undefined Saprotroph\| | Dimorphic Yeast-Yeast | NULL | Probable | As nectar-tap_saprotroph (Põlme et al. 2020); Decay sugar-rich substrates (Põlme et al. 2020); This asexual genus is polyphyletic and contain species that function very differently depend on what they are. Caution should be taken when interpreting guilds for Candida species. | Manolakaki D, et al. 2010. Virulence. 1: 367-375 ((https://doi.org/10.4161/viru.1.5.12796)); Mekbib SB, et al. 2011. Tropical Plant Pathology 36:233-240 ((https://doi.org/10.1590/S1982-56762011000400004)); Põlme S, et al. 2020. Fungal Diversity 105:1-16 ((https://doi.org/10.1007/s13225-020-00466-2)) |
| Otu08331 | 3 | k__Fungi(100);p__Ascomycota(100);c__Saccharomycetes(100);o__Saccharomycetales(100);f__Metschnikowiaceae(100);g__Metschnikowia(100);s__Metschnikowia_pulcherrima(100); | Metschnikowia | 13 | Saprotroph | \|Undefined Saprotroph\| | Yeast | NULL | Probable | As nectar-tap_saprotroph (Põlme et al. 2020); Decay sugar-rich substrates (Põlme et al. 2020) | Põlme S, et al. 2020. Fungal Diversity 105:1-16 ((https://doi.org/10.1007/s13225-020-00466-2)) |
| Otu08332 | 3 | k__Fungi(100);p__Ascomycota(100);c__Saccharomycetes(100);o__Saccharomycetales(100);f__Metschnikowiaceae(100);g__Metschnikowia(100);s__Metschnikowia_pulcherrima(67); | Metschnikowia | 13 | Saprotroph | \|Undefined Saprotroph\| | Yeast | NULL | Probable | As nectar-tap_saprotroph (Põlme et al. 2020); Decay sugar-rich substrates (Põlme et al. 2020) | Põlme S, et al. 2020. Fungal Diversity 105:1-16 ((https://doi.org/10.1007/s13225-020-00466-2)) |
| Otu08333 | 3 | k__Fungi(100);p__Ascomycota(100);c__Saccharomycetes(100);o__Saccharomycetales(100);f__Metschnikowiaceae(100);g__Metschnikowia(100);s__Metschnikowia_pulcherrima(67); | Metschnikowia | 13 | Saprotroph | \|Undefined Saprotroph\| | Yeast | NULL | Probable | As nectar-tap_saprotroph (Põlme et al. 2020); Decay sugar-rich substrates (Põlme et al. 2020) | Põlme S, et al. 2020. Fungal Diversity 105:1-16 ((https://doi.org/10.1007/s13225-020-00466-2)) |
| Otu08335 | 3 | k__Fungi(100);p__Ascomycota(100);c__Saccharomycetes(100);o__Saccharomycetales(100);f__Metschnikowiaceae(100);g__Metschnikowia(100);s__Metschnikowia_pulcherrima(67); | Metschnikowia | 13 | Saprotroph | \|Undefined Saprotroph\| | Yeast | NULL | Probable | As nectar-tap_saprotroph (Põlme et al. 2020); Decay sugar-rich substrates (Põlme et al. 2020) | Põlme S, et al. 2020. Fungal Diversity 105:1-16 ((https://doi.org/10.1007/s13225-020-00466-2)) |
| Otu08336 | 3 | k__Fungi(100);p__Basidiomycota(100);c__Agaricomycetes(100);o__Trechisporales(100);f__Hydnodontaceae(100);g__Trechispora(100);g__Trechispora_unclassified(100); | Trechispora | 13 | Saprotroph | \|Wood Saprotroph\| | Corticioid | White Rot | Highly Probable | As white_rot (Põlme et al. 2020); Decay wood (Põlme et al. 2020) | Gilbertson RL, Ryvarden L. 1987-1987. North American Polypores. Fungiflora, Oslo (ISBN: 978-0945345060); Rinaldi AC, et al. 2008. Fungal Diversity 33:1-45 ((http://www.fungaldiversity.org/fdp/sfdp/33-1.pdf)) (as Uncertain); Tedersoo L, et al. 2010. Mycorrhiza 20:217-263 ((https://doi.org/10.1007/s00572-009-0274-x)) (as not ECM).; Põlme S, et al. 2020. Fungal Diversity 105:1-16 ((https://doi.org/10.1007/s13225-020-00466-2)) |
| Otu08337 | 3 | k__Fungi(100);p__Ascomycota(100);c__Saccharomycetes(100);o__Saccharomycetales(100);f__Metschnikowiaceae(100);g__Metschnikowia(100);s__Metschnikowia_pulcherrima(100); | Metschnikowia | 13 | Saprotroph | \|Undefined Saprotroph\| | Yeast | NULL | Probable | As nectar-tap_saprotroph (Põlme et al. 2020); Decay sugar-rich substrates (Põlme et al. 2020) | Põlme S, et al. 2020. Fungal Diversity 105:1-16 ((https://doi.org/10.1007/s13225-020-00466-2)) |
| Otu08338 | 3 | k__Fungi(100);p__Ascomycota(100);c__Saccharomycetes(100);o__Saccharomycetales(100);f__Metschnikowiaceae(100);g__Metschnikowia(100);s__Metschnikowia_pulcherrima(100); | Metschnikowia | 13 | Saprotroph | \|Undefined Saprotroph\| | Yeast | NULL | Probable | As nectar-tap_saprotroph (Põlme et al. 2020); Decay sugar-rich substrates (Põlme et al. 2020) | Põlme S, et al. 2020. Fungal Diversity 105:1-16 ((https://doi.org/10.1007/s13225-020-00466-2)) |
| Otu08339 | 3 | k__Fungi(100);p__Ascomycota(100);c__Saccharomycetes(100);o__Saccharomycetales(100);f__Metschnikowiaceae(100);g__Metschnikowia(100);s__Metschnikowia_pulcherrima(100); | Metschnikowia | 13 | Saprotroph | \|Undefined Saprotroph\| | Yeast | NULL | Probable | As nectar-tap_saprotroph (Põlme et al. 2020); Decay sugar-rich substrates (Põlme et al. 2020) | Põlme S, et al. 2020. Fungal Diversity 105:1-16 ((https://doi.org/10.1007/s13225-020-00466-2)) |
| Otu08348 | 3 | k__Fungi(100);p__Ascomycota(100);c__Saccharomycetes(100);o__Saccharomycetales(100);f__Metschnikowiaceae(100);g__Metschnikowia(100);s__Metschnikowia_pulcherrima(100); | Metschnikowia | 13 | Saprotroph | \|Undefined Saprotroph\| | Yeast | NULL | Probable | As nectar-tap_saprotroph (Põlme et al. 2020); Decay sugar-rich substrates (Põlme et al. 2020) | Põlme S, et al. 2020. Fungal Diversity 105:1-16 ((https://doi.org/10.1007/s13225-020-00466-2)) |
| Otu08349 | 3 | k__Fungi(100);p__Ascomycota(100);c__Saccharomycetes(100);o__Saccharomycetales(100);f__Metschnikowiaceae(100);g__Metschnikowia(100);s__Metschnikowia_pulcherrima(67); | Metschnikowia | 13 | Saprotroph | \|Undefined Saprotroph\| | Yeast | NULL | Probable | As nectar-tap_saprotroph (Põlme et al. 2020); Decay sugar-rich substrates (Põlme et al. 2020) | Põlme S, et al. 2020. Fungal Diversity 105:1-16 ((https://doi.org/10.1007/s13225-020-00466-2)) |
| Otu08353 | 3 | k__Fungi(100);p__Ascomycota(100);c__Sordariomycetes(100);o__Hypocreales(100);o__Hypocreales_unclassified(100);o__Hypocreales_unclassified(100);o__Hypocreales_unclassified(100); | Hypocreales | 7 | Saprotroph | Undefined Saprotroph | Microfungus | NULL | Possible | NULL | Sterkenburg E, et al. 2015. New Phytologist 207:1145-1158 ((https://doi.org/10.1111/nph.13426)) |
| Otu08354 | 3 | k__Fungi(100);p__Ascomycota(100);c__Saccharomycetes(100);o__Saccharomycetales(100);f__Metschnikowiaceae(100);g__Metschnikowia(100);s__Metschnikowia_pulcherrima(100); | Metschnikowia | 13 | Saprotroph | \|Undefined Saprotroph\| | Yeast | NULL | Probable | As nectar-tap_saprotroph (Põlme et al. 2020); Decay sugar-rich substrates (Põlme et al. 2020) | Põlme S, et al. 2020. Fungal Diversity 105:1-16 ((https://doi.org/10.1007/s13225-020-00466-2)) |
| Otu08357 | 3 | k__Fungi(100);p__Ascomycota(100);c__Sordariomycetes(100);o__Hypocreales(100);f__Ophiocordycipitaceae(100);g__Hirsutella(100);s__Hirsutella_vermicola(100); | Hirsutella | 13 | Pathotroph-Saprotroph | \|Animal Parasite\|-Animal Pathogen-Undefined Saprotroph | Microfungus | NULL | Probable | As animal_decomposer (Põlme et al. 2020); Decay animal material (Põlme et al. 2020) | Tedersoo L, et al. 2014. Science 346:e1256688 ((https://doi.org/10.1126/science.1256688)); Põlme S, et al. 2020. Fungal Diversity 105:1-16 ((https://doi.org/10.1007/s13225-020-00466-2)) |
| Otu08358 | 3 | k__Fungi(100);p__Ascomycota(100);c__Saccharomycetes(100);o__Saccharomycetales(100);f__Metschnikowiaceae(100);g__Metschnikowia(100);s__Metschnikowia_pulcherrima(100); | Metschnikowia | 13 | Saprotroph | \|Undefined Saprotroph\| | Yeast | NULL | Probable | As nectar-tap_saprotroph (Põlme et al. 2020); Decay sugar-rich substrates (Põlme et al. 2020) | Põlme S, et al. 2020. Fungal Diversity 105:1-16 ((https://doi.org/10.1007/s13225-020-00466-2)) |
| Otu08359 | 3 | k__Fungi(100);p__Ascomycota(100);c__Sordariomycetes(100);o__Hypocreales(100);o__Hypocreales_unclassified(100);o__Hypocreales_unclassified(100);o__Hypocreales_unclassified(100); | Hypocreales | 7 | Saprotroph | Undefined Saprotroph | Microfungus | NULL | Possible | NULL | Sterkenburg E, et al. 2015. New Phytologist 207:1145-1158 ((https://doi.org/10.1111/nph.13426)) |
| Otu08361 | 3 | k__Fungi(100);p__Ascomycota(100);c__Saccharomycetes(100);o__Saccharomycetales(100);f__Metschnikowiaceae(100);g__Metschnikowia(100);s__Metschnikowia_pulcherrima(67); | Metschnikowia | 13 | Saprotroph | \|Undefined Saprotroph\| | Yeast | NULL | Probable | As nectar-tap_saprotroph (Põlme et al. 2020); Decay sugar-rich substrates (Põlme et al. 2020) | Põlme S, et al. 2020. Fungal Diversity 105:1-16 ((https://doi.org/10.1007/s13225-020-00466-2)) |
| Otu08364 | 3 | k__Fungi(100);p__Ascomycota(100);c__Saccharomycetes(100);o__Saccharomycetales(100);f__Metschnikowiaceae(100);g__Metschnikowia(100);g__Metschnikowia_unclassified(67); | Metschnikowia | 13 | Saprotroph | \|Undefined Saprotroph\| | Yeast | NULL | Probable | As nectar-tap_saprotroph (Põlme et al. 2020); Decay sugar-rich substrates (Põlme et al. 2020) | Põlme S, et al. 2020. Fungal Diversity 105:1-16 ((https://doi.org/10.1007/s13225-020-00466-2)) |
| Otu08366 | 3 | k__Fungi(100);p__Ascomycota(100);c__Leotiomycetes(100);o__Helotiales(100);f__Vibrisseaceae(100);g__Phialocephala(100);s__Phialocephala_fusca(100); | Phialocephala | 13 | Saprotroph-Symbiotroph | Endophyte-\|Undefined Saprotroph\| | Microfungus | NULL | Highly Probable | As root_endophyte (Põlme et al. 2020); As root_endophyte_dark_septate (Põlme et al. 2020); As soil_saprotroph (Põlme et al. 2020); Dark Septate Endophyte, known from plant roots.; Decay roots, soil (Põlme et al. 2020) | Newsham KK. 2011. New Phytologist 190:783-793 ((https://doi.org/10.1111/j.1469-8137.2010.03611.x)) (as dark septate root endophytes); Põlme S, et al. 2020. Fungal Diversity 105:1-16 ((https://doi.org/10.1007/s13225-020-00466-2)) |
| Otu08368 | 3 | k__Fungi(100);p__Ascomycota(100);c__Saccharomycetes(100);o__Saccharomycetales(100);f__Metschnikowiaceae(100);g__Metschnikowia(100);s__Metschnikowia_pulcherrima(67); | Metschnikowia | 13 | Saprotroph | \|Undefined Saprotroph\| | Yeast | NULL | Probable | As nectar-tap_saprotroph (Põlme et al. 2020); Decay sugar-rich substrates (Põlme et al. 2020) | Põlme S, et al. 2020. Fungal Diversity 105:1-16 ((https://doi.org/10.1007/s13225-020-00466-2)) |
| Otu08369 | 3 | k__Fungi(100);p__Ascomycota(100);c__Sordariomycetes(100);o__Hypocreales(100);o__Hypocreales_unclassified(100);o__Hypocreales_unclassified(100);o__Hypocreales_unclassified(100); | Hypocreales | 7 | Saprotroph | Undefined Saprotroph | Microfungus | NULL | Possible | NULL | Sterkenburg E, et al. 2015. New Phytologist 207:1145-1158 ((https://doi.org/10.1111/nph.13426)) |
| Otu08370 | 3 | k__Fungi(100);p__Ascomycota(100);c__Saccharomycetes(100);o__Saccharomycetales(100);f__Metschnikowiaceae(100);g__Metschnikowia(100);s__Metschnikowia_pulcherrima(100); | Metschnikowia | 13 | Saprotroph | \|Undefined Saprotroph\| | Yeast | NULL | Probable | As nectar-tap_saprotroph (Põlme et al. 2020); Decay sugar-rich substrates (Põlme et al. 2020) | Põlme S, et al. 2020. Fungal Diversity 105:1-16 ((https://doi.org/10.1007/s13225-020-00466-2)) |
| Otu08371 | 3 | k__Fungi(100);p__Ascomycota(100);c__Saccharomycetes(100);o__Saccharomycetales(100);f__Metschnikowiaceae(100);g__Metschnikowia(100);s__Metschnikowia_pulcherrima(100); | Metschnikowia | 13 | Saprotroph | \|Undefined Saprotroph\| | Yeast | NULL | Probable | As nectar-tap_saprotroph (Põlme et al. 2020); Decay sugar-rich substrates (Põlme et al. 2020) | Põlme S, et al. 2020. Fungal Diversity 105:1-16 ((https://doi.org/10.1007/s13225-020-00466-2)) |
| Otu08372 | 3 | k__Fungi(100);p__Basidiomycota(100);c__Agaricomycetes(100);o__Agaricales(100);f__Entolomataceae(100);g__Entoloma(100);s__Entoloma_infula(100); | Entoloma | 13 | Pathotroph-Saprotroph-Symbiotroph | Ectomycorrhizal-Fungal Parasite-\|Undefined Saprotroph\|-Undefined Symbiotroph | Agaricoid | NULL | Possible | As soil_saprotroph (Põlme et al. 2020); Decay roots, soil (Põlme et al. 2020) | Rinaldi AC, et al. 2008. Fungal Diversity 33:1-45 ((http://www.fungaldiversity.org/fdp/sfdp/33-1.pdf)) (as ECM/not ECM); Tedersoo L, et al. 2010. Mycorrhiza 20:217-263 ((https://doi.org/10.1007/s00572-009-0274-x; Põlme S, et al. 2020. Fungal Diversity 105:1-16 ((https://doi.org/10.1007/s13225-020-00466-2)); sensu stricto)); http://mycorrhizas.info/ecmf.html ((http://mycorrhizas.info/ecmf.html)) ((as ECM/not ECM)) |
| Otu08373 | 3 | k__Fungi(100);p__Basidiomycota(100);c__Agaricomycetes(100);o__Agaricales(100);f__Entolomataceae(100);g__unclassified_Entolomataceae(100);s__Entolomataceae_sp(100); | Entolomataceae | 9 | Pathotroph-Saprotroph-Symbiotroph | Ectomycorrhizal-Fungal Parasite-Soil Saprotroph-Undefined Saprotroph | Agaricoid | NULL | Possible | Saprobic or ectomycorrhizal, in both broadleaved and coniferous forests; a few species associated with other substrata such as dung and mosses (Cannon & Kirk 2007) | Cannon PF, Kirk PM. 2007. Fungal Families of the World. CAB International, Cambridge (ISBN: 978-0851998275); List compiled by John Plischke III ((https://goodmushroom.weebly.com/uploads/5/7/1/7/57176535/fungi_on_fungi_pdf_lq.pdf)) |
| Otu08375 | 3 | k__Fungi(100);p__Basidiomycota(100);c__Agaricomycetes(100);o__Agaricales(100);f__Cortinariaceae(100);g__Cortinarius(100);s__Cortinarius_caperatus(100); | Cortinarius | 13 | Symbiotroph | \|Ectomycorrhizal\| | Agaricoid | NULL | Highly Probable | NULL | Rinaldi AC, et al. 2008. Fungal Diversity 33:1-45 ((http://www.fungaldiversity.org/fdp/sfdp/33-1.pdf)); Tedersoo L, et al. 2010. Mycorrhiza 20:217-263 ((https://doi.org/10.1007/s00572-009-0274-x)); Põlme S, et al. 2020. Fungal Diversity 105:1-16 ((https://doi.org/10.1007/s13225-020-00466-2)); http://mycorrhizas.info/ecmf.html ((http://mycorrhizas.info/ecmf.html)) |
| Otu08376 | 3 | k__Fungi(100);p__Ascomycota(100);c__Saccharomycetes(100);o__Saccharomycetales(100);f__Metschnikowiaceae(100);g__Metschnikowia(100);s__Metschnikowia_pulcherrima(100); | Metschnikowia | 13 | Saprotroph | \|Undefined Saprotroph\| | Yeast | NULL | Probable | As nectar-tap_saprotroph (Põlme et al. 2020); Decay sugar-rich substrates (Põlme et al. 2020) | Põlme S, et al. 2020. Fungal Diversity 105:1-16 ((https://doi.org/10.1007/s13225-020-00466-2)) |
| Otu08377 | 3 | k__Fungi(100);p__Ascomycota(100);c__Saccharomycetes(100);o__Saccharomycetales(100);f__Metschnikowiaceae(100);g__Metschnikowia(100);s__Metschnikowia_pulcherrima(67); | Metschnikowia | 13 | Saprotroph | \|Undefined Saprotroph\| | Yeast | NULL | Probable | As nectar-tap_saprotroph (Põlme et al. 2020); Decay sugar-rich substrates (Põlme et al. 2020) | Põlme S, et al. 2020. Fungal Diversity 105:1-16 ((https://doi.org/10.1007/s13225-020-00466-2)) |
| Otu08378 | 3 | k__Fungi(100);p__Basidiomycota(100);c__Agaricomycetes(100);o__Polyporales(100);f__Fomitopsidaceae(100);f__Fomitopsidaceae_unclassified(100);f__Fomitopsidaceae_unclassified(100); | Fomitopsidaceae | 9 | Pathotroph-Saprotroph | Plant Pathogen-Wood Saprotroph | Polyporoid | Brown Rot | Probable | Most species are parasitic on woody plants, causing brown rots (Cannon & Kirk 2007) | Cannon PF, Kirk PM. 2007. Fungal Families of the World. CAB International, Cambridge (ISBN: 978-0851998275) |
| Otu08380 | 3 | k__Fungi(100);p__Ascomycota(100);c__Saccharomycetes(100);o__Saccharomycetales(100);f__Metschnikowiaceae(100);g__Metschnikowia(100);s__Metschnikowia_pulcherrima(67); | Metschnikowia | 13 | Saprotroph | \|Undefined Saprotroph\| | Yeast | NULL | Probable | As nectar-tap_saprotroph (Põlme et al. 2020); Decay sugar-rich substrates (Põlme et al. 2020) | Põlme S, et al. 2020. Fungal Diversity 105:1-16 ((https://doi.org/10.1007/s13225-020-00466-2)) |
| Otu08381 | 3 | k__Fungi(100);p__unclassified_Fungi(100);c__unclassified_Fungi(100);o__unclassified_Fungi(100);f__unclassified_Fungi(100);g__unclassified_Fungi(100);s__Slopeiomyces_cylindrosporus(100); | Slopeiomyces | 13 | Symbiotroph | \|Endophyte\| | Microfungus | NULL | Probable | As root_endophyte (Põlme et al. 2020) | Põlme S, et al. 2020. Fungal Diversity 105:1-16 ((https://doi.org/10.1007/s13225-020-00466-2)) |
| Otu08383 | 3 | k__Fungi(100);p__Ascomycota(100);c__Saccharomycetes(100);o__Saccharomycetales(100);f__Metschnikowiaceae(100);g__Metschnikowia(100);s__Metschnikowia_pulcherrima(100); | Metschnikowia | 13 | Saprotroph | \|Undefined Saprotroph\| | Yeast | NULL | Probable | As nectar-tap_saprotroph (Põlme et al. 2020); Decay sugar-rich substrates (Põlme et al. 2020) | Põlme S, et al. 2020. Fungal Diversity 105:1-16 ((https://doi.org/10.1007/s13225-020-00466-2)) |
| Otu08384 | 3 | k__Fungi(100);p__Ascomycota(100);c__Dothideomycetes(100);o__Pleosporales(100);f__Lophiostomataceae(100);g__Lophiostoma(100);s__Lophiostoma_cf_cynaroidis_A33(100); | Lophiostoma | 13 | Saprotroph | Plant Saprotroph-Undefined Saprotroph-\|Wood Saprotroph\| | Microfungus | NULL | Probable | As litter_saprotroph (Põlme et al. 2020); Decay wood, leaf, fruit, seed (Põlme et al. 2020) | Tedersoo L, et al. 2014. Science 346:e1256688 ((https://doi.org/10.1126/science.1256688)); Põlme S, et al. 2020. Fungal Diversity 105:1-16 ((https://doi.org/10.1007/s13225-020-00466-2)) |
| Otu08385 | 3 | k__Fungi(100);p__Ascomycota(100);c__Saccharomycetes(100);o__Saccharomycetales(100);f__Metschnikowiaceae(100);g__Metschnikowia(100);s__Metschnikowia_pulcherrima(100); | Metschnikowia | 13 | Saprotroph | \|Undefined Saprotroph\| | Yeast | NULL | Probable | As nectar-tap_saprotroph (Põlme et al. 2020); Decay sugar-rich substrates (Põlme et al. 2020) | Põlme S, et al. 2020. Fungal Diversity 105:1-16 ((https://doi.org/10.1007/s13225-020-00466-2)) |
| Otu08388 | 3 | k__Fungi(100);p__Basidiomycota(100);c__Agaricomycetes(100);o__Thelephorales(100);f__Thelephoraceae(100);g__Tomentellopsis(100);g__Tomentellopsis_unclassified(100); | Tomentellopsis | 13 | Symbiotroph | \|Ectomycorrhizal\| | Corticioid | NULL | Highly Probable | NULL | Rinaldi AC, et al. 2008. Fungal Diversity 33:1-45 ((http://www.fungaldiversity.org/fdp/sfdp/33-1.pdf)); Tedersoo L, et al. 2010. Mycorrhiza 20:217-263 ((https://doi.org/10.1007/s00572-009-0274-x)); Põlme S, et al. 2020. Fungal Diversity 105:1-16 ((https://doi.org/10.1007/s13225-020-00466-2)); http://mycorrhizas.info/ecmf.html ((http://mycorrhizas.info/ecmf.html)) |
| Otu08389 | 3 | k__Fungi(100);p__Ascomycota(100);c__Sordariomycetes(100);o__Hypocreales(100);f__Hypocreales_family_Incertae_sedis(100);g__Acremonium(100);s__Acremonium_sp_24_OA_2013(100); | Acremonium | 13 | Pathotroph-Saprotroph-Symbiotroph | Animal Pathogen-Endophyte-Fungal Parasite-Plant Pathogen-\|Undefined Saprotroph\|-Wood Saprotroph | Microfungus | Soft Rot | Possible | As foliar_endophyte (Põlme et al. 2020); culture (Hanada et al. 2010, Del Olmo; Decay leaf, fruit, seed, soil, dung, animal material (Põlme et al. 2020); Endophyte detection method; Ruiz & Arnold 2014); Likely opportunistic human pathogen (Irinyi et al. 2015) | Duncan CG, Eslyn WE. 1966. Mycologia 58:642-645 ((https://doi.org/10.2307/3757045)); Nilsson T. 1973. Studia Forestalia Suecica 104:1-40 ((https://pub.epsilon.slu.se/5711/)); Worrall JJ, et al. 1997. Mycologia 89:199-219 ((https://doi.org/10.2307/3761073)); Hanada RE, et al. 2010. Fungal Biology 114:901-910 ((https://doi.org/10.1016/j.funbio.2010.08.006)); Del Olmo-Ruiz M, Arnold AE. 2014. Mycologia 106:8-21 ((https://doi.org/10.3852/13-098)); Irinyi L, et al. 2015. Medical Mycology 53313-337 ((https://doi.org/10.1093/mmy/myv008)); Põlme S, et al. 2020. Fungal Diversity 105:1-16 ((https://doi.org/10.1007/s13225-020-00466-2)) |
| Otu08391 | 3 | k__Fungi(100);p__Ascomycota(100);c__Sordariomycetes(100);o__Sordariales(100);f__Lasiosphaeriaceae(100);g__Lasiosphaeria(100);s__Lasiosphaeria_ovina(100); | Lasiosphaeria | 13 | Saprotroph | \|Undefined Saprotroph\| | Microfungus | NULL | Probable | NULL | Tedersoo L, et al. 2014. Science 346:e1256688 ((https://doi.org/10.1126/science.1256688)); Põlme S, et al. 2020. Fungal Diversity 105:1-16 ((https://doi.org/10.1007/s13225-020-00466-2)) |
| Otu08392 | 3 | k__Fungi(100);p__Ascomycota(100);c__Sordariomycetes(100);o__Boliniales(100);f__Boliniaceae(100);g__Pseudovalsaria(100);s__Pseudovalsaria_ferruginea(100); | Pseudovalsaria | 13 | Saprotroph | Undefined Saprotroph-\|Wood Saprotroph\| | Microfungus | NULL | Probable | Decay wood (Põlme et al. 2020) | Tedersoo L, et al. 2014. Science 346:e1256688 ((https://doi.org/10.1126/science.1256688)); Põlme S, et al. 2020. Fungal Diversity 105:1-16 ((https://doi.org/10.1007/s13225-020-00466-2)) |
| Otu08397 | 3 | k__Fungi(100);p__Ascomycota(100);c__Sordariomycetes(100);o__Hypocreales(100);f__Nectriaceae(100);f__Nectriaceae_unclassified(100);f__Nectriaceae_unclassified(100); | Nectriaceae | 9 | Pathotroph-Saprotroph-Symbiotroph | Animal Pathogen-Endophyte-Fungal Parasite-Lichen Parasite-Plant Pathogen-Wood Saprotroph | Microfungus | NULL | Possible | Fusarium species are amongst the economically significant plant pathogens (Cannon & Kirk 2007) | Cannon PF, Kirk PM. 2007. Fungal Families of the World. CAB International, Cambridge (ISBN: 978-0851998275); Hirooka Y, et al. 2012. Studies in Mycology 711-210 ((https://doi.org/10.3114/sim0001)); Tedersoo L, et al. 2014. Science 346:e1256688 ((https://doi.org/10.1126/science.1256688)); The American Phytopathological Society Common Names of Plant Diseases ((http://www.apsnet.org/publications/commonnames/Pages/default.aspx)); https://www.cdc.gov ((https://www.cdc.gov)) |
| Otu08399 | 3 | k__Fungi(100);p__Ascomycota(100);c__Sordariomycetes(100);o__Hypocreales(100);o__Hypocreales_unclassified(100);o__Hypocreales_unclassified(100);o__Hypocreales_unclassified(100); | Hypocreales | 7 | Saprotroph | Undefined Saprotroph | Microfungus | NULL | Possible | NULL | Sterkenburg E, et al. 2015. New Phytologist 207:1145-1158 ((https://doi.org/10.1111/nph.13426)) |
| Otu08400 | 3 | k__Fungi(100);p__Ascomycota(100);c__Saccharomycetes(100);o__Saccharomycetales(100);f__Saccharomycetales_family_Incertae_sedis(100);g__Candida(100);g__Candida_unclassified(67); | Candida | 13 | Pathotroph-Saprotroph-Symbiotroph | Animal Pathogen-Endophyte-Endosymbiont-Epiphyte-\|Undefined Saprotroph\| | Dimorphic Yeast-Yeast | NULL | Probable | As nectar-tap_saprotroph (Põlme et al. 2020); Decay sugar-rich substrates (Põlme et al. 2020); This asexual genus is polyphyletic and contain species that function very differently depend on what they are. Caution should be taken when interpreting guilds for Candida species. | Manolakaki D, et al. 2010. Virulence. 1: 367-375 ((https://doi.org/10.4161/viru.1.5.12796)); Mekbib SB, et al. 2011. Tropical Plant Pathology 36:233-240 ((https://doi.org/10.1590/S1982-56762011000400004)); Põlme S, et al. 2020. Fungal Diversity 105:1-16 ((https://doi.org/10.1007/s13225-020-00466-2)) |
| Otu08401 | 3 | k__Fungi(100);p__Ascomycota(100);c__Saccharomycetes(100);o__Saccharomycetales(100);f__Metschnikowiaceae(100);g__Metschnikowia(100);s__Metschnikowia_pulcherrima(100); | Metschnikowia | 13 | Saprotroph | \|Undefined Saprotroph\| | Yeast | NULL | Probable | As nectar-tap_saprotroph (Põlme et al. 2020); Decay sugar-rich substrates (Põlme et al. 2020) | Põlme S, et al. 2020. Fungal Diversity 105:1-16 ((https://doi.org/10.1007/s13225-020-00466-2)) |
| Otu08403 | 3 | k__Fungi(100);p__Ascomycota(100);c__Saccharomycetes(100);o__Saccharomycetales(100);f__Metschnikowiaceae(100);g__Metschnikowia(100);s__Metschnikowia_pulcherrima(100); | Metschnikowia | 13 | Saprotroph | \|Undefined Saprotroph\| | Yeast | NULL | Probable | As nectar-tap_saprotroph (Põlme et al. 2020); Decay sugar-rich substrates (Põlme et al. 2020) | Põlme S, et al. 2020. Fungal Diversity 105:1-16 ((https://doi.org/10.1007/s13225-020-00466-2)) |
| Otu08405 | 3 | k__Fungi(100);p__Basidiomycota(100);c__Agaricomycetes(100);o__Thelephorales(100);f__Thelephoraceae(100);g__Tomentellopsis(100);s__Tomentellopsis_sp(100); | Tomentellopsis | 13 | Symbiotroph | \|Ectomycorrhizal\| | Corticioid | NULL | Highly Probable | NULL | Rinaldi AC, et al. 2008. Fungal Diversity 33:1-45 ((http://www.fungaldiversity.org/fdp/sfdp/33-1.pdf)); Tedersoo L, et al. 2010. Mycorrhiza 20:217-263 ((https://doi.org/10.1007/s00572-009-0274-x)); Põlme S, et al. 2020. Fungal Diversity 105:1-16 ((https://doi.org/10.1007/s13225-020-00466-2)); http://mycorrhizas.info/ecmf.html ((http://mycorrhizas.info/ecmf.html)) |
| Otu08406 | 3 | k__Fungi(100);p__Ascomycota(100);c__Dothideomycetes(100);o__Dothideomycetes_order_Incertae_sedis(100);f__Micropeltidaceae(100);g__Stomiopeltis(100);s__Stomiopeltis_sp_RS52(100); | Stomiopeltis | 13 | Saprotroph-Symbiotroph | \|Epiphyte\|-Plant Saprotroph-Undefined Saprotroph | Microfungus | NULL | Probable | As litter_saprotroph (Põlme et al. 2020); Decay leaf, fruit, seed (Põlme et al. 2020) | Tedersoo L, et al. 2014. Science 346:e1256688 ((https://doi.org/10.1126/science.1256688)); Põlme S, et al. 2020. Fungal Diversity 105:1-16 ((https://doi.org/10.1007/s13225-020-00466-2)) |
| Otu08407 | 3 | k__Fungi(100);p__Ascomycota(100);c__Saccharomycetes(100);o__Saccharomycetales(100);f__Metschnikowiaceae(100);g__Metschnikowia(100);g__Metschnikowia_unclassified(67); | Metschnikowia | 13 | Saprotroph | \|Undefined Saprotroph\| | Yeast | NULL | Probable | As nectar-tap_saprotroph (Põlme et al. 2020); Decay sugar-rich substrates (Põlme et al. 2020) | Põlme S, et al. 2020. Fungal Diversity 105:1-16 ((https://doi.org/10.1007/s13225-020-00466-2)) |
| Otu08408 | 3 | k__Fungi(100);p__Ascomycota(100);c__Sordariomycetes(67);o__Chaetosphaeriales(67);f__Chaetosphaeriaceae(67);f__Chaetosphaeriaceae_unclassified(67);f__Chaetosphaeriaceae_unclassified(67); | Chaetosphaeriaceae | 9 | Saprotroph | Plant Saprotroph-Wood Saprotroph | Microfungus | NULL | Probable | Saprobes associated with decaying woody and herbaceous plant material, including timber and timber products in damp buildings (Cannon & Kirk 2007) | Cannon PF, Kirk PM. 2007. Fungal Families of the World. CAB International, Cambridge (ISBN: 978-0851998275) |
| Otu08409 | 3 | k__Fungi(100);p__Ascomycota(100);c__Saccharomycetes(100);o__Saccharomycetales(100);f__Metschnikowiaceae(100);g__Metschnikowia(100);g__Metschnikowia_unclassified(100); | Metschnikowia | 13 | Saprotroph | \|Undefined Saprotroph\| | Yeast | NULL | Probable | As nectar-tap_saprotroph (Põlme et al. 2020); Decay sugar-rich substrates (Põlme et al. 2020) | Põlme S, et al. 2020. Fungal Diversity 105:1-16 ((https://doi.org/10.1007/s13225-020-00466-2)) |
| Otu08414 | 3 | k__Fungi(100);p__Ascomycota(100);c__Saccharomycetes(100);o__Saccharomycetales(100);f__Metschnikowiaceae(100);g__Metschnikowia(100);s__Metschnikowia_pulcherrima(67); | Metschnikowia | 13 | Saprotroph | \|Undefined Saprotroph\| | Yeast | NULL | Probable | As nectar-tap_saprotroph (Põlme et al. 2020); Decay sugar-rich substrates (Põlme et al. 2020) | Põlme S, et al. 2020. Fungal Diversity 105:1-16 ((https://doi.org/10.1007/s13225-020-00466-2)) |
| Otu08415 | 3 | k__Fungi(100);p__Ascomycota(100);c__Saccharomycetes(100);o__Saccharomycetales(100);f__Metschnikowiaceae(100);g__Metschnikowia(100);s__Metschnikowia_pulcherrima(67); | Metschnikowia | 13 | Saprotroph | \|Undefined Saprotroph\| | Yeast | NULL | Probable | As nectar-tap_saprotroph (Põlme et al. 2020); Decay sugar-rich substrates (Põlme et al. 2020) | Põlme S, et al. 2020. Fungal Diversity 105:1-16 ((https://doi.org/10.1007/s13225-020-00466-2)) |
| Otu08418 | 3 | k__Fungi(100);p__Ascomycota(100);c__Saccharomycetes(100);o__Saccharomycetales(100);f__Metschnikowiaceae(100);g__Metschnikowia(100);s__Metschnikowia_pulcherrima(100); | Metschnikowia | 13 | Saprotroph | \|Undefined Saprotroph\| | Yeast | NULL | Probable | As nectar-tap_saprotroph (Põlme et al. 2020); Decay sugar-rich substrates (Põlme et al. 2020) | Põlme S, et al. 2020. Fungal Diversity 105:1-16 ((https://doi.org/10.1007/s13225-020-00466-2)) |
| Otu08421 | 3 | k__Fungi(100);p__Basidiomycota(100);c__Agaricomycetes(100);o__Corticiales(100);f__Corticiaceae(100);g__Vuilleminia(100);g__Vuilleminia_unclassified(100); | Vuilleminia | 13 | Pathotroph-Saprotroph | Plant Pathogen-Undefined Saprotroph-\|Wood Saprotroph\| | Corticioid | White Rot | Probable | As white_rot (Põlme et al. 2020); As wood_pathogen (Põlme et al. 2020); Decay wood (Põlme et al. 2020) | Tedersoo L, et al. 2014. Science 346:e1256688 ((https://doi.org/10.1126/science.1256688)); Põlme S, et al. 2020. Fungal Diversity 105:1-16 ((https://doi.org/10.1007/s13225-020-00466-2)) |
| Otu08422 | 3 | k__Fungi(100);p__Ascomycota(100);c__Saccharomycetes(100);o__Saccharomycetales(100);f__Metschnikowiaceae(100);g__Metschnikowia(100);s__Metschnikowia_pulcherrima(100); | Metschnikowia | 13 | Saprotroph | \|Undefined Saprotroph\| | Yeast | NULL | Probable | As nectar-tap_saprotroph (Põlme et al. 2020); Decay sugar-rich substrates (Põlme et al. 2020) | Põlme S, et al. 2020. Fungal Diversity 105:1-16 ((https://doi.org/10.1007/s13225-020-00466-2)) |
| Otu08423 | 3 | k__Fungi(100);p__Basidiomycota(100);c__Agaricomycetes(100);o__Agaricales(100);f__Bolbitiaceae(100);g__Bolbitius(100);g__Bolbitius_unclassified(100); | Bolbitius | 13 | Saprotroph | \|Dung Saprotroph\|-Plant Saprotroph | Agaricoid | NULL | Probable | As litter_saprotroph (Põlme et al. 2020); Decay leaf, fruit, seed, dung (Põlme et al. 2020) | Tedersoo L, et al. 2014. Science 346:e1256688 ((https://doi.org/10.1126/science.1256688)); Põlme S, et al. 2020. Fungal Diversity 105:1-16 ((https://doi.org/10.1007/s13225-020-00466-2)) |
| Otu08424 | 3 | k__Fungi(100);p__Ascomycota(100);c__Saccharomycetes(100);o__Saccharomycetales(100);f__Metschnikowiaceae(100);g__Metschnikowia(100);s__Metschnikowia_pulcherrima(100); | Metschnikowia | 13 | Saprotroph | \|Undefined Saprotroph\| | Yeast | NULL | Probable | As nectar-tap_saprotroph (Põlme et al. 2020); Decay sugar-rich substrates (Põlme et al. 2020) | Põlme S, et al. 2020. Fungal Diversity 105:1-16 ((https://doi.org/10.1007/s13225-020-00466-2)) |
| Otu08429 | 3 | k__Fungi(100);p__Ascomycota(100);c__Sordariomycetes(100);o__Hypocreales(100);f__Hypocreales_family_Incertae_sedis(67);f__Hypocreales_family_Incertae_sedis_unclassified(67);f__Hypocreales_family_Incertae_sedis_unclassified(67); | Hypocreales | 7 | Saprotroph | Undefined Saprotroph | Microfungus | NULL | Possible | NULL | Sterkenburg E, et al. 2015. New Phytologist 207:1145-1158 ((https://doi.org/10.1111/nph.13426)) |
| Otu08430 | 3 | k__Fungi(100);p__Ascomycota(100);c__Saccharomycetes(100);o__Saccharomycetales(100);f__Metschnikowiaceae(100);g__Metschnikowia(100);s__Metschnikowia_pulcherrima(100); | Metschnikowia | 13 | Saprotroph | \|Undefined Saprotroph\| | Yeast | NULL | Probable | As nectar-tap_saprotroph (Põlme et al. 2020); Decay sugar-rich substrates (Põlme et al. 2020) | Põlme S, et al. 2020. Fungal Diversity 105:1-16 ((https://doi.org/10.1007/s13225-020-00466-2)) |
| Otu08432 | 3 | k__Fungi(100);p__Basidiomycota(100);c__Agaricomycetes(100);o__Thelephorales(100);f__Bankeraceae(100);g__Hydnellum(100);s__Hydnellum_scrobiculatum(100); | Hydnellum | 13 | Symbiotroph | \|Ectomycorrhizal\| | Agaricoid-Hydnoid | NULL | Highly Probable | NULL | Rinaldi AC, et al. 2008. Fungal Diversity 33:1-45 ((http://www.fungaldiversity.org/fdp/sfdp/33-1.pdf)); Tedersoo L, et al. 2010. Mycorrhiza 20:217-263 ((https://doi.org/10.1007/s00572-009-0274-x)); Põlme S, et al. 2020. Fungal Diversity 105:1-16 ((https://doi.org/10.1007/s13225-020-00466-2)); http://mycorrhizas.info/ecmf.html ((http://mycorrhizas.info/ecmf.html)) |
| Otu08435 | 3 | k__Fungi(100);p__Ascomycota(100);c__Saccharomycetes(100);o__Saccharomycetales(100);f__Metschnikowiaceae(100);g__Metschnikowia(100);g__Metschnikowia_unclassified(100); | Metschnikowia | 13 | Saprotroph | \|Undefined Saprotroph\| | Yeast | NULL | Probable | As nectar-tap_saprotroph (Põlme et al. 2020); Decay sugar-rich substrates (Põlme et al. 2020) | Põlme S, et al. 2020. Fungal Diversity 105:1-16 ((https://doi.org/10.1007/s13225-020-00466-2)) |
| Otu08436 | 3 | k__Fungi(100);p__Ascomycota(100);c__Sordariomycetes(100);o__Hypocreales(100);o__Hypocreales_unclassified(100);o__Hypocreales_unclassified(100);o__Hypocreales_unclassified(100); | Hypocreales | 7 | Saprotroph | Undefined Saprotroph | Microfungus | NULL | Possible | NULL | Sterkenburg E, et al. 2015. New Phytologist 207:1145-1158 ((https://doi.org/10.1111/nph.13426)) |
| Otu08437 | 3 | k__Fungi(100);p__Ascomycota(100);c__Saccharomycetes(100);o__Saccharomycetales(100);f__Metschnikowiaceae(100);g__Metschnikowia(100);s__Metschnikowia_pulcherrima(67); | Metschnikowia | 13 | Saprotroph | \|Undefined Saprotroph\| | Yeast | NULL | Probable | As nectar-tap_saprotroph (Põlme et al. 2020); Decay sugar-rich substrates (Põlme et al. 2020) | Põlme S, et al. 2020. Fungal Diversity 105:1-16 ((https://doi.org/10.1007/s13225-020-00466-2)) |
| Otu08439 | 3 | k__Fungi(100);p__Ascomycota(100);c__Saccharomycetes(100);o__Saccharomycetales(100);f__Metschnikowiaceae(100);g__Metschnikowia(100);s__Metschnikowia_pulcherrima(100); | Metschnikowia | 13 | Saprotroph | \|Undefined Saprotroph\| | Yeast | NULL | Probable | As nectar-tap_saprotroph (Põlme et al. 2020); Decay sugar-rich substrates (Põlme et al. 2020) | Põlme S, et al. 2020. Fungal Diversity 105:1-16 ((https://doi.org/10.1007/s13225-020-00466-2)) |
| Otu08441 | 3 | k__Fungi(100);p__Ascomycota(100);c__Sordariomycetes(100);o__Hypocreales(100);o__Hypocreales_unclassified(100);o__Hypocreales_unclassified(100);o__Hypocreales_unclassified(100); | Hypocreales | 7 | Saprotroph | Undefined Saprotroph | Microfungus | NULL | Possible | NULL | Sterkenburg E, et al. 2015. New Phytologist 207:1145-1158 ((https://doi.org/10.1111/nph.13426)) |
| Otu08442 | 3 | k__Fungi(100);p__Basidiomycota(100);c__Microbotryomycetes(100);o__Sporidiobolales(100);f__Sporidiobolales_family_Incertae_sedis(100);g__Sporobolomyces(100);s__Sporobolomyces_griseoflavus(100); | Sporobolomyces | 13 | Pathotroph-Saprotroph | \|Fungal Parasite\|-Plant Saprotroph | Yeast | NULL | Probable | As litter_saprotroph (Põlme et al. 2020); As mycoparasite (Põlme et al. 2020); Decay fungal material (Põlme et al. 2020) | Tedersoo L, et al. 2014. Science 346:e1256688 ((https://doi.org/10.1126/science.1256688)); Purahong W, et al. 2016. Molecular Ecology 25:4059-4074 ((https://doi.org/10.1111/mec.13739)); Põlme S, et al. 2020. Fungal Diversity 105:1-16 ((https://doi.org/10.1007/s13225-020-00466-2)) |
| Otu08445 | 3 | k__Fungi(100);p__Ascomycota(100);c__Saccharomycetes(100);o__Saccharomycetales(100);f__Metschnikowiaceae(100);g__Metschnikowia(100);s__Metschnikowia_pulcherrima(100); | Metschnikowia | 13 | Saprotroph | \|Undefined Saprotroph\| | Yeast | NULL | Probable | As nectar-tap_saprotroph (Põlme et al. 2020); Decay sugar-rich substrates (Põlme et al. 2020) | Põlme S, et al. 2020. Fungal Diversity 105:1-16 ((https://doi.org/10.1007/s13225-020-00466-2)) |
| Otu08446 | 3 | k__Fungi(100);p__Basidiomycota(100);c__Agaricomycetes(100);o__Agaricales(100);f__Agaricaceae(100);g__Melanophyllum(100);s__Melanophyllum_haematospermum(100); | Melanophyllum | 13 | Saprotroph | \|Undefined Saprotroph\| | Agaricoid | NULL | Probable | As soil_saprotroph (Põlme et al. 2020); Decay soil (Põlme et al. 2020) | Tedersoo L, et al. 2014. Science 346:e1256688 ((https://doi.org/10.1126/science.1256688)); Põlme S, et al. 2020. Fungal Diversity 105:1-16 ((https://doi.org/10.1007/s13225-020-00466-2)) |
| Otu08447 | 3 | k__Fungi(100);p__Basidiomycota(100);c__Agaricomycetes(100);o__Agaricales(100);f__Cortinariaceae(100);g__Phaeocollybia(100);s__Phaeocollybia_olivacea(100); | Phaeocollybia | 13 | Symbiotroph | \|Ectomycorrhizal\| | Agaricoid | NULL | Possible | NULL | Põlme S, et al. 2020. Fungal Diversity 105:1-16 ((https://doi.org/10.1007/s13225-020-00466-2)); http://mycorrhizas.info/ecmf.html ((http://mycorrhizas.info/ecmf.html)) |
| Otu08448 | 3 | k__Fungi(100);p__Basidiomycota(100);c__Agaricomycetes(100);o__Agaricales(100);f__Cortinariaceae(100);g__Phaeocollybia(100);g__Phaeocollybia_unclassified(100); | Phaeocollybia | 13 | Symbiotroph | \|Ectomycorrhizal\| | Agaricoid | NULL | Possible | NULL | Põlme S, et al. 2020. Fungal Diversity 105:1-16 ((https://doi.org/10.1007/s13225-020-00466-2)); http://mycorrhizas.info/ecmf.html ((http://mycorrhizas.info/ecmf.html)) |
| Otu08449 | 3 | k__Fungi(100);p__Ascomycota(100);c__Saccharomycetes(100);o__Saccharomycetales(100);f__Metschnikowiaceae(100);g__Metschnikowia(100);s__Metschnikowia_pulcherrima(100); | Metschnikowia | 13 | Saprotroph | \|Undefined Saprotroph\| | Yeast | NULL | Probable | As nectar-tap_saprotroph (Põlme et al. 2020); Decay sugar-rich substrates (Põlme et al. 2020) | Põlme S, et al. 2020. Fungal Diversity 105:1-16 ((https://doi.org/10.1007/s13225-020-00466-2)) |
| Otu08453 | 3 | k__Fungi(100);p__Ascomycota(100);c__Saccharomycetes(100);o__Saccharomycetales(100);f__Metschnikowiaceae(100);g__Metschnikowia(100);s__Metschnikowia_pulcherrima(100); | Metschnikowia | 13 | Saprotroph | \|Undefined Saprotroph\| | Yeast | NULL | Probable | As nectar-tap_saprotroph (Põlme et al. 2020); Decay sugar-rich substrates (Põlme et al. 2020) | Põlme S, et al. 2020. Fungal Diversity 105:1-16 ((https://doi.org/10.1007/s13225-020-00466-2)) |
| Otu08455 | 3 | k__Fungi(100);p__Ascomycota(100);c__Saccharomycetes(100);o__Saccharomycetales(100);f__Metschnikowiaceae(100);g__Metschnikowia(100);s__Metschnikowia_pulcherrima(67); | Metschnikowia | 13 | Saprotroph | \|Undefined Saprotroph\| | Yeast | NULL | Probable | As nectar-tap_saprotroph (Põlme et al. 2020); Decay sugar-rich substrates (Põlme et al. 2020) | Põlme S, et al. 2020. Fungal Diversity 105:1-16 ((https://doi.org/10.1007/s13225-020-00466-2)) |
| Otu08456 | 3 | k__Fungi(100);p__Basidiomycota(100);c__Agaricomycetes(100);o__Agaricales(100);f__Pluteaceae(100);g__Pluteus(100);s__Pluteus_cf_plautus_UBC_F23908(100); | Pluteus | 13 | Pathotroph-Saprotroph | Bryophyte Parasite-\|Plant Saprotroph\|-Wood Saprotroph | Agaricoid | White Rot | Probable | As litter_saprotroph (Põlme et al. 2020); As white_rot (Põlme et al. 2020); Decay leaf, fruit, seed (Põlme et al. 2020) | Grgurinovic CA, Simpson JA. 2001. Fungal Diversity 8:97-127 ((http://www.fungaldiversity.org/fdp/sfdp/FD_8_97-127.pdf)); James TY, et al. 2006. Nature 443:818-822 ((https://doi.org/10.1038/nature05110)); Tedersoo L, et al. 2014. Science 346:e1256688 ((https://doi.org/10.1126/science.1256688)); Põlme S, et al. 2020. Fungal Diversity 105:1-16 ((https://doi.org/10.1007/s13225-020-00466-2)) |
| Otu08457 | 3 | k__Fungi(100);p__Ascomycota(100);c__Saccharomycetes(100);o__Saccharomycetales(100);f__Metschnikowiaceae(100);g__Metschnikowia(100);s__Metschnikowia_pulcherrima(100); | Metschnikowia | 13 | Saprotroph | \|Undefined Saprotroph\| | Yeast | NULL | Probable | As nectar-tap_saprotroph (Põlme et al. 2020); Decay sugar-rich substrates (Põlme et al. 2020) | Põlme S, et al. 2020. Fungal Diversity 105:1-16 ((https://doi.org/10.1007/s13225-020-00466-2)) |
| Otu08459 | 3 | k__Fungi(100);p__Ascomycota(100);c__Saccharomycetes(100);o__Saccharomycetales(100);f__Metschnikowiaceae(100);g__Metschnikowia(100);s__Metschnikowia_pulcherrima(100); | Metschnikowia | 13 | Saprotroph | \|Undefined Saprotroph\| | Yeast | NULL | Probable | As nectar-tap_saprotroph (Põlme et al. 2020); Decay sugar-rich substrates (Põlme et al. 2020) | Põlme S, et al. 2020. Fungal Diversity 105:1-16 ((https://doi.org/10.1007/s13225-020-00466-2)) |
| Otu08464 | 3 | k__Fungi(100);p__Ascomycota(100);c__Saccharomycetes(100);o__Saccharomycetales(100);f__Metschnikowiaceae(100);g__Metschnikowia(100);s__Metschnikowia_pulcherrima(100); | Metschnikowia | 13 | Saprotroph | \|Undefined Saprotroph\| | Yeast | NULL | Probable | As nectar-tap_saprotroph (Põlme et al. 2020); Decay sugar-rich substrates (Põlme et al. 2020) | Põlme S, et al. 2020. Fungal Diversity 105:1-16 ((https://doi.org/10.1007/s13225-020-00466-2)) |
| Otu08465 | 3 | k__Fungi(100);p__Ascomycota(100);c__Sordariomycetes(100);o__Hypocreales(100);f__Cordycipitaceae(100);g__Lecanicillium(100);s__Lecanicillium_fusisporum(100); | Lecanicillium | 13 | Pathotroph-Saprotroph-Symbiotroph | \|Animal Parasite\|-Animal Pathogen-Endophyte-Undefined Saprotroph | Microfungus | NULL | Probable | As animal_decomposer (Põlme et al. 2020); As foliar_endophyte (Põlme et al. 2020); Decay animal material (Põlme et al. 2020) | Tedersoo L, et al. 2014. Science 346:e1256688 ((https://doi.org/10.1126/science.1256688)); Põlme S, et al. 2020. Fungal Diversity 105:1-16 ((https://doi.org/10.1007/s13225-020-00466-2)) |
| Otu08467 | 3 | k__Fungi(100);p__Basidiomycota(100);c__Agaricomycetes(100);o__Agaricales(100);f__Psathyrellaceae(100);g__Psathyrella(100);g__Psathyrella_unclassified(100); | Psathyrella | 13 | Saprotroph | Plant Saprotroph-\|Wood Saprotroph\| | Agaricoid | NULL | Probable | As litter_saprotroph (Põlme et al. 2020); Decay wood, leaf, fruit, seed (Põlme et al. 2020) | Tedersoo L, et al. 2014. Science 346:e1256688 ((https://doi.org/10.1126/science.1256688)); Põlme S, et al. 2020. Fungal Diversity 105:1-16 ((https://doi.org/10.1007/s13225-020-00466-2)) |
| Otu08468 | 3 | k__Fungi(100);p__Basidiomycota(100);c__Agaricomycetes(100);o__Agaricales(100);o__Agaricales_unclassified(100);o__Agaricales_unclassified(100);o__Agaricales_unclassified(100); | Agaricales | 7 | Pathotroph-Saprotroph-Symbiotroph | Bryophyte Parasite-Dung Saprotroph-Ectomycorrhizal-Fungal Parasite-Leaf Saprotroph-Plant Parasite-Undefined Saprotroph-Wood Saprotroph | Agaricoid-Gasteroid-Microfungus-Secotioid-Yeast | NULL | Possible | Mushrooms and toadstools, Gill fungi, Agarics... lignicolous, sometimes muscicolous or fungicolous, saprobic, mycorrhizal, rarely parasitic on plants or fungi (Kirk et al. 2008) | Kirk PM et al. 2008. Dictionary of the Fungi. Tenth Edition. CAB International, Wallingford (ISBN: 978-0851998268) |
| Otu08469 | 3 | k__Fungi(100);p__Ascomycota(100);c__Saccharomycetes(100);o__Saccharomycetales(100);f__Metschnikowiaceae(100);g__Metschnikowia(100);s__Metschnikowia_pulcherrima(100); | Metschnikowia | 13 | Saprotroph | \|Undefined Saprotroph\| | Yeast | NULL | Probable | As nectar-tap_saprotroph (Põlme et al. 2020); Decay sugar-rich substrates (Põlme et al. 2020) | Põlme S, et al. 2020. Fungal Diversity 105:1-16 ((https://doi.org/10.1007/s13225-020-00466-2)) |
| Otu08470 | 3 | k__Fungi(100);p__Ascomycota(100);c__Saccharomycetes(100);o__Saccharomycetales(100);f__Metschnikowiaceae(100);g__Metschnikowia(100);s__Metschnikowia_pulcherrima(67); | Metschnikowia | 13 | Saprotroph | \|Undefined Saprotroph\| | Yeast | NULL | Probable | As nectar-tap_saprotroph (Põlme et al. 2020); Decay sugar-rich substrates (Põlme et al. 2020) | Põlme S, et al. 2020. Fungal Diversity 105:1-16 ((https://doi.org/10.1007/s13225-020-00466-2)) |
| Otu08472 | 3 | k__Fungi(100);p__Basidiomycota(100);c__Agaricomycetes(100);o__Agaricales(100);f__Agaricaceae(100);g__Lycoperdon(100);g__Lycoperdon_unclassified(100); | Lycoperdon | 13 | Saprotroph | \|Plant Saprotroph\|-Undefined Saprotroph | Gasteroid | NULL | Probable | puffball; As litter_saprotroph (Põlme et al. 2020); Common name ; Decay leaf, fruit, seed (Põlme et al. 2020) | Grgurinovic CA, Simpson JA. 2001. Fungal Diversity 8:97-127 ((http://www.fungaldiversity.org/fdp/sfdp/FD_8_97-127.pdf)); Tedersoo L, et al. 2014. Science 346:e1256688 ((https://doi.org/10.1126/science.1256688)); Põlme S, et al. 2020. Fungal Diversity 105:1-16 ((https://doi.org/10.1007/s13225-020-00466-2)) |
| Otu08476 | 3 | k__Fungi(100);p__Ascomycota(100);c__Saccharomycetes(100);o__Saccharomycetales(100);f__Metschnikowiaceae(100);g__Metschnikowia(100);s__Metschnikowia_pulcherrima(67); | Metschnikowia | 13 | Saprotroph | \|Undefined Saprotroph\| | Yeast | NULL | Probable | As nectar-tap_saprotroph (Põlme et al. 2020); Decay sugar-rich substrates (Põlme et al. 2020) | Põlme S, et al. 2020. Fungal Diversity 105:1-16 ((https://doi.org/10.1007/s13225-020-00466-2)) |
| Otu08478 | 3 | k__Fungi(100);p__Basidiomycota(100);c__Agaricomycetes(100);o__Agaricales(100);f__Agaricaceae(100);g__Morganella(100);s__Morganella_subincarnata(100); | Morganella | 13 | Saprotroph | \|Wood Saprotroph\| | Gasteroid | NULL | Probable | puffball; Common name ; Decay wood (Põlme et al. 2020) | Tedersoo L, et al. 2014. Science 346:e1256688 ((https://doi.org/10.1126/science.1256688)); Põlme S, et al. 2020. Fungal Diversity 105:1-16 ((https://doi.org/10.1007/s13225-020-00466-2)) |
| Otu08479 | 3 | k__Fungi(100);p__Ascomycota(100);c__Saccharomycetes(100);o__Saccharomycetales(100);f__Metschnikowiaceae(100);g__Metschnikowia(100);s__Metschnikowia_pulcherrima(67); | Metschnikowia | 13 | Saprotroph | \|Undefined Saprotroph\| | Yeast | NULL | Probable | As nectar-tap_saprotroph (Põlme et al. 2020); Decay sugar-rich substrates (Põlme et al. 2020) | Põlme S, et al. 2020. Fungal Diversity 105:1-16 ((https://doi.org/10.1007/s13225-020-00466-2)) |
| Otu08481 | 3 | k__Fungi(100);p__Ascomycota(100);c__Saccharomycetes(100);o__Saccharomycetales(100);f__Metschnikowiaceae(100);g__Metschnikowia(100);g__Metschnikowia_unclassified(67); | Metschnikowia | 13 | Saprotroph | \|Undefined Saprotroph\| | Yeast | NULL | Probable | As nectar-tap_saprotroph (Põlme et al. 2020); Decay sugar-rich substrates (Põlme et al. 2020) | Põlme S, et al. 2020. Fungal Diversity 105:1-16 ((https://doi.org/10.1007/s13225-020-00466-2)) |
| Otu08482 | 3 | k__Fungi(100);p__Ascomycota(100);c__Sordariomycetes(100);o__Hypocreales(100);o__Hypocreales_unclassified(100);o__Hypocreales_unclassified(100);o__Hypocreales_unclassified(100); | Hypocreales | 7 | Saprotroph | Undefined Saprotroph | Microfungus | NULL | Possible | NULL | Sterkenburg E, et al. 2015. New Phytologist 207:1145-1158 ((https://doi.org/10.1111/nph.13426)) |
| Otu08483 | 3 | k__Fungi(100);p__Basidiomycota(100);c__Agaricomycetes(100);o__Gloeophyllales(100);f__Gloeophyllaceae(100);g__Gloeophyllum(100);s__Gloeophyllum_sepiarium(100); | Gloeophyllum | 13 | Saprotroph | \|Wood Saprotroph\| | Polyporoid | Brown Rot | Highly Probable | As brown-rot (Põlme et al. 2020); Decay wood (Põlme et al. 2020) | Gilbertson RL, Ryvarden L. 1987-1987. North American Polypores. Fungiflora, Oslo (ISBN: 978-0945345060); Põlme S, et al. 2020. Fungal Diversity 105:1-16 ((https://doi.org/10.1007/s13225-020-00466-2)) |
| Otu08485 | 3 | k__Fungi(100);p__Ascomycota(100);c__Saccharomycetes(100);o__Saccharomycetales(100);f__Metschnikowiaceae(100);g__Metschnikowia(100);s__Metschnikowia_pulcherrima(100); | Metschnikowia | 13 | Saprotroph | \|Undefined Saprotroph\| | Yeast | NULL | Probable | As nectar-tap_saprotroph (Põlme et al. 2020); Decay sugar-rich substrates (Põlme et al. 2020) | Põlme S, et al. 2020. Fungal Diversity 105:1-16 ((https://doi.org/10.1007/s13225-020-00466-2)) |
| Otu08487 | 3 | k__Fungi(100);p__Ascomycota(100);c__Saccharomycetes(100);o__Saccharomycetales(100);f__Metschnikowiaceae(100);g__Metschnikowia(100);s__Metschnikowia_pulcherrima(100); | Metschnikowia | 13 | Saprotroph | \|Undefined Saprotroph\| | Yeast | NULL | Probable | As nectar-tap_saprotroph (Põlme et al. 2020); Decay sugar-rich substrates (Põlme et al. 2020) | Põlme S, et al. 2020. Fungal Diversity 105:1-16 ((https://doi.org/10.1007/s13225-020-00466-2)) |
| Otu08488 | 3 | k__Fungi(100);p__Ascomycota(100);c__Saccharomycetes(100);o__Saccharomycetales(100);f__Metschnikowiaceae(100);g__Metschnikowia(100);s__Metschnikowia_pulcherrima(67); | Metschnikowia | 13 | Saprotroph | \|Undefined Saprotroph\| | Yeast | NULL | Probable | As nectar-tap_saprotroph (Põlme et al. 2020); Decay sugar-rich substrates (Põlme et al. 2020) | Põlme S, et al. 2020. Fungal Diversity 105:1-16 ((https://doi.org/10.1007/s13225-020-00466-2)) |
| Otu08489 | 3 | k__Fungi(100);p__Ascomycota(100);c__Sordariomycetes(100);o__Sordariales(100);f__Chaetomiaceae(100);g__Chaetomium(100);s__Chaetomium_sp_UPML025(100); | Chaetomium | 13 | Pathotroph-Saprotroph-Symbiotroph | Animal Pathogen-Dung Saprotroph-Endophyte-Epiphyte-Plant Pathogen-\|Plant Saprotroph\|-Wood Saprotroph | Microfungus | Soft Rot | Possible | As foliar_endophyte (Põlme et al. 2020); As litter_saprotroph (Põlme et al. 2020); culture (Busby et al. 2016, David et al. 2016); Likely opportunistic human pathogen (Irinyi et al. 2015); culture (Massimo et al. 2015); Endophyte detection method; Decay wood, leaf, fruit, seed, soil, dung (Põlme et al. 2020); derived commodities (Cannon & Kirk 2007); Epiphyte detection method; Saprobic, acting as key decay organisms of plant materials and plant | Seehann G, et al. 1975. List of Fungi in Soft-Rot Tissue Tests. Institute fur Holzbiologie und Holzschutz, Bundesforschungsanstalt fur Forst - und Holzwirtshaft, Hamburg - Lohbrugge, Germany; Cannon PF, Kirk PM. 2007. Fungal Families of the World. CAB International, Cambridge ((https://isbnsearch.org/isbn/9780851998275)); Bills GF, et al. 2013. Current Opinion in Microbiology 16:549-565 ((https://doi.org/10.1016/j.mib.2013.08.001)); Tedersoo L, et al. 2014. Science 346:e1256688 ((https://doi.org/10.1126/science.1256688)); Irinyi L, et al. 2015. Medical Mycology 53:313-337 ((https://doi.org/10.1093/mmy/myv008)); Busby PE, et al. 2016. New Phytologist 209:1681-1692 ((https://doi.org/10.1111/nph.13742)); Põlme S, et al. 2020. Fungal Diversity 105:1-16 ((https://doi.org/10.1007/s13225-020-00466-2)) |
| Otu08490 | 3 | k__Fungi(100);p__Ascomycota(100);c__Dothideomycetes(100);o__Capnodiales(100);f__Mycosphaerellaceae(100);f__Mycosphaerellaceae_unclassified(100);f__Mycosphaerellaceae_unclassified(100); | Mycosphaerellaceae | 9 | Pathotroph-Saprotroph | Plant Pathogen-Undefined Saprotroph | Microfungus | NULL | Probable | Biotrophic, necrotrophic or saprobic on various plant tissues (Cannon & Kirk 2007) | Cannon PF, Kirk PM. 2007. Fungal Families of the World. CAB International, Cambridge (ISBN: 978-0851998275) |
| Otu08491 | 3 | k__Fungi(100);p__Ascomycota(100);c__Saccharomycetes(100);o__Saccharomycetales(100);f__Metschnikowiaceae(100);g__Metschnikowia(100);s__Metschnikowia_pulcherrima(67); | Metschnikowia | 13 | Saprotroph | \|Undefined Saprotroph\| | Yeast | NULL | Probable | As nectar-tap_saprotroph (Põlme et al. 2020); Decay sugar-rich substrates (Põlme et al. 2020) | Põlme S, et al. 2020. Fungal Diversity 105:1-16 ((https://doi.org/10.1007/s13225-020-00466-2)) |
| Otu08492 | 3 | k__Fungi(100);p__Ascomycota(100);c__Sordariomycetes(100);o__Hypocreales(100);o__Hypocreales_unclassified(100);o__Hypocreales_unclassified(100);o__Hypocreales_unclassified(100); | Hypocreales | 7 | Saprotroph | Undefined Saprotroph | Microfungus | NULL | Possible | NULL | Sterkenburg E, et al. 2015. New Phytologist 207:1145-1158 ((https://doi.org/10.1111/nph.13426)) |
| Otu08497 | 3 | k__Fungi(100);p__Basidiomycota(100);c__Agaricomycetes(100);o__Agaricales(100);o__Agaricales_unclassified(100);o__Agaricales_unclassified(100);o__Agaricales_unclassified(100); | Agaricales | 7 | Pathotroph-Saprotroph-Symbiotroph | Bryophyte Parasite-Dung Saprotroph-Ectomycorrhizal-Fungal Parasite-Leaf Saprotroph-Plant Parasite-Undefined Saprotroph-Wood Saprotroph | Agaricoid-Gasteroid-Microfungus-Secotioid-Yeast | NULL | Possible | Mushrooms and toadstools, Gill fungi, Agarics... lignicolous, sometimes muscicolous or fungicolous, saprobic, mycorrhizal, rarely parasitic on plants or fungi (Kirk et al. 2008) | Kirk PM et al. 2008. Dictionary of the Fungi. Tenth Edition. CAB International, Wallingford (ISBN: 978-0851998268) |
| Otu08501 | 3 | k__Fungi(100);p__Ascomycota(100);c__Saccharomycetes(100);o__Saccharomycetales(100);f__Metschnikowiaceae(100);g__Metschnikowia(100);s__Metschnikowia_pulcherrima(67); | Metschnikowia | 13 | Saprotroph | \|Undefined Saprotroph\| | Yeast | NULL | Probable | As nectar-tap_saprotroph (Põlme et al. 2020); Decay sugar-rich substrates (Põlme et al. 2020) | Põlme S, et al. 2020. Fungal Diversity 105:1-16 ((https://doi.org/10.1007/s13225-020-00466-2)) |
| Otu08503 | 3 | k__Fungi(100);p__Basidiomycota(100);c__Agaricomycetes(100);o__Polyporales(100);f__Meripilaceae(100);f__Meripilaceae_unclassified(100);f__Meripilaceae_unclassified(100); | Meripilaceae | 9 | Pathotroph-Saprotroph | Plant Pathogen-Undefined Saprotroph | Polyporoid | NULL | Probable | Saprobic or parasitic on hardwood trees, often growing from buried roots and causing white rot (Cannon & Kirk 2007) | Cannon PF, Kirk PM. 2007. Fungal Families of the World. CAB International, Cambridge (ISBN: 978-0851998275) |
| Otu08504 | 3 | k__Fungi(100);p__Basidiomycota(100);c__Agaricomycetes(100);o__Polyporales(100);f__Fomitopsidaceae(100);g__Antrodia(100);s__Antrodia_pulvinascens(100); | Antrodia | 13 | Saprotroph | \|Wood Saprotroph\| | Corticioid | Brown Rot | Highly Probable | As brown-rot (Põlme et al. 2020); Decay wood (Põlme et al. 2020) | Gilbertson RL, Ryvarden L. 1987-1987. North American Polypores. Fungiflora, Oslo (ISBN: 978-0945345060); Põlme S, et al. 2020. Fungal Diversity 105:1-16 ((https://doi.org/10.1007/s13225-020-00466-2)) |
| Otu08506 | 3 | k__Fungi(100);p__Ascomycota(100);c__Saccharomycetes(100);o__Saccharomycetales(100);f__Metschnikowiaceae(100);g__Metschnikowia(100);g__Metschnikowia_unclassified(100); | Metschnikowia | 13 | Saprotroph | \|Undefined Saprotroph\| | Yeast | NULL | Probable | As nectar-tap_saprotroph (Põlme et al. 2020); Decay sugar-rich substrates (Põlme et al. 2020) | Põlme S, et al. 2020. Fungal Diversity 105:1-16 ((https://doi.org/10.1007/s13225-020-00466-2)) |
| Otu08508 | 3 | k__Fungi(100);p__Basidiomycota(100);c__Exobasidiomycetes(100);o__Entylomatales(100);f__Entylomataceae(100);g__Entyloma(100);s__Entyloma_microsporum(100); | Entyloma | 13 | Pathotroph | \|Plant Pathogen\| | Smut | NULL | Probable | NULL | Tedersoo L, et al. 2014. Science 346:e1256688 ((https://doi.org/10.1126/science.1256688)); Põlme S, et al. 2020. Fungal Diversity 105:1-16 ((https://doi.org/10.1007/s13225-020-00466-2)) |
| Otu08509 | 3 | k__Fungi(100);p__Ascomycota(100);c__Saccharomycetes(100);o__Saccharomycetales(100);f__Metschnikowiaceae(100);g__Metschnikowia(100);s__Metschnikowia_pulcherrima(100); | Metschnikowia | 13 | Saprotroph | \|Undefined Saprotroph\| | Yeast | NULL | Probable | As nectar-tap_saprotroph (Põlme et al. 2020); Decay sugar-rich substrates (Põlme et al. 2020) | Põlme S, et al. 2020. Fungal Diversity 105:1-16 ((https://doi.org/10.1007/s13225-020-00466-2)) |
| Otu08510 | 3 | k__Fungi(100);p__Ascomycota(100);c__Saccharomycetes(100);o__Saccharomycetales(100);f__Metschnikowiaceae(100);g__Metschnikowia(100);s__Metschnikowia_pulcherrima(100); | Metschnikowia | 13 | Saprotroph | \|Undefined Saprotroph\| | Yeast | NULL | Probable | As nectar-tap_saprotroph (Põlme et al. 2020); Decay sugar-rich substrates (Põlme et al. 2020) | Põlme S, et al. 2020. Fungal Diversity 105:1-16 ((https://doi.org/10.1007/s13225-020-00466-2)) |
| Otu08513 | 3 | k__Fungi(100);p__Ascomycota(100);c__Saccharomycetes(100);o__Saccharomycetales(100);f__Metschnikowiaceae(100);g__Metschnikowia(100);s__Metschnikowia_pulcherrima(100); | Metschnikowia | 13 | Saprotroph | \|Undefined Saprotroph\| | Yeast | NULL | Probable | As nectar-tap_saprotroph (Põlme et al. 2020); Decay sugar-rich substrates (Põlme et al. 2020) | Põlme S, et al. 2020. Fungal Diversity 105:1-16 ((https://doi.org/10.1007/s13225-020-00466-2)) |
| Otu08514 | 3 | k__Fungi(100);p__Ascomycota(100);c__Sordariomycetes(100);o__Sordariales(100);f__Lasiosphaeriaceae(100);g__Lasiosphaeris(100);s__Lasiosphaeris_sp_4_ANM_2011(100); | Lasiosphaeris | 13 | Saprotroph | \|Undefined Saprotroph\| | Microfungus | NULL | Probable | NULL | Tedersoo L, et al. 2014. Science 346:e1256688 ((https://doi.org/10.1126/science.1256688)); Põlme S, et al. 2020. Fungal Diversity 105:1-16 ((https://doi.org/10.1007/s13225-020-00466-2)) |
| Otu08515 | 3 | k__Fungi(100);p__Ascomycota(100);c__Saccharomycetes(100);o__Saccharomycetales(100);f__Metschnikowiaceae(100);g__Metschnikowia(100);s__Metschnikowia_pulcherrima(100); | Metschnikowia | 13 | Saprotroph | \|Undefined Saprotroph\| | Yeast | NULL | Probable | As nectar-tap_saprotroph (Põlme et al. 2020); Decay sugar-rich substrates (Põlme et al. 2020) | Põlme S, et al. 2020. Fungal Diversity 105:1-16 ((https://doi.org/10.1007/s13225-020-00466-2)) |
| Otu08517 | 3 | k__Fungi(100);p__Zygomycota(100);c__Zygomycota_class_Incertae_sedis(100);o__Mucorales(100);f__Backusellaceae(100);g__Backusella(100);s__Backusella_lamprospora(100); | Backusella | 13 | Saprotroph | Dung Saprotroph-\|Undefined Saprotroph\| | Microfungus-Sporophores | NULL | Probable | As soil_saprotroph (Põlme et al. 2020); Decay soil, dung (Põlme et al. 2020) | Walther G, et al. 2013. Persoonia 30:11-47 ((https://doi.org/10.3767/003158513X665070)); Põlme S, et al. 2020. Fungal Diversity 105:1-16 ((https://doi.org/10.1007/s13225-020-00466-2)) |
| Otu08518 | 3 | k__Fungi(100);p__Ascomycota(100);c__Saccharomycetes(100);o__Saccharomycetales(100);f__Metschnikowiaceae(100);g__Metschnikowia(100);s__Metschnikowia_pulcherrima(100); | Metschnikowia | 13 | Saprotroph | \|Undefined Saprotroph\| | Yeast | NULL | Probable | As nectar-tap_saprotroph (Põlme et al. 2020); Decay sugar-rich substrates (Põlme et al. 2020) | Põlme S, et al. 2020. Fungal Diversity 105:1-16 ((https://doi.org/10.1007/s13225-020-00466-2)) |
| Otu08519 | 3 | k__Fungi(100);p__Ascomycota(100);c__Saccharomycetes(100);o__Saccharomycetales(100);f__Metschnikowiaceae(100);g__Metschnikowia(100);s__Metschnikowia_pulcherrima(100); | Metschnikowia | 13 | Saprotroph | \|Undefined Saprotroph\| | Yeast | NULL | Probable | As nectar-tap_saprotroph (Põlme et al. 2020); Decay sugar-rich substrates (Põlme et al. 2020) | Põlme S, et al. 2020. Fungal Diversity 105:1-16 ((https://doi.org/10.1007/s13225-020-00466-2)) |
| Otu08521 | 3 | k__Fungi(100);p__Basidiomycota(100);c__Pucciniomycetes(100);o__Pucciniales(100);f__Pucciniaceae(100);g__Puccinia(100);s__Puccinia_graminis_f_sp_tritici(100); | Puccinia | 13 | Pathotroph | \|Plant Pathogen\| | Rust | NULL | Highly Probable | NULL | Tedersoo L, et al. 2014. Science 346:e1256688 ((https://doi.org/10.1126/science.1256688)); Põlme S, et al. 2020. Fungal Diversity 105:1-16 ((https://doi.org/10.1007/s13225-020-00466-2)) |
| Otu08522 | 3 | k__Fungi(100);p__Ascomycota(100);c__Saccharomycetes(100);o__Saccharomycetales(100);f__Metschnikowiaceae(100);g__Metschnikowia(100);s__Metschnikowia_pulcherrima(100); | Metschnikowia | 13 | Saprotroph | \|Undefined Saprotroph\| | Yeast | NULL | Probable | As nectar-tap_saprotroph (Põlme et al. 2020); Decay sugar-rich substrates (Põlme et al. 2020) | Põlme S, et al. 2020. Fungal Diversity 105:1-16 ((https://doi.org/10.1007/s13225-020-00466-2)) |
| Otu08523 | 3 | k__Fungi(100);p__Ascomycota(100);c__Dothideomycetes(100);o__Dothideomycetes_order_Incertae_sedis(100);f__Pseudeurotiaceae(100);g__Pseudeurotium(100);g__Pseudeurotium_unclassified(100); | Pseudeurotium | 13 | Saprotroph | \|Dung Saprotroph\|-Undefined Saprotroph | Microfungus | NULL | Probable | As soil_saprotroph (Põlme et al. 2020); Decay soil, dung (Põlme et al. 2020) | Tedersoo L, et al. 2014. Science 346:e1256688 ((https://doi.org/10.1126/science.1256688)); Põlme S, et al. 2020. Fungal Diversity 105:1-16 ((https://doi.org/10.1007/s13225-020-00466-2)) |
| Otu08526 | 3 | k__Fungi(100);p__Ascomycota(100);c__Saccharomycetes(100);o__Saccharomycetales(100);f__Metschnikowiaceae(100);g__Metschnikowia(100);s__Metschnikowia_pulcherrima(100); | Metschnikowia | 13 | Saprotroph | \|Undefined Saprotroph\| | Yeast | NULL | Probable | As nectar-tap_saprotroph (Põlme et al. 2020); Decay sugar-rich substrates (Põlme et al. 2020) | Põlme S, et al. 2020. Fungal Diversity 105:1-16 ((https://doi.org/10.1007/s13225-020-00466-2)) |
| Otu08529 | 3 | k__Fungi(100);p__Zygomycota(100);c__Zygomycota_class_Incertae_sedis(100);o__Mortierellales(100);f__unclassified_Mortierellales(100);g__unclassified_Mortierellales(100);s__Mortierellales_sp(100); | Mortierellales | 7 | Saprotroph-Symbiotroph | Endophyte-Soil Saprotroph-Undefined Saprotroph | Microfungus | NULL | Possible | Saprobic in soil and thus frequently isolated from soil and material contaminated with soil such as hay or rotten silage and spoil tips (Cannon & Kirk 2007) | Cannon PF, Kirk PM. 2007. Fungal Families of the World. CAB International, Cambridge (ISBN: 978-0851998275); Sterkenburg E, et al. 2015. New Phytologist 207:1145-1158 ((https://doi.org/10.1111/nph.13426)); David AS, et al. 2016. Microbial Ecology 71:912-926 ((https://doi.org/10.1007/s00248-015-0712-6)) |
| Otu08530 | 3 | k__Fungi(100);p__Ascomycota(100);c__Saccharomycetes(100);o__Saccharomycetales(100);f__Metschnikowiaceae(100);g__Metschnikowia(100);s__Metschnikowia_pulcherrima(100); | Metschnikowia | 13 | Saprotroph | \|Undefined Saprotroph\| | Yeast | NULL | Probable | As nectar-tap_saprotroph (Põlme et al. 2020); Decay sugar-rich substrates (Põlme et al. 2020) | Põlme S, et al. 2020. Fungal Diversity 105:1-16 ((https://doi.org/10.1007/s13225-020-00466-2)) |
| Otu08532 | 3 | k__Fungi(100);p__Ascomycota(100);c__Dothideomycetes(100);o__Capnodiales(100);f__Mycosphaerellaceae(100);g__Mycosphaerella(100);s__Mycosphaerella_nyssicola(100); | Mycosphaerella | 13 | Pathotroph-Saprotroph | \|Plant Pathogen\|-Plant Saprotroph | Microfungus | NULL | Probable | As litter_saprotroph (Põlme et al. 2020); Decay leaf, fruit, seed (Põlme et al. 2020) | Tedersoo L, et al. 2014. Science 346:e1256688 ((https://doi.org/10.1126/science.1256688)); Põlme S, et al. 2020. Fungal Diversity 105:1-16 ((https://doi.org/10.1007/s13225-020-00466-2)) |
| Otu08535 | 3 | k__Fungi(100);p__Zygomycota(100);c__Zygomycota_class_Incertae_sedis(100);o__Mucorales(100);f__Umbelopsidaceae(100);g__Umbelopsis(100);s__Umbelopsis_ramanniana(100); | Umbelopsis | 13 | Saprotroph | \|Undefined Saprotroph\| | Microfungus | NULL | Probable | As soil_saprotroph (Põlme et al. 2020); Decay roots, soil (Põlme et al. 2020) | Tedersoo L, et al. 2014. Science 346:e1256688 ((https://doi.org/10.1126/science.1256688)); Põlme S, et al. 2020. Fungal Diversity 105:1-16 ((https://doi.org/10.1007/s13225-020-00466-2)) |
| Otu08541 | 3 | k__Fungi(100);p__Basidiomycota(100);c__Agaricomycetes(100);o__Agaricales(100);f__Inocybaceae(100);g__Crepidotus(100);s__Crepidotus_applanatus(100); | Crepidotus | 13 | Saprotroph | \|Wood Saprotroph\| | Agaricoid | NULL | Highly Probable | Decay wood (Põlme et al. 2020) | Grgurinovic CA, Simpson JA. 2001. Fungal Diversity 8:97-127 ((http://www.fungaldiversity.org/fdp/sfdp/FD_8_97-127.pdf)); Aime MC, et al. 2005. American Journal of Botany 92:74-82 ((https://doi.org/10.3732/ajb.92.1.74)); Tedersoo L, et al. 2014. Science 346:e1256688 ((https://doi.org/10.1126/science.1256688)); Põlme S, et al. 2020. Fungal Diversity 105:1-16 ((https://doi.org/10.1007/s13225-020-00466-2)) |
| Otu08546 | 3 | k__Fungi(100);p__Basidiomycota(100);c__Agaricomycetes(100);o__Auriculariales(100);f__Auriculariales_family_Incertae_sedis(100);g__Auricularia(100);s__Auricularia_sp(100); | Auricularia | 13 | Saprotroph | Undefined Saprotroph-\|Wood Saprotroph\| | Tremelloid | NULL | Probable | Decay wood (Põlme et al. 2020) | Tedersoo L, et al. 2014. Science 346:e1256688 ((https://doi.org/10.1126/science.1256688)); Põlme S, et al. 2020. Fungal Diversity 105:1-16 ((https://doi.org/10.1007/s13225-020-00466-2)) |

<table>
<tr><td>Otu08548</td><td>3</td><td>k__Fungi(100);p__Basidiomycota(100);c__Agaricomycetes(100);o__Auriculariales(100);f__Auriculariales_family_Incertae_sedis(100);g__Auricularia(100);g__Auricularia_unclassified(100);</td><td>Auricularia</td><td>13</td><td>Saprotroph</td><td>Undefined Saprotroph-|Wood Saprotroph|</td><td>Tremelloid</td><td>NULL</td><td>Probable</td><td>Decay wood (Põlme et al. 2020)</td><td>Tedersoo L, et al. 2014. Science 346:e1256688 ((https://doi.org/10.1126/science.1256688)); Põlme S, et al. 2020. Fungal Diversity 105:1-16 ((https://doi.org/10.1007/s13225-020-00466-2))</td></tr>
<tr><td>Otu08550</td><td>3</td><td>k__Fungi(100);p__Ascomycota(100);c__Sordariomycetes(100);o__Chaetosphaeriales(100);f__Chaetosphaeriaceae(100);f__Chaetosphaeriaceae_unclassified(67);f__Chaetosphaeriaceae_unclassified(67);</td><td>Chaetosphaeriaceae</td><td>9</td><td>Saprotroph</td><td>Plant Saprotroph-Wood Saprotroph</td><td>Microfungus</td><td>NULL</td><td>Probable</td><td>Saprobes associated with decaying woody and herbaceous plant material, including timber and timber products in damp buildings (Cannon & Kirk 2007)</td><td>Cannon PF, Kirk PM. 2007. Fungal Families of the World. CAB International, Cambridge (ISBN: 978-0851998275)</td></tr>
<tr><td>Otu08551</td><td>3</td><td>k__Fungi(100);p__Basidiomycota(100);c__Pucciniomycetes(100);o__Pucciniales(100);f__Pucciniaceae(100);g__Puccinia(100);s__Puccinia_brachypodii_var._arrhenatheri(100);</td><td>Puccinia</td><td>13</td><td>Pathotroph</td><td>|Plant Pathogen|</td><td>Rust</td><td>NULL</td><td>Highly Probable</td><td>NULL</td><td>Tedersoo L, et al. 2014. Science 346:e1256688 ((https://doi.org/10.1126/science.1256688)); Põlme S, et al. 2020. Fungal Diversity 105:1-16 ((https://doi.org/10.1007/s13225-020-00466-2))</td></tr>
<tr><td>Otu08552</td><td>3</td><td>k__Fungi(100);p__Ascomycota(100);c__Saccharomycetes(100);o__Saccharomycetales(100);f__Metschnikowiaceae(100);g__Metschnikowia(100);s__Metschnikowia_pulcherrima(67);</td><td>Metschnikowia</td><td>13</td><td>Saprotroph</td><td>|Undefined Saprotroph|</td><td>Yeast</td><td>NULL</td><td>Probable</td><td>As nectar-tap_saprotroph (Põlme et al. 2020); Decay sugar-rich substrates (Põlme et al. 2020)</td><td>Põlme S, et al. 2020. Fungal Diversity 105:1-16 ((https://doi.org/10.1007/s13225-020-00466-2))</td></tr>
<tr><td>Otu08554</td><td>3</td><td>k__Fungi(100);p__Ascomycota(100);c__Saccharomycetes(100);o__Saccharomycetales(100);f__Saccharomycetales_family_Incertae_sedis(100);g__Candida(100);g__Candida_unclassified(67);</td><td>Candida</td><td>13</td><td>Pathotroph-Saprotroph-Symbiotroph</td><td>Animal Pathogen-Endophyte-Endosymbiont-Epiphyte-|Undefined Saprotroph|</td><td>Dimorphic Yeast-Yeast</td><td>NULL</td><td>Probable</td><td>As nectar-tap_saprotroph (Põlme et al. 2020); Decay sugar-rich substrates (Põlme et al. 2020); This asexual genus is polyphyletic and contain species that function very differently depend on what they are. Caution should be taken when interpreting guilds for Candida species.</td><td>Manolakaki D, et al. 2010. Virulence. 1: 367-375 ((https://doi.org/10.4161/viru.1.5.12796)); Mekbib SB, et al. 2011. Tropical Plant Pathology 36:233-240 ((https://doi.org/10.1590/S1982-56762011000400004)); Põlme S, et al. 2020. Fungal Diversity 105:1-16 ((https://doi.org/10.1007/s13225-020-00466-2))</td></tr>
<tr><td>Otu08556</td><td>3</td><td>k__Fungi(100);p__Ascomycota(100);c__Dothideomycetes(100);o__Pleosporales(100);f__Phaeosphaeriaceae(100);f__Phaeosphaeriaceae_unclassified(67);f__Phaeosphaeriaceae_unclassified(67);</td><td>Phaeosphaeriaceae</td><td>9</td><td>Pathotroph-Saprotroph</td><td>Fungal Parasite-Plant Pathogen-Plant Saprotroph</td><td>Microfungus</td><td>NULL</td><td>Probable</td><td>Necrotrophic or saprobic on a wide range of plants, especially monocotyledons; a few species parasitic on other fungi (Cannon & Kirk 2007)</td><td>Cannon PF, Kirk PM. 2007. Fungal Families of the World. CAB International, Cambridge (ISBN: 978-0851998275)</td></tr>
<tr><td>Otu08557</td><td>3</td><td>k__Fungi(100);p__Ascomycota(100);c__Saccharomycetes(100);o__Saccharomycetales(100);f__Metschnikowiaceae(100);g__Metschnikowia(100);s__Metschnikowia_pulcherrima(67);</td><td>Metschnikowia</td><td>13</td><td>Saprotroph</td><td>|Undefined Saprotroph|</td><td>Yeast</td><td>NULL</td><td>Probable</td><td>As nectar-tap_saprotroph (Põlme et al. 2020); Decay sugar-rich substrates (Põlme et al. 2020)</td><td>Põlme S, et al. 2020. Fungal Diversity 105:1-16 ((https://doi.org/10.1007/s13225-020-00466-2))</td></tr>
<tr><td>Otu08558</td><td>3</td><td>k__Fungi(100);p__Ascomycota(100);c__Saccharomycetes(100);o__Saccharomycetales(100);f__Metschnikowiaceae(100);g__Metschnikowia(100);s__Metschnikowia_pulcherrima(100);</td><td>Metschnikowia</td><td>13</td><td>Saprotroph</td><td>|Undefined Saprotroph|</td><td>Yeast</td><td>NULL</td><td>Probable</td><td>As nectar-tap_saprotroph (Põlme et al. 2020); Decay sugar-rich substrates (Põlme et al. 2020)</td><td>Põlme S, et al. 2020. Fungal Diversity 105:1-16 ((https://doi.org/10.1007/s13225-020-00466-2))</td></tr>
<tr><td>Otu08561</td><td>3</td><td>k__Fungi(100);p__Ascomycota(100);c__Saccharomycetes(100);o__Saccharomycetales(100);f__Metschnikowiaceae(100);g__Metschnikowia(100);s__Metschnikowia_pulcherrima(67);</td><td>Metschnikowia</td><td>13</td><td>Saprotroph</td><td>|Undefined Saprotroph|</td><td>Yeast</td><td>NULL</td><td>Probable</td><td>As nectar-tap_saprotroph (Põlme et al. 2020); Decay sugar-rich substrates (Põlme et al. 2020)</td><td>Põlme S, et al. 2020. Fungal Diversity 105:1-16 ((https://doi.org/10.1007/s13225-020-00466-2))</td></tr>
<tr><td>Otu08562</td><td>3</td><td>k__Fungi(100);p__Ascomycota(100);c__Dothideomycetes(100);o__Capnodiales(100);f__Mycosphaerellaceae(100);g__Pseudocercospora(100);s__Pseudocercospora_lyoniae(100);</td><td>Pseudocercospora</td><td>13</td><td>Pathotroph</td><td>|Plant Pathogen|</td><td>Microfungus</td><td>NULL</td><td>Probable</td><td>NULL</td><td>Tedersoo L, et al. 2014. Science 346:e1256688 ((https://doi.org/10.1126/science.1256688)); Põlme S, et al. 2020. Fungal Diversity 105:1-16 ((https://doi.org/10.1007/s13225-020-00466-2))</td></tr>
<tr><td>Otu08563</td><td>3</td><td>k__Fungi(100);p__Ascomycota(100);c__Saccharomycetes(100);o__Saccharomycetales(100);f__Metschnikowiaceae(100);g__Metschnikowia(100);s__Metschnikowia_pulcherrima(100);</td><td>Metschnikowia</td><td>13</td><td>Saprotroph</td><td>|Undefined Saprotroph|</td><td>Yeast</td><td>NULL</td><td>Probable</td><td>As nectar-tap_saprotroph (Põlme et al. 2020); Decay sugar-rich substrates (Põlme et al. 2020)</td><td>Põlme S, et al. 2020. Fungal Diversity 105:1-16 ((https://doi.org/10.1007/s13225-020-00466-2))</td></tr>
<tr><td>Otu08564</td><td>3</td><td>k__Fungi(100);p__Ascomycota(100);c__Saccharomycetes(100);o__Saccharomycetales(100);f__Metschnikowiaceae(100);g__Metschnikowia(100);s__Metschnikowia_pulcherrima(100);</td><td>Metschnikowia</td><td>13</td><td>Saprotroph</td><td>|Undefined Saprotroph|</td><td>Yeast</td><td>NULL</td><td>Probable</td><td>As nectar-tap_saprotroph (Põlme et al. 2020); Decay sugar-rich substrates (Põlme et al. 2020)</td><td>Põlme S, et al. 2020. Fungal Diversity 105:1-16 ((https://doi.org/10.1007/s13225-020-00466-2))</td></tr>
<tr><td>Otu08566</td><td>3</td><td>k__Fungi(100);p__Ascomycota(100);c__Saccharomycetes(100);o__Saccharomycetales(100);f__Metschnikowiaceae(100);g__Metschnikowia(100);s__Metschnikowia_pulcherrima(67);</td><td>Metschnikowia</td><td>13</td><td>Saprotroph</td><td>|Undefined Saprotroph|</td><td>Yeast</td><td>NULL</td><td>Probable</td><td>As nectar-tap_saprotroph (Põlme et al. 2020); Decay sugar-rich substrates (Põlme et al. 2020)</td><td>Põlme S, et al. 2020. Fungal Diversity 105:1-16 ((https://doi.org/10.1007/s13225-020-00466-2))</td></tr>
<tr><td>Otu08567</td><td>3</td><td>k__Fungi(100);p__Ascomycota(100);c__Saccharomycetes(100);o__Saccharomycetales(100);f__Metschnikowiaceae(100);g__Metschnikowia(100);s__Metschnikowia_pulcherrima(100);</td><td>Metschnikowia</td><td>13</td><td>Saprotroph</td><td>|Undefined Saprotroph|</td><td>Yeast</td><td>NULL</td><td>Probable</td><td>As nectar-tap_saprotroph (Põlme et al. 2020); Decay sugar-rich substrates (Põlme et al. 2020)</td><td>Põlme S, et al. 2020. Fungal Diversity 105:1-16 ((https://doi.org/10.1007/s13225-020-00466-2))</td></tr>
<tr><td>Otu08570</td><td>3</td><td>k__Fungi(100);p__Ascomycota(100);c__Dothideomycetes(100);o__Pleosporales(100);f__Montagnulaceae(100);g__unclassified_Montagnulaceae(67);s__Montagnulaceae_sp(67);</td><td>Montagnulaceae</td><td>9</td><td>Pathotroph-Saprotroph</td><td>Fungal Parasite-Plant Pathogen-Undefined Saprotroph</td><td>Microfungus</td><td>NULL</td><td>Probable</td><td>Species in this clade can be saprobic, plant pathogenic, or mycoparasitic (Zhang et al. 2009)</td><td>Zhang Y, et al. 2009. Studies in Mycology 64:85-102 ((https://doi.org/10.3114/sim.2009.64.04))</td></tr>
<tr><td>Otu08571</td><td>3</td><td>k__Fungi(100);p__Ascomycota(100);c__Saccharomycetes(100);o__Saccharomycetales(100);f__Metschnikowiaceae(100);g__Metschnikowia(100);s__Metschnikowia_pulcherrima(67);</td><td>Metschnikowia</td><td>13</td><td>Saprotroph</td><td>|Undefined Saprotroph|</td><td>Yeast</td><td>NULL</td><td>Probable</td><td>As nectar-tap_saprotroph (Põlme et al. 2020); Decay sugar-rich substrates (Põlme et al. 2020)</td><td>Põlme S, et al. 2020. Fungal Diversity 105:1-16 ((https://doi.org/10.1007/s13225-020-00466-2))</td></tr>
<tr><td>Otu08573</td><td>3</td><td>k__Fungi(100);p__Ascomycota(100);c__Saccharomycetes(100);o__Saccharomycetales(100);f__Metschnikowiaceae(100);g__Metschnikowia(100);s__Metschnikowia_pulcherrima(67);</td><td>Metschnikowia</td><td>13</td><td>Saprotroph</td><td>|Undefined Saprotroph|</td><td>Yeast</td><td>NULL</td><td>Probable</td><td>As nectar-tap_saprotroph (Põlme et al. 2020); Decay sugar-rich substrates (Põlme et al. 2020)</td><td>Põlme S, et al. 2020. Fungal Diversity 105:1-16 ((https://doi.org/10.1007/s13225-020-00466-2))</td></tr>
<tr><td>Otu08575</td><td>3</td><td>k__Fungi(100);p__Ascomycota(100);c__Sordariomycetes(100);o__Sordariales(100);f__Lasiosphaeriaceae(100);f__Lasiosphaeriaceae_unclassified(67);f__Lasiosphaeriaceae_unclassified(67);</td><td>Lasiosphaeriaceae</td><td>9</td><td>Saprotroph</td><td>Undefined Saprotroph</td><td>Microfungus</td><td>NULL</td><td>Probable</td><td>Saprobes found dung, soil and rotting vegetation (Cannon & Kirk 2007)</td><td>Cannon PF, Kirk PM. 2007. Fungal Families of the World. CAB International, Cambridge (ISBN: 978-0851998275)</td></tr>
<tr><td>Otu08576</td><td>3</td><td>k__Fungi(100);p__Ascomycota(100);c__Saccharomycetes(100);o__Saccharomycetales(100);f__Metschnikowiaceae(100);g__Metschnikowia(100);s__Metschnikowia_pulcherrima(67);</td><td>Metschnikowia</td><td>13</td><td>Saprotroph</td><td>|Undefined Saprotroph|</td><td>Yeast</td><td>NULL</td><td>Probable</td><td>As nectar-tap_saprotroph (Põlme et al. 2020); Decay sugar-rich substrates (Põlme et al. 2020)</td><td>Põlme S, et al. 2020. Fungal Diversity 105:1-16 ((https://doi.org/10.1007/s13225-020-00466-2))</td></tr>
<tr><td>Otu08579</td><td>3</td><td>k__Fungi(100);p__Ascomycota(100);c__Saccharomycetes(100);o__Saccharomycetales(100);f__Metschnikowiaceae(100);g__Metschnikowia(100);s__Metschnikowia_viticola(100);</td><td>Metschnikowia</td><td>13</td><td>Saprotroph</td><td>|Undefined Saprotroph|</td><td>Yeast</td><td>NULL</td><td>Probable</td><td>As nectar-tap_saprotroph (Põlme et al. 2020); Decay sugar-rich substrates (Põlme et al. 2020)</td><td>Põlme S, et al. 2020. Fungal Diversity 105:1-16 ((https://doi.org/10.1007/s13225-020-00466-2))</td></tr>
<tr><td>Otu08580</td><td>3</td><td>k__Fungi(100);p__Basidiomycota(100);c__Agaricomycetes(100);o__Agaricales(100);o__Agaricales_unclassified(100);o__Agaricales_unclassified(100);o__Agaricales_unclassified(100);</td><td>Agaricales</td><td>7</td><td>Pathotroph-Saprotroph-Symbiotroph</td><td>Bryophyte Parasite-Dung Saprotroph-Ectomycorrhizal-Fungal Parasite-Leaf Saprotroph-Plant Parasite-Undefined Saprotroph-Wood Saprotroph</td><td>Agaricoid-Gasteroid-Microfungus-Secotioid-Yeast</td><td>NULL</td><td>Possible</td><td>Mushrooms and toadstools, Gill fungi, Agarics... lignicolous, sometimes muscicolous or fungicolous, saprobic, mycorrhizal, rarely parasitic on plants or fungi (Kirk et al. 2008)</td><td>Kirk PM et al. 2008. Dictionary of the Fungi. Tenth Edition. CAB International, Wallingford (ISBN: 978-0851998268)</td></tr>
<tr><td>Otu08581</td><td>3</td><td>k__Fungi(100);p__Ascomycota(100);c__Saccharomycetes(100);o__Saccharomycetales(100);f__Metschnikowiaceae(100);g__Metschnikowia(100);s__Metschnikowia_pulcherrima(100);</td><td>Metschnikowia</td><td>13</td><td>Saprotroph</td><td>|Undefined Saprotroph|</td><td>Yeast</td><td>NULL</td><td>Probable</td><td>As nectar-tap_saprotroph (Põlme et al. 2020); Decay sugar-rich substrates (Põlme et al. 2020)</td><td>Põlme S, et al. 2020. Fungal Diversity 105:1-16 ((https://doi.org/10.1007/s13225-020-00466-2))</td></tr>
<tr><td>Otu08583</td><td>3</td><td>k__Fungi(100);p__Ascomycota(100);c__Saccharomycetes(100);o__Saccharomycetales(100);f__Metschnikowiaceae(100);g__Metschnikowia(100);g__Metschnikowia_unclassified(67);</td><td>Metschnikowia</td><td>13</td><td>Saprotroph</td><td>|Undefined Saprotroph|</td><td>Yeast</td><td>NULL</td><td>Probable</td><td>As nectar-tap_saprotroph (Põlme et al. 2020); Decay sugar-rich substrates (Põlme et al. 2020)</td><td>Põlme S, et al. 2020. Fungal Diversity 105:1-16 ((https://doi.org/10.1007/s13225-020-00466-2))</td></tr>
<tr><td>Otu08584</td><td>3</td><td>k__Fungi(100);p__Ascomycota(100);c__Saccharomycetes(100);o__Saccharomycetales(100);f__Metschnikowiaceae(100);g__Metschnikowia(100);s__Metschnikowia_pulcherrima(67);</td><td>Metschnikowia</td><td>13</td><td>Saprotroph</td><td>|Undefined Saprotroph|</td><td>Yeast</td><td>NULL</td><td>Probable</td><td>As nectar-tap_saprotroph (Põlme et al. 2020); Decay sugar-rich substrates (Põlme et al. 2020)</td><td>Põlme S, et al. 2020. Fungal Diversity 105:1-16 ((https://doi.org/10.1007/s13225-020-00466-2))</td></tr>
<tr><td>Otu08586</td><td>3</td><td>k__Fungi(100);p__Ascomycota(100);c__Dothideomycetes(100);o__Capnodiales(100);f__Davidiellaceae(100);g__Cladosporium(100);s__Cladosporium_sphaerospermum(67);</td><td>Cladosporium</td><td>13</td><td>Pathotroph-Saprotroph-Symbiotroph</td><td>Animal Pathogen-Endophyte-Lichen Parasite-Plant Pathogen-|Plant Saprotroph|-Wood Saprotroph</td><td>Microfungus</td><td>NULL</td><td>Possible</td><td>As foliar_endophyte (Põlme et al. 2020); As litter_saprotroph (Põlme et al. 2020); Decay wood, leaf, fruit, seed, soil, dung, animal material (Põlme et al. 2020)</td><td>Seehann G, et al. 1975. List of Fungi in Soft-Rot Tissue Tests. Institute fur Holzbiologie und Holzschutz, Bundesforschungsanstalt fur Forst - und Holzwirtshaft, Hamburg - Lohbrugge, Germany; Lawrey JD, Diederich P. 2016. http://www.lichenicolous.net (accessed 24 May 2017) ((http://www.lichenicolous.net)); Põlme S, et al. 2020. Fungal Diversity 105:1-16 ((https://doi.org/10.1007/s13225-020-00466-2)); The American Phytopathological Society Common Names of Plant Diseases ((http://www.apsnet.org/publications/commonnames/Pages/default.aspx))</td></tr>
<tr><td>Otu08588</td><td>3</td><td>k__Fungi(100);p__Ascomycota(100);c__Saccharomycetes(100);o__Saccharomycetales(100);f__Metschnikowiaceae(100);g__Metschnikowia(100);s__Metschnikowia_pulcherrima(100);</td><td>Metschnikowia</td><td>13</td><td>Saprotroph</td><td>|Undefined Saprotroph|</td><td>Yeast</td><td>NULL</td><td>Probable</td><td>As nectar-tap_saprotroph (Põlme et al. 2020); Decay sugar-rich substrates (Põlme et al. 2020)</td><td>Põlme S, et al. 2020. Fungal Diversity 105:1-16 ((https://doi.org/10.1007/s13225-020-00466-2))</td></tr>
<tr><td>Otu08589</td><td>3</td><td>k__Fungi(100);p__Ascomycota(100);c__Sordariomycetes(100);o__Xylariales(100);f__Amphisphaeriaceae(100);f__Amphisphaeriaceae_unclassified(67);f__Amphisphaeriaceae_unclassified(67);</td><td>Amphisphaeriaceae</td><td>9</td><td>Saprotroph</td><td>Undefined Saprotroph</td><td>Microfungus</td><td>NULL</td><td>Possible</td><td>Saprobes or weak parasites of leaves, stems, etc., often appearing as secondary colonizers following damage to plants by other agents (Cannon & Kirk 2007)</td><td>Cannon PF, Kirk PM. 2007. Fungal Families of the World. CAB International, Cambridge (ISBN: 978-0851998275)</td></tr>
<tr><td>Otu08591</td><td>3</td><td>k__Fungi(100);p__Ascomycota(100);c__Saccharomycetes(100);o__Saccharomycetales(100);f__Metschnikowiaceae(100);g__Metschnikowia(100);g__Metschnikowia_unclassified(100);</td><td>Metschnikowia</td><td>13</td><td>Saprotroph</td><td>|Undefined Saprotroph|</td><td>Yeast</td><td>NULL</td><td>Probable</td><td>As nectar-tap_saprotroph (Põlme et al. 2020); Decay sugar-rich substrates (Põlme et al. 2020)</td><td>Põlme S, et al. 2020. Fungal Diversity 105:1-16 ((https://doi.org/10.1007/s13225-020-00466-2))</td></tr>
<tr><td>Otu08593</td><td>3</td><td>k__Fungi(100);p__Ascomycota(100);c__Saccharomycetes(100);o__Saccharomycetales(100);f__Metschnikowiaceae(100);g__Metschnikowia(100);g__Metschnikowia_unclassified(67);</td><td>Metschnikowia</td><td>13</td><td>Saprotroph</td><td>|Undefined Saprotroph|</td><td>Yeast</td><td>NULL</td><td>Probable</td><td>As nectar-tap_saprotroph (Põlme et al. 2020); Decay sugar-rich substrates (Põlme et al. 2020)</td><td>Põlme S, et al. 2020. Fungal Diversity 105:1-16 ((https://doi.org/10.1007/s13225-020-00466-2))</td></tr>
<tr><td>Otu08595</td><td>3</td><td>k__Fungi(100);p__Ascomycota(100);c__Saccharomycetes(100);o__Saccharomycetales(100);f__Metschnikowiaceae(100);g__Metschnikowia(100);s__Metschnikowia_pulcherrima(67);</td><td>Metschnikowia</td><td>13</td><td>Saprotroph</td><td>|Undefined Saprotroph|</td><td>Yeast</td><td>NULL</td><td>Probable</td><td>As nectar-tap_saprotroph (Põlme et al. 2020); Decay sugar-rich substrates (Põlme et al. 2020)</td><td>Põlme S, et al. 2020. Fungal Diversity 105:1-16 ((https://doi.org/10.1007/s13225-020-00466-2))</td></tr>
<tr><td>Otu08599</td><td>3</td><td>k__Fungi(100);p__Ascomycota(100);c__Saccharomycetes(100);o__Saccharomycetales(100);f__Saccharomycetales_family_Incertae_sedis(100);g__Candida(100);g__Candida_unclassified(100);</td><td>Candida</td><td>13</td><td>Pathotroph-Saprotroph-Symbiotroph</td><td>Animal Pathogen-Endophyte-Endosymbiont-Epiphyte-|Undefined Saprotroph|</td><td>Dimorphic Yeast-Yeast</td><td>NULL</td><td>Probable</td><td>As nectar-tap_saprotroph (Põlme et al. 2020); Decay sugar-rich substrates (Põlme et al. 2020); This asexual genus is polyphyletic and contain species that function very differently depend on what they are. Caution should be taken when interpreting guilds for Candida species.</td><td>Manolakaki D, et al. 2010. Virulence. 1: 367-375 ((https://doi.org/10.4161/viru.1.5.12796)); Mekbib SB, et al. 2011. Tropical Plant Pathology 36:233-240 ((https://doi.org/10.1590/S1982-56762011000400004)); Põlme S, et al. 2020. Fungal Diversity 105:1-16 ((https://doi.org/10.1007/s13225-020-00466-2))</td></tr>
<tr><td>Otu08601</td><td>3</td><td>k__Fungi(100);p__Ascomycota(100);c__Sordariomycetes(100);o__Hypocreales(100);f__Cordycipitaceae(100);g__Beauveria(100);s__Beauveria_virella(100);</td><td>Beauveria</td><td>13</td><td>Pathotroph-Saprotroph-Symbiotroph</td><td>|Animal Parasite|-Animal Pathogen-Endophyte-Undefined Saprotroph</td><td>Microfungus</td><td>NULL</td><td>Probable</td><td>As animal_decomposer (Põlme et al. 2020); As foliar_endophyte (Põlme et al. 2020); Decay animal material (Põlme et al. 2020)</td><td>Tedersoo L, et al. 2014. Science 346:e1256688 ((https://doi.org/10.1126/science.1256688)); Põlme S, et al. 2020. Fungal Diversity 105:1-16 ((https://doi.org/10.1007/s13225-020-00466-2))</td></tr>
<tr><td>Otu08602</td><td>3</td><td>k__Fungi(100);p__Ascomycota(100);c__Saccharomycetes(100);o__Saccharomycetales(100);f__Metschnikowiaceae(100);g__Metschnikowia(100);g__Metschnikowia_unclassified(100);</td><td>Metschnikowia</td><td>13</td><td>Saprotroph</td><td>|Undefined Saprotroph|</td><td>Yeast</td><td>NULL</td><td>Probable</td><td>As nectar-tap_saprotroph (Põlme et al. 2020); Decay sugar-rich substrates (Põlme et al. 2020)</td><td>Põlme S, et al. 2020. Fungal Diversity 105:1-16 ((https://doi.org/10.1007/s13225-020-00466-2))</td></tr>
<tr><td>Otu08604</td><td>3</td><td>k__Fungi(100);p__Ascomycota(100);c__Saccharomycetes(100);o__Saccharomycetales(100);f__Metschnikowiaceae(100);g__Metschnikowia(100);s__Metschnikowia_pulcherrima(100);</td><td>Metschnikowia</td><td>13</td><td>Saprotroph</td><td>|Undefined Saprotroph|</td><td>Yeast</td><td>NULL</td><td>Probable</td><td>As nectar-tap_saprotroph (Põlme et al. 2020); Decay sugar-rich substrates (Põlme et al. 2020)</td><td>Põlme S, et al. 2020. Fungal Diversity 105:1-16 ((https://doi.org/10.1007/s13225-020-00466-2))</td></tr>
<tr><td>Otu08606</td><td>3</td><td>k__Fungi(100);p__Basidiomycota(100);c__Agaricomycetes(100);o__Cantharellales(100);f__Tulasnellaceae(100);g__unclassified_Tulasnellaceae(100);s__Tulasnellaceae_sp(100);</td><td>Tulasnellaceae</td><td>9</td><td>Saprotroph-Symbiotroph</td><td>Ectomycorrhizal-Undefined Saprotroph</td><td>Corticioid</td><td>NULL</td><td>Probable</td><td>Typically found on decaying wood and other rotting vegetation, presumably saprobic in nutrition. Many anamorphic taxa have mycorrhizal associations with orchids. The linkages between teleomorph and anamorph need further study; some Epulorhiza species have associations with the Sebacinaceae rather than Tulasnellaceae (Cannon & Kirk 2007)</td><td>Cannon PF, Kirk PM. 2007. Fungal Families of the World. CAB International, Cambridge (ISBN: 978-0851998275)</td></tr>
<tr><td>Otu08608</td><td>3</td><td>k__Fungi(100);p__Basidiomycota(100);c__Agaricostilbomycetes(100);o__Agaricostilbales(100);f__Chionosphaeraceae(100);f__Chionosphaeraceae_unclassified(100);f__Chionosphaeraceae_unclassified(100);</td><td>Chionosphaeraceae</td><td>9</td><td>Pathotroph-Saprotroph</td><td>Fungal Parasite-Undefined Saprotroph</td><td>Microfungus</td><td>NULL</td><td>Possible</td><td>Species appear to be mycoparasitic, associated with fungi of bark beetle galleries, wood-inhabiting Xylariaceae or lichens. Others are found erumpent from bark or decaying wood and may be sabrobic (Cannon & Kirk 2007)</td><td>Cannon PF, Kirk PM. 2007. Fungal Families of the World. CAB International, Cambridge (ISBN: 978-0851998275)</td></tr>
<tr><td>Otu08610</td><td>3</td><td>k__Fungi(100);p__Ascomycota(100);c__Saccharomycetes(100);o__Saccharomycetales(100);f__Metschnikowiaceae(100);g__Metschnikowia(100);s__Metschnikowia_pulcherrima(100);</td><td>Metschnikowia</td><td>13</td><td>Saprotroph</td><td>|Undefined Saprotroph|</td><td>Yeast</td><td>NULL</td><td>Probable</td><td>As nectar-tap_saprotroph (Põlme et al. 2020); Decay sugar-rich substrates (Põlme et al. 2020)</td><td>Põlme S, et al. 2020. Fungal Diversity 105:1-16 ((https://doi.org/10.1007/s13225-020-00466-2))</td></tr>
<tr><td>Otu08611</td><td>3</td><td>k__Fungi(100);p__Ascomycota(100);c__Saccharomycetes(100);o__Saccharomycetales(100);f__Metschnikowiaceae(100);g__Metschnikowia(100);s__Metschnikowia_pulcherrima(100);</td><td>Metschnikowia</td><td>13</td><td>Saprotroph</td><td>|Undefined Saprotroph|</td><td>Yeast</td><td>NULL</td><td>Probable</td><td>As nectar-tap_saprotroph (Põlme et al. 2020); Decay sugar-rich substrates (Põlme et al. 2020)</td><td>Põlme S, et al. 2020. Fungal Diversity 105:1-16 ((https://doi.org/10.1007/s13225-020-00466-2))</td></tr>
<tr><td>Otu08614</td><td>3</td><td>k__Fungi(100);p__Ascomycota(100);c__Saccharomycetes(100);o__Saccharomycetales(100);f__Metschnikowiaceae(100);g__Metschnikowia(100);s__Metschnikowia_pulcherrima(100);</td><td>Metschnikowia</td><td>13</td><td>Saprotroph</td><td>|Undefined Saprotroph|</td><td>Yeast</td><td>NULL</td><td>Probable</td><td>As nectar-tap_saprotroph (Põlme et al. 2020); Decay sugar-rich substrates (Põlme et al. 2020)</td><td>Põlme S, et al. 2020. Fungal Diversity 105:1-16 ((https://doi.org/10.1007/s13225-020-00466-2))</td></tr>
<tr><td>Otu08616</td><td>3</td><td>k__Fungi(100);p__Ascomycota(100);c__Sordariomycetes(100);o__Xylariales(100);f__Xylariaceae(100);f__Xylariaceae_unclassified(100);f__Xylariaceae_unclassified(100);</td><td>Xylariaceae</td><td>9</td><td>Pathotroph-Saprotroph-Symbiotroph</td><td>Dung Saprotroph-Endophyte-Plant Pathogen-Undefined Saprotroph</td><td>Xylarioid</td><td>NULL</td><td>Probable</td><td>Saprobes or weak pathogens, many endophytes, mainly in wood and bark, occasionally in soil and dung (Cannon & Kirk 2007)</td><td>Cannon PF, Kirk PM. 2007. Fungal Families of the World. CAB International, Cambridge (ISBN: 978-0851998275)</td></tr>
<tr><td>Otu08617</td><td>3</td><td>k__Fungi(100);p__Ascomycota(100);c__Saccharomycetes(100);o__Saccharomycetales(100);f__Metschnikowiaceae(100);g__Metschnikowia(100);s__Metschnikowia_pulcherrima(100);</td><td>Metschnikowia</td><td>13</td><td>Saprotroph</td><td>|Undefined Saprotroph|</td><td>Yeast</td><td>NULL</td><td>Probable</td><td>As nectar-tap_saprotroph (Põlme et al. 2020); Decay sugar-rich substrates (Põlme et al. 2020)</td><td>Põlme S, et al. 2020. Fungal Diversity 105:1-16 ((https://doi.org/10.1007/s13225-020-00466-2))</td></tr>
<tr><td>Otu08621</td><td>3</td><td>k__Fungi(100);p__Ascomycota(100);c__Saccharomycetes(100);o__Saccharomycetales(100);f__Metschnikowiaceae(100);g__Metschnikowia(100);s__Metschnikowia_pulcherrima(100);</td><td>Metschnikowia</td><td>13</td><td>Saprotroph</td><td>|Undefined Saprotroph|</td><td>Yeast</td><td>NULL</td><td>Probable</td><td>As nectar-tap_saprotroph (Põlme et al. 2020); Decay sugar-rich substrates (Põlme et al. 2020)</td><td>Põlme S, et al. 2020. Fungal Diversity 105:1-16 ((https://doi.org/10.1007/s13225-020-00466-2))</td></tr>
<tr><td>Otu08622</td><td>3</td><td>k__Fungi(100);p__Ascomycota(100);c__Saccharomycetes(100);o__Saccharomycetales(100);f__Metschnikowiaceae(100);g__Metschnikowia(100);s__Metschnikowia_pulcherrima(67);</td><td>Metschnikowia</td><td>13</td><td>Saprotroph</td><td>|Undefined Saprotroph|</td><td>Yeast</td><td>NULL</td><td>Probable</td><td>As nectar-tap_saprotroph (Põlme et al. 2020); Decay sugar-rich substrates (Põlme et al. 2020)</td><td>Põlme S, et al. 2020. Fungal Diversity 105:1-16 ((https://doi.org/10.1007/s13225-020-00466-2))</td></tr>
<tr><td>Otu08627</td><td>3</td><td>k__Fungi(100);p__Ascomycota(100);c__Sordariomycetes(100);o__Hypocreales(100);f__Cordycipitaceae(100);g__Beauveria(100);s__Beauveria_virella(100);</td><td>Beauveria</td><td>13</td><td>Pathotroph-Saprotroph-Symbiotroph</td><td>|Animal Parasite|-Animal Pathogen-Endophyte-Undefined Saprotroph</td><td>Microfungus</td><td>NULL</td><td>Probable</td><td>As animal_decomposer (Põlme et al. 2020); As foliar_endophyte (Põlme et al. 2020); Decay animal material (Põlme et al. 2020)</td><td>Tedersoo L, et al. 2014. Science 346:e1256688 ((https://doi.org/10.1126/science.1256688)); Põlme S, et al. 2020. Fungal Diversity 105:1-16 ((https://doi.org/10.1007/s13225-020-00466-2))</td></tr>
<tr><td>Otu08628</td><td>3</td><td>k__Fungi(100);p__Ascomycota(100);c__Saccharomycetes(100);o__Saccharomycetales(100);f__Metschnikowiaceae(100);g__Metschnikowia(100);s__Metschnikowia_pulcherrima(100);</td><td>Metschnikowia</td><td>13</td><td>Saprotroph</td><td>|Undefined Saprotroph|</td><td>Yeast</td><td>NULL</td><td>Probable</td><td>As nectar-tap_saprotroph (Põlme et al. 2020); Decay sugar-rich substrates (Põlme et al. 2020)</td><td>Põlme S, et al. 2020. Fungal Diversity 105:1-16 ((https://doi.org/10.1007/s13225-020-00466-2))</td></tr>
<tr><td>Otu08629</td><td>3</td><td>k__Fungi(100);p__Basidiomycota(100);c__Agaricomycetes(100);o__Gomphales(100);f__Gomphaceae(100);g__Ramaria(100);s__Ramaria_sp(100);</td><td>Ramaria</td><td>13</td><td>Saprotroph-Symbiotroph</td><td>|Ectomycorrhizal|-Plant Saprotroph</td><td>Clavarioid</td><td>NULL</td><td>Possible</td><td>As litter_saprotroph (Põlme et al. 2020)</td><td>Rinaldi AC, et al. 2008. Fungal Diversity 33:1-45 ((http://www.fungaldiversity.org/fdp/sfdp/33-1.pdf)) (as ECM/not ECM); Tedersoo L, et al. 2010. Mycorrhiza 20:217-263 ((10.1007/s00572-009-0274-x)) (pro parte); Põlme S, et al. 2020. Fungal Diversity 105:1-16 ((https://doi.org/10.1007/s13225-020-00466-2)); http://mycorrhizas.info/ecmf.html ((http://mycorrhizas.info/ecmf.html)) (as ECM)</td></tr>
<tr><td>Otu08630</td><td>3</td><td>k__Fungi(100);p__Ascomycota(100);c__Saccharomycetes(100);o__Saccharomycetales(100);f__Metschnikowiaceae(100);g__Metschnikowia(100);s__Metschnikowia_pulcherrima(100);</td><td>Metschnikowia</td><td>13</td><td>Saprotroph</td><td>|Undefined Saprotroph|</td><td>Yeast</td><td>NULL</td><td>Probable</td><td>As nectar-tap_saprotroph (Põlme et al. 2020); Decay sugar-rich substrates (Põlme et al. 2020)</td><td>Põlme S, et al. 2020. Fungal Diversity 105:1-16 ((https://doi.org/10.1007/s13225-020-00466-2))</td></tr>
<tr><td>Otu08632</td><td>3</td><td>k__Fungi(100);p__Basidiomycota(100);c__Agaricomycetes(100);o__Agaricales(100);f__Agaricaceae(100);g__Agaricus(100);s__Agaricus_silvaticus(100);</td><td>Agaricus</td><td>13</td><td>Saprotroph</td><td>|Undefined Saprotroph|</td><td>Agaricoid-Gasteroid-Secotioid</td><td>NULL</td><td>Probable</td><td>As soil_saprotroph (Põlme et al. 2020); Decay soil (Põlme et al. 2020)</td><td>Tedersoo L, et al. 2014. Science 346:e1256688 ((https://doi.org/10.1126/science.1256688)); Põlme S, et al. 2020. Fungal Diversity 105:1-16 ((https://doi.org/10.1007/s13225-020-00466-2))</td></tr>
<tr><td>Otu08634</td><td>3</td><td>k__Fungi(100);p__Ascomycota(100);c__Saccharomycetes(100);o__Saccharomycetales(100);f__Metschnikowiaceae(100);g__Metschnikowia(100);s__Metschnikowia_pulcherrima(100);</td><td>Metschnikowia</td><td>13</td><td>Saprotroph</td><td>|Undefined Saprotroph|</td><td>Yeast</td><td>NULL</td><td>Probable</td><td>As nectar-tap_saprotroph (Põlme et al. 2020); Decay sugar-rich substrates (Põlme et al. 2020)</td><td>Põlme S, et al. 2020. Fungal Diversity 105:1-16 ((https://doi.org/10.1007/s13225-020-00466-2))</td></tr>
<tr><td>Otu08635</td><td>3</td><td>k__Fungi(100);p__Ascomycota(100);c__Saccharomycetes(100);o__Saccharomycetales(100);f__Metschnikowiaceae(100);g__Metschnikowia(100);s__Metschnikowia_pulcherrima(100);</td><td>Metschnikowia</td><td>13</td><td>Saprotroph</td><td>|Undefined Saprotroph|</td><td>Yeast</td><td>NULL</td><td>Probable</td><td>As nectar-tap_saprotroph (Põlme et al. 2020); Decay sugar-rich substrates (Põlme et al. 2020)</td><td>Põlme S, et al. 2020. Fungal Diversity 105:1-16 ((https://doi.org/10.1007/s13225-020-00466-2))</td></tr>
<tr><td>Otu08636</td><td>3</td><td>k__Fungi(100);p__Ascomycota(100);c__Saccharomycetes(100);o__Saccharomycetales(100);f__Metschnikowiaceae(100);g__Metschnikowia(100);s__Metschnikowia_pulcherrima(100);</td><td>Metschnikowia</td><td>13</td><td>Saprotroph</td><td>|Undefined Saprotroph|</td><td>Yeast</td><td>NULL</td><td>Probable</td><td>As nectar-tap_saprotroph (Põlme et al. 2020); Decay sugar-rich substrates (Põlme et al. 2020)</td><td>Põlme S, et al. 2020. Fungal Diversity 105:1-16 ((https://doi.org/10.1007/s13225-020-00466-2))</td></tr>
<tr><td>Otu08638</td><td>3</td><td>k__Fungi(100);p__Basidiomycota(100);c__Agaricomycetes(100);o__Agaricales(100);f__Inocybaceae(100);g__Crepidotus(100);s__Crepidotus_sphaerosporus(100);</td><td>Crepidotus</td><td>13</td><td>Saprotroph</td><td>|Wood Saprotroph|</td><td>Agaricoid</td><td>NULL</td><td>Highly Probable</td><td>Decay wood (Põlme et al. 2020)</td><td>Grgurinovic CA, Simpson JA. 2001. Fungal Diversity 8:97-127 ((http://www.fungaldiversity.org/fdp/sfdp/FD_8_97-127.pdf)); Aime MC, et al. 2005. American Journal of Botany 92:74-82 ((https://doi.org/10.3732/ajb.92.1.74)); Tedersoo L, et al. 2014. Science 346:e1256688 ((https://doi.org/10.1126/science.1256688)); Põlme S, et al. 2020. Fungal Diversity 105:1-16 ((https://doi.org/10.1007/s13225-020-00466-2))</td></tr>
<tr><td>Otu08643</td><td>3</td><td>k__Fungi(100);p__Ascomycota(100);c__Leotiomycetes(100);o__Helotiales(100);f__Vibrisseaceae(100);g__Phialocephala(100);s__Phialocephala_sp_CM16s1(100);</td><td>Phialocephala</td><td>13</td><td>Saprotroph-Symbiotroph</td><td>Endophyte-|Undefined Saprotroph|</td><td>Microfungus</td><td>NULL</td><td>Highly Probable</td><td>As root_endophyte (Põlme et al. 2020); As root_endophyte_dark_septate (Põlme et al. 2020); As soil_saprotroph (Põlme et al. 2020); Dark Septate Endophyte, known from plant roots.; Decay roots, soil (Põlme et al. 2020)</td><td>Newsham KK. 2011. New Phytologist 190:783-793 ((https://doi.org/10.1111/j.1469-8137.2010.03611.x)) (as dark septate root endophytes); Põlme S, et al. 2020. Fungal Diversity 105:1-16 ((https://doi.org/10.1007/s13225-020-00466-2))</td></tr>
<tr><td>Otu08644</td><td>3</td><td>k__Fungi(100);p__Ascomycota(100);c__Saccharomycetes(100);o__Saccharomycetales(100);f__Metschnikowiaceae(100);g__Metschnikowia(100);s__Metschnikowia_pulcherrima(67);</td><td>Metschnikowia</td><td>13</td><td>Saprotroph</td><td>|Undefined Saprotroph|</td><td>Yeast</td><td>NULL</td><td>Probable</td><td>As nectar-tap_saprotroph (Põlme et al. 2020); Decay sugar-rich substrates (Põlme et al. 2020)</td><td>Põlme S, et al. 2020. Fungal Diversity 105:1-16 ((https://doi.org/10.1007/s13225-020-00466-2))</td></tr>
<tr><td>Otu08645</td><td>3</td><td>k__Fungi(100);p__Ascomycota(100);c__Saccharomycetes(100);o__Saccharomycetales(100);f__Saccharomycetales_family_Incertae_sedis(100);g__Candida(100);g__Candida_unclassified(100);</td><td>Candida</td><td>13</td><td>Pathotroph-Saprotroph-Symbiotroph</td><td>Animal Pathogen-Endophyte-Endosymbiont-Epiphyte-|Undefined Saprotroph|</td><td>Dimorphic Yeast-Yeast</td><td>NULL</td><td>Probable</td><td>As nectar-tap_saprotroph (Põlme et al. 2020); Decay sugar-rich substrates (Põlme et al. 2020); This asexual genus is polyphyletic and contain species that function very differently depend on what they are. Caution should be taken when interpreting guilds for Candida species.</td><td>Manolakaki D, et al. 2010. Virulence. 1: 367-375 ((https://doi.org/10.4161/viru.1.5.12796)); Mekbib SB, et al. 2011. Tropical Plant Pathology 36:233-240 ((https://doi.org/10.1590/S1982-56762011000400004)); Põlme S, et al. 2020. Fungal Diversity 105:1-16 ((https://doi.org/10.1007/s13225-020-00466-2))</td></tr>
<tr><td>Otu08646</td><td>3</td><td>k__Fungi(100);p__Basidiomycota(100);c__Agaricostilbomycetes(100);o__Agaricostilbales(100);f__Agaricostilbaceae(100);g__Bensingtonia(100);g__Bensingtonia_unclassified(100);</td><td>Bensingtonia</td><td>13</td><td>Saprotroph</td><td>|Undefined Saprotroph|</td><td>Dimorphic-Facultative Yeast</td><td>NULL</td><td>Highly Probable</td><td>NULL</td><td>Kurtzman CP, et al. (eds.) 2011. The Yeasts, a Taxonomic Study. Fifth Edition. Vols 1-3. Elsevier, San Diego (ISBN: 9780444521491); Põlme S, et al. 2020. Fungal Diversity 105:1-16 ((https://doi.org/10.1007/s13225-020-00466-2))</td></tr>
<tr><td>Otu08649</td><td>3</td><td>k__Fungi(100);p__Ascomycota(100);c__Saccharomycetes(100);o__Saccharomycetales(100);f__Metschnikowiaceae(100);g__Metschnikowia(100);s__Metschnikowia_pulcherrima(100);</td><td>Metschnikowia</td><td>13</td><td>Saprotroph</td><td>|Undefined Saprotroph|</td><td>Yeast</td><td>NULL</td><td>Probable</td><td>As nectar-tap_saprotroph (Põlme et al. 2020); Decay sugar-rich substrates (Põlme et al. 2020)</td><td>Põlme S, et al. 2020. Fungal Diversity 105:1-16 ((https://doi.org/10.1007/s13225-020-00466-2))</td></tr>
<tr><td>Otu08650</td><td>3</td><td>k__Fungi(100);p__Ascomycota(100);c__Saccharomycetes(100);o__Saccharomycetales(100);f__Metschnikowiaceae(100);g__Metschnikowia(100);s__Metschnikowia_pulcherrima(100);</td><td>Metschnikowia</td><td>13</td><td>Saprotroph</td><td>|Undefined Saprotroph|</td><td>Yeast</td><td>NULL</td><td>Probable</td><td>As nectar-tap_saprotroph (Põlme et al. 2020); Decay sugar-rich substrates (Põlme et al. 2020)</td><td>Põlme S, et al. 2020. Fungal Diversity 105:1-16 ((https://doi.org/10.1007/s13225-020-00466-2))</td></tr>
<tr><td>Otu08651</td><td>3</td><td>k__Fungi(100);p__Basidiomycota(100);c__Agaricomycetes(100);o__Corticiales(100);f__Corticiaceae(100);g__Waitea(100);g__Waitea_unclassified(100);</td><td>Waitea</td><td>13</td><td>Pathotroph-Saprotroph</td><td>|Plant Pathogen|-Plant Saprotroph</td><td>Corticioid</td><td>NULL</td><td>Probable</td><td>As litter_saprotroph (Põlme et al. 2020); Decay leaf, fruit, seed (Põlme et al. 2020)</td><td>Tedersoo L, et al. 2014. Science 346:e1256688 ((https://doi.org/10.1126/science.1256688)); Põlme S, et al. 2020. Fungal Diversity 105:1-16 ((https://doi.org/10.1007/s13225-020-00466-2))</td></tr>
<tr><td>Otu08653</td><td>3</td><td>k__Fungi(100);p__Ascomycota(100);c__Saccharomycetes(100);o__Saccharomycetales(100);f__Metschnikowiaceae(100);g__Metschnikowia(100);s__Metschnikowia_pulcherrima(67);</td><td>Metschnikowia</td><td>13</td><td>Saprotroph</td><td>|Undefined Saprotroph|</td><td>Yeast</td><td>NULL</td><td>Probable</td><td>As nectar-tap_saprotroph (Põlme et al. 2020); Decay sugar-rich substrates (Põlme et al. 2020)</td><td>Põlme S, et al. 2020. Fungal Diversity 105:1-16 ((https://doi.org/10.1007/s13225-020-00466-2))</td></tr>
<tr><td>Otu08654</td><td>3</td><td>k__Fungi(100);p__Ascomycota(100);c__Saccharomycetes(100);o__Saccharomycetales(100);f__Metschnikowiaceae(100);g__Metschnikowia(100);s__Metschnikowia_pulcherrima(67);</td><td>Metschnikowia</td><td>13</td><td>Saprotroph</td><td>|Undefined Saprotroph|</td><td>Yeast</td><td>NULL</td><td>Probable</td><td>As nectar-tap_saprotroph (Põlme et al. 2020); Decay sugar-rich substrates (Põlme et al. 2020)</td><td>Põlme S, et al. 2020. Fungal Diversity 105:1-16 ((https://doi.org/10.1007/s13225-020-00466-2))</td></tr>
<tr><td>Otu08655</td><td>3</td><td>k__Fungi(100);p__Ascomycota(100);c__Saccharomycetes(100);o__Saccharomycetales(100);f__Metschnikowiaceae(100);g__Metschnikowia(100);s__Metschnikowia_pulcherrima(67);</td><td>Metschnikowia</td><td>13</td><td>Saprotroph</td><td>|Undefined Saprotroph|</td><td>Yeast</td><td>NULL</td><td>Probable</td><td>As nectar-tap_saprotroph (Põlme et al. 2020); Decay sugar-rich substrates (Põlme et al. 2020)</td><td>Põlme S, et al. 2020. Fungal Diversity 105:1-16 ((https://doi.org/10.1007/s13225-020-00466-2))</td></tr>
<tr><td>Otu08656</td><td>3</td><td>k__Fungi(100);p__Ascomycota(100);c__Saccharomycetes(100);o__Saccharomycetales(100);f__Metschnikowiaceae(100);g__Metschnikowia(100);s__Metschnikowia_pulcherrima(100);</td><td>Metschnikowia</td><td>13</td><td>Saprotroph</td><td>|Undefined Saprotroph|</td><td>Yeast</td><td>NULL</td><td>Probable</td><td>As nectar-tap_saprotroph (Põlme et al. 2020); Decay sugar-rich substrates (Põlme et al. 2020)</td><td>Põlme S, et al. 2020. Fungal Diversity 105:1-16 ((https://doi.org/10.1007/s13225-020-00466-2))</td></tr>
<tr><td>Otu08659</td><td>3</td><td>k__Fungi(100);p__Basidiomycota(100);c__Agaricomycetes(100);o__Agaricales(100);f__Inocybaceae(100);g__Inocybe(100);s__Inocybe_griseoscabrosa(100);</td><td>Inocybe</td><td>13</td><td>Symbiotroph</td><td>|Ectomycorrhizal|</td><td>Agaricoid</td><td>NULL</td><td>Highly Probable</td><td>Sensu stricto (Matheny 2009).</td><td>Rinaldi AC, et al. 2008. Fungal Diversity 33:1-45 ((http://www.fungaldiversity.org/fdp/sfdp/33-1.pdf)); Matheny PB. 2009. McIlvainea 18:11-21 ((https://namyco.org/docs/3_Matheny.pdf)); Tedersoo L, et al. 2010. Mycorrhiza 20:217-263 ((https://doi.org/10.1007/s00572-009-0274-x)); Põlme S, et al. 2020. Fungal Diversity 105:1-16 ((https://doi.org/10.1007/s13225-020-00466-2)); http://mycorrhizas.info/ecmf.html ((http://mycorrhizas.info/ecmf.html))</td></tr>
<tr><td>Otu08662</td><td>3</td><td>k__Fungi(100);p__Basidiomycota(100);c__Agaricomycetes(100);o__Agaricales(100);f__Amanitaceae(100);g__Amanita(100);s__Amanita_volvata(100);</td><td>Amanita</td><td>13</td><td>Saprotroph-Symbiotroph</td><td>|Ectomycorrhizal|-Undefined Saprotroph</td><td>Agaricoid</td><td>NULL</td><td>Probable</td><td>Some members of the genus are deadly poisonous</td><td>Rinaldi AC, et al. 2008. Fungal Diversity 33:1-45 ((http://www.fungaldiversity.org/fdp/sfdp/33-1.pdf)); Tedersoo L, et al. 2010. Mycorrhiza 20:217-263 ((https://doi.org/10.1007/s00572-009-0274-x)); Wolfe BE, et al. 2012. PLoS ONE 7:e39597 ((https://doi.org/10.1371/journal.pone.0039597)); Põlme S, et al. 2020. Fungal Diversity 105:1-16 ((https://doi.org/10.1007/s13225-020-00466-2)); http://mycorrhizas.info/ecmf.html ((http://mycorrhizas.info/ecmf.html)); http://en.wikipedia.org/wiki/List_of_deadly_fungi ((http://en.wikipedia.org/wiki/List_of_deadly_fungi))</td></tr>
<tr><td>Otu08663</td><td>3</td><td>k__Fungi(100);p__Ascomycota(100);c__Saccharomycetes(100);o__Saccharomycetales(100);f__Metschnikowiaceae(100);g__Metschnikowia(100);s__Metschnikowia_pulcherrima(100);</td><td>Metschnikowia</td><td>13</td><td>Saprotroph</td><td>|Undefined Saprotroph|</td><td>Yeast</td><td>NULL</td><td>Probable</td><td>As nectar-tap_saprotroph (Põlme et al. 2020); Decay sugar-rich substrates (Põlme et al. 2020)</td><td>Põlme S, et al. 2020. Fungal Diversity 105:1-16 ((https://doi.org/10.1007/s13225-020-00466-2))</td></tr>
<tr><td>Otu08664</td><td>3</td><td>k__Fungi(100);p__Ascomycota(100);c__Sordariomycetes(100);o__Sordariales(100);f__Chaetomiaceae(100);f__Chaetomiaceae_unclassified(100);f__Chaetomiaceae_unclassified(100);</td><td>Chaetomiaceae</td><td>9</td><td>Pathotroph-Saprotroph-Symbiotroph</td><td>Animal Pathogen-Dung Saprotroph-Endophyte-Epiphyte-Plant Saprotroph-Wood Saprotroph</td><td>Microfungus</td><td>Soft Rot</td><td>Possible</td><td>Saprobic, acting as key decay organisms of plant materials and plant-derived commodities (Cannon & Kirk 2007); Epiphyte detection method-culture (Massimo et al. 2015); Endophyte detection method-culture (Busby et al. 2016, David et al. 2016); Likely opportunistic human pathogen (Irinyi et al. 2015)</td><td>Seehann G, et al. 1975. List of Fungi in Soft-Rot Tissue Tests. Institute fur Holzbiologie und Holzschutz, Bundesforschungsanstalt fur Forst - und Holzwirtshaft, Hamburg - Lohbrugge, Germany; Cannon PF, Kirk PM. 2007. Fungal Families of the World. CAB International, Cambridge ((https://isbnsearch.org/isbn/9780851998275)); Bills GF, et al. 2013. Current Opinion in Microbiology 16:549-565 ((https://doi.org/10.1016/j.mib.2013.08.001)); Tedersoo L, et al. 2014. Science 346:e1256688 ((https://doi.org/10.1126/science.1256688)); Irinyi L, et al. 2015. Medical Mycology 53:313-337 ((https://doi.org/10.1093/mmy/myv008)); Massimo NC, et al. 2015. Microbial Ecology 70:61-76 ((https://doi.org/10.1007/s00248-014-0563-6)); Busby PE, et al. 2016. New Phytologist 209:1681-1692 ((https://doi.org/10.1111/nph.13742)); David AS, et al. 2016. Microbial Ecology 71:912-926 ((https://doi.org/10.1007/s00248-015-0712-6))</td></tr>
<tr><td>Otu08665</td><td>3</td><td>k__Fungi(100);p__Basidiomycota(100);c__Agaricomycetes(100);o__Agaricales(100);o__Agaricales_unclassified(100);o__Agaricales_unclassified(100);o__Agaricales_unclassified(100);</td><td>Agaricales</td><td>7</td><td>Pathotroph-Saprotroph-Symbiotroph</td><td>Bryophyte Parasite-Dung Saprotroph-Ectomycorrhizal-Fungal Parasite-Leaf Saprotroph-Plant Parasite-Undefined Saprotroph-Wood Saprotroph</td><td>Agaricoid-Gasteroid-Microfungus-Secotioid-Yeast</td><td>NULL</td><td>Possible</td><td>Mushrooms and toadstools, Gill fungi, Agarics... lignicolous, sometimes muscicolous or fungicolous, saprobic, mycorrhizal, rarely parasitic on plants or fungi (Kirk et al. 2008)</td><td>Kirk PM et al. 2008. Dictionary of the Fungi. Tenth Edition. CAB International, Wallingford (ISBN: 978-0851998268)</td></tr>
<tr><td>Otu08666</td><td>3</td><td>k__Fungi(100);p__Ascomycota(100);c__Saccharomycetes(100);o__Saccharomycetales(100);f__Metschnikowiaceae(100);g__Metschnikowia(100);g__Metschnikowia_unclassified(100);</td><td>Metschnikowia</td><td>13</td><td>Saprotroph</td><td>|Undefined Saprotroph|</td><td>Yeast</td><td>NULL</td><td>Probable</td><td>As nectar-tap_saprotroph (Põlme et al. 2020); Decay sugar-rich substrates (Põlme et al. 2020)</td><td>Põlme S, et al. 2020. Fungal Diversity 105:1-16 ((https://doi.org/10.1007/s13225-020-00466-2))</td></tr>
<tr><td>Otu08668</td><td>3</td><td>k__Fungi(100);p__Ascomycota(100);c__Saccharomycetes(100);o__Saccharomycetales(100);f__Metschnikowiaceae(100);g__Metschnikowia(100);g__Metschnikowia_unclassified(67);</td><td>Metschnikowia</td><td>13</td><td>Saprotroph</td><td>|Undefined Saprotroph|</td><td>Yeast</td><td>NULL</td><td>Probable</td><td>As nectar-tap_saprotroph (Põlme et al. 2020); Decay sugar-rich substrates (Põlme et al. 2020)</td><td>Põlme S, et al. 2020. Fungal Diversity 105:1-16 ((https://doi.org/10.1007/s13225-020-00466-2))</td></tr>
<tr><td>Otu08670</td><td>3</td><td>k__Fungi(100);p__Ascomycota(100);c__Saccharomycetes(100);o__Saccharomycetales(100);f__Metschnikowiaceae(100);g__Metschnikowia(100);s__Metschnikowia_pulcherrima(67);</td><td>Metschnikowia</td><td>13</td><td>Saprotroph</td><td>|Undefined Saprotroph|</td><td>Yeast</td><td>NULL</td><td>Probable</td><td>As nectar-tap_saprotroph (Põlme et al. 2020); Decay sugar-rich substrates (Põlme et al. 2020)</td><td>Põlme S, et al. 2020. Fungal Diversity 105:1-16 ((https://doi.org/10.1007/s13225-020-00466-2))</td></tr>
<tr><td>Otu08671</td><td>3</td><td>k__Fungi(100);p__Ascomycota(100);c__Saccharomycetes(100);o__Saccharomycetales(100);f__Metschnikowiaceae(100);g__Metschnikowia(100);s__Metschnikowia_pulcherrima(100);</td><td>Metschnikowia</td><td>13</td><td>Saprotroph</td><td>|Undefined Saprotroph|</td><td>Yeast</td><td>NULL</td><td>Probable</td><td>As nectar-tap_saprotroph (Põlme et al. 2020); Decay sugar-rich substrates (Põlme et al. 2020)</td><td>Põlme S, et al. 2020. Fungal Diversity 105:1-16 ((https://doi.org/10.1007/s13225-020-00466-2))</td></tr>
<tr><td>Otu08672</td><td>3</td><td>k__Fungi(100);p__Ascomycota(100);c__Dothideomycetes(100);o__Dothideomycetes_order_Incertae_sedis(100);f__Myxotrichaceae(100);g__Oidiodendron(100);g__Oidiodendron_unclassified(67);</td><td>Oidiodendron</td><td>13</td><td>Pathotroph-Saprotroph-Symbiotroph</td><td>Endophyte-Ericoid Mycorrhizal-|Undefined Saprotroph|</td><td>Microfungus</td><td>NULL</td><td>Probable</td><td>As root_endophyte (Põlme et al. 2020); As soil_saprotroph (Põlme et al. 2020); Dark Septate Endophyte; Decay soil (Põlme et al. 2020)</td><td>Newsham KK. 2011. New Phytologist 190:783-793 ((https://doi.org/10.1111/j.1469-8137.2010.03611.x)) (as dark septate root endophytes); Põlme S, et al. 2020. Fungal Diversity 105:1-16 ((https://doi.org/10.1007/s13225-020-00466-2))</td></tr>
<tr><td>Otu08673</td><td>3</td><td>k__Fungi(100);p__Ascomycota(100);c__Saccharomycetes(100);o__Saccharomycetales(100);f__Metschnikowiaceae(100);g__Metschnikowia(100);s__Metschnikowia_pulcherrima(100);</td><td>Metschnikowia</td><td>13</td><td>Saprotroph</td><td>|Undefined Saprotroph|</td><td>Yeast</td><td>NULL</td><td>Probable</td><td>As nectar-tap_saprotroph (Põlme et al. 2020); Decay sugar-rich substrates (Põlme et al. 2020)</td><td>Põlme S, et al. 2020. Fungal Diversity 105:1-16 ((https://doi.org/10.1007/s13225-020-00466-2))</td></tr>
<tr><td>Otu08674</td><td>3</td><td>k__Fungi(100);p__Ascomycota(100);c__Saccharomycetes(100);o__Saccharomycetales(100);f__Metschnikowiaceae(100);g__Metschnikowia(100);s__Metschnikowia_pulcherrima(100);</td><td>Metschnikowia</td><td>13</td><td>Saprotroph</td><td>|Undefined Saprotroph|</td><td>Yeast</td><td>NULL</td><td>Probable</td><td>As nectar-tap_saprotroph (Põlme et al. 2020); Decay sugar-rich substrates (Põlme et al. 2020)</td><td>Põlme S, et al. 2020. Fungal Diversity 105:1-16 ((https://doi.org/10.1007/s13225-020-00466-2))</td></tr>
<tr><td>Otu08676</td><td>3</td><td>k__Fungi(100);p__Ascomycota(100);c__Saccharomycetes(100);o__Saccharomycetales(100);f__Metschnikowiaceae(100);g__Metschnikowia(100);g__Metschnikowia_unclassified(67);</td><td>Metschnikowia</td><td>13</td><td>Saprotroph</td><td>|Undefined Saprotroph|</td><td>Yeast</td><td>NULL</td><td>Probable</td><td>As nectar-tap_saprotroph (Põlme et al. 2020); Decay sugar-rich substrates (Põlme et al. 2020)</td><td>Põlme S, et al. 2020. Fungal Diversity 105:1-16 ((https://doi.org/10.1007/s13225-020-00466-2))</td></tr>
<tr><td>Otu08677</td><td>3</td><td>k__Fungi(100);p__Ascomycota(100);c__Saccharomycetes(100);o__Saccharomycetales(100);f__Metschnikowiaceae(100);g__Metschnikowia(100);g__Metschnikowia_unclassified(67);</td><td>Metschnikowia</td><td>13</td><td>Saprotroph</td><td>|Undefined Saprotroph|</td><td>Yeast</td><td>NULL</td><td>Probable</td><td>As nectar-tap_saprotroph (Põlme et al. 2020); Decay sugar-rich substrates (Põlme et al. 2020)</td><td>Põlme S, et al. 2020. Fungal Diversity 105:1-16 ((https://doi.org/10.1007/s13225-020-00466-2))</td></tr>
<tr><td>Otu08678</td><td>3</td><td>k__Fungi(100);p__Ascomycota(100);c__Saccharomycetes(100);o__Saccharomycetales(100);f__Metschnikowiaceae(100);g__Metschnikowia(100);s__Metschnikowia_pulcherrima(100);</td><td>Metschnikowia</td><td>13</td><td>Saprotroph</td><td>|Undefined Saprotroph|</td><td>Yeast</td><td>NULL</td><td>Probable</td><td>As nectar-tap_saprotroph (Põlme et al. 2020); Decay sugar-rich substrates (Põlme et al. 2020)</td><td>Põlme S, et al. 2020. Fungal Diversity 105:1-16 ((https://doi.org/10.1007/s13225-020-00466-2))</td></tr>
<tr><td>Otu08679</td><td>3</td><td>k__Fungi(100);p__Ascomycota(100);c__Saccharomycetes(100);o__Saccharomycetales(100);f__Metschnikowiaceae(100);g__Metschnikowia(100);s__Metschnikowia_pulcherrima(67);</td><td>Metschnikowia</td><td>13</td><td>Saprotroph</td><td>|Undefined Saprotroph|</td><td>Yeast</td><td>NULL</td><td>Probable</td><td>As nectar-tap_saprotroph (Põlme et al. 2020); Decay sugar-rich substrates (Põlme et al. 2020)</td><td>Põlme S, et al. 2020. Fungal Diversity 105:1-16 ((https://doi.org/10.1007/s13225-020-00466-2))</td></tr>
<tr><td>Otu08681</td><td>3</td><td>k__Fungi(100);p__Ascomycota(100);c__Saccharomycetes(100);o__Saccharomycetales(100);f__Metschnikowiaceae(100);g__Metschnikowia(100);s__Metschnikowia_pulcherrima(67);</td><td>Metschnikowia</td><td>13</td><td>Saprotroph</td><td>|Undefined Saprotroph|</td><td>Yeast</td><td>NULL</td><td>Probable</td><td>As nectar-tap_saprotroph (Põlme et al. 2020); Decay sugar-rich substrates (Põlme et al. 2020)</td><td>Põlme S, et al. 2020. Fungal Diversity 105:1-16 ((https://doi.org/10.1007/s13225-020-00466-2))</td></tr>
<tr><td>Otu08685</td><td>3</td><td>k__Fungi(100);p__Ascomycota(100);c__Saccharomycetes(100);o__Saccharomycetales(100);f__Metschnikowiaceae(100);g__Metschnikowia(100);s__Metschnikowia_pulcherrima(67);</td><td>Metschnikowia</td><td>13</td><td>Saprotroph</td><td>|Undefined Saprotroph|</td><td>Yeast</td><td>NULL</td><td>Probable</td><td>As nectar-tap_saprotroph (Põlme et al. 2020); Decay sugar-rich substrates (Põlme et al. 2020)</td><td>Põlme S, et al. 2020. Fungal Diversity 105:1-16 ((https://doi.org/10.1007/s13225-020-00466-2))</td></tr>
<tr><td>Otu08686</td><td>3</td><td>k__Fungi(100);p__Basidiomycota(100);c__Agaricomycetes(100);o__Cantharellales(100);f__Tulasnellaceae(100);g__Tulasnella(100);s__Tulasnella_sp_184(100);</td><td>Tulasnella</td><td>13</td><td>Saprotroph-Symbiotroph</td><td>Ectomycorrhizal-|Plant Saprotroph|-Undefined Saprotroph</td><td>Corticioid</td><td>NULL</td><td>Possible</td><td>As litter_saprotroph (Põlme et al. 2020); Decay leaf, fruit, seed (Põlme et al. 2020)</td><td>Rinaldi AC, et al. 2008. Fungal Diversity 33:1-45 ((http://www.fungaldiversity.org/fdp/sfdp/33-1.pdf)); Tedersoo L, et al. 2010. Mycorrhiza 20:217-263 ((https://doi.org/10.1007/s00572-009-0274-x)) (pro parte); Põlme S, et al. 2020. Fungal Diversity 105:1-16 ((https://doi.org/10.1007/s13225-020-00466-2))</td></tr>
<tr><td>Otu08689</td><td>3</td><td>k__Fungi(100);p__Ascomycota(100);c__Saccharomycetes(100);o__Saccharomycetales(100);f__Saccharomycetales_family_Incertae_sedis(100);g__Candida(67);g__Candida_unclassified(67);</td><td>Candida</td><td>13</td><td>Pathotroph-Saprotroph-Symbiotroph</td><td>Animal Pathogen-Endophyte-Endosymbiont-Epiphyte-|Undefined Saprotroph|</td><td>Dimorphic Yeast-Yeast</td><td>NULL</td><td>Probable</td><td>As nectar-tap_saprotroph (Põlme et al. 2020); Decay sugar-rich substrates (Põlme et al. 2020); This asexual genus is polyphyletic and contain species that function very differently depend on what they are. Caution should be taken when interpreting guilds for Candida species.</td><td>Manolakaki D, et al. 2010. Virulence. 1: 367-375 ((https://doi.org/10.4161/viru.1.5.12796)); Mekbib SB, et al. 2011. Tropical Plant Pathology 36:233-240 ((https://doi.org/10.1590/S1982-56762011000400004)); Põlme S, et al. 2020. Fungal Diversity 105:1-16 ((https://doi.org/10.1007/s13225-020-00466-2))</td></tr>
<tr><td>Otu08690</td><td>3</td><td>k__Fungi(100);p__Ascomycota(100);c__Sordariomycetes(100);o__Melanosporales(100);f__Ceratostomataceae(100);g__Harzia(100);s__Harzia_acremonioides(100);</td><td>Harzia</td><td>13</td><td>Saprotroph</td><td>Plant Saprotroph-Undefined Saprotroph-|Wood Saprotroph|</td><td>Microfungus</td><td>NULL</td><td>Probable</td><td>As litter_saprotroph (Põlme et al. 2020); Decay wood, leaf, fruit, seed (Põlme et al. 2020)</td><td>Tedersoo L, et al. 2014. Science 346:e1256688 ((https://doi.org/10.1126/science.1256688)); Põlme S, et al. 2020. Fungal Diversity 105:1-16 ((https://doi.org/10.1007/s13225-020-00466-2))</td></tr>
<tr><td>Otu08692</td><td>3</td><td>k__Fungi(100);p__Basidiomycota(100);c__Agaricomycetes(100);o__Agaricales(100);o__Agaricales_unclassified(100);o__Agaricales_unclassified(100);o__Agaricales_unclassified(100);</td><td>Agaricales</td><td>7</td><td>Pathotroph-Saprotroph-Symbiotroph</td><td>Bryophyte Parasite-Dung Saprotroph-Ectomycorrhizal-Fungal Parasite-Leaf Saprotroph-Plant Parasite-Undefined Saprotroph-Wood Saprotroph</td><td>Agaricoid-Gasteroid-Microfungus-Secotioid-Yeast</td><td>NULL</td><td>Possible</td><td>Mushrooms and toadstools, Gill fungi, Agarics... lignicolous, sometimes muscicolous or fungicolous, saprobic, mycorrhizal, rarely parasitic on plants or fungi (Kirk et al. 2008)</td><td>Kirk PM et al. 2008. Dictionary of the Fungi. Tenth Edition. CAB International, Wallingford (ISBN: 978-0851998268)</td></tr>
<tr><td>Otu08693</td><td>3</td><td>k__Fungi(100);p__Ascomycota(100);c__Dothideomycetes(100);o__Pleosporales(100);f__Venturiaceae(100);f__Venturiaceae_unclassified(100);f__Venturiaceae_unclassified(100);</td><td>Venturiaceae</td><td>9</td><td>Pathotroph-Saprotroph-Symbiotroph</td><td>Endophyte-Plant Pathogen-Undefined Saprotroph</td><td>Microfungus</td><td>NULL</td><td>Probable</td><td>Mostly endophytes and weak parasites of aerial plant parts, including bark and wood, leaf and fruit tissue; rarely biotrophic. Little is known except for the important pathogens (Cannon & Kirk 2007)</td><td>Cannon PF, Kirk PM. 2007. Fungal Families of the World. CAB International, Cambridge (ISBN: 978-0851998275)</td></tr>
<tr><td>Otu08694</td><td>3</td><td>k__Fungi(100);p__Ascomycota(100);c__Saccharomycetes(100);o__Saccharomycetales(100);f__Metschnikowiaceae(100);g__Metschnikowia(100);s__Metschnikowia_pulcherrima(100);</td><td>Metschnikowia</td><td>13</td><td>Saprotroph</td><td>|Undefined Saprotroph|</td><td>Yeast</td><td>NULL</td><td>Probable</td><td>As nectar-tap_saprotroph (Põlme et al. 2020); Decay sugar-rich substrates (Põlme et al. 2020)</td><td>Põlme S, et al. 2020. Fungal Diversity 105:1-16 ((https://doi.org/10.1007/s13225-020-00466-2))</td></tr>
<tr><td>Otu08696</td><td>3</td><td>k__Fungi(100);p__Basidiomycota(100);c__Microbotryomycetes(100);o__Sporidiobolales(100);f__Sporidiobolales_family_Incertae_sedis(100);g__Sporobolomyces(100);s__Sporobolomyces_lactophilus(100);</td><td>Sporobolomyces</td><td>13</td><td>Pathotroph-Saprotroph</td><td>|Fungal Parasite|-Plant Saprotroph</td><td>Yeast</td><td>NULL</td><td>Probable</td><td>As litter_saprotroph (Põlme et al. 2020); As mycoparasite (Põlme et al. 2020); Decay fungal material (Põlme et al. 2020)</td><td>Tedersoo L, et al. 2014. Science 346:e1256688 ((https://doi.org/10.1126/science.1256688)); Purahong W, et al. 2016. Molecular Ecology 25:4059-4074 ((https://doi.org/10.1111/mec.13739)); Põlme S, et al. 2020. Fungal Diversity 105:1-16 ((https://doi.org/10.1007/s13225-020-00466-2))</td></tr>
<tr><td>Otu08697</td><td>3</td><td>k__Fungi(100);p__Ascomycota(100);c__Saccharomycetes(100);o__Saccharomycetales(100);f__Metschnikowiaceae(100);g__Metschnikowia(100);g__Metschnikowia_unclassified(100);</td><td>Metschnikowia</td><td>13</td><td>Saprotroph</td><td>|Undefined Saprotroph|</td><td>Yeast</td><td>NULL</td><td>Probable</td><td>As nectar-tap_saprotroph (Põlme et al. 2020); Decay sugar-rich substrates (Põlme et al. 2020)</td><td>Põlme S, et al. 2020. Fungal Diversity 105:1-16 ((https://doi.org/10.1007/s13225-020-00466-2))</td></tr>
<tr><td>Otu08699</td><td>3</td><td>k__Fungi(100);p__Basidiomycota(100);c__Agaricomycetes(100);o__Agaricales(100);o__Agaricales_unclassified(100);o__Agaricales_unclassified(100);o__Agaricales_unclassified(100);</td><td>Agaricales</td><td>7</td><td>Pathotroph-Saprotroph-Symbiotroph</td><td>Bryophyte Parasite-Dung Saprotroph-Ectomycorrhizal-Fungal Parasite-Leaf Saprotroph-Plant Parasite-Undefined Saprotroph-Wood Saprotroph</td><td>Agaricoid-Gasteroid-Microfungus-Secotioid-Yeast</td><td>NULL</td><td>Possible</td><td>Mushrooms and toadstools, Gill fungi, Agarics... lignicolous, sometimes muscicolous or fungicolous, saprobic, mycorrhizal, rarely parasitic on plants or fungi (Kirk et al. 2008)</td><td>Kirk PM et al. 2008. Dictionary of the Fungi. Tenth Edition. CAB International, Wallingford (ISBN: 978-0851998268)</td></tr>
<tr><td>Otu08700</td><td>3</td><td>k__Fungi(100);p__Basidiomycota(100);c__Agaricomycetes(100);o__Trechisporales(100);f__Hydnodontaceae(100);g__Trechispora(100);s__Trechispora_sp(100);</td><td>Trechispora</td><td>13</td><td>Saprotroph</td><td>|Wood Saprotroph|</td><td>Corticioid</td><td>White Rot</td><td>Highly Probable</td><td>As white_rot (Põlme et al. 2020); Decay wood (Põlme et al. 2020)</td><td>Gilbertson RL, Ryvarden L. 1987-1987. North American Polypores. Fungiflora, Oslo (ISBN: 978-0945345060); Rinaldi AC, et al. 2008. Fungal Diversity 33:1-45 ((http://www.fungaldiversity.org/fdp/sfdp/33-1.pdf)) (as Uncertain); Tedersoo L, et al. 2010. Mycorrhiza 20:217-263 ((https://doi.org/10.1007/s00572-009-0274-x)) (as not ECM).; Põlme S, et al. 2020. Fungal Diversity 105:1-16 ((https://doi.org/10.1007/s13225-020-00466-2))</td></tr>
<tr><td>Otu08702</td><td>3</td><td>k__Fungi(100);p__Ascomycota(100);c__Dothideomycetes(100);o__Botryosphaeriales(100);f__Botryosphaeriaceae(100);g__Endomelanconiopsis(100);g__Endomelanconiopsis_unclassified(100);</td><td>Endomelanconiopsis</td><td>13</td><td>Pathotroph-Saprotroph-Symbiotroph</td><td>|Endophyte|-Plant Pathogen-Undefined Saprotroph</td><td>Microfungus</td><td>NULL</td><td>Probable</td><td>As foliar_endophyte (Põlme et al. 2020); As soil_saprotroph (Põlme et al. 2020); Decay leaf, fruit, seed (Põlme et al. 2020)</td><td>Tedersoo L, et al. 2014. Science 346:e1256688 ((https://doi.org/10.1126/science.1256688)); Põlme S, et al. 2020. Fungal Diversity 105:1-16 ((https://doi.org/10.1007/s13225-020-00466-2))</td></tr>
<tr><td>Otu08703</td><td>3</td><td>k__Fungi(100);p__Ascomycota(100);c__Sordariomycetes(100);o__Hypocreales(100);f__Nectriaceae(100);g__Fusarium(100);g__Fusarium_unclassified(100);</td><td>Fusarium</td><td>13</td><td>Pathotroph-Saprotroph-Symbiotroph</td><td>Animal Pathogen-Endophyte-Lichen Parasite-|Plant Pathogen|-Plant Saprotroph-Undefined Saprotroph-Wood Saprotroph</td><td>Microfungus</td><td>Soft Rot</td><td>Possible</td><td>As foliar_endophyte (Põlme et al. 2020); As litter_saprotroph (Põlme et al. 2020); Causing damping-off disease (Põlme et al. 2020); Decay leaf, fruit, seed, roots, soil, animal material (Põlme et al. 2020); Likely opportunistic human pathogen (Irinyi et al. 2015)</td><td>Duncan CG, Eslyn WE. 1966. Mycologia 58:642-664 ((https://doi.org/10.2307/3757045)); Seehann G, et al. 1975. List of Fungi in Soft-Rot Tissue Tests. Institute fur Holzbiologie und Holzschutz, Bundesforschungsanstalt fur Forst - und Holzwirtshaft, Hamburg - Lohbrugge, Germany; Costa IPMW, et al. 2012. Mycotaxon 119:e26 ((http://mycotaxon.com/resources/checklists/costa_v119_checklist.pdf)); Tedersoo L, et al. 2014. Science 346:e1256688 ((https://doi.org/10.1126/science.1256688)); Irinyi L, et al. 2015. Medical Mycology 53:313-337 ((https://doi.org/10.1093/mmy/myv008)); Schroers, et al. 2016. Mycologia 108:806-819 ((https://doi.org/10.3852/15-255)); Põlme S, et al. 2020. Fungal Diversity 105:1-16 ((https://doi.org/10.1007/s13225-020-00466-2)); The American Phytopathological Society Common Names of Plant Diseases ((http://www.apsnet.org/publications/commonnames/Pages/default.aspx)); Lawrey JD, Diederich P. 2016. http://www.lichenicolous.net (accessed 24 May 2017) ((http://www.lichenicolous.net))</td></tr>
<tr><td>Otu08705</td><td>3</td><td>k__Fungi(100);p__Ascomycota(100);c__Saccharomycetes(100);o__Saccharomycetales(100);f__Metschnikowiaceae(100);g__Metschnikowia(100);s__Metschnikowia_pulcherrima(100);</td><td>Metschnikowia</td><td>13</td><td>Saprotroph</td><td>|Undefined Saprotroph|</td><td>Yeast</td><td>NULL</td><td>Probable</td><td>As nectar-tap_saprotroph (Põlme et al. 2020); Decay sugar-rich substrates (Põlme et al. 2020)</td><td>Põlme S, et al. 2020. Fungal Diversity 105:1-16 ((https://doi.org/10.1007/s13225-020-00466-2))</td></tr>
<tr><td>Otu08706</td><td>3</td><td>k__Fungi(100);p__Ascomycota(100);c__Saccharomycetes(100);o__Saccharomycetales(100);f__Metschnikowiaceae(100);g__Metschnikowia(100);s__Metschnikowia_pulcherrima(100);</td><td>Metschnikowia</td><td>13</td><td>Saprotroph</td><td>|Undefined Saprotroph|</td><td>Yeast</td><td>NULL</td><td>Probable</td><td>As nectar-tap_saprotroph (Põlme et al. 2020); Decay sugar-rich substrates (Põlme et al. 2020)</td><td>Põlme S, et al. 2020. Fungal Diversity 105:1-16 ((https://doi.org/10.1007/s13225-020-00466-2))</td></tr>
<tr><td>Otu08710</td><td>3</td><td>k__Fungi(100);p__Ascomycota(100);c__Saccharomycetes(100);o__Saccharomycetales(100);f__Metschnikowiaceae(100);g__Metschnikowia(100);g__Metschnikowia_unclassified(100);</td><td>Metschnikowia</td><td>13</td><td>Saprotroph</td><td>|Undefined Saprotroph|</td><td>Yeast</td><td>NULL</td><td>Probable</td><td>As nectar-tap_saprotroph (Põlme et al. 2020); Decay sugar-rich substrates (Põlme et al. 2020)</td><td>Põlme S, et al. 2020. Fungal Diversity 105:1-16 ((https://doi.org/10.1007/s13225-020-00466-2))</td></tr>
<tr><td>Otu08713</td><td>3</td><td>k__Fungi(100);p__Ascomycota(100);c__Saccharomycetes(100);o__Saccharomycetales(100);f__Metschnikowiaceae(100);g__Metschnikowia(100);g__Metschnikowia_unclassified(100);</td><td>Metschnikowia</td><td>13</td><td>Saprotroph</td><td>|Undefined Saprotroph|</td><td>Yeast</td><td>NULL</td><td>Probable</td><td>As nectar-tap_saprotroph (Põlme et al. 2020); Decay sugar-rich substrates (Põlme et al. 2020)</td><td>Põlme S, et al. 2020. Fungal Diversity 105:1-16 ((https://doi.org/10.1007/s13225-020-00466-2))</td></tr>
<tr><td>Otu08717</td><td>3</td><td>k__Fungi(100);p__Ascomycota(100);c__Sordariomycetes(100);o__Hypocreales(100);o__Hypocreales_unclassified(100);o__Hypocreales_unclassified(100);o__Hypocreales_unclassified(100);</td><td>Hypocreales</td><td>7</td><td>Saprotroph</td><td>Undefined Saprotroph</td><td>Microfungus</td><td>NULL</td><td>Possible</td><td>NULL</td><td>Sterkenburg E, et al. 2015. New Phytologist 207:1145-1158 ((https://doi.org/10.1111/nph.13426))</td></tr>
<tr><td>Otu08719</td><td>3</td><td>k__Fungi(100);p__Ascomycota(100);c__Saccharomycetes(100);o__Saccharomycetales(100);f__Metschnikowiaceae(100);g__Metschnikowia(100);s__Metschnikowia_pulcherrima(67);</td><td>Metschnikowia</td><td>13</td><td>Saprotroph</td><td>|Undefined Saprotroph|</td><td>Yeast</td><td>NULL</td><td>Probable</td><td>As nectar-tap_saprotroph (Põlme et al. 2020); Decay sugar-rich substrates (Põlme et al. 2020)</td><td>Põlme S, et al. 2020. Fungal Diversity 105:1-16 ((https://doi.org/10.1007/s13225-020-00466-2))</td></tr>
<tr><td>Otu08721</td><td>3</td><td>k__Fungi(100);p__Basidiomycota(100);c__Agaricomycetes(100);o__Agaricales(100);f__Tricholomataceae(100);g__Tricholomopsis(100);s__Tricholomopsis_flammula(100);</td><td>Tricholomopsis</td><td>13</td><td>Saprotroph</td><td>Undefined Saprotroph-|Wood Saprotroph|</td><td>Agaricoid</td><td>White Rot</td><td>Probable</td><td>Decay wood (Põlme et al. 2020)</td><td>Tedersoo L, et al. 2014. Science 346:e1256688 ((https://doi.org/10.1126/science.1256688)); Põlme S, et al. 2020. Fungal Diversity 105:1-16 ((https://doi.org/10.1007/s13225-020-00466-2))</td></tr>
<tr><td>Otu08723</td><td>3</td><td>k__Fungi(100);p__Ascomycota(100);c__Saccharomycetes(100);o__Saccharomycetales(100);f__Metschnikowiaceae(100);g__Metschnikowia(100);s__Metschnikowia_pulcherrima(67);</td><td>Metschnikowia</td><td>13</td><td>Saprotroph</td><td>|Undefined Saprotroph|</td><td>Yeast</td><td>NULL</td><td>Probable</td><td>As nectar-tap_saprotroph (Põlme et al. 2020); Decay sugar-rich substrates (Põlme et al. 2020)</td><td>Põlme S, et al. 2020. Fungal Diversity 105:1-16 ((https://doi.org/10.1007/s13225-020-00466-2))</td></tr>
<tr><td>Otu08724</td><td>3</td><td>k__Fungi(100);p__Ascomycota(100);c__Saccharomycetes(100);o__Saccharomycetales(100);f__Metschnikowiaceae(100);g__Metschnikowia(100);s__Metschnikowia_pulcherrima(100);</td><td>Metschnikowia</td><td>13</td><td>Saprotroph</td><td>|Undefined Saprotroph|</td><td>Yeast</td><td>NULL</td><td>Probable</td><td>As nectar-tap_saprotroph (Põlme et al. 2020); Decay sugar-rich substrates (Põlme et al. 2020)</td><td>Põlme S, et al. 2020. Fungal Diversity 105:1-16 ((https://doi.org/10.1007/s13225-020-00466-2))</td></tr>
<tr><td>Otu08725</td><td>3</td><td>k__Fungi(100);p__Ascomycota(100);c__Saccharomycetes(100);o__Saccharomycetales(100);f__Metschnikowiaceae(100);g__Metschnikowia(100);s__Metschnikowia_pulcherrima(100);</td><td>Metschnikowia</td><td>13</td><td>Saprotroph</td><td>|Undefined Saprotroph|</td><td>Yeast</td><td>NULL</td><td>Probable</td><td>As nectar-tap_saprotroph (Põlme et al. 2020); Decay sugar-rich substrates (Põlme et al. 2020)</td><td>Põlme S, et al. 2020. Fungal Diversity 105:1-16 ((https://doi.org/10.1007/s13225-020-00466-2))</td></tr>
<tr><td>Otu08726</td><td>3</td><td>k__Fungi(100);p__Ascomycota(100);c__Dothideomycetes(100);o__Pleosporales(100);f__Leptosphaeriaceae(100);g__Plenodomus(100);g__Plenodomus_unclassified(100);</td><td>Plenodomus</td><td>13</td><td>Pathotroph-Saprotroph</td><td>|Plant Pathogen|-Plant Saprotroph-Undefined Saprotroph</td><td>Microfungus</td><td>NULL</td><td>Probable</td><td>As litter_saprotroph (Põlme et al. 2020); Decay leaf, fruit, seed (Põlme et al. 2020)</td><td>Tedersoo L, et al. 2014. Science 346:e1256688 ((https://doi.org/10.1126/science.1256688)); Põlme S, et al. 2020. Fungal Diversity 105:1-16 ((https://doi.org/10.1007/s13225-020-00466-2))</td></tr>
<tr><td>Otu08727</td><td>3</td><td>k__Fungi(100);p__Ascomycota(100);c__Saccharomycetes(100);o__Saccharomycetales(100);f__Metschnikowiaceae(100);g__Metschnikowia(100);s__Metschnikowia_pulcherrima(100);</td><td>Metschnikowia</td><td>13</td><td>Saprotroph</td><td>|Undefined Saprotroph|</td><td>Yeast</td><td>NULL</td><td>Probable</td><td>As nectar-tap_saprotroph (Põlme et al. 2020); Decay sugar-rich substrates (Põlme et al. 2020)</td><td>Põlme S, et al. 2020. Fungal Diversity 105:1-16 ((https://doi.org/10.1007/s13225-020-00466-2))</td></tr>
<tr><td>Otu08732</td><td>3</td><td>k__Fungi(100);p__Ascomycota(100);c__Saccharomycetes(100);o__Saccharomycetales(100);f__Metschnikowiaceae(100);g__Metschnikowia(100);s__Metschnikowia_pulcherrima(67);</td><td>Metschnikowia</td><td>13</td><td>Saprotroph</td><td>|Undefined Saprotroph|</td><td>Yeast</td><td>NULL</td><td>Probable</td><td>As nectar-tap_saprotroph (Põlme et al. 2020); Decay sugar-rich substrates (Põlme et al. 2020)</td><td>Põlme S, et al. 2020. Fungal Diversity 105:1-16 ((https://doi.org/10.1007/s13225-020-00466-2))</td></tr>
<tr><td>Otu08734</td><td>3</td><td>k__Fungi(100);p__Ascomycota(100);c__Saccharomycetes(100);o__Saccharomycetales(100);f__Metschnikowiaceae(100);g__Metschnikowia(100);s__Metschnikowia_pulcherrima(67);</td><td>Metschnikowia</td><td>13</td><td>Saprotroph</td><td>|Undefined Saprotroph|</td><td>Yeast</td><td>NULL</td><td>Probable</td><td>As nectar-tap_saprotroph (Põlme et al. 2020); Decay sugar-rich substrates (Põlme et al. 2020)</td><td>Põlme S, et al. 2020. Fungal Diversity 105:1-16 ((https://doi.org/10.1007/s13225-020-00466-2))</td></tr>
<tr><td>Otu08735</td><td>3</td><td>k__Fungi(100);p__Ascomycota(100);c__Saccharomycetes(100);o__Saccharomycetales(100);f__Metschnikowiaceae(100);g__Metschnikowia(100);g__Metschnikowia_unclassified(67);</td><td>Metschnikowia</td><td>13</td><td>Saprotroph</td><td>|Undefined Saprotroph|</td><td>Yeast</td><td>NULL</td><td>Probable</td><td>As nectar-tap_saprotroph (Põlme et al. 2020); Decay sugar-rich substrates (Põlme et al. 2020)</td><td>Põlme S, et al. 2020. Fungal Diversity 105:1-16 ((https://doi.org/10.1007/s13225-020-00466-2))</td></tr>
<tr><td>Otu08737</td><td>3</td><td>k__Fungi(100);p__Basidiomycota(100);c__Agaricomycetes(100);o__Russulales(100);f__Peniophoraceae(100);f__Peniophoraceae_unclassified(100);f__Peniophoraceae_unclassified(100);</td><td>Peniophoraceae</td><td>9</td><td>Pathotroph-Saprotroph</td><td>Plant Pathogen-Wood Saprotroph</td><td>Corticioid</td><td>White Rot</td><td>Probable</td><td>Most species are saprobic, causing rots of standing and fallen wood. Many species are adapted to xeric conditions, being able to grow on rapidly drying dead attached branches (Cannon & Kirk 2007)</td><td>Cannon PF, Kirk PM. 2007. Fungal Families of the World. CAB International, Cambridge (ISBN: 978-0851998275)</td></tr>
<tr><td>Otu08746</td><td>3</td><td>k__Fungi(100);p__Ascomycota(100);c__Dothideomycetes(100);o__Myriangiales(100);f__Elsinoaceae(100);f__Elsinoaceae_unclassified(100);f__Elsinoaceae_unclassified(100);</td><td>Elsinoaceae</td><td>9</td><td>Pathotroph</td><td>Plant Pathogen</td><td>Microfungus</td><td>NULL</td><td>Probable</td><td>Biotrophic or necrotrophic on leaves, stems and fruits of a wide range of plants (Cannon & Kirk 2007)</td><td>Cannon PF, Kirk PM. 2007. Fungal Families of the World. CAB International, Cambridge (ISBN: 978-0851998275)</td></tr>
<tr><td>Otu08748</td><td>3</td><td>k__Fungi(100);p__Ascomycota(100);c__Pezizomycetes(100);o__Pezizales(100);f__Pezizaceae(100);g__Pachyphloeus(100);s__Pachyphloeus_sp_18_RH_2012(100);</td><td>Pachyphloeus</td><td>13</td><td>Symbiotroph</td><td>|Ectomycorrhizal|</td><td>Gasteroid</td><td>Hypogeous</td><td>Highly Probable</td><td>NULL</td><td>Rinaldi AC, et al. 2008. Fungal Diversity 33:1-45 ((http://www.fungaldiversity.org/fdp/sfdp/33-1.pdf)); Tedersoo L, et al. 2010. Mycorrhiza 20:217-263 ((https://doi.org/10.1007/s00572-009-0274-x)); Põlme S, et al. 2020. Fungal Diversity 105:1-16 ((https://doi.org/10.1007/s13225-020-00466-2)); http://mycorrhizas.info/ecmf.html ((http://mycorrhizas.info/ecmf.html))</td></tr>
<tr><td>Otu08749</td><td>3</td><td>k__Fungi(100);p__Ascomycota(100);c__Saccharomycetes(100);o__Saccharomycetales(100);f__Metschnikowiaceae(100);g__Metschnikowia(100);s__Metschnikowia_pulcherrima(100);</td><td>Metschnikowia</td><td>13</td><td>Saprotroph</td><td>|Undefined Saprotroph|</td><td>Yeast</td><td>NULL</td><td>Probable</td><td>As nectar-tap_saprotroph (Põlme et al. 2020); Decay sugar-rich substrates (Põlme et al. 2020)</td><td>Põlme S, et al. 2020. Fungal Diversity 105:1-16 ((https://doi.org/10.1007/s13225-020-00466-2))</td></tr>
<tr><td>Otu08750</td><td>3</td><td>k__Fungi(100);p__Ascomycota(100);c__Saccharomycetes(100);o__Saccharomycetales(100);f__Metschnikowiaceae(100);g__Metschnikowia(100);s__Metschnikowia_pulcherrima(100);</td><td>Metschnikowia</td><td>13</td><td>Saprotroph</td><td>|Undefined Saprotroph|</td><td>Yeast</td><td>NULL</td><td>Probable</td><td>As nectar-tap_saprotroph (Põlme et al. 2020); Decay sugar-rich substrates (Põlme et al. 2020)</td><td>Põlme S, et al. 2020. Fungal Diversity 105:1-16 ((https://doi.org/10.1007/s13225-020-00466-2))</td></tr>
<tr><td>Otu08751</td><td>3</td><td>k__Fungi(100);p__Ascomycota(100);c__Sordariomycetes(100);o__Sordariales(100);f__Cephalothecaceae(100);g__unclassified_Cephalothecaceae(100);s__Cephalothecaceae_sp(100);</td><td>Cephalothecaceae</td><td>9</td><td>Saprotroph</td><td>Fungal Parasite-Wood Saprotroph</td><td>Microfungus</td><td>NULL</td><td>Probable</td><td>Typically saprobic, fruiting on very rotten wood or other fungi (Cannon & Kirk 2007)</td><td>Cannon PF, Kirk PM. 2007. Fungal Families of the World. CAB International, Cambridge (ISBN: 978-0851998275)</td></tr>
<tr><td>Otu08756</td><td>3</td><td>k__Fungi(100);p__Ascomycota(100);c__Saccharomycetes(100);o__Saccharomycetales(100);f__Metschnikowiaceae(100);g__Metschnikowia(100);s__Metschnikowia_pulcherrima(100);</td><td>Metschnikowia</td><td>13</td><td>Saprotroph</td><td>|Undefined Saprotroph|</td><td>Yeast</td><td>NULL</td><td>Probable</td><td>As nectar-tap_saprotroph (Põlme et al. 2020); Decay sugar-rich substrates (Põlme et al. 2020)</td><td>Põlme S, et al. 2020. Fungal Diversity 105:1-16 ((https://doi.org/10.1007/s13225-020-00466-2))</td></tr>
<tr><td>Otu08757</td><td>3</td><td>k__Fungi(100);p__Ascomycota(100);c__Eurotiomycetes(100);o__Eurotiales(100);f__Trichocomaceae(100);g__Penicillium(100);s__Penicillium_sp_CMV_2008c(67);</td><td>Penicillium</td><td>13</td><td>Pathotroph-Saprotroph-Symbiotroph</td><td>Animal Parasite-Dung Saprotroph-Endophyte-Fungal Parasite-Plant Pathogen-|Undefined Saprotroph|-Wood Saprotroph</td><td>Microfungus</td><td>Soft Rot</td><td>Highly Probable</td><td>As foliar_endophyte (Põlme et al. 2020); As mycoparasite (Põlme et al. 2020); Decay soil (Põlme et al. 2020)</td><td>Duncan CG, Eslyn WE. 1966. Mycologia 58:642-664 ((https://doi.org/10.2307/3757045)); Seehann G, et al. 1975. List of Fungi in Soft-Rot Tissue Tests. Institute fur Holzbiologie und Holzschutz, Bundesforschungsanstalt fur Forst - und Holzwirtshaft, Hamburg - Lohbrugge, Germany ((https://www.irg-wp.com/irgdocs/details.php?d6493033-c9ea-40f3-8e3e-5697f32d640d)); Costa IPMW, et al. 2012. Mycotaxon 119:e26 ((http://mycotaxon.com/resources/checklists/costa_v119_checklist.pdf)); Bills GF, et al. 2013. Current Opinion in Microbiology 16:549-565 ((https://doi.org/10.1016/j.mib.2013.08.001)); Tedersoo L, et al. 2014. Science 346:e1256688 ((https://doi.org/10.1126/science.1256688)); Põlme S, et al. 2020. Fungal Diversity 105:1-16 ((https://doi.org/10.1007/s13225-020-00466-2))</td></tr>
<tr><td>Otu08758</td><td>3</td><td>k__Fungi(100);p__Basidiomycota(100);c__Agaricomycetes(100);o__Agaricales(100);f__Agaricaceae(100);g__Bovista(100);s__Bovista_plumbea(100);</td><td>Bovista</td><td>13</td><td>Saprotroph</td><td>|Undefined Saprotroph|</td><td>Gasteroid</td><td>NULL</td><td>Probable</td><td>puffball; As soil_saprotroph (Põlme et al. 2020); Common name ; Decay soil (Põlme et al. 2020)</td><td>Grgurinovic CA, Simpson JA. 2001. Fungal Diversity 8:97-127 ((http://www.fungaldiversity.org/fdp/sfdp/FD_8_97-127.pdf)); Tedersoo L, et al. 2014. Science 346:e1256688 ((https://doi.org/10.1126/science.1256688)); Põlme S, et al. 2020. Fungal Diversity 105:1-16 ((https://doi.org/10.1007/s13225-020-00466-2))</td></tr>
<tr><td>Otu08759</td><td>3</td><td>k__Fungi(100);p__Ascomycota(100);c__Sordariomycetes(100);o__Coniochaetales(100);f__Coniochaetaceae(100);g__Coniochaeta(100);s__Coniochaeta_prunicola(100);</td><td>Coniochaeta</td><td>13</td><td>Pathotroph-Saprotroph-Symbiotroph</td><td>Animal Pathogen-Dung Saprotroph-Endophyte-Lichen Parasite-Plant Pathogen-|Undefined Saprotroph|</td><td>Microfungus</td><td>NULL</td><td>Possible</td><td>As foliar_endophyte (Põlme et al. 2020); Decay leaf, fruit, seed, soil, dung, animal material (Põlme et al. 2020)</td><td>Bell A. 1983. Dung Fungi: An Illustrated Guide to Coprophilous Fungi in New Zealand. Victoria University Press, Wellington (ISBN: 978-0864730015); Damm U, et al. 2010. Persoonia 24:60-80 ((http://doi.org/10.3767/003158510X500705)); Costa IPMW, et al. 2012. Mycotaxon 119:e26 ((http://mycotaxon.com/resources/checklists/costa_v119_checklist.pdf)); Tedersoo L, et al. 2014. Science 346:1256688 ((https://doi.org/10.1126/science.1256688)); Irinyi L, et al. 2015. Medical Mycology 53:313-337 ((https://doi.org/10.1093/mmy/myv008)); Lawrey JD, Diederich P. 2016. http://www.lichenicolous.net (accessed 24 May 2017) ((http://www.lichenicolous.net)); Põlme S, et al. 2020. Fungal Diversity 105:1-16 ((https://doi.org/10.1007/s13225-020-00466-2))</td></tr>
<tr><td>Otu08760</td><td>3</td><td>k__Fungi(100);p__Ascomycota(100);c__Saccharomycetes(100);o__Saccharomycetales(100);f__Metschnikowiaceae(100);g__Metschnikowia(100);s__Metschnikowia_pulcherrima(67);</td><td>Metschnikowia</td><td>13</td><td>Saprotroph</td><td>|Undefined Saprotroph|</td><td>Yeast</td><td>NULL</td><td>Probable</td><td>As nectar-tap_saprotroph (Põlme et al. 2020); Decay sugar-rich substrates (Põlme et al. 2020)</td><td>Põlme S, et al. 2020. Fungal Diversity 105:1-16 ((https://doi.org/10.1007/s13225-020-00466-2))</td></tr>
<tr><td>Otu08765</td><td>3</td><td>k__Fungi(100);p__Ascomycota(100);c__Saccharomycetes(100);o__Saccharomycetales(100);f__Lipomycetaceae(100);g__Lipomyces(100);s__Lipomyces_sp(100);</td><td>Lipomyces</td><td>13</td><td>Saprotroph</td><td>|Undefined Saprotroph|</td><td>Yeast</td><td>NULL</td><td>Probable</td><td>As nectar-tap_saprotroph (Põlme et al. 2020); Decay sugar-rich substrates (Põlme et al. 2020)</td><td>Põlme S, et al. 2020. Fungal Diversity 105:1-16 ((https://doi.org/10.1007/s13225-020-00466-2))</td></tr>
<tr><td>Otu08768</td><td>3</td><td>k__Fungi(100);p__Ascomycota(100);c__Dothideomycetes(100);o__Dothideomycetes_order_Incertae_sedis(100);f__Myxotrichaceae(100);g__Oidiodendron(100);s__Oidiodendron_maius(100);</td><td>Oidiodendron</td><td>13</td><td>Pathotroph-Saprotroph-Symbiotroph</td><td>Endophyte-Ericoid Mycorrhizal-|Undefined Saprotroph|</td><td>Microfungus</td><td>NULL</td><td>Probable</td><td>As root_endophyte (Põlme et al. 2020); As soil_saprotroph (Põlme et al. 2020); Dark Septate Endophyte; Decay soil (Põlme et al. 2020)</td><td>Newsham KK. 2011. New Phytologist 190:783-793 ((https://doi.org/10.1111/j.1469-8137.2010.03611.x)) (as dark septate root endophytes); Põlme S, et al. 2020. Fungal Diversity 105:1-16 ((https://doi.org/10.1007/s13225-020-00466-2))</td></tr>
<tr><td>Otu08769</td><td>3</td><td>k__Fungi(100);p__Ascomycota(100);c__Saccharomycetes(100);o__Saccharomycetales(100);f__Metschnikowiaceae(100);g__Metschnikowia(100);s__Metschnikowia_pulcherrima(100);</td><td>Metschnikowia</td><td>13</td><td>Saprotroph</td><td>|Undefined Saprotroph|</td><td>Yeast</td><td>NULL</td><td>Probable</td><td>As nectar-tap_saprotroph (Põlme et al. 2020); Decay sugar-rich substrates (Põlme et al. 2020)</td><td>Põlme S, et al. 2020. Fungal Diversity 105:1-16 ((https://doi.org/10.1007/s13225-020-00466-2))</td></tr>
<tr><td>Otu08770</td><td>3</td><td>k__Fungi(100);p__Ascomycota(100);c__Saccharomycetes(100);o__Saccharomycetales(100);f__Metschnikowiaceae(100);g__Metschnikowia(100);g__Metschnikowia_unclassified(100);</td><td>Metschnikowia</td><td>13</td><td>Saprotroph</td><td>|Undefined Saprotroph|</td><td>Yeast</td><td>NULL</td><td>Probable</td><td>As nectar-tap_saprotroph (Põlme et al. 2020); Decay sugar-rich substrates (Põlme et al. 2020)</td><td>Põlme S, et al. 2020. Fungal Diversity 105:1-16 ((https://doi.org/10.1007/s13225-020-00466-2))</td></tr>
<tr><td>Otu08771</td><td>3</td><td>k__Fungi(100);p__Ascomycota(100);c__Saccharomycetes(100);o__Saccharomycetales(100);o__Saccharomycetales_unclassified(67);o__Saccharomycetales_unclassified(67);o__Saccharomycetales_unclassified(67);</td><td>Saccharomycetales</td><td>7</td><td>Saprotroph</td><td>Undefined Saprotroph</td><td>Yeast</td><td>NULL</td><td>Possible</td><td>NULL</td><td>Sterkenburg E, et al. 2015. New Phytologist 207:1145-1158 ((https://doi.org/10.1111/nph.13426))</td></tr>
<tr><td>Otu08772</td><td>3</td><td>k__Fungi(100);p__Zygomycota(100);c__Zygomycota_class_Incertae_sedis(100);o__Entomophthorales(100);f__Entomophthoraceae(100);g__Pandora(100);g__Pandora_unclassified(100);</td><td>Pandora</td><td>13</td><td>Pathotroph-Saprotroph</td><td>|Animal Parasite|-Animal Pathogen-Undefined Saprotroph</td><td>Microfungus</td><td>NULL</td><td>Probable</td><td>As animal_decomposer (Põlme et al. 2020); Decay animal material (Põlme et al. 2020)</td><td>Tedersoo L, et al. 2014. Science 346:e1256688 ((https://doi.org/10.1126/science.1256688)); Põlme S, et al. 2020. Fungal Diversity 105:1-16 ((https://doi.org/10.1007/s13225-020-00466-2))</td></tr>
<tr><td>Otu08773</td><td>3</td><td>k__Fungi(100);p__Ascomycota(100);c__Saccharomycetes(100);o__Saccharomycetales(100);f__Metschnikowiaceae(100);g__Metschnikowia(100);g__Metschnikowia_unclassified(100);</td><td>Metschnikowia</td><td>13</td><td>Saprotroph</td><td>|Undefined Saprotroph|</td><td>Yeast</td><td>NULL</td><td>Probable</td><td>As nectar-tap_saprotroph (Põlme et al. 2020); Decay sugar-rich substrates (Põlme et al. 2020)</td><td>Põlme S, et al. 2020. Fungal Diversity 105:1-16 ((https://doi.org/10.1007/s13225-020-00466-2))</td></tr>
<tr><td>Otu08777</td><td>3</td><td>k__Fungi(100);p__Ascomycota(100);c__Saccharomycetes(100);o__Saccharomycetales(100);f__Metschnikowiaceae(100);g__Metschnikowia(100);s__Metschnikowia_pulcherrima(67);</td><td>Metschnikowia</td><td>13</td><td>Saprotroph</td><td>|Undefined Saprotroph|</td><td>Yeast</td><td>NULL</td><td>Probable</td><td>As nectar-tap_saprotroph (Põlme et al. 2020); Decay sugar-rich substrates (Põlme et al. 2020)</td><td>Põlme S, et al. 2020. Fungal Diversity 105:1-16 ((https://doi.org/10.1007/s13225-020-00466-2))</td></tr>
<tr><td>Otu08780</td><td>3</td><td>k__Fungi(100);p__Ascomycota(100);c__Saccharomycetes(100);o__Saccharomycetales(100);f__Metschnikowiaceae(100);g__Metschnikowia(100);s__Metschnikowia_pulcherrima(100);</td><td>Metschnikowia</td><td>13</td><td>Saprotroph</td><td>|Undefined Saprotroph|</td><td>Yeast</td><td>NULL</td><td>Probable</td><td>As nectar-tap_saprotroph (Põlme et al. 2020); Decay sugar-rich substrates (Põlme et al. 2020)</td><td>Põlme S, et al. 2020. Fungal Diversity 105:1-16 ((https://doi.org/10.1007/s13225-020-00466-2))</td></tr>
<tr><td>Otu08782</td><td>3</td><td>k__Fungi(100);p__Ascomycota(100);c__Saccharomycetes(100);o__Saccharomycetales(100);f__Metschnikowiaceae(100);g__Metschnikowia(100);g__Metschnikowia_unclassified(67);</td><td>Metschnikowia</td><td>13</td><td>Saprotroph</td><td>|Undefined Saprotroph|</td><td>Yeast</td><td>NULL</td><td>Probable</td><td>As nectar-tap_saprotroph (Põlme et al. 2020); Decay sugar-rich substrates (Põlme et al. 2020)</td><td>Põlme S, et al. 2020. Fungal Diversity 105:1-16 ((https://doi.org/10.1007/s13225-020-00466-2))</td></tr>
<tr><td>Otu08785</td><td>3</td><td>k__Fungi(100);p__Ascomycota(100);c__Saccharomycetes(100);o__Saccharomycetales(100);f__Metschnikowiaceae(100);g__Metschnikowia(100);s__Metschnikowia_pulcherrima(100);</td><td>Metschnikowia</td><td>13</td><td>Saprotroph</td><td>|Undefined Saprotroph|</td><td>Yeast</td><td>NULL</td><td>Probable</td><td>As nectar-tap_saprotroph (Põlme et al. 2020); Decay sugar-rich substrates (Põlme et al. 2020)</td><td>Põlme S, et al. 2020. Fungal Diversity 105:1-16 ((https://doi.org/10.1007/s13225-020-00466-2))</td></tr>
<tr><td>Otu08789</td><td>3</td><td>k__Fungi(100);p__Ascomycota(100);c__Saccharomycetes(100);o__Saccharomycetales(100);f__Metschnikowiaceae(100);g__Metschnikowia(100);s__Metschnikowia_pulcherrima(67);</td><td>Metschnikowia</td><td>13</td><td>Saprotroph</td><td>|Undefined Saprotroph|</td><td>Yeast</td><td>NULL</td><td>Probable</td><td>As nectar-tap_saprotroph (Põlme et al. 2020); Decay sugar-rich substrates (Põlme et al. 2020)</td><td>Põlme S, et al. 2020. Fungal Diversity 105:1-16 ((https://doi.org/10.1007/s13225-020-00466-2))</td></tr>
<tr><td>Otu08790</td><td>3</td><td>k__Fungi(100);p__Ascomycota(100);c__Saccharomycetes(100);o__Saccharomycetales(100);f__Metschnikowiaceae(100);g__Metschnikowia(100);g__Metschnikowia_unclassified(67);</td><td>Metschnikowia</td><td>13</td><td>Saprotroph</td><td>|Undefined Saprotroph|</td><td>Yeast</td><td>NULL</td><td>Probable</td><td>As nectar-tap_saprotroph (Põlme et al. 2020); Decay sugar-rich substrates (Põlme et al. 2020)</td><td>Põlme S, et al. 2020. Fungal Diversity 105:1-16 ((https://doi.org/10.1007/s13225-020-00466-2))</td></tr>
<tr><td>Otu08791</td><td>3</td><td>k__Fungi(100);p__Ascomycota(100);c__Sordariomycetes(100);o__Sordariales(100);f__Lasiosphaeriaceae(100);g__Zopfiella(67);g__Zopfiella_unclassified(67);</td><td>Zopfiella</td><td>13</td><td>Saprotroph</td><td>|Dung Saprotroph|</td><td>Microfungus</td><td>NULL</td><td>Highly Probable</td><td>Decay dung (Põlme et al. 2020)</td><td>Bills GF, et al. 2013. Current Opinion in Microbiology 16:549-565 ((https://doi.org/10.1016/j.mib.2013.08.001)); Tedersoo L, et al. 2014. Science 346:e1256688 ((https://doi.org/10.1126/science.1256688)); Põlme S, et al. 2020. Fungal Diversity 105:1-16 ((https://doi.org/10.1007/s13225-020-00466-2))</td></tr>
<tr><td>Otu08792</td><td>3</td><td>k__Fungi(100);p__Ascomycota(100);c__Saccharomycetes(100);o__Saccharomycetales(100);f__Metschnikowiaceae(100);g__Metschnikowia(100);g__Metschnikowia_unclassified(100);</td><td>Metschnikowia</td><td>13</td><td>Saprotroph</td><td>|Undefined Saprotroph|</td><td>Yeast</td><td>NULL</td><td>Probable</td><td>As nectar-tap_saprotroph (Põlme et al. 2020); Decay sugar-rich substrates (Põlme et al. 2020)</td><td>Põlme S, et al. 2020. Fungal Diversity 105:1-16 ((https://doi.org/10.1007/s13225-020-00466-2))</td></tr>
<tr><td>Otu06933</td><td>3</td><td>k__Fungi(100);p__Basidiomycota(100);c__Tremellomycetes(100);c__Tremellomycetes_unclassified(100);c__Tremellomycetes_unclassified(100);c__Tremellomycetes_unclassified(100);c__Tremellomycetes_unclassified(100);</td><td>-</td><td>-</td><td>-</td><td>-</td><td>-</td><td>-</td><td>-</td><td>Unassigned</td><td>-</td></tr>
<tr><td>Otu06934</td><td>3</td><td>k__Fungi(100);k__Fungi_unclassified(100);k__Fungi_unclassified(100);k__Fungi_unclassified(100);k__Fungi_unclassified(100);k__Fungi_unclassified(100);k__Fungi_unclassified(100);</td><td>-</td><td>-</td><td>-</td><td>-</td><td>-</td><td>-</td><td>-</td><td>Unassigned</td><td>-</td></tr>
<tr><td>Otu06937</td><td>3</td><td>k__Fungi(100);p__Basidiomycota(100);p__Basidiomycota_unclassified(100);p__Basidiomycota_unclassified(100);p__Basidiomycota_unclassified(100);p__Basidiomycota_unclassified(100);p__Basidiomycota_unclassified(100);</td><td>-</td><td>-</td><td>-</td><td>-</td><td>-</td><td>-</td><td>-</td><td>Unassigned</td><td>-</td></tr>
<tr><td>Otu06938</td><td>3</td><td>k__Fungi(100);p__Ascomycota(100);c__Leotiomycetes(67);o__Helotiales(67);o__Helotiales_unclassified(67);o__Helotiales_unclassified(67);o__Helotiales_unclassified(67);</td><td>-</td><td>-</td><td>-</td><td>-</td><td>-</td><td>-</td><td>-</td><td>Unassigned</td><td>-</td></tr>
<tr><td>Otu06939</td><td>3</td><td>k__Fungi(100);k__Fungi_unclassified(67);k__Fungi_unclassified(67);k__Fungi_unclassified(67);k__Fungi_unclassified(67);k__Fungi_unclassified(67);k__Fungi_unclassified(67);</td><td>-</td><td>-</td><td>-</td><td>-</td><td>-</td><td>-</td><td>-</td><td>Unassigned</td><td>-</td></tr>
<tr><td>Otu06941</td><td>3</td><td>k__Fungi(100);k__Fungi_unclassified(67);k__Fungi_unclassified(67);k__Fungi_unclassified(67);k__Fungi_unclassified(67);k__Fungi_unclassified(67);k__Fungi_unclassified(67);</td><td>-</td><td>-</td><td>-</td><td>-</td><td>-</td><td>-</td><td>-</td><td>Unassigned</td><td>-</td></tr>
<tr><td>Otu06942</td><td>3</td><td>k__Fungi(100);p__Basidiomycota(100);c__Agaricomycetes(100);c__Agaricomycetes_unclassified(100);c__Agaricomycetes_unclassified(100);c__Agaricomycetes_unclassified(100);c__Agaricomycetes_unclassified(100);</td><td>-</td><td>-</td><td>-</td><td>-</td><td>-</td><td>-</td><td>-</td><td>Unassigned</td><td>-</td></tr>
<tr><td>Otu06944</td><td>3</td><td>k__Fungi(100);k__Fungi_unclassified(100);k__Fungi_unclassified(100);k__Fungi_unclassified(100);k__Fungi_unclassified(100);k__Fungi_unclassified(100);k__Fungi_unclassified(100);</td><td>-</td><td>-</td><td>-</td><td>-</td><td>-</td><td>-</td><td>-</td><td>Unassigned</td><td>-</td></tr>
<tr><td>Otu06945</td><td>3</td><td>k__Fungi(100);p__Ascomycota(67);p__Ascomycota_unclassified(67);p__Ascomycota_unclassified(67);p__Ascomycota_unclassified(67);p__Ascomycota_unclassified(67);p__Ascomycota_unclassified(67);</td><td>-</td><td>-</td><td>-</td><td>-</td><td>-</td><td>-</td><td>-</td><td>Unassigned</td><td>-</td></tr>
<tr><td>Otu06947</td><td>3</td><td>k__Fungi(100);p__Ascomycota(100);p__Ascomycota_unclassified(100);p__Ascomycota_unclassified(100);p__Ascomycota_unclassified(100);p__Ascomycota_unclassified(100);p__Ascomycota_unclassified(100);</td><td>-</td><td>-</td><td>-</td><td>-</td><td>-</td><td>-</td><td>-</td><td>Unassigned</td><td>-</td></tr>
<tr><td>Otu06948</td><td>3</td><td>k__Fungi(100);p__Ascomycota(100);p__Ascomycota_unclassified(100);p__Ascomycota_unclassified(100);p__Ascomycota_unclassified(100);p__Ascomycota_unclassified(100);p__Ascomycota_unclassified(100);</td><td>-</td><td>-</td><td>-</td><td>-</td><td>-</td><td>-</td><td>-</td><td>Unassigned</td><td>-</td></tr>
<tr><td>Otu06949</td><td>3</td><td>k__Fungi(100);p__Ascomycota(100);c__Dothideomycetes(100);o__Pleosporales(100);o__Pleosporales_unclassified(100);o__Pleosporales_unclassified(100);o__Pleosporales_unclassified(100);</td><td>-</td><td>-</td><td>-</td><td>-</td><td>-</td><td>-</td><td>-</td><td>Unassigned</td><td>-</td></tr>
<tr><td>Otu06952</td><td>3</td><td>k__Fungi(100);k__Fungi_unclassified(100);k__Fungi_unclassified(100);k__Fungi_unclassified(100);k__Fungi_unclassified(100);k__Fungi_unclassified(100);k__Fungi_unclassified(100);</td><td>-</td><td>-</td><td>-</td><td>-</td><td>-</td><td>-</td><td>-</td><td>Unassigned</td><td>-</td></tr>
<tr><td>Otu06957</td><td>3</td><td>k__Fungi(100);p__Ascomycota(100);p__Ascomycota_unclassified(100);p__Ascomycota_unclassified(100);p__Ascomycota_unclassified(100);p__Ascomycota_unclassified(100);p__Ascomycota_unclassified(100);</td><td>-</td><td>-</td><td>-</td><td>-</td><td>-</td><td>-</td><td>-</td><td>Unassigned</td><td>-</td></tr>
<tr><td>Otu06961</td><td>3</td><td>k__Fungi(100);p__Ascomycota(100);p__Ascomycota_unclassified(100);p__Ascomycota_unclassified(100);p__Ascomycota_unclassified(100);p__Ascomycota_unclassified(100);p__Ascomycota_unclassified(100);</td><td>-</td><td>-</td><td>-</td><td>-</td><td>-</td><td>-</td><td>-</td><td>Unassigned</td><td>-</td></tr>
<tr><td>Otu06963</td><td>3</td><td>k__Fungi(100);p__Basidiomycota(100);c__Agaricomycetes(100);o__Russulales(100);o__Russulales_unclassified(100);o__Russulales_unclassified(100);o__Russulales_unclassified(100);</td><td>-</td><td>-</td><td>-</td><td>-</td><td>-</td><td>-</td><td>-</td><td>Unassigned</td><td>-</td></tr>
<tr><td>Otu06965</td><td>3</td><td>k__Fungi(100);k__Fungi_unclassified(100);k__Fungi_unclassified(100);k__Fungi_unclassified(100);k__Fungi_unclassified(100);k__Fungi_unclassified(100);k__Fungi_unclassified(100);</td><td>-</td><td>-</td><td>-</td><td>-</td><td>-</td><td>-</td><td>-</td><td>Unassigned</td><td>-</td></tr>
<tr><td>Otu06972</td><td>3</td><td>k__Fungi(100);p__Ascomycota(67);p__Ascomycota_unclassified(67);p__Ascomycota_unclassified(67);p__Ascomycota_unclassified(67);p__Ascomycota_unclassified(67);p__Ascomycota_unclassified(67);</td><td>-</td><td>-</td><td>-</td><td>-</td><td>-</td><td>-</td><td>-</td><td>Unassigned</td><td>-</td></tr>
<tr><td>Otu06973</td><td>3</td><td>k__Fungi(100);p__Ascomycota(67);p__Ascomycota_unclassified(67);p__Ascomycota_unclassified(67);p__Ascomycota_unclassified(67);p__Ascomycota_unclassified(67);p__Ascomycota_unclassified(67);</td><td>-</td><td>-</td><td>-</td><td>-</td><td>-</td><td>-</td><td>-</td><td>Unassigned</td><td>-</td></tr>
<tr><td>Otu06976</td><td>3</td><td>k__Fungi(100);p__Ascomycota(100);p__Ascomycota_unclassified(100);p__Ascomycota_unclassified(100);p__Ascomycota_unclassified(100);p__Ascomycota_unclassified(100);p__Ascomycota_unclassified(100);</td><td>-</td><td>-</td><td>-</td><td>-</td><td>-</td><td>-</td><td>-</td><td>Unassigned</td><td>-</td></tr>
<tr><td>Otu06977</td><td>3</td><td>k__Fungi(100);p__Chytridiomycota(100);c__Chytridiomycetes(100);o__Rhizophydiales(100);o__Rhizophydiales_unclassified(100);o__Rhizophydiales_unclassified(100);o__Rhizophydiales_unclassified(100);</td><td>-</td><td>-</td><td>-</td><td>-</td><td>-</td><td>-</td><td>-</td><td>Unassigned</td><td>-</td></tr>
<tr><td>Otu06978</td><td>3</td><td>k__Fungi(100);p__Ascomycota(100);p__Ascomycota_unclassified(67);p__Ascomycota_unclassified(67);p__Ascomycota_unclassified(67);p__Ascomycota_unclassified(67);p__Ascomycota_unclassified(67);</td><td>-</td><td>-</td><td>-</td><td>-</td><td>-</td><td>-</td><td>-</td><td>Unassigned</td><td>-</td></tr>
<tr><td>Otu06979</td><td>3</td><td>k__Fungi(100);p__Ascomycota(67);p__Ascomycota_unclassified(67);p__Ascomycota_unclassified(67);p__Ascomycota_unclassified(67);p__Ascomycota_unclassified(67);p__Ascomycota_unclassified(67);</td><td>-</td><td>-</td><td>-</td><td>-</td><td>-</td><td>-</td><td>-</td><td>Unassigned</td><td>-</td></tr>
<tr><td>Otu06982</td><td>3</td><td>k__Fungi(100);p__Basidiomycota(100);p__Basidiomycota_unclassified(100);p__Basidiomycota_unclassified(100);p__Basidiomycota_unclassified(100);p__Basidiomycota_unclassified(100);p__Basidiomycota_unclassified(100);</td><td>-</td><td>-</td><td>-</td><td>-</td><td>-</td><td>-</td><td>-</td><td>Unassigned</td><td>-</td></tr>
<tr><td>Otu06984</td><td>3</td><td>k__Fungi(100);k__Fungi_unclassified(100);k__Fungi_unclassified(100);k__Fungi_unclassified(100);k__Fungi_unclassified(100);k__Fungi_unclassified(100);k__Fungi_unclassified(100);</td><td>-</td><td>-</td><td>-</td><td>-</td><td>-</td><td>-</td><td>-</td><td>Unassigned</td><td>-</td></tr>
<tr><td>Otu06986</td><td>3</td><td>k__Fungi(100);p__Ascomycota(67);p__Ascomycota_unclassified(67);p__Ascomycota_unclassified(67);p__Ascomycota_unclassified(67);p__Ascomycota_unclassified(67);p__Ascomycota_unclassified(67);</td><td>-</td><td>-</td><td>-</td><td>-</td><td>-</td><td>-</td><td>-</td><td>Unassigned</td><td>-</td></tr>
<tr><td>Otu06987</td><td>3</td><td>k__Fungi(100);p__Basidiomycota(100);c__Agaricomycetes(100);c__Agaricomycetes_unclassified(100);c__Agaricomycetes_unclassified(100);c__Agaricomycetes_unclassified(100);c__Agaricomycetes_unclassified(100);</td><td>-</td><td>-</td><td>-</td><td>-</td><td>-</td><td>-</td><td>-</td><td>Unassigned</td><td>-</td></tr>
<tr><td>Otu06991</td><td>3</td><td>k__Fungi(100);p__Ascomycota(100);p__Ascomycota_unclassified(100);p__Ascomycota_unclassified(100);p__Ascomycota_unclassified(100);p__Ascomycota_unclassified(100);p__Ascomycota_unclassified(100);</td><td>-</td><td>-</td><td>-</td><td>-</td><td>-</td><td>-</td><td>-</td><td>Unassigned</td><td>-</td></tr>
<tr><td>Otu06992</td><td>3</td><td>k__Fungi(100);k__Fungi_unclassified(100);k__Fungi_unclassified(100);k__Fungi_unclassified(100);k__Fungi_unclassified(100);k__Fungi_unclassified(100);k__Fungi_unclassified(100);</td><td>-</td><td>-</td><td>-</td><td>-</td><td>-</td><td>-</td><td>-</td><td>Unassigned</td><td>-</td></tr>
<tr><td>Otu06993</td><td>3</td><td>k__Fungi(100);p__Ascomycota(67);p__Ascomycota_unclassified(67);p__Ascomycota_unclassified(67);p__Ascomycota_unclassified(67);p__Ascomycota_unclassified(67);p__Ascomycota_unclassified(67);</td><td>-</td><td>-</td><td>-</td><td>-</td><td>-</td><td>-</td><td>-</td><td>Unassigned</td><td>-</td></tr>
</table>

| | | | | | | | | | | | |
|---|---|---|---|---|---|---|---|---|---|---|---|
| Otu06994 | 3 | k__Fungi(100);p__Ascomycota(67);p__Ascomycota_unclassified(67);p__Ascomycota_unclassified(67);p__Ascomycota_unclassified(67);p__Ascomycota_unclassified(67);p__Ascomycota_unclassified(67); | - | - | - | - | - | - | - | Unassigned | - |
| Otu07001 | 3 | k__Fungi(100);p__Ascomycota(67);p__Ascomycota_unclassified(67);p__Ascomycota_unclassified(67);p__Ascomycota_unclassified(67);p__Ascomycota_unclassified(67);p__Ascomycota_unclassified(67); | - | - | - | - | - | - | - | Unassigned | - |
| Otu07002 | 3 | k__Fungi(100);k__Fungi_unclassified(100);k__Fungi_unclassified(100);k__Fungi_unclassified(100);k__Fungi_unclassified(100);k__Fungi_unclassified(100);k__Fungi_unclassified(100); | - | - | - | - | - | - | - | Unassigned | - |
| Otu07003 | 3 | k__Fungi(100);k__Fungi_unclassified(100);k__Fungi_unclassified(100);k__Fungi_unclassified(100);k__Fungi_unclassified(100);k__Fungi_unclassified(100);k__Fungi_unclassified(100); | - | - | - | - | - | - | - | Unassigned | - |
| Otu07004 | 3 | k__Fungi(100);p__Ascomycota(67);p__Ascomycota_unclassified(67);p__Ascomycota_unclassified(67);p__Ascomycota_unclassified(67);p__Ascomycota_unclassified(67);p__Ascomycota_unclassified(67); | - | - | - | - | - | - | - | Unassigned | - |
| Otu07011 | 3 | k__Fungi(100);p__Basidiomycota(100);p__Basidiomycota_unclassified(100);p__Basidiomycota_unclassified(100);p__Basidiomycota_unclassified(100);p__Basidiomycota_unclassified(100);p__Basidiomycota_unclassified(100); | - | - | - | - | - | - | - | Unassigned | - |
| Otu07013 | 3 | k__Fungi(100);k__Fungi_unclassified(67);k__Fungi_unclassified(67);k__Fungi_unclassified(67);k__Fungi_unclassified(67);k__Fungi_unclassified(67);k__Fungi_unclassified(67); | - | - | - | - | - | - | - | Unassigned | - |
| Otu07015 | 3 | k__Fungi(100);k__Fungi_unclassified(67);k__Fungi_unclassified(67);k__Fungi_unclassified(67);k__Fungi_unclassified(67);k__Fungi_unclassified(67);k__Fungi_unclassified(67); | - | - | - | - | - | - | - | Unassigned | - |
| Otu07021 | 3 | k__Fungi(100);p__Basidiomycota(100);c__Agaricomycetes(100);o__Polyporales(100);o__Polyporales_unclassified(100);o__Polyporales_unclassified(100);o__Polyporales_unclassified(100); | - | - | - | - | - | - | - | Unassigned | - |
| Otu07023 | 3 | k__Fungi(100);p__Ascomycota(100);p__Ascomycota_unclassified(100);p__Ascomycota_unclassified(100);p__Ascomycota_unclassified(100);p__Ascomycota_unclassified(100);p__Ascomycota_unclassified(100); | - | - | - | - | - | - | - | Unassigned | - |
| Otu07026 | 3 | k__Fungi(100);k__Fungi_unclassified(67);k__Fungi_unclassified(67);k__Fungi_unclassified(67);k__Fungi_unclassified(67);k__Fungi_unclassified(67);k__Fungi_unclassified(67); | - | - | - | - | - | - | - | Unassigned | - |
| Otu07028 | 3 | k__Fungi(100);p__Ascomycota(100);p__Ascomycota_unclassified(100);p__Ascomycota_unclassified(100);p__Ascomycota_unclassified(100);p__Ascomycota_unclassified(100);p__Ascomycota_unclassified(100); | - | - | - | - | - | - | - | Unassigned | - |
| Otu07030 | 3 | k__Fungi(100);p__Basidiomycota(100);c__Agaricomycetes(100);c__Agaricomycetes_unclassified(100);c__Agaricomycetes_unclassified(100);c__Agaricomycetes_unclassified(100);c__Agaricomycetes_unclassified(100); | - | - | - | - | - | - | - | Unassigned | - |
| Otu07031 | 3 | k__Fungi(100);p__Ascomycota(67);p__Ascomycota_unclassified(67);p__Ascomycota_unclassified(67);p__Ascomycota_unclassified(67);p__Ascomycota_unclassified(67);p__Ascomycota_unclassified(67); | - | - | - | - | - | - | - | Unassigned | - |
| Otu07033 | 3 | k__Fungi(100);p__Basidiomycota(100);c__Agaricomycetes(100);c__Agaricomycetes_unclassified(100);c__Agaricomycetes_unclassified(100);c__Agaricomycetes_unclassified(100);c__Agaricomycetes_unclassified(100); | - | - | - | - | - | - | - | Unassigned | - |
| Otu07034 | 3 | k__Fungi(100);p__Ascomycota(67);p__Ascomycota_unclassified(67);p__Ascomycota_unclassified(67);p__Ascomycota_unclassified(67);p__Ascomycota_unclassified(67);p__Ascomycota_unclassified(67); | - | - | - | - | - | - | - | Unassigned | - |
| Otu07038 | 3 | k__Fungi(100);p__Basidiomycota(100);c__Agaricomycetes(100);c__Agaricomycetes_unclassified(100);c__Agaricomycetes_unclassified(100);c__Agaricomycetes_unclassified(100);c__Agaricomycetes_unclassified(100); | - | - | - | - | - | - | - | Unassigned | - |
| Otu07046 | 3 | k__Fungi(100);k__Fungi_unclassified(100);k__Fungi_unclassified(100);k__Fungi_unclassified(100);k__Fungi_unclassified(100);k__Fungi_unclassified(100);k__Fungi_unclassified(100); | - | - | - | - | - | - | - | Unassigned | - |
| Otu07050 | 3 | k__Fungi(100);p__Basidiomycota(100);c__Microbotryomycetes(100);o__Leucosporidiales(100);f__unclassified_Leucosporidiales(100);g__unclassified_Leucosporidiales(100);s__Leucosporidiales_sp(100); | - | - | - | - | - | - | - | Unassigned | - |
| Otu07052 | 3 | k__Fungi(100);k__Fungi_unclassified(100);k__Fungi_unclassified(100);k__Fungi_unclassified(100);k__Fungi_unclassified(100);k__Fungi_unclassified(100);k__Fungi_unclassified(100); | - | - | - | - | - | - | - | Unassigned | - |
| Otu07056 | 3 | k__Fungi(100);p__Basidiomycota(100);p__Basidiomycota_unclassified(100);p__Basidiomycota_unclassified(100);p__Basidiomycota_unclassified(100);p__Basidiomycota_unclassified(100);p__Basidiomycota_unclassified(100); | - | - | - | - | - | - | - | Unassigned | - |
| Otu07060 | 3 | k__Fungi(100);p__Ascomycota(100);p__Ascomycota_unclassified(100);p__Ascomycota_unclassified(100);p__Ascomycota_unclassified(100);p__Ascomycota_unclassified(100);p__Ascomycota_unclassified(100); | - | - | - | - | - | - | - | Unassigned | - |
| Otu07061 | 3 | k__Fungi(100);k__Fungi_unclassified(67);k__Fungi_unclassified(67);k__Fungi_unclassified(67);k__Fungi_unclassified(67);k__Fungi_unclassified(67);k__Fungi_unclassified(67); | - | - | - | - | - | - | - | Unassigned | - |
| Otu07062 | 3 | k__Fungi(100);p__Ascomycota(100);p__Ascomycota_unclassified(67);p__Ascomycota_unclassified(67);p__Ascomycota_unclassified(67);p__Ascomycota_unclassified(67);p__Ascomycota_unclassified(67); | - | - | - | - | - | - | - | Unassigned | - |
| Otu07063 | 3 | k__Fungi(100);k__Fungi_unclassified(67);k__Fungi_unclassified(67);k__Fungi_unclassified(67);k__Fungi_unclassified(67);k__Fungi_unclassified(67);k__Fungi_unclassified(67); | - | - | - | - | - | - | - | Unassigned | - |
| Otu07066 | 3 | k__Fungi(100);p__Ascomycota(100);p__Ascomycota_unclassified(100);p__Ascomycota_unclassified(100);p__Ascomycota_unclassified(100);p__Ascomycota_unclassified(100);p__Ascomycota_unclassified(100); | - | - | - | - | - | - | - | Unassigned | - |
| Otu07067 | 3 | k__Fungi(100);k__Fungi_unclassified(100);k__Fungi_unclassified(100);k__Fungi_unclassified(100);k__Fungi_unclassified(100);k__Fungi_unclassified(100);k__Fungi_unclassified(100); | - | - | - | - | - | - | - | Unassigned | - |
| Otu07071 | 3 | k__Fungi(100);p__Basidiomycota(100);p__Basidiomycota_unclassified(100);p__Basidiomycota_unclassified(100);p__Basidiomycota_unclassified(100);p__Basidiomycota_unclassified(100);p__Basidiomycota_unclassified(100); | - | - | - | - | - | - | - | Unassigned | - |
| Otu07073 | 3 | k__Fungi(100);p__Ascomycota(67);p__Ascomycota_unclassified(67);p__Ascomycota_unclassified(67);p__Ascomycota_unclassified(67);p__Ascomycota_unclassified(67);p__Ascomycota_unclassified(67); | - | - | - | - | - | - | - | Unassigned | - |
| Otu07078 | 3 | k__Fungi(100);p__Basidiomycota(100);c__Agaricomycetes(100);c__Agaricomycetes_unclassified(100);c__Agaricomycetes_unclassified(100);c__Agaricomycetes_unclassified(100);c__Agaricomycetes_unclassified(100); | - | - | - | - | - | - | - | Unassigned | - |
| Otu07081 | 3 | k__Fungi(100);p__Ascomycota(100);p__Ascomycota_unclassified(100);p__Ascomycota_unclassified(100);p__Ascomycota_unclassified(100);p__Ascomycota_unclassified(100);p__Ascomycota_unclassified(100); | - | - | - | - | - | - | - | Unassigned | - |
| Otu07082 | 3 | k__Fungi(100);k__Fungi_unclassified(100);k__Fungi_unclassified(100);k__Fungi_unclassified(100);k__Fungi_unclassified(100);k__Fungi_unclassified(100);k__Fungi_unclassified(100); | - | - | - | - | - | - | - | Unassigned | - |
| Otu07090 | 3 | k__Fungi(100);p__Ascomycota(67);p__Ascomycota_unclassified(67);p__Ascomycota_unclassified(67);p__Ascomycota_unclassified(67);p__Ascomycota_unclassified(67);p__Ascomycota_unclassified(67); | - | - | - | - | - | - | - | Unassigned | - |
| Otu07092 | 3 | k__Fungi(100);p__Ascomycota(67);p__Ascomycota_unclassified(67);p__Ascomycota_unclassified(67);p__Ascomycota_unclassified(67);p__Ascomycota_unclassified(67);p__Ascomycota_unclassified(67); | - | - | - | - | - | - | - | Unassigned | - |
| Otu07100 | 3 | k__Fungi(100);k__Fungi_unclassified(100);k__Fungi_unclassified(100);k__Fungi_unclassified(100);k__Fungi_unclassified(100);k__Fungi_unclassified(100);k__Fungi_unclassified(100); | - | - | - | - | - | - | - | Unassigned | - |
| Otu07101 | 3 | k__Fungi(100);k__Fungi_unclassified(67);k__Fungi_unclassified(67);k__Fungi_unclassified(67);k__Fungi_unclassified(67);k__Fungi_unclassified(67);k__Fungi_unclassified(67); | - | - | - | - | - | - | - | Unassigned | - |
| Otu07102 | 3 | k__Fungi(100);p__Ascomycota(67);p__Ascomycota_unclassified(67);p__Ascomycota_unclassified(67);p__Ascomycota_unclassified(67);p__Ascomycota_unclassified(67);p__Ascomycota_unclassified(67); | - | - | - | - | - | - | - | Unassigned | - |
| Otu07103 | 3 | k__Fungi(100);k__Fungi_unclassified(67);k__Fungi_unclassified(67);k__Fungi_unclassified(67);k__Fungi_unclassified(67);k__Fungi_unclassified(67);k__Fungi_unclassified(67); | - | - | - | - | - | - | - | Unassigned | - |
| Otu07104 | 3 | k__Fungi(100);k__Fungi_unclassified(67);k__Fungi_unclassified(67);k__Fungi_unclassified(67);k__Fungi_unclassified(67);k__Fungi_unclassified(67);k__Fungi_unclassified(67); | - | - | - | - | - | - | - | Unassigned | - |
| Otu07110 | 3 | k__Fungi(100);k__Fungi_unclassified(100);k__Fungi_unclassified(100);k__Fungi_unclassified(100);k__Fungi_unclassified(100);k__Fungi_unclassified(100);k__Fungi_unclassified(100); | - | - | - | - | - | - | - | Unassigned | - |
| Otu07114 | 3 | k__Fungi(100);k__Fungi_unclassified(67);k__Fungi_unclassified(67);k__Fungi_unclassified(67);k__Fungi_unclassified(67);k__Fungi_unclassified(67);k__Fungi_unclassified(67); | - | - | - | - | - | - | - | Unassigned | - |
| Otu07119 | 3 | k__Fungi(100);p__Ascomycota(67);p__Ascomycota_unclassified(67);p__Ascomycota_unclassified(67);p__Ascomycota_unclassified(67);p__Ascomycota_unclassified(67);p__Ascomycota_unclassified(67); | - | - | - | - | - | - | - | Unassigned | - |
| Otu07122 | 3 | k__Fungi(100);k__Fungi_unclassified(100);k__Fungi_unclassified(100);k__Fungi_unclassified(100);k__Fungi_unclassified(100);k__Fungi_unclassified(100);k__Fungi_unclassified(100); | - | - | - | - | - | - | - | Unassigned | - |
| Otu07123 | 3 | k__Fungi(100);p__Ascomycota(67);p__Ascomycota_unclassified(67);p__Ascomycota_unclassified(67);p__Ascomycota_unclassified(67);p__Ascomycota_unclassified(67);p__Ascomycota_unclassified(67); | - | - | - | - | - | - | - | Unassigned | - |
| Otu07124 | 3 | k__Fungi(100);k__Fungi_unclassified(100);k__Fungi_unclassified(100);k__Fungi_unclassified(100);k__Fungi_unclassified(100);k__Fungi_unclassified(100);k__Fungi_unclassified(100); | - | - | - | - | - | - | - | Unassigned | - |
| Otu07128 | 3 | k__Fungi(100);p__Ascomycota(67);p__Ascomycota_unclassified(67);p__Ascomycota_unclassified(67);p__Ascomycota_unclassified(67);p__Ascomycota_unclassified(67);p__Ascomycota_unclassified(67); | - | - | - | - | - | - | - | Unassigned | - |
| Otu07130 | 3 | k__Fungi(100);k__Fungi_unclassified(100);k__Fungi_unclassified(100);k__Fungi_unclassified(100);k__Fungi_unclassified(100);k__Fungi_unclassified(100);k__Fungi_unclassified(100); | - | - | - | - | - | - | - | Unassigned | - |
| Otu07131 | 3 | k__Fungi(100);k__Fungi_unclassified(67);k__Fungi_unclassified(67);k__Fungi_unclassified(67);k__Fungi_unclassified(67);k__Fungi_unclassified(67);k__Fungi_unclassified(67); | - | - | - | - | - | - | - | Unassigned | - |
| Otu07135 | 3 | k__Fungi(100);k__Fungi_unclassified(67);k__Fungi_unclassified(67);k__Fungi_unclassified(67);k__Fungi_unclassified(67);k__Fungi_unclassified(67);k__Fungi_unclassified(67); | - | - | - | - | - | - | - | Unassigned | - |
| Otu07136 | 3 | k__Fungi(100);p__Basidiomycota(100);c__Agaricomycetes(100);c__Agaricomycetes_unclassified(100);c__Agaricomycetes_unclassified(100);c__Agaricomycetes_unclassified(100);c__Agaricomycetes_unclassified(100); | - | - | - | - | - | - | - | Unassigned | - |
| Otu07137 | 3 | k__Fungi(100);p__Basidiomycota(100);c__Agaricostilbomycetes(100);o__Agaricostilbales(100);o__Agaricostilbales_unclassified(100);o__Agaricostilbales_unclassified(100);o__Agaricostilbales_unclassified(100); | - | - | - | - | - | - | - | Unassigned | - |
| Otu07143 | 3 | k__Fungi(100);k__Fungi_unclassified(100);k__Fungi_unclassified(100);k__Fungi_unclassified(100);k__Fungi_unclassified(100);k__Fungi_unclassified(100);k__Fungi_unclassified(100); | - | - | - | - | - | - | - | Unassigned | - |
| Otu07144 | 3 | k__Fungi(100);p__Ascomycota(100);c__Sordariomycetes(100);c__Sordariomycetes_unclassified(67);c__Sordariomycetes_unclassified(67);c__Sordariomycetes_unclassified(67);c__Sordariomycetes_unclassified(67); | - | - | - | - | - | - | - | Unassigned | - |
| Otu07145 | 3 | k__Fungi(100);p__Ascomycota(100);c__Sordariomycetes(100);o__Hypocreales(100);f__Hypocreaceae(100);f__Hypocreaceae_unclassified(100);f__Hypocreaceae_unclassified(100); | - | - | - | - | - | - | - | Unassigned | - |
| Otu07147 | 3 | k__Fungi(100);k__Fungi_unclassified(100);k__Fungi_unclassified(100);k__Fungi_unclassified(100);k__Fungi_unclassified(100);k__Fungi_unclassified(100);k__Fungi_unclassified(100); | - | - | - | - | - | - | - | Unassigned | - |
| Otu07150 | 3 | k__Fungi(100);k__Fungi_unclassified(67);k__Fungi_unclassified(67);k__Fungi_unclassified(67);k__Fungi_unclassified(67);k__Fungi_unclassified(67);k__Fungi_unclassified(67); | - | - | - | - | - | - | - | Unassigned | - |
| Otu07151 | 3 | k__Fungi(100);k__Fungi_unclassified(67);k__Fungi_unclassified(67);k__Fungi_unclassified(67);k__Fungi_unclassified(67);k__Fungi_unclassified(67);k__Fungi_unclassified(67); | - | - | - | - | - | - | - | Unassigned | - |
| Otu07153 | 3 | k__Fungi(100);p__Ascomycota(100);p__Ascomycota_unclassified(100);p__Ascomycota_unclassified(100);p__Ascomycota_unclassified(100);p__Ascomycota_unclassified(100);p__Ascomycota_unclassified(100); | - | - | - | - | - | - | - | Unassigned | - |
| Otu07154 | 3 | k__Fungi(100);p__Basidiomycota(100);c__Agaricomycetes(100);c__Agaricomycetes_unclassified(67);c__Agaricomycetes_unclassified(67);c__Agaricomycetes_unclassified(67);c__Agaricomycetes_unclassified(67); | - | - | - | - | - | - | - | Unassigned | - |
| Otu07156 | 3 | k__Fungi(100);p__Ascomycota(67);p__Ascomycota_unclassified(67);p__Ascomycota_unclassified(67);p__Ascomycota_unclassified(67);p__Ascomycota_unclassified(67);p__Ascomycota_unclassified(67); | - | - | - | - | - | - | - | Unassigned | - |
| Otu07160 | 3 | k__Fungi(100);p__Ascomycota(100);c__Eurotiomycetes(100);o__Chaetothyriales(100);o__Chaetothyriales_unclassified(100);o__Chaetothyriales_unclassified(100);o__Chaetothyriales_unclassified(100); | - | - | - | - | - | - | - | Unassigned | - |
| Otu07162 | 3 | k__Fungi(100);p__Ascomycota(100);p__Ascomycota_unclassified(100);p__Ascomycota_unclassified(100);p__Ascomycota_unclassified(100);p__Ascomycota_unclassified(100);p__Ascomycota_unclassified(100); | - | - | - | - | - | - | - | Unassigned | - |
| Otu07163 | 3 | k__Fungi(100);p__Ascomycota(67);p__Ascomycota_unclassified(67);p__Ascomycota_unclassified(67);p__Ascomycota_unclassified(67);p__Ascomycota_unclassified(67);p__Ascomycota_unclassified(67); | - | - | - | - | - | - | - | Unassigned | - |
| Otu07165 | 3 | k__Fungi(100);k__Fungi_unclassified(100);k__Fungi_unclassified(100);k__Fungi_unclassified(100);k__Fungi_unclassified(100);k__Fungi_unclassified(100);k__Fungi_unclassified(100); | - | - | - | - | - | - | - | Unassigned | - |
| Otu07166 | 3 | k__Fungi(100);p__Ascomycota(67);p__Ascomycota_unclassified(67);p__Ascomycota_unclassified(67);p__Ascomycota_unclassified(67);p__Ascomycota_unclassified(67);p__Ascomycota_unclassified(67); | - | - | - | - | - | - | - | Unassigned | - |
| Otu07167 | 3 | k__Fungi(100);k__Fungi_unclassified(100);k__Fungi_unclassified(100);k__Fungi_unclassified(100);k__Fungi_unclassified(100);k__Fungi_unclassified(100);k__Fungi_unclassified(100); | - | - | - | - | - | - | - | Unassigned | - |
| Otu07169 | 3 | k__Fungi(100);p__Ascomycota(100);p__Ascomycota_unclassified(100);p__Ascomycota_unclassified(100);p__Ascomycota_unclassified(100);p__Ascomycota_unclassified(100);p__Ascomycota_unclassified(100); | - | - | - | - | - | - | - | Unassigned | - |
| Otu07172 | 3 | k__Fungi(100);k__Fungi_unclassified(67);k__Fungi_unclassified(67);k__Fungi_unclassified(67);k__Fungi_unclassified(67);k__Fungi_unclassified(67);k__Fungi_unclassified(67); | - | - | - | - | - | - | - | Unassigned | - |
| Otu07173 | 3 | k__Fungi(100);p__Ascomycota(100);c__Dothideomycetes(67);c__Dothideomycetes_unclassified(67);c__Dothideomycetes_unclassified(67);c__Dothideomycetes_unclassified(67);c__Dothideomycetes_unclassified(67); | - | - | - | - | - | - | - | Unassigned | - |
| Otu07174 | 3 | k__Fungi(100);p__Ascomycota(100);c__Eurotiomycetes(100);c__Eurotiomycetes_unclassified(100);c__Eurotiomycetes_unclassified(100);c__Eurotiomycetes_unclassified(100);c__Eurotiomycetes_unclassified(100); | - | - | - | - | - | - | - | Unassigned | - |
| Otu07175 | 3 | k__Fungi(100);p__Ascomycota(100);p__Ascomycota_unclassified(100);p__Ascomycota_unclassified(100);p__Ascomycota_unclassified(100);p__Ascomycota_unclassified(100);p__Ascomycota_unclassified(100); | - | - | - | - | - | - | - | Unassigned | - |
| Otu07177 | 3 | k__Fungi(100);k__Fungi_unclassified(100);k__Fungi_unclassified(100);k__Fungi_unclassified(100);k__Fungi_unclassified(100);k__Fungi_unclassified(100);k__Fungi_unclassified(100); | - | - | - | - | - | - | - | Unassigned | - |
| Otu07181 | 3 | k__Fungi(100);p__Ascomycota(100);p__Ascomycota_unclassified(100);p__Ascomycota_unclassified(100);p__Ascomycota_unclassified(100);p__Ascomycota_unclassified(100);p__Ascomycota_unclassified(100); | - | - | - | - | - | - | - | Unassigned | - |
| Otu07183 | 3 | k__Fungi(100);p__Ascomycota(100);p__Ascomycota_unclassified(100);p__Ascomycota_unclassified(100);p__Ascomycota_unclassified(100);p__Ascomycota_unclassified(100);p__Ascomycota_unclassified(100); | - | - | - | - | - | - | - | Unassigned | - |
| Otu07185 | 3 | k__Fungi(100);k__Fungi_unclassified(67);k__Fungi_unclassified(67);k__Fungi_unclassified(67);k__Fungi_unclassified(67);k__Fungi_unclassified(67);k__Fungi_unclassified(67); | - | - | - | - | - | - | - | Unassigned | - |
| Otu07189 | 3 | k__Fungi(100);k__Fungi_unclassified(67);k__Fungi_unclassified(67);k__Fungi_unclassified(67);k__Fungi_unclassified(67);k__Fungi_unclassified(67);k__Fungi_unclassified(67); | - | - | - | - | - | - | - | Unassigned | - |
| Otu07190 | 3 | k__Fungi(100);p__Basidiomycota(100);p__Basidiomycota_unclassified(100);p__Basidiomycota_unclassified(100);p__Basidiomycota_unclassified(100);p__Basidiomycota_unclassified(100);p__Basidiomycota_unclassified(100); | - | - | - | - | - | - | - | Unassigned | - |
| Otu07191 | 3 | k__Fungi(100);k__Fungi_unclassified(67);k__Fungi_unclassified(67);k__Fungi_unclassified(67);k__Fungi_unclassified(67);k__Fungi_unclassified(67);k__Fungi_unclassified(67); | - | - | - | - | - | - | - | Unassigned | - |
| Otu07193 | 3 | k__Fungi(100);p__Ascomycota(100);p__Ascomycota_unclassified(100);p__Ascomycota_unclassified(100);p__Ascomycota_unclassified(100);p__Ascomycota_unclassified(100);p__Ascomycota_unclassified(100); | - | - | - | - | - | - | - | Unassigned | - |
| Otu07195 | 3 | k__Fungi(100);k__Fungi_unclassified(67);k__Fungi_unclassified(67);k__Fungi_unclassified(67);k__Fungi_unclassified(67);k__Fungi_unclassified(67);k__Fungi_unclassified(67); | - | - | - | - | - | - | - | Unassigned | - |
| Otu07196 | 3 | k__Fungi(100);k__Fungi_unclassified(100);k__Fungi_unclassified(100);k__Fungi_unclassified(100);k__Fungi_unclassified(100);k__Fungi_unclassified(100);k__Fungi_unclassified(100); | - | - | - | - | - | - | - | Unassigned | - |
| Otu07198 | 3 | k__Fungi(100);k__Fungi_unclassified(67);k__Fungi_unclassified(67);k__Fungi_unclassified(67);k__Fungi_unclassified(67);k__Fungi_unclassified(67);k__Fungi_unclassified(67); | - | - | - | - | - | - | - | Unassigned | - |
| Otu07199 | 3 | k__Fungi(100);p__Ascomycota(100);p__Ascomycota_unclassified(100);p__Ascomycota_unclassified(100);p__Ascomycota_unclassified(100);p__Ascomycota_unclassified(100);p__Ascomycota_unclassified(100); | - | - | - | - | - | - | - | Unassigned | - |
| Otu07201 | 3 | k__Fungi(100);k__Fungi_unclassified(67);k__Fungi_unclassified(67);k__Fungi_unclassified(67);k__Fungi_unclassified(67);k__Fungi_unclassified(67);k__Fungi_unclassified(67); | - | - | - | - | - | - | - | Unassigned | - |
| Otu07202 | 3 | k__Fungi(100);p__Basidiomycota(100);p__Basidiomycota_unclassified(100);p__Basidiomycota_unclassified(100);p__Basidiomycota_unclassified(100);p__Basidiomycota_unclassified(100);p__Basidiomycota_unclassified(100); | - | - | - | - | - | - | - | Unassigned | - |
| Otu07204 | 3 | k__Fungi(100);p__Ascomycota(100);c__Sordariomycetes(100);c__Sordariomycetes_unclassified(100);c__Sordariomycetes_unclassified(100);c__Sordariomycetes_unclassified(100);c__Sordariomycetes_unclassified(100); | - | - | - | - | - | - | - | Unassigned | - |
| Otu07205 | 3 | k__Fungi(100);k__Fungi_unclassified(67);k__Fungi_unclassified(67);k__Fungi_unclassified(67);k__Fungi_unclassified(67);k__Fungi_unclassified(67);k__Fungi_unclassified(67); | - | - | - | - | - | - | - | Unassigned | - |
| Otu07208 | 3 | k__Fungi(100);p__Basidiomycota(100);c__Agaricomycetes(100);o__Trechisporales(100);f__unclassified_Trechisporales(67);g__unclassified_Trechisporales(67);s__Trechisporales_sp(67); | - | - | - | - | - | - | - | Unassigned | - |
| Otu07214 | 3 | k__Fungi(100);p__Ascomycota(67);p__Ascomycota_unclassified(67);p__Ascomycota_unclassified(67);p__Ascomycota_unclassified(67);p__Ascomycota_unclassified(67);p__Ascomycota_unclassified(67); | - | - | - | - | - | - | - | Unassigned | - |
| Otu07215 | 3 | k__Fungi(100);p__Basidiomycota(100);c__Agaricomycetes(100);o__Polyporales(100);f__Phanerochaetaceae(100);f__Phanerochaetaceae_unclassified(100);f__Phanerochaetaceae_unclassified(100); | - | - | - | - | - | - | - | Unassigned | - |
| Otu07216 | 3 | k__Fungi(100);p__Ascomycota(67);p__Ascomycota_unclassified(67);p__Ascomycota_unclassified(67);p__Ascomycota_unclassified(67);p__Ascomycota_unclassified(67);p__Ascomycota_unclassified(67); | - | - | - | - | - | - | - | Unassigned | - |
| Otu07218 | 3 | k__Fungi(100);k__Fungi_unclassified(100);k__Fungi_unclassified(100);k__Fungi_unclassified(100);k__Fungi_unclassified(100);k__Fungi_unclassified(100);k__Fungi_unclassified(100); | - | - | - | - | - | - | - | Unassigned | - |
| Otu07221 | 3 | k__Fungi(100);p__Basidiomycota(100);c__Tremellomycetes(100);c__Tremellomycetes_unclassified(67);c__Tremellomycetes_unclassified(67);c__Tremellomycetes_unclassified(67);c__Tremellomycetes_unclassified(67); | - | - | - | - | - | - | - | Unassigned | - |
| Otu07222 | 3 | k__Fungi(100);p__Ascomycota(100);c__Eurotiomycetes(100);c__Eurotiomycetes_unclassified(100);c__Eurotiomycetes_unclassified(100);c__Eurotiomycetes_unclassified(100);c__Eurotiomycetes_unclassified(100); | - | - | - | - | - | - | - | Unassigned | - |
| Otu07223 | 3 | k__Fungi(100);k__Fungi_unclassified(100);k__Fungi_unclassified(100);k__Fungi_unclassified(100);k__Fungi_unclassified(100);k__Fungi_unclassified(100);k__Fungi_unclassified(100); | - | - | - | - | - | - | - | Unassigned | - |
| Otu07225 | 3 | k__Fungi(100);p__Ascomycota(100);c__Leotiomycetes(100);c__Leotiomycetes_unclassified(67);c__Leotiomycetes_unclassified(67);c__Leotiomycetes_unclassified(67);c__Leotiomycetes_unclassified(67); | - | - | - | - | - | - | - | Unassigned | - |
| Otu07227 | 3 | k__Fungi(100);k__Fungi_unclassified(67);k__Fungi_unclassified(67);k__Fungi_unclassified(67);k__Fungi_unclassified(67);k__Fungi_unclassified(67);k__Fungi_unclassified(67); | - | - | - | - | - | - | - | Unassigned | - |
| Otu07229 | 3 | k__Fungi(100);p__Basidiomycota(100);p__Basidiomycota_unclassified(100);p__Basidiomycota_unclassified(100);p__Basidiomycota_unclassified(100);p__Basidiomycota_unclassified(100);p__Basidiomycota_unclassified(100); | - | - | - | - | - | - | - | Unassigned | - |
| Otu07233 | 3 | k__Fungi(100);p__Ascomycota(100);p__Ascomycota_unclassified(100);p__Ascomycota_unclassified(100);p__Ascomycota_unclassified(100);p__Ascomycota_unclassified(100);p__Ascomycota_unclassified(100); | - | - | - | - | - | - | - | Unassigned | - |
| Otu07235 | 3 | k__Fungi(100);k__Fungi_unclassified(67);k__Fungi_unclassified(67);k__Fungi_unclassified(67);k__Fungi_unclassified(67);k__Fungi_unclassified(67);k__Fungi_unclassified(67); | - | - | - | - | - | - | - | Unassigned | - |
| Otu07242 | 3 | k__Fungi(100);k__Fungi_unclassified(100);k__Fungi_unclassified(100);k__Fungi_unclassified(100);k__Fungi_unclassified(100);k__Fungi_unclassified(100);k__Fungi_unclassified(100); | - | - | - | - | - | - | - | Unassigned | - |
| Otu07244 | 3 | k__Fungi(100);k__Fungi_unclassified(67);k__Fungi_unclassified(67);k__Fungi_unclassified(67);k__Fungi_unclassified(67);k__Fungi_unclassified(67);k__Fungi_unclassified(67); | - | - | - | - | - | - | - | Unassigned | - |
| Otu07245 | 3 | k__Fungi(100);p__Ascomycota(100);p__Ascomycota_unclassified(100);p__Ascomycota_unclassified(100);p__Ascomycota_unclassified(100);p__Ascomycota_unclassified(100);p__Ascomycota_unclassified(100); | - | - | - | - | - | - | - | Unassigned | - |
| Otu07248 | 3 | k__Fungi(100);p__Ascomycota(100);c__Dothideomycetes(100);c__Dothideomycetes_unclassified(67);c__Dothideomycetes_unclassified(67);c__Dothideomycetes_unclassified(67);c__Dothideomycetes_unclassified(67); | - | - | - | - | - | - | - | Unassigned | - |
| Otu07249 | 3 | k__Fungi(100);p__Ascomycota(100);c__Dothideomycetes(67);c__Dothideomycetes_unclassified(67);c__Dothideomycetes_unclassified(67);c__Dothideomycetes_unclassified(67);c__Dothideomycetes_unclassified(67); | - | - | - | - | - | - | - | Unassigned | - |
| Otu07250 | 3 | k__Fungi(100);p__Ascomycota(100);c__Sordariomycetes(100);c__Sordariomycetes_unclassified(100);c__Sordariomycetes_unclassified(100);c__Sordariomycetes_unclassified(100);c__Sordariomycetes_unclassified(100); | - | - | - | - | - | - | - | Unassigned | - |
| Otu07252 | 3 | k__Fungi(100);p__Ascomycota(67);p__Ascomycota_unclassified(67);p__Ascomycota_unclassified(67);p__Ascomycota_unclassified(67);p__Ascomycota_unclassified(67);p__Ascomycota_unclassified(67); | - | - | - | - | - | - | - | Unassigned | - |
| Otu07254 | 3 | k__Fungi(100);p__Ascomycota(100);c__Sordariomycetes(100);c__Sordariomycetes_unclassified(67);c__Sordariomycetes_unclassified(67);c__Sordariomycetes_unclassified(67);c__Sordariomycetes_unclassified(67); | - | - | - | - | - | - | - | Unassigned | - |
| Otu07255 | 3 | k__Fungi(100);k__Fungi_unclassified(67);k__Fungi_unclassified(67);k__Fungi_unclassified(67);k__Fungi_unclassified(67);k__Fungi_unclassified(67);k__Fungi_unclassified(67); | - | - | - | - | - | - | - | Unassigned | - |
| Otu07259 | 3 | k__Fungi(100);p__Ascomycota(67);p__Ascomycota_unclassified(67);p__Ascomycota_unclassified(67);p__Ascomycota_unclassified(67);p__Ascomycota_unclassified(67);p__Ascomycota_unclassified(67); | - | - | - | - | - | - | - | Unassigned | - |
| Otu07260 | 3 | k__Fungi(100);p__Ascomycota(100);p__Ascomycota_unclassified(100);p__Ascomycota_unclassified(100);p__Ascomycota_unclassified(100);p__Ascomycota_unclassified(100);p__Ascomycota_unclassified(100); | - | - | - | - | - | - | - | Unassigned | - |
| Otu07265 | 3 | k__Fungi(100);k__Fungi_unclassified(100);k__Fungi_unclassified(100);k__Fungi_unclassified(100);k__Fungi_unclassified(100);k__Fungi_unclassified(100);k__Fungi_unclassified(100); | - | - | - | - | - | - | - | Unassigned | - |
| Otu07269 | 3 | k__Fungi(100);k__Fungi_unclassified(100);k__Fungi_unclassified(100);k__Fungi_unclassified(100);k__Fungi_unclassified(100);k__Fungi_unclassified(100);k__Fungi_unclassified(100); | - | - | - | - | - | - | - | Unassigned | - |
| Otu07270 | 3 | k__Fungi(100);k__Fungi_unclassified(67);k__Fungi_unclassified(67);k__Fungi_unclassified(67);k__Fungi_unclassified(67);k__Fungi_unclassified(67);k__Fungi_unclassified(67); | - | - | - | - | - | - | - | Unassigned | - |
| Otu07271 | 3 | k__Fungi(100);p__Ascomycota(67);p__Ascomycota_unclassified(67);p__Ascomycota_unclassified(67);p__Ascomycota_unclassified(67);p__Ascomycota_unclassified(67);p__Ascomycota_unclassified(67); | - | - | - | - | - | - | - | Unassigned | - |
| Otu07275 | 3 | k__Fungi(100);p__Ascomycota(67);p__Ascomycota_unclassified(67);p__Ascomycota_unclassified(67);p__Ascomycota_unclassified(67);p__Ascomycota_unclassified(67);p__Ascomycota_unclassified(67); | - | - | - | - | - | - | - | Unassigned | - |
| Otu07276 | 3 | k__Fungi(100);k__Fungi_unclassified(100);k__Fungi_unclassified(100);k__Fungi_unclassified(100);k__Fungi_unclassified(100);k__Fungi_unclassified(100);k__Fungi_unclassified(100); | - | - | - | - | - | - | - | Unassigned | - |
| Otu07278 | 3 | k__Fungi(100);k__Fungi_unclassified(67);k__Fungi_unclassified(67);k__Fungi_unclassified(67);k__Fungi_unclassified(67);k__Fungi_unclassified(67);k__Fungi_unclassified(67); | - | - | - | - | - | - | - | Unassigned | - |
| Otu07280 | 3 | k__Fungi(100);p__Ascomycota(100);p__Ascomycota_unclassified(100);p__Ascomycota_unclassified(100);p__Ascomycota_unclassified(100);p__Ascomycota_unclassified(100);p__Ascomycota_unclassified(100); | - | - | - | - | - | - | - | Unassigned | - |
| Otu07281 | 3 | k__Fungi(100);p__Basidiomycota(100);c__Agaricomycetes(100);c__Agaricomycetes_unclassified(100);c__Agaricomycetes_unclassified(100);c__Agaricomycetes_unclassified(100);c__Agaricomycetes_unclassified(100); | - | - | - | - | - | - | - | Unassigned | - |
| Otu07282 | 3 | k__Fungi(100);k__Fungi_unclassified(67);k__Fungi_unclassified(67);k__Fungi_unclassified(67);k__Fungi_unclassified(67);k__Fungi_unclassified(67);k__Fungi_unclassified(67); | - | - | - | - | - | - | - | Unassigned | - |
| Otu07286 | 3 | k__Fungi(100);p__Basidiomycota(100);c__Agaricomycetes(100);o__Polyporales(100);f__Phanerochaetaceae(100);f__Phanerochaetaceae_unclassified(100);f__Phanerochaetaceae_unclassified(100); | - | - | - | - | - | - | - | Unassigned | - |
| Otu07287 | 3 | k__Fungi(100);p__Ascomycota(100);p__Ascomycota_unclassified(100);p__Ascomycota_unclassified(100);p__Ascomycota_unclassified(100);p__Ascomycota_unclassified(100);p__Ascomycota_unclassified(100); | - | - | - | - | - | - | - | Unassigned | - |
| Otu07288 | 3 | k__Fungi(100);k__Fungi_unclassified(67);k__Fungi_unclassified(67);k__Fungi_unclassified(67);k__Fungi_unclassified(67);k__Fungi_unclassified(67);k__Fungi_unclassified(67); | - | - | - | - | - | - | - | Unassigned | - |
| Otu07290 | 3 | k__Fungi(100);p__Basidiomycota(100);c__Agaricomycetes(100);o__Boletales(100);f__Paxillaceae(67);f__Paxillaceae_unclassified(67);f__Paxillaceae_unclassified(67); | - | - | - | - | - | - | - | Unassigned | - |
| Otu07292 | 3 | k__Fungi(100);p__Ascomycota(100);c__Leotiomycetes(67);o__Helotiales(67);o__Helotiales_unclassified(67);o__Helotiales_unclassified(67);o__Helotiales_unclassified(67); | - | - | - | - | - | - | - | Unassigned | - |
| Otu07293 | 3 | k__Fungi(100);k__Fungi_unclassified(100);k__Fungi_unclassified(100);k__Fungi_unclassified(100);k__Fungi_unclassified(100);k__Fungi_unclassified(100);k__Fungi_unclassified(100); | - | - | - | - | - | - | - | Unassigned | - |
| Otu07295 | 3 | k__Fungi(100);p__Basidiomycota(100);c__Microbotryomycetes(100);o__Sporidiobolales(100);f__Sporidiobolales_family_Incertae_sedis(100);f__Sporidiobolales_family_Incertae_sedis_unclassified(100);f__Sporidiobolales_family_Incertae_sedis_unclassified(100); | - | - | - | - | - | - | - | Unassigned | - |
| Otu07297 | 3 | k__Fungi(100);p__Ascomycota(100);c__Dothideomycetes(100);o__Pleosporales(67);o__Pleosporales_unclassified(67);o__Pleosporales_unclassified(67);o__Pleosporales_unclassified(67); | - | - | - | - | - | - | - | Unassigned | - |
| Otu07298 | 3 | k__Fungi(100);k__Fungi_unclassified(100);k__Fungi_unclassified(100);k__Fungi_unclassified(100);k__Fungi_unclassified(100);k__Fungi_unclassified(100);k__Fungi_unclassified(100); | - | - | - | - | - | - | - | Unassigned | - |
| Otu07299 | 3 | k__Fungi(100);p__Basidiomycota(100);c__Microbotryomycetes(100);o__Sporidiobolales(100);f__Sporidiobolales_family_Incertae_sedis(100);f__Sporidiobolales_family_Incertae_sedis_unclassified(67);f__Sporidiobolales_family_Incertae_sedis_unclassified(67); | - | - | - | - | - | - | - | Unassigned | - |
| Otu07300 | 3 | k__Fungi(100);k__Fungi_unclassified(67);k__Fungi_unclassified(67);k__Fungi_unclassified(67);k__Fungi_unclassified(67);k__Fungi_unclassified(67);k__Fungi_unclassified(67); | - | - | - | - | - | - | - | Unassigned | - |
| Otu07301 | 3 | k__Fungi(100);p__Ascomycota(100);p__Ascomycota_unclassified(100);p__Ascomycota_unclassified(100);p__Ascomycota_unclassified(100);p__Ascomycota_unclassified(100);p__Ascomycota_unclassified(100); | - | - | - | - | - | - | - | Unassigned | - |
| Otu07302 | 3 | k__Fungi(100);p__Ascomycota(100);p__Ascomycota_unclassified(100);p__Ascomycota_unclassified(100);p__Ascomycota_unclassified(100);p__Ascomycota_unclassified(100);p__Ascomycota_unclassified(100); | - | - | - | - | - | - | - | Unassigned | - |
| Otu07307 | 3 | k__Fungi(100);p__Ascomycota(100);p__Ascomycota_unclassified(100);p__Ascomycota_unclassified(100);p__Ascomycota_unclassified(100);p__Ascomycota_unclassified(100);p__Ascomycota_unclassified(100); | - | - | - | - | - | - | - | Unassigned | - |
| Otu07310 | 3 | k__Fungi(100);p__Ascomycota(100);p__Ascomycota_unclassified(100);p__Ascomycota_unclassified(100);p__Ascomycota_unclassified(100);p__Ascomycota_unclassified(100);p__Ascomycota_unclassified(100); | - | - | - | - | - | - | - | Unassigned | - |
| Otu07315 | 3 | k__Fungi(100);p__Ascomycota(100);c__Eurotiomycetes(100);o__Eurotiales(100);f__Trichocomaceae(100);f__Trichocomaceae_unclassified(100);f__Trichocomaceae_unclassified(100); | - | - | - | - | - | - | - | Unassigned | - |
| Otu07318 | 3 | k__Fungi(100);p__Ascomycota(67);p__Ascomycota_unclassified(67);p__Ascomycota_unclassified(67);p__Ascomycota_unclassified(67);p__Ascomycota_unclassified(67);p__Ascomycota_unclassified(67); | - | - | - | - | - | - | - | Unassigned | - |
| Otu07321 | 3 | k__Fungi(100);k__Fungi_unclassified(100);k__Fungi_unclassified(100);k__Fungi_unclassified(100);k__Fungi_unclassified(100);k__Fungi_unclassified(100);k__Fungi_unclassified(100); | - | - | - | - | - | - | - | Unassigned | - |
| Otu07324 | 3 | k__Fungi(100);k__Fungi_unclassified(67);k__Fungi_unclassified(67);k__Fungi_unclassified(67);k__Fungi_unclassified(67);k__Fungi_unclassified(67);k__Fungi_unclassified(67); | - | - | - | - | - | - | - | Unassigned | - |
| Otu07325 | 3 | k__Fungi(100);p__Ascomycota(100);p__Ascomycota_unclassified(100);p__Ascomycota_unclassified(100);p__Ascomycota_unclassified(100);p__Ascomycota_unclassified(100);p__Ascomycota_unclassified(100); | - | - | - | - | - | - | - | Unassigned | - |
| Otu07327 | 3 | k__Fungi(100);k__Fungi_unclassified(67);k__Fungi_unclassified(67);k__Fungi_unclassified(67);k__Fungi_unclassified(67);k__Fungi_unclassified(67);k__Fungi_unclassified(67); | - | - | - | - | - | - | - | Unassigned | - |
| Otu07328 | 3 | k__Fungi(100);p__Ascomycota(100);c__Sordariomycetes(100);c__Sordariomycetes_unclassified(100);c__Sordariomycetes_unclassified(100);c__Sordariomycetes_unclassified(100);c__Sordariomycetes_unclassified(100); | - | - | - | - | - | - | - | Unassigned | - |
| Otu07332 | 3 | k__Fungi(100);p__Ascomycota(100);p__Ascomycota_unclassified(100);p__Ascomycota_unclassified(100);p__Ascomycota_unclassified(100);p__Ascomycota_unclassified(100);p__Ascomycota_unclassified(100); | - | - | - | - | - | - | - | Unassigned | - |
| Otu07339 | 3 | k__Fungi(100);p__Ascomycota(100);c__Lecanoromycetes(100);c__Lecanoromycetes_unclassified(100);c__Lecanoromycetes_unclassified(100);c__Lecanoromycetes_unclassified(100);c__Lecanoromycetes_unclassified(100); | - | - | - | - | - | - | - | Unassigned | - |
| Otu07340 | 3 | k__Fungi(100);p__Ascomycota(100);c__Leotiomycetes(100);o__Helotiales(100);f__Helotiales_family_Incertae_sedis(100);f__Helotiales_family_Incertae_sedis_unclassified(100);f__Helotiales_family_Incertae_sedis_unclassified(100); | - | - | - | - | - | - | - | Unassigned | - |
| Otu07342 | 3 | k__Fungi(100);p__Basidiomycota(100);c__Agaricomycetes(100);o__Polyporales(67);o__Polyporales_unclassified(67);o__Polyporales_unclassified(67);o__Polyporales_unclassified(67); | - | - | - | - | - | - | - | Unassigned | - |
| Otu07344 | 3 | k__Fungi(100);k__Fungi_unclassified(67);k__Fungi_unclassified(67);k__Fungi_unclassified(67);k__Fungi_unclassified(67);k__Fungi_unclassified(67);k__Fungi_unclassified(67); | - | - | - | - | - | - | - | Unassigned | - |
| Otu07346 | 3 | k__Fungi(100);p__Ascomycota(100);c__Dothideomycetes(100);o__Pleosporales(100);o__Pleosporales_unclassified(67);o__Pleosporales_unclassified(67);o__Pleosporales_unclassified(67); | - | - | - | - | - | - | - | Unassigned | - |
| Otu07348 | 3 | k__Fungi(100);p__Ascomycota(67);p__Ascomycota_unclassified(67);p__Ascomycota_unclassified(67);p__Ascomycota_unclassified(67);p__Ascomycota_unclassified(67);p__Ascomycota_unclassified(67); | - | - | - | - | - | - | - | Unassigned | - |
| Otu07350 | 3 | k__Fungi(100);p__Ascomycota(100);p__Ascomycota_unclassified(100);p__Ascomycota_unclassified(100);p__Ascomycota_unclassified(100);p__Ascomycota_unclassified(100);p__Ascomycota_unclassified(100); | - | - | - | - | - | - | - | Unassigned | - |
| Otu07354 | 3 | k__Fungi(100);k__Fungi_unclassified(67);k__Fungi_unclassified(67);k__Fungi_unclassified(67);k__Fungi_unclassified(67);k__Fungi_unclassified(67);k__Fungi_unclassified(67); | - | - | - | - | - | - | - | Unassigned | - |
| Otu07359 | 3 | k__Fungi(100);p__Ascomycota(67);p__Ascomycota_unclassified(67);p__Ascomycota_unclassified(67);p__Ascomycota_unclassified(67);p__Ascomycota_unclassified(67);p__Ascomycota_unclassified(67); | - | - | - | - | - | - | - | Unassigned | - |
| Otu07362 | 3 | k__Fungi(100);p__Ascomycota(100);p__Ascomycota_unclassified(100);p__Ascomycota_unclassified(100);p__Ascomycota_unclassified(100);p__Ascomycota_unclassified(100);p__Ascomycota_unclassified(100); | - | - | - | - | - | - | - | Unassigned | - |
| Otu07363 | 3 | k__Fungi(100);k__Fungi_unclassified(100);k__Fungi_unclassified(100);k__Fungi_unclassified(100);k__Fungi_unclassified(100);k__Fungi_unclassified(100);k__Fungi_unclassified(100); | - | - | - | - | - | - | - | Unassigned | - |
| Otu07365 | 3 | k__Fungi(100);p__Ascomycota(100);p__Ascomycota_unclassified(100);p__Ascomycota_unclassified(100);p__Ascomycota_unclassified(100);p__Ascomycota_unclassified(100);p__Ascomycota_unclassified(100); | - | - | - | - | - | - | - | Unassigned | - |
| Otu07367 | 3 | k__Fungi(100);p__Ascomycota(100);p__Ascomycota_unclassified(100);p__Ascomycota_unclassified(100);p__Ascomycota_unclassified(100);p__Ascomycota_unclassified(100);p__Ascomycota_unclassified(100); | - | - | - | - | - | - | - | Unassigned | - |
| Otu07368 | 3 | k__Fungi(100);p__Basidiomycota(100);c__Agaricomycetes(100);c__Agaricomycetes_unclassified(100);c__Agaricomycetes_unclassified(100);c__Agaricomycetes_unclassified(100);c__Agaricomycetes_unclassified(100); | - | - | - | - | - | - | - | Unassigned | - |
| Otu07370 | 3 | k__Fungi(100);k__Fungi_unclassified(100);k__Fungi_unclassified(100);k__Fungi_unclassified(100);k__Fungi_unclassified(100);k__Fungi_unclassified(100);k__Fungi_unclassified(100); | - | - | - | - | - | - | - | Unassigned | - |
| Otu07371 | 3 | k__Fungi(100);k__Fungi_unclassified(100);k__Fungi_unclassified(100);k__Fungi_unclassified(100);k__Fungi_unclassified(100);k__Fungi_unclassified(100);k__Fungi_unclassified(100); | - | - | - | - | - | - | - | Unassigned | - |
| Otu07375 | 3 | k__Fungi(100);p__Ascomycota(100);c__Sordariomycetes(100);c__Sordariomycetes_unclassified(100);c__Sordariomycetes_unclassified(100);c__Sordariomycetes_unclassified(100);c__Sordariomycetes_unclassified(100); | - | - | - | - | - | - | - | Unassigned | - |
| Otu07376 | 3 | k__Fungi(100);k__Fungi_unclassified(100);k__Fungi_unclassified(100);k__Fungi_unclassified(100);k__Fungi_unclassified(100);k__Fungi_unclassified(100);k__Fungi_unclassified(100); | - | - | - | - | - | - | - | Unassigned | - |
| Otu07379 | 3 | k__Fungi(100);k__Fungi_unclassified(67);k__Fungi_unclassified(67);k__Fungi_unclassified(67);k__Fungi_unclassified(67);k__Fungi_unclassified(67);k__Fungi_unclassified(67); | - | - | - | - | - | - | - | Unassigned | - |
| Otu07383 | 3 | k__Fungi(100);k__Fungi_unclassified(67);k__Fungi_unclassified(67);k__Fungi_unclassified(67);k__Fungi_unclassified(67);k__Fungi_unclassified(67);k__Fungi_unclassified(67); | - | - | - | - | - | - | - | Unassigned | - |
| Otu07384 | 3 | k__Fungi(100);p__Ascomycota(100);c__Eurotiomycetes(100);o__Eurotiales(100);f__Trichocomaceae(100);f__Trichocomaceae_unclassified(67);f__Trichocomaceae_unclassified(67); | - | - | - | - | - | - | - | Unassigned | - |
| Otu07386 | 3 | k__Fungi(100);p__Ascomycota(67);p__Ascomycota_unclassified(67);p__Ascomycota_unclassified(67);p__Ascomycota_unclassified(67);p__Ascomycota_unclassified(67);p__Ascomycota_unclassified(67); | - | - | - | - | - | - | - | Unassigned | - |
| Otu07388 | 3 | k__Fungi(100);k__Fungi_unclassified(100);k__Fungi_unclassified(100);k__Fungi_unclassified(100);k__Fungi_unclassified(100);k__Fungi_unclassified(100);k__Fungi_unclassified(100); | - | - | - | - | - | - | - | Unassigned | - |
| Otu07389 | 3 | k__Fungi(100);k__Fungi_unclassified(67);k__Fungi_unclassified(67);k__Fungi_unclassified(67);k__Fungi_unclassified(67);k__Fungi_unclassified(67);k__Fungi_unclassified(67); | - | - | - | - | - | - | - | Unassigned | - |
| Otu07390 | 3 | k__Fungi(100);k__Fungi_unclassified(67);k__Fungi_unclassified(67);k__Fungi_unclassified(67);k__Fungi_unclassified(67);k__Fungi_unclassified(67);k__Fungi_unclassified(67); | - | - | - | - | - | - | - | Unassigned | - |
| Otu07391 | 3 | k__Fungi(100);k__Fungi_unclassified(67);k__Fungi_unclassified(67);k__Fungi_unclassified(67);k__Fungi_unclassified(67);k__Fungi_unclassified(67);k__Fungi_unclassified(67); | - | - | - | - | - | - | - | Unassigned | - |
| Otu07394 | 3 | k__Fungi(100);k__Fungi_unclassified(67);k__Fungi_unclassified(67);k__Fungi_unclassified(67);k__Fungi_unclassified(67);k__Fungi_unclassified(67);k__Fungi_unclassified(67); | - | - | - | - | - | - | - | Unassigned | - |
| Otu07396 | 3 | k__Fungi(100);k__Fungi_unclassified(100);k__Fungi_unclassified(100);k__Fungi_unclassified(100);k__Fungi_unclassified(100);k__Fungi_unclassified(100);k__Fungi_unclassified(100); | - | - | - | - | - | - | - | Unassigned | - |
| Otu07400 | 3 | k__Fungi(100);p__Basidiomycota(100);c__Agaricomycetes(100);c__Agaricomycetes_unclassified(100);c__Agaricomycetes_unclassified(100);c__Agaricomycetes_unclassified(100);c__Agaricomycetes_unclassified(100); | - | - | - | - | - | - | - | Unassigned | - |
| Otu07401 | 3 | k__Fungi(100);p__Ascomycota(100);p__Ascomycota_unclassified(100);p__Ascomycota_unclassified(100);p__Ascomycota_unclassified(100);p__Ascomycota_unclassified(100);p__Ascomycota_unclassified(100); | - | - | - | - | - | - | - | Unassigned | - |
| Otu07402 | 3 | k__Fungi(100);k__Fungi_unclassified(100);k__Fungi_unclassified(100);k__Fungi_unclassified(100);k__Fungi_unclassified(100);k__Fungi_unclassified(100);k__Fungi_unclassified(100); | - | - | - | - | - | - | - | Unassigned | - |
| Otu07405 | 3 | k__Fungi(100);k__Fungi_unclassified(67);k__Fungi_unclassified(67);k__Fungi_unclassified(67);k__Fungi_unclassified(67);k__Fungi_unclassified(67);k__Fungi_unclassified(67); | - | - | - | - | - | - | - | Unassigned | - |
| Otu07407 | 3 | k__Fungi(100);p__Ascomycota(100);p__Ascomycota_unclassified(100);p__Ascomycota_unclassified(100);p__Ascomycota_unclassified(100);p__Ascomycota_unclassified(100);p__Ascomycota_unclassified(100); | - | - | - | - | - | - | - | Unassigned | - |
| Otu07412 | 3 | k__Fungi(100);p__Basidiomycota(100);p__Basidiomycota_unclassified(100);p__Basidiomycota_unclassified(100);p__Basidiomycota_unclassified(100);p__Basidiomycota_unclassified(100);p__Basidiomycota_unclassified(100); | - | - | - | - | - | - | - | Unassigned | - |
| Otu07414 | 3 | k__Fungi(100);p__Ascomycota(67);p__Ascomycota_unclassified(67);p__Ascomycota_unclassified(67);p__Ascomycota_unclassified(67);p__Ascomycota_unclassified(67);p__Ascomycota_unclassified(67); | - | - | - | - | - | - | - | Unassigned | - |
| Otu07415 | 3 | k__Fungi(100);k__Fungi_unclassified(67);k__Fungi_unclassified(67);k__Fungi_unclassified(67);k__Fungi_unclassified(67);k__Fungi_unclassified(67);k__Fungi_unclassified(67); | - | - | - | - | - | - | - | Unassigned | - |
| Otu07416 | 3 | k__Fungi(100);p__Ascomycota(100);c__Dothideomycetes(100);o__Pleosporales(100);f__unclassified_Pleosporales(67);g__unclassified_Pleosporales(67);s__Pleosporales_sp(67); | - | - | - | - | - | - | - | Unassigned | - |
| Otu07417 | 3 | k__Fungi(100);p__Basidiomycota(100);p__Basidiomycota_unclassified(100);p__Basidiomycota_unclassified(100);p__Basidiomycota_unclassified(100);p__Basidiomycota_unclassified(100);p__Basidiomycota_unclassified(100); | - | - | - | - | - | - | - | Unassigned | - |
| Otu07419 | 3 | k__Fungi(100);p__Basidiomycota(100);p__Basidiomycota_unclassified(100);p__Basidiomycota_unclassified(100);p__Basidiomycota_unclassified(100);p__Basidiomycota_unclassified(100);p__Basidiomycota_unclassified(100); | - | - | - | - | - | - | - | Unassigned | - |
| Otu07422 | 3 | k__Fungi(100);p__Basidiomycota(100);c__Agaricomycetes(100);c__Agaricomycetes_unclassified(100);c__Agaricomycetes_unclassified(100);c__Agaricomycetes_unclassified(100);c__Agaricomycetes_unclassified(100); | - | - | - | - | - | - | - | Unassigned | - |
| Otu07423 | 3 | k__Fungi(100);k__Fungi_unclassified(67);k__Fungi_unclassified(67);k__Fungi_unclassified(67);k__Fungi_unclassified(67);k__Fungi_unclassified(67);k__Fungi_unclassified(67); | - | - | - | - | - | - | - | Unassigned | - |
| Otu07424 | 3 | k__Fungi(100);k__Fungi_unclassified(67);k__Fungi_unclassified(67);k__Fungi_unclassified(67);k__Fungi_unclassified(67);k__Fungi_unclassified(67);k__Fungi_unclassified(67); | - | - | - | - | - | - | - | Unassigned | - |
| Otu07426 | 3 | k__Fungi(100);p__Basidiomycota(100);p__Basidiomycota_unclassified(100);p__Basidiomycota_unclassified(100);p__Basidiomycota_unclassified(100);p__Basidiomycota_unclassified(100);p__Basidiomycota_unclassified(100); | - | - | - | - | - | - | - | Unassigned | - |
| Otu07430 | 3 | k__Fungi(100);k__Fungi_unclassified(67);k__Fungi_unclassified(67);k__Fungi_unclassified(67);k__Fungi_unclassified(67);k__Fungi_unclassified(67);k__Fungi_unclassified(67); | - | - | - | - | - | - | - | Unassigned | - |
| Otu07438 | 3 | k__Fungi(100);p__Chytridiomycota(100);c__unclassified_Chytridiomycota(100);o__unclassified_Chytridiomycota(100);f__unclassified_Chytridiomycota(100);g__unclassified_Chytridiomycota(100);s__Chytridiomycota_sp(100); | - | - | - | - | - | - | - | Unassigned | - |
| Otu07441 | 3 | k__Fungi(100);k__Fungi_unclassified(100);k__Fungi_unclassified(100);k__Fungi_unclassified(100);k__Fungi_unclassified(100);k__Fungi_unclassified(100);k__Fungi_unclassified(100); | - | - | - | - | - | - | - | Unassigned | - |
| Otu07442 | 3 | k__Fungi(100);k__Fungi_unclassified(67);k__Fungi_unclassified(67);k__Fungi_unclassified(67);k__Fungi_unclassified(67);k__Fungi_unclassified(67);k__Fungi_unclassified(67); | - | - | - | - | - | - | - | Unassigned | - |
| Otu07443 | 3 | k__Fungi(100);k__Fungi_unclassified(67);k__Fungi_unclassified(67);k__Fungi_unclassified(67);k__Fungi_unclassified(67);k__Fungi_unclassified(67);k__Fungi_unclassified(67); | - | - | - | - | - | - | - | Unassigned | - |
| Otu07445 | 3 | k__Fungi(100);p__Ascomycota(100);p__Ascomycota_unclassified(100);p__Ascomycota_unclassified(100);p__Ascomycota_unclassified(100);p__Ascomycota_unclassified(100);p__Ascomycota_unclassified(100); | - | - | - | - | - | - | - | Unassigned | - |
| Otu07447 | 3 | k__Fungi(100);p__Ascomycota(100);c__Dothideomycetes(100);c__Dothideomycetes_unclassified(100);c__Dothideomycetes_unclassified(100);c__Dothideomycetes_unclassified(100);c__Dothideomycetes_unclassified(100); | - | - | - | - | - | - | - | Unassigned | - |
| Otu07449 | 3 | k__Fungi(100);p__Basidiomycota(100);c__Agaricomycetes(100);o__Trechisporales(100);f__Hydnodontaceae(100);f__Hydnodontaceae_unclassified(100);f__Hydnodontaceae_unclassified(100); | - | - | - | - | - | - | - | Unassigned | - |
| Otu07450 | 3 | k__Fungi(100);p__Ascomycota(67);p__Ascomycota_unclassified(67);p__Ascomycota_unclassified(67);p__Ascomycota_unclassified(67);p__Ascomycota_unclassified(67);p__Ascomycota_unclassified(67); | - | - | - | - | - | - | - | Unassigned | - |
| Otu07454 | 3 | k__Fungi(100);p__Ascomycota(100);p__Ascomycota_unclassified(100);p__Ascomycota_unclassified(100);p__Ascomycota_unclassified(100);p__Ascomycota_unclassified(100);p__Ascomycota_unclassified(100); | - | - | - | - | - | - | - | Unassigned | - |
| Otu07455 | 3 | k__Fungi(100);p__Ascomycota(67);p__Ascomycota_unclassified(67);p__Ascomycota_unclassified(67);p__Ascomycota_unclassified(67);p__Ascomycota_unclassified(67);p__Ascomycota_unclassified(67); | - | - | - | - | - | - | - | Unassigned | - |
| Otu07457 | 3 | k__Fungi(100);p__Ascomycota(100);p__Ascomycota_unclassified(100);p__Ascomycota_unclassified(100);p__Ascomycota_unclassified(100);p__Ascomycota_unclassified(100);p__Ascomycota_unclassified(100); | - | - | - | - | - | - | - | Unassigned | - |
| Otu07459 | 3 | k__Fungi(100);p__Basidiomycota(100);c__Agaricomycetes(100);c__Agaricomycetes_unclassified(100);c__Agaricomycetes_unclassified(100);c__Agaricomycetes_unclassified(100);c__Agaricomycetes_unclassified(100); | - | - | - | - | - | - | - | Unassigned | - |
| Otu07461 | 3 | k__Fungi(100);k__Fungi_unclassified(67);k__Fungi_unclassified(67);k__Fungi_unclassified(67);k__Fungi_unclassified(67);k__Fungi_unclassified(67);k__Fungi_unclassified(67); | - | - | - | - | - | - | - | Unassigned | - |
| Otu07465 | 3 | k__Fungi(100);k__Fungi_unclassified(67);k__Fungi_unclassified(67);k__Fungi_unclassified(67);k__Fungi_unclassified(67);k__Fungi_unclassified(67);k__Fungi_unclassified(67); | - | - | - | - | - | - | - | Unassigned | - |
| Otu07466 | 3 | k__Fungi(100);k__Fungi_unclassified(67);k__Fungi_unclassified(67);k__Fungi_unclassified(67);k__Fungi_unclassified(67);k__Fungi_unclassified(67);k__Fungi_unclassified(67); | - | - | - | - | - | - | - | Unassigned | - |
| Otu07472 | 3 | k__Fungi(100);p__Ascomycota(67);p__Ascomycota_unclassified(67);p__Ascomycota_unclassified(67);p__Ascomycota_unclassified(67);p__Ascomycota_unclassified(67);p__Ascomycota_unclassified(67); | - | - | - | - | - | - | - | Unassigned | - |
| Otu07473 | 3 | k__Fungi(100);k__Fungi_unclassified(100);k__Fungi_unclassified(100);k__Fungi_unclassified(100);k__Fungi_unclassified(100);k__Fungi_unclassified(100);k__Fungi_unclassified(100); | - | - | - | - | - | - | - | Unassigned | - |
| Otu07475 | 3 | k__Fungi(100);k__Fungi_unclassified(67);k__Fungi_unclassified(67);k__Fungi_unclassified(67);k__Fungi_unclassified(67);k__Fungi_unclassified(67);k__Fungi_unclassified(67); | - | - | - | - | - | - | - | Unassigned | - |
| Otu07476 | 3 | k__Fungi(100);p__Ascomycota(100);p__Ascomycota_unclassified(100);p__Ascomycota_unclassified(100);p__Ascomycota_unclassified(100);p__Ascomycota_unclassified(100);p__Ascomycota_unclassified(100); | - | - | - | - | - | - | - | Unassigned | - |
| Otu07478 | 3 | k__Fungi(100);p__Basidiomycota(100);p__Basidiomycota_unclassified(100);p__Basidiomycota_unclassified(100);p__Basidiomycota_unclassified(100);p__Basidiomycota_unclassified(100);p__Basidiomycota_unclassified(100); | - | - | - | - | - | - | - | Unassigned | - |
| Otu07479 | 3 | k__Fungi(100);p__Ascomycota(100);p__Ascomycota_unclassified(100);p__Ascomycota_unclassified(100);p__Ascomycota_unclassified(100);p__Ascomycota_unclassified(100);p__Ascomycota_unclassified(100); | - | - | - | - | - | - | - | Unassigned | - |
| Otu07484 | 3 | k__Fungi(100);p__Ascomycota(67);p__Ascomycota_unclassified(67);p__Ascomycota_unclassified(67);p__Ascomycota_unclassified(67);p__Ascomycota_unclassified(67);p__Ascomycota_unclassified(67); | - | - | - | - | - | - | - | Unassigned | - |
| Otu07485 | 3 | k__Fungi(100);k__Fungi_unclassified(100);k__Fungi_unclassified(100);k__Fungi_unclassified(100);k__Fungi_unclassified(100);k__Fungi_unclassified(100);k__Fungi_unclassified(100); | - | - | - | - | - | - | - | Unassigned | - |
| Otu07488 | 3 | k__Fungi(100);p__Basidiomycota(100);p__Basidiomycota_unclassified(100);p__Basidiomycota_unclassified(100);p__Basidiomycota_unclassified(100);p__Basidiomycota_unclassified(100);p__Basidiomycota_unclassified(100); | - | - | - | - | - | - | - | Unassigned | - |
| Otu07490 | 3 | k__Fungi(100);p__Ascomycota(100);c__Dothideomycetes(67);c__Dothideomycetes_unclassified(67);c__Dothideomycetes_unclassified(67);c__Dothideomycetes_unclassified(67);c__Dothideomycetes_unclassified(67); | - | - | - | - | - | - | - | Unassigned | - |
| Otu07491 | 3 | k__Fungi(100);k__Fungi_unclassified(67);k__Fungi_unclassified(67);k__Fungi_unclassified(67);k__Fungi_unclassified(67);k__Fungi_unclassified(67);k__Fungi_unclassified(67); | - | - | - | - | - | - | - | Unassigned | - |
| Otu07492 | 3 | k__Fungi(100);p__Ascomycota(100);p__Ascomycota_unclassified(100);p__Ascomycota_unclassified(100);p__Ascomycota_unclassified(100);p__Ascomycota_unclassified(100);p__Ascomycota_unclassified(100); | - | - | - | - | - | - | - | Unassigned | - |
| Otu07496 | 3 | k__Fungi(100);p__Basidiomycota(100);c__Exobasidiomycetes(100);o__Exobasidiales(100);o__Exobasidiales_unclassified(100);o__Exobasidiales_unclassified(100);o__Exobasidiales_unclassified(100); | - | - | - | - | - | - | - | Unassigned | - |

| | | | | | | | | | | |
|---|---|---|---|---|---|---|---|---|---|---|
| Otu07497 | 3 | k__Fungi(100);p__Basidiomycota(100);c__Agaricomycetes(100);c__Agaricomycetes_unclassified(100);c__Agaricomycetes_unclassified(100);c__Agaricomycetes_unclassified(100);c__Agaricomycetes_unclassified(100); | - | - | - | - | - | - | - | Unassigned | - |
| Otu07500 | 3 | k__Fungi(100);k__Fungi_unclassified(100);k__Fungi_unclassified(100);k__Fungi_unclassified(100);k__Fungi_unclassified(100);k__Fungi_unclassified(100);k__Fungi_unclassified(100); | - | - | - | - | - | - | - | Unassigned | - |
| Otu07501 | 3 | k__Fungi(100);p__Ascomycota(100);c__Sordariomycetes(100);c__Sordariomycetes_unclassified(100);c__Sordariomycetes_unclassified(100);c__Sordariomycetes_unclassified(100);c__Sordariomycetes_unclassified(100); | - | - | - | - | - | - | - | Unassigned | - |
| Otu07508 | 3 | k__Fungi(100);k__Fungi_unclassified(67);k__Fungi_unclassified(67);k__Fungi_unclassified(67);k__Fungi_unclassified(67);k__Fungi_unclassified(67);k__Fungi_unclassified(67); | - | - | - | - | - | - | - | Unassigned | - |
| Otu07511 | 3 | k__Fungi(100);k__Fungi_unclassified(100);k__Fungi_unclassified(100);k__Fungi_unclassified(100);k__Fungi_unclassified(100);k__Fungi_unclassified(100);k__Fungi_unclassified(100); | - | - | - | - | - | - | - | Unassigned | - |
| Otu07514 | 3 | k__Fungi(100);p__Ascomycota(100);p__Ascomycota_unclassified(100);p__Ascomycota_unclassified(100);p__Ascomycota_unclassified(100);p__Ascomycota_unclassified(100);p__Ascomycota_unclassified(100); | - | - | - | - | - | - | - | Unassigned | - |
| Otu07515 | 3 | k__Fungi(100);p__Ascomycota(67);p__Ascomycota_unclassified(67);p__Ascomycota_unclassified(67);p__Ascomycota_unclassified(67);p__Ascomycota_unclassified(67);p__Ascomycota_unclassified(67); | - | - | - | - | - | - | - | Unassigned | - |
| Otu07516 | 3 | k__Fungi(100);k__Fungi_unclassified(67);k__Fungi_unclassified(67);k__Fungi_unclassified(67);k__Fungi_unclassified(67);k__Fungi_unclassified(67);k__Fungi_unclassified(67); | - | - | - | - | - | - | - | Unassigned | - |
| Otu07518 | 3 | k__Fungi(100);p__Ascomycota(67);p__Ascomycota_unclassified(67);p__Ascomycota_unclassified(67);p__Ascomycota_unclassified(67);p__Ascomycota_unclassified(67);p__Ascomycota_unclassified(67); | - | - | - | - | - | - | - | Unassigned | - |
| Otu07519 | 3 | k__Fungi(100);p__Basidiomycota(100);c__Agaricomycetes(100);o__Auriculariales(100);f__unclassified_Auriculariales(100);g__unclassified_Auriculariales(100);s__Auriculariales_sp(100); | - | - | - | - | - | - | - | Unassigned | - |
| Otu07520 | 3 | k__Fungi(100);p__Ascomycota(100);c__Pezizomycetes(100);o__Pezizales(100);f__unclassified_Pezizales(100);g__unclassified_Pezizales(100);s__Pezizales_sp(100); | - | - | - | - | - | - | - | Unassigned | - |
| Otu07522 | 3 | k__Fungi(100);p__Ascomycota(67);p__Ascomycota_unclassified(67);p__Ascomycota_unclassified(67);p__Ascomycota_unclassified(67);p__Ascomycota_unclassified(67);p__Ascomycota_unclassified(67); | - | - | - | - | - | - | - | Unassigned | - |
| Otu07524 | 3 | k__Fungi(100);k__Fungi_unclassified(100);k__Fungi_unclassified(100);k__Fungi_unclassified(100);k__Fungi_unclassified(100);k__Fungi_unclassified(100);k__Fungi_unclassified(100); | - | - | - | - | - | - | - | Unassigned | - |
| Otu07527 | 3 | k__Fungi(100);p__Ascomycota(67);p__Ascomycota_unclassified(67);p__Ascomycota_unclassified(67);p__Ascomycota_unclassified(67);p__Ascomycota_unclassified(67);p__Ascomycota_unclassified(67); | - | - | - | - | - | - | - | Unassigned | - |
| Otu07528 | 3 | k__Fungi(100);p__Ascomycota(100);p__Ascomycota_unclassified(100);p__Ascomycota_unclassified(100);p__Ascomycota_unclassified(100);p__Ascomycota_unclassified(100);p__Ascomycota_unclassified(100); | - | - | - | - | - | - | - | Unassigned | - |
| Otu07529 | 3 | k__Fungi(100);k__Fungi_unclassified(100);k__Fungi_unclassified(100);k__Fungi_unclassified(100);k__Fungi_unclassified(100);k__Fungi_unclassified(100);k__Fungi_unclassified(100); | - | - | - | - | - | - | - | Unassigned | - |
| Otu07530 | 3 | k__Fungi(100);p__Ascomycota(100);p__Ascomycota_unclassified(100);p__Ascomycota_unclassified(100);p__Ascomycota_unclassified(100);p__Ascomycota_unclassified(100);p__Ascomycota_unclassified(100); | - | - | - | - | - | - | - | Unassigned | - |
| Otu07533 | 3 | k__Fungi(100);p__Ascomycota(67);p__Ascomycota_unclassified(67);p__Ascomycota_unclassified(67);p__Ascomycota_unclassified(67);p__Ascomycota_unclassified(67);p__Ascomycota_unclassified(67); | - | - | - | - | - | - | - | Unassigned | - |
| Otu07535 | 3 | k__Fungi(100);p__Basidiomycota(100);c__Microbotryomycetes(100);o__Sporidiobolales(100);f__Sporidiobolales_family_Incertae_sedis(100);f__Sporidiobolales_family_Incertae_sedis_unclassified(100);f__Sporidiobolales_family_Incertae_sedis_unclassified(100); | - | - | - | - | - | - | - | Unassigned | - |
| Otu07536 | 3 | k__Fungi(100);k__Fungi_unclassified(100);k__Fungi_unclassified(100);k__Fungi_unclassified(100);k__Fungi_unclassified(100);k__Fungi_unclassified(100);k__Fungi_unclassified(100); | - | - | - | - | - | - | - | Unassigned | - |
| Otu07537 | 3 | k__Fungi(100);p__Basidiomycota(100);c__Microbotryomycetes(100);o__Sporidiobolales(100);f__Sporidiobolales_family_Incertae_sedis(100);f__Sporidiobolales_family_Incertae_sedis_unclassified(67);f__Sporidiobolales_family_Incertae_sedis_unclassified(67); | - | - | - | - | - | - | - | Unassigned | - |
| Otu07541 | 3 | k__Fungi(100);k__Fungi_unclassified(67);k__Fungi_unclassified(67);k__Fungi_unclassified(67);k__Fungi_unclassified(67);k__Fungi_unclassified(67);k__Fungi_unclassified(67); | - | - | - | - | - | - | - | Unassigned | - |
| Otu07544 | 3 | k__Fungi(100);k__Fungi_unclassified(67);k__Fungi_unclassified(67);k__Fungi_unclassified(67);k__Fungi_unclassified(67);k__Fungi_unclassified(67);k__Fungi_unclassified(67); | - | - | - | - | - | - | - | Unassigned | - |
| Otu07545 | 3 | k__Fungi(100);p__Ascomycota(67);p__Ascomycota_unclassified(67);p__Ascomycota_unclassified(67);p__Ascomycota_unclassified(67);p__Ascomycota_unclassified(67);p__Ascomycota_unclassified(67); | - | - | - | - | - | - | - | Unassigned | - |
| Otu07547 | 3 | k__Fungi(100);k__Fungi_unclassified(67);k__Fungi_unclassified(67);k__Fungi_unclassified(67);k__Fungi_unclassified(67);k__Fungi_unclassified(67);k__Fungi_unclassified(67); | - | - | - | - | - | - | - | Unassigned | - |
| Otu07548 | 3 | k__Fungi(100);k__Fungi_unclassified(67);k__Fungi_unclassified(67);k__Fungi_unclassified(67);k__Fungi_unclassified(67);k__Fungi_unclassified(67);k__Fungi_unclassified(67); | - | - | - | - | - | - | - | Unassigned | - |
| Otu07551 | 3 | k__Fungi(100);p__Ascomycota(67);p__Ascomycota_unclassified(67);p__Ascomycota_unclassified(67);p__Ascomycota_unclassified(67);p__Ascomycota_unclassified(67);p__Ascomycota_unclassified(67); | - | - | - | - | - | - | - | Unassigned | - |
| Otu07552 | 3 | k__Fungi(100);k__Fungi_unclassified(67);k__Fungi_unclassified(67);k__Fungi_unclassified(67);k__Fungi_unclassified(67);k__Fungi_unclassified(67);k__Fungi_unclassified(67); | - | - | - | - | - | - | - | Unassigned | - |
| Otu07553 | 3 | k__Fungi(100);k__Fungi_unclassified(67);k__Fungi_unclassified(67);k__Fungi_unclassified(67);k__Fungi_unclassified(67);k__Fungi_unclassified(67);k__Fungi_unclassified(67); | - | - | - | - | - | - | - | Unassigned | - |
| Otu07556 | 3 | k__Fungi(100);k__Fungi_unclassified(67);k__Fungi_unclassified(67);k__Fungi_unclassified(67);k__Fungi_unclassified(67);k__Fungi_unclassified(67);k__Fungi_unclassified(67); | - | - | - | - | - | - | - | Unassigned | - |
| Otu07557 | 3 | k__Fungi(100);k__Fungi_unclassified(100);k__Fungi_unclassified(100);k__Fungi_unclassified(100);k__Fungi_unclassified(100);k__Fungi_unclassified(100);k__Fungi_unclassified(100); | - | - | - | - | - | - | - | Unassigned | - |
| Otu07560 | 3 | k__Fungi(100);p__Ascomycota(100);p__Ascomycota_unclassified(100);p__Ascomycota_unclassified(100);p__Ascomycota_unclassified(100);p__Ascomycota_unclassified(100);p__Ascomycota_unclassified(100); | - | - | - | - | - | - | - | Unassigned | - |
| Otu07563 | 3 | k__Fungi(100);p__Ascomycota(100);p__Ascomycota_unclassified(100);p__Ascomycota_unclassified(100);p__Ascomycota_unclassified(100);p__Ascomycota_unclassified(100);p__Ascomycota_unclassified(100); | - | - | - | - | - | - | - | Unassigned | - |
| Otu07564 | 3 | k__Fungi(100);p__Basidiomycota(100);p__Basidiomycota_unclassified(67);p__Basidiomycota_unclassified(67);p__Basidiomycota_unclassified(67);p__Basidiomycota_unclassified(67);p__Basidiomycota_unclassified(67); | - | - | - | - | - | - | - | Unassigned | - |
| Otu07566 | 3 | k__Fungi(100);p__Ascomycota(67);p__Ascomycota_unclassified(67);p__Ascomycota_unclassified(67);p__Ascomycota_unclassified(67);p__Ascomycota_unclassified(67);p__Ascomycota_unclassified(67); | - | - | - | - | - | - | - | Unassigned | - |
| Otu07567 | 3 | k__Fungi(100);k__Fungi_unclassified(100);k__Fungi_unclassified(100);k__Fungi_unclassified(100);k__Fungi_unclassified(100);k__Fungi_unclassified(100);k__Fungi_unclassified(100); | - | - | - | - | - | - | - | Unassigned | - |
| Otu07568 | 3 | k__Fungi(100);k__Fungi_unclassified(100);k__Fungi_unclassified(100);k__Fungi_unclassified(100);k__Fungi_unclassified(100);k__Fungi_unclassified(100);k__Fungi_unclassified(100); | - | - | - | - | - | - | - | Unassigned | - |
| Otu07569 | 3 | k__Fungi(100);k__Fungi_unclassified(67);k__Fungi_unclassified(67);k__Fungi_unclassified(67);k__Fungi_unclassified(67);k__Fungi_unclassified(67);k__Fungi_unclassified(67); | - | - | - | - | - | - | - | Unassigned | - |
| Otu07570 | 3 | k__Fungi(100);k__Fungi_unclassified(100);k__Fungi_unclassified(100);k__Fungi_unclassified(100);k__Fungi_unclassified(100);k__Fungi_unclassified(100);k__Fungi_unclassified(100); | - | - | - | - | - | - | - | Unassigned | - |
| Otu07571 | 3 | k__Fungi(100);p__Basidiomycota(100);c__Agaricomycetes(100);o__Agaricales(100);f__Niaceae(67);f__Niaceae_unclassified(67);f__Niaceae_unclassified(67); | - | - | - | - | - | - | - | Unassigned | - |
| Otu07572 | 3 | k__Fungi(100);k__Fungi_unclassified(100);k__Fungi_unclassified(100);k__Fungi_unclassified(100);k__Fungi_unclassified(100);k__Fungi_unclassified(100);k__Fungi_unclassified(100); | - | - | - | - | - | - | - | Unassigned | - |
| Otu07576 | 3 | k__Fungi(100);p__Ascomycota(100);p__Ascomycota_unclassified(100);p__Ascomycota_unclassified(100);p__Ascomycota_unclassified(100);p__Ascomycota_unclassified(100);p__Ascomycota_unclassified(100); | - | - | - | - | - | - | - | Unassigned | - |
| Otu07577 | 3 | k__Fungi(100);k__Fungi_unclassified(67);k__Fungi_unclassified(67);k__Fungi_unclassified(67);k__Fungi_unclassified(67);k__Fungi_unclassified(67);k__Fungi_unclassified(67); | - | - | - | - | - | - | - | Unassigned | - |
| Otu07582 | 3 | k__Fungi(100);p__Ascomycota(100);p__Ascomycota_unclassified(100);p__Ascomycota_unclassified(100);p__Ascomycota_unclassified(100);p__Ascomycota_unclassified(100);p__Ascomycota_unclassified(100); | - | - | - | - | - | - | - | Unassigned | - |
| Otu07585 | 3 | k__Fungi(100);p__Ascomycota(100);p__Ascomycota_unclassified(100);p__Ascomycota_unclassified(100);p__Ascomycota_unclassified(100);p__Ascomycota_unclassified(100);p__Ascomycota_unclassified(100); | - | - | - | - | - | - | - | Unassigned | - |
| Otu07586 | 3 | k__Fungi(100);k__Fungi_unclassified(100);k__Fungi_unclassified(100);k__Fungi_unclassified(100);k__Fungi_unclassified(100);k__Fungi_unclassified(100);k__Fungi_unclassified(100); | - | - | - | - | - | - | - | Unassigned | - |
| Otu07588 | 3 | k__Fungi(100);k__Fungi_unclassified(100);k__Fungi_unclassified(100);k__Fungi_unclassified(100);k__Fungi_unclassified(100);k__Fungi_unclassified(100);k__Fungi_unclassified(100); | - | - | - | - | - | - | - | Unassigned | - |
| Otu07589 | 3 | k__Fungi(100);k__Fungi_unclassified(67);k__Fungi_unclassified(67);k__Fungi_unclassified(67);k__Fungi_unclassified(67);k__Fungi_unclassified(67);k__Fungi_unclassified(67); | - | - | - | - | - | - | - | Unassigned | - |
| Otu07590 | 3 | k__Fungi(100);k__Fungi_unclassified(100);k__Fungi_unclassified(100);k__Fungi_unclassified(100);k__Fungi_unclassified(100);k__Fungi_unclassified(100);k__Fungi_unclassified(100); | - | - | - | - | - | - | - | Unassigned | - |
| Otu07591 | 3 | k__Fungi(100);p__Ascomycota(67);p__Ascomycota_unclassified(67);p__Ascomycota_unclassified(67);p__Ascomycota_unclassified(67);p__Ascomycota_unclassified(67);p__Ascomycota_unclassified(67); | - | - | - | - | - | - | - | Unassigned | - |
| Otu07593 | 3 | k__Fungi(100);p__Basidiomycota(100);p__Basidiomycota_unclassified(67);p__Basidiomycota_unclassified(67);p__Basidiomycota_unclassified(67);p__Basidiomycota_unclassified(67);p__Basidiomycota_unclassified(67); | - | - | - | - | - | - | - | Unassigned | - |
| Otu07595 | 3 | k__Fungi(100);k__Fungi_unclassified(67);k__Fungi_unclassified(67);k__Fungi_unclassified(67);k__Fungi_unclassified(67);k__Fungi_unclassified(67);k__Fungi_unclassified(67); | - | - | - | - | - | - | - | Unassigned | - |
| Otu07596 | 3 | k__Fungi(100);k__Fungi_unclassified(100);k__Fungi_unclassified(100);k__Fungi_unclassified(100);k__Fungi_unclassified(100);k__Fungi_unclassified(100);k__Fungi_unclassified(100); | - | - | - | - | - | - | - | Unassigned | - |
| Otu07597 | 3 | k__Fungi(100);k__Fungi_unclassified(100);k__Fungi_unclassified(100);k__Fungi_unclassified(100);k__Fungi_unclassified(100);k__Fungi_unclassified(100);k__Fungi_unclassified(100); | - | - | - | - | - | - | - | Unassigned | - |
| Otu07599 | 3 | k__Fungi(100);p__Ascomycota(100);p__Ascomycota_unclassified(100);p__Ascomycota_unclassified(100);p__Ascomycota_unclassified(100);p__Ascomycota_unclassified(100);p__Ascomycota_unclassified(100); | - | - | - | - | - | - | - | Unassigned | - |
| Otu07601 | 3 | k__Fungi(100);p__Ascomycota(100);p__Ascomycota_unclassified(100);p__Ascomycota_unclassified(100);p__Ascomycota_unclassified(100);p__Ascomycota_unclassified(100);p__Ascomycota_unclassified(100); | - | - | - | - | - | - | - | Unassigned | - |
| Otu07603 | 3 | k__Fungi(100);k__Fungi_unclassified(100);k__Fungi_unclassified(100);k__Fungi_unclassified(100);k__Fungi_unclassified(100);k__Fungi_unclassified(100);k__Fungi_unclassified(100); | - | - | - | - | - | - | - | Unassigned | - |
| Otu07605 | 3 | k__Fungi(100);k__Fungi_unclassified(100);k__Fungi_unclassified(100);k__Fungi_unclassified(100);k__Fungi_unclassified(100);k__Fungi_unclassified(100);k__Fungi_unclassified(100); | - | - | - | - | - | - | - | Unassigned | - |
| Otu07606 | 3 | k__Fungi(100);p__Ascomycota(100);p__Ascomycota_unclassified(100);p__Ascomycota_unclassified(100);p__Ascomycota_unclassified(100);p__Ascomycota_unclassified(100);p__Ascomycota_unclassified(100); | - | - | - | - | - | - | - | Unassigned | - |
| Otu07607 | 3 | k__Fungi(100);p__Ascomycota(100);c__Dothideomycetes(100);o__Pleosporales(100);o__Pleosporales_unclassified(100);o__Pleosporales_unclassified(100);o__Pleosporales_unclassified(100); | - | - | - | - | - | - | - | Unassigned | - |
| Otu07610 | 3 | k__Fungi(100);p__Ascomycota(100);c__Leotiomycetes(100);o__Helotiales(67);o__Helotiales_unclassified(67);o__Helotiales_unclassified(67);o__Helotiales_unclassified(67); | - | - | - | - | - | - | - | Unassigned | - |
| Otu07611 | 3 | k__Fungi(100);p__Chytridiomycota(100);c__unclassified_Chytridiomycota(100);o__unclassified_Chytridiomycota(100);f__unclassified_Chytridiomycota(100);g__unclassified_Chytridiomycota(100);s__Chytridiomycota_sp(100); | - | - | - | - | - | - | - | Unassigned | - |
| Otu07616 | 3 | k__Fungi(100);p__Chytridiomycota(100);c__unclassified_Chytridiomycota(100);o__unclassified_Chytridiomycota(100);f__unclassified_Chytridiomycota(100);g__unclassified_Chytridiomycota(100);s__Chytridiomycota_sp(100); | - | - | - | - | - | - | - | Unassigned | - |
| Otu07618 | 3 | k__Fungi(100);k__Fungi_unclassified(100);k__Fungi_unclassified(100);k__Fungi_unclassified(100);k__Fungi_unclassified(100);k__Fungi_unclassified(100);k__Fungi_unclassified(100); | - | - | - | - | - | - | - | Unassigned | - |
| Otu07621 | 3 | k__Fungi(100);p__Basidiomycota(100);c__Agaricomycetes(100);c__Agaricomycetes_unclassified(100);c__Agaricomycetes_unclassified(100);c__Agaricomycetes_unclassified(100);c__Agaricomycetes_unclassified(100); | - | - | - | - | - | - | - | Unassigned | - |
| Otu07622 | 3 | k__Fungi(100);p__Ascomycota(67);p__Ascomycota_unclassified(67);p__Ascomycota_unclassified(67);p__Ascomycota_unclassified(67);p__Ascomycota_unclassified(67);p__Ascomycota_unclassified(67); | - | - | - | - | - | - | - | Unassigned | - |
| Otu07623 | 3 | k__Fungi(100);k__Fungi_unclassified(100);k__Fungi_unclassified(100);k__Fungi_unclassified(100);k__Fungi_unclassified(100);k__Fungi_unclassified(100);k__Fungi_unclassified(100); | - | - | - | - | - | - | - | Unassigned | - |
| Otu07624 | 3 | k__Fungi(100);k__Fungi_unclassified(100);k__Fungi_unclassified(100);k__Fungi_unclassified(100);k__Fungi_unclassified(100);k__Fungi_unclassified(100);k__Fungi_unclassified(100); | - | - | - | - | - | - | - | Unassigned | - |
| Otu07627 | 3 | k__Fungi(100);p__Ascomycota(67);p__Ascomycota_unclassified(67);p__Ascomycota_unclassified(67);p__Ascomycota_unclassified(67);p__Ascomycota_unclassified(67);p__Ascomycota_unclassified(67); | - | - | - | - | - | - | - | Unassigned | - |
| Otu07629 | 3 | k__Fungi(100);p__Ascomycota(100);p__Ascomycota_unclassified(100);p__Ascomycota_unclassified(100);p__Ascomycota_unclassified(100);p__Ascomycota_unclassified(100);p__Ascomycota_unclassified(100); | - | - | - | - | - | - | - | Unassigned | - |
| Otu07633 | 3 | k__Fungi(100);k__Fungi_unclassified(67);k__Fungi_unclassified(67);k__Fungi_unclassified(67);k__Fungi_unclassified(67);k__Fungi_unclassified(67);k__Fungi_unclassified(67); | - | - | - | - | - | - | - | Unassigned | - |
| Otu07634 | 3 | k__Fungi(100);p__Ascomycota(67);p__Ascomycota_unclassified(67);p__Ascomycota_unclassified(67);p__Ascomycota_unclassified(67);p__Ascomycota_unclassified(67);p__Ascomycota_unclassified(67); | - | - | - | - | - | - | - | Unassigned | - |
| Otu07635 | 3 | k__Fungi(100);k__Fungi_unclassified(100);k__Fungi_unclassified(100);k__Fungi_unclassified(100);k__Fungi_unclassified(100);k__Fungi_unclassified(100);k__Fungi_unclassified(100); | - | - | - | - | - | - | - | Unassigned | - |
| Otu07636 | 3 | k__Fungi(100);p__Ascomycota(67);p__Ascomycota_unclassified(67);p__Ascomycota_unclassified(67);p__Ascomycota_unclassified(67);p__Ascomycota_unclassified(67);p__Ascomycota_unclassified(67); | - | - | - | - | - | - | - | Unassigned | - |
| Otu07637 | 3 | k__Fungi(100);k__Fungi_unclassified(67);k__Fungi_unclassified(67);k__Fungi_unclassified(67);k__Fungi_unclassified(67);k__Fungi_unclassified(67);k__Fungi_unclassified(67); | - | - | - | - | - | - | - | Unassigned | - |
| Otu07639 | 3 | k__Fungi(100);p__Ascomycota(100);p__Ascomycota_unclassified(100);p__Ascomycota_unclassified(100);p__Ascomycota_unclassified(100);p__Ascomycota_unclassified(100);p__Ascomycota_unclassified(100); | - | - | - | - | - | - | - | Unassigned | - |
| Otu07641 | 3 | k__Fungi(100);k__Fungi_unclassified(100);k__Fungi_unclassified(100);k__Fungi_unclassified(100);k__Fungi_unclassified(100);k__Fungi_unclassified(100);k__Fungi_unclassified(100); | - | - | - | - | - | - | - | Unassigned | - |
| Otu07644 | 3 | k__Fungi(100);p__Ascomycota(67);p__Ascomycota_unclassified(67);p__Ascomycota_unclassified(67);p__Ascomycota_unclassified(67);p__Ascomycota_unclassified(67);p__Ascomycota_unclassified(67); | - | - | - | - | - | - | - | Unassigned | - |
| Otu07647 | 3 | k__Fungi(100);p__Ascomycota(100);c__Leotiomycetes(100);o__Helotiales(100);o__Helotiales_unclassified(67);o__Helotiales_unclassified(67);o__Helotiales_unclassified(67); | - | - | - | - | - | - | - | Unassigned | - |
| Otu07649 | 3 | k__Fungi(100);p__Ascomycota(67);p__Ascomycota_unclassified(67);p__Ascomycota_unclassified(67);p__Ascomycota_unclassified(67);p__Ascomycota_unclassified(67);p__Ascomycota_unclassified(67); | - | - | - | - | - | - | - | Unassigned | - |
| Otu07652 | 3 | k__Fungi(100);k__Fungi_unclassified(100);k__Fungi_unclassified(100);k__Fungi_unclassified(100);k__Fungi_unclassified(100);k__Fungi_unclassified(100);k__Fungi_unclassified(100); | - | - | - | - | - | - | - | Unassigned | - |
| Otu07653 | 3 | k__Fungi(100);k__Fungi_unclassified(67);k__Fungi_unclassified(67);k__Fungi_unclassified(67);k__Fungi_unclassified(67);k__Fungi_unclassified(67);k__Fungi_unclassified(67); | - | - | - | - | - | - | - | Unassigned | - |
| Otu07655 | 3 | k__Fungi(100);p__Ascomycota(100);p__Ascomycota_unclassified(100);p__Ascomycota_unclassified(100);p__Ascomycota_unclassified(100);p__Ascomycota_unclassified(100);p__Ascomycota_unclassified(100); | - | - | - | - | - | - | - | Unassigned | - |
| Otu07656 | 3 | k__Fungi(100);k__Fungi_unclassified(100);k__Fungi_unclassified(100);k__Fungi_unclassified(100);k__Fungi_unclassified(100);k__Fungi_unclassified(100);k__Fungi_unclassified(100); | - | - | - | - | - | - | - | Unassigned | - |
| Otu07657 | 3 | k__Fungi(100);p__Ascomycota(100);c__Leotiomycetes(100);o__Helotiales(100);f__Dermateaceae(100);g__unclassified_Dermateaceae(100);s__uncultured_Dermateaceae(100); | - | - | - | - | - | - | - | Unassigned | - |
| Otu07661 | 3 | k__Fungi(100);p__Ascomycota(100);c__Leotiomycetes(100);o__Helotiales(100);f__Helotiales_family_Incertae_sedis(100);f__Helotiales_family_Incertae_sedis_unclassified(100);f__Helotiales_family_Incertae_sedis_unclassified(100); | - | - | - | - | - | - | - | Unassigned | - |
| Otu07663 | 3 | k__Fungi(100);k__Fungi_unclassified(67);k__Fungi_unclassified(67);k__Fungi_unclassified(67);k__Fungi_unclassified(67);k__Fungi_unclassified(67);k__Fungi_unclassified(67); | - | - | - | - | - | - | - | Unassigned | - |
| Otu07665 | 3 | k__Fungi(100);k__Fungi_unclassified(67);k__Fungi_unclassified(67);k__Fungi_unclassified(67);k__Fungi_unclassified(67);k__Fungi_unclassified(67);k__Fungi_unclassified(67); | - | - | - | - | - | - | - | Unassigned | - |
| Otu07668 | 3 | k__Fungi(100);k__Fungi_unclassified(67);k__Fungi_unclassified(67);k__Fungi_unclassified(67);k__Fungi_unclassified(67);k__Fungi_unclassified(67);k__Fungi_unclassified(67); | - | - | - | - | - | - | - | Unassigned | - |
| Otu07671 | 3 | k__Fungi(100);k__Fungi_unclassified(100);k__Fungi_unclassified(100);k__Fungi_unclassified(100);k__Fungi_unclassified(100);k__Fungi_unclassified(100);k__Fungi_unclassified(100); | - | - | - | - | - | - | - | Unassigned | - |
| Otu07677 | 3 | k__Fungi(100);k__Fungi_unclassified(67);k__Fungi_unclassified(67);k__Fungi_unclassified(67);k__Fungi_unclassified(67);k__Fungi_unclassified(67);k__Fungi_unclassified(67); | - | - | - | - | - | - | - | Unassigned | - |
| Otu07680 | 3 | k__Fungi(100);k__Fungi_unclassified(100);k__Fungi_unclassified(100);k__Fungi_unclassified(100);k__Fungi_unclassified(100);k__Fungi_unclassified(100);k__Fungi_unclassified(100); | - | - | - | - | - | - | - | Unassigned | - |
| Otu07681 | 3 | k__Fungi(100);p__Ascomycota(100);c__Leotiomycetes(100);o__Helotiales(100);f__Helotiales_family_Incertae_sedis(100);f__Helotiales_family_Incertae_sedis_unclassified(100);f__Helotiales_family_Incertae_sedis_unclassified(100); | - | - | - | - | - | - | - | Unassigned | - |
| Otu07682 | 3 | k__Fungi(100);k__Fungi_unclassified(100);k__Fungi_unclassified(100);k__Fungi_unclassified(100);k__Fungi_unclassified(100);k__Fungi_unclassified(100);k__Fungi_unclassified(100); | - | - | - | - | - | - | - | Unassigned | - |
| Otu07684 | 3 | k__Fungi(100);k__Fungi_unclassified(67);k__Fungi_unclassified(67);k__Fungi_unclassified(67);k__Fungi_unclassified(67);k__Fungi_unclassified(67);k__Fungi_unclassified(67); | - | - | - | - | - | - | - | Unassigned | - |
| Otu07685 | 3 | k__Fungi(100);p__Ascomycota(67);p__Ascomycota_unclassified(67);p__Ascomycota_unclassified(67);p__Ascomycota_unclassified(67);p__Ascomycota_unclassified(67);p__Ascomycota_unclassified(67); | - | - | - | - | - | - | - | Unassigned | - |
| Otu07692 | 3 | k__Fungi(100);p__Ascomycota(67);p__Ascomycota_unclassified(67);p__Ascomycota_unclassified(67);p__Ascomycota_unclassified(67);p__Ascomycota_unclassified(67);p__Ascomycota_unclassified(67); | - | - | - | - | - | - | - | Unassigned | - |
| Otu07694 | 3 | k__Fungi(100);p__Ascomycota(100);p__Ascomycota_unclassified(100);p__Ascomycota_unclassified(100);p__Ascomycota_unclassified(100);p__Ascomycota_unclassified(100);p__Ascomycota_unclassified(100); | - | - | - | - | - | - | - | Unassigned | - |
| Otu07697 | 3 | k__Fungi(100);k__Fungi_unclassified(67);k__Fungi_unclassified(67);k__Fungi_unclassified(67);k__Fungi_unclassified(67);k__Fungi_unclassified(67);k__Fungi_unclassified(67); | - | - | - | - | - | - | - | Unassigned | - |
| Otu07699 | 3 | k__Fungi(100);k__Fungi_unclassified(100);k__Fungi_unclassified(100);k__Fungi_unclassified(100);k__Fungi_unclassified(100);k__Fungi_unclassified(100);k__Fungi_unclassified(100); | - | - | - | - | - | - | - | Unassigned | - |
| Otu07700 | 3 | k__Fungi(100);k__Fungi_unclassified(67);k__Fungi_unclassified(67);k__Fungi_unclassified(67);k__Fungi_unclassified(67);k__Fungi_unclassified(67);k__Fungi_unclassified(67); | - | - | - | - | - | - | - | Unassigned | - |
| Otu07702 | 3 | k__Fungi(100);k__Fungi_unclassified(100);k__Fungi_unclassified(100);k__Fungi_unclassified(100);k__Fungi_unclassified(100);k__Fungi_unclassified(100);k__Fungi_unclassified(100); | - | - | - | - | - | - | - | Unassigned | - |
| Otu07703 | 3 | k__Fungi(100);k__Fungi_unclassified(67);k__Fungi_unclassified(67);k__Fungi_unclassified(67);k__Fungi_unclassified(67);k__Fungi_unclassified(67);k__Fungi_unclassified(67); | - | - | - | - | - | - | - | Unassigned | - |
| Otu07705 | 3 | k__Fungi(100);p__Ascomycota(100);p__Ascomycota_unclassified(100);p__Ascomycota_unclassified(100);p__Ascomycota_unclassified(100);p__Ascomycota_unclassified(100);p__Ascomycota_unclassified(100); | - | - | - | - | - | - | - | Unassigned | - |
| Otu07708 | 3 | k__Fungi(100);k__Fungi_unclassified(100);k__Fungi_unclassified(100);k__Fungi_unclassified(100);k__Fungi_unclassified(100);k__Fungi_unclassified(100);k__Fungi_unclassified(100); | - | - | - | - | - | - | - | Unassigned | - |
| Otu07710 | 3 | k__Fungi(100);p__Ascomycota(67);p__Ascomycota_unclassified(67);p__Ascomycota_unclassified(67);p__Ascomycota_unclassified(67);p__Ascomycota_unclassified(67);p__Ascomycota_unclassified(67); | - | - | - | - | - | - | - | Unassigned | - |
| Otu07712 | 3 | k__Fungi(100);k__Fungi_unclassified(67);k__Fungi_unclassified(67);k__Fungi_unclassified(67);k__Fungi_unclassified(67);k__Fungi_unclassified(67);k__Fungi_unclassified(67); | - | - | - | - | - | - | - | Unassigned | - |
| Otu07715 | 3 | k__Fungi(100);p__Ascomycota(100);p__Ascomycota_unclassified(100);p__Ascomycota_unclassified(100);p__Ascomycota_unclassified(100);p__Ascomycota_unclassified(100);p__Ascomycota_unclassified(100); | - | - | - | - | - | - | - | Unassigned | - |
| Otu07717 | 3 | k__Fungi(100);p__Ascomycota(100);p__Ascomycota_unclassified(100);p__Ascomycota_unclassified(100);p__Ascomycota_unclassified(100);p__Ascomycota_unclassified(100);p__Ascomycota_unclassified(100); | - | - | - | - | - | - | - | Unassigned | - |
| Otu07723 | 3 | k__Fungi(100);k__Fungi_unclassified(67);k__Fungi_unclassified(67);k__Fungi_unclassified(67);k__Fungi_unclassified(67);k__Fungi_unclassified(67);k__Fungi_unclassified(67); | - | - | - | - | - | - | - | Unassigned | - |
| Otu07724 | 3 | k__Fungi(100);k__Fungi_unclassified(67);k__Fungi_unclassified(67);k__Fungi_unclassified(67);k__Fungi_unclassified(67);k__Fungi_unclassified(67);k__Fungi_unclassified(67); | - | - | - | - | - | - | - | Unassigned | - |
| Otu07725 | 3 | k__Fungi(100);k__Fungi_unclassified(100);k__Fungi_unclassified(100);k__Fungi_unclassified(100);k__Fungi_unclassified(100);k__Fungi_unclassified(100);k__Fungi_unclassified(100); | - | - | - | - | - | - | - | Unassigned | - |
| Otu07728 | 3 | k__Fungi(100);k__Fungi_unclassified(100);k__Fungi_unclassified(100);k__Fungi_unclassified(100);k__Fungi_unclassified(100);k__Fungi_unclassified(100);k__Fungi_unclassified(100); | - | - | - | - | - | - | - | Unassigned | - |
| Otu07730 | 3 | k__Fungi(100);k__Fungi_unclassified(100);k__Fungi_unclassified(100);k__Fungi_unclassified(100);k__Fungi_unclassified(100);k__Fungi_unclassified(100);k__Fungi_unclassified(100); | - | - | - | - | - | - | - | Unassigned | - |
| Otu07731 | 3 | k__Fungi(100);p__Ascomycota(67);p__Ascomycota_unclassified(67);p__Ascomycota_unclassified(67);p__Ascomycota_unclassified(67);p__Ascomycota_unclassified(67);p__Ascomycota_unclassified(67); | - | - | - | - | - | - | - | Unassigned | - |
| Otu07735 | 3 | k__Fungi(100);p__Ascomycota(67);p__Ascomycota_unclassified(67);p__Ascomycota_unclassified(67);p__Ascomycota_unclassified(67);p__Ascomycota_unclassified(67);p__Ascomycota_unclassified(67); | - | - | - | - | - | - | - | Unassigned | - |
| Otu07737 | 3 | k__Fungi(100);p__Ascomycota(100);p__Ascomycota_unclassified(100);p__Ascomycota_unclassified(100);p__Ascomycota_unclassified(100);p__Ascomycota_unclassified(100);p__Ascomycota_unclassified(100); | - | - | - | - | - | - | - | Unassigned | - |
| Otu07742 | 3 | k__Fungi(100);p__Ascomycota(100);p__Ascomycota_unclassified(100);p__Ascomycota_unclassified(100);p__Ascomycota_unclassified(100);p__Ascomycota_unclassified(100);p__Ascomycota_unclassified(100); | - | - | - | - | - | - | - | Unassigned | - |
| Otu07744 | 3 | k__Fungi(100);p__Ascomycota(100);c__Leotiomycetes(67);c__Leotiomycetes_unclassified(67);c__Leotiomycetes_unclassified(67);c__Leotiomycetes_unclassified(67);c__Leotiomycetes_unclassified(67); | - | - | - | - | - | - | - | Unassigned | - |
| Otu07745 | 3 | k__Fungi(100);k__Fungi_unclassified(100);k__Fungi_unclassified(100);k__Fungi_unclassified(100);k__Fungi_unclassified(100);k__Fungi_unclassified(100);k__Fungi_unclassified(100); | - | - | - | - | - | - | - | Unassigned | - |
| Otu07747 | 3 | k__Fungi(100);k__Fungi_unclassified(67);k__Fungi_unclassified(67);k__Fungi_unclassified(67);k__Fungi_unclassified(67);k__Fungi_unclassified(67);k__Fungi_unclassified(67); | - | - | - | - | - | - | - | Unassigned | - |
| Otu07756 | 3 | k__Fungi(100);p__Ascomycota(100);p__Ascomycota_unclassified(100);p__Ascomycota_unclassified(100);p__Ascomycota_unclassified(100);p__Ascomycota_unclassified(100);p__Ascomycota_unclassified(100); | - | - | - | - | - | - | - | Unassigned | - |
| Otu07760 | 3 | k__Fungi(100);p__Ascomycota(67);p__Ascomycota_unclassified(67);p__Ascomycota_unclassified(67);p__Ascomycota_unclassified(67);p__Ascomycota_unclassified(67);p__Ascomycota_unclassified(67); | - | - | - | - | - | - | - | Unassigned | - |
| Otu07761 | 3 | k__Fungi(100);k__Fungi_unclassified(100);k__Fungi_unclassified(100);k__Fungi_unclassified(100);k__Fungi_unclassified(100);k__Fungi_unclassified(100);k__Fungi_unclassified(100); | - | - | - | - | - | - | - | Unassigned | - |
| Otu07763 | 3 | k__Fungi(100);k__Fungi_unclassified(67);k__Fungi_unclassified(67);k__Fungi_unclassified(67);k__Fungi_unclassified(67);k__Fungi_unclassified(67);k__Fungi_unclassified(67); | - | - | - | - | - | - | - | Unassigned | - |
| Otu07764 | 3 | k__Fungi(100);p__Ascomycota(100);p__Ascomycota_unclassified(100);p__Ascomycota_unclassified(100);p__Ascomycota_unclassified(100);p__Ascomycota_unclassified(100);p__Ascomycota_unclassified(100); | - | - | - | - | - | - | - | Unassigned | - |
| Otu07765 | 3 | k__Fungi(100);k__Fungi_unclassified(67);k__Fungi_unclassified(67);k__Fungi_unclassified(67);k__Fungi_unclassified(67);k__Fungi_unclassified(67);k__Fungi_unclassified(67); | - | - | - | - | - | - | - | Unassigned | - |
| Otu07766 | 3 | k__Fungi(100);p__Ascomycota(67);p__Ascomycota_unclassified(67);p__Ascomycota_unclassified(67);p__Ascomycota_unclassified(67);p__Ascomycota_unclassified(67);p__Ascomycota_unclassified(67); | - | - | - | - | - | - | - | Unassigned | - |
| Otu07771 | 3 | k__Fungi(100);k__Fungi_unclassified(100);k__Fungi_unclassified(100);k__Fungi_unclassified(100);k__Fungi_unclassified(100);k__Fungi_unclassified(100);k__Fungi_unclassified(100); | - | - | - | - | - | - | - | Unassigned | - |
| Otu07774 | 3 | k__Fungi(100);p__Ascomycota(100);c__Sordariomycetes(100);o__Sordariales(100);o__Sordariales_unclassified(100);o__Sordariales_unclassified(100);o__Sordariales_unclassified(100); | - | - | - | - | - | - | - | Unassigned | - |
| Otu07775 | 3 | k__Fungi(100);p__Ascomycota(67);p__Ascomycota_unclassified(67);p__Ascomycota_unclassified(67);p__Ascomycota_unclassified(67);p__Ascomycota_unclassified(67);p__Ascomycota_unclassified(67); | - | - | - | - | - | - | - | Unassigned | - |
| Otu07777 | 3 | k__Fungi(100);k__Fungi_unclassified(100);k__Fungi_unclassified(100);k__Fungi_unclassified(100);k__Fungi_unclassified(100);k__Fungi_unclassified(100);k__Fungi_unclassified(100); | - | - | - | - | - | - | - | Unassigned | - |
| Otu07779 | 3 | k__Fungi(100);p__Ascomycota(67);p__Ascomycota_unclassified(67);p__Ascomycota_unclassified(67);p__Ascomycota_unclassified(67);p__Ascomycota_unclassified(67);p__Ascomycota_unclassified(67); | - | - | - | - | - | - | - | Unassigned | - |
| Otu07781 | 3 | k__Fungi(100);p__Ascomycota(67);p__Ascomycota_unclassified(67);p__Ascomycota_unclassified(67);p__Ascomycota_unclassified(67);p__Ascomycota_unclassified(67);p__Ascomycota_unclassified(67); | - | - | - | - | - | - | - | Unassigned | - |
| Otu07782 | 3 | k__Fungi(100);p__Ascomycota(67);p__Ascomycota_unclassified(67);p__Ascomycota_unclassified(67);p__Ascomycota_unclassified(67);p__Ascomycota_unclassified(67);p__Ascomycota_unclassified(67); | - | - | - | - | - | - | - | Unassigned | - |
| Otu07783 | 3 | k__Fungi(100);k__Fungi_unclassified(100);k__Fungi_unclassified(100);k__Fungi_unclassified(100);k__Fungi_unclassified(100);k__Fungi_unclassified(100);k__Fungi_unclassified(100); | - | - | - | - | - | - | - | Unassigned | - |
| Otu07784 | 3 | k__Fungi(100);k__Fungi_unclassified(67);k__Fungi_unclassified(67);k__Fungi_unclassified(67);k__Fungi_unclassified(67);k__Fungi_unclassified(67);k__Fungi_unclassified(67); | - | - | - | - | - | - | - | Unassigned | - |
| Otu07785 | 3 | k__Fungi(100);k__Fungi_unclassified(100);k__Fungi_unclassified(100);k__Fungi_unclassified(100);k__Fungi_unclassified(100);k__Fungi_unclassified(100);k__Fungi_unclassified(100); | - | - | - | - | - | - | - | Unassigned | - |
| Otu07787 | 3 | k__Fungi(100);k__Fungi_unclassified(100);k__Fungi_unclassified(100);k__Fungi_unclassified(100);k__Fungi_unclassified(100);k__Fungi_unclassified(100);k__Fungi_unclassified(100); | - | - | - | - | - | - | - | Unassigned | - |
| Otu07788 | 3 | k__Fungi(100);p__Ascomycota(67);p__Ascomycota_unclassified(67);p__Ascomycota_unclassified(67);p__Ascomycota_unclassified(67);p__Ascomycota_unclassified(67);p__Ascomycota_unclassified(67); | - | - | - | - | - | - | - | Unassigned | - |
| Otu07792 | 3 | k__Fungi(100);p__Ascomycota(100);c__Dothideomycetes(100);o__Pleosporales(100);o__Pleosporales_unclassified(100);o__Pleosporales_unclassified(100);o__Pleosporales_unclassified(100); | - | - | - | - | - | - | - | Unassigned | - |
| Otu07793 | 3 | k__Fungi(100);k__Fungi_unclassified(67);k__Fungi_unclassified(67);k__Fungi_unclassified(67);k__Fungi_unclassified(67);k__Fungi_unclassified(67);k__Fungi_unclassified(67); | - | - | - | - | - | - | - | Unassigned | - |
| Otu07795 | 3 | k__Fungi(100);p__Ascomycota(100);p__Ascomycota_unclassified(100);p__Ascomycota_unclassified(100);p__Ascomycota_unclassified(100);p__Ascomycota_unclassified(100);p__Ascomycota_unclassified(100); | - | - | - | - | - | - | - | Unassigned | - |
| Otu07796 | 3 | k__Fungi(100);k__Fungi_unclassified(100);k__Fungi_unclassified(100);k__Fungi_unclassified(100);k__Fungi_unclassified(100);k__Fungi_unclassified(100);k__Fungi_unclassified(100); | - | - | - | - | - | - | - | Unassigned | - |
| Otu07797 | 3 | k__Fungi(100);p__Ascomycota(100);p__Ascomycota_unclassified(100);p__Ascomycota_unclassified(100);p__Ascomycota_unclassified(100);p__Ascomycota_unclassified(100);p__Ascomycota_unclassified(100); | - | - | - | - | - | - | - | Unassigned | - |
| Otu07798 | 3 | k__Plantae(100);p__unclassified_Plantae(100);c__unclassified_Plantae(100);o__unclassified_Plantae(100);f__unclassified_Plantae(100);g__unclassified_Plantae(100);s__Plantae_sp(100); | - | - | - | - | - | - | - | Unassigned | - |
| Otu07799 | 3 | k__Fungi(100);p__Ascomycota(67);p__Ascomycota_unclassified(67);p__Ascomycota_unclassified(67);p__Ascomycota_unclassified(67);p__Ascomycota_unclassified(67);p__Ascomycota_unclassified(67); | - | - | - | - | - | - | - | Unassigned | - |
| Otu07800 | 3 | k__Fungi(100);p__Ascomycota(100);p__Ascomycota_unclassified(100);p__Ascomycota_unclassified(100);p__Ascomycota_unclassified(100);p__Ascomycota_unclassified(100);p__Ascomycota_unclassified(100); | - | - | - | - | - | - | - | Unassigned | - |
| Otu07802 | 3 | k__Fungi(100);p__Ascomycota(100);c__Leotiomycetes(100);o__Helotiales(100);o__Helotiales_unclassified(100);o__Helotiales_unclassified(100);o__Helotiales_unclassified(100); | - | - | - | - | - | - | - | Unassigned | - |
| Otu07803 | 3 | k__Fungi(100);k__Fungi_unclassified(67);k__Fungi_unclassified(67);k__Fungi_unclassified(67);k__Fungi_unclassified(67);k__Fungi_unclassified(67);k__Fungi_unclassified(67); | - | - | - | - | - | - | - | Unassigned | - |
| Otu07811 | 3 | k__Fungi(100);p__Ascomycota(67);p__Ascomycota_unclassified(67);p__Ascomycota_unclassified(67);p__Ascomycota_unclassified(67);p__Ascomycota_unclassified(67);p__Ascomycota_unclassified(67); | - | - | - | - | - | - | - | Unassigned | - |
| Otu07812 | 3 | k__Fungi(100);p__Ascomycota(100);p__Ascomycota_unclassified(100);p__Ascomycota_unclassified(100);p__Ascomycota_unclassified(100);p__Ascomycota_unclassified(100);p__Ascomycota_unclassified(100); | - | - | - | - | - | - | - | Unassigned | - |
| Otu07813 | 3 | k__Fungi(100);p__Ascomycota(100);p__Ascomycota_unclassified(100);p__Ascomycota_unclassified(100);p__Ascomycota_unclassified(100);p__Ascomycota_unclassified(100);p__Ascomycota_unclassified(100); | - | - | - | - | - | - | - | Unassigned | - |
| Otu07814 | 3 | k__Fungi(100);k__Fungi_unclassified(100);k__Fungi_unclassified(100);k__Fungi_unclassified(100);k__Fungi_unclassified(100);k__Fungi_unclassified(100);k__Fungi_unclassified(100); | - | - | - | - | - | - | - | Unassigned | - |
| Otu07815 | 3 | k__Fungi(100);k__Fungi_unclassified(100);k__Fungi_unclassified(100);k__Fungi_unclassified(100);k__Fungi_unclassified(100);k__Fungi_unclassified(100);k__Fungi_unclassified(100); | - | - | - | - | - | - | - | Unassigned | - |
| Otu07817 | 3 | k__Fungi(100);p__Ascomycota(100);p__Ascomycota_unclassified(67);p__Ascomycota_unclassified(67);p__Ascomycota_unclassified(67);p__Ascomycota_unclassified(67);p__Ascomycota_unclassified(67); | - | - | - | - | - | - | - | Unassigned | - |
| Otu07818 | 3 | k__Fungi(100);p__Ascomycota(100);c__Sordariomycetes(67);c__Sordariomycetes_unclassified(67);c__Sordariomycetes_unclassified(67);c__Sordariomycetes_unclassified(67);c__Sordariomycetes_unclassified(67); | - | - | - | - | - | - | - | Unassigned | - |
| Otu07819 | 3 | k__Fungi(100);p__Ascomycota(100);p__Ascomycota_unclassified(100);p__Ascomycota_unclassified(100);p__Ascomycota_unclassified(100);p__Ascomycota_unclassified(100);p__Ascomycota_unclassified(100); | - | - | - | - | - | - | - | Unassigned | - |
| Otu07820 | 3 | k__Fungi(100);k__Fungi_unclassified(100);k__Fungi_unclassified(100);k__Fungi_unclassified(100);k__Fungi_unclassified(100);k__Fungi_unclassified(100);k__Fungi_unclassified(100); | - | - | - | - | - | - | - | Unassigned | - |
| Otu07821 | 3 | k__Fungi(100);k__Fungi_unclassified(100);k__Fungi_unclassified(100);k__Fungi_unclassified(100);k__Fungi_unclassified(100);k__Fungi_unclassified(100);k__Fungi_unclassified(100); | - | - | - | - | - | - | - | Unassigned | - |
| Otu07822 | 3 | k__Fungi(100);p__Ascomycota(100);p__Ascomycota_unclassified(100);p__Ascomycota_unclassified(100);p__Ascomycota_unclassified(100);p__Ascomycota_unclassified(100);p__Ascomycota_unclassified(100); | - | - | - | - | - | - | - | Unassigned | - |
| Otu07823 | 3 | k__Fungi(100);p__Ascomycota(100);p__Ascomycota_unclassified(100);p__Ascomycota_unclassified(100);p__Ascomycota_unclassified(100);p__Ascomycota_unclassified(100);p__Ascomycota_unclassified(100); | - | - | - | - | - | - | - | Unassigned | - |
| Otu07825 | 3 | k__Plantae(100);p__unclassified_Plantae(100);c__unclassified_Plantae(100);o__unclassified_Plantae(100);f__unclassified_Plantae(100);g__unclassified_Plantae(100);s__Plantae_sp(100); | - | - | - | - | - | - | - | Unassigned | - |
| Otu07826 | 3 | k__Fungi(100);k__Fungi_unclassified(100);k__Fungi_unclassified(100);k__Fungi_unclassified(100);k__Fungi_unclassified(100);k__Fungi_unclassified(100);k__Fungi_unclassified(100); | - | - | - | - | - | - | - | Unassigned | - |
| Otu07827 | 3 | k__Fungi(100);p__Ascomycota(100);p__Ascomycota_unclassified(100);p__Ascomycota_unclassified(100);p__Ascomycota_unclassified(100);p__Ascomycota_unclassified(100);p__Ascomycota_unclassified(100); | - | - | - | - | - | - | - | Unassigned | - |
| Otu07828 | 3 | k__Fungi(100);k__Fungi_unclassified(100);k__Fungi_unclassified(100);k__Fungi_unclassified(100);k__Fungi_unclassified(100);k__Fungi_unclassified(100);k__Fungi_unclassified(100); | - | - | - | - | - | - | - | Unassigned | - |
| Otu07830 | 3 | k__Plantae(100);p__unclassified_Plantae(100);c__unclassified_Plantae(100);o__unclassified_Plantae(100);f__unclassified_Plantae(100);g__unclassified_Plantae(100);s__Plantae_sp(100); | - | - | - | - | - | - | - | Unassigned | - |
| Otu07832 | 3 | k__Fungi(100);k__Fungi_unclassified(67);k__Fungi_unclassified(67);k__Fungi_unclassified(67);k__Fungi_unclassified(67);k__Fungi_unclassified(67);k__Fungi_unclassified(67); | - | - | - | - | - | - | - | Unassigned | - |
| Otu07833 | 3 | k__Fungi(100);p__Ascomycota(100);p__Ascomycota_unclassified(100);p__Ascomycota_unclassified(100);p__Ascomycota_unclassified(100);p__Ascomycota_unclassified(100);p__Ascomycota_unclassified(100); | - | - | - | - | - | - | - | Unassigned | - |
| Otu07834 | 3 | k__Fungi(100);p__Ascomycota(100);p__Ascomycota_unclassified(100);p__Ascomycota_unclassified(100);p__Ascomycota_unclassified(100);p__Ascomycota_unclassified(100);p__Ascomycota_unclassified(100); | - | - | - | - | - | - | - | Unassigned | - |
| Otu07837 | 3 | k__Fungi(100);p__Ascomycota(100);c__Dothideomycetes(100);o__Myriangiales(100);o__Myriangiales_unclassified(100);o__Myriangiales_unclassified(100);o__Myriangiales_unclassified(100); | - | - | - | - | - | - | - | Unassigned | - |
| Otu07839 | 3 | k__Fungi(100);p__Ascomycota(100);p__Ascomycota_unclassified(100);p__Ascomycota_unclassified(100);p__Ascomycota_unclassified(100);p__Ascomycota_unclassified(100);p__Ascomycota_unclassified(100); | - | - | - | - | - | - | - | Unassigned | - |
| Otu07840 | 3 | k__Fungi(100);p__Ascomycota(67);p__Ascomycota_unclassified(67);p__Ascomycota_unclassified(67);p__Ascomycota_unclassified(67);p__Ascomycota_unclassified(67);p__Ascomycota_unclassified(67); | - | - | - | - | - | - | - | Unassigned | - |
| Otu07842 | 3 | k__Fungi(100);k__Fungi_unclassified(67);k__Fungi_unclassified(67);k__Fungi_unclassified(67);k__Fungi_unclassified(67);k__Fungi_unclassified(67);k__Fungi_unclassified(67); | - | - | - | - | - | - | - | Unassigned | - |
| Otu07843 | 3 | k__Fungi(100);p__Ascomycota(100);c__Dothideomycetes(100);o__Pleosporales(100);o__Pleosporales_unclassified(100);o__Pleosporales_unclassified(100);o__Pleosporales_unclassified(100); | - | - | - | - | - | - | - | Unassigned | - |
| Otu07845 | 3 | k__Plantae(67);p__unclassified_Plantae(67);c__unclassified_Plantae(67);o__unclassified_Plantae(67);f__unclassified_Plantae(67);g__unclassified_Plantae(67);s__Plantae_sp(67); | - | - | - | - | - | - | - | Unassigned | - |
| Otu07847 | 3 | k__Plantae(100);p__unclassified_Plantae(100);c__unclassified_Plantae(100);o__unclassified_Plantae(100);f__unclassified_Plantae(100);g__unclassified_Plantae(100);s__Plantae_sp(100); | - | - | - | - | - | - | - | Unassigned | - |
| Otu07848 | 3 | k__Fungi(100);p__Ascomycota(100);c__Sordariomycetes(100);o__Xylariales(100);o__Xylariales_unclassified(100);o__Xylariales_unclassified(100);o__Xylariales_unclassified(100); | - | - | - | - | - | - | - | Unassigned | - |
| Otu07849 | 3 | k__Fungi(100);p__Ascomycota(100);c__Sordariomycetes(100);o__Xylariales(100);o__Xylariales_unclassified(100);o__Xylariales_unclassified(100);o__Xylariales_unclassified(100); | - | - | - | - | - | - | - | Unassigned | - |
| Otu07850 | 3 | k__Plantae(100);p__unclassified_Plantae(100);c__unclassified_Plantae(100);o__unclassified_Plantae(100);f__unclassified_Plantae(100);g__unclassified_Plantae(100);s__Plantae_sp(100); | - | - | - | - | - | - | - | Unassigned | - |
| Otu07851 | 3 | k__Fungi(100);p__Ascomycota(100);p__Ascomycota_unclassified(100);p__Ascomycota_unclassified(100);p__Ascomycota_unclassified(100);p__Ascomycota_unclassified(100);p__Ascomycota_unclassified(100); | - | - | - | - | - | - | - | Unassigned | - |
| Otu07853 | 3 | k__Fungi(100);k__Fungi_unclassified(100);k__Fungi_unclassified(100);k__Fungi_unclassified(100);k__Fungi_unclassified(100);k__Fungi_unclassified(100);k__Fungi_unclassified(100); | - | - | - | - | - | - | - | Unassigned | - |
| Otu07854 | 3 | k__Fungi(100);k__Fungi_unclassified(67);k__Fungi_unclassified(67);k__Fungi_unclassified(67);k__Fungi_unclassified(67);k__Fungi_unclassified(67);k__Fungi_unclassified(67); | - | - | - | - | - | - | - | Unassigned | - |
| Otu07855 | 3 | k__Fungi(100);p__Ascomycota(100);p__Ascomycota_unclassified(100);p__Ascomycota_unclassified(100);p__Ascomycota_unclassified(100);p__Ascomycota_unclassified(100);p__Ascomycota_unclassified(100); | - | - | - | - | - | - | - | Unassigned | - |
| Otu07856 | 3 | k__Fungi(100);k__Fungi_unclassified(67);k__Fungi_unclassified(67);k__Fungi_unclassified(67);k__Fungi_unclassified(67);k__Fungi_unclassified(67);k__Fungi_unclassified(67); | - | - | - | - | - | - | - | Unassigned | - |
| Otu07858 | 3 | k__Fungi(100);p__Ascomycota(67);p__Ascomycota_unclassified(67);p__Ascomycota_unclassified(67);p__Ascomycota_unclassified(67);p__Ascomycota_unclassified(67);p__Ascomycota_unclassified(67); | - | - | - | - | - | - | - | Unassigned | - |
| Otu07861 | 3 | k__Fungi(100);k__Fungi_unclassified(67);k__Fungi_unclassified(67);k__Fungi_unclassified(67);k__Fungi_unclassified(67);k__Fungi_unclassified(67);k__Fungi_unclassified(67); | - | - | - | - | - | - | - | Unassigned | - |
| Otu07865 | 3 | k__Fungi(100);k__Fungi_unclassified(67);k__Fungi_unclassified(67);k__Fungi_unclassified(67);k__Fungi_unclassified(67);k__Fungi_unclassified(67);k__Fungi_unclassified(67); | - | - | - | - | - | - | - | Unassigned | - |
| Otu07866 | 3 | k__Fungi(100);k__Fungi_unclassified(100);k__Fungi_unclassified(100);k__Fungi_unclassified(100);k__Fungi_unclassified(100);k__Fungi_unclassified(100);k__Fungi_unclassified(100); | - | - | - | - | - | - | - | Unassigned | - |
| Otu07868 | 3 | k__Fungi(100);p__Ascomycota(67);p__Ascomycota_unclassified(67);p__Ascomycota_unclassified(67);p__Ascomycota_unclassified(67);p__Ascomycota_unclassified(67);p__Ascomycota_unclassified(67); | - | - | - | - | - | - | - | Unassigned | - |
| Otu07873 | 3 | k__Fungi(100);k__Fungi_unclassified(67);k__Fungi_unclassified(67);k__Fungi_unclassified(67);k__Fungi_unclassified(67);k__Fungi_unclassified(67);k__Fungi_unclassified(67); | - | - | - | - | - | - | - | Unassigned | - |
| Otu07874 | 3 | k__Fungi(100);p__Ascomycota(100);c__Leotiomycetes(100);o__Helotiales(100);o__Helotiales_unclassified(100);o__Helotiales_unclassified(100);o__Helotiales_unclassified(100); | - | - | - | - | - | - | - | Unassigned | - |
| Otu07875 | 3 | k__Fungi(100);p__Ascomycota(67);p__Ascomycota_unclassified(67);p__Ascomycota_unclassified(67);p__Ascomycota_unclassified(67);p__Ascomycota_unclassified(67);p__Ascomycota_unclassified(67); | - | - | - | - | - | - | - | Unassigned | - |
| Otu07878 | 3 | k__Fungi(100);p__Ascomycota(67);p__Ascomycota_unclassified(67);p__Ascomycota_unclassified(67);p__Ascomycota_unclassified(67);p__Ascomycota_unclassified(67);p__Ascomycota_unclassified(67); | - | - | - | - | - | - | - | Unassigned | - |
| Otu07879 | 3 | k__Fungi(100);p__Ascomycota(67);p__Ascomycota_unclassified(67);p__Ascomycota_unclassified(67);p__Ascomycota_unclassified(67);p__Ascomycota_unclassified(67);p__Ascomycota_unclassified(67); | - | - | - | - | - | - | - | Unassigned | - |
| Otu07881 | 3 | k__Fungi(100);k__Fungi_unclassified(67);k__Fungi_unclassified(67);k__Fungi_unclassified(67);k__Fungi_unclassified(67);k__Fungi_unclassified(67);k__Fungi_unclassified(67); | - | - | - | - | - | - | - | Unassigned | - |
| Otu07883 | 3 | k__Fungi(100);k__Fungi_unclassified(67);k__Fungi_unclassified(67);k__Fungi_unclassified(67);k__Fungi_unclassified(67);k__Fungi_unclassified(67);k__Fungi_unclassified(67); | - | - | - | - | - | - | - | Unassigned | - |
| Otu07885 | 3 | k__Fungi(100);k__Fungi_unclassified(100);k__Fungi_unclassified(100);k__Fungi_unclassified(100);k__Fungi_unclassified(100);k__Fungi_unclassified(100);k__Fungi_unclassified(100); | - | - | - | - | - | - | - | Unassigned | - |
| Otu07888 | 3 | k__Fungi(100);k__Fungi_unclassified(67);k__Fungi_unclassified(67);k__Fungi_unclassified(67);k__Fungi_unclassified(67);k__Fungi_unclassified(67);k__Fungi_unclassified(67); | - | - | - | - | - | - | - | Unassigned | - |
| Otu07890 | 3 | k__Fungi(100);k__Fungi_unclassified(100);k__Fungi_unclassified(100);k__Fungi_unclassified(100);k__Fungi_unclassified(100);k__Fungi_unclassified(100);k__Fungi_unclassified(100); | - | - | - | - | - | - | - | Unassigned | - |
| Otu07892 | 3 | k__Fungi(100);k__Fungi_unclassified(67);k__Fungi_unclassified(67);k__Fungi_unclassified(67);k__Fungi_unclassified(67);k__Fungi_unclassified(67);k__Fungi_unclassified(67); | - | - | - | - | - | - | - | Unassigned | - |
| Otu07893 | 3 | k__Fungi(100);k__Fungi_unclassified(67);k__Fungi_unclassified(67);k__Fungi_unclassified(67);k__Fungi_unclassified(67);k__Fungi_unclassified(67);k__Fungi_unclassified(67); | - | - | - | - | - | - | - | Unassigned | - |
| Otu07894 | 3 | k__Fungi(100);k__Fungi_unclassified(67);k__Fungi_unclassified(67);k__Fungi_unclassified(67);k__Fungi_unclassified(67);k__Fungi_unclassified(67);k__Fungi_unclassified(67); | - | - | - | - | - | - | - | Unassigned | - |
| Otu07895 | 3 | k__Fungi(100);p__Ascomycota(100);p__Ascomycota_unclassified(100);p__Ascomycota_unclassified(100);p__Ascomycota_unclassified(100);p__Ascomycota_unclassified(100);p__Ascomycota_unclassified(100); | - | - | - | - | - | - | - | Unassigned | - |
| Otu07898 | 3 | k__Fungi(100);k__Fungi_unclassified(100);k__Fungi_unclassified(100);k__Fungi_unclassified(100);k__Fungi_unclassified(100);k__Fungi_unclassified(100);k__Fungi_unclassified(100); | - | - | - | - | - | - | - | Unassigned | - |
| Otu07903 | 3 | k__Fungi(100);p__Ascomycota(100);p__Ascomycota_unclassified(100);p__Ascomycota_unclassified(100);p__Ascomycota_unclassified(100);p__Ascomycota_unclassified(100);p__Ascomycota_unclassified(100); | - | - | - | - | - | - | - | Unassigned | - |
| Otu07904 | 3 | k__Fungi(100);p__Basidiomycota(100);p__Basidiomycota_unclassified(100);p__Basidiomycota_unclassified(100);p__Basidiomycota_unclassified(100);p__Basidiomycota_unclassified(100);p__Basidiomycota_unclassified(100); | - | - | - | - | - | - | - | Unassigned | - |
| Otu07905 | 3 | k__Fungi(100);k__Fungi_unclassified(67);k__Fungi_unclassified(67);k__Fungi_unclassified(67);k__Fungi_unclassified(67);k__Fungi_unclassified(67);k__Fungi_unclassified(67); | - | - | - | - | - | - | - | Unassigned | - |
| Otu07907 | 3 | k__Fungi(100);p__Ascomycota(67);p__Ascomycota_unclassified(67);p__Ascomycota_unclassified(67);p__Ascomycota_unclassified(67);p__Ascomycota_unclassified(67);p__Ascomycota_unclassified(67); | - | - | - | - | - | - | - | Unassigned | - |
| Otu07908 | 3 | k__Fungi(100);p__Ascomycota(67);p__Ascomycota_unclassified(67);p__Ascomycota_unclassified(67);p__Ascomycota_unclassified(67);p__Ascomycota_unclassified(67);p__Ascomycota_unclassified(67); | - | - | - | - | - | - | - | Unassigned | - |
| Otu07911 | 3 | k__Fungi(100);k__Fungi_unclassified(100);k__Fungi_unclassified(100);k__Fungi_unclassified(100);k__Fungi_unclassified(100);k__Fungi_unclassified(100);k__Fungi_unclassified(100); | - | - | - | - | - | - | - | Unassigned | - |
| Otu07914 | 3 | k__Fungi(100);k__Fungi_unclassified(100);k__Fungi_unclassified(100);k__Fungi_unclassified(100);k__Fungi_unclassified(100);k__Fungi_unclassified(100);k__Fungi_unclassified(100); | - | - | - | - | - | - | - | Unassigned | - |
| Otu07916 | 3 | k__Fungi(100);k__Fungi_unclassified(67);k__Fungi_unclassified(67);k__Fungi_unclassified(67);k__Fungi_unclassified(67);k__Fungi_unclassified(67);k__Fungi_unclassified(67); | - | - | - | - | - | - | - | Unassigned | - |
| Otu07918 | 3 | k__Fungi(100);p__Ascomycota(67);p__Ascomycota_unclassified(67);p__Ascomycota_unclassified(67);p__Ascomycota_unclassified(67);p__Ascomycota_unclassified(67);p__Ascomycota_unclassified(67); | - | - | - | - | - | - | - | Unassigned | - |
| Otu07919 | 3 | k__Fungi(100);k__Fungi_unclassified(67);k__Fungi_unclassified(67);k__Fungi_unclassified(67);k__Fungi_unclassified(67);k__Fungi_unclassified(67);k__Fungi_unclassified(67); | - | - | - | - | - | - | - | Unassigned | - |
| Otu07920 | 3 | k__Fungi(100);p__Ascomycota(67);p__Ascomycota_unclassified(67);p__Ascomycota_unclassified(67);p__Ascomycota_unclassified(67);p__Ascomycota_unclassified(67);p__Ascomycota_unclassified(67); | - | - | - | - | - | - | - | Unassigned | - |

| | | | | | | | | | | | |
|---|---|---|---|---|---|---|---|---|---|---|---|
| Otu07921 | 3 | k__Fungi(100);p__Ascomycota(100);c__Dothideomycetes(100);o__Capnodiales(100);o__Capnodiales_unclassified(100);o__Capnodiales_unclassified(100);o__Capnodiales_unclassified(100); | - | - | - | - | - | - | - | Unassigned | - |
| Otu07922 | 3 | k__Fungi(100);p__Ascomycota(67);p__Ascomycota_unclassified(67);p__Ascomycota_unclassified(67);p__Ascomycota_unclassified(67);p__Ascomycota_unclassified(67);p__Ascomycota_unclassified(67); | - | - | - | - | - | - | - | Unassigned | - |
| Otu07923 | 3 | k__Fungi(100);k__Fungi_unclassified(67);k__Fungi_unclassified(67);k__Fungi_unclassified(67);k__Fungi_unclassified(67);k__Fungi_unclassified(67);k__Fungi_unclassified(67); | - | - | - | - | - | - | - | Unassigned | - |
| Otu07924 | 3 | k__Fungi(100);k__Fungi_unclassified(67);k__Fungi_unclassified(67);k__Fungi_unclassified(67);k__Fungi_unclassified(67);k__Fungi_unclassified(67);k__Fungi_unclassified(67); | - | - | - | - | - | - | - | Unassigned | - |
| Otu07925 | 3 | k__Fungi(100);k__Fungi_unclassified(67);k__Fungi_unclassified(67);k__Fungi_unclassified(67);k__Fungi_unclassified(67);k__Fungi_unclassified(67);k__Fungi_unclassified(67); | - | - | - | - | - | - | - | Unassigned | - |
| Otu07926 | 3 | k__Fungi(100);p__Ascomycota(100);p__Ascomycota_unclassified(100);p__Ascomycota_unclassified(100);p__Ascomycota_unclassified(100);p__Ascomycota_unclassified(100);p__Ascomycota_unclassified(100); | - | - | - | - | - | - | - | Unassigned | - |
| Otu07928 | 3 | k__Fungi(100);k__Fungi_unclassified(100);k__Fungi_unclassified(100);k__Fungi_unclassified(100);k__Fungi_unclassified(100);k__Fungi_unclassified(100);k__Fungi_unclassified(100); | - | - | - | - | - | - | - | Unassigned | - |
| Otu07929 | 3 | k__Fungi(100);p__Ascomycota(100);c__Leotiomycetes(100);o__Helotiales(67);o__Helotiales_unclassified(67);o__Helotiales_unclassified(67);o__Helotiales_unclassified(67); | - | - | - | - | - | - | - | Unassigned | - |
| Otu07931 | 3 | k__Fungi(100);p__Basidiomycota(100);p__Basidiomycota_unclassified(100);p__Basidiomycota_unclassified(100);p__Basidiomycota_unclassified(100);p__Basidiomycota_unclassified(100);p__Basidiomycota_unclassified(100); | - | - | - | - | - | - | - | Unassigned | - |
| Otu07935 | 3 | k__Fungi(100);k__Fungi_unclassified(100);k__Fungi_unclassified(100);k__Fungi_unclassified(100);k__Fungi_unclassified(100);k__Fungi_unclassified(100);k__Fungi_unclassified(100); | - | - | - | - | - | - | - | Unassigned | - |
| Otu07937 | 3 | k__Fungi(100);k__Fungi_unclassified(67);k__Fungi_unclassified(67);k__Fungi_unclassified(67);k__Fungi_unclassified(67);k__Fungi_unclassified(67);k__Fungi_unclassified(67); | - | - | - | - | - | - | - | Unassigned | - |
| Otu07939 | 3 | k__Fungi(100);k__Fungi_unclassified(67);k__Fungi_unclassified(67);k__Fungi_unclassified(67);k__Fungi_unclassified(67);k__Fungi_unclassified(67);k__Fungi_unclassified(67); | - | - | - | - | - | - | - | Unassigned | - |
| Otu07941 | 3 | k__Fungi(100);k__Fungi_unclassified(67);k__Fungi_unclassified(67);k__Fungi_unclassified(67);k__Fungi_unclassified(67);k__Fungi_unclassified(67);k__Fungi_unclassified(67); | - | - | - | - | - | - | - | Unassigned | - |
| Otu07942 | 3 | k__Fungi(100);p__Ascomycota(67);p__Ascomycota_unclassified(67);p__Ascomycota_unclassified(67);p__Ascomycota_unclassified(67);p__Ascomycota_unclassified(67);p__Ascomycota_unclassified(67); | - | - | - | - | - | - | - | Unassigned | - |
| Otu07945 | 3 | k__Fungi(100);p__Ascomycota(67);p__Ascomycota_unclassified(67);p__Ascomycota_unclassified(67);p__Ascomycota_unclassified(67);p__Ascomycota_unclassified(67);p__Ascomycota_unclassified(67); | - | - | - | - | - | - | - | Unassigned | - |
| Otu07946 | 3 | k__Fungi(100);k__Fungi_unclassified(100);k__Fungi_unclassified(100);k__Fungi_unclassified(100);k__Fungi_unclassified(100);k__Fungi_unclassified(100);k__Fungi_unclassified(100); | - | - | - | - | - | - | - | Unassigned | - |
| Otu07948 | 3 | k__Fungi(100);k__Fungi_unclassified(100);k__Fungi_unclassified(100);k__Fungi_unclassified(100);k__Fungi_unclassified(100);k__Fungi_unclassified(100);k__Fungi_unclassified(100); | - | - | - | - | - | - | - | Unassigned | - |
| Otu07949 | 3 | k__Fungi(100);k__Fungi_unclassified(67);k__Fungi_unclassified(67);k__Fungi_unclassified(67);k__Fungi_unclassified(67);k__Fungi_unclassified(67);k__Fungi_unclassified(67); | - | - | - | - | - | - | - | Unassigned | - |
| Otu07952 | 3 | k__Fungi(100);k__Fungi_unclassified(67);k__Fungi_unclassified(67);k__Fungi_unclassified(67);k__Fungi_unclassified(67);k__Fungi_unclassified(67);k__Fungi_unclassified(67); | - | - | - | - | - | - | - | Unassigned | - |
| Otu07953 | 3 | k__Fungi(100);p__Ascomycota(67);p__Ascomycota_unclassified(67);p__Ascomycota_unclassified(67);p__Ascomycota_unclassified(67);p__Ascomycota_unclassified(67);p__Ascomycota_unclassified(67); | - | - | - | - | - | - | - | Unassigned | - |
| Otu07954 | 3 | k__Fungi(100);p__Ascomycota(67);p__Ascomycota_unclassified(67);p__Ascomycota_unclassified(67);p__Ascomycota_unclassified(67);p__Ascomycota_unclassified(67);p__Ascomycota_unclassified(67); | - | - | - | - | - | - | - | Unassigned | - |
| Otu07956 | 3 | k__Fungi(100);k__Fungi_unclassified(67);k__Fungi_unclassified(67);k__Fungi_unclassified(67);k__Fungi_unclassified(67);k__Fungi_unclassified(67);k__Fungi_unclassified(67); | - | - | - | - | - | - | - | Unassigned | - |
| Otu07958 | 3 | k__Fungi(100);p__Ascomycota(100);p__Ascomycota_unclassified(100);p__Ascomycota_unclassified(100);p__Ascomycota_unclassified(100);p__Ascomycota_unclassified(100);p__Ascomycota_unclassified(100); | - | - | - | - | - | - | - | Unassigned | - |
| Otu07959 | 3 | k__Fungi(100);p__Ascomycota(100);p__Ascomycota_unclassified(100);p__Ascomycota_unclassified(100);p__Ascomycota_unclassified(100);p__Ascomycota_unclassified(100);p__Ascomycota_unclassified(100); | - | - | - | - | - | - | - | Unassigned | - |
| Otu07962 | 3 | k__Fungi(100);k__Fungi_unclassified(67);k__Fungi_unclassified(67);k__Fungi_unclassified(67);k__Fungi_unclassified(67);k__Fungi_unclassified(67);k__Fungi_unclassified(67); | - | - | - | - | - | - | - | Unassigned | - |
| Otu07964 | 3 | k__Fungi(100);k__Fungi_unclassified(100);k__Fungi_unclassified(100);k__Fungi_unclassified(100);k__Fungi_unclassified(100);k__Fungi_unclassified(100);k__Fungi_unclassified(100); | - | - | - | - | - | - | - | Unassigned | - |
| Otu07965 | 3 | k__Fungi(100);k__Fungi_unclassified(100);k__Fungi_unclassified(100);k__Fungi_unclassified(100);k__Fungi_unclassified(100);k__Fungi_unclassified(100);k__Fungi_unclassified(100); | - | - | - | - | - | - | - | Unassigned | - |
| Otu07967 | 3 | k__Fungi(100);p__Basidiomycota(100);c__Microbotryomycetes(100);o__Sporidiobolales(100);f__Sporidiobolales_family_Incertae_sedis(100);f__Sporidiobolales_family_Incertae_sedis_unclassified(67);f__Sporidiobolales_family_Incertae_sedis_unclassified(67); | - | - | - | - | - | - | - | Unassigned | - |
| Otu07974 | 3 | k__Fungi(100);p__Ascomycota(100);p__Ascomycota_unclassified(100);p__Ascomycota_unclassified(100);p__Ascomycota_unclassified(100);p__Ascomycota_unclassified(100);p__Ascomycota_unclassified(100); | - | - | - | - | - | - | - | Unassigned | - |
| Otu07975 | 3 | k__Fungi(100);k__Fungi_unclassified(100);k__Fungi_unclassified(100);k__Fungi_unclassified(100);k__Fungi_unclassified(100);k__Fungi_unclassified(100);k__Fungi_unclassified(100); | - | - | - | - | - | - | - | Unassigned | - |
| Otu07976 | 3 | k__Fungi(100);k__Fungi_unclassified(100);k__Fungi_unclassified(100);k__Fungi_unclassified(100);k__Fungi_unclassified(100);k__Fungi_unclassified(100);k__Fungi_unclassified(100); | - | - | - | - | - | - | - | Unassigned | - |
| Otu07977 | 3 | k__Fungi(100);p__Ascomycota(100);c__Dothideomycetes(100);o__Pleosporales(100);o__Pleosporales_unclassified(67);o__Pleosporales_unclassified(67);o__Pleosporales_unclassified(67); | - | - | - | - | - | - | - | Unassigned | - |
| Otu07983 | 3 | k__Fungi(100);k__Fungi_unclassified(67);k__Fungi_unclassified(67);k__Fungi_unclassified(67);k__Fungi_unclassified(67);k__Fungi_unclassified(67);k__Fungi_unclassified(67); | - | - | - | - | - | - | - | Unassigned | - |
| Otu07985 | 3 | k__Fungi(100);p__Ascomycota(100);c__Dothideomycetes(100);o__Pleosporales(100);o__Pleosporales_unclassified(100);o__Pleosporales_unclassified(100);o__Pleosporales_unclassified(100); | - | - | - | - | - | - | - | Unassigned | - |
| Otu07987 | 3 | k__Fungi(100);p__Ascomycota(67);p__Ascomycota_unclassified(67);p__Ascomycota_unclassified(67);p__Ascomycota_unclassified(67);p__Ascomycota_unclassified(67);p__Ascomycota_unclassified(67); | - | - | - | - | - | - | - | Unassigned | - |
| Otu07989 | 3 | k__Fungi(100);p__Ascomycota(100);p__Ascomycota_unclassified(100);p__Ascomycota_unclassified(100);p__Ascomycota_unclassified(100);p__Ascomycota_unclassified(100);p__Ascomycota_unclassified(100); | - | - | - | - | - | - | - | Unassigned | - |
| Otu07992 | 3 | k__Fungi(100);p__Ascomycota(67);p__Ascomycota_unclassified(67);p__Ascomycota_unclassified(67);p__Ascomycota_unclassified(67);p__Ascomycota_unclassified(67);p__Ascomycota_unclassified(67); | - | - | - | - | - | - | - | Unassigned | - |
| Otu07995 | 3 | k__Fungi(100);k__Fungi_unclassified(100);k__Fungi_unclassified(100);k__Fungi_unclassified(100);k__Fungi_unclassified(100);k__Fungi_unclassified(100);k__Fungi_unclassified(100); | - | - | - | - | - | - | - | Unassigned | - |
| Otu07996 | 3 | k__Fungi(100);p__Ascomycota(100);p__Ascomycota_unclassified(100);p__Ascomycota_unclassified(100);p__Ascomycota_unclassified(100);p__Ascomycota_unclassified(100);p__Ascomycota_unclassified(100); | - | - | - | - | - | - | - | Unassigned | - |
| Otu07997 | 3 | k__Fungi(100);k__Fungi_unclassified(67);k__Fungi_unclassified(67);k__Fungi_unclassified(67);k__Fungi_unclassified(67);k__Fungi_unclassified(67);k__Fungi_unclassified(67); | - | - | - | - | - | - | - | Unassigned | - |
| Otu07998 | 3 | k__Fungi(100);k__Fungi_unclassified(100);k__Fungi_unclassified(100);k__Fungi_unclassified(100);k__Fungi_unclassified(100);k__Fungi_unclassified(100);k__Fungi_unclassified(100); | - | - | - | - | - | - | - | Unassigned | - |
| Otu08000 | 3 | k__Fungi(100);p__Ascomycota(100);p__Ascomycota_unclassified(100);p__Ascomycota_unclassified(100);p__Ascomycota_unclassified(100);p__Ascomycota_unclassified(100);p__Ascomycota_unclassified(100); | - | - | - | - | - | - | - | Unassigned | - |
| Otu08001 | 3 | k__Fungi(100);k__Fungi_unclassified(67);k__Fungi_unclassified(67);k__Fungi_unclassified(67);k__Fungi_unclassified(67);k__Fungi_unclassified(67);k__Fungi_unclassified(67); | - | - | - | - | - | - | - | Unassigned | - |
| Otu08007 | 3 | k__Fungi(100);k__Fungi_unclassified(67);k__Fungi_unclassified(67);k__Fungi_unclassified(67);k__Fungi_unclassified(67);k__Fungi_unclassified(67);k__Fungi_unclassified(67); | - | - | - | - | - | - | - | Unassigned | - |
| Otu08009 | 3 | k__Fungi(100);k__Fungi_unclassified(67);k__Fungi_unclassified(67);k__Fungi_unclassified(67);k__Fungi_unclassified(67);k__Fungi_unclassified(67);k__Fungi_unclassified(67); | - | - | - | - | - | - | - | Unassigned | - |
| Otu08010 | 3 | k__Fungi(100);k__Fungi_unclassified(100);k__Fungi_unclassified(100);k__Fungi_unclassified(100);k__Fungi_unclassified(100);k__Fungi_unclassified(100);k__Fungi_unclassified(100); | - | - | - | - | - | - | - | Unassigned | - |
| Otu08018 | 3 | k__Fungi(100);k__Fungi_unclassified(100);k__Fungi_unclassified(100);k__Fungi_unclassified(100);k__Fungi_unclassified(100);k__Fungi_unclassified(100);k__Fungi_unclassified(100); | - | - | - | - | - | - | - | Unassigned | - |
| Otu08019 | 3 | k__Fungi(100);k__Fungi_unclassified(67);k__Fungi_unclassified(67);k__Fungi_unclassified(67);k__Fungi_unclassified(67);k__Fungi_unclassified(67);k__Fungi_unclassified(67); | - | - | - | - | - | - | - | Unassigned | - |
| Otu08023 | 3 | k__Fungi(100);k__Fungi_unclassified(67);k__Fungi_unclassified(67);k__Fungi_unclassified(67);k__Fungi_unclassified(67);k__Fungi_unclassified(67);k__Fungi_unclassified(67); | - | - | - | - | - | - | - | Unassigned | - |
| Otu08026 | 3 | k__Fungi(100);p__Ascomycota(100);c__Dothideomycetes(100);c__Dothideomycetes_unclassified(100);c__Dothideomycetes_unclassified(100);c__Dothideomycetes_unclassified(100);c__Dothideomycetes_unclassified(100); | - | - | - | - | - | - | - | Unassigned | - |
| Otu08028 | 3 | k__Fungi(100);k__Fungi_unclassified(67);k__Fungi_unclassified(67);k__Fungi_unclassified(67);k__Fungi_unclassified(67);k__Fungi_unclassified(67);k__Fungi_unclassified(67); | - | - | - | - | - | - | - | Unassigned | - |
| Otu08029 | 3 | k__Fungi(100);k__Fungi_unclassified(67);k__Fungi_unclassified(67);k__Fungi_unclassified(67);k__Fungi_unclassified(67);k__Fungi_unclassified(67);k__Fungi_unclassified(67); | - | - | - | - | - | - | - | Unassigned | - |
| Otu08032 | 3 | k__Fungi(100);k__Fungi_unclassified(67);k__Fungi_unclassified(67);k__Fungi_unclassified(67);k__Fungi_unclassified(67);k__Fungi_unclassified(67);k__Fungi_unclassified(67); | - | - | - | - | - | - | - | Unassigned | - |
| Otu08033 | 3 | k__Fungi(100);p__Ascomycota(100);c__Leotiomycetes(100);o__Helotiales(100);f__Helotiales_family_Incertae_sedis(100);f__Helotiales_family_Incertae_sedis_unclassified(100);f__Helotiales_family_Incertae_sedis_unclassified(100); | - | - | - | - | - | - | - | Unassigned | - |
| Otu08035 | 3 | k__Fungi(100);p__Ascomycota(67);p__Ascomycota_unclassified(67);p__Ascomycota_unclassified(67);p__Ascomycota_unclassified(67);p__Ascomycota_unclassified(67);p__Ascomycota_unclassified(67); | - | - | - | - | - | - | - | Unassigned | - |
| Otu08037 | 3 | k__Fungi(100);k__Fungi_unclassified(100);k__Fungi_unclassified(100);k__Fungi_unclassified(100);k__Fungi_unclassified(100);k__Fungi_unclassified(100);k__Fungi_unclassified(100); | - | - | - | - | - | - | - | Unassigned | - |
| Otu08041 | 3 | k__Fungi(100);p__Ascomycota(100);p__Ascomycota_unclassified(100);p__Ascomycota_unclassified(100);p__Ascomycota_unclassified(100);p__Ascomycota_unclassified(100);p__Ascomycota_unclassified(100); | - | - | - | - | - | - | - | Unassigned | - |
| Otu08042 | 3 | k__Fungi(100);p__Ascomycota(67);p__Ascomycota_unclassified(67);p__Ascomycota_unclassified(67);p__Ascomycota_unclassified(67);p__Ascomycota_unclassified(67);p__Ascomycota_unclassified(67); | - | - | - | - | - | - | - | Unassigned | - |
| Otu08044 | 3 | k__Fungi(100);p__Ascomycota(100);p__Ascomycota_unclassified(100);p__Ascomycota_unclassified(100);p__Ascomycota_unclassified(100);p__Ascomycota_unclassified(100);p__Ascomycota_unclassified(100); | - | - | - | - | - | - | - | Unassigned | - |
| Otu08045 | 3 | k__Fungi(100);p__Ascomycota(100);p__Ascomycota_unclassified(100);p__Ascomycota_unclassified(100);p__Ascomycota_unclassified(100);p__Ascomycota_unclassified(100);p__Ascomycota_unclassified(100); | - | - | - | - | - | - | - | Unassigned | - |
| Otu08048 | 3 | k__Fungi(100);k__Fungi_unclassified(67);k__Fungi_unclassified(67);k__Fungi_unclassified(67);k__Fungi_unclassified(67);k__Fungi_unclassified(67);k__Fungi_unclassified(67); | - | - | - | - | - | - | - | Unassigned | - |
| Otu08049 | 3 | k__Fungi(100);p__Ascomycota(100);c__Dothideomycetes(100);o__Pleosporales(100);o__Pleosporales_unclassified(100);o__Pleosporales_unclassified(100);o__Pleosporales_unclassified(100); | - | - | - | - | - | - | - | Unassigned | - |
| Otu08053 | 3 | k__Fungi(100);p__Ascomycota(67);p__Ascomycota_unclassified(67);p__Ascomycota_unclassified(67);p__Ascomycota_unclassified(67);p__Ascomycota_unclassified(67);p__Ascomycota_unclassified(67); | - | - | - | - | - | - | - | Unassigned | - |
| Otu08060 | 3 | k__Fungi(100);p__Ascomycota(100);p__Ascomycota_unclassified(100);p__Ascomycota_unclassified(100);p__Ascomycota_unclassified(100);p__Ascomycota_unclassified(100);p__Ascomycota_unclassified(100); | - | - | - | - | - | - | - | Unassigned | - |
| Otu08065 | 3 | k__Fungi(100);p__Ascomycota(100);p__Ascomycota_unclassified(100);p__Ascomycota_unclassified(100);p__Ascomycota_unclassified(100);p__Ascomycota_unclassified(100);p__Ascomycota_unclassified(100); | - | - | - | - | - | - | - | Unassigned | - |
| Otu08066 | 3 | k__Fungi(100);k__Fungi_unclassified(100);k__Fungi_unclassified(100);k__Fungi_unclassified(100);k__Fungi_unclassified(100);k__Fungi_unclassified(100);k__Fungi_unclassified(100); | - | - | - | - | - | - | - | Unassigned | - |
| Otu08069 | 3 | k__Fungi(100);k__Fungi_unclassified(67);k__Fungi_unclassified(67);k__Fungi_unclassified(67);k__Fungi_unclassified(67);k__Fungi_unclassified(67);k__Fungi_unclassified(67); | - | - | - | - | - | - | - | Unassigned | - |
| Otu08070 | 3 | k__Fungi(100);k__Fungi_unclassified(100);k__Fungi_unclassified(100);k__Fungi_unclassified(100);k__Fungi_unclassified(100);k__Fungi_unclassified(100);k__Fungi_unclassified(100); | - | - | - | - | - | - | - | Unassigned | - |
| Otu08071 | 3 | k__Fungi(100);k__Fungi_unclassified(100);k__Fungi_unclassified(100);k__Fungi_unclassified(100);k__Fungi_unclassified(100);k__Fungi_unclassified(100);k__Fungi_unclassified(100); | - | - | - | - | - | - | - | Unassigned | - |
| Otu08073 | 3 | k__Fungi(100);k__Fungi_unclassified(67);k__Fungi_unclassified(67);k__Fungi_unclassified(67);k__Fungi_unclassified(67);k__Fungi_unclassified(67);k__Fungi_unclassified(67); | - | - | - | - | - | - | - | Unassigned | - |
| Otu08076 | 3 | k__Fungi(100);k__Fungi_unclassified(100);k__Fungi_unclassified(100);k__Fungi_unclassified(100);k__Fungi_unclassified(100);k__Fungi_unclassified(100);k__Fungi_unclassified(100); | - | - | - | - | - | - | - | Unassigned | - |
| Otu08077 | 3 | k__Fungi(100);p__Ascomycota(100);c__Dothideomycetes(67);c__Dothideomycetes_unclassified(67);c__Dothideomycetes_unclassified(67);c__Dothideomycetes_unclassified(67);c__Dothideomycetes_unclassified(67); | - | - | - | - | - | - | - | Unassigned | - |
| Otu08079 | 3 | k__Fungi(100);k__Fungi_unclassified(100);k__Fungi_unclassified(100);k__Fungi_unclassified(100);k__Fungi_unclassified(100);k__Fungi_unclassified(100);k__Fungi_unclassified(100); | - | - | - | - | - | - | - | Unassigned | - |
| Otu08081 | 3 | k__Fungi(100);k__Fungi_unclassified(67);k__Fungi_unclassified(67);k__Fungi_unclassified(67);k__Fungi_unclassified(67);k__Fungi_unclassified(67);k__Fungi_unclassified(67); | - | - | - | - | - | - | - | Unassigned | - |
| Otu08084 | 3 | k__Fungi(100);p__Ascomycota(67);p__Ascomycota_unclassified(67);p__Ascomycota_unclassified(67);p__Ascomycota_unclassified(67);p__Ascomycota_unclassified(67);p__Ascomycota_unclassified(67); | - | - | - | - | - | - | - | Unassigned | - |
| Otu08085 | 3 | k__Fungi(100);k__Fungi_unclassified(100);k__Fungi_unclassified(100);k__Fungi_unclassified(100);k__Fungi_unclassified(100);k__Fungi_unclassified(100);k__Fungi_unclassified(100); | - | - | - | - | - | - | - | Unassigned | - |
| Otu08086 | 3 | k__Fungi(100);k__Fungi_unclassified(100);k__Fungi_unclassified(100);k__Fungi_unclassified(100);k__Fungi_unclassified(100);k__Fungi_unclassified(100);k__Fungi_unclassified(100); | - | - | - | - | - | - | - | Unassigned | - |
| Otu08087 | 3 | k__Fungi(100);p__Ascomycota(67);p__Ascomycota_unclassified(67);p__Ascomycota_unclassified(67);p__Ascomycota_unclassified(67);p__Ascomycota_unclassified(67);p__Ascomycota_unclassified(67); | - | - | - | - | - | - | - | Unassigned | - |
| Otu08089 | 3 | k__Fungi(100);p__Ascomycota(67);p__Ascomycota_unclassified(67);p__Ascomycota_unclassified(67);p__Ascomycota_unclassified(67);p__Ascomycota_unclassified(67);p__Ascomycota_unclassified(67); | - | - | - | - | - | - | - | Unassigned | - |
| Otu08091 | 3 | k__Fungi(100);p__unclassified_Fungi(100);c__unclassified_Fungi(100);o__unclassified_Fungi(100);f__unclassified_Fungi(100);g__unclassified_Fungi(100);s__Fungi_sp(100); | - | - | - | - | - | - | - | Unassigned | - |
| Otu08093 | 3 | k__Fungi(100);k__Fungi_unclassified(100);k__Fungi_unclassified(100);k__Fungi_unclassified(100);k__Fungi_unclassified(100);k__Fungi_unclassified(100);k__Fungi_unclassified(100); | - | - | - | - | - | - | - | Unassigned | - |
| Otu08097 | 3 | k__Fungi(100);k__Fungi_unclassified(100);k__Fungi_unclassified(100);k__Fungi_unclassified(100);k__Fungi_unclassified(100);k__Fungi_unclassified(100);k__Fungi_unclassified(100); | - | - | - | - | - | - | - | Unassigned | - |
| Otu08100 | 3 | k__Fungi(100);k__Fungi_unclassified(100);k__Fungi_unclassified(100);k__Fungi_unclassified(100);k__Fungi_unclassified(100);k__Fungi_unclassified(100);k__Fungi_unclassified(100); | - | - | - | - | - | - | - | Unassigned | - |
| Otu08102 | 3 | k__Fungi(100);k__Fungi_unclassified(100);k__Fungi_unclassified(100);k__Fungi_unclassified(100);k__Fungi_unclassified(100);k__Fungi_unclassified(100);k__Fungi_unclassified(100); | - | - | - | - | - | - | - | Unassigned | - |
| Otu08105 | 3 | k__Fungi(100);p__Ascomycota(100);c__Dothideomycetes(100);o__Pleosporales(100);o__Pleosporales_unclassified(100);o__Pleosporales_unclassified(100);o__Pleosporales_unclassified(100); | - | - | - | - | - | - | - | Unassigned | - |
| Otu08106 | 3 | k__Fungi(100);k__Fungi_unclassified(67);k__Fungi_unclassified(67);k__Fungi_unclassified(67);k__Fungi_unclassified(67);k__Fungi_unclassified(67);k__Fungi_unclassified(67); | - | - | - | - | - | - | - | Unassigned | - |
| Otu08112 | 3 | k__Fungi(100);p__Ascomycota(100);p__Ascomycota_unclassified(100);p__Ascomycota_unclassified(100);p__Ascomycota_unclassified(100);p__Ascomycota_unclassified(100);p__Ascomycota_unclassified(100); | - | - | - | - | - | - | - | Unassigned | - |
| Otu08115 | 3 | k__Fungi(100);k__Fungi_unclassified(100);k__Fungi_unclassified(100);k__Fungi_unclassified(100);k__Fungi_unclassified(100);k__Fungi_unclassified(100);k__Fungi_unclassified(100); | - | - | - | - | - | - | - | Unassigned | - |
| Otu08117 | 3 | k__Fungi(100);p__Ascomycota(67);p__Ascomycota_unclassified(67);p__Ascomycota_unclassified(67);p__Ascomycota_unclassified(67);p__Ascomycota_unclassified(67);p__Ascomycota_unclassified(67); | - | - | - | - | - | - | - | Unassigned | - |
| Otu08119 | 3 | k__Fungi(100);k__Fungi_unclassified(67);k__Fungi_unclassified(67);k__Fungi_unclassified(67);k__Fungi_unclassified(67);k__Fungi_unclassified(67);k__Fungi_unclassified(67); | - | - | - | - | - | - | - | Unassigned | - |
| Otu08120 | 3 | k__Fungi(100);k__Fungi_unclassified(67);k__Fungi_unclassified(67);k__Fungi_unclassified(67);k__Fungi_unclassified(67);k__Fungi_unclassified(67);k__Fungi_unclassified(67); | - | - | - | - | - | - | - | Unassigned | - |
| Otu08123 | 3 | k__Fungi(100);p__Ascomycota(100);p__Ascomycota_unclassified(100);p__Ascomycota_unclassified(100);p__Ascomycota_unclassified(100);p__Ascomycota_unclassified(100);p__Ascomycota_unclassified(100); | - | - | - | - | - | - | - | Unassigned | - |
| Otu08124 | 3 | k__Fungi(100);k__Fungi_unclassified(100);k__Fungi_unclassified(100);k__Fungi_unclassified(100);k__Fungi_unclassified(100);k__Fungi_unclassified(100);k__Fungi_unclassified(100); | - | - | - | - | - | - | - | Unassigned | - |
| Otu08125 | 3 | k__Fungi(100);p__Ascomycota(100);p__Ascomycota_unclassified(100);p__Ascomycota_unclassified(100);p__Ascomycota_unclassified(100);p__Ascomycota_unclassified(100);p__Ascomycota_unclassified(100); | - | - | - | - | - | - | - | Unassigned | - |
| Otu08126 | 3 | k__Fungi(100);k__Fungi_unclassified(67);k__Fungi_unclassified(67);k__Fungi_unclassified(67);k__Fungi_unclassified(67);k__Fungi_unclassified(67);k__Fungi_unclassified(67); | - | - | - | - | - | - | - | Unassigned | - |
| Otu08130 | 3 | k__Fungi(100);k__Fungi_unclassified(100);k__Fungi_unclassified(100);k__Fungi_unclassified(100);k__Fungi_unclassified(100);k__Fungi_unclassified(100);k__Fungi_unclassified(100); | - | - | - | - | - | - | - | Unassigned | - |
| Otu08131 | 3 | k__Fungi(100);p__Ascomycota(100);p__Ascomycota_unclassified(100);p__Ascomycota_unclassified(100);p__Ascomycota_unclassified(100);p__Ascomycota_unclassified(100);p__Ascomycota_unclassified(100); | - | - | - | - | - | - | - | Unassigned | - |
| Otu08132 | 3 | k__Fungi(100);p__Ascomycota(67);p__Ascomycota_unclassified(67);p__Ascomycota_unclassified(67);p__Ascomycota_unclassified(67);p__Ascomycota_unclassified(67);p__Ascomycota_unclassified(67); | - | - | - | - | - | - | - | Unassigned | - |
| Otu08135 | 3 | k__Fungi(100);p__Ascomycota(100);p__Ascomycota_unclassified(100);p__Ascomycota_unclassified(100);p__Ascomycota_unclassified(100);p__Ascomycota_unclassified(100);p__Ascomycota_unclassified(100); | - | - | - | - | - | - | - | Unassigned | - |
| Otu08136 | 3 | k__Fungi(100);p__Ascomycota(100);c__Dothideomycetes(67);c__Dothideomycetes_unclassified(67);c__Dothideomycetes_unclassified(67);c__Dothideomycetes_unclassified(67);c__Dothideomycetes_unclassified(67); | - | - | - | - | - | - | - | Unassigned | - |
| Otu08138 | 3 | k__Fungi(100);k__Fungi_unclassified(67);k__Fungi_unclassified(67);k__Fungi_unclassified(67);k__Fungi_unclassified(67);k__Fungi_unclassified(67);k__Fungi_unclassified(67); | - | - | - | - | - | - | - | Unassigned | - |
| Otu08141 | 3 | k__Fungi(100);k__Fungi_unclassified(67);k__Fungi_unclassified(67);k__Fungi_unclassified(67);k__Fungi_unclassified(67);k__Fungi_unclassified(67);k__Fungi_unclassified(67); | - | - | - | - | - | - | - | Unassigned | - |
| Otu08142 | 3 | k__Fungi(100);p__Ascomycota(100);p__Ascomycota_unclassified(100);p__Ascomycota_unclassified(100);p__Ascomycota_unclassified(100);p__Ascomycota_unclassified(100);p__Ascomycota_unclassified(100); | - | - | - | - | - | - | - | Unassigned | - |
| Otu08143 | 3 | k__Fungi(100);p__Ascomycota(100);c__Dothideomycetes(100);o__Hysteriales(100);f__unclassified_Hysteriales(100);g__unclassified_Hysteriales(100);s__Hysteriales_sp(100); | - | - | - | - | - | - | - | Unassigned | - |
| Otu08145 | 3 | k__Fungi(100);p__Ascomycota(67);p__Ascomycota_unclassified(67);p__Ascomycota_unclassified(67);p__Ascomycota_unclassified(67);p__Ascomycota_unclassified(67);p__Ascomycota_unclassified(67); | - | - | - | - | - | - | - | Unassigned | - |
| Otu08146 | 3 | k__Fungi(100);k__Fungi_unclassified(100);k__Fungi_unclassified(100);k__Fungi_unclassified(100);k__Fungi_unclassified(100);k__Fungi_unclassified(100);k__Fungi_unclassified(100); | - | - | - | - | - | - | - | Unassigned | - |
| Otu08149 | 3 | k__Fungi(100);p__Ascomycota(100);c__Lecanoromycetes(100);c__Lecanoromycetes_unclassified(67);c__Lecanoromycetes_unclassified(67);c__Lecanoromycetes_unclassified(67);c__Lecanoromycetes_unclassified(67); | - | - | - | - | - | - | - | Unassigned | - |
| Otu08151 | 3 | k__Fungi(100);k__Fungi_unclassified(100);k__Fungi_unclassified(100);k__Fungi_unclassified(100);k__Fungi_unclassified(100);k__Fungi_unclassified(100);k__Fungi_unclassified(100); | - | - | - | - | - | - | - | Unassigned | - |
| Otu08154 | 3 | k__Fungi(100);p__Ascomycota(67);p__Ascomycota_unclassified(67);p__Ascomycota_unclassified(67);p__Ascomycota_unclassified(67);p__Ascomycota_unclassified(67);p__Ascomycota_unclassified(67); | - | - | - | - | - | - | - | Unassigned | - |
| Otu08155 | 3 | k__Fungi(100);p__Ascomycota(100);p__Ascomycota_unclassified(100);p__Ascomycota_unclassified(100);p__Ascomycota_unclassified(100);p__Ascomycota_unclassified(100);p__Ascomycota_unclassified(100); | - | - | - | - | - | - | - | Unassigned | - |
| Otu08157 | 3 | k__Fungi(100);k__Fungi_unclassified(67);k__Fungi_unclassified(67);k__Fungi_unclassified(67);k__Fungi_unclassified(67);k__Fungi_unclassified(67);k__Fungi_unclassified(67); | - | - | - | - | - | - | - | Unassigned | - |
| Otu08159 | 3 | k__Fungi(100);p__Ascomycota(100);p__Ascomycota_unclassified(100);p__Ascomycota_unclassified(100);p__Ascomycota_unclassified(100);p__Ascomycota_unclassified(100);p__Ascomycota_unclassified(100); | - | - | - | - | - | - | - | Unassigned | - |
| Otu08164 | 3 | k__Fungi(100);p__Ascomycota(100);c__Leotiomycetes(100);c__Leotiomycetes_unclassified(100);c__Leotiomycetes_unclassified(100);c__Leotiomycetes_unclassified(100);c__Leotiomycetes_unclassified(100); | - | - | - | - | - | - | - | Unassigned | - |
| Otu08165 | 3 | k__Fungi(100);p__Ascomycota(100);p__Ascomycota_unclassified(100);p__Ascomycota_unclassified(100);p__Ascomycota_unclassified(100);p__Ascomycota_unclassified(100);p__Ascomycota_unclassified(100); | - | - | - | - | - | - | - | Unassigned | - |
| Otu08166 | 3 | k__Fungi(100);k__Fungi_unclassified(100);k__Fungi_unclassified(100);k__Fungi_unclassified(100);k__Fungi_unclassified(100);k__Fungi_unclassified(100);k__Fungi_unclassified(100); | - | - | - | - | - | - | - | Unassigned | - |
| Otu08168 | 3 | k__Fungi(100);p__Ascomycota(100);p__Ascomycota_unclassified(100);p__Ascomycota_unclassified(100);p__Ascomycota_unclassified(100);p__Ascomycota_unclassified(100);p__Ascomycota_unclassified(100); | - | - | - | - | - | - | - | Unassigned | - |
| Otu08169 | 3 | k__Fungi(100);k__Fungi_unclassified(67);k__Fungi_unclassified(67);k__Fungi_unclassified(67);k__Fungi_unclassified(67);k__Fungi_unclassified(67);k__Fungi_unclassified(67); | - | - | - | - | - | - | - | Unassigned | - |
| Otu08171 | 3 | k__Fungi(100);p__Ascomycota(67);p__Ascomycota_unclassified(67);p__Ascomycota_unclassified(67);p__Ascomycota_unclassified(67);p__Ascomycota_unclassified(67);p__Ascomycota_unclassified(67); | - | - | - | - | - | - | - | Unassigned | - |
| Otu08173 | 3 | k__Fungi(100);p__Ascomycota(100);p__Ascomycota_unclassified(100);p__Ascomycota_unclassified(100);p__Ascomycota_unclassified(100);p__Ascomycota_unclassified(100);p__Ascomycota_unclassified(100); | - | - | - | - | - | - | - | Unassigned | - |
| Otu08175 | 3 | k__Fungi(100);p__Ascomycota(100);p__Ascomycota_unclassified(100);p__Ascomycota_unclassified(100);p__Ascomycota_unclassified(100);p__Ascomycota_unclassified(100);p__Ascomycota_unclassified(100); | - | - | - | - | - | - | - | Unassigned | - |
| Otu08176 | 3 | k__Fungi(100);p__Ascomycota(100);p__Ascomycota_unclassified(100);p__Ascomycota_unclassified(100);p__Ascomycota_unclassified(100);p__Ascomycota_unclassified(100);p__Ascomycota_unclassified(100); | - | - | - | - | - | - | - | Unassigned | - |
| Otu08177 | 3 | k__Fungi(100);p__Ascomycota(67);p__Ascomycota_unclassified(67);p__Ascomycota_unclassified(67);p__Ascomycota_unclassified(67);p__Ascomycota_unclassified(67);p__Ascomycota_unclassified(67); | - | - | - | - | - | - | - | Unassigned | - |
| Otu08180 | 3 | k__Fungi(100);p__Ascomycota(100);p__Ascomycota_unclassified(100);p__Ascomycota_unclassified(100);p__Ascomycota_unclassified(100);p__Ascomycota_unclassified(100);p__Ascomycota_unclassified(100); | - | - | - | - | - | - | - | Unassigned | - |
| Otu08181 | 3 | k__Fungi(100);k__Fungi_unclassified(100);k__Fungi_unclassified(100);k__Fungi_unclassified(100);k__Fungi_unclassified(100);k__Fungi_unclassified(100);k__Fungi_unclassified(100); | - | - | - | - | - | - | - | Unassigned | - |
| Otu08182 | 3 | k__Fungi(100);p__Ascomycota(100);p__Ascomycota_unclassified(100);p__Ascomycota_unclassified(100);p__Ascomycota_unclassified(100);p__Ascomycota_unclassified(100);p__Ascomycota_unclassified(100); | - | - | - | - | - | - | - | Unassigned | - |
| Otu08183 | 3 | k__Fungi(100);p__Ascomycota(67);p__Ascomycota_unclassified(67);p__Ascomycota_unclassified(67);p__Ascomycota_unclassified(67);p__Ascomycota_unclassified(67);p__Ascomycota_unclassified(67); | - | - | - | - | - | - | - | Unassigned | - |
| Otu08184 | 3 | k__Fungi(100);k__Fungi_unclassified(67);k__Fungi_unclassified(67);k__Fungi_unclassified(67);k__Fungi_unclassified(67);k__Fungi_unclassified(67);k__Fungi_unclassified(67); | - | - | - | - | - | - | - | Unassigned | - |
| Otu08187 | 3 | k__Fungi(100);k__Fungi_unclassified(67);k__Fungi_unclassified(67);k__Fungi_unclassified(67);k__Fungi_unclassified(67);k__Fungi_unclassified(67);k__Fungi_unclassified(67); | - | - | - | - | - | - | - | Unassigned | - |
| Otu08193 | 3 | k__Fungi(100);p__Ascomycota(67);p__Ascomycota_unclassified(67);p__Ascomycota_unclassified(67);p__Ascomycota_unclassified(67);p__Ascomycota_unclassified(67);p__Ascomycota_unclassified(67); | - | - | - | - | - | - | - | Unassigned | - |
| Otu08197 | 3 | k__Fungi(100);k__Fungi_unclassified(100);k__Fungi_unclassified(100);k__Fungi_unclassified(100);k__Fungi_unclassified(100);k__Fungi_unclassified(100);k__Fungi_unclassified(100); | - | - | - | - | - | - | - | Unassigned | - |
| Otu08200 | 3 | k__Fungi(100);k__Fungi_unclassified(67);k__Fungi_unclassified(67);k__Fungi_unclassified(67);k__Fungi_unclassified(67);k__Fungi_unclassified(67);k__Fungi_unclassified(67); | - | - | - | - | - | - | - | Unassigned | - |
| Otu08201 | 3 | k__Fungi(100);p__Ascomycota(67);p__Ascomycota_unclassified(67);p__Ascomycota_unclassified(67);p__Ascomycota_unclassified(67);p__Ascomycota_unclassified(67);p__Ascomycota_unclassified(67); | - | - | - | - | - | - | - | Unassigned | - |
| Otu08202 | 3 | k__Fungi(100);p__Ascomycota(100);c__Dothideomycetes(100);o__Capnodiales(100);f__Davidiellaceae(100);f__Davidiellaceae_unclassified(67);f__Davidiellaceae_unclassified(67); | - | - | - | - | - | - | - | Unassigned | - |
| Otu08203 | 3 | k__Fungi(100);p__Ascomycota(67);p__Ascomycota_unclassified(67);p__Ascomycota_unclassified(67);p__Ascomycota_unclassified(67);p__Ascomycota_unclassified(67);p__Ascomycota_unclassified(67); | - | - | - | - | - | - | - | Unassigned | - |
| Otu08204 | 3 | k__Fungi(100);k__Fungi_unclassified(100);k__Fungi_unclassified(100);k__Fungi_unclassified(100);k__Fungi_unclassified(100);k__Fungi_unclassified(100);k__Fungi_unclassified(100); | - | - | - | - | - | - | - | Unassigned | - |
| Otu08205 | 3 | k__Fungi(100);k__Fungi_unclassified(67);k__Fungi_unclassified(67);k__Fungi_unclassified(67);k__Fungi_unclassified(67);k__Fungi_unclassified(67);k__Fungi_unclassified(67); | - | - | - | - | - | - | - | Unassigned | - |
| Otu08206 | 3 | k__Fungi(100);k__Fungi_unclassified(100);k__Fungi_unclassified(100);k__Fungi_unclassified(100);k__Fungi_unclassified(100);k__Fungi_unclassified(100);k__Fungi_unclassified(100); | - | - | - | - | - | - | - | Unassigned | - |
| Otu08210 | 3 | k__Fungi(100);k__Fungi_unclassified(100);k__Fungi_unclassified(100);k__Fungi_unclassified(100);k__Fungi_unclassified(100);k__Fungi_unclassified(100);k__Fungi_unclassified(100); | - | - | - | - | - | - | - | Unassigned | - |
| Otu08211 | 3 | k__Fungi(100);p__Ascomycota(67);p__Ascomycota_unclassified(67);p__Ascomycota_unclassified(67);p__Ascomycota_unclassified(67);p__Ascomycota_unclassified(67);p__Ascomycota_unclassified(67); | - | - | - | - | - | - | - | Unassigned | - |
| Otu08219 | 3 | k__Fungi(100);p__Ascomycota(67);p__Ascomycota_unclassified(67);p__Ascomycota_unclassified(67);p__Ascomycota_unclassified(67);p__Ascomycota_unclassified(67);p__Ascomycota_unclassified(67); | - | - | - | - | - | - | - | Unassigned | - |
| Otu08224 | 3 | k__Fungi(100);k__Fungi_unclassified(67);k__Fungi_unclassified(67);k__Fungi_unclassified(67);k__Fungi_unclassified(67);k__Fungi_unclassified(67);k__Fungi_unclassified(67); | - | - | - | - | - | - | - | Unassigned | - |
| Otu08225 | 3 | k__Fungi(100);k__Fungi_unclassified(100);k__Fungi_unclassified(100);k__Fungi_unclassified(100);k__Fungi_unclassified(100);k__Fungi_unclassified(100);k__Fungi_unclassified(100); | - | - | - | - | - | - | - | Unassigned | - |
| Otu08228 | 3 | k__Fungi(100);k__Fungi_unclassified(67);k__Fungi_unclassified(67);k__Fungi_unclassified(67);k__Fungi_unclassified(67);k__Fungi_unclassified(67);k__Fungi_unclassified(67); | - | - | - | - | - | - | - | Unassigned | - |
| Otu08230 | 3 | k__Fungi(100);k__Fungi_unclassified(67);k__Fungi_unclassified(67);k__Fungi_unclassified(67);k__Fungi_unclassified(67);k__Fungi_unclassified(67);k__Fungi_unclassified(67); | - | - | - | - | - | - | - | Unassigned | - |
| Otu08231 | 3 | k__Fungi(100);p__Ascomycota(67);p__Ascomycota_unclassified(67);p__Ascomycota_unclassified(67);p__Ascomycota_unclassified(67);p__Ascomycota_unclassified(67);p__Ascomycota_unclassified(67); | - | - | - | - | - | - | - | Unassigned | - |
| Otu08236 | 3 | k__Fungi(100);p__Ascomycota(100);p__Ascomycota_unclassified(100);p__Ascomycota_unclassified(100);p__Ascomycota_unclassified(100);p__Ascomycota_unclassified(100);p__Ascomycota_unclassified(100); | - | - | - | - | - | - | - | Unassigned | - |
| Otu08237 | 3 | k__Fungi(100);p__Ascomycota(100);p__Ascomycota_unclassified(100);p__Ascomycota_unclassified(100);p__Ascomycota_unclassified(100);p__Ascomycota_unclassified(100);p__Ascomycota_unclassified(100); | - | - | - | - | - | - | - | Unassigned | - |
| Otu08238 | 3 | k__Fungi(100);p__Ascomycota(100);p__Ascomycota_unclassified(100);p__Ascomycota_unclassified(100);p__Ascomycota_unclassified(100);p__Ascomycota_unclassified(100);p__Ascomycota_unclassified(100); | - | - | - | - | - | - | - | Unassigned | - |
| Otu08241 | 3 | k__Fungi(100);k__Fungi_unclassified(67);k__Fungi_unclassified(67);k__Fungi_unclassified(67);k__Fungi_unclassified(67);k__Fungi_unclassified(67);k__Fungi_unclassified(67); | - | - | - | - | - | - | - | Unassigned | - |
| Otu08242 | 3 | k__Fungi(100);k__Fungi_unclassified(67);k__Fungi_unclassified(67);k__Fungi_unclassified(67);k__Fungi_unclassified(67);k__Fungi_unclassified(67);k__Fungi_unclassified(67); | - | - | - | - | - | - | - | Unassigned | - |
| Otu08244 | 3 | k__Fungi(100);p__Ascomycota(67);p__Ascomycota_unclassified(67);p__Ascomycota_unclassified(67);p__Ascomycota_unclassified(67);p__Ascomycota_unclassified(67);p__Ascomycota_unclassified(67); | - | - | - | - | - | - | - | Unassigned | - |
| Otu08248 | 3 | k__Fungi(100);p__Ascomycota(67);p__Ascomycota_unclassified(67);p__Ascomycota_unclassified(67);p__Ascomycota_unclassified(67);p__Ascomycota_unclassified(67);p__Ascomycota_unclassified(67); | - | - | - | - | - | - | - | Unassigned | - |
| Otu08249 | 3 | k__Fungi(100);k__Fungi_unclassified(67);k__Fungi_unclassified(67);k__Fungi_unclassified(67);k__Fungi_unclassified(67);k__Fungi_unclassified(67);k__Fungi_unclassified(67); | - | - | - | - | - | - | - | Unassigned | - |
| Otu08251 | 3 | k__Fungi(100);k__Fungi_unclassified(67);k__Fungi_unclassified(67);k__Fungi_unclassified(67);k__Fungi_unclassified(67);k__Fungi_unclassified(67);k__Fungi_unclassified(67); | - | - | - | - | - | - | - | Unassigned | - |
| Otu08253 | 3 | k__Fungi(100);k__Fungi_unclassified(67);k__Fungi_unclassified(67);k__Fungi_unclassified(67);k__Fungi_unclassified(67);k__Fungi_unclassified(67);k__Fungi_unclassified(67); | - | - | - | - | - | - | - | Unassigned | - |
| Otu08256 | 3 | k__Fungi(100);k__Fungi_unclassified(67);k__Fungi_unclassified(67);k__Fungi_unclassified(67);k__Fungi_unclassified(67);k__Fungi_unclassified(67);k__Fungi_unclassified(67); | - | - | - | - | - | - | - | Unassigned | - |
| Otu08258 | 3 | k__Fungi(100);k__Fungi_unclassified(67);k__Fungi_unclassified(67);k__Fungi_unclassified(67);k__Fungi_unclassified(67);k__Fungi_unclassified(67);k__Fungi_unclassified(67); | - | - | - | - | - | - | - | Unassigned | - |
| Otu08261 | 3 | k__Fungi(100);k__Fungi_unclassified(100);k__Fungi_unclassified(100);k__Fungi_unclassified(100);k__Fungi_unclassified(100);k__Fungi_unclassified(100);k__Fungi_unclassified(100); | - | - | - | - | - | - | - | Unassigned | - |
| Otu08263 | 3 | k__Fungi(100);p__Ascomycota(100);c__Sordariomycetes(100);o__Hypocreales(100);f__Cordycipitaceae(100);g__unclassified_Cordycipitaceae(100);s__Cordycipitaceae_sp(100); | - | - | - | - | - | - | - | Unassigned | - |
| Otu08265 | 3 | k__Fungi(100);p__Ascomycota(67);p__Ascomycota_unclassified(67);p__Ascomycota_unclassified(67);p__Ascomycota_unclassified(67);p__Ascomycota_unclassified(67);p__Ascomycota_unclassified(67); | - | - | - | - | - | - | - | Unassigned | - |
| Otu08269 | 3 | k__Fungi(100);k__Fungi_unclassified(67);k__Fungi_unclassified(67);k__Fungi_unclassified(67);k__Fungi_unclassified(67);k__Fungi_unclassified(67);k__Fungi_unclassified(67); | - | - | - | - | - | - | - | Unassigned | - |
| Otu08271 | 3 | k__Fungi(100);p__Ascomycota(100);p__Ascomycota_unclassified(100);p__Ascomycota_unclassified(100);p__Ascomycota_unclassified(100);p__Ascomycota_unclassified(100);p__Ascomycota_unclassified(100); | - | - | - | - | - | - | - | Unassigned | - |
| Otu08274 | 3 | k__Fungi(100);p__Ascomycota(100);p__Ascomycota_unclassified(67);p__Ascomycota_unclassified(67);p__Ascomycota_unclassified(67);p__Ascomycota_unclassified(67);p__Ascomycota_unclassified(67); | - | - | - | - | - | - | - | Unassigned | - |
| Otu08275 | 3 | k__Fungi(100);p__Ascomycota(100);p__Ascomycota_unclassified(100);p__Ascomycota_unclassified(100);p__Ascomycota_unclassified(100);p__Ascomycota_unclassified(100);p__Ascomycota_unclassified(100); | - | - | - | - | - | - | - | Unassigned | - |
| Otu08276 | 3 | k__Fungi(100);p__Ascomycota(67);p__Ascomycota_unclassified(67);p__Ascomycota_unclassified(67);p__Ascomycota_unclassified(67);p__Ascomycota_unclassified(67);p__Ascomycota_unclassified(67); | - | - | - | - | - | - | - | Unassigned | - |
| Otu08280 | 3 | k__Fungi(100);p__Ascomycota(100);p__Ascomycota_unclassified(100);p__Ascomycota_unclassified(100);p__Ascomycota_unclassified(100);p__Ascomycota_unclassified(100);p__Ascomycota_unclassified(100); | - | - | - | - | - | - | - | Unassigned | - |
| Otu08281 | 3 | k__Fungi(100);p__Ascomycota(67);p__Ascomycota_unclassified(67);p__Ascomycota_unclassified(67);p__Ascomycota_unclassified(67);p__Ascomycota_unclassified(67);p__Ascomycota_unclassified(67); | - | - | - | - | - | - | - | Unassigned | - |
| Otu08282 | 3 | k__Fungi(100);p__Ascomycota(100);p__Ascomycota_unclassified(100);p__Ascomycota_unclassified(100);p__Ascomycota_unclassified(100);p__Ascomycota_unclassified(100);p__Ascomycota_unclassified(100); | - | - | - | - | - | - | - | Unassigned | - |
| Otu08285 | 3 | k__Fungi(100);p__Ascomycota(100);p__Ascomycota_unclassified(100);p__Ascomycota_unclassified(100);p__Ascomycota_unclassified(100);p__Ascomycota_unclassified(100);p__Ascomycota_unclassified(100); | - | - | - | - | - | - | - | Unassigned | - |
| Otu08293 | 3 | k__Fungi(100);p__Basidiomycota(100);c__Agaricomycetes(100);c__Agaricomycetes_unclassified(100);c__Agaricomycetes_unclassified(100);c__Agaricomycetes_unclassified(100);c__Agaricomycetes_unclassified(100); | - | - | - | - | - | - | - | Unassigned | - |
| Otu08295 | 3 | k__Fungi(100);p__Ascomycota(100);c__Sordariomycetes(100);c__Sordariomycetes_unclassified(100);c__Sordariomycetes_unclassified(100);c__Sordariomycetes_unclassified(100);c__Sordariomycetes_unclassified(100); | - | - | - | - | - | - | - | Unassigned | - |
| Otu08299 | 3 | k__Fungi(100);p__Basidiomycota(100);c__Agaricomycetes(100);o__Cantharellales(100);f__Ceratobasidiaceae(100);g__unclassified_Ceratobasidiaceae(100);s__uncultured_Ceratobasidium(100); | - | - | - | - | - | - | - | Unassigned | - |
| Otu08301 | 3 | k__Fungi(100);k__Fungi_unclassified(100);k__Fungi_unclassified(100);k__Fungi_unclassified(100);k__Fungi_unclassified(100);k__Fungi_unclassified(100);k__Fungi_unclassified(100); | - | - | - | - | - | - | - | Unassigned | - |
| Otu08302 | 3 | k__Fungi(100);p__Basidiomycota(100);c__unclassified_Basidiomycota(100);o__unclassified_Basidiomycota(100);f__unclassified_Basidiomycota(100);g__unclassified_Basidiomycota(100);s__Basidiomycota_sp(100); | - | - | - | - | - | - | - | Unassigned | - |
| Otu08303 | 3 | k__Fungi(100);k__Fungi_unclassified(67);k__Fungi_unclassified(67);k__Fungi_unclassified(67);k__Fungi_unclassified(67);k__Fungi_unclassified(67);k__Fungi_unclassified(67); | - | - | - | - | - | - | - | Unassigned | - |
| Otu08307 | 3 | k__Fungi(100);k__Fungi_unclassified(100);k__Fungi_unclassified(100);k__Fungi_unclassified(100);k__Fungi_unclassified(100);k__Fungi_unclassified(100);k__Fungi_unclassified(100); | - | - | - | - | - | - | - | Unassigned | - |
| Otu08308 | 3 | k__Fungi(100);k__Fungi_unclassified(67);k__Fungi_unclassified(67);k__Fungi_unclassified(67);k__Fungi_unclassified(67);k__Fungi_unclassified(67);k__Fungi_unclassified(67); | - | - | - | - | - | - | - | Unassigned | - |
| Otu08311 | 3 | k__Fungi(100);p__Ascomycota(67);p__Ascomycota_unclassified(67);p__Ascomycota_unclassified(67);p__Ascomycota_unclassified(67);p__Ascomycota_unclassified(67);p__Ascomycota_unclassified(67); | - | - | - | - | - | - | - | Unassigned | - |
| Otu08313 | 3 | k__Fungi(100);k__Fungi_unclassified(67);k__Fungi_unclassified(67);k__Fungi_unclassified(67);k__Fungi_unclassified(67);k__Fungi_unclassified(67);k__Fungi_unclassified(67); | - | - | - | - | - | - | - | Unassigned | - |
| Otu08314 | 3 | k__Fungi(100);k__Fungi_unclassified(67);k__Fungi_unclassified(67);k__Fungi_unclassified(67);k__Fungi_unclassified(67);k__Fungi_unclassified(67);k__Fungi_unclassified(67); | - | - | - | - | - | - | - | Unassigned | - |
| Otu08317 | 3 | k__Fungi(100);p__Ascomycota(100);p__Ascomycota_unclassified(100);p__Ascomycota_unclassified(100);p__Ascomycota_unclassified(100);p__Ascomycota_unclassified(100);p__Ascomycota_unclassified(100); | - | - | - | - | - | - | - | Unassigned | - |
| Otu08322 | 3 | k__Fungi(100);p__Ascomycota(67);p__Ascomycota_unclassified(67);p__Ascomycota_unclassified(67);p__Ascomycota_unclassified(67);p__Ascomycota_unclassified(67);p__Ascomycota_unclassified(67); | - | - | - | - | - | - | - | Unassigned | - |
| Otu08323 | 3 | k__Fungi(100);k__Fungi_unclassified(67);k__Fungi_unclassified(67);k__Fungi_unclassified(67);k__Fungi_unclassified(67);k__Fungi_unclassified(67);k__Fungi_unclassified(67); | - | - | - | - | - | - | - | Unassigned | - |
| Otu08326 | 3 | k__Fungi(100);k__Fungi_unclassified(100);k__Fungi_unclassified(100);k__Fungi_unclassified(100);k__Fungi_unclassified(100);k__Fungi_unclassified(100);k__Fungi_unclassified(100); | - | - | - | - | - | - | - | Unassigned | - |
| Otu08327 | 3 | k__Fungi(100);k__Fungi_unclassified(100);k__Fungi_unclassified(100);k__Fungi_unclassified(100);k__Fungi_unclassified(100);k__Fungi_unclassified(100);k__Fungi_unclassified(100); | - | - | - | - | - | - | - | Unassigned | - |
| Otu08328 | 3 | k__Fungi(100);k__Fungi_unclassified(67);k__Fungi_unclassified(67);k__Fungi_unclassified(67);k__Fungi_unclassified(67);k__Fungi_unclassified(67);k__Fungi_unclassified(67); | - | - | - | - | - | - | - | Unassigned | - |
| Otu08330 | 3 | k__Fungi(100);p__Ascomycota(100);p__Ascomycota_unclassified(67);p__Ascomycota_unclassified(67);p__Ascomycota_unclassified(67);p__Ascomycota_unclassified(67);p__Ascomycota_unclassified(67); | - | - | - | - | - | - | - | Unassigned | - |
| Otu08334 | 3 | k__Fungi(100);p__Ascomycota(100);c__Sordariomycetes(100);c__Sordariomycetes_unclassified(100);c__Sordariomycetes_unclassified(100);c__Sordariomycetes_unclassified(100);c__Sordariomycetes_unclassified(100); | - | - | - | - | - | - | - | Unassigned | - |
| Otu08340 | 3 | k__Fungi(100);p__Ascomycota(67);p__Ascomycota_unclassified(67);p__Ascomycota_unclassified(67);p__Ascomycota_unclassified(67);p__Ascomycota_unclassified(67);p__Ascomycota_unclassified(67); | - | - | - | - | - | - | - | Unassigned | - |
| Otu08341 | 3 | k__Fungi(100);k__Fungi_unclassified(67);k__Fungi_unclassified(67);k__Fungi_unclassified(67);k__Fungi_unclassified(67);k__Fungi_unclassified(67);k__Fungi_unclassified(67); | - | - | - | - | - | - | - | Unassigned | - |
| Otu08342 | 3 | k__Fungi(100);p__Basidiomycota(100);c__Agaricomycetes(100);c__Agaricomycetes_unclassified(100);c__Agaricomycetes_unclassified(100);c__Agaricomycetes_unclassified(100);c__Agaricomycetes_unclassified(100); | - | - | - | - | - | - | - | Unassigned | - |
| Otu08343 | 3 | k__Fungi(100);k__Fungi_unclassified(100);k__Fungi_unclassified(100);k__Fungi_unclassified(100);k__Fungi_unclassified(100);k__Fungi_unclassified(100);k__Fungi_unclassified(100); | - | - | - | - | - | - | - | Unassigned | - |
| Otu08344 | 3 | k__Fungi(100);p__Basidiomycota(100);c__Agaricomycetes(100);o__Trechisporales(100);o__Trechisporales_unclassified(100);o__Trechisporales_unclassified(100);o__Trechisporales_unclassified(100); | - | - | - | - | - | - | - | Unassigned | - |
| Otu08345 | 3 | k__Fungi(100);p__Ascomycota(100);p__Ascomycota_unclassified(100);p__Ascomycota_unclassified(100);p__Ascomycota_unclassified(100);p__Ascomycota_unclassified(100);p__Ascomycota_unclassified(100); | - | - | - | - | - | - | - | Unassigned | - |
| Otu08346 | 3 | k__Fungi(100);p__Ascomycota(67);p__Ascomycota_unclassified(67);p__Ascomycota_unclassified(67);p__Ascomycota_unclassified(67);p__Ascomycota_unclassified(67);p__Ascomycota_unclassified(67); | - | - | - | - | - | - | - | Unassigned | - |
| Otu08347 | 3 | k__Fungi(100);k__Fungi_unclassified(67);k__Fungi_unclassified(67);k__Fungi_unclassified(67);k__Fungi_unclassified(67);k__Fungi_unclassified(67);k__Fungi_unclassified(67); | - | - | - | - | - | - | - | Unassigned | - |
| Otu08350 | 3 | k__Fungi(100);p__Ascomycota(100);p__Ascomycota_unclassified(100);p__Ascomycota_unclassified(100);p__Ascomycota_unclassified(100);p__Ascomycota_unclassified(100);p__Ascomycota_unclassified(100); | - | - | - | - | - | - | - | Unassigned | - |
| Otu08351 | 3 | k__Fungi(100);p__Ascomycota(67);p__Ascomycota_unclassified(67);p__Ascomycota_unclassified(67);p__Ascomycota_unclassified(67);p__Ascomycota_unclassified(67);p__Ascomycota_unclassified(67); | - | - | - | - | - | - | - | Unassigned | - |
| Otu08352 | 3 | k__Fungi(100);k__Fungi_unclassified(100);k__Fungi_unclassified(100);k__Fungi_unclassified(100);k__Fungi_unclassified(100);k__Fungi_unclassified(100);k__Fungi_unclassified(100); | - | - | - | - | - | - | - | Unassigned | - |
| Otu08355 | 3 | k__Fungi(100);k__Fungi_unclassified(100);k__Fungi_unclassified(100);k__Fungi_unclassified(100);k__Fungi_unclassified(100);k__Fungi_unclassified(100);k__Fungi_unclassified(100); | - | - | - | - | - | - | - | Unassigned | - |
| Otu08356 | 3 | k__Fungi(100);k__Fungi_unclassified(67);k__Fungi_unclassified(67);k__Fungi_unclassified(67);k__Fungi_unclassified(67);k__Fungi_unclassified(67);k__Fungi_unclassified(67); | - | - | - | - | - | - | - | Unassigned | - |
| Otu08360 | 3 | k__Fungi(100);p__Ascomycota(67);p__Ascomycota_unclassified(67);p__Ascomycota_unclassified(67);p__Ascomycota_unclassified(67);p__Ascomycota_unclassified(67);p__Ascomycota_unclassified(67); | - | - | - | - | - | - | - | Unassigned | - |
| Otu08362 | 3 | k__Fungi(100);p__Ascomycota(100);p__Ascomycota_unclassified(67);p__Ascomycota_unclassified(67);p__Ascomycota_unclassified(67);p__Ascomycota_unclassified(67);p__Ascomycota_unclassified(67); | - | - | - | - | - | - | - | Unassigned | - |
| Otu08363 | 3 | k__Fungi(100);p__Ascomycota(100);p__Ascomycota_unclassified(100);p__Ascomycota_unclassified(100);p__Ascomycota_unclassified(100);p__Ascomycota_unclassified(100);p__Ascomycota_unclassified(100); | - | - | - | - | - | - | - | Unassigned | - |
| Otu08365 | 3 | k__Fungi(100);k__Fungi_unclassified(100);k__Fungi_unclassified(100);k__Fungi_unclassified(100);k__Fungi_unclassified(100);k__Fungi_unclassified(100);k__Fungi_unclassified(100); | - | - | - | - | - | - | - | Unassigned | - |
| Otu08367 | 3 | k__Fungi(100);k__Fungi_unclassified(100);k__Fungi_unclassified(100);k__Fungi_unclassified(100);k__Fungi_unclassified(100);k__Fungi_unclassified(100);k__Fungi_unclassified(100); | - | - | - | - | - | - | - | Unassigned | - |
| Otu08374 | 3 | k__Fungi(100);p__Ascomycota(100);p__Ascomycota_unclassified(100);p__Ascomycota_unclassified(100);p__Ascomycota_unclassified(100);p__Ascomycota_unclassified(100);p__Ascomycota_unclassified(100); | - | - | - | - | - | - | - | Unassigned | - |
| Otu08379 | 3 | k__Fungi(100);p__Ascomycota(67);p__Ascomycota_unclassified(67);p__Ascomycota_unclassified(67);p__Ascomycota_unclassified(67);p__Ascomycota_unclassified(67);p__Ascomycota_unclassified(67); | - | - | - | - | - | - | - | Unassigned | - |
| Otu08382 | 3 | k__Fungi(100);k__Fungi_unclassified(100);k__Fungi_unclassified(100);k__Fungi_unclassified(100);k__Fungi_unclassified(100);k__Fungi_unclassified(100);k__Fungi_unclassified(100); | - | - | - | - | - | - | - | Unassigned | - |
| Otu08386 | 3 | k__Fungi(100);k__Fungi_unclassified(67);k__Fungi_unclassified(67);k__Fungi_unclassified(67);k__Fungi_unclassified(67);k__Fungi_unclassified(67);k__Fungi_unclassified(67); | - | - | - | - | - | - | - | Unassigned | - |
| Otu08387 | 3 | k__Fungi(100);p__Ascomycota(67);p__Ascomycota_unclassified(67);p__Ascomycota_unclassified(67);p__Ascomycota_unclassified(67);p__Ascomycota_unclassified(67);p__Ascomycota_unclassified(67); | - | - | - | - | - | - | - | Unassigned | - |
| Otu08390 | 3 | k__Fungi(100);k__Fungi_unclassified(100);k__Fungi_unclassified(100);k__Fungi_unclassified(100);k__Fungi_unclassified(100);k__Fungi_unclassified(100);k__Fungi_unclassified(100); | - | - | - | - | - | - | - | Unassigned | - |
| Otu08393 | 3 | k__Fungi(100);p__Ascomycota(100);c__Dothideomycetes(100);o__Pleosporales(100);f__Pleosporales_family_Incertae_sedis(100);f__Pleosporales_family_Incertae_sedis_unclassified(100);f__Pleosporales_family_Incertae_sedis_unclassified(100); | - | - | - | - | - | - | - | Unassigned | - |
| Otu08394 | 3 | k__Fungi(100);p__Ascomycota(100);c__Dothideomycetes(100);o__Pleosporales(100);o__Pleosporales_unclassified(100);o__Pleosporales_unclassified(100);o__Pleosporales_unclassified(100); | - | - | - | - | - | - | - | Unassigned | - |
| Otu08395 | 3 | k__Fungi(100);k__Fungi_unclassified(100);k__Fungi_unclassified(100);k__Fungi_unclassified(100);k__Fungi_unclassified(100);k__Fungi_unclassified(100);k__Fungi_unclassified(100); | - | - | - | - | - | - | - | Unassigned | - |
| Otu08396 | 3 | k__Fungi(100);p__Ascomycota(100);c__Sordariomycetes(100);o__Hypocreales(100);f__Hypocreaceae(100);f__Hypocreaceae_unclassified(100);f__Hypocreaceae_unclassified(100); | - | - | - | - | - | - | - | Unassigned | - |

| | | | | | | | | | | | |
|---|---|---|---|---|---|---|---|---|---|---|---|
| Otu08398 | 3 | k__Fungi(100);p__Ascomycota(100);p__Ascomycota_unclassified(100);p__Ascomycota_unclassified(100);p__Ascomycota_unclassified(100);p__Ascomycota_unclassified(100);p__Ascomycota_unclassified(100); | - | - | - | - | - | - | - | Unassigned | - |
| Otu08402 | 3 | k__Fungi(100);p__Ascomycota(67);p__Ascomycota_unclassified(67);p__Ascomycota_unclassified(67);p__Ascomycota_unclassified(67);p__Ascomycota_unclassified(67);p__Ascomycota_unclassified(67); | - | - | - | - | - | - | - | Unassigned | - |
| Otu08404 | 3 | k__Fungi(100);p__Ascomycota(100);c__Sordariomycetes(100);o__Hypocreales(100);f__Hypocreaceae(100);f__Hypocreaceae_unclassified(100);f__Hypocreaceae_unclassified(100); | - | - | - | - | - | - | - | Unassigned | - |
| Otu08410 | 3 | k__Fungi(100);k__Fungi_unclassified(100);k__Fungi_unclassified(100);k__Fungi_unclassified(100);k__Fungi_unclassified(100);k__Fungi_unclassified(100);k__Fungi_unclassified(100); | - | - | - | - | - | - | - | Unassigned | - |
| Otu08411 | 3 | k__Fungi(100);p__Basidiomycota(100);c__Agaricomycetes(100);o__Boletales(100);f__Suillaceae(100);f__Suillaceae_unclassified(100);f__Suillaceae_unclassified(100); | - | - | - | - | - | - | - | Unassigned | - |
| Otu08412 | 3 | k__Fungi(100);k__Fungi_unclassified(100);k__Fungi_unclassified(100);k__Fungi_unclassified(100);k__Fungi_unclassified(100);k__Fungi_unclassified(100);k__Fungi_unclassified(100); | - | - | - | - | - | - | - | Unassigned | - |
| Otu08413 | 3 | k__Fungi(100);p__Ascomycota(67);p__Ascomycota_unclassified(67);p__Ascomycota_unclassified(67);p__Ascomycota_unclassified(67);p__Ascomycota_unclassified(67);p__Ascomycota_unclassified(67); | - | - | - | - | - | - | - | Unassigned | - |
| Otu08416 | 3 | k__Fungi(100);p__Ascomycota(100);p__Ascomycota_unclassified(100);p__Ascomycota_unclassified(100);p__Ascomycota_unclassified(100);p__Ascomycota_unclassified(100);p__Ascomycota_unclassified(100); | - | - | - | - | - | - | - | Unassigned | - |
| Otu08417 | 3 | k__Fungi(100);p__Basidiomycota(100);c__Agaricomycetes(100);c__Agaricomycetes_unclassified(100);c__Agaricomycetes_unclassified(100);c__Agaricomycetes_unclassified(100);c__Agaricomycetes_unclassified(100); | - | - | - | - | - | - | - | Unassigned | - |
| Otu08419 | 3 | k__Fungi(100);p__Basidiomycota(100);c__Agaricomycetes(100);c__Agaricomycetes_unclassified(100);c__Agaricomycetes_unclassified(100);c__Agaricomycetes_unclassified(100);c__Agaricomycetes_unclassified(100); | - | - | - | - | - | - | - | Unassigned | - |
| Otu08420 | 3 | k__Fungi(100);k__Fungi_unclassified(100);k__Fungi_unclassified(100);k__Fungi_unclassified(100);k__Fungi_unclassified(100);k__Fungi_unclassified(100);k__Fungi_unclassified(100); | - | - | - | - | - | - | - | Unassigned | - |
| Otu08425 | 3 | k__Fungi(100);p__Ascomycota(100);c__Sordariomycetes(100);o__Xylariales(67);o__Xylariales_unclassified(67);o__Xylariales_unclassified(67);o__Xylariales_unclassified(67); | - | - | - | - | - | - | - | Unassigned | - |
| Otu08426 | 3 | k__Fungi(100);k__Fungi_unclassified(100);k__Fungi_unclassified(100);k__Fungi_unclassified(100);k__Fungi_unclassified(100);k__Fungi_unclassified(100);k__Fungi_unclassified(100); | - | - | - | - | - | - | - | Unassigned | - |
| Otu08427 | 3 | k__Fungi(100);p__Ascomycota(100);p__Ascomycota_unclassified(100);p__Ascomycota_unclassified(100);p__Ascomycota_unclassified(100);p__Ascomycota_unclassified(100);p__Ascomycota_unclassified(100); | - | - | - | - | - | - | - | Unassigned | - |
| Otu08428 | 3 | k__Fungi(100);k__Fungi_unclassified(100);k__Fungi_unclassified(100);k__Fungi_unclassified(100);k__Fungi_unclassified(100);k__Fungi_unclassified(100);k__Fungi_unclassified(100); | - | - | - | - | - | - | - | Unassigned | - |
| Otu08431 | 3 | k__Fungi(100);p__Ascomycota(100);p__Ascomycota_unclassified(100);p__Ascomycota_unclassified(100);p__Ascomycota_unclassified(100);p__Ascomycota_unclassified(100);p__Ascomycota_unclassified(100); | - | - | - | - | - | - | - | Unassigned | - |
| Otu08433 | 3 | k__Fungi(100);p__Ascomycota(100);p__Ascomycota_unclassified(100);p__Ascomycota_unclassified(100);p__Ascomycota_unclassified(100);p__Ascomycota_unclassified(100);p__Ascomycota_unclassified(100); | - | - | - | - | - | - | - | Unassigned | - |
| Otu08434 | 3 | k__Fungi(100);p__Ascomycota(100);p__Ascomycota_unclassified(100);p__Ascomycota_unclassified(100);p__Ascomycota_unclassified(100);p__Ascomycota_unclassified(100);p__Ascomycota_unclassified(100); | - | - | - | - | - | - | - | Unassigned | - |
| Otu08438 | 3 | k__Fungi(100);p__Ascomycota(100);p__Ascomycota_unclassified(67);p__Ascomycota_unclassified(67);p__Ascomycota_unclassified(67);p__Ascomycota_unclassified(67);p__Ascomycota_unclassified(67); | - | - | - | - | - | - | - | Unassigned | - |
| Otu08440 | 3 | k__Fungi(100);k__Fungi_unclassified(67);k__Fungi_unclassified(67);k__Fungi_unclassified(67);k__Fungi_unclassified(67);k__Fungi_unclassified(67);k__Fungi_unclassified(67); | - | - | - | - | - | - | - | Unassigned | - |
| Otu08443 | 3 | k__Fungi(100);k__Fungi_unclassified(67);k__Fungi_unclassified(67);k__Fungi_unclassified(67);k__Fungi_unclassified(67);k__Fungi_unclassified(67);k__Fungi_unclassified(67); | - | - | - | - | - | - | - | Unassigned | - |
| Otu08444 | 3 | k__Fungi(100);p__Ascomycota(100);p__Ascomycota_unclassified(100);p__Ascomycota_unclassified(100);p__Ascomycota_unclassified(100);p__Ascomycota_unclassified(100);p__Ascomycota_unclassified(100); | - | - | - | - | - | - | - | Unassigned | - |
| Otu08450 | 3 | k__Fungi(100);p__Ascomycota(100);c__Dothideomycetes(100);o__Pleosporales(100);o__Pleosporales_unclassified(100);o__Pleosporales_unclassified(100);o__Pleosporales_unclassified(100); | - | - | - | - | - | - | - | Unassigned | - |
| Otu08451 | 3 | k__Fungi(100);k__Fungi_unclassified(67);k__Fungi_unclassified(67);k__Fungi_unclassified(67);k__Fungi_unclassified(67);k__Fungi_unclassified(67);k__Fungi_unclassified(67); | - | - | - | - | - | - | - | Unassigned | - |
| Otu08452 | 3 | k__Fungi(100);p__Ascomycota(100);p__Ascomycota_unclassified(100);p__Ascomycota_unclassified(100);p__Ascomycota_unclassified(100);p__Ascomycota_unclassified(100);p__Ascomycota_unclassified(100); | - | - | - | - | - | - | - | Unassigned | - |
| Otu08454 | 3 | k__Fungi(100);p__Ascomycota(100);p__Ascomycota_unclassified(100);p__Ascomycota_unclassified(100);p__Ascomycota_unclassified(100);p__Ascomycota_unclassified(100);p__Ascomycota_unclassified(100); | - | - | - | - | - | - | - | Unassigned | - |
| Otu08458 | 3 | k__Fungi(100);k__Fungi_unclassified(100);k__Fungi_unclassified(100);k__Fungi_unclassified(100);k__Fungi_unclassified(100);k__Fungi_unclassified(100);k__Fungi_unclassified(100); | - | - | - | - | - | - | - | Unassigned | - |
| Otu08460 | 3 | k__Fungi(100);k__Fungi_unclassified(67);k__Fungi_unclassified(67);k__Fungi_unclassified(67);k__Fungi_unclassified(67);k__Fungi_unclassified(67);k__Fungi_unclassified(67); | - | - | - | - | - | - | - | Unassigned | - |
| Otu08461 | 3 | k__Fungi(100);k__Fungi_unclassified(100);k__Fungi_unclassified(100);k__Fungi_unclassified(100);k__Fungi_unclassified(100);k__Fungi_unclassified(100);k__Fungi_unclassified(100); | - | - | - | - | - | - | - | Unassigned | - |
| Otu08462 | 3 | k__Fungi(100);p__Ascomycota(100);p__Ascomycota_unclassified(67);p__Ascomycota_unclassified(67);p__Ascomycota_unclassified(67);p__Ascomycota_unclassified(67);p__Ascomycota_unclassified(67); | - | - | - | - | - | - | - | Unassigned | - |
| Otu08463 | 3 | k__Fungi(100);k__Fungi_unclassified(67);k__Fungi_unclassified(67);k__Fungi_unclassified(67);k__Fungi_unclassified(67);k__Fungi_unclassified(67);k__Fungi_unclassified(67); | - | - | - | - | - | - | - | Unassigned | - |
| Otu08466 | 3 | k__Fungi(100);p__Ascomycota(67);p__Ascomycota_unclassified(67);p__Ascomycota_unclassified(67);p__Ascomycota_unclassified(67);p__Ascomycota_unclassified(67);p__Ascomycota_unclassified(67); | - | - | - | - | - | - | - | Unassigned | - |
| Otu08471 | 3 | k__Fungi(100);p__Ascomycota(67);p__Ascomycota_unclassified(67);p__Ascomycota_unclassified(67);p__Ascomycota_unclassified(67);p__Ascomycota_unclassified(67);p__Ascomycota_unclassified(67); | - | - | - | - | - | - | - | Unassigned | - |
| Otu08473 | 3 | k__Fungi(100);k__Fungi_unclassified(67);k__Fungi_unclassified(67);k__Fungi_unclassified(67);k__Fungi_unclassified(67);k__Fungi_unclassified(67);k__Fungi_unclassified(67); | - | - | - | - | - | - | - | Unassigned | - |
| Otu08474 | 3 | k__Fungi(100);p__Ascomycota(67);p__Ascomycota_unclassified(67);p__Ascomycota_unclassified(67);p__Ascomycota_unclassified(67);p__Ascomycota_unclassified(67);p__Ascomycota_unclassified(67); | - | - | - | - | - | - | - | Unassigned | - |
| Otu08475 | 3 | k__Fungi(100);k__Fungi_unclassified(100);k__Fungi_unclassified(100);k__Fungi_unclassified(100);k__Fungi_unclassified(100);k__Fungi_unclassified(100);k__Fungi_unclassified(100); | - | - | - | - | - | - | - | Unassigned | - |
| Otu08477 | 3 | k__Fungi(100);p__Ascomycota(67);p__Ascomycota_unclassified(67);p__Ascomycota_unclassified(67);p__Ascomycota_unclassified(67);p__Ascomycota_unclassified(67);p__Ascomycota_unclassified(67); | - | - | - | - | - | - | - | Unassigned | - |
| Otu08480 | 3 | k__Fungi(100);p__Ascomycota(100);p__Ascomycota_unclassified(100);p__Ascomycota_unclassified(100);p__Ascomycota_unclassified(100);p__Ascomycota_unclassified(100);p__Ascomycota_unclassified(100); | - | - | - | - | - | - | - | Unassigned | - |
| Otu08484 | 3 | k__Fungi(100);p__Ascomycota(100);p__Ascomycota_unclassified(100);p__Ascomycota_unclassified(100);p__Ascomycota_unclassified(100);p__Ascomycota_unclassified(100);p__Ascomycota_unclassified(100); | - | - | - | - | - | - | - | Unassigned | - |
| Otu08486 | 3 | k__Fungi(100);p__Ascomycota(100);p__Ascomycota_unclassified(100);p__Ascomycota_unclassified(100);p__Ascomycota_unclassified(100);p__Ascomycota_unclassified(100);p__Ascomycota_unclassified(100); | - | - | - | - | - | - | - | Unassigned | - |
| Otu08493 | 3 | k__Fungi(100);p__Ascomycota(100);c__Dothideomycetes(100);o__Myriangiales(100);f__unclassified_Myriangiales(100);g__unclassified_Myriangiales(100);s__Myriangiales_sp(100); | - | - | - | - | - | - | - | Unassigned | - |
| Otu08494 | 3 | k__Fungi(100);p__Ascomycota(100);c__Sordariomycetes(100);o__Sordariales(100);o__Sordariales_unclassified(100);o__Sordariales_unclassified(100);o__Sordariales_unclassified(100); | - | - | - | - | - | - | - | Unassigned | - |
| Otu08495 | 3 | k__Fungi(100);p__Ascomycota(100);c__Sordariomycetes(100);c__Sordariomycetes_unclassified(100);c__Sordariomycetes_unclassified(100);c__Sordariomycetes_unclassified(100);c__Sordariomycetes_unclassified(100); | - | - | - | - | - | - | - | Unassigned | - |
| Otu08496 | 3 | k__Fungi(100);k__Fungi_unclassified(100);k__Fungi_unclassified(100);k__Fungi_unclassified(100);k__Fungi_unclassified(100);k__Fungi_unclassified(100);k__Fungi_unclassified(100); | - | - | - | - | - | - | - | Unassigned | - |
| Otu08498 | 3 | k__Fungi(100);k__Fungi_unclassified(100);k__Fungi_unclassified(100);k__Fungi_unclassified(100);k__Fungi_unclassified(100);k__Fungi_unclassified(100);k__Fungi_unclassified(100); | - | - | - | - | - | - | - | Unassigned | - |
| Otu08499 | 3 | k__Fungi(100);k__Fungi_unclassified(67);k__Fungi_unclassified(67);k__Fungi_unclassified(67);k__Fungi_unclassified(67);k__Fungi_unclassified(67);k__Fungi_unclassified(67); | - | - | - | - | - | - | - | Unassigned | - |
| Otu08500 | 3 | k__Fungi(100);p__Basidiomycota(100);c__Agaricomycetes(100);o__Polyporales(100);o__Polyporales_unclassified(100);o__Polyporales_unclassified(100);o__Polyporales_unclassified(100); | - | - | - | - | - | - | - | Unassigned | - |
| Otu08502 | 3 | k__Fungi(100);k__Fungi_unclassified(100);k__Fungi_unclassified(100);k__Fungi_unclassified(100);k__Fungi_unclassified(100);k__Fungi_unclassified(100);k__Fungi_unclassified(100); | - | - | - | - | - | - | - | Unassigned | - |
| Otu08505 | 3 | k__Fungi(100);p__Basidiomycota(100);c__Agaricomycetes(100);c__Agaricomycetes_unclassified(100);c__Agaricomycetes_unclassified(100);c__Agaricomycetes_unclassified(100);c__Agaricomycetes_unclassified(100); | - | - | - | - | - | - | - | Unassigned | - |
| Otu08507 | 3 | k__Plantae(100);p__unclassified_Plantae(100);c__unclassified_Plantae(100);o__unclassified_Plantae(100);f__unclassified_Plantae(100);g__unclassified_Plantae(100);s__Plantae_sp(100); | - | - | - | - | - | - | - | Unassigned | - |
| Otu08511 | 3 | k__Fungi(100);p__Ascomycota(100);p__Ascomycota_unclassified(100);p__Ascomycota_unclassified(100);p__Ascomycota_unclassified(100);p__Ascomycota_unclassified(100);p__Ascomycota_unclassified(100); | - | - | - | - | - | - | - | Unassigned | - |
| Otu08512 | 3 | k__Fungi(100);p__Ascomycota(100);p__Ascomycota_unclassified(100);p__Ascomycota_unclassified(100);p__Ascomycota_unclassified(100);p__Ascomycota_unclassified(100);p__Ascomycota_unclassified(100); | - | - | - | - | - | - | - | Unassigned | - |
| Otu08516 | 3 | k__Fungi(100);k__Fungi_unclassified(100);k__Fungi_unclassified(100);k__Fungi_unclassified(100);k__Fungi_unclassified(100);k__Fungi_unclassified(100);k__Fungi_unclassified(100); | - | - | - | - | - | - | - | Unassigned | - |
| Otu08520 | 3 | k__Fungi(100);p__Basidiomycota(100);p__Basidiomycota_unclassified(100);p__Basidiomycota_unclassified(100);p__Basidiomycota_unclassified(100);p__Basidiomycota_unclassified(100);p__Basidiomycota_unclassified(100); | - | - | - | - | - | - | - | Unassigned | - |
| Otu08524 | 3 | k__Fungi(100);p__Ascomycota(100);p__Ascomycota_unclassified(67);p__Ascomycota_unclassified(67);p__Ascomycota_unclassified(67);p__Ascomycota_unclassified(67);p__Ascomycota_unclassified(67); | - | - | - | - | - | - | - | Unassigned | - |
| Otu08525 | 3 | k__Fungi(100);k__Fungi_unclassified(67);k__Fungi_unclassified(67);k__Fungi_unclassified(67);k__Fungi_unclassified(67);k__Fungi_unclassified(67);k__Fungi_unclassified(67); | - | - | - | - | - | - | - | Unassigned | - |
| Otu08527 | 3 | k__Fungi(100);p__Ascomycota(100);p__Ascomycota_unclassified(100);p__Ascomycota_unclassified(100);p__Ascomycota_unclassified(100);p__Ascomycota_unclassified(100);p__Ascomycota_unclassified(100); | - | - | - | - | - | - | - | Unassigned | - |
| Otu08528 | 3 | k__Fungi(100);p__Ascomycota(100);p__Ascomycota_unclassified(100);p__Ascomycota_unclassified(100);p__Ascomycota_unclassified(100);p__Ascomycota_unclassified(100);p__Ascomycota_unclassified(100); | - | - | - | - | - | - | - | Unassigned | - |
| Otu08531 | 3 | k__Fungi(100);k__Fungi_unclassified(100);k__Fungi_unclassified(100);k__Fungi_unclassified(100);k__Fungi_unclassified(100);k__Fungi_unclassified(100);k__Fungi_unclassified(100); | - | - | - | - | - | - | - | Unassigned | - |
| Otu08533 | 3 | k__Fungi(100);p__Ascomycota(67);p__Ascomycota_unclassified(67);p__Ascomycota_unclassified(67);p__Ascomycota_unclassified(67);p__Ascomycota_unclassified(67);p__Ascomycota_unclassified(67); | - | - | - | - | - | - | - | Unassigned | - |
| Otu08534 | 3 | k__Fungi(100);k__Fungi_unclassified(100);k__Fungi_unclassified(100);k__Fungi_unclassified(100);k__Fungi_unclassified(100);k__Fungi_unclassified(100);k__Fungi_unclassified(100); | - | - | - | - | - | - | - | Unassigned | - |
| Otu08536 | 3 | k__Fungi(100);p__Ascomycota(100);c__Sordariomycetes(100);c__Sordariomycetes_unclassified(100);c__Sordariomycetes_unclassified(100);c__Sordariomycetes_unclassified(100);c__Sordariomycetes_unclassified(100); | - | - | - | - | - | - | - | Unassigned | - |
| Otu08537 | 3 | k__Fungi(100);k__Fungi_unclassified(67);k__Fungi_unclassified(67);k__Fungi_unclassified(67);k__Fungi_unclassified(67);k__Fungi_unclassified(67);k__Fungi_unclassified(67); | - | - | - | - | - | - | - | Unassigned | - |
| Otu08538 | 3 | k__Fungi(100);k__Fungi_unclassified(100);k__Fungi_unclassified(100);k__Fungi_unclassified(100);k__Fungi_unclassified(100);k__Fungi_unclassified(100);k__Fungi_unclassified(100); | - | - | - | - | - | - | - | Unassigned | - |
| Otu08539 | 3 | k__Fungi(100);k__Fungi_unclassified(100);k__Fungi_unclassified(100);k__Fungi_unclassified(100);k__Fungi_unclassified(100);k__Fungi_unclassified(100);k__Fungi_unclassified(100); | - | - | - | - | - | - | - | Unassigned | - |
| Otu08540 | 3 | k__Fungi(100);k__Fungi_unclassified(67);k__Fungi_unclassified(67);k__Fungi_unclassified(67);k__Fungi_unclassified(67);k__Fungi_unclassified(67);k__Fungi_unclassified(67); | - | - | - | - | - | - | - | Unassigned | - |
| Otu08542 | 3 | k__Fungi(100);p__Ascomycota(100);p__Ascomycota_unclassified(100);p__Ascomycota_unclassified(100);p__Ascomycota_unclassified(100);p__Ascomycota_unclassified(100);p__Ascomycota_unclassified(100); | - | - | - | - | - | - | - | Unassigned | - |
| Otu08543 | 3 | k__Fungi(100);p__Ascomycota(67);p__Ascomycota_unclassified(67);p__Ascomycota_unclassified(67);p__Ascomycota_unclassified(67);p__Ascomycota_unclassified(67);p__Ascomycota_unclassified(67); | - | - | - | - | - | - | - | Unassigned | - |
| Otu08544 | 3 | k__Fungi(100);p__Ascomycota(67);p__Ascomycota_unclassified(67);p__Ascomycota_unclassified(67);p__Ascomycota_unclassified(67);p__Ascomycota_unclassified(67);p__Ascomycota_unclassified(67); | - | - | - | - | - | - | - | Unassigned | - |
| Otu08545 | 3 | k__Fungi(100);k__Fungi_unclassified(100);k__Fungi_unclassified(100);k__Fungi_unclassified(100);k__Fungi_unclassified(100);k__Fungi_unclassified(100);k__Fungi_unclassified(100); | - | - | - | - | - | - | - | Unassigned | - |
| Otu08547 | 3 | k__Fungi(100);p__Ascomycota(100);p__Ascomycota_unclassified(100);p__Ascomycota_unclassified(100);p__Ascomycota_unclassified(100);p__Ascomycota_unclassified(100);p__Ascomycota_unclassified(100); | - | - | - | - | - | - | - | Unassigned | - |
| Otu08549 | 3 | k__Fungi(100);p__Basidiomycota(100);c__Agaricomycetes(100);o__Auriculariales(100);f__Auriculariales_family_Incertae_sedis(100);f__Auriculariales_family_Incertae_sedis_unclassified(100);f__Auriculariales_family_Incertae_sedis_unclassified(100); | - | - | - | - | - | - | - | Unassigned | - |
| Otu08553 | 3 | k__Fungi(100);p__Ascomycota(100);c__Eurotiomycetes(100);c__Eurotiomycetes_unclassified(100);c__Eurotiomycetes_unclassified(100);c__Eurotiomycetes_unclassified(100);c__Eurotiomycetes_unclassified(100); | - | - | - | - | - | - | - | Unassigned | - |
| Otu08555 | 3 | k__Fungi(100);k__Fungi_unclassified(67);k__Fungi_unclassified(67);k__Fungi_unclassified(67);k__Fungi_unclassified(67);k__Fungi_unclassified(67);k__Fungi_unclassified(67); | - | - | - | - | - | - | - | Unassigned | - |
| Otu08559 | 3 | k__Fungi(100);p__Ascomycota(100);p__Ascomycota_unclassified(100);p__Ascomycota_unclassified(100);p__Ascomycota_unclassified(100);p__Ascomycota_unclassified(100);p__Ascomycota_unclassified(100); | - | - | - | - | - | - | - | Unassigned | - |
| Otu08560 | 3 | k__Fungi(100);p__Basidiomycota(67);p__Basidiomycota_unclassified(67);p__Basidiomycota_unclassified(67);p__Basidiomycota_unclassified(67);p__Basidiomycota_unclassified(67);p__Basidiomycota_unclassified(67); | - | - | - | - | - | - | - | Unassigned | - |
| Otu08565 | 3 | k__Fungi(100);p__Ascomycota(100);c__Sordariomycetes(100);o__Xylariales(67);o__Xylariales_unclassified(67);o__Xylariales_unclassified(67);o__Xylariales_unclassified(67); | - | - | - | - | - | - | - | Unassigned | - |
| Otu08568 | 3 | k__Fungi(100);p__Ascomycota(100);c__Dothideomycetes(100);o__Pleosporales(100);f__Pleosporales_family_Incertae_sedis(100);f__Pleosporales_family_Incertae_sedis_unclassified(100);f__Pleosporales_family_Incertae_sedis_unclassified(100); | - | - | - | - | - | - | - | Unassigned | - |
| Otu08569 | 3 | k__Fungi(100);p__Ascomycota(100);p__Ascomycota_unclassified(100);p__Ascomycota_unclassified(100);p__Ascomycota_unclassified(100);p__Ascomycota_unclassified(100);p__Ascomycota_unclassified(100); | - | - | - | - | - | - | - | Unassigned | - |
| Otu08572 | 3 | k__Fungi(100);k__Fungi_unclassified(100);k__Fungi_unclassified(100);k__Fungi_unclassified(100);k__Fungi_unclassified(100);k__Fungi_unclassified(100);k__Fungi_unclassified(100); | - | - | - | - | - | - | - | Unassigned | - |
| Otu08574 | 3 | k__Fungi(100);k__Fungi_unclassified(67);k__Fungi_unclassified(67);k__Fungi_unclassified(67);k__Fungi_unclassified(67);k__Fungi_unclassified(67);k__Fungi_unclassified(67); | - | - | - | - | - | - | - | Unassigned | - |
| Otu08577 | 3 | k__Fungi(100);k__Fungi_unclassified(100);k__Fungi_unclassified(100);k__Fungi_unclassified(100);k__Fungi_unclassified(100);k__Fungi_unclassified(100);k__Fungi_unclassified(100); | - | - | - | - | - | - | - | Unassigned | - |
| Otu08578 | 3 | k__Fungi(100);p__Ascomycota(100);c__Sordariomycetes(100);c__Sordariomycetes_unclassified(100);c__Sordariomycetes_unclassified(100);c__Sordariomycetes_unclassified(100);c__Sordariomycetes_unclassified(100); | - | - | - | - | - | - | - | Unassigned | - |
| Otu08582 | 3 | k__Fungi(100);p__Ascomycota(100);p__Ascomycota_unclassified(100);p__Ascomycota_unclassified(100);p__Ascomycota_unclassified(100);p__Ascomycota_unclassified(100);p__Ascomycota_unclassified(100); | - | - | - | - | - | - | - | Unassigned | - |
| Otu08585 | 3 | k__Fungi(100);p__Ascomycota(67);p__Ascomycota_unclassified(67);p__Ascomycota_unclassified(67);p__Ascomycota_unclassified(67);p__Ascomycota_unclassified(67);p__Ascomycota_unclassified(67); | - | - | - | - | - | - | - | Unassigned | - |
| Otu08587 | 3 | k__Fungi(100);p__Ascomycota(67);p__Ascomycota_unclassified(67);p__Ascomycota_unclassified(67);p__Ascomycota_unclassified(67);p__Ascomycota_unclassified(67);p__Ascomycota_unclassified(67); | - | - | - | - | - | - | - | Unassigned | - |
| Otu08590 | 3 | k__Plantae(100);p__unclassified_Plantae(100);c__unclassified_Plantae(100);o__unclassified_Plantae(100);f__unclassified_Plantae(100);g__unclassified_Plantae(100);s__Plantae_sp(100); | - | - | - | - | - | - | - | Unassigned | - |
| Otu08592 | 3 | k__Fungi(100);p__Ascomycota(100);c__Dothideomycetes(100);o__Pleosporales(100);o__Pleosporales_unclassified(100);o__Pleosporales_unclassified(100);o__Pleosporales_unclassified(100); | - | - | - | - | - | - | - | Unassigned | - |
| Otu08594 | 3 | k__Fungi(100);p__Ascomycota(100);p__Ascomycota_unclassified(100);p__Ascomycota_unclassified(100);p__Ascomycota_unclassified(100);p__Ascomycota_unclassified(100);p__Ascomycota_unclassified(100); | - | - | - | - | - | - | - | Unassigned | - |
| Otu08596 | 3 | k__Fungi(100);p__Ascomycota(100);c__Leotiomycetes(100);o__Helotiales(100);o__Helotiales_unclassified(100);o__Helotiales_unclassified(100);o__Helotiales_unclassified(100); | - | - | - | - | - | - | - | Unassigned | - |
| Otu08597 | 3 | k__Fungi(100);p__Ascomycota(100);p__Ascomycota_unclassified(100);p__Ascomycota_unclassified(100);p__Ascomycota_unclassified(100);p__Ascomycota_unclassified(100);p__Ascomycota_unclassified(100); | - | - | - | - | - | - | - | Unassigned | - |
| Otu08598 | 3 | k__Fungi(100);k__Fungi_unclassified(67);k__Fungi_unclassified(67);k__Fungi_unclassified(67);k__Fungi_unclassified(67);k__Fungi_unclassified(67);k__Fungi_unclassified(67); | - | - | - | - | - | - | - | Unassigned | - |
| Otu08600 | 3 | k__Plantae(100);p__unclassified_Plantae(100);c__unclassified_Plantae(100);o__unclassified_Plantae(100);f__unclassified_Plantae(100);g__unclassified_Plantae(100);s__Plantae_sp(100); | - | - | - | - | - | - | - | Unassigned | - |
| Otu08603 | 3 | k__Fungi(100);p__Ascomycota(100);p__Ascomycota_unclassified(100);p__Ascomycota_unclassified(100);p__Ascomycota_unclassified(100);p__Ascomycota_unclassified(100);p__Ascomycota_unclassified(100); | - | - | - | - | - | - | - | Unassigned | - |
| Otu08605 | 3 | k__Plantae(100);p__unclassified_Plantae(100);c__unclassified_Plantae(100);o__unclassified_Plantae(100);f__unclassified_Plantae(100);g__unclassified_Plantae(100);s__Plantae_sp(100); | - | - | - | - | - | - | - | Unassigned | - |
| Otu08607 | 3 | k__Fungi(100);k__Fungi_unclassified(100);k__Fungi_unclassified(100);k__Fungi_unclassified(100);k__Fungi_unclassified(100);k__Fungi_unclassified(100);k__Fungi_unclassified(100); | - | - | - | - | - | - | - | Unassigned | - |
| Otu08609 | 3 | k__Fungi(100);p__Ascomycota(67);p__Ascomycota_unclassified(67);p__Ascomycota_unclassified(67);p__Ascomycota_unclassified(67);p__Ascomycota_unclassified(67);p__Ascomycota_unclassified(67); | - | - | - | - | - | - | - | Unassigned | - |
| Otu08612 | 3 | k__Fungi(100);p__Basidiomycota(100);p__Basidiomycota_unclassified(100);p__Basidiomycota_unclassified(100);p__Basidiomycota_unclassified(100);p__Basidiomycota_unclassified(100);p__Basidiomycota_unclassified(100); | - | - | - | - | - | - | - | Unassigned | - |
| Otu08613 | 3 | k__Fungi(100);p__Ascomycota(100);p__Ascomycota_unclassified(100);p__Ascomycota_unclassified(100);p__Ascomycota_unclassified(100);p__Ascomycota_unclassified(100);p__Ascomycota_unclassified(100); | - | - | - | - | - | - | - | Unassigned | - |
| Otu08615 | 3 | k__Fungi(100);p__Ascomycota(100);c__Dothideomycetes(100);c__Dothideomycetes_unclassified(100);c__Dothideomycetes_unclassified(100);c__Dothideomycetes_unclassified(100);c__Dothideomycetes_unclassified(100); | - | - | - | - | - | - | - | Unassigned | - |
| Otu08618 | 3 | k__Plantae(100);p__unclassified_Plantae(100);c__unclassified_Plantae(100);o__unclassified_Plantae(100);f__unclassified_Plantae(100);g__unclassified_Plantae(100);s__Plantae_sp(100); | - | - | - | - | - | - | - | Unassigned | - |
| Otu08619 | 3 | k__Plantae(100);p__unclassified_Plantae(100);c__unclassified_Plantae(100);o__unclassified_Plantae(100);f__unclassified_Plantae(100);g__unclassified_Plantae(100);s__Plantae_sp(100); | - | - | - | - | - | - | - | Unassigned | - |
| Otu08620 | 3 | k__Fungi(100);k__Fungi_unclassified(100);k__Fungi_unclassified(100);k__Fungi_unclassified(100);k__Fungi_unclassified(100);k__Fungi_unclassified(100);k__Fungi_unclassified(100); | - | - | - | - | - | - | - | Unassigned | - |
| Otu08623 | 3 | k__Fungi(100);p__Ascomycota(67);p__Ascomycota_unclassified(67);p__Ascomycota_unclassified(67);p__Ascomycota_unclassified(67);p__Ascomycota_unclassified(67);p__Ascomycota_unclassified(67); | - | - | - | - | - | - | - | Unassigned | - |
| Otu08624 | 3 | k__Fungi(100);p__Ascomycota(67);p__Ascomycota_unclassified(67);p__Ascomycota_unclassified(67);p__Ascomycota_unclassified(67);p__Ascomycota_unclassified(67);p__Ascomycota_unclassified(67); | - | - | - | - | - | - | - | Unassigned | - |
| Otu08625 | 3 | k__Fungi(100);k__Fungi_unclassified(67);k__Fungi_unclassified(67);k__Fungi_unclassified(67);k__Fungi_unclassified(67);k__Fungi_unclassified(67);k__Fungi_unclassified(67); | - | - | - | - | - | - | - | Unassigned | - |
| Otu08626 | 3 | k__Fungi(100);p__Ascomycota(67);p__Ascomycota_unclassified(67);p__Ascomycota_unclassified(67);p__Ascomycota_unclassified(67);p__Ascomycota_unclassified(67);p__Ascomycota_unclassified(67); | - | - | - | - | - | - | - | Unassigned | - |
| Otu08631 | 3 | k__Fungi(100);k__Fungi_unclassified(100);k__Fungi_unclassified(100);k__Fungi_unclassified(100);k__Fungi_unclassified(100);k__Fungi_unclassified(100);k__Fungi_unclassified(100); | - | - | - | - | - | - | - | Unassigned | - |
| Otu08633 | 3 | k__Fungi(100);p__Ascomycota(67);p__Ascomycota_unclassified(67);p__Ascomycota_unclassified(67);p__Ascomycota_unclassified(67);p__Ascomycota_unclassified(67);p__Ascomycota_unclassified(67); | - | - | - | - | - | - | - | Unassigned | - |
| Otu08637 | 3 | k__Fungi(100);k__Fungi_unclassified(67);k__Fungi_unclassified(67);k__Fungi_unclassified(67);k__Fungi_unclassified(67);k__Fungi_unclassified(67);k__Fungi_unclassified(67); | - | - | - | - | - | - | - | Unassigned | - |
| Otu08639 | 3 | k__Fungi(100);k__Fungi_unclassified(67);k__Fungi_unclassified(67);k__Fungi_unclassified(67);k__Fungi_unclassified(67);k__Fungi_unclassified(67);k__Fungi_unclassified(67); | - | - | - | - | - | - | - | Unassigned | - |
| Otu08640 | 3 | k__Fungi(100);p__Ascomycota(100);c__Sordariomycetes(100);c__Sordariomycetes_unclassified(100);c__Sordariomycetes_unclassified(100);c__Sordariomycetes_unclassified(100);c__Sordariomycetes_unclassified(100); | - | - | - | - | - | - | - | Unassigned | - |
| Otu08641 | 3 | k__Fungi(100);p__Ascomycota(100);p__Ascomycota_unclassified(100);p__Ascomycota_unclassified(100);p__Ascomycota_unclassified(100);p__Ascomycota_unclassified(100);p__Ascomycota_unclassified(100); | - | - | - | - | - | - | - | Unassigned | - |
| Otu08642 | 3 | k__Fungi(100);p__Ascomycota(100);c__Dothideomycetes(100);o__Pleosporales(100);o__Pleosporales_unclassified(100);o__Pleosporales_unclassified(100);o__Pleosporales_unclassified(100); | - | - | - | - | - | - | - | Unassigned | - |
| Otu08647 | 3 | k__Plantae(100);p__unclassified_Plantae(100);c__unclassified_Plantae(100);o__unclassified_Plantae(100);f__unclassified_Plantae(100);g__unclassified_Plantae(100);s__Plantae_sp(100); | - | - | - | - | - | - | - | Unassigned | - |
| Otu08648 | 3 | k__Fungi(100);k__Fungi_unclassified(100);k__Fungi_unclassified(100);k__Fungi_unclassified(100);k__Fungi_unclassified(100);k__Fungi_unclassified(100);k__Fungi_unclassified(100); | - | - | - | - | - | - | - | Unassigned | - |
| Otu08652 | 3 | k__Fungi(100);p__Basidiomycota(100);c__Agaricomycetes(100);c__Agaricomycetes_unclassified(100);c__Agaricomycetes_unclassified(100);c__Agaricomycetes_unclassified(100);c__Agaricomycetes_unclassified(100); | - | - | - | - | - | - | - | Unassigned | - |
| Otu08657 | 3 | k__Fungi(100);p__Ascomycota(100);c__Dothideomycetes(100);o__Capnodiales(100);f__Davidiellaceae(100);f__Davidiellaceae_unclassified(67);f__Davidiellaceae_unclassified(67); | - | - | - | - | - | - | - | Unassigned | - |
| Otu08658 | 3 | k__Plantae(100);p__unclassified_Plantae(100);c__unclassified_Plantae(100);o__unclassified_Plantae(100);f__unclassified_Plantae(100);g__unclassified_Plantae(100);s__Plantae_sp(100); | - | - | - | - | - | - | - | Unassigned | - |
| Otu08660 | 3 | k__Fungi(100);k__Fungi_unclassified(100);k__Fungi_unclassified(100);k__Fungi_unclassified(100);k__Fungi_unclassified(100);k__Fungi_unclassified(100);k__Fungi_unclassified(100); | - | - | - | - | - | - | - | Unassigned | - |
| Otu08661 | 3 | k__Fungi(100);p__Ascomycota(67);p__Ascomycota_unclassified(67);p__Ascomycota_unclassified(67);p__Ascomycota_unclassified(67);p__Ascomycota_unclassified(67);p__Ascomycota_unclassified(67); | - | - | - | - | - | - | - | Unassigned | - |
| Otu08667 | 3 | k__Fungi(100);k__Fungi_unclassified(67);k__Fungi_unclassified(67);k__Fungi_unclassified(67);k__Fungi_unclassified(67);k__Fungi_unclassified(67);k__Fungi_unclassified(67); | - | - | - | - | - | - | - | Unassigned | - |
| Otu08669 | 3 | k__Fungi(100);k__Fungi_unclassified(100);k__Fungi_unclassified(100);k__Fungi_unclassified(100);k__Fungi_unclassified(100);k__Fungi_unclassified(100);k__Fungi_unclassified(100); | - | - | - | - | - | - | - | Unassigned | - |
| Otu08675 | 3 | k__Fungi(100);p__Ascomycota(100);c__Leotiomycetes(100);o__Helotiales(100);o__Helotiales_unclassified(100);o__Helotiales_unclassified(100);o__Helotiales_unclassified(100); | - | - | - | - | - | - | - | Unassigned | - |
| Otu08680 | 3 | k__Fungi(100);p__Ascomycota(67);p__Ascomycota_unclassified(67);p__Ascomycota_unclassified(67);p__Ascomycota_unclassified(67);p__Ascomycota_unclassified(67);p__Ascomycota_unclassified(67); | - | - | - | - | - | - | - | Unassigned | - |
| Otu08682 | 3 | k__Fungi(100);p__Ascomycota(100);p__Ascomycota_unclassified(100);p__Ascomycota_unclassified(100);p__Ascomycota_unclassified(100);p__Ascomycota_unclassified(100);p__Ascomycota_unclassified(100); | - | - | - | - | - | - | - | Unassigned | - |
| Otu08683 | 3 | k__Fungi(100);p__Ascomycota(67);p__Ascomycota_unclassified(67);p__Ascomycota_unclassified(67);p__Ascomycota_unclassified(67);p__Ascomycota_unclassified(67);p__Ascomycota_unclassified(67); | - | - | - | - | - | - | - | Unassigned | - |
| Otu08684 | 3 | k__Fungi(100);p__Ascomycota(100);p__Ascomycota_unclassified(100);p__Ascomycota_unclassified(100);p__Ascomycota_unclassified(100);p__Ascomycota_unclassified(100);p__Ascomycota_unclassified(100); | - | - | - | - | - | - | - | Unassigned | - |
| Otu08687 | 3 | k__Fungi(100);k__Fungi_unclassified(67);k__Fungi_unclassified(67);k__Fungi_unclassified(67);k__Fungi_unclassified(67);k__Fungi_unclassified(67);k__Fungi_unclassified(67); | - | - | - | - | - | - | - | Unassigned | - |
| Otu08688 | 3 | k__Fungi(100);k__Fungi_unclassified(67);k__Fungi_unclassified(67);k__Fungi_unclassified(67);k__Fungi_unclassified(67);k__Fungi_unclassified(67);k__Fungi_unclassified(67); | - | - | - | - | - | - | - | Unassigned | - |
| Otu08691 | 3 | k__Fungi(100);k__Fungi_unclassified(67);k__Fungi_unclassified(67);k__Fungi_unclassified(67);k__Fungi_unclassified(67);k__Fungi_unclassified(67);k__Fungi_unclassified(67); | - | - | - | - | - | - | - | Unassigned | - |
| Otu08695 | 3 | k__Fungi(100);p__Ascomycota(100);p__Ascomycota_unclassified(100);p__Ascomycota_unclassified(100);p__Ascomycota_unclassified(100);p__Ascomycota_unclassified(100);p__Ascomycota_unclassified(100); | - | - | - | - | - | - | - | Unassigned | - |
| Otu08698 | 3 | k__Fungi(100);k__Fungi_unclassified(100);k__Fungi_unclassified(100);k__Fungi_unclassified(100);k__Fungi_unclassified(100);k__Fungi_unclassified(100);k__Fungi_unclassified(100); | - | - | - | - | - | - | - | Unassigned | - |
| Otu08701 | 3 | k__Fungi(100);p__Ascomycota(67);p__Ascomycota_unclassified(67);p__Ascomycota_unclassified(67);p__Ascomycota_unclassified(67);p__Ascomycota_unclassified(67);p__Ascomycota_unclassified(67); | - | - | - | - | - | - | - | Unassigned | - |
| Otu08704 | 3 | k__Fungi(100);p__Ascomycota(100);p__Ascomycota_unclassified(100);p__Ascomycota_unclassified(100);p__Ascomycota_unclassified(100);p__Ascomycota_unclassified(100);p__Ascomycota_unclassified(100); | - | - | - | - | - | - | - | Unassigned | - |
| Otu08707 | 3 | k__Fungi(100);p__Ascomycota(100);p__Ascomycota_unclassified(100);p__Ascomycota_unclassified(100);p__Ascomycota_unclassified(100);p__Ascomycota_unclassified(100);p__Ascomycota_unclassified(100); | - | - | - | - | - | - | - | Unassigned | - |
| Otu08708 | 3 | k__Fungi(100);k__Fungi_unclassified(100);k__Fungi_unclassified(100);k__Fungi_unclassified(100);k__Fungi_unclassified(100);k__Fungi_unclassified(100);k__Fungi_unclassified(100); | - | - | - | - | - | - | - | Unassigned | - |
| Otu08709 | 3 | k__Fungi(100);p__Ascomycota(67);p__Ascomycota_unclassified(67);p__Ascomycota_unclassified(67);p__Ascomycota_unclassified(67);p__Ascomycota_unclassified(67);p__Ascomycota_unclassified(67); | - | - | - | - | - | - | - | Unassigned | - |
| Otu08711 | 3 | k__Fungi(100);p__Basidiomycota(100);p__Basidiomycota_unclassified(100);p__Basidiomycota_unclassified(100);p__Basidiomycota_unclassified(100);p__Basidiomycota_unclassified(100);p__Basidiomycota_unclassified(100); | - | - | - | - | - | - | - | Unassigned | - |
| Otu08712 | 3 | k__Fungi(100);p__Ascomycota(100);p__Ascomycota_unclassified(100);p__Ascomycota_unclassified(100);p__Ascomycota_unclassified(100);p__Ascomycota_unclassified(100);p__Ascomycota_unclassified(100); | - | - | - | - | - | - | - | Unassigned | - |
| Otu08714 | 3 | k__Fungi(100);p__Basidiomycota(100);c__Microbotryomycetes(100);c__Microbotryomycetes_unclassified(100);c__Microbotryomycetes_unclassified(100);c__Microbotryomycetes_unclassified(100);c__Microbotryomycetes_unclassified(100); | - | - | - | - | - | - | - | Unassigned | - |
| Otu08715 | 3 | k__Fungi(100);p__Ascomycota(67);p__Ascomycota_unclassified(67);p__Ascomycota_unclassified(67);p__Ascomycota_unclassified(67);p__Ascomycota_unclassified(67);p__Ascomycota_unclassified(67); | - | - | - | - | - | - | - | Unassigned | - |
| Otu08716 | 3 | k__Fungi(100);k__Fungi_unclassified(100);k__Fungi_unclassified(100);k__Fungi_unclassified(100);k__Fungi_unclassified(100);k__Fungi_unclassified(100);k__Fungi_unclassified(100); | - | - | - | - | - | - | - | Unassigned | - |
| Otu08718 | 3 | k__Fungi(100);p__Ascomycota(67);p__Ascomycota_unclassified(67);p__Ascomycota_unclassified(67);p__Ascomycota_unclassified(67);p__Ascomycota_unclassified(67);p__Ascomycota_unclassified(67); | - | - | - | - | - | - | - | Unassigned | - |
| Otu08720 | 3 | k__Fungi(100);p__Basidiomycota(100);p__Basidiomycota_unclassified(100);p__Basidiomycota_unclassified(100);p__Basidiomycota_unclassified(100);p__Basidiomycota_unclassified(100);p__Basidiomycota_unclassified(100); | - | - | - | - | - | - | - | Unassigned | - |
| Otu08722 | 3 | k__Fungi(100);p__Basidiomycota(100);p__Basidiomycota_unclassified(100);p__Basidiomycota_unclassified(100);p__Basidiomycota_unclassified(100);p__Basidiomycota_unclassified(100);p__Basidiomycota_unclassified(100); | - | - | - | - | - | - | - | Unassigned | - |
| Otu08728 | 3 | k__Fungi(100);p__Ascomycota(100);p__Ascomycota_unclassified(100);p__Ascomycota_unclassified(100);p__Ascomycota_unclassified(100);p__Ascomycota_unclassified(100);p__Ascomycota_unclassified(100); | - | - | - | - | - | - | - | Unassigned | - |
| Otu08729 | 3 | k__Fungi(100);p__Ascomycota(100);c__Eurotiomycetes(100);o__Eurotiales(100);f__Trichocomaceae(100);f__Trichocomaceae_unclassified(100);f__Trichocomaceae_unclassified(100); | - | - | - | - | - | - | - | Unassigned | - |
| Otu08730 | 3 | k__Fungi(100);p__Ascomycota(100);p__Ascomycota_unclassified(67);p__Ascomycota_unclassified(67);p__Ascomycota_unclassified(67);p__Ascomycota_unclassified(67);p__Ascomycota_unclassified(67); | - | - | - | - | - | - | - | Unassigned | - |
| Otu08731 | 3 | k__Fungi(100);k__Fungi_unclassified(100);k__Fungi_unclassified(100);k__Fungi_unclassified(100);k__Fungi_unclassified(100);k__Fungi_unclassified(100);k__Fungi_unclassified(100); | - | - | - | - | - | - | - | Unassigned | - |
| Otu08733 | 3 | k__Fungi(100);k__Fungi_unclassified(67);k__Fungi_unclassified(67);k__Fungi_unclassified(67);k__Fungi_unclassified(67);k__Fungi_unclassified(67);k__Fungi_unclassified(67); | - | - | - | - | - | - | - | Unassigned | - |
| Otu08736 | 3 | k__Fungi(100);p__Basidiomycota(100);p__Basidiomycota_unclassified(100);p__Basidiomycota_unclassified(100);p__Basidiomycota_unclassified(100);p__Basidiomycota_unclassified(100);p__Basidiomycota_unclassified(100); | - | - | - | - | - | - | - | Unassigned | - |
| Otu08738 | 3 | k__Fungi(100);k__Fungi_unclassified(100);k__Fungi_unclassified(100);k__Fungi_unclassified(100);k__Fungi_unclassified(100);k__Fungi_unclassified(100);k__Fungi_unclassified(100); | - | - | - | - | - | - | - | Unassigned | - |
| Otu08739 | 3 | k__Fungi(100);p__Ascomycota(100);c__Dothideomycetes(100);o__Pleosporales(100);o__Pleosporales_unclassified(100);o__Pleosporales_unclassified(100);o__Pleosporales_unclassified(100); | - | - | - | - | - | - | - | Unassigned | - |
| Otu08740 | 3 | k__Fungi(100);k__Fungi_unclassified(67);k__Fungi_unclassified(67);k__Fungi_unclassified(67);k__Fungi_unclassified(67);k__Fungi_unclassified(67);k__Fungi_unclassified(67); | - | - | - | - | - | - | - | Unassigned | - |
| Otu08741 | 3 | k__Fungi(100);p__Ascomycota(100);p__Ascomycota_unclassified(100);p__Ascomycota_unclassified(100);p__Ascomycota_unclassified(100);p__Ascomycota_unclassified(100);p__Ascomycota_unclassified(100); | - | - | - | - | - | - | - | Unassigned | - |
| Otu08742 | 3 | k__Fungi(100);k__Fungi_unclassified(67);k__Fungi_unclassified(67);k__Fungi_unclassified(67);k__Fungi_unclassified(67);k__Fungi_unclassified(67);k__Fungi_unclassified(67); | - | - | - | - | - | - | - | Unassigned | - |
| Otu08743 | 3 | k__Fungi(100);k__Fungi_unclassified(100);k__Fungi_unclassified(100);k__Fungi_unclassified(100);k__Fungi_unclassified(100);k__Fungi_unclassified(100);k__Fungi_unclassified(100); | - | - | - | - | - | - | - | Unassigned | - |
| Otu08744 | 3 | k__Fungi(100);k__Fungi_unclassified(100);k__Fungi_unclassified(100);k__Fungi_unclassified(100);k__Fungi_unclassified(100);k__Fungi_unclassified(100);k__Fungi_unclassified(100); | - | - | - | - | - | - | - | Unassigned | - |
| Otu08745 | 3 | k__Fungi(100);p__Ascomycota(100);p__Ascomycota_unclassified(100);p__Ascomycota_unclassified(100);p__Ascomycota_unclassified(100);p__Ascomycota_unclassified(100);p__Ascomycota_unclassified(100); | - | - | - | - | - | - | - | Unassigned | - |
| Otu08747 | 3 | k__Fungi(100);k__Fungi_unclassified(100);k__Fungi_unclassified(100);k__Fungi_unclassified(100);k__Fungi_unclassified(100);k__Fungi_unclassified(100);k__Fungi_unclassified(100); | - | - | - | - | - | - | - | Unassigned | - |
| Otu08752 | 3 | k__Fungi(100);k__Fungi_unclassified(67);k__Fungi_unclassified(67);k__Fungi_unclassified(67);k__Fungi_unclassified(67);k__Fungi_unclassified(67);k__Fungi_unclassified(67); | - | - | - | - | - | - | - | Unassigned | - |
| Otu08753 | 3 | k__Fungi(100);p__Ascomycota(100);p__Ascomycota_unclassified(100);p__Ascomycota_unclassified(100);p__Ascomycota_unclassified(100);p__Ascomycota_unclassified(100);p__Ascomycota_unclassified(100); | - | - | - | - | - | - | - | Unassigned | - |
| Otu08754 | 3 | k__Fungi(100);p__Ascomycota(100);p__Ascomycota_unclassified(100);p__Ascomycota_unclassified(100);p__Ascomycota_unclassified(100);p__Ascomycota_unclassified(100);p__Ascomycota_unclassified(100); | - | - | - | - | - | - | - | Unassigned | - |
| Otu08755 | 3 | k__Fungi(100);k__Fungi_unclassified(100);k__Fungi_unclassified(100);k__Fungi_unclassified(100);k__Fungi_unclassified(100);k__Fungi_unclassified(100);k__Fungi_unclassified(100); | - | - | - | - | - | - | - | Unassigned | - |
| Otu08761 | 3 | k__Fungi(100);p__Ascomycota(100);c__Eurotiomycetes(100);o__Chaetothyriales(100);o__Chaetothyriales_unclassified(67);o__Chaetothyriales_unclassified(67);o__Chaetothyriales_unclassified(67); | - | - | - | - | - | - | - | Unassigned | - |
| Otu08762 | 3 | k__Fungi(100);k__Fungi_unclassified(100);k__Fungi_unclassified(100);k__Fungi_unclassified(100);k__Fungi_unclassified(100);k__Fungi_unclassified(100);k__Fungi_unclassified(100); | - | - | - | - | - | - | - | Unassigned | - |
| Otu08763 | 3 | k__Fungi(100);k__Fungi_unclassified(100);k__Fungi_unclassified(100);k__Fungi_unclassified(100);k__Fungi_unclassified(100);k__Fungi_unclassified(100);k__Fungi_unclassified(100); | - | - | - | - | - | - | - | Unassigned | - |
| Otu08764 | 3 | k__Fungi(100);p__Ascomycota(67);p__Ascomycota_unclassified(67);p__Ascomycota_unclassified(67);p__Ascomycota_unclassified(67);p__Ascomycota_unclassified(67);p__Ascomycota_unclassified(67); | - | - | - | - | - | - | - | Unassigned | - |
| Otu08766 | 3 | k__Fungi(100);k__Fungi_unclassified(67);k__Fungi_unclassified(67);k__Fungi_unclassified(67);k__Fungi_unclassified(67);k__Fungi_unclassified(67);k__Fungi_unclassified(67); | - | - | - | - | - | - | - | Unassigned | - |
| Otu08767 | 3 | k__Fungi(100);p__Ascomycota(100);p__Ascomycota_unclassified(100);p__Ascomycota_unclassified(100);p__Ascomycota_unclassified(100);p__Ascomycota_unclassified(100);p__Ascomycota_unclassified(100); | - | - | - | - | - | - | - | Unassigned | - |
| Otu08774 | 3 | k__Fungi(100);p__Ascomycota(67);p__Ascomycota_unclassified(67);p__Ascomycota_unclassified(67);p__Ascomycota_unclassified(67);p__Ascomycota_unclassified(67);p__Ascomycota_unclassified(67); | - | - | - | - | - | - | - | Unassigned | - |
| Otu08775 | 3 | k__Fungi(100);p__Ascomycota(67);p__Ascomycota_unclassified(67);p__Ascomycota_unclassified(67);p__Ascomycota_unclassified(67);p__Ascomycota_unclassified(67);p__Ascomycota_unclassified(67); | - | - | - | - | - | - | - | Unassigned | - |
| Otu08776 | 3 | k__Fungi(100);k__Fungi_unclassified(67);k__Fungi_unclassified(67);k__Fungi_unclassified(67);k__Fungi_unclassified(67);k__Fungi_unclassified(67);k__Fungi_unclassified(67); | - | - | - | - | - | - | - | Unassigned | - |
| Otu08778 | 3 | k__Fungi(100);p__Ascomycota(100);p__Ascomycota_unclassified(100);p__Ascomycota_unclassified(100);p__Ascomycota_unclassified(100);p__Ascomycota_unclassified(100);p__Ascomycota_unclassified(100); | - | - | - | - | - | - | - | Unassigned | - |
| Otu08779 | 3 | k__Fungi(100);k__Fungi_unclassified(100);k__Fungi_unclassified(100);k__Fungi_unclassified(100);k__Fungi_unclassified(100);k__Fungi_unclassified(100);k__Fungi_unclassified(100); | - | - | - | - | - | - | - | Unassigned | - |
| Otu08781 | 3 | k__Fungi(100);p__Ascomycota(100);p__Ascomycota_unclassified(100);p__Ascomycota_unclassified(100);p__Ascomycota_unclassified(100);p__Ascomycota_unclassified(100);p__Ascomycota_unclassified(100); | - | - | - | - | - | - | - | Unassigned | - |
| Otu08783 | 3 | k__Fungi(100);p__Ascomycota(100);p__Ascomycota_unclassified(100);p__Ascomycota_unclassified(100);p__Ascomycota_unclassified(100);p__Ascomycota_unclassified(100);p__Ascomycota_unclassified(100); | - | - | - | - | - | - | - | Unassigned | - |
| Otu08784 | 3 | k__Fungi(100);k__Fungi_unclassified(100);k__Fungi_unclassified(100);k__Fungi_unclassified(100);k__Fungi_unclassified(100);k__Fungi_unclassified(100);k__Fungi_unclassified(100); | - | - | - | - | - | - | - | Unassigned | - |
| Otu08786 | 3 | k__Fungi(100);p__Ascomycota(67);p__Ascomycota_unclassified(67);p__Ascomycota_unclassified(67);p__Ascomycota_unclassified(67);p__Ascomycota_unclassified(67);p__Ascomycota_unclassified(67); | - | - | - | - | - | - | - | Unassigned | - |
| Otu08787 | 3 | k__Fungi(100);p__Ascomycota(100);p__Ascomycota_unclassified(100);p__Ascomycota_unclassified(100);p__Ascomycota_unclassified(100);p__Ascomycota_unclassified(100);p__Ascomycota_unclassified(100); | - | - | - | - | - | - | - | Unassigned | - |
| Otu08788 | 3 | k__Fungi(100);p__Ascomycota(100);p__Ascomycota_unclassified(100);p__Ascomycota_unclassified(100);p__Ascomycota_unclassified(100);p__Ascomycota_unclassified(100);p__Ascomycota_unclassified(100); | - | - | - | - | - | - | - | Unassigned | - |
| Otu08793 | 3 | k__Fungi(100);k__Fungi_unclassified(100);k__Fungi_unclassified(100);k__Fungi_unclassified(100);k__Fungi_unclassified(100);k__Fungi_unclassified(100);k__Fungi_unclassified(100); | - | - | - | - | - | - | - | Unassigned | - |
| Otu08794 | 3 | k__Plantae(100);p__unclassified_Plantae(100);c__unclassified_Plantae(100);o__unclassified_Plantae(100);f__unclassified_Plantae(100);g__unclassified_Plantae(100);s__Plantae_sp(100); | - | - | - | - | - | - | - | Unassigned | - |
| Otu08795 | 3 | k__Fungi(100);k__Fungi_unclassified(100);k__Fungi_unclassified(100);k__Fungi_unclassified(100);k__Fungi_unclassified(100);k__Fungi_unclassified(100);k__Fungi_unclassified(100); | - | - | - | - | - | - | - | Unassigned | - |
| Otu08796 | 2 | k__Fungi(100);p__Ascomycota(100);c__Saccharomycetes(100);o__Saccharomycetales(100);f__Metschnikowiaceae(100);g__Metschnikowia(100);s__Metschnikowia_pulcherrima(100); | Metschnikowia | 13 | Saprotroph | \|Undefined Saprotroph\| | Yeast | NULL | Probable | As nectar-tap_saprotroph (Põlme et al. 2020); Decay sugar-rich substrates (Põlme et al. 2020) | Põlme S, et al. 2020. Fungal Diversity 105:1-16 ((https://doi.org/10.1007/s13225-020-00466-2)) |
| Otu08797 | 2 | k__Fungi(100);p__Ascomycota(100);c__Saccharomycetes(100);o__Saccharomycetales(100);f__Metschnikowiaceae(100);g__Metschnikowia(100);s__Metschnikowia_pulcherrima(100); | Metschnikowia | 13 | Saprotroph | \|Undefined Saprotroph\| | Yeast | NULL | Probable | As nectar-tap_saprotroph (Põlme et al. 2020); Decay sugar-rich substrates (Põlme et al. 2020) | Põlme S, et al. 2020. Fungal Diversity 105:1-16 ((https://doi.org/10.1007/s13225-020-00466-2)) |
| Otu08798 | 2 | k__Fungi(100);p__Ascomycota(100);c__Saccharomycetes(100);o__Saccharomycetales(100);f__Metschnikowiaceae(100);g__Metschnikowia(100);s__Metschnikowia_pulcherrima(100); | Metschnikowia | 13 | Saprotroph | \|Undefined Saprotroph\| | Yeast | NULL | Probable | As nectar-tap_saprotroph (Põlme et al. 2020); Decay sugar-rich substrates (Põlme et al. 2020) | Põlme S, et al. 2020. Fungal Diversity 105:1-16 ((https://doi.org/10.1007/s13225-020-00466-2)) |
| Otu08800 | 2 | k__Fungi(100);p__Ascomycota(100);c__Saccharomycetes(100);o__Saccharomycetales(100);f__Metschnikowiaceae(100);g__Metschnikowia(100);s__Metschnikowia_pulcherrima(100); | Metschnikowia | 13 | Saprotroph | \|Undefined Saprotroph\| | Yeast | NULL | Probable | As nectar-tap_saprotroph (Põlme et al. 2020); Decay sugar-rich substrates (Põlme et al. 2020) | Põlme S, et al. 2020. Fungal Diversity 105:1-16 ((https://doi.org/10.1007/s13225-020-00466-2)) |
| Otu08803 | 2 | k__Fungi(100);p__Ascomycota(100);c__Saccharomycetes(100);o__Saccharomycetales(100);f__Metschnikowiaceae(100);g__Metschnikowia(100);s__Metschnikowia_pulcherrima(100); | Metschnikowia | 13 | Saprotroph | \|Undefined Saprotroph\| | Yeast | NULL | Probable | As nectar-tap_saprotroph (Põlme et al. 2020); Decay sugar-rich substrates (Põlme et al. 2020) | Põlme S, et al. 2020. Fungal Diversity 105:1-16 ((https://doi.org/10.1007/s13225-020-00466-2)) |
| Otu08805 | 2 | k__Fungi(100);p__Ascomycota(100);c__Saccharomycetes(100);o__Saccharomycetales(100);f__Metschnikowiaceae(100);g__Metschnikowia(100);g__Metschnikowia_unclassified(100); | Metschnikowia | 13 | Saprotroph | \|Undefined Saprotroph\| | Yeast | NULL | Probable | As nectar-tap_saprotroph (Põlme et al. 2020); Decay sugar-rich substrates (Põlme et al. 2020) | Põlme S, et al. 2020. Fungal Diversity 105:1-16 ((https://doi.org/10.1007/s13225-020-00466-2)) |
| Otu08808 | 2 | k__Fungi(100);p__Ascomycota(100);c__Saccharomycetes(100);o__Saccharomycetales(100);f__Metschnikowiaceae(100);g__Metschnikowia(100);s__Metschnikowia_pulcherrima(100); | Metschnikowia | 13 | Saprotroph | \|Undefined Saprotroph\| | Yeast | NULL | Probable | As nectar-tap_saprotroph (Põlme et al. 2020); Decay sugar-rich substrates (Põlme et al. 2020) | Põlme S, et al. 2020. Fungal Diversity 105:1-16 ((https://doi.org/10.1007/s13225-020-00466-2)) |
| Otu08812 | 2 | k__Fungi(100);p__Ascomycota(100);c__Saccharomycetes(100);o__Saccharomycetales(100);f__Metschnikowiaceae(100);g__Metschnikowia(100);s__Metschnikowia_pulcherrima(100); | Metschnikowia | 13 | Saprotroph | \|Undefined Saprotroph\| | Yeast | NULL | Probable | As nectar-tap_saprotroph (Põlme et al. 2020); Decay sugar-rich substrates (Põlme et al. 2020) | Põlme S, et al. 2020. Fungal Diversity 105:1-16 ((https://doi.org/10.1007/s13225-020-00466-2)) |
| Otu08814 | 2 | k__Fungi(100);p__Ascomycota(100);c__Saccharomycetes(100);o__Saccharomycetales(100);f__Metschnikowiaceae(100);g__Metschnikowia(100);s__Metschnikowia_pulcherrima(100); | Metschnikowia | 13 | Saprotroph | \|Undefined Saprotroph\| | Yeast | NULL | Probable | As nectar-tap_saprotroph (Põlme et al. 2020); Decay sugar-rich substrates (Põlme et al. 2020) | Põlme S, et al. 2020. Fungal Diversity 105:1-16 ((https://doi.org/10.1007/s13225-020-00466-2)) |
| Otu08815 | 2 | k__Fungi(100);p__Ascomycota(100);c__Saccharomycetes(100);o__Saccharomycetales(100);f__Metschnikowiaceae(100);g__Metschnikowia(100);g__Metschnikowia_unclassified(100); | Metschnikowia | 13 | Saprotroph | \|Undefined Saprotroph\| | Yeast | NULL | Probable | As nectar-tap_saprotroph (Põlme et al. 2020); Decay sugar-rich substrates (Põlme et al. 2020) | Põlme S, et al. 2020. Fungal Diversity 105:1-16 ((https://doi.org/10.1007/s13225-020-00466-2)) |
| Otu08816 | 2 | k__Fungi(100);p__Ascomycota(100);c__Saccharomycetes(100);o__Saccharomycetales(100);f__Metschnikowiaceae(100);g__Metschnikowia(100);g__Metschnikowia_unclassified(100); | Metschnikowia | 13 | Saprotroph | \|Undefined Saprotroph\| | Yeast | NULL | Probable | As nectar-tap_saprotroph (Põlme et al. 2020); Decay sugar-rich substrates (Põlme et al. 2020) | Põlme S, et al. 2020. Fungal Diversity 105:1-16 ((https://doi.org/10.1007/s13225-020-00466-2)) |
| Otu08817 | 2 | k__Fungi(100);p__Ascomycota(100);c__Saccharomycetes(100);o__Saccharomycetales(100);f__Metschnikowiaceae(100);g__Metschnikowia(100);g__Metschnikowia_unclassified(100); | Metschnikowia | 13 | Saprotroph | \|Undefined Saprotroph\| | Yeast | NULL | Probable | As nectar-tap_saprotroph (Põlme et al. 2020); Decay sugar-rich substrates (Põlme et al. 2020) | Põlme S, et al. 2020. Fungal Diversity 105:1-16 ((https://doi.org/10.1007/s13225-020-00466-2)) |
| Otu08820 | 2 | k__Fungi(100);p__Ascomycota(100);c__Saccharomycetes(100);o__Saccharomycetales(100);f__Metschnikowiaceae(100);g__Metschnikowia(100);s__Metschnikowia_pulcherrima(100); | Metschnikowia | 13 | Saprotroph | \|Undefined Saprotroph\| | Yeast | NULL | Probable | As nectar-tap_saprotroph (Põlme et al. 2020); Decay sugar-rich substrates (Põlme et al. 2020) | Põlme S, et al. 2020. Fungal Diversity 105:1-16 ((https://doi.org/10.1007/s13225-020-00466-2)) |
| Otu08822 | 2 | k__Fungi(100);p__Ascomycota(100);c__Saccharomycetes(100);o__Saccharomycetales(100);f__Metschnikowiaceae(100);g__Metschnikowia(100);s__Metschnikowia_pulcherrima(100); | Metschnikowia | 13 | Saprotroph | \|Undefined Saprotroph\| | Yeast | NULL | Probable | As nectar-tap_saprotroph (Põlme et al. 2020); Decay sugar-rich substrates (Põlme et al. 2020) | Põlme S, et al. 2020. Fungal Diversity 105:1-16 ((https://doi.org/10.1007/s13225-020-00466-2)) |
| Otu08823 | 2 | k__Fungi(100);p__Ascomycota(100);c__Sordariomycetes(100);o__Chaetosphaeriales(100);f__Chaetosphaeriaceae(100);g__Chaetosphaeria(100);s__Chaetosphaeria_sp_KRCF710(100); | Chaetosphaeria | 13 | Saprotroph-Symbiotroph | Endophyte-\|Plant Saprotroph\|-Wood Saprotroph | Microfungus | NULL | Probable | As foliar_endophyte (Põlme et al. 2020); As litter_saprotroph (Põlme et al. 2020); Decay wood, leaf, fruit, seed (Põlme et al. 2020) | Réblová M, et al. 2006. Mycological Research 110:104-109 ((https://doi.org/10.1016/j.mycres.2005.09.003)); Tedersoo L, et al. 2014. Science 346:e1256688 ((https://doi.org/10.1126/science.1256688)); Põlme S, et al. 2020. Fungal Diversity 105:1-16 ((https://doi.org/10.1007/s13225-020-00466-2)) |
| Otu08824 | 2 | k__Fungi(100);p__Ascomycota(100);c__Dothideomycetes(100);o__Pleosporales(100);f__Massarinaceae(100);g__Helminthosporium(100);s__Helminthosporium_asterinum(100); | Helminthosporium | 13 | Pathotroph | Fungal Parasite-\|Plant Pathogen\| | Microfungus | NULL | Probable | NULL | Samuels GJ. 2007. Self-published; Tedersoo L, et al. 2014. Science 346:e1256688 ((https://doi.org/10.1126/science.12566)); Põlme S, et al. 2020. Fungal Diversity 105:1-16 ((https://doi.org/10.1007/s13225-020-00466-2)) |
| Otu08827 | 2 | k__Fungi(100);p__Ascomycota(100);c__Saccharomycetes(100);o__Saccharomycetales(100);f__Metschnikowiaceae(100);g__Metschnikowia(100);g__Metschnikowia_unclassified(100); | Metschnikowia | 13 | Saprotroph | \|Undefined Saprotroph\| | Yeast | NULL | Probable | As nectar-tap_saprotroph (Põlme et al. 2020); Decay sugar-rich substrates (Põlme et al. 2020) | Põlme S, et al. 2020. Fungal Diversity 105:1-16 ((https://doi.org/10.1007/s13225-020-00466-2)) |
| Otu08828 | 2 | k__Fungi(100);p__Ascomycota(100);c__Saccharomycetes(100);o__Saccharomycetales(100);f__Metschnikowiaceae(100);g__Metschnikowia(100);s__Metschnikowia_pulcherrima(100); | Metschnikowia | 13 | Saprotroph | \|Undefined Saprotroph\| | Yeast | NULL | Probable | As nectar-tap_saprotroph (Põlme et al. 2020); Decay sugar-rich substrates (Põlme et al. 2020) | Põlme S, et al. 2020. Fungal Diversity 105:1-16 ((https://doi.org/10.1007/s13225-020-00466-2)) |
| Otu08829 | 2 | k__Fungi(100);p__Ascomycota(100);c__Saccharomycetes(100);o__Saccharomycetales(100);f__Metschnikowiaceae(100);g__Metschnikowia(100);g__Metschnikowia_unclassified(100); | Metschnikowia | 13 | Saprotroph | \|Undefined Saprotroph\| | Yeast | NULL | Probable | As nectar-tap_saprotroph (Põlme et al. 2020); Decay sugar-rich substrates (Põlme et al. 2020) | Põlme S, et al. 2020. Fungal Diversity 105:1-16 ((https://doi.org/10.1007/s13225-020-00466-2)) |
| Otu08831 | 2 | k__Fungi(100);p__Ascomycota(100);c__Saccharomycetes(100);o__Saccharomycetales(100);f__Metschnikowiaceae(100);g__Metschnikowia(100);s__Metschnikowia_pulcherrima(100); | Metschnikowia | 13 | Saprotroph | \|Undefined Saprotroph\| | Yeast | NULL | Probable | As nectar-tap_saprotroph (Põlme et al. 2020); Decay sugar-rich substrates (Põlme et al. 2020) | Põlme S, et al. 2020. Fungal Diversity 105:1-16 ((https://doi.org/10.1007/s13225-020-00466-2)) |
| Otu08833 | 2 | k__Fungi(100);p__Ascomycota(100);c__Saccharomycetes(100);o__Saccharomycetales(100);f__Metschnikowiaceae(100);g__Metschnikowia(100);s__Metschnikowia_pulcherrima(100); | Metschnikowia | 13 | Saprotroph | \|Undefined Saprotroph\| | Yeast | NULL | Probable | As nectar-tap_saprotroph (Põlme et al. 2020); Decay sugar-rich substrates (Põlme et al. 2020) | Põlme S, et al. 2020. Fungal Diversity 105:1-16 ((https://doi.org/10.1007/s13225-020-00466-2)) |
| Otu08834 | 2 | k__Fungi(100);p__Ascomycota(100);c__Saccharomycetes(100);o__Saccharomycetales(100);f__Metschnikowiaceae(100);g__Metschnikowia(100);g__Metschnikowia_unclassified(100); | Metschnikowia | 13 | Saprotroph | \|Undefined Saprotroph\| | Yeast | NULL | Probable | As nectar-tap_saprotroph (Põlme et al. 2020); Decay sugar-rich substrates (Põlme et al. 2020) | Põlme S, et al. 2020. Fungal Diversity 105:1-16 ((https://doi.org/10.1007/s13225-020-00466-2)) |
| Otu08835 | 2 | k__Fungi(100);p__Ascomycota(100);c__Saccharomycetes(100);o__Saccharomycetales(100);f__Metschnikowiaceae(100);g__Metschnikowia(100);s__Metschnikowia_pulcherrima(100); | Metschnikowia | 13 | Saprotroph | \|Undefined Saprotroph\| | Yeast | NULL | Probable | As nectar-tap_saprotroph (Põlme et al. 2020); Decay sugar-rich substrates (Põlme et al. 2020) | Põlme S, et al. 2020. Fungal Diversity 105:1-16 ((https://doi.org/10.1007/s13225-020-00466-2)) |
| Otu08836 | 2 | k__Fungi(100);p__Ascomycota(100);c__Saccharomycetes(100);o__Saccharomycetales(100);f__Metschnikowiaceae(100);g__Metschnikowia(100);s__Metschnikowia_pulcherrima(100); | Metschnikowia | 13 | Saprotroph | \|Undefined Saprotroph\| | Yeast | NULL | Probable | As nectar-tap_saprotroph (Põlme et al. 2020); Decay sugar-rich substrates (Põlme et al. 2020) | Põlme S, et al. 2020. Fungal Diversity 105:1-16 ((https://doi.org/10.1007/s13225-020-00466-2)) |
| Otu08837 | 2 | k__Fungi(100);p__Ascomycota(100);c__Saccharomycetes(100);o__Saccharomycetales(100);f__Metschnikowiaceae(100);g__Metschnikowia(100);g__Metschnikowia_unclassified(100); | Metschnikowia | 13 | Saprotroph | \|Undefined Saprotroph\| | Yeast | NULL | Probable | As nectar-tap_saprotroph (Põlme et al. 2020); Decay sugar-rich substrates (Põlme et al. 2020) | Põlme S, et al. 2020. Fungal Diversity 105:1-16 ((https://doi.org/10.1007/s13225-020-00466-2)) |
| Otu08838 | 2 | k__Fungi(100);p__Ascomycota(100);c__Saccharomycetes(100);o__Saccharomycetales(100);f__Metschnikowiaceae(100);g__Metschnikowia(100);g__Metschnikowia_unclassified(100); | Metschnikowia | 13 | Saprotroph | \|Undefined Saprotroph\| | Yeast | NULL | Probable | As nectar-tap_saprotroph (Põlme et al. 2020); Decay sugar-rich substrates (Põlme et al. 2020) | Põlme S, et al. 2020. Fungal Diversity 105:1-16 ((https://doi.org/10.1007/s13225-020-00466-2)) |
| Otu08844 | 2 | k__Fungi(100);p__Ascomycota(100);c__Saccharomycetes(100);o__Saccharomycetales(100);f__Metschnikowiaceae(100);g__Metschnikowia(100);s__Metschnikowia_pulcherrima(100); | Metschnikowia | 13 | Saprotroph | \|Undefined Saprotroph\| | Yeast | NULL | Probable | As nectar-tap_saprotroph (Põlme et al. 2020); Decay sugar-rich substrates (Põlme et al. 2020) | Põlme S, et al. 2020. Fungal Diversity 105:1-16 ((https://doi.org/10.1007/s13225-020-00466-2)) |
| Otu08847 | 2 | k__Fungi(100);p__Ascomycota(100);c__Saccharomycetes(100);o__Saccharomycetales(100);f__Metschnikowiaceae(100);g__Metschnikowia(100);g__Metschnikowia_unclassified(100); | Metschnikowia | 13 | Saprotroph | \|Undefined Saprotroph\| | Yeast | NULL | Probable | As nectar-tap_saprotroph (Põlme et al. 2020); Decay sugar-rich substrates (Põlme et al. 2020) | Põlme S, et al. 2020. Fungal Diversity 105:1-16 ((https://doi.org/10.1007/s13225-020-00466-2)) |
| Otu08849 | 2 | k__Fungi(100);p__Ascomycota(100);c__Saccharomycetes(100);o__Saccharomycetales(100);f__Metschnikowiaceae(100);g__Metschnikowia(100);s__Metschnikowia_pulcherrima(100); | Metschnikowia | 13 | Saprotroph | \|Undefined Saprotroph\| | Yeast | NULL | Probable | As nectar-tap_saprotroph (Põlme et al. 2020); Decay sugar-rich substrates (Põlme et al. 2020) | Põlme S, et al. 2020. Fungal Diversity 105:1-16 ((https://doi.org/10.1007/s13225-020-00466-2)) |

| | | | | | | | | | | | |
|---|---|---|---|---|---|---|---|---|---|---|---|
| Otu08850 | 2 | k__Fungi(100);p__Ascomycota(100);c__Saccharomycetes(100);o__Saccharomycetales(100);f__Saccharomycetales_family_Incertae_sedis(100);f__Saccharomycetales_family_Incertae_sedis_unclassified(100);f__Saccharomycetales_family_Incertae_sedis_unclassified(100); | Saccharomycetales | 7 | Saprotroph | Undefined Saprotroph | Yeast | NULL | Possible | NULL | Sterkenburg E, et al. 2015. New Phytologist 207:1145-1158 ((https://doi.org/10.1111/nph.13426)) |
| Otu08854 | 2 | k__Fungi(100);p__Ascomycota(100);c__Saccharomycetes(100);o__Saccharomycetales(100);f__Metschnikowiaceae(100);g__Metschnikowia(100);s__Metschnikowia_pulcherrima(100); | Metschnikowia | 13 | Saprotroph | \|Undefined Saprotroph\| | Yeast | NULL | Probable | As nectar-tap_saprotroph (Põlme et al. 2020); Decay sugar-rich substrates (Põlme et al. 2020) | Põlme S, et al. 2020. Fungal Diversity 105:1-16 ((https://doi.org/10.1007/s13225-020-00466-2)) |
| Otu08855 | 2 | k__Fungi(100);p__Ascomycota(100);c__Saccharomycetes(100);o__Saccharomycetales(100);f__Metschnikowiaceae(100);g__Metschnikowia(100);g__Metschnikowia_unclassified(100); | Metschnikowia | 13 | Saprotroph | \|Undefined Saprotroph\| | Yeast | NULL | Probable | As nectar-tap_saprotroph (Põlme et al. 2020); Decay sugar-rich substrates (Põlme et al. 2020) | Põlme S, et al. 2020. Fungal Diversity 105:1-16 ((https://doi.org/10.1007/s13225-020-00466-2)) |
| Otu08860 | 2 | k__Fungi(100);p__Ascomycota(100);c__Saccharomycetes(100);o__Saccharomycetales(100);f__Metschnikowiaceae(100);g__Metschnikowia(100);g__Metschnikowia_unclassified(100); | Metschnikowia | 13 | Saprotroph | \|Undefined Saprotroph\| | Yeast | NULL | Probable | As nectar-tap_saprotroph (Põlme et al. 2020); Decay sugar-rich substrates (Põlme et al. 2020) | Põlme S, et al. 2020. Fungal Diversity 105:1-16 ((https://doi.org/10.1007/s13225-020-00466-2)) |
| Otu08861 | 2 | k__Fungi(100);p__Ascomycota(100);c__Saccharomycetes(100);o__Saccharomycetales(100);f__Metschnikowiaceae(100);g__Metschnikowia(100);g__Metschnikowia_unclassified(100); | Metschnikowia | 13 | Saprotroph | \|Undefined Saprotroph\| | Yeast | NULL | Probable | As nectar-tap_saprotroph (Põlme et al. 2020); Decay sugar-rich substrates (Põlme et al. 2020) | Põlme S, et al. 2020. Fungal Diversity 105:1-16 ((https://doi.org/10.1007/s13225-020-00466-2)) |
| Otu08862 | 2 | k__Fungi(100);p__Ascomycota(100);c__Saccharomycetes(100);o__Saccharomycetales(100);f__Metschnikowiaceae(100);g__Metschnikowia(100);g__Metschnikowia_unclassified(100); | Metschnikowia | 13 | Saprotroph | \|Undefined Saprotroph\| | Yeast | NULL | Probable | As nectar-tap_saprotroph (Põlme et al. 2020); Decay sugar-rich substrates (Põlme et al. 2020) | Põlme S, et al. 2020. Fungal Diversity 105:1-16 ((https://doi.org/10.1007/s13225-020-00466-2)) |
| Otu08863 | 2 | k__Fungi(100);p__Ascomycota(100);c__Saccharomycetes(100);o__Saccharomycetales(100);f__Metschnikowiaceae(100);g__Metschnikowia(100);g__Metschnikowia_unclassified(100); | Metschnikowia | 13 | Saprotroph | \|Undefined Saprotroph\| | Yeast | NULL | Probable | As nectar-tap_saprotroph (Põlme et al. 2020); Decay sugar-rich substrates (Põlme et al. 2020) | Põlme S, et al. 2020. Fungal Diversity 105:1-16 ((https://doi.org/10.1007/s13225-020-00466-2)) |
| Otu08867 | 2 | k__Fungi(100);p__Ascomycota(100);c__Saccharomycetes(100);o__Saccharomycetales(100);f__Metschnikowiaceae(100);g__Metschnikowia(100);g__Metschnikowia_unclassified(100); | Metschnikowia | 13 | Saprotroph | \|Undefined Saprotroph\| | Yeast | NULL | Probable | As nectar-tap_saprotroph (Põlme et al. 2020); Decay sugar-rich substrates (Põlme et al. 2020) | Põlme S, et al. 2020. Fungal Diversity 105:1-16 ((https://doi.org/10.1007/s13225-020-00466-2)) |
| Otu08870 | 2 | k__Fungi(100);p__Ascomycota(100);c__Saccharomycetes(100);o__Saccharomycetales(100);f__Metschnikowiaceae(100);g__Metschnikowia(100);s__Metschnikowia_pulcherrima(100); | Metschnikowia | 13 | Saprotroph | \|Undefined Saprotroph\| | Yeast | NULL | Probable | As nectar-tap_saprotroph (Põlme et al. 2020); Decay sugar-rich substrates (Põlme et al. 2020) | Põlme S, et al. 2020. Fungal Diversity 105:1-16 ((https://doi.org/10.1007/s13225-020-00466-2)) |
| Otu08871 | 2 | k__Fungi(100);p__Ascomycota(100);c__Saccharomycetes(100);o__Saccharomycetales(100);f__Metschnikowiaceae(100);g__Metschnikowia(100);s__Metschnikowia_pulcherrima(100); | Metschnikowia | 13 | Saprotroph | \|Undefined Saprotroph\| | Yeast | NULL | Probable | As nectar-tap_saprotroph (Põlme et al. 2020); Decay sugar-rich substrates (Põlme et al. 2020) | Põlme S, et al. 2020. Fungal Diversity 105:1-16 ((https://doi.org/10.1007/s13225-020-00466-2)) |
| Otu08873 | 2 | k__Fungi(100);p__Ascomycota(100);c__Saccharomycetes(100);o__Saccharomycetales(100);f__Metschnikowiaceae(100);g__Metschnikowia(100);s__Metschnikowia_pulcherrima(100); | Metschnikowia | 13 | Saprotroph | \|Undefined Saprotroph\| | Yeast | NULL | Probable | As nectar-tap_saprotroph (Põlme et al. 2020); Decay sugar-rich substrates (Põlme et al. 2020) | Põlme S, et al. 2020. Fungal Diversity 105:1-16 ((https://doi.org/10.1007/s13225-020-00466-2)) |
| Otu08875 | 2 | k__Fungi(100);p__Ascomycota(100);c__Saccharomycetes(100);o__Saccharomycetales(100);f__Metschnikowiaceae(100);g__Metschnikowia(100);s__Metschnikowia_pulcherrima(100); | Metschnikowia | 13 | Saprotroph | \|Undefined Saprotroph\| | Yeast | NULL | Probable | As nectar-tap_saprotroph (Põlme et al. 2020); Decay sugar-rich substrates (Põlme et al. 2020) | Põlme S, et al. 2020. Fungal Diversity 105:1-16 ((https://doi.org/10.1007/s13225-020-00466-2)) |
| Otu08878 | 2 | k__Fungi(100);p__Ascomycota(100);c__Saccharomycetes(100);o__Saccharomycetales(100);f__Metschnikowiaceae(100);g__Metschnikowia(100);s__Metschnikowia_pulcherrima(100); | Metschnikowia | 13 | Saprotroph | \|Undefined Saprotroph\| | Yeast | NULL | Probable | As nectar-tap_saprotroph (Põlme et al. 2020); Decay sugar-rich substrates (Põlme et al. 2020) | Põlme S, et al. 2020. Fungal Diversity 105:1-16 ((https://doi.org/10.1007/s13225-020-00466-2)) |
| Otu08885 | 2 | k__Fungi(100);p__Ascomycota(100);c__Saccharomycetes(100);o__Saccharomycetales(100);f__Metschnikowiaceae(100);g__Metschnikowia(100);g__Metschnikowia_unclassified(100); | Metschnikowia | 13 | Saprotroph | \|Undefined Saprotroph\| | Yeast | NULL | Probable | As nectar-tap_saprotroph (Põlme et al. 2020); Decay sugar-rich substrates (Põlme et al. 2020) | Põlme S, et al. 2020. Fungal Diversity 105:1-16 ((https://doi.org/10.1007/s13225-020-00466-2)) |
| Otu08890 | 2 | k__Fungi(100);p__Ascomycota(100);c__Saccharomycetes(100);o__Saccharomycetales(100);f__Metschnikowiaceae(100);g__Metschnikowia(100);g__Metschnikowia_unclassified(100); | Metschnikowia | 13 | Saprotroph | \|Undefined Saprotroph\| | Yeast | NULL | Probable | As nectar-tap_saprotroph (Põlme et al. 2020); Decay sugar-rich substrates (Põlme et al. 2020) | Põlme S, et al. 2020. Fungal Diversity 105:1-16 ((https://doi.org/10.1007/s13225-020-00466-2)) |
| Otu08894 | 2 | k__Fungi(100);p__Ascomycota(100);c__Saccharomycetes(100);o__Saccharomycetales(100);f__Metschnikowiaceae(100);g__Metschnikowia(100);s__Metschnikowia_pulcherrima(100); | Metschnikowia | 13 | Saprotroph | \|Undefined Saprotroph\| | Yeast | NULL | Probable | As nectar-tap_saprotroph (Põlme et al. 2020); Decay sugar-rich substrates (Põlme et al. 2020) | Põlme S, et al. 2020. Fungal Diversity 105:1-16 ((https://doi.org/10.1007/s13225-020-00466-2)) |
| Otu08896 | 2 | k__Fungi(100);p__Ascomycota(100);c__Eurotiomycetes(100);o__Eurotiales(100);f__Trichocomaceae(100);g__Penicillium(100);s__Penicillium_thiersii(100); | Penicillium | 13 | Pathotroph-Saprotroph-Symbiotroph | Animal Parasite-Dung Saprotroph-Endophyte-Fungal Parasite-Plant Pathogen-\|Undefined Saprotroph\|-Wood Saprotroph | Microfungus | Soft Rot | Highly Probable | As foliar_endophyte (Põlme et al. 2020); As mycoparasite (Põlme et al. 2020); Decay soil (Põlme et al. 2020) | Duncan CG, Eslyn WE. 1966. Mycologia 58:642-664 ((https://doi.org/10.2307/3757045)); Seehann G, et al. 1975. List of Fungi in Soft-Rot Tissue Tests. Institute fur Holzbiologie und Holzschutz, Bundesforschungsanstalt fur Forst - und Holzwirtschaft, Hamburg - Lohbrugge, Germany ((https://www.irg-wp.com/irgdocs/details.php?d6493033-c9ea-40f3-8e3e-5697f32d640d)); Costa IPMW, et al. 2012. Mycotaxon 119:e26 ((http://mycotaxon.com/resources/checklists/costa_v119_checklist.pdf)); Bills GF, et al. 2013. Current Opinion in Microbiology 16: 549-565 ((https://doi.org/10.1016/j.mib.2013.08.001)); Tedersoo L, et al. 2014. Science 346:e1256688 ((https://doi.org/10.1126/science.1256688)); Põlme S, et al. 2020. Fungal Diversity 105:1-16 ((https://doi.org/10.1007/s13225-020-00466-2)) |
| Otu08897 | 2 | k__Fungi(100);p__Ascomycota(100);c__Ascomycota_class_Incertae_sedis(100);o__Ascomycota_order_Incertae_sedis(100);f__Ascomycota_family_Incertae_sedis(100);g__Hansfordia(100);s__Hansfordia_pulvinata(100); | Hansfordia | 13 | Saprotroph | \|Undefined Saprotroph\| | Microfungus | NULL | Probable | NULL | Tedersoo L, et al. 2014. Science 346:e1256688 ((https://doi.org/10.1126/science.1256688)); Põlme S, et al. 2020. Fungal Diversity 105:1-16 ((https://doi.org/10.1007/s13225-020-00466-2)) |
| Otu08898 | 2 | k__Fungi(100);p__Ascomycota(100);c__Saccharomycetes(100);o__Saccharomycetales(100);f__Metschnikowiaceae(100);g__Metschnikowia(100);s__Metschnikowia_pulcherrima(100); | Metschnikowia | 13 | Saprotroph | \|Undefined Saprotroph\| | Yeast | NULL | Probable | As nectar-tap_saprotroph (Põlme et al. 2020); Decay sugar-rich substrates (Põlme et al. 2020) | Põlme S, et al. 2020. Fungal Diversity 105:1-16 ((https://doi.org/10.1007/s13225-020-00466-2)) |
| Otu08899 | 2 | k__Fungi(100);p__Ascomycota(100);c__Saccharomycetes(100);o__Saccharomycetales(100);f__Metschnikowiaceae(100);g__Metschnikowia(100);s__Metschnikowia_pulcherrima(100); | Metschnikowia | 13 | Saprotroph | \|Undefined Saprotroph\| | Yeast | NULL | Probable | As nectar-tap_saprotroph (Põlme et al. 2020); Decay sugar-rich substrates (Põlme et al. 2020) | Põlme S, et al. 2020. Fungal Diversity 105:1-16 ((https://doi.org/10.1007/s13225-020-00466-2)) |
| Otu08901 | 2 | k__Fungi(100);p__Ascomycota(100);c__Saccharomycetes(100);o__Saccharomycetales(100);f__Metschnikowiaceae(100);g__Metschnikowia(100);g__Metschnikowia_unclassified(100); | Metschnikowia | 13 | Saprotroph | \|Undefined Saprotroph\| | Yeast | NULL | Probable | As nectar-tap_saprotroph (Põlme et al. 2020); Decay sugar-rich substrates (Põlme et al. 2020) | Põlme S, et al. 2020. Fungal Diversity 105:1-16 ((https://doi.org/10.1007/s13225-020-00466-2)) |
| Otu08904 | 2 | k__Fungi(100);p__Ascomycota(100);c__Saccharomycetes(100);o__Saccharomycetales(100);f__Metschnikowiaceae(100);g__Metschnikowia(100);s__Metschnikowia_pulcherrima(100); | Metschnikowia | 13 | Saprotroph | \|Undefined Saprotroph\| | Yeast | NULL | Probable | As nectar-tap_saprotroph (Põlme et al. 2020); Decay sugar-rich substrates (Põlme et al. 2020) | Põlme S, et al. 2020. Fungal Diversity 105:1-16 ((https://doi.org/10.1007/s13225-020-00466-2)) |
| Otu08906 | 2 | k__Fungi(100);p__Ascomycota(100);c__Saccharomycetes(100);o__Saccharomycetales(100);f__Metschnikowiaceae(100);g__Metschnikowia(100);s__Metschnikowia_pulcherrima(100); | Metschnikowia | 13 | Saprotroph | \|Undefined Saprotroph\| | Yeast | NULL | Probable | As nectar-tap_saprotroph (Põlme et al. 2020); Decay sugar-rich substrates (Põlme et al. 2020) | Põlme S, et al. 2020. Fungal Diversity 105:1-16 ((https://doi.org/10.1007/s13225-020-00466-2)) |
| Otu08908 | 2 | k__Fungi(100);p__Ascomycota(100);c__Saccharomycetes(100);o__Saccharomycetales(100);f__Metschnikowiaceae(100);g__Metschnikowia(100);s__Metschnikowia_pulcherrima(100); | Metschnikowia | 13 | Saprotroph | \|Undefined Saprotroph\| | Yeast | NULL | Probable | As nectar-tap_saprotroph (Põlme et al. 2020); Decay sugar-rich substrates (Põlme et al. 2020) | Põlme S, et al. 2020. Fungal Diversity 105:1-16 ((https://doi.org/10.1007/s13225-020-00466-2)) |
| Otu08910 | 2 | k__Fungi(100);p__Ascomycota(100);c__Saccharomycetes(100);o__Saccharomycetales(100);f__Metschnikowiaceae(100);g__Metschnikowia(100);g__Metschnikowia_unclassified(100); | Metschnikowia | 13 | Saprotroph | \|Undefined Saprotroph\| | Yeast | NULL | Probable | As nectar-tap_saprotroph (Põlme et al. 2020); Decay sugar-rich substrates (Põlme et al. 2020) | Põlme S, et al. 2020. Fungal Diversity 105:1-16 ((https://doi.org/10.1007/s13225-020-00466-2)) |
| Otu08914 | 2 | k__Fungi(100);p__Ascomycota(100);c__Saccharomycetes(100);o__Saccharomycetales(100);f__Metschnikowiaceae(100);g__Metschnikowia(100);g__Metschnikowia_unclassified(100); | Metschnikowia | 13 | Saprotroph | \|Undefined Saprotroph\| | Yeast | NULL | Probable | As nectar-tap_saprotroph (Põlme et al. 2020); Decay sugar-rich substrates (Põlme et al. 2020) | Põlme S, et al. 2020. Fungal Diversity 105:1-16 ((https://doi.org/10.1007/s13225-020-00466-2)) |
| Otu08916 | 2 | k__Fungi(100);p__Ascomycota(100);c__Saccharomycetes(100);o__Saccharomycetales(100);f__Saccharomycetales_family_Incertae_sedis(100);f__Saccharomycetales_family_Incertae_sedis_unclassified(100);f__Saccharomycetales_family_Incertae_sedis_unclassified(100); | Saccharomycetales | 7 | Saprotroph | Undefined Saprotroph | Yeast | NULL | Possible | NULL | Sterkenburg E, et al. 2015. New Phytologist 207:1145-1158 ((https://doi.org/10.1111/nph.13426)) |
| Otu08917 | 2 | k__Fungi(100);p__Ascomycota(100);c__Saccharomycetes(100);o__Saccharomycetales(100);f__Saccharomycetales_family_Incertae_sedis(100);f__Saccharomycetales_family_Incertae_sedis_unclassified(100);f__Saccharomycetales_family_Incertae_sedis_unclassified(100); | Saccharomycetales | 7 | Saprotroph | Undefined Saprotroph | Yeast | NULL | Possible | NULL | Sterkenburg E, et al. 2015. New Phytologist 207:1145-1158 ((https://doi.org/10.1111/nph.13426)) |
| Otu08918 | 2 | k__Fungi(100);p__Ascomycota(100);c__Saccharomycetes(100);o__Saccharomycetales(100);f__Saccharomycetales_family_Incertae_sedis(100);g__Candida(100);g__Candida_unclassified(100); | Candida | 13 | Pathotroph-Saprotroph-Symbiotroph | Animal Pathogen-Endophyte-Endosymbiont-Epiphyte-\|Undefined Saprotroph\| | Dimorphic Yeast-Yeast | NULL | Probable | As nectar-tap_saprotroph (Põlme et al. 2020); Decay sugar-rich substrates (Põlme et al. 2020); This asexual genus is polyphyletic and contain species that function very differently depend on what they are. Caution should be taken when interpreting guilds for Candida species. | Manolakaki D, et al. 2010. Virulence. 1: 367-375 ((https://doi.org/10.4161/viru.1.5.12796)); Mekbib SB, et al. 2011. Tropical Plant Pathology 36:233-240 ((https://doi.org/10.1590/S1982-56762011000400004)); Põlme S, et al. 2020. Fungal Diversity 105:1-16 ((https://doi.org/10.1007/s13225-020-00466-2)) |
| Otu08919 | 2 | k__Fungi(100);p__Ascomycota(100);c__Saccharomycetes(100);o__Saccharomycetales(100);f__Metschnikowiaceae(100);g__Metschnikowia(100);g__Metschnikowia_unclassified(100); | Metschnikowia | 13 | Saprotroph | \|Undefined Saprotroph\| | Yeast | NULL | Probable | As nectar-tap_saprotroph (Põlme et al. 2020); Decay sugar-rich substrates (Põlme et al. 2020) | Põlme S, et al. 2020. Fungal Diversity 105:1-16 ((https://doi.org/10.1007/s13225-020-00466-2)) |
| Otu08920 | 2 | k__Fungi(100);p__Ascomycota(100);c__Saccharomycetes(100);o__Saccharomycetales(100);f__Metschnikowiaceae(100);g__Metschnikowia(100);g__Metschnikowia_unclassified(100); | Metschnikowia | 13 | Saprotroph | \|Undefined Saprotroph\| | Yeast | NULL | Probable | As nectar-tap_saprotroph (Põlme et al. 2020); Decay sugar-rich substrates (Põlme et al. 2020) | Põlme S, et al. 2020. Fungal Diversity 105:1-16 ((https://doi.org/10.1007/s13225-020-00466-2)) |
| Otu08921 | 2 | k__Fungi(100);p__Ascomycota(100);c__Saccharomycetes(100);o__Saccharomycetales(100);f__Metschnikowiaceae(100);g__Metschnikowia(100);g__Metschnikowia_unclassified(100); | Metschnikowia | 13 | Saprotroph | \|Undefined Saprotroph\| | Yeast | NULL | Probable | As nectar-tap_saprotroph (Põlme et al. 2020); Decay sugar-rich substrates (Põlme et al. 2020) | Põlme S, et al. 2020. Fungal Diversity 105:1-16 ((https://doi.org/10.1007/s13225-020-00466-2)) |
| Otu08922 | 2 | k__Fungi(100);p__Ascomycota(100);c__Saccharomycetes(100);o__Saccharomycetales(100);f__Metschnikowiaceae(100);g__Metschnikowia(100);g__Metschnikowia_unclassified(100); | Metschnikowia | 13 | Saprotroph | \|Undefined Saprotroph\| | Yeast | NULL | Probable | As nectar-tap_saprotroph (Põlme et al. 2020); Decay sugar-rich substrates (Põlme et al. 2020) | Põlme S, et al. 2020. Fungal Diversity 105:1-16 ((https://doi.org/10.1007/s13225-020-00466-2)) |
| Otu08923 | 2 | k__Fungi(100);p__Ascomycota(100);c__Saccharomycetes(100);o__Saccharomycetales(100);f__Saccharomycetales_family_Incertae_sedis(100);g__Candida(100);g__Candida_unclassified(100); | Candida | 13 | Pathotroph-Saprotroph-Symbiotroph | Animal Pathogen-Endophyte-Endosymbiont-Epiphyte-\|Undefined Saprotroph\| | Dimorphic Yeast-Yeast | NULL | Probable | As nectar-tap_saprotroph (Põlme et al. 2020); Decay sugar-rich substrates (Põlme et al. 2020); This asexual genus is polyphyletic and contain species that function very differently depend on what they are. Caution should be taken when interpreting guilds for Candida species. | Manolakaki D, et al. 2010. Virulence. 1: 367-375 ((https://doi.org/10.4161/viru.1.5.12796)); Mekbib SB, et al. 2011. Tropical Plant Pathology 36:233-240 ((https://doi.org/10.1590/S1982-56762011000400004)); Põlme S, et al. 2020. Fungal Diversity 105:1-16 ((https://doi.org/10.1007/s13225-020-00466-2)) |
| Otu08924 | 2 | k__Fungi(100);p__Ascomycota(100);c__Saccharomycetes(100);o__Saccharomycetales(100);f__Metschnikowiaceae(100);g__Metschnikowia(100);g__Metschnikowia_unclassified(100); | Metschnikowia | 13 | Saprotroph | \|Undefined Saprotroph\| | Yeast | NULL | Probable | As nectar-tap_saprotroph (Põlme et al. 2020); Decay sugar-rich substrates (Põlme et al. 2020) | Põlme S, et al. 2020. Fungal Diversity 105:1-16 ((https://doi.org/10.1007/s13225-020-00466-2)) |
| Otu08926 | 2 | k__Fungi(100);p__Ascomycota(100);c__Leotiomycetes(100);o__Helotiales(100);f__Rutstroemiaceae(100);g__Rutstroemia(100);g__Rutstroemia_unclassified(100); | Rutstroemia | 13 | Saprotroph | \|Plant Saprotroph\| | Microfungus | NULL | Probable | As litter_saprotroph (Põlme et al. 2020); Decay leaf, fruit, seed (Põlme et al. 2020); Essentially saprobic on various plant parts, especially seeds and fruits, though initial infection may take place in living tissue (Cannon & Kirk 2007) | Cannon PF, Kirk PM. 2007. Fungal Families of the World. CAB International, Cambridge (ISBN: 978-0851998275); Tedersoo L, et al. 2014. Science 346:e1256688 ((https://doi.org/10.1126/science.1256688)); Põlme S, et al. 2020. Fungal Diversity 105:1-16 ((https://doi.org/10.1007/s13225-020-00466-2)) |
| Otu08927 | 2 | k__Fungi(100);p__Ascomycota(100);c__Saccharomycetes(100);o__Saccharomycetales(100);f__Metschnikowiaceae(100);g__Metschnikowia(100);s__Metschnikowia_pulcherrima(100); | Metschnikowia | 13 | Saprotroph | \|Undefined Saprotroph\| | Yeast | NULL | Probable | As nectar-tap_saprotroph (Põlme et al. 2020); Decay sugar-rich substrates (Põlme et al. 2020) | Põlme S, et al. 2020. Fungal Diversity 105:1-16 ((https://doi.org/10.1007/s13225-020-00466-2)) |
| Otu08928 | 2 | k__Fungi(100);p__Ascomycota(100);c__Saccharomycetes(100);o__Saccharomycetales(100);f__Metschnikowiaceae(100);g__Metschnikowia(100);s__Metschnikowia_pulcherrima(100); | Metschnikowia | 13 | Saprotroph | \|Undefined Saprotroph\| | Yeast | NULL | Probable | As nectar-tap_saprotroph (Põlme et al. 2020); Decay sugar-rich substrates (Põlme et al. 2020) | Põlme S, et al. 2020. Fungal Diversity 105:1-16 ((https://doi.org/10.1007/s13225-020-00466-2)) |
| Otu08931 | 2 | k__Fungi(100);p__Ascomycota(100);c__Saccharomycetes(100);o__Saccharomycetales(100);f__Metschnikowiaceae(100);g__Metschnikowia(100);s__Metschnikowia_pulcherrima(100); | Metschnikowia | 13 | Saprotroph | \|Undefined Saprotroph\| | Yeast | NULL | Probable | As nectar-tap_saprotroph (Põlme et al. 2020); Decay sugar-rich substrates (Põlme et al. 2020) | Põlme S, et al. 2020. Fungal Diversity 105:1-16 ((https://doi.org/10.1007/s13225-020-00466-2)) |
| Otu08933 | 2 | k__Fungi(100);p__Ascomycota(100);c__Saccharomycetes(100);o__Saccharomycetales(100);f__Metschnikowiaceae(100);g__Metschnikowia(100);s__Metschnikowia_pulcherrima(100); | Metschnikowia | 13 | Saprotroph | \|Undefined Saprotroph\| | Yeast | NULL | Probable | As nectar-tap_saprotroph (Põlme et al. 2020); Decay sugar-rich substrates (Põlme et al. 2020) | Põlme S, et al. 2020. Fungal Diversity 105:1-16 ((https://doi.org/10.1007/s13225-020-00466-2)) |
| Otu08934 | 2 | k__Fungi(100);p__Ascomycota(100);c__Saccharomycetes(100);o__Saccharomycetales(100);f__Metschnikowiaceae(100);g__Metschnikowia(100);s__Metschnikowia_pulcherrima(100); | Metschnikowia | 13 | Saprotroph | \|Undefined Saprotroph\| | Yeast | NULL | Probable | As nectar-tap_saprotroph (Põlme et al. 2020); Decay sugar-rich substrates (Põlme et al. 2020) | Põlme S, et al. 2020. Fungal Diversity 105:1-16 ((https://doi.org/10.1007/s13225-020-00466-2)) |
| Otu08938 | 2 | k__Fungi(100);p__Ascomycota(100);c__Saccharomycetes(100);o__Saccharomycetales(100);f__Metschnikowiaceae(100);g__Metschnikowia(100);s__Metschnikowia_pulcherrima(100); | Metschnikowia | 13 | Saprotroph | \|Undefined Saprotroph\| | Yeast | NULL | Probable | As nectar-tap_saprotroph (Põlme et al. 2020); Decay sugar-rich substrates (Põlme et al. 2020) | Põlme S, et al. 2020. Fungal Diversity 105:1-16 ((https://doi.org/10.1007/s13225-020-00466-2)) |
| Otu08939 | 2 | k__Fungi(100);p__Ascomycota(100);c__Saccharomycetes(100);o__Saccharomycetales(100);f__Metschnikowiaceae(100);g__Metschnikowia(100);s__Metschnikowia_pulcherrima(100); | Metschnikowia | 13 | Saprotroph | \|Undefined Saprotroph\| | Yeast | NULL | Probable | As nectar-tap_saprotroph (Põlme et al. 2020); Decay sugar-rich substrates (Põlme et al. 2020) | Põlme S, et al. 2020. Fungal Diversity 105:1-16 ((https://doi.org/10.1007/s13225-020-00466-2)) |
| Otu08940 | 2 | k__Fungi(100);p__Ascomycota(100);c__Saccharomycetes(100);o__Saccharomycetales(100);f__Saccharomycetales_family_Incertae_sedis(100);g__Candida(100);g__Candida_unclassified(100); | Candida | 13 | Pathotroph-Saprotroph-Symbiotroph | Animal Pathogen-Endophyte-Endosymbiont-Epiphyte-\|Undefined Saprotroph\| | Dimorphic Yeast-Yeast | NULL | Probable | As nectar-tap_saprotroph (Põlme et al. 2020); Decay sugar-rich substrates (Põlme et al. 2020); This asexual genus is polyphyletic and contain species that function very differently depend on what they are. Caution should be taken when interpreting guilds for Candida species. | Manolakaki D, et al. 2010. Virulence. 1: 367-375 ((https://doi.org/10.4161/viru.1.5.12796)); Mekbib SB, et al. 2011. Tropical Plant Pathology 36:233-240 ((https://doi.org/10.1590/S1982-56762011000400004)); Põlme S, et al. 2020. Fungal Diversity 105:1-16 ((https://doi.org/10.1007/s13225-020-00466-2)) |
| Otu08942 | 2 | k__Fungi(100);p__Ascomycota(100);c__Saccharomycetes(100);o__Saccharomycetales(100);f__Saccharomycetales_family_Incertae_sedis(100);g__Candida(100);g__Candida_unclassified(100); | Candida | 13 | Pathotroph-Saprotroph-Symbiotroph | Animal Pathogen-Endophyte-Endosymbiont-Epiphyte-\|Undefined Saprotroph\| | Dimorphic Yeast-Yeast | NULL | Probable | As nectar-tap_saprotroph (Põlme et al. 2020); Decay sugar-rich substrates (Põlme et al. 2020); This asexual genus is polyphyletic and contain species that function very differently depend on what they are. Caution should be taken when interpreting guilds for Candida species. | Manolakaki D, et al. 2010. Virulence. 1: 367-375 ((https://doi.org/10.4161/viru.1.5.12796)); Mekbib SB, et al. 2011. Tropical Plant Pathology 36:233-240 ((https://doi.org/10.1590/S1982-56762011000400004)); Põlme S, et al. 2020. Fungal Diversity 105:1-16 ((https://doi.org/10.1007/s13225-020-00466-2)) |
| Otu08944 | 2 | k__Fungi(100);p__Ascomycota(100);c__Saccharomycetes(100);o__Saccharomycetales(100);f__Metschnikowiaceae(100);g__Metschnikowia(100);g__Metschnikowia_unclassified(100); | Metschnikowia | 13 | Saprotroph | \|Undefined Saprotroph\| | Yeast | NULL | Probable | As nectar-tap_saprotroph (Põlme et al. 2020); Decay sugar-rich substrates (Põlme et al. 2020) | Põlme S, et al. 2020. Fungal Diversity 105:1-16 ((https://doi.org/10.1007/s13225-020-00466-2)) |
| Otu08945 | 2 | k__Fungi(100);p__Ascomycota(100);c__Saccharomycetes(100);o__Saccharomycetales(100);f__Metschnikowiaceae(100);g__Metschnikowia(100);g__Metschnikowia_unclassified(100); | Metschnikowia | 13 | Saprotroph | \|Undefined Saprotroph\| | Yeast | NULL | Probable | As nectar-tap_saprotroph (Põlme et al. 2020); Decay sugar-rich substrates (Põlme et al. 2020) | Põlme S, et al. 2020. Fungal Diversity 105:1-16 ((https://doi.org/10.1007/s13225-020-00466-2)) |
| Otu08946 | 2 | k__Fungi(100);p__Ascomycota(100);c__Saccharomycetes(100);o__Saccharomycetales(100);f__Metschnikowiaceae(100);g__Metschnikowia(100);s__Metschnikowia_pulcherrima(100); | Metschnikowia | 13 | Saprotroph | \|Undefined Saprotroph\| | Yeast | NULL | Probable | As nectar-tap_saprotroph (Põlme et al. 2020); Decay sugar-rich substrates (Põlme et al. 2020) | Põlme S, et al. 2020. Fungal Diversity 105:1-16 ((https://doi.org/10.1007/s13225-020-00466-2)) |
| Otu08947 | 2 | k__Fungi(100);p__Ascomycota(100);c__Saccharomycetes(100);o__Saccharomycetales(100);f__Metschnikowiaceae(100);g__Metschnikowia(100);g__Metschnikowia_unclassified(100); | Metschnikowia | 13 | Saprotroph | \|Undefined Saprotroph\| | Yeast | NULL | Probable | As nectar-tap_saprotroph (Põlme et al. 2020); Decay sugar-rich substrates (Põlme et al. 2020) | Põlme S, et al. 2020. Fungal Diversity 105:1-16 ((https://doi.org/10.1007/s13225-020-00466-2)) |
| Otu08949 | 2 | k__Fungi(100);p__Ascomycota(100);c__Saccharomycetes(100);o__Saccharomycetales(100);f__Metschnikowiaceae(100);g__Metschnikowia(100);g__Metschnikowia_unclassified(100); | Metschnikowia | 13 | Saprotroph | \|Undefined Saprotroph\| | Yeast | NULL | Probable | As nectar-tap_saprotroph (Põlme et al. 2020); Decay sugar-rich substrates (Põlme et al. 2020) | Põlme S, et al. 2020. Fungal Diversity 105:1-16 ((https://doi.org/10.1007/s13225-020-00466-2)) |
| Otu08950 | 2 | k__Fungi(100);p__Ascomycota(100);c__Saccharomycetes(100);o__Saccharomycetales(100);f__Metschnikowiaceae(100);g__Metschnikowia(100);s__Metschnikowia_pulcherrima(100); | Metschnikowia | 13 | Saprotroph | \|Undefined Saprotroph\| | Yeast | NULL | Probable | As nectar-tap_saprotroph (Põlme et al. 2020); Decay sugar-rich substrates (Põlme et al. 2020) | Põlme S, et al. 2020. Fungal Diversity 105:1-16 ((https://doi.org/10.1007/s13225-020-00466-2)) |
| Otu08951 | 2 | k__Fungi(100);p__Ascomycota(100);c__Saccharomycetes(100);o__Saccharomycetales(100);f__Metschnikowiaceae(100);g__Metschnikowia(100);g__Metschnikowia_unclassified(100); | Metschnikowia | 13 | Saprotroph | \|Undefined Saprotroph\| | Yeast | NULL | Probable | As nectar-tap_saprotroph (Põlme et al. 2020); Decay sugar-rich substrates (Põlme et al. 2020) | Põlme S, et al. 2020. Fungal Diversity 105:1-16 ((https://doi.org/10.1007/s13225-020-00466-2)) |
| Otu08952 | 2 | k__Fungi(100);p__Basidiomycota(100);c__Microbotryomycetes(100);o__Sporidiobolales(100);f__Sporidiobolales_family_Incertae_sedis(100);g__Sporobolomyces(100);s__Sporobolomyces_lactosus(100); | Sporobolomyces | 13 | Pathotroph-Saprotroph | \|Fungal Parasite\|-Plant Saprotroph | Yeast | NULL | Probable | As litter_saprotroph (Põlme et al. 2020); As mycoparasite (Põlme et al. 2020); Decay fungal material (Põlme et al. 2020) | Tedersoo L, et al. 2014. Science 346:e1256688 ((https://doi.org/10.1126/science.1256688)); Purahong W, et al. 2016. Molecular Ecology 25:4059-4074 ((https://doi.org/10.1111/mec.13739)); Põlme S, et al. 2020. Fungal Diversity 105:1-16 ((https://doi.org/10.1007/s13225-020-00466-2)) |
| Otu08953 | 2 | k__Fungi(100);p__Ascomycota(100);c__Saccharomycetes(100);o__Saccharomycetales(100);f__Saccharomycetales_family_Incertae_sedis(100);g__Candida(100);g__Candida_unclassified(100); | Candida | 13 | Pathotroph-Saprotroph-Symbiotroph | Animal Pathogen-Endophyte-Endosymbiont-Epiphyte-\|Undefined Saprotroph\| | Dimorphic Yeast-Yeast | NULL | Probable | As nectar-tap_saprotroph (Põlme et al. 2020); Decay sugar-rich substrates (Põlme et al. 2020); This asexual genus is polyphyletic and contain species that function very differently depend on what they are. Caution should be taken when interpreting guilds for Candida species. | Manolakaki D, et al. 2010. Virulence. 1: 367-375 ((https://doi.org/10.4161/viru.1.5.12796)); Mekbib SB, et al. 2011. Tropical Plant Pathology 36:233-240 ((https://doi.org/10.1590/S1982-56762011000400004)); Põlme S, et al. 2020. Fungal Diversity 105:1-16 ((https://doi.org/10.1007/s13225-020-00466-2)) |
| Otu08956 | 2 | k__Fungi(100);p__Ascomycota(100);c__Saccharomycetes(100);o__Saccharomycetales(100);f__Metschnikowiaceae(100);g__Metschnikowia(100);s__Metschnikowia_pulcherrima(100); | Metschnikowia | 13 | Saprotroph | \|Undefined Saprotroph\| | Yeast | NULL | Probable | As nectar-tap_saprotroph (Põlme et al. 2020); Decay sugar-rich substrates (Põlme et al. 2020) | Põlme S, et al. 2020. Fungal Diversity 105:1-16 ((https://doi.org/10.1007/s13225-020-00466-2)) |
| Otu08957 | 2 | k__Fungi(100);p__Ascomycota(100);c__Saccharomycetes(100);o__Saccharomycetales(100);f__Metschnikowiaceae(100);g__Metschnikowia(100);s__Metschnikowia_pulcherrima(100); | Metschnikowia | 13 | Saprotroph | \|Undefined Saprotroph\| | Yeast | NULL | Probable | As nectar-tap_saprotroph (Põlme et al. 2020); Decay sugar-rich substrates (Põlme et al. 2020) | Põlme S, et al. 2020. Fungal Diversity 105:1-16 ((https://doi.org/10.1007/s13225-020-00466-2)) |
| Otu08959 | 2 | k__Fungi(100);p__Basidiomycota(100);c__Microbotryomycetes(100);o__Sporidiobolales(100);f__Sporidiobolales_family_Incertae_sedis(100);g__Sporobolomyces(100);s__Sporobolomyces_lactosus(100); | Sporobolomyces | 13 | Pathotroph-Saprotroph | \|Fungal Parasite\|-Plant Saprotroph | Yeast | NULL | Probable | As litter_saprotroph (Põlme et al. 2020); As mycoparasite (Põlme et al. 2020); Decay fungal material (Põlme et al. 2020) | Tedersoo L, et al. 2014. Science 346:e1256688 ((https://doi.org/10.1126/science.1256688)); Purahong W, et al. 2016. Molecular Ecology 25:4059-4074 ((https://doi.org/10.1111/mec.13739)); Põlme S, et al. 2020. Fungal Diversity 105:1-16 ((https://doi.org/10.1007/s13225-020-00466-2)) |
| Otu08966 | 2 | k__Fungi(100);p__Ascomycota(100);c__Saccharomycetes(100);o__Saccharomycetales(100);f__Metschnikowiaceae(100);g__Metschnikowia(100);s__Metschnikowia_pulcherrima(100); | Metschnikowia | 13 | Saprotroph | \|Undefined Saprotroph\| | Yeast | NULL | Probable | As nectar-tap_saprotroph (Põlme et al. 2020); Decay sugar-rich substrates (Põlme et al. 2020) | Põlme S, et al. 2020. Fungal Diversity 105:1-16 ((https://doi.org/10.1007/s13225-020-00466-2)) |
| Otu08967 | 2 | k__Fungi(100);p__Basidiomycota(100);c__Microbotryomycetes(100);o__Sporidiobolales(100);f__Sporidiobolales_family_Incertae_sedis(100);g__Sporobolomyces(100);s__Sporobolomyces_lactosus(100); | Sporobolomyces | 13 | Pathotroph-Saprotroph | \|Fungal Parasite\|-Plant Saprotroph | Yeast | NULL | Probable | As litter_saprotroph (Põlme et al. 2020); As mycoparasite (Põlme et al. 2020); Decay fungal material (Põlme et al. 2020) | Tedersoo L, et al. 2014. Science 346:e1256688 ((https://doi.org/10.1126/science.1256688)); Purahong W, et al. 2016. Molecular Ecology 25:4059-4074 ((https://doi.org/10.1111/mec.13739)); Põlme S, et al. 2020. Fungal Diversity 105:1-16 ((https://doi.org/10.1007/s13225-020-00466-2)) |
| Otu08970 | 2 | k__Fungi(100);p__Ascomycota(100);c__Saccharomycetes(100);o__Saccharomycetales(100);f__Metschnikowiaceae(100);g__Metschnikowia(100);g__Metschnikowia_unclassified(100); | Metschnikowia | 13 | Saprotroph | \|Undefined Saprotroph\| | Yeast | NULL | Probable | As nectar-tap_saprotroph (Põlme et al. 2020); Decay sugar-rich substrates (Põlme et al. 2020) | Põlme S, et al. 2020. Fungal Diversity 105:1-16 ((https://doi.org/10.1007/s13225-020-00466-2)) |
| Otu08972 | 2 | k__Fungi(100);p__Ascomycota(100);c__Saccharomycetes(100);o__Saccharomycetales(100);f__Metschnikowiaceae(100);g__Metschnikowia(100);s__Metschnikowia_pulcherrima(100); | Metschnikowia | 13 | Saprotroph | \|Undefined Saprotroph\| | Yeast | NULL | Probable | As nectar-tap_saprotroph (Põlme et al. 2020); Decay sugar-rich substrates (Põlme et al. 2020) | Põlme S, et al. 2020. Fungal Diversity 105:1-16 ((https://doi.org/10.1007/s13225-020-00466-2)) |
| Otu08973 | 2 | k__Fungi(100);p__Ascomycota(100);c__Saccharomycetes(100);o__Saccharomycetales(100);f__Metschnikowiaceae(100);g__Metschnikowia(100);g__Metschnikowia_unclassified(100); | Metschnikowia | 13 | Saprotroph | \|Undefined Saprotroph\| | Yeast | NULL | Probable | As nectar-tap_saprotroph (Põlme et al. 2020); Decay sugar-rich substrates (Põlme et al. 2020) | Põlme S, et al. 2020. Fungal Diversity 105:1-16 ((https://doi.org/10.1007/s13225-020-00466-2)) |
| Otu08974 | 2 | k__Fungi(100);p__Ascomycota(100);c__Saccharomycetes(100);o__Saccharomycetales(100);f__Metschnikowiaceae(100);g__Metschnikowia(100);g__Metschnikowia_unclassified(100); | Metschnikowia | 13 | Saprotroph | \|Undefined Saprotroph\| | Yeast | NULL | Probable | As nectar-tap_saprotroph (Põlme et al. 2020); Decay sugar-rich substrates (Põlme et al. 2020) | Põlme S, et al. 2020. Fungal Diversity 105:1-16 ((https://doi.org/10.1007/s13225-020-00466-2)) |
| Otu08976 | 2 | k__Fungi(100);p__Ascomycota(100);c__Saccharomycetes(100);o__Saccharomycetales(100);f__Metschnikowiaceae(100);g__Metschnikowia(100);s__Metschnikowia_pulcherrima(100); | Metschnikowia | 13 | Saprotroph | \|Undefined Saprotroph\| | Yeast | NULL | Probable | As nectar-tap_saprotroph (Põlme et al. 2020); Decay sugar-rich substrates (Põlme et al. 2020) | Põlme S, et al. 2020. Fungal Diversity 105:1-16 ((https://doi.org/10.1007/s13225-020-00466-2)) |
| Otu08977 | 2 | k__Fungi(100);p__Ascomycota(100);c__Saccharomycetes(100);o__Saccharomycetales(100);f__Metschnikowiaceae(100);g__Metschnikowia(100);g__Metschnikowia_unclassified(100); | Metschnikowia | 13 | Saprotroph | \|Undefined Saprotroph\| | Yeast | NULL | Probable | As nectar-tap_saprotroph (Põlme et al. 2020); Decay sugar-rich substrates (Põlme et al. 2020) | Põlme S, et al. 2020. Fungal Diversity 105:1-16 ((https://doi.org/10.1007/s13225-020-00466-2)) |
| Otu08978 | 2 | k__Fungi(100);p__Ascomycota(100);c__Saccharomycetes(100);o__Saccharomycetales(100);f__Metschnikowiaceae(100);g__Metschnikowia(100);g__Metschnikowia_unclassified(100); | Metschnikowia | 13 | Saprotroph | \|Undefined Saprotroph\| | Yeast | NULL | Probable | As nectar-tap_saprotroph (Põlme et al. 2020); Decay sugar-rich substrates (Põlme et al. 2020) | Põlme S, et al. 2020. Fungal Diversity 105:1-16 ((https://doi.org/10.1007/s13225-020-00466-2)) |
| Otu08979 | 2 | k__Fungi(100);p__Ascomycota(100);c__Saccharomycetes(100);o__Saccharomycetales(100);f__Metschnikowiaceae(100);g__Metschnikowia(100);s__Metschnikowia_pulcherrima(100); | Metschnikowia | 13 | Saprotroph | \|Undefined Saprotroph\| | Yeast | NULL | Probable | As nectar-tap_saprotroph (Põlme et al. 2020); Decay sugar-rich substrates (Põlme et al. 2020) | Põlme S, et al. 2020. Fungal Diversity 105:1-16 ((https://doi.org/10.1007/s13225-020-00466-2)) |
| Otu08980 | 2 | k__Fungi(100);p__Ascomycota(100);c__Saccharomycetes(100);o__Saccharomycetales(100);f__Metschnikowiaceae(100);g__Metschnikowia(100);g__Metschnikowia_unclassified(100); | Metschnikowia | 13 | Saprotroph | \|Undefined Saprotroph\| | Yeast | NULL | Probable | As nectar-tap_saprotroph (Põlme et al. 2020); Decay sugar-rich substrates (Põlme et al. 2020) | Põlme S, et al. 2020. Fungal Diversity 105:1-16 ((https://doi.org/10.1007/s13225-020-00466-2)) |
| Otu08982 | 2 | k__Fungi(100);p__Ascomycota(100);c__Saccharomycetes(100);o__Saccharomycetales(100);f__Metschnikowiaceae(100);g__Metschnikowia(100);g__Metschnikowia_unclassified(100); | Metschnikowia | 13 | Saprotroph | \|Undefined Saprotroph\| | Yeast | NULL | Probable | As nectar-tap_saprotroph (Põlme et al. 2020); Decay sugar-rich substrates (Põlme et al. 2020) | Põlme S, et al. 2020. Fungal Diversity 105:1-16 ((https://doi.org/10.1007/s13225-020-00466-2)) |
| Otu08988 | 2 | k__Fungi(100);p__Ascomycota(100);c__Saccharomycetes(100);o__Saccharomycetales(100);f__Metschnikowiaceae(100);g__Metschnikowia(100);g__Metschnikowia_unclassified(100); | Metschnikowia | 13 | Saprotroph | \|Undefined Saprotroph\| | Yeast | NULL | Probable | As nectar-tap_saprotroph (Põlme et al. 2020); Decay sugar-rich substrates (Põlme et al. 2020) | Põlme S, et al. 2020. Fungal Diversity 105:1-16 ((https://doi.org/10.1007/s13225-020-00466-2)) |
| Otu08989 | 2 | k__Fungi(100);p__Ascomycota(100);c__Saccharomycetes(100);o__Saccharomycetales(100);f__Metschnikowiaceae(100);g__Metschnikowia(100);s__Metschnikowia_pulcherrima(100); | Metschnikowia | 13 | Saprotroph | \|Undefined Saprotroph\| | Yeast | NULL | Probable | As nectar-tap_saprotroph (Põlme et al. 2020); Decay sugar-rich substrates (Põlme et al. 2020) | Põlme S, et al. 2020. Fungal Diversity 105:1-16 ((https://doi.org/10.1007/s13225-020-00466-2)) |
| Otu08990 | 2 | k__Fungi(100);p__Basidiomycota(100);c__Microbotryomycetes(100);o__Sporidiobolales(100);f__Sporidiobolales_family_Incertae_sedis(100);g__Sporobolomyces(100);s__Sporobolomyces_lactosus(100); | Sporobolomyces | 13 | Pathotroph-Saprotroph | \|Fungal Parasite\|-Plant Saprotroph | Yeast | NULL | Probable | As litter_saprotroph (Põlme et al. 2020); As mycoparasite (Põlme et al. 2020); Decay fungal material (Põlme et al. 2020) | Tedersoo L, et al. 2014. Science 346:e1256688 ((https://doi.org/10.1126/science.1256688)); Purahong W, et al. 2016. Molecular Ecology 25:4059-4074 ((https://doi.org/10.1111/mec.13739)); Põlme S, et al. 2020. Fungal Diversity 105:1-16 ((https://doi.org/10.1007/s13225-020-00466-2)) |
| Otu08991 | 2 | k__Fungi(100);p__Ascomycota(100);c__Saccharomycetes(100);o__Saccharomycetales(100);f__Metschnikowiaceae(100);g__Metschnikowia(100);g__Metschnikowia_unclassified(100); | Metschnikowia | 13 | Saprotroph | \|Undefined Saprotroph\| | Yeast | NULL | Probable | As nectar-tap_saprotroph (Põlme et al. 2020); Decay sugar-rich substrates (Põlme et al. 2020) | Põlme S, et al. 2020. Fungal Diversity 105:1-16 ((https://doi.org/10.1007/s13225-020-00466-2)) |
| Otu08992 | 2 | k__Fungi(100);p__Ascomycota(100);c__Saccharomycetes(100);o__Saccharomycetales(100);f__Metschnikowiaceae(100);g__Metschnikowia(100);g__Metschnikowia_unclassified(100); | Metschnikowia | 13 | Saprotroph | \|Undefined Saprotroph\| | Yeast | NULL | Probable | As nectar-tap_saprotroph (Põlme et al. 2020); Decay sugar-rich substrates (Põlme et al. 2020) | Põlme S, et al. 2020. Fungal Diversity 105:1-16 ((https://doi.org/10.1007/s13225-020-00466-2)) |
| Otu08993 | 2 | k__Fungi(100);p__Ascomycota(100);c__Saccharomycetes(100);o__Saccharomycetales(100);f__Metschnikowiaceae(100);g__Metschnikowia(100);s__Metschnikowia_pulcherrima(100); | Metschnikowia | 13 | Saprotroph | \|Undefined Saprotroph\| | Yeast | NULL | Probable | As nectar-tap_saprotroph (Põlme et al. 2020); Decay sugar-rich substrates (Põlme et al. 2020) | Põlme S, et al. 2020. Fungal Diversity 105:1-16 ((https://doi.org/10.1007/s13225-020-00466-2)) |
| Otu09000 | 2 | k__Fungi(100);p__Ascomycota(100);c__Saccharomycetes(100);o__Saccharomycetales(100);f__Metschnikowiaceae(100);g__Metschnikowia(100);s__Metschnikowia_pulcherrima(100); | Metschnikowia | 13 | Saprotroph | \|Undefined Saprotroph\| | Yeast | NULL | Probable | As nectar-tap_saprotroph (Põlme et al. 2020); Decay sugar-rich substrates (Põlme et al. 2020) | Põlme S, et al. 2020. Fungal Diversity 105:1-16 ((https://doi.org/10.1007/s13225-020-00466-2)) |
| Otu09001 | 2 | k__Fungi(100);p__Ascomycota(100);c__Saccharomycetes(100);o__Saccharomycetales(100);f__Metschnikowiaceae(100);g__Metschnikowia(100);g__Metschnikowia_unclassified(100); | Metschnikowia | 13 | Saprotroph | \|Undefined Saprotroph\| | Yeast | NULL | Probable | As nectar-tap_saprotroph (Põlme et al. 2020); Decay sugar-rich substrates (Põlme et al. 2020) | Põlme S, et al. 2020. Fungal Diversity 105:1-16 ((https://doi.org/10.1007/s13225-020-00466-2)) |
| Otu09002 | 2 | k__Fungi(100);p__Ascomycota(100);c__Saccharomycetes(100);o__Saccharomycetales(100);f__Metschnikowiaceae(100);g__Metschnikowia(100);g__Metschnikowia_unclassified(100); | Metschnikowia | 13 | Saprotroph | \|Undefined Saprotroph\| | Yeast | NULL | Probable | As nectar-tap_saprotroph (Põlme et al. 2020); Decay sugar-rich substrates (Põlme et al. 2020) | Põlme S, et al. 2020. Fungal Diversity 105:1-16 ((https://doi.org/10.1007/s13225-020-00466-2)) |
| Otu09004 | 2 | k__Fungi(100);p__Ascomycota(100);c__Saccharomycetes(100);o__Saccharomycetales(100);f__Metschnikowiaceae(100);g__Metschnikowia(100);g__Metschnikowia_unclassified(100); | Metschnikowia | 13 | Saprotroph | \|Undefined Saprotroph\| | Yeast | NULL | Probable | As nectar-tap_saprotroph (Põlme et al. 2020); Decay sugar-rich substrates (Põlme et al. 2020) | Põlme S, et al. 2020. Fungal Diversity 105:1-16 ((https://doi.org/10.1007/s13225-020-00466-2)) |
| Otu09005 | 2 | k__Fungi(100);p__Ascomycota(100);c__Saccharomycetes(100);o__Saccharomycetales(100);f__Metschnikowiaceae(100);g__Metschnikowia(100);g__Metschnikowia_unclassified(100); | Metschnikowia | 13 | Saprotroph | \|Undefined Saprotroph\| | Yeast | NULL | Probable | As nectar-tap_saprotroph (Põlme et al. 2020); Decay sugar-rich substrates (Põlme et al. 2020) | Põlme S, et al. 2020. Fungal Diversity 105:1-16 ((https://doi.org/10.1007/s13225-020-00466-2)) |
| Otu09006 | 2 | k__Fungi(100);p__Ascomycota(100);c__Saccharomycetes(100);o__Saccharomycetales(100);f__Metschnikowiaceae(100);g__Metschnikowia(100);s__Metschnikowia_pulcherrima(100); | Metschnikowia | 13 | Saprotroph | \|Undefined Saprotroph\| | Yeast | NULL | Probable | As nectar-tap_saprotroph (Põlme et al. 2020); Decay sugar-rich substrates (Põlme et al. 2020) | Põlme S, et al. 2020. Fungal Diversity 105:1-16 ((https://doi.org/10.1007/s13225-020-00466-2)) |
| Otu09009 | 2 | k__Fungi(100);p__Ascomycota(100);c__Saccharomycetes(100);o__Saccharomycetales(100);f__Metschnikowiaceae(100);g__Metschnikowia(100);s__Metschnikowia_pulcherrima(100); | Metschnikowia | 13 | Saprotroph | \|Undefined Saprotroph\| | Yeast | NULL | Probable | As nectar-tap_saprotroph (Põlme et al. 2020); Decay sugar-rich substrates (Põlme et al. 2020) | Põlme S, et al. 2020. Fungal Diversity 105:1-16 ((https://doi.org/10.1007/s13225-020-00466-2)) |
| Otu09012 | 2 | k__Fungi(100);p__Ascomycota(100);c__Saccharomycetes(100);o__Saccharomycetales(100);f__Metschnikowiaceae(100);g__Metschnikowia(100);g__Metschnikowia_unclassified(100); | Metschnikowia | 13 | Saprotroph | \|Undefined Saprotroph\| | Yeast | NULL | Probable | As nectar-tap_saprotroph (Põlme et al. 2020); Decay sugar-rich substrates (Põlme et al. 2020) | Põlme S, et al. 2020. Fungal Diversity 105:1-16 ((https://doi.org/10.1007/s13225-020-00466-2)) |
| Otu09014 | 2 | k__Fungi(100);p__Ascomycota(100);c__Saccharomycetes(100);o__Saccharomycetales(100);f__Metschnikowiaceae(100);g__Metschnikowia(100);g__Metschnikowia_unclassified(100); | Metschnikowia | 13 | Saprotroph | \|Undefined Saprotroph\| | Yeast | NULL | Probable | As nectar-tap_saprotroph (Põlme et al. 2020); Decay sugar-rich substrates (Põlme et al. 2020) | Põlme S, et al. 2020. Fungal Diversity 105:1-16 ((https://doi.org/10.1007/s13225-020-00466-2)) |
| Otu09016 | 2 | k__Fungi(100);p__Ascomycota(100);c__Saccharomycetes(100);o__Saccharomycetales(100);f__Metschnikowiaceae(100);g__Metschnikowia(100);s__Metschnikowia_pulcherrima(100); | Metschnikowia | 13 | Saprotroph | \|Undefined Saprotroph\| | Yeast | NULL | Probable | As nectar-tap_saprotroph (Põlme et al. 2020); Decay sugar-rich substrates (Põlme et al. 2020) | Põlme S, et al. 2020. Fungal Diversity 105:1-16 ((https://doi.org/10.1007/s13225-020-00466-2)) |
| Otu09018 | 2 | k__Fungi(100);p__Ascomycota(100);c__Saccharomycetes(100);o__Saccharomycetales(100);f__Metschnikowiaceae(100);g__Metschnikowia(100);g__Metschnikowia_unclassified(100); | Metschnikowia | 13 | Saprotroph | \|Undefined Saprotroph\| | Yeast | NULL | Probable | As nectar-tap_saprotroph (Põlme et al. 2020); Decay sugar-rich substrates (Põlme et al. 2020) | Põlme S, et al. 2020. Fungal Diversity 105:1-16 ((https://doi.org/10.1007/s13225-020-00466-2)) |
| Otu09019 | 2 | k__Fungi(100);p__Ascomycota(100);c__Saccharomycetes(100);o__Saccharomycetales(100);f__Metschnikowiaceae(100);g__Metschnikowia(100);g__Metschnikowia_unclassified(100); | Metschnikowia | 13 | Saprotroph | \|Undefined Saprotroph\| | Yeast | NULL | Probable | As nectar-tap_saprotroph (Põlme et al. 2020); Decay sugar-rich substrates (Põlme et al. 2020) | Põlme S, et al. 2020. Fungal Diversity 105:1-16 ((https://doi.org/10.1007/s13225-020-00466-2)) |
| Otu09020 | 2 | k__Fungi(100);p__Ascomycota(100);c__Saccharomycetes(100);o__Saccharomycetales(100);f__Metschnikowiaceae(100);g__Metschnikowia(100);s__Metschnikowia_pulcherrima(100); | Metschnikowia | 13 | Saprotroph | \|Undefined Saprotroph\| | Yeast | NULL | Probable | As nectar-tap_saprotroph (Põlme et al. 2020); Decay sugar-rich substrates (Põlme et al. 2020) | Põlme S, et al. 2020. Fungal Diversity 105:1-16 ((https://doi.org/10.1007/s13225-020-00466-2)) |
| Otu09021 | 2 | k__Fungi(100);p__Ascomycota(100);c__Saccharomycetes(100);o__Saccharomycetales(100);f__Metschnikowiaceae(100);g__Metschnikowia(100);s__Metschnikowia_pulcherrima(100); | Metschnikowia | 13 | Saprotroph | \|Undefined Saprotroph\| | Yeast | NULL | Probable | As nectar-tap_saprotroph (Põlme et al. 2020); Decay sugar-rich substrates (Põlme et al. 2020) | Põlme S, et al. 2020. Fungal Diversity 105:1-16 ((https://doi.org/10.1007/s13225-020-00466-2)) |
| Otu09024 | 2 | k__Fungi(100);p__Ascomycota(100);c__Saccharomycetes(100);o__Saccharomycetales(100);f__Metschnikowiaceae(100);g__Metschnikowia(100);s__Metschnikowia_pulcherrima(100); | Metschnikowia | 13 | Saprotroph | \|Undefined Saprotroph\| | Yeast | NULL | Probable | As nectar-tap_saprotroph (Põlme et al. 2020); Decay sugar-rich substrates (Põlme et al. 2020) | Põlme S, et al. 2020. Fungal Diversity 105:1-16 ((https://doi.org/10.1007/s13225-020-00466-2)) |
| Otu09025 | 2 | k__Fungi(100);p__Ascomycota(100);c__Saccharomycetes(100);o__Saccharomycetales(100);f__Metschnikowiaceae(100);g__Metschnikowia(100);g__Metschnikowia_unclassified(100); | Metschnikowia | 13 | Saprotroph | \|Undefined Saprotroph\| | Yeast | NULL | Probable | As nectar-tap_saprotroph (Põlme et al. 2020); Decay sugar-rich substrates (Põlme et al. 2020) | Põlme S, et al. 2020. Fungal Diversity 105:1-16 ((https://doi.org/10.1007/s13225-020-00466-2)) |
| Otu09027 | 2 | k__Fungi(100);p__Ascomycota(100);c__Saccharomycetes(100);o__Saccharomycetales(100);f__Metschnikowiaceae(100);g__Metschnikowia(100);g__Metschnikowia_unclassified(100); | Metschnikowia | 13 | Saprotroph | \|Undefined Saprotroph\| | Yeast | NULL | Probable | As nectar-tap_saprotroph (Põlme et al. 2020); Decay sugar-rich substrates (Põlme et al. 2020) | Põlme S, et al. 2020. Fungal Diversity 105:1-16 ((https://doi.org/10.1007/s13225-020-00466-2)) |
| Otu09029 | 2 | k__Fungi(100);p__Ascomycota(100);c__Saccharomycetes(100);o__Saccharomycetales(100);f__Metschnikowiaceae(100);g__Metschnikowia(100);s__Metschnikowia_pulcherrima(100); | Metschnikowia | 13 | Saprotroph | \|Undefined Saprotroph\| | Yeast | NULL | Probable | As nectar-tap_saprotroph (Põlme et al. 2020); Decay sugar-rich substrates (Põlme et al. 2020) | Põlme S, et al. 2020. Fungal Diversity 105:1-16 ((https://doi.org/10.1007/s13225-020-00466-2)) |
| Otu09032 | 2 | k__Fungi(100);p__Ascomycota(100);c__Saccharomycetes(100);o__Saccharomycetales(100);f__Metschnikowiaceae(100);g__Metschnikowia(100);g__Metschnikowia_unclassified(100); | Metschnikowia | 13 | Saprotroph | \|Undefined Saprotroph\| | Yeast | NULL | Probable | As nectar-tap_saprotroph (Põlme et al. 2020); Decay sugar-rich substrates (Põlme et al. 2020) | Põlme S, et al. 2020. Fungal Diversity 105:1-16 ((https://doi.org/10.1007/s13225-020-00466-2)) |
| Otu09034 | 2 | k__Fungi(100);p__Ascomycota(100);c__Saccharomycetes(100);o__Saccharomycetales(100);f__Metschnikowiaceae(100);g__Metschnikowia(100);s__Metschnikowia_pulcherrima(100); | Metschnikowia | 13 | Saprotroph | \|Undefined Saprotroph\| | Yeast | NULL | Probable | As nectar-tap_saprotroph (Põlme et al. 2020); Decay sugar-rich substrates (Põlme et al. 2020) | Põlme S, et al. 2020. Fungal Diversity 105:1-16 ((https://doi.org/10.1007/s13225-020-00466-2)) |
| Otu09035 | 2 | k__Fungi(100);p__Ascomycota(100);c__Saccharomycetes(100);o__Saccharomycetales(100);f__Metschnikowiaceae(100);g__Metschnikowia(100);s__Metschnikowia_pulcherrima(100); | Metschnikowia | 13 | Saprotroph | \|Undefined Saprotroph\| | Yeast | NULL | Probable | As nectar-tap_saprotroph (Põlme et al. 2020); Decay sugar-rich substrates (Põlme et al. 2020) | Põlme S, et al. 2020. Fungal Diversity 105:1-16 ((https://doi.org/10.1007/s13225-020-00466-2)) |
| Otu09036 | 2 | k__Fungi(100);p__Ascomycota(100);c__Saccharomycetes(100);o__Saccharomycetales(100);f__Metschnikowiaceae(100);g__Metschnikowia(100);g__Metschnikowia_unclassified(100); | Metschnikowia | 13 | Saprotroph | \|Undefined Saprotroph\| | Yeast | NULL | Probable | As nectar-tap_saprotroph (Põlme et al. 2020); Decay sugar-rich substrates (Põlme et al. 2020) | Põlme S, et al. 2020. Fungal Diversity 105:1-16 ((https://doi.org/10.1007/s13225-020-00466-2)) |
| Otu09037 | 2 | k__Fungi(100);p__Ascomycota(100);c__Saccharomycetes(100);o__Saccharomycetales(100);f__Metschnikowiaceae(100);g__Metschnikowia(100);s__Metschnikowia_pulcherrima(100); | Metschnikowia | 13 | Saprotroph | \|Undefined Saprotroph\| | Yeast | NULL | Probable | As nectar-tap_saprotroph (Põlme et al. 2020); Decay sugar-rich substrates (Põlme et al. 2020) | Põlme S, et al. 2020. Fungal Diversity 105:1-16 ((https://doi.org/10.1007/s13225-020-00466-2)) |
| Otu09040 | 2 | k__Fungi(100);p__Ascomycota(100);c__Saccharomycetes(100);o__Saccharomycetales(100);f__Metschnikowiaceae(100);g__Metschnikowia(100);g__Metschnikowia_unclassified(100); | Metschnikowia | 13 | Saprotroph | \|Undefined Saprotroph\| | Yeast | NULL | Probable | As nectar-tap_saprotroph (Põlme et al. 2020); Decay sugar-rich substrates (Põlme et al. 2020) | Põlme S, et al. 2020. Fungal Diversity 105:1-16 ((https://doi.org/10.1007/s13225-020-00466-2)) |
| Otu09041 | 2 | k__Fungi(100);p__Ascomycota(100);c__Saccharomycetes(100);o__Saccharomycetales(100);f__Metschnikowiaceae(100);g__Metschnikowia(100);s__Metschnikowia_pulcherrima(100); | Metschnikowia | 13 | Saprotroph | \|Undefined Saprotroph\| | Yeast | NULL | Probable | As nectar-tap_saprotroph (Põlme et al. 2020); Decay sugar-rich substrates (Põlme et al. 2020) | Põlme S, et al. 2020. Fungal Diversity 105:1-16 ((https://doi.org/10.1007/s13225-020-00466-2)) |
| Otu09043 | 2 | k__Fungi(100);p__Ascomycota(100);c__Saccharomycetes(100);o__Saccharomycetales(100);f__Metschnikowiaceae(100);g__Metschnikowia(100);g__Metschnikowia_unclassified(100); | Metschnikowia | 13 | Saprotroph | \|Undefined Saprotroph\| | Yeast | NULL | Probable | As nectar-tap_saprotroph (Põlme et al. 2020); Decay sugar-rich substrates (Põlme et al. 2020) | Põlme S, et al. 2020. Fungal Diversity 105:1-16 ((https://doi.org/10.1007/s13225-020-00466-2)) |
| Otu09047 | 2 | k__Fungi(100);p__Ascomycota(100);c__Saccharomycetes(100);o__Saccharomycetales(100);f__Metschnikowiaceae(100);g__Metschnikowia(100);s__Metschnikowia_pulcherrima(100); | Metschnikowia | 13 | Saprotroph | \|Undefined Saprotroph\| | Yeast | NULL | Probable | As nectar-tap_saprotroph (Põlme et al. 2020); Decay sugar-rich substrates (Põlme et al. 2020) | Põlme S, et al. 2020. Fungal Diversity 105:1-16 ((https://doi.org/10.1007/s13225-020-00466-2)) |
| Otu09048 | 2 | k__Fungi(100);p__Ascomycota(100);c__Saccharomycetes(100);o__Saccharomycetales(100);f__Metschnikowiaceae(100);g__Metschnikowia(100);g__Metschnikowia_unclassified(100); | Metschnikowia | 13 | Saprotroph | \|Undefined Saprotroph\| | Yeast | NULL | Probable | As nectar-tap_saprotroph (Põlme et al. 2020); Decay sugar-rich substrates (Põlme et al. 2020) | Põlme S, et al. 2020. Fungal Diversity 105:1-16 ((https://doi.org/10.1007/s13225-020-00466-2)) |
| Otu09051 | 2 | k__Fungi(100);p__Ascomycota(100);c__Saccharomycetes(100);o__Saccharomycetales(100);f__Metschnikowiaceae(100);g__Metschnikowia(100);s__Metschnikowia_pulcherrima(100); | Metschnikowia | 13 | Saprotroph | \|Undefined Saprotroph\| | Yeast | NULL | Probable | As nectar-tap_saprotroph (Põlme et al. 2020); Decay sugar-rich substrates (Põlme et al. 2020) | Põlme S, et al. 2020. Fungal Diversity 105:1-16 ((https://doi.org/10.1007/s13225-020-00466-2)) |
| Otu09052 | 2 | k__Fungi(100);p__Ascomycota(100);c__Saccharomycetes(100);o__Saccharomycetales(100);f__Metschnikowiaceae(100);g__Metschnikowia(100);g__Metschnikowia_unclassified(100); | Metschnikowia | 13 | Saprotroph | \|Undefined Saprotroph\| | Yeast | NULL | Probable | As nectar-tap_saprotroph (Põlme et al. 2020); Decay sugar-rich substrates (Põlme et al. 2020) | Põlme S, et al. 2020. Fungal Diversity 105:1-16 ((https://doi.org/10.1007/s13225-020-00466-2)) |
| Otu09053 | 2 | k__Fungi(100);p__Ascomycota(100);c__Saccharomycetes(100);o__Saccharomycetales(100);f__Metschnikowiaceae(100);g__Metschnikowia(100);s__Metschnikowia_pulcherrima(100); | Metschnikowia | 13 | Saprotroph | \|Undefined Saprotroph\| | Yeast | NULL | Probable | As nectar-tap_saprotroph (Põlme et al. 2020); Decay sugar-rich substrates (Põlme et al. 2020) | Põlme S, et al. 2020. Fungal Diversity 105:1-16 ((https://doi.org/10.1007/s13225-020-00466-2)) |
| Otu09054 | 2 | k__Fungi(100);p__Ascomycota(100);c__Saccharomycetes(100);o__Saccharomycetales(100);f__Metschnikowiaceae(100);g__Metschnikowia(100);g__Metschnikowia_unclassified(100); | Metschnikowia | 13 | Saprotroph | \|Undefined Saprotroph\| | Yeast | NULL | Probable | As nectar-tap_saprotroph (Põlme et al. 2020); Decay sugar-rich substrates (Põlme et al. 2020) | Põlme S, et al. 2020. Fungal Diversity 105:1-16 ((https://doi.org/10.1007/s13225-020-00466-2)) |
| Otu09055 | 2 | k__Fungi(100);p__Ascomycota(100);c__Saccharomycetes(100);o__Saccharomycetales(100);f__Metschnikowiaceae(100);g__Metschnikowia(100);s__Metschnikowia_pulcherrima(100); | Metschnikowia | 13 | Saprotroph | \|Undefined Saprotroph\| | Yeast | NULL | Probable | As nectar-tap_saprotroph (Põlme et al. 2020); Decay sugar-rich substrates (Põlme et al. 2020) | Põlme S, et al. 2020. Fungal Diversity 105:1-16 ((https://doi.org/10.1007/s13225-020-00466-2)) |
| Otu09057 | 2 | k__Fungi(100);p__Ascomycota(100);c__Saccharomycetes(100);o__Saccharomycetales(100);f__Metschnikowiaceae(100);g__Metschnikowia(100);s__Metschnikowia_pulcherrima(100); | Metschnikowia | 13 | Saprotroph | \|Undefined Saprotroph\| | Yeast | NULL | Probable | As nectar-tap_saprotroph (Põlme et al. 2020); Decay sugar-rich substrates (Põlme et al. 2020) | Põlme S, et al. 2020. Fungal Diversity 105:1-16 ((https://doi.org/10.1007/s13225-020-00466-2)) |
| Otu09058 | 2 | k__Fungi(100);p__Ascomycota(100);c__Saccharomycetes(100);o__Saccharomycetales(100);f__Saccharomycetales_family_Incertae_sedis(100);g__Candida(100);g__Candida_unclassified(100); | Candida | 13 | Pathotroph-Saprotroph-Symbiotroph | Animal Pathogen-Endophyte-Endosymbiont-Epiphyte-\|Undefined Saprotroph\| | Dimorphic Yeast-Yeast | NULL | Probable | As nectar-tap_saprotroph (Põlme et al. 2020); Decay sugar-rich substrates (Põlme et al. 2020); This asexual genus is polyphyletic and contain species that function very differently depend on what they are. Caution should be taken when interpreting guilds for Candida species. | Manolakaki D, et al. 2010. Virulence. 1: 367-375 ((https://doi.org/10.4161/viru.1.5.12796)); Mekbib SB, et al. 2011. Tropical Plant Pathology 36:233-240 ((https://doi.org/10.1590/S1982-56762011000400004)); Põlme S, et al. 2020. Fungal Diversity 105:1-16 ((https://doi.org/10.1007/s13225-020-00466-2)) |
| Otu09059 | 2 | k__Fungi(100);p__Ascomycota(100);c__Saccharomycetes(100);o__Saccharomycetales(100);f__Metschnikowiaceae(100);g__Metschnikowia(100);g__Metschnikowia_unclassified(100); | Metschnikowia | 13 | Saprotroph | \|Undefined Saprotroph\| | Yeast | NULL | Probable | As nectar-tap_saprotroph (Põlme et al. 2020); Decay sugar-rich substrates (Põlme et al. 2020) | Põlme S, et al. 2020. Fungal Diversity 105:1-16 ((https://doi.org/10.1007/s13225-020-00466-2)) |
| Otu09060 | 2 | k__Fungi(100);p__Basidiomycota(100);c__Tremellomycetes(100);o__Tremellales(100);f__Tremellales_family_Incertae_sedis(100);g__Dioszegia(100);s__Dioszegia_changbaiensis(100); | Dioszegia | 13 | Saprotroph | \|Plant Saprotroph\|-Undefined Saprotroph | Yeast | NULL | Probable | As litter_saprotroph (Põlme et al. 2020); Decay leaf, fruit, seed (Põlme et al. 2020); The Dioszegia subclade sensu Inacio (Inacio et al. 20015) is characterized by orange; yellow or orange to deep orange colonies. Isolated from plant leaves and soil. | Põlme S, et al. 2020. Fungal Diversity 105:1-16 ((https://doi.org/10.1007/s13225-020-00466-2)); Takashima M, et al. 2011 International Journal of Systematic and Evolutionary Microbiology 61:1736-1739 ((https://doi.org/10.1099/ijs.0.025254-0)) |
| Otu09061 | 2 | k__Fungi(100);p__Ascomycota(100);c__Saccharomycetes(100);o__Saccharomycetales(100);f__Metschnikowiaceae(100);g__Metschnikowia(100);s__Metschnikowia_pulcherrima(100); | Metschnikowia | 13 | Saprotroph | \|Undefined Saprotroph\| | Yeast | NULL | Probable | As nectar-tap_saprotroph (Põlme et al. 2020); Decay sugar-rich substrates (Põlme et al. 2020) | Põlme S, et al. 2020. Fungal Diversity 105:1-16 ((https://doi.org/10.1007/s13225-020-00466-2)) |
| Otu09067 | 2 | k__Fungi(100);p__Ascomycota(100);c__Saccharomycetes(100);o__Saccharomycetales(100);f__Metschnikowiaceae(100);g__Metschnikowia(100);g__Metschnikowia_unclassified(100); | Metschnikowia | 13 | Saprotroph | \|Undefined Saprotroph\| | Yeast | NULL | Probable | As nectar-tap_saprotroph (Põlme et al. 2020); Decay sugar-rich substrates (Põlme et al. 2020) | Põlme S, et al. 2020. Fungal Diversity 105:1-16 ((https://doi.org/10.1007/s13225-020-00466-2)) |
| Otu09070 | 2 | k__Fungi(100);p__Ascomycota(100);c__Saccharomycetes(100);o__Saccharomycetales(100);f__Metschnikowiaceae(100);g__Metschnikowia(100);s__Metschnikowia_pulcherrima(100); | Metschnikowia | 13 | Saprotroph | \|Undefined Saprotroph\| | Yeast | NULL | Probable | As nectar-tap_saprotroph (Põlme et al. 2020); Decay sugar-rich substrates (Põlme et al. 2020) | Põlme S, et al. 2020. Fungal Diversity 105:1-16 ((https://doi.org/10.1007/s13225-020-00466-2)) |
| Otu09071 | 2 | k__Fungi(100);p__Ascomycota(100);c__Saccharomycetes(100);o__Saccharomycetales(100);f__Metschnikowiaceae(100);g__Metschnikowia(100);s__Metschnikowia_pulcherrima(100); | Metschnikowia | 13 | Saprotroph | \|Undefined Saprotroph\| | Yeast | NULL | Probable | As nectar-tap_saprotroph (Põlme et al. 2020); Decay sugar-rich substrates (Põlme et al. 2020) | Põlme S, et al. 2020. Fungal Diversity 105:1-16 ((https://doi.org/10.1007/s13225-020-00466-2)) |
| Otu09072 | 2 | k__Fungi(100);p__Ascomycota(100);c__Saccharomycetes(100);o__Saccharomycetales(100);f__Metschnikowiaceae(100);g__Metschnikowia(100);g__Metschnikowia_unclassified(100); | Metschnikowia | 13 | Saprotroph | \|Undefined Saprotroph\| | Yeast | NULL | Probable | As nectar-tap_saprotroph (Põlme et al. 2020); Decay sugar-rich substrates (Põlme et al. 2020) | Põlme S, et al. 2020. Fungal Diversity 105:1-16 ((https://doi.org/10.1007/s13225-020-00466-2)) |
| Otu09074 | 2 | k__Fungi(100);p__Ascomycota(100);c__Saccharomycetes(100);o__Saccharomycetales(100);f__Metschnikowiaceae(100);g__Metschnikowia(100);s__Metschnikowia_pulcherrima(100); | Metschnikowia | 13 | Saprotroph | \|Undefined Saprotroph\| | Yeast | NULL | Probable | As nectar-tap_saprotroph (Põlme et al. 2020); Decay sugar-rich substrates (Põlme et al. 2020) | Põlme S, et al. 2020. Fungal Diversity 105:1-16 ((https://doi.org/10.1007/s13225-020-00466-2)) |
| Otu09078 | 2 | k__Fungi(100);p__Ascomycota(100);c__Saccharomycetes(100);o__Saccharomycetales(100);f__Metschnikowiaceae(100);g__Metschnikowia(100);g__Metschnikowia_unclassified(100); | Metschnikowia | 13 | Saprotroph | \|Undefined Saprotroph\| | Yeast | NULL | Probable | As nectar-tap_saprotroph (Põlme et al. 2020); Decay sugar-rich substrates (Põlme et al. 2020) | Põlme S, et al. 2020. Fungal Diversity 105:1-16 ((https://doi.org/10.1007/s13225-020-00466-2)) |
| Otu09079 | 2 | k__Fungi(100);p__Ascomycota(100);c__Saccharomycetes(100);o__Saccharomycetales(100);f__Metschnikowiaceae(100);g__Metschnikowia(100);s__Metschnikowia_pulcherrima(100); | Metschnikowia | 13 | Saprotroph | \|Undefined Saprotroph\| | Yeast | NULL | Probable | As nectar-tap_saprotroph (Põlme et al. 2020); Decay sugar-rich substrates (Põlme et al. 2020) | Põlme S, et al. 2020. Fungal Diversity 105:1-16 ((https://doi.org/10.1007/s13225-020-00466-2)) |
| Otu09080 | 2 | k__Fungi(100);p__Ascomycota(100);c__Saccharomycetes(100);o__Saccharomycetales(100);f__Metschnikowiaceae(100);g__Metschnikowia(100);s__Metschnikowia_pulcherrima(100); | Metschnikowia | 13 | Saprotroph | \|Undefined Saprotroph\| | Yeast | NULL | Probable | As nectar-tap_saprotroph (Põlme et al. 2020); Decay sugar-rich substrates (Põlme et al. 2020) | Põlme S, et al. 2020. Fungal Diversity 105:1-16 ((https://doi.org/10.1007/s13225-020-00466-2)) |
| Otu09082 | 2 | k__Fungi(100);p__Ascomycota(100);c__Saccharomycetes(100);o__Saccharomycetales(100);f__Saccharomycetales_family_Incertae_sedis(100);g__Candida(100);g__Candida_unclassified(100); | Candida | 13 | Pathotroph-Saprotroph-Symbiotroph | Animal Pathogen-Endophyte-Endosymbiont-Epiphyte-\|Undefined Saprotroph\| | Dimorphic Yeast-Yeast | NULL | Probable | As nectar-tap_saprotroph (Põlme et al. 2020); Decay sugar-rich substrates (Põlme et al. 2020); This asexual genus is polyphyletic and contain species that function very differently depend on what they are. Caution should be taken when interpreting guilds for Candida species. | Manolakaki D, et al. 2010. Virulence. 1: 367-375 ((https://doi.org/10.4161/viru.1.5.12796)); Mekbib SB, et al. 2011. Tropical Plant Pathology 36:233-240 ((https://doi.org/10.1590/S1982-56762011000400004)); Põlme S, et al. 2020. Fungal Diversity 105:1-16 ((https://doi.org/10.1007/s13225-020-00466-2)) |
| Otu09084 | 2 | k__Fungi(100);p__Ascomycota(100);c__Saccharomycetes(100);o__Saccharomycetales(100);f__Metschnikowiaceae(100);g__Metschnikowia(100);s__Metschnikowia_pulcherrima(100); | Metschnikowia | 13 | Saprotroph | \|Undefined Saprotroph\| | Yeast | NULL | Probable | As nectar-tap_saprotroph (Põlme et al. 2020); Decay sugar-rich substrates (Põlme et al. 2020) | Põlme S, et al. 2020. Fungal Diversity 105:1-16 ((https://doi.org/10.1007/s13225-020-00466-2)) |
| Otu09086 | 2 | k__Fungi(100);p__Ascomycota(100);c__Saccharomycetes(100);o__Saccharomycetales(100);f__Metschnikowiaceae(100);g__Metschnikowia(100);s__Metschnikowia_pulcherrima(100); | Metschnikowia | 13 | Saprotroph | \|Undefined Saprotroph\| | Yeast | NULL | Probable | As nectar-tap_saprotroph (Põlme et al. 2020); Decay sugar-rich substrates (Põlme et al. 2020) | Põlme S, et al. 2020. Fungal Diversity 105:1-16 ((https://doi.org/10.1007/s13225-020-00466-2)) |
| Otu09090 | 2 | k__Fungi(100);p__Ascomycota(100);c__Saccharomycetes(100);o__Saccharomycetales(100);f__Metschnikowiaceae(100);g__Metschnikowia(100);g__Metschnikowia_unclassified(100); | Metschnikowia | 13 | Saprotroph | \|Undefined Saprotroph\| | Yeast | NULL | Probable | As nectar-tap_saprotroph (Põlme et al. 2020); Decay sugar-rich substrates (Põlme et al. 2020) | Põlme S, et al. 2020. Fungal Diversity 105:1-16 ((https://doi.org/10.1007/s13225-020-00466-2)) |
| Otu09093 | 2 | k__Fungi(100);p__Ascomycota(100);c__Saccharomycetes(100);o__Saccharomycetales(100);f__Metschnikowiaceae(100);g__Metschnikowia(100);s__Metschnikowia_pulcherrima(100); | Metschnikowia | 13 | Saprotroph | \|Undefined Saprotroph\| | Yeast | NULL | Probable | As nectar-tap_saprotroph (Põlme et al. 2020); Decay sugar-rich substrates (Põlme et al. 2020) | Põlme S, et al. 2020. Fungal Diversity 105:1-16 ((https://doi.org/10.1007/s13225-020-00466-2)) |
| Otu09094 | 2 | k__Fungi(100);p__Ascomycota(100);c__Saccharomycetes(100);o__Saccharomycetales(100);f__Metschnikowiaceae(100);g__Metschnikowia(100);s__Metschnikowia_pulcherrima(100); | Metschnikowia | 13 | Saprotroph | \|Undefined Saprotroph\| | Yeast | NULL | Probable | As nectar-tap_saprotroph (Põlme et al. 2020); Decay sugar-rich substrates (Põlme et al. 2020) | Põlme S, et al. 2020. Fungal Diversity 105:1-16 ((https://doi.org/10.1007/s13225-020-00466-2)) |
| Otu09095 | 2 | k__Fungi(100);p__Ascomycota(100);c__Saccharomycetes(100);o__Saccharomycetales(100);f__Metschnikowiaceae(100);g__Metschnikowia(100);s__Metschnikowia_pulcherrima(100); | Metschnikowia | 13 | Saprotroph | \|Undefined Saprotroph\| | Yeast | NULL | Probable | As nectar-tap_saprotroph (Põlme et al. 2020); Decay sugar-rich substrates (Põlme et al. 2020) | Põlme S, et al. 2020. Fungal Diversity 105:1-16 ((https://doi.org/10.1007/s13225-020-00466-2)) |
| Otu09096 | 2 | k__Fungi(100);p__Ascomycota(100);c__Saccharomycetes(100);o__Saccharomycetales(100);f__Metschnikowiaceae(100);g__Metschnikowia(100);s__Metschnikowia_pulcherrima(100); | Metschnikowia | 13 | Saprotroph | \|Undefined Saprotroph\| | Yeast | NULL | Probable | As nectar-tap_saprotroph (Põlme et al. 2020); Decay sugar-rich substrates (Põlme et al. 2020) | Põlme S, et al. 2020. Fungal Diversity 105:1-16 ((https://doi.org/10.1007/s13225-020-00466-2)) |
| Otu09099 | 2 | k__Fungi(100);p__Ascomycota(100);c__Saccharomycetes(100);o__Saccharomycetales(100);f__Saccharomycetales_family_Incertae_sedis(100);g__Candida(100);g__Candida_unclassified(100); | Candida | 13 | Pathotroph-Saprotroph-Symbiotroph | Animal Pathogen-Endophyte-Endosymbiont-Epiphyte-\|Undefined Saprotroph\| | Dimorphic Yeast-Yeast | NULL | Probable | As nectar-tap_saprotroph (Põlme et al. 2020); Decay sugar-rich substrates (Põlme et al. 2020); This asexual genus is polyphyletic and contain species that function very differently depend on what they are. Caution should be taken when interpreting guilds for Candida species. | Manolakaki D, et al. 2010. Virulence. 1: 367-375 ((https://doi.org/10.4161/viru.1.5.12796)); Mekbib SB, et al. 2011. Tropical Plant Pathology 36:233-240 ((https://doi.org/10.1590/S1982-56762011000400004)); Põlme S, et al. 2020. Fungal Diversity 105:1-16 ((https://doi.org/10.1007/s13225-020-00466-2)) |
| Otu09100 | 2 | k__Fungi(100);p__Ascomycota(100);c__Saccharomycetes(100);o__Saccharomycetales(100);f__Metschnikowiaceae(100);g__Metschnikowia(100);s__Metschnikowia_pulcherrima(100); | Metschnikowia | 13 | Saprotroph | \|Undefined Saprotroph\| | Yeast | NULL | Probable | As nectar-tap_saprotroph (Põlme et al. 2020); Decay sugar-rich substrates (Põlme et al. 2020) | Põlme S, et al. 2020. Fungal Diversity 105:1-16 ((https://doi.org/10.1007/s13225-020-00466-2)) |
| Otu09101 | 2 | k__Fungi(100);p__Ascomycota(100);c__Saccharomycetes(100);o__Saccharomycetales(100);f__Saccharomycetales_family_Incertae_sedis(100);g__Candida(100);g__Candida_unclassified(100); | Candida | 13 | Pathotroph-Saprotroph-Symbiotroph | Animal Pathogen-Endophyte-Endosymbiont-Epiphyte-\|Undefined Saprotroph\| | Dimorphic Yeast-Yeast | NULL | Probable | As nectar-tap_saprotroph (Põlme et al. 2020); Decay sugar-rich substrates (Põlme et al. 2020); This asexual genus is polyphyletic and contain species that function very differently depend on what they are. Caution should be taken when interpreting guilds for Candida species. | Manolakaki D, et al. 2010. Virulence. 1: 367-375 ((https://doi.org/10.4161/viru.1.5.12796)); Mekbib SB, et al. 2011. Tropical Plant Pathology 36:233-240 ((https://doi.org/10.1590/S1982-56762011000400004)); Põlme S, et al. 2020. Fungal Diversity 105:1-16 ((https://doi.org/10.1007/s13225-020-00466-2)) |
| Otu09104 | 2 | k__Fungi(100);p__Ascomycota(100);c__Saccharomycetes(100);o__Saccharomycetales(100);f__Metschnikowiaceae(100);g__Metschnikowia(100);g__Metschnikowia_unclassified(100); | Metschnikowia | 13 | Saprotroph | \|Undefined Saprotroph\| | Yeast | NULL | Probable | As nectar-tap_saprotroph (Põlme et al. 2020); Decay sugar-rich substrates (Põlme et al. 2020) | Põlme S, et al. 2020. Fungal Diversity 105:1-16 ((https://doi.org/10.1007/s13225-020-00466-2)) |
| Otu09106 | 2 | k__Fungi(100);p__Ascomycota(100);c__Saccharomycetes(100);o__Saccharomycetales(100);f__Metschnikowiaceae(100);g__Metschnikowia(100);s__Metschnikowia_pulcherrima(100); | Metschnikowia | 13 | Saprotroph | \|Undefined Saprotroph\| | Yeast | NULL | Probable | As nectar-tap_saprotroph (Põlme et al. 2020); Decay sugar-rich substrates (Põlme et al. 2020) | Põlme S, et al. 2020. Fungal Diversity 105:1-16 ((https://doi.org/10.1007/s13225-020-00466-2)) |
| Otu09107 | 2 | k__Fungi(100);p__Ascomycota(100);c__Saccharomycetes(100);o__Saccharomycetales(100);f__Metschnikowiaceae(100);g__Metschnikowia(100);g__Metschnikowia_unclassified(100); | Metschnikowia | 13 | Saprotroph | \|Undefined Saprotroph\| | Yeast | NULL | Probable | As nectar-tap_saprotroph (Põlme et al. 2020); Decay sugar-rich substrates (Põlme et al. 2020) | Põlme S, et al. 2020. Fungal Diversity 105:1-16 ((https://doi.org/10.1007/s13225-020-00466-2)) |
| Otu09109 | 2 | k__Fungi(100);p__Ascomycota(100);c__Saccharomycetes(100);o__Saccharomycetales(100);f__Metschnikowiaceae(100);g__Metschnikowia(100);s__Metschnikowia_pulcherrima(100); | Metschnikowia | 13 | Saprotroph | \|Undefined Saprotroph\| | Yeast | NULL | Probable | As nectar-tap_saprotroph (Põlme et al. 2020); Decay sugar-rich substrates (Põlme et al. 2020) | Põlme S, et al. 2020. Fungal Diversity 105:1-16 ((https://doi.org/10.1007/s13225-020-00466-2)) |
| Otu09110 | 2 | k__Fungi(100);p__Ascomycota(100);c__Saccharomycetes(100);o__Saccharomycetales(100);f__Metschnikowiaceae(100);g__Metschnikowia(100);s__Metschnikowia_pulcherrima(100); | Metschnikowia | 13 | Saprotroph | \|Undefined Saprotroph\| | Yeast | NULL | Probable | As nectar-tap_saprotroph (Põlme et al. 2020); Decay sugar-rich substrates (Põlme et al. 2020) | Põlme S, et al. 2020. Fungal Diversity 105:1-16 ((https://doi.org/10.1007/s13225-020-00466-2)) |
| Otu09112 | 2 | k__Fungi(100);p__Ascomycota(100);c__Saccharomycetes(100);o__Saccharomycetales(100);f__Metschnikowiaceae(100);g__Metschnikowia(100);s__Metschnikowia_pulcherrima(100); | Metschnikowia | 13 | Saprotroph | \|Undefined Saprotroph\| | Yeast | NULL | Probable | As nectar-tap_saprotroph (Põlme et al. 2020); Decay sugar-rich substrates (Põlme et al. 2020) | Põlme S, et al. 2020. Fungal Diversity 105:1-16 ((https://doi.org/10.1007/s13225-020-00466-2)) |
| Otu09113 | 2 | k__Fungi(100);p__Ascomycota(100);c__Saccharomycetes(100);o__Saccharomycetales(100);f__Metschnikowiaceae(100);g__Metschnikowia(100);s__Metschnikowia_pulcherrima(100); | Metschnikowia | 13 | Saprotroph | \|Undefined Saprotroph\| | Yeast | NULL | Probable | As nectar-tap_saprotroph (Põlme et al. 2020); Decay sugar-rich substrates (Põlme et al. 2020) | Põlme S, et al. 2020. Fungal Diversity 105:1-16 ((https://doi.org/10.1007/s13225-020-00466-2)) |
| Otu09114 | 2 | k__Fungi(100);p__Ascomycota(100);c__Saccharomycetes(100);o__Saccharomycetales(100);f__Metschnikowiaceae(100);g__Metschnikowia(100);s__Metschnikowia_pulcherrima(100); | Metschnikowia | 13 | Saprotroph | \|Undefined Saprotroph\| | Yeast | NULL | Probable | As nectar-tap_saprotroph (Põlme et al. 2020); Decay sugar-rich substrates (Põlme et al. 2020) | Põlme S, et al. 2020. Fungal Diversity 105:1-16 ((https://doi.org/10.1007/s13225-020-00466-2)) |
| Otu09119 | 2 | k__Fungi(100);p__Ascomycota(100);c__Saccharomycetes(100);o__Saccharomycetales(100);f__Metschnikowiaceae(100);g__Metschnikowia(100);s__Metschnikowia_pulcherrima(100); | Metschnikowia | 13 | Saprotroph | \|Undefined Saprotroph\| | Yeast | NULL | Probable | As nectar-tap_saprotroph (Põlme et al. 2020); Decay sugar-rich substrates (Põlme et al. 2020) | Põlme S, et al. 2020. Fungal Diversity 105:1-16 ((https://doi.org/10.1007/s13225-020-00466-2)) |
| Otu09120 | 2 | k__Fungi(100);p__Ascomycota(100);c__Saccharomycetes(100);o__Saccharomycetales(100);f__Metschnikowiaceae(100);g__Metschnikowia(100);s__Metschnikowia_pulcherrima(100); | Metschnikowia | 13 | Saprotroph | \|Undefined Saprotroph\| | Yeast | NULL | Probable | As nectar-tap_saprotroph (Põlme et al. 2020); Decay sugar-rich substrates (Põlme et al. 2020) | Põlme S, et al. 2020. Fungal Diversity 105:1-16 ((https://doi.org/10.1007/s13225-020-00466-2)) |
| Otu09123 | 2 | k__Fungi(100);p__Ascomycota(100);c__Saccharomycetes(100);o__Saccharomycetales(100);f__Metschnikowiaceae(100);g__Metschnikowia(100);s__Metschnikowia_pulcherrima(100); | Metschnikowia | 13 | Saprotroph | \|Undefined Saprotroph\| | Yeast | NULL | Probable | As nectar-tap_saprotroph (Põlme et al. 2020); Decay sugar-rich substrates (Põlme et al. 2020) | Põlme S, et al. 2020. Fungal Diversity 105:1-16 ((https://doi.org/10.1007/s13225-020-00466-2)) |
| Otu09124 | 2 | k__Fungi(100);p__Ascomycota(100);c__Saccharomycetes(100);o__Saccharomycetales(100);f__Metschnikowiaceae(100);g__Metschnikowia(100);g__Metschnikowia_unclassified(100); | Metschnikowia | 13 | Saprotroph | \|Undefined Saprotroph\| | Yeast | NULL | Probable | As nectar-tap_saprotroph (Põlme et al. 2020); Decay sugar-rich substrates (Põlme et al. 2020) | Põlme S, et al. 2020. Fungal Diversity 105:1-16 ((https://doi.org/10.1007/s13225-020-00466-2)) |
| Otu09126 | 2 | k__Fungi(100);p__Ascomycota(100);c__Saccharomycetes(100);o__Saccharomycetales(100);f__Metschnikowiaceae(100);g__Metschnikowia(100);g__Metschnikowia_unclassified(100); | Metschnikowia | 13 | Saprotroph | \|Undefined Saprotroph\| | Yeast | NULL | Probable | As nectar-tap_saprotroph (Põlme et al. 2020); Decay sugar-rich substrates (Põlme et al. 2020) | Põlme S, et al. 2020. Fungal Diversity 105:1-16 ((https://doi.org/10.1007/s13225-020-00466-2)) |
| Otu09127 | 2 | k__Fungi(100);p__Ascomycota(100);c__Saccharomycetes(100);o__Saccharomycetales(100);f__Metschnikowiaceae(100);g__Metschnikowia(100);g__Metschnikowia_unclassified(100); | Metschnikowia | 13 | Saprotroph | \|Undefined Saprotroph\| | Yeast | NULL | Probable | As nectar-tap_saprotroph (Põlme et al. 2020); Decay sugar-rich substrates (Põlme et al. 2020) | Põlme S, et al. 2020. Fungal Diversity 105:1-16 ((https://doi.org/10.1007/s13225-020-00466-2)) |
| Otu09128 | 2 | k__Fungi(100);p__Ascomycota(100);c__Saccharomycetes(100);o__Saccharomycetales(100);f__Metschnikowiaceae(100);g__Metschnikowia(100);s__Metschnikowia_pulcherrima(100); | Metschnikowia | 13 | Saprotroph | \|Undefined Saprotroph\| | Yeast | NULL | Probable | As nectar-tap_saprotroph (Põlme et al. 2020); Decay sugar-rich substrates (Põlme et al. 2020) | Põlme S, et al. 2020. Fungal Diversity 105:1-16 ((https://doi.org/10.1007/s13225-020-00466-2)) |
| Otu09129 | 2 | k__Fungi(100);p__Ascomycota(100);c__Saccharomycetes(100);o__Saccharomycetales(100);f__Metschnikowiaceae(100);g__Metschnikowia(100);g__Metschnikowia_unclassified(100); | Metschnikowia | 13 | Saprotroph | \|Undefined Saprotroph\| | Yeast | NULL | Probable | As nectar-tap_saprotroph (Põlme et al. 2020); Decay sugar-rich substrates (Põlme et al. 2020) | Põlme S, et al. 2020. Fungal Diversity 105:1-16 ((https://doi.org/10.1007/s13225-020-00466-2)) |
| Otu09132 | 2 | k__Fungi(100);p__Ascomycota(100);c__Saccharomycetes(100);o__Saccharomycetales(100);f__Metschnikowiaceae(100);g__Metschnikowia(100);g__Metschnikowia_unclassified(100); | Metschnikowia | 13 | Saprotroph | \|Undefined Saprotroph\| | Yeast | NULL | Probable | As nectar-tap_saprotroph (Põlme et al. 2020); Decay sugar-rich substrates (Põlme et al. 2020) | Põlme S, et al. 2020. Fungal Diversity 105:1-16 ((https://doi.org/10.1007/s13225-020-00466-2)) |
| Otu09133 | 2 | k__Fungi(100);p__Ascomycota(100);c__Saccharomycetes(100);o__Saccharomycetales(100);f__Metschnikowiaceae(100);g__Metschnikowia(100);s__Metschnikowia_pulcherrima(100); | Metschnikowia | 13 | Saprotroph | \|Undefined Saprotroph\| | Yeast | NULL | Probable | As nectar-tap_saprotroph (Põlme et al. 2020); Decay sugar-rich substrates (Põlme et al. 2020) | Põlme S, et al. 2020. Fungal Diversity 105:1-16 ((https://doi.org/10.1007/s13225-020-00466-2)) |
| Otu09135 | 2 | k__Fungi(100);p__Ascomycota(100);c__Saccharomycetes(100);o__Saccharomycetales(100);f__Metschnikowiaceae(100);g__Metschnikowia(100);g__Metschnikowia_unclassified(100); | Metschnikowia | 13 | Saprotroph | \|Undefined Saprotroph\| | Yeast | NULL | Probable | As nectar-tap_saprotroph (Põlme et al. 2020); Decay sugar-rich substrates (Põlme et al. 2020) | Põlme S, et al. 2020. Fungal Diversity 105:1-16 ((https://doi.org/10.1007/s13225-020-00466-2)) |
| Otu09137 | 2 | k__Fungi(100);p__Ascomycota(100);c__Saccharomycetes(100);o__Saccharomycetales(100);f__Metschnikowiaceae(100);g__Metschnikowia(100);s__Metschnikowia_pulcherrima(100); | Metschnikowia | 13 | Saprotroph | \|Undefined Saprotroph\| | Yeast | NULL | Probable | As nectar-tap_saprotroph (Põlme et al. 2020); Decay sugar-rich substrates (Põlme et al. 2020) | Põlme S, et al. 2020. Fungal Diversity 105:1-16 ((https://doi.org/10.1007/s13225-020-00466-2)) |
| Otu09138 | 2 | k__Fungi(100);p__Ascomycota(100);c__Saccharomycetes(100);o__Saccharomycetales(100);f__Metschnikowiaceae(100);g__Metschnikowia(100);g__Metschnikowia_unclassified(100); | Metschnikowia | 13 | Saprotroph | \|Undefined Saprotroph\| | Yeast | NULL | Probable | As nectar-tap_saprotroph (Põlme et al. 2020); Decay sugar-rich substrates (Põlme et al. 2020) | Põlme S, et al. 2020. Fungal Diversity 105:1-16 ((https://doi.org/10.1007/s13225-020-00466-2)) |
| Otu09139 | 2 | k__Fungi(100);p__Ascomycota(100);c__Saccharomycetes(100);o__Saccharomycetales(100);f__Metschnikowiaceae(100);g__Metschnikowia(100);s__Metschnikowia_pulcherrima(100); | Metschnikowia | 13 | Saprotroph | \|Undefined Saprotroph\| | Yeast | NULL | Probable | As nectar-tap_saprotroph (Põlme et al. 2020); Decay sugar-rich substrates (Põlme et al. 2020) | Põlme S, et al. 2020. Fungal Diversity 105:1-16 ((https://doi.org/10.1007/s13225-020-00466-2)) |
| Otu09147 | 2 | k__Fungi(100);p__Ascomycota(100);c__Saccharomycetes(100);o__Saccharomycetales(100);f__Metschnikowiaceae(100);g__Metschnikowia(100);s__Metschnikowia_pulcherrima(100); | Metschnikowia | 13 | Saprotroph | \|Undefined Saprotroph\| | Yeast | NULL | Probable | As nectar-tap_saprotroph (Põlme et al. 2020); Decay sugar-rich substrates (Põlme et al. 2020) | Põlme S, et al. 2020. Fungal Diversity 105:1-16 ((https://doi.org/10.1007/s13225-020-00466-2)) |
| Otu09151 | 2 | k__Fungi(100);p__Ascomycota(100);c__Saccharomycetes(100);o__Saccharomycetales(100);f__Metschnikowiaceae(100);g__Metschnikowia(100);g__Metschnikowia_unclassified(100); | Metschnikowia | 13 | Saprotroph | \|Undefined Saprotroph\| | Yeast | NULL | Probable | As nectar-tap_saprotroph (Põlme et al. 2020); Decay sugar-rich substrates (Põlme et al. 2020) | Põlme S, et al. 2020. Fungal Diversity 105:1-16 ((https://doi.org/10.1007/s13225-020-00466-2)) |
| Otu09154 | 2 | k__Fungi(100);p__Ascomycota(100);c__Saccharomycetes(100);o__Saccharomycetales(100);f__Metschnikowiaceae(100);g__Metschnikowia(100);g__Metschnikowia_unclassified(100); | Metschnikowia | 13 | Saprotroph | \|Undefined Saprotroph\| | Yeast | NULL | Probable | As nectar-tap_saprotroph (Põlme et al. 2020); Decay sugar-rich substrates (Põlme et al. 2020) | Põlme S, et al. 2020. Fungal Diversity 105:1-16 ((https://doi.org/10.1007/s13225-020-00466-2)) |
| Otu09156 | 2 | k__Fungi(100);p__Ascomycota(100);c__Saccharomycetes(100);o__Saccharomycetales(100);f__Metschnikowiaceae(100);g__Metschnikowia(100);s__Metschnikowia_pulcherrima(100); | Metschnikowia | 13 | Saprotroph | \|Undefined Saprotroph\| | Yeast | NULL | Probable | As nectar-tap_saprotroph (Põlme et al. 2020); Decay sugar-rich substrates (Põlme et al. 2020) | Põlme S, et al. 2020. Fungal Diversity 105:1-16 ((https://doi.org/10.1007/s13225-020-00466-2)) |
| Otu09158 | 2 | k__Fungi(100);p__Ascomycota(100);c__Saccharomycetes(100);o__Saccharomycetales(100);f__Metschnikowiaceae(100);g__Metschnikowia(100);s__Metschnikowia_pulcherrima(100); | Metschnikowia | 13 | Saprotroph | \|Undefined Saprotroph\| | Yeast | NULL | Probable | As nectar-tap_saprotroph (Põlme et al. 2020); Decay sugar-rich substrates (Põlme et al. 2020) | Põlme S, et al. 2020. Fungal Diversity 105:1-16 ((https://doi.org/10.1007/s13225-020-00466-2)) |
| Otu09159 | 2 | k__Fungi(100);p__Ascomycota(100);c__Saccharomycetes(100);o__Saccharomycetales(100);f__Metschnikowiaceae(100);g__Metschnikowia(100);g__Metschnikowia_unclassified(100); | Metschnikowia | 13 | Saprotroph | \|Undefined Saprotroph\| | Yeast | NULL | Probable | As nectar-tap_saprotroph (Põlme et al. 2020); Decay sugar-rich substrates (Põlme et al. 2020) | Põlme S, et al. 2020. Fungal Diversity 105:1-16 ((https://doi.org/10.1007/s13225-020-00466-2)) |
| Otu09161 | 2 | k__Fungi(100);p__Ascomycota(100);c__Saccharomycetes(100);o__Saccharomycetales(100);f__Metschnikowiaceae(100);g__Metschnikowia(100);s__Metschnikowia_pulcherrima(100); | Metschnikowia | 13 | Saprotroph | \|Undefined Saprotroph\| | Yeast | NULL | Probable | As nectar-tap_saprotroph (Põlme et al. 2020); Decay sugar-rich substrates (Põlme et al. 2020) | Põlme S, et al. 2020. Fungal Diversity 105:1-16 ((https://doi.org/10.1007/s13225-020-00466-2)) |
| Otu09164 | 2 | k__Fungi(100);p__Ascomycota(100);c__Saccharomycetes(100);o__Saccharomycetales(100);f__Metschnikowiaceae(100);g__Metschnikowia(100);g__Metschnikowia_unclassified(100); | Metschnikowia | 13 | Saprotroph | \|Undefined Saprotroph\| | Yeast | NULL | Probable | As nectar-tap_saprotroph (Põlme et al. 2020); Decay sugar-rich substrates (Põlme et al. 2020) | Põlme S, et al. 2020. Fungal Diversity 105:1-16 ((https://doi.org/10.1007/s13225-020-00466-2)) |
| Otu09166 | 2 | k__Fungi(100);p__Ascomycota(100);c__Saccharomycetes(100);o__Saccharomycetales(100);f__Metschnikowiaceae(100);g__Metschnikowia(100);s__Metschnikowia_pulcherrima(100); | Metschnikowia | 13 | Saprotroph | \|Undefined Saprotroph\| | Yeast | NULL | Probable | As nectar-tap_saprotroph (Põlme et al. 2020); Decay sugar-rich substrates (Põlme et al. 2020) | Põlme S, et al. 2020. Fungal Diversity 105:1-16 ((https://doi.org/10.1007/s13225-020-00466-2)) |
| Otu09167 | 2 | k__Fungi(100);p__Ascomycota(100);c__Saccharomycetes(100);o__Saccharomycetales(100);f__Metschnikowiaceae(100);g__Metschnikowia(100);s__Metschnikowia_pulcherrima(100); | Metschnikowia | 13 | Saprotroph | \|Undefined Saprotroph\| | Yeast | NULL | Probable | As nectar-tap_saprotroph (Põlme et al. 2020); Decay sugar-rich substrates (Põlme et al. 2020) | Põlme S, et al. 2020. Fungal Diversity 105:1-16 ((https://doi.org/10.1007/s13225-020-00466-2)) |
| Otu09171 | 2 | k__Fungi(100);p__Ascomycota(100);c__Sordariomycetes(100);o__Hypocreales(100);f__Ophiocordycipitaceae(100);g__Elaphocordyceps(100);g__Elaphocordyceps_unclassified(100); | Elaphocordyceps | 13 | Pathotroph | Fungal Parasite | Secotioid-Spathuloid | NULL | Highly Probable | NULL | Samuels GJ. 2007. Self-published; Sung G-H, et al. 2007. Studies in Mycology 57:5-59 ((https://www.studiesinmycology.org/index.php/issue/59-studies-in-mycology-no-57)); List compiled by John Plischke III ((https://goodmushroom.weebly.com/uploads/5/7/1/7/57176535/fungi_on_fungi_pdf_lq.pdf)) |
| Otu09172 | 2 | k__Fungi(100);p__Ascomycota(100);c__Saccharomycetes(100);o__Saccharomycetales(100);f__Metschnikowiaceae(100);g__Metschnikowia(100);s__Metschnikowia_pulcherrima(100); | Metschnikowia | 13 | Saprotroph | \|Undefined Saprotroph\| | Yeast | NULL | Probable | As nectar-tap_saprotroph (Põlme et al. 2020); Decay sugar-rich substrates (Põlme et al. 2020) | Põlme S, et al. 2020. Fungal Diversity 105:1-16 ((https://doi.org/10.1007/s13225-020-00466-2)) |
| Otu09173 | 2 | k__Fungi(100);p__Ascomycota(100);c__Saccharomycetes(100);o__Saccharomycetales(100);f__Metschnikowiaceae(100);g__Metschnikowia(100);s__Metschnikowia_pulcherrima(100); | Metschnikowia | 13 | Saprotroph | \|Undefined Saprotroph\| | Yeast | NULL | Probable | As nectar-tap_saprotroph (Põlme et al. 2020); Decay sugar-rich substrates (Põlme et al. 2020) | Põlme S, et al. 2020. Fungal Diversity 105:1-16 ((https://doi.org/10.1007/s13225-020-00466-2)) |
| Otu09177 | 2 | k__Fungi(100);p__Ascomycota(100);c__Saccharomycetes(100);o__Saccharomycetales(100);f__Metschnikowiaceae(100);g__Metschnikowia(100);s__Metschnikowia_pulcherrima(100); | Metschnikowia | 13 | Saprotroph | \|Undefined Saprotroph\| | Yeast | NULL | Probable | As nectar-tap_saprotroph (Põlme et al. 2020); Decay sugar-rich substrates (Põlme et al. 2020) | Põlme S, et al. 2020. Fungal Diversity 105:1-16 ((https://doi.org/10.1007/s13225-020-00466-2)) |
| Otu09181 | 2 | k__Fungi(100);p__Ascomycota(100);c__Saccharomycetes(100);o__Saccharomycetales(100);f__Metschnikowiaceae(100);g__Metschnikowia(100);g__Metschnikowia_unclassified(100); | Metschnikowia | 13 | Saprotroph | \|Undefined Saprotroph\| | Yeast | NULL | Probable | As nectar-tap_saprotroph (Põlme et al. 2020); Decay sugar-rich substrates (Põlme et al. 2020) | Põlme S, et al. 2020. Fungal Diversity 105:1-16 ((https://doi.org/10.1007/s13225-020-00466-2)) |
| Otu09184 | 2 | k__Fungi(100);p__Ascomycota(100);c__Saccharomycetes(100);o__Saccharomycetales(100);f__Metschnikowiaceae(100);g__Metschnikowia(100);s__Metschnikowia_pulcherrima(100); | Metschnikowia | 13 | Saprotroph | \|Undefined Saprotroph\| | Yeast | NULL | Probable | As nectar-tap_saprotroph (Põlme et al. 2020); Decay sugar-rich substrates (Põlme et al. 2020) | Põlme S, et al. 2020. Fungal Diversity 105:1-16 ((https://doi.org/10.1007/s13225-020-00466-2)) |
| Otu09185 | 2 | k__Fungi(100);p__Ascomycota(100);c__Saccharomycetes(100);o__Saccharomycetales(100);f__Metschnikowiaceae(100);g__Metschnikowia(100);s__Metschnikowia_pulcherrima(100); | Metschnikowia | 13 | Saprotroph | \|Undefined Saprotroph\| | Yeast | NULL | Probable | As nectar-tap_saprotroph (Põlme et al. 2020); Decay sugar-rich substrates (Põlme et al. 2020) | Põlme S, et al. 2020. Fungal Diversity 105:1-16 ((https://doi.org/10.1007/s13225-020-00466-2)) |
| Otu09186 | 2 | k__Fungi(100);p__Ascomycota(100);c__Saccharomycetes(100);o__Saccharomycetales(100);f__Metschnikowiaceae(100);g__Metschnikowia(100);s__Metschnikowia_pulcherrima(100); | Metschnikowia | 13 | Saprotroph | \|Undefined Saprotroph\| | Yeast | NULL | Probable | As nectar-tap_saprotroph (Põlme et al. 2020); Decay sugar-rich substrates (Põlme et al. 2020) | Põlme S, et al. 2020. Fungal Diversity 105:1-16 ((https://doi.org/10.1007/s13225-020-00466-2)) |
| Otu09187 | 2 | k__Fungi(100);p__Ascomycota(100);c__Saccharomycetes(100);o__Saccharomycetales(100);f__Metschnikowiaceae(100);g__Metschnikowia(100);s__Metschnikowia_pulcherrima(100); | Metschnikowia | 13 | Saprotroph | \|Undefined Saprotroph\| | Yeast | NULL | Probable | As nectar-tap_saprotroph (Põlme et al. 2020); Decay sugar-rich substrates (Põlme et al. 2020) | Põlme S, et al. 2020. Fungal Diversity 105:1-16 ((https://doi.org/10.1007/s13225-020-00466-2)) |
| Otu09188 | 2 | k__Fungi(100);p__Ascomycota(100);c__Saccharomycetes(100);o__Saccharomycetales(100);f__Metschnikowiaceae(100);g__Metschnikowia(100);s__Metschnikowia_pulcherrima(100); | Metschnikowia | 13 | Saprotroph | \|Undefined Saprotroph\| | Yeast | NULL | Probable | As nectar-tap_saprotroph (Põlme et al. 2020); Decay sugar-rich substrates (Põlme et al. 2020) | Põlme S, et al. 2020. Fungal Diversity 105:1-16 ((https://doi.org/10.1007/s13225-020-00466-2)) |
| Otu09195 | 2 | k__Fungi(100);p__Ascomycota(100);c__Saccharomycetes(100);o__Saccharomycetales(100);f__Metschnikowiaceae(100);g__Metschnikowia(100);s__Metschnikowia_pulcherrima(100); | Metschnikowia | 13 | Saprotroph | \|Undefined Saprotroph\| | Yeast | NULL | Probable | As nectar-tap_saprotroph (Põlme et al. 2020); Decay sugar-rich substrates (Põlme et al. 2020) | Põlme S, et al. 2020. Fungal Diversity 105:1-16 ((https://doi.org/10.1007/s13225-020-00466-2)) |
| Otu09199 | 2 | k__Fungi(100);p__Ascomycota(100);c__Saccharomycetes(100);o__Saccharomycetales(100);f__Metschnikowiaceae(100);g__Metschnikowia(100);s__Metschnikowia_pulcherrima(100); | Metschnikowia | 13 | Saprotroph | \|Undefined Saprotroph\| | Yeast | NULL | Probable | As nectar-tap_saprotroph (Põlme et al. 2020); Decay sugar-rich substrates (Põlme et al. 2020) | Põlme S, et al. 2020. Fungal Diversity 105:1-16 ((https://doi.org/10.1007/s13225-020-00466-2)) |
| Otu09200 | 2 | k__Fungi(100);p__Ascomycota(100);c__Saccharomycetes(100);o__Saccharomycetales(100);f__Metschnikowiaceae(100);g__Metschnikowia(100);s__Metschnikowia_pulcherrima(100); | Metschnikowia | 13 | Saprotroph | \|Undefined Saprotroph\| | Yeast | NULL | Probable | As nectar-tap_saprotroph (Põlme et al. 2020); Decay sugar-rich substrates (Põlme et al. 2020) | Põlme S, et al. 2020. Fungal Diversity 105:1-16 ((https://doi.org/10.1007/s13225-020-00466-2)) |
| Otu09201 | 2 | k__Fungi(100);p__Ascomycota(100);c__Saccharomycetes(100);o__Saccharomycetales(100);f__Metschnikowiaceae(100);g__Metschnikowia(100);g__Metschnikowia_unclassified(100); | Metschnikowia | 13 | Saprotroph | \|Undefined Saprotroph\| | Yeast | NULL | Probable | As nectar-tap_saprotroph (Põlme et al. 2020); Decay sugar-rich substrates (Põlme et al. 2020) | Põlme S, et al. 2020. Fungal Diversity 105:1-16 ((https://doi.org/10.1007/s13225-020-00466-2)) |
| Otu09202 | 2 | k__Fungi(100);p__Ascomycota(100);c__Saccharomycetes(100);o__Saccharomycetales(100);f__Metschnikowiaceae(100);g__Metschnikowia(100);g__Metschnikowia_unclassified(100); | Metschnikowia | 13 | Saprotroph | \|Undefined Saprotroph\| | Yeast | NULL | Probable | As nectar-tap_saprotroph (Põlme et al. 2020); Decay sugar-rich substrates (Põlme et al. 2020) | Põlme S, et al. 2020. Fungal Diversity 105:1-16 ((https://doi.org/10.1007/s13225-020-00466-2)) |
| Otu09205 | 2 | k__Fungi(100);p__Ascomycota(100);c__Saccharomycetes(100);o__Saccharomycetales(100);f__Metschnikowiaceae(100);g__Metschnikowia(100);s__Metschnikowia_pulcherrima(100); | Metschnikowia | 13 | Saprotroph | \|Undefined Saprotroph\| | Yeast | NULL | Probable | As nectar-tap_saprotroph (Põlme et al. 2020); Decay sugar-rich substrates (Põlme et al. 2020) | Põlme S, et al. 2020. Fungal Diversity 105:1-16 ((https://doi.org/10.1007/s13225-020-00466-2)) |
| Otu09206 | 2 | k__Fungi(100);p__Ascomycota(100);c__Saccharomycetes(100);o__Saccharomycetales(100);f__Metschnikowiaceae(100);g__Metschnikowia(100);s__Metschnikowia_pulcherrima(100); | Metschnikowia | 13 | Saprotroph | \|Undefined Saprotroph\| | Yeast | NULL | Probable | As nectar-tap_saprotroph (Põlme et al. 2020); Decay sugar-rich substrates (Põlme et al. 2020) | Põlme S, et al. 2020. Fungal Diversity 105:1-16 ((https://doi.org/10.1007/s13225-020-00466-2)) |
| Otu09207 | 2 | k__Fungi(100);p__Ascomycota(100);c__Saccharomycetes(100);o__Saccharomycetales(100);f__Metschnikowiaceae(100);g__Metschnikowia(100);s__Metschnikowia_pulcherrima(100); | Metschnikowia | 13 | Saprotroph | \|Undefined Saprotroph\| | Yeast | NULL | Probable | As nectar-tap_saprotroph (Põlme et al. 2020); Decay sugar-rich substrates (Põlme et al. 2020) | Põlme S, et al. 2020. Fungal Diversity 105:1-16 ((https://doi.org/10.1007/s13225-020-00466-2)) |
| Otu09211 | 2 | k__Fungi(100);p__Ascomycota(100);c__Saccharomycetes(100);o__Saccharomycetales(100);f__Metschnikowiaceae(100);g__Metschnikowia(100);g__Metschnikowia_unclassified(100); | Metschnikowia | 13 | Saprotroph | \|Undefined Saprotroph\| | Yeast | NULL | Probable | As nectar-tap_saprotroph (Põlme et al. 2020); Decay sugar-rich substrates (Põlme et al. 2020) | Põlme S, et al. 2020. Fungal Diversity 105:1-16 ((https://doi.org/10.1007/s13225-020-00466-2)) |
| Otu09212 | 2 | k__Fungi(100);p__Ascomycota(100);c__Saccharomycetes(100);o__Saccharomycetales(100);f__Metschnikowiaceae(100);g__Metschnikowia(100);s__Metschnikowia_pulcherrima(100); | Metschnikowia | 13 | Saprotroph | \|Undefined Saprotroph\| | Yeast | NULL | Probable | As nectar-tap_saprotroph (Põlme et al. 2020); Decay sugar-rich substrates (Põlme et al. 2020) | Põlme S, et al. 2020. Fungal Diversity 105:1-16 ((https://doi.org/10.1007/s13225-020-00466-2)) |
| Otu09213 | 2 | k__Fungi(100);p__Ascomycota(100);c__Saccharomycetes(100);o__Saccharomycetales(100);f__Metschnikowiaceae(100);g__Metschnikowia(100);s__Metschnikowia_pulcherrima(100); | Metschnikowia | 13 | Saprotroph | \|Undefined Saprotroph\| | Yeast | NULL | Probable | As nectar-tap_saprotroph (Põlme et al. 2020); Decay sugar-rich substrates (Põlme et al. 2020) | Põlme S, et al. 2020. Fungal Diversity 105:1-16 ((https://doi.org/10.1007/s13225-020-00466-2)) |
| Otu09214 | 2 | k__Fungi(100);p__Ascomycota(100);c__Dothideomycetes(100);o__Capnodiales(100);f__Capnodiales_family_Incertae_sedis(100);g__Toxicocladosporium(100);s__Toxicocladosporium_sp(100); | Toxicocladosporium | 13 | Pathotroph-Symbiotroph | Endophyte-\|Plant Pathogen\| | Microfungus | NULL | Probable | As foliar_endophyte (Põlme et al. 2020); Decay leaf, fruit, seed (Põlme et al. 2020); Isolated from leaves of Posoqueria and Eucalyptus camaldulensis (occurring within a single small lesion) as well as moldy paint | Crous PW, et al. 2009. Persoonia 22:83-94 ((https://doi.org/10.3767/003158509X439364)); Põlme S, et al. 2020. Fungal Diversity 105:1-16 ((https://doi.org/10.1007/s13225-020-00466-2)) |
| Otu09215 | 2 | k__Fungi(100);p__Ascomycota(100);c__Saccharomycetes(100);o__Saccharomycetales(100);o__Saccharomycetales_unclassified(100);o__Saccharomycetales_unclassified(100);o__Saccharomycetales_unclassified(100); | Saccharomycetales | 7 | Saprotroph | Undefined Saprotroph | Yeast | NULL | Possible | NULL | Sterkenburg E, et al. 2015. New Phytologist 207:1145-1158 ((https://doi.org/10.1111/nph.13426)) |
| Otu09216 | 2 | k__Fungi(100);p__Ascomycota(100);c__Saccharomycetes(100);o__Saccharomycetales(100);f__Metschnikowiaceae(100);g__Metschnikowia(100);s__Metschnikowia_pulcherrima(100); | Metschnikowia | 13 | Saprotroph | \|Undefined Saprotroph\| | Yeast | NULL | Probable | As nectar-tap_saprotroph (Põlme et al. 2020); Decay sugar-rich substrates (Põlme et al. 2020) | Põlme S, et al. 2020. Fungal Diversity 105:1-16 ((https://doi.org/10.1007/s13225-020-00466-2)) |
| Otu09217 | 2 | k__Fungi(100);p__Ascomycota(100);c__Saccharomycetes(100);o__Saccharomycetales(100);f__Metschnikowiaceae(100);g__Metschnikowia(100);s__Metschnikowia_pulcherrima(100); | Metschnikowia | 13 | Saprotroph | \|Undefined Saprotroph\| | Yeast | NULL | Probable | As nectar-tap_saprotroph (Põlme et al. 2020); Decay sugar-rich substrates (Põlme et al. 2020) | Põlme S, et al. 2020. Fungal Diversity 105:1-16 ((https://doi.org/10.1007/s13225-020-00466-2)) |
| Otu09218 | 2 | k__Fungi(100);p__Ascomycota(100);c__Saccharomycetes(100);o__Saccharomycetales(100);f__Metschnikowiaceae(100);g__Metschnikowia(100);g__Metschnikowia_unclassified(100); | Metschnikowia | 13 | Saprotroph | \|Undefined Saprotroph\| | Yeast | NULL | Probable | As nectar-tap_saprotroph (Põlme et al. 2020); Decay sugar-rich substrates (Põlme et al. 2020) | Põlme S, et al. 2020. Fungal Diversity 105:1-16 ((https://doi.org/10.1007/s13225-020-00466-2)) |

| | | | | | | | | | | | |
|---|---|---|---|---|---|---|---|---|---|---|---|
| Otu09219 | 2 | k__Fungi(100);p__Ascomycota(100);c__Saccharomycetes(100);o__Saccharomycetales(100);f__Metschnikowiaceae(100);g__Metschnikowia(100);g__Metschnikowia_unclassified(100); | Metschnikowia | 13 | Saprotroph | \|Undefined Saprotroph\| | Yeast | NULL | Probable | As nectar-tap_saprotroph (Põlme et al. 2020); Decay sugar-rich substrates (Põlme et al. 2020) | Põlme S, et al. 2020. Fungal Diversity 105:1-16 ((https://doi.org/10.1007/s13225-020-00466-2)) |
| Otu09220 | 2 | k__Fungi(100);p__Ascomycota(100);c__Saccharomycetes(100);o__Saccharomycetales(100);f__Metschnikowiaceae(100);g__Metschnikowia(100);s__Metschnikowia_pulcherrima(100); | Metschnikowia | 13 | Saprotroph | \|Undefined Saprotroph\| | Yeast | NULL | Probable | As nectar-tap_saprotroph (Põlme et al. 2020); Decay sugar-rich substrates (Põlme et al. 2020) | Põlme S, et al. 2020. Fungal Diversity 105:1-16 ((https://doi.org/10.1007/s13225-020-00466-2)) |
| Otu09223 | 2 | k__Fungi(100);p__Ascomycota(100);c__Saccharomycetes(100);o__Saccharomycetales(100);f__Metschnikowiaceae(100);g__Metschnikowia(100);g__Metschnikowia_unclassified(100); | Metschnikowia | 13 | Saprotroph | \|Undefined Saprotroph\| | Yeast | NULL | Probable | As nectar-tap_saprotroph (Põlme et al. 2020); Decay sugar-rich substrates (Põlme et al. 2020) | Põlme S, et al. 2020. Fungal Diversity 105:1-16 ((https://doi.org/10.1007/s13225-020-00466-2)) |
| Otu09224 | 2 | k__Fungi(100);p__Ascomycota(100);c__Saccharomycetes(100);o__Saccharomycetales(100);f__Metschnikowiaceae(100);g__Metschnikowia(100);s__Metschnikowia_pulcherrima(100); | Metschnikowia | 13 | Saprotroph | \|Undefined Saprotroph\| | Yeast | NULL | Probable | As nectar-tap_saprotroph (Põlme et al. 2020); Decay sugar-rich substrates (Põlme et al. 2020) | Põlme S, et al. 2020. Fungal Diversity 105:1-16 ((https://doi.org/10.1007/s13225-020-00466-2)) |
| Otu09226 | 2 | k__Fungi(100);p__Ascomycota(100);c__Saccharomycetes(100);o__Saccharomycetales(100);f__Metschnikowiaceae(100);g__Metschnikowia(100);g__Metschnikowia_unclassified(100); | Metschnikowia | 13 | Saprotroph | \|Undefined Saprotroph\| | Yeast | NULL | Probable | As nectar-tap_saprotroph (Põlme et al. 2020); Decay sugar-rich substrates (Põlme et al. 2020) | Põlme S, et al. 2020. Fungal Diversity 105:1-16 ((https://doi.org/10.1007/s13225-020-00466-2)) |
| Otu09227 | 2 | k__Fungi(100);p__Ascomycota(100);c__Saccharomycetes(100);o__Saccharomycetales(100);f__Metschnikowiaceae(100);g__Metschnikowia(100);s__Metschnikowia_pulcherrima(100); | Metschnikowia | 13 | Saprotroph | \|Undefined Saprotroph\| | Yeast | NULL | Probable | As nectar-tap_saprotroph (Põlme et al. 2020); Decay sugar-rich substrates (Põlme et al. 2020) | Põlme S, et al. 2020. Fungal Diversity 105:1-16 ((https://doi.org/10.1007/s13225-020-00466-2)) |
| Otu09229 | 2 | k__Fungi(100);p__Ascomycota(100);c__Saccharomycetes(100);o__Saccharomycetales(100);f__Metschnikowiaceae(100);g__Metschnikowia(100);g__Metschnikowia_unclassified(100); | Metschnikowia | 13 | Saprotroph | \|Undefined Saprotroph\| | Yeast | NULL | Probable | As nectar-tap_saprotroph (Põlme et al. 2020); Decay sugar-rich substrates (Põlme et al. 2020) | Põlme S, et al. 2020. Fungal Diversity 105:1-16 ((https://doi.org/10.1007/s13225-020-00466-2)) |
| Otu09232 | 2 | k__Fungi(100);p__Ascomycota(100);c__Saccharomycetes(100);o__Saccharomycetales(100);f__Metschnikowiaceae(100);g__Metschnikowia(100);s__Metschnikowia_pulcherrima(100); | Metschnikowia | 13 | Saprotroph | \|Undefined Saprotroph\| | Yeast | NULL | Probable | As nectar-tap_saprotroph (Põlme et al. 2020); Decay sugar-rich substrates (Põlme et al. 2020) | Põlme S, et al. 2020. Fungal Diversity 105:1-16 ((https://doi.org/10.1007/s13225-020-00466-2)) |
| Otu09235 | 2 | k__Fungi(100);p__Ascomycota(100);c__Saccharomycetes(100);o__Saccharomycetales(100);f__Metschnikowiaceae(100);g__Metschnikowia(100);g__Metschnikowia_unclassified(100); | Metschnikowia | 13 | Saprotroph | \|Undefined Saprotroph\| | Yeast | NULL | Probable | As nectar-tap_saprotroph (Põlme et al. 2020); Decay sugar-rich substrates (Põlme et al. 2020) | Põlme S, et al. 2020. Fungal Diversity 105:1-16 ((https://doi.org/10.1007/s13225-020-00466-2)) |
| Otu09236 | 2 | k__Fungi(100);p__Ascomycota(100);c__Dothideomycetes(100);o__Pleosporales(100);f__Phaeosphaeriaceae(100);g__Phaeosphaeria(100);s__Phaeosphaeria_triglochinicola(100); | Phaeosphaeria | 13 | Pathotroph-Saprotroph-Symbiotroph | Endophyte-Plant Pathogen-\|Plant Saprotroph\| | Microfungus | NULL | Probable | As litter_saprotroph (Põlme et al. 2020); Decay leaf, fruit, seed (Põlme et al. 2020) | Tedersoo L, et al. 2014. Science 346:e1256688 ((https://doi.org/10.1126/science.1256688));David AS, et al. 2016. Microbial Ecology 71: 912-926 ((https://doi.org/10.1007/s00248-015-0712-6)); Põlme S, et al. 2020. Fungal Diversity 105:1-16 ((https://doi.org/10.1007/s13225-020-00466-2)) |
| Otu09237 | 2 | k__Fungi(100);p__Ascomycota(100);c__Saccharomycetes(100);o__Saccharomycetales(100);f__Metschnikowiaceae(100);g__Metschnikowia(100);g__Metschnikowia_unclassified(100); | Metschnikowia | 13 | Saprotroph | \|Undefined Saprotroph\| | Yeast | NULL | Probable | As nectar-tap_saprotroph (Põlme et al. 2020); Decay sugar-rich substrates (Põlme et al. 2020) | Põlme S, et al. 2020. Fungal Diversity 105:1-16 ((https://doi.org/10.1007/s13225-020-00466-2)) |
| Otu09240 | 2 | k__Fungi(100);p__Ascomycota(100);c__Saccharomycetes(100);o__Saccharomycetales(100);f__Metschnikowiaceae(100);g__Metschnikowia(100);g__Metschnikowia_unclassified(100); | Metschnikowia | 13 | Saprotroph | \|Undefined Saprotroph\| | Yeast | NULL | Probable | As nectar-tap_saprotroph (Põlme et al. 2020); Decay sugar-rich substrates (Põlme et al. 2020) | Põlme S, et al. 2020. Fungal Diversity 105:1-16 ((https://doi.org/10.1007/s13225-020-00466-2)) |
| Otu09242 | 2 | k__Fungi(100);p__Ascomycota(100);c__Saccharomycetes(100);o__Saccharomycetales(100);f__Metschnikowiaceae(100);g__Metschnikowia(100);g__Metschnikowia_unclassified(100); | Metschnikowia | 13 | Saprotroph | \|Undefined Saprotroph\| | Yeast | NULL | Probable | As nectar-tap_saprotroph (Põlme et al. 2020); Decay sugar-rich substrates (Põlme et al. 2020) | Põlme S, et al. 2020. Fungal Diversity 105:1-16 ((https://doi.org/10.1007/s13225-020-00466-2)) |
| Otu09245 | 2 | k__Fungi(100);p__Ascomycota(100);c__Saccharomycetes(100);o__Saccharomycetales(100);f__Metschnikowiaceae(100);g__Metschnikowia(100);s__Metschnikowia_pulcherrima(100); | Metschnikowia | 13 | Saprotroph | \|Undefined Saprotroph\| | Yeast | NULL | Probable | As nectar-tap_saprotroph (Põlme et al. 2020); Decay sugar-rich substrates (Põlme et al. 2020) | Põlme S, et al. 2020. Fungal Diversity 105:1-16 ((https://doi.org/10.1007/s13225-020-00466-2)) |
| Otu09246 | 2 | k__Fungi(100);p__Ascomycota(100);c__Saccharomycetes(100);o__Saccharomycetales(100);f__Metschnikowiaceae(100);g__Metschnikowia(100);g__Metschnikowia_unclassified(100); | Metschnikowia | 13 | Saprotroph | \|Undefined Saprotroph\| | Yeast | NULL | Probable | As nectar-tap_saprotroph (Põlme et al. 2020); Decay sugar-rich substrates (Põlme et al. 2020) | Põlme S, et al. 2020. Fungal Diversity 105:1-16 ((https://doi.org/10.1007/s13225-020-00466-2)) |
| Otu09251 | 2 | k__Fungi(100);p__Ascomycota(100);c__Saccharomycetes(100);o__Saccharomycetales(100);f__Metschnikowiaceae(100);g__Metschnikowia(100);s__Metschnikowia_pulcherrima(100); | Metschnikowia | 13 | Saprotroph | \|Undefined Saprotroph\| | Yeast | NULL | Probable | As nectar-tap_saprotroph (Põlme et al. 2020); Decay sugar-rich substrates (Põlme et al. 2020) | Põlme S, et al. 2020. Fungal Diversity 105:1-16 ((https://doi.org/10.1007/s13225-020-00466-2)) |
| Otu09254 | 2 | k__Fungi(100);p__Ascomycota(100);c__Saccharomycetes(100);o__Saccharomycetales(100);f__Metschnikowiaceae(100);g__Metschnikowia(100);s__Metschnikowia_pulcherrima(100); | Metschnikowia | 13 | Saprotroph | \|Undefined Saprotroph\| | Yeast | NULL | Probable | As nectar-tap_saprotroph (Põlme et al. 2020); Decay sugar-rich substrates (Põlme et al. 2020) | Põlme S, et al. 2020. Fungal Diversity 105:1-16 ((https://doi.org/10.1007/s13225-020-00466-2)) |
| Otu09257 | 2 | k__Fungi(100);p__Ascomycota(100);c__Leotiomycetes(100);o__Helotiales(100);f__Vibrisseaceae(100);f__Vibrisseaceae_unclassified(100);f__Vibrisseaceae_unclassified(100); | Vibrisseaceae | 9 | Saprotroph-Symbiotroph | Endophyte-Undefined Saprotroph-Wood Saprotroph | Microfungus | NULL | Possible | Saprobic, usually on wood and bark immersed in fresh water. Some species may be root endophytes (Cannon & Kirk 2007) | Cannon PF, Kirk PM. 2007. Fungal Families of the World. CAB International, Cambridge (ISBN: 978-0851998275) |
| Otu09258 | 2 | k__Fungi(100);p__Ascomycota(100);c__Saccharomycetes(100);o__Saccharomycetales(100);f__Metschnikowiaceae(100);g__Metschnikowia(100);g__Metschnikowia_unclassified(100); | Metschnikowia | 13 | Saprotroph | \|Undefined Saprotroph\| | Yeast | NULL | Probable | As nectar-tap_saprotroph (Põlme et al. 2020); Decay sugar-rich substrates (Põlme et al. 2020) | Põlme S, et al. 2020. Fungal Diversity 105:1-16 ((https://doi.org/10.1007/s13225-020-00466-2)) |
| Otu09259 | 2 | k__Fungi(100);p__Ascomycota(100);c__Saccharomycetes(100);o__Saccharomycetales(100);f__Metschnikowiaceae(100);g__Metschnikowia(100);s__Metschnikowia_pulcherrima(100); | Metschnikowia | 13 | Saprotroph | \|Undefined Saprotroph\| | Yeast | NULL | Probable | As nectar-tap_saprotroph (Põlme et al. 2020); Decay sugar-rich substrates (Põlme et al. 2020) | Põlme S, et al. 2020. Fungal Diversity 105:1-16 ((https://doi.org/10.1007/s13225-020-00466-2)) |
| Otu09261 | 2 | k__Fungi(100);p__Ascomycota(100);c__Saccharomycetes(100);o__Saccharomycetales(100);f__Metschnikowiaceae(100);g__Metschnikowia(100);g__Metschnikowia_unclassified(100); | Metschnikowia | 13 | Saprotroph | \|Undefined Saprotroph\| | Yeast | NULL | Probable | As nectar-tap_saprotroph (Põlme et al. 2020); Decay sugar-rich substrates (Põlme et al. 2020) | Põlme S, et al. 2020. Fungal Diversity 105:1-16 ((https://doi.org/10.1007/s13225-020-00466-2)) |
| Otu09262 | 2 | k__Fungi(100);p__Ascomycota(100);c__Saccharomycetes(100);o__Saccharomycetales(100);f__Metschnikowiaceae(100);g__Metschnikowia(100);s__Metschnikowia_pulcherrima(100); | Metschnikowia | 13 | Saprotroph | \|Undefined Saprotroph\| | Yeast | NULL | Probable | As nectar-tap_saprotroph (Põlme et al. 2020); Decay sugar-rich substrates (Põlme et al. 2020) | Põlme S, et al. 2020. Fungal Diversity 105:1-16 ((https://doi.org/10.1007/s13225-020-00466-2)) |
| Otu09263 | 2 | k__Fungi(100);p__Ascomycota(100);c__Saccharomycetes(100);o__Saccharomycetales(100);f__Metschnikowiaceae(100);g__Metschnikowia(100);s__Metschnikowia_pulcherrima(100); | Metschnikowia | 13 | Saprotroph | \|Undefined Saprotroph\| | Yeast | NULL | Probable | As nectar-tap_saprotroph (Põlme et al. 2020); Decay sugar-rich substrates (Põlme et al. 2020) | Põlme S, et al. 2020. Fungal Diversity 105:1-16 ((https://doi.org/10.1007/s13225-020-00466-2)) |
| Otu09264 | 2 | k__Fungi(100);p__Ascomycota(100);c__Saccharomycetes(100);o__Saccharomycetales(100);f__Metschnikowiaceae(100);g__Metschnikowia(100);s__Metschnikowia_pulcherrima(100); | Metschnikowia | 13 | Saprotroph | \|Undefined Saprotroph\| | Yeast | NULL | Probable | As nectar-tap_saprotroph (Põlme et al. 2020); Decay sugar-rich substrates (Põlme et al. 2020) | Põlme S, et al. 2020. Fungal Diversity 105:1-16 ((https://doi.org/10.1007/s13225-020-00466-2)) |
| Otu09267 | 2 | k__Fungi(100);p__Ascomycota(100);c__Saccharomycetes(100);o__Saccharomycetales(100);f__Metschnikowiaceae(100);g__Metschnikowia(100);g__Metschnikowia_unclassified(100); | Metschnikowia | 13 | Saprotroph | \|Undefined Saprotroph\| | Yeast | NULL | Probable | As nectar-tap_saprotroph (Põlme et al. 2020); Decay sugar-rich substrates (Põlme et al. 2020) | Põlme S, et al. 2020. Fungal Diversity 105:1-16 ((https://doi.org/10.1007/s13225-020-00466-2)) |
| Otu09270 | 2 | k__Fungi(100);p__Ascomycota(100);c__Saccharomycetes(100);o__Saccharomycetales(100);f__Metschnikowiaceae(100);g__Metschnikowia(100);s__Metschnikowia_pulcherrima(100); | Metschnikowia | 13 | Saprotroph | \|Undefined Saprotroph\| | Yeast | NULL | Probable | As nectar-tap_saprotroph (Põlme et al. 2020); Decay sugar-rich substrates (Põlme et al. 2020) | Põlme S, et al. 2020. Fungal Diversity 105:1-16 ((https://doi.org/10.1007/s13225-020-00466-2)) |
| Otu09272 | 2 | k__Fungi(100);p__Ascomycota(100);c__Saccharomycetes(100);o__Saccharomycetales(100);f__Metschnikowiaceae(100);g__Metschnikowia(100);g__Metschnikowia_unclassified(100); | Metschnikowia | 13 | Saprotroph | \|Undefined Saprotroph\| | Yeast | NULL | Probable | As nectar-tap_saprotroph (Põlme et al. 2020); Decay sugar-rich substrates (Põlme et al. 2020) | Põlme S, et al. 2020. Fungal Diversity 105:1-16 ((https://doi.org/10.1007/s13225-020-00466-2)) |
| Otu09273 | 2 | k__Fungi(100);p__Ascomycota(100);c__Saccharomycetes(100);o__Saccharomycetales(100);f__Metschnikowiaceae(100);g__Metschnikowia(100);g__Metschnikowia_unclassified(100); | Metschnikowia | 13 | Saprotroph | \|Undefined Saprotroph\| | Yeast | NULL | Probable | As nectar-tap_saprotroph (Põlme et al. 2020); Decay sugar-rich substrates (Põlme et al. 2020) | Põlme S, et al. 2020. Fungal Diversity 105:1-16 ((https://doi.org/10.1007/s13225-020-00466-2)) |
| Otu09274 | 2 | k__Fungi(100);p__Ascomycota(100);c__Saccharomycetes(100);o__Saccharomycetales(100);f__Metschnikowiaceae(100);g__Metschnikowia(100);s__Metschnikowia_pulcherrima(100); | Metschnikowia | 13 | Saprotroph | \|Undefined Saprotroph\| | Yeast | NULL | Probable | As nectar-tap_saprotroph (Põlme et al. 2020); Decay sugar-rich substrates (Põlme et al. 2020) | Põlme S, et al. 2020. Fungal Diversity 105:1-16 ((https://doi.org/10.1007/s13225-020-00466-2)) |
| Otu09276 | 2 | k__Fungi(100);p__Ascomycota(100);c__Saccharomycetes(100);o__Saccharomycetales(100);f__Metschnikowiaceae(100);g__Metschnikowia(100);s__Metschnikowia_pulcherrima(100); | Metschnikowia | 13 | Saprotroph | \|Undefined Saprotroph\| | Yeast | NULL | Probable | As nectar-tap_saprotroph (Põlme et al. 2020); Decay sugar-rich substrates (Põlme et al. 2020) | Põlme S, et al. 2020. Fungal Diversity 105:1-16 ((https://doi.org/10.1007/s13225-020-00466-2)) |
| Otu09283 | 2 | k__Fungi(100);p__Ascomycota(100);c__Saccharomycetes(100);o__Saccharomycetales(100);f__Metschnikowiaceae(100);g__Metschnikowia(100);s__Metschnikowia_pulcherrima(100); | Metschnikowia | 13 | Saprotroph | \|Undefined Saprotroph\| | Yeast | NULL | Probable | As nectar-tap_saprotroph (Põlme et al. 2020); Decay sugar-rich substrates (Põlme et al. 2020) | Põlme S, et al. 2020. Fungal Diversity 105:1-16 ((https://doi.org/10.1007/s13225-020-00466-2)) |
| Otu09285 | 2 | k__Fungi(100);p__Ascomycota(100);c__Saccharomycetes(100);o__Saccharomycetales(100);f__Metschnikowiaceae(100);g__Metschnikowia(100);s__Metschnikowia_pulcherrima(100); | Metschnikowia | 13 | Saprotroph | \|Undefined Saprotroph\| | Yeast | NULL | Probable | As nectar-tap_saprotroph (Põlme et al. 2020); Decay sugar-rich substrates (Põlme et al. 2020) | Põlme S, et al. 2020. Fungal Diversity 105:1-16 ((https://doi.org/10.1007/s13225-020-00466-2)) |
| Otu09286 | 2 | k__Fungi(100);p__Ascomycota(100);c__Saccharomycetes(100);o__Saccharomycetales(100);f__Metschnikowiaceae(100);g__Metschnikowia(100);s__Metschnikowia_pulcherrima(100); | Metschnikowia | 13 | Saprotroph | \|Undefined Saprotroph\| | Yeast | NULL | Probable | As nectar-tap_saprotroph (Põlme et al. 2020); Decay sugar-rich substrates (Põlme et al. 2020) | Põlme S, et al. 2020. Fungal Diversity 105:1-16 ((https://doi.org/10.1007/s13225-020-00466-2)) |
| Otu09287 | 2 | k__Fungi(100);p__Ascomycota(100);c__Saccharomycetes(100);o__Saccharomycetales(100);f__Metschnikowiaceae(100);g__Metschnikowia(100);g__Metschnikowia_unclassified(100); | Metschnikowia | 13 | Saprotroph | \|Undefined Saprotroph\| | Yeast | NULL | Probable | As nectar-tap_saprotroph (Põlme et al. 2020); Decay sugar-rich substrates (Põlme et al. 2020) | Põlme S, et al. 2020. Fungal Diversity 105:1-16 ((https://doi.org/10.1007/s13225-020-00466-2)) |
| Otu09288 | 2 | k__Fungi(100);p__Ascomycota(100);c__Saccharomycetes(100);o__Saccharomycetales(100);f__Metschnikowiaceae(100);g__Metschnikowia(100);s__Metschnikowia_pulcherrima(100); | Metschnikowia | 13 | Saprotroph | \|Undefined Saprotroph\| | Yeast | NULL | Probable | As nectar-tap_saprotroph (Põlme et al. 2020); Decay sugar-rich substrates (Põlme et al. 2020) | Põlme S, et al. 2020. Fungal Diversity 105:1-16 ((https://doi.org/10.1007/s13225-020-00466-2)) |
| Otu09291 | 2 | k__Fungi(100);p__Ascomycota(100);c__Saccharomycetes(100);o__Saccharomycetales(100);f__Metschnikowiaceae(100);g__Metschnikowia(100);s__Metschnikowia_pulcherrima(100); | Metschnikowia | 13 | Saprotroph | \|Undefined Saprotroph\| | Yeast | NULL | Probable | As nectar-tap_saprotroph (Põlme et al. 2020); Decay sugar-rich substrates (Põlme et al. 2020) | Põlme S, et al. 2020. Fungal Diversity 105:1-16 ((https://doi.org/10.1007/s13225-020-00466-2)) |
| Otu09296 | 2 | k__Fungi(100);p__Ascomycota(100);c__Saccharomycetes(100);o__Saccharomycetales(100);f__Metschnikowiaceae(100);g__Metschnikowia(100);s__Metschnikowia_pulcherrima(100); | Metschnikowia | 13 | Saprotroph | \|Undefined Saprotroph\| | Yeast | NULL | Probable | As nectar-tap_saprotroph (Põlme et al. 2020); Decay sugar-rich substrates (Põlme et al. 2020) | Põlme S, et al. 2020. Fungal Diversity 105:1-16 ((https://doi.org/10.1007/s13225-020-00466-2)) |
| Otu09297 | 2 | k__Fungi(100);p__Ascomycota(100);c__Saccharomycetes(100);o__Saccharomycetales(100);f__Metschnikowiaceae(100);g__Metschnikowia(100);s__Metschnikowia_pulcherrima(100); | Metschnikowia | 13 | Saprotroph | \|Undefined Saprotroph\| | Yeast | NULL | Probable | As nectar-tap_saprotroph (Põlme et al. 2020); Decay sugar-rich substrates (Põlme et al. 2020) | Põlme S, et al. 2020. Fungal Diversity 105:1-16 ((https://doi.org/10.1007/s13225-020-00466-2)) |
| Otu09302 | 2 | k__Fungi(100);p__Ascomycota(100);c__Saccharomycetes(100);o__Saccharomycetales(100);f__Metschnikowiaceae(100);g__Metschnikowia(100);s__Metschnikowia_pulcherrima(100); | Metschnikowia | 13 | Saprotroph | \|Undefined Saprotroph\| | Yeast | NULL | Probable | As nectar-tap_saprotroph (Põlme et al. 2020); Decay sugar-rich substrates (Põlme et al. 2020) | Põlme S, et al. 2020. Fungal Diversity 105:1-16 ((https://doi.org/10.1007/s13225-020-00466-2)) |
| Otu09303 | 2 | k__Fungi(100);p__Ascomycota(100);c__Saccharomycetes(100);o__Saccharomycetales(100);f__Metschnikowiaceae(100);g__Metschnikowia(100);s__Metschnikowia_pulcherrima(100); | Metschnikowia | 13 | Saprotroph | \|Undefined Saprotroph\| | Yeast | NULL | Probable | As nectar-tap_saprotroph (Põlme et al. 2020); Decay sugar-rich substrates (Põlme et al. 2020) | Põlme S, et al. 2020. Fungal Diversity 105:1-16 ((https://doi.org/10.1007/s13225-020-00466-2)) |
| Otu09306 | 2 | k__Fungi(100);p__Ascomycota(100);c__Saccharomycetes(100);o__Saccharomycetales(100);f__Metschnikowiaceae(100);g__Metschnikowia(100);s__Metschnikowia_pulcherrima(100); | Metschnikowia | 13 | Saprotroph | \|Undefined Saprotroph\| | Yeast | NULL | Probable | As nectar-tap_saprotroph (Põlme et al. 2020); Decay sugar-rich substrates (Põlme et al. 2020) | Põlme S, et al. 2020. Fungal Diversity 105:1-16 ((https://doi.org/10.1007/s13225-020-00466-2)) |
| Otu09311 | 2 | k__Fungi(100);p__Ascomycota(100);c__Saccharomycetes(100);o__Saccharomycetales(100);f__Metschnikowiaceae(100);g__Metschnikowia(100);g__Metschnikowia_unclassified(100); | Metschnikowia | 13 | Saprotroph | \|Undefined Saprotroph\| | Yeast | NULL | Probable | As nectar-tap_saprotroph (Põlme et al. 2020); Decay sugar-rich substrates (Põlme et al. 2020) | Põlme S, et al. 2020. Fungal Diversity 105:1-16 ((https://doi.org/10.1007/s13225-020-00466-2)) |
| Otu09312 | 2 | k__Fungi(100);p__Ascomycota(100);c__Saccharomycetes(100);o__Saccharomycetales(100);f__Metschnikowiaceae(100);g__Metschnikowia(100);s__Metschnikowia_pulcherrima(100); | Metschnikowia | 13 | Saprotroph | \|Undefined Saprotroph\| | Yeast | NULL | Probable | As nectar-tap_saprotroph (Põlme et al. 2020); Decay sugar-rich substrates (Põlme et al. 2020) | Põlme S, et al. 2020. Fungal Diversity 105:1-16 ((https://doi.org/10.1007/s13225-020-00466-2)) |
| Otu09313 | 2 | k__Fungi(100);p__Ascomycota(100);c__Saccharomycetes(100);o__Saccharomycetales(100);f__Metschnikowiaceae(100);g__Metschnikowia(100);s__Metschnikowia_pulcherrima(100); | Metschnikowia | 13 | Saprotroph | \|Undefined Saprotroph\| | Yeast | NULL | Probable | As nectar-tap_saprotroph (Põlme et al. 2020); Decay sugar-rich substrates (Põlme et al. 2020) | Põlme S, et al. 2020. Fungal Diversity 105:1-16 ((https://doi.org/10.1007/s13225-020-00466-2)) |
| Otu09316 | 2 | k__Fungi(100);p__Ascomycota(100);c__Saccharomycetes(100);o__Saccharomycetales(100);f__Metschnikowiaceae(100);g__Metschnikowia(100);s__Metschnikowia_pulcherrima(100); | Metschnikowia | 13 | Saprotroph | \|Undefined Saprotroph\| | Yeast | NULL | Probable | As nectar-tap_saprotroph (Põlme et al. 2020); Decay sugar-rich substrates (Põlme et al. 2020) | Põlme S, et al. 2020. Fungal Diversity 105:1-16 ((https://doi.org/10.1007/s13225-020-00466-2)) |
| Otu09318 | 2 | k__Fungi(100);p__Ascomycota(100);c__Saccharomycetes(100);o__Saccharomycetales(100);f__Metschnikowiaceae(100);g__Metschnikowia(100);g__Metschnikowia_unclassified(100); | Metschnikowia | 13 | Saprotroph | \|Undefined Saprotroph\| | Yeast | NULL | Probable | As nectar-tap_saprotroph (Põlme et al. 2020); Decay sugar-rich substrates (Põlme et al. 2020) | Põlme S, et al. 2020. Fungal Diversity 105:1-16 ((https://doi.org/10.1007/s13225-020-00466-2)) |
| Otu09319 | 2 | k__Fungi(100);p__Ascomycota(100);c__Saccharomycetes(100);o__Saccharomycetales(100);f__Metschnikowiaceae(100);g__Metschnikowia(100);s__Metschnikowia_pulcherrima(100); | Metschnikowia | 13 | Saprotroph | \|Undefined Saprotroph\| | Yeast | NULL | Probable | As nectar-tap_saprotroph (Põlme et al. 2020); Decay sugar-rich substrates (Põlme et al. 2020) | Põlme S, et al. 2020. Fungal Diversity 105:1-16 ((https://doi.org/10.1007/s13225-020-00466-2)) |
| Otu09320 | 2 | k__Fungi(100);p__Ascomycota(100);c__Saccharomycetes(100);o__Saccharomycetales(100);f__Metschnikowiaceae(100);g__Metschnikowia(100);s__Metschnikowia_pulcherrima(100); | Metschnikowia | 13 | Saprotroph | \|Undefined Saprotroph\| | Yeast | NULL | Probable | As nectar-tap_saprotroph (Põlme et al. 2020); Decay sugar-rich substrates (Põlme et al. 2020) | Põlme S, et al. 2020. Fungal Diversity 105:1-16 ((https://doi.org/10.1007/s13225-020-00466-2)) |
| Otu09321 | 2 | k__Fungi(100);p__Ascomycota(100);c__Saccharomycetes(100);o__Saccharomycetales(100);f__Metschnikowiaceae(100);g__Metschnikowia(100);s__Metschnikowia_pulcherrima(100); | Metschnikowia | 13 | Saprotroph | \|Undefined Saprotroph\| | Yeast | NULL | Probable | As nectar-tap_saprotroph (Põlme et al. 2020); Decay sugar-rich substrates (Põlme et al. 2020) | Põlme S, et al. 2020. Fungal Diversity 105:1-16 ((https://doi.org/10.1007/s13225-020-00466-2)) |
| Otu09323 | 2 | k__Fungi(100);p__Ascomycota(100);c__Saccharomycetes(100);o__Saccharomycetales(100);f__Metschnikowiaceae(100);g__Metschnikowia(100);g__Metschnikowia_unclassified(100); | Metschnikowia | 13 | Saprotroph | \|Undefined Saprotroph\| | Yeast | NULL | Probable | As nectar-tap_saprotroph (Põlme et al. 2020); Decay sugar-rich substrates (Põlme et al. 2020) | Põlme S, et al. 2020. Fungal Diversity 105:1-16 ((https://doi.org/10.1007/s13225-020-00466-2)) |
| Otu09324 | 2 | k__Fungi(100);p__Ascomycota(100);c__Saccharomycetes(100);o__Saccharomycetales(100);f__Metschnikowiaceae(100);g__Metschnikowia(100);g__Metschnikowia_unclassified(100); | Metschnikowia | 13 | Saprotroph | \|Undefined Saprotroph\| | Yeast | NULL | Probable | As nectar-tap_saprotroph (Põlme et al. 2020); Decay sugar-rich substrates (Põlme et al. 2020) | Põlme S, et al. 2020. Fungal Diversity 105:1-16 ((https://doi.org/10.1007/s13225-020-00466-2)) |
| Otu09331 | 2 | k__Fungi(100);p__Ascomycota(100);c__Saccharomycetes(100);o__Saccharomycetales(100);f__Metschnikowiaceae(100);g__Metschnikowia(100);s__Metschnikowia_pulcherrima(100); | Metschnikowia | 13 | Saprotroph | \|Undefined Saprotroph\| | Yeast | NULL | Probable | As nectar-tap_saprotroph (Põlme et al. 2020); Decay sugar-rich substrates (Põlme et al. 2020) | Põlme S, et al. 2020. Fungal Diversity 105:1-16 ((https://doi.org/10.1007/s13225-020-00466-2)) |
| Otu09334 | 2 | k__Fungi(100);p__Ascomycota(100);c__Saccharomycetes(100);o__Saccharomycetales(100);f__Metschnikowiaceae(100);g__Metschnikowia(100);s__Metschnikowia_pulcherrima(100); | Metschnikowia | 13 | Saprotroph | \|Undefined Saprotroph\| | Yeast | NULL | Probable | As nectar-tap_saprotroph (Põlme et al. 2020); Decay sugar-rich substrates (Põlme et al. 2020) | Põlme S, et al. 2020. Fungal Diversity 105:1-16 ((https://doi.org/10.1007/s13225-020-00466-2)) |
| Otu09338 | 2 | k__Fungi(100);p__Ascomycota(100);c__Saccharomycetes(100);o__Saccharomycetales(100);f__Metschnikowiaceae(100);g__Metschnikowia(100);g__Metschnikowia_unclassified(100); | Metschnikowia | 13 | Saprotroph | \|Undefined Saprotroph\| | Yeast | NULL | Probable | As nectar-tap_saprotroph (Põlme et al. 2020); Decay sugar-rich substrates (Põlme et al. 2020) | Põlme S, et al. 2020. Fungal Diversity 105:1-16 ((https://doi.org/10.1007/s13225-020-00466-2)) |
| Otu09339 | 2 | k__Fungi(100);p__Ascomycota(100);c__Saccharomycetes(100);o__Saccharomycetales(100);f__Metschnikowiaceae(100);g__Metschnikowia(100);s__Metschnikowia_pulcherrima(100); | Metschnikowia | 13 | Saprotroph | \|Undefined Saprotroph\| | Yeast | NULL | Probable | As nectar-tap_saprotroph (Põlme et al. 2020); Decay sugar-rich substrates (Põlme et al. 2020) | Põlme S, et al. 2020. Fungal Diversity 105:1-16 ((https://doi.org/10.1007/s13225-020-00466-2)) |
| Otu09341 | 2 | k__Fungi(100);p__Ascomycota(100);c__Saccharomycetes(100);o__Saccharomycetales(100);f__Metschnikowiaceae(100);g__Metschnikowia(100);g__Metschnikowia_unclassified(100); | Metschnikowia | 13 | Saprotroph | \|Undefined Saprotroph\| | Yeast | NULL | Probable | As nectar-tap_saprotroph (Põlme et al. 2020); Decay sugar-rich substrates (Põlme et al. 2020) | Põlme S, et al. 2020. Fungal Diversity 105:1-16 ((https://doi.org/10.1007/s13225-020-00466-2)) |
| Otu09343 | 2 | k__Fungi(100);p__Ascomycota(100);c__Saccharomycetes(100);o__Saccharomycetales(100);f__Metschnikowiaceae(100);g__Metschnikowia(100);s__Metschnikowia_pulcherrima(100); | Metschnikowia | 13 | Saprotroph | \|Undefined Saprotroph\| | Yeast | NULL | Probable | As nectar-tap_saprotroph (Põlme et al. 2020); Decay sugar-rich substrates (Põlme et al. 2020) | Põlme S, et al. 2020. Fungal Diversity 105:1-16 ((https://doi.org/10.1007/s13225-020-00466-2)) |
| Otu09344 | 2 | k__Fungi(100);p__Ascomycota(100);c__Saccharomycetes(100);o__Saccharomycetales(100);f__Metschnikowiaceae(100);g__Metschnikowia(100);s__Metschnikowia_pulcherrima(100); | Metschnikowia | 13 | Saprotroph | \|Undefined Saprotroph\| | Yeast | NULL | Probable | As nectar-tap_saprotroph (Põlme et al. 2020); Decay sugar-rich substrates (Põlme et al. 2020) | Põlme S, et al. 2020. Fungal Diversity 105:1-16 ((https://doi.org/10.1007/s13225-020-00466-2)) |
| Otu09345 | 2 | k__Fungi(100);p__Ascomycota(100);c__Saccharomycetes(100);o__Saccharomycetales(100);f__Metschnikowiaceae(100);g__Metschnikowia(100);s__Metschnikowia_pulcherrima(100); | Metschnikowia | 13 | Saprotroph | \|Undefined Saprotroph\| | Yeast | NULL | Probable | As nectar-tap_saprotroph (Põlme et al. 2020); Decay sugar-rich substrates (Põlme et al. 2020) | Põlme S, et al. 2020. Fungal Diversity 105:1-16 ((https://doi.org/10.1007/s13225-020-00466-2)) |
| Otu09346 | 2 | k__Fungi(100);p__Ascomycota(100);c__Saccharomycetes(100);o__Saccharomycetales(100);f__Metschnikowiaceae(100);g__Metschnikowia(100);g__Metschnikowia_unclassified(100); | Metschnikowia | 13 | Saprotroph | \|Undefined Saprotroph\| | Yeast | NULL | Probable | As nectar-tap_saprotroph (Põlme et al. 2020); Decay sugar-rich substrates (Põlme et al. 2020) | Põlme S, et al. 2020. Fungal Diversity 105:1-16 ((https://doi.org/10.1007/s13225-020-00466-2)) |
| Otu09347 | 2 | k__Fungi(100);p__Ascomycota(100);c__Saccharomycetes(100);o__Saccharomycetales(100);f__Metschnikowiaceae(100);g__Metschnikowia(100);s__Metschnikowia_pulcherrima(100); | Metschnikowia | 13 | Saprotroph | \|Undefined Saprotroph\| | Yeast | NULL | Probable | As nectar-tap_saprotroph (Põlme et al. 2020); Decay sugar-rich substrates (Põlme et al. 2020) | Põlme S, et al. 2020. Fungal Diversity 105:1-16 ((https://doi.org/10.1007/s13225-020-00466-2)) |
| Otu09348 | 2 | k__Fungi(100);p__Ascomycota(100);c__Saccharomycetes(100);o__Saccharomycetales(100);f__Metschnikowiaceae(100);g__Metschnikowia(100);s__Metschnikowia_pulcherrima(100); | Metschnikowia | 13 | Saprotroph | \|Undefined Saprotroph\| | Yeast | NULL | Probable | As nectar-tap_saprotroph (Põlme et al. 2020); Decay sugar-rich substrates (Põlme et al. 2020) | Põlme S, et al. 2020. Fungal Diversity 105:1-16 ((https://doi.org/10.1007/s13225-020-00466-2)) |
| Otu09349 | 2 | k__Fungi(100);p__Ascomycota(100);c__Saccharomycetes(100);o__Saccharomycetales(100);f__Metschnikowiaceae(100);g__Metschnikowia(100);s__Metschnikowia_pulcherrima(100); | Metschnikowia | 13 | Saprotroph | \|Undefined Saprotroph\| | Yeast | NULL | Probable | As nectar-tap_saprotroph (Põlme et al. 2020); Decay sugar-rich substrates (Põlme et al. 2020) | Põlme S, et al. 2020. Fungal Diversity 105:1-16 ((https://doi.org/10.1007/s13225-020-00466-2)) |
| Otu09352 | 2 | k__Fungi(100);p__Ascomycota(100);c__Saccharomycetes(100);o__Saccharomycetales(100);f__Metschnikowiaceae(100);g__Metschnikowia(100);s__Metschnikowia_pulcherrima(100); | Metschnikowia | 13 | Saprotroph | \|Undefined Saprotroph\| | Yeast | NULL | Probable | As nectar-tap_saprotroph (Põlme et al. 2020); Decay sugar-rich substrates (Põlme et al. 2020) | Põlme S, et al. 2020. Fungal Diversity 105:1-16 ((https://doi.org/10.1007/s13225-020-00466-2)) |
| Otu09353 | 2 | k__Fungi(100);p__Ascomycota(100);c__Saccharomycetes(100);o__Saccharomycetales(100);f__Metschnikowiaceae(100);g__Metschnikowia(100);s__Metschnikowia_pulcherrima(100); | Metschnikowia | 13 | Saprotroph | \|Undefined Saprotroph\| | Yeast | NULL | Probable | As nectar-tap_saprotroph (Põlme et al. 2020); Decay sugar-rich substrates (Põlme et al. 2020) | Põlme S, et al. 2020. Fungal Diversity 105:1-16 ((https://doi.org/10.1007/s13225-020-00466-2)) |
| Otu09354 | 2 | k__Fungi(100);p__Ascomycota(100);c__Saccharomycetes(100);o__Saccharomycetales(100);f__Metschnikowiaceae(100);g__Metschnikowia(100);g__Metschnikowia_unclassified(100); | Metschnikowia | 13 | Saprotroph | \|Undefined Saprotroph\| | Yeast | NULL | Probable | As nectar-tap_saprotroph (Põlme et al. 2020); Decay sugar-rich substrates (Põlme et al. 2020) | Põlme S, et al. 2020. Fungal Diversity 105:1-16 ((https://doi.org/10.1007/s13225-020-00466-2)) |
| Otu09355 | 2 | k__Fungi(100);p__Ascomycota(100);c__Saccharomycetes(100);o__Saccharomycetales(100);f__Metschnikowiaceae(100);g__Metschnikowia(100);g__Metschnikowia_unclassified(100); | Metschnikowia | 13 | Saprotroph | \|Undefined Saprotroph\| | Yeast | NULL | Probable | As nectar-tap_saprotroph (Põlme et al. 2020); Decay sugar-rich substrates (Põlme et al. 2020) | Põlme S, et al. 2020. Fungal Diversity 105:1-16 ((https://doi.org/10.1007/s13225-020-00466-2)) |
| Otu09356 | 2 | k__Fungi(100);p__Ascomycota(100);c__Saccharomycetes(100);o__Saccharomycetales(100);f__Metschnikowiaceae(100);g__Metschnikowia(100);s__Metschnikowia_pulcherrima(100); | Metschnikowia | 13 | Saprotroph | \|Undefined Saprotroph\| | Yeast | NULL | Probable | As nectar-tap_saprotroph (Põlme et al. 2020); Decay sugar-rich substrates (Põlme et al. 2020) | Põlme S, et al. 2020. Fungal Diversity 105:1-16 ((https://doi.org/10.1007/s13225-020-00466-2)) |
| Otu09357 | 2 | k__Fungi(100);p__Ascomycota(100);c__Saccharomycetes(100);o__Saccharomycetales(100);f__Metschnikowiaceae(100);g__Metschnikowia(100);g__Metschnikowia_unclassified(100); | Metschnikowia | 13 | Saprotroph | \|Undefined Saprotroph\| | Yeast | NULL | Probable | As nectar-tap_saprotroph (Põlme et al. 2020); Decay sugar-rich substrates (Põlme et al. 2020) | Põlme S, et al. 2020. Fungal Diversity 105:1-16 ((https://doi.org/10.1007/s13225-020-00466-2)) |
| Otu09358 | 2 | k__Fungi(100);p__Ascomycota(100);c__Saccharomycetes(100);o__Saccharomycetales(100);f__Metschnikowiaceae(100);g__Metschnikowia(100);s__Metschnikowia_pulcherrima(100); | Metschnikowia | 13 | Saprotroph | \|Undefined Saprotroph\| | Yeast | NULL | Probable | As nectar-tap_saprotroph (Põlme et al. 2020); Decay sugar-rich substrates (Põlme et al. 2020) | Põlme S, et al. 2020. Fungal Diversity 105:1-16 ((https://doi.org/10.1007/s13225-020-00466-2)) |
| Otu09362 | 2 | k__Fungi(100);p__Ascomycota(100);c__Saccharomycetes(100);o__Saccharomycetales(100);f__Metschnikowiaceae(100);g__Metschnikowia(100);g__Metschnikowia_unclassified(100); | Metschnikowia | 13 | Saprotroph | \|Undefined Saprotroph\| | Yeast | NULL | Probable | As nectar-tap_saprotroph (Põlme et al. 2020); Decay sugar-rich substrates (Põlme et al. 2020) | Põlme S, et al. 2020. Fungal Diversity 105:1-16 ((https://doi.org/10.1007/s13225-020-00466-2)) |
| Otu09364 | 2 | k__Fungi(100);p__Ascomycota(100);c__Saccharomycetes(100);o__Saccharomycetales(100);f__Metschnikowiaceae(100);g__Metschnikowia(100);s__Metschnikowia_pulcherrima(100); | Metschnikowia | 13 | Saprotroph | \|Undefined Saprotroph\| | Yeast | NULL | Probable | As nectar-tap_saprotroph (Põlme et al. 2020); Decay sugar-rich substrates (Põlme et al. 2020) | Põlme S, et al. 2020. Fungal Diversity 105:1-16 ((https://doi.org/10.1007/s13225-020-00466-2)) |
| Otu09365 | 2 | k__Fungi(100);p__Ascomycota(100);c__Saccharomycetes(100);o__Saccharomycetales(100);f__Metschnikowiaceae(100);g__Metschnikowia(100);s__Metschnikowia_pulcherrima(100); | Metschnikowia | 13 | Saprotroph | \|Undefined Saprotroph\| | Yeast | NULL | Probable | As nectar-tap_saprotroph (Põlme et al. 2020); Decay sugar-rich substrates (Põlme et al. 2020) | Põlme S, et al. 2020. Fungal Diversity 105:1-16 ((https://doi.org/10.1007/s13225-020-00466-2)) |
| Otu09367 | 2 | k__Fungi(100);p__Ascomycota(100);c__Saccharomycetes(100);o__Saccharomycetales(100);f__Metschnikowiaceae(100);g__Metschnikowia(100);s__Metschnikowia_pulcherrima(100); | Metschnikowia | 13 | Saprotroph | \|Undefined Saprotroph\| | Yeast | NULL | Probable | As nectar-tap_saprotroph (Põlme et al. 2020); Decay sugar-rich substrates (Põlme et al. 2020) | Põlme S, et al. 2020. Fungal Diversity 105:1-16 ((https://doi.org/10.1007/s13225-020-00466-2)) |
| Otu09368 | 2 | k__Fungi(100);p__Ascomycota(100);c__Leotiomycetes(100);o__Helotiales(100);f__Dermateaceae(100);g__Cryptosporiopsis(100);s__Cryptosporiopsis_actinidiae(100); | Cryptosporiopsis | 13 | Saprotroph | \|Undefined Saprotroph\| | Microfungus | NULL | Probable | NULL | Tedersoo L, et al. 2014. Science 346:e1256688 ((https://doi.org/10.1126/science.1256688)); Põlme S, et al. 2020. Fungal Diversity 105:1-16 ((https://doi.org/10.1007/s13225-020-00466-2)) |
| Otu09369 | 2 | k__Fungi(100);p__Ascomycota(100);c__Saccharomycetes(100);o__Saccharomycetales(100);f__Metschnikowiaceae(100);g__Metschnikowia(100);s__Metschnikowia_pulcherrima(100); | Metschnikowia | 13 | Saprotroph | \|Undefined Saprotroph\| | Yeast | NULL | Probable | As nectar-tap_saprotroph (Põlme et al. 2020); Decay sugar-rich substrates (Põlme et al. 2020) | Põlme S, et al. 2020. Fungal Diversity 105:1-16 ((https://doi.org/10.1007/s13225-020-00466-2)) |
| Otu09371 | 2 | k__Fungi(100);p__Ascomycota(100);c__Saccharomycetes(100);o__Saccharomycetales(100);f__Metschnikowiaceae(100);g__Metschnikowia(100);s__Metschnikowia_pulcherrima(100); | Metschnikowia | 13 | Saprotroph | \|Undefined Saprotroph\| | Yeast | NULL | Probable | As nectar-tap_saprotroph (Põlme et al. 2020); Decay sugar-rich substrates (Põlme et al. 2020) | Põlme S, et al. 2020. Fungal Diversity 105:1-16 ((https://doi.org/10.1007/s13225-020-00466-2)) |
| Otu09374 | 2 | k__Fungi(100);p__Ascomycota(100);c__Saccharomycetes(100);o__Saccharomycetales(100);f__Metschnikowiaceae(100);g__Metschnikowia(100);s__Metschnikowia_pulcherrima(100); | Metschnikowia | 13 | Saprotroph | \|Undefined Saprotroph\| | Yeast | NULL | Probable | As nectar-tap_saprotroph (Põlme et al. 2020); Decay sugar-rich substrates (Põlme et al. 2020) | Põlme S, et al. 2020. Fungal Diversity 105:1-16 ((https://doi.org/10.1007/s13225-020-00466-2)) |
| Otu09376 | 2 | k__Fungi(100);p__Ascomycota(100);c__Saccharomycetes(100);o__Saccharomycetales(100);f__Metschnikowiaceae(100);g__Metschnikowia(100);s__Metschnikowia_pulcherrima(100); | Metschnikowia | 13 | Saprotroph | \|Undefined Saprotroph\| | Yeast | NULL | Probable | As nectar-tap_saprotroph (Põlme et al. 2020); Decay sugar-rich substrates (Põlme et al. 2020) | Põlme S, et al. 2020. Fungal Diversity 105:1-16 ((https://doi.org/10.1007/s13225-020-00466-2)) |
| Otu09377 | 2 | k__Fungi(100);p__Ascomycota(100);c__Saccharomycetes(100);o__Saccharomycetales(100);f__Metschnikowiaceae(100);g__Metschnikowia(100);g__Metschnikowia_unclassified(100); | Metschnikowia | 13 | Saprotroph | \|Undefined Saprotroph\| | Yeast | NULL | Probable | As nectar-tap_saprotroph (Põlme et al. 2020); Decay sugar-rich substrates (Põlme et al. 2020) | Põlme S, et al. 2020. Fungal Diversity 105:1-16 ((https://doi.org/10.1007/s13225-020-00466-2)) |
| Otu09378 | 2 | k__Fungi(100);p__Ascomycota(100);c__Saccharomycetes(100);o__Saccharomycetales(100);f__Metschnikowiaceae(100);g__Metschnikowia(100);g__Metschnikowia_unclassified(100); | Metschnikowia | 13 | Saprotroph | \|Undefined Saprotroph\| | Yeast | NULL | Probable | As nectar-tap_saprotroph (Põlme et al. 2020); Decay sugar-rich substrates (Põlme et al. 2020) | Põlme S, et al. 2020. Fungal Diversity 105:1-16 ((https://doi.org/10.1007/s13225-020-00466-2)) |
| Otu09379 | 2 | k__Fungi(100);p__Ascomycota(100);c__Saccharomycetes(100);o__Saccharomycetales(100);f__Metschnikowiaceae(100);g__Metschnikowia(100);s__Metschnikowia_pulcherrima(100); | Metschnikowia | 13 | Saprotroph | \|Undefined Saprotroph\| | Yeast | NULL | Probable | As nectar-tap_saprotroph (Põlme et al. 2020); Decay sugar-rich substrates (Põlme et al. 2020) | Põlme S, et al. 2020. Fungal Diversity 105:1-16 ((https://doi.org/10.1007/s13225-020-00466-2)) |
| Otu09380 | 2 | k__Fungi(100);p__Ascomycota(100);c__Saccharomycetes(100);o__Saccharomycetales(100);f__Metschnikowiaceae(100);g__Metschnikowia(100);s__Metschnikowia_pulcherrima(100); | Metschnikowia | 13 | Saprotroph | \|Undefined Saprotroph\| | Yeast | NULL | Probable | As nectar-tap_saprotroph (Põlme et al. 2020); Decay sugar-rich substrates (Põlme et al. 2020) | Põlme S, et al. 2020. Fungal Diversity 105:1-16 ((https://doi.org/10.1007/s13225-020-00466-2)) |
| Otu09383 | 2 | k__Fungi(100);p__Ascomycota(100);c__Saccharomycetes(100);o__Saccharomycetales(100);f__Metschnikowiaceae(100);g__Metschnikowia(100);s__Metschnikowia_pulcherrima(100); | Metschnikowia | 13 | Saprotroph | \|Undefined Saprotroph\| | Yeast | NULL | Probable | As nectar-tap_saprotroph (Põlme et al. 2020); Decay sugar-rich substrates (Põlme et al. 2020) | Põlme S, et al. 2020. Fungal Diversity 105:1-16 ((https://doi.org/10.1007/s13225-020-00466-2)) |
| Otu09391 | 2 | k__Fungi(100);p__Ascomycota(100);c__Saccharomycetes(100);o__Saccharomycetales(100);f__Metschnikowiaceae(100);g__Metschnikowia(100);s__Metschnikowia_pulcherrima(100); | Metschnikowia | 13 | Saprotroph | \|Undefined Saprotroph\| | Yeast | NULL | Probable | As nectar-tap_saprotroph (Põlme et al. 2020); Decay sugar-rich substrates (Põlme et al. 2020) | Põlme S, et al. 2020. Fungal Diversity 105:1-16 ((https://doi.org/10.1007/s13225-020-00466-2)) |
| Otu09392 | 2 | k__Fungi(100);p__Ascomycota(100);c__Saccharomycetes(100);o__Saccharomycetales(100);f__Metschnikowiaceae(100);g__Metschnikowia(100);g__Metschnikowia_unclassified(100); | Metschnikowia | 13 | Saprotroph | \|Undefined Saprotroph\| | Yeast | NULL | Probable | As nectar-tap_saprotroph (Põlme et al. 2020); Decay sugar-rich substrates (Põlme et al. 2020) | Põlme S, et al. 2020. Fungal Diversity 105:1-16 ((https://doi.org/10.1007/s13225-020-00466-2)) |
| Otu09394 | 2 | k__Fungi(100);p__Ascomycota(100);c__Saccharomycetes(100);o__Saccharomycetales(100);f__Metschnikowiaceae(100);g__Metschnikowia(100);g__Metschnikowia_unclassified(100); | Metschnikowia | 13 | Saprotroph | \|Undefined Saprotroph\| | Yeast | NULL | Probable | As nectar-tap_saprotroph (Põlme et al. 2020); Decay sugar-rich substrates (Põlme et al. 2020) | Põlme S, et al. 2020. Fungal Diversity 105:1-16 ((https://doi.org/10.1007/s13225-020-00466-2)) |
| Otu09396 | 2 | k__Fungi(100);p__Ascomycota(100);c__Saccharomycetes(100);o__Saccharomycetales(100);f__Metschnikowiaceae(100);g__Metschnikowia(100);s__Metschnikowia_pulcherrima(100); | Metschnikowia | 13 | Saprotroph | \|Undefined Saprotroph\| | Yeast | NULL | Probable | As nectar-tap_saprotroph (Põlme et al. 2020); Decay sugar-rich substrates (Põlme et al. 2020) | Põlme S, et al. 2020. Fungal Diversity 105:1-16 ((https://doi.org/10.1007/s13225-020-00466-2)) |
| Otu09398 | 2 | k__Fungi(100);p__Ascomycota(100);c__Saccharomycetes(100);o__Saccharomycetales(100);f__Metschnikowiaceae(100);g__Metschnikowia(100);s__Metschnikowia_pulcherrima(100); | Metschnikowia | 13 | Saprotroph | \|Undefined Saprotroph\| | Yeast | NULL | Probable | As nectar-tap_saprotroph (Põlme et al. 2020); Decay sugar-rich substrates (Põlme et al. 2020) | Põlme S, et al. 2020. Fungal Diversity 105:1-16 ((https://doi.org/10.1007/s13225-020-00466-2)) |
| Otu09399 | 2 | k__Fungi(100);p__Ascomycota(100);c__Saccharomycetes(100);o__Saccharomycetales(100);f__Metschnikowiaceae(100);g__Metschnikowia(100);s__Metschnikowia_pulcherrima(100); | Metschnikowia | 13 | Saprotroph | \|Undefined Saprotroph\| | Yeast | NULL | Probable | As nectar-tap_saprotroph (Põlme et al. 2020); Decay sugar-rich substrates (Põlme et al. 2020) | Põlme S, et al. 2020. Fungal Diversity 105:1-16 ((https://doi.org/10.1007/s13225-020-00466-2)) |
| Otu09400 | 2 | k__Fungi(100);p__Ascomycota(100);c__Saccharomycetes(100);o__Saccharomycetales(100);f__Metschnikowiaceae(100);g__Metschnikowia(100);s__Metschnikowia_pulcherrima(100); | Metschnikowia | 13 | Saprotroph | \|Undefined Saprotroph\| | Yeast | NULL | Probable | As nectar-tap_saprotroph (Põlme et al. 2020); Decay sugar-rich substrates (Põlme et al. 2020) | Põlme S, et al. 2020. Fungal Diversity 105:1-16 ((https://doi.org/10.1007/s13225-020-00466-2)) |
| Otu09405 | 2 | k__Fungi(100);p__Ascomycota(100);c__Saccharomycetes(100);o__Saccharomycetales(100);f__Metschnikowiaceae(100);g__Metschnikowia(100);g__Metschnikowia_unclassified(100); | Metschnikowia | 13 | Saprotroph | \|Undefined Saprotroph\| | Yeast | NULL | Probable | As nectar-tap_saprotroph (Põlme et al. 2020); Decay sugar-rich substrates (Põlme et al. 2020) | Põlme S, et al. 2020. Fungal Diversity 105:1-16 ((https://doi.org/10.1007/s13225-020-00466-2)) |
| Otu09406 | 2 | k__Fungi(100);p__Ascomycota(100);c__Saccharomycetes(100);o__Saccharomycetales(100);f__Metschnikowiaceae(100);g__Metschnikowia(100);s__Metschnikowia_pulcherrima(100); | Metschnikowia | 13 | Saprotroph | \|Undefined Saprotroph\| | Yeast | NULL | Probable | As nectar-tap_saprotroph (Põlme et al. 2020); Decay sugar-rich substrates (Põlme et al. 2020) | Põlme S, et al. 2020. Fungal Diversity 105:1-16 ((https://doi.org/10.1007/s13225-020-00466-2)) |
| Otu09408 | 2 | k__Fungi(100);p__Ascomycota(100);c__Saccharomycetes(100);o__Saccharomycetales(100);f__Metschnikowiaceae(100);g__Metschnikowia(100);g__Metschnikowia_unclassified(100); | Metschnikowia | 13 | Saprotroph | \|Undefined Saprotroph\| | Yeast | NULL | Probable | As nectar-tap_saprotroph (Põlme et al. 2020); Decay sugar-rich substrates (Põlme et al. 2020) | Põlme S, et al. 2020. Fungal Diversity 105:1-16 ((https://doi.org/10.1007/s13225-020-00466-2)) |
| Otu09410 | 2 | k__Fungi(100);p__Ascomycota(100);c__Saccharomycetes(100);o__Saccharomycetales(100);f__Metschnikowiaceae(100);g__Metschnikowia(100);g__Metschnikowia_unclassified(100); | Metschnikowia | 13 | Saprotroph | \|Undefined Saprotroph\| | Yeast | NULL | Probable | As nectar-tap_saprotroph (Põlme et al. 2020); Decay sugar-rich substrates (Põlme et al. 2020) | Põlme S, et al. 2020. Fungal Diversity 105:1-16 ((https://doi.org/10.1007/s13225-020-00466-2)) |
| Otu09413 | 2 | k__Fungi(100);p__Ascomycota(100);c__Saccharomycetes(100);o__Saccharomycetales(100);f__Metschnikowiaceae(100);g__Metschnikowia(100);s__Metschnikowia_pulcherrima(100); | Metschnikowia | 13 | Saprotroph | \|Undefined Saprotroph\| | Yeast | NULL | Probable | As nectar-tap_saprotroph (Põlme et al. 2020); Decay sugar-rich substrates (Põlme et al. 2020) | Põlme S, et al. 2020. Fungal Diversity 105:1-16 ((https://doi.org/10.1007/s13225-020-00466-2)) |
| Otu09414 | 2 | k__Fungi(100);p__Ascomycota(100);c__Saccharomycetes(100);o__Saccharomycetales(100);f__Metschnikowiaceae(100);g__Metschnikowia(100);g__Metschnikowia_unclassified(100); | Metschnikowia | 13 | Saprotroph | \|Undefined Saprotroph\| | Yeast | NULL | Probable | As nectar-tap_saprotroph (Põlme et al. 2020); Decay sugar-rich substrates (Põlme et al. 2020) | Põlme S, et al. 2020. Fungal Diversity 105:1-16 ((https://doi.org/10.1007/s13225-020-00466-2)) |
| Otu09415 | 2 | k__Fungi(100);p__Ascomycota(100);c__Saccharomycetes(100);o__Saccharomycetales(100);f__Metschnikowiaceae(100);g__Metschnikowia(100);s__Metschnikowia_pulcherrima(100); | Metschnikowia | 13 | Saprotroph | \|Undefined Saprotroph\| | Yeast | NULL | Probable | As nectar-tap_saprotroph (Põlme et al. 2020); Decay sugar-rich substrates (Põlme et al. 2020) | Põlme S, et al. 2020. Fungal Diversity 105:1-16 ((https://doi.org/10.1007/s13225-020-00466-2)) |
| Otu09416 | 2 | k__Fungi(100);p__Ascomycota(100);c__Saccharomycetes(100);o__Saccharomycetales(100);f__Metschnikowiaceae(100);g__Metschnikowia(100);g__Metschnikowia_unclassified(100); | Metschnikowia | 13 | Saprotroph | \|Undefined Saprotroph\| | Yeast | NULL | Probable | As nectar-tap_saprotroph (Põlme et al. 2020); Decay sugar-rich substrates (Põlme et al. 2020) | Põlme S, et al. 2020. Fungal Diversity 105:1-16 ((https://doi.org/10.1007/s13225-020-00466-2)) |
| Otu09418 | 2 | k__Fungi(100);p__Ascomycota(100);c__Saccharomycetes(100);o__Saccharomycetales(100);f__Metschnikowiaceae(100);g__Metschnikowia(100);g__Metschnikowia_unclassified(100); | Metschnikowia | 13 | Saprotroph | \|Undefined Saprotroph\| | Yeast | NULL | Probable | As nectar-tap_saprotroph (Põlme et al. 2020); Decay sugar-rich substrates (Põlme et al. 2020) | Põlme S, et al. 2020. Fungal Diversity 105:1-16 ((https://doi.org/10.1007/s13225-020-00466-2)) |
| Otu09424 | 2 | k__Fungi(100);p__Ascomycota(100);c__Saccharomycetes(100);o__Saccharomycetales(100);f__Metschnikowiaceae(100);g__Metschnikowia(100);s__Metschnikowia_pulcherrima(100); | Metschnikowia | 13 | Saprotroph | \|Undefined Saprotroph\| | Yeast | NULL | Probable | As nectar-tap_saprotroph (Põlme et al. 2020); Decay sugar-rich substrates (Põlme et al. 2020) | Põlme S, et al. 2020. Fungal Diversity 105:1-16 ((https://doi.org/10.1007/s13225-020-00466-2)) |
| Otu09425 | 2 | k__Fungi(100);p__Ascomycota(100);c__Saccharomycetes(100);o__Saccharomycetales(100);f__Metschnikowiaceae(100);g__Metschnikowia(100);s__Metschnikowia_pulcherrima(100); | Metschnikowia | 13 | Saprotroph | \|Undefined Saprotroph\| | Yeast | NULL | Probable | As nectar-tap_saprotroph (Põlme et al. 2020); Decay sugar-rich substrates (Põlme et al. 2020) | Põlme S, et al. 2020. Fungal Diversity 105:1-16 ((https://doi.org/10.1007/s13225-020-00466-2)) |
| Otu09427 | 2 | k__Fungi(100);p__Ascomycota(100);c__Saccharomycetes(100);o__Saccharomycetales(100);f__Metschnikowiaceae(100);g__Metschnikowia(100);s__Metschnikowia_pulcherrima(100); | Metschnikowia | 13 | Saprotroph | \|Undefined Saprotroph\| | Yeast | NULL | Probable | As nectar-tap_saprotroph (Põlme et al. 2020); Decay sugar-rich substrates (Põlme et al. 2020) | Põlme S, et al. 2020. Fungal Diversity 105:1-16 ((https://doi.org/10.1007/s13225-020-00466-2)) |
| Otu09428 | 2 | k__Fungi(100);p__Ascomycota(100);c__Saccharomycetes(100);o__Saccharomycetales(100);f__Metschnikowiaceae(100);g__Metschnikowia(100);g__Metschnikowia_unclassified(100); | Metschnikowia | 13 | Saprotroph | \|Undefined Saprotroph\| | Yeast | NULL | Probable | As nectar-tap_saprotroph (Põlme et al. 2020); Decay sugar-rich substrates (Põlme et al. 2020) | Põlme S, et al. 2020. Fungal Diversity 105:1-16 ((https://doi.org/10.1007/s13225-020-00466-2)) |
| Otu09431 | 2 | k__Fungi(100);p__Ascomycota(100);c__Saccharomycetes(100);o__Saccharomycetales(100);f__Metschnikowiaceae(100);g__Metschnikowia(100);g__Metschnikowia_unclassified(100); | Metschnikowia | 13 | Saprotroph | \|Undefined Saprotroph\| | Yeast | NULL | Probable | As nectar-tap_saprotroph (Põlme et al. 2020); Decay sugar-rich substrates (Põlme et al. 2020) | Põlme S, et al. 2020. Fungal Diversity 105:1-16 ((https://doi.org/10.1007/s13225-020-00466-2)) |
| Otu09432 | 2 | k__Fungi(100);p__Ascomycota(100);c__Saccharomycetes(100);o__Saccharomycetales(100);f__Metschnikowiaceae(100);g__Metschnikowia(100);g__Metschnikowia_unclassified(100); | Metschnikowia | 13 | Saprotroph | \|Undefined Saprotroph\| | Yeast | NULL | Probable | As nectar-tap_saprotroph (Põlme et al. 2020); Decay sugar-rich substrates (Põlme et al. 2020) | Põlme S, et al. 2020. Fungal Diversity 105:1-16 ((https://doi.org/10.1007/s13225-020-00466-2)) |
| Otu09433 | 2 | k__Fungi(100);p__Ascomycota(100);c__Saccharomycetes(100);o__Saccharomycetales(100);f__Metschnikowiaceae(100);g__Metschnikowia(100);g__Metschnikowia_unclassified(100); | Metschnikowia | 13 | Saprotroph | \|Undefined Saprotroph\| | Yeast | NULL | Probable | As nectar-tap_saprotroph (Põlme et al. 2020); Decay sugar-rich substrates (Põlme et al. 2020) | Põlme S, et al. 2020. Fungal Diversity 105:1-16 ((https://doi.org/10.1007/s13225-020-00466-2)) |
| Otu09434 | 2 | k__Fungi(100);p__Ascomycota(100);c__Saccharomycetes(100);o__Saccharomycetales(100);f__Metschnikowiaceae(100);g__Metschnikowia(100);g__Metschnikowia_unclassified(100); | Metschnikowia | 13 | Saprotroph | \|Undefined Saprotroph\| | Yeast | NULL | Probable | As nectar-tap_saprotroph (Põlme et al. 2020); Decay sugar-rich substrates (Põlme et al. 2020) | Põlme S, et al. 2020. Fungal Diversity 105:1-16 ((https://doi.org/10.1007/s13225-020-00466-2)) |
| Otu09436 | 2 | k__Fungi(100);p__Ascomycota(100);c__Saccharomycetes(100);o__Saccharomycetales(100);f__Metschnikowiaceae(100);g__Metschnikowia(100);g__Metschnikowia_unclassified(100); | Metschnikowia | 13 | Saprotroph | \|Undefined Saprotroph\| | Yeast | NULL | Probable | As nectar-tap_saprotroph (Põlme et al. 2020); Decay sugar-rich substrates (Põlme et al. 2020) | Põlme S, et al. 2020. Fungal Diversity 105:1-16 ((https://doi.org/10.1007/s13225-020-00466-2)) |
| Otu09437 | 2 | k__Fungi(100);p__Ascomycota(100);c__Saccharomycetes(100);o__Saccharomycetales(100);f__Metschnikowiaceae(100);g__Metschnikowia(100);g__Metschnikowia_unclassified(100); | Metschnikowia | 13 | Saprotroph | \|Undefined Saprotroph\| | Yeast | NULL | Probable | As nectar-tap_saprotroph (Põlme et al. 2020); Decay sugar-rich substrates (Põlme et al. 2020) | Põlme S, et al. 2020. Fungal Diversity 105:1-16 ((https://doi.org/10.1007/s13225-020-00466-2)) |
| Otu09443 | 2 | k__Fungi(100);p__Ascomycota(100);c__Saccharomycetes(100);o__Saccharomycetales(100);f__Metschnikowiaceae(100);g__Metschnikowia(100);s__Metschnikowia_pulcherrima(100); | Metschnikowia | 13 | Saprotroph | \|Undefined Saprotroph\| | Yeast | NULL | Probable | As nectar-tap_saprotroph (Põlme et al. 2020); Decay sugar-rich substrates (Põlme et al. 2020) | Põlme S, et al. 2020. Fungal Diversity 105:1-16 ((https://doi.org/10.1007/s13225-020-00466-2)) |
| Otu09445 | 2 | k__Fungi(100);p__Ascomycota(100);c__Saccharomycetes(100);o__Saccharomycetales(100);f__Metschnikowiaceae(100);g__Metschnikowia(100);s__Metschnikowia_pulcherrima(100); | Metschnikowia | 13 | Saprotroph | \|Undefined Saprotroph\| | Yeast | NULL | Probable | As nectar-tap_saprotroph (Põlme et al. 2020); Decay sugar-rich substrates (Põlme et al. 2020) | Põlme S, et al. 2020. Fungal Diversity 105:1-16 ((https://doi.org/10.1007/s13225-020-00466-2)) |
| Otu09446 | 2 | k__Fungi(100);p__Ascomycota(100);c__Saccharomycetes(100);o__Saccharomycetales(100);f__Metschnikowiaceae(100);g__Metschnikowia(100);g__Metschnikowia_unclassified(100); | Metschnikowia | 13 | Saprotroph | \|Undefined Saprotroph\| | Yeast | NULL | Probable | As nectar-tap_saprotroph (Põlme et al. 2020); Decay sugar-rich substrates (Põlme et al. 2020) | Põlme S, et al. 2020. Fungal Diversity 105:1-16 ((https://doi.org/10.1007/s13225-020-00466-2)) |
| Otu09447 | 2 | k__Fungi(100);p__Ascomycota(100);c__Saccharomycetes(100);o__Saccharomycetales(100);f__Metschnikowiaceae(100);g__Metschnikowia(100);s__Metschnikowia_pulcherrima(100); | Metschnikowia | 13 | Saprotroph | \|Undefined Saprotroph\| | Yeast | NULL | Probable | As nectar-tap_saprotroph (Põlme et al. 2020); Decay sugar-rich substrates (Põlme et al. 2020) | Põlme S, et al. 2020. Fungal Diversity 105:1-16 ((https://doi.org/10.1007/s13225-020-00466-2)) |
| Otu09454 | 2 | k__Fungi(100);p__Ascomycota(100);c__Saccharomycetes(100);o__Saccharomycetales(100);f__Metschnikowiaceae(100);g__Metschnikowia(100);s__Metschnikowia_pulcherrima(100); | Metschnikowia | 13 | Saprotroph | \|Undefined Saprotroph\| | Yeast | NULL | Probable | As nectar-tap_saprotroph (Põlme et al. 2020); Decay sugar-rich substrates (Põlme et al. 2020) | Põlme S, et al. 2020. Fungal Diversity 105:1-16 ((https://doi.org/10.1007/s13225-020-00466-2)) |
| Otu09455 | 2 | k__Fungi(100);p__Ascomycota(100);c__Saccharomycetes(100);o__Saccharomycetales(100);f__Metschnikowiaceae(100);g__Metschnikowia(100);s__Metschnikowia_pulcherrima(100); | Metschnikowia | 13 | Saprotroph | \|Undefined Saprotroph\| | Yeast | NULL | Probable | As nectar-tap_saprotroph (Põlme et al. 2020); Decay sugar-rich substrates (Põlme et al. 2020) | Põlme S, et al. 2020. Fungal Diversity 105:1-16 ((https://doi.org/10.1007/s13225-020-00466-2)) |
| Otu09456 | 2 | k__Fungi(100);p__Ascomycota(100);c__Saccharomycetes(100);o__Saccharomycetales(100);f__Metschnikowiaceae(100);g__Metschnikowia(100);s__Metschnikowia_pulcherrima(100); | Metschnikowia | 13 | Saprotroph | \|Undefined Saprotroph\| | Yeast | NULL | Probable | As nectar-tap_saprotroph (Põlme et al. 2020); Decay sugar-rich substrates (Põlme et al. 2020) | Põlme S, et al. 2020. Fungal Diversity 105:1-16 ((https://doi.org/10.1007/s13225-020-00466-2)) |
| Otu09464 | 2 | k__Fungi(100);p__Ascomycota(100);c__Saccharomycetes(100);o__Saccharomycetales(100);f__Metschnikowiaceae(100);g__Metschnikowia(100);s__Metschnikowia_pulcherrima(100); | Metschnikowia | 13 | Saprotroph | \|Undefined Saprotroph\| | Yeast | NULL | Probable | As nectar-tap_saprotroph (Põlme et al. 2020); Decay sugar-rich substrates (Põlme et al. 2020) | Põlme S, et al. 2020. Fungal Diversity 105:1-16 ((https://doi.org/10.1007/s13225-020-00466-2)) |
| Otu09465 | 2 | k__Fungi(100);p__Ascomycota(100);c__Saccharomycetes(100);o__Saccharomycetales(100);f__Metschnikowiaceae(100);g__Metschnikowia(100);g__Metschnikowia_unclassified(100); | Metschnikowia | 13 | Saprotroph | \|Undefined Saprotroph\| | Yeast | NULL | Probable | As nectar-tap_saprotroph (Põlme et al. 2020); Decay sugar-rich substrates (Põlme et al. 2020) | Põlme S, et al. 2020. Fungal Diversity 105:1-16 ((https://doi.org/10.1007/s13225-020-00466-2)) |
| Otu09467 | 2 | k__Fungi(100);p__Ascomycota(100);c__Saccharomycetes(100);o__Saccharomycetales(100);f__Metschnikowiaceae(100);g__Metschnikowia(100);s__Metschnikowia_pulcherrima(100); | Metschnikowia | 13 | Saprotroph | \|Undefined Saprotroph\| | Yeast | NULL | Probable | As nectar-tap_saprotroph (Põlme et al. 2020); Decay sugar-rich substrates (Põlme et al. 2020) | Põlme S, et al. 2020. Fungal Diversity 105:1-16 ((https://doi.org/10.1007/s13225-020-00466-2)) |
| Otu09470 | 2 | k__Fungi(100);p__Ascomycota(100);c__Saccharomycetes(100);o__Saccharomycetales(100);f__Metschnikowiaceae(100);g__Metschnikowia(100);g__Metschnikowia_unclassified(100); | Metschnikowia | 13 | Saprotroph | \|Undefined Saprotroph\| | Yeast | NULL | Probable | As nectar-tap_saprotroph (Põlme et al. 2020); Decay sugar-rich substrates (Põlme et al. 2020) | Põlme S, et al. 2020. Fungal Diversity 105:1-16 ((https://doi.org/10.1007/s13225-020-00466-2)) |
| Otu09471 | 2 | k__Fungi(100);p__Ascomycota(100);c__Dothideomycetes(100);o__Pleosporales(100);f__Tubeufiaceae(100);f__Tubeufiaceae_unclassified(100);f__Tubeufiaceae_unclassified(100); | Tubeufiaceae | 9 | Pathotroph-Saprotroph | Animal Pathogen-Undefined Saprotroph | NULL | NULL | Probable | Biotrophic or saprobic, especially on scale insects or other fungi. The anamorphic forms are frequently aero-aquatic (Cannon & Kirk 2007) | Cannon PF, Kirk PM. 2007. Fungal Families of the World. CAB International, Cambridge (ISBN: 978-0851998275) |
| Otu09478 | 2 | k__Fungi(100);p__Ascomycota(100);c__Saccharomycetes(100);o__Saccharomycetales(100);f__Metschnikowiaceae(100);g__Metschnikowia(100);g__Metschnikowia_unclassified(100); | Metschnikowia | 13 | Saprotroph | \|Undefined Saprotroph\| | Yeast | NULL | Probable | As nectar-tap_saprotroph (Põlme et al. 2020); Decay sugar-rich substrates (Põlme et al. 2020) | Põlme S, et al. 2020. Fungal Diversity 105:1-16 ((https://doi.org/10.1007/s13225-020-00466-2)) |
| Otu09480 | 2 | k__Fungi(100);p__Ascomycota(100);c__Dothideomycetes(100);o__Botryosphaeriales(100);f__Botryosphaeriaceae(100);f__Botryosphaeriaceae_unclassified(100);f__Botryosphaeriaceae_unclassified(100); | Botryosphaeriaceae | 9 | Pathotroph-Saprotroph-Symbiotroph | Endophyte-Plant Pathogen-Wood Saprotroph | Microfungus | NULL | Probable | Biotrophic, necrotrophic, or saprobic, especially associated with woody tissue (Cannon & Kirk 2007) | Cannon PF, Kirk PM. 2007. Fungal Families of the World. CAB International, Cambridge (ISBN: 978-0851998275) |
| Otu09481 | 2 | k__Fungi(100);p__Ascomycota(100);c__Dothideomycetes(100);o__Capnodiales(100);f__Teratosphaeriaceae(100);g__Teratosphaeria(100);g__Teratosphaeria_unclassified(100); | Teratosphaeria | 13 | Pathotroph-Saprotroph | \|Plant Pathogen\|-Plant Saprotroph-Wood Saprotroph | Microfungus | NULL | Probable | As litter_saprotroph (Põlme et al. 2020); As wood_pathogen (Põlme et al. 2020); Decay wood, leaf, fruit, seed (Põlme et al. 2020) | Yamazaki A, et al. 2010. Mycoscience 51:443-455 ((https://www.jstage.jst.go.jp/article/mycosci/51/6/51_MYC51443/_pdf)); Tedersoo L, et al. 2014. Science 346:e1256688 ((https://doi.org/10.1126/science.1256688)); Põlme S, et al. 2020. Fungal Diversity 105:1-16 ((https://doi.org/10.1007/s13225-020-00466-2)) |
| Otu09482 | 2 | k__Fungi(100);p__Ascomycota(100);c__Saccharomycetes(100);o__Saccharomycetales(100);f__Metschnikowiaceae(100);g__Metschnikowia(100);s__Metschnikowia_pulcherrima(100); | Metschnikowia | 13 | Saprotroph | \|Undefined Saprotroph\| | Yeast | NULL | Probable | As nectar-tap_saprotroph (Põlme et al. 2020); Decay sugar-rich substrates (Põlme et al. 2020) | Põlme S, et al. 2020. Fungal Diversity 105:1-16 ((https://doi.org/10.1007/s13225-020-00466-2)) |
| Otu09483 | 2 | k__Fungi(100);p__Ascomycota(100);c__Saccharomycetes(100);o__Saccharomycetales(100);f__Metschnikowiaceae(100);g__Metschnikowia(100);g__Metschnikowia_unclassified(100); | Metschnikowia | 13 | Saprotroph | \|Undefined Saprotroph\| | Yeast | NULL | Probable | As nectar-tap_saprotroph (Põlme et al. 2020); Decay sugar-rich substrates (Põlme et al. 2020) | Põlme S, et al. 2020. Fungal Diversity 105:1-16 ((https://doi.org/10.1007/s13225-020-00466-2)) |
| Otu09484 | 2 | k__Fungi(100);p__Ascomycota(100);c__Saccharomycetes(100);o__Saccharomycetales(100);f__Metschnikowiaceae(100);g__Metschnikowia(100);s__Metschnikowia_pulcherrima(100); | Metschnikowia | 13 | Saprotroph | \|Undefined Saprotroph\| | Yeast | NULL | Probable | As nectar-tap_saprotroph (Põlme et al. 2020); Decay sugar-rich substrates (Põlme et al. 2020) | Põlme S, et al. 2020. Fungal Diversity 105:1-16 ((https://doi.org/10.1007/s13225-020-00466-2)) |
| Otu09486 | 2 | k__Fungi(100);p__Ascomycota(100);c__Saccharomycetes(100);o__Saccharomycetales(100);f__Metschnikowiaceae(100);g__Metschnikowia(100);s__Metschnikowia_pulcherrima(100); | Metschnikowia | 13 | Saprotroph | \|Undefined Saprotroph\| | Yeast | NULL | Probable | As nectar-tap_saprotroph (Põlme et al. 2020); Decay sugar-rich substrates (Põlme et al. 2020) | Põlme S, et al. 2020. Fungal Diversity 105:1-16 ((https://doi.org/10.1007/s13225-020-00466-2)) |
| Otu09487 | 2 | k__Fungi(100);p__Ascomycota(100);c__Saccharomycetes(100);o__Saccharomycetales(100);f__Metschnikowiaceae(100);g__Metschnikowia(100);s__Metschnikowia_pulcherrima(100); | Metschnikowia | 13 | Saprotroph | \|Undefined Saprotroph\| | Yeast | NULL | Probable | As nectar-tap_saprotroph (Põlme et al. 2020); Decay sugar-rich substrates (Põlme et al. 2020) | Põlme S, et al. 2020. Fungal Diversity 105:1-16 ((https://doi.org/10.1007/s13225-020-00466-2)) |
| Otu09488 | 2 | k__Fungi(100);p__Ascomycota(100);c__Saccharomycetes(100);o__Saccharomycetales(100);f__Metschnikowiaceae(100);g__Metschnikowia(100);s__Metschnikowia_pulcherrima(100); | Metschnikowia | 13 | Saprotroph | \|Undefined Saprotroph\| | Yeast | NULL | Probable | As nectar-tap_saprotroph (Põlme et al. 2020); Decay sugar-rich substrates (Põlme et al. 2020) | Põlme S, et al. 2020. Fungal Diversity 105:1-16 ((https://doi.org/10.1007/s13225-020-00466-2)) |
| Otu09489 | 2 | k__Fungi(100);p__Ascomycota(100);c__Dothideomycetes(100);o__Capnodiales(100);f__Teratosphaeriaceae(100);g__Devriesia(100);s__Devriesia_sp(100); | Devriesia | 13 | Pathotroph-Saprotroph | \|Plant Pathogen\|-Plant Saprotroph | Microfungus | NULL | Probable | As litter_saprotroph (Põlme et al. 2020); Decay leaf, fruit, seed (Põlme et al. 2020) | Tedersoo L, et al. 2014. Science 346:e1256688 ((https://doi.org/10.1126/science.1256688)); Põlme S, et al. 2020. Fungal Diversity 105:1-16 ((https://doi.org/10.1007/s13225-020-00466-2)) |
| Otu09490 | 2 | k__Fungi(100);p__Ascomycota(100);c__Saccharomycetes(100);o__Saccharomycetales(100);f__Metschnikowiaceae(100);g__Metschnikowia(100);s__Metschnikowia_pulcherrima(100); | Metschnikowia | 13 | Saprotroph | \|Undefined Saprotroph\| | Yeast | NULL | Probable | As nectar-tap_saprotroph (Põlme et al. 2020); Decay sugar-rich substrates (Põlme et al. 2020) | Põlme S, et al. 2020. Fungal Diversity 105:1-16 ((https://doi.org/10.1007/s13225-020-00466-2)) |
| Otu09491 | 2 | k__Fungi(100);p__Ascomycota(100);c__Saccharomycetes(100);o__Saccharomycetales(100);f__Metschnikowiaceae(100);g__Metschnikowia(100);g__Metschnikowia_unclassified(100); | Metschnikowia | 13 | Saprotroph | \|Undefined Saprotroph\| | Yeast | NULL | Probable | As nectar-tap_saprotroph (Põlme et al. 2020); Decay sugar-rich substrates (Põlme et al. 2020) | Põlme S, et al. 2020. Fungal Diversity 105:1-16 ((https://doi.org/10.1007/s13225-020-00466-2)) |
| Otu09492 | 2 | k__Fungi(100);p__Ascomycota(100);c__Saccharomycetes(100);o__Saccharomycetales(100);f__Metschnikowiaceae(100);g__Metschnikowia(100);s__Metschnikowia_pulcherrima(100); | Metschnikowia | 13 | Saprotroph | \|Undefined Saprotroph\| | Yeast | NULL | Probable | As nectar-tap_saprotroph (Põlme et al. 2020); Decay sugar-rich substrates (Põlme et al. 2020) | Põlme S, et al. 2020. Fungal Diversity 105:1-16 ((https://doi.org/10.1007/s13225-020-00466-2)) |
| Otu09494 | 2 | k__Fungi(100);p__Ascomycota(100);c__Saccharomycetes(100);o__Saccharomycetales(100);f__Metschnikowiaceae(100);g__Metschnikowia(100);g__Metschnikowia_unclassified(100); | Metschnikowia | 13 | Saprotroph | \|Undefined Saprotroph\| | Yeast | NULL | Probable | As nectar-tap_saprotroph (Põlme et al. 2020); Decay sugar-rich substrates (Põlme et al. 2020) | Põlme S, et al. 2020. Fungal Diversity 105:1-16 ((https://doi.org/10.1007/s13225-020-00466-2)) |
| Otu09496 | 2 | k__Fungi(100);p__Ascomycota(100);c__Saccharomycetes(100);o__Saccharomycetales(100);f__Metschnikowiaceae(100);g__Metschnikowia(100);s__Metschnikowia_pulcherrima(100); | Metschnikowia | 13 | Saprotroph | \|Undefined Saprotroph\| | Yeast | NULL | Probable | As nectar-tap_saprotroph (Põlme et al. 2020); Decay sugar-rich substrates (Põlme et al. 2020) | Põlme S, et al. 2020. Fungal Diversity 105:1-16 ((https://doi.org/10.1007/s13225-020-00466-2)) |
| Otu09498 | 2 | k__Fungi(100);p__Ascomycota(100);c__Saccharomycetes(100);o__Saccharomycetales(100);f__Metschnikowiaceae(100);g__Metschnikowia(100);g__Metschnikowia_unclassified(100); | Metschnikowia | 13 | Saprotroph | \|Undefined Saprotroph\| | Yeast | NULL | Probable | As nectar-tap_saprotroph (Põlme et al. 2020); Decay sugar-rich substrates (Põlme et al. 2020) | Põlme S, et al. 2020. Fungal Diversity 105:1-16 ((https://doi.org/10.1007/s13225-020-00466-2)) |
| Otu09500 | 2 | k__Fungi(100);p__Ascomycota(100);c__Saccharomycetes(100);o__Saccharomycetales(100);f__Metschnikowiaceae(100);g__Metschnikowia(100);s__Metschnikowia_pulcherrima(100); | Metschnikowia | 13 | Saprotroph | \|Undefined Saprotroph\| | Yeast | NULL | Probable | As nectar-tap_saprotroph (Põlme et al. 2020); Decay sugar-rich substrates (Põlme et al. 2020) | Põlme S, et al. 2020. Fungal Diversity 105:1-16 ((https://doi.org/10.1007/s13225-020-00466-2)) |
| Otu09501 | 2 | k__Fungi(100);p__Ascomycota(100);c__Saccharomycetes(100);o__Saccharomycetales(100);f__Metschnikowiaceae(100);g__Metschnikowia(100);s__Metschnikowia_pulcherrima(100); | Metschnikowia | 13 | Saprotroph | \|Undefined Saprotroph\| | Yeast | NULL | Probable | As nectar-tap_saprotroph (Põlme et al. 2020); Decay sugar-rich substrates (Põlme et al. 2020) | Põlme S, et al. 2020. Fungal Diversity 105:1-16 ((https://doi.org/10.1007/s13225-020-00466-2)) |
| Otu09502 | 2 | k__Fungi(100);p__Ascomycota(100);c__Saccharomycetes(100);o__Saccharomycetales(100);f__Metschnikowiaceae(100);g__Metschnikowia(100);g__Metschnikowia_unclassified(100); | Metschnikowia | 13 | Saprotroph | \|Undefined Saprotroph\| | Yeast | NULL | Probable | As nectar-tap_saprotroph (Põlme et al. 2020); Decay sugar-rich substrates (Põlme et al. 2020) | Põlme S, et al. 2020. Fungal Diversity 105:1-16 ((https://doi.org/10.1007/s13225-020-00466-2)) |
| Otu09503 | 2 | k__Fungi(100);p__Ascomycota(100);c__Leotiomycetes(100);o__Helotiales(100);f__Vibrisseaceae(100);g__Phialocephala(100);s__Phialocephala_fusca(100); | Phialocephala | 13 | Saprotroph-Symbiotroph | Endophyte-\|Undefined Saprotroph\| | Microfungus | NULL | Highly Probable | As root_endophyte (Põlme et al. 2020); As root_endophyte_dark_septate (Põlme et al. 2020); As soil_saprotroph (Põlme et al. 2020); Dark Septate Endophyte, known from plant roots.; Decay roots, soil (Põlme et al. 2020) | Newsham KK. 2011. New Phytologist 190:783-793 ((https://doi.org/10.1111/j.1469-8137.2010.03611.x)) (as dark septate root endophytes); Põlme S, et al. 2020. Fungal Diversity 105:1-16 ((https://doi.org/10.1007/s13225-020-00466-2)) |
| Otu09504 | 2 | k__Fungi(100);p__Ascomycota(100);c__Saccharomycetes(100);o__Saccharomycetales(100);f__Metschnikowiaceae(100);g__Metschnikowia(100);s__Metschnikowia_pulcherrima(100); | Metschnikowia | 13 | Saprotroph | \|Undefined Saprotroph\| | Yeast | NULL | Probable | As nectar-tap_saprotroph (Põlme et al. 2020); Decay sugar-rich substrates (Põlme et al. 2020) | Põlme S, et al. 2020. Fungal Diversity 105:1-16 ((https://doi.org/10.1007/s13225-020-00466-2)) |
| Otu09506 | 2 | k__Fungi(100);p__Ascomycota(100);c__Dothideomycetes(100);o__Pleosporales(100);f__Leptosphaeriaceae(100);f__Leptosphaeriaceae_unclassified(100);f__Leptosphaeriaceae_unclassified(100); | Leptosphaeriaceae | 9 | Pathotroph-Saprotroph | Plant Pathogen-Wood Saprotroph | Microfungus | NULL | Probable | Saprobic or weakly necrotrophic on stems or leaves (Cannon & Kirk 2007) | Cannon PF, Kirk PM. 2007. Fungal Families of the World. CAB International, Cambridge (ISBN: 978-0851998275) |
| Otu09508 | 2 | k__Fungi(100);p__Ascomycota(100);c__Saccharomycetes(100);o__Saccharomycetales(100);f__Metschnikowiaceae(100);g__Metschnikowia(100);s__Metschnikowia_pulcherrima(100); | Metschnikowia | 13 | Saprotroph | \|Undefined Saprotroph\| | Yeast | NULL | Probable | As nectar-tap_saprotroph (Põlme et al. 2020); Decay sugar-rich substrates (Põlme et al. 2020) | Põlme S, et al. 2020. Fungal Diversity 105:1-16 ((https://doi.org/10.1007/s13225-020-00466-2)) |
| Otu09509 | 2 | k__Fungi(100);p__Ascomycota(100);c__Saccharomycetes(100);o__Saccharomycetales(100);f__Metschnikowiaceae(100);g__Metschnikowia(100);s__Metschnikowia_pulcherrima(100); | Metschnikowia | 13 | Saprotroph | \|Undefined Saprotroph\| | Yeast | NULL | Probable | As nectar-tap_saprotroph (Põlme et al. 2020); Decay sugar-rich substrates (Põlme et al. 2020) | Põlme S, et al. 2020. Fungal Diversity 105:1-16 ((https://doi.org/10.1007/s13225-020-00466-2)) |
| Otu09513 | 2 | k__Fungi(100);p__Ascomycota(100);c__Sordariomycetes(100);o__Coniochaetales(100);f__Coniochaetaceae(100);g__unclassified_Coniochaetaceae(100);s__Coniochaetaceae_sp(100); | Coniochaetaceae | 9 | Pathotroph-Saprotroph-Symbiotroph | Animal Pathogen-Dung Saprotroph-Endophyte-Lichen Parasite-Plant Pathogen-Undefined Saprotroph | Microfungus | NULL | Possible | Saprobic on wood and bark (especially in dry habitats), coprophilous or isolated from soil and similar substrata (Cannon & Kirk 2007) | Cannon PF, Kirk PM. 2007. Fungal Families of the World. CAB International, Cambridge (ISBN: 978-0851998275); Damm U, et al. 2010. Persoonia 24:60-80 ((http://doi.org/10.3767/003158510X500705)); Irinyi L, et al. 2015. Medical Mycology 53:313-337 ((https://doi.org/10.1093/mmy/myv008)); Lawrey JD, Diederich P. 2016. http://www.lichenicolous.net (accessed 24 May 2017) ((http://www.lichenicolous.net)) |
| Otu09514 | 2 | k__Fungi(100);p__Ascomycota(100);c__Saccharomycetes(100);o__Saccharomycetales(100);f__Metschnikowiaceae(100);g__Metschnikowia(100);s__Metschnikowia_pulcherrima(100); | Metschnikowia | 13 | Saprotroph | \|Undefined Saprotroph\| | Yeast | NULL | Probable | As nectar-tap_saprotroph (Põlme et al. 2020); Decay sugar-rich substrates (Põlme et al. 2020) | Põlme S, et al. 2020. Fungal Diversity 105:1-16 ((https://doi.org/10.1007/s13225-020-00466-2)) |
| Otu09515 | 2 | k__Fungi(100);p__Ascomycota(100);c__Saccharomycetes(100);o__Saccharomycetales(100);f__Metschnikowiaceae(100);g__Metschnikowia(100);g__Metschnikowia_unclassified(100); | Metschnikowia | 13 | Saprotroph | \|Undefined Saprotroph\| | Yeast | NULL | Probable | As nectar-tap_saprotroph (Põlme et al. 2020); Decay sugar-rich substrates (Põlme et al. 2020) | Põlme S, et al. 2020. Fungal Diversity 105:1-16 ((https://doi.org/10.1007/s13225-020-00466-2)) |
| Otu09516 | 2 | k__Fungi(100);p__Ascomycota(100);c__Saccharomycetes(100);o__Saccharomycetales(100);f__Metschnikowiaceae(100);g__Metschnikowia(100);s__Metschnikowia_pulcherrima(100); | Metschnikowia | 13 | Saprotroph | \|Undefined Saprotroph\| | Yeast | NULL | Probable | As nectar-tap_saprotroph (Põlme et al. 2020); Decay sugar-rich substrates (Põlme et al. 2020) | Põlme S, et al. 2020. Fungal Diversity 105:1-16 ((https://doi.org/10.1007/s13225-020-00466-2)) |
| Otu09517 | 2 | k__Fungi(100);p__Ascomycota(100);c__Saccharomycetes(100);o__Saccharomycetales(100);f__Metschnikowiaceae(100);g__Metschnikowia(100);g__Metschnikowia_unclassified(100); | Metschnikowia | 13 | Saprotroph | \|Undefined Saprotroph\| | Yeast | NULL | Probable | As nectar-tap_saprotroph (Põlme et al. 2020); Decay sugar-rich substrates (Põlme et al. 2020) | Põlme S, et al. 2020. Fungal Diversity 105:1-16 ((https://doi.org/10.1007/s13225-020-00466-2)) |
| Otu09518 | 2 | k__Fungi(100);p__Ascomycota(100);c__Saccharomycetes(100);o__Saccharomycetales(100);f__Metschnikowiaceae(100);g__Metschnikowia(100);g__Metschnikowia_unclassified(100); | Metschnikowia | 13 | Saprotroph | \|Undefined Saprotroph\| | Yeast | NULL | Probable | As nectar-tap_saprotroph (Põlme et al. 2020); Decay sugar-rich substrates (Põlme et al. 2020) | Põlme S, et al. 2020. Fungal Diversity 105:1-16 ((https://doi.org/10.1007/s13225-020-00466-2)) |
| Otu09519 | 2 | k__Fungi(100);p__Ascomycota(100);c__Saccharomycetes(100);o__Saccharomycetales(100);f__Metschnikowiaceae(100);g__Metschnikowia(100);g__Metschnikowia_unclassified(100); | Metschnikowia | 13 | Saprotroph | \|Undefined Saprotroph\| | Yeast | NULL | Probable | As nectar-tap_saprotroph (Põlme et al. 2020); Decay sugar-rich substrates (Põlme et al. 2020) | Põlme S, et al. 2020. Fungal Diversity 105:1-16 ((https://doi.org/10.1007/s13225-020-00466-2)) |
| Otu09521 | 2 | k__Fungi(100);p__Ascomycota(100);c__Saccharomycetes(100);o__Saccharomycetales(100);f__Metschnikowiaceae(100);g__Metschnikowia(100);s__Metschnikowia_pulcherrima(100); | Metschnikowia | 13 | Saprotroph | \|Undefined Saprotroph\| | Yeast | NULL | Probable | As nectar-tap_saprotroph (Põlme et al. 2020); Decay sugar-rich substrates (Põlme et al. 2020) | Põlme S, et al. 2020. Fungal Diversity 105:1-16 ((https://doi.org/10.1007/s13225-020-00466-2)) |
| Otu09524 | 2 | k__Fungi(100);p__Ascomycota(100);c__Saccharomycetes(100);o__Saccharomycetales(100);f__Metschnikowiaceae(100);g__Metschnikowia(100);s__Metschnikowia_pulcherrima(100); | Metschnikowia | 13 | Saprotroph | \|Undefined Saprotroph\| | Yeast | NULL | Probable | As nectar-tap_saprotroph (Põlme et al. 2020); Decay sugar-rich substrates (Põlme et al. 2020) | Põlme S, et al. 2020. Fungal Diversity 105:1-16 ((https://doi.org/10.1007/s13225-020-00466-2)) |
| Otu09525 | 2 | k__Fungi(100);p__Ascomycota(100);c__Saccharomycetes(100);o__Saccharomycetales(100);f__Metschnikowiaceae(100);g__Metschnikowia(100);g__Metschnikowia_unclassified(100); | Metschnikowia | 13 | Saprotroph | \|Undefined Saprotroph\| | Yeast | NULL | Probable | As nectar-tap_saprotroph (Põlme et al. 2020); Decay sugar-rich substrates (Põlme et al. 2020) | Põlme S, et al. 2020. Fungal Diversity 105:1-16 ((https://doi.org/10.1007/s13225-020-00466-2)) |
| Otu09526 | 2 | k__Fungi(100);p__Ascomycota(100);c__Saccharomycetes(100);o__Saccharomycetales(100);f__Metschnikowiaceae(100);g__Metschnikowia(100);g__Metschnikowia_unclassified(100); | Metschnikowia | 13 | Saprotroph | \|Undefined Saprotroph\| | Yeast | NULL | Probable | As nectar-tap_saprotroph (Põlme et al. 2020); Decay sugar-rich substrates (Põlme et al. 2020) | Põlme S, et al. 2020. Fungal Diversity 105:1-16 ((https://doi.org/10.1007/s13225-020-00466-2)) |
| Otu09527 | 2 | k__Fungi(100);p__Ascomycota(100);c__Saccharomycetes(100);o__Saccharomycetales(100);f__Metschnikowiaceae(100);g__Metschnikowia(100);g__Metschnikowia_unclassified(100); | Metschnikowia | 13 | Saprotroph | \|Undefined Saprotroph\| | Yeast | NULL | Probable | As nectar-tap_saprotroph (Põlme et al. 2020); Decay sugar-rich substrates (Põlme et al. 2020) | Põlme S, et al. 2020. Fungal Diversity 105:1-16 ((https://doi.org/10.1007/s13225-020-00466-2)) |
| Otu09529 | 2 | k__Fungi(100);p__Ascomycota(100);c__Saccharomycetes(100);o__Saccharomycetales(100);f__Metschnikowiaceae(100);g__Metschnikowia(100);s__Metschnikowia_pulcherrima(100); | Metschnikowia | 13 | Saprotroph | \|Undefined Saprotroph\| | Yeast | NULL | Probable | As nectar-tap_saprotroph (Põlme et al. 2020); Decay sugar-rich substrates (Põlme et al. 2020) | Põlme S, et al. 2020. Fungal Diversity 105:1-16 ((https://doi.org/10.1007/s13225-020-00466-2)) |
| Otu09530 | 2 | k__Fungi(100);p__Ascomycota(100);c__Saccharomycetes(100);o__Saccharomycetales(100);f__Metschnikowiaceae(100);g__Metschnikowia(100);s__Metschnikowia_pulcherrima(100); | Metschnikowia | 13 | Saprotroph | \|Undefined Saprotroph\| | Yeast | NULL | Probable | As nectar-tap_saprotroph (Põlme et al. 2020); Decay sugar-rich substrates (Põlme et al. 2020) | Põlme S, et al. 2020. Fungal Diversity 105:1-16 ((https://doi.org/10.1007/s13225-020-00466-2)) |
| Otu09536 | 2 | k__Fungi(100);p__Ascomycota(100);c__Saccharomycetes(100);o__Saccharomycetales(100);f__Metschnikowiaceae(100);g__Metschnikowia(100);s__Metschnikowia_pulcherrima(100); | Metschnikowia | 13 | Saprotroph | \|Undefined Saprotroph\| | Yeast | NULL | Probable | As nectar-tap_saprotroph (Põlme et al. 2020); Decay sugar-rich substrates (Põlme et al. 2020) | Põlme S, et al. 2020. Fungal Diversity 105:1-16 ((https://doi.org/10.1007/s13225-020-00466-2)) |
| Otu09541 | 2 | k__Fungi(100);p__Ascomycota(100);c__Saccharomycetes(100);o__Saccharomycetales(100);f__Metschnikowiaceae(100);g__Metschnikowia(100);g__Metschnikowia_unclassified(100); | Metschnikowia | 13 | Saprotroph | \|Undefined Saprotroph\| | Yeast | NULL | Probable | As nectar-tap_saprotroph (Põlme et al. 2020); Decay sugar-rich substrates (Põlme et al. 2020) | Põlme S, et al. 2020. Fungal Diversity 105:1-16 ((https://doi.org/10.1007/s13225-020-00466-2)) |
| Otu09543 | 2 | k__Fungi(100);p__Ascomycota(100);c__Saccharomycetes(100);o__Saccharomycetales(100);f__Saccharomycetales_family_Incertae_sedis(100);g__Candida(100);g__Candida_unclassified(100); | Candida | 13 | Pathotroph-Saprotroph-Symbiotroph | Animal Pathogen-Endophyte-Endosymbiont-Epiphyte-\|Undefined Saprotroph\| | Dimorphic Yeast-Yeast | NULL | Probable | As nectar-tap_saprotroph (Põlme et al. 2020); Decay sugar-rich substrates (Põlme et al. 2020); This asexual genus is polyphyletic and contain species that function very differently depend on what they are. Caution should be taken when interpreting guilds for Candida species. | Manolakaki D, et al. 2010. Virulence. 1: 367-375 ((https://doi.org/10.4161/viru.1.5.12796)); Mekbib SB, et al. 2011. Tropical Plant Pathology 36:233-240 ((https://doi.org/10.1590/S1982-56762011000400004)); Põlme S, et al. 2020. Fungal Diversity 105:1-16 ((https://doi.org/10.1007/s13225-020-00466-2)) |
| Otu09546 | 2 | k__Fungi(100);p__Ascomycota(100);c__Leotiomycetes(100);o__Helotiales(100);f__Helotiaceae(100);g__Varicosporium(100);s__Varicosporium_delicatum(100); | Varicosporium | 13 | Saprotroph | \|Plant Saprotroph\|-Undefined Saprotroph | Microfungus | NULL | Probable | As litter_saprotroph (Põlme et al. 2020); Decay leaf, fruit, seed (Põlme et al. 2020) | Tedersoo L, et al. 2014. Science 346:e1256688 ((https://doi.org/10.1126/science.1256688)); Põlme S, et al. 2020. Fungal Diversity 105:1-16 ((https://doi.org/10.1007/s13225-020-00466-2)) |
| Otu09548 | 2 | k__Fungi(100);p__Ascomycota(100);c__Saccharomycetes(100);o__Saccharomycetales(100);f__Metschnikowiaceae(100);g__Metschnikowia(100);s__Metschnikowia_pulcherrima(100); | Metschnikowia | 13 | Saprotroph | \|Undefined Saprotroph\| | Yeast | NULL | Probable | As nectar-tap_saprotroph (Põlme et al. 2020); Decay sugar-rich substrates (Põlme et al. 2020) | Põlme S, et al. 2020. Fungal Diversity 105:1-16 ((https://doi.org/10.1007/s13225-020-00466-2)) |
| Otu09549 | 2 | k__Fungi(100);p__Ascomycota(100);c__Saccharomycetes(100);o__Saccharomycetales(100);f__Metschnikowiaceae(100);g__Metschnikowia(100);s__Metschnikowia_pulcherrima(100); | Metschnikowia | 13 | Saprotroph | \|Undefined Saprotroph\| | Yeast | NULL | Probable | As nectar-tap_saprotroph (Põlme et al. 2020); Decay sugar-rich substrates (Põlme et al. 2020) | Põlme S, et al. 2020. Fungal Diversity 105:1-16 ((https://doi.org/10.1007/s13225-020-00466-2)) |
| Otu09551 | 2 | k__Fungi(100);p__Ascomycota(100);c__Saccharomycetes(100);o__Saccharomycetales(100);f__Metschnikowiaceae(100);g__Metschnikowia(100);g__Metschnikowia_unclassified(100); | Metschnikowia | 13 | Saprotroph | \|Undefined Saprotroph\| | Yeast | NULL | Probable | As nectar-tap_saprotroph (Põlme et al. 2020); Decay sugar-rich substrates (Põlme et al. 2020) | Põlme S, et al. 2020. Fungal Diversity 105:1-16 ((https://doi.org/10.1007/s13225-020-00466-2)) |
| Otu09552 | 2 | k__Fungi(100);p__Ascomycota(100);c__Saccharomycetes(100);o__Saccharomycetales(100);f__Metschnikowiaceae(100);g__Metschnikowia(100);s__Metschnikowia_pulcherrima(100); | Metschnikowia | 13 | Saprotroph | \|Undefined Saprotroph\| | Yeast | NULL | Probable | As nectar-tap_saprotroph (Põlme et al. 2020); Decay sugar-rich substrates (Põlme et al. 2020) | Põlme S, et al. 2020. Fungal Diversity 105:1-16 ((https://doi.org/10.1007/s13225-020-00466-2)) |
| Otu09555 | 2 | k__Fungi(100);p__Ascomycota(100);c__Saccharomycetes(100);o__Saccharomycetales(100);f__Metschnikowiaceae(100);g__Metschnikowia(100);g__Metschnikowia_unclassified(100); | Metschnikowia | 13 | Saprotroph | \|Undefined Saprotroph\| | Yeast | NULL | Probable | As nectar-tap_saprotroph (Põlme et al. 2020); Decay sugar-rich substrates (Põlme et al. 2020) | Põlme S, et al. 2020. Fungal Diversity 105:1-16 ((https://doi.org/10.1007/s13225-020-00466-2)) |
| Otu09556 | 2 | k__Fungi(100);p__Ascomycota(100);c__Saccharomycetes(100);o__Saccharomycetales(100);f__Metschnikowiaceae(100);g__Metschnikowia(100);s__Metschnikowia_pulcherrima(100); | Metschnikowia | 13 | Saprotroph | \|Undefined Saprotroph\| | Yeast | NULL | Probable | As nectar-tap_saprotroph (Põlme et al. 2020); Decay sugar-rich substrates (Põlme et al. 2020) | Põlme S, et al. 2020. Fungal Diversity 105:1-16 ((https://doi.org/10.1007/s13225-020-00466-2)) |
| Otu09557 | 2 | k__Fungi(100);p__Ascomycota(100);c__Saccharomycetes(100);o__Saccharomycetales(100);f__Metschnikowiaceae(100);g__Metschnikowia(100);s__Metschnikowia_pulcherrima(100); | Metschnikowia | 13 | Saprotroph | \|Undefined Saprotroph\| | Yeast | NULL | Probable | As nectar-tap_saprotroph (Põlme et al. 2020); Decay sugar-rich substrates (Põlme et al. 2020) | Põlme S, et al. 2020. Fungal Diversity 105:1-16 ((https://doi.org/10.1007/s13225-020-00466-2)) |
| Otu09558 | 2 | k__Fungi(100);p__Ascomycota(100);c__Dothideomycetes(100);o__Pleosporales(100);f__Phaeosphaeriaceae(100);f__Phaeosphaeriaceae_unclassified(100);f__Phaeosphaeriaceae_unclassified(100); | Phaeosphaeriaceae | 9 | Pathotroph-Saprotroph | Fungal Parasite-Plant Pathogen-Plant Saprotroph | Microfungus | NULL | Probable | Necrotrophic or saprobic on a wide range of plants, especially monocotyledons; a few species parasitic on other fungi (Cannon & Kirk 2007) | Cannon PF, Kirk PM. 2007. Fungal Families of the World. CAB International, Cambridge (ISBN: 978-0851998275) |
| Otu09559 | 2 | k__Fungi(100);p__Ascomycota(100);c__Saccharomycetes(100);o__Saccharomycetales(100);f__Metschnikowiaceae(100);g__Metschnikowia(100);s__Metschnikowia_pulcherrima(100); | Metschnikowia | 13 | Saprotroph | \|Undefined Saprotroph\| | Yeast | NULL | Probable | As nectar-tap_saprotroph (Põlme et al. 2020); Decay sugar-rich substrates (Põlme et al. 2020) | Põlme S, et al. 2020. Fungal Diversity 105:1-16 ((https://doi.org/10.1007/s13225-020-00466-2)) |
| Otu09561 | 2 | k__Fungi(100);p__Ascomycota(100);c__Saccharomycetes(100);o__Saccharomycetales(100);f__Metschnikowiaceae(100);g__Metschnikowia(100);s__Metschnikowia_pulcherrima(100); | Metschnikowia | 13 | Saprotroph | \|Undefined Saprotroph\| | Yeast | NULL | Probable | As nectar-tap_saprotroph (Põlme et al. 2020); Decay sugar-rich substrates (Põlme et al. 2020) | Põlme S, et al. 2020. Fungal Diversity 105:1-16 ((https://doi.org/10.1007/s13225-020-00466-2)) |
| Otu09564 | 2 | k__Fungi(100);p__Ascomycota(100);c__Saccharomycetes(100);o__Saccharomycetales(100);f__Metschnikowiaceae(100);g__Metschnikowia(100);g__Metschnikowia_unclassified(100); | Metschnikowia | 13 | Saprotroph | \|Undefined Saprotroph\| | Yeast | NULL | Probable | As nectar-tap_saprotroph (Põlme et al. 2020); Decay sugar-rich substrates (Põlme et al. 2020) | Põlme S, et al. 2020. Fungal Diversity 105:1-16 ((https://doi.org/10.1007/s13225-020-00466-2)) |
| Otu09568 | 2 | k__Fungi(100);p__Ascomycota(100);c__Saccharomycetes(100);o__Saccharomycetales(100);f__Metschnikowiaceae(100);g__Metschnikowia(100);s__Metschnikowia_pulcherrima(100); | Metschnikowia | 13 | Saprotroph | \|Undefined Saprotroph\| | Yeast | NULL | Probable | As nectar-tap_saprotroph (Põlme et al. 2020); Decay sugar-rich substrates (Põlme et al. 2020) | Põlme S, et al. 2020. Fungal Diversity 105:1-16 ((https://doi.org/10.1007/s13225-020-00466-2)) |
| Otu09571 | 2 | k__Fungi(100);p__Ascomycota(100);c__Saccharomycetes(100);o__Saccharomycetales(100);f__Saccharomycetales_family_Incertae_sedis(100);g__Candida(100);g__Candida_unclassified(100); | Candida | 13 | Pathotroph-Saprotroph-Symbiotroph | Animal Pathogen-Endophyte-Endosymbiont-Epiphyte-\|Undefined Saprotroph\| | Dimorphic Yeast-Yeast | NULL | Probable | As nectar-tap_saprotroph (Põlme et al. 2020); Decay sugar-rich substrates (Põlme et al. 2020); This asexual genus is polyphyletic and contain species that function very differently depend on what they are. Caution should be taken when interpreting guilds for Candida species. | Manolakaki D, et al. 2010. Virulence. 1: 367-375 ((https://doi.org/10.4161/viru.1.5.12796)); Mekbib SB, et al. 2011. Tropical Plant Pathology 36:233-240 ((https://doi.org/10.1590/S1982-56762011000400004)); Põlme S, et al. 2020. Fungal Diversity 105:1-16 ((https://doi.org/10.1007/s13225-020-00466-2)) |
| Otu09572 | 2 | k__Fungi(100);p__Ascomycota(100);c__Saccharomycetes(100);o__Saccharomycetales(100);f__Metschnikowiaceae(100);g__Metschnikowia(100);s__Metschnikowia_pulcherrima(100); | Metschnikowia | 13 | Saprotroph | \|Undefined Saprotroph\| | Yeast | NULL | Probable | As nectar-tap_saprotroph (Põlme et al. 2020); Decay sugar-rich substrates (Põlme et al. 2020) | Põlme S, et al. 2020. Fungal Diversity 105:1-16 ((https://doi.org/10.1007/s13225-020-00466-2)) |
| Otu09573 | 2 | k__Fungi(100);p__Ascomycota(100);c__Saccharomycetes(100);o__Saccharomycetales(100);f__Metschnikowiaceae(100);g__Metschnikowia(100);g__Metschnikowia_unclassified(100); | Metschnikowia | 13 | Saprotroph | \|Undefined Saprotroph\| | Yeast | NULL | Probable | As nectar-tap_saprotroph (Põlme et al. 2020); Decay sugar-rich substrates (Põlme et al. 2020) | Põlme S, et al. 2020. Fungal Diversity 105:1-16 ((https://doi.org/10.1007/s13225-020-00466-2)) |
| Otu09574 | 2 | k__Fungi(100);p__Ascomycota(100);c__Saccharomycetes(100);o__Saccharomycetales(100);f__Metschnikowiaceae(100);g__Metschnikowia(100);s__Metschnikowia_pulcherrima(100); | Metschnikowia | 13 | Saprotroph | \|Undefined Saprotroph\| | Yeast | NULL | Probable | As nectar-tap_saprotroph (Põlme et al. 2020); Decay sugar-rich substrates (Põlme et al. 2020) | Põlme S, et al. 2020. Fungal Diversity 105:1-16 ((https://doi.org/10.1007/s13225-020-00466-2)) |
| Otu09575 | 2 | k__Fungi(100);p__Ascomycota(100);c__Saccharomycetes(100);o__Saccharomycetales(100);f__Metschnikowiaceae(100);g__Metschnikowia(100);s__Metschnikowia_pulcherrima(100); | Metschnikowia | 13 | Saprotroph | \|Undefined Saprotroph\| | Yeast | NULL | Probable | As nectar-tap_saprotroph (Põlme et al. 2020); Decay sugar-rich substrates (Põlme et al. 2020) | Põlme S, et al. 2020. Fungal Diversity 105:1-16 ((https://doi.org/10.1007/s13225-020-00466-2)) |
| Otu09576 | 2 | k__Fungi(100);p__Ascomycota(100);c__Saccharomycetes(100);o__Saccharomycetales(100);f__Metschnikowiaceae(100);g__Metschnikowia(100);g__Metschnikowia_unclassified(100); | Metschnikowia | 13 | Saprotroph | \|Undefined Saprotroph\| | Yeast | NULL | Probable | As nectar-tap_saprotroph (Põlme et al. 2020); Decay sugar-rich substrates (Põlme et al. 2020) | Põlme S, et al. 2020. Fungal Diversity 105:1-16 ((https://doi.org/10.1007/s13225-020-00466-2)) |
| Otu09577 | 2 | k__Fungi(100);p__Ascomycota(100);c__Saccharomycetes(100);o__Saccharomycetales(100);f__Metschnikowiaceae(100);g__Metschnikowia(100);s__Metschnikowia_pulcherrima(100); | Metschnikowia | 13 | Saprotroph | \|Undefined Saprotroph\| | Yeast | NULL | Probable | As nectar-tap_saprotroph (Põlme et al. 2020); Decay sugar-rich substrates (Põlme et al. 2020) | Põlme S, et al. 2020. Fungal Diversity 105:1-16 ((https://doi.org/10.1007/s13225-020-00466-2)) |
| Otu09579 | 2 | k__Fungi(100);p__Ascomycota(100);c__Saccharomycetes(100);o__Saccharomycetales(100);f__Metschnikowiaceae(100);g__Metschnikowia(100);s__Metschnikowia_pulcherrima(100); | Metschnikowia | 13 | Saprotroph | \|Undefined Saprotroph\| | Yeast | NULL | Probable | As nectar-tap_saprotroph (Põlme et al. 2020); Decay sugar-rich substrates (Põlme et al. 2020) | Põlme S, et al. 2020. Fungal Diversity 105:1-16 ((https://doi.org/10.1007/s13225-020-00466-2)) |
| Otu09582 | 2 | k__Fungi(100);p__Ascomycota(100);c__Saccharomycetes(100);o__Saccharomycetales(100);f__Metschnikowiaceae(100);g__Metschnikowia(100);s__Metschnikowia_pulcherrima(100); | Metschnikowia | 13 | Saprotroph | \|Undefined Saprotroph\| | Yeast | NULL | Probable | As nectar-tap_saprotroph (Põlme et al. 2020); Decay sugar-rich substrates (Põlme et al. 2020) | Põlme S, et al. 2020. Fungal Diversity 105:1-16 ((https://doi.org/10.1007/s13225-020-00466-2)) |
| Otu09584 | 2 | k__Fungi(100);p__Ascomycota(100);c__Saccharomycetes(100);o__Saccharomycetales(100);f__Metschnikowiaceae(100);g__Metschnikowia(100);g__Metschnikowia_unclassified(100); | Metschnikowia | 13 | Saprotroph | \|Undefined Saprotroph\| | Yeast | NULL | Probable | As nectar-tap_saprotroph (Põlme et al. 2020); Decay sugar-rich substrates (Põlme et al. 2020) | Põlme S, et al. 2020. Fungal Diversity 105:1-16 ((https://doi.org/10.1007/s13225-020-00466-2)) |
| Otu09585 | 2 | k__Fungi(100);p__Ascomycota(100);c__Saccharomycetes(100);o__Saccharomycetales(100);f__Metschnikowiaceae(100);g__Metschnikowia(100);s__Metschnikowia_pulcherrima(100); | Metschnikowia | 13 | Saprotroph | \|Undefined Saprotroph\| | Yeast | NULL | Probable | As nectar-tap_saprotroph (Põlme et al. 2020); Decay sugar-rich substrates (Põlme et al. 2020) | Põlme S, et al. 2020. Fungal Diversity 105:1-16 ((https://doi.org/10.1007/s13225-020-00466-2)) |
| Otu09586 | 2 | k__Fungi(100);p__Ascomycota(100);c__Sordariomycetes(100);o__Xylariales(100);f__Diatrypaceae(100);g__Libertella(100);s__Libertella_sp(100); | Libertella | 13 | Pathotroph | \|Plant Pathogen\| | Microfungus | NULL | Probable | NULL | Tedersoo L, et al. 2014. Science 346:e1256688 ((https://doi.org/10.1126/science.1256688)); Põlme S, et al. 2020. Fungal Diversity 105:1-16 ((https://doi.org/10.1007/s13225-020-00466-2)) |
| Otu09587 | 2 | k__Fungi(100);p__Ascomycota(100);c__Saccharomycetes(100);o__Saccharomycetales(100);f__Metschnikowiaceae(100);g__Metschnikowia(100);g__Metschnikowia_unclassified(100); | Metschnikowia | 13 | Saprotroph | \|Undefined Saprotroph\| | Yeast | NULL | Probable | As nectar-tap_saprotroph (Põlme et al. 2020); Decay sugar-rich substrates (Põlme et al. 2020) | Põlme S, et al. 2020. Fungal Diversity 105:1-16 ((https://doi.org/10.1007/s13225-020-00466-2)) |
| Otu09588 | 2 | k__Fungi(100);p__Ascomycota(100);c__Saccharomycetes(100);o__Saccharomycetales(100);f__Metschnikowiaceae(100);g__Metschnikowia(100);s__Metschnikowia_pulcherrima(100); | Metschnikowia | 13 | Saprotroph | \|Undefined Saprotroph\| | Yeast | NULL | Probable | As nectar-tap_saprotroph (Põlme et al. 2020); Decay sugar-rich substrates (Põlme et al. 2020) | Põlme S, et al. 2020. Fungal Diversity 105:1-16 ((https://doi.org/10.1007/s13225-020-00466-2)) |
| Otu09589 | 2 | k__Fungi(100);p__Ascomycota(100);c__Saccharomycetes(100);o__Saccharomycetales(100);f__Metschnikowiaceae(100);g__Metschnikowia(100);s__Metschnikowia_pulcherrima(100); | Metschnikowia | 13 | Saprotroph | \|Undefined Saprotroph\| | Yeast | NULL | Probable | As nectar-tap_saprotroph (Põlme et al. 2020); Decay sugar-rich substrates (Põlme et al. 2020) | Põlme S, et al. 2020. Fungal Diversity 105:1-16 ((https://doi.org/10.1007/s13225-020-00466-2)) |
| Otu09592 | 2 | k__Fungi(100);p__Ascomycota(100);c__Saccharomycetes(100);o__Saccharomycetales(100);f__Saccharomycetales_family_Incertae_sedis(100);g__Candida(100);g__Candida_unclassified(100); | Candida | 13 | Pathotroph-Saprotroph-Symbiotroph | Animal Pathogen-Endophyte-Endosymbiont-Epiphyte-\|Undefined Saprotroph\| | Dimorphic Yeast-Yeast | NULL | Probable | As nectar-tap_saprotroph (Põlme et al. 2020); Decay sugar-rich substrates (Põlme et al. 2020); This asexual genus is polyphyletic and contain species that function very differently depend on what they are. Caution should be taken when interpreting guilds for Candida species. | Manolakaki D, et al. 2010. Virulence. 1: 367-375 ((https://doi.org/10.4161/viru.1.5.12796)); Mekbib SB, et al. 2011. Tropical Plant Pathology 36:233-240 ((https://doi.org/10.1590/S1982-56762011000400004)); Põlme S, et al. 2020. Fungal Diversity 105:1-16 ((https://doi.org/10.1007/s13225-020-00466-2)) |
| Otu09593 | 2 | k__Fungi(100);p__Ascomycota(100);c__Saccharomycetes(100);o__Saccharomycetales(100);f__Metschnikowiaceae(100);g__Metschnikowia(100);g__Metschnikowia_unclassified(100); | Metschnikowia | 13 | Saprotroph | \|Undefined Saprotroph\| | Yeast | NULL | Probable | As nectar-tap_saprotroph (Põlme et al. 2020); Decay sugar-rich substrates (Põlme et al. 2020) | Põlme S, et al. 2020. Fungal Diversity 105:1-16 ((https://doi.org/10.1007/s13225-020-00466-2)) |
| Otu09594 | 2 | k__Fungi(100);p__Ascomycota(100);c__Saccharomycetes(100);o__Saccharomycetales(100);f__Metschnikowiaceae(100);g__Metschnikowia(100);s__Metschnikowia_pulcherrima(100); | Metschnikowia | 13 | Saprotroph | \|Undefined Saprotroph\| | Yeast | NULL | Probable | As nectar-tap_saprotroph (Põlme et al. 2020); Decay sugar-rich substrates (Põlme et al. 2020) | Põlme S, et al. 2020. Fungal Diversity 105:1-16 ((https://doi.org/10.1007/s13225-020-00466-2)) |
| Otu09595 | 2 | k__Fungi(100);p__Ascomycota(100);c__Dothideomycetes(100);o__Pleosporales(100);f__Lophiostomataceae(100);g__Lophiostoma(100);s__Lophiostoma_sp_CBS_109932(100); | Lophiostoma | 13 | Saprotroph | Plant Saprotroph-Undefined Saprotroph-\|Wood Saprotroph\| | Microfungus | NULL | Probable | As litter_saprotroph (Põlme et al. 2020); Decay wood, leaf, fruit, seed (Põlme et al. 2020) | Tedersoo L, et al. 2014. Science 346:e1256688 ((https://doi.org/10.1126/science.1256688)); Põlme S, et al. 2020. Fungal Diversity 105:1-16 ((https://doi.org/10.1007/s13225-020-00466-2)) |
| Otu09596 | 2 | k__Fungi(100);p__Ascomycota(100);c__Saccharomycetes(100);o__Saccharomycetales(100);f__Metschnikowiaceae(100);g__Metschnikowia(100);g__Metschnikowia_unclassified(100); | Metschnikowia | 13 | Saprotroph | \|Undefined Saprotroph\| | Yeast | NULL | Probable | As nectar-tap_saprotroph (Põlme et al. 2020); Decay sugar-rich substrates (Põlme et al. 2020) | Põlme S, et al. 2020. Fungal Diversity 105:1-16 ((https://doi.org/10.1007/s13225-020-00466-2)) |
| Otu09600 | 2 | k__Fungi(100);p__Ascomycota(100);c__Saccharomycetes(100);o__Saccharomycetales(100);f__Metschnikowiaceae(100);g__Metschnikowia(100);g__Metschnikowia_unclassified(100); | Metschnikowia | 13 | Saprotroph | \|Undefined Saprotroph\| | Yeast | NULL | Probable | As nectar-tap_saprotroph (Põlme et al. 2020); Decay sugar-rich substrates (Põlme et al. 2020) | Põlme S, et al. 2020. Fungal Diversity 105:1-16 ((https://doi.org/10.1007/s13225-020-00466-2)) |
| Otu09602 | 2 | k__Fungi(100);p__Ascomycota(100);c__Saccharomycetes(100);o__Saccharomycetales(100);f__Metschnikowiaceae(100);g__Metschnikowia(100);s__Metschnikowia_pulcherrima(100); | Metschnikowia | 13 | Saprotroph | \|Undefined Saprotroph\| | Yeast | NULL | Probable | As nectar-tap_saprotroph (Põlme et al. 2020); Decay sugar-rich substrates (Põlme et al. 2020) | Põlme S, et al. 2020. Fungal Diversity 105:1-16 ((https://doi.org/10.1007/s13225-020-00466-2)) |
| Otu09605 | 2 | k__Fungi(100);p__Ascomycota(100);c__Saccharomycetes(100);o__Saccharomycetales(100);f__Metschnikowiaceae(100);g__Metschnikowia(100);g__Metschnikowia_unclassified(100); | Metschnikowia | 13 | Saprotroph | \|Undefined Saprotroph\| | Yeast | NULL | Probable | As nectar-tap_saprotroph (Põlme et al. 2020); Decay sugar-rich substrates (Põlme et al. 2020) | Põlme S, et al. 2020. Fungal Diversity 105:1-16 ((https://doi.org/10.1007/s13225-020-00466-2)) |
| Otu09608 | 2 | k__Fungi(100);p__Ascomycota(100);c__Saccharomycetes(100);o__Saccharomycetales(100);f__Metschnikowiaceae(100);g__Metschnikowia(100);s__Metschnikowia_pulcherrima(100); | Metschnikowia | 13 | Saprotroph | \|Undefined Saprotroph\| | Yeast | NULL | Probable | As nectar-tap_saprotroph (Põlme et al. 2020); Decay sugar-rich substrates (Põlme et al. 2020) | Põlme S, et al. 2020. Fungal Diversity 105:1-16 ((https://doi.org/10.1007/s13225-020-00466-2)) |
| Otu09609 | 2 | k__Fungi(100);p__Ascomycota(100);c__Saccharomycetes(100);o__Saccharomycetales(100);f__Saccharomycetales_family_Incertae_sedis(100);g__Candida(100);g__Candida_unclassified(100); | Candida | 13 | Pathotroph-Saprotroph-Symbiotroph | Animal Pathogen-Endophyte-Endosymbiont-Epiphyte-\|Undefined Saprotroph\| | Dimorphic Yeast-Yeast | NULL | Probable | As nectar-tap_saprotroph (Põlme et al. 2020); Decay sugar-rich substrates (Põlme et al. 2020); This asexual genus is polyphyletic and contain species that function very differently depend on what they are. Caution should be taken when interpreting guilds for Candida species. | Manolakaki D, et al. 2010. Virulence. 1: 367-375 ((https://doi.org/10.4161/viru.1.5.12796)); Mekbib SB, et al. 2011. Tropical Plant Pathology 36:233-240 ((https://doi.org/10.1590/S1982-56762011000400004)); Põlme S, et al. 2020. Fungal Diversity 105:1-16 ((https://doi.org/10.1007/s13225-020-00466-2)) |
| Otu09611 | 2 | k__Fungi(100);p__Ascomycota(100);c__Saccharomycetes(100);o__Saccharomycetales(100);f__Metschnikowiaceae(100);g__Metschnikowia(100);s__Metschnikowia_pulcherrima(100); | Metschnikowia | 13 | Saprotroph | \|Undefined Saprotroph\| | Yeast | NULL | Probable | As nectar-tap_saprotroph (Põlme et al. 2020); Decay sugar-rich substrates (Põlme et al. 2020) | Põlme S, et al. 2020. Fungal Diversity 105:1-16 ((https://doi.org/10.1007/s13225-020-00466-2)) |
| Otu09612 | 2 | k__Fungi(100);p__Ascomycota(100);c__Saccharomycetes(100);o__Saccharomycetales(100);f__Metschnikowiaceae(100);g__Metschnikowia(100);g__Metschnikowia_unclassified(100); | Metschnikowia | 13 | Saprotroph | \|Undefined Saprotroph\| | Yeast | NULL | Probable | As nectar-tap_saprotroph (Põlme et al. 2020); Decay sugar-rich substrates (Põlme et al. 2020) | Põlme S, et al. 2020. Fungal Diversity 105:1-16 ((https://doi.org/10.1007/s13225-020-00466-2)) |

| | | | | | | | | | | | |
|---|---|---|---|---|---|---|---|---|---|---|---|
| Otu09613 | 2 | k__Fungi(100);p__Ascomycota(100);c__Saccharomycetes(100);o__Saccharomycetales(100);f__Metschnikowiaceae(100);g__Metschnikowia(100);s__Metschnikowia_pulcherrima(100); | Metschnikowia | 13 | Saprotroph | \|Undefined Saprotroph\| | Yeast | NULL | Probable | As nectar-tap_saprotroph (Põlme et al. 2020); Decay sugar-rich substrates (Põlme et al. 2020) | Põlme S, et al. 2020. Fungal Diversity 105:1-16 ((https://doi.org/10.1007/s13225-020-00466-2)) |
| Otu09614 | 2 | k__Fungi(100);p__Ascomycota(100);c__Saccharomycetes(100);o__Saccharomycetales(100);f__Metschnikowiaceae(100);g__Metschnikowia(100);s__Metschnikowia_pulcherrima(100); | Metschnikowia | 13 | Saprotroph | \|Undefined Saprotroph\| | Yeast | NULL | Probable | As nectar-tap_saprotroph (Põlme et al. 2020); Decay sugar-rich substrates (Põlme et al. 2020) | Põlme S, et al. 2020. Fungal Diversity 105:1-16 ((https://doi.org/10.1007/s13225-020-00466-2)) |
| Otu09616 | 2 | k__Fungi(100);p__Ascomycota(100);c__Saccharomycetes(100);o__Saccharomycetales(100);f__Metschnikowiaceae(100);g__Metschnikowia(100);s__Metschnikowia_pulcherrima(100); | Metschnikowia | 13 | Saprotroph | \|Undefined Saprotroph\| | Yeast | NULL | Probable | As nectar-tap_saprotroph (Põlme et al. 2020); Decay sugar-rich substrates (Põlme et al. 2020) | Põlme S, et al. 2020. Fungal Diversity 105:1-16 ((https://doi.org/10.1007/s13225-020-00466-2)) |
| Otu09617 | 2 | k__Fungi(100);p__Ascomycota(100);c__Saccharomycetes(100);o__Saccharomycetales(100);f__Metschnikowiaceae(100);g__Metschnikowia(100);g__Metschnikowia_unclassified(100); | Metschnikowia | 13 | Saprotroph | \|Undefined Saprotroph\| | Yeast | NULL | Probable | As nectar-tap_saprotroph (Põlme et al. 2020); Decay sugar-rich substrates (Põlme et al. 2020) | Põlme S, et al. 2020. Fungal Diversity 105:1-16 ((https://doi.org/10.1007/s13225-020-00466-2)) |
| Otu09618 | 2 | k__Fungi(100);p__Ascomycota(100);c__Saccharomycetes(100);o__Saccharomycetales(100);f__Saccharomycetales_family_Incertae_sedis(100);g__Candida(100);g__Candida_unclassified(100); | Candida | 13 | Pathotroph-Saprotroph-Symbiotroph | Animal Pathogen-Endophyte-Endosymbiont-Epiphyte-\|Undefined Saprotroph\| | Dimorphic Yeast-Yeast | NULL | Probable | As nectar-tap_saprotroph (Põlme et al. 2020); Decay sugar-rich substrates (Põlme et al. 2020); This asexual genus is polyphyletic and contain species that function very differently depend on what they are. Caution should be taken when interpreting guilds for Candida species. | Manolakaki D, et al. 2010. Virulence. 1: 367-375 ((https://doi.org/10.4161/viru.1.5.12796)); Mekbib SB, et al. 2011. Tropical Plant Pathology 36:233-240 ((https://doi.org/10.1590/S1982-56762011000400004)); Põlme S, et al. 2020. Fungal Diversity 105:1-16 ((https://doi.org/10.1007/s13225-020-00466-2)) |
| Otu09619 | 2 | k__Fungi(100);p__Ascomycota(100);c__Saccharomycetes(100);o__Saccharomycetales(100);f__Metschnikowiaceae(100);g__Metschnikowia(100);g__Metschnikowia_unclassified(100); | Metschnikowia | 13 | Saprotroph | \|Undefined Saprotroph\| | Yeast | NULL | Probable | As nectar-tap_saprotroph (Põlme et al. 2020); Decay sugar-rich substrates (Põlme et al. 2020) | Põlme S, et al. 2020. Fungal Diversity 105:1-16 ((https://doi.org/10.1007/s13225-020-00466-2)) |
| Otu09621 | 2 | k__Fungi(100);p__Ascomycota(100);c__Saccharomycetes(100);o__Saccharomycetales(100);f__Metschnikowiaceae(100);g__Metschnikowia(100);s__Metschnikowia_pulcherrima(100); | Metschnikowia | 13 | Saprotroph | \|Undefined Saprotroph\| | Yeast | NULL | Probable | As nectar-tap_saprotroph (Põlme et al. 2020); Decay sugar-rich substrates (Põlme et al. 2020) | Põlme S, et al. 2020. Fungal Diversity 105:1-16 ((https://doi.org/10.1007/s13225-020-00466-2)) |
| Otu09622 | 2 | k__Fungi(100);p__Ascomycota(100);c__Saccharomycetes(100);o__Saccharomycetales(100);f__Metschnikowiaceae(100);g__Metschnikowia(100);s__Metschnikowia_pulcherrima(100); | Metschnikowia | 13 | Saprotroph | \|Undefined Saprotroph\| | Yeast | NULL | Probable | As nectar-tap_saprotroph (Põlme et al. 2020); Decay sugar-rich substrates (Põlme et al. 2020) | Põlme S, et al. 2020. Fungal Diversity 105:1-16 ((https://doi.org/10.1007/s13225-020-00466-2)) |
| Otu09624 | 2 | k__Fungi(100);p__Ascomycota(100);c__Saccharomycetes(100);o__Saccharomycetales(100);f__Metschnikowiaceae(100);g__Metschnikowia(100);s__Metschnikowia_pulcherrima(100); | Metschnikowia | 13 | Saprotroph | \|Undefined Saprotroph\| | Yeast | NULL | Probable | As nectar-tap_saprotroph (Põlme et al. 2020); Decay sugar-rich substrates (Põlme et al. 2020) | Põlme S, et al. 2020. Fungal Diversity 105:1-16 ((https://doi.org/10.1007/s13225-020-00466-2)) |
| Otu09625 | 2 | k__Fungi(100);p__Ascomycota(100);c__Saccharomycetes(100);o__Saccharomycetales(100);f__Metschnikowiaceae(100);g__Metschnikowia(100);s__Metschnikowia_pulcherrima(100); | Metschnikowia | 13 | Saprotroph | \|Undefined Saprotroph\| | Yeast | NULL | Probable | As nectar-tap_saprotroph (Põlme et al. 2020); Decay sugar-rich substrates (Põlme et al. 2020) | Põlme S, et al. 2020. Fungal Diversity 105:1-16 ((https://doi.org/10.1007/s13225-020-00466-2)) |
| Otu09629 | 2 | k__Fungi(100);p__Ascomycota(100);c__Saccharomycetes(100);o__Saccharomycetales(100);f__Metschnikowiaceae(100);g__Metschnikowia(100);s__Metschnikowia_pulcherrima(100); | Metschnikowia | 13 | Saprotroph | \|Undefined Saprotroph\| | Yeast | NULL | Probable | As nectar-tap_saprotroph (Põlme et al. 2020); Decay sugar-rich substrates (Põlme et al. 2020) | Põlme S, et al. 2020. Fungal Diversity 105:1-16 ((https://doi.org/10.1007/s13225-020-00466-2)) |
| Otu09633 | 2 | k__Fungi(100);p__Ascomycota(100);c__Saccharomycetes(100);o__Saccharomycetales(100);f__Metschnikowiaceae(100);g__Metschnikowia(100);s__Metschnikowia_pulcherrima(100); | Metschnikowia | 13 | Saprotroph | \|Undefined Saprotroph\| | Yeast | NULL | Probable | As nectar-tap_saprotroph (Põlme et al. 2020); Decay sugar-rich substrates (Põlme et al. 2020) | Põlme S, et al. 2020. Fungal Diversity 105:1-16 ((https://doi.org/10.1007/s13225-020-00466-2)) |
| Otu09634 | 2 | k__Fungi(100);p__Ascomycota(100);c__Saccharomycetes(100);o__Saccharomycetales(100);f__Metschnikowiaceae(100);g__Metschnikowia(100);s__Metschnikowia_pulcherrima(100); | Metschnikowia | 13 | Saprotroph | \|Undefined Saprotroph\| | Yeast | NULL | Probable | As nectar-tap_saprotroph (Põlme et al. 2020); Decay sugar-rich substrates (Põlme et al. 2020) | Põlme S, et al. 2020. Fungal Diversity 105:1-16 ((https://doi.org/10.1007/s13225-020-00466-2)) |
| Otu09638 | 2 | k__Fungi(100);p__Ascomycota(100);c__Saccharomycetes(100);o__Saccharomycetales(100);f__Metschnikowiaceae(100);g__Metschnikowia(100);s__Metschnikowia_pulcherrima(100); | Metschnikowia | 13 | Saprotroph | \|Undefined Saprotroph\| | Yeast | NULL | Probable | As nectar-tap_saprotroph (Põlme et al. 2020); Decay sugar-rich substrates (Põlme et al. 2020) | Põlme S, et al. 2020. Fungal Diversity 105:1-16 ((https://doi.org/10.1007/s13225-020-00466-2)) |
| Otu09640 | 2 | k__Fungi(100);p__Ascomycota(100);c__Saccharomycetes(100);o__Saccharomycetales(100);f__Metschnikowiaceae(100);g__Metschnikowia(100);g__Metschnikowia_unclassified(100); | Metschnikowia | 13 | Saprotroph | \|Undefined Saprotroph\| | Yeast | NULL | Probable | As nectar-tap_saprotroph (Põlme et al. 2020); Decay sugar-rich substrates (Põlme et al. 2020) | Põlme S, et al. 2020. Fungal Diversity 105:1-16 ((https://doi.org/10.1007/s13225-020-00466-2)) |
| Otu09641 | 2 | k__Fungi(100);p__Ascomycota(100);c__Saccharomycetes(100);o__Saccharomycetales(100);f__Metschnikowiaceae(100);g__Metschnikowia(100);g__Metschnikowia_unclassified(100); | Metschnikowia | 13 | Saprotroph | \|Undefined Saprotroph\| | Yeast | NULL | Probable | As nectar-tap_saprotroph (Põlme et al. 2020); Decay sugar-rich substrates (Põlme et al. 2020) | Põlme S, et al. 2020. Fungal Diversity 105:1-16 ((https://doi.org/10.1007/s13225-020-00466-2)) |
| Otu09642 | 2 | k__Fungi(100);p__Ascomycota(100);c__Saccharomycetes(100);o__Saccharomycetales(100);f__Metschnikowiaceae(100);g__Metschnikowia(100);s__Metschnikowia_pulcherrima(100); | Metschnikowia | 13 | Saprotroph | \|Undefined Saprotroph\| | Yeast | NULL | Probable | As nectar-tap_saprotroph (Põlme et al. 2020); Decay sugar-rich substrates (Põlme et al. 2020) | Põlme S, et al. 2020. Fungal Diversity 105:1-16 ((https://doi.org/10.1007/s13225-020-00466-2)) |
| Otu09643 | 2 | k__Fungi(100);p__Ascomycota(100);c__Dothideomycetes(100);o__Botryosphaeriales(100);f__Botryosphaeriaceae(100);g__Diplodia(100);g__Diplodia_unclassified(100); | Diplodia | 13 | Pathotroph-Saprotroph-Symbiotroph | Epiphyte-\|Plant Pathogen\|-Plant Saprotroph-Wood Saprotroph | Microfungus | NULL | Possible | As litter_saprotroph (Põlme et al. 2020); Cause of soft; culture (Carisse et al. 2000); Decay leaf, fruit, seed (Põlme et al. 2020); rot; Epiphyte detection method | Duncan CG, Eslyn WE. 1966. Mycologia 58:642-645 ((https://doi.org/10.2307/3757045)); Carisse O, et al. 2000.Phytopathology 90:31-37 ((https://doi.org/10.1094/PHYTO.2000.90.1.31)); Põlme S, et al. 2020. Fungal Diversity 105:1-16 ((https://doi.org/10.1007/s13225-020-00466-2)) |
| Otu09650 | 2 | k__Fungi(100);p__Ascomycota(100);c__Saccharomycetes(100);o__Saccharomycetales(100);f__Metschnikowiaceae(100);g__Metschnikowia(100);s__Metschnikowia_pulcherrima(100); | Metschnikowia | 13 | Saprotroph | \|Undefined Saprotroph\| | Yeast | NULL | Probable | As nectar-tap_saprotroph (Põlme et al. 2020); Decay sugar-rich substrates (Põlme et al. 2020) | Põlme S, et al. 2020. Fungal Diversity 105:1-16 ((https://doi.org/10.1007/s13225-020-00466-2)) |
| Otu09652 | 2 | k__Fungi(100);p__Ascomycota(100);c__Saccharomycetes(100);o__Saccharomycetales(100);f__Metschnikowiaceae(100);g__Metschnikowia(100);g__Metschnikowia_unclassified(100); | Metschnikowia | 13 | Saprotroph | \|Undefined Saprotroph\| | Yeast | NULL | Probable | As nectar-tap_saprotroph (Põlme et al. 2020); Decay sugar-rich substrates (Põlme et al. 2020) | Põlme S, et al. 2020. Fungal Diversity 105:1-16 ((https://doi.org/10.1007/s13225-020-00466-2)) |
| Otu09654 | 2 | k__Fungi(100);p__Ascomycota(100);c__Saccharomycetes(100);o__Saccharomycetales(100);f__Metschnikowiaceae(100);g__Metschnikowia(100);s__Metschnikowia_pulcherrima(100); | Metschnikowia | 13 | Saprotroph | \|Undefined Saprotroph\| | Yeast | NULL | Probable | As nectar-tap_saprotroph (Põlme et al. 2020); Decay sugar-rich substrates (Põlme et al. 2020) | Põlme S, et al. 2020. Fungal Diversity 105:1-16 ((https://doi.org/10.1007/s13225-020-00466-2)) |
| Otu09656 | 2 | k__Fungi(100);p__Ascomycota(100);c__Saccharomycetes(100);o__Saccharomycetales(100);f__Metschnikowiaceae(100);g__Metschnikowia(100);s__Metschnikowia_pulcherrima(100); | Metschnikowia | 13 | Saprotroph | \|Undefined Saprotroph\| | Yeast | NULL | Probable | As nectar-tap_saprotroph (Põlme et al. 2020); Decay sugar-rich substrates (Põlme et al. 2020) | Põlme S, et al. 2020. Fungal Diversity 105:1-16 ((https://doi.org/10.1007/s13225-020-00466-2)) |
| Otu09657 | 2 | k__Fungi(100);p__Ascomycota(100);c__Saccharomycetes(100);o__Saccharomycetales(100);f__Metschnikowiaceae(100);g__Metschnikowia(100);s__Metschnikowia_pulcherrima(100); | Metschnikowia | 13 | Saprotroph | \|Undefined Saprotroph\| | Yeast | NULL | Probable | As nectar-tap_saprotroph (Põlme et al. 2020); Decay sugar-rich substrates (Põlme et al. 2020) | Põlme S, et al. 2020. Fungal Diversity 105:1-16 ((https://doi.org/10.1007/s13225-020-00466-2)) |
| Otu09661 | 2 | k__Fungi(100);p__Ascomycota(100);c__Saccharomycetes(100);o__Saccharomycetales(100);f__Metschnikowiaceae(100);g__Metschnikowia(100);s__Metschnikowia_pulcherrima(100); | Metschnikowia | 13 | Saprotroph | \|Undefined Saprotroph\| | Yeast | NULL | Probable | As nectar-tap_saprotroph (Põlme et al. 2020); Decay sugar-rich substrates (Põlme et al. 2020) | Põlme S, et al. 2020. Fungal Diversity 105:1-16 ((https://doi.org/10.1007/s13225-020-00466-2)) |
| Otu09662 | 2 | k__Fungi(100);p__Ascomycota(100);c__Saccharomycetes(100);o__Saccharomycetales(100);f__Metschnikowiaceae(100);g__Metschnikowia(100);g__Metschnikowia_unclassified(100); | Metschnikowia | 13 | Saprotroph | \|Undefined Saprotroph\| | Yeast | NULL | Probable | As nectar-tap_saprotroph (Põlme et al. 2020); Decay sugar-rich substrates (Põlme et al. 2020) | Põlme S, et al. 2020. Fungal Diversity 105:1-16 ((https://doi.org/10.1007/s13225-020-00466-2)) |
| Otu09663 | 2 | k__Fungi(100);p__Ascomycota(100);c__Saccharomycetes(100);o__Saccharomycetales(100);f__Metschnikowiaceae(100);g__Metschnikowia(100);g__Metschnikowia_unclassified(100); | Metschnikowia | 13 | Saprotroph | \|Undefined Saprotroph\| | Yeast | NULL | Probable | As nectar-tap_saprotroph (Põlme et al. 2020); Decay sugar-rich substrates (Põlme et al. 2020) | Põlme S, et al. 2020. Fungal Diversity 105:1-16 ((https://doi.org/10.1007/s13225-020-00466-2)) |
| Otu09664 | 2 | k__Fungi(100);p__Ascomycota(100);c__Saccharomycetes(100);o__Saccharomycetales(100);f__Metschnikowiaceae(100);g__Metschnikowia(100);g__Metschnikowia_unclassified(100); | Metschnikowia | 13 | Saprotroph | \|Undefined Saprotroph\| | Yeast | NULL | Probable | As nectar-tap_saprotroph (Põlme et al. 2020); Decay sugar-rich substrates (Põlme et al. 2020) | Põlme S, et al. 2020. Fungal Diversity 105:1-16 ((https://doi.org/10.1007/s13225-020-00466-2)) |
| Otu09665 | 2 | k__Fungi(100);p__Ascomycota(100);c__Saccharomycetes(100);o__Saccharomycetales(100);f__Metschnikowiaceae(100);g__Metschnikowia(100);g__Metschnikowia_unclassified(100); | Metschnikowia | 13 | Saprotroph | \|Undefined Saprotroph\| | Yeast | NULL | Probable | As nectar-tap_saprotroph (Põlme et al. 2020); Decay sugar-rich substrates (Põlme et al. 2020) | Põlme S, et al. 2020. Fungal Diversity 105:1-16 ((https://doi.org/10.1007/s13225-020-00466-2)) |
| Otu09673 | 2 | k__Fungi(100);p__Ascomycota(100);c__Saccharomycetes(100);o__Saccharomycetales(100);f__Metschnikowiaceae(100);g__Metschnikowia(100);s__Metschnikowia_pulcherrima(100); | Metschnikowia | 13 | Saprotroph | \|Undefined Saprotroph\| | Yeast | NULL | Probable | As nectar-tap_saprotroph (Põlme et al. 2020); Decay sugar-rich substrates (Põlme et al. 2020) | Põlme S, et al. 2020. Fungal Diversity 105:1-16 ((https://doi.org/10.1007/s13225-020-00466-2)) |
| Otu09675 | 2 | k__Fungi(100);p__Ascomycota(100);c__Saccharomycetes(100);o__Saccharomycetales(100);f__Metschnikowiaceae(100);g__Metschnikowia(100);g__Metschnikowia_unclassified(100); | Metschnikowia | 13 | Saprotroph | \|Undefined Saprotroph\| | Yeast | NULL | Probable | As nectar-tap_saprotroph (Põlme et al. 2020); Decay sugar-rich substrates (Põlme et al. 2020) | Põlme S, et al. 2020. Fungal Diversity 105:1-16 ((https://doi.org/10.1007/s13225-020-00466-2)) |
| Otu09676 | 2 | k__Fungi(100);p__Ascomycota(100);c__Saccharomycetes(100);o__Saccharomycetales(100);f__Metschnikowiaceae(100);g__Metschnikowia(100);g__Metschnikowia_unclassified(100); | Metschnikowia | 13 | Saprotroph | \|Undefined Saprotroph\| | Yeast | NULL | Probable | As nectar-tap_saprotroph (Põlme et al. 2020); Decay sugar-rich substrates (Põlme et al. 2020) | Põlme S, et al. 2020. Fungal Diversity 105:1-16 ((https://doi.org/10.1007/s13225-020-00466-2)) |
| Otu09678 | 2 | k__Fungi(100);p__Ascomycota(100);c__Saccharomycetes(100);o__Saccharomycetales(100);f__Metschnikowiaceae(100);g__Metschnikowia(100);s__Metschnikowia_pulcherrima(100); | Metschnikowia | 13 | Saprotroph | \|Undefined Saprotroph\| | Yeast | NULL | Probable | As nectar-tap_saprotroph (Põlme et al. 2020); Decay sugar-rich substrates (Põlme et al. 2020) | Põlme S, et al. 2020. Fungal Diversity 105:1-16 ((https://doi.org/10.1007/s13225-020-00466-2)) |
| Otu09680 | 2 | k__Fungi(100);p__Ascomycota(100);c__Saccharomycetes(100);o__Saccharomycetales(100);f__Metschnikowiaceae(100);g__Metschnikowia(100);g__Metschnikowia_unclassified(100); | Metschnikowia | 13 | Saprotroph | \|Undefined Saprotroph\| | Yeast | NULL | Probable | As nectar-tap_saprotroph (Põlme et al. 2020); Decay sugar-rich substrates (Põlme et al. 2020) | Põlme S, et al. 2020. Fungal Diversity 105:1-16 ((https://doi.org/10.1007/s13225-020-00466-2)) |
| Otu09681 | 2 | k__Fungi(100);p__Ascomycota(100);c__Saccharomycetes(100);o__Saccharomycetales(100);f__Metschnikowiaceae(100);g__Metschnikowia(100);s__Metschnikowia_pulcherrima(100); | Metschnikowia | 13 | Saprotroph | \|Undefined Saprotroph\| | Yeast | NULL | Probable | As nectar-tap_saprotroph (Põlme et al. 2020); Decay sugar-rich substrates (Põlme et al. 2020) | Põlme S, et al. 2020. Fungal Diversity 105:1-16 ((https://doi.org/10.1007/s13225-020-00466-2)) |
| Otu09682 | 2 | k__Fungi(100);p__Ascomycota(100);c__Saccharomycetes(100);o__Saccharomycetales(100);f__Metschnikowiaceae(100);g__Metschnikowia(100);s__Metschnikowia_pulcherrima(100); | Metschnikowia | 13 | Saprotroph | \|Undefined Saprotroph\| | Yeast | NULL | Probable | As nectar-tap_saprotroph (Põlme et al. 2020); Decay sugar-rich substrates (Põlme et al. 2020) | Põlme S, et al. 2020. Fungal Diversity 105:1-16 ((https://doi.org/10.1007/s13225-020-00466-2)) |
| Otu09683 | 2 | k__Fungi(100);p__Ascomycota(100);c__Saccharomycetes(100);o__Saccharomycetales(100);f__Metschnikowiaceae(100);g__Metschnikowia(100);g__Metschnikowia_unclassified(100); | Metschnikowia | 13 | Saprotroph | \|Undefined Saprotroph\| | Yeast | NULL | Probable | As nectar-tap_saprotroph (Põlme et al. 2020); Decay sugar-rich substrates (Põlme et al. 2020) | Põlme S, et al. 2020. Fungal Diversity 105:1-16 ((https://doi.org/10.1007/s13225-020-00466-2)) |
| Otu09684 | 2 | k__Fungi(100);p__Ascomycota(100);c__Dothideomycetes(100);o__Pleosporales(100);f__Venturiaceae(100);g__Venturia(100);g__Venturia_unclassified(100); | Venturia | 13 | Pathotroph-Pathotroph-Saprotroph | \|Plant Pathogen\|-Plant Saprotroph | Microfungus | NULL | Probable | As litter_saprotroph (Põlme et al. 2020); Decay leaf, fruit, seed (Põlme et al. 2020) | Tedersoo L, et al. 2014. Science 346:e1256688 ((https://doi.org/10.1126/science.1256688)); Põlme S, et al. 2020. Fungal Diversity 105:1-16 ((https://doi.org/10.1007/s13225-020-00466-2)) |
| Otu09685 | 2 | k__Fungi(100);p__Ascomycota(100);c__Saccharomycetes(100);o__Saccharomycetales(100);f__Metschnikowiaceae(100);g__Metschnikowia(100);s__Metschnikowia_pulcherrima(100); | Metschnikowia | 13 | Saprotroph | \|Undefined Saprotroph\| | Yeast | NULL | Probable | As nectar-tap_saprotroph (Põlme et al. 2020); Decay sugar-rich substrates (Põlme et al. 2020) | Põlme S, et al. 2020. Fungal Diversity 105:1-16 ((https://doi.org/10.1007/s13225-020-00466-2)) |
| Otu09686 | 2 | k__Fungi(100);p__Ascomycota(100);c__Saccharomycetes(100);o__Saccharomycetales(100);f__Metschnikowiaceae(100);g__Metschnikowia(100);s__Metschnikowia_pulcherrima(100); | Metschnikowia | 13 | Saprotroph | \|Undefined Saprotroph\| | Yeast | NULL | Probable | As nectar-tap_saprotroph (Põlme et al. 2020); Decay sugar-rich substrates (Põlme et al. 2020) | Põlme S, et al. 2020. Fungal Diversity 105:1-16 ((https://doi.org/10.1007/s13225-020-00466-2)) |
| Otu09687 | 2 | k__Fungi(100);p__Ascomycota(100);c__Saccharomycetes(100);o__Saccharomycetales(100);f__Metschnikowiaceae(100);g__Metschnikowia(100);g__Metschnikowia_unclassified(100); | Metschnikowia | 13 | Saprotroph | \|Undefined Saprotroph\| | Yeast | NULL | Probable | As nectar-tap_saprotroph (Põlme et al. 2020); Decay sugar-rich substrates (Põlme et al. 2020) | Põlme S, et al. 2020. Fungal Diversity 105:1-16 ((https://doi.org/10.1007/s13225-020-00466-2)) |
| Otu09688 | 2 | k__Fungi(100);p__Ascomycota(100);c__Saccharomycetes(100);o__Saccharomycetales(100);f__Metschnikowiaceae(100);g__Metschnikowia(100);g__Metschnikowia_unclassified(100); | Metschnikowia | 13 | Saprotroph | \|Undefined Saprotroph\| | Yeast | NULL | Probable | As nectar-tap_saprotroph (Põlme et al. 2020); Decay sugar-rich substrates (Põlme et al. 2020) | Põlme S, et al. 2020. Fungal Diversity 105:1-16 ((https://doi.org/10.1007/s13225-020-00466-2)) |
| Otu09690 | 2 | k__Fungi(100);p__Ascomycota(100);c__Saccharomycetes(100);o__Saccharomycetales(100);f__Metschnikowiaceae(100);g__Metschnikowia(100);s__Metschnikowia_pulcherrima(100); | Metschnikowia | 13 | Saprotroph | \|Undefined Saprotroph\| | Yeast | NULL | Probable | As nectar-tap_saprotroph (Põlme et al. 2020); Decay sugar-rich substrates (Põlme et al. 2020) | Põlme S, et al. 2020. Fungal Diversity 105:1-16 ((https://doi.org/10.1007/s13225-020-00466-2)) |
| Otu09692 | 2 | k__Fungi(100);p__Ascomycota(100);c__Saccharomycetes(100);o__Saccharomycetales(100);f__Metschnikowiaceae(100);g__Metschnikowia(100);g__Metschnikowia_unclassified(100); | Metschnikowia | 13 | Saprotroph | \|Undefined Saprotroph\| | Yeast | NULL | Probable | As nectar-tap_saprotroph (Põlme et al. 2020); Decay sugar-rich substrates (Põlme et al. 2020) | Põlme S, et al. 2020. Fungal Diversity 105:1-16 ((https://doi.org/10.1007/s13225-020-00466-2)) |
| Otu09697 | 2 | k__Fungi(100);p__Ascomycota(100);c__Saccharomycetes(100);o__Saccharomycetales(100);f__Metschnikowiaceae(100);g__Metschnikowia(100);g__Metschnikowia_unclassified(100); | Metschnikowia | 13 | Saprotroph | \|Undefined Saprotroph\| | Yeast | NULL | Probable | As nectar-tap_saprotroph (Põlme et al. 2020); Decay sugar-rich substrates (Põlme et al. 2020) | Põlme S, et al. 2020. Fungal Diversity 105:1-16 ((https://doi.org/10.1007/s13225-020-00466-2)) |
| Otu09699 | 2 | k__Fungi(100);p__Ascomycota(100);c__Saccharomycetes(100);o__Saccharomycetales(100);f__Metschnikowiaceae(100);g__Metschnikowia(100);g__Metschnikowia_unclassified(100); | Metschnikowia | 13 | Saprotroph | \|Undefined Saprotroph\| | Yeast | NULL | Probable | As nectar-tap_saprotroph (Põlme et al. 2020); Decay sugar-rich substrates (Põlme et al. 2020) | Põlme S, et al. 2020. Fungal Diversity 105:1-16 ((https://doi.org/10.1007/s13225-020-00466-2)) |
| Otu09700 | 2 | k__Fungi(100);p__Ascomycota(100);c__Saccharomycetes(100);o__Saccharomycetales(100);f__Metschnikowiaceae(100);g__Metschnikowia(100);g__Metschnikowia_unclassified(100); | Metschnikowia | 13 | Saprotroph | \|Undefined Saprotroph\| | Yeast | NULL | Probable | As nectar-tap_saprotroph (Põlme et al. 2020); Decay sugar-rich substrates (Põlme et al. 2020) | Põlme S, et al. 2020. Fungal Diversity 105:1-16 ((https://doi.org/10.1007/s13225-020-00466-2)) |
| Otu09702 | 2 | k__Fungi(100);p__Ascomycota(100);c__Saccharomycetes(100);o__Saccharomycetales(100);f__Metschnikowiaceae(100);g__Metschnikowia(100);g__Metschnikowia_unclassified(100); | Metschnikowia | 13 | Saprotroph | \|Undefined Saprotroph\| | Yeast | NULL | Probable | As nectar-tap_saprotroph (Põlme et al. 2020); Decay sugar-rich substrates (Põlme et al. 2020) | Põlme S, et al. 2020. Fungal Diversity 105:1-16 ((https://doi.org/10.1007/s13225-020-00466-2)) |
| Otu09703 | 2 | k__Fungi(100);p__Ascomycota(100);c__Saccharomycetes(100);o__Saccharomycetales(100);f__Metschnikowiaceae(100);g__Metschnikowia(100);g__Metschnikowia_unclassified(100); | Metschnikowia | 13 | Saprotroph | \|Undefined Saprotroph\| | Yeast | NULL | Probable | As nectar-tap_saprotroph (Põlme et al. 2020); Decay sugar-rich substrates (Põlme et al. 2020) | Põlme S, et al. 2020. Fungal Diversity 105:1-16 ((https://doi.org/10.1007/s13225-020-00466-2)) |
| Otu09708 | 2 | k__Fungi(100);p__Ascomycota(100);c__Saccharomycetes(100);o__Saccharomycetales(100);f__Metschnikowiaceae(100);g__Metschnikowia(100);g__Metschnikowia_unclassified(100); | Metschnikowia | 13 | Saprotroph | \|Undefined Saprotroph\| | Yeast | NULL | Probable | As nectar-tap_saprotroph (Põlme et al. 2020); Decay sugar-rich substrates (Põlme et al. 2020) | Põlme S, et al. 2020. Fungal Diversity 105:1-16 ((https://doi.org/10.1007/s13225-020-00466-2)) |
| Otu09710 | 2 | k__Fungi(100);p__Ascomycota(100);c__Saccharomycetes(100);o__Saccharomycetales(100);f__Metschnikowiaceae(100);g__Metschnikowia(100);s__Metschnikowia_pulcherrima(100); | Metschnikowia | 13 | Saprotroph | \|Undefined Saprotroph\| | Yeast | NULL | Probable | As nectar-tap_saprotroph (Põlme et al. 2020); Decay sugar-rich substrates (Põlme et al. 2020) | Põlme S, et al. 2020. Fungal Diversity 105:1-16 ((https://doi.org/10.1007/s13225-020-00466-2)) |
| Otu09711 | 2 | k__Fungi(100);p__Ascomycota(100);c__Saccharomycetes(100);o__Saccharomycetales(100);f__Metschnikowiaceae(100);g__Metschnikowia(100);s__Metschnikowia_pulcherrima(100); | Metschnikowia | 13 | Saprotroph | \|Undefined Saprotroph\| | Yeast | NULL | Probable | As nectar-tap_saprotroph (Põlme et al. 2020); Decay sugar-rich substrates (Põlme et al. 2020) | Põlme S, et al. 2020. Fungal Diversity 105:1-16 ((https://doi.org/10.1007/s13225-020-00466-2)) |
| Otu09713 | 2 | k__Fungi(100);p__Ascomycota(100);c__Saccharomycetes(100);o__Saccharomycetales(100);f__Metschnikowiaceae(100);g__Metschnikowia(100);s__Metschnikowia_pulcherrima(100); | Metschnikowia | 13 | Saprotroph | \|Undefined Saprotroph\| | Yeast | NULL | Probable | As nectar-tap_saprotroph (Põlme et al. 2020); Decay sugar-rich substrates (Põlme et al. 2020) | Põlme S, et al. 2020. Fungal Diversity 105:1-16 ((https://doi.org/10.1007/s13225-020-00466-2)) |
| Otu09714 | 2 | k__Fungi(100);p__Ascomycota(100);c__Saccharomycetes(100);o__Saccharomycetales(100);f__Metschnikowiaceae(100);g__Metschnikowia(100);s__Metschnikowia_pulcherrima(100); | Metschnikowia | 13 | Saprotroph | \|Undefined Saprotroph\| | Yeast | NULL | Probable | As nectar-tap_saprotroph (Põlme et al. 2020); Decay sugar-rich substrates (Põlme et al. 2020) | Põlme S, et al. 2020. Fungal Diversity 105:1-16 ((https://doi.org/10.1007/s13225-020-00466-2)) |
| Otu09716 | 2 | k__Fungi(100);p__Ascomycota(100);c__Saccharomycetes(100);o__Saccharomycetales(100);f__Metschnikowiaceae(100);g__Metschnikowia(100);s__Metschnikowia_pulcherrima(100); | Metschnikowia | 13 | Saprotroph | \|Undefined Saprotroph\| | Yeast | NULL | Probable | As nectar-tap_saprotroph (Põlme et al. 2020); Decay sugar-rich substrates (Põlme et al. 2020) | Põlme S, et al. 2020. Fungal Diversity 105:1-16 ((https://doi.org/10.1007/s13225-020-00466-2)) |
| Otu09717 | 2 | k__Fungi(100);p__Ascomycota(100);c__Saccharomycetes(100);o__Saccharomycetales(100);f__Metschnikowiaceae(100);g__Metschnikowia(100);s__Metschnikowia_pulcherrima(100); | Metschnikowia | 13 | Saprotroph | \|Undefined Saprotroph\| | Yeast | NULL | Probable | As nectar-tap_saprotroph (Põlme et al. 2020); Decay sugar-rich substrates (Põlme et al. 2020) | Põlme S, et al. 2020. Fungal Diversity 105:1-16 ((https://doi.org/10.1007/s13225-020-00466-2)) |
| Otu09718 | 2 | k__Fungi(100);p__Ascomycota(100);c__Saccharomycetes(100);o__Saccharomycetales(100);f__Metschnikowiaceae(100);g__Metschnikowia(100);g__Metschnikowia_unclassified(100); | Metschnikowia | 13 | Saprotroph | \|Undefined Saprotroph\| | Yeast | NULL | Probable | As nectar-tap_saprotroph (Põlme et al. 2020); Decay sugar-rich substrates (Põlme et al. 2020) | Põlme S, et al. 2020. Fungal Diversity 105:1-16 ((https://doi.org/10.1007/s13225-020-00466-2)) |
| Otu09719 | 2 | k__Fungi(100);p__Ascomycota(100);c__Saccharomycetes(100);o__Saccharomycetales(100);f__Metschnikowiaceae(100);g__Metschnikowia(100);s__Metschnikowia_pulcherrima(100); | Metschnikowia | 13 | Saprotroph | \|Undefined Saprotroph\| | Yeast | NULL | Probable | As nectar-tap_saprotroph (Põlme et al. 2020); Decay sugar-rich substrates (Põlme et al. 2020) | Põlme S, et al. 2020. Fungal Diversity 105:1-16 ((https://doi.org/10.1007/s13225-020-00466-2)) |
| Otu09720 | 2 | k__Fungi(100);p__Ascomycota(100);c__Saccharomycetes(100);o__Saccharomycetales(100);f__Metschnikowiaceae(100);g__Metschnikowia(100);g__Metschnikowia_unclassified(100); | Metschnikowia | 13 | Saprotroph | \|Undefined Saprotroph\| | Yeast | NULL | Probable | As nectar-tap_saprotroph (Põlme et al. 2020); Decay sugar-rich substrates (Põlme et al. 2020) | Põlme S, et al. 2020. Fungal Diversity 105:1-16 ((https://doi.org/10.1007/s13225-020-00466-2)) |
| Otu09721 | 2 | k__Fungi(100);p__Ascomycota(100);c__Saccharomycetes(100);o__Saccharomycetales(100);f__Metschnikowiaceae(100);g__Metschnikowia(100);g__Metschnikowia_unclassified(100); | Metschnikowia | 13 | Saprotroph | \|Undefined Saprotroph\| | Yeast | NULL | Probable | As nectar-tap_saprotroph (Põlme et al. 2020); Decay sugar-rich substrates (Põlme et al. 2020) | Põlme S, et al. 2020. Fungal Diversity 105:1-16 ((https://doi.org/10.1007/s13225-020-00466-2)) |
| Otu09722 | 2 | k__Fungi(100);p__Ascomycota(100);c__Saccharomycetes(100);o__Saccharomycetales(100);f__Saccharomycetales_family_Incertae_sedis(100);f__Saccharomycetales_family_Incertae_sedis_unclassified(100);f__Saccharomycetales_family_Incertae_sedis_unclassified(100); | Saccharomycetales | 7 | Saprotroph | Undefined Saprotroph | Yeast | NULL | Possible | NULL | Sterkenburg E, et al. 2015. New Phytologist 207:1145-1158 ((https://doi.org/10.1111/nph.13426)) |
| Otu09728 | 2 | k__Fungi(100);p__Ascomycota(100);c__Saccharomycetes(100);o__Saccharomycetales(100);f__Metschnikowiaceae(100);g__Metschnikowia(100);g__Metschnikowia_unclassified(100); | Metschnikowia | 13 | Saprotroph | \|Undefined Saprotroph\| | Yeast | NULL | Probable | As nectar-tap_saprotroph (Põlme et al. 2020); Decay sugar-rich substrates (Põlme et al. 2020) | Põlme S, et al. 2020. Fungal Diversity 105:1-16 ((https://doi.org/10.1007/s13225-020-00466-2)) |
| Otu09730 | 2 | k__Fungi(100);p__Ascomycota(100);c__Saccharomycetes(100);o__Saccharomycetales(100);f__Metschnikowiaceae(100);g__Metschnikowia(100);s__Metschnikowia_pulcherrima(100); | Metschnikowia | 13 | Saprotroph | \|Undefined Saprotroph\| | Yeast | NULL | Probable | As nectar-tap_saprotroph (Põlme et al. 2020); Decay sugar-rich substrates (Põlme et al. 2020) | Põlme S, et al. 2020. Fungal Diversity 105:1-16 ((https://doi.org/10.1007/s13225-020-00466-2)) |
| Otu09731 | 2 | k__Fungi(100);p__Ascomycota(100);c__Saccharomycetes(100);o__Saccharomycetales(100);f__Metschnikowiaceae(100);g__Metschnikowia(100);s__Metschnikowia_pulcherrima(100); | Metschnikowia | 13 | Saprotroph | \|Undefined Saprotroph\| | Yeast | NULL | Probable | As nectar-tap_saprotroph (Põlme et al. 2020); Decay sugar-rich substrates (Põlme et al. 2020) | Põlme S, et al. 2020. Fungal Diversity 105:1-16 ((https://doi.org/10.1007/s13225-020-00466-2)) |
| Otu09733 | 2 | k__Fungi(100);p__Ascomycota(100);c__Saccharomycetes(100);o__Saccharomycetales(100);f__Metschnikowiaceae(100);g__Metschnikowia(100);s__Metschnikowia_pulcherrima(100); | Metschnikowia | 13 | Saprotroph | \|Undefined Saprotroph\| | Yeast | NULL | Probable | As nectar-tap_saprotroph (Põlme et al. 2020); Decay sugar-rich substrates (Põlme et al. 2020) | Põlme S, et al. 2020. Fungal Diversity 105:1-16 ((https://doi.org/10.1007/s13225-020-00466-2)) |
| Otu09734 | 2 | k__Fungi(100);p__Ascomycota(100);c__Saccharomycetes(100);o__Saccharomycetales(100);f__Metschnikowiaceae(100);g__Metschnikowia(100);s__Metschnikowia_pulcherrima(100); | Metschnikowia | 13 | Saprotroph | \|Undefined Saprotroph\| | Yeast | NULL | Probable | As nectar-tap_saprotroph (Põlme et al. 2020); Decay sugar-rich substrates (Põlme et al. 2020) | Põlme S, et al. 2020. Fungal Diversity 105:1-16 ((https://doi.org/10.1007/s13225-020-00466-2)) |
| Otu09735 | 2 | k__Fungi(100);p__Ascomycota(100);c__Saccharomycetes(100);o__Saccharomycetales(100);f__Metschnikowiaceae(100);g__Metschnikowia(100);g__Metschnikowia_unclassified(100); | Metschnikowia | 13 | Saprotroph | \|Undefined Saprotroph\| | Yeast | NULL | Probable | As nectar-tap_saprotroph (Põlme et al. 2020); Decay sugar-rich substrates (Põlme et al. 2020) | Põlme S, et al. 2020. Fungal Diversity 105:1-16 ((https://doi.org/10.1007/s13225-020-00466-2)) |
| Otu09736 | 2 | k__Fungi(100);p__Ascomycota(100);c__Saccharomycetes(100);o__Saccharomycetales(100);o__Saccharomycetales_unclassified(100);o__Saccharomycetales_unclassified(100);o__Saccharomycetales_unclassified(100); | Saccharomycetales | 7 | Saprotroph | Undefined Saprotroph | Yeast | NULL | Possible | NULL | Sterkenburg E, et al. 2015. New Phytologist 207:1145-1158 ((https://doi.org/10.1111/nph.13426)) |
| Otu09737 | 2 | k__Fungi(100);p__Ascomycota(100);c__Saccharomycetes(100);o__Saccharomycetales(100);f__Metschnikowiaceae(100);g__Metschnikowia(100);s__Metschnikowia_pulcherrima(100); | Metschnikowia | 13 | Saprotroph | \|Undefined Saprotroph\| | Yeast | NULL | Probable | As nectar-tap_saprotroph (Põlme et al. 2020); Decay sugar-rich substrates (Põlme et al. 2020) | Põlme S, et al. 2020. Fungal Diversity 105:1-16 ((https://doi.org/10.1007/s13225-020-00466-2)) |
| Otu09739 | 2 | k__Fungi(100);p__Ascomycota(100);c__Saccharomycetes(100);o__Saccharomycetales(100);f__Metschnikowiaceae(100);g__Metschnikowia(100);g__Metschnikowia_unclassified(100); | Metschnikowia | 13 | Saprotroph | \|Undefined Saprotroph\| | Yeast | NULL | Probable | As nectar-tap_saprotroph (Põlme et al. 2020); Decay sugar-rich substrates (Põlme et al. 2020) | Põlme S, et al. 2020. Fungal Diversity 105:1-16 ((https://doi.org/10.1007/s13225-020-00466-2)) |
| Otu09740 | 2 | k__Fungi(100);p__Ascomycota(100);c__Saccharomycetes(100);o__Saccharomycetales(100);f__Metschnikowiaceae(100);g__Metschnikowia(100);g__Metschnikowia_unclassified(100); | Metschnikowia | 13 | Saprotroph | \|Undefined Saprotroph\| | Yeast | NULL | Probable | As nectar-tap_saprotroph (Põlme et al. 2020); Decay sugar-rich substrates (Põlme et al. 2020) | Põlme S, et al. 2020. Fungal Diversity 105:1-16 ((https://doi.org/10.1007/s13225-020-00466-2)) |
| Otu09742 | 2 | k__Fungi(100);p__Ascomycota(100);c__Saccharomycetes(100);o__Saccharomycetales(100);f__Metschnikowiaceae(100);g__Metschnikowia(100);s__Metschnikowia_pulcherrima(100); | Metschnikowia | 13 | Saprotroph | \|Undefined Saprotroph\| | Yeast | NULL | Probable | As nectar-tap_saprotroph (Põlme et al. 2020); Decay sugar-rich substrates (Põlme et al. 2020) | Põlme S, et al. 2020. Fungal Diversity 105:1-16 ((https://doi.org/10.1007/s13225-020-00466-2)) |
| Otu09743 | 2 | k__Fungi(100);p__Ascomycota(100);c__Saccharomycetes(100);o__Saccharomycetales(100);f__Metschnikowiaceae(100);g__Metschnikowia(100);s__Metschnikowia_pulcherrima(100); | Metschnikowia | 13 | Saprotroph | \|Undefined Saprotroph\| | Yeast | NULL | Probable | As nectar-tap_saprotroph (Põlme et al. 2020); Decay sugar-rich substrates (Põlme et al. 2020) | Põlme S, et al. 2020. Fungal Diversity 105:1-16 ((https://doi.org/10.1007/s13225-020-00466-2)) |
| Otu09744 | 2 | k__Fungi(100);p__Ascomycota(100);c__Saccharomycetes(100);o__Saccharomycetales(100);f__Metschnikowiaceae(100);g__Metschnikowia(100);g__Metschnikowia_unclassified(100); | Metschnikowia | 13 | Saprotroph | \|Undefined Saprotroph\| | Yeast | NULL | Probable | As nectar-tap_saprotroph (Põlme et al. 2020); Decay sugar-rich substrates (Põlme et al. 2020) | Põlme S, et al. 2020. Fungal Diversity 105:1-16 ((https://doi.org/10.1007/s13225-020-00466-2)) |
| Otu09746 | 2 | k__Fungi(100);p__Ascomycota(100);c__Saccharomycetes(100);o__Saccharomycetales(100);f__Metschnikowiaceae(100);g__Metschnikowia(100);s__Metschnikowia_pulcherrima(100); | Metschnikowia | 13 | Saprotroph | \|Undefined Saprotroph\| | Yeast | NULL | Probable | As nectar-tap_saprotroph (Põlme et al. 2020); Decay sugar-rich substrates (Põlme et al. 2020) | Põlme S, et al. 2020. Fungal Diversity 105:1-16 ((https://doi.org/10.1007/s13225-020-00466-2)) |
| Otu09748 | 2 | k__Fungi(100);p__Ascomycota(100);c__Dothideomycetes(100);o__Dothideomycetes_order_Incertae_sedis(100);f__Dothideomycetes_family_Incertae_sedis(100);g__Pseudoramichloridium(100);s__Pseudoramichloridium_henryi(100); | Pseudoramichloridium | 13 | Pathotroph-Saprotroph | Plant Pathogen-\|Undefined Saprotroph\| | Microfungus | NULL | Probable | As soil_saprotroph (Põlme et al. 2020); Decay leaf, fruit, seed, soil (Põlme et al. 2020) | Põlme S, et al. 2020. Fungal Diversity 105:1-16 ((https://doi.org/10.1007/s13225-020-00466-2)) |
| Otu09749 | 2 | k__Fungi(100);p__Ascomycota(100);c__Saccharomycetes(100);o__Saccharomycetales(100);f__Metschnikowiaceae(100);g__Metschnikowia(100);g__Metschnikowia_unclassified(100); | Metschnikowia | 13 | Saprotroph | \|Undefined Saprotroph\| | Yeast | NULL | Probable | As nectar-tap_saprotroph (Põlme et al. 2020); Decay sugar-rich substrates (Põlme et al. 2020) | Põlme S, et al. 2020. Fungal Diversity 105:1-16 ((https://doi.org/10.1007/s13225-020-00466-2)) |
| Otu09751 | 2 | k__Fungi(100);p__Basidiomycota(100);c__Tremellomycetes(100);o__Tremellales(100);f__Tremellales_family_Incertae_sedis(100);g__Dioszegia(100);s__Dioszegia_zsoltii_var._yunnanensis(100); | Dioszegia | 13 | Saprotroph | \|Plant Saprotroph\|-Undefined Saprotroph | Yeast | NULL | Probable | As litter_saprotroph (Põlme et al. 2020); Decay leaf, fruit, seed (Põlme et al. 2020); The Dioszegia subclade sensu Inacio (Inacio et al. 20015) is characterized by orange; yellow or orange to deep orange colonies. Isolated from plant leaves and soil. | Põlme S, et al. 2020. Fungal Diversity 105:1-16 ((https://doi.org/10.1007/s13225-020-00466-2)); Takashima M, et al. 2011 International Journal of Systematic and Evolutionary Microbiology 61:1736-1739 ((https://doi.org/10.1099/ijs.0.025254-0)) |
| Otu09753 | 2 | k__Fungi(100);p__Ascomycota(100);c__Saccharomycetes(100);o__Saccharomycetales(100);f__Saccharomycetales_family_Incertae_sedis(100);g__Candida(100);g__Candida_unclassified(100); | Candida | 13 | Pathotroph-Saprotroph-Symbiotroph | Animal Pathogen-Endophyte-Endosymbiont-Epiphyte-\|Undefined Saprotroph\| | Dimorphic Yeast-Yeast | NULL | Probable | As nectar-tap_saprotroph (Põlme et al. 2020); Decay sugar-rich substrates (Põlme et al. 2020); This asexual genus is polyphyletic and contain species that function very differently depend on what they are. Caution should be taken when interpreting guilds for Candida species. | Manolakaki D, et al. 2010. Virulence. 1: 367-375 ((https://doi.org/10.4161/viru.1.5.12796)); Mekbib SB, et al. 2011. Tropical Plant Pathology 36:233-240 ((https://doi.org/10.1590/S1982-56762011000400004)); Põlme S, et al. 2020. Fungal Diversity 105:1-16 ((https://doi.org/10.1007/s13225-020-00466-2)) |
| Otu09756 | 2 | k__Fungi(100);p__Ascomycota(100);c__Saccharomycetes(100);o__Saccharomycetales(100);f__Metschnikowiaceae(100);g__Metschnikowia(100);g__Metschnikowia_unclassified(100); | Metschnikowia | 13 | Saprotroph | \|Undefined Saprotroph\| | Yeast | NULL | Probable | As nectar-tap_saprotroph (Põlme et al. 2020); Decay sugar-rich substrates (Põlme et al. 2020) | Põlme S, et al. 2020. Fungal Diversity 105:1-16 ((https://doi.org/10.1007/s13225-020-00466-2)) |
| Otu09757 | 2 | k__Fungi(100);p__Ascomycota(100);c__Saccharomycetes(100);o__Saccharomycetales(100);f__Metschnikowiaceae(100);g__Metschnikowia(100);s__Metschnikowia_pulcherrima(100); | Metschnikowia | 13 | Saprotroph | \|Undefined Saprotroph\| | Yeast | NULL | Probable | As nectar-tap_saprotroph (Põlme et al. 2020); Decay sugar-rich substrates (Põlme et al. 2020) | Põlme S, et al. 2020. Fungal Diversity 105:1-16 ((https://doi.org/10.1007/s13225-020-00466-2)) |
| Otu09758 | 2 | k__Fungi(100);p__Ascomycota(100);c__Saccharomycetes(100);o__Saccharomycetales(100);f__Metschnikowiaceae(100);g__Metschnikowia(100);s__Metschnikowia_pulcherrima(100); | Metschnikowia | 13 | Saprotroph | \|Undefined Saprotroph\| | Yeast | NULL | Probable | As nectar-tap_saprotroph (Põlme et al. 2020); Decay sugar-rich substrates (Põlme et al. 2020) | Põlme S, et al. 2020. Fungal Diversity 105:1-16 ((https://doi.org/10.1007/s13225-020-00466-2)) |
| Otu09760 | 2 | k__Fungi(100);p__Ascomycota(100);c__Saccharomycetes(100);o__Saccharomycetales(100);f__Metschnikowiaceae(100);g__Metschnikowia(100);s__Metschnikowia_pulcherrima(100); | Metschnikowia | 13 | Saprotroph | \|Undefined Saprotroph\| | Yeast | NULL | Probable | As nectar-tap_saprotroph (Põlme et al. 2020); Decay sugar-rich substrates (Põlme et al. 2020) | Põlme S, et al. 2020. Fungal Diversity 105:1-16 ((https://doi.org/10.1007/s13225-020-00466-2)) |
| Otu09762 | 2 | k__Fungi(100);p__Ascomycota(100);c__Saccharomycetes(100);o__Saccharomycetales(100);f__Saccharomycetales_family_Incertae_sedis(100);f__Saccharomycetales_family_Incertae_sedis_unclassified(100);f__Saccharomycetales_family_Incertae_sedis_unclassified(100); | Saccharomycetales | 7 | Saprotroph | Undefined Saprotroph | Yeast | NULL | Possible | NULL | Sterkenburg E, et al. 2015. New Phytologist 207:1145-1158 ((https://doi.org/10.1111/nph.13426)) |
| Otu09763 | 2 | k__Fungi(100);p__Ascomycota(100);c__Saccharomycetes(100);o__Saccharomycetales(100);f__Metschnikowiaceae(100);g__Metschnikowia(100);s__Metschnikowia_pulcherrima(100); | Metschnikowia | 13 | Saprotroph | \|Undefined Saprotroph\| | Yeast | NULL | Probable | As nectar-tap_saprotroph (Põlme et al. 2020); Decay sugar-rich substrates (Põlme et al. 2020) | Põlme S, et al. 2020. Fungal Diversity 105:1-16 ((https://doi.org/10.1007/s13225-020-00466-2)) |
| Otu09765 | 2 | k__Fungi(100);p__Ascomycota(100);c__Saccharomycetes(100);o__Saccharomycetales(100);f__Metschnikowiaceae(100);g__Metschnikowia(100);g__Metschnikowia_unclassified(100); | Metschnikowia | 13 | Saprotroph | \|Undefined Saprotroph\| | Yeast | NULL | Probable | As nectar-tap_saprotroph (Põlme et al. 2020); Decay sugar-rich substrates (Põlme et al. 2020) | Põlme S, et al. 2020. Fungal Diversity 105:1-16 ((https://doi.org/10.1007/s13225-020-00466-2)) |
| Otu09767 | 2 | k__Fungi(100);p__Ascomycota(100);c__Saccharomycetes(100);o__Saccharomycetales(100);f__Metschnikowiaceae(100);g__Metschnikowia(100);g__Metschnikowia_unclassified(100); | Metschnikowia | 13 | Saprotroph | \|Undefined Saprotroph\| | Yeast | NULL | Probable | As nectar-tap_saprotroph (Põlme et al. 2020); Decay sugar-rich substrates (Põlme et al. 2020) | Põlme S, et al. 2020. Fungal Diversity 105:1-16 ((https://doi.org/10.1007/s13225-020-00466-2)) |
| Otu09768 | 2 | k__Fungi(100);p__Ascomycota(100);c__Saccharomycetes(100);o__Saccharomycetales(100);f__Metschnikowiaceae(100);g__Metschnikowia(100);s__Metschnikowia_pulcherrima(100); | Metschnikowia | 13 | Saprotroph | \|Undefined Saprotroph\| | Yeast | NULL | Probable | As nectar-tap_saprotroph (Põlme et al. 2020); Decay sugar-rich substrates (Põlme et al. 2020) | Põlme S, et al. 2020. Fungal Diversity 105:1-16 ((https://doi.org/10.1007/s13225-020-00466-2)) |
| Otu09769 | 2 | k__Fungi(100);p__Ascomycota(100);c__Saccharomycetes(100);o__Saccharomycetales(100);f__Metschnikowiaceae(100);g__Metschnikowia(100);g__Metschnikowia_unclassified(100); | Metschnikowia | 13 | Saprotroph | \|Undefined Saprotroph\| | Yeast | NULL | Probable | As nectar-tap_saprotroph (Põlme et al. 2020); Decay sugar-rich substrates (Põlme et al. 2020) | Põlme S, et al. 2020. Fungal Diversity 105:1-16 ((https://doi.org/10.1007/s13225-020-00466-2)) |
| Otu09770 | 2 | k__Fungi(100);p__Ascomycota(100);c__Saccharomycetes(100);o__Saccharomycetales(100);f__Metschnikowiaceae(100);g__Metschnikowia(100);g__Metschnikowia_unclassified(100); | Metschnikowia | 13 | Saprotroph | \|Undefined Saprotroph\| | Yeast | NULL | Probable | As nectar-tap_saprotroph (Põlme et al. 2020); Decay sugar-rich substrates (Põlme et al. 2020) | Põlme S, et al. 2020. Fungal Diversity 105:1-16 ((https://doi.org/10.1007/s13225-020-00466-2)) |
| Otu09771 | 2 | k__Fungi(100);p__Ascomycota(100);c__Saccharomycetes(100);o__Saccharomycetales(100);f__Metschnikowiaceae(100);g__Metschnikowia(100);g__Metschnikowia_unclassified(100); | Metschnikowia | 13 | Saprotroph | \|Undefined Saprotroph\| | Yeast | NULL | Probable | As nectar-tap_saprotroph (Põlme et al. 2020); Decay sugar-rich substrates (Põlme et al. 2020) | Põlme S, et al. 2020. Fungal Diversity 105:1-16 ((https://doi.org/10.1007/s13225-020-00466-2)) |
| Otu09772 | 2 | k__Fungi(100);p__Ascomycota(100);c__Saccharomycetes(100);o__Saccharomycetales(100);f__Metschnikowiaceae(100);g__Metschnikowia(100);s__Metschnikowia_pulcherrima(100); | Metschnikowia | 13 | Saprotroph | \|Undefined Saprotroph\| | Yeast | NULL | Probable | As nectar-tap_saprotroph (Põlme et al. 2020); Decay sugar-rich substrates (Põlme et al. 2020) | Põlme S, et al. 2020. Fungal Diversity 105:1-16 ((https://doi.org/10.1007/s13225-020-00466-2)) |
| Otu09774 | 2 | k__Fungi(100);p__Ascomycota(100);c__Saccharomycetes(100);o__Saccharomycetales(100);f__Metschnikowiaceae(100);g__Metschnikowia(100);g__Metschnikowia_unclassified(100); | Metschnikowia | 13 | Saprotroph | \|Undefined Saprotroph\| | Yeast | NULL | Probable | As nectar-tap_saprotroph (Põlme et al. 2020); Decay sugar-rich substrates (Põlme et al. 2020) | Põlme S, et al. 2020. Fungal Diversity 105:1-16 ((https://doi.org/10.1007/s13225-020-00466-2)) |
| Otu09777 | 2 | k__Fungi(100);p__Ascomycota(100);c__Saccharomycetes(100);o__Saccharomycetales(100);f__Saccharomycetales_family_Incertae_sedis(100);g__Candida(100);g__Candida_unclassified(100); | Candida | 13 | Pathotroph-Saprotroph-Symbiotroph | Animal Pathogen-Endophyte-Endosymbiont-Epiphyte-\|Undefined Saprotroph\| | Dimorphic Yeast-Yeast | NULL | Probable | As nectar-tap_saprotroph (Põlme et al. 2020); Decay sugar-rich substrates (Põlme et al. 2020); This asexual genus is polyphyletic and contain species that function very differently depend on what they are. Caution should be taken when interpreting guilds for Candida species. | Manolakaki D, et al. 2010. Virulence. 1: 367-375 ((https://doi.org/10.4161/viru.1.5.12796)); Mekbib SB, et al. 2011. Tropical Plant Pathology 36:233-240 ((https://doi.org/10.1590/S1982-56762011000400004)); Põlme S, et al. 2020. Fungal Diversity 105:1-16 ((https://doi.org/10.1007/s13225-020-00466-2)) |
| Otu09780 | 2 | k__Fungi(100);p__Ascomycota(100);c__Saccharomycetes(100);o__Saccharomycetales(100);f__Metschnikowiaceae(100);g__Metschnikowia(100);s__Metschnikowia_pulcherrima(100); | Metschnikowia | 13 | Saprotroph | \|Undefined Saprotroph\| | Yeast | NULL | Probable | As nectar-tap_saprotroph (Põlme et al. 2020); Decay sugar-rich substrates (Põlme et al. 2020) | Põlme S, et al. 2020. Fungal Diversity 105:1-16 ((https://doi.org/10.1007/s13225-020-00466-2)) |
| Otu09781 | 2 | k__Fungi(100);p__Ascomycota(100);c__Saccharomycetes(100);o__Saccharomycetales(100);f__Metschnikowiaceae(100);g__Metschnikowia(100);g__Metschnikowia_unclassified(100); | Metschnikowia | 13 | Saprotroph | \|Undefined Saprotroph\| | Yeast | NULL | Probable | As nectar-tap_saprotroph (Põlme et al. 2020); Decay sugar-rich substrates (Põlme et al. 2020) | Põlme S, et al. 2020. Fungal Diversity 105:1-16 ((https://doi.org/10.1007/s13225-020-00466-2)) |
| Otu09782 | 2 | k__Fungi(100);p__Ascomycota(100);c__Saccharomycetes(100);o__Saccharomycetales(100);o__Saccharomycetales_unclassified(100);o__Saccharomycetales_unclassified(100);o__Saccharomycetales_unclassified(100); | Saccharomycetales | 7 | Saprotroph | Undefined Saprotroph | Yeast | NULL | Possible | NULL | Sterkenburg E, et al. 2015. New Phytologist 207:1145-1158 ((https://doi.org/10.1111/nph.13426)) |
| Otu09785 | 2 | k__Fungi(100);p__Ascomycota(100);c__Saccharomycetes(100);o__Saccharomycetales(100);f__Metschnikowiaceae(100);g__Metschnikowia(100);g__Metschnikowia_unclassified(100); | Metschnikowia | 13 | Saprotroph | \|Undefined Saprotroph\| | Yeast | NULL | Probable | As nectar-tap_saprotroph (Põlme et al. 2020); Decay sugar-rich substrates (Põlme et al. 2020) | Põlme S, et al. 2020. Fungal Diversity 105:1-16 ((https://doi.org/10.1007/s13225-020-00466-2)) |
| Otu09786 | 2 | k__Fungi(100);p__Ascomycota(100);c__Saccharomycetes(100);o__Saccharomycetales(100);f__Metschnikowiaceae(100);g__Metschnikowia(100);g__Metschnikowia_unclassified(100); | Metschnikowia | 13 | Saprotroph | \|Undefined Saprotroph\| | Yeast | NULL | Probable | As nectar-tap_saprotroph (Põlme et al. 2020); Decay sugar-rich substrates (Põlme et al. 2020) | Põlme S, et al. 2020. Fungal Diversity 105:1-16 ((https://doi.org/10.1007/s13225-020-00466-2)) |
| Otu09788 | 2 | k__Fungi(100);p__Ascomycota(100);c__Saccharomycetes(100);o__Saccharomycetales(100);f__Metschnikowiaceae(100);g__Metschnikowia(100);s__Metschnikowia_pulcherrima(100); | Metschnikowia | 13 | Saprotroph | \|Undefined Saprotroph\| | Yeast | NULL | Probable | As nectar-tap_saprotroph (Põlme et al. 2020); Decay sugar-rich substrates (Põlme et al. 2020) | Põlme S, et al. 2020. Fungal Diversity 105:1-16 ((https://doi.org/10.1007/s13225-020-00466-2)) |
| Otu09790 | 2 | k__Fungi(100);p__Ascomycota(100);c__Saccharomycetes(100);o__Saccharomycetales(100);f__Metschnikowiaceae(100);g__Metschnikowia(100);s__Metschnikowia_pulcherrima(100); | Metschnikowia | 13 | Saprotroph | \|Undefined Saprotroph\| | Yeast | NULL | Probable | As nectar-tap_saprotroph (Põlme et al. 2020); Decay sugar-rich substrates (Põlme et al. 2020) | Põlme S, et al. 2020. Fungal Diversity 105:1-16 ((https://doi.org/10.1007/s13225-020-00466-2)) |
| Otu09791 | 2 | k__Fungi(100);p__Ascomycota(100);c__Saccharomycetes(100);o__Saccharomycetales(100);f__Metschnikowiaceae(100);g__Metschnikowia(100);g__Metschnikowia_unclassified(100); | Metschnikowia | 13 | Saprotroph | \|Undefined Saprotroph\| | Yeast | NULL | Probable | As nectar-tap_saprotroph (Põlme et al. 2020); Decay sugar-rich substrates (Põlme et al. 2020) | Põlme S, et al. 2020. Fungal Diversity 105:1-16 ((https://doi.org/10.1007/s13225-020-00466-2)) |
| Otu09792 | 2 | k__Fungi(100);p__Ascomycota(100);c__Saccharomycetes(100);o__Saccharomycetales(100);f__Saccharomycetales_family_Incertae_sedis(100);g__Candida(100);g__Candida_unclassified(100); | Candida | 13 | Pathotroph-Saprotroph-Symbiotroph | Animal Pathogen-Endophyte-Endosymbiont-Epiphyte-\|Undefined Saprotroph\| | Dimorphic Yeast-Yeast | NULL | Probable | As nectar-tap_saprotroph (Põlme et al. 2020); Decay sugar-rich substrates (Põlme et al. 2020); This asexual genus is polyphyletic and contain species that function very differently depend on what they are. Caution should be taken when interpreting guilds for Candida species. | Manolakaki D, et al. 2010. Virulence. 1: 367-375 ((https://doi.org/10.4161/viru.1.5.12796)); Mekbib SB, et al. 2011. Tropical Plant Pathology 36:233-240 ((https://doi.org/10.1590/S1982-56762011000400004)); Põlme S, et al. 2020. Fungal Diversity 105:1-16 ((https://doi.org/10.1007/s13225-020-00466-2)) |
| Otu09793 | 2 | k__Fungi(100);p__Ascomycota(100);c__Sordariomycetes(100);o__Hypocreales(100);f__Nectriaceae(100);g__Fusarium(100);s__Fusarium_tricinctum(100); | Fusarium | 13 | Pathotroph-Saprotroph-Symbiotroph | Animal Pathogen-Endophyte-Lichen Parasite-\|Plant Pathogen\|-Plant Saprotroph-Undefined Saprotroph-Wood Saprotroph | Microfungus | Soft Rot | Possible | As foliar_endophyte (Põlme et al. 2020); As litter_saprotroph (Põlme et al. 2020); Causing damping-off disease (Põlme et al. 2020); Decay leaf, fruit, seed, roots, soil, animal material (Põlme et al. 2020); Likely opportunistic human pathogen (Irinyi et al. 2015) | Duncan CG, Eslyn WE. 1966. Mycologia 58:642-664 ((https://doi.org/10.2307/3757045)); Seehann G, et al. 1975. List of Fungi in Soft-Rot Tissue Tests. Institute fur Holzbiologie und Holzschutz, Bundesforschungsanstalt fur Forst - und Holzwirtshaft, Hamburg - Lohbrugge, Germany; Costa IPMW, et al. 2012. Mycotaxon 119:e26 ((http://mycotaxon.com/resources/checklists/costa_v119_checklist.pdf)); Tedersoo L, et al. 2014. Science 346:e1256688 ((https://doi.org/10.1126/science.1256688)); Irinyi L, et al. 2015. Medical Mycology 53:313-337 ((https://doi.org/10.1093/mmy/myv008)); Schroers, et al. 2016. Mycologia 108:806-819 ((https://doi.org/10.3852/15-255)); Põlme S, et al. 2020. Fungal Diversity 105:1-16 ((https://doi.org/10.1007/s13225-020-00466-2)); The American Phytopathological Society Common Names of Plant Diseases ((http://www.apsnet.org/publications/commonnames/Pages/default.aspx)); Lawrey JD, Diederich P. 2016. http://www.lichenicolous.net (accessed 24 May 2017) ((http://www.lichenicolous.net)) |
| Otu09794 | 2 | k__Fungi(100);p__Ascomycota(100);c__Saccharomycetes(100);o__Saccharomycetales(100);f__Metschnikowiaceae(100);g__Metschnikowia(100);s__Metschnikowia_pulcherrima(100); | Metschnikowia | 13 | Saprotroph | \|Undefined Saprotroph\| | Yeast | NULL | Probable | As nectar-tap_saprotroph (Põlme et al. 2020); Decay sugar-rich substrates (Põlme et al. 2020) | Põlme S, et al. 2020. Fungal Diversity 105:1-16 ((https://doi.org/10.1007/s13225-020-00466-2)) |
| Otu09796 | 2 | k__Fungi(100);p__Ascomycota(100);c__Saccharomycetes(100);o__Saccharomycetales(100);f__Metschnikowiaceae(100);g__Metschnikowia(100);g__Metschnikowia_unclassified(100); | Metschnikowia | 13 | Saprotroph | \|Undefined Saprotroph\| | Yeast | NULL | Probable | As nectar-tap_saprotroph (Põlme et al. 2020); Decay sugar-rich substrates (Põlme et al. 2020) | Põlme S, et al. 2020. Fungal Diversity 105:1-16 ((https://doi.org/10.1007/s13225-020-00466-2)) |
| Otu09798 | 2 | k__Fungi(100);p__Ascomycota(100);c__Saccharomycetes(100);o__Saccharomycetales(100);f__Metschnikowiaceae(100);g__Metschnikowia(100);s__Metschnikowia_pulcherrima(100); | Metschnikowia | 13 | Saprotroph | \|Undefined Saprotroph\| | Yeast | NULL | Probable | As nectar-tap_saprotroph (Põlme et al. 2020); Decay sugar-rich substrates (Põlme et al. 2020) | Põlme S, et al. 2020. Fungal Diversity 105:1-16 ((https://doi.org/10.1007/s13225-020-00466-2)) |
| Otu09799 | 2 | k__Fungi(100);p__Ascomycota(100);c__Saccharomycetes(100);o__Saccharomycetales(100);f__Metschnikowiaceae(100);g__Metschnikowia(100);g__Metschnikowia_unclassified(100); | Metschnikowia | 13 | Saprotroph | \|Undefined Saprotroph\| | Yeast | NULL | Probable | As nectar-tap_saprotroph (Põlme et al. 2020); Decay sugar-rich substrates (Põlme et al. 2020) | Põlme S, et al. 2020. Fungal Diversity 105:1-16 ((https://doi.org/10.1007/s13225-020-00466-2)) |
| Otu09800 | 2 | k__Fungi(100);p__Ascomycota(100);c__Saccharomycetes(100);o__Saccharomycetales(100);f__Metschnikowiaceae(100);g__Metschnikowia(100);g__Metschnikowia_unclassified(100); | Metschnikowia | 13 | Saprotroph | \|Undefined Saprotroph\| | Yeast | NULL | Probable | As nectar-tap_saprotroph (Põlme et al. 2020); Decay sugar-rich substrates (Põlme et al. 2020) | Põlme S, et al. 2020. Fungal Diversity 105:1-16 ((https://doi.org/10.1007/s13225-020-00466-2)) |
| Otu09802 | 2 | k__Fungi(100);p__Ascomycota(100);c__Saccharomycetes(100);o__Saccharomycetales(100);f__Metschnikowiaceae(100);g__Metschnikowia(100);s__Metschnikowia_pulcherrima(100); | Metschnikowia | 13 | Saprotroph | \|Undefined Saprotroph\| | Yeast | NULL | Probable | As nectar-tap_saprotroph (Põlme et al. 2020); Decay sugar-rich substrates (Põlme et al. 2020) | Põlme S, et al. 2020. Fungal Diversity 105:1-16 ((https://doi.org/10.1007/s13225-020-00466-2)) |
| Otu09804 | 2 | k__Fungi(100);p__Basidiomycota(100);c__Tremellomycetes(100);o__Tremellales(100);f__Tremellales_family_Incertae_sedis(100);g__Cryptococcus(100);s__Cryptococcus_sp_JCM_11356(100); | Cryptococcus | 13 | Pathotroph-Saprotroph-Symbiotroph | Animal Pathogen-Endophyte-Epiphyte-\|Undefined Saprotroph\| | Dimorphic Yeast-Yeast | NULL | Possible | As foliar_endophyte (Põlme et al. 2020); Decay leaf, fruit, seed, soil, dung, animal material (Põlme et al. 2020) | Andrews JH, et al. 1983. Phytopathology 73:228-234 ((https://doi.org/10.1094/Phyto-73-228)); Kurtzman CP, et al. (eds.) 2011. The Yeasts, a Taxonomic Study. Fifth Edition. Vols 1-3. Elsevier, San Diego (ISBN: 9780444521491); Põlme S, et al. 2020. Fungal Diversity 105:1-16 ((https://doi.org/10.1007/s13225-020-00466-2)) |
| Otu09805 | 2 | k__Fungi(100);p__Ascomycota(100);c__Saccharomycetes(100);o__Saccharomycetales(100);f__Metschnikowiaceae(100);g__Metschnikowia(100);g__Metschnikowia_unclassified(100); | Metschnikowia | 13 | Saprotroph | \|Undefined Saprotroph\| | Yeast | NULL | Probable | As nectar-tap_saprotroph (Põlme et al. 2020); Decay sugar-rich substrates (Põlme et al. 2020) | Põlme S, et al. 2020. Fungal Diversity 105:1-16 ((https://doi.org/10.1007/s13225-020-00466-2)) |
| Otu09807 | 2 | k__Fungi(100);p__Ascomycota(100);c__Saccharomycetes(100);o__Saccharomycetales(100);f__Metschnikowiaceae(100);g__Metschnikowia(100);g__Metschnikowia_unclassified(100); | Metschnikowia | 13 | Saprotroph | \|Undefined Saprotroph\| | Yeast | NULL | Probable | As nectar-tap_saprotroph (Põlme et al. 2020); Decay sugar-rich substrates (Põlme et al. 2020) | Põlme S, et al. 2020. Fungal Diversity 105:1-16 ((https://doi.org/10.1007/s13225-020-00466-2)) |
| Otu09808 | 2 | k__Fungi(100);p__Ascomycota(100);c__Saccharomycetes(100);o__Saccharomycetales(100);f__Metschnikowiaceae(100);g__Metschnikowia(100);s__Metschnikowia_pulcherrima(100); | Metschnikowia | 13 | Saprotroph | \|Undefined Saprotroph\| | Yeast | NULL | Probable | As nectar-tap_saprotroph (Põlme et al. 2020); Decay sugar-rich substrates (Põlme et al. 2020) | Põlme S, et al. 2020. Fungal Diversity 105:1-16 ((https://doi.org/10.1007/s13225-020-00466-2)) |
| Otu09809 | 2 | k__Fungi(100);p__Ascomycota(100);c__Saccharomycetes(100);o__Saccharomycetales(100);f__Metschnikowiaceae(100);g__Metschnikowia(100);s__Metschnikowia_pulcherrima(100); | Metschnikowia | 13 | Saprotroph | \|Undefined Saprotroph\| | Yeast | NULL | Probable | As nectar-tap_saprotroph (Põlme et al. 2020); Decay sugar-rich substrates (Põlme et al. 2020) | Põlme S, et al. 2020. Fungal Diversity 105:1-16 ((https://doi.org/10.1007/s13225-020-00466-2)) |
| Otu09810 | 2 | k__Fungi(100);p__Ascomycota(100);c__Saccharomycetes(100);o__Saccharomycetales(100);f__Metschnikowiaceae(100);g__Metschnikowia(100);s__Metschnikowia_pulcherrima(100); | Metschnikowia | 13 | Saprotroph | \|Undefined Saprotroph\| | Yeast | NULL | Probable | As nectar-tap_saprotroph (Põlme et al. 2020); Decay sugar-rich substrates (Põlme et al. 2020) | Põlme S, et al. 2020. Fungal Diversity 105:1-16 ((https://doi.org/10.1007/s13225-020-00466-2)) |
| Otu09811 | 2 | k__Fungi(100);p__Ascomycota(100);c__Saccharomycetes(100);o__Saccharomycetales(100);f__Metschnikowiaceae(100);g__Metschnikowia(100);s__Metschnikowia_pulcherrima(100); | Metschnikowia | 13 | Saprotroph | \|Undefined Saprotroph\| | Yeast | NULL | Probable | As nectar-tap_saprotroph (Põlme et al. 2020); Decay sugar-rich substrates (Põlme et al. 2020) | Põlme S, et al. 2020. Fungal Diversity 105:1-16 ((https://doi.org/10.1007/s13225-020-00466-2)) |
| Otu09813 | 2 | k__Fungi(100);p__Ascomycota(100);c__Saccharomycetes(100);o__Saccharomycetales(100);o__Saccharomycetales_unclassified(100);o__Saccharomycetales_unclassified(100);o__Saccharomycetales_unclassified(100); | Saccharomycetales | 7 | Saprotroph | Undefined Saprotroph | Yeast | NULL | Possible | NULL | Sterkenburg E, et al. 2015. New Phytologist 207:1145-1158 ((https://doi.org/10.1111/nph.13426)) |
| Otu09814 | 2 | k__Fungi(100);p__Ascomycota(100);c__Saccharomycetes(100);o__Saccharomycetales(100);f__Metschnikowiaceae(100);g__Metschnikowia(100);g__Metschnikowia_unclassified(100); | Metschnikowia | 13 | Saprotroph | \|Undefined Saprotroph\| | Yeast | NULL | Probable | As nectar-tap_saprotroph (Põlme et al. 2020); Decay sugar-rich substrates (Põlme et al. 2020) | Põlme S, et al. 2020. Fungal Diversity 105:1-16 ((https://doi.org/10.1007/s13225-020-00466-2)) |
| Otu09815 | 2 | k__Fungi(100);p__Ascomycota(100);c__Saccharomycetes(100);o__Saccharomycetales(100);f__Metschnikowiaceae(100);g__Metschnikowia(100);s__Metschnikowia_pulcherrima(100); | Metschnikowia | 13 | Saprotroph | \|Undefined Saprotroph\| | Yeast | NULL | Probable | As nectar-tap_saprotroph (Põlme et al. 2020); Decay sugar-rich substrates (Põlme et al. 2020) | Põlme S, et al. 2020. Fungal Diversity 105:1-16 ((https://doi.org/10.1007/s13225-020-00466-2)) |
| Otu09817 | 2 | k__Fungi(100);p__Ascomycota(100);c__Saccharomycetes(100);o__Saccharomycetales(100);f__Metschnikowiaceae(100);g__Metschnikowia(100);s__Metschnikowia_pulcherrima(100); | Metschnikowia | 13 | Saprotroph | \|Undefined Saprotroph\| | Yeast | NULL | Probable | As nectar-tap_saprotroph (Põlme et al. 2020); Decay sugar-rich substrates (Põlme et al. 2020) | Põlme S, et al. 2020. Fungal Diversity 105:1-16 ((https://doi.org/10.1007/s13225-020-00466-2)) |
| Otu09818 | 2 | k__Fungi(100);p__Ascomycota(100);c__Saccharomycetes(100);o__Saccharomycetales(100);f__Metschnikowiaceae(100);g__Metschnikowia(100);s__Metschnikowia_pulcherrima(100); | Metschnikowia | 13 | Saprotroph | \|Undefined Saprotroph\| | Yeast | NULL | Probable | As nectar-tap_saprotroph (Põlme et al. 2020); Decay sugar-rich substrates (Põlme et al. 2020) | Põlme S, et al. 2020. Fungal Diversity 105:1-16 ((https://doi.org/10.1007/s13225-020-00466-2)) |
| Otu09819 | 2 | k__Fungi(100);p__Ascomycota(100);c__Saccharomycetes(100);o__Saccharomycetales(100);f__Metschnikowiaceae(100);g__Metschnikowia(100);s__Metschnikowia_pulcherrima(100); | Metschnikowia | 13 | Saprotroph | \|Undefined Saprotroph\| | Yeast | NULL | Probable | As nectar-tap_saprotroph (Põlme et al. 2020); Decay sugar-rich substrates (Põlme et al. 2020) | Põlme S, et al. 2020. Fungal Diversity 105:1-16 ((https://doi.org/10.1007/s13225-020-00466-2)) |
| Otu09820 | 2 | k__Fungi(100);p__Ascomycota(100);c__Saccharomycetes(100);o__Saccharomycetales(100);f__Metschnikowiaceae(100);g__Metschnikowia(100);s__Metschnikowia_pulcherrima(100); | Metschnikowia | 13 | Saprotroph | \|Undefined Saprotroph\| | Yeast | NULL | Probable | As nectar-tap_saprotroph (Põlme et al. 2020); Decay sugar-rich substrates (Põlme et al. 2020) | Põlme S, et al. 2020. Fungal Diversity 105:1-16 ((https://doi.org/10.1007/s13225-020-00466-2)) |
| Otu09821 | 2 | k__Fungi(100);p__Ascomycota(100);c__Saccharomycetes(100);o__Saccharomycetales(100);f__Metschnikowiaceae(100);g__Metschnikowia(100);g__Metschnikowia_unclassified(100); | Metschnikowia | 13 | Saprotroph | \|Undefined Saprotroph\| | Yeast | NULL | Probable | As nectar-tap_saprotroph (Põlme et al. 2020); Decay sugar-rich substrates (Põlme et al. 2020) | Põlme S, et al. 2020. Fungal Diversity 105:1-16 ((https://doi.org/10.1007/s13225-020-00466-2)) |
| Otu09822 | 2 | k__Fungi(100);p__Ascomycota(100);c__Saccharomycetes(100);o__Saccharomycetales(100);f__Metschnikowiaceae(100);g__Metschnikowia(100);s__Metschnikowia_pulcherrima(100); | Metschnikowia | 13 | Saprotroph | \|Undefined Saprotroph\| | Yeast | NULL | Probable | As nectar-tap_saprotroph (Põlme et al. 2020); Decay sugar-rich substrates (Põlme et al. 2020) | Põlme S, et al. 2020. Fungal Diversity 105:1-16 ((https://doi.org/10.1007/s13225-020-00466-2)) |
| Otu09824 | 2 | k__Fungi(100);p__Ascomycota(100);c__Saccharomycetes(100);o__Saccharomycetales(100);f__Metschnikowiaceae(100);g__Metschnikowia(100);g__Metschnikowia_unclassified(100); | Metschnikowia | 13 | Saprotroph | \|Undefined Saprotroph\| | Yeast | NULL | Probable | As nectar-tap_saprotroph (Põlme et al. 2020); Decay sugar-rich substrates (Põlme et al. 2020) | Põlme S, et al. 2020. Fungal Diversity 105:1-16 ((https://doi.org/10.1007/s13225-020-00466-2)) |
| Otu09826 | 2 | k__Fungi(100);p__Ascomycota(100);c__Saccharomycetes(100);o__Saccharomycetales(100);f__Metschnikowiaceae(100);g__Metschnikowia(100);g__Metschnikowia_unclassified(100); | Metschnikowia | 13 | Saprotroph | \|Undefined Saprotroph\| | Yeast | NULL | Probable | As nectar-tap_saprotroph (Põlme et al. 2020); Decay sugar-rich substrates (Põlme et al. 2020) | Põlme S, et al. 2020. Fungal Diversity 105:1-16 ((https://doi.org/10.1007/s13225-020-00466-2)) |
| Otu09827 | 2 | k__Fungi(100);p__Ascomycota(100);c__Saccharomycetes(100);o__Saccharomycetales(100);f__Metschnikowiaceae(100);g__Metschnikowia(100);s__Metschnikowia_pulcherrima(100); | Metschnikowia | 13 | Saprotroph | \|Undefined Saprotroph\| | Yeast | NULL | Probable | As nectar-tap_saprotroph (Põlme et al. 2020); Decay sugar-rich substrates (Põlme et al. 2020) | Põlme S, et al. 2020. Fungal Diversity 105:1-16 ((https://doi.org/10.1007/s13225-020-00466-2)) |
| Otu09830 | 2 | k__Fungi(100);p__Ascomycota(100);c__Saccharomycetes(100);o__Saccharomycetales(100);f__Metschnikowiaceae(100);g__Metschnikowia(100);s__Metschnikowia_pulcherrima(100); | Metschnikowia | 13 | Saprotroph | \|Undefined Saprotroph\| | Yeast | NULL | Probable | As nectar-tap_saprotroph (Põlme et al. 2020); Decay sugar-rich substrates (Põlme et al. 2020) | Põlme S, et al. 2020. Fungal Diversity 105:1-16 ((https://doi.org/10.1007/s13225-020-00466-2)) |
| Otu09832 | 2 | k__Fungi(100);p__Ascomycota(100);c__Saccharomycetes(100);o__Saccharomycetales(100);f__Metschnikowiaceae(100);g__Metschnikowia(100);g__Metschnikowia_unclassified(100); | Metschnikowia | 13 | Saprotroph | \|Undefined Saprotroph\| | Yeast | NULL | Probable | As nectar-tap_saprotroph (Põlme et al. 2020); Decay sugar-rich substrates (Põlme et al. 2020) | Põlme S, et al. 2020. Fungal Diversity 105:1-16 ((https://doi.org/10.1007/s13225-020-00466-2)) |
| Otu09833 | 2 | k__Fungi(100);p__Ascomycota(100);c__Saccharomycetes(100);o__Saccharomycetales(100);f__Metschnikowiaceae(100);g__Metschnikowia(100);s__Metschnikowia_pulcherrima(100); | Metschnikowia | 13 | Saprotroph | \|Undefined Saprotroph\| | Yeast | NULL | Probable | As nectar-tap_saprotroph (Põlme et al. 2020); Decay sugar-rich substrates (Põlme et al. 2020) | Põlme S, et al. 2020. Fungal Diversity 105:1-16 ((https://doi.org/10.1007/s13225-020-00466-2)) |
| Otu09834 | 2 | k__Fungi(100);p__Ascomycota(100);c__Saccharomycetes(100);o__Saccharomycetales(100);f__Metschnikowiaceae(100);g__Metschnikowia(100);s__Metschnikowia_pulcherrima(100); | Metschnikowia | 13 | Saprotroph | \|Undefined Saprotroph\| | Yeast | NULL | Probable | As nectar-tap_saprotroph (Põlme et al. 2020); Decay sugar-rich substrates (Põlme et al. 2020) | Põlme S, et al. 2020. Fungal Diversity 105:1-16 ((https://doi.org/10.1007/s13225-020-00466-2)) |
| Otu09835 | 2 | k__Fungi(100);p__Ascomycota(100);c__Saccharomycetes(100);o__Saccharomycetales(100);f__Metschnikowiaceae(100);g__Metschnikowia(100);s__Metschnikowia_pulcherrima(100); | Metschnikowia | 13 | Saprotroph | \|Undefined Saprotroph\| | Yeast | NULL | Probable | As nectar-tap_saprotroph (Põlme et al. 2020); Decay sugar-rich substrates (Põlme et al. 2020) | Põlme S, et al. 2020. Fungal Diversity 105:1-16 ((https://doi.org/10.1007/s13225-020-00466-2)) |
| Otu09836 | 2 | k__Fungi(100);p__Ascomycota(100);c__Saccharomycetes(100);o__Saccharomycetales(100);f__Metschnikowiaceae(100);g__Metschnikowia(100);s__Metschnikowia_pulcherrima(100); | Metschnikowia | 13 | Saprotroph | \|Undefined Saprotroph\| | Yeast | NULL | Probable | As nectar-tap_saprotroph (Põlme et al. 2020); Decay sugar-rich substrates (Põlme et al. 2020) | Põlme S, et al. 2020. Fungal Diversity 105:1-16 ((https://doi.org/10.1007/s13225-020-00466-2)) |
| Otu09837 | 2 | k__Fungi(100);p__Ascomycota(100);c__Saccharomycetes(100);o__Saccharomycetales(100);f__Metschnikowiaceae(100);g__Metschnikowia(100);s__Metschnikowia_pulcherrima(100); | Metschnikowia | 13 | Saprotroph | \|Undefined Saprotroph\| | Yeast | NULL | Probable | As nectar-tap_saprotroph (Põlme et al. 2020); Decay sugar-rich substrates (Põlme et al. 2020) | Põlme S, et al. 2020. Fungal Diversity 105:1-16 ((https://doi.org/10.1007/s13225-020-00466-2)) |
| Otu09838 | 2 | k__Fungi(100);p__Ascomycota(100);c__Saccharomycetes(100);o__Saccharomycetales(100);f__Metschnikowiaceae(100);g__Metschnikowia(100);s__Metschnikowia_pulcherrima(100); | Metschnikowia | 13 | Saprotroph | \|Undefined Saprotroph\| | Yeast | NULL | Probable | As nectar-tap_saprotroph (Põlme et al. 2020); Decay sugar-rich substrates (Põlme et al. 2020) | Põlme S, et al. 2020. Fungal Diversity 105:1-16 ((https://doi.org/10.1007/s13225-020-00466-2)) |
| Otu09839 | 2 | k__Fungi(100);p__Ascomycota(100);c__Saccharomycetes(100);o__Saccharomycetales(100);f__Metschnikowiaceae(100);g__Metschnikowia(100);s__Metschnikowia_pulcherrima(100); | Metschnikowia | 13 | Saprotroph | \|Undefined Saprotroph\| | Yeast | NULL | Probable | As nectar-tap_saprotroph (Põlme et al. 2020); Decay sugar-rich substrates (Põlme et al. 2020) | Põlme S, et al. 2020. Fungal Diversity 105:1-16 ((https://doi.org/10.1007/s13225-020-00466-2)) |
| Otu09841 | 2 | k__Fungi(100);p__Ascomycota(100);c__Saccharomycetes(100);o__Saccharomycetales(100);f__Metschnikowiaceae(100);g__Metschnikowia(100);g__Metschnikowia_unclassified(100); | Metschnikowia | 13 | Saprotroph | \|Undefined Saprotroph\| | Yeast | NULL | Probable | As nectar-tap_saprotroph (Põlme et al. 2020); Decay sugar-rich substrates (Põlme et al. 2020) | Põlme S, et al. 2020. Fungal Diversity 105:1-16 ((https://doi.org/10.1007/s13225-020-00466-2)) |
| Otu09842 | 2 | k__Fungi(100);p__Ascomycota(100);c__Saccharomycetes(100);o__Saccharomycetales(100);f__Metschnikowiaceae(100);g__Metschnikowia(100);g__Metschnikowia_unclassified(100); | Metschnikowia | 13 | Saprotroph | \|Undefined Saprotroph\| | Yeast | NULL | Probable | As nectar-tap_saprotroph (Põlme et al. 2020); Decay sugar-rich substrates (Põlme et al. 2020) | Põlme S, et al. 2020. Fungal Diversity 105:1-16 ((https://doi.org/10.1007/s13225-020-00466-2)) |
| Otu09843 | 2 | k__Fungi(100);p__Ascomycota(100);c__Saccharomycetes(100);o__Saccharomycetales(100);f__Metschnikowiaceae(100);g__Metschnikowia(100);s__Metschnikowia_pulcherrima(100); | Metschnikowia | 13 | Saprotroph | \|Undefined Saprotroph\| | Yeast | NULL | Probable | As nectar-tap_saprotroph (Põlme et al. 2020); Decay sugar-rich substrates (Põlme et al. 2020) | Põlme S, et al. 2020. Fungal Diversity 105:1-16 ((https://doi.org/10.1007/s13225-020-00466-2)) |
| Otu09844 | 2 | k__Fungi(100);p__Ascomycota(100);c__Saccharomycetes(100);o__Saccharomycetales(100);f__Metschnikowiaceae(100);g__Metschnikowia(100);g__Metschnikowia_unclassified(100); | Metschnikowia | 13 | Saprotroph | \|Undefined Saprotroph\| | Yeast | NULL | Probable | As nectar-tap_saprotroph (Põlme et al. 2020); Decay sugar-rich substrates (Põlme et al. 2020) | Põlme S, et al. 2020. Fungal Diversity 105:1-16 ((https://doi.org/10.1007/s13225-020-00466-2)) |
| Otu09846 | 2 | k__Fungi(100);p__Basidiomycota(100);c__Microbotryomycetes(100);o__Sporidiobolales(100);f__Sporidiobolales_family_Incertae_sedis(100);g__Rhodotorula(100);s__Rhodotorula_mucilaginosa(100); | Rhodotorula | 13 | Pathotroph-Saprotroph-Symbiotroph | Animal Pathogen-Animal Symbiotroph-Endophyte-Plant Pathogen-\|Undefined Saprotroph\| | Yeast | NULL | Probable | As foliar_endophyte (Põlme et al. 2020); Opportunistic human pathogen | Põlme S, et al. 2020. Fungal Diversity 105:1-16 ((https://doi.org/10.1007/s13225-020-00466-2)); https://www.cdc.gov ((https://www.cdc.gov)) |
| Otu09847 | 2 | k__Fungi(100);p__Ascomycota(100);c__Saccharomycetes(100);o__Saccharomycetales(100);f__Metschnikowiaceae(100);g__Metschnikowia(100);s__Metschnikowia_pulcherrima(100); | Metschnikowia | 13 | Saprotroph | \|Undefined Saprotroph\| | Yeast | NULL | Probable | As nectar-tap_saprotroph (Põlme et al. 2020); Decay sugar-rich substrates (Põlme et al. 2020) | Põlme S, et al. 2020. Fungal Diversity 105:1-16 ((https://doi.org/10.1007/s13225-020-00466-2)) |
| Otu09848 | 2 | k__Fungi(100);p__Ascomycota(100);c__Saccharomycetes(100);o__Saccharomycetales(100);f__Metschnikowiaceae(100);g__Metschnikowia(100);g__Metschnikowia_unclassified(100); | Metschnikowia | 13 | Saprotroph | \|Undefined Saprotroph\| | Yeast | NULL | Probable | As nectar-tap_saprotroph (Põlme et al. 2020); Decay sugar-rich substrates (Põlme et al. 2020) | Põlme S, et al. 2020. Fungal Diversity 105:1-16 ((https://doi.org/10.1007/s13225-020-00466-2)) |
| Otu09850 | 2 | k__Fungi(100);p__Ascomycota(100);c__Saccharomycetes(100);o__Saccharomycetales(100);f__Metschnikowiaceae(100);g__Metschnikowia(100);s__Metschnikowia_pulcherrima(100); | Metschnikowia | 13 | Saprotroph | \|Undefined Saprotroph\| | Yeast | NULL | Probable | As nectar-tap_saprotroph (Põlme et al. 2020); Decay sugar-rich substrates (Põlme et al. 2020) | Põlme S, et al. 2020. Fungal Diversity 105:1-16 ((https://doi.org/10.1007/s13225-020-00466-2)) |
| Otu09851 | 2 | k__Fungi(100);p__Ascomycota(100);c__Saccharomycetes(100);o__Saccharomycetales(100);f__Metschnikowiaceae(100);g__Metschnikowia(100);s__Metschnikowia_pulcherrima(100); | Metschnikowia | 13 | Saprotroph | \|Undefined Saprotroph\| | Yeast | NULL | Probable | As nectar-tap_saprotroph (Põlme et al. 2020); Decay sugar-rich substrates (Põlme et al. 2020) | Põlme S, et al. 2020. Fungal Diversity 105:1-16 ((https://doi.org/10.1007/s13225-020-00466-2)) |
| Otu09852 | 2 | k__Fungi(100);p__Ascomycota(100);c__Saccharomycetes(100);o__Saccharomycetales(100);f__Metschnikowiaceae(100);g__Metschnikowia(100);s__Metschnikowia_pulcherrima(100); | Metschnikowia | 13 | Saprotroph | \|Undefined Saprotroph\| | Yeast | NULL | Probable | As nectar-tap_saprotroph (Põlme et al. 2020); Decay sugar-rich substrates (Põlme et al. 2020) | Põlme S, et al. 2020. Fungal Diversity 105:1-16 ((https://doi.org/10.1007/s13225-020-00466-2)) |
| Otu09853 | 2 | k__Fungi(100);p__Ascomycota(100);c__Saccharomycetes(100);o__Saccharomycetales(100);f__Metschnikowiaceae(100);g__Metschnikowia(100);g__Metschnikowia_unclassified(100); | Metschnikowia | 13 | Saprotroph | \|Undefined Saprotroph\| | Yeast | NULL | Probable | As nectar-tap_saprotroph (Põlme et al. 2020); Decay sugar-rich substrates (Põlme et al. 2020) | Põlme S, et al. 2020. Fungal Diversity 105:1-16 ((https://doi.org/10.1007/s13225-020-00466-2)) |
| Otu09855 | 2 | k__Fungi(100);p__Ascomycota(100);c__Saccharomycetes(100);o__Saccharomycetales(100);f__Metschnikowiaceae(100);g__Metschnikowia(100);s__Metschnikowia_pulcherrima(100); | Metschnikowia | 13 | Saprotroph | \|Undefined Saprotroph\| | Yeast | NULL | Probable | As nectar-tap_saprotroph (Põlme et al. 2020); Decay sugar-rich substrates (Põlme et al. 2020) | Põlme S, et al. 2020. Fungal Diversity 105:1-16 ((https://doi.org/10.1007/s13225-020-00466-2)) |
| Otu09856 | 2 | k__Fungi(100);p__Ascomycota(100);c__Saccharomycetes(100);o__Saccharomycetales(100);f__Metschnikowiaceae(100);g__Metschnikowia(100);g__Metschnikowia_unclassified(100); | Metschnikowia | 13 | Saprotroph | \|Undefined Saprotroph\| | Yeast | NULL | Probable | As nectar-tap_saprotroph (Põlme et al. 2020); Decay sugar-rich substrates (Põlme et al. 2020) | Põlme S, et al. 2020. Fungal Diversity 105:1-16 ((https://doi.org/10.1007/s13225-020-00466-2)) |
| Otu09857 | 2 | k__Fungi(100);p__Ascomycota(100);c__Saccharomycetes(100);o__Saccharomycetales(100);f__Metschnikowiaceae(100);g__Metschnikowia(100);s__Metschnikowia_pulcherrima(100); | Metschnikowia | 13 | Saprotroph | \|Undefined Saprotroph\| | Yeast | NULL | Probable | As nectar-tap_saprotroph (Põlme et al. 2020); Decay sugar-rich substrates (Põlme et al. 2020) | Põlme S, et al. 2020. Fungal Diversity 105:1-16 ((https://doi.org/10.1007/s13225-020-00466-2)) |
| Otu09858 | 2 | k__Fungi(100);p__Ascomycota(100);c__Saccharomycetes(100);o__Saccharomycetales(100);f__Metschnikowiaceae(100);g__Metschnikowia(100);s__Metschnikowia_pulcherrima(100); | Metschnikowia | 13 | Saprotroph | \|Undefined Saprotroph\| | Yeast | NULL | Probable | As nectar-tap_saprotroph (Põlme et al. 2020); Decay sugar-rich substrates (Põlme et al. 2020) | Põlme S, et al. 2020. Fungal Diversity 105:1-16 ((https://doi.org/10.1007/s13225-020-00466-2)) |
| Otu09860 | 2 | k__Fungi(100);p__Ascomycota(100);c__Saccharomycetes(100);o__Saccharomycetales(100);f__Metschnikowiaceae(100);g__Metschnikowia(100);g__Metschnikowia_unclassified(100); | Metschnikowia | 13 | Saprotroph | \|Undefined Saprotroph\| | Yeast | NULL | Probable | As nectar-tap_saprotroph (Põlme et al. 2020); Decay sugar-rich substrates (Põlme et al. 2020) | Põlme S, et al. 2020. Fungal Diversity 105:1-16 ((https://doi.org/10.1007/s13225-020-00466-2)) |
| Otu09862 | 2 | k__Fungi(100);p__Ascomycota(100);c__Saccharomycetes(100);o__Saccharomycetales(100);f__Metschnikowiaceae(100);g__Metschnikowia(100);g__Metschnikowia_unclassified(100); | Metschnikowia | 13 | Saprotroph | \|Undefined Saprotroph\| | Yeast | NULL | Probable | As nectar-tap_saprotroph (Põlme et al. 2020); Decay sugar-rich substrates (Põlme et al. 2020) | Põlme S, et al. 2020. Fungal Diversity 105:1-16 ((https://doi.org/10.1007/s13225-020-00466-2)) |
| Otu09863 | 2 | k__Fungi(100);p__Ascomycota(100);c__Saccharomycetes(100);o__Saccharomycetales(100);f__Metschnikowiaceae(100);g__Metschnikowia(100);g__Metschnikowia_unclassified(100); | Metschnikowia | 13 | Saprotroph | \|Undefined Saprotroph\| | Yeast | NULL | Probable | As nectar-tap_saprotroph (Põlme et al. 2020); Decay sugar-rich substrates (Põlme et al. 2020) | Põlme S, et al. 2020. Fungal Diversity 105:1-16 ((https://doi.org/10.1007/s13225-020-00466-2)) |
| Otu09864 | 2 | k__Fungi(100);p__Ascomycota(100);c__Saccharomycetes(100);o__Saccharomycetales(100);f__Metschnikowiaceae(100);g__Metschnikowia(100);g__Metschnikowia_unclassified(100); | Metschnikowia | 13 | Saprotroph | \|Undefined Saprotroph\| | Yeast | NULL | Probable | As nectar-tap_saprotroph (Põlme et al. 2020); Decay sugar-rich substrates (Põlme et al. 2020) | Põlme S, et al. 2020. Fungal Diversity 105:1-16 ((https://doi.org/10.1007/s13225-020-00466-2)) |
| Otu09865 | 2 | k__Fungi(100);p__Ascomycota(100);c__Saccharomycetes(100);o__Saccharomycetales(100);f__Metschnikowiaceae(100);g__Metschnikowia(100);s__Metschnikowia_pulcherrima(100); | Metschnikowia | 13 | Saprotroph | \|Undefined Saprotroph\| | Yeast | NULL | Probable | As nectar-tap_saprotroph (Põlme et al. 2020); Decay sugar-rich substrates (Põlme et al. 2020) | Põlme S, et al. 2020. Fungal Diversity 105:1-16 ((https://doi.org/10.1007/s13225-020-00466-2)) |
| Otu09867 | 2 | k__Fungi(100);p__Ascomycota(100);c__Saccharomycetes(100);o__Saccharomycetales(100);f__Metschnikowiaceae(100);g__Metschnikowia(100);s__Metschnikowia_pulcherrima(100); | Metschnikowia | 13 | Saprotroph | \|Undefined Saprotroph\| | Yeast | NULL | Probable | As nectar-tap_saprotroph (Põlme et al. 2020); Decay sugar-rich substrates (Põlme et al. 2020) | Põlme S, et al. 2020. Fungal Diversity 105:1-16 ((https://doi.org/10.1007/s13225-020-00466-2)) |
| Otu09868 | 2 | k__Fungi(100);p__Ascomycota(100);c__Saccharomycetes(100);o__Saccharomycetales(100);f__Metschnikowiaceae(100);g__Metschnikowia(100);s__Metschnikowia_pulcherrima(100); | Metschnikowia | 13 | Saprotroph | \|Undefined Saprotroph\| | Yeast | NULL | Probable | As nectar-tap_saprotroph (Põlme et al. 2020); Decay sugar-rich substrates (Põlme et al. 2020) | Põlme S, et al. 2020. Fungal Diversity 105:1-16 ((https://doi.org/10.1007/s13225-020-00466-2)) |
| Otu09871 | 2 | k__Fungi(100);p__Ascomycota(100);c__Leotiomycetes(100);o__Helotiales(100);f__Helotiales_family_Incertae_sedis(100);g__Ascocoryne(100);s__Ascocoryne_sp(100); | Ascocoryne | 13 | Saprotroph | Undefined Saprotroph-\|Wood Saprotroph\| | Tremelloid | NULL | Probable | Decay wood (Põlme et al. 2020) | Tedersoo L, et al. 2014. Science 346:e1256688 ((https://doi.org/10.1126/science.1256688)); Põlme S, et al. 2020. Fungal Diversity 105:1-16 ((https://doi.org/10.1007/s13225-020-00466-2)) |
| Otu09873 | 2 | k__Fungi(100);p__Ascomycota(100);c__Leotiomycetes(100);o__Helotiales(100);f__Hyaloscyphaceae(100);g__Lachnum(100);s__Lachnum_rachidicola(100); | Lachnum | 13 | Saprotroph-Symbiotroph | Endophyte-\|Plant Saprotroph\|-Undefined Saprotroph-Wood Saprotroph | Microfungus | NULL | Probable | As litter_saprotroph (Põlme et al. 2020); As root_endophyte (Põlme et al. 2020); Decay wood, leaf, fruit, seed (Põlme et al. 2020) | Tedersoo L, et al. 2014. Science 346:e1256688 ((https://doi.org/10.1126/science.1256688)); Põlme S, et al. 2020. Fungal Diversity 105:1-16 ((https://doi.org/10.1007/s13225-020-00466-2)) |
| Otu09874 | 2 | k__Fungi(100);p__Ascomycota(100);c__Saccharomycetes(100);o__Saccharomycetales(100);f__Saccharomycetales_family_Incertae_sedis(100);f__Saccharomycetales_family_Incertae_sedis_unclassified(100);f__Saccharomycetales_family_Incertae_sedis_unclassified(100); | Saccharomycetales | 7 | Saprotroph | Undefined Saprotroph | Yeast | NULL | Possible | NULL | Sterkenburg E, et al. 2015. New Phytologist 207:1145-1158 ((https://doi.org/10.1111/nph.13426)) |
| Otu09875 | 2 | k__Fungi(100);p__Ascomycota(100);c__Leotiomycetes(100);o__Leotiales(100);f__Bulgariaceae(100);g__Holwaya(100);s__Holwaya_sp(100); | Holwaya | 13 | Saprotroph | Undefined Saprotroph-\|Wood Saprotroph\| | Microfungus | NULL | Probable | Decay wood (Põlme et al. 2020) | Tedersoo L, et al. 2014. Science 346:e1256688 ((https://doi.org/10.1126/science.1256688)); Põlme S, et al. 2020. Fungal Diversity 105:1-16 ((https://doi.org/10.1007/s13225-020-00466-2)) |
| Otu09877 | 2 | k__Fungi(100);p__Ascomycota(100);c__Saccharomycetes(100);o__Saccharomycetales(100);f__Metschnikowiaceae(100);g__Metschnikowia(100);g__Metschnikowia_unclassified(100); | Metschnikowia | 13 | Saprotroph | \|Undefined Saprotroph\| | Yeast | NULL | Probable | As nectar-tap_saprotroph (Põlme et al. 2020); Decay sugar-rich substrates (Põlme et al. 2020) | Põlme S, et al. 2020. Fungal Diversity 105:1-16 ((https://doi.org/10.1007/s13225-020-00466-2)) |
| Otu09880 | 2 | k__Fungi(100);p__Ascomycota(100);c__Saccharomycetes(100);o__Saccharomycetales(100);f__Metschnikowiaceae(100);g__Metschnikowia(100);g__Metschnikowia_unclassified(100); | Metschnikowia | 13 | Saprotroph | \|Undefined Saprotroph\| | Yeast | NULL | Probable | As nectar-tap_saprotroph (Põlme et al. 2020); Decay sugar-rich substrates (Põlme et al. 2020) | Põlme S, et al. 2020. Fungal Diversity 105:1-16 ((https://doi.org/10.1007/s13225-020-00466-2)) |
| Otu09883 | 2 | k__Fungi(100);p__Ascomycota(100);c__Eurotiomycetes(100);o__Chaetothyriales(100);f__Herpotrichiellaceae(100);g__Phaeomoniella(100);g__Phaeomoniella_unclassified(100); | Phaeomoniella | 13 | Pathotroph | \|Plant Pathogen\| | Dimorphic-Microfungus; Facultative Yeast (Tedersoo et | NULL | Highly Probable | Pathotroph isolated from necrosis in Prunus wood (Damm et al. 2010) | Damm U, et al. 2010. Persoonia 24:60-80 ((http://doi.org/10.3767/003158510X500705)); Tedersoo L, et al. 2014. Science 346:e1256688 ((https://doi.org/10.1126/science.1256688)); Põlme S, et al. 2020. Fungal Diversity 105:1-16 ((https://doi.org/10.1007/s13225-020-00466-2)) |
| Otu09884 | 2 | k__Fungi(100);p__Ascomycota(100);c__Saccharomycetes(100);o__Saccharomycetales(100);f__Metschnikowiaceae(100);g__Metschnikowia(100);s__Metschnikowia_pulcherrima(100); | Metschnikowia | 13 | Saprotroph | \|Undefined Saprotroph\| | Yeast | NULL | Probable | As nectar-tap_saprotroph (Põlme et al. 2020); Decay sugar-rich substrates (Põlme et al. 2020) | Põlme S, et al. 2020. Fungal Diversity 105:1-16 ((https://doi.org/10.1007/s13225-020-00466-2)) |
| Otu09885 | 2 | k__Fungi(100);p__Ascomycota(100);c__Saccharomycetes(100);o__Saccharomycetales(100);f__Metschnikowiaceae(100);g__Metschnikowia(100);s__Metschnikowia_pulcherrima(100); | Metschnikowia | 13 | Saprotroph | \|Undefined Saprotroph\| | Yeast | NULL | Probable | As nectar-tap_saprotroph (Põlme et al. 2020); Decay sugar-rich substrates (Põlme et al. 2020) | Põlme S, et al. 2020. Fungal Diversity 105:1-16 ((https://doi.org/10.1007/s13225-020-00466-2)) |
| Otu09886 | 2 | k__Fungi(100);p__Ascomycota(100);c__Dothideomycetes(100);o__Capnodiales(100);f__Mycosphaerellaceae(100);g__Mycosphaerella(100);s__Mycosphaerella_nyssicola(100); | Mycosphaerella | 13 | Pathotroph-Saprotroph | \|Plant Pathogen\|-Plant Saprotroph | Microfungus | NULL | Probable | As litter_saprotroph (Põlme et al. 2020); Decay leaf, fruit, seed (Põlme et al. 2020) | Tedersoo L, et al. 2014. Science 346:e1256688 ((https://doi.org/10.1126/science.1256688)); Põlme S, et al. 2020. Fungal Diversity 105:1-16 ((https://doi.org/10.1007/s13225-020-00466-2)) |
| Otu09890 | 2 | k__Fungi(100);p__Ascomycota(100);c__Saccharomycetes(100);o__Saccharomycetales(100);f__Metschnikowiaceae(100);g__Metschnikowia(100);s__Metschnikowia_pulcherrima(100); | Metschnikowia | 13 | Saprotroph | \|Undefined Saprotroph\| | Yeast | NULL | Probable | As nectar-tap_saprotroph (Põlme et al. 2020); Decay sugar-rich substrates (Põlme et al. 2020) | Põlme S, et al. 2020. Fungal Diversity 105:1-16 ((https://doi.org/10.1007/s13225-020-00466-2)) |
| Otu09896 | 2 | k__Fungi(100);p__Ascomycota(100);c__Saccharomycetes(100);o__Saccharomycetales(100);f__Metschnikowiaceae(100);g__Metschnikowia(100);g__Metschnikowia_unclassified(100); | Metschnikowia | 13 | Saprotroph | \|Undefined Saprotroph\| | Yeast | NULL | Probable | As nectar-tap_saprotroph (Põlme et al. 2020); Decay sugar-rich substrates (Põlme et al. 2020) | Põlme S, et al. 2020. Fungal Diversity 105:1-16 ((https://doi.org/10.1007/s13225-020-00466-2)) |
| Otu09899 | 2 | k__Fungi(100);p__Ascomycota(100);c__Saccharomycetes(100);o__Saccharomycetales(100);f__Metschnikowiaceae(100);g__Metschnikowia(100);g__Metschnikowia_unclassified(100); | Metschnikowia | 13 | Saprotroph | \|Undefined Saprotroph\| | Yeast | NULL | Probable | As nectar-tap_saprotroph (Põlme et al. 2020); Decay sugar-rich substrates (Põlme et al. 2020) | Põlme S, et al. 2020. Fungal Diversity 105:1-16 ((https://doi.org/10.1007/s13225-020-00466-2)) |
| Otu09900 | 2 | k__Fungi(100);p__Ascomycota(100);c__Dothideomycetes(100);o__Capnodiales(100);f__Capnodiales_family_Incertae_sedis(100);g__Rachicladosporium(100);s__Rachicladosporium_sp_HX6(100); | Rachicladosporium | 13 | Saprotroph | \|Plant Saprotroph\| | Microfungus | NULL | Probable | As litter_saprotroph (Põlme et al. 2020); Decay leaf, fruit, seed (Põlme et al. 2020) | Põlme S, et al. 2020. Fungal Diversity 105:1-16 ((https://doi.org/10.1007/s13225-020-00466-2)) |
| Otu09904 | 2 | k__Fungi(100);p__Ascomycota(100);c__Saccharomycetes(100);o__Saccharomycetales(100);f__Metschnikowiaceae(100);g__Metschnikowia(100);g__Metschnikowia_unclassified(100); | Metschnikowia | 13 | Saprotroph | \|Undefined Saprotroph\| | Yeast | NULL | Probable | As nectar-tap_saprotroph (Põlme et al. 2020); Decay sugar-rich substrates (Põlme et al. 2020) | Põlme S, et al. 2020. Fungal Diversity 105:1-16 ((https://doi.org/10.1007/s13225-020-00466-2)) |
| Otu09905 | 2 | k__Fungi(100);p__Ascomycota(100);c__Saccharomycetes(100);o__Saccharomycetales(100);f__Metschnikowiaceae(100);g__Metschnikowia(100);s__Metschnikowia_pulcherrima(100); | Metschnikowia | 13 | Saprotroph | \|Undefined Saprotroph\| | Yeast | NULL | Probable | As nectar-tap_saprotroph (Põlme et al. 2020); Decay sugar-rich substrates (Põlme et al. 2020) | Põlme S, et al. 2020. Fungal Diversity 105:1-16 ((https://doi.org/10.1007/s13225-020-00466-2)) |
| Otu09908 | 2 | k__Fungi(100);p__Ascomycota(100);c__Saccharomycetes(100);o__Saccharomycetales(100);f__Metschnikowiaceae(100);g__Metschnikowia(100);g__Metschnikowia_unclassified(100); | Metschnikowia | 13 | Saprotroph | \|Undefined Saprotroph\| | Yeast | NULL | Probable | As nectar-tap_saprotroph (Põlme et al. 2020); Decay sugar-rich substrates (Põlme et al. 2020) | Põlme S, et al. 2020. Fungal Diversity 105:1-16 ((https://doi.org/10.1007/s13225-020-00466-2)) |
| Otu09909 | 2 | k__Fungi(100);p__Ascomycota(100);c__Saccharomycetes(100);o__Saccharomycetales(100);f__Metschnikowiaceae(100);g__Metschnikowia(100);s__Metschnikowia_pulcherrima(100); | Metschnikowia | 13 | Saprotroph | \|Undefined Saprotroph\| | Yeast | NULL | Probable | As nectar-tap_saprotroph (Põlme et al. 2020); Decay sugar-rich substrates (Põlme et al. 2020) | Põlme S, et al. 2020. Fungal Diversity 105:1-16 ((https://doi.org/10.1007/s13225-020-00466-2)) |
| Otu09910 | 2 | k__Fungi(100);p__Ascomycota(100);c__Saccharomycetes(100);o__Saccharomycetales(100);f__Metschnikowiaceae(100);g__Metschnikowia(100);s__Metschnikowia_pulcherrima(100); | Metschnikowia | 13 | Saprotroph | \|Undefined Saprotroph\| | Yeast | NULL | Probable | As nectar-tap_saprotroph (Põlme et al. 2020); Decay sugar-rich substrates (Põlme et al. 2020) | Põlme S, et al. 2020. Fungal Diversity 105:1-16 ((https://doi.org/10.1007/s13225-020-00466-2)) |
| Otu09916 | 2 | k__Fungi(100);p__Ascomycota(100);c__Saccharomycetes(100);o__Saccharomycetales(100);f__Metschnikowiaceae(100);g__Metschnikowia(100);s__Metschnikowia_pulcherrima(100); | Metschnikowia | 13 | Saprotroph | \|Undefined Saprotroph\| | Yeast | NULL | Probable | As nectar-tap_saprotroph (Põlme et al. 2020); Decay sugar-rich substrates (Põlme et al. 2020) | Põlme S, et al. 2020. Fungal Diversity 105:1-16 ((https://doi.org/10.1007/s13225-020-00466-2)) |
| Otu09917 | 2 | k__Fungi(100);p__Ascomycota(100);c__Saccharomycetes(100);o__Saccharomycetales(100);f__Metschnikowiaceae(100);g__Metschnikowia(100);s__Metschnikowia_pulcherrima(100); | Metschnikowia | 13 | Saprotroph | \|Undefined Saprotroph\| | Yeast | NULL | Probable | As nectar-tap_saprotroph (Põlme et al. 2020); Decay sugar-rich substrates (Põlme et al. 2020) | Põlme S, et al. 2020. Fungal Diversity 105:1-16 ((https://doi.org/10.1007/s13225-020-00466-2)) |
| Otu09920 | 2 | k__Fungi(100);p__Ascomycota(100);c__Saccharomycetes(100);o__Saccharomycetales(100);f__Metschnikowiaceae(100);g__Metschnikowia(100);s__Metschnikowia_pulcherrima(100); | Metschnikowia | 13 | Saprotroph | \|Undefined Saprotroph\| | Yeast | NULL | Probable | As nectar-tap_saprotroph (Põlme et al. 2020); Decay sugar-rich substrates (Põlme et al. 2020) | Põlme S, et al. 2020. Fungal Diversity 105:1-16 ((https://doi.org/10.1007/s13225-020-00466-2)) |
| Otu09923 | 2 | k__Fungi(100);p__Ascomycota(100);c__Saccharomycetes(100);o__Saccharomycetales(100);f__Metschnikowiaceae(100);g__Metschnikowia(100);g__Metschnikowia_unclassified(100); | Metschnikowia | 13 | Saprotroph | \|Undefined Saprotroph\| | Yeast | NULL | Probable | As nectar-tap_saprotroph (Põlme et al. 2020); Decay sugar-rich substrates (Põlme et al. 2020) | Põlme S, et al. 2020. Fungal Diversity 105:1-16 ((https://doi.org/10.1007/s13225-020-00466-2)) |
| Otu09924 | 2 | k__Fungi(100);p__Ascomycota(100);c__Saccharomycetes(100);o__Saccharomycetales(100);f__Metschnikowiaceae(100);g__Metschnikowia(100);s__Metschnikowia_pulcherrima(100); | Metschnikowia | 13 | Saprotroph | \|Undefined Saprotroph\| | Yeast | NULL | Probable | As nectar-tap_saprotroph (Põlme et al. 2020); Decay sugar-rich substrates (Põlme et al. 2020) | Põlme S, et al. 2020. Fungal Diversity 105:1-16 ((https://doi.org/10.1007/s13225-020-00466-2)) |
| Otu09926 | 2 | k__Fungi(100);p__Ascomycota(100);c__Saccharomycetes(100);o__Saccharomycetales(100);f__Metschnikowiaceae(100);g__Metschnikowia(100);s__Metschnikowia_pulcherrima(100); | Metschnikowia | 13 | Saprotroph | \|Undefined Saprotroph\| | Yeast | NULL | Probable | As nectar-tap_saprotroph (Põlme et al. 2020); Decay sugar-rich substrates (Põlme et al. 2020) | Põlme S, et al. 2020. Fungal Diversity 105:1-16 ((https://doi.org/10.1007/s13225-020-00466-2)) |
| Otu09928 | 2 | k__Fungi(100);p__Ascomycota(100);c__Saccharomycetes(100);o__Saccharomycetales(100);f__Metschnikowiaceae(100);g__Metschnikowia(100);g__Metschnikowia_unclassified(100); | Metschnikowia | 13 | Saprotroph | \|Undefined Saprotroph\| | Yeast | NULL | Probable | As nectar-tap_saprotroph (Põlme et al. 2020); Decay sugar-rich substrates (Põlme et al. 2020) | Põlme S, et al. 2020. Fungal Diversity 105:1-16 ((https://doi.org/10.1007/s13225-020-00466-2)) |
| Otu09930 | 2 | k__Fungi(100);p__Ascomycota(100);c__Saccharomycetes(100);o__Saccharomycetales(100);f__Metschnikowiaceae(100);g__Metschnikowia(100);s__Metschnikowia_pulcherrima(100); | Metschnikowia | 13 | Saprotroph | \|Undefined Saprotroph\| | Yeast | NULL | Probable | As nectar-tap_saprotroph (Põlme et al. 2020); Decay sugar-rich substrates (Põlme et al. 2020) | Põlme S, et al. 2020. Fungal Diversity 105:1-16 ((https://doi.org/10.1007/s13225-020-00466-2)) |
| Otu09931 | 2 | k__Fungi(100);p__Ascomycota(100);c__Saccharomycetes(100);o__Saccharomycetales(100);f__Metschnikowiaceae(100);g__Metschnikowia(100);s__Metschnikowia_pulcherrima(100); | Metschnikowia | 13 | Saprotroph | \|Undefined Saprotroph\| | Yeast | NULL | Probable | As nectar-tap_saprotroph (Põlme et al. 2020); Decay sugar-rich substrates (Põlme et al. 2020) | Põlme S, et al. 2020. Fungal Diversity 105:1-16 ((https://doi.org/10.1007/s13225-020-00466-2)) |
| Otu09934 | 2 | k__Fungi(100);p__Ascomycota(100);c__Saccharomycetes(100);o__Saccharomycetales(100);f__Metschnikowiaceae(100);g__Metschnikowia(100);g__Metschnikowia_unclassified(100); | Metschnikowia | 13 | Saprotroph | \|Undefined Saprotroph\| | Yeast | NULL | Probable | As nectar-tap_saprotroph (Põlme et al. 2020); Decay sugar-rich substrates (Põlme et al. 2020) | Põlme S, et al. 2020. Fungal Diversity 105:1-16 ((https://doi.org/10.1007/s13225-020-00466-2)) |
| Otu09935 | 2 | k__Fungi(100);p__Ascomycota(100);c__Saccharomycetes(100);o__Saccharomycetales(100);f__Metschnikowiaceae(100);g__Metschnikowia(100);s__Metschnikowia_pulcherrima(100); | Metschnikowia | 13 | Saprotroph | \|Undefined Saprotroph\| | Yeast | NULL | Probable | As nectar-tap_saprotroph (Põlme et al. 2020); Decay sugar-rich substrates (Põlme et al. 2020) | Põlme S, et al. 2020. Fungal Diversity 105:1-16 ((https://doi.org/10.1007/s13225-020-00466-2)) |
| Otu09939 | 2 | k__Fungi(100);p__Ascomycota(100);c__Saccharomycetes(100);o__Saccharomycetales(100);f__Metschnikowiaceae(100);g__Metschnikowia(100);s__Metschnikowia_pulcherrima(100); | Metschnikowia | 13 | Saprotroph | \|Undefined Saprotroph\| | Yeast | NULL | Probable | As nectar-tap_saprotroph (Põlme et al. 2020); Decay sugar-rich substrates (Põlme et al. 2020) | Põlme S, et al. 2020. Fungal Diversity 105:1-16 ((https://doi.org/10.1007/s13225-020-00466-2)) |
| Otu09941 | 2 | k__Fungi(100);p__Ascomycota(100);c__Saccharomycetes(100);o__Saccharomycetales(100);f__Metschnikowiaceae(100);g__Metschnikowia(100);g__Metschnikowia_unclassified(100); | Metschnikowia | 13 | Saprotroph | \|Undefined Saprotroph\| | Yeast | NULL | Probable | As nectar-tap_saprotroph (Põlme et al. 2020); Decay sugar-rich substrates (Põlme et al. 2020) | Põlme S, et al. 2020. Fungal Diversity 105:1-16 ((https://doi.org/10.1007/s13225-020-00466-2)) |
| Otu09943 | 2 | k__Fungi(100);p__Ascomycota(100);c__Saccharomycetes(100);o__Saccharomycetales(100);f__Saccharomycetales_family_Incertae_sedis(100);g__Candida(100);g__Candida_unclassified(100); | Candida | 13 | Pathotroph-Saprotroph-Symbiotroph | Animal Pathogen-Endophyte-Endosymbiont-Epiphyte-\|Undefined Saprotroph\| | Dimorphic Yeast-Yeast | NULL | Probable | As nectar-tap_saprotroph (Põlme et al. 2020); Decay sugar-rich substrates (Põlme et al. 2020); This asexual genus is polyphyletic and contain species that function very differently depend on what they are. Caution should be taken when interpreting guilds for Candida species. | Manolakaki D, et al. 2010. Virulence. 1: 367-375 ((https://doi.org/10.4161/viru.1.5.12796)); Mekbib SB, et al. 2011. Tropical Plant Pathology 36:233-240 ((https://doi.org/10.1590/S1982-56762011000400004)); Põlme S, et al. 2020. Fungal Diversity 105:1-16 ((https://doi.org/10.1007/s13225-020-00466-2)) |
| Otu09949 | 2 | k__Fungi(100);p__Ascomycota(100);c__Saccharomycetes(100);o__Saccharomycetales(100);f__Metschnikowiaceae(100);g__Metschnikowia(100);s__Metschnikowia_pulcherrima(100); | Metschnikowia | 13 | Saprotroph | \|Undefined Saprotroph\| | Yeast | NULL | Probable | As nectar-tap_saprotroph (Põlme et al. 2020); Decay sugar-rich substrates (Põlme et al. 2020) | Põlme S, et al. 2020. Fungal Diversity 105:1-16 ((https://doi.org/10.1007/s13225-020-00466-2)) |
| Otu09950 | 2 | k__Fungi(100);p__Ascomycota(100);c__Saccharomycetes(100);o__Saccharomycetales(100);f__Saccharomycetales_family_Incertae_sedis(100);g__Candida(100);g__Candida_unclassified(100); | Candida | 13 | Pathotroph-Saprotroph-Symbiotroph | Animal Pathogen-Endophyte-Endosymbiont-Epiphyte-\|Undefined Saprotroph\| | Dimorphic Yeast-Yeast | NULL | Probable | As nectar-tap_saprotroph (Põlme et al. 2020); Decay sugar-rich substrates (Põlme et al. 2020); This asexual genus is polyphyletic and contain species that function very differently depend on what they are. Caution should be taken when interpreting guilds for Candida species. | Manolakaki D, et al. 2010. Virulence. 1: 367-375 ((https://doi.org/10.4161/viru.1.5.12796)); Mekbib SB, et al. 2011. Tropical Plant Pathology 36:233-240 ((https://doi.org/10.1590/S1982-56762011000400004)); Põlme S, et al. 2020. Fungal Diversity 105:1-16 ((https://doi.org/10.1007/s13225-020-00466-2)) |

| | | | | | | | | | | | |
|---|---|---|---|---|---|---|---|---|---|---|---|
| Otu09951 | 2 | k__Fungi(100);p__Ascomycota(100);c__Sordariomycetes(100);o__Sordariomycetes_order_Incertae_sedis(100);f__Sordariomycetes_family_Incertae_sedis(100);g__Myrmecridium(100);s__Myrmecridium_schulzeri(100); | Myrmecridium | 13 | Saprotroph | \|Undefined Saprotroph\| | Microfungus | NULL | Probable | As soil_saprotroph (Põlme et al. 2020); Decay soil (Põlme et al. 2020) | Tedersoo L, et al. 2014. Science 346:e1256688 ((https://doi.org/10.1126/science.1256688)); Põlme S, et al. 2020. Fungal Diversity 105:1-16 ((https://doi.org/10.1007/s13225-020-00466-2)) |
| Otu09952 | 2 | k__Fungi(100);p__Ascomycota(100);c__Leotiomycetes(100);o__Helotiales(100);f__Hyaloscyphaceae(100);g__Cistella(100);s__Cistella_sp(100); | Cistella | 13 | Saprotroph-Symbiotroph | Endophyte-Plant Saprotroph-Undefined Saprotroph-Wood Saprotroph | Microfungus | NULL | Probable | As foliar_endophyte (Põlme et al. 2020); As litter_saprotroph (Põlme et al. 2020); Decay wood, leaf, fruit, seed (Põlme et al. 2020) | Tedersoo L, et al. 2014. Science 346:e1256688 ((https://doi.org/10.1126/science.1256688)); Põlme S, et al. 2020. Fungal Diversity 105:1-16 ((https://doi.org/10.1007/s13225-020-00466-2)) |
| Otu09953 | 2 | k__Fungi(100);p__Ascomycota(100);c__Saccharomycetes(100);o__Saccharomycetales(100);f__Metschnikowiaceae(100);g__Metschnikowia(100);g__Metschnikowia_unclassified(100); | Metschnikowia | 13 | Saprotroph | \|Undefined Saprotroph\| | Yeast | NULL | Probable | As nectar-tap_saprotroph (Põlme et al. 2020); Decay sugar-rich substrates (Põlme et al. 2020) | Põlme S, et al. 2020. Fungal Diversity 105:1-16 ((https://doi.org/10.1007/s13225-020-00466-2)) |
| Otu09955 | 2 | k__Fungi(100);p__Ascomycota(100);c__Saccharomycetes(100);o__Saccharomycetales(100);f__Metschnikowiaceae(100);g__Metschnikowia(100);g__Metschnikowia_unclassified(100); | Metschnikowia | 13 | Saprotroph | \|Undefined Saprotroph\| | Yeast | NULL | Probable | As nectar-tap_saprotroph (Põlme et al. 2020); Decay sugar-rich substrates (Põlme et al. 2020) | Põlme S, et al. 2020. Fungal Diversity 105:1-16 ((https://doi.org/10.1007/s13225-020-00466-2)) |
| Otu09956 | 2 | k__Fungi(100);p__Ascomycota(100);c__Dothideomycetes(100);o__Pleosporales(100);f__Venturiaceae(100);f__Venturiaceae_unclassified(100);f__Venturiaceae_unclassified(100); | Venturiaceae | 9 | Pathotroph-Saprotroph-Symbiotroph | Endophyte-Plant Pathogen-Undefined Saprotroph | Microfungus | NULL | Probable | Mostly endophytes and weak parasites of aerial plant parts, including bark and wood, leaf and fruit tissue; rarely biotrophic. Little is known except for the important pathogens (Cannon & Kirk 2007) | Cannon PF, Kirk PM. 2007. Fungal Families of the World. CAB International, Cambridge (ISBN: 978-0851998275) |
| Otu09957 | 2 | k__Fungi(100);p__Ascomycota(100);c__Saccharomycetes(100);o__Saccharomycetales(100);f__Metschnikowiaceae(100);g__Metschnikowia(100);s__Metschnikowia_pulcherrima(100); | Metschnikowia | 13 | Saprotroph | \|Undefined Saprotroph\| | Yeast | NULL | Probable | As nectar-tap_saprotroph (Põlme et al. 2020); Decay sugar-rich substrates (Põlme et al. 2020) | Põlme S, et al. 2020. Fungal Diversity 105:1-16 ((https://doi.org/10.1007/s13225-020-00466-2)) |
| Otu09958 | 2 | k__Fungi(100);p__Ascomycota(100);c__Saccharomycetes(100);o__Saccharomycetales(100);f__Metschnikowiaceae(100);g__Metschnikowia(100);s__Metschnikowia_pulcherrima(100); | Metschnikowia | 13 | Saprotroph | \|Undefined Saprotroph\| | Yeast | NULL | Probable | As nectar-tap_saprotroph (Põlme et al. 2020); Decay sugar-rich substrates (Põlme et al. 2020) | Põlme S, et al. 2020. Fungal Diversity 105:1-16 ((https://doi.org/10.1007/s13225-020-00466-2)) |
| Otu09962 | 2 | k__Fungi(100);p__Ascomycota(100);c__Saccharomycetes(100);o__Saccharomycetales(100);f__Metschnikowiaceae(100);g__Metschnikowia(100);g__Metschnikowia_unclassified(100); | Metschnikowia | 13 | Saprotroph | \|Undefined Saprotroph\| | Yeast | NULL | Probable | As nectar-tap_saprotroph (Põlme et al. 2020); Decay sugar-rich substrates (Põlme et al. 2020) | Põlme S, et al. 2020. Fungal Diversity 105:1-16 ((https://doi.org/10.1007/s13225-020-00466-2)) |
| Otu09963 | 2 | k__Fungi(100);p__Ascomycota(100);c__Saccharomycetes(100);o__Saccharomycetales(100);f__Metschnikowiaceae(100);g__Metschnikowia(100);s__Metschnikowia_pulcherrima(100); | Metschnikowia | 13 | Saprotroph | \|Undefined Saprotroph\| | Yeast | NULL | Probable | As nectar-tap_saprotroph (Põlme et al. 2020); Decay sugar-rich substrates (Põlme et al. 2020) | Põlme S, et al. 2020. Fungal Diversity 105:1-16 ((https://doi.org/10.1007/s13225-020-00466-2)) |
| Otu09966 | 2 | k__Fungi(100);p__Ascomycota(100);c__Saccharomycetes(100);o__Saccharomycetales(100);f__Metschnikowiaceae(100);g__Metschnikowia(100);s__Metschnikowia_pulcherrima(100); | Metschnikowia | 13 | Saprotroph | \|Undefined Saprotroph\| | Yeast | NULL | Probable | As nectar-tap_saprotroph (Põlme et al. 2020); Decay sugar-rich substrates (Põlme et al. 2020) | Põlme S, et al. 2020. Fungal Diversity 105:1-16 ((https://doi.org/10.1007/s13225-020-00466-2)) |
| Otu09969 | 2 | k__Fungi(100);p__Ascomycota(100);c__Saccharomycetes(100);o__Saccharomycetales(100);f__Metschnikowiaceae(100);g__Metschnikowia(100);g__Metschnikowia_unclassified(100); | Metschnikowia | 13 | Saprotroph | \|Undefined Saprotroph\| | Yeast | NULL | Probable | As nectar-tap_saprotroph (Põlme et al. 2020); Decay sugar-rich substrates (Põlme et al. 2020) | Põlme S, et al. 2020. Fungal Diversity 105:1-16 ((https://doi.org/10.1007/s13225-020-00466-2)) |
| Otu09970 | 2 | k__Fungi(100);p__Ascomycota(100);c__Saccharomycetes(100);o__Saccharomycetales(100);f__Metschnikowiaceae(100);g__Metschnikowia(100);s__Metschnikowia_pulcherrima(100); | Metschnikowia | 13 | Saprotroph | \|Undefined Saprotroph\| | Yeast | NULL | Probable | As nectar-tap_saprotroph (Põlme et al. 2020); Decay sugar-rich substrates (Põlme et al. 2020) | Põlme S, et al. 2020. Fungal Diversity 105:1-16 ((https://doi.org/10.1007/s13225-020-00466-2)) |
| Otu09975 | 2 | k__Fungi(100);p__Ascomycota(100);c__Saccharomycetes(100);o__Saccharomycetales(100);f__Saccharomycetales_family_Incertae_sedis(100);g__Candida(100);g__Candida_unclassified(100); | Candida | 13 | Pathotroph-Saprotroph-Symbiotroph | Animal Pathogen-Endophyte-Endosymbiont-Epiphyte-\|Undefined Saprotroph\| | Dimorphic Yeast-Yeast | NULL | Probable | As nectar-tap_saprotroph (Põlme et al. 2020); Decay sugar-rich substrates (Põlme et al. 2020); This asexual genus is polyphyletic and contain species that function very differently depend on what they are. Caution should be taken when interpreting guilds for Candida species. | Manolakaki D, et al. 2010. Virulence. 1: 367-375 ((https://doi.org/10.4161/viru.1.5.12796)); Mekbib SB, et al. 2011. Tropical Plant Pathology 36:233-240 ((https://doi.org/10.1590/S1982-56762011000400004)); Põlme S, et al. 2020. Fungal Diversity 105:1-16 ((https://doi.org/10.1007/s13225-020-00466-2)) |
| Otu09979 | 2 | k__Fungi(100);p__Ascomycota(100);c__Saccharomycetes(100);o__Saccharomycetales(100);f__Metschnikowiaceae(100);g__Metschnikowia(100);s__Metschnikowia_pulcherrima(100); | Metschnikowia | 13 | Saprotroph | \|Undefined Saprotroph\| | Yeast | NULL | Probable | As nectar-tap_saprotroph (Põlme et al. 2020); Decay sugar-rich substrates (Põlme et al. 2020) | Põlme S, et al. 2020. Fungal Diversity 105:1-16 ((https://doi.org/10.1007/s13225-020-00466-2)) |
| Otu09980 | 2 | k__Fungi(100);p__Ascomycota(100);c__Saccharomycetes(100);o__Saccharomycetales(100);f__Saccharomycetales_family_Incertae_sedis(100);g__Candida(100);g__Candida_unclassified(100); | Candida | 13 | Pathotroph-Saprotroph-Symbiotroph | Animal Pathogen-Endophyte-Endosymbiont-Epiphyte-\|Undefined Saprotroph\| | Dimorphic Yeast-Yeast | NULL | Probable | As nectar-tap_saprotroph (Põlme et al. 2020); Decay sugar-rich substrates (Põlme et al. 2020); This asexual genus is polyphyletic and contain species that function very differently depend on what they are. Caution should be taken when interpreting guilds for Candida species. | Manolakaki D, et al. 2010. Virulence. 1: 367-375 ((https://doi.org/10.4161/viru.1.5.12796)); Mekbib SB, et al. 2011. Tropical Plant Pathology 36:233-240 ((https://doi.org/10.1590/S1982-56762011000400004)); Põlme S, et al. 2020. Fungal Diversity 105:1-16 ((https://doi.org/10.1007/s13225-020-00466-2)) |
| Otu09982 | 2 | k__Fungi(100);p__Ascomycota(100);c__Saccharomycetes(100);o__Saccharomycetales(100);f__Metschnikowiaceae(100);g__Metschnikowia(100);s__Metschnikowia_pulcherrima(100); | Metschnikowia | 13 | Saprotroph | \|Undefined Saprotroph\| | Yeast | NULL | Probable | As nectar-tap_saprotroph (Põlme et al. 2020); Decay sugar-rich substrates (Põlme et al. 2020) | Põlme S, et al. 2020. Fungal Diversity 105:1-16 ((https://doi.org/10.1007/s13225-020-00466-2)) |
| Otu09983 | 2 | k__Fungi(100);p__Ascomycota(100);c__Saccharomycetes(100);o__Saccharomycetales(100);f__Metschnikowiaceae(100);g__Metschnikowia(100);s__Metschnikowia_pulcherrima(100); | Metschnikowia | 13 | Saprotroph | \|Undefined Saprotroph\| | Yeast | NULL | Probable | As nectar-tap_saprotroph (Põlme et al. 2020); Decay sugar-rich substrates (Põlme et al. 2020) | Põlme S, et al. 2020. Fungal Diversity 105:1-16 ((https://doi.org/10.1007/s13225-020-00466-2)) |
| Otu09984 | 2 | k__Fungi(100);p__Ascomycota(100);c__Saccharomycetes(100);o__Saccharomycetales(100);f__Metschnikowiaceae(100);g__Metschnikowia(100);g__Metschnikowia_unclassified(100); | Metschnikowia | 13 | Saprotroph | \|Undefined Saprotroph\| | Yeast | NULL | Probable | As nectar-tap_saprotroph (Põlme et al. 2020); Decay sugar-rich substrates (Põlme et al. 2020) | Põlme S, et al. 2020. Fungal Diversity 105:1-16 ((https://doi.org/10.1007/s13225-020-00466-2)) |
| Otu09986 | 2 | k__Fungi(100);p__Ascomycota(100);c__Saccharomycetes(100);o__Saccharomycetales(100);f__Metschnikowiaceae(100);g__Metschnikowia(100);g__Metschnikowia_unclassified(100); | Metschnikowia | 13 | Saprotroph | \|Undefined Saprotroph\| | Yeast | NULL | Probable | As nectar-tap_saprotroph (Põlme et al. 2020); Decay sugar-rich substrates (Põlme et al. 2020) | Põlme S, et al. 2020. Fungal Diversity 105:1-16 ((https://doi.org/10.1007/s13225-020-00466-2)) |
| Otu09989 | 2 | k__Fungi(100);p__Ascomycota(100);c__Saccharomycetes(100);o__Saccharomycetales(100);f__Metschnikowiaceae(100);g__Metschnikowia(100);s__Metschnikowia_pulcherrima(100); | Metschnikowia | 13 | Saprotroph | \|Undefined Saprotroph\| | Yeast | NULL | Probable | As nectar-tap_saprotroph (Põlme et al. 2020); Decay sugar-rich substrates (Põlme et al. 2020) | Põlme S, et al. 2020. Fungal Diversity 105:1-16 ((https://doi.org/10.1007/s13225-020-00466-2)) |
| Otu09990 | 2 | k__Fungi(100);p__Ascomycota(100);c__Saccharomycetes(100);o__Saccharomycetales(100);f__Metschnikowiaceae(100);g__Metschnikowia(100);s__Metschnikowia_pulcherrima(100); | Metschnikowia | 13 | Saprotroph | \|Undefined Saprotroph\| | Yeast | NULL | Probable | As nectar-tap_saprotroph (Põlme et al. 2020); Decay sugar-rich substrates (Põlme et al. 2020) | Põlme S, et al. 2020. Fungal Diversity 105:1-16 ((https://doi.org/10.1007/s13225-020-00466-2)) |
| Otu09991 | 2 | k__Fungi(100);p__Ascomycota(100);c__Saccharomycetes(100);o__Saccharomycetales(100);f__Metschnikowiaceae(100);g__Metschnikowia(100);s__Metschnikowia_pulcherrima(100); | Metschnikowia | 13 | Saprotroph | \|Undefined Saprotroph\| | Yeast | NULL | Probable | As nectar-tap_saprotroph (Põlme et al. 2020); Decay sugar-rich substrates (Põlme et al. 2020) | Põlme S, et al. 2020. Fungal Diversity 105:1-16 ((https://doi.org/10.1007/s13225-020-00466-2)) |
| Otu09992 | 2 | k__Fungi(100);p__Ascomycota(100);c__Saccharomycetes(100);o__Saccharomycetales(100);f__Metschnikowiaceae(100);g__Metschnikowia(100);s__Metschnikowia_pulcherrima(100); | Metschnikowia | 13 | Saprotroph | \|Undefined Saprotroph\| | Yeast | NULL | Probable | As nectar-tap_saprotroph (Põlme et al. 2020); Decay sugar-rich substrates (Põlme et al. 2020) | Põlme S, et al. 2020. Fungal Diversity 105:1-16 ((https://doi.org/10.1007/s13225-020-00466-2)) |
| Otu09993 | 2 | k__Fungi(100);p__Ascomycota(100);c__Saccharomycetes(100);o__Saccharomycetales(100);f__Metschnikowiaceae(100);g__Metschnikowia(100);s__Metschnikowia_pulcherrima(100); | Metschnikowia | 13 | Saprotroph | \|Undefined Saprotroph\| | Yeast | NULL | Probable | As nectar-tap_saprotroph (Põlme et al. 2020); Decay sugar-rich substrates (Põlme et al. 2020) | Põlme S, et al. 2020. Fungal Diversity 105:1-16 ((https://doi.org/10.1007/s13225-020-00466-2)) |
| Otu09994 | 2 | k__Fungi(100);p__Ascomycota(100);c__Saccharomycetes(100);o__Saccharomycetales(100);f__Metschnikowiaceae(100);g__Metschnikowia(100);s__Metschnikowia_pulcherrima(100); | Metschnikowia | 13 | Saprotroph | \|Undefined Saprotroph\| | Yeast | NULL | Probable | As nectar-tap_saprotroph (Põlme et al. 2020); Decay sugar-rich substrates (Põlme et al. 2020) | Põlme S, et al. 2020. Fungal Diversity 105:1-16 ((https://doi.org/10.1007/s13225-020-00466-2)) |
| Otu09997 | 2 | k__Fungi(100);p__Ascomycota(100);c__Saccharomycetes(100);o__Saccharomycetales(100);f__Metschnikowiaceae(100);g__Metschnikowia(100);g__Metschnikowia_unclassified(100); | Metschnikowia | 13 | Saprotroph | \|Undefined Saprotroph\| | Yeast | NULL | Probable | As nectar-tap_saprotroph (Põlme et al. 2020); Decay sugar-rich substrates (Põlme et al. 2020) | Põlme S, et al. 2020. Fungal Diversity 105:1-16 ((https://doi.org/10.1007/s13225-020-00466-2)) |
| Otu10002 | 2 | k__Fungi(100);p__Ascomycota(100);c__Saccharomycetes(100);o__Saccharomycetales(100);f__Metschnikowiaceae(100);g__Metschnikowia(100);s__Metschnikowia_pulcherrima(100); | Metschnikowia | 13 | Saprotroph | \|Undefined Saprotroph\| | Yeast | NULL | Probable | As nectar-tap_saprotroph (Põlme et al. 2020); Decay sugar-rich substrates (Põlme et al. 2020) | Põlme S, et al. 2020. Fungal Diversity 105:1-16 ((https://doi.org/10.1007/s13225-020-00466-2)) |
| Otu10003 | 2 | k__Fungi(100);p__Ascomycota(100);c__Saccharomycetes(100);o__Saccharomycetales(100);f__Metschnikowiaceae(100);g__Metschnikowia(100);s__Metschnikowia_pulcherrima(100); | Metschnikowia | 13 | Saprotroph | \|Undefined Saprotroph\| | Yeast | NULL | Probable | As nectar-tap_saprotroph (Põlme et al. 2020); Decay sugar-rich substrates (Põlme et al. 2020) | Põlme S, et al. 2020. Fungal Diversity 105:1-16 ((https://doi.org/10.1007/s13225-020-00466-2)) |
| Otu10004 | 2 | k__Fungi(100);p__Ascomycota(100);c__Saccharomycetes(100);o__Saccharomycetales(100);f__Metschnikowiaceae(100);g__Metschnikowia(100);g__Metschnikowia_unclassified(100); | Metschnikowia | 13 | Saprotroph | \|Undefined Saprotroph\| | Yeast | NULL | Probable | As nectar-tap_saprotroph (Põlme et al. 2020); Decay sugar-rich substrates (Põlme et al. 2020) | Põlme S, et al. 2020. Fungal Diversity 105:1-16 ((https://doi.org/10.1007/s13225-020-00466-2)) |
| Otu10006 | 2 | k__Fungi(100);p__Ascomycota(100);c__Saccharomycetes(100);o__Saccharomycetales(100);f__Metschnikowiaceae(100);g__Metschnikowia(100);s__Metschnikowia_pulcherrima(100); | Metschnikowia | 13 | Saprotroph | \|Undefined Saprotroph\| | Yeast | NULL | Probable | As nectar-tap_saprotroph (Põlme et al. 2020); Decay sugar-rich substrates (Põlme et al. 2020) | Põlme S, et al. 2020. Fungal Diversity 105:1-16 ((https://doi.org/10.1007/s13225-020-00466-2)) |
| Otu10010 | 2 | k__Fungi(100);p__Ascomycota(100);c__Saccharomycetes(100);o__Saccharomycetales(100);f__Metschnikowiaceae(100);g__Metschnikowia(100);s__Metschnikowia_pulcherrima(100); | Metschnikowia | 13 | Saprotroph | \|Undefined Saprotroph\| | Yeast | NULL | Probable | As nectar-tap_saprotroph (Põlme et al. 2020); Decay sugar-rich substrates (Põlme et al. 2020) | Põlme S, et al. 2020. Fungal Diversity 105:1-16 ((https://doi.org/10.1007/s13225-020-00466-2)) |
| Otu10011 | 2 | k__Fungi(100);p__Ascomycota(100);c__Saccharomycetes(100);o__Saccharomycetales(100);f__Metschnikowiaceae(100);g__Metschnikowia(100);g__Metschnikowia_unclassified(100); | Metschnikowia | 13 | Saprotroph | \|Undefined Saprotroph\| | Yeast | NULL | Probable | As nectar-tap_saprotroph (Põlme et al. 2020); Decay sugar-rich substrates (Põlme et al. 2020) | Põlme S, et al. 2020. Fungal Diversity 105:1-16 ((https://doi.org/10.1007/s13225-020-00466-2)) |
| Otu10012 | 2 | k__Fungi(100);p__Ascomycota(100);c__Saccharomycetes(100);o__Saccharomycetales(100);f__Metschnikowiaceae(100);g__Metschnikowia(100);s__Metschnikowia_pulcherrima(100); | Metschnikowia | 13 | Saprotroph | \|Undefined Saprotroph\| | Yeast | NULL | Probable | As nectar-tap_saprotroph (Põlme et al. 2020); Decay sugar-rich substrates (Põlme et al. 2020) | Põlme S, et al. 2020. Fungal Diversity 105:1-16 ((https://doi.org/10.1007/s13225-020-00466-2)) |
| Otu10013 | 2 | k__Fungi(100);p__Ascomycota(100);c__Saccharomycetes(100);o__Saccharomycetales(100);f__Metschnikowiaceae(100);g__Metschnikowia(100);g__Metschnikowia_unclassified(100); | Metschnikowia | 13 | Saprotroph | \|Undefined Saprotroph\| | Yeast | NULL | Probable | As nectar-tap_saprotroph (Põlme et al. 2020); Decay sugar-rich substrates (Põlme et al. 2020) | Põlme S, et al. 2020. Fungal Diversity 105:1-16 ((https://doi.org/10.1007/s13225-020-00466-2)) |
| Otu10014 | 2 | k__Fungi(100);p__Ascomycota(100);c__Saccharomycetes(100);o__Saccharomycetales(100);f__Metschnikowiaceae(100);g__Metschnikowia(100);g__Metschnikowia_unclassified(100); | Metschnikowia | 13 | Saprotroph | \|Undefined Saprotroph\| | Yeast | NULL | Probable | As nectar-tap_saprotroph (Põlme et al. 2020); Decay sugar-rich substrates (Põlme et al. 2020) | Põlme S, et al. 2020. Fungal Diversity 105:1-16 ((https://doi.org/10.1007/s13225-020-00466-2)) |
| Otu10015 | 2 | k__Fungi(100);p__Ascomycota(100);c__Saccharomycetes(100);o__Saccharomycetales(100);f__Metschnikowiaceae(100);g__Metschnikowia(100);s__Metschnikowia_pulcherrima(100); | Metschnikowia | 13 | Saprotroph | \|Undefined Saprotroph\| | Yeast | NULL | Probable | As nectar-tap_saprotroph (Põlme et al. 2020); Decay sugar-rich substrates (Põlme et al. 2020) | Põlme S, et al. 2020. Fungal Diversity 105:1-16 ((https://doi.org/10.1007/s13225-020-00466-2)) |
| Otu10016 | 2 | k__Fungi(100);p__Ascomycota(100);c__Saccharomycetes(100);o__Saccharomycetales(100);f__Metschnikowiaceae(100);g__Metschnikowia(100);s__Metschnikowia_pulcherrima(100); | Metschnikowia | 13 | Saprotroph | \|Undefined Saprotroph\| | Yeast | NULL | Probable | As nectar-tap_saprotroph (Põlme et al. 2020); Decay sugar-rich substrates (Põlme et al. 2020) | Põlme S, et al. 2020. Fungal Diversity 105:1-16 ((https://doi.org/10.1007/s13225-020-00466-2)) |
| Otu10017 | 2 | k__Fungi(100);p__Ascomycota(100);c__Saccharomycetes(100);o__Saccharomycetales(100);f__Metschnikowiaceae(100);g__Metschnikowia(100);s__Metschnikowia_pulcherrima(100); | Metschnikowia | 13 | Saprotroph | \|Undefined Saprotroph\| | Yeast | NULL | Probable | As nectar-tap_saprotroph (Põlme et al. 2020); Decay sugar-rich substrates (Põlme et al. 2020) | Põlme S, et al. 2020. Fungal Diversity 105:1-16 ((https://doi.org/10.1007/s13225-020-00466-2)) |
| Otu10019 | 2 | k__Fungi(100);p__Ascomycota(100);c__Saccharomycetes(100);o__Saccharomycetales(100);f__Metschnikowiaceae(100);g__Metschnikowia(100);s__Metschnikowia_pulcherrima(100); | Metschnikowia | 13 | Saprotroph | \|Undefined Saprotroph\| | Yeast | NULL | Probable | As nectar-tap_saprotroph (Põlme et al. 2020); Decay sugar-rich substrates (Põlme et al. 2020) | Põlme S, et al. 2020. Fungal Diversity 105:1-16 ((https://doi.org/10.1007/s13225-020-00466-2)) |
| Otu10020 | 2 | k__Fungi(100);p__Ascomycota(100);c__Saccharomycetes(100);o__Saccharomycetales(100);f__Metschnikowiaceae(100);g__Metschnikowia(100);s__Metschnikowia_pulcherrima(100); | Metschnikowia | 13 | Saprotroph | \|Undefined Saprotroph\| | Yeast | NULL | Probable | As nectar-tap_saprotroph (Põlme et al. 2020); Decay sugar-rich substrates (Põlme et al. 2020) | Põlme S, et al. 2020. Fungal Diversity 105:1-16 ((https://doi.org/10.1007/s13225-020-00466-2)) |
| Otu10023 | 2 | k__Fungi(100);p__Ascomycota(100);c__Saccharomycetes(100);o__Saccharomycetales(100);f__Metschnikowiaceae(100);g__Metschnikowia(100);s__Metschnikowia_pulcherrima(100); | Metschnikowia | 13 | Saprotroph | \|Undefined Saprotroph\| | Yeast | NULL | Probable | As nectar-tap_saprotroph (Põlme et al. 2020); Decay sugar-rich substrates (Põlme et al. 2020) | Põlme S, et al. 2020. Fungal Diversity 105:1-16 ((https://doi.org/10.1007/s13225-020-00466-2)) |
| Otu10024 | 2 | k__Fungi(100);p__Ascomycota(100);c__Saccharomycetes(100);o__Saccharomycetales(100);f__Metschnikowiaceae(100);g__Metschnikowia(100);s__Metschnikowia_pulcherrima(100); | Metschnikowia | 13 | Saprotroph | \|Undefined Saprotroph\| | Yeast | NULL | Probable | As nectar-tap_saprotroph (Põlme et al. 2020); Decay sugar-rich substrates (Põlme et al. 2020) | Põlme S, et al. 2020. Fungal Diversity 105:1-16 ((https://doi.org/10.1007/s13225-020-00466-2)) |
| Otu10026 | 2 | k__Fungi(100);p__Ascomycota(100);c__Saccharomycetes(100);o__Saccharomycetales(100);f__Metschnikowiaceae(100);g__Metschnikowia(100);s__Metschnikowia_pulcherrima(100); | Metschnikowia | 13 | Saprotroph | \|Undefined Saprotroph\| | Yeast | NULL | Probable | As nectar-tap_saprotroph (Põlme et al. 2020); Decay sugar-rich substrates (Põlme et al. 2020) | Põlme S, et al. 2020. Fungal Diversity 105:1-16 ((https://doi.org/10.1007/s13225-020-00466-2)) |
| Otu10028 | 2 | k__Fungi(100);p__Ascomycota(100);c__Saccharomycetes(100);o__Saccharomycetales(100);f__Metschnikowiaceae(100);g__Metschnikowia(100);s__Metschnikowia_pulcherrima(100); | Metschnikowia | 13 | Saprotroph | \|Undefined Saprotroph\| | Yeast | NULL | Probable | As nectar-tap_saprotroph (Põlme et al. 2020); Decay sugar-rich substrates (Põlme et al. 2020) | Põlme S, et al. 2020. Fungal Diversity 105:1-16 ((https://doi.org/10.1007/s13225-020-00466-2)) |
| Otu10030 | 2 | k__Fungi(100);p__Ascomycota(100);c__Saccharomycetes(100);o__Saccharomycetales(100);f__Metschnikowiaceae(100);g__Metschnikowia(100);s__Metschnikowia_pulcherrima(100); | Metschnikowia | 13 | Saprotroph | \|Undefined Saprotroph\| | Yeast | NULL | Probable | As nectar-tap_saprotroph (Põlme et al. 2020); Decay sugar-rich substrates (Põlme et al. 2020) | Põlme S, et al. 2020. Fungal Diversity 105:1-16 ((https://doi.org/10.1007/s13225-020-00466-2)) |
| Otu10031 | 2 | k__Fungi(100);p__Ascomycota(100);c__Saccharomycetes(100);o__Saccharomycetales(100);f__Metschnikowiaceae(100);g__Metschnikowia(100);s__Metschnikowia_pulcherrima(100); | Metschnikowia | 13 | Saprotroph | \|Undefined Saprotroph\| | Yeast | NULL | Probable | As nectar-tap_saprotroph (Põlme et al. 2020); Decay sugar-rich substrates (Põlme et al. 2020) | Põlme S, et al. 2020. Fungal Diversity 105:1-16 ((https://doi.org/10.1007/s13225-020-00466-2)) |
| Otu10033 | 2 | k__Fungi(100);p__Ascomycota(100);c__Saccharomycetes(100);o__Saccharomycetales(100);f__Metschnikowiaceae(100);g__Metschnikowia(100);s__Metschnikowia_pulcherrima(100); | Metschnikowia | 13 | Saprotroph | \|Undefined Saprotroph\| | Yeast | NULL | Probable | As nectar-tap_saprotroph (Põlme et al. 2020); Decay sugar-rich substrates (Põlme et al. 2020) | Põlme S, et al. 2020. Fungal Diversity 105:1-16 ((https://doi.org/10.1007/s13225-020-00466-2)) |
| Otu10035 | 2 | k__Fungi(100);p__Ascomycota(100);c__Saccharomycetes(100);o__Saccharomycetales(100);f__Metschnikowiaceae(100);g__Metschnikowia(100);g__Metschnikowia_unclassified(100); | Metschnikowia | 13 | Saprotroph | \|Undefined Saprotroph\| | Yeast | NULL | Probable | As nectar-tap_saprotroph (Põlme et al. 2020); Decay sugar-rich substrates (Põlme et al. 2020) | Põlme S, et al. 2020. Fungal Diversity 105:1-16 ((https://doi.org/10.1007/s13225-020-00466-2)) |
| Otu10039 | 2 | k__Fungi(100);p__Ascomycota(100);c__Sordariomycetes(100);o__Coniochaetales(100);f__Coniochaetaceae(100);f__Coniochaetaceae_unclassified(100);f__Coniochaetaceae_unclassified(100); | Coniochaetaceae | 9 | Pathotroph-Saprotroph-Symbiotroph | Animal Pathogen-Dung Saprotroph-Endophyte-Lichen Parasite-Plant Pathogen-Undefined Saprotroph | Microfungus | NULL | Possible | Saprobic on wood and bark (especially in dry habitats), coprophilous or isolated from soil and similar substrata (Cannon & Kirk 2007) | Cannon PF, Kirk PM. 2007. Fungal Families of the World. CAB International, Cambridge (ISBN: 978-0851998275); Damm U, et al. 2010. Persoonia 24:60-80 ((http://doi.org/10.3767/003158510X500705)); Irinyi L, et al. 2015. Medical Mycology 53:313-337 ((https://doi.org/10.1093/mmy/myv008)); Lawrey JD, Diederich P. 2016. http://www.lichenicolous.net (accessed 24 May 2017) ((http://www.lichenicolous.net)) |
| Otu10040 | 2 | k__Fungi(100);p__Ascomycota(100);c__Saccharomycetes(100);o__Saccharomycetales(100);f__Saccharomycetales_family_Incertae_sedis(100);g__Candida(100);g__Candida_unclassified(100); | Candida | 13 | Pathotroph-Saprotroph-Symbiotroph | Animal Pathogen-Endophyte-Endosymbiont-Epiphyte-\|Undefined Saprotroph\| | Dimorphic Yeast-Yeast | NULL | Probable | As nectar-tap_saprotroph (Põlme et al. 2020); Decay sugar-rich substrates (Põlme et al. 2020); This asexual genus is polyphyletic and contain species that function very differently depend on what they are. Caution should be taken when interpreting guilds for Candida species. | Manolakaki D, et al. 2010. Virulence. 1: 367-375 ((https://doi.org/10.4161/viru.1.5.12796)); Mekbib SB, et al. 2011. Tropical Plant Pathology 36:233-240 ((https://doi.org/10.1590/S1982-56762011000400004)); Põlme S, et al. 2020. Fungal Diversity 105:1-16 ((https://doi.org/10.1007/s13225-020-00466-2)) |
| Otu10044 | 2 | k__Fungi(100);p__Ascomycota(100);c__Saccharomycetes(100);o__Saccharomycetales(100);f__Metschnikowiaceae(100);g__Metschnikowia(100);s__Metschnikowia_pulcherrima(100); | Metschnikowia | 13 | Saprotroph | \|Undefined Saprotroph\| | Yeast | NULL | Probable | As nectar-tap_saprotroph (Põlme et al. 2020); Decay sugar-rich substrates (Põlme et al. 2020) | Põlme S, et al. 2020. Fungal Diversity 105:1-16 ((https://doi.org/10.1007/s13225-020-00466-2)) |
| Otu10046 | 2 | k__Fungi(100);p__Ascomycota(100);c__Saccharomycetes(100);o__Saccharomycetales(100);f__Metschnikowiaceae(100);g__Metschnikowia(100);s__Metschnikowia_pulcherrima(100); | Metschnikowia | 13 | Saprotroph | \|Undefined Saprotroph\| | Yeast | NULL | Probable | As nectar-tap_saprotroph (Põlme et al. 2020); Decay sugar-rich substrates (Põlme et al. 2020) | Põlme S, et al. 2020. Fungal Diversity 105:1-16 ((https://doi.org/10.1007/s13225-020-00466-2)) |
| Otu10047 | 2 | k__Fungi(100);p__Ascomycota(100);c__Saccharomycetes(100);o__Saccharomycetales(100);f__Metschnikowiaceae(100);g__Metschnikowia(100);g__Metschnikowia_unclassified(100); | Metschnikowia | 13 | Saprotroph | \|Undefined Saprotroph\| | Yeast | NULL | Probable | As nectar-tap_saprotroph (Põlme et al. 2020); Decay sugar-rich substrates (Põlme et al. 2020) | Põlme S, et al. 2020. Fungal Diversity 105:1-16 ((https://doi.org/10.1007/s13225-020-00466-2)) |
| Otu10049 | 2 | k__Fungi(100);p__Ascomycota(100);c__Saccharomycetes(100);o__Saccharomycetales(100);f__Metschnikowiaceae(100);g__Metschnikowia(100);g__Metschnikowia_unclassified(100); | Metschnikowia | 13 | Saprotroph | \|Undefined Saprotroph\| | Yeast | NULL | Probable | As nectar-tap_saprotroph (Põlme et al. 2020); Decay sugar-rich substrates (Põlme et al. 2020) | Põlme S, et al. 2020. Fungal Diversity 105:1-16 ((https://doi.org/10.1007/s13225-020-00466-2)) |
| Otu10051 | 2 | k__Fungi(100);p__Ascomycota(100);c__Saccharomycetes(100);o__Saccharomycetales(100);f__Metschnikowiaceae(100);g__Metschnikowia(100);s__Metschnikowia_pulcherrima(100); | Metschnikowia | 13 | Saprotroph | \|Undefined Saprotroph\| | Yeast | NULL | Probable | As nectar-tap_saprotroph (Põlme et al. 2020); Decay sugar-rich substrates (Põlme et al. 2020) | Põlme S, et al. 2020. Fungal Diversity 105:1-16 ((https://doi.org/10.1007/s13225-020-00466-2)) |
| Otu10052 | 2 | k__Fungi(100);p__Ascomycota(100);c__Saccharomycetes(100);o__Saccharomycetales(100);f__Metschnikowiaceae(100);g__Metschnikowia(100);s__Metschnikowia_pulcherrima(100); | Metschnikowia | 13 | Saprotroph | \|Undefined Saprotroph\| | Yeast | NULL | Probable | As nectar-tap_saprotroph (Põlme et al. 2020); Decay sugar-rich substrates (Põlme et al. 2020) | Põlme S, et al. 2020. Fungal Diversity 105:1-16 ((https://doi.org/10.1007/s13225-020-00466-2)) |
| Otu10053 | 2 | k__Fungi(100);p__Ascomycota(100);c__Saccharomycetes(100);o__Saccharomycetales(100);f__Saccharomycetales_family_Incertae_sedis(100);g__Candida(100);g__Candida_unclassified(100); | Candida | 13 | Pathotroph-Saprotroph-Symbiotroph | Animal Pathogen-Endophyte-Endosymbiont-Epiphyte-\|Undefined Saprotroph\| | Dimorphic Yeast-Yeast | NULL | Probable | As nectar-tap_saprotroph (Põlme et al. 2020); Decay sugar-rich substrates (Põlme et al. 2020); This asexual genus is polyphyletic and contain species that function very differently depend on what they are. Caution should be taken when interpreting guilds for Candida species. | Manolakaki D, et al. 2010. Virulence. 1: 367-375 ((https://doi.org/10.4161/viru.1.5.12796)); Mekbib SB, et al. 2011. Tropical Plant Pathology 36:233-240 ((https://doi.org/10.1590/S1982-56762011000400004)); Põlme S, et al. 2020. Fungal Diversity 105:1-16 ((https://doi.org/10.1007/s13225-020-00466-2)) |
| Otu10054 | 2 | k__Fungi(100);p__Ascomycota(100);c__Saccharomycetes(100);o__Saccharomycetales(100);f__Metschnikowiaceae(100);g__Metschnikowia(100);g__Metschnikowia_unclassified(100); | Metschnikowia | 13 | Saprotroph | \|Undefined Saprotroph\| | Yeast | NULL | Probable | As nectar-tap_saprotroph (Põlme et al. 2020); Decay sugar-rich substrates (Põlme et al. 2020) | Põlme S, et al. 2020. Fungal Diversity 105:1-16 ((https://doi.org/10.1007/s13225-020-00466-2)) |
| Otu10055 | 2 | k__Fungi(100);p__Ascomycota(100);c__Saccharomycetes(100);o__Saccharomycetales(100);f__Metschnikowiaceae(100);g__Metschnikowia(100);s__Metschnikowia_pulcherrima(100); | Metschnikowia | 13 | Saprotroph | \|Undefined Saprotroph\| | Yeast | NULL | Probable | As nectar-tap_saprotroph (Põlme et al. 2020); Decay sugar-rich substrates (Põlme et al. 2020) | Põlme S, et al. 2020. Fungal Diversity 105:1-16 ((https://doi.org/10.1007/s13225-020-00466-2)) |
| Otu10056 | 2 | k__Fungi(100);p__Ascomycota(100);c__Saccharomycetes(100);o__Saccharomycetales(100);f__Metschnikowiaceae(100);g__Metschnikowia(100);s__Metschnikowia_pulcherrima(100); | Metschnikowia | 13 | Saprotroph | \|Undefined Saprotroph\| | Yeast | NULL | Probable | As nectar-tap_saprotroph (Põlme et al. 2020); Decay sugar-rich substrates (Põlme et al. 2020) | Põlme S, et al. 2020. Fungal Diversity 105:1-16 ((https://doi.org/10.1007/s13225-020-00466-2)) |
| Otu10058 | 2 | k__Fungi(100);p__Ascomycota(100);c__Saccharomycetes(100);o__Saccharomycetales(100);f__Metschnikowiaceae(100);g__Metschnikowia(100);s__Metschnikowia_pulcherrima(100); | Metschnikowia | 13 | Saprotroph | \|Undefined Saprotroph\| | Yeast | NULL | Probable | As nectar-tap_saprotroph (Põlme et al. 2020); Decay sugar-rich substrates (Põlme et al. 2020) | Põlme S, et al. 2020. Fungal Diversity 105:1-16 ((https://doi.org/10.1007/s13225-020-00466-2)) |
| Otu10065 | 2 | k__Fungi(100);p__Ascomycota(100);c__Saccharomycetes(100);o__Saccharomycetales(100);f__Metschnikowiaceae(100);g__Metschnikowia(100);s__Metschnikowia_pulcherrima(100); | Metschnikowia | 13 | Saprotroph | \|Undefined Saprotroph\| | Yeast | NULL | Probable | As nectar-tap_saprotroph (Põlme et al. 2020); Decay sugar-rich substrates (Põlme et al. 2020) | Põlme S, et al. 2020. Fungal Diversity 105:1-16 ((https://doi.org/10.1007/s13225-020-00466-2)) |
| Otu10066 | 2 | k__Fungi(100);p__Ascomycota(100);c__Leotiomycetes(100);o__Helotiales(100);f__Helotiales_family_Incertae_sedis(100);g__Infundichalara(100);s__Infundichalara_sp(100); | Infundichalara | 13 | Saprotroph | \|Plant Saprotroph\|-Undefined Saprotroph | Microfungus | NULL | Probable | As litter_saprotroph (Põlme et al. 2020); Decay leaf, fruit, seed (Põlme et al. 2020); Host - Pinus (Põlme et al. 2020) | Tedersoo L, et al. 2014. Science 346:e1256688 ((https://doi.org/10.1126/science.1256688)); Põlme S, et al. 2020. Fungal Diversity 105:1-16 ((https://doi.org/10.1007/s13225-020-00466-2)) |
| Otu10067 | 2 | k__Fungi(100);p__Ascomycota(100);c__Saccharomycetes(100);o__Saccharomycetales(100);f__Metschnikowiaceae(100);g__Metschnikowia(100);s__Metschnikowia_pulcherrima(100); | Metschnikowia | 13 | Saprotroph | \|Undefined Saprotroph\| | Yeast | NULL | Probable | As nectar-tap_saprotroph (Põlme et al. 2020); Decay sugar-rich substrates (Põlme et al. 2020) | Põlme S, et al. 2020. Fungal Diversity 105:1-16 ((https://doi.org/10.1007/s13225-020-00466-2)) |
| Otu10071 | 2 | k__Fungi(100);p__Ascomycota(100);c__Sordariomycetes(100);o__Hypocreales(100);f__Ophiocordycipitaceae(100);f__Ophiocordycipitaceae_unclassified(100);f__Ophiocordycipitaceae_unclassified(100); | Ophiocordycipitaceae | 9 | Pathotroph-Symbiotroph | Endophyte-Insect Pathogen | Microfungus | NULL | Probable | A family of parasitic fungi (Sung et al. 2007) | Sung GH, et al. 2007. Studies in Mycology 57:5-59 ((https://doi.org/10.3114/sim.2007.57.01)); Costa IPMW, et al. 2012. Mycotaxon 119: e26 ((http://mycotaxon.com/resources/checklists/costa_v119_checklist.pdf)) |
| Otu10076 | 2 | k__Fungi(100);p__Ascomycota(100);c__Saccharomycetes(100);o__Saccharomycetales(100);f__Saccharomycetales_family_Incertae_sedis(100);f__Saccharomycetales_family_Incertae_sedis_unclassified(100);f__Saccharomycetales_family_Incertae_sedis_unclassified(100); | Saccharomycetales | 7 | Saprotroph | Undefined Saprotroph | Yeast | NULL | Possible | NULL | Sterkenburg E, et al. 2015. New Phytologist 207:1145-1158 ((https://doi.org/10.1111/nph.13426)) |
| Otu10079 | 2 | k__Fungi(100);p__Ascomycota(100);c__Saccharomycetes(100);o__Saccharomycetales(100);f__Metschnikowiaceae(100);g__Metschnikowia(100);s__Metschnikowia_pulcherrima(100); | Metschnikowia | 13 | Saprotroph | \|Undefined Saprotroph\| | Yeast | NULL | Probable | As nectar-tap_saprotroph (Põlme et al. 2020); Decay sugar-rich substrates (Põlme et al. 2020) | Põlme S, et al. 2020. Fungal Diversity 105:1-16 ((https://doi.org/10.1007/s13225-020-00466-2)) |
| Otu10084 | 2 | k__Fungi(100);p__Ascomycota(100);c__Saccharomycetes(100);o__Saccharomycetales(100);f__Metschnikowiaceae(100);g__Metschnikowia(100);g__Metschnikowia_unclassified(100); | Metschnikowia | 13 | Saprotroph | \|Undefined Saprotroph\| | Yeast | NULL | Probable | As nectar-tap_saprotroph (Põlme et al. 2020); Decay sugar-rich substrates (Põlme et al. 2020) | Põlme S, et al. 2020. Fungal Diversity 105:1-16 ((https://doi.org/10.1007/s13225-020-00466-2)) |
| Otu10086 | 2 | k__Fungi(100);p__Ascomycota(100);c__Saccharomycetes(100);o__Saccharomycetales(100);f__Metschnikowiaceae(100);g__Metschnikowia(100);s__Metschnikowia_pulcherrima(100); | Metschnikowia | 13 | Saprotroph | \|Undefined Saprotroph\| | Yeast | NULL | Probable | As nectar-tap_saprotroph (Põlme et al. 2020); Decay sugar-rich substrates (Põlme et al. 2020) | Põlme S, et al. 2020. Fungal Diversity 105:1-16 ((https://doi.org/10.1007/s13225-020-00466-2)) |
| Otu10087 | 2 | k__Fungi(100);p__Ascomycota(100);c__Saccharomycetes(100);o__Saccharomycetales(100);f__Metschnikowiaceae(100);g__Metschnikowia(100);s__Metschnikowia_pulcherrima(100); | Metschnikowia | 13 | Saprotroph | \|Undefined Saprotroph\| | Yeast | NULL | Probable | As nectar-tap_saprotroph (Põlme et al. 2020); Decay sugar-rich substrates (Põlme et al. 2020) | Põlme S, et al. 2020. Fungal Diversity 105:1-16 ((https://doi.org/10.1007/s13225-020-00466-2)) |
| Otu10088 | 2 | k__Fungi(100);p__Ascomycota(100);c__Saccharomycetes(100);o__Saccharomycetales(100);f__Metschnikowiaceae(100);g__Metschnikowia(100);s__Metschnikowia_pulcherrima(100); | Metschnikowia | 13 | Saprotroph | \|Undefined Saprotroph\| | Yeast | NULL | Probable | As nectar-tap_saprotroph (Põlme et al. 2020); Decay sugar-rich substrates (Põlme et al. 2020) | Põlme S, et al. 2020. Fungal Diversity 105:1-16 ((https://doi.org/10.1007/s13225-020-00466-2)) |
| Otu10090 | 2 | k__Fungi(100);p__Ascomycota(100);c__Saccharomycetes(100);o__Saccharomycetales(100);f__Metschnikowiaceae(100);g__Metschnikowia(100);g__Metschnikowia_unclassified(100); | Metschnikowia | 13 | Saprotroph | \|Undefined Saprotroph\| | Yeast | NULL | Probable | As nectar-tap_saprotroph (Põlme et al. 2020); Decay sugar-rich substrates (Põlme et al. 2020) | Põlme S, et al. 2020. Fungal Diversity 105:1-16 ((https://doi.org/10.1007/s13225-020-00466-2)) |
| Otu10091 | 2 | k__Fungi(100);p__Ascomycota(100);c__Saccharomycetes(100);o__Saccharomycetales(100);o__Saccharomycetales_unclassified(100);o__Saccharomycetales_unclassified(100);o__Saccharomycetales_unclassified(100); | Saccharomycetales | 7 | Saprotroph | Undefined Saprotroph | Yeast | NULL | Possible | NULL | Sterkenburg E, et al. 2015. New Phytologist 207:1145-1158 ((https://doi.org/10.1111/nph.13426)) |
| Otu10092 | 2 | k__Fungi(100);p__Ascomycota(100);c__Saccharomycetes(100);o__Saccharomycetales(100);f__Metschnikowiaceae(100);g__Metschnikowia(100);g__Metschnikowia_unclassified(100); | Metschnikowia | 13 | Saprotroph | \|Undefined Saprotroph\| | Yeast | NULL | Probable | As nectar-tap_saprotroph (Põlme et al. 2020); Decay sugar-rich substrates (Põlme et al. 2020) | Põlme S, et al. 2020. Fungal Diversity 105:1-16 ((https://doi.org/10.1007/s13225-020-00466-2)) |
| Otu10096 | 2 | k__Fungi(100);p__Ascomycota(100);c__Dothideomycetes(100);o__Capnodiales(100);f__Davidiellaceae(100);g__Cladosporium(100);s__Cladosporium_sphaerospermum(100); | Cladosporium | 13 | Pathotroph-Saprotroph-Symbiotroph | Animal Pathogen-Endophyte-Lichen Parasite-Plant Pathogen-\|Plant Saprotroph\|-Wood Saprotroph | Microfungus | NULL | Possible | As foliar_endophyte (Põlme et al. 2020); As litter_saprotroph (Põlme et al. 2020); Decay wood, leaf, fruit, seed, soil, dung, animal material (Põlme et al. 2020) | Seehann G, et al. 1975. List of Fungi in Soft-Rot Tissue Tests. Institute fur Holzbiologie und Holzschutz, Bundesforschungsanstalt fur Forst - und Holzwirtshaft, Hamburg - Lohbrugge, Germany; Lawrey JD, Diederich P. 2016. http://www.lichenicolous.net (accessed 24 May 2017) ((http://www.lichenicolous.net)); Põlme S, et al. 2020. Fungal Diversity 105:1-16 ((https://doi.org/10.1007/s13225-020-00466-2)); The American Phytopathological Society Common Names of Plant Diseases ((http://www.apsnet.org/publications/commonnames/Pages/default.aspx)) |
| Otu10097 | 2 | k__Fungi(100);p__Ascomycota(100);c__Saccharomycetes(100);o__Saccharomycetales(100);f__Metschnikowiaceae(100);g__Metschnikowia(100);s__Metschnikowia_pulcherrima(100); | Metschnikowia | 13 | Saprotroph | \|Undefined Saprotroph\| | Yeast | NULL | Probable | As nectar-tap_saprotroph (Põlme et al. 2020); Decay sugar-rich substrates (Põlme et al. 2020) | Põlme S, et al. 2020. Fungal Diversity 105:1-16 ((https://doi.org/10.1007/s13225-020-00466-2)) |
| Otu10100 | 2 | k__Fungi(100);p__Ascomycota(100);c__Saccharomycetes(100);o__Saccharomycetales(100);f__Metschnikowiaceae(100);g__Metschnikowia(100);s__Metschnikowia_pulcherrima(100); | Metschnikowia | 13 | Saprotroph | \|Undefined Saprotroph\| | Yeast | NULL | Probable | As nectar-tap_saprotroph (Põlme et al. 2020); Decay sugar-rich substrates (Põlme et al. 2020) | Põlme S, et al. 2020. Fungal Diversity 105:1-16 ((https://doi.org/10.1007/s13225-020-00466-2)) |
| Otu10101 | 2 | k__Fungi(100);p__Ascomycota(100);c__Saccharomycetes(100);o__Saccharomycetales(100);f__Metschnikowiaceae(100);g__Metschnikowia(100);g__Metschnikowia_unclassified(100); | Metschnikowia | 13 | Saprotroph | \|Undefined Saprotroph\| | Yeast | NULL | Probable | As nectar-tap_saprotroph (Põlme et al. 2020); Decay sugar-rich substrates (Põlme et al. 2020) | Põlme S, et al. 2020. Fungal Diversity 105:1-16 ((https://doi.org/10.1007/s13225-020-00466-2)) |
| Otu10102 | 2 | k__Fungi(100);p__Ascomycota(100);c__Saccharomycetes(100);o__Saccharomycetales(100);f__Metschnikowiaceae(100);g__Metschnikowia(100);s__Metschnikowia_pulcherrima(100); | Metschnikowia | 13 | Saprotroph | \|Undefined Saprotroph\| | Yeast | NULL | Probable | As nectar-tap_saprotroph (Põlme et al. 2020); Decay sugar-rich substrates (Põlme et al. 2020) | Põlme S, et al. 2020. Fungal Diversity 105:1-16 ((https://doi.org/10.1007/s13225-020-00466-2)) |
| Otu10103 | 2 | k__Fungi(100);p__Ascomycota(100);c__Saccharomycetes(100);o__Saccharomycetales(100);f__Metschnikowiaceae(100);g__Metschnikowia(100);g__Metschnikowia_unclassified(100); | Metschnikowia | 13 | Saprotroph | \|Undefined Saprotroph\| | Yeast | NULL | Probable | As nectar-tap_saprotroph (Põlme et al. 2020); Decay sugar-rich substrates (Põlme et al. 2020) | Põlme S, et al. 2020. Fungal Diversity 105:1-16 ((https://doi.org/10.1007/s13225-020-00466-2)) |
| Otu10107 | 2 | k__Fungi(100);p__Ascomycota(100);c__Saccharomycetes(100);o__Saccharomycetales(100);f__Metschnikowiaceae(100);g__Metschnikowia(100);s__Metschnikowia_pulcherrima(100); | Metschnikowia | 13 | Saprotroph | \|Undefined Saprotroph\| | Yeast | NULL | Probable | As nectar-tap_saprotroph (Põlme et al. 2020); Decay sugar-rich substrates (Põlme et al. 2020) | Põlme S, et al. 2020. Fungal Diversity 105:1-16 ((https://doi.org/10.1007/s13225-020-00466-2)) |
| Otu10112 | 2 | k__Fungi(100);p__Ascomycota(100);c__Saccharomycetes(100);o__Saccharomycetales(100);f__Saccharomycetaceae(100);g__Kazachstania(100);s__Kazachstania_martiniae(100); | Kazachstania | 13 | Saprotroph | \|Undefined Saprotroph\| | Yeast | NULL | Probable | As nectar-tap_saprotroph (Põlme et al. 2020); Decay sugar-rich substrates (Põlme et al. 2020) | Põlme S, et al. 2020. Fungal Diversity 105:1-16 ((https://doi.org/10.1007/s13225-020-00466-2)) |
| Otu10114 | 2 | k__Fungi(100);p__Ascomycota(100);c__Saccharomycetes(100);o__Saccharomycetales(100);f__Metschnikowiaceae(100);g__Metschnikowia(100);s__Metschnikowia_pulcherrima(100); | Metschnikowia | 13 | Saprotroph | \|Undefined Saprotroph\| | Yeast | NULL | Probable | As nectar-tap_saprotroph (Põlme et al. 2020); Decay sugar-rich substrates (Põlme et al. 2020) | Põlme S, et al. 2020. Fungal Diversity 105:1-16 ((https://doi.org/10.1007/s13225-020-00466-2)) |
| Otu10115 | 2 | k__Fungi(100);p__Ascomycota(100);c__Saccharomycetes(100);o__Saccharomycetales(100);f__Metschnikowiaceae(100);g__Metschnikowia(100);s__Metschnikowia_pulcherrima(100); | Metschnikowia | 13 | Saprotroph | \|Undefined Saprotroph\| | Yeast | NULL | Probable | As nectar-tap_saprotroph (Põlme et al. 2020); Decay sugar-rich substrates (Põlme et al. 2020) | Põlme S, et al. 2020. Fungal Diversity 105:1-16 ((https://doi.org/10.1007/s13225-020-00466-2)) |
| Otu10116 | 2 | k__Fungi(100);p__Ascomycota(100);c__Saccharomycetes(100);o__Saccharomycetales(100);f__Saccharomycetales_family_Incertae_sedis(100);g__Candida(100);g__Candida_unclassified(100); | Candida | 13 | Pathotroph-Saprotroph-Symbiotroph | Animal Pathogen-Endophyte-Endosymbiont-Epiphyte-\|Undefined Saprotroph\| | Dimorphic Yeast-Yeast | NULL | Probable | As nectar-tap_saprotroph (Põlme et al. 2020); Decay sugar-rich substrates (Põlme et al. 2020); This asexual genus is polyphyletic and contain species that function very differently depend on what they are. Caution should be taken when interpreting guilds for Candida species. | Manolakaki D, et al. 2010. Virulence. 1: 367-375 ((https://doi.org/10.4161/viru.1.5.12796)); Mekbib SB, et al. 2011. Tropical Plant Pathology 36:233-240 ((https://doi.org/10.1590/S1982-56762011000400004)); Põlme S, et al. 2020. Fungal Diversity 105:1-16 ((https://doi.org/10.1007/s13225-020-00466-2)) |
| Otu10117 | 2 | k__Fungi(100);p__Ascomycota(100);c__Saccharomycetes(100);o__Saccharomycetales(100);f__Metschnikowiaceae(100);g__Metschnikowia(100);g__Metschnikowia_unclassified(100); | Metschnikowia | 13 | Saprotroph | \|Undefined Saprotroph\| | Yeast | NULL | Probable | As nectar-tap_saprotroph (Põlme et al. 2020); Decay sugar-rich substrates (Põlme et al. 2020) | Põlme S, et al. 2020. Fungal Diversity 105:1-16 ((https://doi.org/10.1007/s13225-020-00466-2)) |
| Otu10119 | 2 | k__Fungi(100);p__Ascomycota(100);c__Saccharomycetes(100);o__Saccharomycetales(100);f__Metschnikowiaceae(100);g__Metschnikowia(100);s__Metschnikowia_pulcherrima(100); | Metschnikowia | 13 | Saprotroph | \|Undefined Saprotroph\| | Yeast | NULL | Probable | As nectar-tap_saprotroph (Põlme et al. 2020); Decay sugar-rich substrates (Põlme et al. 2020) | Põlme S, et al. 2020. Fungal Diversity 105:1-16 ((https://doi.org/10.1007/s13225-020-00466-2)) |
| Otu10120 | 2 | k__Fungi(100);p__Ascomycota(100);c__Saccharomycetes(100);o__Saccharomycetales(100);f__Saccharomycetales_family_Incertae_sedis(100);g__Candida(100);g__Candida_unclassified(100); | Candida | 13 | Pathotroph-Saprotroph-Symbiotroph | Animal Pathogen-Endophyte-Endosymbiont-Epiphyte-\|Undefined Saprotroph\| | Dimorphic Yeast-Yeast | NULL | Probable | As nectar-tap_saprotroph (Põlme et al. 2020); Decay sugar-rich substrates (Põlme et al. 2020); This asexual genus is polyphyletic and contain species that function very differently depend on what they are. Caution should be taken when interpreting guilds for Candida species. | Manolakaki D, et al. 2010. Virulence. 1: 367-375 ((https://doi.org/10.4161/viru.1.5.12796)); Mekbib SB, et al. 2011. Tropical Plant Pathology 36:233-240 ((https://doi.org/10.1590/S1982-56762011000400004)); Põlme S, et al. 2020. Fungal Diversity 105:1-16 ((https://doi.org/10.1007/s13225-020-00466-2)) |
| Otu10122 | 2 | k__Fungi(100);p__Ascomycota(100);c__Saccharomycetes(100);o__Saccharomycetales(100);f__Metschnikowiaceae(100);g__Metschnikowia(100);s__Metschnikowia_pulcherrima(100); | Metschnikowia | 13 | Saprotroph | \|Undefined Saprotroph\| | Yeast | NULL | Probable | As nectar-tap_saprotroph (Põlme et al. 2020); Decay sugar-rich substrates (Põlme et al. 2020) | Põlme S, et al. 2020. Fungal Diversity 105:1-16 ((https://doi.org/10.1007/s13225-020-00466-2)) |
| Otu10124 | 2 | k__Fungi(100);p__Ascomycota(100);c__Saccharomycetes(100);o__Saccharomycetales(100);f__Metschnikowiaceae(100);g__Metschnikowia(100);s__Metschnikowia_pulcherrima(100); | Metschnikowia | 13 | Saprotroph | \|Undefined Saprotroph\| | Yeast | NULL | Probable | As nectar-tap_saprotroph (Põlme et al. 2020); Decay sugar-rich substrates (Põlme et al. 2020) | Põlme S, et al. 2020. Fungal Diversity 105:1-16 ((https://doi.org/10.1007/s13225-020-00466-2)) |
| Otu10125 | 2 | k__Fungi(100);p__Ascomycota(100);c__Saccharomycetes(100);o__Saccharomycetales(100);f__Metschnikowiaceae(100);g__Metschnikowia(100);s__Metschnikowia_pulcherrima(100); | Metschnikowia | 13 | Saprotroph | \|Undefined Saprotroph\| | Yeast | NULL | Probable | As nectar-tap_saprotroph (Põlme et al. 2020); Decay sugar-rich substrates (Põlme et al. 2020) | Põlme S, et al. 2020. Fungal Diversity 105:1-16 ((https://doi.org/10.1007/s13225-020-00466-2)) |
| Otu10126 | 2 | k__Fungi(100);p__Ascomycota(100);c__Saccharomycetes(100);o__Saccharomycetales(100);f__Metschnikowiaceae(100);g__Metschnikowia(100);s__Metschnikowia_pulcherrima(100); | Metschnikowia | 13 | Saprotroph | \|Undefined Saprotroph\| | Yeast | NULL | Probable | As nectar-tap_saprotroph (Põlme et al. 2020); Decay sugar-rich substrates (Põlme et al. 2020) | Põlme S, et al. 2020. Fungal Diversity 105:1-16 ((https://doi.org/10.1007/s13225-020-00466-2)) |
| Otu10127 | 2 | k__Fungi(100);p__Ascomycota(100);c__Saccharomycetes(100);o__Saccharomycetales(100);f__Metschnikowiaceae(100);g__Metschnikowia(100);s__Metschnikowia_pulcherrima(100); | Metschnikowia | 13 | Saprotroph | \|Undefined Saprotroph\| | Yeast | NULL | Probable | As nectar-tap_saprotroph (Põlme et al. 2020); Decay sugar-rich substrates (Põlme et al. 2020) | Põlme S, et al. 2020. Fungal Diversity 105:1-16 ((https://doi.org/10.1007/s13225-020-00466-2)) |
| Otu10128 | 2 | k__Fungi(100);p__Ascomycota(100);c__Dothideomycetes(100);o__Capnodiales(100);f__Mycosphaerellaceae(100);g__Zasmidium(100);s__Zasmidium_nocoxi(100); | Zasmidium | 13 | Pathotroph-Saprotroph | \|Plant Pathogen\|-Plant Saprotroph-Undefined Saprotroph | Microfungus | NULL | Probable | As litter_saprotroph (Põlme et al. 2020); Decay leaf, fruit, seed (Põlme et al. 2020) | Tedersoo L, et al. 2014. Science 346:e1256688 ((https://doi.org/10.1126/science.1256688)); Põlme S, et al. 2020. Fungal Diversity 105:1-16 ((https://doi.org/10.1007/s13225-020-00466-2)) |
| Otu10132 | 2 | k__Fungi(100);p__Ascomycota(100);c__Saccharomycetes(100);o__Saccharomycetales(100);f__Metschnikowiaceae(100);g__Metschnikowia(100);s__Metschnikowia_pulcherrima(100); | Metschnikowia | 13 | Saprotroph | \|Undefined Saprotroph\| | Yeast | NULL | Probable | As nectar-tap_saprotroph (Põlme et al. 2020); Decay sugar-rich substrates (Põlme et al. 2020) | Põlme S, et al. 2020. Fungal Diversity 105:1-16 ((https://doi.org/10.1007/s13225-020-00466-2)) |
| Otu10133 | 2 | k__Fungi(100);p__Ascomycota(100);c__Saccharomycetes(100);o__Saccharomycetales(100);f__Metschnikowiaceae(100);g__Metschnikowia(100);s__Metschnikowia_pulcherrima(100); | Metschnikowia | 13 | Saprotroph | \|Undefined Saprotroph\| | Yeast | NULL | Probable | As nectar-tap_saprotroph (Põlme et al. 2020); Decay sugar-rich substrates (Põlme et al. 2020) | Põlme S, et al. 2020. Fungal Diversity 105:1-16 ((https://doi.org/10.1007/s13225-020-00466-2)) |
| Otu10134 | 2 | k__Fungi(100);p__Ascomycota(100);c__Saccharomycetes(100);o__Saccharomycetales(100);f__Metschnikowiaceae(100);g__Metschnikowia(100);s__Metschnikowia_pulcherrima(100); | Metschnikowia | 13 | Saprotroph | \|Undefined Saprotroph\| | Yeast | NULL | Probable | As nectar-tap_saprotroph (Põlme et al. 2020); Decay sugar-rich substrates (Põlme et al. 2020) | Põlme S, et al. 2020. Fungal Diversity 105:1-16 ((https://doi.org/10.1007/s13225-020-00466-2)) |
| Otu10138 | 2 | k__Fungi(100);p__Ascomycota(100);c__Saccharomycetes(100);o__Saccharomycetales(100);f__Metschnikowiaceae(100);g__Metschnikowia(100);s__Metschnikowia_pulcherrima(100); | Metschnikowia | 13 | Saprotroph | \|Undefined Saprotroph\| | Yeast | NULL | Probable | As nectar-tap_saprotroph (Põlme et al. 2020); Decay sugar-rich substrates (Põlme et al. 2020) | Põlme S, et al. 2020. Fungal Diversity 105:1-16 ((https://doi.org/10.1007/s13225-020-00466-2)) |
| Otu10139 | 2 | k__Fungi(100);p__Ascomycota(100);c__Saccharomycetes(100);o__Saccharomycetales(100);f__Metschnikowiaceae(100);g__Metschnikowia(100);g__Metschnikowia_unclassified(100); | Metschnikowia | 13 | Saprotroph | \|Undefined Saprotroph\| | Yeast | NULL | Probable | As nectar-tap_saprotroph (Põlme et al. 2020); Decay sugar-rich substrates (Põlme et al. 2020) | Põlme S, et al. 2020. Fungal Diversity 105:1-16 ((https://doi.org/10.1007/s13225-020-00466-2)) |
| Otu10140 | 2 | k__Fungi(100);p__Ascomycota(100);c__Saccharomycetes(100);o__Saccharomycetales(100);f__Metschnikowiaceae(100);g__Metschnikowia(100);s__Metschnikowia_pulcherrima(100); | Metschnikowia | 13 | Saprotroph | \|Undefined Saprotroph\| | Yeast | NULL | Probable | As nectar-tap_saprotroph (Põlme et al. 2020); Decay sugar-rich substrates (Põlme et al. 2020) | Põlme S, et al. 2020. Fungal Diversity 105:1-16 ((https://doi.org/10.1007/s13225-020-00466-2)) |
| Otu10143 | 2 | k__Fungi(100);p__Ascomycota(100);c__Saccharomycetes(100);o__Saccharomycetales(100);f__Metschnikowiaceae(100);g__Metschnikowia(100);s__Metschnikowia_pulcherrima(100); | Metschnikowia | 13 | Saprotroph | \|Undefined Saprotroph\| | Yeast | NULL | Probable | As nectar-tap_saprotroph (Põlme et al. 2020); Decay sugar-rich substrates (Põlme et al. 2020) | Põlme S, et al. 2020. Fungal Diversity 105:1-16 ((https://doi.org/10.1007/s13225-020-00466-2)) |
| Otu10144 | 2 | k__Fungi(100);p__Ascomycota(100);c__Saccharomycetes(100);o__Saccharomycetales(100);f__Metschnikowiaceae(100);g__Metschnikowia(100);s__Metschnikowia_pulcherrima(100); | Metschnikowia | 13 | Saprotroph | \|Undefined Saprotroph\| | Yeast | NULL | Probable | As nectar-tap_saprotroph (Põlme et al. 2020); Decay sugar-rich substrates (Põlme et al. 2020) | Põlme S, et al. 2020. Fungal Diversity 105:1-16 ((https://doi.org/10.1007/s13225-020-00466-2)) |
| Otu10146 | 2 | k__Fungi(100);p__Ascomycota(100);c__Saccharomycetes(100);o__Saccharomycetales(100);f__Metschnikowiaceae(100);g__Metschnikowia(100);s__Metschnikowia_pulcherrima(100); | Metschnikowia | 13 | Saprotroph | \|Undefined Saprotroph\| | Yeast | NULL | Probable | As nectar-tap_saprotroph (Põlme et al. 2020); Decay sugar-rich substrates (Põlme et al. 2020) | Põlme S, et al. 2020. Fungal Diversity 105:1-16 ((https://doi.org/10.1007/s13225-020-00466-2)) |
| Otu10147 | 2 | k__Fungi(100);p__Ascomycota(100);c__Sordariomycetes(100);o__Hypocreales(100);f__Nectriaceae(100);f__Nectriaceae_unclassified(100);f__Nectriaceae_unclassified(100); | Nectriaceae | 9 | Pathotroph-Saprotroph-Symbiotroph | Animal Pathogen-Endophyte-Fungal Parasite-Lichen Parasite-Plant Pathogen-Wood Saprotroph | Microfungus | NULL | Possible | Fusarium species are amongst the economically significant plant pathogens (Cannon & Kirk 2007) | Cannon PF, Kirk PM. 2007. Fungal Families of the World. CAB International, Cambridge (ISBN: 978-0851998275); Hirooka Y, et al. 2012. Studies in Mycology 711-210 ((https://doi.org/10.3114/sim0001)); Tedersoo L, et al. 2014. Science 346:e1256688 ((https://doi.org/10.1126/science.1256688)); The American Phytopathological Society Common Names of Plant Diseases ((http://www.apsnet.org/publications/commonnames/Pages/default.aspx)); https://www.cdc.gov ((https://www.cdc.gov)) |
| Otu10148 | 2 | k__Fungi(100);p__Ascomycota(100);c__Saccharomycetes(100);o__Saccharomycetales(100);f__Metschnikowiaceae(100);g__Metschnikowia(100);g__Metschnikowia_unclassified(100); | Metschnikowia | 13 | Saprotroph | \|Undefined Saprotroph\| | Yeast | NULL | Probable | As nectar-tap_saprotroph (Põlme et al. 2020); Decay sugar-rich substrates (Põlme et al. 2020) | Põlme S, et al. 2020. Fungal Diversity 105:1-16 ((https://doi.org/10.1007/s13225-020-00466-2)) |
| Otu10149 | 2 | k__Fungi(100);p__Ascomycota(100);c__Dothideomycetes(100);o__Capnodiales(100);f__Davidiellaceae(100);g__Cladosporium(100);g__Cladosporium_unclassified(100); | Cladosporium | 13 | Pathotroph-Saprotroph-Symbiotroph | Animal Pathogen-Endophyte-Lichen Parasite-Plant Pathogen-\|Plant Saprotroph\|-Wood Saprotroph | Microfungus | NULL | Possible | As foliar_endophyte (Põlme et al. 2020); As litter_saprotroph (Põlme et al. 2020); Decay wood, leaf, fruit, seed, soil, dung, animal material (Põlme et al. 2020) | Seehann G, et al. 1975. List of Fungi in Soft-Rot Tissue Tests. Institute fur Holzbiologie und Holzschutz, Bundesforschungsanstalt fur Forst - und Holzwirtshaft, Hamburg - Lohbrugge, Germany; Lawrey JD, Diederich P. 2016. http://www.lichenicolous.net (accessed 24 May 2017) ((http://www.lichenicolous.net)); Põlme S, et al. 2020. Fungal Diversity 105:1-16 ((https://doi.org/10.1007/s13225-020-00466-2)); The American Phytopathological Society Common Names of Plant Diseases ((http://www.apsnet.org/publications/commonnames/Pages/default.aspx)) |
| Otu10152 | 2 | k__Fungi(100);p__Ascomycota(100);c__Saccharomycetes(100);o__Saccharomycetales(100);f__Metschnikowiaceae(100);g__Metschnikowia(100);g__Metschnikowia_unclassified(100); | Metschnikowia | 13 | Saprotroph | \|Undefined Saprotroph\| | Yeast | NULL | Probable | As nectar-tap_saprotroph (Põlme et al. 2020); Decay sugar-rich substrates (Põlme et al. 2020) | Põlme S, et al. 2020. Fungal Diversity 105:1-16 ((https://doi.org/10.1007/s13225-020-00466-2)) |
| Otu10159 | 2 | k__Fungi(100);p__Ascomycota(100);c__Saccharomycetes(100);o__Saccharomycetales(100);f__Metschnikowiaceae(100);g__Metschnikowia(100);g__Metschnikowia_unclassified(100); | Metschnikowia | 13 | Saprotroph | \|Undefined Saprotroph\| | Yeast | NULL | Probable | As nectar-tap_saprotroph (Põlme et al. 2020); Decay sugar-rich substrates (Põlme et al. 2020) | Põlme S, et al. 2020. Fungal Diversity 105:1-16 ((https://doi.org/10.1007/s13225-020-00466-2)) |
| Otu10160 | 2 | k__Fungi(100);p__Ascomycota(100);c__Saccharomycetes(100);o__Saccharomycetales(100);f__Metschnikowiaceae(100);g__Metschnikowia(100);s__Metschnikowia_pulcherrima(100); | Metschnikowia | 13 | Saprotroph | \|Undefined Saprotroph\| | Yeast | NULL | Probable | As nectar-tap_saprotroph (Põlme et al. 2020); Decay sugar-rich substrates (Põlme et al. 2020) | Põlme S, et al. 2020. Fungal Diversity 105:1-16 ((https://doi.org/10.1007/s13225-020-00466-2)) |
| Otu10161 | 2 | k__Fungi(100);p__Ascomycota(100);c__Saccharomycetes(100);o__Saccharomycetales(100);f__Metschnikowiaceae(100);g__Metschnikowia(100);g__Metschnikowia_unclassified(100); | Metschnikowia | 13 | Saprotroph | \|Undefined Saprotroph\| | Yeast | NULL | Probable | As nectar-tap_saprotroph (Põlme et al. 2020); Decay sugar-rich substrates (Põlme et al. 2020) | Põlme S, et al. 2020. Fungal Diversity 105:1-16 ((https://doi.org/10.1007/s13225-020-00466-2)) |
| Otu10162 | 2 | k__Fungi(100);p__Ascomycota(100);c__Saccharomycetes(100);o__Saccharomycetales(100);f__Metschnikowiaceae(100);g__Metschnikowia(100);s__Metschnikowia_pulcherrima(100); | Metschnikowia | 13 | Saprotroph | \|Undefined Saprotroph\| | Yeast | NULL | Probable | As nectar-tap_saprotroph (Põlme et al. 2020); Decay sugar-rich substrates (Põlme et al. 2020) | Põlme S, et al. 2020. Fungal Diversity 105:1-16 ((https://doi.org/10.1007/s13225-020-00466-2)) |
| Otu10163 | 2 | k__Fungi(100);p__Ascomycota(100);c__Saccharomycetes(100);o__Saccharomycetales(100);f__Metschnikowiaceae(100);g__Metschnikowia(100);g__Metschnikowia_unclassified(100); | Metschnikowia | 13 | Saprotroph | \|Undefined Saprotroph\| | Yeast | NULL | Probable | As nectar-tap_saprotroph (Põlme et al. 2020); Decay sugar-rich substrates (Põlme et al. 2020) | Põlme S, et al. 2020. Fungal Diversity 105:1-16 ((https://doi.org/10.1007/s13225-020-00466-2)) |
| Otu10164 | 2 | k__Fungi(100);p__Ascomycota(100);c__Saccharomycetes(100);o__Saccharomycetales(100);f__Metschnikowiaceae(100);g__Metschnikowia(100);s__Metschnikowia_pulcherrima(100); | Metschnikowia | 13 | Saprotroph | \|Undefined Saprotroph\| | Yeast | NULL | Probable | As nectar-tap_saprotroph (Põlme et al. 2020); Decay sugar-rich substrates (Põlme et al. 2020) | Põlme S, et al. 2020. Fungal Diversity 105:1-16 ((https://doi.org/10.1007/s13225-020-00466-2)) |
| Otu10167 | 2 | k__Fungi(100);p__Ascomycota(100);c__Saccharomycetes(100);o__Saccharomycetales(100);f__Metschnikowiaceae(100);g__Metschnikowia(100);s__Metschnikowia_pulcherrima(100); | Metschnikowia | 13 | Saprotroph | \|Undefined Saprotroph\| | Yeast | NULL | Probable | As nectar-tap_saprotroph (Põlme et al. 2020); Decay sugar-rich substrates (Põlme et al. 2020) | Põlme S, et al. 2020. Fungal Diversity 105:1-16 ((https://doi.org/10.1007/s13225-020-00466-2)) |
| Otu10170 | 2 | k__Fungi(100);p__Ascomycota(100);c__Saccharomycetes(100);o__Saccharomycetales(100);f__Metschnikowiaceae(100);g__Metschnikowia(100);s__Metschnikowia_pulcherrima(100); | Metschnikowia | 13 | Saprotroph | \|Undefined Saprotroph\| | Yeast | NULL | Probable | As nectar-tap_saprotroph (Põlme et al. 2020); Decay sugar-rich substrates (Põlme et al. 2020) | Põlme S, et al. 2020. Fungal Diversity 105:1-16 ((https://doi.org/10.1007/s13225-020-00466-2)) |
| Otu10172 | 2 | k__Fungi(100);p__Ascomycota(100);c__Saccharomycetes(100);o__Saccharomycetales(100);f__Metschnikowiaceae(100);g__Metschnikowia(100);s__Metschnikowia_pulcherrima(100); | Metschnikowia | 13 | Saprotroph | \|Undefined Saprotroph\| | Yeast | NULL | Probable | As nectar-tap_saprotroph (Põlme et al. 2020); Decay sugar-rich substrates (Põlme et al. 2020) | Põlme S, et al. 2020. Fungal Diversity 105:1-16 ((https://doi.org/10.1007/s13225-020-00466-2)) |
| Otu10174 | 2 | k__Fungi(100);p__Ascomycota(100);c__Leotiomycetes(100);o__Helotiales(100);f__Hyaloscyphaceae(100);g__Lachnum(100);g__Lachnum_unclassified(100); | Lachnum | 13 | Saprotroph-Symbiotroph | Endophyte-\|Plant Saprotroph\|-Undefined Saprotroph-Wood Saprotroph | Microfungus | NULL | Probable | As litter_saprotroph (Põlme et al. 2020); As root_endophyte (Põlme et al. 2020); Decay wood, leaf, fruit, seed (Põlme et al. 2020) | Tedersoo L, et al. 2014. Science 346:e1256688 ((https://doi.org/10.1126/science.1256688)); Põlme S, et al. 2020. Fungal Diversity 105:1-16 ((https://doi.org/10.1007/s13225-020-00466-2)) |
| Otu10175 | 2 | k__Fungi(100);p__Ascomycota(100);c__Saccharomycetes(100);o__Saccharomycetales(100);f__Metschnikowiaceae(100);g__Metschnikowia(100);g__Metschnikowia_unclassified(100); | Metschnikowia | 13 | Saprotroph | \|Undefined Saprotroph\| | Yeast | NULL | Probable | As nectar-tap_saprotroph (Põlme et al. 2020); Decay sugar-rich substrates (Põlme et al. 2020) | Põlme S, et al. 2020. Fungal Diversity 105:1-16 ((https://doi.org/10.1007/s13225-020-00466-2)) |
| Otu10176 | 2 | k__Fungi(100);p__Ascomycota(100);c__Saccharomycetes(100);o__Saccharomycetales(100);f__Metschnikowiaceae(100);g__Metschnikowia(100);s__Metschnikowia_pulcherrima(100); | Metschnikowia | 13 | Saprotroph | \|Undefined Saprotroph\| | Yeast | NULL | Probable | As nectar-tap_saprotroph (Põlme et al. 2020); Decay sugar-rich substrates (Põlme et al. 2020) | Põlme S, et al. 2020. Fungal Diversity 105:1-16 ((https://doi.org/10.1007/s13225-020-00466-2)) |
| Otu10177 | 2 | k__Fungi(100);p__Ascomycota(100);c__Saccharomycetes(100);o__Saccharomycetales(100);f__Metschnikowiaceae(100);g__Metschnikowia(100);g__Metschnikowia_unclassified(100); | Metschnikowia | 13 | Saprotroph | \|Undefined Saprotroph\| | Yeast | NULL | Probable | As nectar-tap_saprotroph (Põlme et al. 2020); Decay sugar-rich substrates (Põlme et al. 2020) | Põlme S, et al. 2020. Fungal Diversity 105:1-16 ((https://doi.org/10.1007/s13225-020-00466-2)) |
| Otu10178 | 2 | k__Fungi(100);p__Ascomycota(100);c__Saccharomycetes(100);o__Saccharomycetales(100);f__Metschnikowiaceae(100);g__Metschnikowia(100);s__Metschnikowia_pulcherrima(100); | Metschnikowia | 13 | Saprotroph | \|Undefined Saprotroph\| | Yeast | NULL | Probable | As nectar-tap_saprotroph (Põlme et al. 2020); Decay sugar-rich substrates (Põlme et al. 2020) | Põlme S, et al. 2020. Fungal Diversity 105:1-16 ((https://doi.org/10.1007/s13225-020-00466-2)) |
| Otu10179 | 2 | k__Fungi(100);p__Ascomycota(100);c__Saccharomycetes(100);o__Saccharomycetales(100);f__Metschnikowiaceae(100);g__Metschnikowia(100);g__Metschnikowia_unclassified(100); | Metschnikowia | 13 | Saprotroph | \|Undefined Saprotroph\| | Yeast | NULL | Probable | As nectar-tap_saprotroph (Põlme et al. 2020); Decay sugar-rich substrates (Põlme et al. 2020) | Põlme S, et al. 2020. Fungal Diversity 105:1-16 ((https://doi.org/10.1007/s13225-020-00466-2)) |
| Otu10181 | 2 | k__Fungi(100);p__Ascomycota(100);c__Saccharomycetes(100);o__Saccharomycetales(100);f__Metschnikowiaceae(100);g__Metschnikowia(100);g__Metschnikowia_unclassified(100); | Metschnikowia | 13 | Saprotroph | \|Undefined Saprotroph\| | Yeast | NULL | Probable | As nectar-tap_saprotroph (Põlme et al. 2020); Decay sugar-rich substrates (Põlme et al. 2020) | Põlme S, et al. 2020. Fungal Diversity 105:1-16 ((https://doi.org/10.1007/s13225-020-00466-2)) |
| Otu10183 | 2 | k__Fungi(100);p__Ascomycota(100);c__Dothideomycetes(100);o__Dothideomycetes_order_Incertae_sedis(100);f__Myxotrichaceae(100);g__Oidiodendron(100);s__Oidiodendron_maius(100); | Oidiodendron | 13 | Pathotroph-Saprotroph-Symbiotroph | Endophyte-Ericoid Mycorrhizal-\|Undefined Saprotroph\| | Microfungus | NULL | Probable | As root_endophyte (Põlme et al. 2020); As soil_saprotroph (Põlme et al. 2020); Dark Septate Endophyte; Decay soil (Põlme et al. 2020) | Newsham KK. 2011. New Phytologist 190:783-793 ((https://doi.org/10.1111/j.1469-8137.2010.03611.x)) (as dark septate root endophytes); Põlme S, et al. 2020. Fungal Diversity 105:1-16 ((https://doi.org/10.1007/s13225-020-00466-2)) |
| Otu10184 | 2 | k__Fungi(100);p__Ascomycota(100);c__Saccharomycetes(100);o__Saccharomycetales(100);f__Metschnikowiaceae(100);g__Metschnikowia(100);s__Metschnikowia_pulcherrima(100); | Metschnikowia | 13 | Saprotroph | \|Undefined Saprotroph\| | Yeast | NULL | Probable | As nectar-tap_saprotroph (Põlme et al. 2020); Decay sugar-rich substrates (Põlme et al. 2020) | Põlme S, et al. 2020. Fungal Diversity 105:1-16 ((https://doi.org/10.1007/s13225-020-00466-2)) |
| Otu10185 | 2 | k__Fungi(100);p__Ascomycota(100);c__Saccharomycetes(100);o__Saccharomycetales(100);f__Metschnikowiaceae(100);g__Metschnikowia(100);s__Metschnikowia_pulcherrima(100); | Metschnikowia | 13 | Saprotroph | \|Undefined Saprotroph\| | Yeast | NULL | Probable | As nectar-tap_saprotroph (Põlme et al. 2020); Decay sugar-rich substrates (Põlme et al. 2020) | Põlme S, et al. 2020. Fungal Diversity 105:1-16 ((https://doi.org/10.1007/s13225-020-00466-2)) |
| Otu10186 | 2 | k__Fungi(100);p__Ascomycota(100);c__Leotiomycetes(100);o__Leotiomycetes_order_Incertae_sedis(100);f__Leotiomycetes_family_Incertae_sedis(100);g__Geomyces(100);g__Geomyces_unclassified(100); | Geomyces | 13 | Saprotroph | \|Undefined Saprotroph\| | Microfungus | NULL | Probable | As soil_saprotroph (Põlme et al. 2020); Decay soil (Põlme et al. 2020); Nose Bat Syndrome causing Geomyces destructans, are now placed in the genus Pseudogymnoascus (see Minnis & Lindner 2013); This genus is now restricted to one species Geomyces auratus; however, it is important to note that the majority of species that once comprised this genus, including the White | Minnis AM, Lindner DL. 2013. Fungal Biology 117:638-649 ((https://doi.org/10.1016/j.funbio.2013.07.001)); Tedersoo L, et al. 2014. Science 346:e1256688 ((https://doi.org/10.1126/science.1256688)); Põlme S, et al. 2020. Fungal Diversity 105:1-16 ((https://doi.org/10.1007/s13225-020-00466-2)) |
| Otu10187 | 2 | k__Fungi(100);p__Ascomycota(100);c__Saccharomycetes(100);o__Saccharomycetales(100);f__Metschnikowiaceae(100);g__Metschnikowia(100);g__Metschnikowia_unclassified(100); | Metschnikowia | 13 | Saprotroph | \|Undefined Saprotroph\| | Yeast | NULL | Probable | As nectar-tap_saprotroph (Põlme et al. 2020); Decay sugar-rich substrates (Põlme et al. 2020) | Põlme S, et al. 2020. Fungal Diversity 105:1-16 ((https://doi.org/10.1007/s13225-020-00466-2)) |
| Otu10188 | 2 | k__Fungi(100);p__Ascomycota(100);c__Saccharomycetes(100);o__Saccharomycetales(100);f__Metschnikowiaceae(100);g__Metschnikowia(100);s__Metschnikowia_pulcherrima(100); | Metschnikowia | 13 | Saprotroph | \|Undefined Saprotroph\| | Yeast | NULL | Probable | As nectar-tap_saprotroph (Põlme et al. 2020); Decay sugar-rich substrates (Põlme et al. 2020) | Põlme S, et al. 2020. Fungal Diversity 105:1-16 ((https://doi.org/10.1007/s13225-020-00466-2)) |
| Otu10190 | 2 | k__Fungi(100);p__Ascomycota(100);c__Saccharomycetes(100);o__Saccharomycetales(100);f__Metschnikowiaceae(100);g__Metschnikowia(100);s__Metschnikowia_pulcherrima(100); | Metschnikowia | 13 | Saprotroph | \|Undefined Saprotroph\| | Yeast | NULL | Probable | As nectar-tap_saprotroph (Põlme et al. 2020); Decay sugar-rich substrates (Põlme et al. 2020) | Põlme S, et al. 2020. Fungal Diversity 105:1-16 ((https://doi.org/10.1007/s13225-020-00466-2)) |
| Otu10193 | 2 | k__Fungi(100);p__Ascomycota(100);c__Saccharomycetes(100);o__Saccharomycetales(100);f__Metschnikowiaceae(100);g__Metschnikowia(100);s__Metschnikowia_pulcherrima(100); | Metschnikowia | 13 | Saprotroph | \|Undefined Saprotroph\| | Yeast | NULL | Probable | As nectar-tap_saprotroph (Põlme et al. 2020); Decay sugar-rich substrates (Põlme et al. 2020) | Põlme S, et al. 2020. Fungal Diversity 105:1-16 ((https://doi.org/10.1007/s13225-020-00466-2)) |
| Otu10195 | 2 | k__Fungi(100);p__Ascomycota(100);c__Leotiomycetes(100);o__Helotiales(100);f__Helotiales_family_Incertae_sedis(100);g__Glarea(100);s__Glarea_lozoyensis(100); | Glarea | 13 | Saprotroph | \|Undefined Saprotroph\| | Microfungus | NULL | Probable | As soil_saprotroph (Põlme et al. 2020); Decay soil (Põlme et al. 2020) | Tedersoo L, et al. 2014. Science 346:e1256688 ((https://doi.org/10.1126/science.1256688)); Põlme S, et al. 2020. Fungal Diversity 105:1-16 ((https://doi.org/10.1007/s13225-020-00466-2)) |
| Otu10196 | 2 | k__Fungi(100);p__Ascomycota(100);c__Saccharomycetes(100);o__Saccharomycetales(100);f__Metschnikowiaceae(100);g__Metschnikowia(100);s__Metschnikowia_pulcherrima(100); | Metschnikowia | 13 | Saprotroph | \|Undefined Saprotroph\| | Yeast | NULL | Probable | As nectar-tap_saprotroph (Põlme et al. 2020); Decay sugar-rich substrates (Põlme et al. 2020) | Põlme S, et al. 2020. Fungal Diversity 105:1-16 ((https://doi.org/10.1007/s13225-020-00466-2)) |
| Otu10198 | 2 | k__Fungi(100);p__Ascomycota(100);c__Saccharomycetes(100);o__Saccharomycetales(100);f__Saccharomycetales_family_Incertae_sedis(100);f__Saccharomycetales_family_Incertae_sedis_unclassified(100);f__Saccharomycetales_family_Incertae_sedis_unclassified(100); | Saccharomycetales | 7 | Saprotroph | Undefined Saprotroph | Yeast | NULL | Possible | NULL | Sterkenburg E, et al. 2015. New Phytologist 207:1145-1158 ((https://doi.org/10.1111/nph.13426)) |
| Otu10200 | 2 | k__Fungi(100);p__Ascomycota(100);c__Saccharomycetes(100);o__Saccharomycetales(100);f__Metschnikowiaceae(100);g__Metschnikowia(100);s__Metschnikowia_pulcherrima(100); | Metschnikowia | 13 | Saprotroph | \|Undefined Saprotroph\| | Yeast | NULL | Probable | As nectar-tap_saprotroph (Põlme et al. 2020); Decay sugar-rich substrates (Põlme et al. 2020) | Põlme S, et al. 2020. Fungal Diversity 105:1-16 ((https://doi.org/10.1007/s13225-020-00466-2)) |
| Otu10204 | 2 | k__Fungi(100);p__Ascomycota(100);c__Saccharomycetes(100);o__Saccharomycetales(100);f__Metschnikowiaceae(100);g__Metschnikowia(100);s__Metschnikowia_pulcherrima(100); | Metschnikowia | 13 | Saprotroph | \|Undefined Saprotroph\| | Yeast | NULL | Probable | As nectar-tap_saprotroph (Põlme et al. 2020); Decay sugar-rich substrates (Põlme et al. 2020) | Põlme S, et al. 2020. Fungal Diversity 105:1-16 ((https://doi.org/10.1007/s13225-020-00466-2)) |
| Otu10205 | 2 | k__Fungi(100);p__Ascomycota(100);c__Saccharomycetes(100);o__Saccharomycetales(100);f__Metschnikowiaceae(100);g__Metschnikowia(100);s__Metschnikowia_pulcherrima(100); | Metschnikowia | 13 | Saprotroph | \|Undefined Saprotroph\| | Yeast | NULL | Probable | As nectar-tap_saprotroph (Põlme et al. 2020); Decay sugar-rich substrates (Põlme et al. 2020) | Põlme S, et al. 2020. Fungal Diversity 105:1-16 ((https://doi.org/10.1007/s13225-020-00466-2)) |
| Otu10207 | 2 | k__Fungi(100);p__Ascomycota(100);c__Saccharomycetes(100);o__Saccharomycetales(100);f__Metschnikowiaceae(100);g__Metschnikowia(100);g__Metschnikowia_unclassified(100); | Metschnikowia | 13 | Saprotroph | \|Undefined Saprotroph\| | Yeast | NULL | Probable | As nectar-tap_saprotroph (Põlme et al. 2020); Decay sugar-rich substrates (Põlme et al. 2020) | Põlme S, et al. 2020. Fungal Diversity 105:1-16 ((https://doi.org/10.1007/s13225-020-00466-2)) |
| Otu10209 | 2 | k__Fungi(100);p__Ascomycota(100);c__Saccharomycetes(100);o__Saccharomycetales(100);f__Metschnikowiaceae(100);g__Metschnikowia(100);g__Metschnikowia_unclassified(100); | Metschnikowia | 13 | Saprotroph | \|Undefined Saprotroph\| | Yeast | NULL | Probable | As nectar-tap_saprotroph (Põlme et al. 2020); Decay sugar-rich substrates (Põlme et al. 2020) | Põlme S, et al. 2020. Fungal Diversity 105:1-16 ((https://doi.org/10.1007/s13225-020-00466-2)) |
| Otu10214 | 2 | k__Fungi(100);p__Ascomycota(100);c__Saccharomycetes(100);o__Saccharomycetales(100);f__Metschnikowiaceae(100);g__Metschnikowia(100);s__Metschnikowia_pulcherrima(100); | Metschnikowia | 13 | Saprotroph | \|Undefined Saprotroph\| | Yeast | NULL | Probable | As nectar-tap_saprotroph (Põlme et al. 2020); Decay sugar-rich substrates (Põlme et al. 2020) | Põlme S, et al. 2020. Fungal Diversity 105:1-16 ((https://doi.org/10.1007/s13225-020-00466-2)) |
| Otu10215 | 2 | k__Fungi(100);p__Ascomycota(100);c__Saccharomycetes(100);o__Saccharomycetales(100);f__Metschnikowiaceae(100);g__Metschnikowia(100);s__Metschnikowia_pulcherrima(100); | Metschnikowia | 13 | Saprotroph | \|Undefined Saprotroph\| | Yeast | NULL | Probable | As nectar-tap_saprotroph (Põlme et al. 2020); Decay sugar-rich substrates (Põlme et al. 2020) | Põlme S, et al. 2020. Fungal Diversity 105:1-16 ((https://doi.org/10.1007/s13225-020-00466-2)) |
| Otu10216 | 2 | k__Fungi(100);p__Ascomycota(100);c__Dothideomycetes(100);o__Capnodiales(100);f__Teratosphaeriaceae(100);f__Teratosphaeriaceae_unclassified(100);f__Teratosphaeriaceae_unclassified(100); | Teratosphaeriaceae | 9 | Pathotroph-Saprotroph | Animal Pathogen-Plant Pathogen-Undefined Saprotroph | NULL | NULL | Probable | Includes numerous saprobic, extremophilic, human opportunistic, and plant pathogenic fungi (Quaedvlieg et al. 2014) | Quaedvlieg W, et al. 2014. Persoonia 3:31-40 ((https://doi.org/10.3767/003158514X681981)) |
| Otu10218 | 2 | k__Fungi(100);p__Ascomycota(100);c__Saccharomycetes(100);o__Saccharomycetales(100);f__Metschnikowiaceae(100);g__Metschnikowia(100);s__Metschnikowia_pulcherrima(100); | Metschnikowia | 13 | Saprotroph | \|Undefined Saprotroph\| | Yeast | NULL | Probable | As nectar-tap_saprotroph (Põlme et al. 2020); Decay sugar-rich substrates (Põlme et al. 2020) | Põlme S, et al. 2020. Fungal Diversity 105:1-16 ((https://doi.org/10.1007/s13225-020-00466-2)) |
| Otu10220 | 2 | k__Fungi(100);p__Ascomycota(100);c__Saccharomycetes(100);o__Saccharomycetales(100);f__Metschnikowiaceae(100);g__Metschnikowia(100);g__Metschnikowia_unclassified(100); | Metschnikowia | 13 | Saprotroph | \|Undefined Saprotroph\| | Yeast | NULL | Probable | As nectar-tap_saprotroph (Põlme et al. 2020); Decay sugar-rich substrates (Põlme et al. 2020) | Põlme S, et al. 2020. Fungal Diversity 105:1-16 ((https://doi.org/10.1007/s13225-020-00466-2)) |
| Otu10223 | 2 | k__Fungi(100);p__Ascomycota(100);c__Saccharomycetes(100);o__Saccharomycetales(100);f__Metschnikowiaceae(100);g__Metschnikowia(100);s__Metschnikowia_pulcherrima(100); | Metschnikowia | 13 | Saprotroph | \|Undefined Saprotroph\| | Yeast | NULL | Probable | As nectar-tap_saprotroph (Põlme et al. 2020); Decay sugar-rich substrates (Põlme et al. 2020) | Põlme S, et al. 2020. Fungal Diversity 105:1-16 ((https://doi.org/10.1007/s13225-020-00466-2)) |
| Otu10224 | 2 | k__Fungi(100);p__Ascomycota(100);c__Saccharomycetes(100);o__Saccharomycetales(100);f__Metschnikowiaceae(100);g__Metschnikowia(100);g__Metschnikowia_unclassified(100); | Metschnikowia | 13 | Saprotroph | \|Undefined Saprotroph\| | Yeast | NULL | Probable | As nectar-tap_saprotroph (Põlme et al. 2020); Decay sugar-rich substrates (Põlme et al. 2020) | Põlme S, et al. 2020. Fungal Diversity 105:1-16 ((https://doi.org/10.1007/s13225-020-00466-2)) |
| Otu10225 | 2 | k__Fungi(100);p__Ascomycota(100);c__Saccharomycetes(100);o__Saccharomycetales(100);f__Metschnikowiaceae(100);g__Metschnikowia(100);s__Metschnikowia_pulcherrima(100); | Metschnikowia | 13 | Saprotroph | \|Undefined Saprotroph\| | Yeast | NULL | Probable | As nectar-tap_saprotroph (Põlme et al. 2020); Decay sugar-rich substrates (Põlme et al. 2020) | Põlme S, et al. 2020. Fungal Diversity 105:1-16 ((https://doi.org/10.1007/s13225-020-00466-2)) |
| Otu10226 | 2 | k__Fungi(100);p__Ascomycota(100);c__Saccharomycetes(100);o__Saccharomycetales(100);f__Saccharomycetales_family_Incertae_sedis(100);g__Candida(100);g__Candida_unclassified(100); | Candida | 13 | Pathotroph-Saprotroph-Symbiotroph | Animal Pathogen-Endophyte-Endosymbiont-Epiphyte-\|Undefined Saprotroph\| | Dimorphic Yeast-Yeast | NULL | Probable | As nectar-tap_saprotroph (Põlme et al. 2020); Decay sugar-rich substrates (Põlme et al. 2020); This asexual genus is polyphyletic and contain species that function very differently depend on what they are. Caution should be taken when interpreting guilds for Candida species. | Manolakaki D, et al. 2010. Virulence. 1: 367-375 ((https://doi.org/10.4161/viru.1.5.12796)); Mekbib SB, et al. 2011. Tropical Plant Pathology 36:233-240 ((https://doi.org/10.1590/S1982-56762011000400004)); Põlme S, et al. 2020. Fungal Diversity 105:1-16 ((https://doi.org/10.1007/s13225-020-00466-2)) |
| Otu10229 | 2 | k__Fungi(100);p__Ascomycota(100);c__Saccharomycetes(100);o__Saccharomycetales(100);f__Metschnikowiaceae(100);g__Metschnikowia(100);s__Metschnikowia_pulcherrima(100); | Metschnikowia | 13 | Saprotroph | \|Undefined Saprotroph\| | Yeast | NULL | Probable | As nectar-tap_saprotroph (Põlme et al. 2020); Decay sugar-rich substrates (Põlme et al. 2020) | Põlme S, et al. 2020. Fungal Diversity 105:1-16 ((https://doi.org/10.1007/s13225-020-00466-2)) |
| Otu10232 | 2 | k__Fungi(100);p__Ascomycota(100);c__Saccharomycetes(100);o__Saccharomycetales(100);f__Metschnikowiaceae(100);g__Metschnikowia(100);s__Metschnikowia_pulcherrima(100); | Metschnikowia | 13 | Saprotroph | \|Undefined Saprotroph\| | Yeast | NULL | Probable | As nectar-tap_saprotroph (Põlme et al. 2020); Decay sugar-rich substrates (Põlme et al. 2020) | Põlme S, et al. 2020. Fungal Diversity 105:1-16 ((https://doi.org/10.1007/s13225-020-00466-2)) |
| Otu10233 | 2 | k__Fungi(100);p__Ascomycota(100);c__Saccharomycetes(100);o__Saccharomycetales(100);f__Saccharomycetales_family_Incertae_sedis(100);g__Candida(100);g__Candida_unclassified(100); | Candida | 13 | Pathotroph-Saprotroph-Symbiotroph | Animal Pathogen-Endophyte-Endosymbiont-Epiphyte-\|Undefined Saprotroph\| | Dimorphic Yeast-Yeast | NULL | Probable | As nectar-tap_saprotroph (Põlme et al. 2020); Decay sugar-rich substrates (Põlme et al. 2020); This asexual genus is polyphyletic and contain species that function very differently depend on what they are. Caution should be taken when interpreting guilds for Candida species. | Manolakaki D, et al. 2010. Virulence. 1: 367-375 ((https://doi.org/10.4161/viru.1.5.12796)); Mekbib SB, et al. 2011. Tropical Plant Pathology 36:233-240 ((https://doi.org/10.1590/S1982-56762011000400004)); Põlme S, et al. 2020. Fungal Diversity 105:1-16 ((https://doi.org/10.1007/s13225-020-00466-2)) |
| Otu10234 | 2 | k__Fungi(100);p__Ascomycota(100);c__Saccharomycetes(100);o__Saccharomycetales(100);f__Metschnikowiaceae(100);g__Metschnikowia(100);s__Metschnikowia_pulcherrima(100); | Metschnikowia | 13 | Saprotroph | \|Undefined Saprotroph\| | Yeast | NULL | Probable | As nectar-tap_saprotroph (Põlme et al. 2020); Decay sugar-rich substrates (Põlme et al. 2020) | Põlme S, et al. 2020. Fungal Diversity 105:1-16 ((https://doi.org/10.1007/s13225-020-00466-2)) |
| Otu10235 | 2 | k__Fungi(100);p__Ascomycota(100);c__Dothideomycetes(100);o__Dothideomycetes_order_Incertae_sedis(100);f__Myxotrichaceae(100);g__Oidiodendron(100);g__Oidiodendron_unclassified(100); | Oidiodendron | 13 | Pathotroph-Saprotroph-Symbiotroph | Endophyte-Ericoid Mycorrhizal-\|Undefined Saprotroph\| | Microfungus | NULL | Probable | As root_endophyte (Põlme et al. 2020); As soil_saprotroph (Põlme et al. 2020); Dark Septate Endophyte; Decay soil (Põlme et al. 2020) | Newsham KK. 2011. New Phytologist 190:783-793 ((https://doi.org/10.1111/j.1469-8137.2010.03611.x)) (as dark septate root endophytes); Põlme S, et al. 2020. Fungal Diversity 105:1-16 ((https://doi.org/10.1007/s13225-020-00466-2)) |
| Otu10239 | 2 | k__Fungi(100);p__Ascomycota(100);c__Saccharomycetes(100);o__Saccharomycetales(100);f__Metschnikowiaceae(100);g__Metschnikowia(100);g__Metschnikowia_unclassified(100); | Metschnikowia | 13 | Saprotroph | \|Undefined Saprotroph\| | Yeast | NULL | Probable | As nectar-tap_saprotroph (Põlme et al. 2020); Decay sugar-rich substrates (Põlme et al. 2020) | Põlme S, et al. 2020. Fungal Diversity 105:1-16 ((https://doi.org/10.1007/s13225-020-00466-2)) |
| Otu10240 | 2 | k__Fungi(100);p__Ascomycota(100);c__Saccharomycetes(100);o__Saccharomycetales(100);f__Metschnikowiaceae(100);g__Metschnikowia(100);s__Metschnikowia_pulcherrima(100); | Metschnikowia | 13 | Saprotroph | \|Undefined Saprotroph\| | Yeast | NULL | Probable | As nectar-tap_saprotroph (Põlme et al. 2020); Decay sugar-rich substrates (Põlme et al. 2020) | Põlme S, et al. 2020. Fungal Diversity 105:1-16 ((https://doi.org/10.1007/s13225-020-00466-2)) |
| Otu10241 | 2 | k__Fungi(100);p__Ascomycota(100);c__Saccharomycetes(100);o__Saccharomycetales(100);f__Metschnikowiaceae(100);g__Metschnikowia(100);g__Metschnikowia_unclassified(100); | Metschnikowia | 13 | Saprotroph | \|Undefined Saprotroph\| | Yeast | NULL | Probable | As nectar-tap_saprotroph (Põlme et al. 2020); Decay sugar-rich substrates (Põlme et al. 2020) | Põlme S, et al. 2020. Fungal Diversity 105:1-16 ((https://doi.org/10.1007/s13225-020-00466-2)) |
| Otu10244 | 2 | k__Fungi(100);p__Ascomycota(100);c__Saccharomycetes(100);o__Saccharomycetales(100);f__Metschnikowiaceae(100);g__Metschnikowia(100);s__Metschnikowia_pulcherrima(100); | Metschnikowia | 13 | Saprotroph | \|Undefined Saprotroph\| | Yeast | NULL | Probable | As nectar-tap_saprotroph (Põlme et al. 2020); Decay sugar-rich substrates (Põlme et al. 2020) | Põlme S, et al. 2020. Fungal Diversity 105:1-16 ((https://doi.org/10.1007/s13225-020-00466-2)) |
| Otu10245 | 2 | k__Fungi(100);p__Ascomycota(100);c__Saccharomycetes(100);o__Saccharomycetales(100);f__Metschnikowiaceae(100);g__Metschnikowia(100);g__Metschnikowia_unclassified(100); | Metschnikowia | 13 | Saprotroph | \|Undefined Saprotroph\| | Yeast | NULL | Probable | As nectar-tap_saprotroph (Põlme et al. 2020); Decay sugar-rich substrates (Põlme et al. 2020) | Põlme S, et al. 2020. Fungal Diversity 105:1-16 ((https://doi.org/10.1007/s13225-020-00466-2)) |
| Otu10246 | 2 | k__Fungi(100);p__Ascomycota(100);c__Saccharomycetes(100);o__Saccharomycetales(100);f__Metschnikowiaceae(100);g__Metschnikowia(100);g__Metschnikowia_unclassified(100); | Metschnikowia | 13 | Saprotroph | \|Undefined Saprotroph\| | Yeast | NULL | Probable | As nectar-tap_saprotroph (Põlme et al. 2020); Decay sugar-rich substrates (Põlme et al. 2020) | Põlme S, et al. 2020. Fungal Diversity 105:1-16 ((https://doi.org/10.1007/s13225-020-00466-2)) |
| Otu10247 | 2 | k__Fungi(100);p__Ascomycota(100);c__Saccharomycetes(100);o__Saccharomycetales(100);f__Metschnikowiaceae(100);g__Metschnikowia(100);g__Metschnikowia_unclassified(100); | Metschnikowia | 13 | Saprotroph | \|Undefined Saprotroph\| | Yeast | NULL | Probable | As nectar-tap_saprotroph (Põlme et al. 2020); Decay sugar-rich substrates (Põlme et al. 2020) | Põlme S, et al. 2020. Fungal Diversity 105:1-16 ((https://doi.org/10.1007/s13225-020-00466-2)) |
| Otu10248 | 2 | k__Fungi(100);p__Ascomycota(100);c__Saccharomycetes(100);o__Saccharomycetales(100);f__Metschnikowiaceae(100);g__Metschnikowia(100);g__Metschnikowia_unclassified(100); | Metschnikowia | 13 | Saprotroph | \|Undefined Saprotroph\| | Yeast | NULL | Probable | As nectar-tap_saprotroph (Põlme et al. 2020); Decay sugar-rich substrates (Põlme et al. 2020) | Põlme S, et al. 2020. Fungal Diversity 105:1-16 ((https://doi.org/10.1007/s13225-020-00466-2)) |
| Otu10249 | 2 | k__Fungi(100);p__Ascomycota(100);c__Saccharomycetes(100);o__Saccharomycetales(100);f__Metschnikowiaceae(100);g__Metschnikowia(100);s__Metschnikowia_pulcherrima(100); | Metschnikowia | 13 | Saprotroph | \|Undefined Saprotroph\| | Yeast | NULL | Probable | As nectar-tap_saprotroph (Põlme et al. 2020); Decay sugar-rich substrates (Põlme et al. 2020) | Põlme S, et al. 2020. Fungal Diversity 105:1-16 ((https://doi.org/10.1007/s13225-020-00466-2)) |
| Otu10250 | 2 | k__Fungi(100);p__Ascomycota(100);c__Saccharomycetes(100);o__Saccharomycetales(100);f__Metschnikowiaceae(100);g__Metschnikowia(100);g__Metschnikowia_unclassified(100); | Metschnikowia | 13 | Saprotroph | \|Undefined Saprotroph\| | Yeast | NULL | Probable | As nectar-tap_saprotroph (Põlme et al. 2020); Decay sugar-rich substrates (Põlme et al. 2020) | Põlme S, et al. 2020. Fungal Diversity 105:1-16 ((https://doi.org/10.1007/s13225-020-00466-2)) |
| Otu10253 | 2 | k__Fungi(100);p__Ascomycota(100);c__Saccharomycetes(100);o__Saccharomycetales(100);f__Metschnikowiaceae(100);g__Metschnikowia(100);s__Metschnikowia_pulcherrima(100); | Metschnikowia | 13 | Saprotroph | \|Undefined Saprotroph\| | Yeast | NULL | Probable | As nectar-tap_saprotroph (Põlme et al. 2020); Decay sugar-rich substrates (Põlme et al. 2020) | Põlme S, et al. 2020. Fungal Diversity 105:1-16 ((https://doi.org/10.1007/s13225-020-00466-2)) |
| Otu10254 | 2 | k__Fungi(100);p__Ascomycota(100);c__Saccharomycetes(100);o__Saccharomycetales(100);f__Metschnikowiaceae(100);g__Metschnikowia(100);g__Metschnikowia_unclassified(100); | Metschnikowia | 13 | Saprotroph | \|Undefined Saprotroph\| | Yeast | NULL | Probable | As nectar-tap_saprotroph (Põlme et al. 2020); Decay sugar-rich substrates (Põlme et al. 2020) | Põlme S, et al. 2020. Fungal Diversity 105:1-16 ((https://doi.org/10.1007/s13225-020-00466-2)) |
| Otu10255 | 2 | k__Fungi(100);p__Ascomycota(100);c__Saccharomycetes(100);o__Saccharomycetales(100);f__Metschnikowiaceae(100);g__Metschnikowia(100);s__Metschnikowia_pulcherrima(100); | Metschnikowia | 13 | Saprotroph | \|Undefined Saprotroph\| | Yeast | NULL | Probable | As nectar-tap_saprotroph (Põlme et al. 2020); Decay sugar-rich substrates (Põlme et al. 2020) | Põlme S, et al. 2020. Fungal Diversity 105:1-16 ((https://doi.org/10.1007/s13225-020-00466-2)) |
| Otu10258 | 2 | k__Fungi(100);p__Ascomycota(100);c__Saccharomycetes(100);o__Saccharomycetales(100);f__Metschnikowiaceae(100);g__Metschnikowia(100);g__Metschnikowia_unclassified(100); | Metschnikowia | 13 | Saprotroph | \|Undefined Saprotroph\| | Yeast | NULL | Probable | As nectar-tap_saprotroph (Põlme et al. 2020); Decay sugar-rich substrates (Põlme et al. 2020) | Põlme S, et al. 2020. Fungal Diversity 105:1-16 ((https://doi.org/10.1007/s13225-020-00466-2)) |
| Otu10259 | 2 | k__Fungi(100);p__Ascomycota(100);c__Saccharomycetes(100);o__Saccharomycetales(100);f__Metschnikowiaceae(100);g__Metschnikowia(100);g__Metschnikowia_unclassified(100); | Metschnikowia | 13 | Saprotroph | \|Undefined Saprotroph\| | Yeast | NULL | Probable | As nectar-tap_saprotroph (Põlme et al. 2020); Decay sugar-rich substrates (Põlme et al. 2020) | Põlme S, et al. 2020. Fungal Diversity 105:1-16 ((https://doi.org/10.1007/s13225-020-00466-2)) |
| Otu10260 | 2 | k__Fungi(100);p__Ascomycota(100);c__Saccharomycetes(100);o__Saccharomycetales(100);f__Metschnikowiaceae(100);g__Metschnikowia(100);s__Metschnikowia_pulcherrima(100); | Metschnikowia | 13 | Saprotroph | \|Undefined Saprotroph\| | Yeast | NULL | Probable | As nectar-tap_saprotroph (Põlme et al. 2020); Decay sugar-rich substrates (Põlme et al. 2020) | Põlme S, et al. 2020. Fungal Diversity 105:1-16 ((https://doi.org/10.1007/s13225-020-00466-2)) |
| Otu10261 | 2 | k__Fungi(100);p__Ascomycota(100);c__Saccharomycetes(100);o__Saccharomycetales(100);f__Metschnikowiaceae(100);g__Metschnikowia(100);s__Metschnikowia_pulcherrima(100); | Metschnikowia | 13 | Saprotroph | \|Undefined Saprotroph\| | Yeast | NULL | Probable | As nectar-tap_saprotroph (Põlme et al. 2020); Decay sugar-rich substrates (Põlme et al. 2020) | Põlme S, et al. 2020. Fungal Diversity 105:1-16 ((https://doi.org/10.1007/s13225-020-00466-2)) |
| Otu10262 | 2 | k__Fungi(100);p__Ascomycota(100);c__Saccharomycetes(100);o__Saccharomycetales(100);f__Metschnikowiaceae(100);g__Metschnikowia(100);g__Metschnikowia_unclassified(100); | Metschnikowia | 13 | Saprotroph | \|Undefined Saprotroph\| | Yeast | NULL | Probable | As nectar-tap_saprotroph (Põlme et al. 2020); Decay sugar-rich substrates (Põlme et al. 2020) | Põlme S, et al. 2020. Fungal Diversity 105:1-16 ((https://doi.org/10.1007/s13225-020-00466-2)) |
| Otu10263 | 2 | k__Fungi(100);p__Ascomycota(100);c__Saccharomycetes(100);o__Saccharomycetales(100);f__Metschnikowiaceae(100);g__Metschnikowia(100);g__Metschnikowia_unclassified(100); | Metschnikowia | 13 | Saprotroph | \|Undefined Saprotroph\| | Yeast | NULL | Probable | As nectar-tap_saprotroph (Põlme et al. 2020); Decay sugar-rich substrates (Põlme et al. 2020) | Põlme S, et al. 2020. Fungal Diversity 105:1-16 ((https://doi.org/10.1007/s13225-020-00466-2)) |
| Otu10265 | 2 | k__Fungi(100);p__Ascomycota(100);c__Saccharomycetes(100);o__Saccharomycetales(100);f__Metschnikowiaceae(100);g__Metschnikowia(100);g__Metschnikowia_unclassified(100); | Metschnikowia | 13 | Saprotroph | \|Undefined Saprotroph\| | Yeast | NULL | Probable | As nectar-tap_saprotroph (Põlme et al. 2020); Decay sugar-rich substrates (Põlme et al. 2020) | Põlme S, et al. 2020. Fungal Diversity 105:1-16 ((https://doi.org/10.1007/s13225-020-00466-2)) |
| Otu10266 | 2 | k__Fungi(100);p__Ascomycota(100);c__Saccharomycetes(100);o__Saccharomycetales(100);f__Metschnikowiaceae(100);g__Metschnikowia(100);g__Metschnikowia_unclassified(100); | Metschnikowia | 13 | Saprotroph | \|Undefined Saprotroph\| | Yeast | NULL | Probable | As nectar-tap_saprotroph (Põlme et al. 2020); Decay sugar-rich substrates (Põlme et al. 2020) | Põlme S, et al. 2020. Fungal Diversity 105:1-16 ((https://doi.org/10.1007/s13225-020-00466-2)) |
| Otu10267 | 2 | k__Fungi(100);p__Ascomycota(100);c__Saccharomycetes(100);o__Saccharomycetales(100);f__Metschnikowiaceae(100);g__Metschnikowia(100);s__Metschnikowia_pulcherrima(100); | Metschnikowia | 13 | Saprotroph | \|Undefined Saprotroph\| | Yeast | NULL | Probable | As nectar-tap_saprotroph (Põlme et al. 2020); Decay sugar-rich substrates (Põlme et al. 2020) | Põlme S, et al. 2020. Fungal Diversity 105:1-16 ((https://doi.org/10.1007/s13225-020-00466-2)) |
| Otu10270 | 2 | k__Fungi(100);p__Ascomycota(100);c__Saccharomycetes(100);o__Saccharomycetales(100);f__Metschnikowiaceae(100);g__Metschnikowia(100);g__Metschnikowia_unclassified(100); | Metschnikowia | 13 | Saprotroph | \|Undefined Saprotroph\| | Yeast | NULL | Probable | As nectar-tap_saprotroph (Põlme et al. 2020); Decay sugar-rich substrates (Põlme et al. 2020) | Põlme S, et al. 2020. Fungal Diversity 105:1-16 ((https://doi.org/10.1007/s13225-020-00466-2)) |
| Otu10271 | 2 | k__Fungi(100);p__Ascomycota(100);c__Saccharomycetes(100);o__Saccharomycetales(100);f__Metschnikowiaceae(100);g__Metschnikowia(100);s__Metschnikowia_pulcherrima(100); | Metschnikowia | 13 | Saprotroph | \|Undefined Saprotroph\| | Yeast | NULL | Probable | As nectar-tap_saprotroph (Põlme et al. 2020); Decay sugar-rich substrates (Põlme et al. 2020) | Põlme S, et al. 2020. Fungal Diversity 105:1-16 ((https://doi.org/10.1007/s13225-020-00466-2)) |
| Otu10273 | 2 | k__Fungi(100);p__Ascomycota(100);c__Saccharomycetes(100);o__Saccharomycetales(100);f__Metschnikowiaceae(100);g__Metschnikowia(100);g__Metschnikowia_unclassified(100); | Metschnikowia | 13 | Saprotroph | \|Undefined Saprotroph\| | Yeast | NULL | Probable | As nectar-tap_saprotroph (Põlme et al. 2020); Decay sugar-rich substrates (Põlme et al. 2020) | Põlme S, et al. 2020. Fungal Diversity 105:1-16 ((https://doi.org/10.1007/s13225-020-00466-2)) |
| Otu10274 | 2 | k__Fungi(100);p__Ascomycota(100);c__Saccharomycetes(100);o__Saccharomycetales(100);f__Metschnikowiaceae(100);g__Metschnikowia(100);s__Metschnikowia_pulcherrima(100); | Metschnikowia | 13 | Saprotroph | \|Undefined Saprotroph\| | Yeast | NULL | Probable | As nectar-tap_saprotroph (Põlme et al. 2020); Decay sugar-rich substrates (Põlme et al. 2020) | Põlme S, et al. 2020. Fungal Diversity 105:1-16 ((https://doi.org/10.1007/s13225-020-00466-2)) |
| Otu10278 | 2 | k__Fungi(100);p__Ascomycota(100);c__Saccharomycetes(100);o__Saccharomycetales(100);f__Metschnikowiaceae(100);g__Metschnikowia(100);g__Metschnikowia_unclassified(100); | Metschnikowia | 13 | Saprotroph | \|Undefined Saprotroph\| | Yeast | NULL | Probable | As nectar-tap_saprotroph (Põlme et al. 2020); Decay sugar-rich substrates (Põlme et al. 2020) | Põlme S, et al. 2020. Fungal Diversity 105:1-16 ((https://doi.org/10.1007/s13225-020-00466-2)) |
| Otu10279 | 2 | k__Fungi(100);p__Ascomycota(100);c__Saccharomycetes(100);o__Saccharomycetales(100);f__Metschnikowiaceae(100);g__Metschnikowia(100);g__Metschnikowia_unclassified(100); | Metschnikowia | 13 | Saprotroph | \|Undefined Saprotroph\| | Yeast | NULL | Probable | As nectar-tap_saprotroph (Põlme et al. 2020); Decay sugar-rich substrates (Põlme et al. 2020) | Põlme S, et al. 2020. Fungal Diversity 105:1-16 ((https://doi.org/10.1007/s13225-020-00466-2)) |
| Otu10280 | 2 | k__Fungi(100);p__Ascomycota(100);c__Saccharomycetes(100);o__Saccharomycetales(100);f__Metschnikowiaceae(100);g__Metschnikowia(100);s__Metschnikowia_pulcherrima(100); | Metschnikowia | 13 | Saprotroph | \|Undefined Saprotroph\| | Yeast | NULL | Probable | As nectar-tap_saprotroph (Põlme et al. 2020); Decay sugar-rich substrates (Põlme et al. 2020) | Põlme S, et al. 2020. Fungal Diversity 105:1-16 ((https://doi.org/10.1007/s13225-020-00466-2)) |
| Otu10281 | 2 | k__Fungi(100);p__Ascomycota(100);c__Saccharomycetes(100);o__Saccharomycetales(100);f__Metschnikowiaceae(100);g__Metschnikowia(100);s__Metschnikowia_pulcherrima(100); | Metschnikowia | 13 | Saprotroph | \|Undefined Saprotroph\| | Yeast | NULL | Probable | As nectar-tap_saprotroph (Põlme et al. 2020); Decay sugar-rich substrates (Põlme et al. 2020) | Põlme S, et al. 2020. Fungal Diversity 105:1-16 ((https://doi.org/10.1007/s13225-020-00466-2)) |
| Otu10283 | 2 | k__Fungi(100);p__Basidiomycota(100);c__Microbotryomycetes(100);o__Sporidiobolales(100);f__Sporidiobolales_family_Incertae_sedis(100);g__Sporobolomyces(100);s__Sporobolomyces_lactosus(100); | Sporobolomyces | 13 | Pathotroph-Saprotroph | \|Fungal Parasite\|-Plant Saprotroph | Yeast | NULL | Probable | As litter_saprotroph (Põlme et al. 2020); As mycoparasite (Põlme et al. 2020); Decay fungal material (Põlme et al. 2020) | Tedersoo L, et al. 2014. Science 346:e1256688 ((https://doi.org/10.1126/science.1256688)); Purahong W, et al. 2016. Molecular Ecology 25:4059-4074 ((https://doi.org/10.1111/mec.13739)); Põlme S, et al. 2020. Fungal Diversity 105:1-16 ((https://doi.org/10.1007/s13225-020-00466-2)) |
| Otu10284 | 2 | k__Fungi(100);p__Ascomycota(100);c__Saccharomycetes(100);o__Saccharomycetales(100);f__Metschnikowiaceae(100);g__Metschnikowia(100);s__Metschnikowia_pulcherrima(100); | Metschnikowia | 13 | Saprotroph | \|Undefined Saprotroph\| | Yeast | NULL | Probable | As nectar-tap_saprotroph (Põlme et al. 2020); Decay sugar-rich substrates (Põlme et al. 2020) | Põlme S, et al. 2020. Fungal Diversity 105:1-16 ((https://doi.org/10.1007/s13225-020-00466-2)) |
| Otu10285 | 2 | k__Fungi(100);p__Ascomycota(100);c__Saccharomycetes(100);o__Saccharomycetales(100);f__Metschnikowiaceae(100);g__Metschnikowia(100);s__Metschnikowia_pulcherrima(100); | Metschnikowia | 13 | Saprotroph | \|Undefined Saprotroph\| | Yeast | NULL | Probable | As nectar-tap_saprotroph (Põlme et al. 2020); Decay sugar-rich substrates (Põlme et al. 2020) | Põlme S, et al. 2020. Fungal Diversity 105:1-16 ((https://doi.org/10.1007/s13225-020-00466-2)) |
| Otu10286 | 2 | k__Fungi(100);p__Ascomycota(100);c__Saccharomycetes(100);o__Saccharomycetales(100);f__Metschnikowiaceae(100);g__Metschnikowia(100);g__Metschnikowia_unclassified(100); | Metschnikowia | 13 | Saprotroph | \|Undefined Saprotroph\| | Yeast | NULL | Probable | As nectar-tap_saprotroph (Põlme et al. 2020); Decay sugar-rich substrates (Põlme et al. 2020) | Põlme S, et al. 2020. Fungal Diversity 105:1-16 ((https://doi.org/10.1007/s13225-020-00466-2)) |

| | | | | | | | | | | | |
|---|---|---|---|---|---|---|---|---|---|---|---|
| Otu10288 | 2 | k__Fungi(100);p__Ascomycota(100);c__Saccharomycetes(100);o__Saccharomycetales(100);f__Saccharomycetales_family_Incertae_sedis(100);g__Candida(100);g__Candida_unclassified(100); | Candida | 13 | Pathotroph-Saprotroph-Symbiotroph | Animal Pathogen-Endophyte-Endosymbiont-Epiphyte-\|Undefined Saprotroph\| | Dimorphic Yeast-Yeast | NULL | Probable | As nectar-tap_saprotroph (Põlme et al. 2020); Decay sugar-rich substrates (Põlme et al. 2020); This asexual genus is polyphyletic and contain species that function very differently depend on what they are. Caution should be taken when interpreting guilds for Candida species. | Manolakaki D, et al. 2010. Virulence. 1: 367-375 ((https://doi.org/10.4161/viru.1.5.12796)); Mekbib SB, et al. 2011. Tropical Plant Pathology 36:233-240 ((https://doi.org/10.1590/S1982-56762011000400004)); Põlme S, et al. 2020. Fungal Diversity 105:1-16 ((https://doi.org/10.1007/s13225-020-00466-2)) |
| Otu10292 | 2 | k__Fungi(100);p__Ascomycota(100);c__Saccharomycetes(100);o__Saccharomycetales(100);f__Metschnikowiaceae(100);g__Metschnikowia(100);s__Metschnikowia_pulcherrima(100); | Metschnikowia | 13 | Saprotroph | \|Undefined Saprotroph\| | Yeast | NULL | Probable | As nectar-tap_saprotroph (Põlme et al. 2020); Decay sugar-rich substrates (Põlme et al. 2020) | Põlme S, et al. 2020. Fungal Diversity 105:1-16 ((https://doi.org/10.1007/s13225-020-00466-2)) |
| Otu10295 | 2 | k__Fungi(100);p__Ascomycota(100);c__Saccharomycetes(100);o__Saccharomycetales(100);f__Metschnikowiaceae(100);g__Metschnikowia(100);g__Metschnikowia_unclassified(100); | Metschnikowia | 13 | Saprotroph | \|Undefined Saprotroph\| | Yeast | NULL | Probable | As nectar-tap_saprotroph (Põlme et al. 2020); Decay sugar-rich substrates (Põlme et al. 2020) | Põlme S, et al. 2020. Fungal Diversity 105:1-16 ((https://doi.org/10.1007/s13225-020-00466-2)) |
| Otu10296 | 2 | k__Fungi(100);p__Ascomycota(100);c__Saccharomycetes(100);o__Saccharomycetales(100);f__Metschnikowiaceae(100);g__Metschnikowia(100);s__Metschnikowia_pulcherrima(100); | Metschnikowia | 13 | Saprotroph | \|Undefined Saprotroph\| | Yeast | NULL | Probable | As nectar-tap_saprotroph (Põlme et al. 2020); Decay sugar-rich substrates (Põlme et al. 2020) | Põlme S, et al. 2020. Fungal Diversity 105:1-16 ((https://doi.org/10.1007/s13225-020-00466-2)) |
| Otu10300 | 2 | k__Fungi(100);p__Ascomycota(100);c__Saccharomycetes(100);o__Saccharomycetales(100);f__Metschnikowiaceae(100);g__Metschnikowia(100);s__Metschnikowia_pulcherrima(100); | Metschnikowia | 13 | Saprotroph | \|Undefined Saprotroph\| | Yeast | NULL | Probable | As nectar-tap_saprotroph (Põlme et al. 2020); Decay sugar-rich substrates (Põlme et al. 2020) | Põlme S, et al. 2020. Fungal Diversity 105:1-16 ((https://doi.org/10.1007/s13225-020-00466-2)) |
| Otu10303 | 2 | k__Fungi(100);p__Ascomycota(100);c__Saccharomycetes(100);o__Saccharomycetales(100);f__Metschnikowiaceae(100);g__Metschnikowia(100);s__Metschnikowia_pulcherrima(100); | Metschnikowia | 13 | Saprotroph | \|Undefined Saprotroph\| | Yeast | NULL | Probable | As nectar-tap_saprotroph (Põlme et al. 2020); Decay sugar-rich substrates (Põlme et al. 2020) | Põlme S, et al. 2020. Fungal Diversity 105:1-16 ((https://doi.org/10.1007/s13225-020-00466-2)) |
| Otu10306 | 2 | k__Fungi(100);p__Ascomycota(100);c__Saccharomycetes(100);o__Saccharomycetales(100);f__Metschnikowiaceae(100);g__Metschnikowia(100);g__Metschnikowia_unclassified(100); | Metschnikowia | 13 | Saprotroph | \|Undefined Saprotroph\| | Yeast | NULL | Probable | As nectar-tap_saprotroph (Põlme et al. 2020); Decay sugar-rich substrates (Põlme et al. 2020) | Põlme S, et al. 2020. Fungal Diversity 105:1-16 ((https://doi.org/10.1007/s13225-020-00466-2)) |
| Otu10307 | 2 | k__Fungi(100);p__Ascomycota(100);c__Saccharomycetes(100);o__Saccharomycetales(100);f__Metschnikowiaceae(100);g__Metschnikowia(100);s__Metschnikowia_pulcherrima(100); | Metschnikowia | 13 | Saprotroph | \|Undefined Saprotroph\| | Yeast | NULL | Probable | As nectar-tap_saprotroph (Põlme et al. 2020); Decay sugar-rich substrates (Põlme et al. 2020) | Põlme S, et al. 2020. Fungal Diversity 105:1-16 ((https://doi.org/10.1007/s13225-020-00466-2)) |
| Otu10309 | 2 | k__Fungi(100);p__Ascomycota(100);c__Saccharomycetes(100);o__Saccharomycetales(100);f__Metschnikowiaceae(100);g__Metschnikowia(100);s__Metschnikowia_pulcherrima(100); | Metschnikowia | 13 | Saprotroph | \|Undefined Saprotroph\| | Yeast | NULL | Probable | As nectar-tap_saprotroph (Põlme et al. 2020); Decay sugar-rich substrates (Põlme et al. 2020) | Põlme S, et al. 2020. Fungal Diversity 105:1-16 ((https://doi.org/10.1007/s13225-020-00466-2)) |
| Otu10310 | 2 | k__Fungi(100);p__Ascomycota(100);c__Saccharomycetes(100);o__Saccharomycetales(100);f__Metschnikowiaceae(100);g__Metschnikowia(100);s__Metschnikowia_pulcherrima(100); | Metschnikowia | 13 | Saprotroph | \|Undefined Saprotroph\| | Yeast | NULL | Probable | As nectar-tap_saprotroph (Põlme et al. 2020); Decay sugar-rich substrates (Põlme et al. 2020) | Põlme S, et al. 2020. Fungal Diversity 105:1-16 ((https://doi.org/10.1007/s13225-020-00466-2)) |
| Otu10316 | 2 | k__Fungi(100);p__Ascomycota(100);c__Saccharomycetes(100);o__Saccharomycetales(100);f__Metschnikowiaceae(100);g__Metschnikowia(100);g__Metschnikowia_unclassified(100); | Metschnikowia | 13 | Saprotroph | \|Undefined Saprotroph\| | Yeast | NULL | Probable | As nectar-tap_saprotroph (Põlme et al. 2020); Decay sugar-rich substrates (Põlme et al. 2020) | Põlme S, et al. 2020. Fungal Diversity 105:1-16 ((https://doi.org/10.1007/s13225-020-00466-2)) |
| Otu10317 | 2 | k__Fungi(100);p__Ascomycota(100);c__Saccharomycetes(100);o__Saccharomycetales(100);f__Metschnikowiaceae(100);g__Metschnikowia(100);s__Metschnikowia_pulcherrima(100); | Metschnikowia | 13 | Saprotroph | \|Undefined Saprotroph\| | Yeast | NULL | Probable | As nectar-tap_saprotroph (Põlme et al. 2020); Decay sugar-rich substrates (Põlme et al. 2020) | Põlme S, et al. 2020. Fungal Diversity 105:1-16 ((https://doi.org/10.1007/s13225-020-00466-2)) |
| Otu10318 | 2 | k__Fungi(100);p__Ascomycota(100);c__Saccharomycetes(100);o__Saccharomycetales(100);f__Metschnikowiaceae(100);g__Metschnikowia(100);s__Metschnikowia_pulcherrima(100); | Metschnikowia | 13 | Saprotroph | \|Undefined Saprotroph\| | Yeast | NULL | Probable | As nectar-tap_saprotroph (Põlme et al. 2020); Decay sugar-rich substrates (Põlme et al. 2020) | Põlme S, et al. 2020. Fungal Diversity 105:1-16 ((https://doi.org/10.1007/s13225-020-00466-2)) |
| Otu10320 | 2 | k__Fungi(100);p__Ascomycota(100);c__Saccharomycetes(100);o__Saccharomycetales(100);f__Metschnikowiaceae(100);g__Metschnikowia(100);g__Metschnikowia_unclassified(100); | Metschnikowia | 13 | Saprotroph | \|Undefined Saprotroph\| | Yeast | NULL | Probable | As nectar-tap_saprotroph (Põlme et al. 2020); Decay sugar-rich substrates (Põlme et al. 2020) | Põlme S, et al. 2020. Fungal Diversity 105:1-16 ((https://doi.org/10.1007/s13225-020-00466-2)) |
| Otu10322 | 2 | k__Fungi(100);p__Ascomycota(100);c__Saccharomycetes(100);o__Saccharomycetales(100);f__Metschnikowiaceae(100);g__Metschnikowia(100);g__Metschnikowia_unclassified(100); | Metschnikowia | 13 | Saprotroph | \|Undefined Saprotroph\| | Yeast | NULL | Probable | As nectar-tap_saprotroph (Põlme et al. 2020); Decay sugar-rich substrates (Põlme et al. 2020) | Põlme S, et al. 2020. Fungal Diversity 105:1-16 ((https://doi.org/10.1007/s13225-020-00466-2)) |
| Otu10323 | 2 | k__Fungi(100);p__Ascomycota(100);c__Saccharomycetes(100);o__Saccharomycetales(100);f__Metschnikowiaceae(100);g__Metschnikowia(100);g__Metschnikowia_unclassified(100); | Metschnikowia | 13 | Saprotroph | \|Undefined Saprotroph\| | Yeast | NULL | Probable | As nectar-tap_saprotroph (Põlme et al. 2020); Decay sugar-rich substrates (Põlme et al. 2020) | Põlme S, et al. 2020. Fungal Diversity 105:1-16 ((https://doi.org/10.1007/s13225-020-00466-2)) |
| Otu10324 | 2 | k__Fungi(100);p__Ascomycota(100);c__Saccharomycetes(100);o__Saccharomycetales(100);f__Metschnikowiaceae(100);g__Metschnikowia(100);s__Metschnikowia_pulcherrima(100); | Metschnikowia | 13 | Saprotroph | \|Undefined Saprotroph\| | Yeast | NULL | Probable | As nectar-tap_saprotroph (Põlme et al. 2020); Decay sugar-rich substrates (Põlme et al. 2020) | Põlme S, et al. 2020. Fungal Diversity 105:1-16 ((https://doi.org/10.1007/s13225-020-00466-2)) |
| Otu10325 | 2 | k__Fungi(100);p__Ascomycota(100);c__Saccharomycetes(100);o__Saccharomycetales(100);f__Metschnikowiaceae(100);g__Metschnikowia(100);g__Metschnikowia_unclassified(100); | Metschnikowia | 13 | Saprotroph | \|Undefined Saprotroph\| | Yeast | NULL | Probable | As nectar-tap_saprotroph (Põlme et al. 2020); Decay sugar-rich substrates (Põlme et al. 2020) | Põlme S, et al. 2020. Fungal Diversity 105:1-16 ((https://doi.org/10.1007/s13225-020-00466-2)) |
| Otu10326 | 2 | k__Fungi(100);p__Ascomycota(100);c__Saccharomycetes(100);o__Saccharomycetales(100);f__Metschnikowiaceae(100);g__Metschnikowia(100);s__Metschnikowia_pulcherrima(100); | Metschnikowia | 13 | Saprotroph | \|Undefined Saprotroph\| | Yeast | NULL | Probable | As nectar-tap_saprotroph (Põlme et al. 2020); Decay sugar-rich substrates (Põlme et al. 2020) | Põlme S, et al. 2020. Fungal Diversity 105:1-16 ((https://doi.org/10.1007/s13225-020-00466-2)) |
| Otu10327 | 2 | k__Fungi(100);p__Ascomycota(100);c__Saccharomycetes(100);o__Saccharomycetales(100);f__Metschnikowiaceae(100);g__Metschnikowia(100);s__Metschnikowia_pulcherrima(100); | Metschnikowia | 13 | Saprotroph | \|Undefined Saprotroph\| | Yeast | NULL | Probable | As nectar-tap_saprotroph (Põlme et al. 2020); Decay sugar-rich substrates (Põlme et al. 2020) | Põlme S, et al. 2020. Fungal Diversity 105:1-16 ((https://doi.org/10.1007/s13225-020-00466-2)) |
| Otu10328 | 2 | k__Fungi(100);p__Ascomycota(100);c__Saccharomycetes(100);o__Saccharomycetales(100);f__Metschnikowiaceae(100);g__Metschnikowia(100);s__Metschnikowia_pulcherrima(100); | Metschnikowia | 13 | Saprotroph | \|Undefined Saprotroph\| | Yeast | NULL | Probable | As nectar-tap_saprotroph (Põlme et al. 2020); Decay sugar-rich substrates (Põlme et al. 2020) | Põlme S, et al. 2020. Fungal Diversity 105:1-16 ((https://doi.org/10.1007/s13225-020-00466-2)) |
| Otu10329 | 2 | k__Fungi(100);p__Ascomycota(100);c__Saccharomycetes(100);o__Saccharomycetales(100);f__Saccharomycetales_family_Incertae_sedis(100);f__Saccharomycetales_family_Incertae_sedis_unclassified(100);f__Saccharomycetales_family_Incertae_sedis_unclassified(100); | Saccharomycetales | 7 | Saprotroph | Undefined Saprotroph | Yeast | NULL | Possible | NULL | Sterkenburg E, et al. 2015. New Phytologist 207:1145-1158 ((https://doi.org/10.1111/nph.13426)) |
| Otu10331 | 2 | k__Fungi(100);p__Ascomycota(100);c__Saccharomycetes(100);o__Saccharomycetales(100);f__Metschnikowiaceae(100);g__Metschnikowia(100);g__Metschnikowia_unclassified(100); | Metschnikowia | 13 | Saprotroph | \|Undefined Saprotroph\| | Yeast | NULL | Probable | As nectar-tap_saprotroph (Põlme et al. 2020); Decay sugar-rich substrates (Põlme et al. 2020) | Põlme S, et al. 2020. Fungal Diversity 105:1-16 ((https://doi.org/10.1007/s13225-020-00466-2)) |
| Otu10334 | 2 | k__Fungi(100);p__Ascomycota(100);c__Saccharomycetes(100);o__Saccharomycetales(100);f__Metschnikowiaceae(100);g__Metschnikowia(100);s__Metschnikowia_pulcherrima(100); | Metschnikowia | 13 | Saprotroph | \|Undefined Saprotroph\| | Yeast | NULL | Probable | As nectar-tap_saprotroph (Põlme et al. 2020); Decay sugar-rich substrates (Põlme et al. 2020) | Põlme S, et al. 2020. Fungal Diversity 105:1-16 ((https://doi.org/10.1007/s13225-020-00466-2)) |
| Otu10335 | 2 | k__Fungi(100);p__Ascomycota(100);c__Saccharomycetes(100);o__Saccharomycetales(100);f__Metschnikowiaceae(100);g__Metschnikowia(100);g__Metschnikowia_unclassified(100); | Metschnikowia | 13 | Saprotroph | \|Undefined Saprotroph\| | Yeast | NULL | Probable | As nectar-tap_saprotroph (Põlme et al. 2020); Decay sugar-rich substrates (Põlme et al. 2020) | Põlme S, et al. 2020. Fungal Diversity 105:1-16 ((https://doi.org/10.1007/s13225-020-00466-2)) |
| Otu10338 | 2 | k__Fungi(100);p__Ascomycota(100);c__Saccharomycetes(100);o__Saccharomycetales(100);f__Saccharomycetales_family_Incertae_sedis(100);g__Candida(100);g__Candida_unclassified(100); | Candida | 13 | Pathotroph-Saprotroph-Symbiotroph | Animal Pathogen-Endophyte-Endosymbiont-Epiphyte-\|Undefined Saprotroph\| | Dimorphic Yeast-Yeast | NULL | Probable | As nectar-tap_saprotroph (Põlme et al. 2020); Decay sugar-rich substrates (Põlme et al. 2020); This asexual genus is polyphyletic and contain species that function very differently depend on what they are. Caution should be taken when interpreting guilds for Candida species. | Manolakaki D, et al. 2010. Virulence. 1: 367-375 ((https://doi.org/10.4161/viru.1.5.12796)); Mekbib SB, et al. 2011. Tropical Plant Pathology 36:233-240 ((https://doi.org/10.1590/S1982-56762011000400004)); Põlme S, et al. 2020. Fungal Diversity 105:1-16 ((https://doi.org/10.1007/s13225-020-00466-2)) |
| Otu10347 | 2 | k__Fungi(100);p__Ascomycota(100);c__Saccharomycetes(100);o__Saccharomycetales(100);f__Metschnikowiaceae(100);g__Metschnikowia(100);s__Metschnikowia_pulcherrima(100); | Metschnikowia | 13 | Saprotroph | \|Undefined Saprotroph\| | Yeast | NULL | Probable | As nectar-tap_saprotroph (Põlme et al. 2020); Decay sugar-rich substrates (Põlme et al. 2020) | Põlme S, et al. 2020. Fungal Diversity 105:1-16 ((https://doi.org/10.1007/s13225-020-00466-2)) |
| Otu10348 | 2 | k__Fungi(100);p__Ascomycota(100);c__Saccharomycetes(100);o__Saccharomycetales(100);f__Metschnikowiaceae(100);g__Metschnikowia(100);g__Metschnikowia_unclassified(100); | Metschnikowia | 13 | Saprotroph | \|Undefined Saprotroph\| | Yeast | NULL | Probable | As nectar-tap_saprotroph (Põlme et al. 2020); Decay sugar-rich substrates (Põlme et al. 2020) | Põlme S, et al. 2020. Fungal Diversity 105:1-16 ((https://doi.org/10.1007/s13225-020-00466-2)) |
| Otu10349 | 2 | k__Fungi(100);p__Ascomycota(100);c__Saccharomycetes(100);o__Saccharomycetales(100);f__Metschnikowiaceae(100);g__Metschnikowia(100);s__Metschnikowia_pulcherrima(100); | Metschnikowia | 13 | Saprotroph | \|Undefined Saprotroph\| | Yeast | NULL | Probable | As nectar-tap_saprotroph (Põlme et al. 2020); Decay sugar-rich substrates (Põlme et al. 2020) | Põlme S, et al. 2020. Fungal Diversity 105:1-16 ((https://doi.org/10.1007/s13225-020-00466-2)) |
| Otu10350 | 2 | k__Fungi(100);p__Ascomycota(100);c__Saccharomycetes(100);o__Saccharomycetales(100);f__Metschnikowiaceae(100);g__Metschnikowia(100);s__Metschnikowia_pulcherrima(100); | Metschnikowia | 13 | Saprotroph | \|Undefined Saprotroph\| | Yeast | NULL | Probable | As nectar-tap_saprotroph (Põlme et al. 2020); Decay sugar-rich substrates (Põlme et al. 2020) | Põlme S, et al. 2020. Fungal Diversity 105:1-16 ((https://doi.org/10.1007/s13225-020-00466-2)) |
| Otu10351 | 2 | k__Fungi(100);p__Ascomycota(100);c__Saccharomycetes(100);o__Saccharomycetales(100);f__Metschnikowiaceae(100);g__Metschnikowia(100);g__Metschnikowia_unclassified(100); | Metschnikowia | 13 | Saprotroph | \|Undefined Saprotroph\| | Yeast | NULL | Probable | As nectar-tap_saprotroph (Põlme et al. 2020); Decay sugar-rich substrates (Põlme et al. 2020) | Põlme S, et al. 2020. Fungal Diversity 105:1-16 ((https://doi.org/10.1007/s13225-020-00466-2)) |
| Otu10355 | 2 | k__Fungi(100);p__Ascomycota(100);c__Saccharomycetes(100);o__Saccharomycetales(100);f__Metschnikowiaceae(100);g__Metschnikowia(100);s__Metschnikowia_pulcherrima(100); | Metschnikowia | 13 | Saprotroph | \|Undefined Saprotroph\| | Yeast | NULL | Probable | As nectar-tap_saprotroph (Põlme et al. 2020); Decay sugar-rich substrates (Põlme et al. 2020) | Põlme S, et al. 2020. Fungal Diversity 105:1-16 ((https://doi.org/10.1007/s13225-020-00466-2)) |
| Otu10356 | 2 | k__Fungi(100);p__Ascomycota(100);c__Saccharomycetes(100);o__Saccharomycetales(100);f__Metschnikowiaceae(100);g__Metschnikowia(100);s__Metschnikowia_pulcherrima(100); | Metschnikowia | 13 | Saprotroph | \|Undefined Saprotroph\| | Yeast | NULL | Probable | As nectar-tap_saprotroph (Põlme et al. 2020); Decay sugar-rich substrates (Põlme et al. 2020) | Põlme S, et al. 2020. Fungal Diversity 105:1-16 ((https://doi.org/10.1007/s13225-020-00466-2)) |
| Otu10357 | 2 | k__Fungi(100);p__Ascomycota(100);c__Saccharomycetes(100);o__Saccharomycetales(100);f__Metschnikowiaceae(100);g__Metschnikowia(100);s__Metschnikowia_pulcherrima(100); | Metschnikowia | 13 | Saprotroph | \|Undefined Saprotroph\| | Yeast | NULL | Probable | As nectar-tap_saprotroph (Põlme et al. 2020); Decay sugar-rich substrates (Põlme et al. 2020) | Põlme S, et al. 2020. Fungal Diversity 105:1-16 ((https://doi.org/10.1007/s13225-020-00466-2)) |
| Otu10358 | 2 | k__Fungi(100);p__Ascomycota(100);c__Saccharomycetes(100);o__Saccharomycetales(100);f__Metschnikowiaceae(100);g__Metschnikowia(100);s__Metschnikowia_pulcherrima(100); | Metschnikowia | 13 | Saprotroph | \|Undefined Saprotroph\| | Yeast | NULL | Probable | As nectar-tap_saprotroph (Põlme et al. 2020); Decay sugar-rich substrates (Põlme et al. 2020) | Põlme S, et al. 2020. Fungal Diversity 105:1-16 ((https://doi.org/10.1007/s13225-020-00466-2)) |
| Otu10359 | 2 | k__Fungi(100);p__Ascomycota(100);c__Saccharomycetes(100);o__Saccharomycetales(100);f__Metschnikowiaceae(100);g__Metschnikowia(100);s__Metschnikowia_pulcherrima(100); | Metschnikowia | 13 | Saprotroph | \|Undefined Saprotroph\| | Yeast | NULL | Probable | As nectar-tap_saprotroph (Põlme et al. 2020); Decay sugar-rich substrates (Põlme et al. 2020) | Põlme S, et al. 2020. Fungal Diversity 105:1-16 ((https://doi.org/10.1007/s13225-020-00466-2)) |
| Otu10360 | 2 | k__Fungi(100);p__Ascomycota(100);c__Saccharomycetes(100);o__Saccharomycetales(100);f__Saccharomycetales_family_Incertae_sedis(100);f__Saccharomycetales_family_Incertae_sedis_unclassified(100);f__Saccharomycetales_family_Incertae_sedis_unclassified(100); | Saccharomycetales | 7 | Saprotroph | Undefined Saprotroph | Yeast | NULL | Possible | NULL | Sterkenburg E, et al. 2015. New Phytologist 207:1145-1158 ((https://doi.org/10.1111/nph.13426)) |
| Otu10362 | 2 | k__Fungi(100);p__Ascomycota(100);c__Saccharomycetes(100);o__Saccharomycetales(100);f__Metschnikowiaceae(100);g__Metschnikowia(100);g__Metschnikowia_unclassified(100); | Metschnikowia | 13 | Saprotroph | \|Undefined Saprotroph\| | Yeast | NULL | Probable | As nectar-tap_saprotroph (Põlme et al. 2020); Decay sugar-rich substrates (Põlme et al. 2020) | Põlme S, et al. 2020. Fungal Diversity 105:1-16 ((https://doi.org/10.1007/s13225-020-00466-2)) |
| Otu10363 | 2 | k__Fungi(100);p__Ascomycota(100);c__Saccharomycetes(100);o__Saccharomycetales(100);f__Metschnikowiaceae(100);g__Metschnikowia(100);s__Metschnikowia_pulcherrima(100); | Metschnikowia | 13 | Saprotroph | \|Undefined Saprotroph\| | Yeast | NULL | Probable | As nectar-tap_saprotroph (Põlme et al. 2020); Decay sugar-rich substrates (Põlme et al. 2020) | Põlme S, et al. 2020. Fungal Diversity 105:1-16 ((https://doi.org/10.1007/s13225-020-00466-2)) |
| Otu10364 | 2 | k__Fungi(100);p__Ascomycota(100);c__Saccharomycetes(100);o__Saccharomycetales(100);f__Metschnikowiaceae(100);g__Metschnikowia(100);g__Metschnikowia_unclassified(100); | Metschnikowia | 13 | Saprotroph | \|Undefined Saprotroph\| | Yeast | NULL | Probable | As nectar-tap_saprotroph (Põlme et al. 2020); Decay sugar-rich substrates (Põlme et al. 2020) | Põlme S, et al. 2020. Fungal Diversity 105:1-16 ((https://doi.org/10.1007/s13225-020-00466-2)) |
| Otu10366 | 2 | k__Fungi(100);p__Ascomycota(100);c__Saccharomycetes(100);o__Saccharomycetales(100);f__Metschnikowiaceae(100);g__Metschnikowia(100);g__Metschnikowia_unclassified(100); | Metschnikowia | 13 | Saprotroph | \|Undefined Saprotroph\| | Yeast | NULL | Probable | As nectar-tap_saprotroph (Põlme et al. 2020); Decay sugar-rich substrates (Põlme et al. 2020) | Põlme S, et al. 2020. Fungal Diversity 105:1-16 ((https://doi.org/10.1007/s13225-020-00466-2)) |
| Otu10369 | 2 | k__Fungi(100);p__Ascomycota(100);c__Saccharomycetes(100);o__Saccharomycetales(100);f__Metschnikowiaceae(100);g__Metschnikowia(100);s__Metschnikowia_pulcherrima(100); | Metschnikowia | 13 | Saprotroph | \|Undefined Saprotroph\| | Yeast | NULL | Probable | As nectar-tap_saprotroph (Põlme et al. 2020); Decay sugar-rich substrates (Põlme et al. 2020) | Põlme S, et al. 2020. Fungal Diversity 105:1-16 ((https://doi.org/10.1007/s13225-020-00466-2)) |
| Otu10370 | 2 | k__Fungi(100);p__Ascomycota(100);c__Saccharomycetes(100);o__Saccharomycetales(100);f__Metschnikowiaceae(100);g__Metschnikowia(100);g__Metschnikowia_unclassified(100); | Metschnikowia | 13 | Saprotroph | \|Undefined Saprotroph\| | Yeast | NULL | Probable | As nectar-tap_saprotroph (Põlme et al. 2020); Decay sugar-rich substrates (Põlme et al. 2020) | Põlme S, et al. 2020. Fungal Diversity 105:1-16 ((https://doi.org/10.1007/s13225-020-00466-2)) |
| Otu10372 | 2 | k__Fungi(100);p__Ascomycota(100);c__Saccharomycetes(100);o__Saccharomycetales(100);f__Metschnikowiaceae(100);g__Metschnikowia(100);g__Metschnikowia_unclassified(100); | Metschnikowia | 13 | Saprotroph | \|Undefined Saprotroph\| | Yeast | NULL | Probable | As nectar-tap_saprotroph (Põlme et al. 2020); Decay sugar-rich substrates (Põlme et al. 2020) | Põlme S, et al. 2020. Fungal Diversity 105:1-16 ((https://doi.org/10.1007/s13225-020-00466-2)) |
| Otu10374 | 2 | k__Fungi(100);p__Ascomycota(100);c__Saccharomycetes(100);o__Saccharomycetales(100);f__Metschnikowiaceae(100);g__Metschnikowia(100);s__Metschnikowia_pulcherrima(100); | Metschnikowia | 13 | Saprotroph | \|Undefined Saprotroph\| | Yeast | NULL | Probable | As nectar-tap_saprotroph (Põlme et al. 2020); Decay sugar-rich substrates (Põlme et al. 2020) | Põlme S, et al. 2020. Fungal Diversity 105:1-16 ((https://doi.org/10.1007/s13225-020-00466-2)) |
| Otu10376 | 2 | k__Fungi(100);p__Ascomycota(100);c__Saccharomycetes(100);o__Saccharomycetales(100);f__Metschnikowiaceae(100);g__Metschnikowia(100);s__Metschnikowia_pulcherrima(100); | Metschnikowia | 13 | Saprotroph | \|Undefined Saprotroph\| | Yeast | NULL | Probable | As nectar-tap_saprotroph (Põlme et al. 2020); Decay sugar-rich substrates (Põlme et al. 2020) | Põlme S, et al. 2020. Fungal Diversity 105:1-16 ((https://doi.org/10.1007/s13225-020-00466-2)) |
| Otu10377 | 2 | k__Fungi(100);p__Ascomycota(100);c__Saccharomycetes(100);o__Saccharomycetales(100);f__Metschnikowiaceae(100);g__Metschnikowia(100);s__Metschnikowia_pulcherrima(100); | Metschnikowia | 13 | Saprotroph | \|Undefined Saprotroph\| | Yeast | NULL | Probable | As nectar-tap_saprotroph (Põlme et al. 2020); Decay sugar-rich substrates (Põlme et al. 2020) | Põlme S, et al. 2020. Fungal Diversity 105:1-16 ((https://doi.org/10.1007/s13225-020-00466-2)) |
| Otu10379 | 2 | k__Fungi(100);p__Ascomycota(100);c__Saccharomycetes(100);o__Saccharomycetales(100);f__Metschnikowiaceae(100);g__Metschnikowia(100);s__Metschnikowia_pulcherrima(100); | Metschnikowia | 13 | Saprotroph | \|Undefined Saprotroph\| | Yeast | NULL | Probable | As nectar-tap_saprotroph (Põlme et al. 2020); Decay sugar-rich substrates (Põlme et al. 2020) | Põlme S, et al. 2020. Fungal Diversity 105:1-16 ((https://doi.org/10.1007/s13225-020-00466-2)) |
| Otu10382 | 2 | k__Fungi(100);p__Ascomycota(100);c__Saccharomycetes(100);o__Saccharomycetales(100);f__Metschnikowiaceae(100);g__Metschnikowia(100);s__Metschnikowia_pulcherrima(100); | Metschnikowia | 13 | Saprotroph | \|Undefined Saprotroph\| | Yeast | NULL | Probable | As nectar-tap_saprotroph (Põlme et al. 2020); Decay sugar-rich substrates (Põlme et al. 2020) | Põlme S, et al. 2020. Fungal Diversity 105:1-16 ((https://doi.org/10.1007/s13225-020-00466-2)) |
| Otu10386 | 2 | k__Fungi(100);p__Ascomycota(100);c__Saccharomycetes(100);o__Saccharomycetales(100);f__Metschnikowiaceae(100);g__Metschnikowia(100);s__Metschnikowia_pulcherrima(100); | Metschnikowia | 13 | Saprotroph | \|Undefined Saprotroph\| | Yeast | NULL | Probable | As nectar-tap_saprotroph (Põlme et al. 2020); Decay sugar-rich substrates (Põlme et al. 2020) | Põlme S, et al. 2020. Fungal Diversity 105:1-16 ((https://doi.org/10.1007/s13225-020-00466-2)) |
| Otu10392 | 2 | k__Fungi(100);p__Ascomycota(100);c__Saccharomycetes(100);o__Saccharomycetales(100);f__Metschnikowiaceae(100);g__Metschnikowia(100);s__Metschnikowia_pulcherrima(100); | Metschnikowia | 13 | Saprotroph | \|Undefined Saprotroph\| | Yeast | NULL | Probable | As nectar-tap_saprotroph (Põlme et al. 2020); Decay sugar-rich substrates (Põlme et al. 2020) | Põlme S, et al. 2020. Fungal Diversity 105:1-16 ((https://doi.org/10.1007/s13225-020-00466-2)) |
| Otu10396 | 2 | k__Fungi(100);p__Ascomycota(100);c__Saccharomycetes(100);o__Saccharomycetales(100);f__Metschnikowiaceae(100);g__Metschnikowia(100);g__Metschnikowia_unclassified(100); | Metschnikowia | 13 | Saprotroph | \|Undefined Saprotroph\| | Yeast | NULL | Probable | As nectar-tap_saprotroph (Põlme et al. 2020); Decay sugar-rich substrates (Põlme et al. 2020) | Põlme S, et al. 2020. Fungal Diversity 105:1-16 ((https://doi.org/10.1007/s13225-020-00466-2)) |
| Otu10397 | 2 | k__Fungi(100);p__Ascomycota(100);c__Dothideomycetes(100);o__Pleosporales(100);f__Venturiaceae(100);g__Venturia(100);g__Venturia_unclassified(100); | Venturia | 13 | Pathotroph-Pathotroph-Saprotroph | \|Plant Pathogen\|-Plant Saprotroph | Microfungus | NULL | Probable | As litter_saprotroph (Põlme et al. 2020); Decay leaf, fruit, seed (Põlme et al. 2020) | Tedersoo L, et al. 2014. Science 346:e1256688 ((https://doi.org/10.1126/science.1256688)); Põlme S, et al. 2020. Fungal Diversity 105:1-16 ((https://doi.org/10.1007/s13225-020-00466-2)) |
| Otu10399 | 2 | k__Fungi(100);p__Ascomycota(100);c__Dothideomycetes(100);o__Pleosporales(100);f__Venturiaceae(100);g__Venturia(100);s__Venturia_asperata(100); | Venturia | 13 | Pathotroph-Pathotroph-Saprotroph | \|Plant Pathogen\|-Plant Saprotroph | Microfungus | NULL | Probable | As litter_saprotroph (Põlme et al. 2020); Decay leaf, fruit, seed (Põlme et al. 2020) | Tedersoo L, et al. 2014. Science 346:e1256688 ((https://doi.org/10.1126/science.1256688)); Põlme S, et al. 2020. Fungal Diversity 105:1-16 ((https://doi.org/10.1007/s13225-020-00466-2)) |
| Otu10402 | 2 | k__Fungi(100);p__Ascomycota(100);c__Dothideomycetes(100);o__Capnodiales(100);f__Mycosphaerellaceae(100);g__Pseudocercosporella(100);s__Pseudocercosporella_sp(100); | Pseudocercosporella | 13 | Pathotroph | \|Plant Pathogen\| | Microfungus | NULL | Probable | NULL | Tedersoo L, et al. 2014. Science 346:e1256688 ((https://doi.org/10.1126/science.1256688)); Põlme S, et al. 2020. Fungal Diversity 105:1-16 ((https://doi.org/10.1007/s13225-020-00466-2)) |
| Otu10403 | 2 | k__Fungi(100);p__Ascomycota(100);c__Saccharomycetes(100);o__Saccharomycetales(100);f__Metschnikowiaceae(100);g__Metschnikowia(100);g__Metschnikowia_unclassified(100); | Metschnikowia | 13 | Saprotroph | \|Undefined Saprotroph\| | Yeast | NULL | Probable | As nectar-tap_saprotroph (Põlme et al. 2020); Decay sugar-rich substrates (Põlme et al. 2020) | Põlme S, et al. 2020. Fungal Diversity 105:1-16 ((https://doi.org/10.1007/s13225-020-00466-2)) |
| Otu10404 | 2 | k__Fungi(100);p__Ascomycota(100);c__Saccharomycetes(100);o__Saccharomycetales(100);f__Metschnikowiaceae(100);g__Metschnikowia(100);g__Metschnikowia_unclassified(100); | Metschnikowia | 13 | Saprotroph | \|Undefined Saprotroph\| | Yeast | NULL | Probable | As nectar-tap_saprotroph (Põlme et al. 2020); Decay sugar-rich substrates (Põlme et al. 2020) | Põlme S, et al. 2020. Fungal Diversity 105:1-16 ((https://doi.org/10.1007/s13225-020-00466-2)) |
| Otu10405 | 2 | k__Fungi(100);p__Ascomycota(100);c__Saccharomycetes(100);o__Saccharomycetales(100);f__Metschnikowiaceae(100);g__Metschnikowia(100);s__Metschnikowia_pulcherrima(100); | Metschnikowia | 13 | Saprotroph | \|Undefined Saprotroph\| | Yeast | NULL | Probable | As nectar-tap_saprotroph (Põlme et al. 2020); Decay sugar-rich substrates (Põlme et al. 2020) | Põlme S, et al. 2020. Fungal Diversity 105:1-16 ((https://doi.org/10.1007/s13225-020-00466-2)) |
| Otu10407 | 2 | k__Fungi(100);p__Ascomycota(100);c__Saccharomycetes(100);o__Saccharomycetales(100);f__Metschnikowiaceae(100);g__Metschnikowia(100);s__Metschnikowia_pulcherrima(100); | Metschnikowia | 13 | Saprotroph | \|Undefined Saprotroph\| | Yeast | NULL | Probable | As nectar-tap_saprotroph (Põlme et al. 2020); Decay sugar-rich substrates (Põlme et al. 2020) | Põlme S, et al. 2020. Fungal Diversity 105:1-16 ((https://doi.org/10.1007/s13225-020-00466-2)) |
| Otu10409 | 2 | k__Fungi(100);p__Ascomycota(100);c__Saccharomycetes(100);o__Saccharomycetales(100);f__Metschnikowiaceae(100);g__Metschnikowia(100);s__Metschnikowia_pulcherrima(100); | Metschnikowia | 13 | Saprotroph | \|Undefined Saprotroph\| | Yeast | NULL | Probable | As nectar-tap_saprotroph (Põlme et al. 2020); Decay sugar-rich substrates (Põlme et al. 2020) | Põlme S, et al. 2020. Fungal Diversity 105:1-16 ((https://doi.org/10.1007/s13225-020-00466-2)) |
| Otu10411 | 2 | k__Fungi(100);p__Ascomycota(100);c__Saccharomycetes(100);o__Saccharomycetales(100);f__Saccharomycetales_family_Incertae_sedis(100);f__Saccharomycetales_family_Incertae_sedis_unclassified(100);f__Saccharomycetales_family_Incertae_sedis_unclassified(100); | Saccharomycetales | 7 | Saprotroph | Undefined Saprotroph | Yeast | NULL | Possible | NULL | Sterkenburg E, et al. 2015. New Phytologist 207:1145-1158 ((https://doi.org/10.1111/nph.13426)) |
| Otu10414 | 2 | k__Fungi(100);p__Ascomycota(100);c__Saccharomycetes(100);o__Saccharomycetales(100);f__Metschnikowiaceae(100);g__Metschnikowia(100);s__Metschnikowia_pulcherrima(100); | Metschnikowia | 13 | Saprotroph | \|Undefined Saprotroph\| | Yeast | NULL | Probable | As nectar-tap_saprotroph (Põlme et al. 2020); Decay sugar-rich substrates (Põlme et al. 2020) | Põlme S, et al. 2020. Fungal Diversity 105:1-16 ((https://doi.org/10.1007/s13225-020-00466-2)) |
| Otu10416 | 2 | k__Fungi(100);p__Ascomycota(100);c__Saccharomycetes(100);o__Saccharomycetales(100);f__Saccharomycetales_family_Incertae_sedis(100);g__Candida(100);g__Candida_unclassified(100); | Candida | 13 | Pathotroph-Saprotroph-Symbiotroph | Animal Pathogen-Endophyte-Endosymbiont-Epiphyte-\|Undefined Saprotroph\| | Dimorphic Yeast-Yeast | NULL | Probable | As nectar-tap_saprotroph (Põlme et al. 2020); Decay sugar-rich substrates (Põlme et al. 2020); This asexual genus is polyphyletic and contain species that function very differently depend on what they are. Caution should be taken when interpreting guilds for Candida species. | Manolakaki D, et al. 2010. Virulence. 1: 367-375 ((https://doi.org/10.4161/viru.1.5.12796)); Mekbib SB, et al. 2011. Tropical Plant Pathology 36:233-240 ((https://doi.org/10.1590/S1982-56762011000400004)); Põlme S, et al. 2020. Fungal Diversity 105:1-16 ((https://doi.org/10.1007/s13225-020-00466-2)) |
| Otu10419 | 2 | k__Fungi(100);p__Ascomycota(100);c__Saccharomycetes(100);o__Saccharomycetales(100);f__Metschnikowiaceae(100);g__Metschnikowia(100);g__Metschnikowia_unclassified(100); | Metschnikowia | 13 | Saprotroph | \|Undefined Saprotroph\| | Yeast | NULL | Probable | As nectar-tap_saprotroph (Põlme et al. 2020); Decay sugar-rich substrates (Põlme et al. 2020) | Põlme S, et al. 2020. Fungal Diversity 105:1-16 ((https://doi.org/10.1007/s13225-020-00466-2)) |
| Otu10420 | 2 | k__Fungi(100);p__Ascomycota(100);c__Saccharomycetes(100);o__Saccharomycetales(100);f__Metschnikowiaceae(100);g__Metschnikowia(100);s__Metschnikowia_pulcherrima(100); | Metschnikowia | 13 | Saprotroph | \|Undefined Saprotroph\| | Yeast | NULL | Probable | As nectar-tap_saprotroph (Põlme et al. 2020); Decay sugar-rich substrates (Põlme et al. 2020) | Põlme S, et al. 2020. Fungal Diversity 105:1-16 ((https://doi.org/10.1007/s13225-020-00466-2)) |
| Otu10427 | 2 | k__Fungi(100);p__Ascomycota(100);c__Saccharomycetes(100);o__Saccharomycetales(100);f__Metschnikowiaceae(100);g__Metschnikowia(100);s__Metschnikowia_pulcherrima(100); | Metschnikowia | 13 | Saprotroph | \|Undefined Saprotroph\| | Yeast | NULL | Probable | As nectar-tap_saprotroph (Põlme et al. 2020); Decay sugar-rich substrates (Põlme et al. 2020) | Põlme S, et al. 2020. Fungal Diversity 105:1-16 ((https://doi.org/10.1007/s13225-020-00466-2)) |
| Otu10432 | 2 | k__Fungi(100);p__Ascomycota(100);c__Saccharomycetes(100);o__Saccharomycetales(100);f__Metschnikowiaceae(100);g__Metschnikowia(100);s__Metschnikowia_pulcherrima(100); | Metschnikowia | 13 | Saprotroph | \|Undefined Saprotroph\| | Yeast | NULL | Probable | As nectar-tap_saprotroph (Põlme et al. 2020); Decay sugar-rich substrates (Põlme et al. 2020) | Põlme S, et al. 2020. Fungal Diversity 105:1-16 ((https://doi.org/10.1007/s13225-020-00466-2)) |
| Otu10433 | 2 | k__Fungi(100);p__Ascomycota(100);c__Saccharomycetes(100);o__Saccharomycetales(100);f__Metschnikowiaceae(100);g__Metschnikowia(100);g__Metschnikowia_unclassified(100); | Metschnikowia | 13 | Saprotroph | \|Undefined Saprotroph\| | Yeast | NULL | Probable | As nectar-tap_saprotroph (Põlme et al. 2020); Decay sugar-rich substrates (Põlme et al. 2020) | Põlme S, et al. 2020. Fungal Diversity 105:1-16 ((https://doi.org/10.1007/s13225-020-00466-2)) |
| Otu10434 | 2 | k__Fungi(100);p__Ascomycota(100);c__Saccharomycetes(100);o__Saccharomycetales(100);f__Metschnikowiaceae(100);g__Metschnikowia(100);s__Metschnikowia_pulcherrima(100); | Metschnikowia | 13 | Saprotroph | \|Undefined Saprotroph\| | Yeast | NULL | Probable | As nectar-tap_saprotroph (Põlme et al. 2020); Decay sugar-rich substrates (Põlme et al. 2020) | Põlme S, et al. 2020. Fungal Diversity 105:1-16 ((https://doi.org/10.1007/s13225-020-00466-2)) |
| Otu10435 | 2 | k__Fungi(100);p__Ascomycota(100);c__Ascomycota_class_Incertae_sedis(100);o__Ascomycota_order_Incertae_sedis(100);f__Ascomycota_family_Incertae_sedis(100);g__Microcyclospora(100);s__Microcyclospora_quercina(100); | Microcyclospora | 13 | Pathotroph | \|Plant Pathogen\| | Microfungus | NULL | Probable | NULL | Põlme S, et al. 2020. Fungal Diversity 105:1-16 ((https://doi.org/10.1007/s13225-020-00466-2)) |
| Otu10436 | 2 | k__Fungi(100);p__Ascomycota(100);c__Saccharomycetes(100);o__Saccharomycetales(100);f__Metschnikowiaceae(100);g__Metschnikowia(100);g__Metschnikowia_unclassified(100); | Metschnikowia | 13 | Saprotroph | \|Undefined Saprotroph\| | Yeast | NULL | Probable | As nectar-tap_saprotroph (Põlme et al. 2020); Decay sugar-rich substrates (Põlme et al. 2020) | Põlme S, et al. 2020. Fungal Diversity 105:1-16 ((https://doi.org/10.1007/s13225-020-00466-2)) |
| Otu10437 | 2 | k__Fungi(100);p__Ascomycota(100);c__Saccharomycetes(100);o__Saccharomycetales(100);f__Metschnikowiaceae(100);g__Metschnikowia(100);s__Metschnikowia_pulcherrima(100); | Metschnikowia | 13 | Saprotroph | \|Undefined Saprotroph\| | Yeast | NULL | Probable | As nectar-tap_saprotroph (Põlme et al. 2020); Decay sugar-rich substrates (Põlme et al. 2020) | Põlme S, et al. 2020. Fungal Diversity 105:1-16 ((https://doi.org/10.1007/s13225-020-00466-2)) |
| Otu10438 | 2 | k__Fungi(100);p__Ascomycota(100);c__Saccharomycetes(100);o__Saccharomycetales(100);f__Metschnikowiaceae(100);g__Metschnikowia(100);s__Metschnikowia_pulcherrima(100); | Metschnikowia | 13 | Saprotroph | \|Undefined Saprotroph\| | Yeast | NULL | Probable | As nectar-tap_saprotroph (Põlme et al. 2020); Decay sugar-rich substrates (Põlme et al. 2020) | Põlme S, et al. 2020. Fungal Diversity 105:1-16 ((https://doi.org/10.1007/s13225-020-00466-2)) |
| Otu10439 | 2 | k__Fungi(100);p__Ascomycota(100);c__Saccharomycetes(100);o__Saccharomycetales(100);f__Metschnikowiaceae(100);g__Metschnikowia(100);g__Metschnikowia_unclassified(100); | Metschnikowia | 13 | Saprotroph | \|Undefined Saprotroph\| | Yeast | NULL | Probable | As nectar-tap_saprotroph (Põlme et al. 2020); Decay sugar-rich substrates (Põlme et al. 2020) | Põlme S, et al. 2020. Fungal Diversity 105:1-16 ((https://doi.org/10.1007/s13225-020-00466-2)) |
| Otu10440 | 2 | k__Fungi(100);p__Ascomycota(100);c__Saccharomycetes(100);o__Saccharomycetales(100);f__Metschnikowiaceae(100);g__Metschnikowia(100);s__Metschnikowia_pulcherrima(100); | Metschnikowia | 13 | Saprotroph | \|Undefined Saprotroph\| | Yeast | NULL | Probable | As nectar-tap_saprotroph (Põlme et al. 2020); Decay sugar-rich substrates (Põlme et al. 2020) | Põlme S, et al. 2020. Fungal Diversity 105:1-16 ((https://doi.org/10.1007/s13225-020-00466-2)) |
| Otu10441 | 2 | k__Fungi(100);p__Ascomycota(100);c__Saccharomycetes(100);o__Saccharomycetales(100);f__Metschnikowiaceae(100);g__Metschnikowia(100);g__Metschnikowia_unclassified(100); | Metschnikowia | 13 | Saprotroph | \|Undefined Saprotroph\| | Yeast | NULL | Probable | As nectar-tap_saprotroph (Põlme et al. 2020); Decay sugar-rich substrates (Põlme et al. 2020) | Põlme S, et al. 2020. Fungal Diversity 105:1-16 ((https://doi.org/10.1007/s13225-020-00466-2)) |
| Otu10442 | 2 | k__Fungi(100);p__Basidiomycota(100);c__Microbotryomycetes(100);o__Sporidiobolales(100);f__Sporidiobolales_family_Incertae_sedis(100);g__Sporobolomyces(100);s__Sporobolomyces_lactosus(100); | Sporobolomyces | 13 | Pathotroph-Saprotroph | \|Fungal Parasite\|-Plant Saprotroph | Yeast | NULL | Probable | As litter_saprotroph (Põlme et al. 2020); As mycoparasite (Põlme et al. 2020); Decay fungal material (Põlme et al. 2020) | Tedersoo L, et al. 2014. Science 346:e1256688 ((https://doi.org/10.1126/science.1256688)); Purahong W, et al. 2016. Molecular Ecology 25:4059-4074 ((https://doi.org/10.1111/mec.13739)); Põlme S, et al. 2020. Fungal Diversity 105:1-16 ((https://doi.org/10.1007/s13225-020-00466-2)) |
| Otu10445 | 2 | k__Fungi(100);p__Ascomycota(100);c__Dothideomycetes(100);o__Capnodiales(100);f__Mycosphaerellaceae(100);f__Mycosphaerellaceae_unclassified(100);f__Mycosphaerellaceae_unclassified(100); | Mycosphaerellaceae | 9 | Pathotroph-Saprotroph | Plant Pathogen-Undefined Saprotroph | Microfungus | NULL | Probable | Biotrophic, necrotrophic or saprobic on various plant tissues (Cannon & Kirk 2007) | Cannon PF, Kirk PM. 2007. Fungal Families of the World. CAB International, Cambridge (ISBN: 978-0851998275) |
| Otu10449 | 2 | k__Fungi(100);p__Ascomycota(100);c__Saccharomycetes(100);o__Saccharomycetales(100);f__Metschnikowiaceae(100);g__Metschnikowia(100);g__Metschnikowia_unclassified(100); | Metschnikowia | 13 | Saprotroph | \|Undefined Saprotroph\| | Yeast | NULL | Probable | As nectar-tap_saprotroph (Põlme et al. 2020); Decay sugar-rich substrates (Põlme et al. 2020) | Põlme S, et al. 2020. Fungal Diversity 105:1-16 ((https://doi.org/10.1007/s13225-020-00466-2)) |
| Otu10450 | 2 | k__Fungi(100);p__Ascomycota(100);c__Saccharomycetes(100);o__Saccharomycetales(100);f__Metschnikowiaceae(100);g__Metschnikowia(100);g__Metschnikowia_unclassified(100); | Metschnikowia | 13 | Saprotroph | \|Undefined Saprotroph\| | Yeast | NULL | Probable | As nectar-tap_saprotroph (Põlme et al. 2020); Decay sugar-rich substrates (Põlme et al. 2020) | Põlme S, et al. 2020. Fungal Diversity 105:1-16 ((https://doi.org/10.1007/s13225-020-00466-2)) |
| Otu10451 | 2 | k__Fungi(100);p__Ascomycota(100);c__Saccharomycetes(100);o__Saccharomycetales(100);f__Metschnikowiaceae(100);g__Metschnikowia(100);s__Metschnikowia_pulcherrima(100); | Metschnikowia | 13 | Saprotroph | \|Undefined Saprotroph\| | Yeast | NULL | Probable | As nectar-tap_saprotroph (Põlme et al. 2020); Decay sugar-rich substrates (Põlme et al. 2020) | Põlme S, et al. 2020. Fungal Diversity 105:1-16 ((https://doi.org/10.1007/s13225-020-00466-2)) |
| Otu10452 | 2 | k__Fungi(100);p__Ascomycota(100);c__Saccharomycetes(100);o__Saccharomycetales(100);f__Metschnikowiaceae(100);g__Metschnikowia(100);g__Metschnikowia_unclassified(100); | Metschnikowia | 13 | Saprotroph | \|Undefined Saprotroph\| | Yeast | NULL | Probable | As nectar-tap_saprotroph (Põlme et al. 2020); Decay sugar-rich substrates (Põlme et al. 2020) | Põlme S, et al. 2020. Fungal Diversity 105:1-16 ((https://doi.org/10.1007/s13225-020-00466-2)) |
| Otu10453 | 2 | k__Fungi(100);p__Ascomycota(100);c__Dothideomycetes(100);o__Dothideomycetes_order_Incertae_sedis(100);f__Pseudeurotiaceae(100);g__Pseudeurotium(100);g__Pseudeurotium_unclassified(100); | Pseudeurotium | 13 | Saprotroph | \|Dung Saprotroph\|-Undefined Saprotroph | Microfungus | NULL | Probable | As soil_saprotroph (Põlme et al. 2020); Decay soil, dung (Põlme et al. 2020) | Tedersoo L, et al. 2014. Science 346:e1256688 ((https://doi.org/10.1126/science.1256688)); Põlme S, et al. 2020. Fungal Diversity 105:1-16 ((https://doi.org/10.1007/s13225-020-00466-2)) |
| Otu10454 | 2 | k__Fungi(100);p__Ascomycota(100);c__Saccharomycetes(100);o__Saccharomycetales(100);f__Metschnikowiaceae(100);g__Metschnikowia(100);s__Metschnikowia_pulcherrima(100); | Metschnikowia | 13 | Saprotroph | \|Undefined Saprotroph\| | Yeast | NULL | Probable | As nectar-tap_saprotroph (Põlme et al. 2020); Decay sugar-rich substrates (Põlme et al. 2020) | Põlme S, et al. 2020. Fungal Diversity 105:1-16 ((https://doi.org/10.1007/s13225-020-00466-2)) |
| Otu10455 | 2 | k__Fungi(100);p__Ascomycota(100);c__Saccharomycetes(100);o__Saccharomycetales(100);f__Metschnikowiaceae(100);g__Metschnikowia(100);g__Metschnikowia_unclassified(100); | Metschnikowia | 13 | Saprotroph | \|Undefined Saprotroph\| | Yeast | NULL | Probable | As nectar-tap_saprotroph (Põlme et al. 2020); Decay sugar-rich substrates (Põlme et al. 2020) | Põlme S, et al. 2020. Fungal Diversity 105:1-16 ((https://doi.org/10.1007/s13225-020-00466-2)) |
| Otu10458 | 2 | k__Fungi(100);p__Ascomycota(100);c__Saccharomycetes(100);o__Saccharomycetales(100);f__Metschnikowiaceae(100);g__Metschnikowia(100);s__Metschnikowia_pulcherrima(100); | Metschnikowia | 13 | Saprotroph | \|Undefined Saprotroph\| | Yeast | NULL | Probable | As nectar-tap_saprotroph (Põlme et al. 2020); Decay sugar-rich substrates (Põlme et al. 2020) | Põlme S, et al. 2020. Fungal Diversity 105:1-16 ((https://doi.org/10.1007/s13225-020-00466-2)) |
| Otu10459 | 2 | k__Fungi(100);p__Ascomycota(100);c__Saccharomycetes(100);o__Saccharomycetales(100);f__Metschnikowiaceae(100);g__Metschnikowia(100);g__Metschnikowia_unclassified(100); | Metschnikowia | 13 | Saprotroph | \|Undefined Saprotroph\| | Yeast | NULL | Probable | As nectar-tap_saprotroph (Põlme et al. 2020); Decay sugar-rich substrates (Põlme et al. 2020) | Põlme S, et al. 2020. Fungal Diversity 105:1-16 ((https://doi.org/10.1007/s13225-020-00466-2)) |
| Otu10462 | 2 | k__Fungi(100);p__Ascomycota(100);c__Saccharomycetes(100);o__Saccharomycetales(100);f__Metschnikowiaceae(100);g__Metschnikowia(100);g__Metschnikowia_unclassified(100); | Metschnikowia | 13 | Saprotroph | \|Undefined Saprotroph\| | Yeast | NULL | Probable | As nectar-tap_saprotroph (Põlme et al. 2020); Decay sugar-rich substrates (Põlme et al. 2020) | Põlme S, et al. 2020. Fungal Diversity 105:1-16 ((https://doi.org/10.1007/s13225-020-00466-2)) |
| Otu10463 | 2 | k__Fungi(100);p__Ascomycota(100);c__Saccharomycetes(100);o__Saccharomycetales(100);f__Metschnikowiaceae(100);g__Metschnikowia(100);g__Metschnikowia_unclassified(100); | Metschnikowia | 13 | Saprotroph | \|Undefined Saprotroph\| | Yeast | NULL | Probable | As nectar-tap_saprotroph (Põlme et al. 2020); Decay sugar-rich substrates (Põlme et al. 2020) | Põlme S, et al. 2020. Fungal Diversity 105:1-16 ((https://doi.org/10.1007/s13225-020-00466-2)) |
| Otu10464 | 2 | k__Fungi(100);p__Ascomycota(100);c__Saccharomycetes(100);o__Saccharomycetales(100);f__Metschnikowiaceae(100);g__Metschnikowia(100);s__Metschnikowia_pulcherrima(100); | Metschnikowia | 13 | Saprotroph | \|Undefined Saprotroph\| | Yeast | NULL | Probable | As nectar-tap_saprotroph (Põlme et al. 2020); Decay sugar-rich substrates (Põlme et al. 2020) | Põlme S, et al. 2020. Fungal Diversity 105:1-16 ((https://doi.org/10.1007/s13225-020-00466-2)) |
| Otu10465 | 2 | k__Fungi(100);p__Basidiomycota(100);c__Microbotryomycetes(100);o__Sporidiobolales(100);f__Sporidiobolales_family_Incertae_sedis(100);g__Sporobolomyces(100);s__Sporobolomyces_lactosus(100); | Sporobolomyces | 13 | Pathotroph-Saprotroph | \|Fungal Parasite\|-Plant Saprotroph | Yeast | NULL | Probable | As litter_saprotroph (Põlme et al. 2020); As mycoparasite (Põlme et al. 2020); Decay fungal material (Põlme et al. 2020) | Tedersoo L, et al. 2014. Science 346:e1256688 ((https://doi.org/10.1126/science.1256688)); Purahong W, et al. 2016. Molecular Ecology 25:4059-4074 ((https://doi.org/10.1111/mec.13739)); Põlme S, et al. 2020. Fungal Diversity 105:1-16 ((https://doi.org/10.1007/s13225-020-00466-2)) |
| Otu10467 | 2 | k__Fungi(100);p__Ascomycota(100);c__Saccharomycetes(100);o__Saccharomycetales(100);f__Saccharomycetales_family_Incertae_sedis(100);g__Candida(100);g__Candida_unclassified(100); | Candida | 13 | Pathotroph-Saprotroph-Symbiotroph | Animal Pathogen-Endophyte-Endosymbiont-Epiphyte-\|Undefined Saprotroph\| | Dimorphic Yeast-Yeast | NULL | Probable | As nectar-tap_saprotroph (Põlme et al. 2020); Decay sugar-rich substrates (Põlme et al. 2020); This asexual genus is polyphyletic and contain species that function very differently depend on what they are. Caution should be taken when interpreting guilds for Candida species. | Manolakaki D, et al. 2010. Virulence. 1: 367-375 ((https://doi.org/10.4161/viru.1.5.12796)); Mekbib SB, et al. 2011. Tropical Plant Pathology 36:233-240 ((https://doi.org/10.1590/S1982-56762011000400004)); Põlme S, et al. 2020. Fungal Diversity 105:1-16 ((https://doi.org/10.1007/s13225-020-00466-2)) |
| Otu10468 | 2 | k__Fungi(100);p__Ascomycota(100);c__Dothideomycetes(100);o__Capnodiales(100);f__Teratosphaeriaceae(100);g__Catenulostroma(100);s__Catenulostroma_hermanusense(100); | Catenulostroma | 13 | Pathotroph | \|Plant Pathogen\| | Microfungus | NULL | Probable | NULL | Tedersoo L, et al. 2014. Science 346:e1256688 ((https://doi.org/10.1126/science.1256688)); Põlme S, et al. 2020. Fungal Diversity 105:1-16 ((https://doi.org/10.1007/s13225-020-00466-2)) |
| Otu10469 | 2 | k__Fungi(100);p__Ascomycota(100);c__Saccharomycetes(100);o__Saccharomycetales(100);f__Saccharomycetales_family_Incertae_sedis(100);f__Saccharomycetales_family_Incertae_sedis_unclassified(100);f__Saccharomycetales_family_Incertae_sedis_unclassified(100); | Saccharomycetales | 7 | Saprotroph | Undefined Saprotroph | Yeast | NULL | Possible | NULL | Sterkenburg E, et al. 2015. New Phytologist 207:1145-1158 ((https://doi.org/10.1111/nph.13426)) |
| Otu10470 | 2 | k__Fungi(100);p__Ascomycota(100);c__Saccharomycetes(100);o__Saccharomycetales(100);f__Metschnikowiaceae(100);g__Metschnikowia(100);g__Metschnikowia_unclassified(100); | Metschnikowia | 13 | Saprotroph | \|Undefined Saprotroph\| | Yeast | NULL | Probable | As nectar-tap_saprotroph (Põlme et al. 2020); Decay sugar-rich substrates (Põlme et al. 2020) | Põlme S, et al. 2020. Fungal Diversity 105:1-16 ((https://doi.org/10.1007/s13225-020-00466-2)) |
| Otu10473 | 2 | k__Fungi(100);p__Ascomycota(100);c__Saccharomycetes(100);o__Saccharomycetales(100);f__Saccharomycetales_family_Incertae_sedis(100);g__Candida(100);g__Candida_unclassified(100); | Candida | 13 | Pathotroph-Saprotroph-Symbiotroph | Animal Pathogen-Endophyte-Endosymbiont-Epiphyte-\|Undefined Saprotroph\| | Dimorphic Yeast-Yeast | NULL | Probable | As nectar-tap_saprotroph (Põlme et al. 2020); Decay sugar-rich substrates (Põlme et al. 2020); This asexual genus is polyphyletic and contain species that function very differently depend on what they are. Caution should be taken when interpreting guilds for Candida species. | Manolakaki D, et al. 2010. Virulence. 1: 367-375 ((https://doi.org/10.4161/viru.1.5.12796)); Mekbib SB, et al. 2011. Tropical Plant Pathology 36:233-240 ((https://doi.org/10.1590/S1982-56762011000400004)); Põlme S, et al. 2020. Fungal Diversity 105:1-16 ((https://doi.org/10.1007/s13225-020-00466-2)) |
| Otu10474 | 2 | k__Fungi(100);p__Ascomycota(100);c__Saccharomycetes(100);o__Saccharomycetales(100);f__Metschnikowiaceae(100);g__Metschnikowia(100);g__Metschnikowia_unclassified(100); | Metschnikowia | 13 | Saprotroph | \|Undefined Saprotroph\| | Yeast | NULL | Probable | As nectar-tap_saprotroph (Põlme et al. 2020); Decay sugar-rich substrates (Põlme et al. 2020) | Põlme S, et al. 2020. Fungal Diversity 105:1-16 ((https://doi.org/10.1007/s13225-020-00466-2)) |
| Otu10476 | 2 | k__Fungi(100);p__Ascomycota(100);c__Saccharomycetes(100);o__Saccharomycetales(100);f__Metschnikowiaceae(100);g__Metschnikowia(100);g__Metschnikowia_unclassified(100); | Metschnikowia | 13 | Saprotroph | \|Undefined Saprotroph\| | Yeast | NULL | Probable | As nectar-tap_saprotroph (Põlme et al. 2020); Decay sugar-rich substrates (Põlme et al. 2020) | Põlme S, et al. 2020. Fungal Diversity 105:1-16 ((https://doi.org/10.1007/s13225-020-00466-2)) |
| Otu10477 | 2 | k__Fungi(100);p__Ascomycota(100);c__Saccharomycetes(100);o__Saccharomycetales(100);f__Metschnikowiaceae(100);g__Metschnikowia(100);g__Metschnikowia_unclassified(100); | Metschnikowia | 13 | Saprotroph | \|Undefined Saprotroph\| | Yeast | NULL | Probable | As nectar-tap_saprotroph (Põlme et al. 2020); Decay sugar-rich substrates (Põlme et al. 2020) | Põlme S, et al. 2020. Fungal Diversity 105:1-16 ((https://doi.org/10.1007/s13225-020-00466-2)) |
| Otu10478 | 2 | k__Fungi(100);p__Ascomycota(100);c__Saccharomycetes(100);o__Saccharomycetales(100);f__Metschnikowiaceae(100);g__Metschnikowia(100);s__Metschnikowia_pulcherrima(100); | Metschnikowia | 13 | Saprotroph | \|Undefined Saprotroph\| | Yeast | NULL | Probable | As nectar-tap_saprotroph (Põlme et al. 2020); Decay sugar-rich substrates (Põlme et al. 2020) | Põlme S, et al. 2020. Fungal Diversity 105:1-16 ((https://doi.org/10.1007/s13225-020-00466-2)) |
| Otu10479 | 2 | k__Fungi(100);p__Basidiomycota(100);c__Microbotryomycetes(100);o__Sporidiobolales(100);f__Sporidiobolales_family_Incertae_sedis(100);g__Sporobolomyces(100);s__Sporobolomyces_lactosus(100); | Sporobolomyces | 13 | Pathotroph-Saprotroph | \|Fungal Parasite\|-Plant Saprotroph | Yeast | NULL | Probable | As litter_saprotroph (Põlme et al. 2020); As mycoparasite (Põlme et al. 2020); Decay fungal material (Põlme et al. 2020) | Tedersoo L, et al. 2014. Science 346:e1256688 ((https://doi.org/10.1126/science.1256688)); Purahong W, et al. 2016. Molecular Ecology 25:4059-4074 ((https://doi.org/10.1111/mec.13739)); Põlme S, et al. 2020. Fungal Diversity 105:1-16 ((https://doi.org/10.1007/s13225-020-00466-2)) |
| Otu10481 | 2 | k__Fungi(100);p__Ascomycota(100);c__Saccharomycetes(100);o__Saccharomycetales(100);f__Metschnikowiaceae(100);g__Metschnikowia(100);g__Metschnikowia_unclassified(100); | Metschnikowia | 13 | Saprotroph | \|Undefined Saprotroph\| | Yeast | NULL | Probable | As nectar-tap_saprotroph (Põlme et al. 2020); Decay sugar-rich substrates (Põlme et al. 2020) | Põlme S, et al. 2020. Fungal Diversity 105:1-16 ((https://doi.org/10.1007/s13225-020-00466-2)) |
| Otu10483 | 2 | k__Fungi(100);p__Ascomycota(100);c__Saccharomycetes(100);o__Saccharomycetales(100);f__Metschnikowiaceae(100);g__Metschnikowia(100);s__Metschnikowia_pulcherrima(100); | Metschnikowia | 13 | Saprotroph | \|Undefined Saprotroph\| | Yeast | NULL | Probable | As nectar-tap_saprotroph (Põlme et al. 2020); Decay sugar-rich substrates (Põlme et al. 2020) | Põlme S, et al. 2020. Fungal Diversity 105:1-16 ((https://doi.org/10.1007/s13225-020-00466-2)) |
| Otu10485 | 2 | k__Fungi(100);p__Ascomycota(100);c__Saccharomycetes(100);o__Saccharomycetales(100);f__Metschnikowiaceae(100);g__Metschnikowia(100);g__Metschnikowia_unclassified(100); | Metschnikowia | 13 | Saprotroph | \|Undefined Saprotroph\| | Yeast | NULL | Probable | As nectar-tap_saprotroph (Põlme et al. 2020); Decay sugar-rich substrates (Põlme et al. 2020) | Põlme S, et al. 2020. Fungal Diversity 105:1-16 ((https://doi.org/10.1007/s13225-020-00466-2)) |
| Otu10486 | 2 | k__Fungi(100);p__Ascomycota(100);c__Saccharomycetes(100);o__Saccharomycetales(100);f__Metschnikowiaceae(100);g__Metschnikowia(100);s__Metschnikowia_pulcherrima(100); | Metschnikowia | 13 | Saprotroph | \|Undefined Saprotroph\| | Yeast | NULL | Probable | As nectar-tap_saprotroph (Põlme et al. 2020); Decay sugar-rich substrates (Põlme et al. 2020) | Põlme S, et al. 2020. Fungal Diversity 105:1-16 ((https://doi.org/10.1007/s13225-020-00466-2)) |
| Otu10487 | 2 | k__Fungi(100);p__Ascomycota(100);c__Saccharomycetes(100);o__Saccharomycetales(100);f__Metschnikowiaceae(100);g__Metschnikowia(100);s__Metschnikowia_pulcherrima(100); | Metschnikowia | 13 | Saprotroph | \|Undefined Saprotroph\| | Yeast | NULL | Probable | As nectar-tap_saprotroph (Põlme et al. 2020); Decay sugar-rich substrates (Põlme et al. 2020) | Põlme S, et al. 2020. Fungal Diversity 105:1-16 ((https://doi.org/10.1007/s13225-020-00466-2)) |
| Otu10490 | 2 | k__Fungi(100);p__Ascomycota(100);c__Saccharomycetes(100);o__Saccharomycetales(100);f__Metschnikowiaceae(100);g__Metschnikowia(100);g__Metschnikowia_unclassified(100); | Metschnikowia | 13 | Saprotroph | \|Undefined Saprotroph\| | Yeast | NULL | Probable | As nectar-tap_saprotroph (Põlme et al. 2020); Decay sugar-rich substrates (Põlme et al. 2020) | Põlme S, et al. 2020. Fungal Diversity 105:1-16 ((https://doi.org/10.1007/s13225-020-00466-2)) |
| Otu10491 | 2 | k__Fungi(100);p__Ascomycota(100);c__Saccharomycetes(100);o__Saccharomycetales(100);f__Metschnikowiaceae(100);g__Metschnikowia(100);s__Metschnikowia_pulcherrima(100); | Metschnikowia | 13 | Saprotroph | \|Undefined Saprotroph\| | Yeast | NULL | Probable | As nectar-tap_saprotroph (Põlme et al. 2020); Decay sugar-rich substrates (Põlme et al. 2020) | Põlme S, et al. 2020. Fungal Diversity 105:1-16 ((https://doi.org/10.1007/s13225-020-00466-2)) |
| Otu10492 | 2 | k__Fungi(100);p__Ascomycota(100);c__Saccharomycetes(100);o__Saccharomycetales(100);f__Metschnikowiaceae(100);g__Metschnikowia(100);g__Metschnikowia_unclassified(100); | Metschnikowia | 13 | Saprotroph | \|Undefined Saprotroph\| | Yeast | NULL | Probable | As nectar-tap_saprotroph (Põlme et al. 2020); Decay sugar-rich substrates (Põlme et al. 2020) | Põlme S, et al. 2020. Fungal Diversity 105:1-16 ((https://doi.org/10.1007/s13225-020-00466-2)) |
| Otu10493 | 2 | k__Fungi(100);p__Ascomycota(100);c__Saccharomycetes(100);o__Saccharomycetales(100);f__Metschnikowiaceae(100);g__Metschnikowia(100);g__Metschnikowia_unclassified(100); | Metschnikowia | 13 | Saprotroph | \|Undefined Saprotroph\| | Yeast | NULL | Probable | As nectar-tap_saprotroph (Põlme et al. 2020); Decay sugar-rich substrates (Põlme et al. 2020) | Põlme S, et al. 2020. Fungal Diversity 105:1-16 ((https://doi.org/10.1007/s13225-020-00466-2)) |
| Otu10494 | 2 | k__Fungi(100);p__Ascomycota(100);c__Saccharomycetes(100);o__Saccharomycetales(100);f__Metschnikowiaceae(100);g__Metschnikowia(100);g__Metschnikowia_unclassified(100); | Metschnikowia | 13 | Saprotroph | \|Undefined Saprotroph\| | Yeast | NULL | Probable | As nectar-tap_saprotroph (Põlme et al. 2020); Decay sugar-rich substrates (Põlme et al. 2020) | Põlme S, et al. 2020. Fungal Diversity 105:1-16 ((https://doi.org/10.1007/s13225-020-00466-2)) |
| Otu10495 | 2 | k__Fungi(100);p__Ascomycota(100);c__Saccharomycetes(100);o__Saccharomycetales(100);f__Metschnikowiaceae(100);g__Metschnikowia(100);g__Metschnikowia_unclassified(100); | Metschnikowia | 13 | Saprotroph | \|Undefined Saprotroph\| | Yeast | NULL | Probable | As nectar-tap_saprotroph (Põlme et al. 2020); Decay sugar-rich substrates (Põlme et al. 2020) | Põlme S, et al. 2020. Fungal Diversity 105:1-16 ((https://doi.org/10.1007/s13225-020-00466-2)) |
| Otu10496 | 2 | k__Fungi(100);p__Ascomycota(100);c__Saccharomycetes(100);o__Saccharomycetales(100);f__Metschnikowiaceae(100);g__Metschnikowia(100);g__Metschnikowia_unclassified(100); | Metschnikowia | 13 | Saprotroph | \|Undefined Saprotroph\| | Yeast | NULL | Probable | As nectar-tap_saprotroph (Põlme et al. 2020); Decay sugar-rich substrates (Põlme et al. 2020) | Põlme S, et al. 2020. Fungal Diversity 105:1-16 ((https://doi.org/10.1007/s13225-020-00466-2)) |
| Otu10500 | 2 | k__Fungi(100);p__Ascomycota(100);c__Saccharomycetes(100);o__Saccharomycetales(100);f__Metschnikowiaceae(100);g__Metschnikowia(100);g__Metschnikowia_unclassified(100); | Metschnikowia | 13 | Saprotroph | \|Undefined Saprotroph\| | Yeast | NULL | Probable | As nectar-tap_saprotroph (Põlme et al. 2020); Decay sugar-rich substrates (Põlme et al. 2020) | Põlme S, et al. 2020. Fungal Diversity 105:1-16 ((https://doi.org/10.1007/s13225-020-00466-2)) |
| Otu10501 | 2 | k__Fungi(100);p__Ascomycota(100);c__Saccharomycetes(100);o__Saccharomycetales(100);f__Metschnikowiaceae(100);g__Metschnikowia(100);s__Metschnikowia_pulcherrima(100); | Metschnikowia | 13 | Saprotroph | \|Undefined Saprotroph\| | Yeast | NULL | Probable | As nectar-tap_saprotroph (Põlme et al. 2020); Decay sugar-rich substrates (Põlme et al. 2020) | Põlme S, et al. 2020. Fungal Diversity 105:1-16 ((https://doi.org/10.1007/s13225-020-00466-2)) |
| Otu10507 | 2 | k__Fungi(100);p__Ascomycota(100);c__Saccharomycetes(100);o__Saccharomycetales(100);f__Metschnikowiaceae(100);g__Metschnikowia(100);s__Metschnikowia_pulcherrima(100); | Metschnikowia | 13 | Saprotroph | \|Undefined Saprotroph\| | Yeast | NULL | Probable | As nectar-tap_saprotroph (Põlme et al. 2020); Decay sugar-rich substrates (Põlme et al. 2020) | Põlme S, et al. 2020. Fungal Diversity 105:1-16 ((https://doi.org/10.1007/s13225-020-00466-2)) |
| Otu10508 | 2 | k__Fungi(100);p__Ascomycota(100);c__Saccharomycetes(100);o__Saccharomycetales(100);f__Metschnikowiaceae(100);g__Metschnikowia(100);g__Metschnikowia_unclassified(100); | Metschnikowia | 13 | Saprotroph | \|Undefined Saprotroph\| | Yeast | NULL | Probable | As nectar-tap_saprotroph (Põlme et al. 2020); Decay sugar-rich substrates (Põlme et al. 2020) | Põlme S, et al. 2020. Fungal Diversity 105:1-16 ((https://doi.org/10.1007/s13225-020-00466-2)) |
| Otu10509 | 2 | k__Fungi(100);p__Ascomycota(100);c__Saccharomycetes(100);o__Saccharomycetales(100);f__Metschnikowiaceae(100);g__Metschnikowia(100);s__Metschnikowia_pulcherrima(100); | Metschnikowia | 13 | Saprotroph | \|Undefined Saprotroph\| | Yeast | NULL | Probable | As nectar-tap_saprotroph (Põlme et al. 2020); Decay sugar-rich substrates (Põlme et al. 2020) | Põlme S, et al. 2020. Fungal Diversity 105:1-16 ((https://doi.org/10.1007/s13225-020-00466-2)) |
| Otu10510 | 2 | k__Fungi(100);p__Ascomycota(100);c__Saccharomycetes(100);o__Saccharomycetales(100);f__Metschnikowiaceae(100);g__Metschnikowia(100);g__Metschnikowia_unclassified(100); | Metschnikowia | 13 | Saprotroph | \|Undefined Saprotroph\| | Yeast | NULL | Probable | As nectar-tap_saprotroph (Põlme et al. 2020); Decay sugar-rich substrates (Põlme et al. 2020) | Põlme S, et al. 2020. Fungal Diversity 105:1-16 ((https://doi.org/10.1007/s13225-020-00466-2)) |
| Otu10511 | 2 | k__Fungi(100);p__Ascomycota(100);c__Saccharomycetes(100);o__Saccharomycetales(100);f__Metschnikowiaceae(100);g__Metschnikowia(100);g__Metschnikowia_unclassified(100); | Metschnikowia | 13 | Saprotroph | \|Undefined Saprotroph\| | Yeast | NULL | Probable | As nectar-tap_saprotroph (Põlme et al. 2020); Decay sugar-rich substrates (Põlme et al. 2020) | Põlme S, et al. 2020. Fungal Diversity 105:1-16 ((https://doi.org/10.1007/s13225-020-00466-2)) |
| Otu10512 | 2 | k__Fungi(100);p__Ascomycota(100);c__Saccharomycetes(100);o__Saccharomycetales(100);f__Metschnikowiaceae(100);g__Metschnikowia(100);g__Metschnikowia_unclassified(100); | Metschnikowia | 13 | Saprotroph | \|Undefined Saprotroph\| | Yeast | NULL | Probable | As nectar-tap_saprotroph (Põlme et al. 2020); Decay sugar-rich substrates (Põlme et al. 2020) | Põlme S, et al. 2020. Fungal Diversity 105:1-16 ((https://doi.org/10.1007/s13225-020-00466-2)) |
| Otu10513 | 2 | k__Fungi(100);p__Ascomycota(100);c__Saccharomycetes(100);o__Saccharomycetales(100);f__Metschnikowiaceae(100);g__Metschnikowia(100);s__Metschnikowia_pulcherrima(100); | Metschnikowia | 13 | Saprotroph | \|Undefined Saprotroph\| | Yeast | NULL | Probable | As nectar-tap_saprotroph (Põlme et al. 2020); Decay sugar-rich substrates (Põlme et al. 2020) | Põlme S, et al. 2020. Fungal Diversity 105:1-16 ((https://doi.org/10.1007/s13225-020-00466-2)) |
| Otu10515 | 2 | k__Fungi(100);p__Ascomycota(100);c__Saccharomycetes(100);o__Saccharomycetales(100);f__Metschnikowiaceae(100);g__Metschnikowia(100);g__Metschnikowia_unclassified(100); | Metschnikowia | 13 | Saprotroph | \|Undefined Saprotroph\| | Yeast | NULL | Probable | As nectar-tap_saprotroph (Põlme et al. 2020); Decay sugar-rich substrates (Põlme et al. 2020) | Põlme S, et al. 2020. Fungal Diversity 105:1-16 ((https://doi.org/10.1007/s13225-020-00466-2)) |
| Otu10519 | 2 | k__Fungi(100);p__Ascomycota(100);c__Saccharomycetes(100);o__Saccharomycetales(100);f__Metschnikowiaceae(100);g__Metschnikowia(100);g__Metschnikowia_unclassified(100); | Metschnikowia | 13 | Saprotroph | \|Undefined Saprotroph\| | Yeast | NULL | Probable | As nectar-tap_saprotroph (Põlme et al. 2020); Decay sugar-rich substrates (Põlme et al. 2020) | Põlme S, et al. 2020. Fungal Diversity 105:1-16 ((https://doi.org/10.1007/s13225-020-00466-2)) |
| Otu10520 | 2 | k__Fungi(100);p__Ascomycota(100);c__Saccharomycetes(100);o__Saccharomycetales(100);f__Metschnikowiaceae(100);g__Metschnikowia(100);g__Metschnikowia_unclassified(100); | Metschnikowia | 13 | Saprotroph | \|Undefined Saprotroph\| | Yeast | NULL | Probable | As nectar-tap_saprotroph (Põlme et al. 2020); Decay sugar-rich substrates (Põlme et al. 2020) | Põlme S, et al. 2020. Fungal Diversity 105:1-16 ((https://doi.org/10.1007/s13225-020-00466-2)) |
| Otu10521 | 2 | k__Fungi(100);p__Ascomycota(100);c__Saccharomycetes(100);o__Saccharomycetales(100);f__Metschnikowiaceae(100);g__Metschnikowia(100);g__Metschnikowia_unclassified(100); | Metschnikowia | 13 | Saprotroph | \|Undefined Saprotroph\| | Yeast | NULL | Probable | As nectar-tap_saprotroph (Põlme et al. 2020); Decay sugar-rich substrates (Põlme et al. 2020) | Põlme S, et al. 2020. Fungal Diversity 105:1-16 ((https://doi.org/10.1007/s13225-020-00466-2)) |
| Otu10522 | 2 | k__Fungi(100);p__Ascomycota(100);c__Saccharomycetes(100);o__Saccharomycetales(100);f__Metschnikowiaceae(100);g__Metschnikowia(100);g__Metschnikowia_unclassified(100); | Metschnikowia | 13 | Saprotroph | \|Undefined Saprotroph\| | Yeast | NULL | Probable | As nectar-tap_saprotroph (Põlme et al. 2020); Decay sugar-rich substrates (Põlme et al. 2020) | Põlme S, et al. 2020. Fungal Diversity 105:1-16 ((https://doi.org/10.1007/s13225-020-00466-2)) |
| Otu10523 | 2 | k__Fungi(100);p__Ascomycota(100);c__Saccharomycetes(100);o__Saccharomycetales(100);f__Metschnikowiaceae(100);g__Metschnikowia(100);s__Metschnikowia_pulcherrima(100); | Metschnikowia | 13 | Saprotroph | \|Undefined Saprotroph\| | Yeast | NULL | Probable | As nectar-tap_saprotroph (Põlme et al. 2020); Decay sugar-rich substrates (Põlme et al. 2020) | Põlme S, et al. 2020. Fungal Diversity 105:1-16 ((https://doi.org/10.1007/s13225-020-00466-2)) |
| Otu10524 | 2 | k__Fungi(100);p__Ascomycota(100);c__Saccharomycetes(100);o__Saccharomycetales(100);f__Metschnikowiaceae(100);g__Metschnikowia(100);g__Metschnikowia_unclassified(100); | Metschnikowia | 13 | Saprotroph | \|Undefined Saprotroph\| | Yeast | NULL | Probable | As nectar-tap_saprotroph (Põlme et al. 2020); Decay sugar-rich substrates (Põlme et al. 2020) | Põlme S, et al. 2020. Fungal Diversity 105:1-16 ((https://doi.org/10.1007/s13225-020-00466-2)) |
| Otu10525 | 2 | k__Fungi(100);p__Ascomycota(100);c__Saccharomycetes(100);o__Saccharomycetales(100);f__Metschnikowiaceae(100);g__Metschnikowia(100);s__Metschnikowia_pulcherrima(100); | Metschnikowia | 13 | Saprotroph | \|Undefined Saprotroph\| | Yeast | NULL | Probable | As nectar-tap_saprotroph (Põlme et al. 2020); Decay sugar-rich substrates (Põlme et al. 2020) | Põlme S, et al. 2020. Fungal Diversity 105:1-16 ((https://doi.org/10.1007/s13225-020-00466-2)) |
| Otu10526 | 2 | k__Fungi(100);p__Ascomycota(100);c__Leotiomycetes(100);o__Helotiales(100);f__Vibrisseaceae(100);g__Phialocephala(100);s__Phialocephala_fusca(100); | Phialocephala | 13 | Saprotroph-Symbiotroph | Endophyte-\|Undefined Saprotroph\| | Microfungus | NULL | Highly Probable | As root_endophyte (Põlme et al. 2020); As root_endophyte_dark_septate (Põlme et al. 2020); As soil_saprotroph (Põlme et al. 2020); Dark Septate Endophyte, known from plant roots.; Decay roots, soil (Põlme et al. 2020) | Newsham KK. 2011. New Phytologist 190:783-793 ((https://doi.org/10.1111/j.1469-8137.2010.03611.x)) (as dark septate root endophytes); Põlme S, et al. 2020. Fungal Diversity 105:1-16 ((https://doi.org/10.1007/s13225-020-00466-2)) |
| Otu10527 | 2 | k__Fungi(100);p__Ascomycota(100);c__Saccharomycetes(100);o__Saccharomycetales(100);f__Metschnikowiaceae(100);g__Metschnikowia(100);g__Metschnikowia_unclassified(100); | Metschnikowia | 13 | Saprotroph | \|Undefined Saprotroph\| | Yeast | NULL | Probable | As nectar-tap_saprotroph (Põlme et al. 2020); Decay sugar-rich substrates (Põlme et al. 2020) | Põlme S, et al. 2020. Fungal Diversity 105:1-16 ((https://doi.org/10.1007/s13225-020-00466-2)) |
| Otu10528 | 2 | k__Fungi(100);p__Ascomycota(100);c__Saccharomycetes(100);o__Saccharomycetales(100);f__Metschnikowiaceae(100);g__Metschnikowia(100);s__Metschnikowia_pulcherrima(100); | Metschnikowia | 13 | Saprotroph | \|Undefined Saprotroph\| | Yeast | NULL | Probable | As nectar-tap_saprotroph (Põlme et al. 2020); Decay sugar-rich substrates (Põlme et al. 2020) | Põlme S, et al. 2020. Fungal Diversity 105:1-16 ((https://doi.org/10.1007/s13225-020-00466-2)) |
| Otu10532 | 2 | k__Fungi(100);p__Ascomycota(100);c__Saccharomycetes(100);o__Saccharomycetales(100);f__Metschnikowiaceae(100);g__Metschnikowia(100);s__Metschnikowia_pulcherrima(100); | Metschnikowia | 13 | Saprotroph | \|Undefined Saprotroph\| | Yeast | NULL | Probable | As nectar-tap_saprotroph (Põlme et al. 2020); Decay sugar-rich substrates (Põlme et al. 2020) | Põlme S, et al. 2020. Fungal Diversity 105:1-16 ((https://doi.org/10.1007/s13225-020-00466-2)) |
| Otu10533 | 2 | k__Fungi(100);p__Basidiomycota(100);c__Microbotryomycetes(100);o__Sporidiobolales(100);f__Sporidiobolales_family_Incertae_sedis(100);g__Rhodotorula(100);g__Rhodotorula_unclassified(100); | Rhodotorula | 13 | Pathotroph-Saprotroph-Symbiotroph | Animal Pathogen-Animal Symbiotroph-Endophyte-Plant Pathogen-\|Undefined Saprotroph\| | Yeast | NULL | Probable | As foliar_endophyte (Põlme et al. 2020); Opportunistic human pathogen | Põlme S, et al. 2020. Fungal Diversity 105:1-16 ((https://doi.org/10.1007/s13225-020-00466-2)); https://www.cdc.gov ((https://www.cdc.gov)) |
| Otu10536 | 2 | k__Fungi(100);p__Ascomycota(100);c__Saccharomycetes(100);o__Saccharomycetales(100);f__Metschnikowiaceae(100);g__Metschnikowia(100);s__Metschnikowia_pulcherrima(100); | Metschnikowia | 13 | Saprotroph | \|Undefined Saprotroph\| | Yeast | NULL | Probable | As nectar-tap_saprotroph (Põlme et al. 2020); Decay sugar-rich substrates (Põlme et al. 2020) | Põlme S, et al. 2020. Fungal Diversity 105:1-16 ((https://doi.org/10.1007/s13225-020-00466-2)) |
| Otu10537 | 2 | k__Fungi(100);p__Ascomycota(100);c__Saccharomycetes(100);o__Saccharomycetales(100);f__Metschnikowiaceae(100);g__Metschnikowia(100);g__Metschnikowia_unclassified(100); | Metschnikowia | 13 | Saprotroph | \|Undefined Saprotroph\| | Yeast | NULL | Probable | As nectar-tap_saprotroph (Põlme et al. 2020); Decay sugar-rich substrates (Põlme et al. 2020) | Põlme S, et al. 2020. Fungal Diversity 105:1-16 ((https://doi.org/10.1007/s13225-020-00466-2)) |
| Otu10539 | 2 | k__Fungi(100);p__Ascomycota(100);c__Saccharomycetes(100);o__Saccharomycetales(100);f__Metschnikowiaceae(100);g__Metschnikowia(100);s__Metschnikowia_pulcherrima(100); | Metschnikowia | 13 | Saprotroph | \|Undefined Saprotroph\| | Yeast | NULL | Probable | As nectar-tap_saprotroph (Põlme et al. 2020); Decay sugar-rich substrates (Põlme et al. 2020) | Põlme S, et al. 2020. Fungal Diversity 105:1-16 ((https://doi.org/10.1007/s13225-020-00466-2)) |
| Otu10545 | 2 | k__Fungi(100);p__Ascomycota(100);c__Saccharomycetes(100);o__Saccharomycetales(100);f__Metschnikowiaceae(100);g__Metschnikowia(100);g__Metschnikowia_unclassified(100); | Metschnikowia | 13 | Saprotroph | \|Undefined Saprotroph\| | Yeast | NULL | Probable | As nectar-tap_saprotroph (Põlme et al. 2020); Decay sugar-rich substrates (Põlme et al. 2020) | Põlme S, et al. 2020. Fungal Diversity 105:1-16 ((https://doi.org/10.1007/s13225-020-00466-2)) |
| Otu10548 | 2 | k__Fungi(100);p__Ascomycota(100);c__Saccharomycetes(100);o__Saccharomycetales(100);f__Metschnikowiaceae(100);g__Metschnikowia(100);g__Metschnikowia_unclassified(100); | Metschnikowia | 13 | Saprotroph | \|Undefined Saprotroph\| | Yeast | NULL | Probable | As nectar-tap_saprotroph (Põlme et al. 2020); Decay sugar-rich substrates (Põlme et al. 2020) | Põlme S, et al. 2020. Fungal Diversity 105:1-16 ((https://doi.org/10.1007/s13225-020-00466-2)) |
| Otu10549 | 2 | k__Fungi(100);p__Ascomycota(100);c__Saccharomycetes(100);o__Saccharomycetales(100);f__Metschnikowiaceae(100);g__Metschnikowia(100);g__Metschnikowia_unclassified(100); | Metschnikowia | 13 | Saprotroph | \|Undefined Saprotroph\| | Yeast | NULL | Probable | As nectar-tap_saprotroph (Põlme et al. 2020); Decay sugar-rich substrates (Põlme et al. 2020) | Põlme S, et al. 2020. Fungal Diversity 105:1-16 ((https://doi.org/10.1007/s13225-020-00466-2)) |
| Otu10550 | 2 | k__Fungi(100);p__Ascomycota(100);c__Saccharomycetes(100);o__Saccharomycetales(100);f__Saccharomycetales_family_Incertae_sedis(100);g__Candida(100);g__Candida_unclassified(100); | Candida | 13 | Pathotroph-Saprotroph-Symbiotroph | Animal Pathogen-Endophyte-Endosymbiont-Epiphyte-\|Undefined Saprotroph\| | Dimorphic Yeast-Yeast | NULL | Probable | As nectar-tap_saprotroph (Põlme et al. 2020); Decay sugar-rich substrates (Põlme et al. 2020); This asexual genus is polyphyletic and contain species that function very differently depend on what they are. Caution should be taken when interpreting guilds for Candida species. | Manolakaki D, et al. 2010. Virulence. 1: 367-375 ((https://doi.org/10.4161/viru.1.5.12796)); Mekbib SB, et al. 2011. Tropical Plant Pathology 36:233-240 ((https://doi.org/10.1590/S1982-56762011000400004)); Põlme S, et al. 2020. Fungal Diversity 105:1-16 ((https://doi.org/10.1007/s13225-020-00466-2)) |
| Otu10551 | 2 | k__Fungi(100);p__Ascomycota(100);c__Saccharomycetes(100);o__Saccharomycetales(100);o__Saccharomycetales_unclassified(100);o__Saccharomycetales_unclassified(100);o__Saccharomycetales_unclassified(100); | Saccharomycetales | 7 | Saprotroph | Undefined Saprotroph | Yeast | NULL | Possible | NULL | Sterkenburg E, et al. 2015. New Phytologist 207:1145-1158 ((https://doi.org/10.1111/nph.13426)) |
| Otu10556 | 2 | k__Fungi(100);p__Ascomycota(100);c__Leotiomycetes(100);o__Helotiales(100);f__Hyaloscyphaceae(100);f__Hyaloscyphaceae_unclassified(100);f__Hyaloscyphaceae_unclassified(100); | Hyaloscyphaceae | 9 | Saprotroph | Plant Saprotroph-Wood Saprotroph | Microfungus | NULL | Probable | Saprobic on woody and herbaceous material (Cannon & Kirk 2007) | Cannon PF, Kirk PM. 2007. Fungal Families of the World. CAB International, Cambridge (ISBN: 978-0851998275) |
| Otu10557 | 2 | k__Fungi(100);p__Ascomycota(100);c__Saccharomycetes(100);o__Saccharomycetales(100);f__Metschnikowiaceae(100);g__Metschnikowia(100);g__Metschnikowia_unclassified(100); | Metschnikowia | 13 | Saprotroph | \|Undefined Saprotroph\| | Yeast | NULL | Probable | As nectar-tap_saprotroph (Põlme et al. 2020); Decay sugar-rich substrates (Põlme et al. 2020) | Põlme S, et al. 2020. Fungal Diversity 105:1-16 ((https://doi.org/10.1007/s13225-020-00466-2)) |
| Otu10558 | 2 | k__Fungi(100);p__Ascomycota(100);c__Saccharomycetes(100);o__Saccharomycetales(100);f__Metschnikowiaceae(100);g__Metschnikowia(100);s__Metschnikowia_pulcherrima(100); | Metschnikowia | 13 | Saprotroph | \|Undefined Saprotroph\| | Yeast | NULL | Probable | As nectar-tap_saprotroph (Põlme et al. 2020); Decay sugar-rich substrates (Põlme et al. 2020) | Põlme S, et al. 2020. Fungal Diversity 105:1-16 ((https://doi.org/10.1007/s13225-020-00466-2)) |
| Otu10559 | 2 | k__Fungi(100);p__Ascomycota(100);c__Leotiomycetes(100);o__Helotiales(100);f__Helotiales_family_Incertae_sedis(100);g__Cadophora(100);s__Cadophora_melinii(100); | Cadophora | 13 | Pathotroph-Saprotroph-Symbiotroph | Ectomycorrhizal-Endophyte-Plant Pathogen-\|Plant Saprotroph\| | Microfungus | NULL | Highly Probable | As foliar_endophyte (Põlme et al. 2020); As litter_saprotroph (Põlme et al. 2020); Dark Septate Endophyte, known from plant roots.; Decay leaf, fruit, seed (Põlme et al. 2020) | Tedersoo L, et al. 2010. Mycorrhiza 20:217-263 ((https://doi.org/10.1007/s00572-009-0274-x)); Newsham KK. 2011. New Phytologist 190: 783-793 ((https://doi.org/10.1111/j.1469-8137.2010.03611.x)) (as DSE); Põlme S, et al. 2020. Fungal Diversity 105:1-16 ((https://doi.org/10.1007/s13225-020-00466-2)) |
| Otu10560 | 2 | k__Fungi(100);p__Ascomycota(100);c__Saccharomycetes(100);o__Saccharomycetales(100);f__Metschnikowiaceae(100);g__Metschnikowia(100);s__Metschnikowia_pulcherrima(100); | Metschnikowia | 13 | Saprotroph | \|Undefined Saprotroph\| | Yeast | NULL | Probable | As nectar-tap_saprotroph (Põlme et al. 2020); Decay sugar-rich substrates (Põlme et al. 2020) | Põlme S, et al. 2020. Fungal Diversity 105:1-16 ((https://doi.org/10.1007/s13225-020-00466-2)) |
| Otu10563 | 2 | k__Fungi(100);p__Ascomycota(100);c__Saccharomycetes(100);o__Saccharomycetales(100);f__Metschnikowiaceae(100);g__Metschnikowia(100);s__Metschnikowia_pulcherrima(100); | Metschnikowia | 13 | Saprotroph | \|Undefined Saprotroph\| | Yeast | NULL | Probable | As nectar-tap_saprotroph (Põlme et al. 2020); Decay sugar-rich substrates (Põlme et al. 2020) | Põlme S, et al. 2020. Fungal Diversity 105:1-16 ((https://doi.org/10.1007/s13225-020-00466-2)) |
| Otu10565 | 2 | k__Fungi(100);p__Ascomycota(100);c__Saccharomycetes(100);o__Saccharomycetales(100);f__Metschnikowiaceae(100);g__Metschnikowia(100);s__Metschnikowia_pulcherrima(100); | Metschnikowia | 13 | Saprotroph | \|Undefined Saprotroph\| | Yeast | NULL | Probable | As nectar-tap_saprotroph (Põlme et al. 2020); Decay sugar-rich substrates (Põlme et al. 2020) | Põlme S, et al. 2020. Fungal Diversity 105:1-16 ((https://doi.org/10.1007/s13225-020-00466-2)) |
| Otu10566 | 2 | k__Fungi(100);p__Ascomycota(100);c__Saccharomycetes(100);o__Saccharomycetales(100);f__Saccharomycetales_family_Incertae_sedis(100);g__Candida(100);g__Candida_unclassified(100); | Candida | 13 | Pathotroph-Saprotroph-Symbiotroph | Animal Pathogen-Endophyte-Endosymbiont-Epiphyte-\|Undefined Saprotroph\| | Dimorphic Yeast-Yeast | NULL | Probable | As nectar-tap_saprotroph (Põlme et al. 2020); Decay sugar-rich substrates (Põlme et al. 2020); This asexual genus is polyphyletic and contain species that function very differently depend on what they are. Caution should be taken when interpreting guilds for Candida species. | Manolakaki D, et al. 2010. Virulence. 1: 367-375 ((https://doi.org/10.4161/viru.1.5.12796)); Mekbib SB, et al. 2011. Tropical Plant Pathology 36:233-240 ((https://doi.org/10.1590/S1982-56762011000400004)); Põlme S, et al. 2020. Fungal Diversity 105:1-16 ((https://doi.org/10.1007/s13225-020-00466-2)) |
| Otu10567 | 2 | k__Fungi(100);p__Ascomycota(100);c__Dothideomycetes(100);o__Capnodiales(100);f__Mycosphaerellaceae(100);f__Mycosphaerellaceae_unclassified(100);f__Mycosphaerellaceae_unclassified(100); | Mycosphaerellaceae | 9 | Pathotroph-Saprotroph | Plant Pathogen-Undefined Saprotroph | Microfungus | NULL | Probable | Biotrophic, necrotrophic or saprobic on various plant tissues (Cannon & Kirk 2007) | Cannon PF, Kirk PM. 2007. Fungal Families of the World. CAB International, Cambridge (ISBN: 978-0851998275) |
| Otu10570 | 2 | k__Fungi(100);p__Ascomycota(100);c__Saccharomycetes(100);o__Saccharomycetales(100);f__Metschnikowiaceae(100);g__Metschnikowia(100);g__Metschnikowia_unclassified(100); | Metschnikowia | 13 | Saprotroph | \|Undefined Saprotroph\| | Yeast | NULL | Probable | As nectar-tap_saprotroph (Põlme et al. 2020); Decay sugar-rich substrates (Põlme et al. 2020) | Põlme S, et al. 2020. Fungal Diversity 105:1-16 ((https://doi.org/10.1007/s13225-020-00466-2)) |
| Otu10573 | 2 | k__Fungi(100);p__Ascomycota(100);c__Saccharomycetes(100);o__Saccharomycetales(100);f__Metschnikowiaceae(100);g__Metschnikowia(100);g__Metschnikowia_unclassified(100); | Metschnikowia | 13 | Saprotroph | \|Undefined Saprotroph\| | Yeast | NULL | Probable | As nectar-tap_saprotroph (Põlme et al. 2020); Decay sugar-rich substrates (Põlme et al. 2020) | Põlme S, et al. 2020. Fungal Diversity 105:1-16 ((https://doi.org/10.1007/s13225-020-00466-2)) |
| Otu10576 | 2 | k__Fungi(100);p__Ascomycota(100);c__Saccharomycetes(100);o__Saccharomycetales(100);f__Metschnikowiaceae(100);g__Metschnikowia(100);s__Metschnikowia_pulcherrima(100); | Metschnikowia | 13 | Saprotroph | \|Undefined Saprotroph\| | Yeast | NULL | Probable | As nectar-tap_saprotroph (Põlme et al. 2020); Decay sugar-rich substrates (Põlme et al. 2020) | Põlme S, et al. 2020. Fungal Diversity 105:1-16 ((https://doi.org/10.1007/s13225-020-00466-2)) |
| Otu10577 | 2 | k__Fungi(100);p__Ascomycota(100);c__Sordariomycetes(100);o__Xylariales(100);f__Amphisphaeriaceae(100);f__Amphisphaeriaceae_unclassified(100);f__Amphisphaeriaceae_unclassified(100); | Amphisphaeriaceae | 9 | Saprotroph | Undefined Saprotroph | Microfungus | NULL | Possible | Saprobes or weak parasites of leaves, stems, etc., often appearing as secondary colonizers following damage to plants by other agents (Cannon & Kirk 2007) | Cannon PF, Kirk PM. 2007. Fungal Families of the World. CAB International, Cambridge (ISBN: 978-0851998275) |
| Otu10580 | 2 | k__Fungi(100);p__Ascomycota(100);c__Sordariomycetes(100);o__Xylariales(100);f__Amphisphaeriaceae(100);g__Monochaetia(100);s__Monochaetia_sp_PSH2004I_1031(100); | Monochaetia | 13 | Pathotroph | \|Plant Pathogen\| | Microfungus | NULL | Probable | NULL | Tedersoo L, et al. 2014. Science 346:e1256688 ((https://doi.org/10.1126/science.1256688)); Põlme S, et al. 2020. Fungal Diversity 105:1-16 ((https://doi.org/10.1007/s13225-020-00466-2)) |
| Otu10581 | 2 | k__Fungi(100);p__Ascomycota(100);c__Saccharomycetes(100);o__Saccharomycetales(100);f__Metschnikowiaceae(100);g__Metschnikowia(100);g__Metschnikowia_unclassified(100); | Metschnikowia | 13 | Saprotroph | \|Undefined Saprotroph\| | Yeast | NULL | Probable | As nectar-tap_saprotroph (Põlme et al. 2020); Decay sugar-rich substrates (Põlme et al. 2020) | Põlme S, et al. 2020. Fungal Diversity 105:1-16 ((https://doi.org/10.1007/s13225-020-00466-2)) |
| Otu10582 | 2 | k__Fungi(100);p__Ascomycota(100);c__Saccharomycetes(100);o__Saccharomycetales(100);f__Metschnikowiaceae(100);g__Metschnikowia(100);s__Metschnikowia_pulcherrima(100); | Metschnikowia | 13 | Saprotroph | \|Undefined Saprotroph\| | Yeast | NULL | Probable | As nectar-tap_saprotroph (Põlme et al. 2020); Decay sugar-rich substrates (Põlme et al. 2020) | Põlme S, et al. 2020. Fungal Diversity 105:1-16 ((https://doi.org/10.1007/s13225-020-00466-2)) |
| Otu10584 | 2 | k__Fungi(100);p__Ascomycota(100);c__Saccharomycetes(100);o__Saccharomycetales(100);f__Metschnikowiaceae(100);g__Metschnikowia(100);s__Metschnikowia_pulcherrima(100); | Metschnikowia | 13 | Saprotroph | \|Undefined Saprotroph\| | Yeast | NULL | Probable | As nectar-tap_saprotroph (Põlme et al. 2020); Decay sugar-rich substrates (Põlme et al. 2020) | Põlme S, et al. 2020. Fungal Diversity 105:1-16 ((https://doi.org/10.1007/s13225-020-00466-2)) |
| Otu10585 | 2 | k__Fungi(100);p__Ascomycota(100);c__Saccharomycetes(100);o__Saccharomycetales(100);f__Metschnikowiaceae(100);g__Metschnikowia(100);s__Metschnikowia_pulcherrima(100); | Metschnikowia | 13 | Saprotroph | \|Undefined Saprotroph\| | Yeast | NULL | Probable | As nectar-tap_saprotroph (Põlme et al. 2020); Decay sugar-rich substrates (Põlme et al. 2020) | Põlme S, et al. 2020. Fungal Diversity 105:1-16 ((https://doi.org/10.1007/s13225-020-00466-2)) |
| Otu10586 | 2 | k__Fungi(100);p__Ascomycota(100);c__Saccharomycetes(100);o__Saccharomycetales(100);f__Metschnikowiaceae(100);g__Metschnikowia(100);s__Metschnikowia_pulcherrima(100); | Metschnikowia | 13 | Saprotroph | \|Undefined Saprotroph\| | Yeast | NULL | Probable | As nectar-tap_saprotroph (Põlme et al. 2020); Decay sugar-rich substrates (Põlme et al. 2020) | Põlme S, et al. 2020. Fungal Diversity 105:1-16 ((https://doi.org/10.1007/s13225-020-00466-2)) |
| Otu10588 | 2 | k__Fungi(100);p__Ascomycota(100);c__Sordariomycetes(100);o__Xylariales(100);f__Xylariaceae(100);f__Xylariaceae_unclassified(100);f__Xylariaceae_unclassified(100); | Xylariaceae | 9 | Pathotroph-Saprotroph-Symbiotroph | Dung Saprotroph-Endophyte-Plant Pathogen-Undefined Saprotroph | Xylarioid | NULL | Probable | Saprobes or weak pathogens, many endophytes, mainly in wood and bark, occasionally in soil and dung (Cannon & Kirk 2007) | Cannon PF, Kirk PM. 2007. Fungal Families of the World. CAB International, Cambridge (ISBN: 978-0851998275) |
| Otu10590 | 2 | k__Fungi(100);p__Ascomycota(100);c__Saccharomycetes(100);o__Saccharomycetales(100);f__Metschnikowiaceae(100);g__Metschnikowia(100);g__Metschnikowia_unclassified(100); | Metschnikowia | 13 | Saprotroph | \|Undefined Saprotroph\| | Yeast | NULL | Probable | As nectar-tap_saprotroph (Põlme et al. 2020); Decay sugar-rich substrates (Põlme et al. 2020) | Põlme S, et al. 2020. Fungal Diversity 105:1-16 ((https://doi.org/10.1007/s13225-020-00466-2)) |
| Otu10591 | 2 | k__Fungi(100);p__Ascomycota(100);c__Orbiliomycetes(100);o__Orbiliales(100);f__Orbiliaceae(100);g__unclassified_Orbiliaceae(100);s__Orbiliaceae_sp(100); | Orbiliaceae | 9 | Saprotroph | Wood Saprotroph | Microfungus | NULL | Probable | Saprobic, found especially on wood in both wet and dry habitats. Many species are nematophagous, forming adhesive knobs, nets or ring-like traps in the presence of a food source; perhaps a strategy for life in low-nitrogen environments. There have been reports of associations with algae for a few species, but an established lichen association appears not to exist (Cannon & Kirk 2007) | Cannon PF, Kirk PM. 2007. Fungal Families of the World. CAB International, Cambridge (ISBN: 978-0851998275) |
| Otu10593 | 2 | k__Fungi(100);p__Ascomycota(100);c__Saccharomycetes(100);o__Saccharomycetales(100);f__Metschnikowiaceae(100);g__Metschnikowia(100);s__Metschnikowia_pulcherrima(100); | Metschnikowia | 13 | Saprotroph | \|Undefined Saprotroph\| | Yeast | NULL | Probable | As nectar-tap_saprotroph (Põlme et al. 2020); Decay sugar-rich substrates (Põlme et al. 2020) | Põlme S, et al. 2020. Fungal Diversity 105:1-16 ((https://doi.org/10.1007/s13225-020-00466-2)) |
| Otu10595 | 2 | k__Fungi(100);p__Ascomycota(100);c__Saccharomycetes(100);o__Saccharomycetales(100);f__Metschnikowiaceae(100);g__Metschnikowia(100);s__Metschnikowia_pulcherrima(100); | Metschnikowia | 13 | Saprotroph | \|Undefined Saprotroph\| | Yeast | NULL | Probable | As nectar-tap_saprotroph (Põlme et al. 2020); Decay sugar-rich substrates (Põlme et al. 2020) | Põlme S, et al. 2020. Fungal Diversity 105:1-16 ((https://doi.org/10.1007/s13225-020-00466-2)) |
| Otu10596 | 2 | k__Fungi(100);p__Ascomycota(100);c__Sordariomycetes(100);o__Sordariomycetes_order_Incertae_sedis(100);f__Sordariomycetes_family_Incertae_sedis(100);g__Xylomelasma(100);s__Xylomelasma_sp_ZK29/08(100); | Xylomelasma | 13 | Saprotroph | Undefined Saprotroph-\|Wood Saprotroph\| | Microfungus | NULL | Probable | Decay wood (Põlme et al. 2020) | Tedersoo L, et al. 2014. Science 346:e1256688 ((https://doi.org/10.1126/science.1256688)); Põlme S, et al. 2020. Fungal Diversity 105:1-16 ((https://doi.org/10.1007/s13225-020-00466-2)) |
| Otu10598 | 2 | k__Fungi(100);p__Ascomycota(100);c__Saccharomycetes(100);o__Saccharomycetales(100);f__Metschnikowiaceae(100);g__Metschnikowia(100);g__Metschnikowia_unclassified(100); | Metschnikowia | 13 | Saprotroph | \|Undefined Saprotroph\| | Yeast | NULL | Probable | As nectar-tap_saprotroph (Põlme et al. 2020); Decay sugar-rich substrates (Põlme et al. 2020) | Põlme S, et al. 2020. Fungal Diversity 105:1-16 ((https://doi.org/10.1007/s13225-020-00466-2)) |
| Otu10605 | 2 | k__Fungi(100);p__Ascomycota(100);c__Saccharomycetes(100);o__Saccharomycetales(100);f__Metschnikowiaceae(100);g__Metschnikowia(100);g__Metschnikowia_unclassified(100); | Metschnikowia | 13 | Saprotroph | \|Undefined Saprotroph\| | Yeast | NULL | Probable | As nectar-tap_saprotroph (Põlme et al. 2020); Decay sugar-rich substrates (Põlme et al. 2020) | Põlme S, et al. 2020. Fungal Diversity 105:1-16 ((https://doi.org/10.1007/s13225-020-00466-2)) |
| Otu10608 | 2 | k__Fungi(100);p__Ascomycota(100);c__Saccharomycetes(100);o__Saccharomycetales(100);f__Metschnikowiaceae(100);g__Metschnikowia(100);s__Metschnikowia_pulcherrima(100); | Metschnikowia | 13 | Saprotroph | \|Undefined Saprotroph\| | Yeast | NULL | Probable | As nectar-tap_saprotroph (Põlme et al. 2020); Decay sugar-rich substrates (Põlme et al. 2020) | Põlme S, et al. 2020. Fungal Diversity 105:1-16 ((https://doi.org/10.1007/s13225-020-00466-2)) |
| Otu10610 | 2 | k__Fungi(100);p__Ascomycota(100);c__Eurotiomycetes(100);o__Mycocaliciales(100);f__Mycocaliciaceae(100);g__Mycocalicium(100);s__Mycocalicium_subtile(100); | Mycocalicium | 13 | Pathotroph-Saprotroph-Symbiotroph | Lichen Parasite-Lichenized-\|Undefined Saprotroph\| | Calicioid-Thallus | NULL | Possible | As resin_saprotroph (Põlme et al. 2020); licenized taxa; Lichenized and non | Esslinger TL. 2014. North Dakota State University ((http://www.ndsu.edu/pubweb/~esslinge/chcklst/chcklst7.htm)); Põlme S, et al. 2020. Fungal Diversity 105:1-16 ((https://doi.org/10.1007/s13225-020-00466-2)) |
| Otu10611 | 2 | k__Fungi(100);p__Ascomycota(100);c__Sordariomycetes(100);o__Hypocreales(100);o__Hypocreales_unclassified(100);o__Hypocreales_unclassified(100);o__Hypocreales_unclassified(100); | Hypocreales | 7 | Saprotroph | Undefined Saprotroph | Microfungus | NULL | Possible | NULL | Sterkenburg E, et al. 2015. New Phytologist 207:1145-1158 ((https://doi.org/10.1111/nph.13426)) |
| Otu10612 | 2 | k__Fungi(100);p__Ascomycota(100);c__Eurotiomycetes(100);o__Eurotiales(100);f__Trichocomaceae(100);g__Penicillium(100);g__Penicillium_unclassified(100); | Penicillium | 13 | Pathotroph-Saprotroph-Symbiotroph | Animal Parasite-Dung Saprotroph-Endophyte-Fungal Parasite-Plant Pathogen-\|Undefined Saprotroph\|-Wood Saprotroph | Microfungus | Soft Rot | Highly Probable | As foliar_endophyte (Põlme et al. 2020); As mycoparasite (Põlme et al. 2020); Decay soil (Põlme et al. 2020) | Duncan CG, Eslyn WE. 1966. Mycologia 58:642-664 ((https://doi.org/10.2307/3757045)); Seehann G, et al. 1975. List of Fungi in Soft-Rot Tissue Tests. Institute fur Holzbiologie und Holzschutz, Bundesforschungsanstalt fur Forst - und Holzwirtshaft, Hamburg - Lohbrugge, Germany ((https://www.irg-wp.com/irgdocs/details.php?d6493033-c9ea-40f3-8e3e-5697f32d640d)); Costa IPMW, et al. 2012. Mycotaxon 119:e26 ((http://mycotaxon.com/resources/checklists/costa_v119_checklist.pdf)); Bills GF, et al. 2013. Current Opinion in Microbiology 16: 549-565 ((https://doi.org/10.1016/j.mib.2013.08.001)); Tedersoo L, et al. 2014. Science 346:e1256688 ((https://doi.org/10.1126/science.1256688)); Põlme S, et al. 2020. Fungal Diversity 105:1-16 ((https://doi.org/10.1007/s13225-020-00466-2)) |
| Otu10615 | 2 | k__Fungi(100);p__Ascomycota(100);c__Eurotiomycetes(100);o__Eurotiales(100);f__Trichocomaceae(100);g__Penicillium(100);s__Penicillium_daleae(100); | Penicillium | 13 | Pathotroph-Saprotroph-Symbiotroph | Animal Parasite-Dung Saprotroph-Endophyte-Fungal Parasite-Plant Pathogen-\|Undefined Saprotroph\|-Wood Saprotroph | Microfungus | Soft Rot | Highly Probable | As foliar_endophyte (Põlme et al. 2020); As mycoparasite (Põlme et al. 2020); Decay soil (Põlme et al. 2020) | Duncan CG, Eslyn WE. 1966. Mycologia 58:642-664 ((https://doi.org/10.2307/3757045)); Seehann G, et al. 1975. List of Fungi in Soft-Rot Tissue Tests. Institute fur Holzbiologie und Holzschutz, Bundesforschungsanstalt fur Forst - und Holzwirtshaft, Hamburg - Lohbrugge, Germany ((https://www.irg-wp.com/irgdocs/details.php?d6493033-c9ea-40f3-8e3e-5697f32d640d)); Costa IPMW, et al. 2012. Mycotaxon 119:e26 ((http://mycotaxon.com/resources/checklists/costa_v119_checklist.pdf)); Bills GF, et al. 2013. Current Opinion in Microbiology 16: 549-565 ((https://doi.org/10.1016/j.mib.2013.08.001)); Tedersoo L, et al. 2014. Science 346:e1256688 ((https://doi.org/10.1126/science.1256688)); Põlme S, et al. 2020. Fungal Diversity 105:1-16 ((https://doi.org/10.1007/s13225-020-00466-2)) |
| Otu10619 | 2 | k__Fungi(100);p__Ascomycota(100);c__Saccharomycetes(100);o__Saccharomycetales(100);f__Metschnikowiaceae(100);g__Metschnikowia(100);s__Metschnikowia_pulcherrima(100); | Metschnikowia | 13 | Saprotroph | \|Undefined Saprotroph\| | Yeast | NULL | Probable | As nectar-tap_saprotroph (Põlme et al. 2020); Decay sugar-rich substrates (Põlme et al. 2020) | Põlme S, et al. 2020. Fungal Diversity 105:1-16 ((https://doi.org/10.1007/s13225-020-00466-2)) |
| Otu10622 | 2 | k__Fungi(100);p__Ascomycota(100);c__Saccharomycetes(100);o__Saccharomycetales(100);f__Metschnikowiaceae(100);g__Metschnikowia(100);s__Metschnikowia_pulcherrima(100); | Metschnikowia | 13 | Saprotroph | \|Undefined Saprotroph\| | Yeast | NULL | Probable | As nectar-tap_saprotroph (Põlme et al. 2020); Decay sugar-rich substrates (Põlme et al. 2020) | Põlme S, et al. 2020. Fungal Diversity 105:1-16 ((https://doi.org/10.1007/s13225-020-00466-2)) |
| Otu10624 | 2 | k__Fungi(100);p__Ascomycota(100);c__Saccharomycetes(100);o__Saccharomycetales(100);f__Metschnikowiaceae(100);g__Metschnikowia(100);s__Metschnikowia_pulcherrima(100); | Metschnikowia | 13 | Saprotroph | \|Undefined Saprotroph\| | Yeast | NULL | Probable | As nectar-tap_saprotroph (Põlme et al. 2020); Decay sugar-rich substrates (Põlme et al. 2020) | Põlme S, et al. 2020. Fungal Diversity 105:1-16 ((https://doi.org/10.1007/s13225-020-00466-2)) |
| Otu10628 | 2 | k__Fungi(100);p__Ascomycota(100);c__Saccharomycetes(100);o__Saccharomycetales(100);f__Metschnikowiaceae(100);g__Metschnikowia(100);s__Metschnikowia_pulcherrima(100); | Metschnikowia | 13 | Saprotroph | \|Undefined Saprotroph\| | Yeast | NULL | Probable | As nectar-tap_saprotroph (Põlme et al. 2020); Decay sugar-rich substrates (Põlme et al. 2020) | Põlme S, et al. 2020. Fungal Diversity 105:1-16 ((https://doi.org/10.1007/s13225-020-00466-2)) |
| Otu10632 | 2 | k__Fungi(100);p__Ascomycota(100);c__Saccharomycetes(100);o__Saccharomycetales(100);f__Metschnikowiaceae(100);g__Metschnikowia(100);s__Metschnikowia_pulcherrima(100); | Metschnikowia | 13 | Saprotroph | \|Undefined Saprotroph\| | Yeast | NULL | Probable | As nectar-tap_saprotroph (Põlme et al. 2020); Decay sugar-rich substrates (Põlme et al. 2020) | Põlme S, et al. 2020. Fungal Diversity 105:1-16 ((https://doi.org/10.1007/s13225-020-00466-2)) |
| Otu10633 | 2 | k__Fungi(100);p__Ascomycota(100);c__Saccharomycetes(100);o__Saccharomycetales(100);f__Metschnikowiaceae(100);g__Metschnikowia(100);g__Metschnikowia_unclassified(100); | Metschnikowia | 13 | Saprotroph | \|Undefined Saprotroph\| | Yeast | NULL | Probable | As nectar-tap_saprotroph (Põlme et al. 2020); Decay sugar-rich substrates (Põlme et al. 2020) | Põlme S, et al. 2020. Fungal Diversity 105:1-16 ((https://doi.org/10.1007/s13225-020-00466-2)) |
| Otu10634 | 2 | k__Fungi(100);p__Ascomycota(100);c__Saccharomycetes(100);o__Saccharomycetales(100);f__Metschnikowiaceae(100);g__Metschnikowia(100);g__Metschnikowia_unclassified(100); | Metschnikowia | 13 | Saprotroph | \|Undefined Saprotroph\| | Yeast | NULL | Probable | As nectar-tap_saprotroph (Põlme et al. 2020); Decay sugar-rich substrates (Põlme et al. 2020) | Põlme S, et al. 2020. Fungal Diversity 105:1-16 ((https://doi.org/10.1007/s13225-020-00466-2)) |
| Otu10635 | 2 | k__Fungi(100);p__Ascomycota(100);c__Saccharomycetes(100);o__Saccharomycetales(100);f__Metschnikowiaceae(100);g__Metschnikowia(100);g__Metschnikowia_unclassified(100); | Metschnikowia | 13 | Saprotroph | \|Undefined Saprotroph\| | Yeast | NULL | Probable | As nectar-tap_saprotroph (Põlme et al. 2020); Decay sugar-rich substrates (Põlme et al. 2020) | Põlme S, et al. 2020. Fungal Diversity 105:1-16 ((https://doi.org/10.1007/s13225-020-00466-2)) |
| Otu10636 | 2 | k__Fungi(100);p__Basidiomycota(100);c__Tremellomycetes(100);o__Tremellales(100);f__Tremellales_family_Incertae_sedis(100);g__Dioszegia(100);s__Dioszegia_zsoltii_var._zsoltii(100); | Dioszegia | 13 | Saprotroph | \|Plant Saprotroph\|-Undefined Saprotroph | Yeast | NULL | Probable | As litter_saprotroph (Põlme et al. 2020); Decay leaf, fruit, seed (Põlme et al. 2020); The Dioszegia subclade sensu Inacio (Inacio et al. 20015) is characterized by orange; yellow or orange to deep orange colonies. Isolated from plant leaves and soil. | Põlme S, et al. 2020. Fungal Diversity 105:1-16 ((https://doi.org/10.1007/s13225-020-00466-2)); Takashima M, et al. 2011 International Journal of Systematic and Evolutionary Microbiology 61:1736-1739 ((https://doi.org/10.1099/ijs.0.025254-0)) |
| Otu10637 | 2 | k__Fungi(100);p__Ascomycota(100);c__Saccharomycetes(100);o__Saccharomycetales(100);f__Metschnikowiaceae(100);g__Metschnikowia(100);s__Metschnikowia_pulcherrima(100); | Metschnikowia | 13 | Saprotroph | \|Undefined Saprotroph\| | Yeast | NULL | Probable | As nectar-tap_saprotroph (Põlme et al. 2020); Decay sugar-rich substrates (Põlme et al. 2020) | Põlme S, et al. 2020. Fungal Diversity 105:1-16 ((https://doi.org/10.1007/s13225-020-00466-2)) |
| Otu10638 | 2 | k__Fungi(100);p__Ascomycota(100);c__Saccharomycetes(100);o__Saccharomycetales(100);f__Metschnikowiaceae(100);g__Metschnikowia(100);g__Metschnikowia_unclassified(100); | Metschnikowia | 13 | Saprotroph | \|Undefined Saprotroph\| | Yeast | NULL | Probable | As nectar-tap_saprotroph (Põlme et al. 2020); Decay sugar-rich substrates (Põlme et al. 2020) | Põlme S, et al. 2020. Fungal Diversity 105:1-16 ((https://doi.org/10.1007/s13225-020-00466-2)) |

| | | | | | | | | | | | |
|---|---|---|---|---|---|---|---|---|---|---|---|
| Otu10640 | 2 | k__Fungi(100);p__Ascomycota(100);c__Saccharomycetes(100);o__Saccharomycetales(100);f__Metschnikowiaceae(100);g__Metschnikowia(100);s__Metschnikowia_pulcherrima(100); | Metschnikowia | 13 | Saprotroph | \|Undefined Saprotroph\| | Yeast | NULL | Probable | As nectar-tap_saprotroph (Põlme et al. 2020); Decay sugar-rich substrates (Põlme et al. 2020) | Põlme S, et al. 2020. Fungal Diversity 105:1-16 ((https://doi.org/10.1007/s13225-020-00466-2)) |
| Otu10647 | 2 | k__Fungi(100);p__Ascomycota(100);c__Saccharomycetes(100);o__Saccharomycetales(100);f__Metschnikowiaceae(100);g__Metschnikowia(100);g__Metschnikowia_unclassified(100); | Metschnikowia | 13 | Saprotroph | \|Undefined Saprotroph\| | Yeast | NULL | Probable | As nectar-tap_saprotroph (Põlme et al. 2020); Decay sugar-rich substrates (Põlme et al. 2020) | Põlme S, et al. 2020. Fungal Diversity 105:1-16 ((https://doi.org/10.1007/s13225-020-00466-2)) |
| Otu10649 | 2 | k__Fungi(100);p__Ascomycota(100);c__Saccharomycetes(100);o__Saccharomycetales(100);f__Metschnikowiaceae(100);g__Metschnikowia(100);s__Metschnikowia_pulcherrima(100); | Metschnikowia | 13 | Saprotroph | \|Undefined Saprotroph\| | Yeast | NULL | Probable | As nectar-tap_saprotroph (Põlme et al. 2020); Decay sugar-rich substrates (Põlme et al. 2020) | Põlme S, et al. 2020. Fungal Diversity 105:1-16 ((https://doi.org/10.1007/s13225-020-00466-2)) |
| Otu10650 | 2 | k__Fungi(100);p__Ascomycota(100);c__Saccharomycetes(100);o__Saccharomycetales(100);f__Metschnikowiaceae(100);g__Metschnikowia(100);g__Metschnikowia_unclassified(100); | Metschnikowia | 13 | Saprotroph | \|Undefined Saprotroph\| | Yeast | NULL | Probable | As nectar-tap_saprotroph (Põlme et al. 2020); Decay sugar-rich substrates (Põlme et al. 2020) | Põlme S, et al. 2020. Fungal Diversity 105:1-16 ((https://doi.org/10.1007/s13225-020-00466-2)) |
| Otu10651 | 2 | k__Fungi(100);p__Ascomycota(100);c__Sordariomycetes(100);o__Hypocreales(100);f__Hypocreales_family_Incertae_sedis(100);g__Acremonium(100);g__Acremonium_unclassified(100); | Acremonium | 13 | Pathotroph-Saprotroph-Symbiotroph | Animal Pathogen-Endophyte-Fungal Parasite-Plant Pathogen-\|Undefined Saprotroph\|-Wood Saprotroph | Microfungus | Soft Rot | Possible | As foliar_endophyte (Põlme et al. 2020); culture (Hanada et al. 2010, Del Olmo; Decay leaf, fruit, seed, soil, dung, animal material (Põlme et al. 2020); Endophyte detection method; Ruiz & Arnold 2014); Likely opportunistic human pathogen (Irinyi et al. 2015) | Duncan CG, Eslyn WE. 1966. Mycologia 58:642-645 ((https://doi.org/10.2307/3757045)); Nilsson T. 1973. Studia Forestalia Suecica 104: 1-40 ((https://pub.epsilon.slu.se/5711/)); Worrall JJ, et al. 1997. Mycologia 89:199-219 ((https://doi.org/10.2307/3761073)); Hanada RE, et al. 2010. Fungal Biology 114:901-910 ((https://doi.org/10.1016/j.funbio.2010.08.006)); Del Olmo-Ruiz M, Arnold AE. 2014. Mycologia 106: 8-21 ((https://doi.org/10.3852/13-098)); Irinyi L, et al. 2015. Medical Mycology 53313-337 ((https://doi.org/10.1093/mmy/myv008)); Põlme S, et al. 2020. Fungal Diversity 105:1-16 ((https://doi.org/10.1007/s13225-020-00466-2)) |
| Otu10653 | 2 | k__Fungi(100);p__Ascomycota(100);c__Saccharomycetes(100);o__Saccharomycetales(100);f__Metschnikowiaceae(100);g__Metschnikowia(100);s__Metschnikowia_pulcherrima(100); | Metschnikowia | 13 | Saprotroph | \|Undefined Saprotroph\| | Yeast | NULL | Probable | As nectar-tap_saprotroph (Põlme et al. 2020); Decay sugar-rich substrates (Põlme et al. 2020) | Põlme S, et al. 2020. Fungal Diversity 105:1-16 ((https://doi.org/10.1007/s13225-020-00466-2)) |
| Otu10654 | 2 | k__Fungi(100);p__Ascomycota(100);c__Dothideomycetes(100);o__Dothideales(100);f__Dothideaceae(100);g__Endoconidioma(100);s__Endoconidioma_populi(100); | Endoconidioma | 13 | Saprotroph | Undefined Saprotroph-\|Wood Saprotroph\| | Microfungus | NULL | Probable | Decay: wood (Põlme et al. 2020) | Tedersoo L, et al. 2014. Science 346:e1256688 ((https://doi.org/10.1126/science.1256688)); Põlme S, et al. 2020. Fungal Diversity 105:1-16 ((https://doi.org/10.1007/s13225-020-00466-2)) |
| Otu10656 | 2 | k__Fungi(100);p__Zygomycota(100);c__Zygomycota_class_Incertae_sedis(100);o__Mortierellales(100);f__Mortierellaceae(100);g__Mortierella(100);g__Mortierella_unclassified(100); | Mortierella | 13 | Saprotroph-Symbiotroph | Endophyte-Plant Saprotroph-\|Undefined Saprotroph\| | Microfungus | NULL | Probable | As chitinolytic (Põlme et al. 2020); As soil_saprotroph (Põlme et al. 2020); Decay roots, soil (Põlme et al. 2020); Saprobic in soil and thus frequenctly isolated from soil and material contaminated with soil such as hay or rotten silage and spoil tips (Cannon & Kirk 2007) | Cannon PF, Kirk PM. 2007. Fungal Families of the World. CAB International, Cambridge (ISBN: 978-0851998275); Tedersoo L, et al. 2014. Science 346:e1256688 ((https://doi.org/10.1126/science.1256688)); Purahong W, et al. 2016. Molecular Ecology 25:4059-4074 ((https://doi.org/10.1111/mec.13739)); Põlme S, et al. 2020. Fungal Diversity 105:1-16 ((https://doi.org/10.1007/s13225-020-00466-2)) |
| Otu10658 | 2 | k__Fungi(100);p__Zygomycota(100);c__Zygomycota_class_Incertae_sedis(100);o__Mucorales(100);f__Umbelopsidaceae(100);g__Umbelopsis(100);s__Umbelopsis_ramanniana(100); | Umbelopsis | 13 | Saprotroph | \|Undefined Saprotroph\| | Microfungus | NULL | Probable | As soil_saprotroph (Põlme et al. 2020); Decay roots, soil (Põlme et al. 2020) | Tedersoo L, et al. 2014. Science 346:e1256688 ((https://doi.org/10.1126/science.1256688)); Põlme S, et al. 2020. Fungal Diversity 105:1-16 ((https://doi.org/10.1007/s13225-020-00466-2)) |
| Otu10661 | 2 | k__Fungi(100);p__Zygomycota(100);c__Zygomycota_class_Incertae_sedis(100);o__Mucorales(100);f__Umbelopsidaceae(100);g__Umbelopsis(100);g__Umbelopsis_unclassified(100); | Umbelopsis | 13 | Saprotroph | \|Undefined Saprotroph\| | Microfungus | NULL | Probable | As soil_saprotroph (Põlme et al. 2020); Decay roots, soil (Põlme et al. 2020) | Tedersoo L, et al. 2014. Science 346:e1256688 ((https://doi.org/10.1126/science.1256688)); Põlme S, et al. 2020. Fungal Diversity 105:1-16 ((https://doi.org/10.1007/s13225-020-00466-2)) |
| Otu10662 | 2 | k__Fungi(100);p__Ascomycota(100);c__Saccharomycetes(100);o__Saccharomycetales(100);f__Metschnikowiaceae(100);g__Metschnikowia(100);s__Metschnikowia_pulcherrima(100); | Metschnikowia | 13 | Saprotroph | \|Undefined Saprotroph\| | Yeast | NULL | Probable | As nectar-tap_saprotroph (Põlme et al. 2020); Decay sugar-rich substrates (Põlme et al. 2020) | Põlme S, et al. 2020. Fungal Diversity 105:1-16 ((https://doi.org/10.1007/s13225-020-00466-2)) |
| Otu10665 | 2 | k__Fungi(100);p__Ascomycota(100);c__Dothideomycetes(100);o__Pleosporales(100);f__Pleosporales_family_Incertae_sedis(100);g__Periconia(100);s__Periconia_sp_CY137(100); | Periconia | 13 | Pathotroph-Saprotroph-Symbiotroph | Endophyte-\|Plant Pathogen\|-Wood Saprotroph | Microfungus | Soft Rot | Possible | Asteraceae, Musaceae (Costa et al. 2012); As foliar_endophyte (Põlme et al. 2020); Dark Septate Endophyte, known from plant roots (Mandyam et al. 2010); Endophyte Host ; Decay leaf, fruit, seed (Põlme et al. 2020) | Duncan CG, Eslyn WE. 1966. Mycologia 58:642-645 ((https://doi.org/10.2307/3757045)); Costa IPMW, et al. 2012. Mycotaxon 119:e26 ((http://mycotaxon.com/resources/checklists/costa_v119_checklist.pdf)); Põlme S, et al. 2020. Fungal Diversity 105:1-16 ((https://doi.org/10.1007/s13225-020-00466-2)); Mandyam K, et al. 2012. Mycorrhiza 22:109-119 ((https://doi.org/10.1007/s00572-011-0386-y)); Knapp DG, et al. 2012. PLOS ONE 7:e32570 (( https://doi.org/10.1371/journal.pone.0032570)); The American Phytopathological Society Common Names of Plant Diseases ((http://www.apsnet.org/publications/commonnames/Pages/default.aspx)) |
| Otu10666 | 2 | k__Fungi(100);p__Ascomycota(100);c__Saccharomycetes(100);o__Saccharomycetales(100);f__Saccharomycetaceae(100);g__Kazachstania(100);s__Kazachstania_intestinalis(100); | Kazachstania | 13 | Saprotroph | \|Undefined Saprotroph\| | Yeast | NULL | Probable | As nectar-tap_saprotroph (Põlme et al. 2020); Decay sugar-rich substrates (Põlme et al. 2020) | Põlme S, et al. 2020. Fungal Diversity 105:1-16 ((https://doi.org/10.1007/s13225-020-00466-2)) |
| Otu10667 | 2 | k__Fungi(100);p__Ascomycota(100);c__Sordariomycetes(100);o__Hypocreales(100);f__Nectriaceae(100);f__Nectriaceae_unclassified(100);f__Nectriaceae_unclassified(100); | Nectriaceae | 9 | Pathotroph-Saprotroph-Symbiotroph | Animal Pathogen-Endophyte-Fungal Parasite-Lichen Parasite-Plant Pathogen-Wood Saprotroph | Microfungus | NULL | Possible | Fusarium species are amongst the economically significant plant pathogens (Cannon & Kirk 2007) | Cannon PF, Kirk PM. 2007. Fungal Families of the World. CAB International, Cambridge (ISBN: 978-0851998275); Hirooka Y, et al. 2012. Studies in Mycology 711-210 ((https://doi.org/10.3114/sim0001)); Tedersoo L, et al. 2014. Science 346:e1256688 ((https://doi.org/10.1126/science.1256688)); The American Phytopathological Society Common Names of Plant Diseases ((http://www.apsnet.org/publications/commonnames/Pages/default.aspx)); https://www.cdc.gov ((https://www.cdc.gov)) |
| Otu10668 | 2 | k__Fungi(100);p__Ascomycota(100);c__Saccharomycetes(100);o__Saccharomycetales(100);f__Metschnikowiaceae(100);g__Metschnikowia(100);g__Metschnikowia_unclassified(100); | Metschnikowia | 13 | Saprotroph | \|Undefined Saprotroph\| | Yeast | NULL | Probable | As nectar-tap_saprotroph (Põlme et al. 2020); Decay sugar-rich substrates (Põlme et al. 2020) | Põlme S, et al. 2020. Fungal Diversity 105:1-16 ((https://doi.org/10.1007/s13225-020-00466-2)) |
| Otu10672 | 2 | k__Fungi(100);p__Ascomycota(100);c__Saccharomycetes(100);o__Saccharomycetales(100);f__Metschnikowiaceae(100);g__Metschnikowia(100);s__Metschnikowia_pulcherrima(100); | Metschnikowia | 13 | Saprotroph | \|Undefined Saprotroph\| | Yeast | NULL | Probable | As nectar-tap_saprotroph (Põlme et al. 2020); Decay sugar-rich substrates (Põlme et al. 2020) | Põlme S, et al. 2020. Fungal Diversity 105:1-16 ((https://doi.org/10.1007/s13225-020-00466-2)) |
| Otu10673 | 2 | k__Fungi(100);p__Ascomycota(100);c__Saccharomycetes(100);o__Saccharomycetales(100);f__Metschnikowiaceae(100);g__Metschnikowia(100);g__Metschnikowia_unclassified(100); | Metschnikowia | 13 | Saprotroph | \|Undefined Saprotroph\| | Yeast | NULL | Probable | As nectar-tap_saprotroph (Põlme et al. 2020); Decay sugar-rich substrates (Põlme et al. 2020) | Põlme S, et al. 2020. Fungal Diversity 105:1-16 ((https://doi.org/10.1007/s13225-020-00466-2)) |
| Otu10678 | 2 | k__Fungi(100);p__Ascomycota(100);c__Saccharomycetes(100);o__Saccharomycetales(100);f__Metschnikowiaceae(100);g__Metschnikowia(100);g__Metschnikowia_unclassified(100); | Metschnikowia | 13 | Saprotroph | \|Undefined Saprotroph\| | Yeast | NULL | Probable | As nectar-tap_saprotroph (Põlme et al. 2020); Decay sugar-rich substrates (Põlme et al. 2020) | Põlme S, et al. 2020. Fungal Diversity 105:1-16 ((https://doi.org/10.1007/s13225-020-00466-2)) |
| Otu10679 | 2 | k__Fungi(100);p__Ascomycota(100);c__Saccharomycetes(100);o__Saccharomycetales(100);f__Metschnikowiaceae(100);g__Metschnikowia(100);s__Metschnikowia_pulcherrima(100); | Metschnikowia | 13 | Saprotroph | \|Undefined Saprotroph\| | Yeast | NULL | Probable | As nectar-tap_saprotroph (Põlme et al. 2020); Decay sugar-rich substrates (Põlme et al. 2020) | Põlme S, et al. 2020. Fungal Diversity 105:1-16 ((https://doi.org/10.1007/s13225-020-00466-2)) |
| Otu10681 | 2 | k__Fungi(100);p__Ascomycota(100);c__Saccharomycetes(100);o__Saccharomycetales(100);f__Metschnikowiaceae(100);g__Metschnikowia(100);g__Metschnikowia_unclassified(100); | Metschnikowia | 13 | Saprotroph | \|Undefined Saprotroph\| | Yeast | NULL | Probable | As nectar-tap_saprotroph (Põlme et al. 2020); Decay sugar-rich substrates (Põlme et al. 2020) | Põlme S, et al. 2020. Fungal Diversity 105:1-16 ((https://doi.org/10.1007/s13225-020-00466-2)) |
| Otu10682 | 2 | k__Fungi(100);p__Ascomycota(100);c__Saccharomycetes(100);o__Saccharomycetales(100);f__Metschnikowiaceae(100);g__Metschnikowia(100);s__Metschnikowia_pulcherrima(100); | Metschnikowia | 13 | Saprotroph | \|Undefined Saprotroph\| | Yeast | NULL | Probable | As nectar-tap_saprotroph (Põlme et al. 2020); Decay sugar-rich substrates (Põlme et al. 2020) | Põlme S, et al. 2020. Fungal Diversity 105:1-16 ((https://doi.org/10.1007/s13225-020-00466-2)) |
| Otu10687 | 2 | k__Fungi(100);p__Ascomycota(100);c__Saccharomycetes(100);o__Saccharomycetales(100);f__Saccharomycetales_family_Incertae_sedis(100);g__Candida(100);g__Candida_unclassified(100); | Candida | 13 | Pathotroph-Saprotroph-Symbiotroph | Animal Pathogen-Endophyte-Endosymbiont-Epiphyte-\|Undefined Saprotroph\| | Dimorphic Yeast-Yeast | NULL | Probable | As nectar-tap_saprotroph (Põlme et al. 2020); Decay sugar-rich substrates (Põlme et al. 2020); This asexual genus is polyphyletic and contain species that function very differently depend on what they are. Caution should be taken when interpreting guilds for Candida species. | Manolakaki D, et al. 2010. Virulence. 1: 367-375 ((https://doi.org/10.4161/viru.1.5.12796)); Mekbib SB, et al. 2011. Tropical Plant Pathology 36:233-240 ((https://doi.org/10.1590/S1982-56762011000400004)); Põlme S, et al. 2020. Fungal Diversity 105:1-16 ((https://doi.org/10.1007/s13225-020-00466-2)) |
| Otu10690 | 2 | k__Fungi(100);p__Ascomycota(100);c__Sordariomycetes(100);o__Hypocreales(100);f__Ophiocordycipitaceae(100);f__Ophiocordycipitaceae_unclassified(100);f__Ophiocordycipitaceae_unclassified(100); | Ophiocordycipitaceae | 9 | Pathotroph-Symbiotroph | Endophyte-Insect Pathogen | Microfungus | NULL | Probable | A family of parasitic fungi (Sung et al. 2007) | Sung GH, et al. 2007. Studies in Mycology 57:5-59 ((https://doi.org/10.3114/sim.2007.57.01)); Costa IPMW, et al. 2012. Mycotaxon 119: e26 ((http://mycotaxon.com/resources/checklists/costa_v119_checklist.pdf)) |
| Otu10692 | 2 | k__Fungi(100);p__Ascomycota(100);c__Saccharomycetes(100);o__Saccharomycetales(100);f__Metschnikowiaceae(100);g__Metschnikowia(100);s__Metschnikowia_pulcherrima(100); | Metschnikowia | 13 | Saprotroph | \|Undefined Saprotroph\| | Yeast | NULL | Probable | As nectar-tap_saprotroph (Põlme et al. 2020); Decay sugar-rich substrates (Põlme et al. 2020) | Põlme S, et al. 2020. Fungal Diversity 105:1-16 ((https://doi.org/10.1007/s13225-020-00466-2)) |
| Otu10693 | 2 | k__Fungi(100);p__Ascomycota(100);c__Sordariomycetes(100);o__Hypocreales(100);f__Ophiocordycipitaceae(100);g__Ophiocordyceps(100);s__Ophiocordyceps_heteropoda_var_langyashanensis(100); | Ophiocordyceps | 13 | Pathotroph-Saprotroph-Symbiotroph | Endophyte-\|Insect Pathogen\|-Undefined Saprotroph | Microfungus | NULL | Probable | As animal_decomposer (Põlme et al. 2020); Contains about 140 species that grow on insects (Kirk et al. 2008); Decay animal material (Põlme et al. 2020) | Kirk PM et al. 2008. Dictionary of the Fungi. Tenth Edition. CAB International, Wallingford (ISBN: 978-0851998268); Costa IPMW, et al. 2012. Mycotaxon 119:e26 ((http://mycotaxon.com/resources/checklists/costa_v119_checklist.pdf)); Tedersoo L, et al. 2014. Science 346: e1256688 ((https://doi.org/10.1126/science.1256688)); Põlme S, et al. 2020. Fungal Diversity 105:1-16 ((https://doi.org/10.1007/s13225-020-00466-2)) |
| Otu10695 | 2 | k__Fungi(100);p__Ascomycota(100);c__Saccharomycetes(100);o__Saccharomycetales(100);f__Metschnikowiaceae(100);g__Metschnikowia(100);s__Metschnikowia_pulcherrima(100); | Metschnikowia | 13 | Saprotroph | \|Undefined Saprotroph\| | Yeast | NULL | Probable | As nectar-tap_saprotroph (Põlme et al. 2020); Decay sugar-rich substrates (Põlme et al. 2020) | Põlme S, et al. 2020. Fungal Diversity 105:1-16 ((https://doi.org/10.1007/s13225-020-00466-2)) |
| Otu10698 | 2 | k__Fungi(100);p__Ascomycota(100);c__Saccharomycetes(100);o__Saccharomycetales(100);f__Metschnikowiaceae(100);g__Metschnikowia(100);s__Metschnikowia_pulcherrima(100); | Metschnikowia | 13 | Saprotroph | \|Undefined Saprotroph\| | Yeast | NULL | Probable | As nectar-tap_saprotroph (Põlme et al. 2020); Decay sugar-rich substrates (Põlme et al. 2020) | Põlme S, et al. 2020. Fungal Diversity 105:1-16 ((https://doi.org/10.1007/s13225-020-00466-2)) |
| Otu10701 | 2 | k__Fungi(100);p__Ascomycota(100);c__Saccharomycetes(100);o__Saccharomycetales(100);f__Metschnikowiaceae(100);g__Metschnikowia(100);s__Metschnikowia_pulcherrima(100); | Metschnikowia | 13 | Saprotroph | \|Undefined Saprotroph\| | Yeast | NULL | Probable | As nectar-tap_saprotroph (Põlme et al. 2020); Decay sugar-rich substrates (Põlme et al. 2020) | Põlme S, et al. 2020. Fungal Diversity 105:1-16 ((https://doi.org/10.1007/s13225-020-00466-2)) |
| Otu10703 | 2 | k__Fungi(100);p__Ascomycota(100);c__Saccharomycetes(100);o__Saccharomycetales(100);f__Saccharomycetales_family_Incertae_sedis(100);g__Candida(100);g__Candida_unclassified(100); | Candida | 13 | Pathotroph-Saprotroph-Symbiotroph | Animal Pathogen-Endophyte-Endosymbiont-Epiphyte-\|Undefined Saprotroph\| | Dimorphic Yeast-Yeast | NULL | Probable | As nectar-tap_saprotroph (Põlme et al. 2020); Decay sugar-rich substrates (Põlme et al. 2020); This asexual genus is polyphyletic and contain species that function very differently depend on what they are. Caution should be taken when interpreting guilds for Candida species. | Manolakaki D, et al. 2010. Virulence. 1: 367-375 ((https://doi.org/10.4161/viru.1.5.12796)); Mekbib SB, et al. 2011. Tropical Plant Pathology 36:233-240 ((https://doi.org/10.1590/S1982-56762011000400004)); Põlme S, et al. 2020. Fungal Diversity 105:1-16 ((https://doi.org/10.1007/s13225-020-00466-2)) |
| Otu10704 | 2 | k__Fungi(100);p__Ascomycota(100);c__Saccharomycetes(100);o__Saccharomycetales(100);f__Metschnikowiaceae(100);g__Metschnikowia(100);s__Metschnikowia_pulcherrima(100); | Metschnikowia | 13 | Saprotroph | \|Undefined Saprotroph\| | Yeast | NULL | Probable | As nectar-tap_saprotroph (Põlme et al. 2020); Decay sugar-rich substrates (Põlme et al. 2020) | Põlme S, et al. 2020. Fungal Diversity 105:1-16 ((https://doi.org/10.1007/s13225-020-00466-2)) |
| Otu10705 | 2 | k__Fungi(100);p__Ascomycota(100);c__Saccharomycetes(100);o__Saccharomycetales(100);f__Metschnikowiaceae(100);g__Metschnikowia(100);g__Metschnikowia_unclassified(100); | Metschnikowia | 13 | Saprotroph | \|Undefined Saprotroph\| | Yeast | NULL | Probable | As nectar-tap_saprotroph (Põlme et al. 2020); Decay sugar-rich substrates (Põlme et al. 2020) | Põlme S, et al. 2020. Fungal Diversity 105:1-16 ((https://doi.org/10.1007/s13225-020-00466-2)) |
| Otu10708 | 2 | k__Fungi(100);p__Ascomycota(100);c__Sordariomycetes(100);o__Hypocreales(100);f__Hypocreales_family_Incertae_sedis(100);g__Stachybotrys(100);s__Stachybotrys_eucylindrospora(100); | Stachybotrys | 13 | Pathotroph-Saprotroph | Plant Pathogen-Plant Saprotroph-\|Wood Saprotroph\| | Microfungus | Soft Rot | Highly Probable | As litter_saprotroph (Põlme et al. 2020); Decay wood, leaf, fruit, seed (Põlme et al. 2020) | Seehann G, et al. 1975. List of Fungi in Soft-Rot Tissue Tests. Institute fur Holzbiologie und Holzschutz, Bundesforschungsanstalt fur Forst - und Holzwirtshaft, Hamburg - Lohbrugge, Germany; Costa IPMW, et al. 2012. Mycotaxon 119:e26 ((http://mycotaxon.com/resources/checklists/costa_v119_checklist.pdf)); Tedersoo L, et al. 2014. Science 346:e1256688 ((https://doi.org/10.1126/science.1256688)); Põlme S, et al. 2020. Fungal Diversity 105:1-16 ((https://doi.org/10.1007/s13225-020-00466-2)); The American Phytopathological Society Common Names of Plant Diseases ((http://www.apsnet.org/publications/commonnames/Pages/default.aspx)) |
| Otu10709 | 2 | k__Fungi(100);p__Ascomycota(100);c__Sordariomycetes(100);o__Microascales(100);f__Halosphaeriaceae(100);g__unclassified_Halosphaeriaceae(100);s__Halosphaeriaceae_sp(100); | Halosphaeriaceae | 9 | Saprotroph | Undefined Saprotroph-Wood Saprotroph | Microfungus | NULL | Probable | Found in aquatic habitats, mostly in marine environments; saprobic on submerged or intertidal wood, occasionally associated with sand grains and similar substrata (Cannon & Kirk 2007) | Cannon PF, Kirk PM. 2007. Fungal Families of the World. CAB International, Cambridge (ISBN: 978-0851998275) |
| Otu10711 | 2 | k__Fungi(100);p__Ascomycota(100);c__Sordariomycetes(100);o__Sordariomycetes_order_Incertae_sedis(100);f__Sordariomycetes_family_Incertae_sedis(100);g__Eucasphaeria(100);s__Eucasphaeria_capensis(100); | Eucasphaeria | 13 | Pathotroph-Saprotroph | \|Plant Pathogen\|-Plant Saprotroph-Undefined Saprotroph | Microfungus | NULL | Probable | As litter_saprotroph (Põlme et al. 2020); Decay leaf, fruit, seed (Põlme et al. 2020) | Tedersoo L, et al. 2014. Science 346:e1256688 ((https://doi.org/10.1126/science.1256688)); Põlme S, et al. 2020. Fungal Diversity 105:1-16 ((https://doi.org/10.1007/s13225-020-00466-2)) |
| Otu10713 | 2 | k__Fungi(100);p__Ascomycota(100);c__Saccharomycetes(100);o__Saccharomycetales(100);f__Metschnikowiaceae(100);g__Metschnikowia(100);s__Metschnikowia_pulcherrima(100); | Metschnikowia | 13 | Saprotroph | \|Undefined Saprotroph\| | Yeast | NULL | Probable | As nectar-tap_saprotroph (Põlme et al. 2020); Decay sugar-rich substrates (Põlme et al. 2020) | Põlme S, et al. 2020. Fungal Diversity 105:1-16 ((https://doi.org/10.1007/s13225-020-00466-2)) |
| Otu10714 | 2 | k__Fungi(100);p__Ascomycota(100);c__Saccharomycetes(100);o__Saccharomycetales(100);f__Metschnikowiaceae(100);g__Metschnikowia(100);s__Metschnikowia_pulcherrima(100); | Metschnikowia | 13 | Saprotroph | \|Undefined Saprotroph\| | Yeast | NULL | Probable | As nectar-tap_saprotroph (Põlme et al. 2020); Decay sugar-rich substrates (Põlme et al. 2020) | Põlme S, et al. 2020. Fungal Diversity 105:1-16 ((https://doi.org/10.1007/s13225-020-00466-2)) |
| Otu10715 | 2 | k__Fungi(100);p__Ascomycota(100);c__Sordariomycetes(100);o__Hypocreales(100);o__Hypocreales_unclassified(100);o__Hypocreales_unclassified(100);o__Hypocreales_unclassified(100); | Hypocreales | 7 | Saprotroph | Undefined Saprotroph | Microfungus | NULL | Possible | NULL | Sterkenburg E, et al. 2015. New Phytologist 207:1145-1158 ((https://doi.org/10.1111/nph.13426)) |
| Otu10716 | 2 | k__Fungi(100);p__Ascomycota(100);c__Sordariomycetes(100);o__Sordariomycetes_order_Incertae_sedis(100);f__Glomerellaceae(100);g__Colletotrichum(100);g__Colletotrichum_unclassified(100); | Colletotrichum | 13 | Pathotroph-Saprotroph-Symbiotroph | Endophyte-\|Plant Pathogen\|-Plant Saprotroph | Microfungus | NULL | Probable | As foliar_endophyte (Põlme et al. 2020); As litter_saprotroph (Põlme et al. 2020); Decay leaf, fruit, seed (Põlme et al. 2020) | Costa IPMW, et al. 2012. Mycotaxon 119:e26 ((http://mycotaxon.com/resources/checklists/costa_v119_checklist.pdf)); Tedersoo L, et al. 2014. Science 346:e1256688 ((https://doi.org/10.1126/science.1256688)); Põlme S, et al. 2020. Fungal Diversity 105:1-16 ((https://doi.org/10.1007/s13225-020-00466-2)); The American Phytopathological Society Common Names of Plant Diseases ((http://www.apsnet.org/publications/commonnames/Pages/default.aspx)) |
| Otu10718 | 2 | k__Fungi(100);p__Ascomycota(100);c__Saccharomycetes(100);o__Saccharomycetales(100);f__Metschnikowiaceae(100);g__Metschnikowia(100);g__Metschnikowia_unclassified(100); | Metschnikowia | 13 | Saprotroph | \|Undefined Saprotroph\| | Yeast | NULL | Probable | As nectar-tap_saprotroph (Põlme et al. 2020); Decay sugar-rich substrates (Põlme et al. 2020) | Põlme S, et al. 2020. Fungal Diversity 105:1-16 ((https://doi.org/10.1007/s13225-020-00466-2)) |
| Otu10719 | 2 | k__Fungi(100);p__Ascomycota(100);c__Saccharomycetes(100);o__Saccharomycetales(100);f__Metschnikowiaceae(100);g__Metschnikowia(100);s__Metschnikowia_pulcherrima(100); | Metschnikowia | 13 | Saprotroph | \|Undefined Saprotroph\| | Yeast | NULL | Probable | As nectar-tap_saprotroph (Põlme et al. 2020); Decay sugar-rich substrates (Põlme et al. 2020) | Põlme S, et al. 2020. Fungal Diversity 105:1-16 ((https://doi.org/10.1007/s13225-020-00466-2)) |
| Otu10720 | 2 | k__Fungi(100);p__Ascomycota(100);c__Saccharomycetes(100);o__Saccharomycetales(100);f__Metschnikowiaceae(100);g__Metschnikowia(100);g__Metschnikowia_unclassified(100); | Metschnikowia | 13 | Saprotroph | \|Undefined Saprotroph\| | Yeast | NULL | Probable | As nectar-tap_saprotroph (Põlme et al. 2020); Decay sugar-rich substrates (Põlme et al. 2020) | Põlme S, et al. 2020. Fungal Diversity 105:1-16 ((https://doi.org/10.1007/s13225-020-00466-2)) |
| Otu10723 | 2 | k__Fungi(100);p__Ascomycota(100);c__Saccharomycetes(100);o__Saccharomycetales(100);f__Metschnikowiaceae(100);g__Metschnikowia(100);g__Metschnikowia_unclassified(100); | Metschnikowia | 13 | Saprotroph | \|Undefined Saprotroph\| | Yeast | NULL | Probable | As nectar-tap_saprotroph (Põlme et al. 2020); Decay sugar-rich substrates (Põlme et al. 2020) | Põlme S, et al. 2020. Fungal Diversity 105:1-16 ((https://doi.org/10.1007/s13225-020-00466-2)) |
| Otu10732 | 2 | k__Fungi(100);p__Ascomycota(100);c__Saccharomycetes(100);o__Saccharomycetales(100);f__Metschnikowiaceae(100);g__Metschnikowia(100);g__Metschnikowia_unclassified(100); | Metschnikowia | 13 | Saprotroph | \|Undefined Saprotroph\| | Yeast | NULL | Probable | As nectar-tap_saprotroph (Põlme et al. 2020); Decay sugar-rich substrates (Põlme et al. 2020) | Põlme S, et al. 2020. Fungal Diversity 105:1-16 ((https://doi.org/10.1007/s13225-020-00466-2)) |
| Otu10733 | 2 | k__Fungi(100);p__Ascomycota(100);c__Saccharomycetes(100);o__Saccharomycetales(100);f__Metschnikowiaceae(100);g__Metschnikowia(100);s__Metschnikowia_pulcherrima(100); | Metschnikowia | 13 | Saprotroph | \|Undefined Saprotroph\| | Yeast | NULL | Probable | As nectar-tap_saprotroph (Põlme et al. 2020); Decay sugar-rich substrates (Põlme et al. 2020) | Põlme S, et al. 2020. Fungal Diversity 105:1-16 ((https://doi.org/10.1007/s13225-020-00466-2)) |
| Otu10734 | 2 | k__Fungi(100);p__Ascomycota(100);c__Saccharomycetes(100);o__Saccharomycetales(100);f__Metschnikowiaceae(100);g__Metschnikowia(100);s__Metschnikowia_pulcherrima(100); | Metschnikowia | 13 | Saprotroph | \|Undefined Saprotroph\| | Yeast | NULL | Probable | As nectar-tap_saprotroph (Põlme et al. 2020); Decay sugar-rich substrates (Põlme et al. 2020) | Põlme S, et al. 2020. Fungal Diversity 105:1-16 ((https://doi.org/10.1007/s13225-020-00466-2)) |
| Otu10735 | 2 | k__Fungi(100);p__Ascomycota(100);c__Saccharomycetes(100);o__Saccharomycetales(100);f__Saccharomycetaceae(100);g__Kazachstania(100);s__Kazachstania_martiniae(100); | Kazachstania | 13 | Saprotroph | \|Undefined Saprotroph\| | Yeast | NULL | Probable | As nectar-tap_saprotroph (Põlme et al. 2020); Decay animal material, sugar-rich substrates (Põlme et al. 2020) | Põlme S, et al. 2020. Fungal Diversity 105:1-16 ((https://doi.org/10.1007/s13225-020-00466-2)) |
| Otu10736 | 2 | k__Fungi(100);p__Ascomycota(100);c__Saccharomycetes(100);o__Saccharomycetales(100);f__Saccharomycetaceae(100);g__Torulaspora(100);s__Torulaspora_delbrueckii(100); | Torulaspora | 13 | Pathotroph-Saprotroph | Animal Parasite-\|Undefined Saprotroph\| | Yeast | NULL | Probable | As nectar-tap_saprotroph (Põlme et al. 2020); Decay sugar-rich substrates (Põlme et al. 2020) | Põlme S, et al. 2020. Fungal Diversity 105:1-16 ((https://doi.org/10.1007/s13225-020-00466-2)) |
| Otu10737 | 2 | k__Fungi(100);p__Ascomycota(100);c__Saccharomycetes(100);o__Saccharomycetales(100);f__Metschnikowiaceae(100);g__Metschnikowia(100);s__Metschnikowia_pulcherrima(100); | Metschnikowia | 13 | Saprotroph | \|Undefined Saprotroph\| | Yeast | NULL | Probable | As nectar-tap_saprotroph (Põlme et al. 2020); Decay sugar-rich substrates (Põlme et al. 2020) | Põlme S, et al. 2020. Fungal Diversity 105:1-16 ((https://doi.org/10.1007/s13225-020-00466-2)) |
| Otu10738 | 2 | k__Fungi(100);p__Ascomycota(100);c__Saccharomycetes(100);o__Saccharomycetales(100);f__Saccharomycetaceae(100);g__Saccharomyces(100);g__Saccharomyces_unclassified(100); | Saccharomyces | 13 | Saprotroph | \|Undefined Saprotroph\| | Yeast | NULL | Probable | As nectar-tap_saprotroph (Põlme et al. 2020); Decay sugar-rich substrates (Põlme et al. 2020) | James TY, et al. 2006. Nature 443:818-822 ((https://doi.org/10.1038/nature05110)); Põlme S, et al. 2020. Fungal Diversity 105:1-16 ((https://doi.org/10.1007/s13225-020-00466-2)) |
| Otu10739 | 2 | k__Fungi(100);p__Ascomycota(100);c__Saccharomycetes(100);o__Saccharomycetales(100);f__Saccharomycetaceae(100);g__Saccharomyces(100);g__Saccharomyces_unclassified(100); | Saccharomyces | 13 | Saprotroph | \|Undefined Saprotroph\| | Yeast | NULL | Probable | As nectar-tap_saprotroph (Põlme et al. 2020); Decay sugar-rich substrates (Põlme et al. 2020) | James TY, et al. 2006. Nature 443:818-822 ((https://doi.org/10.1038/nature05110)); Põlme S, et al. 2020. Fungal Diversity 105:1-16 ((https://doi.org/10.1007/s13225-020-00466-2)) |
| Otu10740 | 2 | k__Fungi(100);p__Ascomycota(100);c__Saccharomycetes(100);o__Saccharomycetales(100);f__Saccharomycetaceae(100);g__Saccharomyces(100);g__Saccharomyces_unclassified(100); | Saccharomyces | 13 | Saprotroph | \|Undefined Saprotroph\| | Yeast | NULL | Probable | As nectar-tap_saprotroph (Põlme et al. 2020); Decay sugar-rich substrates (Põlme et al. 2020) | James TY, et al. 2006. Nature 443:818-822 ((https://doi.org/10.1038/nature05110)); Põlme S, et al. 2020. Fungal Diversity 105:1-16 ((https://doi.org/10.1007/s13225-020-00466-2)) |
| Otu10741 | 2 | k__Fungi(100);p__Ascomycota(100);c__Saccharomycetes(100);o__Saccharomycetales(100);f__Saccharomycetaceae(100);g__Saccharomyces(100);g__Saccharomyces_unclassified(100); | Saccharomyces | 13 | Saprotroph | \|Undefined Saprotroph\| | Yeast | NULL | Probable | As nectar-tap_saprotroph (Põlme et al. 2020); Decay sugar-rich substrates (Põlme et al. 2020) | James TY, et al. 2006. Nature 443:818-822 ((https://doi.org/10.1038/nature05110)); Põlme S, et al. 2020. Fungal Diversity 105:1-16 ((https://doi.org/10.1007/s13225-020-00466-2)) |
| Otu10742 | 2 | k__Fungi(100);p__Ascomycota(100);c__Saccharomycetes(100);o__Saccharomycetales(100);f__Saccharomycetaceae(100);g__Saccharomyces(100);g__Saccharomyces_unclassified(100); | Saccharomyces | 13 | Saprotroph | \|Undefined Saprotroph\| | Yeast | NULL | Probable | As nectar-tap_saprotroph (Põlme et al. 2020); Decay sugar-rich substrates (Põlme et al. 2020) | James TY, et al. 2006. Nature 443:818-822 ((https://doi.org/10.1038/nature05110)); Põlme S, et al. 2020. Fungal Diversity 105:1-16 ((https://doi.org/10.1007/s13225-020-00466-2)) |
| Otu10746 | 2 | k__Fungi(100);p__Ascomycota(100);c__Saccharomycetes(100);o__Saccharomycetales(100);f__Metschnikowiaceae(100);g__Metschnikowia(100);s__Metschnikowia_pulcherrima(100); | Metschnikowia | 13 | Saprotroph | \|Undefined Saprotroph\| | Yeast | NULL | Probable | As nectar-tap_saprotroph (Põlme et al. 2020); Decay sugar-rich substrates (Põlme et al. 2020) | Põlme S, et al. 2020. Fungal Diversity 105:1-16 ((https://doi.org/10.1007/s13225-020-00466-2)) |
| Otu10747 | 2 | k__Fungi(100);p__Ascomycota(100);c__Saccharomycetes(100);o__Saccharomycetales(100);f__Metschnikowiaceae(100);g__Metschnikowia(100);g__Metschnikowia_unclassified(100); | Metschnikowia | 13 | Saprotroph | \|Undefined Saprotroph\| | Yeast | NULL | Probable | As nectar-tap_saprotroph (Põlme et al. 2020); Decay sugar-rich substrates (Põlme et al. 2020) | Põlme S, et al. 2020. Fungal Diversity 105:1-16 ((https://doi.org/10.1007/s13225-020-00466-2)) |
| Otu10749 | 2 | k__Fungi(100);p__Ascomycota(100);c__Saccharomycetes(100);o__Saccharomycetales(100);o__Saccharomycetales_unclassified(100);o__Saccharomycetales_unclassified(100);o__Saccharomycetales_unclassified(100); | Saccharomycetales | 7 | Saprotroph | Undefined Saprotroph | Yeast | NULL | Possible | NULL | Sterkenburg E, et al. 2015. New Phytologist 207:1145-1158 ((https://doi.org/10.1111/nph.13426)) |
| Otu10750 | 2 | k__Fungi(100);p__Ascomycota(100);c__Saccharomycetes(100);o__Saccharomycetales(100);f__Metschnikowiaceae(100);g__Metschnikowia(100);s__Metschnikowia_pulcherrima(100); | Metschnikowia | 13 | Saprotroph | \|Undefined Saprotroph\| | Yeast | NULL | Probable | As nectar-tap_saprotroph (Põlme et al. 2020); Decay sugar-rich substrates (Põlme et al. 2020) | Põlme S, et al. 2020. Fungal Diversity 105:1-16 ((https://doi.org/10.1007/s13225-020-00466-2)) |
| Otu10751 | 2 | k__Fungi(100);p__Ascomycota(100);c__Saccharomycetes(100);o__Saccharomycetales(100);f__Metschnikowiaceae(100);g__Metschnikowia(100);s__Metschnikowia_pulcherrima(100); | Metschnikowia | 13 | Saprotroph | \|Undefined Saprotroph\| | Yeast | NULL | Probable | As nectar-tap_saprotroph (Põlme et al. 2020); Decay sugar-rich substrates (Põlme et al. 2020) | Põlme S, et al. 2020. Fungal Diversity 105:1-16 ((https://doi.org/10.1007/s13225-020-00466-2)) |
| Otu10752 | 2 | k__Fungi(100);p__Ascomycota(100);c__Saccharomycetes(100);o__Saccharomycetales(100);f__Metschnikowiaceae(100);g__Metschnikowia(100);s__Metschnikowia_pulcherrima(100); | Metschnikowia | 13 | Saprotroph | \|Undefined Saprotroph\| | Yeast | NULL | Probable | As nectar-tap_saprotroph (Põlme et al. 2020); Decay sugar-rich substrates (Põlme et al. 2020) | Põlme S, et al. 2020. Fungal Diversity 105:1-16 ((https://doi.org/10.1007/s13225-020-00466-2)) |
| Otu10753 | 2 | k__Fungi(100);p__Ascomycota(100);c__Saccharomycetes(100);o__Saccharomycetales(100);f__Metschnikowiaceae(100);g__Metschnikowia(100);g__Metschnikowia_unclassified(100); | Metschnikowia | 13 | Saprotroph | \|Undefined Saprotroph\| | Yeast | NULL | Probable | As nectar-tap_saprotroph (Põlme et al. 2020); Decay sugar-rich substrates (Põlme et al. 2020) | Põlme S, et al. 2020. Fungal Diversity 105:1-16 ((https://doi.org/10.1007/s13225-020-00466-2)) |
| Otu10755 | 2 | k__Fungi(100);p__Ascomycota(100);c__Saccharomycetes(100);o__Saccharomycetales(100);f__Metschnikowiaceae(100);g__Metschnikowia(100);g__Metschnikowia_unclassified(100); | Metschnikowia | 13 | Saprotroph | \|Undefined Saprotroph\| | Yeast | NULL | Probable | As nectar-tap_saprotroph (Põlme et al. 2020); Decay sugar-rich substrates (Põlme et al. 2020) | Põlme S, et al. 2020. Fungal Diversity 105:1-16 ((https://doi.org/10.1007/s13225-020-00466-2)) |
| Otu10762 | 2 | k__Fungi(100);p__Ascomycota(100);c__Saccharomycetes(100);o__Saccharomycetales(100);f__Metschnikowiaceae(100);g__Metschnikowia(100);s__Metschnikowia_pulcherrima(100); | Metschnikowia | 13 | Saprotroph | \|Undefined Saprotroph\| | Yeast | NULL | Probable | As nectar-tap_saprotroph (Põlme et al. 2020); Decay sugar-rich substrates (Põlme et al. 2020) | Põlme S, et al. 2020. Fungal Diversity 105:1-16 ((https://doi.org/10.1007/s13225-020-00466-2)) |
| Otu10764 | 2 | k__Fungi(100);p__Ascomycota(100);c__Saccharomycetes(100);o__Saccharomycetales(100);f__Metschnikowiaceae(100);g__Metschnikowia(100);s__Metschnikowia_pulcherrima(100); | Metschnikowia | 13 | Saprotroph | \|Undefined Saprotroph\| | Yeast | NULL | Probable | As nectar-tap_saprotroph (Põlme et al. 2020); Decay sugar-rich substrates (Põlme et al. 2020) | Põlme S, et al. 2020. Fungal Diversity 105:1-16 ((https://doi.org/10.1007/s13225-020-00466-2)) |
| Otu10766 | 2 | k__Fungi(100);p__Ascomycota(100);c__Saccharomycetes(100);o__Saccharomycetales(100);f__Metschnikowiaceae(100);g__Metschnikowia(100);s__Metschnikowia_pulcherrima(100); | Metschnikowia | 13 | Saprotroph | \|Undefined Saprotroph\| | Yeast | NULL | Probable | As nectar-tap_saprotroph (Põlme et al. 2020); Decay sugar-rich substrates (Põlme et al. 2020) | Põlme S, et al. 2020. Fungal Diversity 105:1-16 ((https://doi.org/10.1007/s13225-020-00466-2)) |
| Otu10767 | 2 | k__Fungi(100);p__Ascomycota(100);c__Saccharomycetes(100);o__Saccharomycetales(100);f__Metschnikowiaceae(100);g__Metschnikowia(100);s__Metschnikowia_pulcherrima(100); | Metschnikowia | 13 | Saprotroph | \|Undefined Saprotroph\| | Yeast | NULL | Probable | As nectar-tap_saprotroph (Põlme et al. 2020); Decay sugar-rich substrates (Põlme et al. 2020) | Põlme S, et al. 2020. Fungal Diversity 105:1-16 ((https://doi.org/10.1007/s13225-020-00466-2)) |
| Otu10768 | 2 | k__Fungi(100);p__Ascomycota(100);c__Saccharomycetes(100);o__Saccharomycetales(100);f__Metschnikowiaceae(100);g__Metschnikowia(100);s__Metschnikowia_pulcherrima(100); | Metschnikowia | 13 | Saprotroph | \|Undefined Saprotroph\| | Yeast | NULL | Probable | As nectar-tap_saprotroph (Põlme et al. 2020); Decay sugar-rich substrates (Põlme et al. 2020) | Põlme S, et al. 2020. Fungal Diversity 105:1-16 ((https://doi.org/10.1007/s13225-020-00466-2)) |
| Otu10769 | 2 | k__Fungi(100);p__Ascomycota(100);c__Saccharomycetes(100);o__Saccharomycetales(100);f__Metschnikowiaceae(100);g__Metschnikowia(100);s__Metschnikowia_pulcherrima(100); | Metschnikowia | 13 | Saprotroph | \|Undefined Saprotroph\| | Yeast | NULL | Probable | As nectar-tap_saprotroph (Põlme et al. 2020); Decay sugar-rich substrates (Põlme et al. 2020) | Põlme S, et al. 2020. Fungal Diversity 105:1-16 ((https://doi.org/10.1007/s13225-020-00466-2)) |
| Otu10771 | 2 | k__Fungi(100);p__Basidiomycota(100);c__Microbotryomycetes(100);o__Sporidiobolales(100);f__Sporidiobolales_family_Incertae_sedis(100);g__Sporobolomyces(100);s__Sporobolomyces_ruberrimus(100); | Sporobolomyces | 13 | Pathotroph-Saprotroph | \|Fungal Parasite\|-Plant Saprotroph | Yeast | NULL | Probable | As litter_saprotroph (Põlme et al. 2020); As mycoparasite (Põlme et al. 2020); Decay fungal material (Põlme et al. 2020) | Tedersoo L, et al. 2014. Science 346:e1256688 ((https://doi.org/10.1126/science.1256688)); Purahong W, et al. 2016. Molecular Ecology 25:4059-4074 ((https://doi.org/10.1111/mec.13739)); Põlme S, et al. 2020. Fungal Diversity 105:1-16 ((https://doi.org/10.1007/s13225-020-00466-2)) |
| Otu10772 | 2 | k__Fungi(100);p__Ascomycota(100);c__Saccharomycetes(100);o__Saccharomycetales(100);f__Metschnikowiaceae(100);g__Metschnikowia(100);g__Metschnikowia_unclassified(100); | Metschnikowia | 13 | Saprotroph | \|Undefined Saprotroph\| | Yeast | NULL | Probable | As nectar-tap_saprotroph (Põlme et al. 2020); Decay sugar-rich substrates (Põlme et al. 2020) | Põlme S, et al. 2020. Fungal Diversity 105:1-16 ((https://doi.org/10.1007/s13225-020-00466-2)) |
| Otu10776 | 2 | k__Fungi(100);p__Ascomycota(100);c__Saccharomycetes(100);o__Saccharomycetales(100);f__Metschnikowiaceae(100);g__Metschnikowia(100);s__Metschnikowia_pulcherrima(100); | Metschnikowia | 13 | Saprotroph | \|Undefined Saprotroph\| | Yeast | NULL | Probable | As nectar-tap_saprotroph (Põlme et al. 2020); Decay sugar-rich substrates (Põlme et al. 2020) | Põlme S, et al. 2020. Fungal Diversity 105:1-16 ((https://doi.org/10.1007/s13225-020-00466-2)) |
| Otu10778 | 2 | k__Fungi(100);p__Ascomycota(100);c__Leotiomycetes(100);o__Erysiphales(100);f__Erysiphaceae(100);g__Erysiphe(100);s__Erysiphe_pulchra(100); | Erysiphe | 13 | Pathotroph | \|Plant Pathogen\| | Microfungus | NULL | Probable | Biotrophic, with affected plant tissues damaged but usually not killing outright (Cannon & Kirk 2007); Common name - powdery mildew | Tedersoo L, et al. 2014. Science 346:e1256688 ((https://doi.org/10.1126/science.1256688)); Põlme S, et al. 2020. Fungal Diversity 105:1-16 ((https://doi.org/10.1007/s13225-020-00466-2)) |
| Otu10779 | 2 | k__Fungi(100);p__Ascomycota(100);c__Saccharomycetes(100);o__Saccharomycetales(100);o__Saccharomycetales_unclassified(100);o__Saccharomycetales_unclassified(100);o__Saccharomycetales_unclassified(100); | Saccharomycetales | 7 | Saprotroph | Undefined Saprotroph | Yeast | NULL | Possible | NULL | Sterkenburg E, et al. 2015. New Phytologist 207:1145-1158 ((https://doi.org/10.1111/nph.13426)) |
| Otu10781 | 2 | k__Fungi(100);p__Ascomycota(100);c__Dothideomycetes(100);o__Botryosphaeriales(100);f__Botryosphaeriaceae(100);f__Botryosphaeriaceae_unclassified(100);f__Botryosphaeriaceae_unclassified(100); | Botryosphaeriaceae | 9 | Pathotroph-Saprotroph-Symbiotroph | Endophyte-Plant Pathogen-Wood Saprotroph | Microfungus | NULL | Probable | Biotrophic, necrotrophic, or saprobic, especially associated with woody tissue (Cannon & Kirk 2007) | Cannon PF, Kirk PM. 2007. Fungal Families of the World. CAB International, Cambridge (ISBN: 978-0851998275) |
| Otu10782 | 2 | k__Fungi(100);p__Ascomycota(100);c__Dothideomycetes(100);o__Pleosporales(100);f__Pleomassariaceae(100);g__Trematosphaeria(100);s__Trematosphaeria_pertusa(100); | Trematosphaeria | 13 | Saprotroph | Undefined Saprotroph-\|Wood Saprotroph\| | Microfungus | NULL | Probable | Decay wood (Põlme et al. 2020) | Tedersoo L, et al. 2014. Science 346:e1256688 ((https://doi.org/10.1126/science.1256688)); Põlme S, et al. 2020. Fungal Diversity 105:1-16 ((https://doi.org/10.1007/s13225-020-00466-2)) |
| Otu10783 | 2 | k__Fungi(100);p__Ascomycota(100);c__Saccharomycetes(100);o__Saccharomycetales(100);f__Metschnikowiaceae(100);g__Metschnikowia(100);s__Metschnikowia_pulcherrima(100); | Metschnikowia | 13 | Saprotroph | \|Undefined Saprotroph\| | Yeast | NULL | Probable | As nectar-tap_saprotroph (Põlme et al. 2020); Decay sugar-rich substrates (Põlme et al. 2020) | Põlme S, et al. 2020. Fungal Diversity 105:1-16 ((https://doi.org/10.1007/s13225-020-00466-2)) |
| Otu10784 | 2 | k__Fungi(100);p__Ascomycota(100);c__Saccharomycetes(100);o__Saccharomycetales(100);f__Metschnikowiaceae(100);g__Metschnikowia(100);s__Metschnikowia_pulcherrima(100); | Metschnikowia | 13 | Saprotroph | \|Undefined Saprotroph\| | Yeast | NULL | Probable | As nectar-tap_saprotroph (Põlme et al. 2020); Decay sugar-rich substrates (Põlme et al. 2020) | Põlme S, et al. 2020. Fungal Diversity 105:1-16 ((https://doi.org/10.1007/s13225-020-00466-2)) |
| Otu10785 | 2 | k__Fungi(100);p__Ascomycota(100);c__Saccharomycetes(100);o__Saccharomycetales(100);f__Metschnikowiaceae(100);g__Metschnikowia(100);s__Metschnikowia_pulcherrima(100); | Metschnikowia | 13 | Saprotroph | \|Undefined Saprotroph\| | Yeast | NULL | Probable | As nectar-tap_saprotroph (Põlme et al. 2020); Decay sugar-rich substrates (Põlme et al. 2020) | Põlme S, et al. 2020. Fungal Diversity 105:1-16 ((https://doi.org/10.1007/s13225-020-00466-2)) |
| Otu10787 | 2 | k__Fungi(100);p__Ascomycota(100);c__Saccharomycetes(100);o__Saccharomycetales(100);f__Metschnikowiaceae(100);g__Metschnikowia(100);s__Metschnikowia_pulcherrima(100); | Metschnikowia | 13 | Saprotroph | \|Undefined Saprotroph\| | Yeast | NULL | Probable | As nectar-tap_saprotroph (Põlme et al. 2020); Decay sugar-rich substrates (Põlme et al. 2020) | Põlme S, et al. 2020. Fungal Diversity 105:1-16 ((https://doi.org/10.1007/s13225-020-00466-2)) |
| Otu10789 | 2 | k__Fungi(100);p__Ascomycota(100);c__Saccharomycetes(100);o__Saccharomycetales(100);f__Metschnikowiaceae(100);g__Metschnikowia(100);s__Metschnikowia_pulcherrima(100); | Metschnikowia | 13 | Saprotroph | \|Undefined Saprotroph\| | Yeast | NULL | Probable | As nectar-tap_saprotroph (Põlme et al. 2020); Decay sugar-rich substrates (Põlme et al. 2020) | Põlme S, et al. 2020. Fungal Diversity 105:1-16 ((https://doi.org/10.1007/s13225-020-00466-2)) |
| Otu10790 | 2 | k__Fungi(100);p__Ascomycota(100);c__Saccharomycetes(100);o__Saccharomycetales(100);f__Saccharomycetales_family_Incertae_sedis(100);g__Candida(100);g__Candida_unclassified(100); | Candida | 13 | Pathotroph-Saprotroph-Symbiotroph | Animal Pathogen-Endophyte-Endosymbiont-Epiphyte-\|Undefined Saprotroph\| | Dimorphic Yeast-Yeast | NULL | Probable | As nectar-tap_saprotroph (Põlme et al. 2020); Decay sugar-rich substrates (Põlme et al. 2020); This asexual genus is polyphyletic and contain species that function very differently depend on what they are. Caution should be taken when interpreting guilds for Candida species. | Manolakaki D, et al. 2010. Virulence. 1: 367-375 ((https://doi.org/10.4161/viru.1.5.12796)); Mekbib SB, et al. 2011. Tropical Plant Pathology 36:233-240 ((https://doi.org/10.1590/S1982-56762011000400004)); Põlme S, et al. 2020. Fungal Diversity 105:1-16 ((https://doi.org/10.1007/s13225-020-00466-2)) |
| Otu10794 | 2 | k__Fungi(100);p__Ascomycota(100);c__Saccharomycetes(100);o__Saccharomycetales(100);f__Metschnikowiaceae(100);g__Metschnikowia(100);g__Metschnikowia_unclassified(100); | Metschnikowia | 13 | Saprotroph | \|Undefined Saprotroph\| | Yeast | NULL | Probable | As nectar-tap_saprotroph (Põlme et al. 2020); Decay sugar-rich substrates (Põlme et al. 2020) | Põlme S, et al. 2020. Fungal Diversity 105:1-16 ((https://doi.org/10.1007/s13225-020-00466-2)) |
| Otu10796 | 2 | k__Fungi(100);p__Ascomycota(100);c__Saccharomycetes(100);o__Saccharomycetales(100);f__Metschnikowiaceae(100);g__Metschnikowia(100);g__Metschnikowia_unclassified(100); | Metschnikowia | 13 | Saprotroph | \|Undefined Saprotroph\| | Yeast | NULL | Probable | As nectar-tap_saprotroph (Põlme et al. 2020); Decay sugar-rich substrates (Põlme et al. 2020) | Põlme S, et al. 2020. Fungal Diversity 105:1-16 ((https://doi.org/10.1007/s13225-020-00466-2)) |
| Otu10797 | 2 | k__Fungi(100);p__Ascomycota(100);c__Saccharomycetes(100);o__Saccharomycetales(100);f__Metschnikowiaceae(100);g__Metschnikowia(100);s__Metschnikowia_pulcherrima(100); | Metschnikowia | 13 | Saprotroph | \|Undefined Saprotroph\| | Yeast | NULL | Probable | As nectar-tap_saprotroph (Põlme et al. 2020); Decay sugar-rich substrates (Põlme et al. 2020) | Põlme S, et al. 2020. Fungal Diversity 105:1-16 ((https://doi.org/10.1007/s13225-020-00466-2)) |
| Otu10800 | 2 | k__Fungi(100);p__Ascomycota(100);c__Saccharomycetes(100);o__Saccharomycetales(100);f__Metschnikowiaceae(100);g__Metschnikowia(100);g__Metschnikowia_unclassified(100); | Metschnikowia | 13 | Saprotroph | \|Undefined Saprotroph\| | Yeast | NULL | Probable | As nectar-tap_saprotroph (Põlme et al. 2020); Decay sugar-rich substrates (Põlme et al. 2020) | Põlme S, et al. 2020. Fungal Diversity 105:1-16 ((https://doi.org/10.1007/s13225-020-00466-2)) |
| Otu10801 | 2 | k__Fungi(100);p__Ascomycota(100);c__Leotiomycetes(100);o__Rhytismatales(100);f__Rhytismataceae(100);f__Rhytismataceae_unclassified(100);f__Rhytismataceae_unclassified(100); | Rhytismataceae | 9 | Pathotroph-Saprotroph-Symbiotroph | Endophyte-Leaf Saprotroph-Plant Pathogen | Microfungus | NULL | Probable | Biotrophic, necrotrophic or saprobic on leaves and bark, often initially as endophytes, causing tar spots, premature leaf fall and occasionally witches broom symptoms (Cannon & Kirk 2007) | Cannon PF, Kirk PM. 2007. Fungal Families of the World. CAB International, Cambridge (ISBN: 978-0851998275) |
| Otu10804 | 2 | k__Fungi(100);p__Ascomycota(100);c__Saccharomycetes(100);o__Saccharomycetales(100);f__Saccharomycetales_family_Incertae_sedis(100);f__Saccharomycetales_family_Incertae_sedis_unclassified(100);f__Saccharomycetales_family_Incertae_sedis_unclassified(100); | Saccharomycetales | 7 | Saprotroph | Undefined Saprotroph | Yeast | NULL | Possible | NULL | Sterkenburg E, et al. 2015. New Phytologist 207:1145-1158 ((https://doi.org/10.1111/nph.13426)) |
| Otu10805 | 2 | k__Fungi(100);p__Ascomycota(100);c__Saccharomycetes(100);o__Saccharomycetales(100);f__Metschnikowiaceae(100);g__Metschnikowia(100);g__Metschnikowia_unclassified(100); | Metschnikowia | 13 | Saprotroph | \|Undefined Saprotroph\| | Yeast | NULL | Probable | As nectar-tap_saprotroph (Põlme et al. 2020); Decay sugar-rich substrates (Põlme et al. 2020) | Põlme S, et al. 2020. Fungal Diversity 105:1-16 ((https://doi.org/10.1007/s13225-020-00466-2)) |
| Otu10806 | 2 | k__Fungi(100);p__Ascomycota(100);c__Saccharomycetes(100);o__Saccharomycetales(100);f__Metschnikowiaceae(100);g__Metschnikowia(100);g__Metschnikowia_unclassified(100); | Metschnikowia | 13 | Saprotroph | \|Undefined Saprotroph\| | Yeast | NULL | Probable | As nectar-tap_saprotroph (Põlme et al. 2020); Decay sugar-rich substrates (Põlme et al. 2020) | Põlme S, et al. 2020. Fungal Diversity 105:1-16 ((https://doi.org/10.1007/s13225-020-00466-2)) |
| Otu10807 | 2 | k__Fungi(100);p__Ascomycota(100);c__Saccharomycetes(100);o__Saccharomycetales(100);f__Metschnikowiaceae(100);g__Metschnikowia(100);s__Metschnikowia_pulcherrima(100); | Metschnikowia | 13 | Saprotroph | \|Undefined Saprotroph\| | Yeast | NULL | Probable | As nectar-tap_saprotroph (Põlme et al. 2020); Decay sugar-rich substrates (Põlme et al. 2020) | Põlme S, et al. 2020. Fungal Diversity 105:1-16 ((https://doi.org/10.1007/s13225-020-00466-2)) |
| Otu10808 | 2 | k__Fungi(100);p__Ascomycota(100);c__Saccharomycetes(100);o__Saccharomycetales(100);f__Metschnikowiaceae(100);g__Metschnikowia(100);g__Metschnikowia_unclassified(100); | Metschnikowia | 13 | Saprotroph | \|Undefined Saprotroph\| | Yeast | NULL | Probable | As nectar-tap_saprotroph (Põlme et al. 2020); Decay sugar-rich substrates (Põlme et al. 2020) | Põlme S, et al. 2020. Fungal Diversity 105:1-16 ((https://doi.org/10.1007/s13225-020-00466-2)) |
| Otu10809 | 2 | k__Fungi(100);p__Ascomycota(100);c__Saccharomycetes(100);o__Saccharomycetales(100);f__Metschnikowiaceae(100);g__Metschnikowia(100);s__Metschnikowia_pulcherrima(100); | Metschnikowia | 13 | Saprotroph | \|Undefined Saprotroph\| | Yeast | NULL | Probable | As nectar-tap_saprotroph (Põlme et al. 2020); Decay sugar-rich substrates (Põlme et al. 2020) | Põlme S, et al. 2020. Fungal Diversity 105:1-16 ((https://doi.org/10.1007/s13225-020-00466-2)) |
| Otu10810 | 2 | k__Fungi(100);p__Ascomycota(100);c__Saccharomycetes(100);o__Saccharomycetales(100);f__Metschnikowiaceae(100);g__Metschnikowia(100);s__Metschnikowia_pulcherrima(100); | Metschnikowia | 13 | Saprotroph | \|Undefined Saprotroph\| | Yeast | NULL | Probable | As nectar-tap_saprotroph (Põlme et al. 2020); Decay sugar-rich substrates (Põlme et al. 2020) | Põlme S, et al. 2020. Fungal Diversity 105:1-16 ((https://doi.org/10.1007/s13225-020-00466-2)) |
| Otu10811 | 2 | k__Fungi(100);p__Ascomycota(100);c__Saccharomycetes(100);o__Saccharomycetales(100);f__Metschnikowiaceae(100);g__Metschnikowia(100);g__Metschnikowia_unclassified(100); | Metschnikowia | 13 | Saprotroph | \|Undefined Saprotroph\| | Yeast | NULL | Probable | As nectar-tap_saprotroph (Põlme et al. 2020); Decay sugar-rich substrates (Põlme et al. 2020) | Põlme S, et al. 2020. Fungal Diversity 105:1-16 ((https://doi.org/10.1007/s13225-020-00466-2)) |
| Otu10813 | 2 | k__Fungi(100);p__Ascomycota(100);c__Saccharomycetes(100);o__Saccharomycetales(100);f__Metschnikowiaceae(100);g__Metschnikowia(100);s__Metschnikowia_pulcherrima(100); | Metschnikowia | 13 | Saprotroph | \|Undefined Saprotroph\| | Yeast | NULL | Probable | As nectar-tap_saprotroph (Põlme et al. 2020); Decay sugar-rich substrates (Põlme et al. 2020) | Põlme S, et al. 2020. Fungal Diversity 105:1-16 ((https://doi.org/10.1007/s13225-020-00466-2)) |
| Otu10815 | 2 | k__Fungi(100);p__Ascomycota(100);c__Saccharomycetes(100);o__Saccharomycetales(100);f__Metschnikowiaceae(100);g__Metschnikowia(100);s__Metschnikowia_pulcherrima(100); | Metschnikowia | 13 | Saprotroph | \|Undefined Saprotroph\| | Yeast | NULL | Probable | As nectar-tap_saprotroph (Põlme et al. 2020); Decay sugar-rich substrates (Põlme et al. 2020) | Põlme S, et al. 2020. Fungal Diversity 105:1-16 ((https://doi.org/10.1007/s13225-020-00466-2)) |
| Otu10816 | 2 | k__Fungi(100);p__Ascomycota(100);c__Saccharomycetes(100);o__Saccharomycetales(100);f__Metschnikowiaceae(100);g__Metschnikowia(100);s__Metschnikowia_pulcherrima(100); | Metschnikowia | 13 | Saprotroph | \|Undefined Saprotroph\| | Yeast | NULL | Probable | As nectar-tap_saprotroph (Põlme et al. 2020); Decay sugar-rich substrates (Põlme et al. 2020) | Põlme S, et al. 2020. Fungal Diversity 105:1-16 ((https://doi.org/10.1007/s13225-020-00466-2)) |
| Otu10818 | 2 | k__Fungi(100);p__Ascomycota(100);c__Saccharomycetes(100);o__Saccharomycetales(100);f__Metschnikowiaceae(100);g__Metschnikowia(100);g__Metschnikowia_unclassified(100); | Metschnikowia | 13 | Saprotroph | \|Undefined Saprotroph\| | Yeast | NULL | Probable | As nectar-tap_saprotroph (Põlme et al. 2020); Decay sugar-rich substrates (Põlme et al. 2020) | Põlme S, et al. 2020. Fungal Diversity 105:1-16 ((https://doi.org/10.1007/s13225-020-00466-2)) |
| Otu10819 | 2 | k__Fungi(100);p__Ascomycota(100);c__Saccharomycetes(100);o__Saccharomycetales(100);f__Metschnikowiaceae(100);g__Metschnikowia(100);s__Metschnikowia_pulcherrima(100); | Metschnikowia | 13 | Saprotroph | \|Undefined Saprotroph\| | Yeast | NULL | Probable | As nectar-tap_saprotroph (Põlme et al. 2020); Decay sugar-rich substrates (Põlme et al. 2020) | Põlme S, et al. 2020. Fungal Diversity 105:1-16 ((https://doi.org/10.1007/s13225-020-00466-2)) |
| Otu10820 | 2 | k__Fungi(100);p__Ascomycota(100);c__Saccharomycetes(100);o__Saccharomycetales(100);f__Metschnikowiaceae(100);g__Metschnikowia(100);g__Metschnikowia_unclassified(100); | Metschnikowia | 13 | Saprotroph | \|Undefined Saprotroph\| | Yeast | NULL | Probable | As nectar-tap_saprotroph (Põlme et al. 2020); Decay sugar-rich substrates (Põlme et al. 2020) | Põlme S, et al. 2020. Fungal Diversity 105:1-16 ((https://doi.org/10.1007/s13225-020-00466-2)) |
| Otu10822 | 2 | k__Fungi(100);p__Ascomycota(100);c__Saccharomycetes(100);o__Saccharomycetales(100);f__Metschnikowiaceae(100);g__Metschnikowia(100);s__Metschnikowia_pulcherrima(100); | Metschnikowia | 13 | Saprotroph | \|Undefined Saprotroph\| | Yeast | NULL | Probable | As nectar-tap_saprotroph (Põlme et al. 2020); Decay sugar-rich substrates (Põlme et al. 2020) | Põlme S, et al. 2020. Fungal Diversity 105:1-16 ((https://doi.org/10.1007/s13225-020-00466-2)) |
| Otu10824 | 2 | k__Fungi(100);p__Ascomycota(100);c__Saccharomycetes(100);o__Saccharomycetales(100);f__Metschnikowiaceae(100);g__Metschnikowia(100);g__Metschnikowia_unclassified(100); | Metschnikowia | 13 | Saprotroph | \|Undefined Saprotroph\| | Yeast | NULL | Probable | As nectar-tap_saprotroph (Põlme et al. 2020); Decay sugar-rich substrates (Põlme et al. 2020) | Põlme S, et al. 2020. Fungal Diversity 105:1-16 ((https://doi.org/10.1007/s13225-020-00466-2)) |
| Otu10825 | 2 | k__Fungi(100);p__Ascomycota(100);c__Saccharomycetes(100);o__Saccharomycetales(100);f__Metschnikowiaceae(100);g__Metschnikowia(100);g__Metschnikowia_unclassified(100); | Metschnikowia | 13 | Saprotroph | \|Undefined Saprotroph\| | Yeast | NULL | Probable | As nectar-tap_saprotroph (Põlme et al. 2020); Decay sugar-rich substrates (Põlme et al. 2020) | Põlme S, et al. 2020. Fungal Diversity 105:1-16 ((https://doi.org/10.1007/s13225-020-00466-2)) |
| Otu10827 | 2 | k__Fungi(100);p__Ascomycota(100);c__Saccharomycetes(100);o__Saccharomycetales(100);f__Metschnikowiaceae(100);g__Metschnikowia(100);s__Metschnikowia_pulcherrima(100); | Metschnikowia | 13 | Saprotroph | \|Undefined Saprotroph\| | Yeast | NULL | Probable | As nectar-tap_saprotroph (Põlme et al. 2020); Decay sugar-rich substrates (Põlme et al. 2020) | Põlme S, et al. 2020. Fungal Diversity 105:1-16 ((https://doi.org/10.1007/s13225-020-00466-2)) |
| Otu10829 | 2 | k__Fungi(100);p__Ascomycota(100);c__Saccharomycetes(100);o__Saccharomycetales(100);f__Metschnikowiaceae(100);g__Metschnikowia(100);g__Metschnikowia_unclassified(100); | Metschnikowia | 13 | Saprotroph | \|Undefined Saprotroph\| | Yeast | NULL | Probable | As nectar-tap_saprotroph (Põlme et al. 2020); Decay sugar-rich substrates (Põlme et al. 2020) | Põlme S, et al. 2020. Fungal Diversity 105:1-16 ((https://doi.org/10.1007/s13225-020-00466-2)) |
| Otu10832 | 2 | k__Fungi(100);p__Ascomycota(100);c__Saccharomycetes(100);o__Saccharomycetales(100);f__Metschnikowiaceae(100);g__Metschnikowia(100);s__Metschnikowia_pulcherrima(100); | Metschnikowia | 13 | Saprotroph | \|Undefined Saprotroph\| | Yeast | NULL | Probable | As nectar-tap_saprotroph (Põlme et al. 2020); Decay sugar-rich substrates (Põlme et al. 2020) | Põlme S, et al. 2020. Fungal Diversity 105:1-16 ((https://doi.org/10.1007/s13225-020-00466-2)) |
| Otu10833 | 2 | k__Fungi(100);p__Ascomycota(100);c__Sordariomycetes(100);o__Sordariales(100);f__Lasiosphaeriaceae(100);g__Podospora(100);g__Podospora_unclassified(100); | Podospora | 13 | Saprotroph-Symbiotroph | \|Dung Saprotroph\|-Endophyte-Plant Saprotroph-Undefined Saprotroph | Microfungus | NULL | Possible | As foliar_endophyte (Põlme et al. 2020); As litter_saprotroph (Põlme et al. 2020); culture (David et al. 2016); Decay leaf, fruit, seed, dung (Põlme et al. 2020); Endophyte detection method | Bell A. 1983. Dung Fungi: An Illustrated Guide to Coprophilous Fungi in New Zealand. Victoria University Press, Wellington (ISBN: 978-0864730015); Tedersoo L, et al. 2014. Science 346:e1256688 ((https://doi.org/10.1126/science.1256688)); David AS, et al. 2016. Microbial Ecology 71:912-926 ((https://doi.org/10.1007/s00248-015-0712-6)); Põlme S, et al. 2020. Fungal Diversity 105:1-16 ((https://doi.org/10.1007/s13225-020-00466-2)) |
| Otu10835 | 2 | k__Fungi(100);p__Ascomycota(100);c__Saccharomycetes(100);o__Saccharomycetales(100);f__Metschnikowiaceae(100);g__Metschnikowia(100);g__Metschnikowia_unclassified(100); | Metschnikowia | 13 | Saprotroph | \|Undefined Saprotroph\| | Yeast | NULL | Probable | As nectar-tap_saprotroph (Põlme et al. 2020); Decay sugar-rich substrates (Põlme et al. 2020) | Põlme S, et al. 2020. Fungal Diversity 105:1-16 ((https://doi.org/10.1007/s13225-020-00466-2)) |
| Otu10836 | 2 | k__Fungi(100);p__Ascomycota(100);c__Saccharomycetes(100);o__Saccharomycetales(100);f__Metschnikowiaceae(100);g__Metschnikowia(100);s__Metschnikowia_pulcherrima(100); | Metschnikowia | 13 | Saprotroph | \|Undefined Saprotroph\| | Yeast | NULL | Probable | As nectar-tap_saprotroph (Põlme et al. 2020); Decay sugar-rich substrates (Põlme et al. 2020) | Põlme S, et al. 2020. Fungal Diversity 105:1-16 ((https://doi.org/10.1007/s13225-020-00466-2)) |
| Otu10837 | 2 | k__Fungi(100);p__Ascomycota(100);c__Saccharomycetes(100);o__Saccharomycetales(100);f__Metschnikowiaceae(100);g__Metschnikowia(100);s__Metschnikowia_pulcherrima(100); | Metschnikowia | 13 | Saprotroph | \|Undefined Saprotroph\| | Yeast | NULL | Probable | As nectar-tap_saprotroph (Põlme et al. 2020); Decay sugar-rich substrates (Põlme et al. 2020) | Põlme S, et al. 2020. Fungal Diversity 105:1-16 ((https://doi.org/10.1007/s13225-020-00466-2)) |
| Otu10839 | 2 | k__Fungi(100);p__Ascomycota(100);c__Saccharomycetes(100);o__Saccharomycetales(100);f__Metschnikowiaceae(100);g__Metschnikowia(100);s__Metschnikowia_pulcherrima(100); | Metschnikowia | 13 | Saprotroph | \|Undefined Saprotroph\| | Yeast | NULL | Probable | As nectar-tap_saprotroph (Põlme et al. 2020); Decay sugar-rich substrates (Põlme et al. 2020) | Põlme S, et al. 2020. Fungal Diversity 105:1-16 ((https://doi.org/10.1007/s13225-020-00466-2)) |
| Otu10840 | 2 | k__Fungi(100);p__Ascomycota(100);c__Saccharomycetes(100);o__Saccharomycetales(100);f__Metschnikowiaceae(100);g__Metschnikowia(100);s__Metschnikowia_pulcherrima(100); | Metschnikowia | 13 | Saprotroph | \|Undefined Saprotroph\| | Yeast | NULL | Probable | As nectar-tap_saprotroph (Põlme et al. 2020); Decay sugar-rich substrates (Põlme et al. 2020) | Põlme S, et al. 2020. Fungal Diversity 105:1-16 ((https://doi.org/10.1007/s13225-020-00466-2)) |
| Otu10841 | 2 | k__Fungi(100);p__Ascomycota(100);c__Saccharomycetes(100);o__Saccharomycetales(100);f__Metschnikowiaceae(100);g__Metschnikowia(100);g__Metschnikowia_unclassified(100); | Metschnikowia | 13 | Saprotroph | \|Undefined Saprotroph\| | Yeast | NULL | Probable | As nectar-tap_saprotroph (Põlme et al. 2020); Decay sugar-rich substrates (Põlme et al. 2020) | Põlme S, et al. 2020. Fungal Diversity 105:1-16 ((https://doi.org/10.1007/s13225-020-00466-2)) |
| Otu10842 | 2 | k__Fungi(100);p__Ascomycota(100);c__Saccharomycetes(100);o__Saccharomycetales(100);f__Metschnikowiaceae(100);g__Metschnikowia(100);s__Metschnikowia_pulcherrima(100); | Metschnikowia | 13 | Saprotroph | \|Undefined Saprotroph\| | Yeast | NULL | Probable | As nectar-tap_saprotroph (Põlme et al. 2020); Decay sugar-rich substrates (Põlme et al. 2020) | Põlme S, et al. 2020. Fungal Diversity 105:1-16 ((https://doi.org/10.1007/s13225-020-00466-2)) |
| Otu10843 | 2 | k__Fungi(100);p__Ascomycota(100);c__Saccharomycetes(100);o__Saccharomycetales(100);f__Metschnikowiaceae(100);g__Metschnikowia(100);s__Metschnikowia_pulcherrima(100); | Metschnikowia | 13 | Saprotroph | \|Undefined Saprotroph\| | Yeast | NULL | Probable | As nectar-tap_saprotroph (Põlme et al. 2020); Decay sugar-rich substrates (Põlme et al. 2020) | Põlme S, et al. 2020. Fungal Diversity 105:1-16 ((https://doi.org/10.1007/s13225-020-00466-2)) |
| Otu10844 | 2 | k__Fungi(100);p__Ascomycota(100);c__Saccharomycetes(100);o__Saccharomycetales(100);f__Metschnikowiaceae(100);g__Metschnikowia(100);g__Metschnikowia_unclassified(100); | Metschnikowia | 13 | Saprotroph | \|Undefined Saprotroph\| | Yeast | NULL | Probable | As nectar-tap_saprotroph (Põlme et al. 2020); Decay sugar-rich substrates (Põlme et al. 2020) | Põlme S, et al. 2020. Fungal Diversity 105:1-16 ((https://doi.org/10.1007/s13225-020-00466-2)) |
| Otu10845 | 2 | k__Fungi(100);p__Ascomycota(100);c__Saccharomycetes(100);o__Saccharomycetales(100);f__Saccharomycetales_family_Incertae_sedis(100);g__Candida(100);g__Candida_unclassified(100); | Candida | 13 | Pathotroph-Saprotroph-Symbiotroph | Animal Pathogen-Endophyte-Endosymbiont-Epiphyte-\|Undefined Saprotroph\| | Dimorphic Yeast-Yeast | NULL | Probable | As nectar-tap_saprotroph (Põlme et al. 2020); Decay sugar-rich substrates (Põlme et al. 2020); This asexual genus is polyphyletic and contain species that function very differently depend on what they are. Caution should be taken when interpreting guilds for Candida species. | Manolakaki D, et al. 2010. Virulence. 1: 367-375 ((https://doi.org/10.4161/viru.1.5.12796)); Mekbib SB, et al. 2011. Tropical Plant Pathology 36:233-240 ((https://doi.org/10.1590/S1982-56762011000400004)); Põlme S, et al. 2020. Fungal Diversity 105:1-16 ((https://doi.org/10.1007/s13225-020-00466-2)) |
| Otu10846 | 2 | k__Fungi(100);p__Ascomycota(100);c__Saccharomycetes(100);o__Saccharomycetales(100);f__Metschnikowiaceae(100);g__Metschnikowia(100);g__Metschnikowia_unclassified(100); | Metschnikowia | 13 | Saprotroph | \|Undefined Saprotroph\| | Yeast | NULL | Probable | As nectar-tap_saprotroph (Põlme et al. 2020); Decay sugar-rich substrates (Põlme et al. 2020) | Põlme S, et al. 2020. Fungal Diversity 105:1-16 ((https://doi.org/10.1007/s13225-020-00466-2)) |
| Otu10850 | 2 | k__Fungi(100);p__Ascomycota(100);c__Dothideomycetes(100);o__Dothideomycetes_order_Incertae_sedis(100);f__Myxotrichaceae(100);g__Oidiodendron(100);s__Oidiodendron_griseum(100); | Oidiodendron | 13 | Pathotroph-Saprotroph-Symbiotroph | Endophyte-Ericoid Mycorrhizal-\|Undefined Saprotroph\| | Microfungus | NULL | Probable | As root_endophyte (Põlme et al. 2020); As soil_saprotroph (Põlme et al. 2020); Dark Septate Endophyte; Decay soil (Põlme et al. 2020) | Newsham KK. 2011. New Phytologist 190:783-793 ((https://doi.org/10.1111/j.1469-8137.2010.03611.x)) (as dark septate root endophytes); Põlme S, et al. 2020. Fungal Diversity 105:1-16 ((https://doi.org/10.1007/s13225-020-00466-2)) |
| Otu10856 | 2 | k__Fungi(100);p__Ascomycota(100);c__Saccharomycetes(100);o__Saccharomycetales(100);f__Metschnikowiaceae(100);g__Metschnikowia(100);s__Metschnikowia_pulcherrima(100); | Metschnikowia | 13 | Saprotroph | \|Undefined Saprotroph\| | Yeast | NULL | Probable | As nectar-tap_saprotroph (Põlme et al. 2020); Decay sugar-rich substrates (Põlme et al. 2020) | Põlme S, et al. 2020. Fungal Diversity 105:1-16 ((https://doi.org/10.1007/s13225-020-00466-2)) |
| Otu10857 | 2 | k__Fungi(100);p__Ascomycota(100);c__Saccharomycetes(100);o__Saccharomycetales(100);f__Metschnikowiaceae(100);g__Metschnikowia(100);g__Metschnikowia_unclassified(100); | Metschnikowia | 13 | Saprotroph | \|Undefined Saprotroph\| | Yeast | NULL | Probable | As nectar-tap_saprotroph (Põlme et al. 2020); Decay sugar-rich substrates (Põlme et al. 2020) | Põlme S, et al. 2020. Fungal Diversity 105:1-16 ((https://doi.org/10.1007/s13225-020-00466-2)) |
| Otu10858 | 2 | k__Fungi(100);p__Ascomycota(100);c__Saccharomycetes(100);o__Saccharomycetales(100);f__Metschnikowiaceae(100);g__Metschnikowia(100);s__Metschnikowia_pulcherrima(100); | Metschnikowia | 13 | Saprotroph | \|Undefined Saprotroph\| | Yeast | NULL | Probable | As nectar-tap_saprotroph (Põlme et al. 2020); Decay sugar-rich substrates (Põlme et al. 2020) | Põlme S, et al. 2020. Fungal Diversity 105:1-16 ((https://doi.org/10.1007/s13225-020-00466-2)) |
| Otu10859 | 2 | k__Fungi(100);p__Ascomycota(100);c__Saccharomycetes(100);o__Saccharomycetales(100);f__Metschnikowiaceae(100);g__Metschnikowia(100);s__Metschnikowia_pulcherrima(100); | Metschnikowia | 13 | Saprotroph | \|Undefined Saprotroph\| | Yeast | NULL | Probable | As nectar-tap_saprotroph (Põlme et al. 2020); Decay sugar-rich substrates (Põlme et al. 2020) | Põlme S, et al. 2020. Fungal Diversity 105:1-16 ((https://doi.org/10.1007/s13225-020-00466-2)) |
| Otu10860 | 2 | k__Fungi(100);p__Ascomycota(100);c__Saccharomycetes(100);o__Saccharomycetales(100);f__Metschnikowiaceae(100);g__Metschnikowia(100);s__Metschnikowia_pulcherrima(100); | Metschnikowia | 13 | Saprotroph | \|Undefined Saprotroph\| | Yeast | NULL | Probable | As nectar-tap_saprotroph (Põlme et al. 2020); Decay sugar-rich substrates (Põlme et al. 2020) | Põlme S, et al. 2020. Fungal Diversity 105:1-16 ((https://doi.org/10.1007/s13225-020-00466-2)) |
| Otu10861 | 2 | k__Fungi(100);p__Ascomycota(100);c__Saccharomycetes(100);o__Saccharomycetales(100);f__Metschnikowiaceae(100);g__Metschnikowia(100);s__Metschnikowia_pulcherrima(100); | Metschnikowia | 13 | Saprotroph | \|Undefined Saprotroph\| | Yeast | NULL | Probable | As nectar-tap_saprotroph (Põlme et al. 2020); Decay sugar-rich substrates (Põlme et al. 2020) | Põlme S, et al. 2020. Fungal Diversity 105:1-16 ((https://doi.org/10.1007/s13225-020-00466-2)) |
| Otu10865 | 2 | k__Fungi(100);p__Ascomycota(100);c__Saccharomycetes(100);o__Saccharomycetales(100);f__Metschnikowiaceae(100);g__Metschnikowia(100);s__Metschnikowia_pulcherrima(100); | Metschnikowia | 13 | Saprotroph | \|Undefined Saprotroph\| | Yeast | NULL | Probable | As nectar-tap_saprotroph (Põlme et al. 2020); Decay sugar-rich substrates (Põlme et al. 2020) | Põlme S, et al. 2020. Fungal Diversity 105:1-16 ((https://doi.org/10.1007/s13225-020-00466-2)) |
| Otu10866 | 2 | k__Fungi(100);p__Ascomycota(100);c__Saccharomycetes(100);o__Saccharomycetales(100);f__Metschnikowiaceae(100);g__Metschnikowia(100);s__Metschnikowia_pulcherrima(100); | Metschnikowia | 13 | Saprotroph | \|Undefined Saprotroph\| | Yeast | NULL | Probable | As nectar-tap_saprotroph (Põlme et al. 2020); Decay sugar-rich substrates (Põlme et al. 2020) | Põlme S, et al. 2020. Fungal Diversity 105:1-16 ((https://doi.org/10.1007/s13225-020-00466-2)) |
| Otu10867 | 2 | k__Fungi(100);p__Ascomycota(100);c__Saccharomycetes(100);o__Saccharomycetales(100);f__Metschnikowiaceae(100);g__Metschnikowia(100);g__Metschnikowia_unclassified(100); | Metschnikowia | 13 | Saprotroph | \|Undefined Saprotroph\| | Yeast | NULL | Probable | As nectar-tap_saprotroph (Põlme et al. 2020); Decay sugar-rich substrates (Põlme et al. 2020) | Põlme S, et al. 2020. Fungal Diversity 105:1-16 ((https://doi.org/10.1007/s13225-020-00466-2)) |
| Otu10868 | 2 | k__Fungi(100);p__Ascomycota(100);c__Saccharomycetes(100);o__Saccharomycetales(100);f__Metschnikowiaceae(100);g__Metschnikowia(100);s__Metschnikowia_pulcherrima(100); | Metschnikowia | 13 | Saprotroph | \|Undefined Saprotroph\| | Yeast | NULL | Probable | As nectar-tap_saprotroph (Põlme et al. 2020); Decay sugar-rich substrates (Põlme et al. 2020) | Põlme S, et al. 2020. Fungal Diversity 105:1-16 ((https://doi.org/10.1007/s13225-020-00466-2)) |
| Otu10871 | 2 | k__Fungi(100);p__Ascomycota(100);c__Saccharomycetes(100);o__Saccharomycetales(100);f__Metschnikowiaceae(100);g__Metschnikowia(100);s__Metschnikowia_pulcherrima(100); | Metschnikowia | 13 | Saprotroph | \|Undefined Saprotroph\| | Yeast | NULL | Probable | As nectar-tap_saprotroph (Põlme et al. 2020); Decay sugar-rich substrates (Põlme et al. 2020) | Põlme S, et al. 2020. Fungal Diversity 105:1-16 ((https://doi.org/10.1007/s13225-020-00466-2)) |
| Otu10872 | 2 | k__Fungi(100);p__Ascomycota(100);c__Saccharomycetes(100);o__Saccharomycetales(100);f__Saccharomycetales_family_Incertae_sedis(100);g__Candida(100);g__Candida_unclassified(100); | Candida | 13 | Pathotroph-Saprotroph-Symbiotroph | Animal Pathogen-Endophyte-Endosymbiont-Epiphyte-\|Undefined Saprotroph\| | Dimorphic Yeast-Yeast | NULL | Probable | As nectar-tap_saprotroph (Põlme et al. 2020); Decay sugar-rich substrates (Põlme et al. 2020); This asexual genus is polyphyletic and contain species that function very differently depend on what they are. Caution should be taken when interpreting guilds for Candida species. | Manolakaki D, et al. 2010. Virulence. 1: 367-375 ((https://doi.org/10.4161/viru.1.5.12796)); Mekbib SB, et al. 2011. Tropical Plant Pathology 36:233-240 ((https://doi.org/10.1590/S1982-56762011000400004)); Põlme S, et al. 2020. Fungal Diversity 105:1-16 ((https://doi.org/10.1007/s13225-020-00466-2)) |
| Otu10873 | 2 | k__Fungi(100);p__Ascomycota(100);c__Saccharomycetes(100);o__Saccharomycetales(100);f__Metschnikowiaceae(100);g__Metschnikowia(100);g__Metschnikowia_unclassified(100); | Metschnikowia | 13 | Saprotroph | \|Undefined Saprotroph\| | Yeast | NULL | Probable | As nectar-tap_saprotroph (Põlme et al. 2020); Decay sugar-rich substrates (Põlme et al. 2020) | Põlme S, et al. 2020. Fungal Diversity 105:1-16 ((https://doi.org/10.1007/s13225-020-00466-2)) |
| Otu10875 | 2 | k__Fungi(100);p__Ascomycota(100);c__Saccharomycetes(100);o__Saccharomycetales(100);f__Metschnikowiaceae(100);g__Metschnikowia(100);s__Metschnikowia_pulcherrima(100); | Metschnikowia | 13 | Saprotroph | \|Undefined Saprotroph\| | Yeast | NULL | Probable | As nectar-tap_saprotroph (Põlme et al. 2020); Decay sugar-rich substrates (Põlme et al. 2020) | Põlme S, et al. 2020. Fungal Diversity 105:1-16 ((https://doi.org/10.1007/s13225-020-00466-2)) |
| Otu10878 | 2 | k__Fungi(100);p__Ascomycota(100);c__Saccharomycetes(100);o__Saccharomycetales(100);f__Metschnikowiaceae(100);g__Metschnikowia(100);s__Metschnikowia_pulcherrima(100); | Metschnikowia | 13 | Saprotroph | \|Undefined Saprotroph\| | Yeast | NULL | Probable | As nectar-tap_saprotroph (Põlme et al. 2020); Decay sugar-rich substrates (Põlme et al. 2020) | Põlme S, et al. 2020. Fungal Diversity 105:1-16 ((https://doi.org/10.1007/s13225-020-00466-2)) |
| Otu10879 | 2 | k__Fungi(100);p__Ascomycota(100);c__Saccharomycetes(100);o__Saccharomycetales(100);f__Metschnikowiaceae(100);g__Metschnikowia(100);s__Metschnikowia_pulcherrima(100); | Metschnikowia | 13 | Saprotroph | \|Undefined Saprotroph\| | Yeast | NULL | Probable | As nectar-tap_saprotroph (Põlme et al. 2020); Decay sugar-rich substrates (Põlme et al. 2020) | Põlme S, et al. 2020. Fungal Diversity 105:1-16 ((https://doi.org/10.1007/s13225-020-00466-2)) |
| Otu10884 | 2 | k__Fungi(100);p__Ascomycota(100);c__Saccharomycetes(100);o__Saccharomycetales(100);f__Metschnikowiaceae(100);g__Metschnikowia(100);s__Metschnikowia_pulcherrima(100); | Metschnikowia | 13 | Saprotroph | \|Undefined Saprotroph\| | Yeast | NULL | Probable | As nectar-tap_saprotroph (Põlme et al. 2020); Decay sugar-rich substrates (Põlme et al. 2020) | Põlme S, et al. 2020. Fungal Diversity 105:1-16 ((https://doi.org/10.1007/s13225-020-00466-2)) |
| Otu10887 | 2 | k__Fungi(100);p__Ascomycota(100);c__Saccharomycetes(100);o__Saccharomycetales(100);f__Metschnikowiaceae(100);g__Metschnikowia(100);s__Metschnikowia_pulcherrima(100); | Metschnikowia | 13 | Saprotroph | \|Undefined Saprotroph\| | Yeast | NULL | Probable | As nectar-tap_saprotroph (Põlme et al. 2020); Decay sugar-rich substrates (Põlme et al. 2020) | Põlme S, et al. 2020. Fungal Diversity 105:1-16 ((https://doi.org/10.1007/s13225-020-00466-2)) |
| Otu10889 | 2 | k__Fungi(100);p__Ascomycota(100);c__Saccharomycetes(100);o__Saccharomycetales(100);f__Metschnikowiaceae(100);g__Metschnikowia(100);s__Metschnikowia_pulcherrima(100); | Metschnikowia | 13 | Saprotroph | \|Undefined Saprotroph\| | Yeast | NULL | Probable | As nectar-tap_saprotroph (Põlme et al. 2020); Decay sugar-rich substrates (Põlme et al. 2020) | Põlme S, et al. 2020. Fungal Diversity 105:1-16 ((https://doi.org/10.1007/s13225-020-00466-2)) |
| Otu10890 | 2 | k__Fungi(100);p__Ascomycota(100);c__Saccharomycetes(100);o__Saccharomycetales(100);f__Metschnikowiaceae(100);g__Metschnikowia(100);g__Metschnikowia_unclassified(100); | Metschnikowia | 13 | Saprotroph | \|Undefined Saprotroph\| | Yeast | NULL | Probable | As nectar-tap_saprotroph (Põlme et al. 2020); Decay sugar-rich substrates (Põlme et al. 2020) | Põlme S, et al. 2020. Fungal Diversity 105:1-16 ((https://doi.org/10.1007/s13225-020-00466-2)) |
| Otu10895 | 2 | k__Fungi(100);p__Ascomycota(100);c__Saccharomycetes(100);o__Saccharomycetales(100);f__Metschnikowiaceae(100);g__Metschnikowia(100);s__Metschnikowia_pulcherrima(100); | Metschnikowia | 13 | Saprotroph | \|Undefined Saprotroph\| | Yeast | NULL | Probable | As nectar-tap_saprotroph (Põlme et al. 2020); Decay sugar-rich substrates (Põlme et al. 2020) | Põlme S, et al. 2020. Fungal Diversity 105:1-16 ((https://doi.org/10.1007/s13225-020-00466-2)) |
| Otu10897 | 2 | k__Fungi(100);p__Ascomycota(100);c__Saccharomycetes(100);o__Saccharomycetales(100);f__Metschnikowiaceae(100);g__Metschnikowia(100);s__Metschnikowia_pulcherrima(100); | Metschnikowia | 13 | Saprotroph | \|Undefined Saprotroph\| | Yeast | NULL | Probable | As nectar-tap_saprotroph (Põlme et al. 2020); Decay sugar-rich substrates (Põlme et al. 2020) | Põlme S, et al. 2020. Fungal Diversity 105:1-16 ((https://doi.org/10.1007/s13225-020-00466-2)) |
| Otu10898 | 2 | k__Fungi(100);p__Ascomycota(100);c__Saccharomycetes(100);o__Saccharomycetales(100);f__Metschnikowiaceae(100);g__Metschnikowia(100);s__Metschnikowia_pulcherrima(100); | Metschnikowia | 13 | Saprotroph | \|Undefined Saprotroph\| | Yeast | NULL | Probable | As nectar-tap_saprotroph (Põlme et al. 2020); Decay sugar-rich substrates (Põlme et al. 2020) | Põlme S, et al. 2020. Fungal Diversity 105:1-16 ((https://doi.org/10.1007/s13225-020-00466-2)) |
| Otu10900 | 2 | k__Fungi(100);p__Ascomycota(100);c__Saccharomycetes(100);o__Saccharomycetales(100);f__Metschnikowiaceae(100);g__Metschnikowia(100);g__Metschnikowia_unclassified(100); | Metschnikowia | 13 | Saprotroph | \|Undefined Saprotroph\| | Yeast | NULL | Probable | As nectar-tap_saprotroph (Põlme et al. 2020); Decay sugar-rich substrates (Põlme et al. 2020) | Põlme S, et al. 2020. Fungal Diversity 105:1-16 ((https://doi.org/10.1007/s13225-020-00466-2)) |
| Otu10901 | 2 | k__Fungi(100);p__Ascomycota(100);c__Saccharomycetes(100);o__Saccharomycetales(100);f__Metschnikowiaceae(100);g__Metschnikowia(100);s__Metschnikowia_pulcherrima(100); | Metschnikowia | 13 | Saprotroph | \|Undefined Saprotroph\| | Yeast | NULL | Probable | As nectar-tap_saprotroph (Põlme et al. 2020); Decay sugar-rich substrates (Põlme et al. 2020) | Põlme S, et al. 2020. Fungal Diversity 105:1-16 ((https://doi.org/10.1007/s13225-020-00466-2)) |
| Otu10902 | 2 | k__Fungi(100);p__Ascomycota(100);c__Saccharomycetes(100);o__Saccharomycetales(100);f__Metschnikowiaceae(100);g__Metschnikowia(100);s__Metschnikowia_pulcherrima(100); | Metschnikowia | 13 | Saprotroph | \|Undefined Saprotroph\| | Yeast | NULL | Probable | As nectar-tap_saprotroph (Põlme et al. 2020); Decay sugar-rich substrates (Põlme et al. 2020) | Põlme S, et al. 2020. Fungal Diversity 105:1-16 ((https://doi.org/10.1007/s13225-020-00466-2)) |
| Otu10903 | 2 | k__Fungi(100);p__Ascomycota(100);c__Saccharomycetes(100);o__Saccharomycetales(100);f__Metschnikowiaceae(100);g__Metschnikowia(100);s__Metschnikowia_pulcherrima(100); | Metschnikowia | 13 | Saprotroph | \|Undefined Saprotroph\| | Yeast | NULL | Probable | As nectar-tap_saprotroph (Põlme et al. 2020); Decay sugar-rich substrates (Põlme et al. 2020) | Põlme S, et al. 2020. Fungal Diversity 105:1-16 ((https://doi.org/10.1007/s13225-020-00466-2)) |
| Otu10904 | 2 | k__Fungi(100);p__Ascomycota(100);c__Saccharomycetes(100);o__Saccharomycetales(100);f__Saccharomycetales_family_Incertae_sedis(100);g__Candida(100);g__Candida_unclassified(100); | Candida | 13 | Pathotroph-Saprotroph-Symbiotroph | Animal Pathogen-Endophyte-Endosymbiont-Epiphyte-\|Undefined Saprotroph\| | Dimorphic Yeast-Yeast | NULL | Probable | As nectar-tap_saprotroph (Põlme et al. 2020); Decay sugar-rich substrates (Põlme et al. 2020); This asexual genus is polyphyletic and contain species that function very differently depend on what they are. Caution should be taken when interpreting guilds for Candida species. | Manolakaki D, et al. 2010. Virulence. 1: 367-375 ((https://doi.org/10.4161/viru.1.5.12796)); Mekbib SB, et al. 2011. Tropical Plant Pathology 36:233-240 ((https://doi.org/10.1590/S1982-56762011000400004)); Põlme S, et al. 2020. Fungal Diversity 105:1-16 ((https://doi.org/10.1007/s13225-020-00466-2)) |
| Otu10907 | 2 | k__Fungi(100);p__Ascomycota(100);c__Saccharomycetes(100);o__Saccharomycetales(100);f__Metschnikowiaceae(100);g__Metschnikowia(100);s__Metschnikowia_pulcherrima(100); | Metschnikowia | 13 | Saprotroph | \|Undefined Saprotroph\| | Yeast | NULL | Probable | As nectar-tap_saprotroph (Põlme et al. 2020); Decay sugar-rich substrates (Põlme et al. 2020) | Põlme S, et al. 2020. Fungal Diversity 105:1-16 ((https://doi.org/10.1007/s13225-020-00466-2)) |
| Otu10908 | 2 | k__Fungi(100);p__Ascomycota(100);c__Saccharomycetes(100);o__Saccharomycetales(100);f__Metschnikowiaceae(100);g__Metschnikowia(100);s__Metschnikowia_pulcherrima(100); | Metschnikowia | 13 | Saprotroph | \|Undefined Saprotroph\| | Yeast | NULL | Probable | As nectar-tap_saprotroph (Põlme et al. 2020); Decay sugar-rich substrates (Põlme et al. 2020) | Põlme S, et al. 2020. Fungal Diversity 105:1-16 ((https://doi.org/10.1007/s13225-020-00466-2)) |
| Otu10909 | 2 | k__Fungi(100);p__Ascomycota(100);c__Saccharomycetes(100);o__Saccharomycetales(100);f__Metschnikowiaceae(100);g__Metschnikowia(100);s__Metschnikowia_pulcherrima(100); | Metschnikowia | 13 | Saprotroph | \|Undefined Saprotroph\| | Yeast | NULL | Probable | As nectar-tap_saprotroph (Põlme et al. 2020); Decay sugar-rich substrates (Põlme et al. 2020) | Põlme S, et al. 2020. Fungal Diversity 105:1-16 ((https://doi.org/10.1007/s13225-020-00466-2)) |
| Otu10911 | 2 | k__Fungi(100);p__Ascomycota(100);c__Saccharomycetes(100);o__Saccharomycetales(100);f__Metschnikowiaceae(100);g__Metschnikowia(100);g__Metschnikowia_unclassified(100); | Metschnikowia | 13 | Saprotroph | \|Undefined Saprotroph\| | Yeast | NULL | Probable | As nectar-tap_saprotroph (Põlme et al. 2020); Decay sugar-rich substrates (Põlme et al. 2020) | Põlme S, et al. 2020. Fungal Diversity 105:1-16 ((https://doi.org/10.1007/s13225-020-00466-2)) |
| Otu10916 | 2 | k__Fungi(100);p__Zygomycota(100);c__Zygomycota_class_Incertae_sedis(100);o__Mucorales(100);f__Mucoraceae(100);g__Mucor(100);s__Mucor_genevensis(100); | Mucor | 13 | Saprotroph | \|Undefined Saprotroph\| | Microfungus | NULL | Probable | As soil_saprotroph (Põlme et al. 2020); Decay soil (Põlme et al. 2020) | Tedersoo L, et al. 2014. Science 346:e1256688 ((https://doi.org/10.1126/science.1256688)); Põlme S, et al. 2020. Fungal Diversity 105:1-16 ((https://doi.org/10.1007/s13225-020-00466-2)) |
| Otu10917 | 2 | k__Fungi(100);p__Zygomycota(100);c__Zygomycota_class_Incertae_sedis(100);o__Mucorales(100);f__Mucoraceae(100);g__Mucor(100);s__Mucor_genevensis(100); | Mucor | 13 | Saprotroph | \|Undefined Saprotroph\| | Microfungus | NULL | Probable | As soil_saprotroph (Põlme et al. 2020); Decay soil (Põlme et al. 2020) | Tedersoo L, et al. 2014. Science 346:e1256688 ((https://doi.org/10.1126/science.1256688)); Põlme S, et al. 2020. Fungal Diversity 105:1-16 ((https://doi.org/10.1007/s13225-020-00466-2)) |
| Otu10918 | 2 | k__Fungi(100);p__Zygomycota(100);c__Zygomycota_class_Incertae_sedis(100);o__Mucorales(100);f__Mucoraceae(100);g__Mucor(100);s__Mucor_genevensis(100); | Mucor | 13 | Saprotroph | \|Undefined Saprotroph\| | Microfungus | NULL | Probable | As soil_saprotroph (Põlme et al. 2020); Decay soil (Põlme et al. 2020) | Tedersoo L, et al. 2014. Science 346:e1256688 ((https://doi.org/10.1126/science.1256688)); Põlme S, et al. 2020. Fungal Diversity 105:1-16 ((https://doi.org/10.1007/s13225-020-00466-2)) |
| Otu10921 | 2 | k__Fungi(100);p__Ascomycota(100);c__Saccharomycetes(100);o__Saccharomycetales(100);f__Metschnikowiaceae(100);g__Metschnikowia(100);s__Metschnikowia_pulcherrima(100); | Metschnikowia | 13 | Saprotroph | \|Undefined Saprotroph\| | Yeast | NULL | Probable | As nectar-tap_saprotroph (Põlme et al. 2020); Decay sugar-rich substrates (Põlme et al. 2020) | Põlme S, et al. 2020. Fungal Diversity 105:1-16 ((https://doi.org/10.1007/s13225-020-00466-2)) |
| Otu10922 | 2 | k__Fungi(100);p__Ascomycota(100);c__Saccharomycetes(100);o__Saccharomycetales(100);f__Metschnikowiaceae(100);g__Metschnikowia(100);g__Metschnikowia_unclassified(100); | Metschnikowia | 13 | Saprotroph | \|Undefined Saprotroph\| | Yeast | NULL | Probable | As nectar-tap_saprotroph (Põlme et al. 2020); Decay sugar-rich substrates (Põlme et al. 2020) | Põlme S, et al. 2020. Fungal Diversity 105:1-16 ((https://doi.org/10.1007/s13225-020-00466-2)) |
| Otu10927 | 2 | k__Fungi(100);p__Ascomycota(100);c__Saccharomycetes(100);o__Saccharomycetales(100);f__Metschnikowiaceae(100);g__Metschnikowia(100);s__Metschnikowia_pulcherrima(100); | Metschnikowia | 13 | Saprotroph | \|Undefined Saprotroph\| | Yeast | NULL | Probable | As nectar-tap_saprotroph (Põlme et al. 2020); Decay sugar-rich substrates (Põlme et al. 2020) | Põlme S, et al. 2020. Fungal Diversity 105:1-16 ((https://doi.org/10.1007/s13225-020-00466-2)) |
| Otu10929 | 2 | k__Fungi(100);p__Ascomycota(100);c__Saccharomycetes(100);o__Saccharomycetales(100);f__Saccharomycetales_family_Incertae_sedis(100);g__Candida(100);g__Candida_unclassified(100); | Candida | 13 | Pathotroph-Saprotroph-Symbiotroph | Animal Pathogen-Endophyte-Endosymbiont-Epiphyte-\|Undefined Saprotroph\| | Dimorphic Yeast-Yeast | NULL | Probable | As nectar-tap_saprotroph (Põlme et al. 2020); Decay sugar-rich substrates (Põlme et al. 2020); This asexual genus is polyphyletic and contain species that function very differently depend on what they are. Caution should be taken when interpreting guilds for Candida species. | Manolakaki D, et al. 2010. Virulence. 1: 367-375 ((https://doi.org/10.4161/viru.1.5.12796)); Mekbib SB, et al. 2011. Tropical Plant Pathology 36:233-240 ((https://doi.org/10.1590/S1982-56762011000400004)); Põlme S, et al. 2020. Fungal Diversity 105:1-16 ((https://doi.org/10.1007/s13225-020-00466-2)) |
| Otu10933 | 2 | k__Fungi(100);p__Zygomycota(100);c__Zygomycota_class_Incertae_sedis(100);o__Mucorales(100);f__Mucoraceae(100);g__Mucor(100);s__Mucor_genevensis(100); | Mucor | 13 | Saprotroph | \|Undefined Saprotroph\| | Microfungus | NULL | Probable | As soil_saprotroph (Põlme et al. 2020); Decay soil (Põlme et al. 2020) | Tedersoo L, et al. 2014. Science 346:e1256688 ((https://doi.org/10.1126/science.1256688)); Põlme S, et al. 2020. Fungal Diversity 105:1-16 ((https://doi.org/10.1007/s13225-020-00466-2)) |
| Otu10936 | 2 | k__Fungi(100);p__Ascomycota(100);c__Saccharomycetes(100);o__Saccharomycetales(100);f__Metschnikowiaceae(100);g__Metschnikowia(100);s__Metschnikowia_pulcherrima(100); | Metschnikowia | 13 | Saprotroph | \|Undefined Saprotroph\| | Yeast | NULL | Probable | As nectar-tap_saprotroph (Põlme et al. 2020); Decay sugar-rich substrates (Põlme et al. 2020) | Põlme S, et al. 2020. Fungal Diversity 105:1-16 ((https://doi.org/10.1007/s13225-020-00466-2)) |
| Otu10940 | 2 | k__Fungi(100);p__Ascomycota(100);c__Saccharomycetes(100);o__Saccharomycetales(100);f__Saccharomycetales_family_Incertae_sedis(100);g__Candida(100);g__Candida_unclassified(100); | Candida | 13 | Pathotroph-Saprotroph-Symbiotroph | Animal Pathogen-Endophyte-Endosymbiont-Epiphyte-\|Undefined Saprotroph\| | Dimorphic Yeast-Yeast | NULL | Probable | As nectar-tap_saprotroph (Põlme et al. 2020); Decay sugar-rich substrates (Põlme et al. 2020); This asexual genus is polyphyletic and contain species that function very differently depend on what they are. Caution should be taken when interpreting guilds for Candida species. | Manolakaki D, et al. 2010. Virulence. 1: 367-375 ((https://doi.org/10.4161/viru.1.5.12796)); Mekbib SB, et al. 2011. Tropical Plant Pathology 36:233-240 ((https://doi.org/10.1590/S1982-56762011000400004)); Põlme S, et al. 2020. Fungal Diversity 105:1-16 ((https://doi.org/10.1007/s13225-020-00466-2)) |
| Otu10944 | 2 | k__Fungi(100);p__Ascomycota(100);c__Saccharomycetes(100);o__Saccharomycetales(100);f__Metschnikowiaceae(100);g__Metschnikowia(100);s__Metschnikowia_pulcherrima(100); | Metschnikowia | 13 | Saprotroph | \|Undefined Saprotroph\| | Yeast | NULL | Probable | As nectar-tap_saprotroph (Põlme et al. 2020); Decay sugar-rich substrates (Põlme et al. 2020) | Põlme S, et al. 2020. Fungal Diversity 105:1-16 ((https://doi.org/10.1007/s13225-020-00466-2)) |
| Otu10946 | 2 | k__Fungi(100);p__Ascomycota(100);c__Saccharomycetes(100);o__Saccharomycetales(100);f__Metschnikowiaceae(100);g__Metschnikowia(100);s__Metschnikowia_pulcherrima(100); | Metschnikowia | 13 | Saprotroph | \|Undefined Saprotroph\| | Yeast | NULL | Probable | As nectar-tap_saprotroph (Põlme et al. 2020); Decay sugar-rich substrates (Põlme et al. 2020) | Põlme S, et al. 2020. Fungal Diversity 105:1-16 ((https://doi.org/10.1007/s13225-020-00466-2)) |
| Otu10948 | 2 | k__Fungi(100);p__Ascomycota(100);c__Saccharomycetes(100);o__Saccharomycetales(100);f__Metschnikowiaceae(100);g__Metschnikowia(100);g__Metschnikowia_unclassified(100); | Metschnikowia | 13 | Saprotroph | \|Undefined Saprotroph\| | Yeast | NULL | Probable | As nectar-tap_saprotroph (Põlme et al. 2020); Decay sugar-rich substrates (Põlme et al. 2020) | Põlme S, et al. 2020. Fungal Diversity 105:1-16 ((https://doi.org/10.1007/s13225-020-00466-2)) |
| Otu10957 | 2 | k__Fungi(100);p__Ascomycota(100);c__Saccharomycetes(100);o__Saccharomycetales(100);f__Metschnikowiaceae(100);g__Metschnikowia(100);g__Metschnikowia_unclassified(100); | Metschnikowia | 13 | Saprotroph | \|Undefined Saprotroph\| | Yeast | NULL | Probable | As nectar-tap_saprotroph (Põlme et al. 2020); Decay sugar-rich substrates (Põlme et al. 2020) | Põlme S, et al. 2020. Fungal Diversity 105:1-16 ((https://doi.org/10.1007/s13225-020-00466-2)) |
| Otu10961 | 2 | k__Fungi(100);p__Ascomycota(100);c__Saccharomycetes(100);o__Saccharomycetales(100);f__Metschnikowiaceae(100);g__Metschnikowia(100);s__Metschnikowia_pulcherrima(100); | Metschnikowia | 13 | Saprotroph | \|Undefined Saprotroph\| | Yeast | NULL | Probable | As nectar-tap_saprotroph (Põlme et al. 2020); Decay sugar-rich substrates (Põlme et al. 2020) | Põlme S, et al. 2020. Fungal Diversity 105:1-16 ((https://doi.org/10.1007/s13225-020-00466-2)) |
| Otu10962 | 2 | k__Fungi(100);p__Basidiomycota(100);c__Wallemiomycetes(100);o__Wallemiales(100);f__Wallemiaceae(100);g__Wallemia(100);s__Wallemia_sebi(100); | Wallemia | 13 | Saprotroph | \|Undefined Saprotroph\| | Microfungus | NULL | Probable | Decay sugar-rich substrates (Põlme et al. 2020) | Tedersoo L, et al. 2014. Science 346:e1256688 ((https://doi.org/10.1126/science.1256688)); Põlme S, et al. 2020. Fungal Diversity 105:1-16 ((https://doi.org/10.1007/s13225-020-00466-2)) |
| Otu10963 | 2 | k__Fungi(100);p__Ascomycota(100);c__Saccharomycetes(100);o__Saccharomycetales(100);f__Metschnikowiaceae(100);g__Metschnikowia(100);s__Metschnikowia_pulcherrima(100); | Metschnikowia | 13 | Saprotroph | \|Undefined Saprotroph\| | Yeast | NULL | Probable | As nectar-tap_saprotroph (Põlme et al. 2020); Decay sugar-rich substrates (Põlme et al. 2020) | Põlme S, et al. 2020. Fungal Diversity 105:1-16 ((https://doi.org/10.1007/s13225-020-00466-2)) |
| Otu10965 | 2 | k__Fungi(100);p__Basidiomycota(100);c__Wallemiomycetes(100);o__Wallemiales(100);f__Wallemiaceae(100);g__Wallemia(100);s__Wallemia_sebi(100); | Wallemia | 13 | Saprotroph | \|Undefined Saprotroph\| | Microfungus | NULL | Probable | Decay sugar-rich substrates (Põlme et al. 2020) | Tedersoo L, et al. 2014. Science 346:e1256688 ((https://doi.org/10.1126/science.1256688)); Põlme S, et al. 2020. Fungal Diversity 105:1-16 ((https://doi.org/10.1007/s13225-020-00466-2)) |
| Otu10966 | 2 | k__Fungi(100);p__Basidiomycota(100);c__Wallemiomycetes(100);o__Wallemiales(100);f__Wallemiaceae(100);g__Wallemia(100);s__Wallemia_sebi(100); | Wallemia | 13 | Saprotroph | \|Undefined Saprotroph\| | Microfungus | NULL | Probable | Decay sugar-rich substrates (Põlme et al. 2020) | Tedersoo L, et al. 2014. Science 346:e1256688 ((https://doi.org/10.1126/science.1256688)); Põlme S, et al. 2020. Fungal Diversity 105:1-16 ((https://doi.org/10.1007/s13225-020-00466-2)) |
| Otu10967 | 2 | k__Fungi(100);p__Basidiomycota(100);c__Wallemiomycetes(100);o__Wallemiales(100);f__Wallemiaceae(100);g__Wallemia(100);s__Wallemia_sebi(100); | Wallemia | 13 | Saprotroph | \|Undefined Saprotroph\| | Microfungus | NULL | Probable | Decay sugar-rich substrates (Põlme et al. 2020) | Tedersoo L, et al. 2014. Science 346:e1256688 ((https://doi.org/10.1126/science.1256688)); Põlme S, et al. 2020. Fungal Diversity 105:1-16 ((https://doi.org/10.1007/s13225-020-00466-2)) |
| Otu10968 | 2 | k__Fungi(100);p__Ascomycota(100);c__Saccharomycetes(100);o__Saccharomycetales(100);f__Metschnikowiaceae(100);g__Metschnikowia(100);g__Metschnikowia_unclassified(100); | Metschnikowia | 13 | Saprotroph | \|Undefined Saprotroph\| | Yeast | NULL | Probable | As nectar-tap_saprotroph (Põlme et al. 2020); Decay sugar-rich substrates (Põlme et al. 2020) | Põlme S, et al. 2020. Fungal Diversity 105:1-16 ((https://doi.org/10.1007/s13225-020-00466-2)) |
| Otu10969 | 2 | k__Fungi(100);p__Ascomycota(100);c__Saccharomycetes(100);o__Saccharomycetales(100);f__Metschnikowiaceae(100);g__Metschnikowia(100);g__Metschnikowia_unclassified(100); | Metschnikowia | 13 | Saprotroph | \|Undefined Saprotroph\| | Yeast | NULL | Probable | As nectar-tap_saprotroph (Põlme et al. 2020); Decay sugar-rich substrates (Põlme et al. 2020) | Põlme S, et al. 2020. Fungal Diversity 105:1-16 ((https://doi.org/10.1007/s13225-020-00466-2)) |

| | | | | | | | | | | | |
|---|---|---|---|---|---|---|---|---|---|---|---|
| Otu10971 | 2 | k__Fungi(100);p__Ascomycota(100);c__Saccharomycetes(100);o__Saccharomycetales(100);f__Metschnikowiaceae(100);g__Metschnikowia(100);s__Metschnikowia_pulcherrima(100); | Metschnikowia | 13 | Saprotroph | \|Undefined Saprotroph\| | Yeast | NULL | Probable | [illegible] | [illegible] |
| Otu10972 | 2 | k__Fungi(100);p__Basidiomycota(100);c__Microbotryomycetes(100);o__Sporidiobolales(100);f__Sporidiobolales_family_Incertae_sedis(100);g__Rhodotorula(100);s__Rhodotorula_taiwanensis(100); | Rhodotorula | 13 | Pathotroph-Saprotroph-Symbiotroph | Animal Pathogen-Animal Symbiotroph-Endophyte-Plant Pathogen-\|Undefined Saprotroph\| | Yeast | NULL | Probable | [illegible] | [illegible] |
| Otu10973 | 2 | k__Fungi(100);p__Zygomycota(100);c__Zygomycota_class_Incertae_sedis(100);o__Mucorales(100);f__Mucoraceae(100);g__Mucor(100);s__Mucor_hiemalis(100); | Mucor | 13 | Saprotroph | \|Undefined Saprotroph\| | Microfungus | NULL | Probable | [illegible] | [illegible] |
| Otu10974 | 2 | k__Fungi(100);p__Zygomycota(100);c__Zygomycota_class_Incertae_sedis(100);o__Mucorales(100);f__Mucoraceae(100);g__Mucor(100);s__Mucor_genevensis(100); | Mucor | 13 | Saprotroph | \|Undefined Saprotroph\| | Microfungus | NULL | Probable | [illegible] | [illegible] |
| Otu10975 | 2 | k__Fungi(100);p__Ascomycota(100);c__Saccharomycetes(100);o__Saccharomycetales(100);f__Metschnikowiaceae(100);g__Metschnikowia(100);s__Metschnikowia_pulcherrima(100); | Metschnikowia | 13 | Saprotroph | \|Undefined Saprotroph\| | Yeast | NULL | Probable | [illegible] | [illegible] |
| Otu10980 | 2 | k__Fungi(100);p__Zygomycota(100);c__Zygomycota_class_Incertae_sedis(100);o__Mucorales(100);f__Mucoraceae(100);g__Mucor(100);s__Mucor_genevensis(100); | Mucor | 13 | Saprotroph | \|Undefined Saprotroph\| | Microfungus | NULL | Probable | [illegible] | [illegible] |
| Otu10981 | 2 | k__Fungi(100);p__Ascomycota(100);c__Saccharomycetes(100);o__Saccharomycetales(100);f__Metschnikowiaceae(100);g__Metschnikowia(100);s__Metschnikowia_pulcherrima(100); | Metschnikowia | 13 | Saprotroph | \|Undefined Saprotroph\| | Yeast | NULL | Probable | [illegible] | [illegible] |
| Otu10982 | 2 | k__Fungi(100);p__Ascomycota(100);c__Saccharomycetes(100);o__Saccharomycetales(100);f__Metschnikowiaceae(100);g__Metschnikowia(100);s__Metschnikowia_pulcherrima(100); | Metschnikowia | 13 | Saprotroph | \|Undefined Saprotroph\| | Yeast | NULL | Probable | [illegible] | [illegible] |
| Otu10983 | 2 | k__Fungi(100);p__Ascomycota(100);c__Saccharomycetes(100);o__Saccharomycetales(100);f__Saccharomycetales_family_Incertae_sedis(100);g__Candida(100);g__Candida_unclassified(100); | Candida | 13 | Pathotroph-Saprotroph-Symbiotroph | Animal Pathogen-Endophyte-Endosymbiont-Epiphyte-\|Undefined Saprotroph\| | Dimorphic Yeast-Yeast | NULL | Probable | [illegible] | [illegible] |
| Otu10984 | 2 | k__Fungi(100);p__Zygomycota(100);c__Zygomycota_class_Incertae_sedis(100);o__Mucorales(100);f__Mucoraceae(100);g__Mucor(100);g__Mucor_unclassified(100); | Mucor | 13 | Saprotroph | \|Undefined Saprotroph\| | Microfungus | NULL | Probable | [illegible] | [illegible] |
| Otu10985 | 2 | k__Fungi(100);p__Ascomycota(100);c__Saccharomycetes(100);o__Saccharomycetales(100);f__Metschnikowiaceae(100);g__Metschnikowia(100);s__Metschnikowia_pulcherrima(100); | Metschnikowia | 13 | Saprotroph | \|Undefined Saprotroph\| | Yeast | NULL | Probable | [illegible] | [illegible] |
| Otu10986 | 2 | k__Fungi(100);p__Ascomycota(100);c__Dothideomycetes(100);o__Capnodiales(100);f__Mycosphaerellaceae(100);g__Passalora(100);g__Passalora_unclassified(100); | Passalora | 13 | Pathotroph | \|Plant Pathogen\| | Microfungus | NULL | Probable | NULL | [illegible] |
| Otu10987 | 2 | k__Fungi(100);p__Basidiomycota(100);c__Tremellomycetes(100);o__Tremellales(100);f__Tremellales_family_Incertae_sedis(100);g__Fellomyces(100);s__Fellomyces_horovitziae(100); | Fellomyces | 13 | Pathotroph-Saprotroph | Fungal Parasite-\|Undefined Saprotroph\| | Yeast | NULL | Probable | [illegible] | [illegible] |
| Otu10990 | 2 | k__Fungi(100);p__Ascomycota(100);c__Saccharomycetes(100);o__Saccharomycetales(100);f__Metschnikowiaceae(100);g__Metschnikowia(100);g__Metschnikowia_unclassified(100); | Metschnikowia | 13 | Saprotroph | \|Undefined Saprotroph\| | Yeast | NULL | Probable | [illegible] | [illegible] |
| Otu10992 | 2 | k__Fungi(100);p__Zygomycota(100);c__Zygomycota_class_Incertae_sedis(100);o__Mucorales(100);f__Mucoraceae(100);g__Mucor(100);g__Mucor_unclassified(100); | Mucor | 13 | Saprotroph | \|Undefined Saprotroph\| | Microfungus | NULL | Probable | [illegible] | [illegible] |
| Otu10994 | 2 | k__Fungi(100);p__Zygomycota(100);c__Zygomycota_class_Incertae_sedis(100);o__Mucorales(100);f__Mucoraceae(100);g__Mucor(100);s__Mucor_ucrainicus(100); | Mucor | 13 | Saprotroph | \|Undefined Saprotroph\| | Microfungus | NULL | Probable | [illegible] | [illegible] |
| Otu10996 | 2 | k__Fungi(100);p__Ascomycota(100);c__Saccharomycetes(100);o__Saccharomycetales(100);f__Metschnikowiaceae(100);g__Metschnikowia(100);g__Metschnikowia_unclassified(100); | Metschnikowia | 13 | Saprotroph | \|Undefined Saprotroph\| | Yeast | NULL | Probable | [illegible] | [illegible] |
| Otu10997 | 2 | k__Fungi(100);p__Ascomycota(100);c__Saccharomycetes(100);o__Saccharomycetales(100);f__Metschnikowiaceae(100);g__Metschnikowia(100);g__Metschnikowia_unclassified(100); | Metschnikowia | 13 | Saprotroph | \|Undefined Saprotroph\| | Yeast | NULL | Probable | [illegible] | [illegible] |
| Otu10998 | 2 | k__Fungi(100);p__Ascomycota(100);c__Saccharomycetes(100);o__Saccharomycetales(100);f__Metschnikowiaceae(100);g__Metschnikowia(100);s__Metschnikowia_pulcherrima(100); | Metschnikowia | 13 | Saprotroph | \|Undefined Saprotroph\| | Yeast | NULL | Probable | [illegible] | [illegible] |
| Otu10999 | 2 | k__Fungi(100);p__Ascomycota(100);c__Saccharomycetes(100);o__Saccharomycetales(100);f__Metschnikowiaceae(100);g__Metschnikowia(100);s__Metschnikowia_pulcherrima(100); | Metschnikowia | 13 | Saprotroph | \|Undefined Saprotroph\| | Yeast | NULL | Probable | [illegible] | [illegible] |
| Otu11000 | 2 | k__Fungi(100);p__Basidiomycota(100);c__Agaricomycetes(100);o__Russulales(100);f__Russulaceae(100);g__Russula(100);g__Russula_unclassified(100); | Russula | 13 | Symbiotroph | \|Ectomycorrhizal\| | Agaricoid | NULL | Highly Probable | NULL | [illegible] |
| Otu11001 | 2 | k__Fungi(100);p__Ascomycota(100);c__Saccharomycetes(100);o__Saccharomycetales(100);f__Metschnikowiaceae(100);g__Metschnikowia(100);s__Metschnikowia_pulcherrima(100); | Metschnikowia | 13 | Saprotroph | \|Undefined Saprotroph\| | Yeast | NULL | Probable | [illegible] | [illegible] |
| Otu11002 | 2 | k__Fungi(100);p__Ascomycota(100);c__Saccharomycetes(100);o__Saccharomycetales(100);f__Saccharomycetales_family_Incertae_sedis(100);g__Candida(100);g__Candida_unclassified(100); | Candida | 13 | Pathotroph-Saprotroph-Symbiotroph | Animal Pathogen-Endophyte-Endosymbiont-Epiphyte-\|Undefined Saprotroph\| | Dimorphic Yeast-Yeast | NULL | Probable | [illegible] | [illegible] |
| Otu11003 | 2 | k__Fungi(100);p__Ascomycota(100);c__Saccharomycetes(100);o__Saccharomycetales(100);f__Metschnikowiaceae(100);g__Metschnikowia(100);g__Metschnikowia_unclassified(100); | Metschnikowia | 13 | Saprotroph | \|Undefined Saprotroph\| | Yeast | NULL | Probable | [illegible] | [illegible] |
| Otu11004 | 2 | k__Fungi(100);p__Ascomycota(100);c__Saccharomycetes(100);o__Saccharomycetales(100);f__Metschnikowiaceae(100);g__Metschnikowia(100);g__Metschnikowia_unclassified(100); | Metschnikowia | 13 | Saprotroph | \|Undefined Saprotroph\| | Yeast | NULL | Probable | [illegible] | [illegible] |
| Otu11006 | 2 | k__Fungi(100);p__Ascomycota(100);c__Sordariomycetes(100);o__Hypocreales(100);f__Nectriaceae(100);g__Cosmospora(100);s__Cosmospora_coccinea(100); | Cosmospora | 13 | Pathotroph | \|Fungal Parasite\| | Microfungus | NULL | Highly Probable | [illegible] | [illegible] |
| Otu11007 | 2 | k__Fungi(100);p__Ascomycota(100);c__Saccharomycetes(100);o__Saccharomycetales(100);f__Metschnikowiaceae(100);g__Metschnikowia(100);s__Metschnikowia_pulcherrima(100); | Metschnikowia | 13 | Saprotroph | \|Undefined Saprotroph\| | Yeast | NULL | Probable | [illegible] | [illegible] |
| Otu11008 | 2 | k__Fungi(100);p__Ascomycota(100);c__Saccharomycetes(100);o__Saccharomycetales(100);f__Metschnikowiaceae(100);g__Metschnikowia(100);s__Metschnikowia_pulcherrima(100); | Metschnikowia | 13 | Saprotroph | \|Undefined Saprotroph\| | Yeast | NULL | Probable | [illegible] | [illegible] |
| Otu11010 | 2 | k__Fungi(100);p__Ascomycota(100);c__Eurotiomycetes(100);o__Eurotiales(100);f__Trichocomaceae(100);g__Penicillium(100);g__Penicillium_unclassified(100); | Penicillium | 13 | Pathotroph-Saprotroph-Symbiotroph | Animal Parasite-Dung Saprotroph-Endophyte-Fungal Parasite-Plant Pathogen-\|Undefined Saprotroph\|-Wood Saprotroph | Microfungus | Soft Rot | Highly Probable | [illegible] | [illegible] |
| Otu11012 | 2 | k__Fungi(100);p__Ascomycota(100);c__Eurotiomycetes(100);o__Onygenales(100);f__Arthrodermataceae(100);f__Arthrodermataceae_unclassified(100);f__Arthrodermataceae_unclassified(100); | Arthrodermataceae | 9 | Pathotroph-Saprotroph | Animal Pathogen-Undefined Saprotroph | NULL | NULL | Probable | [illegible] | [illegible] |
| Otu11013 | 2 | k__Fungi(100);p__Ascomycota(100);c__Saccharomycetes(100);o__Saccharomycetales(100);f__Metschnikowiaceae(100);g__Metschnikowia(100);g__Metschnikowia_unclassified(100); | Metschnikowia | 13 | Saprotroph | \|Undefined Saprotroph\| | Yeast | NULL | Probable | [illegible] | [illegible] |
| Otu11014 | 2 | k__Fungi(100);p__Ascomycota(100);c__Saccharomycetes(100);o__Saccharomycetales(100);f__Metschnikowiaceae(100);g__Metschnikowia(100);g__Metschnikowia_unclassified(100); | Metschnikowia | 13 | Saprotroph | \|Undefined Saprotroph\| | Yeast | NULL | Probable | [illegible] | [illegible] |
| Otu11016 | 2 | k__Fungi(100);p__Ascomycota(100);c__Saccharomycetes(100);o__Saccharomycetales(100);o__Saccharomycetales_unclassified(100);o__Saccharomycetales_unclassified(100);o__Saccharomycetales_unclassified(100); | Saccharomycetales | 7 | Saprotroph | Undefined Saprotroph | Yeast | NULL | Possible | NULL | [illegible] |
| Otu11017 | 2 | k__Fungi(100);p__Zygomycota(100);c__Zygomycota_class_Incertae_sedis(100);o__Basidiobolales(100);f__Basidiobolaceae(100);g__Basidiobolus(100);g__Basidiobolus_unclassified(100); | Basidiobolus | 13 | Pathotroph-Saprotroph-Symbiotroph | Animal Pathogen-Animal Symbiotroph-\|Undefined Saprotroph\| | Microfungus | NULL | Probable | [illegible] | [illegible] |
| Otu11019 | 2 | k__Fungi(100);p__Ascomycota(100);c__Saccharomycetes(100);o__Saccharomycetales(100);f__Metschnikowiaceae(100);g__Metschnikowia(100);g__Metschnikowia_unclassified(100); | Metschnikowia | 13 | Saprotroph | \|Undefined Saprotroph\| | Yeast | NULL | Probable | [illegible] | [illegible] |
| Otu11020 | 2 | k__Fungi(100);p__Ascomycota(100);c__Saccharomycetes(100);o__Saccharomycetales(100);f__Metschnikowiaceae(100);g__Metschnikowia(100);s__Metschnikowia_pulcherrima(100); | Metschnikowia | 13 | Saprotroph | \|Undefined Saprotroph\| | Yeast | NULL | Probable | [illegible] | [illegible] |
| Otu11021 | 2 | k__Fungi(100);p__Zygomycota(100);c__Zygomycota_class_Incertae_sedis(100);o__Mortierellales(100);f__Mortierellaceae(100);g__unclassified_Mortierellaceae(100);s__Mortierellaceae_sp(100); | Mortierellaceae | 9 | Saprotroph-Symbiotroph | Endophyte-Litter Saprotroph-Soil Saprotroph-Undefined Saprotroph | Microfungus | NULL | Possible | [illegible] | [illegible] |
| Otu11022 | 2 | k__Fungi(100);p__Zygomycota(100);c__Zygomycota_class_Incertae_sedis(100);o__Mortierellales(100);f__Mortierellaceae(100);g__unclassified_Mortierellaceae(100);s__Mortierellaceae_sp(100); | Mortierellaceae | 9 | Saprotroph-Symbiotroph | Endophyte-Litter Saprotroph-Soil Saprotroph-Undefined Saprotroph | Microfungus | NULL | Possible | [illegible] | [illegible] |
| Otu11023 | 2 | k__Fungi(100);p__Ascomycota(100);c__Saccharomycetes(100);o__Saccharomycetales(100);f__Metschnikowiaceae(100);g__Metschnikowia(100);g__Metschnikowia_unclassified(100); | Metschnikowia | 13 | Saprotroph | \|Undefined Saprotroph\| | Yeast | NULL | Probable | [illegible] | [illegible] |
| Otu11025 | 2 | k__Fungi(100);p__Zygomycota(100);c__Zygomycota_class_Incertae_sedis(100);o__Mortierellales(100);f__unclassified_Mortierellales(100);g__unclassified_Mortierellales(100);s__Mortierellales_sp(100); | Mortierellales | 7 | Saprotroph-Symbiotroph | Endophyte-Soil Saprotroph-Undefined Saprotroph | Microfungus | NULL | Possible | [illegible] | [illegible] |
| Otu11026 | 2 | k__Fungi(100);p__Basidiomycota(100);c__Agaricomycetes(100);o__Agaricales(100);o__Agaricales_unclassified(100);o__Agaricales_unclassified(100);o__Agaricales_unclassified(100); | Agaricales | 7 | Pathotroph-Saprotroph-Symbiotroph | Bryophyte Parasite-Dung Saprotroph-Ectomycorrhizal-Fungal Parasite-Leaf Saprotroph-Plant Parasite-Undefined Saprotroph-Wood Saprotroph | Agaricoid-Gasteroid-Microfungus-Secotioid-Yeast | NULL | Possible | [illegible] | [illegible] |
| Otu11029 | 2 | k__Fungi(100);p__Ascomycota(100);c__Saccharomycetes(100);o__Saccharomycetales(100);f__Metschnikowiaceae(100);g__Metschnikowia(100);s__Metschnikowia_pulcherrima(100); | Metschnikowia | 13 | Saprotroph | \|Undefined Saprotroph\| | Yeast | NULL | Probable | [illegible] | [illegible] |
| Otu11030 | 2 | k__Fungi(100);p__Ascomycota(100);c__Saccharomycetes(100);o__Saccharomycetales(100);f__Metschnikowiaceae(100);g__Metschnikowia(100);g__Metschnikowia_unclassified(100); | Metschnikowia | 13 | Saprotroph | \|Undefined Saprotroph\| | Yeast | NULL | Probable | [illegible] | [illegible] |
| Otu11032 | 2 | k__Fungi(100);p__Ascomycota(100);c__Saccharomycetes(100);o__Saccharomycetales(100);f__Metschnikowiaceae(100);g__Metschnikowia(100);g__Metschnikowia_unclassified(100); | Metschnikowia | 13 | Saprotroph | \|Undefined Saprotroph\| | Yeast | NULL | Probable | [illegible] | [illegible] |
| Otu11033 | 2 | k__Fungi(100);p__Basidiomycota(100);c__Agaricomycetes(100);o__Agaricales(100);f__Niaceae(100);g__Lachnella(100);s__Lachnella_alboviolascens(100); | Lachnella | 13 | Saprotroph | Undefined Saprotroph-\|Wood Saprotroph\| | Cyphelloid | NULL | Probable | [illegible] | [illegible] |
| Otu11034 | 2 | k__Fungi(100);p__Basidiomycota(100);c__Agaricomycetes(100);o__Agaricales(100);o__Agaricales_unclassified(100);o__Agaricales_unclassified(100);o__Agaricales_unclassified(100); | Agaricales | 7 | Pathotroph-Saprotroph-Symbiotroph | Bryophyte Parasite-Dung Saprotroph-Ectomycorrhizal-Fungal Parasite-Leaf Saprotroph-Plant Parasite-Undefined Saprotroph-Wood Saprotroph | Agaricoid-Gasteroid-Microfungus-Secotioid-Yeast | NULL | Possible | [illegible] | [illegible] |
| Otu11035 | 2 | k__Fungi(100);p__Ascomycota(100);c__Saccharomycetes(100);o__Saccharomycetales(100);f__Metschnikowiaceae(100);g__Metschnikowia(100);s__Metschnikowia_pulcherrima(100); | Metschnikowia | 13 | Saprotroph | \|Undefined Saprotroph\| | Yeast | NULL | Probable | [illegible] | [illegible] |
| Otu11037 | 2 | k__Fungi(100);p__Ascomycota(100);c__Saccharomycetes(100);o__Saccharomycetales(100);f__Metschnikowiaceae(100);g__Metschnikowia(100);g__Metschnikowia_unclassified(100); | Metschnikowia | 13 | Saprotroph | \|Undefined Saprotroph\| | Yeast | NULL | Probable | [illegible] | [illegible] |
| Otu11040 | 2 | k__Fungi(100);p__Ascomycota(100);c__Saccharomycetes(100);o__Saccharomycetales(100);f__Metschnikowiaceae(100);g__Metschnikowia(100);s__Metschnikowia_pulcherrima(100); | Metschnikowia | 13 | Saprotroph | \|Undefined Saprotroph\| | Yeast | NULL | Probable | [illegible] | [illegible] |
| Otu11041 | 2 | k__Fungi(100);p__Ascomycota(100);c__Saccharomycetes(100);o__Saccharomycetales(100);f__Metschnikowiaceae(100);g__Metschnikowia(100);g__Metschnikowia_unclassified(100); | Metschnikowia | 13 | Saprotroph | \|Undefined Saprotroph\| | Yeast | NULL | Probable | [illegible] | [illegible] |
| Otu11042 | 2 | k__Fungi(100);p__Ascomycota(100);c__Dothideomycetes(100);o__Dothideomycetes_order_Incertae_sedis(100);f__Myxotrichaceae(100);g__Oidiodendron(100);g__Oidiodendron_unclassified(100); | Oidiodendron | 13 | Pathotroph-Saprotroph-Symbiotroph | Endophyte-Ericoid Mycorrhizal-\|Undefined Saprotroph\| | Microfungus | NULL | Probable | [illegible] | [illegible] |
| Otu11049 | 2 | k__Fungi(100);p__Basidiomycota(100);c__Agaricomycetes(100);o__Russulales(100);f__Russulaceae(100);g__Russula(100);s__Russula_ionochlora(100); | Russula | 13 | Symbiotroph | \|Ectomycorrhizal\| | Agaricoid | NULL | Highly Probable | NULL | [illegible] |
| Otu11050 | 2 | k__Fungi(100);p__Basidiomycota(100);c__Tremellomycetes(100);o__Tremellales(100);f__Tremellales_family_Incertae_sedis(100);g__Dioszegia(100);s__Dioszegia_zsoltii_var._yunnanensis(100); | Dioszegia | 13 | Saprotroph | \|Plant Saprotroph\|-Undefined Saprotroph | Yeast | NULL | Probable | [illegible] | [illegible] |
| Otu11052 | 2 | k__Fungi(100);p__Basidiomycota(100);c__Agaricomycetes(100);o__Russulales(100);f__Auriscalpiaceae(100);g__Lentinellus(100);s__Lentinellus_cochleatus(100); | Lentinellus | 13 | Pathotroph-Saprotroph | Plant Pathogen-Undefined Saprotroph-\|Wood Saprotroph\| | Agaricoid | White Rot | Probable | [illegible] | [illegible] |
| Otu11054 | 2 | k__Fungi(100);p__Ascomycota(100);c__Sordariomycetes(100);o__Hypocreales(100);f__Hypocreales_family_Incertae_sedis(100);g__Calcarisporium(100);s__Calcarisporium_sp(100); | Calcarisporium | 13 | Pathotroph | \|Fungal Parasite\| | Microfungus | NULL | Probable | [illegible] | [illegible] |
| Otu11055 | 2 | k__Fungi(100);p__Ascomycota(100);c__Sordariomycetes(100);o__Sordariales(100);f__Cephalothecaceae(100);g__Cryptendoxyla(100);s__Cryptendoxyla_hypophloia(100); | Cryptendoxyla | 13 | Saprotroph | Undefined Saprotroph-\|Wood Saprotroph\| | Microfungus | NULL | Probable | [illegible] | [illegible] |
| Otu11058 | 2 | k__Fungi(100);p__Ascomycota(100);c__Pezizomycetes(100);o__Pezizales(100);f__Pyronemataceae(100);g__Pyronema(100);s__Pyronema_sp_GSL5_6_2(100); | Pyronema | 13 | Saprotroph-Symbiotroph | Endophyte-Plant Saprotroph-\|Undefined Saprotroph\| | Microfungus | NULL | Probable | [illegible] | [illegible] |
| Otu11059 | 2 | k__Fungi(100);p__Ascomycota(100);c__Saccharomycetes(100);o__Saccharomycetales(100);f__Metschnikowiaceae(100);g__Metschnikowia(100);s__Metschnikowia_pulcherrima(100); | Metschnikowia | 13 | Saprotroph | \|Undefined Saprotroph\| | Yeast | NULL | Probable | [illegible] | [illegible] |
| Otu11060 | 2 | k__Fungi(100);p__Ascomycota(100);c__Saccharomycetes(100);o__Saccharomycetales(100);f__Metschnikowiaceae(100);g__Metschnikowia(100);s__Metschnikowia_pulcherrima(100); | Metschnikowia | 13 | Saprotroph | \|Undefined Saprotroph\| | Yeast | NULL | Probable | [illegible] | [illegible] |
| Otu11061 | 2 | k__Fungi(100);p__Ascomycota(100);c__Saccharomycetes(100);o__Saccharomycetales(100);f__Metschnikowiaceae(100);g__Metschnikowia(100);s__Metschnikowia_pulcherrima(100); | Metschnikowia | 13 | Saprotroph | \|Undefined Saprotroph\| | Yeast | NULL | Probable | [illegible] | [illegible] |
| Otu11062 | 2 | k__Fungi(100);p__Ascomycota(100);c__Saccharomycetes(100);o__Saccharomycetales(100);f__Metschnikowiaceae(100);g__Metschnikowia(100);s__Metschnikowia_pulcherrima(100); | Metschnikowia | 13 | Saprotroph | \|Undefined Saprotroph\| | Yeast | NULL | Probable | [illegible] | [illegible] |
| Otu11064 | 2 | k__Fungi(100);p__Ascomycota(100);c__Eurotiomycetes(100);o__Chaetothyriales(100);f__Herpotrichiellaceae(100);f__Herpotrichiellaceae_unclassified(100);f__Herpotrichiellaceae_unclassified(100); | Herpotrichiellaceae | 9 | Pathotroph-Saprotroph | Animal Pathogen-Fungal Parasite-Undefined Saprotroph | Facultative Yeast-Microfungus | NULL | Probable | [illegible] | [illegible] |
| Otu11065 | 2 | k__Fungi(100);p__Ascomycota(100);c__Dothideomycetes(100);o__Pleosporales(100);f__Lophiostomataceae(100);g__Lophiostoma(100);g__Lophiostoma_unclassified(100); | Lophiostoma | 13 | Saprotroph | Plant Saprotroph-Undefined Saprotroph-\|Wood Saprotroph\| | Microfungus | NULL | Probable | [illegible] | [illegible] |
| Otu11067 | 2 | k__Fungi(100);p__Ascomycota(100);c__Saccharomycetes(100);o__Saccharomycetales(100);f__Metschnikowiaceae(100);g__Metschnikowia(100);g__Metschnikowia_unclassified(100); | Metschnikowia | 13 | Saprotroph | \|Undefined Saprotroph\| | Yeast | NULL | Probable | [illegible] | [illegible] |
| Otu11068 | 2 | k__Fungi(100);p__Ascomycota(100);c__Saccharomycetes(100);o__Saccharomycetales(100);f__Metschnikowiaceae(100);g__Metschnikowia(100);g__Metschnikowia_unclassified(100); | Metschnikowia | 13 | Saprotroph | \|Undefined Saprotroph\| | Yeast | NULL | Probable | [illegible] | [illegible] |
| Otu11069 | 2 | k__Fungi(100);p__Ascomycota(100);c__Eurotiomycetes(100);o__Chaetothyriales(100);f__Herpotrichiellaceae(100);f__Herpotrichiellaceae_unclassified(100);f__Herpotrichiellaceae_unclassified(100); | Herpotrichiellaceae | 9 | Pathotroph-Saprotroph | Animal Pathogen-Fungal Parasite-Undefined Saprotroph | Facultative Yeast-Microfungus | NULL | Probable | [illegible] | [illegible] |
| Otu11070 | 2 | k__Fungi(100);p__Ascomycota(100);c__Eurotiomycetes(100);o__Chaetothyriales(100);f__Herpotrichiellaceae(100);f__Herpotrichiellaceae_unclassified(100);f__Herpotrichiellaceae_unclassified(100); | Herpotrichiellaceae | 9 | Pathotroph-Saprotroph | Animal Pathogen-Fungal Parasite-Undefined Saprotroph | Facultative Yeast-Microfungus | NULL | Probable | [illegible] | [illegible] |
| Otu11071 | 2 | k__Fungi(100);p__Ascomycota(100);c__Saccharomycetes(100);o__Saccharomycetales(100);f__Metschnikowiaceae(100);g__Metschnikowia(100);g__Metschnikowia_unclassified(100); | Metschnikowia | 13 | Saprotroph | \|Undefined Saprotroph\| | Yeast | NULL | Probable | [illegible] | [illegible] |
| Otu11072 | 2 | k__Fungi(100);p__Ascomycota(100);c__Eurotiomycetes(100);o__Chaetothyriales(100);f__Herpotrichiellaceae(100);g__Exophiala(100);g__Exophiala_unclassified(100); | Exophiala | 13 | Pathotroph-Saprotroph-Symbiotroph | \|Animal Parasite\|-Animal Pathogen-Endophyte-Plant Saprotroph-Undefined Saprotroph | Dimorphic-Facultative Yeast | NULL | Probable | [illegible] | [illegible] |
| Otu11074 | 2 | k__Fungi(100);p__Ascomycota(100);c__Eurotiomycetes(100);o__Chaetothyriales(100);f__Herpotrichiellaceae(100);g__Exophiala(100);g__Exophiala_unclassified(100); | Exophiala | 13 | Pathotroph-Saprotroph-Symbiotroph | \|Animal Parasite\|-Animal Pathogen-Endophyte-Plant Saprotroph-Undefined Saprotroph | Dimorphic-Facultative Yeast | NULL | Probable | [illegible] | [illegible] |
| Otu11077 | 2 | k__Fungi(100);p__Ascomycota(100);c__Sordariomycetes(100);o__Sordariomycetes_order_Incertae_sedis(100);f__Magnaporthaceae(100);g__Pyricularia(100);s__Pyricularia_sp_ASR_31(100); | Pyricularia | 13 | Pathotroph | \|Plant Pathogen\| | Microfungus | NULL | Probable | NULL | [illegible] |
| Otu11078 | 2 | k__Fungi(100);p__Ascomycota(100);c__Eurotiomycetes(100);o__Chaetothyriales(100);f__Herpotrichiellaceae(100);f__Herpotrichiellaceae_unclassified(100);f__Herpotrichiellaceae_unclassified(100); | Herpotrichiellaceae | 9 | Pathotroph-Saprotroph | Animal Pathogen-Fungal Parasite-Undefined Saprotroph | Facultative Yeast-Microfungus | NULL | Probable | [illegible] | [illegible] |
| Otu11080 | 2 | k__Fungi(100);p__Ascomycota(100);c__Saccharomycetes(100);o__Saccharomycetales(100);f__Metschnikowiaceae(100);g__Metschnikowia(100);s__Metschnikowia_pulcherrima(100); | Metschnikowia | 13 | Saprotroph | \|Undefined Saprotroph\| | Yeast | NULL | Probable | [illegible] | [illegible] |
| Otu11081 | 2 | k__Fungi(100);p__Ascomycota(100);c__Sordariomycetes(100);o__Sordariales(100);f__Chaetomiaceae(100);f__Chaetomiaceae_unclassified(100);f__Chaetomiaceae_unclassified(100); | Chaetomiaceae | 9 | Pathotroph-Saprotroph-Symbiotroph | Animal Pathogen-Dung Saprotroph-Endophyte-Epiphyte-Plant Saprotroph-Wood Saprotroph | Microfungus | Soft Rot | Possible | [illegible] | [illegible] |
| Otu11085 | 2 | k__Fungi(100);p__Ascomycota(100);c__Saccharomycetes(100);o__Saccharomycetales(100);f__Metschnikowiaceae(100);g__Metschnikowia(100);s__Metschnikowia_pulcherrima(100); | Metschnikowia | 13 | Saprotroph | \|Undefined Saprotroph\| | Yeast | NULL | Probable | [illegible] | [illegible] |
| Otu11089 | 2 | k__Fungi(100);p__Basidiomycota(100);c__Agaricomycetes(100);o__Boletales(100);f__Gomphidiaceae(100);g__Gomphidius(100);g__Gomphidius_unclassified(100); | Gomphidius | 13 | Pathotroph-Symbiotroph | \|Ectomycorrhizal\|-Fungal Parasite | Agaricoid | NULL | Probable | [illegible] | [illegible] |
| Otu11091 | 2 | k__Fungi(100);p__Basidiomycota(100);c__Tremellomycetes(100);o__Tremellales(100);f__Tremellales_family_Incertae_sedis(100);g__Cryptococcus(100);s__Cryptococcus_sp_JCM_11356(100); | Cryptococcus | 13 | Pathotroph-Saprotroph-Symbiotroph | Animal Pathogen-Endophyte-Epiphyte-\|Undefined Saprotroph\| | Dimorphic Yeast-Yeast | NULL | Possible | [illegible] | [illegible] |
| Otu11092 | 2 | k__Fungi(100);p__Ascomycota(100);c__Saccharomycetes(100);o__Saccharomycetales(100);f__Metschnikowiaceae(100);g__Metschnikowia(100);g__Metschnikowia_unclassified(100); | Metschnikowia | 13 | Saprotroph | \|Undefined Saprotroph\| | Yeast | NULL | Probable | [illegible] | [illegible] |
| Otu11093 | 2 | k__Fungi(100);p__Basidiomycota(100);c__Microbotryomycetes(100);o__Sporidiobolales(100);f__Sporidiobolales_family_Incertae_sedis(100);g__Sporobolomyces(100);s__Sporobolomyces_lactosus(100); | Sporobolomyces | 13 | Pathotroph-Saprotroph | \|Fungal Parasite\|-Plant Saprotroph | Yeast | NULL | Probable | [illegible] | [illegible] |
| Otu11095 | 2 | k__Fungi(100);p__Ascomycota(100);c__Saccharomycetes(100);o__Saccharomycetales(100);f__Metschnikowiaceae(100);g__Metschnikowia(100);g__Metschnikowia_unclassified(100); | Metschnikowia | 13 | Saprotroph | \|Undefined Saprotroph\| | Yeast | NULL | Probable | [illegible] | [illegible] |
| Otu11097 | 2 | k__Fungi(100);p__Ascomycota(100);c__Saccharomycetes(100);o__Saccharomycetales(100);f__Metschnikowiaceae(100);g__Metschnikowia(100);g__Metschnikowia_unclassified(100); | Metschnikowia | 13 | Saprotroph | \|Undefined Saprotroph\| | Yeast | NULL | Probable | [illegible] | [illegible] |
| Otu11098 | 2 | k__Fungi(100);p__Ascomycota(100);c__Saccharomycetes(100);o__Saccharomycetales(100);f__Metschnikowiaceae(100);g__Metschnikowia(100);g__Metschnikowia_unclassified(100); | Metschnikowia | 13 | Saprotroph | \|Undefined Saprotroph\| | Yeast | NULL | Probable | [illegible] | [illegible] |
| Otu11099 | 2 | k__Fungi(100);p__Ascomycota(100);c__Saccharomycetes(100);o__Saccharomycetales(100);f__Metschnikowiaceae(100);g__Metschnikowia(100);s__Metschnikowia_pulcherrima(100); | Metschnikowia | 13 | Saprotroph | \|Undefined Saprotroph\| | Yeast | NULL | Probable | [illegible] | [illegible] |
| Otu11100 | 2 | k__Fungi(100);p__Ascomycota(100);c__Saccharomycetes(100);o__Saccharomycetales(100);f__Metschnikowiaceae(100);g__Metschnikowia(100);s__Metschnikowia_pulcherrima(100); | Metschnikowia | 13 | Saprotroph | \|Undefined Saprotroph\| | Yeast | NULL | Probable | [illegible] | [illegible] |
| Otu11101 | 2 | k__Fungi(100);p__Basidiomycota(100);c__Microbotryomycetes(100);o__Sporidiobolales(100);f__Sporidiobolales_family_Incertae_sedis(100);g__Sporobolomyces(100);s__Sporobolomyces_lactosus(100); | Sporobolomyces | 13 | Pathotroph-Saprotroph | \|Fungal Parasite\|-Plant Saprotroph | Yeast | NULL | Probable | [illegible] | [illegible] |
| Otu11103 | 2 | k__Fungi(100);p__Ascomycota(100);c__Saccharomycetes(100);o__Saccharomycetales(100);f__Saccharomycetales_family_Incertae_sedis(100);g__Candida(100);g__Candida_unclassified(100); | Candida | 13 | Pathotroph-Saprotroph-Symbiotroph | Animal Pathogen-Endophyte-Endosymbiont-Epiphyte-\|Undefined Saprotroph\| | Dimorphic Yeast-Yeast | NULL | Probable | [illegible] | [illegible] |
| Otu11107 | 2 | k__Fungi(100);p__Basidiomycota(100);c__Agaricomycetes(100);o__Hymenochaetales(100);f__Hymenochaetaceae(100);g__Hymenochaete(100);g__Hymenochaete_unclassified(100); | Hymenochaete | 13 | Pathotroph-Saprotroph | Plant Pathogen-Undefined Saprotroph-\|Wood Saprotroph\| | Corticioid-Polyporoid | White Rot | Probable | [illegible] | [illegible] |
| Otu11108 | 2 | k__Fungi(100);p__Ascomycota(100);c__Saccharomycetes(100);o__Saccharomycetales(100);f__Metschnikowiaceae(100);g__Metschnikowia(100);g__Metschnikowia_unclassified(100); | Metschnikowia | 13 | Saprotroph | \|Undefined Saprotroph\| | Yeast | NULL | Probable | [illegible] | [illegible] |
| Otu11109 | 2 | k__Fungi(100);p__Basidiomycota(100);c__Tremellomycetes(100);o__Tremellales(100);f__Tremellales_family_Incertae_sedis(100);f__Tremellales_family_Incertae_sedis_unclassified(100);f__Tremellales_family_Incertae_sedis_unclassified(100); | Tremellales | 7 | Pathotroph-Saprotroph-Symbiotroph | Fungal Parasite-Undefined Saprotroph | Tremelloid-Yeast | NULL | Possible | [illegible] | [illegible] |
| Otu11110 | 2 | k__Fungi(100);p__Basidiomycota(100);c__Tremellomycetes(100);o__Tremellales(100);o__Tremellales_unclassified(100);o__Tremellales_unclassified(100);o__Tremellales_unclassified(100); | Tremellales | 7 | Pathotroph-Saprotroph-Symbiotroph | Fungal Parasite-Undefined Saprotroph | Tremelloid-Yeast | NULL | Possible | [illegible] | [illegible] |
| Otu11112 | 2 | k__Fungi(100);p__Ascomycota(100);c__Saccharomycetes(100);o__Saccharomycetales(100);f__Metschnikowiaceae(100);g__Metschnikowia(100);s__Metschnikowia_pulcherrima(100); | Metschnikowia | 13 | Saprotroph | \|Undefined Saprotroph\| | Yeast | NULL | Probable | [illegible] | [illegible] |
| Otu11115 | 2 | k__Fungi(100);p__Ascomycota(100);c__Saccharomycetes(100);o__Saccharomycetales(100);f__Metschnikowiaceae(100);g__Metschnikowia(100);g__Metschnikowia_unclassified(100); | Metschnikowia | 13 | Saprotroph | \|Undefined Saprotroph\| | Yeast | NULL | Probable | [illegible] | [illegible] |
| Otu11116 | 2 | k__Fungi(100);p__Basidiomycota(100);c__Tremellomycetes(100);o__Tremellales(100);f__Tremellales_family_Incertae_sedis(100);g__Cryptococcus(100);s__Cryptococcus_aff_amylolyticus_AS_22501(100); | Cryptococcus | 13 | Pathotroph-Saprotroph-Symbiotroph | Animal Pathogen-Endophyte-Epiphyte-\|Undefined Saprotroph\| | Dimorphic Yeast-Yeast | NULL | Possible | [illegible] | [illegible] |
| Otu11118 | 2 | k__Fungi(100);p__Ascomycota(100);c__Saccharomycetes(100);o__Saccharomycetales(100);f__Saccharomycetales_family_Incertae_sedis(100);g__Candida(100);g__Candida_unclassified(100); | Candida | 13 | Pathotroph-Saprotroph-Symbiotroph | Animal Pathogen-Endophyte-Endosymbiont-Epiphyte-\|Undefined Saprotroph\| | Dimorphic Yeast-Yeast | NULL | Probable | [illegible] | [illegible] |
| Otu11122 | 2 | k__Fungi(100);p__Basidiomycota(100);c__Agaricomycetes(100);o__Agaricales(100);f__Mycenaceae(100);g__Mycena(100);s__Mycena_sp(100); | Mycena | 13 | Pathotroph-Saprotroph | Plant Pathogen-\|Plant Saprotroph\|-Undefined Saprotroph-Wood Saprotroph | Agaricoid | NULL | Probable | [illegible] | [illegible] |
| Otu11123 | 2 | k__Fungi(100);p__Basidiomycota(100);c__Agaricomycetes(100);o__Agaricales(100);f__Mycenaceae(100);g__Mycena(100);g__Mycena_unclassified(100); | Mycena | 13 | Pathotroph-Saprotroph | Plant Pathogen-\|Plant Saprotroph\|-Undefined Saprotroph-Wood Saprotroph | Agaricoid | NULL | Probable | [illegible] | [illegible] |
| Otu11127 | 2 | k__Fungi(100);p__Basidiomycota(100);c__Agaricomycetes(100);o__Russulales(100);f__Stereaceae(100);f__Stereaceae_unclassified(100);f__Stereaceae_unclassified(100); | Stereaceae | 9 | Saprotroph | Undefined Saprotroph-Wood Saprotroph | Corticioid-Polyporoid | NULL | Probable | [illegible] | [illegible] |
| Otu11128 | 2 | k__Fungi(100);p__Ascomycota(100);c__Saccharomycetes(100);o__Saccharomycetales(100);f__Metschnikowiaceae(100);g__Metschnikowia(100);s__Metschnikowia_pulcherrima(100); | Metschnikowia | 13 | Saprotroph | \|Undefined Saprotroph\| | Yeast | NULL | Probable | [illegible] | [illegible] |
| Otu11129 | 2 | k__Fungi(100);p__Basidiomycota(100);c__Agaricomycetes(100);o__Thelephorales(100);f__Thelephoraceae(100);g__Tomentella(100);g__Tomentella_unclassified(100); | Tomentella | 13 | Symbiotroph | \|Ectomycorrhizal\| | Corticioid | NULL | Highly Probable | NULL | [illegible] |
| Otu11130 | 2 | k__Fungi(100);p__Basidiomycota(100);c__Agaricomycetes(100);o__Thelephorales(100);f__Thelephoraceae(100);g__Tomentella(100);s__Tomentella_lateritia(100); | Tomentella | 13 | Symbiotroph | \|Ectomycorrhizal\| | Corticioid | NULL | Highly Probable | NULL | [illegible] |
| Otu11131 | 2 | k__Fungi(100);p__Ascomycota(100);c__Ascomycota_class_Incertae_sedis(100);o__Ascomycota_order_Incertae_sedis(100);f__Ascomycota_family_Incertae_sedis(100);g__Sympodiella(100);s__Sympodiella_acicola(100); | Sympodiella | 13 | Saprotroph | \|Plant Saprotroph\|-Undefined Saprotroph | Microfungus | NULL | Probable | [illegible] | [illegible] |
| Otu11132 | 2 | k__Fungi(100);p__Basidiomycota(100);c__Agaricomycetes(100);o__Thelephorales(100);f__Thelephoraceae(100);g__Tomentella(100);s__Tomentella_sp(100); | Tomentella | 13 | Symbiotroph | \|Ectomycorrhizal\| | Corticioid | NULL | Highly Probable | NULL | [illegible] |
| Otu11133 | 2 | k__Fungi(100);p__Ascomycota(100);c__Dothideomycetes(100);o__Pleosporales(100);f__Cucurbitariaceae(100);g__Pyrenochaetopsis(100);s__Pyrenochaetopsis_pratorum(100); | Pyrenochaetopsis | 13 | Pathotroph-Saprotroph-Symbiotroph | \|Endophyte\|-Lichen Parasite-Plant Saprotroph-Undefined Saprotroph | Microfungus | NULL | Possible | [illegible] | [illegible] |
| Otu11134 | 2 | k__Fungi(100);p__Ascomycota(100);c__Saccharomycetes(100);o__Saccharomycetales(100);f__Metschnikowiaceae(100);g__Metschnikowia(100);g__Metschnikowia_unclassified(100); | Metschnikowia | 13 | Saprotroph | \|Undefined Saprotroph\| | Yeast | NULL | Probable | [illegible] | [illegible] |
| Otu11135 | 2 | k__Fungi(100);p__Ascomycota(100);c__Saccharomycetes(100);o__Saccharomycetales(100);f__Metschnikowiaceae(100);g__Metschnikowia(100);s__Metschnikowia_pulcherrima(100); | Metschnikowia | 13 | Saprotroph | \|Undefined Saprotroph\| | Yeast | NULL | Probable | [illegible] | [illegible] |
| Otu11136 | 2 | k__Fungi(100);p__Ascomycota(100);c__Saccharomycetes(100);o__Saccharomycetales(100);f__Metschnikowiaceae(100);g__Metschnikowia(100);s__Metschnikowia_pulcherrima(100); | Metschnikowia | 13 | Saprotroph | \|Undefined Saprotroph\| | Yeast | NULL | Probable | [illegible] | [illegible] |
| Otu11138 | 2 | k__Fungi(100);p__Ascomycota(100);c__Saccharomycetes(100);o__Saccharomycetales(100);f__Metschnikowiaceae(100);g__Metschnikowia(100);g__Metschnikowia_unclassified(100); | Metschnikowia | 13 | Saprotroph | \|Undefined Saprotroph\| | Yeast | NULL | Probable | [illegible] | [illegible] |
| Otu11139 | 2 | k__Fungi(100);p__Ascomycota(100);c__Saccharomycetes(100);o__Saccharomycetales(100);f__Metschnikowiaceae(100);g__Metschnikowia(100);s__Metschnikowia_pulcherrima(100); | Metschnikowia | 13 | Saprotroph | \|Undefined Saprotroph\| | Yeast | NULL | Probable | [illegible] | [illegible] |
| Otu11140 | 2 | k__Fungi(100);p__Ascomycota(100);c__Saccharomycetes(100);o__Saccharomycetales(100);f__Metschnikowiaceae(100);g__Metschnikowia(100);g__Metschnikowia_unclassified(100); | Metschnikowia | 13 | Saprotroph | \|Undefined Saprotroph\| | Yeast | NULL | Probable | [illegible] | [illegible] |
| Otu11143 | 2 | k__Fungi(100);p__Ascomycota(100);c__Sordariomycetes(100);o__Sordariales(100);f__Cephalothecaceae(100);g__Phialemonium(100);s__Phialemonium_atrogriseum(100); | Phialemonium | 13 | Pathotroph-Saprotroph | \|Animal Parasite\|-Undefined Saprotroph | Microfungus | NULL | Probable | [illegible] | [illegible] |
| Otu11144 | 2 | k__Fungi(100);p__Ascomycota(100);c__Saccharomycetes(100);o__Saccharomycetales(100);f__Metschnikowiaceae(100);g__Metschnikowia(100);s__Metschnikowia_pulcherrima(100); | Metschnikowia | 13 | Saprotroph | \|Undefined Saprotroph\| | Yeast | NULL | Probable | [illegible] | [illegible] |
| Otu11145 | 2 | k__Fungi(100);p__Basidiomycota(100);c__Microbotryomycetes(100);o__Sporidiobolales(100);f__Sporidiobolales_family_Incertae_sedis(100);g__Rhodotorula(100);s__Rhodotorula_mucilaginosa(100); | Rhodotorula | 13 | Pathotroph-Saprotroph-Symbiotroph | Animal Pathogen-Animal Symbiotroph-Endophyte-Plant Pathogen-\|Undefined Saprotroph\| | Yeast | NULL | Probable | [illegible] | [illegible] |
| Otu11147 | 2 | k__Fungi(100);p__Ascomycota(100);c__Saccharomycetes(100);o__Saccharomycetales(100);f__Metschnikowiaceae(100);g__Metschnikowia(100);g__Metschnikowia_unclassified(100); | Metschnikowia | 13 | Saprotroph | \|Undefined Saprotroph\| | Yeast | NULL | Probable | [illegible] | [illegible] |
| Otu11148 | 2 | k__Fungi(100);p__Ascomycota(100);c__Saccharomycetes(100);o__Saccharomycetales(100);f__Metschnikowiaceae(100);g__Metschnikowia(100);g__Metschnikowia_unclassified(100); | Metschnikowia | 13 | Saprotroph | \|Undefined Saprotroph\| | Yeast | NULL | Probable | [illegible] | [illegible] |
| Otu11149 | 2 | k__Fungi(100);p__Ascomycota(100);c__Saccharomycetes(100);o__Saccharomycetales(100);f__Metschnikowiaceae(100);g__Metschnikowia(100);g__Metschnikowia_unclassified(100); | Metschnikowia | 13 | Saprotroph | \|Undefined Saprotroph\| | Yeast | NULL | Probable | [illegible] | [illegible] |
| Otu11151 | 2 | k__Fungi(100);p__Ascomycota(100);c__Saccharomycetes(100);o__Saccharomycetales(100);f__Metschnikowiaceae(100);g__Metschnikowia(100);s__Metschnikowia_pulcherrima(100); | Metschnikowia | 13 | Saprotroph | \|Undefined Saprotroph\| | Yeast | NULL | Probable | [illegible] | [illegible] |
| Otu11152 | 2 | k__Fungi(100);p__Ascomycota(100);c__Saccharomycetes(100);o__Saccharomycetales(100);f__Metschnikowiaceae(100);g__Metschnikowia(100);s__Metschnikowia_pulcherrima(100); | Metschnikowia | 13 | Saprotroph | \|Undefined Saprotroph\| | Yeast | NULL | Probable | [illegible] | [illegible] |
| Otu11154 | 2 | k__Fungi(100);p__Basidiomycota(100);c__Microbotryomycetes(100);o__Sporidiobolales(100);f__Sporidiobolales_family_Incertae_sedis(100);g__Rhodotorula(100);s__Rhodotorula_taiwanensis(100); | Rhodotorula | 13 | Pathotroph-Saprotroph-Symbiotroph | Animal Pathogen-Animal Symbiotroph-Endophyte-Plant Pathogen-\|Undefined Saprotroph\| | Yeast | NULL | Probable | [illegible] | [illegible] |
| Otu11155 | 2 | k__Fungi(100);p__Basidiomycota(100);c__Microbotryomycetes(100);o__Sporidiobolales(100);f__Sporidiobolales_family_Incertae_sedis(100);g__Rhodotorula(100);s__Rhodotorula_taiwanensis(100); | Rhodotorula | 13 | Pathotroph-Saprotroph-Symbiotroph | Animal Pathogen-Animal Symbiotroph-Endophyte-Plant Pathogen-\|Undefined Saprotroph\| | Yeast | NULL | Probable | [illegible] | [illegible] |
| Otu11156 | 2 | k__Fungi(100);p__Basidiomycota(100);c__Microbotryomycetes(100);o__Sporidiobolales(100);f__Sporidiobolales_family_Incertae_sedis(100);g__Rhodotorula(100);s__Rhodotorula_mucilaginosa(100); | Rhodotorula | 13 | Pathotroph-Saprotroph-Symbiotroph | Animal Pathogen-Animal Symbiotroph-Endophyte-Plant Pathogen-\|Undefined Saprotroph\| | Yeast | NULL | Probable | [illegible] | [illegible] |
| Otu11158 | 2 | k__Fungi(100);p__Ascomycota(100);c__Saccharomycetes(100);o__Saccharomycetales(100);f__Metschnikowiaceae(100);g__Metschnikowia(100);s__Metschnikowia_pulcherrima(100); | Metschnikowia | 13 | Saprotroph | \|Undefined Saprotroph\| | Yeast | NULL | Probable | [illegible] | [illegible] |
| Otu11159 | 2 | k__Fungi(100);p__Basidiomycota(100);c__Tremellomycetes(100);o__Tremellales(100);f__Tremellales_family_Incertae_sedis(100);g__Auriculibuller(100);s__Auriculibuller_fuscus(100); | Auriculibuller | 13 | Saprotroph | \|Undefined Saprotroph\| | Yeast | NULL | Probable | NULL | [illegible] |
| Otu11160 | 2 | k__Fungi(100);p__Basidiomycota(100);c__Microbotryomycetes(100);o__Sporidiobolales(100);f__Sporidiobolales_family_Incertae_sedis(100);g__Rhodotorula(100);s__Rhodotorula_taiwanensis(100); | Rhodotorula | 13 | Pathotroph-Saprotroph-Symbiotroph | Animal Pathogen-Animal Symbiotroph-Endophyte-Plant Pathogen-\|Undefined Saprotroph\| | Yeast | NULL | Probable | [illegible] | [illegible] |
| Otu11161 | 2 | k__Fungi(100);p__Ascomycota(100);c__Saccharomycetes(100);o__Saccharomycetales(100);f__Metschnikowiaceae(100);g__Metschnikowia(100);s__Metschnikowia_pulcherrima(100); | Metschnikowia | 13 | Saprotroph | \|Undefined Saprotroph\| | Yeast | NULL | Probable | [illegible] | [illegible] |
| Otu11162 | 2 | k__Fungi(100);p__Ascomycota(100);c__Saccharomycetes(100);o__Saccharomycetales(100);f__Metschnikowiaceae(100);g__Metschnikowia(100);s__Metschnikowia_pulcherrima(100); | Metschnikowia | 13 | Saprotroph | \|Undefined Saprotroph\| | Yeast | NULL | Probable | [illegible] | [illegible] |
| Otu11163 | 2 | k__Fungi(100);p__Basidiomycota(100);c__Agaricomycetes(100);o__Russulales(100);f__Peniophoraceae(100);g__Peniophora(100);g__Peniophora_unclassified(100); | Peniophora | 13 | Pathotroph-Saprotroph | Plant Pathogen-\|Wood Saprotroph\| | Corticioid | White Rot | Probable | [illegible] | [illegible] |
| Otu11164 | 2 | k__Fungi(100);p__Basidiomycota(100);c__Agaricomycetes(100);o__Russulales(100);f__Peniophoraceae(100);f__Peniophoraceae_unclassified(100);f__Peniophoraceae_unclassified(100); | Peniophoraceae | 9 | Pathotroph-Saprotroph | Plant Pathogen-Wood Saprotroph | Corticioid | White Rot | Probable | [illegible] | [illegible] |
| Otu11166 | 2 | k__Fungi(100);p__Ascomycota(100);c__Saccharomycetes(100);o__Saccharomycetales(100);f__Metschnikowiaceae(100);g__Metschnikowia(100);g__Metschnikowia_unclassified(100); | Metschnikowia | 13 | Saprotroph | \|Undefined Saprotroph\| | Yeast | NULL | Probable | [illegible] | [illegible] |
| Otu11170 | 2 | k__Fungi(100);p__Ascomycota(100);c__Saccharomycetes(100);o__Saccharomycetales(100);f__Metschnikowiaceae(100);g__Metschnikowia(100);s__Metschnikowia_pulcherrima(100); | Metschnikowia | 13 | Saprotroph | \|Undefined Saprotroph\| | Yeast | NULL | Probable | [illegible] | [illegible] |
| Otu11171 | 2 | k__Fungi(100);p__Basidiomycota(100);c__Agaricomycetes(100);o__Russulales(100);f__Russulaceae(100);g__Russula(100);s__Russula_sp(100); | Russula | 13 | Symbiotroph | \|Ectomycorrhizal\| | Agaricoid | NULL | Highly Probable | NULL | [illegible] |
| Otu11174 | 2 | k__Fungi(100);p__Ascomycota(100);c__Saccharomycetes(100);o__Saccharomycetales(100);f__Metschnikowiaceae(100);g__Metschnikowia(100);g__Metschnikowia_unclassified(100); | Metschnikowia | 13 | Saprotroph | \|Undefined Saprotroph\| | Yeast | NULL | Probable | [illegible] | [illegible] |
| Otu11175 | 2 | k__Fungi(100);p__Basidiomycota(100);c__Agaricomycetes(100);o__Russulales(100);f__Russulaceae(100);f__Russulaceae_unclassified(100);f__Russulaceae_unclassified(100); | Russulaceae | 9 | Symbiotroph | Ectomycorrhizal | Agaricoid-Corticioid-Gasteroid-Secotioid | NULL | Probable | [illegible] | [illegible] |
| Otu11177 | 2 | k__Fungi(100);p__Basidiomycota(100);c__Cystobasidiomycetes(100);o__Cystobasidiomycetes_order_Incertae_sedis(100);f__Cystobasidiomycetes_family_Incertae_sedis(100);g__Sakaguchia(100);s__Sakaguchia_dacryoidea(100); | Sakaguchia | 13 | Saprotroph | \|Undefined Saprotroph\| | Dimorphic-Yeast | NULL | Probable | [illegible] | [illegible] |
| Otu11180 | 2 | k__Fungi(100);p__Ascomycota(100);c__Saccharomycetes(100);o__Saccharomycetales(100);f__Metschnikowiaceae(100);g__Metschnikowia(100);g__Metschnikowia_unclassified(100); | Metschnikowia | 13 | Saprotroph | \|Undefined Saprotroph\| | Yeast | NULL | Probable | [illegible] | [illegible] |
| Otu11181 | 2 | k__Fungi(100);p__Ascomycota(100);c__Saccharomycetes(100);o__Saccharomycetales(100);f__Metschnikowiaceae(100);g__Metschnikowia(100);s__Metschnikowia_pulcherrima(100); | Metschnikowia | 13 | Saprotroph | \|Undefined Saprotroph\| | Yeast | NULL | Probable | [illegible] | [illegible] |
| Otu11182 | 2 | k__Fungi(100);p__Ascomycota(100);c__Saccharomycetes(100);o__Saccharomycetales(100);f__Metschnikowiaceae(100);g__Metschnikowia(100);s__Metschnikowia_pulcherrima(100); | Metschnikowia | 13 | Saprotroph | \|Undefined Saprotroph\| | Yeast | NULL | Probable | [illegible] | [illegible] |
| Otu11183 | 2 | k__Fungi(100);p__Ascomycota(100);c__Saccharomycetes(100);o__Saccharomycetales(100);f__Metschnikowiaceae(100);g__Metschnikowia(100);g__Metschnikowia_unclassified(100); | Metschnikowia | 13 | Saprotroph | \|Undefined Saprotroph\| | Yeast | NULL | Probable | [illegible] | [illegible] |
| Otu11184 | 2 | k__Fungi(100);p__Basidiomycota(100);c__Agaricomycetes(100);o__Agaricales(100);f__Psathyrellaceae(100);g__Psathyrella(100);g__Psathyrella_unclassified(100); | Psathyrella | 13 | Saprotroph | Plant Saprotroph-\|Wood Saprotroph\| | Agaricoid | NULL | Probable | [illegible] | [illegible] |
| Otu11185 | 2 | k__Fungi(100);p__Basidiomycota(100);c__Agaricomycetes(100);o__Polyporales(100);f__Meruliaceae(100);g__Scopuloides(100);s__Scopuloides_hydnoides(100); | Scopuloides | 13 | Saprotroph | Undefined Saprotroph-\|Wood Saprotroph\| | Corticioid | White Rot | Probable | [illegible] | [illegible] |
| Otu11186 | 2 | k__Fungi(100);p__Basidiomycota(100);c__Agaricomycetes(100);o__Polyporales(100);f__Meruliaceae(100);g__Irpex(100);s__Irpex_sp_JSL_2009a(100); | Irpex | 13 | Saprotroph | \|Wood Saprotroph\| | Corticioid | White Rot | Highly Probable | [illegible] | [illegible] |
| Otu11188 | 2 | k__Fungi(100);p__Basidiomycota(100);c__Agaricomycetes(100);o__Polyporales(100);f__Polyporaceae(100);f__Polyporaceae_unclassified(100);f__Polyporaceae_unclassified(100); | Polyporaceae | 9 | Saprotroph | Wood Saprotroph | Polyporoid | Brown Rot-White Rot | Probable | [illegible] | [illegible] |
| Otu11193 | 2 | k__Fungi(100);p__Basidiomycota(100);c__Agaricomycetes(100);o__Thelephorales(100);f__Thelephoraceae(100);g__Tomentella(100);s__Tomentella_ellisii(100); | Tomentella | 13 | Symbiotroph | \|Ectomycorrhizal\| | Corticioid | NULL | Highly Probable | NULL | [illegible] |
| Otu11194 | 2 | k__Fungi(100);p__Ascomycota(100);c__Saccharomycetes(100);o__Saccharomycetales(100);f__Metschnikowiaceae(100);g__Metschnikowia(100);s__Metschnikowia_pulcherrima(100); | Metschnikowia | 13 | Saprotroph | \|Undefined Saprotroph\| | Yeast | NULL | Probable | [illegible] | [illegible] |
| Otu11195 | 2 | k__Fungi(100);p__Ascomycota(100);c__Saccharomycetes(100);o__Saccharomycetales(100);f__Saccharomycetales_family_Incertae_sedis(100);g__Candida(100);g__Candida_unclassified(100); | Candida | 13 | Pathotroph-Saprotroph-Symbiotroph | Animal Pathogen-Endophyte-Endosymbiont-Epiphyte-\|Undefined Saprotroph\| | Dimorphic Yeast-Yeast | NULL | Probable | [illegible] | [illegible] |
| Otu11196 | 2 | k__Fungi(100);p__Basidiomycota(100);c__Agaricomycetes(100);o__Hymenochaetales(100);f__Hymenochaetaceae(100);g__Inonotus(100);s__Inonotus_hispidus(100); | Inonotus | 13 | Pathotroph-Saprotroph | Plant Pathogen-\|Wood Saprotroph\| | Polyporoid | White Rot | Highly Probable | [illegible] | [illegible] |
| Otu11197 | 2 | k__Fungi(100);p__Ascomycota(100);c__Saccharomycetes(100);o__Saccharomycetales(100);f__Metschnikowiaceae(100);g__Metschnikowia(100);s__Metschnikowia_pulcherrima(100); | Metschnikowia | 13 | Saprotroph | \|Undefined Saprotroph\| | Yeast | NULL | Probable | [illegible] | [illegible] |
| Otu11198 | 2 | k__Fungi(100);p__Basidiomycota(100);c__Agaricomycetes(100);o__Hymenochaetales(100);f__Hymenochaetaceae(100);g__Inonotus(100);g__Inonotus_unclassified(100); | Inonotus | 13 | Pathotroph-Saprotroph | Plant Pathogen-\|Wood Saprotroph\| | Polyporoid | White Rot | Highly Probable | [illegible] | [illegible] |

| | | | | | | | | | | | |
|---|---|---|---|---|---|---|---|---|---|---|---|
| Otu11199 | 2 | k__Fungi(100);p__Ascomycota(100);c__Saccharomycetes(100);o__Saccharomycetales(100);f__Metschnikowiaceae(100);g__Metschnikowia(100);s__Metschnikowia_pulcherrima(100); | Metschnikowia | 13 | Saprotroph | \|Undefined Saprotroph\| | Yeast | NULL | Probable | As nectar-tap_saprotroph (Põlme et al. 2020); Decay sugar-rich substrates (Põlme et al. 2020) | Põlme S, et al. 2020. Fungal Diversity 105:1-16 ((https://doi.org/10.1007/s13225-020-00466-2)) |
| Otu11201 | 2 | k__Fungi(100);p__Ascomycota(100);c__Saccharomycetes(100);o__Saccharomycetales(100);f__Saccharomycetales_family_Incertae_sedis(100);g__Candida(100);g__Candida_unclassified(100); | Candida | 13 | Pathotroph-Saprotroph-Symbiotroph | Animal Pathogen-Endophyte-Endosymbiont-Epiphyte-\|Undefined Saprotroph\| | Dimorphic Yeast-Yeast | NULL | Probable | As nectar-tap_saprotroph (Põlme et al. 2020); Decay sugar-rich substrates (Põlme et al. 2020); This asexual genus is polyphyletic and contain species that function very differently depend on what they are. Caution should be taken when interpreting guilds for Candida species. | Manolakaki D, et al. 2010. Virulence. 1: 367-375 ((https://doi.org/10.4161/viru.1.5.12796)); Mekbib SB, et al. 2011. Tropical Plant Pathology 36:233-240 ((https://doi.org/10.1590/S1982-56762011000400004)); Põlme S, et al. 2020. Fungal Diversity 105:1-16 ((https://doi.org/10.1007/s13225-020-00466-2)) |
| Otu11202 | 2 | k__Fungi(100);p__Basidiomycota(100);c__Agaricomycetes(100);o__Russulales(100);f__Peniophoraceae(100);f__Peniophoraceae_unclassified(100);f__Peniophoraceae_unclassified(100); | Peniophoraceae | 9 | Pathotroph-Saprotroph | Plant Pathogen-Wood Saprotroph | Corticioid | White Rot | Probable | Most species are saprobic, causing rots of standing and fallen wood. Many species are adapted to xeric conditions, being able to grow on rapidly drying dead attached branches (Cannon & Kirk 2007) | Cannon PF, Kirk PM. 2007. Fungal Families of the World. CAB International, Cambridge (ISBN: 978-0851998275) |
| Otu11203 | 2 | k__Fungi(100);p__Ascomycota(100);c__Sordariomycetes(100);o__Hypocreales(100);f__Nectriaceae(100);g__Fusarium(100);g__Fusarium_unclassified(100); | Fusarium | 13 | Pathotroph-Saprotroph-Symbiotroph | Animal Pathogen-Endophyte-Lichen Parasite-\|Plant Pathogen\|-Plant Saprotroph-Undefined Saprotroph-Wood Saprotroph | Microfungus | Soft Rot | Possible | As foliar_endophyte (Põlme et al. 2020); As litter_saprotroph (Põlme et al. 2020); Causing damping-off disease (Põlme et al. 2020); Decay leaf, fruit, seed, roots, soil, animal material (Põlme et al. 2020); Likely opportunistic human pathogen (Irinyi et al. 2015) | Duncan CG, Eslyn WE. 1966. Mycologia 58:642-664 ((https://doi.org/10.2307/3757045)); Seehann G, et al. 1975. List of Fungi in Soft-Rot Tissue Tests. Institute fur Holzbiologie und Holzschutz, Bundesforschungsanstalt fur Forst - und Holzwirtshaft, Hamburg - Lohbrugge, Germany; Costa IPMW, et al. 2012. Mycotaxon 119:e26 ((http://mycotaxon.com/resources/checklists/costa_v119_checklist.pdf)); Tedersoo L, et al. 2014. Science 346:e1256688 ((https://doi.org/10.1126/science.1256688)); Irinyi L, et al. 2015. Medical Mycology 53:313-337 ((https://doi.org/10.1093/mmy/myv008)); Schroers, et al. 2016. Mycologia 108:806-819 ((https://doi.org/10.3852/15-255)); Põlme S, et al. 2020. Fungal Diversity 105:1-16 ((https://doi.org/10.1007/s13225-020-00466-2)); The American Phytopathological Society Common Names of Plant Diseases ((http://www.apsnet.org/publications/commonnames/Pages/default.aspx)); Lawrey JD, Diederich P. 2016. http://www.lichenicolous.net (accessed 24 May 2017) ((http://www.lichenicolous.net)) |
| Otu11204 | 2 | k__Fungi(100);p__Ascomycota(100);c__Saccharomycetes(100);o__Saccharomycetales(100);f__Metschnikowiaceae(100);g__Metschnikowia(100);s__Metschnikowia_pulcherrima(100); | Metschnikowia | 13 | Saprotroph | \|Undefined Saprotroph\| | Yeast | NULL | Probable | As nectar-tap_saprotroph (Põlme et al. 2020); Decay sugar-rich substrates (Põlme et al. 2020) | Põlme S, et al. 2020. Fungal Diversity 105:1-16 ((https://doi.org/10.1007/s13225-020-00466-2)) |
| Otu11205 | 2 | k__Fungi(100);p__Basidiomycota(100);c__Microbotryomycetes(100);o__Sporidiobolales(100);f__Sporidiobolales_family_Incertae_sedis(100);g__Sporobolomyces(100);s__Sporobolomyces_lactosus(100); | Sporobolomyces | 13 | Pathotroph-Saprotroph | \|Fungal Parasite\|-Plant Saprotroph | Yeast | NULL | Probable | As litter_saprotroph (Põlme et al. 2020); As mycoparasite (Põlme et al. 2020); Decay fungal material (Põlme et al. 2020) | Tedersoo L, et al. 2014. Science 346:e1256688 ((https://doi.org/10.1126/science.1256688)); Purahong W, et al. 2016. Molecular Ecology 25:4059-4074 ((https://doi.org/10.1111/mec.13739)); Põlme S, et al. 2020. Fungal Diversity 105:1-16 ((https://doi.org/10.1007/s13225-020-00466-2)) |
| Otu11206 | 2 | k__Fungi(100);p__Ascomycota(100);c__Saccharomycetes(100);o__Saccharomycetales(100);f__Metschnikowiaceae(100);g__Metschnikowia(100);g__Metschnikowia_unclassified(100); | Metschnikowia | 13 | Saprotroph | \|Undefined Saprotroph\| | Yeast | NULL | Probable | As nectar-tap_saprotroph (Põlme et al. 2020); Decay sugar-rich substrates (Põlme et al. 2020) | Põlme S, et al. 2020. Fungal Diversity 105:1-16 ((https://doi.org/10.1007/s13225-020-00466-2)) |
| Otu11207 | 2 | k__Fungi(100);p__Ascomycota(100);c__Saccharomycetes(100);o__Saccharomycetales(100);f__Metschnikowiaceae(100);g__Metschnikowia(100);s__Metschnikowia_pulcherrima(100); | Metschnikowia | 13 | Saprotroph | \|Undefined Saprotroph\| | Yeast | NULL | Probable | As nectar-tap_saprotroph (Põlme et al. 2020); Decay sugar-rich substrates (Põlme et al. 2020) | Põlme S, et al. 2020. Fungal Diversity 105:1-16 ((https://doi.org/10.1007/s13225-020-00466-2)) |
| Otu11208 | 2 | k__Fungi(100);p__Basidiomycota(100);c__Agaricomycetes(100);o__Russulales(100);f__Russulaceae(100);g__Russula(100);g__Russula_unclassified(100); | Russula | 13 | Symbiotroph | \|Ectomycorrhizal\| | Agaricoid | NULL | Highly Probable | NULL | Rinaldi AC, et al. 2008. Fungal Diversity 33:1-45 ((http://www.fungaldiversity.org/fdp/sfdp/33-1.pdf)); Tedersoo L, et al. 2010. Mycorrhiza 20:217-263 ((https://doi.org/10.1007/s00572-009-0274-x)); Põlme S, et al. 2020. Fungal Diversity 105:1-16 ((https://doi.org/10.1007/s13225-020-00466-2)); http://mycorrhizas.info/ecmf.html ((http://mycorrhizas.info/ecmf.html)) |
| Otu11210 | 2 | k__Fungi(100);p__Ascomycota(100);c__Saccharomycetes(100);o__Saccharomycetales(100);f__Metschnikowiaceae(100);g__Metschnikowia(100);s__Metschnikowia_pulcherrima(100); | Metschnikowia | 13 | Saprotroph | \|Undefined Saprotroph\| | Yeast | NULL | Probable | As nectar-tap_saprotroph (Põlme et al. 2020); Decay sugar-rich substrates (Põlme et al. 2020) | Põlme S, et al. 2020. Fungal Diversity 105:1-16 ((https://doi.org/10.1007/s13225-020-00466-2)) |
| Otu11211 | 2 | k__Fungi(100);p__Basidiomycota(100);c__Agaricomycetes(100);o__Thelephorales(100);f__Thelephoraceae(100);f__Thelephoraceae_unclassified(100);f__Thelephoraceae_unclassified(100); | Thelephoraceae | 9 | Saprotroph-Symbiotroph | Ectomycorrhizal-Undefined Saprotroph | Clavarioid | NULL | Probable | On the ground, on soil, a few on wood. At least some speices ectomycorrhizal (Cannon & Kirk 2007) | Cannon PF, Kirk PM. 2007. Fungal Families of the World. CAB International, Cambridge (ISBN: 978-0851998275) |
| Otu11213 | 2 | k__Fungi(100);p__Basidiomycota(100);c__Microbotryomycetes(100);o__Sporidiobolales(100);f__Sporidiobolales_family_Incertae_sedis(100);g__Sporobolomyces(100);s__Sporobolomyces_lactosus(100); | Sporobolomyces | 13 | Pathotroph-Saprotroph | \|Fungal Parasite\|-Plant Saprotroph | Yeast | NULL | Probable | As litter_saprotroph (Põlme et al. 2020); As mycoparasite (Põlme et al. 2020); Decay fungal material (Põlme et al. 2020) | Tedersoo L, et al. 2014. Science 346:e1256688 ((https://doi.org/10.1126/science.1256688)); Purahong W, et al. 2016. Molecular Ecology 25:4059-4074 ((https://doi.org/10.1111/mec.13739)); Põlme S, et al. 2020. Fungal Diversity 105:1-16 ((https://doi.org/10.1007/s13225-020-00466-2)) |
| Otu11215 | 2 | k__Fungi(100);p__Ascomycota(100);c__Saccharomycetes(100);o__Saccharomycetales(100);f__Metschnikowiaceae(100);g__Metschnikowia(100);s__Metschnikowia_pulcherrima(100); | Metschnikowia | 13 | Saprotroph | \|Undefined Saprotroph\| | Yeast | NULL | Probable | As nectar-tap_saprotroph (Põlme et al. 2020); Decay sugar-rich substrates (Põlme et al. 2020) | Põlme S, et al. 2020. Fungal Diversity 105:1-16 ((https://doi.org/10.1007/s13225-020-00466-2)) |
| Otu11216 | 2 | k__Fungi(100);p__Ascomycota(100);c__Saccharomycetes(100);o__Saccharomycetales(100);f__Metschnikowiaceae(100);g__Metschnikowia(100);s__Metschnikowia_pulcherrima(100); | Metschnikowia | 13 | Saprotroph | \|Undefined Saprotroph\| | Yeast | NULL | Probable | As nectar-tap_saprotroph (Põlme et al. 2020); Decay sugar-rich substrates (Põlme et al. 2020) | Põlme S, et al. 2020. Fungal Diversity 105:1-16 ((https://doi.org/10.1007/s13225-020-00466-2)) |
| Otu11217 | 2 | k__Fungi(100);p__Ascomycota(100);c__Sordariomycetes(100);o__Hypocreales(100);f__Hypocreales_family_Incertae_sedis(100);f__Hypocreales_family_Incertae_sedis_unclassified(100);f__Hypocreales_family_Incertae_sedis_unclassified(100); | Hypocreales | 7 | Saprotroph | Undefined Saprotroph | Microfungus | NULL | Possible | NULL | Sterkenburg E, et al. 2015. New Phytologist 207:1145-1158 ((https://doi.org/10.1111/nph.13426)) |
| Otu11219 | 2 | k__Fungi(100);p__Ascomycota(100);c__Saccharomycetes(100);o__Saccharomycetales(100);f__Saccharomycetales_family_Incertae_sedis(100);g__Candida(100);g__Candida_unclassified(100); | Candida | 13 | Pathotroph-Saprotroph-Symbiotroph | Animal Pathogen-Endophyte-Endosymbiont-Epiphyte-\|Undefined Saprotroph\| | Dimorphic Yeast-Yeast | NULL | Probable | As nectar-tap_saprotroph (Põlme et al. 2020); Decay sugar-rich substrates (Põlme et al. 2020); This asexual genus is polyphyletic and contain species that function very differently depend on what they are. Caution should be taken when interpreting guilds for Candida species. | Manolakaki D, et al. 2010. Virulence. 1: 367-375 ((https://doi.org/10.4161/viru.1.5.12796)); Mekbib SB, et al. 2011. Tropical Plant Pathology 36:233-240 ((https://doi.org/10.1590/S1982-56762011000400004)); Põlme S, et al. 2020. Fungal Diversity 105:1-16 ((https://doi.org/10.1007/s13225-020-00466-2)) |
| Otu11226 | 2 | k__Fungi(100);p__Ascomycota(100);c__Saccharomycetes(100);o__Saccharomycetales(100);f__Metschnikowiaceae(100);g__Metschnikowia(100);g__Metschnikowia_unclassified(100); | Metschnikowia | 13 | Saprotroph | \|Undefined Saprotroph\| | Yeast | NULL | Probable | As nectar-tap_saprotroph (Põlme et al. 2020); Decay sugar-rich substrates (Põlme et al. 2020) | Põlme S, et al. 2020. Fungal Diversity 105:1-16 ((https://doi.org/10.1007/s13225-020-00466-2)) |
| Otu11228 | 2 | k__Fungi(100);p__Ascomycota(100);c__Sordariomycetes(100);o__Hypocreales(100);f__Nectriaceae(100);f__Nectriaceae_unclassified(100);f__Nectriaceae_unclassified(100); | Nectriaceae | 9 | Pathotroph-Saprotroph-Symbiotroph | Animal Pathogen-Endophyte-Fungal Parasite-Lichen Parasite-Plant Pathogen-Wood Saprotroph | Microfungus | NULL | Possible | Fusarium species are amongst the economically significant plant pathogens (Cannon & Kirk 2007) | Cannon PF, Kirk PM. 2007. Fungal Families of the World. CAB International, Cambridge (ISBN: 978-0851998275); Hirooka Y, et al. 2012. Studies in Mycology 711-210 ((https://doi.org/10.3114/sim0001)); Tedersoo L, et al. 2014. Science 346:e1256688 ((https://doi.org/10.1126/science.1256688)); The American Phytopathological Society Common Names of Plant Diseases ((http://www.apsnet.org/publications/commonnames/Pages/default.aspx)); https://www.cdc.gov ((https://www.cdc.gov)) |
| Otu11229 | 2 | k__Fungi(100);p__Ascomycota(100);c__Saccharomycetes(100);o__Saccharomycetales(100);f__Metschnikowiaceae(100);g__Metschnikowia(100);g__Metschnikowia_unclassified(100); | Metschnikowia | 13 | Saprotroph | \|Undefined Saprotroph\| | Yeast | NULL | Probable | As nectar-tap_saprotroph (Põlme et al. 2020); Decay sugar-rich substrates (Põlme et al. 2020) | Põlme S, et al. 2020. Fungal Diversity 105:1-16 ((https://doi.org/10.1007/s13225-020-00466-2)) |
| Otu11230 | 2 | k__Fungi(100);p__Ascomycota(100);c__Saccharomycetes(100);o__Saccharomycetales(100);f__Metschnikowiaceae(100);g__Metschnikowia(100);g__Metschnikowia_unclassified(100); | Metschnikowia | 13 | Saprotroph | \|Undefined Saprotroph\| | Yeast | NULL | Probable | As nectar-tap_saprotroph (Põlme et al. 2020); Decay sugar-rich substrates (Põlme et al. 2020) | Põlme S, et al. 2020. Fungal Diversity 105:1-16 ((https://doi.org/10.1007/s13225-020-00466-2)) |
| Otu11231 | 2 | k__Fungi(100);p__Ascomycota(100);c__Sordariomycetes(100);o__Hypocreales(100);o__Hypocreales_unclassified(100);o__Hypocreales_unclassified(100);o__Hypocreales_unclassified(100); | Hypocreales | 7 | Saprotroph | Undefined Saprotroph | Microfungus | NULL | Possible | NULL | Sterkenburg E, et al. 2015. New Phytologist 207:1145-1158 ((https://doi.org/10.1111/nph.13426)) |
| Otu11232 | 2 | k__Fungi(100);p__Ascomycota(100);c__Sordariomycetes(100);o__Diaporthales(100);f__Gnomoniaceae(100);g__Gnomonia(100);s__Gnomonia_rostellata(100); | Gnomonia | 13 | Pathotroph-Saprotroph-Symbiotroph | \|Endophyte\|-Plant Pathogen-Plant Saprotroph-Undefined Saprotroph | Microfungus | NULL | Probable | As foliar_endophyte (Põlme et al. 2020); As litter_saprotroph (Põlme et al. 2020); Decay leaf, fruit, seed (Põlme et al. 2020); Most species are saprobes or endophytes in leaves and bark, but a few are neurotrophic pathogens (Cannon & Kirk 2007) | Cannon PF, Kirk PM. 2007. Fungal Families of the World. CAB International, Cambridge (ISBN: 978-0851998275); Tedersoo L, et al. 2014. Science 346:e1256688 ((https://doi.org/10.1126/science.1256688)); Põlme S, et al. 2020. Fungal Diversity 105:1-16 ((https://doi.org/10.1007/s13225-020-00466-2)) |
| Otu11233 | 2 | k__Fungi(100);p__Ascomycota(100);c__Sordariomycetes(100);o__Hypocreales(100);o__Hypocreales_unclassified(100);o__Hypocreales_unclassified(100);o__Hypocreales_unclassified(100); | Hypocreales | 7 | Saprotroph | Undefined Saprotroph | Microfungus | NULL | Possible | NULL | Sterkenburg E, et al. 2015. New Phytologist 207:1145-1158 ((https://doi.org/10.1111/nph.13426)) |
| Otu11234 | 2 | k__Fungi(100);p__Ascomycota(100);c__Sordariomycetes(100);o__Xylariales(100);f__Amphisphaeriaceae(100);f__Amphisphaeriaceae_unclassified(100);f__Amphisphaeriaceae_unclassified(100); | Amphisphaeriaceae | 9 | Saprotroph | Undefined Saprotroph | Microfungus | NULL | Possible | Saprobes or weak parasites of leaves, stems, etc., often appearing as secondary colonizers following damage to plants by other agents (Cannon & Kirk 2007) | Cannon PF, Kirk PM. 2007. Fungal Families of the World. CAB International, Cambridge (ISBN: 978-0851998275) |
| Otu11235 | 2 | k__Fungi(100);p__Ascomycota(100);c__Ascomycota_class_Incertae_sedis(100);o__Ascomycota_order_Incertae_sedis(100);f__Ascomycota_family_Incertae_sedis(100);g__Phialemoniopsis(100);s__Phialemoniopsis_cornearis(100); | Phialemoniopsis | 13 | Pathotroph-Saprotroph | \|Animal Parasite\|-Undefined Saprotroph | Microfungus | NULL | Probable | As animal_decomposer (Põlme et al. 2020); Decay animal material (Põlme et al. 2020) | Põlme S, et al. 2020. Fungal Diversity 105:1-16 ((https://doi.org/10.1007/s13225-020-00466-2)) |
| Otu11236 | 2 | k__Fungi(100);p__Ascomycota(100);c__Sordariomycetes(100);o__Microascales(100);f__Halosphaeriaceae(100);g__unclassified_Halosphaeriaceae(100);s__Halosphaeriaceae_sp(100); | Halosphaeriaceae | 9 | Saprotroph | Undefined Saprotroph-Wood Saprotroph | Microfungus | NULL | Probable | Found in aquatic habitats, mostly in marine environments; saprobic on submerged or intertidal wood, occasionally associated with sand grains and similar substrata (Cannon & Kirk 2007) | Cannon PF, Kirk PM. 2007. Fungal Families of the World. CAB International, Cambridge (ISBN: 978-0851998275) |
| Otu11237 | 2 | k__Fungi(100);p__Ascomycota(100);c__Dothideomycetes(100);o__Botryosphaeriales(100);f__Botryosphaeriaceae(100);f__Botryosphaeriaceae_unclassified(100);f__Botryosphaeriaceae_unclassified(100); | Botryosphaeriaceae | 9 | Pathotroph-Saprotroph-Symbiotroph | Endophyte-Plant Pathogen-Wood Saprotroph | Microfungus | NULL | Probable | Biotrophic, necrotrophic, or saprobic, especially associated with woody tissue (Cannon & Kirk 2007) | Cannon PF, Kirk PM. 2007. Fungal Families of the World. CAB International, Cambridge (ISBN: 978-0851998275) |
| Otu11239 | 2 | k__Fungi(100);p__Ascomycota(100);c__Dothideomycetes(100);o__Pleosporales(100);f__Sporormiaceae(100);f__Sporormiaceae_unclassified(100);f__Sporormiaceae_unclassified(100); | Sporormiaceae | 9 | Saprotroph | Dung Saprotroph-Plant Saprotroph | Microfungus | NULL | Probable | Saprobic, associated with dung and rotting vegetation (Cannon & Kirk 2007) | Cannon PF, Kirk PM. 2007. Fungal Families of the World. CAB International, Cambridge (ISBN: 978-0851998275) |
| Otu11240 | 2 | k__Fungi(100);p__Ascomycota(100);c__Dothideomycetes(100);o__Pleosporales(100);f__Pleosporaceae(100);g__Stemphylium(100);s__Stemphylium_sp_1662(100); | Stemphylium | 13 | Pathotroph-Saprotroph | \|Plant Pathogen\|-Plant Saprotroph-Wood Saprotroph | Microfungus | Soft Rot | Highly Probable | As litter_saprotroph (Põlme et al. 2020); Decay leaf, fruit, seed (Põlme et al. 2020) | Duncan CG, Eslyn WE. 1966. Mycologia 58:642-664 ((https://doi.org/10.2307/3757045)); Seehann G, et al. 1975. List of Fungi in Soft-Rot Tissue Tests. Institute fur Holzbiologie und Holzschutz, Bundesforschungsanstalt fur Forst - und Holzwirtshaft, Hamburg - Lohbrugge, Germany ((https://www.irg-wp.com/irgdocs/details.php?d6493033-c9ea-40f3-8e3e-5697f32d640d)); Costa IPMW, et al. 2012. Mycotaxon 119:e26 ((http://mycotaxon.com/resources/checklists/costa_v119_checklist.pdf)); Tedersoo L, et al. 2014. Science 346:e1256688 ((https://doi.org/10.1126/science.1256688)); Põlme S, et al. 2020. Fungal Diversity 105:1-16 ((https://doi.org/10.1007/s13225-020-00466-2)); The American Phytopathological Society Common Names of Plant Diseases ((https://www.apsnet.org/edcenter/resources/commonnames/Pages/default.aspx)) |
| Otu11241 | 2 | k__Fungi(100);p__Ascomycota(100);c__Sordariomycetes(100);o__Microascales(100);f__Microascaceae(100);g__Microascus(100);s__Microascus_sp_r249(100); | Microascus | 13 | Pathotroph-Saprotroph-Symbiotroph | Animal Pathogen-Dung Saprotroph-Endophyte-Plant Pathogen-Plant Saprotroph-\|Undefined Saprotroph\| | Microfungus | NULL | Possible | As foliar_endophyte (Põlme et al. 2020); As litter_saprotroph (Põlme et al. 2020); As soil_saprotroph (Põlme et al. 2020); Decay leaf, fruit, seed, soil (Põlme et al. 2020) | James TY, et al. 2006. Nature 443:818-822 ((https://doi.org/10.1038/nature05110)); Costa IPMW, et al. 2012. Mycotaxon 119:e26 (as Endophyte) ((http://mycotaxon.com/resources/checklists/costa_v119_checklist.pdf)); Bills GF, et al. 2013. Current Opinion in Microbiology 16:549-565 ((https://doi.org/10.1016/j.mib.2013.08.001)); Irinyi L, et al. 2015. Medical Mycology 53:313-337 ((https://doi.org/10.1093/mmy/myv008)); Põlme S, et al. 2020. Fungal Diversity 105:1-16 ((https://doi.org/10.1007/s13225-020-00466-2)) |
| Otu11242 | 2 | k__Fungi(100);p__Ascomycota(100);c__Saccharomycetes(100);o__Saccharomycetales(100);f__Metschnikowiaceae(100);g__Metschnikowia(100);s__Metschnikowia_pulcherrima(100); | Metschnikowia | 13 | Saprotroph | \|Undefined Saprotroph\| | Yeast | NULL | Probable | As nectar-tap_saprotroph (Põlme et al. 2020); Decay sugar-rich substrates (Põlme et al. 2020) | Põlme S, et al. 2020. Fungal Diversity 105:1-16 ((https://doi.org/10.1007/s13225-020-00466-2)) |
| Otu11244 | 2 | k__Fungi(100);p__Ascomycota(100);c__Saccharomycetes(100);o__Saccharomycetales(100);f__Metschnikowiaceae(100);g__Metschnikowia(100);s__Metschnikowia_pulcherrima(100); | Metschnikowia | 13 | Saprotroph | \|Undefined Saprotroph\| | Yeast | NULL | Probable | As nectar-tap_saprotroph (Põlme et al. 2020); Decay sugar-rich substrates (Põlme et al. 2020) | Põlme S, et al. 2020. Fungal Diversity 105:1-16 ((https://doi.org/10.1007/s13225-020-00466-2)) |
| Otu11247 | 2 | k__Fungi(100);p__Ascomycota(100);c__Saccharomycetes(100);o__Saccharomycetales(100);f__Metschnikowiaceae(100);g__Metschnikowia(100);g__Metschnikowia_unclassified(100); | Metschnikowia | 13 | Saprotroph | \|Undefined Saprotroph\| | Yeast | NULL | Probable | As nectar-tap_saprotroph (Põlme et al. 2020); Decay sugar-rich substrates (Põlme et al. 2020) | Põlme S, et al. 2020. Fungal Diversity 105:1-16 ((https://doi.org/10.1007/s13225-020-00466-2)) |
| Otu11248 | 2 | k__Fungi(100);p__Ascomycota(100);c__Sordariomycetes(100);o__Diaporthales(100);f__Diaporthales_family_Incertae_sedis(100);g__Ophiognomonia(100);s__Ophiognomonia_melanostyla(100); | Ophiognomonia | 13 | Pathotroph | \|Plant Pathogen\| | Microfungus | NULL | Highly Probable | See Marssoniella juglandis, Ophiognomonia leptostyla, and Syn. Gnomonia leptostyla; Causes leaf blotch and spots, also known as walnut anthracnose or walnut black spot (Sogonov et al. 2008) | Sogonov MV, et al. 2008. Studies in Mycology 62:1-79 ((https://www.studiesinmycology.org/sim/Sim62/03_Leaf-inhabiting_genera_of_the_Gnomoniaceae,_Diaporthales.pdf)); Põlme S, et al. 2020. Fungal Diversity 105:1-16 ((https://doi.org/10.1007/s13225-020-00466-2)) |
| Otu11249 | 2 | k__Fungi(100);p__Ascomycota(100);c__Dothideomycetes(100);o__Pleosporales(100);f__Phaeosphaeriaceae(100);g__Phaeosphaeria(100);s__Phaeosphaeria_triglochinicola(100); | Phaeosphaeria | 13 | Pathotroph-Saprotroph-Symbiotroph | Endophyte-Plant Pathogen-\|Plant Saprotroph\| | Microfungus | NULL | Probable | As litter_saprotroph (Põlme et al. 2020); Decay leaf, fruit, seed (Põlme et al. 2020) | Tedersoo L, et al. 2014. Science 346:e1256688 ((https://doi.org/10.1126/science.1256688));David AS, et al. 2016. Microbial Ecology 71: 912-926 ((https://doi.org/10.1007/s00248-015-0712-6)); Põlme S, et al. 2020. Fungal Diversity 105:1-16 ((https://doi.org/10.1007/s13225-020-00466-2)) |
| Otu11252 | 2 | k__Fungi(100);p__Ascomycota(100);c__Saccharomycetes(100);o__Saccharomycetales(100);f__Metschnikowiaceae(100);g__Metschnikowia(100);s__Metschnikowia_pulcherrima(100); | Metschnikowia | 13 | Saprotroph | \|Undefined Saprotroph\| | Yeast | NULL | Probable | As nectar-tap_saprotroph (Põlme et al. 2020); Decay sugar-rich substrates (Põlme et al. 2020) | Põlme S, et al. 2020. Fungal Diversity 105:1-16 ((https://doi.org/10.1007/s13225-020-00466-2)) |
| Otu11253 | 2 | k__Fungi(100);p__Ascomycota(100);c__Saccharomycetes(100);o__Saccharomycetales(100);f__Metschnikowiaceae(100);g__Metschnikowia(100);g__Metschnikowia_unclassified(100); | Metschnikowia | 13 | Saprotroph | \|Undefined Saprotroph\| | Yeast | NULL | Probable | As nectar-tap_saprotroph (Põlme et al. 2020); Decay sugar-rich substrates (Põlme et al. 2020) | Põlme S, et al. 2020. Fungal Diversity 105:1-16 ((https://doi.org/10.1007/s13225-020-00466-2)) |
| Otu11254 | 2 | k__Fungi(100);p__Ascomycota(100);c__Sordariomycetes(100);o__Microascales(100);f__Microascaceae(100);g__Microascus(100);g__Microascus_unclassified(100); | Microascus | 13 | Pathotroph-Saprotroph-Symbiotroph | Animal Pathogen-Dung Saprotroph-Endophyte-Plant Pathogen-Plant Saprotroph-\|Undefined Saprotroph\| | Microfungus | NULL | Possible | As foliar_endophyte (Põlme et al. 2020); As litter_saprotroph (Põlme et al. 2020); As soil_saprotroph (Põlme et al. 2020); Decay leaf, fruit, seed, soil (Põlme et al. 2020) | James TY, et al. 2006. Nature 443:818-822 ((https://doi.org/10.1038/nature05110)); Costa IPMW, et al. 2012. Mycotaxon 119:e26 (as Endophyte) ((http://mycotaxon.com/resources/checklists/costa_v119_checklist.pdf)); Bills GF, et al. 2013. Current Opinion in Microbiology 16:549-565 ((https://doi.org/10.1016/j.mib.2013.08.001)); Irinyi L, et al. 2015. Medical Mycology 53:313-337 ((https://doi.org/10.1093/mmy/myv008)); Põlme S, et al. 2020. Fungal Diversity 105:1-16 ((https://doi.org/10.1007/s13225-020-00466-2)) |
| Otu11255 | 2 | k__Fungi(100);p__Ascomycota(100);c__Sordariomycetes(100);o__Hypocreales(100);o__Hypocreales_unclassified(100);o__Hypocreales_unclassified(100);o__Hypocreales_unclassified(100); | Hypocreales | 7 | Saprotroph | Undefined Saprotroph | Microfungus | NULL | Possible | NULL | Sterkenburg E, et al. 2015. New Phytologist 207:1145-1158 ((https://doi.org/10.1111/nph.13426)) |
| Otu11256 | 2 | k__Fungi(100);p__Ascomycota(100);c__Saccharomycetes(100);o__Saccharomycetales(100);f__Metschnikowiaceae(100);g__Metschnikowia(100);s__Metschnikowia_pulcherrima(100); | Metschnikowia | 13 | Saprotroph | \|Undefined Saprotroph\| | Yeast | NULL | Probable | As nectar-tap_saprotroph (Põlme et al. 2020); Decay sugar-rich substrates (Põlme et al. 2020) | Põlme S, et al. 2020. Fungal Diversity 105:1-16 ((https://doi.org/10.1007/s13225-020-00466-2)) |
| Otu11257 | 2 | k__Fungi(100);p__Ascomycota(100);c__Sordariomycetes(100);o__Xylariales(100);f__Xylariaceae(100);g__unclassified_Xylariaceae(100);s__Xylariaceae_sp(100); | Xylariaceae | 9 | Pathotroph-Saprotroph-Symbiotroph | Dung Saprotroph-Endophyte-Plant Pathogen-Undefined Saprotroph | Xylarioid | NULL | Probable | Saprobes or weak pathogens, many endophytes, mainly in wood and bark, occasionally in soil and dung (Cannon & Kirk 2007) | Cannon PF, Kirk PM. 2007. Fungal Families of the World. CAB International, Cambridge (ISBN: 978-0851998275) |
| Otu11258 | 2 | k__Fungi(100);p__Ascomycota(100);c__Saccharomycetes(100);o__Saccharomycetales(100);f__Metschnikowiaceae(100);g__Metschnikowia(100);s__Metschnikowia_pulcherrima(100); | Metschnikowia | 13 | Saprotroph | \|Undefined Saprotroph\| | Yeast | NULL | Probable | As nectar-tap_saprotroph (Põlme et al. 2020); Decay sugar-rich substrates (Põlme et al. 2020) | Põlme S, et al. 2020. Fungal Diversity 105:1-16 ((https://doi.org/10.1007/s13225-020-00466-2)) |
| Otu11259 | 2 | k__Fungi(100);p__Ascomycota(100);c__Saccharomycetes(100);o__Saccharomycetales(100);f__Metschnikowiaceae(100);g__Metschnikowia(100);g__Metschnikowia_unclassified(100); | Metschnikowia | 13 | Saprotroph | \|Undefined Saprotroph\| | Yeast | NULL | Probable | As nectar-tap_saprotroph (Põlme et al. 2020); Decay sugar-rich substrates (Põlme et al. 2020) | Põlme S, et al. 2020. Fungal Diversity 105:1-16 ((https://doi.org/10.1007/s13225-020-00466-2)) |
| Otu11261 | 2 | k__Fungi(100);p__Ascomycota(100);c__Saccharomycetes(100);o__Saccharomycetales(100);f__Metschnikowiaceae(100);g__Metschnikowia(100);s__Metschnikowia_pulcherrima(100); | Metschnikowia | 13 | Saprotroph | \|Undefined Saprotroph\| | Yeast | NULL | Probable | As nectar-tap_saprotroph (Põlme et al. 2020); Decay sugar-rich substrates (Põlme et al. 2020) | Põlme S, et al. 2020. Fungal Diversity 105:1-16 ((https://doi.org/10.1007/s13225-020-00466-2)) |
| Otu11262 | 2 | k__Fungi(100);p__Ascomycota(100);c__Sordariomycetes(100);o__Hypocreales(100);f__Hypocreales_family_Incertae_sedis(100);g__Stilbella(100);s__Stilbella_sp(100); | Stilbella | 13 | Pathotroph-Saprotroph-Symbiotroph | \|Dung Saprotroph\|-Endophyte-Plant Pathogen-Plant Saprotroph-Wood Saprotroph | Microfungus | Soft Rot | Possible | As foliar_endophyte (Põlme et al. 2020); As litter_saprotroph (Põlme et al. 2020); Decay leaf, fruit, seed, dung (Põlme et al. 2020) | Duncan CG, Eslyn WE. 1966. Mycologia 58:642-645 ((https://doi.org/10.2307/3757045)); Tedersoo L, et al. 2014. Science 346:e1256688 ((https://doi.org/10.1126/science.1256688)); Põlme S, et al. 2020. Fungal Diversity 105:1-16 ((https://doi.org/10.1007/s13225-020-00466-2)) |
| Otu11263 | 2 | k__Fungi(100);p__Ascomycota(100);c__Saccharomycetes(100);o__Saccharomycetales(100);f__Metschnikowiaceae(100);g__Metschnikowia(100);g__Metschnikowia_unclassified(100); | Metschnikowia | 13 | Saprotroph | \|Undefined Saprotroph\| | Yeast | NULL | Probable | As nectar-tap_saprotroph (Põlme et al. 2020); Decay sugar-rich substrates (Põlme et al. 2020) | Põlme S, et al. 2020. Fungal Diversity 105:1-16 ((https://doi.org/10.1007/s13225-020-00466-2)) |
| Otu11265 | 2 | k__Fungi(100);p__Ascomycota(100);c__Saccharomycetes(100);o__Saccharomycetales(100);f__Metschnikowiaceae(100);g__Metschnikowia(100);s__Metschnikowia_pulcherrima(100); | Metschnikowia | 13 | Saprotroph | \|Undefined Saprotroph\| | Yeast | NULL | Probable | As nectar-tap_saprotroph (Põlme et al. 2020); Decay sugar-rich substrates (Põlme et al. 2020) | Põlme S, et al. 2020. Fungal Diversity 105:1-16 ((https://doi.org/10.1007/s13225-020-00466-2)) |
| Otu11266 | 2 | k__Fungi(100);p__Ascomycota(100);c__Saccharomycetes(100);o__Saccharomycetales(100);f__Metschnikowiaceae(100);g__Metschnikowia(100);g__Metschnikowia_unclassified(100); | Metschnikowia | 13 | Saprotroph | \|Undefined Saprotroph\| | Yeast | NULL | Probable | As nectar-tap_saprotroph (Põlme et al. 2020); Decay sugar-rich substrates (Põlme et al. 2020) | Põlme S, et al. 2020. Fungal Diversity 105:1-16 ((https://doi.org/10.1007/s13225-020-00466-2)) |
| Otu11268 | 2 | k__Fungi(100);p__Ascomycota(100);c__Saccharomycetes(100);o__Saccharomycetales(100);f__Metschnikowiaceae(100);g__Metschnikowia(100);s__Metschnikowia_pulcherrima(100); | Metschnikowia | 13 | Saprotroph | \|Undefined Saprotroph\| | Yeast | NULL | Probable | As nectar-tap_saprotroph (Põlme et al. 2020); Decay sugar-rich substrates (Põlme et al. 2020) | Põlme S, et al. 2020. Fungal Diversity 105:1-16 ((https://doi.org/10.1007/s13225-020-00466-2)) |
| Otu11269 | 2 | k__Fungi(100);p__Ascomycota(100);c__Sordariomycetes(100);o__Hypocreales(100);f__Nectriaceae(100);g__Nectria(100);s__Nectria_cinnabarina(100); | Nectria | 13 | Pathotroph-Saprotroph-Symbiotroph | Animal Pathogen-Endophyte-Fungal Parasite-Lichen Parasite-\|Plant Pathogen\|-Plant Saprotroph-Wood Saprotroph | Microfungus | NULL | Possible | As foliar_endophyte (Põlme et al. 2020); As litter_saprotroph (Põlme et al. 2020); Decay wood, leaf, fruit, seed, roots, soil (Põlme et al. 2020) | Hirooka Y, et al. 2012. Studies in Mycology 71:1-210 ((https://doi.org/10.3114/sim0001)); Tedersoo L, et al. 2014. Science 346:e1256688 ((https://doi.org/10.1126/science.1256688)); Põlme S, et al. 2020. Fungal Diversity 105:1-16 ((https://doi.org/10.1007/s13225-020-00466-2)) |
| Otu11270 | 2 | k__Fungi(100);p__Ascomycota(100);c__Saccharomycetes(100);o__Saccharomycetales(100);f__Metschnikowiaceae(100);g__Metschnikowia(100);s__Metschnikowia_pulcherrima(100); | Metschnikowia | 13 | Saprotroph | \|Undefined Saprotroph\| | Yeast | NULL | Probable | As nectar-tap_saprotroph (Põlme et al. 2020); Decay sugar-rich substrates (Põlme et al. 2020) | Põlme S, et al. 2020. Fungal Diversity 105:1-16 ((https://doi.org/10.1007/s13225-020-00466-2)) |
| Otu11272 | 2 | k__Fungi(100);p__Ascomycota(100);c__Sordariomycetes(100);o__Hypocreales(100);o__Hypocreales_unclassified(100);o__Hypocreales_unclassified(100);o__Hypocreales_unclassified(100); | Hypocreales | 7 | Saprotroph | Undefined Saprotroph | Microfungus | NULL | Possible | NULL | Sterkenburg E, et al. 2015. New Phytologist 207:1145-1158 ((https://doi.org/10.1111/nph.13426)) |
| Otu11273 | 2 | k__Fungi(100);p__Ascomycota(100);c__Saccharomycetes(100);o__Saccharomycetales(100);f__Metschnikowiaceae(100);g__Metschnikowia(100);g__Metschnikowia_unclassified(100); | Metschnikowia | 13 | Saprotroph | \|Undefined Saprotroph\| | Yeast | NULL | Probable | As nectar-tap_saprotroph (Põlme et al. 2020); Decay sugar-rich substrates (Põlme et al. 2020) | Põlme S, et al. 2020. Fungal Diversity 105:1-16 ((https://doi.org/10.1007/s13225-020-00466-2)) |
| Otu11274 | 2 | k__Fungi(100);p__Ascomycota(100);c__Saccharomycetes(100);o__Saccharomycetales(100);f__Saccharomycetales_family_Incertae_sedis(100);g__Candida(100);s__Candida_sp_AS23486(100); | Candida | 13 | Pathotroph-Saprotroph-Symbiotroph | Animal Pathogen-Endophyte-Endosymbiont-Epiphyte-\|Undefined Saprotroph\| | Dimorphic Yeast-Yeast | NULL | Probable | As nectar-tap_saprotroph (Põlme et al. 2020); Decay sugar-rich substrates (Põlme et al. 2020); This asexual genus is polyphyletic and contain species that function very differently depend on what they are. Caution should be taken when interpreting guilds for Candida species. | Manolakaki D, et al. 2010. Virulence. 1: 367-375 ((https://doi.org/10.4161/viru.1.5.12796)); Mekbib SB, et al. 2011. Tropical Plant Pathology 36:233-240 ((https://doi.org/10.1590/S1982-56762011000400004)); Põlme S, et al. 2020. Fungal Diversity 105:1-16 ((https://doi.org/10.1007/s13225-020-00466-2)) |
| Otu11275 | 2 | k__Fungi(100);p__Ascomycota(100);c__Saccharomycetes(100);o__Saccharomycetales(100);f__Metschnikowiaceae(100);g__Metschnikowia(100);g__Metschnikowia_unclassified(100); | Metschnikowia | 13 | Saprotroph | \|Undefined Saprotroph\| | Yeast | NULL | Probable | As nectar-tap_saprotroph (Põlme et al. 2020); Decay sugar-rich substrates (Põlme et al. 2020) | Põlme S, et al. 2020. Fungal Diversity 105:1-16 ((https://doi.org/10.1007/s13225-020-00466-2)) |
| Otu11278 | 2 | k__Fungi(100);p__Ascomycota(100);c__Saccharomycetes(100);o__Saccharomycetales(100);f__Metschnikowiaceae(100);g__Metschnikowia(100);s__Metschnikowia_pulcherrima(100); | Metschnikowia | 13 | Saprotroph | \|Undefined Saprotroph\| | Yeast | NULL | Probable | As nectar-tap_saprotroph (Põlme et al. 2020); Decay sugar-rich substrates (Põlme et al. 2020) | Põlme S, et al. 2020. Fungal Diversity 105:1-16 ((https://doi.org/10.1007/s13225-020-00466-2)) |
| Otu11279 | 2 | k__Fungi(100);p__Ascomycota(100);c__Sordariomycetes(100);o__Hypocreales(100);f__Hypocreales_family_Incertae_sedis(100);g__Acremonium(100);s__Acremonium_cavaraeanum(100); | Acremonium | 13 | Pathotroph-Saprotroph-Symbiotroph | Animal Pathogen-Endophyte-Fungal Parasite-Plant Pathogen-\|Undefined Saprotroph\|-Wood Saprotroph | Microfungus | Soft Rot | Possible | As foliar_endophyte (Põlme et al. 2020); culture (Hanada et al. 2010, Del Olmo; Decay leaf, fruit, seed, soil, dung, animal material (Põlme et al. 2020); Endophyte detection method; Ruiz & Arnold 2014); Likely opportunistic human pathogen (Irinyi et al. 2015) | Duncan CG, Eslyn WE. 1966. Mycologia 58:642-645 ((https://doi.org/10.2307/3757045)); Nilsson T. 1973. Studia Forestalia Suecica 104: 1-40 ((https://pub.epsilon.slu.se/5711/)); Worrall JJ, et al. 1997. Mycologia 89:199-219 ((https://doi.org/10.2307/3761073)); Hanada RE, et al. 2010. Fungal Biology 114:901-910 ((https://doi.org/10.1016/j.funbio.2010.08.006)); Del Olmo-Ruiz M, Arnold AE. 2014. Mycologia 106: 8-21 ((https://doi.org/10.3852/13-098)); Irinyi L, et al. 2015. Medical Mycology 53313-337 ((https://doi.org/10.1093/mmy/myv008)); Põlme S, et al. 2020. Fungal Diversity 105:1-16 ((https://doi.org/10.1007/s13225-020-00466-2)) |
| Otu11281 | 2 | k__Fungi(100);p__Ascomycota(100);c__Saccharomycetes(100);o__Saccharomycetales(100);f__Metschnikowiaceae(100);g__Metschnikowia(100);g__Metschnikowia_unclassified(100); | Metschnikowia | 13 | Saprotroph | \|Undefined Saprotroph\| | Yeast | NULL | Probable | As nectar-tap_saprotroph (Põlme et al. 2020); Decay sugar-rich substrates (Põlme et al. 2020) | Põlme S, et al. 2020. Fungal Diversity 105:1-16 ((https://doi.org/10.1007/s13225-020-00466-2)) |
| Otu11282 | 2 | k__Fungi(100);p__Ascomycota(100);c__Sordariomycetes(100);o__Hypocreales(100);o__Hypocreales_unclassified(100);o__Hypocreales_unclassified(100);o__Hypocreales_unclassified(100); | Hypocreales | 7 | Saprotroph | Undefined Saprotroph | Microfungus | NULL | Possible | NULL | Sterkenburg E, et al. 2015. New Phytologist 207:1145-1158 ((https://doi.org/10.1111/nph.13426)) |
| Otu11286 | 2 | k__Fungi(100);p__Ascomycota(100);c__Saccharomycetes(100);o__Saccharomycetales(100);f__Metschnikowiaceae(100);g__Metschnikowia(100);s__Metschnikowia_pulcherrima(100); | Metschnikowia | 13 | Saprotroph | \|Undefined Saprotroph\| | Yeast | NULL | Probable | As nectar-tap_saprotroph (Põlme et al. 2020); Decay sugar-rich substrates (Põlme et al. 2020) | Põlme S, et al. 2020. Fungal Diversity 105:1-16 ((https://doi.org/10.1007/s13225-020-00466-2)) |
| Otu11287 | 2 | k__Fungi(100);p__Ascomycota(100);c__Saccharomycetes(100);o__Saccharomycetales(100);f__Metschnikowiaceae(100);g__Metschnikowia(100);s__Metschnikowia_pulcherrima(100); | Metschnikowia | 13 | Saprotroph | \|Undefined Saprotroph\| | Yeast | NULL | Probable | As nectar-tap_saprotroph (Põlme et al. 2020); Decay sugar-rich substrates (Põlme et al. 2020) | Põlme S, et al. 2020. Fungal Diversity 105:1-16 ((https://doi.org/10.1007/s13225-020-00466-2)) |
| Otu11290 | 2 | k__Fungi(100);p__Ascomycota(100);c__Leotiomycetes(100);o__Helotiales(100);f__Hyaloscyphaceae(100);f__Hyaloscyphaceae_unclassified(100);f__Hyaloscyphaceae_unclassified(100); | Hyaloscyphaceae | 9 | Saprotroph | Plant Saprotroph-Wood Saprotroph | Microfungus | NULL | Probable | Saprobic on woody and herbaceous material (Cannon & Kirk 2007) | Cannon PF, Kirk PM. 2007. Fungal Families of the World. CAB International, Cambridge (ISBN: 978-0851998275) |
| Otu11292 | 2 | k__Fungi(100);p__Ascomycota(100);c__Saccharomycetes(100);o__Saccharomycetales(100);f__Metschnikowiaceae(100);g__Metschnikowia(100);s__Metschnikowia_pulcherrima(100); | Metschnikowia | 13 | Saprotroph | \|Undefined Saprotroph\| | Yeast | NULL | Probable | As nectar-tap_saprotroph (Põlme et al. 2020); Decay sugar-rich substrates (Põlme et al. 2020) | Põlme S, et al. 2020. Fungal Diversity 105:1-16 ((https://doi.org/10.1007/s13225-020-00466-2)) |
| Otu11294 | 2 | k__Fungi(100);p__Ascomycota(100);c__Saccharomycetes(100);o__Saccharomycetales(100);f__Metschnikowiaceae(100);g__Metschnikowia(100);s__Metschnikowia_pulcherrima(100); | Metschnikowia | 13 | Saprotroph | \|Undefined Saprotroph\| | Yeast | NULL | Probable | As nectar-tap_saprotroph (Põlme et al. 2020); Decay sugar-rich substrates (Põlme et al. 2020) | Põlme S, et al. 2020. Fungal Diversity 105:1-16 ((https://doi.org/10.1007/s13225-020-00466-2)) |
| Otu11295 | 2 | k__Fungi(100);p__Ascomycota(100);c__Sordariomycetes(100);o__Microascales(100);f__Halosphaeriaceae(100);g__unclassified_Halosphaeriaceae(100);s__Halosphaeriaceae_sp(100); | Halosphaeriaceae | 9 | Saprotroph | Undefined Saprotroph-Wood Saprotroph | Microfungus | NULL | Probable | Found in aquatic habitats, mostly in marine environments; saprobic on submerged or intertidal wood, occasionally associated with sand grains and similar substrata (Cannon & Kirk 2007) | Cannon PF, Kirk PM. 2007. Fungal Families of the World. CAB International, Cambridge (ISBN: 978-0851998275) |
| Otu11296 | 2 | k__Fungi(100);p__Ascomycota(100);c__Saccharomycetes(100);o__Saccharomycetales(100);f__Metschnikowiaceae(100);g__Metschnikowia(100);g__Metschnikowia_unclassified(100); | Metschnikowia | 13 | Saprotroph | \|Undefined Saprotroph\| | Yeast | NULL | Probable | As nectar-tap_saprotroph (Põlme et al. 2020); Decay sugar-rich substrates (Põlme et al. 2020) | Põlme S, et al. 2020. Fungal Diversity 105:1-16 ((https://doi.org/10.1007/s13225-020-00466-2)) |
| Otu11297 | 2 | k__Fungi(100);p__Ascomycota(100);c__Saccharomycetes(100);o__Saccharomycetales(100);f__Metschnikowiaceae(100);g__Metschnikowia(100);s__Metschnikowia_pulcherrima(100); | Metschnikowia | 13 | Saprotroph | \|Undefined Saprotroph\| | Yeast | NULL | Probable | As nectar-tap_saprotroph (Põlme et al. 2020); Decay sugar-rich substrates (Põlme et al. 2020) | Põlme S, et al. 2020. Fungal Diversity 105:1-16 ((https://doi.org/10.1007/s13225-020-00466-2)) |
| Otu11298 | 2 | k__Fungi(100);p__Ascomycota(100);c__Sordariomycetes(100);o__Hypocreales(100);f__Hypocreales_family_Incertae_sedis(100);g__Myrothecium(100);g__Myrothecium_unclassified(100); | Myrothecium | 13 | Pathotroph-Saprotroph | Fungal Parasite-\|Plant Saprotroph\|-Undefined Saprotroph | Microfungus | NULL | Probable | As litter_saprotroph (Põlme et al. 2020); As mycoparasite (Põlme et al. 2020); Decay leaf, fruit, seed (Põlme et al. 2020) | Tedersoo L, et al. 2014. Science 346:e1256688 ((https://doi.org/10.1126/science.1256688)); Põlme S, et al. 2020. Fungal Diversity 105:1-16 ((https://doi.org/10.1007/s13225-020-00466-2)) |
| Otu11300 | 2 | k__Fungi(100);p__Ascomycota(100);c__Sordariomycetes(100);o__Hypocreales(100);f__Nectriaceae(100);g__unclassified_Nectriaceae(100);s__Nectriaceae_sp(100); | Nectriaceae | 9 | Pathotroph-Saprotroph-Symbiotroph | Animal Pathogen-Endophyte-Fungal Parasite-Lichen Parasite-Plant Pathogen-Wood Saprotroph | Microfungus | NULL | Possible | Fusarium species are amongst the economically significant plant pathogens (Cannon & Kirk 2007) | Cannon PF, Kirk PM. 2007. Fungal Families of the World. CAB International, Cambridge (ISBN: 978-0851998275); Hirooka Y, et al. 2012. Studies in Mycology 711-210 ((https://doi.org/10.3114/sim0001)); Tedersoo L, et al. 2014. Science 346:e1256688 ((https://doi.org/10.1126/science.1256688)); The American Phytopathological Society Common Names of Plant Diseases ((http://www.apsnet.org/publications/commonnames/Pages/default.aspx)); https://www.cdc.gov ((https://www.cdc.gov)) |
| Otu11301 | 2 | k__Fungi(100);p__Ascomycota(100);c__Sordariomycetes(100);o__Hypocreales(100);f__Nectriaceae(100);g__Fusicolla(100);s__Fusicolla_violacea(100); | Fusicolla | 13 | Pathotroph | \|Fungal Parasite\| | Microfungus | NULL | Probable | As mycoparasite (Põlme et al. 2020); Decay fungal material (Põlme et al. 2020) | Põlme S, et al. 2020. Fungal Diversity 105:1-16 ((https://doi.org/10.1007/s13225-020-00466-2)) |
| Otu11302 | 2 | k__Fungi(100);p__Ascomycota(100);c__Sordariomycetes(100);o__Hypocreales(100);f__Clavicipitaceae(100);g__Pochonia(100);s__Pochonia_sp(100); | Pochonia | 13 | Pathotroph-Saprotroph | \|Animal Parasite\|-Animal Pathogen-Undefined Saprotroph | Microfungus | NULL | Probable | As animal_decomposer (Põlme et al. 2020); Decay animal material (Põlme et al. 2020) | Tedersoo L, et al. 2014. Science 346:e1256688 ((https://doi.org/10.1126/science.1256688)); Põlme S, et al. 2020. Fungal Diversity 105:1-16 ((https://doi.org/10.1007/s13225-020-00466-2)) |
| Otu11305 | 2 | k__Fungi(100);p__Ascomycota(100);c__Sordariomycetes(100);o__Hypocreales(100);f__Nectriaceae(100);g__unclassified_Nectriaceae(100);s__Nectriaceae_sp(100); | Nectriaceae | 9 | Pathotroph-Saprotroph-Symbiotroph | Animal Pathogen-Endophyte-Fungal Parasite-Lichen Parasite-Plant Pathogen-Wood Saprotroph | Microfungus | NULL | Possible | Fusarium species are amongst the economically significant plant pathogens (Cannon & Kirk 2007) | Cannon PF, Kirk PM. 2007. Fungal Families of the World. CAB International, Cambridge (ISBN: 978-0851998275); Hirooka Y, et al. 2012. Studies in Mycology 711-210 ((https://doi.org/10.3114/sim0001)); Tedersoo L, et al. 2014. Science 346:e1256688 ((https://doi.org/10.1126/science.1256688)); The American Phytopathological Society Common Names of Plant Diseases ((http://www.apsnet.org/publications/commonnames/Pages/default.aspx)); https://www.cdc.gov ((https://www.cdc.gov)) |
| Otu11306 | 2 | k__Fungi(100);p__Ascomycota(100);c__Sordariomycetes(100);o__Microascales(100);f__Halosphaeriaceae(100);g__unclassified_Halosphaeriaceae(100);s__Halosphaeriaceae_sp(100); | Halosphaeriaceae | 9 | Saprotroph | Undefined Saprotroph-Wood Saprotroph | Microfungus | NULL | Probable | Found in aquatic habitats, mostly in marine environments; saprobic on submerged or intertidal wood, occasionally associated with sand grains and similar substrata (Cannon & Kirk 2007) | Cannon PF, Kirk PM. 2007. Fungal Families of the World. CAB International, Cambridge (ISBN: 978-0851998275) |
| Otu11307 | 2 | k__Fungi(100);p__Ascomycota(100);c__Sordariomycetes(100);o__Microascales(100);f__Halosphaeriaceae(100);g__unclassified_Halosphaeriaceae(100);s__Halosphaeriaceae_sp(100); | Halosphaeriaceae | 9 | Saprotroph | Undefined Saprotroph-Wood Saprotroph | Microfungus | NULL | Probable | Found in aquatic habitats, mostly in marine environments; saprobic on submerged or intertidal wood, occasionally associated with sand grains and similar substrata (Cannon & Kirk 2007) | Cannon PF, Kirk PM. 2007. Fungal Families of the World. CAB International, Cambridge (ISBN: 978-0851998275) |
| Otu11308 | 2 | k__Fungi(100);p__Ascomycota(100);c__Saccharomycetes(100);o__Saccharomycetales(100);f__Metschnikowiaceae(100);g__Metschnikowia(100);g__Metschnikowia_unclassified(100); | Metschnikowia | 13 | Saprotroph | \|Undefined Saprotroph\| | Yeast | NULL | Probable | As nectar-tap_saprotroph (Põlme et al. 2020); Decay sugar-rich substrates (Põlme et al. 2020) | Põlme S, et al. 2020. Fungal Diversity 105:1-16 ((https://doi.org/10.1007/s13225-020-00466-2)) |
| Otu11309 | 2 | k__Fungi(100);p__Ascomycota(100);c__Sordariomycetes(100);o__Boliniales(100);f__Boliniaceae(100);g__Endoxyla(100);s__Endoxyla_sp(100); | Endoxyla | 13 | Saprotroph | Undefined Saprotroph-\|Wood Saprotroph\| | Microfungus | NULL | Probable | Decay wood (Põlme et al. 2020) | Tedersoo L, et al. 2014. Science 346:e1256688 ((https://doi.org/10.1126/science.1256688)); Põlme S, et al. 2020. Fungal Diversity 105:1-16 ((https://doi.org/10.1007/s13225-020-00466-2)) |
| Otu11310 | 2 | k__Fungi(100);p__Ascomycota(100);c__Saccharomycetes(100);o__Saccharomycetales(100);f__Metschnikowiaceae(100);g__Metschnikowia(100);s__Metschnikowia_pulcherrima(100); | Metschnikowia | 13 | Saprotroph | \|Undefined Saprotroph\| | Yeast | NULL | Probable | As nectar-tap_saprotroph (Põlme et al. 2020); Decay sugar-rich substrates (Põlme et al. 2020) | Põlme S, et al. 2020. Fungal Diversity 105:1-16 ((https://doi.org/10.1007/s13225-020-00466-2)) |
| Otu11311 | 2 | k__Fungi(100);p__Ascomycota(100);c__Saccharomycetes(100);o__Saccharomycetales(100);f__Metschnikowiaceae(100);g__Metschnikowia(100);s__Metschnikowia_pulcherrima(100); | Metschnikowia | 13 | Saprotroph | \|Undefined Saprotroph\| | Yeast | NULL | Probable | As nectar-tap_saprotroph (Põlme et al. 2020); Decay sugar-rich substrates (Põlme et al. 2020) | Põlme S, et al. 2020. Fungal Diversity 105:1-16 ((https://doi.org/10.1007/s13225-020-00466-2)) |
| Otu11312 | 2 | k__Fungi(100);p__Ascomycota(100);c__Sordariomycetes(100);o__Hypocreales(100);f__Hypocreales_family_Incertae_sedis(100);g__Acremonium(100);s__Acremonium_spinosum(100); | Acremonium | 13 | Pathotroph-Saprotroph-Symbiotroph | Animal Pathogen-Endophyte-Fungal Parasite-Plant Pathogen-\|Undefined Saprotroph\|-Wood Saprotroph | Microfungus | Soft Rot | Possible | As foliar_endophyte (Põlme et al. 2020); culture (Hanada et al. 2010, Del Olmo; Decay leaf, fruit, seed, soil, dung, animal material (Põlme et al. 2020); Endophyte detection method; Ruiz & Arnold 2014); Likely opportunistic human pathogen (Irinyi et al. 2015) | Duncan CG, Eslyn WE. 1966. Mycologia 58:642-645 ((https://doi.org/10.2307/3757045)); Nilsson T. 1973. Studia Forestalia Suecica 104: 1-40 ((https://pub.epsilon.slu.se/5711/)); Worrall JJ, et al. 1997. Mycologia 89:199-219 ((https://doi.org/10.2307/3761073)); Hanada RE, et al. 2010. Fungal Biology 114:901-910 ((https://doi.org/10.1016/j.funbio.2010.08.006)); Del Olmo-Ruiz M, Arnold AE. 2014. Mycologia 106: 8-21 ((https://doi.org/10.3852/13-098)); Irinyi L, et al. 2015. Medical Mycology 53313-337 ((https://doi.org/10.1093/mmy/myv008)); Põlme S, et al. 2020. Fungal Diversity 105:1-16 ((https://doi.org/10.1007/s13225-020-00466-2)) |
| Otu11313 | 2 | k__Fungi(100);p__Ascomycota(100);c__Sordariomycetes(100);o__Hypocreales(100);f__Hypocreales_family_Incertae_sedis(100);g__Acremonium(100);s__Acremonium_spinosum(100); | Acremonium | 13 | Pathotroph-Saprotroph-Symbiotroph | Animal Pathogen-Endophyte-Fungal Parasite-Plant Pathogen-\|Undefined Saprotroph\|-Wood Saprotroph | Microfungus | Soft Rot | Possible | As foliar_endophyte (Põlme et al. 2020); culture (Hanada et al. 2010, Del Olmo; Decay leaf, fruit, seed, soil, dung, animal material (Põlme et al. 2020); Endophyte detection method; Ruiz & Arnold 2014); Likely opportunistic human pathogen (Irinyi et al. 2015) | Duncan CG, Eslyn WE. 1966. Mycologia 58:642-645 ((https://doi.org/10.2307/3757045)); Nilsson T. 1973. Studia Forestalia Suecica 104: 1-40 ((https://pub.epsilon.slu.se/5711/)); Worrall JJ, et al. 1997. Mycologia 89:199-219 ((https://doi.org/10.2307/3761073)); Hanada RE, et al. 2010. Fungal Biology 114:901-910 ((https://doi.org/10.1016/j.funbio.2010.08.006)); Del Olmo-Ruiz M, Arnold AE. 2014. Mycologia 106: 8-21 ((https://doi.org/10.3852/13-098)); Irinyi L, et al. 2015. Medical Mycology 53313-337 ((https://doi.org/10.1093/mmy/myv008)); Põlme S, et al. 2020. Fungal Diversity 105:1-16 ((https://doi.org/10.1007/s13225-020-00466-2)) |
| Otu11315 | 2 | k__Fungi(100);p__Ascomycota(100);c__Saccharomycetes(100);o__Saccharomycetales(100);f__Metschnikowiaceae(100);g__Metschnikowia(100);g__Metschnikowia_unclassified(100); | Metschnikowia | 13 | Saprotroph | \|Undefined Saprotroph\| | Yeast | NULL | Probable | As nectar-tap_saprotroph (Põlme et al. 2020); Decay sugar-rich substrates (Põlme et al. 2020) | Põlme S, et al. 2020. Fungal Diversity 105:1-16 ((https://doi.org/10.1007/s13225-020-00466-2)) |
| Otu11316 | 2 | k__Fungi(100);p__Ascomycota(100);c__Saccharomycetes(100);o__Saccharomycetales(100);f__Metschnikowiaceae(100);g__Metschnikowia(100);g__Metschnikowia_unclassified(100); | Metschnikowia | 13 | Saprotroph | \|Undefined Saprotroph\| | Yeast | NULL | Probable | As nectar-tap_saprotroph (Põlme et al. 2020); Decay sugar-rich substrates (Põlme et al. 2020) | Põlme S, et al. 2020. Fungal Diversity 105:1-16 ((https://doi.org/10.1007/s13225-020-00466-2)) |
| Otu11317 | 2 | k__Fungi(100);p__Ascomycota(100);c__Saccharomycetes(100);o__Saccharomycetales(100);f__Metschnikowiaceae(100);g__Metschnikowia(100);s__Metschnikowia_pulcherrima(100); | Metschnikowia | 13 | Saprotroph | \|Undefined Saprotroph\| | Yeast | NULL | Probable | As nectar-tap_saprotroph (Põlme et al. 2020); Decay sugar-rich substrates (Põlme et al. 2020) | Põlme S, et al. 2020. Fungal Diversity 105:1-16 ((https://doi.org/10.1007/s13225-020-00466-2)) |
| Otu11318 | 2 | k__Fungi(100);p__Ascomycota(100);c__Sordariomycetes(100);o__Hypocreales(100);f__Hypocreaceae(100);g__Sphaerostilbella(100);s__Sphaerostilbella_novae_zelandiae(100); | Sphaerostilbella | 13 | Pathotroph | \|Fungal Parasite\| | Microfungus | NULL | Highly Probable | As mycoparasite (Põlme et al. 2020); Decay fungal material (Põlme et al. 2020); Host - Stereum (Põlme et al. 2020); Occurs on Stereum species | Tedersoo L, et al. 2014. Science 346:e1256688 ((https://doi.org/10.1126/science.1256688)); Põlme S, et al. 2020. Fungal Diversity 105:1-16 ((https://doi.org/10.1007/s13225-020-00466-2)); Samuels GJ. 2007 self-published. |
| Otu11320 | 2 | k__Fungi(100);p__Ascomycota(100);c__Sordariomycetes(100);o__Hypocreales(100);f__Nectriaceae(100);f__Nectriaceae_unclassified(100);f__Nectriaceae_unclassified(100); | Nectriaceae | 9 | Pathotroph-Saprotroph-Symbiotroph | Animal Pathogen-Endophyte-Fungal Parasite-Lichen Parasite-Plant Pathogen-Wood Saprotroph | Microfungus | NULL | Possible | Fusarium species are amongst the economically significant plant pathogens (Cannon & Kirk 2007) | Cannon PF, Kirk PM. 2007. Fungal Families of the World. CAB International, Cambridge (ISBN: 978-0851998275); Hirooka Y, et al. 2012. Studies in Mycology 711-210 ((https://doi.org/10.3114/sim0001)); Tedersoo L, et al. 2014. Science 346:e1256688 ((https://doi.org/10.1126/science.1256688)); The American Phytopathological Society Common Names of Plant Diseases ((http://www.apsnet.org/publications/commonnames/Pages/default.aspx)); https://www.cdc.gov ((https://www.cdc.gov)) |
| Otu11321 | 2 | k__Fungi(100);p__Ascomycota(100);c__Saccharomycetes(100);o__Saccharomycetales(100);f__Metschnikowiaceae(100);g__Metschnikowia(100);s__Metschnikowia_pulcherrima(100); | Metschnikowia | 13 | Saprotroph | \|Undefined Saprotroph\| | Yeast | NULL | Probable | As nectar-tap_saprotroph (Põlme et al. 2020); Decay sugar-rich substrates (Põlme et al. 2020) | Põlme S, et al. 2020. Fungal Diversity 105:1-16 ((https://doi.org/10.1007/s13225-020-00466-2)) |
| Otu11322 | 2 | k__Fungi(100);p__Ascomycota(100);c__Saccharomycetes(100);o__Saccharomycetales(100);f__Metschnikowiaceae(100);g__Metschnikowia(100);g__Metschnikowia_unclassified(100); | Metschnikowia | 13 | Saprotroph | \|Undefined Saprotroph\| | Yeast | NULL | Probable | As nectar-tap_saprotroph (Põlme et al. 2020); Decay sugar-rich substrates (Põlme et al. 2020) | Põlme S, et al. 2020. Fungal Diversity 105:1-16 ((https://doi.org/10.1007/s13225-020-00466-2)) |
| Otu11323 | 2 | k__Fungi(100);p__Ascomycota(100);c__Saccharomycetes(100);o__Saccharomycetales(100);f__Metschnikowiaceae(100);g__Metschnikowia(100);s__Metschnikowia_pulcherrima(100); | Metschnikowia | 13 | Saprotroph | \|Undefined Saprotroph\| | Yeast | NULL | Probable | As nectar-tap_saprotroph (Põlme et al. 2020); Decay sugar-rich substrates (Põlme et al. 2020) | Põlme S, et al. 2020. Fungal Diversity 105:1-16 ((https://doi.org/10.1007/s13225-020-00466-2)) |
| Otu11326 | 2 | k__Fungi(100);p__Ascomycota(100);c__Lecanoromycetes(100);o__Ostropales(100);f__Stictidaceae(100);f__Stictidaceae_unclassified(100);f__Stictidaceae_unclassified(100); | Stictidaceae | 9 | Symbiotroph | Lichenized-Wood Saprotroph | Microfungus-Thallus | NULL | Probable | Saprobic on wood and stems, a few species lichenized or lignicolous (Cannon & Kirk 2007) | Cannon PF, Kirk PM. 2007. Fungal Families of the World. CAB International, Cambridge (ISBN: 978-0851998275) |
| Otu11329 | 2 | k__Fungi(100);p__Ascomycota(100);c__Saccharomycetes(100);o__Saccharomycetales(100);f__Saccharomycetales_family_Incertae_sedis(100);g__Candida(100);g__Candida_unclassified(100); | Candida | 13 | Pathotroph-Saprotroph-Symbiotroph | Animal Pathogen-Endophyte-Endosymbiont-Epiphyte-\|Undefined Saprotroph\| | Dimorphic Yeast-Yeast | NULL | Probable | As nectar-tap_saprotroph (Põlme et al. 2020); Decay sugar-rich substrates (Põlme et al. 2020); This asexual genus is polyphyletic and contain species that function very differently depend on what they are. Caution should be taken when interpreting guilds for Candida species. | Manolakaki D, et al. 2010. Virulence. 1: 367-375 ((https://doi.org/10.4161/viru.1.5.12796)); Mekbib SB, et al. 2011. Tropical Plant Pathology 36:233-240 ((https://doi.org/10.1590/S1982-56762011000400004)); Põlme S, et al. 2020. Fungal Diversity 105:1-16 ((https://doi.org/10.1007/s13225-020-00466-2)) |
| Otu11330 | 2 | k__Fungi(100);p__Ascomycota(100);c__Eurotiomycetes(100);o__Eurotiales(100);f__Trichocomaceae(100);g__Rasamsonia(100);s__Rasamsonia_brevistipitata(100); | Rasamsonia | 13 | Pathotroph-Saprotroph | Animal Parasite-\|Undefined Saprotroph\| | Microfungus | NULL | Probable | As soil_saprotroph (Põlme et al. 2020); Decay soil, animal material (Põlme et al. 2020) | Tedersoo L, et al. 2014. Science 346:e1256688 ((https://doi.org/10.1126/science.1256688)); Põlme S, et al. 2020. Fungal Diversity 105:1-16 ((https://doi.org/10.1007/s13225-020-00466-2)) |
| Otu11332 | 2 | k__Fungi(100);p__Ascomycota(100);c__Taphrinomycetes(100);o__Taphrinales(100);f__Taphrinaceae(100);f__Taphrinaceae_unclassified(100);f__Taphrinaceae_unclassified(100); | Taphrinaceae | 9 | Pathotroph | Plant Pathogen | Microfungus | NULL | Probable | Biotrophic on plants, usually causing hyperplasia (galls, witches broom) or lesions (Cannon & Kirk 2007) | Cannon PF, Kirk PM. 2007. Fungal Families of the World. CAB International, Cambridge (ISBN: 978-0851998275) |
| Otu11336 | 2 | k__Fungi(100);p__Ascomycota(100);c__Saccharomycetes(100);o__Saccharomycetales(100);f__Metschnikowiaceae(100);g__Metschnikowia(100);s__Metschnikowia_pulcherrima(100); | Metschnikowia | 13 | Saprotroph | \|Undefined Saprotroph\| | Yeast | NULL | Probable | As nectar-tap_saprotroph (Põlme et al. 2020); Decay sugar-rich substrates (Põlme et al. 2020) | Põlme S, et al. 2020. Fungal Diversity 105:1-16 ((https://doi.org/10.1007/s13225-020-00466-2)) |
| Otu11337 | 2 | k__Fungi(100);p__Ascomycota(100);c__Taphrinomycetes(100);o__Taphrinales(100);f__Taphrinaceae(100);g__Taphrina(100);g__Taphrina_unclassified(100); | Taphrina | 13 | Pathotroph | \|Plant Pathogen\| | Dimorphic-Microfungus | NULL | Probable | NULL | Põlme S, et al. 2020. Fungal Diversity 105:1-16 ((https://doi.org/10.1007/s13225-020-00466-2)); The American Phytopathological Society Common Names of Plant Diseases ((http://www.apsnet.org/publications/commonnames/Pages/default.aspx)) |
| Otu11340 | 2 | k__Fungi(100);p__Ascomycota(100);c__Taphrinomycetes(100);o__Taphrinales(100);f__Taphrinaceae(100);g__Taphrina(100);s__Taphrina_populina(100); | Taphrina | 13 | Pathotroph | \|Plant Pathogen\| | Dimorphic-Microfungus | NULL | Probable | NULL | Põlme S, et al. 2020. Fungal Diversity 105:1-16 ((https://doi.org/10.1007/s13225-020-00466-2)); The American Phytopathological Society Common Names of Plant Diseases ((http://www.apsnet.org/publications/commonnames/Pages/default.aspx)) |
| Otu11341 | 2 | k__Fungi(100);p__Ascomycota(100);c__Taphrinomycetes(100);o__Taphrinales(100);f__Taphrinaceae(100);g__Taphrina(100);s__Taphrina_sp(100); | Taphrina | 13 | Pathotroph | \|Plant Pathogen\| | Dimorphic-Microfungus | NULL | Probable | NULL | Põlme S, et al. 2020. Fungal Diversity 105:1-16 ((https://doi.org/10.1007/s13225-020-00466-2)); The American Phytopathological Society Common Names of Plant Diseases ((http://www.apsnet.org/publications/commonnames/Pages/default.aspx)) |
| Otu11343 | 2 | k__Fungi(100);p__Ascomycota(100);c__Saccharomycetes(100);o__Saccharomycetales(100);f__Metschnikowiaceae(100);g__Metschnikowia(100);s__Metschnikowia_pulcherrima(100); | Metschnikowia | 13 | Saprotroph | \|Undefined Saprotroph\| | Yeast | NULL | Probable | As nectar-tap_saprotroph (Põlme et al. 2020); Decay sugar-rich substrates (Põlme et al. 2020) | Põlme S, et al. 2020. Fungal Diversity 105:1-16 ((https://doi.org/10.1007/s13225-020-00466-2)) |
| Otu11345 | 2 | k__Fungi(100);p__Ascomycota(100);c__Saccharomycetes(100);o__Saccharomycetales(100);f__Metschnikowiaceae(100);g__Metschnikowia(100);g__Metschnikowia_unclassified(100); | Metschnikowia | 13 | Saprotroph | \|Undefined Saprotroph\| | Yeast | NULL | Probable | As nectar-tap_saprotroph (Põlme et al. 2020); Decay sugar-rich substrates (Põlme et al. 2020) | Põlme S, et al. 2020. Fungal Diversity 105:1-16 ((https://doi.org/10.1007/s13225-020-00466-2)) |
| Otu11346 | 2 | k__Fungi(100);p__Ascomycota(100);c__Saccharomycetes(100);o__Saccharomycetales(100);f__Metschnikowiaceae(100);g__Metschnikowia(100);s__Metschnikowia_pulcherrima(100); | Metschnikowia | 13 | Saprotroph | \|Undefined Saprotroph\| | Yeast | NULL | Probable | As nectar-tap_saprotroph (Põlme et al. 2020); Decay sugar-rich substrates (Põlme et al. 2020) | Põlme S, et al. 2020. Fungal Diversity 105:1-16 ((https://doi.org/10.1007/s13225-020-00466-2)) |
| Otu11347 | 2 | k__Fungi(100);p__Ascomycota(100);c__Saccharomycetes(100);o__Saccharomycetales(100);f__Metschnikowiaceae(100);g__Metschnikowia(100);s__Metschnikowia_pulcherrima(100); | Metschnikowia | 13 | Saprotroph | \|Undefined Saprotroph\| | Yeast | NULL | Probable | As nectar-tap_saprotroph (Põlme et al. 2020); Decay sugar-rich substrates (Põlme et al. 2020) | Põlme S, et al. 2020. Fungal Diversity 105:1-16 ((https://doi.org/10.1007/s13225-020-00466-2)) |
| Otu11349 | 2 | k__Fungi(100);p__Ascomycota(100);c__Saccharomycetes(100);o__Saccharomycetales(100);f__Metschnikowiaceae(100);g__Metschnikowia(100);s__Metschnikowia_pulcherrima(100); | Metschnikowia | 13 | Saprotroph | \|Undefined Saprotroph\| | Yeast | NULL | Probable | As nectar-tap_saprotroph (Põlme et al. 2020); Decay sugar-rich substrates (Põlme et al. 2020) | Põlme S, et al. 2020. Fungal Diversity 105:1-16 ((https://doi.org/10.1007/s13225-020-00466-2)) |
| Otu11350 | 2 | k__Fungi(100);p__Ascomycota(100);c__Eurotiomycetes(100);o__Chaetothyriales(100);f__Herpotrichiellaceae(100);g__Cladophialophora(100);g__Cladophialophora_unclassified(100); | Cladophialophora | 13 | Pathotroph-Saprotroph-Symbiotroph | Animal Pathogen-Endophyte-Fungal Parasite-\|Undefined Saprotroph\| | Dimorphic-Microfungus | NULL | Probable | As mycoparasite (Põlme et al. 2020); As root_endophyte_dark_septate (Põlme et al. 2020); As soil_saprotroph (Põlme et al. 2020); Decay soil (Põlme et al. 2020) | James TY, et al. 2006. Nature 443:818-822 ((https://doi.org/10.1038/nature05110)); Põlme S, et al. 2020. Fungal Diversity 105:1-16 ((https://doi.org/10.1007/s13225-020-00466-2)) |
| Otu11351 | 2 | k__Fungi(100);p__Ascomycota(100);c__Eurotiomycetes(100);o__Chaetothyriales(100);f__Herpotrichiellaceae(100);g__Cladophialophora(100);g__Cladophialophora_unclassified(100); | Cladophialophora | 13 | Pathotroph-Saprotroph-Symbiotroph | Animal Pathogen-Endophyte-Fungal Parasite-\|Undefined Saprotroph\| | Dimorphic-Microfungus | NULL | Probable | As mycoparasite (Põlme et al. 2020); As root_endophyte_dark_septate (Põlme et al. 2020); As soil_saprotroph (Põlme et al. 2020); Decay soil (Põlme et al. 2020) | James TY, et al. 2006. Nature 443:818-822 ((https://doi.org/10.1038/nature05110)); Põlme S, et al. 2020. Fungal Diversity 105:1-16 ((https://doi.org/10.1007/s13225-020-00466-2)) |
| Otu11352 | 2 | k__Fungi(100);p__Ascomycota(100);c__Eurotiomycetes(100);o__Chaetothyriales(100);f__Herpotrichiellaceae(100);f__Herpotrichiellaceae_unclassified(100);f__Herpotrichiellaceae_unclassified(100); | Herpotrichiellaceae | 9 | Pathotroph-Saprotroph | Animal Pathogen-Fungal Parasite-Undefined Saprotroph | Facultative Yeast-Microfungus | NULL | Probable | Mostly saprobic on plants or other fungi, isolated from soil etc., and from medical sources (Cannon & Kirk 2007) | Cannon PF, Kirk PM. 2007. Fungal Families of the World. CAB International, Cambridge (ISBN: 978-0851998275) |
| Otu11353 | 2 | k__Fungi(100);p__Ascomycota(100);c__Eurotiomycetes(100);o__Chaetothyriales(100);f__Herpotrichiellaceae(100);g__Rhinocladiella(100);g__Rhinocladiella_unclassified(100); | Rhinocladiella | 13 | Pathotroph-Saprotroph-Symbiotroph | Animal Parasite-Endophyte-\|Undefined Saprotroph\|-Wood Saprotroph | Dimorphic-Facultative Yeast | Soft Rot | Probable | As root_endophyte_dark_septate (Põlme et al. 2020); As soil_saprotroph (Põlme et al. 2020); Decay soil, animal material (Põlme et al. 2020) | Nilsson T. 1973. Studia Forestalia Suecica 104:1-40 ((https://pub.epsilon.slu.se/5711/)); Seehann G, et al. 1975. List of Fungi in Soft-Rot Tissue Tests. Institute fur Holzbiologie und Holzschutz, Bundesforschungsanstalt fur Forst - und Holzwirtshaft, Hamburg - Lohbrugge, Germany; Costa IPMW, et al. 2012. Mycotaxon 119:e26 ((http://mycotaxon.com/resources/checklists/costa_v119_checklist.pdf)); Tedersoo L, et al. 2014. Science 346:e1256688 ((https://doi.org/10.1126/science.1256688)); Põlme S, et al. 2020. Fungal Diversity 105:1-16 ((https://doi.org/10.1007/s13225-020-00466-2)) |
| Otu11355 | 2 | k__Fungi(100);p__Ascomycota(100);c__Saccharomycetes(100);o__Saccharomycetales(100);f__Metschnikowiaceae(100);g__Metschnikowia(100);g__Metschnikowia_unclassified(100); | Metschnikowia | 13 | Saprotroph | \|Undefined Saprotroph\| | Yeast | NULL | Probable | As nectar-tap_saprotroph (Põlme et al. 2020); Decay sugar-rich substrates (Põlme et al. 2020) | Põlme S, et al. 2020. Fungal Diversity 105:1-16 ((https://doi.org/10.1007/s13225-020-00466-2)) |
| Otu11356 | 2 | k__Fungi(100);p__Ascomycota(100);c__Saccharomycetes(100);o__Saccharomycetales(100);f__Metschnikowiaceae(100);g__Metschnikowia(100);g__Metschnikowia_unclassified(100); | Metschnikowia | 13 | Saprotroph | \|Undefined Saprotroph\| | Yeast | NULL | Probable | As nectar-tap_saprotroph (Põlme et al. 2020); Decay sugar-rich substrates (Põlme et al. 2020) | Põlme S, et al. 2020. Fungal Diversity 105:1-16 ((https://doi.org/10.1007/s13225-020-00466-2)) |
| Otu11357 | 2 | k__Fungi(100);p__Ascomycota(100);c__Saccharomycetes(100);o__Saccharomycetales(100);f__Debaryomycetaceae(100);g__Priceomyces(100);g__Priceomyces_unclassified(100); | Priceomyces | 13 | Saprotroph | \|Undefined Saprotroph\| | Yeast | NULL | Probable | As nectar-tap_saprotroph (Põlme et al. 2020); Decay sugar-rich substrates (Põlme et al. 2020) | Põlme S, et al. 2020. Fungal Diversity 105:1-16 ((https://doi.org/10.1007/s13225-020-00466-2)) |
| Otu11358 | 2 | k__Fungi(100);p__Ascomycota(100);c__Eurotiomycetes(100);o__Chaetothyriales(100);f__Herpotrichiellaceae(100);g__Capronia(100);s__Capronia_semi-immersa(100); | Capronia | 13 | Pathotroph-Saprotroph-Symbiotroph | Animal Pathogen-Endophyte-Fungal Parasite-\|Undefined Saprotroph\| | Dimorphic-Facultative Yeast | NULL | Highly Probable | As mycoparasite (Põlme et al. 2020); As root_endophyte_dark_septate (Põlme et al. 2020); As soil_saprotroph (Põlme et al. 2020); Dark Septate Endophyte, known from plant roots.; Decay soil (Põlme et al. 2020) | Tedersoo L, et al. 2014. Science 346:e1256688 ((https://doi.org/10.1126/science.1256688)); Põlme S, et al. 2020. Fungal Diversity 105:1-16 ((https://doi.org/10.1007/s13225-020-00466-2)) |
| Otu11359 | 2 | k__Fungi(100);p__Ascomycota(100);c__Saccharomycetes(100);o__Saccharomycetales(100);f__Metschnikowiaceae(100);g__Metschnikowia(100);s__Metschnikowia_pulcherrima(100); | Metschnikowia | 13 | Saprotroph | \|Undefined Saprotroph\| | Yeast | NULL | Probable | As nectar-tap_saprotroph (Põlme et al. 2020); Decay sugar-rich substrates (Põlme et al. 2020) | Põlme S, et al. 2020. Fungal Diversity 105:1-16 ((https://doi.org/10.1007/s13225-020-00466-2)) |
| Otu11360 | 2 | k__Fungi(100);p__Ascomycota(100);c__Dothideomycetes(100);o__Pleosporales(100);f__Pleosporaceae(100);f__Pleosporaceae_unclassified(100);f__Pleosporaceae_unclassified(100); | Pleosporaceae | 9 | Pathotroph-Saprotroph | Endophyte-Lichen Parasite-Plant Pathogen-Undefined Saprotroph | Microfungus | NULL | Probable | Necrotrophic pathogens and saprobes, especially associated with grasses. Many species are seed-borne (Cannon & Kirk 2007) | Cannon PF, Kirk PM. 2007. Fungal Families of the World. CAB International, Cambridge (ISBN: 978-0851998275); David AS, et al. 2016. Microbial Ecology 71:912-926 ((https://doi.org/10.1007/s00248-015-0712-6)); Lawrey JD, Diederich P. 2016. http://www.lichenicolous.net (accessed 24 May 2017) ((http://www.lichenicolous.net)) |
| Otu11361 | 2 | k__Fungi(100);p__Ascomycota(100);c__Saccharomycetes(100);o__Saccharomycetales(100);f__Metschnikowiaceae(100);g__Metschnikowia(100);s__Metschnikowia_pulcherrima(100); | Metschnikowia | 13 | Saprotroph | \|Undefined Saprotroph\| | Yeast | NULL | Probable | As nectar-tap_saprotroph (Põlme et al. 2020); Decay sugar-rich substrates (Põlme et al. 2020) | Põlme S, et al. 2020. Fungal Diversity 105:1-16 ((https://doi.org/10.1007/s13225-020-00466-2)) |
| Otu11362 | 2 | k__Fungi(100);p__Ascomycota(100);c__Sordariomycetes(100);o__Diaporthales(100);f__Diaporthaceae(100);g__Diaporthe(100);g__Diaporthe_unclassified(100); | Diaporthe | 13 | Pathotroph-Saprotroph-Symbiotroph | Endophyte-\|Plant Pathogen\|-Plant Saprotroph | Microfungus | NULL | Possible | As foliar_endophyte (Põlme et al. 2020); As litter_saprotroph (Põlme et al. 2020); Decay leaf, fruit, seed (Põlme et al. 2020) | Tedersoo L, et al. 2014. Science 346:e1256688 ((https://doi.org/10.1126/science.1256688)); Põlme S, et al. 2020. Fungal Diversity 105:1-16 ((https://doi.org/10.1007/s13225-020-00466-2)) |
| Otu11363 | 2 | k__Fungi(100);p__Ascomycota(100);c__Saccharomycetes(100);o__Saccharomycetales(100);f__Metschnikowiaceae(100);g__Metschnikowia(100);g__Metschnikowia_unclassified(100); | Metschnikowia | 13 | Saprotroph | \|Undefined Saprotroph\| | Yeast | NULL | Probable | As nectar-tap_saprotroph (Põlme et al. 2020); Decay sugar-rich substrates (Põlme et al. 2020) | Põlme S, et al. 2020. Fungal Diversity 105:1-16 ((https://doi.org/10.1007/s13225-020-00466-2)) |
| Otu11364 | 2 | k__Fungi(100);p__Ascomycota(100);c__Saccharomycetes(100);o__Saccharomycetales(100);f__Metschnikowiaceae(100);g__Metschnikowia(100);s__Metschnikowia_pulcherrima(100); | Metschnikowia | 13 | Saprotroph | \|Undefined Saprotroph\| | Yeast | NULL | Probable | As nectar-tap_saprotroph (Põlme et al. 2020); Decay sugar-rich substrates (Põlme et al. 2020) | Põlme S, et al. 2020. Fungal Diversity 105:1-16 ((https://doi.org/10.1007/s13225-020-00466-2)) |
| Otu11365 | 2 | k__Fungi(100);p__Ascomycota(100);c__Eurotiomycetes(100);o__Verrucariales(100);f__Verrucariaceae(100);f__Verrucariaceae_unclassified(100);f__Verrucariaceae_unclassified(100); | Verrucariaceae | 9 | Symbiotroph | Lichenized | Thallus | NULL | Highly Probable | Lichenized with green algae, on rocks, soil, etc., more rarely on wood and bark (Cannon & Kirk 2007) | Cannon PF, Kirk PM. 2007. Fungal Families of the World. CAB International, Cambridge (ISBN: 978-0851998275) |
| Otu11366 | 2 | k__Fungi(100);p__Ascomycota(100);c__Saccharomycetes(100);o__Saccharomycetales(100);f__Metschnikowiaceae(100);g__Metschnikowia(100);g__Metschnikowia_unclassified(100); | Metschnikowia | 13 | Saprotroph | \|Undefined Saprotroph\| | Yeast | NULL | Probable | As nectar-tap_saprotroph (Põlme et al. 2020); Decay sugar-rich substrates (Põlme et al. 2020) | Põlme S, et al. 2020. Fungal Diversity 105:1-16 ((https://doi.org/10.1007/s13225-020-00466-2)) |
| Otu11367 | 2 | k__Fungi(100);p__Ascomycota(100);c__Saccharomycetes(100);o__Saccharomycetales(100);f__Metschnikowiaceae(100);g__Metschnikowia(100);g__Metschnikowia_unclassified(100); | Metschnikowia | 13 | Saprotroph | \|Undefined Saprotroph\| | Yeast | NULL | Probable | As nectar-tap_saprotroph (Põlme et al. 2020); Decay sugar-rich substrates (Põlme et al. 2020) | Põlme S, et al. 2020. Fungal Diversity 105:1-16 ((https://doi.org/10.1007/s13225-020-00466-2)) |
| Otu11369 | 2 | k__Fungi(100);p__Ascomycota(100);c__Saccharomycetes(100);o__Saccharomycetales(100);f__Metschnikowiaceae(100);g__Metschnikowia(100);g__Metschnikowia_unclassified(100); | Metschnikowia | 13 | Saprotroph | \|Undefined Saprotroph\| | Yeast | NULL | Probable | As nectar-tap_saprotroph (Põlme et al. 2020); Decay sugar-rich substrates (Põlme et al. 2020) | Põlme S, et al. 2020. Fungal Diversity 105:1-16 ((https://doi.org/10.1007/s13225-020-00466-2)) |
| Otu11371 | 2 | k__Fungi(100);p__Ascomycota(100);c__Saccharomycetes(100);o__Saccharomycetales(100);f__Saccharomycetaceae(100);g__Williopsis(100);s__Williopsis_sp(100); | Williopsis | 13 | Saprotroph-Symbiotroph | Endophyte-Plant Saprotroph-\|Undefined Saprotroph\| | Microfungus-Yeast | NULL | Possible | As foliar_endophyte (Põlme et al. 2020); As litter_saprotroph (Põlme et al. 2020); As soil_saprotroph (Põlme et al. 2020); culture (Del Olmo; Decay leaf, fruit, seed, soil (Põlme et al. 2020); Endophyte detection method; Ruiz & Arnold 2014) | Del Olmo-Ruiz M, Arnold AE. 2014. Mycologia 106:8-21 ((https://doi.org/10.3852/13-098)); Põlme S, et al. 2020. Fungal Diversity 105:1-16 ((https://doi.org/10.1007/s13225-020-00466-2)) |
| Otu11372 | 2 | k__Fungi(100);p__Ascomycota(100);c__Saccharomycetes(100);o__Saccharomycetales(100);f__Metschnikowiaceae(100);g__Metschnikowia(100);g__Metschnikowia_unclassified(100); | Metschnikowia | 13 | Saprotroph | \|Undefined Saprotroph\| | Yeast | NULL | Probable | As nectar-tap_saprotroph (Põlme et al. 2020); Decay sugar-rich substrates (Põlme et al. 2020) | Põlme S, et al. 2020. Fungal Diversity 105:1-16 ((https://doi.org/10.1007/s13225-020-00466-2)) |
| Otu11373 | 2 | k__Fungi(100);p__Ascomycota(100);c__Saccharomycetes(100);o__Saccharomycetales(100);f__Saccharomycetaceae(100);g__Kazachstania(100);s__Kazachstania_intestinalis(100); | Kazachstania | 13 | Saprotroph | \|Undefined Saprotroph\| | Yeast | NULL | Probable | As nectar-tap_saprotroph (Põlme et al. 2020); Decay sugar-rich substrates (Põlme et al. 2020) | Põlme S, et al. 2020. Fungal Diversity 105:1-16 ((https://doi.org/10.1007/s13225-020-00466-2)) |
| Otu11376 | 2 | k__Fungi(100);p__Ascomycota(100);c__Saccharomycetes(100);o__Saccharomycetales(100);f__Metschnikowiaceae(100);g__Metschnikowia(100);g__Metschnikowia_unclassified(100); | Metschnikowia | 13 | Saprotroph | \|Undefined Saprotroph\| | Yeast | NULL | Probable | As nectar-tap_saprotroph (Põlme et al. 2020); Decay sugar-rich substrates (Põlme et al. 2020) | Põlme S, et al. 2020. Fungal Diversity 105:1-16 ((https://doi.org/10.1007/s13225-020-00466-2)) |
| Otu11378 | 2 | k__Fungi(100);p__Ascomycota(100);c__Saccharomycetes(100);o__Saccharomycetales(100);f__Metschnikowiaceae(100);g__Metschnikowia(100);s__Metschnikowia_pulcherrima(100); | Metschnikowia | 13 | Saprotroph | \|Undefined Saprotroph\| | Yeast | NULL | Probable | As nectar-tap_saprotroph (Põlme et al. 2020); Decay sugar-rich substrates (Põlme et al. 2020) | Põlme S, et al. 2020. Fungal Diversity 105:1-16 ((https://doi.org/10.1007/s13225-020-00466-2)) |
| Otu11379 | 2 | k__Fungi(100);p__Ascomycota(100);c__Saccharomycetes(100);o__Saccharomycetales(100);f__Metschnikowiaceae(100);g__Metschnikowia(100);s__Metschnikowia_pulcherrima(100); | Metschnikowia | 13 | Saprotroph | \|Undefined Saprotroph\| | Yeast | NULL | Probable | As nectar-tap_saprotroph (Põlme et al. 2020); Decay sugar-rich substrates (Põlme et al. 2020) | Põlme S, et al. 2020. Fungal Diversity 105:1-16 ((https://doi.org/10.1007/s13225-020-00466-2)) |
| Otu11380 | 2 | k__Fungi(100);p__Ascomycota(100);c__Saccharomycetes(100);o__Saccharomycetales(100);f__Metschnikowiaceae(100);g__Metschnikowia(100);s__Metschnikowia_pulcherrima(100); | Metschnikowia | 13 | Saprotroph | \|Undefined Saprotroph\| | Yeast | NULL | Probable | As nectar-tap_saprotroph (Põlme et al. 2020); Decay sugar-rich substrates (Põlme et al. 2020) | Põlme S, et al. 2020. Fungal Diversity 105:1-16 ((https://doi.org/10.1007/s13225-020-00466-2)) |
| Otu11381 | 2 | k__Fungi(100);p__Ascomycota(100);c__Saccharomycetes(100);o__Saccharomycetales(100);f__Metschnikowiaceae(100);g__Metschnikowia(100);g__Metschnikowia_unclassified(100); | Metschnikowia | 13 | Saprotroph | \|Undefined Saprotroph\| | Yeast | NULL | Probable | As nectar-tap_saprotroph (Põlme et al. 2020); Decay sugar-rich substrates (Põlme et al. 2020) | Põlme S, et al. 2020. Fungal Diversity 105:1-16 ((https://doi.org/10.1007/s13225-020-00466-2)) |
| Otu11382 | 2 | k__Fungi(100);p__Ascomycota(100);c__Saccharomycetes(100);o__Saccharomycetales(100);f__Metschnikowiaceae(100);g__Metschnikowia(100);s__Metschnikowia_pulcherrima(100); | Metschnikowia | 13 | Saprotroph | \|Undefined Saprotroph\| | Yeast | NULL | Probable | As nectar-tap_saprotroph (Põlme et al. 2020); Decay sugar-rich substrates (Põlme et al. 2020) | Põlme S, et al. 2020. Fungal Diversity 105:1-16 ((https://doi.org/10.1007/s13225-020-00466-2)) |
| Otu11383 | 2 | k__Fungi(100);p__Ascomycota(100);c__Saccharomycetes(100);o__Saccharomycetales(100);f__Metschnikowiaceae(100);g__Metschnikowia(100);s__Metschnikowia_pulcherrima(100); | Metschnikowia | 13 | Saprotroph | \|Undefined Saprotroph\| | Yeast | NULL | Probable | As nectar-tap_saprotroph (Põlme et al. 2020); Decay sugar-rich substrates (Põlme et al. 2020) | Põlme S, et al. 2020. Fungal Diversity 105:1-16 ((https://doi.org/10.1007/s13225-020-00466-2)) |
| Otu11384 | 2 | k__Fungi(100);p__Ascomycota(100);c__Saccharomycetes(100);o__Saccharomycetales(100);f__Saccharomycetaceae(100);g__Kazachstania(100);s__Kazachstania_intestinalis(100); | Kazachstania | 13 | Saprotroph | \|Undefined Saprotroph\| | Yeast | NULL | Probable | As nectar-tap_saprotroph (Põlme et al. 2020); Decay sugar-rich substrates (Põlme et al. 2020) | Põlme S, et al. 2020. Fungal Diversity 105:1-16 ((https://doi.org/10.1007/s13225-020-00466-2)) |
| Otu11385 | 2 | k__Fungi(100);p__Ascomycota(100);c__Saccharomycetes(100);o__Saccharomycetales(100);f__Metschnikowiaceae(100);g__Metschnikowia(100);s__Metschnikowia_pulcherrima(100); | Metschnikowia | 13 | Saprotroph | \|Undefined Saprotroph\| | Yeast | NULL | Probable | As nectar-tap_saprotroph (Põlme et al. 2020); Decay sugar-rich substrates (Põlme et al. 2020) | Põlme S, et al. 2020. Fungal Diversity 105:1-16 ((https://doi.org/10.1007/s13225-020-00466-2)) |
| Otu11386 | 2 | k__Fungi(100);p__Ascomycota(100);c__Saccharomycetes(100);o__Saccharomycetales(100);f__Saccharomycetaceae(100);g__Lachancea(100);s__Lachancea_thermotolerans(100); | Lachancea | 13 | Pathotroph-Saprotroph | Animal Parasite-Animal Pathogen-\|Undefined Saprotroph\| | Yeast | NULL | Possible | As nectar-tap_saprotroph (Põlme et al. 2020); Decay animal material, sugar-rich substrates (Põlme et al. 2020); Likely opportunistic human pathogen (Irinyi et al. 2015; see L. fermentati) | Irinyi L, et al. 2015. Medical Mycology 53:313-337 ((https://doi.org/10.1093/mmy/myv008)); Põlme S, et al. 2020. Fungal Diversity 105:1-16 ((https://doi.org/10.1007/s13225-020-00466-2)) |
| Otu11387 | 2 | k__Fungi(100);p__Ascomycota(100);c__Saccharomycetes(100);o__Saccharomycetales(100);f__Metschnikowiaceae(100);g__Metschnikowia(100);g__Metschnikowia_unclassified(100); | Metschnikowia | 13 | Saprotroph | \|Undefined Saprotroph\| | Yeast | NULL | Probable | As nectar-tap_saprotroph (Põlme et al. 2020); Decay sugar-rich substrates (Põlme et al. 2020) | Põlme S, et al. 2020. Fungal Diversity 105:1-16 ((https://doi.org/10.1007/s13225-020-00466-2)) |
| Otu11388 | 2 | k__Fungi(100);p__Ascomycota(100);c__Saccharomycetes(100);o__Saccharomycetales(100);f__Saccharomycetaceae(100);g__Lachancea(100);s__Lachancea_thermotolerans(100); | Lachancea | 13 | Pathotroph-Saprotroph | Animal Parasite-Animal Pathogen-\|Undefined Saprotroph\| | Yeast | NULL | Possible | As nectar-tap_saprotroph (Põlme et al. 2020); Decay animal material, sugar-rich substrates (Põlme et al. 2020); Likely opportunistic human pathogen (Irinyi et al. 2015; see L. fermentati) | Irinyi L, et al. 2015. Medical Mycology 53:313-337 ((https://doi.org/10.1093/mmy/myv008)); Põlme S, et al. 2020. Fungal Diversity 105:1-16 ((https://doi.org/10.1007/s13225-020-00466-2)) |
| Otu11389 | 2 | k__Fungi(100);p__Ascomycota(100);c__Saccharomycetes(100);o__Saccharomycetales(100);f__Metschnikowiaceae(100);g__Metschnikowia(100);s__Metschnikowia_pulcherrima(100); | Metschnikowia | 13 | Saprotroph | \|Undefined Saprotroph\| | Yeast | NULL | Probable | As nectar-tap_saprotroph (Põlme et al. 2020); Decay sugar-rich substrates (Põlme et al. 2020) | Põlme S, et al. 2020. Fungal Diversity 105:1-16 ((https://doi.org/10.1007/s13225-020-00466-2)) |
| Otu11390 | 2 | k__Fungi(100);p__Ascomycota(100);c__Saccharomycetes(100);o__Saccharomycetales(100);f__Saccharomycetaceae(100);g__Kazachstania(100);s__Kazachstania_intestinalis(100); | Kazachstania | 13 | Saprotroph | \|Undefined Saprotroph\| | Yeast | NULL | Probable | As nectar-tap_saprotroph (Põlme et al. 2020); Decay sugar-rich substrates (Põlme et al. 2020) | Põlme S, et al. 2020. Fungal Diversity 105:1-16 ((https://doi.org/10.1007/s13225-020-00466-2)) |
| Otu11391 | 2 | k__Fungi(100);p__Ascomycota(100);c__Dothideomycetes(100);o__Capnodiales(100);f__Capnodiales_family_Incertae_sedis(100);g__Capnobotryella(100);s__Capnobotryella_renispora(100); | Capnobotryella | 13 | Saprotroph | \|Plant Saprotroph\|-Undefined Saprotroph | Microfungus | NULL | Probable | As litter_saprotroph (Põlme et al. 2020); Decay leaf, fruit, seed, soil (Põlme et al. 2020) | Tedersoo L, et al. 2014. Science 346:e1256688 ((https://doi.org/10.1126/science.1256688)); Põlme S, et al. 2020. Fungal Diversity 105:1-16 ((https://doi.org/10.1007/s13225-020-00466-2)) |
| Otu11392 | 2 | k__Fungi(100);p__Ascomycota(100);c__Saccharomycetes(100);o__Saccharomycetales(100);f__Metschnikowiaceae(100);g__Metschnikowia(100);s__Metschnikowia_pulcherrima(100); | Metschnikowia | 13 | Saprotroph | \|Undefined Saprotroph\| | Yeast | NULL | Probable | As nectar-tap_saprotroph (Põlme et al. 2020); Decay sugar-rich substrates (Põlme et al. 2020) | Põlme S, et al. 2020. Fungal Diversity 105:1-16 ((https://doi.org/10.1007/s13225-020-00466-2)) |
| Otu11393 | 2 | k__Fungi(100);p__Ascomycota(100);c__Saccharomycetes(100);o__Saccharomycetales(100);f__Metschnikowiaceae(100);g__Metschnikowia(100);s__Metschnikowia_pulcherrima(100); | Metschnikowia | 13 | Saprotroph | \|Undefined Saprotroph\| | Yeast | NULL | Probable | As nectar-tap_saprotroph (Põlme et al. 2020); Decay sugar-rich substrates (Põlme et al. 2020) | Põlme S, et al. 2020. Fungal Diversity 105:1-16 ((https://doi.org/10.1007/s13225-020-00466-2)) |
| Otu11394 | 2 | k__Fungi(100);p__Ascomycota(100);c__Saccharomycetes(100);o__Saccharomycetales(100);f__Metschnikowiaceae(100);g__Metschnikowia(100);g__Metschnikowia_unclassified(100); | Metschnikowia | 13 | Saprotroph | \|Undefined Saprotroph\| | Yeast | NULL | Probable | As nectar-tap_saprotroph (Põlme et al. 2020); Decay sugar-rich substrates (Põlme et al. 2020) | Põlme S, et al. 2020. Fungal Diversity 105:1-16 ((https://doi.org/10.1007/s13225-020-00466-2)) |
| Otu11396 | 2 | k__Fungi(100);p__Ascomycota(100);c__Saccharomycetes(100);o__Saccharomycetales(100);f__Metschnikowiaceae(100);g__Metschnikowia(100);s__Metschnikowia_pulcherrima(100); | Metschnikowia | 13 | Saprotroph | \|Undefined Saprotroph\| | Yeast | NULL | Probable | As nectar-tap_saprotroph (Põlme et al. 2020); Decay sugar-rich substrates (Põlme et al. 2020) | Põlme S, et al. 2020. Fungal Diversity 105:1-16 ((https://doi.org/10.1007/s13225-020-00466-2)) |
| Otu11397 | 2 | k__Fungi(100);p__Ascomycota(100);c__Saccharomycetes(100);o__Saccharomycetales(100);f__Saccharomycetaceae(100);g__Kazachstania(100);s__Kazachstania_intestinalis(100); | Kazachstania | 13 | Saprotroph | \|Undefined Saprotroph\| | Yeast | NULL | Probable | As nectar-tap_saprotroph (Põlme et al. 2020); Decay sugar-rich substrates (Põlme et al. 2020) | Põlme S, et al. 2020. Fungal Diversity 105:1-16 ((https://doi.org/10.1007/s13225-020-00466-2)) |
| Otu11399 | 2 | k__Fungi(100);p__Ascomycota(100);c__Sordariomycetes(100);o__Diaporthales(100);f__Valsaceae(100);g__Cytospora(100);g__Cytospora_unclassified(100); | Cytospora | 13 | Pathotroph-Saprotroph-Symbiotroph | Endophyte-\|Plant Pathogen\|-Plant Saprotroph-Wood Saprotroph | Microfungus | Soft Rot | Probable | As foliar_endophyte (Põlme et al. 2020); As litter_saprotroph (Põlme et al. 2020); Decay leaf, fruit, seed (Põlme et al. 2020) | Duncan CG, Eslyn WE. 1966. Mycologia 58:642-645 ((https://doi.org/10.2307/3757045)); Tedersoo L, et al. 2014. Science 346:e1256688 ((https://doi.org/10.1126/science.1256688)); Põlme S, et al. 2020. Fungal Diversity 105:1-16 ((https://doi.org/10.1007/s13225-020-00466-2)) |
| Otu11400 | 2 | k__Fungi(100);p__Ascomycota(100);c__Saccharomycetes(100);o__Saccharomycetales(100);f__Metschnikowiaceae(100);g__Metschnikowia(100);g__Metschnikowia_unclassified(100); | Metschnikowia | 13 | Saprotroph | \|Undefined Saprotroph\| | Yeast | NULL | Probable | As nectar-tap_saprotroph (Põlme et al. 2020); Decay sugar-rich substrates (Põlme et al. 2020) | Põlme S, et al. 2020. Fungal Diversity 105:1-16 ((https://doi.org/10.1007/s13225-020-00466-2)) |
| Otu11402 | 2 | k__Fungi(100);p__Ascomycota(100);c__Saccharomycetes(100);o__Saccharomycetales(100);f__Metschnikowiaceae(100);g__Metschnikowia(100);g__Metschnikowia_unclassified(100); | Metschnikowia | 13 | Saprotroph | \|Undefined Saprotroph\| | Yeast | NULL | Probable | As nectar-tap_saprotroph (Põlme et al. 2020); Decay sugar-rich substrates (Põlme et al. 2020) | Põlme S, et al. 2020. Fungal Diversity 105:1-16 ((https://doi.org/10.1007/s13225-020-00466-2)) |
| Otu11403 | 2 | k__Fungi(100);p__Ascomycota(100);c__Saccharomycetes(100);o__Saccharomycetales(100);f__Metschnikowiaceae(100);g__Metschnikowia(100);g__Metschnikowia_unclassified(100); | Metschnikowia | 13 | Saprotroph | \|Undefined Saprotroph\| | Yeast | NULL | Probable | As nectar-tap_saprotroph (Põlme et al. 2020); Decay sugar-rich substrates (Põlme et al. 2020) | Põlme S, et al. 2020. Fungal Diversity 105:1-16 ((https://doi.org/10.1007/s13225-020-00466-2)) |
| Otu11404 | 2 | k__Fungi(100);p__Ascomycota(100);c__Saccharomycetes(100);o__Saccharomycetales(100);f__Metschnikowiaceae(100);g__Metschnikowia(100);g__Metschnikowia_unclassified(100); | Metschnikowia | 13 | Saprotroph | \|Undefined Saprotroph\| | Yeast | NULL | Probable | As nectar-tap_saprotroph (Põlme et al. 2020); Decay sugar-rich substrates (Põlme et al. 2020) | Põlme S, et al. 2020. Fungal Diversity 105:1-16 ((https://doi.org/10.1007/s13225-020-00466-2)) |
| Otu11405 | 2 | k__Fungi(100);p__Ascomycota(100);c__Saccharomycetes(100);o__Saccharomycetales(100);f__Metschnikowiaceae(100);g__Metschnikowia(100);g__Metschnikowia_unclassified(100); | Metschnikowia | 13 | Saprotroph | \|Undefined Saprotroph\| | Yeast | NULL | Probable | As nectar-tap_saprotroph (Põlme et al. 2020); Decay sugar-rich substrates (Põlme et al. 2020) | Põlme S, et al. 2020. Fungal Diversity 105:1-16 ((https://doi.org/10.1007/s13225-020-00466-2)) |
| Otu11407 | 2 | k__Fungi(100);p__Ascomycota(100);c__Saccharomycetes(100);o__Saccharomycetales(100);f__Metschnikowiaceae(100);g__Metschnikowia(100);s__Metschnikowia_pulcherrima(100); | Metschnikowia | 13 | Saprotroph | \|Undefined Saprotroph\| | Yeast | NULL | Probable | As nectar-tap_saprotroph (Põlme et al. 2020); Decay sugar-rich substrates (Põlme et al. 2020) | Põlme S, et al. 2020. Fungal Diversity 105:1-16 ((https://doi.org/10.1007/s13225-020-00466-2)) |
| Otu11409 | 2 | k__Fungi(100);p__Ascomycota(100);c__Saccharomycetes(100);o__Saccharomycetales(100);f__Metschnikowiaceae(100);g__Metschnikowia(100);s__Metschnikowia_pulcherrima(100); | Metschnikowia | 13 | Saprotroph | \|Undefined Saprotroph\| | Yeast | NULL | Probable | As nectar-tap_saprotroph (Põlme et al. 2020); Decay sugar-rich substrates (Põlme et al. 2020) | Põlme S, et al. 2020. Fungal Diversity 105:1-16 ((https://doi.org/10.1007/s13225-020-00466-2)) |
| Otu11410 | 2 | k__Fungi(100);p__Ascomycota(100);c__Saccharomycetes(100);o__Saccharomycetales(100);f__Metschnikowiaceae(100);g__Metschnikowia(100);g__Metschnikowia_unclassified(100); | Metschnikowia | 13 | Saprotroph | \|Undefined Saprotroph\| | Yeast | NULL | Probable | As nectar-tap_saprotroph (Põlme et al. 2020); Decay sugar-rich substrates (Põlme et al. 2020) | Põlme S, et al. 2020. Fungal Diversity 105:1-16 ((https://doi.org/10.1007/s13225-020-00466-2)) |
| Otu11411 | 2 | k__Fungi(100);p__Ascomycota(100);c__Saccharomycetes(100);o__Saccharomycetales(100);f__Metschnikowiaceae(100);g__Metschnikowia(100);s__Metschnikowia_pulcherrima(100); | Metschnikowia | 13 | Saprotroph | \|Undefined Saprotroph\| | Yeast | NULL | Probable | As nectar-tap_saprotroph (Põlme et al. 2020); Decay sugar-rich substrates (Põlme et al. 2020) | Põlme S, et al. 2020. Fungal Diversity 105:1-16 ((https://doi.org/10.1007/s13225-020-00466-2)) |
| Otu11412 | 2 | k__Fungi(100);p__Basidiomycota(100);c__Tremellomycetes(100);o__Tremellales(100);f__Tremellales_family_Incertae_sedis(100);g__Hannaella(100);s__Hannaella_luteola(100); | Hannaella | 13 | Pathotroph-Saprotroph | \|Animal Parasite\|-Undefined Saprotroph | Yeast | NULL | Probable | As animal_decomposer (Põlme et al. 2020); Decay animal material (Põlme et al. 2020) | Põlme S, et al. 2020. Fungal Diversity 105:1-16 ((https://doi.org/10.1007/s13225-020-00466-2)) |

| | | | | | | | | | | | |
|---|---|---|---|---|---|---|---|---|---|---|---|
| Otu11413 | 2 | k__Fungi(100);p__Ascomycota(100);c__Saccharomycetes(100);o__Saccharomycetales(100);f__Saccharomycetaceae(100);g__Kazachstania(100);s__Kazachstania_intestinalis(100); | Kazachstania | 13 | Saprotroph | \|Undefined Saprotroph\| | Yeast | NULL | Probable | As nectar-tap_saprotroph (Põlme et al. 2020); Decay sugar-rich substrates (Põlme et al. 2020) | Põlme S, et al. 2020. Fungal Diversity 105:1-16 ((https://doi.org/10.1007/s13225-020-00466-2)) |
| Otu11414 | 2 | k__Fungi(100);p__Ascomycota(100);c__Saccharomycetes(100);o__Saccharomycetales(100);f__Metschnikowiaceae(100);g__Metschnikowia(100);g__Metschnikowia_unclassified(100); | Metschnikowia | 13 | Saprotroph | \|Undefined Saprotroph\| | Yeast | NULL | Probable | As nectar-tap_saprotroph (Põlme et al. 2020); Decay sugar-rich substrates (Põlme et al. 2020) | Põlme S, et al. 2020. Fungal Diversity 105:1-16 ((https://doi.org/10.1007/s13225-020-00466-2)) |
| Otu11415 | 2 | k__Fungi(100);p__Ascomycota(100);c__Saccharomycetes(100);o__Saccharomycetales(100);f__Metschnikowiaceae(100);g__Metschnikowia(100);g__Metschnikowia_unclassified(100); | Metschnikowia | 13 | Saprotroph | \|Undefined Saprotroph\| | Yeast | NULL | Probable | As nectar-tap_saprotroph (Põlme et al. 2020); Decay sugar-rich substrates (Põlme et al. 2020) | Põlme S, et al. 2020. Fungal Diversity 105:1-16 ((https://doi.org/10.1007/s13225-020-00466-2)) |
| Otu11416 | 2 | k__Fungi(100);p__Ascomycota(100);c__Saccharomycetes(100);o__Saccharomycetales(100);f__Metschnikowiaceae(100);g__Metschnikowia(100);s__Metschnikowia_pulcherrima(100); | Metschnikowia | 13 | Saprotroph | \|Undefined Saprotroph\| | Yeast | NULL | Probable | As nectar-tap_saprotroph (Põlme et al. 2020); Decay sugar-rich substrates (Põlme et al. 2020) | Põlme S, et al. 2020. Fungal Diversity 105:1-16 ((https://doi.org/10.1007/s13225-020-00466-2)) |
| Otu11417 | 2 | k__Fungi(100);p__Ascomycota(100);c__Saccharomycetes(100);o__Saccharomycetales(100);f__Metschnikowiaceae(100);g__Metschnikowia(100);s__Metschnikowia_pulcherrima(100); | Metschnikowia | 13 | Saprotroph | \|Undefined Saprotroph\| | Yeast | NULL | Probable | As nectar-tap_saprotroph (Põlme et al. 2020); Decay sugar-rich substrates (Põlme et al. 2020) | Põlme S, et al. 2020. Fungal Diversity 105:1-16 ((https://doi.org/10.1007/s13225-020-00466-2)) |
| Otu11418 | 2 | k__Fungi(100);p__Ascomycota(100);c__Saccharomycetes(100);o__Saccharomycetales(100);f__Metschnikowiaceae(100);g__Metschnikowia(100);g__Metschnikowia_unclassified(100); | Metschnikowia | 13 | Saprotroph | \|Undefined Saprotroph\| | Yeast | NULL | Probable | As nectar-tap_saprotroph (Põlme et al. 2020); Decay sugar-rich substrates (Põlme et al. 2020) | Põlme S, et al. 2020. Fungal Diversity 105:1-16 ((https://doi.org/10.1007/s13225-020-00466-2)) |
| Otu11419 | 2 | k__Fungi(100);p__Ascomycota(100);c__Sordariomycetes(100);o__Sordariales(100);f__Chaetomiaceae(100);g__Humicola(100);g__Humicola_unclassified(100); | Humicola | 13 | Pathotroph-Saprotroph | Plant Pathogen-Undefined Saprotroph-\|Wood Saprotroph\| | Microfungus | Soft Rot | Probable | Decay wood (Põlme et al. 2020) | Seehann G, et al. 1975. List of Fungi in Soft-Rot Tissue Tests. Institute fur Holzbiologie und Holzschutz, Bundesforschungsanstalt fur Forst - und Holzwirtshaft, Hamburg - Lohbrugge, Germany; Costa IPMW, et al. 2012. Mycotaxon 119:e26 ((http://mycotaxon.com/resources/checklists/costa_v119_checklist.pdf)); Tedersoo L, et al. 2014. Science 346:e1256688 ((https://doi.org/10.1126/science.1256688)); Põlme S, et al. 2020. Fungal Diversity 105:1-16 ((https://doi.org/10.1007/s13225-020-00466-2)) |
| Otu11420 | 2 | k__Fungi(100);p__Ascomycota(100);c__Saccharomycetes(100);o__Saccharomycetales(100);f__Metschnikowiaceae(100);g__Metschnikowia(100);g__Metschnikowia_unclassified(100); | Metschnikowia | 13 | Saprotroph | \|Undefined Saprotroph\| | Yeast | NULL | Probable | As nectar-tap_saprotroph (Põlme et al. 2020); Decay sugar-rich substrates (Põlme et al. 2020) | Põlme S, et al. 2020. Fungal Diversity 105:1-16 ((https://doi.org/10.1007/s13225-020-00466-2)) |
| Otu11421 | 2 | k__Fungi(100);p__Ascomycota(100);c__Saccharomycetes(100);o__Saccharomycetales(100);f__Metschnikowiaceae(100);g__Metschnikowia(100);s__Metschnikowia_pulcherrima(100); | Metschnikowia | 13 | Saprotroph | \|Undefined Saprotroph\| | Yeast | NULL | Probable | As nectar-tap_saprotroph (Põlme et al. 2020); Decay sugar-rich substrates (Põlme et al. 2020) | Põlme S, et al. 2020. Fungal Diversity 105:1-16 ((https://doi.org/10.1007/s13225-020-00466-2)) |
| Otu11423 | 2 | k__Fungi(100);p__Ascomycota(100);c__Saccharomycetes(100);o__Saccharomycetales(100);f__Saccharomycetaceae(100);g__Kazachstania(100);s__Kazachstania_intestinalis(100); | Kazachstania | 13 | Saprotroph | \|Undefined Saprotroph\| | Yeast | NULL | Probable | As nectar-tap_saprotroph (Põlme et al. 2020); Decay sugar-rich substrates (Põlme et al. 2020) | Põlme S, et al. 2020. Fungal Diversity 105:1-16 ((https://doi.org/10.1007/s13225-020-00466-2)) |
| Otu11424 | 2 | k__Fungi(100);p__Ascomycota(100);c__Saccharomycetes(100);o__Saccharomycetales(100);f__Saccharomycetaceae(100);g__Kazachstania(100);s__Kazachstania_intestinalis(100); | Kazachstania | 13 | Saprotroph | \|Undefined Saprotroph\| | Yeast | NULL | Probable | As nectar-tap_saprotroph (Põlme et al. 2020); Decay sugar-rich substrates (Põlme et al. 2020) | Põlme S, et al. 2020. Fungal Diversity 105:1-16 ((https://doi.org/10.1007/s13225-020-00466-2)) |
| Otu11425 | 2 | k__Fungi(100);p__Ascomycota(100);c__Saccharomycetes(100);o__Saccharomycetales(100);f__Saccharomycetaceae(100);g__Kazachstania(100);s__Kazachstania_intestinalis(100); | Kazachstania | 13 | Saprotroph | \|Undefined Saprotroph\| | Yeast | NULL | Probable | As nectar-tap_saprotroph (Põlme et al. 2020); Decay sugar-rich substrates (Põlme et al. 2020) | Põlme S, et al. 2020. Fungal Diversity 105:1-16 ((https://doi.org/10.1007/s13225-020-00466-2)) |
| Otu11426 | 2 | k__Fungi(100);p__Ascomycota(100);c__Saccharomycetes(100);o__Saccharomycetales(100);f__Saccharomycetales_family_Incertae_sedis(100);g__Candida(100);g__Candida_unclassified(100); | Candida | 13 | Pathotroph-Saprotroph-Symbiotroph | Animal Pathogen-Endophyte-Endosymbiont-Epiphyte-\|Undefined Saprotroph\| | Dimorphic Yeast-Yeast | NULL | Probable | As nectar-tap_saprotroph (Põlme et al. 2020); Decay sugar-rich substrates (Põlme et al. 2020); This asexual genus is polyphyletic and contain species that function very differently depend on what they are. Caution should be taken when interpreting guilds for Candida species. | Manolakaki D, et al. 2010. Virulence. 1: 367-375 ((https://doi.org/10.4161/viru.1.5.12796)); Mekbib SB, et al. 2011. Tropical Plant Pathology 36:233-240 ((https://doi.org/10.1590/S1982-56762011000400004)); Põlme S, et al. 2020. Fungal Diversity 105:1-16 ((https://doi.org/10.1007/s13225-020-00466-2)) |
| Otu11427 | 2 | k__Fungi(100);p__Ascomycota(100);c__Saccharomycetes(100);o__Saccharomycetales(100);f__Metschnikowiaceae(100);g__Metschnikowia(100);s__Metschnikowia_pulcherrima(100); | Metschnikowia | 13 | Saprotroph | \|Undefined Saprotroph\| | Yeast | NULL | Probable | As nectar-tap_saprotroph (Põlme et al. 2020); Decay sugar-rich substrates (Põlme et al. 2020) | Põlme S, et al. 2020. Fungal Diversity 105:1-16 ((https://doi.org/10.1007/s13225-020-00466-2)) |
| Otu11430 | 2 | k__Fungi(100);p__Ascomycota(100);c__Saccharomycetes(100);o__Saccharomycetales(100);f__Metschnikowiaceae(100);g__Metschnikowia(100);g__Metschnikowia_unclassified(100); | Metschnikowia | 13 | Saprotroph | \|Undefined Saprotroph\| | Yeast | NULL | Probable | As nectar-tap_saprotroph (Põlme et al. 2020); Decay sugar-rich substrates (Põlme et al. 2020) | Põlme S, et al. 2020. Fungal Diversity 105:1-16 ((https://doi.org/10.1007/s13225-020-00466-2)) |
| Otu11431 | 2 | k__Fungi(100);p__Ascomycota(100);c__Saccharomycetes(100);o__Saccharomycetales(100);f__Metschnikowiaceae(100);g__Metschnikowia(100);s__Metschnikowia_pulcherrima(100); | Metschnikowia | 13 | Saprotroph | \|Undefined Saprotroph\| | Yeast | NULL | Probable | As nectar-tap_saprotroph (Põlme et al. 2020); Decay sugar-rich substrates (Põlme et al. 2020) | Põlme S, et al. 2020. Fungal Diversity 105:1-16 ((https://doi.org/10.1007/s13225-020-00466-2)) |
| Otu11433 | 2 | k__Fungi(100);p__Ascomycota(100);c__Saccharomycetes(100);o__Saccharomycetales(100);f__Metschnikowiaceae(100);g__Metschnikowia(100);s__Metschnikowia_pulcherrima(100); | Metschnikowia | 13 | Saprotroph | \|Undefined Saprotroph\| | Yeast | NULL | Probable | As nectar-tap_saprotroph (Põlme et al. 2020); Decay sugar-rich substrates (Põlme et al. 2020) | Põlme S, et al. 2020. Fungal Diversity 105:1-16 ((https://doi.org/10.1007/s13225-020-00466-2)) |
| Otu11434 | 2 | k__Fungi(100);p__Ascomycota(100);c__Saccharomycetes(100);o__Saccharomycetales(100);f__Metschnikowiaceae(100);g__Metschnikowia(100);s__Metschnikowia_pulcherrima(100); | Metschnikowia | 13 | Saprotroph | \|Undefined Saprotroph\| | Yeast | NULL | Probable | As nectar-tap_saprotroph (Põlme et al. 2020); Decay sugar-rich substrates (Põlme et al. 2020) | Põlme S, et al. 2020. Fungal Diversity 105:1-16 ((https://doi.org/10.1007/s13225-020-00466-2)) |
| Otu11436 | 2 | k__Fungi(100);p__Ascomycota(100);c__Saccharomycetes(100);o__Saccharomycetales(100);f__Metschnikowiaceae(100);g__Metschnikowia(100);s__Metschnikowia_pulcherrima(100); | Metschnikowia | 13 | Saprotroph | \|Undefined Saprotroph\| | Yeast | NULL | Probable | As nectar-tap_saprotroph (Põlme et al. 2020); Decay sugar-rich substrates (Põlme et al. 2020) | Põlme S, et al. 2020. Fungal Diversity 105:1-16 ((https://doi.org/10.1007/s13225-020-00466-2)) |
| Otu11437 | 2 | k__Fungi(100);p__Ascomycota(100);c__Saccharomycetes(100);o__Saccharomycetales(100);f__Metschnikowiaceae(100);g__Metschnikowia(100);s__Metschnikowia_pulcherrima(100); | Metschnikowia | 13 | Saprotroph | \|Undefined Saprotroph\| | Yeast | NULL | Probable | As nectar-tap_saprotroph (Põlme et al. 2020); Decay sugar-rich substrates (Põlme et al. 2020) | Põlme S, et al. 2020. Fungal Diversity 105:1-16 ((https://doi.org/10.1007/s13225-020-00466-2)) |
| Otu11438 | 2 | k__Fungi(100);p__Ascomycota(100);c__Saccharomycetes(100);o__Saccharomycetales(100);f__Metschnikowiaceae(100);g__Metschnikowia(100);s__Metschnikowia_pulcherrima(100); | Metschnikowia | 13 | Saprotroph | \|Undefined Saprotroph\| | Yeast | NULL | Probable | As nectar-tap_saprotroph (Põlme et al. 2020); Decay sugar-rich substrates (Põlme et al. 2020) | Põlme S, et al. 2020. Fungal Diversity 105:1-16 ((https://doi.org/10.1007/s13225-020-00466-2)) |
| Otu11441 | 2 | k__Fungi(100);p__Ascomycota(100);c__Saccharomycetes(100);o__Saccharomycetales(100);f__Metschnikowiaceae(100);g__Metschnikowia(100);s__Metschnikowia_pulcherrima(100); | Metschnikowia | 13 | Saprotroph | \|Undefined Saprotroph\| | Yeast | NULL | Probable | As nectar-tap_saprotroph (Põlme et al. 2020); Decay sugar-rich substrates (Põlme et al. 2020) | Põlme S, et al. 2020. Fungal Diversity 105:1-16 ((https://doi.org/10.1007/s13225-020-00466-2)) |
| Otu11442 | 2 | k__Fungi(100);p__Ascomycota(100);c__Saccharomycetes(100);o__Saccharomycetales(100);f__Metschnikowiaceae(100);g__Metschnikowia(100);g__Metschnikowia_unclassified(100); | Metschnikowia | 13 | Saprotroph | \|Undefined Saprotroph\| | Yeast | NULL | Probable | As nectar-tap_saprotroph (Põlme et al. 2020); Decay sugar-rich substrates (Põlme et al. 2020) | Põlme S, et al. 2020. Fungal Diversity 105:1-16 ((https://doi.org/10.1007/s13225-020-00466-2)) |
| Otu11444 | 2 | k__Fungi(100);p__Ascomycota(100);c__Saccharomycetes(100);o__Saccharomycetales(100);f__Metschnikowiaceae(100);g__Metschnikowia(100);s__Metschnikowia_pulcherrima(100); | Metschnikowia | 13 | Saprotroph | \|Undefined Saprotroph\| | Yeast | NULL | Probable | As nectar-tap_saprotroph (Põlme et al. 2020); Decay sugar-rich substrates (Põlme et al. 2020) | Põlme S, et al. 2020. Fungal Diversity 105:1-16 ((https://doi.org/10.1007/s13225-020-00466-2)) |
| Otu11445 | 2 | k__Fungi(100);p__Ascomycota(100);c__Saccharomycetes(100);o__Saccharomycetales(100);f__Metschnikowiaceae(100);g__Metschnikowia(100);s__Metschnikowia_pulcherrima(100); | Metschnikowia | 13 | Saprotroph | \|Undefined Saprotroph\| | Yeast | NULL | Probable | As nectar-tap_saprotroph (Põlme et al. 2020); Decay sugar-rich substrates (Põlme et al. 2020) | Põlme S, et al. 2020. Fungal Diversity 105:1-16 ((https://doi.org/10.1007/s13225-020-00466-2)) |
| Otu11447 | 2 | k__Fungi(100);p__Ascomycota(100);c__Saccharomycetes(100);o__Saccharomycetales(100);f__Metschnikowiaceae(100);g__Metschnikowia(100);g__Metschnikowia_unclassified(100); | Metschnikowia | 13 | Saprotroph | \|Undefined Saprotroph\| | Yeast | NULL | Probable | As nectar-tap_saprotroph (Põlme et al. 2020); Decay sugar-rich substrates (Põlme et al. 2020) | Põlme S, et al. 2020. Fungal Diversity 105:1-16 ((https://doi.org/10.1007/s13225-020-00466-2)) |
| Otu11449 | 2 | k__Fungi(100);p__Ascomycota(100);c__Saccharomycetes(100);o__Saccharomycetales(100);f__Metschnikowiaceae(100);g__Metschnikowia(100);g__Metschnikowia_unclassified(100); | Metschnikowia | 13 | Saprotroph | \|Undefined Saprotroph\| | Yeast | NULL | Probable | As nectar-tap_saprotroph (Põlme et al. 2020); Decay sugar-rich substrates (Põlme et al. 2020) | Põlme S, et al. 2020. Fungal Diversity 105:1-16 ((https://doi.org/10.1007/s13225-020-00466-2)) |
| Otu11450 | 2 | k__Fungi(100);p__Ascomycota(100);c__Saccharomycetes(100);o__Saccharomycetales(100);f__Metschnikowiaceae(100);g__Metschnikowia(100);s__Metschnikowia_pulcherrima(100); | Metschnikowia | 13 | Saprotroph | \|Undefined Saprotroph\| | Yeast | NULL | Probable | As nectar-tap_saprotroph (Põlme et al. 2020); Decay sugar-rich substrates (Põlme et al. 2020) | Põlme S, et al. 2020. Fungal Diversity 105:1-16 ((https://doi.org/10.1007/s13225-020-00466-2)) |
| Otu11452 | 2 | k__Fungi(100);p__Ascomycota(100);c__Saccharomycetes(100);o__Saccharomycetales(100);f__Metschnikowiaceae(100);g__Metschnikowia(100);g__Metschnikowia_unclassified(100); | Metschnikowia | 13 | Saprotroph | \|Undefined Saprotroph\| | Yeast | NULL | Probable | As nectar-tap_saprotroph (Põlme et al. 2020); Decay sugar-rich substrates (Põlme et al. 2020) | Põlme S, et al. 2020. Fungal Diversity 105:1-16 ((https://doi.org/10.1007/s13225-020-00466-2)) |
| Otu11454 | 2 | k__Fungi(100);p__Ascomycota(100);c__Saccharomycetes(100);o__Saccharomycetales(100);f__Metschnikowiaceae(100);g__Metschnikowia(100);s__Metschnikowia_pulcherrima(100); | Metschnikowia | 13 | Saprotroph | \|Undefined Saprotroph\| | Yeast | NULL | Probable | As nectar-tap_saprotroph (Põlme et al. 2020); Decay sugar-rich substrates (Põlme et al. 2020) | Põlme S, et al. 2020. Fungal Diversity 105:1-16 ((https://doi.org/10.1007/s13225-020-00466-2)) |
| Otu11455 | 2 | k__Fungi(100);p__Ascomycota(100);c__Saccharomycetes(100);o__Saccharomycetales(100);f__Metschnikowiaceae(100);g__Metschnikowia(100);s__Metschnikowia_pulcherrima(100); | Metschnikowia | 13 | Saprotroph | \|Undefined Saprotroph\| | Yeast | NULL | Probable | As nectar-tap_saprotroph (Põlme et al. 2020); Decay sugar-rich substrates (Põlme et al. 2020) | Põlme S, et al. 2020. Fungal Diversity 105:1-16 ((https://doi.org/10.1007/s13225-020-00466-2)) |
| Otu11458 | 2 | k__Fungi(100);p__Ascomycota(100);c__Saccharomycetes(100);o__Saccharomycetales(100);f__Metschnikowiaceae(100);g__Metschnikowia(100);s__Metschnikowia_pulcherrima(100); | Metschnikowia | 13 | Saprotroph | \|Undefined Saprotroph\| | Yeast | NULL | Probable | As nectar-tap_saprotroph (Põlme et al. 2020); Decay sugar-rich substrates (Põlme et al. 2020) | Põlme S, et al. 2020. Fungal Diversity 105:1-16 ((https://doi.org/10.1007/s13225-020-00466-2)) |
| Otu11459 | 2 | k__Fungi(100);p__Ascomycota(100);c__Sordariomycetes(100);o__Coniochaetales(100);f__Coniochaetaceae(100);g__Coniochaeta(100);s__Coniochaeta_prunicola(100); | Coniochaeta | 13 | Pathotroph-Saprotroph-Symbiotroph | Animal Pathogen-Dung Saprotroph-Endophyte-Lichen Parasite-Plant Pathogen-\|Undefined Saprotroph\| | Microfungus | NULL | Possible | As foliar_endophyte (Põlme et al. 2020); Decay leaf, fruit, seed, soil, dung, animal material (Põlme et al. 2020) | Bell A. 1983. Dung Fungi: An Illustrated Guide to Coprophilous Fungi in New Zealand. Victoria University Press, Wellington (ISBN: 978-0864730015); Damm U, et al. 2010. Persoonia 24:60-80 ((http://doi.org/10.3767/003158510X500705)); Costa IPMW, et al. 2012. Mycotaxon 119:e26 ((http://mycotaxon.com/resources/checklists/costa_v119_checklist.pdf)); Tedersoo L, et al. 2014. Science 346:1256688 ((https://doi.org/:10.1126/science.1256688)); Irinyi L, et al. 2015. Medical Mycology 53:313-337 ((https://doi.org/10.1093/mmy/myv008)); Lawrey JD, Diederich P. 2016. http://www.lichenicolous.net (accessed 24 May 2017) ((http://www.lichenicolous.net)); Põlme S, et al. 2020. Fungal Diversity 105:1-16 ((https://doi.org/10.1007/s13225-020-00466-2)) |
| Otu11463 | 2 | k__Fungi(100);p__Ascomycota(100);c__Saccharomycetes(100);o__Saccharomycetales(100);f__Metschnikowiaceae(100);g__Metschnikowia(100);g__Metschnikowia_unclassified(100); | Metschnikowia | 13 | Saprotroph | \|Undefined Saprotroph\| | Yeast | NULL | Probable | As nectar-tap_saprotroph (Põlme et al. 2020); Decay sugar-rich substrates (Põlme et al. 2020) | Põlme S, et al. 2020. Fungal Diversity 105:1-16 ((https://doi.org/10.1007/s13225-020-00466-2)) |
| Otu11464 | 2 | k__Fungi(100);p__Ascomycota(100);c__Saccharomycetes(100);o__Saccharomycetales(100);f__Metschnikowiaceae(100);g__Metschnikowia(100);s__Metschnikowia_pulcherrima(100); | Metschnikowia | 13 | Saprotroph | \|Undefined Saprotroph\| | Yeast | NULL | Probable | As nectar-tap_saprotroph (Põlme et al. 2020); Decay sugar-rich substrates (Põlme et al. 2020) | Põlme S, et al. 2020. Fungal Diversity 105:1-16 ((https://doi.org/10.1007/s13225-020-00466-2)) |
| Otu11465 | 2 | k__Fungi(100);p__Ascomycota(100);c__Saccharomycetes(100);o__Saccharomycetales(100);f__Metschnikowiaceae(100);g__Metschnikowia(100);s__Metschnikowia_pulcherrima(100); | Metschnikowia | 13 | Saprotroph | \|Undefined Saprotroph\| | Yeast | NULL | Probable | As nectar-tap_saprotroph (Põlme et al. 2020); Decay sugar-rich substrates (Põlme et al. 2020) | Põlme S, et al. 2020. Fungal Diversity 105:1-16 ((https://doi.org/10.1007/s13225-020-00466-2)) |
| Otu11467 | 2 | k__Fungi(100);p__Ascomycota(100);c__Saccharomycetes(100);o__Saccharomycetales(100);f__Metschnikowiaceae(100);g__Metschnikowia(100);s__Metschnikowia_pulcherrima(100); | Metschnikowia | 13 | Saprotroph | \|Undefined Saprotroph\| | Yeast | NULL | Probable | As nectar-tap_saprotroph (Põlme et al. 2020); Decay sugar-rich substrates (Põlme et al. 2020) | Põlme S, et al. 2020. Fungal Diversity 105:1-16 ((https://doi.org/10.1007/s13225-020-00466-2)) |
| Otu11468 | 2 | k__Fungi(100);p__Ascomycota(100);c__Saccharomycetes(100);o__Saccharomycetales(100);f__Metschnikowiaceae(100);g__Metschnikowia(100);s__Metschnikowia_pulcherrima(100); | Metschnikowia | 13 | Saprotroph | \|Undefined Saprotroph\| | Yeast | NULL | Probable | As nectar-tap_saprotroph (Põlme et al. 2020); Decay sugar-rich substrates (Põlme et al. 2020) | Põlme S, et al. 2020. Fungal Diversity 105:1-16 ((https://doi.org/10.1007/s13225-020-00466-2)) |
| Otu11471 | 2 | k__Fungi(100);p__Ascomycota(100);c__Sordariomycetes(100);o__Hypocreales(100);o__Hypocreales_unclassified(100);o__Hypocreales_unclassified(100);o__Hypocreales_unclassified(100); | Hypocreales | 7 | Saprotroph | Undefined Saprotroph | Microfungus | NULL | Possible | NULL | Sterkenburg E, et al. 2015. New Phytologist 207:1145-1158 ((https://doi.org/10.1111/nph.13426)) |
| Otu11473 | 2 | k__Fungi(100);p__Ascomycota(100);c__Saccharomycetes(100);o__Saccharomycetales(100);f__Metschnikowiaceae(100);g__Metschnikowia(100);s__Metschnikowia_pulcherrima(100); | Metschnikowia | 13 | Saprotroph | \|Undefined Saprotroph\| | Yeast | NULL | Probable | As nectar-tap_saprotroph (Põlme et al. 2020); Decay sugar-rich substrates (Põlme et al. 2020) | Põlme S, et al. 2020. Fungal Diversity 105:1-16 ((https://doi.org/10.1007/s13225-020-00466-2)) |
| Otu11474 | 2 | k__Fungi(100);p__Ascomycota(100);c__Saccharomycetes(100);o__Saccharomycetales(100);f__Metschnikowiaceae(100);g__Metschnikowia(100);s__Metschnikowia_pulcherrima(100); | Metschnikowia | 13 | Saprotroph | \|Undefined Saprotroph\| | Yeast | NULL | Probable | As nectar-tap_saprotroph (Põlme et al. 2020); Decay sugar-rich substrates (Põlme et al. 2020) | Põlme S, et al. 2020. Fungal Diversity 105:1-16 ((https://doi.org/10.1007/s13225-020-00466-2)) |
| Otu11475 | 2 | k__Fungi(100);p__Ascomycota(100);c__Saccharomycetes(100);o__Saccharomycetales(100);f__Metschnikowiaceae(100);g__Metschnikowia(100);s__Metschnikowia_pulcherrima(100); | Metschnikowia | 13 | Saprotroph | \|Undefined Saprotroph\| | Yeast | NULL | Probable | As nectar-tap_saprotroph (Põlme et al. 2020); Decay sugar-rich substrates (Põlme et al. 2020) | Põlme S, et al. 2020. Fungal Diversity 105:1-16 ((https://doi.org/10.1007/s13225-020-00466-2)) |
| Otu11477 | 2 | k__Fungi(100);p__Ascomycota(100);c__Saccharomycetes(100);o__Saccharomycetales(100);f__Metschnikowiaceae(100);g__Metschnikowia(100);s__Metschnikowia_pulcherrima(100); | Metschnikowia | 13 | Saprotroph | \|Undefined Saprotroph\| | Yeast | NULL | Probable | As nectar-tap_saprotroph (Põlme et al. 2020); Decay sugar-rich substrates (Põlme et al. 2020) | Põlme S, et al. 2020. Fungal Diversity 105:1-16 ((https://doi.org/10.1007/s13225-020-00466-2)) |
| Otu11478 | 2 | k__Fungi(100);p__Ascomycota(100);c__Saccharomycetes(100);o__Saccharomycetales(100);f__Metschnikowiaceae(100);g__Metschnikowia(100);s__Metschnikowia_pulcherrima(100); | Metschnikowia | 13 | Saprotroph | \|Undefined Saprotroph\| | Yeast | NULL | Probable | As nectar-tap_saprotroph (Põlme et al. 2020); Decay sugar-rich substrates (Põlme et al. 2020) | Põlme S, et al. 2020. Fungal Diversity 105:1-16 ((https://doi.org/10.1007/s13225-020-00466-2)) |
| Otu11480 | 2 | k__Fungi(100);p__Ascomycota(100);c__Saccharomycetes(100);o__Saccharomycetales(100);f__Metschnikowiaceae(100);g__Metschnikowia(100);s__Metschnikowia_pulcherrima(100); | Metschnikowia | 13 | Saprotroph | \|Undefined Saprotroph\| | Yeast | NULL | Probable | As nectar-tap_saprotroph (Põlme et al. 2020); Decay sugar-rich substrates (Põlme et al. 2020) | Põlme S, et al. 2020. Fungal Diversity 105:1-16 ((https://doi.org/10.1007/s13225-020-00466-2)) |
| Otu11483 | 2 | k__Fungi(100);p__Ascomycota(100);c__Saccharomycetes(100);o__Saccharomycetales(100);f__Metschnikowiaceae(100);g__Metschnikowia(100);g__Metschnikowia_unclassified(100); | Metschnikowia | 13 | Saprotroph | \|Undefined Saprotroph\| | Yeast | NULL | Probable | As nectar-tap_saprotroph (Põlme et al. 2020); Decay sugar-rich substrates (Põlme et al. 2020) | Põlme S, et al. 2020. Fungal Diversity 105:1-16 ((https://doi.org/10.1007/s13225-020-00466-2)) |
| Otu11484 | 2 | k__Fungi(100);p__Ascomycota(100);c__Saccharomycetes(100);o__Saccharomycetales(100);f__Metschnikowiaceae(100);g__Metschnikowia(100);g__Metschnikowia_unclassified(100); | Metschnikowia | 13 | Saprotroph | \|Undefined Saprotroph\| | Yeast | NULL | Probable | As nectar-tap_saprotroph (Põlme et al. 2020); Decay sugar-rich substrates (Põlme et al. 2020) | Põlme S, et al. 2020. Fungal Diversity 105:1-16 ((https://doi.org/10.1007/s13225-020-00466-2)) |
| Otu11485 | 2 | k__Fungi(100);p__Ascomycota(100);c__Saccharomycetes(100);o__Saccharomycetales(100);f__Metschnikowiaceae(100);g__Metschnikowia(100);g__Metschnikowia_unclassified(100); | Metschnikowia | 13 | Saprotroph | \|Undefined Saprotroph\| | Yeast | NULL | Probable | As nectar-tap_saprotroph (Põlme et al. 2020); Decay sugar-rich substrates (Põlme et al. 2020) | Põlme S, et al. 2020. Fungal Diversity 105:1-16 ((https://doi.org/10.1007/s13225-020-00466-2)) |
| Otu11486 | 2 | k__Fungi(100);p__Ascomycota(100);c__Saccharomycetes(100);o__Saccharomycetales(100);f__Metschnikowiaceae(100);g__Metschnikowia(100);g__Metschnikowia_unclassified(100); | Metschnikowia | 13 | Saprotroph | \|Undefined Saprotroph\| | Yeast | NULL | Probable | As nectar-tap_saprotroph (Põlme et al. 2020); Decay sugar-rich substrates (Põlme et al. 2020) | Põlme S, et al. 2020. Fungal Diversity 105:1-16 ((https://doi.org/10.1007/s13225-020-00466-2)) |
| Otu11487 | 2 | k__Fungi(100);p__Ascomycota(100);c__Saccharomycetes(100);o__Saccharomycetales(100);f__Metschnikowiaceae(100);g__Metschnikowia(100);g__Metschnikowia_unclassified(100); | Metschnikowia | 13 | Saprotroph | \|Undefined Saprotroph\| | Yeast | NULL | Probable | As nectar-tap_saprotroph (Põlme et al. 2020); Decay sugar-rich substrates (Põlme et al. 2020) | Põlme S, et al. 2020. Fungal Diversity 105:1-16 ((https://doi.org/10.1007/s13225-020-00466-2)) |
| Otu11488 | 2 | k__Fungi(100);p__Ascomycota(100);c__Saccharomycetes(100);o__Saccharomycetales(100);f__Metschnikowiaceae(100);g__Metschnikowia(100);g__Metschnikowia_unclassified(100); | Metschnikowia | 13 | Saprotroph | \|Undefined Saprotroph\| | Yeast | NULL | Probable | As nectar-tap_saprotroph (Põlme et al. 2020); Decay sugar-rich substrates (Põlme et al. 2020) | Põlme S, et al. 2020. Fungal Diversity 105:1-16 ((https://doi.org/10.1007/s13225-020-00466-2)) |
| Otu11490 | 2 | k__Fungi(100);p__Basidiomycota(100);c__Wallemiomycetes(100);o__Wallemiales(100);f__Wallemiaceae(100);g__Wallemia(100);s__Wallemia_sp_SJ_2014(100); | Wallemia | 13 | Saprotroph | \|Undefined Saprotroph\| | Microfungus | NULL | Probable | Decay sugar-rich substrates (Põlme et al. 2020) | Tedersoo L, et al. 2014. Science 346:e1256688 ((https://doi.org/10.1126/science.1256688)); Põlme S, et al. 2020. Fungal Diversity 105:1-16 ((https://doi.org/10.1007/s13225-020-00466-2)) |
| Otu11496 | 2 | k__Fungi(100);p__Ascomycota(100);c__Saccharomycetes(100);o__Saccharomycetales(100);f__Metschnikowiaceae(100);g__Metschnikowia(100);g__Metschnikowia_unclassified(100); | Metschnikowia | 13 | Saprotroph | \|Undefined Saprotroph\| | Yeast | NULL | Probable | As nectar-tap_saprotroph (Põlme et al. 2020); Decay sugar-rich substrates (Põlme et al. 2020) | Põlme S, et al. 2020. Fungal Diversity 105:1-16 ((https://doi.org/10.1007/s13225-020-00466-2)) |
| Otu11503 | 2 | k__Fungi(100);p__Ascomycota(100);c__Sordariomycetes(100);o__Hypocreales(100);f__Hypocreales_family_Incertae_sedis(100);g__Acremonium(100);s__Acremonium_sp_24_OA_2013(100); | Acremonium | 13 | Pathotroph-Saprotroph-Symbiotroph | Animal Pathogen-Endophyte-Fungal Parasite-Plant Pathogen-\|Undefined Saprotroph\|-Wood Saprotroph | Microfungus | Soft Rot | Possible | As foliar_endophyte (Põlme et al. 2020); culture (Hanada et al. 2010, Del Olmo; Decay leaf, fruit, seed, soil, dung, animal material (Põlme et al. 2020); Endophyte detection method; Ruiz & Arnold 2014); Likely opportunistic human pathogen (Irinyi et al. 2015) | Duncan CG, Eslyn WE. 1966. Mycologia 58:642-645 ((https://doi.org/10.2307/3757045)); Nilsson T. 1973. Studia Forestalia Suecica 104: 1-40 ((https://pub.epsilon.slu.se/5711/)); Worrall JJ, et al. 1997. Mycologia 89:199-219 ((https://doi.org/10.2307/3761073)); Hanada RE, et al. 2010. Fungal Biology 114:901-910 ((https://doi.org/10.1016/j.funbio.2010.08.006)); Del Olmo-Ruiz M, Arnold AE. 2014. Mycologia 106: 8-21 ((https://doi.org/10.3852/13-098)); Irinyi L, et al. 2015. Medical Mycology 53:313-337 ((https://doi.org/10.1093/mmy/myv008)); Põlme S, et al. 2020. Fungal Diversity 105:1-16 ((https://doi.org/10.1007/s13225-020-00466-2)) |
| Otu11506 | 2 | k__Fungi(100);p__Basidiomycota(100);c__Agaricomycetes(100);o__Agaricales(100);f__Pleurotaceae(100);g__Pleurotus(100);s__Pleurotus_dryinus(100); | Pleurotus | 13 | Pathotroph-Saprotroph | Endophyte-Nematophagous-Plant Pathogen-\|Wood Saprotroph\| | Agaricoid | White Rot | Probable | As white_rot (Põlme et al. 2020); As wood_pathogen (Põlme et al. 2020); Decay wood, animal material (Põlme et al. 2020); This genus is known to be nematophagous, capturing and consuming nematodes. | James TY, et al. 2006. Nature 443:818-822 ((https://doi.org/10.1038/nature05110)); Costa IPMW, et al. 2012. Mycotaxon 119:e26 ((http://mycotaxon.com/resources/checklists/costa_v119_checklist.pdf)); Tedersoo L, et al. 2014. Science 346:e1256688 ((https://doi.org/10.1126/science.1256688)); Põlme S, et al. 2020. Fungal Diversity 105:1-16 ((https://doi.org/10.1007/s13225-020-00466-2)); The American Phytopathological Society Common Names of Plant Diseases ((http://www.apsnet.org/publications/commonnames/Pages/default.aspx)) |
| Otu11510 | 2 | k__Fungi(100);p__Ascomycota(100);c__Saccharomycetes(100);o__Saccharomycetales(100);f__Metschnikowiaceae(100);g__Metschnikowia(100);g__Metschnikowia_unclassified(100); | Metschnikowia | 13 | Saprotroph | \|Undefined Saprotroph\| | Yeast | NULL | Probable | As nectar-tap_saprotroph (Põlme et al. 2020); Decay sugar-rich substrates (Põlme et al. 2020) | Põlme S, et al. 2020. Fungal Diversity 105:1-16 ((https://doi.org/10.1007/s13225-020-00466-2)) |
| Otu11511 | 2 | k__Fungi(100);p__Ascomycota(100);c__Saccharomycetes(100);o__Saccharomycetales(100);f__Metschnikowiaceae(100);g__Metschnikowia(100);g__Metschnikowia_unclassified(100); | Metschnikowia | 13 | Saprotroph | \|Undefined Saprotroph\| | Yeast | NULL | Probable | As nectar-tap_saprotroph (Põlme et al. 2020); Decay sugar-rich substrates (Põlme et al. 2020) | Põlme S, et al. 2020. Fungal Diversity 105:1-16 ((https://doi.org/10.1007/s13225-020-00466-2)) |
| Otu11512 | 2 | k__Fungi(100);p__Ascomycota(100);c__Saccharomycetes(100);o__Saccharomycetales(100);f__Metschnikowiaceae(100);g__Metschnikowia(100);s__Metschnikowia_pulcherrima(100); | Metschnikowia | 13 | Saprotroph | \|Undefined Saprotroph\| | Yeast | NULL | Probable | As nectar-tap_saprotroph (Põlme et al. 2020); Decay sugar-rich substrates (Põlme et al. 2020) | Põlme S, et al. 2020. Fungal Diversity 105:1-16 ((https://doi.org/10.1007/s13225-020-00466-2)) |
| Otu11513 | 2 | k__Fungi(100);p__Ascomycota(100);c__Saccharomycetes(100);o__Saccharomycetales(100);f__Metschnikowiaceae(100);g__Metschnikowia(100);s__Metschnikowia_viticola(100); | Metschnikowia | 13 | Saprotroph | \|Undefined Saprotroph\| | Yeast | NULL | Probable | As nectar-tap_saprotroph (Põlme et al. 2020); Decay sugar-rich substrates (Põlme et al. 2020) | Põlme S, et al. 2020. Fungal Diversity 105:1-16 ((https://doi.org/10.1007/s13225-020-00466-2)) |
| Otu11515 | 2 | k__Fungi(100);p__Ascomycota(100);c__Saccharomycetes(100);o__Saccharomycetales(100);f__Metschnikowiaceae(100);g__Metschnikowia(100);s__Metschnikowia_pulcherrima(100); | Metschnikowia | 13 | Saprotroph | \|Undefined Saprotroph\| | Yeast | NULL | Probable | As nectar-tap_saprotroph (Põlme et al. 2020); Decay sugar-rich substrates (Põlme et al. 2020) | Põlme S, et al. 2020. Fungal Diversity 105:1-16 ((https://doi.org/10.1007/s13225-020-00466-2)) |
| Otu11516 | 2 | k__Fungi(100);p__Ascomycota(100);c__Saccharomycetes(100);o__Saccharomycetales(100);f__Saccharomycetales_family_Incertae_sedis(100);g__Candida(100);g__Candida_unclassified(100); | Candida | 13 | Pathotroph-Saprotroph-Symbiotroph | Animal Pathogen-Endophyte-Endosymbiont-Epiphyte-\|Undefined Saprotroph\| | Dimorphic Yeast-Yeast | NULL | Probable | As nectar-tap_saprotroph (Põlme et al. 2020); Decay sugar-rich substrates (Põlme et al. 2020); This asexual genus is polyphyletic and contain species that function very differently depend on what they are. Caution should be taken when interpreting guilds for Candida species. | Manolakaki D, et al. 2010. Virulence. 1: 367-375 ((https://doi.org/10.4161/viru.1.5.12796)); Mekbib SB, et al. 2011. Tropical Plant Pathology 36:233-240 ((https://doi.org/10.1590/S1982-56762011000400004)); Põlme S, et al. 2020. Fungal Diversity 105:1-16 ((https://doi.org/10.1007/s13225-020-00466-2)) |
| Otu11517 | 2 | k__Fungi(100);p__Ascomycota(100);c__Saccharomycetes(100);o__Saccharomycetales(100);f__Metschnikowiaceae(100);g__Metschnikowia(100);g__Metschnikowia_unclassified(100); | Metschnikowia | 13 | Saprotroph | \|Undefined Saprotroph\| | Yeast | NULL | Probable | As nectar-tap_saprotroph (Põlme et al. 2020); Decay sugar-rich substrates (Põlme et al. 2020) | Põlme S, et al. 2020. Fungal Diversity 105:1-16 ((https://doi.org/10.1007/s13225-020-00466-2)) |
| Otu11518 | 2 | k__Fungi(100);p__Ascomycota(100);c__Saccharomycetes(100);o__Saccharomycetales(100);f__Metschnikowiaceae(100);g__Metschnikowia(100);s__Metschnikowia_pulcherrima(100); | Metschnikowia | 13 | Saprotroph | \|Undefined Saprotroph\| | Yeast | NULL | Probable | As nectar-tap_saprotroph (Põlme et al. 2020); Decay sugar-rich substrates (Põlme et al. 2020) | Põlme S, et al. 2020. Fungal Diversity 105:1-16 ((https://doi.org/10.1007/s13225-020-00466-2)) |
| Otu11519 | 2 | k__Fungi(100);p__Ascomycota(100);c__Saccharomycetes(100);o__Saccharomycetales(100);f__Metschnikowiaceae(100);g__Metschnikowia(100);g__Metschnikowia_unclassified(100); | Metschnikowia | 13 | Saprotroph | \|Undefined Saprotroph\| | Yeast | NULL | Probable | As nectar-tap_saprotroph (Põlme et al. 2020); Decay sugar-rich substrates (Põlme et al. 2020) | Põlme S, et al. 2020. Fungal Diversity 105:1-16 ((https://doi.org/10.1007/s13225-020-00466-2)) |
| Otu11520 | 2 | k__Fungi(100);p__Ascomycota(100);c__Saccharomycetes(100);o__Saccharomycetales(100);f__Metschnikowiaceae(100);g__Metschnikowia(100);g__Metschnikowia_unclassified(100); | Metschnikowia | 13 | Saprotroph | \|Undefined Saprotroph\| | Yeast | NULL | Probable | As nectar-tap_saprotroph (Põlme et al. 2020); Decay sugar-rich substrates (Põlme et al. 2020) | Põlme S, et al. 2020. Fungal Diversity 105:1-16 ((https://doi.org/10.1007/s13225-020-00466-2)) |
| Otu11521 | 2 | k__Fungi(100);p__Ascomycota(100);c__Saccharomycetes(100);o__Saccharomycetales(100);f__Metschnikowiaceae(100);g__Metschnikowia(100);g__Metschnikowia_unclassified(100); | Metschnikowia | 13 | Saprotroph | \|Undefined Saprotroph\| | Yeast | NULL | Probable | As nectar-tap_saprotroph (Põlme et al. 2020); Decay sugar-rich substrates (Põlme et al. 2020) | Põlme S, et al. 2020. Fungal Diversity 105:1-16 ((https://doi.org/10.1007/s13225-020-00466-2)) |
| Otu11525 | 2 | k__Fungi(100);p__Ascomycota(100);c__Saccharomycetes(100);o__Saccharomycetales(100);f__Metschnikowiaceae(100);g__Metschnikowia(100);g__Metschnikowia_unclassified(100); | Metschnikowia | 13 | Saprotroph | \|Undefined Saprotroph\| | Yeast | NULL | Probable | As nectar-tap_saprotroph (Põlme et al. 2020); Decay sugar-rich substrates (Põlme et al. 2020) | Põlme S, et al. 2020. Fungal Diversity 105:1-16 ((https://doi.org/10.1007/s13225-020-00466-2)) |
| Otu11527 | 2 | k__Fungi(100);p__Ascomycota(100);c__Saccharomycetes(100);o__Saccharomycetales(100);f__Metschnikowiaceae(100);g__Metschnikowia(100);s__Metschnikowia_pulcherrima(100); | Metschnikowia | 13 | Saprotroph | \|Undefined Saprotroph\| | Yeast | NULL | Probable | As nectar-tap_saprotroph (Põlme et al. 2020); Decay sugar-rich substrates (Põlme et al. 2020) | Põlme S, et al. 2020. Fungal Diversity 105:1-16 ((https://doi.org/10.1007/s13225-020-00466-2)) |
| Otu11528 | 2 | k__Fungi(100);p__Ascomycota(100);c__Saccharomycetes(100);o__Saccharomycetales(100);f__Metschnikowiaceae(100);g__Metschnikowia(100);g__Metschnikowia_unclassified(100); | Metschnikowia | 13 | Saprotroph | \|Undefined Saprotroph\| | Yeast | NULL | Probable | As nectar-tap_saprotroph (Põlme et al. 2020); Decay sugar-rich substrates (Põlme et al. 2020) | Põlme S, et al. 2020. Fungal Diversity 105:1-16 ((https://doi.org/10.1007/s13225-020-00466-2)) |
| Otu11529 | 2 | k__Fungi(100);p__Ascomycota(100);c__Saccharomycetes(100);o__Saccharomycetales(100);f__Metschnikowiaceae(100);g__Metschnikowia(100);s__Metschnikowia_pulcherrima(100); | Metschnikowia | 13 | Saprotroph | \|Undefined Saprotroph\| | Yeast | NULL | Probable | As nectar-tap_saprotroph (Põlme et al. 2020); Decay sugar-rich substrates (Põlme et al. 2020) | Põlme S, et al. 2020. Fungal Diversity 105:1-16 ((https://doi.org/10.1007/s13225-020-00466-2)) |
| Otu11531 | 2 | k__Fungi(100);p__Ascomycota(100);c__Saccharomycetes(100);o__Saccharomycetales(100);f__Metschnikowiaceae(100);g__Metschnikowia(100);g__Metschnikowia_unclassified(100); | Metschnikowia | 13 | Saprotroph | \|Undefined Saprotroph\| | Yeast | NULL | Probable | As nectar-tap_saprotroph (Põlme et al. 2020); Decay sugar-rich substrates (Põlme et al. 2020) | Põlme S, et al. 2020. Fungal Diversity 105:1-16 ((https://doi.org/10.1007/s13225-020-00466-2)) |
| Otu11533 | 2 | k__Fungi(100);p__Ascomycota(100);c__Saccharomycetes(100);o__Saccharomycetales(100);f__Metschnikowiaceae(100);g__Metschnikowia(100);g__Metschnikowia_unclassified(100); | Metschnikowia | 13 | Saprotroph | \|Undefined Saprotroph\| | Yeast | NULL | Probable | As nectar-tap_saprotroph (Põlme et al. 2020); Decay sugar-rich substrates (Põlme et al. 2020) | Põlme S, et al. 2020. Fungal Diversity 105:1-16 ((https://doi.org/10.1007/s13225-020-00466-2)) |
| Otu11534 | 2 | k__Fungi(100);p__Ascomycota(100);c__Saccharomycetes(100);o__Saccharomycetales(100);f__Metschnikowiaceae(100);g__Metschnikowia(100);g__Metschnikowia_unclassified(100); | Metschnikowia | 13 | Saprotroph | \|Undefined Saprotroph\| | Yeast | NULL | Probable | As nectar-tap_saprotroph (Põlme et al. 2020); Decay sugar-rich substrates (Põlme et al. 2020) | Põlme S, et al. 2020. Fungal Diversity 105:1-16 ((https://doi.org/10.1007/s13225-020-00466-2)) |
| Otu11535 | 2 | k__Fungi(100);p__Ascomycota(100);c__Saccharomycetes(100);o__Saccharomycetales(100);f__Metschnikowiaceae(100);g__Metschnikowia(100);g__Metschnikowia_unclassified(100); | Metschnikowia | 13 | Saprotroph | \|Undefined Saprotroph\| | Yeast | NULL | Probable | As nectar-tap_saprotroph (Põlme et al. 2020); Decay sugar-rich substrates (Põlme et al. 2020) | Põlme S, et al. 2020. Fungal Diversity 105:1-16 ((https://doi.org/10.1007/s13225-020-00466-2)) |
| Otu11537 | 2 | k__Fungi(100);p__Ascomycota(100);c__Saccharomycetes(100);o__Saccharomycetales(100);f__Metschnikowiaceae(100);g__Metschnikowia(100);g__Metschnikowia_unclassified(100); | Metschnikowia | 13 | Saprotroph | \|Undefined Saprotroph\| | Yeast | NULL | Probable | As nectar-tap_saprotroph (Põlme et al. 2020); Decay sugar-rich substrates (Põlme et al. 2020) | Põlme S, et al. 2020. Fungal Diversity 105:1-16 ((https://doi.org/10.1007/s13225-020-00466-2)) |
| Otu11539 | 2 | k__Fungi(100);p__Ascomycota(100);c__Saccharomycetes(100);o__Saccharomycetales(100);f__Metschnikowiaceae(100);g__Metschnikowia(100);g__Metschnikowia_unclassified(100); | Metschnikowia | 13 | Saprotroph | \|Undefined Saprotroph\| | Yeast | NULL | Probable | As nectar-tap_saprotroph (Põlme et al. 2020); Decay sugar-rich substrates (Põlme et al. 2020) | Põlme S, et al. 2020. Fungal Diversity 105:1-16 ((https://doi.org/10.1007/s13225-020-00466-2)) |
| Otu11540 | 2 | k__Fungi(100);p__Ascomycota(100);c__Saccharomycetes(100);o__Saccharomycetales(100);f__Metschnikowiaceae(100);g__Metschnikowia(100);s__Metschnikowia_pulcherrima(100); | Metschnikowia | 13 | Saprotroph | \|Undefined Saprotroph\| | Yeast | NULL | Probable | As nectar-tap_saprotroph (Põlme et al. 2020); Decay sugar-rich substrates (Põlme et al. 2020) | Põlme S, et al. 2020. Fungal Diversity 105:1-16 ((https://doi.org/10.1007/s13225-020-00466-2)) |
| Otu11545 | 2 | k__Fungi(100);p__Ascomycota(100);c__Saccharomycetes(100);o__Saccharomycetales(100);f__Metschnikowiaceae(100);g__Metschnikowia(100);s__Metschnikowia_pulcherrima(100); | Metschnikowia | 13 | Saprotroph | \|Undefined Saprotroph\| | Yeast | NULL | Probable | As nectar-tap_saprotroph (Põlme et al. 2020); Decay sugar-rich substrates (Põlme et al. 2020) | Põlme S, et al. 2020. Fungal Diversity 105:1-16 ((https://doi.org/10.1007/s13225-020-00466-2)) |
| Otu11546 | 2 | k__Fungi(100);p__Ascomycota(100);c__Sordariomycetes(100);o__Xylariales(100);f__Xylariaceae(100);g__Anthostomella(100);s__Anthostomella_sp(100); | Anthostomella | 13 | Saprotroph-Symbiotroph | \|Endophyte\|-Plant Saprotroph-Undefined Saprotroph | Microfungus | NULL | Probable | As foliar_endophyte (Põlme et al. 2020); As litter_saprotroph (Põlme et al. 2020); Decay leaf, fruit, seed (Põlme et al. 2020) | Tedersoo L, et al. 2014. Science 346:e1256688 ((https://doi.org/10.1126/science.1256688)); Põlme S, et al. 2020. Fungal Diversity 105:1-16 ((https://doi.org/10.1007/s13225-020-00466-2)) |
| Otu11548 | 2 | k__Fungi(100);p__Ascomycota(100);c__Sordariomycetes(100);o__Sordariales(100);f__Lasiosphaeriaceae(100);g__Apodus(100);s__Apodus_sp(100); | Apodus | 13 | Saprotroph | \|Dung Saprotroph\|-Undefined Saprotroph | Microfungus | NULL | Probable | Decay dung (Põlme et al. 2020) | Tedersoo L, et al. 2014. Science 346:e1256688 ((https://doi.org/10.1126/science.1256688)); Põlme S, et al. 2020. Fungal Diversity 105:1-16 ((https://doi.org/10.1007/s13225-020-00466-2)) |
| Otu11550 | 2 | k__Fungi(100);p__Ascomycota(100);c__Saccharomycetes(100);o__Saccharomycetales(100);f__Metschnikowiaceae(100);g__Metschnikowia(100);s__Metschnikowia_pulcherrima(100); | Metschnikowia | 13 | Saprotroph | \|Undefined Saprotroph\| | Yeast | NULL | Probable | As nectar-tap_saprotroph (Põlme et al. 2020); Decay sugar-rich substrates (Põlme et al. 2020) | Põlme S, et al. 2020. Fungal Diversity 105:1-16 ((https://doi.org/10.1007/s13225-020-00466-2)) |
| Otu11551 | 2 | k__Fungi(100);p__Ascomycota(100);c__Dothideomycetes(100);o__Pleosporales(100);f__Pleosporaceae(100);g__Chalastospora(100);s__Chalastospora_ellipsoidea(100); | Chalastospora | 13 | Pathotroph-Saprotroph | \|Plant Pathogen\|-Plant Saprotroph | Microfungus | NULL | Probable | As litter_saprotroph (Põlme et al. 2020); Decay leaf, fruit, seed (Põlme et al. 2020) | Põlme S, et al. 2020. Fungal Diversity 105:1-16 ((https://doi.org/10.1007/s13225-020-00466-2)); The American Phytopathological Society Common Names of Plant Diseases ((http://www.apsnet.org/publications/commonnames/Pages/default.aspx)) |
| Otu11552 | 2 | k__Fungi(100);p__Ascomycota(100);c__Dothideomycetes(100);o__Pleosporales(100);f__Tubeufiaceae(100);g__unclassified_Tubeufiaceae(100);s__Tubeufiaceae_sp(100); | Tubeufiaceae | 9 | Pathotroph-Saprotroph | Animal Pathogen-Undefined Saprotroph | NULL | NULL | Probable | Biotrophic or saprobic, especially on scale insects or other fungi. The anamorphic forms are fequently aero-aquatic (Cannon & Kirk 2007) | Cannon PF, Kirk PM. 2007. Fungal Families of the World. CAB International, Cambridge (ISBN: 978-0851998275) |
| Otu11554 | 2 | k__Fungi(100);p__Ascomycota(100);c__Saccharomycetes(100);o__Saccharomycetales(100);f__Metschnikowiaceae(100);g__Metschnikowia(100);s__Metschnikowia_pulcherrima(100); | Metschnikowia | 13 | Saprotroph | \|Undefined Saprotroph\| | Yeast | NULL | Probable | As nectar-tap_saprotroph (Põlme et al. 2020); Decay sugar-rich substrates (Põlme et al. 2020) | Põlme S, et al. 2020. Fungal Diversity 105:1-16 ((https://doi.org/10.1007/s13225-020-00466-2)) |
| Otu11556 | 2 | k__Fungi(100);p__Ascomycota(100);c__Saccharomycetes(100);o__Saccharomycetales(100);f__Metschnikowiaceae(100);g__Metschnikowia(100);s__Metschnikowia_pulcherrima(100); | Metschnikowia | 13 | Saprotroph | \|Undefined Saprotroph\| | Yeast | NULL | Probable | As nectar-tap_saprotroph (Põlme et al. 2020); Decay sugar-rich substrates (Põlme et al. 2020) | Põlme S, et al. 2020. Fungal Diversity 105:1-16 ((https://doi.org/10.1007/s13225-020-00466-2)) |
| Otu11557 | 2 | k__Fungi(100);p__Ascomycota(100);c__Saccharomycetes(100);o__Saccharomycetales(100);f__Metschnikowiaceae(100);g__Metschnikowia(100);g__Metschnikowia_unclassified(100); | Metschnikowia | 13 | Saprotroph | \|Undefined Saprotroph\| | Yeast | NULL | Probable | As nectar-tap_saprotroph (Põlme et al. 2020); Decay sugar-rich substrates (Põlme et al. 2020) | Põlme S, et al. 2020. Fungal Diversity 105:1-16 ((https://doi.org/10.1007/s13225-020-00466-2)) |
| Otu11558 | 2 | k__Fungi(100);p__Ascomycota(100);c__Saccharomycetes(100);o__Saccharomycetales(100);f__Saccharomycetales_family_Incertae_sedis(100);g__Candida(100);g__Candida_unclassified(100); | Candida | 13 | Pathotroph-Saprotroph-Symbiotroph | Animal Pathogen-Endophyte-Endosymbiont-Epiphyte-\|Undefined Saprotroph\| | Dimorphic Yeast-Yeast | NULL | Probable | As nectar-tap_saprotroph (Põlme et al. 2020); Decay sugar-rich substrates (Põlme et al. 2020); This asexual genus is polyphyletic and contain species that function very differently depend on what they are. Caution should be taken when interpreting guilds for Candida species. | Manolakaki D, et al. 2010. Virulence. 1: 367-375 ((https://doi.org/10.4161/viru.1.5.12796)); Mekbib SB, et al. 2011. Tropical Plant Pathology 36:233-240 ((https://doi.org/10.1590/S1982-56762011000400004)); Põlme S, et al. 2020. Fungal Diversity 105:1-16 ((https://doi.org/10.1007/s13225-020-00466-2)) |
| Otu11559 | 2 | k__Fungi(100);p__Ascomycota(100);c__Saccharomycetes(100);o__Saccharomycetales(100);f__Metschnikowiaceae(100);g__Metschnikowia(100);s__Metschnikowia_pulcherrima(100); | Metschnikowia | 13 | Saprotroph | \|Undefined Saprotroph\| | Yeast | NULL | Probable | As nectar-tap_saprotroph (Põlme et al. 2020); Decay sugar-rich substrates (Põlme et al. 2020) | Põlme S, et al. 2020. Fungal Diversity 105:1-16 ((https://doi.org/10.1007/s13225-020-00466-2)) |
| Otu11563 | 2 | k__Fungi(100);p__Ascomycota(100);c__Saccharomycetes(100);o__Saccharomycetales(100);f__Metschnikowiaceae(100);g__Metschnikowia(100);g__Metschnikowia_unclassified(100); | Metschnikowia | 13 | Saprotroph | \|Undefined Saprotroph\| | Yeast | NULL | Probable | As nectar-tap_saprotroph (Põlme et al. 2020); Decay sugar-rich substrates (Põlme et al. 2020) | Põlme S, et al. 2020. Fungal Diversity 105:1-16 ((https://doi.org/10.1007/s13225-020-00466-2)) |
| Otu11564 | 2 | k__Fungi(100);p__Ascomycota(100);c__Saccharomycetes(100);o__Saccharomycetales(100);f__Metschnikowiaceae(100);g__Metschnikowia(100);s__Metschnikowia_pulcherrima(100); | Metschnikowia | 13 | Saprotroph | \|Undefined Saprotroph\| | Yeast | NULL | Probable | As nectar-tap_saprotroph (Põlme et al. 2020); Decay sugar-rich substrates (Põlme et al. 2020) | Põlme S, et al. 2020. Fungal Diversity 105:1-16 ((https://doi.org/10.1007/s13225-020-00466-2)) |
| Otu11565 | 2 | k__Fungi(100);p__Ascomycota(100);c__Saccharomycetes(100);o__Saccharomycetales(100);f__Metschnikowiaceae(100);g__Metschnikowia(100);s__Metschnikowia_pulcherrima(100); | Metschnikowia | 13 | Saprotroph | \|Undefined Saprotroph\| | Yeast | NULL | Probable | As nectar-tap_saprotroph (Põlme et al. 2020); Decay sugar-rich substrates (Põlme et al. 2020) | Põlme S, et al. 2020. Fungal Diversity 105:1-16 ((https://doi.org/10.1007/s13225-020-00466-2)) |
| Otu11566 | 2 | k__Fungi(100);p__Ascomycota(100);c__Saccharomycetes(100);o__Saccharomycetales(100);f__Metschnikowiaceae(100);g__Metschnikowia(100);s__Metschnikowia_pulcherrima(100); | Metschnikowia | 13 | Saprotroph | \|Undefined Saprotroph\| | Yeast | NULL | Probable | As nectar-tap_saprotroph (Põlme et al. 2020); Decay sugar-rich substrates (Põlme et al. 2020) | Põlme S, et al. 2020. Fungal Diversity 105:1-16 ((https://doi.org/10.1007/s13225-020-00466-2)) |
| Otu11567 | 2 | k__Fungi(100);p__Ascomycota(100);c__Saccharomycetes(100);o__Saccharomycetales(100);f__Metschnikowiaceae(100);g__Metschnikowia(100);s__Metschnikowia_pulcherrima(100); | Metschnikowia | 13 | Saprotroph | \|Undefined Saprotroph\| | Yeast | NULL | Probable | As nectar-tap_saprotroph (Põlme et al. 2020); Decay sugar-rich substrates (Põlme et al. 2020) | Põlme S, et al. 2020. Fungal Diversity 105:1-16 ((https://doi.org/10.1007/s13225-020-00466-2)) |
| Otu11570 | 2 | k__Fungi(100);p__Ascomycota(100);c__Saccharomycetes(100);o__Saccharomycetales(100);f__Metschnikowiaceae(100);g__Metschnikowia(100);g__Metschnikowia_unclassified(100); | Metschnikowia | 13 | Saprotroph | \|Undefined Saprotroph\| | Yeast | NULL | Probable | As nectar-tap_saprotroph (Põlme et al. 2020); Decay sugar-rich substrates (Põlme et al. 2020) | Põlme S, et al. 2020. Fungal Diversity 105:1-16 ((https://doi.org/10.1007/s13225-020-00466-2)) |
| Otu11571 | 2 | k__Fungi(100);p__Ascomycota(100);c__Saccharomycetes(100);o__Saccharomycetales(100);f__Metschnikowiaceae(100);g__Metschnikowia(100);s__Metschnikowia_pulcherrima(100); | Metschnikowia | 13 | Saprotroph | \|Undefined Saprotroph\| | Yeast | NULL | Probable | As nectar-tap_saprotroph (Põlme et al. 2020); Decay sugar-rich substrates (Põlme et al. 2020) | Põlme S, et al. 2020. Fungal Diversity 105:1-16 ((https://doi.org/10.1007/s13225-020-00466-2)) |
| Otu11572 | 2 | k__Fungi(100);p__Ascomycota(100);c__Saccharomycetes(100);o__Saccharomycetales(100);f__Metschnikowiaceae(100);g__Metschnikowia(100);s__Metschnikowia_pulcherrima(100); | Metschnikowia | 13 | Saprotroph | \|Undefined Saprotroph\| | Yeast | NULL | Probable | As nectar-tap_saprotroph (Põlme et al. 2020); Decay sugar-rich substrates (Põlme et al. 2020) | Põlme S, et al. 2020. Fungal Diversity 105:1-16 ((https://doi.org/10.1007/s13225-020-00466-2)) |
| Otu11573 | 2 | k__Fungi(100);p__Ascomycota(100);c__Saccharomycetes(100);o__Saccharomycetales(100);f__Metschnikowiaceae(100);g__Metschnikowia(100);s__Metschnikowia_pulcherrima(100); | Metschnikowia | 13 | Saprotroph | \|Undefined Saprotroph\| | Yeast | NULL | Probable | As nectar-tap_saprotroph (Põlme et al. 2020); Decay sugar-rich substrates (Põlme et al. 2020) | Põlme S, et al. 2020. Fungal Diversity 105:1-16 ((https://doi.org/10.1007/s13225-020-00466-2)) |
| Otu11574 | 2 | k__Fungi(100);p__Ascomycota(100);c__Saccharomycetes(100);o__Saccharomycetales(100);f__Metschnikowiaceae(100);g__Metschnikowia(100);s__Metschnikowia_viticola(100); | Metschnikowia | 13 | Saprotroph | \|Undefined Saprotroph\| | Yeast | NULL | Probable | As nectar-tap_saprotroph (Põlme et al. 2020); Decay sugar-rich substrates (Põlme et al. 2020) | Põlme S, et al. 2020. Fungal Diversity 105:1-16 ((https://doi.org/10.1007/s13225-020-00466-2)) |
| Otu11575 | 2 | k__Fungi(100);p__Ascomycota(100);c__Saccharomycetes(100);o__Saccharomycetales(100);f__Metschnikowiaceae(100);g__Metschnikowia(100);g__Metschnikowia_unclassified(100); | Metschnikowia | 13 | Saprotroph | \|Undefined Saprotroph\| | Yeast | NULL | Probable | As nectar-tap_saprotroph (Põlme et al. 2020); Decay sugar-rich substrates (Põlme et al. 2020) | Põlme S, et al. 2020. Fungal Diversity 105:1-16 ((https://doi.org/10.1007/s13225-020-00466-2)) |
| Otu11576 | 2 | k__Fungi(100);p__Ascomycota(100);c__Saccharomycetes(100);o__Saccharomycetales(100);f__Metschnikowiaceae(100);g__Metschnikowia(100);s__Metschnikowia_pulcherrima(100); | Metschnikowia | 13 | Saprotroph | \|Undefined Saprotroph\| | Yeast | NULL | Probable | As nectar-tap_saprotroph (Põlme et al. 2020); Decay sugar-rich substrates (Põlme et al. 2020) | Põlme S, et al. 2020. Fungal Diversity 105:1-16 ((https://doi.org/10.1007/s13225-020-00466-2)) |
| Otu11577 | 2 | k__Fungi(100);p__Ascomycota(100);c__Saccharomycetes(100);o__Saccharomycetales(100);f__Metschnikowiaceae(100);g__Metschnikowia(100);s__Metschnikowia_pulcherrima(100); | Metschnikowia | 13 | Saprotroph | \|Undefined Saprotroph\| | Yeast | NULL | Probable | As nectar-tap_saprotroph (Põlme et al. 2020); Decay sugar-rich substrates (Põlme et al. 2020) | Põlme S, et al. 2020. Fungal Diversity 105:1-16 ((https://doi.org/10.1007/s13225-020-00466-2)) |
| Otu11578 | 2 | k__Fungi(100);p__Ascomycota(100);c__Saccharomycetes(100);o__Saccharomycetales(100);f__Metschnikowiaceae(100);g__Metschnikowia(100);g__Metschnikowia_unclassified(100); | Metschnikowia | 13 | Saprotroph | \|Undefined Saprotroph\| | Yeast | NULL | Probable | As nectar-tap_saprotroph (Põlme et al. 2020); Decay sugar-rich substrates (Põlme et al. 2020) | Põlme S, et al. 2020. Fungal Diversity 105:1-16 ((https://doi.org/10.1007/s13225-020-00466-2)) |
| Otu11579 | 2 | k__Fungi(100);p__Ascomycota(100);c__Saccharomycetes(100);o__Saccharomycetales(100);f__Metschnikowiaceae(100);g__Metschnikowia(100);s__Metschnikowia_pulcherrima(100); | Metschnikowia | 13 | Saprotroph | \|Undefined Saprotroph\| | Yeast | NULL | Probable | As nectar-tap_saprotroph (Põlme et al. 2020); Decay sugar-rich substrates (Põlme et al. 2020) | Põlme S, et al. 2020. Fungal Diversity 105:1-16 ((https://doi.org/10.1007/s13225-020-00466-2)) |
| Otu11580 | 2 | k__Fungi(100);p__Ascomycota(100);c__Saccharomycetes(100);o__Saccharomycetales(100);f__Metschnikowiaceae(100);g__Metschnikowia(100);s__Metschnikowia_pulcherrima(100); | Metschnikowia | 13 | Saprotroph | \|Undefined Saprotroph\| | Yeast | NULL | Probable | As nectar-tap_saprotroph (Põlme et al. 2020); Decay sugar-rich substrates (Põlme et al. 2020) | Põlme S, et al. 2020. Fungal Diversity 105:1-16 ((https://doi.org/10.1007/s13225-020-00466-2)) |
| Otu11582 | 2 | k__Fungi(100);p__Ascomycota(100);c__Saccharomycetes(100);o__Saccharomycetales(100);f__Metschnikowiaceae(100);g__Metschnikowia(100);s__Metschnikowia_pulcherrima(100); | Metschnikowia | 13 | Saprotroph | \|Undefined Saprotroph\| | Yeast | NULL | Probable | As nectar-tap_saprotroph (Põlme et al. 2020); Decay sugar-rich substrates (Põlme et al. 2020) | Põlme S, et al. 2020. Fungal Diversity 105:1-16 ((https://doi.org/10.1007/s13225-020-00466-2)) |
| Otu11583 | 2 | k__Fungi(100);p__Ascomycota(100);c__Saccharomycetes(100);o__Saccharomycetales(100);f__Saccharomycetales_family_Incertae_sedis(100);g__Candida(100);g__Candida_unclassified(100); | Candida | 13 | Pathotroph-Saprotroph-Symbiotroph | Animal Pathogen-Endophyte-Endosymbiont-Epiphyte-\|Undefined Saprotroph\| | Dimorphic Yeast-Yeast | NULL | Probable | As nectar-tap_saprotroph (Põlme et al. 2020); Decay sugar-rich substrates (Põlme et al. 2020); This asexual genus is polyphyletic and contain species that function very differently depend on what they are. Caution should be taken when interpreting guilds for Candida species. | Manolakaki D, et al. 2010. Virulence. 1: 367-375 ((https://doi.org/10.4161/viru.1.5.12796)); Mekbib SB, et al. 2011. Tropical Plant Pathology 36:233-240 ((https://doi.org/10.1590/S1982-56762011000400004)); Põlme S, et al. 2020. Fungal Diversity 105:1-16 ((https://doi.org/10.1007/s13225-020-00466-2)) |
| Otu11585 | 2 | k__Fungi(100);p__Ascomycota(100);c__Saccharomycetes(100);o__Saccharomycetales(100);f__Metschnikowiaceae(100);g__Metschnikowia(100);s__Metschnikowia_pulcherrima(100); | Metschnikowia | 13 | Saprotroph | \|Undefined Saprotroph\| | Yeast | NULL | Probable | As nectar-tap_saprotroph (Põlme et al. 2020); Decay sugar-rich substrates (Põlme et al. 2020) | Põlme S, et al. 2020. Fungal Diversity 105:1-16 ((https://doi.org/10.1007/s13225-020-00466-2)) |
| Otu11586 | 2 | k__Fungi(100);p__Ascomycota(100);c__Saccharomycetes(100);o__Saccharomycetales(100);f__Metschnikowiaceae(100);g__Metschnikowia(100);s__Metschnikowia_pulcherrima(100); | Metschnikowia | 13 | Saprotroph | \|Undefined Saprotroph\| | Yeast | NULL | Probable | As nectar-tap_saprotroph (Põlme et al. 2020); Decay sugar-rich substrates (Põlme et al. 2020) | Põlme S, et al. 2020. Fungal Diversity 105:1-16 ((https://doi.org/10.1007/s13225-020-00466-2)) |
| Otu11587 | 2 | k__Fungi(100);p__Ascomycota(100);c__Saccharomycetes(100);o__Saccharomycetales(100);f__Metschnikowiaceae(100);g__Metschnikowia(100);s__Metschnikowia_pulcherrima(100); | Metschnikowia | 13 | Saprotroph | \|Undefined Saprotroph\| | Yeast | NULL | Probable | As nectar-tap_saprotroph (Põlme et al. 2020); Decay sugar-rich substrates (Põlme et al. 2020) | Põlme S, et al. 2020. Fungal Diversity 105:1-16 ((https://doi.org/10.1007/s13225-020-00466-2)) |
| Otu11588 | 2 | k__Fungi(100);p__Ascomycota(100);c__Saccharomycetes(100);o__Saccharomycetales(100);f__Metschnikowiaceae(100);g__Metschnikowia(100);s__Metschnikowia_pulcherrima(100); | Metschnikowia | 13 | Saprotroph | \|Undefined Saprotroph\| | Yeast | NULL | Probable | As nectar-tap_saprotroph (Põlme et al. 2020); Decay sugar-rich substrates (Põlme et al. 2020) | Põlme S, et al. 2020. Fungal Diversity 105:1-16 ((https://doi.org/10.1007/s13225-020-00466-2)) |
| Otu11589 | 2 | k__Fungi(100);p__Ascomycota(100);c__Saccharomycetes(100);o__Saccharomycetales(100);f__Metschnikowiaceae(100);g__Metschnikowia(100);s__Metschnikowia_pulcherrima(100); | Metschnikowia | 13 | Saprotroph | \|Undefined Saprotroph\| | Yeast | NULL | Probable | As nectar-tap_saprotroph (Põlme et al. 2020); Decay sugar-rich substrates (Põlme et al. 2020) | Põlme S, et al. 2020. Fungal Diversity 105:1-16 ((https://doi.org/10.1007/s13225-020-00466-2)) |
| Otu11590 | 2 | k__Fungi(100);p__Ascomycota(100);c__Saccharomycetes(100);o__Saccharomycetales(100);f__Metschnikowiaceae(100);g__Metschnikowia(100);s__Metschnikowia_pulcherrima(100); | Metschnikowia | 13 | Saprotroph | \|Undefined Saprotroph\| | Yeast | NULL | Probable | As nectar-tap_saprotroph (Põlme et al. 2020); Decay sugar-rich substrates (Põlme et al. 2020) | Põlme S, et al. 2020. Fungal Diversity 105:1-16 ((https://doi.org/10.1007/s13225-020-00466-2)) |
| Otu11591 | 2 | k__Fungi(100);p__Ascomycota(100);c__Saccharomycetes(100);o__Saccharomycetales(100);f__Metschnikowiaceae(100);g__Metschnikowia(100);s__Metschnikowia_pulcherrima(100); | Metschnikowia | 13 | Saprotroph | \|Undefined Saprotroph\| | Yeast | NULL | Probable | As nectar-tap_saprotroph (Põlme et al. 2020); Decay sugar-rich substrates (Põlme et al. 2020) | Põlme S, et al. 2020. Fungal Diversity 105:1-16 ((https://doi.org/10.1007/s13225-020-00466-2)) |
| Otu11592 | 2 | k__Fungi(100);p__Ascomycota(100);c__Saccharomycetes(100);o__Saccharomycetales(100);f__Metschnikowiaceae(100);g__Metschnikowia(100);s__Metschnikowia_pulcherrima(100); | Metschnikowia | 13 | Saprotroph | \|Undefined Saprotroph\| | Yeast | NULL | Probable | As nectar-tap_saprotroph (Põlme et al. 2020); Decay sugar-rich substrates (Põlme et al. 2020) | Põlme S, et al. 2020. Fungal Diversity 105:1-16 ((https://doi.org/10.1007/s13225-020-00466-2)) |
| Otu11593 | 2 | k__Fungi(100);p__Ascomycota(100);c__Saccharomycetes(100);o__Saccharomycetales(100);f__Metschnikowiaceae(100);g__Metschnikowia(100);s__Metschnikowia_pulcherrima(100); | Metschnikowia | 13 | Saprotroph | \|Undefined Saprotroph\| | Yeast | NULL | Probable | As nectar-tap_saprotroph (Põlme et al. 2020); Decay sugar-rich substrates (Põlme et al. 2020) | Põlme S, et al. 2020. Fungal Diversity 105:1-16 ((https://doi.org/10.1007/s13225-020-00466-2)) |
| Otu11595 | 2 | k__Fungi(100);p__Ascomycota(100);c__Saccharomycetes(100);o__Saccharomycetales(100);f__Metschnikowiaceae(100);g__Metschnikowia(100);s__Metschnikowia_pulcherrima(100); | Metschnikowia | 13 | Saprotroph | \|Undefined Saprotroph\| | Yeast | NULL | Probable | As nectar-tap_saprotroph (Põlme et al. 2020); Decay sugar-rich substrates (Põlme et al. 2020) | Põlme S, et al. 2020. Fungal Diversity 105:1-16 ((https://doi.org/10.1007/s13225-020-00466-2)) |
| Otu11596 | 2 | k__Fungi(100);p__Ascomycota(100);c__Saccharomycetes(100);o__Saccharomycetales(100);f__Metschnikowiaceae(100);g__Metschnikowia(100);g__Metschnikowia_unclassified(100); | Metschnikowia | 13 | Saprotroph | \|Undefined Saprotroph\| | Yeast | NULL | Probable | As nectar-tap_saprotroph (Põlme et al. 2020); Decay sugar-rich substrates (Põlme et al. 2020) | Põlme S, et al. 2020. Fungal Diversity 105:1-16 ((https://doi.org/10.1007/s13225-020-00466-2)) |
| Otu11597 | 2 | k__Fungi(100);p__Ascomycota(100);c__Saccharomycetes(100);o__Saccharomycetales(100);f__Metschnikowiaceae(100);g__Metschnikowia(100);s__Metschnikowia_pulcherrima(100); | Metschnikowia | 13 | Saprotroph | \|Undefined Saprotroph\| | Yeast | NULL | Probable | As nectar-tap_saprotroph (Põlme et al. 2020); Decay sugar-rich substrates (Põlme et al. 2020) | Põlme S, et al. 2020. Fungal Diversity 105:1-16 ((https://doi.org/10.1007/s13225-020-00466-2)) |
| Otu11598 | 2 | k__Fungi(100);p__Ascomycota(100);c__Saccharomycetes(100);o__Saccharomycetales(100);f__Metschnikowiaceae(100);g__Metschnikowia(100);s__Metschnikowia_pulcherrima(100); | Metschnikowia | 13 | Saprotroph | \|Undefined Saprotroph\| | Yeast | NULL | Probable | As nectar-tap_saprotroph (Põlme et al. 2020); Decay sugar-rich substrates (Põlme et al. 2020) | Põlme S, et al. 2020. Fungal Diversity 105:1-16 ((https://doi.org/10.1007/s13225-020-00466-2)) |
| Otu11600 | 2 | k__Fungi(100);p__Ascomycota(100);c__Saccharomycetes(100);o__Saccharomycetales(100);f__Metschnikowiaceae(100);g__Metschnikowia(100);g__Metschnikowia_unclassified(100); | Metschnikowia | 13 | Saprotroph | \|Undefined Saprotroph\| | Yeast | NULL | Probable | As nectar-tap_saprotroph (Põlme et al. 2020); Decay sugar-rich substrates (Põlme et al. 2020) | Põlme S, et al. 2020. Fungal Diversity 105:1-16 ((https://doi.org/10.1007/s13225-020-00466-2)) |
| Otu11601 | 2 | k__Fungi(100);p__Ascomycota(100);c__Saccharomycetes(100);o__Saccharomycetales(100);f__Metschnikowiaceae(100);g__Metschnikowia(100);g__Metschnikowia_unclassified(100); | Metschnikowia | 13 | Saprotroph | \|Undefined Saprotroph\| | Yeast | NULL | Probable | As nectar-tap_saprotroph (Põlme et al. 2020); Decay sugar-rich substrates (Põlme et al. 2020) | Põlme S, et al. 2020. Fungal Diversity 105:1-16 ((https://doi.org/10.1007/s13225-020-00466-2)) |
| Otu11603 | 2 | k__Fungi(100);p__Ascomycota(100);c__Saccharomycetes(100);o__Saccharomycetales(100);f__Metschnikowiaceae(100);g__Metschnikowia(100);s__Metschnikowia_pulcherrima(100); | Metschnikowia | 13 | Saprotroph | \|Undefined Saprotroph\| | Yeast | NULL | Probable | As nectar-tap_saprotroph (Põlme et al. 2020); Decay sugar-rich substrates (Põlme et al. 2020) | Põlme S, et al. 2020. Fungal Diversity 105:1-16 ((https://doi.org/10.1007/s13225-020-00466-2)) |
| Otu11605 | 2 | k__Fungi(100);p__Ascomycota(100);c__Saccharomycetes(100);o__Saccharomycetales(100);f__Metschnikowiaceae(100);g__Metschnikowia(100);s__Metschnikowia_pulcherrima(100); | Metschnikowia | 13 | Saprotroph | \|Undefined Saprotroph\| | Yeast | NULL | Probable | As nectar-tap_saprotroph (Põlme et al. 2020); Decay sugar-rich substrates (Põlme et al. 2020) | Põlme S, et al. 2020. Fungal Diversity 105:1-16 ((https://doi.org/10.1007/s13225-020-00466-2)) |
| Otu11606 | 2 | k__Fungi(100);p__Ascomycota(100);c__Saccharomycetes(100);o__Saccharomycetales(100);f__Metschnikowiaceae(100);g__Metschnikowia(100);s__Metschnikowia_pulcherrima(100); | Metschnikowia | 13 | Saprotroph | \|Undefined Saprotroph\| | Yeast | NULL | Probable | As nectar-tap_saprotroph (Põlme et al. 2020); Decay sugar-rich substrates (Põlme et al. 2020) | Põlme S, et al. 2020. Fungal Diversity 105:1-16 ((https://doi.org/10.1007/s13225-020-00466-2)) |
| Otu11607 | 2 | k__Fungi(100);p__Ascomycota(100);c__Saccharomycetes(100);o__Saccharomycetales(100);f__Metschnikowiaceae(100);g__Metschnikowia(100);g__Metschnikowia_unclassified(100); | Metschnikowia | 13 | Saprotroph | \|Undefined Saprotroph\| | Yeast | NULL | Probable | As nectar-tap_saprotroph (Põlme et al. 2020); Decay sugar-rich substrates (Põlme et al. 2020) | Põlme S, et al. 2020. Fungal Diversity 105:1-16 ((https://doi.org/10.1007/s13225-020-00466-2)) |
| Otu11608 | 2 | k__Fungi(100);p__Ascomycota(100);c__Saccharomycetes(100);o__Saccharomycetales(100);f__Metschnikowiaceae(100);g__Metschnikowia(100);g__Metschnikowia_unclassified(100); | Metschnikowia | 13 | Saprotroph | \|Undefined Saprotroph\| | Yeast | NULL | Probable | As nectar-tap_saprotroph (Põlme et al. 2020); Decay sugar-rich substrates (Põlme et al. 2020) | Põlme S, et al. 2020. Fungal Diversity 105:1-16 ((https://doi.org/10.1007/s13225-020-00466-2)) |
| Otu11609 | 2 | k__Fungi(100);p__Ascomycota(100);c__Saccharomycetes(100);o__Saccharomycetales(100);f__Metschnikowiaceae(100);g__Metschnikowia(100);s__Metschnikowia_viticola(100); | Metschnikowia | 13 | Saprotroph | \|Undefined Saprotroph\| | Yeast | NULL | Probable | As nectar-tap_saprotroph (Põlme et al. 2020); Decay sugar-rich substrates (Põlme et al. 2020) | Põlme S, et al. 2020. Fungal Diversity 105:1-16 ((https://doi.org/10.1007/s13225-020-00466-2)) |
| Otu11611 | 2 | k__Fungi(100);p__Ascomycota(100);c__Saccharomycetes(100);o__Saccharomycetales(100);f__Metschnikowiaceae(100);g__Metschnikowia(100);s__Metschnikowia_pulcherrima(100); | Metschnikowia | 13 | Saprotroph | \|Undefined Saprotroph\| | Yeast | NULL | Probable | As nectar-tap_saprotroph (Põlme et al. 2020); Decay sugar-rich substrates (Põlme et al. 2020) | Põlme S, et al. 2020. Fungal Diversity 105:1-16 ((https://doi.org/10.1007/s13225-020-00466-2)) |
| Otu11612 | 2 | k__Fungi(100);p__Ascomycota(100);c__Saccharomycetes(100);o__Saccharomycetales(100);f__Metschnikowiaceae(100);g__Metschnikowia(100);s__Metschnikowia_pulcherrima(100); | Metschnikowia | 13 | Saprotroph | \|Undefined Saprotroph\| | Yeast | NULL | Probable | As nectar-tap_saprotroph (Põlme et al. 2020); Decay sugar-rich substrates (Põlme et al. 2020) | Põlme S, et al. 2020. Fungal Diversity 105:1-16 ((https://doi.org/10.1007/s13225-020-00466-2)) |
| Otu11615 | 2 | k__Fungi(100);p__Ascomycota(100);c__Saccharomycetes(100);o__Saccharomycetales(100);f__Metschnikowiaceae(100);g__Metschnikowia(100);g__Metschnikowia_unclassified(100); | Metschnikowia | 13 | Saprotroph | \|Undefined Saprotroph\| | Yeast | NULL | Probable | As nectar-tap_saprotroph (Põlme et al. 2020); Decay sugar-rich substrates (Põlme et al. 2020) | Põlme S, et al. 2020. Fungal Diversity 105:1-16 ((https://doi.org/10.1007/s13225-020-00466-2)) |
| Otu11616 | 2 | k__Fungi(100);p__Ascomycota(100);c__Saccharomycetes(100);o__Saccharomycetales(100);f__Metschnikowiaceae(100);g__Metschnikowia(100);g__Metschnikowia_unclassified(100); | Metschnikowia | 13 | Saprotroph | \|Undefined Saprotroph\| | Yeast | NULL | Probable | As nectar-tap_saprotroph (Põlme et al. 2020); Decay sugar-rich substrates (Põlme et al. 2020) | Põlme S, et al. 2020. Fungal Diversity 105:1-16 ((https://doi.org/10.1007/s13225-020-00466-2)) |
| Otu11619 | 2 | k__Fungi(100);p__Ascomycota(100);c__Saccharomycetes(100);o__Saccharomycetales(100);f__Metschnikowiaceae(100);g__Metschnikowia(100);s__Metschnikowia_pulcherrima(100); | Metschnikowia | 13 | Saprotroph | \|Undefined Saprotroph\| | Yeast | NULL | Probable | As nectar-tap_saprotroph (Põlme et al. 2020); Decay sugar-rich substrates (Põlme et al. 2020) | Põlme S, et al. 2020. Fungal Diversity 105:1-16 ((https://doi.org/10.1007/s13225-020-00466-2)) |
| Otu11620 | 2 | k__Fungi(100);p__Ascomycota(100);c__Saccharomycetes(100);o__Saccharomycetales(100);f__Metschnikowiaceae(100);g__Metschnikowia(100);g__Metschnikowia_unclassified(100); | Metschnikowia | 13 | Saprotroph | \|Undefined Saprotroph\| | Yeast | NULL | Probable | As nectar-tap_saprotroph (Põlme et al. 2020); Decay sugar-rich substrates (Põlme et al. 2020) | Põlme S, et al. 2020. Fungal Diversity 105:1-16 ((https://doi.org/10.1007/s13225-020-00466-2)) |
| Otu11621 | 2 | k__Fungi(100);p__Ascomycota(100);c__Saccharomycetes(100);o__Saccharomycetales(100);f__Metschnikowiaceae(100);g__Metschnikowia(100);s__Metschnikowia_pulcherrima(100); | Metschnikowia | 13 | Saprotroph | \|Undefined Saprotroph\| | Yeast | NULL | Probable | As nectar-tap_saprotroph (Põlme et al. 2020); Decay sugar-rich substrates (Põlme et al. 2020) | Põlme S, et al. 2020. Fungal Diversity 105:1-16 ((https://doi.org/10.1007/s13225-020-00466-2)) |
| Otu11622 | 2 | k__Fungi(100);p__Ascomycota(100);c__Saccharomycetes(100);o__Saccharomycetales(100);f__Metschnikowiaceae(100);g__Metschnikowia(100);s__Metschnikowia_pulcherrima(100); | Metschnikowia | 13 | Saprotroph | \|Undefined Saprotroph\| | Yeast | NULL | Probable | As nectar-tap_saprotroph (Põlme et al. 2020); Decay sugar-rich substrates (Põlme et al. 2020) | Põlme S, et al. 2020. Fungal Diversity 105:1-16 ((https://doi.org/10.1007/s13225-020-00466-2)) |
| Otu11623 | 2 | k__Fungi(100);p__Ascomycota(100);c__Saccharomycetes(100);o__Saccharomycetales(100);f__Metschnikowiaceae(100);g__Metschnikowia(100);s__Metschnikowia_pulcherrima(100); | Metschnikowia | 13 | Saprotroph | \|Undefined Saprotroph\| | Yeast | NULL | Probable | As nectar-tap_saprotroph (Põlme et al. 2020); Decay sugar-rich substrates (Põlme et al. 2020) | Põlme S, et al. 2020. Fungal Diversity 105:1-16 ((https://doi.org/10.1007/s13225-020-00466-2)) |
| Otu11625 | 2 | k__Fungi(100);p__Ascomycota(100);c__Saccharomycetes(100);o__Saccharomycetales(100);f__Metschnikowiaceae(100);g__Metschnikowia(100);s__Metschnikowia_pulcherrima(100); | Metschnikowia | 13 | Saprotroph | \|Undefined Saprotroph\| | Yeast | NULL | Probable | As nectar-tap_saprotroph (Põlme et al. 2020); Decay sugar-rich substrates (Põlme et al. 2020) | Põlme S, et al. 2020. Fungal Diversity 105:1-16 ((https://doi.org/10.1007/s13225-020-00466-2)) |
| Otu11627 | 2 | k__Fungi(100);p__Ascomycota(100);c__Saccharomycetes(100);o__Saccharomycetales(100);f__Metschnikowiaceae(100);g__Metschnikowia(100);s__Metschnikowia_pulcherrima(100); | Metschnikowia | 13 | Saprotroph | \|Undefined Saprotroph\| | Yeast | NULL | Probable | As nectar-tap_saprotroph (Põlme et al. 2020); Decay sugar-rich substrates (Põlme et al. 2020) | Põlme S, et al. 2020. Fungal Diversity 105:1-16 ((https://doi.org/10.1007/s13225-020-00466-2)) |
| Otu11630 | 2 | k__Fungi(100);p__Chytridiomycota(100);c__Chytridiomycetes(100);o__Spizellomycetales(100);f__Spizellomycetaceae(100);f__Spizellomycetaceae_unclassified(100);f__Spizellomycetaceae_unclassified(100); | Spizellomycetaceae | 9 | Pathotroph | Plant Pathogen-Undefined Parasite-Undefined Saprotroph | Microfungus | NULL | Probable | Common in the soil and also as parasites of a range of soil organisms and plants (Cannon & Kirk 2007) | Cannon PF, Kirk PM. 2007. Fungal Families of the World. CAB International, Cambridge (ISBN: 978-0851998275) |
| Otu11633 | 2 | k__Fungi(100);p__Ascomycota(100);c__Saccharomycetes(100);o__Saccharomycetales(100);f__Metschnikowiaceae(100);g__Metschnikowia(100);g__Metschnikowia_unclassified(100); | Metschnikowia | 13 | Saprotroph | \|Undefined Saprotroph\| | Yeast | NULL | Probable | As nectar-tap_saprotroph (Põlme et al. 2020); Decay sugar-rich substrates (Põlme et al. 2020) | Põlme S, et al. 2020. Fungal Diversity 105:1-16 ((https://doi.org/10.1007/s13225-020-00466-2)) |
| Otu11634 | 2 | k__Fungi(100);p__Ascomycota(100);c__Saccharomycetes(100);o__Saccharomycetales(100);f__Metschnikowiaceae(100);g__Metschnikowia(100);s__Metschnikowia_pulcherrima(100); | Metschnikowia | 13 | Saprotroph | \|Undefined Saprotroph\| | Yeast | NULL | Probable | As nectar-tap_saprotroph (Põlme et al. 2020); Decay sugar-rich substrates (Põlme et al. 2020) | Põlme S, et al. 2020. Fungal Diversity 105:1-16 ((https://doi.org/10.1007/s13225-020-00466-2)) |
| Otu11635 | 2 | k__Fungi(100);p__Ascomycota(100);c__Saccharomycetes(100);o__Saccharomycetales(100);f__Metschnikowiaceae(100);g__Metschnikowia(100);s__Metschnikowia_pulcherrima(100); | Metschnikowia | 13 | Saprotroph | \|Undefined Saprotroph\| | Yeast | NULL | Probable | As nectar-tap_saprotroph (Põlme et al. 2020); Decay sugar-rich substrates (Põlme et al. 2020) | Põlme S, et al. 2020. Fungal Diversity 105:1-16 ((https://doi.org/10.1007/s13225-020-00466-2)) |
| Otu11636 | 2 | k__Fungi(100);p__Ascomycota(100);c__Saccharomycetes(100);o__Saccharomycetales(100);f__Metschnikowiaceae(100);g__Metschnikowia(100);s__Metschnikowia_pulcherrima(100); | Metschnikowia | 13 | Saprotroph | \|Undefined Saprotroph\| | Yeast | NULL | Probable | As nectar-tap_saprotroph (Põlme et al. 2020); Decay sugar-rich substrates (Põlme et al. 2020) | Põlme S, et al. 2020. Fungal Diversity 105:1-16 ((https://doi.org/10.1007/s13225-020-00466-2)) |
| Otu11637 | 2 | k__Fungi(100);p__Ascomycota(100);c__Saccharomycetes(100);o__Saccharomycetales(100);f__Metschnikowiaceae(100);g__Metschnikowia(100);s__Metschnikowia_pulcherrima(100); | Metschnikowia | 13 | Saprotroph | \|Undefined Saprotroph\| | Yeast | NULL | Probable | As nectar-tap_saprotroph (Põlme et al. 2020); Decay sugar-rich substrates (Põlme et al. 2020) | Põlme S, et al. 2020. Fungal Diversity 105:1-16 ((https://doi.org/10.1007/s13225-020-00466-2)) |
| Otu11639 | 2 | k__Fungi(100);p__Ascomycota(100);c__Saccharomycetes(100);o__Saccharomycetales(100);f__Metschnikowiaceae(100);g__Metschnikowia(100);g__Metschnikowia_unclassified(100); | Metschnikowia | 13 | Saprotroph | \|Undefined Saprotroph\| | Yeast | NULL | Probable | As nectar-tap_saprotroph (Põlme et al. 2020); Decay sugar-rich substrates (Põlme et al. 2020) | Põlme S, et al. 2020. Fungal Diversity 105:1-16 ((https://doi.org/10.1007/s13225-020-00466-2)) |
| Otu11640 | 2 | k__Fungi(100);p__Ascomycota(100);c__Saccharomycetes(100);o__Saccharomycetales(100);f__Metschnikowiaceae(100);g__Metschnikowia(100);g__Metschnikowia_unclassified(100); | Metschnikowia | 13 | Saprotroph | \|Undefined Saprotroph\| | Yeast | NULL | Probable | As nectar-tap_saprotroph (Põlme et al. 2020); Decay sugar-rich substrates (Põlme et al. 2020) | Põlme S, et al. 2020. Fungal Diversity 105:1-16 ((https://doi.org/10.1007/s13225-020-00466-2)) |
| Otu11641 | 2 | k__Fungi(100);p__Ascomycota(100);c__Saccharomycetes(100);o__Saccharomycetales(100);f__Metschnikowiaceae(100);g__Metschnikowia(100);s__Metschnikowia_pulcherrima(100); | Metschnikowia | 13 | Saprotroph | \|Undefined Saprotroph\| | Yeast | NULL | Probable | As nectar-tap_saprotroph (Põlme et al. 2020); Decay sugar-rich substrates (Põlme et al. 2020) | Põlme S, et al. 2020. Fungal Diversity 105:1-16 ((https://doi.org/10.1007/s13225-020-00466-2)) |
| Otu11642 | 2 | k__Fungi(100);p__Ascomycota(100);c__Saccharomycetes(100);o__Saccharomycetales(100);f__Metschnikowiaceae(100);g__Metschnikowia(100);g__Metschnikowia_unclassified(100); | Metschnikowia | 13 | Saprotroph | \|Undefined Saprotroph\| | Yeast | NULL | Probable | As nectar-tap_saprotroph (Põlme et al. 2020); Decay sugar-rich substrates (Põlme et al. 2020) | Põlme S, et al. 2020. Fungal Diversity 105:1-16 ((https://doi.org/10.1007/s13225-020-00466-2)) |
| Otu11643 | 2 | k__Fungi(100);p__Basidiomycota(100);c__Agaricostilbomycetes(100);o__Agaricostilbales(100);f__Chionosphaeraceae(100);g__Kurtzmanomyces(100);s__Kurtzmanomyces_nectairei(100); | Kurtzmanomyces | 13 | Saprotroph | \|Undefined Saprotroph\| | Dimorphic | NULL | Probable | NULL | Põlme S, et al. 2020. Fungal Diversity 105:1-16 ((https://doi.org/10.1007/s13225-020-00466-2)) |
| Otu11644 | 2 | k__Fungi(100);p__Basidiomycota(100);c__Agaricomycetes(100);o__Agaricales(100);f__Cortinariaceae(100);g__Cortinarius(100);g__Cortinarius_unclassified(100); | Cortinarius | 13 | Symbiotroph | \|Ectomycorrhizal\| | Agaricoid | NULL | Highly Probable | NULL | Rinaldi AC, et al. 2008. Fungal Diversity 33:1-45 ((http://www.fungaldiversity.org/fdp/sfdp/33-1.pdf)); Tedersoo L, et al. 2010. Mycorrhiza 20:217-263 ((https://doi.org/10.1007/s00572-009-0274-x)); Põlme S, et al. 2020. Fungal Diversity 105:1-16 ((https://doi.org/10.1007/s13225-020-00466-2)); http://mycorrhizas.info/ecmf.html ((http://mycorrhizas.info/ecmf.html)) |
| Otu11645 | 2 | k__Fungi(100);p__Basidiomycota(100);c__Agaricomycetes(100);o__Agaricales(100);f__Cortinariaceae(100);g__Cortinarius(100);s__Cortinarius_bivelus(100); | Cortinarius | 13 | Symbiotroph | \|Ectomycorrhizal\| | Agaricoid | NULL | Highly Probable | NULL | Rinaldi AC, et al. 2008. Fungal Diversity 33:1-45 ((http://www.fungaldiversity.org/fdp/sfdp/33-1.pdf)); Tedersoo L, et al. 2010. Mycorrhiza 20:217-263 ((https://doi.org/10.1007/s00572-009-0274-x)); Põlme S, et al. 2020. Fungal Diversity 105:1-16 ((https://doi.org/10.1007/s13225-020-00466-2)); http://mycorrhizas.info/ecmf.html ((http://mycorrhizas.info/ecmf.html)) |
| Otu11646 | 2 | k__Fungi(100);p__Basidiomycota(100);c__Agaricomycetes(100);o__Boletales(100);f__Boletaceae(100);g__Boletus(100);g__Boletus_unclassified(100); | Boletus | 13 | Pathotroph-Symbiotroph | \|Ectomycorrhizal\|-Fungal Parasite | Agaricoid-Boletoid | NULL | Highly Probable | NULL | Rinaldi AC, et al. 2008. Fungal Diversity 33:1-45 ((http://www.fungaldiversity.org/fdp/sfdp/33-1.pdf)); Tedersoo L, et al. 2010. Mycorrhiza 20:217-263 ((https://doi.org/10.1007/s00572-009-0274-x)); Põlme S, et al. 2020. Fungal Diversity 105:1-16 ((https://doi.org/10.1007/s13225-020-00466-2)); http://mycorrhizas.info/ecmf.html ((http://mycorrhizas.info/ecmf.html)); List compiled by John Plischke III ((https://goodmushroom.weebly.com/uploads/5/7/1/7/57176535/fungi_on_fungi_pdf_lq.pdf)) |
| Otu11647 | 2 | k__Fungi(100);p__Basidiomycota(100);c__Agaricomycetes(100);o__Boletales(100);f__Boletaceae(100);f__Boletaceae_unclassified(100);f__Boletaceae_unclassified(100); | Boletaceae | 9 | Pathotroph-Saprotroph-Symbiotroph | Ectomycorrhizal-Fungal Parasite-Plant Saprotroph-Wood Saprotroph | Boletoid | NULL | Probable | Most species form ectomycorrhizal associations with gymnosperm or angiosperm trees. Some are saprobes on decaying wood and leaf litter, a few are parasitic on other fungi (Cannon & Kirk 2007) | Cannon PF, Kirk PM. 2007. Fungal Families of the World. CAB International, Cambridge (ISBN: 978-0851998275) |
| Otu11648 | 2 | k__Fungi(100);p__Basidiomycota(100);c__Agaricomycetes(100);o__Agaricales(100);f__Entolomataceae(100);g__Entoloma(100);s__Entoloma_insidiosum(100); | Entoloma | 13 | Pathotroph-Saprotroph-Symbiotroph | Ectomycorrhizal-Fungal Parasite-\|Undefined Saprotroph\|-Undefined Symbiotroph | Agaricoid | NULL | Possible | As soil_saprotroph (Põlme et al. 2020); Decay roots, soil (Põlme et al. 2020) | Rinaldi AC, et al. 2008. Fungal Diversity 33:1-45 ((http://www.fungaldiversity.org/fdp/sfdp/33-1.pdf)) (as ECM/not ECM); Tedersoo L, et al. 2010. Mycorrhiza 20:217-263 ((https://doi.org/10.1007/s00572-009-0274-x; Põlme S, et al. 2020. Fungal Diversity 105:1-16 ((https://doi.org/10.1007/s13225-020-00466-2)); sensu stricto)); http://mycorrhizas.info/ecmf.html ((http://mycorrhizas.info/ecmf.html)) ((as ECM/not ECM)) |
| Otu11649 | 2 | k__Fungi(100);p__Basidiomycota(100);c__Agaricomycetes(100);o__Polyporales(100);f__Phanerochaetaceae(100);g__Ceriporiopsis(100);s__Ceriporiopsis_sp_PRM_899300(100); | Ceriporiopsis | 13 | Saprotroph | \|Wood Saprotroph\| | Corticioid | White Rot | Highly Probable | As white_rot (Põlme et al. 2020); Decay wood (Põlme et al. 2020) | Gilbertson RL, Ryvarden L. 1987-1987. North American Polypores. Fungiflora, Oslo (ISBN: 978-0945345060); Tedersoo L, et al. 2014. Science 346:e1256688 ((https://doi.org/10.1126/science.1256688)); Põlme S, et al. 2020. Fungal Diversity 105:1-16 ((https://doi.org/10.1007/s13225-020-00466-2)) |
| Otu11651 | 2 | k__Fungi(100);p__Basidiomycota(100);c__Agaricomycetes(100);o__Cantharellales(100);f__Ceratobasidiaceae(100);g__Ceratobasidium(100);s__Ceratobasidium_sp_AG_H(100); | Ceratobasidium | 13 | Pathotroph-Saprotroph-Symbiotroph | Ectomycorrhizal-Endomycorrhizal-Orchid Mycorrhizal-\|Plant Pathogen\|-Plant Saprotroph-Undefined Saprotroph | Corticioid-Microfungus | NULL | Possible | especially of Orchidaceae but also even bryophytes (Cannon & Kirk 2007); As litter_saprotroph (Põlme et al. 2020); Best known as plant pathogens, but also saprobic. Species are endomycorrhizal with a wide range of plant roots ; Decay leaf, fruit, seed, roots, soil (Põlme et al. 2020) | Cannon PF, Kirk PM. 2007. Fungal Families of the World. CAB International, Cambridge (ISBN: 978-0851998275); Tedersoo L, et al. 2014. Science 346:e1256688 ((https://doi.org/10.1126/science.1256688)); Põlme S, et al. 2020. Fungal Diversity 105:1-16 ((https://doi.org/10.1007/s13225-020-00466-2)) |
| Otu11652 | 2 | k__Fungi(100);p__Basidiomycota(100);c__Agaricomycetes(100);o__Cantharellales(100);f__Ceratobasidiaceae(100);g__Ceratobasidium(100);g__Ceratobasidium_unclassified(100); | Ceratobasidium | 13 | Pathotroph-Saprotroph-Symbiotroph | Ectomycorrhizal-Endomycorrhizal-Orchid Mycorrhizal-\|Plant Pathogen\|-Plant Saprotroph-Undefined Saprotroph | Corticioid-Microfungus | NULL | Possible | especially of Orchidaceae but also even bryophytes (Cannon & Kirk 2007); As litter_saprotroph (Põlme et al. 2020); Best known as plant pathogens, but also saprobic. Species are endomycorrhizal with a wide range of plant roots ; Decay leaf, fruit, seed, roots, soil (Põlme et al. 2020) | Cannon PF, Kirk PM. 2007. Fungal Families of the World. CAB International, Cambridge (ISBN: 978-0851998275); Tedersoo L, et al. 2014. Science 346:e1256688 ((https://doi.org/10.1126/science.1256688)); Põlme S, et al. 2020. Fungal Diversity 105:1-16 ((https://doi.org/10.1007/s13225-020-00466-2)) |
| Otu11653 | 2 | k__Fungi(100);p__Basidiomycota(100);c__Exobasidiomycetes(100);o__Exobasidiales(100);f__Exobasidiaceae(100);g__Exobasidium(100);g__Exobasidium_unclassified(100); | Exobasidium | 13 | Pathotroph | \|Plant Pathogen\| | Leaf Blister-Leaf Gall-Leaf Spot | NULL | Probable | NULL | Tedersoo L, et al. 2014. Science 346:e1256688 ((https://doi.org/10.1126/science.1256688)); Põlme S, et al. 2020. Fungal Diversity 105:1-16 ((https://doi.org/10.1007/s13225-020-00466-2)) |
| Otu11657 | 2 | k__Fungi(100);p__Basidiomycota(100);c__Agaricomycetes(100);o__Agaricales(100);f__Cortinariaceae(100);g__Cortinarius(100);g__Cortinarius_unclassified(100); | Cortinarius | 13 | Symbiotroph | \|Ectomycorrhizal\| | Agaricoid | NULL | Highly Probable | NULL | Rinaldi AC, et al. 2008. Fungal Diversity 33:1-45 ((http://www.fungaldiversity.org/fdp/sfdp/33-1.pdf)); Tedersoo L, et al. 2010. Mycorrhiza 20:217-263 ((https://doi.org/10.1007/s00572-009-0274-x)); Põlme S, et al. 2020. Fungal Diversity 105:1-16 ((https://doi.org/10.1007/s13225-020-00466-2)); http://mycorrhizas.info/ecmf.html ((http://mycorrhizas.info/ecmf.html)) |
| Otu11659 | 2 | k__Fungi(100);p__Basidiomycota(100);c__Agaricomycetes(100);o__Agaricales(100);f__Pluteaceae(100);g__Pluteus(100);s__Pluteus_thomsonii(100); | Pluteus | 13 | Pathotroph-Saprotroph | Bryophyte Parasite-\|Plant Saprotroph\|-Wood Saprotroph | Agaricoid | White Rot | Probable | As litter_saprotroph (Põlme et al. 2020); As white_rot (Põlme et al. 2020); Decay leaf, fruit, seed (Põlme et al. 2020) | Grgurinovic CA, Simpson JA. 2001. Fungal Diversity 8:97-127 ((http://www.fungaldiversity.org/fdp/sfdp/FD_8_97-127.pdf)); James TY, et al. 2006. Nature 443:818-822 ((https://doi.org/10.1038/nature05110)); Tedersoo L, et al. 2014. Science 346:e1256688 ((https://doi.org/10.1126/science.1256688)); Põlme S, et al. 2020. Fungal Diversity 105:1-16 ((https://doi.org/10.1007/s13225-020-00466-2)) |
| Otu11660 | 2 | k__Fungi(100);p__Basidiomycota(100);c__Agaricostilbomycetes(100);o__Agaricostilbales(100);f__Chionosphaeraceae(100);g__Kurtzmanomyces(100);s__Kurtzmanomyces_tardus(100); | Kurtzmanomyces | 13 | Saprotroph | \|Undefined Saprotroph\| | Dimorphic | NULL | Probable | NULL | Põlme S, et al. 2020. Fungal Diversity 105:1-16 ((https://doi.org/10.1007/s13225-020-00466-2)) |
| Otu11670 | 2 | k__Fungi(100);p__Basidiomycota(100);c__Agaricomycetes(100);o__Russulales(100);f__Albatrellaceae(100);g__Albatrellus(100);s__Albatrellus_higanensis(100); | Albatrellus | 13 | Symbiotroph | \|Ectomycorrhizal\| | Agaricoid-Polyporoid | NULL | Highly Probable | Common name - sheeps polypore (Gilbertson & Ryvarden 1987-1987) | Gilbertson RL, Ryvarden L. 1987-1987. North American Polypores. Fungiflora, Oslo (ISBN: 978-0945345060); Rinaldi AC, et al. 2008. Fungal Diversity 33:1-45 ((http://www.fungaldiversity.org/fdp/sfdp/33-1.pdf)); Tedersoo L, et al. 2010. Mycorrhiza 20:217-263 ((https://doi.org/10.1007/s00572-009-0274-x)); Põlme S, et al. 2020. Fungal Diversity 105:1-16 ((https://doi.org/10.1007/s13225-020-00466-2)); http://mycorrhizas.info/ecmf.html ((http://mycorrhizas.info/ecmf.html)) |
| Otu11671 | 2 | k__Fungi(100);p__Basidiomycota(100);c__Agaricomycetes(100);o__Boletales(100);f__Boletaceae(100);g__Boletus(100);g__Boletus_unclassified(100); | Boletus | 13 | Pathotroph-Symbiotroph | \|Ectomycorrhizal\|-Fungal Parasite | Agaricoid-Boletoid | NULL | Highly Probable | NULL | Rinaldi AC, et al. 2008. Fungal Diversity 33:1-45 ((http://www.fungaldiversity.org/fdp/sfdp/33-1.pdf)); Tedersoo L, et al. 2010. Mycorrhiza 20:217-263 ((https://doi.org/10.1007/s00572-009-0274-x)); Põlme S, et al. 2020. Fungal Diversity 105:1-16 ((https://doi.org/10.1007/s13225-020-00466-2)); http://mycorrhizas.info/ecmf.html ((http://mycorrhizas.info/ecmf.html)); List compiled by John Plischke III ((https://goodmushroom.weebly.com/uploads/5/7/1/7/57176535/fungi_on_fungi_pdf_lq.pdf)) |
| Otu11672 | 2 | k__Fungi(100);p__Basidiomycota(100);c__Agaricomycetes(100);o__Boletales(100);f__Suillaceae(100);g__Suillus(100);s__Suillus_sp(100); | Suillus | 13 | Symbiotroph | \|Ectomycorrhizal\| | Agaricoid-Boletoid | NULL | Highly Probable | Host - Pinaceae (Põlme et al. 2020) | Rinaldi AC, et al. 2008. Fungal Diversity 33:1-45 ((http://www.fungaldiversity.org/fdp/sfdp/33-1.pdf)); Tedersoo L, et al. 2010. Mycorrhiza 20:217-263 ((https://doi.org/10.1007/s00572-009-0274-x)); Põlme S, et al. 2020. Fungal Diversity 105:1-16 ((https://doi.org/10.1007/s13225-020-00466-2)); http://mycorrhizas.info/ecmf.html ((http://mycorrhizas.info/ecmf.html)) |
| Otu11674 | 2 | k__Fungi(100);p__Basidiomycota(100);c__Agaricomycetes(100);o__Auriculariales(100);f__Auriculariales_family_Incertae_sedis(100);g__Exidia(100);g__Exidia_unclassified(100); | Exidia | 13 | Saprotroph-Symbiotroph | Endophyte-Plant Saprotroph-Undefined Saprotroph-\|Wood Saprotroph\| | Tremelloid | NULL | Probable | As foliar_endophyte (Põlme et al. 2020); As litter_saprotroph (Põlme et al. 2020); Decay wood, leaf, fruit, seed (Põlme et al. 2020) | Tedersoo L, et al. 2014. Science 346:e1256688 ((https://doi.org/10.1126/science.1256688)); David AS, et al. 2016. Microbial Ecology 71: 912-926 ((https://doi.org/10.1007/s00248-015-0712-6)); Põlme S, et al. 2020. Fungal Diversity 105:1-16 ((https://doi.org/10.1007/s13225-020-00466-2)) |
| Otu11677 | 2 | k__Fungi(100);p__Basidiomycota(100);c__Agaricomycetes(100);o__Polyporales(100);f__Polyporaceae(100);g__Perenniporiella(100);s__Perenniporiella_tepeitensis(100); | Perenniporiella | 13 | Saprotroph | Undefined Saprotroph-\|Wood Saprotroph\| | Polyporoid | White Rot | Probable | As white_rot (Põlme et al. 2020); Decay wood (Põlme et al. 2020) | Tedersoo L, et al. 2014. Science 346:e1256688 ((https://doi.org/10.1126/science.1256688)); Põlme S, et al. 2020. Fungal Diversity 105:1-16 ((https://doi.org/10.1007/s13225-020-00466-2)) |
| Otu11679 | 2 | k__Fungi(100);p__Basidiomycota(100);c__Agaricomycetes(100);o__Agaricales(100);f__Tricholomataceae(100);g__Tricholoma(100);g__Tricholoma_unclassified(100); | Tricholoma | 13 | Pathotroph-Symbiotroph | \|Ectomycorrhizal\|-Fungal Parasite | Agaricoid | NULL | Highly Probable | Tricholoma species are ectomycorrhizal, occurring in coniferous and broadleaved forests (Cannon & Kirk 2007); T. sclerotoideum is parasitic on Helvella spp. | Cannon PF, Kirk PM. 2007. Fungal Families of the World. CAB International, Cambridge (ISBN: 978-0851998275); Rinaldi AC, et al. 2008. Fungal Diversity 33:1-45 ((http://www.fungaldiversity.org/fdp/sfdp/33-1.pdf)); Tedersoo L, et al. 2010. Mycorrhiza 20:217-263 ((https://doi.org/10.1007/s00572-009-0274-x)); Põlme S, et al. 2020. Fungal Diversity 105:1-16 ((https://doi.org/10.1007/s13225-020-00466-2)); http://mycorrhizas.info/ecmf.html ((http://mycorrhizas.info/ecmf.html)) |
| Otu11680 | 2 | k__Fungi(100);p__Basidiomycota(100);c__Agaricomycetes(100);o__Thelephorales(100);f__Bankeraceae(100);g__Sarcodon(100);g__Sarcodon_unclassified(100); | Sarcodon | 13 | Symbiotroph | \|Ectomycorrhizal\| | Agaricoid-Hydnoid | NULL | Highly Probable | NULL | Rinaldi AC, et al. 2008. Fungal Diversity 33:1-45 ((http://www.fungaldiversity.org/fdp/sfdp/33-1.pdf)); Tedersoo L, et al. 2010. Mycorrhiza 20:217-263 ((https://doi.org/10.1007/s00572-009-0274-x)); Põlme S, et al. 2020. Fungal Diversity 105:1-16 ((https://doi.org/10.1007/s13225-020-00466-2)); http://mycorrhizas.info/ecmf.html ((http://mycorrhizas.info/ecmf.html)) |
| Otu11681 | 2 | k__Fungi(100);p__Basidiomycota(100);c__Agaricomycetes(100);o__Polyporales(100);f__Fomitopsidaceae(100);g__Fibroporia(100);s__Fibroporia_gossypium(100); | Fibroporia | 13 | Saprotroph | \|Wood Saprotroph\| | Corticioid | Brown Rot | Probable | As brown-rot (Põlme et al. 2020); Decay wood (Põlme et al. 2020) | Tedersoo L, et al. 2014. Science 346:e1256688 ((https://doi.org/10.1126/science.1256688)); Põlme S, et al. 2020. Fungal Diversity 105:1-16 ((https://doi.org/10.1007/s13225-020-00466-2)) |
| Otu11685 | 2 | k__Fungi(100);p__Basidiomycota(100);c__Agaricomycetes(100);o__Polyporales(100);f__Phanerochaetaceae(100);g__Antrodiella(100);s__Antrodiella_romellii(100); | Antrodiella | 13 | Saprotroph | \|Wood Saprotroph\| | Corticioid-Polyporoid | White Rot | Highly Probable | As white_rot (Põlme et al. 2020); Decay wood (Põlme et al. 2020) | Gilbertson RL, Ryvarden L. 1987-1987. North American Polypores. Fungiflora, Oslo (ISBN: 978-0945345060); Põlme S, et al. 2020. Fungal Diversity 105:1-16 ((https://doi.org/10.1007/s13225-020-00466-2)) |

| | | | | | | | | | | | |
|---|---|---|---|---|---|---|---|---|---|---|---|
| Otu11686 | 2 | k__Fungi(100);p__Basidiomycota(100);c__Agaricomycetes(100);o__Atheliales(100);f__Atheliaceae(100);g__Tylospora(100);s__Tylospora_sp(100); | Tylospora | 13 | Symbiotroph | \|Ectomycorrhizal\| | Corticioid | NULL | Highly Probable | NULL | Rinaldi AC, et al. 2008. Fungal Diversity 33:1-45 ((http://www.fungaldiversity.org/fdp/sfdp/33-1.pdf)); Tedersoo L, et al. 2010. Mycorrhiza 20:217-263 ((https://doi.org/10.1007/s00572-009-0274-x)); Põlme S, et al. 2020. Fungal Diversity 105:1-16 ((https://doi.org/10.1007/s13225-020-00466-2)); http://mycorrhizas.info/ecmf.html ((http://mycorrhizas.info/ecmf.html)) |
| Otu11687 | 2 | k__Fungi(100);p__Basidiomycota(100);c__Agaricomycetes(100);o__Agaricales(100);f__Inocybaceae(100);g__Crepidotus(100);s__Crepidotus_variabilis(100); | Crepidotus | 13 | Saprotroph | \|Wood Saprotroph\| | Agaricoid | NULL | Highly Probable | Decay wood (Põlme et al. 2020) | Grgurinovic CA, Simpson JA. 2001. Fungal Diversity 8:97-127 ((http://www.fungaldiversity.org/fdp/sfdp/FD_8_97-127.pdf)); Aime MC, et al. 2005. American Journal of Botany 92:74-82 ((https://doi.org/10.3732/ajb.92.1.74)); Tedersoo L, et al. 2014. Science 346:e1256688 ((https://doi.org/10.1126/science.1256688)); Põlme S, et al. 2020. Fungal Diversity 105:1-16 ((https://doi.org/10.1007/s13225-020-00466-2)) |
| Otu11688 | 2 | k__Fungi(100);p__Basidiomycota(100);c__Agaricomycetes(100);o__Agaricales(100);f__Inocybaceae(100);g__Tubaria(100);g__Tubaria_unclassified(100); | Tubaria | 13 | Saprotroph | \|Plant Saprotroph\|-Wood Saprotroph | Agaricoid | NULL | Probable | As litter_saprotroph (Põlme et al. 2020); Decay leaf, fruit, seed (Põlme et al. 2020) | Tedersoo L, et al. 2014. Science 346:e1256688 ((https://doi.org/10.1126/science.1256688)); Põlme S, et al. 2020. Fungal Diversity 105:1-16 ((https://doi.org/10.1007/s13225-020-00466-2)) |
| Otu11689 | 2 | k__Fungi(100);p__Basidiomycota(100);c__Agaricomycetes(100);o__Agaricales(100);f__Agaricaceae(100);g__Calvatia(100);s__Calvatia_cyathiformis(100); | Calvatia | 13 | Saprotroph | \|Undefined Saprotroph\| | Gasteroid | NULL | Probable | puffball; As soil_saprotroph (Põlme et al. 2020); Common name ; Decay soil (Põlme et al. 2020) | Grgurinovic CA, Simpson JA. 2001. Fungal Diversity 8:97-127 ((http://www.fungaldiversity.org/fdp/sfdp/FD_8_97-127.pdf)); Tedersoo L, et al. 2014. Science 346:e1256688 ((https://doi.org/10.1126/science.1256688)); Põlme S, et al. 2020. Fungal Diversity 105:1-16 ((https://doi.org/10.1007/s13225-020-00466-2)) |
| Otu11690 | 2 | k__Fungi(100);p__Basidiomycota(100);c__Agaricomycetes(100);o__Agaricales(100);f__Inocybaceae(100);g__Crepidotus(100);s__Crepidotus_variabilis(100); | Crepidotus | 13 | Saprotroph | \|Wood Saprotroph\| | Agaricoid | NULL | Highly Probable | Decay wood (Põlme et al. 2020) | Grgurinovic CA, Simpson JA. 2001. Fungal Diversity 8:97-127 ((http://www.fungaldiversity.org/fdp/sfdp/FD_8_97-127.pdf)); Aime MC, et al. 2005. American Journal of Botany 92:74-82 ((https://doi.org/10.3732/ajb.92.1.74)); Tedersoo L, et al. 2014. Science 346:e1256688 ((https://doi.org/10.1126/science.1256688)); Põlme S, et al. 2020. Fungal Diversity 105:1-16 ((https://doi.org/10.1007/s13225-020-00466-2)) |
| Otu11691 | 2 | k__Fungi(100);p__Basidiomycota(100);c__Agaricomycetes(100);o__Agaricales(100);f__Psathyrellaceae(100);g__Coprinopsis(100);s__Coprinopsis_villosa(100); | Coprinopsis | 13 | Saprotroph | \|Undefined Saprotroph\| | Agaricoid | NULL | Probable | As soil_saprotroph (Põlme et al. 2020); Decay soil (Põlme et al. 2020) | Tedersoo L, et al. 2014. Science 346:e1256688 ((https://doi.org/10.1126/science.1256688)); Põlme S, et al. 2020. Fungal Diversity 105:1-16 ((https://doi.org/10.1007/s13225-020-00466-2)) |
| Otu11692 | 2 | k__Fungi(100);p__Basidiomycota(100);c__Agaricomycetes(100);o__Agaricales(100);f__Strophariaceae(100);f__Strophariaceae_unclassified(100);f__Strophariaceae_unclassified(100); | Strophariaceae | 9 | Pathotroph-Saprotroph | Dung Saprotroph-Plant Parasite-Soil Saprotroph-Undefined Saprotroph-Wood Saprotroph | Agaricoid | NULL | Probable | Very variable ecology; Pholiota species are lignicolous and can act as weak parasites while most other taxa are either sparobes in grassy habitats or are found in dung. Species are not ectomycorrhizal (Cannon & Kirk 2007) | Cannon PF, Kirk PM. 2007. Fungal Families of the World. CAB International, Cambridge (ISBN: 978-0851998275) |
| Otu11693 | 2 | k__Fungi(100);p__Basidiomycota(100);c__Agaricomycetes(100);o__Atheliales(100);f__Atheliaceae(100);g__Byssocorticium(100);s__Byssocorticium_sp_1827_CG_2012(100); | Byssocorticium | 13 | Saprotroph-Symbiotroph | \|Ectomycorrhizal\|-Undefined Saprotroph | Corticioid | NULL | Highly Probable | NULL | Rinaldi AC, et al. 2008. Fungal Diversity 33:1-45 ((http://www.fungaldiversity.org/fdp/sfdp/33-1.pdf)); Tedersoo L, et al. 2010. Mycorrhiza 20:217-263 ((https://doi.org/10.1007/s00572-009-0274-x)); Taylor DL, et al. 2014. Ecological Monographs 84:3-20 ((https://doi.org/10.1890/12-1693.1)); Põlme S, et al. 2020. Fungal Diversity 105:1-16 ((https://doi.org/10.1007/s13225-020-00466-2)); http://mycorrhizas.info/ecmf.html ((http://mycorrhizas.info/ecmf.html)) |
| Otu11694 | 2 | k__Fungi(100);p__Basidiomycota(100);c__Agaricomycetes(100);o__Atheliales(100);f__Atheliaceae(100);g__Byssocorticium(100);s__Byssocorticium_caeruleum(100); | Byssocorticium | 13 | Saprotroph-Symbiotroph | \|Ectomycorrhizal\|-Undefined Saprotroph | Corticioid | NULL | Highly Probable | NULL | Rinaldi AC, et al. 2008. Fungal Diversity 33:1-45 ((http://www.fungaldiversity.org/fdp/sfdp/33-1.pdf)); Tedersoo L, et al. 2010. Mycorrhiza 20:217-263 ((https://doi.org/10.1007/s00572-009-0274-x)); Taylor DL, et al. 2014. Ecological Monographs 84:3-20 ((https://doi.org/10.1890/12-1693.1)); Põlme S, et al. 2020. Fungal Diversity 105:1-16 ((https://doi.org/10.1007/s13225-020-00466-2)); http://mycorrhizas.info/ecmf.html ((http://mycorrhizas.info/ecmf.html)) |
| Otu11695 | 2 | k__Fungi(100);p__Basidiomycota(100);c__Agaricomycetes(100);o__Agaricales(100);f__Pluteaceae(100);g__Pluteus(100);s__Pluteus_salicinus(100); | Pluteus | 13 | Pathotroph-Saprotroph | Bryophyte Parasite-\|Plant Saprotroph\|-Wood Saprotroph | Agaricoid | White Rot | Probable | As litter_saprotroph (Põlme et al. 2020); As white_rot (Põlme et al. 2020); Decay leaf, fruit, seed (Põlme et al. 2020) | Grgurinovic CA, Simpson JA. 2001. Fungal Diversity 8:97-127 ((http://www.fungaldiversity.org/fdp/sfdp/FD_8_97-127.pdf)); James TY, et al. 2006. Nature 443:818-822 ((https://doi.org/10.1038/nature05110)); Tedersoo L, et al. 2014. Science 346:e1256688 ((https://doi.org/10.1126/science.1256688)); Põlme S, et al. 2020. Fungal Diversity 105:1-16 ((https://doi.org/10.1007/s13225-020-00466-2)) |
| Otu11696 | 2 | k__Fungi(100);p__Basidiomycota(100);c__Agaricomycetes(100);o__Agaricales(100);f__Bolbitiaceae(100);g__Conocybe(100);s__Conocybe_aff_ochrostriata_NL_0830(100); | Conocybe | 13 | Saprotroph | Dung Saprotroph-Plant Saprotroph-\|Undefined Saprotroph\| | Agaricoid | NULL | Probable | As litter_saprotroph (Põlme et al. 2020); As soil_saprotroph (Põlme et al. 2020); Decay soil (Põlme et al. 2020) | Tedersoo L, et al. 2014. Science 346:e1256688 ((https://doi.org/10.1126/science.1256688)); Põlme S, et al. 2020. Fungal Diversity 105:1-16 ((https://doi.org/10.1007/s13225-020-00466-2)) |
| Otu11698 | 2 | k__Fungi(100);p__Basidiomycota(100);c__Microbotryomycetes(100);o__Sporidiobolales(100);f__Sporidiobolales_family_Incertae_sedis(100);g__Sporobolomyces(100);s__Sporobolomyces_coprosmicola(100); | Sporobolomyces | 13 | Pathotroph-Saprotroph | \|Fungal Parasite\|-Plant Saprotroph | Yeast | NULL | Probable | As litter_saprotroph (Põlme et al. 2020); As mycoparasite (Põlme et al. 2020); Decay fungal material (Põlme et al. 2020) | Tedersoo L, et al. 2014. Science 346:e1256688 ((https://doi.org/10.1126/science.1256688)); Purahong W, et al. 2016. Molecular Ecology 25:4059-4074 ((https://doi.org/10.1111/mec.13739)); Põlme S, et al. 2020. Fungal Diversity 105:1-16 ((https://doi.org/10.1007/s13225-020-00466-2)) |
| Otu11699 | 2 | k__Fungi(100);p__Basidiomycota(100);c__Agaricomycetes(100);o__Corticiales(100);f__Corticiaceae(100);g__Waitea(100);s__Rhizoctonia_oryzae(100); | Rhizoctonia | 13 | Pathotroph-Saprotroph-Symbiotroph | Ectomycorrhizal-Orchid Mycorrhizal-\|Plant Pathogen\|-Plant Saprotroph-Wood Saprotroph | Corticioid-Microfungus | Soft Rot | Probable | As litter_saprotroph (Põlme et al. 2020); Decay leaf, fruit, seed, roots, soil (Põlme et al. 2020); Only symbiotic (perhaps leaning parasitic) in special cases with orchids, otherwise typically a plant pathogen. | Seehann G, et al. 1975. List of Fungi in Soft-Rot Tissue Tests. Institute fur Holzbiologie und Holzschutz, Bundesforschungsanstalt fur Forst - und Holzwirtshaft, Hamburg - Lohbrugge, Germany; Costa IPMW, et al. 2012. Mycotaxon 119:e26 ((http://mycotaxon.com/resources/checklists/costa_v119_checklist.pdf)); Tedersoo L, et al. 2014. Science 346:e1256688 ((https://doi.org/10.1126/science.1256688)); Põlme S, et al. 2020. Fungal Diversity 105:1-16 ((https://doi.org/10.1007/s13225-020-00466-2)); List compiled by Sara Branco with input from Nhu Nguyen and Else Vellinga; The American Phytopathological Society Common Names of Plant Diseases ((http://www.apsnet.org/publications/commonnames/Pages/default.aspx)) |
| Otu11700 | 2 | k__Fungi(100);p__Basidiomycota(100);c__Agaricostilbomycetes(100);o__Agaricostilbales(100);f__Agaricostilbaceae(100);g__Bensingtonia(100);g__Bensingtonia_unclassified(100); | Bensingtonia | 13 | Saprotroph | \|Undefined Saprotroph\| | Dimorphic-Facultative Yeast | NULL | Highly Probable | NULL | Kurtzman CP, et al. (eds.) 2011. The Yeasts, a Taxonomic Study. Fifth Edition. Vols 1-3. Elsevier, San Diego (ISBN: 9780444521491); Põlme S, et al. 2020. Fungal Diversity 105:1-16 ((https://doi.org/10.1007/s13225-020-00466-2)) |
| Otu11702 | 2 | k__Fungi(100);p__Basidiomycota(100);c__Agaricomycetes(100);o__Cantharellales(100);f__Botryobasidiaceae(100);g__Haplotrichum(100);s__Haplotrichum_conspersum(100); | Haplotrichum | 13 | Saprotroph | \|Wood Saprotroph\| | Corticioid | White Rot | Probable | As white_rot (Põlme et al. 2020); Decay wood (Põlme et al. 2020) | Põlme S, et al. 2020. Fungal Diversity 105:1-16 ((https://doi.org/10.1007/s13225-020-00466-2)) |
| Otu11709 | 2 | k__Fungi(100);p__Ascomycota(100);c__Dothideomycetes(100);o__Pleosporales(100);f__Pleosporales_family_Incertae_sedis(100);g__Ochrocladosporium(100);s__Ochrocladosporium_sp(100); | Ochrocladosporium | 13 | Saprotroph | Undefined Saprotroph-\|Wood Saprotroph\| | Microfungus | NULL | Probable | Decay wood (Põlme et al. 2020) | Tedersoo L, et al. 2014. Science 346:e1256688 ((https://doi.org/10.1126/science.1256688)); Põlme S, et al. 2020. Fungal Diversity 105:1-16 ((https://doi.org/10.1007/s13225-020-00466-2)) |
| Otu11710 | 2 | k__Fungi(100);p__Ascomycota(100);c__Dothideomycetes(100);o__Capnodiales(100);f__Teratosphaeriaceae(100);g__Teratosphaeria(100);g__Teratosphaeria_unclassified(100); | Teratosphaeria | 13 | Pathotroph-Saprotroph | \|Plant Pathogen\|-Plant Saprotroph-Wood Saprotroph | Microfungus | NULL | Probable | As litter_saprotroph (Põlme et al. 2020); As wood_pathogen (Põlme et al. 2020); Decay wood, leaf, fruit, seed (Põlme et al. 2020) | Yamazaki A, et al. 2010. Mycoscience 51:443-455 ((https://www.jstage.jst.go.jp/article/mycosci/51/6/51_MYC51443/_pdf)); Tedersoo L, et al. 2014. Science 346:e1256688 ((https://doi.org/10.1126/science.1256688)); Põlme S, et al. 2020. Fungal Diversity 105:1-16 ((https://doi.org/10.1007/s13225-020-00466-2)) |
| Otu11711 | 2 | k__Fungi(100);p__Basidiomycota(100);c__Agaricomycetes(100);o__Agaricales(100);f__Inocybaceae(100);g__Inocybe(100);g__Inocybe_unclassified(100); | Inocybe | 13 | Symbiotroph | \|Ectomycorrhizal\| | Agaricoid | NULL | Highly Probable | Sensu stricto (Matheny 2009). | Rinaldi AC, et al. 2008. Fungal Diversity 33:1-45 ((http://www.fungaldiversity.org/fdp/sfdp/33-1.pdf)); Matheny PB. 2009. McIlvainea 18:11-21 ((https://namyco.org/docs/3_Matheny.pdf)); Tedersoo L, et al. 2010. Mycorrhiza 20:217-263 ((https://doi.org/10.1007/s00572-009-0274-x)); Põlme S, et al. 2020. Fungal Diversity 105:1-16 ((https://doi.org/10.1007/s13225-020-00466-2)); http://mycorrhizas.info/ecmf.html ((http://mycorrhizas.info/ecmf.html)) |
| Otu11714 | 2 | k__Fungi(100);p__Ascomycota(100);c__Lecanoromycetes(100);o__Lecanorales(100);o__Lecanorales_unclassified(100);o__Lecanorales_unclassified(100);o__Lecanorales_unclassified(100); | Lecanorales | 7 | Pathotroph-Symbiotroph | Lichen Parasite-Lichenized | Microfungus-Thallus | NULL | Probable | Lichenized with green algae, occasionally lichenicolous, on rocks, mosses and wood (Cannon & Kirk 2007); ...an order of mostly lichen-forming fungi (Kirk et al. 2008) | Cannon PF, Kirk PM. 2007. Fungal Families of the World. CAB International, Cambridge (ISBN: 978-0851998275); Kirk PM et al. 2008. Dictionary of the Fungi. Tenth Edition. CAB International, Wallingford (ISBN: 978-0851998268); Esslinger TL. 2014. North Dakota State University ((http://www.ndsu.edu/pubweb/~esslinge/chcklst/chcklst7.htm)) |
| Otu11719 | 2 | k__Fungi(100);p__Ascomycota(100);c__Lecanoromycetes(100);o__Lecanorales(100);o__Lecanorales_unclassified(100);o__Lecanorales_unclassified(100);o__Lecanorales_unclassified(100); | Lecanorales | 7 | Pathotroph-Symbiotroph | Lichen Parasite-Lichenized | Microfungus-Thallus | NULL | Probable | Lichenized with green algae, occasionally lichenicolous, on rocks, mosses and wood (Cannon & Kirk 2007); ...an order of mostly lichen-forming fungi (Kirk et al. 2008) | Cannon PF, Kirk PM. 2007. Fungal Families of the World. CAB International, Cambridge (ISBN: 978-0851998275); Kirk PM et al. 2008. Dictionary of the Fungi. Tenth Edition. CAB International, Wallingford (ISBN: 978-0851998268); Esslinger TL. 2014. North Dakota State University ((http://www.ndsu.edu/pubweb/~esslinge/chcklst/chcklst7.htm)) |
| Otu11722 | 2 | k__Fungi(100);p__Ascomycota(100);c__Leotiomycetes(100);o__Helotiales(100);f__Helotiales_family_Incertae_sedis(100);g__Spirosphaera(100);s__Spirosphaera_cupreorufescens(100); | Spirosphaera | 13 | Saprotroph | \|Plant Saprotroph\|-Undefined Saprotroph | Microfungus | NULL | Probable | As litter_saprotroph (Põlme et al. 2020); Decay leaf, fruit, seed (Põlme et al. 2020) | Tedersoo L, et al. 2014. Science 346:e1256688 ((https://doi.org/10.1126/science.1256688)); Põlme S, et al. 2020. Fungal Diversity 105:1-16 ((https://doi.org/10.1007/s13225-020-00466-2)) |
| Otu11723 | 2 | k__Fungi(100);p__Ascomycota(100);c__Dothideomycetes(100);o__Pleosporales(100);f__Phaeosphaeriaceae(100);g__Stagonospora(100);s__Stagonospora_pseudovitensis(100); | Stagonospora | 13 | Pathotroph | \|Plant Pathogen\| | Microfungus | NULL | Probable | NULL | Tedersoo L, et al. 2014. Science 346:e1256688 ((https://doi.org/10.1126/science.1256688)); Põlme S, et al. 2020. Fungal Diversity 105:1-16 ((https://doi.org/10.1007/s13225-020-00466-2)) |
| Otu11724 | 2 | k__Fungi(100);p__Basidiomycota(100);c__Agaricomycetes(100);o__Agaricales(100);f__Agaricaceae(100);g__Agaricus(100);g__Agaricus_unclassified(100); | Agaricus | 13 | Saprotroph | \|Undefined Saprotroph\| | Agaricoid-Gasteroid-Secotioid | NULL | Probable | As soil_saprotroph (Põlme et al. 2020); Decay soil (Põlme et al. 2020) | Tedersoo L, et al. 2014. Science 346:e1256688 ((https://doi.org/10.1126/science.1256688)); Põlme S, et al. 2020. Fungal Diversity 105:1-16 ((https://doi.org/10.1007/s13225-020-00466-2)) |
| Otu11728 | 2 | k__Fungi(100);p__Zygomycota(100);c__Zygomycota_class_Incertae_sedis(100);o__Mortierellales(100);f__Mortierellaceae(100);g__Mortierella(100);s__Mortierella_exigua(100); | Mortierella | 13 | Saprotroph-Symbiotroph | Endophyte-Plant Saprotroph-\|Undefined Saprotroph\| | Microfungus | NULL | Probable | As chitinolytic (Põlme et al. 2020); As soil_saprotroph (Põlme et al. 2020); Decay roots, soil (Põlme et al. 2020); Saprobic in soil and thus frequenctly isolated from soil and material contaminated with soil such as hay or rotten silage and spoil tips (Cannon & Kirk 2007) | Cannon PF, Kirk PM. 2007. Fungal Families of the World. CAB International, Cambridge (ISBN: 978-0851998275); Tedersoo L, et al. 2014. Science 346:e1256688 ((https://doi.org/10.1126/science.1256688)); Purahong W, et al. 2016. Molecular Ecology 25:4059-4074 ((https://doi.org/10.1111/mec.13739)); Põlme S, et al. 2020. Fungal Diversity 105:1-16 ((https://doi.org/10.1007/s13225-020-00466-2)) |
| Otu11730 | 2 | k__Fungi(100);p__Basidiomycota(100);c__Agaricostilbomycetes(100);o__Agaricostilbales(100);f__Agaricostilbaceae(100);f__Agaricostilbaceae_unclassified(100);f__Agaricostilbaceae_unclassified(100); | Agaricostilbaceae | 9 | Saprotroph | Undefined Saprotroph | NULL | NULL | Possible | Probably saprobic, usually associated with dead palm tissues (Cannon & Kirk 2007) | Cannon PF, Kirk PM. 2007. Fungal Families of the World. CAB International, Cambridge (ISBN: 978-0851998275) |
| Otu11732 | 2 | k__Fungi(100);p__Basidiomycota(100);c__Tremellomycetes(100);o__Filobasidiales(100);f__Filobasidiaceae(100);g__Cryptococcus(100);s__Cryptococcus_skinneri(100); | Cryptococcus | 13 | Pathotroph-Saprotroph-Symbiotroph | Animal Pathogen-Endophyte-Epiphyte-\|Undefined Saprotroph\| | Dimorphic Yeast-Yeast | NULL | Possible | As foliar_endophyte (Põlme et al. 2020); Decay leaf, fruit, seed, soil, dung, animal material (Põlme et al. 2020) | Andrews JH, et al. 1983. Phytopathology 73:228-234 ((https://doi.org/10.1094/Phyto-73-228)); Kurtzman CP, et al. (eds.) 2011. The Yeasts, a Taxonomic Study. Fifth Edition. Vols 1-3. Elsevier, San Diego (ISBN: 9780444521491); Põlme S, et al. 2020. Fungal Diversity 105:1-16 ((https://doi.org/10.1007/s13225-020-00466-2)) |
| Otu11733 | 2 | k__Fungi(100);p__Basidiomycota(100);c__Agaricomycetes(100);o__Agaricales(100);o__Agaricales_unclassified(100);o__Agaricales_unclassified(100);o__Agaricales_unclassified(100); | Agaricales | 7 | Pathotroph-Saprotroph-Symbiotroph | Bryophyte Parasite-Dung Saprotroph-Ectomycorrhizal-Fungal Parasite-Leaf Saprotroph-Plant Parasite-Undefined Saprotroph-Wood Saprotroph | Agaricoid-Gasteroid-Microfungus-Secotioid-Yeast | NULL | Possible | Mushrooms and toadstools, Gill fungi, Agarics... lignicolous, sometimes muscicolous or fungicolous, saprobic, mycorrhizal, rarely parasitic on plants or fungi (Kirk et al. 2008) | Kirk PM et al. 2008. Dictionary of the Fungi. Tenth Edition. CAB International, Wallingford (ISBN: 978-0851998268) |
| Otu11735 | 2 | k__Fungi(100);p__Zygomycota(100);c__Zygomycota_class_Incertae_sedis(100);o__Mucorales(100);f__Mucoraceae(100);g__Mucor(100);s__Mucor_flavus(100); | Mucor | 13 | Saprotroph | \|Undefined Saprotroph\| | Microfungus | NULL | Probable | As soil_saprotroph (Põlme et al. 2020); Decay soil (Põlme et al. 2020) | Tedersoo L, et al. 2014. Science 346:e1256688 ((https://doi.org/10.1126/science.1256688)); Põlme S, et al. 2020. Fungal Diversity 105:1-16 ((https://doi.org/10.1007/s13225-020-00466-2)) |
| Otu11736 | 2 | k__Fungi(100);p__Basidiomycota(100);c__Agaricomycetes(100);o__Agaricales(100);f__Amanitaceae(100);g__Amanita(100);s__Amanita_vaginata(100); | Amanita | 13 | Saprotroph-Symbiotroph | \|Ectomycorrhizal\|-Undefined Saprotroph | Agaricoid | NULL | Probable | Some members of the genus are deadly poisonous | Rinaldi AC, et al. 2008. Fungal Diversity 33:1-45 ((http://www.fungaldiversity.org/fdp/sfdp/33-1.pdf)); Tedersoo L, et al. 2010. Mycorrhiza 20:217-263 ((https://doi.org/10.1007/s00572-009-0274-x)); Wolfe BE, et al. 2012. PLoS ONE 7:e39597 ((https://doi.org/10.1371/journal.pone.0039597)); Põlme S, et al. 2020. Fungal Diversity 105:1-16 ((https://doi.org/10.1007/s13225-020-00466-2)); http://mycorrhizas.info/ecmf.html ((http://mycorrhizas.info/ecmf.html)); http://en.wikipedia.org/wiki/List_of_deadly_fungi ((http://en.wikipedia.org/wiki/List_of_deadly_fungi)) |
| Otu11737 | 2 | k__Fungi(100);p__Basidiomycota(100);c__Agaricomycetes(100);o__Boletales(100);f__Gyroporaceae(100);g__Gyroporus(100);s__Gyroporus_castaneus(100); | Gyroporus | 13 | Symbiotroph | \|Ectomycorrhizal\| | Agaricoid-Boletoid | NULL | Highly Probable | NULL | Rinaldi AC, et al. 2008. Fungal Diversity 33:1-45 ((http://www.fungaldiversity.org/fdp/sfdp/33-1.pdf)); Tedersoo L, et al. 2010. Mycorrhiza 20:217-263 ((https://doi.org/10.1007/s00572-009-0274-x)); Põlme S, et al. 2020. Fungal Diversity 105:1-16 ((https://doi.org/10.1007/s13225-020-00466-2)); http://mycorrhizas.info/ecmf.html ((http://mycorrhizas.info/ecmf.html)) |
| Otu11738 | 2 | k__Fungi(100);p__Basidiomycota(100);c__Agaricomycetes(100);o__Agaricales(100);f__Inocybaceae(100);g__Inocybe(100);g__Inocybe_unclassified(100); | Inocybe | 13 | Symbiotroph | \|Ectomycorrhizal\| | Agaricoid | NULL | Highly Probable | Sensu stricto (Matheny 2009). | Rinaldi AC, et al. 2008. Fungal Diversity 33:1-45 ((http://www.fungaldiversity.org/fdp/sfdp/33-1.pdf)); Matheny PB. 2009. McIlvainea 18:11-21 ((https://namyco.org/docs/3_Matheny.pdf)); Tedersoo L, et al. 2010. Mycorrhiza 20:217-263 ((https://doi.org/10.1007/s00572-009-0274-x)); Põlme S, et al. 2020. Fungal Diversity 105:1-16 ((https://doi.org/10.1007/s13225-020-00466-2)); http://mycorrhizas.info/ecmf.html ((http://mycorrhizas.info/ecmf.html)) |
| Otu11739 | 2 | k__Fungi(100);p__Basidiomycota(100);c__Agaricomycetes(100);o__Agaricales(100);f__Inocybaceae(100);g__Inocybe(100);s__Inocybe_cf_subochracea_PBM2615(100); | Inocybe | 13 | Symbiotroph | \|Ectomycorrhizal\| | Agaricoid | NULL | Highly Probable | Sensu stricto (Matheny 2009). | Rinaldi AC, et al. 2008. Fungal Diversity 33:1-45 ((http://www.fungaldiversity.org/fdp/sfdp/33-1.pdf)); Matheny PB. 2009. McIlvainea 18:11-21 ((https://namyco.org/docs/3_Matheny.pdf)); Tedersoo L, et al. 2010. Mycorrhiza 20:217-263 ((https://doi.org/10.1007/s00572-009-0274-x)); Põlme S, et al. 2020. Fungal Diversity 105:1-16 ((https://doi.org/10.1007/s13225-020-00466-2)); http://mycorrhizas.info/ecmf.html ((http://mycorrhizas.info/ecmf.html)) |
| Otu11740 | 2 | k__Fungi(100);p__Basidiomycota(100);c__Agaricomycetes(100);o__Agaricales(100);f__Inocybaceae(100);g__Crepidotus(100);s__Crepidotus_mollis(100); | Crepidotus | 13 | Saprotroph | \|Wood Saprotroph\| | Agaricoid | NULL | Highly Probable | Decay wood (Põlme et al. 2020) | Grgurinovic CA, Simpson JA. 2001. Fungal Diversity 8:97-127 ((http://www.fungaldiversity.org/fdp/sfdp/FD_8_97-127.pdf)); Aime MC, et al. 2005. American Journal of Botany 92:74-82 ((https://doi.org/10.3732/ajb.92.1.74)); Tedersoo L, et al. 2014. Science 346:e1256688 ((https://doi.org/10.1126/science.1256688)); Põlme S, et al. 2020. Fungal Diversity 105:1-16 ((https://doi.org/10.1007/s13225-020-00466-2)) |
| Otu11741 | 2 | k__Fungi(100);p__Basidiomycota(100);c__Agaricomycetes(100);o__Agaricales(100);o__Agaricales_unclassified(100);o__Agaricales_unclassified(100);o__Agaricales_unclassified(100); | Agaricales | 7 | Pathotroph-Saprotroph-Symbiotroph | Bryophyte Parasite-Dung Saprotroph-Ectomycorrhizal-Fungal Parasite-Leaf Saprotroph-Plant Parasite-Undefined Saprotroph-Wood Saprotroph | Agaricoid-Gasteroid-Microfungus-Secotioid-Yeast | NULL | Possible | Mushrooms and toadstools, Gill fungi, Agarics... lignicolous, sometimes muscicolous or fungicolous, saprobic, mycorrhizal, rarely parasitic on plants or fungi (Kirk et al. 2008) | Kirk PM et al. 2008. Dictionary of the Fungi. Tenth Edition. CAB International, Wallingford (ISBN: 978-0851998268) |
| Otu11742 | 2 | k__Fungi(100);p__Basidiomycota(100);c__Agaricomycetes(100);o__Agaricales(100);f__Inocybaceae(100);g__Inocybe(100);s__Inocybe_lacera_var._helobia(100); | Inocybe | 13 | Symbiotroph | \|Ectomycorrhizal\| | Agaricoid | NULL | Highly Probable | Sensu stricto (Matheny 2009). | Rinaldi AC, et al. 2008. Fungal Diversity 33:1-45 ((http://www.fungaldiversity.org/fdp/sfdp/33-1.pdf)); Matheny PB. 2009. McIlvainea 18:11-21 ((https://namyco.org/docs/3_Matheny.pdf)); Tedersoo L, et al. 2010. Mycorrhiza 20:217-263 ((https://doi.org/10.1007/s00572-009-0274-x)); Põlme S, et al. 2020. Fungal Diversity 105:1-16 ((https://doi.org/10.1007/s13225-020-00466-2)); http://mycorrhizas.info/ecmf.html ((http://mycorrhizas.info/ecmf.html)) |
| Otu11743 | 2 | k__Fungi(100);p__Basidiomycota(100);c__Agaricomycetes(100);o__Agaricales(100);f__Inocybaceae(100);g__Inocybe(100);s__Inocybe_geophylla(100); | Inocybe | 13 | Symbiotroph | \|Ectomycorrhizal\| | Agaricoid | NULL | Highly Probable | Sensu stricto (Matheny 2009). | Rinaldi AC, et al. 2008. Fungal Diversity 33:1-45 ((http://www.fungaldiversity.org/fdp/sfdp/33-1.pdf)); Matheny PB. 2009. McIlvainea 18:11-21 ((https://namyco.org/docs/3_Matheny.pdf)); Tedersoo L, et al. 2010. Mycorrhiza 20:217-263 ((https://doi.org/10.1007/s00572-009-0274-x)); Põlme S, et al. 2020. Fungal Diversity 105:1-16 ((https://doi.org/10.1007/s13225-020-00466-2)); http://mycorrhizas.info/ecmf.html ((http://mycorrhizas.info/ecmf.html)) |
| Otu11744 | 2 | k__Fungi(100);p__Basidiomycota(100);c__Agaricomycetes(100);o__Agaricales(100);f__Inocybaceae(100);g__Inocybe(100);g__Inocybe_unclassified(100); | Inocybe | 13 | Symbiotroph | \|Ectomycorrhizal\| | Agaricoid | NULL | Highly Probable | Sensu stricto (Matheny 2009). | Rinaldi AC, et al. 2008. Fungal Diversity 33:1-45 ((http://www.fungaldiversity.org/fdp/sfdp/33-1.pdf)); Matheny PB. 2009. McIlvainea 18:11-21 ((https://namyco.org/docs/3_Matheny.pdf)); Tedersoo L, et al. 2010. Mycorrhiza 20:217-263 ((https://doi.org/10.1007/s00572-009-0274-x)); Põlme S, et al. 2020. Fungal Diversity 105:1-16 ((https://doi.org/10.1007/s13225-020-00466-2)); http://mycorrhizas.info/ecmf.html ((http://mycorrhizas.info/ecmf.html)) |
| Otu11745 | 2 | k__Fungi(100);p__Basidiomycota(100);c__Agaricomycetes(100);o__Gomphales(100);f__Gomphaceae(100);f__Gomphaceae_unclassified(100);f__Gomphaceae_unclassified(100); | Gomphaceae | 9 | Saprotroph-Symbiotroph | Ectomycorrhizal-Undefined Saprotroph-Wood Saprotroph | Clavarioid-Gomphoid | NULL | Probable | Terrestrial or lignicolous, either saprobic or forming ectomycorrhizal relationships with woody plants (Cannon & Kirk 2007) | Cannon PF, Kirk PM. 2007. Fungal Families of the World. CAB International, Cambridge (ISBN: 978-0851998275) |
| Otu11746 | 2 | k__Fungi(100);p__Basidiomycota(100);c__Agaricomycetes(100);o__Gomphales(100);f__Gomphaceae(100);g__Ramaria(100);s__Ramaria_stricta(100); | Ramaria | 13 | Saprotroph-Symbiotroph | \|Ectomycorrhizal\|-Plant Saprotroph | Clavarioid | NULL | Possible | As litter_saprotroph (Põlme et al. 2020) | Rinaldi AC, et al. 2008. Fungal Diversity 33:1-45 ((http://www.fungaldiversity.org/fdp/sfdp/33-1.pdf)) (as ECM/not ECM); Tedersoo L, et al. 2010. Mycorrhiza 20:217-263 ((10.1007/s00572-009-0274-x)) (pro parte); Põlme S, et al. 2020. Fungal Diversity 105:1-16 ((https://doi.org/10.1007/s13225-020-00466-2)); http://mycorrhizas.info/ecmf.html ((http://mycorrhizas.info/ecmf.html)) (as ECM) |
| Otu11748 | 2 | k__Fungi(100);p__Basidiomycota(100);c__Agaricomycetes(100);o__Agaricales(100);f__Inocybaceae(100);g__Inocybe(100);g__Inocybe_unclassified(100); | Inocybe | 13 | Symbiotroph | \|Ectomycorrhizal\| | Agaricoid | NULL | Highly Probable | Sensu stricto (Matheny 2009). | Rinaldi AC, et al. 2008. Fungal Diversity 33:1-45 ((http://www.fungaldiversity.org/fdp/sfdp/33-1.pdf)); Matheny PB. 2009. McIlvainea 18:11-21 ((https://namyco.org/docs/3_Matheny.pdf)); Tedersoo L, et al. 2010. Mycorrhiza 20:217-263 ((https://doi.org/10.1007/s00572-009-0274-x)); Põlme S, et al. 2020. Fungal Diversity 105:1-16 ((https://doi.org/10.1007/s13225-020-00466-2)); http://mycorrhizas.info/ecmf.html ((http://mycorrhizas.info/ecmf.html)) |
| Otu11749 | 2 | k__Fungi(100);p__Basidiomycota(100);c__Agaricomycetes(100);o__Agaricales(100);f__Inocybaceae(100);g__Inocybe(100);g__Inocybe_unclassified(100); | Inocybe | 13 | Symbiotroph | \|Ectomycorrhizal\| | Agaricoid | NULL | Highly Probable | Sensu stricto (Matheny 2009). | Rinaldi AC, et al. 2008. Fungal Diversity 33:1-45 ((http://www.fungaldiversity.org/fdp/sfdp/33-1.pdf)); Matheny PB. 2009. McIlvainea 18:11-21 ((https://namyco.org/docs/3_Matheny.pdf)); Tedersoo L, et al. 2010. Mycorrhiza 20:217-263 ((https://doi.org/10.1007/s00572-009-0274-x)); Põlme S, et al. 2020. Fungal Diversity 105:1-16 ((https://doi.org/10.1007/s13225-020-00466-2)); http://mycorrhizas.info/ecmf.html ((http://mycorrhizas.info/ecmf.html)) |
| Otu11750 | 2 | k__Fungi(100);p__Ascomycota(100);c__Pezizomycetes(100);o__Pezizales(100);f__Discinaceae(100);g__Hydnotrya(100);s__Hydnotrya_sp(100); | Hydnotrya | 13 | Symbiotroph | \|Ectomycorrhizal\| | Gasteroid | Hypogeous | Highly Probable | Common name - false truffle; Syn. Geoporella | Rinaldi AC, et al. 2008. Fungal Diversity 33:1-45 ((http://www.fungaldiversity.org/fdp/sfdp/33-1.pdf)); Tedersoo L, et al. 2010. Mycorrhiza 20:217-263 ((https://doi.org/10.1007/s00572-009-0274-x)); Põlme S, et al. 2020. Fungal Diversity 105:1-16 ((https://doi.org/10.1007/s13225-020-00466-2)); http://mycorrhizas.info/ecmf.html ((http://mycorrhizas.info/ecmf.html)) |
| Otu11751 | 2 | k__Fungi(100);p__Ascomycota(100);c__Sordariomycetes(100);o__Hypocreales(100);o__Hypocreales_unclassified(100);o__Hypocreales_unclassified(100);o__Hypocreales_unclassified(100); | Hypocreales | 7 | Saprotroph | Undefined Saprotroph | Microfungus | NULL | Possible | NULL | Sterkenburg E, et al. 2015. New Phytologist 207:1145-1158 ((https://doi.org/10.1111/nph.13426)) |
| Otu11752 | 2 | k__Fungi(100);p__Ascomycota(100);c__Sordariomycetes(100);o__Sordariomycetes_order_Incertae_sedis(100);f__Glomerellaceae(100);g__Colletotrichum(100);g__Colletotrichum_unclassified(100); | Colletotrichum | 13 | Pathotroph-Saprotroph-Symbiotroph | Endophyte-\|Plant Pathogen\|-Plant Saprotroph | Microfungus | NULL | Probable | As foliar_endophyte (Põlme et al. 2020); As litter_saprotroph (Põlme et al. 2020); Decay leaf, fruit, seed (Põlme et al. 2020) | Costa IPMW, et al. 2012. Mycotaxon 119:e26 ((http://mycotaxon.com/resources/checklists/costa_v119_checklist.pdf)); Tedersoo L, et al. 2014. Science 346:e1256688 ((https://doi.org/10.1126/science.1256688)); Põlme S, et al. 2020. Fungal Diversity 105:1-16 ((https://doi.org/10.1007/s13225-020-00466-2)); The American Phytopathological Society Common Names of Plant Diseases ((http://www.apsnet.org/publications/commonnames/Pages/default.aspx)) |
| Otu11753 | 2 | k__Fungi(100);p__Ascomycota(100);c__Sordariomycetes(100);o__Hypocreales(100);f__Cordycipitaceae(100);g__Cordyceps(100);s__Cordyceps_sp_JQY_2013(100); | Cordyceps | 13 | Pathotroph-Symbiotroph | Clavicipitaceous Endophyte-Fungal Parasite-\|Insect Pathogen\| | Cordyceptoid | NULL | Possible | As foliar_endophyte (Põlme et al. 2020); Decay animal material (Põlme et al. 2020) | Samuels GJ. 2007. Self-published; Costa IPMW, et al. 2012. Mycotaxon 119:e26 ((http://mycotaxon.com/resources/checklists/costa_v119_checklist.pdf)); Tedersoo L, et al. 2014. Science 346:e1256688 ((https://doi.org/10.1126/science.1256688)); Põlme S, et al. 2020. Fungal Diversity 105:1-16 ((https://doi.org/10.1007/s13225-020-00466-2)) |
| Otu11754 | 2 | k__Fungi(100);p__Ascomycota(100);c__Sordariomycetes(100);o__Hypocreales(100);f__Hypocreales_family_Incertae_sedis(100);f__Hypocreales_family_Incertae_sedis_unclassified(100);f__Hypocreales_family_Incertae_sedis_unclassified(100); | Hypocreales | 7 | Saprotroph | Undefined Saprotroph | Microfungus | NULL | Possible | NULL | Sterkenburg E, et al. 2015. New Phytologist 207:1145-1158 ((https://doi.org/10.1111/nph.13426)) |
| Otu11756 | 2 | k__Fungi(100);p__Ascomycota(100);c__Sordariomycetes(100);o__Hypocreales(100);f__Clavicipitaceae(100);f__Clavicipitaceae_unclassified(100);f__Clavicipitaceae_unclassified(100); | Clavicipitaceae | 9 | Pathotroph-Symbiotroph | Endophyte-Epiphyte-Fungal Parasite-Insect Parasite | Microfungus | NULL | Possible | Parasitic, endophytic or epiphytic on Insecta, other fungi or Gamineae (Cannon & Kirk 2007) | Cannon PF, Kirk PM. 2007. Fungal Families of the World. CAB International, Cambridge (ISBN: 978-0851998275) |
| Otu11757 | 2 | k__Fungi(100);p__Ascomycota(100);c__Sordariomycetes(100);o__Hypocreales(100);o__Hypocreales_unclassified(100);o__Hypocreales_unclassified(100);o__Hypocreales_unclassified(100); | Hypocreales | 7 | Saprotroph | Undefined Saprotroph | Microfungus | NULL | Possible | NULL | Sterkenburg E, et al. 2015. New Phytologist 207:1145-1158 ((https://doi.org/10.1111/nph.13426)) |
| Otu11758 | 2 | k__Fungi(100);p__Ascomycota(100);c__Sordariomycetes(100);o__Hypocreales(100);o__Hypocreales_unclassified(100);o__Hypocreales_unclassified(100);o__Hypocreales_unclassified(100); | Hypocreales | 7 | Saprotroph | Undefined Saprotroph | Microfungus | NULL | Possible | NULL | Sterkenburg E, et al. 2015. New Phytologist 207:1145-1158 ((https://doi.org/10.1111/nph.13426)) |
| Otu11760 | 2 | k__Fungi(100);p__Ascomycota(100);c__Sordariomycetes(100);o__Hypocreales(100);o__Hypocreales_unclassified(100);o__Hypocreales_unclassified(100);o__Hypocreales_unclassified(100); | Hypocreales | 7 | Saprotroph | Undefined Saprotroph | Microfungus | NULL | Possible | NULL | Sterkenburg E, et al. 2015. New Phytologist 207:1145-1158 ((https://doi.org/10.1111/nph.13426)) |
| Otu11761 | 2 | k__Fungi(100);p__Ascomycota(100);c__Sordariomycetes(100);o__Diaporthales(100);f__Gnomoniaceae(100);f__Gnomoniaceae_unclassified(100);f__Gnomoniaceae_unclassified(100); | Gnomoniaceae | 9 | Pathotroph-Saprotroph-Symbiotroph | Endophyte-Plant Pathogen-Undefined Saprotroph | Microfungus | NULL | Probable | Most species are saprobes or endophytes in leaves and bark, but a few are neurotrophic pathogens (Cannon & Kirk 2007) | Cannon PF, Kirk PM. 2007. Fungal Families of the World. CAB International, Cambridge (ISBN: 978-0851998275) |
| Otu11762 | 2 | k__Fungi(100);p__Ascomycota(100);c__Sordariomycetes(100);o__Diaporthales(100);f__Diaporthaceae(100);g__Phomopsis(100);s__Phomopsis_sp_STAM_32(100); | Phomopsis | 13 | Pathotroph-Pathotroph-Saprotroph-Symbio | Endophyte-\|Plant Pathogen\|-Wood Saprotroph | Microfungus | Soft Rot-Soft Rot | Probable | As foliar_endophyte (Põlme et al. 2020); Decay leaf, fruit, seed (Põlme et al. 2020) | Seehann G, et al. 1975. List of Fungi in Soft-Rot Tissue Tests. Institute fur Holzbiologie und Holzschutz, Bundesforschungsanstalt fur Forst - und Holzwirtshaft, Hamburg - Lohbrugge, Germany; Costa IPMW, et al. 2012. Mycotaxon 119:e26 ((http://mycotaxon.com/resources/checklists/costa_v119_checklist.pdf)); Tedersoo L, et al. 2014. Science 346:e1256688 ((https://doi.org/10.1126/science.1256688)); Põlme S, et al. 2020. Fungal Diversity 105:1-16 ((https://doi.org/10.1007/s13225-020-00466-2)) |
| Otu11763 | 2 | k__Fungi(100);p__Ascomycota(100);c__Eurotiomycetes(100);o__Eurotiales(100);f__Trichocomaceae(100);g__Penicillium(100);s__Penicillium_roseopurpureum(100); | Penicillium | 13 | Pathotroph-Saprotroph-Symbiotroph | Animal Parasite-Dung Saprotroph-Endophyte-Fungal Parasite-Plant Pathogen-\|Undefined Saprotroph\|-Wood Saprotroph | Microfungus | Soft Rot | Highly Probable | As foliar_endophyte (Põlme et al. 2020); As mycoparasite (Põlme et al. 2020); Decay soil (Põlme et al. 2020) | Duncan CG, Eslyn WE. 1966. Mycologia 58:642-664 ((https://doi.org/10.2307/3757045)); Seehann G, et al. 1975. List of Fungi in Soft-Rot Tissue Tests. Institute fur Holzbiologie und Holzschutz, Bundesforschungsanstalt fur Forst - und Holzwirtshaft, Hamburg - Lohbrugge, Germany ((https://www.irg-wp.com/irgdocs/details.php?d6493033-c9ea-40f3-8e3e-5697f32d640d)); Costa IPMW, et al. 2012. Mycotaxon 119:e26 ((http://mycotaxon.com/resources/checklists/costa_v119_checklist.pdf)); Bills GF, et al. 2013. Current Opinion in Microbiology 16: 549-565 ((https://doi.org/10.1016/j.mib.2013.08.001)); Tedersoo L, et al. 2014. Science 346:e1256688 ((https://doi.org/10.1126/science.1256688)); Põlme S, et al. 2020. Fungal Diversity 105:1-16 ((https://doi.org/10.1007/s13225-020-00466-2)) |
| Otu11764 | 2 | k__Fungi(100);p__Ascomycota(100);c__Sordariomycetes(100);o__Hypocreales(100);f__Hypocreaceae(100);g__Hypocrea(100);s__Hypocrea_tremelloides(100); | Hypocrea | 13 | Pathotroph | Endophyte-Fungal Parasite | Microfungus | NULL | Probable | NULL | Tedersoo L, et al. 2014. Science 346:e1256688 ((https://doi.org/10.1126/science.1256688)); Christian N, et al. 2016. Microbial Ecology 72: 621-632 ((https://doi.org/10.1007/s00248-016-0804-y)) |
| Otu11765 | 2 | k__Fungi(100);p__Ascomycota(100);c__Lecanoromycetes(100);o__Ostropales(100);f__Stictidaceae(100);g__Stictis(100);s__Stictis_populorum(100); | Stictis | 13 | Symbiotroph | \|Lichenized\| | Thallus | NULL | Highly Probable | NULL | Esslinger TL. 2014. North Dakota State University ((http://www.ndsu.edu/pubweb/~esslinge/chcklst/chcklst7.htm)); Põlme S, et al. 2020. Fungal Diversity 105:1-16 ((https://doi.org/10.1007/s13225-020-00466-2)) |
| Otu11767 | 2 | k__Fungi(100);p__Ascomycota(100);c__Ascomycota_class_Incertae_sedis(100);o__Ascomycota_order_Incertae_sedis(100);f__Ascomycota_family_Incertae_sedis(100);g__Veronaea(100);g__Veronaea_unclassified(100); | Veronaea | 13 | Pathotroph-Saprotroph | Dung Saprotroph-\|Plant Pathogen\|-Plant Saprotroph | Dimorphic | NULL | Probable | As litter_saprotroph (Põlme et al. 2020); Decay leaf, fruit, seed (Põlme et al. 2020) | Põlme S, et al. 2020. Fungal Diversity 105:1-16 ((https://doi.org/10.1007/s13225-020-00466-2)); The American Phytopathological Society Common Names of Plant Diseases ((http://www.apsnet.org/publications/commonnames/Pages/default.aspx)) |
| Otu11768 | 2 | k__Fungi(100);p__Ascomycota(100);c__Eurotiomycetes(100);o__Chaetothyriales(100);f__Herpotrichiellaceae(100);g__Cladophialophora(100);s__Cladophialophora_sp_LS_2013(100); | Cladophialophora | 13 | Pathotroph-Saprotroph-Symbiotroph | Animal Pathogen-Endophyte-Fungal Parasite-\|Undefined Saprotroph\| | Dimorphic-Microfungus | NULL | Probable | As mycoparasite (Põlme et al. 2020); As root_endophyte_dark_septate (Põlme et al. 2020); As soil_saprotroph (Põlme et al. 2020); Decay soil (Põlme et al. 2020) | James TY, et al. 2006. Nature 443:818-822 ((https://doi.org/10.1038/nature05110)); Põlme S, et al. 2020. Fungal Diversity 105:1-16 ((https://doi.org/10.1007/s13225-020-00466-2)) |
| Otu11769 | 2 | k__Fungi(100);p__Ascomycota(100);c__Lecanoromycetes(100);o__Teloschistales(100);f__Physciaceae(100);g__Physcia(100);s__Physcia_sp(100); | Physcia | 13 | Symbiotroph | \|Lichenized\| | Thallus | NULL | Highly Probable | NULL | Esslinger TL. 2014. North Dakota State University ((http://www.ndsu.edu/pubweb/~esslinge/chcklst/chcklst7.htm)); Põlme S, et al. 2020. Fungal Diversity 105:1-16 ((https://doi.org/10.1007/s13225-020-00466-2)) |
| Otu11772 | 2 | k__Fungi(100);p__Ascomycota(100);c__Eurotiomycetes(100);o__Pyrenulales(100);f__Massariaceae(100);g__Massaria(100);s__Massaria_anomia(100); | Massaria | 13 | Saprotroph | Undefined Saprotroph-\|Wood Saprotroph\| | Microfungus | NULL | Probable | Decay wood (Põlme et al. 2020) | Tedersoo L, et al. 2014. Science 346:e1256688 ((https://doi.org/10.1126/science.1256688)); Põlme S, et al. 2020. Fungal Diversity 105:1-16 ((https://doi.org/10.1007/s13225-020-00466-2)) |
| Otu11773 | 2 | k__Fungi(100);p__Ascomycota(100);c__Ascomycota_class_Incertae_sedis(100);o__Ascomycota_order_Incertae_sedis(100);f__Ascomycota_family_Incertae_sedis(100);g__Veronaea(100);s__Veronaea_compacta(100); | Veronaea | 13 | Pathotroph-Saprotroph | Dung Saprotroph-\|Plant Pathogen\|-Plant Saprotroph | Dimorphic | NULL | Probable | As litter_saprotroph (Põlme et al. 2020); Decay leaf, fruit, seed (Põlme et al. 2020) | Põlme S, et al. 2020. Fungal Diversity 105:1-16 ((https://doi.org/10.1007/s13225-020-00466-2)); The American Phytopathological Society Common Names of Plant Diseases ((http://www.apsnet.org/publications/commonnames/Pages/default.aspx)) |
| Otu11774 | 2 | k__Fungi(100);p__Ascomycota(100);c__Eurotiomycetes(100);o__Chaetothyriales(100);f__Herpotrichiellaceae(100);g__Cladophialophora(100);g__Cladophialophora_unclassified(100); | Cladophialophora | 13 | Pathotroph-Saprotroph-Symbiotroph | Animal Pathogen-Endophyte-Fungal Parasite-\|Undefined Saprotroph\| | Dimorphic-Microfungus | NULL | Probable | As mycoparasite (Põlme et al. 2020); As root_endophyte_dark_septate (Põlme et al. 2020); As soil_saprotroph (Põlme et al. 2020); Decay soil (Põlme et al. 2020) | James TY, et al. 2006. Nature 443:818-822 ((https://doi.org/10.1038/nature05110)); Põlme S, et al. 2020. Fungal Diversity 105:1-16 ((https://doi.org/10.1007/s13225-020-00466-2)) |
| Otu11775 | 2 | k__Fungi(100);p__Ascomycota(100);c__Eurotiomycetes(100);o__Chaetothyriales(100);f__Herpotrichiellaceae(100);f__Herpotrichiellaceae_unclassified(100);f__Herpotrichiellaceae_unclassified(100); | Herpotrichiellaceae | 9 | Pathotroph-Saprotroph | Animal Pathogen-Fungal Parasite-Undefined Saprotroph | Facultative Yeast-Microfungus | NULL | Probable | Mostly saprobic on plants or other fungi, isolated from soil etc., and from medical sources (Cannon & Kirk 2007) | Cannon PF, Kirk PM. 2007. Fungal Families of the World. CAB International, Cambridge (ISBN: 978-0851998275) |
| Otu11776 | 2 | k__Fungi(100);p__Ascomycota(100);c__Eurotiomycetes(100);o__Chaetothyriales(100);f__Herpotrichiellaceae(100);g__Cladophialophora(100);g__Cladophialophora_unclassified(100); | Cladophialophora | 13 | Pathotroph-Saprotroph-Symbiotroph | Animal Pathogen-Endophyte-Fungal Parasite-\|Undefined Saprotroph\| | Dimorphic-Microfungus | NULL | Probable | As mycoparasite (Põlme et al. 2020); As root_endophyte_dark_septate (Põlme et al. 2020); As soil_saprotroph (Põlme et al. 2020); Decay soil (Põlme et al. 2020) | James TY, et al. 2006. Nature 443:818-822 ((https://doi.org/10.1038/nature05110)); Põlme S, et al. 2020. Fungal Diversity 105:1-16 ((https://doi.org/10.1007/s13225-020-00466-2)) |
| Otu11778 | 2 | k__Fungi(100);p__Ascomycota(100);c__Pezizomycetes(100);o__Pezizales(100);f__Pezizaceae(100);g__Pachyphloeus(100);s__Pachyphloeus_sp_15_RH_2012(100); | Pachyphloeus | 13 | Symbiotroph | \|Ectomycorrhizal\| | Gasteroid | Hypogeous | Highly Probable | NULL | Rinaldi AC, et al. 2008. Fungal Diversity 33:1-45 ((http://www.fungaldiversity.org/fdp/sfdp/33-1.pdf)); Tedersoo L, et al. 2010. Mycorrhiza 20:217-263 ((https://doi.org/10.1007/s00572-009-0274-x)); Põlme S, et al. 2020. Fungal Diversity 105:1-16 ((https://doi.org/10.1007/s13225-020-00466-2)); http://mycorrhizas.info/ecmf.html ((http://mycorrhizas.info/ecmf.html)) |
| Otu11783 | 2 | k__Fungi(100);p__Ascomycota(100);c__Sordariomycetes(100);o__Xylariales(100);f__Xylariaceae(100);g__Hypoxylon(100);g__Hypoxylon_unclassified(100); | Hypoxylon | 13 | Saprotroph-Saprotroph-Symbiotroph | Endophyte-Undefined Saprotroph-\|Wood Saprotroph\| | Corticioid-Microfungus-Xylarioid | NULL | Probable | As foliar_endophyte (Põlme et al. 2020); Decay wood, leaf, fruit, seed (Põlme et al. 2020); Host - angiosperms (Põlme et al. 2020) | Tedersoo L, et al. 2014. Science 346:e1256688 ((https://doi.org/10.1126/science.1256688)); Põlme S, et al. 2020. Fungal Diversity 105:1-16 ((https://doi.org/10.1007/s13225-020-00466-2)) |
| Otu11785 | 2 | k__Fungi(100);p__Ascomycota(100);c__Sordariomycetes(100);o__Xylariales(100);f__Diatrypaceae(100);g__Eutypa(100);g__Eutypa_unclassified(100); | Eutypa | 13 | Pathotroph | \|Plant Pathogen\| | Microfungus | NULL | Probable | NULL | Tedersoo L, et al. 2014. Science 346:e1256688 ((https://doi.org/10.1126/science.1256688)); Põlme S, et al. 2020. Fungal Diversity 105:1-16 ((https://doi.org/10.1007/s13225-020-00466-2)) |
| Otu11786 | 2 | k__Fungi(100);p__Ascomycota(100);c__Sordariomycetes(100);o__Xylariales(100);f__Diatrypaceae(100);g__Eutypa(100);s__Eutypa_spinosa(100); | Eutypa | 13 | Pathotroph | \|Plant Pathogen\| | Microfungus | NULL | Probable | NULL | Tedersoo L, et al. 2014. Science 346:e1256688 ((https://doi.org/10.1126/science.1256688)); Põlme S, et al. 2020. Fungal Diversity 105:1-16 ((https://doi.org/10.1007/s13225-020-00466-2)) |
| Otu11788 | 2 | k__Fungi(100);p__Ascomycota(100);c__Orbiliomycetes(100);o__Orbiliales(100);f__Orbiliaceae(100);g__Arthrobotrys(100);s__Arthrobotrys_conoides(100); | Arthrobotrys | 13 | Pathotroph-Saprotroph | \|Animal Parasite\|-Dung Saprotroph-Nematophagous-Wood Saprotroph | Microfungus | NULL | Probable | Decay wood, leaf, fruit, seed, soil, animal material (Põlme et al. 2020); Nematode-trapping fungus (Yang et al. 2020) | Seifert KA, Kendrick WB, Murase G. 1983. A Key to Hyphomycetes on Dung. University of Waterloo Press, Waterloo; Tedersoo L, et al. 2014. Science 346:e1256688 ((https://doi.org/10.1126/science.1256688)); Yang C-T, et al. 2020. Proceedings of the National Academy of Sciences 117:6762-6770 ((https://doi.org/10.1073/pnas.1919726117)) |
| Otu11791 | 2 | k__Fungi(100);p__Ascomycota(100);c__Eurotiomycetes(100);o__Chaetothyriales(100);f__Herpotrichiellaceae(100);g__Cladophialophora(100);s__Cladophialophora_sp_99003d(100); | Cladophialophora | 13 | Pathotroph-Saprotroph-Symbiotroph | Animal Pathogen-Endophyte-Fungal Parasite-\|Undefined Saprotroph\| | Dimorphic-Microfungus | NULL | Probable | As mycoparasite (Põlme et al. 2020); As root_endophyte_dark_septate (Põlme et al. 2020); As soil_saprotroph (Põlme et al. 2020); Decay soil (Põlme et al. 2020) | James TY, et al. 2006. Nature 443:818-822 ((https://doi.org/10.1038/nature05110)); Põlme S, et al. 2020. Fungal Diversity 105:1-16 ((https://doi.org/10.1007/s13225-020-00466-2)) |
| Otu11792 | 2 | k__Fungi(100);p__Ascomycota(100);c__Eurotiomycetes(100);o__Eurotiales(100);f__Trichocomaceae(100);g__Penicillium(100);g__Penicillium_unclassified(100); | Penicillium | 13 | Pathotroph-Saprotroph-Symbiotroph | Animal Parasite-Dung Saprotroph-Endophyte-Fungal Parasite-Plant Pathogen-\|Undefined Saprotroph\|-Wood Saprotroph | Microfungus | Soft Rot | Highly Probable | As foliar_endophyte (Põlme et al. 2020); As mycoparasite (Põlme et al. 2020); Decay soil (Põlme et al. 2020) | Duncan CG, Eslyn WE. 1966. Mycologia 58:642-664 ((https://doi.org/10.2307/3757045)); Seehann G, et al. 1975. List of Fungi in Soft-Rot Tissue Tests. Institute fur Holzbiologie und Holzschutz, Bundesforschungsanstalt fur Forst - und Holzwirtshaft, Hamburg - Lohbrugge, Germany ((https://www.irg-wp.com/irgdocs/details.php?d6493033-c9ea-40f3-8e3e-5697f32d640d)); Costa IPMW, et al. 2012. Mycotaxon 119:e26 ((http://mycotaxon.com/resources/checklists/costa_v119_checklist.pdf)); Bills GF, et al. 2013. Current Opinion in Microbiology 16: 549-565 ((https://doi.org/10.1016/j.mib.2013.08.001)); Tedersoo L, et al. 2014. Science 346:e1256688 ((https://doi.org/10.1126/science.1256688)); Põlme S, et al. 2020. Fungal Diversity 105:1-16 ((https://doi.org/10.1007/s13225-020-00466-2)) |
| Otu11793 | 2 | k__Fungi(100);p__Ascomycota(100);c__Eurotiomycetes(100);o__Eurotiales(100);f__Trichocomaceae(100);g__Penicillium(100);s__Penicillium_sp_NRRL_28159(100); | Penicillium | 13 | Pathotroph-Saprotroph-Symbiotroph | Animal Parasite-Dung Saprotroph-Endophyte-Fungal Parasite-Plant Pathogen-\|Undefined Saprotroph\|-Wood Saprotroph | Microfungus | Soft Rot | Highly Probable | As foliar_endophyte (Põlme et al. 2020); As mycoparasite (Põlme et al. 2020); Decay soil (Põlme et al. 2020) | Duncan CG, Eslyn WE. 1966. Mycologia 58:642-664 ((https://doi.org/10.2307/3757045)); Seehann G, et al. 1975. List of Fungi in Soft-Rot Tissue Tests. Institute fur Holzbiologie und Holzschutz, Bundesforschungsanstalt fur Forst - und Holzwirtshaft, Hamburg - Lohbrugge, Germany ((https://www.irg-wp.com/irgdocs/details.php?d6493033-c9ea-40f3-8e3e-5697f32d640d)); Costa IPMW, et al. 2012. Mycotaxon 119:e26 ((http://mycotaxon.com/resources/checklists/costa_v119_checklist.pdf)); Bills GF, et al. 2013. Current Opinion in Microbiology 16: 549-565 ((https://doi.org/10.1016/j.mib.2013.08.001)); Tedersoo L, et al. 2014. Science 346:e1256688 ((https://doi.org/10.1126/science.1256688)); Põlme S, et al. 2020. Fungal Diversity 105:1-16 ((https://doi.org/10.1007/s13225-020-00466-2)) |
| Otu11795 | 2 | k__Fungi(100);p__Ascomycota(100);c__Lecanoromycetes(100);o__Lecanorales(100);f__Cladoniaceae(100);g__Cladonia(100);g__Cladonia_unclassified(100); | Cladonia | 13 | Symbiotroph | \|Lichenized\| | Thallus | NULL | Highly Probable | NULL | James TY, et al. 2006. Nature 443:818-822 ((https://doi.org/10.1038/nature05110)); Põlme S, et al. 2020. Fungal Diversity 105:1-16 ((https://doi.org/10.1007/s13225-020-00466-2)) |
| Otu11797 | 2 | k__Fungi(100);p__Ascomycota(100);c__Saccharomycetes(100);o__Saccharomycetales(100);f__Pichiaceae(100);g__Yamadazyma(100);s__Yamadazyma_phyllophila(100); | Yamadazyma | 13 | Saprotroph | \|Undefined Saprotroph\| | Yeast | NULL | Probable | As nectar-tap_saprotroph (Põlme et al. 2020); Decay sugar-rich substrates (Põlme et al. 2020) | Põlme S, et al. 2020. Fungal Diversity 105:1-16 ((https://doi.org/10.1007/s13225-020-00466-2)) |
| Otu11801 | 2 | k__Fungi(100);p__Ascomycota(100);c__Dothideomycetes(100);o__Myriangiales(100);f__Elsinoaceae(100);f__Elsinoaceae_unclassified(100);f__Elsinoaceae_unclassified(100); | Elsinoaceae | 9 | Pathotroph | Plant Pathogen | Microfungus | NULL | Probable | Biotrophic or necrotrophic on leaves, stems and fruits of a wide range of plants (Cannon & Kirk 2007) | Cannon PF, Kirk PM. 2007. Fungal Families of the World. CAB International, Cambridge (ISBN: 978-0851998275) |
| Otu11803 | 2 | k__Fungi(100);p__Ascomycota(100);c__Sordariomycetes(100);o__Hypocreales(100);o__Hypocreales_unclassified(100);o__Hypocreales_unclassified(100);o__Hypocreales_unclassified(100); | Hypocreales | 7 | Saprotroph | Undefined Saprotroph | Microfungus | NULL | Possible | NULL | Sterkenburg E, et al. 2015. New Phytologist 207:1145-1158 ((https://doi.org/10.1111/nph.13426)) |
| Otu11804 | 2 | k__Fungi(100);p__Basidiomycota(100);c__Agaricomycetes(100);o__Agaricales(100);f__Mycenaceae(100);g__Mycena(100);g__Mycena_unclassified(100); | Mycena | 13 | Pathotroph-Saprotroph | Plant Pathogen-\|Plant Saprotroph\|-Undefined Saprotroph-Wood Saprotroph | Agaricoid | NULL | Probable | As litter_saprotroph (Põlme et al. 2020); Decay leaf, fruit, seed (Põlme et al. 2020) | Grgurinovic CA, Simpson JA. 2001. Fungal Diversity 8:97-127 ((http://www.fungaldiversity.org/fdp/sfdp/FD_8_97-127.pdf)); Tedersoo L, et al. 2014. Science 346:e1256688 ((https://doi.org/10.1126/science.1256688)); Põlme S, et al. 2020. Fungal Diversity 105:1-16 ((https://doi.org/10.1007/s13225-020-00466-2)); The American Phytopathological Society Common Names of Plant Diseases ((http://www.apsnet.org/publications/commonnames/Pages/default.aspx)) |
| Otu11805 | 2 | k__Fungi(100);p__Basidiomycota(100);c__Agaricomycetes(100);o__Cantharellales(100);f__Botryobasidiaceae(100);f__Botryobasidiaceae_unclassified(100);f__Botryobasidiaceae_unclassified(100); | Botryobasidiaceae | 9 | Saprotroph | Undefined Saprotroph-Wood Saprotroph | Corticioid | NULL | Probable | Saprobic, typically on rotten wood (Cannon & Kirk 2007) | Cannon PF, Kirk PM. 2007. Fungal Families of the World. CAB International, Cambridge (ISBN: 978-0851998275) |
| Otu11807 | 2 | k__Fungi(100);p__Basidiomycota(100);c__Agaricomycetes(100);o__Russulales(100);f__Russulaceae(100);g__Russula(100);g__Russula_unclassified(100); | Russula | 13 | Symbiotroph | \|Ectomycorrhizal\| | Agaricoid | NULL | Highly Probable | NULL | Rinaldi AC, et al. 2008. Fungal Diversity 33:1-45 ((http://www.fungaldiversity.org/fdp/sfdp/33-1.pdf)); Tedersoo L, et al. 2010. Mycorrhiza 20:217-263 ((https://doi.org/10.1007/s00572-009-0274-x)); Põlme S, et al. 2020. Fungal Diversity 105:1-16 ((https://doi.org/10.1007/s13225-020-00466-2)); http://mycorrhizas.info/ecmf.html ((http://mycorrhizas.info/ecmf.html)) |
| Otu11809 | 2 | k__Fungi(100);p__Basidiomycota(100);c__Exobasidiomycetes(100);o__Tilletiales(100);f__Tilletiaceae(100);g__Tilletia(100);s__Tilletia_indica(100); | Tilletia | 13 | Pathotroph | \|Plant Pathogen\| | Smut | NULL | Probable | Host - Poaceae (Põlme et al. 2020) | Tedersoo L, et al. 2014. Science 346:e1256688 ((https://doi.org/10.1126/science.1256688)); Põlme S, et al. 2020. Fungal Diversity 105:1-16 ((https://doi.org/10.1007/s13225-020-00466-2)) |
| Otu11811 | 2 | k__Fungi(100);p__Basidiomycota(100);c__Agaricomycetes(100);o__Thelephorales(100);f__Thelephoraceae(100);g__Thelephora(100);s__Thelephora_ganbajun(100); | Thelephora | 13 | Symbiotroph | \|Ectomycorrhizal\| | Clavarioid | NULL | Highly Probable | NULL | Rinaldi AC, et al. 2008. Fungal Diversity 33:1-45 ((http://www.fungaldiversity.org/fdp/sfdp/33-1.pdf)); Tedersoo L, et al. 2010. Mycorrhiza 20:217-263 ((https://doi.org/10.1007/s00572-009-0274-x)); Põlme S, et al. 2020. Fungal Diversity 105:1-16 ((https://doi.org/10.1007/s13225-020-00466-2)); http://mycorrhizas.info/ecmf.html ((http://mycorrhizas.info/ecmf.html)) |
| Otu11812 | 2 | k__Fungi(100);p__Basidiomycota(100);c__Agaricomycetes(100);o__Thelephorales(100);f__Thelephoraceae(100);g__Tomentella(100);g__Tomentella_unclassified(100); | Tomentella | 13 | Symbiotroph | \|Ectomycorrhizal\| | Corticioid | NULL | Highly Probable | NULL | Rinaldi AC, et al. 2008. Fungal Diversity 33:1-45 ((http://www.fungaldiversity.org/fdp/sfdp/33-1.pdf)); Tedersoo L, et al. 2010. Mycorrhiza 20:217-263 ((https://doi.org/10.1007/s00572-009-0274-x)); Põlme S, et al. 2020. Fungal Diversity 105:1-16 ((https://doi.org/10.1007/s13225-020-00466-2)); http://mycorrhizas.info/ecmf.html ((http://mycorrhizas.info/ecmf.html)) |
| Otu11813 | 2 | k__Fungi(100);p__Basidiomycota(100);c__Agaricomycetes(100);o__Thelephorales(100);f__Thelephoraceae(100);g__Tomentella(100);s__Tomentella_sp(100); | Tomentella | 13 | Symbiotroph | \|Ectomycorrhizal\| | Corticioid | NULL | Highly Probable | NULL | Rinaldi AC, et al. 2008. Fungal Diversity 33:1-45 ((http://www.fungaldiversity.org/fdp/sfdp/33-1.pdf)); Tedersoo L, et al. 2010. Mycorrhiza 20:217-263 ((https://doi.org/10.1007/s00572-009-0274-x)); Põlme S, et al. 2020. Fungal Diversity 105:1-16 ((https://doi.org/10.1007/s13225-020-00466-2)); http://mycorrhizas.info/ecmf.html ((http://mycorrhizas.info/ecmf.html)) |
| Otu11814 | 2 | k__Fungi(100);p__Basidiomycota(100);c__Agaricomycetes(100);o__Thelephorales(100);f__Thelephoraceae(100);g__Tomentella(100);g__Tomentella_unclassified(100); | Tomentella | 13 | Symbiotroph | \|Ectomycorrhizal\| | Corticioid | NULL | Highly Probable | NULL | Rinaldi AC, et al. 2008. Fungal Diversity 33:1-45 ((http://www.fungaldiversity.org/fdp/sfdp/33-1.pdf)); Tedersoo L, et al. 2010. Mycorrhiza 20:217-263 ((https://doi.org/10.1007/s00572-009-0274-x)); Põlme S, et al. 2020. Fungal Diversity 105:1-16 ((https://doi.org/10.1007/s13225-020-00466-2)); http://mycorrhizas.info/ecmf.html ((http://mycorrhizas.info/ecmf.html)) |
| Otu11816 | 2 | k__Fungi(100);p__Basidiomycota(100);c__Tremellomycetes(100);o__Tremellales(100);f__Tremellales_family_Incertae_sedis(100);g__Cryptococcus(100);s__Cryptococcus_sp_TC2_092(100); | Cryptococcus | 13 | Pathotroph-Saprotroph-Symbiotroph | Animal Pathogen-Endophyte-Epiphyte-\|Undefined Saprotroph\| | Dimorphic Yeast-Yeast | NULL | Possible | As foliar_endophyte (Põlme et al. 2020); Decay leaf, fruit, seed, soil, dung, animal material (Põlme et al. 2020) | Andrews JH, et al. 1983. Phytopathology 73:228-234 ((https://doi.org/10.1094/Phyto-73-228)); Kurtzman CP, et al. (eds.) 2011. The Yeasts, a Taxonomic Study. Fifth Edition. Vols 1-3. Elsevier, San Diego (ISBN: 9780444521491); Põlme S, et al. 2020. Fungal Diversity 105:1-16 ((https://doi.org/10.1007/s13225-020-00466-2)) |
| Otu11818 | 2 | k__Fungi(100);p__Basidiomycota(100);c__Agaricomycetes(100);o__Russulales(100);f__Russulaceae(100);g__Russula(100);s__Russula_compacta(100); | Russula | 13 | Symbiotroph | \|Ectomycorrhizal\| | Agaricoid | NULL | Highly Probable | NULL | Rinaldi AC, et al. 2008. Fungal Diversity 33:1-45 ((http://www.fungaldiversity.org/fdp/sfdp/33-1.pdf)); Tedersoo L, et al. 2010. Mycorrhiza 20:217-263 ((https://doi.org/10.1007/s00572-009-0274-x)); Põlme S, et al. 2020. Fungal Diversity 105:1-16 ((https://doi.org/10.1007/s13225-020-00466-2)); http://mycorrhizas.info/ecmf.html ((http://mycorrhizas.info/ecmf.html)) |
| Otu11819 | 2 | k__Fungi(100);p__Basidiomycota(100);c__Agaricomycetes(100);o__Agaricales(100);f__Pleurotaceae(100);g__Hohenbuehelia(100);s__Hohenbuehelia_unguicularis(100); | Hohenbuehelia | 13 | Pathotroph-Saprotroph | Nematophagous-Undefined Saprotroph-\|Wood Saprotroph\| | Agaricoid | White Rot | Probable | As white_rot (Põlme et al. 2020); Decay wood, animal material (Põlme et al. 2020) | Tedersoo L, et al. 2014. Science 346:e1256688 ((https://doi.org/10.1126/science.1256688)); Põlme S, et al. 2020. Fungal Diversity 105:1-16 ((https://doi.org/10.1007/s13225-020-00466-2)) |
| Otu11820 | 2 | k__Fungi(100);p__Basidiomycota(100);c__Agaricomycetes(100);o__Agaricales(100);o__Agaricales_unclassified(100);o__Agaricales_unclassified(100);o__Agaricales_unclassified(100); | Agaricales | 7 | Pathotroph-Saprotroph-Symbiotroph | Bryophyte Parasite-Dung Saprotroph-Ectomycorrhizal-Fungal Parasite-Leaf Saprotroph-Plant Parasite-Undefined Saprotroph-Wood Saprotroph | Agaricoid-Gasteroid-Microfungus-Secotioid-Yeast | NULL | Possible | Mushrooms and toadstools, Gill fungi, Agarics... lignicolous, sometimes muscicolous or fungicolous, saprobic, mycorrhizal, rarely parasitic on plants or fungi (Kirk et al. 2008) | Kirk PM et al. 2008. Dictionary of the Fungi. Tenth Edition. CAB International, Wallingford (ISBN: 978-0851998268) |
| Otu11821 | 2 | k__Fungi(100);p__Basidiomycota(100);c__Tremellomycetes(100);o__Tremellales(100);f__Tremellales_family_Incertae_sedis(100);f__Tremellales_family_Incertae_sedis_unclassified(100);f__Tremellales_family_Incertae_sedis_unclassified(100); | Tremellales | 7 | Pathotroph-Saprotroph-Symbiotroph | Fungal Parasite-Undefined Saprotroph | Tremelloid-Yeast | NULL | Possible | The order contains both teleomorphic and anamorphic species, most of the latter being yeasts | Sterkenburg E, et al. 2015. New Phytologist 207:1145-1158 ((https://doi.org/10.1111/nph.13426)) |
| Otu11822 | 2 | k__Fungi(100);p__Basidiomycota(100);c__Agaricomycetes(100);o__Agaricales(100);f__Marasmiaceae(100);g__Marasmius(100);s__Marasmius_pulcherripes(100); | Marasmius | 13 | Pathotroph-Saprotroph-Symbiotroph | Endophyte-Plant Pathogen-\|Plant Saprotroph\|-Undefined Saprotroph | Agaricoid | NULL | Probable | As litter_saprotroph (Põlme et al. 2020); Decay leaf, fruit, seed (Põlme et al. 2020); Often one of the most prominent fungal components of tropical forests, playing a major role in nutrient cycling (Cannon & Kirk 2007) | Cannon PF, Kirk PM. 2007. Fungal Families of the World. CAB International, Cambridge (ISBN: 978-0851998275); Tedersoo L, et al. 2014. Science 346:e1256688 ((https://doi.org/10.1126/science.1256688)); David AS, et al. 2016. Microbial Ecology 71:912-926 ((https://doi.org/10.1007/s00248-015-0712-6)); Põlme S, et al. 2020. Fungal Diversity 105:1-16 ((https://doi.org/10.1007/s13225-020-00466-2)) |
| Otu11823 | 2 | k__Fungi(100);p__Basidiomycota(100);c__Tremellomycetes(100);o__Tremellales(100);f__Tremellales_family_Incertae_sedis(100);f__Tremellales_family_Incertae_sedis_unclassified(100);f__Tremellales_family_Incertae_sedis_unclassified(100); | Tremellales | 7 | Pathotroph-Saprotroph-Symbiotroph | Fungal Parasite-Undefined Saprotroph | Tremelloid-Yeast | NULL | Possible | The order contains both teleomorphic and anamorphic species, most of the latter being yeasts | Sterkenburg E, et al. 2015. New Phytologist 207:1145-1158 ((https://doi.org/10.1111/nph.13426)) |
| Otu11826 | 2 | k__Fungi(100);p__Basidiomycota(100);c__Agaricomycetes(100);o__Cantharellales(100);f__Hydnaceae(100);g__Sistotrema(100);s__Sistotrema_sp(100); | Sistotrema | 13 | Saprotroph-Symbiotroph | Ectomycorrhizal-\|Plant Saprotroph\|-Wood Saprotroph | Corticioid | White Rot | Possible | As litter_saprotroph (Põlme et al. 2020); Decay leaf, fruit, seed (Põlme et al. 2020) | Gilbertson RL, Ryvarden L. 1987-1987. North American Polypores. Fungiflora, Oslo (ISBN: 978-0945345060); Rinaldi AC, et al. 2008. Fungal Diversity 33:1-45 ((http://www.fungaldiversity.org/fdp/sfdp/33-1.pdf)) (as ECM/not ECM); Tedersoo L, et al. 2010. Mycorrhiza 20:217-263 ((https://doi.org/10.1007/s00572-009-0274-x)) (pro parte); Põlme S, et al. 2020. Fungal Diversity 105:1-16 ((https://doi.org/10.1007/s13225-020-00466-2)); http://mycorrhizas.info/ecmf.html ((http://mycorrhizas.info/ecmf.html)) (as ECM/not ECM). |
| Otu11827 | 2 | k__Fungi(100);p__Basidiomycota(100);c__Agaricomycetes(100);o__Cantharellales(100);f__Clavulinaceae(100);g__Clavulina(100);g__Clavulina_unclassified(100); | Clavulina | 13 | Symbiotroph | \|Ectomycorrhizal\| | Clavarioid | NULL | Highly Probable | NULL | Tedersoo L, et al. 2014. Science 346:e1256688 ((https://doi.org/10.1126/science.1256688)); Põlme S, et al. 2020. Fungal Diversity 105:1-16 ((https://doi.org/10.1007/s13225-020-00466-2)) |
| Otu11828 | 2 | k__Fungi(100);p__Basidiomycota(100);c__Microbotryomycetes(100);o__Sporidiobolales(100);f__Sporidiobolales_family_Incertae_sedis(100);g__Rhodotorula(100);s__Rhodotorula_yarrowii(100); | Rhodotorula | 13 | Pathotroph-Saprotroph-Symbiotroph | Animal Pathogen-Animal Symbiotroph-Endophyte-Plant Pathogen-\|Undefined Saprotroph\| | Yeast | NULL | Probable | As foliar_endophyte (Põlme et al. 2020); Opportunistic human pathogen | Põlme S, et al. 2020. Fungal Diversity 105:1-16 ((https://doi.org/10.1007/s13225-020-00466-2)); https://www.cdc.gov ((https://www.cdc.gov)) |
| Otu11830 | 2 | k__Fungi(100);p__Basidiomycota(100);c__Agaricomycetes(100);o__Agaricales(100);f__Pluteaceae(100);g__Pluteus(100);s__Pluteus_aurantiorugosus(100); | Pluteus | 13 | Pathotroph-Saprotroph | Bryophyte Parasite-\|Plant Saprotroph\|-Wood Saprotroph | Agaricoid | White Rot | Probable | As litter_saprotroph (Põlme et al. 2020); As white_rot (Põlme et al. 2020); Decay leaf, fruit, seed (Põlme et al. 2020) | Grgurinovic CA, Simpson JA. 2001. Fungal Diversity 8:97-127 ((http://www.fungaldiversity.org/fdp/sfdp/FD_8_97-127.pdf)); James TY, et al. 2006. Nature 443:818-822 ((https://doi.org/10.1038/nature05110)); Tedersoo L, et al. 2014. Science 346:e1256688 ((https://doi.org/10.1126/science.1256688)); Põlme S, et al. 2020. Fungal Diversity 105:1-16 ((https://doi.org/10.1007/s13225-020-00466-2)) |
| Otu11831 | 2 | k__Fungi(100);p__Basidiomycota(100);c__Agaricomycetes(100);o__Agaricales(100);f__Mycenaceae(100);g__Mycena(100);s__Mycena_acicula(100); | Mycena | 13 | Pathotroph-Saprotroph | Plant Pathogen-\|Plant Saprotroph\|-Undefined Saprotroph-Wood Saprotroph | Agaricoid | NULL | Probable | As litter_saprotroph (Põlme et al. 2020); Decay leaf, fruit, seed (Põlme et al. 2020) | Grgurinovic CA, Simpson JA. 2001. Fungal Diversity 8:97-127 ((http://www.fungaldiversity.org/fdp/sfdp/FD_8_97-127.pdf)); Tedersoo L, et al. 2014. Science 346:e1256688 ((https://doi.org/10.1126/science.1256688)); Põlme S, et al. 2020. Fungal Diversity 105:1-16 ((https://doi.org/10.1007/s13225-020-00466-2)); The American Phytopathological Society Common Names of Plant Diseases ((http://www.apsnet.org/publications/commonnames/Pages/default.aspx)) |
| Otu11832 | 2 | k__Fungi(100);p__Ascomycota(100);c__Eurotiomycetes(100);o__Chaetothyriales(100);f__Herpotrichiellaceae(100);g__Exophiala(100);s__Exophiala_moniliae(100); | Exophiala | 13 | Pathotroph-Saprotroph-Symbiotroph | \|Animal Parasite\|-Animal Pathogen-Endophyte-Plant Saprotroph-Undefined Saprotroph | Dimorphic-Facultative Yeast | NULL | Probable | As litter_saprotroph (Põlme et al. 2020); As root_endophyte_dark_septate (Põlme et al. 2020); Decay leaf, fruit, seed, animal material (Põlme et al. 2020) | Tedersoo L, et al. 2014. Science 346:e1256688 ((https://doi.org/10.1126/science.1256688)), Nagy LG, et al. 2014. Nature Communications 5:e4471 ((https://doi.org/10.1038/ncomms5471)); Põlme S, et al. 2020. Fungal Diversity 105:1-16 ((https://doi.org/10.1007/s13225-020-00466-2)) |
| Otu11834 | 2 | k__Fungi(100);p__Basidiomycota(100);c__Agaricomycetes(100);o__Trechisporales(100);f__Hydnodontaceae(100);g__Trechispora(100);s__Trechispora_sp(100); | Trechispora | 13 | Saprotroph | \|Wood Saprotroph\| | Corticioid | White Rot | Highly Probable | As white_rot (Põlme et al. 2020); Decay wood (Põlme et al. 2020) | Gilbertson RL, Ryvarden L. 1987-1987. North American Polypores. Fungiflora, Oslo (ISBN: 978-0945345060); Rinaldi AC, et al. 2008. Fungal Diversity 33:1-45 ((http://www.fungaldiversity.org/fdp/sfdp/33-1.pdf)) (as Uncertain); Tedersoo L, et al. 2010. Mycorrhiza 20:217-263 ((https://doi.org/10.1007/s00572-009-0274-x)) (as not ECM).; Põlme S, et al. 2020. Fungal Diversity 105:1-16 ((https://doi.org/10.1007/s13225-020-00466-2)) |
| Otu11835 | 2 | k__Fungi(100);p__Ascomycota(100);c__Pezizomycetes(100);o__Pezizales(100);f__Sarcosomataceae(100);g__Pseudoplectania(100);s__Pseudoplectania_nigrella(100); | Pseudoplectania | 13 | Saprotroph | \|Plant Saprotroph\|-Undefined Saprotroph | Microfungus-Pezizoid | NULL | Probable | As litter_saprotroph (Põlme et al. 2020); Decay leaf, fruit, seed (Põlme et al. 2020) | Tedersoo L, et al. 2014. Science 346:e1256688 ((https://doi.org/10.1126/science.1256688)); Põlme S, et al. 2020. Fungal Diversity 105:1-16 ((https://doi.org/10.1007/s13225-020-00466-2)) |
| Otu11839 | 2 | k__Fungi(100);p__Ascomycota(100);c__Saccharomycetes(100);o__Saccharomycetales(100);f__Saccharomycetales_family_Incertae_sedis(100);g__Wickerhamomyces(100);s__Wickerhamomyces_sp_ES22S03(100); | Wickerhamomyces | 13 | Saprotroph | \|Plant Saprotroph\|-Undefined Saprotroph | Yeast | NULL | Probable | As litter_saprotroph (Põlme et al. 2020); As nectar-tap_saprotroph (Põlme et al. 2020); Decay leaf, fruit, seed, sugar-rich substrates (Põlme et al. 2020) | Kurtzman CP, et al. 2008. FEMS Yeast Research 8:939-954 ((https://doi.org/10.1111/j.1567-1364.2008.00419.x)); Põlme S, et al. 2020. Fungal Diversity 105:1-16 ((https://doi.org/10.1007/s13225-020-00466-2)) |
| Otu11842 | 2 | k__Fungi(100);p__Basidiomycota(100);c__Pucciniomycetes(100);o__Pucciniales(100);f__Pucciniastraceae(100);g__Pucciniastrum(100);s__Pucciniastrum_epilobii(100); | Pucciniastrum | 13 | Pathotroph | \|Plant Pathogen\| | Rust | NULL | Highly Probable | NULL | Tedersoo L, et al. 2014. Science 346:e1256688 ((https://doi.org/10.1126/science.1256688)); Põlme S, et al. 2020. Fungal Diversity 105:1-16 ((https://doi.org/10.1007/s13225-020-00466-2)) |
| Otu11844 | 2 | k__Fungi(100);p__Ascomycota(100);c__Saccharomycetes(100);o__Saccharomycetales(100);f__Metschnikowiaceae(100);g__Metschnikowia(100);g__Metschnikowia_unclassified(100); | Metschnikowia | 13 | Saprotroph | \|Undefined Saprotroph\| | Yeast | NULL | Probable | As nectar-tap_saprotroph (Põlme et al. 2020); Decay sugar-rich substrates (Põlme et al. 2020) | Põlme S, et al. 2020. Fungal Diversity 105:1-16 ((https://doi.org/10.1007/s13225-020-00466-2)) |
| Otu11845 | 2 | k__Fungi(100);p__Basidiomycota(100);c__Agaricomycetes(100);o__Agaricales(100);f__unclassified_Agaricales(100);g__unclassified_Agaricales(100);s__Agaricales_sp(100); | Agaricales | 7 | Pathotroph-Saprotroph-Symbiotroph | Bryophyte Parasite-Dung Saprotroph-Ectomycorrhizal-Fungal Parasite-Leaf Saprotroph-Plant Parasite-Undefined Saprotroph-Wood Saprotroph | Agaricoid-Gasteroid-Microfungus-Secotioid-Yeast | NULL | Possible | Mushrooms and toadstools, Gill fungi, Agarics... lignicolous, sometimes muscicolous or fungicolous, saprobic, mycorrhizal, rarely parasitic on plants or fungi (Kirk et al. 2008) | Kirk PM et al. 2008. Dictionary of the Fungi. Tenth Edition. CAB International, Wallingford (ISBN: 978-0851998268) |
| Otu11846 | 2 | k__Fungi(100);p__Basidiomycota(100);c__Agaricomycetes(100);o__Sebacinales(100);f__Sebacinaceae(100);g__Tremellodendron(100);g__Tremellodendron_unclassified(100); | Tremellodendron | 13 | Symbiotroph | \|Ectomycorrhizal\| | Clavarioid | NULL | Highly Probable | NULL | Rinaldi AC, et al. 2008. Fungal Diversity 33:1-45 ((http://www.fungaldiversity.org/fdp/sfdp/33-1.pdf)); Tedersoo L, et al. 2010. Mycorrhiza 20:217-263 ((https://doi.org/10.1007/s00572-009-0274-x)); Põlme S, et al. 2020. Fungal Diversity 105:1-16 ((https://doi.org/10.1007/s13225-020-00466-2)) |
| Otu11847 | 2 | k__Fungi(100);p__Basidiomycota(100);c__Agaricomycetes(100);o__Russulales(100);f__Lachnocladiaceae(100);g__Asterostroma(100);s__Asterostroma_macrosporum(100); | Asterostroma | 13 | Saprotroph | Undefined Saprotroph-\|Wood Saprotroph\| | Corticioid | White Rot | Probable | As white_rot (Põlme et al. 2020); Decay wood (Põlme et al. 2020) | Tedersoo L, et al. 2014. Science 346:e1256688 ((https://doi.org/10.1126/science.1256688)); Põlme S, et al. 2020. Fungal Diversity 105:1-16 ((https://doi.org/10.1007/s13225-020-00466-2)) |
| Otu11848 | 2 | k__Fungi(100);p__Basidiomycota(100);c__Agaricomycetes(100);o__Russulales(100);f__Lachnocladiaceae(100);g__Asterostroma(100);g__Asterostroma_unclassified(100); | Asterostroma | 13 | Saprotroph | Undefined Saprotroph-\|Wood Saprotroph\| | Corticioid | White Rot | Probable | As white_rot (Põlme et al. 2020); Decay wood (Põlme et al. 2020) | Tedersoo L, et al. 2014. Science 346:e1256688 ((https://doi.org/10.1126/science.1256688)); Põlme S, et al. 2020. Fungal Diversity 105:1-16 ((https://doi.org/10.1007/s13225-020-00466-2)) |

| | | | | | | | | | | | |
|---|---|---|---|---|---|---|---|---|---|---|---|
| Otu11850 | 2 | k__Fungi(100);p__Basidiomycota(100);c__Microbotryomycetes(100);o__Sporidiobolales(100);f__Sporidiobolales_family_Incertae_sedis(100);g__Rhodotorula(100);g__Rhodotorula_unclassified(100); | Rhodotorula | 13 | Pathotroph-Saprotroph-Symbiotroph | Animal Pathogen-Animal Symbiotroph-Endophyte-Plant Pathogen-\|Undefined Saprotroph\| | Yeast | NULL | Probable | As foliar_endophyte (Põlme et al. 2020); Opportunistic human pathogen | Põlme S, et al. 2020. Fungal Diversity 105:1-16 ((https://doi.org/10.1007/s13225-020-00466-2)); https://www.cdc.gov ((https://www.cdc.gov)) |
| Otu11851 | 2 | k__Fungi(100);p__Basidiomycota(100);c__Cystobasidiomycetes(100);o__Erythrobasidiales(100);f__Erythrobasidiales_family_Incertae_sedis(100);g__Erythrobasidium(100);s__Erythrobasidium_sp(100); | Erythrobasidium | 13 | Pathotroph-Saprotroph | \|Fungal Parasite\|-Undefined Saprotroph | Yeast | NULL | Possible | As mycoparasite (Põlme et al. 2020); Decay fungal material (Põlme et al. 2020) | Sugiyama J, Hamamoto M. 1998. The Yeasts 4:654-655 ((https://doi.org/10.1016/B978-044481312-1/50086-1)); Põlme S, et al. 2020. Fungal Diversity 105:1-16 ((https://doi.org/10.1007/s13225-020-00466-2)) |
| Otu11853 | 2 | k__Fungi(100);p__Basidiomycota(100);c__Microbotryomycetes(100);o__Sporidiobolales(100);f__Sporidiobolales_family_Incertae_sedis(100);g__Rhodotorula(100);s__Rhodotorula_oryzicola(100); | Rhodotorula | 13 | Pathotroph-Saprotroph-Symbiotroph | Animal Pathogen-Animal Symbiotroph-Endophyte-Plant Pathogen-\|Undefined Saprotroph\| | Yeast | NULL | Probable | As foliar_endophyte (Põlme et al. 2020); Opportunistic human pathogen | Põlme S, et al. 2020. Fungal Diversity 105:1-16 ((https://doi.org/10.1007/s13225-020-00466-2)); https://www.cdc.gov ((https://www.cdc.gov)) |
| Otu11855 | 2 | k__Fungi(100);p__Basidiomycota(100);c__Agaricomycetes(100);o__Russulales(100);f__Russulaceae(100);g__Russula(100);s__Russula_lilacea(100); | Russula | 13 | Symbiotroph | \|Ectomycorrhizal\| | Agaricoid | NULL | Highly Probable | NULL | Rinaldi AC, et al. 2008. Fungal Diversity 33:1-45 ((http://www.fungaldiversity.org/fdp/sfdp/33-1.pdf)); Tedersoo L, et al. 2010. Mycorrhiza 20:217-263 ((https://doi.org/10.1007/s00572-009-0274-x)); Põlme S, et al. 2020. Fungal Diversity 105:1-16 ((https://doi.org/10.1007/s13225-020-00466-2)); http://mycorrhizas.info/ecmf.html ((http://mycorrhizas.info/ecmf.html)) |
| Otu11856 | 2 | k__Fungi(100);p__Basidiomycota(100);c__Agaricomycetes(100);o__Russulales(100);f__Russulaceae(100);g__Russula(100);g__Russula_unclassified(100); | Russula | 13 | Symbiotroph | \|Ectomycorrhizal\| | Agaricoid | NULL | Highly Probable | NULL | Rinaldi AC, et al. 2008. Fungal Diversity 33:1-45 ((http://www.fungaldiversity.org/fdp/sfdp/33-1.pdf)); Tedersoo L, et al. 2010. Mycorrhiza 20:217-263 ((https://doi.org/10.1007/s00572-009-0274-x)); Põlme S, et al. 2020. Fungal Diversity 105:1-16 ((https://doi.org/10.1007/s13225-020-00466-2)); http://mycorrhizas.info/ecmf.html ((http://mycorrhizas.info/ecmf.html)) |
| Otu11857 | 2 | k__Fungi(100);p__Ascomycota(100);c__Saccharomycetes(100);o__Saccharomycetales(100);f__Metschnikowiaceae(100);g__Metschnikowia(100);s__Metschnikowia_pulcherrima(100); | Metschnikowia | 13 | Saprotroph | \|Undefined Saprotroph\| | Yeast | NULL | Probable | As nectar-tap_saprotroph (Põlme et al. 2020); Decay sugar-rich substrates (Põlme et al. 2020) | Põlme S, et al. 2020. Fungal Diversity 105:1-16 ((https://doi.org/10.1007/s13225-020-00466-2)) |
| Otu11860 | 2 | k__Fungi(100);p__Ascomycota(100);c__Saccharomycetes(100);o__Saccharomycetales(100);f__Saccharomycetales_family_Incertae_sedis(100);g__Candida(100);g__Candida_unclassified(100); | Candida | 13 | Pathotroph-Saprotroph-Symbiotroph | Animal Pathogen-Endophyte-Endosymbiont-Epiphyte-\|Undefined Saprotroph\| | Dimorphic Yeast-Yeast | NULL | Probable | As nectar-tap_saprotroph (Põlme et al. 2020); Decay sugar-rich substrates (Põlme et al. 2020); This asexual genus is polyphyletic and contain species that function very differently depend on what they are. Caution should be taken when interpreting guilds for Candida species. | Manolakaki D, et al. 2010. Virulence. 1: 367-375 ((https://doi.org/10.4161/viru.1.5.12796)); Mekbib SB, et al. 2011. Tropical Plant Pathology 36:233-240 ((https://doi.org/10.1590/S1982-56762011000400004)); Põlme S, et al. 2020. Fungal Diversity 105:1-16 ((https://doi.org/10.1007/s13225-020-00466-2)) |
| Otu11861 | 2 | k__Fungi(100);p__Ascomycota(100);c__Saccharomycetes(100);o__Saccharomycetales(100);f__Metschnikowiaceae(100);g__Metschnikowia(100);s__Metschnikowia_pulcherrima(100); | Metschnikowia | 13 | Saprotroph | \|Undefined Saprotroph\| | Yeast | NULL | Probable | As nectar-tap_saprotroph (Põlme et al. 2020); Decay sugar-rich substrates (Põlme et al. 2020) | Põlme S, et al. 2020. Fungal Diversity 105:1-16 ((https://doi.org/10.1007/s13225-020-00466-2)) |
| Otu11864 | 2 | k__Fungi(100);p__Ascomycota(100);c__Saccharomycetes(100);o__Saccharomycetales(100);f__Metschnikowiaceae(100);g__Metschnikowia(100);g__Metschnikowia_unclassified(100); | Metschnikowia | 13 | Saprotroph | \|Undefined Saprotroph\| | Yeast | NULL | Probable | As nectar-tap_saprotroph (Põlme et al. 2020); Decay sugar-rich substrates (Põlme et al. 2020) | Põlme S, et al. 2020. Fungal Diversity 105:1-16 ((https://doi.org/10.1007/s13225-020-00466-2)) |
| Otu11865 | 2 | k__Fungi(100);p__Ascomycota(100);c__Saccharomycetes(100);o__Saccharomycetales(100);f__Saccharomycetales_family_Incertae_sedis(100);g__Candida(100);g__Candida_unclassified(100); | Candida | 13 | Pathotroph-Saprotroph-Symbiotroph | Animal Pathogen-Endophyte-Endosymbiont-Epiphyte-\|Undefined Saprotroph\| | Dimorphic Yeast-Yeast | NULL | Probable | As nectar-tap_saprotroph (Põlme et al. 2020); Decay sugar-rich substrates (Põlme et al. 2020); This asexual genus is polyphyletic and contain species that function very differently depend on what they are. Caution should be taken when interpreting guilds for Candida species. | Manolakaki D, et al. 2010. Virulence. 1: 367-375 ((https://doi.org/10.4161/viru.1.5.12796)); Mekbib SB, et al. 2011. Tropical Plant Pathology 36:233-240 ((https://doi.org/10.1590/S1982-56762011000400004)); Põlme S, et al. 2020. Fungal Diversity 105:1-16 ((https://doi.org/10.1007/s13225-020-00466-2)) |
| Otu11866 | 2 | k__Fungi(100);p__Ascomycota(100);c__Saccharomycetes(100);o__Saccharomycetales(100);f__Metschnikowiaceae(100);g__Metschnikowia(100);s__Metschnikowia_pulcherrima(100); | Metschnikowia | 13 | Saprotroph | \|Undefined Saprotroph\| | Yeast | NULL | Probable | As nectar-tap_saprotroph (Põlme et al. 2020); Decay sugar-rich substrates (Põlme et al. 2020) | Põlme S, et al. 2020. Fungal Diversity 105:1-16 ((https://doi.org/10.1007/s13225-020-00466-2)) |
| Otu11867 | 2 | k__Fungi(100);p__Ascomycota(100);c__Saccharomycetes(100);o__Saccharomycetales(100);f__Metschnikowiaceae(100);g__Metschnikowia(100);s__Metschnikowia_pulcherrima(100); | Metschnikowia | 13 | Saprotroph | \|Undefined Saprotroph\| | Yeast | NULL | Probable | As nectar-tap_saprotroph (Põlme et al. 2020); Decay sugar-rich substrates (Põlme et al. 2020) | Põlme S, et al. 2020. Fungal Diversity 105:1-16 ((https://doi.org/10.1007/s13225-020-00466-2)) |
| Otu11869 | 2 | k__Fungi(100);p__Ascomycota(100);c__Saccharomycetes(100);o__Saccharomycetales(100);f__Saccharomycetales_family_Incertae_sedis(100);g__Candida(100);g__Candida_unclassified(100); | Candida | 13 | Pathotroph-Saprotroph-Symbiotroph | Animal Pathogen-Endophyte-Endosymbiont-Epiphyte-\|Undefined Saprotroph\| | Dimorphic Yeast-Yeast | NULL | Probable | As nectar-tap_saprotroph (Põlme et al. 2020); Decay sugar-rich substrates (Põlme et al. 2020); This asexual genus is polyphyletic and contain species that function very differently depend on what they are. Caution should be taken when interpreting guilds for Candida species. | Manolakaki D, et al. 2010. Virulence. 1: 367-375 ((https://doi.org/10.4161/viru.1.5.12796)); Mekbib SB, et al. 2011. Tropical Plant Pathology 36:233-240 ((https://doi.org/10.1590/S1982-56762011000400004)); Põlme S, et al. 2020. Fungal Diversity 105:1-16 ((https://doi.org/10.1007/s13225-020-00466-2)) |
| Otu11871 | 2 | k__Fungi(100);p__Ascomycota(100);c__Saccharomycetes(100);o__Saccharomycetales(100);f__Metschnikowiaceae(100);g__Metschnikowia(100);s__Metschnikowia_pulcherrima(100); | Metschnikowia | 13 | Saprotroph | \|Undefined Saprotroph\| | Yeast | NULL | Probable | As nectar-tap_saprotroph (Põlme et al. 2020); Decay sugar-rich substrates (Põlme et al. 2020) | Põlme S, et al. 2020. Fungal Diversity 105:1-16 ((https://doi.org/10.1007/s13225-020-00466-2)) |
| Otu11872 | 2 | k__Fungi(100);p__Ascomycota(100);c__Saccharomycetes(100);o__Saccharomycetales(100);f__Metschnikowiaceae(100);g__Metschnikowia(100);s__Metschnikowia_pulcherrima(100); | Metschnikowia | 13 | Saprotroph | \|Undefined Saprotroph\| | Yeast | NULL | Probable | As nectar-tap_saprotroph (Põlme et al. 2020); Decay sugar-rich substrates (Põlme et al. 2020) | Põlme S, et al. 2020. Fungal Diversity 105:1-16 ((https://doi.org/10.1007/s13225-020-00466-2)) |
| Otu11873 | 2 | k__Fungi(100);p__Ascomycota(100);c__Saccharomycetes(100);o__Saccharomycetales(100);f__Metschnikowiaceae(100);g__Metschnikowia(100);s__Metschnikowia_pulcherrima(100); | Metschnikowia | 13 | Saprotroph | \|Undefined Saprotroph\| | Yeast | NULL | Probable | As nectar-tap_saprotroph (Põlme et al. 2020); Decay sugar-rich substrates (Põlme et al. 2020) | Põlme S, et al. 2020. Fungal Diversity 105:1-16 ((https://doi.org/10.1007/s13225-020-00466-2)) |
| Otu11874 | 2 | k__Fungi(100);p__Ascomycota(100);c__Saccharomycetes(100);o__Saccharomycetales(100);f__Metschnikowiaceae(100);g__Metschnikowia(100);g__Metschnikowia_unclassified(100); | Metschnikowia | 13 | Saprotroph | \|Undefined Saprotroph\| | Yeast | NULL | Probable | As nectar-tap_saprotroph (Põlme et al. 2020); Decay sugar-rich substrates (Põlme et al. 2020) | Põlme S, et al. 2020. Fungal Diversity 105:1-16 ((https://doi.org/10.1007/s13225-020-00466-2)) |
| Otu11875 | 2 | k__Fungi(100);p__Ascomycota(100);c__Saccharomycetes(100);o__Saccharomycetales(100);f__Metschnikowiaceae(100);g__Metschnikowia(100);s__Metschnikowia_pulcherrima(100); | Metschnikowia | 13 | Saprotroph | \|Undefined Saprotroph\| | Yeast | NULL | Probable | As nectar-tap_saprotroph (Põlme et al. 2020); Decay sugar-rich substrates (Põlme et al. 2020) | Põlme S, et al. 2020. Fungal Diversity 105:1-16 ((https://doi.org/10.1007/s13225-020-00466-2)) |
| Otu11878 | 2 | k__Fungi(100);p__Ascomycota(100);c__Saccharomycetes(100);o__Saccharomycetales(100);f__Metschnikowiaceae(100);g__Metschnikowia(100);s__Metschnikowia_pulcherrima(100); | Metschnikowia | 13 | Saprotroph | \|Undefined Saprotroph\| | Yeast | NULL | Probable | As nectar-tap_saprotroph (Põlme et al. 2020); Decay sugar-rich substrates (Põlme et al. 2020) | Põlme S, et al. 2020. Fungal Diversity 105:1-16 ((https://doi.org/10.1007/s13225-020-00466-2)) |
| Otu11879 | 2 | k__Fungi(100);p__Ascomycota(100);c__Saccharomycetes(100);o__Saccharomycetales(100);f__Metschnikowiaceae(100);g__Metschnikowia(100);g__Metschnikowia_unclassified(100); | Metschnikowia | 13 | Saprotroph | \|Undefined Saprotroph\| | Yeast | NULL | Probable | As nectar-tap_saprotroph (Põlme et al. 2020); Decay sugar-rich substrates (Põlme et al. 2020) | Põlme S, et al. 2020. Fungal Diversity 105:1-16 ((https://doi.org/10.1007/s13225-020-00466-2)) |
| Otu11881 | 2 | k__Fungi(100);p__Ascomycota(100);c__Saccharomycetes(100);o__Saccharomycetales(100);f__Metschnikowiaceae(100);g__Metschnikowia(100);s__Metschnikowia_pulcherrima(100); | Metschnikowia | 13 | Saprotroph | \|Undefined Saprotroph\| | Yeast | NULL | Probable | As nectar-tap_saprotroph (Põlme et al. 2020); Decay sugar-rich substrates (Põlme et al. 2020) | Põlme S, et al. 2020. Fungal Diversity 105:1-16 ((https://doi.org/10.1007/s13225-020-00466-2)) |
| Otu11883 | 2 | k__Fungi(100);p__Ascomycota(100);c__Saccharomycetes(100);o__Saccharomycetales(100);f__Saccharomycetales_family_Incertae_sedis(100);g__Candida(100);g__Candida_unclassified(100); | Candida | 13 | Pathotroph-Saprotroph-Symbiotroph | Animal Pathogen-Endophyte-Endosymbiont-Epiphyte-\|Undefined Saprotroph\| | Dimorphic Yeast-Yeast | NULL | Probable | As nectar-tap_saprotroph (Põlme et al. 2020); Decay sugar-rich substrates (Põlme et al. 2020); This asexual genus is polyphyletic and contain species that function very differently depend on what they are. Caution should be taken when interpreting guilds for Candida species. | Manolakaki D, et al. 2010. Virulence. 1: 367-375 ((https://doi.org/10.4161/viru.1.5.12796)); Mekbib SB, et al. 2011. Tropical Plant Pathology 36:233-240 ((https://doi.org/10.1590/S1982-56762011000400004)); Põlme S, et al. 2020. Fungal Diversity 105:1-16 ((https://doi.org/10.1007/s13225-020-00466-2)) |
| Otu11884 | 2 | k__Fungi(100);p__Ascomycota(100);c__Saccharomycetes(100);o__Saccharomycetales(100);f__Saccharomycetales_family_Incertae_sedis(100);g__Candida(100);g__Candida_unclassified(100); | Candida | 13 | Pathotroph-Saprotroph-Symbiotroph | Animal Pathogen-Endophyte-Endosymbiont-Epiphyte-\|Undefined Saprotroph\| | Dimorphic Yeast-Yeast | NULL | Probable | As nectar-tap_saprotroph (Põlme et al. 2020); Decay sugar-rich substrates (Põlme et al. 2020); This asexual genus is polyphyletic and contain species that function very differently depend on what they are. Caution should be taken when interpreting guilds for Candida species. | Manolakaki D, et al. 2010. Virulence. 1: 367-375 ((https://doi.org/10.4161/viru.1.5.12796)); Mekbib SB, et al. 2011. Tropical Plant Pathology 36:233-240 ((https://doi.org/10.1590/S1982-56762011000400004)); Põlme S, et al. 2020. Fungal Diversity 105:1-16 ((https://doi.org/10.1007/s13225-020-00466-2)) |
| Otu11885 | 2 | k__Fungi(100);p__Ascomycota(100);c__Saccharomycetes(100);o__Saccharomycetales(100);f__Metschnikowiaceae(100);g__Metschnikowia(100);s__Metschnikowia_pulcherrima(100); | Metschnikowia | 13 | Saprotroph | \|Undefined Saprotroph\| | Yeast | NULL | Probable | As nectar-tap_saprotroph (Põlme et al. 2020); Decay sugar-rich substrates (Põlme et al. 2020) | Põlme S, et al. 2020. Fungal Diversity 105:1-16 ((https://doi.org/10.1007/s13225-020-00466-2)) |
| Otu11888 | 2 | k__Fungi(100);p__Ascomycota(100);c__Saccharomycetes(100);o__Saccharomycetales(100);f__Metschnikowiaceae(100);g__Metschnikowia(100);s__Metschnikowia_pulcherrima(100); | Metschnikowia | 13 | Saprotroph | \|Undefined Saprotroph\| | Yeast | NULL | Probable | As nectar-tap_saprotroph (Põlme et al. 2020); Decay sugar-rich substrates (Põlme et al. 2020) | Põlme S, et al. 2020. Fungal Diversity 105:1-16 ((https://doi.org/10.1007/s13225-020-00466-2)) |
| Otu11889 | 2 | k__Fungi(100);p__Ascomycota(100);c__Saccharomycetes(100);o__Saccharomycetales(100);f__Metschnikowiaceae(100);g__Metschnikowia(100);s__Metschnikowia_pulcherrima(100); | Metschnikowia | 13 | Saprotroph | \|Undefined Saprotroph\| | Yeast | NULL | Probable | As nectar-tap_saprotroph (Põlme et al. 2020); Decay sugar-rich substrates (Põlme et al. 2020) | Põlme S, et al. 2020. Fungal Diversity 105:1-16 ((https://doi.org/10.1007/s13225-020-00466-2)) |
| Otu11891 | 2 | k__Fungi(100);p__Ascomycota(100);c__Saccharomycetes(100);o__Saccharomycetales(100);f__Saccharomycetales_family_Incertae_sedis(100);f__Saccharomycetales_family_Incertae_sedis_unclassified(100);f__Saccharomycetales_family_Incertae_sedis_unclassified(100); | Saccharomycetales | 7 | Saprotroph | Undefined Saprotroph | Yeast | NULL | Possible | NULL | Sterkenburg E, et al. 2015. New Phytologist 207:1145-1158 ((https://doi.org/10.1111/nph.13426)) |
| Otu11894 | 2 | k__Fungi(100);p__Ascomycota(100);c__Saccharomycetes(100);o__Saccharomycetales(100);f__Saccharomycetales_family_Incertae_sedis(100);g__Candida(100);g__Candida_unclassified(100); | Candida | 13 | Pathotroph-Saprotroph-Symbiotroph | Animal Pathogen-Endophyte-Endosymbiont-Epiphyte-\|Undefined Saprotroph\| | Dimorphic Yeast-Yeast | NULL | Probable | As nectar-tap_saprotroph (Põlme et al. 2020); Decay sugar-rich substrates (Põlme et al. 2020); This asexual genus is polyphyletic and contain species that function very differently depend on what they are. Caution should be taken when interpreting guilds for Candida species. | Manolakaki D, et al. 2010. Virulence. 1: 367-375 ((https://doi.org/10.4161/viru.1.5.12796)); Mekbib SB, et al. 2011. Tropical Plant Pathology 36:233-240 ((https://doi.org/10.1590/S1982-56762011000400004)); Põlme S, et al. 2020. Fungal Diversity 105:1-16 ((https://doi.org/10.1007/s13225-020-00466-2)) |
| Otu11898 | 2 | k__Fungi(100);p__Ascomycota(100);c__Saccharomycetes(100);o__Saccharomycetales(100);f__Metschnikowiaceae(100);g__Metschnikowia(100);s__Metschnikowia_pulcherrima(100); | Metschnikowia | 13 | Saprotroph | \|Undefined Saprotroph\| | Yeast | NULL | Probable | As nectar-tap_saprotroph (Põlme et al. 2020); Decay sugar-rich substrates (Põlme et al. 2020) | Põlme S, et al. 2020. Fungal Diversity 105:1-16 ((https://doi.org/10.1007/s13225-020-00466-2)) |
| Otu11900 | 2 | k__Fungi(100);p__Ascomycota(100);c__Saccharomycetes(100);o__Saccharomycetales(100);f__Metschnikowiaceae(100);g__Metschnikowia(100);s__Metschnikowia_pulcherrima(100); | Metschnikowia | 13 | Saprotroph | \|Undefined Saprotroph\| | Yeast | NULL | Probable | As nectar-tap_saprotroph (Põlme et al. 2020); Decay sugar-rich substrates (Põlme et al. 2020) | Põlme S, et al. 2020. Fungal Diversity 105:1-16 ((https://doi.org/10.1007/s13225-020-00466-2)) |
| Otu11904 | 2 | k__Fungi(100);p__Ascomycota(100);c__Saccharomycetes(100);o__Saccharomycetales(100);f__Saccharomycetales_family_Incertae_sedis(100);g__Candida(100);g__Candida_unclassified(100); | Candida | 13 | Pathotroph-Saprotroph-Symbiotroph | Animal Pathogen-Endophyte-Endosymbiont-Epiphyte-\|Undefined Saprotroph\| | Dimorphic Yeast-Yeast | NULL | Probable | As nectar-tap_saprotroph (Põlme et al. 2020); Decay sugar-rich substrates (Põlme et al. 2020); This asexual genus is polyphyletic and contain species that function very differently depend on what they are. Caution should be taken when interpreting guilds for Candida species. | Manolakaki D, et al. 2010. Virulence. 1: 367-375 ((https://doi.org/10.4161/viru.1.5.12796)); Mekbib SB, et al. 2011. Tropical Plant Pathology 36:233-240 ((https://doi.org/10.1590/S1982-56762011000400004)); Põlme S, et al. 2020. Fungal Diversity 105:1-16 ((https://doi.org/10.1007/s13225-020-00466-2)) |
| Otu11908 | 2 | k__Fungi(100);p__Ascomycota(100);c__Dothideomycetes(100);o__Dothideomycetes_order_Incertae_sedis(100);f__Myxotrichaceae(100);g__Oidiodendron(100);s__Oidiodendron_maius(100); | Oidiodendron | 13 | Pathotroph-Saprotroph-Symbiotroph | Endophyte-Ericoid Mycorrhizal-\|Undefined Saprotroph\| | Microfungus | NULL | Probable | As root_endophyte (Põlme et al. 2020); As soil_saprotroph (Põlme et al. 2020); Dark Septate Endophyte; Decay soil (Põlme et al. 2020) | Newsham KK. 2011. New Phytologist 190:783-793 ((https://doi.org/10.1111/j.1469-8137.2010.03611.x)) (as dark septate root endophytes); Põlme S, et al. 2020. Fungal Diversity 105:1-16 ((https://doi.org/10.1007/s13225-020-00466-2)) |
| Otu11909 | 2 | k__Fungi(100);p__Ascomycota(100);c__Saccharomycetes(100);o__Saccharomycetales(100);f__Metschnikowiaceae(100);g__Metschnikowia(100);g__Metschnikowia_unclassified(100); | Metschnikowia | 13 | Saprotroph | \|Undefined Saprotroph\| | Yeast | NULL | Probable | As nectar-tap_saprotroph (Põlme et al. 2020); Decay sugar-rich substrates (Põlme et al. 2020) | Põlme S, et al. 2020. Fungal Diversity 105:1-16 ((https://doi.org/10.1007/s13225-020-00466-2)) |
| Otu11911 | 2 | k__Fungi(100);p__Ascomycota(100);c__Saccharomycetes(100);o__Saccharomycetales(100);f__Metschnikowiaceae(100);g__Metschnikowia(100);s__Metschnikowia_pulcherrima(100); | Metschnikowia | 13 | Saprotroph | \|Undefined Saprotroph\| | Yeast | NULL | Probable | As nectar-tap_saprotroph (Põlme et al. 2020); Decay sugar-rich substrates (Põlme et al. 2020) | Põlme S, et al. 2020. Fungal Diversity 105:1-16 ((https://doi.org/10.1007/s13225-020-00466-2)) |
| Otu11912 | 2 | k__Fungi(100);p__Ascomycota(100);c__Saccharomycetes(100);o__Saccharomycetales(100);f__Metschnikowiaceae(100);g__Metschnikowia(100);g__Metschnikowia_unclassified(100); | Metschnikowia | 13 | Saprotroph | \|Undefined Saprotroph\| | Yeast | NULL | Probable | As nectar-tap_saprotroph (Põlme et al. 2020); Decay sugar-rich substrates (Põlme et al. 2020) | Põlme S, et al. 2020. Fungal Diversity 105:1-16 ((https://doi.org/10.1007/s13225-020-00466-2)) |
| Otu11913 | 2 | k__Fungi(100);p__Ascomycota(100);c__Saccharomycetes(100);o__Saccharomycetales(100);f__Metschnikowiaceae(100);g__Metschnikowia(100);g__Metschnikowia_unclassified(100); | Metschnikowia | 13 | Saprotroph | \|Undefined Saprotroph\| | Yeast | NULL | Probable | As nectar-tap_saprotroph (Põlme et al. 2020); Decay sugar-rich substrates (Põlme et al. 2020) | Põlme S, et al. 2020. Fungal Diversity 105:1-16 ((https://doi.org/10.1007/s13225-020-00466-2)) |
| Otu11914 | 2 | k__Fungi(100);p__Ascomycota(100);c__Saccharomycetes(100);o__Saccharomycetales(100);f__Metschnikowiaceae(100);g__Metschnikowia(100);s__Metschnikowia_pulcherrima(100); | Metschnikowia | 13 | Saprotroph | \|Undefined Saprotroph\| | Yeast | NULL | Probable | As nectar-tap_saprotroph (Põlme et al. 2020); Decay sugar-rich substrates (Põlme et al. 2020) | Põlme S, et al. 2020. Fungal Diversity 105:1-16 ((https://doi.org/10.1007/s13225-020-00466-2)) |
| Otu11915 | 2 | k__Fungi(100);p__Ascomycota(100);c__Saccharomycetes(100);o__Saccharomycetales(100);f__Metschnikowiaceae(100);g__Metschnikowia(100);s__Metschnikowia_pulcherrima(100); | Metschnikowia | 13 | Saprotroph | \|Undefined Saprotroph\| | Yeast | NULL | Probable | As nectar-tap_saprotroph (Põlme et al. 2020); Decay sugar-rich substrates (Põlme et al. 2020) | Põlme S, et al. 2020. Fungal Diversity 105:1-16 ((https://doi.org/10.1007/s13225-020-00466-2)) |
| Otu11916 | 2 | k__Fungi(100);p__Ascomycota(100);c__Saccharomycetes(100);o__Saccharomycetales(100);f__Metschnikowiaceae(100);g__Metschnikowia(100);g__Metschnikowia_unclassified(100); | Metschnikowia | 13 | Saprotroph | \|Undefined Saprotroph\| | Yeast | NULL | Probable | As nectar-tap_saprotroph (Põlme et al. 2020); Decay sugar-rich substrates (Põlme et al. 2020) | Põlme S, et al. 2020. Fungal Diversity 105:1-16 ((https://doi.org/10.1007/s13225-020-00466-2)) |
| Otu11917 | 2 | k__Fungi(100);p__Ascomycota(100);c__Saccharomycetes(100);o__Saccharomycetales(100);f__Metschnikowiaceae(100);g__Metschnikowia(100);g__Metschnikowia_unclassified(100); | Metschnikowia | 13 | Saprotroph | \|Undefined Saprotroph\| | Yeast | NULL | Probable | As nectar-tap_saprotroph (Põlme et al. 2020); Decay sugar-rich substrates (Põlme et al. 2020) | Põlme S, et al. 2020. Fungal Diversity 105:1-16 ((https://doi.org/10.1007/s13225-020-00466-2)) |
| Otu11918 | 2 | k__Fungi(100);p__Ascomycota(100);c__Dothideomycetes(100);o__Pleosporales(100);f__Venturiaceae(100);g__Fusicladium(100);s__Fusicladium_caryigenum(100); | Fusicladium | 13 | Pathotroph-Symbiotroph | Endophyte-Lichen Parasite-\|Plant Pathogen\| | Microfungus | NULL | Probable | NULL | Tedersoo L, et al. 2014. Science 346:e1256688 ((https://doi.org/10.1126/science.1256688)); Põlme S, et al. 2020. Fungal Diversity 105:1-16 ((https://doi.org/10.1007/s13225-020-00466-2)) |
| Otu11919 | 2 | k__Fungi(100);p__Ascomycota(100);c__Saccharomycetes(100);o__Saccharomycetales(100);f__Metschnikowiaceae(100);g__Metschnikowia(100);g__Metschnikowia_unclassified(100); | Metschnikowia | 13 | Saprotroph | \|Undefined Saprotroph\| | Yeast | NULL | Probable | As nectar-tap_saprotroph (Põlme et al. 2020); Decay sugar-rich substrates (Põlme et al. 2020) | Põlme S, et al. 2020. Fungal Diversity 105:1-16 ((https://doi.org/10.1007/s13225-020-00466-2)) |
| Otu11920 | 2 | k__Fungi(100);p__Ascomycota(100);c__Saccharomycetes(100);o__Saccharomycetales(100);f__Metschnikowiaceae(100);g__Metschnikowia(100);g__Metschnikowia_unclassified(100); | Metschnikowia | 13 | Saprotroph | \|Undefined Saprotroph\| | Yeast | NULL | Probable | As nectar-tap_saprotroph (Põlme et al. 2020); Decay sugar-rich substrates (Põlme et al. 2020) | Põlme S, et al. 2020. Fungal Diversity 105:1-16 ((https://doi.org/10.1007/s13225-020-00466-2)) |
| Otu11922 | 2 | k__Fungi(100);p__Ascomycota(100);c__Saccharomycetes(100);o__Saccharomycetales(100);f__Metschnikowiaceae(100);g__Metschnikowia(100);s__Metschnikowia_pulcherrima(100); | Metschnikowia | 13 | Saprotroph | \|Undefined Saprotroph\| | Yeast | NULL | Probable | As nectar-tap_saprotroph (Põlme et al. 2020); Decay sugar-rich substrates (Põlme et al. 2020) | Põlme S, et al. 2020. Fungal Diversity 105:1-16 ((https://doi.org/10.1007/s13225-020-00466-2)) |
| Otu11923 | 2 | k__Fungi(100);p__Ascomycota(100);c__Saccharomycetes(100);o__Saccharomycetales(100);f__Metschnikowiaceae(100);g__Metschnikowia(100);s__Metschnikowia_pulcherrima(100); | Metschnikowia | 13 | Saprotroph | \|Undefined Saprotroph\| | Yeast | NULL | Probable | As nectar-tap_saprotroph (Põlme et al. 2020); Decay sugar-rich substrates (Põlme et al. 2020) | Põlme S, et al. 2020. Fungal Diversity 105:1-16 ((https://doi.org/10.1007/s13225-020-00466-2)) |
| Otu11926 | 2 | k__Fungi(100);p__Ascomycota(100);c__Saccharomycetes(100);o__Saccharomycetales(100);f__Metschnikowiaceae(100);g__Metschnikowia(100);s__Metschnikowia_pulcherrima(100); | Metschnikowia | 13 | Saprotroph | \|Undefined Saprotroph\| | Yeast | NULL | Probable | As nectar-tap_saprotroph (Põlme et al. 2020); Decay sugar-rich substrates (Põlme et al. 2020) | Põlme S, et al. 2020. Fungal Diversity 105:1-16 ((https://doi.org/10.1007/s13225-020-00466-2)) |
| Otu11928 | 2 | k__Fungi(100);p__Ascomycota(100);c__Saccharomycetes(100);o__Saccharomycetales(100);f__Metschnikowiaceae(100);g__Metschnikowia(100);s__Metschnikowia_pulcherrima(100); | Metschnikowia | 13 | Saprotroph | \|Undefined Saprotroph\| | Yeast | NULL | Probable | As nectar-tap_saprotroph (Põlme et al. 2020); Decay sugar-rich substrates (Põlme et al. 2020) | Põlme S, et al. 2020. Fungal Diversity 105:1-16 ((https://doi.org/10.1007/s13225-020-00466-2)) |
| Otu11929 | 2 | k__Fungi(100);p__Ascomycota(100);c__Saccharomycetes(100);o__Saccharomycetales(100);o__Saccharomycetales_unclassified(100);o__Saccharomycetales_unclassified(100);o__Saccharomycetales_unclassified(100); | Saccharomycetales | 7 | Saprotroph | Undefined Saprotroph | Yeast | NULL | Possible | NULL | Sterkenburg E, et al. 2015. New Phytologist 207:1145-1158 ((https://doi.org/10.1111/nph.13426)) |
| Otu11930 | 2 | k__Fungi(100);p__Ascomycota(100);c__Saccharomycetes(100);o__Saccharomycetales(100);f__Saccharomycetales_family_Incertae_sedis(100);g__Candida(100);g__Candida_unclassified(100); | Candida | 13 | Pathotroph-Saprotroph-Symbiotroph | Animal Pathogen-Endophyte-Endosymbiont-Epiphyte-\|Undefined Saprotroph\| | Dimorphic Yeast-Yeast | NULL | Probable | As nectar-tap_saprotroph (Põlme et al. 2020); Decay sugar-rich substrates (Põlme et al. 2020); This asexual genus is polyphyletic and contain species that function very differently depend on what they are. Caution should be taken when interpreting guilds for Candida species. | Manolakaki D, et al. 2010. Virulence. 1: 367-375 ((https://doi.org/10.4161/viru.1.5.12796)); Mekbib SB, et al. 2011. Tropical Plant Pathology 36:233-240 ((https://doi.org/10.1590/S1982-56762011000400004)); Põlme S, et al. 2020. Fungal Diversity 105:1-16 ((https://doi.org/10.1007/s13225-020-00466-2)) |
| Otu11935 | 2 | k__Fungi(100);p__Basidiomycota(100);c__Microbotryomycetes(100);o__Sporidiobolales(100);f__Sporidiobolales_family_Incertae_sedis(100);g__Sporobolomyces(100);s__Sporobolomyces_lactosus(100); | Sporobolomyces | 13 | Pathotroph-Saprotroph | \|Fungal Parasite\|-Plant Saprotroph | Yeast | NULL | Probable | As litter_saprotroph (Põlme et al. 2020); As mycoparasite (Põlme et al. 2020); Decay fungal material (Põlme et al. 2020) | Tedersoo L, et al. 2014. Science 346:e1256688 ((https://doi.org/10.1126/science.1256688)); Purahong W, et al. 2016. Molecular Ecology 25:4059-4074 ((https://doi.org/10.1111/mec.13739)); Põlme S, et al. 2020. Fungal Diversity 105:1-16 ((https://doi.org/10.1007/s13225-020-00466-2)) |
| Otu11936 | 2 | k__Fungi(100);p__Ascomycota(100);c__Saccharomycetes(100);o__Saccharomycetales(100);f__Metschnikowiaceae(100);g__Metschnikowia(100);g__Metschnikowia_unclassified(100); | Metschnikowia | 13 | Saprotroph | \|Undefined Saprotroph\| | Yeast | NULL | Probable | As nectar-tap_saprotroph (Põlme et al. 2020); Decay sugar-rich substrates (Põlme et al. 2020) | Põlme S, et al. 2020. Fungal Diversity 105:1-16 ((https://doi.org/10.1007/s13225-020-00466-2)) |
| Otu11939 | 2 | k__Fungi(100);p__Ascomycota(100);c__Saccharomycetes(100);o__Saccharomycetales(100);f__Metschnikowiaceae(100);g__Metschnikowia(100);g__Metschnikowia_unclassified(100); | Metschnikowia | 13 | Saprotroph | \|Undefined Saprotroph\| | Yeast | NULL | Probable | As nectar-tap_saprotroph (Põlme et al. 2020); Decay sugar-rich substrates (Põlme et al. 2020) | Põlme S, et al. 2020. Fungal Diversity 105:1-16 ((https://doi.org/10.1007/s13225-020-00466-2)) |
| Otu11940 | 2 | k__Fungi(100);p__Ascomycota(100);c__Saccharomycetes(100);o__Saccharomycetales(100);f__Metschnikowiaceae(100);g__Metschnikowia(100);s__Metschnikowia_pulcherrima(100); | Metschnikowia | 13 | Saprotroph | \|Undefined Saprotroph\| | Yeast | NULL | Probable | As nectar-tap_saprotroph (Põlme et al. 2020); Decay sugar-rich substrates (Põlme et al. 2020) | Põlme S, et al. 2020. Fungal Diversity 105:1-16 ((https://doi.org/10.1007/s13225-020-00466-2)) |
| Otu11941 | 2 | k__Fungi(100);p__Ascomycota(100);c__Saccharomycetes(100);o__Saccharomycetales(100);f__Metschnikowiaceae(100);g__Metschnikowia(100);s__Metschnikowia_pulcherrima(100); | Metschnikowia | 13 | Saprotroph | \|Undefined Saprotroph\| | Yeast | NULL | Probable | As nectar-tap_saprotroph (Põlme et al. 2020); Decay sugar-rich substrates (Põlme et al. 2020) | Põlme S, et al. 2020. Fungal Diversity 105:1-16 ((https://doi.org/10.1007/s13225-020-00466-2)) |
| Otu11943 | 2 | k__Fungi(100);p__Ascomycota(100);c__Saccharomycetes(100);o__Saccharomycetales(100);f__Metschnikowiaceae(100);g__Metschnikowia(100);s__Metschnikowia_pulcherrima(100); | Metschnikowia | 13 | Saprotroph | \|Undefined Saprotroph\| | Yeast | NULL | Probable | As nectar-tap_saprotroph (Põlme et al. 2020); Decay sugar-rich substrates (Põlme et al. 2020) | Põlme S, et al. 2020. Fungal Diversity 105:1-16 ((https://doi.org/10.1007/s13225-020-00466-2)) |
| Otu11944 | 2 | k__Fungi(100);p__Ascomycota(100);c__Saccharomycetes(100);o__Saccharomycetales(100);f__Metschnikowiaceae(100);g__Metschnikowia(100);g__Metschnikowia_unclassified(100); | Metschnikowia | 13 | Saprotroph | \|Undefined Saprotroph\| | Yeast | NULL | Probable | As nectar-tap_saprotroph (Põlme et al. 2020); Decay sugar-rich substrates (Põlme et al. 2020) | Põlme S, et al. 2020. Fungal Diversity 105:1-16 ((https://doi.org/10.1007/s13225-020-00466-2)) |
| Otu11945 | 2 | k__Fungi(100);p__Ascomycota(100);c__Saccharomycetes(100);o__Saccharomycetales(100);f__Metschnikowiaceae(100);g__Metschnikowia(100);g__Metschnikowia_unclassified(100); | Metschnikowia | 13 | Saprotroph | \|Undefined Saprotroph\| | Yeast | NULL | Probable | As nectar-tap_saprotroph (Põlme et al. 2020); Decay sugar-rich substrates (Põlme et al. 2020) | Põlme S, et al. 2020. Fungal Diversity 105:1-16 ((https://doi.org/10.1007/s13225-020-00466-2)) |
| Otu11946 | 2 | k__Fungi(100);p__Ascomycota(100);c__Saccharomycetes(100);o__Saccharomycetales(100);f__Metschnikowiaceae(100);g__Metschnikowia(100);s__Metschnikowia_pulcherrima(100); | Metschnikowia | 13 | Saprotroph | \|Undefined Saprotroph\| | Yeast | NULL | Probable | As nectar-tap_saprotroph (Põlme et al. 2020); Decay sugar-rich substrates (Põlme et al. 2020) | Põlme S, et al. 2020. Fungal Diversity 105:1-16 ((https://doi.org/10.1007/s13225-020-00466-2)) |
| Otu11947 | 2 | k__Fungi(100);p__Ascomycota(100);c__Saccharomycetes(100);o__Saccharomycetales(100);f__Metschnikowiaceae(100);g__Metschnikowia(100);s__Metschnikowia_pulcherrima(100); | Metschnikowia | 13 | Saprotroph | \|Undefined Saprotroph\| | Yeast | NULL | Probable | As nectar-tap_saprotroph (Põlme et al. 2020); Decay sugar-rich substrates (Põlme et al. 2020) | Põlme S, et al. 2020. Fungal Diversity 105:1-16 ((https://doi.org/10.1007/s13225-020-00466-2)) |
| Otu11949 | 2 | k__Fungi(100);p__Ascomycota(100);c__Saccharomycetes(100);o__Saccharomycetales(100);f__Metschnikowiaceae(100);g__Metschnikowia(100);s__Metschnikowia_pulcherrima(100); | Metschnikowia | 13 | Saprotroph | \|Undefined Saprotroph\| | Yeast | NULL | Probable | As nectar-tap_saprotroph (Põlme et al. 2020); Decay sugar-rich substrates (Põlme et al. 2020) | Põlme S, et al. 2020. Fungal Diversity 105:1-16 ((https://doi.org/10.1007/s13225-020-00466-2)) |
| Otu11950 | 2 | k__Fungi(100);p__Ascomycota(100);c__Saccharomycetes(100);o__Saccharomycetales(100);f__Metschnikowiaceae(100);g__Metschnikowia(100);s__Metschnikowia_pulcherrima(100); | Metschnikowia | 13 | Saprotroph | \|Undefined Saprotroph\| | Yeast | NULL | Probable | As nectar-tap_saprotroph (Põlme et al. 2020); Decay sugar-rich substrates (Põlme et al. 2020) | Põlme S, et al. 2020. Fungal Diversity 105:1-16 ((https://doi.org/10.1007/s13225-020-00466-2)) |
| Otu11951 | 2 | k__Fungi(100);p__Ascomycota(100);c__Saccharomycetes(100);o__Saccharomycetales(100);f__Metschnikowiaceae(100);g__Metschnikowia(100);s__Metschnikowia_pulcherrima(100); | Metschnikowia | 13 | Saprotroph | \|Undefined Saprotroph\| | Yeast | NULL | Probable | As nectar-tap_saprotroph (Põlme et al. 2020); Decay sugar-rich substrates (Põlme et al. 2020) | Põlme S, et al. 2020. Fungal Diversity 105:1-16 ((https://doi.org/10.1007/s13225-020-00466-2)) |
| Otu11954 | 2 | k__Fungi(100);p__Ascomycota(100);c__Saccharomycetes(100);o__Saccharomycetales(100);f__Metschnikowiaceae(100);g__Metschnikowia(100);g__Metschnikowia_unclassified(100); | Metschnikowia | 13 | Saprotroph | \|Undefined Saprotroph\| | Yeast | NULL | Probable | As nectar-tap_saprotroph (Põlme et al. 2020); Decay sugar-rich substrates (Põlme et al. 2020) | Põlme S, et al. 2020. Fungal Diversity 105:1-16 ((https://doi.org/10.1007/s13225-020-00466-2)) |
| Otu11959 | 2 | k__Fungi(100);p__Ascomycota(100);c__Saccharomycetes(100);o__Saccharomycetales(100);f__Saccharomycetales_family_Incertae_sedis(100);f__Saccharomycetales_family_Incertae_sedis_unclassified(100);f__Saccharomycetales_family_Incertae_sedis_unclassified(100); | Saccharomycetales | 7 | Saprotroph | Undefined Saprotroph | Yeast | NULL | Possible | NULL | Sterkenburg E, et al. 2015. New Phytologist 207:1145-1158 ((https://doi.org/10.1111/nph.13426)) |
| Otu11961 | 2 | k__Fungi(100);p__Ascomycota(100);c__Saccharomycetes(100);o__Saccharomycetales(100);f__Metschnikowiaceae(100);g__Metschnikowia(100);g__Metschnikowia_unclassified(100); | Metschnikowia | 13 | Saprotroph | \|Undefined Saprotroph\| | Yeast | NULL | Probable | As nectar-tap_saprotroph (Põlme et al. 2020); Decay sugar-rich substrates (Põlme et al. 2020) | Põlme S, et al. 2020. Fungal Diversity 105:1-16 ((https://doi.org/10.1007/s13225-020-00466-2)) |
| Otu11962 | 2 | k__Fungi(100);p__Ascomycota(100);c__Saccharomycetes(100);o__Saccharomycetales(100);f__Metschnikowiaceae(100);g__Metschnikowia(100);s__Metschnikowia_pulcherrima(100); | Metschnikowia | 13 | Saprotroph | \|Undefined Saprotroph\| | Yeast | NULL | Probable | As nectar-tap_saprotroph (Põlme et al. 2020); Decay sugar-rich substrates (Põlme et al. 2020) | Põlme S, et al. 2020. Fungal Diversity 105:1-16 ((https://doi.org/10.1007/s13225-020-00466-2)) |
| Otu11964 | 2 | k__Fungi(100);p__Ascomycota(100);c__Saccharomycetes(100);o__Saccharomycetales(100);f__Metschnikowiaceae(100);g__Metschnikowia(100);g__Metschnikowia_unclassified(100); | Metschnikowia | 13 | Saprotroph | \|Undefined Saprotroph\| | Yeast | NULL | Probable | As nectar-tap_saprotroph (Põlme et al. 2020); Decay sugar-rich substrates (Põlme et al. 2020) | Põlme S, et al. 2020. Fungal Diversity 105:1-16 ((https://doi.org/10.1007/s13225-020-00466-2)) |
| Otu11965 | 2 | k__Fungi(100);p__Ascomycota(100);c__Saccharomycetes(100);o__Saccharomycetales(100);f__Metschnikowiaceae(100);g__Metschnikowia(100);s__Metschnikowia_pulcherrima(100); | Metschnikowia | 13 | Saprotroph | \|Undefined Saprotroph\| | Yeast | NULL | Probable | As nectar-tap_saprotroph (Põlme et al. 2020); Decay sugar-rich substrates (Põlme et al. 2020) | Põlme S, et al. 2020. Fungal Diversity 105:1-16 ((https://doi.org/10.1007/s13225-020-00466-2)) |
| Otu11967 | 2 | k__Fungi(100);p__Ascomycota(100);c__Saccharomycetes(100);o__Saccharomycetales(100);f__Metschnikowiaceae(100);g__Metschnikowia(100);s__Metschnikowia_pulcherrima(100); | Metschnikowia | 13 | Saprotroph | \|Undefined Saprotroph\| | Yeast | NULL | Probable | As nectar-tap_saprotroph (Põlme et al. 2020); Decay sugar-rich substrates (Põlme et al. 2020) | Põlme S, et al. 2020. Fungal Diversity 105:1-16 ((https://doi.org/10.1007/s13225-020-00466-2)) |
| Otu11968 | 2 | k__Fungi(100);p__Ascomycota(100);c__Saccharomycetes(100);o__Saccharomycetales(100);f__Metschnikowiaceae(100);g__Metschnikowia(100);s__Metschnikowia_pulcherrima(100); | Metschnikowia | 13 | Saprotroph | \|Undefined Saprotroph\| | Yeast | NULL | Probable | As nectar-tap_saprotroph (Põlme et al. 2020); Decay sugar-rich substrates (Põlme et al. 2020) | Põlme S, et al. 2020. Fungal Diversity 105:1-16 ((https://doi.org/10.1007/s13225-020-00466-2)) |
| Otu11969 | 2 | k__Fungi(100);p__Ascomycota(100);c__Saccharomycetes(100);o__Saccharomycetales(100);f__Metschnikowiaceae(100);g__Metschnikowia(100);s__Metschnikowia_pulcherrima(100); | Metschnikowia | 13 | Saprotroph | \|Undefined Saprotroph\| | Yeast | NULL | Probable | As nectar-tap_saprotroph (Põlme et al. 2020); Decay sugar-rich substrates (Põlme et al. 2020) | Põlme S, et al. 2020. Fungal Diversity 105:1-16 ((https://doi.org/10.1007/s13225-020-00466-2)) |
| Otu11971 | 2 | k__Fungi(100);p__Ascomycota(100);c__Saccharomycetes(100);o__Saccharomycetales(100);f__Saccharomycetales_family_Incertae_sedis(100);g__Candida(100);g__Candida_unclassified(100); | Candida | 13 | Pathotroph-Saprotroph-Symbiotroph | Animal Pathogen-Endophyte-Endosymbiont-Epiphyte-\|Undefined Saprotroph\| | Dimorphic Yeast-Yeast | NULL | Probable | As nectar-tap_saprotroph (Põlme et al. 2020); Decay sugar-rich substrates (Põlme et al. 2020); This asexual genus is polyphyletic and contain species that function very differently depend on what they are. Caution should be taken when interpreting guilds for Candida species. | Manolakaki D, et al. 2010. Virulence. 1: 367-375 ((https://doi.org/10.4161/viru.1.5.12796)); Mekbib SB, et al. 2011. Tropical Plant Pathology 36:233-240 ((https://doi.org/10.1590/S1982-56762011000400004)); Põlme S, et al. 2020. Fungal Diversity 105:1-16 ((https://doi.org/10.1007/s13225-020-00466-2)) |
| Otu11972 | 2 | k__Fungi(100);p__Ascomycota(100);c__Saccharomycetes(100);o__Saccharomycetales(100);f__Metschnikowiaceae(100);g__Metschnikowia(100);g__Metschnikowia_unclassified(100); | Metschnikowia | 13 | Saprotroph | \|Undefined Saprotroph\| | Yeast | NULL | Probable | As nectar-tap_saprotroph (Põlme et al. 2020); Decay sugar-rich substrates (Põlme et al. 2020) | Põlme S, et al. 2020. Fungal Diversity 105:1-16 ((https://doi.org/10.1007/s13225-020-00466-2)) |
| Otu11973 | 2 | k__Fungi(100);p__Ascomycota(100);c__Saccharomycetes(100);o__Saccharomycetales(100);f__Metschnikowiaceae(100);g__Metschnikowia(100);g__Metschnikowia_unclassified(100); | Metschnikowia | 13 | Saprotroph | \|Undefined Saprotroph\| | Yeast | NULL | Probable | As nectar-tap_saprotroph (Põlme et al. 2020); Decay sugar-rich substrates (Põlme et al. 2020) | Põlme S, et al. 2020. Fungal Diversity 105:1-16 ((https://doi.org/10.1007/s13225-020-00466-2)) |
| Otu11974 | 2 | k__Fungi(100);p__Ascomycota(100);c__Saccharomycetes(100);o__Saccharomycetales(100);f__Metschnikowiaceae(100);g__Metschnikowia(100);s__Metschnikowia_pulcherrima(100); | Metschnikowia | 13 | Saprotroph | \|Undefined Saprotroph\| | Yeast | NULL | Probable | As nectar-tap_saprotroph (Põlme et al. 2020); Decay sugar-rich substrates (Põlme et al. 2020) | Põlme S, et al. 2020. Fungal Diversity 105:1-16 ((https://doi.org/10.1007/s13225-020-00466-2)) |
| Otu11975 | 2 | k__Fungi(100);p__Ascomycota(100);c__Saccharomycetes(100);o__Saccharomycetales(100);f__Metschnikowiaceae(100);g__Metschnikowia(100);s__Metschnikowia_pulcherrima(100); | Metschnikowia | 13 | Saprotroph | \|Undefined Saprotroph\| | Yeast | NULL | Probable | As nectar-tap_saprotroph (Põlme et al. 2020); Decay sugar-rich substrates (Põlme et al. 2020) | Põlme S, et al. 2020. Fungal Diversity 105:1-16 ((https://doi.org/10.1007/s13225-020-00466-2)) |
| Otu11976 | 2 | k__Fungi(100);p__Ascomycota(100);c__Saccharomycetes(100);o__Saccharomycetales(100);f__Metschnikowiaceae(100);g__Metschnikowia(100);s__Metschnikowia_pulcherrima(100); | Metschnikowia | 13 | Saprotroph | \|Undefined Saprotroph\| | Yeast | NULL | Probable | As nectar-tap_saprotroph (Põlme et al. 2020); Decay sugar-rich substrates (Põlme et al. 2020) | Põlme S, et al. 2020. Fungal Diversity 105:1-16 ((https://doi.org/10.1007/s13225-020-00466-2)) |
| Otu11977 | 2 | k__Fungi(100);p__Ascomycota(100);c__Saccharomycetes(100);o__Saccharomycetales(100);f__Metschnikowiaceae(100);g__Metschnikowia(100);g__Metschnikowia_unclassified(100); | Metschnikowia | 13 | Saprotroph | \|Undefined Saprotroph\| | Yeast | NULL | Probable | As nectar-tap_saprotroph (Põlme et al. 2020); Decay sugar-rich substrates (Põlme et al. 2020) | Põlme S, et al. 2020. Fungal Diversity 105:1-16 ((https://doi.org/10.1007/s13225-020-00466-2)) |
| Otu11978 | 2 | k__Fungi(100);p__Ascomycota(100);c__Saccharomycetes(100);o__Saccharomycetales(100);f__Metschnikowiaceae(100);g__Metschnikowia(100);g__Metschnikowia_unclassified(100); | Metschnikowia | 13 | Saprotroph | \|Undefined Saprotroph\| | Yeast | NULL | Probable | As nectar-tap_saprotroph (Põlme et al. 2020); Decay sugar-rich substrates (Põlme et al. 2020) | Põlme S, et al. 2020. Fungal Diversity 105:1-16 ((https://doi.org/10.1007/s13225-020-00466-2)) |
| Otu11979 | 2 | k__Fungi(100);p__Ascomycota(100);c__Saccharomycetes(100);o__Saccharomycetales(100);f__Metschnikowiaceae(100);g__Metschnikowia(100);g__Metschnikowia_unclassified(100); | Metschnikowia | 13 | Saprotroph | \|Undefined Saprotroph\| | Yeast | NULL | Probable | As nectar-tap_saprotroph (Põlme et al. 2020); Decay sugar-rich substrates (Põlme et al. 2020) | Põlme S, et al. 2020. Fungal Diversity 105:1-16 ((https://doi.org/10.1007/s13225-020-00466-2)) |
| Otu11980 | 2 | k__Fungi(100);p__Ascomycota(100);c__Saccharomycetes(100);o__Saccharomycetales(100);f__Metschnikowiaceae(100);g__Metschnikowia(100);g__Metschnikowia_unclassified(100); | Metschnikowia | 13 | Saprotroph | \|Undefined Saprotroph\| | Yeast | NULL | Probable | As nectar-tap_saprotroph (Põlme et al. 2020); Decay sugar-rich substrates (Põlme et al. 2020) | Põlme S, et al. 2020. Fungal Diversity 105:1-16 ((https://doi.org/10.1007/s13225-020-00466-2)) |
| Otu11981 | 2 | k__Fungi(100);p__Ascomycota(100);c__Saccharomycetes(100);o__Saccharomycetales(100);f__Metschnikowiaceae(100);g__Metschnikowia(100);s__Metschnikowia_pulcherrima(100); | Metschnikowia | 13 | Saprotroph | \|Undefined Saprotroph\| | Yeast | NULL | Probable | As nectar-tap_saprotroph (Põlme et al. 2020); Decay sugar-rich substrates (Põlme et al. 2020) | Põlme S, et al. 2020. Fungal Diversity 105:1-16 ((https://doi.org/10.1007/s13225-020-00466-2)) |
| Otu11982 | 2 | k__Fungi(100);p__Ascomycota(100);c__Saccharomycetes(100);o__Saccharomycetales(100);f__Metschnikowiaceae(100);g__Metschnikowia(100);s__Metschnikowia_pulcherrima(100); | Metschnikowia | 13 | Saprotroph | \|Undefined Saprotroph\| | Yeast | NULL | Probable | As nectar-tap_saprotroph (Põlme et al. 2020); Decay sugar-rich substrates (Põlme et al. 2020) | Põlme S, et al. 2020. Fungal Diversity 105:1-16 ((https://doi.org/10.1007/s13225-020-00466-2)) |
| Otu11986 | 2 | k__Fungi(100);p__Ascomycota(100);c__Saccharomycetes(100);o__Saccharomycetales(100);f__Metschnikowiaceae(100);g__Metschnikowia(100);g__Metschnikowia_unclassified(100); | Metschnikowia | 13 | Saprotroph | \|Undefined Saprotroph\| | Yeast | NULL | Probable | As nectar-tap_saprotroph (Põlme et al. 2020); Decay sugar-rich substrates (Põlme et al. 2020) | Põlme S, et al. 2020. Fungal Diversity 105:1-16 ((https://doi.org/10.1007/s13225-020-00466-2)) |
| Otu11987 | 2 | k__Fungi(100);p__Ascomycota(100);c__Saccharomycetes(100);o__Saccharomycetales(100);f__Metschnikowiaceae(100);g__Metschnikowia(100);g__Metschnikowia_unclassified(100); | Metschnikowia | 13 | Saprotroph | \|Undefined Saprotroph\| | Yeast | NULL | Probable | As nectar-tap_saprotroph (Põlme et al. 2020); Decay sugar-rich substrates (Põlme et al. 2020) | Põlme S, et al. 2020. Fungal Diversity 105:1-16 ((https://doi.org/10.1007/s13225-020-00466-2)) |
| Otu11990 | 2 | k__Fungi(100);p__Ascomycota(100);c__Saccharomycetes(100);o__Saccharomycetales(100);f__Metschnikowiaceae(100);g__Metschnikowia(100);s__Metschnikowia_pulcherrima(100); | Metschnikowia | 13 | Saprotroph | \|Undefined Saprotroph\| | Yeast | NULL | Probable | As nectar-tap_saprotroph (Põlme et al. 2020); Decay sugar-rich substrates (Põlme et al. 2020) | Põlme S, et al. 2020. Fungal Diversity 105:1-16 ((https://doi.org/10.1007/s13225-020-00466-2)) |
| Otu11992 | 2 | k__Fungi(100);p__Ascomycota(100);c__Saccharomycetes(100);o__Saccharomycetales(100);f__Metschnikowiaceae(100);g__Metschnikowia(100);s__Metschnikowia_pulcherrima(100); | Metschnikowia | 13 | Saprotroph | \|Undefined Saprotroph\| | Yeast | NULL | Probable | As nectar-tap_saprotroph (Põlme et al. 2020); Decay sugar-rich substrates (Põlme et al. 2020) | Põlme S, et al. 2020. Fungal Diversity 105:1-16 ((https://doi.org/10.1007/s13225-020-00466-2)) |
| Otu11993 | 2 | k__Fungi(100);p__Basidiomycota(100);c__Microbotryomycetes(100);o__Sporidiobolales(100);f__Sporidiobolales_family_Incertae_sedis(100);g__Sporobolomyces(100);s__Sporobolomyces_lactosus(100); | Sporobolomyces | 13 | Pathotroph-Saprotroph | \|Fungal Parasite\|-Plant Saprotroph | Yeast | NULL | Probable | As litter_saprotroph (Põlme et al. 2020); As mycoparasite (Põlme et al. 2020); Decay fungal material (Põlme et al. 2020) | Tedersoo L, et al. 2014. Science 346:e1256688 ((https://doi.org/10.1126/science.1256688)); Purahong W, et al. 2016. Molecular Ecology 25:4059-4074 ((https://doi.org/10.1111/mec.13739)); Põlme S, et al. 2020. Fungal Diversity 105:1-16 ((https://doi.org/10.1007/s13225-020-00466-2)) |
| Otu11995 | 2 | k__Fungi(100);p__Ascomycota(100);c__Saccharomycetes(100);o__Saccharomycetales(100);f__Metschnikowiaceae(100);g__Metschnikowia(100);s__Metschnikowia_pulcherrima(100); | Metschnikowia | 13 | Saprotroph | \|Undefined Saprotroph\| | Yeast | NULL | Probable | As nectar-tap_saprotroph (Põlme et al. 2020); Decay sugar-rich substrates (Põlme et al. 2020) | Põlme S, et al. 2020. Fungal Diversity 105:1-16 ((https://doi.org/10.1007/s13225-020-00466-2)) |
| Otu11997 | 2 | k__Fungi(100);p__Ascomycota(100);c__Saccharomycetes(100);o__Saccharomycetales(100);f__Metschnikowiaceae(100);g__Metschnikowia(100);s__Metschnikowia_pulcherrima(100); | Metschnikowia | 13 | Saprotroph | \|Undefined Saprotroph\| | Yeast | NULL | Probable | As nectar-tap_saprotroph (Põlme et al. 2020); Decay sugar-rich substrates (Põlme et al. 2020) | Põlme S, et al. 2020. Fungal Diversity 105:1-16 ((https://doi.org/10.1007/s13225-020-00466-2)) |
| Otu11999 | 2 | k__Fungi(100);p__Ascomycota(100);c__Saccharomycetes(100);o__Saccharomycetales(100);f__Metschnikowiaceae(100);g__Metschnikowia(100);s__Metschnikowia_pulcherrima(100); | Metschnikowia | 13 | Saprotroph | \|Undefined Saprotroph\| | Yeast | NULL | Probable | As nectar-tap_saprotroph (Põlme et al. 2020); Decay sugar-rich substrates (Põlme et al. 2020) | Põlme S, et al. 2020. Fungal Diversity 105:1-16 ((https://doi.org/10.1007/s13225-020-00466-2)) |
| Otu12000 | 2 | k__Fungi(100);p__Ascomycota(100);c__Saccharomycetes(100);o__Saccharomycetales(100);f__Metschnikowiaceae(100);g__Metschnikowia(100);s__Metschnikowia_pulcherrima(100); | Metschnikowia | 13 | Saprotroph | \|Undefined Saprotroph\| | Yeast | NULL | Probable | As nectar-tap_saprotroph (Põlme et al. 2020); Decay sugar-rich substrates (Põlme et al. 2020) | Põlme S, et al. 2020. Fungal Diversity 105:1-16 ((https://doi.org/10.1007/s13225-020-00466-2)) |
| Otu12001 | 2 | k__Fungi(100);p__Ascomycota(100);c__Saccharomycetes(100);o__Saccharomycetales(100);f__Saccharomycetales_family_Incertae_sedis(100);f__Saccharomycetales_family_Incertae_sedis_unclassified(100);f__Saccharomycetales_family_Incertae_sedis_unclassified(100); | Saccharomycetales | 7 | Saprotroph | Undefined Saprotroph | Yeast | NULL | Possible | NULL | Sterkenburg E, et al. 2015. New Phytologist 207:1145-1158 ((https://doi.org/10.1111/nph.13426)) |
| Otu12002 | 2 | k__Fungi(100);p__Ascomycota(100);c__Saccharomycetes(100);o__Saccharomycetales(100);f__Metschnikowiaceae(100);g__Metschnikowia(100);g__Metschnikowia_unclassified(100); | Metschnikowia | 13 | Saprotroph | \|Undefined Saprotroph\| | Yeast | NULL | Probable | As nectar-tap_saprotroph (Põlme et al. 2020); Decay sugar-rich substrates (Põlme et al. 2020) | Põlme S, et al. 2020. Fungal Diversity 105:1-16 ((https://doi.org/10.1007/s13225-020-00466-2)) |
| Otu12003 | 2 | k__Fungi(100);p__Ascomycota(100);c__Saccharomycetes(100);o__Saccharomycetales(100);f__Metschnikowiaceae(100);g__Metschnikowia(100);s__Metschnikowia_viticola(100); | Metschnikowia | 13 | Saprotroph | \|Undefined Saprotroph\| | Yeast | NULL | Probable | As nectar-tap_saprotroph (Põlme et al. 2020); Decay sugar-rich substrates (Põlme et al. 2020) | Põlme S, et al. 2020. Fungal Diversity 105:1-16 ((https://doi.org/10.1007/s13225-020-00466-2)) |
| Otu12004 | 2 | k__Fungi(100);p__Ascomycota(100);c__Saccharomycetes(100);o__Saccharomycetales(100);f__Metschnikowiaceae(100);g__Metschnikowia(100);g__Metschnikowia_unclassified(100); | Metschnikowia | 13 | Saprotroph | \|Undefined Saprotroph\| | Yeast | NULL | Probable | As nectar-tap_saprotroph (Põlme et al. 2020); Decay sugar-rich substrates (Põlme et al. 2020) | Põlme S, et al. 2020. Fungal Diversity 105:1-16 ((https://doi.org/10.1007/s13225-020-00466-2)) |
| Otu12005 | 2 | k__Fungi(100);p__Ascomycota(100);c__Saccharomycetes(100);o__Saccharomycetales(100);f__Metschnikowiaceae(100);g__Metschnikowia(100);g__Metschnikowia_unclassified(100); | Metschnikowia | 13 | Saprotroph | \|Undefined Saprotroph\| | Yeast | NULL | Probable | As nectar-tap_saprotroph (Põlme et al. 2020); Decay sugar-rich substrates (Põlme et al. 2020) | Põlme S, et al. 2020. Fungal Diversity 105:1-16 ((https://doi.org/10.1007/s13225-020-00466-2)) |
| Otu12006 | 2 | k__Fungi(100);p__Ascomycota(100);c__Saccharomycetes(100);o__Saccharomycetales(100);f__Saccharomycetales_family_Incertae_sedis(100);g__Candida(100);g__Candida_unclassified(100); | Candida | 13 | Pathotroph-Saprotroph-Symbiotroph | Animal Pathogen-Endophyte-Endosymbiont-Epiphyte-\|Undefined Saprotroph\| | Dimorphic Yeast-Yeast | NULL | Probable | As nectar-tap_saprotroph (Põlme et al. 2020); Decay sugar-rich substrates (Põlme et al. 2020); This asexual genus is polyphyletic and contain species that function very differently depend on what they are. Caution should be taken when interpreting guilds for Candida species. | Manolakaki D, et al. 2010. Virulence. 1: 367-375 ((https://doi.org/10.4161/viru.1.5.12796)); Mekbib SB, et al. 2011. Tropical Plant Pathology 36:233-240 ((https://doi.org/10.1590/S1982-56762011000400004)); Põlme S, et al. 2020. Fungal Diversity 105:1-16 ((https://doi.org/10.1007/s13225-020-00466-2)) |
| Otu12007 | 2 | k__Fungi(100);p__Ascomycota(100);c__Saccharomycetes(100);o__Saccharomycetales(100);f__Metschnikowiaceae(100);g__Metschnikowia(100);s__Metschnikowia_pulcherrima(100); | Metschnikowia | 13 | Saprotroph | \|Undefined Saprotroph\| | Yeast | NULL | Probable | As nectar-tap_saprotroph (Põlme et al. 2020); Decay sugar-rich substrates (Põlme et al. 2020) | Põlme S, et al. 2020. Fungal Diversity 105:1-16 ((https://doi.org/10.1007/s13225-020-00466-2)) |
| Otu12008 | 2 | k__Fungi(100);p__Ascomycota(100);c__Saccharomycetes(100);o__Saccharomycetales(100);f__Metschnikowiaceae(100);g__Metschnikowia(100);s__Metschnikowia_pulcherrima(100); | Metschnikowia | 13 | Saprotroph | \|Undefined Saprotroph\| | Yeast | NULL | Probable | As nectar-tap_saprotroph (Põlme et al. 2020); Decay sugar-rich substrates (Põlme et al. 2020) | Põlme S, et al. 2020. Fungal Diversity 105:1-16 ((https://doi.org/10.1007/s13225-020-00466-2)) |
| Otu12009 | 2 | k__Fungi(100);p__Ascomycota(100);c__Saccharomycetes(100);o__Saccharomycetales(100);f__Saccharomycetales_family_Incertae_sedis(100);f__Saccharomycetales_family_Incertae_sedis_unclassified(100);f__Saccharomycetales_family_Incertae_sedis_unclassified(100); | Saccharomycetales | 7 | Saprotroph | Undefined Saprotroph | Yeast | NULL | Possible | NULL | Sterkenburg E, et al. 2015. New Phytologist 207:1145-1158 ((https://doi.org/10.1111/nph.13426)) |
| Otu12010 | 2 | k__Fungi(100);p__Ascomycota(100);c__Saccharomycetes(100);o__Saccharomycetales(100);f__Metschnikowiaceae(100);g__Metschnikowia(100);g__Metschnikowia_unclassified(100); | Metschnikowia | 13 | Saprotroph | \|Undefined Saprotroph\| | Yeast | NULL | Probable | As nectar-tap_saprotroph (Põlme et al. 2020); Decay sugar-rich substrates (Põlme et al. 2020) | Põlme S, et al. 2020. Fungal Diversity 105:1-16 ((https://doi.org/10.1007/s13225-020-00466-2)) |
| Otu12011 | 2 | k__Fungi(100);p__Ascomycota(100);c__Saccharomycetes(100);o__Saccharomycetales(100);f__Saccharomycetales_family_Incertae_sedis(100);g__Candida(100);g__Candida_unclassified(100); | Candida | 13 | Pathotroph-Saprotroph-Symbiotroph | Animal Pathogen-Endophyte-Endosymbiont-Epiphyte-\|Undefined Saprotroph\| | Dimorphic Yeast-Yeast | NULL | Probable | As nectar-tap_saprotroph (Põlme et al. 2020); Decay sugar-rich substrates (Põlme et al. 2020); This asexual genus is polyphyletic and contain species that function very differently depend on what they are. Caution should be taken when interpreting guilds for Candida species. | Manolakaki D, et al. 2010. Virulence. 1: 367-375 ((https://doi.org/10.4161/viru.1.5.12796)); Mekbib SB, et al. 2011. Tropical Plant Pathology 36:233-240 ((https://doi.org/10.1590/S1982-56762011000400004)); Põlme S, et al. 2020. Fungal Diversity 105:1-16 ((https://doi.org/10.1007/s13225-020-00466-2)) |
| Otu12012 | 2 | k__Fungi(100);p__Ascomycota(100);c__Saccharomycetes(100);o__Saccharomycetales(100);f__Metschnikowiaceae(100);g__Metschnikowia(100);s__Metschnikowia_pulcherrima(100); | Metschnikowia | 13 | Saprotroph | \|Undefined Saprotroph\| | Yeast | NULL | Probable | As nectar-tap_saprotroph (Põlme et al. 2020); Decay sugar-rich substrates (Põlme et al. 2020) | Põlme S, et al. 2020. Fungal Diversity 105:1-16 ((https://doi.org/10.1007/s13225-020-00466-2)) |
| Otu12013 | 2 | k__Fungi(100);p__Ascomycota(100);c__Saccharomycetes(100);o__Saccharomycetales(100);f__Metschnikowiaceae(100);g__Metschnikowia(100);g__Metschnikowia_unclassified(100); | Metschnikowia | 13 | Saprotroph | \|Undefined Saprotroph\| | Yeast | NULL | Probable | As nectar-tap_saprotroph (Põlme et al. 2020); Decay sugar-rich substrates (Põlme et al. 2020) | Põlme S, et al. 2020. Fungal Diversity 105:1-16 ((https://doi.org/10.1007/s13225-020-00466-2)) |
| Otu12015 | 2 | k__Fungi(100);p__Ascomycota(100);c__Saccharomycetes(100);o__Saccharomycetales(100);f__Metschnikowiaceae(100);g__Metschnikowia(100);s__Metschnikowia_pulcherrima(100); | Metschnikowia | 13 | Saprotroph | \|Undefined Saprotroph\| | Yeast | NULL | Probable | As nectar-tap_saprotroph (Põlme et al. 2020); Decay sugar-rich substrates (Põlme et al. 2020) | Põlme S, et al. 2020. Fungal Diversity 105:1-16 ((https://doi.org/10.1007/s13225-020-00466-2)) |
| Otu12016 | 2 | k__Fungi(100);p__Ascomycota(100);c__Saccharomycetes(100);o__Saccharomycetales(100);f__Metschnikowiaceae(100);g__Metschnikowia(100);g__Metschnikowia_unclassified(100); | Metschnikowia | 13 | Saprotroph | \|Undefined Saprotroph\| | Yeast | NULL | Probable | As nectar-tap_saprotroph (Põlme et al. 2020); Decay sugar-rich substrates (Põlme et al. 2020) | Põlme S, et al. 2020. Fungal Diversity 105:1-16 ((https://doi.org/10.1007/s13225-020-00466-2)) |
| Otu12017 | 2 | k__Fungi(100);p__Ascomycota(100);c__Saccharomycetes(100);o__Saccharomycetales(100);f__Metschnikowiaceae(100);g__Metschnikowia(100);g__Metschnikowia_unclassified(100); | Metschnikowia | 13 | Saprotroph | \|Undefined Saprotroph\| | Yeast | NULL | Probable | As nectar-tap_saprotroph (Põlme et al. 2020); Decay sugar-rich substrates (Põlme et al. 2020) | Põlme S, et al. 2020. Fungal Diversity 105:1-16 ((https://doi.org/10.1007/s13225-020-00466-2)) |
| Otu12018 | 2 | k__Fungi(100);p__Ascomycota(100);c__Saccharomycetes(100);o__Saccharomycetales(100);f__Metschnikowiaceae(100);g__Metschnikowia(100);s__Metschnikowia_pulcherrima(100); | Metschnikowia | 13 | Saprotroph | \|Undefined Saprotroph\| | Yeast | NULL | Probable | As nectar-tap_saprotroph (Põlme et al. 2020); Decay sugar-rich substrates (Põlme et al. 2020) | Põlme S, et al. 2020. Fungal Diversity 105:1-16 ((https://doi.org/10.1007/s13225-020-00466-2)) |
| Otu12019 | 2 | k__Fungi(100);p__Ascomycota(100);c__Saccharomycetes(100);o__Saccharomycetales(100);f__Metschnikowiaceae(100);g__Metschnikowia(100);s__Metschnikowia_pulcherrima(100); | Metschnikowia | 13 | Saprotroph | \|Undefined Saprotroph\| | Yeast | NULL | Probable | As nectar-tap_saprotroph (Põlme et al. 2020); Decay sugar-rich substrates (Põlme et al. 2020) | Põlme S, et al. 2020. Fungal Diversity 105:1-16 ((https://doi.org/10.1007/s13225-020-00466-2)) |
| Otu12022 | 2 | k__Fungi(100);p__Ascomycota(100);c__Saccharomycetes(100);o__Saccharomycetales(100);f__Metschnikowiaceae(100);g__Metschnikowia(100);s__Metschnikowia_pulcherrima(100); | Metschnikowia | 13 | Saprotroph | \|Undefined Saprotroph\| | Yeast | NULL | Probable | As nectar-tap_saprotroph (Põlme et al. 2020); Decay sugar-rich substrates (Põlme et al. 2020) | Põlme S, et al. 2020. Fungal Diversity 105:1-16 ((https://doi.org/10.1007/s13225-020-00466-2)) |
| Otu12028 | 2 | k__Fungi(100);p__Ascomycota(100);c__Saccharomycetes(100);o__Saccharomycetales(100);f__Metschnikowiaceae(100);g__Metschnikowia(100);s__Metschnikowia_pulcherrima(100); | Metschnikowia | 13 | Saprotroph | \|Undefined Saprotroph\| | Yeast | NULL | Probable | As nectar-tap_saprotroph (Põlme et al. 2020); Decay sugar-rich substrates (Põlme et al. 2020) | Põlme S, et al. 2020. Fungal Diversity 105:1-16 ((https://doi.org/10.1007/s13225-020-00466-2)) |
| Otu12029 | 2 | k__Fungi(100);p__Ascomycota(100);c__Saccharomycetes(100);o__Saccharomycetales(100);f__Metschnikowiaceae(100);g__Metschnikowia(100);s__Metschnikowia_pulcherrima(100); | Metschnikowia | 13 | Saprotroph | \|Undefined Saprotroph\| | Yeast | NULL | Probable | As nectar-tap_saprotroph (Põlme et al. 2020); Decay sugar-rich substrates (Põlme et al. 2020) | Põlme S, et al. 2020. Fungal Diversity 105:1-16 ((https://doi.org/10.1007/s13225-020-00466-2)) |
| Otu12030 | 2 | k__Fungi(100);p__Ascomycota(100);c__Saccharomycetes(100);o__Saccharomycetales(100);f__Metschnikowiaceae(100);g__Metschnikowia(100);s__Metschnikowia_pulcherrima(100); | Metschnikowia | 13 | Saprotroph | \|Undefined Saprotroph\| | Yeast | NULL | Probable | As nectar-tap_saprotroph (Põlme et al. 2020); Decay sugar-rich substrates (Põlme et al. 2020) | Põlme S, et al. 2020. Fungal Diversity 105:1-16 ((https://doi.org/10.1007/s13225-020-00466-2)) |
| Otu12034 | 2 | k__Fungi(100);p__Ascomycota(100);c__Saccharomycetes(100);o__Saccharomycetales(100);f__Metschnikowiaceae(100);g__Metschnikowia(100);s__Metschnikowia_pulcherrima(100); | Metschnikowia | 13 | Saprotroph | \|Undefined Saprotroph\| | Yeast | NULL | Probable | As nectar-tap_saprotroph (Põlme et al. 2020); Decay sugar-rich substrates (Põlme et al. 2020) | Põlme S, et al. 2020. Fungal Diversity 105:1-16 ((https://doi.org/10.1007/s13225-020-00466-2)) |
| Otu12037 | 2 | k__Fungi(100);p__Ascomycota(100);c__Saccharomycetes(100);o__Saccharomycetales(100);f__Saccharomycetales_family_Incertae_sedis(100);f__Saccharomycetales_family_Incertae_sedis_unclassified(100);f__Saccharomycetales_family_Incertae_sedis_unclassified(100); | Saccharomycetales | 7 | Saprotroph | Undefined Saprotroph | Yeast | NULL | Possible | NULL | Sterkenburg E, et al. 2015. New Phytologist 207:1145-1158 ((https://doi.org/10.1111/nph.13426)) |
| Otu12039 | 2 | k__Fungi(100);p__Ascomycota(100);c__Saccharomycetes(100);o__Saccharomycetales(100);f__Metschnikowiaceae(100);g__Metschnikowia(100);s__Metschnikowia_pulcherrima(100); | Metschnikowia | 13 | Saprotroph | \|Undefined Saprotroph\| | Yeast | NULL | Probable | As nectar-tap_saprotroph (Põlme et al. 2020); Decay sugar-rich substrates (Põlme et al. 2020) | Põlme S, et al. 2020. Fungal Diversity 105:1-16 ((https://doi.org/10.1007/s13225-020-00466-2)) |
| Otu12040 | 2 | k__Fungi(100);p__Ascomycota(100);c__Saccharomycetes(100);o__Saccharomycetales(100);o__Saccharomycetales_unclassified(100);o__Saccharomycetales_unclassified(100);o__Saccharomycetales_unclassified(100); | Saccharomycetales | 7 | Saprotroph | Undefined Saprotroph | Yeast | NULL | Possible | NULL | Sterkenburg E, et al. 2015. New Phytologist 207:1145-1158 ((https://doi.org/10.1111/nph.13426)) |
| Otu12042 | 2 | k__Fungi(100);p__Ascomycota(100);c__Saccharomycetes(100);o__Saccharomycetales(100);f__Metschnikowiaceae(100);g__Metschnikowia(100);g__Metschnikowia_unclassified(100); | Metschnikowia | 13 | Saprotroph | \|Undefined Saprotroph\| | Yeast | NULL | Probable | As nectar-tap_saprotroph (Põlme et al. 2020); Decay sugar-rich substrates (Põlme et al. 2020) | Põlme S, et al. 2020. Fungal Diversity 105:1-16 ((https://doi.org/10.1007/s13225-020-00466-2)) |
| Otu12043 | 2 | k__Fungi(100);p__Ascomycota(100);c__Saccharomycetes(100);o__Saccharomycetales(100);f__Metschnikowiaceae(100);g__Metschnikowia(100);s__Metschnikowia_pulcherrima(100); | Metschnikowia | 13 | Saprotroph | \|Undefined Saprotroph\| | Yeast | NULL | Probable | As nectar-tap_saprotroph (Põlme et al. 2020); Decay sugar-rich substrates (Põlme et al. 2020) | Põlme S, et al. 2020. Fungal Diversity 105:1-16 ((https://doi.org/10.1007/s13225-020-00466-2)) |
| Otu12045 | 2 | k__Fungi(100);p__Ascomycota(100);c__Saccharomycetes(100);o__Saccharomycetales(100);f__Metschnikowiaceae(100);g__Metschnikowia(100);s__Metschnikowia_pulcherrima(100); | Metschnikowia | 13 | Saprotroph | \|Undefined Saprotroph\| | Yeast | NULL | Probable | As nectar-tap_saprotroph (Põlme et al. 2020); Decay sugar-rich substrates (Põlme et al. 2020) | Põlme S, et al. 2020. Fungal Diversity 105:1-16 ((https://doi.org/10.1007/s13225-020-00466-2)) |
| Otu12046 | 2 | k__Fungi(100);p__Ascomycota(100);c__Saccharomycetes(100);o__Saccharomycetales(100);f__Metschnikowiaceae(100);g__Metschnikowia(100);s__Metschnikowia_pulcherrima(100); | Metschnikowia | 13 | Saprotroph | \|Undefined Saprotroph\| | Yeast | NULL | Probable | As nectar-tap_saprotroph (Põlme et al. 2020); Decay sugar-rich substrates (Põlme et al. 2020) | Põlme S, et al. 2020. Fungal Diversity 105:1-16 ((https://doi.org/10.1007/s13225-020-00466-2)) |
| Otu12047 | 2 | k__Fungi(100);p__Ascomycota(100);c__Saccharomycetes(100);o__Saccharomycetales(100);f__Metschnikowiaceae(100);g__Metschnikowia(100);g__Metschnikowia_unclassified(100); | Metschnikowia | 13 | Saprotroph | \|Undefined Saprotroph\| | Yeast | NULL | Probable | As nectar-tap_saprotroph (Põlme et al. 2020); Decay sugar-rich substrates (Põlme et al. 2020) | Põlme S, et al. 2020. Fungal Diversity 105:1-16 ((https://doi.org/10.1007/s13225-020-00466-2)) |
| Otu12049 | 2 | k__Fungi(100);p__Ascomycota(100);c__Saccharomycetes(100);o__Saccharomycetales(100);f__Metschnikowiaceae(100);g__Metschnikowia(100);s__Metschnikowia_pulcherrima(100); | Metschnikowia | 13 | Saprotroph | \|Undefined Saprotroph\| | Yeast | NULL | Probable | As nectar-tap_saprotroph (Põlme et al. 2020); Decay sugar-rich substrates (Põlme et al. 2020) | Põlme S, et al. 2020. Fungal Diversity 105:1-16 ((https://doi.org/10.1007/s13225-020-00466-2)) |
| Otu12050 | 2 | k__Fungi(100);p__Basidiomycota(100);c__Microbotryomycetes(100);o__Sporidiobolales(100);f__Sporidiobolales_family_Incertae_sedis(100);g__Sporobolomyces(100);s__Sporobolomyces_lactosus(100); | Sporobolomyces | 13 | Pathotroph-Saprotroph | \|Fungal Parasite\|-Plant Saprotroph | Yeast | NULL | Probable | As litter_saprotroph (Põlme et al. 2020); As mycoparasite (Põlme et al. 2020); Decay fungal material (Põlme et al. 2020) | Tedersoo L, et al. 2014. Science 346:e1256688 ((https://doi.org/10.1126/science.1256688)); Purahong W, et al. 2016. Molecular Ecology 25:4059-4074 ((https://doi.org/10.1111/mec.13739)); Põlme S, et al. 2020. Fungal Diversity 105:1-16 ((https://doi.org/10.1007/s13225-020-00466-2)) |
| Otu12051 | 2 | k__Fungi(100);p__Ascomycota(100);c__Saccharomycetes(100);o__Saccharomycetales(100);f__Metschnikowiaceae(100);g__Metschnikowia(100);s__Metschnikowia_pulcherrima(100); | Metschnikowia | 13 | Saprotroph | \|Undefined Saprotroph\| | Yeast | NULL | Probable | As nectar-tap_saprotroph (Põlme et al. 2020); Decay sugar-rich substrates (Põlme et al. 2020) | Põlme S, et al. 2020. Fungal Diversity 105:1-16 ((https://doi.org/10.1007/s13225-020-00466-2)) |
| Otu12053 | 2 | k__Fungi(100);p__Ascomycota(100);c__Saccharomycetes(100);o__Saccharomycetales(100);f__Metschnikowiaceae(100);g__Metschnikowia(100);g__Metschnikowia_unclassified(100); | Metschnikowia | 13 | Saprotroph | \|Undefined Saprotroph\| | Yeast | NULL | Probable | As nectar-tap_saprotroph (Põlme et al. 2020); Decay sugar-rich substrates (Põlme et al. 2020) | Põlme S, et al. 2020. Fungal Diversity 105:1-16 ((https://doi.org/10.1007/s13225-020-00466-2)) |
| Otu12054 | 2 | k__Fungi(100);p__Ascomycota(100);c__Saccharomycetes(100);o__Saccharomycetales(100);f__Metschnikowiaceae(100);g__Metschnikowia(100);g__Metschnikowia_unclassified(100); | Metschnikowia | 13 | Saprotroph | \|Undefined Saprotroph\| | Yeast | NULL | Probable | As nectar-tap_saprotroph (Põlme et al. 2020); Decay sugar-rich substrates (Põlme et al. 2020) | Põlme S, et al. 2020. Fungal Diversity 105:1-16 ((https://doi.org/10.1007/s13225-020-00466-2)) |
| Otu12056 | 2 | k__Fungi(100);p__Ascomycota(100);c__Saccharomycetes(100);o__Saccharomycetales(100);f__Metschnikowiaceae(100);g__Metschnikowia(100);s__Metschnikowia_pulcherrima(100); | Metschnikowia | 13 | Saprotroph | \|Undefined Saprotroph\| | Yeast | NULL | Probable | As nectar-tap_saprotroph (Põlme et al. 2020); Decay sugar-rich substrates (Põlme et al. 2020) | Põlme S, et al. 2020. Fungal Diversity 105:1-16 ((https://doi.org/10.1007/s13225-020-00466-2)) |
| Otu12057 | 2 | k__Fungi(100);p__Ascomycota(100);c__Saccharomycetes(100);o__Saccharomycetales(100);f__Metschnikowiaceae(100);g__Metschnikowia(100);s__Metschnikowia_viticola(100); | Metschnikowia | 13 | Saprotroph | \|Undefined Saprotroph\| | Yeast | NULL | Probable | As nectar-tap_saprotroph (Põlme et al. 2020); Decay sugar-rich substrates (Põlme et al. 2020) | Põlme S, et al. 2020. Fungal Diversity 105:1-16 ((https://doi.org/10.1007/s13225-020-00466-2)) |
| Otu12058 | 2 | k__Fungi(100);p__Ascomycota(100);c__Saccharomycetes(100);o__Saccharomycetales(100);f__Metschnikowiaceae(100);g__Metschnikowia(100);s__Metschnikowia_pulcherrima(100); | Metschnikowia | 13 | Saprotroph | \|Undefined Saprotroph\| | Yeast | NULL | Probable | As nectar-tap_saprotroph (Põlme et al. 2020); Decay sugar-rich substrates (Põlme et al. 2020) | Põlme S, et al. 2020. Fungal Diversity 105:1-16 ((https://doi.org/10.1007/s13225-020-00466-2)) |
| Otu12059 | 2 | k__Fungi(100);p__Ascomycota(100);c__Saccharomycetes(100);o__Saccharomycetales(100);f__Metschnikowiaceae(100);g__Metschnikowia(100);s__Metschnikowia_pulcherrima(100); | Metschnikowia | 13 | Saprotroph | \|Undefined Saprotroph\| | Yeast | NULL | Probable | As nectar-tap_saprotroph (Põlme et al. 2020); Decay sugar-rich substrates (Põlme et al. 2020) | Põlme S, et al. 2020. Fungal Diversity 105:1-16 ((https://doi.org/10.1007/s13225-020-00466-2)) |
| Otu12060 | 2 | k__Fungi(100);p__Ascomycota(100);c__Saccharomycetes(100);o__Saccharomycetales(100);f__Metschnikowiaceae(100);g__Metschnikowia(100);s__Metschnikowia_pulcherrima(100); | Metschnikowia | 13 | Saprotroph | \|Undefined Saprotroph\| | Yeast | NULL | Probable | As nectar-tap_saprotroph (Põlme et al. 2020); Decay sugar-rich substrates (Põlme et al. 2020) | Põlme S, et al. 2020. Fungal Diversity 105:1-16 ((https://doi.org/10.1007/s13225-020-00466-2)) |
| Otu12061 | 2 | k__Fungi(100);p__Ascomycota(100);c__Saccharomycetes(100);o__Saccharomycetales(100);f__Metschnikowiaceae(100);g__Metschnikowia(100);g__Metschnikowia_unclassified(100); | Metschnikowia | 13 | Saprotroph | \|Undefined Saprotroph\| | Yeast | NULL | Probable | As nectar-tap_saprotroph (Põlme et al. 2020); Decay sugar-rich substrates (Põlme et al. 2020) | Põlme S, et al. 2020. Fungal Diversity 105:1-16 ((https://doi.org/10.1007/s13225-020-00466-2)) |
| Otu12062 | 2 | k__Fungi(100);p__Ascomycota(100);c__Saccharomycetes(100);o__Saccharomycetales(100);f__Metschnikowiaceae(100);g__Metschnikowia(100);g__Metschnikowia_unclassified(100); | Metschnikowia | 13 | Saprotroph | \|Undefined Saprotroph\| | Yeast | NULL | Probable | As nectar-tap_saprotroph (Põlme et al. 2020); Decay sugar-rich substrates (Põlme et al. 2020) | Põlme S, et al. 2020. Fungal Diversity 105:1-16 ((https://doi.org/10.1007/s13225-020-00466-2)) |
| Otu12063 | 2 | k__Fungi(100);p__Basidiomycota(100);c__Microbotryomycetes(100);o__Sporidiobolales(100);f__Sporidiobolales_family_Incertae_sedis(100);g__Sporobolomyces(100);s__Sporobolomyces_lactosus(100); | Sporobolomyces | 13 | Pathotroph-Saprotroph | \|Fungal Parasite\|-Plant Saprotroph | Yeast | NULL | Probable | As litter_saprotroph (Põlme et al. 2020); As mycoparasite (Põlme et al. 2020); Decay fungal material (Põlme et al. 2020) | Tedersoo L, et al. 2014. Science 346:e1256688 ((https://doi.org/10.1126/science.1256688)); Purahong W, et al. 2016. Molecular Ecology 25:4059-4074 ((https://doi.org/10.1111/mec.13739)); Põlme S, et al. 2020. Fungal Diversity 105:1-16 ((https://doi.org/10.1007/s13225-020-00466-2)) |
| Otu12064 | 2 | k__Fungi(100);p__Ascomycota(100);c__Saccharomycetes(100);o__Saccharomycetales(100);f__Metschnikowiaceae(100);g__Metschnikowia(100);g__Metschnikowia_unclassified(100); | Metschnikowia | 13 | Saprotroph | \|Undefined Saprotroph\| | Yeast | NULL | Probable | As nectar-tap_saprotroph (Põlme et al. 2020); Decay sugar-rich substrates (Põlme et al. 2020) | Põlme S, et al. 2020. Fungal Diversity 105:1-16 ((https://doi.org/10.1007/s13225-020-00466-2)) |
| Otu12066 | 2 | k__Fungi(100);p__Ascomycota(100);c__Saccharomycetes(100);o__Saccharomycetales(100);f__Metschnikowiaceae(100);g__Metschnikowia(100);g__Metschnikowia_unclassified(100); | Metschnikowia | 13 | Saprotroph | \|Undefined Saprotroph\| | Yeast | NULL | Probable | As nectar-tap_saprotroph (Põlme et al. 2020); Decay sugar-rich substrates (Põlme et al. 2020) | Põlme S, et al. 2020. Fungal Diversity 105:1-16 ((https://doi.org/10.1007/s13225-020-00466-2)) |
| Otu12067 | 2 | k__Fungi(100);p__Ascomycota(100);c__Saccharomycetes(100);o__Saccharomycetales(100);f__Metschnikowiaceae(100);g__Metschnikowia(100);g__Metschnikowia_unclassified(100); | Metschnikowia | 13 | Saprotroph | \|Undefined Saprotroph\| | Yeast | NULL | Probable | As nectar-tap_saprotroph (Põlme et al. 2020); Decay sugar-rich substrates (Põlme et al. 2020) | Põlme S, et al. 2020. Fungal Diversity 105:1-16 ((https://doi.org/10.1007/s13225-020-00466-2)) |
| Otu12068 | 2 | k__Fungi(100);p__Ascomycota(100);c__Saccharomycetes(100);o__Saccharomycetales(100);f__Metschnikowiaceae(100);g__Metschnikowia(100);g__Metschnikowia_unclassified(100); | Metschnikowia | 13 | Saprotroph | \|Undefined Saprotroph\| | Yeast | NULL | Probable | As nectar-tap_saprotroph (Põlme et al. 2020); Decay sugar-rich substrates (Põlme et al. 2020) | Põlme S, et al. 2020. Fungal Diversity 105:1-16 ((https://doi.org/10.1007/s13225-020-00466-2)) |
| Otu12069 | 2 | k__Fungi(100);p__Ascomycota(100);c__Saccharomycetes(100);o__Saccharomycetales(100);f__Metschnikowiaceae(100);g__Metschnikowia(100);g__Metschnikowia_unclassified(100); | Metschnikowia | 13 | Saprotroph | \|Undefined Saprotroph\| | Yeast | NULL | Probable | As nectar-tap_saprotroph (Põlme et al. 2020); Decay sugar-rich substrates (Põlme et al. 2020) | Põlme S, et al. 2020. Fungal Diversity 105:1-16 ((https://doi.org/10.1007/s13225-020-00466-2)) |
| Otu12072 | 2 | k__Fungi(100);p__Ascomycota(100);c__Saccharomycetes(100);o__Saccharomycetales(100);f__Metschnikowiaceae(100);g__Metschnikowia(100);g__Metschnikowia_unclassified(100); | Metschnikowia | 13 | Saprotroph | \|Undefined Saprotroph\| | Yeast | NULL | Probable | As nectar-tap_saprotroph (Põlme et al. 2020); Decay sugar-rich substrates (Põlme et al. 2020) | Põlme S, et al. 2020. Fungal Diversity 105:1-16 ((https://doi.org/10.1007/s13225-020-00466-2)) |
| Otu12073 | 2 | k__Fungi(100);p__Ascomycota(100);c__Saccharomycetes(100);o__Saccharomycetales(100);f__Metschnikowiaceae(100);g__Metschnikowia(100);g__Metschnikowia_unclassified(100); | Metschnikowia | 13 | Saprotroph | \|Undefined Saprotroph\| | Yeast | NULL | Probable | As nectar-tap_saprotroph (Põlme et al. 2020); Decay sugar-rich substrates (Põlme et al. 2020) | Põlme S, et al. 2020. Fungal Diversity 105:1-16 ((https://doi.org/10.1007/s13225-020-00466-2)) |
| Otu12074 | 2 | k__Fungi(100);p__Ascomycota(100);c__Saccharomycetes(100);o__Saccharomycetales(100);f__Metschnikowiaceae(100);g__Metschnikowia(100);s__Metschnikowia_pulcherrima(100); | Metschnikowia | 13 | Saprotroph | \|Undefined Saprotroph\| | Yeast | NULL | Probable | As nectar-tap_saprotroph (Põlme et al. 2020); Decay sugar-rich substrates (Põlme et al. 2020) | Põlme S, et al. 2020. Fungal Diversity 105:1-16 ((https://doi.org/10.1007/s13225-020-00466-2)) |
| Otu12075 | 2 | k__Fungi(100);p__Ascomycota(100);c__Saccharomycetes(100);o__Saccharomycetales(100);f__Metschnikowiaceae(100);g__Metschnikowia(100);g__Metschnikowia_unclassified(100); | Metschnikowia | 13 | Saprotroph | \|Undefined Saprotroph\| | Yeast | NULL | Probable | As nectar-tap_saprotroph (Põlme et al. 2020); Decay sugar-rich substrates (Põlme et al. 2020) | Põlme S, et al. 2020. Fungal Diversity 105:1-16 ((https://doi.org/10.1007/s13225-020-00466-2)) |
| Otu12076 | 2 | k__Fungi(100);p__Ascomycota(100);c__Saccharomycetes(100);o__Saccharomycetales(100);f__Metschnikowiaceae(100);g__Metschnikowia(100);s__Metschnikowia_pulcherrima(100); | Metschnikowia | 13 | Saprotroph | \|Undefined Saprotroph\| | Yeast | NULL | Probable | As nectar-tap_saprotroph (Põlme et al. 2020); Decay sugar-rich substrates (Põlme et al. 2020) | Põlme S, et al. 2020. Fungal Diversity 105:1-16 ((https://doi.org/10.1007/s13225-020-00466-2)) |
| Otu12077 | 2 | k__Fungi(100);p__Ascomycota(100);c__Saccharomycetes(100);o__Saccharomycetales(100);f__Metschnikowiaceae(100);g__Metschnikowia(100);g__Metschnikowia_unclassified(100); | Metschnikowia | 13 | Saprotroph | \|Undefined Saprotroph\| | Yeast | NULL | Probable | As nectar-tap_saprotroph (Põlme et al. 2020); Decay sugar-rich substrates (Põlme et al. 2020) | Põlme S, et al. 2020. Fungal Diversity 105:1-16 ((https://doi.org/10.1007/s13225-020-00466-2)) |
| Otu12078 | 2 | k__Fungi(100);p__Ascomycota(100);c__Saccharomycetes(100);o__Saccharomycetales(100);f__Metschnikowiaceae(100);g__Metschnikowia(100);g__Metschnikowia_unclassified(100); | Metschnikowia | 13 | Saprotroph | \|Undefined Saprotroph\| | Yeast | NULL | Probable | As nectar-tap_saprotroph (Põlme et al. 2020); Decay sugar-rich substrates (Põlme et al. 2020) | Põlme S, et al. 2020. Fungal Diversity 105:1-16 ((https://doi.org/10.1007/s13225-020-00466-2)) |
| Otu12081 | 2 | k__Fungi(100);p__Ascomycota(100);c__Saccharomycetes(100);o__Saccharomycetales(100);f__Saccharomycetales_family_Incertae_sedis(100);g__Candida(100);g__Candida_unclassified(100); | Candida | 13 | Pathotroph-Saprotroph-Symbiotroph | Animal Pathogen-Endophyte-Endosymbiont-Epiphyte-\|Undefined Saprotroph\| | Dimorphic Yeast-Yeast | NULL | Probable | As nectar-tap_saprotroph (Põlme et al. 2020); Decay sugar-rich substrates (Põlme et al. 2020); This asexual genus is polyphyletic and contain species that function very differently depend on what they are. Caution should be taken when interpreting guilds for Candida species. | Manolakaki D, et al. 2010. Virulence. 1: 367-375 ((https://doi.org/10.4161/viru.1.5.12796)); Mekbib SB, et al. 2011. Tropical Plant Pathology 36:233-240 ((https://doi.org/10.1590/S1982-56762011000400004)); Põlme S, et al. 2020. Fungal Diversity 105:1-16 ((https://doi.org/10.1007/s13225-020-00466-2)) |
| Otu12082 | 2 | k__Fungi(100);p__Ascomycota(100);c__Saccharomycetes(100);o__Saccharomycetales(100);f__Metschnikowiaceae(100);g__Metschnikowia(100);s__Metschnikowia_pulcherrima(100); | Metschnikowia | 13 | Saprotroph | \|Undefined Saprotroph\| | Yeast | NULL | Probable | As nectar-tap_saprotroph (Põlme et al. 2020); Decay sugar-rich substrates (Põlme et al. 2020) | Põlme S, et al. 2020. Fungal Diversity 105:1-16 ((https://doi.org/10.1007/s13225-020-00466-2)) |
| Otu12083 | 2 | k__Fungi(100);p__Ascomycota(100);c__Saccharomycetes(100);o__Saccharomycetales(100);f__Metschnikowiaceae(100);g__Metschnikowia(100);g__Metschnikowia_unclassified(100); | Metschnikowia | 13 | Saprotroph | \|Undefined Saprotroph\| | Yeast | NULL | Probable | As nectar-tap_saprotroph (Põlme et al. 2020); Decay sugar-rich substrates (Põlme et al. 2020) | Põlme S, et al. 2020. Fungal Diversity 105:1-16 ((https://doi.org/10.1007/s13225-020-00466-2)) |
| Otu12085 | 2 | k__Fungi(100);p__Ascomycota(100);c__Saccharomycetes(100);o__Saccharomycetales(100);o__Saccharomycetales_unclassified(100);o__Saccharomycetales_unclassified(100);o__Saccharomycetales_unclassified(100); | Saccharomycetales | 7 | Saprotroph | Undefined Saprotroph | Yeast | NULL | Possible | NULL | Sterkenburg E, et al. 2015. New Phytologist 207:1145-1158 ((https://doi.org/10.1111/nph.13426)) |
| Otu12086 | 2 | k__Fungi(100);p__Ascomycota(100);c__Saccharomycetes(100);o__Saccharomycetales(100);f__Metschnikowiaceae(100);g__Metschnikowia(100);g__Metschnikowia_unclassified(100); | Metschnikowia | 13 | Saprotroph | \|Undefined Saprotroph\| | Yeast | NULL | Probable | As nectar-tap_saprotroph (Põlme et al. 2020); Decay sugar-rich substrates (Põlme et al. 2020) | Põlme S, et al. 2020. Fungal Diversity 105:1-16 ((https://doi.org/10.1007/s13225-020-00466-2)) |
| Otu12088 | 2 | k__Fungi(100);p__Ascomycota(100);c__Saccharomycetes(100);o__Saccharomycetales(100);f__Metschnikowiaceae(100);g__Metschnikowia(100);g__Metschnikowia_unclassified(100); | Metschnikowia | 13 | Saprotroph | \|Undefined Saprotroph\| | Yeast | NULL | Probable | As nectar-tap_saprotroph (Põlme et al. 2020); Decay sugar-rich substrates (Põlme et al. 2020) | Põlme S, et al. 2020. Fungal Diversity 105:1-16 ((https://doi.org/10.1007/s13225-020-00466-2)) |
| Otu12089 | 2 | k__Fungi(100);p__Ascomycota(100);c__Saccharomycetes(100);o__Saccharomycetales(100);f__Metschnikowiaceae(100);g__Metschnikowia(100);s__Metschnikowia_pulcherrima(100); | Metschnikowia | 13 | Saprotroph | \|Undefined Saprotroph\| | Yeast | NULL | Probable | As nectar-tap_saprotroph (Põlme et al. 2020); Decay sugar-rich substrates (Põlme et al. 2020) | Põlme S, et al. 2020. Fungal Diversity 105:1-16 ((https://doi.org/10.1007/s13225-020-00466-2)) |
| Otu12092 | 2 | k__Fungi(100);p__Ascomycota(100);c__Leotiomycetes(100);o__Helotiales(100);f__Hyaloscyphaceae(100);g__Lachnum(100);s__Lachnum_spartinae(100); | Lachnum | 13 | Saprotroph-Symbiotroph | Endophyte-\|Plant Saprotroph\|-Undefined Saprotroph-Wood Saprotroph | Microfungus | NULL | Probable | As litter_saprotroph (Põlme et al. 2020); As root_endophyte (Põlme et al. 2020); Decay wood, leaf, fruit, seed (Põlme et al. 2020) | Tedersoo L, et al. 2014. Science 346:e1256688 ((https://doi.org/10.1126/science.1256688)); Põlme S, et al. 2020. Fungal Diversity 105:1-16 ((https://doi.org/10.1007/s13225-020-00466-2)) |
| Otu12093 | 2 | k__Fungi(100);p__Ascomycota(100);c__Saccharomycetes(100);o__Saccharomycetales(100);f__Metschnikowiaceae(100);g__Metschnikowia(100);s__Metschnikowia_pulcherrima(100); | Metschnikowia | 13 | Saprotroph | \|Undefined Saprotroph\| | Yeast | NULL | Probable | As nectar-tap_saprotroph (Põlme et al. 2020); Decay sugar-rich substrates (Põlme et al. 2020) | Põlme S, et al. 2020. Fungal Diversity 105:1-16 ((https://doi.org/10.1007/s13225-020-00466-2)) |
| Otu12097 | 2 | k__Fungi(100);p__Ascomycota(100);c__Saccharomycetes(100);o__Saccharomycetales(100);f__Metschnikowiaceae(100);g__Metschnikowia(100);g__Metschnikowia_unclassified(100); | Metschnikowia | 13 | Saprotroph | \|Undefined Saprotroph\| | Yeast | NULL | Probable | As nectar-tap_saprotroph (Põlme et al. 2020); Decay sugar-rich substrates (Põlme et al. 2020) | Põlme S, et al. 2020. Fungal Diversity 105:1-16 ((https://doi.org/10.1007/s13225-020-00466-2)) |
| Otu12098 | 2 | k__Fungi(100);p__Ascomycota(100);c__Saccharomycetes(100);o__Saccharomycetales(100);f__Metschnikowiaceae(100);g__Metschnikowia(100);g__Metschnikowia_unclassified(100); | Metschnikowia | 13 | Saprotroph | \|Undefined Saprotroph\| | Yeast | NULL | Probable | As nectar-tap_saprotroph (Põlme et al. 2020); Decay sugar-rich substrates (Põlme et al. 2020) | Põlme S, et al. 2020. Fungal Diversity 105:1-16 ((https://doi.org/10.1007/s13225-020-00466-2)) |
| Otu12099 | 2 | k__Fungi(100);p__Ascomycota(100);c__Saccharomycetes(100);o__Saccharomycetales(100);f__Metschnikowiaceae(100);g__Metschnikowia(100);g__Metschnikowia_unclassified(100); | Metschnikowia | 13 | Saprotroph | \|Undefined Saprotroph\| | Yeast | NULL | Probable | As nectar-tap_saprotroph (Põlme et al. 2020); Decay sugar-rich substrates (Põlme et al. 2020) | Põlme S, et al. 2020. Fungal Diversity 105:1-16 ((https://doi.org/10.1007/s13225-020-00466-2)) |
| Otu12100 | 2 | k__Fungi(100);p__Ascomycota(100);c__Saccharomycetes(100);o__Saccharomycetales(100);f__Metschnikowiaceae(100);g__Metschnikowia(100);s__Metschnikowia_pulcherrima(100); | Metschnikowia | 13 | Saprotroph | \|Undefined Saprotroph\| | Yeast | NULL | Probable | As nectar-tap_saprotroph (Põlme et al. 2020); Decay sugar-rich substrates (Põlme et al. 2020) | Põlme S, et al. 2020. Fungal Diversity 105:1-16 ((https://doi.org/10.1007/s13225-020-00466-2)) |
| Otu12102 | 2 | k__Fungi(100);p__Ascomycota(100);c__Saccharomycetes(100);o__Saccharomycetales(100);f__Metschnikowiaceae(100);g__Metschnikowia(100);g__Metschnikowia_unclassified(100); | Metschnikowia | 13 | Saprotroph | \|Undefined Saprotroph\| | Yeast | NULL | Probable | As nectar-tap_saprotroph (Põlme et al. 2020); Decay sugar-rich substrates (Põlme et al. 2020) | Põlme S, et al. 2020. Fungal Diversity 105:1-16 ((https://doi.org/10.1007/s13225-020-00466-2)) |
| Otu12104 | 2 | k__Fungi(100);p__Ascomycota(100);c__Saccharomycetes(100);o__Saccharomycetales(100);f__Metschnikowiaceae(100);g__Metschnikowia(100);s__Metschnikowia_pulcherrima(100); | Metschnikowia | 13 | Saprotroph | \|Undefined Saprotroph\| | Yeast | NULL | Probable | As nectar-tap_saprotroph (Põlme et al. 2020); Decay sugar-rich substrates (Põlme et al. 2020) | Põlme S, et al. 2020. Fungal Diversity 105:1-16 ((https://doi.org/10.1007/s13225-020-00466-2)) |
| Otu12106 | 2 | k__Fungi(100);p__Ascomycota(100);c__Saccharomycetes(100);o__Saccharomycetales(100);f__Metschnikowiaceae(100);g__Metschnikowia(100);s__Metschnikowia_pulcherrima(100); | Metschnikowia | 13 | Saprotroph | \|Undefined Saprotroph\| | Yeast | NULL | Probable | As nectar-tap_saprotroph (Põlme et al. 2020); Decay sugar-rich substrates (Põlme et al. 2020) | Põlme S, et al. 2020. Fungal Diversity 105:1-16 ((https://doi.org/10.1007/s13225-020-00466-2)) |
| Otu12108 | 2 | k__Fungi(100);p__Ascomycota(100);c__Saccharomycetes(100);o__Saccharomycetales(100);f__Metschnikowiaceae(100);g__Metschnikowia(100);s__Metschnikowia_pulcherrima(100); | Metschnikowia | 13 | Saprotroph | \|Undefined Saprotroph\| | Yeast | NULL | Probable | As nectar-tap_saprotroph (Põlme et al. 2020); Decay sugar-rich substrates (Põlme et al. 2020) | Põlme S, et al. 2020. Fungal Diversity 105:1-16 ((https://doi.org/10.1007/s13225-020-00466-2)) |
| Otu12109 | 2 | k__Fungi(100);p__Ascomycota(100);c__Saccharomycetes(100);o__Saccharomycetales(100);f__Metschnikowiaceae(100);g__Metschnikowia(100);g__Metschnikowia_unclassified(100); | Metschnikowia | 13 | Saprotroph | \|Undefined Saprotroph\| | Yeast | NULL | Probable | As nectar-tap_saprotroph (Põlme et al. 2020); Decay sugar-rich substrates (Põlme et al. 2020) | Põlme S, et al. 2020. Fungal Diversity 105:1-16 ((https://doi.org/10.1007/s13225-020-00466-2)) |
| Otu12111 | 2 | k__Fungi(100);p__Ascomycota(100);c__Saccharomycetes(100);o__Saccharomycetales(100);f__Metschnikowiaceae(100);g__Metschnikowia(100);g__Metschnikowia_unclassified(100); | Metschnikowia | 13 | Saprotroph | \|Undefined Saprotroph\| | Yeast | NULL | Probable | As nectar-tap_saprotroph (Põlme et al. 2020); Decay sugar-rich substrates (Põlme et al. 2020) | Põlme S, et al. 2020. Fungal Diversity 105:1-16 ((https://doi.org/10.1007/s13225-020-00466-2)) |
| Otu12112 | 2 | k__Fungi(100);p__Ascomycota(100);c__Saccharomycetes(100);o__Saccharomycetales(100);f__Metschnikowiaceae(100);g__Metschnikowia(100);s__Metschnikowia_pulcherrima(100); | Metschnikowia | 13 | Saprotroph | \|Undefined Saprotroph\| | Yeast | NULL | Probable | As nectar-tap_saprotroph (Põlme et al. 2020); Decay sugar-rich substrates (Põlme et al. 2020) | Põlme S, et al. 2020. Fungal Diversity 105:1-16 ((https://doi.org/10.1007/s13225-020-00466-2)) |
| Otu12114 | 2 | k__Fungi(100);p__Ascomycota(100);c__Saccharomycetes(100);o__Saccharomycetales(100);f__Metschnikowiaceae(100);g__Metschnikowia(100);s__Metschnikowia_pulcherrima(100); | Metschnikowia | 13 | Saprotroph | \|Undefined Saprotroph\| | Yeast | NULL | Probable | As nectar-tap_saprotroph (Põlme et al. 2020); Decay sugar-rich substrates (Põlme et al. 2020) | Põlme S, et al. 2020. Fungal Diversity 105:1-16 ((https://doi.org/10.1007/s13225-020-00466-2)) |
| Otu12115 | 2 | k__Fungi(100);p__Ascomycota(100);c__Saccharomycetes(100);o__Saccharomycetales(100);f__Metschnikowiaceae(100);g__Metschnikowia(100);g__Metschnikowia_unclassified(100); | Metschnikowia | 13 | Saprotroph | \|Undefined Saprotroph\| | Yeast | NULL | Probable | As nectar-tap_saprotroph (Põlme et al. 2020); Decay sugar-rich substrates (Põlme et al. 2020) | Põlme S, et al. 2020. Fungal Diversity 105:1-16 ((https://doi.org/10.1007/s13225-020-00466-2)) |
| Otu12116 | 2 | k__Fungi(100);p__Ascomycota(100);c__Saccharomycetes(100);o__Saccharomycetales(100);f__Metschnikowiaceae(100);g__Metschnikowia(100);s__Metschnikowia_pulcherrima(100); | Metschnikowia | 13 | Saprotroph | \|Undefined Saprotroph\| | Yeast | NULL | Probable | As nectar-tap_saprotroph (Põlme et al. 2020); Decay sugar-rich substrates (Põlme et al. 2020) | Põlme S, et al. 2020. Fungal Diversity 105:1-16 ((https://doi.org/10.1007/s13225-020-00466-2)) |
| Otu12117 | 2 | k__Fungi(100);p__Zygomycota(100);c__Zygomycota_class_Incertae_sedis(100);o__Mucorales(100);f__Mucoraceae(100);g__Mucor(100);g__Mucor_unclassified(100); | Mucor | 13 | Saprotroph | \|Undefined Saprotroph\| | Microfungus | NULL | Probable | As soil_saprotroph (Põlme et al. 2020); Decay soil (Põlme et al. 2020) | Tedersoo L, et al. 2014. Science 346:e1256688 ((https://doi.org/10.1126/science.1256688)); Põlme S, et al. 2020. Fungal Diversity 105:1-16 ((https://doi.org/10.1007/s13225-020-00466-2)) |
| Otu12118 | 2 | k__Fungi(100);p__Ascomycota(100);c__Saccharomycetes(100);o__Saccharomycetales(100);f__Metschnikowiaceae(100);g__Metschnikowia(100);g__Metschnikowia_unclassified(100); | Metschnikowia | 13 | Saprotroph | \|Undefined Saprotroph\| | Yeast | NULL | Probable | As nectar-tap_saprotroph (Põlme et al. 2020); Decay sugar-rich substrates (Põlme et al. 2020) | Põlme S, et al. 2020. Fungal Diversity 105:1-16 ((https://doi.org/10.1007/s13225-020-00466-2)) |
| Otu12119 | 2 | k__Fungi(100);p__Ascomycota(100);c__Saccharomycetes(100);o__Saccharomycetales(100);f__Metschnikowiaceae(100);g__Metschnikowia(100);g__Metschnikowia_unclassified(100); | Metschnikowia | 13 | Saprotroph | \|Undefined Saprotroph\| | Yeast | NULL | Probable | As nectar-tap_saprotroph (Põlme et al. 2020); Decay sugar-rich substrates (Põlme et al. 2020) | Põlme S, et al. 2020. Fungal Diversity 105:1-16 ((https://doi.org/10.1007/s13225-020-00466-2)) |
| Otu12120 | 2 | k__Fungi(100);p__Ascomycota(100);c__Saccharomycetes(100);o__Saccharomycetales(100);f__Metschnikowiaceae(100);g__Metschnikowia(100);s__Metschnikowia_pulcherrima(100); | Metschnikowia | 13 | Saprotroph | \|Undefined Saprotroph\| | Yeast | NULL | Probable | As nectar-tap_saprotroph (Põlme et al. 2020); Decay sugar-rich substrates (Põlme et al. 2020) | Põlme S, et al. 2020. Fungal Diversity 105:1-16 ((https://doi.org/10.1007/s13225-020-00466-2)) |
| Otu12124 | 2 | k__Fungi(100);p__Ascomycota(100);c__Saccharomycetes(100);o__Saccharomycetales(100);f__Metschnikowiaceae(100);g__Metschnikowia(100);g__Metschnikowia_unclassified(100); | Metschnikowia | 13 | Saprotroph | \|Undefined Saprotroph\| | Yeast | NULL | Probable | As nectar-tap_saprotroph (Põlme et al. 2020); Decay sugar-rich substrates (Põlme et al. 2020) | Põlme S, et al. 2020. Fungal Diversity 105:1-16 ((https://doi.org/10.1007/s13225-020-00466-2)) |
| Otu12125 | 2 | k__Fungi(100);p__Ascomycota(100);c__Saccharomycetes(100);o__Saccharomycetales(100);f__Metschnikowiaceae(100);g__Metschnikowia(100);g__Metschnikowia_unclassified(100); | Metschnikowia | 13 | Saprotroph | \|Undefined Saprotroph\| | Yeast | NULL | Probable | As nectar-tap_saprotroph (Põlme et al. 2020); Decay sugar-rich substrates (Põlme et al. 2020) | Põlme S, et al. 2020. Fungal Diversity 105:1-16 ((https://doi.org/10.1007/s13225-020-00466-2)) |
| Otu12126 | 2 | k__Fungi(100);p__Ascomycota(100);c__Saccharomycetes(100);o__Saccharomycetales(100);f__Metschnikowiaceae(100);g__Metschnikowia(100);s__Metschnikowia_pulcherrima(100); | Metschnikowia | 13 | Saprotroph | \|Undefined Saprotroph\| | Yeast | NULL | Probable | As nectar-tap_saprotroph (Põlme et al. 2020); Decay sugar-rich substrates (Põlme et al. 2020) | Põlme S, et al. 2020. Fungal Diversity 105:1-16 ((https://doi.org/10.1007/s13225-020-00466-2)) |
| Otu12127 | 2 | k__Fungi(100);p__Ascomycota(100);c__Saccharomycetes(100);o__Saccharomycetales(100);f__Metschnikowiaceae(100);g__Metschnikowia(100);g__Metschnikowia_unclassified(100); | Metschnikowia | 13 | Saprotroph | \|Undefined Saprotroph\| | Yeast | NULL | Probable | As nectar-tap_saprotroph (Põlme et al. 2020); Decay sugar-rich substrates (Põlme et al. 2020) | Põlme S, et al. 2020. Fungal Diversity 105:1-16 ((https://doi.org/10.1007/s13225-020-00466-2)) |
| Otu12128 | 2 | k__Fungi(100);p__Ascomycota(100);c__Saccharomycetes(100);o__Saccharomycetales(100);f__Metschnikowiaceae(100);g__Metschnikowia(100);s__Metschnikowia_pulcherrima(100); | Metschnikowia | 13 | Saprotroph | \|Undefined Saprotroph\| | Yeast | NULL | Probable | As nectar-tap_saprotroph (Põlme et al. 2020); Decay sugar-rich substrates (Põlme et al. 2020) | Põlme S, et al. 2020. Fungal Diversity 105:1-16 ((https://doi.org/10.1007/s13225-020-00466-2)) |
| Otu12129 | 2 | k__Fungi(100);p__Ascomycota(100);c__Saccharomycetes(100);o__Saccharomycetales(100);f__Metschnikowiaceae(100);g__Metschnikowia(100);g__Metschnikowia_unclassified(100); | Metschnikowia | 13 | Saprotroph | \|Undefined Saprotroph\| | Yeast | NULL | Probable | As nectar-tap_saprotroph (Põlme et al. 2020); Decay sugar-rich substrates (Põlme et al. 2020) | Põlme S, et al. 2020. Fungal Diversity 105:1-16 ((https://doi.org/10.1007/s13225-020-00466-2)) |
| Otu12130 | 2 | k__Fungi(100);p__Ascomycota(100);c__Saccharomycetes(100);o__Saccharomycetales(100);f__Metschnikowiaceae(100);g__Metschnikowia(100);s__Metschnikowia_pulcherrima(100); | Metschnikowia | 13 | Saprotroph | \|Undefined Saprotroph\| | Yeast | NULL | Probable | As nectar-tap_saprotroph (Põlme et al. 2020); Decay sugar-rich substrates (Põlme et al. 2020) | Põlme S, et al. 2020. Fungal Diversity 105:1-16 ((https://doi.org/10.1007/s13225-020-00466-2)) |
| Otu12131 | 2 | k__Fungi(100);p__Ascomycota(100);c__Saccharomycetes(100);o__Saccharomycetales(100);f__Metschnikowiaceae(100);g__Metschnikowia(100);s__Metschnikowia_pulcherrima(100); | Metschnikowia | 13 | Saprotroph | \|Undefined Saprotroph\| | Yeast | NULL | Probable | As nectar-tap_saprotroph (Põlme et al. 2020); Decay sugar-rich substrates (Põlme et al. 2020) | Põlme S, et al. 2020. Fungal Diversity 105:1-16 ((https://doi.org/10.1007/s13225-020-00466-2)) |
| Otu12132 | 2 | k__Fungi(100);p__Ascomycota(100);c__Saccharomycetes(100);o__Saccharomycetales(100);f__Metschnikowiaceae(100);g__Metschnikowia(100);g__Metschnikowia_unclassified(100); | Metschnikowia | 13 | Saprotroph | \|Undefined Saprotroph\| | Yeast | NULL | Probable | As nectar-tap_saprotroph (Põlme et al. 2020); Decay sugar-rich substrates (Põlme et al. 2020) | Põlme S, et al. 2020. Fungal Diversity 105:1-16 ((https://doi.org/10.1007/s13225-020-00466-2)) |
| Otu12134 | 2 | k__Fungi(100);p__Ascomycota(100);c__Saccharomycetes(100);o__Saccharomycetales(100);f__Metschnikowiaceae(100);g__Metschnikowia(100);s__Metschnikowia_pulcherrima(100); | Metschnikowia | 13 | Saprotroph | \|Undefined Saprotroph\| | Yeast | NULL | Probable | As nectar-tap_saprotroph (Põlme et al. 2020); Decay sugar-rich substrates (Põlme et al. 2020) | Põlme S, et al. 2020. Fungal Diversity 105:1-16 ((https://doi.org/10.1007/s13225-020-00466-2)) |
| Otu12136 | 2 | k__Fungi(100);p__Ascomycota(100);c__Saccharomycetes(100);o__Saccharomycetales(100);f__Metschnikowiaceae(100);g__Metschnikowia(100);g__Metschnikowia_unclassified(100); | Metschnikowia | 13 | Saprotroph | \|Undefined Saprotroph\| | Yeast | NULL | Probable | As nectar-tap_saprotroph (Põlme et al. 2020); Decay sugar-rich substrates (Põlme et al. 2020) | Põlme S, et al. 2020. Fungal Diversity 105:1-16 ((https://doi.org/10.1007/s13225-020-00466-2)) |
| Otu12137 | 2 | k__Fungi(100);p__Ascomycota(100);c__Saccharomycetes(100);o__Saccharomycetales(100);f__Metschnikowiaceae(100);g__Metschnikowia(100);g__Metschnikowia_unclassified(100); | Metschnikowia | 13 | Saprotroph | \|Undefined Saprotroph\| | Yeast | NULL | Probable | As nectar-tap_saprotroph (Põlme et al. 2020); Decay sugar-rich substrates (Põlme et al. 2020) | Põlme S, et al. 2020. Fungal Diversity 105:1-16 ((https://doi.org/10.1007/s13225-020-00466-2)) |

| | | | | | | | | | | | |
|---|---|---|---|---|---|---|---|---|---|---|---|
| Otu12138 | 2 | k__Fungi(100);p__Ascomycota(100);c__Saccharomycetes(100);o__Saccharomycetales(100);f__Metschnikowiaceae(100);g__Metschnikowia(100);s__Metschnikowia_pulcherrima(100); | Metschnikowia | 13 | Saprotroph | \|Undefined Saprotroph\| | Yeast | NULL | Probable | As nectar-tap_saprotroph (Põlme et al. 2020); Decay sugar-rich substrates (Põlme et al. 2020) | Põlme S, et al. 2020. Fungal Diversity 105:1-16 ((https://doi.org/10.1007/s13225-020-00466-2)) |
| Otu12139 | 2 | k__Fungi(100);p__Ascomycota(100);c__Saccharomycetes(100);o__Saccharomycetales(100);f__Metschnikowiaceae(100);g__Metschnikowia(100);s__Metschnikowia_pulcherrima(100); | Metschnikowia | 13 | Saprotroph | \|Undefined Saprotroph\| | Yeast | NULL | Probable | As nectar-tap_saprotroph (Põlme et al. 2020); Decay sugar-rich substrates (Põlme et al. 2020) | Põlme S, et al. 2020. Fungal Diversity 105:1-16 ((https://doi.org/10.1007/s13225-020-00466-2)) |
| Otu12140 | 2 | k__Fungi(100);p__Ascomycota(100);c__Saccharomycetes(100);o__Saccharomycetales(100);f__Metschnikowiaceae(100);g__Metschnikowia(100);s__Metschnikowia_pulcherrima(100); | Metschnikowia | 13 | Saprotroph | \|Undefined Saprotroph\| | Yeast | NULL | Probable | As nectar-tap_saprotroph (Põlme et al. 2020); Decay sugar-rich substrates (Põlme et al. 2020) | Põlme S, et al. 2020. Fungal Diversity 105:1-16 ((https://doi.org/10.1007/s13225-020-00466-2)) |
| Otu12141 | 2 | k__Fungi(100);p__Ascomycota(100);c__Dothideomycetes(100);o__Dothideomycetes_order_Incertae_sedis(100);f__Myxotrichaceae(100);g__Oidiodendron(100);s__Oidiodendron_myxotrichoides(100); | Oidiodendron | 13 | Pathotroph-Saprotroph-Symbiotroph | Endophyte-Ericoid Mycorrhizal-\|Undefined Saprotroph\| | Microfungus | NULL | Probable | As root_endophyte (Põlme et al. 2020); As soil_saprotroph (Põlme et al. 2020); Dark Septate Endophyte; Decay soil (Põlme et al. 2020) | Newsham KK. 2011. New Phytologist 190:783-793 ((https://doi.org/10.1111/j.1469-8137.2010.03611.x)) (as dark septate root endophytes); Põlme S, et al. 2020. Fungal Diversity 105:1-16 ((https://doi.org/10.1007/s13225-020-00466-2)) |
| Otu12142 | 2 | k__Fungi(100);p__Ascomycota(100);c__Saccharomycetes(100);o__Saccharomycetales(100);f__Metschnikowiaceae(100);g__Metschnikowia(100);s__Metschnikowia_pulcherrima(100); | Metschnikowia | 13 | Saprotroph | \|Undefined Saprotroph\| | Yeast | NULL | Probable | As nectar-tap_saprotroph (Põlme et al. 2020); Decay sugar-rich substrates (Põlme et al. 2020) | Põlme S, et al. 2020. Fungal Diversity 105:1-16 ((https://doi.org/10.1007/s13225-020-00466-2)) |
| Otu12144 | 2 | k__Fungi(100);p__Ascomycota(100);c__Saccharomycetes(100);o__Saccharomycetales(100);f__Metschnikowiaceae(100);g__Metschnikowia(100);g__Metschnikowia_unclassified(100); | Metschnikowia | 13 | Saprotroph | \|Undefined Saprotroph\| | Yeast | NULL | Probable | As nectar-tap_saprotroph (Põlme et al. 2020); Decay sugar-rich substrates (Põlme et al. 2020) | Põlme S, et al. 2020. Fungal Diversity 105:1-16 ((https://doi.org/10.1007/s13225-020-00466-2)) |
| Otu12145 | 2 | k__Fungi(100);p__Ascomycota(100);c__Saccharomycetes(100);o__Saccharomycetales(100);f__Metschnikowiaceae(100);g__Metschnikowia(100);s__Metschnikowia_pulcherrima(100); | Metschnikowia | 13 | Saprotroph | \|Undefined Saprotroph\| | Yeast | NULL | Probable | As nectar-tap_saprotroph (Põlme et al. 2020); Decay sugar-rich substrates (Põlme et al. 2020) | Põlme S, et al. 2020. Fungal Diversity 105:1-16 ((https://doi.org/10.1007/s13225-020-00466-2)) |
| Otu12146 | 2 | k__Fungi(100);p__Ascomycota(100);c__Saccharomycetes(100);o__Saccharomycetales(100);f__Metschnikowiaceae(100);g__Metschnikowia(100);s__Metschnikowia_pulcherrima(100); | Metschnikowia | 13 | Saprotroph | \|Undefined Saprotroph\| | Yeast | NULL | Probable | As nectar-tap_saprotroph (Põlme et al. 2020); Decay sugar-rich substrates (Põlme et al. 2020) | Põlme S, et al. 2020. Fungal Diversity 105:1-16 ((https://doi.org/10.1007/s13225-020-00466-2)) |
| Otu12147 | 2 | k__Fungi(100);p__Ascomycota(100);c__Saccharomycetes(100);o__Saccharomycetales(100);f__Metschnikowiaceae(100);g__Metschnikowia(100);s__Metschnikowia_pulcherrima(100); | Metschnikowia | 13 | Saprotroph | \|Undefined Saprotroph\| | Yeast | NULL | Probable | As nectar-tap_saprotroph (Põlme et al. 2020); Decay sugar-rich substrates (Põlme et al. 2020) | Põlme S, et al. 2020. Fungal Diversity 105:1-16 ((https://doi.org/10.1007/s13225-020-00466-2)) |
| Otu12150 | 2 | k__Fungi(100);p__Ascomycota(100);c__Saccharomycetes(100);o__Saccharomycetales(100);f__Metschnikowiaceae(100);g__Metschnikowia(100);g__Metschnikowia_unclassified(100); | Metschnikowia | 13 | Saprotroph | \|Undefined Saprotroph\| | Yeast | NULL | Probable | As nectar-tap_saprotroph (Põlme et al. 2020); Decay sugar-rich substrates (Põlme et al. 2020) | Põlme S, et al. 2020. Fungal Diversity 105:1-16 ((https://doi.org/10.1007/s13225-020-00466-2)) |
| Otu12151 | 2 | k__Fungi(100);p__Ascomycota(100);c__Saccharomycetes(100);o__Saccharomycetales(100);f__Metschnikowiaceae(100);g__Metschnikowia(100);g__Metschnikowia_unclassified(100); | Metschnikowia | 13 | Saprotroph | \|Undefined Saprotroph\| | Yeast | NULL | Probable | As nectar-tap_saprotroph (Põlme et al. 2020); Decay sugar-rich substrates (Põlme et al. 2020) | Põlme S, et al. 2020. Fungal Diversity 105:1-16 ((https://doi.org/10.1007/s13225-020-00466-2)) |
| Otu12152 | 2 | k__Fungi(100);p__Ascomycota(100);c__Saccharomycetes(100);o__Saccharomycetales(100);f__Metschnikowiaceae(100);g__Metschnikowia(100);g__Metschnikowia_unclassified(100); | Metschnikowia | 13 | Saprotroph | \|Undefined Saprotroph\| | Yeast | NULL | Probable | As nectar-tap_saprotroph (Põlme et al. 2020); Decay sugar-rich substrates (Põlme et al. 2020) | Põlme S, et al. 2020. Fungal Diversity 105:1-16 ((https://doi.org/10.1007/s13225-020-00466-2)) |
| Otu12153 | 2 | k__Fungi(100);p__Ascomycota(100);c__Saccharomycetes(100);o__Saccharomycetales(100);f__Metschnikowiaceae(100);g__Metschnikowia(100);g__Metschnikowia_unclassified(100); | Metschnikowia | 13 | Saprotroph | \|Undefined Saprotroph\| | Yeast | NULL | Probable | As nectar-tap_saprotroph (Põlme et al. 2020); Decay sugar-rich substrates (Põlme et al. 2020) | Põlme S, et al. 2020. Fungal Diversity 105:1-16 ((https://doi.org/10.1007/s13225-020-00466-2)) |
| Otu12157 | 2 | k__Fungi(100);p__Ascomycota(100);c__Saccharomycetes(100);o__Saccharomycetales(100);f__Metschnikowiaceae(100);g__Metschnikowia(100);g__Metschnikowia_unclassified(100); | Metschnikowia | 13 | Saprotroph | \|Undefined Saprotroph\| | Yeast | NULL | Probable | As nectar-tap_saprotroph (Põlme et al. 2020); Decay sugar-rich substrates (Põlme et al. 2020) | Põlme S, et al. 2020. Fungal Diversity 105:1-16 ((https://doi.org/10.1007/s13225-020-00466-2)) |
| Otu12162 | 2 | k__Fungi(100);p__Ascomycota(100);c__Saccharomycetes(100);o__Saccharomycetales(100);f__Metschnikowiaceae(100);g__Metschnikowia(100);s__Metschnikowia_pulcherrima(100); | Metschnikowia | 13 | Saprotroph | \|Undefined Saprotroph\| | Yeast | NULL | Probable | As nectar-tap_saprotroph (Põlme et al. 2020); Decay sugar-rich substrates (Põlme et al. 2020) | Põlme S, et al. 2020. Fungal Diversity 105:1-16 ((https://doi.org/10.1007/s13225-020-00466-2)) |
| Otu12163 | 2 | k__Fungi(100);p__Ascomycota(100);c__Saccharomycetes(100);o__Saccharomycetales(100);f__Metschnikowiaceae(100);g__Metschnikowia(100);g__Metschnikowia_unclassified(100); | Metschnikowia | 13 | Saprotroph | \|Undefined Saprotroph\| | Yeast | NULL | Probable | As nectar-tap_saprotroph (Põlme et al. 2020); Decay sugar-rich substrates (Põlme et al. 2020) | Põlme S, et al. 2020. Fungal Diversity 105:1-16 ((https://doi.org/10.1007/s13225-020-00466-2)) |
| Otu12166 | 2 | k__Fungi(100);p__Ascomycota(100);c__Saccharomycetes(100);o__Saccharomycetales(100);f__Metschnikowiaceae(100);g__Metschnikowia(100);s__Metschnikowia_pulcherrima(100); | Metschnikowia | 13 | Saprotroph | \|Undefined Saprotroph\| | Yeast | NULL | Probable | As nectar-tap_saprotroph (Põlme et al. 2020); Decay sugar-rich substrates (Põlme et al. 2020) | Põlme S, et al. 2020. Fungal Diversity 105:1-16 ((https://doi.org/10.1007/s13225-020-00466-2)) |
| Otu12167 | 2 | k__Fungi(100);p__Ascomycota(100);c__Saccharomycetes(100);o__Saccharomycetales(100);f__Metschnikowiaceae(100);g__Metschnikowia(100);g__Metschnikowia_unclassified(100); | Metschnikowia | 13 | Saprotroph | \|Undefined Saprotroph\| | Yeast | NULL | Probable | As nectar-tap_saprotroph (Põlme et al. 2020); Decay sugar-rich substrates (Põlme et al. 2020) | Põlme S, et al. 2020. Fungal Diversity 105:1-16 ((https://doi.org/10.1007/s13225-020-00466-2)) |
| Otu12168 | 2 | k__Fungi(100);p__Ascomycota(100);c__Saccharomycetes(100);o__Saccharomycetales(100);f__Metschnikowiaceae(100);g__Metschnikowia(100);g__Metschnikowia_unclassified(100); | Metschnikowia | 13 | Saprotroph | \|Undefined Saprotroph\| | Yeast | NULL | Probable | As nectar-tap_saprotroph (Põlme et al. 2020); Decay sugar-rich substrates (Põlme et al. 2020) | Põlme S, et al. 2020. Fungal Diversity 105:1-16 ((https://doi.org/10.1007/s13225-020-00466-2)) |
| Otu12169 | 2 | k__Fungi(100);p__Ascomycota(100);c__Saccharomycetes(100);o__Saccharomycetales(100);f__Metschnikowiaceae(100);g__Metschnikowia(100);g__Metschnikowia_unclassified(100); | Metschnikowia | 13 | Saprotroph | \|Undefined Saprotroph\| | Yeast | NULL | Probable | As nectar-tap_saprotroph (Põlme et al. 2020); Decay sugar-rich substrates (Põlme et al. 2020) | Põlme S, et al. 2020. Fungal Diversity 105:1-16 ((https://doi.org/10.1007/s13225-020-00466-2)) |
| Otu12171 | 2 | k__Fungi(100);p__Ascomycota(100);c__Saccharomycetes(100);o__Saccharomycetales(100);f__Saccharomycetales_family_Incertae_sedis(100);g__Candida(100);g__Candida_unclassified(100); | Candida | 13 | Pathotroph-Saprotroph-Symbiotroph | Animal Pathogen-Endophyte-Endosymbiont-Epiphyte-\|Undefined Saprotroph\| | Dimorphic Yeast-Yeast | NULL | Probable | As nectar-tap_saprotroph (Põlme et al. 2020); Decay sugar-rich substrates (Põlme et al. 2020); This asexual genus is polyphyletic and contain species that function very differently depend on what they are. Caution should be taken when interpreting guilds for Candida species. | Manolakaki D, et al. 2010. Virulence. 1: 367-375 ((https://doi.org/10.4161/viru.1.5.12796)); Mekbib SB, et al. 2011. Tropical Plant Pathology 36:233-240 ((https://doi.org/10.1590/S1982-56762011000400004)); Põlme S, et al. 2020. Fungal Diversity 105:1-16 ((https://doi.org/10.1007/s13225-020-00466-2)) |
| Otu12172 | 2 | k__Fungi(100);p__Ascomycota(100);c__Saccharomycetes(100);o__Saccharomycetales(100);f__Metschnikowiaceae(100);g__Metschnikowia(100);s__Metschnikowia_pulcherrima(100); | Metschnikowia | 13 | Saprotroph | \|Undefined Saprotroph\| | Yeast | NULL | Probable | As nectar-tap_saprotroph (Põlme et al. 2020); Decay sugar-rich substrates (Põlme et al. 2020) | Põlme S, et al. 2020. Fungal Diversity 105:1-16 ((https://doi.org/10.1007/s13225-020-00466-2)) |
| Otu12175 | 2 | k__Fungi(100);p__Ascomycota(100);c__Saccharomycetes(100);o__Saccharomycetales(100);f__Metschnikowiaceae(100);g__Metschnikowia(100);g__Metschnikowia_unclassified(100); | Metschnikowia | 13 | Saprotroph | \|Undefined Saprotroph\| | Yeast | NULL | Probable | As nectar-tap_saprotroph (Põlme et al. 2020); Decay sugar-rich substrates (Põlme et al. 2020) | Põlme S, et al. 2020. Fungal Diversity 105:1-16 ((https://doi.org/10.1007/s13225-020-00466-2)) |
| Otu12177 | 2 | k__Fungi(100);p__Ascomycota(100);c__Saccharomycetes(100);o__Saccharomycetales(100);f__Metschnikowiaceae(100);g__Metschnikowia(100);s__Metschnikowia_pulcherrima(100); | Metschnikowia | 13 | Saprotroph | \|Undefined Saprotroph\| | Yeast | NULL | Probable | As nectar-tap_saprotroph (Põlme et al. 2020); Decay sugar-rich substrates (Põlme et al. 2020) | Põlme S, et al. 2020. Fungal Diversity 105:1-16 ((https://doi.org/10.1007/s13225-020-00466-2)) |
| Otu12179 | 2 | k__Fungi(100);p__Basidiomycota(100);c__Microbotryomycetes(100);o__Sporidiobolales(100);f__Sporidiobolales_family_Incertae_sedis(100);g__Sporobolomyces(100);s__Sporobolomyces_lactosus(100); | Sporobolomyces | 13 | Pathotroph-Saprotroph | \|Fungal Parasite\|-Plant Saprotroph | Yeast | NULL | Probable | As litter_saprotroph (Põlme et al. 2020); As mycoparasite (Põlme et al. 2020); Decay fungal material (Põlme et al. 2020) | Tedersoo L, et al. 2014. Science 346:e1256688 ((https://doi.org/10.1126/science.1256688)); Purahong W, et al. 2016. Molecular Ecology 25:4059-4074 ((https://doi.org/10.1111/mec.13739)); Põlme S, et al. 2020. Fungal Diversity 105:1-16 ((https://doi.org/10.1007/s13225-020-00466-2)) |
| Otu12180 | 2 | k__Fungi(100);p__Ascomycota(100);c__Dothideomycetes(100);o__Pleosporales(100);f__Tubeufiaceae(100);f__Tubeufiaceae_unclassified(100);f__Tubeufiaceae_unclassified(100); | Tubeufiaceae | 9 | Pathotroph-Saprotroph | Animal Pathogen-Undefined Saprotroph | NULL | NULL | Probable | Biotrophic or saprobic, especially on scale insects or other fungi. The anamorphic forms are fequently aero-aquatic (Cannon & Kirk 2007) | Cannon PF, Kirk PM. 2007. Fungal Families of the World. CAB International, Cambridge (ISBN: 978-0851998275) |
| Otu12184 | 2 | k__Fungi(100);p__Ascomycota(100);c__Sordariomycetes(100);o__Sordariales(100);f__Chaetomiaceae(100);g__Chaetomium(100);g__Chaetomium_unclassified(100); | Chaetomium | 13 | Pathotroph-Saprotroph-Symbiotroph | Animal Pathogen-Dung Saprotroph-Endophyte-Epiphyte-Plant Pathogen-\|Plant Saprotroph\|-Wood Saprotroph | Microfungus | Soft Rot | Possible | As foliar_endophyte (Põlme et al. 2020); As litter_saprotroph (Põlme et al. 2020); culture (Busby et al. 2016, David et al. 2016); Likely opportunistic human pathogen (Irinyi et al. 2015); culture (Massimo et al. 2015); Endophyte detection method; Decay wood, leaf, fruit, seed, soil, dung (Põlme et al. 2020); derived commodities (Cannon & Kirk 2007); Epiphyte detection method; Saprobic, acting as key decay organisms of plant materials and plant | Seehann G, et al. 1975. List of Fungi in Soft-Rot Tissue Tests. Institute fur Holzbiologie und Holzschutz, Bundesforschungsanstalt fur Forst - und Holzwirtshaft, Hamburg - Lohbrugge, Germany; Cannon PF, Kirk PM. 2007. Fungal Families of the World. CAB International, Cambridge ((https://isbnsearch.org/isbn/9780851998275)); Bills GF, et al. 2013. Current Opinion in Microbiology 16:549-565 ((https://doi.org/10.1016/j.mib.2013.08.001)); Tedersoo L, et al. 2014. Science 346:e1256688 ((https://doi.org/10.1126/science.1256688)); Irinyi L, et al. 2015. Medical Mycology 53:313-337 ((https://doi.org/10.1093/mmy/myv008)); Busby PE, et al. 2016. New Phytologist 209:1681-1692 ((https://doi.org/10.1111/nph.13742)); Põlme S, et al. 2020. Fungal Diversity 105:1-16 ((https://doi.org/10.1007/s13225-020-00466-2)) |
| Otu12185 | 2 | k__Fungi(100);p__Ascomycota(100);c__Dothideomycetes(100);o__Pleosporales(100);f__Pleomassariaceae(100);g__unclassified_Pleomassariaceae(100);s__Pleomassariaceae_sp(100); | Pleomassariaceae | 9 | Pathotroph-Saprotroph | Plant Pathogen-Plant Saprotroph | Microfungus | NULL | Probable | Saprobic or necrotrophic on wood, bark and herbaceous material (Cannon & Kirk 2007) | Cannon PF, Kirk PM. 2007. Fungal Families of the World. CAB International, Cambridge (ISBN: 978-0851998275) |
| Otu12186 | 2 | k__Fungi(100);p__Ascomycota(100);c__Eurotiomycetes(100);o__Chaetothyriales(100);f__Herpotrichiellaceae(100);g__Rhinocladiella(100);s__Rhinocladiella_sp_VL271(100); | Rhinocladiella | 13 | Pathotroph-Saprotroph-Symbiotroph | Animal Parasite-Endophyte-\|Undefined Saprotroph\|-Wood Saprotroph | Dimorphic-Facultative Yeast | Soft Rot | Probable | As root_endophyte_dark_septate (Põlme et al. 2020); As soil_saprotroph (Põlme et al. 2020); Decay soil, animal material (Põlme et al. 2020) | Nilsson T. 1973. Studia Forestalia Suecica 104:1-40 ((https://pub.epsilon.slu.se/5711/)); Seehann G, et al. 1975. List of Fungi in Soft-Rot Tissue Tests. Institute fur Holzbiologie und Holzschutz, Bundesforschungsanstalt fur Forst - und Holzwirtshaft, Hamburg - Lohbrugge, Germany; Costa IPMW, et al. 2012. Mycotaxon 119:e26 ((http://mycotaxon.com/resources/checklists/costa_v119_checklist.pdf)); Tedersoo L, et al. 2014. Science 346:e1256688 ((https://doi.org/10.1126/science.1256688)); Põlme S, et al. 2020. Fungal Diversity 105:1-16 ((https://doi.org/10.1007/s13225-020-00466-2)) |
| Otu12189 | 2 | k__Fungi(100);p__Ascomycota(100);c__Saccharomycetes(100);o__Saccharomycetales(100);f__Metschnikowiaceae(100);g__Metschnikowia(100);g__Metschnikowia_unclassified(100); | Metschnikowia | 13 | Saprotroph | \|Undefined Saprotroph\| | Yeast | NULL | Probable | As nectar-tap_saprotroph (Põlme et al. 2020); Decay sugar-rich substrates (Põlme et al. 2020) | Põlme S, et al. 2020. Fungal Diversity 105:1-16 ((https://doi.org/10.1007/s13225-020-00466-2)) |
| Otu12190 | 2 | k__Fungi(100);p__Ascomycota(100);c__Leotiomycetes(100);o__Helotiales(100);f__Helotiaceae(100);g__Crocicreas(100);s__Crocicreas_cyathoideum_var._cyathoideum(100); | Crocicreas | 13 | Saprotroph | \|Plant Saprotroph\|-Undefined Saprotroph | Microfungus-Pezizoid | NULL | Probable | As litter_saprotroph (Põlme et al. 2020); Decay leaf, fruit, seed (Põlme et al. 2020) | Carpenter SE, 1983. Memoirs of The New York Botanical Garden 33:1-290 (ISBN: 978-0893272302); Põlme S, et al. 2020. Fungal Diversity 105:1-16 ((https://doi.org/10.1007/s13225-020-00466-2)) |
| Otu12191 | 2 | k__Fungi(100);p__Basidiomycota(100);c__Microbotryomycetes(100);o__Sporidiobolales(100);f__Sporidiobolales_family_Incertae_sedis(100);g__Sporobolomyces(100);s__Sporobolomyces_lactosus(100); | Sporobolomyces | 13 | Pathotroph-Saprotroph | \|Fungal Parasite\|-Plant Saprotroph | Yeast | NULL | Probable | As litter_saprotroph (Põlme et al. 2020); As mycoparasite (Põlme et al. 2020); Decay fungal material (Põlme et al. 2020) | Tedersoo L, et al. 2014. Science 346:e1256688 ((https://doi.org/10.1126/science.1256688)); Purahong W, et al. 2016. Molecular Ecology 25:4059-4074 ((https://doi.org/10.1111/mec.13739)); Põlme S, et al. 2020. Fungal Diversity 105:1-16 ((https://doi.org/10.1007/s13225-020-00466-2)) |
| Otu12193 | 2 | k__Fungi(100);p__Basidiomycota(100);c__Microbotryomycetes(100);o__Sporidiobolales(100);f__Sporidiobolales_family_Incertae_sedis(100);g__Sporobolomyces(100);s__Sporobolomyces_lactosus(100); | Sporobolomyces | 13 | Pathotroph-Saprotroph | \|Fungal Parasite\|-Plant Saprotroph | Yeast | NULL | Probable | As litter_saprotroph (Põlme et al. 2020); As mycoparasite (Põlme et al. 2020); Decay fungal material (Põlme et al. 2020) | Tedersoo L, et al. 2014. Science 346:e1256688 ((https://doi.org/10.1126/science.1256688)); Purahong W, et al. 2016. Molecular Ecology 25:4059-4074 ((https://doi.org/10.1111/mec.13739)); Põlme S, et al. 2020. Fungal Diversity 105:1-16 ((https://doi.org/10.1007/s13225-020-00466-2)) |
| Otu12194 | 2 | k__Fungi(100);p__Ascomycota(100);c__Saccharomycetes(100);o__Saccharomycetales(100);f__Metschnikowiaceae(100);g__Metschnikowia(100);g__Metschnikowia_unclassified(100); | Metschnikowia | 13 | Saprotroph | \|Undefined Saprotroph\| | Yeast | NULL | Probable | As nectar-tap_saprotroph (Põlme et al. 2020); Decay sugar-rich substrates (Põlme et al. 2020) | Põlme S, et al. 2020. Fungal Diversity 105:1-16 ((https://doi.org/10.1007/s13225-020-00466-2)) |
| Otu12195 | 2 | k__Fungi(100);p__Ascomycota(100);c__Saccharomycetes(100);o__Saccharomycetales(100);f__Metschnikowiaceae(100);g__Metschnikowia(100);s__Metschnikowia_pulcherrima(100); | Metschnikowia | 13 | Saprotroph | \|Undefined Saprotroph\| | Yeast | NULL | Probable | As nectar-tap_saprotroph (Põlme et al. 2020); Decay sugar-rich substrates (Põlme et al. 2020) | Põlme S, et al. 2020. Fungal Diversity 105:1-16 ((https://doi.org/10.1007/s13225-020-00466-2)) |
| Otu12196 | 2 | k__Fungi(100);p__Ascomycota(100);c__Saccharomycetes(100);o__Saccharomycetales(100);f__Metschnikowiaceae(100);g__Metschnikowia(100);s__Metschnikowia_pulcherrima(100); | Metschnikowia | 13 | Saprotroph | \|Undefined Saprotroph\| | Yeast | NULL | Probable | As nectar-tap_saprotroph (Põlme et al. 2020); Decay sugar-rich substrates (Põlme et al. 2020) | Põlme S, et al. 2020. Fungal Diversity 105:1-16 ((https://doi.org/10.1007/s13225-020-00466-2)) |
| Otu12197 | 2 | k__Fungi(100);p__Ascomycota(100);c__Saccharomycetes(100);o__Saccharomycetales(100);f__Metschnikowiaceae(100);g__Metschnikowia(100);s__Metschnikowia_pulcherrima(100); | Metschnikowia | 13 | Saprotroph | \|Undefined Saprotroph\| | Yeast | NULL | Probable | As nectar-tap_saprotroph (Põlme et al. 2020); Decay sugar-rich substrates (Põlme et al. 2020) | Põlme S, et al. 2020. Fungal Diversity 105:1-16 ((https://doi.org/10.1007/s13225-020-00466-2)) |
| Otu12198 | 2 | k__Fungi(100);p__Ascomycota(100);c__Saccharomycetes(100);o__Saccharomycetales(100);f__Metschnikowiaceae(100);g__Metschnikowia(100);g__Metschnikowia_unclassified(100); | Metschnikowia | 13 | Saprotroph | \|Undefined Saprotroph\| | Yeast | NULL | Probable | As nectar-tap_saprotroph (Põlme et al. 2020); Decay sugar-rich substrates (Põlme et al. 2020) | Põlme S, et al. 2020. Fungal Diversity 105:1-16 ((https://doi.org/10.1007/s13225-020-00466-2)) |
| Otu12200 | 2 | k__Fungi(100);p__Ascomycota(100);c__Eurotiomycetes(100);o__Chaetothyriales(100);f__Herpotrichiellaceae(100);g__Phialophora(100);s__Phialophora_alba(100); | Phialophora | 13 | Pathotroph-Saprotroph-Symbiotroph | Animal Pathogen-Endophyte-Ericoid Mycorrhizal-Plant Pathogen-\|Plant Saprotroph\|-Wood Saprotroph | Dimorphic-Facultative Yeast-Microfungus | NULL | Possible | As foliar_endophyte (Põlme et al. 2020); As litter_saprotroph (Põlme et al. 2020); Consult the current literature for updated names on this paraphyletic grouping that included the Black Yeasts, Dark Septate Endophytes, etc.; Decay wood, leaf, fruit, seed, soil, dung, animal material (Põlme et al. 2020) | Nilsson T. 1973. Studia Forestalia Suecica 104:1-40 ((https://pub.epsilon.slu.se/5711/)); de Hoog GS, et al. 2000. Mycoses 43:409-416 ((https://doi.org/10.1111/j.1439-0507.2000.00601.x)); Rinaldi AC, et al. 2008. Fungal Diversity 3:31-45 ((https://www.fungaldiversity.org/fdp/sfdp/33-1.pdf)); Newsham KK. 2011. New Phytologist 190:783-793 ((https://doi.org/10.1111/j.1469-8137.2010.03611.x)) (as DSE); Irinyi L, et al. 2015. Medical Mycology 53:313-337 ((https://doi.org/10.1093/mmy/myv008)); Põlme S, et al. 2020. Fungal Diversity 105:1-16 ((https://doi.org/10.1007/s13225-020-00466-2)) |
| Otu12201 | 2 | k__Fungi(100);p__Ascomycota(100);c__Saccharomycetes(100);o__Saccharomycetales(100);f__Metschnikowiaceae(100);g__Metschnikowia(100);s__Metschnikowia_pulcherrima(100); | Metschnikowia | 13 | Saprotroph | \|Undefined Saprotroph\| | Yeast | NULL | Probable | As nectar-tap_saprotroph (Põlme et al. 2020); Decay sugar-rich substrates (Põlme et al. 2020) | Põlme S, et al. 2020. Fungal Diversity 105:1-16 ((https://doi.org/10.1007/s13225-020-00466-2)) |
| Otu12202 | 2 | k__Fungi(100);p__Ascomycota(100);c__Leotiomycetes(100);o__Helotiales(100);f__Vibrisseaceae(100);f__Vibrisseaceae_unclassified(100);f__Vibrisseaceae_unclassified(100); | Vibrisseaceae | 9 | Saprotroph-Symbiotroph | Endophyte-Undefined Saprotroph-Wood Saprotroph | Microfungus | NULL | Possible | Saprobic, usually on wood and bark immersed in fresh water. Some species may be root endophytes (Cannon & Kirk 2007) | Cannon PF, Kirk PM. 2007. Fungal Families of the World. CAB International, Cambridge (ISBN: 978-0851998275) |
| Otu12204 | 2 | k__Fungi(100);p__Ascomycota(100);c__Ascomycota_class_Incertae_sedis(100);o__Ascomycota_order_Incertae_sedis(100);f__Ascomycota_family_Incertae_sedis(100);g__Chalara(100);s__Chalara_hyalocuspica(100); | Chalara | 13 | Pathotroph-Saprotroph-Symbiotroph | Endophyte-Plant Pathogen-\|Plant Saprotroph\|-Wood Saprotroph | Microfungus | Soft Rot | Probable | As foliar_endophyte (Põlme et al. 2020); As litter_saprotroph (Põlme et al. 2020); As wood_pathogen (Põlme et al. 2020); Decay wood, leaf, fruit, seed (Põlme et al. 2020) | Duncan CG, Eslyn WE. 1966. Mycologia 58:642-645 ((https://doi.org/10.2307/3757045)); Tedersoo L, et al. 2014. Science 346:e1256688 ((https://doi.org/10.1126/science.1256688)); Põlme S, et al. 2020. Fungal Diversity 105:1-16 ((https://doi.org/10.1007/s13225-020-00466-2)) |
| Otu12205 | 2 | k__Fungi(100);p__Ascomycota(100);c__Leotiomycetes(100);o__Rhytismatales(100);f__Rhytismataceae(100);g__unclassified_Rhytismataceae(100);s__Rhytismataceae_sp(100); | Rhytismataceae | 9 | Pathotroph-Saprotroph-Symbiotroph | Endophyte-Leaf Saprotroph-Plant Pathogen | Microfungus | NULL | Probable | Biotrophic, necrotrophic or saprobic on leaves and bark, often initially as endophytes, causing tar spots, premature leaf fall and occasionally witches broom symptoms (Cannon & Kirk 2007) | Cannon PF, Kirk PM. 2007. Fungal Families of the World. CAB International, Cambridge (ISBN: 978-0851998275) |
| Otu12206 | 2 | k__Fungi(100);p__Ascomycota(100);c__Saccharomycetes(100);o__Saccharomycetales(100);f__Metschnikowiaceae(100);g__Metschnikowia(100);g__Metschnikowia_unclassified(100); | Metschnikowia | 13 | Saprotroph | \|Undefined Saprotroph\| | Yeast | NULL | Probable | As nectar-tap_saprotroph (Põlme et al. 2020); Decay sugar-rich substrates (Põlme et al. 2020) | Põlme S, et al. 2020. Fungal Diversity 105:1-16 ((https://doi.org/10.1007/s13225-020-00466-2)) |
| Otu12207 | 2 | k__Fungi(100);p__Ascomycota(100);c__Saccharomycetes(100);o__Saccharomycetales(100);f__Metschnikowiaceae(100);g__Metschnikowia(100);g__Metschnikowia_unclassified(100); | Metschnikowia | 13 | Saprotroph | \|Undefined Saprotroph\| | Yeast | NULL | Probable | As nectar-tap_saprotroph (Põlme et al. 2020); Decay sugar-rich substrates (Põlme et al. 2020) | Põlme S, et al. 2020. Fungal Diversity 105:1-16 ((https://doi.org/10.1007/s13225-020-00466-2)) |
| Otu12209 | 2 | k__Fungi(100);p__Ascomycota(100);c__Saccharomycetes(100);o__Saccharomycetales(100);f__Saccharomycetales_family_Incertae_sedis(100);g__Candida(100);g__Candida_unclassified(100); | Candida | 13 | Pathotroph-Saprotroph-Symbiotroph | Animal Pathogen-Endophyte-Endosymbiont-Epiphyte-\|Undefined Saprotroph\| | Dimorphic Yeast-Yeast | NULL | Probable | As nectar-tap_saprotroph (Põlme et al. 2020); Decay sugar-rich substrates (Põlme et al. 2020); This asexual genus is polyphyletic and contain species that function very differently depend on what they are. Caution should be taken when interpreting guilds for Candida species. | Manolakaki D, et al. 2010. Virulence. 1: 367-375 ((https://doi.org/10.4161/viru.1.5.12796)); Mekbib SB, et al. 2011. Tropical Plant Pathology 36:233-240 ((https://doi.org/10.1590/S1982-56762011000400004)); Põlme S, et al. 2020. Fungal Diversity 105:1-16 ((https://doi.org/10.1007/s13225-020-00466-2)) |
| Otu12214 | 2 | k__Fungi(100);p__Ascomycota(100);c__Saccharomycetes(100);o__Saccharomycetales(100);f__Metschnikowiaceae(100);g__Metschnikowia(100);s__Metschnikowia_pulcherrima(100); | Metschnikowia | 13 | Saprotroph | \|Undefined Saprotroph\| | Yeast | NULL | Probable | As nectar-tap_saprotroph (Põlme et al. 2020); Decay sugar-rich substrates (Põlme et al. 2020) | Põlme S, et al. 2020. Fungal Diversity 105:1-16 ((https://doi.org/10.1007/s13225-020-00466-2)) |
| Otu12215 | 2 | k__Fungi(100);p__Ascomycota(100);c__Saccharomycetes(100);o__Saccharomycetales(100);f__Metschnikowiaceae(100);g__Metschnikowia(100);g__Metschnikowia_unclassified(100); | Metschnikowia | 13 | Saprotroph | \|Undefined Saprotroph\| | Yeast | NULL | Probable | As nectar-tap_saprotroph (Põlme et al. 2020); Decay sugar-rich substrates (Põlme et al. 2020) | Põlme S, et al. 2020. Fungal Diversity 105:1-16 ((https://doi.org/10.1007/s13225-020-00466-2)) |
| Otu12216 | 2 | k__Fungi(100);p__Ascomycota(100);c__Saccharomycetes(100);o__Saccharomycetales(100);f__Metschnikowiaceae(100);g__Metschnikowia(100);s__Metschnikowia_pulcherrima(100); | Metschnikowia | 13 | Saprotroph | \|Undefined Saprotroph\| | Yeast | NULL | Probable | As nectar-tap_saprotroph (Põlme et al. 2020); Decay sugar-rich substrates (Põlme et al. 2020) | Põlme S, et al. 2020. Fungal Diversity 105:1-16 ((https://doi.org/10.1007/s13225-020-00466-2)) |
| Otu12217 | 2 | k__Fungi(100);p__Ascomycota(100);c__Saccharomycetes(100);o__Saccharomycetales(100);f__Metschnikowiaceae(100);g__Metschnikowia(100);g__Metschnikowia_unclassified(100); | Metschnikowia | 13 | Saprotroph | \|Undefined Saprotroph\| | Yeast | NULL | Probable | As nectar-tap_saprotroph (Põlme et al. 2020); Decay sugar-rich substrates (Põlme et al. 2020) | Põlme S, et al. 2020. Fungal Diversity 105:1-16 ((https://doi.org/10.1007/s13225-020-00466-2)) |
| Otu12218 | 2 | k__Fungi(100);p__Ascomycota(100);c__Saccharomycetes(100);o__Saccharomycetales(100);f__Metschnikowiaceae(100);g__Metschnikowia(100);s__Metschnikowia_pulcherrima(100); | Metschnikowia | 13 | Saprotroph | \|Undefined Saprotroph\| | Yeast | NULL | Probable | As nectar-tap_saprotroph (Põlme et al. 2020); Decay sugar-rich substrates (Põlme et al. 2020) | Põlme S, et al. 2020. Fungal Diversity 105:1-16 ((https://doi.org/10.1007/s13225-020-00466-2)) |
| Otu12219 | 2 | k__Fungi(100);p__Ascomycota(100);c__Saccharomycetes(100);o__Saccharomycetales(100);f__Metschnikowiaceae(100);g__Metschnikowia(100);g__Metschnikowia_unclassified(100); | Metschnikowia | 13 | Saprotroph | \|Undefined Saprotroph\| | Yeast | NULL | Probable | As nectar-tap_saprotroph (Põlme et al. 2020); Decay sugar-rich substrates (Põlme et al. 2020) | Põlme S, et al. 2020. Fungal Diversity 105:1-16 ((https://doi.org/10.1007/s13225-020-00466-2)) |
| Otu12220 | 2 | k__Fungi(100);p__Ascomycota(100);c__Saccharomycetes(100);o__Saccharomycetales(100);f__Metschnikowiaceae(100);g__Metschnikowia(100);g__Metschnikowia_unclassified(100); | Metschnikowia | 13 | Saprotroph | \|Undefined Saprotroph\| | Yeast | NULL | Probable | As nectar-tap_saprotroph (Põlme et al. 2020); Decay sugar-rich substrates (Põlme et al. 2020) | Põlme S, et al. 2020. Fungal Diversity 105:1-16 ((https://doi.org/10.1007/s13225-020-00466-2)) |
| Otu12221 | 2 | k__Fungi(100);p__Ascomycota(100);c__Saccharomycetes(100);o__Saccharomycetales(100);f__Saccharomycetaceae(100);g__Saccharomyces(100);g__Saccharomyces_unclassified(100); | Saccharomyces | 13 | Saprotroph | \|Undefined Saprotroph\| | Yeast | NULL | Probable | As nectar-tap_saprotroph (Põlme et al. 2020); Decay sugar-rich substrates (Põlme et al. 2020) | James TY, et al. 2006. Nature 443:818-822 ((https://doi.org/10.1038/nature05110)); Põlme S, et al. 2020. Fungal Diversity 105:1-16 ((https://doi.org/10.1007/s13225-020-00466-2)) |
| Otu12222 | 2 | k__Fungi(100);p__Ascomycota(100);c__Saccharomycetes(100);o__Saccharomycetales(100);f__Metschnikowiaceae(100);g__Metschnikowia(100);g__Metschnikowia_unclassified(100); | Metschnikowia | 13 | Saprotroph | \|Undefined Saprotroph\| | Yeast | NULL | Probable | As nectar-tap_saprotroph (Põlme et al. 2020); Decay sugar-rich substrates (Põlme et al. 2020) | Põlme S, et al. 2020. Fungal Diversity 105:1-16 ((https://doi.org/10.1007/s13225-020-00466-2)) |
| Otu12223 | 2 | k__Fungi(100);p__Ascomycota(100);c__Saccharomycetes(100);o__Saccharomycetales(100);f__Metschnikowiaceae(100);g__Metschnikowia(100);g__Metschnikowia_unclassified(100); | Metschnikowia | 13 | Saprotroph | \|Undefined Saprotroph\| | Yeast | NULL | Probable | As nectar-tap_saprotroph (Põlme et al. 2020); Decay sugar-rich substrates (Põlme et al. 2020) | Põlme S, et al. 2020. Fungal Diversity 105:1-16 ((https://doi.org/10.1007/s13225-020-00466-2)) |
| Otu12225 | 2 | k__Fungi(100);p__Ascomycota(100);c__Saccharomycetes(100);o__Saccharomycetales(100);f__Metschnikowiaceae(100);g__Metschnikowia(100);g__Metschnikowia_unclassified(100); | Metschnikowia | 13 | Saprotroph | \|Undefined Saprotroph\| | Yeast | NULL | Probable | As nectar-tap_saprotroph (Põlme et al. 2020); Decay sugar-rich substrates (Põlme et al. 2020) | Põlme S, et al. 2020. Fungal Diversity 105:1-16 ((https://doi.org/10.1007/s13225-020-00466-2)) |
| Otu12227 | 2 | k__Fungi(100);p__Ascomycota(100);c__Sordariomycetes(100);o__Sordariomycetes_order_Incertae_sedis(100);f__Sordariomycetes_family_Incertae_sedis(100);g__Myrmecridium(100);s__Myrmecridium_schulzeri(100); | Myrmecridium | 13 | Saprotroph | \|Undefined Saprotroph\| | Microfungus | NULL | Probable | As soil_saprotroph (Põlme et al. 2020); Decay soil (Põlme et al. 2020) | Tedersoo L, et al. 2014. Science 346:e1256688 ((https://doi.org/10.1126/science.1256688)); Põlme S, et al. 2020. Fungal Diversity 105:1-16 ((https://doi.org/10.1007/s13225-020-00466-2)) |
| Otu12228 | 2 | k__Fungi(100);p__Ascomycota(100);c__Saccharomycetes(100);o__Saccharomycetales(100);f__Metschnikowiaceae(100);g__Metschnikowia(100);g__Metschnikowia_unclassified(100); | Metschnikowia | 13 | Saprotroph | \|Undefined Saprotroph\| | Yeast | NULL | Probable | As nectar-tap_saprotroph (Põlme et al. 2020); Decay sugar-rich substrates (Põlme et al. 2020) | Põlme S, et al. 2020. Fungal Diversity 105:1-16 ((https://doi.org/10.1007/s13225-020-00466-2)) |
| Otu12230 | 2 | k__Fungi(100);p__Ascomycota(100);c__Dothideomycetes(100);o__Pleosporales(100);f__Sporormiaceae(100);g__Preussia(100);s__Preussia_sp_GP_2006a(100); | Preussia | 13 | Saprotroph | \|Dung Saprotroph\|-Undefined Saprotroph | Microfungus | NULL | Probable | Decay dung (Põlme et al. 2020) | Tedersoo L, et al. 2014. Science 346:e1256688 ((https://doi.org/10.1126/science.1256688)); Põlme S, et al. 2020. Fungal Diversity 105:1-16 ((https://doi.org/10.1007/s13225-020-00466-2)) |
| Otu12231 | 2 | k__Fungi(100);p__Ascomycota(100);c__Dothideomycetes(100);o__Capnodiales(100);f__Mycosphaerellaceae(100);f__Mycosphaerellaceae_unclassified(100);f__Mycosphaerellaceae_unclassified(100); | Mycosphaerellaceae | 9 | Pathotroph-Saprotroph | Plant Pathogen-Undefined Saprotroph | Microfungus | NULL | Probable | Biotrophic, necrotrophic or saprobic on various plant tissues (Cannon & Kirk 2007) | Cannon PF, Kirk PM. 2007. Fungal Families of the World. CAB International, Cambridge (ISBN: 978-0851998275) |
| Otu12241 | 2 | k__Fungi(100);p__Ascomycota(100);c__Saccharomycetes(100);o__Saccharomycetales(100);f__Metschnikowiaceae(100);g__Metschnikowia(100);s__Metschnikowia_pulcherrima(100); | Metschnikowia | 13 | Saprotroph | \|Undefined Saprotroph\| | Yeast | NULL | Probable | As nectar-tap_saprotroph (Põlme et al. 2020); Decay sugar-rich substrates (Põlme et al. 2020) | Põlme S, et al. 2020. Fungal Diversity 105:1-16 ((https://doi.org/10.1007/s13225-020-00466-2)) |
| Otu12243 | 2 | k__Fungi(100);p__Ascomycota(100);c__Saccharomycetes(100);o__Saccharomycetales(100);f__Metschnikowiaceae(100);g__Metschnikowia(100);s__Metschnikowia_pulcherrima(100); | Metschnikowia | 13 | Saprotroph | \|Undefined Saprotroph\| | Yeast | NULL | Probable | As nectar-tap_saprotroph (Põlme et al. 2020); Decay sugar-rich substrates (Põlme et al. 2020) | Põlme S, et al. 2020. Fungal Diversity 105:1-16 ((https://doi.org/10.1007/s13225-020-00466-2)) |
| Otu12244 | 2 | k__Fungi(100);p__Ascomycota(100);c__Saccharomycetes(100);o__Saccharomycetales(100);f__Metschnikowiaceae(100);g__Metschnikowia(100);s__Metschnikowia_pulcherrima(100); | Metschnikowia | 13 | Saprotroph | \|Undefined Saprotroph\| | Yeast | NULL | Probable | As nectar-tap_saprotroph (Põlme et al. 2020); Decay sugar-rich substrates (Põlme et al. 2020) | Põlme S, et al. 2020. Fungal Diversity 105:1-16 ((https://doi.org/10.1007/s13225-020-00466-2)) |
| Otu12248 | 2 | k__Fungi(100);p__Ascomycota(100);c__Saccharomycetes(100);o__Saccharomycetales(100);f__Metschnikowiaceae(100);g__Metschnikowia(100);s__Metschnikowia_pulcherrima(100); | Metschnikowia | 13 | Saprotroph | \|Undefined Saprotroph\| | Yeast | NULL | Probable | As nectar-tap_saprotroph (Põlme et al. 2020); Decay sugar-rich substrates (Põlme et al. 2020) | Põlme S, et al. 2020. Fungal Diversity 105:1-16 ((https://doi.org/10.1007/s13225-020-00466-2)) |
| Otu12250 | 2 | k__Fungi(100);p__Ascomycota(100);c__Saccharomycetes(100);o__Saccharomycetales(100);f__Metschnikowiaceae(100);g__Metschnikowia(100);s__Metschnikowia_pulcherrima(100); | Metschnikowia | 13 | Saprotroph | \|Undefined Saprotroph\| | Yeast | NULL | Probable | As nectar-tap_saprotroph (Põlme et al. 2020); Decay sugar-rich substrates (Põlme et al. 2020) | Põlme S, et al. 2020. Fungal Diversity 105:1-16 ((https://doi.org/10.1007/s13225-020-00466-2)) |
| Otu12251 | 2 | k__Fungi(100);p__Ascomycota(100);c__Saccharomycetes(100);o__Saccharomycetales(100);f__Metschnikowiaceae(100);g__Metschnikowia(100);s__Metschnikowia_pulcherrima(100); | Metschnikowia | 13 | Saprotroph | \|Undefined Saprotroph\| | Yeast | NULL | Probable | As nectar-tap_saprotroph (Põlme et al. 2020); Decay sugar-rich substrates (Põlme et al. 2020) | Põlme S, et al. 2020. Fungal Diversity 105:1-16 ((https://doi.org/10.1007/s13225-020-00466-2)) |
| Otu12252 | 2 | k__Fungi(100);p__Ascomycota(100);c__Saccharomycetes(100);o__Saccharomycetales(100);f__Metschnikowiaceae(100);g__Metschnikowia(100);g__Metschnikowia_unclassified(100); | Metschnikowia | 13 | Saprotroph | \|Undefined Saprotroph\| | Yeast | NULL | Probable | As nectar-tap_saprotroph (Põlme et al. 2020); Decay sugar-rich substrates (Põlme et al. 2020) | Põlme S, et al. 2020. Fungal Diversity 105:1-16 ((https://doi.org/10.1007/s13225-020-00466-2)) |
| Otu12254 | 2 | k__Fungi(100);p__Ascomycota(100);c__Saccharomycetes(100);o__Saccharomycetales(100);f__Metschnikowiaceae(100);g__Metschnikowia(100);s__Metschnikowia_pulcherrima(100); | Metschnikowia | 13 | Saprotroph | \|Undefined Saprotroph\| | Yeast | NULL | Probable | As nectar-tap_saprotroph (Põlme et al. 2020); Decay sugar-rich substrates (Põlme et al. 2020) | Põlme S, et al. 2020. Fungal Diversity 105:1-16 ((https://doi.org/10.1007/s13225-020-00466-2)) |
| Otu12259 | 2 | k__Fungi(100);p__Ascomycota(100);c__Saccharomycetes(100);o__Saccharomycetales(100);f__Metschnikowiaceae(100);g__Metschnikowia(100);g__Metschnikowia_unclassified(100); | Metschnikowia | 13 | Saprotroph | \|Undefined Saprotroph\| | Yeast | NULL | Probable | As nectar-tap_saprotroph (Põlme et al. 2020); Decay sugar-rich substrates (Põlme et al. 2020) | Põlme S, et al. 2020. Fungal Diversity 105:1-16 ((https://doi.org/10.1007/s13225-020-00466-2)) |
| Otu12260 | 2 | k__Fungi(100);p__Ascomycota(100);c__Saccharomycetes(100);o__Saccharomycetales(100);f__Metschnikowiaceae(100);g__Metschnikowia(100);s__Metschnikowia_pulcherrima(100); | Metschnikowia | 13 | Saprotroph | \|Undefined Saprotroph\| | Yeast | NULL | Probable | As nectar-tap_saprotroph (Põlme et al. 2020); Decay sugar-rich substrates (Põlme et al. 2020) | Põlme S, et al. 2020. Fungal Diversity 105:1-16 ((https://doi.org/10.1007/s13225-020-00466-2)) |
| Otu12261 | 2 | k__Fungi(100);p__Ascomycota(100);c__Leotiomycetes(100);o__Helotiales(100);f__Hyaloscyphaceae(100);g__Dasyscyphella(100);s__Dasyscyphella_sp_FC_2209(100); | Dasyscyphella | 13 | Saprotroph | Plant Saprotroph-Undefined Saprotroph-\|Wood Saprotroph\| | Microfungus-Pezizoid | NULL | Probable | As litter_saprotroph (Põlme et al. 2020); Decay wood, leaf, fruit, seed (Põlme et al. 2020) | Tedersoo L, et al. 2014. Science 346:e1256688 ((https://doi.org/10.1126/science.1256688)); Põlme S, et al. 2020. Fungal Diversity 105:1-16 ((https://doi.org/10.1007/s13225-020-00466-2)) |
| Otu12262 | 2 | k__Fungi(100);p__Ascomycota(100);c__Leotiomycetes(100);o__Helotiales(100);f__Helotiaceae(100);g__Hymenoscyphus(100);g__Hymenoscyphus_unclassified(100); | Hymenoscyphus | 13 | Pathotroph-Saprotroph-Symbiotroph | Bryophyte Parasite-Ectomycorrhizal-Ericoid Mycorrhizal-Plant Pathogen-\|Plant Saprotroph\|-Undefined Saprotroph | Microfungus | NULL | Possible | As litter_saprotroph (Põlme et al. 2020); Dark Septate Microfungus; Syn. Pezoloma among other (see e.g., Pezoloma ericae); Decay leaf, fruit, seed (Põlme et al. 2020) | Baral HO, Krieglsteiner L. 2006. Acta Mycologica 41:11-20 ((https://doi.org/10.5586/am.2006.003)) (as a bryophyte parasite); Huhtinen S, et al. 2010. Nova Hedwigia 90:413-431 ((https://doi.org/10.1127/0029-5035/2010/0090-0413)); Bent E, et al. 2011. Applied and Environmental Microbiology 77:3351-3359 ((https://doi.org/10.1128/AEM.02575-10)) (as ectomycorrhizae); Tedersoo L, et al. 2014. Science 346:e1256688 ((https://doi.org/10.1126/science.1256688)) (as saprotroph); Põlme S, et al. 2020. Fungal Diversity 105:1-16 ((https://doi.org/10.1007/s13225-020-00466-2)) |
| Otu12263 | 2 | k__Fungi(100);p__Ascomycota(100);c__Sordariomycetes(100);o__Sordariales(100);f__Chaetomiaceae(100);g__Humicola(100);s__Humicola_grisea_var._grisea(100); | Humicola | 13 | Pathotroph-Saprotroph | Plant Pathogen-Undefined Saprotroph-\|Wood Saprotroph\| | Microfungus | Soft Rot | Probable | Decay wood (Põlme et al. 2020) | Seehann G, et al. 1975. List of Fungi in Soft-Rot Tissue Tests. Institute fur Holzbiologie und Holzschutz, Bundesforschungsanstalt fur Forst - und Holzwirtshaft, Hamburg - Lohbrugge, Germany; Costa IPMW, et al. 2012. Mycotaxon 119:e26 ((http://mycotaxon.com/resources/checklists/costa_v119_checklist.pdf)); Tedersoo L, et al. 2014. Science 346:e1256688 ((https://doi.org/10.1126/science.1256688)); Põlme S, et al. 2020. Fungal Diversity 105:1-16 ((https://doi.org/10.1007/s13225-020-00466-2)) |
| Otu12265 | 2 | k__Fungi(100);p__Basidiomycota(100);c__Microbotryomycetes(100);o__Sporidiobolales(100);f__Sporidiobolales_family_Incertae_sedis(100);g__Sporobolomyces(100);s__Sporobolomyces_lactosus(100); | Sporobolomyces | 13 | Pathotroph-Saprotroph | \|Fungal Parasite\|-Plant Saprotroph | Yeast | NULL | Probable | As litter_saprotroph (Põlme et al. 2020); As mycoparasite (Põlme et al. 2020); Decay fungal material (Põlme et al. 2020) | Tedersoo L, et al. 2014. Science 346:e1256688 ((https://doi.org/10.1126/science.1256688)); Purahong W, et al. 2016. Molecular Ecology 25:4059-4074 ((https://doi.org/10.1111/mec.13739)); Põlme S, et al. 2020. Fungal Diversity 105:1-16 ((https://doi.org/10.1007/s13225-020-00466-2)) |
| Otu12267 | 2 | k__Fungi(100);p__Ascomycota(100);c__Saccharomycetes(100);o__Saccharomycetales(100);f__Metschnikowiaceae(100);g__Metschnikowia(100);s__Metschnikowia_pulcherrima(100); | Metschnikowia | 13 | Saprotroph | \|Undefined Saprotroph\| | Yeast | NULL | Probable | As nectar-tap_saprotroph (Põlme et al. 2020); Decay sugar-rich substrates (Põlme et al. 2020) | Põlme S, et al. 2020. Fungal Diversity 105:1-16 ((https://doi.org/10.1007/s13225-020-00466-2)) |
| Otu12268 | 2 | k__Fungi(100);p__Ascomycota(100);c__Dothideomycetes(100);o__Pleosporales(100);f__Venturiaceae(100);g__Venturia(100);g__Venturia_unclassified(100); | Venturia | 13 | Pathotroph-Pathotroph-Saprotroph | \|Plant Pathogen\|-Plant Saprotroph | Microfungus | NULL | Probable | As litter_saprotroph (Põlme et al. 2020); Decay leaf, fruit, seed (Põlme et al. 2020) | Tedersoo L, et al. 2014. Science 346:e1256688 ((https://doi.org/10.1126/science.1256688)); Põlme S, et al. 2020. Fungal Diversity 105:1-16 ((https://doi.org/10.1007/s13225-020-00466-2)) |
| Otu12269 | 2 | k__Fungi(100);p__Ascomycota(100);c__Saccharomycetes(100);o__Saccharomycetales(100);f__Metschnikowiaceae(100);g__Metschnikowia(100);s__Metschnikowia_pulcherrima(100); | Metschnikowia | 13 | Saprotroph | \|Undefined Saprotroph\| | Yeast | NULL | Probable | As nectar-tap_saprotroph (Põlme et al. 2020); Decay sugar-rich substrates (Põlme et al. 2020) | Põlme S, et al. 2020. Fungal Diversity 105:1-16 ((https://doi.org/10.1007/s13225-020-00466-2)) |
| Otu12270 | 2 | k__Fungi(100);p__Ascomycota(100);c__Dothideomycetes(100);o__Pleosporales(100);f__Sporormiaceae(100);g__Preussia(100);s__Preussia_sp_GP_2006a(100); | Preussia | 13 | Saprotroph | \|Dung Saprotroph\|-Undefined Saprotroph | Microfungus | NULL | Probable | Decay dung (Põlme et al. 2020) | Tedersoo L, et al. 2014. Science 346:e1256688 ((https://doi.org/10.1126/science.1256688)); Põlme S, et al. 2020. Fungal Diversity 105:1-16 ((https://doi.org/10.1007/s13225-020-00466-2)) |
| Otu12271 | 2 | k__Fungi(100);p__Basidiomycota(100);c__Microbotryomycetes(100);o__Sporidiobolales(100);f__Sporidiobolales_family_Incertae_sedis(100);g__Sporobolomyces(100);s__Sporobolomyces_lactosus(100); | Sporobolomyces | 13 | Pathotroph-Saprotroph | \|Fungal Parasite\|-Plant Saprotroph | Yeast | NULL | Probable | As litter_saprotroph (Põlme et al. 2020); As mycoparasite (Põlme et al. 2020); Decay fungal material (Põlme et al. 2020) | Tedersoo L, et al. 2014. Science 346:e1256688 ((https://doi.org/10.1126/science.1256688)); Purahong W, et al. 2016. Molecular Ecology 25:4059-4074 ((https://doi.org/10.1111/mec.13739)); Põlme S, et al. 2020. Fungal Diversity 105:1-16 ((https://doi.org/10.1007/s13225-020-00466-2)) |
| Otu12272 | 2 | k__Fungi(100);p__Ascomycota(100);c__Saccharomycetes(100);o__Saccharomycetales(100);f__Metschnikowiaceae(100);g__Metschnikowia(100);s__Metschnikowia_pulcherrima(100); | Metschnikowia | 13 | Saprotroph | \|Undefined Saprotroph\| | Yeast | NULL | Probable | As nectar-tap_saprotroph (Põlme et al. 2020); Decay sugar-rich substrates (Põlme et al. 2020) | Põlme S, et al. 2020. Fungal Diversity 105:1-16 ((https://doi.org/10.1007/s13225-020-00466-2)) |
| Otu12273 | 2 | k__Fungi(100);p__Ascomycota(100);c__Saccharomycetes(100);o__Saccharomycetales(100);f__Metschnikowiaceae(100);g__Metschnikowia(100);s__Metschnikowia_pulcherrima(100); | Metschnikowia | 13 | Saprotroph | \|Undefined Saprotroph\| | Yeast | NULL | Probable | As nectar-tap_saprotroph (Põlme et al. 2020); Decay sugar-rich substrates (Põlme et al. 2020) | Põlme S, et al. 2020. Fungal Diversity 105:1-16 ((https://doi.org/10.1007/s13225-020-00466-2)) |
| Otu12274 | 2 | k__Fungi(100);p__Ascomycota(100);c__Saccharomycetes(100);o__Saccharomycetales(100);f__Metschnikowiaceae(100);g__Metschnikowia(100);s__Metschnikowia_pulcherrima(100); | Metschnikowia | 13 | Saprotroph | \|Undefined Saprotroph\| | Yeast | NULL | Probable | As nectar-tap_saprotroph (Põlme et al. 2020); Decay sugar-rich substrates (Põlme et al. 2020) | Põlme S, et al. 2020. Fungal Diversity 105:1-16 ((https://doi.org/10.1007/s13225-020-00466-2)) |
| Otu12276 | 2 | k__Fungi(100);p__Basidiomycota(100);c__Microbotryomycetes(100);o__Sporidiobolales(100);f__Sporidiobolales_family_Incertae_sedis(100);g__Sporobolomyces(100);s__Sporobolomyces_lactosus(100); | Sporobolomyces | 13 | Pathotroph-Saprotroph | \|Fungal Parasite\|-Plant Saprotroph | Yeast | NULL | Probable | As litter_saprotroph (Põlme et al. 2020); As mycoparasite (Põlme et al. 2020); Decay fungal material (Põlme et al. 2020) | Tedersoo L, et al. 2014. Science 346:e1256688 ((https://doi.org/10.1126/science.1256688)); Purahong W, et al. 2016. Molecular Ecology 25:4059-4074 ((https://doi.org/10.1111/mec.13739)); Põlme S, et al. 2020. Fungal Diversity 105:1-16 ((https://doi.org/10.1007/s13225-020-00466-2)) |
| Otu12277 | 2 | k__Fungi(100);p__Basidiomycota(100);c__Tremellomycetes(100);o__Tremellales(100);f__Tremellales_family_Incertae_sedis(100);g__Hannaella(100);s__Hannaella_luteola(100); | Hannaella | 13 | Pathotroph-Saprotroph | \|Animal Parasite\|-Undefined Saprotroph | Yeast | NULL | Probable | As animal_decomposer (Põlme et al. 2020); Decay animal material (Põlme et al. 2020) | Põlme S, et al. 2020. Fungal Diversity 105:1-16 ((https://doi.org/10.1007/s13225-020-00466-2)) |
| Otu12278 | 2 | k__Fungi(100);p__Basidiomycota(100);c__Microbotryomycetes(100);o__Sporidiobolales(100);f__Sporidiobolales_family_Incertae_sedis(100);g__Rhodotorula(100);s__Rhodotorula_mucilaginosa(100); | Rhodotorula | 13 | Pathotroph-Saprotroph-Symbiotroph | Animal Pathogen-Animal Symbiotroph-Endophyte-Plant Pathogen-\|Undefined Saprotroph\| | Yeast | NULL | Probable | As foliar_endophyte (Põlme et al. 2020); Opportunistic human pathogen | Põlme S, et al. 2020. Fungal Diversity 105:1-16 ((https://doi.org/10.1007/s13225-020-00466-2)); https://www.cdc.gov ((https://www.cdc.gov)) |
| Otu12280 | 2 | k__Fungi(100);p__Ascomycota(100);c__Saccharomycetes(100);o__Saccharomycetales(100);f__Metschnikowiaceae(100);g__Metschnikowia(100);g__Metschnikowia_unclassified(100); | Metschnikowia | 13 | Saprotroph | \|Undefined Saprotroph\| | Yeast | NULL | Probable | As nectar-tap_saprotroph (Põlme et al. 2020); Decay sugar-rich substrates (Põlme et al. 2020) | Põlme S, et al. 2020. Fungal Diversity 105:1-16 ((https://doi.org/10.1007/s13225-020-00466-2)) |
| Otu12281 | 2 | k__Fungi(100);p__Basidiomycota(100);c__Microbotryomycetes(100);o__Sporidiobolales(100);f__Sporidiobolales_family_Incertae_sedis(100);g__Sporobolomyces(100);s__Sporobolomyces_lactosus(100); | Sporobolomyces | 13 | Pathotroph-Saprotroph | \|Fungal Parasite\|-Plant Saprotroph | Yeast | NULL | Probable | As litter_saprotroph (Põlme et al. 2020); As mycoparasite (Põlme et al. 2020); Decay fungal material (Põlme et al. 2020) | Tedersoo L, et al. 2014. Science 346:e1256688 ((https://doi.org/10.1126/science.1256688)); Purahong W, et al. 2016. Molecular Ecology 25:4059-4074 ((https://doi.org/10.1111/mec.13739)); Põlme S, et al. 2020. Fungal Diversity 105:1-16 ((https://doi.org/10.1007/s13225-020-00466-2)) |
| Otu12282 | 2 | k__Fungi(100);p__Ascomycota(100);c__Saccharomycetes(100);o__Saccharomycetales(100);f__Metschnikowiaceae(100);g__Metschnikowia(100);s__Metschnikowia_pulcherrima(100); | Metschnikowia | 13 | Saprotroph | \|Undefined Saprotroph\| | Yeast | NULL | Probable | As nectar-tap_saprotroph (Põlme et al. 2020); Decay sugar-rich substrates (Põlme et al. 2020) | Põlme S, et al. 2020. Fungal Diversity 105:1-16 ((https://doi.org/10.1007/s13225-020-00466-2)) |
| Otu12283 | 2 | k__Fungi(100);p__Basidiomycota(100);c__Microbotryomycetes(100);o__Sporidiobolales(100);f__Sporidiobolales_family_Incertae_sedis(100);g__Sporobolomyces(100);s__Sporobolomyces_lactosus(100); | Sporobolomyces | 13 | Pathotroph-Saprotroph | \|Fungal Parasite\|-Plant Saprotroph | Yeast | NULL | Probable | As litter_saprotroph (Põlme et al. 2020); As mycoparasite (Põlme et al. 2020); Decay fungal material (Põlme et al. 2020) | Tedersoo L, et al. 2014. Science 346:e1256688 ((https://doi.org/10.1126/science.1256688)); Purahong W, et al. 2016. Molecular Ecology 25:4059-4074 ((https://doi.org/10.1111/mec.13739)); Põlme S, et al. 2020. Fungal Diversity 105:1-16 ((https://doi.org/10.1007/s13225-020-00466-2)) |
| Otu12284 | 2 | k__Fungi(100);p__Basidiomycota(100);c__Microbotryomycetes(100);o__Sporidiobolales(100);f__Sporidiobolales_family_Incertae_sedis(100);g__Sporobolomyces(100);s__Sporobolomyces_lactosus(100); | Sporobolomyces | 13 | Pathotroph-Saprotroph | \|Fungal Parasite\|-Plant Saprotroph | Yeast | NULL | Probable | As litter_saprotroph (Põlme et al. 2020); As mycoparasite (Põlme et al. 2020); Decay fungal material (Põlme et al. 2020) | Tedersoo L, et al. 2014. Science 346:e1256688 ((https://doi.org/10.1126/science.1256688)); Purahong W, et al. 2016. Molecular Ecology 25:4059-4074 ((https://doi.org/10.1111/mec.13739)); Põlme S, et al. 2020. Fungal Diversity 105:1-16 ((https://doi.org/10.1007/s13225-020-00466-2)) |
| Otu12285 | 2 | k__Fungi(100);p__Ascomycota(100);c__Saccharomycetes(100);o__Saccharomycetales(100);f__Metschnikowiaceae(100);g__Metschnikowia(100);g__Metschnikowia_unclassified(100); | Metschnikowia | 13 | Saprotroph | \|Undefined Saprotroph\| | Yeast | NULL | Probable | As nectar-tap_saprotroph (Põlme et al. 2020); Decay sugar-rich substrates (Põlme et al. 2020) | Põlme S, et al. 2020. Fungal Diversity 105:1-16 ((https://doi.org/10.1007/s13225-020-00466-2)) |
| Otu12286 | 2 | k__Fungi(100);p__Ascomycota(100);c__Saccharomycetes(100);o__Saccharomycetales(100);f__Metschnikowiaceae(100);g__Metschnikowia(100);s__Metschnikowia_pulcherrima(100); | Metschnikowia | 13 | Saprotroph | \|Undefined Saprotroph\| | Yeast | NULL | Probable | As nectar-tap_saprotroph (Põlme et al. 2020); Decay sugar-rich substrates (Põlme et al. 2020) | Põlme S, et al. 2020. Fungal Diversity 105:1-16 ((https://doi.org/10.1007/s13225-020-00466-2)) |
| Otu12288 | 2 | k__Fungi(100);p__Basidiomycota(100);c__Microbotryomycetes(100);o__Sporidiobolales(100);f__Sporidiobolales_family_Incertae_sedis(100);g__Sporobolomyces(100);s__Sporobolomyces_lactosus(100); | Sporobolomyces | 13 | Pathotroph-Saprotroph | \|Fungal Parasite\|-Plant Saprotroph | Yeast | NULL | Probable | As litter_saprotroph (Põlme et al. 2020); As mycoparasite (Põlme et al. 2020); Decay fungal material (Põlme et al. 2020) | Tedersoo L, et al. 2014. Science 346:e1256688 ((https://doi.org/10.1126/science.1256688)); Purahong W, et al. 2016. Molecular Ecology 25:4059-4074 ((https://doi.org/10.1111/mec.13739)); Põlme S, et al. 2020. Fungal Diversity 105:1-16 ((https://doi.org/10.1007/s13225-020-00466-2)) |
| Otu12290 | 2 | k__Fungi(100);p__Ascomycota(100);c__Saccharomycetes(100);o__Saccharomycetales(100);f__Saccharomycetales_family_Incertae_sedis(100);g__Candida(100);g__Candida_unclassified(100); | Candida | 13 | Pathotroph-Saprotroph-Symbiotroph | Animal Pathogen-Endophyte-Endosymbiont-Epiphyte-\|Undefined Saprotroph\| | Dimorphic Yeast-Yeast | NULL | Probable | As nectar-tap_saprotroph (Põlme et al. 2020); Decay sugar-rich substrates (Põlme et al. 2020); This asexual genus is polyphyletic and contain species that function very differently depend on what they are. Caution should be taken when interpreting guilds for Candida species. | Manolakaki D, et al. 2010. Virulence. 1: 367-375 ((https://doi.org/10.4161/viru.1.5.12796)); Mekbib SB, et al. 2011. Tropical Plant Pathology 36:233-240 ((https://doi.org/10.1590/S1982-56762011000400004)); Põlme S, et al. 2020. Fungal Diversity 105:1-16 ((https://doi.org/10.1007/s13225-020-00466-2)) |
| Otu12293 | 2 | k__Fungi(100);p__Basidiomycota(100);c__Agaricomycetes(100);o__Cantharellales(100);f__Ceratobasidiaceae(100);g__Thanatephorus(100);s__Thanatephorus_cucumeris(100); | Thanatephorus | 13 | Pathotroph-Saprotroph | \|Plant Pathogen\|-Plant Saprotroph | Corticioid | NULL | Probable | As litter_saprotroph (Põlme et al. 2020); Decay leaf, fruit, seed (Põlme et al. 2020) | Tedersoo L, et al. 2014. Science 346:e1256688 ((https://doi.org/10.1126/science.1256688)); Põlme S, et al. 2020. Fungal Diversity 105:1-16 ((https://doi.org/10.1007/s13225-020-00466-2)) |
| Otu12295 | 2 | k__Fungi(100);p__Basidiomycota(100);c__Agaricomycetes(100);o__Polyporales(100);f__Fomitopsidaceae(100);g__Antrodia(100);s__Antrodia_minuta(100); | Antrodia | 13 | Saprotroph | \|Wood Saprotroph\| | Corticioid | Brown Rot | Highly Probable | As brown-rot (Põlme et al. 2020); Decay wood (Põlme et al. 2020) | Gilbertson RL, Ryvarden L. 1987-1987. North American Polypores. Fungiflora, Oslo (ISBN: 978-0945345060); Põlme S, et al. 2020. Fungal Diversity 105:1-16 ((https://doi.org/10.1007/s13225-020-00466-2)) |
| Otu12296 | 2 | k__Fungi(100);p__Ascomycota(100);c__Saccharomycetes(100);o__Saccharomycetales(100);f__Metschnikowiaceae(100);g__Metschnikowia(100);g__Metschnikowia_unclassified(100); | Metschnikowia | 13 | Saprotroph | \|Undefined Saprotroph\| | Yeast | NULL | Probable | As nectar-tap_saprotroph (Põlme et al. 2020); Decay sugar-rich substrates (Põlme et al. 2020) | Põlme S, et al. 2020. Fungal Diversity 105:1-16 ((https://doi.org/10.1007/s13225-020-00466-2)) |
| Otu12297 | 2 | k__Fungi(100);p__Basidiomycota(100);c__Agaricomycetes(100);o__Polyporales(100);f__Polyporaceae(100);g__Trametes(100);s__Trametes_versicolor(100); | Trametes | 13 | Pathotroph-Saprotroph | Plant Pathogen-\|Wood Saprotroph\| | Polyporoid | White Rot | Highly Probable | As white_rot (Põlme et al. 2020); As wood_pathogen (Põlme et al. 2020); Decay wood (Põlme et al. 2020) | Gilbertson RL, Ryvarden L. 1987-1987. North American Polypores. Fungiflora, Oslo (ISBN: 978-0945345060); Põlme S, et al. 2020. Fungal Diversity 105:1-16 ((https://doi.org/10.1007/s13225-020-00466-2)) |
| Otu12298 | 2 | k__Fungi(100);p__Basidiomycota(100);c__Wallemiomycetes(100);o__Wallemiales(100);f__Wallemiaceae(100);g__Wallemia(100);s__Wallemia_sebi(100); | Wallemia | 13 | Saprotroph | \|Undefined Saprotroph\| | Microfungus | NULL | Probable | Decay sugar-rich substrates (Põlme et al. 2020) | Tedersoo L, et al. 2014. Science 346:e1256688 ((https://doi.org/10.1126/science.1256688)); Põlme S, et al. 2020. Fungal Diversity 105:1-16 ((https://doi.org/10.1007/s13225-020-00466-2)) |
| Otu12300 | 2 | k__Fungi(100);p__Ascomycota(100);c__Saccharomycetes(100);o__Saccharomycetales(100);f__Metschnikowiaceae(100);g__Metschnikowia(100);s__Metschnikowia_pulcherrima(100); | Metschnikowia | 13 | Saprotroph | \|Undefined Saprotroph\| | Yeast | NULL | Probable | As nectar-tap_saprotroph (Põlme et al. 2020); Decay sugar-rich substrates (Põlme et al. 2020) | Põlme S, et al. 2020. Fungal Diversity 105:1-16 ((https://doi.org/10.1007/s13225-020-00466-2)) |
| Otu12303 | 2 | k__Fungi(100);p__Ascomycota(100);c__Saccharomycetes(100);o__Saccharomycetales(100);f__Metschnikowiaceae(100);g__Metschnikowia(100);s__Metschnikowia_pulcherrima(100); | Metschnikowia | 13 | Saprotroph | \|Undefined Saprotroph\| | Yeast | NULL | Probable | As nectar-tap_saprotroph (Põlme et al. 2020); Decay sugar-rich substrates (Põlme et al. 2020) | Põlme S, et al. 2020. Fungal Diversity 105:1-16 ((https://doi.org/10.1007/s13225-020-00466-2)) |
| Otu12304 | 2 | k__Fungi(100);p__Ascomycota(100);c__Saccharomycetes(100);o__Saccharomycetales(100);f__Metschnikowiaceae(100);g__Metschnikowia(100);s__Metschnikowia_pulcherrima(100); | Metschnikowia | 13 | Saprotroph | \|Undefined Saprotroph\| | Yeast | NULL | Probable | As nectar-tap_saprotroph (Põlme et al. 2020); Decay sugar-rich substrates (Põlme et al. 2020) | Põlme S, et al. 2020. Fungal Diversity 105:1-16 ((https://doi.org/10.1007/s13225-020-00466-2)) |
| Otu12305 | 2 | k__Fungi(100);p__Basidiomycota(100);c__Agaricomycetes(100);o__Cantharellales(100);f__Ceratobasidiaceae(100);g__Ceratobasidium(100);g__Ceratobasidium_unclassified(100); | Ceratobasidium | 13 | Pathotroph-Saprotroph-Symbiotroph | Ectomycorrhizal-Endomycorrhizal-Orchid Mycorrhizal-\|Plant Pathogen\|-Plant Saprotroph-Undefined Saprotroph | Corticioid-Microfungus | NULL | Possible | especially of Orchidaceae but also even bryophytes (Cannon & Kirk 2007); As litter_saprotroph (Põlme et al. 2020); Best known as plant pathogens, but also saprobic. Species are endomycorrhizal with a wide range of plant roots ; Decay leaf, fruit, seed, roots, soil (Põlme et al. 2020) | Cannon PF, Kirk PM. 2007. Fungal Families of the World. CAB International, Cambridge (ISBN: 978-0851998275); Tedersoo L, et al. 2014. Science 346:e1256688 ((https://doi.org/10.1126/science.1256688)); Põlme S, et al. 2020. Fungal Diversity 105:1-16 ((https://doi.org/10.1007/s13225-020-00466-2)) |
| Otu12306 | 2 | k__Fungi(100);p__Basidiomycota(100);c__Agaricomycetes(100);o__Polyporales(100);f__Meripilaceae(100);g__Rigidoporus(100);s__Rigidoporus_sp_JV0511_23(100); | Rigidoporus | 13 | Saprotroph | \|Wood Saprotroph\| | Polyporoid | White Rot | Highly Probable | As white_rot (Põlme et al. 2020); Decay wood (Põlme et al. 2020) | Gilbertson RL, Ryvarden L. 1987-1987. North American Polypores. Fungiflora, Oslo (ISBN: 978-0945345060); Põlme S, et al. 2020. Fungal Diversity 105:1-16 ((https://doi.org/10.1007/s13225-020-00466-2)) |
| Otu12307 | 2 | k__Fungi(100);p__Basidiomycota(100);c__Agaricomycetes(100);o__Polyporales(100);f__Meruliaceae(100);g__Hyphoderma(100);s__Hyphoderma_setigerum(100); | Hyphoderma | 13 | Saprotroph | Undefined Saprotroph-\|Wood Saprotroph\| | Corticioid | White Rot | Probable | As white_rot (Põlme et al. 2020); Decay wood (Põlme et al. 2020) | Tedersoo L, et al. 2014. Science 346:e1256688 ((https://doi.org/10.1126/science.1256688)); Põlme S, et al. 2020. Fungal Diversity 105:1-16 ((https://doi.org/10.1007/s13225-020-00466-2)) |
| Otu12310 | 2 | k__Fungi(100);p__Basidiomycota(100);c__Agaricomycetes(100);o__Cantharellales(100);f__Ceratobasidiaceae(100);g__Rhizoctonia(100);s__Rhizoctonia_sp_KW214(100); | Rhizoctonia | 13 | Pathotroph-Saprotroph-Symbiotroph | Ectomycorrhizal-Orchid Mycorrhizal-\|Plant Pathogen\|-Plant Saprotroph-Wood Saprotroph | Corticioid-Microfungus | Soft Rot | Probable | As litter_saprotroph (Põlme et al. 2020); Decay leaf, fruit, seed, roots, soil (Põlme et al. 2020); Only symbiotic (perhaps leaning parasitic) in special cases with orchids, otherwise typically a plant pathogen. | Seehann G, et al. 1975. List of Fungi in Soft-Rot Tissue Tests. Institute fur Holzbiologie und Holzschutz, Bundesforschungsanstalt fur Forst - und Holzwirtshaft, Hamburg - Lohbrugge, Germany; Costa IPMW, et al. 2012. Mycotaxon 119:e26 ((http://mycotaxon.com/resources/checklists/costa_v119_checklist.pdf)); Tedersoo L, et al. 2014. Science 346:e1256688 ((https://doi.org/10.1126/science.1256688)); Põlme S, et al. 2020. Fungal Diversity 105:1-16 ((https://doi.org/10.1007/s13225-020-00466-2)); List compiled by Sara Branco with input from Nhu Nguyen and Else Vellinga; The American Phytopathological Society Common Names of Plant Diseases ((http://www.apsnet.org/publications/commonnames/Pages/default.aspx)) |
| Otu12311 | 2 | k__Fungi(100);p__Ascomycota(100);c__Saccharomycetes(100);o__Saccharomycetales(100);f__Metschnikowiaceae(100);g__Metschnikowia(100);g__Metschnikowia_unclassified(100); | Metschnikowia | 13 | Saprotroph | \|Undefined Saprotroph\| | Yeast | NULL | Probable | As nectar-tap_saprotroph (Põlme et al. 2020); Decay sugar-rich substrates (Põlme et al. 2020) | Põlme S, et al. 2020. Fungal Diversity 105:1-16 ((https://doi.org/10.1007/s13225-020-00466-2)) |
| Otu12317 | 2 | k__Fungi(100);p__Zygomycota(100);c__Zygomycota_class_Incertae_sedis(100);o__Kickxellales(100);f__Kickxellaceae(100);g__Ramicandelaber(100);s__Ramicandelaber_sp(100); | Ramicandelaber | 13 | Saprotroph | \|Undefined Saprotroph\| | Microfungus | NULL | Probable | As soil_saprotroph (Põlme et al. 2020); Decay soil (Põlme et al. 2020) | Tedersoo L, et al. 2014. Science 346:e1256688 ((https://doi.org/10.1126/science.1256688)); Põlme S, et al. 2020. Fungal Diversity 105:1-16 ((https://doi.org/10.1007/s13225-020-00466-2)) |
| Otu12321 | 2 | k__Fungi(100);p__Basidiomycota(100);c__Agaricomycetes(100);o__Agaricales(100);f__Tricholomataceae(100);g__Tricholoma(100);s__Tricholoma_fulvum(100); | Tricholoma | 13 | Pathotroph-Symbiotroph | \|Ectomycorrhizal\|-Fungal Parasite | Agaricoid | NULL | Highly Probable | Tricholoma species are ectomycorrhizal, occurring in coniferous and broadleaved forests (Cannon & Kirk 2007); T. sclerotoideum is parasitic on Helvella spp. | Cannon PF, Kirk PM. 2007. Fungal Families of the World. CAB International, Cambridge (ISBN: 978-0851998275); Rinaldi AC, et al. 2008. Fungal Diversity 33:1-45 ((http://www.fungaldiversity.org/fdp/sfdp/33-1.pdf)); Tedersoo L, et al. 2010. Mycorrhiza 20:217-263 ((https://doi.org/10.1007/s00572-009-0274-x)); Põlme S, et al. 2020. Fungal Diversity 105:1-16 ((https://doi.org/10.1007/s13225-020-00466-2)); http://mycorrhizas.info/ecmf.html ((http://mycorrhizas.info/ecmf.html)) |
| Otu12322 | 2 | k__Fungi(100);p__Basidiomycota(100);c__Agaricomycetes(100);o__Corticiales(100);f__Corticiaceae(100);f__Corticiaceae_unclassified(100);f__Corticiaceae_unclassified(100); | Corticiaceae | 9 | Pathotroph-Saprotroph | Lichen Parasite-Plant Pathogen-Wood Saprotroph | Corticioid-Microfungus | NULL | Possible | Saprobes or pathogens on a wide range of mainly woody substrata; a few species are lichenicolous and some are grass pathogens (Cannon & Kirk 2007) | Cannon PF, Kirk PM. 2007. Fungal Families of the World. CAB International, Cambridge (ISBN: 978-0851998275) |
| Otu12323 | 2 | k__Fungi(100);p__Basidiomycota(100);c__Microbotryomycetes(100);o__Sporidiobolales(100);f__Sporidiobolales_family_Incertae_sedis(100);g__Rhodotorula(100);s__Rhodotorula_taiwanensis(100); | Rhodotorula | 13 | Pathotroph-Saprotroph-Symbiotroph | Animal Pathogen-Animal Symbiotroph-Endophyte-Plant Pathogen-\|Undefined Saprotroph\| | Yeast | NULL | Probable | As foliar_endophyte (Põlme et al. 2020); Opportunistic human pathogen | Põlme S, et al. 2020. Fungal Diversity 105:1-16 ((https://doi.org/10.1007/s13225-020-00466-2)); https://www.cdc.gov ((https://www.cdc.gov)) |
| Otu12325 | 2 | k__Fungi(100);p__Ascomycota(100);c__Saccharomycetes(100);o__Saccharomycetales(100);f__Metschnikowiaceae(100);g__Metschnikowia(100);s__Metschnikowia_pulcherrima(100); | Metschnikowia | 13 | Saprotroph | \|Undefined Saprotroph\| | Yeast | NULL | Probable | As nectar-tap_saprotroph (Põlme et al. 2020); Decay sugar-rich substrates (Põlme et al. 2020) | Põlme S, et al. 2020. Fungal Diversity 105:1-16 ((https://doi.org/10.1007/s13225-020-00466-2)) |
| Otu12327 | 2 | k__Fungi(100);p__Ascomycota(100);c__Saccharomycetes(100);o__Saccharomycetales(100);f__Metschnikowiaceae(100);g__Metschnikowia(100);g__Metschnikowia_unclassified(100); | Metschnikowia | 13 | Saprotroph | \|Undefined Saprotroph\| | Yeast | NULL | Probable | As nectar-tap_saprotroph (Põlme et al. 2020); Decay sugar-rich substrates (Põlme et al. 2020) | Põlme S, et al. 2020. Fungal Diversity 105:1-16 ((https://doi.org/10.1007/s13225-020-00466-2)) |
| Otu12329 | 2 | k__Fungi(100);p__Ascomycota(100);c__Saccharomycetes(100);o__Saccharomycetales(100);f__Metschnikowiaceae(100);g__Metschnikowia(100);s__Metschnikowia_pulcherrima(100); | Metschnikowia | 13 | Saprotroph | \|Undefined Saprotroph\| | Yeast | NULL | Probable | As nectar-tap_saprotroph (Põlme et al. 2020); Decay sugar-rich substrates (Põlme et al. 2020) | Põlme S, et al. 2020. Fungal Diversity 105:1-16 ((https://doi.org/10.1007/s13225-020-00466-2)) |
| Otu12330 | 2 | k__Fungi(100);p__Basidiomycota(100);c__Agaricomycetes(100);o__Trechisporales(100);f__Hydnodontaceae(100);g__Sistotremastrum(100);s__Sistotremastrum_guttuliferum(100); | Sistotremastrum | 13 | Saprotroph | Undefined Saprotroph-\|Wood Saprotroph\| | Corticioid | White Rot | Probable | As white_rot (Põlme et al. 2020); Decay wood (Põlme et al. 2020) | Tedersoo L, et al. 2014. Science 346:e1256688 ((https://doi.org/10.1126/science.1256688)); Põlme S, et al. 2020. Fungal Diversity 105:1-16 ((https://doi.org/10.1007/s13225-020-00466-2)) |
| Otu12331 | 2 | k__Fungi(100);p__Ascomycota(100);c__Saccharomycetes(100);o__Saccharomycetales(100);f__Metschnikowiaceae(100);g__Metschnikowia(100);g__Metschnikowia_unclassified(100); | Metschnikowia | 13 | Saprotroph | \|Undefined Saprotroph\| | Yeast | NULL | Probable | As nectar-tap_saprotroph (Põlme et al. 2020); Decay sugar-rich substrates (Põlme et al. 2020) | Põlme S, et al. 2020. Fungal Diversity 105:1-16 ((https://doi.org/10.1007/s13225-020-00466-2)) |
| Otu12332 | 2 | k__Fungi(100);p__Ascomycota(100);c__Saccharomycetes(100);o__Saccharomycetales(100);f__Metschnikowiaceae(100);g__Metschnikowia(100);g__Metschnikowia_unclassified(100); | Metschnikowia | 13 | Saprotroph | \|Undefined Saprotroph\| | Yeast | NULL | Probable | As nectar-tap_saprotroph (Põlme et al. 2020); Decay sugar-rich substrates (Põlme et al. 2020) | Põlme S, et al. 2020. Fungal Diversity 105:1-16 ((https://doi.org/10.1007/s13225-020-00466-2)) |
| Otu12333 | 2 | k__Fungi(100);p__Basidiomycota(100);c__Agaricomycetes(100);o__Polyporales(100);f__Fomitopsidaceae(100);g__Postia(100);s__Postia_tephroleuca(100); | Postia | 13 | Saprotroph | \|Wood Saprotroph\| | Polyporoid | Brown Rot | Highly Probable | As brown-rot (Põlme et al. 2020); Decay wood (Põlme et al. 2020) | Jordan CR, et al. 1996. Holzforschung - International Journal of the Biology, Chemistry, Physics and Technology of Wood 50:312-318 ((https://doi.org/10.1515/hfsg.1996.50.4.312)); Põlme S, et al. 2020. Fungal Diversity 105:1-16 ((https://doi.org/10.1007/s13225-020-00466-2)) |
| Otu12334 | 2 | k__Fungi(100);p__Basidiomycota(100);c__Agaricomycetes(100);o__Polyporales(100);f__Fomitopsidaceae(100);g__Postia(100);s__Postia_ceriflua(100); | Postia | 13 | Saprotroph | \|Wood Saprotroph\| | Polyporoid | Brown Rot | Highly Probable | As brown-rot (Põlme et al. 2020); Decay wood (Põlme et al. 2020) | Jordan CR, et al. 1996. Holzforschung - International Journal of the Biology, Chemistry, Physics and Technology of Wood 50:312-318 ((https://doi.org/10.1515/hfsg.1996.50.4.312)); Põlme S, et al. 2020. Fungal Diversity 105:1-16 ((https://doi.org/10.1007/s13225-020-00466-2)) |
| Otu12336 | 2 | k__Fungi(100);p__Basidiomycota(100);c__Agaricomycetes(100);o__Agaricales(100);f__Tricholomataceae(100);g__Clitocybe(100);s__Clitocybe_sp_MRD022_08(100); | Clitocybe | 13 | Pathotroph-Saprotroph | \|Plant Saprotroph\| | Agaricoid | NULL | Probable | As litter_saprotroph (Põlme et al. 2020); Decay leaf, fruit, seed (Põlme et al. 2020) | Tedersoo L, et al. 2014. Science 346:e1256688 ((https://doi.org/10.1126/science.1256688)); Põlme S, et al. 2020. Fungal Diversity 105:1-16 ((https://doi.org/10.1007/s13225-020-00466-2)) |
| Otu12337 | 2 | k__Fungi(100);p__Zygomycota(100);c__Zygomycota_class_Incertae_sedis(100);o__Mucorales(100);f__Backusellaceae(100);g__Backusella(100);s__Backusella_lamprospora(100); | Backusella | 13 | Saprotroph | Dung Saprotroph-\|Undefined Saprotroph\| | Microfungus-Sporophores | NULL | Probable | As soil_saprotroph (Põlme et al. 2020); Decay soil, dung (Põlme et al. 2020) | Walther G, et al. 2013. Persoonia 30:11-47 ((https://doi.org/10.3767/003158513X665070)); Põlme S, et al. 2020. Fungal Diversity 105:1-16 ((https://doi.org/10.1007/s13225-020-00466-2)) |
| Otu12338 | 2 | k__Fungi(100);p__Ascomycota(100);c__Saccharomycetes(100);o__Saccharomycetales(100);f__Metschnikowiaceae(100);g__Metschnikowia(100);s__Metschnikowia_pulcherrima(100); | Metschnikowia | 13 | Saprotroph | \|Undefined Saprotroph\| | Yeast | NULL | Probable | As nectar-tap_saprotroph (Põlme et al. 2020); Decay sugar-rich substrates (Põlme et al. 2020) | Põlme S, et al. 2020. Fungal Diversity 105:1-16 ((https://doi.org/10.1007/s13225-020-00466-2)) |
| Otu12339 | 2 | k__Fungi(100);p__Ascomycota(100);c__Saccharomycetes(100);o__Saccharomycetales(100);f__Metschnikowiaceae(100);g__Metschnikowia(100);g__Metschnikowia_unclassified(100); | Metschnikowia | 13 | Saprotroph | \|Undefined Saprotroph\| | Yeast | NULL | Probable | As nectar-tap_saprotroph (Põlme et al. 2020); Decay sugar-rich substrates (Põlme et al. 2020) | Põlme S, et al. 2020. Fungal Diversity 105:1-16 ((https://doi.org/10.1007/s13225-020-00466-2)) |
| Otu12340 | 2 | k__Fungi(100);p__Ascomycota(100);c__Saccharomycetes(100);o__Saccharomycetales(100);f__Metschnikowiaceae(100);g__Metschnikowia(100);g__Metschnikowia_unclassified(100); | Metschnikowia | 13 | Saprotroph | \|Undefined Saprotroph\| | Yeast | NULL | Probable | As nectar-tap_saprotroph (Põlme et al. 2020); Decay sugar-rich substrates (Põlme et al. 2020) | Põlme S, et al. 2020. Fungal Diversity 105:1-16 ((https://doi.org/10.1007/s13225-020-00466-2)) |
| Otu12341 | 2 | k__Fungi(100);p__Ascomycota(100);c__Saccharomycetes(100);o__Saccharomycetales(100);f__Saccharomycetales_family_Incertae_sedis(100);f__Saccharomycetales_family_Incertae_sedis_unclassified(100);f__Saccharomycetales_family_Incertae_sedis_unclassified(100); | Saccharomycetales | 7 | Saprotroph | Undefined Saprotroph | Yeast | NULL | Possible | NULL | Sterkenburg E, et al. 2015. New Phytologist 207:1145-1158 ((https://doi.org/10.1111/nph.13426)) |
| Otu12342 | 2 | k__Fungi(100);p__Ascomycota(100);c__Saccharomycetes(100);o__Saccharomycetales(100);f__Metschnikowiaceae(100);g__Metschnikowia(100);g__Metschnikowia_unclassified(100); | Metschnikowia | 13 | Saprotroph | \|Undefined Saprotroph\| | Yeast | NULL | Probable | As nectar-tap_saprotroph (Põlme et al. 2020); Decay sugar-rich substrates (Põlme et al. 2020) | Põlme S, et al. 2020. Fungal Diversity 105:1-16 ((https://doi.org/10.1007/s13225-020-00466-2)) |
| Otu12343 | 2 | k__Fungi(100);p__Ascomycota(100);c__Saccharomycetes(100);o__Saccharomycetales(100);f__Metschnikowiaceae(100);g__Metschnikowia(100);g__Metschnikowia_unclassified(100); | Metschnikowia | 13 | Saprotroph | \|Undefined Saprotroph\| | Yeast | NULL | Probable | As nectar-tap_saprotroph (Põlme et al. 2020); Decay sugar-rich substrates (Põlme et al. 2020) | Põlme S, et al. 2020. Fungal Diversity 105:1-16 ((https://doi.org/10.1007/s13225-020-00466-2)) |
| Otu12344 | 2 | k__Fungi(100);p__Basidiomycota(100);c__Microbotryomycetes(100);o__Sporidiobolales(100);f__Sporidiobolales_family_Incertae_sedis(100);g__Rhodotorula(100);s__Rhodotorula_graminis(100); | Rhodotorula | 13 | Pathotroph-Saprotroph-Symbiotroph | Animal Pathogen-Animal Symbiotroph-Endophyte-Plant Pathogen-\|Undefined Saprotroph\| | Yeast | NULL | Probable | As foliar_endophyte (Põlme et al. 2020); Opportunistic human pathogen | Põlme S, et al. 2020. Fungal Diversity 105:1-16 ((https://doi.org/10.1007/s13225-020-00466-2)); https://www.cdc.gov ((https://www.cdc.gov)) |
| Otu12346 | 2 | k__Fungi(100);p__Ascomycota(100);c__Saccharomycetes(100);o__Saccharomycetales(100);f__Metschnikowiaceae(100);g__Metschnikowia(100);g__Metschnikowia_unclassified(100); | Metschnikowia | 13 | Saprotroph | \|Undefined Saprotroph\| | Yeast | NULL | Probable | As nectar-tap_saprotroph (Põlme et al. 2020); Decay sugar-rich substrates (Põlme et al. 2020) | Põlme S, et al. 2020. Fungal Diversity 105:1-16 ((https://doi.org/10.1007/s13225-020-00466-2)) |
| Otu12348 | 2 | k__Fungi(100);p__Basidiomycota(100);c__Agaricomycetes(100);o__Agaricales(100);o__Agaricales_unclassified(100);o__Agaricales_unclassified(100);o__Agaricales_unclassified(100); | Agaricales | 7 | Pathotroph-Saprotroph-Symbiotroph | Bryophyte Parasite-Dung Saprotroph-Ectomycorrhizal-Fungal Parasite-Leaf Saprotroph-Plant Parasite-Undefined Saprotroph-Wood Saprotroph | Agaricoid-Gasteroid-Microfungus-Secotioid-Yeast | NULL | Possible | Mushrooms and toadstools, Gill fungi, Agarics... lignicolous, sometimes muscicolous or fungicolous, saprobic, mycorrhizal, rarely parasitic on plants or fungi (Kirk et al. 2008) | Kirk PM et al. 2008. Dictionary of the Fungi. Tenth Edition. CAB International, Wallingford (ISBN: 978-0851998268) |
| Otu12351 | 2 | k__Fungi(100);p__Ascomycota(100);c__Saccharomycetes(100);o__Saccharomycetales(100);f__Metschnikowiaceae(100);g__Metschnikowia(100);g__Metschnikowia_unclassified(100); | Metschnikowia | 13 | Saprotroph | \|Undefined Saprotroph\| | Yeast | NULL | Probable | As nectar-tap_saprotroph (Põlme et al. 2020); Decay sugar-rich substrates (Põlme et al. 2020) | Põlme S, et al. 2020. Fungal Diversity 105:1-16 ((https://doi.org/10.1007/s13225-020-00466-2)) |
| Otu12354 | 2 | k__Fungi(100);p__Ascomycota(100);c__Saccharomycetes(100);o__Saccharomycetales(100);f__Metschnikowiaceae(100);g__Metschnikowia(100);g__Metschnikowia_unclassified(100); | Metschnikowia | 13 | Saprotroph | \|Undefined Saprotroph\| | Yeast | NULL | Probable | As nectar-tap_saprotroph (Põlme et al. 2020); Decay sugar-rich substrates (Põlme et al. 2020) | Põlme S, et al. 2020. Fungal Diversity 105:1-16 ((https://doi.org/10.1007/s13225-020-00466-2)) |
| Otu12356 | 2 | k__Fungi(100);p__Ascomycota(100);c__Saccharomycetes(100);o__Saccharomycetales(100);f__Metschnikowiaceae(100);g__Metschnikowia(100);s__Metschnikowia_pulcherrima(100); | Metschnikowia | 13 | Saprotroph | \|Undefined Saprotroph\| | Yeast | NULL | Probable | As nectar-tap_saprotroph (Põlme et al. 2020); Decay sugar-rich substrates (Põlme et al. 2020) | Põlme S, et al. 2020. Fungal Diversity 105:1-16 ((https://doi.org/10.1007/s13225-020-00466-2)) |
| Otu12358 | 2 | k__Fungi(100);p__Basidiomycota(100);c__Agaricomycetes(100);o__Boletales(100);f__Coniophoraceae(100);g__Coniophora(100);s__Coniophora_puteana(100); | Coniophora | 13 | Saprotroph | \|Wood Saprotroph\| | Corticioid | Brown Rot | Probable | As brown-rot (Põlme et al. 2020); Decay wood (Põlme et al. 2020) | Tedersoo L, et al. 2014. Science 346:e1256688 ((https://doi.org/10.1126/science.1256688)); Põlme S, et al. 2020. Fungal Diversity 105:1-16 ((https://doi.org/10.1007/s13225-020-00466-2)) |
| Otu12361 | 2 | k__Fungi(100);p__Basidiomycota(100);c__Microbotryomycetes(100);o__Sporidiobolales(100);f__Sporidiobolales_family_Incertae_sedis(100);g__Rhodotorula(100);g__Rhodotorula_unclassified(100); | Rhodotorula | 13 | Pathotroph-Saprotroph-Symbiotroph | Animal Pathogen-Animal Symbiotroph-Endophyte-Plant Pathogen-\|Undefined Saprotroph\| | Yeast | NULL | Probable | As foliar_endophyte (Põlme et al. 2020); Opportunistic human pathogen | Põlme S, et al. 2020. Fungal Diversity 105:1-16 ((https://doi.org/10.1007/s13225-020-00466-2)); https://www.cdc.gov ((https://www.cdc.gov)) |
| Otu12363 | 2 | k__Fungi(100);p__Ascomycota(100);c__Saccharomycetes(100);o__Saccharomycetales(100);f__Metschnikowiaceae(100);g__Metschnikowia(100);s__Metschnikowia_pulcherrima(100); | Metschnikowia | 13 | Saprotroph | \|Undefined Saprotroph\| | Yeast | NULL | Probable | As nectar-tap_saprotroph (Põlme et al. 2020); Decay sugar-rich substrates (Põlme et al. 2020) | Põlme S, et al. 2020. Fungal Diversity 105:1-16 ((https://doi.org/10.1007/s13225-020-00466-2)) |
| Otu12364 | 2 | k__Fungi(100);p__Basidiomycota(100);c__Agaricomycetes(100);o__Cantharellales(100);f__Ceratobasidiaceae(100);g__Ceratobasidium(100);s__Ceratobasidium_sp(100); | Ceratobasidium | 13 | Pathotroph-Saprotroph-Symbiotroph | Ectomycorrhizal-Endomycorrhizal-Orchid Mycorrhizal-\|Plant Pathogen\|-Plant Saprotroph-Undefined Saprotroph | Corticioid-Microfungus | NULL | Possible | especially of Orchidaceae but also even bryophytes (Cannon & Kirk 2007); As litter_saprotroph (Põlme et al. 2020); Best known as plant pathogens, but also saprobic. Species are endomycorrhizal with a wide range of plant roots ; Decay leaf, fruit, seed, roots, soil (Põlme et al. 2020) | Cannon PF, Kirk PM. 2007. Fungal Families of the World. CAB International, Cambridge (ISBN: 978-0851998275); Tedersoo L, et al. 2014. Science 346:e1256688 ((https://doi.org/10.1126/science.1256688)); Põlme S, et al. 2020. Fungal Diversity 105:1-16 ((https://doi.org/10.1007/s13225-020-00466-2)) |
| Otu12365 | 2 | k__Fungi(100);p__Ascomycota(100);c__Saccharomycetes(100);o__Saccharomycetales(100);f__Metschnikowiaceae(100);g__Metschnikowia(100);g__Metschnikowia_unclassified(100); | Metschnikowia | 13 | Saprotroph | \|Undefined Saprotroph\| | Yeast | NULL | Probable | As nectar-tap_saprotroph (Põlme et al. 2020); Decay sugar-rich substrates (Põlme et al. 2020) | Põlme S, et al. 2020. Fungal Diversity 105:1-16 ((https://doi.org/10.1007/s13225-020-00466-2)) |
| Otu12369 | 2 | k__Fungi(100);p__Ascomycota(100);c__Saccharomycetes(100);o__Saccharomycetales(100);f__Metschnikowiaceae(100);g__Metschnikowia(100);g__Metschnikowia_unclassified(100); | Metschnikowia | 13 | Saprotroph | \|Undefined Saprotroph\| | Yeast | NULL | Probable | As nectar-tap_saprotroph (Põlme et al. 2020); Decay sugar-rich substrates (Põlme et al. 2020) | Põlme S, et al. 2020. Fungal Diversity 105:1-16 ((https://doi.org/10.1007/s13225-020-00466-2)) |
| Otu12370 | 2 | k__Fungi(100);p__Ascomycota(100);c__Saccharomycetes(100);o__Saccharomycetales(100);f__Metschnikowiaceae(100);g__Metschnikowia(100);g__Metschnikowia_unclassified(100); | Metschnikowia | 13 | Saprotroph | \|Undefined Saprotroph\| | Yeast | NULL | Probable | As nectar-tap_saprotroph (Põlme et al. 2020); Decay sugar-rich substrates (Põlme et al. 2020) | Põlme S, et al. 2020. Fungal Diversity 105:1-16 ((https://doi.org/10.1007/s13225-020-00466-2)) |
| Otu12371 | 2 | k__Fungi(100);p__Basidiomycota(100);c__Microbotryomycetes(100);o__Sporidiobolales(100);f__Sporidiobolales_family_Incertae_sedis(100);g__Sporobolomyces(100);s__Sporobolomyces_lactosus(100); | Sporobolomyces | 13 | Pathotroph-Saprotroph | \|Fungal Parasite\|-Plant Saprotroph | Yeast | NULL | Probable | As litter_saprotroph (Põlme et al. 2020); As mycoparasite (Põlme et al. 2020); Decay fungal material (Põlme et al. 2020) | Tedersoo L, et al. 2014. Science 346:e1256688 ((https://doi.org/10.1126/science.1256688)); Purahong W, et al. 2016. Molecular Ecology 25:4059-4074 ((https://doi.org/10.1111/mec.13739)); Põlme S, et al. 2020. Fungal Diversity 105:1-16 ((https://doi.org/10.1007/s13225-020-00466-2)) |
| Otu12372 | 2 | k__Fungi(100);p__Ascomycota(100);c__Saccharomycetes(100);o__Saccharomycetales(100);f__Metschnikowiaceae(100);g__Metschnikowia(100);s__Metschnikowia_pulcherrima(100); | Metschnikowia | 13 | Saprotroph | \|Undefined Saprotroph\| | Yeast | NULL | Probable | As nectar-tap_saprotroph (Põlme et al. 2020); Decay sugar-rich substrates (Põlme et al. 2020) | Põlme S, et al. 2020. Fungal Diversity 105:1-16 ((https://doi.org/10.1007/s13225-020-00466-2)) |
| Otu12373 | 2 | k__Fungi(100);p__Basidiomycota(100);c__Agaricomycetes(100);o__Boletales(100);f__Boletaceae(100);g__Chalciporus(100);g__Chalciporus_unclassified(100); | Chalciporus | 13 | Pathotroph-Symbiotroph | Ectomycorrhizal-\|Fungal Parasite\| | Agaricoid-Boletoid | NULL | Possible | As mycoparasite (Põlme et al. 2020); Decay fungal material (Põlme et al. 2020); Host - Amanita (Põlme et al. 2020); Putatively mycoparasitic (Nuhn et al. 2013); Can usually form erratic mycorrhizal associations with seed plants (Zhang et al. 2015) | Nuhn ME et al. 2013. Fungal Biology 117:7-8 ((https://doi.org/10.1016/j.funbio.2013.04.008)); Tedersoo L, et al. 2014. Science 346:e1256688 ((https://doi.org/10.1126/science.1256688)); Zhang M et al. 2015. Mycoscience 57:20-25 ((https://doi.org/10.1016/j.myc.2015.07.004)); Põlme S, et al. 2020. Fungal Diversity 105:1-16 ((https://doi.org/10.1007/s13225-020-00466-2)) |
| Otu12374 | 2 | k__Fungi(100);p__Basidiomycota(100);c__Agaricomycetes(100);o__Agaricales(100);f__Inocybaceae(100);g__Flammulaster(100);s__Flammulaster_limulatus_var._limulatus(100); | Flammulaster | 13 | Saprotroph | \|Plant Saprotroph\|-Undefined Saprotroph-Wood Saprotroph | Agaricoid | NULL | Probable | As litter_saprotroph (Põlme et al. 2020); Decay leaf, fruit, seed (Põlme et al. 2020) | Tedersoo L, et al. 2014. Science 346:e1256688 ((https://doi.org/10.1126/science.1256688)); Põlme S, et al. 2020. Fungal Diversity 105:1-16 ((https://doi.org/10.1007/s13225-020-00466-2)) |
| Otu12375 | 2 | k__Fungi(100);p__Ascomycota(100);c__Saccharomycetes(100);o__Saccharomycetales(100);f__Metschnikowiaceae(100);g__Metschnikowia(100);s__Metschnikowia_pulcherrima(100); | Metschnikowia | 13 | Saprotroph | \|Undefined Saprotroph\| | Yeast | NULL | Probable | As nectar-tap_saprotroph (Põlme et al. 2020); Decay sugar-rich substrates (Põlme et al. 2020) | Põlme S, et al. 2020. Fungal Diversity 105:1-16 ((https://doi.org/10.1007/s13225-020-00466-2)) |
| Otu12376 | 2 | k__Fungi(100);p__Basidiomycota(100);c__Agaricomycetes(100);o__Cantharellales(100);f__Ceratobasidiaceae(100);g__Ceratobasidium(100);s__Ceratobasidium_sp_AG_I(100); | Ceratobasidium | 13 | Pathotroph-Saprotroph-Symbiotroph | Ectomycorrhizal-Endomycorrhizal-Orchid Mycorrhizal-\|Plant Pathogen\|-Plant Saprotroph-Undefined Saprotroph | Corticioid-Microfungus | NULL | Possible | especially of Orchidaceae but also even bryophytes (Cannon & Kirk 2007); As litter_saprotroph (Põlme et al. 2020); Best known as plant pathogens, but also saprobic. Species are endomycorrhizal with a wide range of plant roots ; Decay leaf, fruit, seed, roots, soil (Põlme et al. 2020) | Cannon PF, Kirk PM. 2007. Fungal Families of the World. CAB International, Cambridge (ISBN: 978-0851998275); Tedersoo L, et al. 2014. Science 346:e1256688 ((https://doi.org/10.1126/science.1256688)); Põlme S, et al. 2020. Fungal Diversity 105:1-16 ((https://doi.org/10.1007/s13225-020-00466-2)) |
| Otu12378 | 2 | k__Fungi(100);p__Ascomycota(100);c__Saccharomycetes(100);o__Saccharomycetales(100);f__Metschnikowiaceae(100);g__Metschnikowia(100);s__Metschnikowia_pulcherrima(100); | Metschnikowia | 13 | Saprotroph | \|Undefined Saprotroph\| | Yeast | NULL | Probable | As nectar-tap_saprotroph (Põlme et al. 2020); Decay sugar-rich substrates (Põlme et al. 2020) | Põlme S, et al. 2020. Fungal Diversity 105:1-16 ((https://doi.org/10.1007/s13225-020-00466-2)) |
| Otu12380 | 2 | k__Fungi(100);p__Ascomycota(100);c__Saccharomycetes(100);o__Saccharomycetales(100);f__Metschnikowiaceae(100);g__Metschnikowia(100);g__Metschnikowia_unclassified(100); | Metschnikowia | 13 | Saprotroph | \|Undefined Saprotroph\| | Yeast | NULL | Probable | As nectar-tap_saprotroph (Põlme et al. 2020); Decay sugar-rich substrates (Põlme et al. 2020) | Põlme S, et al. 2020. Fungal Diversity 105:1-16 ((https://doi.org/10.1007/s13225-020-00466-2)) |
| Otu12381 | 2 | k__Fungi(100);p__Ascomycota(100);c__Saccharomycetes(100);o__Saccharomycetales(100);f__Metschnikowiaceae(100);g__Metschnikowia(100);g__Metschnikowia_unclassified(100); | Metschnikowia | 13 | Saprotroph | \|Undefined Saprotroph\| | Yeast | NULL | Probable | As nectar-tap_saprotroph (Põlme et al. 2020); Decay sugar-rich substrates (Põlme et al. 2020) | Põlme S, et al. 2020. Fungal Diversity 105:1-16 ((https://doi.org/10.1007/s13225-020-00466-2)) |
| Otu12382 | 2 | k__Fungi(100);p__Basidiomycota(100);c__Agaricomycetes(100);o__Polyporales(100);f__Meruliaceae(100);g__Hypochnicium(100);s__Hypochnicium_punctulatum(100); | Hypochnicium | 13 | Saprotroph | Undefined Saprotroph-\|Wood Saprotroph\| | Corticioid | White Rot | Probable | As white_rot (Põlme et al. 2020); Decay wood (Põlme et al. 2020) | Tedersoo L, et al. 2014. Science 346:e1256688 ((https://doi.org/10.1126/science.1256688)); Põlme S, et al. 2020. Fungal Diversity 105:1-16 ((https://doi.org/10.1007/s13225-020-00466-2)) |

| | | | | | | | | | | | | |
|---|---|---|---|---|---|---|---|---|---|---|---|---|
| Otu12388 | | 2 | k__Fungi(100);p__Basidiomycota(100);c__Agaricomycetes(100);o__Hymenochaetales(100);f__Hymenochaetales_family_Incertae_sedis(100);g__Trichaptum(100);s__Trichaptum_biforme(100); | Trichaptum | 13 | Saprotroph | \|Wood Saprotroph\| | Polyporoid | White Rot | Highly Probable | As white_rot (Põlme et al. 2020); Decay wood (Põlme et al. 2020) | Gilbertson RL, Ryvarden L. 1987-1987. North American Polypores. Fungiflora, Oslo (ISBN: 978-0945345060); Põlme S, et al. 2020. Fungal Diversity 105:1-16 ((https://doi.org/10.1007/s13225-020-00466-2)) |
| Otu12391 | | 2 | k__Fungi(100);p__Basidiomycota(100);c__Agaricomycetes(100);o__Trechisporales(100);f__Hydnodontaceae(100);g__Trechispora(100);g__Trechispora_unclassified(100); | Trechispora | 13 | Saprotroph | \|Wood Saprotroph\| | Corticioid | White Rot | Highly Probable | As white_rot (Põlme et al. 2020); Decay wood (Põlme et al. 2020) | Gilbertson RL, Ryvarden L. 1987-1987. North American Polypores. Fungiflora, Oslo (ISBN: 978-0945345060); Rinaldi AC, et al. 2008. Fungal Diversity 33:1-45 ((http://www.fungaldiversity.org/fdp/sfdp/33-1.pdf)) (as Uncertain); Tedersoo L, et al. 2010. Mycorrhiza 20:217-263 ((https://doi.org/10.1007/s00572-009-0274-x)) (as not ECM).; Põlme S, et al. 2020. Fungal Diversity 105:1-16 ((https://doi.org/10.1007/s13225-020-00466-2)) |
| Otu12393 | | 2 | k__Fungi(100);p__Basidiomycota(100);c__Agaricomycetes(100);o__Polyporales(100);f__Fomitopsidaceae(100);g__Antrodia(100);g__Antrodia_unclassified(100); | Antrodia | 13 | Saprotroph | \|Wood Saprotroph\| | Corticioid | Brown Rot | Highly Probable | As brown-rot (Põlme et al. 2020); Decay wood (Põlme et al. 2020) | Gilbertson RL, Ryvarden L. 1987-1987. North American Polypores. Fungiflora, Oslo (ISBN: 978-0945345060); Põlme S, et al. 2020. Fungal Diversity 105:1-16 ((https://doi.org/10.1007/s13225-020-00466-2)) |
| Otu12395 | | 2 | k__Fungi(100);p__Basidiomycota(100);c__Agaricomycetes(100);o__Polyporales(100);f__Meruliaceae(100);g__Hyphoderma(100);s__Hyphoderma_praetermissum(100); | Hyphoderma | 13 | Saprotroph | Undefined Saprotroph-\|Wood Saprotroph\| | Corticioid | White Rot | Probable | As white_rot (Põlme et al. 2020); Decay wood (Põlme et al. 2020) | Tedersoo L, et al. 2014. Science 346:e1256688 ((https://doi.org/10.1126/science.1256688)); Põlme S, et al. 2020. Fungal Diversity 105:1-16 ((https://doi.org/10.1007/s13225-020-00466-2)) |
| Otu12397 | | 2 | k__Fungi(100);p__Ascomycota(100);c__Saccharomycetes(100);o__Saccharomycetales(100);f__Metschnikowiaceae(100);g__Metschnikowia(100);g__Metschnikowia_unclassified(100); | Metschnikowia | 13 | Saprotroph | \|Undefined Saprotroph\| | Yeast | NULL | Probable | As nectar-tap_saprotroph (Põlme et al. 2020); Decay sugar-rich substrates (Põlme et al. 2020) | Põlme S, et al. 2020. Fungal Diversity 105:1-16 ((https://doi.org/10.1007/s13225-020-00466-2)) |
| Otu12398 | | 2 | k__Fungi(100);p__Ascomycota(100);c__Dothideomycetes(100);o__Dothideomycetes_order_Incertae_sedis(100);f__Myxotrichaceae(100);f__Myxotrichaceae_unclassified(100);f__Myxotrichaceae_unclassified(100); | Myxotrichaceae | 9 | Saprotroph-Symbiotroph | Ectomycorrhizal-Undefined Saprotroph | Microfungus | NULL | Probable | Many species are cellulolytic, occurring on paper, straw etc. Others are mycorrhizal associated with roots of Ericaceae and Epacridaceae (Cannon & Kirk 2007) | Cannon PF, Kirk PM. 2007. Fungal Families of the World. CAB International, Cambridge (ISBN: 978-0851998275) |
| Otu12399 | | 2 | k__Fungi(100);p__Ascomycota(100);c__Saccharomycetes(100);o__Saccharomycetales(100);f__Metschnikowiaceae(100);g__Metschnikowia(100);g__Metschnikowia_unclassified(100); | Metschnikowia | 13 | Saprotroph | \|Undefined Saprotroph\| | Yeast | NULL | Probable | As nectar-tap_saprotroph (Põlme et al. 2020); Decay sugar-rich substrates (Põlme et al. 2020) | Põlme S, et al. 2020. Fungal Diversity 105:1-16 ((https://doi.org/10.1007/s13225-020-00466-2)) |
| Otu12400 | | 2 | k__Fungi(100);p__Ascomycota(100);c__Saccharomycetes(100);o__Saccharomycetales(100);f__Metschnikowiaceae(100);g__Metschnikowia(100);s__Metschnikowia_pulcherrima(100); | Metschnikowia | 13 | Saprotroph | \|Undefined Saprotroph\| | Yeast | NULL | Probable | As nectar-tap_saprotroph (Põlme et al. 2020); Decay sugar-rich substrates (Põlme et al. 2020) | Põlme S, et al. 2020. Fungal Diversity 105:1-16 ((https://doi.org/10.1007/s13225-020-00466-2)) |
| Otu12401 | | 2 | k__Fungi(100);p__Ascomycota(100);c__Saccharomycetes(100);o__Saccharomycetales(100);f__Metschnikowiaceae(100);g__Metschnikowia(100);g__Metschnikowia_unclassified(100); | Metschnikowia | 13 | Saprotroph | \|Undefined Saprotroph\| | Yeast | NULL | Probable | As nectar-tap_saprotroph (Põlme et al. 2020); Decay sugar-rich substrates (Põlme et al. 2020) | Põlme S, et al. 2020. Fungal Diversity 105:1-16 ((https://doi.org/10.1007/s13225-020-00466-2)) |
| Otu12402 | | 2 | k__Fungi(100);p__Ascomycota(100);c__Saccharomycetes(100);o__Saccharomycetales(100);f__Metschnikowiaceae(100);g__Metschnikowia(100);s__Metschnikowia_pulcherrima(100); | Metschnikowia | 13 | Saprotroph | \|Undefined Saprotroph\| | Yeast | NULL | Probable | As nectar-tap_saprotroph (Põlme et al. 2020); Decay sugar-rich substrates (Põlme et al. 2020) | Põlme S, et al. 2020. Fungal Diversity 105:1-16 ((https://doi.org/10.1007/s13225-020-00466-2)) |
| Otu12403 | | 2 | k__Fungi(100);p__Ascomycota(100);c__Saccharomycetes(100);o__Saccharomycetales(100);f__Metschnikowiaceae(100);g__Metschnikowia(100);g__Metschnikowia_unclassified(100); | Metschnikowia | 13 | Saprotroph | \|Undefined Saprotroph\| | Yeast | NULL | Probable | As nectar-tap_saprotroph (Põlme et al. 2020); Decay sugar-rich substrates (Põlme et al. 2020) | Põlme S, et al. 2020. Fungal Diversity 105:1-16 ((https://doi.org/10.1007/s13225-020-00466-2)) |
| Otu12407 | | 2 | k__Fungi(100);p__Ascomycota(100);c__Saccharomycetes(100);o__Saccharomycetales(100);f__Metschnikowiaceae(100);g__Metschnikowia(100);s__Metschnikowia_pulcherrima(100); | Metschnikowia | 13 | Saprotroph | \|Undefined Saprotroph\| | Yeast | NULL | Probable | As nectar-tap_saprotroph (Põlme et al. 2020); Decay sugar-rich substrates (Põlme et al. 2020) | Põlme S, et al. 2020. Fungal Diversity 105:1-16 ((https://doi.org/10.1007/s13225-020-00466-2)) |
| Otu12408 | | 2 | k__Fungi(100);p__Basidiomycota(100);c__Agaricomycetes(100);o__Boletales(100);f__Boletaceae(100);g__Boletus(100);s__Boletus_sp(100); | Boletus | 13 | Pathotroph-Symbiotroph | \|Ectomycorrhizal\|-Fungal Parasite | Agaricoid-Boletoid | NULL | Highly Probable | NULL | Rinaldi AC, et al. 2008. Fungal Diversity 33:1-45 ((http://www.fungaldiversity.org/fdp/sfdp/33-1.pdf)); Tedersoo L, et al. 2010. Mycorrhiza 20:217-263 ((https://doi.org/10.1007/s00572-009-0274-x)); Põlme S, et al. 2020. Fungal Diversity 105:1-16 ((https://doi.org/10.1007/s13225-020-00466-2)); http://mycorrhizas.info/ecmf.html ((http://mycorrhizas.info/ecmf.html)); List compiled by John Plischke III ((https://goodmushroom.weebly.com/uploads/5/7/1/7/57176535/fungi_on_fungi_pdf_lq.pdf)) |
| Otu12409 | | 2 | k__Fungi(100);p__Ascomycota(100);c__Saccharomycetes(100);o__Saccharomycetales(100);f__Metschnikowiaceae(100);g__Metschnikowia(100);g__Metschnikowia_unclassified(100); | Metschnikowia | 13 | Saprotroph | \|Undefined Saprotroph\| | Yeast | NULL | Probable | As nectar-tap_saprotroph (Põlme et al. 2020); Decay sugar-rich substrates (Põlme et al. 2020) | Põlme S, et al. 2020. Fungal Diversity 105:1-16 ((https://doi.org/10.1007/s13225-020-00466-2)) |
| Otu12410 | | 2 | k__Fungi(100);p__Ascomycota(100);c__Saccharomycetes(100);o__Saccharomycetales(100);f__Metschnikowiaceae(100);g__Metschnikowia(100);g__Metschnikowia_unclassified(100); | Metschnikowia | 13 | Saprotroph | \|Undefined Saprotroph\| | Yeast | NULL | Probable | As nectar-tap_saprotroph (Põlme et al. 2020); Decay sugar-rich substrates (Põlme et al. 2020) | Põlme S, et al. 2020. Fungal Diversity 105:1-16 ((https://doi.org/10.1007/s13225-020-00466-2)) |
| Otu12412 | | 2 | k__Fungi(100);p__Ascomycota(100);c__Saccharomycetes(100);o__Saccharomycetales(100);f__Metschnikowiaceae(100);g__Metschnikowia(100);g__Metschnikowia_unclassified(100); | Metschnikowia | 13 | Saprotroph | \|Undefined Saprotroph\| | Yeast | NULL | Probable | As nectar-tap_saprotroph (Põlme et al. 2020); Decay sugar-rich substrates (Põlme et al. 2020) | Põlme S, et al. 2020. Fungal Diversity 105:1-16 ((https://doi.org/10.1007/s13225-020-00466-2)) |
| Otu12413 | | 2 | k__Fungi(100);p__Basidiomycota(100);c__Agaricomycetes(100);o__Boletales(100);f__Suillaceae(100);g__Suillus(100);s__Suillus_spectabilis(100); | Suillus | 13 | Symbiotroph | \|Ectomycorrhizal\| | Agaricoid-Boletoid | NULL | Highly Probable | Host - Pinaceae (Põlme et al. 2020) | Rinaldi AC, et al. 2008. Fungal Diversity 33:1-45 ((http://www.fungaldiversity.org/fdp/sfdp/33-1.pdf)); Tedersoo L, et al. 2010. Mycorrhiza 20:217-263 ((https://doi.org/10.1007/s00572-009-0274-x)); Põlme S, et al. 2020. Fungal Diversity 105:1-16 ((https://doi.org/10.1007/s13225-020-00466-2)); http://mycorrhizas.info/ecmf.html ((http://mycorrhizas.info/ecmf.html)) |
| Otu12414 | | 2 | k__Fungi(100);p__Ascomycota(100);c__Saccharomycetes(100);o__Saccharomycetales(100);f__Metschnikowiaceae(100);g__Metschnikowia(100);g__Metschnikowia_unclassified(100); | Metschnikowia | 13 | Saprotroph | \|Undefined Saprotroph\| | Yeast | NULL | Probable | As nectar-tap_saprotroph (Põlme et al. 2020); Decay sugar-rich substrates (Põlme et al. 2020) | Põlme S, et al. 2020. Fungal Diversity 105:1-16 ((https://doi.org/10.1007/s13225-020-00466-2)) |
| Otu12416 | | 2 | k__Fungi(100);p__Basidiomycota(100);c__Microbotryomycetes(100);o__Sporidiobolales(100);f__Sporidiobolales_family_Incertae_sedis(100);g__Rhodotorula(100);s__Rhodotorula_sp(100); | Rhodotorula | 13 | Pathotroph-Saprotroph-Symbiotroph | Animal Pathogen-Animal Symbiotroph-Endophyte-Plant Pathogen-\|Undefined Saprotroph\| | Yeast | NULL | Probable | As foliar_endophyte (Põlme et al. 2020); Opportunistic human pathogen | Põlme S, et al. 2020. Fungal Diversity 105:1-16 ((https://doi.org/10.1007/s13225-020-00466-2)); https://www.cdc.gov ((https://www.cdc.gov)) |
| Otu12420 | | 2 | k__Fungi(100);p__Ascomycota(100);c__Saccharomycetes(100);o__Saccharomycetales(100);f__Metschnikowiaceae(100);g__Metschnikowia(100);s__Metschnikowia_pulcherrima(100); | Metschnikowia | 13 | Saprotroph | \|Undefined Saprotroph\| | Yeast | NULL | Probable | As nectar-tap_saprotroph (Põlme et al. 2020); Decay sugar-rich substrates (Põlme et al. 2020) | Põlme S, et al. 2020. Fungal Diversity 105:1-16 ((https://doi.org/10.1007/s13225-020-00466-2)) |
| Otu12421 | | 2 | k__Fungi(100);p__Ascomycota(100);c__Saccharomycetes(100);o__Saccharomycetales(100);f__Metschnikowiaceae(100);g__Metschnikowia(100);s__Metschnikowia_pulcherrima(100); | Metschnikowia | 13 | Saprotroph | \|Undefined Saprotroph\| | Yeast | NULL | Probable | As nectar-tap_saprotroph (Põlme et al. 2020); Decay sugar-rich substrates (Põlme et al. 2020) | Põlme S, et al. 2020. Fungal Diversity 105:1-16 ((https://doi.org/10.1007/s13225-020-00466-2)) |
| Otu12422 | | 2 | k__Fungi(100);p__Ascomycota(100);c__Saccharomycetes(100);o__Saccharomycetales(100);f__Metschnikowiaceae(100);g__Metschnikowia(100);g__Metschnikowia_unclassified(100); | Metschnikowia | 13 | Saprotroph | \|Undefined Saprotroph\| | Yeast | NULL | Probable | As nectar-tap_saprotroph (Põlme et al. 2020); Decay sugar-rich substrates (Põlme et al. 2020) | Põlme S, et al. 2020. Fungal Diversity 105:1-16 ((https://doi.org/10.1007/s13225-020-00466-2)) |
| Otu12423 | | 2 | k__Fungi(100);p__Basidiomycota(100);c__Microbotryomycetes(100);o__Sporidiobolales(100);f__Sporidiobolales_family_Incertae_sedis(100);g__Rhodotorula(100);s__Rhodotorula_nothofagi(100); | Rhodotorula | 13 | Pathotroph-Saprotroph-Symbiotroph | Animal Pathogen-Animal Symbiotroph-Endophyte-Plant Pathogen-\|Undefined Saprotroph\| | Yeast | NULL | Probable | As foliar_endophyte (Põlme et al. 2020); Opportunistic human pathogen | Põlme S, et al. 2020. Fungal Diversity 105:1-16 ((https://doi.org/10.1007/s13225-020-00466-2)); https://www.cdc.gov ((https://www.cdc.gov)) |
| Otu12425 | | 2 | k__Fungi(100);p__Ascomycota(100);c__Saccharomycetes(100);o__Saccharomycetales(100);f__Saccharomycetales_family_Incertae_sedis(100);g__Candida(100);g__Candida_unclassified(100); | Candida | 13 | Pathotroph-Saprotroph-Symbiotroph | Animal Pathogen-Endophyte-Endosymbiont-Epiphyte-\|Undefined Saprotroph\| | Dimorphic Yeast-Yeast | NULL | Probable | As nectar-tap_saprotroph (Põlme et al. 2020); Decay sugar-rich substrates (Põlme et al. 2020); This asexual genus is polyphyletic and contain species that function very differently depend on what they are. Caution should be taken when interpreting guilds for Candida species. | Manolakaki D, et al. 2010. Virulence. 1: 367-375 ((https://doi.org/10.4161/viru.1.5.12796)); Mekbib SB, et al. 2011. Tropical Plant Pathology 36:233-240 ((https://doi.org/10.1590/S1982-56762011000400004)); Põlme S, et al. 2020. Fungal Diversity 105:1-16 ((https://doi.org/10.1007/s13225-020-00466-2)) |
| Otu12426 | | 2 | k__Fungi(100);p__Ascomycota(100);c__Saccharomycetes(100);o__Saccharomycetales(100);f__Metschnikowiaceae(100);g__Metschnikowia(100);s__Metschnikowia_pulcherrima(100); | Metschnikowia | 13 | Saprotroph | \|Undefined Saprotroph\| | Yeast | NULL | Probable | As nectar-tap_saprotroph (Põlme et al. 2020); Decay sugar-rich substrates (Põlme et al. 2020) | Põlme S, et al. 2020. Fungal Diversity 105:1-16 ((https://doi.org/10.1007/s13225-020-00466-2)) |
| Otu12428 | | 2 | k__Fungi(100);p__Ascomycota(100);c__Saccharomycetes(100);o__Saccharomycetales(100);f__Metschnikowiaceae(100);g__Metschnikowia(100);s__Metschnikowia_pulcherrima(100); | Metschnikowia | 13 | Saprotroph | \|Undefined Saprotroph\| | Yeast | NULL | Probable | As nectar-tap_saprotroph (Põlme et al. 2020); Decay sugar-rich substrates (Põlme et al. 2020) | Põlme S, et al. 2020. Fungal Diversity 105:1-16 ((https://doi.org/10.1007/s13225-020-00466-2)) |
| Otu12431 | | 2 | k__Fungi(100);p__Ascomycota(100);c__Saccharomycetes(100);o__Saccharomycetales(100);f__Metschnikowiaceae(100);g__Metschnikowia(100);s__Metschnikowia_viticola(100); | Metschnikowia | 13 | Saprotroph | \|Undefined Saprotroph\| | Yeast | NULL | Probable | As nectar-tap_saprotroph (Põlme et al. 2020); Decay sugar-rich substrates (Põlme et al. 2020) | Põlme S, et al. 2020. Fungal Diversity 105:1-16 ((https://doi.org/10.1007/s13225-020-00466-2)) |
| Otu12433 | | 2 | k__Fungi(100);p__Basidiomycota(100);c__Microbotryomycetes(100);o__Sporidiobolales(100);f__Sporidiobolales_family_Incertae_sedis(100);g__Rhodotorula(100);g__Rhodotorula_unclassified(100); | Rhodotorula | 13 | Pathotroph-Saprotroph-Symbiotroph | Animal Pathogen-Animal Symbiotroph-Endophyte-Plant Pathogen-\|Undefined Saprotroph\| | Yeast | NULL | Probable | As foliar_endophyte (Põlme et al. 2020); Opportunistic human pathogen | Põlme S, et al. 2020. Fungal Diversity 105:1-16 ((https://doi.org/10.1007/s13225-020-00466-2)); https://www.cdc.gov ((https://www.cdc.gov)) |
| Otu12435 | | 2 | k__Fungi(100);p__Ascomycota(100);c__Saccharomycetes(100);o__Saccharomycetales(100);f__Metschnikowiaceae(100);g__Metschnikowia(100);g__Metschnikowia_unclassified(100); | Metschnikowia | 13 | Saprotroph | \|Undefined Saprotroph\| | Yeast | NULL | Probable | As nectar-tap_saprotroph (Põlme et al. 2020); Decay sugar-rich substrates (Põlme et al. 2020) | Põlme S, et al. 2020. Fungal Diversity 105:1-16 ((https://doi.org/10.1007/s13225-020-00466-2)) |
| Otu12436 | | 2 | k__Fungi(100);p__Ascomycota(100);c__Saccharomycetes(100);o__Saccharomycetales(100);f__Metschnikowiaceae(100);g__Metschnikowia(100);s__Metschnikowia_pulcherrima(100); | Metschnikowia | 13 | Saprotroph | \|Undefined Saprotroph\| | Yeast | NULL | Probable | As nectar-tap_saprotroph (Põlme et al. 2020); Decay sugar-rich substrates (Põlme et al. 2020) | Põlme S, et al. 2020. Fungal Diversity 105:1-16 ((https://doi.org/10.1007/s13225-020-00466-2)) |
| Otu12437 | | 2 | k__Fungi(100);p__Ascomycota(100);c__Saccharomycetes(100);o__Saccharomycetales(100);f__Metschnikowiaceae(100);g__Metschnikowia(100);g__Metschnikowia_unclassified(100); | Metschnikowia | 13 | Saprotroph | \|Undefined Saprotroph\| | Yeast | NULL | Probable | As nectar-tap_saprotroph (Põlme et al. 2020); Decay sugar-rich substrates (Põlme et al. 2020) | Põlme S, et al. 2020. Fungal Diversity 105:1-16 ((https://doi.org/10.1007/s13225-020-00466-2)) |
| Otu12438 | | 2 | k__Fungi(100);p__Ascomycota(100);c__Saccharomycetes(100);o__Saccharomycetales(100);f__Metschnikowiaceae(100);g__Metschnikowia(100);s__Metschnikowia_pulcherrima(100); | Metschnikowia | 13 | Saprotroph | \|Undefined Saprotroph\| | Yeast | NULL | Probable | As nectar-tap_saprotroph (Põlme et al. 2020); Decay sugar-rich substrates (Põlme et al. 2020) | Põlme S, et al. 2020. Fungal Diversity 105:1-16 ((https://doi.org/10.1007/s13225-020-00466-2)) |
| Otu12439 | | 2 | k__Fungi(100);p__Ascomycota(100);c__Saccharomycetes(100);o__Saccharomycetales(100);f__Metschnikowiaceae(100);g__Metschnikowia(100);s__Metschnikowia_pulcherrima(100); | Metschnikowia | 13 | Saprotroph | \|Undefined Saprotroph\| | Yeast | NULL | Probable | As nectar-tap_saprotroph (Põlme et al. 2020); Decay sugar-rich substrates (Põlme et al. 2020) | Põlme S, et al. 2020. Fungal Diversity 105:1-16 ((https://doi.org/10.1007/s13225-020-00466-2)) |
| Otu12442 | | 2 | k__Fungi(100);p__Basidiomycota(100);c__Agaricomycetes(100);o__Boletales(100);f__Boletaceae(100);g__Boletus(100);g__Boletus_unclassified(100); | Boletus | 13 | Pathotroph-Symbiotroph | \|Ectomycorrhizal\|-Fungal Parasite | Agaricoid-Boletoid | NULL | Highly Probable | NULL | Rinaldi AC, et al. 2008. Fungal Diversity 33:1-45 ((http://www.fungaldiversity.org/fdp/sfdp/33-1.pdf)); Tedersoo L, et al. 2010. Mycorrhiza 20:217-263 ((https://doi.org/10.1007/s00572-009-0274-x)); Põlme S, et al. 2020. Fungal Diversity 105:1-16 ((https://doi.org/10.1007/s13225-020-00466-2)); http://mycorrhizas.info/ecmf.html ((http://mycorrhizas.info/ecmf.html)); List compiled by John Plischke III ((https://goodmushroom.weebly.com/uploads/5/7/1/7/57176535/fungi_on_fungi_pdf_lq.pdf)) |
| Otu12443 | | 2 | k__Fungi(100);p__Ascomycota(100);c__Saccharomycetes(100);o__Saccharomycetales(100);f__Metschnikowiaceae(100);g__Metschnikowia(100);g__Metschnikowia_unclassified(100); | Metschnikowia | 13 | Saprotroph | \|Undefined Saprotroph\| | Yeast | NULL | Probable | As nectar-tap_saprotroph (Põlme et al. 2020); Decay sugar-rich substrates (Põlme et al. 2020) | Põlme S, et al. 2020. Fungal Diversity 105:1-16 ((https://doi.org/10.1007/s13225-020-00466-2)) |
| Otu12444 | | 2 | k__Fungi(100);p__Ascomycota(100);c__Saccharomycetes(100);o__Saccharomycetales(100);f__Metschnikowiaceae(100);g__Metschnikowia(100);s__Metschnikowia_pulcherrima(100); | Metschnikowia | 13 | Saprotroph | \|Undefined Saprotroph\| | Yeast | NULL | Probable | As nectar-tap_saprotroph (Põlme et al. 2020); Decay sugar-rich substrates (Põlme et al. 2020) | Põlme S, et al. 2020. Fungal Diversity 105:1-16 ((https://doi.org/10.1007/s13225-020-00466-2)) |
| Otu12445 | | 2 | k__Fungi(100);p__Ascomycota(100);c__Saccharomycetes(100);o__Saccharomycetales(100);f__Metschnikowiaceae(100);g__Metschnikowia(100);s__Metschnikowia_pulcherrima(100); | Metschnikowia | 13 | Saprotroph | \|Undefined Saprotroph\| | Yeast | NULL | Probable | As nectar-tap_saprotroph (Põlme et al. 2020); Decay sugar-rich substrates (Põlme et al. 2020) | Põlme S, et al. 2020. Fungal Diversity 105:1-16 ((https://doi.org/10.1007/s13225-020-00466-2)) |
| Otu12446 | | 2 | k__Fungi(100);p__Ascomycota(100);c__Saccharomycetes(100);o__Saccharomycetales(100);f__Metschnikowiaceae(100);g__Metschnikowia(100);s__Metschnikowia_pulcherrima(100); | Metschnikowia | 13 | Saprotroph | \|Undefined Saprotroph\| | Yeast | NULL | Probable | As nectar-tap_saprotroph (Põlme et al. 2020); Decay sugar-rich substrates (Põlme et al. 2020) | Põlme S, et al. 2020. Fungal Diversity 105:1-16 ((https://doi.org/10.1007/s13225-020-00466-2)) |
| Otu12447 | | 2 | k__Fungi(100);p__Ascomycota(100);c__Saccharomycetes(100);o__Saccharomycetales(100);f__Metschnikowiaceae(100);g__Metschnikowia(100);g__Metschnikowia_unclassified(100); | Metschnikowia | 13 | Saprotroph | \|Undefined Saprotroph\| | Yeast | NULL | Probable | As nectar-tap_saprotroph (Põlme et al. 2020); Decay sugar-rich substrates (Põlme et al. 2020) | Põlme S, et al. 2020. Fungal Diversity 105:1-16 ((https://doi.org/10.1007/s13225-020-00466-2)) |
| Otu12448 | | 2 | k__Fungi(100);p__Ascomycota(100);c__Saccharomycetes(100);o__Saccharomycetales(100);f__Metschnikowiaceae(100);g__Metschnikowia(100);s__Metschnikowia_pulcherrima(100); | Metschnikowia | 13 | Saprotroph | \|Undefined Saprotroph\| | Yeast | NULL | Probable | As nectar-tap_saprotroph (Põlme et al. 2020); Decay sugar-rich substrates (Põlme et al. 2020) | Põlme S, et al. 2020. Fungal Diversity 105:1-16 ((https://doi.org/10.1007/s13225-020-00466-2)) |
| Otu12451 | | 2 | k__Fungi(100);p__Basidiomycota(100);c__Tremellomycetes(100);o__Tremellales(100);o__Tremellales_unclassified(100);o__Tremellales_unclassified(100);o__Tremellales_unclassified(100); | Tremellales | 7 | Pathotroph-Saprotroph-Symbiotroph | Fungal Parasite-Undefined Saprotroph | Tremelloid-Yeast | NULL | Possible | The order contains both teleomorphic and anamorphic species, most of the latter being yeasts | Sterkenburg E, et al. 2015. New Phytologist 207:1145-1158 ((https://doi.org/10.1111/nph.13426)) |
| Otu12452 | | 2 | k__Fungi(100);p__Basidiomycota(100);c__Tremellomycetes(100);o__Filobasidiales(100);f__Filobasidiaceae(100);g__Cryptococcus(100);s__Cryptococcus_friedmannii(100); | Cryptococcus | 13 | Pathotroph-Saprotroph-Symbiotroph | Animal Pathogen-Endophyte-Epiphyte-\|Undefined Saprotroph\| | Dimorphic Yeast-Yeast | NULL | Possible | As foliar_endophyte (Põlme et al. 2020); Decay leaf, fruit, seed, soil, dung, animal material (Põlme et al. 2020) | Andrews JH, et al. 1983. Phytopathology 73:228-234 ((https://doi.org/10.1094/Phyto-73-228)); Kurtzman CP, et al. (eds.) 2011. The Yeasts, a Taxonomic Study. Fifth Edition. Vols 1-3. Elsevier, San Diego (ISBN: 9780444521491); Põlme S, et al. 2020. Fungal Diversity 105:1-16 ((https://doi.org/10.1007/s13225-020-00466-2)) |
| Otu12453 | | 2 | k__Fungi(100);p__Ascomycota(100);c__Saccharomycetes(100);o__Saccharomycetales(100);f__Metschnikowiaceae(100);g__Metschnikowia(100);g__Metschnikowia_unclassified(100); | Metschnikowia | 13 | Saprotroph | \|Undefined Saprotroph\| | Yeast | NULL | Probable | As nectar-tap_saprotroph (Põlme et al. 2020); Decay sugar-rich substrates (Põlme et al. 2020) | Põlme S, et al. 2020. Fungal Diversity 105:1-16 ((https://doi.org/10.1007/s13225-020-00466-2)) |
| Otu12455 | | 2 | k__Fungi(100);p__Basidiomycota(100);c__Agaricomycetes(100);o__Agaricales(100);f__Agaricaceae(100);f__Agaricaceae_unclassified(100);f__Agaricaceae_unclassified(100); | Agaricaceae | 9 | Saprotroph | Undefined Saprotroph | Agaricoid-Gasteroid-Secotioid | NULL | Probable | Primarily saprobes in grassland and woodland situations (Cannon & Kirk 2007) | Cannon PF, Kirk PM. 2007. Fungal Families of the World. CAB International, Cambridge (ISBN: 978-0851998275) |
| Otu12458 | | 2 | k__Fungi(100);p__Ascomycota(100);c__Saccharomycetes(100);o__Saccharomycetales(100);f__Metschnikowiaceae(100);g__Metschnikowia(100);s__Metschnikowia_pulcherrima(100); | Metschnikowia | 13 | Saprotroph | \|Undefined Saprotroph\| | Yeast | NULL | Probable | As nectar-tap_saprotroph (Põlme et al. 2020); Decay sugar-rich substrates (Põlme et al. 2020) | Põlme S, et al. 2020. Fungal Diversity 105:1-16 ((https://doi.org/10.1007/s13225-020-00466-2)) |
| Otu12459 | | 2 | k__Fungi(100);p__Ascomycota(100);c__Saccharomycetes(100);o__Saccharomycetales(100);f__Metschnikowiaceae(100);g__Metschnikowia(100);g__Metschnikowia_unclassified(100); | Metschnikowia | 13 | Saprotroph | \|Undefined Saprotroph\| | Yeast | NULL | Probable | As nectar-tap_saprotroph (Põlme et al. 2020); Decay sugar-rich substrates (Põlme et al. 2020) | Põlme S, et al. 2020. Fungal Diversity 105:1-16 ((https://doi.org/10.1007/s13225-020-00466-2)) |
| Otu12460 | | 2 | k__Fungi(100);p__Ascomycota(100);c__Saccharomycetes(100);o__Saccharomycetales(100);f__Metschnikowiaceae(100);g__Metschnikowia(100);s__Metschnikowia_pulcherrima(100); | Metschnikowia | 13 | Saprotroph | \|Undefined Saprotroph\| | Yeast | NULL | Probable | As nectar-tap_saprotroph (Põlme et al. 2020); Decay sugar-rich substrates (Põlme et al. 2020) | Põlme S, et al. 2020. Fungal Diversity 105:1-16 ((https://doi.org/10.1007/s13225-020-00466-2)) |
| Otu12461 | | 2 | k__Fungi(100);p__Ascomycota(100);c__Saccharomycetes(100);o__Saccharomycetales(100);f__Metschnikowiaceae(100);g__Metschnikowia(100);s__Metschnikowia_pulcherrima(100); | Metschnikowia | 13 | Saprotroph | \|Undefined Saprotroph\| | Yeast | NULL | Probable | As nectar-tap_saprotroph (Põlme et al. 2020); Decay sugar-rich substrates (Põlme et al. 2020) | Põlme S, et al. 2020. Fungal Diversity 105:1-16 ((https://doi.org/10.1007/s13225-020-00466-2)) |
| Otu12463 | | 2 | k__Fungi(100);p__Ascomycota(100);c__Saccharomycetes(100);o__Saccharomycetales(100);f__Metschnikowiaceae(100);g__Metschnikowia(100);s__Metschnikowia_pulcherrima(100); | Metschnikowia | 13 | Saprotroph | \|Undefined Saprotroph\| | Yeast | NULL | Probable | As nectar-tap_saprotroph (Põlme et al. 2020); Decay sugar-rich substrates (Põlme et al. 2020) | Põlme S, et al. 2020. Fungal Diversity 105:1-16 ((https://doi.org/10.1007/s13225-020-00466-2)) |
| Otu12465 | | 2 | k__Fungi(100);p__Ascomycota(100);c__Saccharomycetes(100);o__Saccharomycetales(100);f__Metschnikowiaceae(100);g__Metschnikowia(100);g__Metschnikowia_unclassified(100); | Metschnikowia | 13 | Saprotroph | \|Undefined Saprotroph\| | Yeast | NULL | Probable | As nectar-tap_saprotroph (Põlme et al. 2020); Decay sugar-rich substrates (Põlme et al. 2020) | Põlme S, et al. 2020. Fungal Diversity 105:1-16 ((https://doi.org/10.1007/s13225-020-00466-2)) |
| Otu12466 | | 2 | k__Fungi(100);p__Ascomycota(100);c__Saccharomycetes(100);o__Saccharomycetales(100);f__Metschnikowiaceae(100);g__Metschnikowia(100);s__Metschnikowia_pulcherrima(100); | Metschnikowia | 13 | Saprotroph | \|Undefined Saprotroph\| | Yeast | NULL | Probable | As nectar-tap_saprotroph (Põlme et al. 2020); Decay sugar-rich substrates (Põlme et al. 2020) | Põlme S, et al. 2020. Fungal Diversity 105:1-16 ((https://doi.org/10.1007/s13225-020-00466-2)) |
| Otu12467 | | 2 | k__Fungi(100);p__Ascomycota(100);c__Saccharomycetes(100);o__Saccharomycetales(100);f__Metschnikowiaceae(100);g__Metschnikowia(100);s__Metschnikowia_pulcherrima(100); | Metschnikowia | 13 | Saprotroph | \|Undefined Saprotroph\| | Yeast | NULL | Probable | As nectar-tap_saprotroph (Põlme et al. 2020); Decay sugar-rich substrates (Põlme et al. 2020) | Põlme S, et al. 2020. Fungal Diversity 105:1-16 ((https://doi.org/10.1007/s13225-020-00466-2)) |
| Otu12470 | | 2 | k__Fungi(100);p__Basidiomycota(100);c__Agaricomycetes(100);o__Polyporales(100);f__Phanerochaetaceae(100);g__Ceriporia(100);g__Ceriporia_unclassified(100); | Ceriporia | 13 | Saprotroph | \|Wood Saprotroph\| | Corticioid | White Rot | Highly Probable | As white_rot (Põlme et al. 2020); Decay wood (Põlme et al. 2020) | Tedersoo L, et al. 2014. Science 346:e1256688 ((https://doi.org/10.1126/science.1256688)); Põlme S, et al. 2020. Fungal Diversity 105:1-16 ((https://doi.org/10.1007/s13225-020-00466-2)) |
| Otu12471 | | 2 | k__Fungi(100);p__Ascomycota(100);c__Saccharomycetes(100);o__Saccharomycetales(100);f__Metschnikowiaceae(100);g__Metschnikowia(100);g__Metschnikowia_unclassified(100); | Metschnikowia | 13 | Saprotroph | \|Undefined Saprotroph\| | Yeast | NULL | Probable | As nectar-tap_saprotroph (Põlme et al. 2020); Decay sugar-rich substrates (Põlme et al. 2020) | Põlme S, et al. 2020. Fungal Diversity 105:1-16 ((https://doi.org/10.1007/s13225-020-00466-2)) |
| Otu12473 | | 2 | k__Fungi(100);p__Ascomycota(100);c__Saccharomycetes(100);o__Saccharomycetales(100);f__Metschnikowiaceae(100);g__Metschnikowia(100);g__Metschnikowia_unclassified(100); | Metschnikowia | 13 | Saprotroph | \|Undefined Saprotroph\| | Yeast | NULL | Probable | As nectar-tap_saprotroph (Põlme et al. 2020); Decay sugar-rich substrates (Põlme et al. 2020) | Põlme S, et al. 2020. Fungal Diversity 105:1-16 ((https://doi.org/10.1007/s13225-020-00466-2)) |
| Otu12474 | | 2 | k__Fungi(100);p__Ascomycota(100);c__Saccharomycetes(100);o__Saccharomycetales(100);f__Metschnikowiaceae(100);g__Metschnikowia(100);g__Metschnikowia_unclassified(100); | Metschnikowia | 13 | Saprotroph | \|Undefined Saprotroph\| | Yeast | NULL | Probable | As nectar-tap_saprotroph (Põlme et al. 2020); Decay sugar-rich substrates (Põlme et al. 2020) | Põlme S, et al. 2020. Fungal Diversity 105:1-16 ((https://doi.org/10.1007/s13225-020-00466-2)) |
| Otu12475 | | 2 | k__Fungi(100);p__Ascomycota(100);c__Saccharomycetes(100);o__Saccharomycetales(100);f__Metschnikowiaceae(100);g__Metschnikowia(100);s__Metschnikowia_pulcherrima(100); | Metschnikowia | 13 | Saprotroph | \|Undefined Saprotroph\| | Yeast | NULL | Probable | As nectar-tap_saprotroph (Põlme et al. 2020); Decay sugar-rich substrates (Põlme et al. 2020) | Põlme S, et al. 2020. Fungal Diversity 105:1-16 ((https://doi.org/10.1007/s13225-020-00466-2)) |
| Otu12477 | | 2 | k__Fungi(100);p__Basidiomycota(100);c__Agaricomycetes(100);o__Polyporales(100);f__Xenasmataceae(100);g__Xenasma(100);s__Xenasma_rimicola(100); | Xenasma | 13 | Saprotroph | Undefined Saprotroph-\|Wood Saprotroph\| | Corticioid | White Rot | Probable | As white_rot (Põlme et al. 2020); Decay wood (Põlme et al. 2020) | Tedersoo L, et al. 2014. Science 346:e1256688 ((https://doi.org/10.1126/science.1256688)); Põlme S, et al. 2020. Fungal Diversity 105:1-16 ((https://doi.org/10.1007/s13225-020-00466-2)) |
| Otu12478 | | 2 | k__Fungi(100);p__Ascomycota(100);c__Saccharomycetes(100);o__Saccharomycetales(100);f__Metschnikowiaceae(100);g__Metschnikowia(100);s__Metschnikowia_pulcherrima(100); | Metschnikowia | 13 | Saprotroph | \|Undefined Saprotroph\| | Yeast | NULL | Probable | As nectar-tap_saprotroph (Põlme et al. 2020); Decay sugar-rich substrates (Põlme et al. 2020) | Põlme S, et al. 2020. Fungal Diversity 105:1-16 ((https://doi.org/10.1007/s13225-020-00466-2)) |
| Otu12480 | | 2 | k__Fungi(100);p__Ascomycota(100);c__Leotiomycetes(100);o__Helotiales(100);f__Vibrisseaceae(100);g__Phialocephala(100);s__Phialocephala_fusca(100); | Phialocephala | 13 | Saprotroph-Symbiotroph | Endophyte-\|Undefined Saprotroph\| | Microfungus | NULL | Highly Probable | As root_endophyte (Põlme et al. 2020); As root_endophyte_dark_septate (Põlme et al. 2020); As soil_saprotroph (Põlme et al. 2020); Dark Septate Endophyte, known from plant roots.; Decay roots, soil (Põlme et al. 2020) | Newsham KK. 2011. New Phytologist 190:783-793 ((https://doi.org/10.1111/j.1469-8137.2010.03611.x)) (as dark septate root endophytes); Põlme S, et al. 2020. Fungal Diversity 105:1-16 ((https://doi.org/10.1007/s13225-020-00466-2)) |
| Otu12481 | | 2 | k__Fungi(100);p__Ascomycota(100);c__Saccharomycetes(100);o__Saccharomycetales(100);f__Metschnikowiaceae(100);g__Metschnikowia(100);s__Metschnikowia_pulcherrima(100); | Metschnikowia | 13 | Saprotroph | \|Undefined Saprotroph\| | Yeast | NULL | Probable | As nectar-tap_saprotroph (Põlme et al. 2020); Decay sugar-rich substrates (Põlme et al. 2020) | Põlme S, et al. 2020. Fungal Diversity 105:1-16 ((https://doi.org/10.1007/s13225-020-00466-2)) |
| Otu12484 | | 2 | k__Fungi(100);p__Ascomycota(100);c__Saccharomycetes(100);o__Saccharomycetales(100);f__Metschnikowiaceae(100);g__Metschnikowia(100);s__Metschnikowia_pulcherrima(100); | Metschnikowia | 13 | Saprotroph | \|Undefined Saprotroph\| | Yeast | NULL | Probable | As nectar-tap_saprotroph (Põlme et al. 2020); Decay sugar-rich substrates (Põlme et al. 2020) | Põlme S, et al. 2020. Fungal Diversity 105:1-16 ((https://doi.org/10.1007/s13225-020-00466-2)) |
| Otu12485 | | 2 | k__Fungi(100);p__Ascomycota(100);c__Saccharomycetes(100);o__Saccharomycetales(100);f__Metschnikowiaceae(100);g__Metschnikowia(100);g__Metschnikowia_unclassified(100); | Metschnikowia | 13 | Saprotroph | \|Undefined Saprotroph\| | Yeast | NULL | Probable | As nectar-tap_saprotroph (Põlme et al. 2020); Decay sugar-rich substrates (Põlme et al. 2020) | Põlme S, et al. 2020. Fungal Diversity 105:1-16 ((https://doi.org/10.1007/s13225-020-00466-2)) |
| Otu12486 | | 2 | k__Fungi(100);p__Ascomycota(100);c__Saccharomycetes(100);o__Saccharomycetales(100);f__Metschnikowiaceae(100);g__Metschnikowia(100);s__Metschnikowia_pulcherrima(100); | Metschnikowia | 13 | Saprotroph | \|Undefined Saprotroph\| | Yeast | NULL | Probable | As nectar-tap_saprotroph (Põlme et al. 2020); Decay sugar-rich substrates (Põlme et al. 2020) | Põlme S, et al. 2020. Fungal Diversity 105:1-16 ((https://doi.org/10.1007/s13225-020-00466-2)) |
| Otu12487 | | 2 | k__Fungi(100);p__Ascomycota(100);c__Saccharomycetes(100);o__Saccharomycetales(100);f__Metschnikowiaceae(100);g__Metschnikowia(100);s__Metschnikowia_pulcherrima(100); | Metschnikowia | 13 | Saprotroph | \|Undefined Saprotroph\| | Yeast | NULL | Probable | As nectar-tap_saprotroph (Põlme et al. 2020); Decay sugar-rich substrates (Põlme et al. 2020) | Põlme S, et al. 2020. Fungal Diversity 105:1-16 ((https://doi.org/10.1007/s13225-020-00466-2)) |
| Otu12489 | | 2 | k__Fungi(100);p__Basidiomycota(100);c__Agaricomycetes(100);o__Trechisporales(100);f__Hydnodontaceae(100);g__Trechispora(100);g__Trechispora_unclassified(100); | Trechispora | 13 | Saprotroph | \|Wood Saprotroph\| | Corticioid | White Rot | Highly Probable | As white_rot (Põlme et al. 2020); Decay wood (Põlme et al. 2020) | Gilbertson RL, Ryvarden L. 1987-1987. North American Polypores. Fungiflora, Oslo (ISBN: 978-0945345060); Rinaldi AC, et al. 2008. Fungal Diversity 33:1-45 ((http://www.fungaldiversity.org/fdp/sfdp/33-1.pdf)) (as Uncertain); Tedersoo L, et al. 2010. Mycorrhiza 20:217-263 ((https://doi.org/10.1007/s00572-009-0274-x)) (as not ECM).; Põlme S, et al. 2020. Fungal Diversity 105:1-16 ((https://doi.org/10.1007/s13225-020-00466-2)) |
| Otu12490 | | 2 | k__Fungi(100);p__Basidiomycota(100);c__Agaricomycetes(100);o__Trechisporales(100);f__Hydnodontaceae(100);g__Trechispora(100);s__Trechispora_cohaerens(100); | Trechispora | 13 | Saprotroph | \|Wood Saprotroph\| | Corticioid | White Rot | Highly Probable | As white_rot (Põlme et al. 2020); Decay wood (Põlme et al. 2020) | Gilbertson RL, Ryvarden L. 1987-1987. North American Polypores. Fungiflora, Oslo (ISBN: 978-0945345060); Rinaldi AC, et al. 2008. Fungal Diversity 33:1-45 ((http://www.fungaldiversity.org/fdp/sfdp/33-1.pdf)) (as Uncertain); Tedersoo L, et al. 2010. Mycorrhiza 20:217-263 ((https://doi.org/10.1007/s00572-009-0274-x)) (as not ECM).; Põlme S, et al. 2020. Fungal Diversity 105:1-16 ((https://doi.org/10.1007/s13225-020-00466-2)) |
| Otu12491 | | 2 | k__Fungi(100);p__Basidiomycota(100);c__Tremellomycetes(100);o__Filobasidiales(100);f__Filobasidiaceae(100);g__Cryptococcus(100);s__Cryptococcus_sp(100); | Cryptococcus | 13 | Pathotroph-Saprotroph-Symbiotroph | Animal Pathogen-Endophyte-Epiphyte-\|Undefined Saprotroph\| | Dimorphic Yeast-Yeast | NULL | Possible | As foliar_endophyte (Põlme et al. 2020); Decay leaf, fruit, seed, soil, dung, animal material (Põlme et al. 2020) | Andrews JH, et al. 1983. Phytopathology 73:228-234 ((https://doi.org/10.1094/Phyto-73-228)); Kurtzman CP, et al. (eds.) 2011. The Yeasts, a Taxonomic Study. Fifth Edition. Vols 1-3. Elsevier, San Diego (ISBN: 9780444521491); Põlme S, et al. 2020. Fungal Diversity 105:1-16 ((https://doi.org/10.1007/s13225-020-00466-2)) |
| Otu12493 | | 2 | k__Fungi(100);p__Basidiomycota(100);c__Agaricomycetes(100);o__Trechisporales(100);f__Hydnodontaceae(100);g__Trechispora(100);s__Trechispora_sp(100); | Trechispora | 13 | Saprotroph | \|Wood Saprotroph\| | Corticioid | White Rot | Highly Probable | As white_rot (Põlme et al. 2020); Decay wood (Põlme et al. 2020) | Gilbertson RL, Ryvarden L. 1987-1987. North American Polypores. Fungiflora, Oslo (ISBN: 978-0945345060); Rinaldi AC, et al. 2008. Fungal Diversity 33:1-45 ((http://www.fungaldiversity.org/fdp/sfdp/33-1.pdf)) (as Uncertain); Tedersoo L, et al. 2010. Mycorrhiza 20:217-263 ((https://doi.org/10.1007/s00572-009-0274-x)) (as not ECM).; Põlme S, et al. 2020. Fungal Diversity 105:1-16 ((https://doi.org/10.1007/s13225-020-00466-2)) |
| Otu12494 | | 2 | k__Fungi(100);p__Basidiomycota(100);c__Agaricomycetes(100);o__Gomphales(100);f__Gomphaceae(100);g__Ramaria(100);s__Ramaria_araiospora(100); | Ramaria | 13 | Saprotroph-Symbiotroph | \|Ectomycorrhizal\|-Plant Saprotroph | Clavarioid | NULL | Possible | As litter_saprotroph (Põlme et al. 2020) | Rinaldi AC, et al. 2008. Fungal Diversity 33:1-45 ((http://www.fungaldiversity.org/fdp/sfdp/33-1.pdf)) (as ECM/not ECM); Tedersoo L, et al. 2010. Mycorrhiza 20:217-263 ((10.1007/s00572-009-0274-x)) (pro parte); Põlme S, et al. 2020. Fungal Diversity 105:1-16 ((https://doi.org/10.1007/s13225-020-00466-2)); http://mycorrhizas.info/ecmf.html ((http://mycorrhizas.info/ecmf.html)) (as ECM) |
| Otu12495 | | 2 | k__Fungi(100);p__Basidiomycota(100);c__Agaricomycetes(100);o__Hymenochaetales(100);f__Hymenochaetaceae(100);g__Botryodontia(100);s__Botryodontia_millavensis(100); | Botryodontia | 13 | Saprotroph | Undefined Saprotroph-\|Wood Saprotroph\| | Corticioid | White Rot | Probable | As white_rot (Põlme et al. 2020); Decay wood (Põlme et al. 2020) | Tedersoo L, et al. 2014. Science 346:e1256688 ((https://doi.org/10.1126/science.1256688)); Põlme S, et al. 2020. Fungal Diversity 105:1-16 ((https://doi.org/10.1007/s13225-020-00466-2)) |
| Otu12496 | | 2 | k__Fungi(100);p__Basidiomycota(100);c__Agaricomycetes(100);o__Polyporales(100);f__Meruliaceae(100);g__Steccherinum(100);g__Steccherinum_unclassified(100); | Steccherinum | 13 | Saprotroph | Undefined Saprotroph-\|Wood Saprotroph\| | Corticioid | White Rot | Probable | As white_rot (Põlme et al. 2020); Decay wood (Põlme et al. 2020) | Tedersoo L, et al. 2014. Science 346:e1256688 ((https://doi.org/10.1126/science.1256688)); Põlme S, et al. 2020. Fungal Diversity 105:1-16 ((https://doi.org/10.1007/s13225-020-00466-2)) |
| Otu12497 | | 2 | k__Fungi(100);p__Basidiomycota(100);c__Pucciniomycetes(100);o__Pucciniales(100);f__Pucciniaceae(100);g__Puccinia(100);s__Puccinia_bartholomaei(100); | Puccinia | 13 | Pathotroph | \|Plant Pathogen\| | Rust | NULL | Highly Probable | NULL | Tedersoo L, et al. 2014. Science 346:e1256688 ((https://doi.org/10.1126/science.1256688)); Põlme S, et al. 2020. Fungal Diversity 105:1-16 ((https://doi.org/10.1007/s13225-020-00466-2)) |
| Otu12498 | | 2 | k__Fungi(100);p__Ascomycota(100);c__Saccharomycetes(100);o__Saccharomycetales(100);f__Metschnikowiaceae(100);g__Metschnikowia(100);g__Metschnikowia_unclassified(100); | Metschnikowia | 13 | Saprotroph | \|Undefined Saprotroph\| | Yeast | NULL | Probable | As nectar-tap_saprotroph (Põlme et al. 2020); Decay sugar-rich substrates (Põlme et al. 2020) | Põlme S, et al. 2020. Fungal Diversity 105:1-16 ((https://doi.org/10.1007/s13225-020-00466-2)) |
| Otu12499 | | 2 | k__Fungi(100);p__Basidiomycota(100);c__Agaricomycetes(100);o__Polyporales(100);f__Polyporaceae(100);f__Polyporaceae_unclassified(100);f__Polyporaceae_unclassified(100); | Polyporaceae | 9 | Saprotroph | Wood Saprotroph | Polyporoid | Brown Rot-White | Probable | Lignicolous, causing brown or white rot, mostly of broadleaved trees (Cannon & Kirk 2007) | Cannon PF, Kirk PM. 2007. Fungal Families of the World. CAB International, Cambridge (ISBN: 978-0851998275) |
| Otu12500 | | 2 | k__Fungi(100);p__Ascomycota(100);c__Saccharomycetes(100);o__Saccharomycetales(100);f__Metschnikowiaceae(100);g__Metschnikowia(100);s__Metschnikowia_pulcherrima(100); | Metschnikowia | 13 | Saprotroph | \|Undefined Saprotroph\| | Yeast | NULL | Probable | As nectar-tap_saprotroph (Põlme et al. 2020); Decay sugar-rich substrates (Põlme et al. 2020) | Põlme S, et al. 2020. Fungal Diversity 105:1-16 ((https://doi.org/10.1007/s13225-020-00466-2)) |
| Otu12501 | | 2 | k__Fungi(100);p__Basidiomycota(100);c__Agaricomycetes(100);o__Hymenochaetales(100);f__Schizoporaceae(100);f__Schizoporaceae_unclassified(100);f__Schizoporaceae_unclassified(100); | Schizoporaceae | 9 | Saprotroph | Wood Saprotroph | Corticioid | White Rot | Probable | Saprobic, causing white rot of standing and fallen wood of coniferous and broadleaved trees, rarely on herbaceous stems (Cannon & Kirk 2007) | Cannon PF, Kirk PM. 2007. Fungal Families of the World. CAB International, Cambridge (ISBN: 978-0851998275) |
| Otu12503 | | 2 | k__Fungi(100);p__Basidiomycota(100);c__Microbotryomycetes(100);o__Sporidiobolales(100);f__Sporidiobolales_family_Incertae_sedis(100);g__Sporobolomyces(100);s__Sporobolomyces_subbrunneus(100); | Sporobolomyces | 13 | Pathotroph-Saprotroph | \|Fungal Parasite\|-Plant Saprotroph | Yeast | NULL | Probable | As litter_saprotroph (Põlme et al. 2020); As mycoparasite (Põlme et al. 2020); Decay fungal material (Põlme et al. 2020) | Tedersoo L, et al. 2014. Science 346:e1256688 ((https://doi.org/10.1126/science.1256688)); Purahong W, et al. 2016. Molecular Ecology 25:4059-4074 ((https://doi.org/10.1111/mec.13739)); Põlme S, et al. 2020. Fungal Diversity 105:1-16 ((https://doi.org/10.1007/s13225-020-00466-2)) |
| Otu12504 | | 2 | k__Fungi(100);p__Ascomycota(100);c__Saccharomycetes(100);o__Saccharomycetales(100);f__Metschnikowiaceae(100);g__Metschnikowia(100);g__Metschnikowia_unclassified(100); | Metschnikowia | 13 | Saprotroph | \|Undefined Saprotroph\| | Yeast | NULL | Probable | As nectar-tap_saprotroph (Põlme et al. 2020); Decay sugar-rich substrates (Põlme et al. 2020) | Põlme S, et al. 2020. Fungal Diversity 105:1-16 ((https://doi.org/10.1007/s13225-020-00466-2)) |
| Otu12506 | | 2 | k__Fungi(100);p__Ascomycota(100);c__Saccharomycetes(100);o__Saccharomycetales(100);f__Metschnikowiaceae(100);g__Metschnikowia(100);g__Metschnikowia_unclassified(100); | Metschnikowia | 13 | Saprotroph | \|Undefined Saprotroph\| | Yeast | NULL | Probable | As nectar-tap_saprotroph (Põlme et al. 2020); Decay sugar-rich substrates (Põlme et al. 2020) | Põlme S, et al. 2020. Fungal Diversity 105:1-16 ((https://doi.org/10.1007/s13225-020-00466-2)) |
| Otu12509 | | 2 | k__Fungi(100);p__Ascomycota(100);c__Saccharomycetes(100);o__Saccharomycetales(100);f__Metschnikowiaceae(100);g__Metschnikowia(100);g__Metschnikowia_unclassified(100); | Metschnikowia | 13 | Saprotroph | \|Undefined Saprotroph\| | Yeast | NULL | Probable | As nectar-tap_saprotroph (Põlme et al. 2020); Decay sugar-rich substrates (Põlme et al. 2020) | Põlme S, et al. 2020. Fungal Diversity 105:1-16 ((https://doi.org/10.1007/s13225-020-00466-2)) |
| Otu12510 | | 2 | k__Fungi(100);p__Ascomycota(100);c__Saccharomycetes(100);o__Saccharomycetales(100);f__Metschnikowiaceae(100);g__Metschnikowia(100);s__Metschnikowia_pulcherrima(100); | Metschnikowia | 13 | Saprotroph | \|Undefined Saprotroph\| | Yeast | NULL | Probable | As nectar-tap_saprotroph (Põlme et al. 2020); Decay sugar-rich substrates (Põlme et al. 2020) | Põlme S, et al. 2020. Fungal Diversity 105:1-16 ((https://doi.org/10.1007/s13225-020-00466-2)) |
| Otu12511 | | 2 | k__Fungi(100);p__Ascomycota(100);c__Saccharomycetes(100);o__Saccharomycetales(100);f__Metschnikowiaceae(100);g__Metschnikowia(100);g__Metschnikowia_unclassified(100); | Metschnikowia | 13 | Saprotroph | \|Undefined Saprotroph\| | Yeast | NULL | Probable | As nectar-tap_saprotroph (Põlme et al. 2020); Decay sugar-rich substrates (Põlme et al. 2020) | Põlme S, et al. 2020. Fungal Diversity 105:1-16 ((https://doi.org/10.1007/s13225-020-00466-2)) |
| Otu12512 | | 2 | k__Fungi(100);p__Ascomycota(100);c__Saccharomycetes(100);o__Saccharomycetales(100);f__Metschnikowiaceae(100);g__Metschnikowia(100);g__Metschnikowia_unclassified(100); | Metschnikowia | 13 | Saprotroph | \|Undefined Saprotroph\| | Yeast | NULL | Probable | As nectar-tap_saprotroph (Põlme et al. 2020); Decay sugar-rich substrates (Põlme et al. 2020) | Põlme S, et al. 2020. Fungal Diversity 105:1-16 ((https://doi.org/10.1007/s13225-020-00466-2)) |
| Otu12514 | | 2 | k__Fungi(100);p__Ascomycota(100);c__Saccharomycetes(100);o__Saccharomycetales(100);f__Metschnikowiaceae(100);g__Metschnikowia(100);s__Metschnikowia_pulcherrima(100); | Metschnikowia | 13 | Saprotroph | \|Undefined Saprotroph\| | Yeast | NULL | Probable | As nectar-tap_saprotroph (Põlme et al. 2020); Decay sugar-rich substrates (Põlme et al. 2020) | Põlme S, et al. 2020. Fungal Diversity 105:1-16 ((https://doi.org/10.1007/s13225-020-00466-2)) |
| Otu12515 | | 2 | k__Fungi(100);p__Ascomycota(100);c__Saccharomycetes(100);o__Saccharomycetales(100);f__Metschnikowiaceae(100);g__Metschnikowia(100);g__Metschnikowia_unclassified(100); | Metschnikowia | 13 | Saprotroph | \|Undefined Saprotroph\| | Yeast | NULL | Probable | As nectar-tap_saprotroph (Põlme et al. 2020); Decay sugar-rich substrates (Põlme et al. 2020) | Põlme S, et al. 2020. Fungal Diversity 105:1-16 ((https://doi.org/10.1007/s13225-020-00466-2)) |
| Otu12518 | | 2 | k__Fungi(100);p__Ascomycota(100);c__Saccharomycetes(100);o__Saccharomycetales(100);f__Metschnikowiaceae(100);g__Metschnikowia(100);s__Metschnikowia_pulcherrima(100); | Metschnikowia | 13 | Saprotroph | \|Undefined Saprotroph\| | Yeast | NULL | Probable | As nectar-tap_saprotroph (Põlme et al. 2020); Decay sugar-rich substrates (Põlme et al. 2020) | Põlme S, et al. 2020. Fungal Diversity 105:1-16 ((https://doi.org/10.1007/s13225-020-00466-2)) |
| Otu12521 | | 2 | k__Fungi(100);p__Basidiomycota(100);c__Agaricomycetes(100);o__Polyporales(100);f__Meripilaceae(100);g__Physisporinus(100);s__Physisporinus_vitreus(100); | Physisporinus | 13 | Saprotroph | \|Wood Saprotroph\| | Corticioid | White Rot | Highly Probable | As white_rot (Põlme et al. 2020); Decay wood (Põlme et al. 2020) | Gilbertson RL, Ryvarden L. 1987-1987. North American Polypores. Fungiflora, Oslo (ISBN: 978-0945345060); Põlme S, et al. 2020. Fungal Diversity 105:1-16 ((https://doi.org/10.1007/s13225-020-00466-2)) |
| Otu12526 | | 2 | k__Fungi(100);p__Ascomycota(100);c__Saccharomycetes(100);o__Saccharomycetales(100);f__Metschnikowiaceae(100);g__Metschnikowia(100);g__Metschnikowia_unclassified(100); | Metschnikowia | 13 | Saprotroph | \|Undefined Saprotroph\| | Yeast | NULL | Probable | As nectar-tap_saprotroph (Põlme et al. 2020); Decay sugar-rich substrates (Põlme et al. 2020) | Põlme S, et al. 2020. Fungal Diversity 105:1-16 ((https://doi.org/10.1007/s13225-020-00466-2)) |
| Otu12528 | | 2 | k__Fungi(100);p__Ascomycota(100);c__Saccharomycetes(100);o__Saccharomycetales(100);f__Metschnikowiaceae(100);g__Metschnikowia(100);s__Metschnikowia_pulcherrima(100); | Metschnikowia | 13 | Saprotroph | \|Undefined Saprotroph\| | Yeast | NULL | Probable | As nectar-tap_saprotroph (Põlme et al. 2020); Decay sugar-rich substrates (Põlme et al. 2020) | Põlme S, et al. 2020. Fungal Diversity 105:1-16 ((https://doi.org/10.1007/s13225-020-00466-2)) |
| Otu12529 | | 2 | k__Fungi(100);p__Basidiomycota(100);c__Agaricomycetes(100);o__Agaricales(100);f__Psathyrellaceae(100);g__Coprinopsis(100);s__Coprinopsis_echinospora(100); | Coprinopsis | 13 | Saprotroph | \|Undefined Saprotroph\| | Agaricoid | NULL | Probable | As soil_saprotroph (Põlme et al. 2020); Decay soil (Põlme et al. 2020) | Tedersoo L, et al. 2014. Science 346:e1256688 ((https://doi.org/10.1126/science.1256688)); Põlme S, et al. 2020. Fungal Diversity 105:1-16 ((https://doi.org/10.1007/s13225-020-00466-2)) |
| Otu12530 | | 2 | k__Fungi(100);p__Ascomycota(100);c__Saccharomycetes(100);o__Saccharomycetales(100);f__Metschnikowiaceae(100);g__Metschnikowia(100);g__Metschnikowia_unclassified(100); | Metschnikowia | 13 | Saprotroph | \|Undefined Saprotroph\| | Yeast | NULL | Probable | As nectar-tap_saprotroph (Põlme et al. 2020); Decay sugar-rich substrates (Põlme et al. 2020) | Põlme S, et al. 2020. Fungal Diversity 105:1-16 ((https://doi.org/10.1007/s13225-020-00466-2)) |
| Otu12531 | | 2 | k__Fungi(100);p__Ascomycota(100);c__Saccharomycetes(100);o__Saccharomycetales(100);f__Metschnikowiaceae(100);g__Metschnikowia(100);g__Metschnikowia_unclassified(100); | Metschnikowia | 13 | Saprotroph | \|Undefined Saprotroph\| | Yeast | NULL | Probable | As nectar-tap_saprotroph (Põlme et al. 2020); Decay sugar-rich substrates (Põlme et al. 2020) | Põlme S, et al. 2020. Fungal Diversity 105:1-16 ((https://doi.org/10.1007/s13225-020-00466-2)) |
| Otu12532 | | 2 | k__Fungi(100);p__Basidiomycota(100);c__Agaricomycetes(100);o__Agaricales(100);f__Psathyrellaceae(100);g__Coprinopsis(100);s__Coprinopsis_strossmayeri(100); | Coprinopsis | 13 | Saprotroph | \|Undefined Saprotroph\| | Agaricoid | NULL | Probable | As soil_saprotroph (Põlme et al. 2020); Decay soil (Põlme et al. 2020) | Tedersoo L, et al. 2014. Science 346:e1256688 ((https://doi.org/10.1126/science.1256688)); Põlme S, et al. 2020. Fungal Diversity 105:1-16 ((https://doi.org/10.1007/s13225-020-00466-2)) |
| Otu12533 | | 2 | k__Fungi(100);p__Ascomycota(100);c__Saccharomycetes(100);o__Saccharomycetales(100);f__Metschnikowiaceae(100);g__Metschnikowia(100);s__Metschnikowia_pulcherrima(100); | Metschnikowia | 13 | Saprotroph | \|Undefined Saprotroph\| | Yeast | NULL | Probable | As nectar-tap_saprotroph (Põlme et al. 2020); Decay sugar-rich substrates (Põlme et al. 2020) | Põlme S, et al. 2020. Fungal Diversity 105:1-16 ((https://doi.org/10.1007/s13225-020-00466-2)) |
| Otu12534 | | 2 | k__Fungi(100);p__Ascomycota(100);c__Saccharomycetes(100);o__Saccharomycetales(100);f__Metschnikowiaceae(100);g__Metschnikowia(100);s__Metschnikowia_pulcherrima(100); | Metschnikowia | 13 | Saprotroph | \|Undefined Saprotroph\| | Yeast | NULL | Probable | As nectar-tap_saprotroph (Põlme et al. 2020); Decay sugar-rich substrates (Põlme et al. 2020) | Põlme S, et al. 2020. Fungal Diversity 105:1-16 ((https://doi.org/10.1007/s13225-020-00466-2)) |
| Otu12540 | | 2 | k__Fungi(100);p__Ascomycota(100);c__Saccharomycetes(100);o__Saccharomycetales(100);f__Metschnikowiaceae(100);g__Metschnikowia(100);s__Metschnikowia_pulcherrima(100); | Metschnikowia | 13 | Saprotroph | \|Undefined Saprotroph\| | Yeast | NULL | Probable | As nectar-tap_saprotroph (Põlme et al. 2020); Decay sugar-rich substrates (Põlme et al. 2020) | Põlme S, et al. 2020. Fungal Diversity 105:1-16 ((https://doi.org/10.1007/s13225-020-00466-2)) |
| Otu12541 | | 2 | k__Fungi(100);p__Basidiomycota(100);c__Agaricomycetes(100);o__Agaricales(100);f__Strophariaceae(100);f__Strophariaceae_unclassified(100);f__Strophariaceae_unclassified(100); | Strophariaceae | 9 | Pathotroph-Saprotroph | Dung Saprotroph-Plant Parasite-Soil Saprotroph-Undefined Saprotroph-Wood Saprotroph | Agaricoid | NULL | Probable | Very variable ecology; Pholiota species are lignicolous and can act as weak parasites while most other taxa are either saprobes in grassy habitats or are found in dung. Species are not ectomycorrhizal (Cannon & Kirk 2007) | Cannon PF, Kirk PM. 2007. Fungal Families of the World. CAB International, Cambridge (ISBN: 978-0851998275) |
| Otu12546 | | 2 | k__Fungi(100);p__Basidiomycota(100);c__Agaricomycetes(100);o__Polyporales(100);f__Phanerochaetaceae(100);g__Ceriporiopsis(100);s__Ceriporiopsis_sp_DLL2011_1(100); | Ceriporiopsis | 13 | Saprotroph | \|Wood Saprotroph\| | Corticioid | White Rot | Highly Probable | As white_rot (Põlme et al. 2020); Decay wood (Põlme et al. 2020) | Gilbertson RL, Ryvarden L. 1987-1987. North American Polypores. Fungiflora, Oslo (ISBN: 978-0945345060); Tedersoo L, et al. 2014. Science 346:e1256688 ((https://doi.org/10.1126/science.1256688)); Põlme S, et al. 2020. Fungal Diversity 105:1-16 ((https://doi.org/10.1007/s13225-020-00466-2)) |
| Otu12547 | | 2 | k__Fungi(100);p__Basidiomycota(100);c__Agaricomycetes(100);o__Agaricales(100);f__Amanitaceae(100);g__Amanita(100);s__Amanita_sp(100); | Amanita | 13 | Saprotroph-Symbiotroph | \|Ectomycorrhizal\|-Undefined Saprotroph | Agaricoid | NULL | Probable | Some members of the genus are deadly poisonous | Rinaldi AC, et al. 2008. Fungal Diversity 33:1-45 ((http://www.fungaldiversity.org/fdp/sfdp/33-1.pdf)); Tedersoo L, et al. 2010. Mycorrhiza 20:217-263 ((https://doi.org/10.1007/s00572-009-0274-x)); Wolfe BE, et al. 2012. PLoS ONE 7:e39597 ((https://doi.org/10.1371/journal.pone.0039597)); Põlme S, et al. 2020. Fungal Diversity 105:1-16 ((https://doi.org/10.1007/s13225-020-00466-2)); http://mycorrhizas.info/ecmf.html ((http://mycorrhizas.info/ecmf.html)); http://en.wikipedia.org/wiki/List_of_deadly_fungi ((http://en.wikipedia.org/wiki/List_of_deadly_fungi)) |
| Otu12548 | | 2 | k__Fungi(100);p__Basidiomycota(100);c__Agaricomycetes(100);o__Agaricales(100);f__Inocybaceae(100);g__Crepidotus(100);s__Crepidotus_crocophyllus(100); | Crepidotus | 13 | Saprotroph | \|Wood Saprotroph\| | Agaricoid | NULL | Highly Probable | Decay wood (Põlme et al. 2020) | Grgurinovic CA, Simpson JA. 2001. Fungal Diversity 8:97-127 ((http://www.fungaldiversity.org/fdp/sfdp/FD_8_97-127.pdf)); Aime MC, et al. 2005. American Journal of Botany 92:74-82 ((https://doi.org/10.3732/ajb.92.1.74)); Tedersoo L, et al. 2014. Science 346:e1256688 ((https://doi.org/10.1126/science.1256688)); Põlme S, et al. 2020. Fungal Diversity 105:1-16 ((https://doi.org/10.1007/s13225-020-00466-2)) |
| Otu12550 | | 2 | k__Fungi(100);p__Ascomycota(100);c__Saccharomycetes(100);o__Saccharomycetales(100);f__Metschnikowiaceae(100);g__Metschnikowia(100);s__Metschnikowia_pulcherrima(100); | Metschnikowia | 13 | Saprotroph | \|Undefined Saprotroph\| | Yeast | NULL | Probable | As nectar-tap_saprotroph (Põlme et al. 2020); Decay sugar-rich substrates (Põlme et al. 2020) | Põlme S, et al. 2020. Fungal Diversity 105:1-16 ((https://doi.org/10.1007/s13225-020-00466-2)) |
| Otu12551 | | 2 | k__Fungi(100);p__Ascomycota(100);c__Saccharomycetes(100);o__Saccharomycetales(100);f__Metschnikowiaceae(100);g__Metschnikowia(100);s__Metschnikowia_pulcherrima(100); | Metschnikowia | 13 | Saprotroph | \|Undefined Saprotroph\| | Yeast | NULL | Probable | As nectar-tap_saprotroph (Põlme et al. 2020); Decay sugar-rich substrates (Põlme et al. 2020) | Põlme S, et al. 2020. Fungal Diversity 105:1-16 ((https://doi.org/10.1007/s13225-020-00466-2)) |
| Otu12552 | | 2 | k__Fungi(100);p__Basidiomycota(100);c__Microbotryomycetes(100);o__Sporidiobolales(100);f__Sporidiobolales_family_Incertae_sedis(100);g__Sporobolomyces(100);s__Sporobolomyces_ruberrimus(100); | Sporobolomyces | 13 | Pathotroph-Saprotroph | \|Fungal Parasite\|-Plant Saprotroph | Yeast | NULL | Probable | As litter_saprotroph (Põlme et al. 2020); As mycoparasite (Põlme et al. 2020); Decay fungal material (Põlme et al. 2020) | Tedersoo L, et al. 2014. Science 346:e1256688 ((https://doi.org/10.1126/science.1256688)); Purahong W, et al. 2016. Molecular Ecology 25:4059-4074 ((https://doi.org/10.1111/mec.13739)); Põlme S, et al. 2020. Fungal Diversity 105:1-16 ((https://doi.org/10.1007/s13225-020-00466-2)) |
| Otu12553 | | 2 | k__Fungi(100);p__Basidiomycota(100);c__Microbotryomycetes(100);o__Sporidiobolales(100);f__Sporidiobolales_family_Incertae_sedis(100);g__Sporobolomyces(100);s__Sporobolomyces_sp_TY_223(100); | Sporobolomyces | 13 | Pathotroph-Saprotroph | \|Fungal Parasite\|-Plant Saprotroph | Yeast | NULL | Probable | As litter_saprotroph (Põlme et al. 2020); As mycoparasite (Põlme et al. 2020); Decay fungal material (Põlme et al. 2020) | Tedersoo L, et al. 2014. Science 346:e1256688 ((https://doi.org/10.1126/science.1256688)); Purahong W, et al. 2016. Molecular Ecology 25:4059-4074 ((https://doi.org/10.1111/mec.13739)); Põlme S, et al. 2020. Fungal Diversity 105:1-16 ((https://doi.org/10.1007/s13225-020-00466-2)) |
| Otu12555 | | 2 | k__Fungi(100);p__Basidiomycota(100);c__Microbotryomycetes(100);o__Sporidiobolales(100);f__Sporidiobolales_family_Incertae_sedis(100);g__Sporobolomyces(100);g__Sporobolomyces_unclassified(100); | Sporobolomyces | 13 | Pathotroph-Saprotroph | \|Fungal Parasite\|-Plant Saprotroph | Yeast | NULL | Probable | As litter_saprotroph (Põlme et al. 2020); As mycoparasite (Põlme et al. 2020); Decay fungal material (Põlme et al. 2020) | Tedersoo L, et al. 2014. Science 346:e1256688 ((https://doi.org/10.1126/science.1256688)); Purahong W, et al. 2016. Molecular Ecology 25:4059-4074 ((https://doi.org/10.1111/mec.13739)); Põlme S, et al. 2020. Fungal Diversity 105:1-16 ((https://doi.org/10.1007/s13225-020-00466-2)) |
| Otu12556 | | 2 | k__Fungi(100);p__Ascomycota(100);c__Saccharomycetes(100);o__Saccharomycetales(100);o__Saccharomycetales_unclassified(100);o__Saccharomycetales_unclassified(100);o__Saccharomycetales_unclassified(100); | Saccharomycetales | 7 | Saprotroph | Undefined Saprotroph | Yeast | NULL | Possible | NULL | Sterkenburg E, et al. 2015. New Phytologist 207:1145-1158 ((https://doi.org/10.1111/nph.13426)) |
| Otu12557 | | 2 | k__Fungi(100);p__Ascomycota(100);c__Saccharomycetes(100);o__Saccharomycetales(100);f__Metschnikowiaceae(100);g__Metschnikowia(100);g__Metschnikowia_unclassified(100); | Metschnikowia | 13 | Saprotroph | \|Undefined Saprotroph\| | Yeast | NULL | Probable | As nectar-tap_saprotroph (Põlme et al. 2020); Decay sugar-rich substrates (Põlme et al. 2020) | Põlme S, et al. 2020. Fungal Diversity 105:1-16 ((https://doi.org/10.1007/s13225-020-00466-2)) |
| Otu12558 | | 2 | k__Fungi(100);p__Ascomycota(100);c__Saccharomycetes(100);o__Saccharomycetales(100);f__Metschnikowiaceae(100);g__Metschnikowia(100);g__Metschnikowia_unclassified(100); | Metschnikowia | 13 | Saprotroph | \|Undefined Saprotroph\| | Yeast | NULL | Probable | As nectar-tap_saprotroph (Põlme et al. 2020); Decay sugar-rich substrates (Põlme et al. 2020) | Põlme S, et al. 2020. Fungal Diversity 105:1-16 ((https://doi.org/10.1007/s13225-020-00466-2)) |
| Otu12559 | | 2 | k__Fungi(100);p__Ascomycota(100);c__Saccharomycetes(100);o__Saccharomycetales(100);f__Metschnikowiaceae(100);g__Metschnikowia(100);s__Metschnikowia_pulcherrima(100); | Metschnikowia | 13 | Saprotroph | \|Undefined Saprotroph\| | Yeast | NULL | Probable | As nectar-tap_saprotroph (Põlme et al. 2020); Decay sugar-rich substrates (Põlme et al. 2020) | Põlme S, et al. 2020. Fungal Diversity 105:1-16 ((https://doi.org/10.1007/s13225-020-00466-2)) |
| Otu12561 | | 2 | k__Fungi(100);p__Ascomycota(100);c__Saccharomycetes(100);o__Saccharomycetales(100);f__Metschnikowiaceae(100);g__Metschnikowia(100);g__Metschnikowia_unclassified(100); | Metschnikowia | 13 | Saprotroph | \|Undefined Saprotroph\| | Yeast | NULL | Probable | As nectar-tap_saprotroph (Põlme et al. 2020); Decay sugar-rich substrates (Põlme et al. 2020) | Põlme S, et al. 2020. Fungal Diversity 105:1-16 ((https://doi.org/10.1007/s13225-020-00466-2)) |
| Otu12564 | | 2 | k__Fungi(100);p__Ascomycota(100);c__Saccharomycetes(100);o__Saccharomycetales(100);f__Metschnikowiaceae(100);g__Metschnikowia(100);g__Metschnikowia_unclassified(100); | Metschnikowia | 13 | Saprotroph | \|Undefined Saprotroph\| | Yeast | NULL | Probable | As nectar-tap_saprotroph (Põlme et al. 2020); Decay sugar-rich substrates (Põlme et al. 2020) | Põlme S, et al. 2020. Fungal Diversity 105:1-16 ((https://doi.org/10.1007/s13225-020-00466-2)) |
| Otu12565 | | 2 | k__Fungi(100);p__Basidiomycota(100);c__Agaricomycetes(100);o__Agaricales(100);f__Lyophyllaceae(100);f__Lyophyllaceae_unclassified(100);f__Lyophyllaceae_unclassified(100); | Lyophyllaceae | 9 | Pathotroph-Saprotroph-Symbiotroph | Ectomycorrhizal-Fungal Pathogen-Undefined Saprotroph | Agaricoid | NULL | Possible | NULL | Kirk PM et al. 2008. Dictionary of the Fungi. Tenth Edition. CAB International, Wallingford (ISBN: 978-0851998268) |
| Otu12566 | | 2 | k__Fungi(100);p__Ascomycota(100);c__Saccharomycetes(100);o__Saccharomycetales(100);f__Metschnikowiaceae(100);g__Metschnikowia(100);g__Metschnikowia_unclassified(100); | Metschnikowia | 13 | Saprotroph | \|Undefined Saprotroph\| | Yeast | NULL | Probable | As nectar-tap_saprotroph (Põlme et al. 2020); Decay sugar-rich substrates (Põlme et al. 2020) | Põlme S, et al. 2020. Fungal Diversity 105:1-16 ((https://doi.org/10.1007/s13225-020-00466-2)) |
| Otu12567 | | 2 | k__Fungi(100);p__Basidiomycota(100);c__Agaricomycetes(100);o__Agaricales(100);f__Psathyrellaceae(100);g__Coprinellus(100);s__Coprinellus_sp(100); | Coprinellus | 13 | Saprotroph | \|Undefined Saprotroph\| | Agaricoid | NULL | Probable | As soil_saprotroph (Põlme et al. 2020); Decay soil (Põlme et al. 2020) | Tedersoo L, et al. 2014. Science 346:e1256688 ((https://doi.org/10.1126/science.1256688)); Põlme S, et al. 2020. Fungal Diversity 105:1-16 ((https://doi.org/10.1007/s13225-020-00466-2)) |
| Otu12568 | | 2 | k__Fungi(100);p__Ascomycota(100);c__Saccharomycetes(100);o__Saccharomycetales(100);f__Metschnikowiaceae(100);g__Metschnikowia(100);s__Metschnikowia_pulcherrima(100); | Metschnikowia | 13 | Saprotroph | \|Undefined Saprotroph\| | Yeast | NULL | Probable | As nectar-tap_saprotroph (Põlme et al. 2020); Decay sugar-rich substrates (Põlme et al. 2020) | Põlme S, et al. 2020. Fungal Diversity 105:1-16 ((https://doi.org/10.1007/s13225-020-00466-2)) |
| Otu12569 | | 2 | k__Fungi(100);p__Ascomycota(100);c__Saccharomycetes(100);o__Saccharomycetales(100);f__Saccharomycetales_family_Incertae_sedis(100);g__Candida(100);g__Candida_unclassified(100); | Candida | 13 | Pathotroph-Saprotroph-Symbiotroph | Animal Pathogen-Endophyte-Endosymbiont-Epiphyte-\|Undefined Saprotroph\| | Dimorphic Yeast-Yeast | NULL | Probable | As nectar-tap_saprotroph (Põlme et al. 2020); Decay sugar-rich substrates (Põlme et al. 2020); This asexual genus is polyphyletic and contain species that function very differently depend on what they are. Caution should be taken when interpreting guilds for Candida species. | Manolakaki D, et al. 2010. Virulence. 1: 367-375 ((https://doi.org/10.4161/viru.1.5.12796)); Mekbib SB, et al. 2011. Tropical Plant Pathology 36:233-240 ((https://doi.org/10.1590/S1982-56762011000400004)); Põlme S, et al. 2020. Fungal Diversity 105:1-16 ((https://doi.org/10.1007/s13225-020-00466-2)) |
| Otu12570 | | 2 | k__Fungi(100);p__Ascomycota(100);c__Saccharomycetes(100);o__Saccharomycetales(100);f__Metschnikowiaceae(100);g__Metschnikowia(100);g__Metschnikowia_unclassified(100); | Metschnikowia | 13 | Saprotroph | \|Undefined Saprotroph\| | Yeast | NULL | Probable | As nectar-tap_saprotroph (Põlme et al. 2020); Decay sugar-rich substrates (Põlme et al. 2020) | Põlme S, et al. 2020. Fungal Diversity 105:1-16 ((https://doi.org/10.1007/s13225-020-00466-2)) |
| Otu12573 | | 2 | k__Fungi(100);p__Ascomycota(100);c__Saccharomycetes(100);o__Saccharomycetales(100);f__Metschnikowiaceae(100);g__Metschnikowia(100);g__Metschnikowia_unclassified(100); | Metschnikowia | 13 | Saprotroph | \|Undefined Saprotroph\| | Yeast | NULL | Probable | As nectar-tap_saprotroph (Põlme et al. 2020); Decay sugar-rich substrates (Põlme et al. 2020) | Põlme S, et al. 2020. Fungal Diversity 105:1-16 ((https://doi.org/10.1007/s13225-020-00466-2)) |
| Otu12575 | | 2 | k__Fungi(100);p__Ascomycota(100);c__Saccharomycetes(100);o__Saccharomycetales(100);f__Metschnikowiaceae(100);g__Metschnikowia(100);s__Metschnikowia_pulcherrima(100); | Metschnikowia | 13 | Saprotroph | \|Undefined Saprotroph\| | Yeast | NULL | Probable | As nectar-tap_saprotroph (Põlme et al. 2020); Decay sugar-rich substrates (Põlme et al. 2020) | Põlme S, et al. 2020. Fungal Diversity 105:1-16 ((https://doi.org/10.1007/s13225-020-00466-2)) |
| Otu12576 | | 2 | k__Fungi(100);p__Ascomycota(100);c__Saccharomycetes(100);o__Saccharomycetales(100);f__Metschnikowiaceae(100);g__Metschnikowia(100);g__Metschnikowia_unclassified(100); | Metschnikowia | 13 | Saprotroph | \|Undefined Saprotroph\| | Yeast | NULL | Probable | As nectar-tap_saprotroph (Põlme et al. 2020); Decay sugar-rich substrates (Põlme et al. 2020) | Põlme S, et al. 2020. Fungal Diversity 105:1-16 ((https://doi.org/10.1007/s13225-020-00466-2)) |
| Otu12577 | | 2 | k__Fungi(100);p__Ascomycota(100);c__Saccharomycetes(100);o__Saccharomycetales(100);f__Metschnikowiaceae(100);g__Metschnikowia(100);g__Metschnikowia_unclassified(100); | Metschnikowia | 13 | Saprotroph | \|Undefined Saprotroph\| | Yeast | NULL | Probable | As nectar-tap_saprotroph (Põlme et al. 2020); Decay sugar-rich substrates (Põlme et al. 2020) | Põlme S, et al. 2020. Fungal Diversity 105:1-16 ((https://doi.org/10.1007/s13225-020-00466-2)) |
| Otu12581 | | 2 | k__Fungi(100);p__Basidiomycota(100);c__Agaricomycetes(100);o__Agaricales(100);f__Cortinariaceae(100);g__Cortinarius(100);g__Cortinarius_unclassified(100); | Cortinarius | 13 | Symbiotroph | \|Ectomycorrhizal\| | Agaricoid | NULL | Highly Probable | NULL | Rinaldi AC, et al. 2008. Fungal Diversity 33:1-45 ((http://www.fungaldiversity.org/fdp/sfdp/33-1.pdf)); Tedersoo L, et al. 2010. Mycorrhiza 20:217-263 ((https://doi.org/10.1007/s00572-009-0274-x)); Põlme S, et al. 2020. Fungal Diversity 105:1-16 ((https://doi.org/10.1007/s13225-020-00466-2)); http://mycorrhizas.info/ecmf.html ((http://mycorrhizas.info/ecmf.html)) |
| Otu12583 | | 2 | k__Fungi(100);p__Ascomycota(100);c__Saccharomycetes(100);o__Saccharomycetales(100);f__Metschnikowiaceae(100);g__Metschnikowia(100);s__Metschnikowia_pulcherrima(100); | Metschnikowia | 13 | Saprotroph | \|Undefined Saprotroph\| | Yeast | NULL | Probable | As nectar-tap_saprotroph (Põlme et al. 2020); Decay sugar-rich substrates (Põlme et al. 2020) | Põlme S, et al. 2020. Fungal Diversity 105:1-16 ((https://doi.org/10.1007/s13225-020-00466-2)) |
| Otu12584 | | 2 | k__Fungi(100);p__Ascomycota(100);c__Saccharomycetes(100);o__Saccharomycetales(100);f__Metschnikowiaceae(100);g__Metschnikowia(100);s__Metschnikowia_pulcherrima(100); | Metschnikowia | 13 | Saprotroph | \|Undefined Saprotroph\| | Yeast | NULL | Probable | As nectar-tap_saprotroph (Põlme et al. 2020); Decay sugar-rich substrates (Põlme et al. 2020) | Põlme S, et al. 2020. Fungal Diversity 105:1-16 ((https://doi.org/10.1007/s13225-020-00466-2)) |
| Otu12585 | | 2 | k__Fungi(100);p__Ascomycota(100);c__Dothideomycetes(100);o__Pleosporales(100);f__Pleosporaceae(100);g__Alternaria(100);s__Alternaria_porri(100); | Alternaria | 13 | Pathotroph-Saprotroph-Symbiotroph | Animal Pathogen-Endophyte-\|Plant Pathogen\|-Plant Saprotroph-Wood Saprotroph | Microfungus | Soft Rot | Possible | As foliar_endophyte (Põlme et al. 2020); As litter_saprotroph (Põlme et al. 2020); Decay leaf, fruit, seed (Põlme et al. 2020) | Duncan CG, Eslyn WE. 1966. Mycologia 58:642-664 ((https://doi.org/10.2307/3757045)); Seehann G, et al. 1975. List of Fungi in Soft-Rot Tissue Tests. Institute fur Holzbiologie und Holzschutz, Bundesforschungsanstalt fur Forst - und Holzwirtshaft, Hamburg - Lohbrugge, Germany ((https://www.irg-wp.com/irgdocs/details.php?d6493033-c9ea-40f3-8e3e-5697f32d640d)); Tedersoo L, et al. 2014. Science 346:e1256688 ((https://doi.org/10.1126/science.1256688)); Põlme S, et al. 2020. Fungal Diversity 105:1-16 ((https://doi.org/10.1007/s13225-020-00466-2)) |
| Otu12587 | | 2 | k__Fungi(100);p__Ascomycota(100);c__Saccharomycetes(100);o__Saccharomycetales(100);f__Saccharomycetales_family_Incertae_sedis(100);g__Candida(100);g__Candida_unclassified(100); | Candida | 13 | Pathotroph-Saprotroph-Symbiotroph | Animal Pathogen-Endophyte-Endosymbiont-Epiphyte-\|Undefined Saprotroph\| | Dimorphic Yeast-Yeast | NULL | Probable | As nectar-tap_saprotroph (Põlme et al. 2020); Decay sugar-rich substrates (Põlme et al. 2020); This asexual genus is polyphyletic and contain species that function very differently depend on what they are. Caution should be taken when interpreting guilds for Candida species. | Manolakaki D, et al. 2010. Virulence. 1: 367-375 ((https://doi.org/10.4161/viru.1.5.12796)); Mekbib SB, et al. 2011. Tropical Plant Pathology 36:233-240 ((https://doi.org/10.1590/S1982-56762011000400004)); Põlme S, et al. 2020. Fungal Diversity 105:1-16 ((https://doi.org/10.1007/s13225-020-00466-2)) |
| Otu12589 | | 2 | k__Fungi(100);p__Basidiomycota(100);c__Agaricomycetes(100);o__Agaricales(100);f__Cortinariaceae(100);g__Cortinarius(100);g__Cortinarius_unclassified(100); | Cortinarius | 13 | Symbiotroph | \|Ectomycorrhizal\| | Agaricoid | NULL | Highly Probable | NULL | Rinaldi AC, et al. 2008. Fungal Diversity 33:1-45 ((http://www.fungaldiversity.org/fdp/sfdp/33-1.pdf)); Tedersoo L, et al. 2010. Mycorrhiza 20:217-263 ((https://doi.org/10.1007/s00572-009-0274-x)); Põlme S, et al. 2020. Fungal Diversity 105:1-16 ((https://doi.org/10.1007/s13225-020-00466-2)); http://mycorrhizas.info/ecmf.html ((http://mycorrhizas.info/ecmf.html)) |
| Otu12590 | | 2 | k__Fungi(100);p__Ascomycota(100);c__Saccharomycetes(100);o__Saccharomycetales(100);f__Metschnikowiaceae(100);g__Metschnikowia(100);s__Metschnikowia_pulcherrima(100); | Metschnikowia | 13 | Saprotroph | \|Undefined Saprotroph\| | Yeast | NULL | Probable | As nectar-tap_saprotroph (Põlme et al. 2020); Decay sugar-rich substrates (Põlme et al. 2020) | Põlme S, et al. 2020. Fungal Diversity 105:1-16 ((https://doi.org/10.1007/s13225-020-00466-2)) |
| Otu12591 | | 2 | k__Fungi(100);p__Ascomycota(100);c__Saccharomycetes(100);o__Saccharomycetales(100);f__Metschnikowiaceae(100);g__Metschnikowia(100);g__Metschnikowia_unclassified(100); | Metschnikowia | 13 | Saprotroph | \|Undefined Saprotroph\| | Yeast | NULL | Probable | As nectar-tap_saprotroph (Põlme et al. 2020); Decay sugar-rich substrates (Põlme et al. 2020) | Põlme S, et al. 2020. Fungal Diversity 105:1-16 ((https://doi.org/10.1007/s13225-020-00466-2)) |
| Otu12592 | | 2 | k__Fungi(100);p__Ascomycota(100);c__Saccharomycetes(100);o__Saccharomycetales(100);f__Metschnikowiaceae(100);g__Metschnikowia(100);g__Metschnikowia_unclassified(100); | Metschnikowia | 13 | Saprotroph | \|Undefined Saprotroph\| | Yeast | NULL | Probable | As nectar-tap_saprotroph (Põlme et al. 2020); Decay sugar-rich substrates (Põlme et al. 2020) | Põlme S, et al. 2020. Fungal Diversity 105:1-16 ((https://doi.org/10.1007/s13225-020-00466-2)) |
| Otu12596 | | 2 | k__Fungi(100);p__Ascomycota(100);c__Saccharomycetes(100);o__Saccharomycetales(100);f__Metschnikowiaceae(100);g__Metschnikowia(100);s__Metschnikowia_pulcherrima(100); | Metschnikowia | 13 | Saprotroph | \|Undefined Saprotroph\| | Yeast | NULL | Probable | As nectar-tap_saprotroph (Põlme et al. 2020); Decay sugar-rich substrates (Põlme et al. 2020) | Põlme S, et al. 2020. Fungal Diversity 105:1-16 ((https://doi.org/10.1007/s13225-020-00466-2)) |
| Otu12597 | | 2 | k__Fungi(100);p__Basidiomycota(100);c__Tremellomycetes(100);o__Filobasidiales(100);f__Filobasidiaceae(100);g__Cryptococcus(100);s__Cryptococcus_friedmannii(100); | Cryptococcus | 13 | Pathotroph-Saprotroph-Symbiotroph | Animal Pathogen-Endophyte-Epiphyte-\|Undefined Saprotroph\| | Dimorphic Yeast-Yeast | NULL | Possible | As foliar_endophyte (Põlme et al. 2020); Decay leaf, fruit, seed, soil, dung, animal material (Põlme et al. 2020) | Andrews JH, et al. 1983. Phytopathology 73:228-234 ((https://doi.org/10.1094/Phyto-73-228)); Kurtzman CP, et al. (eds.) 2011. The Yeasts, a Taxonomic Study. Fifth Edition. Vols 1-3. Elsevier, San Diego (ISBN: 9780444521491); Põlme S, et al. 2020. Fungal Diversity 105:1-16 ((https://doi.org/10.1007/s13225-020-00466-2)) |
| Otu12598 | | 2 | k__Fungi(100);p__Ascomycota(100);c__Saccharomycetes(100);o__Saccharomycetales(100);f__Metschnikowiaceae(100);g__Metschnikowia(100);g__Metschnikowia_unclassified(100); | Metschnikowia | 13 | Saprotroph | \|Undefined Saprotroph\| | Yeast | NULL | Probable | As nectar-tap_saprotroph (Põlme et al. 2020); Decay sugar-rich substrates (Põlme et al. 2020) | Põlme S, et al. 2020. Fungal Diversity 105:1-16 ((https://doi.org/10.1007/s13225-020-00466-2)) |
| Otu12601 | | 2 | k__Fungi(100);p__Ascomycota(100);c__Saccharomycetes(100);o__Saccharomycetales(100);f__Metschnikowiaceae(100);g__Metschnikowia(100);s__Metschnikowia_pulcherrima(100); | Metschnikowia | 13 | Saprotroph | \|Undefined Saprotroph\| | Yeast | NULL | Probable | As nectar-tap_saprotroph (Põlme et al. 2020); Decay sugar-rich substrates (Põlme et al. 2020) | Põlme S, et al. 2020. Fungal Diversity 105:1-16 ((https://doi.org/10.1007/s13225-020-00466-2)) |
| Otu12602 | | 2 | k__Fungi(100);p__Ascomycota(100);c__Saccharomycetes(100);o__Saccharomycetales(100);f__Saccharomycetaceae(100);g__Saccharomyces(100);g__Saccharomyces_unclassified(100); | Saccharomyces | 13 | Saprotroph | \|Undefined Saprotroph\| | Yeast | NULL | Probable | As nectar-tap_saprotroph (Põlme et al. 2020); Decay sugar-rich substrates (Põlme et al. 2020) | James TY, et al. 2006. Nature 443:818-822 ((https://doi.org/10.1038/nature05110)); Põlme S, et al. 2020. Fungal Diversity 105:1-16 ((https://doi.org/10.1007/s13225-020-00466-2)) |
| Otu12604 | | 2 | k__Fungi(100);p__Ascomycota(100);c__Saccharomycetes(100);o__Saccharomycetales(100);f__Metschnikowiaceae(100);g__Metschnikowia(100);g__Metschnikowia_unclassified(100); | Metschnikowia | 13 | Saprotroph | \|Undefined Saprotroph\| | Yeast | NULL | Probable | As nectar-tap_saprotroph (Põlme et al. 2020); Decay sugar-rich substrates (Põlme et al. 2020) | Põlme S, et al. 2020. Fungal Diversity 105:1-16 ((https://doi.org/10.1007/s13225-020-00466-2)) |
| Otu12605 | | 2 | k__Fungi(100);p__Basidiomycota(100);c__Agaricomycetes(100);o__Trechisporales(100);f__Hydnodontaceae(100);g__Trechispora(100);g__Trechispora_unclassified(100); | Trechispora | 13 | Saprotroph | \|Wood Saprotroph\| | Corticioid | White Rot | Highly Probable | As white_rot (Põlme et al. 2020); Decay wood (Põlme et al. 2020) | Gilbertson RL, Ryvarden L. 1987-1987. North American Polypores. Fungiflora, Oslo (ISBN: 978-0945345060); Rinaldi AC, et al. 2008. Fungal Diversity 33:1-45 ((http://www.fungaldiversity.org/fdp/sfdp/33-1.pdf)) (as Uncertain); Tedersoo L, et al. 2010. Mycorrhiza 20:217-263 ((https://doi.org/10.1007/s00572-009-0274-x)) (as not ECM).; Põlme S, et al. 2020. Fungal Diversity 105:1-16 ((https://doi.org/10.1007/s13225-020-00466-2)) |
| Otu12606 | | 2 | k__Fungi(100);p__Ascomycota(100);c__Saccharomycetes(100);o__Saccharomycetales(100);f__Metschnikowiaceae(100);g__Metschnikowia(100);s__Metschnikowia_pulcherrima(100); | Metschnikowia | 13 | Saprotroph | \|Undefined Saprotroph\| | Yeast | NULL | Probable | As nectar-tap_saprotroph (Põlme et al. 2020); Decay sugar-rich substrates (Põlme et al. 2020) | Põlme S, et al. 2020. Fungal Diversity 105:1-16 ((https://doi.org/10.1007/s13225-020-00466-2)) |
| Otu12607 | | 2 | k__Fungi(100);p__Ascomycota(100);c__Saccharomycetes(100);o__Saccharomycetales(100);f__Metschnikowiaceae(100);g__Metschnikowia(100);s__Metschnikowia_pulcherrima(100); | Metschnikowia | 13 | Saprotroph | \|Undefined Saprotroph\| | Yeast | NULL | Probable | As nectar-tap_saprotroph (Põlme et al. 2020); Decay sugar-rich substrates (Põlme et al. 2020) | Põlme S, et al. 2020. Fungal Diversity 105:1-16 ((https://doi.org/10.1007/s13225-020-00466-2)) |
| Otu12609 | | 2 | k__Fungi(100);p__Ascomycota(100);c__Saccharomycetes(100);o__Saccharomycetales(100);f__Metschnikowiaceae(100);g__Metschnikowia(100);s__Metschnikowia_pulcherrima(100); | Metschnikowia | 13 | Saprotroph | \|Undefined Saprotroph\| | Yeast | NULL | Probable | As nectar-tap_saprotroph (Põlme et al. 2020); Decay sugar-rich substrates (Põlme et al. 2020) | Põlme S, et al. 2020. Fungal Diversity 105:1-16 ((https://doi.org/10.1007/s13225-020-00466-2)) |
| Otu12610 | | 2 | k__Fungi(100);p__Ascomycota(100);c__Saccharomycetes(100);o__Saccharomycetales(100);f__Metschnikowiaceae(100);g__Metschnikowia(100);s__Metschnikowia_pulcherrima(100); | Metschnikowia | 13 | Saprotroph | \|Undefined Saprotroph\| | Yeast | NULL | Probable | As nectar-tap_saprotroph (Põlme et al. 2020); Decay sugar-rich substrates (Põlme et al. 2020) | Põlme S, et al. 2020. Fungal Diversity 105:1-16 ((https://doi.org/10.1007/s13225-020-00466-2)) |
| Otu12611 | | 2 | k__Fungi(100);p__Basidiomycota(100);c__Agaricomycetes(100);o__Russulales(100);f__Peniophoraceae(100);g__Dendrophora(100);s__Dendrophora_sp(100); | Dendrophora | 13 | Saprotroph | Undefined Saprotroph-\|Wood Saprotroph\| | Corticioid | White Rot | Probable | As white_rot (Põlme et al. 2020); Decay wood (Põlme et al. 2020) | Tedersoo L, et al. 2014. Science 346:e1256688 ((https://doi.org/10.1126/science.1256688)); Põlme S, et al. 2020. Fungal Diversity 105:1-16 ((https://doi.org/10.1007/s13225-020-00466-2)) |
| Otu12612 | | 2 | k__Fungi(100);p__Ascomycota(100);c__Saccharomycetes(100);o__Saccharomycetales(100);f__Metschnikowiaceae(100);g__Metschnikowia(100);g__Metschnikowia_unclassified(100); | Metschnikowia | 13 | Saprotroph | \|Undefined Saprotroph\| | Yeast | NULL | Probable | As nectar-tap_saprotroph (Põlme et al. 2020); Decay sugar-rich substrates (Põlme et al. 2020) | Põlme S, et al. 2020. Fungal Diversity 105:1-16 ((https://doi.org/10.1007/s13225-020-00466-2)) |
| Otu12613 | | 2 | k__Fungi(100);p__Ascomycota(100);c__Saccharomycetes(100);o__Saccharomycetales(100);f__Metschnikowiaceae(100);g__Metschnikowia(100);s__Metschnikowia_pulcherrima(100); | Metschnikowia | 13 | Saprotroph | \|Undefined Saprotroph\| | Yeast | NULL | Probable | As nectar-tap_saprotroph (Põlme et al. 2020); Decay sugar-rich substrates (Põlme et al. 2020) | Põlme S, et al. 2020. Fungal Diversity 105:1-16 ((https://doi.org/10.1007/s13225-020-00466-2)) |
| Otu12614 | | 2 | k__Fungi(100);p__Ascomycota(100);c__Sordariomycetes(100);o__Xylariales(100);f__Amphisphaeriaceae(100);g__Pestalotiopsis(100);g__Pestalotiopsis_unclassified(100); | Pestalotiopsis | 13 | Pathotroph | \|Plant Pathogen\| | Microfungus | NULL | Probable | NULL | Tedersoo L, et al. 2014. Science 346:e1256688 ((https://doi.org/10.1126/science.1256688)); Põlme S, et al. 2020. Fungal Diversity 105:1-16 ((https://doi.org/10.1007/s13225-020-00466-2)) |
| Otu12615 | | 2 | k__Fungi(100);p__Ascomycota(100);c__Saccharomycetes(100);o__Saccharomycetales(100);f__Saccharomycetaceae(100);g__Lachancea(100);s__Lachancea_thermotolerans(100); | Lachancea | 13 | Pathotroph-Saprotroph | Animal Parasite-Animal Pathogen-\|Undefined Saprotroph\| | Yeast | NULL | Possible | As nectar-tap_saprotroph (Põlme et al. 2020); Decay animal material, sugar-rich substrates (Põlme et al. 2020); Likely opportunistic human pathogen (Irinyi et al. 2015; see L. fermentati) | Irinyi L, et al. 2015. Medical Mycology 53:313-337 ((https://doi.org/10.1093/mmy/myv008)); Põlme S, et al. 2020. Fungal Diversity 105:1-16 ((https://doi.org/10.1007/s13225-020-00466-2)) |
| Otu12616 | | 2 | k__Fungi(100);p__Ascomycota(100);c__Saccharomycetes(100);o__Saccharomycetales(100);f__Metschnikowiaceae(100);g__Metschnikowia(100);g__Metschnikowia_unclassified(100); | Metschnikowia | 13 | Saprotroph | \|Undefined Saprotroph\| | Yeast | NULL | Probable | As nectar-tap_saprotroph (Põlme et al. 2020); Decay sugar-rich substrates (Põlme et al. 2020) | Põlme S, et al. 2020. Fungal Diversity 105:1-16 ((https://doi.org/10.1007/s13225-020-00466-2)) |
| Otu12617 | | 2 | k__Fungi(100);p__Ascomycota(100);c__Saccharomycetes(100);o__Saccharomycetales(100);f__Metschnikowiaceae(100);g__Metschnikowia(100);g__Metschnikowia_unclassified(100); | Metschnikowia | 13 | Saprotroph | \|Undefined Saprotroph\| | Yeast | NULL | Probable | As nectar-tap_saprotroph (Põlme et al. 2020); Decay sugar-rich substrates (Põlme et al. 2020) | Põlme S, et al. 2020. Fungal Diversity 105:1-16 ((https://doi.org/10.1007/s13225-020-00466-2)) |
| Otu12620 | | 2 | k__Fungi(100);p__Basidiomycota(100);c__Agaricomycetes(100);o__Agaricales(100);f__Bolbitiaceae(100);g__Conocybe(100);g__Conocybe_unclassified(100); | Conocybe | 13 | Saprotroph | Dung Saprotroph-Plant Saprotroph-\|Undefined Saprotroph\| | Agaricoid | NULL | Probable | As litter_saprotroph (Põlme et al. 2020); As soil_saprotroph (Põlme et al. 2020); Decay soil (Põlme et al. 2020) | Tedersoo L, et al. 2014. Science 346:e1256688 ((https://doi.org/10.1126/science.1256688)); Põlme S, et al. 2020. Fungal Diversity 105:1-16 ((https://doi.org/10.1007/s13225-020-00466-2)) |
| Otu12621 | | 2 | k__Fungi(100);p__Basidiomycota(100);c__Agaricomycetes(100);o__Agaricales(100);f__Cortinariaceae(100);g__Cortinarius(100);g__Cortinarius_unclassified(100); | Cortinarius | 13 | Symbiotroph | \|Ectomycorrhizal\| | Agaricoid | NULL | Highly Probable | NULL | Rinaldi AC, et al. 2008. Fungal Diversity 33:1-45 ((http://www.fungaldiversity.org/fdp/sfdp/33-1.pdf)); Tedersoo L, et al. 2010. Mycorrhiza 20:217-263 ((https://doi.org/10.1007/s00572-009-0274-x)); Põlme S, et al. 2020. Fungal Diversity 105:1-16 ((https://doi.org/10.1007/s13225-020-00466-2)); http://mycorrhizas.info/ecmf.html ((http://mycorrhizas.info/ecmf.html)) |
| Otu12622 | | 2 | k__Fungi(100);p__Ascomycota(100);c__Saccharomycetes(100);o__Saccharomycetales(100);f__Metschnikowiaceae(100);g__Metschnikowia(100);s__Metschnikowia_pulcherrima(100); | Metschnikowia | 13 | Saprotroph | \|Undefined Saprotroph\| | Yeast | NULL | Probable | As nectar-tap_saprotroph (Põlme et al. 2020); Decay sugar-rich substrates (Põlme et al. 2020) | Põlme S, et al. 2020. Fungal Diversity 105:1-16 ((https://doi.org/10.1007/s13225-020-00466-2)) |
| Otu12623 | | 2 | k__Fungi(100);p__Basidiomycota(100);c__Agaricomycetes(100);o__Agaricales(100);f__Tricholomataceae(100);g__Collybia(100);s__Collybia_cookei(100); | Collybia | 13 | Pathotroph-Saprotroph | \|Fungal Parasite\|-Plant Pathogen | Agaricoid | NULL | Highly Probable | As mycoparasite (Põlme et al. 2020); Decay fungal material (Põlme et al. 2020) | Tedersoo L, et al. 2014. Science 346:e1256688 ((https://doi.org/10.1126/science.1256688)); Põlme S, et al. 2020. Fungal Diversity 105:1-16 ((https://doi.org/10.1007/s13225-020-00466-2)) |
| Otu12625 | | 2 | k__Fungi(100);p__Ascomycota(100);c__Saccharomycetes(100);o__Saccharomycetales(100);f__Metschnikowiaceae(100);g__Metschnikowia(100);s__Metschnikowia_pulcherrima(100); | Metschnikowia | 13 | Saprotroph | \|Undefined Saprotroph\| | Yeast | NULL | Probable | As nectar-tap_saprotroph (Põlme et al. 2020); Decay sugar-rich substrates (Põlme et al. 2020) | Põlme S, et al. 2020. Fungal Diversity 105:1-16 ((https://doi.org/10.1007/s13225-020-00466-2)) |
| Otu12626 | | 2 | k__Fungi(100);p__Ascomycota(100);c__Saccharomycetes(100);o__Saccharomycetales(100);f__Metschnikowiaceae(100);g__Metschnikowia(100);g__Metschnikowia_unclassified(100); | Metschnikowia | 13 | Saprotroph | \|Undefined Saprotroph\| | Yeast | NULL | Probable | As nectar-tap_saprotroph (Põlme et al. 2020); Decay sugar-rich substrates (Põlme et al. 2020) | Põlme S, et al. 2020. Fungal Diversity 105:1-16 ((https://doi.org/10.1007/s13225-020-00466-2)) |
| Otu12627 | | 2 | k__Fungi(100);p__Basidiomycota(100);c__Agaricomycetes(100);o__Agaricales(100);o__Agaricales_unclassified(100);o__Agaricales_unclassified(100);o__Agaricales_unclassified(100); | Agaricales | 7 | Pathotroph-Saprotroph-Symbiotroph | Bryophyte Parasite-Dung Saprotroph-Ectomycorrhizal-Fungal Parasite-Leaf Saprotroph-Plant Parasite-Undefined Saprotroph-Wood Saprotroph | Agaricoid-Gasteroid-Microfungus-Secotioid-Yeast | NULL | Possible | Mushrooms and toadstools, Gill fungi, Agarics... lignicolous, sometimes muscicolous or fungicolous, saprobic, mycorrhizal, rarely parasitic on plants or fungi (Kirk et al. 2008) | Kirk PM et al. 2008. Dictionary of the Fungi. Tenth Edition. CAB International, Wallingford (ISBN: 978-0851998268) |
| Otu12631 | | 2 | k__Fungi(100);p__Ascomycota(100);c__Saccharomycetes(100);o__Saccharomycetales(100);f__Metschnikowiaceae(100);g__Metschnikowia(100);s__Metschnikowia_pulcherrima(100); | Metschnikowia | 13 | Saprotroph | \|Undefined Saprotroph\| | Yeast | NULL | Probable | As nectar-tap_saprotroph (Põlme et al. 2020); Decay sugar-rich substrates (Põlme et al. 2020) | Põlme S, et al. 2020. Fungal Diversity 105:1-16 ((https://doi.org/10.1007/s13225-020-00466-2)) |

| | | | | | | | | | | | |
|---|---|---|---|---|---|---|---|---|---|---|---|
| Otu12632 | 2 | k__Fungi(100);p__Basidiomycota(100);c__Microbotryomycetes(100);o__Sporidiobolales(100);f__Sporidiobolales_family_Incertae_sedis(100);g__Rhodotorula(100);s__Rhodotorula_graminis(100); | Rhodotorula | 13 | Pathotroph-Saprotroph-Symbiotroph | Animal Pathogen-Animal Symbiotroph-Endophyte-Plant Pathogen-\|Undefined Saprotroph\| | Yeast | NULL | Probable | As foliar_endophyte (Põlme et al. 2020); Opportunistic human pathogen | Põlme S, et al. 2020. Fungal Diversity 105:1-16 ((https://doi.org/10.1007/s13225-020-00466-2)); https://www.cdc.gov ((https://www.cdc.gov)) |
| Otu12633 | 2 | k__Fungi(100);p__Ascomycota(100);c__Saccharomycetes(100);o__Saccharomycetales(100);f__Metschnikowiaceae(100);g__Metschnikowia(100);s__Metschnikowia_pulcherrima(100); | Metschnikowia | 13 | Saprotroph | \|Undefined Saprotroph\| | Yeast | NULL | Probable | As nectar-tap_saprotroph (Põlme et al. 2020); Decay sugar-rich substrates (Põlme et al. 2020) | Põlme S, et al. 2020. Fungal Diversity 105:1-16 ((https://doi.org/10.1007/s13225-020-00466-2)) |
| Otu12637 | 2 | k__Fungi(100);p__Basidiomycota(100);c__Agaricomycetes(100);o__Agaricomycetes_order_Incertae_sedis(100);f__Agaricomycetes_family_Incertae_sedis(100);g__Resinicium(100);s__Resinicium_sp(100); | Resinicium | 13 | Saprotroph | Undefined Saprotroph-\|Wood Saprotroph\| | Corticioid | White Rot | Probable | As white_rot (Põlme et al. 2020); Decay wood (Põlme et al. 2020) | Tedersoo L, et al. 2014. Science 346:e1256688 ((https://doi.org/10.1126/science.1256688)); Põlme S, et al. 2020. Fungal Diversity 105:1-16 ((https://doi.org/10.1007/s13225-020-00466-2)) |
| Otu12640 | 2 | k__Fungi(100);p__Ascomycota(100);c__Saccharomycetes(100);o__Saccharomycetales(100);f__Metschnikowiaceae(100);g__Metschnikowia(100);g__Metschnikowia_unclassified(100); | Metschnikowia | 13 | Saprotroph | \|Undefined Saprotroph\| | Yeast | NULL | Probable | As nectar-tap_saprotroph (Põlme et al. 2020); Decay sugar-rich substrates (Põlme et al. 2020) | Põlme S, et al. 2020. Fungal Diversity 105:1-16 ((https://doi.org/10.1007/s13225-020-00466-2)) |
| Otu12641 | 2 | k__Fungi(100);p__Ascomycota(100);c__Saccharomycetes(100);o__Saccharomycetales(100);f__Saccharomycetales_family_Incertae_sedis(100);f__Saccharomycetales_family_Incertae_sedis_unclassified(100);f__Saccharomycetales_family_Incertae_sedis_unclassified(100); | Saccharomycetales | 7 | Saprotroph | Undefined Saprotroph | Yeast | NULL | Possible | NULL | Sterkenburg E, et al. 2015. New Phytologist 207:1145-1158 ((https://doi.org/10.1111/nph.13426)) |
| Otu12643 | 2 | k__Fungi(100);p__Ascomycota(100);c__Saccharomycetes(100);o__Saccharomycetales(100);f__Metschnikowiaceae(100);g__Metschnikowia(100);g__Metschnikowia_unclassified(100); | Metschnikowia | 13 | Saprotroph | \|Undefined Saprotroph\| | Yeast | NULL | Probable | As nectar-tap_saprotroph (Põlme et al. 2020); Decay sugar-rich substrates (Põlme et al. 2020) | Põlme S, et al. 2020. Fungal Diversity 105:1-16 ((https://doi.org/10.1007/s13225-020-00466-2)) |
| Otu12644 | 2 | k__Fungi(100);p__Ascomycota(100);c__Saccharomycetes(100);o__Saccharomycetales(100);f__Metschnikowiaceae(100);g__Metschnikowia(100);s__Metschnikowia_pulcherrima(100); | Metschnikowia | 13 | Saprotroph | \|Undefined Saprotroph\| | Yeast | NULL | Probable | As nectar-tap_saprotroph (Põlme et al. 2020); Decay sugar-rich substrates (Põlme et al. 2020) | Põlme S, et al. 2020. Fungal Diversity 105:1-16 ((https://doi.org/10.1007/s13225-020-00466-2)) |
| Otu12645 | 2 | k__Fungi(100);p__Ascomycota(100);c__Saccharomycetes(100);o__Saccharomycetales(100);f__Metschnikowiaceae(100);g__Metschnikowia(100);g__Metschnikowia_unclassified(100); | Metschnikowia | 13 | Saprotroph | \|Undefined Saprotroph\| | Yeast | NULL | Probable | As nectar-tap_saprotroph (Põlme et al. 2020); Decay sugar-rich substrates (Põlme et al. 2020) | Põlme S, et al. 2020. Fungal Diversity 105:1-16 ((https://doi.org/10.1007/s13225-020-00466-2)) |
| Otu12646 | 2 | k__Fungi(100);p__Ascomycota(100);c__Saccharomycetes(100);o__Saccharomycetales(100);f__Metschnikowiaceae(100);g__Metschnikowia(100);s__Metschnikowia_pulcherrima(100); | Metschnikowia | 13 | Saprotroph | \|Undefined Saprotroph\| | Yeast | NULL | Probable | As nectar-tap_saprotroph (Põlme et al. 2020); Decay sugar-rich substrates (Põlme et al. 2020) | Põlme S, et al. 2020. Fungal Diversity 105:1-16 ((https://doi.org/10.1007/s13225-020-00466-2)) |
| Otu12647 | 2 | k__Fungi(100);p__Ascomycota(100);c__Saccharomycetes(100);o__Saccharomycetales(100);f__Saccharomycetales_family_Incertae_sedis(100);g__Candida(100);g__Candida_unclassified(100); | Candida | 13 | Pathotroph-Saprotroph-Symbiotroph | Animal Pathogen-Endophyte-Endosymbiont-Epiphyte-\|Undefined Saprotroph\| | Dimorphic Yeast-Yeast | NULL | Probable | As nectar-tap_saprotroph (Põlme et al. 2020); Decay sugar-rich substrates (Põlme et al. 2020); This asexual genus is polyphyletic and contain species that function very differently depend on what they are. Caution should be taken when interpreting guilds for Candida species. | Manolakaki D, et al. 2010. Virulence. 1: 367-375 ((https://doi.org/10.4161/viru.1.5.12796)); Mekbib SB, et al. 2011. Tropical Plant Pathology 36:233-240 ((https://doi.org/10.1590/S1982-56762011000400004)); Põlme S, et al. 2020. Fungal Diversity 105:1-16 ((https://doi.org/10.1007/s13225-020-00466-2)) |
| Otu12648 | 2 | k__Fungi(100);p__Ascomycota(100);c__Saccharomycetes(100);o__Saccharomycetales(100);f__Metschnikowiaceae(100);g__Metschnikowia(100);s__Metschnikowia_pulcherrima(100); | Metschnikowia | 13 | Saprotroph | \|Undefined Saprotroph\| | Yeast | NULL | Probable | As nectar-tap_saprotroph (Põlme et al. 2020); Decay sugar-rich substrates (Põlme et al. 2020) | Põlme S, et al. 2020. Fungal Diversity 105:1-16 ((https://doi.org/10.1007/s13225-020-00466-2)) |
| Otu12650 | 2 | k__Fungi(100);p__Ascomycota(100);c__Saccharomycetes(100);o__Saccharomycetales(100);f__Metschnikowiaceae(100);g__Metschnikowia(100);s__Metschnikowia_pulcherrima(100); | Metschnikowia | 13 | Saprotroph | \|Undefined Saprotroph\| | Yeast | NULL | Probable | As nectar-tap_saprotroph (Põlme et al. 2020); Decay sugar-rich substrates (Põlme et al. 2020) | Põlme S, et al. 2020. Fungal Diversity 105:1-16 ((https://doi.org/10.1007/s13225-020-00466-2)) |
| Otu12652 | 2 | k__Fungi(100);p__Basidiomycota(100);c__Microbotryomycetes(100);o__Sporidiobolales(100);f__Sporidiobolales_family_Incertae_sedis(100);g__Sporobolomyces(100);s__Sporobolomyces_lactosus(100); | Sporobolomyces | 13 | Pathotroph-Saprotroph | \|Fungal Parasite\|-Plant Saprotroph | Yeast | NULL | Probable | As litter_saprotroph (Põlme et al. 2020); As mycoparasite (Põlme et al. 2020); Decay fungal material (Põlme et al. 2020) | Tedersoo L, et al. 2014. Science 346:e1256688 ((https://doi.org/10.1126/science.1256688)); Purahong W, et al. 2016. Molecular Ecology 25:4059-4074 ((https://doi.org/10.1111/mec.13739)); Põlme S, et al. 2020. Fungal Diversity 105:1-16 ((https://doi.org/10.1007/s13225-020-00466-2)) |
| Otu12653 | 2 | k__Fungi(100);p__Ascomycota(100);c__Saccharomycetes(100);o__Saccharomycetales(100);f__Metschnikowiaceae(100);g__Metschnikowia(100);s__Metschnikowia_pulcherrima(100); | Metschnikowia | 13 | Saprotroph | \|Undefined Saprotroph\| | Yeast | NULL | Probable | As nectar-tap_saprotroph (Põlme et al. 2020); Decay sugar-rich substrates (Põlme et al. 2020) | Põlme S, et al. 2020. Fungal Diversity 105:1-16 ((https://doi.org/10.1007/s13225-020-00466-2)) |
| Otu12655 | 2 | k__Fungi(100);p__Ascomycota(100);c__Saccharomycetes(100);o__Saccharomycetales(100);f__Metschnikowiaceae(100);g__Metschnikowia(100);s__Metschnikowia_pulcherrima(100); | Metschnikowia | 13 | Saprotroph | \|Undefined Saprotroph\| | Yeast | NULL | Probable | As nectar-tap_saprotroph (Põlme et al. 2020); Decay sugar-rich substrates (Põlme et al. 2020) | Põlme S, et al. 2020. Fungal Diversity 105:1-16 ((https://doi.org/10.1007/s13225-020-00466-2)) |
| Otu12656 | 2 | k__Fungi(100);p__Ascomycota(100);c__Saccharomycetes(100);o__Saccharomycetales(100);f__Metschnikowiaceae(100);g__Metschnikowia(100);g__Metschnikowia_unclassified(100); | Metschnikowia | 13 | Saprotroph | \|Undefined Saprotroph\| | Yeast | NULL | Probable | As nectar-tap_saprotroph (Põlme et al. 2020); Decay sugar-rich substrates (Põlme et al. 2020) | Põlme S, et al. 2020. Fungal Diversity 105:1-16 ((https://doi.org/10.1007/s13225-020-00466-2)) |
| Otu12657 | 2 | k__Fungi(100);p__Ascomycota(100);c__Saccharomycetes(100);o__Saccharomycetales(100);f__Metschnikowiaceae(100);g__Metschnikowia(100);g__Metschnikowia_unclassified(100); | Metschnikowia | 13 | Saprotroph | \|Undefined Saprotroph\| | Yeast | NULL | Probable | As nectar-tap_saprotroph (Põlme et al. 2020); Decay sugar-rich substrates (Põlme et al. 2020) | Põlme S, et al. 2020. Fungal Diversity 105:1-16 ((https://doi.org/10.1007/s13225-020-00466-2)) |
| Otu12659 | 2 | k__Fungi(100);p__Zygomycota(100);c__Zygomycota_class_Incertae_sedis(100);o__Mucorales(100);f__Mucoraceae(100);g__Mucor(100);s__Mucor_circinelloides_f._lusitanicus(100); | Mucor | 13 | Saprotroph | \|Undefined Saprotroph\| | Microfungus | NULL | Probable | As soil_saprotroph (Põlme et al. 2020); Decay soil (Põlme et al. 2020) | Tedersoo L, et al. 2014. Science 346:e1256688 ((https://doi.org/10.1126/science.1256688)); Põlme S, et al. 2020. Fungal Diversity 105:1-16 ((https://doi.org/10.1007/s13225-020-00466-2)) |
| Otu12661 | 2 | k__Fungi(100);p__Basidiomycota(100);c__Agaricomycetes(100);o__Agaricales(100);f__Cortinariaceae(100);g__Cortinarius(100);g__Cortinarius_unclassified(100); | Cortinarius | 13 | Symbiotroph | \|Ectomycorrhizal\| | Agaricoid | NULL | Highly Probable | NULL | Rinaldi AC, et al. 2008. Fungal Diversity 33:1-45 ((http://www.fungaldiversity.org/fdp/sfdp/33-1.pdf)); Tedersoo L, et al. 2010. Mycorrhiza 20:217-263 ((https://doi.org/10.1007/s00572-009-0274-x)); Põlme S, et al. 2020. Fungal Diversity 105:1-16 ((https://doi.org/10.1007/s13225-020-00466-2)); http://mycorrhizas.info/ecmf.html ((http://mycorrhizas.info/ecmf.html)) |
| Otu12664 | 2 | k__Fungi(100);p__Ascomycota(100);c__Saccharomycetes(100);o__Saccharomycetales(100);f__Metschnikowiaceae(100);g__Metschnikowia(100);g__Metschnikowia_unclassified(100); | Metschnikowia | 13 | Saprotroph | \|Undefined Saprotroph\| | Yeast | NULL | Probable | As nectar-tap_saprotroph (Põlme et al. 2020); Decay sugar-rich substrates (Põlme et al. 2020) | Põlme S, et al. 2020. Fungal Diversity 105:1-16 ((https://doi.org/10.1007/s13225-020-00466-2)) |
| Otu12665 | 2 | k__Fungi(100);p__Basidiomycota(100);c__Agaricomycetes(100);o__Agaricales(100);f__Cortinariaceae(100);g__Cortinarius(100);g__Cortinarius_unclassified(100); | Cortinarius | 13 | Symbiotroph | \|Ectomycorrhizal\| | Agaricoid | NULL | Highly Probable | NULL | Rinaldi AC, et al. 2008. Fungal Diversity 33:1-45 ((http://www.fungaldiversity.org/fdp/sfdp/33-1.pdf)); Tedersoo L, et al. 2010. Mycorrhiza 20:217-263 ((https://doi.org/10.1007/s00572-009-0274-x)); Põlme S, et al. 2020. Fungal Diversity 105:1-16 ((https://doi.org/10.1007/s13225-020-00466-2)); http://mycorrhizas.info/ecmf.html ((http://mycorrhizas.info/ecmf.html)) |
| Otu12666 | 2 | k__Fungi(100);p__Basidiomycota(100);c__Agaricomycetes(100);o__Agaricales(100);f__Cortinariaceae(100);g__Cortinarius(100);g__Cortinarius_unclassified(100); | Cortinarius | 13 | Symbiotroph | \|Ectomycorrhizal\| | Agaricoid | NULL | Highly Probable | NULL | Rinaldi AC, et al. 2008. Fungal Diversity 33:1-45 ((http://www.fungaldiversity.org/fdp/sfdp/33-1.pdf)); Tedersoo L, et al. 2010. Mycorrhiza 20:217-263 ((https://doi.org/10.1007/s00572-009-0274-x)); Põlme S, et al. 2020. Fungal Diversity 105:1-16 ((https://doi.org/10.1007/s13225-020-00466-2)); http://mycorrhizas.info/ecmf.html ((http://mycorrhizas.info/ecmf.html)) |
| Otu12667 | 2 | k__Fungi(100);p__Basidiomycota(100);c__Agaricomycetes(100);o__Agaricales(100);f__Cortinariaceae(100);g__Cortinarius(100);g__Cortinarius_unclassified(100); | Cortinarius | 13 | Symbiotroph | \|Ectomycorrhizal\| | Agaricoid | NULL | Highly Probable | NULL | Rinaldi AC, et al. 2008. Fungal Diversity 33:1-45 ((http://www.fungaldiversity.org/fdp/sfdp/33-1.pdf)); Tedersoo L, et al. 2010. Mycorrhiza 20:217-263 ((https://doi.org/10.1007/s00572-009-0274-x)); Põlme S, et al. 2020. Fungal Diversity 105:1-16 ((https://doi.org/10.1007/s13225-020-00466-2)); http://mycorrhizas.info/ecmf.html ((http://mycorrhizas.info/ecmf.html)) |
| Otu12668 | 2 | k__Fungi(100);p__Basidiomycota(100);c__Agaricomycetes(100);o__Agaricales(100);f__Cortinariaceae(100);g__Cortinarius(100);g__Cortinarius_unclassified(100); | Cortinarius | 13 | Symbiotroph | \|Ectomycorrhizal\| | Agaricoid | NULL | Highly Probable | NULL | Rinaldi AC, et al. 2008. Fungal Diversity 33:1-45 ((http://www.fungaldiversity.org/fdp/sfdp/33-1.pdf)); Tedersoo L, et al. 2010. Mycorrhiza 20:217-263 ((https://doi.org/10.1007/s00572-009-0274-x)); Põlme S, et al. 2020. Fungal Diversity 105:1-16 ((https://doi.org/10.1007/s13225-020-00466-2)); http://mycorrhizas.info/ecmf.html ((http://mycorrhizas.info/ecmf.html)) |
| Otu12669 | 2 | k__Fungi(100);p__Basidiomycota(100);c__Agaricomycetes(100);o__Agaricales(100);f__Cortinariaceae(100);g__Cortinarius(100);g__Cortinarius_unclassified(100); | Cortinarius | 13 | Symbiotroph | \|Ectomycorrhizal\| | Agaricoid | NULL | Highly Probable | NULL | Rinaldi AC, et al. 2008. Fungal Diversity 33:1-45 ((http://www.fungaldiversity.org/fdp/sfdp/33-1.pdf)); Tedersoo L, et al. 2010. Mycorrhiza 20:217-263 ((https://doi.org/10.1007/s00572-009-0274-x)); Põlme S, et al. 2020. Fungal Diversity 105:1-16 ((https://doi.org/10.1007/s13225-020-00466-2)); http://mycorrhizas.info/ecmf.html ((http://mycorrhizas.info/ecmf.html)) |
| Otu12670 | 2 | k__Fungi(100);p__Ascomycota(100);c__Saccharomycetes(100);o__Saccharomycetales(100);f__Saccharomycetales_family_Incertae_sedis(100);g__Candida(100);g__Candida_unclassified(100); | Candida | 13 | Pathotroph-Saprotroph-Symbiotroph | Animal Pathogen-Endophyte-Endosymbiont-Epiphyte-\|Undefined Saprotroph\| | Dimorphic Yeast-Yeast | NULL | Probable | As nectar-tap_saprotroph (Põlme et al. 2020); Decay sugar-rich substrates (Põlme et al. 2020); This asexual genus is polyphyletic and contain species that function very differently depend on what they are. Caution should be taken when interpreting guilds for Candida species. | Manolakaki D, et al. 2010. Virulence. 1: 367-375 ((https://doi.org/10.4161/viru.1.5.12796)); Mekbib SB, et al. 2011. Tropical Plant Pathology 36:233-240 ((https://doi.org/10.1590/S1982-56762011000400004)); Põlme S, et al. 2020. Fungal Diversity 105:1-16 ((https://doi.org/10.1007/s13225-020-00466-2)) |
| Otu12673 | 2 | k__Fungi(100);p__Ascomycota(100);c__Saccharomycetes(100);o__Saccharomycetales(100);f__Metschnikowiaceae(100);g__Metschnikowia(100);s__Metschnikowia_pulcherrima(100); | Metschnikowia | 13 | Saprotroph | \|Undefined Saprotroph\| | Yeast | NULL | Probable | As nectar-tap_saprotroph (Põlme et al. 2020); Decay sugar-rich substrates (Põlme et al. 2020) | Põlme S, et al. 2020. Fungal Diversity 105:1-16 ((https://doi.org/10.1007/s13225-020-00466-2)) |
| Otu12677 | 2 | k__Fungi(100);p__Ascomycota(100);c__Saccharomycetes(100);o__Saccharomycetales(100);f__Metschnikowiaceae(100);g__Metschnikowia(100);g__Metschnikowia_unclassified(100); | Metschnikowia | 13 | Saprotroph | \|Undefined Saprotroph\| | Yeast | NULL | Probable | As nectar-tap_saprotroph (Põlme et al. 2020); Decay sugar-rich substrates (Põlme et al. 2020) | Põlme S, et al. 2020. Fungal Diversity 105:1-16 ((https://doi.org/10.1007/s13225-020-00466-2)) |
| Otu12681 | 2 | k__Fungi(100);p__Basidiomycota(100);c__Agaricomycetes(100);o__Polyporales(100);f__Meruliaceae(100);g__Phlebia(100);s__Phlebia_subserialis(100); | Phlebia | 13 | Saprotroph | \|Wood Saprotroph\| | Corticioid | White Rot | Highly Probable | As white_rot (Põlme et al. 2020); Decay wood (Põlme et al. 2020) | Kirk PM et al. 2008. Dictionary of the Fungi. Tenth Edition. CAB International, Wallingford (ISBN: 978-0851998268); Põlme S, et al. 2020. Fungal Diversity 105:1-16 ((https://doi.org/10.1007/s13225-020-00466-2)) |
| Otu12682 | 2 | k__Fungi(100);p__Ascomycota(100);c__Saccharomycetes(100);o__Saccharomycetales(100);f__Metschnikowiaceae(100);g__Metschnikowia(100);s__Metschnikowia_pulcherrima(100); | Metschnikowia | 13 | Saprotroph | \|Undefined Saprotroph\| | Yeast | NULL | Probable | As nectar-tap_saprotroph (Põlme et al. 2020); Decay sugar-rich substrates (Põlme et al. 2020) | Põlme S, et al. 2020. Fungal Diversity 105:1-16 ((https://doi.org/10.1007/s13225-020-00466-2)) |
| Otu12685 | 2 | k__Fungi(100);p__Ascomycota(100);c__Saccharomycetes(100);o__Saccharomycetales(100);f__Metschnikowiaceae(100);g__Metschnikowia(100);s__Metschnikowia_pulcherrima(100); | Metschnikowia | 13 | Saprotroph | \|Undefined Saprotroph\| | Yeast | NULL | Probable | As nectar-tap_saprotroph (Põlme et al. 2020); Decay sugar-rich substrates (Põlme et al. 2020) | Põlme S, et al. 2020. Fungal Diversity 105:1-16 ((https://doi.org/10.1007/s13225-020-00466-2)) |
| Otu12689 | 2 | k__Fungi(100);p__Ascomycota(100);c__Saccharomycetes(100);o__Saccharomycetales(100);f__Metschnikowiaceae(100);g__Metschnikowia(100);s__Metschnikowia_pulcherrima(100); | Metschnikowia | 13 | Saprotroph | \|Undefined Saprotroph\| | Yeast | NULL | Probable | As nectar-tap_saprotroph (Põlme et al. 2020); Decay sugar-rich substrates (Põlme et al. 2020) | Põlme S, et al. 2020. Fungal Diversity 105:1-16 ((https://doi.org/10.1007/s13225-020-00466-2)) |
| Otu12691 | 2 | k__Fungi(100);p__Ascomycota(100);c__Saccharomycetes(100);o__Saccharomycetales(100);f__Metschnikowiaceae(100);g__Metschnikowia(100);s__Metschnikowia_pulcherrima(100); | Metschnikowia | 13 | Saprotroph | \|Undefined Saprotroph\| | Yeast | NULL | Probable | As nectar-tap_saprotroph (Põlme et al. 2020); Decay sugar-rich substrates (Põlme et al. 2020) | Põlme S, et al. 2020. Fungal Diversity 105:1-16 ((https://doi.org/10.1007/s13225-020-00466-2)) |
| Otu12694 | 2 | k__Fungi(100);p__Ascomycota(100);c__Saccharomycetes(100);o__Saccharomycetales(100);f__Saccharomycetales_family_Incertae_sedis(100);g__Candida(100);g__Candida_unclassified(100); | Candida | 13 | Pathotroph-Saprotroph-Symbiotroph | Animal Pathogen-Endophyte-Endosymbiont-Epiphyte-\|Undefined Saprotroph\| | Dimorphic Yeast-Yeast | NULL | Probable | As nectar-tap_saprotroph (Põlme et al. 2020); Decay sugar-rich substrates (Põlme et al. 2020); This asexual genus is polyphyletic and contain species that function very differently depend on what they are. Caution should be taken when interpreting guilds for Candida species. | Manolakaki D, et al. 2010. Virulence. 1: 367-375 ((https://doi.org/10.4161/viru.1.5.12796)); Mekbib SB, et al. 2011. Tropical Plant Pathology 36:233-240 ((https://doi.org/10.1590/S1982-56762011000400004)); Põlme S, et al. 2020. Fungal Diversity 105:1-16 ((https://doi.org/10.1007/s13225-020-00466-2)) |
| Otu12695 | 2 | k__Fungi(100);p__Ascomycota(100);c__Saccharomycetes(100);o__Saccharomycetales(100);f__Metschnikowiaceae(100);g__Metschnikowia(100);g__Metschnikowia_unclassified(100); | Metschnikowia | 13 | Saprotroph | \|Undefined Saprotroph\| | Yeast | NULL | Probable | As nectar-tap_saprotroph (Põlme et al. 2020); Decay sugar-rich substrates (Põlme et al. 2020) | Põlme S, et al. 2020. Fungal Diversity 105:1-16 ((https://doi.org/10.1007/s13225-020-00466-2)) |
| Otu12698 | 2 | k__Fungi(100);p__Ascomycota(100);c__Saccharomycetes(100);o__Saccharomycetales(100);f__Metschnikowiaceae(100);g__Metschnikowia(100);g__Metschnikowia_unclassified(100); | Metschnikowia | 13 | Saprotroph | \|Undefined Saprotroph\| | Yeast | NULL | Probable | As nectar-tap_saprotroph (Põlme et al. 2020); Decay sugar-rich substrates (Põlme et al. 2020) | Põlme S, et al. 2020. Fungal Diversity 105:1-16 ((https://doi.org/10.1007/s13225-020-00466-2)) |
| Otu12700 | 2 | k__Fungi(100);p__Ascomycota(100);c__Saccharomycetes(100);o__Saccharomycetales(100);f__Metschnikowiaceae(100);g__Metschnikowia(100);s__Metschnikowia_pulcherrima(100); | Metschnikowia | 13 | Saprotroph | \|Undefined Saprotroph\| | Yeast | NULL | Probable | As nectar-tap_saprotroph (Põlme et al. 2020); Decay sugar-rich substrates (Põlme et al. 2020) | Põlme S, et al. 2020. Fungal Diversity 105:1-16 ((https://doi.org/10.1007/s13225-020-00466-2)) |
| Otu12701 | 2 | k__Fungi(100);p__Basidiomycota(100);c__Ustilaginomycetes(100);o__Ustilaginales(100);f__Ustilaginaceae(100);g__unclassified_Ustilaginaceae(100);s__Ustilaginaceae_sp(100); | Ustilaginaceae | 9 | Pathotroph | Plant Pathogen | Smut | NULL | Probable | Biotrophic in living tissue of Gramineae and Cyperaceae (Cannon & Kirk 2007) | Cannon PF, Kirk PM. 2007. Fungal Families of the World. CAB International, Cambridge (ISBN: 978-0851998275) |
| Otu12702 | 2 | k__Fungi(100);p__Ascomycota(100);c__Saccharomycetes(100);o__Saccharomycetales(100);f__Metschnikowiaceae(100);g__Metschnikowia(100);s__Metschnikowia_pulcherrima(100); | Metschnikowia | 13 | Saprotroph | \|Undefined Saprotroph\| | Yeast | NULL | Probable | As nectar-tap_saprotroph (Põlme et al. 2020); Decay sugar-rich substrates (Põlme et al. 2020) | Põlme S, et al. 2020. Fungal Diversity 105:1-16 ((https://doi.org/10.1007/s13225-020-00466-2)) |
| Otu12705 | 2 | k__Fungi(100);p__Ascomycota(100);c__Dothideomycetes(100);o__Dothideomycetes_order_Incertae_sedis(100);f__Myxotrichaceae(100);g__Oidiodendron(100);s__Oidiodendron_maius(100); | Oidiodendron | 13 | Pathotroph-Saprotroph-Symbiotroph | Endophyte-Ericoid Mycorrhizal-\|Undefined Saprotroph\| | Microfungus | NULL | Probable | As root_endophyte (Põlme et al. 2020); As soil_saprotroph (Põlme et al. 2020); Dark Septate Endophyte; Decay soil (Põlme et al. 2020) | Newsham KK. 2011. New Phytologist 190:783-793 ((https://doi.org/10.1111/j.1469-8137.2010.03611.x)) (as dark septate root endophytes); Põlme S, et al. 2020. Fungal Diversity 105:1-16 ((https://doi.org/10.1007/s13225-020-00466-2)) |
| Otu12707 | 2 | k__Fungi(100);p__Basidiomycota(100);c__Agaricomycetes(100);o__Polyporales(100);f__Phanerochaetaceae(100);g__Phanerochaete(100);g__Phanerochaete_unclassified(100); | Phanerochaete | 13 | Saprotroph | \|Wood Saprotroph\| | Corticioid | White Rot | Highly Probable | As white_rot (Põlme et al. 2020); Decay wood (Põlme et al. 2020) | Glenn JK, et al. 1983. Biochemical and Biophysical Research Communications 114:1077-1083 ((https://doi.org/10.1016/0006-291x((83))90672-1)); Põlme S, et al. 2020. Fungal Diversity 105:1-16 ((https://doi.org/10.1007/s13225-020-00466-2)) |
| Otu12709 | 2 | k__Fungi(100);p__Ascomycota(100);c__Saccharomycetes(100);o__Saccharomycetales(100);f__Metschnikowiaceae(100);g__Metschnikowia(100);s__Metschnikowia_pulcherrima(100); | Metschnikowia | 13 | Saprotroph | \|Undefined Saprotroph\| | Yeast | NULL | Probable | As nectar-tap_saprotroph (Põlme et al. 2020); Decay sugar-rich substrates (Põlme et al. 2020) | Põlme S, et al. 2020. Fungal Diversity 105:1-16 ((https://doi.org/10.1007/s13225-020-00466-2)) |
| Otu12710 | 2 | k__Fungi(100);p__Ascomycota(100);c__Saccharomycetes(100);o__Saccharomycetales(100);f__Metschnikowiaceae(100);g__Metschnikowia(100);s__Metschnikowia_pulcherrima(100); | Metschnikowia | 13 | Saprotroph | \|Undefined Saprotroph\| | Yeast | NULL | Probable | As nectar-tap_saprotroph (Põlme et al. 2020); Decay sugar-rich substrates (Põlme et al. 2020) | Põlme S, et al. 2020. Fungal Diversity 105:1-16 ((https://doi.org/10.1007/s13225-020-00466-2)) |
| Otu12711 | 2 | k__Fungi(100);p__Ascomycota(100);c__Saccharomycetes(100);o__Saccharomycetales(100);f__Metschnikowiaceae(100);g__Metschnikowia(100);s__Metschnikowia_pulcherrima(100); | Metschnikowia | 13 | Saprotroph | \|Undefined Saprotroph\| | Yeast | NULL | Probable | As nectar-tap_saprotroph (Põlme et al. 2020); Decay sugar-rich substrates (Põlme et al. 2020) | Põlme S, et al. 2020. Fungal Diversity 105:1-16 ((https://doi.org/10.1007/s13225-020-00466-2)) |
| Otu12712 | 2 | k__Fungi(100);p__Ascomycota(100);c__Saccharomycetes(100);o__Saccharomycetales(100);f__Metschnikowiaceae(100);g__Metschnikowia(100);g__Metschnikowia_unclassified(100); | Metschnikowia | 13 | Saprotroph | \|Undefined Saprotroph\| | Yeast | NULL | Probable | As nectar-tap_saprotroph (Põlme et al. 2020); Decay sugar-rich substrates (Põlme et al. 2020) | Põlme S, et al. 2020. Fungal Diversity 105:1-16 ((https://doi.org/10.1007/s13225-020-00466-2)) |
| Otu12713 | 2 | k__Fungi(100);p__Ascomycota(100);c__Saccharomycetes(100);o__Saccharomycetales(100);f__Metschnikowiaceae(100);g__Metschnikowia(100);s__Metschnikowia_pulcherrima(100); | Metschnikowia | 13 | Saprotroph | \|Undefined Saprotroph\| | Yeast | NULL | Probable | As nectar-tap_saprotroph (Põlme et al. 2020); Decay sugar-rich substrates (Põlme et al. 2020) | Põlme S, et al. 2020. Fungal Diversity 105:1-16 ((https://doi.org/10.1007/s13225-020-00466-2)) |
| Otu12715 | 2 | k__Fungi(100);p__Ascomycota(100);c__Saccharomycetes(100);o__Saccharomycetales(100);f__Metschnikowiaceae(100);g__Metschnikowia(100);s__Metschnikowia_pulcherrima(100); | Metschnikowia | 13 | Saprotroph | \|Undefined Saprotroph\| | Yeast | NULL | Probable | As nectar-tap_saprotroph (Põlme et al. 2020); Decay sugar-rich substrates (Põlme et al. 2020) | Põlme S, et al. 2020. Fungal Diversity 105:1-16 ((https://doi.org/10.1007/s13225-020-00466-2)) |
| Otu12716 | 2 | k__Fungi(100);p__Ascomycota(100);c__Saccharomycetes(100);o__Saccharomycetales(100);f__Metschnikowiaceae(100);g__Metschnikowia(100);s__Metschnikowia_pulcherrima(100); | Metschnikowia | 13 | Saprotroph | \|Undefined Saprotroph\| | Yeast | NULL | Probable | As nectar-tap_saprotroph (Põlme et al. 2020); Decay sugar-rich substrates (Põlme et al. 2020) | Põlme S, et al. 2020. Fungal Diversity 105:1-16 ((https://doi.org/10.1007/s13225-020-00466-2)) |
| Otu12717 | 2 | k__Fungi(100);p__Basidiomycota(100);c__Agaricomycetes(100);o__Cantharellales(100);f__Clavulinaceae(100);g__Clavulina(100);g__Clavulina_unclassified(100); | Clavulina | 13 | Symbiotroph | \|Ectomycorrhizal\| | Clavarioid | NULL | Highly Probable | NULL | Tedersoo L, et al. 2014. Science 346:e1256688 ((https://doi.org/10.1126/science.1256688)); Põlme S, et al. 2020. Fungal Diversity 105:1-16 ((https://doi.org/10.1007/s13225-020-00466-2)) |
| Otu12720 | 2 | k__Fungi(100);p__Ascomycota(100);c__Saccharomycetes(100);o__Saccharomycetales(100);f__Metschnikowiaceae(100);g__Metschnikowia(100);s__Metschnikowia_pulcherrima(100); | Metschnikowia | 13 | Saprotroph | \|Undefined Saprotroph\| | Yeast | NULL | Probable | As nectar-tap_saprotroph (Põlme et al. 2020); Decay sugar-rich substrates (Põlme et al. 2020) | Põlme S, et al. 2020. Fungal Diversity 105:1-16 ((https://doi.org/10.1007/s13225-020-00466-2)) |
| Otu12721 | 2 | k__Fungi(100);p__Ascomycota(100);c__Saccharomycetes(100);o__Saccharomycetales(100);f__Metschnikowiaceae(100);g__Metschnikowia(100);s__Metschnikowia_pulcherrima(100); | Metschnikowia | 13 | Saprotroph | \|Undefined Saprotroph\| | Yeast | NULL | Probable | As nectar-tap_saprotroph (Põlme et al. 2020); Decay sugar-rich substrates (Põlme et al. 2020) | Põlme S, et al. 2020. Fungal Diversity 105:1-16 ((https://doi.org/10.1007/s13225-020-00466-2)) |
| Otu12723 | 2 | k__Fungi(100);p__Ascomycota(100);c__Saccharomycetes(100);o__Saccharomycetales(100);f__Metschnikowiaceae(100);g__Metschnikowia(100);s__Metschnikowia_pulcherrima(100); | Metschnikowia | 13 | Saprotroph | \|Undefined Saprotroph\| | Yeast | NULL | Probable | As nectar-tap_saprotroph (Põlme et al. 2020); Decay sugar-rich substrates (Põlme et al. 2020) | Põlme S, et al. 2020. Fungal Diversity 105:1-16 ((https://doi.org/10.1007/s13225-020-00466-2)) |
| Otu12724 | 2 | k__Fungi(100);p__Ascomycota(100);c__Saccharomycetes(100);o__Saccharomycetales(100);f__Metschnikowiaceae(100);g__Metschnikowia(100);g__Metschnikowia_unclassified(100); | Metschnikowia | 13 | Saprotroph | \|Undefined Saprotroph\| | Yeast | NULL | Probable | As nectar-tap_saprotroph (Põlme et al. 2020); Decay sugar-rich substrates (Põlme et al. 2020) | Põlme S, et al. 2020. Fungal Diversity 105:1-16 ((https://doi.org/10.1007/s13225-020-00466-2)) |
| Otu12726 | 2 | k__Fungi(100);p__Basidiomycota(100);c__Agaricomycetes(100);o__Cantharellales(100);f__Hydnaceae(100);g__Sistotrema(100);s__Sistotrema_muscicola(100); | Sistotrema | 13 | Saprotroph-Symbiotroph | Ectomycorrhizal-\|Plant Saprotroph\|-Wood Saprotroph | Corticioid | White Rot | Possible | As litter_saprotroph (Põlme et al. 2020); Decay leaf, fruit, seed (Põlme et al. 2020) | Gilbertson RL, Ryvarden L. 1987-1987. North American Polypores. Fungiflora, Oslo (ISBN: 978-0945345060); Rinaldi AC, et al. 2008. Fungal Diversity 33:1-45 ((http://www.fungaldiversity.org/fdp/sfdp/33-1.pdf)) (as ECM/not ECM); Tedersoo L, et al. 2010. Mycorrhiza 20:217-263 ((https://doi.org/10.1007/s00572-009-0274-x)) (pro parte); Põlme S, et al. 2020. Fungal Diversity 105:1-16 ((https://doi.org/10.1007/s13225-020-00466-2)); http://mycorrhizas.info/ecmf.html ((http://mycorrhizas.info/ecmf.html)) (as ECM/not ECM). |
| Otu12727 | 2 | k__Fungi(100);p__Ascomycota(100);c__Saccharomycetes(100);o__Saccharomycetales(100);f__Metschnikowiaceae(100);g__Metschnikowia(100);g__Metschnikowia_unclassified(100); | Metschnikowia | 13 | Saprotroph | \|Undefined Saprotroph\| | Yeast | NULL | Probable | As nectar-tap_saprotroph (Põlme et al. 2020); Decay sugar-rich substrates (Põlme et al. 2020) | Põlme S, et al. 2020. Fungal Diversity 105:1-16 ((https://doi.org/10.1007/s13225-020-00466-2)) |
| Otu12728 | 2 | k__Fungi(100);p__Ascomycota(100);c__Saccharomycetes(100);o__Saccharomycetales(100);f__Metschnikowiaceae(100);g__Metschnikowia(100);s__Metschnikowia_pulcherrima(100); | Metschnikowia | 13 | Saprotroph | \|Undefined Saprotroph\| | Yeast | NULL | Probable | As nectar-tap_saprotroph (Põlme et al. 2020); Decay sugar-rich substrates (Põlme et al. 2020) | Põlme S, et al. 2020. Fungal Diversity 105:1-16 ((https://doi.org/10.1007/s13225-020-00466-2)) |
| Otu12729 | 2 | k__Fungi(100);p__Ascomycota(100);c__Saccharomycetes(100);o__Saccharomycetales(100);f__Metschnikowiaceae(100);g__Metschnikowia(100);s__Metschnikowia_pulcherrima(100); | Metschnikowia | 13 | Saprotroph | \|Undefined Saprotroph\| | Yeast | NULL | Probable | As nectar-tap_saprotroph (Põlme et al. 2020); Decay sugar-rich substrates (Põlme et al. 2020) | Põlme S, et al. 2020. Fungal Diversity 105:1-16 ((https://doi.org/10.1007/s13225-020-00466-2)) |
| Otu12731 | 2 | k__Fungi(100);p__Ascomycota(100);c__Sordariomycetes(100);o__Hypocreales(100);f__Nectriaceae(100);g__Fusarium(100);g__Fusarium_unclassified(100); | Fusarium | 13 | Pathotroph-Saprotroph-Symbiotroph | Animal Pathogen-Endophyte-Lichen Parasite-\|Plant Pathogen\|-Plant Saprotroph-Undefined Saprotroph-Wood Saprotroph | Microfungus | Soft Rot | Possible | As foliar_endophyte (Põlme et al. 2020); As litter_saprotroph (Põlme et al. 2020); Causing damping-off disease (Põlme et al. 2020); Decay leaf, fruit, seed, roots, soil, animal material (Põlme et al. 2020); Likely opportunistic human pathogen (Irinyi et al. 2015) | Duncan CG, Eslyn WE. 1966. Mycologia 58:642-664 ((https://doi.org/10.2307/3757045)); Seehann G, et al. 1975. List of Fungi in Soft-Rot Tissue Tests. Institute fur Holzbiologie und Holzschutz, Bundesforschungsanstalt fur Forst - und Holzwirtshaft, Hamburg - Lohbrugge, Germany; Costa IPMW, et al. 2012. Mycotaxon 119:e26 ((http://mycotaxon.com/resources/checklists/costa_v119_checklist.pdf)); Tedersoo L, et al. 2014. Science 346:e1256688 ((https://doi.org/10.1126/science.1256688)); Irinyi L, et al. 2015. Medical Mycology 53:313-337 ((https://doi.org/10.1093/mmy/myv008)); Schroers, et al. 2016. Mycologia 108:806-819 ((https://doi.org/10.3852/15-255)); Põlme S, et al. 2020. Fungal Diversity 105:1-16 ((https://doi.org/10.1007/s13225-020-00466-2)); The American Phytopathological Society Common Names of Plant Diseases (http://www.apsnet.org/publications/commonnames/Pages/default.aspx)); Lawrey JD, Diederich P. 2016. http://www.lichenicolous.net (accessed 24 May 2017) ((http://www.lichenicolous.net)) |
| Otu12732 | 2 | k__Fungi(100);p__Basidiomycota(100);c__Agaricostilbomycetes(100);o__Agaricostilbales(100);f__Chionosphaeraceae(100);g__Kurtzmanomyces(100);g__Kurtzmanomyces_unclassified(100); | Kurtzmanomyces | 13 | Saprotroph | \|Undefined Saprotroph\| | Dimorphic | NULL | Probable | NULL | Põlme S, et al. 2020. Fungal Diversity 105:1-16 ((https://doi.org/10.1007/s13225-020-00466-2)) |
| Otu12733 | 2 | k__Fungi(100);p__Ascomycota(100);c__Saccharomycetes(100);o__Saccharomycetales(100);f__Metschnikowiaceae(100);g__Metschnikowia(100);g__Metschnikowia_unclassified(100); | Metschnikowia | 13 | Saprotroph | \|Undefined Saprotroph\| | Yeast | NULL | Probable | As nectar-tap_saprotroph (Põlme et al. 2020); Decay sugar-rich substrates (Põlme et al. 2020) | Põlme S, et al. 2020. Fungal Diversity 105:1-16 ((https://doi.org/10.1007/s13225-020-00466-2)) |
| Otu12737 | 2 | k__Fungi(100);p__Basidiomycota(100);c__Exobasidiomycetes(100);o__Microstromatales(100);f__Microstromataceae(100);g__Microstroma(100);s__Microstroma_juglandis(100); | Microstroma | 13 | Pathotroph | \|Plant Pathogen\| | Dimorphic | NULL | Probable | NULL | Tedersoo L, et al. 2014. Science 346:e1256688 ((https://doi.org/10.1126/science.1256688)); Põlme S, et al. 2020. Fungal Diversity 105:1-16 ((https://doi.org/10.1007/s13225-020-00466-2)) |
| Otu12740 | 2 | k__Fungi(100);p__Ascomycota(100);c__Saccharomycetes(100);o__Saccharomycetales(100);f__Metschnikowiaceae(100);g__Metschnikowia(100);s__Metschnikowia_pulcherrima(100); | Metschnikowia | 13 | Saprotroph | \|Undefined Saprotroph\| | Yeast | NULL | Probable | As nectar-tap_saprotroph (Põlme et al. 2020); Decay sugar-rich substrates (Põlme et al. 2020) | Põlme S, et al. 2020. Fungal Diversity 105:1-16 ((https://doi.org/10.1007/s13225-020-00466-2)) |
| Otu12741 | 2 | k__Fungi(100);p__Basidiomycota(100);c__Microbotryomycetes(100);o__Sporidiobolales(100);f__Sporidiobolales_family_Incertae_sedis(100);g__Rhodotorula(100);s__Rhodotorula_sp(100); | Rhodotorula | 13 | Pathotroph-Saprotroph-Symbiotroph | Animal Pathogen-Animal Symbiotroph-Endophyte-Plant Pathogen-\|Undefined Saprotroph\| | Yeast | NULL | Probable | As foliar_endophyte (Põlme et al. 2020); Opportunistic human pathogen | Põlme S, et al. 2020. Fungal Diversity 105:1-16 ((https://doi.org/10.1007/s13225-020-00466-2)); https://www.cdc.gov ((https://www.cdc.gov)) |
| Otu12744 | 2 | k__Fungi(100);p__Ascomycota(100);c__Saccharomycetes(100);o__Saccharomycetales(100);f__Metschnikowiaceae(100);g__Metschnikowia(100);s__Metschnikowia_pulcherrima(100); | Metschnikowia | 13 | Saprotroph | \|Undefined Saprotroph\| | Yeast | NULL | Probable | As nectar-tap_saprotroph (Põlme et al. 2020); Decay sugar-rich substrates (Põlme et al. 2020) | Põlme S, et al. 2020. Fungal Diversity 105:1-16 ((https://doi.org/10.1007/s13225-020-00466-2)) |
| Otu12745 | 2 | k__Fungi(100);p__Ascomycota(100);c__Saccharomycetes(100);o__Saccharomycetales(100);f__Metschnikowiaceae(100);g__Metschnikowia(100);s__Metschnikowia_pulcherrima(100); | Metschnikowia | 13 | Saprotroph | \|Undefined Saprotroph\| | Yeast | NULL | Probable | As nectar-tap_saprotroph (Põlme et al. 2020); Decay sugar-rich substrates (Põlme et al. 2020) | Põlme S, et al. 2020. Fungal Diversity 105:1-16 ((https://doi.org/10.1007/s13225-020-00466-2)) |
| Otu12746 | 2 | k__Fungi(100);p__Ascomycota(100);c__Saccharomycetes(100);o__Saccharomycetales(100);f__Metschnikowiaceae(100);g__Metschnikowia(100);g__Metschnikowia_unclassified(100); | Metschnikowia | 13 | Saprotroph | \|Undefined Saprotroph\| | Yeast | NULL | Probable | As nectar-tap_saprotroph (Põlme et al. 2020); Decay sugar-rich substrates (Põlme et al. 2020) | Põlme S, et al. 2020. Fungal Diversity 105:1-16 ((https://doi.org/10.1007/s13225-020-00466-2)) |
| Otu12747 | 2 | k__Fungi(100);p__Ascomycota(100);c__Saccharomycetes(100);o__Saccharomycetales(100);f__Metschnikowiaceae(100);g__Metschnikowia(100);g__Metschnikowia_unclassified(100); | Metschnikowia | 13 | Saprotroph | \|Undefined Saprotroph\| | Yeast | NULL | Probable | As nectar-tap_saprotroph (Põlme et al. 2020); Decay sugar-rich substrates (Põlme et al. 2020) | Põlme S, et al. 2020. Fungal Diversity 105:1-16 ((https://doi.org/10.1007/s13225-020-00466-2)) |
| Otu12750 | 2 | k__Fungi(100);p__Ascomycota(100);c__Saccharomycetes(100);o__Saccharomycetales(100);f__Metschnikowiaceae(100);g__Metschnikowia(100);s__Metschnikowia_pulcherrima(100); | Metschnikowia | 13 | Saprotroph | \|Undefined Saprotroph\| | Yeast | NULL | Probable | As nectar-tap_saprotroph (Põlme et al. 2020); Decay sugar-rich substrates (Põlme et al. 2020) | Põlme S, et al. 2020. Fungal Diversity 105:1-16 ((https://doi.org/10.1007/s13225-020-00466-2)) |
| Otu12751 | 2 | k__Fungi(100);p__Ascomycota(100);c__Saccharomycetes(100);o__Saccharomycetales(100);f__Metschnikowiaceae(100);g__Metschnikowia(100);g__Metschnikowia_unclassified(100); | Metschnikowia | 13 | Saprotroph | \|Undefined Saprotroph\| | Yeast | NULL | Probable | As nectar-tap_saprotroph (Põlme et al. 2020); Decay sugar-rich substrates (Põlme et al. 2020) | Põlme S, et al. 2020. Fungal Diversity 105:1-16 ((https://doi.org/10.1007/s13225-020-00466-2)) |
| Otu12753 | 2 | k__Fungi(100);p__Ascomycota(100);c__Saccharomycetes(100);o__Saccharomycetales(100);f__Metschnikowiaceae(100);g__Metschnikowia(100);s__Metschnikowia_pulcherrima(100); | Metschnikowia | 13 | Saprotroph | \|Undefined Saprotroph\| | Yeast | NULL | Probable | As nectar-tap_saprotroph (Põlme et al. 2020); Decay sugar-rich substrates (Põlme et al. 2020) | Põlme S, et al. 2020. Fungal Diversity 105:1-16 ((https://doi.org/10.1007/s13225-020-00466-2)) |
| Otu12755 | 2 | k__Fungi(100);p__Ascomycota(100);c__Saccharomycetes(100);o__Saccharomycetales(100);f__Metschnikowiaceae(100);g__Metschnikowia(100);s__Metschnikowia_pulcherrima(100); | Metschnikowia | 13 | Saprotroph | \|Undefined Saprotroph\| | Yeast | NULL | Probable | As nectar-tap_saprotroph (Põlme et al. 2020); Decay sugar-rich substrates (Põlme et al. 2020) | Põlme S, et al. 2020. Fungal Diversity 105:1-16 ((https://doi.org/10.1007/s13225-020-00466-2)) |
| Otu12756 | 2 | k__Fungi(100);p__Ascomycota(100);c__Saccharomycetes(100);o__Saccharomycetales(100);f__Metschnikowiaceae(100);g__Metschnikowia(100);s__Metschnikowia_pulcherrima(100); | Metschnikowia | 13 | Saprotroph | \|Undefined Saprotroph\| | Yeast | NULL | Probable | As nectar-tap_saprotroph (Põlme et al. 2020); Decay sugar-rich substrates (Põlme et al. 2020) | Põlme S, et al. 2020. Fungal Diversity 105:1-16 ((https://doi.org/10.1007/s13225-020-00466-2)) |
| Otu12758 | 2 | k__Fungi(100);p__Ascomycota(100);c__Dothideomycetes(100);o__Pleosporales(100);f__Phaeosphaeriaceae(100);g__Sclerostagonospora(100);s__Sclerostagonospora_sp(100); | Sclerostagonospora | 13 | Pathotroph-Saprotroph | Plant Pathogen-\|Plant Saprotroph\|-Undefined Saprotroph | Microfungus | NULL | Probable | As litter_saprotroph (Põlme et al. 2020); Decay leaf, fruit, seed (Põlme et al. 2020) | Tedersoo L, et al. 2014. Science 346:e1256688 ((https://doi.org/10.1126/science.1256688)); Põlme S, et al. 2020. Fungal Diversity 105:1-16 ((https://doi.org/10.1007/s13225-020-00466-2)) |
| Otu12763 | 2 | k__Fungi(100);p__Ascomycota(100);c__Saccharomycetes(100);o__Saccharomycetales(100);f__Metschnikowiaceae(100);g__Metschnikowia(100);s__Metschnikowia_pulcherrima(100); | Metschnikowia | 13 | Saprotroph | \|Undefined Saprotroph\| | Yeast | NULL | Probable | As nectar-tap_saprotroph (Põlme et al. 2020); Decay sugar-rich substrates (Põlme et al. 2020) | Põlme S, et al. 2020. Fungal Diversity 105:1-16 ((https://doi.org/10.1007/s13225-020-00466-2)) |
| Otu12768 | 2 | k__Fungi(100);p__Ascomycota(100);c__Saccharomycetes(100);o__Saccharomycetales(100);f__Metschnikowiaceae(100);g__Metschnikowia(100);s__Metschnikowia_pulcherrima(100); | Metschnikowia | 13 | Saprotroph | \|Undefined Saprotroph\| | Yeast | NULL | Probable | As nectar-tap_saprotroph (Põlme et al. 2020); Decay sugar-rich substrates (Põlme et al. 2020) | Põlme S, et al. 2020. Fungal Diversity 105:1-16 ((https://doi.org/10.1007/s13225-020-00466-2)) |
| Otu12770 | 2 | k__Fungi(100);p__Ascomycota(100);c__Saccharomycetes(100);o__Saccharomycetales(100);f__Metschnikowiaceae(100);g__Metschnikowia(100);s__Metschnikowia_pulcherrima(100); | Metschnikowia | 13 | Saprotroph | \|Undefined Saprotroph\| | Yeast | NULL | Probable | As nectar-tap_saprotroph (Põlme et al. 2020); Decay sugar-rich substrates (Põlme et al. 2020) | Põlme S, et al. 2020. Fungal Diversity 105:1-16 ((https://doi.org/10.1007/s13225-020-00466-2)) |
| Otu12772 | 2 | k__Fungi(100);p__Ascomycota(100);c__Saccharomycetes(100);o__Saccharomycetales(100);f__Metschnikowiaceae(100);g__Metschnikowia(100);s__Metschnikowia_pulcherrima(100); | Metschnikowia | 13 | Saprotroph | \|Undefined Saprotroph\| | Yeast | NULL | Probable | As nectar-tap_saprotroph (Põlme et al. 2020); Decay sugar-rich substrates (Põlme et al. 2020) | Põlme S, et al. 2020. Fungal Diversity 105:1-16 ((https://doi.org/10.1007/s13225-020-00466-2)) |
| Otu12775 | 2 | k__Fungi(100);p__Ascomycota(100);c__Saccharomycetes(100);o__Saccharomycetales(100);f__Metschnikowiaceae(100);g__Metschnikowia(100);s__Metschnikowia_pulcherrima(100); | Metschnikowia | 13 | Saprotroph | \|Undefined Saprotroph\| | Yeast | NULL | Probable | As nectar-tap_saprotroph (Põlme et al. 2020); Decay sugar-rich substrates (Põlme et al. 2020) | Põlme S, et al. 2020. Fungal Diversity 105:1-16 ((https://doi.org/10.1007/s13225-020-00466-2)) |
| Otu12778 | 2 | k__Fungi(100);p__Ascomycota(100);c__Saccharomycetes(100);o__Saccharomycetales(100);f__Metschnikowiaceae(100);g__Metschnikowia(100);g__Metschnikowia_unclassified(100); | Metschnikowia | 13 | Saprotroph | \|Undefined Saprotroph\| | Yeast | NULL | Probable | As nectar-tap_saprotroph (Põlme et al. 2020); Decay sugar-rich substrates (Põlme et al. 2020) | Põlme S, et al. 2020. Fungal Diversity 105:1-16 ((https://doi.org/10.1007/s13225-020-00466-2)) |
| Otu12779 | 2 | k__Fungi(100);p__Ascomycota(100);c__Sordariomycetes(100);o__Hypocreales(100);f__Nectriaceae(100);g__Gibberella(100);s__Gibberella_intricans(100); | Gibberella | 13 | Pathotroph-Saprotroph-Symbiotroph | Endophyte-\|Plant Pathogen\|-Plant Saprotroph-Undefined Saprotroph | Microfungus | Soft Rot | Probable | As foliar_endophyte (Põlme et al. 2020); As litter_saprotroph (Põlme et al. 2020); Causing damping-off disease (Põlme et al. 2020); Decay leaf, fruit, seed, roots, soil, animal material (Põlme et al. 2020) | Tedersoo L, et al. 2014. Science 346:e1256688 ((https://doi.org/10.1126/science.1256688)); Põlme S, et al. 2020. Fungal Diversity 105:1-16 ((https://doi.org/10.1007/s13225-020-00466-2)) |
| Otu12784 | 2 | k__Fungi(100);p__Ascomycota(100);c__Saccharomycetes(100);o__Saccharomycetales(100);f__Saccharomycetales_family_Incertae_sedis(100);f__Saccharomycetales_family_Incertae_sedis_unclassified(100);f__Saccharomycetales_family_Incertae_sedis_unclassified(100); | Saccharomycetales | 7 | Saprotroph | Undefined Saprotroph | Yeast | NULL | Possible | NULL | Sterkenburg E, et al. 2015. New Phytologist 207:1145-1158 ((https://doi.org/10.1111/nph.13426)) |
| Otu12786 | 2 | k__Fungi(100);p__Ascomycota(100);c__Saccharomycetes(100);o__Saccharomycetales(100);f__Metschnikowiaceae(100);g__Metschnikowia(100);g__Metschnikowia_unclassified(100); | Metschnikowia | 13 | Saprotroph | \|Undefined Saprotroph\| | Yeast | NULL | Probable | As nectar-tap_saprotroph (Põlme et al. 2020); Decay sugar-rich substrates (Põlme et al. 2020) | Põlme S, et al. 2020. Fungal Diversity 105:1-16 ((https://doi.org/10.1007/s13225-020-00466-2)) |
| Otu12787 | 2 | k__Fungi(100);p__Ascomycota(100);c__Saccharomycetes(100);o__Saccharomycetales(100);f__Metschnikowiaceae(100);g__Metschnikowia(100);s__Metschnikowia_pulcherrima(100); | Metschnikowia | 13 | Saprotroph | \|Undefined Saprotroph\| | Yeast | NULL | Probable | As nectar-tap_saprotroph (Põlme et al. 2020); Decay sugar-rich substrates (Põlme et al. 2020) | Põlme S, et al. 2020. Fungal Diversity 105:1-16 ((https://doi.org/10.1007/s13225-020-00466-2)) |
| Otu12788 | 2 | k__Fungi(100);p__Ascomycota(100);c__Saccharomycetes(100);o__Saccharomycetales(100);f__Metschnikowiaceae(100);g__Metschnikowia(100);g__Metschnikowia_unclassified(100); | Metschnikowia | 13 | Saprotroph | \|Undefined Saprotroph\| | Yeast | NULL | Probable | As nectar-tap_saprotroph (Põlme et al. 2020); Decay sugar-rich substrates (Põlme et al. 2020) | Põlme S, et al. 2020. Fungal Diversity 105:1-16 ((https://doi.org/10.1007/s13225-020-00466-2)) |
| Otu12792 | 2 | k__Fungi(100);p__Ascomycota(100);c__Saccharomycetes(100);o__Saccharomycetales(100);f__Metschnikowiaceae(100);g__Metschnikowia(100);s__Metschnikowia_pulcherrima(100); | Metschnikowia | 13 | Saprotroph | \|Undefined Saprotroph\| | Yeast | NULL | Probable | As nectar-tap_saprotroph (Põlme et al. 2020); Decay sugar-rich substrates (Põlme et al. 2020) | Põlme S, et al. 2020. Fungal Diversity 105:1-16 ((https://doi.org/10.1007/s13225-020-00466-2)) |
| Otu12793 | 2 | k__Fungi(100);p__Ascomycota(100);c__Saccharomycetes(100);o__Saccharomycetales(100);f__Metschnikowiaceae(100);g__Metschnikowia(100);s__Metschnikowia_pulcherrima(100); | Metschnikowia | 13 | Saprotroph | \|Undefined Saprotroph\| | Yeast | NULL | Probable | As nectar-tap_saprotroph (Põlme et al. 2020); Decay sugar-rich substrates (Põlme et al. 2020) | Põlme S, et al. 2020. Fungal Diversity 105:1-16 ((https://doi.org/10.1007/s13225-020-00466-2)) |
| Otu12796 | 2 | k__Fungi(100);p__Ascomycota(100);c__Saccharomycetes(100);o__Saccharomycetales(100);f__Metschnikowiaceae(100);g__Metschnikowia(100);s__Metschnikowia_pulcherrima(100); | Metschnikowia | 13 | Saprotroph | \|Undefined Saprotroph\| | Yeast | NULL | Probable | As nectar-tap_saprotroph (Põlme et al. 2020); Decay sugar-rich substrates (Põlme et al. 2020) | Põlme S, et al. 2020. Fungal Diversity 105:1-16 ((https://doi.org/10.1007/s13225-020-00466-2)) |
| Otu12797 | 2 | k__Fungi(100);p__Ascomycota(100);c__Saccharomycetes(100);o__Saccharomycetales(100);f__Metschnikowiaceae(100);g__Metschnikowia(100);g__Metschnikowia_unclassified(100); | Metschnikowia | 13 | Saprotroph | \|Undefined Saprotroph\| | Yeast | NULL | Probable | As nectar-tap_saprotroph (Põlme et al. 2020); Decay sugar-rich substrates (Põlme et al. 2020) | Põlme S, et al. 2020. Fungal Diversity 105:1-16 ((https://doi.org/10.1007/s13225-020-00466-2)) |
| Otu12798 | 2 | k__Fungi(100);p__Ascomycota(100);c__Saccharomycetes(100);o__Saccharomycetales(100);f__Metschnikowiaceae(100);g__Metschnikowia(100);s__Metschnikowia_pulcherrima(100); | Metschnikowia | 13 | Saprotroph | \|Undefined Saprotroph\| | Yeast | NULL | Probable | As nectar-tap_saprotroph (Põlme et al. 2020); Decay sugar-rich substrates (Põlme et al. 2020) | Põlme S, et al. 2020. Fungal Diversity 105:1-16 ((https://doi.org/10.1007/s13225-020-00466-2)) |
| Otu12800 | 2 | k__Fungi(100);p__Ascomycota(100);c__Saccharomycetes(100);o__Saccharomycetales(100);f__Metschnikowiaceae(100);g__Metschnikowia(100);g__Metschnikowia_unclassified(100); | Metschnikowia | 13 | Saprotroph | \|Undefined Saprotroph\| | Yeast | NULL | Probable | As nectar-tap_saprotroph (Põlme et al. 2020); Decay sugar-rich substrates (Põlme et al. 2020) | Põlme S, et al. 2020. Fungal Diversity 105:1-16 ((https://doi.org/10.1007/s13225-020-00466-2)) |
| Otu12801 | 2 | k__Fungi(100);p__Ascomycota(100);c__Saccharomycetes(100);o__Saccharomycetales(100);f__Metschnikowiaceae(100);g__Metschnikowia(100);g__Metschnikowia_unclassified(100); | Metschnikowia | 13 | Saprotroph | \|Undefined Saprotroph\| | Yeast | NULL | Probable | As nectar-tap_saprotroph (Põlme et al. 2020); Decay sugar-rich substrates (Põlme et al. 2020) | Põlme S, et al. 2020. Fungal Diversity 105:1-16 ((https://doi.org/10.1007/s13225-020-00466-2)) |
| Otu12802 | 2 | k__Fungi(100);p__Ascomycota(100);c__Dothideomycetes(100);o__Capnodiales(100);f__Mycosphaerellaceae(100);g__Mycosphaerella(100);g__Mycosphaerella_unclassified(100); | Mycosphaerella | 13 | Pathotroph-Saprotroph | \|Plant Pathogen\|-Plant Saprotroph | Microfungus | NULL | Probable | As litter_saprotroph (Põlme et al. 2020); Decay leaf, fruit, seed (Põlme et al. 2020) | Tedersoo L, et al. 2014. Science 346:e1256688 ((https://doi.org/10.1126/science.1256688)); Põlme S, et al. 2020. Fungal Diversity 105:1-16 ((https://doi.org/10.1007/s13225-020-00466-2)) |
| Otu12803 | 2 | k__Fungi(100);p__Ascomycota(100);c__Saccharomycetes(100);o__Saccharomycetales(100);f__Metschnikowiaceae(100);g__Metschnikowia(100);s__Metschnikowia_pulcherrima(100); | Metschnikowia | 13 | Saprotroph | \|Undefined Saprotroph\| | Yeast | NULL | Probable | As nectar-tap_saprotroph (Põlme et al. 2020); Decay sugar-rich substrates (Põlme et al. 2020) | Põlme S, et al. 2020. Fungal Diversity 105:1-16 ((https://doi.org/10.1007/s13225-020-00466-2)) |
| Otu12804 | 2 | k__Fungi(100);p__Ascomycota(100);c__Saccharomycetes(100);o__Saccharomycetales(100);f__Metschnikowiaceae(100);g__Metschnikowia(100);g__Metschnikowia_unclassified(100); | Metschnikowia | 13 | Saprotroph | \|Undefined Saprotroph\| | Yeast | NULL | Probable | As nectar-tap_saprotroph (Põlme et al. 2020); Decay sugar-rich substrates (Põlme et al. 2020) | Põlme S, et al. 2020. Fungal Diversity 105:1-16 ((https://doi.org/10.1007/s13225-020-00466-2)) |
| Otu12808 | 2 | k__Fungi(100);p__Chytridiomycota(100);c__Chytridiomycetes(100);o__Rhizophydiales(100);f__Protrudomycetaceae(100);g__Protrudomyces(100);s__Protrudomyces_lateralis(100); | Protrudomyces | 13 | Saprotroph | \|Pollen Saprotroph\| | Chytrid-Microfungus | NULL | Probable | Decay pollen (Põlme et al. 2020) | Põlme S, et al. 2020. Fungal Diversity 105:1-16 ((https://doi.org/10.1007/s13225-020-00466-2)) |
| Otu12809 | 2 | k__Fungi(100);p__Ascomycota(100);c__Dothideomycetes(100);o__Dothideomycetes_order_Incertae_sedis(100);f__Pseudeurotiaceae(100);g__Pseudeurotium(100);g__Pseudeurotium_unclassified(100); | Pseudeurotium | 13 | Saprotroph | \|Dung Saprotroph\|-Undefined Saprotroph | Microfungus | NULL | Probable | As soil_saprotroph (Põlme et al. 2020); Decay soil, dung (Põlme et al. 2020) | Tedersoo L, et al. 2014. Science 346:e1256688 ((https://doi.org/10.1126/science.1256688)); Põlme S, et al. 2020. Fungal Diversity 105:1-16 ((https://doi.org/10.1007/s13225-020-00466-2)) |
| Otu12810 | 2 | k__Fungi(100);p__Ascomycota(100);c__Saccharomycetes(100);o__Saccharomycetales(100);f__Metschnikowiaceae(100);g__Metschnikowia(100);g__Metschnikowia_unclassified(100); | Metschnikowia | 13 | Saprotroph | \|Undefined Saprotroph\| | Yeast | NULL | Probable | As nectar-tap_saprotroph (Põlme et al. 2020); Decay sugar-rich substrates (Põlme et al. 2020) | Põlme S, et al. 2020. Fungal Diversity 105:1-16 ((https://doi.org/10.1007/s13225-020-00466-2)) |
| Otu12811 | 2 | k__Fungi(100);p__Ascomycota(100);c__Leotiomycetes(100);o__Helotiales(100);f__Helotiaceae(100);g__Articulospora(100);s__Articulospora_atra(100); | Articulospora | 13 | Saprotroph | \|Plant Saprotroph\|-Undefined Saprotroph | Microfungus | NULL | Probable | As litter_saprotroph (Põlme et al. 2020); Decay leaf, fruit, seed (Põlme et al. 2020) | Tedersoo L, et al. 2014. Science 346:e1256688 ((https://doi.org/10.1126/science.1256688)); Põlme S, et al. 2020. Fungal Diversity 105:1-16 ((https://doi.org/10.1007/s13225-020-00466-2)) |
| Otu12812 | 2 | k__Fungi(100);p__Ascomycota(100);c__Saccharomycetes(100);o__Saccharomycetales(100);f__Metschnikowiaceae(100);g__Metschnikowia(100);g__Metschnikowia_unclassified(100); | Metschnikowia | 13 | Saprotroph | \|Undefined Saprotroph\| | Yeast | NULL | Probable | As nectar-tap_saprotroph (Põlme et al. 2020); Decay sugar-rich substrates (Põlme et al. 2020) | Põlme S, et al. 2020. Fungal Diversity 105:1-16 ((https://doi.org/10.1007/s13225-020-00466-2)) |
| Otu12813 | 2 | k__Fungi(100);p__Ascomycota(100);c__Saccharomycetes(100);o__Saccharomycetales(100);f__Metschnikowiaceae(100);g__Metschnikowia(100);s__Metschnikowia_pulcherrima(100); | Metschnikowia | 13 | Saprotroph | \|Undefined Saprotroph\| | Yeast | NULL | Probable | As nectar-tap_saprotroph (Põlme et al. 2020); Decay sugar-rich substrates (Põlme et al. 2020) | Põlme S, et al. 2020. Fungal Diversity 105:1-16 ((https://doi.org/10.1007/s13225-020-00466-2)) |
| Otu12815 | 2 | k__Fungi(100);p__Ascomycota(100);c__Saccharomycetes(100);o__Saccharomycetales(100);f__Metschnikowiaceae(100);g__Metschnikowia(100);g__Metschnikowia_unclassified(100); | Metschnikowia | 13 | Saprotroph | \|Undefined Saprotroph\| | Yeast | NULL | Probable | As nectar-tap_saprotroph (Põlme et al. 2020); Decay sugar-rich substrates (Põlme et al. 2020) | Põlme S, et al. 2020. Fungal Diversity 105:1-16 ((https://doi.org/10.1007/s13225-020-00466-2)) |
| Otu12818 | 2 | k__Fungi(100);p__Ascomycota(100);c__Saccharomycetes(100);o__Saccharomycetales(100);f__Metschnikowiaceae(100);g__Metschnikowia(100);s__Metschnikowia_pulcherrima(100); | Metschnikowia | 13 | Saprotroph | \|Undefined Saprotroph\| | Yeast | NULL | Probable | As nectar-tap_saprotroph (Põlme et al. 2020); Decay sugar-rich substrates (Põlme et al. 2020) | Põlme S, et al. 2020. Fungal Diversity 105:1-16 ((https://doi.org/10.1007/s13225-020-00466-2)) |
| Otu12819 | 2 | k__Fungi(100);p__Basidiomycota(100);c__Microbotryomycetes(100);o__Sporidiobolales(100);f__Sporidiobolales_family_Incertae_sedis(100);g__Rhodotorula(100);s__Rhodotorula_taiwanensis(100); | Rhodotorula | 13 | Pathotroph-Saprotroph-Symbiotroph | Animal Pathogen-Animal Symbiotroph-Endophyte-Plant Pathogen-\|Undefined Saprotroph\| | Yeast | NULL | Probable | As foliar_endophyte (Põlme et al. 2020); Opportunistic human pathogen | Põlme S, et al. 2020. Fungal Diversity 105:1-16 ((https://doi.org/10.1007/s13225-020-00466-2)); https://www.cdc.gov ((https://www.cdc.gov)) |
| Otu12821 | 2 | k__Fungi(100);p__Ascomycota(100);c__Saccharomycetes(100);o__Saccharomycetales(100);o__Saccharomycetales_unclassified(100);o__Saccharomycetales_unclassified(100);o__Saccharomycetales_unclassified(100); | Saccharomycetales | 7 | Saprotroph | Undefined Saprotroph | Yeast | NULL | Possible | NULL | Sterkenburg E, et al. 2015. New Phytologist 207:1145-1158 ((https://doi.org/10.1111/nph.13426)) |
| Otu12824 | 2 | k__Fungi(100);p__Basidiomycota(100);c__Microbotryomycetes(100);o__Sporidiobolales(100);f__Sporidiobolales_family_Incertae_sedis(100);g__Rhodotorula(100);g__Rhodotorula_unclassified(100); | Rhodotorula | 13 | Pathotroph-Saprotroph-Symbiotroph | Animal Pathogen-Animal Symbiotroph-Endophyte-Plant Pathogen-\|Undefined Saprotroph\| | Yeast | NULL | Probable | As foliar_endophyte (Põlme et al. 2020); Opportunistic human pathogen | Põlme S, et al. 2020. Fungal Diversity 105:1-16 ((https://doi.org/10.1007/s13225-020-00466-2)); https://www.cdc.gov ((https://www.cdc.gov)) |
| Otu12826 | 2 | k__Fungi(100);p__Ascomycota(100);c__Saccharomycetes(100);o__Saccharomycetales(100);f__Metschnikowiaceae(100);g__Metschnikowia(100);g__Metschnikowia_unclassified(100); | Metschnikowia | 13 | Saprotroph | \|Undefined Saprotroph\| | Yeast | NULL | Probable | As nectar-tap_saprotroph (Põlme et al. 2020); Decay sugar-rich substrates (Põlme et al. 2020) | Põlme S, et al. 2020. Fungal Diversity 105:1-16 ((https://doi.org/10.1007/s13225-020-00466-2)) |
| Otu12827 | 2 | k__Fungi(100);p__Ascomycota(100);c__Saccharomycetes(100);o__Saccharomycetales(100);f__Metschnikowiaceae(100);g__Metschnikowia(100);s__Metschnikowia_pulcherrima(100); | Metschnikowia | 13 | Saprotroph | \|Undefined Saprotroph\| | Yeast | NULL | Probable | As nectar-tap_saprotroph (Põlme et al. 2020); Decay sugar-rich substrates (Põlme et al. 2020) | Põlme S, et al. 2020. Fungal Diversity 105:1-16 ((https://doi.org/10.1007/s13225-020-00466-2)) |
| Otu12828 | 2 | k__Fungi(100);p__Ascomycota(100);c__Saccharomycetes(100);o__Saccharomycetales(100);f__Metschnikowiaceae(100);g__Metschnikowia(100);g__Metschnikowia_unclassified(100); | Metschnikowia | 13 | Saprotroph | \|Undefined Saprotroph\| | Yeast | NULL | Probable | As nectar-tap_saprotroph (Põlme et al. 2020); Decay sugar-rich substrates (Põlme et al. 2020) | Põlme S, et al. 2020. Fungal Diversity 105:1-16 ((https://doi.org/10.1007/s13225-020-00466-2)) |
| Otu12829 | 2 | k__Fungi(100);p__Basidiomycota(100);c__Microbotryomycetes(100);o__Sporidiobolales(100);f__Sporidiobolales_family_Incertae_sedis(100);g__Rhodotorula(100);s__Rhodotorula_taiwanensis(100); | Rhodotorula | 13 | Pathotroph-Saprotroph-Symbiotroph | Animal Pathogen-Animal Symbiotroph-Endophyte-Plant Pathogen-\|Undefined Saprotroph\| | Yeast | NULL | Probable | As foliar_endophyte (Põlme et al. 2020); Opportunistic human pathogen | Põlme S, et al. 2020. Fungal Diversity 105:1-16 ((https://doi.org/10.1007/s13225-020-00466-2)); https://www.cdc.gov ((https://www.cdc.gov)) |
| Otu12830 | 2 | k__Fungi(100);p__Ascomycota(100);c__Saccharomycetes(100);o__Saccharomycetales(100);f__Metschnikowiaceae(100);g__Metschnikowia(100);s__Metschnikowia_pulcherrima(100); | Metschnikowia | 13 | Saprotroph | \|Undefined Saprotroph\| | Yeast | NULL | Probable | As nectar-tap_saprotroph (Põlme et al. 2020); Decay sugar-rich substrates (Põlme et al. 2020) | Põlme S, et al. 2020. Fungal Diversity 105:1-16 ((https://doi.org/10.1007/s13225-020-00466-2)) |
| Otu12831 | 2 | k__Fungi(100);p__Ascomycota(100);c__Saccharomycetes(100);o__Saccharomycetales(100);f__Saccharomycetales_family_Incertae_sedis(100);g__Candida(100);g__Candida_unclassified(100); | Candida | 13 | Pathotroph-Saprotroph-Symbiotroph | Animal Pathogen-Endophyte-Endosymbiont-Epiphyte-\|Undefined Saprotroph\| | Dimorphic Yeast-Yeast | NULL | Probable | As nectar-tap_saprotroph (Põlme et al. 2020); Decay sugar-rich substrates (Põlme et al. 2020); This asexual genus is polyphyletic and contain species that function very differently depend on what they are. Caution should be taken when interpreting guilds for Candida species. | Manolakaki D, et al. 2010. Virulence. 1: 367-375 ((https://doi.org/10.4161/viru.1.5.12796)); Mekbib SB, et al. 2011. Tropical Plant Pathology 36:233-240 ((https://doi.org/10.1590/S1982-56762011000400004)); Põlme S, et al. 2020. Fungal Diversity 105:1-16 ((https://doi.org/10.1007/s13225-020-00466-2)) |
| Otu12832 | 2 | k__Fungi(100);p__Ascomycota(100);c__Saccharomycetes(100);o__Saccharomycetales(100);f__Metschnikowiaceae(100);g__Metschnikowia(100);s__Metschnikowia_pulcherrima(100); | Metschnikowia | 13 | Saprotroph | \|Undefined Saprotroph\| | Yeast | NULL | Probable | As nectar-tap_saprotroph (Põlme et al. 2020); Decay sugar-rich substrates (Põlme et al. 2020) | Põlme S, et al. 2020. Fungal Diversity 105:1-16 ((https://doi.org/10.1007/s13225-020-00466-2)) |
| Otu12836 | 2 | k__Fungi(100);p__Ascomycota(100);c__Saccharomycetes(100);o__Saccharomycetales(100);f__Metschnikowiaceae(100);g__Metschnikowia(100);s__Metschnikowia_pulcherrima(100); | Metschnikowia | 13 | Saprotroph | \|Undefined Saprotroph\| | Yeast | NULL | Probable | As nectar-tap_saprotroph (Põlme et al. 2020); Decay sugar-rich substrates (Põlme et al. 2020) | Põlme S, et al. 2020. Fungal Diversity 105:1-16 ((https://doi.org/10.1007/s13225-020-00466-2)) |
| Otu12837 | 2 | k__Fungi(100);p__Ascomycota(100);c__Saccharomycetes(100);o__Saccharomycetales(100);f__Saccharomycetales_family_Incertae_sedis(100);f__Saccharomycetales_family_Incertae_sedis_unclassified(100);f__Saccharomycetales_family_Incertae_sedis_unclassified(100); | Saccharomycetales | 7 | Saprotroph | Undefined Saprotroph | Yeast | NULL | Possible | NULL | Sterkenburg E, et al. 2015. New Phytologist 207:1145-1158 ((https://doi.org/10.1111/nph.13426)) |
| Otu12841 | 2 | k__Fungi(100);p__Basidiomycota(100);c__Ustilaginomycetes(100);o__Ustilaginales(100);f__Ustilaginaceae(100);g__Pseudozyma(100);s__Pseudozyma_aphidis(100); | Pseudozyma | 13 | Pathotroph-Symbiotroph | Endophyte-\|Epiphyte\|-Fungal Parasite | Yeast | NULL | Probable | As foliar_endophyte (Põlme et al. 2020); As mycoparasite (Põlme et al. 2020); Decay leaf, fruit, seed, fungal material (Põlme et al. 2020) | Põlme S, et al. 2020. Fungal Diversity 105:1-16 ((https://doi.org/10.1007/s13225-020-00466-2)) |
| Otu12842 | 2 | k__Fungi(100);p__Ascomycota(100);c__Saccharomycetes(100);o__Saccharomycetales(100);f__Metschnikowiaceae(100);g__Metschnikowia(100);s__Metschnikowia_pulcherrima(100); | Metschnikowia | 13 | Saprotroph | \|Undefined Saprotroph\| | Yeast | NULL | Probable | As nectar-tap_saprotroph (Põlme et al. 2020); Decay sugar-rich substrates (Põlme et al. 2020) | Põlme S, et al. 2020. Fungal Diversity 105:1-16 ((https://doi.org/10.1007/s13225-020-00466-2)) |
| Otu12843 | 2 | k__Fungi(100);p__Ascomycota(100);c__Saccharomycetes(100);o__Saccharomycetales(100);f__Metschnikowiaceae(100);g__Metschnikowia(100);s__Metschnikowia_pulcherrima(100); | Metschnikowia | 13 | Saprotroph | \|Undefined Saprotroph\| | Yeast | NULL | Probable | As nectar-tap_saprotroph (Põlme et al. 2020); Decay sugar-rich substrates (Põlme et al. 2020) | Põlme S, et al. 2020. Fungal Diversity 105:1-16 ((https://doi.org/10.1007/s13225-020-00466-2)) |
| Otu12844 | 2 | k__Fungi(100);p__Basidiomycota(100);c__Agaricomycetes(100);o__Polyporales(100);f__Meruliaceae(100);g__Phlebia(100);s__Phlebia_tremellosa(100); | Phlebia | 13 | Saprotroph | \|Wood Saprotroph\| | Corticioid | White Rot | Highly Probable | As white_rot (Põlme et al. 2020); Decay wood (Põlme et al. 2020) | Kirk PM et al. 2008. Dictionary of the Fungi. Tenth Edition. CAB International, Wallingford (ISBN: 978-0851998268); Põlme S, et al. 2020. Fungal Diversity 105:1-16 ((https://doi.org/10.1007/s13225-020-00466-2)) |
| Otu12846 | 2 | k__Fungi(100);p__Ascomycota(100);c__Saccharomycetes(100);o__Saccharomycetales(100);f__Metschnikowiaceae(100);g__Metschnikowia(100);g__Metschnikowia_unclassified(100); | Metschnikowia | 13 | Saprotroph | \|Undefined Saprotroph\| | Yeast | NULL | Probable | As nectar-tap_saprotroph (Põlme et al. 2020); Decay sugar-rich substrates (Põlme et al. 2020) | Põlme S, et al. 2020. Fungal Diversity 105:1-16 ((https://doi.org/10.1007/s13225-020-00466-2)) |
| Otu12847 | 2 | k__Fungi(100);p__Ascomycota(100);c__Saccharomycetes(100);o__Saccharomycetales(100);f__Metschnikowiaceae(100);g__Metschnikowia(100);g__Metschnikowia_unclassified(100); | Metschnikowia | 13 | Saprotroph | \|Undefined Saprotroph\| | Yeast | NULL | Probable | As nectar-tap_saprotroph (Põlme et al. 2020); Decay sugar-rich substrates (Põlme et al. 2020) | Põlme S, et al. 2020. Fungal Diversity 105:1-16 ((https://doi.org/10.1007/s13225-020-00466-2)) |
| Otu12848 | 2 | k__Fungi(100);p__Ascomycota(100);c__Saccharomycetes(100);o__Saccharomycetales(100);f__Metschnikowiaceae(100);g__Metschnikowia(100);s__Metschnikowia_pulcherrima(100); | Metschnikowia | 13 | Saprotroph | \|Undefined Saprotroph\| | Yeast | NULL | Probable | As nectar-tap_saprotroph (Põlme et al. 2020); Decay sugar-rich substrates (Põlme et al. 2020) | Põlme S, et al. 2020. Fungal Diversity 105:1-16 ((https://doi.org/10.1007/s13225-020-00466-2)) |
| Otu12849 | 2 | k__Fungi(100);p__Ascomycota(100);c__Saccharomycetes(100);o__Saccharomycetales(100);f__Metschnikowiaceae(100);g__Metschnikowia(100);s__Metschnikowia_pulcherrima(100); | Metschnikowia | 13 | Saprotroph | \|Undefined Saprotroph\| | Yeast | NULL | Probable | As nectar-tap_saprotroph (Põlme et al. 2020); Decay sugar-rich substrates (Põlme et al. 2020) | Põlme S, et al. 2020. Fungal Diversity 105:1-16 ((https://doi.org/10.1007/s13225-020-00466-2)) |
| Otu12852 | 2 | k__Fungi(100);p__Ascomycota(100);c__Saccharomycetes(100);o__Saccharomycetales(100);f__Metschnikowiaceae(100);g__Metschnikowia(100);s__Metschnikowia_pulcherrima(100); | Metschnikowia | 13 | Saprotroph | \|Undefined Saprotroph\| | Yeast | NULL | Probable | As nectar-tap_saprotroph (Põlme et al. 2020); Decay sugar-rich substrates (Põlme et al. 2020) | Põlme S, et al. 2020. Fungal Diversity 105:1-16 ((https://doi.org/10.1007/s13225-020-00466-2)) |
| Otu12854 | 2 | k__Fungi(100);p__Ascomycota(100);c__Saccharomycetes(100);o__Saccharomycetales(100);f__Metschnikowiaceae(100);g__Metschnikowia(100);s__Metschnikowia_pulcherrima(100); | Metschnikowia | 13 | Saprotroph | \|Undefined Saprotroph\| | Yeast | NULL | Probable | As nectar-tap_saprotroph (Põlme et al. 2020); Decay sugar-rich substrates (Põlme et al. 2020) | Põlme S, et al. 2020. Fungal Diversity 105:1-16 ((https://doi.org/10.1007/s13225-020-00466-2)) |
| Otu12855 | 2 | k__Fungi(100);p__Ascomycota(100);c__Saccharomycetes(100);o__Saccharomycetales(100);f__Metschnikowiaceae(100);g__Metschnikowia(100);g__Metschnikowia_unclassified(100); | Metschnikowia | 13 | Saprotroph | \|Undefined Saprotroph\| | Yeast | NULL | Probable | As nectar-tap_saprotroph (Põlme et al. 2020); Decay sugar-rich substrates (Põlme et al. 2020) | Põlme S, et al. 2020. Fungal Diversity 105:1-16 ((https://doi.org/10.1007/s13225-020-00466-2)) |
| Otu12856 | 2 | k__Fungi(100);p__Ascomycota(100);c__Saccharomycetes(100);o__Saccharomycetales(100);f__Metschnikowiaceae(100);g__Metschnikowia(100);s__Metschnikowia_pulcherrima(100); | Metschnikowia | 13 | Saprotroph | \|Undefined Saprotroph\| | Yeast | NULL | Probable | As nectar-tap_saprotroph (Põlme et al. 2020); Decay sugar-rich substrates (Põlme et al. 2020) | Põlme S, et al. 2020. Fungal Diversity 105:1-16 ((https://doi.org/10.1007/s13225-020-00466-2)) |
| Otu12857 | 2 | k__Fungi(100);p__Ascomycota(100);c__Saccharomycetes(100);o__Saccharomycetales(100);f__Metschnikowiaceae(100);g__Metschnikowia(100);g__Metschnikowia_unclassified(100); | Metschnikowia | 13 | Saprotroph | \|Undefined Saprotroph\| | Yeast | NULL | Probable | As nectar-tap_saprotroph (Põlme et al. 2020); Decay sugar-rich substrates (Põlme et al. 2020) | Põlme S, et al. 2020. Fungal Diversity 105:1-16 ((https://doi.org/10.1007/s13225-020-00466-2)) |
| Otu12859 | 2 | k__Fungi(100);p__Ascomycota(100);c__Saccharomycetes(100);o__Saccharomycetales(100);f__Metschnikowiaceae(100);g__Metschnikowia(100);s__Metschnikowia_pulcherrima(100); | Metschnikowia | 13 | Saprotroph | \|Undefined Saprotroph\| | Yeast | NULL | Probable | As nectar-tap_saprotroph (Põlme et al. 2020); Decay sugar-rich substrates (Põlme et al. 2020) | Põlme S, et al. 2020. Fungal Diversity 105:1-16 ((https://doi.org/10.1007/s13225-020-00466-2)) |
| Otu12860 | 2 | k__Fungi(100);p__Ascomycota(100);c__Saccharomycetes(100);o__Saccharomycetales(100);f__Metschnikowiaceae(100);g__Metschnikowia(100);g__Metschnikowia_unclassified(100); | Metschnikowia | 13 | Saprotroph | \|Undefined Saprotroph\| | Yeast | NULL | Probable | As nectar-tap_saprotroph (Põlme et al. 2020); Decay sugar-rich substrates (Põlme et al. 2020) | Põlme S, et al. 2020. Fungal Diversity 105:1-16 ((https://doi.org/10.1007/s13225-020-00466-2)) |
| Otu12864 | 2 | k__Fungi(100);p__Ascomycota(100);c__Saccharomycetes(100);o__Saccharomycetales(100);f__Metschnikowiaceae(100);g__Metschnikowia(100);s__Metschnikowia_pulcherrima(100); | Metschnikowia | 13 | Saprotroph | \|Undefined Saprotroph\| | Yeast | NULL | Probable | As nectar-tap_saprotroph (Põlme et al. 2020); Decay sugar-rich substrates (Põlme et al. 2020) | Põlme S, et al. 2020. Fungal Diversity 105:1-16 ((https://doi.org/10.1007/s13225-020-00466-2)) |
| Otu12865 | 2 | k__Fungi(100);p__Basidiomycota(100);c__Tremellomycetes(100);o__Tremellales(100);f__Tremellales_family_Incertae_sedis(100);g__Hannaella(100);s__Hannaella_luteola(100); | Hannaella | 13 | Pathotroph-Saprotroph | \|Animal Parasite\|-Undefined Saprotroph | Yeast | NULL | Probable | As animal_decomposer (Põlme et al. 2020); Decay animal material (Põlme et al. 2020) | Põlme S, et al. 2020. Fungal Diversity 105:1-16 ((https://doi.org/10.1007/s13225-020-00466-2)) |
| Otu12867 | 2 | k__Fungi(100);p__Zygomycota(100);c__Zygomycota_class_Incertae_sedis(100);o__Mucorales(100);f__Mucoraceae(100);g__Mucor(100);s__Mucor_sp_JJP_2009a(100); | Mucor | 13 | Saprotroph | \|Undefined Saprotroph\| | Microfungus | NULL | Probable | As soil_saprotroph (Põlme et al. 2020); Decay soil (Põlme et al. 2020) | Tedersoo L, et al. 2014. Science 346:e1256688 ((https://doi.org/10.1126/science.1256688)); Põlme S, et al. 2020. Fungal Diversity 105:1-16 ((https://doi.org/10.1007/s13225-020-00466-2)) |
| Otu12868 | 2 | k__Fungi(100);p__Ascomycota(100);c__Saccharomycetes(100);o__Saccharomycetales(100);f__Metschnikowiaceae(100);g__Metschnikowia(100);s__Metschnikowia_pulcherrima(100); | Metschnikowia | 13 | Saprotroph | \|Undefined Saprotroph\| | Yeast | NULL | Probable | As nectar-tap_saprotroph (Põlme et al. 2020); Decay sugar-rich substrates (Põlme et al. 2020) | Põlme S, et al. 2020. Fungal Diversity 105:1-16 ((https://doi.org/10.1007/s13225-020-00466-2)) |
| Otu12870 | 2 | k__Fungi(100);p__Ascomycota(100);c__Saccharomycetes(100);o__Saccharomycetales(100);f__Metschnikowiaceae(100);g__Metschnikowia(100);g__Metschnikowia_unclassified(100); | Metschnikowia | 13 | Saprotroph | \|Undefined Saprotroph\| | Yeast | NULL | Probable | As nectar-tap_saprotroph (Põlme et al. 2020); Decay sugar-rich substrates (Põlme et al. 2020) | Põlme S, et al. 2020. Fungal Diversity 105:1-16 ((https://doi.org/10.1007/s13225-020-00466-2)) |
| Otu12872 | 2 | k__Fungi(100);p__Ascomycota(100);c__Saccharomycetes(100);o__Saccharomycetales(100);f__Metschnikowiaceae(100);g__Metschnikowia(100);s__Metschnikowia_pulcherrima(100); | Metschnikowia | 13 | Saprotroph | \|Undefined Saprotroph\| | Yeast | NULL | Probable | As nectar-tap_saprotroph (Põlme et al. 2020); Decay sugar-rich substrates (Põlme et al. 2020) | Põlme S, et al. 2020. Fungal Diversity 105:1-16 ((https://doi.org/10.1007/s13225-020-00466-2)) |
| Otu12873 | 2 | k__Fungi(100);p__Ascomycota(100);c__Saccharomycetes(100);o__Saccharomycetales(100);f__Metschnikowiaceae(100);g__Metschnikowia(100);s__Metschnikowia_pulcherrima(100); | Metschnikowia | 13 | Saprotroph | \|Undefined Saprotroph\| | Yeast | NULL | Probable | As nectar-tap_saprotroph (Põlme et al. 2020); Decay sugar-rich substrates (Põlme et al. 2020) | Põlme S, et al. 2020. Fungal Diversity 105:1-16 ((https://doi.org/10.1007/s13225-020-00466-2)) |
| Otu12874 | 2 | k__Fungi(100);p__Ascomycota(100);c__Saccharomycetes(100);o__Saccharomycetales(100);f__Metschnikowiaceae(100);g__Metschnikowia(100);s__Metschnikowia_pulcherrima(100); | Metschnikowia | 13 | Saprotroph | \|Undefined Saprotroph\| | Yeast | NULL | Probable | As nectar-tap_saprotroph (Põlme et al. 2020); Decay sugar-rich substrates (Põlme et al. 2020) | Põlme S, et al. 2020. Fungal Diversity 105:1-16 ((https://doi.org/10.1007/s13225-020-00466-2)) |
| Otu12875 | 2 | k__Fungi(100);p__Ascomycota(100);c__Saccharomycetes(100);o__Saccharomycetales(100);f__Saccharomycetales_family_Incertae_sedis(100);g__Candida(100);g__Candida_unclassified(100); | Candida | 13 | Pathotroph-Saprotroph-Symbiotroph | Animal Pathogen-Endophyte-Endosymbiont-Epiphyte-\|Undefined Saprotroph\| | Dimorphic Yeast-Yeast | NULL | Probable | As nectar-tap_saprotroph (Põlme et al. 2020); Decay sugar-rich substrates (Põlme et al. 2020); This asexual genus is polyphyletic and contain species that function very differently depend on what they are. Caution should be taken when interpreting guilds for Candida species. | Manolakaki D, et al. 2010. Virulence. 1: 367-375 ((https://doi.org/10.4161/viru.1.5.12796)); Mekbib SB, et al. 2011. Tropical Plant Pathology 36:233-240 ((https://doi.org/10.1590/S1982-56762011000400004)); Põlme S, et al. 2020. Fungal Diversity 105:1-16 ((https://doi.org/10.1007/s13225-020-00466-2)) |
| Otu12878 | 2 | k__Fungi(100);p__Ascomycota(100);c__Saccharomycetes(100);o__Saccharomycetales(100);f__Metschnikowiaceae(100);g__Metschnikowia(100);s__Metschnikowia_pulcherrima(100); | Metschnikowia | 13 | Saprotroph | \|Undefined Saprotroph\| | Yeast | NULL | Probable | As nectar-tap_saprotroph (Põlme et al. 2020); Decay sugar-rich substrates (Põlme et al. 2020) | Põlme S, et al. 2020. Fungal Diversity 105:1-16 ((https://doi.org/10.1007/s13225-020-00466-2)) |
| Otu12879 | 2 | k__Fungi(100);p__Ascomycota(100);c__Saccharomycetes(100);o__Saccharomycetales(100);f__Metschnikowiaceae(100);g__Metschnikowia(100);s__Metschnikowia_pulcherrima(100); | Metschnikowia | 13 | Saprotroph | \|Undefined Saprotroph\| | Yeast | NULL | Probable | As nectar-tap_saprotroph (Põlme et al. 2020); Decay sugar-rich substrates (Põlme et al. 2020) | Põlme S, et al. 2020. Fungal Diversity 105:1-16 ((https://doi.org/10.1007/s13225-020-00466-2)) |
| Otu12881 | 2 | k__Fungi(100);p__Basidiomycota(100);c__Exobasidiomycetes(100);o__Exobasidiomycetes_order_Incertae_sedis(100);f__Exobasidiomycetes_family_Incertae_sedis(100);g__Tilletiopsis(100);s__Tilletiopsis_minor(100); | Tilletiopsis | 13 | Pathotroph-Saprotroph | Endophyte-Fungal Parasite-\|Plant Pathogen\|-Plant Saprotroph-Undefined Saprotroph | Microfungus-Yeast | NULL | Probable | As litter_saprotroph (Põlme et al. 2020); As mycoparasite (Põlme et al. 2020); Decay leaf, fruit, seed (Põlme et al. 2020); Isolated from leaves...dead leaves (Nyland 1950) | Nyland G. 1950. Mycologia 42:487-496 ((ttps://doi.org/10.1080/00275514.1950.12017853)); Põlme S, et al. 2020. Fungal Diversity 105:1-16 ((https://doi.org/10.1007/s13225-020-00466-2)) |
| Otu12883 | 2 | k__Fungi(100);p__Ascomycota(100);c__Saccharomycetes(100);o__Saccharomycetales(100);f__Metschnikowiaceae(100);g__Metschnikowia(100);s__Metschnikowia_pulcherrima(100); | Metschnikowia | 13 | Saprotroph | \|Undefined Saprotroph\| | Yeast | NULL | Probable | As nectar-tap_saprotroph (Põlme et al. 2020); Decay sugar-rich substrates (Põlme et al. 2020) | Põlme S, et al. 2020. Fungal Diversity 105:1-16 ((https://doi.org/10.1007/s13225-020-00466-2)) |
| Otu12884 | 2 | k__Fungi(100);p__Ascomycota(100);c__Saccharomycetes(100);o__Saccharomycetales(100);f__Metschnikowiaceae(100);g__Metschnikowia(100);s__Metschnikowia_pulcherrima(100); | Metschnikowia | 13 | Saprotroph | \|Undefined Saprotroph\| | Yeast | NULL | Probable | As nectar-tap_saprotroph (Põlme et al. 2020); Decay sugar-rich substrates (Põlme et al. 2020) | Põlme S, et al. 2020. Fungal Diversity 105:1-16 ((https://doi.org/10.1007/s13225-020-00466-2)) |
| Otu12885 | 2 | k__Fungi(100);p__Basidiomycota(100);c__Agaricomycetes(100);o__Agaricomycetes_order_Incertae_sedis(100);f__Agaricomycetes_family_Incertae_sedis(100);g__Peniophorella(100);s__Peniophorella_odontiiformis(100); | Peniophorella | 13 | Saprotroph | Undefined Saprotroph-\|Wood Saprotroph\| | Corticioid | White Rot | Probable | As white_rot (Põlme et al. 2020); Decay wood (Põlme et al. 2020) | Tedersoo L, et al. 2014. Science 346:e1256688 ((https://doi.org/10.1126/science.1256688)); Põlme S, et al. 2020. Fungal Diversity 105:1-16 ((https://doi.org/10.1007/s13225-020-00466-2)) |
| Otu12890 | 2 | k__Fungi(100);p__Basidiomycota(100);c__Microbotryomycetes(100);o__Sporidiobolales(100);f__Sporidiobolales_family_Incertae_sedis(100);g__Rhodotorula(100);s__Rhodotorula_crocea(100); | Rhodotorula | 13 | Pathotroph-Saprotroph-Symbiotroph | Animal Pathogen-Animal Symbiotroph-Endophyte-Plant Pathogen-\|Undefined Saprotroph\| | Yeast | NULL | Probable | As foliar_endophyte (Põlme et al. 2020); Opportunistic human pathogen | Põlme S, et al. 2020. Fungal Diversity 105:1-16 ((https://doi.org/10.1007/s13225-020-00466-2)); https://www.cdc.gov ((https://www.cdc.gov)) |
| Otu12892 | 2 | k__Fungi(100);p__Ascomycota(100);c__Saccharomycetes(100);o__Saccharomycetales(100);f__Saccharomycetales_family_Incertae_sedis(100);f__Saccharomycetales_family_Incertae_sedis_unclassified(100);f__Saccharomycetales_family_Incertae_sedis_unclassified(100); | Saccharomycetales | 7 | Saprotroph | Undefined Saprotroph | Yeast | NULL | Possible | NULL | Sterkenburg E, et al. 2015. New Phytologist 207:1145-1158 ((https://doi.org/10.1111/nph.13426)) |
| Otu12893 | 2 | k__Fungi(100);p__Zygomycota(100);c__Zygomycota_class_Incertae_sedis(100);o__Mortierellales(100);f__Mortierellaceae(100);g__Mortierella(100);g__Mortierella_unclassified(100); | Mortierella | 13 | Saprotroph-Symbiotroph | Endophyte-Plant Saprotroph-\|Undefined Saprotroph\| | Microfungus | NULL | Probable | As chitinolytic (Põlme et al. 2020); As soil_saprotroph (Põlme et al. 2020); Decay roots, soil (Põlme et al. 2020); Saprobic in soil and thus frequently isolated from soil and material contaminated with soil such as hay or rotten silage and spoil tips (Cannon & Kirk 2007) | Cannon PF, Kirk PM. 2007. Fungal Families of the World. CAB International, Cambridge (ISBN: 978-0851998275); Tedersoo L, et al. 2014. Science 346:e1256688 ((https://doi.org/10.1126/science.1256688)); Purahong W, et al. 2016. Molecular Ecology 25:4059-4074 ((https://doi.org/10.1111/mec.13739)); Põlme S, et al. 2020. Fungal Diversity 105:1-16 ((https://doi.org/10.1007/s13225-020-00466-2)) |
| Otu12894 | 2 | k__Fungi(100);p__Ascomycota(100);c__Saccharomycetes(100);o__Saccharomycetales(100);f__Metschnikowiaceae(100);g__Metschnikowia(100);g__Metschnikowia_unclassified(100); | Metschnikowia | 13 | Saprotroph | \|Undefined Saprotroph\| | Yeast | NULL | Probable | As nectar-tap_saprotroph (Põlme et al. 2020); Decay sugar-rich substrates (Põlme et al. 2020) | Põlme S, et al. 2020. Fungal Diversity 105:1-16 ((https://doi.org/10.1007/s13225-020-00466-2)) |
| Otu12897 | 2 | k__Fungi(100);p__Ascomycota(100);c__Saccharomycetes(100);o__Saccharomycetales(100);f__Metschnikowiaceae(100);g__Metschnikowia(100);g__Metschnikowia_unclassified(100); | Metschnikowia | 13 | Saprotroph | \|Undefined Saprotroph\| | Yeast | NULL | Probable | As nectar-tap_saprotroph (Põlme et al. 2020); Decay sugar-rich substrates (Põlme et al. 2020) | Põlme S, et al. 2020. Fungal Diversity 105:1-16 ((https://doi.org/10.1007/s13225-020-00466-2)) |
| Otu12898 | 2 | k__Fungi(100);p__Basidiomycota(100);c__Exobasidiomycetes(100);o__Microstromatales(100);f__Microstromataceae(100);g__Microstroma(100);s__Microstroma_juglandis(100); | Microstroma | 13 | Pathotroph | \|Plant Pathogen\| | Dimorphic | NULL | Probable | NULL | Tedersoo L, et al. 2014. Science 346:e1256688 ((https://doi.org/10.1126/science.1256688)); Põlme S, et al. 2020. Fungal Diversity 105:1-16 ((https://doi.org/10.1007/s13225-020-00466-2)) |
| Otu12902 | 2 | k__Fungi(100);p__Ascomycota(100);c__Saccharomycetes(100);o__Saccharomycetales(100);f__Metschnikowiaceae(100);g__Metschnikowia(100);s__Metschnikowia_pulcherrima(100); | Metschnikowia | 13 | Saprotroph | \|Undefined Saprotroph\| | Yeast | NULL | Probable | As nectar-tap_saprotroph (Põlme et al. 2020); Decay sugar-rich substrates (Põlme et al. 2020) | Põlme S, et al. 2020. Fungal Diversity 105:1-16 ((https://doi.org/10.1007/s13225-020-00466-2)) |
| Otu12904 | 2 | k__Fungi(100);p__Ascomycota(100);c__Saccharomycetes(100);o__Saccharomycetales(100);f__Metschnikowiaceae(100);g__Metschnikowia(100);s__Metschnikowia_pulcherrima(100); | Metschnikowia | 13 | Saprotroph | \|Undefined Saprotroph\| | Yeast | NULL | Probable | As nectar-tap_saprotroph (Põlme et al. 2020); Decay sugar-rich substrates (Põlme et al. 2020) | Põlme S, et al. 2020. Fungal Diversity 105:1-16 ((https://doi.org/10.1007/s13225-020-00466-2)) |
| Otu12906 | 2 | k__Fungi(100);p__Ascomycota(100);c__Saccharomycetes(100);o__Saccharomycetales(100);o__Saccharomycetales_unclassified(100);o__Saccharomycetales_unclassified(100);o__Saccharomycetales_unclassified(100); | Saccharomycetales | 7 | Saprotroph | Undefined Saprotroph | Yeast | NULL | Possible | NULL | Sterkenburg E, et al. 2015. New Phytologist 207:1145-1158 ((https://doi.org/10.1111/nph.13426)) |
| Otu12907 | 2 | k__Fungi(100);p__Ascomycota(100);c__Saccharomycetes(100);o__Saccharomycetales(100);f__Metschnikowiaceae(100);g__Metschnikowia(100);s__Metschnikowia_pulcherrima(100); | Metschnikowia | 13 | Saprotroph | \|Undefined Saprotroph\| | Yeast | NULL | Probable | As nectar-tap_saprotroph (Põlme et al. 2020); Decay sugar-rich substrates (Põlme et al. 2020) | Põlme S, et al. 2020. Fungal Diversity 105:1-16 ((https://doi.org/10.1007/s13225-020-00466-2)) |
| Otu12909 | 2 | k__Fungi(100);p__Ascomycota(100);c__Saccharomycetes(100);o__Saccharomycetales(100);f__Metschnikowiaceae(100);g__Metschnikowia(100);s__Metschnikowia_pulcherrima(100); | Metschnikowia | 13 | Saprotroph | \|Undefined Saprotroph\| | Yeast | NULL | Probable | As nectar-tap_saprotroph (Põlme et al. 2020); Decay sugar-rich substrates (Põlme et al. 2020) | Põlme S, et al. 2020. Fungal Diversity 105:1-16 ((https://doi.org/10.1007/s13225-020-00466-2)) |
| Otu12910 | 2 | k__Fungi(100);p__Ascomycota(100);c__Saccharomycetes(100);o__Saccharomycetales(100);f__Metschnikowiaceae(100);g__Metschnikowia(100);s__Metschnikowia_pulcherrima(100); | Metschnikowia | 13 | Saprotroph | \|Undefined Saprotroph\| | Yeast | NULL | Probable | As nectar-tap_saprotroph (Põlme et al. 2020); Decay sugar-rich substrates (Põlme et al. 2020) | Põlme S, et al. 2020. Fungal Diversity 105:1-16 ((https://doi.org/10.1007/s13225-020-00466-2)) |
| Otu12913 | 2 | k__Fungi(100);p__Ascomycota(100);c__Saccharomycetes(100);o__Saccharomycetales(100);f__Metschnikowiaceae(100);g__Metschnikowia(100);s__Metschnikowia_pulcherrima(100); | Metschnikowia | 13 | Saprotroph | \|Undefined Saprotroph\| | Yeast | NULL | Probable | As nectar-tap_saprotroph (Põlme et al. 2020); Decay sugar-rich substrates (Põlme et al. 2020) | Põlme S, et al. 2020. Fungal Diversity 105:1-16 ((https://doi.org/10.1007/s13225-020-00466-2)) |
| Otu12915 | 2 | k__Fungi(100);p__Ascomycota(100);c__Sordariomycetes(100);o__Diaporthales(100);f__Diaporthales_family_Incertae_sedis(100);g__Harknessia(100);g__Harknessia_unclassified(100); | Harknessia | 13 | Pathotroph-Saprotroph-Symbiotroph | \|Endophyte\|-Plant Pathogen-Plant Saprotroph-Undefined Saprotroph | Microfungus | NULL | Probable | As foliar_endophyte (Põlme et al. 2020); As litter_saprotroph (Põlme et al. 2020); Decay leaf, fruit, seed (Põlme et al. 2020); The genus includes around 60 species that range ecologically from being endophytes, to saprobes, or foliar pathogens (Lee et al. 2004) | Lee S, et al. 2004. Studies in Mycology 50:235-252 ((https://www.studiesinmycology.org/index.php/issue/52-studies-in-mycology-no-50)); Tedersoo L, et al. 2014. Science 346:e1256688 ((https://doi.org/10.1126/science.1256688)); Põlme S, et al. 2020. Fungal Diversity 105:1-16 ((https://doi.org/10.1007/s13225-020-00466-2)) |
| Otu12916 | 2 | k__Fungi(100);p__Ascomycota(100);c__Saccharomycetes(100);o__Saccharomycetales(100);f__Metschnikowiaceae(100);g__Metschnikowia(100);s__Metschnikowia_pulcherrima(100); | Metschnikowia | 13 | Saprotroph | \|Undefined Saprotroph\| | Yeast | NULL | Probable | As nectar-tap_saprotroph (Põlme et al. 2020); Decay sugar-rich substrates (Põlme et al. 2020) | Põlme S, et al. 2020. Fungal Diversity 105:1-16 ((https://doi.org/10.1007/s13225-020-00466-2)) |
| Otu12917 | 2 | k__Fungi(100);p__Ascomycota(100);c__Saccharomycetes(100);o__Saccharomycetales(100);f__Saccharomycetales_family_Incertae_sedis(100);g__Candida(100);g__Candida_unclassified(100); | Candida | 13 | Pathotroph-Saprotroph-Symbiotroph | Animal Pathogen-Endophyte-Endosymbiont-Epiphyte-\|Undefined Saprotroph\| | Dimorphic Yeast-Yeast | NULL | Probable | As nectar-tap_saprotroph (Põlme et al. 2020); Decay sugar-rich substrates (Põlme et al. 2020); This asexual genus is polyphyletic and contain species that function very differently depend on what they are. Caution should be taken when interpreting guilds for Candida species. | Manolakaki D, et al. 2010. Virulence. 1: 367-375 ((https://doi.org/10.4161/viru.1.5.12796)); Mekbib SB, et al. 2011. Tropical Plant Pathology 36:233-240 ((https://doi.org/10.1590/S1982-56762011000400004)); Põlme S, et al. 2020. Fungal Diversity 105:1-16 ((https://doi.org/10.1007/s13225-020-00466-2)) |
| Otu12918 | 2 | k__Fungi(100);p__Ascomycota(100);c__Saccharomycetes(100);o__Saccharomycetales(100);f__Metschnikowiaceae(100);g__Metschnikowia(100);s__Metschnikowia_pulcherrima(100); | Metschnikowia | 13 | Saprotroph | \|Undefined Saprotroph\| | Yeast | NULL | Probable | As nectar-tap_saprotroph (Põlme et al. 2020); Decay sugar-rich substrates (Põlme et al. 2020) | Põlme S, et al. 2020. Fungal Diversity 105:1-16 ((https://doi.org/10.1007/s13225-020-00466-2)) |
| Otu12919 | 2 | k__Fungi(100);p__Ascomycota(100);c__Saccharomycetes(100);o__Saccharomycetales(100);f__Metschnikowiaceae(100);g__Metschnikowia(100);s__Metschnikowia_pulcherrima(100); | Metschnikowia | 13 | Saprotroph | \|Undefined Saprotroph\| | Yeast | NULL | Probable | As nectar-tap_saprotroph (Põlme et al. 2020); Decay sugar-rich substrates (Põlme et al. 2020) | Põlme S, et al. 2020. Fungal Diversity 105:1-16 ((https://doi.org/10.1007/s13225-020-00466-2)) |
| Otu12920 | 2 | k__Fungi(100);p__Ascomycota(100);c__Saccharomycetes(100);o__Saccharomycetales(100);f__Metschnikowiaceae(100);g__Metschnikowia(100);s__Metschnikowia_pulcherrima(100); | Metschnikowia | 13 | Saprotroph | \|Undefined Saprotroph\| | Yeast | NULL | Probable | As nectar-tap_saprotroph (Põlme et al. 2020); Decay sugar-rich substrates (Põlme et al. 2020) | Põlme S, et al. 2020. Fungal Diversity 105:1-16 ((https://doi.org/10.1007/s13225-020-00466-2)) |
| Otu12921 | 2 | k__Fungi(100);p__Ascomycota(100);c__Saccharomycetes(100);o__Saccharomycetales(100);f__Metschnikowiaceae(100);g__Metschnikowia(100);g__Metschnikowia_unclassified(100); | Metschnikowia | 13 | Saprotroph | \|Undefined Saprotroph\| | Yeast | NULL | Probable | As nectar-tap_saprotroph (Põlme et al. 2020); Decay sugar-rich substrates (Põlme et al. 2020) | Põlme S, et al. 2020. Fungal Diversity 105:1-16 ((https://doi.org/10.1007/s13225-020-00466-2)) |
| Otu12922 | 2 | k__Fungi(100);p__Ascomycota(100);c__Saccharomycetes(100);o__Saccharomycetales(100);f__Metschnikowiaceae(100);g__Metschnikowia(100);g__Metschnikowia_unclassified(100); | Metschnikowia | 13 | Saprotroph | \|Undefined Saprotroph\| | Yeast | NULL | Probable | As nectar-tap_saprotroph (Põlme et al. 2020); Decay sugar-rich substrates (Põlme et al. 2020) | Põlme S, et al. 2020. Fungal Diversity 105:1-16 ((https://doi.org/10.1007/s13225-020-00466-2)) |
| Otu12925 | 2 | k__Fungi(100);p__Ascomycota(100);c__Saccharomycetes(100);o__Saccharomycetales(100);f__Metschnikowiaceae(100);g__Metschnikowia(100);g__Metschnikowia_unclassified(100); | Metschnikowia | 13 | Saprotroph | \|Undefined Saprotroph\| | Yeast | NULL | Probable | As nectar-tap_saprotroph (Põlme et al. 2020); Decay sugar-rich substrates (Põlme et al. 2020) | Põlme S, et al. 2020. Fungal Diversity 105:1-16 ((https://doi.org/10.1007/s13225-020-00466-2)) |
| Otu12926 | 2 | k__Fungi(100);p__Ascomycota(100);c__Saccharomycetes(100);o__Saccharomycetales(100);f__Metschnikowiaceae(100);g__Metschnikowia(100);s__Metschnikowia_pulcherrima(100); | Metschnikowia | 13 | Saprotroph | \|Undefined Saprotroph\| | Yeast | NULL | Probable | As nectar-tap_saprotroph (Põlme et al. 2020); Decay sugar-rich substrates (Põlme et al. 2020) | Põlme S, et al. 2020. Fungal Diversity 105:1-16 ((https://doi.org/10.1007/s13225-020-00466-2)) |
| Otu12933 | 2 | k__Fungi(100);p__Ascomycota(100);c__Saccharomycetes(100);o__Saccharomycetales(100);f__Metschnikowiaceae(100);g__Metschnikowia(100);g__Metschnikowia_unclassified(100); | Metschnikowia | 13 | Saprotroph | \|Undefined Saprotroph\| | Yeast | NULL | Probable | As nectar-tap_saprotroph (Põlme et al. 2020); Decay sugar-rich substrates (Põlme et al. 2020) | Põlme S, et al. 2020. Fungal Diversity 105:1-16 ((https://doi.org/10.1007/s13225-020-00466-2)) |
| Otu12934 | 2 | k__Fungi(100);p__Ascomycota(100);c__Saccharomycetes(100);o__Saccharomycetales(100);f__Metschnikowiaceae(100);g__Metschnikowia(100);g__Metschnikowia_unclassified(100); | Metschnikowia | 13 | Saprotroph | \|Undefined Saprotroph\| | Yeast | NULL | Probable | As nectar-tap_saprotroph (Põlme et al. 2020); Decay sugar-rich substrates (Põlme et al. 2020) | Põlme S, et al. 2020. Fungal Diversity 105:1-16 ((https://doi.org/10.1007/s13225-020-00466-2)) |
| Otu12935 | 2 | k__Fungi(100);p__Ascomycota(100);c__Saccharomycetes(100);o__Saccharomycetales(100);f__Metschnikowiaceae(100);g__Metschnikowia(100);s__Metschnikowia_pulcherrima(100); | Metschnikowia | 13 | Saprotroph | \|Undefined Saprotroph\| | Yeast | NULL | Probable | As nectar-tap_saprotroph (Põlme et al. 2020); Decay sugar-rich substrates (Põlme et al. 2020) | Põlme S, et al. 2020. Fungal Diversity 105:1-16 ((https://doi.org/10.1007/s13225-020-00466-2)) |
| Otu12936 | 2 | k__Fungi(100);p__Ascomycota(100);c__Saccharomycetes(100);o__Saccharomycetales(100);f__Metschnikowiaceae(100);g__Metschnikowia(100);s__Metschnikowia_pulcherrima(100); | Metschnikowia | 13 | Saprotroph | \|Undefined Saprotroph\| | Yeast | NULL | Probable | As nectar-tap_saprotroph (Põlme et al. 2020); Decay sugar-rich substrates (Põlme et al. 2020) | Põlme S, et al. 2020. Fungal Diversity 105:1-16 ((https://doi.org/10.1007/s13225-020-00466-2)) |
| Otu12937 | 2 | k__Fungi(100);p__Ascomycota(100);c__Saccharomycetes(100);o__Saccharomycetales(100);f__Metschnikowiaceae(100);g__Metschnikowia(100);s__Metschnikowia_pulcherrima(100); | Metschnikowia | 13 | Saprotroph | \|Undefined Saprotroph\| | Yeast | NULL | Probable | As nectar-tap_saprotroph (Põlme et al. 2020); Decay sugar-rich substrates (Põlme et al. 2020) | Põlme S, et al. 2020. Fungal Diversity 105:1-16 ((https://doi.org/10.1007/s13225-020-00466-2)) |

| | | | | | | | | | | | |
|---|---|---|---|---|---|---|---|---|---|---|---|
| Otu12938 | 2 | k__Fungi(100);p__Basidiomycota(100);c__Agaricomycetes(100);o__Boletales(100);f__Gyroporaceae(100);g__Gyroporus(100);s__Gyroporus_castaneus(100); | Gyroporus | 13 | Symbiotroph | \|Ectomycorrhizal\| | Agaricoid-Boletoid | NULL | Highly Probable | NULL | Rinaldi AC, et al. 2008. Fungal Diversity 33:1-45 ((http://www.fungaldiversity.org/fdp/sfdp/33-1.pdf)); Tedersoo L, et al. 2010. Mycorrhiza 20:217-263 ((https://doi.org/10.1007/s00572-009-0274-x)); Põlme S, et al. 2020. Fungal Diversity 105:1-16 ((https://doi.org/10.1007/s13225-020-00466-2)); http://mycorrhizas.info/ecmf.html ((http://mycorrhizas.info/ecmf.html)) |
| Otu12939 | 2 | k__Fungi(100);p__Ascomycota(100);c__Saccharomycetes(100);o__Saccharomycetales(100);f__Metschnikowiaceae(100);g__Metschnikowia(100);s__Metschnikowia_pulcherrima(100); | Metschnikowia | 13 | Saprotroph | \|Undefined Saprotroph\| | Yeast | NULL | Probable | As nectar-tap_saprotroph (Põlme et al. 2020); Decay sugar-rich substrates (Põlme et al. 2020) | Põlme S, et al. 2020. Fungal Diversity 105:1-16 ((https://doi.org/10.1007/s13225-020-00466-2)) |
| Otu12946 | 2 | k__Fungi(100);p__Basidiomycota(100);c__Microbotryomycetes(100);o__Microbotryomycetes_order_Incertae_sedis(100);f__Microbotryomycetes_family_Incertae_sedis(100);g__Curvibasidium(100);s__Curvibasidium_cygneicollum(100); | Curvibasidium | 13 | Saprotroph-Symbiotroph | Epiphyte-Plant Saprotroph-Undefined Saprotroph-\|Wood Saprotroph\| | Dimorphic-Microfungus | NULL | Possible | As litter_saprotroph (Põlme et al. 2020); Decay wood, leaf, fruit, seed, soil (Põlme et al. 2020) | Sampaio JP, et al. 2004. International Journal of Systematic and Evolutionary Microbiology 54:1401-1407 ((https://doi.org/10.1099/ijs.0.03037-0)); Põlme S, et al. 2020. Fungal Diversity 105:1-16 ((https://doi.org/10.1007/s13225-020-00466-2)) |
| Otu12951 | 2 | k__Fungi(100);p__Ascomycota(100);c__Saccharomycetes(100);o__Saccharomycetales(100);f__Metschnikowiaceae(100);g__Metschnikowia(100);s__Metschnikowia_pulcherrima(100); | Metschnikowia | 13 | Saprotroph | \|Undefined Saprotroph\| | Yeast | NULL | Probable | As nectar-tap_saprotroph (Põlme et al. 2020); Decay sugar-rich substrates (Põlme et al. 2020) | Põlme S, et al. 2020. Fungal Diversity 105:1-16 ((https://doi.org/10.1007/s13225-020-00466-2)) |
| Otu12953 | 2 | k__Fungi(100);p__Ascomycota(100);c__Saccharomycetes(100);o__Saccharomycetales(100);f__Metschnikowiaceae(100);g__Metschnikowia(100);s__Metschnikowia_pulcherrima(100); | Metschnikowia | 13 | Saprotroph | \|Undefined Saprotroph\| | Yeast | NULL | Probable | As nectar-tap_saprotroph (Põlme et al. 2020); Decay sugar-rich substrates (Põlme et al. 2020) | Põlme S, et al. 2020. Fungal Diversity 105:1-16 ((https://doi.org/10.1007/s13225-020-00466-2)) |
| Otu12954 | 2 | k__Fungi(100);p__Ascomycota(100);c__Saccharomycetes(100);o__Saccharomycetales(100);f__Metschnikowiaceae(100);g__Metschnikowia(100);s__Metschnikowia_pulcherrima(100); | Metschnikowia | 13 | Saprotroph | \|Undefined Saprotroph\| | Yeast | NULL | Probable | As nectar-tap_saprotroph (Põlme et al. 2020); Decay sugar-rich substrates (Põlme et al. 2020) | Põlme S, et al. 2020. Fungal Diversity 105:1-16 ((https://doi.org/10.1007/s13225-020-00466-2)) |
| Otu12955 | 2 | k__Fungi(100);p__Ascomycota(100);c__Saccharomycetes(100);o__Saccharomycetales(100);f__Metschnikowiaceae(100);g__Metschnikowia(100);g__Metschnikowia_unclassified(100); | Metschnikowia | 13 | Saprotroph | \|Undefined Saprotroph\| | Yeast | NULL | Probable | As nectar-tap_saprotroph (Põlme et al. 2020); Decay sugar-rich substrates (Põlme et al. 2020) | Põlme S, et al. 2020. Fungal Diversity 105:1-16 ((https://doi.org/10.1007/s13225-020-00466-2)) |
| Otu12956 | 2 | k__Fungi(100);p__Ascomycota(100);c__Saccharomycetes(100);o__Saccharomycetales(100);f__Metschnikowiaceae(100);g__Metschnikowia(100);g__Metschnikowia_unclassified(100); | Metschnikowia | 13 | Saprotroph | \|Undefined Saprotroph\| | Yeast | NULL | Probable | As nectar-tap_saprotroph (Põlme et al. 2020); Decay sugar-rich substrates (Põlme et al. 2020) | Põlme S, et al. 2020. Fungal Diversity 105:1-16 ((https://doi.org/10.1007/s13225-020-00466-2)) |
| Otu12957 | 2 | k__Fungi(100);p__Ascomycota(100);c__Saccharomycetes(100);o__Saccharomycetales(100);f__Metschnikowiaceae(100);g__Metschnikowia(100);s__Metschnikowia_pulcherrima(100); | Metschnikowia | 13 | Saprotroph | \|Undefined Saprotroph\| | Yeast | NULL | Probable | As nectar-tap_saprotroph (Põlme et al. 2020); Decay sugar-rich substrates (Põlme et al. 2020) | Põlme S, et al. 2020. Fungal Diversity 105:1-16 ((https://doi.org/10.1007/s13225-020-00466-2)) |
| Otu12958 | 2 | k__Fungi(100);p__Ascomycota(100);c__Saccharomycetes(100);o__Saccharomycetales(100);f__Metschnikowiaceae(100);g__Metschnikowia(100);g__Metschnikowia_unclassified(100); | Metschnikowia | 13 | Saprotroph | \|Undefined Saprotroph\| | Yeast | NULL | Probable | As nectar-tap_saprotroph (Põlme et al. 2020); Decay sugar-rich substrates (Põlme et al. 2020) | Põlme S, et al. 2020. Fungal Diversity 105:1-16 ((https://doi.org/10.1007/s13225-020-00466-2)) |
| Otu12959 | 2 | k__Fungi(100);p__Basidiomycota(100);c__Agaricomycetes(100);o__Agaricales(100);o__Agaricales_unclassified(100);o__Agaricales_unclassified(100);o__Agaricales_unclassified(100); | Agaricales | 7 | Pathotroph-Saprotroph-Symbiotroph | Bryophyte Parasite-Dung Saprotroph-Ectomycorrhizal-Fungal Parasite-Leaf Saprotroph-Plant Parasite-Undefined Saprotroph-Wood Saprotroph | Agaricoid-Gasteroid-Microfungus-Secotioid-Yeast | NULL | Possible | Mushrooms and toadstools, Gill fungi, Agarics... lignicolous, sometimes muscicolous or fungicolous, saprobic, mycorrhizal, rarely parasitic on plants or fungi (Kirk et al. 2008) | Kirk PM et al. 2008. Dictionary of the Fungi. Tenth Edition. CAB International, Wallingford (ISBN: 978-0851998268) |
| Otu12960 | 2 | k__Fungi(100);p__Ascomycota(100);c__Saccharomycetes(100);o__Saccharomycetales(100);f__Metschnikowiaceae(100);g__Metschnikowia(100);g__Metschnikowia_unclassified(100); | Metschnikowia | 13 | Saprotroph | \|Undefined Saprotroph\| | Yeast | NULL | Probable | As nectar-tap_saprotroph (Põlme et al. 2020); Decay sugar-rich substrates (Põlme et al. 2020) | Põlme S, et al. 2020. Fungal Diversity 105:1-16 ((https://doi.org/10.1007/s13225-020-00466-2)) |
| Otu12962 | 2 | k__Fungi(100);p__Ascomycota(100);c__Saccharomycetes(100);o__Saccharomycetales(100);f__Metschnikowiaceae(100);g__Metschnikowia(100);s__Metschnikowia_pulcherrima(100); | Metschnikowia | 13 | Saprotroph | \|Undefined Saprotroph\| | Yeast | NULL | Probable | As nectar-tap_saprotroph (Põlme et al. 2020); Decay sugar-rich substrates (Põlme et al. 2020) | Põlme S, et al. 2020. Fungal Diversity 105:1-16 ((https://doi.org/10.1007/s13225-020-00466-2)) |
| Otu08799 | 2 | k__Fungi(100);p__Ascomycota(100);p__Ascomycota_unclassified(100);p__Ascomycota_unclassified(100);p__Ascomycota_unclassified(100);p__Ascomycota_unclassified(100);p__Ascomycota_unclassified(100); | - | - | - | - | - | - | - | Unassigned | - |
| Otu08801 | 2 | k__Fungi(100);p__Ascomycota(100);p__Ascomycota_unclassified(100);p__Ascomycota_unclassified(100);p__Ascomycota_unclassified(100);p__Ascomycota_unclassified(100);p__Ascomycota_unclassified(100); | - | - | - | - | - | - | - | Unassigned | - |
| Otu08802 | 2 | k__Fungi(100);k__Fungi_unclassified(100);k__Fungi_unclassified(100);k__Fungi_unclassified(100);k__Fungi_unclassified(100);k__Fungi_unclassified(100);k__Fungi_unclassified(100); | - | - | - | - | - | - | - | Unassigned | - |
| Otu08804 | 2 | k__Fungi(100);k__Fungi_unclassified(100);k__Fungi_unclassified(100);k__Fungi_unclassified(100);k__Fungi_unclassified(100);k__Fungi_unclassified(100);k__Fungi_unclassified(100); | - | - | - | - | - | - | - | Unassigned | - |
| Otu08806 | 2 | k__Fungi(100);p__Ascomycota(100);p__Ascomycota_unclassified(100);p__Ascomycota_unclassified(100);p__Ascomycota_unclassified(100);p__Ascomycota_unclassified(100);p__Ascomycota_unclassified(100); | - | - | - | - | - | - | - | Unassigned | - |
| Otu08807 | 2 | k__Fungi(100);k__Fungi_unclassified(100);k__Fungi_unclassified(100);k__Fungi_unclassified(100);k__Fungi_unclassified(100);k__Fungi_unclassified(100);k__Fungi_unclassified(100); | - | - | - | - | - | - | - | Unassigned | - |
| Otu08809 | 2 | k__Fungi(100);k__Fungi_unclassified(100);k__Fungi_unclassified(100);k__Fungi_unclassified(100);k__Fungi_unclassified(100);k__Fungi_unclassified(100);k__Fungi_unclassified(100); | - | - | - | - | - | - | - | Unassigned | - |
| Otu08810 | 2 | k__Fungi(100);k__Fungi_unclassified(100);k__Fungi_unclassified(100);k__Fungi_unclassified(100);k__Fungi_unclassified(100);k__Fungi_unclassified(100);k__Fungi_unclassified(100); | - | - | - | - | - | - | - | Unassigned | - |
| Otu08811 | 2 | k__Fungi(100);k__Fungi_unclassified(100);k__Fungi_unclassified(100);k__Fungi_unclassified(100);k__Fungi_unclassified(100);k__Fungi_unclassified(100);k__Fungi_unclassified(100); | - | - | - | - | - | - | - | Unassigned | - |
| Otu08813 | 2 | k__Fungi(100);k__Fungi_unclassified(100);k__Fungi_unclassified(100);k__Fungi_unclassified(100);k__Fungi_unclassified(100);k__Fungi_unclassified(100);k__Fungi_unclassified(100); | - | - | - | - | - | - | - | Unassigned | - |
| Otu08818 | 2 | k__Fungi(100);p__Ascomycota(100);p__Ascomycota_unclassified(100);p__Ascomycota_unclassified(100);p__Ascomycota_unclassified(100);p__Ascomycota_unclassified(100);p__Ascomycota_unclassified(100); | - | - | - | - | - | - | - | Unassigned | - |
| Otu08819 | 2 | k__Fungi(100);k__Fungi_unclassified(100);k__Fungi_unclassified(100);k__Fungi_unclassified(100);k__Fungi_unclassified(100);k__Fungi_unclassified(100);k__Fungi_unclassified(100); | - | - | - | - | - | - | - | Unassigned | - |
| Otu08821 | 2 | k__Fungi(100);k__Fungi_unclassified(100);k__Fungi_unclassified(100);k__Fungi_unclassified(100);k__Fungi_unclassified(100);k__Fungi_unclassified(100);k__Fungi_unclassified(100); | - | - | - | - | - | - | - | Unassigned | - |
| Otu08825 | 2 | k__Fungi(100);k__Fungi_unclassified(100);k__Fungi_unclassified(100);k__Fungi_unclassified(100);k__Fungi_unclassified(100);k__Fungi_unclassified(100);k__Fungi_unclassified(100); | - | - | - | - | - | - | - | Unassigned | - |
| Otu08826 | 2 | k__Fungi(100);k__Fungi_unclassified(100);k__Fungi_unclassified(100);k__Fungi_unclassified(100);k__Fungi_unclassified(100);k__Fungi_unclassified(100);k__Fungi_unclassified(100); | - | - | - | - | - | - | - | Unassigned | - |
| Otu08830 | 2 | k__Fungi(100);k__Fungi_unclassified(100);k__Fungi_unclassified(100);k__Fungi_unclassified(100);k__Fungi_unclassified(100);k__Fungi_unclassified(100);k__Fungi_unclassified(100); | - | - | - | - | - | - | - | Unassigned | - |
| Otu08832 | 2 | k__Fungi(100);k__Fungi_unclassified(100);k__Fungi_unclassified(100);k__Fungi_unclassified(100);k__Fungi_unclassified(100);k__Fungi_unclassified(100);k__Fungi_unclassified(100); | - | - | - | - | - | - | - | Unassigned | - |
| Otu08839 | 2 | k__Fungi(100);p__Ascomycota(100);p__Ascomycota_unclassified(100);p__Ascomycota_unclassified(100);p__Ascomycota_unclassified(100);p__Ascomycota_unclassified(100);p__Ascomycota_unclassified(100); | - | - | - | - | - | - | - | Unassigned | - |
| Otu08840 | 2 | k__Fungi(100);k__Fungi_unclassified(100);k__Fungi_unclassified(100);k__Fungi_unclassified(100);k__Fungi_unclassified(100);k__Fungi_unclassified(100);k__Fungi_unclassified(100); | - | - | - | - | - | - | - | Unassigned | - |
| Otu08841 | 2 | k__Fungi(100);k__Fungi_unclassified(100);k__Fungi_unclassified(100);k__Fungi_unclassified(100);k__Fungi_unclassified(100);k__Fungi_unclassified(100);k__Fungi_unclassified(100); | - | - | - | - | - | - | - | Unassigned | - |
| Otu08842 | 2 | k__Fungi(100);k__Fungi_unclassified(100);k__Fungi_unclassified(100);k__Fungi_unclassified(100);k__Fungi_unclassified(100);k__Fungi_unclassified(100);k__Fungi_unclassified(100); | - | - | - | - | - | - | - | Unassigned | - |
| Otu08843 | 2 | k__Fungi(100);k__Fungi_unclassified(100);k__Fungi_unclassified(100);k__Fungi_unclassified(100);k__Fungi_unclassified(100);k__Fungi_unclassified(100);k__Fungi_unclassified(100); | - | - | - | - | - | - | - | Unassigned | - |
| Otu08845 | 2 | k__Fungi(100);k__Fungi_unclassified(100);k__Fungi_unclassified(100);k__Fungi_unclassified(100);k__Fungi_unclassified(100);k__Fungi_unclassified(100);k__Fungi_unclassified(100); | - | - | - | - | - | - | - | Unassigned | - |
| Otu08846 | 2 | k__Fungi(100);k__Fungi_unclassified(100);k__Fungi_unclassified(100);k__Fungi_unclassified(100);k__Fungi_unclassified(100);k__Fungi_unclassified(100);k__Fungi_unclassified(100); | - | - | - | - | - | - | - | Unassigned | - |
| Otu08848 | 2 | k__Fungi(100);p__Ascomycota(100);p__Ascomycota_unclassified(100);p__Ascomycota_unclassified(100);p__Ascomycota_unclassified(100);p__Ascomycota_unclassified(100);p__Ascomycota_unclassified(100); | - | - | - | - | - | - | - | Unassigned | - |
| Otu08851 | 2 | k__Fungi(100);k__Fungi_unclassified(100);k__Fungi_unclassified(100);k__Fungi_unclassified(100);k__Fungi_unclassified(100);k__Fungi_unclassified(100);k__Fungi_unclassified(100); | - | - | - | - | - | - | - | Unassigned | - |
| Otu08852 | 2 | k__Fungi(100);k__Fungi_unclassified(100);k__Fungi_unclassified(100);k__Fungi_unclassified(100);k__Fungi_unclassified(100);k__Fungi_unclassified(100);k__Fungi_unclassified(100); | - | - | - | - | - | - | - | Unassigned | - |
| Otu08853 | 2 | k__Fungi(100);k__Fungi_unclassified(100);k__Fungi_unclassified(100);k__Fungi_unclassified(100);k__Fungi_unclassified(100);k__Fungi_unclassified(100);k__Fungi_unclassified(100); | - | - | - | - | - | - | - | Unassigned | - |
| Otu08856 | 2 | k__Fungi(100);k__Fungi_unclassified(100);k__Fungi_unclassified(100);k__Fungi_unclassified(100);k__Fungi_unclassified(100);k__Fungi_unclassified(100);k__Fungi_unclassified(100); | - | - | - | - | - | - | - | Unassigned | - |
| Otu08857 | 2 | k__Fungi(100);p__Ascomycota(100);p__Ascomycota_unclassified(100);p__Ascomycota_unclassified(100);p__Ascomycota_unclassified(100);p__Ascomycota_unclassified(100);p__Ascomycota_unclassified(100); | - | - | - | - | - | - | - | Unassigned | - |
| Otu08858 | 2 | k__Fungi(100);p__Ascomycota(100);p__Ascomycota_unclassified(100);p__Ascomycota_unclassified(100);p__Ascomycota_unclassified(100);p__Ascomycota_unclassified(100);p__Ascomycota_unclassified(100); | - | - | - | - | - | - | - | Unassigned | - |
| Otu08859 | 2 | k__Fungi(100);k__Fungi_unclassified(100);k__Fungi_unclassified(100);k__Fungi_unclassified(100);k__Fungi_unclassified(100);k__Fungi_unclassified(100);k__Fungi_unclassified(100); | - | - | - | - | - | - | - | Unassigned | - |
| Otu08864 | 2 | k__Fungi(100);k__Fungi_unclassified(100);k__Fungi_unclassified(100);k__Fungi_unclassified(100);k__Fungi_unclassified(100);k__Fungi_unclassified(100);k__Fungi_unclassified(100); | - | - | - | - | - | - | - | Unassigned | - |
| Otu08865 | 2 | k__Fungi(100);k__Fungi_unclassified(100);k__Fungi_unclassified(100);k__Fungi_unclassified(100);k__Fungi_unclassified(100);k__Fungi_unclassified(100);k__Fungi_unclassified(100); | - | - | - | - | - | - | - | Unassigned | - |
| Otu08866 | 2 | k__Fungi(100);k__Fungi_unclassified(100);k__Fungi_unclassified(100);k__Fungi_unclassified(100);k__Fungi_unclassified(100);k__Fungi_unclassified(100);k__Fungi_unclassified(100); | - | - | - | - | - | - | - | Unassigned | - |
| Otu08868 | 2 | k__Fungi(100);k__Fungi_unclassified(100);k__Fungi_unclassified(100);k__Fungi_unclassified(100);k__Fungi_unclassified(100);k__Fungi_unclassified(100);k__Fungi_unclassified(100); | - | - | - | - | - | - | - | Unassigned | - |
| Otu08869 | 2 | k__Fungi(100);k__Fungi_unclassified(100);k__Fungi_unclassified(100);k__Fungi_unclassified(100);k__Fungi_unclassified(100);k__Fungi_unclassified(100);k__Fungi_unclassified(100); | - | - | - | - | - | - | - | Unassigned | - |
| Otu08872 | 2 | k__Fungi(100);k__Fungi_unclassified(100);k__Fungi_unclassified(100);k__Fungi_unclassified(100);k__Fungi_unclassified(100);k__Fungi_unclassified(100);k__Fungi_unclassified(100); | - | - | - | - | - | - | - | Unassigned | - |
| Otu08874 | 2 | k__Fungi(100);k__Fungi_unclassified(100);k__Fungi_unclassified(100);k__Fungi_unclassified(100);k__Fungi_unclassified(100);k__Fungi_unclassified(100);k__Fungi_unclassified(100); | - | - | - | - | - | - | - | Unassigned | - |
| Otu08876 | 2 | k__Fungi(100);k__Fungi_unclassified(100);k__Fungi_unclassified(100);k__Fungi_unclassified(100);k__Fungi_unclassified(100);k__Fungi_unclassified(100);k__Fungi_unclassified(100); | - | - | - | - | - | - | - | Unassigned | - |
| Otu08877 | 2 | k__Fungi(100);k__Fungi_unclassified(100);k__Fungi_unclassified(100);k__Fungi_unclassified(100);k__Fungi_unclassified(100);k__Fungi_unclassified(100);k__Fungi_unclassified(100); | - | - | - | - | - | - | - | Unassigned | - |
| Otu08879 | 2 | k__Fungi(100);k__Fungi_unclassified(100);k__Fungi_unclassified(100);k__Fungi_unclassified(100);k__Fungi_unclassified(100);k__Fungi_unclassified(100);k__Fungi_unclassified(100); | - | - | - | - | - | - | - | Unassigned | - |
| Otu08880 | 2 | k__Fungi(100);k__Fungi_unclassified(100);k__Fungi_unclassified(100);k__Fungi_unclassified(100);k__Fungi_unclassified(100);k__Fungi_unclassified(100);k__Fungi_unclassified(100); | - | - | - | - | - | - | - | Unassigned | - |
| Otu08881 | 2 | k__Fungi(100);k__Fungi_unclassified(100);k__Fungi_unclassified(100);k__Fungi_unclassified(100);k__Fungi_unclassified(100);k__Fungi_unclassified(100);k__Fungi_unclassified(100); | - | - | - | - | - | - | - | Unassigned | - |
| Otu08882 | 2 | k__Fungi(100);p__Ascomycota(100);p__Ascomycota_unclassified(100);p__Ascomycota_unclassified(100);p__Ascomycota_unclassified(100);p__Ascomycota_unclassified(100);p__Ascomycota_unclassified(100); | - | - | - | - | - | - | - | Unassigned | - |
| Otu08883 | 2 | k__Fungi(100);k__Fungi_unclassified(100);k__Fungi_unclassified(100);k__Fungi_unclassified(100);k__Fungi_unclassified(100);k__Fungi_unclassified(100);k__Fungi_unclassified(100); | - | - | - | - | - | - | - | Unassigned | - |
| Otu08884 | 2 | k__Fungi(100);k__Fungi_unclassified(100);k__Fungi_unclassified(100);k__Fungi_unclassified(100);k__Fungi_unclassified(100);k__Fungi_unclassified(100);k__Fungi_unclassified(100); | - | - | - | - | - | - | - | Unassigned | - |
| Otu08886 | 2 | k__Fungi(100);p__Ascomycota(100);p__Ascomycota_unclassified(100);p__Ascomycota_unclassified(100);p__Ascomycota_unclassified(100);p__Ascomycota_unclassified(100);p__Ascomycota_unclassified(100); | - | - | - | - | - | - | - | Unassigned | - |
| Otu08887 | 2 | k__Fungi(100);k__Fungi_unclassified(100);k__Fungi_unclassified(100);k__Fungi_unclassified(100);k__Fungi_unclassified(100);k__Fungi_unclassified(100);k__Fungi_unclassified(100); | - | - | - | - | - | - | - | Unassigned | - |
| Otu08888 | 2 | k__Fungi(100);k__Fungi_unclassified(100);k__Fungi_unclassified(100);k__Fungi_unclassified(100);k__Fungi_unclassified(100);k__Fungi_unclassified(100);k__Fungi_unclassified(100); | - | - | - | - | - | - | - | Unassigned | - |
| Otu08889 | 2 | k__Fungi(100);k__Fungi_unclassified(100);k__Fungi_unclassified(100);k__Fungi_unclassified(100);k__Fungi_unclassified(100);k__Fungi_unclassified(100);k__Fungi_unclassified(100); | - | - | - | - | - | - | - | Unassigned | - |
| Otu08891 | 2 | k__Fungi(100);k__Fungi_unclassified(100);k__Fungi_unclassified(100);k__Fungi_unclassified(100);k__Fungi_unclassified(100);k__Fungi_unclassified(100);k__Fungi_unclassified(100); | - | - | - | - | - | - | - | Unassigned | - |
| Otu08892 | 2 | k__Fungi(100);k__Fungi_unclassified(100);k__Fungi_unclassified(100);k__Fungi_unclassified(100);k__Fungi_unclassified(100);k__Fungi_unclassified(100);k__Fungi_unclassified(100); | - | - | - | - | - | - | - | Unassigned | - |
| Otu08893 | 2 | k__Fungi(100);p__Ascomycota(100);c__Dothideomycetes(100);o__Pleosporales(100);o__Pleosporales_unclassified(100);o__Pleosporales_unclassified(100);o__Pleosporales_unclassified(100); | - | - | - | - | - | - | - | Unassigned | - |
| Otu08895 | 2 | k__Fungi(100);k__Fungi_unclassified(100);k__Fungi_unclassified(100);k__Fungi_unclassified(100);k__Fungi_unclassified(100);k__Fungi_unclassified(100);k__Fungi_unclassified(100); | - | - | - | - | - | - | - | Unassigned | - |
| Otu08900 | 2 | k__Fungi(100);k__Fungi_unclassified(100);k__Fungi_unclassified(100);k__Fungi_unclassified(100);k__Fungi_unclassified(100);k__Fungi_unclassified(100);k__Fungi_unclassified(100); | - | - | - | - | - | - | - | Unassigned | - |
| Otu08902 | 2 | k__Fungi(100);k__Fungi_unclassified(100);k__Fungi_unclassified(100);k__Fungi_unclassified(100);k__Fungi_unclassified(100);k__Fungi_unclassified(100);k__Fungi_unclassified(100); | - | - | - | - | - | - | - | Unassigned | - |
| Otu08903 | 2 | k__Fungi(100);k__Fungi_unclassified(100);k__Fungi_unclassified(100);k__Fungi_unclassified(100);k__Fungi_unclassified(100);k__Fungi_unclassified(100);k__Fungi_unclassified(100); | - | - | - | - | - | - | - | Unassigned | - |
| Otu08905 | 2 | k__Fungi(100);p__Ascomycota(100);p__Ascomycota_unclassified(100);p__Ascomycota_unclassified(100);p__Ascomycota_unclassified(100);p__Ascomycota_unclassified(100);p__Ascomycota_unclassified(100); | - | - | - | - | - | - | - | Unassigned | - |
| Otu08907 | 2 | k__Fungi(100);k__Fungi_unclassified(100);k__Fungi_unclassified(100);k__Fungi_unclassified(100);k__Fungi_unclassified(100);k__Fungi_unclassified(100);k__Fungi_unclassified(100); | - | - | - | - | - | - | - | Unassigned | - |
| Otu08909 | 2 | k__Fungi(100);k__Fungi_unclassified(100);k__Fungi_unclassified(100);k__Fungi_unclassified(100);k__Fungi_unclassified(100);k__Fungi_unclassified(100);k__Fungi_unclassified(100); | - | - | - | - | - | - | - | Unassigned | - |
| Otu08911 | 2 | k__Fungi(100);k__Fungi_unclassified(100);k__Fungi_unclassified(100);k__Fungi_unclassified(100);k__Fungi_unclassified(100);k__Fungi_unclassified(100);k__Fungi_unclassified(100); | - | - | - | - | - | - | - | Unassigned | - |
| Otu08912 | 2 | k__Fungi(100);p__Ascomycota(100);p__Ascomycota_unclassified(100);p__Ascomycota_unclassified(100);p__Ascomycota_unclassified(100);p__Ascomycota_unclassified(100);p__Ascomycota_unclassified(100); | - | - | - | - | - | - | - | Unassigned | - |
| Otu08913 | 2 | k__Fungi(100);k__Fungi_unclassified(100);k__Fungi_unclassified(100);k__Fungi_unclassified(100);k__Fungi_unclassified(100);k__Fungi_unclassified(100);k__Fungi_unclassified(100); | - | - | - | - | - | - | - | Unassigned | - |
| Otu08915 | 2 | k__Fungi(100);k__Fungi_unclassified(100);k__Fungi_unclassified(100);k__Fungi_unclassified(100);k__Fungi_unclassified(100);k__Fungi_unclassified(100);k__Fungi_unclassified(100); | - | - | - | - | - | - | - | Unassigned | - |
| Otu08925 | 2 | k__Fungi(100);p__Ascomycota(100);c__Leotiomycetes(100);o__Helotiales(100);o__Helotiales_unclassified(100);o__Helotiales_unclassified(100);o__Helotiales_unclassified(100); | - | - | - | - | - | - | - | Unassigned | - |
| Otu08929 | 2 | k__Fungi(100);k__Fungi_unclassified(100);k__Fungi_unclassified(100);k__Fungi_unclassified(100);k__Fungi_unclassified(100);k__Fungi_unclassified(100);k__Fungi_unclassified(100); | - | - | - | - | - | - | - | Unassigned | - |
| Otu08930 | 2 | k__Fungi(100);k__Fungi_unclassified(100);k__Fungi_unclassified(100);k__Fungi_unclassified(100);k__Fungi_unclassified(100);k__Fungi_unclassified(100);k__Fungi_unclassified(100); | - | - | - | - | - | - | - | Unassigned | - |
| Otu08932 | 2 | k__Fungi(100);k__Fungi_unclassified(100);k__Fungi_unclassified(100);k__Fungi_unclassified(100);k__Fungi_unclassified(100);k__Fungi_unclassified(100);k__Fungi_unclassified(100); | - | - | - | - | - | - | - | Unassigned | - |
| Otu08935 | 2 | k__Fungi(100);k__Fungi_unclassified(100);k__Fungi_unclassified(100);k__Fungi_unclassified(100);k__Fungi_unclassified(100);k__Fungi_unclassified(100);k__Fungi_unclassified(100); | - | - | - | - | - | - | - | Unassigned | - |
| Otu08936 | 2 | k__Fungi(100);k__Fungi_unclassified(100);k__Fungi_unclassified(100);k__Fungi_unclassified(100);k__Fungi_unclassified(100);k__Fungi_unclassified(100);k__Fungi_unclassified(100); | - | - | - | - | - | - | - | Unassigned | - |
| Otu08937 | 2 | k__Fungi(100);p__Ascomycota(100);p__Ascomycota_unclassified(100);p__Ascomycota_unclassified(100);p__Ascomycota_unclassified(100);p__Ascomycota_unclassified(100);p__Ascomycota_unclassified(100); | - | - | - | - | - | - | - | Unassigned | - |
| Otu08941 | 2 | k__Fungi(100);k__Fungi_unclassified(100);k__Fungi_unclassified(100);k__Fungi_unclassified(100);k__Fungi_unclassified(100);k__Fungi_unclassified(100);k__Fungi_unclassified(100); | - | - | - | - | - | - | - | Unassigned | - |
| Otu08943 | 2 | k__Fungi(100);k__Fungi_unclassified(100);k__Fungi_unclassified(100);k__Fungi_unclassified(100);k__Fungi_unclassified(100);k__Fungi_unclassified(100);k__Fungi_unclassified(100); | - | - | - | - | - | - | - | Unassigned | - |
| Otu08948 | 2 | k__Fungi(100);p__Ascomycota(100);p__Ascomycota_unclassified(100);p__Ascomycota_unclassified(100);p__Ascomycota_unclassified(100);p__Ascomycota_unclassified(100);p__Ascomycota_unclassified(100); | - | - | - | - | - | - | - | Unassigned | - |
| Otu08954 | 2 | k__Fungi(100);k__Fungi_unclassified(100);k__Fungi_unclassified(100);k__Fungi_unclassified(100);k__Fungi_unclassified(100);k__Fungi_unclassified(100);k__Fungi_unclassified(100); | - | - | - | - | - | - | - | Unassigned | - |
| Otu08955 | 2 | k__Fungi(100);k__Fungi_unclassified(100);k__Fungi_unclassified(100);k__Fungi_unclassified(100);k__Fungi_unclassified(100);k__Fungi_unclassified(100);k__Fungi_unclassified(100); | - | - | - | - | - | - | - | Unassigned | - |
| Otu08958 | 2 | k__Fungi(100);p__Ascomycota(100);p__Ascomycota_unclassified(100);p__Ascomycota_unclassified(100);p__Ascomycota_unclassified(100);p__Ascomycota_unclassified(100);p__Ascomycota_unclassified(100); | - | - | - | - | - | - | - | Unassigned | - |
| Otu08960 | 2 | k__Fungi(100);k__Fungi_unclassified(100);k__Fungi_unclassified(100);k__Fungi_unclassified(100);k__Fungi_unclassified(100);k__Fungi_unclassified(100);k__Fungi_unclassified(100); | - | - | - | - | - | - | - | Unassigned | - |
| Otu08961 | 2 | k__Fungi(100);p__Ascomycota(100);p__Ascomycota_unclassified(100);p__Ascomycota_unclassified(100);p__Ascomycota_unclassified(100);p__Ascomycota_unclassified(100);p__Ascomycota_unclassified(100); | - | - | - | - | - | - | - | Unassigned | - |
| Otu08962 | 2 | k__Fungi(100);p__Ascomycota(100);p__Ascomycota_unclassified(100);p__Ascomycota_unclassified(100);p__Ascomycota_unclassified(100);p__Ascomycota_unclassified(100);p__Ascomycota_unclassified(100); | - | - | - | - | - | - | - | Unassigned | - |
| Otu08963 | 2 | k__Fungi(100);k__Fungi_unclassified(100);k__Fungi_unclassified(100);k__Fungi_unclassified(100);k__Fungi_unclassified(100);k__Fungi_unclassified(100);k__Fungi_unclassified(100); | - | - | - | - | - | - | - | Unassigned | - |
| Otu08964 | 2 | k__Fungi(100);p__Ascomycota(100);p__Ascomycota_unclassified(100);p__Ascomycota_unclassified(100);p__Ascomycota_unclassified(100);p__Ascomycota_unclassified(100);p__Ascomycota_unclassified(100); | - | - | - | - | - | - | - | Unassigned | - |
| Otu08965 | 2 | k__Fungi(100);p__Ascomycota(100);p__Ascomycota_unclassified(100);p__Ascomycota_unclassified(100);p__Ascomycota_unclassified(100);p__Ascomycota_unclassified(100);p__Ascomycota_unclassified(100); | - | - | - | - | - | - | - | Unassigned | - |
| Otu08968 | 2 | k__Fungi(100);k__Fungi_unclassified(100);k__Fungi_unclassified(100);k__Fungi_unclassified(100);k__Fungi_unclassified(100);k__Fungi_unclassified(100);k__Fungi_unclassified(100); | - | - | - | - | - | - | - | Unassigned | - |
| Otu08969 | 2 | k__Fungi(100);k__Fungi_unclassified(100);k__Fungi_unclassified(100);k__Fungi_unclassified(100);k__Fungi_unclassified(100);k__Fungi_unclassified(100);k__Fungi_unclassified(100); | - | - | - | - | - | - | - | Unassigned | - |
| Otu08971 | 2 | k__Fungi(100);k__Fungi_unclassified(100);k__Fungi_unclassified(100);k__Fungi_unclassified(100);k__Fungi_unclassified(100);k__Fungi_unclassified(100);k__Fungi_unclassified(100); | - | - | - | - | - | - | - | Unassigned | - |
| Otu08975 | 2 | k__Fungi(100);k__Fungi_unclassified(100);k__Fungi_unclassified(100);k__Fungi_unclassified(100);k__Fungi_unclassified(100);k__Fungi_unclassified(100);k__Fungi_unclassified(100); | - | - | - | - | - | - | - | Unassigned | - |
| Otu08981 | 2 | k__Fungi(100);k__Fungi_unclassified(100);k__Fungi_unclassified(100);k__Fungi_unclassified(100);k__Fungi_unclassified(100);k__Fungi_unclassified(100);k__Fungi_unclassified(100); | - | - | - | - | - | - | - | Unassigned | - |
| Otu08983 | 2 | k__Fungi(100);k__Fungi_unclassified(100);k__Fungi_unclassified(100);k__Fungi_unclassified(100);k__Fungi_unclassified(100);k__Fungi_unclassified(100);k__Fungi_unclassified(100); | - | - | - | - | - | - | - | Unassigned | - |
| Otu08984 | 2 | k__Fungi(100);p__Ascomycota(100);p__Ascomycota_unclassified(100);p__Ascomycota_unclassified(100);p__Ascomycota_unclassified(100);p__Ascomycota_unclassified(100);p__Ascomycota_unclassified(100); | - | - | - | - | - | - | - | Unassigned | - |
| Otu08985 | 2 | k__Fungi(100);k__Fungi_unclassified(100);k__Fungi_unclassified(100);k__Fungi_unclassified(100);k__Fungi_unclassified(100);k__Fungi_unclassified(100);k__Fungi_unclassified(100); | - | - | - | - | - | - | - | Unassigned | - |
| Otu08986 | 2 | k__Fungi(100);p__Ascomycota(100);p__Ascomycota_unclassified(100);p__Ascomycota_unclassified(100);p__Ascomycota_unclassified(100);p__Ascomycota_unclassified(100);p__Ascomycota_unclassified(100); | - | - | - | - | - | - | - | Unassigned | - |
| Otu08987 | 2 | k__Fungi(100);k__Fungi_unclassified(100);k__Fungi_unclassified(100);k__Fungi_unclassified(100);k__Fungi_unclassified(100);k__Fungi_unclassified(100);k__Fungi_unclassified(100); | - | - | - | - | - | - | - | Unassigned | - |
| Otu08994 | 2 | k__Fungi(100);k__Fungi_unclassified(100);k__Fungi_unclassified(100);k__Fungi_unclassified(100);k__Fungi_unclassified(100);k__Fungi_unclassified(100);k__Fungi_unclassified(100); | - | - | - | - | - | - | - | Unassigned | - |
| Otu08995 | 2 | k__Fungi(100);k__Fungi_unclassified(100);k__Fungi_unclassified(100);k__Fungi_unclassified(100);k__Fungi_unclassified(100);k__Fungi_unclassified(100);k__Fungi_unclassified(100); | - | - | - | - | - | - | - | Unassigned | - |
| Otu08996 | 2 | k__Fungi(100);k__Fungi_unclassified(100);k__Fungi_unclassified(100);k__Fungi_unclassified(100);k__Fungi_unclassified(100);k__Fungi_unclassified(100);k__Fungi_unclassified(100); | - | - | - | - | - | - | - | Unassigned | - |
| Otu08997 | 2 | k__Fungi(100);p__Ascomycota(100);p__Ascomycota_unclassified(100);p__Ascomycota_unclassified(100);p__Ascomycota_unclassified(100);p__Ascomycota_unclassified(100);p__Ascomycota_unclassified(100); | - | - | - | - | - | - | - | Unassigned | - |
| Otu08998 | 2 | k__Fungi(100);k__Fungi_unclassified(100);k__Fungi_unclassified(100);k__Fungi_unclassified(100);k__Fungi_unclassified(100);k__Fungi_unclassified(100);k__Fungi_unclassified(100); | - | - | - | - | - | - | - | Unassigned | - |
| Otu08999 | 2 | k__Fungi(100);k__Fungi_unclassified(100);k__Fungi_unclassified(100);k__Fungi_unclassified(100);k__Fungi_unclassified(100);k__Fungi_unclassified(100);k__Fungi_unclassified(100); | - | - | - | - | - | - | - | Unassigned | - |
| Otu09003 | 2 | k__Fungi(100);p__Ascomycota(100);p__Ascomycota_unclassified(100);p__Ascomycota_unclassified(100);p__Ascomycota_unclassified(100);p__Ascomycota_unclassified(100);p__Ascomycota_unclassified(100); | - | - | - | - | - | - | - | Unassigned | - |
| Otu09007 | 2 | k__Fungi(100);k__Fungi_unclassified(100);k__Fungi_unclassified(100);k__Fungi_unclassified(100);k__Fungi_unclassified(100);k__Fungi_unclassified(100);k__Fungi_unclassified(100); | - | - | - | - | - | - | - | Unassigned | - |
| Otu09008 | 2 | k__Fungi(100);k__Fungi_unclassified(100);k__Fungi_unclassified(100);k__Fungi_unclassified(100);k__Fungi_unclassified(100);k__Fungi_unclassified(100);k__Fungi_unclassified(100); | - | - | - | - | - | - | - | Unassigned | - |
| Otu09010 | 2 | k__Fungi(100);k__Fungi_unclassified(100);k__Fungi_unclassified(100);k__Fungi_unclassified(100);k__Fungi_unclassified(100);k__Fungi_unclassified(100);k__Fungi_unclassified(100); | - | - | - | - | - | - | - | Unassigned | - |
| Otu09011 | 2 | k__Fungi(100);k__Fungi_unclassified(100);k__Fungi_unclassified(100);k__Fungi_unclassified(100);k__Fungi_unclassified(100);k__Fungi_unclassified(100);k__Fungi_unclassified(100); | - | - | - | - | - | - | - | Unassigned | - |
| Otu09013 | 2 | k__Fungi(100);k__Fungi_unclassified(100);k__Fungi_unclassified(100);k__Fungi_unclassified(100);k__Fungi_unclassified(100);k__Fungi_unclassified(100);k__Fungi_unclassified(100); | - | - | - | - | - | - | - | Unassigned | - |
| Otu09015 | 2 | k__Fungi(100);k__Fungi_unclassified(100);k__Fungi_unclassified(100);k__Fungi_unclassified(100);k__Fungi_unclassified(100);k__Fungi_unclassified(100);k__Fungi_unclassified(100); | - | - | - | - | - | - | - | Unassigned | - |
| Otu09017 | 2 | k__Fungi(100);k__Fungi_unclassified(100);k__Fungi_unclassified(100);k__Fungi_unclassified(100);k__Fungi_unclassified(100);k__Fungi_unclassified(100);k__Fungi_unclassified(100); | - | - | - | - | - | - | - | Unassigned | - |
| Otu09022 | 2 | k__Fungi(100);p__Ascomycota(100);p__Ascomycota_unclassified(100);p__Ascomycota_unclassified(100);p__Ascomycota_unclassified(100);p__Ascomycota_unclassified(100);p__Ascomycota_unclassified(100); | - | - | - | - | - | - | - | Unassigned | - |
| Otu09023 | 2 | k__Fungi(100);k__Fungi_unclassified(100);k__Fungi_unclassified(100);k__Fungi_unclassified(100);k__Fungi_unclassified(100);k__Fungi_unclassified(100);k__Fungi_unclassified(100); | - | - | - | - | - | - | - | Unassigned | - |
| Otu09026 | 2 | k__Fungi(100);k__Fungi_unclassified(100);k__Fungi_unclassified(100);k__Fungi_unclassified(100);k__Fungi_unclassified(100);k__Fungi_unclassified(100);k__Fungi_unclassified(100); | - | - | - | - | - | - | - | Unassigned | - |
| Otu09028 | 2 | k__Fungi(100);k__Fungi_unclassified(100);k__Fungi_unclassified(100);k__Fungi_unclassified(100);k__Fungi_unclassified(100);k__Fungi_unclassified(100);k__Fungi_unclassified(100); | - | - | - | - | - | - | - | Unassigned | - |
| Otu09030 | 2 | k__Fungi(100);k__Fungi_unclassified(100);k__Fungi_unclassified(100);k__Fungi_unclassified(100);k__Fungi_unclassified(100);k__Fungi_unclassified(100);k__Fungi_unclassified(100); | - | - | - | - | - | - | - | Unassigned | - |
| Otu09031 | 2 | k__Fungi(100);k__Fungi_unclassified(100);k__Fungi_unclassified(100);k__Fungi_unclassified(100);k__Fungi_unclassified(100);k__Fungi_unclassified(100);k__Fungi_unclassified(100); | - | - | - | - | - | - | - | Unassigned | - |
| Otu09033 | 2 | k__Fungi(100);k__Fungi_unclassified(100);k__Fungi_unclassified(100);k__Fungi_unclassified(100);k__Fungi_unclassified(100);k__Fungi_unclassified(100);k__Fungi_unclassified(100); | - | - | - | - | - | - | - | Unassigned | - |
| Otu09038 | 2 | k__Fungi(100);k__Fungi_unclassified(100);k__Fungi_unclassified(100);k__Fungi_unclassified(100);k__Fungi_unclassified(100);k__Fungi_unclassified(100);k__Fungi_unclassified(100); | - | - | - | - | - | - | - | Unassigned | - |
| Otu09039 | 2 | k__Fungi(100);k__Fungi_unclassified(100);k__Fungi_unclassified(100);k__Fungi_unclassified(100);k__Fungi_unclassified(100);k__Fungi_unclassified(100);k__Fungi_unclassified(100); | - | - | - | - | - | - | - | Unassigned | - |
| Otu09042 | 2 | k__Fungi(100);p__Ascomycota(100);p__Ascomycota_unclassified(100);p__Ascomycota_unclassified(100);p__Ascomycota_unclassified(100);p__Ascomycota_unclassified(100);p__Ascomycota_unclassified(100); | - | - | - | - | - | - | - | Unassigned | - |
| Otu09044 | 2 | k__Fungi(100);k__Fungi_unclassified(100);k__Fungi_unclassified(100);k__Fungi_unclassified(100);k__Fungi_unclassified(100);k__Fungi_unclassified(100);k__Fungi_unclassified(100); | - | - | - | - | - | - | - | Unassigned | - |
| Otu09045 | 2 | k__Fungi(100);k__Fungi_unclassified(100);k__Fungi_unclassified(100);k__Fungi_unclassified(100);k__Fungi_unclassified(100);k__Fungi_unclassified(100);k__Fungi_unclassified(100); | - | - | - | - | - | - | - | Unassigned | - |
| Otu09046 | 2 | k__Fungi(100);p__Ascomycota(100);p__Ascomycota_unclassified(100);p__Ascomycota_unclassified(100);p__Ascomycota_unclassified(100);p__Ascomycota_unclassified(100);p__Ascomycota_unclassified(100); | - | - | - | - | - | - | - | Unassigned | - |
| Otu09049 | 2 | k__Fungi(100);k__Fungi_unclassified(100);k__Fungi_unclassified(100);k__Fungi_unclassified(100);k__Fungi_unclassified(100);k__Fungi_unclassified(100);k__Fungi_unclassified(100); | - | - | - | - | - | - | - | Unassigned | - |
| Otu09050 | 2 | k__Fungi(100);k__Fungi_unclassified(100);k__Fungi_unclassified(100);k__Fungi_unclassified(100);k__Fungi_unclassified(100);k__Fungi_unclassified(100);k__Fungi_unclassified(100); | - | - | - | - | - | - | - | Unassigned | - |
| Otu09056 | 2 | k__Fungi(100);k__Fungi_unclassified(100);k__Fungi_unclassified(100);k__Fungi_unclassified(100);k__Fungi_unclassified(100);k__Fungi_unclassified(100);k__Fungi_unclassified(100); | - | - | - | - | - | - | - | Unassigned | - |
| Otu09062 | 2 | k__Fungi(100);k__Fungi_unclassified(100);k__Fungi_unclassified(100);k__Fungi_unclassified(100);k__Fungi_unclassified(100);k__Fungi_unclassified(100);k__Fungi_unclassified(100); | - | - | - | - | - | - | - | Unassigned | - |
| Otu09063 | 2 | k__Fungi(100);k__Fungi_unclassified(100);k__Fungi_unclassified(100);k__Fungi_unclassified(100);k__Fungi_unclassified(100);k__Fungi_unclassified(100);k__Fungi_unclassified(100); | - | - | - | - | - | - | - | Unassigned | - |
| Otu09064 | 2 | k__Fungi(100);k__Fungi_unclassified(100);k__Fungi_unclassified(100);k__Fungi_unclassified(100);k__Fungi_unclassified(100);k__Fungi_unclassified(100);k__Fungi_unclassified(100); | - | - | - | - | - | - | - | Unassigned | - |
| Otu09065 | 2 | k__Fungi(100);k__Fungi_unclassified(100);k__Fungi_unclassified(100);k__Fungi_unclassified(100);k__Fungi_unclassified(100);k__Fungi_unclassified(100);k__Fungi_unclassified(100); | - | - | - | - | - | - | - | Unassigned | - |
| Otu09066 | 2 | k__Fungi(100);k__Fungi_unclassified(100);k__Fungi_unclassified(100);k__Fungi_unclassified(100);k__Fungi_unclassified(100);k__Fungi_unclassified(100);k__Fungi_unclassified(100); | - | - | - | - | - | - | - | Unassigned | - |
| Otu09068 | 2 | k__Fungi(100);p__Ascomycota(100);p__Ascomycota_unclassified(100);p__Ascomycota_unclassified(100);p__Ascomycota_unclassified(100);p__Ascomycota_unclassified(100);p__Ascomycota_unclassified(100); | - | - | - | - | - | - | - | Unassigned | - |
| Otu09069 | 2 | k__Fungi(100);k__Fungi_unclassified(100);k__Fungi_unclassified(100);k__Fungi_unclassified(100);k__Fungi_unclassified(100);k__Fungi_unclassified(100);k__Fungi_unclassified(100); | - | - | - | - | - | - | - | Unassigned | - |
| Otu09073 | 2 | k__Fungi(100);k__Fungi_unclassified(100);k__Fungi_unclassified(100);k__Fungi_unclassified(100);k__Fungi_unclassified(100);k__Fungi_unclassified(100);k__Fungi_unclassified(100); | - | - | - | - | - | - | - | Unassigned | - |
| Otu09075 | 2 | k__Fungi(100);k__Fungi_unclassified(100);k__Fungi_unclassified(100);k__Fungi_unclassified(100);k__Fungi_unclassified(100);k__Fungi_unclassified(100);k__Fungi_unclassified(100); | - | - | - | - | - | - | - | Unassigned | - |
| Otu09076 | 2 | k__Fungi(100);k__Fungi_unclassified(100);k__Fungi_unclassified(100);k__Fungi_unclassified(100);k__Fungi_unclassified(100);k__Fungi_unclassified(100);k__Fungi_unclassified(100); | - | - | - | - | - | - | - | Unassigned | - |
| Otu09077 | 2 | k__Fungi(100);k__Fungi_unclassified(100);k__Fungi_unclassified(100);k__Fungi_unclassified(100);k__Fungi_unclassified(100);k__Fungi_unclassified(100);k__Fungi_unclassified(100); | - | - | - | - | - | - | - | Unassigned | - |
| Otu09081 | 2 | k__Fungi(100);k__Fungi_unclassified(100);k__Fungi_unclassified(100);k__Fungi_unclassified(100);k__Fungi_unclassified(100);k__Fungi_unclassified(100);k__Fungi_unclassified(100); | - | - | - | - | - | - | - | Unassigned | - |
| Otu09083 | 2 | k__Fungi(100);k__Fungi_unclassified(100);k__Fungi_unclassified(100);k__Fungi_unclassified(100);k__Fungi_unclassified(100);k__Fungi_unclassified(100);k__Fungi_unclassified(100); | - | - | - | - | - | - | - | Unassigned | - |
| Otu09085 | 2 | k__Fungi(100);k__Fungi_unclassified(100);k__Fungi_unclassified(100);k__Fungi_unclassified(100);k__Fungi_unclassified(100);k__Fungi_unclassified(100);k__Fungi_unclassified(100); | - | - | - | - | - | - | - | Unassigned | - |
| Otu09087 | 2 | k__Fungi(100);p__Ascomycota(100);p__Ascomycota_unclassified(100);p__Ascomycota_unclassified(100);p__Ascomycota_unclassified(100);p__Ascomycota_unclassified(100);p__Ascomycota_unclassified(100); | - | - | - | - | - | - | - | Unassigned | - |
| Otu09088 | 2 | k__Fungi(100);p__Ascomycota(100);p__Ascomycota_unclassified(100);p__Ascomycota_unclassified(100);p__Ascomycota_unclassified(100);p__Ascomycota_unclassified(100);p__Ascomycota_unclassified(100); | - | - | - | - | - | - | - | Unassigned | - |
| Otu09089 | 2 | k__Fungi(100);k__Fungi_unclassified(100);k__Fungi_unclassified(100);k__Fungi_unclassified(100);k__Fungi_unclassified(100);k__Fungi_unclassified(100);k__Fungi_unclassified(100); | - | - | - | - | - | - | - | Unassigned | - |
| Otu09091 | 2 | k__Fungi(100);p__Ascomycota(100);p__Ascomycota_unclassified(100);p__Ascomycota_unclassified(100);p__Ascomycota_unclassified(100);p__Ascomycota_unclassified(100);p__Ascomycota_unclassified(100); | - | - | - | - | - | - | - | Unassigned | - |
| Otu09092 | 2 | k__Fungi(100);k__Fungi_unclassified(100);k__Fungi_unclassified(100);k__Fungi_unclassified(100);k__Fungi_unclassified(100);k__Fungi_unclassified(100);k__Fungi_unclassified(100); | - | - | - | - | - | - | - | Unassigned | - |
| Otu09097 | 2 | k__Fungi(100);p__Ascomycota(100);p__Ascomycota_unclassified(100);p__Ascomycota_unclassified(100);p__Ascomycota_unclassified(100);p__Ascomycota_unclassified(100);p__Ascomycota_unclassified(100); | - | - | - | - | - | - | - | Unassigned | - |
| Otu09098 | 2 | k__Fungi(100);k__Fungi_unclassified(100);k__Fungi_unclassified(100);k__Fungi_unclassified(100);k__Fungi_unclassified(100);k__Fungi_unclassified(100);k__Fungi_unclassified(100); | - | - | - | - | - | - | - | Unassigned | - |
| Otu09102 | 2 | k__Fungi(100);k__Fungi_unclassified(100);k__Fungi_unclassified(100);k__Fungi_unclassified(100);k__Fungi_unclassified(100);k__Fungi_unclassified(100);k__Fungi_unclassified(100); | - | - | - | - | - | - | - | Unassigned | - |
| Otu09103 | 2 | k__Fungi(100);k__Fungi_unclassified(100);k__Fungi_unclassified(100);k__Fungi_unclassified(100);k__Fungi_unclassified(100);k__Fungi_unclassified(100);k__Fungi_unclassified(100); | - | - | - | - | - | - | - | Unassigned | - |
| Otu09105 | 2 | k__Fungi(100);p__Ascomycota(100);p__Ascomycota_unclassified(100);p__Ascomycota_unclassified(100);p__Ascomycota_unclassified(100);p__Ascomycota_unclassified(100);p__Ascomycota_unclassified(100); | - | - | - | - | - | - | - | Unassigned | - |
| Otu09108 | 2 | k__Fungi(100);k__Fungi_unclassified(100);k__Fungi_unclassified(100);k__Fungi_unclassified(100);k__Fungi_unclassified(100);k__Fungi_unclassified(100);k__Fungi_unclassified(100); | - | - | - | - | - | - | - | Unassigned | - |
| Otu09111 | 2 | k__Fungi(100);p__Ascomycota(100);p__Ascomycota_unclassified(100);p__Ascomycota_unclassified(100);p__Ascomycota_unclassified(100);p__Ascomycota_unclassified(100);p__Ascomycota_unclassified(100); | - | - | - | - | - | - | - | Unassigned | - |
| Otu09115 | 2 | k__Fungi(100);k__Fungi_unclassified(100);k__Fungi_unclassified(100);k__Fungi_unclassified(100);k__Fungi_unclassified(100);k__Fungi_unclassified(100);k__Fungi_unclassified(100); | - | - | - | - | - | - | - | Unassigned | - |
| Otu09116 | 2 | k__Fungi(100);p__Ascomycota(100);p__Ascomycota_unclassified(100);p__Ascomycota_unclassified(100);p__Ascomycota_unclassified(100);p__Ascomycota_unclassified(100);p__Ascomycota_unclassified(100); | - | - | - | - | - | - | - | Unassigned | - |
| Otu09117 | 2 | k__Fungi(100);p__Ascomycota(100);p__Ascomycota_unclassified(100);p__Ascomycota_unclassified(100);p__Ascomycota_unclassified(100);p__Ascomycota_unclassified(100);p__Ascomycota_unclassified(100); | - | - | - | - | - | - | - | Unassigned | - |
| Otu09118 | 2 | k__Fungi(100);k__Fungi_unclassified(100);k__Fungi_unclassified(100);k__Fungi_unclassified(100);k__Fungi_unclassified(100);k__Fungi_unclassified(100);k__Fungi_unclassified(100); | - | - | - | - | - | - | - | Unassigned | - |
| Otu09121 | 2 | k__Fungi(100);k__Fungi_unclassified(100);k__Fungi_unclassified(100);k__Fungi_unclassified(100);k__Fungi_unclassified(100);k__Fungi_unclassified(100);k__Fungi_unclassified(100); | - | - | - | - | - | - | - | Unassigned | - |
| Otu09122 | 2 | k__Fungi(100);k__Fungi_unclassified(100);k__Fungi_unclassified(100);k__Fungi_unclassified(100);k__Fungi_unclassified(100);k__Fungi_unclassified(100);k__Fungi_unclassified(100); | - | - | - | - | - | - | - | Unassigned | - |
| Otu09125 | 2 | k__Fungi(100);p__Ascomycota(100);p__Ascomycota_unclassified(100);p__Ascomycota_unclassified(100);p__Ascomycota_unclassified(100);p__Ascomycota_unclassified(100);p__Ascomycota_unclassified(100); | - | - | - | - | - | - | - | Unassigned | - |
| Otu09130 | 2 | k__Fungi(100);p__Ascomycota(100);p__Ascomycota_unclassified(100);p__Ascomycota_unclassified(100);p__Ascomycota_unclassified(100);p__Ascomycota_unclassified(100);p__Ascomycota_unclassified(100); | - | - | - | - | - | - | - | Unassigned | - |
| Otu09131 | 2 | k__Fungi(100);k__Fungi_unclassified(100);k__Fungi_unclassified(100);k__Fungi_unclassified(100);k__Fungi_unclassified(100);k__Fungi_unclassified(100);k__Fungi_unclassified(100); | - | - | - | - | - | - | - | Unassigned | - |
| Otu09134 | 2 | k__Fungi(100);k__Fungi_unclassified(100);k__Fungi_unclassified(100);k__Fungi_unclassified(100);k__Fungi_unclassified(100);k__Fungi_unclassified(100);k__Fungi_unclassified(100); | - | - | - | - | - | - | - | Unassigned | - |
| Otu09136 | 2 | k__Fungi(100);k__Fungi_unclassified(100);k__Fungi_unclassified(100);k__Fungi_unclassified(100);k__Fungi_unclassified(100);k__Fungi_unclassified(100);k__Fungi_unclassified(100); | - | - | - | - | - | - | - | Unassigned | - |
| Otu09140 | 2 | k__Fungi(100);k__Fungi_unclassified(100);k__Fungi_unclassified(100);k__Fungi_unclassified(100);k__Fungi_unclassified(100);k__Fungi_unclassified(100);k__Fungi_unclassified(100); | - | - | - | - | - | - | - | Unassigned | - |
| Otu09141 | 2 | k__Fungi(100);k__Fungi_unclassified(100);k__Fungi_unclassified(100);k__Fungi_unclassified(100);k__Fungi_unclassified(100);k__Fungi_unclassified(100);k__Fungi_unclassified(100); | - | - | - | - | - | - | - | Unassigned | - |
| Otu09142 | 2 | k__Fungi(100);k__Fungi_unclassified(100);k__Fungi_unclassified(100);k__Fungi_unclassified(100);k__Fungi_unclassified(100);k__Fungi_unclassified(100);k__Fungi_unclassified(100); | - | - | - | - | - | - | - | Unassigned | - |
| Otu09143 | 2 | k__Fungi(100);p__Ascomycota(100);p__Ascomycota_unclassified(100);p__Ascomycota_unclassified(100);p__Ascomycota_unclassified(100);p__Ascomycota_unclassified(100);p__Ascomycota_unclassified(100); | - | - | - | - | - | - | - | Unassigned | - |
| Otu09144 | 2 | k__Fungi(100);k__Fungi_unclassified(100);k__Fungi_unclassified(100);k__Fungi_unclassified(100);k__Fungi_unclassified(100);k__Fungi_unclassified(100);k__Fungi_unclassified(100); | - | - | - | - | - | - | - | Unassigned | - |
| Otu09145 | 2 | k__Fungi(100);k__Fungi_unclassified(100);k__Fungi_unclassified(100);k__Fungi_unclassified(100);k__Fungi_unclassified(100);k__Fungi_unclassified(100);k__Fungi_unclassified(100); | - | - | - | - | - | - | - | Unassigned | - |
| Otu09146 | 2 | k__Fungi(100);p__Ascomycota(100);p__Ascomycota_unclassified(100);p__Ascomycota_unclassified(100);p__Ascomycota_unclassified(100);p__Ascomycota_unclassified(100);p__Ascomycota_unclassified(100); | - | - | - | - | - | - | - | Unassigned | - |
| Otu09148 | 2 | k__Fungi(100);p__Ascomycota(100);p__Ascomycota_unclassified(100);p__Ascomycota_unclassified(100);p__Ascomycota_unclassified(100);p__Ascomycota_unclassified(100);p__Ascomycota_unclassified(100); | - | - | - | - | - | - | - | Unassigned | - |
| Otu09149 | 2 | k__Fungi(100);k__Fungi_unclassified(100);k__Fungi_unclassified(100);k__Fungi_unclassified(100);k__Fungi_unclassified(100);k__Fungi_unclassified(100);k__Fungi_unclassified(100); | - | - | - | - | - | - | - | Unassigned | - |
| Otu09150 | 2 | k__Fungi(100);k__Fungi_unclassified(100);k__Fungi_unclassified(100);k__Fungi_unclassified(100);k__Fungi_unclassified(100);k__Fungi_unclassified(100);k__Fungi_unclassified(100); | - | - | - | - | - | - | - | Unassigned | - |
| Otu09152 | 2 | k__Fungi(100);k__Fungi_unclassified(100);k__Fungi_unclassified(100);k__Fungi_unclassified(100);k__Fungi_unclassified(100);k__Fungi_unclassified(100);k__Fungi_unclassified(100); | - | - | - | - | - | - | - | Unassigned | - |
| Otu09153 | 2 | k__Fungi(100);k__Fungi_unclassified(100);k__Fungi_unclassified(100);k__Fungi_unclassified(100);k__Fungi_unclassified(100);k__Fungi_unclassified(100);k__Fungi_unclassified(100); | - | - | - | - | - | - | - | Unassigned | - |
| Otu09155 | 2 | k__Fungi(100);k__Fungi_unclassified(100);k__Fungi_unclassified(100);k__Fungi_unclassified(100);k__Fungi_unclassified(100);k__Fungi_unclassified(100);k__Fungi_unclassified(100); | - | - | - | - | - | - | - | Unassigned | - |
| Otu09157 | 2 | k__Fungi(100);p__Ascomycota(100);p__Ascomycota_unclassified(100);p__Ascomycota_unclassified(100);p__Ascomycota_unclassified(100);p__Ascomycota_unclassified(100);p__Ascomycota_unclassified(100); | - | - | - | - | - | - | - | Unassigned | - |
| Otu09160 | 2 | k__Fungi(100);k__Fungi_unclassified(100);k__Fungi_unclassified(100);k__Fungi_unclassified(100);k__Fungi_unclassified(100);k__Fungi_unclassified(100);k__Fungi_unclassified(100); | - | - | - | - | - | - | - | Unassigned | - |
| Otu09162 | 2 | k__Fungi(100);k__Fungi_unclassified(100);k__Fungi_unclassified(100);k__Fungi_unclassified(100);k__Fungi_unclassified(100);k__Fungi_unclassified(100);k__Fungi_unclassified(100); | - | - | - | - | - | - | - | Unassigned | - |
| Otu09163 | 2 | k__Fungi(100);k__Fungi_unclassified(100);k__Fungi_unclassified(100);k__Fungi_unclassified(100);k__Fungi_unclassified(100);k__Fungi_unclassified(100);k__Fungi_unclassified(100); | - | - | - | - | - | - | - | Unassigned | - |
| Otu09165 | 2 | k__Fungi(100);p__Ascomycota(100);p__Ascomycota_unclassified(100);p__Ascomycota_unclassified(100);p__Ascomycota_unclassified(100);p__Ascomycota_unclassified(100);p__Ascomycota_unclassified(100); | - | - | - | - | - | - | - | Unassigned | - |
| Otu09168 | 2 | k__Fungi(100);k__Fungi_unclassified(100);k__Fungi_unclassified(100);k__Fungi_unclassified(100);k__Fungi_unclassified(100);k__Fungi_unclassified(100);k__Fungi_unclassified(100); | - | - | - | - | - | - | - | Unassigned | - |
| Otu09169 | 2 | k__Fungi(100);k__Fungi_unclassified(100);k__Fungi_unclassified(100);k__Fungi_unclassified(100);k__Fungi_unclassified(100);k__Fungi_unclassified(100);k__Fungi_unclassified(100); | - | - | - | - | - | - | - | Unassigned | - |
| Otu09170 | 2 | k__Fungi(100);k__Fungi_unclassified(100);k__Fungi_unclassified(100);k__Fungi_unclassified(100);k__Fungi_unclassified(100);k__Fungi_unclassified(100);k__Fungi_unclassified(100); | - | - | - | - | - | - | - | Unassigned | - |
| Otu09174 | 2 | k__Fungi(100);k__Fungi_unclassified(100);k__Fungi_unclassified(100);k__Fungi_unclassified(100);k__Fungi_unclassified(100);k__Fungi_unclassified(100);k__Fungi_unclassified(100); | - | - | - | - | - | - | - | Unassigned | - |
| Otu09175 | 2 | k__Fungi(100);k__Fungi_unclassified(100);k__Fungi_unclassified(100);k__Fungi_unclassified(100);k__Fungi_unclassified(100);k__Fungi_unclassified(100);k__Fungi_unclassified(100); | - | - | - | - | - | - | - | Unassigned | - |
| Otu09176 | 2 | k__Fungi(100);p__Ascomycota(100);p__Ascomycota_unclassified(100);p__Ascomycota_unclassified(100);p__Ascomycota_unclassified(100);p__Ascomycota_unclassified(100);p__Ascomycota_unclassified(100); | - | - | - | - | - | - | - | Unassigned | - |
| Otu09178 | 2 | k__Fungi(100);k__Fungi_unclassified(100);k__Fungi_unclassified(100);k__Fungi_unclassified(100);k__Fungi_unclassified(100);k__Fungi_unclassified(100);k__Fungi_unclassified(100); | - | - | - | - | - | - | - | Unassigned | - |
| Otu09179 | 2 | k__Fungi(100);p__Ascomycota(100);c__Leotiomycetes(100);c__Leotiomycetes_unclassified(100);c__Leotiomycetes_unclassified(100);c__Leotiomycetes_unclassified(100);c__Leotiomycetes_unclassified(100); | - | - | - | - | - | - | - | Unassigned | - |
| Otu09180 | 2 | k__Fungi(100);k__Fungi_unclassified(100);k__Fungi_unclassified(100);k__Fungi_unclassified(100);k__Fungi_unclassified(100);k__Fungi_unclassified(100);k__Fungi_unclassified(100); | - | - | - | - | - | - | - | Unassigned | - |
| Otu09182 | 2 | k__Fungi(100);k__Fungi_unclassified(100);k__Fungi_unclassified(100);k__Fungi_unclassified(100);k__Fungi_unclassified(100);k__Fungi_unclassified(100);k__Fungi_unclassified(100); | - | - | - | - | - | - | - | Unassigned | - |
| Otu09183 | 2 | k__Fungi(100);k__Fungi_unclassified(100);k__Fungi_unclassified(100);k__Fungi_unclassified(100);k__Fungi_unclassified(100);k__Fungi_unclassified(100);k__Fungi_unclassified(100); | - | - | - | - | - | - | - | Unassigned | - |
| Otu09189 | 2 | k__Fungi(100);k__Fungi_unclassified(100);k__Fungi_unclassified(100);k__Fungi_unclassified(100);k__Fungi_unclassified(100);k__Fungi_unclassified(100);k__Fungi_unclassified(100); | - | - | - | - | - | - | - | Unassigned | - |
| Otu09190 | 2 | k__Fungi(100);p__Ascomycota(100);p__Ascomycota_unclassified(100);p__Ascomycota_unclassified(100);p__Ascomycota_unclassified(100);p__Ascomycota_unclassified(100);p__Ascomycota_unclassified(100); | - | - | - | - | - | - | - | Unassigned | - |
| Otu09191 | 2 | k__Fungi(100);k__Fungi_unclassified(100);k__Fungi_unclassified(100);k__Fungi_unclassified(100);k__Fungi_unclassified(100);k__Fungi_unclassified(100);k__Fungi_unclassified(100); | - | - | - | - | - | - | - | Unassigned | - |
| Otu09192 | 2 | k__Fungi(100);k__Fungi_unclassified(100);k__Fungi_unclassified(100);k__Fungi_unclassified(100);k__Fungi_unclassified(100);k__Fungi_unclassified(100);k__Fungi_unclassified(100); | - | - | - | - | - | - | - | Unassigned | - |
| Otu09193 | 2 | k__Fungi(100);k__Fungi_unclassified(100);k__Fungi_unclassified(100);k__Fungi_unclassified(100);k__Fungi_unclassified(100);k__Fungi_unclassified(100);k__Fungi_unclassified(100); | - | - | - | - | - | - | - | Unassigned | - |
| Otu09194 | 2 | k__Fungi(100);k__Fungi_unclassified(100);k__Fungi_unclassified(100);k__Fungi_unclassified(100);k__Fungi_unclassified(100);k__Fungi_unclassified(100);k__Fungi_unclassified(100); | - | - | - | - | - | - | - | Unassigned | - |
| Otu09196 | 2 | k__Fungi(100);k__Fungi_unclassified(100);k__Fungi_unclassified(100);k__Fungi_unclassified(100);k__Fungi_unclassified(100);k__Fungi_unclassified(100);k__Fungi_unclassified(100); | - | - | - | - | - | - | - | Unassigned | - |
| Otu09197 | 2 | k__Fungi(100);p__Ascomycota(100);p__Ascomycota_unclassified(100);p__Ascomycota_unclassified(100);p__Ascomycota_unclassified(100);p__Ascomycota_unclassified(100);p__Ascomycota_unclassified(100); | - | - | - | - | - | - | - | Unassigned | - |
| Otu09198 | 2 | k__Fungi(100);k__Fungi_unclassified(100);k__Fungi_unclassified(100);k__Fungi_unclassified(100);k__Fungi_unclassified(100);k__Fungi_unclassified(100);k__Fungi_unclassified(100); | - | - | - | - | - | - | - | Unassigned | - |
| Otu09203 | 2 | k__Fungi(100);k__Fungi_unclassified(100);k__Fungi_unclassified(100);k__Fungi_unclassified(100);k__Fungi_unclassified(100);k__Fungi_unclassified(100);k__Fungi_unclassified(100); | - | - | - | - | - | - | - | Unassigned | - |
| Otu09204 | 2 | k__Fungi(100);k__Fungi_unclassified(100);k__Fungi_unclassified(100);k__Fungi_unclassified(100);k__Fungi_unclassified(100);k__Fungi_unclassified(100);k__Fungi_unclassified(100); | - | - | - | - | - | - | - | Unassigned | - |

| | | | | | | | | | | | |
|---|---|---|---|---|---|---|---|---|---|---|---|
| Otu09208 | 2 | k__Fungi(100);k__Fungi_unclassified(100);k__Fungi_unclassified(100);k__Fungi_unclassified(100);k__Fungi_unclassified(100);k__Fungi_unclassified(100);k__Fungi_unclassified(100); | - | - | - | - | - | - | - | Unassigned | - |
| Otu09209 | 2 | k__Fungi(100);k__Fungi_unclassified(100);k__Fungi_unclassified(100);k__Fungi_unclassified(100);k__Fungi_unclassified(100);k__Fungi_unclassified(100);k__Fungi_unclassified(100); | - | - | - | - | - | - | - | Unassigned | - |
| Otu09210 | 2 | k__Fungi(100);k__Fungi_unclassified(100);k__Fungi_unclassified(100);k__Fungi_unclassified(100);k__Fungi_unclassified(100);k__Fungi_unclassified(100);k__Fungi_unclassified(100); | - | - | - | - | - | - | - | Unassigned | - |
| Otu09221 | 2 | k__Fungi(100);k__Fungi_unclassified(100);k__Fungi_unclassified(100);k__Fungi_unclassified(100);k__Fungi_unclassified(100);k__Fungi_unclassified(100);k__Fungi_unclassified(100); | - | - | - | - | - | - | - | Unassigned | - |
| Otu09222 | 2 | k__Fungi(100);k__Fungi_unclassified(100);k__Fungi_unclassified(100);k__Fungi_unclassified(100);k__Fungi_unclassified(100);k__Fungi_unclassified(100);k__Fungi_unclassified(100); | - | - | - | - | - | - | - | Unassigned | - |
| Otu09225 | 2 | k__Fungi(100);k__Fungi_unclassified(100);k__Fungi_unclassified(100);k__Fungi_unclassified(100);k__Fungi_unclassified(100);k__Fungi_unclassified(100);k__Fungi_unclassified(100); | - | - | - | - | - | - | - | Unassigned | - |
| Otu09228 | 2 | k__Fungi(100);k__Fungi_unclassified(100);k__Fungi_unclassified(100);k__Fungi_unclassified(100);k__Fungi_unclassified(100);k__Fungi_unclassified(100);k__Fungi_unclassified(100); | - | - | - | - | - | - | - | Unassigned | - |
| Otu09230 | 2 | k__Fungi(100);p__Ascomycota(100);p__Ascomycota_unclassified(100);p__Ascomycota_unclassified(100);p__Ascomycota_unclassified(100);p__Ascomycota_unclassified(100);p__Ascomycota_unclassified(100); | - | - | - | - | - | - | - | Unassigned | - |
| Otu09231 | 2 | k__Fungi(100);k__Fungi_unclassified(100);k__Fungi_unclassified(100);k__Fungi_unclassified(100);k__Fungi_unclassified(100);k__Fungi_unclassified(100);k__Fungi_unclassified(100); | - | - | - | - | - | - | - | Unassigned | - |
| Otu09233 | 2 | k__Fungi(100);k__Fungi_unclassified(100);k__Fungi_unclassified(100);k__Fungi_unclassified(100);k__Fungi_unclassified(100);k__Fungi_unclassified(100);k__Fungi_unclassified(100); | - | - | - | - | - | - | - | Unassigned | - |
| Otu09234 | 2 | k__Fungi(100);p__Ascomycota(100);c__Dothideomycetes(100);o__Pleosporales(100);f__unclassified_Pleosporales(100);g__unclassified_Pleosporales(100);s__Pleosporales_sp(100); | - | - | - | - | - | - | - | Unassigned | - |
| Otu09238 | 2 | k__Fungi(100);k__Fungi_unclassified(100);k__Fungi_unclassified(100);k__Fungi_unclassified(100);k__Fungi_unclassified(100);k__Fungi_unclassified(100);k__Fungi_unclassified(100); | - | - | - | - | - | - | - | Unassigned | - |
| Otu09239 | 2 | k__Fungi(100);k__Fungi_unclassified(100);k__Fungi_unclassified(100);k__Fungi_unclassified(100);k__Fungi_unclassified(100);k__Fungi_unclassified(100);k__Fungi_unclassified(100); | - | - | - | - | - | - | - | Unassigned | - |
| Otu09241 | 2 | k__Fungi(100);k__Fungi_unclassified(100);k__Fungi_unclassified(100);k__Fungi_unclassified(100);k__Fungi_unclassified(100);k__Fungi_unclassified(100);k__Fungi_unclassified(100); | - | - | - | - | - | - | - | Unassigned | - |
| Otu09243 | 2 | k__Fungi(100);p__Ascomycota(100);p__Ascomycota_unclassified(100);p__Ascomycota_unclassified(100);p__Ascomycota_unclassified(100);p__Ascomycota_unclassified(100);p__Ascomycota_unclassified(100); | - | - | - | - | - | - | - | Unassigned | - |
| Otu09244 | 2 | k__Fungi(100);p__Ascomycota(100);p__Ascomycota_unclassified(100);p__Ascomycota_unclassified(100);p__Ascomycota_unclassified(100);p__Ascomycota_unclassified(100);p__Ascomycota_unclassified(100); | - | - | - | - | - | - | - | Unassigned | - |
| Otu09247 | 2 | k__Fungi(100);k__Fungi_unclassified(100);k__Fungi_unclassified(100);k__Fungi_unclassified(100);k__Fungi_unclassified(100);k__Fungi_unclassified(100);k__Fungi_unclassified(100); | - | - | - | - | - | - | - | Unassigned | - |
| Otu09248 | 2 | k__Fungi(100);k__Fungi_unclassified(100);k__Fungi_unclassified(100);k__Fungi_unclassified(100);k__Fungi_unclassified(100);k__Fungi_unclassified(100);k__Fungi_unclassified(100); | - | - | - | - | - | - | - | Unassigned | - |
| Otu09249 | 2 | k__Fungi(100);k__Fungi_unclassified(100);k__Fungi_unclassified(100);k__Fungi_unclassified(100);k__Fungi_unclassified(100);k__Fungi_unclassified(100);k__Fungi_unclassified(100); | - | - | - | - | - | - | - | Unassigned | - |
| Otu09250 | 2 | k__Fungi(100);p__Ascomycota(100);c__Dothideomycetes(100);c__Dothideomycetes_unclassified(100);c__Dothideomycetes_unclassified(100);c__Dothideomycetes_unclassified(100);c__Dothideomycetes_unclassified(100); | - | - | - | - | - | - | - | Unassigned | - |
| Otu09252 | 2 | k__Fungi(100);p__Ascomycota(100);c__Leotiomycetes(100);c__Leotiomycetes_unclassified(100);c__Leotiomycetes_unclassified(100);c__Leotiomycetes_unclassified(100);c__Leotiomycetes_unclassified(100); | - | - | - | - | - | - | - | Unassigned | - |
| Otu09253 | 2 | k__Fungi(100);k__Fungi_unclassified(100);k__Fungi_unclassified(100);k__Fungi_unclassified(100);k__Fungi_unclassified(100);k__Fungi_unclassified(100);k__Fungi_unclassified(100); | - | - | - | - | - | - | - | Unassigned | - |
| Otu09255 | 2 | k__Fungi(100);p__Ascomycota(100);p__Ascomycota_unclassified(100);p__Ascomycota_unclassified(100);p__Ascomycota_unclassified(100);p__Ascomycota_unclassified(100);p__Ascomycota_unclassified(100); | - | - | - | - | - | - | - | Unassigned | - |
| Otu09256 | 2 | k__Fungi(100);k__Fungi_unclassified(100);k__Fungi_unclassified(100);k__Fungi_unclassified(100);k__Fungi_unclassified(100);k__Fungi_unclassified(100);k__Fungi_unclassified(100); | - | - | - | - | - | - | - | Unassigned | - |
| Otu09260 | 2 | k__Fungi(100);k__Fungi_unclassified(100);k__Fungi_unclassified(100);k__Fungi_unclassified(100);k__Fungi_unclassified(100);k__Fungi_unclassified(100);k__Fungi_unclassified(100); | - | - | - | - | - | - | - | Unassigned | - |
| Otu09265 | 2 | k__Fungi(100);p__Ascomycota(100);p__Ascomycota_unclassified(100);p__Ascomycota_unclassified(100);p__Ascomycota_unclassified(100);p__Ascomycota_unclassified(100);p__Ascomycota_unclassified(100); | - | - | - | - | - | - | - | Unassigned | - |
| Otu09266 | 2 | k__Fungi(100);p__Ascomycota(100);p__Ascomycota_unclassified(100);p__Ascomycota_unclassified(100);p__Ascomycota_unclassified(100);p__Ascomycota_unclassified(100);p__Ascomycota_unclassified(100); | - | - | - | - | - | - | - | Unassigned | - |
| Otu09268 | 2 | k__Fungi(100);k__Fungi_unclassified(100);k__Fungi_unclassified(100);k__Fungi_unclassified(100);k__Fungi_unclassified(100);k__Fungi_unclassified(100);k__Fungi_unclassified(100); | - | - | - | - | - | - | - | Unassigned | - |
| Otu09269 | 2 | k__Fungi(100);k__Fungi_unclassified(100);k__Fungi_unclassified(100);k__Fungi_unclassified(100);k__Fungi_unclassified(100);k__Fungi_unclassified(100);k__Fungi_unclassified(100); | - | - | - | - | - | - | - | Unassigned | - |
| Otu09271 | 2 | k__Fungi(100);p__Ascomycota(100);p__Ascomycota_unclassified(100);p__Ascomycota_unclassified(100);p__Ascomycota_unclassified(100);p__Ascomycota_unclassified(100);p__Ascomycota_unclassified(100); | - | - | - | - | - | - | - | Unassigned | - |
| Otu09275 | 2 | k__Fungi(100);k__Fungi_unclassified(100);k__Fungi_unclassified(100);k__Fungi_unclassified(100);k__Fungi_unclassified(100);k__Fungi_unclassified(100);k__Fungi_unclassified(100); | - | - | - | - | - | - | - | Unassigned | - |
| Otu09277 | 2 | k__Fungi(100);p__Ascomycota(100);c__Leotiomycetes(100);c__Leotiomycetes_unclassified(100);c__Leotiomycetes_unclassified(100);c__Leotiomycetes_unclassified(100);c__Leotiomycetes_unclassified(100); | - | - | - | - | - | - | - | Unassigned | - |
| Otu09278 | 2 | k__Fungi(100);p__Ascomycota(100);c__Leotiomycetes(100);c__Leotiomycetes_unclassified(100);c__Leotiomycetes_unclassified(100);c__Leotiomycetes_unclassified(100);c__Leotiomycetes_unclassified(100); | - | - | - | - | - | - | - | Unassigned | - |
| Otu09279 | 2 | k__Fungi(100);k__Fungi_unclassified(100);k__Fungi_unclassified(100);k__Fungi_unclassified(100);k__Fungi_unclassified(100);k__Fungi_unclassified(100);k__Fungi_unclassified(100); | - | - | - | - | - | - | - | Unassigned | - |
| Otu09280 | 2 | k__Fungi(100);p__Ascomycota(100);p__Ascomycota_unclassified(100);p__Ascomycota_unclassified(100);p__Ascomycota_unclassified(100);p__Ascomycota_unclassified(100);p__Ascomycota_unclassified(100); | - | - | - | - | - | - | - | Unassigned | - |
| Otu09281 | 2 | k__Fungi(100);k__Fungi_unclassified(100);k__Fungi_unclassified(100);k__Fungi_unclassified(100);k__Fungi_unclassified(100);k__Fungi_unclassified(100);k__Fungi_unclassified(100); | - | - | - | - | - | - | - | Unassigned | - |
| Otu09282 | 2 | k__Fungi(100);p__Ascomycota(100);c__Leotiomycetes(100);o__Helotiales(100);o__Helotiales_unclassified(100);o__Helotiales_unclassified(100);o__Helotiales_unclassified(100); | - | - | - | - | - | - | - | Unassigned | - |
| Otu09284 | 2 | k__Fungi(100);p__Ascomycota(100);p__Ascomycota_unclassified(100);p__Ascomycota_unclassified(100);p__Ascomycota_unclassified(100);p__Ascomycota_unclassified(100);p__Ascomycota_unclassified(100); | - | - | - | - | - | - | - | Unassigned | - |
| Otu09289 | 2 | k__Fungi(100);p__Ascomycota(100);p__Ascomycota_unclassified(100);p__Ascomycota_unclassified(100);p__Ascomycota_unclassified(100);p__Ascomycota_unclassified(100);p__Ascomycota_unclassified(100); | - | - | - | - | - | - | - | Unassigned | - |
| Otu09290 | 2 | k__Fungi(100);p__Ascomycota(100);c__Dothideomycetes(100);o__Capnodiales(100);o__Capnodiales_unclassified(100);o__Capnodiales_unclassified(100);o__Capnodiales_unclassified(100); | - | - | - | - | - | - | - | Unassigned | - |
| Otu09292 | 2 | k__Fungi(100);k__Fungi_unclassified(100);k__Fungi_unclassified(100);k__Fungi_unclassified(100);k__Fungi_unclassified(100);k__Fungi_unclassified(100);k__Fungi_unclassified(100); | - | - | - | - | - | - | - | Unassigned | - |
| Otu09293 | 2 | k__Fungi(100);k__Fungi_unclassified(100);k__Fungi_unclassified(100);k__Fungi_unclassified(100);k__Fungi_unclassified(100);k__Fungi_unclassified(100);k__Fungi_unclassified(100); | - | - | - | - | - | - | - | Unassigned | - |
| Otu09294 | 2 | k__Fungi(100);k__Fungi_unclassified(100);k__Fungi_unclassified(100);k__Fungi_unclassified(100);k__Fungi_unclassified(100);k__Fungi_unclassified(100);k__Fungi_unclassified(100); | - | - | - | - | - | - | - | Unassigned | - |
| Otu09295 | 2 | k__Fungi(100);k__Fungi_unclassified(100);k__Fungi_unclassified(100);k__Fungi_unclassified(100);k__Fungi_unclassified(100);k__Fungi_unclassified(100);k__Fungi_unclassified(100); | - | - | - | - | - | - | - | Unassigned | - |
| Otu09298 | 2 | k__Fungi(100);k__Fungi_unclassified(100);k__Fungi_unclassified(100);k__Fungi_unclassified(100);k__Fungi_unclassified(100);k__Fungi_unclassified(100);k__Fungi_unclassified(100); | - | - | - | - | - | - | - | Unassigned | - |
| Otu09299 | 2 | k__Fungi(100);k__Fungi_unclassified(100);k__Fungi_unclassified(100);k__Fungi_unclassified(100);k__Fungi_unclassified(100);k__Fungi_unclassified(100);k__Fungi_unclassified(100); | - | - | - | - | - | - | - | Unassigned | - |
| Otu09300 | 2 | k__Fungi(100);k__Fungi_unclassified(100);k__Fungi_unclassified(100);k__Fungi_unclassified(100);k__Fungi_unclassified(100);k__Fungi_unclassified(100);k__Fungi_unclassified(100); | - | - | - | - | - | - | - | Unassigned | - |
| Otu09301 | 2 | k__Fungi(100);k__Fungi_unclassified(100);k__Fungi_unclassified(100);k__Fungi_unclassified(100);k__Fungi_unclassified(100);k__Fungi_unclassified(100);k__Fungi_unclassified(100); | - | - | - | - | - | - | - | Unassigned | - |
| Otu09304 | 2 | k__Fungi(100);k__Fungi_unclassified(100);k__Fungi_unclassified(100);k__Fungi_unclassified(100);k__Fungi_unclassified(100);k__Fungi_unclassified(100);k__Fungi_unclassified(100); | - | - | - | - | - | - | - | Unassigned | - |
| Otu09305 | 2 | k__Fungi(100);k__Fungi_unclassified(100);k__Fungi_unclassified(100);k__Fungi_unclassified(100);k__Fungi_unclassified(100);k__Fungi_unclassified(100);k__Fungi_unclassified(100); | - | - | - | - | - | - | - | Unassigned | - |
| Otu09307 | 2 | k__Fungi(100);k__Fungi_unclassified(100);k__Fungi_unclassified(100);k__Fungi_unclassified(100);k__Fungi_unclassified(100);k__Fungi_unclassified(100);k__Fungi_unclassified(100); | - | - | - | - | - | - | - | Unassigned | - |
| Otu09308 | 2 | k__Fungi(100);k__Fungi_unclassified(100);k__Fungi_unclassified(100);k__Fungi_unclassified(100);k__Fungi_unclassified(100);k__Fungi_unclassified(100);k__Fungi_unclassified(100); | - | - | - | - | - | - | - | Unassigned | - |
| Otu09309 | 2 | k__Fungi(100);k__Fungi_unclassified(100);k__Fungi_unclassified(100);k__Fungi_unclassified(100);k__Fungi_unclassified(100);k__Fungi_unclassified(100);k__Fungi_unclassified(100); | - | - | - | - | - | - | - | Unassigned | - |
| Otu09310 | 2 | k__Fungi(100);k__Fungi_unclassified(100);k__Fungi_unclassified(100);k__Fungi_unclassified(100);k__Fungi_unclassified(100);k__Fungi_unclassified(100);k__Fungi_unclassified(100); | - | - | - | - | - | - | - | Unassigned | - |
| Otu09314 | 2 | k__Fungi(100);k__Fungi_unclassified(100);k__Fungi_unclassified(100);k__Fungi_unclassified(100);k__Fungi_unclassified(100);k__Fungi_unclassified(100);k__Fungi_unclassified(100); | - | - | - | - | - | - | - | Unassigned | - |
| Otu09315 | 2 | k__Fungi(100);k__Fungi_unclassified(100);k__Fungi_unclassified(100);k__Fungi_unclassified(100);k__Fungi_unclassified(100);k__Fungi_unclassified(100);k__Fungi_unclassified(100); | - | - | - | - | - | - | - | Unassigned | - |
| Otu09317 | 2 | k__Fungi(100);p__Ascomycota(100);p__Ascomycota_unclassified(100);p__Ascomycota_unclassified(100);p__Ascomycota_unclassified(100);p__Ascomycota_unclassified(100);p__Ascomycota_unclassified(100); | - | - | - | - | - | - | - | Unassigned | - |
| Otu09322 | 2 | k__Fungi(100);p__Ascomycota(100);p__Ascomycota_unclassified(100);p__Ascomycota_unclassified(100);p__Ascomycota_unclassified(100);p__Ascomycota_unclassified(100);p__Ascomycota_unclassified(100); | - | - | - | - | - | - | - | Unassigned | - |
| Otu09325 | 2 | k__Fungi(100);p__Ascomycota(100);p__Ascomycota_unclassified(100);p__Ascomycota_unclassified(100);p__Ascomycota_unclassified(100);p__Ascomycota_unclassified(100);p__Ascomycota_unclassified(100); | - | - | - | - | - | - | - | Unassigned | - |
| Otu09326 | 2 | k__Fungi(100);k__Fungi_unclassified(100);k__Fungi_unclassified(100);k__Fungi_unclassified(100);k__Fungi_unclassified(100);k__Fungi_unclassified(100);k__Fungi_unclassified(100); | - | - | - | - | - | - | - | Unassigned | - |
| Otu09327 | 2 | k__Fungi(100);k__Fungi_unclassified(100);k__Fungi_unclassified(100);k__Fungi_unclassified(100);k__Fungi_unclassified(100);k__Fungi_unclassified(100);k__Fungi_unclassified(100); | - | - | - | - | - | - | - | Unassigned | - |
| Otu09328 | 2 | k__Fungi(100);k__Fungi_unclassified(100);k__Fungi_unclassified(100);k__Fungi_unclassified(100);k__Fungi_unclassified(100);k__Fungi_unclassified(100);k__Fungi_unclassified(100); | - | - | - | - | - | - | - | Unassigned | - |
| Otu09329 | 2 | k__Fungi(100);k__Fungi_unclassified(100);k__Fungi_unclassified(100);k__Fungi_unclassified(100);k__Fungi_unclassified(100);k__Fungi_unclassified(100);k__Fungi_unclassified(100); | - | - | - | - | - | - | - | Unassigned | - |
| Otu09330 | 2 | k__Fungi(100);k__Fungi_unclassified(100);k__Fungi_unclassified(100);k__Fungi_unclassified(100);k__Fungi_unclassified(100);k__Fungi_unclassified(100);k__Fungi_unclassified(100); | - | - | - | - | - | - | - | Unassigned | - |
| Otu09332 | 2 | k__Fungi(100);k__Fungi_unclassified(100);k__Fungi_unclassified(100);k__Fungi_unclassified(100);k__Fungi_unclassified(100);k__Fungi_unclassified(100);k__Fungi_unclassified(100); | - | - | - | - | - | - | - | Unassigned | - |
| Otu09333 | 2 | k__Fungi(100);k__Fungi_unclassified(100);k__Fungi_unclassified(100);k__Fungi_unclassified(100);k__Fungi_unclassified(100);k__Fungi_unclassified(100);k__Fungi_unclassified(100); | - | - | - | - | - | - | - | Unassigned | - |
| Otu09335 | 2 | k__Fungi(100);k__Fungi_unclassified(100);k__Fungi_unclassified(100);k__Fungi_unclassified(100);k__Fungi_unclassified(100);k__Fungi_unclassified(100);k__Fungi_unclassified(100); | - | - | - | - | - | - | - | Unassigned | - |
| Otu09336 | 2 | k__Fungi(100);p__Ascomycota(100);p__Ascomycota_unclassified(100);p__Ascomycota_unclassified(100);p__Ascomycota_unclassified(100);p__Ascomycota_unclassified(100);p__Ascomycota_unclassified(100); | - | - | - | - | - | - | - | Unassigned | - |
| Otu09337 | 2 | k__Fungi(100);p__Ascomycota(100);p__Ascomycota_unclassified(100);p__Ascomycota_unclassified(100);p__Ascomycota_unclassified(100);p__Ascomycota_unclassified(100);p__Ascomycota_unclassified(100); | - | - | - | - | - | - | - | Unassigned | - |
| Otu09340 | 2 | k__Fungi(100);k__Fungi_unclassified(100);k__Fungi_unclassified(100);k__Fungi_unclassified(100);k__Fungi_unclassified(100);k__Fungi_unclassified(100);k__Fungi_unclassified(100); | - | - | - | - | - | - | - | Unassigned | - |
| Otu09342 | 2 | k__Fungi(100);k__Fungi_unclassified(100);k__Fungi_unclassified(100);k__Fungi_unclassified(100);k__Fungi_unclassified(100);k__Fungi_unclassified(100);k__Fungi_unclassified(100); | - | - | - | - | - | - | - | Unassigned | - |
| Otu09350 | 2 | k__Fungi(100);k__Fungi_unclassified(100);k__Fungi_unclassified(100);k__Fungi_unclassified(100);k__Fungi_unclassified(100);k__Fungi_unclassified(100);k__Fungi_unclassified(100); | - | - | - | - | - | - | - | Unassigned | - |
| Otu09351 | 2 | k__Fungi(100);k__Fungi_unclassified(100);k__Fungi_unclassified(100);k__Fungi_unclassified(100);k__Fungi_unclassified(100);k__Fungi_unclassified(100);k__Fungi_unclassified(100); | - | - | - | - | - | - | - | Unassigned | - |
| Otu09359 | 2 | k__Fungi(100);p__Ascomycota(100);p__Ascomycota_unclassified(100);p__Ascomycota_unclassified(100);p__Ascomycota_unclassified(100);p__Ascomycota_unclassified(100);p__Ascomycota_unclassified(100); | - | - | - | - | - | - | - | Unassigned | - |
| Otu09360 | 2 | k__Fungi(100);k__Fungi_unclassified(100);k__Fungi_unclassified(100);k__Fungi_unclassified(100);k__Fungi_unclassified(100);k__Fungi_unclassified(100);k__Fungi_unclassified(100); | - | - | - | - | - | - | - | Unassigned | - |
| Otu09361 | 2 | k__Fungi(100);k__Fungi_unclassified(100);k__Fungi_unclassified(100);k__Fungi_unclassified(100);k__Fungi_unclassified(100);k__Fungi_unclassified(100);k__Fungi_unclassified(100); | - | - | - | - | - | - | - | Unassigned | - |
| Otu09363 | 2 | k__Fungi(100);k__Fungi_unclassified(100);k__Fungi_unclassified(100);k__Fungi_unclassified(100);k__Fungi_unclassified(100);k__Fungi_unclassified(100);k__Fungi_unclassified(100); | - | - | - | - | - | - | - | Unassigned | - |
| Otu09366 | 2 | k__Fungi(100);k__Fungi_unclassified(100);k__Fungi_unclassified(100);k__Fungi_unclassified(100);k__Fungi_unclassified(100);k__Fungi_unclassified(100);k__Fungi_unclassified(100); | - | - | - | - | - | - | - | Unassigned | - |
| Otu09370 | 2 | k__Fungi(100);p__Ascomycota(100);p__Ascomycota_unclassified(100);p__Ascomycota_unclassified(100);p__Ascomycota_unclassified(100);p__Ascomycota_unclassified(100);p__Ascomycota_unclassified(100); | - | - | - | - | - | - | - | Unassigned | - |
| Otu09372 | 2 | k__Fungi(100);k__Fungi_unclassified(100);k__Fungi_unclassified(100);k__Fungi_unclassified(100);k__Fungi_unclassified(100);k__Fungi_unclassified(100);k__Fungi_unclassified(100); | - | - | - | - | - | - | - | Unassigned | - |
| Otu09373 | 2 | k__Fungi(100);k__Fungi_unclassified(100);k__Fungi_unclassified(100);k__Fungi_unclassified(100);k__Fungi_unclassified(100);k__Fungi_unclassified(100);k__Fungi_unclassified(100); | - | - | - | - | - | - | - | Unassigned | - |
| Otu09375 | 2 | k__Fungi(100);k__Fungi_unclassified(100);k__Fungi_unclassified(100);k__Fungi_unclassified(100);k__Fungi_unclassified(100);k__Fungi_unclassified(100);k__Fungi_unclassified(100); | - | - | - | - | - | - | - | Unassigned | - |
| Otu09381 | 2 | k__Fungi(100);p__Ascomycota(100);p__Ascomycota_unclassified(100);p__Ascomycota_unclassified(100);p__Ascomycota_unclassified(100);p__Ascomycota_unclassified(100);p__Ascomycota_unclassified(100); | - | - | - | - | - | - | - | Unassigned | - |
| Otu09382 | 2 | k__Fungi(100);k__Fungi_unclassified(100);k__Fungi_unclassified(100);k__Fungi_unclassified(100);k__Fungi_unclassified(100);k__Fungi_unclassified(100);k__Fungi_unclassified(100); | - | - | - | - | - | - | - | Unassigned | - |
| Otu09384 | 2 | k__Fungi(100);p__Ascomycota(100);p__Ascomycota_unclassified(100);p__Ascomycota_unclassified(100);p__Ascomycota_unclassified(100);p__Ascomycota_unclassified(100);p__Ascomycota_unclassified(100); | - | - | - | - | - | - | - | Unassigned | - |
| Otu09385 | 2 | k__Fungi(100);p__Basidiomycota(100);p__Basidiomycota_unclassified(100);p__Basidiomycota_unclassified(100);p__Basidiomycota_unclassified(100);p__Basidiomycota_unclassified(100);p__Basidiomycota_unclassified(100); | - | - | - | - | - | - | - | Unassigned | - |
| Otu09386 | 2 | k__Fungi(100);p__Ascomycota(100);p__Ascomycota_unclassified(100);p__Ascomycota_unclassified(100);p__Ascomycota_unclassified(100);p__Ascomycota_unclassified(100);p__Ascomycota_unclassified(100); | - | - | - | - | - | - | - | Unassigned | - |
| Otu09387 | 2 | k__Fungi(100);k__Fungi_unclassified(100);k__Fungi_unclassified(100);k__Fungi_unclassified(100);k__Fungi_unclassified(100);k__Fungi_unclassified(100);k__Fungi_unclassified(100); | - | - | - | - | - | - | - | Unassigned | - |
| Otu09388 | 2 | k__Fungi(100);k__Fungi_unclassified(100);k__Fungi_unclassified(100);k__Fungi_unclassified(100);k__Fungi_unclassified(100);k__Fungi_unclassified(100);k__Fungi_unclassified(100); | - | - | - | - | - | - | - | Unassigned | - |
| Otu09389 | 2 | k__Fungi(100);k__Fungi_unclassified(100);k__Fungi_unclassified(100);k__Fungi_unclassified(100);k__Fungi_unclassified(100);k__Fungi_unclassified(100);k__Fungi_unclassified(100); | - | - | - | - | - | - | - | Unassigned | - |
| Otu09390 | 2 | k__Fungi(100);k__Fungi_unclassified(100);k__Fungi_unclassified(100);k__Fungi_unclassified(100);k__Fungi_unclassified(100);k__Fungi_unclassified(100);k__Fungi_unclassified(100); | - | - | - | - | - | - | - | Unassigned | - |
| Otu09393 | 2 | k__Fungi(100);k__Fungi_unclassified(100);k__Fungi_unclassified(100);k__Fungi_unclassified(100);k__Fungi_unclassified(100);k__Fungi_unclassified(100);k__Fungi_unclassified(100); | - | - | - | - | - | - | - | Unassigned | - |
| Otu09395 | 2 | k__Fungi(100);p__Ascomycota(100);p__Ascomycota_unclassified(100);p__Ascomycota_unclassified(100);p__Ascomycota_unclassified(100);p__Ascomycota_unclassified(100);p__Ascomycota_unclassified(100); | - | - | - | - | - | - | - | Unassigned | - |
| Otu09397 | 2 | k__Fungi(100);k__Fungi_unclassified(100);k__Fungi_unclassified(100);k__Fungi_unclassified(100);k__Fungi_unclassified(100);k__Fungi_unclassified(100);k__Fungi_unclassified(100); | - | - | - | - | - | - | - | Unassigned | - |
| Otu09401 | 2 | k__Fungi(100);k__Fungi_unclassified(100);k__Fungi_unclassified(100);k__Fungi_unclassified(100);k__Fungi_unclassified(100);k__Fungi_unclassified(100);k__Fungi_unclassified(100); | - | - | - | - | - | - | - | Unassigned | - |
| Otu09402 | 2 | k__Fungi(100);k__Fungi_unclassified(100);k__Fungi_unclassified(100);k__Fungi_unclassified(100);k__Fungi_unclassified(100);k__Fungi_unclassified(100);k__Fungi_unclassified(100); | - | - | - | - | - | - | - | Unassigned | - |
| Otu09403 | 2 | k__Fungi(100);k__Fungi_unclassified(100);k__Fungi_unclassified(100);k__Fungi_unclassified(100);k__Fungi_unclassified(100);k__Fungi_unclassified(100);k__Fungi_unclassified(100); | - | - | - | - | - | - | - | Unassigned | - |
| Otu09404 | 2 | k__Fungi(100);k__Fungi_unclassified(100);k__Fungi_unclassified(100);k__Fungi_unclassified(100);k__Fungi_unclassified(100);k__Fungi_unclassified(100);k__Fungi_unclassified(100); | - | - | - | - | - | - | - | Unassigned | - |
| Otu09407 | 2 | k__Fungi(100);k__Fungi_unclassified(100);k__Fungi_unclassified(100);k__Fungi_unclassified(100);k__Fungi_unclassified(100);k__Fungi_unclassified(100);k__Fungi_unclassified(100); | - | - | - | - | - | - | - | Unassigned | - |
| Otu09409 | 2 | k__Fungi(100);k__Fungi_unclassified(100);k__Fungi_unclassified(100);k__Fungi_unclassified(100);k__Fungi_unclassified(100);k__Fungi_unclassified(100);k__Fungi_unclassified(100); | - | - | - | - | - | - | - | Unassigned | - |
| Otu09411 | 2 | k__Fungi(100);k__Fungi_unclassified(100);k__Fungi_unclassified(100);k__Fungi_unclassified(100);k__Fungi_unclassified(100);k__Fungi_unclassified(100);k__Fungi_unclassified(100); | - | - | - | - | - | - | - | Unassigned | - |
| Otu09412 | 2 | k__Fungi(100);k__Fungi_unclassified(100);k__Fungi_unclassified(100);k__Fungi_unclassified(100);k__Fungi_unclassified(100);k__Fungi_unclassified(100);k__Fungi_unclassified(100); | - | - | - | - | - | - | - | Unassigned | - |
| Otu09417 | 2 | k__Fungi(100);k__Fungi_unclassified(100);k__Fungi_unclassified(100);k__Fungi_unclassified(100);k__Fungi_unclassified(100);k__Fungi_unclassified(100);k__Fungi_unclassified(100); | - | - | - | - | - | - | - | Unassigned | - |
| Otu09419 | 2 | k__Fungi(100);k__Fungi_unclassified(100);k__Fungi_unclassified(100);k__Fungi_unclassified(100);k__Fungi_unclassified(100);k__Fungi_unclassified(100);k__Fungi_unclassified(100); | - | - | - | - | - | - | - | Unassigned | - |
| Otu09420 | 2 | k__Fungi(100);k__Fungi_unclassified(100);k__Fungi_unclassified(100);k__Fungi_unclassified(100);k__Fungi_unclassified(100);k__Fungi_unclassified(100);k__Fungi_unclassified(100); | - | - | - | - | - | - | - | Unassigned | - |
| Otu09421 | 2 | k__Fungi(100);k__Fungi_unclassified(100);k__Fungi_unclassified(100);k__Fungi_unclassified(100);k__Fungi_unclassified(100);k__Fungi_unclassified(100);k__Fungi_unclassified(100); | - | - | - | - | - | - | - | Unassigned | - |
| Otu09422 | 2 | k__Fungi(100);k__Fungi_unclassified(100);k__Fungi_unclassified(100);k__Fungi_unclassified(100);k__Fungi_unclassified(100);k__Fungi_unclassified(100);k__Fungi_unclassified(100); | - | - | - | - | - | - | - | Unassigned | - |
| Otu09423 | 2 | k__Fungi(100);k__Fungi_unclassified(100);k__Fungi_unclassified(100);k__Fungi_unclassified(100);k__Fungi_unclassified(100);k__Fungi_unclassified(100);k__Fungi_unclassified(100); | - | - | - | - | - | - | - | Unassigned | - |
| Otu09426 | 2 | k__Fungi(100);k__Fungi_unclassified(100);k__Fungi_unclassified(100);k__Fungi_unclassified(100);k__Fungi_unclassified(100);k__Fungi_unclassified(100);k__Fungi_unclassified(100); | - | - | - | - | - | - | - | Unassigned | - |
| Otu09429 | 2 | k__Fungi(100);p__Ascomycota(100);p__Ascomycota_unclassified(100);p__Ascomycota_unclassified(100);p__Ascomycota_unclassified(100);p__Ascomycota_unclassified(100);p__Ascomycota_unclassified(100); | - | - | - | - | - | - | - | Unassigned | - |
| Otu09430 | 2 | k__Fungi(100);k__Fungi_unclassified(100);k__Fungi_unclassified(100);k__Fungi_unclassified(100);k__Fungi_unclassified(100);k__Fungi_unclassified(100);k__Fungi_unclassified(100); | - | - | - | - | - | - | - | Unassigned | - |
| Otu09435 | 2 | k__Fungi(100);k__Fungi_unclassified(100);k__Fungi_unclassified(100);k__Fungi_unclassified(100);k__Fungi_unclassified(100);k__Fungi_unclassified(100);k__Fungi_unclassified(100); | - | - | - | - | - | - | - | Unassigned | - |
| Otu09438 | 2 | k__Fungi(100);p__Ascomycota(100);p__Ascomycota_unclassified(100);p__Ascomycota_unclassified(100);p__Ascomycota_unclassified(100);p__Ascomycota_unclassified(100);p__Ascomycota_unclassified(100); | - | - | - | - | - | - | - | Unassigned | - |
| Otu09439 | 2 | k__Fungi(100);k__Fungi_unclassified(100);k__Fungi_unclassified(100);k__Fungi_unclassified(100);k__Fungi_unclassified(100);k__Fungi_unclassified(100);k__Fungi_unclassified(100); | - | - | - | - | - | - | - | Unassigned | - |
| Otu09440 | 2 | k__Fungi(100);k__Fungi_unclassified(100);k__Fungi_unclassified(100);k__Fungi_unclassified(100);k__Fungi_unclassified(100);k__Fungi_unclassified(100);k__Fungi_unclassified(100); | - | - | - | - | - | - | - | Unassigned | - |
| Otu09441 | 2 | k__Fungi(100);p__Ascomycota(100);p__Ascomycota_unclassified(100);p__Ascomycota_unclassified(100);p__Ascomycota_unclassified(100);p__Ascomycota_unclassified(100);p__Ascomycota_unclassified(100); | - | - | - | - | - | - | - | Unassigned | - |
| Otu09442 | 2 | k__Fungi(100);k__Fungi_unclassified(100);k__Fungi_unclassified(100);k__Fungi_unclassified(100);k__Fungi_unclassified(100);k__Fungi_unclassified(100);k__Fungi_unclassified(100); | - | - | - | - | - | - | - | Unassigned | - |
| Otu09444 | 2 | k__Fungi(100);k__Fungi_unclassified(100);k__Fungi_unclassified(100);k__Fungi_unclassified(100);k__Fungi_unclassified(100);k__Fungi_unclassified(100);k__Fungi_unclassified(100); | - | - | - | - | - | - | - | Unassigned | - |
| Otu09448 | 2 | k__Fungi(100);k__Fungi_unclassified(100);k__Fungi_unclassified(100);k__Fungi_unclassified(100);k__Fungi_unclassified(100);k__Fungi_unclassified(100);k__Fungi_unclassified(100); | - | - | - | - | - | - | - | Unassigned | - |
| Otu09449 | 2 | k__Fungi(100);k__Fungi_unclassified(100);k__Fungi_unclassified(100);k__Fungi_unclassified(100);k__Fungi_unclassified(100);k__Fungi_unclassified(100);k__Fungi_unclassified(100); | - | - | - | - | - | - | - | Unassigned | - |
| Otu09450 | 2 | k__Fungi(100);k__Fungi_unclassified(100);k__Fungi_unclassified(100);k__Fungi_unclassified(100);k__Fungi_unclassified(100);k__Fungi_unclassified(100);k__Fungi_unclassified(100); | - | - | - | - | - | - | - | Unassigned | - |
| Otu09451 | 2 | k__Fungi(100);k__Fungi_unclassified(100);k__Fungi_unclassified(100);k__Fungi_unclassified(100);k__Fungi_unclassified(100);k__Fungi_unclassified(100);k__Fungi_unclassified(100); | - | - | - | - | - | - | - | Unassigned | - |
| Otu09452 | 2 | k__Fungi(100);k__Fungi_unclassified(100);k__Fungi_unclassified(100);k__Fungi_unclassified(100);k__Fungi_unclassified(100);k__Fungi_unclassified(100);k__Fungi_unclassified(100); | - | - | - | - | - | - | - | Unassigned | - |
| Otu09453 | 2 | k__Fungi(100);k__Fungi_unclassified(100);k__Fungi_unclassified(100);k__Fungi_unclassified(100);k__Fungi_unclassified(100);k__Fungi_unclassified(100);k__Fungi_unclassified(100); | - | - | - | - | - | - | - | Unassigned | - |
| Otu09457 | 2 | k__Fungi(100);k__Fungi_unclassified(100);k__Fungi_unclassified(100);k__Fungi_unclassified(100);k__Fungi_unclassified(100);k__Fungi_unclassified(100);k__Fungi_unclassified(100); | - | - | - | - | - | - | - | Unassigned | - |
| Otu09458 | 2 | k__Fungi(100);k__Fungi_unclassified(100);k__Fungi_unclassified(100);k__Fungi_unclassified(100);k__Fungi_unclassified(100);k__Fungi_unclassified(100);k__Fungi_unclassified(100); | - | - | - | - | - | - | - | Unassigned | - |
| Otu09459 | 2 | k__Fungi(100);k__Fungi_unclassified(100);k__Fungi_unclassified(100);k__Fungi_unclassified(100);k__Fungi_unclassified(100);k__Fungi_unclassified(100);k__Fungi_unclassified(100); | - | - | - | - | - | - | - | Unassigned | - |
| Otu09460 | 2 | k__Fungi(100);k__Fungi_unclassified(100);k__Fungi_unclassified(100);k__Fungi_unclassified(100);k__Fungi_unclassified(100);k__Fungi_unclassified(100);k__Fungi_unclassified(100); | - | - | - | - | - | - | - | Unassigned | - |
| Otu09461 | 2 | k__Fungi(100);k__Fungi_unclassified(100);k__Fungi_unclassified(100);k__Fungi_unclassified(100);k__Fungi_unclassified(100);k__Fungi_unclassified(100);k__Fungi_unclassified(100); | - | - | - | - | - | - | - | Unassigned | - |
| Otu09462 | 2 | k__Fungi(100);p__Ascomycota(100);p__Ascomycota_unclassified(100);p__Ascomycota_unclassified(100);p__Ascomycota_unclassified(100);p__Ascomycota_unclassified(100);p__Ascomycota_unclassified(100); | - | - | - | - | - | - | - | Unassigned | - |
| Otu09463 | 2 | k__Fungi(100);p__Ascomycota(100);p__Ascomycota_unclassified(100);p__Ascomycota_unclassified(100);p__Ascomycota_unclassified(100);p__Ascomycota_unclassified(100);p__Ascomycota_unclassified(100); | - | - | - | - | - | - | - | Unassigned | - |
| Otu09466 | 2 | k__Fungi(100);k__Fungi_unclassified(100);k__Fungi_unclassified(100);k__Fungi_unclassified(100);k__Fungi_unclassified(100);k__Fungi_unclassified(100);k__Fungi_unclassified(100); | - | - | - | - | - | - | - | Unassigned | - |
| Otu09468 | 2 | k__Fungi(100);k__Fungi_unclassified(100);k__Fungi_unclassified(100);k__Fungi_unclassified(100);k__Fungi_unclassified(100);k__Fungi_unclassified(100);k__Fungi_unclassified(100); | - | - | - | - | - | - | - | Unassigned | - |
| Otu09469 | 2 | k__Fungi(100);k__Fungi_unclassified(100);k__Fungi_unclassified(100);k__Fungi_unclassified(100);k__Fungi_unclassified(100);k__Fungi_unclassified(100);k__Fungi_unclassified(100); | - | - | - | - | - | - | - | Unassigned | - |
| Otu09472 | 2 | k__Fungi(100);k__Fungi_unclassified(100);k__Fungi_unclassified(100);k__Fungi_unclassified(100);k__Fungi_unclassified(100);k__Fungi_unclassified(100);k__Fungi_unclassified(100); | - | - | - | - | - | - | - | Unassigned | - |
| Otu09473 | 2 | k__Fungi(100);p__Ascomycota(100);p__Ascomycota_unclassified(100);p__Ascomycota_unclassified(100);p__Ascomycota_unclassified(100);p__Ascomycota_unclassified(100);p__Ascomycota_unclassified(100); | - | - | - | - | - | - | - | Unassigned | - |
| Otu09474 | 2 | k__Fungi(100);p__Ascomycota(100);p__Ascomycota_unclassified(100);p__Ascomycota_unclassified(100);p__Ascomycota_unclassified(100);p__Ascomycota_unclassified(100);p__Ascomycota_unclassified(100); | - | - | - | - | - | - | - | Unassigned | - |
| Otu09475 | 2 | k__Fungi(100);k__Fungi_unclassified(100);k__Fungi_unclassified(100);k__Fungi_unclassified(100);k__Fungi_unclassified(100);k__Fungi_unclassified(100);k__Fungi_unclassified(100); | - | - | - | - | - | - | - | Unassigned | - |
| Otu09476 | 2 | k__Fungi(100);p__Ascomycota(100);c__Dothideomycetes(100);o__Pleosporales(100);o__Pleosporales_unclassified(100);o__Pleosporales_unclassified(100);o__Pleosporales_unclassified(100); | - | - | - | - | - | - | - | Unassigned | - |
| Otu09477 | 2 | k__Fungi(100);p__Ascomycota(100);p__Ascomycota_unclassified(100);p__Ascomycota_unclassified(100);p__Ascomycota_unclassified(100);p__Ascomycota_unclassified(100);p__Ascomycota_unclassified(100); | - | - | - | - | - | - | - | Unassigned | - |
| Otu09479 | 2 | k__Fungi(100);k__Fungi_unclassified(100);k__Fungi_unclassified(100);k__Fungi_unclassified(100);k__Fungi_unclassified(100);k__Fungi_unclassified(100);k__Fungi_unclassified(100); | - | - | - | - | - | - | - | Unassigned | - |
| Otu09485 | 2 | k__Fungi(100);p__Ascomycota(100);p__Ascomycota_unclassified(100);p__Ascomycota_unclassified(100);p__Ascomycota_unclassified(100);p__Ascomycota_unclassified(100);p__Ascomycota_unclassified(100); | - | - | - | - | - | - | - | Unassigned | - |
| Otu09493 | 2 | k__Fungi(100);k__Fungi_unclassified(100);k__Fungi_unclassified(100);k__Fungi_unclassified(100);k__Fungi_unclassified(100);k__Fungi_unclassified(100);k__Fungi_unclassified(100); | - | - | - | - | - | - | - | Unassigned | - |
| Otu09495 | 2 | k__Fungi(100);p__Ascomycota(100);p__Ascomycota_unclassified(100);p__Ascomycota_unclassified(100);p__Ascomycota_unclassified(100);p__Ascomycota_unclassified(100);p__Ascomycota_unclassified(100); | - | - | - | - | - | - | - | Unassigned | - |
| Otu09497 | 2 | k__Fungi(100);k__Fungi_unclassified(100);k__Fungi_unclassified(100);k__Fungi_unclassified(100);k__Fungi_unclassified(100);k__Fungi_unclassified(100);k__Fungi_unclassified(100); | - | - | - | - | - | - | - | Unassigned | - |
| Otu09499 | 2 | k__Fungi(100);p__Ascomycota(100);p__Ascomycota_unclassified(100);p__Ascomycota_unclassified(100);p__Ascomycota_unclassified(100);p__Ascomycota_unclassified(100);p__Ascomycota_unclassified(100); | - | - | - | - | - | - | - | Unassigned | - |
| Otu09505 | 2 | k__Fungi(100);k__Fungi_unclassified(100);k__Fungi_unclassified(100);k__Fungi_unclassified(100);k__Fungi_unclassified(100);k__Fungi_unclassified(100);k__Fungi_unclassified(100); | - | - | - | - | - | - | - | Unassigned | - |
| Otu09507 | 2 | k__Fungi(100);k__Fungi_unclassified(100);k__Fungi_unclassified(100);k__Fungi_unclassified(100);k__Fungi_unclassified(100);k__Fungi_unclassified(100);k__Fungi_unclassified(100); | - | - | - | - | - | - | - | Unassigned | - |
| Otu09510 | 2 | k__Fungi(100);k__Fungi_unclassified(100);k__Fungi_unclassified(100);k__Fungi_unclassified(100);k__Fungi_unclassified(100);k__Fungi_unclassified(100);k__Fungi_unclassified(100); | - | - | - | - | - | - | - | Unassigned | - |
| Otu09511 | 2 | k__Fungi(100);p__Ascomycota(100);p__Ascomycota_unclassified(100);p__Ascomycota_unclassified(100);p__Ascomycota_unclassified(100);p__Ascomycota_unclassified(100);p__Ascomycota_unclassified(100); | - | - | - | - | - | - | - | Unassigned | - |
| Otu09512 | 2 | k__Fungi(100);k__Fungi_unclassified(100);k__Fungi_unclassified(100);k__Fungi_unclassified(100);k__Fungi_unclassified(100);k__Fungi_unclassified(100);k__Fungi_unclassified(100); | - | - | - | - | - | - | - | Unassigned | - |
| Otu09520 | 2 | k__Fungi(100);k__Fungi_unclassified(100);k__Fungi_unclassified(100);k__Fungi_unclassified(100);k__Fungi_unclassified(100);k__Fungi_unclassified(100);k__Fungi_unclassified(100); | - | - | - | - | - | - | - | Unassigned | - |
| Otu09522 | 2 | k__Fungi(100);p__Ascomycota(100);p__Ascomycota_unclassified(100);p__Ascomycota_unclassified(100);p__Ascomycota_unclassified(100);p__Ascomycota_unclassified(100);p__Ascomycota_unclassified(100); | - | - | - | - | - | - | - | Unassigned | - |
| Otu09523 | 2 | k__Fungi(100);p__Ascomycota(100);p__Ascomycota_unclassified(100);p__Ascomycota_unclassified(100);p__Ascomycota_unclassified(100);p__Ascomycota_unclassified(100);p__Ascomycota_unclassified(100); | - | - | - | - | - | - | - | Unassigned | - |
| Otu09528 | 2 | k__Fungi(100);k__Fungi_unclassified(100);k__Fungi_unclassified(100);k__Fungi_unclassified(100);k__Fungi_unclassified(100);k__Fungi_unclassified(100);k__Fungi_unclassified(100); | - | - | - | - | - | - | - | Unassigned | - |
| Otu09531 | 2 | k__Fungi(100);k__Fungi_unclassified(100);k__Fungi_unclassified(100);k__Fungi_unclassified(100);k__Fungi_unclassified(100);k__Fungi_unclassified(100);k__Fungi_unclassified(100); | - | - | - | - | - | - | - | Unassigned | - |
| Otu09532 | 2 | k__Fungi(100);k__Fungi_unclassified(100);k__Fungi_unclassified(100);k__Fungi_unclassified(100);k__Fungi_unclassified(100);k__Fungi_unclassified(100);k__Fungi_unclassified(100); | - | - | - | - | - | - | - | Unassigned | - |
| Otu09533 | 2 | k__Fungi(100);k__Fungi_unclassified(100);k__Fungi_unclassified(100);k__Fungi_unclassified(100);k__Fungi_unclassified(100);k__Fungi_unclassified(100);k__Fungi_unclassified(100); | - | - | - | - | - | - | - | Unassigned | - |
| Otu09534 | 2 | k__Fungi(100);k__Fungi_unclassified(100);k__Fungi_unclassified(100);k__Fungi_unclassified(100);k__Fungi_unclassified(100);k__Fungi_unclassified(100);k__Fungi_unclassified(100); | - | - | - | - | - | - | - | Unassigned | - |
| Otu09535 | 2 | k__Fungi(100);k__Fungi_unclassified(100);k__Fungi_unclassified(100);k__Fungi_unclassified(100);k__Fungi_unclassified(100);k__Fungi_unclassified(100);k__Fungi_unclassified(100); | - | - | - | - | - | - | - | Unassigned | - |
| Otu09537 | 2 | k__Fungi(100);p__Ascomycota(100);c__Dothideomycetes(100);c__Dothideomycetes_unclassified(100);c__Dothideomycetes_unclassified(100);c__Dothideomycetes_unclassified(100);c__Dothideomycetes_unclassified(100); | - | - | - | - | - | - | - | Unassigned | - |
| Otu09538 | 2 | k__Fungi(100);p__Ascomycota(100);p__Ascomycota_unclassified(100);p__Ascomycota_unclassified(100);p__Ascomycota_unclassified(100);p__Ascomycota_unclassified(100);p__Ascomycota_unclassified(100); | - | - | - | - | - | - | - | Unassigned | - |
| Otu09539 | 2 | k__Fungi(100);k__Fungi_unclassified(100);k__Fungi_unclassified(100);k__Fungi_unclassified(100);k__Fungi_unclassified(100);k__Fungi_unclassified(100);k__Fungi_unclassified(100); | - | - | - | - | - | - | - | Unassigned | - |
| Otu09540 | 2 | k__Fungi(100);k__Fungi_unclassified(100);k__Fungi_unclassified(100);k__Fungi_unclassified(100);k__Fungi_unclassified(100);k__Fungi_unclassified(100);k__Fungi_unclassified(100); | - | - | - | - | - | - | - | Unassigned | - |
| Otu09542 | 2 | k__Fungi(100);k__Fungi_unclassified(100);k__Fungi_unclassified(100);k__Fungi_unclassified(100);k__Fungi_unclassified(100);k__Fungi_unclassified(100);k__Fungi_unclassified(100); | - | - | - | - | - | - | - | Unassigned | - |
| Otu09544 | 2 | k__Fungi(100);k__Fungi_unclassified(100);k__Fungi_unclassified(100);k__Fungi_unclassified(100);k__Fungi_unclassified(100);k__Fungi_unclassified(100);k__Fungi_unclassified(100); | - | - | - | - | - | - | - | Unassigned | - |
| Otu09545 | 2 | k__Fungi(100);k__Fungi_unclassified(100);k__Fungi_unclassified(100);k__Fungi_unclassified(100);k__Fungi_unclassified(100);k__Fungi_unclassified(100);k__Fungi_unclassified(100); | - | - | - | - | - | - | - | Unassigned | - |
| Otu09547 | 2 | k__Fungi(100);k__Fungi_unclassified(100);k__Fungi_unclassified(100);k__Fungi_unclassified(100);k__Fungi_unclassified(100);k__Fungi_unclassified(100);k__Fungi_unclassified(100); | - | - | - | - | - | - | - | Unassigned | - |
| Otu09550 | 2 | k__Fungi(100);p__Ascomycota(100);p__Ascomycota_unclassified(100);p__Ascomycota_unclassified(100);p__Ascomycota_unclassified(100);p__Ascomycota_unclassified(100);p__Ascomycota_unclassified(100); | - | - | - | - | - | - | - | Unassigned | - |
| Otu09553 | 2 | k__Fungi(100);k__Fungi_unclassified(100);k__Fungi_unclassified(100);k__Fungi_unclassified(100);k__Fungi_unclassified(100);k__Fungi_unclassified(100);k__Fungi_unclassified(100); | - | - | - | - | - | - | - | Unassigned | - |
| Otu09554 | 2 | k__Fungi(100);k__Fungi_unclassified(100);k__Fungi_unclassified(100);k__Fungi_unclassified(100);k__Fungi_unclassified(100);k__Fungi_unclassified(100);k__Fungi_unclassified(100); | - | - | - | - | - | - | - | Unassigned | - |
| Otu09560 | 2 | k__Fungi(100);k__Fungi_unclassified(100);k__Fungi_unclassified(100);k__Fungi_unclassified(100);k__Fungi_unclassified(100);k__Fungi_unclassified(100);k__Fungi_unclassified(100); | - | - | - | - | - | - | - | Unassigned | - |
| Otu09562 | 2 | k__Fungi(100);p__Ascomycota(100);p__Ascomycota_unclassified(100);p__Ascomycota_unclassified(100);p__Ascomycota_unclassified(100);p__Ascomycota_unclassified(100);p__Ascomycota_unclassified(100); | - | - | - | - | - | - | - | Unassigned | - |
| Otu09563 | 2 | k__Fungi(100);p__Ascomycota(100);c__Leotiomycetes(100);o__Helotiales(100);o__Helotiales_unclassified(100);o__Helotiales_unclassified(100);o__Helotiales_unclassified(100); | - | - | - | - | - | - | - | Unassigned | - |
| Otu09565 | 2 | k__Fungi(100);k__Fungi_unclassified(100);k__Fungi_unclassified(100);k__Fungi_unclassified(100);k__Fungi_unclassified(100);k__Fungi_unclassified(100);k__Fungi_unclassified(100); | - | - | - | - | - | - | - | Unassigned | - |
| Otu09566 | 2 | k__Fungi(100);p__Ascomycota(100);p__Ascomycota_unclassified(100);p__Ascomycota_unclassified(100);p__Ascomycota_unclassified(100);p__Ascomycota_unclassified(100);p__Ascomycota_unclassified(100); | - | - | - | - | - | - | - | Unassigned | - |
| Otu09567 | 2 | k__Fungi(100);p__Ascomycota(100);p__Ascomycota_unclassified(100);p__Ascomycota_unclassified(100);p__Ascomycota_unclassified(100);p__Ascomycota_unclassified(100);p__Ascomycota_unclassified(100); | - | - | - | - | - | - | - | Unassigned | - |
| Otu09569 | 2 | k__Fungi(100);k__Fungi_unclassified(100);k__Fungi_unclassified(100);k__Fungi_unclassified(100);k__Fungi_unclassified(100);k__Fungi_unclassified(100);k__Fungi_unclassified(100); | - | - | - | - | - | - | - | Unassigned | - |
| Otu09570 | 2 | k__Fungi(100);p__Ascomycota(100);p__Ascomycota_unclassified(100);p__Ascomycota_unclassified(100);p__Ascomycota_unclassified(100);p__Ascomycota_unclassified(100);p__Ascomycota_unclassified(100); | - | - | - | - | - | - | - | Unassigned | - |
| Otu09578 | 2 | k__Fungi(100);k__Fungi_unclassified(100);k__Fungi_unclassified(100);k__Fungi_unclassified(100);k__Fungi_unclassified(100);k__Fungi_unclassified(100);k__Fungi_unclassified(100); | - | - | - | - | - | - | - | Unassigned | - |
| Otu09580 | 2 | k__Fungi(100);k__Fungi_unclassified(100);k__Fungi_unclassified(100);k__Fungi_unclassified(100);k__Fungi_unclassified(100);k__Fungi_unclassified(100);k__Fungi_unclassified(100); | - | - | - | - | - | - | - | Unassigned | - |
| Otu09581 | 2 | k__Fungi(100);k__Fungi_unclassified(100);k__Fungi_unclassified(100);k__Fungi_unclassified(100);k__Fungi_unclassified(100);k__Fungi_unclassified(100);k__Fungi_unclassified(100); | - | - | - | - | - | - | - | Unassigned | - |
| Otu09583 | 2 | k__Fungi(100);k__Fungi_unclassified(100);k__Fungi_unclassified(100);k__Fungi_unclassified(100);k__Fungi_unclassified(100);k__Fungi_unclassified(100);k__Fungi_unclassified(100); | - | - | - | - | - | - | - | Unassigned | - |
| Otu09590 | 2 | k__Fungi(100);k__Fungi_unclassified(100);k__Fungi_unclassified(100);k__Fungi_unclassified(100);k__Fungi_unclassified(100);k__Fungi_unclassified(100);k__Fungi_unclassified(100); | - | - | - | - | - | - | - | Unassigned | - |
| Otu09591 | 2 | k__Fungi(100);k__Fungi_unclassified(100);k__Fungi_unclassified(100);k__Fungi_unclassified(100);k__Fungi_unclassified(100);k__Fungi_unclassified(100);k__Fungi_unclassified(100); | - | - | - | - | - | - | - | Unassigned | - |
| Otu09597 | 2 | k__Fungi(100);k__Fungi_unclassified(100);k__Fungi_unclassified(100);k__Fungi_unclassified(100);k__Fungi_unclassified(100);k__Fungi_unclassified(100);k__Fungi_unclassified(100); | - | - | - | - | - | - | - | Unassigned | - |
| Otu09598 | 2 | k__Fungi(100);p__Ascomycota(100);p__Ascomycota_unclassified(100);p__Ascomycota_unclassified(100);p__Ascomycota_unclassified(100);p__Ascomycota_unclassified(100);p__Ascomycota_unclassified(100); | - | - | - | - | - | - | - | Unassigned | - |
| Otu09599 | 2 | k__Fungi(100);p__Ascomycota(100);p__Ascomycota_unclassified(100);p__Ascomycota_unclassified(100);p__Ascomycota_unclassified(100);p__Ascomycota_unclassified(100);p__Ascomycota_unclassified(100); | - | - | - | - | - | - | - | Unassigned | - |
| Otu09601 | 2 | k__Fungi(100);p__Basidiomycota(100);p__Basidiomycota_unclassified(100);p__Basidiomycota_unclassified(100);p__Basidiomycota_unclassified(100);p__Basidiomycota_unclassified(100);p__Basidiomycota_unclassified(100); | - | - | - | - | - | - | - | Unassigned | - |
| Otu09603 | 2 | k__Fungi(100);k__Fungi_unclassified(100);k__Fungi_unclassified(100);k__Fungi_unclassified(100);k__Fungi_unclassified(100);k__Fungi_unclassified(100);k__Fungi_unclassified(100); | - | - | - | - | - | - | - | Unassigned | - |
| Otu09604 | 2 | k__Fungi(100);k__Fungi_unclassified(100);k__Fungi_unclassified(100);k__Fungi_unclassified(100);k__Fungi_unclassified(100);k__Fungi_unclassified(100);k__Fungi_unclassified(100); | - | - | - | - | - | - | - | Unassigned | - |
| Otu09606 | 2 | k__Fungi(100);p__Ascomycota(100);p__Ascomycota_unclassified(100);p__Ascomycota_unclassified(100);p__Ascomycota_unclassified(100);p__Ascomycota_unclassified(100);p__Ascomycota_unclassified(100); | - | - | - | - | - | - | - | Unassigned | - |
| Otu09607 | 2 | k__Fungi(100);k__Fungi_unclassified(100);k__Fungi_unclassified(100);k__Fungi_unclassified(100);k__Fungi_unclassified(100);k__Fungi_unclassified(100);k__Fungi_unclassified(100); | - | - | - | - | - | - | - | Unassigned | - |
| Otu09610 | 2 | k__Fungi(100);p__Ascomycota(100);p__Ascomycota_unclassified(100);p__Ascomycota_unclassified(100);p__Ascomycota_unclassified(100);p__Ascomycota_unclassified(100);p__Ascomycota_unclassified(100); | - | - | - | - | - | - | - | Unassigned | - |
| Otu09615 | 2 | k__Fungi(100);k__Fungi_unclassified(100);k__Fungi_unclassified(100);k__Fungi_unclassified(100);k__Fungi_unclassified(100);k__Fungi_unclassified(100);k__Fungi_unclassified(100); | - | - | - | - | - | - | - | Unassigned | - |
| Otu09620 | 2 | k__Fungi(100);k__Fungi_unclassified(100);k__Fungi_unclassified(100);k__Fungi_unclassified(100);k__Fungi_unclassified(100);k__Fungi_unclassified(100);k__Fungi_unclassified(100); | - | - | - | - | - | - | - | Unassigned | - |
| Otu09623 | 2 | k__Fungi(100);k__Fungi_unclassified(100);k__Fungi_unclassified(100);k__Fungi_unclassified(100);k__Fungi_unclassified(100);k__Fungi_unclassified(100);k__Fungi_unclassified(100); | - | - | - | - | - | - | - | Unassigned | - |
| Otu09626 | 2 | k__Fungi(100);k__Fungi_unclassified(100);k__Fungi_unclassified(100);k__Fungi_unclassified(100);k__Fungi_unclassified(100);k__Fungi_unclassified(100);k__Fungi_unclassified(100); | - | - | - | - | - | - | - | Unassigned | - |
| Otu09627 | 2 | k__Fungi(100);k__Fungi_unclassified(100);k__Fungi_unclassified(100);k__Fungi_unclassified(100);k__Fungi_unclassified(100);k__Fungi_unclassified(100);k__Fungi_unclassified(100); | - | - | - | - | - | - | - | Unassigned | - |
| Otu09628 | 2 | k__Fungi(100);p__Ascomycota(100);p__Ascomycota_unclassified(100);p__Ascomycota_unclassified(100);p__Ascomycota_unclassified(100);p__Ascomycota_unclassified(100);p__Ascomycota_unclassified(100); | - | - | - | - | - | - | - | Unassigned | - |
| Otu09630 | 2 | k__Fungi(100);p__Ascomycota(100);p__Ascomycota_unclassified(100);p__Ascomycota_unclassified(100);p__Ascomycota_unclassified(100);p__Ascomycota_unclassified(100);p__Ascomycota_unclassified(100); | - | - | - | - | - | - | - | Unassigned | - |
| Otu09631 | 2 | k__Fungi(100);p__Ascomycota(100);p__Ascomycota_unclassified(100);p__Ascomycota_unclassified(100);p__Ascomycota_unclassified(100);p__Ascomycota_unclassified(100);p__Ascomycota_unclassified(100); | - | - | - | - | - | - | - | Unassigned | - |
| Otu09632 | 2 | k__Fungi(100);k__Fungi_unclassified(100);k__Fungi_unclassified(100);k__Fungi_unclassified(100);k__Fungi_unclassified(100);k__Fungi_unclassified(100);k__Fungi_unclassified(100); | - | - | - | - | - | - | - | Unassigned | - |
| Otu09635 | 2 | k__Fungi(100);p__Ascomycota(100);p__Ascomycota_unclassified(100);p__Ascomycota_unclassified(100);p__Ascomycota_unclassified(100);p__Ascomycota_unclassified(100);p__Ascomycota_unclassified(100); | - | - | - | - | - | - | - | Unassigned | - |
| Otu09636 | 2 | k__Fungi(100);k__Fungi_unclassified(100);k__Fungi_unclassified(100);k__Fungi_unclassified(100);k__Fungi_unclassified(100);k__Fungi_unclassified(100);k__Fungi_unclassified(100); | - | - | - | - | - | - | - | Unassigned | - |
| Otu09637 | 2 | k__Fungi(100);p__Ascomycota(100);p__Ascomycota_unclassified(100);p__Ascomycota_unclassified(100);p__Ascomycota_unclassified(100);p__Ascomycota_unclassified(100);p__Ascomycota_unclassified(100); | - | - | - | - | - | - | - | Unassigned | - |
| Otu09639 | 2 | k__Fungi(100);k__Fungi_unclassified(100);k__Fungi_unclassified(100);k__Fungi_unclassified(100);k__Fungi_unclassified(100);k__Fungi_unclassified(100);k__Fungi_unclassified(100); | - | - | - | - | - | - | - | Unassigned | - |
| Otu09644 | 2 | k__Fungi(100);k__Fungi_unclassified(100);k__Fungi_unclassified(100);k__Fungi_unclassified(100);k__Fungi_unclassified(100);k__Fungi_unclassified(100);k__Fungi_unclassified(100); | - | - | - | - | - | - | - | Unassigned | - |
| Otu09645 | 2 | k__Fungi(100);k__Fungi_unclassified(100);k__Fungi_unclassified(100);k__Fungi_unclassified(100);k__Fungi_unclassified(100);k__Fungi_unclassified(100);k__Fungi_unclassified(100); | - | - | - | - | - | - | - | Unassigned | - |
| Otu09646 | 2 | k__Fungi(100);k__Fungi_unclassified(100);k__Fungi_unclassified(100);k__Fungi_unclassified(100);k__Fungi_unclassified(100);k__Fungi_unclassified(100);k__Fungi_unclassified(100); | - | - | - | - | - | - | - | Unassigned | - |
| Otu09647 | 2 | k__Fungi(100);k__Fungi_unclassified(100);k__Fungi_unclassified(100);k__Fungi_unclassified(100);k__Fungi_unclassified(100);k__Fungi_unclassified(100);k__Fungi_unclassified(100); | - | - | - | - | - | - | - | Unassigned | - |
| Otu09648 | 2 | k__Fungi(100);k__Fungi_unclassified(100);k__Fungi_unclassified(100);k__Fungi_unclassified(100);k__Fungi_unclassified(100);k__Fungi_unclassified(100);k__Fungi_unclassified(100); | - | - | - | - | - | - | - | Unassigned | - |
| Otu09649 | 2 | k__Fungi(100);p__Ascomycota(100);p__Ascomycota_unclassified(100);p__Ascomycota_unclassified(100);p__Ascomycota_unclassified(100);p__Ascomycota_unclassified(100);p__Ascomycota_unclassified(100); | - | - | - | - | - | - | - | Unassigned | - |
| Otu09651 | 2 | k__Fungi(100);k__Fungi_unclassified(100);k__Fungi_unclassified(100);k__Fungi_unclassified(100);k__Fungi_unclassified(100);k__Fungi_unclassified(100);k__Fungi_unclassified(100); | - | - | - | - | - | - | - | Unassigned | - |
| Otu09653 | 2 | k__Fungi(100);k__Fungi_unclassified(100);k__Fungi_unclassified(100);k__Fungi_unclassified(100);k__Fungi_unclassified(100);k__Fungi_unclassified(100);k__Fungi_unclassified(100); | - | - | - | - | - | - | - | Unassigned | - |
| Otu09655 | 2 | k__Fungi(100);k__Fungi_unclassified(100);k__Fungi_unclassified(100);k__Fungi_unclassified(100);k__Fungi_unclassified(100);k__Fungi_unclassified(100);k__Fungi_unclassified(100); | - | - | - | - | - | - | - | Unassigned | - |

| | | | | | | | | | | | |
|---|---|---|---|---|---|---|---|---|---|---|---|
| Otu09658 | 2 | k__Fungi(100);k__Fungi_unclassified(100);k__Fungi_unclassified(100);k__Fungi_unclassified(100);k__Fungi_unclassified(100);k__Fungi_unclassified(100); | - | - | - | - | - | - | - | Unassigned | - |
| Otu09659 | 2 | k__Fungi(100);k__Fungi_unclassified(100);k__Fungi_unclassified(100);k__Fungi_unclassified(100);k__Fungi_unclassified(100);k__Fungi_unclassified(100); | - | - | - | - | - | - | - | Unassigned | - |
| Otu09660 | 2 | k__Fungi(100);p__Ascomycota(100);p__Ascomycota_unclassified(100);p__Ascomycota_unclassified(100);p__Ascomycota_unclassified(100);p__Ascomycota_unclassified(100);p__Ascomycota_unclassified(100); | - | - | - | - | - | - | - | Unassigned | - |
| Otu09666 | 2 | k__Fungi(100);k__Fungi_unclassified(100);k__Fungi_unclassified(100);k__Fungi_unclassified(100);k__Fungi_unclassified(100);k__Fungi_unclassified(100); | - | - | - | - | - | - | - | Unassigned | - |
| Otu09667 | 2 | k__Fungi(100);k__Fungi_unclassified(100);k__Fungi_unclassified(100);k__Fungi_unclassified(100);k__Fungi_unclassified(100);k__Fungi_unclassified(100); | - | - | - | - | - | - | - | Unassigned | - |
| Otu09668 | 2 | k__Fungi(100);k__Fungi_unclassified(100);k__Fungi_unclassified(100);k__Fungi_unclassified(100);k__Fungi_unclassified(100);k__Fungi_unclassified(100); | - | - | - | - | - | - | - | Unassigned | - |
| Otu09669 | 2 | k__Fungi(100);p__Ascomycota(100);p__Ascomycota_unclassified(100);p__Ascomycota_unclassified(100);p__Ascomycota_unclassified(100);p__Ascomycota_unclassified(100);p__Ascomycota_unclassified(100); | - | - | - | - | - | - | - | Unassigned | - |
| Otu09670 | 2 | k__Fungi(100);k__Fungi_unclassified(100);k__Fungi_unclassified(100);k__Fungi_unclassified(100);k__Fungi_unclassified(100);k__Fungi_unclassified(100); | - | - | - | - | - | - | - | Unassigned | - |
| Otu09671 | 2 | k__Fungi(100);p__Ascomycota(100);p__Ascomycota_unclassified(100);p__Ascomycota_unclassified(100);p__Ascomycota_unclassified(100);p__Ascomycota_unclassified(100);p__Ascomycota_unclassified(100); | - | - | - | - | - | - | - | Unassigned | - |
| Otu09672 | 2 | k__Fungi(100);p__Ascomycota(100);p__Ascomycota_unclassified(100);p__Ascomycota_unclassified(100);p__Ascomycota_unclassified(100);p__Ascomycota_unclassified(100);p__Ascomycota_unclassified(100); | - | - | - | - | - | - | - | Unassigned | - |
| Otu09674 | 2 | k__Fungi(100);k__Fungi_unclassified(100);k__Fungi_unclassified(100);k__Fungi_unclassified(100);k__Fungi_unclassified(100);k__Fungi_unclassified(100); | - | - | - | - | - | - | - | Unassigned | - |
| Otu09677 | 2 | k__Fungi(100);k__Fungi_unclassified(100);k__Fungi_unclassified(100);k__Fungi_unclassified(100);k__Fungi_unclassified(100);k__Fungi_unclassified(100); | - | - | - | - | - | - | - | Unassigned | - |
| Otu09679 | 2 | k__Fungi(100);k__Fungi_unclassified(100);k__Fungi_unclassified(100);k__Fungi_unclassified(100);k__Fungi_unclassified(100);k__Fungi_unclassified(100); | - | - | - | - | - | - | - | Unassigned | - |
| Otu09689 | 2 | k__Fungi(100);k__Fungi_unclassified(100);k__Fungi_unclassified(100);k__Fungi_unclassified(100);k__Fungi_unclassified(100);k__Fungi_unclassified(100); | - | - | - | - | - | - | - | Unassigned | - |
| Otu09691 | 2 | k__Fungi(100);k__Fungi_unclassified(100);k__Fungi_unclassified(100);k__Fungi_unclassified(100);k__Fungi_unclassified(100);k__Fungi_unclassified(100); | - | - | - | - | - | - | - | Unassigned | - |
| Otu09693 | 2 | k__Fungi(100);k__Fungi_unclassified(100);k__Fungi_unclassified(100);k__Fungi_unclassified(100);k__Fungi_unclassified(100);k__Fungi_unclassified(100); | - | - | - | - | - | - | - | Unassigned | - |
| Otu09694 | 2 | k__Fungi(100);k__Fungi_unclassified(100);k__Fungi_unclassified(100);k__Fungi_unclassified(100);k__Fungi_unclassified(100);k__Fungi_unclassified(100); | - | - | - | - | - | - | - | Unassigned | - |
| Otu09695 | 2 | k__Fungi(100);p__Ascomycota(100);p__Ascomycota_unclassified(100);p__Ascomycota_unclassified(100);p__Ascomycota_unclassified(100);p__Ascomycota_unclassified(100);p__Ascomycota_unclassified(100); | - | - | - | - | - | - | - | Unassigned | - |
| Otu09696 | 2 | k__Fungi(100);k__Fungi_unclassified(100);k__Fungi_unclassified(100);k__Fungi_unclassified(100);k__Fungi_unclassified(100);k__Fungi_unclassified(100); | - | - | - | - | - | - | - | Unassigned | - |
| Otu09698 | 2 | k__Fungi(100);k__Fungi_unclassified(100);k__Fungi_unclassified(100);k__Fungi_unclassified(100);k__Fungi_unclassified(100);k__Fungi_unclassified(100); | - | - | - | - | - | - | - | Unassigned | - |
| Otu09701 | 2 | k__Fungi(100);k__Fungi_unclassified(100);k__Fungi_unclassified(100);k__Fungi_unclassified(100);k__Fungi_unclassified(100);k__Fungi_unclassified(100); | - | - | - | - | - | - | - | Unassigned | - |
| Otu09704 | 2 | k__Fungi(100);k__Fungi_unclassified(100);k__Fungi_unclassified(100);k__Fungi_unclassified(100);k__Fungi_unclassified(100);k__Fungi_unclassified(100); | - | - | - | - | - | - | - | Unassigned | - |
| Otu09705 | 2 | k__Fungi(100);p__Ascomycota(100);p__Ascomycota_unclassified(100);p__Ascomycota_unclassified(100);p__Ascomycota_unclassified(100);p__Ascomycota_unclassified(100);p__Ascomycota_unclassified(100); | - | - | - | - | - | - | - | Unassigned | - |
| Otu09706 | 2 | k__Fungi(100);k__Fungi_unclassified(100);k__Fungi_unclassified(100);k__Fungi_unclassified(100);k__Fungi_unclassified(100);k__Fungi_unclassified(100); | - | - | - | - | - | - | - | Unassigned | - |
| Otu09707 | 2 | k__Fungi(100);k__Fungi_unclassified(100);k__Fungi_unclassified(100);k__Fungi_unclassified(100);k__Fungi_unclassified(100);k__Fungi_unclassified(100); | - | - | - | - | - | - | - | Unassigned | - |
| Otu09709 | 2 | k__Fungi(100);k__Fungi_unclassified(100);k__Fungi_unclassified(100);k__Fungi_unclassified(100);k__Fungi_unclassified(100);k__Fungi_unclassified(100); | - | - | - | - | - | - | - | Unassigned | - |
| Otu09712 | 2 | k__Fungi(100);k__Fungi_unclassified(100);k__Fungi_unclassified(100);k__Fungi_unclassified(100);k__Fungi_unclassified(100);k__Fungi_unclassified(100); | - | - | - | - | - | - | - | Unassigned | - |
| Otu09715 | 2 | k__Fungi(100);k__Fungi_unclassified(100);k__Fungi_unclassified(100);k__Fungi_unclassified(100);k__Fungi_unclassified(100);k__Fungi_unclassified(100); | - | - | - | - | - | - | - | Unassigned | - |
| Otu09723 | 2 | k__Fungi(100);k__Fungi_unclassified(100);k__Fungi_unclassified(100);k__Fungi_unclassified(100);k__Fungi_unclassified(100);k__Fungi_unclassified(100); | - | - | - | - | - | - | - | Unassigned | - |
| Otu09724 | 2 | k__Fungi(100);k__Fungi_unclassified(100);k__Fungi_unclassified(100);k__Fungi_unclassified(100);k__Fungi_unclassified(100);k__Fungi_unclassified(100); | - | - | - | - | - | - | - | Unassigned | - |
| Otu09725 | 2 | k__Fungi(100);k__Fungi_unclassified(100);k__Fungi_unclassified(100);k__Fungi_unclassified(100);k__Fungi_unclassified(100);k__Fungi_unclassified(100); | - | - | - | - | - | - | - | Unassigned | - |
| Otu09726 | 2 | k__Fungi(100);k__Fungi_unclassified(100);k__Fungi_unclassified(100);k__Fungi_unclassified(100);k__Fungi_unclassified(100);k__Fungi_unclassified(100); | - | - | - | - | - | - | - | Unassigned | - |
| Otu09727 | 2 | k__Fungi(100);k__Fungi_unclassified(100);k__Fungi_unclassified(100);k__Fungi_unclassified(100);k__Fungi_unclassified(100);k__Fungi_unclassified(100); | - | - | - | - | - | - | - | Unassigned | - |
| Otu09729 | 2 | k__Fungi(100);k__Fungi_unclassified(100);k__Fungi_unclassified(100);k__Fungi_unclassified(100);k__Fungi_unclassified(100);k__Fungi_unclassified(100); | - | - | - | - | - | - | - | Unassigned | - |
| Otu09732 | 2 | k__Fungi(100);p__Ascomycota(100);p__Ascomycota_unclassified(100);p__Ascomycota_unclassified(100);p__Ascomycota_unclassified(100);p__Ascomycota_unclassified(100);p__Ascomycota_unclassified(100); | - | - | - | - | - | - | - | Unassigned | - |
| Otu09738 | 2 | k__Fungi(100);k__Fungi_unclassified(100);k__Fungi_unclassified(100);k__Fungi_unclassified(100);k__Fungi_unclassified(100);k__Fungi_unclassified(100); | - | - | - | - | - | - | - | Unassigned | - |
| Otu09741 | 2 | k__Fungi(100);k__Fungi_unclassified(100);k__Fungi_unclassified(100);k__Fungi_unclassified(100);k__Fungi_unclassified(100);k__Fungi_unclassified(100); | - | - | - | - | - | - | - | Unassigned | - |
| Otu09745 | 2 | k__Fungi(100);k__Fungi_unclassified(100);k__Fungi_unclassified(100);k__Fungi_unclassified(100);k__Fungi_unclassified(100);k__Fungi_unclassified(100); | - | - | - | - | - | - | - | Unassigned | - |
| Otu09747 | 2 | k__Fungi(100);k__Fungi_unclassified(100);k__Fungi_unclassified(100);k__Fungi_unclassified(100);k__Fungi_unclassified(100);k__Fungi_unclassified(100); | - | - | - | - | - | - | - | Unassigned | - |
| Otu09750 | 2 | k__Fungi(100);k__Fungi_unclassified(100);k__Fungi_unclassified(100);k__Fungi_unclassified(100);k__Fungi_unclassified(100);k__Fungi_unclassified(100); | - | - | - | - | - | - | - | Unassigned | - |
| Otu09752 | 2 | k__Fungi(100);k__Fungi_unclassified(100);k__Fungi_unclassified(100);k__Fungi_unclassified(100);k__Fungi_unclassified(100);k__Fungi_unclassified(100); | - | - | - | - | - | - | - | Unassigned | - |
| Otu09754 | 2 | k__Fungi(100);k__Fungi_unclassified(100);k__Fungi_unclassified(100);k__Fungi_unclassified(100);k__Fungi_unclassified(100);k__Fungi_unclassified(100); | - | - | - | - | - | - | - | Unassigned | - |
| Otu09755 | 2 | k__Fungi(100);p__Ascomycota(100);p__Ascomycota_unclassified(100);p__Ascomycota_unclassified(100);p__Ascomycota_unclassified(100);p__Ascomycota_unclassified(100);p__Ascomycota_unclassified(100); | - | - | - | - | - | - | - | Unassigned | - |
| Otu09759 | 2 | k__Fungi(100);p__Ascomycota(100);p__Ascomycota_unclassified(100);p__Ascomycota_unclassified(100);p__Ascomycota_unclassified(100);p__Ascomycota_unclassified(100);p__Ascomycota_unclassified(100); | - | - | - | - | - | - | - | Unassigned | - |
| Otu09761 | 2 | k__Fungi(100);p__Ascomycota(100);p__Ascomycota_unclassified(100);p__Ascomycota_unclassified(100);p__Ascomycota_unclassified(100);p__Ascomycota_unclassified(100);p__Ascomycota_unclassified(100); | - | - | - | - | - | - | - | Unassigned | - |
| Otu09764 | 2 | k__Fungi(100);k__Fungi_unclassified(100);k__Fungi_unclassified(100);k__Fungi_unclassified(100);k__Fungi_unclassified(100);k__Fungi_unclassified(100); | - | - | - | - | - | - | - | Unassigned | - |
| Otu09766 | 2 | k__Fungi(100);p__Ascomycota(100);p__Ascomycota_unclassified(100);p__Ascomycota_unclassified(100);p__Ascomycota_unclassified(100);p__Ascomycota_unclassified(100);p__Ascomycota_unclassified(100); | - | - | - | - | - | - | - | Unassigned | - |
| Otu09773 | 2 | k__Fungi(100);k__Fungi_unclassified(100);k__Fungi_unclassified(100);k__Fungi_unclassified(100);k__Fungi_unclassified(100);k__Fungi_unclassified(100); | - | - | - | - | - | - | - | Unassigned | - |
| Otu09775 | 2 | k__Fungi(100);k__Fungi_unclassified(100);k__Fungi_unclassified(100);k__Fungi_unclassified(100);k__Fungi_unclassified(100);k__Fungi_unclassified(100); | - | - | - | - | - | - | - | Unassigned | - |
| Otu09776 | 2 | k__Fungi(100);k__Fungi_unclassified(100);k__Fungi_unclassified(100);k__Fungi_unclassified(100);k__Fungi_unclassified(100);k__Fungi_unclassified(100); | - | - | - | - | - | - | - | Unassigned | - |
| Otu09778 | 2 | k__Fungi(100);p__Ascomycota(100);p__Ascomycota_unclassified(100);p__Ascomycota_unclassified(100);p__Ascomycota_unclassified(100);p__Ascomycota_unclassified(100);p__Ascomycota_unclassified(100); | - | - | - | - | - | - | - | Unassigned | - |
| Otu09779 | 2 | k__Fungi(100);k__Fungi_unclassified(100);k__Fungi_unclassified(100);k__Fungi_unclassified(100);k__Fungi_unclassified(100);k__Fungi_unclassified(100); | - | - | - | - | - | - | - | Unassigned | - |
| Otu09783 | 2 | k__Fungi(100);k__Fungi_unclassified(100);k__Fungi_unclassified(100);k__Fungi_unclassified(100);k__Fungi_unclassified(100);k__Fungi_unclassified(100); | - | - | - | - | - | - | - | Unassigned | - |
| Otu09784 | 2 | k__Fungi(100);p__Ascomycota(100);p__Ascomycota_unclassified(100);p__Ascomycota_unclassified(100);p__Ascomycota_unclassified(100);p__Ascomycota_unclassified(100);p__Ascomycota_unclassified(100); | - | - | - | - | - | - | - | Unassigned | - |
| Otu09787 | 2 | k__Fungi(100);p__Ascomycota(100);p__Ascomycota_unclassified(100);p__Ascomycota_unclassified(100);p__Ascomycota_unclassified(100);p__Ascomycota_unclassified(100);p__Ascomycota_unclassified(100); | - | - | - | - | - | - | - | Unassigned | - |
| Otu09789 | 2 | k__Fungi(100);k__Fungi_unclassified(100);k__Fungi_unclassified(100);k__Fungi_unclassified(100);k__Fungi_unclassified(100);k__Fungi_unclassified(100); | - | - | - | - | - | - | - | Unassigned | - |
| Otu09795 | 2 | k__Fungi(100);k__Fungi_unclassified(100);k__Fungi_unclassified(100);k__Fungi_unclassified(100);k__Fungi_unclassified(100);k__Fungi_unclassified(100); | - | - | - | - | - | - | - | Unassigned | - |
| Otu09797 | 2 | k__Fungi(100);k__Fungi_unclassified(100);k__Fungi_unclassified(100);k__Fungi_unclassified(100);k__Fungi_unclassified(100);k__Fungi_unclassified(100); | - | - | - | - | - | - | - | Unassigned | - |
| Otu09801 | 2 | k__Fungi(100);k__Fungi_unclassified(100);k__Fungi_unclassified(100);k__Fungi_unclassified(100);k__Fungi_unclassified(100);k__Fungi_unclassified(100); | - | - | - | - | - | - | - | Unassigned | - |
| Otu09803 | 2 | k__Fungi(100);k__Fungi_unclassified(100);k__Fungi_unclassified(100);k__Fungi_unclassified(100);k__Fungi_unclassified(100);k__Fungi_unclassified(100); | - | - | - | - | - | - | - | Unassigned | - |
| Otu09806 | 2 | k__Fungi(100);k__Fungi_unclassified(100);k__Fungi_unclassified(100);k__Fungi_unclassified(100);k__Fungi_unclassified(100);k__Fungi_unclassified(100); | - | - | - | - | - | - | - | Unassigned | - |
| Otu09812 | 2 | k__Fungi(100);k__Fungi_unclassified(100);k__Fungi_unclassified(100);k__Fungi_unclassified(100);k__Fungi_unclassified(100);k__Fungi_unclassified(100); | - | - | - | - | - | - | - | Unassigned | - |
| Otu09816 | 2 | k__Fungi(100);p__Ascomycota(100);p__Ascomycota_unclassified(100);p__Ascomycota_unclassified(100);p__Ascomycota_unclassified(100);p__Ascomycota_unclassified(100);p__Ascomycota_unclassified(100); | - | - | - | - | - | - | - | Unassigned | - |
| Otu09823 | 2 | k__Fungi(100);k__Fungi_unclassified(100);k__Fungi_unclassified(100);k__Fungi_unclassified(100);k__Fungi_unclassified(100);k__Fungi_unclassified(100); | - | - | - | - | - | - | - | Unassigned | - |
| Otu09825 | 2 | k__Fungi(100);k__Fungi_unclassified(100);k__Fungi_unclassified(100);k__Fungi_unclassified(100);k__Fungi_unclassified(100);k__Fungi_unclassified(100); | - | - | - | - | - | - | - | Unassigned | - |
| Otu09828 | 2 | k__Fungi(100);k__Fungi_unclassified(100);k__Fungi_unclassified(100);k__Fungi_unclassified(100);k__Fungi_unclassified(100);k__Fungi_unclassified(100); | - | - | - | - | - | - | - | Unassigned | - |
| Otu09829 | 2 | k__Fungi(100);p__Ascomycota(100);p__Ascomycota_unclassified(100);p__Ascomycota_unclassified(100);p__Ascomycota_unclassified(100);p__Ascomycota_unclassified(100);p__Ascomycota_unclassified(100); | - | - | - | - | - | - | - | Unassigned | - |
| Otu09831 | 2 | k__Fungi(100);k__Fungi_unclassified(100);k__Fungi_unclassified(100);k__Fungi_unclassified(100);k__Fungi_unclassified(100);k__Fungi_unclassified(100); | - | - | - | - | - | - | - | Unassigned | - |
| Otu09840 | 2 | k__Fungi(100);k__Fungi_unclassified(100);k__Fungi_unclassified(100);k__Fungi_unclassified(100);k__Fungi_unclassified(100);k__Fungi_unclassified(100); | - | - | - | - | - | - | - | Unassigned | - |
| Otu09845 | 2 | k__Fungi(100);p__Basidiomycota(100);c__Agaricomycetes(100);o__Polyporales(100);o__Polyporales_unclassified(100);o__Polyporales_unclassified(100);o__Polyporales_unclassified(100); | - | - | - | - | - | - | - | Unassigned | - |
| Otu09849 | 2 | k__Fungi(100);k__Fungi_unclassified(100);k__Fungi_unclassified(100);k__Fungi_unclassified(100);k__Fungi_unclassified(100);k__Fungi_unclassified(100); | - | - | - | - | - | - | - | Unassigned | - |
| Otu09854 | 2 | k__Fungi(100);p__Ascomycota(100);p__Ascomycota_unclassified(100);p__Ascomycota_unclassified(100);p__Ascomycota_unclassified(100);p__Ascomycota_unclassified(100);p__Ascomycota_unclassified(100); | - | - | - | - | - | - | - | Unassigned | - |
| Otu09859 | 2 | k__Fungi(100);k__Fungi_unclassified(100);k__Fungi_unclassified(100);k__Fungi_unclassified(100);k__Fungi_unclassified(100);k__Fungi_unclassified(100); | - | - | - | - | - | - | - | Unassigned | - |
| Otu09861 | 2 | k__Fungi(100);k__Fungi_unclassified(100);k__Fungi_unclassified(100);k__Fungi_unclassified(100);k__Fungi_unclassified(100);k__Fungi_unclassified(100); | - | - | - | - | - | - | - | Unassigned | - |
| Otu09866 | 2 | k__Fungi(100);k__Fungi_unclassified(100);k__Fungi_unclassified(100);k__Fungi_unclassified(100);k__Fungi_unclassified(100);k__Fungi_unclassified(100); | - | - | - | - | - | - | - | Unassigned | - |
| Otu09869 | 2 | k__Fungi(100);k__Fungi_unclassified(100);k__Fungi_unclassified(100);k__Fungi_unclassified(100);k__Fungi_unclassified(100);k__Fungi_unclassified(100); | - | - | - | - | - | - | - | Unassigned | - |
| Otu09870 | 2 | k__Fungi(100);k__Fungi_unclassified(100);k__Fungi_unclassified(100);k__Fungi_unclassified(100);k__Fungi_unclassified(100);k__Fungi_unclassified(100); | - | - | - | - | - | - | - | Unassigned | - |
| Otu09872 | 2 | k__Fungi(100);k__Fungi_unclassified(100);k__Fungi_unclassified(100);k__Fungi_unclassified(100);k__Fungi_unclassified(100);k__Fungi_unclassified(100); | - | - | - | - | - | - | - | Unassigned | - |
| Otu09876 | 2 | k__Fungi(100);p__Ascomycota(100);c__Dothideomycetes(100);o__Capnodiales(100);o__Capnodiales_unclassified(100);o__Capnodiales_unclassified(100);o__Capnodiales_unclassified(100); | - | - | - | - | - | - | - | Unassigned | - |
| Otu09878 | 2 | k__Fungi(100);k__Fungi_unclassified(100);k__Fungi_unclassified(100);k__Fungi_unclassified(100);k__Fungi_unclassified(100);k__Fungi_unclassified(100); | - | - | - | - | - | - | - | Unassigned | - |
| Otu09879 | 2 | k__Fungi(100);p__Ascomycota(100);c__Dothideomycetes(100);o__Pleosporales(100);f__Pleosporales_family_Incertae_sedis(100);f__Pleosporales_family_Incertae_sedis_unclassified(100);f__Pleosporales_family_Incertae_sedis_unclassified(100); | - | - | - | - | - | - | - | Unassigned | - |
| Otu09881 | 2 | k__Fungi(100);p__Ascomycota(100);p__Ascomycota_unclassified(100);p__Ascomycota_unclassified(100);p__Ascomycota_unclassified(100);p__Ascomycota_unclassified(100);p__Ascomycota_unclassified(100); | - | - | - | - | - | - | - | Unassigned | - |
| Otu09882 | 2 | k__Fungi(100);p__Ascomycota(100);c__Leotiomycetes(100);c__Leotiomycetes_unclassified(100);c__Leotiomycetes_unclassified(100);c__Leotiomycetes_unclassified(100);c__Leotiomycetes_unclassified(100); | - | - | - | - | - | - | - | Unassigned | - |
| Otu09887 | 2 | k__Fungi(100);p__Ascomycota(100);p__Ascomycota_unclassified(100);p__Ascomycota_unclassified(100);p__Ascomycota_unclassified(100);p__Ascomycota_unclassified(100);p__Ascomycota_unclassified(100); | - | - | - | - | - | - | - | Unassigned | - |
| Otu09888 | 2 | k__Fungi(100);k__Fungi_unclassified(100);k__Fungi_unclassified(100);k__Fungi_unclassified(100);k__Fungi_unclassified(100);k__Fungi_unclassified(100); | - | - | - | - | - | - | - | Unassigned | - |
| Otu09889 | 2 | k__Fungi(100);p__Ascomycota(100);p__Ascomycota_unclassified(100);p__Ascomycota_unclassified(100);p__Ascomycota_unclassified(100);p__Ascomycota_unclassified(100);p__Ascomycota_unclassified(100); | - | - | - | - | - | - | - | Unassigned | - |
| Otu09891 | 2 | k__Fungi(100);p__Ascomycota(100);p__Ascomycota_unclassified(100);p__Ascomycota_unclassified(100);p__Ascomycota_unclassified(100);p__Ascomycota_unclassified(100);p__Ascomycota_unclassified(100); | - | - | - | - | - | - | - | Unassigned | - |
| Otu09892 | 2 | k__Fungi(100);k__Fungi_unclassified(100);k__Fungi_unclassified(100);k__Fungi_unclassified(100);k__Fungi_unclassified(100);k__Fungi_unclassified(100); | - | - | - | - | - | - | - | Unassigned | - |
| Otu09893 | 2 | k__Fungi(100);p__Ascomycota(100);p__Ascomycota_unclassified(100);p__Ascomycota_unclassified(100);p__Ascomycota_unclassified(100);p__Ascomycota_unclassified(100);p__Ascomycota_unclassified(100); | - | - | - | - | - | - | - | Unassigned | - |
| Otu09894 | 2 | k__Fungi(100);k__Fungi_unclassified(100);k__Fungi_unclassified(100);k__Fungi_unclassified(100);k__Fungi_unclassified(100);k__Fungi_unclassified(100); | - | - | - | - | - | - | - | Unassigned | - |
| Otu09895 | 2 | k__Fungi(100);k__Fungi_unclassified(100);k__Fungi_unclassified(100);k__Fungi_unclassified(100);k__Fungi_unclassified(100);k__Fungi_unclassified(100); | - | - | - | - | - | - | - | Unassigned | - |
| Otu09897 | 2 | k__Fungi(100);k__Fungi_unclassified(100);k__Fungi_unclassified(100);k__Fungi_unclassified(100);k__Fungi_unclassified(100);k__Fungi_unclassified(100); | - | - | - | - | - | - | - | Unassigned | - |
| Otu09898 | 2 | k__Fungi(100);p__Ascomycota(100);p__Ascomycota_unclassified(100);p__Ascomycota_unclassified(100);p__Ascomycota_unclassified(100);p__Ascomycota_unclassified(100);p__Ascomycota_unclassified(100); | - | - | - | - | - | - | - | Unassigned | - |
| Otu09901 | 2 | k__Fungi(100);p__Ascomycota(100);p__Ascomycota_unclassified(100);p__Ascomycota_unclassified(100);p__Ascomycota_unclassified(100);p__Ascomycota_unclassified(100);p__Ascomycota_unclassified(100); | - | - | - | - | - | - | - | Unassigned | - |
| Otu09902 | 2 | k__Fungi(100);k__Fungi_unclassified(100);k__Fungi_unclassified(100);k__Fungi_unclassified(100);k__Fungi_unclassified(100);k__Fungi_unclassified(100); | - | - | - | - | - | - | - | Unassigned | - |
| Otu09903 | 2 | k__Fungi(100);k__Fungi_unclassified(100);k__Fungi_unclassified(100);k__Fungi_unclassified(100);k__Fungi_unclassified(100);k__Fungi_unclassified(100); | - | - | - | - | - | - | - | Unassigned | - |
| Otu09906 | 2 | k__Fungi(100);k__Fungi_unclassified(100);k__Fungi_unclassified(100);k__Fungi_unclassified(100);k__Fungi_unclassified(100);k__Fungi_unclassified(100); | - | - | - | - | - | - | - | Unassigned | - |
| Otu09907 | 2 | k__Fungi(100);p__Ascomycota(100);p__Ascomycota_unclassified(100);p__Ascomycota_unclassified(100);p__Ascomycota_unclassified(100);p__Ascomycota_unclassified(100);p__Ascomycota_unclassified(100); | - | - | - | - | - | - | - | Unassigned | - |
| Otu09911 | 2 | k__Fungi(100);k__Fungi_unclassified(100);k__Fungi_unclassified(100);k__Fungi_unclassified(100);k__Fungi_unclassified(100);k__Fungi_unclassified(100); | - | - | - | - | - | - | - | Unassigned | - |
| Otu09912 | 2 | k__Fungi(100);k__Fungi_unclassified(100);k__Fungi_unclassified(100);k__Fungi_unclassified(100);k__Fungi_unclassified(100);k__Fungi_unclassified(100); | - | - | - | - | - | - | - | Unassigned | - |
| Otu09913 | 2 | k__Fungi(100);k__Fungi_unclassified(100);k__Fungi_unclassified(100);k__Fungi_unclassified(100);k__Fungi_unclassified(100);k__Fungi_unclassified(100); | - | - | - | - | - | - | - | Unassigned | - |
| Otu09914 | 2 | k__Fungi(100);k__Fungi_unclassified(100);k__Fungi_unclassified(100);k__Fungi_unclassified(100);k__Fungi_unclassified(100);k__Fungi_unclassified(100); | - | - | - | - | - | - | - | Unassigned | - |
| Otu09915 | 2 | k__Fungi(100);k__Fungi_unclassified(100);k__Fungi_unclassified(100);k__Fungi_unclassified(100);k__Fungi_unclassified(100);k__Fungi_unclassified(100); | - | - | - | - | - | - | - | Unassigned | - |
| Otu09918 | 2 | k__Fungi(100);k__Fungi_unclassified(100);k__Fungi_unclassified(100);k__Fungi_unclassified(100);k__Fungi_unclassified(100);k__Fungi_unclassified(100); | - | - | - | - | - | - | - | Unassigned | - |
| Otu09919 | 2 | k__Fungi(100);p__Ascomycota(100);p__Ascomycota_unclassified(100);p__Ascomycota_unclassified(100);p__Ascomycota_unclassified(100);p__Ascomycota_unclassified(100);p__Ascomycota_unclassified(100); | - | - | - | - | - | - | - | Unassigned | - |
| Otu09921 | 2 | k__Fungi(100);k__Fungi_unclassified(100);k__Fungi_unclassified(100);k__Fungi_unclassified(100);k__Fungi_unclassified(100);k__Fungi_unclassified(100); | - | - | - | - | - | - | - | Unassigned | - |
| Otu09922 | 2 | k__Fungi(100);p__Ascomycota(100);c__Dothideomycetes(100);o__Pleosporales(100);o__Pleosporales_unclassified(100);o__Pleosporales_unclassified(100);o__Pleosporales_unclassified(100); | - | - | - | - | - | - | - | Unassigned | - |
| Otu09925 | 2 | k__Fungi(100);k__Fungi_unclassified(100);k__Fungi_unclassified(100);k__Fungi_unclassified(100);k__Fungi_unclassified(100);k__Fungi_unclassified(100); | - | - | - | - | - | - | - | Unassigned | - |
| Otu09927 | 2 | k__Fungi(100);p__Ascomycota(100);p__Ascomycota_unclassified(100);p__Ascomycota_unclassified(100);p__Ascomycota_unclassified(100);p__Ascomycota_unclassified(100);p__Ascomycota_unclassified(100); | - | - | - | - | - | - | - | Unassigned | - |
| Otu09929 | 2 | k__Fungi(100);k__Fungi_unclassified(100);k__Fungi_unclassified(100);k__Fungi_unclassified(100);k__Fungi_unclassified(100);k__Fungi_unclassified(100); | - | - | - | - | - | - | - | Unassigned | - |
| Otu09932 | 2 | k__Fungi(100);p__Ascomycota(100);p__Ascomycota_unclassified(100);p__Ascomycota_unclassified(100);p__Ascomycota_unclassified(100);p__Ascomycota_unclassified(100);p__Ascomycota_unclassified(100); | - | - | - | - | - | - | - | Unassigned | - |
| Otu09933 | 2 | k__Fungi(100);k__Fungi_unclassified(100);k__Fungi_unclassified(100);k__Fungi_unclassified(100);k__Fungi_unclassified(100);k__Fungi_unclassified(100); | - | - | - | - | - | - | - | Unassigned | - |
| Otu09936 | 2 | k__Fungi(100);k__Fungi_unclassified(100);k__Fungi_unclassified(100);k__Fungi_unclassified(100);k__Fungi_unclassified(100);k__Fungi_unclassified(100); | - | - | - | - | - | - | - | Unassigned | - |
| Otu09937 | 2 | k__Fungi(100);p__Ascomycota(100);c__Sordariomycetes(100);o__Sordariomycetes_order_Incertae_sedis(100);o__Sordariomycetes_order_Incertae_sedis_unclassified(100);o__Sordariomycetes_order_Incertae_sedis_unclassified(100);o__Sordariomycetes_order_Incertae_sedis_uncl | - | - | - | - | - | - | - | Unassigned | - |
| Otu09938 | 2 | k__Fungi(100);k__Fungi_unclassified(100);k__Fungi_unclassified(100);k__Fungi_unclassified(100);k__Fungi_unclassified(100);k__Fungi_unclassified(100); | - | - | - | - | - | - | - | Unassigned | - |
| Otu09940 | 2 | k__Fungi(100);k__Fungi_unclassified(100);k__Fungi_unclassified(100);k__Fungi_unclassified(100);k__Fungi_unclassified(100);k__Fungi_unclassified(100); | - | - | - | - | - | - | - | Unassigned | - |
| Otu09942 | 2 | k__Fungi(100);k__Fungi_unclassified(100);k__Fungi_unclassified(100);k__Fungi_unclassified(100);k__Fungi_unclassified(100);k__Fungi_unclassified(100); | - | - | - | - | - | - | - | Unassigned | - |
| Otu09944 | 2 | k__Fungi(100);k__Fungi_unclassified(100);k__Fungi_unclassified(100);k__Fungi_unclassified(100);k__Fungi_unclassified(100);k__Fungi_unclassified(100); | - | - | - | - | - | - | - | Unassigned | - |
| Otu09945 | 2 | k__Fungi(100);p__Ascomycota(100);p__Ascomycota_unclassified(100);p__Ascomycota_unclassified(100);p__Ascomycota_unclassified(100);p__Ascomycota_unclassified(100);p__Ascomycota_unclassified(100); | - | - | - | - | - | - | - | Unassigned | - |
| Otu09946 | 2 | k__Fungi(100);k__Fungi_unclassified(100);k__Fungi_unclassified(100);k__Fungi_unclassified(100);k__Fungi_unclassified(100);k__Fungi_unclassified(100); | - | - | - | - | - | - | - | Unassigned | - |
| Otu09947 | 2 | k__Fungi(100);k__Fungi_unclassified(100);k__Fungi_unclassified(100);k__Fungi_unclassified(100);k__Fungi_unclassified(100);k__Fungi_unclassified(100); | - | - | - | - | - | - | - | Unassigned | - |
| Otu09948 | 2 | k__Fungi(100);p__Ascomycota(100);p__Ascomycota_unclassified(100);p__Ascomycota_unclassified(100);p__Ascomycota_unclassified(100);p__Ascomycota_unclassified(100);p__Ascomycota_unclassified(100); | - | - | - | - | - | - | - | Unassigned | - |
| Otu09954 | 2 | k__Fungi(100);k__Fungi_unclassified(100);k__Fungi_unclassified(100);k__Fungi_unclassified(100);k__Fungi_unclassified(100);k__Fungi_unclassified(100); | - | - | - | - | - | - | - | Unassigned | - |
| Otu09959 | 2 | k__Fungi(100);p__Ascomycota(100);p__Ascomycota_unclassified(100);p__Ascomycota_unclassified(100);p__Ascomycota_unclassified(100);p__Ascomycota_unclassified(100);p__Ascomycota_unclassified(100); | - | - | - | - | - | - | - | Unassigned | - |
| Otu09960 | 2 | k__Fungi(100);k__Fungi_unclassified(100);k__Fungi_unclassified(100);k__Fungi_unclassified(100);k__Fungi_unclassified(100);k__Fungi_unclassified(100); | - | - | - | - | - | - | - | Unassigned | - |
| Otu09961 | 2 | k__Fungi(100);p__Ascomycota(100);p__Ascomycota_unclassified(100);p__Ascomycota_unclassified(100);p__Ascomycota_unclassified(100);p__Ascomycota_unclassified(100);p__Ascomycota_unclassified(100); | - | - | - | - | - | - | - | Unassigned | - |
| Otu09964 | 2 | k__Fungi(100);k__Fungi_unclassified(100);k__Fungi_unclassified(100);k__Fungi_unclassified(100);k__Fungi_unclassified(100);k__Fungi_unclassified(100); | - | - | - | - | - | - | - | Unassigned | - |
| Otu09965 | 2 | k__Fungi(100);k__Fungi_unclassified(100);k__Fungi_unclassified(100);k__Fungi_unclassified(100);k__Fungi_unclassified(100);k__Fungi_unclassified(100); | - | - | - | - | - | - | - | Unassigned | - |
| Otu09967 | 2 | k__Fungi(100);k__Fungi_unclassified(100);k__Fungi_unclassified(100);k__Fungi_unclassified(100);k__Fungi_unclassified(100);k__Fungi_unclassified(100); | - | - | - | - | - | - | - | Unassigned | - |
| Otu09968 | 2 | k__Fungi(100);k__Fungi_unclassified(100);k__Fungi_unclassified(100);k__Fungi_unclassified(100);k__Fungi_unclassified(100);k__Fungi_unclassified(100); | - | - | - | - | - | - | - | Unassigned | - |
| Otu09971 | 2 | k__Fungi(100);p__Ascomycota(100);p__Ascomycota_unclassified(100);p__Ascomycota_unclassified(100);p__Ascomycota_unclassified(100);p__Ascomycota_unclassified(100);p__Ascomycota_unclassified(100); | - | - | - | - | - | - | - | Unassigned | - |
| Otu09972 | 2 | k__Fungi(100);k__Fungi_unclassified(100);k__Fungi_unclassified(100);k__Fungi_unclassified(100);k__Fungi_unclassified(100);k__Fungi_unclassified(100); | - | - | - | - | - | - | - | Unassigned | - |
| Otu09973 | 2 | k__Fungi(100);k__Fungi_unclassified(100);k__Fungi_unclassified(100);k__Fungi_unclassified(100);k__Fungi_unclassified(100);k__Fungi_unclassified(100); | - | - | - | - | - | - | - | Unassigned | - |
| Otu09974 | 2 | k__Fungi(100);k__Fungi_unclassified(100);k__Fungi_unclassified(100);k__Fungi_unclassified(100);k__Fungi_unclassified(100);k__Fungi_unclassified(100); | - | - | - | - | - | - | - | Unassigned | - |
| Otu09976 | 2 | k__Fungi(100);p__Ascomycota(100);c__Leotiomycetes(100);c__Leotiomycetes_unclassified(100);c__Leotiomycetes_unclassified(100);c__Leotiomycetes_unclassified(100);c__Leotiomycetes_unclassified(100); | - | - | - | - | - | - | - | Unassigned | - |
| Otu09977 | 2 | k__Fungi(100);k__Fungi_unclassified(100);k__Fungi_unclassified(100);k__Fungi_unclassified(100);k__Fungi_unclassified(100);k__Fungi_unclassified(100); | - | - | - | - | - | - | - | Unassigned | - |
| Otu09978 | 2 | k__Fungi(100);k__Fungi_unclassified(100);k__Fungi_unclassified(100);k__Fungi_unclassified(100);k__Fungi_unclassified(100);k__Fungi_unclassified(100); | - | - | - | - | - | - | - | Unassigned | - |
| Otu09981 | 2 | k__Fungi(100);k__Fungi_unclassified(100);k__Fungi_unclassified(100);k__Fungi_unclassified(100);k__Fungi_unclassified(100);k__Fungi_unclassified(100); | - | - | - | - | - | - | - | Unassigned | - |
| Otu09985 | 2 | k__Fungi(100);k__Fungi_unclassified(100);k__Fungi_unclassified(100);k__Fungi_unclassified(100);k__Fungi_unclassified(100);k__Fungi_unclassified(100); | - | - | - | - | - | - | - | Unassigned | - |
| Otu09987 | 2 | k__Fungi(100);k__Fungi_unclassified(100);k__Fungi_unclassified(100);k__Fungi_unclassified(100);k__Fungi_unclassified(100);k__Fungi_unclassified(100); | - | - | - | - | - | - | - | Unassigned | - |
| Otu09988 | 2 | k__Fungi(100);k__Fungi_unclassified(100);k__Fungi_unclassified(100);k__Fungi_unclassified(100);k__Fungi_unclassified(100);k__Fungi_unclassified(100); | - | - | - | - | - | - | - | Unassigned | - |
| Otu09995 | 2 | k__Fungi(100);k__Fungi_unclassified(100);k__Fungi_unclassified(100);k__Fungi_unclassified(100);k__Fungi_unclassified(100);k__Fungi_unclassified(100); | - | - | - | - | - | - | - | Unassigned | - |
| Otu09996 | 2 | k__Fungi(100);k__Fungi_unclassified(100);k__Fungi_unclassified(100);k__Fungi_unclassified(100);k__Fungi_unclassified(100);k__Fungi_unclassified(100); | - | - | - | - | - | - | - | Unassigned | - |
| Otu09998 | 2 | k__Fungi(100);p__Ascomycota(100);p__Ascomycota_unclassified(100);p__Ascomycota_unclassified(100);p__Ascomycota_unclassified(100);p__Ascomycota_unclassified(100);p__Ascomycota_unclassified(100); | - | - | - | - | - | - | - | Unassigned | - |
| Otu09999 | 2 | k__Fungi(100);p__Ascomycota(100);p__Ascomycota_unclassified(100);p__Ascomycota_unclassified(100);p__Ascomycota_unclassified(100);p__Ascomycota_unclassified(100);p__Ascomycota_unclassified(100); | - | - | - | - | - | - | - | Unassigned | - |
| Otu10000 | 2 | k__Fungi(100);p__Ascomycota(100);p__Ascomycota_unclassified(100);p__Ascomycota_unclassified(100);p__Ascomycota_unclassified(100);p__Ascomycota_unclassified(100);p__Ascomycota_unclassified(100); | - | - | - | - | - | - | - | Unassigned | - |
| Otu10001 | 2 | k__Fungi(100);k__Fungi_unclassified(100);k__Fungi_unclassified(100);k__Fungi_unclassified(100);k__Fungi_unclassified(100);k__Fungi_unclassified(100); | - | - | - | - | - | - | - | Unassigned | - |
| Otu10005 | 2 | k__Fungi(100);k__Fungi_unclassified(100);k__Fungi_unclassified(100);k__Fungi_unclassified(100);k__Fungi_unclassified(100);k__Fungi_unclassified(100); | - | - | - | - | - | - | - | Unassigned | - |
| Otu10007 | 2 | k__Fungi(100);p__Ascomycota(100);p__Ascomycota_unclassified(100);p__Ascomycota_unclassified(100);p__Ascomycota_unclassified(100);p__Ascomycota_unclassified(100);p__Ascomycota_unclassified(100); | - | - | - | - | - | - | - | Unassigned | - |
| Otu10008 | 2 | k__Fungi(100);k__Fungi_unclassified(100);k__Fungi_unclassified(100);k__Fungi_unclassified(100);k__Fungi_unclassified(100);k__Fungi_unclassified(100); | - | - | - | - | - | - | - | Unassigned | - |
| Otu10009 | 2 | k__Fungi(100);k__Fungi_unclassified(100);k__Fungi_unclassified(100);k__Fungi_unclassified(100);k__Fungi_unclassified(100);k__Fungi_unclassified(100); | - | - | - | - | - | - | - | Unassigned | - |
| Otu10018 | 2 | k__Fungi(100);p__Ascomycota(100);p__Ascomycota_unclassified(100);p__Ascomycota_unclassified(100);p__Ascomycota_unclassified(100);p__Ascomycota_unclassified(100);p__Ascomycota_unclassified(100); | - | - | - | - | - | - | - | Unassigned | - |
| Otu10021 | 2 | k__Fungi(100);k__Fungi_unclassified(100);k__Fungi_unclassified(100);k__Fungi_unclassified(100);k__Fungi_unclassified(100);k__Fungi_unclassified(100); | - | - | - | - | - | - | - | Unassigned | - |
| Otu10022 | 2 | k__Fungi(100);k__Fungi_unclassified(100);k__Fungi_unclassified(100);k__Fungi_unclassified(100);k__Fungi_unclassified(100);k__Fungi_unclassified(100); | - | - | - | - | - | - | - | Unassigned | - |
| Otu10025 | 2 | k__Fungi(100);k__Fungi_unclassified(100);k__Fungi_unclassified(100);k__Fungi_unclassified(100);k__Fungi_unclassified(100);k__Fungi_unclassified(100); | - | - | - | - | - | - | - | Unassigned | - |
| Otu10027 | 2 | k__Fungi(100);k__Fungi_unclassified(100);k__Fungi_unclassified(100);k__Fungi_unclassified(100);k__Fungi_unclassified(100);k__Fungi_unclassified(100); | - | - | - | - | - | - | - | Unassigned | - |
| Otu10029 | 2 | k__Fungi(100);k__Fungi_unclassified(100);k__Fungi_unclassified(100);k__Fungi_unclassified(100);k__Fungi_unclassified(100);k__Fungi_unclassified(100); | - | - | - | - | - | - | - | Unassigned | - |
| Otu10032 | 2 | k__Fungi(100);p__Ascomycota(100);p__Ascomycota_unclassified(100);p__Ascomycota_unclassified(100);p__Ascomycota_unclassified(100);p__Ascomycota_unclassified(100);p__Ascomycota_unclassified(100); | - | - | - | - | - | - | - | Unassigned | - |
| Otu10034 | 2 | k__Fungi(100);p__Ascomycota(100);p__Ascomycota_unclassified(100);p__Ascomycota_unclassified(100);p__Ascomycota_unclassified(100);p__Ascomycota_unclassified(100);p__Ascomycota_unclassified(100); | - | - | - | - | - | - | - | Unassigned | - |
| Otu10036 | 2 | k__Fungi(100);k__Fungi_unclassified(100);k__Fungi_unclassified(100);k__Fungi_unclassified(100);k__Fungi_unclassified(100);k__Fungi_unclassified(100); | - | - | - | - | - | - | - | Unassigned | - |
| Otu10037 | 2 | k__Fungi(100);k__Fungi_unclassified(100);k__Fungi_unclassified(100);k__Fungi_unclassified(100);k__Fungi_unclassified(100);k__Fungi_unclassified(100); | - | - | - | - | - | - | - | Unassigned | - |
| Otu10038 | 2 | k__Fungi(100);k__Fungi_unclassified(100);k__Fungi_unclassified(100);k__Fungi_unclassified(100);k__Fungi_unclassified(100);k__Fungi_unclassified(100); | - | - | - | - | - | - | - | Unassigned | - |
| Otu10041 | 2 | k__Fungi(100);k__Fungi_unclassified(100);k__Fungi_unclassified(100);k__Fungi_unclassified(100);k__Fungi_unclassified(100);k__Fungi_unclassified(100); | - | - | - | - | - | - | - | Unassigned | - |
| Otu10042 | 2 | k__Fungi(100);p__Ascomycota(100);p__Ascomycota_unclassified(100);p__Ascomycota_unclassified(100);p__Ascomycota_unclassified(100);p__Ascomycota_unclassified(100);p__Ascomycota_unclassified(100); | - | - | - | - | - | - | - | Unassigned | - |
| Otu10043 | 2 | k__Fungi(100);p__Ascomycota(100);p__Ascomycota_unclassified(100);p__Ascomycota_unclassified(100);p__Ascomycota_unclassified(100);p__Ascomycota_unclassified(100);p__Ascomycota_unclassified(100); | - | - | - | - | - | - | - | Unassigned | - |
| Otu10045 | 2 | k__Fungi(100);k__Fungi_unclassified(100);k__Fungi_unclassified(100);k__Fungi_unclassified(100);k__Fungi_unclassified(100);k__Fungi_unclassified(100); | - | - | - | - | - | - | - | Unassigned | - |
| Otu10048 | 2 | k__Fungi(100);p__Ascomycota(100);p__Ascomycota_unclassified(100);p__Ascomycota_unclassified(100);p__Ascomycota_unclassified(100);p__Ascomycota_unclassified(100);p__Ascomycota_unclassified(100); | - | - | - | - | - | - | - | Unassigned | - |
| Otu10050 | 2 | k__Fungi(100);p__Ascomycota(100);p__Ascomycota_unclassified(100);p__Ascomycota_unclassified(100);p__Ascomycota_unclassified(100);p__Ascomycota_unclassified(100);p__Ascomycota_unclassified(100); | - | - | - | - | - | - | - | Unassigned | - |
| Otu10057 | 2 | k__Fungi(100);k__Fungi_unclassified(100);k__Fungi_unclassified(100);k__Fungi_unclassified(100);k__Fungi_unclassified(100);k__Fungi_unclassified(100); | - | - | - | - | - | - | - | Unassigned | - |
| Otu10059 | 2 | k__Fungi(100);p__Ascomycota(100);c__Sordariomycetes(100);o__Sordariales(100);o__Sordariales_unclassified(100);o__Sordariales_unclassified(100);o__Sordariales_unclassified(100); | - | - | - | - | - | - | - | Unassigned | - |
| Otu10060 | 2 | k__Fungi(100);k__Fungi_unclassified(100);k__Fungi_unclassified(100);k__Fungi_unclassified(100);k__Fungi_unclassified(100);k__Fungi_unclassified(100); | - | - | - | - | - | - | - | Unassigned | - |
| Otu10061 | 2 | k__Fungi(100);p__Ascomycota(100);p__Ascomycota_unclassified(100);p__Ascomycota_unclassified(100);p__Ascomycota_unclassified(100);p__Ascomycota_unclassified(100);p__Ascomycota_unclassified(100); | - | - | - | - | - | - | - | Unassigned | - |
| Otu10062 | 2 | k__Fungi(100);k__Fungi_unclassified(100);k__Fungi_unclassified(100);k__Fungi_unclassified(100);k__Fungi_unclassified(100);k__Fungi_unclassified(100); | - | - | - | - | - | - | - | Unassigned | - |
| Otu10063 | 2 | k__Fungi(100);k__Fungi_unclassified(100);k__Fungi_unclassified(100);k__Fungi_unclassified(100);k__Fungi_unclassified(100);k__Fungi_unclassified(100); | - | - | - | - | - | - | - | Unassigned | - |
| Otu10064 | 2 | k__Fungi(100);k__Fungi_unclassified(100);k__Fungi_unclassified(100);k__Fungi_unclassified(100);k__Fungi_unclassified(100);k__Fungi_unclassified(100); | - | - | - | - | - | - | - | Unassigned | - |
| Otu10068 | 2 | k__Fungi(100);k__Fungi_unclassified(100);k__Fungi_unclassified(100);k__Fungi_unclassified(100);k__Fungi_unclassified(100);k__Fungi_unclassified(100); | - | - | - | - | - | - | - | Unassigned | - |
| Otu10069 | 2 | k__Fungi(100);k__Fungi_unclassified(100);k__Fungi_unclassified(100);k__Fungi_unclassified(100);k__Fungi_unclassified(100);k__Fungi_unclassified(100); | - | - | - | - | - | - | - | Unassigned | - |
| Otu10070 | 2 | k__Fungi(100);p__Ascomycota(100);p__Ascomycota_unclassified(100);p__Ascomycota_unclassified(100);p__Ascomycota_unclassified(100);p__Ascomycota_unclassified(100);p__Ascomycota_unclassified(100); | - | - | - | - | - | - | - | Unassigned | - |
| Otu10072 | 2 | k__Fungi(100);k__Fungi_unclassified(100);k__Fungi_unclassified(100);k__Fungi_unclassified(100);k__Fungi_unclassified(100);k__Fungi_unclassified(100); | - | - | - | - | - | - | - | Unassigned | - |
| Otu10073 | 2 | k__Fungi(100);k__Fungi_unclassified(100);k__Fungi_unclassified(100);k__Fungi_unclassified(100);k__Fungi_unclassified(100);k__Fungi_unclassified(100); | - | - | - | - | - | - | - | Unassigned | - |
| Otu10074 | 2 | k__Fungi(100);k__Fungi_unclassified(100);k__Fungi_unclassified(100);k__Fungi_unclassified(100);k__Fungi_unclassified(100);k__Fungi_unclassified(100); | - | - | - | - | - | - | - | Unassigned | - |
| Otu10075 | 2 | k__Fungi(100);k__Fungi_unclassified(100);k__Fungi_unclassified(100);k__Fungi_unclassified(100);k__Fungi_unclassified(100);k__Fungi_unclassified(100); | - | - | - | - | - | - | - | Unassigned | - |
| Otu10077 | 2 | k__Fungi(100);k__Fungi_unclassified(100);k__Fungi_unclassified(100);k__Fungi_unclassified(100);k__Fungi_unclassified(100);k__Fungi_unclassified(100); | - | - | - | - | - | - | - | Unassigned | - |
| Otu10078 | 2 | k__Fungi(100);k__Fungi_unclassified(100);k__Fungi_unclassified(100);k__Fungi_unclassified(100);k__Fungi_unclassified(100);k__Fungi_unclassified(100); | - | - | - | - | - | - | - | Unassigned | - |
| Otu10080 | 2 | k__Fungi(100);k__Fungi_unclassified(100);k__Fungi_unclassified(100);k__Fungi_unclassified(100);k__Fungi_unclassified(100);k__Fungi_unclassified(100); | - | - | - | - | - | - | - | Unassigned | - |
| Otu10081 | 2 | k__Fungi(100);p__Ascomycota(100);p__Ascomycota_unclassified(100);p__Ascomycota_unclassified(100);p__Ascomycota_unclassified(100);p__Ascomycota_unclassified(100);p__Ascomycota_unclassified(100); | - | - | - | - | - | - | - | Unassigned | - |
| Otu10082 | 2 | k__Fungi(100);k__Fungi_unclassified(100);k__Fungi_unclassified(100);k__Fungi_unclassified(100);k__Fungi_unclassified(100);k__Fungi_unclassified(100); | - | - | - | - | - | - | - | Unassigned | - |
| Otu10083 | 2 | k__Fungi(100);k__Fungi_unclassified(100);k__Fungi_unclassified(100);k__Fungi_unclassified(100);k__Fungi_unclassified(100);k__Fungi_unclassified(100); | - | - | - | - | - | - | - | Unassigned | - |
| Otu10085 | 2 | k__Fungi(100);k__Fungi_unclassified(100);k__Fungi_unclassified(100);k__Fungi_unclassified(100);k__Fungi_unclassified(100);k__Fungi_unclassified(100); | - | - | - | - | - | - | - | Unassigned | - |
| Otu10089 | 2 | k__Fungi(100);k__Fungi_unclassified(100);k__Fungi_unclassified(100);k__Fungi_unclassified(100);k__Fungi_unclassified(100);k__Fungi_unclassified(100); | - | - | - | - | - | - | - | Unassigned | - |
| Otu10093 | 2 | k__Fungi(100);p__Ascomycota(100);p__Ascomycota_unclassified(100);p__Ascomycota_unclassified(100);p__Ascomycota_unclassified(100);p__Ascomycota_unclassified(100);p__Ascomycota_unclassified(100); | - | - | - | - | - | - | - | Unassigned | - |
| Otu10094 | 2 | k__Fungi(100);k__Fungi_unclassified(100);k__Fungi_unclassified(100);k__Fungi_unclassified(100);k__Fungi_unclassified(100);k__Fungi_unclassified(100); | - | - | - | - | - | - | - | Unassigned | - |
| Otu10095 | 2 | k__Fungi(100);p__Ascomycota(100);c__Leotiomycetes(100);o__Helotiales(100);o__Helotiales_unclassified(100);o__Helotiales_unclassified(100);o__Helotiales_unclassified(100); | - | - | - | - | - | - | - | Unassigned | - |
| Otu10098 | 2 | k__Fungi(100);p__Ascomycota(100);p__Ascomycota_unclassified(100);p__Ascomycota_unclassified(100);p__Ascomycota_unclassified(100);p__Ascomycota_unclassified(100);p__Ascomycota_unclassified(100); | - | - | - | - | - | - | - | Unassigned | - |
| Otu10099 | 2 | k__Fungi(100);k__Fungi_unclassified(100);k__Fungi_unclassified(100);k__Fungi_unclassified(100);k__Fungi_unclassified(100);k__Fungi_unclassified(100); | - | - | - | - | - | - | - | Unassigned | - |
| Otu10104 | 2 | k__Fungi(100);k__Fungi_unclassified(100);k__Fungi_unclassified(100);k__Fungi_unclassified(100);k__Fungi_unclassified(100);k__Fungi_unclassified(100); | - | - | - | - | - | - | - | Unassigned | - |
| Otu10105 | 2 | k__Fungi(100);p__Ascomycota(100);c__Dothideomycetes(100);o__Pleosporales(100);o__Pleosporales_unclassified(100);o__Pleosporales_unclassified(100);o__Pleosporales_unclassified(100); | - | - | - | - | - | - | - | Unassigned | - |
| Otu10106 | 2 | k__Fungi(100);k__Fungi_unclassified(100);k__Fungi_unclassified(100);k__Fungi_unclassified(100);k__Fungi_unclassified(100);k__Fungi_unclassified(100); | - | - | - | - | - | - | - | Unassigned | - |
| Otu10108 | 2 | k__Fungi(100);p__Ascomycota(100);p__Ascomycota_unclassified(100);p__Ascomycota_unclassified(100);p__Ascomycota_unclassified(100);p__Ascomycota_unclassified(100);p__Ascomycota_unclassified(100); | - | - | - | - | - | - | - | Unassigned | - |
| Otu10109 | 2 | k__Fungi(100);k__Fungi_unclassified(100);k__Fungi_unclassified(100);k__Fungi_unclassified(100);k__Fungi_unclassified(100);k__Fungi_unclassified(100); | - | - | - | - | - | - | - | Unassigned | - |
| Otu10110 | 2 | k__Fungi(100);p__Ascomycota(100);p__Ascomycota_unclassified(100);p__Ascomycota_unclassified(100);p__Ascomycota_unclassified(100);p__Ascomycota_unclassified(100);p__Ascomycota_unclassified(100); | - | - | - | - | - | - | - | Unassigned | - |
| Otu10111 | 2 | k__Fungi(100);k__Fungi_unclassified(100);k__Fungi_unclassified(100);k__Fungi_unclassified(100);k__Fungi_unclassified(100);k__Fungi_unclassified(100); | - | - | - | - | - | - | - | Unassigned | - |
| Otu10113 | 2 | k__Fungi(100);k__Fungi_unclassified(100);k__Fungi_unclassified(100);k__Fungi_unclassified(100);k__Fungi_unclassified(100);k__Fungi_unclassified(100); | - | - | - | - | - | - | - | Unassigned | - |
| Otu10118 | 2 | k__Fungi(100);p__Ascomycota(100);p__Ascomycota_unclassified(100);p__Ascomycota_unclassified(100);p__Ascomycota_unclassified(100);p__Ascomycota_unclassified(100);p__Ascomycota_unclassified(100); | - | - | - | - | - | - | - | Unassigned | - |
| Otu10121 | 2 | k__Fungi(100);p__Ascomycota(100);p__Ascomycota_unclassified(100);p__Ascomycota_unclassified(100);p__Ascomycota_unclassified(100);p__Ascomycota_unclassified(100);p__Ascomycota_unclassified(100); | - | - | - | - | - | - | - | Unassigned | - |
| Otu10123 | 2 | k__Fungi(100);k__Fungi_unclassified(100);k__Fungi_unclassified(100);k__Fungi_unclassified(100);k__Fungi_unclassified(100);k__Fungi_unclassified(100); | - | - | - | - | - | - | - | Unassigned | - |
| Otu10129 | 2 | k__Fungi(100);k__Fungi_unclassified(100);k__Fungi_unclassified(100);k__Fungi_unclassified(100);k__Fungi_unclassified(100);k__Fungi_unclassified(100); | - | - | - | - | - | - | - | Unassigned | - |
| Otu10130 | 2 | k__Fungi(100);k__Fungi_unclassified(100);k__Fungi_unclassified(100);k__Fungi_unclassified(100);k__Fungi_unclassified(100);k__Fungi_unclassified(100); | - | - | - | - | - | - | - | Unassigned | - |
| Otu10131 | 2 | k__Fungi(100);k__Fungi_unclassified(100);k__Fungi_unclassified(100);k__Fungi_unclassified(100);k__Fungi_unclassified(100);k__Fungi_unclassified(100); | - | - | - | - | - | - | - | Unassigned | - |
| Otu10135 | 2 | k__Fungi(100);p__Ascomycota(100);p__Ascomycota_unclassified(100);p__Ascomycota_unclassified(100);p__Ascomycota_unclassified(100);p__Ascomycota_unclassified(100);p__Ascomycota_unclassified(100); | - | - | - | - | - | - | - | Unassigned | - |
| Otu10136 | 2 | k__Fungi(100);k__Fungi_unclassified(100);k__Fungi_unclassified(100);k__Fungi_unclassified(100);k__Fungi_unclassified(100);k__Fungi_unclassified(100); | - | - | - | - | - | - | - | Unassigned | - |
| Otu10137 | 2 | k__Fungi(100);p__Ascomycota(100);p__Ascomycota_unclassified(100);p__Ascomycota_unclassified(100);p__Ascomycota_unclassified(100);p__Ascomycota_unclassified(100);p__Ascomycota_unclassified(100); | - | - | - | - | - | - | - | Unassigned | - |
| Otu10141 | 2 | k__Fungi(100);k__Fungi_unclassified(100);k__Fungi_unclassified(100);k__Fungi_unclassified(100);k__Fungi_unclassified(100);k__Fungi_unclassified(100); | - | - | - | - | - | - | - | Unassigned | - |
| Otu10142 | 2 | k__Fungi(100);p__Ascomycota(100);p__Ascomycota_unclassified(100);p__Ascomycota_unclassified(100);p__Ascomycota_unclassified(100);p__Ascomycota_unclassified(100);p__Ascomycota_unclassified(100); | - | - | - | - | - | - | - | Unassigned | - |
| Otu10145 | 2 | k__Fungi(100);k__Fungi_unclassified(100);k__Fungi_unclassified(100);k__Fungi_unclassified(100);k__Fungi_unclassified(100);k__Fungi_unclassified(100); | - | - | - | - | - | - | - | Unassigned | - |
| Otu10150 | 2 | k__Fungi(100);k__Fungi_unclassified(100);k__Fungi_unclassified(100);k__Fungi_unclassified(100);k__Fungi_unclassified(100);k__Fungi_unclassified(100); | - | - | - | - | - | - | - | Unassigned | - |

| | | | | | | | | | | | |
|---|---|---|---|---|---|---|---|---|---|---|---|
| Otu10151 | 2 | k__Fungi(100);k__Fungi_unclassified(100);k__Fungi_unclassified(100);k__Fungi_unclassified(100);k__Fungi_unclassified(100);k__Fungi_unclassified(100);k__Fungi_unclassified(100); | - | - | - | - | - | - | - | Unassigned | - |
| Otu10153 | 2 | k__Fungi(100);p__Ascomycota(100);p__Ascomycota_unclassified(100);p__Ascomycota_unclassified(100);p__Ascomycota_unclassified(100);p__Ascomycota_unclassified(100);p__Ascomycota_unclassified(100); | - | - | - | - | - | - | - | Unassigned | - |
| Otu10154 | 2 | k__Fungi(100);p__Ascomycota(100);c__Dothideomycetes(100);o__Pleosporales(100);o__Pleosporales_unclassified(100);o__Pleosporales_unclassified(100);o__Pleosporales_unclassified(100); | - | - | - | - | - | - | - | Unassigned | - |
| Otu10155 | 2 | k__Fungi(100);k__Fungi_unclassified(100);k__Fungi_unclassified(100);k__Fungi_unclassified(100);k__Fungi_unclassified(100);k__Fungi_unclassified(100);k__Fungi_unclassified(100); | - | - | - | - | - | - | - | Unassigned | - |
| Otu10156 | 2 | k__Fungi(100);k__Fungi_unclassified(100);k__Fungi_unclassified(100);k__Fungi_unclassified(100);k__Fungi_unclassified(100);k__Fungi_unclassified(100);k__Fungi_unclassified(100); | - | - | - | - | - | - | - | Unassigned | - |
| Otu10157 | 2 | k__Fungi(100);k__Fungi_unclassified(100);k__Fungi_unclassified(100);k__Fungi_unclassified(100);k__Fungi_unclassified(100);k__Fungi_unclassified(100);k__Fungi_unclassified(100); | - | - | - | - | - | - | - | Unassigned | - |
| Otu10158 | 2 | k__Fungi(100);k__Fungi_unclassified(100);k__Fungi_unclassified(100);k__Fungi_unclassified(100);k__Fungi_unclassified(100);k__Fungi_unclassified(100);k__Fungi_unclassified(100); | - | - | - | - | - | - | - | Unassigned | - |
| Otu10165 | 2 | k__Fungi(100);k__Fungi_unclassified(100);k__Fungi_unclassified(100);k__Fungi_unclassified(100);k__Fungi_unclassified(100);k__Fungi_unclassified(100);k__Fungi_unclassified(100); | - | - | - | - | - | - | - | Unassigned | - |
| Otu10166 | 2 | k__Fungi(100);k__Fungi_unclassified(100);k__Fungi_unclassified(100);k__Fungi_unclassified(100);k__Fungi_unclassified(100);k__Fungi_unclassified(100);k__Fungi_unclassified(100); | - | - | - | - | - | - | - | Unassigned | - |
| Otu10168 | 2 | k__Fungi(100);k__Fungi_unclassified(100);k__Fungi_unclassified(100);k__Fungi_unclassified(100);k__Fungi_unclassified(100);k__Fungi_unclassified(100);k__Fungi_unclassified(100); | - | - | - | - | - | - | - | Unassigned | - |
| Otu10169 | 2 | k__Fungi(100);k__Fungi_unclassified(100);k__Fungi_unclassified(100);k__Fungi_unclassified(100);k__Fungi_unclassified(100);k__Fungi_unclassified(100);k__Fungi_unclassified(100); | - | - | - | - | - | - | - | Unassigned | - |
| Otu10171 | 2 | k__Fungi(100);p__Ascomycota(100);c__Dothideomycetes(100);o__Pleosporales(100);o__Pleosporales_unclassified(100);o__Pleosporales_unclassified(100);o__Pleosporales_unclassified(100); | - | - | - | - | - | - | - | Unassigned | - |
| Otu10173 | 2 | k__Fungi(100);p__Ascomycota(100);p__Ascomycota_unclassified(100);p__Ascomycota_unclassified(100);p__Ascomycota_unclassified(100);p__Ascomycota_unclassified(100);p__Ascomycota_unclassified(100); | - | - | - | - | - | - | - | Unassigned | - |
| Otu10180 | 2 | k__Fungi(100);k__Fungi_unclassified(100);k__Fungi_unclassified(100);k__Fungi_unclassified(100);k__Fungi_unclassified(100);k__Fungi_unclassified(100);k__Fungi_unclassified(100); | - | - | - | - | - | - | - | Unassigned | - |
| Otu10182 | 2 | k__Fungi(100);p__Ascomycota(100);c__Leotiomycetes(100);o__Helotiales(100);o__Helotiales_unclassified(100);o__Helotiales_unclassified(100);o__Helotiales_unclassified(100); | - | - | - | - | - | - | - | Unassigned | - |
| Otu10189 | 2 | k__Fungi(100);k__Fungi_unclassified(100);k__Fungi_unclassified(100);k__Fungi_unclassified(100);k__Fungi_unclassified(100);k__Fungi_unclassified(100);k__Fungi_unclassified(100); | - | - | - | - | - | - | - | Unassigned | - |
| Otu10191 | 2 | k__Fungi(100);k__Fungi_unclassified(100);k__Fungi_unclassified(100);k__Fungi_unclassified(100);k__Fungi_unclassified(100);k__Fungi_unclassified(100);k__Fungi_unclassified(100); | - | - | - | - | - | - | - | Unassigned | - |
| Otu10192 | 2 | k__Fungi(100);p__Ascomycota(100);p__Ascomycota_unclassified(100);p__Ascomycota_unclassified(100);p__Ascomycota_unclassified(100);p__Ascomycota_unclassified(100);p__Ascomycota_unclassified(100); | - | - | - | - | - | - | - | Unassigned | - |
| Otu10194 | 2 | k__Fungi(100);k__Fungi_unclassified(100);k__Fungi_unclassified(100);k__Fungi_unclassified(100);k__Fungi_unclassified(100);k__Fungi_unclassified(100);k__Fungi_unclassified(100); | - | - | - | - | - | - | - | Unassigned | - |
| Otu10197 | 2 | k__Fungi(100);k__Fungi_unclassified(100);k__Fungi_unclassified(100);k__Fungi_unclassified(100);k__Fungi_unclassified(100);k__Fungi_unclassified(100);k__Fungi_unclassified(100); | - | - | - | - | - | - | - | Unassigned | - |
| Otu10199 | 2 | k__Fungi(100);k__Fungi_unclassified(100);k__Fungi_unclassified(100);k__Fungi_unclassified(100);k__Fungi_unclassified(100);k__Fungi_unclassified(100);k__Fungi_unclassified(100); | - | - | - | - | - | - | - | Unassigned | - |
| Otu10201 | 2 | k__Fungi(100);k__Fungi_unclassified(100);k__Fungi_unclassified(100);k__Fungi_unclassified(100);k__Fungi_unclassified(100);k__Fungi_unclassified(100);k__Fungi_unclassified(100); | - | - | - | - | - | - | - | Unassigned | - |
| Otu10202 | 2 | k__Fungi(100);k__Fungi_unclassified(100);k__Fungi_unclassified(100);k__Fungi_unclassified(100);k__Fungi_unclassified(100);k__Fungi_unclassified(100);k__Fungi_unclassified(100); | - | - | - | - | - | - | - | Unassigned | - |
| Otu10203 | 2 | k__Fungi(100);p__Ascomycota(100);p__Ascomycota_unclassified(100);p__Ascomycota_unclassified(100);p__Ascomycota_unclassified(100);p__Ascomycota_unclassified(100);p__Ascomycota_unclassified(100); | - | - | - | - | - | - | - | Unassigned | - |
| Otu10206 | 2 | k__Fungi(100);k__Fungi_unclassified(100);k__Fungi_unclassified(100);k__Fungi_unclassified(100);k__Fungi_unclassified(100);k__Fungi_unclassified(100);k__Fungi_unclassified(100); | - | - | - | - | - | - | - | Unassigned | - |
| Otu10208 | 2 | k__Fungi(100);p__Ascomycota(100);p__Ascomycota_unclassified(100);p__Ascomycota_unclassified(100);p__Ascomycota_unclassified(100);p__Ascomycota_unclassified(100);p__Ascomycota_unclassified(100); | - | - | - | - | - | - | - | Unassigned | - |
| Otu10210 | 2 | k__Fungi(100);k__Fungi_unclassified(100);k__Fungi_unclassified(100);k__Fungi_unclassified(100);k__Fungi_unclassified(100);k__Fungi_unclassified(100);k__Fungi_unclassified(100); | - | - | - | - | - | - | - | Unassigned | - |
| Otu10211 | 2 | k__Fungi(100);k__Fungi_unclassified(100);k__Fungi_unclassified(100);k__Fungi_unclassified(100);k__Fungi_unclassified(100);k__Fungi_unclassified(100);k__Fungi_unclassified(100); | - | - | - | - | - | - | - | Unassigned | - |
| Otu10212 | 2 | k__Fungi(100);p__Ascomycota(100);p__Ascomycota_unclassified(100);p__Ascomycota_unclassified(100);p__Ascomycota_unclassified(100);p__Ascomycota_unclassified(100);p__Ascomycota_unclassified(100); | - | - | - | - | - | - | - | Unassigned | - |
| Otu10213 | 2 | k__Fungi(100);p__Ascomycota(100);c__Leotiomycetes(100);o__Helotiales(100);o__Helotiales_unclassified(100);o__Helotiales_unclassified(100);o__Helotiales_unclassified(100); | - | - | - | - | - | - | - | Unassigned | - |
| Otu10217 | 2 | k__Fungi(100);p__Ascomycota(100);p__Ascomycota_unclassified(100);p__Ascomycota_unclassified(100);p__Ascomycota_unclassified(100);p__Ascomycota_unclassified(100);p__Ascomycota_unclassified(100); | - | - | - | - | - | - | - | Unassigned | - |
| Otu10219 | 2 | k__Fungi(100);k__Fungi_unclassified(100);k__Fungi_unclassified(100);k__Fungi_unclassified(100);k__Fungi_unclassified(100);k__Fungi_unclassified(100);k__Fungi_unclassified(100); | - | - | - | - | - | - | - | Unassigned | - |
| Otu10221 | 2 | k__Fungi(100);k__Fungi_unclassified(100);k__Fungi_unclassified(100);k__Fungi_unclassified(100);k__Fungi_unclassified(100);k__Fungi_unclassified(100);k__Fungi_unclassified(100); | - | - | - | - | - | - | - | Unassigned | - |
| Otu10222 | 2 | k__Fungi(100);k__Fungi_unclassified(100);k__Fungi_unclassified(100);k__Fungi_unclassified(100);k__Fungi_unclassified(100);k__Fungi_unclassified(100);k__Fungi_unclassified(100); | - | - | - | - | - | - | - | Unassigned | - |
| Otu10227 | 2 | k__Fungi(100);k__Fungi_unclassified(100);k__Fungi_unclassified(100);k__Fungi_unclassified(100);k__Fungi_unclassified(100);k__Fungi_unclassified(100);k__Fungi_unclassified(100); | - | - | - | - | - | - | - | Unassigned | - |
| Otu10228 | 2 | k__Fungi(100);k__Fungi_unclassified(100);k__Fungi_unclassified(100);k__Fungi_unclassified(100);k__Fungi_unclassified(100);k__Fungi_unclassified(100);k__Fungi_unclassified(100); | - | - | - | - | - | - | - | Unassigned | - |
| Otu10230 | 2 | k__Fungi(100);k__Fungi_unclassified(100);k__Fungi_unclassified(100);k__Fungi_unclassified(100);k__Fungi_unclassified(100);k__Fungi_unclassified(100);k__Fungi_unclassified(100); | - | - | - | - | - | - | - | Unassigned | - |
| Otu10231 | 2 | k__Fungi(100);p__Ascomycota(100);c__Leotiomycetes(100);c__Leotiomycetes_unclassified(100);c__Leotiomycetes_unclassified(100);c__Leotiomycetes_unclassified(100);c__Leotiomycetes_unclassified(100); | - | - | - | - | - | - | - | Unassigned | - |
| Otu10236 | 2 | k__Fungi(100);k__Fungi_unclassified(100);k__Fungi_unclassified(100);k__Fungi_unclassified(100);k__Fungi_unclassified(100);k__Fungi_unclassified(100);k__Fungi_unclassified(100); | - | - | - | - | - | - | - | Unassigned | - |
| Otu10237 | 2 | k__Fungi(100);k__Fungi_unclassified(100);k__Fungi_unclassified(100);k__Fungi_unclassified(100);k__Fungi_unclassified(100);k__Fungi_unclassified(100);k__Fungi_unclassified(100); | - | - | - | - | - | - | - | Unassigned | - |
| Otu10238 | 2 | k__Fungi(100);k__Fungi_unclassified(100);k__Fungi_unclassified(100);k__Fungi_unclassified(100);k__Fungi_unclassified(100);k__Fungi_unclassified(100);k__Fungi_unclassified(100); | - | - | - | - | - | - | - | Unassigned | - |
| Otu10242 | 2 | k__Fungi(100);k__Fungi_unclassified(100);k__Fungi_unclassified(100);k__Fungi_unclassified(100);k__Fungi_unclassified(100);k__Fungi_unclassified(100);k__Fungi_unclassified(100); | - | - | - | - | - | - | - | Unassigned | - |
| Otu10243 | 2 | k__Fungi(100);k__Fungi_unclassified(100);k__Fungi_unclassified(100);k__Fungi_unclassified(100);k__Fungi_unclassified(100);k__Fungi_unclassified(100);k__Fungi_unclassified(100); | - | - | - | - | - | - | - | Unassigned | - |
| Otu10251 | 2 | k__Fungi(100);k__Fungi_unclassified(100);k__Fungi_unclassified(100);k__Fungi_unclassified(100);k__Fungi_unclassified(100);k__Fungi_unclassified(100);k__Fungi_unclassified(100); | - | - | - | - | - | - | - | Unassigned | - |
| Otu10252 | 2 | k__Fungi(100);p__Ascomycota(100);p__Ascomycota_unclassified(100);p__Ascomycota_unclassified(100);p__Ascomycota_unclassified(100);p__Ascomycota_unclassified(100);p__Ascomycota_unclassified(100); | - | - | - | - | - | - | - | Unassigned | - |
| Otu10256 | 2 | k__Fungi(100);k__Fungi_unclassified(100);k__Fungi_unclassified(100);k__Fungi_unclassified(100);k__Fungi_unclassified(100);k__Fungi_unclassified(100);k__Fungi_unclassified(100); | - | - | - | - | - | - | - | Unassigned | - |
| Otu10257 | 2 | k__Fungi(100);k__Fungi_unclassified(100);k__Fungi_unclassified(100);k__Fungi_unclassified(100);k__Fungi_unclassified(100);k__Fungi_unclassified(100);k__Fungi_unclassified(100); | - | - | - | - | - | - | - | Unassigned | - |
| Otu10264 | 2 | k__Fungi(100);p__Ascomycota(100);c__Sordariomycetes(100);c__Sordariomycetes_unclassified(100);c__Sordariomycetes_unclassified(100);c__Sordariomycetes_unclassified(100);c__Sordariomycetes_unclassified(100); | - | - | - | - | - | - | - | Unassigned | - |
| Otu10268 | 2 | k__Fungi(100);k__Fungi_unclassified(100);k__Fungi_unclassified(100);k__Fungi_unclassified(100);k__Fungi_unclassified(100);k__Fungi_unclassified(100);k__Fungi_unclassified(100); | - | - | - | - | - | - | - | Unassigned | - |
| Otu10269 | 2 | k__Fungi(100);p__Ascomycota(100);p__Ascomycota_unclassified(100);p__Ascomycota_unclassified(100);p__Ascomycota_unclassified(100);p__Ascomycota_unclassified(100);p__Ascomycota_unclassified(100); | - | - | - | - | - | - | - | Unassigned | - |
| Otu10272 | 2 | k__Fungi(100);p__Ascomycota(100);p__Ascomycota_unclassified(100);p__Ascomycota_unclassified(100);p__Ascomycota_unclassified(100);p__Ascomycota_unclassified(100);p__Ascomycota_unclassified(100); | - | - | - | - | - | - | - | Unassigned | - |
| Otu10275 | 2 | k__Fungi(100);p__Ascomycota(100);p__Ascomycota_unclassified(100);p__Ascomycota_unclassified(100);p__Ascomycota_unclassified(100);p__Ascomycota_unclassified(100);p__Ascomycota_unclassified(100); | - | - | - | - | - | - | - | Unassigned | - |
| Otu10276 | 2 | k__Fungi(100);k__Fungi_unclassified(100);k__Fungi_unclassified(100);k__Fungi_unclassified(100);k__Fungi_unclassified(100);k__Fungi_unclassified(100);k__Fungi_unclassified(100); | - | - | - | - | - | - | - | Unassigned | - |
| Otu10277 | 2 | k__Fungi(100);p__Ascomycota(100);c__Dothideomycetes(100);c__Dothideomycetes_unclassified(100);c__Dothideomycetes_unclassified(100);c__Dothideomycetes_unclassified(100);c__Dothideomycetes_unclassified(100); | - | - | - | - | - | - | - | Unassigned | - |
| Otu10282 | 2 | k__Fungi(100);p__Ascomycota(100);p__Ascomycota_unclassified(100);p__Ascomycota_unclassified(100);p__Ascomycota_unclassified(100);p__Ascomycota_unclassified(100);p__Ascomycota_unclassified(100); | - | - | - | - | - | - | - | Unassigned | - |
| Otu10287 | 2 | k__Fungi(100);k__Fungi_unclassified(100);k__Fungi_unclassified(100);k__Fungi_unclassified(100);k__Fungi_unclassified(100);k__Fungi_unclassified(100);k__Fungi_unclassified(100); | - | - | - | - | - | - | - | Unassigned | - |
| Otu10289 | 2 | k__Fungi(100);k__Fungi_unclassified(100);k__Fungi_unclassified(100);k__Fungi_unclassified(100);k__Fungi_unclassified(100);k__Fungi_unclassified(100);k__Fungi_unclassified(100); | - | - | - | - | - | - | - | Unassigned | - |
| Otu10290 | 2 | k__Fungi(100);k__Fungi_unclassified(100);k__Fungi_unclassified(100);k__Fungi_unclassified(100);k__Fungi_unclassified(100);k__Fungi_unclassified(100);k__Fungi_unclassified(100); | - | - | - | - | - | - | - | Unassigned | - |
| Otu10291 | 2 | k__Fungi(100);k__Fungi_unclassified(100);k__Fungi_unclassified(100);k__Fungi_unclassified(100);k__Fungi_unclassified(100);k__Fungi_unclassified(100);k__Fungi_unclassified(100); | - | - | - | - | - | - | - | Unassigned | - |
| Otu10293 | 2 | k__Fungi(100);k__Fungi_unclassified(100);k__Fungi_unclassified(100);k__Fungi_unclassified(100);k__Fungi_unclassified(100);k__Fungi_unclassified(100);k__Fungi_unclassified(100); | - | - | - | - | - | - | - | Unassigned | - |
| Otu10294 | 2 | k__Fungi(100);k__Fungi_unclassified(100);k__Fungi_unclassified(100);k__Fungi_unclassified(100);k__Fungi_unclassified(100);k__Fungi_unclassified(100);k__Fungi_unclassified(100); | - | - | - | - | - | - | - | Unassigned | - |
| Otu10297 | 2 | k__Fungi(100);k__Fungi_unclassified(100);k__Fungi_unclassified(100);k__Fungi_unclassified(100);k__Fungi_unclassified(100);k__Fungi_unclassified(100);k__Fungi_unclassified(100); | - | - | - | - | - | - | - | Unassigned | - |
| Otu10298 | 2 | k__Fungi(100);k__Fungi_unclassified(100);k__Fungi_unclassified(100);k__Fungi_unclassified(100);k__Fungi_unclassified(100);k__Fungi_unclassified(100);k__Fungi_unclassified(100); | - | - | - | - | - | - | - | Unassigned | - |
| Otu10299 | 2 | k__Fungi(100);k__Fungi_unclassified(100);k__Fungi_unclassified(100);k__Fungi_unclassified(100);k__Fungi_unclassified(100);k__Fungi_unclassified(100);k__Fungi_unclassified(100); | - | - | - | - | - | - | - | Unassigned | - |
| Otu10301 | 2 | k__Fungi(100);p__Ascomycota(100);p__Ascomycota_unclassified(100);p__Ascomycota_unclassified(100);p__Ascomycota_unclassified(100);p__Ascomycota_unclassified(100);p__Ascomycota_unclassified(100); | - | - | - | - | - | - | - | Unassigned | - |
| Otu10302 | 2 | k__Fungi(100);k__Fungi_unclassified(100);k__Fungi_unclassified(100);k__Fungi_unclassified(100);k__Fungi_unclassified(100);k__Fungi_unclassified(100);k__Fungi_unclassified(100); | - | - | - | - | - | - | - | Unassigned | - |
| Otu10304 | 2 | k__Fungi(100);k__Fungi_unclassified(100);k__Fungi_unclassified(100);k__Fungi_unclassified(100);k__Fungi_unclassified(100);k__Fungi_unclassified(100);k__Fungi_unclassified(100); | - | - | - | - | - | - | - | Unassigned | - |
| Otu10305 | 2 | k__Fungi(100);k__Fungi_unclassified(100);k__Fungi_unclassified(100);k__Fungi_unclassified(100);k__Fungi_unclassified(100);k__Fungi_unclassified(100);k__Fungi_unclassified(100); | - | - | - | - | - | - | - | Unassigned | - |
| Otu10308 | 2 | k__Fungi(100);p__Ascomycota(100);p__Ascomycota_unclassified(100);p__Ascomycota_unclassified(100);p__Ascomycota_unclassified(100);p__Ascomycota_unclassified(100);p__Ascomycota_unclassified(100); | - | - | - | - | - | - | - | Unassigned | - |
| Otu10311 | 2 | k__Fungi(100);p__Ascomycota(100);p__Ascomycota_unclassified(100);p__Ascomycota_unclassified(100);p__Ascomycota_unclassified(100);p__Ascomycota_unclassified(100);p__Ascomycota_unclassified(100); | - | - | - | - | - | - | - | Unassigned | - |
| Otu10312 | 2 | k__Fungi(100);k__Fungi_unclassified(100);k__Fungi_unclassified(100);k__Fungi_unclassified(100);k__Fungi_unclassified(100);k__Fungi_unclassified(100);k__Fungi_unclassified(100); | - | - | - | - | - | - | - | Unassigned | - |
| Otu10313 | 2 | k__Fungi(100);p__Ascomycota(100);p__Ascomycota_unclassified(100);p__Ascomycota_unclassified(100);p__Ascomycota_unclassified(100);p__Ascomycota_unclassified(100);p__Ascomycota_unclassified(100); | - | - | - | - | - | - | - | Unassigned | - |
| Otu10314 | 2 | k__Fungi(100);p__Ascomycota(100);p__Ascomycota_unclassified(100);p__Ascomycota_unclassified(100);p__Ascomycota_unclassified(100);p__Ascomycota_unclassified(100);p__Ascomycota_unclassified(100); | - | - | - | - | - | - | - | Unassigned | - |
| Otu10315 | 2 | k__Fungi(100);k__Fungi_unclassified(100);k__Fungi_unclassified(100);k__Fungi_unclassified(100);k__Fungi_unclassified(100);k__Fungi_unclassified(100);k__Fungi_unclassified(100); | - | - | - | - | - | - | - | Unassigned | - |
| Otu10319 | 2 | k__Fungi(100);k__Fungi_unclassified(100);k__Fungi_unclassified(100);k__Fungi_unclassified(100);k__Fungi_unclassified(100);k__Fungi_unclassified(100);k__Fungi_unclassified(100); | - | - | - | - | - | - | - | Unassigned | - |
| Otu10321 | 2 | k__Fungi(100);k__Fungi_unclassified(100);k__Fungi_unclassified(100);k__Fungi_unclassified(100);k__Fungi_unclassified(100);k__Fungi_unclassified(100);k__Fungi_unclassified(100); | - | - | - | - | - | - | - | Unassigned | - |
| Otu10330 | 2 | k__Fungi(100);k__Fungi_unclassified(100);k__Fungi_unclassified(100);k__Fungi_unclassified(100);k__Fungi_unclassified(100);k__Fungi_unclassified(100);k__Fungi_unclassified(100); | - | - | - | - | - | - | - | Unassigned | - |
| Otu10332 | 2 | k__Fungi(100);k__Fungi_unclassified(100);k__Fungi_unclassified(100);k__Fungi_unclassified(100);k__Fungi_unclassified(100);k__Fungi_unclassified(100);k__Fungi_unclassified(100); | - | - | - | - | - | - | - | Unassigned | - |
| Otu10333 | 2 | k__Fungi(100);p__Ascomycota(100);p__Ascomycota_unclassified(100);p__Ascomycota_unclassified(100);p__Ascomycota_unclassified(100);p__Ascomycota_unclassified(100);p__Ascomycota_unclassified(100); | - | - | - | - | - | - | - | Unassigned | - |
| Otu10336 | 2 | k__Fungi(100);k__Fungi_unclassified(100);k__Fungi_unclassified(100);k__Fungi_unclassified(100);k__Fungi_unclassified(100);k__Fungi_unclassified(100);k__Fungi_unclassified(100); | - | - | - | - | - | - | - | Unassigned | - |
| Otu10337 | 2 | k__Fungi(100);k__Fungi_unclassified(100);k__Fungi_unclassified(100);k__Fungi_unclassified(100);k__Fungi_unclassified(100);k__Fungi_unclassified(100);k__Fungi_unclassified(100); | - | - | - | - | - | - | - | Unassigned | - |
| Otu10339 | 2 | k__Fungi(100);p__Ascomycota(100);p__Ascomycota_unclassified(100);p__Ascomycota_unclassified(100);p__Ascomycota_unclassified(100);p__Ascomycota_unclassified(100);p__Ascomycota_unclassified(100); | - | - | - | - | - | - | - | Unassigned | - |
| Otu10340 | 2 | k__Fungi(100);k__Fungi_unclassified(100);k__Fungi_unclassified(100);k__Fungi_unclassified(100);k__Fungi_unclassified(100);k__Fungi_unclassified(100);k__Fungi_unclassified(100); | - | - | - | - | - | - | - | Unassigned | - |
| Otu10341 | 2 | k__Fungi(100);k__Fungi_unclassified(100);k__Fungi_unclassified(100);k__Fungi_unclassified(100);k__Fungi_unclassified(100);k__Fungi_unclassified(100);k__Fungi_unclassified(100); | - | - | - | - | - | - | - | Unassigned | - |
| Otu10342 | 2 | k__Fungi(100);k__Fungi_unclassified(100);k__Fungi_unclassified(100);k__Fungi_unclassified(100);k__Fungi_unclassified(100);k__Fungi_unclassified(100);k__Fungi_unclassified(100); | - | - | - | - | - | - | - | Unassigned | - |
| Otu10343 | 2 | k__Fungi(100);k__Fungi_unclassified(100);k__Fungi_unclassified(100);k__Fungi_unclassified(100);k__Fungi_unclassified(100);k__Fungi_unclassified(100);k__Fungi_unclassified(100); | - | - | - | - | - | - | - | Unassigned | - |
| Otu10344 | 2 | k__Fungi(100);p__Ascomycota(100);p__Ascomycota_unclassified(100);p__Ascomycota_unclassified(100);p__Ascomycota_unclassified(100);p__Ascomycota_unclassified(100);p__Ascomycota_unclassified(100); | - | - | - | - | - | - | - | Unassigned | - |
| Otu10345 | 2 | k__Fungi(100);k__Fungi_unclassified(100);k__Fungi_unclassified(100);k__Fungi_unclassified(100);k__Fungi_unclassified(100);k__Fungi_unclassified(100);k__Fungi_unclassified(100); | - | - | - | - | - | - | - | Unassigned | - |
| Otu10346 | 2 | k__Fungi(100);k__Fungi_unclassified(100);k__Fungi_unclassified(100);k__Fungi_unclassified(100);k__Fungi_unclassified(100);k__Fungi_unclassified(100);k__Fungi_unclassified(100); | - | - | - | - | - | - | - | Unassigned | - |
| Otu10352 | 2 | k__Fungi(100);k__Fungi_unclassified(100);k__Fungi_unclassified(100);k__Fungi_unclassified(100);k__Fungi_unclassified(100);k__Fungi_unclassified(100);k__Fungi_unclassified(100); | - | - | - | - | - | - | - | Unassigned | - |
| Otu10353 | 2 | k__Fungi(100);p__Ascomycota(100);p__Ascomycota_unclassified(100);p__Ascomycota_unclassified(100);p__Ascomycota_unclassified(100);p__Ascomycota_unclassified(100);p__Ascomycota_unclassified(100); | - | - | - | - | - | - | - | Unassigned | - |
| Otu10354 | 2 | k__Fungi(100);k__Fungi_unclassified(100);k__Fungi_unclassified(100);k__Fungi_unclassified(100);k__Fungi_unclassified(100);k__Fungi_unclassified(100);k__Fungi_unclassified(100); | - | - | - | - | - | - | - | Unassigned | - |
| Otu10361 | 2 | k__Fungi(100);k__Fungi_unclassified(100);k__Fungi_unclassified(100);k__Fungi_unclassified(100);k__Fungi_unclassified(100);k__Fungi_unclassified(100);k__Fungi_unclassified(100); | - | - | - | - | - | - | - | Unassigned | - |
| Otu10365 | 2 | k__Fungi(100);k__Fungi_unclassified(100);k__Fungi_unclassified(100);k__Fungi_unclassified(100);k__Fungi_unclassified(100);k__Fungi_unclassified(100);k__Fungi_unclassified(100); | - | - | - | - | - | - | - | Unassigned | - |
| Otu10367 | 2 | k__Fungi(100);k__Fungi_unclassified(100);k__Fungi_unclassified(100);k__Fungi_unclassified(100);k__Fungi_unclassified(100);k__Fungi_unclassified(100);k__Fungi_unclassified(100); | - | - | - | - | - | - | - | Unassigned | - |
| Otu10368 | 2 | k__Fungi(100);k__Fungi_unclassified(100);k__Fungi_unclassified(100);k__Fungi_unclassified(100);k__Fungi_unclassified(100);k__Fungi_unclassified(100);k__Fungi_unclassified(100); | - | - | - | - | - | - | - | Unassigned | - |
| Otu10371 | 2 | k__Fungi(100);p__Ascomycota(100);p__Ascomycota_unclassified(100);p__Ascomycota_unclassified(100);p__Ascomycota_unclassified(100);p__Ascomycota_unclassified(100);p__Ascomycota_unclassified(100); | - | - | - | - | - | - | - | Unassigned | - |
| Otu10373 | 2 | k__Fungi(100);p__Ascomycota(100);p__Ascomycota_unclassified(100);p__Ascomycota_unclassified(100);p__Ascomycota_unclassified(100);p__Ascomycota_unclassified(100);p__Ascomycota_unclassified(100); | - | - | - | - | - | - | - | Unassigned | - |
| Otu10375 | 2 | k__Fungi(100);k__Fungi_unclassified(100);k__Fungi_unclassified(100);k__Fungi_unclassified(100);k__Fungi_unclassified(100);k__Fungi_unclassified(100);k__Fungi_unclassified(100); | - | - | - | - | - | - | - | Unassigned | - |
| Otu10378 | 2 | k__Fungi(100);k__Fungi_unclassified(100);k__Fungi_unclassified(100);k__Fungi_unclassified(100);k__Fungi_unclassified(100);k__Fungi_unclassified(100);k__Fungi_unclassified(100); | - | - | - | - | - | - | - | Unassigned | - |
| Otu10380 | 2 | k__Fungi(100);k__Fungi_unclassified(100);k__Fungi_unclassified(100);k__Fungi_unclassified(100);k__Fungi_unclassified(100);k__Fungi_unclassified(100);k__Fungi_unclassified(100); | - | - | - | - | - | - | - | Unassigned | - |
| Otu10381 | 2 | k__Fungi(100);k__Fungi_unclassified(100);k__Fungi_unclassified(100);k__Fungi_unclassified(100);k__Fungi_unclassified(100);k__Fungi_unclassified(100);k__Fungi_unclassified(100); | - | - | - | - | - | - | - | Unassigned | - |
| Otu10383 | 2 | k__Fungi(100);p__Ascomycota(100);p__Ascomycota_unclassified(100);p__Ascomycota_unclassified(100);p__Ascomycota_unclassified(100);p__Ascomycota_unclassified(100);p__Ascomycota_unclassified(100); | - | - | - | - | - | - | - | Unassigned | - |
| Otu10384 | 2 | k__Fungi(100);p__Ascomycota(100);p__Ascomycota_unclassified(100);p__Ascomycota_unclassified(100);p__Ascomycota_unclassified(100);p__Ascomycota_unclassified(100);p__Ascomycota_unclassified(100); | - | - | - | - | - | - | - | Unassigned | - |
| Otu10385 | 2 | k__Fungi(100);p__Ascomycota(100);p__Ascomycota_unclassified(100);p__Ascomycota_unclassified(100);p__Ascomycota_unclassified(100);p__Ascomycota_unclassified(100);p__Ascomycota_unclassified(100); | - | - | - | - | - | - | - | Unassigned | - |
| Otu10387 | 2 | k__Fungi(100);p__Ascomycota(100);c__Sordariomycetes(100);c__Sordariomycetes_unclassified(100);c__Sordariomycetes_unclassified(100);c__Sordariomycetes_unclassified(100);c__Sordariomycetes_unclassified(100); | - | - | - | - | - | - | - | Unassigned | - |
| Otu10388 | 2 | k__Fungi(100);k__Fungi_unclassified(100);k__Fungi_unclassified(100);k__Fungi_unclassified(100);k__Fungi_unclassified(100);k__Fungi_unclassified(100);k__Fungi_unclassified(100); | - | - | - | - | - | - | - | Unassigned | - |
| Otu10389 | 2 | k__Fungi(100);k__Fungi_unclassified(100);k__Fungi_unclassified(100);k__Fungi_unclassified(100);k__Fungi_unclassified(100);k__Fungi_unclassified(100);k__Fungi_unclassified(100); | - | - | - | - | - | - | - | Unassigned | - |
| Otu10390 | 2 | k__Fungi(100);k__Fungi_unclassified(100);k__Fungi_unclassified(100);k__Fungi_unclassified(100);k__Fungi_unclassified(100);k__Fungi_unclassified(100);k__Fungi_unclassified(100); | - | - | - | - | - | - | - | Unassigned | - |
| Otu10391 | 2 | k__Fungi(100);k__Fungi_unclassified(100);k__Fungi_unclassified(100);k__Fungi_unclassified(100);k__Fungi_unclassified(100);k__Fungi_unclassified(100);k__Fungi_unclassified(100); | - | - | - | - | - | - | - | Unassigned | - |
| Otu10393 | 2 | k__Fungi(100);p__Ascomycota(100);p__Ascomycota_unclassified(100);p__Ascomycota_unclassified(100);p__Ascomycota_unclassified(100);p__Ascomycota_unclassified(100);p__Ascomycota_unclassified(100); | - | - | - | - | - | - | - | Unassigned | - |
| Otu10394 | 2 | k__Fungi(100);k__Fungi_unclassified(100);k__Fungi_unclassified(100);k__Fungi_unclassified(100);k__Fungi_unclassified(100);k__Fungi_unclassified(100);k__Fungi_unclassified(100); | - | - | - | - | - | - | - | Unassigned | - |
| Otu10395 | 2 | k__Fungi(100);p__Ascomycota(100);p__Ascomycota_unclassified(100);p__Ascomycota_unclassified(100);p__Ascomycota_unclassified(100);p__Ascomycota_unclassified(100);p__Ascomycota_unclassified(100); | - | - | - | - | - | - | - | Unassigned | - |
| Otu10398 | 2 | k__Fungi(100);k__Fungi_unclassified(100);k__Fungi_unclassified(100);k__Fungi_unclassified(100);k__Fungi_unclassified(100);k__Fungi_unclassified(100);k__Fungi_unclassified(100); | - | - | - | - | - | - | - | Unassigned | - |
| Otu10400 | 2 | k__Fungi(100);p__Ascomycota(100);c__Leotiomycetes(100);o__Helotiales(100);o__Helotiales_unclassified(100);o__Helotiales_unclassified(100);o__Helotiales_unclassified(100); | - | - | - | - | - | - | - | Unassigned | - |
| Otu10401 | 2 | k__Fungi(100);p__Ascomycota(100);p__Ascomycota_unclassified(100);p__Ascomycota_unclassified(100);p__Ascomycota_unclassified(100);p__Ascomycota_unclassified(100);p__Ascomycota_unclassified(100); | - | - | - | - | - | - | - | Unassigned | - |
| Otu10406 | 2 | k__Fungi(100);k__Fungi_unclassified(100);k__Fungi_unclassified(100);k__Fungi_unclassified(100);k__Fungi_unclassified(100);k__Fungi_unclassified(100);k__Fungi_unclassified(100); | - | - | - | - | - | - | - | Unassigned | - |
| Otu10408 | 2 | k__Fungi(100);k__Fungi_unclassified(100);k__Fungi_unclassified(100);k__Fungi_unclassified(100);k__Fungi_unclassified(100);k__Fungi_unclassified(100);k__Fungi_unclassified(100); | - | - | - | - | - | - | - | Unassigned | - |
| Otu10410 | 2 | k__Fungi(100);k__Fungi_unclassified(100);k__Fungi_unclassified(100);k__Fungi_unclassified(100);k__Fungi_unclassified(100);k__Fungi_unclassified(100);k__Fungi_unclassified(100); | - | - | - | - | - | - | - | Unassigned | - |
| Otu10412 | 2 | k__Fungi(100);k__Fungi_unclassified(100);k__Fungi_unclassified(100);k__Fungi_unclassified(100);k__Fungi_unclassified(100);k__Fungi_unclassified(100);k__Fungi_unclassified(100); | - | - | - | - | - | - | - | Unassigned | - |
| Otu10413 | 2 | k__Fungi(100);k__Fungi_unclassified(100);k__Fungi_unclassified(100);k__Fungi_unclassified(100);k__Fungi_unclassified(100);k__Fungi_unclassified(100);k__Fungi_unclassified(100); | - | - | - | - | - | - | - | Unassigned | - |
| Otu10415 | 2 | k__Fungi(100);k__Fungi_unclassified(100);k__Fungi_unclassified(100);k__Fungi_unclassified(100);k__Fungi_unclassified(100);k__Fungi_unclassified(100);k__Fungi_unclassified(100); | - | - | - | - | - | - | - | Unassigned | - |
| Otu10417 | 2 | k__Fungi(100);k__Fungi_unclassified(100);k__Fungi_unclassified(100);k__Fungi_unclassified(100);k__Fungi_unclassified(100);k__Fungi_unclassified(100);k__Fungi_unclassified(100); | - | - | - | - | - | - | - | Unassigned | - |
| Otu10418 | 2 | k__Fungi(100);k__Fungi_unclassified(100);k__Fungi_unclassified(100);k__Fungi_unclassified(100);k__Fungi_unclassified(100);k__Fungi_unclassified(100);k__Fungi_unclassified(100); | - | - | - | - | - | - | - | Unassigned | - |
| Otu10421 | 2 | k__Fungi(100);k__Fungi_unclassified(100);k__Fungi_unclassified(100);k__Fungi_unclassified(100);k__Fungi_unclassified(100);k__Fungi_unclassified(100);k__Fungi_unclassified(100); | - | - | - | - | - | - | - | Unassigned | - |
| Otu10422 | 2 | k__Fungi(100);k__Fungi_unclassified(100);k__Fungi_unclassified(100);k__Fungi_unclassified(100);k__Fungi_unclassified(100);k__Fungi_unclassified(100);k__Fungi_unclassified(100); | - | - | - | - | - | - | - | Unassigned | - |
| Otu10423 | 2 | k__Fungi(100);k__Fungi_unclassified(100);k__Fungi_unclassified(100);k__Fungi_unclassified(100);k__Fungi_unclassified(100);k__Fungi_unclassified(100);k__Fungi_unclassified(100); | - | - | - | - | - | - | - | Unassigned | - |
| Otu10424 | 2 | k__Fungi(100);p__Ascomycota(100);p__Ascomycota_unclassified(100);p__Ascomycota_unclassified(100);p__Ascomycota_unclassified(100);p__Ascomycota_unclassified(100);p__Ascomycota_unclassified(100); | - | - | - | - | - | - | - | Unassigned | - |
| Otu10425 | 2 | k__Fungi(100);k__Fungi_unclassified(100);k__Fungi_unclassified(100);k__Fungi_unclassified(100);k__Fungi_unclassified(100);k__Fungi_unclassified(100);k__Fungi_unclassified(100); | - | - | - | - | - | - | - | Unassigned | - |
| Otu10426 | 2 | k__Fungi(100);k__Fungi_unclassified(100);k__Fungi_unclassified(100);k__Fungi_unclassified(100);k__Fungi_unclassified(100);k__Fungi_unclassified(100);k__Fungi_unclassified(100); | - | - | - | - | - | - | - | Unassigned | - |
| Otu10428 | 2 | k__Fungi(100);k__Fungi_unclassified(100);k__Fungi_unclassified(100);k__Fungi_unclassified(100);k__Fungi_unclassified(100);k__Fungi_unclassified(100);k__Fungi_unclassified(100); | - | - | - | - | - | - | - | Unassigned | - |
| Otu10429 | 2 | k__Fungi(100);p__Ascomycota(100);p__Ascomycota_unclassified(100);p__Ascomycota_unclassified(100);p__Ascomycota_unclassified(100);p__Ascomycota_unclassified(100);p__Ascomycota_unclassified(100); | - | - | - | - | - | - | - | Unassigned | - |
| Otu10430 | 2 | k__Fungi(100);k__Fungi_unclassified(100);k__Fungi_unclassified(100);k__Fungi_unclassified(100);k__Fungi_unclassified(100);k__Fungi_unclassified(100);k__Fungi_unclassified(100); | - | - | - | - | - | - | - | Unassigned | - |
| Otu10431 | 2 | k__Fungi(100);k__Fungi_unclassified(100);k__Fungi_unclassified(100);k__Fungi_unclassified(100);k__Fungi_unclassified(100);k__Fungi_unclassified(100);k__Fungi_unclassified(100); | - | - | - | - | - | - | - | Unassigned | - |
| Otu10443 | 2 | k__Fungi(100);k__Fungi_unclassified(100);k__Fungi_unclassified(100);k__Fungi_unclassified(100);k__Fungi_unclassified(100);k__Fungi_unclassified(100);k__Fungi_unclassified(100); | - | - | - | - | - | - | - | Unassigned | - |
| Otu10444 | 2 | k__Fungi(100);k__Fungi_unclassified(100);k__Fungi_unclassified(100);k__Fungi_unclassified(100);k__Fungi_unclassified(100);k__Fungi_unclassified(100);k__Fungi_unclassified(100); | - | - | - | - | - | - | - | Unassigned | - |
| Otu10446 | 2 | k__Fungi(100);k__Fungi_unclassified(100);k__Fungi_unclassified(100);k__Fungi_unclassified(100);k__Fungi_unclassified(100);k__Fungi_unclassified(100);k__Fungi_unclassified(100); | - | - | - | - | - | - | - | Unassigned | - |
| Otu10447 | 2 | k__Fungi(100);p__Ascomycota(100);p__Ascomycota_unclassified(100);p__Ascomycota_unclassified(100);p__Ascomycota_unclassified(100);p__Ascomycota_unclassified(100);p__Ascomycota_unclassified(100); | - | - | - | - | - | - | - | Unassigned | - |
| Otu10448 | 2 | k__Fungi(100);k__Fungi_unclassified(100);k__Fungi_unclassified(100);k__Fungi_unclassified(100);k__Fungi_unclassified(100);k__Fungi_unclassified(100);k__Fungi_unclassified(100); | - | - | - | - | - | - | - | Unassigned | - |
| Otu10456 | 2 | k__Fungi(100);k__Fungi_unclassified(100);k__Fungi_unclassified(100);k__Fungi_unclassified(100);k__Fungi_unclassified(100);k__Fungi_unclassified(100);k__Fungi_unclassified(100); | - | - | - | - | - | - | - | Unassigned | - |
| Otu10457 | 2 | k__Fungi(100);k__Fungi_unclassified(100);k__Fungi_unclassified(100);k__Fungi_unclassified(100);k__Fungi_unclassified(100);k__Fungi_unclassified(100);k__Fungi_unclassified(100); | - | - | - | - | - | - | - | Unassigned | - |
| Otu10460 | 2 | k__Fungi(100);k__Fungi_unclassified(100);k__Fungi_unclassified(100);k__Fungi_unclassified(100);k__Fungi_unclassified(100);k__Fungi_unclassified(100);k__Fungi_unclassified(100); | - | - | - | - | - | - | - | Unassigned | - |
| Otu10461 | 2 | k__Fungi(100);p__Ascomycota(100);p__Ascomycota_unclassified(100);p__Ascomycota_unclassified(100);p__Ascomycota_unclassified(100);p__Ascomycota_unclassified(100);p__Ascomycota_unclassified(100); | - | - | - | - | - | - | - | Unassigned | - |
| Otu10466 | 2 | k__Fungi(100);k__Fungi_unclassified(100);k__Fungi_unclassified(100);k__Fungi_unclassified(100);k__Fungi_unclassified(100);k__Fungi_unclassified(100);k__Fungi_unclassified(100); | - | - | - | - | - | - | - | Unassigned | - |
| Otu10471 | 2 | k__Fungi(100);k__Fungi_unclassified(100);k__Fungi_unclassified(100);k__Fungi_unclassified(100);k__Fungi_unclassified(100);k__Fungi_unclassified(100);k__Fungi_unclassified(100); | - | - | - | - | - | - | - | Unassigned | - |
| Otu10472 | 2 | k__Fungi(100);p__Ascomycota(100);p__Ascomycota_unclassified(100);p__Ascomycota_unclassified(100);p__Ascomycota_unclassified(100);p__Ascomycota_unclassified(100);p__Ascomycota_unclassified(100); | - | - | - | - | - | - | - | Unassigned | - |
| Otu10475 | 2 | k__Fungi(100);k__Fungi_unclassified(100);k__Fungi_unclassified(100);k__Fungi_unclassified(100);k__Fungi_unclassified(100);k__Fungi_unclassified(100);k__Fungi_unclassified(100); | - | - | - | - | - | - | - | Unassigned | - |
| Otu10480 | 2 | k__Fungi(100);k__Fungi_unclassified(100);k__Fungi_unclassified(100);k__Fungi_unclassified(100);k__Fungi_unclassified(100);k__Fungi_unclassified(100);k__Fungi_unclassified(100); | - | - | - | - | - | - | - | Unassigned | - |
| Otu10482 | 2 | k__Fungi(100);p__Ascomycota(100);p__Ascomycota_unclassified(100);p__Ascomycota_unclassified(100);p__Ascomycota_unclassified(100);p__Ascomycota_unclassified(100);p__Ascomycota_unclassified(100); | - | - | - | - | - | - | - | Unassigned | - |
| Otu10484 | 2 | k__Fungi(100);p__Ascomycota(100);p__Ascomycota_unclassified(100);p__Ascomycota_unclassified(100);p__Ascomycota_unclassified(100);p__Ascomycota_unclassified(100);p__Ascomycota_unclassified(100); | - | - | - | - | - | - | - | Unassigned | - |
| Otu10488 | 2 | k__Fungi(100);p__Ascomycota(100);p__Ascomycota_unclassified(100);p__Ascomycota_unclassified(100);p__Ascomycota_unclassified(100);p__Ascomycota_unclassified(100);p__Ascomycota_unclassified(100); | - | - | - | - | - | - | - | Unassigned | - |
| Otu10489 | 2 | k__Fungi(100);k__Fungi_unclassified(100);k__Fungi_unclassified(100);k__Fungi_unclassified(100);k__Fungi_unclassified(100);k__Fungi_unclassified(100);k__Fungi_unclassified(100); | - | - | - | - | - | - | - | Unassigned | - |
| Otu10497 | 2 | k__Fungi(100);k__Fungi_unclassified(100);k__Fungi_unclassified(100);k__Fungi_unclassified(100);k__Fungi_unclassified(100);k__Fungi_unclassified(100);k__Fungi_unclassified(100); | - | - | - | - | - | - | - | Unassigned | - |
| Otu10498 | 2 | k__Fungi(100);p__Ascomycota(100);p__Ascomycota_unclassified(100);p__Ascomycota_unclassified(100);p__Ascomycota_unclassified(100);p__Ascomycota_unclassified(100);p__Ascomycota_unclassified(100); | - | - | - | - | - | - | - | Unassigned | - |
| Otu10499 | 2 | k__Fungi(100);p__Ascomycota(100);c__Leotiomycetes(100);o__Helotiales(100);o__Helotiales_unclassified(100);o__Helotiales_unclassified(100);o__Helotiales_unclassified(100); | - | - | - | - | - | - | - | Unassigned | - |
| Otu10502 | 2 | k__Fungi(100);k__Fungi_unclassified(100);k__Fungi_unclassified(100);k__Fungi_unclassified(100);k__Fungi_unclassified(100);k__Fungi_unclassified(100);k__Fungi_unclassified(100); | - | - | - | - | - | - | - | Unassigned | - |
| Otu10503 | 2 | k__Fungi(100);k__Fungi_unclassified(100);k__Fungi_unclassified(100);k__Fungi_unclassified(100);k__Fungi_unclassified(100);k__Fungi_unclassified(100);k__Fungi_unclassified(100); | - | - | - | - | - | - | - | Unassigned | - |
| Otu10504 | 2 | k__Fungi(100);k__Fungi_unclassified(100);k__Fungi_unclassified(100);k__Fungi_unclassified(100);k__Fungi_unclassified(100);k__Fungi_unclassified(100);k__Fungi_unclassified(100); | - | - | - | - | - | - | - | Unassigned | - |
| Otu10505 | 2 | k__Fungi(100);k__Fungi_unclassified(100);k__Fungi_unclassified(100);k__Fungi_unclassified(100);k__Fungi_unclassified(100);k__Fungi_unclassified(100);k__Fungi_unclassified(100); | - | - | - | - | - | - | - | Unassigned | - |
| Otu10506 | 2 | k__Fungi(100);p__Ascomycota(100);p__Ascomycota_unclassified(100);p__Ascomycota_unclassified(100);p__Ascomycota_unclassified(100);p__Ascomycota_unclassified(100);p__Ascomycota_unclassified(100); | - | - | - | - | - | - | - | Unassigned | - |
| Otu10514 | 2 | k__Fungi(100);k__Fungi_unclassified(100);k__Fungi_unclassified(100);k__Fungi_unclassified(100);k__Fungi_unclassified(100);k__Fungi_unclassified(100);k__Fungi_unclassified(100); | - | - | - | - | - | - | - | Unassigned | - |
| Otu10516 | 2 | k__Fungi(100);k__Fungi_unclassified(100);k__Fungi_unclassified(100);k__Fungi_unclassified(100);k__Fungi_unclassified(100);k__Fungi_unclassified(100);k__Fungi_unclassified(100); | - | - | - | - | - | - | - | Unassigned | - |
| Otu10517 | 2 | k__Fungi(100);k__Fungi_unclassified(100);k__Fungi_unclassified(100);k__Fungi_unclassified(100);k__Fungi_unclassified(100);k__Fungi_unclassified(100);k__Fungi_unclassified(100); | - | - | - | - | - | - | - | Unassigned | - |
| Otu10518 | 2 | k__Fungi(100);k__Fungi_unclassified(100);k__Fungi_unclassified(100);k__Fungi_unclassified(100);k__Fungi_unclassified(100);k__Fungi_unclassified(100);k__Fungi_unclassified(100); | - | - | - | - | - | - | - | Unassigned | - |
| Otu10529 | 2 | k__Fungi(100);k__Fungi_unclassified(100);k__Fungi_unclassified(100);k__Fungi_unclassified(100);k__Fungi_unclassified(100);k__Fungi_unclassified(100);k__Fungi_unclassified(100); | - | - | - | - | - | - | - | Unassigned | - |
| Otu10530 | 2 | k__Fungi(100);k__Fungi_unclassified(100);k__Fungi_unclassified(100);k__Fungi_unclassified(100);k__Fungi_unclassified(100);k__Fungi_unclassified(100);k__Fungi_unclassified(100); | - | - | - | - | - | - | - | Unassigned | - |
| Otu10531 | 2 | k__Fungi(100);k__Fungi_unclassified(100);k__Fungi_unclassified(100);k__Fungi_unclassified(100);k__Fungi_unclassified(100);k__Fungi_unclassified(100);k__Fungi_unclassified(100); | - | - | - | - | - | - | - | Unassigned | - |
| Otu10534 | 2 | k__Fungi(100);p__Ascomycota(100);c__Dothideomycetes(100);c__Dothideomycetes_unclassified(100);c__Dothideomycetes_unclassified(100);c__Dothideomycetes_unclassified(100);c__Dothideomycetes_unclassified(100); | - | - | - | - | - | - | - | Unassigned | - |
| Otu10535 | 2 | k__Fungi(100);k__Fungi_unclassified(100);k__Fungi_unclassified(100);k__Fungi_unclassified(100);k__Fungi_unclassified(100);k__Fungi_unclassified(100);k__Fungi_unclassified(100); | - | - | - | - | - | - | - | Unassigned | - |
| Otu10538 | 2 | k__Fungi(100);p__Ascomycota(100);c__Leotiomycetes(100);c__Leotiomycetes_unclassified(100);c__Leotiomycetes_unclassified(100);c__Leotiomycetes_unclassified(100);c__Leotiomycetes_unclassified(100); | - | - | - | - | - | - | - | Unassigned | - |
| Otu10540 | 2 | k__Fungi(100);k__Fungi_unclassified(100);k__Fungi_unclassified(100);k__Fungi_unclassified(100);k__Fungi_unclassified(100);k__Fungi_unclassified(100);k__Fungi_unclassified(100); | - | - | - | - | - | - | - | Unassigned | - |
| Otu10541 | 2 | k__Fungi(100);k__Fungi_unclassified(100);k__Fungi_unclassified(100);k__Fungi_unclassified(100);k__Fungi_unclassified(100);k__Fungi_unclassified(100);k__Fungi_unclassified(100); | - | - | - | - | - | - | - | Unassigned | - |
| Otu10542 | 2 | k__Fungi(100);k__Fungi_unclassified(100);k__Fungi_unclassified(100);k__Fungi_unclassified(100);k__Fungi_unclassified(100);k__Fungi_unclassified(100);k__Fungi_unclassified(100); | - | - | - | - | - | - | - | Unassigned | - |
| Otu10543 | 2 | k__Fungi(100);k__Fungi_unclassified(100);k__Fungi_unclassified(100);k__Fungi_unclassified(100);k__Fungi_unclassified(100);k__Fungi_unclassified(100);k__Fungi_unclassified(100); | - | - | - | - | - | - | - | Unassigned | - |
| Otu10544 | 2 | k__Fungi(100);k__Fungi_unclassified(100);k__Fungi_unclassified(100);k__Fungi_unclassified(100);k__Fungi_unclassified(100);k__Fungi_unclassified(100);k__Fungi_unclassified(100); | - | - | - | - | - | - | - | Unassigned | - |
| Otu10546 | 2 | k__Fungi(100);k__Fungi_unclassified(100);k__Fungi_unclassified(100);k__Fungi_unclassified(100);k__Fungi_unclassified(100);k__Fungi_unclassified(100);k__Fungi_unclassified(100); | - | - | - | - | - | - | - | Unassigned | - |
| Otu10547 | 2 | k__Fungi(100);k__Fungi_unclassified(100);k__Fungi_unclassified(100);k__Fungi_unclassified(100);k__Fungi_unclassified(100);k__Fungi_unclassified(100);k__Fungi_unclassified(100); | - | - | - | - | - | - | - | Unassigned | - |
| Otu10552 | 2 | k__Fungi(100);k__Fungi_unclassified(100);k__Fungi_unclassified(100);k__Fungi_unclassified(100);k__Fungi_unclassified(100);k__Fungi_unclassified(100);k__Fungi_unclassified(100); | - | - | - | - | - | - | - | Unassigned | - |
| Otu10553 | 2 | k__Fungi(100);p__Ascomycota(100);p__Ascomycota_unclassified(100);p__Ascomycota_unclassified(100);p__Ascomycota_unclassified(100);p__Ascomycota_unclassified(100);p__Ascomycota_unclassified(100); | - | - | - | - | - | - | - | Unassigned | - |
| Otu10554 | 2 | k__Fungi(100);k__Fungi_unclassified(100);k__Fungi_unclassified(100);k__Fungi_unclassified(100);k__Fungi_unclassified(100);k__Fungi_unclassified(100);k__Fungi_unclassified(100); | - | - | - | - | - | - | - | Unassigned | - |
| Otu10555 | 2 | k__Fungi(100);p__Ascomycota(100);p__Ascomycota_unclassified(100);p__Ascomycota_unclassified(100);p__Ascomycota_unclassified(100);p__Ascomycota_unclassified(100);p__Ascomycota_unclassified(100); | - | - | - | - | - | - | - | Unassigned | - |
| Otu10561 | 2 | k__Fungi(100);k__Fungi_unclassified(100);k__Fungi_unclassified(100);k__Fungi_unclassified(100);k__Fungi_unclassified(100);k__Fungi_unclassified(100);k__Fungi_unclassified(100); | - | - | - | - | - | - | - | Unassigned | - |
| Otu10562 | 2 | k__Fungi(100);p__Ascomycota(100);p__Ascomycota_unclassified(100);p__Ascomycota_unclassified(100);p__Ascomycota_unclassified(100);p__Ascomycota_unclassified(100);p__Ascomycota_unclassified(100); | - | - | - | - | - | - | - | Unassigned | - |
| Otu10564 | 2 | k__Fungi(100);k__Fungi_unclassified(100);k__Fungi_unclassified(100);k__Fungi_unclassified(100);k__Fungi_unclassified(100);k__Fungi_unclassified(100);k__Fungi_unclassified(100); | - | - | - | - | - | - | - | Unassigned | - |
| Otu10568 | 2 | k__Fungi(100);k__Fungi_unclassified(100);k__Fungi_unclassified(100);k__Fungi_unclassified(100);k__Fungi_unclassified(100);k__Fungi_unclassified(100);k__Fungi_unclassified(100); | - | - | - | - | - | - | - | Unassigned | - |
| Otu10569 | 2 | k__Fungi(100);k__Fungi_unclassified(100);k__Fungi_unclassified(100);k__Fungi_unclassified(100);k__Fungi_unclassified(100);k__Fungi_unclassified(100);k__Fungi_unclassified(100); | - | - | - | - | - | - | - | Unassigned | - |
| Otu10571 | 2 | k__Fungi(100);p__Ascomycota(100);p__Ascomycota_unclassified(100);p__Ascomycota_unclassified(100);p__Ascomycota_unclassified(100);p__Ascomycota_unclassified(100);p__Ascomycota_unclassified(100); | - | - | - | - | - | - | - | Unassigned | - |
| Otu10572 | 2 | k__Fungi(100);k__Fungi_unclassified(100);k__Fungi_unclassified(100);k__Fungi_unclassified(100);k__Fungi_unclassified(100);k__Fungi_unclassified(100);k__Fungi_unclassified(100); | - | - | - | - | - | - | - | Unassigned | - |
| Otu10574 | 2 | k__Fungi(100);p__Ascomycota(100);p__Ascomycota_unclassified(100);p__Ascomycota_unclassified(100);p__Ascomycota_unclassified(100);p__Ascomycota_unclassified(100);p__Ascomycota_unclassified(100); | - | - | - | - | - | - | - | Unassigned | - |
| Otu10575 | 2 | k__Fungi(100);p__Ascomycota(100);p__Ascomycota_unclassified(100);p__Ascomycota_unclassified(100);p__Ascomycota_unclassified(100);p__Ascomycota_unclassified(100);p__Ascomycota_unclassified(100); | - | - | - | - | - | - | - | Unassigned | - |
| Otu10578 | 2 | k__Fungi(100);k__Fungi_unclassified(100);k__Fungi_unclassified(100);k__Fungi_unclassified(100);k__Fungi_unclassified(100);k__Fungi_unclassified(100);k__Fungi_unclassified(100); | - | - | - | - | - | - | - | Unassigned | - |
| Otu10579 | 2 | k__Fungi(100);k__Fungi_unclassified(100);k__Fungi_unclassified(100);k__Fungi_unclassified(100);k__Fungi_unclassified(100);k__Fungi_unclassified(100);k__Fungi_unclassified(100); | - | - | - | - | - | - | - | Unassigned | - |
| Otu10583 | 2 | k__Fungi(100);p__Ascomycota(100);p__Ascomycota_unclassified(100);p__Ascomycota_unclassified(100);p__Ascomycota_unclassified(100);p__Ascomycota_unclassified(100);p__Ascomycota_unclassified(100); | - | - | - | - | - | - | - | Unassigned | - |
| Otu10587 | 2 | k__Fungi(100);p__Ascomycota(100);p__Ascomycota_unclassified(100);p__Ascomycota_unclassified(100);p__Ascomycota_unclassified(100);p__Ascomycota_unclassified(100);p__Ascomycota_unclassified(100); | - | - | - | - | - | - | - | Unassigned | - |
| Otu10589 | 2 | k__Fungi(100);p__Ascomycota(100);p__Ascomycota_unclassified(100);p__Ascomycota_unclassified(100);p__Ascomycota_unclassified(100);p__Ascomycota_unclassified(100);p__Ascomycota_unclassified(100); | - | - | - | - | - | - | - | Unassigned | - |
| Otu10592 | 2 | k__Fungi(100);p__Ascomycota(100);c__Sordariomycetes(100);c__Sordariomycetes_unclassified(100);c__Sordariomycetes_unclassified(100);c__Sordariomycetes_unclassified(100);c__Sordariomycetes_unclassified(100); | - | - | - | - | - | - | - | Unassigned | - |
| Otu10594 | 2 | k__Fungi(100);p__Ascomycota(100);p__Ascomycota_unclassified(100);p__Ascomycota_unclassified(100);p__Ascomycota_unclassified(100);p__Ascomycota_unclassified(100);p__Ascomycota_unclassified(100); | - | - | - | - | - | - | - | Unassigned | - |
| Otu10597 | 2 | k__Fungi(100);k__Fungi_unclassified(100);k__Fungi_unclassified(100);k__Fungi_unclassified(100);k__Fungi_unclassified(100);k__Fungi_unclassified(100);k__Fungi_unclassified(100); | - | - | - | - | - | - | - | Unassigned | - |
| Otu10599 | 2 | k__Fungi(100);p__Ascomycota(100);p__Ascomycota_unclassified(100);p__Ascomycota_unclassified(100);p__Ascomycota_unclassified(100);p__Ascomycota_unclassified(100);p__Ascomycota_unclassified(100); | - | - | - | - | - | - | - | Unassigned | - |
| Otu10600 | 2 | k__Fungi(100);p__Ascomycota(100);p__Ascomycota_unclassified(100);p__Ascomycota_unclassified(100);p__Ascomycota_unclassified(100);p__Ascomycota_unclassified(100);p__Ascomycota_unclassified(100); | - | - | - | - | - | - | - | Unassigned | - |
| Otu10601 | 2 | k__Fungi(100);p__Ascomycota(100);p__Ascomycota_unclassified(100);p__Ascomycota_unclassified(100);p__Ascomycota_unclassified(100);p__Ascomycota_unclassified(100);p__Ascomycota_unclassified(100); | - | - | - | - | - | - | - | Unassigned | - |
| Otu10602 | 2 | k__Fungi(100);p__Ascomycota(100);p__Ascomycota_unclassified(100);p__Ascomycota_unclassified(100);p__Ascomycota_unclassified(100);p__Ascomycota_unclassified(100);p__Ascomycota_unclassified(100); | - | - | - | - | - | - | - | Unassigned | - |
| Otu10603 | 2 | k__Fungi(100);p__Ascomycota(100);p__Ascomycota_unclassified(100);p__Ascomycota_unclassified(100);p__Ascomycota_unclassified(100);p__Ascomycota_unclassified(100);p__Ascomycota_unclassified(100); | - | - | - | - | - | - | - | Unassigned | - |
| Otu10604 | 2 | k__Fungi(100);p__Ascomycota(100);p__Ascomycota_unclassified(100);p__Ascomycota_unclassified(100);p__Ascomycota_unclassified(100);p__Ascomycota_unclassified(100);p__Ascomycota_unclassified(100); | - | - | - | - | - | - | - | Unassigned | - |
| Otu10606 | 2 | k__Fungi(100);k__Fungi_unclassified(100);k__Fungi_unclassified(100);k__Fungi_unclassified(100);k__Fungi_unclassified(100);k__Fungi_unclassified(100);k__Fungi_unclassified(100); | - | - | - | - | - | - | - | Unassigned | - |
| Otu10607 | 2 | k__Fungi(100);p__Ascomycota(100);c__Sordariomycetes(100);o__Hypocreales(100);f__Cordycipitaceae(100);f__Cordycipitaceae_unclassified(100);f__Cordycipitaceae_unclassified(100); | - | - | - | - | - | - | - | Unassigned | - |
| Otu10609 | 2 | k__Fungi(100);p__Ascomycota(100);p__Ascomycota_unclassified(100);p__Ascomycota_unclassified(100);p__Ascomycota_unclassified(100);p__Ascomycota_unclassified(100);p__Ascomycota_unclassified(100); | - | - | - | - | - | - | - | Unassigned | - |
| Otu10613 | 2 | k__Fungi(100);p__Ascomycota(100);c__Eurotiomycetes(100);o__Eurotiales(100);f__Trichocomaceae(100);f__Trichocomaceae_unclassified(100);f__Trichocomaceae_unclassified(100); | - | - | - | - | - | - | - | Unassigned | - |
| Otu10614 | 2 | k__Fungi(100);k__Fungi_unclassified(100);k__Fungi_unclassified(100);k__Fungi_unclassified(100);k__Fungi_unclassified(100);k__Fungi_unclassified(100);k__Fungi_unclassified(100); | - | - | - | - | - | - | - | Unassigned | - |
| Otu10616 | 2 | k__Fungi(100);p__Ascomycota(100);p__Ascomycota_unclassified(100);p__Ascomycota_unclassified(100);p__Ascomycota_unclassified(100);p__Ascomycota_unclassified(100);p__Ascomycota_unclassified(100); | - | - | - | - | - | - | - | Unassigned | - |
| Otu10617 | 2 | k__Fungi(100);p__Ascomycota(100);c__Eurotiomycetes(100);o__Onygenales(100);o__Onygenales_unclassified(100);o__Onygenales_unclassified(100);o__Onygenales_unclassified(100); | - | - | - | - | - | - | - | Unassigned | - |
| Otu10618 | 2 | k__Fungi(100);p__Ascomycota(100);c__Eurotiomycetes(100);c__Eurotiomycetes_unclassified(100);c__Eurotiomycetes_unclassified(100);c__Eurotiomycetes_unclassified(100);c__Eurotiomycetes_unclassified(100); | - | - | - | - | - | - | - | Unassigned | - |
| Otu10620 | 2 | k__Fungi(100);p__Ascomycota(100);p__Ascomycota_unclassified(100);p__Ascomycota_unclassified(100);p__Ascomycota_unclassified(100);p__Ascomycota_unclassified(100);p__Ascomycota_unclassified(100); | - | - | - | - | - | - | - | Unassigned | - |
| Otu10621 | 2 | k__Fungi(100);p__Ascomycota(100);p__Ascomycota_unclassified(100);p__Ascomycota_unclassified(100);p__Ascomycota_unclassified(100);p__Ascomycota_unclassified(100);p__Ascomycota_unclassified(100); | - | - | - | - | - | - | - | Unassigned | - |
| Otu10623 | 2 | k__Fungi(100);p__Ascomycota(100);c__Eurotiomycetes(100);o__Eurotiales(100);f__Trichocomaceae(100);f__Trichocomaceae_unclassified(100);f__Trichocomaceae_unclassified(100); | - | - | - | - | - | - | - | Unassigned | - |
| Otu10625 | 2 | k__Plantae(100);p__unclassified_Plantae(100);c__unclassified_Plantae(100);o__unclassified_Plantae(100);f__unclassified_Plantae(100);g__unclassified_Plantae(100);s__Plantae_sp(100); | - | - | - | - | - | - | - | Unassigned | - |

| | | | | | | | | | | | |
|---|---|---|---|---|---|---|---|---|---|---|---|
| Otu10626 | 2 | k__Plantae(100);p__unclassified_Plantae(100);c__unclassified_Plantae(100);o__unclassified_Plantae(100);f__unclassified_Plantae(100);g__unclassified_Plantae(100);s__Plantae_sp(100); | - | - | - | - | - | - | - | Unassigned | - |
| Otu10627 | 2 | k__Fungi(100);k__Fungi_unclassified(100);k__Fungi_unclassified(100);k__Fungi_unclassified(100);k__Fungi_unclassified(100);k__Fungi_unclassified(100);k__Fungi_unclassified(100); | - | - | - | - | - | - | - | Unassigned | - |
| Otu10629 | 2 | k__Plantae(100);p__unclassified_Plantae(100);c__unclassified_Plantae(100);o__unclassified_Plantae(100);f__unclassified_Plantae(100);g__unclassified_Plantae(100);s__Plantae_sp(100); | - | - | - | - | - | - | - | Unassigned | - |
| Otu10630 | 2 | k__Plantae(100);p__unclassified_Plantae(100);c__unclassified_Plantae(100);o__unclassified_Plantae(100);f__unclassified_Plantae(100);g__unclassified_Plantae(100);s__Plantae_sp(100); | - | - | - | - | - | - | - | Unassigned | - |
| Otu10631 | 2 | k__Plantae(100);p__unclassified_Plantae(100);c__unclassified_Plantae(100);o__unclassified_Plantae(100);f__unclassified_Plantae(100);g__unclassified_Plantae(100);s__Plantae_sp(100); | - | - | - | - | - | - | - | Unassigned | - |
| Otu10639 | 2 | unknown;unknown_unclassified(0);unknown_unclassified(0);unknown_unclassified(0);unknown_unclassified(0);unknown_unclassified(0);unknown_unclassified(0); | - | - | - | - | - | - | - | Unassigned | - |
| Otu10641 | 2 | k__Fungi(100);p__Ascomycota(100);p__Ascomycota_unclassified(100);p__Ascomycota_unclassified(100);p__Ascomycota_unclassified(100);p__Ascomycota_unclassified(100);p__Ascomycota_unclassified(100); | - | - | - | - | - | - | - | Unassigned | - |
| Otu10642 | 2 | k__Plantae(100);p__unclassified_Plantae(100);c__unclassified_Plantae(100);o__unclassified_Plantae(100);f__unclassified_Plantae(100);g__unclassified_Plantae(100);s__Plantae_sp(100); | - | - | - | - | - | - | - | Unassigned | - |
| Otu10643 | 2 | unknown;unknown_unclassified(0);unknown_unclassified(0);unknown_unclassified(0);unknown_unclassified(0);unknown_unclassified(0);unknown_unclassified(0); | - | - | - | - | - | - | - | Unassigned | - |
| Otu10644 | 2 | unknown;unknown_unclassified(0);unknown_unclassified(0);unknown_unclassified(0);unknown_unclassified(0);unknown_unclassified(0);unknown_unclassified(0); | - | - | - | - | - | - | - | Unassigned | - |
| Otu10645 | 2 | k__Plantae(100);p__unclassified_Plantae(100);c__unclassified_Plantae(100);o__unclassified_Plantae(100);f__unclassified_Plantae(100);g__unclassified_Plantae(100);s__Plantae_sp(100); | - | - | - | - | - | - | - | Unassigned | - |
| Otu10646 | 2 | k__Plantae(100);p__unclassified_Plantae(100);c__unclassified_Plantae(100);o__unclassified_Plantae(100);f__unclassified_Plantae(100);g__unclassified_Plantae(100);s__Plantae_sp(100); | - | - | - | - | - | - | - | Unassigned | - |
| Otu10648 | 2 | k__Fungi(100);k__Fungi_unclassified(100);k__Fungi_unclassified(100);k__Fungi_unclassified(100);k__Fungi_unclassified(100);k__Fungi_unclassified(100);k__Fungi_unclassified(100); | - | - | - | - | - | - | - | Unassigned | - |
| Otu10652 | 2 | k__Fungi(100);k__Fungi_unclassified(100);k__Fungi_unclassified(100);k__Fungi_unclassified(100);k__Fungi_unclassified(100);k__Fungi_unclassified(100);k__Fungi_unclassified(100); | - | - | - | - | - | - | - | Unassigned | - |
| Otu10655 | 2 | k__Fungi(100);k__Fungi_unclassified(100);k__Fungi_unclassified(100);k__Fungi_unclassified(100);k__Fungi_unclassified(100);k__Fungi_unclassified(100);k__Fungi_unclassified(100); | - | - | - | - | - | - | - | Unassigned | - |
| Otu10657 | 2 | k__Fungi(100);p__Ascomycota(100);p__Ascomycota_unclassified(100);p__Ascomycota_unclassified(100);p__Ascomycota_unclassified(100);p__Ascomycota_unclassified(100);p__Ascomycota_unclassified(100); | - | - | - | - | - | - | - | Unassigned | - |
| Otu10659 | 2 | k__Fungi(100);k__Fungi_unclassified(100);k__Fungi_unclassified(100);k__Fungi_unclassified(100);k__Fungi_unclassified(100);k__Fungi_unclassified(100);k__Fungi_unclassified(100); | - | - | - | - | - | - | - | Unassigned | - |
| Otu10660 | 2 | k__Fungi(100);p__Ascomycota(100);p__Ascomycota_unclassified(100);p__Ascomycota_unclassified(100);p__Ascomycota_unclassified(100);p__Ascomycota_unclassified(100);p__Ascomycota_unclassified(100); | - | - | - | - | - | - | - | Unassigned | - |
| Otu10663 | 2 | k__Fungi(100);p__Ascomycota(100);p__Ascomycota_unclassified(100);p__Ascomycota_unclassified(100);p__Ascomycota_unclassified(100);p__Ascomycota_unclassified(100);p__Ascomycota_unclassified(100); | - | - | - | - | - | - | - | Unassigned | - |
| Otu10664 | 2 | k__Fungi(100);p__Ascomycota(100);p__Ascomycota_unclassified(100);p__Ascomycota_unclassified(100);p__Ascomycota_unclassified(100);p__Ascomycota_unclassified(100);p__Ascomycota_unclassified(100); | - | - | - | - | - | - | - | Unassigned | - |
| Otu10669 | 2 | k__Fungi(100);p__Ascomycota(100);p__Ascomycota_unclassified(100);p__Ascomycota_unclassified(100);p__Ascomycota_unclassified(100);p__Ascomycota_unclassified(100);p__Ascomycota_unclassified(100); | - | - | - | - | - | - | - | Unassigned | - |
| Otu10670 | 2 | k__Fungi(100);k__Fungi_unclassified(100);k__Fungi_unclassified(100);k__Fungi_unclassified(100);k__Fungi_unclassified(100);k__Fungi_unclassified(100);k__Fungi_unclassified(100); | - | - | - | - | - | - | - | Unassigned | - |
| Otu10671 | 2 | k__Fungi(100);k__Fungi_unclassified(100);k__Fungi_unclassified(100);k__Fungi_unclassified(100);k__Fungi_unclassified(100);k__Fungi_unclassified(100);k__Fungi_unclassified(100); | - | - | - | - | - | - | - | Unassigned | - |
| Otu10674 | 2 | k__Fungi(100);p__Ascomycota(100);p__Ascomycota_unclassified(100);p__Ascomycota_unclassified(100);p__Ascomycota_unclassified(100);p__Ascomycota_unclassified(100);p__Ascomycota_unclassified(100); | - | - | - | - | - | - | - | Unassigned | - |
| Otu10675 | 2 | k__Plantae(100);p__unclassified_Plantae(100);c__unclassified_Plantae(100);o__unclassified_Plantae(100);f__unclassified_Plantae(100);g__unclassified_Plantae(100);s__Plantae_sp(100); | - | - | - | - | - | - | - | Unassigned | - |
| Otu10676 | 2 | k__Fungi(100);k__Fungi_unclassified(100);k__Fungi_unclassified(100);k__Fungi_unclassified(100);k__Fungi_unclassified(100);k__Fungi_unclassified(100);k__Fungi_unclassified(100); | - | - | - | - | - | - | - | Unassigned | - |
| Otu10677 | 2 | k__Fungi(100);k__Fungi_unclassified(100);k__Fungi_unclassified(100);k__Fungi_unclassified(100);k__Fungi_unclassified(100);k__Fungi_unclassified(100);k__Fungi_unclassified(100); | - | - | - | - | - | - | - | Unassigned | - |
| Otu10680 | 2 | k__Fungi(100);k__Fungi_unclassified(100);k__Fungi_unclassified(100);k__Fungi_unclassified(100);k__Fungi_unclassified(100);k__Fungi_unclassified(100);k__Fungi_unclassified(100); | - | - | - | - | - | - | - | Unassigned | - |
| Otu10683 | 2 | k__Fungi(100);p__Ascomycota(100);p__Ascomycota_unclassified(100);p__Ascomycota_unclassified(100);p__Ascomycota_unclassified(100);p__Ascomycota_unclassified(100);p__Ascomycota_unclassified(100); | - | - | - | - | - | - | - | Unassigned | - |
| Otu10684 | 2 | k__Fungi(100);p__Ascomycota(100);p__Ascomycota_unclassified(100);p__Ascomycota_unclassified(100);p__Ascomycota_unclassified(100);p__Ascomycota_unclassified(100);p__Ascomycota_unclassified(100); | - | - | - | - | - | - | - | Unassigned | - |
| Otu10685 | 2 | k__Fungi(100);p__Ascomycota(100);p__Ascomycota_unclassified(100);p__Ascomycota_unclassified(100);p__Ascomycota_unclassified(100);p__Ascomycota_unclassified(100);p__Ascomycota_unclassified(100); | - | - | - | - | - | - | - | Unassigned | - |
| Otu10686 | 2 | k__Fungi(100);p__Ascomycota(100);p__Ascomycota_unclassified(100);p__Ascomycota_unclassified(100);p__Ascomycota_unclassified(100);p__Ascomycota_unclassified(100);p__Ascomycota_unclassified(100); | - | - | - | - | - | - | - | Unassigned | - |
| Otu10688 | 2 | k__Fungi(100);p__Ascomycota(100);p__Ascomycota_unclassified(100);p__Ascomycota_unclassified(100);p__Ascomycota_unclassified(100);p__Ascomycota_unclassified(100);p__Ascomycota_unclassified(100); | - | - | - | - | - | - | - | Unassigned | - |
| Otu10689 | 2 | k__Fungi(100);p__Ascomycota(100);p__Ascomycota_unclassified(100);p__Ascomycota_unclassified(100);p__Ascomycota_unclassified(100);p__Ascomycota_unclassified(100);p__Ascomycota_unclassified(100); | - | - | - | - | - | - | - | Unassigned | - |
| Otu10691 | 2 | k__Fungi(100);p__Ascomycota(100);p__Ascomycota_unclassified(100);p__Ascomycota_unclassified(100);p__Ascomycota_unclassified(100);p__Ascomycota_unclassified(100);p__Ascomycota_unclassified(100); | - | - | - | - | - | - | - | Unassigned | - |
| Otu10694 | 2 | k__Fungi(100);p__Ascomycota(100);c__Sordariomycetes(100);c__Sordariomycetes_unclassified(100);c__Sordariomycetes_unclassified(100);c__Sordariomycetes_unclassified(100);c__Sordariomycetes_unclassified(100); | - | - | - | - | - | - | - | Unassigned | - |
| Otu10696 | 2 | k__Fungi(100);k__Fungi_unclassified(100);k__Fungi_unclassified(100);k__Fungi_unclassified(100);k__Fungi_unclassified(100);k__Fungi_unclassified(100);k__Fungi_unclassified(100); | - | - | - | - | - | - | - | Unassigned | - |
| Otu10697 | 2 | k__Fungi(100);k__Fungi_unclassified(100);k__Fungi_unclassified(100);k__Fungi_unclassified(100);k__Fungi_unclassified(100);k__Fungi_unclassified(100);k__Fungi_unclassified(100); | - | - | - | - | - | - | - | Unassigned | - |
| Otu10699 | 2 | k__Fungi(100);p__Ascomycota(100);p__Ascomycota_unclassified(100);p__Ascomycota_unclassified(100);p__Ascomycota_unclassified(100);p__Ascomycota_unclassified(100);p__Ascomycota_unclassified(100); | - | - | - | - | - | - | - | Unassigned | - |
| Otu10700 | 2 | k__Fungi(100);k__Fungi_unclassified(100);k__Fungi_unclassified(100);k__Fungi_unclassified(100);k__Fungi_unclassified(100);k__Fungi_unclassified(100);k__Fungi_unclassified(100); | - | - | - | - | - | - | - | Unassigned | - |
| Otu10702 | 2 | k__Fungi(100);k__Fungi_unclassified(100);k__Fungi_unclassified(100);k__Fungi_unclassified(100);k__Fungi_unclassified(100);k__Fungi_unclassified(100);k__Fungi_unclassified(100); | - | - | - | - | - | - | - | Unassigned | - |
| Otu10706 | 2 | k__Fungi(100);k__Fungi_unclassified(100);k__Fungi_unclassified(100);k__Fungi_unclassified(100);k__Fungi_unclassified(100);k__Fungi_unclassified(100);k__Fungi_unclassified(100); | - | - | - | - | - | - | - | Unassigned | - |
| Otu10707 | 2 | k__Fungi(100);p__Ascomycota(100);p__Ascomycota_unclassified(100);p__Ascomycota_unclassified(100);p__Ascomycota_unclassified(100);p__Ascomycota_unclassified(100);p__Ascomycota_unclassified(100); | - | - | - | - | - | - | - | Unassigned | - |
| Otu10710 | 2 | k__Fungi(100);p__Ascomycota(100);c__Sordariomycetes(100);c__Sordariomycetes_unclassified(100);c__Sordariomycetes_unclassified(100);c__Sordariomycetes_unclassified(100);c__Sordariomycetes_unclassified(100); | - | - | - | - | - | - | - | Unassigned | - |
| Otu10712 | 2 | k__Fungi(100);p__Ascomycota(100);c__Sordariomycetes(100);c__Sordariomycetes_unclassified(100);c__Sordariomycetes_unclassified(100);c__Sordariomycetes_unclassified(100);c__Sordariomycetes_unclassified(100); | - | - | - | - | - | - | - | Unassigned | - |
| Otu10717 | 2 | k__Fungi(100);p__Ascomycota(100);c__Sordariomycetes(100);c__Sordariomycetes_unclassified(100);c__Sordariomycetes_unclassified(100);c__Sordariomycetes_unclassified(100);c__Sordariomycetes_unclassified(100); | - | - | - | - | - | - | - | Unassigned | - |
| Otu10721 | 2 | k__Fungi(100);p__Ascomycota(100);p__Ascomycota_unclassified(100);p__Ascomycota_unclassified(100);p__Ascomycota_unclassified(100);p__Ascomycota_unclassified(100);p__Ascomycota_unclassified(100); | - | - | - | - | - | - | - | Unassigned | - |
| Otu10722 | 2 | k__Fungi(100);p__Ascomycota(100);p__Ascomycota_unclassified(100);p__Ascomycota_unclassified(100);p__Ascomycota_unclassified(100);p__Ascomycota_unclassified(100);p__Ascomycota_unclassified(100); | - | - | - | - | - | - | - | Unassigned | - |
| Otu10724 | 2 | k__Fungi(100);p__Ascomycota(100);p__Ascomycota_unclassified(100);p__Ascomycota_unclassified(100);p__Ascomycota_unclassified(100);p__Ascomycota_unclassified(100);p__Ascomycota_unclassified(100); | - | - | - | - | - | - | - | Unassigned | - |
| Otu10725 | 2 | k__Fungi(100);p__Ascomycota(100);p__Ascomycota_unclassified(100);p__Ascomycota_unclassified(100);p__Ascomycota_unclassified(100);p__Ascomycota_unclassified(100);p__Ascomycota_unclassified(100); | - | - | - | - | - | - | - | Unassigned | - |
| Otu10726 | 2 | k__Fungi(100);k__Fungi_unclassified(100);k__Fungi_unclassified(100);k__Fungi_unclassified(100);k__Fungi_unclassified(100);k__Fungi_unclassified(100);k__Fungi_unclassified(100); | - | - | - | - | - | - | - | Unassigned | - |
| Otu10727 | 2 | k__Fungi(100);p__Ascomycota(100);p__Ascomycota_unclassified(100);p__Ascomycota_unclassified(100);p__Ascomycota_unclassified(100);p__Ascomycota_unclassified(100);p__Ascomycota_unclassified(100); | - | - | - | - | - | - | - | Unassigned | - |
| Otu10728 | 2 | k__Fungi(100);k__Fungi_unclassified(100);k__Fungi_unclassified(100);k__Fungi_unclassified(100);k__Fungi_unclassified(100);k__Fungi_unclassified(100);k__Fungi_unclassified(100); | - | - | - | - | - | - | - | Unassigned | - |
| Otu10729 | 2 | k__Fungi(100);k__Fungi_unclassified(100);k__Fungi_unclassified(100);k__Fungi_unclassified(100);k__Fungi_unclassified(100);k__Fungi_unclassified(100);k__Fungi_unclassified(100); | - | - | - | - | - | - | - | Unassigned | - |
| Otu10730 | 2 | k__Fungi(100);k__Fungi_unclassified(100);k__Fungi_unclassified(100);k__Fungi_unclassified(100);k__Fungi_unclassified(100);k__Fungi_unclassified(100);k__Fungi_unclassified(100); | - | - | - | - | - | - | - | Unassigned | - |
| Otu10731 | 2 | k__Fungi(100);p__Ascomycota(100);p__Ascomycota_unclassified(100);p__Ascomycota_unclassified(100);p__Ascomycota_unclassified(100);p__Ascomycota_unclassified(100);p__Ascomycota_unclassified(100); | - | - | - | - | - | - | - | Unassigned | - |
| Otu10743 | 2 | k__Fungi(100);p__Ascomycota(100);p__Ascomycota_unclassified(100);p__Ascomycota_unclassified(100);p__Ascomycota_unclassified(100);p__Ascomycota_unclassified(100);p__Ascomycota_unclassified(100); | - | - | - | - | - | - | - | Unassigned | - |
| Otu10744 | 2 | k__Fungi(100);k__Fungi_unclassified(100);k__Fungi_unclassified(100);k__Fungi_unclassified(100);k__Fungi_unclassified(100);k__Fungi_unclassified(100);k__Fungi_unclassified(100); | - | - | - | - | - | - | - | Unassigned | - |
| Otu10745 | 2 | k__Fungi(100);k__Fungi_unclassified(100);k__Fungi_unclassified(100);k__Fungi_unclassified(100);k__Fungi_unclassified(100);k__Fungi_unclassified(100);k__Fungi_unclassified(100); | - | - | - | - | - | - | - | Unassigned | - |
| Otu10748 | 2 | k__Fungi(100);k__Fungi_unclassified(100);k__Fungi_unclassified(100);k__Fungi_unclassified(100);k__Fungi_unclassified(100);k__Fungi_unclassified(100);k__Fungi_unclassified(100); | - | - | - | - | - | - | - | Unassigned | - |
| Otu10754 | 2 | k__Fungi(100);p__Ascomycota(100);p__Ascomycota_unclassified(100);p__Ascomycota_unclassified(100);p__Ascomycota_unclassified(100);p__Ascomycota_unclassified(100);p__Ascomycota_unclassified(100); | - | - | - | - | - | - | - | Unassigned | - |
| Otu10756 | 2 | k__Fungi(100);k__Fungi_unclassified(100);k__Fungi_unclassified(100);k__Fungi_unclassified(100);k__Fungi_unclassified(100);k__Fungi_unclassified(100);k__Fungi_unclassified(100); | - | - | - | - | - | - | - | Unassigned | - |
| Otu10757 | 2 | k__Fungi(100);p__Ascomycota(100);p__Ascomycota_unclassified(100);p__Ascomycota_unclassified(100);p__Ascomycota_unclassified(100);p__Ascomycota_unclassified(100);p__Ascomycota_unclassified(100); | - | - | - | - | - | - | - | Unassigned | - |
| Otu10758 | 2 | k__Fungi(100);p__Ascomycota(100);p__Ascomycota_unclassified(100);p__Ascomycota_unclassified(100);p__Ascomycota_unclassified(100);p__Ascomycota_unclassified(100);p__Ascomycota_unclassified(100); | - | - | - | - | - | - | - | Unassigned | - |
| Otu10759 | 2 | k__Fungi(100);k__Fungi_unclassified(100);k__Fungi_unclassified(100);k__Fungi_unclassified(100);k__Fungi_unclassified(100);k__Fungi_unclassified(100);k__Fungi_unclassified(100); | - | - | - | - | - | - | - | Unassigned | - |
| Otu10760 | 2 | k__Fungi(100);k__Fungi_unclassified(100);k__Fungi_unclassified(100);k__Fungi_unclassified(100);k__Fungi_unclassified(100);k__Fungi_unclassified(100);k__Fungi_unclassified(100); | - | - | - | - | - | - | - | Unassigned | - |
| Otu10761 | 2 | k__Fungi(100);p__Ascomycota(100);p__Ascomycota_unclassified(100);p__Ascomycota_unclassified(100);p__Ascomycota_unclassified(100);p__Ascomycota_unclassified(100);p__Ascomycota_unclassified(100); | - | - | - | - | - | - | - | Unassigned | - |
| Otu10763 | 2 | k__Fungi(100);p__Ascomycota(100);p__Ascomycota_unclassified(100);p__Ascomycota_unclassified(100);p__Ascomycota_unclassified(100);p__Ascomycota_unclassified(100);p__Ascomycota_unclassified(100); | - | - | - | - | - | - | - | Unassigned | - |
| Otu10765 | 2 | k__Fungi(100);p__Ascomycota(100);p__Ascomycota_unclassified(100);p__Ascomycota_unclassified(100);p__Ascomycota_unclassified(100);p__Ascomycota_unclassified(100);p__Ascomycota_unclassified(100); | - | - | - | - | - | - | - | Unassigned | - |
| Otu10770 | 2 | k__Fungi(100);p__Ascomycota(100);p__Ascomycota_unclassified(100);p__Ascomycota_unclassified(100);p__Ascomycota_unclassified(100);p__Ascomycota_unclassified(100);p__Ascomycota_unclassified(100); | - | - | - | - | - | - | - | Unassigned | - |
| Otu10773 | 2 | k__Fungi(100);p__Ascomycota(100);c__Dothideomycetes(100);c__Dothideomycetes_unclassified(100);c__Dothideomycetes_unclassified(100);c__Dothideomycetes_unclassified(100);c__Dothideomycetes_unclassified(100); | - | - | - | - | - | - | - | Unassigned | - |
| Otu10774 | 2 | k__Fungi(100);p__Ascomycota(100);p__Ascomycota_unclassified(100);p__Ascomycota_unclassified(100);p__Ascomycota_unclassified(100);p__Ascomycota_unclassified(100);p__Ascomycota_unclassified(100); | - | - | - | - | - | - | - | Unassigned | - |
| Otu10775 | 2 | k__Fungi(100);p__Ascomycota(100);p__Ascomycota_unclassified(100);p__Ascomycota_unclassified(100);p__Ascomycota_unclassified(100);p__Ascomycota_unclassified(100);p__Ascomycota_unclassified(100); | - | - | - | - | - | - | - | Unassigned | - |
| Otu10777 | 2 | k__Fungi(100);p__Ascomycota(100);c__Dothideomycetes(100);o__Myriangiales(100);o__Myriangiales_unclassified(100);o__Myriangiales_unclassified(100);o__Myriangiales_unclassified(100); | - | - | - | - | - | - | - | Unassigned | - |
| Otu10780 | 2 | k__Fungi(100);k__Fungi_unclassified(100);k__Fungi_unclassified(100);k__Fungi_unclassified(100);k__Fungi_unclassified(100);k__Fungi_unclassified(100);k__Fungi_unclassified(100); | - | - | - | - | - | - | - | Unassigned | - |
| Otu10786 | 2 | k__Fungi(100);k__Fungi_unclassified(100);k__Fungi_unclassified(100);k__Fungi_unclassified(100);k__Fungi_unclassified(100);k__Fungi_unclassified(100);k__Fungi_unclassified(100); | - | - | - | - | - | - | - | Unassigned | - |
| Otu10788 | 2 | k__Fungi(100);k__Fungi_unclassified(100);k__Fungi_unclassified(100);k__Fungi_unclassified(100);k__Fungi_unclassified(100);k__Fungi_unclassified(100);k__Fungi_unclassified(100); | - | - | - | - | - | - | - | Unassigned | - |
| Otu10791 | 2 | k__Fungi(100);p__Ascomycota(100);p__Ascomycota_unclassified(100);p__Ascomycota_unclassified(100);p__Ascomycota_unclassified(100);p__Ascomycota_unclassified(100);p__Ascomycota_unclassified(100); | - | - | - | - | - | - | - | Unassigned | - |
| Otu10792 | 2 | k__Fungi(100);k__Fungi_unclassified(100);k__Fungi_unclassified(100);k__Fungi_unclassified(100);k__Fungi_unclassified(100);k__Fungi_unclassified(100);k__Fungi_unclassified(100); | - | - | - | - | - | - | - | Unassigned | - |
| Otu10793 | 2 | k__Fungi(100);k__Fungi_unclassified(100);k__Fungi_unclassified(100);k__Fungi_unclassified(100);k__Fungi_unclassified(100);k__Fungi_unclassified(100);k__Fungi_unclassified(100); | - | - | - | - | - | - | - | Unassigned | - |
| Otu10795 | 2 | k__Fungi(100);p__Ascomycota(100);p__Ascomycota_unclassified(100);p__Ascomycota_unclassified(100);p__Ascomycota_unclassified(100);p__Ascomycota_unclassified(100);p__Ascomycota_unclassified(100); | - | - | - | - | - | - | - | Unassigned | - |
| Otu10798 | 2 | k__Fungi(100);p__Ascomycota(100);c__Dothideomycetes(100);o__Capnodiales(100);f__Davidiellaceae(100);f__Davidiellaceae_unclassified(100);f__Davidiellaceae_unclassified(100); | - | - | - | - | - | - | - | Unassigned | - |
| Otu10799 | 2 | k__Fungi(100);k__Fungi_unclassified(100);k__Fungi_unclassified(100);k__Fungi_unclassified(100);k__Fungi_unclassified(100);k__Fungi_unclassified(100);k__Fungi_unclassified(100); | - | - | - | - | - | - | - | Unassigned | - |
| Otu10802 | 2 | k__Fungi(100);k__Fungi_unclassified(100);k__Fungi_unclassified(100);k__Fungi_unclassified(100);k__Fungi_unclassified(100);k__Fungi_unclassified(100);k__Fungi_unclassified(100); | - | - | - | - | - | - | - | Unassigned | - |
| Otu10803 | 2 | k__Fungi(100);p__Ascomycota(100);p__Ascomycota_unclassified(100);p__Ascomycota_unclassified(100);p__Ascomycota_unclassified(100);p__Ascomycota_unclassified(100);p__Ascomycota_unclassified(100); | - | - | - | - | - | - | - | Unassigned | - |
| Otu10812 | 2 | k__Fungi(100);p__Ascomycota(100);p__Ascomycota_unclassified(100);p__Ascomycota_unclassified(100);p__Ascomycota_unclassified(100);p__Ascomycota_unclassified(100);p__Ascomycota_unclassified(100); | - | - | - | - | - | - | - | Unassigned | - |
| Otu10814 | 2 | k__Fungi(100);k__Fungi_unclassified(100);k__Fungi_unclassified(100);k__Fungi_unclassified(100);k__Fungi_unclassified(100);k__Fungi_unclassified(100);k__Fungi_unclassified(100); | - | - | - | - | - | - | - | Unassigned | - |
| Otu10817 | 2 | k__Fungi(100);p__Ascomycota(100);p__Ascomycota_unclassified(100);p__Ascomycota_unclassified(100);p__Ascomycota_unclassified(100);p__Ascomycota_unclassified(100);p__Ascomycota_unclassified(100); | - | - | - | - | - | - | - | Unassigned | - |
| Otu10821 | 2 | k__Fungi(100);p__Ascomycota(100);p__Ascomycota_unclassified(100);p__Ascomycota_unclassified(100);p__Ascomycota_unclassified(100);p__Ascomycota_unclassified(100);p__Ascomycota_unclassified(100); | - | - | - | - | - | - | - | Unassigned | - |
| Otu10823 | 2 | k__Fungi(100);p__Ascomycota(100);p__Ascomycota_unclassified(100);p__Ascomycota_unclassified(100);p__Ascomycota_unclassified(100);p__Ascomycota_unclassified(100);p__Ascomycota_unclassified(100); | - | - | - | - | - | - | - | Unassigned | - |
| Otu10826 | 2 | k__Fungi(100);k__Fungi_unclassified(100);k__Fungi_unclassified(100);k__Fungi_unclassified(100);k__Fungi_unclassified(100);k__Fungi_unclassified(100);k__Fungi_unclassified(100); | - | - | - | - | - | - | - | Unassigned | - |
| Otu10828 | 2 | k__Fungi(100);k__Fungi_unclassified(100);k__Fungi_unclassified(100);k__Fungi_unclassified(100);k__Fungi_unclassified(100);k__Fungi_unclassified(100);k__Fungi_unclassified(100); | - | - | - | - | - | - | - | Unassigned | - |
| Otu10830 | 2 | k__Fungi(100);k__Fungi_unclassified(100);k__Fungi_unclassified(100);k__Fungi_unclassified(100);k__Fungi_unclassified(100);k__Fungi_unclassified(100);k__Fungi_unclassified(100); | - | - | - | - | - | - | - | Unassigned | - |
| Otu10831 | 2 | k__Fungi(100);k__Fungi_unclassified(100);k__Fungi_unclassified(100);k__Fungi_unclassified(100);k__Fungi_unclassified(100);k__Fungi_unclassified(100);k__Fungi_unclassified(100); | - | - | - | - | - | - | - | Unassigned | - |
| Otu10834 | 2 | k__Fungi(100);k__Fungi_unclassified(100);k__Fungi_unclassified(100);k__Fungi_unclassified(100);k__Fungi_unclassified(100);k__Fungi_unclassified(100);k__Fungi_unclassified(100); | - | - | - | - | - | - | - | Unassigned | - |
| Otu10838 | 2 | k__Fungi(100);p__Ascomycota(100);p__Ascomycota_unclassified(100);p__Ascomycota_unclassified(100);p__Ascomycota_unclassified(100);p__Ascomycota_unclassified(100);p__Ascomycota_unclassified(100); | - | - | - | - | - | - | - | Unassigned | - |
| Otu10847 | 2 | k__Fungi(100);k__Fungi_unclassified(100);k__Fungi_unclassified(100);k__Fungi_unclassified(100);k__Fungi_unclassified(100);k__Fungi_unclassified(100);k__Fungi_unclassified(100); | - | - | - | - | - | - | - | Unassigned | - |
| Otu10848 | 2 | k__Fungi(100);k__Fungi_unclassified(100);k__Fungi_unclassified(100);k__Fungi_unclassified(100);k__Fungi_unclassified(100);k__Fungi_unclassified(100);k__Fungi_unclassified(100); | - | - | - | - | - | - | - | Unassigned | - |
| Otu10849 | 2 | k__Fungi(100);k__Fungi_unclassified(100);k__Fungi_unclassified(100);k__Fungi_unclassified(100);k__Fungi_unclassified(100);k__Fungi_unclassified(100);k__Fungi_unclassified(100); | - | - | - | - | - | - | - | Unassigned | - |
| Otu10851 | 2 | k__Fungi(100);k__Fungi_unclassified(100);k__Fungi_unclassified(100);k__Fungi_unclassified(100);k__Fungi_unclassified(100);k__Fungi_unclassified(100);k__Fungi_unclassified(100); | - | - | - | - | - | - | - | Unassigned | - |
| Otu10852 | 2 | k__Fungi(100);k__Fungi_unclassified(100);k__Fungi_unclassified(100);k__Fungi_unclassified(100);k__Fungi_unclassified(100);k__Fungi_unclassified(100);k__Fungi_unclassified(100); | - | - | - | - | - | - | - | Unassigned | - |
| Otu10853 | 2 | k__Fungi(100);k__Fungi_unclassified(100);k__Fungi_unclassified(100);k__Fungi_unclassified(100);k__Fungi_unclassified(100);k__Fungi_unclassified(100);k__Fungi_unclassified(100); | - | - | - | - | - | - | - | Unassigned | - |
| Otu10854 | 2 | k__Fungi(100);k__Fungi_unclassified(100);k__Fungi_unclassified(100);k__Fungi_unclassified(100);k__Fungi_unclassified(100);k__Fungi_unclassified(100);k__Fungi_unclassified(100); | - | - | - | - | - | - | - | Unassigned | - |
| Otu10855 | 2 | k__Fungi(100);k__Fungi_unclassified(100);k__Fungi_unclassified(100);k__Fungi_unclassified(100);k__Fungi_unclassified(100);k__Fungi_unclassified(100);k__Fungi_unclassified(100); | - | - | - | - | - | - | - | Unassigned | - |
| Otu10862 | 2 | k__Fungi(100);k__Fungi_unclassified(100);k__Fungi_unclassified(100);k__Fungi_unclassified(100);k__Fungi_unclassified(100);k__Fungi_unclassified(100);k__Fungi_unclassified(100); | - | - | - | - | - | - | - | Unassigned | - |
| Otu10863 | 2 | k__Fungi(100);k__Fungi_unclassified(100);k__Fungi_unclassified(100);k__Fungi_unclassified(100);k__Fungi_unclassified(100);k__Fungi_unclassified(100);k__Fungi_unclassified(100); | - | - | - | - | - | - | - | Unassigned | - |
| Otu10864 | 2 | k__Fungi(100);k__Fungi_unclassified(100);k__Fungi_unclassified(100);k__Fungi_unclassified(100);k__Fungi_unclassified(100);k__Fungi_unclassified(100);k__Fungi_unclassified(100); | - | - | - | - | - | - | - | Unassigned | - |
| Otu10869 | 2 | k__Fungi(100);p__Ascomycota(100);p__Ascomycota_unclassified(100);p__Ascomycota_unclassified(100);p__Ascomycota_unclassified(100);p__Ascomycota_unclassified(100);p__Ascomycota_unclassified(100); | - | - | - | - | - | - | - | Unassigned | - |
| Otu10870 | 2 | k__Fungi(100);k__Fungi_unclassified(100);k__Fungi_unclassified(100);k__Fungi_unclassified(100);k__Fungi_unclassified(100);k__Fungi_unclassified(100);k__Fungi_unclassified(100); | - | - | - | - | - | - | - | Unassigned | - |
| Otu10874 | 2 | k__Fungi(100);k__Fungi_unclassified(100);k__Fungi_unclassified(100);k__Fungi_unclassified(100);k__Fungi_unclassified(100);k__Fungi_unclassified(100);k__Fungi_unclassified(100); | - | - | - | - | - | - | - | Unassigned | - |
| Otu10876 | 2 | k__Fungi(100);p__Ascomycota(100);p__Ascomycota_unclassified(100);p__Ascomycota_unclassified(100);p__Ascomycota_unclassified(100);p__Ascomycota_unclassified(100);p__Ascomycota_unclassified(100); | - | - | - | - | - | - | - | Unassigned | - |
| Otu10877 | 2 | k__Fungi(100);p__Ascomycota(100);p__Ascomycota_unclassified(100);p__Ascomycota_unclassified(100);p__Ascomycota_unclassified(100);p__Ascomycota_unclassified(100);p__Ascomycota_unclassified(100); | - | - | - | - | - | - | - | Unassigned | - |
| Otu10880 | 2 | k__Fungi(100);k__Fungi_unclassified(100);k__Fungi_unclassified(100);k__Fungi_unclassified(100);k__Fungi_unclassified(100);k__Fungi_unclassified(100);k__Fungi_unclassified(100); | - | - | - | - | - | - | - | Unassigned | - |
| Otu10881 | 2 | k__Fungi(100);k__Fungi_unclassified(100);k__Fungi_unclassified(100);k__Fungi_unclassified(100);k__Fungi_unclassified(100);k__Fungi_unclassified(100);k__Fungi_unclassified(100); | - | - | - | - | - | - | - | Unassigned | - |
| Otu10882 | 2 | k__Fungi(100);k__Fungi_unclassified(100);k__Fungi_unclassified(100);k__Fungi_unclassified(100);k__Fungi_unclassified(100);k__Fungi_unclassified(100);k__Fungi_unclassified(100); | - | - | - | - | - | - | - | Unassigned | - |
| Otu10883 | 2 | k__Fungi(100);k__Fungi_unclassified(100);k__Fungi_unclassified(100);k__Fungi_unclassified(100);k__Fungi_unclassified(100);k__Fungi_unclassified(100);k__Fungi_unclassified(100); | - | - | - | - | - | - | - | Unassigned | - |
| Otu10885 | 2 | k__Fungi(100);p__Ascomycota(100);p__Ascomycota_unclassified(100);p__Ascomycota_unclassified(100);p__Ascomycota_unclassified(100);p__Ascomycota_unclassified(100);p__Ascomycota_unclassified(100); | - | - | - | - | - | - | - | Unassigned | - |
| Otu10886 | 2 | k__Fungi(100);p__Ascomycota(100);p__Ascomycota_unclassified(100);p__Ascomycota_unclassified(100);p__Ascomycota_unclassified(100);p__Ascomycota_unclassified(100);p__Ascomycota_unclassified(100); | - | - | - | - | - | - | - | Unassigned | - |
| Otu10888 | 2 | k__Fungi(100);k__Fungi_unclassified(100);k__Fungi_unclassified(100);k__Fungi_unclassified(100);k__Fungi_unclassified(100);k__Fungi_unclassified(100);k__Fungi_unclassified(100); | - | - | - | - | - | - | - | Unassigned | - |
| Otu10891 | 2 | k__Fungi(100);p__Ascomycota(100);c__Dothideomycetes(100);o__Capnodiales(100);o__Capnodiales_unclassified(100);o__Capnodiales_unclassified(100);o__Capnodiales_unclassified(100); | - | - | - | - | - | - | - | Unassigned | - |
| Otu10892 | 2 | k__Fungi(100);k__Fungi_unclassified(100);k__Fungi_unclassified(100);k__Fungi_unclassified(100);k__Fungi_unclassified(100);k__Fungi_unclassified(100);k__Fungi_unclassified(100); | - | - | - | - | - | - | - | Unassigned | - |
| Otu10893 | 2 | k__Fungi(100);k__Fungi_unclassified(100);k__Fungi_unclassified(100);k__Fungi_unclassified(100);k__Fungi_unclassified(100);k__Fungi_unclassified(100);k__Fungi_unclassified(100); | - | - | - | - | - | - | - | Unassigned | - |
| Otu10894 | 2 | k__Plantae(100);p__unclassified_Plantae(100);c__unclassified_Plantae(100);o__unclassified_Plantae(100);f__unclassified_Plantae(100);g__unclassified_Plantae(100);s__Plantae_sp(100); | - | - | - | - | - | - | - | Unassigned | - |
| Otu10896 | 2 | k__Fungi(100);k__Fungi_unclassified(100);k__Fungi_unclassified(100);k__Fungi_unclassified(100);k__Fungi_unclassified(100);k__Fungi_unclassified(100);k__Fungi_unclassified(100); | - | - | - | - | - | - | - | Unassigned | - |
| Otu10899 | 2 | k__Fungi(100);p__Ascomycota(100);p__Ascomycota_unclassified(100);p__Ascomycota_unclassified(100);p__Ascomycota_unclassified(100);p__Ascomycota_unclassified(100);p__Ascomycota_unclassified(100); | - | - | - | - | - | - | - | Unassigned | - |
| Otu10905 | 2 | k__Fungi(100);k__Fungi_unclassified(100);k__Fungi_unclassified(100);k__Fungi_unclassified(100);k__Fungi_unclassified(100);k__Fungi_unclassified(100);k__Fungi_unclassified(100); | - | - | - | - | - | - | - | Unassigned | - |
| Otu10906 | 2 | k__Plantae(100);p__unclassified_Plantae(100);c__unclassified_Plantae(100);o__unclassified_Plantae(100);f__unclassified_Plantae(100);g__unclassified_Plantae(100);s__Plantae_sp(100); | - | - | - | - | - | - | - | Unassigned | - |
| Otu10910 | 2 | k__Plantae(100);p__unclassified_Plantae(100);c__unclassified_Plantae(100);o__unclassified_Plantae(100);f__unclassified_Plantae(100);g__unclassified_Plantae(100);s__Plantae_sp(100); | - | - | - | - | - | - | - | Unassigned | - |
| Otu10912 | 2 | k__Fungi(100);k__Fungi_unclassified(100);k__Fungi_unclassified(100);k__Fungi_unclassified(100);k__Fungi_unclassified(100);k__Fungi_unclassified(100);k__Fungi_unclassified(100); | - | - | - | - | - | - | - | Unassigned | - |
| Otu10913 | 2 | k__Fungi(100);k__Fungi_unclassified(100);k__Fungi_unclassified(100);k__Fungi_unclassified(100);k__Fungi_unclassified(100);k__Fungi_unclassified(100);k__Fungi_unclassified(100); | - | - | - | - | - | - | - | Unassigned | - |
| Otu10914 | 2 | k__Fungi(100);k__Fungi_unclassified(100);k__Fungi_unclassified(100);k__Fungi_unclassified(100);k__Fungi_unclassified(100);k__Fungi_unclassified(100);k__Fungi_unclassified(100); | - | - | - | - | - | - | - | Unassigned | - |
| Otu10915 | 2 | k__Fungi(100);k__Fungi_unclassified(100);k__Fungi_unclassified(100);k__Fungi_unclassified(100);k__Fungi_unclassified(100);k__Fungi_unclassified(100);k__Fungi_unclassified(100); | - | - | - | - | - | - | - | Unassigned | - |
| Otu10919 | 2 | k__Plantae(100);p__unclassified_Plantae(100);c__unclassified_Plantae(100);o__unclassified_Plantae(100);f__unclassified_Plantae(100);g__unclassified_Plantae(100);s__Plantae_sp(100); | - | - | - | - | - | - | - | Unassigned | - |
| Otu10920 | 2 | k__Fungi(100);p__Ascomycota(100);p__Ascomycota_unclassified(100);p__Ascomycota_unclassified(100);p__Ascomycota_unclassified(100);p__Ascomycota_unclassified(100);p__Ascomycota_unclassified(100); | - | - | - | - | - | - | - | Unassigned | - |
| Otu10923 | 2 | k__Fungi(100);k__Fungi_unclassified(100);k__Fungi_unclassified(100);k__Fungi_unclassified(100);k__Fungi_unclassified(100);k__Fungi_unclassified(100);k__Fungi_unclassified(100); | - | - | - | - | - | - | - | Unassigned | - |
| Otu10924 | 2 | k__Plantae(100);p__unclassified_Plantae(100);c__unclassified_Plantae(100);o__unclassified_Plantae(100);f__unclassified_Plantae(100);g__unclassified_Plantae(100);s__Plantae_sp(100); | - | - | - | - | - | - | - | Unassigned | - |
| Otu10925 | 2 | k__Plantae(100);p__unclassified_Plantae(100);c__unclassified_Plantae(100);o__unclassified_Plantae(100);f__unclassified_Plantae(100);g__unclassified_Plantae(100);s__Plantae_sp(100); | - | - | - | - | - | - | - | Unassigned | - |
| Otu10926 | 2 | k__Fungi(100);k__Fungi_unclassified(100);k__Fungi_unclassified(100);k__Fungi_unclassified(100);k__Fungi_unclassified(100);k__Fungi_unclassified(100);k__Fungi_unclassified(100); | - | - | - | - | - | - | - | Unassigned | - |
| Otu10928 | 2 | k__Plantae(100);p__unclassified_Plantae(100);c__unclassified_Plantae(100);o__unclassified_Plantae(100);f__unclassified_Plantae(100);g__unclassified_Plantae(100);s__Plantae_sp(100); | - | - | - | - | - | - | - | Unassigned | - |
| Otu10930 | 2 | k__Plantae(100);p__unclassified_Plantae(100);c__unclassified_Plantae(100);o__unclassified_Plantae(100);f__unclassified_Plantae(100);g__unclassified_Plantae(100);s__Plantae_sp(100); | - | - | - | - | - | - | - | Unassigned | - |
| Otu10931 | 2 | k__Plantae(100);p__unclassified_Plantae(100);c__unclassified_Plantae(100);o__unclassified_Plantae(100);f__unclassified_Plantae(100);g__unclassified_Plantae(100);s__Plantae_sp(100); | - | - | - | - | - | - | - | Unassigned | - |
| Otu10932 | 2 | k__Plantae(100);p__unclassified_Plantae(100);c__unclassified_Plantae(100);o__unclassified_Plantae(100);f__unclassified_Plantae(100);g__unclassified_Plantae(100);s__Plantae_sp(100); | - | - | - | - | - | - | - | Unassigned | - |
| Otu10934 | 2 | k__Plantae(100);p__unclassified_Plantae(100);c__unclassified_Plantae(100);o__unclassified_Plantae(100);f__unclassified_Plantae(100);g__unclassified_Plantae(100);s__Plantae_sp(100); | - | - | - | - | - | - | - | Unassigned | - |
| Otu10935 | 2 | k__Plantae(100);p__unclassified_Plantae(100);c__unclassified_Plantae(100);o__unclassified_Plantae(100);f__unclassified_Plantae(100);g__unclassified_Plantae(100);s__Plantae_sp(100); | - | - | - | - | - | - | - | Unassigned | - |
| Otu10937 | 2 | k__Plantae(100);p__unclassified_Plantae(100);c__unclassified_Plantae(100);o__unclassified_Plantae(100);f__unclassified_Plantae(100);g__unclassified_Plantae(100);s__Plantae_sp(100); | - | - | - | - | - | - | - | Unassigned | - |
| Otu10938 | 2 | k__Plantae(100);p__unclassified_Plantae(100);c__unclassified_Plantae(100);o__unclassified_Plantae(100);f__unclassified_Plantae(100);g__unclassified_Plantae(100);s__Plantae_sp(100); | - | - | - | - | - | - | - | Unassigned | - |
| Otu10939 | 2 | k__Plantae(100);p__unclassified_Plantae(100);c__unclassified_Plantae(100);o__unclassified_Plantae(100);f__unclassified_Plantae(100);g__unclassified_Plantae(100);s__Plantae_sp(100); | - | - | - | - | - | - | - | Unassigned | - |
| Otu10941 | 2 | k__Fungi(100);k__Fungi_unclassified(100);k__Fungi_unclassified(100);k__Fungi_unclassified(100);k__Fungi_unclassified(100);k__Fungi_unclassified(100);k__Fungi_unclassified(100); | - | - | - | - | - | - | - | Unassigned | - |
| Otu10942 | 2 | k__Plantae(100);p__unclassified_Plantae(100);c__unclassified_Plantae(100);o__unclassified_Plantae(100);f__unclassified_Plantae(100);g__unclassified_Plantae(100);s__Plantae_sp(100); | - | - | - | - | - | - | - | Unassigned | - |
| Otu10943 | 2 | k__Plantae(100);p__unclassified_Plantae(100);c__unclassified_Plantae(100);o__unclassified_Plantae(100);f__unclassified_Plantae(100);g__unclassified_Plantae(100);s__Plantae_sp(100); | - | - | - | - | - | - | - | Unassigned | - |
| Otu10945 | 2 | k__Plantae(100);p__unclassified_Plantae(100);c__unclassified_Plantae(100);o__unclassified_Plantae(100);f__unclassified_Plantae(100);g__unclassified_Plantae(100);s__Plantae_sp(100); | - | - | - | - | - | - | - | Unassigned | - |
| Otu10947 | 2 | k__Fungi(100);k__Fungi_unclassified(100);k__Fungi_unclassified(100);k__Fungi_unclassified(100);k__Fungi_unclassified(100);k__Fungi_unclassified(100);k__Fungi_unclassified(100); | - | - | - | - | - | - | - | Unassigned | - |
| Otu10949 | 2 | k__Fungi(100);k__Fungi_unclassified(100);k__Fungi_unclassified(100);k__Fungi_unclassified(100);k__Fungi_unclassified(100);k__Fungi_unclassified(100);k__Fungi_unclassified(100); | - | - | - | - | - | - | - | Unassigned | - |
| Otu10950 | 2 | k__Fungi(100);k__Fungi_unclassified(100);k__Fungi_unclassified(100);k__Fungi_unclassified(100);k__Fungi_unclassified(100);k__Fungi_unclassified(100);k__Fungi_unclassified(100); | - | - | - | - | - | - | - | Unassigned | - |
| Otu10951 | 2 | k__Fungi(100);k__Fungi_unclassified(100);k__Fungi_unclassified(100);k__Fungi_unclassified(100);k__Fungi_unclassified(100);k__Fungi_unclassified(100);k__Fungi_unclassified(100); | - | - | - | - | - | - | - | Unassigned | - |
| Otu10952 | 2 | k__Fungi(100);k__Fungi_unclassified(100);k__Fungi_unclassified(100);k__Fungi_unclassified(100);k__Fungi_unclassified(100);k__Fungi_unclassified(100);k__Fungi_unclassified(100); | - | - | - | - | - | - | - | Unassigned | - |
| Otu10953 | 2 | k__Fungi(100);k__Fungi_unclassified(100);k__Fungi_unclassified(100);k__Fungi_unclassified(100);k__Fungi_unclassified(100);k__Fungi_unclassified(100);k__Fungi_unclassified(100); | - | - | - | - | - | - | - | Unassigned | - |
| Otu10954 | 2 | k__Fungi(100);k__Fungi_unclassified(100);k__Fungi_unclassified(100);k__Fungi_unclassified(100);k__Fungi_unclassified(100);k__Fungi_unclassified(100);k__Fungi_unclassified(100); | - | - | - | - | - | - | - | Unassigned | - |
| Otu10955 | 2 | k__Fungi(100);k__Fungi_unclassified(100);k__Fungi_unclassified(100);k__Fungi_unclassified(100);k__Fungi_unclassified(100);k__Fungi_unclassified(100);k__Fungi_unclassified(100); | - | - | - | - | - | - | - | Unassigned | - |
| Otu10956 | 2 | k__Fungi(100);k__Fungi_unclassified(100);k__Fungi_unclassified(100);k__Fungi_unclassified(100);k__Fungi_unclassified(100);k__Fungi_unclassified(100);k__Fungi_unclassified(100); | - | - | - | - | - | - | - | Unassigned | - |
| Otu10958 | 2 | k__Fungi(100);k__Fungi_unclassified(100);k__Fungi_unclassified(100);k__Fungi_unclassified(100);k__Fungi_unclassified(100);k__Fungi_unclassified(100);k__Fungi_unclassified(100); | - | - | - | - | - | - | - | Unassigned | - |
| Otu10959 | 2 | k__Fungi(100);k__Fungi_unclassified(100);k__Fungi_unclassified(100);k__Fungi_unclassified(100);k__Fungi_unclassified(100);k__Fungi_unclassified(100);k__Fungi_unclassified(100); | - | - | - | - | - | - | - | Unassigned | - |
| Otu10960 | 2 | k__Fungi(100);k__Fungi_unclassified(100);k__Fungi_unclassified(100);k__Fungi_unclassified(100);k__Fungi_unclassified(100);k__Fungi_unclassified(100);k__Fungi_unclassified(100); | - | - | - | - | - | - | - | Unassigned | - |
| Otu10964 | 2 | k__Fungi(100);k__Fungi_unclassified(100);k__Fungi_unclassified(100);k__Fungi_unclassified(100);k__Fungi_unclassified(100);k__Fungi_unclassified(100);k__Fungi_unclassified(100); | - | - | - | - | - | - | - | Unassigned | - |
| Otu10970 | 2 | k__Fungi(100);k__Fungi_unclassified(100);k__Fungi_unclassified(100);k__Fungi_unclassified(100);k__Fungi_unclassified(100);k__Fungi_unclassified(100);k__Fungi_unclassified(100); | - | - | - | - | - | - | - | Unassigned | - |
| Otu10976 | 2 | k__Fungi(100);k__Fungi_unclassified(100);k__Fungi_unclassified(100);k__Fungi_unclassified(100);k__Fungi_unclassified(100);k__Fungi_unclassified(100);k__Fungi_unclassified(100); | - | - | - | - | - | - | - | Unassigned | - |
| Otu10977 | 2 | k__Fungi(100);p__Ascomycota(100);p__Ascomycota_unclassified(100);p__Ascomycota_unclassified(100);p__Ascomycota_unclassified(100);p__Ascomycota_unclassified(100);p__Ascomycota_unclassified(100); | - | - | - | - | - | - | - | Unassigned | - |
| Otu10978 | 2 | k__Fungi(100);k__Fungi_unclassified(100);k__Fungi_unclassified(100);k__Fungi_unclassified(100);k__Fungi_unclassified(100);k__Fungi_unclassified(100);k__Fungi_unclassified(100); | - | - | - | - | - | - | - | Unassigned | - |
| Otu10979 | 2 | k__Fungi(100);k__Fungi_unclassified(100);k__Fungi_unclassified(100);k__Fungi_unclassified(100);k__Fungi_unclassified(100);k__Fungi_unclassified(100);k__Fungi_unclassified(100); | - | - | - | - | - | - | - | Unassigned | - |
| Otu10988 | 2 | k__Fungi(100);p__Basidiomycota(100);c__Agaricomycetes(100);o__Russulales(100);o__Russulales_unclassified(100);o__Russulales_unclassified(100);o__Russulales_unclassified(100); | - | - | - | - | - | - | - | Unassigned | - |
| Otu10989 | 2 | k__Fungi(100);k__Fungi_unclassified(100);k__Fungi_unclassified(100);k__Fungi_unclassified(100);k__Fungi_unclassified(100);k__Fungi_unclassified(100);k__Fungi_unclassified(100); | - | - | - | - | - | - | - | Unassigned | - |
| Otu10991 | 2 | k__Fungi(100);k__Fungi_unclassified(100);k__Fungi_unclassified(100);k__Fungi_unclassified(100);k__Fungi_unclassified(100);k__Fungi_unclassified(100);k__Fungi_unclassified(100); | - | - | - | - | - | - | - | Unassigned | - |
| Otu10993 | 2 | k__Fungi(100);k__Fungi_unclassified(100);k__Fungi_unclassified(100);k__Fungi_unclassified(100);k__Fungi_unclassified(100);k__Fungi_unclassified(100);k__Fungi_unclassified(100); | - | - | - | - | - | - | - | Unassigned | - |
| Otu10995 | 2 | k__Fungi(100);p__Basidiomycota(100);p__Basidiomycota_unclassified(100);p__Basidiomycota_unclassified(100);p__Basidiomycota_unclassified(100);p__Basidiomycota_unclassified(100);p__Basidiomycota_unclassified(100); | - | - | - | - | - | - | - | Unassigned | - |
| Otu11005 | 2 | k__Fungi(100);k__Fungi_unclassified(100);k__Fungi_unclassified(100);k__Fungi_unclassified(100);k__Fungi_unclassified(100);k__Fungi_unclassified(100);k__Fungi_unclassified(100); | - | - | - | - | - | - | - | Unassigned | - |
| Otu11009 | 2 | k__Fungi(100);p__Ascomycota(100);c__Sordariomycetes(100);o__Hypocreales(100);f__Hypocreaceae(100);f__Hypocreaceae_unclassified(100);f__Hypocreaceae_unclassified(100); | - | - | - | - | - | - | - | Unassigned | - |
| Otu11011 | 2 | k__Fungi(100);p__Ascomycota(100);p__Ascomycota_unclassified(100);p__Ascomycota_unclassified(100);p__Ascomycota_unclassified(100);p__Ascomycota_unclassified(100);p__Ascomycota_unclassified(100); | - | - | - | - | - | - | - | Unassigned | - |
| Otu11015 | 2 | k__Fungi(100);k__Fungi_unclassified(100);k__Fungi_unclassified(100);k__Fungi_unclassified(100);k__Fungi_unclassified(100);k__Fungi_unclassified(100);k__Fungi_unclassified(100); | - | - | - | - | - | - | - | Unassigned | - |
| Otu11018 | 2 | k__Fungi(100);k__Fungi_unclassified(100);k__Fungi_unclassified(100);k__Fungi_unclassified(100);k__Fungi_unclassified(100);k__Fungi_unclassified(100);k__Fungi_unclassified(100); | - | - | - | - | - | - | - | Unassigned | - |
| Otu11024 | 2 | k__Fungi(100);k__Fungi_unclassified(100);k__Fungi_unclassified(100);k__Fungi_unclassified(100);k__Fungi_unclassified(100);k__Fungi_unclassified(100);k__Fungi_unclassified(100); | - | - | - | - | - | - | - | Unassigned | - |
| Otu11027 | 2 | k__Fungi(100);k__Fungi_unclassified(100);k__Fungi_unclassified(100);k__Fungi_unclassified(100);k__Fungi_unclassified(100);k__Fungi_unclassified(100);k__Fungi_unclassified(100); | - | - | - | - | - | - | - | Unassigned | - |
| Otu11028 | 2 | k__Fungi(100);p__Ascomycota(100);p__Ascomycota_unclassified(100);p__Ascomycota_unclassified(100);p__Ascomycota_unclassified(100);p__Ascomycota_unclassified(100);p__Ascomycota_unclassified(100); | - | - | - | - | - | - | - | Unassigned | - |
| Otu11031 | 2 | k__Fungi(100);k__Fungi_unclassified(100);k__Fungi_unclassified(100);k__Fungi_unclassified(100);k__Fungi_unclassified(100);k__Fungi_unclassified(100);k__Fungi_unclassified(100); | - | - | - | - | - | - | - | Unassigned | - |
| Otu11036 | 2 | k__Fungi(100);p__Ascomycota(100);p__Ascomycota_unclassified(100);p__Ascomycota_unclassified(100);p__Ascomycota_unclassified(100);p__Ascomycota_unclassified(100);p__Ascomycota_unclassified(100); | - | - | - | - | - | - | - | Unassigned | - |
| Otu11038 | 2 | k__Fungi(100);k__Fungi_unclassified(100);k__Fungi_unclassified(100);k__Fungi_unclassified(100);k__Fungi_unclassified(100);k__Fungi_unclassified(100);k__Fungi_unclassified(100); | - | - | - | - | - | - | - | Unassigned | - |
| Otu11039 | 2 | k__Fungi(100);k__Fungi_unclassified(100);k__Fungi_unclassified(100);k__Fungi_unclassified(100);k__Fungi_unclassified(100);k__Fungi_unclassified(100);k__Fungi_unclassified(100); | - | - | - | - | - | - | - | Unassigned | - |
| Otu11043 | 2 | k__Fungi(100);k__Fungi_unclassified(100);k__Fungi_unclassified(100);k__Fungi_unclassified(100);k__Fungi_unclassified(100);k__Fungi_unclassified(100);k__Fungi_unclassified(100); | - | - | - | - | - | - | - | Unassigned | - |
| Otu11044 | 2 | k__Fungi(100);p__Basidiomycota(100);c__Agaricomycetes(100);c__Agaricomycetes_unclassified(100);c__Agaricomycetes_unclassified(100);c__Agaricomycetes_unclassified(100);c__Agaricomycetes_unclassified(100); | - | - | - | - | - | - | - | Unassigned | - |
| Otu11045 | 2 | k__Fungi(100);p__Basidiomycota(100);c__Agaricomycetes(100);c__Agaricomycetes_unclassified(100);c__Agaricomycetes_unclassified(100);c__Agaricomycetes_unclassified(100);c__Agaricomycetes_unclassified(100); | - | - | - | - | - | - | - | Unassigned | - |
| Otu11046 | 2 | k__Fungi(100);k__Fungi_unclassified(100);k__Fungi_unclassified(100);k__Fungi_unclassified(100);k__Fungi_unclassified(100);k__Fungi_unclassified(100);k__Fungi_unclassified(100); | - | - | - | - | - | - | - | Unassigned | - |
| Otu11047 | 2 | k__Fungi(100);p__Basidiomycota(100);p__Basidiomycota_unclassified(100);p__Basidiomycota_unclassified(100);p__Basidiomycota_unclassified(100);p__Basidiomycota_unclassified(100);p__Basidiomycota_unclassified(100); | - | - | - | - | - | - | - | Unassigned | - |
| Otu11048 | 2 | k__Fungi(100);p__Basidiomycota(100);c__Agaricomycetes(100);c__Agaricomycetes_unclassified(100);c__Agaricomycetes_unclassified(100);c__Agaricomycetes_unclassified(100);c__Agaricomycetes_unclassified(100); | - | - | - | - | - | - | - | Unassigned | - |
| Otu11051 | 2 | k__Fungi(100);p__Ascomycota(100);p__Ascomycota_unclassified(100);p__Ascomycota_unclassified(100);p__Ascomycota_unclassified(100);p__Ascomycota_unclassified(100);p__Ascomycota_unclassified(100); | - | - | - | - | - | - | - | Unassigned | - |
| Otu11053 | 2 | k__Fungi(100);k__Fungi_unclassified(100);k__Fungi_unclassified(100);k__Fungi_unclassified(100);k__Fungi_unclassified(100);k__Fungi_unclassified(100);k__Fungi_unclassified(100); | - | - | - | - | - | - | - | Unassigned | - |
| Otu11056 | 2 | k__Fungi(100);p__Ascomycota(100);p__Ascomycota_unclassified(100);p__Ascomycota_unclassified(100);p__Ascomycota_unclassified(100);p__Ascomycota_unclassified(100);p__Ascomycota_unclassified(100); | - | - | - | - | - | - | - | Unassigned | - |
| Otu11057 | 2 | k__Fungi(100);k__Fungi_unclassified(100);k__Fungi_unclassified(100);k__Fungi_unclassified(100);k__Fungi_unclassified(100);k__Fungi_unclassified(100);k__Fungi_unclassified(100); | - | - | - | - | - | - | - | Unassigned | - |
| Otu11063 | 2 | k__Fungi(100);p__Ascomycota(100);p__Ascomycota_unclassified(100);p__Ascomycota_unclassified(100);p__Ascomycota_unclassified(100);p__Ascomycota_unclassified(100);p__Ascomycota_unclassified(100); | - | - | - | - | - | - | - | Unassigned | - |
| Otu11066 | 2 | k__Fungi(100);k__Fungi_unclassified(100);k__Fungi_unclassified(100);k__Fungi_unclassified(100);k__Fungi_unclassified(100);k__Fungi_unclassified(100);k__Fungi_unclassified(100); | - | - | - | - | - | - | - | Unassigned | - |
| Otu11073 | 2 | k__Fungi(100);p__Basidiomycota(100);p__Basidiomycota_unclassified(100);p__Basidiomycota_unclassified(100);p__Basidiomycota_unclassified(100);p__Basidiomycota_unclassified(100);p__Basidiomycota_unclassified(100); | - | - | - | - | - | - | - | Unassigned | - |
| Otu11075 | 2 | k__Fungi(100);k__Fungi_unclassified(100);k__Fungi_unclassified(100);k__Fungi_unclassified(100);k__Fungi_unclassified(100);k__Fungi_unclassified(100);k__Fungi_unclassified(100); | - | - | - | - | - | - | - | Unassigned | - |
| Otu11076 | 2 | k__Fungi(100);p__Ascomycota(100);c__Eurotiomycetes(100);c__Eurotiomycetes_unclassified(100);c__Eurotiomycetes_unclassified(100);c__Eurotiomycetes_unclassified(100);c__Eurotiomycetes_unclassified(100); | - | - | - | - | - | - | - | Unassigned | - |
| Otu11079 | 2 | k__Fungi(100);k__Fungi_unclassified(100);k__Fungi_unclassified(100);k__Fungi_unclassified(100);k__Fungi_unclassified(100);k__Fungi_unclassified(100);k__Fungi_unclassified(100); | - | - | - | - | - | - | - | Unassigned | - |
| Otu11082 | 2 | k__Fungi(100);p__Ascomycota(100);c__Sordariomycetes(100);c__Sordariomycetes_unclassified(100);c__Sordariomycetes_unclassified(100);c__Sordariomycetes_unclassified(100);c__Sordariomycetes_unclassified(100); | - | - | - | - | - | - | - | Unassigned | - |
| Otu11083 | 2 | k__Fungi(100);p__Ascomycota(100);p__Ascomycota_unclassified(100);p__Ascomycota_unclassified(100);p__Ascomycota_unclassified(100);p__Ascomycota_unclassified(100);p__Ascomycota_unclassified(100); | - | - | - | - | - | - | - | Unassigned | - |
| Otu11084 | 2 | k__Fungi(100);p__Ascomycota(100);p__Ascomycota_unclassified(100);p__Ascomycota_unclassified(100);p__Ascomycota_unclassified(100);p__Ascomycota_unclassified(100);p__Ascomycota_unclassified(100); | - | - | - | - | - | - | - | Unassigned | - |
| Otu11086 | 2 | k__Fungi(100);p__Basidiomycota(100);p__Basidiomycota_unclassified(100);p__Basidiomycota_unclassified(100);p__Basidiomycota_unclassified(100);p__Basidiomycota_unclassified(100);p__Basidiomycota_unclassified(100); | - | - | - | - | - | - | - | Unassigned | - |
| Otu11087 | 2 | k__Fungi(100);p__Ascomycota(100);p__Ascomycota_unclassified(100);p__Ascomycota_unclassified(100);p__Ascomycota_unclassified(100);p__Ascomycota_unclassified(100);p__Ascomycota_unclassified(100); | - | - | - | - | - | - | - | Unassigned | - |
| Otu11088 | 2 | k__Fungi(100);k__Fungi_unclassified(100);k__Fungi_unclassified(100);k__Fungi_unclassified(100);k__Fungi_unclassified(100);k__Fungi_unclassified(100);k__Fungi_unclassified(100); | - | - | - | - | - | - | - | Unassigned | - |
| Otu11090 | 2 | k__Fungi(100);p__Basidiomycota(100);p__Basidiomycota_unclassified(100);p__Basidiomycota_unclassified(100);p__Basidiomycota_unclassified(100);p__Basidiomycota_unclassified(100);p__Basidiomycota_unclassified(100); | - | - | - | - | - | - | - | Unassigned | - |
| Otu11094 | 2 | k__Fungi(100);p__Basidiomycota(100);c__Agaricomycetes(100);o__Cantharellales(100);o__Cantharellales_unclassified(100);o__Cantharellales_unclassified(100);o__Cantharellales_unclassified(100); | - | - | - | - | - | - | - | Unassigned | - |
| Otu11096 | 2 | k__Fungi(100);k__Fungi_unclassified(100);k__Fungi_unclassified(100);k__Fungi_unclassified(100);k__Fungi_unclassified(100);k__Fungi_unclassified(100);k__Fungi_unclassified(100); | - | - | - | - | - | - | - | Unassigned | - |
| Otu11102 | 2 | k__Fungi(100);k__Fungi_unclassified(100);k__Fungi_unclassified(100);k__Fungi_unclassified(100);k__Fungi_unclassified(100);k__Fungi_unclassified(100);k__Fungi_unclassified(100); | - | - | - | - | - | - | - | Unassigned | - |
| Otu11104 | 2 | k__Fungi(100);k__Fungi_unclassified(100);k__Fungi_unclassified(100);k__Fungi_unclassified(100);k__Fungi_unclassified(100);k__Fungi_unclassified(100);k__Fungi_unclassified(100); | - | - | - | - | - | - | - | Unassigned | - |
| Otu11105 | 2 | k__Fungi(100);p__Ascomycota(100);p__Ascomycota_unclassified(100);p__Ascomycota_unclassified(100);p__Ascomycota_unclassified(100);p__Ascomycota_unclassified(100);p__Ascomycota_unclassified(100); | - | - | - | - | - | - | - | Unassigned | - |
| Otu11106 | 2 | k__Fungi(100);p__Ascomycota(100);p__Ascomycota_unclassified(100);p__Ascomycota_unclassified(100);p__Ascomycota_unclassified(100);p__Ascomycota_unclassified(100);p__Ascomycota_unclassified(100); | - | - | - | - | - | - | - | Unassigned | - |
| Otu11111 | 2 | k__Fungi(100);k__Fungi_unclassified(100);k__Fungi_unclassified(100);k__Fungi_unclassified(100);k__Fungi_unclassified(100);k__Fungi_unclassified(100);k__Fungi_unclassified(100); | - | - | - | - | - | - | - | Unassigned | - |

| | | | | | | | | | | |
|---|---|---|---|---|---|---|---|---|---|---|
| Otu11113 | 2 | k__Fungi(100);k__Fungi_unclassified(100);k__Fungi_unclassified(100);k__Fungi_unclassified(100);k__Fungi_unclassified(100);k__Fungi_unclassified(100);k__Fungi_unclassified(100); | - | - | - | - | - | - | - | Unassigned | - |
| Otu11114 | 2 | k__Fungi(100);k__Fungi_unclassified(100);k__Fungi_unclassified(100);k__Fungi_unclassified(100);k__Fungi_unclassified(100);k__Fungi_unclassified(100);k__Fungi_unclassified(100); | - | - | - | - | - | - | - | Unassigned | - |
| Otu11117 | 2 | k__Fungi(100);p__Basidiomycota(100);c__Agaricomycetes(100);c__Agaricomycetes_unclassified(100);c__Agaricomycetes_unclassified(100);c__Agaricomycetes_unclassified(100);c__Agaricomycetes_unclassified(100); | - | - | - | - | - | - | - | Unassigned | - |
| Otu11119 | 2 | k__Fungi(100);k__Fungi_unclassified(100);k__Fungi_unclassified(100);k__Fungi_unclassified(100);k__Fungi_unclassified(100);k__Fungi_unclassified(100);k__Fungi_unclassified(100); | - | - | - | - | - | - | - | Unassigned | - |
| Otu11120 | 2 | k__Fungi(100);p__Ascomycota(100);p__Ascomycota_unclassified(100);p__Ascomycota_unclassified(100);p__Ascomycota_unclassified(100);p__Ascomycota_unclassified(100);p__Ascomycota_unclassified(100); | - | - | - | - | - | - | - | Unassigned | - |
| Otu11121 | 2 | k__Fungi(100);k__Fungi_unclassified(100);k__Fungi_unclassified(100);k__Fungi_unclassified(100);k__Fungi_unclassified(100);k__Fungi_unclassified(100);k__Fungi_unclassified(100); | - | - | - | - | - | - | - | Unassigned | - |
| Otu11124 | 2 | k__Fungi(100);p__Ascomycota(100);p__Ascomycota_unclassified(100);p__Ascomycota_unclassified(100);p__Ascomycota_unclassified(100);p__Ascomycota_unclassified(100);p__Ascomycota_unclassified(100); | - | - | - | - | - | - | - | Unassigned | - |
| Otu11125 | 2 | k__Fungi(100);p__Ascomycota(100);p__Ascomycota_unclassified(100);p__Ascomycota_unclassified(100);p__Ascomycota_unclassified(100);p__Ascomycota_unclassified(100);p__Ascomycota_unclassified(100); | - | - | - | - | - | - | - | Unassigned | - |
| Otu11126 | 2 | k__Fungi(100);p__Ascomycota(100);p__Ascomycota_unclassified(100);p__Ascomycota_unclassified(100);p__Ascomycota_unclassified(100);p__Ascomycota_unclassified(100);p__Ascomycota_unclassified(100); | - | - | - | - | - | - | - | Unassigned | - |
| Otu11137 | 2 | k__Fungi(100);p__Ascomycota(100);p__Ascomycota_unclassified(100);p__Ascomycota_unclassified(100);p__Ascomycota_unclassified(100);p__Ascomycota_unclassified(100);p__Ascomycota_unclassified(100); | - | - | - | - | - | - | - | Unassigned | - |
| Otu11141 | 2 | k__Fungi(100);k__Fungi_unclassified(100);k__Fungi_unclassified(100);k__Fungi_unclassified(100);k__Fungi_unclassified(100);k__Fungi_unclassified(100);k__Fungi_unclassified(100); | - | - | - | - | - | - | - | Unassigned | - |
| Otu11142 | 2 | k__Fungi(100);p__Ascomycota(100);p__Ascomycota_unclassified(100);p__Ascomycota_unclassified(100);p__Ascomycota_unclassified(100);p__Ascomycota_unclassified(100);p__Ascomycota_unclassified(100); | - | - | - | - | - | - | - | Unassigned | - |
| Otu11146 | 2 | k__Fungi(100);p__Ascomycota(100);c__Dothideomycetes(100);o__Capnodiales(100);o__Capnodiales_unclassified(100);o__Capnodiales_unclassified(100);o__Capnodiales_unclassified(100); | - | - | - | - | - | - | - | Unassigned | - |
| Otu11150 | 2 | k__Fungi(100);k__Fungi_unclassified(100);k__Fungi_unclassified(100);k__Fungi_unclassified(100);k__Fungi_unclassified(100);k__Fungi_unclassified(100);k__Fungi_unclassified(100); | - | - | - | - | - | - | - | Unassigned | - |
| Otu11153 | 2 | k__Fungi(100);k__Fungi_unclassified(100);k__Fungi_unclassified(100);k__Fungi_unclassified(100);k__Fungi_unclassified(100);k__Fungi_unclassified(100);k__Fungi_unclassified(100); | - | - | - | - | - | - | - | Unassigned | - |
| Otu11157 | 2 | k__Fungi(100);p__Ascomycota(100);p__Ascomycota_unclassified(100);p__Ascomycota_unclassified(100);p__Ascomycota_unclassified(100);p__Ascomycota_unclassified(100);p__Ascomycota_unclassified(100); | - | - | - | - | - | - | - | Unassigned | - |
| Otu11165 | 2 | k__Fungi(100);k__Fungi_unclassified(100);k__Fungi_unclassified(100);k__Fungi_unclassified(100);k__Fungi_unclassified(100);k__Fungi_unclassified(100);k__Fungi_unclassified(100); | - | - | - | - | - | - | - | Unassigned | - |
| Otu11167 | 2 | k__Fungi(100);p__Basidiomycota(100);c__Agaricomycetes(100);o__Russulales(100);o__Russulales_unclassified(100);o__Russulales_unclassified(100);o__Russulales_unclassified(100); | - | - | - | - | - | - | - | Unassigned | - |
| Otu11168 | 2 | k__Fungi(100);k__Fungi_unclassified(100);k__Fungi_unclassified(100);k__Fungi_unclassified(100);k__Fungi_unclassified(100);k__Fungi_unclassified(100);k__Fungi_unclassified(100); | - | - | - | - | - | - | - | Unassigned | - |
| Otu11169 | 2 | k__Fungi(100);p__Ascomycota(100);p__Ascomycota_unclassified(100);p__Ascomycota_unclassified(100);p__Ascomycota_unclassified(100);p__Ascomycota_unclassified(100);p__Ascomycota_unclassified(100); | - | - | - | - | - | - | - | Unassigned | - |
| Otu11172 | 2 | k__Fungi(100);k__Fungi_unclassified(100);k__Fungi_unclassified(100);k__Fungi_unclassified(100);k__Fungi_unclassified(100);k__Fungi_unclassified(100);k__Fungi_unclassified(100); | - | - | - | - | - | - | - | Unassigned | - |
| Otu11173 | 2 | k__Fungi(100);k__Fungi_unclassified(100);k__Fungi_unclassified(100);k__Fungi_unclassified(100);k__Fungi_unclassified(100);k__Fungi_unclassified(100);k__Fungi_unclassified(100); | - | - | - | - | - | - | - | Unassigned | - |
| Otu11176 | 2 | k__Fungi(100);p__Basidiomycota(100);c__Agaricomycetes(100);c__Agaricomycetes_unclassified(100);c__Agaricomycetes_unclassified(100);c__Agaricomycetes_unclassified(100);c__Agaricomycetes_unclassified(100); | - | - | - | - | - | - | - | Unassigned | - |
| Otu11178 | 2 | k__Fungi(100);k__Fungi_unclassified(100);k__Fungi_unclassified(100);k__Fungi_unclassified(100);k__Fungi_unclassified(100);k__Fungi_unclassified(100);k__Fungi_unclassified(100); | - | - | - | - | - | - | - | Unassigned | - |
| Otu11179 | 2 | k__Fungi(100);k__Fungi_unclassified(100);k__Fungi_unclassified(100);k__Fungi_unclassified(100);k__Fungi_unclassified(100);k__Fungi_unclassified(100);k__Fungi_unclassified(100); | - | - | - | - | - | - | - | Unassigned | - |
| Otu11187 | 2 | k__Fungi(100);p__Ascomycota(100);p__Ascomycota_unclassified(100);p__Ascomycota_unclassified(100);p__Ascomycota_unclassified(100);p__Ascomycota_unclassified(100);p__Ascomycota_unclassified(100); | - | - | - | - | - | - | - | Unassigned | - |
| Otu11189 | 2 | k__Fungi(100);k__Fungi_unclassified(100);k__Fungi_unclassified(100);k__Fungi_unclassified(100);k__Fungi_unclassified(100);k__Fungi_unclassified(100);k__Fungi_unclassified(100); | - | - | - | - | - | - | - | Unassigned | - |
| Otu11190 | 2 | k__Fungi(100);k__Fungi_unclassified(100);k__Fungi_unclassified(100);k__Fungi_unclassified(100);k__Fungi_unclassified(100);k__Fungi_unclassified(100);k__Fungi_unclassified(100); | - | - | - | - | - | - | - | Unassigned | - |
| Otu11191 | 2 | k__Fungi(100);p__Ascomycota(100);p__Ascomycota_unclassified(100);p__Ascomycota_unclassified(100);p__Ascomycota_unclassified(100);p__Ascomycota_unclassified(100);p__Ascomycota_unclassified(100); | - | - | - | - | - | - | - | Unassigned | - |
| Otu11192 | 2 | k__Fungi(100);p__Ascomycota(100);c__Sordariomycetes(100);o__Diaporthales(100);o__Diaporthales_unclassified(100);o__Diaporthales_unclassified(100);o__Diaporthales_unclassified(100); | - | - | - | - | - | - | - | Unassigned | - |
| Otu11200 | 2 | k__Fungi(100);p__Basidiomycota(100);c__Agaricomycetes(100);o__Trechisporales(100);f__unclassified_Trechisporales(100);g__unclassified_Trechisporales(100);s__Trechisporales_sp(100); | - | - | - | - | - | - | - | Unassigned | - |
| Otu11209 | 2 | k__Fungi(100);k__Fungi_unclassified(100);k__Fungi_unclassified(100);k__Fungi_unclassified(100);k__Fungi_unclassified(100);k__Fungi_unclassified(100);k__Fungi_unclassified(100); | - | - | - | - | - | - | - | Unassigned | - |
| Otu11212 | 2 | k__Fungi(100);k__Fungi_unclassified(100);k__Fungi_unclassified(100);k__Fungi_unclassified(100);k__Fungi_unclassified(100);k__Fungi_unclassified(100);k__Fungi_unclassified(100); | - | - | - | - | - | - | - | Unassigned | - |
| Otu11214 | 2 | k__Fungi(100);k__Fungi_unclassified(100);k__Fungi_unclassified(100);k__Fungi_unclassified(100);k__Fungi_unclassified(100);k__Fungi_unclassified(100);k__Fungi_unclassified(100); | - | - | - | - | - | - | - | Unassigned | - |
| Otu11218 | 2 | k__Fungi(100);p__Ascomycota(100);p__Ascomycota_unclassified(100);p__Ascomycota_unclassified(100);p__Ascomycota_unclassified(100);p__Ascomycota_unclassified(100);p__Ascomycota_unclassified(100); | - | - | - | - | - | - | - | Unassigned | - |
| Otu11220 | 2 | k__Fungi(100);k__Fungi_unclassified(100);k__Fungi_unclassified(100);k__Fungi_unclassified(100);k__Fungi_unclassified(100);k__Fungi_unclassified(100);k__Fungi_unclassified(100); | - | - | - | - | - | - | - | Unassigned | - |
| Otu11221 | 2 | k__Fungi(100);k__Fungi_unclassified(100);k__Fungi_unclassified(100);k__Fungi_unclassified(100);k__Fungi_unclassified(100);k__Fungi_unclassified(100);k__Fungi_unclassified(100); | - | - | - | - | - | - | - | Unassigned | - |
| Otu11222 | 2 | k__Fungi(100);p__Ascomycota(100);p__Ascomycota_unclassified(100);p__Ascomycota_unclassified(100);p__Ascomycota_unclassified(100);p__Ascomycota_unclassified(100);p__Ascomycota_unclassified(100); | - | - | - | - | - | - | - | Unassigned | - |
| Otu11223 | 2 | k__Fungi(100);p__Ascomycota(100);c__Sordariomycetes(100);o__Hypocreales(100);f__Hypocreaceae(100);f__Hypocreaceae_unclassified(100);f__Hypocreaceae_unclassified(100); | - | - | - | - | - | - | - | Unassigned | - |
| Otu11224 | 2 | k__Fungi(100);k__Fungi_unclassified(100);k__Fungi_unclassified(100);k__Fungi_unclassified(100);k__Fungi_unclassified(100);k__Fungi_unclassified(100);k__Fungi_unclassified(100); | - | - | - | - | - | - | - | Unassigned | - |
| Otu11225 | 2 | k__Fungi(100);p__Ascomycota(100);p__Ascomycota_unclassified(100);p__Ascomycota_unclassified(100);p__Ascomycota_unclassified(100);p__Ascomycota_unclassified(100);p__Ascomycota_unclassified(100); | - | - | - | - | - | - | - | Unassigned | - |
| Otu11227 | 2 | k__Fungi(100);p__Ascomycota(100);p__Ascomycota_unclassified(100);p__Ascomycota_unclassified(100);p__Ascomycota_unclassified(100);p__Ascomycota_unclassified(100);p__Ascomycota_unclassified(100); | - | - | - | - | - | - | - | Unassigned | - |
| Otu11238 | 2 | k__Fungi(100);p__Ascomycota(100);p__Ascomycota_unclassified(100);p__Ascomycota_unclassified(100);p__Ascomycota_unclassified(100);p__Ascomycota_unclassified(100);p__Ascomycota_unclassified(100); | - | - | - | - | - | - | - | Unassigned | - |
| Otu11243 | 2 | k__Fungi(100);p__Ascomycota(100);c__Taphrinomycetes(100);o__Taphrinales(100);f__Taphrinaceae(100);g__Lalaria(100);s__Lalaria_inositophila(100); | - | - | - | - | - | - | - | Unassigned | - |
| Otu11245 | 2 | k__Fungi(100);p__Ascomycota(100);c__Sordariomycetes(100);o__Hypocreales(100);f__Cordycipitaceae(100);f__Cordycipitaceae_unclassified(100);f__Cordycipitaceae_unclassified(100); | - | - | - | - | - | - | - | Unassigned | - |
| Otu11246 | 2 | k__Fungi(100);p__Ascomycota(100);p__Ascomycota_unclassified(100);p__Ascomycota_unclassified(100);p__Ascomycota_unclassified(100);p__Ascomycota_unclassified(100);p__Ascomycota_unclassified(100); | - | - | - | - | - | - | - | Unassigned | - |
| Otu11250 | 2 | k__Fungi(100);k__Fungi_unclassified(100);k__Fungi_unclassified(100);k__Fungi_unclassified(100);k__Fungi_unclassified(100);k__Fungi_unclassified(100);k__Fungi_unclassified(100); | - | - | - | - | - | - | - | Unassigned | - |
| Otu11251 | 2 | k__Fungi(100);p__Ascomycota(100);p__Ascomycota_unclassified(100);p__Ascomycota_unclassified(100);p__Ascomycota_unclassified(100);p__Ascomycota_unclassified(100);p__Ascomycota_unclassified(100); | - | - | - | - | - | - | - | Unassigned | - |
| Otu11260 | 2 | k__Fungi(100);p__Ascomycota(100);c__Sordariomycetes(100);c__Sordariomycetes_unclassified(100);c__Sordariomycetes_unclassified(100);c__Sordariomycetes_unclassified(100);c__Sordariomycetes_unclassified(100); | - | - | - | - | - | - | - | Unassigned | - |
| Otu11264 | 2 | k__Fungi(100);p__Ascomycota(100);c__Sordariomycetes(100);c__Sordariomycetes_unclassified(100);c__Sordariomycetes_unclassified(100);c__Sordariomycetes_unclassified(100);c__Sordariomycetes_unclassified(100); | - | - | - | - | - | - | - | Unassigned | - |
| Otu11267 | 2 | k__Fungi(100);p__Ascomycota(100);c__Sordariomycetes(100);c__Sordariomycetes_unclassified(100);c__Sordariomycetes_unclassified(100);c__Sordariomycetes_unclassified(100);c__Sordariomycetes_unclassified(100); | - | - | - | - | - | - | - | Unassigned | - |
| Otu11271 | 2 | k__Fungi(100);k__Fungi_unclassified(100);k__Fungi_unclassified(100);k__Fungi_unclassified(100);k__Fungi_unclassified(100);k__Fungi_unclassified(100);k__Fungi_unclassified(100); | - | - | - | - | - | - | - | Unassigned | - |
| Otu11276 | 2 | k__Fungi(100);k__Fungi_unclassified(100);k__Fungi_unclassified(100);k__Fungi_unclassified(100);k__Fungi_unclassified(100);k__Fungi_unclassified(100);k__Fungi_unclassified(100); | - | - | - | - | - | - | - | Unassigned | - |
| Otu11277 | 2 | k__Fungi(100);p__Ascomycota(100);p__Ascomycota_unclassified(100);p__Ascomycota_unclassified(100);p__Ascomycota_unclassified(100);p__Ascomycota_unclassified(100);p__Ascomycota_unclassified(100); | - | - | - | - | - | - | - | Unassigned | - |
| Otu11280 | 2 | k__Fungi(100);p__Ascomycota(100);p__Ascomycota_unclassified(100);p__Ascomycota_unclassified(100);p__Ascomycota_unclassified(100);p__Ascomycota_unclassified(100);p__Ascomycota_unclassified(100); | - | - | - | - | - | - | - | Unassigned | - |
| Otu11283 | 2 | k__Fungi(100);k__Fungi_unclassified(100);k__Fungi_unclassified(100);k__Fungi_unclassified(100);k__Fungi_unclassified(100);k__Fungi_unclassified(100);k__Fungi_unclassified(100); | - | - | - | - | - | - | - | Unassigned | - |
| Otu11284 | 2 | k__Fungi(100);k__Fungi_unclassified(100);k__Fungi_unclassified(100);k__Fungi_unclassified(100);k__Fungi_unclassified(100);k__Fungi_unclassified(100);k__Fungi_unclassified(100); | - | - | - | - | - | - | - | Unassigned | - |
| Otu11285 | 2 | k__Fungi(100);k__Fungi_unclassified(100);k__Fungi_unclassified(100);k__Fungi_unclassified(100);k__Fungi_unclassified(100);k__Fungi_unclassified(100);k__Fungi_unclassified(100); | - | - | - | - | - | - | - | Unassigned | - |
| Otu11288 | 2 | k__Fungi(100);k__Fungi_unclassified(100);k__Fungi_unclassified(100);k__Fungi_unclassified(100);k__Fungi_unclassified(100);k__Fungi_unclassified(100);k__Fungi_unclassified(100); | - | - | - | - | - | - | - | Unassigned | - |
| Otu11289 | 2 | k__Fungi(100);p__Ascomycota(100);p__Ascomycota_unclassified(100);p__Ascomycota_unclassified(100);p__Ascomycota_unclassified(100);p__Ascomycota_unclassified(100);p__Ascomycota_unclassified(100); | - | - | - | - | - | - | - | Unassigned | - |
| Otu11291 | 2 | k__Fungi(100);k__Fungi_unclassified(100);k__Fungi_unclassified(100);k__Fungi_unclassified(100);k__Fungi_unclassified(100);k__Fungi_unclassified(100);k__Fungi_unclassified(100); | - | - | - | - | - | - | - | Unassigned | - |
| Otu11293 | 2 | k__Fungi(100);p__Ascomycota(100);p__Ascomycota_unclassified(100);p__Ascomycota_unclassified(100);p__Ascomycota_unclassified(100);p__Ascomycota_unclassified(100);p__Ascomycota_unclassified(100); | - | - | - | - | - | - | - | Unassigned | - |
| Otu11299 | 2 | k__Fungi(100);p__Ascomycota(100);p__Ascomycota_unclassified(100);p__Ascomycota_unclassified(100);p__Ascomycota_unclassified(100);p__Ascomycota_unclassified(100);p__Ascomycota_unclassified(100); | - | - | - | - | - | - | - | Unassigned | - |
| Otu11303 | 2 | k__Fungi(100);p__Ascomycota(100);c__Sordariomycetes(100);o__Hypocreales(100);f__Hypocreaceae(100);f__Hypocreaceae_unclassified(100);f__Hypocreaceae_unclassified(100); | - | - | - | - | - | - | - | Unassigned | - |
| Otu11304 | 2 | k__Fungi(100);p__Ascomycota(100);c__Sordariomycetes(100);o__Hypocreales(100);f__Hypocreaceae(100);f__Hypocreaceae_unclassified(100);f__Hypocreaceae_unclassified(100); | - | - | - | - | - | - | - | Unassigned | - |
| Otu11314 | 2 | k__Fungi(100);p__Ascomycota(100);c__Sordariomycetes(100);c__Sordariomycetes_unclassified(100);c__Sordariomycetes_unclassified(100);c__Sordariomycetes_unclassified(100);c__Sordariomycetes_unclassified(100); | - | - | - | - | - | - | - | Unassigned | - |
| Otu11319 | 2 | k__Fungi(100);k__Fungi_unclassified(100);k__Fungi_unclassified(100);k__Fungi_unclassified(100);k__Fungi_unclassified(100);k__Fungi_unclassified(100);k__Fungi_unclassified(100); | - | - | - | - | - | - | - | Unassigned | - |
| Otu11324 | 2 | k__Fungi(100);p__Ascomycota(100);p__Ascomycota_unclassified(100);p__Ascomycota_unclassified(100);p__Ascomycota_unclassified(100);p__Ascomycota_unclassified(100);p__Ascomycota_unclassified(100); | - | - | - | - | - | - | - | Unassigned | - |
| Otu11325 | 2 | k__Fungi(100);k__Fungi_unclassified(100);k__Fungi_unclassified(100);k__Fungi_unclassified(100);k__Fungi_unclassified(100);k__Fungi_unclassified(100);k__Fungi_unclassified(100); | - | - | - | - | - | - | - | Unassigned | - |
| Otu11327 | 2 | k__Fungi(100);p__Ascomycota(100);p__Ascomycota_unclassified(100);p__Ascomycota_unclassified(100);p__Ascomycota_unclassified(100);p__Ascomycota_unclassified(100);p__Ascomycota_unclassified(100); | - | - | - | - | - | - | - | Unassigned | - |
| Otu11328 | 2 | k__Fungi(100);p__Ascomycota(100);c__Eurotiomycetes(100);c__Eurotiomycetes_unclassified(100);c__Eurotiomycetes_unclassified(100);c__Eurotiomycetes_unclassified(100);c__Eurotiomycetes_unclassified(100); | - | - | - | - | - | - | - | Unassigned | - |
| Otu11331 | 2 | k__Fungi(100);p__Ascomycota(100);c__Eurotiomycetes(100);c__Eurotiomycetes_unclassified(100);c__Eurotiomycetes_unclassified(100);c__Eurotiomycetes_unclassified(100);c__Eurotiomycetes_unclassified(100); | - | - | - | - | - | - | - | Unassigned | - |
| Otu11333 | 2 | k__Fungi(100);p__Ascomycota(100);p__Ascomycota_unclassified(100);p__Ascomycota_unclassified(100);p__Ascomycota_unclassified(100);p__Ascomycota_unclassified(100);p__Ascomycota_unclassified(100); | - | - | - | - | - | - | - | Unassigned | - |
| Otu11334 | 2 | k__Fungi(100);p__Ascomycota(100);p__Ascomycota_unclassified(100);p__Ascomycota_unclassified(100);p__Ascomycota_unclassified(100);p__Ascomycota_unclassified(100);p__Ascomycota_unclassified(100); | - | - | - | - | - | - | - | Unassigned | - |
| Otu11335 | 2 | k__Fungi(100);k__Fungi_unclassified(100);k__Fungi_unclassified(100);k__Fungi_unclassified(100);k__Fungi_unclassified(100);k__Fungi_unclassified(100);k__Fungi_unclassified(100); | - | - | - | - | - | - | - | Unassigned | - |
| Otu11338 | 2 | k__Fungi(100);k__Fungi_unclassified(100);k__Fungi_unclassified(100);k__Fungi_unclassified(100);k__Fungi_unclassified(100);k__Fungi_unclassified(100);k__Fungi_unclassified(100); | - | - | - | - | - | - | - | Unassigned | - |
| Otu11339 | 2 | k__Fungi(100);k__Fungi_unclassified(100);k__Fungi_unclassified(100);k__Fungi_unclassified(100);k__Fungi_unclassified(100);k__Fungi_unclassified(100);k__Fungi_unclassified(100); | - | - | - | - | - | - | - | Unassigned | - |
| Otu11342 | 2 | k__Fungi(100);k__Fungi_unclassified(100);k__Fungi_unclassified(100);k__Fungi_unclassified(100);k__Fungi_unclassified(100);k__Fungi_unclassified(100);k__Fungi_unclassified(100); | - | - | - | - | - | - | - | Unassigned | - |
| Otu11344 | 2 | k__Fungi(100);p__Ascomycota(100);c__Taphrinomycetes(100);o__Taphrinales(100);f__Taphrinaceae(100);g__Lalaria(100);s__Lalaria_inositophila(100); | - | - | - | - | - | - | - | Unassigned | - |
| Otu11348 | 2 | k__Fungi(100);p__Ascomycota(100);p__Ascomycota_unclassified(100);p__Ascomycota_unclassified(100);p__Ascomycota_unclassified(100);p__Ascomycota_unclassified(100);p__Ascomycota_unclassified(100); | - | - | - | - | - | - | - | Unassigned | - |
| Otu11354 | 2 | k__Fungi(100);p__Ascomycota(100);c__Eurotiomycetes(100);c__Eurotiomycetes_unclassified(100);c__Eurotiomycetes_unclassified(100);c__Eurotiomycetes_unclassified(100);c__Eurotiomycetes_unclassified(100); | - | - | - | - | - | - | - | Unassigned | - |
| Otu11368 | 2 | k__Fungi(100);p__Ascomycota(100);p__Ascomycota_unclassified(100);p__Ascomycota_unclassified(100);p__Ascomycota_unclassified(100);p__Ascomycota_unclassified(100);p__Ascomycota_unclassified(100); | - | - | - | - | - | - | - | Unassigned | - |
| Otu11370 | 2 | k__Fungi(100);p__Ascomycota(100);c__Dothideomycetes(100);c__Dothideomycetes_unclassified(100);c__Dothideomycetes_unclassified(100);c__Dothideomycetes_unclassified(100);c__Dothideomycetes_unclassified(100); | - | - | - | - | - | - | - | Unassigned | - |
| Otu11374 | 2 | k__Fungi(100);k__Fungi_unclassified(100);k__Fungi_unclassified(100);k__Fungi_unclassified(100);k__Fungi_unclassified(100);k__Fungi_unclassified(100);k__Fungi_unclassified(100); | - | - | - | - | - | - | - | Unassigned | - |
| Otu11375 | 2 | k__Fungi(100);k__Fungi_unclassified(100);k__Fungi_unclassified(100);k__Fungi_unclassified(100);k__Fungi_unclassified(100);k__Fungi_unclassified(100);k__Fungi_unclassified(100); | - | - | - | - | - | - | - | Unassigned | - |
| Otu11377 | 2 | k__Fungi(100);k__Fungi_unclassified(100);k__Fungi_unclassified(100);k__Fungi_unclassified(100);k__Fungi_unclassified(100);k__Fungi_unclassified(100);k__Fungi_unclassified(100); | - | - | - | - | - | - | - | Unassigned | - |
| Otu11395 | 2 | k__Fungi(100);p__Ascomycota(100);p__Ascomycota_unclassified(100);p__Ascomycota_unclassified(100);p__Ascomycota_unclassified(100);p__Ascomycota_unclassified(100);p__Ascomycota_unclassified(100); | - | - | - | - | - | - | - | Unassigned | - |
| Otu11398 | 2 | k__Fungi(100);k__Fungi_unclassified(100);k__Fungi_unclassified(100);k__Fungi_unclassified(100);k__Fungi_unclassified(100);k__Fungi_unclassified(100);k__Fungi_unclassified(100); | - | - | - | - | - | - | - | Unassigned | - |
| Otu11401 | 2 | k__Fungi(100);k__Fungi_unclassified(100);k__Fungi_unclassified(100);k__Fungi_unclassified(100);k__Fungi_unclassified(100);k__Fungi_unclassified(100);k__Fungi_unclassified(100); | - | - | - | - | - | - | - | Unassigned | - |
| Otu11406 | 2 | k__Fungi(100);p__Ascomycota(100);p__Ascomycota_unclassified(100);p__Ascomycota_unclassified(100);p__Ascomycota_unclassified(100);p__Ascomycota_unclassified(100);p__Ascomycota_unclassified(100); | - | - | - | - | - | - | - | Unassigned | - |
| Otu11408 | 2 | k__Fungi(100);p__Ascomycota(100);p__Ascomycota_unclassified(100);p__Ascomycota_unclassified(100);p__Ascomycota_unclassified(100);p__Ascomycota_unclassified(100);p__Ascomycota_unclassified(100); | - | - | - | - | - | - | - | Unassigned | - |
| Otu11422 | 2 | k__Fungi(100);k__Fungi_unclassified(100);k__Fungi_unclassified(100);k__Fungi_unclassified(100);k__Fungi_unclassified(100);k__Fungi_unclassified(100);k__Fungi_unclassified(100); | - | - | - | - | - | - | - | Unassigned | - |
| Otu11428 | 2 | k__Fungi(100);k__Fungi_unclassified(100);k__Fungi_unclassified(100);k__Fungi_unclassified(100);k__Fungi_unclassified(100);k__Fungi_unclassified(100);k__Fungi_unclassified(100); | - | - | - | - | - | - | - | Unassigned | - |
| Otu11429 | 2 | k__Fungi(100);p__Ascomycota(100);p__Ascomycota_unclassified(100);p__Ascomycota_unclassified(100);p__Ascomycota_unclassified(100);p__Ascomycota_unclassified(100);p__Ascomycota_unclassified(100); | - | - | - | - | - | - | - | Unassigned | - |
| Otu11432 | 2 | k__Fungi(100);k__Fungi_unclassified(100);k__Fungi_unclassified(100);k__Fungi_unclassified(100);k__Fungi_unclassified(100);k__Fungi_unclassified(100);k__Fungi_unclassified(100); | - | - | - | - | - | - | - | Unassigned | - |
| Otu11435 | 2 | k__Fungi(100);k__Fungi_unclassified(100);k__Fungi_unclassified(100);k__Fungi_unclassified(100);k__Fungi_unclassified(100);k__Fungi_unclassified(100);k__Fungi_unclassified(100); | - | - | - | - | - | - | - | Unassigned | - |
| Otu11439 | 2 | k__Fungi(100);k__Fungi_unclassified(100);k__Fungi_unclassified(100);k__Fungi_unclassified(100);k__Fungi_unclassified(100);k__Fungi_unclassified(100);k__Fungi_unclassified(100); | - | - | - | - | - | - | - | Unassigned | - |
| Otu11440 | 2 | k__Fungi(100);k__Fungi_unclassified(100);k__Fungi_unclassified(100);k__Fungi_unclassified(100);k__Fungi_unclassified(100);k__Fungi_unclassified(100);k__Fungi_unclassified(100); | - | - | - | - | - | - | - | Unassigned | - |
| Otu11443 | 2 | k__Fungi(100);k__Fungi_unclassified(100);k__Fungi_unclassified(100);k__Fungi_unclassified(100);k__Fungi_unclassified(100);k__Fungi_unclassified(100);k__Fungi_unclassified(100); | - | - | - | - | - | - | - | Unassigned | - |
| Otu11446 | 2 | k__Fungi(100);k__Fungi_unclassified(100);k__Fungi_unclassified(100);k__Fungi_unclassified(100);k__Fungi_unclassified(100);k__Fungi_unclassified(100);k__Fungi_unclassified(100); | - | - | - | - | - | - | - | Unassigned | - |
| Otu11448 | 2 | k__Fungi(100);k__Fungi_unclassified(100);k__Fungi_unclassified(100);k__Fungi_unclassified(100);k__Fungi_unclassified(100);k__Fungi_unclassified(100);k__Fungi_unclassified(100); | - | - | - | - | - | - | - | Unassigned | - |
| Otu11451 | 2 | k__Fungi(100);p__Ascomycota(100);p__Ascomycota_unclassified(100);p__Ascomycota_unclassified(100);p__Ascomycota_unclassified(100);p__Ascomycota_unclassified(100);p__Ascomycota_unclassified(100); | - | - | - | - | - | - | - | Unassigned | - |
| Otu11453 | 2 | k__Fungi(100);k__Fungi_unclassified(100);k__Fungi_unclassified(100);k__Fungi_unclassified(100);k__Fungi_unclassified(100);k__Fungi_unclassified(100);k__Fungi_unclassified(100); | - | - | - | - | - | - | - | Unassigned | - |
| Otu11456 | 2 | k__Fungi(100);p__Ascomycota(100);p__Ascomycota_unclassified(100);p__Ascomycota_unclassified(100);p__Ascomycota_unclassified(100);p__Ascomycota_unclassified(100);p__Ascomycota_unclassified(100); | - | - | - | - | - | - | - | Unassigned | - |
| Otu11457 | 2 | k__Fungi(100);p__Ascomycota(100);p__Ascomycota_unclassified(100);p__Ascomycota_unclassified(100);p__Ascomycota_unclassified(100);p__Ascomycota_unclassified(100);p__Ascomycota_unclassified(100); | - | - | - | - | - | - | - | Unassigned | - |
| Otu11460 | 2 | k__Fungi(100);k__Fungi_unclassified(100);k__Fungi_unclassified(100);k__Fungi_unclassified(100);k__Fungi_unclassified(100);k__Fungi_unclassified(100);k__Fungi_unclassified(100); | - | - | - | - | - | - | - | Unassigned | - |
| Otu11461 | 2 | k__Fungi(100);p__Ascomycota(100);p__Ascomycota_unclassified(100);p__Ascomycota_unclassified(100);p__Ascomycota_unclassified(100);p__Ascomycota_unclassified(100);p__Ascomycota_unclassified(100); | - | - | - | - | - | - | - | Unassigned | - |
| Otu11462 | 2 | k__Fungi(100);k__Fungi_unclassified(100);k__Fungi_unclassified(100);k__Fungi_unclassified(100);k__Fungi_unclassified(100);k__Fungi_unclassified(100);k__Fungi_unclassified(100); | - | - | - | - | - | - | - | Unassigned | - |
| Otu11466 | 2 | k__Fungi(100);k__Fungi_unclassified(100);k__Fungi_unclassified(100);k__Fungi_unclassified(100);k__Fungi_unclassified(100);k__Fungi_unclassified(100);k__Fungi_unclassified(100); | - | - | - | - | - | - | - | Unassigned | - |
| Otu11469 | 2 | k__Fungi(100);k__Fungi_unclassified(100);k__Fungi_unclassified(100);k__Fungi_unclassified(100);k__Fungi_unclassified(100);k__Fungi_unclassified(100);k__Fungi_unclassified(100); | - | - | - | - | - | - | - | Unassigned | - |
| Otu11470 | 2 | k__Fungi(100);k__Fungi_unclassified(100);k__Fungi_unclassified(100);k__Fungi_unclassified(100);k__Fungi_unclassified(100);k__Fungi_unclassified(100);k__Fungi_unclassified(100); | - | - | - | - | - | - | - | Unassigned | - |
| Otu11472 | 2 | k__Fungi(100);p__Ascomycota(100);p__Ascomycota_unclassified(100);p__Ascomycota_unclassified(100);p__Ascomycota_unclassified(100);p__Ascomycota_unclassified(100);p__Ascomycota_unclassified(100); | - | - | - | - | - | - | - | Unassigned | - |
| Otu11476 | 2 | k__Fungi(100);k__Fungi_unclassified(100);k__Fungi_unclassified(100);k__Fungi_unclassified(100);k__Fungi_unclassified(100);k__Fungi_unclassified(100);k__Fungi_unclassified(100); | - | - | - | - | - | - | - | Unassigned | - |
| Otu11479 | 2 | k__Fungi(100);p__Ascomycota(100);p__Ascomycota_unclassified(100);p__Ascomycota_unclassified(100);p__Ascomycota_unclassified(100);p__Ascomycota_unclassified(100);p__Ascomycota_unclassified(100); | - | - | - | - | - | - | - | Unassigned | - |
| Otu11481 | 2 | k__Fungi(100);p__Ascomycota(100);p__Ascomycota_unclassified(100);p__Ascomycota_unclassified(100);p__Ascomycota_unclassified(100);p__Ascomycota_unclassified(100);p__Ascomycota_unclassified(100); | - | - | - | - | - | - | - | Unassigned | - |
| Otu11482 | 2 | k__Fungi(100);k__Fungi_unclassified(100);k__Fungi_unclassified(100);k__Fungi_unclassified(100);k__Fungi_unclassified(100);k__Fungi_unclassified(100);k__Fungi_unclassified(100); | - | - | - | - | - | - | - | Unassigned | - |
| Otu11489 | 2 | k__Fungi(100);p__Ascomycota(100);c__Dothideomycetes(100);o__Pleosporales(100);o__Pleosporales_unclassified(100);o__Pleosporales_unclassified(100);o__Pleosporales_unclassified(100); | - | - | - | - | - | - | - | Unassigned | - |
| Otu11491 | 2 | k__Fungi(100);k__Fungi_unclassified(100);k__Fungi_unclassified(100);k__Fungi_unclassified(100);k__Fungi_unclassified(100);k__Fungi_unclassified(100);k__Fungi_unclassified(100); | - | - | - | - | - | - | - | Unassigned | - |
| Otu11492 | 2 | k__Fungi(100);p__Basidiomycota(100);c__Tremellomycetes(100);c__Tremellomycetes_unclassified(100);c__Tremellomycetes_unclassified(100);c__Tremellomycetes_unclassified(100);c__Tremellomycetes_unclassified(100); | - | - | - | - | - | - | - | Unassigned | - |
| Otu11493 | 2 | k__Fungi(100);k__Fungi_unclassified(100);k__Fungi_unclassified(100);k__Fungi_unclassified(100);k__Fungi_unclassified(100);k__Fungi_unclassified(100);k__Fungi_unclassified(100); | - | - | - | - | - | - | - | Unassigned | - |
| Otu11494 | 2 | k__Fungi(100);k__Fungi_unclassified(100);k__Fungi_unclassified(100);k__Fungi_unclassified(100);k__Fungi_unclassified(100);k__Fungi_unclassified(100);k__Fungi_unclassified(100); | - | - | - | - | - | - | - | Unassigned | - |
| Otu11495 | 2 | k__Fungi(100);k__Fungi_unclassified(100);k__Fungi_unclassified(100);k__Fungi_unclassified(100);k__Fungi_unclassified(100);k__Fungi_unclassified(100);k__Fungi_unclassified(100); | - | - | - | - | - | - | - | Unassigned | - |
| Otu11497 | 2 | k__Fungi(100);k__Fungi_unclassified(100);k__Fungi_unclassified(100);k__Fungi_unclassified(100);k__Fungi_unclassified(100);k__Fungi_unclassified(100);k__Fungi_unclassified(100); | - | - | - | - | - | - | - | Unassigned | - |
| Otu11498 | 2 | k__Fungi(100);k__Fungi_unclassified(100);k__Fungi_unclassified(100);k__Fungi_unclassified(100);k__Fungi_unclassified(100);k__Fungi_unclassified(100);k__Fungi_unclassified(100); | - | - | - | - | - | - | - | Unassigned | - |
| Otu11499 | 2 | k__Fungi(100);k__Fungi_unclassified(100);k__Fungi_unclassified(100);k__Fungi_unclassified(100);k__Fungi_unclassified(100);k__Fungi_unclassified(100);k__Fungi_unclassified(100); | - | - | - | - | - | - | - | Unassigned | - |
| Otu11500 | 2 | k__Fungi(100);k__Fungi_unclassified(100);k__Fungi_unclassified(100);k__Fungi_unclassified(100);k__Fungi_unclassified(100);k__Fungi_unclassified(100);k__Fungi_unclassified(100); | - | - | - | - | - | - | - | Unassigned | - |
| Otu11501 | 2 | k__Plantae(100);p__unclassified_Plantae(100);c__unclassified_Plantae(100);o__unclassified_Plantae(100);f__unclassified_Plantae(100);g__unclassified_Plantae(100);s__Plantae_sp(100); | - | - | - | - | - | - | - | Unassigned | - |
| Otu11502 | 2 | k__Plantae(100);p__unclassified_Plantae(100);c__unclassified_Plantae(100);o__unclassified_Plantae(100);f__unclassified_Plantae(100);g__unclassified_Plantae(100);s__Plantae_sp(100); | - | - | - | - | - | - | - | Unassigned | - |
| Otu11504 | 2 | k__Fungi(100);p__Ascomycota(100);c__Sordariomycetes(100);c__Sordariomycetes_unclassified(100);c__Sordariomycetes_unclassified(100);c__Sordariomycetes_unclassified(100);c__Sordariomycetes_unclassified(100); | - | - | - | - | - | - | - | Unassigned | - |
| Otu11505 | 2 | k__Fungi(100);p__Ascomycota(100);c__Leotiomycetes(100);o__Helotiales(100);o__Helotiales_unclassified(100);o__Helotiales_unclassified(100);o__Helotiales_unclassified(100); | - | - | - | - | - | - | - | Unassigned | - |
| Otu11507 | 2 | k__Fungi(100);k__Fungi_unclassified(100);k__Fungi_unclassified(100);k__Fungi_unclassified(100);k__Fungi_unclassified(100);k__Fungi_unclassified(100);k__Fungi_unclassified(100); | - | - | - | - | - | - | - | Unassigned | - |
| Otu11508 | 2 | k__Fungi(100);k__Fungi_unclassified(100);k__Fungi_unclassified(100);k__Fungi_unclassified(100);k__Fungi_unclassified(100);k__Fungi_unclassified(100);k__Fungi_unclassified(100); | - | - | - | - | - | - | - | Unassigned | - |
| Otu11509 | 2 | k__Fungi(100);p__Ascomycota(100);p__Ascomycota_unclassified(100);p__Ascomycota_unclassified(100);p__Ascomycota_unclassified(100);p__Ascomycota_unclassified(100);p__Ascomycota_unclassified(100); | - | - | - | - | - | - | - | Unassigned | - |
| Otu11514 | 2 | k__Fungi(100);p__Ascomycota(100);p__Ascomycota_unclassified(100);p__Ascomycota_unclassified(100);p__Ascomycota_unclassified(100);p__Ascomycota_unclassified(100);p__Ascomycota_unclassified(100); | - | - | - | - | - | - | - | Unassigned | - |
| Otu11522 | 2 | k__Fungi(100);k__Fungi_unclassified(100);k__Fungi_unclassified(100);k__Fungi_unclassified(100);k__Fungi_unclassified(100);k__Fungi_unclassified(100);k__Fungi_unclassified(100); | - | - | - | - | - | - | - | Unassigned | - |
| Otu11523 | 2 | k__Fungi(100);k__Fungi_unclassified(100);k__Fungi_unclassified(100);k__Fungi_unclassified(100);k__Fungi_unclassified(100);k__Fungi_unclassified(100);k__Fungi_unclassified(100); | - | - | - | - | - | - | - | Unassigned | - |
| Otu11524 | 2 | k__Fungi(100);k__Fungi_unclassified(100);k__Fungi_unclassified(100);k__Fungi_unclassified(100);k__Fungi_unclassified(100);k__Fungi_unclassified(100);k__Fungi_unclassified(100); | - | - | - | - | - | - | - | Unassigned | - |
| Otu11526 | 2 | k__Fungi(100);k__Fungi_unclassified(100);k__Fungi_unclassified(100);k__Fungi_unclassified(100);k__Fungi_unclassified(100);k__Fungi_unclassified(100);k__Fungi_unclassified(100); | - | - | - | - | - | - | - | Unassigned | - |
| Otu11530 | 2 | k__Fungi(100);k__Fungi_unclassified(100);k__Fungi_unclassified(100);k__Fungi_unclassified(100);k__Fungi_unclassified(100);k__Fungi_unclassified(100);k__Fungi_unclassified(100); | - | - | - | - | - | - | - | Unassigned | - |
| Otu11532 | 2 | k__Fungi(100);k__Fungi_unclassified(100);k__Fungi_unclassified(100);k__Fungi_unclassified(100);k__Fungi_unclassified(100);k__Fungi_unclassified(100);k__Fungi_unclassified(100); | - | - | - | - | - | - | - | Unassigned | - |
| Otu11536 | 2 | k__Fungi(100);p__Ascomycota(100);c__Eurotiomycetes(100);c__Eurotiomycetes_unclassified(100);c__Eurotiomycetes_unclassified(100);c__Eurotiomycetes_unclassified(100);c__Eurotiomycetes_unclassified(100); | - | - | - | - | - | - | - | Unassigned | - |
| Otu11538 | 2 | k__Fungi(100);k__Fungi_unclassified(100);k__Fungi_unclassified(100);k__Fungi_unclassified(100);k__Fungi_unclassified(100);k__Fungi_unclassified(100);k__Fungi_unclassified(100); | - | - | - | - | - | - | - | Unassigned | - |
| Otu11541 | 2 | k__Fungi(100);k__Fungi_unclassified(100);k__Fungi_unclassified(100);k__Fungi_unclassified(100);k__Fungi_unclassified(100);k__Fungi_unclassified(100);k__Fungi_unclassified(100); | - | - | - | - | - | - | - | Unassigned | - |
| Otu11542 | 2 | k__Fungi(100);k__Fungi_unclassified(100);k__Fungi_unclassified(100);k__Fungi_unclassified(100);k__Fungi_unclassified(100);k__Fungi_unclassified(100);k__Fungi_unclassified(100); | - | - | - | - | - | - | - | Unassigned | - |
| Otu11543 | 2 | k__Fungi(100);k__Fungi_unclassified(100);k__Fungi_unclassified(100);k__Fungi_unclassified(100);k__Fungi_unclassified(100);k__Fungi_unclassified(100);k__Fungi_unclassified(100); | - | - | - | - | - | - | - | Unassigned | - |
| Otu11544 | 2 | k__Fungi(100);k__Fungi_unclassified(100);k__Fungi_unclassified(100);k__Fungi_unclassified(100);k__Fungi_unclassified(100);k__Fungi_unclassified(100);k__Fungi_unclassified(100); | - | - | - | - | - | - | - | Unassigned | - |
| Otu11547 | 2 | k__Fungi(100);p__Ascomycota(100);c__Dothideomycetes(100);o__Pleosporales(100);o__Pleosporales_unclassified(100);o__Pleosporales_unclassified(100);o__Pleosporales_unclassified(100); | - | - | - | - | - | - | - | Unassigned | - |
| Otu11549 | 2 | k__Fungi(100);p__Ascomycota(100);p__Ascomycota_unclassified(100);p__Ascomycota_unclassified(100);p__Ascomycota_unclassified(100);p__Ascomycota_unclassified(100);p__Ascomycota_unclassified(100); | - | - | - | - | - | - | - | Unassigned | - |
| Otu11553 | 2 | k__Fungi(100);p__Ascomycota(100);p__Ascomycota_unclassified(100);p__Ascomycota_unclassified(100);p__Ascomycota_unclassified(100);p__Ascomycota_unclassified(100);p__Ascomycota_unclassified(100); | - | - | - | - | - | - | - | Unassigned | - |
| Otu11555 | 2 | k__Fungi(100);p__Ascomycota(100);c__Dothideomycetes(100);o__Pleosporales(100);o__Pleosporales_unclassified(100);o__Pleosporales_unclassified(100);o__Pleosporales_unclassified(100); | - | - | - | - | - | - | - | Unassigned | - |
| Otu11560 | 2 | k__Fungi(100);k__Fungi_unclassified(100);k__Fungi_unclassified(100);k__Fungi_unclassified(100);k__Fungi_unclassified(100);k__Fungi_unclassified(100);k__Fungi_unclassified(100); | - | - | - | - | - | - | - | Unassigned | - |
| Otu11561 | 2 | k__Fungi(100);k__Fungi_unclassified(100);k__Fungi_unclassified(100);k__Fungi_unclassified(100);k__Fungi_unclassified(100);k__Fungi_unclassified(100);k__Fungi_unclassified(100); | - | - | - | - | - | - | - | Unassigned | - |
| Otu11562 | 2 | k__Fungi(100);p__Ascomycota(100);c__Sordariomycetes(100);o__Sordariales(100);o__Sordariales_unclassified(100);o__Sordariales_unclassified(100);o__Sordariales_unclassified(100); | - | - | - | - | - | - | - | Unassigned | - |
| Otu11568 | 2 | k__Fungi(100);k__Fungi_unclassified(100);k__Fungi_unclassified(100);k__Fungi_unclassified(100);k__Fungi_unclassified(100);k__Fungi_unclassified(100);k__Fungi_unclassified(100); | - | - | - | - | - | - | - | Unassigned | - |
| Otu11569 | 2 | k__Fungi(100);k__Fungi_unclassified(100);k__Fungi_unclassified(100);k__Fungi_unclassified(100);k__Fungi_unclassified(100);k__Fungi_unclassified(100);k__Fungi_unclassified(100); | - | - | - | - | - | - | - | Unassigned | - |
| Otu11581 | 2 | k__Fungi(100);p__Ascomycota(100);p__Ascomycota_unclassified(100);p__Ascomycota_unclassified(100);p__Ascomycota_unclassified(100);p__Ascomycota_unclassified(100);p__Ascomycota_unclassified(100); | - | - | - | - | - | - | - | Unassigned | - |
| Otu11584 | 2 | k__Fungi(100);k__Fungi_unclassified(100);k__Fungi_unclassified(100);k__Fungi_unclassified(100);k__Fungi_unclassified(100);k__Fungi_unclassified(100);k__Fungi_unclassified(100); | - | - | - | - | - | - | - | Unassigned | - |
| Otu11594 | 2 | k__Fungi(100);p__Ascomycota(100);p__Ascomycota_unclassified(100);p__Ascomycota_unclassified(100);p__Ascomycota_unclassified(100);p__Ascomycota_unclassified(100);p__Ascomycota_unclassified(100); | - | - | - | - | - | - | - | Unassigned | - |
| Otu11599 | 2 | k__Fungi(100);k__Fungi_unclassified(100);k__Fungi_unclassified(100);k__Fungi_unclassified(100);k__Fungi_unclassified(100);k__Fungi_unclassified(100);k__Fungi_unclassified(100); | - | - | - | - | - | - | - | Unassigned | - |
| Otu11602 | 2 | k__Fungi(100);k__Fungi_unclassified(100);k__Fungi_unclassified(100);k__Fungi_unclassified(100);k__Fungi_unclassified(100);k__Fungi_unclassified(100);k__Fungi_unclassified(100); | - | - | - | - | - | - | - | Unassigned | - |
| Otu11604 | 2 | k__Fungi(100);k__Fungi_unclassified(100);k__Fungi_unclassified(100);k__Fungi_unclassified(100);k__Fungi_unclassified(100);k__Fungi_unclassified(100);k__Fungi_unclassified(100); | - | - | - | - | - | - | - | Unassigned | - |
| Otu11610 | 2 | k__Fungi(100);k__Fungi_unclassified(100);k__Fungi_unclassified(100);k__Fungi_unclassified(100);k__Fungi_unclassified(100);k__Fungi_unclassified(100);k__Fungi_unclassified(100); | - | - | - | - | - | - | - | Unassigned | - |
| Otu11613 | 2 | k__Fungi(100);k__Fungi_unclassified(100);k__Fungi_unclassified(100);k__Fungi_unclassified(100);k__Fungi_unclassified(100);k__Fungi_unclassified(100);k__Fungi_unclassified(100); | - | - | - | - | - | - | - | Unassigned | - |
| Otu11614 | 2 | k__Fungi(100);p__Ascomycota(100);p__Ascomycota_unclassified(100);p__Ascomycota_unclassified(100);p__Ascomycota_unclassified(100);p__Ascomycota_unclassified(100);p__Ascomycota_unclassified(100); | - | - | - | - | - | - | - | Unassigned | - |
| Otu11617 | 2 | k__Fungi(100);k__Fungi_unclassified(100);k__Fungi_unclassified(100);k__Fungi_unclassified(100);k__Fungi_unclassified(100);k__Fungi_unclassified(100);k__Fungi_unclassified(100); | - | - | - | - | - | - | - | Unassigned | - |
| Otu11618 | 2 | k__Fungi(100);p__Ascomycota(100);p__Ascomycota_unclassified(100);p__Ascomycota_unclassified(100);p__Ascomycota_unclassified(100);p__Ascomycota_unclassified(100);p__Ascomycota_unclassified(100); | - | - | - | - | - | - | - | Unassigned | - |
| Otu11624 | 2 | k__Fungi(100);k__Fungi_unclassified(100);k__Fungi_unclassified(100);k__Fungi_unclassified(100);k__Fungi_unclassified(100);k__Fungi_unclassified(100);k__Fungi_unclassified(100); | - | - | - | - | - | - | - | Unassigned | - |
| Otu11626 | 2 | k__Fungi(100);p__Ascomycota(100);p__Ascomycota_unclassified(100);p__Ascomycota_unclassified(100);p__Ascomycota_unclassified(100);p__Ascomycota_unclassified(100);p__Ascomycota_unclassified(100); | - | - | - | - | - | - | - | Unassigned | - |
| Otu11628 | 2 | k__Fungi(100);k__Fungi_unclassified(100);k__Fungi_unclassified(100);k__Fungi_unclassified(100);k__Fungi_unclassified(100);k__Fungi_unclassified(100);k__Fungi_unclassified(100); | - | - | - | - | - | - | - | Unassigned | - |
| Otu11629 | 2 | k__Fungi(100);p__Ascomycota(100);p__Ascomycota_unclassified(100);p__Ascomycota_unclassified(100);p__Ascomycota_unclassified(100);p__Ascomycota_unclassified(100);p__Ascomycota_unclassified(100); | - | - | - | - | - | - | - | Unassigned | - |
| Otu11631 | 2 | k__Fungi(100);k__Fungi_unclassified(100);k__Fungi_unclassified(100);k__Fungi_unclassified(100);k__Fungi_unclassified(100);k__Fungi_unclassified(100);k__Fungi_unclassified(100); | - | - | - | - | - | - | - | Unassigned | - |
| Otu11632 | 2 | k__Fungi(100);k__Fungi_unclassified(100);k__Fungi_unclassified(100);k__Fungi_unclassified(100);k__Fungi_unclassified(100);k__Fungi_unclassified(100);k__Fungi_unclassified(100); | - | - | - | - | - | - | - | Unassigned | - |
| Otu11638 | 2 | k__Fungi(100);p__Ascomycota(100);p__Ascomycota_unclassified(100);p__Ascomycota_unclassified(100);p__Ascomycota_unclassified(100);p__Ascomycota_unclassified(100);p__Ascomycota_unclassified(100); | - | - | - | - | - | - | - | Unassigned | - |
| Otu11650 | 2 | k__Fungi(100);p__Basidiomycota(100);c__Agaricomycetes(100);c__Agaricomycetes_unclassified(100);c__Agaricomycetes_unclassified(100);c__Agaricomycetes_unclassified(100);c__Agaricomycetes_unclassified(100); | - | - | - | - | - | - | - | Unassigned | - |
| Otu11654 | 2 | k__Fungi(100);p__Basidiomycota(100);p__Basidiomycota_unclassified(100);p__Basidiomycota_unclassified(100);p__Basidiomycota_unclassified(100);p__Basidiomycota_unclassified(100);p__Basidiomycota_unclassified(100); | - | - | - | - | - | - | - | Unassigned | - |
| Otu11655 | 2 | k__Fungi(100);p__Basidiomycota(100);c__Microbotryomycetes(100);c__Microbotryomycetes_unclassified(100);c__Microbotryomycetes_unclassified(100);c__Microbotryomycetes_unclassified(100);c__Microbotryomycetes_unclassified(100); | - | - | - | - | - | - | - | Unassigned | - |
| Otu11656 | 2 | k__Fungi(100);p__Basidiomycota(100);p__Basidiomycota_unclassified(100);p__Basidiomycota_unclassified(100);p__Basidiomycota_unclassified(100);p__Basidiomycota_unclassified(100);p__Basidiomycota_unclassified(100); | - | - | - | - | - | - | - | Unassigned | - |
| Otu11658 | 2 | k__Fungi(100);p__Basidiomycota(100);c__Agaricomycetes(100);c__Agaricomycetes_unclassified(100);c__Agaricomycetes_unclassified(100);c__Agaricomycetes_unclassified(100);c__Agaricomycetes_unclassified(100); | - | - | - | - | - | - | - | Unassigned | - |
| Otu11661 | 2 | k__Fungi(100);k__Fungi_unclassified(100);k__Fungi_unclassified(100);k__Fungi_unclassified(100);k__Fungi_unclassified(100);k__Fungi_unclassified(100);k__Fungi_unclassified(100); | - | - | - | - | - | - | - | Unassigned | - |
| Otu11662 | 2 | k__Fungi(100);k__Fungi_unclassified(100);k__Fungi_unclassified(100);k__Fungi_unclassified(100);k__Fungi_unclassified(100);k__Fungi_unclassified(100);k__Fungi_unclassified(100); | - | - | - | - | - | - | - | Unassigned | - |
| Otu11663 | 2 | k__Fungi(100);p__Ascomycota(100);p__Ascomycota_unclassified(100);p__Ascomycota_unclassified(100);p__Ascomycota_unclassified(100);p__Ascomycota_unclassified(100);p__Ascomycota_unclassified(100); | - | - | - | - | - | - | - | Unassigned | - |
| Otu11664 | 2 | k__Fungi(100);k__Fungi_unclassified(100);k__Fungi_unclassified(100);k__Fungi_unclassified(100);k__Fungi_unclassified(100);k__Fungi_unclassified(100);k__Fungi_unclassified(100); | - | - | - | - | - | - | - | Unassigned | - |
| Otu11665 | 2 | k__Fungi(100);k__Fungi_unclassified(100);k__Fungi_unclassified(100);k__Fungi_unclassified(100);k__Fungi_unclassified(100);k__Fungi_unclassified(100);k__Fungi_unclassified(100); | - | - | - | - | - | - | - | Unassigned | - |
| Otu11666 | 2 | k__Fungi(100);p__Ascomycota(100);p__Ascomycota_unclassified(100);p__Ascomycota_unclassified(100);p__Ascomycota_unclassified(100);p__Ascomycota_unclassified(100);p__Ascomycota_unclassified(100); | - | - | - | - | - | - | - | Unassigned | - |
| Otu11667 | 2 | k__Fungi(100);k__Fungi_unclassified(100);k__Fungi_unclassified(100);k__Fungi_unclassified(100);k__Fungi_unclassified(100);k__Fungi_unclassified(100);k__Fungi_unclassified(100); | - | - | - | - | - | - | - | Unassigned | - |
| Otu11668 | 2 | k__Fungi(100);p__Ascomycota(100);p__Ascomycota_unclassified(100);p__Ascomycota_unclassified(100);p__Ascomycota_unclassified(100);p__Ascomycota_unclassified(100);p__Ascomycota_unclassified(100); | - | - | - | - | - | - | - | Unassigned | - |
| Otu11669 | 2 | k__Fungi(100);p__Ascomycota(100);p__Ascomycota_unclassified(100);p__Ascomycota_unclassified(100);p__Ascomycota_unclassified(100);p__Ascomycota_unclassified(100);p__Ascomycota_unclassified(100); | - | - | - | - | - | - | - | Unassigned | - |
| Otu11673 | 2 | k__Fungi(100);p__Basidiomycota(100);c__Agaricomycetes(100);c__Agaricomycetes_unclassified(100);c__Agaricomycetes_unclassified(100);c__Agaricomycetes_unclassified(100);c__Agaricomycetes_unclassified(100); | - | - | - | - | - | - | - | Unassigned | - |
| Otu11675 | 2 | k__Fungi(100);p__Basidiomycota(100);c__Agaricomycetes(100);o__Polyporales(100);o__Polyporales_unclassified(100);o__Polyporales_unclassified(100);o__Polyporales_unclassified(100); | - | - | - | - | - | - | - | Unassigned | - |
| Otu11676 | 2 | k__Fungi(100);p__Basidiomycota(100);c__Agaricomycetes(100);o__Boletales(100);o__Boletales_unclassified(100);o__Boletales_unclassified(100);o__Boletales_unclassified(100); | - | - | - | - | - | - | - | Unassigned | - |
| Otu11678 | 2 | k__Fungi(100);p__Basidiomycota(100);c__Agaricomycetes(100);o__Polyporales(100);o__Polyporales_unclassified(100);o__Polyporales_unclassified(100);o__Polyporales_unclassified(100); | - | - | - | - | - | - | - | Unassigned | - |
| Otu11682 | 2 | k__Fungi(100);p__Basidiomycota(100);c__Agaricomycetes(100);o__Polyporales(100);o__Polyporales_unclassified(100);o__Polyporales_unclassified(100);o__Polyporales_unclassified(100); | - | - | - | - | - | - | - | Unassigned | - |
| Otu11683 | 2 | k__Fungi(100);p__Basidiomycota(100);c__Agaricomycetes(100);o__Polyporales(100);o__Polyporales_unclassified(100);o__Polyporales_unclassified(100);o__Polyporales_unclassified(100); | - | - | - | - | - | - | - | Unassigned | - |
| Otu11684 | 2 | k__Fungi(100);k__Fungi_unclassified(100);k__Fungi_unclassified(100);k__Fungi_unclassified(100);k__Fungi_unclassified(100);k__Fungi_unclassified(100);k__Fungi_unclassified(100); | - | - | - | - | - | - | - | Unassigned | - |
| Otu11697 | 2 | k__Fungi(100);p__Ascomycota(100);c__Dothideomycetes(100);o__Pleosporales(100);o__Pleosporales_unclassified(100);o__Pleosporales_unclassified(100);o__Pleosporales_unclassified(100); | - | - | - | - | - | - | - | Unassigned | - |
| Otu11701 | 2 | k__Fungi(100);p__Basidiomycota(100);p__Basidiomycota_unclassified(100);p__Basidiomycota_unclassified(100);p__Basidiomycota_unclassified(100);p__Basidiomycota_unclassified(100);p__Basidiomycota_unclassified(100); | - | - | - | - | - | - | - | Unassigned | - |
| Otu11703 | 2 | k__Plantae(100);p__unclassified_Plantae(100);c__unclassified_Plantae(100);o__unclassified_Plantae(100);f__unclassified_Plantae(100);g__unclassified_Plantae(100);s__Plantae_sp(100); | - | - | - | - | - | - | - | Unassigned | - |
| Otu11704 | 2 | k__Plantae(100);p__unclassified_Plantae(100);c__unclassified_Plantae(100);o__unclassified_Plantae(100);f__unclassified_Plantae(100);g__unclassified_Plantae(100);s__Plantae_sp(100); | - | - | - | - | - | - | - | Unassigned | - |
| Otu11705 | 2 | k__Fungi(100);p__Ascomycota(100);c__Dothideomycetes(100);o__Capnodiales(100);o__Capnodiales_unclassified(100);o__Capnodiales_unclassified(100);o__Capnodiales_unclassified(100); | - | - | - | - | - | - | - | Unassigned | - |
| Otu11706 | 2 | k__Fungi(100);p__Ascomycota(100);c__Dothideomycetes(100);o__Capnodiales(100);o__Capnodiales_unclassified(100);o__Capnodiales_unclassified(100);o__Capnodiales_unclassified(100); | - | - | - | - | - | - | - | Unassigned | - |
| Otu11707 | 2 | k__Fungi(100);p__Ascomycota(100);p__Ascomycota_unclassified(100);p__Ascomycota_unclassified(100);p__Ascomycota_unclassified(100);p__Ascomycota_unclassified(100);p__Ascomycota_unclassified(100); | - | - | - | - | - | - | - | Unassigned | - |
| Otu11708 | 2 | k__Fungi(100);p__Ascomycota(100);p__Ascomycota_unclassified(100);p__Ascomycota_unclassified(100);p__Ascomycota_unclassified(100);p__Ascomycota_unclassified(100);p__Ascomycota_unclassified(100); | - | - | - | - | - | - | - | Unassigned | - |
| Otu11712 | 2 | k__Fungi(100);p__Ascomycota(100);p__Ascomycota_unclassified(100);p__Ascomycota_unclassified(100);p__Ascomycota_unclassified(100);p__Ascomycota_unclassified(100);p__Ascomycota_unclassified(100); | - | - | - | - | - | - | - | Unassigned | - |
| Otu11713 | 2 | k__Fungi(100);p__Ascomycota(100);c__Sordariomycetes(100);c__Sordariomycetes_unclassified(100);c__Sordariomycetes_unclassified(100);c__Sordariomycetes_unclassified(100);c__Sordariomycetes_unclassified(100); | - | - | - | - | - | - | - | Unassigned | - |
| Otu11715 | 2 | k__Fungi(100);p__Ascomycota(100);c__Dothideomycetes(100);o__Pleosporales(100);o__Pleosporales_unclassified(100);o__Pleosporales_unclassified(100);o__Pleosporales_unclassified(100); | - | - | - | - | - | - | - | Unassigned | - |
| Otu11716 | 2 | k__Fungi(100);p__Ascomycota(100);c__Dothideomycetes(100);o__Pleosporales(100);f__unclassified_Pleosporales(100);g__unclassified_Pleosporales(100);s__Pleosporales_sp(100); | - | - | - | - | - | - | - | Unassigned | - |
| Otu11717 | 2 | k__Fungi(100);p__Ascomycota(100);c__Dothideomycetes(100);o__Capnodiales(100);f__Davidiellaceae(100);f__Davidiellaceae_unclassified(100);f__Davidiellaceae_unclassified(100); | - | - | - | - | - | - | - | Unassigned | - |
| Otu11718 | 2 | k__Fungi(100);p__Ascomycota(100);p__Ascomycota_unclassified(100);p__Ascomycota_unclassified(100);p__Ascomycota_unclassified(100);p__Ascomycota_unclassified(100);p__Ascomycota_unclassified(100); | - | - | - | - | - | - | - | Unassigned | - |
| Otu11720 | 2 | k__Fungi(100);p__Ascomycota(100);c__Dothideomycetes(100);o__Pleosporales(100);o__Pleosporales_unclassified(100);o__Pleosporales_unclassified(100);o__Pleosporales_unclassified(100); | - | - | - | - | - | - | - | Unassigned | - |
| Otu11721 | 2 | k__Fungi(100);p__Ascomycota(100);p__Ascomycota_unclassified(100);p__Ascomycota_unclassified(100);p__Ascomycota_unclassified(100);p__Ascomycota_unclassified(100);p__Ascomycota_unclassified(100); | - | - | - | - | - | - | - | Unassigned | - |
| Otu11725 | 2 | k__Fungi(100);k__Fungi_unclassified(100);k__Fungi_unclassified(100);k__Fungi_unclassified(100);k__Fungi_unclassified(100);k__Fungi_unclassified(100);k__Fungi_unclassified(100); | - | - | - | - | - | - | - | Unassigned | - |
| Otu11726 | 2 | k__Fungi(100);k__Fungi_unclassified(100);k__Fungi_unclassified(100);k__Fungi_unclassified(100);k__Fungi_unclassified(100);k__Fungi_unclassified(100);k__Fungi_unclassified(100); | - | - | - | - | - | - | - | Unassigned | - |
| Otu11727 | 2 | k__Fungi(100);k__Fungi_unclassified(100);k__Fungi_unclassified(100);k__Fungi_unclassified(100);k__Fungi_unclassified(100);k__Fungi_unclassified(100);k__Fungi_unclassified(100); | - | - | - | - | - | - | - | Unassigned | - |
| Otu11729 | 2 | k__Fungi(100);p__Basidiomycota(100);p__Basidiomycota_unclassified(100);p__Basidiomycota_unclassified(100);p__Basidiomycota_unclassified(100);p__Basidiomycota_unclassified(100);p__Basidiomycota_unclassified(100); | - | - | - | - | - | - | - | Unassigned | - |
| Otu11731 | 2 | k__Fungi(100);p__Basidiomycota(100);c__Agaricomycetes(100);c__Agaricomycetes_unclassified(100);c__Agaricomycetes_unclassified(100);c__Agaricomycetes_unclassified(100);c__Agaricomycetes_unclassified(100); | - | - | - | - | - | - | - | Unassigned | - |
| Otu11734 | 2 | k__Fungi(100);p__Basidiomycota(100);p__Basidiomycota_unclassified(100);p__Basidiomycota_unclassified(100);p__Basidiomycota_unclassified(100);p__Basidiomycota_unclassified(100);p__Basidiomycota_unclassified(100); | - | - | - | - | - | - | - | Unassigned | - |

| | | | | | | | | | | Unassigned | |
|---|---|---|---|---|---|---|---|---|---|---|---|
| Otu11747 | 2 | k__Fungi(100);p__Basidiomycota(100);c__Agaricomycetes(100);c__Agaricomycetes_unclassified(100);c__Agaricomycetes_unclassified(100);c__Agaricomycetes_unclassified(100);c__Agaricomycetes_unclassified(100); | - | - | - | - | - | - | - | Unassigned | - |
| Otu11755 | 2 | k__Fungi(100);p__Ascomycota(100);c__Sordariomycetes(100);c__Sordariomycetes_unclassified(100);c__Sordariomycetes_unclassified(100);c__Sordariomycetes_unclassified(100);c__Sordariomycetes_unclassified(100); | - | - | - | - | - | - | - | Unassigned | - |
| Otu11759 | 2 | k__Fungi(100);p__Ascomycota(100);c__Sordariomycetes(100);o__Microascales(100);o__Microascales_unclassified(100);o__Microascales_unclassified(100);o__Microascales_unclassified(100); | - | - | - | - | - | - | - | Unassigned | - |
| Otu11766 | 2 | k__Fungi(100);p__Ascomycota(100);p__Ascomycota_unclassified(100);p__Ascomycota_unclassified(100);p__Ascomycota_unclassified(100);p__Ascomycota_unclassified(100);p__Ascomycota_unclassified(100); | - | - | - | - | - | - | - | Unassigned | - |
| Otu11770 | 2 | k__Fungi(100);p__Ascomycota(100);c__Eurotiomycetes(100);c__Eurotiomycetes_unclassified(100);c__Eurotiomycetes_unclassified(100);c__Eurotiomycetes_unclassified(100);c__Eurotiomycetes_unclassified(100); | - | - | - | - | - | - | - | Unassigned | - |
| Otu11771 | 2 | k__Fungi(100);p__Ascomycota(100);c__Eurotiomycetes(100);c__Eurotiomycetes_unclassified(100);c__Eurotiomycetes_unclassified(100);c__Eurotiomycetes_unclassified(100);c__Eurotiomycetes_unclassified(100); | - | - | - | - | - | - | - | Unassigned | - |
| Otu11777 | 2 | k__Fungi(100);p__Ascomycota(100);c__Eurotiomycetes(100);o__Eurotiales(100);f__Trichocomaceae(100);f__Trichocomaceae_unclassified(100);f__Trichocomaceae_unclassified(100); | - | - | - | - | - | - | - | Unassigned | - |
| Otu11779 | 2 | k__Fungi(100);p__Ascomycota(100);c__Sordariomycetes(100);c__Sordariomycetes_unclassified(100);c__Sordariomycetes_unclassified(100);c__Sordariomycetes_unclassified(100);c__Sordariomycetes_unclassified(100); | - | - | - | - | - | - | - | Unassigned | - |
| Otu11780 | 2 | k__Fungi(100);p__Ascomycota(100);c__Sordariomycetes(100);c__Sordariomycetes_unclassified(100);c__Sordariomycetes_unclassified(100);c__Sordariomycetes_unclassified(100);c__Sordariomycetes_unclassified(100); | - | - | - | - | - | - | - | Unassigned | - |
| Otu11781 | 2 | k__Fungi(100);p__Ascomycota(100);c__Sordariomycetes(100);c__Sordariomycetes_unclassified(100);c__Sordariomycetes_unclassified(100);c__Sordariomycetes_unclassified(100);c__Sordariomycetes_unclassified(100); | - | - | - | - | - | - | - | Unassigned | - |
| Otu11782 | 2 | k__Fungi(100);p__Ascomycota(100);c__Sordariomycetes(100);c__Sordariomycetes_unclassified(100);c__Sordariomycetes_unclassified(100);c__Sordariomycetes_unclassified(100);c__Sordariomycetes_unclassified(100); | - | - | - | - | - | - | - | Unassigned | - |
| Otu11784 | 2 | k__Fungi(100);p__Ascomycota(100);c__Sordariomycetes(100);o__Xylariales(100);o__Xylariales_unclassified(100);o__Xylariales_unclassified(100);o__Xylariales_unclassified(100); | - | - | - | - | - | - | - | Unassigned | - |
| Otu11787 | 2 | k__Fungi(100);p__Ascomycota(100);p__Ascomycota_unclassified(100);p__Ascomycota_unclassified(100);p__Ascomycota_unclassified(100);p__Ascomycota_unclassified(100);p__Ascomycota_unclassified(100); | - | - | - | - | - | - | - | Unassigned | - |
| Otu11789 | 2 | k__Fungi(100);p__Ascomycota(100);p__Ascomycota_unclassified(100);p__Ascomycota_unclassified(100);p__Ascomycota_unclassified(100);p__Ascomycota_unclassified(100);p__Ascomycota_unclassified(100); | - | - | - | - | - | - | - | Unassigned | - |
| Otu11790 | 2 | k__Fungi(100);p__Ascomycota(100);p__Ascomycota_unclassified(100);p__Ascomycota_unclassified(100);p__Ascomycota_unclassified(100);p__Ascomycota_unclassified(100);p__Ascomycota_unclassified(100); | - | - | - | - | - | - | - | Unassigned | - |
| Otu11794 | 2 | k__Fungi(100);p__Ascomycota(100);p__Ascomycota_unclassified(100);p__Ascomycota_unclassified(100);p__Ascomycota_unclassified(100);p__Ascomycota_unclassified(100);p__Ascomycota_unclassified(100); | - | - | - | - | - | - | - | Unassigned | - |
| Otu11796 | 2 | k__Fungi(100);p__Ascomycota(100);p__Ascomycota_unclassified(100);p__Ascomycota_unclassified(100);p__Ascomycota_unclassified(100);p__Ascomycota_unclassified(100);p__Ascomycota_unclassified(100); | - | - | - | - | - | - | - | Unassigned | - |
| Otu11798 | 2 | k__Fungi(100);p__Ascomycota(100);c__Dothideomycetes(100);o__Pleosporales(100);o__Pleosporales_unclassified(100);o__Pleosporales_unclassified(100);o__Pleosporales_unclassified(100); | - | - | - | - | - | - | - | Unassigned | - |
| Otu11799 | 2 | k__Fungi(100);p__Ascomycota(100);c__Dothideomycetes(100);o__Pleosporales(100);o__Pleosporales_unclassified(100);o__Pleosporales_unclassified(100);o__Pleosporales_unclassified(100); | - | - | - | - | - | - | - | Unassigned | - |
| Otu11800 | 2 | k__Fungi(100);p__Ascomycota(100);c__Dothideomycetes(100);o__Pleosporales(100);o__Pleosporales_unclassified(100);o__Pleosporales_unclassified(100);o__Pleosporales_unclassified(100); | - | - | - | - | - | - | - | Unassigned | - |
| Otu11802 | 2 | k__Fungi(100);p__Ascomycota(100);p__Ascomycota_unclassified(100);p__Ascomycota_unclassified(100);p__Ascomycota_unclassified(100);p__Ascomycota_unclassified(100);p__Ascomycota_unclassified(100); | - | - | - | - | - | - | - | Unassigned | - |
| Otu11806 | 2 | k__Fungi(100);p__Basidiomycota(100);c__Agaricomycetes(100);c__Agaricomycetes_unclassified(100);c__Agaricomycetes_unclassified(100);c__Agaricomycetes_unclassified(100);c__Agaricomycetes_unclassified(100); | - | - | - | - | - | - | - | Unassigned | - |
| Otu11808 | 2 | k__Fungi(100);p__Basidiomycota(100);p__Basidiomycota_unclassified(100);p__Basidiomycota_unclassified(100);p__Basidiomycota_unclassified(100);p__Basidiomycota_unclassified(100);p__Basidiomycota_unclassified(100); | - | - | - | - | - | - | - | Unassigned | - |
| Otu11810 | 2 | k__Fungi(100);p__Basidiomycota(100);c__Agaricostilbomycetes(100);o__Agaricostilbales(100);o__Agaricostilbales_unclassified(100);o__Agaricostilbales_unclassified(100);o__Agaricostilbales_unclassified(100); | - | - | - | - | - | - | - | Unassigned | - |
| Otu11815 | 2 | k__Fungi(100);p__Basidiomycota(100);c__Agaricomycetes(100);c__Agaricomycetes_unclassified(100);c__Agaricomycetes_unclassified(100);c__Agaricomycetes_unclassified(100);c__Agaricomycetes_unclassified(100); | - | - | - | - | - | - | - | Unassigned | - |
| Otu11817 | 2 | k__Fungi(100);k__Fungi_unclassified(100);k__Fungi_unclassified(100);k__Fungi_unclassified(100);k__Fungi_unclassified(100);k__Fungi_unclassified(100);k__Fungi_unclassified(100); | - | - | - | - | - | - | - | Unassigned | - |
| Otu11824 | 2 | k__Fungi(100);p__Basidiomycota(100);c__Agaricomycetes(100);c__Agaricomycetes_unclassified(100);c__Agaricomycetes_unclassified(100);c__Agaricomycetes_unclassified(100);c__Agaricomycetes_unclassified(100); | - | - | - | - | - | - | - | Unassigned | - |
| Otu11825 | 2 | k__Fungi(100);p__Basidiomycota(100);c__Agaricomycetes(100);o__Cantharellales(100);o__Cantharellales_unclassified(100);o__Cantharellales_unclassified(100);o__Cantharellales_unclassified(100); | - | - | - | - | - | - | - | Unassigned | - |
| Otu11829 | 2 | k__Fungi(100);p__Basidiomycota(100);p__Basidiomycota_unclassified(100);p__Basidiomycota_unclassified(100);p__Basidiomycota_unclassified(100);p__Basidiomycota_unclassified(100);p__Basidiomycota_unclassified(100); | - | - | - | - | - | - | - | Unassigned | - |
| Otu11833 | 2 | k__Fungi(100);p__Ascomycota(100);p__Ascomycota_unclassified(100);p__Ascomycota_unclassified(100);p__Ascomycota_unclassified(100);p__Ascomycota_unclassified(100);p__Ascomycota_unclassified(100); | - | - | - | - | - | - | - | Unassigned | - |
| Otu11836 | 2 | k__Fungi(100);p__Ascomycota(100);p__Ascomycota_unclassified(100);p__Ascomycota_unclassified(100);p__Ascomycota_unclassified(100);p__Ascomycota_unclassified(100);p__Ascomycota_unclassified(100); | - | - | - | - | - | - | - | Unassigned | - |
| Otu11837 | 2 | k__Fungi(100);p__Ascomycota(100);p__Ascomycota_unclassified(100);p__Ascomycota_unclassified(100);p__Ascomycota_unclassified(100);p__Ascomycota_unclassified(100);p__Ascomycota_unclassified(100); | - | - | - | - | - | - | - | Unassigned | - |
| Otu11838 | 2 | k__Fungi(100);k__Fungi_unclassified(100);k__Fungi_unclassified(100);k__Fungi_unclassified(100);k__Fungi_unclassified(100);k__Fungi_unclassified(100);k__Fungi_unclassified(100); | - | - | - | - | - | - | - | Unassigned | - |
| Otu11840 | 2 | k__Fungi(100);k__Fungi_unclassified(100);k__Fungi_unclassified(100);k__Fungi_unclassified(100);k__Fungi_unclassified(100);k__Fungi_unclassified(100);k__Fungi_unclassified(100); | - | - | - | - | - | - | - | Unassigned | - |
| Otu11841 | 2 | k__Fungi(100);p__Ascomycota(100);p__Ascomycota_unclassified(100);p__Ascomycota_unclassified(100);p__Ascomycota_unclassified(100);p__Ascomycota_unclassified(100);p__Ascomycota_unclassified(100); | - | - | - | - | - | - | - | Unassigned | - |
| Otu11843 | 2 | k__Fungi(100);p__Basidiomycota(100);c__Agaricomycetes(100);o__Sebacinales(100);o__Sebacinales_unclassified(100);o__Sebacinales_unclassified(100);o__Sebacinales_unclassified(100); | - | - | - | - | - | - | - | Unassigned | - |
| Otu11849 | 2 | k__Fungi(100);p__Basidiomycota(100);c__Agaricomycetes(100);c__Agaricomycetes_unclassified(100);c__Agaricomycetes_unclassified(100);c__Agaricomycetes_unclassified(100);c__Agaricomycetes_unclassified(100); | - | - | - | - | - | - | - | Unassigned | - |
| Otu11852 | 2 | k__Fungi(100);p__Basidiomycota(100);c__Microbotryomycetes(100);o__Sporidiobolales(100);f__Sporidiobolales_family_Incertae_sedis(100);f__Sporidiobolales_family_Incertae_sedis_unclassified(100);f__Sporidiobolales_family_Incertae_sedis_unclassified(100); | - | - | - | - | - | - | - | Unassigned | - |
| Otu11854 | 2 | k__Fungi(100);p__Basidiomycota(100);p__Basidiomycota_unclassified(100);p__Basidiomycota_unclassified(100);p__Basidiomycota_unclassified(100);p__Basidiomycota_unclassified(100);p__Basidiomycota_unclassified(100); | - | - | - | - | - | - | - | Unassigned | - |
| Otu11858 | 2 | k__Fungi(100);k__Fungi_unclassified(100);k__Fungi_unclassified(100);k__Fungi_unclassified(100);k__Fungi_unclassified(100);k__Fungi_unclassified(100);k__Fungi_unclassified(100); | - | - | - | - | - | - | - | Unassigned | - |
| Otu11859 | 2 | k__Fungi(100);p__Ascomycota(100);p__Ascomycota_unclassified(100);p__Ascomycota_unclassified(100);p__Ascomycota_unclassified(100);p__Ascomycota_unclassified(100);p__Ascomycota_unclassified(100); | - | - | - | - | - | - | - | Unassigned | - |
| Otu11862 | 2 | k__Fungi(100);k__Fungi_unclassified(100);k__Fungi_unclassified(100);k__Fungi_unclassified(100);k__Fungi_unclassified(100);k__Fungi_unclassified(100);k__Fungi_unclassified(100); | - | - | - | - | - | - | - | Unassigned | - |
| Otu11863 | 2 | k__Fungi(100);k__Fungi_unclassified(100);k__Fungi_unclassified(100);k__Fungi_unclassified(100);k__Fungi_unclassified(100);k__Fungi_unclassified(100);k__Fungi_unclassified(100); | - | - | - | - | - | - | - | Unassigned | - |
| Otu11868 | 2 | k__Fungi(100);k__Fungi_unclassified(100);k__Fungi_unclassified(100);k__Fungi_unclassified(100);k__Fungi_unclassified(100);k__Fungi_unclassified(100);k__Fungi_unclassified(100); | - | - | - | - | - | - | - | Unassigned | - |
| Otu11870 | 2 | k__Fungi(100);k__Fungi_unclassified(100);k__Fungi_unclassified(100);k__Fungi_unclassified(100);k__Fungi_unclassified(100);k__Fungi_unclassified(100);k__Fungi_unclassified(100); | - | - | - | - | - | - | - | Unassigned | - |
| Otu11876 | 2 | k__Fungi(100);k__Fungi_unclassified(100);k__Fungi_unclassified(100);k__Fungi_unclassified(100);k__Fungi_unclassified(100);k__Fungi_unclassified(100);k__Fungi_unclassified(100); | - | - | - | - | - | - | - | Unassigned | - |
| Otu11877 | 2 | k__Fungi(100);k__Fungi_unclassified(100);k__Fungi_unclassified(100);k__Fungi_unclassified(100);k__Fungi_unclassified(100);k__Fungi_unclassified(100);k__Fungi_unclassified(100); | - | - | - | - | - | - | - | Unassigned | - |
| Otu11880 | 2 | k__Fungi(100);k__Fungi_unclassified(100);k__Fungi_unclassified(100);k__Fungi_unclassified(100);k__Fungi_unclassified(100);k__Fungi_unclassified(100);k__Fungi_unclassified(100); | - | - | - | - | - | - | - | Unassigned | - |
| Otu11882 | 2 | k__Fungi(100);p__Ascomycota(100);p__Ascomycota_unclassified(100);p__Ascomycota_unclassified(100);p__Ascomycota_unclassified(100);p__Ascomycota_unclassified(100);p__Ascomycota_unclassified(100); | - | - | - | - | - | - | - | Unassigned | - |
| Otu11886 | 2 | k__Fungi(100);k__Fungi_unclassified(100);k__Fungi_unclassified(100);k__Fungi_unclassified(100);k__Fungi_unclassified(100);k__Fungi_unclassified(100);k__Fungi_unclassified(100); | - | - | - | - | - | - | - | Unassigned | - |
| Otu11887 | 2 | k__Fungi(100);k__Fungi_unclassified(100);k__Fungi_unclassified(100);k__Fungi_unclassified(100);k__Fungi_unclassified(100);k__Fungi_unclassified(100);k__Fungi_unclassified(100); | - | - | - | - | - | - | - | Unassigned | - |
| Otu11890 | 2 | k__Fungi(100);k__Fungi_unclassified(100);k__Fungi_unclassified(100);k__Fungi_unclassified(100);k__Fungi_unclassified(100);k__Fungi_unclassified(100);k__Fungi_unclassified(100); | - | - | - | - | - | - | - | Unassigned | - |
| Otu11892 | 2 | k__Fungi(100);k__Fungi_unclassified(100);k__Fungi_unclassified(100);k__Fungi_unclassified(100);k__Fungi_unclassified(100);k__Fungi_unclassified(100);k__Fungi_unclassified(100); | - | - | - | - | - | - | - | Unassigned | - |
| Otu11893 | 2 | k__Fungi(100);p__Ascomycota(100);c__Dothideomycetes(100);o__Pleosporales(100);o__Pleosporales_unclassified(100);o__Pleosporales_unclassified(100);o__Pleosporales_unclassified(100); | - | - | - | - | - | - | - | Unassigned | - |
| Otu11895 | 2 | k__Fungi(100);k__Fungi_unclassified(100);k__Fungi_unclassified(100);k__Fungi_unclassified(100);k__Fungi_unclassified(100);k__Fungi_unclassified(100);k__Fungi_unclassified(100); | - | - | - | - | - | - | - | Unassigned | - |
| Otu11896 | 2 | k__Fungi(100);k__Fungi_unclassified(100);k__Fungi_unclassified(100);k__Fungi_unclassified(100);k__Fungi_unclassified(100);k__Fungi_unclassified(100);k__Fungi_unclassified(100); | - | - | - | - | - | - | - | Unassigned | - |
| Otu11897 | 2 | k__Fungi(100);k__Fungi_unclassified(100);k__Fungi_unclassified(100);k__Fungi_unclassified(100);k__Fungi_unclassified(100);k__Fungi_unclassified(100);k__Fungi_unclassified(100); | - | - | - | - | - | - | - | Unassigned | - |
| Otu11899 | 2 | k__Fungi(100);k__Fungi_unclassified(100);k__Fungi_unclassified(100);k__Fungi_unclassified(100);k__Fungi_unclassified(100);k__Fungi_unclassified(100);k__Fungi_unclassified(100); | - | - | - | - | - | - | - | Unassigned | - |
| Otu11901 | 2 | k__Fungi(100);k__Fungi_unclassified(100);k__Fungi_unclassified(100);k__Fungi_unclassified(100);k__Fungi_unclassified(100);k__Fungi_unclassified(100);k__Fungi_unclassified(100); | - | - | - | - | - | - | - | Unassigned | - |
| Otu11902 | 2 | k__Fungi(100);k__Fungi_unclassified(100);k__Fungi_unclassified(100);k__Fungi_unclassified(100);k__Fungi_unclassified(100);k__Fungi_unclassified(100);k__Fungi_unclassified(100); | - | - | - | - | - | - | - | Unassigned | - |
| Otu11903 | 2 | k__Fungi(100);p__Ascomycota(100);c__Leotiomycetes(100);o__Helotiales(100);o__Helotiales_unclassified(100);o__Helotiales_unclassified(100);o__Helotiales_unclassified(100); | - | - | - | - | - | - | - | Unassigned | - |
| Otu11905 | 2 | k__Fungi(100);k__Fungi_unclassified(100);k__Fungi_unclassified(100);k__Fungi_unclassified(100);k__Fungi_unclassified(100);k__Fungi_unclassified(100);k__Fungi_unclassified(100); | - | - | - | - | - | - | - | Unassigned | - |
| Otu11906 | 2 | k__Fungi(100);k__Fungi_unclassified(100);k__Fungi_unclassified(100);k__Fungi_unclassified(100);k__Fungi_unclassified(100);k__Fungi_unclassified(100);k__Fungi_unclassified(100); | - | - | - | - | - | - | - | Unassigned | - |
| Otu11907 | 2 | k__Fungi(100);k__Fungi_unclassified(100);k__Fungi_unclassified(100);k__Fungi_unclassified(100);k__Fungi_unclassified(100);k__Fungi_unclassified(100);k__Fungi_unclassified(100); | - | - | - | - | - | - | - | Unassigned | - |
| Otu11910 | 2 | k__Fungi(100);k__Fungi_unclassified(100);k__Fungi_unclassified(100);k__Fungi_unclassified(100);k__Fungi_unclassified(100);k__Fungi_unclassified(100);k__Fungi_unclassified(100); | - | - | - | - | - | - | - | Unassigned | - |
| Otu11921 | 2 | k__Fungi(100);k__Fungi_unclassified(100);k__Fungi_unclassified(100);k__Fungi_unclassified(100);k__Fungi_unclassified(100);k__Fungi_unclassified(100);k__Fungi_unclassified(100); | - | - | - | - | - | - | - | Unassigned | - |
| Otu11924 | 2 | k__Fungi(100);k__Fungi_unclassified(100);k__Fungi_unclassified(100);k__Fungi_unclassified(100);k__Fungi_unclassified(100);k__Fungi_unclassified(100);k__Fungi_unclassified(100); | - | - | - | - | - | - | - | Unassigned | - |
| Otu11925 | 2 | k__Fungi(100);k__Fungi_unclassified(100);k__Fungi_unclassified(100);k__Fungi_unclassified(100);k__Fungi_unclassified(100);k__Fungi_unclassified(100);k__Fungi_unclassified(100); | - | - | - | - | - | - | - | Unassigned | - |
| Otu11927 | 2 | k__Fungi(100);k__Fungi_unclassified(100);k__Fungi_unclassified(100);k__Fungi_unclassified(100);k__Fungi_unclassified(100);k__Fungi_unclassified(100);k__Fungi_unclassified(100); | - | - | - | - | - | - | - | Unassigned | - |
| Otu11931 | 2 | k__Fungi(100);p__Ascomycota(100);p__Ascomycota_unclassified(100);p__Ascomycota_unclassified(100);p__Ascomycota_unclassified(100);p__Ascomycota_unclassified(100);p__Ascomycota_unclassified(100); | - | - | - | - | - | - | - | Unassigned | - |
| Otu11932 | 2 | k__Fungi(100);p__Ascomycota(100);c__Leotiomycetes(100);o__Helotiales(100);f__unclassified_Helotiales(100);g__unclassified_Helotiales(100);g__unclassified_Helotiales_unclassified(100); | - | - | - | - | - | - | - | Unassigned | - |
| Otu11933 | 2 | k__Fungi(100);k__Fungi_unclassified(100);k__Fungi_unclassified(100);k__Fungi_unclassified(100);k__Fungi_unclassified(100);k__Fungi_unclassified(100);k__Fungi_unclassified(100); | - | - | - | - | - | - | - | Unassigned | - |
| Otu11934 | 2 | k__Fungi(100);k__Fungi_unclassified(100);k__Fungi_unclassified(100);k__Fungi_unclassified(100);k__Fungi_unclassified(100);k__Fungi_unclassified(100);k__Fungi_unclassified(100); | - | - | - | - | - | - | - | Unassigned | - |
| Otu11937 | 2 | k__Fungi(100);k__Fungi_unclassified(100);k__Fungi_unclassified(100);k__Fungi_unclassified(100);k__Fungi_unclassified(100);k__Fungi_unclassified(100);k__Fungi_unclassified(100); | - | - | - | - | - | - | - | Unassigned | - |
| Otu11938 | 2 | k__Fungi(100);k__Fungi_unclassified(100);k__Fungi_unclassified(100);k__Fungi_unclassified(100);k__Fungi_unclassified(100);k__Fungi_unclassified(100);k__Fungi_unclassified(100); | - | - | - | - | - | - | - | Unassigned | - |
| Otu11942 | 2 | k__Fungi(100);k__Fungi_unclassified(100);k__Fungi_unclassified(100);k__Fungi_unclassified(100);k__Fungi_unclassified(100);k__Fungi_unclassified(100);k__Fungi_unclassified(100); | - | - | - | - | - | - | - | Unassigned | - |
| Otu11948 | 2 | k__Fungi(100);p__Ascomycota(100);p__Ascomycota_unclassified(100);p__Ascomycota_unclassified(100);p__Ascomycota_unclassified(100);p__Ascomycota_unclassified(100);p__Ascomycota_unclassified(100); | - | - | - | - | - | - | - | Unassigned | - |
| Otu11952 | 2 | k__Fungi(100);k__Fungi_unclassified(100);k__Fungi_unclassified(100);k__Fungi_unclassified(100);k__Fungi_unclassified(100);k__Fungi_unclassified(100);k__Fungi_unclassified(100); | - | - | - | - | - | - | - | Unassigned | - |
| Otu11953 | 2 | k__Fungi(100);k__Fungi_unclassified(100);k__Fungi_unclassified(100);k__Fungi_unclassified(100);k__Fungi_unclassified(100);k__Fungi_unclassified(100);k__Fungi_unclassified(100); | - | - | - | - | - | - | - | Unassigned | - |
| Otu11955 | 2 | k__Fungi(100);k__Fungi_unclassified(100);k__Fungi_unclassified(100);k__Fungi_unclassified(100);k__Fungi_unclassified(100);k__Fungi_unclassified(100);k__Fungi_unclassified(100); | - | - | - | - | - | - | - | Unassigned | - |
| Otu11956 | 2 | k__Fungi(100);k__Fungi_unclassified(100);k__Fungi_unclassified(100);k__Fungi_unclassified(100);k__Fungi_unclassified(100);k__Fungi_unclassified(100);k__Fungi_unclassified(100); | - | - | - | - | - | - | - | Unassigned | - |
| Otu11957 | 2 | k__Fungi(100);k__Fungi_unclassified(100);k__Fungi_unclassified(100);k__Fungi_unclassified(100);k__Fungi_unclassified(100);k__Fungi_unclassified(100);k__Fungi_unclassified(100); | - | - | - | - | - | - | - | Unassigned | - |
| Otu11958 | 2 | k__Fungi(100);k__Fungi_unclassified(100);k__Fungi_unclassified(100);k__Fungi_unclassified(100);k__Fungi_unclassified(100);k__Fungi_unclassified(100);k__Fungi_unclassified(100); | - | - | - | - | - | - | - | Unassigned | - |
| Otu11960 | 2 | k__Fungi(100);k__Fungi_unclassified(100);k__Fungi_unclassified(100);k__Fungi_unclassified(100);k__Fungi_unclassified(100);k__Fungi_unclassified(100);k__Fungi_unclassified(100); | - | - | - | - | - | - | - | Unassigned | - |
| Otu11963 | 2 | k__Fungi(100);k__Fungi_unclassified(100);k__Fungi_unclassified(100);k__Fungi_unclassified(100);k__Fungi_unclassified(100);k__Fungi_unclassified(100);k__Fungi_unclassified(100); | - | - | - | - | - | - | - | Unassigned | - |
| Otu11966 | 2 | k__Fungi(100);k__Fungi_unclassified(100);k__Fungi_unclassified(100);k__Fungi_unclassified(100);k__Fungi_unclassified(100);k__Fungi_unclassified(100);k__Fungi_unclassified(100); | - | - | - | - | - | - | - | Unassigned | - |
| Otu11970 | 2 | k__Fungi(100);p__Ascomycota(100);p__Ascomycota_unclassified(100);p__Ascomycota_unclassified(100);p__Ascomycota_unclassified(100);p__Ascomycota_unclassified(100);p__Ascomycota_unclassified(100); | - | - | - | - | - | - | - | Unassigned | - |
| Otu11983 | 2 | k__Fungi(100);k__Fungi_unclassified(100);k__Fungi_unclassified(100);k__Fungi_unclassified(100);k__Fungi_unclassified(100);k__Fungi_unclassified(100);k__Fungi_unclassified(100); | - | - | - | - | - | - | - | Unassigned | - |
| Otu11984 | 2 | k__Fungi(100);p__Ascomycota(100);p__Ascomycota_unclassified(100);p__Ascomycota_unclassified(100);p__Ascomycota_unclassified(100);p__Ascomycota_unclassified(100);p__Ascomycota_unclassified(100); | - | - | - | - | - | - | - | Unassigned | - |
| Otu11985 | 2 | k__Fungi(100);k__Fungi_unclassified(100);k__Fungi_unclassified(100);k__Fungi_unclassified(100);k__Fungi_unclassified(100);k__Fungi_unclassified(100);k__Fungi_unclassified(100); | - | - | - | - | - | - | - | Unassigned | - |
| Otu11988 | 2 | k__Fungi(100);k__Fungi_unclassified(100);k__Fungi_unclassified(100);k__Fungi_unclassified(100);k__Fungi_unclassified(100);k__Fungi_unclassified(100);k__Fungi_unclassified(100); | - | - | - | - | - | - | - | Unassigned | - |
| Otu11989 | 2 | k__Fungi(100);p__Ascomycota(100);p__Ascomycota_unclassified(100);p__Ascomycota_unclassified(100);p__Ascomycota_unclassified(100);p__Ascomycota_unclassified(100);p__Ascomycota_unclassified(100); | - | - | - | - | - | - | - | Unassigned | - |
| Otu11991 | 2 | k__Fungi(100);k__Fungi_unclassified(100);k__Fungi_unclassified(100);k__Fungi_unclassified(100);k__Fungi_unclassified(100);k__Fungi_unclassified(100);k__Fungi_unclassified(100); | - | - | - | - | - | - | - | Unassigned | - |
| Otu11994 | 2 | k__Fungi(100);p__Ascomycota(100);c__Sordariomycetes(100);c__Sordariomycetes_unclassified(100);c__Sordariomycetes_unclassified(100);c__Sordariomycetes_unclassified(100);c__Sordariomycetes_unclassified(100); | - | - | - | - | - | - | - | Unassigned | - |
| Otu11996 | 2 | k__Fungi(100);k__Fungi_unclassified(100);k__Fungi_unclassified(100);k__Fungi_unclassified(100);k__Fungi_unclassified(100);k__Fungi_unclassified(100);k__Fungi_unclassified(100); | - | - | - | - | - | - | - | Unassigned | - |
| Otu11998 | 2 | k__Fungi(100);k__Fungi_unclassified(100);k__Fungi_unclassified(100);k__Fungi_unclassified(100);k__Fungi_unclassified(100);k__Fungi_unclassified(100);k__Fungi_unclassified(100); | - | - | - | - | - | - | - | Unassigned | - |
| Otu12014 | 2 | k__Fungi(100);p__Ascomycota(100);p__Ascomycota_unclassified(100);p__Ascomycota_unclassified(100);p__Ascomycota_unclassified(100);p__Ascomycota_unclassified(100);p__Ascomycota_unclassified(100); | - | - | - | - | - | - | - | Unassigned | - |
| Otu12020 | 2 | k__Fungi(100);p__Ascomycota(100);p__Ascomycota_unclassified(100);p__Ascomycota_unclassified(100);p__Ascomycota_unclassified(100);p__Ascomycota_unclassified(100);p__Ascomycota_unclassified(100); | - | - | - | - | - | - | - | Unassigned | - |
| Otu12021 | 2 | k__Fungi(100);p__Ascomycota(100);p__Ascomycota_unclassified(100);p__Ascomycota_unclassified(100);p__Ascomycota_unclassified(100);p__Ascomycota_unclassified(100);p__Ascomycota_unclassified(100); | - | - | - | - | - | - | - | Unassigned | - |
| Otu12023 | 2 | k__Fungi(100);k__Fungi_unclassified(100);k__Fungi_unclassified(100);k__Fungi_unclassified(100);k__Fungi_unclassified(100);k__Fungi_unclassified(100);k__Fungi_unclassified(100); | - | - | - | - | - | - | - | Unassigned | - |
| Otu12024 | 2 | k__Fungi(100);k__Fungi_unclassified(100);k__Fungi_unclassified(100);k__Fungi_unclassified(100);k__Fungi_unclassified(100);k__Fungi_unclassified(100);k__Fungi_unclassified(100); | - | - | - | - | - | - | - | Unassigned | - |
| Otu12025 | 2 | k__Fungi(100);p__Ascomycota(100);p__Ascomycota_unclassified(100);p__Ascomycota_unclassified(100);p__Ascomycota_unclassified(100);p__Ascomycota_unclassified(100);p__Ascomycota_unclassified(100); | - | - | - | - | - | - | - | Unassigned | - |
| Otu12026 | 2 | k__Fungi(100);k__Fungi_unclassified(100);k__Fungi_unclassified(100);k__Fungi_unclassified(100);k__Fungi_unclassified(100);k__Fungi_unclassified(100);k__Fungi_unclassified(100); | - | - | - | - | - | - | - | Unassigned | - |
| Otu12027 | 2 | k__Fungi(100);k__Fungi_unclassified(100);k__Fungi_unclassified(100);k__Fungi_unclassified(100);k__Fungi_unclassified(100);k__Fungi_unclassified(100);k__Fungi_unclassified(100); | - | - | - | - | - | - | - | Unassigned | - |
| Otu12031 | 2 | k__Fungi(100);k__Fungi_unclassified(100);k__Fungi_unclassified(100);k__Fungi_unclassified(100);k__Fungi_unclassified(100);k__Fungi_unclassified(100);k__Fungi_unclassified(100); | - | - | - | - | - | - | - | Unassigned | - |
| Otu12032 | 2 | k__Fungi(100);k__Fungi_unclassified(100);k__Fungi_unclassified(100);k__Fungi_unclassified(100);k__Fungi_unclassified(100);k__Fungi_unclassified(100);k__Fungi_unclassified(100); | - | - | - | - | - | - | - | Unassigned | - |
| Otu12033 | 2 | k__Fungi(100);k__Fungi_unclassified(100);k__Fungi_unclassified(100);k__Fungi_unclassified(100);k__Fungi_unclassified(100);k__Fungi_unclassified(100);k__Fungi_unclassified(100); | - | - | - | - | - | - | - | Unassigned | - |
| Otu12035 | 2 | k__Fungi(100);k__Fungi_unclassified(100);k__Fungi_unclassified(100);k__Fungi_unclassified(100);k__Fungi_unclassified(100);k__Fungi_unclassified(100);k__Fungi_unclassified(100); | - | - | - | - | - | - | - | Unassigned | - |
| Otu12036 | 2 | k__Fungi(100);k__Fungi_unclassified(100);k__Fungi_unclassified(100);k__Fungi_unclassified(100);k__Fungi_unclassified(100);k__Fungi_unclassified(100);k__Fungi_unclassified(100); | - | - | - | - | - | - | - | Unassigned | - |
| Otu12038 | 2 | k__Fungi(100);k__Fungi_unclassified(100);k__Fungi_unclassified(100);k__Fungi_unclassified(100);k__Fungi_unclassified(100);k__Fungi_unclassified(100);k__Fungi_unclassified(100); | - | - | - | - | - | - | - | Unassigned | - |
| Otu12041 | 2 | k__Fungi(100);k__Fungi_unclassified(100);k__Fungi_unclassified(100);k__Fungi_unclassified(100);k__Fungi_unclassified(100);k__Fungi_unclassified(100);k__Fungi_unclassified(100); | - | - | - | - | - | - | - | Unassigned | - |
| Otu12044 | 2 | k__Fungi(100);k__Fungi_unclassified(100);k__Fungi_unclassified(100);k__Fungi_unclassified(100);k__Fungi_unclassified(100);k__Fungi_unclassified(100);k__Fungi_unclassified(100); | - | - | - | - | - | - | - | Unassigned | - |
| Otu12048 | 2 | k__Fungi(100);p__Ascomycota(100);p__Ascomycota_unclassified(100);p__Ascomycota_unclassified(100);p__Ascomycota_unclassified(100);p__Ascomycota_unclassified(100);p__Ascomycota_unclassified(100); | - | - | - | - | - | - | - | Unassigned | - |
| Otu12052 | 2 | k__Fungi(100);p__Ascomycota(100);p__Ascomycota_unclassified(100);p__Ascomycota_unclassified(100);p__Ascomycota_unclassified(100);p__Ascomycota_unclassified(100);p__Ascomycota_unclassified(100); | - | - | - | - | - | - | - | Unassigned | - |
| Otu12055 | 2 | k__Fungi(100);k__Fungi_unclassified(100);k__Fungi_unclassified(100);k__Fungi_unclassified(100);k__Fungi_unclassified(100);k__Fungi_unclassified(100);k__Fungi_unclassified(100); | - | - | - | - | - | - | - | Unassigned | - |
| Otu12065 | 2 | k__Fungi(100);k__Fungi_unclassified(100);k__Fungi_unclassified(100);k__Fungi_unclassified(100);k__Fungi_unclassified(100);k__Fungi_unclassified(100);k__Fungi_unclassified(100); | - | - | - | - | - | - | - | Unassigned | - |
| Otu12070 | 2 | k__Fungi(100);k__Fungi_unclassified(100);k__Fungi_unclassified(100);k__Fungi_unclassified(100);k__Fungi_unclassified(100);k__Fungi_unclassified(100);k__Fungi_unclassified(100); | - | - | - | - | - | - | - | Unassigned | - |
| Otu12071 | 2 | k__Fungi(100);k__Fungi_unclassified(100);k__Fungi_unclassified(100);k__Fungi_unclassified(100);k__Fungi_unclassified(100);k__Fungi_unclassified(100);k__Fungi_unclassified(100); | - | - | - | - | - | - | - | Unassigned | - |
| Otu12079 | 2 | k__Fungi(100);k__Fungi_unclassified(100);k__Fungi_unclassified(100);k__Fungi_unclassified(100);k__Fungi_unclassified(100);k__Fungi_unclassified(100);k__Fungi_unclassified(100); | - | - | - | - | - | - | - | Unassigned | - |
| Otu12080 | 2 | k__Fungi(100);k__Fungi_unclassified(100);k__Fungi_unclassified(100);k__Fungi_unclassified(100);k__Fungi_unclassified(100);k__Fungi_unclassified(100);k__Fungi_unclassified(100); | - | - | - | - | - | - | - | Unassigned | - |
| Otu12084 | 2 | k__Fungi(100);k__Fungi_unclassified(100);k__Fungi_unclassified(100);k__Fungi_unclassified(100);k__Fungi_unclassified(100);k__Fungi_unclassified(100);k__Fungi_unclassified(100); | - | - | - | - | - | - | - | Unassigned | - |
| Otu12087 | 2 | k__Fungi(100);k__Fungi_unclassified(100);k__Fungi_unclassified(100);k__Fungi_unclassified(100);k__Fungi_unclassified(100);k__Fungi_unclassified(100);k__Fungi_unclassified(100); | - | - | - | - | - | - | - | Unassigned | - |
| Otu12090 | 2 | k__Fungi(100);k__Fungi_unclassified(100);k__Fungi_unclassified(100);k__Fungi_unclassified(100);k__Fungi_unclassified(100);k__Fungi_unclassified(100);k__Fungi_unclassified(100); | - | - | - | - | - | - | - | Unassigned | - |
| Otu12091 | 2 | k__Fungi(100);p__Ascomycota(100);p__Ascomycota_unclassified(100);p__Ascomycota_unclassified(100);p__Ascomycota_unclassified(100);p__Ascomycota_unclassified(100);p__Ascomycota_unclassified(100); | - | - | - | - | - | - | - | Unassigned | - |
| Otu12094 | 2 | k__Fungi(100);p__Ascomycota(100);p__Ascomycota_unclassified(100);p__Ascomycota_unclassified(100);p__Ascomycota_unclassified(100);p__Ascomycota_unclassified(100);p__Ascomycota_unclassified(100); | - | - | - | - | - | - | - | Unassigned | - |
| Otu12095 | 2 | k__Fungi(100);k__Fungi_unclassified(100);k__Fungi_unclassified(100);k__Fungi_unclassified(100);k__Fungi_unclassified(100);k__Fungi_unclassified(100);k__Fungi_unclassified(100); | - | - | - | - | - | - | - | Unassigned | - |
| Otu12096 | 2 | k__Fungi(100);p__Ascomycota(100);p__Ascomycota_unclassified(100);p__Ascomycota_unclassified(100);p__Ascomycota_unclassified(100);p__Ascomycota_unclassified(100);p__Ascomycota_unclassified(100); | - | - | - | - | - | - | - | Unassigned | - |
| Otu12101 | 2 | k__Fungi(100);k__Fungi_unclassified(100);k__Fungi_unclassified(100);k__Fungi_unclassified(100);k__Fungi_unclassified(100);k__Fungi_unclassified(100);k__Fungi_unclassified(100); | - | - | - | - | - | - | - | Unassigned | - |
| Otu12103 | 2 | k__Fungi(100);k__Fungi_unclassified(100);k__Fungi_unclassified(100);k__Fungi_unclassified(100);k__Fungi_unclassified(100);k__Fungi_unclassified(100);k__Fungi_unclassified(100); | - | - | - | - | - | - | - | Unassigned | - |
| Otu12105 | 2 | k__Fungi(100);k__Fungi_unclassified(100);k__Fungi_unclassified(100);k__Fungi_unclassified(100);k__Fungi_unclassified(100);k__Fungi_unclassified(100);k__Fungi_unclassified(100); | - | - | - | - | - | - | - | Unassigned | - |
| Otu12107 | 2 | k__Fungi(100);p__Ascomycota(100);c__Leotiomycetes(100);o__Helotiales(100);o__Helotiales_unclassified(100);o__Helotiales_unclassified(100);o__Helotiales_unclassified(100); | - | - | - | - | - | - | - | Unassigned | - |
| Otu12110 | 2 | k__Fungi(100);k__Fungi_unclassified(100);k__Fungi_unclassified(100);k__Fungi_unclassified(100);k__Fungi_unclassified(100);k__Fungi_unclassified(100);k__Fungi_unclassified(100); | - | - | - | - | - | - | - | Unassigned | - |
| Otu12113 | 2 | k__Fungi(100);k__Fungi_unclassified(100);k__Fungi_unclassified(100);k__Fungi_unclassified(100);k__Fungi_unclassified(100);k__Fungi_unclassified(100);k__Fungi_unclassified(100); | - | - | - | - | - | - | - | Unassigned | - |
| Otu12121 | 2 | k__Fungi(100);k__Fungi_unclassified(100);k__Fungi_unclassified(100);k__Fungi_unclassified(100);k__Fungi_unclassified(100);k__Fungi_unclassified(100);k__Fungi_unclassified(100); | - | - | - | - | - | - | - | Unassigned | - |
| Otu12122 | 2 | k__Plantae(100);p__unclassified_Plantae(100);c__unclassified_Plantae(100);o__unclassified_Plantae(100);f__unclassified_Plantae(100);g__unclassified_Plantae(100);s__Plantae_sp(100); | - | - | - | - | - | - | - | Unassigned | - |
| Otu12123 | 2 | k__Fungi(100);k__Fungi_unclassified(100);k__Fungi_unclassified(100);k__Fungi_unclassified(100);k__Fungi_unclassified(100);k__Fungi_unclassified(100);k__Fungi_unclassified(100); | - | - | - | - | - | - | - | Unassigned | - |
| Otu12133 | 2 | k__Fungi(100);k__Fungi_unclassified(100);k__Fungi_unclassified(100);k__Fungi_unclassified(100);k__Fungi_unclassified(100);k__Fungi_unclassified(100);k__Fungi_unclassified(100); | - | - | - | - | - | - | - | Unassigned | - |
| Otu12135 | 2 | k__Fungi(100);p__Ascomycota(100);p__Ascomycota_unclassified(100);p__Ascomycota_unclassified(100);p__Ascomycota_unclassified(100);p__Ascomycota_unclassified(100);p__Ascomycota_unclassified(100); | - | - | - | - | - | - | - | Unassigned | - |
| Otu12143 | 2 | k__Fungi(100);k__Fungi_unclassified(100);k__Fungi_unclassified(100);k__Fungi_unclassified(100);k__Fungi_unclassified(100);k__Fungi_unclassified(100);k__Fungi_unclassified(100); | - | - | - | - | - | - | - | Unassigned | - |
| Otu12148 | 2 | k__Fungi(100);p__Ascomycota(100);p__Ascomycota_unclassified(100);p__Ascomycota_unclassified(100);p__Ascomycota_unclassified(100);p__Ascomycota_unclassified(100);p__Ascomycota_unclassified(100); | - | - | - | - | - | - | - | Unassigned | - |
| Otu12149 | 2 | k__Fungi(100);p__Ascomycota(100);p__Ascomycota_unclassified(100);p__Ascomycota_unclassified(100);p__Ascomycota_unclassified(100);p__Ascomycota_unclassified(100);p__Ascomycota_unclassified(100); | - | - | - | - | - | - | - | Unassigned | - |
| Otu12154 | 2 | k__Fungi(100);p__Ascomycota(100);p__Ascomycota_unclassified(100);p__Ascomycota_unclassified(100);p__Ascomycota_unclassified(100);p__Ascomycota_unclassified(100);p__Ascomycota_unclassified(100); | - | - | - | - | - | - | - | Unassigned | - |
| Otu12155 | 2 | k__Fungi(100);k__Fungi_unclassified(100);k__Fungi_unclassified(100);k__Fungi_unclassified(100);k__Fungi_unclassified(100);k__Fungi_unclassified(100);k__Fungi_unclassified(100); | - | - | - | - | - | - | - | Unassigned | - |
| Otu12156 | 2 | k__Fungi(100);k__Fungi_unclassified(100);k__Fungi_unclassified(100);k__Fungi_unclassified(100);k__Fungi_unclassified(100);k__Fungi_unclassified(100);k__Fungi_unclassified(100); | - | - | - | - | - | - | - | Unassigned | - |
| Otu12158 | 2 | k__Fungi(100);k__Fungi_unclassified(100);k__Fungi_unclassified(100);k__Fungi_unclassified(100);k__Fungi_unclassified(100);k__Fungi_unclassified(100);k__Fungi_unclassified(100); | - | - | - | - | - | - | - | Unassigned | - |
| Otu12159 | 2 | k__Fungi(100);k__Fungi_unclassified(100);k__Fungi_unclassified(100);k__Fungi_unclassified(100);k__Fungi_unclassified(100);k__Fungi_unclassified(100);k__Fungi_unclassified(100); | - | - | - | - | - | - | - | Unassigned | - |
| Otu12160 | 2 | k__Fungi(100);k__Fungi_unclassified(100);k__Fungi_unclassified(100);k__Fungi_unclassified(100);k__Fungi_unclassified(100);k__Fungi_unclassified(100);k__Fungi_unclassified(100); | - | - | - | - | - | - | - | Unassigned | - |
| Otu12161 | 2 | k__Fungi(100);p__Ascomycota(100);p__Ascomycota_unclassified(100);p__Ascomycota_unclassified(100);p__Ascomycota_unclassified(100);p__Ascomycota_unclassified(100);p__Ascomycota_unclassified(100); | - | - | - | - | - | - | - | Unassigned | - |
| Otu12164 | 2 | k__Fungi(100);k__Fungi_unclassified(100);k__Fungi_unclassified(100);k__Fungi_unclassified(100);k__Fungi_unclassified(100);k__Fungi_unclassified(100);k__Fungi_unclassified(100); | - | - | - | - | - | - | - | Unassigned | - |
| Otu12165 | 2 | k__Fungi(100);p__Ascomycota(100);p__Ascomycota_unclassified(100);p__Ascomycota_unclassified(100);p__Ascomycota_unclassified(100);p__Ascomycota_unclassified(100);p__Ascomycota_unclassified(100); | - | - | - | - | - | - | - | Unassigned | - |
| Otu12170 | 2 | k__Fungi(100);k__Fungi_unclassified(100);k__Fungi_unclassified(100);k__Fungi_unclassified(100);k__Fungi_unclassified(100);k__Fungi_unclassified(100);k__Fungi_unclassified(100); | - | - | - | - | - | - | - | Unassigned | - |
| Otu12173 | 2 | k__Fungi(100);p__Ascomycota(100);p__Ascomycota_unclassified(100);p__Ascomycota_unclassified(100);p__Ascomycota_unclassified(100);p__Ascomycota_unclassified(100);p__Ascomycota_unclassified(100); | - | - | - | - | - | - | - | Unassigned | - |
| Otu12174 | 2 | k__Fungi(100);p__Ascomycota(100);c__Sordariomycetes(100);c__Sordariomycetes_unclassified(100);c__Sordariomycetes_unclassified(100);c__Sordariomycetes_unclassified(100);c__Sordariomycetes_unclassified(100); | - | - | - | - | - | - | - | Unassigned | - |
| Otu12176 | 2 | k__Fungi(100);k__Fungi_unclassified(100);k__Fungi_unclassified(100);k__Fungi_unclassified(100);k__Fungi_unclassified(100);k__Fungi_unclassified(100);k__Fungi_unclassified(100); | - | - | - | - | - | - | - | Unassigned | - |
| Otu12178 | 2 | k__Fungi(100);k__Fungi_unclassified(100);k__Fungi_unclassified(100);k__Fungi_unclassified(100);k__Fungi_unclassified(100);k__Fungi_unclassified(100);k__Fungi_unclassified(100); | - | - | - | - | - | - | - | Unassigned | - |
| Otu12181 | 2 | k__Fungi(100);p__Ascomycota(100);p__Ascomycota_unclassified(100);p__Ascomycota_unclassified(100);p__Ascomycota_unclassified(100);p__Ascomycota_unclassified(100);p__Ascomycota_unclassified(100); | - | - | - | - | - | - | - | Unassigned | - |
| Otu12182 | 2 | k__Fungi(100);p__Ascomycota(100);p__Ascomycota_unclassified(100);p__Ascomycota_unclassified(100);p__Ascomycota_unclassified(100);p__Ascomycota_unclassified(100);p__Ascomycota_unclassified(100); | - | - | - | - | - | - | - | Unassigned | - |
| Otu12183 | 2 | k__Fungi(100);p__Ascomycota(100);c__Sordariomycetes(100);o__Sordariales(100);o__Sordariales_unclassified(100);o__Sordariales_unclassified(100);o__Sordariales_unclassified(100); | - | - | - | - | - | - | - | Unassigned | - |
| Otu12187 | 2 | k__Fungi(100);p__Ascomycota(100);c__Dothideomycetes(100);o__Pleosporales(100);o__Pleosporales_unclassified(100);o__Pleosporales_unclassified(100);o__Pleosporales_unclassified(100); | - | - | - | - | - | - | - | Unassigned | - |
| Otu12188 | 2 | k__Fungi(100);k__Fungi_unclassified(100);k__Fungi_unclassified(100);k__Fungi_unclassified(100);k__Fungi_unclassified(100);k__Fungi_unclassified(100);k__Fungi_unclassified(100); | - | - | - | - | - | - | - | Unassigned | - |
| Otu12192 | 2 | k__Fungi(100);p__Ascomycota(100);p__Ascomycota_unclassified(100);p__Ascomycota_unclassified(100);p__Ascomycota_unclassified(100);p__Ascomycota_unclassified(100);p__Ascomycota_unclassified(100); | - | - | - | - | - | - | - | Unassigned | - |
| Otu12199 | 2 | k__Fungi(100);k__Fungi_unclassified(100);k__Fungi_unclassified(100);k__Fungi_unclassified(100);k__Fungi_unclassified(100);k__Fungi_unclassified(100);k__Fungi_unclassified(100); | - | - | - | - | - | - | - | Unassigned | - |
| Otu12203 | 2 | k__Fungi(100);k__Fungi_unclassified(100);k__Fungi_unclassified(100);k__Fungi_unclassified(100);k__Fungi_unclassified(100);k__Fungi_unclassified(100);k__Fungi_unclassified(100); | - | - | - | - | - | - | - | Unassigned | - |
| Otu12208 | 2 | k__Fungi(100);k__Fungi_unclassified(100);k__Fungi_unclassified(100);k__Fungi_unclassified(100);k__Fungi_unclassified(100);k__Fungi_unclassified(100);k__Fungi_unclassified(100); | - | - | - | - | - | - | - | Unassigned | - |
| Otu12210 | 2 | k__Fungi(100);p__Ascomycota(100);p__Ascomycota_unclassified(100);p__Ascomycota_unclassified(100);p__Ascomycota_unclassified(100);p__Ascomycota_unclassified(100);p__Ascomycota_unclassified(100); | - | - | - | - | - | - | - | Unassigned | - |
| Otu12211 | 2 | k__Fungi(100);k__Fungi_unclassified(100);k__Fungi_unclassified(100);k__Fungi_unclassified(100);k__Fungi_unclassified(100);k__Fungi_unclassified(100);k__Fungi_unclassified(100); | - | - | - | - | - | - | - | Unassigned | - |
| Otu12212 | 2 | k__Fungi(100);k__Fungi_unclassified(100);k__Fungi_unclassified(100);k__Fungi_unclassified(100);k__Fungi_unclassified(100);k__Fungi_unclassified(100);k__Fungi_unclassified(100); | - | - | - | - | - | - | - | Unassigned | - |
| Otu12213 | 2 | k__Fungi(100);k__Fungi_unclassified(100);k__Fungi_unclassified(100);k__Fungi_unclassified(100);k__Fungi_unclassified(100);k__Fungi_unclassified(100);k__Fungi_unclassified(100); | - | - | - | - | - | - | - | Unassigned | - |
| Otu12224 | 2 | k__Fungi(100);p__Ascomycota(100);p__Ascomycota_unclassified(100);p__Ascomycota_unclassified(100);p__Ascomycota_unclassified(100);p__Ascomycota_unclassified(100);p__Ascomycota_unclassified(100); | - | - | - | - | - | - | - | Unassigned | - |
| Otu12226 | 2 | k__Fungi(100);p__Ascomycota(100);p__Ascomycota_unclassified(100);p__Ascomycota_unclassified(100);p__Ascomycota_unclassified(100);p__Ascomycota_unclassified(100);p__Ascomycota_unclassified(100); | - | - | - | - | - | - | - | Unassigned | - |
| Otu12229 | 2 | k__Fungi(100);k__Fungi_unclassified(100);k__Fungi_unclassified(100);k__Fungi_unclassified(100);k__Fungi_unclassified(100);k__Fungi_unclassified(100);k__Fungi_unclassified(100); | - | - | - | - | - | - | - | Unassigned | - |
| Otu12232 | 2 | k__Fungi(100);p__Ascomycota(100);p__Ascomycota_unclassified(100);p__Ascomycota_unclassified(100);p__Ascomycota_unclassified(100);p__Ascomycota_unclassified(100);p__Ascomycota_unclassified(100); | - | - | - | - | - | - | - | Unassigned | - |
| Otu12233 | 2 | k__Fungi(100);k__Fungi_unclassified(100);k__Fungi_unclassified(100);k__Fungi_unclassified(100);k__Fungi_unclassified(100);k__Fungi_unclassified(100);k__Fungi_unclassified(100); | - | - | - | - | - | - | - | Unassigned | - |
| Otu12234 | 2 | k__Fungi(100);k__Fungi_unclassified(100);k__Fungi_unclassified(100);k__Fungi_unclassified(100);k__Fungi_unclassified(100);k__Fungi_unclassified(100);k__Fungi_unclassified(100); | - | - | - | - | - | - | - | Unassigned | - |
| Otu12235 | 2 | k__Fungi(100);k__Fungi_unclassified(100);k__Fungi_unclassified(100);k__Fungi_unclassified(100);k__Fungi_unclassified(100);k__Fungi_unclassified(100);k__Fungi_unclassified(100); | - | - | - | - | - | - | - | Unassigned | - |
| Otu12236 | 2 | k__Fungi(100);p__Ascomycota(100);p__Ascomycota_unclassified(100);p__Ascomycota_unclassified(100);p__Ascomycota_unclassified(100);p__Ascomycota_unclassified(100);p__Ascomycota_unclassified(100); | - | - | - | - | - | - | - | Unassigned | - |
| Otu12237 | 2 | k__Fungi(100);k__Fungi_unclassified(100);k__Fungi_unclassified(100);k__Fungi_unclassified(100);k__Fungi_unclassified(100);k__Fungi_unclassified(100);k__Fungi_unclassified(100); | - | - | - | - | - | - | - | Unassigned | - |
| Otu12238 | 2 | k__Fungi(100);k__Fungi_unclassified(100);k__Fungi_unclassified(100);k__Fungi_unclassified(100);k__Fungi_unclassified(100);k__Fungi_unclassified(100);k__Fungi_unclassified(100); | - | - | - | - | - | - | - | Unassigned | - |
| Otu12239 | 2 | k__Fungi(100);k__Fungi_unclassified(100);k__Fungi_unclassified(100);k__Fungi_unclassified(100);k__Fungi_unclassified(100);k__Fungi_unclassified(100);k__Fungi_unclassified(100); | - | - | - | - | - | - | - | Unassigned | - |
| Otu12240 | 2 | k__Fungi(100);k__Fungi_unclassified(100);k__Fungi_unclassified(100);k__Fungi_unclassified(100);k__Fungi_unclassified(100);k__Fungi_unclassified(100);k__Fungi_unclassified(100); | - | - | - | - | - | - | - | Unassigned | - |
| Otu12242 | 2 | k__Fungi(100);p__Ascomycota(100);p__Ascomycota_unclassified(100);p__Ascomycota_unclassified(100);p__Ascomycota_unclassified(100);p__Ascomycota_unclassified(100);p__Ascomycota_unclassified(100); | - | - | - | - | - | - | - | Unassigned | - |
| Otu12245 | 2 | k__Fungi(100);k__Fungi_unclassified(100);k__Fungi_unclassified(100);k__Fungi_unclassified(100);k__Fungi_unclassified(100);k__Fungi_unclassified(100);k__Fungi_unclassified(100); | - | - | - | - | - | - | - | Unassigned | - |
| Otu12246 | 2 | k__Fungi(100);k__Fungi_unclassified(100);k__Fungi_unclassified(100);k__Fungi_unclassified(100);k__Fungi_unclassified(100);k__Fungi_unclassified(100);k__Fungi_unclassified(100); | - | - | - | - | - | - | - | Unassigned | - |
| Otu12247 | 2 | k__Fungi(100);p__Ascomycota(100);p__Ascomycota_unclassified(100);p__Ascomycota_unclassified(100);p__Ascomycota_unclassified(100);p__Ascomycota_unclassified(100);p__Ascomycota_unclassified(100); | - | - | - | - | - | - | - | Unassigned | - |
| Otu12249 | 2 | k__Fungi(100);k__Fungi_unclassified(100);k__Fungi_unclassified(100);k__Fungi_unclassified(100);k__Fungi_unclassified(100);k__Fungi_unclassified(100);k__Fungi_unclassified(100); | - | - | - | - | - | - | - | Unassigned | - |
| Otu12253 | 2 | k__Fungi(100);k__Fungi_unclassified(100);k__Fungi_unclassified(100);k__Fungi_unclassified(100);k__Fungi_unclassified(100);k__Fungi_unclassified(100);k__Fungi_unclassified(100); | - | - | - | - | - | - | - | Unassigned | - |
| Otu12255 | 2 | k__Fungi(100);k__Fungi_unclassified(100);k__Fungi_unclassified(100);k__Fungi_unclassified(100);k__Fungi_unclassified(100);k__Fungi_unclassified(100);k__Fungi_unclassified(100); | - | - | - | - | - | - | - | Unassigned | - |
| Otu12256 | 2 | k__Fungi(100);k__Fungi_unclassified(100);k__Fungi_unclassified(100);k__Fungi_unclassified(100);k__Fungi_unclassified(100);k__Fungi_unclassified(100);k__Fungi_unclassified(100); | - | - | - | - | - | - | - | Unassigned | - |
| Otu12257 | 2 | k__Fungi(100);k__Fungi_unclassified(100);k__Fungi_unclassified(100);k__Fungi_unclassified(100);k__Fungi_unclassified(100);k__Fungi_unclassified(100);k__Fungi_unclassified(100); | - | - | - | - | - | - | - | Unassigned | - |
| Otu12258 | 2 | k__Fungi(100);k__Fungi_unclassified(100);k__Fungi_unclassified(100);k__Fungi_unclassified(100);k__Fungi_unclassified(100);k__Fungi_unclassified(100);k__Fungi_unclassified(100); | - | - | - | - | - | - | - | Unassigned | - |
| Otu12264 | 2 | k__Fungi(100);p__Ascomycota(100);c__Leotiomycetes(100);o__Helotiales(100);o__Helotiales_unclassified(100);o__Helotiales_unclassified(100);o__Helotiales_unclassified(100); | - | - | - | - | - | - | - | Unassigned | - |
| Otu12266 | 2 | k__Fungi(100);p__Ascomycota(100);p__Ascomycota_unclassified(100);p__Ascomycota_unclassified(100);p__Ascomycota_unclassified(100);p__Ascomycota_unclassified(100);p__Ascomycota_unclassified(100); | - | - | - | - | - | - | - | Unassigned | - |
| Otu12275 | 2 | k__Fungi(100);p__Ascomycota(100);p__Ascomycota_unclassified(100);p__Ascomycota_unclassified(100);p__Ascomycota_unclassified(100);p__Ascomycota_unclassified(100);p__Ascomycota_unclassified(100); | - | - | - | - | - | - | - | Unassigned | - |
| Otu12279 | 2 | k__Fungi(100);k__Fungi_unclassified(100);k__Fungi_unclassified(100);k__Fungi_unclassified(100);k__Fungi_unclassified(100);k__Fungi_unclassified(100);k__Fungi_unclassified(100); | - | - | - | - | - | - | - | Unassigned | - |
| Otu12287 | 2 | k__Fungi(100);p__Ascomycota(100);p__Ascomycota_unclassified(100);p__Ascomycota_unclassified(100);p__Ascomycota_unclassified(100);p__Ascomycota_unclassified(100);p__Ascomycota_unclassified(100); | - | - | - | - | - | - | - | Unassigned | - |
| Otu12289 | 2 | k__Fungi(100);k__Fungi_unclassified(100);k__Fungi_unclassified(100);k__Fungi_unclassified(100);k__Fungi_unclassified(100);k__Fungi_unclassified(100);k__Fungi_unclassified(100); | - | - | - | - | - | - | - | Unassigned | - |
| Otu12291 | 2 | k__Fungi(100);p__Ascomycota(100);p__Ascomycota_unclassified(100);p__Ascomycota_unclassified(100);p__Ascomycota_unclassified(100);p__Ascomycota_unclassified(100);p__Ascomycota_unclassified(100); | - | - | - | - | - | - | - | Unassigned | - |
| Otu12292 | 2 | k__Fungi(100);k__Fungi_unclassified(100);k__Fungi_unclassified(100);k__Fungi_unclassified(100);k__Fungi_unclassified(100);k__Fungi_unclassified(100);k__Fungi_unclassified(100); | - | - | - | - | - | - | - | Unassigned | - |
| Otu12294 | 2 | k__Fungi(100);k__Fungi_unclassified(100);k__Fungi_unclassified(100);k__Fungi_unclassified(100);k__Fungi_unclassified(100);k__Fungi_unclassified(100);k__Fungi_unclassified(100); | - | - | - | - | - | - | - | Unassigned | - |
| Otu12299 | 2 | k__Fungi(100);k__Fungi_unclassified(100);k__Fungi_unclassified(100);k__Fungi_unclassified(100);k__Fungi_unclassified(100);k__Fungi_unclassified(100);k__Fungi_unclassified(100); | - | - | - | - | - | - | - | Unassigned | - |
| Otu12301 | 2 | k__Fungi(100);p__Basidiomycota(100);c__Microbotryomycetes(100);o__Sporidiobolales(100);f__Sporidiobolales_family_Incertae_sedis(100);f__Sporidiobolales_family_Incertae_sedis_unclassified(100);f__Sporidiobolales_family_Incertae_sedis_unclassified(100); | - | - | - | - | - | - | - | Unassigned | - |
| Otu12302 | 2 | k__Fungi(100);p__Basidiomycota(100);c__Agaricomycetes(100);o__Cantharellales(100);f__Ceratobasidiaceae(100);g__unclassified_Ceratobasidiaceae(100);s__uncultured_Ceratobasidium(100); | - | - | - | - | - | - | - | Unassigned | - |
| Otu12308 | 2 | k__Fungi(100);p__Ascomycota(100);c__Dothideomycetes(100);o__Pleosporales(100);f__Pleosporales_family_Incertae_sedis(100);f__Pleosporales_family_Incertae_sedis_unclassified(100);f__Pleosporales_family_Incertae_sedis_unclassified(100); | - | - | - | - | - | - | - | Unassigned | - |
| Otu12309 | 2 | k__Fungi(100);k__Fungi_unclassified(100);k__Fungi_unclassified(100);k__Fungi_unclassified(100);k__Fungi_unclassified(100);k__Fungi_unclassified(100);k__Fungi_unclassified(100); | - | - | - | - | - | - | - | Unassigned | - |
| Otu12312 | 2 | k__Fungi(100);k__Fungi_unclassified(100);k__Fungi_unclassified(100);k__Fungi_unclassified(100);k__Fungi_unclassified(100);k__Fungi_unclassified(100);k__Fungi_unclassified(100); | - | - | - | - | - | - | - | Unassigned | - |
| Otu12313 | 2 | k__Fungi(100);p__Basidiomycota(100);c__Microbotryomycetes(100);o__Sporidiobolales(100);f__Sporidiobolales_family_Incertae_sedis(100);f__Sporidiobolales_family_Incertae_sedis_unclassified(100);f__Sporidiobolales_family_Incertae_sedis_unclassified(100); | - | - | - | - | - | - | - | Unassigned | - |
| Otu12314 | 2 | k__Fungi(100);k__Fungi_unclassified(100);k__Fungi_unclassified(100);k__Fungi_unclassified(100);k__Fungi_unclassified(100);k__Fungi_unclassified(100);k__Fungi_unclassified(100); | - | - | - | - | - | - | - | Unassigned | - |
| Otu12315 | 2 | k__Fungi(100);p__Ascomycota(100);p__Ascomycota_unclassified(100);p__Ascomycota_unclassified(100);p__Ascomycota_unclassified(100);p__Ascomycota_unclassified(100);p__Ascomycota_unclassified(100); | - | - | - | - | - | - | - | Unassigned | - |
| Otu12316 | 2 | k__Fungi(100);k__Fungi_unclassified(100);k__Fungi_unclassified(100);k__Fungi_unclassified(100);k__Fungi_unclassified(100);k__Fungi_unclassified(100);k__Fungi_unclassified(100); | - | - | - | - | - | - | - | Unassigned | - |
| Otu12318 | 2 | k__Fungi(100);k__Fungi_unclassified(100);k__Fungi_unclassified(100);k__Fungi_unclassified(100);k__Fungi_unclassified(100);k__Fungi_unclassified(100);k__Fungi_unclassified(100); | - | - | - | - | - | - | - | Unassigned | - |
| Otu12319 | 2 | k__Fungi(100);k__Fungi_unclassified(100);k__Fungi_unclassified(100);k__Fungi_unclassified(100);k__Fungi_unclassified(100);k__Fungi_unclassified(100);k__Fungi_unclassified(100); | - | - | - | - | - | - | - | Unassigned | - |
| Otu12320 | 2 | k__Fungi(100);p__Ascomycota(100);p__Ascomycota_unclassified(100);p__Ascomycota_unclassified(100);p__Ascomycota_unclassified(100);p__Ascomycota_unclassified(100);p__Ascomycota_unclassified(100); | - | - | - | - | - | - | - | Unassigned | - |
| Otu12324 | 2 | k__Plantae(100);p__unclassified_Plantae(100);c__unclassified_Plantae(100);o__unclassified_Plantae(100);f__unclassified_Plantae(100);g__unclassified_Plantae(100);s__Plantae_sp(100); | - | - | - | - | - | - | - | Unassigned | - |
| Otu12326 | 2 | k__Fungi(100);k__Fungi_unclassified(100);k__Fungi_unclassified(100);k__Fungi_unclassified(100);k__Fungi_unclassified(100);k__Fungi_unclassified(100);k__Fungi_unclassified(100); | - | - | - | - | - | - | - | Unassigned | - |
| Otu12328 | 2 | k__Fungi(100);k__Fungi_unclassified(100);k__Fungi_unclassified(100);k__Fungi_unclassified(100);k__Fungi_unclassified(100);k__Fungi_unclassified(100);k__Fungi_unclassified(100); | - | - | - | - | - | - | - | Unassigned | - |
| Otu12335 | 2 | k__Fungi(100);k__Fungi_unclassified(100);k__Fungi_unclassified(100);k__Fungi_unclassified(100);k__Fungi_unclassified(100);k__Fungi_unclassified(100);k__Fungi_unclassified(100); | - | - | - | - | - | - | - | Unassigned | - |

| | | | | | | | | | | | |
|---|---|---|---|---|---|---|---|---|---|---|---|
| Otu12345 | 2 | k__Fungi(100);k__Fungi_unclassified(100);k__Fungi_unclassified(100);k__Fungi_unclassified(100);k__Fungi_unclassified(100);k__Fungi_unclassified(100); | - | - | - | - | - | - | - | Unassigned | - |
| Otu12347 | 2 | k__Fungi(100);p__Ascomycota(100);p__Ascomycota_unclassified(100);p__Ascomycota_unclassified(100);p__Ascomycota_unclassified(100);p__Ascomycota_unclassified(100); | - | - | - | - | - | - | - | Unassigned | - |
| Otu12349 | 2 | k__Fungi(100);k__Fungi_unclassified(100);k__Fungi_unclassified(100);k__Fungi_unclassified(100);k__Fungi_unclassified(100);k__Fungi_unclassified(100); | - | - | - | - | - | - | - | Unassigned | - |
| Otu12350 | 2 | k__Fungi(100);k__Fungi_unclassified(100);k__Fungi_unclassified(100);k__Fungi_unclassified(100);k__Fungi_unclassified(100);k__Fungi_unclassified(100); | - | - | - | - | - | - | - | Unassigned | - |
| Otu12352 | 2 | k__Fungi(100);k__Fungi_unclassified(100);k__Fungi_unclassified(100);k__Fungi_unclassified(100);k__Fungi_unclassified(100);k__Fungi_unclassified(100); | - | - | - | - | - | - | - | Unassigned | - |
| Otu12353 | 2 | k__Fungi(100);p__Ascomycota(100);p__Ascomycota_unclassified(100);p__Ascomycota_unclassified(100);p__Ascomycota_unclassified(100);p__Ascomycota_unclassified(100); | - | - | - | - | - | - | - | Unassigned | - |
| Otu12355 | 2 | k__Fungi(100);k__Fungi_unclassified(100);k__Fungi_unclassified(100);k__Fungi_unclassified(100);k__Fungi_unclassified(100);k__Fungi_unclassified(100); | - | - | - | - | - | - | - | Unassigned | - |
| Otu12357 | 2 | k__Fungi(100);k__Fungi_unclassified(100);k__Fungi_unclassified(100);k__Fungi_unclassified(100);k__Fungi_unclassified(100);k__Fungi_unclassified(100); | - | - | - | - | - | - | - | Unassigned | - |
| Otu12359 | 2 | k__Fungi(100);k__Fungi_unclassified(100);k__Fungi_unclassified(100);k__Fungi_unclassified(100);k__Fungi_unclassified(100);k__Fungi_unclassified(100); | - | - | - | - | - | - | - | Unassigned | - |
| Otu12360 | 2 | k__Fungi(100);p__Ascomycota(100);p__Ascomycota_unclassified(100);p__Ascomycota_unclassified(100);p__Ascomycota_unclassified(100);p__Ascomycota_unclassified(100); | - | - | - | - | - | - | - | Unassigned | - |
| Otu12362 | 2 | k__Fungi(100);p__Ascomycota(100);p__Ascomycota_unclassified(100);p__Ascomycota_unclassified(100);p__Ascomycota_unclassified(100);p__Ascomycota_unclassified(100); | - | - | - | - | - | - | - | Unassigned | - |
| Otu12366 | 2 | k__Fungi(100);p__Ascomycota(100);c__Taphrinomycetes(100);o__Taphrinales(100);f__Taphrinaceae(100);g__Lalaria(100);s__Lalaria_inositophila(100); | - | - | - | - | - | - | - | Unassigned | - |
| Otu12367 | 2 | k__Fungi(100);p__Ascomycota(100);p__Ascomycota_unclassified(100);p__Ascomycota_unclassified(100);p__Ascomycota_unclassified(100);p__Ascomycota_unclassified(100); | - | - | - | - | - | - | - | Unassigned | - |
| Otu12368 | 2 | k__Fungi(100);k__Fungi_unclassified(100);k__Fungi_unclassified(100);k__Fungi_unclassified(100);k__Fungi_unclassified(100);k__Fungi_unclassified(100); | - | - | - | - | - | - | - | Unassigned | - |
| Otu12377 | 2 | k__Fungi(100);k__Fungi_unclassified(100);k__Fungi_unclassified(100);k__Fungi_unclassified(100);k__Fungi_unclassified(100);k__Fungi_unclassified(100); | - | - | - | - | - | - | - | Unassigned | - |
| Otu12379 | 2 | k__Fungi(100);p__Ascomycota(100);p__Ascomycota_unclassified(100);p__Ascomycota_unclassified(100);p__Ascomycota_unclassified(100);p__Ascomycota_unclassified(100); | - | - | - | - | - | - | - | Unassigned | - |
| Otu12383 | 2 | k__Fungi(100);p__Basidiomycota(100);c__Agaricomycetes(100);o__Auriculariales(100);o__Auriculariales_unclassified(100);o__Auriculariales_unclassified(100);o__Auriculariales_unclassified(100); | - | - | - | - | - | - | - | Unassigned | - |
| Otu12384 | 2 | k__Fungi(100);k__Fungi_unclassified(100);k__Fungi_unclassified(100);k__Fungi_unclassified(100);k__Fungi_unclassified(100);k__Fungi_unclassified(100); | - | - | - | - | - | - | - | Unassigned | - |
| Otu12385 | 2 | k__Fungi(100);k__Fungi_unclassified(100);k__Fungi_unclassified(100);k__Fungi_unclassified(100);k__Fungi_unclassified(100);k__Fungi_unclassified(100); | - | - | - | - | - | - | - | Unassigned | - |
| Otu12386 | 2 | k__Fungi(100);p__Basidiomycota(100);c__Agaricomycetes(100);c__Agaricomycetes_unclassified(100);c__Agaricomycetes_unclassified(100);c__Agaricomycetes_unclassified(100);c__Agaricomycetes_unclassified(100); | - | - | - | - | - | - | - | Unassigned | - |
| Otu12387 | 2 | k__Fungi(100);p__Basidiomycota(100);c__Agaricomycetes(100);c__Agaricomycetes_unclassified(100);c__Agaricomycetes_unclassified(100);c__Agaricomycetes_unclassified(100);c__Agaricomycetes_unclassified(100); | - | - | - | - | - | - | - | Unassigned | - |
| Otu12389 | 2 | k__Fungi(100);k__Fungi_unclassified(100);k__Fungi_unclassified(100);k__Fungi_unclassified(100);k__Fungi_unclassified(100);k__Fungi_unclassified(100); | - | - | - | - | - | - | - | Unassigned | - |
| Otu12390 | 2 | k__Fungi(100);k__Fungi_unclassified(100);k__Fungi_unclassified(100);k__Fungi_unclassified(100);k__Fungi_unclassified(100);k__Fungi_unclassified(100); | - | - | - | - | - | - | - | Unassigned | - |
| Otu12392 | 2 | k__Fungi(100);k__Fungi_unclassified(100);k__Fungi_unclassified(100);k__Fungi_unclassified(100);k__Fungi_unclassified(100);k__Fungi_unclassified(100); | - | - | - | - | - | - | - | Unassigned | - |
| Otu12394 | 2 | k__Fungi(100);p__Basidiomycota(100);p__Basidiomycota_unclassified(100);p__Basidiomycota_unclassified(100);p__Basidiomycota_unclassified(100);p__Basidiomycota_unclassified(100);p__Basidiomycota_unclassified(100); | - | - | - | - | - | - | - | Unassigned | - |
| Otu12396 | 2 | k__Fungi(100);p__Ascomycota(100);c__Leotiomycetes(100);o__Helotiales(100);o__Helotiales_unclassified(100);o__Helotiales_unclassified(100);o__Helotiales_unclassified(100); | - | - | - | - | - | - | - | Unassigned | - |
| Otu12404 | 2 | k__Fungi(100);p__Ascomycota(100);p__Ascomycota_unclassified(100);p__Ascomycota_unclassified(100);p__Ascomycota_unclassified(100);p__Ascomycota_unclassified(100); | - | - | - | - | - | - | - | Unassigned | - |
| Otu12405 | 2 | k__Fungi(100);p__Ascomycota(100);p__Ascomycota_unclassified(100);p__Ascomycota_unclassified(100);p__Ascomycota_unclassified(100);p__Ascomycota_unclassified(100); | - | - | - | - | - | - | - | Unassigned | - |
| Otu12406 | 2 | k__Fungi(100);p__Ascomycota(100);p__Ascomycota_unclassified(100);p__Ascomycota_unclassified(100);p__Ascomycota_unclassified(100);p__Ascomycota_unclassified(100); | - | - | - | - | - | - | - | Unassigned | - |
| Otu12411 | 2 | k__Fungi(100);k__Fungi_unclassified(100);k__Fungi_unclassified(100);k__Fungi_unclassified(100);k__Fungi_unclassified(100);k__Fungi_unclassified(100); | - | - | - | - | - | - | - | Unassigned | - |
| Otu12415 | 2 | k__Fungi(100);p__Ascomycota(100);p__Ascomycota_unclassified(100);p__Ascomycota_unclassified(100);p__Ascomycota_unclassified(100);p__Ascomycota_unclassified(100); | - | - | - | - | - | - | - | Unassigned | - |
| Otu12417 | 2 | k__Fungi(100);p__Ascomycota(100);p__Ascomycota_unclassified(100);p__Ascomycota_unclassified(100);p__Ascomycota_unclassified(100);p__Ascomycota_unclassified(100); | - | - | - | - | - | - | - | Unassigned | - |
| Otu12418 | 2 | k__Fungi(100);k__Fungi_unclassified(100);k__Fungi_unclassified(100);k__Fungi_unclassified(100);k__Fungi_unclassified(100);k__Fungi_unclassified(100); | - | - | - | - | - | - | - | Unassigned | - |
| Otu12419 | 2 | k__Fungi(100);k__Fungi_unclassified(100);k__Fungi_unclassified(100);k__Fungi_unclassified(100);k__Fungi_unclassified(100);k__Fungi_unclassified(100); | - | - | - | - | - | - | - | Unassigned | - |
| Otu12424 | 2 | k__Fungi(100);p__Ascomycota(100);p__Ascomycota_unclassified(100);p__Ascomycota_unclassified(100);p__Ascomycota_unclassified(100);p__Ascomycota_unclassified(100); | - | - | - | - | - | - | - | Unassigned | - |
| Otu12427 | 2 | k__Fungi(100);k__Fungi_unclassified(100);k__Fungi_unclassified(100);k__Fungi_unclassified(100);k__Fungi_unclassified(100);k__Fungi_unclassified(100); | - | - | - | - | - | - | - | Unassigned | - |
| Otu12429 | 2 | k__Fungi(100);k__Fungi_unclassified(100);k__Fungi_unclassified(100);k__Fungi_unclassified(100);k__Fungi_unclassified(100);k__Fungi_unclassified(100); | - | - | - | - | - | - | - | Unassigned | - |
| Otu12430 | 2 | k__Fungi(100);k__Fungi_unclassified(100);k__Fungi_unclassified(100);k__Fungi_unclassified(100);k__Fungi_unclassified(100);k__Fungi_unclassified(100); | - | - | - | - | - | - | - | Unassigned | - |
| Otu12432 | 2 | k__Fungi(100);p__Ascomycota(100);p__Ascomycota_unclassified(100);p__Ascomycota_unclassified(100);p__Ascomycota_unclassified(100);p__Ascomycota_unclassified(100); | - | - | - | - | - | - | - | Unassigned | - |
| Otu12434 | 2 | k__Fungi(100);p__Ascomycota(100);c__Sordariomycetes(100);c__Sordariomycetes_unclassified(100);c__Sordariomycetes_unclassified(100);c__Sordariomycetes_unclassified(100);c__Sordariomycetes_unclassified(100); | - | - | - | - | - | - | - | Unassigned | - |
| Otu12440 | 2 | k__Fungi(100);p__Ascomycota(100);p__Ascomycota_unclassified(100);p__Ascomycota_unclassified(100);p__Ascomycota_unclassified(100);p__Ascomycota_unclassified(100); | - | - | - | - | - | - | - | Unassigned | - |
| Otu12441 | 2 | k__Fungi(100);p__Ascomycota(100);p__Ascomycota_unclassified(100);p__Ascomycota_unclassified(100);p__Ascomycota_unclassified(100);p__Ascomycota_unclassified(100); | - | - | - | - | - | - | - | Unassigned | - |
| Otu12449 | 2 | k__Fungi(100);k__Fungi_unclassified(100);k__Fungi_unclassified(100);k__Fungi_unclassified(100);k__Fungi_unclassified(100);k__Fungi_unclassified(100); | - | - | - | - | - | - | - | Unassigned | - |
| Otu12450 | 2 | k__Fungi(100);k__Fungi_unclassified(100);k__Fungi_unclassified(100);k__Fungi_unclassified(100);k__Fungi_unclassified(100);k__Fungi_unclassified(100); | - | - | - | - | - | - | - | Unassigned | - |
| Otu12454 | 2 | k__Fungi(100);p__Ascomycota(100);p__Ascomycota_unclassified(100);p__Ascomycota_unclassified(100);p__Ascomycota_unclassified(100);p__Ascomycota_unclassified(100); | - | - | - | - | - | - | - | Unassigned | - |
| Otu12456 | 2 | k__Fungi(100);k__Fungi_unclassified(100);k__Fungi_unclassified(100);k__Fungi_unclassified(100);k__Fungi_unclassified(100);k__Fungi_unclassified(100); | - | - | - | - | - | - | - | Unassigned | - |
| Otu12457 | 2 | k__Fungi(100);p__Basidiomycota(100);c__Microbotryomycetes(100);o__Sporidiobolales(100);f__Sporidiobolales_family_Incertae_sedis(100);f__Sporidiobolales_family_Incertae_sedis_unclassified(100);f__Sporidiobolales_family_Incertae_sedis_unclassified(100); | - | - | - | - | - | - | - | Unassigned | - |
| Otu12462 | 2 | k__Fungi(100);p__Ascomycota(100);p__Ascomycota_unclassified(100);p__Ascomycota_unclassified(100);p__Ascomycota_unclassified(100);p__Ascomycota_unclassified(100); | - | - | - | - | - | - | - | Unassigned | - |
| Otu12464 | 2 | k__Fungi(100);p__Ascomycota(100);p__Ascomycota_unclassified(100);p__Ascomycota_unclassified(100);p__Ascomycota_unclassified(100);p__Ascomycota_unclassified(100); | - | - | - | - | - | - | - | Unassigned | - |
| Otu12468 | 2 | k__Fungi(100);p__Zygomycota(100);c__Zygomycota_class_Incertae_sedis(100);o__Kickxellales(100);o__Kickxellales_unclassified(100);o__Kickxellales_unclassified(100);o__Kickxellales_unclassified(100); | - | - | - | - | - | - | - | Unassigned | - |
| Otu12469 | 2 | k__Fungi(100);p__Ascomycota(100);p__Ascomycota_unclassified(100);p__Ascomycota_unclassified(100);p__Ascomycota_unclassified(100);p__Ascomycota_unclassified(100); | - | - | - | - | - | - | - | Unassigned | - |
| Otu12472 | 2 | k__Fungi(100);p__Ascomycota(100);c__Dothideomycetes(100);o__Capnodiales(100);o__Capnodiales_unclassified(100);o__Capnodiales_unclassified(100);o__Capnodiales_unclassified(100); | - | - | - | - | - | - | - | Unassigned | - |
| Otu12476 | 2 | k__Fungi(100);k__Fungi_unclassified(100);k__Fungi_unclassified(100);k__Fungi_unclassified(100);k__Fungi_unclassified(100);k__Fungi_unclassified(100); | - | - | - | - | - | - | - | Unassigned | - |
| Otu12479 | 2 | k__Fungi(100);k__Fungi_unclassified(100);k__Fungi_unclassified(100);k__Fungi_unclassified(100);k__Fungi_unclassified(100);k__Fungi_unclassified(100); | - | - | - | - | - | - | - | Unassigned | - |
| Otu12482 | 2 | k__Fungi(100);p__Ascomycota(100);p__Ascomycota_unclassified(100);p__Ascomycota_unclassified(100);p__Ascomycota_unclassified(100);p__Ascomycota_unclassified(100); | - | - | - | - | - | - | - | Unassigned | - |
| Otu12483 | 2 | k__Fungi(100);k__Fungi_unclassified(100);k__Fungi_unclassified(100);k__Fungi_unclassified(100);k__Fungi_unclassified(100);k__Fungi_unclassified(100); | - | - | - | - | - | - | - | Unassigned | - |
| Otu12488 | 2 | k__Fungi(100);k__Fungi_unclassified(100);k__Fungi_unclassified(100);k__Fungi_unclassified(100);k__Fungi_unclassified(100);k__Fungi_unclassified(100); | - | - | - | - | - | - | - | Unassigned | - |
| Otu12492 | 2 | k__Fungi(100);k__Fungi_unclassified(100);k__Fungi_unclassified(100);k__Fungi_unclassified(100);k__Fungi_unclassified(100);k__Fungi_unclassified(100); | - | - | - | - | - | - | - | Unassigned | - |
| Otu12502 | 2 | k__Fungi(100);k__Fungi_unclassified(100);k__Fungi_unclassified(100);k__Fungi_unclassified(100);k__Fungi_unclassified(100);k__Fungi_unclassified(100); | - | - | - | - | - | - | - | Unassigned | - |
| Otu12505 | 2 | k__Fungi(100);k__Fungi_unclassified(100);k__Fungi_unclassified(100);k__Fungi_unclassified(100);k__Fungi_unclassified(100);k__Fungi_unclassified(100); | - | - | - | - | - | - | - | Unassigned | - |
| Otu12507 | 2 | k__Fungi(100);p__Ascomycota(100);p__Ascomycota_unclassified(100);p__Ascomycota_unclassified(100);p__Ascomycota_unclassified(100);p__Ascomycota_unclassified(100); | - | - | - | - | - | - | - | Unassigned | - |
| Otu12508 | 2 | k__Fungi(100);p__Ascomycota(100);p__Ascomycota_unclassified(100);p__Ascomycota_unclassified(100);p__Ascomycota_unclassified(100);p__Ascomycota_unclassified(100); | - | - | - | - | - | - | - | Unassigned | - |
| Otu12513 | 2 | k__Fungi(100);k__Fungi_unclassified(100);k__Fungi_unclassified(100);k__Fungi_unclassified(100);k__Fungi_unclassified(100);k__Fungi_unclassified(100); | - | - | - | - | - | - | - | Unassigned | - |
| Otu12516 | 2 | k__Fungi(100);p__Basidiomycota(100);c__Agaricomycetes(100);o__Polyporales(100);f__Phanerochaetaceae(100);f__Phanerochaetaceae_unclassified(100);f__Phanerochaetaceae_unclassified(100); | - | - | - | - | - | - | - | Unassigned | - |
| Otu12517 | 2 | k__Fungi(100);p__Basidiomycota(100);c__Agaricomycetes(100);c__Agaricomycetes_unclassified(100);c__Agaricomycetes_unclassified(100);c__Agaricomycetes_unclassified(100);c__Agaricomycetes_unclassified(100); | - | - | - | - | - | - | - | Unassigned | - |
| Otu12519 | 2 | k__Fungi(100);k__Fungi_unclassified(100);k__Fungi_unclassified(100);k__Fungi_unclassified(100);k__Fungi_unclassified(100);k__Fungi_unclassified(100); | - | - | - | - | - | - | - | Unassigned | - |
| Otu12520 | 2 | k__Fungi(100);k__Fungi_unclassified(100);k__Fungi_unclassified(100);k__Fungi_unclassified(100);k__Fungi_unclassified(100);k__Fungi_unclassified(100); | - | - | - | - | - | - | - | Unassigned | - |
| Otu12522 | 2 | k__Fungi(100);p__Basidiomycota(100);c__Agaricomycetes(100);o__Cantharellales(100);o__Cantharellales_unclassified(100);o__Cantharellales_unclassified(100);o__Cantharellales_unclassified(100); | - | - | - | - | - | - | - | Unassigned | - |
| Otu12523 | 2 | k__Fungi(100);p__Ascomycota(100);p__Ascomycota_unclassified(100);p__Ascomycota_unclassified(100);p__Ascomycota_unclassified(100);p__Ascomycota_unclassified(100); | - | - | - | - | - | - | - | Unassigned | - |
| Otu12524 | 2 | k__Fungi(100);k__Fungi_unclassified(100);k__Fungi_unclassified(100);k__Fungi_unclassified(100);k__Fungi_unclassified(100);k__Fungi_unclassified(100); | - | - | - | - | - | - | - | Unassigned | - |
| Otu12525 | 2 | k__Fungi(100);k__Fungi_unclassified(100);k__Fungi_unclassified(100);k__Fungi_unclassified(100);k__Fungi_unclassified(100);k__Fungi_unclassified(100); | - | - | - | - | - | - | - | Unassigned | - |
| Otu12527 | 2 | k__Fungi(100);k__Fungi_unclassified(100);k__Fungi_unclassified(100);k__Fungi_unclassified(100);k__Fungi_unclassified(100);k__Fungi_unclassified(100); | - | - | - | - | - | - | - | Unassigned | - |
| Otu12535 | 2 | k__Fungi(100);p__Ascomycota(100);p__Ascomycota_unclassified(100);p__Ascomycota_unclassified(100);p__Ascomycota_unclassified(100);p__Ascomycota_unclassified(100); | - | - | - | - | - | - | - | Unassigned | - |
| Otu12536 | 2 | k__Fungi(100);p__Basidiomycota(100);c__Agaricomycetes(100);o__Polyporales(100);o__Polyporales_unclassified(100);o__Polyporales_unclassified(100);o__Polyporales_unclassified(100); | - | - | - | - | - | - | - | Unassigned | - |
| Otu12537 | 2 | k__Fungi(100);p__Basidiomycota(100);p__Basidiomycota_unclassified(100);p__Basidiomycota_unclassified(100);p__Basidiomycota_unclassified(100);p__Basidiomycota_unclassified(100);p__Basidiomycota_unclassified(100); | - | - | - | - | - | - | - | Unassigned | - |
| Otu12538 | 2 | k__Fungi(100);k__Fungi_unclassified(100);k__Fungi_unclassified(100);k__Fungi_unclassified(100);k__Fungi_unclassified(100);k__Fungi_unclassified(100); | - | - | - | - | - | - | - | Unassigned | - |
| Otu12539 | 2 | k__Fungi(100);k__Fungi_unclassified(100);k__Fungi_unclassified(100);k__Fungi_unclassified(100);k__Fungi_unclassified(100);k__Fungi_unclassified(100); | - | - | - | - | - | - | - | Unassigned | - |
| Otu12542 | 2 | k__Fungi(100);k__Fungi_unclassified(100);k__Fungi_unclassified(100);k__Fungi_unclassified(100);k__Fungi_unclassified(100);k__Fungi_unclassified(100); | - | - | - | - | - | - | - | Unassigned | - |
| Otu12543 | 2 | k__Fungi(100);k__Fungi_unclassified(100);k__Fungi_unclassified(100);k__Fungi_unclassified(100);k__Fungi_unclassified(100);k__Fungi_unclassified(100); | - | - | - | - | - | - | - | Unassigned | - |
| Otu12544 | 2 | k__Fungi(100);k__Fungi_unclassified(100);k__Fungi_unclassified(100);k__Fungi_unclassified(100);k__Fungi_unclassified(100);k__Fungi_unclassified(100); | - | - | - | - | - | - | - | Unassigned | - |
| Otu12545 | 2 | k__Fungi(100);p__Basidiomycota(100);c__Microbotryomycetes(100);o__Sporidiobolales(100);f__Sporidiobolales_family_Incertae_sedis(100);f__Sporidiobolales_family_Incertae_sedis_unclassified(100);f__Sporidiobolales_family_Incertae_sedis_unclassified(100); | - | - | - | - | - | - | - | Unassigned | - |
| Otu12549 | 2 | k__Fungi(100);k__Fungi_unclassified(100);k__Fungi_unclassified(100);k__Fungi_unclassified(100);k__Fungi_unclassified(100);k__Fungi_unclassified(100); | - | - | - | - | - | - | - | Unassigned | - |
| Otu12554 | 2 | k__Fungi(100);k__Fungi_unclassified(100);k__Fungi_unclassified(100);k__Fungi_unclassified(100);k__Fungi_unclassified(100);k__Fungi_unclassified(100); | - | - | - | - | - | - | - | Unassigned | - |
| Otu12560 | 2 | k__Fungi(100);p__Ascomycota(100);p__Ascomycota_unclassified(100);p__Ascomycota_unclassified(100);p__Ascomycota_unclassified(100);p__Ascomycota_unclassified(100); | - | - | - | - | - | - | - | Unassigned | - |
| Otu12562 | 2 | k__Fungi(100);k__Fungi_unclassified(100);k__Fungi_unclassified(100);k__Fungi_unclassified(100);k__Fungi_unclassified(100);k__Fungi_unclassified(100); | - | - | - | - | - | - | - | Unassigned | - |
| Otu12563 | 2 | k__Fungi(100);p__Ascomycota(100);p__Ascomycota_unclassified(100);p__Ascomycota_unclassified(100);p__Ascomycota_unclassified(100);p__Ascomycota_unclassified(100); | - | - | - | - | - | - | - | Unassigned | - |
| Otu12571 | 2 | k__Fungi(100);p__Ascomycota(100);p__Ascomycota_unclassified(100);p__Ascomycota_unclassified(100);p__Ascomycota_unclassified(100);p__Ascomycota_unclassified(100); | - | - | - | - | - | - | - | Unassigned | - |
| Otu12572 | 2 | k__Fungi(100);p__Ascomycota(100);p__Ascomycota_unclassified(100);p__Ascomycota_unclassified(100);p__Ascomycota_unclassified(100);p__Ascomycota_unclassified(100); | - | - | - | - | - | - | - | Unassigned | - |
| Otu12574 | 2 | k__Fungi(100);k__Fungi_unclassified(100);k__Fungi_unclassified(100);k__Fungi_unclassified(100);k__Fungi_unclassified(100);k__Fungi_unclassified(100); | - | - | - | - | - | - | - | Unassigned | - |
| Otu12578 | 2 | k__Fungi(100);k__Fungi_unclassified(100);k__Fungi_unclassified(100);k__Fungi_unclassified(100);k__Fungi_unclassified(100);k__Fungi_unclassified(100); | - | - | - | - | - | - | - | Unassigned | - |
| Otu12579 | 2 | k__Fungi(100);p__Basidiomycota(100);c__Agaricomycetes(100);o__Polyporales(100);o__Polyporales_unclassified(100);o__Polyporales_unclassified(100);o__Polyporales_unclassified(100); | - | - | - | - | - | - | - | Unassigned | - |
| Otu12580 | 2 | k__Fungi(100);k__Fungi_unclassified(100);k__Fungi_unclassified(100);k__Fungi_unclassified(100);k__Fungi_unclassified(100);k__Fungi_unclassified(100); | - | - | - | - | - | - | - | Unassigned | - |
| Otu12582 | 2 | k__Fungi(100);p__Ascomycota(100);p__Ascomycota_unclassified(100);p__Ascomycota_unclassified(100);p__Ascomycota_unclassified(100);p__Ascomycota_unclassified(100); | - | - | - | - | - | - | - | Unassigned | - |
| Otu12586 | 2 | k__Fungi(100);k__Fungi_unclassified(100);k__Fungi_unclassified(100);k__Fungi_unclassified(100);k__Fungi_unclassified(100);k__Fungi_unclassified(100); | - | - | - | - | - | - | - | Unassigned | - |
| Otu12588 | 2 | k__Fungi(100);p__Basidiomycota(100);c__Microbotryomycetes(100);o__Sporidiobolales(100);f__Sporidiobolales_family_Incertae_sedis(100);f__Sporidiobolales_family_Incertae_sedis_unclassified(100);f__Sporidiobolales_family_Incertae_sedis_unclassified(100); | - | - | - | - | - | - | - | Unassigned | - |
| Otu12593 | 2 | k__Fungi(100);p__Basidiomycota(100);c__Agaricomycetes(100);c__Agaricomycetes_unclassified(100);c__Agaricomycetes_unclassified(100);c__Agaricomycetes_unclassified(100);c__Agaricomycetes_unclassified(100); | - | - | - | - | - | - | - | Unassigned | - |
| Otu12594 | 2 | k__Fungi(100);p__Ascomycota(100);p__Ascomycota_unclassified(100);p__Ascomycota_unclassified(100);p__Ascomycota_unclassified(100);p__Ascomycota_unclassified(100); | - | - | - | - | - | - | - | Unassigned | - |
| Otu12595 | 2 | k__Fungi(100);k__Fungi_unclassified(100);k__Fungi_unclassified(100);k__Fungi_unclassified(100);k__Fungi_unclassified(100);k__Fungi_unclassified(100); | - | - | - | - | - | - | - | Unassigned | - |
| Otu12599 | 2 | k__Fungi(100);p__Ascomycota(100);p__Ascomycota_unclassified(100);p__Ascomycota_unclassified(100);p__Ascomycota_unclassified(100);p__Ascomycota_unclassified(100); | - | - | - | - | - | - | - | Unassigned | - |
| Otu12600 | 2 | k__Fungi(100);k__Fungi_unclassified(100);k__Fungi_unclassified(100);k__Fungi_unclassified(100);k__Fungi_unclassified(100);k__Fungi_unclassified(100); | - | - | - | - | - | - | - | Unassigned | - |
| Otu12603 | 2 | k__Fungi(100);k__Fungi_unclassified(100);k__Fungi_unclassified(100);k__Fungi_unclassified(100);k__Fungi_unclassified(100);k__Fungi_unclassified(100); | - | - | - | - | - | - | - | Unassigned | - |
| Otu12608 | 2 | k__Fungi(100);k__Fungi_unclassified(100);k__Fungi_unclassified(100);k__Fungi_unclassified(100);k__Fungi_unclassified(100);k__Fungi_unclassified(100); | - | - | - | - | - | - | - | Unassigned | - |
| Otu12618 | 2 | k__Fungi(100);k__Fungi_unclassified(100);k__Fungi_unclassified(100);k__Fungi_unclassified(100);k__Fungi_unclassified(100);k__Fungi_unclassified(100); | - | - | - | - | - | - | - | Unassigned | - |
| Otu12619 | 2 | k__Fungi(100);k__Fungi_unclassified(100);k__Fungi_unclassified(100);k__Fungi_unclassified(100);k__Fungi_unclassified(100);k__Fungi_unclassified(100); | - | - | - | - | - | - | - | Unassigned | - |
| Otu12624 | 2 | k__Fungi(100);p__Ascomycota(100);p__Ascomycota_unclassified(100);p__Ascomycota_unclassified(100);p__Ascomycota_unclassified(100);p__Ascomycota_unclassified(100); | - | - | - | - | - | - | - | Unassigned | - |
| Otu12628 | 2 | k__Fungi(100);k__Fungi_unclassified(100);k__Fungi_unclassified(100);k__Fungi_unclassified(100);k__Fungi_unclassified(100);k__Fungi_unclassified(100); | - | - | - | - | - | - | - | Unassigned | - |
| Otu12629 | 2 | k__Fungi(100);k__Fungi_unclassified(100);k__Fungi_unclassified(100);k__Fungi_unclassified(100);k__Fungi_unclassified(100);k__Fungi_unclassified(100); | - | - | - | - | - | - | - | Unassigned | - |
| Otu12630 | 2 | k__Fungi(100);p__Basidiomycota(100);c__Agaricomycetes(100);o__Boletales(100);o__Boletales_unclassified(100);o__Boletales_unclassified(100);o__Boletales_unclassified(100); | - | - | - | - | - | - | - | Unassigned | - |
| Otu12634 | 2 | k__Fungi(100);k__Fungi_unclassified(100);k__Fungi_unclassified(100);k__Fungi_unclassified(100);k__Fungi_unclassified(100);k__Fungi_unclassified(100); | - | - | - | - | - | - | - | Unassigned | - |
| Otu12635 | 2 | k__Fungi(100);k__Fungi_unclassified(100);k__Fungi_unclassified(100);k__Fungi_unclassified(100);k__Fungi_unclassified(100);k__Fungi_unclassified(100); | - | - | - | - | - | - | - | Unassigned | - |
| Otu12636 | 2 | k__Fungi(100);p__Ascomycota(100);p__Ascomycota_unclassified(100);p__Ascomycota_unclassified(100);p__Ascomycota_unclassified(100);p__Ascomycota_unclassified(100); | - | - | - | - | - | - | - | Unassigned | - |
| Otu12638 | 2 | k__Fungi(100);p__Basidiomycota(100);p__Basidiomycota_unclassified(100);p__Basidiomycota_unclassified(100);p__Basidiomycota_unclassified(100);p__Basidiomycota_unclassified(100);p__Basidiomycota_unclassified(100); | - | - | - | - | - | - | - | Unassigned | - |
| Otu12639 | 2 | k__Fungi(100);k__Fungi_unclassified(100);k__Fungi_unclassified(100);k__Fungi_unclassified(100);k__Fungi_unclassified(100);k__Fungi_unclassified(100); | - | - | - | - | - | - | - | Unassigned | - |
| Otu12642 | 2 | k__Fungi(100);k__Fungi_unclassified(100);k__Fungi_unclassified(100);k__Fungi_unclassified(100);k__Fungi_unclassified(100);k__Fungi_unclassified(100); | - | - | - | - | - | - | - | Unassigned | - |
| Otu12649 | 2 | k__Fungi(100);p__Ascomycota(100);p__Ascomycota_unclassified(100);p__Ascomycota_unclassified(100);p__Ascomycota_unclassified(100);p__Ascomycota_unclassified(100); | - | - | - | - | - | - | - | Unassigned | - |
| Otu12651 | 2 | k__Fungi(100);k__Fungi_unclassified(100);k__Fungi_unclassified(100);k__Fungi_unclassified(100);k__Fungi_unclassified(100);k__Fungi_unclassified(100); | - | - | - | - | - | - | - | Unassigned | - |
| Otu12654 | 2 | k__Fungi(100);p__Ascomycota(100);p__Ascomycota_unclassified(100);p__Ascomycota_unclassified(100);p__Ascomycota_unclassified(100);p__Ascomycota_unclassified(100); | - | - | - | - | - | - | - | Unassigned | - |
| Otu12658 | 2 | k__Fungi(100);k__Fungi_unclassified(100);k__Fungi_unclassified(100);k__Fungi_unclassified(100);k__Fungi_unclassified(100);k__Fungi_unclassified(100); | - | - | - | - | - | - | - | Unassigned | - |
| Otu12660 | 2 | k__Fungi(100);k__Fungi_unclassified(100);k__Fungi_unclassified(100);k__Fungi_unclassified(100);k__Fungi_unclassified(100);k__Fungi_unclassified(100); | - | - | - | - | - | - | - | Unassigned | - |
| Otu12662 | 2 | k__Fungi(100);k__Fungi_unclassified(100);k__Fungi_unclassified(100);k__Fungi_unclassified(100);k__Fungi_unclassified(100);k__Fungi_unclassified(100); | - | - | - | - | - | - | - | Unassigned | - |
| Otu12663 | 2 | k__Fungi(100);p__Ascomycota(100);p__Ascomycota_unclassified(100);p__Ascomycota_unclassified(100);p__Ascomycota_unclassified(100);p__Ascomycota_unclassified(100); | - | - | - | - | - | - | - | Unassigned | - |
| Otu12671 | 2 | k__Fungi(100);k__Fungi_unclassified(100);k__Fungi_unclassified(100);k__Fungi_unclassified(100);k__Fungi_unclassified(100);k__Fungi_unclassified(100); | - | - | - | - | - | - | - | Unassigned | - |
| Otu12672 | 2 | k__Fungi(100);k__Fungi_unclassified(100);k__Fungi_unclassified(100);k__Fungi_unclassified(100);k__Fungi_unclassified(100);k__Fungi_unclassified(100); | - | - | - | - | - | - | - | Unassigned | - |
| Otu12674 | 2 | k__Fungi(100);p__Ascomycota(100);p__Ascomycota_unclassified(100);p__Ascomycota_unclassified(100);p__Ascomycota_unclassified(100);p__Ascomycota_unclassified(100); | - | - | - | - | - | - | - | Unassigned | - |
| Otu12675 | 2 | k__Fungi(100);p__Basidiomycota(100);c__Agaricomycetes(100);c__Agaricomycetes_unclassified(100);c__Agaricomycetes_unclassified(100);c__Agaricomycetes_unclassified(100);c__Agaricomycetes_unclassified(100); | - | - | - | - | - | - | - | Unassigned | - |
| Otu12676 | 2 | k__Fungi(100);k__Fungi_unclassified(100);k__Fungi_unclassified(100);k__Fungi_unclassified(100);k__Fungi_unclassified(100);k__Fungi_unclassified(100); | - | - | - | - | - | - | - | Unassigned | - |
| Otu12678 | 2 | k__Fungi(100);p__Basidiomycota(100);c__Agaricomycetes(100);o__Corticiales(100);f__unclassified_Corticiales(100);g__unclassified_Corticiales(100);s__Corticiales_sp(100); | - | - | - | - | - | - | - | Unassigned | - |
| Otu12679 | 2 | k__Fungi(100);k__Fungi_unclassified(100);k__Fungi_unclassified(100);k__Fungi_unclassified(100);k__Fungi_unclassified(100);k__Fungi_unclassified(100); | - | - | - | - | - | - | - | Unassigned | - |
| Otu12680 | 2 | k__Fungi(100);k__Fungi_unclassified(100);k__Fungi_unclassified(100);k__Fungi_unclassified(100);k__Fungi_unclassified(100);k__Fungi_unclassified(100); | - | - | - | - | - | - | - | Unassigned | - |
| Otu12683 | 2 | k__Fungi(100);k__Fungi_unclassified(100);k__Fungi_unclassified(100);k__Fungi_unclassified(100);k__Fungi_unclassified(100);k__Fungi_unclassified(100); | - | - | - | - | - | - | - | Unassigned | - |
| Otu12684 | 2 | k__Fungi(100);p__Basidiomycota(100);p__Basidiomycota_unclassified(100);p__Basidiomycota_unclassified(100);p__Basidiomycota_unclassified(100);p__Basidiomycota_unclassified(100);p__Basidiomycota_unclassified(100); | - | - | - | - | - | - | - | Unassigned | - |
| Otu12686 | 2 | k__Fungi(100);k__Fungi_unclassified(100);k__Fungi_unclassified(100);k__Fungi_unclassified(100);k__Fungi_unclassified(100);k__Fungi_unclassified(100); | - | - | - | - | - | - | - | Unassigned | - |
| Otu12687 | 2 | k__Fungi(100);p__Basidiomycota(100);c__Agaricomycetes(100);c__Agaricomycetes_unclassified(100);c__Agaricomycetes_unclassified(100);c__Agaricomycetes_unclassified(100);c__Agaricomycetes_unclassified(100); | - | - | - | - | - | - | - | Unassigned | - |
| Otu12688 | 2 | k__Fungi(100);p__Basidiomycota(100);c__Agaricomycetes(100);c__Agaricomycetes_unclassified(100);c__Agaricomycetes_unclassified(100);c__Agaricomycetes_unclassified(100);c__Agaricomycetes_unclassified(100); | - | - | - | - | - | - | - | Unassigned | - |
| Otu12690 | 2 | k__Fungi(100);k__Fungi_unclassified(100);k__Fungi_unclassified(100);k__Fungi_unclassified(100);k__Fungi_unclassified(100);k__Fungi_unclassified(100); | - | - | - | - | - | - | - | Unassigned | - |
| Otu12692 | 2 | k__Fungi(100);k__Fungi_unclassified(100);k__Fungi_unclassified(100);k__Fungi_unclassified(100);k__Fungi_unclassified(100);k__Fungi_unclassified(100); | - | - | - | - | - | - | - | Unassigned | - |
| Otu12693 | 2 | k__Fungi(100);k__Fungi_unclassified(100);k__Fungi_unclassified(100);k__Fungi_unclassified(100);k__Fungi_unclassified(100);k__Fungi_unclassified(100); | - | - | - | - | - | - | - | Unassigned | - |
| Otu12696 | 2 | k__Fungi(100);k__Fungi_unclassified(100);k__Fungi_unclassified(100);k__Fungi_unclassified(100);k__Fungi_unclassified(100);k__Fungi_unclassified(100); | - | - | - | - | - | - | - | Unassigned | - |
| Otu12697 | 2 | k__Fungi(100);p__Ascomycota(100);p__Ascomycota_unclassified(100);p__Ascomycota_unclassified(100);p__Ascomycota_unclassified(100);p__Ascomycota_unclassified(100); | - | - | - | - | - | - | - | Unassigned | - |
| Otu12699 | 2 | k__Fungi(100);k__Fungi_unclassified(100);k__Fungi_unclassified(100);k__Fungi_unclassified(100);k__Fungi_unclassified(100);k__Fungi_unclassified(100); | - | - | - | - | - | - | - | Unassigned | - |
| Otu12703 | 2 | k__Fungi(100);p__Basidiomycota(100);p__Basidiomycota_unclassified(100);p__Basidiomycota_unclassified(100);p__Basidiomycota_unclassified(100);p__Basidiomycota_unclassified(100);p__Basidiomycota_unclassified(100); | - | - | - | - | - | - | - | Unassigned | - |
| Otu12704 | 2 | k__Fungi(100);k__Fungi_unclassified(100);k__Fungi_unclassified(100);k__Fungi_unclassified(100);k__Fungi_unclassified(100);k__Fungi_unclassified(100); | - | - | - | - | - | - | - | Unassigned | - |
| Otu12706 | 2 | k__Fungi(100);p__Ascomycota(100);p__Ascomycota_unclassified(100);p__Ascomycota_unclassified(100);p__Ascomycota_unclassified(100);p__Ascomycota_unclassified(100); | - | - | - | - | - | - | - | Unassigned | - |
| Otu12708 | 2 | k__Fungi(100);k__Fungi_unclassified(100);k__Fungi_unclassified(100);k__Fungi_unclassified(100);k__Fungi_unclassified(100);k__Fungi_unclassified(100); | - | - | - | - | - | - | - | Unassigned | - |
| Otu12714 | 2 | k__Fungi(100);k__Fungi_unclassified(100);k__Fungi_unclassified(100);k__Fungi_unclassified(100);k__Fungi_unclassified(100);k__Fungi_unclassified(100); | - | - | - | - | - | - | - | Unassigned | - |
| Otu12718 | 2 | k__Fungi(100);k__Fungi_unclassified(100);k__Fungi_unclassified(100);k__Fungi_unclassified(100);k__Fungi_unclassified(100);k__Fungi_unclassified(100); | - | - | - | - | - | - | - | Unassigned | - |
| Otu12719 | 2 | k__Fungi(100);k__Fungi_unclassified(100);k__Fungi_unclassified(100);k__Fungi_unclassified(100);k__Fungi_unclassified(100);k__Fungi_unclassified(100); | - | - | - | - | - | - | - | Unassigned | - |
| Otu12722 | 2 | k__Fungi(100);k__Fungi_unclassified(100);k__Fungi_unclassified(100);k__Fungi_unclassified(100);k__Fungi_unclassified(100);k__Fungi_unclassified(100); | - | - | - | - | - | - | - | Unassigned | - |
| Otu12725 | 2 | k__Fungi(100);k__Fungi_unclassified(100);k__Fungi_unclassified(100);k__Fungi_unclassified(100);k__Fungi_unclassified(100);k__Fungi_unclassified(100); | - | - | - | - | - | - | - | Unassigned | - |
| Otu12730 | 2 | k__Fungi(100);k__Fungi_unclassified(100);k__Fungi_unclassified(100);k__Fungi_unclassified(100);k__Fungi_unclassified(100);k__Fungi_unclassified(100); | - | - | - | - | - | - | - | Unassigned | - |
| Otu12734 | 2 | k__Fungi(100);k__Fungi_unclassified(100);k__Fungi_unclassified(100);k__Fungi_unclassified(100);k__Fungi_unclassified(100);k__Fungi_unclassified(100); | - | - | - | - | - | - | - | Unassigned | - |
| Otu12735 | 2 | k__Fungi(100);k__Fungi_unclassified(100);k__Fungi_unclassified(100);k__Fungi_unclassified(100);k__Fungi_unclassified(100);k__Fungi_unclassified(100); | - | - | - | - | - | - | - | Unassigned | - |
| Otu12736 | 2 | k__Fungi(100);k__Fungi_unclassified(100);k__Fungi_unclassified(100);k__Fungi_unclassified(100);k__Fungi_unclassified(100);k__Fungi_unclassified(100); | - | - | - | - | - | - | - | Unassigned | - |
| Otu12738 | 2 | k__Fungi(100);p__Basidiomycota(100);p__Basidiomycota_unclassified(100);p__Basidiomycota_unclassified(100);p__Basidiomycota_unclassified(100);p__Basidiomycota_unclassified(100);p__Basidiomycota_unclassified(100); | - | - | - | - | - | - | - | Unassigned | - |
| Otu12739 | 2 | k__Fungi(100);p__Ascomycota(100);p__Ascomycota_unclassified(100);p__Ascomycota_unclassified(100);p__Ascomycota_unclassified(100);p__Ascomycota_unclassified(100); | - | - | - | - | - | - | - | Unassigned | - |
| Otu12742 | 2 | k__Fungi(100);k__Fungi_unclassified(100);k__Fungi_unclassified(100);k__Fungi_unclassified(100);k__Fungi_unclassified(100);k__Fungi_unclassified(100); | - | - | - | - | - | - | - | Unassigned | - |
| Otu12743 | 2 | k__Fungi(100);p__Ascomycota(100);p__Ascomycota_unclassified(100);p__Ascomycota_unclassified(100);p__Ascomycota_unclassified(100);p__Ascomycota_unclassified(100); | - | - | - | - | - | - | - | Unassigned | - |
| Otu12748 | 2 | k__Fungi(100);p__Basidiomycota(100);p__Basidiomycota_unclassified(100);p__Basidiomycota_unclassified(100);p__Basidiomycota_unclassified(100);p__Basidiomycota_unclassified(100);p__Basidiomycota_unclassified(100); | - | - | - | - | - | - | - | Unassigned | - |
| Otu12749 | 2 | k__Fungi(100);k__Fungi_unclassified(100);k__Fungi_unclassified(100);k__Fungi_unclassified(100);k__Fungi_unclassified(100);k__Fungi_unclassified(100); | - | - | - | - | - | - | - | Unassigned | - |
| Otu12752 | 2 | k__Fungi(100);k__Fungi_unclassified(100);k__Fungi_unclassified(100);k__Fungi_unclassified(100);k__Fungi_unclassified(100);k__Fungi_unclassified(100); | - | - | - | - | - | - | - | Unassigned | - |
| Otu12754 | 2 | k__Fungi(100);p__Ascomycota(100);p__Ascomycota_unclassified(100);p__Ascomycota_unclassified(100);p__Ascomycota_unclassified(100);p__Ascomycota_unclassified(100); | - | - | - | - | - | - | - | Unassigned | - |
| Otu12757 | 2 | k__Fungi(100);p__Ascomycota(100);p__Ascomycota_unclassified(100);p__Ascomycota_unclassified(100);p__Ascomycota_unclassified(100);p__Ascomycota_unclassified(100); | - | - | - | - | - | - | - | Unassigned | - |
| Otu12759 | 2 | k__Fungi(100);k__Fungi_unclassified(100);k__Fungi_unclassified(100);k__Fungi_unclassified(100);k__Fungi_unclassified(100);k__Fungi_unclassified(100); | - | - | - | - | - | - | - | Unassigned | - |
| Otu12760 | 2 | k__Fungi(100);p__Ascomycota(100);p__Ascomycota_unclassified(100);p__Ascomycota_unclassified(100);p__Ascomycota_unclassified(100);p__Ascomycota_unclassified(100); | - | - | - | - | - | - | - | Unassigned | - |
| Otu12761 | 2 | k__Fungi(100);k__Fungi_unclassified(100);k__Fungi_unclassified(100);k__Fungi_unclassified(100);k__Fungi_unclassified(100);k__Fungi_unclassified(100); | - | - | - | - | - | - | - | Unassigned | - |
| Otu12762 | 2 | k__Fungi(100);p__Ascomycota(100);p__Ascomycota_unclassified(100);p__Ascomycota_unclassified(100);p__Ascomycota_unclassified(100);p__Ascomycota_unclassified(100); | - | - | - | - | - | - | - | Unassigned | - |
| Otu12764 | 2 | k__Fungi(100);p__Basidiomycota(100);c__Agaricomycetes(100);c__Agaricomycetes_unclassified(100);c__Agaricomycetes_unclassified(100);c__Agaricomycetes_unclassified(100);c__Agaricomycetes_unclassified(100); | - | - | - | - | - | - | - | Unassigned | - |
| Otu12765 | 2 | k__Fungi(100);p__Ascomycota(100);p__Ascomycota_unclassified(100);p__Ascomycota_unclassified(100);p__Ascomycota_unclassified(100);p__Ascomycota_unclassified(100); | - | - | - | - | - | - | - | Unassigned | - |
| Otu12766 | 2 | k__Fungi(100);k__Fungi_unclassified(100);k__Fungi_unclassified(100);k__Fungi_unclassified(100);k__Fungi_unclassified(100);k__Fungi_unclassified(100); | - | - | - | - | - | - | - | Unassigned | - |
| Otu12767 | 2 | k__Fungi(100);k__Fungi_unclassified(100);k__Fungi_unclassified(100);k__Fungi_unclassified(100);k__Fungi_unclassified(100);k__Fungi_unclassified(100); | - | - | - | - | - | - | - | Unassigned | - |
| Otu12769 | 2 | k__Fungi(100);k__Fungi_unclassified(100);k__Fungi_unclassified(100);k__Fungi_unclassified(100);k__Fungi_unclassified(100);k__Fungi_unclassified(100); | - | - | - | - | - | - | - | Unassigned | - |
| Otu12771 | 2 | k__Fungi(100);k__Fungi_unclassified(100);k__Fungi_unclassified(100);k__Fungi_unclassified(100);k__Fungi_unclassified(100);k__Fungi_unclassified(100); | - | - | - | - | - | - | - | Unassigned | - |
| Otu12773 | 2 | k__Fungi(100);k__Fungi_unclassified(100);k__Fungi_unclassified(100);k__Fungi_unclassified(100);k__Fungi_unclassified(100);k__Fungi_unclassified(100); | - | - | - | - | - | - | - | Unassigned | - |
| Otu12774 | 2 | k__Fungi(100);p__Ascomycota(100);p__Ascomycota_unclassified(100);p__Ascomycota_unclassified(100);p__Ascomycota_unclassified(100);p__Ascomycota_unclassified(100); | - | - | - | - | - | - | - | Unassigned | - |
| Otu12776 | 2 | k__Fungi(100);k__Fungi_unclassified(100);k__Fungi_unclassified(100);k__Fungi_unclassified(100);k__Fungi_unclassified(100);k__Fungi_unclassified(100); | - | - | - | - | - | - | - | Unassigned | - |
| Otu12777 | 2 | k__Fungi(100);k__Fungi_unclassified(100);k__Fungi_unclassified(100);k__Fungi_unclassified(100);k__Fungi_unclassified(100);k__Fungi_unclassified(100); | - | - | - | - | - | - | - | Unassigned | - |
| Otu12780 | 2 | k__Fungi(100);k__Fungi_unclassified(100);k__Fungi_unclassified(100);k__Fungi_unclassified(100);k__Fungi_unclassified(100);k__Fungi_unclassified(100); | - | - | - | - | - | - | - | Unassigned | - |
| Otu12781 | 2 | k__Fungi(100);k__Fungi_unclassified(100);k__Fungi_unclassified(100);k__Fungi_unclassified(100);k__Fungi_unclassified(100);k__Fungi_unclassified(100); | - | - | - | - | - | - | - | Unassigned | - |
| Otu12782 | 2 | k__Fungi(100);p__Ascomycota(100);c__Leotiomycetes(100);c__Leotiomycetes_unclassified(100);c__Leotiomycetes_unclassified(100);c__Leotiomycetes_unclassified(100);c__Leotiomycetes_unclassified(100); | - | - | - | - | - | - | - | Unassigned | - |
| Otu12783 | 2 | k__Fungi(100);p__Ascomycota(100);p__Ascomycota_unclassified(100);p__Ascomycota_unclassified(100);p__Ascomycota_unclassified(100);p__Ascomycota_unclassified(100); | - | - | - | - | - | - | - | Unassigned | - |
| Otu12785 | 2 | k__Fungi(100);k__Fungi_unclassified(100);k__Fungi_unclassified(100);k__Fungi_unclassified(100);k__Fungi_unclassified(100);k__Fungi_unclassified(100); | - | - | - | - | - | - | - | Unassigned | - |
| Otu12789 | 2 | k__Fungi(100);p__Ascomycota(100);p__Ascomycota_unclassified(100);p__Ascomycota_unclassified(100);p__Ascomycota_unclassified(100);p__Ascomycota_unclassified(100); | - | - | - | - | - | - | - | Unassigned | - |
| Otu12790 | 2 | k__Fungi(100);k__Fungi_unclassified(100);k__Fungi_unclassified(100);k__Fungi_unclassified(100);k__Fungi_unclassified(100);k__Fungi_unclassified(100); | - | - | - | - | - | - | - | Unassigned | - |
| Otu12791 | 2 | k__Fungi(100);k__Fungi_unclassified(100);k__Fungi_unclassified(100);k__Fungi_unclassified(100);k__Fungi_unclassified(100);k__Fungi_unclassified(100); | - | - | - | - | - | - | - | Unassigned | - |
| Otu12794 | 2 | k__Fungi(100);p__Ascomycota(100);c__Dothideomycetes(100);o__Capnodiales(100);o__Capnodiales_unclassified(100);o__Capnodiales_unclassified(100);o__Capnodiales_unclassified(100); | - | - | - | - | - | - | - | Unassigned | - |
| Otu12795 | 2 | k__Fungi(100);p__Ascomycota(100);c__Eurotiomycetes(100);o__Verrucariales(100);f__unclassified_Verrucariales(100);g__unclassified_Verrucariales(100);s__Verrucariales_sp(100); | - | - | - | - | - | - | - | Unassigned | - |
| Otu12799 | 2 | k__Fungi(100);k__Fungi_unclassified(100);k__Fungi_unclassified(100);k__Fungi_unclassified(100);k__Fungi_unclassified(100);k__Fungi_unclassified(100); | - | - | - | - | - | - | - | Unassigned | - |
| Otu12805 | 2 | k__Fungi(100);p__Basidiomycota(100);c__Microbotryomycetes(100);o__Sporidiobolales(100);f__Sporidiobolales_family_Incertae_sedis(100);g__unclassified_Sporidiobolales(100);s__Sporidiobolales_sp(100); | - | - | - | - | - | - | - | Unassigned | - |
| Otu12806 | 2 | k__Fungi(100);p__Ascomycota(100);c__Dothideomycetes(100);o__Capnodiales(100);f__Davidiellaceae(100);f__Davidiellaceae_unclassified(100);f__Davidiellaceae_unclassified(100); | - | - | - | - | - | - | - | Unassigned | - |
| Otu12807 | 2 | k__Fungi(100);k__Fungi_unclassified(100);k__Fungi_unclassified(100);k__Fungi_unclassified(100);k__Fungi_unclassified(100);k__Fungi_unclassified(100); | - | - | - | - | - | - | - | Unassigned | - |
| Otu12814 | 2 | k__Fungi(100);k__Fungi_unclassified(100);k__Fungi_unclassified(100);k__Fungi_unclassified(100);k__Fungi_unclassified(100);k__Fungi_unclassified(100); | - | - | - | - | - | - | - | Unassigned | - |
| Otu12816 | 2 | k__Fungi(100);k__Fungi_unclassified(100);k__Fungi_unclassified(100);k__Fungi_unclassified(100);k__Fungi_unclassified(100);k__Fungi_unclassified(100); | - | - | - | - | - | - | - | Unassigned | - |
| Otu12817 | 2 | k__Fungi(100);k__Fungi_unclassified(100);k__Fungi_unclassified(100);k__Fungi_unclassified(100);k__Fungi_unclassified(100);k__Fungi_unclassified(100); | - | - | - | - | - | - | - | Unassigned | - |
| Otu12820 | 2 | k__Fungi(100);p__Ascomycota(100);p__Ascomycota_unclassified(100);p__Ascomycota_unclassified(100);p__Ascomycota_unclassified(100);p__Ascomycota_unclassified(100); | - | - | - | - | - | - | - | Unassigned | - |
| Otu12822 | 2 | k__Fungi(100);k__Fungi_unclassified(100);k__Fungi_unclassified(100);k__Fungi_unclassified(100);k__Fungi_unclassified(100);k__Fungi_unclassified(100); | - | - | - | - | - | - | - | Unassigned | - |
| Otu12823 | 2 | k__Fungi(100);p__Ascomycota(100);p__Ascomycota_unclassified(100);p__Ascomycota_unclassified(100);p__Ascomycota_unclassified(100);p__Ascomycota_unclassified(100); | - | - | - | - | - | - | - | Unassigned | - |
| Otu12825 | 2 | k__Fungi(100);p__Ascomycota(100);c__Sordariomycetes(100);c__Sordariomycetes_unclassified(100);c__Sordariomycetes_unclassified(100);c__Sordariomycetes_unclassified(100);c__Sordariomycetes_unclassified(100); | - | - | - | - | - | - | - | Unassigned | - |
| Otu12833 | 2 | k__Fungi(100);k__Fungi_unclassified(100);k__Fungi_unclassified(100);k__Fungi_unclassified(100);k__Fungi_unclassified(100);k__Fungi_unclassified(100); | - | - | - | - | - | - | - | Unassigned | - |
| Otu12834 | 2 | k__Fungi(100);k__Fungi_unclassified(100);k__Fungi_unclassified(100);k__Fungi_unclassified(100);k__Fungi_unclassified(100);k__Fungi_unclassified(100); | - | - | - | - | - | - | - | Unassigned | - |
| Otu12835 | 2 | k__Fungi(100);k__Fungi_unclassified(100);k__Fungi_unclassified(100);k__Fungi_unclassified(100);k__Fungi_unclassified(100);k__Fungi_unclassified(100); | - | - | - | - | - | - | - | Unassigned | - |
| Otu12838 | 2 | k__Fungi(100);k__Fungi_unclassified(100);k__Fungi_unclassified(100);k__Fungi_unclassified(100);k__Fungi_unclassified(100);k__Fungi_unclassified(100); | - | - | - | - | - | - | - | Unassigned | - |
| Otu12839 | 2 | k__Fungi(100);p__Ascomycota(100);p__Ascomycota_unclassified(100);p__Ascomycota_unclassified(100);p__Ascomycota_unclassified(100);p__Ascomycota_unclassified(100); | - | - | - | - | - | - | - | Unassigned | - |
| Otu12840 | 2 | k__Fungi(100);k__Fungi_unclassified(100);k__Fungi_unclassified(100);k__Fungi_unclassified(100);k__Fungi_unclassified(100);k__Fungi_unclassified(100); | - | - | - | - | - | - | - | Unassigned | - |
| Otu12845 | 2 | k__Fungi(100);k__Fungi_unclassified(100);k__Fungi_unclassified(100);k__Fungi_unclassified(100);k__Fungi_unclassified(100);k__Fungi_unclassified(100); | - | - | - | - | - | - | - | Unassigned | - |
| Otu12850 | 2 | k__Fungi(100);k__Fungi_unclassified(100);k__Fungi_unclassified(100);k__Fungi_unclassified(100);k__Fungi_unclassified(100);k__Fungi_unclassified(100); | - | - | - | - | - | - | - | Unassigned | - |
| Otu12851 | 2 | k__Fungi(100);p__Ascomycota(100);p__Ascomycota_unclassified(100);p__Ascomycota_unclassified(100);p__Ascomycota_unclassified(100);p__Ascomycota_unclassified(100); | - | - | - | - | - | - | - | Unassigned | - |
| Otu12853 | 2 | k__Fungi(100);k__Fungi_unclassified(100);k__Fungi_unclassified(100);k__Fungi_unclassified(100);k__Fungi_unclassified(100);k__Fungi_unclassified(100); | - | - | - | - | - | - | - | Unassigned | - |
| Otu12858 | 2 | k__Fungi(100);k__Fungi_unclassified(100);k__Fungi_unclassified(100);k__Fungi_unclassified(100);k__Fungi_unclassified(100);k__Fungi_unclassified(100); | - | - | - | - | - | - | - | Unassigned | - |
| Otu12861 | 2 | k__Fungi(100);p__Ascomycota(100);p__Ascomycota_unclassified(100);p__Ascomycota_unclassified(100);p__Ascomycota_unclassified(100);p__Ascomycota_unclassified(100); | - | - | - | - | - | - | - | Unassigned | - |
| Otu12862 | 2 | k__Fungi(100);k__Fungi_unclassified(100);k__Fungi_unclassified(100);k__Fungi_unclassified(100);k__Fungi_unclassified(100);k__Fungi_unclassified(100); | - | - | - | - | - | - | - | Unassigned | - |
| Otu12863 | 2 | k__Fungi(100);p__Ascomycota(100);c__Eurotiomycetes(100);o__Chaetothyriales(100);o__Chaetothyriales_unclassified(100);o__Chaetothyriales_unclassified(100);o__Chaetothyriales_unclassified(100); | - | - | - | - | - | - | - | Unassigned | - |
| Otu12866 | 2 | k__Fungi(100);p__Ascomycota(100);p__Ascomycota_unclassified(100);p__Ascomycota_unclassified(100);p__Ascomycota_unclassified(100);p__Ascomycota_unclassified(100); | - | - | - | - | - | - | - | Unassigned | - |
| Otu12869 | 2 | k__Fungi(100);p__Ascomycota(100);p__Ascomycota_unclassified(100);p__Ascomycota_unclassified(100);p__Ascomycota_unclassified(100);p__Ascomycota_unclassified(100); | - | - | - | - | - | - | - | Unassigned | - |
| Otu12871 | 2 | k__Fungi(100);k__Fungi_unclassified(100);k__Fungi_unclassified(100);k__Fungi_unclassified(100);k__Fungi_unclassified(100);k__Fungi_unclassified(100); | - | - | - | - | - | - | - | Unassigned | - |
| Otu12876 | 2 | k__Fungi(100);p__Ascomycota(100);p__Ascomycota_unclassified(100);p__Ascomycota_unclassified(100);p__Ascomycota_unclassified(100);p__Ascomycota_unclassified(100); | - | - | - | - | - | - | - | Unassigned | - |
| Otu12877 | 2 | k__Fungi(100);k__Fungi_unclassified(100);k__Fungi_unclassified(100);k__Fungi_unclassified(100);k__Fungi_unclassified(100);k__Fungi_unclassified(100); | - | - | - | - | - | - | - | Unassigned | - |

| | | | | | | | | | | | |
|---|---|---|---|---|---|---|---|---|---|---|---|
| Otu12880 | 2 | k__Fungi(100);k__Fungi_unclassified(100);k__Fungi_unclassified(100);k__Fungi_unclassified(100);k__Fungi_unclassified(100);k__Fungi_unclassified(100);k__Fungi_unclassified(100); | - | - | - | - | - | - | - | Unassigned | - |
| Otu12882 | 2 | k__Fungi(100);p__Basidiomycota(100);p__Basidiomycota_unclassified(100);p__Basidiomycota_unclassified(100);p__Basidiomycota_unclassified(100);p__Basidiomycota_unclassified(100);p__Basidiomycota_unclassified(100); | - | - | - | - | - | - | - | Unassigned | - |
| Otu12886 | 2 | k__Fungi(100);p__Ascomycota(100);p__Ascomycota_unclassified(100);p__Ascomycota_unclassified(100);p__Ascomycota_unclassified(100);p__Ascomycota_unclassified(100);p__Ascomycota_unclassified(100); | - | - | - | - | - | - | - | Unassigned | - |
| Otu12887 | 2 | k__Fungi(100);k__Fungi_unclassified(100);k__Fungi_unclassified(100);k__Fungi_unclassified(100);k__Fungi_unclassified(100);k__Fungi_unclassified(100);k__Fungi_unclassified(100); | - | - | - | - | - | - | - | Unassigned | - |
| Otu12888 | 2 | k__Fungi(100);p__Ascomycota(100);p__Ascomycota_unclassified(100);p__Ascomycota_unclassified(100);p__Ascomycota_unclassified(100);p__Ascomycota_unclassified(100);p__Ascomycota_unclassified(100); | - | - | - | - | - | - | - | Unassigned | - |
| Otu12889 | 2 | k__Fungi(100);p__Ascomycota(100);p__Ascomycota_unclassified(100);p__Ascomycota_unclassified(100);p__Ascomycota_unclassified(100);p__Ascomycota_unclassified(100);p__Ascomycota_unclassified(100); | - | - | - | - | - | - | - | Unassigned | - |
| Otu12891 | 2 | k__Fungi(100);k__Fungi_unclassified(100);k__Fungi_unclassified(100);k__Fungi_unclassified(100);k__Fungi_unclassified(100);k__Fungi_unclassified(100);k__Fungi_unclassified(100); | - | - | - | - | - | - | - | Unassigned | - |
| Otu12895 | 2 | k__Fungi(100);p__Ascomycota(100);p__Ascomycota_unclassified(100);p__Ascomycota_unclassified(100);p__Ascomycota_unclassified(100);p__Ascomycota_unclassified(100);p__Ascomycota_unclassified(100); | - | - | - | - | - | - | - | Unassigned | - |
| Otu12896 | 2 | k__Fungi(100);k__Fungi_unclassified(100);k__Fungi_unclassified(100);k__Fungi_unclassified(100);k__Fungi_unclassified(100);k__Fungi_unclassified(100);k__Fungi_unclassified(100); | - | - | - | - | - | - | - | Unassigned | - |
| Otu12899 | 2 | k__Fungi(100);p__Ascomycota(100);p__Ascomycota_unclassified(100);p__Ascomycota_unclassified(100);p__Ascomycota_unclassified(100);p__Ascomycota_unclassified(100);p__Ascomycota_unclassified(100); | - | - | - | - | - | - | - | Unassigned | - |
| Otu12900 | 2 | k__Fungi(100);p__Basidiomycota(100);p__Basidiomycota_unclassified(100);p__Basidiomycota_unclassified(100);p__Basidiomycota_unclassified(100);p__Basidiomycota_unclassified(100);p__Basidiomycota_unclassified(100); | - | - | - | - | - | - | - | Unassigned | - |
| Otu12901 | 2 | k__Fungi(100);p__Basidiomycota(100);p__Basidiomycota_unclassified(100);p__Basidiomycota_unclassified(100);p__Basidiomycota_unclassified(100);p__Basidiomycota_unclassified(100);p__Basidiomycota_unclassified(100); | - | - | - | - | - | - | - | Unassigned | - |
| Otu12903 | 2 | k__Fungi(100);k__Fungi_unclassified(100);k__Fungi_unclassified(100);k__Fungi_unclassified(100);k__Fungi_unclassified(100);k__Fungi_unclassified(100);k__Fungi_unclassified(100); | - | - | - | - | - | - | - | Unassigned | - |
| Otu12905 | 2 | k__Fungi(100);p__Ascomycota(100);p__Ascomycota_unclassified(100);p__Ascomycota_unclassified(100);p__Ascomycota_unclassified(100);p__Ascomycota_unclassified(100);p__Ascomycota_unclassified(100); | - | - | - | - | - | - | - | Unassigned | - |
| Otu12908 | 2 | k__Fungi(100);k__Fungi_unclassified(100);k__Fungi_unclassified(100);k__Fungi_unclassified(100);k__Fungi_unclassified(100);k__Fungi_unclassified(100);k__Fungi_unclassified(100); | - | - | - | - | - | - | - | Unassigned | - |
| Otu12911 | 2 | k__Fungi(100);p__Ascomycota(100);p__Ascomycota_unclassified(100);p__Ascomycota_unclassified(100);p__Ascomycota_unclassified(100);p__Ascomycota_unclassified(100);p__Ascomycota_unclassified(100); | - | - | - | - | - | - | - | Unassigned | - |
| Otu12912 | 2 | k__Fungi(100);k__Fungi_unclassified(100);k__Fungi_unclassified(100);k__Fungi_unclassified(100);k__Fungi_unclassified(100);k__Fungi_unclassified(100);k__Fungi_unclassified(100); | - | - | - | - | - | - | - | Unassigned | - |
| Otu12914 | 2 | k__Fungi(100);p__Ascomycota(100);p__Ascomycota_unclassified(100);p__Ascomycota_unclassified(100);p__Ascomycota_unclassified(100);p__Ascomycota_unclassified(100);p__Ascomycota_unclassified(100); | - | - | - | - | - | - | - | Unassigned | - |
| Otu12923 | 2 | k__Fungi(100);k__Fungi_unclassified(100);k__Fungi_unclassified(100);k__Fungi_unclassified(100);k__Fungi_unclassified(100);k__Fungi_unclassified(100);k__Fungi_unclassified(100); | - | - | - | - | - | - | - | Unassigned | - |
| Otu12924 | 2 | k__Fungi(100);k__Fungi_unclassified(100);k__Fungi_unclassified(100);k__Fungi_unclassified(100);k__Fungi_unclassified(100);k__Fungi_unclassified(100);k__Fungi_unclassified(100); | - | - | - | - | - | - | - | Unassigned | - |
| Otu12927 | 2 | k__Fungi(100);p__Basidiomycota(100);p__Basidiomycota_unclassified(100);p__Basidiomycota_unclassified(100);p__Basidiomycota_unclassified(100);p__Basidiomycota_unclassified(100);p__Basidiomycota_unclassified(100); | - | - | - | - | - | - | - | Unassigned | - |
| Otu12928 | 2 | k__Fungi(100);k__Fungi_unclassified(100);k__Fungi_unclassified(100);k__Fungi_unclassified(100);k__Fungi_unclassified(100);k__Fungi_unclassified(100);k__Fungi_unclassified(100); | - | - | - | - | - | - | - | Unassigned | - |
| Otu12929 | 2 | k__Fungi(100);k__Fungi_unclassified(100);k__Fungi_unclassified(100);k__Fungi_unclassified(100);k__Fungi_unclassified(100);k__Fungi_unclassified(100);k__Fungi_unclassified(100); | - | - | - | - | - | - | - | Unassigned | - |
| Otu12930 | 2 | k__Fungi(100);k__Fungi_unclassified(100);k__Fungi_unclassified(100);k__Fungi_unclassified(100);k__Fungi_unclassified(100);k__Fungi_unclassified(100);k__Fungi_unclassified(100); | - | - | - | - | - | - | - | Unassigned | - |
| Otu12931 | 2 | k__Fungi(100);p__Ascomycota(100);c__Sordariomycetes(100);c__Sordariomycetes_unclassified(100);c__Sordariomycetes_unclassified(100);c__Sordariomycetes_unclassified(100);c__Sordariomycetes_unclassified(100); | - | - | - | - | - | - | - | Unassigned | - |
| Otu12932 | 2 | k__Fungi(100);k__Fungi_unclassified(100);k__Fungi_unclassified(100);k__Fungi_unclassified(100);k__Fungi_unclassified(100);k__Fungi_unclassified(100);k__Fungi_unclassified(100); | - | - | - | - | - | - | - | Unassigned | - |
| Otu12940 | 2 | k__Fungi(100);k__Fungi_unclassified(100);k__Fungi_unclassified(100);k__Fungi_unclassified(100);k__Fungi_unclassified(100);k__Fungi_unclassified(100);k__Fungi_unclassified(100); | - | - | - | - | - | - | - | Unassigned | - |
| Otu12941 | 2 | k__Fungi(100);p__Basidiomycota(100);c__Agaricomycetes(100);c__Agaricomycetes_unclassified(100);c__Agaricomycetes_unclassified(100);c__Agaricomycetes_unclassified(100);c__Agaricomycetes_unclassified(100); | - | - | - | - | - | - | - | Unassigned | - |
| Otu12942 | 2 | k__Fungi(100);p__Basidiomycota(100);c__Agaricomycetes(100);o__Polyporales(100);o__Polyporales_unclassified(100);o__Polyporales_unclassified(100);o__Polyporales_unclassified(100); | - | - | - | - | - | - | - | Unassigned | - |
| Otu12943 | 2 | k__Fungi(100);k__Fungi_unclassified(100);k__Fungi_unclassified(100);k__Fungi_unclassified(100);k__Fungi_unclassified(100);k__Fungi_unclassified(100);k__Fungi_unclassified(100); | - | - | - | - | - | - | - | Unassigned | - |
| Otu12944 | 2 | k__Fungi(100);k__Fungi_unclassified(100);k__Fungi_unclassified(100);k__Fungi_unclassified(100);k__Fungi_unclassified(100);k__Fungi_unclassified(100);k__Fungi_unclassified(100); | - | - | - | - | - | - | - | Unassigned | - |
| Otu12945 | 2 | k__Fungi(100);k__Fungi_unclassified(100);k__Fungi_unclassified(100);k__Fungi_unclassified(100);k__Fungi_unclassified(100);k__Fungi_unclassified(100);k__Fungi_unclassified(100); | - | - | - | - | - | - | - | Unassigned | - |
| Otu12947 | 2 | k__Fungi(100);p__Basidiomycota(100);p__Basidiomycota_unclassified(100);p__Basidiomycota_unclassified(100);p__Basidiomycota_unclassified(100);p__Basidiomycota_unclassified(100);p__Basidiomycota_unclassified(100); | - | - | - | - | - | - | - | Unassigned | - |
| Otu12948 | 2 | k__Fungi(100);p__Ascomycota(100);p__Ascomycota_unclassified(100);p__Ascomycota_unclassified(100);p__Ascomycota_unclassified(100);p__Ascomycota_unclassified(100);p__Ascomycota_unclassified(100); | - | - | - | - | - | - | - | Unassigned | - |
| Otu12949 | 2 | k__Fungi(100);k__Fungi_unclassified(100);k__Fungi_unclassified(100);k__Fungi_unclassified(100);k__Fungi_unclassified(100);k__Fungi_unclassified(100);k__Fungi_unclassified(100); | - | - | - | - | - | - | - | Unassigned | - |
| Otu12950 | 2 | k__Fungi(100);k__Fungi_unclassified(100);k__Fungi_unclassified(100);k__Fungi_unclassified(100);k__Fungi_unclassified(100);k__Fungi_unclassified(100);k__Fungi_unclassified(100); | - | - | - | - | - | - | - | Unassigned | - |
| Otu12952 | 2 | k__Fungi(100);k__Fungi_unclassified(100);k__Fungi_unclassified(100);k__Fungi_unclassified(100);k__Fungi_unclassified(100);k__Fungi_unclassified(100);k__Fungi_unclassified(100); | - | - | - | - | - | - | - | Unassigned | - |
| Otu12961 | 2 | k__Fungi(100);k__Fungi_unclassified(100);k__Fungi_unclassified(100);k__Fungi_unclassified(100);k__Fungi_unclassified(100);k__Fungi_unclassified(100);k__Fungi_unclassified(100); | - | - | - | - | - | - | - | Unassigned | - |
| Otu12963 | 2 | k__Fungi(100);k__Fungi_unclassified(100);k__Fungi_unclassified(100);k__Fungi_unclassified(100);k__Fungi_unclassified(100);k__Fungi_unclassified(100);k__Fungi_unclassified(100); | - | - | - | - | - | - | - | Unassigned | - |
| Otu12964 | 2 | k__Fungi(100);k__Fungi_unclassified(100);k__Fungi_unclassified(100);k__Fungi_unclassified(100);k__Fungi_unclassified(100);k__Fungi_unclassified(100);k__Fungi_unclassified(100); | - | - | - | - | - | - | - | Unassigned | - |
| Otu12965 | 2 | k__Fungi(100);p__Basidiomycota(100);c__Agaricostilbomycetes(100);o__Agaricostilbales(100);o__Agaricostilbales_unclassified(100);o__Agaricostilbales_unclassified(100);o__Agaricostilbales_unclassified(100); | - | - | - | - | - | - | - | Unassigned | - |
| Otu12966 | 1 | k__Fungi(100);p__Ascomycota(100);c__Saccharomycetes(100);o__Saccharomycetales(100);f__Metschnikowiaceae(100);g__Metschnikowia(100);s__Metschnikowia_pulcherrima(100); | Metschnikowia | 13 | Saprotroph | \|Undefined Saprotroph\| | Yeast | NULL | Probable | As nectar-tap_saprotroph (Põlme et al. 2020); Decay sugar-rich substrates (Põlme et al. 2020) | Põlme S, et al. 2020. Fungal Diversity 105:1-16 ((https://doi.org/10.1007/s13225-020-00466-2)) |
| Otu12967 | 1 | k__Fungi(100);p__Ascomycota(100);c__Saccharomycetes(100);o__Saccharomycetales(100);f__Metschnikowiaceae(100);g__Metschnikowia(100);s__Metschnikowia_pulcherrima(100); | Metschnikowia | 13 | Saprotroph | \|Undefined Saprotroph\| | Yeast | NULL | Probable | As nectar-tap_saprotroph (Põlme et al. 2020); Decay sugar-rich substrates (Põlme et al. 2020) | Põlme S, et al. 2020. Fungal Diversity 105:1-16 ((https://doi.org/10.1007/s13225-020-00466-2)) |
| Otu12970 | 1 | k__Fungi(100);p__Ascomycota(100);c__Saccharomycetes(100);o__Saccharomycetales(100);f__Metschnikowiaceae(100);g__Metschnikowia(100);g__Metschnikowia_unclassified(100); | Metschnikowia | 13 | Saprotroph | \|Undefined Saprotroph\| | Yeast | NULL | Probable | As nectar-tap_saprotroph (Põlme et al. 2020); Decay sugar-rich substrates (Põlme et al. 2020) | Põlme S, et al. 2020. Fungal Diversity 105:1-16 ((https://doi.org/10.1007/s13225-020-00466-2)) |
| Otu12972 | 1 | k__Fungi(100);p__Ascomycota(100);c__Saccharomycetes(100);o__Saccharomycetales(100);f__Metschnikowiaceae(100);g__Metschnikowia(100);s__Metschnikowia_pulcherrima(100); | Metschnikowia | 13 | Saprotroph | \|Undefined Saprotroph\| | Yeast | NULL | Probable | As nectar-tap_saprotroph (Põlme et al. 2020); Decay sugar-rich substrates (Põlme et al. 2020) | Põlme S, et al. 2020. Fungal Diversity 105:1-16 ((https://doi.org/10.1007/s13225-020-00466-2)) |
| Otu12973 | 1 | k__Fungi(100);p__Ascomycota(100);c__Saccharomycetes(100);o__Saccharomycetales(100);f__Metschnikowiaceae(100);g__Metschnikowia(100);s__Metschnikowia_pulcherrima(100); | Metschnikowia | 13 | Saprotroph | \|Undefined Saprotroph\| | Yeast | NULL | Probable | As nectar-tap_saprotroph (Põlme et al. 2020); Decay sugar-rich substrates (Põlme et al. 2020) | Põlme S, et al. 2020. Fungal Diversity 105:1-16 ((https://doi.org/10.1007/s13225-020-00466-2)) |
| Otu12976 | 1 | k__Fungi(100);p__Ascomycota(100);c__Saccharomycetes(100);o__Saccharomycetales(100);f__Metschnikowiaceae(100);g__Metschnikowia(100);s__Metschnikowia_pulcherrima(100); | Metschnikowia | 13 | Saprotroph | \|Undefined Saprotroph\| | Yeast | NULL | Probable | As nectar-tap_saprotroph (Põlme et al. 2020); Decay sugar-rich substrates (Põlme et al. 2020) | Põlme S, et al. 2020. Fungal Diversity 105:1-16 ((https://doi.org/10.1007/s13225-020-00466-2)) |
| Otu12977 | 1 | k__Fungi(100);p__Ascomycota(100);c__Saccharomycetes(100);o__Saccharomycetales(100);f__Metschnikowiaceae(100);g__Metschnikowia(100);s__Metschnikowia_pulcherrima(100); | Metschnikowia | 13 | Saprotroph | \|Undefined Saprotroph\| | Yeast | NULL | Probable | As nectar-tap_saprotroph (Põlme et al. 2020); Decay sugar-rich substrates (Põlme et al. 2020) | Põlme S, et al. 2020. Fungal Diversity 105:1-16 ((https://doi.org/10.1007/s13225-020-00466-2)) |
| Otu12978 | 1 | k__Fungi(100);p__Ascomycota(100);c__Saccharomycetes(100);o__Saccharomycetales(100);f__Metschnikowiaceae(100);g__Metschnikowia(100);s__Metschnikowia_pulcherrima(100); | Metschnikowia | 13 | Saprotroph | \|Undefined Saprotroph\| | Yeast | NULL | Probable | As nectar-tap_saprotroph (Põlme et al. 2020); Decay sugar-rich substrates (Põlme et al. 2020) | Põlme S, et al. 2020. Fungal Diversity 105:1-16 ((https://doi.org/10.1007/s13225-020-00466-2)) |
| Otu12983 | 1 | k__Fungi(100);p__Ascomycota(100);c__Saccharomycetes(100);o__Saccharomycetales(100);f__Metschnikowiaceae(100);g__Metschnikowia(100);s__Metschnikowia_pulcherrima(100); | Metschnikowia | 13 | Saprotroph | \|Undefined Saprotroph\| | Yeast | NULL | Probable | As nectar-tap_saprotroph (Põlme et al. 2020); Decay sugar-rich substrates (Põlme et al. 2020) | Põlme S, et al. 2020. Fungal Diversity 105:1-16 ((https://doi.org/10.1007/s13225-020-00466-2)) |
| Otu12984 | 1 | k__Fungi(100);p__Ascomycota(100);c__Saccharomycetes(100);o__Saccharomycetales(100);f__Metschnikowiaceae(100);g__Metschnikowia(100);s__Metschnikowia_pulcherrima(100); | Metschnikowia | 13 | Saprotroph | \|Undefined Saprotroph\| | Yeast | NULL | Probable | As nectar-tap_saprotroph (Põlme et al. 2020); Decay sugar-rich substrates (Põlme et al. 2020) | Põlme S, et al. 2020. Fungal Diversity 105:1-16 ((https://doi.org/10.1007/s13225-020-00466-2)) |
| Otu12987 | 1 | k__Fungi(100);p__Ascomycota(100);c__Saccharomycetes(100);o__Saccharomycetales(100);f__Metschnikowiaceae(100);g__Metschnikowia(100);s__Metschnikowia_pulcherrima(100); | Metschnikowia | 13 | Saprotroph | \|Undefined Saprotroph\| | Yeast | NULL | Probable | As nectar-tap_saprotroph (Põlme et al. 2020); Decay sugar-rich substrates (Põlme et al. 2020) | Põlme S, et al. 2020. Fungal Diversity 105:1-16 ((https://doi.org/10.1007/s13225-020-00466-2)) |
| Otu12991 | 1 | k__Fungi(100);p__Ascomycota(100);c__Saccharomycetes(100);o__Saccharomycetales(100);f__Metschnikowiaceae(100);g__Metschnikowia(100);g__Metschnikowia_unclassified(100); | Metschnikowia | 13 | Saprotroph | \|Undefined Saprotroph\| | Yeast | NULL | Probable | As nectar-tap_saprotroph (Põlme et al. 2020); Decay sugar-rich substrates (Põlme et al. 2020) | Põlme S, et al. 2020. Fungal Diversity 105:1-16 ((https://doi.org/10.1007/s13225-020-00466-2)) |
| Otu12993 | 1 | k__Fungi(100);p__Ascomycota(100);c__Saccharomycetes(100);o__Saccharomycetales(100);f__Metschnikowiaceae(100);g__Metschnikowia(100);s__Metschnikowia_pulcherrima(100); | Metschnikowia | 13 | Saprotroph | \|Undefined Saprotroph\| | Yeast | NULL | Probable | As nectar-tap_saprotroph (Põlme et al. 2020); Decay sugar-rich substrates (Põlme et al. 2020) | Põlme S, et al. 2020. Fungal Diversity 105:1-16 ((https://doi.org/10.1007/s13225-020-00466-2)) |
| Otu12994 | 1 | k__Fungi(100);p__Ascomycota(100);c__Saccharomycetes(100);o__Saccharomycetales(100);f__Metschnikowiaceae(100);g__Metschnikowia(100);s__Metschnikowia_pulcherrima(100); | Metschnikowia | 13 | Saprotroph | \|Undefined Saprotroph\| | Yeast | NULL | Probable | As nectar-tap_saprotroph (Põlme et al. 2020); Decay sugar-rich substrates (Põlme et al. 2020) | Põlme S, et al. 2020. Fungal Diversity 105:1-16 ((https://doi.org/10.1007/s13225-020-00466-2)) |
| Otu12995 | 1 | k__Fungi(100);p__Ascomycota(100);c__Saccharomycetes(100);o__Saccharomycetales(100);f__Metschnikowiaceae(100);g__Metschnikowia(100);s__Metschnikowia_pulcherrima(100); | Metschnikowia | 13 | Saprotroph | \|Undefined Saprotroph\| | Yeast | NULL | Probable | As nectar-tap_saprotroph (Põlme et al. 2020); Decay sugar-rich substrates (Põlme et al. 2020) | Põlme S, et al. 2020. Fungal Diversity 105:1-16 ((https://doi.org/10.1007/s13225-020-00466-2)) |
| Otu12997 | 1 | k__Fungi(100);p__Ascomycota(100);c__Saccharomycetes(100);o__Saccharomycetales(100);f__Metschnikowiaceae(100);g__Metschnikowia(100);s__Metschnikowia_pulcherrima(100); | Metschnikowia | 13 | Saprotroph | \|Undefined Saprotroph\| | Yeast | NULL | Probable | As nectar-tap_saprotroph (Põlme et al. 2020); Decay sugar-rich substrates (Põlme et al. 2020) | Põlme S, et al. 2020. Fungal Diversity 105:1-16 ((https://doi.org/10.1007/s13225-020-00466-2)) |
| Otu12998 | 1 | k__Fungi(100);p__Ascomycota(100);c__Saccharomycetes(100);o__Saccharomycetales(100);f__Metschnikowiaceae(100);g__Metschnikowia(100);g__Metschnikowia_unclassified(100); | Metschnikowia | 13 | Saprotroph | \|Undefined Saprotroph\| | Yeast | NULL | Probable | As nectar-tap_saprotroph (Põlme et al. 2020); Decay sugar-rich substrates (Põlme et al. 2020) | Põlme S, et al. 2020. Fungal Diversity 105:1-16 ((https://doi.org/10.1007/s13225-020-00466-2)) |
| Otu12999 | 1 | k__Fungi(100);p__Ascomycota(100);c__Saccharomycetes(100);o__Saccharomycetales(100);f__Metschnikowiaceae(100);g__Metschnikowia(100);s__Metschnikowia_pulcherrima(100); | Metschnikowia | 13 | Saprotroph | \|Undefined Saprotroph\| | Yeast | NULL | Probable | As nectar-tap_saprotroph (Põlme et al. 2020); Decay sugar-rich substrates (Põlme et al. 2020) | Põlme S, et al. 2020. Fungal Diversity 105:1-16 ((https://doi.org/10.1007/s13225-020-00466-2)) |
| Otu13000 | 1 | k__Fungi(100);p__Ascomycota(100);c__Saccharomycetes(100);o__Saccharomycetales(100);f__Metschnikowiaceae(100);g__Metschnikowia(100);s__Metschnikowia_pulcherrima(100); | Metschnikowia | 13 | Saprotroph | \|Undefined Saprotroph\| | Yeast | NULL | Probable | As nectar-tap_saprotroph (Põlme et al. 2020); Decay sugar-rich substrates (Põlme et al. 2020) | Põlme S, et al. 2020. Fungal Diversity 105:1-16 ((https://doi.org/10.1007/s13225-020-00466-2)) |
| Otu13001 | 1 | k__Fungi(100);p__Ascomycota(100);c__Saccharomycetes(100);o__Saccharomycetales(100);f__Metschnikowiaceae(100);g__Metschnikowia(100);g__Metschnikowia_unclassified(100); | Metschnikowia | 13 | Saprotroph | \|Undefined Saprotroph\| | Yeast | NULL | Probable | As nectar-tap_saprotroph (Põlme et al. 2020); Decay sugar-rich substrates (Põlme et al. 2020) | Põlme S, et al. 2020. Fungal Diversity 105:1-16 ((https://doi.org/10.1007/s13225-020-00466-2)) |
| Otu13002 | 1 | k__Fungi(100);p__Ascomycota(100);c__Saccharomycetes(100);o__Saccharomycetales(100);f__Metschnikowiaceae(100);g__Metschnikowia(100);s__Metschnikowia_pulcherrima(100); | Metschnikowia | 13 | Saprotroph | \|Undefined Saprotroph\| | Yeast | NULL | Probable | As nectar-tap_saprotroph (Põlme et al. 2020); Decay sugar-rich substrates (Põlme et al. 2020) | Põlme S, et al. 2020. Fungal Diversity 105:1-16 ((https://doi.org/10.1007/s13225-020-00466-2)) |
| Otu13003 | 1 | k__Fungi(100);p__Ascomycota(100);c__Saccharomycetes(100);o__Saccharomycetales(100);f__Metschnikowiaceae(100);g__Metschnikowia(100);s__Metschnikowia_pulcherrima(100); | Metschnikowia | 13 | Saprotroph | \|Undefined Saprotroph\| | Yeast | NULL | Probable | As nectar-tap_saprotroph (Põlme et al. 2020); Decay sugar-rich substrates (Põlme et al. 2020) | Põlme S, et al. 2020. Fungal Diversity 105:1-16 ((https://doi.org/10.1007/s13225-020-00466-2)) |
| Otu13005 | 1 | k__Fungi(100);p__Ascomycota(100);c__Saccharomycetes(100);o__Saccharomycetales(100);f__Metschnikowiaceae(100);g__Metschnikowia(100);s__Metschnikowia_pulcherrima(100); | Metschnikowia | 13 | Saprotroph | \|Undefined Saprotroph\| | Yeast | NULL | Probable | As nectar-tap_saprotroph (Põlme et al. 2020); Decay sugar-rich substrates (Põlme et al. 2020) | Põlme S, et al. 2020. Fungal Diversity 105:1-16 ((https://doi.org/10.1007/s13225-020-00466-2)) |
| Otu13006 | 1 | k__Fungi(100);p__Ascomycota(100);c__Saccharomycetes(100);o__Saccharomycetales(100);f__Metschnikowiaceae(100);g__Metschnikowia(100);s__Metschnikowia_pulcherrima(100); | Metschnikowia | 13 | Saprotroph | \|Undefined Saprotroph\| | Yeast | NULL | Probable | As nectar-tap_saprotroph (Põlme et al. 2020); Decay sugar-rich substrates (Põlme et al. 2020) | Põlme S, et al. 2020. Fungal Diversity 105:1-16 ((https://doi.org/10.1007/s13225-020-00466-2)) |
| Otu13008 | 1 | k__Fungi(100);p__Ascomycota(100);c__Saccharomycetes(100);o__Saccharomycetales(100);f__Metschnikowiaceae(100);g__Metschnikowia(100);g__Metschnikowia_unclassified(100); | Metschnikowia | 13 | Saprotroph | \|Undefined Saprotroph\| | Yeast | NULL | Probable | As nectar-tap_saprotroph (Põlme et al. 2020); Decay sugar-rich substrates (Põlme et al. 2020) | Põlme S, et al. 2020. Fungal Diversity 105:1-16 ((https://doi.org/10.1007/s13225-020-00466-2)) |
| Otu13011 | 1 | k__Fungi(100);p__Ascomycota(100);c__Saccharomycetes(100);o__Saccharomycetales(100);f__Metschnikowiaceae(100);g__Metschnikowia(100);s__Metschnikowia_pulcherrima(100); | Metschnikowia | 13 | Saprotroph | \|Undefined Saprotroph\| | Yeast | NULL | Probable | As nectar-tap_saprotroph (Põlme et al. 2020); Decay sugar-rich substrates (Põlme et al. 2020) | Põlme S, et al. 2020. Fungal Diversity 105:1-16 ((https://doi.org/10.1007/s13225-020-00466-2)) |
| Otu13012 | 1 | k__Fungi(100);p__Ascomycota(100);c__Saccharomycetes(100);o__Saccharomycetales(100);f__Metschnikowiaceae(100);g__Metschnikowia(100);s__Metschnikowia_pulcherrima(100); | Metschnikowia | 13 | Saprotroph | \|Undefined Saprotroph\| | Yeast | NULL | Probable | As nectar-tap_saprotroph (Põlme et al. 2020); Decay sugar-rich substrates (Põlme et al. 2020) | Põlme S, et al. 2020. Fungal Diversity 105:1-16 ((https://doi.org/10.1007/s13225-020-00466-2)) |
| Otu13015 | 1 | k__Fungi(100);p__Ascomycota(100);c__Saccharomycetes(100);o__Saccharomycetales(100);f__Metschnikowiaceae(100);g__Metschnikowia(100);s__Metschnikowia_pulcherrima(100); | Metschnikowia | 13 | Saprotroph | \|Undefined Saprotroph\| | Yeast | NULL | Probable | As nectar-tap_saprotroph (Põlme et al. 2020); Decay sugar-rich substrates (Põlme et al. 2020) | Põlme S, et al. 2020. Fungal Diversity 105:1-16 ((https://doi.org/10.1007/s13225-020-00466-2)) |
| Otu13018 | 1 | k__Fungi(100);p__Ascomycota(100);c__Saccharomycetes(100);o__Saccharomycetales(100);f__Metschnikowiaceae(100);g__Metschnikowia(100);g__Metschnikowia_unclassified(100); | Metschnikowia | 13 | Saprotroph | \|Undefined Saprotroph\| | Yeast | NULL | Probable | As nectar-tap_saprotroph (Põlme et al. 2020); Decay sugar-rich substrates (Põlme et al. 2020) | Põlme S, et al. 2020. Fungal Diversity 105:1-16 ((https://doi.org/10.1007/s13225-020-00466-2)) |
| Otu13020 | 1 | k__Fungi(100);p__Ascomycota(100);c__Saccharomycetes(100);o__Saccharomycetales(100);f__Metschnikowiaceae(100);g__Metschnikowia(100);s__Metschnikowia_pulcherrima(100); | Metschnikowia | 13 | Saprotroph | \|Undefined Saprotroph\| | Yeast | NULL | Probable | As nectar-tap_saprotroph (Põlme et al. 2020); Decay sugar-rich substrates (Põlme et al. 2020) | Põlme S, et al. 2020. Fungal Diversity 105:1-16 ((https://doi.org/10.1007/s13225-020-00466-2)) |
| Otu13021 | 1 | k__Fungi(100);p__Ascomycota(100);c__Saccharomycetes(100);o__Saccharomycetales(100);f__Metschnikowiaceae(100);g__Metschnikowia(100);s__Metschnikowia_pulcherrima(100); | Metschnikowia | 13 | Saprotroph | \|Undefined Saprotroph\| | Yeast | NULL | Probable | As nectar-tap_saprotroph (Põlme et al. 2020); Decay sugar-rich substrates (Põlme et al. 2020) | Põlme S, et al. 2020. Fungal Diversity 105:1-16 ((https://doi.org/10.1007/s13225-020-00466-2)) |
| Otu13024 | 1 | k__Fungi(100);p__Ascomycota(100);c__Saccharomycetes(100);o__Saccharomycetales(100);f__Metschnikowiaceae(100);g__Metschnikowia(100);s__Metschnikowia_pulcherrima(100); | Metschnikowia | 13 | Saprotroph | \|Undefined Saprotroph\| | Yeast | NULL | Probable | As nectar-tap_saprotroph (Põlme et al. 2020); Decay sugar-rich substrates (Põlme et al. 2020) | Põlme S, et al. 2020. Fungal Diversity 105:1-16 ((https://doi.org/10.1007/s13225-020-00466-2)) |
| Otu13033 | 1 | k__Fungi(100);p__Ascomycota(100);c__Saccharomycetes(100);o__Saccharomycetales(100);f__Metschnikowiaceae(100);g__Metschnikowia(100);s__Metschnikowia_pulcherrima(100); | Metschnikowia | 13 | Saprotroph | \|Undefined Saprotroph\| | Yeast | NULL | Probable | As nectar-tap_saprotroph (Põlme et al. 2020); Decay sugar-rich substrates (Põlme et al. 2020) | Põlme S, et al. 2020. Fungal Diversity 105:1-16 ((https://doi.org/10.1007/s13225-020-00466-2)) |
| Otu13034 | 1 | k__Fungi(100);p__Ascomycota(100);c__Saccharomycetes(100);o__Saccharomycetales(100);f__Metschnikowiaceae(100);g__Metschnikowia(100);s__Metschnikowia_pulcherrima(100); | Metschnikowia | 13 | Saprotroph | \|Undefined Saprotroph\| | Yeast | NULL | Probable | As nectar-tap_saprotroph (Põlme et al. 2020); Decay sugar-rich substrates (Põlme et al. 2020) | Põlme S, et al. 2020. Fungal Diversity 105:1-16 ((https://doi.org/10.1007/s13225-020-00466-2)) |
| Otu13035 | 1 | k__Fungi(100);p__Ascomycota(100);c__Saccharomycetes(100);o__Saccharomycetales(100);f__Metschnikowiaceae(100);g__Metschnikowia(100);g__Metschnikowia_unclassified(100); | Metschnikowia | 13 | Saprotroph | \|Undefined Saprotroph\| | Yeast | NULL | Probable | As nectar-tap_saprotroph (Põlme et al. 2020); Decay sugar-rich substrates (Põlme et al. 2020) | Põlme S, et al. 2020. Fungal Diversity 105:1-16 ((https://doi.org/10.1007/s13225-020-00466-2)) |
| Otu13036 | 1 | k__Fungi(100);p__Ascomycota(100);c__Saccharomycetes(100);o__Saccharomycetales(100);f__Metschnikowiaceae(100);g__Metschnikowia(100);g__Metschnikowia_unclassified(100); | Metschnikowia | 13 | Saprotroph | \|Undefined Saprotroph\| | Yeast | NULL | Probable | As nectar-tap_saprotroph (Põlme et al. 2020); Decay sugar-rich substrates (Põlme et al. 2020) | Põlme S, et al. 2020. Fungal Diversity 105:1-16 ((https://doi.org/10.1007/s13225-020-00466-2)) |
| Otu13037 | 1 | k__Fungi(100);p__Ascomycota(100);c__Saccharomycetes(100);o__Saccharomycetales(100);f__Metschnikowiaceae(100);g__Metschnikowia(100);s__Metschnikowia_pulcherrima(100); | Metschnikowia | 13 | Saprotroph | \|Undefined Saprotroph\| | Yeast | NULL | Probable | As nectar-tap_saprotroph (Põlme et al. 2020); Decay sugar-rich substrates (Põlme et al. 2020) | Põlme S, et al. 2020. Fungal Diversity 105:1-16 ((https://doi.org/10.1007/s13225-020-00466-2)) |
| Otu13040 | 1 | k__Fungi(100);p__Ascomycota(100);c__Saccharomycetes(100);o__Saccharomycetales(100);f__Metschnikowiaceae(100);g__Metschnikowia(100);s__Metschnikowia_pulcherrima(100); | Metschnikowia | 13 | Saprotroph | \|Undefined Saprotroph\| | Yeast | NULL | Probable | As nectar-tap_saprotroph (Põlme et al. 2020); Decay sugar-rich substrates (Põlme et al. 2020) | Põlme S, et al. 2020. Fungal Diversity 105:1-16 ((https://doi.org/10.1007/s13225-020-00466-2)) |
| Otu13041 | 1 | k__Fungi(100);p__Ascomycota(100);c__Saccharomycetes(100);o__Saccharomycetales(100);f__Metschnikowiaceae(100);g__Metschnikowia(100);g__Metschnikowia_unclassified(100); | Metschnikowia | 13 | Saprotroph | \|Undefined Saprotroph\| | Yeast | NULL | Probable | As nectar-tap_saprotroph (Põlme et al. 2020); Decay sugar-rich substrates (Põlme et al. 2020) | Põlme S, et al. 2020. Fungal Diversity 105:1-16 ((https://doi.org/10.1007/s13225-020-00466-2)) |
| Otu13044 | 1 | k__Fungi(100);p__Ascomycota(100);c__Saccharomycetes(100);o__Saccharomycetales(100);f__Metschnikowiaceae(100);g__Metschnikowia(100);g__Metschnikowia_unclassified(100); | Metschnikowia | 13 | Saprotroph | \|Undefined Saprotroph\| | Yeast | NULL | Probable | As nectar-tap_saprotroph (Põlme et al. 2020); Decay sugar-rich substrates (Põlme et al. 2020) | Põlme S, et al. 2020. Fungal Diversity 105:1-16 ((https://doi.org/10.1007/s13225-020-00466-2)) |
| Otu13048 | 1 | k__Fungi(100);p__Ascomycota(100);c__Saccharomycetes(100);o__Saccharomycetales(100);f__Metschnikowiaceae(100);g__Metschnikowia(100);g__Metschnikowia_unclassified(100); | Metschnikowia | 13 | Saprotroph | \|Undefined Saprotroph\| | Yeast | NULL | Probable | As nectar-tap_saprotroph (Põlme et al. 2020); Decay sugar-rich substrates (Põlme et al. 2020) | Põlme S, et al. 2020. Fungal Diversity 105:1-16 ((https://doi.org/10.1007/s13225-020-00466-2)) |
| Otu13049 | 1 | k__Fungi(100);p__Ascomycota(100);c__Saccharomycetes(100);o__Saccharomycetales(100);f__Metschnikowiaceae(100);g__Metschnikowia(100);s__Metschnikowia_pulcherrima(100); | Metschnikowia | 13 | Saprotroph | \|Undefined Saprotroph\| | Yeast | NULL | Probable | As nectar-tap_saprotroph (Põlme et al. 2020); Decay sugar-rich substrates (Põlme et al. 2020) | Põlme S, et al. 2020. Fungal Diversity 105:1-16 ((https://doi.org/10.1007/s13225-020-00466-2)) |
| Otu13050 | 1 | k__Fungi(100);p__Ascomycota(100);c__Saccharomycetes(100);o__Saccharomycetales(100);f__Metschnikowiaceae(100);g__Metschnikowia(100);g__Metschnikowia_unclassified(100); | Metschnikowia | 13 | Saprotroph | \|Undefined Saprotroph\| | Yeast | NULL | Probable | As nectar-tap_saprotroph (Põlme et al. 2020); Decay sugar-rich substrates (Põlme et al. 2020) | Põlme S, et al. 2020. Fungal Diversity 105:1-16 ((https://doi.org/10.1007/s13225-020-00466-2)) |
| Otu13051 | 1 | k__Fungi(100);p__Ascomycota(100);c__Saccharomycetes(100);o__Saccharomycetales(100);f__Metschnikowiaceae(100);g__Metschnikowia(100);g__Metschnikowia_unclassified(100); | Metschnikowia | 13 | Saprotroph | \|Undefined Saprotroph\| | Yeast | NULL | Probable | As nectar-tap_saprotroph (Põlme et al. 2020); Decay sugar-rich substrates (Põlme et al. 2020) | Põlme S, et al. 2020. Fungal Diversity 105:1-16 ((https://doi.org/10.1007/s13225-020-00466-2)) |
| Otu13052 | 1 | k__Fungi(100);p__Ascomycota(100);c__Saccharomycetes(100);o__Saccharomycetales(100);f__Metschnikowiaceae(100);g__Metschnikowia(100);g__Metschnikowia_unclassified(100); | Metschnikowia | 13 | Saprotroph | \|Undefined Saprotroph\| | Yeast | NULL | Probable | As nectar-tap_saprotroph (Põlme et al. 2020); Decay sugar-rich substrates (Põlme et al. 2020) | Põlme S, et al. 2020. Fungal Diversity 105:1-16 ((https://doi.org/10.1007/s13225-020-00466-2)) |
| Otu13055 | 1 | k__Fungi(100);p__Ascomycota(100);c__Saccharomycetes(100);o__Saccharomycetales(100);f__Metschnikowiaceae(100);g__Metschnikowia(100);g__Metschnikowia_unclassified(100); | Metschnikowia | 13 | Saprotroph | \|Undefined Saprotroph\| | Yeast | NULL | Probable | As nectar-tap_saprotroph (Põlme et al. 2020); Decay sugar-rich substrates (Põlme et al. 2020) | Põlme S, et al. 2020. Fungal Diversity 105:1-16 ((https://doi.org/10.1007/s13225-020-00466-2)) |
| Otu13056 | 1 | k__Fungi(100);p__Ascomycota(100);c__Saccharomycetes(100);o__Saccharomycetales(100);f__Metschnikowiaceae(100);g__Metschnikowia(100);g__Metschnikowia_unclassified(100); | Metschnikowia | 13 | Saprotroph | \|Undefined Saprotroph\| | Yeast | NULL | Probable | As nectar-tap_saprotroph (Põlme et al. 2020); Decay sugar-rich substrates (Põlme et al. 2020) | Põlme S, et al. 2020. Fungal Diversity 105:1-16 ((https://doi.org/10.1007/s13225-020-00466-2)) |
| Otu13058 | 1 | k__Fungi(100);p__Ascomycota(100);c__Saccharomycetes(100);o__Saccharomycetales(100);f__Metschnikowiaceae(100);g__Metschnikowia(100);s__Metschnikowia_pulcherrima(100); | Metschnikowia | 13 | Saprotroph | \|Undefined Saprotroph\| | Yeast | NULL | Probable | As nectar-tap_saprotroph (Põlme et al. 2020); Decay sugar-rich substrates (Põlme et al. 2020) | Põlme S, et al. 2020. Fungal Diversity 105:1-16 ((https://doi.org/10.1007/s13225-020-00466-2)) |
| Otu13059 | 1 | k__Fungi(100);p__Ascomycota(100);c__Saccharomycetes(100);o__Saccharomycetales(100);f__Metschnikowiaceae(100);g__Metschnikowia(100);s__Metschnikowia_pulcherrima(100); | Metschnikowia | 13 | Saprotroph | \|Undefined Saprotroph\| | Yeast | NULL | Probable | As nectar-tap_saprotroph (Põlme et al. 2020); Decay sugar-rich substrates (Põlme et al. 2020) | Põlme S, et al. 2020. Fungal Diversity 105:1-16 ((https://doi.org/10.1007/s13225-020-00466-2)) |
| Otu13061 | 1 | k__Fungi(100);p__Ascomycota(100);c__Saccharomycetes(100);o__Saccharomycetales(100);f__Metschnikowiaceae(100);g__Metschnikowia(100);s__Metschnikowia_pulcherrima(100); | Metschnikowia | 13 | Saprotroph | \|Undefined Saprotroph\| | Yeast | NULL | Probable | As nectar-tap_saprotroph (Põlme et al. 2020); Decay sugar-rich substrates (Põlme et al. 2020) | Põlme S, et al. 2020. Fungal Diversity 105:1-16 ((https://doi.org/10.1007/s13225-020-00466-2)) |
| Otu13062 | 1 | k__Fungi(100);p__Ascomycota(100);c__Saccharomycetes(100);o__Saccharomycetales(100);f__Metschnikowiaceae(100);g__Metschnikowia(100);s__Metschnikowia_pulcherrima(100); | Metschnikowia | 13 | Saprotroph | \|Undefined Saprotroph\| | Yeast | NULL | Probable | As nectar-tap_saprotroph (Põlme et al. 2020); Decay sugar-rich substrates (Põlme et al. 2020) | Põlme S, et al. 2020. Fungal Diversity 105:1-16 ((https://doi.org/10.1007/s13225-020-00466-2)) |
| Otu13063 | 1 | k__Fungi(100);p__Ascomycota(100);c__Saccharomycetes(100);o__Saccharomycetales(100);f__Metschnikowiaceae(100);g__Metschnikowia(100);s__Metschnikowia_pulcherrima(100); | Metschnikowia | 13 | Saprotroph | \|Undefined Saprotroph\| | Yeast | NULL | Probable | As nectar-tap_saprotroph (Põlme et al. 2020); Decay sugar-rich substrates (Põlme et al. 2020) | Põlme S, et al. 2020. Fungal Diversity 105:1-16 ((https://doi.org/10.1007/s13225-020-00466-2)) |
| Otu13064 | 1 | k__Fungi(100);p__Ascomycota(100);c__Saccharomycetes(100);o__Saccharomycetales(100);f__Metschnikowiaceae(100);g__Metschnikowia(100);s__Metschnikowia_pulcherrima(100); | Metschnikowia | 13 | Saprotroph | \|Undefined Saprotroph\| | Yeast | NULL | Probable | As nectar-tap_saprotroph (Põlme et al. 2020); Decay sugar-rich substrates (Põlme et al. 2020) | Põlme S, et al. 2020. Fungal Diversity 105:1-16 ((https://doi.org/10.1007/s13225-020-00466-2)) |
| Otu13065 | 1 | k__Fungi(100);p__Ascomycota(100);c__Saccharomycetes(100);o__Saccharomycetales(100);f__Metschnikowiaceae(100);g__Metschnikowia(100);s__Metschnikowia_pulcherrima(100); | Metschnikowia | 13 | Saprotroph | \|Undefined Saprotroph\| | Yeast | NULL | Probable | As nectar-tap_saprotroph (Põlme et al. 2020); Decay sugar-rich substrates (Põlme et al. 2020) | Põlme S, et al. 2020. Fungal Diversity 105:1-16 ((https://doi.org/10.1007/s13225-020-00466-2)) |
| Otu13066 | 1 | k__Fungi(100);p__Ascomycota(100);c__Saccharomycetes(100);o__Saccharomycetales(100);f__Metschnikowiaceae(100);g__Metschnikowia(100);s__Metschnikowia_pulcherrima(100); | Metschnikowia | 13 | Saprotroph | \|Undefined Saprotroph\| | Yeast | NULL | Probable | As nectar-tap_saprotroph (Põlme et al. 2020); Decay sugar-rich substrates (Põlme et al. 2020) | Põlme S, et al. 2020. Fungal Diversity 105:1-16 ((https://doi.org/10.1007/s13225-020-00466-2)) |
| Otu13069 | 1 | k__Fungi(100);p__Ascomycota(100);c__Saccharomycetes(100);o__Saccharomycetales(100);f__Metschnikowiaceae(100);g__Metschnikowia(100);s__Metschnikowia_pulcherrima(100); | Metschnikowia | 13 | Saprotroph | \|Undefined Saprotroph\| | Yeast | NULL | Probable | As nectar-tap_saprotroph (Põlme et al. 2020); Decay sugar-rich substrates (Põlme et al. 2020) | Põlme S, et al. 2020. Fungal Diversity 105:1-16 ((https://doi.org/10.1007/s13225-020-00466-2)) |
| Otu13070 | 1 | k__Fungi(100);p__Ascomycota(100);c__Saccharomycetes(100);o__Saccharomycetales(100);f__Metschnikowiaceae(100);g__Metschnikowia(100);s__Metschnikowia_pulcherrima(100); | Metschnikowia | 13 | Saprotroph | \|Undefined Saprotroph\| | Yeast | NULL | Probable | As nectar-tap_saprotroph (Põlme et al. 2020); Decay sugar-rich substrates (Põlme et al. 2020) | Põlme S, et al. 2020. Fungal Diversity 105:1-16 ((https://doi.org/10.1007/s13225-020-00466-2)) |
| Otu13071 | 1 | k__Fungi(100);p__Ascomycota(100);c__Saccharomycetes(100);o__Saccharomycetales(100);f__Metschnikowiaceae(100);g__Metschnikowia(100);s__Metschnikowia_pulcherrima(100); | Metschnikowia | 13 | Saprotroph | \|Undefined Saprotroph\| | Yeast | NULL | Probable | As nectar-tap_saprotroph (Põlme et al. 2020); Decay sugar-rich substrates (Põlme et al. 2020) | Põlme S, et al. 2020. Fungal Diversity 105:1-16 ((https://doi.org/10.1007/s13225-020-00466-2)) |
| Otu13074 | 1 | k__Fungi(100);p__Ascomycota(100);c__Saccharomycetes(100);o__Saccharomycetales(100);f__Metschnikowiaceae(100);g__Metschnikowia(100);s__Metschnikowia_pulcherrima(100); | Metschnikowia | 13 | Saprotroph | \|Undefined Saprotroph\| | Yeast | NULL | Probable | As nectar-tap_saprotroph (Põlme et al. 2020); Decay sugar-rich substrates (Põlme et al. 2020) | Põlme S, et al. 2020. Fungal Diversity 105:1-16 ((https://doi.org/10.1007/s13225-020-00466-2)) |
| Otu13076 | 1 | k__Fungi(100);p__Ascomycota(100);c__Saccharomycetes(100);o__Saccharomycetales(100);f__Metschnikowiaceae(100);g__Metschnikowia(100);g__Metschnikowia_unclassified(100); | Metschnikowia | 13 | Saprotroph | \|Undefined Saprotroph\| | Yeast | NULL | Probable | As nectar-tap_saprotroph (Põlme et al. 2020); Decay sugar-rich substrates (Põlme et al. 2020) | Põlme S, et al. 2020. Fungal Diversity 105:1-16 ((https://doi.org/10.1007/s13225-020-00466-2)) |
| Otu13077 | 1 | k__Fungi(100);p__Ascomycota(100);c__Saccharomycetes(100);o__Saccharomycetales(100);f__Metschnikowiaceae(100);g__Metschnikowia(100);s__Metschnikowia_pulcherrima(100); | Metschnikowia | 13 | Saprotroph | \|Undefined Saprotroph\| | Yeast | NULL | Probable | As nectar-tap_saprotroph (Põlme et al. 2020); Decay sugar-rich substrates (Põlme et al. 2020) | Põlme S, et al. 2020. Fungal Diversity 105:1-16 ((https://doi.org/10.1007/s13225-020-00466-2)) |
| Otu13079 | 1 | k__Fungi(100);p__Ascomycota(100);c__Saccharomycetes(100);o__Saccharomycetales(100);f__Metschnikowiaceae(100);g__Metschnikowia(100);s__Metschnikowia_pulcherrima(100); | Metschnikowia | 13 | Saprotroph | \|Undefined Saprotroph\| | Yeast | NULL | Probable | As nectar-tap_saprotroph (Põlme et al. 2020); Decay sugar-rich substrates (Põlme et al. 2020) | Põlme S, et al. 2020. Fungal Diversity 105:1-16 ((https://doi.org/10.1007/s13225-020-00466-2)) |
| Otu13080 | 1 | k__Fungi(100);p__Ascomycota(100);c__Saccharomycetes(100);o__Saccharomycetales(100);f__Metschnikowiaceae(100);g__Metschnikowia(100);s__Metschnikowia_pulcherrima(100); | Metschnikowia | 13 | Saprotroph | \|Undefined Saprotroph\| | Yeast | NULL | Probable | As nectar-tap_saprotroph (Põlme et al. 2020); Decay sugar-rich substrates (Põlme et al. 2020) | Põlme S, et al. 2020. Fungal Diversity 105:1-16 ((https://doi.org/10.1007/s13225-020-00466-2)) |
| Otu13081 | 1 | k__Fungi(100);p__Ascomycota(100);c__Saccharomycetes(100);o__Saccharomycetales(100);f__Metschnikowiaceae(100);g__Metschnikowia(100);s__Metschnikowia_pulcherrima(100); | Metschnikowia | 13 | Saprotroph | \|Undefined Saprotroph\| | Yeast | NULL | Probable | As nectar-tap_saprotroph (Põlme et al. 2020); Decay sugar-rich substrates (Põlme et al. 2020) | Põlme S, et al. 2020. Fungal Diversity 105:1-16 ((https://doi.org/10.1007/s13225-020-00466-2)) |
| Otu13082 | 1 | k__Fungi(100);p__Ascomycota(100);c__Saccharomycetes(100);o__Saccharomycetales(100);f__Metschnikowiaceae(100);g__Metschnikowia(100);s__Metschnikowia_pulcherrima(100); | Metschnikowia | 13 | Saprotroph | \|Undefined Saprotroph\| | Yeast | NULL | Probable | As nectar-tap_saprotroph (Põlme et al. 2020); Decay sugar-rich substrates (Põlme et al. 2020) | Põlme S, et al. 2020. Fungal Diversity 105:1-16 ((https://doi.org/10.1007/s13225-020-00466-2)) |
| Otu13084 | 1 | k__Fungi(100);p__Ascomycota(100);c__Saccharomycetes(100);o__Saccharomycetales(100);f__Metschnikowiaceae(100);g__Metschnikowia(100);s__Metschnikowia_pulcherrima(100); | Metschnikowia | 13 | Saprotroph | \|Undefined Saprotroph\| | Yeast | NULL | Probable | As nectar-tap_saprotroph (Põlme et al. 2020); Decay sugar-rich substrates (Põlme et al. 2020) | Põlme S, et al. 2020. Fungal Diversity 105:1-16 ((https://doi.org/10.1007/s13225-020-00466-2)) |
| Otu13085 | 1 | k__Fungi(100);p__Ascomycota(100);c__Saccharomycetes(100);o__Saccharomycetales(100);f__Metschnikowiaceae(100);g__Metschnikowia(100);s__Metschnikowia_pulcherrima(100); | Metschnikowia | 13 | Saprotroph | \|Undefined Saprotroph\| | Yeast | NULL | Probable | As nectar-tap_saprotroph (Põlme et al. 2020); Decay sugar-rich substrates (Põlme et al. 2020) | Põlme S, et al. 2020. Fungal Diversity 105:1-16 ((https://doi.org/10.1007/s13225-020-00466-2)) |
| Otu13087 | 1 | k__Fungi(100);p__Ascomycota(100);c__Saccharomycetes(100);o__Saccharomycetales(100);f__Metschnikowiaceae(100);g__Metschnikowia(100);g__Metschnikowia_unclassified(100); | Metschnikowia | 13 | Saprotroph | \|Undefined Saprotroph\| | Yeast | NULL | Probable | As nectar-tap_saprotroph (Põlme et al. 2020); Decay sugar-rich substrates (Põlme et al. 2020) | Põlme S, et al. 2020. Fungal Diversity 105:1-16 ((https://doi.org/10.1007/s13225-020-00466-2)) |
| Otu13088 | 1 | k__Fungi(100);p__Ascomycota(100);c__Saccharomycetes(100);o__Saccharomycetales(100);f__Metschnikowiaceae(100);g__Metschnikowia(100);g__Metschnikowia_unclassified(100); | Metschnikowia | 13 | Saprotroph | \|Undefined Saprotroph\| | Yeast | NULL | Probable | As nectar-tap_saprotroph (Põlme et al. 2020); Decay sugar-rich substrates (Põlme et al. 2020) | Põlme S, et al. 2020. Fungal Diversity 105:1-16 ((https://doi.org/10.1007/s13225-020-00466-2)) |
| Otu13089 | 1 | k__Fungi(100);p__Ascomycota(100);c__Saccharomycetes(100);o__Saccharomycetales(100);f__Metschnikowiaceae(100);g__Metschnikowia(100);s__Metschnikowia_pulcherrima(100); | Metschnikowia | 13 | Saprotroph | \|Undefined Saprotroph\| | Yeast | NULL | Probable | As nectar-tap_saprotroph (Põlme et al. 2020); Decay sugar-rich substrates (Põlme et al. 2020) | Põlme S, et al. 2020. Fungal Diversity 105:1-16 ((https://doi.org/10.1007/s13225-020-00466-2)) |
| Otu13090 | 1 | k__Fungi(100);p__Ascomycota(100);c__Saccharomycetes(100);o__Saccharomycetales(100);f__Metschnikowiaceae(100);g__Metschnikowia(100);g__Metschnikowia_unclassified(100); | Metschnikowia | 13 | Saprotroph | \|Undefined Saprotroph\| | Yeast | NULL | Probable | As nectar-tap_saprotroph (Põlme et al. 2020); Decay sugar-rich substrates (Põlme et al. 2020) | Põlme S, et al. 2020. Fungal Diversity 105:1-16 ((https://doi.org/10.1007/s13225-020-00466-2)) |
| Otu13091 | 1 | k__Fungi(100);p__Ascomycota(100);c__Saccharomycetes(100);o__Saccharomycetales(100);f__Metschnikowiaceae(100);g__Metschnikowia(100);s__Metschnikowia_pulcherrima(100); | Metschnikowia | 13 | Saprotroph | \|Undefined Saprotroph\| | Yeast | NULL | Probable | As nectar-tap_saprotroph (Põlme et al. 2020); Decay sugar-rich substrates (Põlme et al. 2020) | Põlme S, et al. 2020. Fungal Diversity 105:1-16 ((https://doi.org/10.1007/s13225-020-00466-2)) |
| Otu13095 | 1 | k__Fungi(100);p__Ascomycota(100);c__Saccharomycetes(100);o__Saccharomycetales(100);f__Metschnikowiaceae(100);g__Metschnikowia(100);s__Metschnikowia_pulcherrima(100); | Metschnikowia | 13 | Saprotroph | \|Undefined Saprotroph\| | Yeast | NULL | Probable | As nectar-tap_saprotroph (Põlme et al. 2020); Decay sugar-rich substrates (Põlme et al. 2020) | Põlme S, et al. 2020. Fungal Diversity 105:1-16 ((https://doi.org/10.1007/s13225-020-00466-2)) |
| Otu13096 | 1 | k__Fungi(100);p__Ascomycota(100);c__Saccharomycetes(100);o__Saccharomycetales(100);f__Metschnikowiaceae(100);g__Metschnikowia(100);s__Metschnikowia_pulcherrima(100); | Metschnikowia | 13 | Saprotroph | \|Undefined Saprotroph\| | Yeast | NULL | Probable | As nectar-tap_saprotroph (Põlme et al. 2020); Decay sugar-rich substrates (Põlme et al. 2020) | Põlme S, et al. 2020. Fungal Diversity 105:1-16 ((https://doi.org/10.1007/s13225-020-00466-2)) |
| Otu13098 | 1 | k__Fungi(100);p__Ascomycota(100);c__Saccharomycetes(100);o__Saccharomycetales(100);f__Metschnikowiaceae(100);g__Metschnikowia(100);s__Metschnikowia_pulcherrima(100); | Metschnikowia | 13 | Saprotroph | \|Undefined Saprotroph\| | Yeast | NULL | Probable | As nectar-tap_saprotroph (Põlme et al. 2020); Decay sugar-rich substrates (Põlme et al. 2020) | Põlme S, et al. 2020. Fungal Diversity 105:1-16 ((https://doi.org/10.1007/s13225-020-00466-2)) |
| Otu13099 | 1 | k__Fungi(100);p__Ascomycota(100);c__Saccharomycetes(100);o__Saccharomycetales(100);f__Metschnikowiaceae(100);g__Metschnikowia(100);s__Metschnikowia_pulcherrima(100); | Metschnikowia | 13 | Saprotroph | \|Undefined Saprotroph\| | Yeast | NULL | Probable | As nectar-tap_saprotroph (Põlme et al. 2020); Decay sugar-rich substrates (Põlme et al. 2020) | Põlme S, et al. 2020. Fungal Diversity 105:1-16 ((https://doi.org/10.1007/s13225-020-00466-2)) |
| Otu13101 | 1 | k__Fungi(100);p__Ascomycota(100);c__Saccharomycetes(100);o__Saccharomycetales(100);f__Metschnikowiaceae(100);g__Metschnikowia(100);s__Metschnikowia_pulcherrima(100); | Metschnikowia | 13 | Saprotroph | \|Undefined Saprotroph\| | Yeast | NULL | Probable | As nectar-tap_saprotroph (Põlme et al. 2020); Decay sugar-rich substrates (Põlme et al. 2020) | Põlme S, et al. 2020. Fungal Diversity 105:1-16 ((https://doi.org/10.1007/s13225-020-00466-2)) |
| Otu13103 | 1 | k__Fungi(100);p__Ascomycota(100);c__Saccharomycetes(100);o__Saccharomycetales(100);f__Metschnikowiaceae(100);g__Metschnikowia(100);s__Metschnikowia_pulcherrima(100); | Metschnikowia | 13 | Saprotroph | \|Undefined Saprotroph\| | Yeast | NULL | Probable | As nectar-tap_saprotroph (Põlme et al. 2020); Decay sugar-rich substrates (Põlme et al. 2020) | Põlme S, et al. 2020. Fungal Diversity 105:1-16 ((https://doi.org/10.1007/s13225-020-00466-2)) |
| Otu13106 | 1 | k__Fungi(100);p__Ascomycota(100);c__Saccharomycetes(100);o__Saccharomycetales(100);f__Metschnikowiaceae(100);g__Metschnikowia(100);s__Metschnikowia_pulcherrima(100); | Metschnikowia | 13 | Saprotroph | \|Undefined Saprotroph\| | Yeast | NULL | Probable | As nectar-tap_saprotroph (Põlme et al. 2020); Decay sugar-rich substrates (Põlme et al. 2020) | Põlme S, et al. 2020. Fungal Diversity 105:1-16 ((https://doi.org/10.1007/s13225-020-00466-2)) |
| Otu13107 | 1 | k__Fungi(100);p__Ascomycota(100);c__Saccharomycetes(100);o__Saccharomycetales(100);f__Metschnikowiaceae(100);g__Metschnikowia(100);s__Metschnikowia_pulcherrima(100); | Metschnikowia | 13 | Saprotroph | \|Undefined Saprotroph\| | Yeast | NULL | Probable | As nectar-tap_saprotroph (Põlme et al. 2020); Decay sugar-rich substrates (Põlme et al. 2020) | Põlme S, et al. 2020. Fungal Diversity 105:1-16 ((https://doi.org/10.1007/s13225-020-00466-2)) |
| Otu13108 | 1 | k__Fungi(100);p__Ascomycota(100);c__Saccharomycetes(100);o__Saccharomycetales(100);f__Metschnikowiaceae(100);g__Metschnikowia(100);s__Metschnikowia_pulcherrima(100); | Metschnikowia | 13 | Saprotroph | \|Undefined Saprotroph\| | Yeast | NULL | Probable | As nectar-tap_saprotroph (Põlme et al. 2020); Decay sugar-rich substrates (Põlme et al. 2020) | Põlme S, et al. 2020. Fungal Diversity 105:1-16 ((https://doi.org/10.1007/s13225-020-00466-2)) |
| Otu13109 | 1 | k__Fungi(100);p__Ascomycota(100);c__Saccharomycetes(100);o__Saccharomycetales(100);f__Metschnikowiaceae(100);g__Metschnikowia(100);s__Metschnikowia_pulcherrima(100); | Metschnikowia | 13 | Saprotroph | \|Undefined Saprotroph\| | Yeast | NULL | Probable | As nectar-tap_saprotroph (Põlme et al. 2020); Decay sugar-rich substrates (Põlme et al. 2020) | Põlme S, et al. 2020. Fungal Diversity 105:1-16 ((https://doi.org/10.1007/s13225-020-00466-2)) |
| Otu13111 | 1 | k__Fungi(100);p__Ascomycota(100);c__Saccharomycetes(100);o__Saccharomycetales(100);f__Metschnikowiaceae(100);g__Metschnikowia(100);s__Metschnikowia_pulcherrima(100); | Metschnikowia | 13 | Saprotroph | \|Undefined Saprotroph\| | Yeast | NULL | Probable | As nectar-tap_saprotroph (Põlme et al. 2020); Decay sugar-rich substrates (Põlme et al. 2020) | Põlme S, et al. 2020. Fungal Diversity 105:1-16 ((https://doi.org/10.1007/s13225-020-00466-2)) |
| Otu13112 | 1 | k__Fungi(100);p__Ascomycota(100);c__Saccharomycetes(100);o__Saccharomycetales(100);f__Metschnikowiaceae(100);g__Metschnikowia(100);s__Metschnikowia_pulcherrima(100); | Metschnikowia | 13 | Saprotroph | \|Undefined Saprotroph\| | Yeast | NULL | Probable | As nectar-tap_saprotroph (Põlme et al. 2020); Decay sugar-rich substrates (Põlme et al. 2020) | Põlme S, et al. 2020. Fungal Diversity 105:1-16 ((https://doi.org/10.1007/s13225-020-00466-2)) |
| Otu13114 | 1 | k__Fungi(100);p__Ascomycota(100);c__Saccharomycetes(100);o__Saccharomycetales(100);f__Metschnikowiaceae(100);g__Metschnikowia(100);g__Metschnikowia_unclassified(100); | Metschnikowia | 13 | Saprotroph | \|Undefined Saprotroph\| | Yeast | NULL | Probable | As nectar-tap_saprotroph (Põlme et al. 2020); Decay sugar-rich substrates (Põlme et al. 2020) | Põlme S, et al. 2020. Fungal Diversity 105:1-16 ((https://doi.org/10.1007/s13225-020-00466-2)) |
| Otu13116 | 1 | k__Fungi(100);p__Ascomycota(100);c__Saccharomycetes(100);o__Saccharomycetales(100);f__Metschnikowiaceae(100);g__Metschnikowia(100);s__Metschnikowia_pulcherrima(100); | Metschnikowia | 13 | Saprotroph | \|Undefined Saprotroph\| | Yeast | NULL | Probable | As nectar-tap_saprotroph (Põlme et al. 2020); Decay sugar-rich substrates (Põlme et al. 2020) | Põlme S, et al. 2020. Fungal Diversity 105:1-16 ((https://doi.org/10.1007/s13225-020-00466-2)) |
| Otu13118 | 1 | k__Fungi(100);p__Ascomycota(100);c__Saccharomycetes(100);o__Saccharomycetales(100);f__Metschnikowiaceae(100);g__Metschnikowia(100);g__Metschnikowia_unclassified(100); | Metschnikowia | 13 | Saprotroph | \|Undefined Saprotroph\| | Yeast | NULL | Probable | As nectar-tap_saprotroph (Põlme et al. 2020); Decay sugar-rich substrates (Põlme et al. 2020) | Põlme S, et al. 2020. Fungal Diversity 105:1-16 ((https://doi.org/10.1007/s13225-020-00466-2)) |
| Otu13120 | 1 | k__Fungi(100);p__Ascomycota(100);c__Saccharomycetes(100);o__Saccharomycetales(100);f__Metschnikowiaceae(100);g__Metschnikowia(100);g__Metschnikowia_unclassified(100); | Metschnikowia | 13 | Saprotroph | \|Undefined Saprotroph\| | Yeast | NULL | Probable | As nectar-tap_saprotroph (Põlme et al. 2020); Decay sugar-rich substrates (Põlme et al. 2020) | Põlme S, et al. 2020. Fungal Diversity 105:1-16 ((https://doi.org/10.1007/s13225-020-00466-2)) |
| Otu13121 | 1 | k__Fungi(100);p__Ascomycota(100);c__Saccharomycetes(100);o__Saccharomycetales(100);f__Metschnikowiaceae(100);g__Metschnikowia(100);g__Metschnikowia_unclassified(100); | Metschnikowia | 13 | Saprotroph | \|Undefined Saprotroph\| | Yeast | NULL | Probable | As nectar-tap_saprotroph (Põlme et al. 2020); Decay sugar-rich substrates (Põlme et al. 2020) | Põlme S, et al. 2020. Fungal Diversity 105:1-16 ((https://doi.org/10.1007/s13225-020-00466-2)) |
| Otu13122 | 1 | k__Fungi(100);p__Ascomycota(100);c__Saccharomycetes(100);o__Saccharomycetales(100);f__Metschnikowiaceae(100);g__Metschnikowia(100);s__Metschnikowia_pulcherrima(100); | Metschnikowia | 13 | Saprotroph | \|Undefined Saprotroph\| | Yeast | NULL | Probable | As nectar-tap_saprotroph (Põlme et al. 2020); Decay sugar-rich substrates (Põlme et al. 2020) | Põlme S, et al. 2020. Fungal Diversity 105:1-16 ((https://doi.org/10.1007/s13225-020-00466-2)) |
| Otu13123 | 1 | k__Fungi(100);p__Ascomycota(100);c__Saccharomycetes(100);o__Saccharomycetales(100);f__Metschnikowiaceae(100);g__Metschnikowia(100);s__Metschnikowia_pulcherrima(100); | Metschnikowia | 13 | Saprotroph | \|Undefined Saprotroph\| | Yeast | NULL | Probable | As nectar-tap_saprotroph (Põlme et al. 2020); Decay sugar-rich substrates (Põlme et al. 2020) | Põlme S, et al. 2020. Fungal Diversity 105:1-16 ((https://doi.org/10.1007/s13225-020-00466-2)) |
| Otu13125 | 1 | k__Fungi(100);p__Ascomycota(100);c__Saccharomycetes(100);o__Saccharomycetales(100);f__Metschnikowiaceae(100);g__Metschnikowia(100);s__Metschnikowia_pulcherrima(100); | Metschnikowia | 13 | Saprotroph | \|Undefined Saprotroph\| | Yeast | NULL | Probable | As nectar-tap_saprotroph (Põlme et al. 2020); Decay sugar-rich substrates (Põlme et al. 2020) | Põlme S, et al. 2020. Fungal Diversity 105:1-16 ((https://doi.org/10.1007/s13225-020-00466-2)) |
| Otu13126 | 1 | k__Fungi(100);p__Ascomycota(100);c__Saccharomycetes(100);o__Saccharomycetales(100);f__Metschnikowiaceae(100);g__Metschnikowia(100);s__Metschnikowia_pulcherrima(100); | Metschnikowia | 13 | Saprotroph | \|Undefined Saprotroph\| | Yeast | NULL | Probable | As nectar-tap_saprotroph (Põlme et al. 2020); Decay sugar-rich substrates (Põlme et al. 2020) | Põlme S, et al. 2020. Fungal Diversity 105:1-16 ((https://doi.org/10.1007/s13225-020-00466-2)) |
| Otu13127 | 1 | k__Fungi(100);p__Ascomycota(100);c__Saccharomycetes(100);o__Saccharomycetales(100);f__Metschnikowiaceae(100);g__Metschnikowia(100);s__Metschnikowia_pulcherrima(100); | Metschnikowia | 13 | Saprotroph | \|Undefined Saprotroph\| | Yeast | NULL | Probable | As nectar-tap_saprotroph (Põlme et al. 2020); Decay sugar-rich substrates (Põlme et al. 2020) | Põlme S, et al. 2020. Fungal Diversity 105:1-16 ((https://doi.org/10.1007/s13225-020-00466-2)) |
| Otu13128 | 1 | k__Fungi(100);p__Ascomycota(100);c__Saccharomycetes(100);o__Saccharomycetales(100);f__Metschnikowiaceae(100);g__Metschnikowia(100);s__Metschnikowia_pulcherrima(100); | Metschnikowia | 13 | Saprotroph | \|Undefined Saprotroph\| | Yeast | NULL | Probable | As nectar-tap_saprotroph (Põlme et al. 2020); Decay sugar-rich substrates (Põlme et al. 2020) | Põlme S, et al. 2020. Fungal Diversity 105:1-16 ((https://doi.org/10.1007/s13225-020-00466-2)) |
| Otu13129 | 1 | k__Fungi(100);p__Ascomycota(100);c__Saccharomycetes(100);o__Saccharomycetales(100);f__Metschnikowiaceae(100);g__Metschnikowia(100);s__Metschnikowia_pulcherrima(100); | Metschnikowia | 13 | Saprotroph | \|Undefined Saprotroph\| | Yeast | NULL | Probable | As nectar-tap_saprotroph (Põlme et al. 2020); Decay sugar-rich substrates (Põlme et al. 2020) | Põlme S, et al. 2020. Fungal Diversity 105:1-16 ((https://doi.org/10.1007/s13225-020-00466-2)) |
| Otu13130 | 1 | k__Fungi(100);p__Ascomycota(100);c__Saccharomycetes(100);o__Saccharomycetales(100);f__Metschnikowiaceae(100);g__Metschnikowia(100);s__Metschnikowia_pulcherrima(100); | Metschnikowia | 13 | Saprotroph | \|Undefined Saprotroph\| | Yeast | NULL | Probable | As nectar-tap_saprotroph (Põlme et al. 2020); Decay sugar-rich substrates (Põlme et al. 2020) | Põlme S, et al. 2020. Fungal Diversity 105:1-16 ((https://doi.org/10.1007/s13225-020-00466-2)) |
| Otu13131 | 1 | k__Fungi(100);p__Ascomycota(100);c__Saccharomycetes(100);o__Saccharomycetales(100);f__Metschnikowiaceae(100);g__Metschnikowia(100);s__Metschnikowia_pulcherrima(100); | Metschnikowia | 13 | Saprotroph | \|Undefined Saprotroph\| | Yeast | NULL | Probable | As nectar-tap_saprotroph (Põlme et al. 2020); Decay sugar-rich substrates (Põlme et al. 2020) | Põlme S, et al. 2020. Fungal Diversity 105:1-16 ((https://doi.org/10.1007/s13225-020-00466-2)) |
| Otu13133 | 1 | k__Fungi(100);p__Ascomycota(100);c__Saccharomycetes(100);o__Saccharomycetales(100);f__Metschnikowiaceae(100);g__Metschnikowia(100);s__Metschnikowia_pulcherrima(100); | Metschnikowia | 13 | Saprotroph | \|Undefined Saprotroph\| | Yeast | NULL | Probable | As nectar-tap_saprotroph (Põlme et al. 2020); Decay sugar-rich substrates (Põlme et al. 2020) | Põlme S, et al. 2020. Fungal Diversity 105:1-16 ((https://doi.org/10.1007/s13225-020-00466-2)) |
| Otu13134 | 1 | k__Fungi(100);p__Ascomycota(100);c__Saccharomycetes(100);o__Saccharomycetales(100);f__Metschnikowiaceae(100);g__Metschnikowia(100);s__Metschnikowia_pulcherrima(100); | Metschnikowia | 13 | Saprotroph | \|Undefined Saprotroph\| | Yeast | NULL | Probable | As nectar-tap_saprotroph (Põlme et al. 2020); Decay sugar-rich substrates (Põlme et al. 2020) | Põlme S, et al. 2020. Fungal Diversity 105:1-16 ((https://doi.org/10.1007/s13225-020-00466-2)) |
| Otu13136 | 1 | k__Fungi(100);p__Ascomycota(100);c__Saccharomycetes(100);o__Saccharomycetales(100);f__Metschnikowiaceae(100);g__Metschnikowia(100);s__Metschnikowia_pulcherrima(100); | Metschnikowia | 13 | Saprotroph | \|Undefined Saprotroph\| | Yeast | NULL | Probable | As nectar-tap_saprotroph (Põlme et al. 2020); Decay sugar-rich substrates (Põlme et al. 2020) | Põlme S, et al. 2020. Fungal Diversity 105:1-16 ((https://doi.org/10.1007/s13225-020-00466-2)) |
| Otu13137 | 1 | k__Fungi(100);p__Ascomycota(100);c__Saccharomycetes(100);o__Saccharomycetales(100);f__Metschnikowiaceae(100);g__Metschnikowia(100);g__Metschnikowia_unclassified(100); | Metschnikowia | 13 | Saprotroph | \|Undefined Saprotroph\| | Yeast | NULL | Probable | As nectar-tap_saprotroph (Põlme et al. 2020); Decay sugar-rich substrates (Põlme et al. 2020) | Põlme S, et al. 2020. Fungal Diversity 105:1-16 ((https://doi.org/10.1007/s13225-020-00466-2)) |
| Otu13138 | 1 | k__Fungi(100);p__Ascomycota(100);c__Saccharomycetes(100);o__Saccharomycetales(100);f__Metschnikowiaceae(100);g__Metschnikowia(100);s__Metschnikowia_pulcherrima(100); | Metschnikowia | 13 | Saprotroph | \|Undefined Saprotroph\| | Yeast | NULL | Probable | As nectar-tap_saprotroph (Põlme et al. 2020); Decay sugar-rich substrates (Põlme et al. 2020) | Põlme S, et al. 2020. Fungal Diversity 105:1-16 ((https://doi.org/10.1007/s13225-020-00466-2)) |
| Otu13139 | 1 | k__Fungi(100);p__Ascomycota(100);c__Saccharomycetes(100);o__Saccharomycetales(100);f__Metschnikowiaceae(100);g__Metschnikowia(100);g__Metschnikowia_unclassified(100); | Metschnikowia | 13 | Saprotroph | \|Undefined Saprotroph\| | Yeast | NULL | Probable | As nectar-tap_saprotroph (Põlme et al. 2020); Decay sugar-rich substrates (Põlme et al. 2020) | Põlme S, et al. 2020. Fungal Diversity 105:1-16 ((https://doi.org/10.1007/s13225-020-00466-2)) |
| Otu13140 | 1 | k__Fungi(100);p__Ascomycota(100);c__Saccharomycetes(100);o__Saccharomycetales(100);f__Metschnikowiaceae(100);g__Metschnikowia(100);g__Metschnikowia_unclassified(100); | Metschnikowia | 13 | Saprotroph | \|Undefined Saprotroph\| | Yeast | NULL | Probable | As nectar-tap_saprotroph (Põlme et al. 2020); Decay sugar-rich substrates (Põlme et al. 2020) | Põlme S, et al. 2020. Fungal Diversity 105:1-16 ((https://doi.org/10.1007/s13225-020-00466-2)) |
| Otu13143 | 1 | k__Fungi(100);p__Ascomycota(100);c__Saccharomycetes(100);o__Saccharomycetales(100);f__Metschnikowiaceae(100);g__Metschnikowia(100);s__Metschnikowia_pulcherrima(100); | Metschnikowia | 13 | Saprotroph | \|Undefined Saprotroph\| | Yeast | NULL | Probable | As nectar-tap_saprotroph (Põlme et al. 2020); Decay sugar-rich substrates (Põlme et al. 2020) | Põlme S, et al. 2020. Fungal Diversity 105:1-16 ((https://doi.org/10.1007/s13225-020-00466-2)) |
| Otu13144 | 1 | k__Fungi(100);p__Ascomycota(100);c__Saccharomycetes(100);o__Saccharomycetales(100);f__Metschnikowiaceae(100);g__Metschnikowia(100);s__Metschnikowia_pulcherrima(100); | Metschnikowia | 13 | Saprotroph | \|Undefined Saprotroph\| | Yeast | NULL | Probable | As nectar-tap_saprotroph (Põlme et al. 2020); Decay sugar-rich substrates (Põlme et al. 2020) | Põlme S, et al. 2020. Fungal Diversity 105:1-16 ((https://doi.org/10.1007/s13225-020-00466-2)) |
| Otu13146 | 1 | k__Fungi(100);p__Ascomycota(100);c__Saccharomycetes(100);o__Saccharomycetales(100);f__Metschnikowiaceae(100);g__Metschnikowia(100);s__Metschnikowia_pulcherrima(100); | Metschnikowia | 13 | Saprotroph | \|Undefined Saprotroph\| | Yeast | NULL | Probable | As nectar-tap_saprotroph (Põlme et al. 2020); Decay sugar-rich substrates (Põlme et al. 2020) | Põlme S, et al. 2020. Fungal Diversity 105:1-16 ((https://doi.org/10.1007/s13225-020-00466-2)) |
| Otu13147 | 1 | k__Fungi(100);p__Ascomycota(100);c__Saccharomycetes(100);o__Saccharomycetales(100);f__Metschnikowiaceae(100);g__Metschnikowia(100);s__Metschnikowia_pulcherrima(100); | Metschnikowia | 13 | Saprotroph | \|Undefined Saprotroph\| | Yeast | NULL | Probable | As nectar-tap_saprotroph (Põlme et al. 2020); Decay sugar-rich substrates (Põlme et al. 2020) | Põlme S, et al. 2020. Fungal Diversity 105:1-16 ((https://doi.org/10.1007/s13225-020-00466-2)) |
| Otu13148 | 1 | k__Fungi(100);p__Ascomycota(100);c__Saccharomycetes(100);o__Saccharomycetales(100);f__Metschnikowiaceae(100);g__Metschnikowia(100);s__Metschnikowia_pulcherrima(100); | Metschnikowia | 13 | Saprotroph | \|Undefined Saprotroph\| | Yeast | NULL | Probable | As nectar-tap_saprotroph (Põlme et al. 2020); Decay sugar-rich substrates (Põlme et al. 2020) | Põlme S, et al. 2020. Fungal Diversity 105:1-16 ((https://doi.org/10.1007/s13225-020-00466-2)) |
| Otu13149 | 1 | k__Fungi(100);p__Ascomycota(100);c__Saccharomycetes(100);o__Saccharomycetales(100);f__Metschnikowiaceae(100);g__Metschnikowia(100);s__Metschnikowia_pulcherrima(100); | Metschnikowia | 13 | Saprotroph | \|Undefined Saprotroph\| | Yeast | NULL | Probable | As nectar-tap_saprotroph (Põlme et al. 2020); Decay sugar-rich substrates (Põlme et al. 2020) | Põlme S, et al. 2020. Fungal Diversity 105:1-16 ((https://doi.org/10.1007/s13225-020-00466-2)) |
| Otu13152 | 1 | k__Fungi(100);p__Ascomycota(100);c__Saccharomycetes(100);o__Saccharomycetales(100);f__Metschnikowiaceae(100);g__Metschnikowia(100);s__Metschnikowia_pulcherrima(100); | Metschnikowia | 13 | Saprotroph | \|Undefined Saprotroph\| | Yeast | NULL | Probable | As nectar-tap_saprotroph (Põlme et al. 2020); Decay sugar-rich substrates (Põlme et al. 2020) | Põlme S, et al. 2020. Fungal Diversity 105:1-16 ((https://doi.org/10.1007/s13225-020-00466-2)) |
| Otu13153 | 1 | k__Fungi(100);p__Ascomycota(100);c__Saccharomycetes(100);o__Saccharomycetales(100);f__Metschnikowiaceae(100);g__Metschnikowia(100);s__Metschnikowia_pulcherrima(100); | Metschnikowia | 13 | Saprotroph | \|Undefined Saprotroph\| | Yeast | NULL | Probable | As nectar-tap_saprotroph (Põlme et al. 2020); Decay sugar-rich substrates (Põlme et al. 2020) | Põlme S, et al. 2020. Fungal Diversity 105:1-16 ((https://doi.org/10.1007/s13225-020-00466-2)) |
| Otu13156 | 1 | k__Fungi(100);p__Ascomycota(100);c__Saccharomycetes(100);o__Saccharomycetales(100);f__Metschnikowiaceae(100);g__Metschnikowia(100);s__Metschnikowia_pulcherrima(100); | Metschnikowia | 13 | Saprotroph | \|Undefined Saprotroph\| | Yeast | NULL | Probable | As nectar-tap_saprotroph (Põlme et al. 2020); Decay sugar-rich substrates (Põlme et al. 2020) | Põlme S, et al. 2020. Fungal Diversity 105:1-16 ((https://doi.org/10.1007/s13225-020-00466-2)) |
| Otu13157 | 1 | k__Fungi(100);p__Ascomycota(100);c__Saccharomycetes(100);o__Saccharomycetales(100);f__Metschnikowiaceae(100);g__Metschnikowia(100);s__Metschnikowia_pulcherrima(100); | Metschnikowia | 13 | Saprotroph | \|Undefined Saprotroph\| | Yeast | NULL | Probable | As nectar-tap_saprotroph (Põlme et al. 2020); Decay sugar-rich substrates (Põlme et al. 2020) | Põlme S, et al. 2020. Fungal Diversity 105:1-16 ((https://doi.org/10.1007/s13225-020-00466-2)) |
| Otu13158 | 1 | k__Fungi(100);p__Basidiomycota(100);c__Microbotryomycetes(100);o__Sporidiobolales(100);f__Sporidiobolales_family_Incertae_sedis(100);g__Sporobolomyces(100);s__Sporobolomyces_lactosus(100); | Sporobolomyces | 13 | Pathotroph-Saprotroph | \|Fungal Parasite\|-Plant Saprotroph | Yeast | NULL | Probable | As litter_saprotroph (Põlme et al. 2020); As mycoparasite (Põlme et al. 2020); Decay fungal material (Põlme et al. 2020) | Tedersoo L, et al. 2014. Science 346:e1256688 ((https://doi.org/10.1126/science.1256688)); Purahong W, et al. 2016. Molecular Ecology 25:4059-4074 ((https://doi.org/10.1111/mec.13739)); Põlme S, et al. 2020. Fungal Diversity 105:1-16 ((https://doi.org/10.1007/s13225-020-00466-2)) |
| Otu13160 | 1 | k__Fungi(100);p__Ascomycota(100);c__Saccharomycetes(100);o__Saccharomycetales(100);f__Metschnikowiaceae(100);g__Metschnikowia(100);g__Metschnikowia_unclassified(100); | Metschnikowia | 13 | Saprotroph | \|Undefined Saprotroph\| | Yeast | NULL | Probable | As nectar-tap_saprotroph (Põlme et al. 2020); Decay sugar-rich substrates (Põlme et al. 2020) | Põlme S, et al. 2020. Fungal Diversity 105:1-16 ((https://doi.org/10.1007/s13225-020-00466-2)) |
| Otu13161 | 1 | k__Fungi(100);p__Ascomycota(100);c__Saccharomycetes(100);o__Saccharomycetales(100);f__Metschnikowiaceae(100);g__Metschnikowia(100);s__Metschnikowia_pulcherrima(100); | Metschnikowia | 13 | Saprotroph | \|Undefined Saprotroph\| | Yeast | NULL | Probable | As nectar-tap_saprotroph (Põlme et al. 2020); Decay sugar-rich substrates (Põlme et al. 2020) | Põlme S, et al. 2020. Fungal Diversity 105:1-16 ((https://doi.org/10.1007/s13225-020-00466-2)) |
| Otu13165 | 1 | k__Fungi(100);p__Ascomycota(100);c__Saccharomycetes(100);o__Saccharomycetales(100);f__Metschnikowiaceae(100);g__Metschnikowia(100);s__Metschnikowia_pulcherrima(100); | Metschnikowia | 13 | Saprotroph | \|Undefined Saprotroph\| | Yeast | NULL | Probable | As nectar-tap_saprotroph (Põlme et al. 2020); Decay sugar-rich substrates (Põlme et al. 2020) | Põlme S, et al. 2020. Fungal Diversity 105:1-16 ((https://doi.org/10.1007/s13225-020-00466-2)) |
| Otu13167 | 1 | k__Fungi(100);p__Ascomycota(100);c__Saccharomycetes(100);o__Saccharomycetales(100);f__Metschnikowiaceae(100);g__Metschnikowia(100);s__Metschnikowia_pulcherrima(100); | Metschnikowia | 13 | Saprotroph | \|Undefined Saprotroph\| | Yeast | NULL | Probable | As nectar-tap_saprotroph (Põlme et al. 2020); Decay sugar-rich substrates (Põlme et al. 2020) | Põlme S, et al. 2020. Fungal Diversity 105:1-16 ((https://doi.org/10.1007/s13225-020-00466-2)) |
| Otu13169 | 1 | k__Fungi(100);p__Ascomycota(100);c__Saccharomycetes(100);o__Saccharomycetales(100);f__Metschnikowiaceae(100);g__Metschnikowia(100);g__Metschnikowia_unclassified(100); | Metschnikowia | 13 | Saprotroph | \|Undefined Saprotroph\| | Yeast | NULL | Probable | As nectar-tap_saprotroph (Põlme et al. 2020); Decay sugar-rich substrates (Põlme et al. 2020) | Põlme S, et al. 2020. Fungal Diversity 105:1-16 ((https://doi.org/10.1007/s13225-020-00466-2)) |
| Otu13171 | 1 | k__Fungi(100);p__Ascomycota(100);c__Saccharomycetes(100);o__Saccharomycetales(100);f__Metschnikowiaceae(100);g__Metschnikowia(100);s__Metschnikowia_pulcherrima(100); | Metschnikowia | 13 | Saprotroph | \|Undefined Saprotroph\| | Yeast | NULL | Probable | As nectar-tap_saprotroph (Põlme et al. 2020); Decay sugar-rich substrates (Põlme et al. 2020) | Põlme S, et al. 2020. Fungal Diversity 105:1-16 ((https://doi.org/10.1007/s13225-020-00466-2)) |
| Otu13172 | 1 | k__Fungi(100);p__Ascomycota(100);c__Saccharomycetes(100);o__Saccharomycetales(100);f__Metschnikowiaceae(100);g__Metschnikowia(100);s__Metschnikowia_pulcherrima(100); | Metschnikowia | 13 | Saprotroph | \|Undefined Saprotroph\| | Yeast | NULL | Probable | As nectar-tap_saprotroph (Põlme et al. 2020); Decay sugar-rich substrates (Põlme et al. 2020) | Põlme S, et al. 2020. Fungal Diversity 105:1-16 ((https://doi.org/10.1007/s13225-020-00466-2)) |
| Otu13173 | 1 | k__Fungi(100);p__Ascomycota(100);c__Saccharomycetes(100);o__Saccharomycetales(100);f__Metschnikowiaceae(100);g__Metschnikowia(100);s__Metschnikowia_pulcherrima(100); | Metschnikowia | 13 | Saprotroph | \|Undefined Saprotroph\| | Yeast | NULL | Probable | As nectar-tap_saprotroph (Põlme et al. 2020); Decay sugar-rich substrates (Põlme et al. 2020) | Põlme S, et al. 2020. Fungal Diversity 105:1-16 ((https://doi.org/10.1007/s13225-020-00466-2)) |
| Otu13174 | 1 | k__Fungi(100);p__Ascomycota(100);c__Saccharomycetes(100);o__Saccharomycetales(100);f__Metschnikowiaceae(100);g__Metschnikowia(100);s__Metschnikowia_pulcherrima(100); | Metschnikowia | 13 | Saprotroph | \|Undefined Saprotroph\| | Yeast | NULL | Probable | As nectar-tap_saprotroph (Põlme et al. 2020); Decay sugar-rich substrates (Põlme et al. 2020) | Põlme S, et al. 2020. Fungal Diversity 105:1-16 ((https://doi.org/10.1007/s13225-020-00466-2)) |
| Otu13175 | 1 | k__Fungi(100);p__Ascomycota(100);c__Saccharomycetes(100);o__Saccharomycetales(100);f__Metschnikowiaceae(100);g__Metschnikowia(100);s__Metschnikowia_pulcherrima(100); | Metschnikowia | 13 | Saprotroph | \|Undefined Saprotroph\| | Yeast | NULL | Probable | As nectar-tap_saprotroph (Põlme et al. 2020); Decay sugar-rich substrates (Põlme et al. 2020) | Põlme S, et al. 2020. Fungal Diversity 105:1-16 ((https://doi.org/10.1007/s13225-020-00466-2)) |
| Otu13176 | 1 | k__Fungi(100);p__Ascomycota(100);c__Saccharomycetes(100);o__Saccharomycetales(100);f__Metschnikowiaceae(100);g__Metschnikowia(100);s__Metschnikowia_pulcherrima(100); | Metschnikowia | 13 | Saprotroph | \|Undefined Saprotroph\| | Yeast | NULL | Probable | As nectar-tap_saprotroph (Põlme et al. 2020); Decay sugar-rich substrates (Põlme et al. 2020) | Põlme S, et al. 2020. Fungal Diversity 105:1-16 ((https://doi.org/10.1007/s13225-020-00466-2)) |
| Otu13177 | 1 | k__Fungi(100);p__Ascomycota(100);c__Saccharomycetes(100);o__Saccharomycetales(100);f__Metschnikowiaceae(100);g__Metschnikowia(100);s__Metschnikowia_pulcherrima(100); | Metschnikowia | 13 | Saprotroph | \|Undefined Saprotroph\| | Yeast | NULL | Probable | As nectar-tap_saprotroph (Põlme et al. 2020); Decay sugar-rich substrates (Põlme et al. 2020) | Põlme S, et al. 2020. Fungal Diversity 105:1-16 ((https://doi.org/10.1007/s13225-020-00466-2)) |
| Otu13179 | 1 | k__Fungi(100);p__Ascomycota(100);c__Saccharomycetes(100);o__Saccharomycetales(100);f__Metschnikowiaceae(100);g__Metschnikowia(100);s__Metschnikowia_pulcherrima(100); | Metschnikowia | 13 | Saprotroph | \|Undefined Saprotroph\| | Yeast | NULL | Probable | As nectar-tap_saprotroph (Põlme et al. 2020); Decay sugar-rich substrates (Põlme et al. 2020) | Põlme S, et al. 2020. Fungal Diversity 105:1-16 ((https://doi.org/10.1007/s13225-020-00466-2)) |
| Otu13180 | 1 | k__Fungi(100);p__Ascomycota(100);c__Saccharomycetes(100);o__Saccharomycetales(100);f__Metschnikowiaceae(100);g__Metschnikowia(100);g__Metschnikowia_unclassified(100); | Metschnikowia | 13 | Saprotroph | \|Undefined Saprotroph\| | Yeast | NULL | Probable | As nectar-tap_saprotroph (Põlme et al. 2020); Decay sugar-rich substrates (Põlme et al. 2020) | Põlme S, et al. 2020. Fungal Diversity 105:1-16 ((https://doi.org/10.1007/s13225-020-00466-2)) |
| Otu13181 | 1 | k__Fungi(100);p__Ascomycota(100);c__Saccharomycetes(100);o__Saccharomycetales(100);f__Metschnikowiaceae(100);g__Metschnikowia(100);s__Metschnikowia_pulcherrima(100); | Metschnikowia | 13 | Saprotroph | \|Undefined Saprotroph\| | Yeast | NULL | Probable | As nectar-tap_saprotroph (Põlme et al. 2020); Decay sugar-rich substrates (Põlme et al. 2020) | Põlme S, et al. 2020. Fungal Diversity 105:1-16 ((https://doi.org/10.1007/s13225-020-00466-2)) |
| Otu13182 | 1 | k__Fungi(100);p__Ascomycota(100);c__Saccharomycetes(100);o__Saccharomycetales(100);f__Metschnikowiaceae(100);g__Metschnikowia(100);s__Metschnikowia_pulcherrima(100); | Metschnikowia | 13 | Saprotroph | \|Undefined Saprotroph\| | Yeast | NULL | Probable | As nectar-tap_saprotroph (Põlme et al. 2020); Decay sugar-rich substrates (Põlme et al. 2020) | Põlme S, et al. 2020. Fungal Diversity 105:1-16 ((https://doi.org/10.1007/s13225-020-00466-2)) |
| Otu13184 | 1 | k__Fungi(100);p__Ascomycota(100);c__Saccharomycetes(100);o__Saccharomycetales(100);f__Metschnikowiaceae(100);g__Metschnikowia(100);s__Metschnikowia_pulcherrima(100); | Metschnikowia | 13 | Saprotroph | \|Undefined Saprotroph\| | Yeast | NULL | Probable | As nectar-tap_saprotroph (Põlme et al. 2020); Decay sugar-rich substrates (Põlme et al. 2020) | Põlme S, et al. 2020. Fungal Diversity 105:1-16 ((https://doi.org/10.1007/s13225-020-00466-2)) |
| Otu13185 | 1 | k__Fungi(100);p__Ascomycota(100);c__Saccharomycetes(100);o__Saccharomycetales(100);f__Metschnikowiaceae(100);g__Metschnikowia(100);s__Metschnikowia_pulcherrima(100); | Metschnikowia | 13 | Saprotroph | \|Undefined Saprotroph\| | Yeast | NULL | Probable | As nectar-tap_saprotroph (Põlme et al. 2020); Decay sugar-rich substrates (Põlme et al. 2020) | Põlme S, et al. 2020. Fungal Diversity 105:1-16 ((https://doi.org/10.1007/s13225-020-00466-2)) |
| Otu13186 | 1 | k__Fungi(100);p__Ascomycota(100);c__Saccharomycetes(100);o__Saccharomycetales(100);f__Metschnikowiaceae(100);g__Metschnikowia(100);s__Metschnikowia_pulcherrima(100); | Metschnikowia | 13 | Saprotroph | \|Undefined Saprotroph\| | Yeast | NULL | Probable | As nectar-tap_saprotroph (Põlme et al. 2020); Decay sugar-rich substrates (Põlme et al. 2020) | Põlme S, et al. 2020. Fungal Diversity 105:1-16 ((https://doi.org/10.1007/s13225-020-00466-2)) |
| Otu13188 | 1 | k__Fungi(100);p__Ascomycota(100);c__Saccharomycetes(100);o__Saccharomycetales(100);f__Metschnikowiaceae(100);g__Metschnikowia(100);s__Metschnikowia_pulcherrima(100); | Metschnikowia | 13 | Saprotroph | \|Undefined Saprotroph\| | Yeast | NULL | Probable | As nectar-tap_saprotroph (Põlme et al. 2020); Decay sugar-rich substrates (Põlme et al. 2020) | Põlme S, et al. 2020. Fungal Diversity 105:1-16 ((https://doi.org/10.1007/s13225-020-00466-2)) |
| Otu13189 | 1 | k__Fungi(100);p__Ascomycota(100);c__Saccharomycetes(100);o__Saccharomycetales(100);f__Metschnikowiaceae(100);g__Metschnikowia(100);g__Metschnikowia_unclassified(100); | Metschnikowia | 13 | Saprotroph | \|Undefined Saprotroph\| | Yeast | NULL | Probable | As nectar-tap_saprotroph (Põlme et al. 2020); Decay sugar-rich substrates (Põlme et al. 2020) | Põlme S, et al. 2020. Fungal Diversity 105:1-16 ((https://doi.org/10.1007/s13225-020-00466-2)) |
| Otu13191 | 1 | k__Fungi(100);p__Ascomycota(100);c__Saccharomycetes(100);o__Saccharomycetales(100);f__Metschnikowiaceae(100);g__Metschnikowia(100);s__Metschnikowia_pulcherrima(100); | Metschnikowia | 13 | Saprotroph | \|Undefined Saprotroph\| | Yeast | NULL | Probable | As nectar-tap_saprotroph (Põlme et al. 2020); Decay sugar-rich substrates (Põlme et al. 2020) | Põlme S, et al. 2020. Fungal Diversity 105:1-16 ((https://doi.org/10.1007/s13225-020-00466-2)) |
| Otu13194 | 1 | k__Fungi(100);p__Ascomycota(100);c__Saccharomycetes(100);o__Saccharomycetales(100);f__Metschnikowiaceae(100);g__Metschnikowia(100);s__Metschnikowia_pulcherrima(100); | Metschnikowia | 13 | Saprotroph | \|Undefined Saprotroph\| | Yeast | NULL | Probable | As nectar-tap_saprotroph (Põlme et al. 2020); Decay sugar-rich substrates (Põlme et al. 2020) | Põlme S, et al. 2020. Fungal Diversity 105:1-16 ((https://doi.org/10.1007/s13225-020-00466-2)) |
| Otu13195 | 1 | k__Fungi(100);p__Ascomycota(100);c__Saccharomycetes(100);o__Saccharomycetales(100);f__Metschnikowiaceae(100);g__Metschnikowia(100);s__Metschnikowia_pulcherrima(100); | Metschnikowia | 13 | Saprotroph | \|Undefined Saprotroph\| | Yeast | NULL | Probable | As nectar-tap_saprotroph (Põlme et al. 2020); Decay sugar-rich substrates (Põlme et al. 2020) | Põlme S, et al. 2020. Fungal Diversity 105:1-16 ((https://doi.org/10.1007/s13225-020-00466-2)) |
| Otu13196 | 1 | k__Fungi(100);p__Ascomycota(100);c__Saccharomycetes(100);o__Saccharomycetales(100);f__Metschnikowiaceae(100);g__Metschnikowia(100);s__Metschnikowia_pulcherrima(100); | Metschnikowia | 13 | Saprotroph | \|Undefined Saprotroph\| | Yeast | NULL | Probable | As nectar-tap_saprotroph (Põlme et al. 2020); Decay sugar-rich substrates (Põlme et al. 2020) | Põlme S, et al. 2020. Fungal Diversity 105:1-16 ((https://doi.org/10.1007/s13225-020-00466-2)) |
| Otu13197 | 1 | k__Fungi(100);p__Ascomycota(100);c__Saccharomycetes(100);o__Saccharomycetales(100);f__Saccharomycetales_family_Incertae_sedis(100);g__Candida(100);g__Candida_unclassified(100); | Candida | 13 | Pathotroph-Saprotroph-Symbiotroph | Animal Pathogen-Endophyte-Endosymbiont-Epiphyte-\|Undefined Saprotroph\| | Dimorphic Yeast-Yeast | NULL | Probable | As nectar-tap_saprotroph (Põlme et al. 2020); Decay sugar-rich substrates (Põlme et al. 2020); This asexual genus is polyphyletic and contain species that function very differently depend on what they are. Caution should be taken when interpreting guilds for Candida species. | Manolakaki D, et al. 2010. Virulence. 1: 367-375 ((https://doi.org/10.4161/viru.1.5.12796)); Mekbib SB, et al. 2011. Tropical Plant Pathology 36:233-240 ((https://doi.org/10.1590/S1982-56762011000400004)); Põlme S, et al. 2020. Fungal Diversity 105:1-16 ((https://doi.org/10.1007/s13225-020-00466-2)) |
| Otu13199 | 1 | k__Fungi(100);p__Ascomycota(100);c__Saccharomycetes(100);o__Saccharomycetales(100);f__Metschnikowiaceae(100);g__Metschnikowia(100);g__Metschnikowia_unclassified(100); | Metschnikowia | 13 | Saprotroph | \|Undefined Saprotroph\| | Yeast | NULL | Probable | As nectar-tap_saprotroph (Põlme et al. 2020); Decay sugar-rich substrates (Põlme et al. 2020) | Põlme S, et al. 2020. Fungal Diversity 105:1-16 ((https://doi.org/10.1007/s13225-020-00466-2)) |
| Otu13203 | 1 | k__Fungi(100);p__Ascomycota(100);c__Saccharomycetes(100);o__Saccharomycetales(100);f__Metschnikowiaceae(100);g__Metschnikowia(100);s__Metschnikowia_pulcherrima(100); | Metschnikowia | 13 | Saprotroph | \|Undefined Saprotroph\| | Yeast | NULL | Probable | As nectar-tap_saprotroph (Põlme et al. 2020); Decay sugar-rich substrates (Põlme et al. 2020) | Põlme S, et al. 2020. Fungal Diversity 105:1-16 ((https://doi.org/10.1007/s13225-020-00466-2)) |
| Otu13204 | 1 | k__Fungi(100);p__Ascomycota(100);c__Saccharomycetes(100);o__Saccharomycetales(100);f__Metschnikowiaceae(100);g__Metschnikowia(100);s__Metschnikowia_pulcherrima(100); | Metschnikowia | 13 | Saprotroph | \|Undefined Saprotroph\| | Yeast | NULL | Probable | As nectar-tap_saprotroph (Põlme et al. 2020); Decay sugar-rich substrates (Põlme et al. 2020) | Põlme S, et al. 2020. Fungal Diversity 105:1-16 ((https://doi.org/10.1007/s13225-020-00466-2)) |
| Otu13205 | 1 | k__Fungi(100);p__Ascomycota(100);c__Saccharomycetes(100);o__Saccharomycetales(100);f__Saccharomycetales_family_Incertae_sedis(100);g__Candida(100);g__Candida_unclassified(100); | Candida | 13 | Pathotroph-Saprotroph-Symbiotroph | Animal Pathogen-Endophyte-Endosymbiont-Epiphyte-\|Undefined Saprotroph\| | Dimorphic Yeast-Yeast | NULL | Probable | As nectar-tap_saprotroph (Põlme et al. 2020); Decay sugar-rich substrates (Põlme et al. 2020); This asexual genus is polyphyletic and contain species that function very differently depend on what they are. Caution should be taken when interpreting guilds for Candida species. | Manolakaki D, et al. 2010. Virulence. 1: 367-375 ((https://doi.org/10.4161/viru.1.5.12796)); Mekbib SB, et al. 2011. Tropical Plant Pathology 36:233-240 ((https://doi.org/10.1590/S1982-56762011000400004)); Põlme S, et al. 2020. Fungal Diversity 105:1-16 ((https://doi.org/10.1007/s13225-020-00466-2)) |
| Otu13207 | 1 | k__Fungi(100);p__Ascomycota(100);c__Saccharomycetes(100);o__Saccharomycetales(100);f__Metschnikowiaceae(100);g__Metschnikowia(100);s__Metschnikowia_pulcherrima(100); | Metschnikowia | 13 | Saprotroph | \|Undefined Saprotroph\| | Yeast | NULL | Probable | As nectar-tap_saprotroph (Põlme et al. 2020); Decay sugar-rich substrates (Põlme et al. 2020) | Põlme S, et al. 2020. Fungal Diversity 105:1-16 ((https://doi.org/10.1007/s13225-020-00466-2)) |
| Otu13209 | 1 | k__Fungi(100);p__Ascomycota(100);c__Saccharomycetes(100);o__Saccharomycetales(100);f__Metschnikowiaceae(100);g__Metschnikowia(100);s__Metschnikowia_pulcherrima(100); | Metschnikowia | 13 | Saprotroph | \|Undefined Saprotroph\| | Yeast | NULL | Probable | As nectar-tap_saprotroph (Põlme et al. 2020); Decay sugar-rich substrates (Põlme et al. 2020) | Põlme S, et al. 2020. Fungal Diversity 105:1-16 ((https://doi.org/10.1007/s13225-020-00466-2)) |
| Otu13210 | 1 | k__Fungi(100);p__Ascomycota(100);c__Saccharomycetes(100);o__Saccharomycetales(100);f__Metschnikowiaceae(100);g__Metschnikowia(100);g__Metschnikowia_unclassified(100); | Metschnikowia | 13 | Saprotroph | \|Undefined Saprotroph\| | Yeast | NULL | Probable | As nectar-tap_saprotroph (Põlme et al. 2020); Decay sugar-rich substrates (Põlme et al. 2020) | Põlme S, et al. 2020. Fungal Diversity 105:1-16 ((https://doi.org/10.1007/s13225-020-00466-2)) |
| Otu13211 | 1 | k__Fungi(100);p__Ascomycota(100);c__Saccharomycetes(100);o__Saccharomycetales(100);f__Metschnikowiaceae(100);g__Metschnikowia(100);s__Metschnikowia_pulcherrima(100); | Metschnikowia | 13 | Saprotroph | \|Undefined Saprotroph\| | Yeast | NULL | Probable | As nectar-tap_saprotroph (Põlme et al. 2020); Decay sugar-rich substrates (Põlme et al. 2020) | Põlme S, et al. 2020. Fungal Diversity 105:1-16 ((https://doi.org/10.1007/s13225-020-00466-2)) |
| Otu13212 | 1 | k__Fungi(100);p__Ascomycota(100);c__Saccharomycetes(100);o__Saccharomycetales(100);f__Metschnikowiaceae(100);g__Metschnikowia(100);s__Metschnikowia_pulcherrima(100); | Metschnikowia | 13 | Saprotroph | \|Undefined Saprotroph\| | Yeast | NULL | Probable | As nectar-tap_saprotroph (Põlme et al. 2020); Decay sugar-rich substrates (Põlme et al. 2020) | Põlme S, et al. 2020. Fungal Diversity 105:1-16 ((https://doi.org/10.1007/s13225-020-00466-2)) |
| Otu13213 | 1 | k__Fungi(100);p__Ascomycota(100);c__Saccharomycetes(100);o__Saccharomycetales(100);f__Saccharomycetales_family_Incertae_sedis(100);f__Saccharomycetales_family_Incertae_sedis_unclassified(100);f__Saccharomycetales_family_Incertae_sedis_unclassified(100); | Saccharomycetales | 7 | Saprotroph | Undefined Saprotroph | Yeast | NULL | Possible | NULL | Sterkenburg E, et al. 2015. New Phytologist 207:1145-1158 ((https://doi.org/10.1111/nph.13426)) |
| Otu13214 | 1 | k__Fungi(100);p__Ascomycota(100);c__Saccharomycetes(100);o__Saccharomycetales(100);f__Metschnikowiaceae(100);g__Metschnikowia(100);g__Metschnikowia_unclassified(100); | Metschnikowia | 13 | Saprotroph | \|Undefined Saprotroph\| | Yeast | NULL | Probable | As nectar-tap_saprotroph (Põlme et al. 2020); Decay sugar-rich substrates (Põlme et al. 2020) | Põlme S, et al. 2020. Fungal Diversity 105:1-16 ((https://doi.org/10.1007/s13225-020-00466-2)) |
| Otu13220 | 1 | k__Fungi(100);p__Ascomycota(100);c__Saccharomycetes(100);o__Saccharomycetales(100);f__Metschnikowiaceae(100);g__Metschnikowia(100);s__Metschnikowia_pulcherrima(100); | Metschnikowia | 13 | Saprotroph | \|Undefined Saprotroph\| | Yeast | NULL | Probable | As nectar-tap_saprotroph (Põlme et al. 2020); Decay sugar-rich substrates (Põlme et al. 2020) | Põlme S, et al. 2020. Fungal Diversity 105:1-16 ((https://doi.org/10.1007/s13225-020-00466-2)) |
| Otu13222 | 1 | k__Fungi(100);p__Ascomycota(100);c__Saccharomycetes(100);o__Saccharomycetales(100);f__Metschnikowiaceae(100);g__Metschnikowia(100);g__Metschnikowia_unclassified(100); | Metschnikowia | 13 | Saprotroph | \|Undefined Saprotroph\| | Yeast | NULL | Probable | As nectar-tap_saprotroph (Põlme et al. 2020); Decay sugar-rich substrates (Põlme et al. 2020) | Põlme S, et al. 2020. Fungal Diversity 105:1-16 ((https://doi.org/10.1007/s13225-020-00466-2)) |
| Otu13224 | 1 | k__Fungi(100);p__Ascomycota(100);c__Saccharomycetes(100);o__Saccharomycetales(100);f__Metschnikowiaceae(100);g__Metschnikowia(100);s__Metschnikowia_pulcherrima(100); | Metschnikowia | 13 | Saprotroph | \|Undefined Saprotroph\| | Yeast | NULL | Probable | As nectar-tap_saprotroph (Põlme et al. 2020); Decay sugar-rich substrates (Põlme et al. 2020) | Põlme S, et al. 2020. Fungal Diversity 105:1-16 ((https://doi.org/10.1007/s13225-020-00466-2)) |
| Otu13225 | 1 | k__Fungi(100);p__Ascomycota(100);c__Saccharomycetes(100);o__Saccharomycetales(100);f__Metschnikowiaceae(100);g__Metschnikowia(100);g__Metschnikowia_unclassified(100); | Metschnikowia | 13 | Saprotroph | \|Undefined Saprotroph\| | Yeast | NULL | Probable | As nectar-tap_saprotroph (Põlme et al. 2020); Decay sugar-rich substrates (Põlme et al. 2020) | Põlme S, et al. 2020. Fungal Diversity 105:1-16 ((https://doi.org/10.1007/s13225-020-00466-2)) |
| Otu13226 | 1 | k__Fungi(100);p__Ascomycota(100);c__Saccharomycetes(100);o__Saccharomycetales(100);f__Metschnikowiaceae(100);g__Metschnikowia(100);s__Metschnikowia_pulcherrima(100); | Metschnikowia | 13 | Saprotroph | \|Undefined Saprotroph\| | Yeast | NULL | Probable | As nectar-tap_saprotroph (Põlme et al. 2020); Decay sugar-rich substrates (Põlme et al. 2020) | Põlme S, et al. 2020. Fungal Diversity 105:1-16 ((https://doi.org/10.1007/s13225-020-00466-2)) |
| Otu13227 | 1 | k__Fungi(100);p__Ascomycota(100);c__Saccharomycetes(100);o__Saccharomycetales(100);f__Metschnikowiaceae(100);g__Metschnikowia(100);s__Metschnikowia_pulcherrima(100); | Metschnikowia | 13 | Saprotroph | \|Undefined Saprotroph\| | Yeast | NULL | Probable | As nectar-tap_saprotroph (Põlme et al. 2020); Decay sugar-rich substrates (Põlme et al. 2020) | Põlme S, et al. 2020. Fungal Diversity 105:1-16 ((https://doi.org/10.1007/s13225-020-00466-2)) |
| Otu13228 | 1 | k__Fungi(100);p__Ascomycota(100);c__Saccharomycetes(100);o__Saccharomycetales(100);f__Metschnikowiaceae(100);g__Metschnikowia(100);s__Metschnikowia_pulcherrima(100); | Metschnikowia | 13 | Saprotroph | \|Undefined Saprotroph\| | Yeast | NULL | Probable | As nectar-tap_saprotroph (Põlme et al. 2020); Decay sugar-rich substrates (Põlme et al. 2020) | Põlme S, et al. 2020. Fungal Diversity 105:1-16 ((https://doi.org/10.1007/s13225-020-00466-2)) |
| Otu13229 | 1 | k__Fungi(100);p__Ascomycota(100);c__Saccharomycetes(100);o__Saccharomycetales(100);f__Metschnikowiaceae(100);g__Metschnikowia(100);s__Metschnikowia_pulcherrima(100); | Metschnikowia | 13 | Saprotroph | \|Undefined Saprotroph\| | Yeast | NULL | Probable | As nectar-tap_saprotroph (Põlme et al. 2020); Decay sugar-rich substrates (Põlme et al. 2020) | Põlme S, et al. 2020. Fungal Diversity 105:1-16 ((https://doi.org/10.1007/s13225-020-00466-2)) |
| Otu13231 | 1 | k__Fungi(100);p__Ascomycota(100);c__Saccharomycetes(100);o__Saccharomycetales(100);f__Metschnikowiaceae(100);g__Metschnikowia(100);s__Metschnikowia_pulcherrima(100); | Metschnikowia | 13 | Saprotroph | \|Undefined Saprotroph\| | Yeast | NULL | Probable | As nectar-tap_saprotroph (Põlme et al. 2020); Decay sugar-rich substrates (Põlme et al. 2020) | Põlme S, et al. 2020. Fungal Diversity 105:1-16 ((https://doi.org/10.1007/s13225-020-00466-2)) |
| Otu13232 | 1 | k__Fungi(100);p__Ascomycota(100);c__Saccharomycetes(100);o__Saccharomycetales(100);f__Metschnikowiaceae(100);g__Metschnikowia(100);s__Metschnikowia_pulcherrima(100); | Metschnikowia | 13 | Saprotroph | \|Undefined Saprotroph\| | Yeast | NULL | Probable | As nectar-tap_saprotroph (Põlme et al. 2020); Decay sugar-rich substrates (Põlme et al. 2020) | Põlme S, et al. 2020. Fungal Diversity 105:1-16 ((https://doi.org/10.1007/s13225-020-00466-2)) |
| Otu13234 | 1 | k__Fungi(100);p__Ascomycota(100);c__Saccharomycetes(100);o__Saccharomycetales(100);f__Metschnikowiaceae(100);g__Metschnikowia(100);s__Metschnikowia_pulcherrima(100); | Metschnikowia | 13 | Saprotroph | \|Undefined Saprotroph\| | Yeast | NULL | Probable | As nectar-tap_saprotroph (Põlme et al. 2020); Decay sugar-rich substrates (Põlme et al. 2020) | Põlme S, et al. 2020. Fungal Diversity 105:1-16 ((https://doi.org/10.1007/s13225-020-00466-2)) |
| Otu13235 | 1 | k__Fungi(100);p__Ascomycota(100);c__Saccharomycetes(100);o__Saccharomycetales(100);f__Metschnikowiaceae(100);g__Metschnikowia(100);g__Metschnikowia_unclassified(100); | Metschnikowia | 13 | Saprotroph | \|Undefined Saprotroph\| | Yeast | NULL | Probable | As nectar-tap_saprotroph (Põlme et al. 2020); Decay sugar-rich substrates (Põlme et al. 2020) | Põlme S, et al. 2020. Fungal Diversity 105:1-16 ((https://doi.org/10.1007/s13225-020-00466-2)) |
| Otu13237 | 1 | k__Fungi(100);p__Ascomycota(100);c__Saccharomycetes(100);o__Saccharomycetales(100);f__Metschnikowiaceae(100);g__Metschnikowia(100);g__Metschnikowia_unclassified(100); | Metschnikowia | 13 | Saprotroph | \|Undefined Saprotroph\| | Yeast | NULL | Probable | As nectar-tap_saprotroph (Põlme et al. 2020); Decay sugar-rich substrates (Põlme et al. 2020) | Põlme S, et al. 2020. Fungal Diversity 105:1-16 ((https://doi.org/10.1007/s13225-020-00466-2)) |
| Otu13238 | 1 | k__Fungi(100);p__Ascomycota(100);c__Saccharomycetes(100);o__Saccharomycetales(100);f__Metschnikowiaceae(100);g__Metschnikowia(100);g__Metschnikowia_unclassified(100); | Metschnikowia | 13 | Saprotroph | \|Undefined Saprotroph\| | Yeast | NULL | Probable | As nectar-tap_saprotroph (Põlme et al. 2020); Decay sugar-rich substrates (Põlme et al. 2020) | Põlme S, et al. 2020. Fungal Diversity 105:1-16 ((https://doi.org/10.1007/s13225-020-00466-2)) |
| Otu13239 | 1 | k__Fungi(100);p__Ascomycota(100);c__Saccharomycetes(100);o__Saccharomycetales(100);f__Metschnikowiaceae(100);g__Metschnikowia(100);s__Metschnikowia_pulcherrima(100); | Metschnikowia | 13 | Saprotroph | \|Undefined Saprotroph\| | Yeast | NULL | Probable | As nectar-tap_saprotroph (Põlme et al. 2020); Decay sugar-rich substrates (Põlme et al. 2020) | Põlme S, et al. 2020. Fungal Diversity 105:1-16 ((https://doi.org/10.1007/s13225-020-00466-2)) |
| Otu13240 | 1 | k__Fungi(100);p__Ascomycota(100);c__Saccharomycetes(100);o__Saccharomycetales(100);f__Metschnikowiaceae(100);g__Metschnikowia(100);g__Metschnikowia_unclassified(100); | Metschnikowia | 13 | Saprotroph | \|Undefined Saprotroph\| | Yeast | NULL | Probable | As nectar-tap_saprotroph (Põlme et al. 2020); Decay sugar-rich substrates (Põlme et al. 2020) | Põlme S, et al. 2020. Fungal Diversity 105:1-16 ((https://doi.org/10.1007/s13225-020-00466-2)) |
| Otu13241 | 1 | k__Fungi(100);p__Ascomycota(100);c__Saccharomycetes(100);o__Saccharomycetales(100);f__Metschnikowiaceae(100);g__Metschnikowia(100);g__Metschnikowia_unclassified(100); | Metschnikowia | 13 | Saprotroph | \|Undefined Saprotroph\| | Yeast | NULL | Probable | As nectar-tap_saprotroph (Põlme et al. 2020); Decay sugar-rich substrates (Põlme et al. 2020) | Põlme S, et al. 2020. Fungal Diversity 105:1-16 ((https://doi.org/10.1007/s13225-020-00466-2)) |
| Otu13242 | 1 | k__Fungi(100);p__Ascomycota(100);c__Saccharomycetes(100);o__Saccharomycetales(100);f__Metschnikowiaceae(100);g__Metschnikowia(100);s__Metschnikowia_pulcherrima(100); | Metschnikowia | 13 | Saprotroph | \|Undefined Saprotroph\| | Yeast | NULL | Probable | As nectar-tap_saprotroph (Põlme et al. 2020); Decay sugar-rich substrates (Põlme et al. 2020) | Põlme S, et al. 2020. Fungal Diversity 105:1-16 ((https://doi.org/10.1007/s13225-020-00466-2)) |

| | | | | | | | | | | | |
|---|---|---|---|---|---|---|---|---|---|---|---|
| Otu13243 | 1 | k__Fungi(100);p__Ascomycota(100);c__Saccharomycetes(100);o__Saccharomycetales(100);f__Metschnikowiaceae(100);g__Metschnikowia(100);s__Metschnikowia_pulcherrima(100); | Metschnikowia | 13 | Saprotroph | \|Undefined Saprotroph\| | Yeast | NULL | Probable | As nectar-tap_saprotroph (Põlme et al. 2020); Decay sugar-rich substrates (Põlme et al. 2020) | Põlme S, et al. 2020. Fungal Diversity 105:1-16 ((https://doi.org/10.1007/s13225-020-00466-2)) |
| Otu13244 | 1 | k__Fungi(100);p__Ascomycota(100);c__Saccharomycetes(100);o__Saccharomycetales(100);f__Metschnikowiaceae(100);g__Metschnikowia(100);s__Metschnikowia_pulcherrima(100); | Metschnikowia | 13 | Saprotroph | \|Undefined Saprotroph\| | Yeast | NULL | Probable | As nectar-tap_saprotroph (Põlme et al. 2020); Decay sugar-rich substrates (Põlme et al. 2020) | Põlme S, et al. 2020. Fungal Diversity 105:1-16 ((https://doi.org/10.1007/s13225-020-00466-2)) |
| Otu13246 | 1 | k__Fungi(100);p__Ascomycota(100);c__Saccharomycetes(100);o__Saccharomycetales(100);o__Saccharomycetales_unclassified(100);o__Saccharomycetales_unclassified(100);o__Saccharomycetales_unclassified(100); | Saccharomycetales | 7 | Saprotroph | Undefined Saprotroph | Yeast | NULL | Possible | NULL | Sterkenburg E, et al. 2015. New Phytologist 207:1145-1158 ((https://doi.org/10.1111/nph.13426)) |
| Otu13247 | 1 | k__Fungi(100);p__Ascomycota(100);c__Saccharomycetes(100);o__Saccharomycetales(100);f__Metschnikowiaceae(100);g__Metschnikowia(100);s__Metschnikowia_pulcherrima(100); | Metschnikowia | 13 | Saprotroph | \|Undefined Saprotroph\| | Yeast | NULL | Probable | As nectar-tap_saprotroph (Põlme et al. 2020); Decay sugar-rich substrates (Põlme et al. 2020) | Põlme S, et al. 2020. Fungal Diversity 105:1-16 ((https://doi.org/10.1007/s13225-020-00466-2)) |
| Otu13248 | 1 | k__Fungi(100);p__Ascomycota(100);c__Saccharomycetes(100);o__Saccharomycetales(100);f__Metschnikowiaceae(100);g__Metschnikowia(100);g__Metschnikowia_unclassified(100); | Metschnikowia | 13 | Saprotroph | \|Undefined Saprotroph\| | Yeast | NULL | Probable | As nectar-tap_saprotroph (Põlme et al. 2020); Decay sugar-rich substrates (Põlme et al. 2020) | Põlme S, et al. 2020. Fungal Diversity 105:1-16 ((https://doi.org/10.1007/s13225-020-00466-2)) |
| Otu13251 | 1 | k__Fungi(100);p__Ascomycota(100);c__Saccharomycetes(100);o__Saccharomycetales(100);f__Metschnikowiaceae(100);g__Metschnikowia(100);s__Metschnikowia_pulcherrima(100); | Metschnikowia | 13 | Saprotroph | \|Undefined Saprotroph\| | Yeast | NULL | Probable | As nectar-tap_saprotroph (Põlme et al. 2020); Decay sugar-rich substrates (Põlme et al. 2020) | Põlme S, et al. 2020. Fungal Diversity 105:1-16 ((https://doi.org/10.1007/s13225-020-00466-2)) |
| Otu13254 | 1 | k__Fungi(100);p__Ascomycota(100);c__Saccharomycetes(100);o__Saccharomycetales(100);f__Metschnikowiaceae(100);g__Metschnikowia(100);s__Metschnikowia_pulcherrima(100); | Metschnikowia | 13 | Saprotroph | \|Undefined Saprotroph\| | Yeast | NULL | Probable | As nectar-tap_saprotroph (Põlme et al. 2020); Decay sugar-rich substrates (Põlme et al. 2020) | Põlme S, et al. 2020. Fungal Diversity 105:1-16 ((https://doi.org/10.1007/s13225-020-00466-2)) |
| Otu13255 | 1 | k__Fungi(100);p__Ascomycota(100);c__Saccharomycetes(100);o__Saccharomycetales(100);f__Metschnikowiaceae(100);g__Metschnikowia(100);s__Metschnikowia_pulcherrima(100); | Metschnikowia | 13 | Saprotroph | \|Undefined Saprotroph\| | Yeast | NULL | Probable | As nectar-tap_saprotroph (Põlme et al. 2020); Decay sugar-rich substrates (Põlme et al. 2020) | Põlme S, et al. 2020. Fungal Diversity 105:1-16 ((https://doi.org/10.1007/s13225-020-00466-2)) |
| Otu13257 | 1 | k__Fungi(100);p__Ascomycota(100);c__Saccharomycetes(100);o__Saccharomycetales(100);f__Metschnikowiaceae(100);g__Metschnikowia(100);g__Metschnikowia_unclassified(100); | Metschnikowia | 13 | Saprotroph | \|Undefined Saprotroph\| | Yeast | NULL | Probable | As nectar-tap_saprotroph (Põlme et al. 2020); Decay sugar-rich substrates (Põlme et al. 2020) | Põlme S, et al. 2020. Fungal Diversity 105:1-16 ((https://doi.org/10.1007/s13225-020-00466-2)) |
| Otu13260 | 1 | k__Fungi(100);p__Ascomycota(100);c__Saccharomycetes(100);o__Saccharomycetales(100);f__Metschnikowiaceae(100);g__Metschnikowia(100);s__Metschnikowia_pulcherrima(100); | Metschnikowia | 13 | Saprotroph | \|Undefined Saprotroph\| | Yeast | NULL | Probable | As nectar-tap_saprotroph (Põlme et al. 2020); Decay sugar-rich substrates (Põlme et al. 2020) | Põlme S, et al. 2020. Fungal Diversity 105:1-16 ((https://doi.org/10.1007/s13225-020-00466-2)) |
| Otu13261 | 1 | k__Fungi(100);p__Ascomycota(100);c__Saccharomycetes(100);o__Saccharomycetales(100);f__Metschnikowiaceae(100);g__Metschnikowia(100);s__Metschnikowia_pulcherrima(100); | Metschnikowia | 13 | Saprotroph | \|Undefined Saprotroph\| | Yeast | NULL | Probable | As nectar-tap_saprotroph (Põlme et al. 2020); Decay sugar-rich substrates (Põlme et al. 2020) | Põlme S, et al. 2020. Fungal Diversity 105:1-16 ((https://doi.org/10.1007/s13225-020-00466-2)) |
| Otu13262 | 1 | k__Fungi(100);p__Ascomycota(100);c__Saccharomycetes(100);o__Saccharomycetales(100);f__Saccharomycetales_family_Incertae_sedis(100);g__Candida(100);g__Candida_unclassified(100); | Candida | 13 | Pathotroph-Saprotroph-Symbiotroph | Animal Pathogen-Endophyte-Endosymbiont-Epiphyte-\|Undefined Saprotroph\| | Dimorphic Yeast-Yeast | NULL | Probable | As nectar-tap_saprotroph (Põlme et al. 2020); Decay sugar-rich substrates (Põlme et al. 2020); This asexual genus is polyphyletic and contain species that function very differently depend on what they are. Caution should be taken when interpreting guilds for Candida species. | Manolakaki D, et al. 2010. Virulence. 1: 367-375 ((https://doi.org/10.4161/viru.1.5.12796)); Mekbib SB, et al. 2011. Tropical Plant Pathology 36:233-240 ((https://doi.org/10.1590/S1982-56762011000400004)); Põlme S, et al. 2020. Fungal Diversity 105:1-16 ((https://doi.org/10.1007/s13225-020-00466-2)) |
| Otu13265 | 1 | k__Fungi(100);p__Ascomycota(100);c__Saccharomycetes(100);o__Saccharomycetales(100);f__Metschnikowiaceae(100);g__Metschnikowia(100);s__Metschnikowia_pulcherrima(100); | Metschnikowia | 13 | Saprotroph | \|Undefined Saprotroph\| | Yeast | NULL | Probable | As nectar-tap_saprotroph (Põlme et al. 2020); Decay sugar-rich substrates (Põlme et al. 2020) | Põlme S, et al. 2020. Fungal Diversity 105:1-16 ((https://doi.org/10.1007/s13225-020-00466-2)) |
| Otu13266 | 1 | k__Fungi(100);p__Ascomycota(100);c__Saccharomycetes(100);o__Saccharomycetales(100);f__Metschnikowiaceae(100);g__Metschnikowia(100);s__Metschnikowia_pulcherrima(100); | Metschnikowia | 13 | Saprotroph | \|Undefined Saprotroph\| | Yeast | NULL | Probable | As nectar-tap_saprotroph (Põlme et al. 2020); Decay sugar-rich substrates (Põlme et al. 2020) | Põlme S, et al. 2020. Fungal Diversity 105:1-16 ((https://doi.org/10.1007/s13225-020-00466-2)) |
| Otu13267 | 1 | k__Fungi(100);p__Ascomycota(100);c__Saccharomycetes(100);o__Saccharomycetales(100);f__Metschnikowiaceae(100);g__Metschnikowia(100);s__Metschnikowia_pulcherrima(100); | Metschnikowia | 13 | Saprotroph | \|Undefined Saprotroph\| | Yeast | NULL | Probable | As nectar-tap_saprotroph (Põlme et al. 2020); Decay sugar-rich substrates (Põlme et al. 2020) | Põlme S, et al. 2020. Fungal Diversity 105:1-16 ((https://doi.org/10.1007/s13225-020-00466-2)) |
| Otu13269 | 1 | k__Fungi(100);p__Ascomycota(100);c__Saccharomycetes(100);o__Saccharomycetales(100);f__Metschnikowiaceae(100);g__Metschnikowia(100);s__Metschnikowia_pulcherrima(100); | Metschnikowia | 13 | Saprotroph | \|Undefined Saprotroph\| | Yeast | NULL | Probable | As nectar-tap_saprotroph (Põlme et al. 2020); Decay sugar-rich substrates (Põlme et al. 2020) | Põlme S, et al. 2020. Fungal Diversity 105:1-16 ((https://doi.org/10.1007/s13225-020-00466-2)) |
| Otu13270 | 1 | k__Fungi(100);p__Ascomycota(100);c__Saccharomycetes(100);o__Saccharomycetales(100);f__Metschnikowiaceae(100);g__Metschnikowia(100);s__Metschnikowia_pulcherrima(100); | Metschnikowia | 13 | Saprotroph | \|Undefined Saprotroph\| | Yeast | NULL | Probable | As nectar-tap_saprotroph (Põlme et al. 2020); Decay sugar-rich substrates (Põlme et al. 2020) | Põlme S, et al. 2020. Fungal Diversity 105:1-16 ((https://doi.org/10.1007/s13225-020-00466-2)) |
| Otu13271 | 1 | k__Fungi(100);p__Ascomycota(100);c__Saccharomycetes(100);o__Saccharomycetales(100);f__Metschnikowiaceae(100);g__Metschnikowia(100);s__Metschnikowia_pulcherrima(100); | Metschnikowia | 13 | Saprotroph | \|Undefined Saprotroph\| | Yeast | NULL | Probable | As nectar-tap_saprotroph (Põlme et al. 2020); Decay sugar-rich substrates (Põlme et al. 2020) | Põlme S, et al. 2020. Fungal Diversity 105:1-16 ((https://doi.org/10.1007/s13225-020-00466-2)) |
| Otu13272 | 1 | k__Fungi(100);p__Ascomycota(100);c__Saccharomycetes(100);o__Saccharomycetales(100);f__Metschnikowiaceae(100);g__Metschnikowia(100);s__Metschnikowia_pulcherrima(100); | Metschnikowia | 13 | Saprotroph | \|Undefined Saprotroph\| | Yeast | NULL | Probable | As nectar-tap_saprotroph (Põlme et al. 2020); Decay sugar-rich substrates (Põlme et al. 2020) | Põlme S, et al. 2020. Fungal Diversity 105:1-16 ((https://doi.org/10.1007/s13225-020-00466-2)) |
| Otu13273 | 1 | k__Fungi(100);p__Ascomycota(100);c__Saccharomycetes(100);o__Saccharomycetales(100);f__Metschnikowiaceae(100);g__Metschnikowia(100);s__Metschnikowia_pulcherrima(100); | Metschnikowia | 13 | Saprotroph | \|Undefined Saprotroph\| | Yeast | NULL | Probable | As nectar-tap_saprotroph (Põlme et al. 2020); Decay sugar-rich substrates (Põlme et al. 2020) | Põlme S, et al. 2020. Fungal Diversity 105:1-16 ((https://doi.org/10.1007/s13225-020-00466-2)) |
| Otu13275 | 1 | k__Fungi(100);p__Ascomycota(100);c__Saccharomycetes(100);o__Saccharomycetales(100);f__Metschnikowiaceae(100);g__Metschnikowia(100);s__Metschnikowia_pulcherrima(100); | Metschnikowia | 13 | Saprotroph | \|Undefined Saprotroph\| | Yeast | NULL | Probable | As nectar-tap_saprotroph (Põlme et al. 2020); Decay sugar-rich substrates (Põlme et al. 2020) | Põlme S, et al. 2020. Fungal Diversity 105:1-16 ((https://doi.org/10.1007/s13225-020-00466-2)) |
| Otu13276 | 1 | k__Fungi(100);p__Ascomycota(100);c__Saccharomycetes(100);o__Saccharomycetales(100);f__Metschnikowiaceae(100);g__Metschnikowia(100);g__Metschnikowia_unclassified(100); | Metschnikowia | 13 | Saprotroph | \|Undefined Saprotroph\| | Yeast | NULL | Probable | As nectar-tap_saprotroph (Põlme et al. 2020); Decay sugar-rich substrates (Põlme et al. 2020) | Põlme S, et al. 2020. Fungal Diversity 105:1-16 ((https://doi.org/10.1007/s13225-020-00466-2)) |
| Otu13277 | 1 | k__Fungi(100);p__Ascomycota(100);c__Saccharomycetes(100);o__Saccharomycetales(100);f__Metschnikowiaceae(100);g__Metschnikowia(100);s__Metschnikowia_pulcherrima(100); | Metschnikowia | 13 | Saprotroph | \|Undefined Saprotroph\| | Yeast | NULL | Probable | As nectar-tap_saprotroph (Põlme et al. 2020); Decay sugar-rich substrates (Põlme et al. 2020) | Põlme S, et al. 2020. Fungal Diversity 105:1-16 ((https://doi.org/10.1007/s13225-020-00466-2)) |
| Otu13280 | 1 | k__Fungi(100);p__Ascomycota(100);c__Saccharomycetes(100);o__Saccharomycetales(100);f__Metschnikowiaceae(100);g__Metschnikowia(100);g__Metschnikowia_unclassified(100); | Metschnikowia | 13 | Saprotroph | \|Undefined Saprotroph\| | Yeast | NULL | Probable | As nectar-tap_saprotroph (Põlme et al. 2020); Decay sugar-rich substrates (Põlme et al. 2020) | Põlme S, et al. 2020. Fungal Diversity 105:1-16 ((https://doi.org/10.1007/s13225-020-00466-2)) |
| Otu13283 | 1 | k__Fungi(100);p__Ascomycota(100);c__Saccharomycetes(100);o__Saccharomycetales(100);f__Metschnikowiaceae(100);g__Metschnikowia(100);g__Metschnikowia_unclassified(100); | Metschnikowia | 13 | Saprotroph | \|Undefined Saprotroph\| | Yeast | NULL | Probable | As nectar-tap_saprotroph (Põlme et al. 2020); Decay sugar-rich substrates (Põlme et al. 2020) | Põlme S, et al. 2020. Fungal Diversity 105:1-16 ((https://doi.org/10.1007/s13225-020-00466-2)) |
| Otu13285 | 1 | k__Fungi(100);p__Ascomycota(100);c__Saccharomycetes(100);o__Saccharomycetales(100);f__Metschnikowiaceae(100);g__Metschnikowia(100);s__Metschnikowia_pulcherrima(100); | Metschnikowia | 13 | Saprotroph | \|Undefined Saprotroph\| | Yeast | NULL | Probable | As nectar-tap_saprotroph (Põlme et al. 2020); Decay sugar-rich substrates (Põlme et al. 2020) | Põlme S, et al. 2020. Fungal Diversity 105:1-16 ((https://doi.org/10.1007/s13225-020-00466-2)) |
| Otu13288 | 1 | k__Fungi(100);p__Ascomycota(100);c__Saccharomycetes(100);o__Saccharomycetales(100);f__Metschnikowiaceae(100);g__Metschnikowia(100);s__Metschnikowia_pulcherrima(100); | Metschnikowia | 13 | Saprotroph | \|Undefined Saprotroph\| | Yeast | NULL | Probable | As nectar-tap_saprotroph (Põlme et al. 2020); Decay sugar-rich substrates (Põlme et al. 2020) | Põlme S, et al. 2020. Fungal Diversity 105:1-16 ((https://doi.org/10.1007/s13225-020-00466-2)) |
| Otu13289 | 1 | k__Fungi(100);p__Ascomycota(100);c__Saccharomycetes(100);o__Saccharomycetales(100);f__Metschnikowiaceae(100);g__Metschnikowia(100);g__Metschnikowia_unclassified(100); | Metschnikowia | 13 | Saprotroph | \|Undefined Saprotroph\| | Yeast | NULL | Probable | As nectar-tap_saprotroph (Põlme et al. 2020); Decay sugar-rich substrates (Põlme et al. 2020) | Põlme S, et al. 2020. Fungal Diversity 105:1-16 ((https://doi.org/10.1007/s13225-020-00466-2)) |
| Otu13291 | 1 | k__Fungi(100);p__Ascomycota(100);c__Saccharomycetes(100);o__Saccharomycetales(100);f__Metschnikowiaceae(100);g__Metschnikowia(100);s__Metschnikowia_pulcherrima(100); | Metschnikowia | 13 | Saprotroph | \|Undefined Saprotroph\| | Yeast | NULL | Probable | As nectar-tap_saprotroph (Põlme et al. 2020); Decay sugar-rich substrates (Põlme et al. 2020) | Põlme S, et al. 2020. Fungal Diversity 105:1-16 ((https://doi.org/10.1007/s13225-020-00466-2)) |
| Otu13292 | 1 | k__Fungi(100);p__Ascomycota(100);c__Saccharomycetes(100);o__Saccharomycetales(100);f__Metschnikowiaceae(100);g__Metschnikowia(100);s__Metschnikowia_pulcherrima(100); | Metschnikowia | 13 | Saprotroph | \|Undefined Saprotroph\| | Yeast | NULL | Probable | As nectar-tap_saprotroph (Põlme et al. 2020); Decay sugar-rich substrates (Põlme et al. 2020) | Põlme S, et al. 2020. Fungal Diversity 105:1-16 ((https://doi.org/10.1007/s13225-020-00466-2)) |
| Otu13295 | 1 | k__Fungi(100);p__Ascomycota(100);c__Saccharomycetes(100);o__Saccharomycetales(100);f__Metschnikowiaceae(100);g__Metschnikowia(100);s__Metschnikowia_pulcherrima(100); | Metschnikowia | 13 | Saprotroph | \|Undefined Saprotroph\| | Yeast | NULL | Probable | As nectar-tap_saprotroph (Põlme et al. 2020); Decay sugar-rich substrates (Põlme et al. 2020) | Põlme S, et al. 2020. Fungal Diversity 105:1-16 ((https://doi.org/10.1007/s13225-020-00466-2)) |
| Otu13296 | 1 | k__Fungi(100);p__Ascomycota(100);c__Saccharomycetes(100);o__Saccharomycetales(100);f__Metschnikowiaceae(100);g__Metschnikowia(100);s__Metschnikowia_pulcherrima(100); | Metschnikowia | 13 | Saprotroph | \|Undefined Saprotroph\| | Yeast | NULL | Probable | As nectar-tap_saprotroph (Põlme et al. 2020); Decay sugar-rich substrates (Põlme et al. 2020) | Põlme S, et al. 2020. Fungal Diversity 105:1-16 ((https://doi.org/10.1007/s13225-020-00466-2)) |
| Otu13297 | 1 | k__Fungi(100);p__Ascomycota(100);c__Saccharomycetes(100);o__Saccharomycetales(100);f__Metschnikowiaceae(100);g__Metschnikowia(100);g__Metschnikowia_unclassified(100); | Metschnikowia | 13 | Saprotroph | \|Undefined Saprotroph\| | Yeast | NULL | Probable | As nectar-tap_saprotroph (Põlme et al. 2020); Decay sugar-rich substrates (Põlme et al. 2020) | Põlme S, et al. 2020. Fungal Diversity 105:1-16 ((https://doi.org/10.1007/s13225-020-00466-2)) |
| Otu13298 | 1 | k__Fungi(100);p__Ascomycota(100);c__Saccharomycetes(100);o__Saccharomycetales(100);f__Metschnikowiaceae(100);g__Metschnikowia(100);s__Metschnikowia_pulcherrima(100); | Metschnikowia | 13 | Saprotroph | \|Undefined Saprotroph\| | Yeast | NULL | Probable | As nectar-tap_saprotroph (Põlme et al. 2020); Decay sugar-rich substrates (Põlme et al. 2020) | Põlme S, et al. 2020. Fungal Diversity 105:1-16 ((https://doi.org/10.1007/s13225-020-00466-2)) |
| Otu13299 | 1 | k__Fungi(100);p__Ascomycota(100);c__Saccharomycetes(100);o__Saccharomycetales(100);f__Metschnikowiaceae(100);g__Metschnikowia(100);s__Metschnikowia_pulcherrima(100); | Metschnikowia | 13 | Saprotroph | \|Undefined Saprotroph\| | Yeast | NULL | Probable | As nectar-tap_saprotroph (Põlme et al. 2020); Decay sugar-rich substrates (Põlme et al. 2020) | Põlme S, et al. 2020. Fungal Diversity 105:1-16 ((https://doi.org/10.1007/s13225-020-00466-2)) |
| Otu13303 | 1 | k__Fungi(100);p__Ascomycota(100);c__Saccharomycetes(100);o__Saccharomycetales(100);f__Metschnikowiaceae(100);g__Metschnikowia(100);g__Metschnikowia_unclassified(100); | Metschnikowia | 13 | Saprotroph | \|Undefined Saprotroph\| | Yeast | NULL | Probable | As nectar-tap_saprotroph (Põlme et al. 2020); Decay sugar-rich substrates (Põlme et al. 2020) | Põlme S, et al. 2020. Fungal Diversity 105:1-16 ((https://doi.org/10.1007/s13225-020-00466-2)) |
| Otu13304 | 1 | k__Fungi(100);p__Ascomycota(100);c__Saccharomycetes(100);o__Saccharomycetales(100);f__Metschnikowiaceae(100);g__Metschnikowia(100);g__Metschnikowia_unclassified(100); | Metschnikowia | 13 | Saprotroph | \|Undefined Saprotroph\| | Yeast | NULL | Probable | As nectar-tap_saprotroph (Põlme et al. 2020); Decay sugar-rich substrates (Põlme et al. 2020) | Põlme S, et al. 2020. Fungal Diversity 105:1-16 ((https://doi.org/10.1007/s13225-020-00466-2)) |
| Otu13305 | 1 | k__Fungi(100);p__Ascomycota(100);c__Saccharomycetes(100);o__Saccharomycetales(100);f__Metschnikowiaceae(100);g__Metschnikowia(100);s__Metschnikowia_pulcherrima(100); | Metschnikowia | 13 | Saprotroph | \|Undefined Saprotroph\| | Yeast | NULL | Probable | As nectar-tap_saprotroph (Põlme et al. 2020); Decay sugar-rich substrates (Põlme et al. 2020) | Põlme S, et al. 2020. Fungal Diversity 105:1-16 ((https://doi.org/10.1007/s13225-020-00466-2)) |
| Otu13306 | 1 | k__Fungi(100);p__Ascomycota(100);c__Saccharomycetes(100);o__Saccharomycetales(100);f__Metschnikowiaceae(100);g__Metschnikowia(100);g__Metschnikowia_unclassified(100); | Metschnikowia | 13 | Saprotroph | \|Undefined Saprotroph\| | Yeast | NULL | Probable | As nectar-tap_saprotroph (Põlme et al. 2020); Decay sugar-rich substrates (Põlme et al. 2020) | Põlme S, et al. 2020. Fungal Diversity 105:1-16 ((https://doi.org/10.1007/s13225-020-00466-2)) |
| Otu13307 | 1 | k__Fungi(100);p__Ascomycota(100);c__Saccharomycetes(100);o__Saccharomycetales(100);f__Metschnikowiaceae(100);g__Metschnikowia(100);s__Metschnikowia_pulcherrima(100); | Metschnikowia | 13 | Saprotroph | \|Undefined Saprotroph\| | Yeast | NULL | Probable | As nectar-tap_saprotroph (Põlme et al. 2020); Decay sugar-rich substrates (Põlme et al. 2020) | Põlme S, et al. 2020. Fungal Diversity 105:1-16 ((https://doi.org/10.1007/s13225-020-00466-2)) |
| Otu13308 | 1 | k__Fungi(100);p__Ascomycota(100);c__Saccharomycetes(100);o__Saccharomycetales(100);f__Metschnikowiaceae(100);g__Metschnikowia(100);s__Metschnikowia_pulcherrima(100); | Metschnikowia | 13 | Saprotroph | \|Undefined Saprotroph\| | Yeast | NULL | Probable | As nectar-tap_saprotroph (Põlme et al. 2020); Decay sugar-rich substrates (Põlme et al. 2020) | Põlme S, et al. 2020. Fungal Diversity 105:1-16 ((https://doi.org/10.1007/s13225-020-00466-2)) |
| Otu13309 | 1 | k__Fungi(100);p__Ascomycota(100);c__Saccharomycetes(100);o__Saccharomycetales(100);f__Metschnikowiaceae(100);g__Metschnikowia(100);s__Metschnikowia_pulcherrima(100); | Metschnikowia | 13 | Saprotroph | \|Undefined Saprotroph\| | Yeast | NULL | Probable | As nectar-tap_saprotroph (Põlme et al. 2020); Decay sugar-rich substrates (Põlme et al. 2020) | Põlme S, et al. 2020. Fungal Diversity 105:1-16 ((https://doi.org/10.1007/s13225-020-00466-2)) |
| Otu13310 | 1 | k__Fungi(100);p__Ascomycota(100);c__Saccharomycetes(100);o__Saccharomycetales(100);f__Metschnikowiaceae(100);g__Metschnikowia(100);s__Metschnikowia_pulcherrima(100); | Metschnikowia | 13 | Saprotroph | \|Undefined Saprotroph\| | Yeast | NULL | Probable | As nectar-tap_saprotroph (Põlme et al. 2020); Decay sugar-rich substrates (Põlme et al. 2020) | Põlme S, et al. 2020. Fungal Diversity 105:1-16 ((https://doi.org/10.1007/s13225-020-00466-2)) |
| Otu13312 | 1 | k__Fungi(100);p__Ascomycota(100);c__Saccharomycetes(100);o__Saccharomycetales(100);f__Metschnikowiaceae(100);g__Metschnikowia(100);s__Metschnikowia_pulcherrima(100); | Metschnikowia | 13 | Saprotroph | \|Undefined Saprotroph\| | Yeast | NULL | Probable | As nectar-tap_saprotroph (Põlme et al. 2020); Decay sugar-rich substrates (Põlme et al. 2020) | Põlme S, et al. 2020. Fungal Diversity 105:1-16 ((https://doi.org/10.1007/s13225-020-00466-2)) |
| Otu13313 | 1 | k__Fungi(100);p__Ascomycota(100);c__Saccharomycetes(100);o__Saccharomycetales(100);f__Metschnikowiaceae(100);g__Metschnikowia(100);s__Metschnikowia_pulcherrima(100); | Metschnikowia | 13 | Saprotroph | \|Undefined Saprotroph\| | Yeast | NULL | Probable | As nectar-tap_saprotroph (Põlme et al. 2020); Decay sugar-rich substrates (Põlme et al. 2020) | Põlme S, et al. 2020. Fungal Diversity 105:1-16 ((https://doi.org/10.1007/s13225-020-00466-2)) |
| Otu13314 | 1 | k__Fungi(100);p__Ascomycota(100);c__Saccharomycetes(100);o__Saccharomycetales(100);f__Metschnikowiaceae(100);g__Metschnikowia(100);s__Metschnikowia_pulcherrima(100); | Metschnikowia | 13 | Saprotroph | \|Undefined Saprotroph\| | Yeast | NULL | Probable | As nectar-tap_saprotroph (Põlme et al. 2020); Decay sugar-rich substrates (Põlme et al. 2020) | Põlme S, et al. 2020. Fungal Diversity 105:1-16 ((https://doi.org/10.1007/s13225-020-00466-2)) |
| Otu13317 | 1 | k__Fungi(100);p__Ascomycota(100);c__Saccharomycetes(100);o__Saccharomycetales(100);f__Metschnikowiaceae(100);g__Metschnikowia(100);s__Metschnikowia_pulcherrima(100); | Metschnikowia | 13 | Saprotroph | \|Undefined Saprotroph\| | Yeast | NULL | Probable | As nectar-tap_saprotroph (Põlme et al. 2020); Decay sugar-rich substrates (Põlme et al. 2020) | Põlme S, et al. 2020. Fungal Diversity 105:1-16 ((https://doi.org/10.1007/s13225-020-00466-2)) |
| Otu13318 | 1 | k__Fungi(100);p__Ascomycota(100);c__Saccharomycetes(100);o__Saccharomycetales(100);f__Metschnikowiaceae(100);g__Metschnikowia(100);s__Metschnikowia_pulcherrima(100); | Metschnikowia | 13 | Saprotroph | \|Undefined Saprotroph\| | Yeast | NULL | Probable | As nectar-tap_saprotroph (Põlme et al. 2020); Decay sugar-rich substrates (Põlme et al. 2020) | Põlme S, et al. 2020. Fungal Diversity 105:1-16 ((https://doi.org/10.1007/s13225-020-00466-2)) |
| Otu13319 | 1 | k__Fungi(100);p__Ascomycota(100);c__Saccharomycetes(100);o__Saccharomycetales(100);f__Metschnikowiaceae(100);g__Metschnikowia(100);s__Metschnikowia_pulcherrima(100); | Metschnikowia | 13 | Saprotroph | \|Undefined Saprotroph\| | Yeast | NULL | Probable | As nectar-tap_saprotroph (Põlme et al. 2020); Decay sugar-rich substrates (Põlme et al. 2020) | Põlme S, et al. 2020. Fungal Diversity 105:1-16 ((https://doi.org/10.1007/s13225-020-00466-2)) |
| Otu13320 | 1 | k__Fungi(100);p__Ascomycota(100);c__Saccharomycetes(100);o__Saccharomycetales(100);f__Metschnikowiaceae(100);g__Metschnikowia(100);s__Metschnikowia_pulcherrima(100); | Metschnikowia | 13 | Saprotroph | \|Undefined Saprotroph\| | Yeast | NULL | Probable | As nectar-tap_saprotroph (Põlme et al. 2020); Decay sugar-rich substrates (Põlme et al. 2020) | Põlme S, et al. 2020. Fungal Diversity 105:1-16 ((https://doi.org/10.1007/s13225-020-00466-2)) |
| Otu13321 | 1 | k__Fungi(100);p__Ascomycota(100);c__Saccharomycetes(100);o__Saccharomycetales(100);f__Metschnikowiaceae(100);g__Metschnikowia(100);s__Metschnikowia_pulcherrima(100); | Metschnikowia | 13 | Saprotroph | \|Undefined Saprotroph\| | Yeast | NULL | Probable | As nectar-tap_saprotroph (Põlme et al. 2020); Decay sugar-rich substrates (Põlme et al. 2020) | Põlme S, et al. 2020. Fungal Diversity 105:1-16 ((https://doi.org/10.1007/s13225-020-00466-2)) |
| Otu13322 | 1 | k__Fungi(100);p__Ascomycota(100);c__Saccharomycetes(100);o__Saccharomycetales(100);f__Metschnikowiaceae(100);g__Metschnikowia(100);g__Metschnikowia_unclassified(100); | Metschnikowia | 13 | Saprotroph | \|Undefined Saprotroph\| | Yeast | NULL | Probable | As nectar-tap_saprotroph (Põlme et al. 2020); Decay sugar-rich substrates (Põlme et al. 2020) | Põlme S, et al. 2020. Fungal Diversity 105:1-16 ((https://doi.org/10.1007/s13225-020-00466-2)) |
| Otu13323 | 1 | k__Fungi(100);p__Ascomycota(100);c__Saccharomycetes(100);o__Saccharomycetales(100);f__Metschnikowiaceae(100);g__Metschnikowia(100);g__Metschnikowia_unclassified(100); | Metschnikowia | 13 | Saprotroph | \|Undefined Saprotroph\| | Yeast | NULL | Probable | As nectar-tap_saprotroph (Põlme et al. 2020); Decay sugar-rich substrates (Põlme et al. 2020) | Põlme S, et al. 2020. Fungal Diversity 105:1-16 ((https://doi.org/10.1007/s13225-020-00466-2)) |
| Otu13325 | 1 | k__Fungi(100);p__Ascomycota(100);c__Saccharomycetes(100);o__Saccharomycetales(100);f__Metschnikowiaceae(100);g__Metschnikowia(100);g__Metschnikowia_unclassified(100); | Metschnikowia | 13 | Saprotroph | \|Undefined Saprotroph\| | Yeast | NULL | Probable | As nectar-tap_saprotroph (Põlme et al. 2020); Decay sugar-rich substrates (Põlme et al. 2020) | Põlme S, et al. 2020. Fungal Diversity 105:1-16 ((https://doi.org/10.1007/s13225-020-00466-2)) |
| Otu13326 | 1 | k__Fungi(100);p__Ascomycota(100);c__Saccharomycetes(100);o__Saccharomycetales(100);f__Metschnikowiaceae(100);g__Metschnikowia(100);g__Metschnikowia_unclassified(100); | Metschnikowia | 13 | Saprotroph | \|Undefined Saprotroph\| | Yeast | NULL | Probable | As nectar-tap_saprotroph (Põlme et al. 2020); Decay sugar-rich substrates (Põlme et al. 2020) | Põlme S, et al. 2020. Fungal Diversity 105:1-16 ((https://doi.org/10.1007/s13225-020-00466-2)) |
| Otu13327 | 1 | k__Fungi(100);p__Ascomycota(100);c__Saccharomycetes(100);o__Saccharomycetales(100);f__Metschnikowiaceae(100);g__Metschnikowia(100);s__Metschnikowia_pulcherrima(100); | Metschnikowia | 13 | Saprotroph | \|Undefined Saprotroph\| | Yeast | NULL | Probable | As nectar-tap_saprotroph (Põlme et al. 2020); Decay sugar-rich substrates (Põlme et al. 2020) | Põlme S, et al. 2020. Fungal Diversity 105:1-16 ((https://doi.org/10.1007/s13225-020-00466-2)) |
| Otu13328 | 1 | k__Fungi(100);p__Ascomycota(100);c__Saccharomycetes(100);o__Saccharomycetales(100);f__Metschnikowiaceae(100);g__Metschnikowia(100);s__Metschnikowia_pulcherrima(100); | Metschnikowia | 13 | Saprotroph | \|Undefined Saprotroph\| | Yeast | NULL | Probable | As nectar-tap_saprotroph (Põlme et al. 2020); Decay sugar-rich substrates (Põlme et al. 2020) | Põlme S, et al. 2020. Fungal Diversity 105:1-16 ((https://doi.org/10.1007/s13225-020-00466-2)) |
| Otu13331 | 1 | k__Fungi(100);p__Ascomycota(100);c__Saccharomycetes(100);o__Saccharomycetales(100);f__Metschnikowiaceae(100);g__Metschnikowia(100);g__Metschnikowia_unclassified(100); | Metschnikowia | 13 | Saprotroph | \|Undefined Saprotroph\| | Yeast | NULL | Probable | As nectar-tap_saprotroph (Põlme et al. 2020); Decay sugar-rich substrates (Põlme et al. 2020) | Põlme S, et al. 2020. Fungal Diversity 105:1-16 ((https://doi.org/10.1007/s13225-020-00466-2)) |
| Otu13333 | 1 | k__Fungi(100);p__Ascomycota(100);c__Saccharomycetes(100);o__Saccharomycetales(100);f__Metschnikowiaceae(100);g__Metschnikowia(100);s__Metschnikowia_pulcherrima(100); | Metschnikowia | 13 | Saprotroph | \|Undefined Saprotroph\| | Yeast | NULL | Probable | As nectar-tap_saprotroph (Põlme et al. 2020); Decay sugar-rich substrates (Põlme et al. 2020) | Põlme S, et al. 2020. Fungal Diversity 105:1-16 ((https://doi.org/10.1007/s13225-020-00466-2)) |
| Otu13334 | 1 | k__Fungi(100);p__Ascomycota(100);c__Saccharomycetes(100);o__Saccharomycetales(100);f__Metschnikowiaceae(100);g__Metschnikowia(100);s__Metschnikowia_pulcherrima(100); | Metschnikowia | 13 | Saprotroph | \|Undefined Saprotroph\| | Yeast | NULL | Probable | As nectar-tap_saprotroph (Põlme et al. 2020); Decay sugar-rich substrates (Põlme et al. 2020) | Põlme S, et al. 2020. Fungal Diversity 105:1-16 ((https://doi.org/10.1007/s13225-020-00466-2)) |
| Otu13336 | 1 | k__Fungi(100);p__Ascomycota(100);c__Saccharomycetes(100);o__Saccharomycetales(100);f__Metschnikowiaceae(100);g__Metschnikowia(100);g__Metschnikowia_unclassified(100); | Metschnikowia | 13 | Saprotroph | \|Undefined Saprotroph\| | Yeast | NULL | Probable | As nectar-tap_saprotroph (Põlme et al. 2020); Decay sugar-rich substrates (Põlme et al. 2020) | Põlme S, et al. 2020. Fungal Diversity 105:1-16 ((https://doi.org/10.1007/s13225-020-00466-2)) |
| Otu13337 | 1 | k__Fungi(100);p__Ascomycota(100);c__Saccharomycetes(100);o__Saccharomycetales(100);f__Metschnikowiaceae(100);g__Metschnikowia(100);s__Metschnikowia_pulcherrima(100); | Metschnikowia | 13 | Saprotroph | \|Undefined Saprotroph\| | Yeast | NULL | Probable | As nectar-tap_saprotroph (Põlme et al. 2020); Decay sugar-rich substrates (Põlme et al. 2020) | Põlme S, et al. 2020. Fungal Diversity 105:1-16 ((https://doi.org/10.1007/s13225-020-00466-2)) |
| Otu13338 | 1 | k__Fungi(100);p__Ascomycota(100);c__Saccharomycetes(100);o__Saccharomycetales(100);f__Metschnikowiaceae(100);g__Metschnikowia(100);g__Metschnikowia_unclassified(100); | Metschnikowia | 13 | Saprotroph | \|Undefined Saprotroph\| | Yeast | NULL | Probable | As nectar-tap_saprotroph (Põlme et al. 2020); Decay sugar-rich substrates (Põlme et al. 2020) | Põlme S, et al. 2020. Fungal Diversity 105:1-16 ((https://doi.org/10.1007/s13225-020-00466-2)) |
| Otu13339 | 1 | k__Fungi(100);p__Ascomycota(100);c__Saccharomycetes(100);o__Saccharomycetales(100);f__Metschnikowiaceae(100);g__Metschnikowia(100);g__Metschnikowia_unclassified(100); | Metschnikowia | 13 | Saprotroph | \|Undefined Saprotroph\| | Yeast | NULL | Probable | As nectar-tap_saprotroph (Põlme et al. 2020); Decay sugar-rich substrates (Põlme et al. 2020) | Põlme S, et al. 2020. Fungal Diversity 105:1-16 ((https://doi.org/10.1007/s13225-020-00466-2)) |
| Otu13340 | 1 | k__Fungi(100);p__Ascomycota(100);c__Saccharomycetes(100);o__Saccharomycetales(100);f__Saccharomycetales_family_Incertae_sedis(100);g__Candida(100);g__Candida_unclassified(100); | Candida | 13 | Pathotroph-Saprotroph-Symbiotroph | Animal Pathogen-Endophyte-Endosymbiont-Epiphyte-\|Undefined Saprotroph\| | Dimorphic Yeast-Yeast | NULL | Probable | As nectar-tap_saprotroph (Põlme et al. 2020); Decay sugar-rich substrates (Põlme et al. 2020); This asexual genus is polyphyletic and contain species that function very differently depend on what they are. Caution should be taken when interpreting guilds for Candida species. | Manolakaki D, et al. 2010. Virulence. 1: 367-375 ((https://doi.org/10.4161/viru.1.5.12796)); Mekbib SB, et al. 2011. Tropical Plant Pathology 36:233-240 ((https://doi.org/10.1590/S1982-56762011000400004)); Põlme S, et al. 2020. Fungal Diversity 105:1-16 ((https://doi.org/10.1007/s13225-020-00466-2)) |
| Otu13341 | 1 | k__Fungi(100);p__Ascomycota(100);c__Saccharomycetes(100);o__Saccharomycetales(100);f__Metschnikowiaceae(100);g__Metschnikowia(100);s__Metschnikowia_pulcherrima(100); | Metschnikowia | 13 | Saprotroph | \|Undefined Saprotroph\| | Yeast | NULL | Probable | As nectar-tap_saprotroph (Põlme et al. 2020); Decay sugar-rich substrates (Põlme et al. 2020) | Põlme S, et al. 2020. Fungal Diversity 105:1-16 ((https://doi.org/10.1007/s13225-020-00466-2)) |
| Otu13342 | 1 | k__Fungi(100);p__Ascomycota(100);c__Saccharomycetes(100);o__Saccharomycetales(100);f__Metschnikowiaceae(100);g__Metschnikowia(100);s__Metschnikowia_pulcherrima(100); | Metschnikowia | 13 | Saprotroph | \|Undefined Saprotroph\| | Yeast | NULL | Probable | As nectar-tap_saprotroph (Põlme et al. 2020); Decay sugar-rich substrates (Põlme et al. 2020) | Põlme S, et al. 2020. Fungal Diversity 105:1-16 ((https://doi.org/10.1007/s13225-020-00466-2)) |
| Otu13343 | 1 | k__Fungi(100);p__Ascomycota(100);c__Saccharomycetes(100);o__Saccharomycetales(100);f__Metschnikowiaceae(100);g__Metschnikowia(100);s__Metschnikowia_pulcherrima(100); | Metschnikowia | 13 | Saprotroph | \|Undefined Saprotroph\| | Yeast | NULL | Probable | As nectar-tap_saprotroph (Põlme et al. 2020); Decay sugar-rich substrates (Põlme et al. 2020) | Põlme S, et al. 2020. Fungal Diversity 105:1-16 ((https://doi.org/10.1007/s13225-020-00466-2)) |
| Otu13344 | 1 | k__Fungi(100);p__Ascomycota(100);c__Saccharomycetes(100);o__Saccharomycetales(100);f__Metschnikowiaceae(100);g__Metschnikowia(100);s__Metschnikowia_pulcherrima(100); | Metschnikowia | 13 | Saprotroph | \|Undefined Saprotroph\| | Yeast | NULL | Probable | As nectar-tap_saprotroph (Põlme et al. 2020); Decay sugar-rich substrates (Põlme et al. 2020) | Põlme S, et al. 2020. Fungal Diversity 105:1-16 ((https://doi.org/10.1007/s13225-020-00466-2)) |
| Otu13345 | 1 | k__Fungi(100);p__Ascomycota(100);c__Saccharomycetes(100);o__Saccharomycetales(100);f__Metschnikowiaceae(100);g__Metschnikowia(100);s__Metschnikowia_pulcherrima(100); | Metschnikowia | 13 | Saprotroph | \|Undefined Saprotroph\| | Yeast | NULL | Probable | As nectar-tap_saprotroph (Põlme et al. 2020); Decay sugar-rich substrates (Põlme et al. 2020) | Põlme S, et al. 2020. Fungal Diversity 105:1-16 ((https://doi.org/10.1007/s13225-020-00466-2)) |
| Otu13348 | 1 | k__Fungi(100);p__Ascomycota(100);c__Saccharomycetes(100);o__Saccharomycetales(100);f__Metschnikowiaceae(100);g__Metschnikowia(100);s__Metschnikowia_pulcherrima(100); | Metschnikowia | 13 | Saprotroph | \|Undefined Saprotroph\| | Yeast | NULL | Probable | As nectar-tap_saprotroph (Põlme et al. 2020); Decay sugar-rich substrates (Põlme et al. 2020) | Põlme S, et al. 2020. Fungal Diversity 105:1-16 ((https://doi.org/10.1007/s13225-020-00466-2)) |
| Otu13349 | 1 | k__Fungi(100);p__Ascomycota(100);c__Saccharomycetes(100);o__Saccharomycetales(100);f__Metschnikowiaceae(100);g__Metschnikowia(100);g__Metschnikowia_unclassified(100); | Metschnikowia | 13 | Saprotroph | \|Undefined Saprotroph\| | Yeast | NULL | Probable | As nectar-tap_saprotroph (Põlme et al. 2020); Decay sugar-rich substrates (Põlme et al. 2020) | Põlme S, et al. 2020. Fungal Diversity 105:1-16 ((https://doi.org/10.1007/s13225-020-00466-2)) |
| Otu13350 | 1 | k__Fungi(100);p__Ascomycota(100);c__Saccharomycetes(100);o__Saccharomycetales(100);f__Metschnikowiaceae(100);g__Metschnikowia(100);s__Metschnikowia_pulcherrima(100); | Metschnikowia | 13 | Saprotroph | \|Undefined Saprotroph\| | Yeast | NULL | Probable | As nectar-tap_saprotroph (Põlme et al. 2020); Decay sugar-rich substrates (Põlme et al. 2020) | Põlme S, et al. 2020. Fungal Diversity 105:1-16 ((https://doi.org/10.1007/s13225-020-00466-2)) |
| Otu13352 | 1 | k__Fungi(100);p__Ascomycota(100);c__Saccharomycetes(100);o__Saccharomycetales(100);f__Metschnikowiaceae(100);g__Metschnikowia(100);s__Metschnikowia_pulcherrima(100); | Metschnikowia | 13 | Saprotroph | \|Undefined Saprotroph\| | Yeast | NULL | Probable | As nectar-tap_saprotroph (Põlme et al. 2020); Decay sugar-rich substrates (Põlme et al. 2020) | Põlme S, et al. 2020. Fungal Diversity 105:1-16 ((https://doi.org/10.1007/s13225-020-00466-2)) |
| Otu13356 | 1 | k__Fungi(100);p__Ascomycota(100);c__Saccharomycetes(100);o__Saccharomycetales(100);f__Metschnikowiaceae(100);g__Metschnikowia(100);s__Metschnikowia_pulcherrima(100); | Metschnikowia | 13 | Saprotroph | \|Undefined Saprotroph\| | Yeast | NULL | Probable | As nectar-tap_saprotroph (Põlme et al. 2020); Decay sugar-rich substrates (Põlme et al. 2020) | Põlme S, et al. 2020. Fungal Diversity 105:1-16 ((https://doi.org/10.1007/s13225-020-00466-2)) |
| Otu13358 | 1 | k__Fungi(100);p__Ascomycota(100);c__Saccharomycetes(100);o__Saccharomycetales(100);f__Metschnikowiaceae(100);g__Metschnikowia(100);s__Metschnikowia_pulcherrima(100); | Metschnikowia | 13 | Saprotroph | \|Undefined Saprotroph\| | Yeast | NULL | Probable | As nectar-tap_saprotroph (Põlme et al. 2020); Decay sugar-rich substrates (Põlme et al. 2020) | Põlme S, et al. 2020. Fungal Diversity 105:1-16 ((https://doi.org/10.1007/s13225-020-00466-2)) |
| Otu13359 | 1 | k__Fungi(100);p__Ascomycota(100);c__Saccharomycetes(100);o__Saccharomycetales(100);f__Metschnikowiaceae(100);g__Metschnikowia(100);s__Metschnikowia_pulcherrima(100); | Metschnikowia | 13 | Saprotroph | \|Undefined Saprotroph\| | Yeast | NULL | Probable | As nectar-tap_saprotroph (Põlme et al. 2020); Decay sugar-rich substrates (Põlme et al. 2020) | Põlme S, et al. 2020. Fungal Diversity 105:1-16 ((https://doi.org/10.1007/s13225-020-00466-2)) |
| Otu13360 | 1 | k__Fungi(100);p__Ascomycota(100);c__Saccharomycetes(100);o__Saccharomycetales(100);f__Metschnikowiaceae(100);g__Metschnikowia(100);s__Metschnikowia_pulcherrima(100); | Metschnikowia | 13 | Saprotroph | \|Undefined Saprotroph\| | Yeast | NULL | Probable | As nectar-tap_saprotroph (Põlme et al. 2020); Decay sugar-rich substrates (Põlme et al. 2020) | Põlme S, et al. 2020. Fungal Diversity 105:1-16 ((https://doi.org/10.1007/s13225-020-00466-2)) |
| Otu13361 | 1 | k__Fungi(100);p__Ascomycota(100);c__Saccharomycetes(100);o__Saccharomycetales(100);f__Metschnikowiaceae(100);g__Metschnikowia(100);g__Metschnikowia_unclassified(100); | Metschnikowia | 13 | Saprotroph | \|Undefined Saprotroph\| | Yeast | NULL | Probable | As nectar-tap_saprotroph (Põlme et al. 2020); Decay sugar-rich substrates (Põlme et al. 2020) | Põlme S, et al. 2020. Fungal Diversity 105:1-16 ((https://doi.org/10.1007/s13225-020-00466-2)) |
| Otu13362 | 1 | k__Fungi(100);p__Ascomycota(100);c__Saccharomycetes(100);o__Saccharomycetales(100);f__Metschnikowiaceae(100);g__Metschnikowia(100);s__Metschnikowia_pulcherrima(100); | Metschnikowia | 13 | Saprotroph | \|Undefined Saprotroph\| | Yeast | NULL | Probable | As nectar-tap_saprotroph (Põlme et al. 2020); Decay sugar-rich substrates (Põlme et al. 2020) | Põlme S, et al. 2020. Fungal Diversity 105:1-16 ((https://doi.org/10.1007/s13225-020-00466-2)) |
| Otu13364 | 1 | k__Fungi(100);p__Ascomycota(100);c__Saccharomycetes(100);o__Saccharomycetales(100);f__Metschnikowiaceae(100);g__Metschnikowia(100);g__Metschnikowia_unclassified(100); | Metschnikowia | 13 | Saprotroph | \|Undefined Saprotroph\| | Yeast | NULL | Probable | As nectar-tap_saprotroph (Põlme et al. 2020); Decay sugar-rich substrates (Põlme et al. 2020) | Põlme S, et al. 2020. Fungal Diversity 105:1-16 ((https://doi.org/10.1007/s13225-020-00466-2)) |
| Otu13366 | 1 | k__Fungi(100);p__Ascomycota(100);c__Saccharomycetes(100);o__Saccharomycetales(100);f__Metschnikowiaceae(100);g__Metschnikowia(100);g__Metschnikowia_unclassified(100); | Metschnikowia | 13 | Saprotroph | \|Undefined Saprotroph\| | Yeast | NULL | Probable | As nectar-tap_saprotroph (Põlme et al. 2020); Decay sugar-rich substrates (Põlme et al. 2020) | Põlme S, et al. 2020. Fungal Diversity 105:1-16 ((https://doi.org/10.1007/s13225-020-00466-2)) |
| Otu13367 | 1 | k__Fungi(100);p__Ascomycota(100);c__Saccharomycetes(100);o__Saccharomycetales(100);f__Metschnikowiaceae(100);g__Metschnikowia(100);s__Metschnikowia_pulcherrima(100); | Metschnikowia | 13 | Saprotroph | \|Undefined Saprotroph\| | Yeast | NULL | Probable | As nectar-tap_saprotroph (Põlme et al. 2020); Decay sugar-rich substrates (Põlme et al. 2020) | Põlme S, et al. 2020. Fungal Diversity 105:1-16 ((https://doi.org/10.1007/s13225-020-00466-2)) |
| Otu13369 | 1 | k__Fungi(100);p__Ascomycota(100);c__Saccharomycetes(100);o__Saccharomycetales(100);f__Metschnikowiaceae(100);g__Metschnikowia(100);g__Metschnikowia_unclassified(100); | Metschnikowia | 13 | Saprotroph | \|Undefined Saprotroph\| | Yeast | NULL | Probable | As nectar-tap_saprotroph (Põlme et al. 2020); Decay sugar-rich substrates (Põlme et al. 2020) | Põlme S, et al. 2020. Fungal Diversity 105:1-16 ((https://doi.org/10.1007/s13225-020-00466-2)) |
| Otu13370 | 1 | k__Fungi(100);p__Ascomycota(100);c__Saccharomycetes(100);o__Saccharomycetales(100);f__Metschnikowiaceae(100);g__Metschnikowia(100);s__Metschnikowia_pulcherrima(100); | Metschnikowia | 13 | Saprotroph | \|Undefined Saprotroph\| | Yeast | NULL | Probable | As nectar-tap_saprotroph (Põlme et al. 2020); Decay sugar-rich substrates (Põlme et al. 2020) | Põlme S, et al. 2020. Fungal Diversity 105:1-16 ((https://doi.org/10.1007/s13225-020-00466-2)) |
| Otu13371 | 1 | k__Fungi(100);p__Ascomycota(100);c__Saccharomycetes(100);o__Saccharomycetales(100);f__Metschnikowiaceae(100);g__Metschnikowia(100);s__Metschnikowia_pulcherrima(100); | Metschnikowia | 13 | Saprotroph | \|Undefined Saprotroph\| | Yeast | NULL | Probable | As nectar-tap_saprotroph (Põlme et al. 2020); Decay sugar-rich substrates (Põlme et al. 2020) | Põlme S, et al. 2020. Fungal Diversity 105:1-16 ((https://doi.org/10.1007/s13225-020-00466-2)) |
| Otu13372 | 1 | k__Fungi(100);p__Ascomycota(100);c__Saccharomycetes(100);o__Saccharomycetales(100);f__Metschnikowiaceae(100);g__Metschnikowia(100);s__Metschnikowia_pulcherrima(100); | Metschnikowia | 13 | Saprotroph | \|Undefined Saprotroph\| | Yeast | NULL | Probable | As nectar-tap_saprotroph (Põlme et al. 2020); Decay sugar-rich substrates (Põlme et al. 2020) | Põlme S, et al. 2020. Fungal Diversity 105:1-16 ((https://doi.org/10.1007/s13225-020-00466-2)) |
| Otu13374 | 1 | k__Fungi(100);p__Ascomycota(100);c__Saccharomycetes(100);o__Saccharomycetales(100);f__Metschnikowiaceae(100);g__Metschnikowia(100);g__Metschnikowia_unclassified(100); | Metschnikowia | 13 | Saprotroph | \|Undefined Saprotroph\| | Yeast | NULL | Probable | As nectar-tap_saprotroph (Põlme et al. 2020); Decay sugar-rich substrates (Põlme et al. 2020) | Põlme S, et al. 2020. Fungal Diversity 105:1-16 ((https://doi.org/10.1007/s13225-020-00466-2)) |
| Otu13375 | 1 | k__Fungi(100);p__Ascomycota(100);c__Saccharomycetes(100);o__Saccharomycetales(100);f__Metschnikowiaceae(100);g__Metschnikowia(100);s__Metschnikowia_pulcherrima(100); | Metschnikowia | 13 | Saprotroph | \|Undefined Saprotroph\| | Yeast | NULL | Probable | As nectar-tap_saprotroph (Põlme et al. 2020); Decay sugar-rich substrates (Põlme et al. 2020) | Põlme S, et al. 2020. Fungal Diversity 105:1-16 ((https://doi.org/10.1007/s13225-020-00466-2)) |
| Otu13376 | 1 | k__Fungi(100);p__Ascomycota(100);c__Saccharomycetes(100);o__Saccharomycetales(100);f__Metschnikowiaceae(100);g__Metschnikowia(100);g__Metschnikowia_unclassified(100); | Metschnikowia | 13 | Saprotroph | \|Undefined Saprotroph\| | Yeast | NULL | Probable | As nectar-tap_saprotroph (Põlme et al. 2020); Decay sugar-rich substrates (Põlme et al. 2020) | Põlme S, et al. 2020. Fungal Diversity 105:1-16 ((https://doi.org/10.1007/s13225-020-00466-2)) |
| Otu13377 | 1 | k__Fungi(100);p__Ascomycota(100);c__Saccharomycetes(100);o__Saccharomycetales(100);f__Metschnikowiaceae(100);g__Metschnikowia(100);s__Metschnikowia_pulcherrima(100); | Metschnikowia | 13 | Saprotroph | \|Undefined Saprotroph\| | Yeast | NULL | Probable | As nectar-tap_saprotroph (Põlme et al. 2020); Decay sugar-rich substrates (Põlme et al. 2020) | Põlme S, et al. 2020. Fungal Diversity 105:1-16 ((https://doi.org/10.1007/s13225-020-00466-2)) |
| Otu13378 | 1 | k__Fungi(100);p__Ascomycota(100);c__Saccharomycetes(100);o__Saccharomycetales(100);f__Metschnikowiaceae(100);g__Metschnikowia(100);s__Metschnikowia_pulcherrima(100); | Metschnikowia | 13 | Saprotroph | \|Undefined Saprotroph\| | Yeast | NULL | Probable | As nectar-tap_saprotroph (Põlme et al. 2020); Decay sugar-rich substrates (Põlme et al. 2020) | Põlme S, et al. 2020. Fungal Diversity 105:1-16 ((https://doi.org/10.1007/s13225-020-00466-2)) |
| Otu13380 | 1 | k__Fungi(100);p__Ascomycota(100);c__Saccharomycetes(100);o__Saccharomycetales(100);f__Metschnikowiaceae(100);g__Metschnikowia(100);g__Metschnikowia_unclassified(100); | Metschnikowia | 13 | Saprotroph | \|Undefined Saprotroph\| | Yeast | NULL | Probable | As nectar-tap_saprotroph (Põlme et al. 2020); Decay sugar-rich substrates (Põlme et al. 2020) | Põlme S, et al. 2020. Fungal Diversity 105:1-16 ((https://doi.org/10.1007/s13225-020-00466-2)) |
| Otu13382 | 1 | k__Fungi(100);p__Ascomycota(100);c__Saccharomycetes(100);o__Saccharomycetales(100);f__Metschnikowiaceae(100);g__Metschnikowia(100);s__Metschnikowia_pulcherrima(100); | Metschnikowia | 13 | Saprotroph | \|Undefined Saprotroph\| | Yeast | NULL | Probable | As nectar-tap_saprotroph (Põlme et al. 2020); Decay sugar-rich substrates (Põlme et al. 2020) | Põlme S, et al. 2020. Fungal Diversity 105:1-16 ((https://doi.org/10.1007/s13225-020-00466-2)) |
| Otu13383 | 1 | k__Fungi(100);p__Ascomycota(100);c__Saccharomycetes(100);o__Saccharomycetales(100);f__Metschnikowiaceae(100);g__Metschnikowia(100);g__Metschnikowia_unclassified(100); | Metschnikowia | 13 | Saprotroph | \|Undefined Saprotroph\| | Yeast | NULL | Probable | As nectar-tap_saprotroph (Põlme et al. 2020); Decay sugar-rich substrates (Põlme et al. 2020) | Põlme S, et al. 2020. Fungal Diversity 105:1-16 ((https://doi.org/10.1007/s13225-020-00466-2)) |
| Otu13384 | 1 | k__Fungi(100);p__Ascomycota(100);c__Saccharomycetes(100);o__Saccharomycetales(100);f__Metschnikowiaceae(100);g__Metschnikowia(100);s__Metschnikowia_pulcherrima(100); | Metschnikowia | 13 | Saprotroph | \|Undefined Saprotroph\| | Yeast | NULL | Probable | As nectar-tap_saprotroph (Põlme et al. 2020); Decay sugar-rich substrates (Põlme et al. 2020) | Põlme S, et al. 2020. Fungal Diversity 105:1-16 ((https://doi.org/10.1007/s13225-020-00466-2)) |
| Otu13385 | 1 | k__Fungi(100);p__Ascomycota(100);c__Saccharomycetes(100);o__Saccharomycetales(100);f__Metschnikowiaceae(100);g__Metschnikowia(100);g__Metschnikowia_unclassified(100); | Metschnikowia | 13 | Saprotroph | \|Undefined Saprotroph\| | Yeast | NULL | Probable | As nectar-tap_saprotroph (Põlme et al. 2020); Decay sugar-rich substrates (Põlme et al. 2020) | Põlme S, et al. 2020. Fungal Diversity 105:1-16 ((https://doi.org/10.1007/s13225-020-00466-2)) |
| Otu13386 | 1 | k__Fungi(100);p__Ascomycota(100);c__Saccharomycetes(100);o__Saccharomycetales(100);f__Metschnikowiaceae(100);g__Metschnikowia(100);s__Metschnikowia_pulcherrima(100); | Metschnikowia | 13 | Saprotroph | \|Undefined Saprotroph\| | Yeast | NULL | Probable | As nectar-tap_saprotroph (Põlme et al. 2020); Decay sugar-rich substrates (Põlme et al. 2020) | Põlme S, et al. 2020. Fungal Diversity 105:1-16 ((https://doi.org/10.1007/s13225-020-00466-2)) |
| Otu13388 | 1 | k__Fungi(100);p__Ascomycota(100);c__Saccharomycetes(100);o__Saccharomycetales(100);f__Metschnikowiaceae(100);g__Metschnikowia(100);s__Metschnikowia_pulcherrima(100); | Metschnikowia | 13 | Saprotroph | \|Undefined Saprotroph\| | Yeast | NULL | Probable | As nectar-tap_saprotroph (Põlme et al. 2020); Decay sugar-rich substrates (Põlme et al. 2020) | Põlme S, et al. 2020. Fungal Diversity 105:1-16 ((https://doi.org/10.1007/s13225-020-00466-2)) |
| Otu13389 | 1 | k__Fungi(100);p__Ascomycota(100);c__Saccharomycetes(100);o__Saccharomycetales(100);f__Metschnikowiaceae(100);g__Metschnikowia(100);s__Metschnikowia_pulcherrima(100); | Metschnikowia | 13 | Saprotroph | \|Undefined Saprotroph\| | Yeast | NULL | Probable | As nectar-tap_saprotroph (Põlme et al. 2020); Decay sugar-rich substrates (Põlme et al. 2020) | Põlme S, et al. 2020. Fungal Diversity 105:1-16 ((https://doi.org/10.1007/s13225-020-00466-2)) |
| Otu13391 | 1 | k__Fungi(100);p__Ascomycota(100);c__Saccharomycetes(100);o__Saccharomycetales(100);f__Metschnikowiaceae(100);g__Metschnikowia(100);s__Metschnikowia_pulcherrima(100); | Metschnikowia | 13 | Saprotroph | \|Undefined Saprotroph\| | Yeast | NULL | Probable | As nectar-tap_saprotroph (Põlme et al. 2020); Decay sugar-rich substrates (Põlme et al. 2020) | Põlme S, et al. 2020. Fungal Diversity 105:1-16 ((https://doi.org/10.1007/s13225-020-00466-2)) |
| Otu13392 | 1 | k__Fungi(100);p__Ascomycota(100);c__Saccharomycetes(100);o__Saccharomycetales(100);f__Metschnikowiaceae(100);g__Metschnikowia(100);s__Metschnikowia_pulcherrima(100); | Metschnikowia | 13 | Saprotroph | \|Undefined Saprotroph\| | Yeast | NULL | Probable | As nectar-tap_saprotroph (Põlme et al. 2020); Decay sugar-rich substrates (Põlme et al. 2020) | Põlme S, et al. 2020. Fungal Diversity 105:1-16 ((https://doi.org/10.1007/s13225-020-00466-2)) |
| Otu13393 | 1 | k__Fungi(100);p__Ascomycota(100);c__Saccharomycetes(100);o__Saccharomycetales(100);f__Metschnikowiaceae(100);g__Metschnikowia(100);s__Metschnikowia_pulcherrima(100); | Metschnikowia | 13 | Saprotroph | \|Undefined Saprotroph\| | Yeast | NULL | Probable | As nectar-tap_saprotroph (Põlme et al. 2020); Decay sugar-rich substrates (Põlme et al. 2020) | Põlme S, et al. 2020. Fungal Diversity 105:1-16 ((https://doi.org/10.1007/s13225-020-00466-2)) |
| Otu13394 | 1 | k__Fungi(100);p__Ascomycota(100);c__Saccharomycetes(100);o__Saccharomycetales(100);f__Metschnikowiaceae(100);g__Metschnikowia(100);s__Metschnikowia_pulcherrima(100); | Metschnikowia | 13 | Saprotroph | \|Undefined Saprotroph\| | Yeast | NULL | Probable | As nectar-tap_saprotroph (Põlme et al. 2020); Decay sugar-rich substrates (Põlme et al. 2020) | Põlme S, et al. 2020. Fungal Diversity 105:1-16 ((https://doi.org/10.1007/s13225-020-00466-2)) |
| Otu13395 | 1 | k__Fungi(100);p__Ascomycota(100);c__Saccharomycetes(100);o__Saccharomycetales(100);f__Saccharomycetales_family_Incertae_sedis(100);g__Candida(100);g__Candida_unclassified(100); | Candida | 13 | Pathotroph-Saprotroph-Symbiotroph | Animal Pathogen-Endophyte-Endosymbiont-Epiphyte-\|Undefined Saprotroph\| | Dimorphic Yeast-Yeast | NULL | Probable | As nectar-tap_saprotroph (Põlme et al. 2020); Decay sugar-rich substrates (Põlme et al. 2020); This asexual genus is polyphyletic and contain species that function very differently depend on what they are. Caution should be taken when interpreting guilds for Candida species. | Manolakaki D, et al. 2010. Virulence. 1: 367-375 ((https://doi.org/10.4161/viru.1.5.12796)); Mekbib SB, et al. 2011. Tropical Plant Pathology 36:233-240 ((https://doi.org/10.1590/S1982-56762011000400004)); Põlme S, et al. 2020. Fungal Diversity 105:1-16 ((https://doi.org/10.1007/s13225-020-00466-2)) |
| Otu13396 | 1 | k__Fungi(100);p__Ascomycota(100);c__Saccharomycetes(100);o__Saccharomycetales(100);f__Metschnikowiaceae(100);g__Metschnikowia(100);g__Metschnikowia_unclassified(100); | Metschnikowia | 13 | Saprotroph | \|Undefined Saprotroph\| | Yeast | NULL | Probable | As nectar-tap_saprotroph (Põlme et al. 2020); Decay sugar-rich substrates (Põlme et al. 2020) | Põlme S, et al. 2020. Fungal Diversity 105:1-16 ((https://doi.org/10.1007/s13225-020-00466-2)) |
| Otu13397 | 1 | k__Fungi(100);p__Ascomycota(100);c__Saccharomycetes(100);o__Saccharomycetales(100);f__Metschnikowiaceae(100);g__Metschnikowia(100);s__Metschnikowia_pulcherrima(100); | Metschnikowia | 13 | Saprotroph | \|Undefined Saprotroph\| | Yeast | NULL | Probable | As nectar-tap_saprotroph (Põlme et al. 2020); Decay sugar-rich substrates (Põlme et al. 2020) | Põlme S, et al. 2020. Fungal Diversity 105:1-16 ((https://doi.org/10.1007/s13225-020-00466-2)) |
| Otu13399 | 1 | k__Fungi(100);p__Ascomycota(100);c__Saccharomycetes(100);o__Saccharomycetales(100);f__Metschnikowiaceae(100);g__Metschnikowia(100);s__Metschnikowia_pulcherrima(100); | Metschnikowia | 13 | Saprotroph | \|Undefined Saprotroph\| | Yeast | NULL | Probable | As nectar-tap_saprotroph (Põlme et al. 2020); Decay sugar-rich substrates (Põlme et al. 2020) | Põlme S, et al. 2020. Fungal Diversity 105:1-16 ((https://doi.org/10.1007/s13225-020-00466-2)) |
| Otu13400 | 1 | k__Fungi(100);p__Ascomycota(100);c__Saccharomycetes(100);o__Saccharomycetales(100);f__Metschnikowiaceae(100);g__Metschnikowia(100);s__Metschnikowia_pulcherrima(100); | Metschnikowia | 13 | Saprotroph | \|Undefined Saprotroph\| | Yeast | NULL | Probable | As nectar-tap_saprotroph (Põlme et al. 2020); Decay sugar-rich substrates (Põlme et al. 2020) | Põlme S, et al. 2020. Fungal Diversity 105:1-16 ((https://doi.org/10.1007/s13225-020-00466-2)) |
| Otu13401 | 1 | k__Fungi(100);p__Ascomycota(100);c__Saccharomycetes(100);o__Saccharomycetales(100);f__Metschnikowiaceae(100);g__Metschnikowia(100);s__Metschnikowia_pulcherrima(100); | Metschnikowia | 13 | Saprotroph | \|Undefined Saprotroph\| | Yeast | NULL | Probable | As nectar-tap_saprotroph (Põlme et al. 2020); Decay sugar-rich substrates (Põlme et al. 2020) | Põlme S, et al. 2020. Fungal Diversity 105:1-16 ((https://doi.org/10.1007/s13225-020-00466-2)) |
| Otu13403 | 1 | k__Fungi(100);p__Ascomycota(100);c__Saccharomycetes(100);o__Saccharomycetales(100);f__Metschnikowiaceae(100);g__Metschnikowia(100);s__Metschnikowia_pulcherrima(100); | Metschnikowia | 13 | Saprotroph | \|Undefined Saprotroph\| | Yeast | NULL | Probable | As nectar-tap_saprotroph (Põlme et al. 2020); Decay sugar-rich substrates (Põlme et al. 2020) | Põlme S, et al. 2020. Fungal Diversity 105:1-16 ((https://doi.org/10.1007/s13225-020-00466-2)) |
| Otu13404 | 1 | k__Fungi(100);p__Ascomycota(100);c__Saccharomycetes(100);o__Saccharomycetales(100);f__Metschnikowiaceae(100);g__Metschnikowia(100);s__Metschnikowia_pulcherrima(100); | Metschnikowia | 13 | Saprotroph | \|Undefined Saprotroph\| | Yeast | NULL | Probable | As nectar-tap_saprotroph (Põlme et al. 2020); Decay sugar-rich substrates (Põlme et al. 2020) | Põlme S, et al. 2020. Fungal Diversity 105:1-16 ((https://doi.org/10.1007/s13225-020-00466-2)) |
| Otu13405 | 1 | k__Fungi(100);p__Ascomycota(100);c__Saccharomycetes(100);o__Saccharomycetales(100);f__Metschnikowiaceae(100);g__Metschnikowia(100);s__Metschnikowia_pulcherrima(100); | Metschnikowia | 13 | Saprotroph | \|Undefined Saprotroph\| | Yeast | NULL | Probable | As nectar-tap_saprotroph (Põlme et al. 2020); Decay sugar-rich substrates (Põlme et al. 2020) | Põlme S, et al. 2020. Fungal Diversity 105:1-16 ((https://doi.org/10.1007/s13225-020-00466-2)) |
| Otu13406 | 1 | k__Fungi(100);p__Ascomycota(100);c__Saccharomycetes(100);o__Saccharomycetales(100);f__Metschnikowiaceae(100);g__Metschnikowia(100);s__Metschnikowia_pulcherrima(100); | Metschnikowia | 13 | Saprotroph | \|Undefined Saprotroph\| | Yeast | NULL | Probable | As nectar-tap_saprotroph (Põlme et al. 2020); Decay sugar-rich substrates (Põlme et al. 2020) | Põlme S, et al. 2020. Fungal Diversity 105:1-16 ((https://doi.org/10.1007/s13225-020-00466-2)) |
| Otu13408 | 1 | k__Fungi(100);p__Ascomycota(100);c__Saccharomycetes(100);o__Saccharomycetales(100);f__Metschnikowiaceae(100);g__Metschnikowia(100);s__Metschnikowia_pulcherrima(100); | Metschnikowia | 13 | Saprotroph | \|Undefined Saprotroph\| | Yeast | NULL | Probable | As nectar-tap_saprotroph (Põlme et al. 2020); Decay sugar-rich substrates (Põlme et al. 2020) | Põlme S, et al. 2020. Fungal Diversity 105:1-16 ((https://doi.org/10.1007/s13225-020-00466-2)) |
| Otu13409 | 1 | k__Fungi(100);p__Ascomycota(100);c__Saccharomycetes(100);o__Saccharomycetales(100);f__Metschnikowiaceae(100);g__Metschnikowia(100);s__Metschnikowia_pulcherrima(100); | Metschnikowia | 13 | Saprotroph | \|Undefined Saprotroph\| | Yeast | NULL | Probable | As nectar-tap_saprotroph (Põlme et al. 2020); Decay sugar-rich substrates (Põlme et al. 2020) | Põlme S, et al. 2020. Fungal Diversity 105:1-16 ((https://doi.org/10.1007/s13225-020-00466-2)) |
| Otu13411 | 1 | k__Fungi(100);p__Ascomycota(100);c__Saccharomycetes(100);o__Saccharomycetales(100);f__Metschnikowiaceae(100);g__Metschnikowia(100);s__Metschnikowia_pulcherrima(100); | Metschnikowia | 13 | Saprotroph | \|Undefined Saprotroph\| | Yeast | NULL | Probable | As nectar-tap_saprotroph (Põlme et al. 2020); Decay sugar-rich substrates (Põlme et al. 2020) | Põlme S, et al. 2020. Fungal Diversity 105:1-16 ((https://doi.org/10.1007/s13225-020-00466-2)) |
| Otu13413 | 1 | k__Fungi(100);p__Ascomycota(100);c__Saccharomycetes(100);o__Saccharomycetales(100);f__Metschnikowiaceae(100);g__Metschnikowia(100);s__Metschnikowia_pulcherrima(100); | Metschnikowia | 13 | Saprotroph | \|Undefined Saprotroph\| | Yeast | NULL | Probable | As nectar-tap_saprotroph (Põlme et al. 2020); Decay sugar-rich substrates (Põlme et al. 2020) | Põlme S, et al. 2020. Fungal Diversity 105:1-16 ((https://doi.org/10.1007/s13225-020-00466-2)) |
| Otu13414 | 1 | k__Fungi(100);p__Ascomycota(100);c__Saccharomycetes(100);o__Saccharomycetales(100);f__Metschnikowiaceae(100);g__Metschnikowia(100);s__Metschnikowia_pulcherrima(100); | Metschnikowia | 13 | Saprotroph | \|Undefined Saprotroph\| | Yeast | NULL | Probable | As nectar-tap_saprotroph (Põlme et al. 2020); Decay sugar-rich substrates (Põlme et al. 2020) | Põlme S, et al. 2020. Fungal Diversity 105:1-16 ((https://doi.org/10.1007/s13225-020-00466-2)) |
| Otu13415 | 1 | k__Fungi(100);p__Ascomycota(100);c__Saccharomycetes(100);o__Saccharomycetales(100);f__Metschnikowiaceae(100);g__Metschnikowia(100);s__Metschnikowia_pulcherrima(100); | Metschnikowia | 13 | Saprotroph | \|Undefined Saprotroph\| | Yeast | NULL | Probable | As nectar-tap_saprotroph (Põlme et al. 2020); Decay sugar-rich substrates (Põlme et al. 2020) | Põlme S, et al. 2020. Fungal Diversity 105:1-16 ((https://doi.org/10.1007/s13225-020-00466-2)) |
| Otu13416 | 1 | k__Fungi(100);p__Ascomycota(100);c__Saccharomycetes(100);o__Saccharomycetales(100);f__Metschnikowiaceae(100);g__Metschnikowia(100);s__Metschnikowia_pulcherrima(100); | Metschnikowia | 13 | Saprotroph | \|Undefined Saprotroph\| | Yeast | NULL | Probable | As nectar-tap_saprotroph (Põlme et al. 2020); Decay sugar-rich substrates (Põlme et al. 2020) | Põlme S, et al. 2020. Fungal Diversity 105:1-16 ((https://doi.org/10.1007/s13225-020-00466-2)) |
| Otu13417 | 1 | k__Fungi(100);p__Ascomycota(100);c__Saccharomycetes(100);o__Saccharomycetales(100);f__Metschnikowiaceae(100);g__Metschnikowia(100);s__Metschnikowia_pulcherrima(100); | Metschnikowia | 13 | Saprotroph | \|Undefined Saprotroph\| | Yeast | NULL | Probable | As nectar-tap_saprotroph (Põlme et al. 2020); Decay sugar-rich substrates (Põlme et al. 2020) | Põlme S, et al. 2020. Fungal Diversity 105:1-16 ((https://doi.org/10.1007/s13225-020-00466-2)) |
| Otu13418 | 1 | k__Fungi(100);p__Ascomycota(100);c__Saccharomycetes(100);o__Saccharomycetales(100);f__Metschnikowiaceae(100);g__Metschnikowia(100);g__Metschnikowia_unclassified(100); | Metschnikowia | 13 | Saprotroph | \|Undefined Saprotroph\| | Yeast | NULL | Probable | As nectar-tap_saprotroph (Põlme et al. 2020); Decay sugar-rich substrates (Põlme et al. 2020) | Põlme S, et al. 2020. Fungal Diversity 105:1-16 ((https://doi.org/10.1007/s13225-020-00466-2)) |
| Otu13419 | 1 | k__Fungi(100);p__Ascomycota(100);c__Saccharomycetes(100);o__Saccharomycetales(100);f__Metschnikowiaceae(100);g__Metschnikowia(100);s__Metschnikowia_pulcherrima(100); | Metschnikowia | 13 | Saprotroph | \|Undefined Saprotroph\| | Yeast | NULL | Probable | As nectar-tap_saprotroph (Põlme et al. 2020); Decay sugar-rich substrates (Põlme et al. 2020) | Põlme S, et al. 2020. Fungal Diversity 105:1-16 ((https://doi.org/10.1007/s13225-020-00466-2)) |
| Otu13420 | 1 | k__Fungi(100);p__Ascomycota(100);c__Saccharomycetes(100);o__Saccharomycetales(100);f__Metschnikowiaceae(100);g__Metschnikowia(100);s__Metschnikowia_pulcherrima(100); | Metschnikowia | 13 | Saprotroph | \|Undefined Saprotroph\| | Yeast | NULL | Probable | As nectar-tap_saprotroph (Põlme et al. 2020); Decay sugar-rich substrates (Põlme et al. 2020) | Põlme S, et al. 2020. Fungal Diversity 105:1-16 ((https://doi.org/10.1007/s13225-020-00466-2)) |
| Otu13422 | 1 | k__Fungi(100);p__Ascomycota(100);c__Saccharomycetes(100);o__Saccharomycetales(100);f__Metschnikowiaceae(100);g__Metschnikowia(100);g__Metschnikowia_unclassified(100); | Metschnikowia | 13 | Saprotroph | \|Undefined Saprotroph\| | Yeast | NULL | Probable | As nectar-tap_saprotroph (Põlme et al. 2020); Decay sugar-rich substrates (Põlme et al. 2020) | Põlme S, et al. 2020. Fungal Diversity 105:1-16 ((https://doi.org/10.1007/s13225-020-00466-2)) |
| Otu13423 | 1 | k__Fungi(100);p__Ascomycota(100);c__Saccharomycetes(100);o__Saccharomycetales(100);f__Metschnikowiaceae(100);g__Metschnikowia(100);s__Metschnikowia_pulcherrima(100); | Metschnikowia | 13 | Saprotroph | \|Undefined Saprotroph\| | Yeast | NULL | Probable | As nectar-tap_saprotroph (Põlme et al. 2020); Decay sugar-rich substrates (Põlme et al. 2020) | Põlme S, et al. 2020. Fungal Diversity 105:1-16 ((https://doi.org/10.1007/s13225-020-00466-2)) |
| Otu13424 | 1 | k__Fungi(100);p__Ascomycota(100);c__Saccharomycetes(100);o__Saccharomycetales(100);f__Metschnikowiaceae(100);g__Metschnikowia(100);s__Metschnikowia_pulcherrima(100); | Metschnikowia | 13 | Saprotroph | \|Undefined Saprotroph\| | Yeast | NULL | Probable | As nectar-tap_saprotroph (Põlme et al. 2020); Decay sugar-rich substrates (Põlme et al. 2020) | Põlme S, et al. 2020. Fungal Diversity 105:1-16 ((https://doi.org/10.1007/s13225-020-00466-2)) |
| Otu13425 | 1 | k__Fungi(100);p__Ascomycota(100);c__Saccharomycetes(100);o__Saccharomycetales(100);f__Metschnikowiaceae(100);g__Metschnikowia(100);s__Metschnikowia_pulcherrima(100); | Metschnikowia | 13 | Saprotroph | \|Undefined Saprotroph\| | Yeast | NULL | Probable | As nectar-tap_saprotroph (Põlme et al. 2020); Decay sugar-rich substrates (Põlme et al. 2020) | Põlme S, et al. 2020. Fungal Diversity 105:1-16 ((https://doi.org/10.1007/s13225-020-00466-2)) |
| Otu13426 | 1 | k__Fungi(100);p__Ascomycota(100);c__Saccharomycetes(100);o__Saccharomycetales(100);f__Metschnikowiaceae(100);g__Metschnikowia(100);s__Metschnikowia_pulcherrima(100); | Metschnikowia | 13 | Saprotroph | \|Undefined Saprotroph\| | Yeast | NULL | Probable | As nectar-tap_saprotroph (Põlme et al. 2020); Decay sugar-rich substrates (Põlme et al. 2020) | Põlme S, et al. 2020. Fungal Diversity 105:1-16 ((https://doi.org/10.1007/s13225-020-00466-2)) |
| Otu13427 | 1 | k__Fungi(100);p__Ascomycota(100);c__Saccharomycetes(100);o__Saccharomycetales(100);f__Metschnikowiaceae(100);g__Metschnikowia(100);s__Metschnikowia_pulcherrima(100); | Metschnikowia | 13 | Saprotroph | \|Undefined Saprotroph\| | Yeast | NULL | Probable | As nectar-tap_saprotroph (Põlme et al. 2020); Decay sugar-rich substrates (Põlme et al. 2020) | Põlme S, et al. 2020. Fungal Diversity 105:1-16 ((https://doi.org/10.1007/s13225-020-00466-2)) |
| Otu13432 | 1 | k__Fungi(100);p__Ascomycota(100);c__Saccharomycetes(100);o__Saccharomycetales(100);f__Metschnikowiaceae(100);g__Metschnikowia(100);g__Metschnikowia_unclassified(100); | Metschnikowia | 13 | Saprotroph | \|Undefined Saprotroph\| | Yeast | NULL | Probable | As nectar-tap_saprotroph (Põlme et al. 2020); Decay sugar-rich substrates (Põlme et al. 2020) | Põlme S, et al. 2020. Fungal Diversity 105:1-16 ((https://doi.org/10.1007/s13225-020-00466-2)) |
| Otu13433 | 1 | k__Fungi(100);p__Ascomycota(100);c__Saccharomycetes(100);o__Saccharomycetales(100);f__Metschnikowiaceae(100);g__Metschnikowia(100);s__Metschnikowia_pulcherrima(100); | Metschnikowia | 13 | Saprotroph | \|Undefined Saprotroph\| | Yeast | NULL | Probable | As nectar-tap_saprotroph (Põlme et al. 2020); Decay sugar-rich substrates (Põlme et al. 2020) | Põlme S, et al. 2020. Fungal Diversity 105:1-16 ((https://doi.org/10.1007/s13225-020-00466-2)) |
| Otu13435 | 1 | k__Fungi(100);p__Ascomycota(100);c__Saccharomycetes(100);o__Saccharomycetales(100);f__Metschnikowiaceae(100);g__Metschnikowia(100);g__Metschnikowia_unclassified(100); | Metschnikowia | 13 | Saprotroph | \|Undefined Saprotroph\| | Yeast | NULL | Probable | As nectar-tap_saprotroph (Põlme et al. 2020); Decay sugar-rich substrates (Põlme et al. 2020) | Põlme S, et al. 2020. Fungal Diversity 105:1-16 ((https://doi.org/10.1007/s13225-020-00466-2)) |
| Otu13436 | 1 | k__Fungi(100);p__Ascomycota(100);c__Saccharomycetes(100);o__Saccharomycetales(100);f__Metschnikowiaceae(100);g__Metschnikowia(100);s__Metschnikowia_pulcherrima(100); | Metschnikowia | 13 | Saprotroph | \|Undefined Saprotroph\| | Yeast | NULL | Probable | As nectar-tap_saprotroph (Põlme et al. 2020); Decay sugar-rich substrates (Põlme et al. 2020) | Põlme S, et al. 2020. Fungal Diversity 105:1-16 ((https://doi.org/10.1007/s13225-020-00466-2)) |
| Otu13441 | 1 | k__Fungi(100);p__Ascomycota(100);c__Saccharomycetes(100);o__Saccharomycetales(100);f__Metschnikowiaceae(100);g__Metschnikowia(100);s__Metschnikowia_pulcherrima(100); | Metschnikowia | 13 | Saprotroph | \|Undefined Saprotroph\| | Yeast | NULL | Probable | As nectar-tap_saprotroph (Põlme et al. 2020); Decay sugar-rich substrates (Põlme et al. 2020) | Põlme S, et al. 2020. Fungal Diversity 105:1-16 ((https://doi.org/10.1007/s13225-020-00466-2)) |
| Otu13442 | 1 | k__Fungi(100);p__Ascomycota(100);c__Saccharomycetes(100);o__Saccharomycetales(100);f__Metschnikowiaceae(100);g__Metschnikowia(100);s__Metschnikowia_pulcherrima(100); | Metschnikowia | 13 | Saprotroph | \|Undefined Saprotroph\| | Yeast | NULL | Probable | As nectar-tap_saprotroph (Põlme et al. 2020); Decay sugar-rich substrates (Põlme et al. 2020) | Põlme S, et al. 2020. Fungal Diversity 105:1-16 ((https://doi.org/10.1007/s13225-020-00466-2)) |
| Otu13444 | 1 | k__Fungi(100);p__Ascomycota(100);c__Saccharomycetes(100);o__Saccharomycetales(100);f__Metschnikowiaceae(100);g__Metschnikowia(100);s__Metschnikowia_pulcherrima(100); | Metschnikowia | 13 | Saprotroph | \|Undefined Saprotroph\| | Yeast | NULL | Probable | As nectar-tap_saprotroph (Põlme et al. 2020); Decay sugar-rich substrates (Põlme et al. 2020) | Põlme S, et al. 2020. Fungal Diversity 105:1-16 ((https://doi.org/10.1007/s13225-020-00466-2)) |
| Otu13449 | 1 | k__Fungi(100);p__Ascomycota(100);c__Saccharomycetes(100);o__Saccharomycetales(100);f__Metschnikowiaceae(100);g__Metschnikowia(100);s__Metschnikowia_pulcherrima(100); | Metschnikowia | 13 | Saprotroph | \|Undefined Saprotroph\| | Yeast | NULL | Probable | As nectar-tap_saprotroph (Põlme et al. 2020); Decay sugar-rich substrates (Põlme et al. 2020) | Põlme S, et al. 2020. Fungal Diversity 105:1-16 ((https://doi.org/10.1007/s13225-020-00466-2)) |
| Otu13451 | 1 | k__Fungi(100);p__Ascomycota(100);c__Saccharomycetes(100);o__Saccharomycetales(100);f__Metschnikowiaceae(100);g__Metschnikowia(100);s__Metschnikowia_pulcherrima(100); | Metschnikowia | 13 | Saprotroph | \|Undefined Saprotroph\| | Yeast | NULL | Probable | As nectar-tap_saprotroph (Põlme et al. 2020); Decay sugar-rich substrates (Põlme et al. 2020) | Põlme S, et al. 2020. Fungal Diversity 105:1-16 ((https://doi.org/10.1007/s13225-020-00466-2)) |
| Otu13454 | 1 | k__Fungi(100);p__Ascomycota(100);c__Saccharomycetes(100);o__Saccharomycetales(100);f__Metschnikowiaceae(100);g__Metschnikowia(100);g__Metschnikowia_unclassified(100); | Metschnikowia | 13 | Saprotroph | \|Undefined Saprotroph\| | Yeast | NULL | Probable | As nectar-tap_saprotroph (Põlme et al. 2020); Decay sugar-rich substrates (Põlme et al. 2020) | Põlme S, et al. 2020. Fungal Diversity 105:1-16 ((https://doi.org/10.1007/s13225-020-00466-2)) |
| Otu13457 | 1 | k__Fungi(100);p__Ascomycota(100);c__Saccharomycetes(100);o__Saccharomycetales(100);f__Metschnikowiaceae(100);g__Metschnikowia(100);s__Metschnikowia_pulcherrima(100); | Metschnikowia | 13 | Saprotroph | \|Undefined Saprotroph\| | Yeast | NULL | Probable | As nectar-tap_saprotroph (Põlme et al. 2020); Decay sugar-rich substrates (Põlme et al. 2020) | Põlme S, et al. 2020. Fungal Diversity 105:1-16 ((https://doi.org/10.1007/s13225-020-00466-2)) |
| Otu13459 | 1 | k__Fungi(100);p__Ascomycota(100);c__Saccharomycetes(100);o__Saccharomycetales(100);f__Metschnikowiaceae(100);g__Metschnikowia(100);s__Metschnikowia_pulcherrima(100); | Metschnikowia | 13 | Saprotroph | \|Undefined Saprotroph\| | Yeast | NULL | Probable | As nectar-tap_saprotroph (Põlme et al. 2020); Decay sugar-rich substrates (Põlme et al. 2020) | Põlme S, et al. 2020. Fungal Diversity 105:1-16 ((https://doi.org/10.1007/s13225-020-00466-2)) |
| Otu13460 | 1 | k__Fungi(100);p__Ascomycota(100);c__Saccharomycetes(100);o__Saccharomycetales(100);f__Metschnikowiaceae(100);g__Metschnikowia(100);s__Metschnikowia_pulcherrima(100); | Metschnikowia | 13 | Saprotroph | \|Undefined Saprotroph\| | Yeast | NULL | Probable | As nectar-tap_saprotroph (Põlme et al. 2020); Decay sugar-rich substrates (Põlme et al. 2020) | Põlme S, et al. 2020. Fungal Diversity 105:1-16 ((https://doi.org/10.1007/s13225-020-00466-2)) |
| Otu13462 | 1 | k__Fungi(100);p__Ascomycota(100);c__Saccharomycetes(100);o__Saccharomycetales(100);f__Metschnikowiaceae(100);g__Metschnikowia(100);s__Metschnikowia_pulcherrima(100); | Metschnikowia | 13 | Saprotroph | \|Undefined Saprotroph\| | Yeast | NULL | Probable | As nectar-tap_saprotroph (Põlme et al. 2020); Decay sugar-rich substrates (Põlme et al. 2020) | Põlme S, et al. 2020. Fungal Diversity 105:1-16 ((https://doi.org/10.1007/s13225-020-00466-2)) |
| Otu13463 | 1 | k__Fungi(100);p__Ascomycota(100);c__Saccharomycetes(100);o__Saccharomycetales(100);f__Metschnikowiaceae(100);g__Metschnikowia(100);s__Metschnikowia_pulcherrima(100); | Metschnikowia | 13 | Saprotroph | \|Undefined Saprotroph\| | Yeast | NULL | Probable | As nectar-tap_saprotroph (Põlme et al. 2020); Decay sugar-rich substrates (Põlme et al. 2020) | Põlme S, et al. 2020. Fungal Diversity 105:1-16 ((https://doi.org/10.1007/s13225-020-00466-2)) |
| Otu13464 | 1 | k__Fungi(100);p__Ascomycota(100);c__Saccharomycetes(100);o__Saccharomycetales(100);f__Metschnikowiaceae(100);g__Metschnikowia(100);s__Metschnikowia_pulcherrima(100); | Metschnikowia | 13 | Saprotroph | \|Undefined Saprotroph\| | Yeast | NULL | Probable | As nectar-tap_saprotroph (Põlme et al. 2020); Decay sugar-rich substrates (Põlme et al. 2020) | Põlme S, et al. 2020. Fungal Diversity 105:1-16 ((https://doi.org/10.1007/s13225-020-00466-2)) |
| Otu13469 | 1 | k__Fungi(100);p__Ascomycota(100);c__Saccharomycetes(100);o__Saccharomycetales(100);f__Metschnikowiaceae(100);g__Metschnikowia(100);g__Metschnikowia_unclassified(100); | Metschnikowia | 13 | Saprotroph | \|Undefined Saprotroph\| | Yeast | NULL | Probable | As nectar-tap_saprotroph (Põlme et al. 2020); Decay sugar-rich substrates (Põlme et al. 2020) | Põlme S, et al. 2020. Fungal Diversity 105:1-16 ((https://doi.org/10.1007/s13225-020-00466-2)) |
| Otu13477 | 1 | k__Fungi(100);p__Ascomycota(100);c__Saccharomycetes(100);o__Saccharomycetales(100);f__Metschnikowiaceae(100);g__Metschnikowia(100);s__Metschnikowia_pulcherrima(100); | Metschnikowia | 13 | Saprotroph | \|Undefined Saprotroph\| | Yeast | NULL | Probable | As nectar-tap_saprotroph (Põlme et al. 2020); Decay sugar-rich substrates (Põlme et al. 2020) | Põlme S, et al. 2020. Fungal Diversity 105:1-16 ((https://doi.org/10.1007/s13225-020-00466-2)) |
| Otu13478 | 1 | k__Fungi(100);p__Ascomycota(100);c__Saccharomycetes(100);o__Saccharomycetales(100);f__Metschnikowiaceae(100);g__Metschnikowia(100);s__Metschnikowia_pulcherrima(100); | Metschnikowia | 13 | Saprotroph | \|Undefined Saprotroph\| | Yeast | NULL | Probable | As nectar-tap_saprotroph (Põlme et al. 2020); Decay sugar-rich substrates (Põlme et al. 2020) | Põlme S, et al. 2020. Fungal Diversity 105:1-16 ((https://doi.org/10.1007/s13225-020-00466-2)) |
| Otu13481 | 1 | k__Fungi(100);p__Ascomycota(100);c__Saccharomycetes(100);o__Saccharomycetales(100);f__Metschnikowiaceae(100);g__Metschnikowia(100);s__Metschnikowia_pulcherrima(100); | Metschnikowia | 13 | Saprotroph | \|Undefined Saprotroph\| | Yeast | NULL | Probable | As nectar-tap_saprotroph (Põlme et al. 2020); Decay sugar-rich substrates (Põlme et al. 2020) | Põlme S, et al. 2020. Fungal Diversity 105:1-16 ((https://doi.org/10.1007/s13225-020-00466-2)) |
| Otu13484 | 1 | k__Fungi(100);p__Ascomycota(100);c__Saccharomycetes(100);o__Saccharomycetales(100);f__Metschnikowiaceae(100);g__Metschnikowia(100);s__Metschnikowia_pulcherrima(100); | Metschnikowia | 13 | Saprotroph | \|Undefined Saprotroph\| | Yeast | NULL | Probable | As nectar-tap_saprotroph (Põlme et al. 2020); Decay sugar-rich substrates (Põlme et al. 2020) | Põlme S, et al. 2020. Fungal Diversity 105:1-16 ((https://doi.org/10.1007/s13225-020-00466-2)) |
| Otu13485 | 1 | k__Fungi(100);p__Ascomycota(100);c__Saccharomycetes(100);o__Saccharomycetales(100);f__Metschnikowiaceae(100);g__Metschnikowia(100);s__Metschnikowia_pulcherrima(100); | Metschnikowia | 13 | Saprotroph | \|Undefined Saprotroph\| | Yeast | NULL | Probable | As nectar-tap_saprotroph (Põlme et al. 2020); Decay sugar-rich substrates (Põlme et al. 2020) | Põlme S, et al. 2020. Fungal Diversity 105:1-16 ((https://doi.org/10.1007/s13225-020-00466-2)) |
| Otu13487 | 1 | k__Fungi(100);p__Ascomycota(100);c__Saccharomycetes(100);o__Saccharomycetales(100);f__Metschnikowiaceae(100);g__Metschnikowia(100);s__Metschnikowia_pulcherrima(100); | Metschnikowia | 13 | Saprotroph | \|Undefined Saprotroph\| | Yeast | NULL | Probable | As nectar-tap_saprotroph (Põlme et al. 2020); Decay sugar-rich substrates (Põlme et al. 2020) | Põlme S, et al. 2020. Fungal Diversity 105:1-16 ((https://doi.org/10.1007/s13225-020-00466-2)) |
| Otu13488 | 1 | k__Fungi(100);p__Ascomycota(100);c__Saccharomycetes(100);o__Saccharomycetales(100);f__Metschnikowiaceae(100);g__Metschnikowia(100);g__Metschnikowia_unclassified(100); | Metschnikowia | 13 | Saprotroph | \|Undefined Saprotroph\| | Yeast | NULL | Probable | As nectar-tap_saprotroph (Põlme et al. 2020); Decay sugar-rich substrates (Põlme et al. 2020) | Põlme S, et al. 2020. Fungal Diversity 105:1-16 ((https://doi.org/10.1007/s13225-020-00466-2)) |
| Otu13490 | 1 | k__Fungi(100);p__Ascomycota(100);c__Saccharomycetes(100);o__Saccharomycetales(100);f__Metschnikowiaceae(100);g__Metschnikowia(100);s__Metschnikowia_pulcherrima(100); | Metschnikowia | 13 | Saprotroph | \|Undefined Saprotroph\| | Yeast | NULL | Probable | As nectar-tap_saprotroph (Põlme et al. 2020); Decay sugar-rich substrates (Põlme et al. 2020) | Põlme S, et al. 2020. Fungal Diversity 105:1-16 ((https://doi.org/10.1007/s13225-020-00466-2)) |
| Otu13491 | 1 | k__Fungi(100);p__Ascomycota(100);c__Saccharomycetes(100);o__Saccharomycetales(100);f__Metschnikowiaceae(100);g__Metschnikowia(100);s__Metschnikowia_pulcherrima(100); | Metschnikowia | 13 | Saprotroph | \|Undefined Saprotroph\| | Yeast | NULL | Probable | As nectar-tap_saprotroph (Põlme et al. 2020); Decay sugar-rich substrates (Põlme et al. 2020) | Põlme S, et al. 2020. Fungal Diversity 105:1-16 ((https://doi.org/10.1007/s13225-020-00466-2)) |
| Otu13498 | 1 | k__Fungi(100);p__Ascomycota(100);c__Saccharomycetes(100);o__Saccharomycetales(100);f__Metschnikowiaceae(100);g__Metschnikowia(100);s__Metschnikowia_pulcherrima(100); | Metschnikowia | 13 | Saprotroph | \|Undefined Saprotroph\| | Yeast | NULL | Probable | As nectar-tap_saprotroph (Põlme et al. 2020); Decay sugar-rich substrates (Põlme et al. 2020) | Põlme S, et al. 2020. Fungal Diversity 105:1-16 ((https://doi.org/10.1007/s13225-020-00466-2)) |
| Otu13501 | 1 | k__Fungi(100);p__Ascomycota(100);c__Saccharomycetes(100);o__Saccharomycetales(100);f__Metschnikowiaceae(100);g__Metschnikowia(100);g__Metschnikowia_unclassified(100); | Metschnikowia | 13 | Saprotroph | \|Undefined Saprotroph\| | Yeast | NULL | Probable | As nectar-tap_saprotroph (Põlme et al. 2020); Decay sugar-rich substrates (Põlme et al. 2020) | Põlme S, et al. 2020. Fungal Diversity 105:1-16 ((https://doi.org/10.1007/s13225-020-00466-2)) |
| Otu13503 | 1 | k__Fungi(100);p__Ascomycota(100);c__Saccharomycetes(100);o__Saccharomycetales(100);f__Metschnikowiaceae(100);g__Metschnikowia(100);s__Metschnikowia_pulcherrima(100); | Metschnikowia | 13 | Saprotroph | \|Undefined Saprotroph\| | Yeast | NULL | Probable | As nectar-tap_saprotroph (Põlme et al. 2020); Decay sugar-rich substrates (Põlme et al. 2020) | Põlme S, et al. 2020. Fungal Diversity 105:1-16 ((https://doi.org/10.1007/s13225-020-00466-2)) |
| Otu13507 | 1 | k__Fungi(100);p__Ascomycota(100);c__Saccharomycetes(100);o__Saccharomycetales(100);f__Metschnikowiaceae(100);g__Metschnikowia(100);g__Metschnikowia_unclassified(100); | Metschnikowia | 13 | Saprotroph | \|Undefined Saprotroph\| | Yeast | NULL | Probable | As nectar-tap_saprotroph (Põlme et al. 2020); Decay sugar-rich substrates (Põlme et al. 2020) | Põlme S, et al. 2020. Fungal Diversity 105:1-16 ((https://doi.org/10.1007/s13225-020-00466-2)) |
| Otu13508 | 1 | k__Fungi(100);p__Ascomycota(100);c__Saccharomycetes(100);o__Saccharomycetales(100);f__Metschnikowiaceae(100);g__Metschnikowia(100);g__Metschnikowia_unclassified(100); | Metschnikowia | 13 | Saprotroph | \|Undefined Saprotroph\| | Yeast | NULL | Probable | As nectar-tap_saprotroph (Põlme et al. 2020); Decay sugar-rich substrates (Põlme et al. 2020) | Põlme S, et al. 2020. Fungal Diversity 105:1-16 ((https://doi.org/10.1007/s13225-020-00466-2)) |
| Otu13509 | 1 | k__Fungi(100);p__Ascomycota(100);c__Sordariomycetes(100);o__Hypocreales(100);f__Nectriaceae(100);g__Fusarium(100);g__Fusarium_unclassified(100); | Fusarium | 13 | Pathotroph-Saprotroph-Symbiotroph | Animal Pathogen-Endophyte-Lichen Parasite-\|Plant Pathogen\|-Plant Saprotroph-Undefined Saprotroph-Wood Saprotroph | Microfungus | Soft Rot | Possible | As foliar_endophyte (Põlme et al. 2020); As litter_saprotroph (Põlme et al. 2020); Causing damping-off disease (Põlme et al. 2020); Decay leaf, fruit, seed, roots, soil, animal material (Põlme et al. 2020); Likely opportunistic human pathogen (Irinyi et al. 2015) | Duncan CG, Eslyn WE. 1966. Mycologia 58:642-664 ((https://doi.org/10.2307/3757045)); Seehann G, et al. 1975. List of Fungi in Soft-Rot Tissue Tests. Institute fur Holzbiologie und Holzschutz, Bundesforschungsanstalt fur Forst - und Holzwirtshaft, Hamburg - Lohbrugge, Germany; Costa IPMW, et al. 2012. Mycotaxon 119:e26 ((http://mycotaxon.com/resources/checklists/costa_v119_checklist.pdf)); Tedersoo L, et al. 2014. Science 346:e1256688 ((https://doi.org/10.1126/science.1256688)); Irinyi L, et al. 2015. Medical Mycology 53:313-337 ((https://doi.org/10.1093/mmy/myv008)); Schroers, et al. 2016. Mycologia 108:806-819 ((https://doi.org/10.3852/15-255)); Põlme S, et al. 2020. Fungal Diversity 105:1-16 ((https://doi.org/10.1007/s13225-020-00466-2)); The American Phytopathological Society Common Names of Plant Diseases ((http://www.apsnet.org/publications/commonnames/Pages/default.aspx)); Lawrey JD, Diederich P. 2016. http://www.lichenicolous.net (accessed 24 May 2017) ((http://www.lichenicolous.net)) |
| Otu13512 | 1 | k__Fungi(100);p__Ascomycota(100);c__Saccharomycetes(100);o__Saccharomycetales(100);f__Metschnikowiaceae(100);g__Metschnikowia(100);s__Metschnikowia_pulcherrima(100); | Metschnikowia | 13 | Saprotroph | \|Undefined Saprotroph\| | Yeast | NULL | Probable | As nectar-tap_saprotroph (Põlme et al. 2020); Decay sugar-rich substrates (Põlme et al. 2020) | Põlme S, et al. 2020. Fungal Diversity 105:1-16 ((https://doi.org/10.1007/s13225-020-00466-2)) |
| Otu13513 | 1 | k__Fungi(100);p__Ascomycota(100);c__Saccharomycetes(100);o__Saccharomycetales(100);f__Metschnikowiaceae(100);g__Metschnikowia(100);s__Metschnikowia_pulcherrima(100); | Metschnikowia | 13 | Saprotroph | \|Undefined Saprotroph\| | Yeast | NULL | Probable | As nectar-tap_saprotroph (Põlme et al. 2020); Decay sugar-rich substrates (Põlme et al. 2020) | Põlme S, et al. 2020. Fungal Diversity 105:1-16 ((https://doi.org/10.1007/s13225-020-00466-2)) |
| Otu13515 | 1 | k__Fungi(100);p__Ascomycota(100);c__Saccharomycetes(100);o__Saccharomycetales(100);f__Metschnikowiaceae(100);g__Metschnikowia(100);s__Metschnikowia_pulcherrima(100); | Metschnikowia | 13 | Saprotroph | \|Undefined Saprotroph\| | Yeast | NULL | Probable | As nectar-tap_saprotroph (Põlme et al. 2020); Decay sugar-rich substrates (Põlme et al. 2020) | Põlme S, et al. 2020. Fungal Diversity 105:1-16 ((https://doi.org/10.1007/s13225-020-00466-2)) |
| Otu13517 | 1 | k__Fungi(100);p__Ascomycota(100);c__Saccharomycetes(100);o__Saccharomycetales(100);f__Metschnikowiaceae(100);g__Metschnikowia(100);s__Metschnikowia_pulcherrima(100); | Metschnikowia | 13 | Saprotroph | \|Undefined Saprotroph\| | Yeast | NULL | Probable | As nectar-tap_saprotroph (Põlme et al. 2020); Decay sugar-rich substrates (Põlme et al. 2020) | Põlme S, et al. 2020. Fungal Diversity 105:1-16 ((https://doi.org/10.1007/s13225-020-00466-2)) |
| Otu13519 | 1 | k__Fungi(100);p__Ascomycota(100);c__Saccharomycetes(100);o__Saccharomycetales(100);f__Metschnikowiaceae(100);g__Metschnikowia(100);g__Metschnikowia_unclassified(100); | Metschnikowia | 13 | Saprotroph | \|Undefined Saprotroph\| | Yeast | NULL | Probable | As nectar-tap_saprotroph (Põlme et al. 2020); Decay sugar-rich substrates (Põlme et al. 2020) | Põlme S, et al. 2020. Fungal Diversity 105:1-16 ((https://doi.org/10.1007/s13225-020-00466-2)) |
| Otu13520 | 1 | k__Fungi(100);p__Ascomycota(100);c__Saccharomycetes(100);o__Saccharomycetales(100);f__Metschnikowiaceae(100);g__Metschnikowia(100);s__Metschnikowia_pulcherrima(100); | Metschnikowia | 13 | Saprotroph | \|Undefined Saprotroph\| | Yeast | NULL | Probable | As nectar-tap_saprotroph (Põlme et al. 2020); Decay sugar-rich substrates (Põlme et al. 2020) | Põlme S, et al. 2020. Fungal Diversity 105:1-16 ((https://doi.org/10.1007/s13225-020-00466-2)) |
| Otu13523 | 1 | k__Fungi(100);p__Ascomycota(100);c__Saccharomycetes(100);o__Saccharomycetales(100);f__Metschnikowiaceae(100);g__Metschnikowia(100);g__Metschnikowia_unclassified(100); | Metschnikowia | 13 | Saprotroph | \|Undefined Saprotroph\| | Yeast | NULL | Probable | As nectar-tap_saprotroph (Põlme et al. 2020); Decay sugar-rich substrates (Põlme et al. 2020) | Põlme S, et al. 2020. Fungal Diversity 105:1-16 ((https://doi.org/10.1007/s13225-020-00466-2)) |
| Otu13525 | 1 | k__Fungi(100);p__Ascomycota(100);c__Saccharomycetes(100);o__Saccharomycetales(100);f__Metschnikowiaceae(100);g__Metschnikowia(100);s__Metschnikowia_pulcherrima(100); | Metschnikowia | 13 | Saprotroph | \|Undefined Saprotroph\| | Yeast | NULL | Probable | As nectar-tap_saprotroph (Põlme et al. 2020); Decay sugar-rich substrates (Põlme et al. 2020) | Põlme S, et al. 2020. Fungal Diversity 105:1-16 ((https://doi.org/10.1007/s13225-020-00466-2)) |
| Otu13527 | 1 | k__Fungi(100);p__Ascomycota(100);c__Saccharomycetes(100);o__Saccharomycetales(100);f__Metschnikowiaceae(100);g__Metschnikowia(100);s__Metschnikowia_pulcherrima(100); | Metschnikowia | 13 | Saprotroph | \|Undefined Saprotroph\| | Yeast | NULL | Probable | As nectar-tap_saprotroph (Põlme et al. 2020); Decay sugar-rich substrates (Põlme et al. 2020) | Põlme S, et al. 2020. Fungal Diversity 105:1-16 ((https://doi.org/10.1007/s13225-020-00466-2)) |
| Otu13529 | 1 | k__Fungi(100);p__Ascomycota(100);c__Saccharomycetes(100);o__Saccharomycetales(100);f__Metschnikowiaceae(100);g__Metschnikowia(100);g__Metschnikowia_unclassified(100); | Metschnikowia | 13 | Saprotroph | \|Undefined Saprotroph\| | Yeast | NULL | Probable | As nectar-tap_saprotroph (Põlme et al. 2020); Decay sugar-rich substrates (Põlme et al. 2020) | Põlme S, et al. 2020. Fungal Diversity 105:1-16 ((https://doi.org/10.1007/s13225-020-00466-2)) |
| Otu13530 | 1 | k__Fungi(100);p__Ascomycota(100);c__Saccharomycetes(100);o__Saccharomycetales(100);f__Metschnikowiaceae(100);g__Metschnikowia(100);s__Metschnikowia_pulcherrima(100); | Metschnikowia | 13 | Saprotroph | \|Undefined Saprotroph\| | Yeast | NULL | Probable | As nectar-tap_saprotroph (Põlme et al. 2020); Decay sugar-rich substrates (Põlme et al. 2020) | Põlme S, et al. 2020. Fungal Diversity 105:1-16 ((https://doi.org/10.1007/s13225-020-00466-2)) |
| Otu13531 | 1 | k__Fungi(100);p__Ascomycota(100);c__Saccharomycetes(100);o__Saccharomycetales(100);f__Metschnikowiaceae(100);g__Metschnikowia(100);s__Metschnikowia_pulcherrima(100); | Metschnikowia | 13 | Saprotroph | \|Undefined Saprotroph\| | Yeast | NULL | Probable | As nectar-tap_saprotroph (Põlme et al. 2020); Decay sugar-rich substrates (Põlme et al. 2020) | Põlme S, et al. 2020. Fungal Diversity 105:1-16 ((https://doi.org/10.1007/s13225-020-00466-2)) |
| Otu13532 | 1 | k__Fungi(100);p__Ascomycota(100);c__Saccharomycetes(100);o__Saccharomycetales(100);f__Metschnikowiaceae(100);g__Metschnikowia(100);s__Metschnikowia_pulcherrima(100); | Metschnikowia | 13 | Saprotroph | \|Undefined Saprotroph\| | Yeast | NULL | Probable | As nectar-tap_saprotroph (Põlme et al. 2020); Decay sugar-rich substrates (Põlme et al. 2020) | Põlme S, et al. 2020. Fungal Diversity 105:1-16 ((https://doi.org/10.1007/s13225-020-00466-2)) |
| Otu13534 | 1 | k__Fungi(100);p__Ascomycota(100);c__Saccharomycetes(100);o__Saccharomycetales(100);f__Metschnikowiaceae(100);g__Metschnikowia(100);s__Metschnikowia_pulcherrima(100); | Metschnikowia | 13 | Saprotroph | \|Undefined Saprotroph\| | Yeast | NULL | Probable | As nectar-tap_saprotroph (Põlme et al. 2020); Decay sugar-rich substrates (Põlme et al. 2020) | Põlme S, et al. 2020. Fungal Diversity 105:1-16 ((https://doi.org/10.1007/s13225-020-00466-2)) |
| Otu13538 | 1 | k__Fungi(100);p__Ascomycota(100);c__Saccharomycetes(100);o__Saccharomycetales(100);f__Metschnikowiaceae(100);g__Metschnikowia(100);s__Metschnikowia_pulcherrima(100); | Metschnikowia | 13 | Saprotroph | \|Undefined Saprotroph\| | Yeast | NULL | Probable | As nectar-tap_saprotroph (Põlme et al. 2020); Decay sugar-rich substrates (Põlme et al. 2020) | Põlme S, et al. 2020. Fungal Diversity 105:1-16 ((https://doi.org/10.1007/s13225-020-00466-2)) |
| Otu13545 | 1 | k__Fungi(100);p__Ascomycota(100);c__Saccharomycetes(100);o__Saccharomycetales(100);f__Metschnikowiaceae(100);g__Metschnikowia(100);g__Metschnikowia_unclassified(100); | Metschnikowia | 13 | Saprotroph | \|Undefined Saprotroph\| | Yeast | NULL | Probable | As nectar-tap_saprotroph (Põlme et al. 2020); Decay sugar-rich substrates (Põlme et al. 2020) | Põlme S, et al. 2020. Fungal Diversity 105:1-16 ((https://doi.org/10.1007/s13225-020-00466-2)) |
| Otu13546 | 1 | k__Fungi(100);p__Ascomycota(100);c__Saccharomycetes(100);o__Saccharomycetales(100);f__Metschnikowiaceae(100);g__Metschnikowia(100);s__Metschnikowia_pulcherrima(100); | Metschnikowia | 13 | Saprotroph | \|Undefined Saprotroph\| | Yeast | NULL | Probable | As nectar-tap_saprotroph (Põlme et al. 2020); Decay sugar-rich substrates (Põlme et al. 2020) | Põlme S, et al. 2020. Fungal Diversity 105:1-16 ((https://doi.org/10.1007/s13225-020-00466-2)) |
| Otu13548 | 1 | k__Fungi(100);p__Ascomycota(100);c__Saccharomycetes(100);o__Saccharomycetales(100);f__Metschnikowiaceae(100);g__Metschnikowia(100);s__Metschnikowia_pulcherrima(100); | Metschnikowia | 13 | Saprotroph | \|Undefined Saprotroph\| | Yeast | NULL | Probable | As nectar-tap_saprotroph (Põlme et al. 2020); Decay sugar-rich substrates (Põlme et al. 2020) | Põlme S, et al. 2020. Fungal Diversity 105:1-16 ((https://doi.org/10.1007/s13225-020-00466-2)) |
| Otu13549 | 1 | k__Fungi(100);p__Ascomycota(100);c__Saccharomycetes(100);o__Saccharomycetales(100);f__Metschnikowiaceae(100);g__Metschnikowia(100);s__Metschnikowia_pulcherrima(100); | Metschnikowia | 13 | Saprotroph | \|Undefined Saprotroph\| | Yeast | NULL | Probable | As nectar-tap_saprotroph (Põlme et al. 2020); Decay sugar-rich substrates (Põlme et al. 2020) | Põlme S, et al. 2020. Fungal Diversity 105:1-16 ((https://doi.org/10.1007/s13225-020-00466-2)) |
| Otu13550 | 1 | k__Fungi(100);p__Ascomycota(100);c__Saccharomycetes(100);o__Saccharomycetales(100);f__Saccharomycetales_family_Incertae_sedis(100);g__Candida(100);g__Candida_unclassified(100); | Candida | 13 | Pathotroph-Saprotroph-Symbiotroph | Animal Pathogen-Endophyte-Endosymbiont-Epiphyte-\|Undefined Saprotroph\| | Dimorphic Yeast-Yeast | NULL | Probable | As nectar-tap_saprotroph (Põlme et al. 2020); Decay sugar-rich substrates (Põlme et al. 2020); This asexual genus is polyphyletic and contain species that function very differently depend on what they are. Caution should be taken when interpreting guilds for Candida species. | Manolakaki D, et al. 2010. Virulence. 1: 367-375 ((https://doi.org/10.4161/viru.1.5.12796)); Mekbib SB, et al. 2011. Tropical Plant Pathology 36:233-240 ((https://doi.org/10.1590/S1982-56762011000400004)); Põlme S, et al. 2020. Fungal Diversity 105:1-16 ((https://doi.org/10.1007/s13225-020-00466-2)) |
| Otu13553 | 1 | k__Fungi(100);p__Ascomycota(100);c__Dothideomycetes(100);o__Dothideomycetes_order_Incertae_sedis(100);f__Myxotrichaceae(100);g__Oidiodendron(100);s__Oidiodendron_maius(100); | Oidiodendron | 13 | Pathotroph-Saprotroph-Symbiotroph | Endophyte-Ericoid Mycorrhizal-\|Undefined Saprotroph\| | Microfungus | NULL | Probable | As root_endophyte (Põlme et al. 2020); As soil_saprotroph (Põlme et al. 2020); Dark Septate Endophyte; Decay soil (Põlme et al. 2020) | Newsham KK. 2011. New Phytologist 190:783-793 ((https://doi.org/10.1111/j.1469-8137.2010.03611.x)) (as dark septate root endophytes); Põlme S, et al. 2020. Fungal Diversity 105:1-16 ((https://doi.org/10.1007/s13225-020-00466-2)) |
| Otu13557 | 1 | k__Fungi(100);p__Ascomycota(100);c__Saccharomycetes(100);o__Saccharomycetales(100);f__Metschnikowiaceae(100);g__Metschnikowia(100);s__Metschnikowia_pulcherrima(100); | Metschnikowia | 13 | Saprotroph | \|Undefined Saprotroph\| | Yeast | NULL | Probable | As nectar-tap_saprotroph (Põlme et al. 2020); Decay sugar-rich substrates (Põlme et al. 2020) | Põlme S, et al. 2020. Fungal Diversity 105:1-16 ((https://doi.org/10.1007/s13225-020-00466-2)) |
| Otu13560 | 1 | k__Fungi(100);p__Ascomycota(100);c__Saccharomycetes(100);o__Saccharomycetales(100);f__Metschnikowiaceae(100);g__Metschnikowia(100);s__Metschnikowia_pulcherrima(100); | Metschnikowia | 13 | Saprotroph | \|Undefined Saprotroph\| | Yeast | NULL | Probable | As nectar-tap_saprotroph (Põlme et al. 2020); Decay sugar-rich substrates (Põlme et al. 2020) | Põlme S, et al. 2020. Fungal Diversity 105:1-16 ((https://doi.org/10.1007/s13225-020-00466-2)) |
| Otu13561 | 1 | k__Fungi(100);p__Ascomycota(100);c__Saccharomycetes(100);o__Saccharomycetales(100);f__Metschnikowiaceae(100);g__Metschnikowia(100);g__Metschnikowia_unclassified(100); | Metschnikowia | 13 | Saprotroph | \|Undefined Saprotroph\| | Yeast | NULL | Probable | As nectar-tap_saprotroph (Põlme et al. 2020); Decay sugar-rich substrates (Põlme et al. 2020) | Põlme S, et al. 2020. Fungal Diversity 105:1-16 ((https://doi.org/10.1007/s13225-020-00466-2)) |
| Otu13563 | 1 | k__Fungi(100);p__Ascomycota(100);c__Saccharomycetes(100);o__Saccharomycetales(100);f__Metschnikowiaceae(100);g__Metschnikowia(100);s__Metschnikowia_pulcherrima(100); | Metschnikowia | 13 | Saprotroph | \|Undefined Saprotroph\| | Yeast | NULL | Probable | As nectar-tap_saprotroph (Põlme et al. 2020); Decay sugar-rich substrates (Põlme et al. 2020) | Põlme S, et al. 2020. Fungal Diversity 105:1-16 ((https://doi.org/10.1007/s13225-020-00466-2)) |
| Otu13564 | 1 | k__Fungi(100);p__Ascomycota(100);c__Saccharomycetes(100);o__Saccharomycetales(100);f__Metschnikowiaceae(100);g__Metschnikowia(100);s__Metschnikowia_pulcherrima(100); | Metschnikowia | 13 | Saprotroph | \|Undefined Saprotroph\| | Yeast | NULL | Probable | As nectar-tap_saprotroph (Põlme et al. 2020); Decay sugar-rich substrates (Põlme et al. 2020) | Põlme S, et al. 2020. Fungal Diversity 105:1-16 ((https://doi.org/10.1007/s13225-020-00466-2)) |
| Otu13565 | 1 | k__Fungi(100);p__Ascomycota(100);c__Saccharomycetes(100);o__Saccharomycetales(100);f__Metschnikowiaceae(100);g__Metschnikowia(100);g__Metschnikowia_unclassified(100); | Metschnikowia | 13 | Saprotroph | \|Undefined Saprotroph\| | Yeast | NULL | Probable | As nectar-tap_saprotroph (Põlme et al. 2020); Decay sugar-rich substrates (Põlme et al. 2020) | Põlme S, et al. 2020. Fungal Diversity 105:1-16 ((https://doi.org/10.1007/s13225-020-00466-2)) |
| Otu13566 | 1 | k__Fungi(100);p__Ascomycota(100);c__Saccharomycetes(100);o__Saccharomycetales(100);f__Metschnikowiaceae(100);g__Metschnikowia(100);s__Metschnikowia_pulcherrima(100); | Metschnikowia | 13 | Saprotroph | \|Undefined Saprotroph\| | Yeast | NULL | Probable | As nectar-tap_saprotroph (Põlme et al. 2020); Decay sugar-rich substrates (Põlme et al. 2020) | Põlme S, et al. 2020. Fungal Diversity 105:1-16 ((https://doi.org/10.1007/s13225-020-00466-2)) |
| Otu13568 | 1 | k__Fungi(100);p__Ascomycota(100);c__Saccharomycetes(100);o__Saccharomycetales(100);f__Metschnikowiaceae(100);g__Metschnikowia(100);s__Metschnikowia_pulcherrima(100); | Metschnikowia | 13 | Saprotroph | \|Undefined Saprotroph\| | Yeast | NULL | Probable | As nectar-tap_saprotroph (Põlme et al. 2020); Decay sugar-rich substrates (Põlme et al. 2020) | Põlme S, et al. 2020. Fungal Diversity 105:1-16 ((https://doi.org/10.1007/s13225-020-00466-2)) |
| Otu13569 | 1 | k__Fungi(100);p__Ascomycota(100);c__Saccharomycetes(100);o__Saccharomycetales(100);f__Metschnikowiaceae(100);g__Metschnikowia(100);s__Metschnikowia_pulcherrima(100); | Metschnikowia | 13 | Saprotroph | \|Undefined Saprotroph\| | Yeast | NULL | Probable | As nectar-tap_saprotroph (Põlme et al. 2020); Decay sugar-rich substrates (Põlme et al. 2020) | Põlme S, et al. 2020. Fungal Diversity 105:1-16 ((https://doi.org/10.1007/s13225-020-00466-2)) |
| Otu13570 | 1 | k__Fungi(100);p__Ascomycota(100);c__Saccharomycetes(100);o__Saccharomycetales(100);f__Metschnikowiaceae(100);g__Metschnikowia(100);s__Metschnikowia_pulcherrima(100); | Metschnikowia | 13 | Saprotroph | \|Undefined Saprotroph\| | Yeast | NULL | Probable | As nectar-tap_saprotroph (Põlme et al. 2020); Decay sugar-rich substrates (Põlme et al. 2020) | Põlme S, et al. 2020. Fungal Diversity 105:1-16 ((https://doi.org/10.1007/s13225-020-00466-2)) |
| Otu13571 | 1 | k__Fungi(100);p__Ascomycota(100);c__Saccharomycetes(100);o__Saccharomycetales(100);f__Metschnikowiaceae(100);g__Metschnikowia(100);s__Metschnikowia_pulcherrima(100); | Metschnikowia | 13 | Saprotroph | \|Undefined Saprotroph\| | Yeast | NULL | Probable | As nectar-tap_saprotroph (Põlme et al. 2020); Decay sugar-rich substrates (Põlme et al. 2020) | Põlme S, et al. 2020. Fungal Diversity 105:1-16 ((https://doi.org/10.1007/s13225-020-00466-2)) |
| Otu13576 | 1 | k__Fungi(100);p__Ascomycota(100);c__Saccharomycetes(100);o__Saccharomycetales(100);f__Metschnikowiaceae(100);g__Metschnikowia(100);s__Metschnikowia_pulcherrima(100); | Metschnikowia | 13 | Saprotroph | \|Undefined Saprotroph\| | Yeast | NULL | Probable | As nectar-tap_saprotroph (Põlme et al. 2020); Decay sugar-rich substrates (Põlme et al. 2020) | Põlme S, et al. 2020. Fungal Diversity 105:1-16 ((https://doi.org/10.1007/s13225-020-00466-2)) |
| Otu13577 | 1 | k__Fungi(100);p__Ascomycota(100);c__Saccharomycetes(100);o__Saccharomycetales(100);f__Metschnikowiaceae(100);g__Metschnikowia(100);s__Metschnikowia_pulcherrima(100); | Metschnikowia | 13 | Saprotroph | \|Undefined Saprotroph\| | Yeast | NULL | Probable | As nectar-tap_saprotroph (Põlme et al. 2020); Decay sugar-rich substrates (Põlme et al. 2020) | Põlme S, et al. 2020. Fungal Diversity 105:1-16 ((https://doi.org/10.1007/s13225-020-00466-2)) |
| Otu13578 | 1 | k__Fungi(100);p__Ascomycota(100);c__Saccharomycetes(100);o__Saccharomycetales(100);f__Metschnikowiaceae(100);g__Metschnikowia(100);g__Metschnikowia_unclassified(100); | Metschnikowia | 13 | Saprotroph | \|Undefined Saprotroph\| | Yeast | NULL | Probable | As nectar-tap_saprotroph (Põlme et al. 2020); Decay sugar-rich substrates (Põlme et al. 2020) | Põlme S, et al. 2020. Fungal Diversity 105:1-16 ((https://doi.org/10.1007/s13225-020-00466-2)) |
| Otu13581 | 1 | k__Fungi(100);p__Ascomycota(100);c__Saccharomycetes(100);o__Saccharomycetales(100);f__Metschnikowiaceae(100);g__Metschnikowia(100);g__Metschnikowia_unclassified(100); | Metschnikowia | 13 | Saprotroph | \|Undefined Saprotroph\| | Yeast | NULL | Probable | As nectar-tap_saprotroph (Põlme et al. 2020); Decay sugar-rich substrates (Põlme et al. 2020) | Põlme S, et al. 2020. Fungal Diversity 105:1-16 ((https://doi.org/10.1007/s13225-020-00466-2)) |
| Otu13582 | 1 | k__Fungi(100);p__Ascomycota(100);c__Saccharomycetes(100);o__Saccharomycetales(100);f__Metschnikowiaceae(100);g__Metschnikowia(100);g__Metschnikowia_unclassified(100); | Metschnikowia | 13 | Saprotroph | \|Undefined Saprotroph\| | Yeast | NULL | Probable | As nectar-tap_saprotroph (Põlme et al. 2020); Decay sugar-rich substrates (Põlme et al. 2020) | Põlme S, et al. 2020. Fungal Diversity 105:1-16 ((https://doi.org/10.1007/s13225-020-00466-2)) |
| Otu13584 | 1 | k__Fungi(100);p__Ascomycota(100);c__Saccharomycetes(100);o__Saccharomycetales(100);f__Metschnikowiaceae(100);g__Metschnikowia(100);s__Metschnikowia_pulcherrima(100); | Metschnikowia | 13 | Saprotroph | \|Undefined Saprotroph\| | Yeast | NULL | Probable | As nectar-tap_saprotroph (Põlme et al. 2020); Decay sugar-rich substrates (Põlme et al. 2020) | Põlme S, et al. 2020. Fungal Diversity 105:1-16 ((https://doi.org/10.1007/s13225-020-00466-2)) |
| Otu13587 | 1 | k__Fungi(100);p__Ascomycota(100);c__Saccharomycetes(100);o__Saccharomycetales(100);f__Metschnikowiaceae(100);g__Metschnikowia(100);g__Metschnikowia_unclassified(100); | Metschnikowia | 13 | Saprotroph | \|Undefined Saprotroph\| | Yeast | NULL | Probable | As nectar-tap_saprotroph (Põlme et al. 2020); Decay sugar-rich substrates (Põlme et al. 2020) | Põlme S, et al. 2020. Fungal Diversity 105:1-16 ((https://doi.org/10.1007/s13225-020-00466-2)) |
| Otu13588 | 1 | k__Fungi(100);p__Ascomycota(100);c__Saccharomycetes(100);o__Saccharomycetales(100);f__Metschnikowiaceae(100);g__Metschnikowia(100);s__Metschnikowia_pulcherrima(100); | Metschnikowia | 13 | Saprotroph | \|Undefined Saprotroph\| | Yeast | NULL | Probable | As nectar-tap_saprotroph (Põlme et al. 2020); Decay sugar-rich substrates (Põlme et al. 2020) | Põlme S, et al. 2020. Fungal Diversity 105:1-16 ((https://doi.org/10.1007/s13225-020-00466-2)) |
| Otu13590 | 1 | k__Fungi(100);p__Ascomycota(100);c__Saccharomycetes(100);o__Saccharomycetales(100);f__Metschnikowiaceae(100);g__Metschnikowia(100);g__Metschnikowia_unclassified(100); | Metschnikowia | 13 | Saprotroph | \|Undefined Saprotroph\| | Yeast | NULL | Probable | As nectar-tap_saprotroph (Põlme et al. 2020); Decay sugar-rich substrates (Põlme et al. 2020) | Põlme S, et al. 2020. Fungal Diversity 105:1-16 ((https://doi.org/10.1007/s13225-020-00466-2)) |
| Otu13591 | 1 | k__Fungi(100);p__Ascomycota(100);c__Saccharomycetes(100);o__Saccharomycetales(100);f__Metschnikowiaceae(100);g__Metschnikowia(100);s__Metschnikowia_pulcherrima(100); | Metschnikowia | 13 | Saprotroph | \|Undefined Saprotroph\| | Yeast | NULL | Probable | As nectar-tap_saprotroph (Põlme et al. 2020); Decay sugar-rich substrates (Põlme et al. 2020) | Põlme S, et al. 2020. Fungal Diversity 105:1-16 ((https://doi.org/10.1007/s13225-020-00466-2)) |

| | | | | | | | | | | | |
|---|---|---|---|---|---|---|---|---|---|---|---|
| Otu13592 | 1 | k__Fungi(100);p__Ascomycota(100);c__Saccharomycetes(100);o__Saccharomycetales(100);f__Metschnikowiaceae(100);g__Metschnikowia(100);s__Metschnikowia_pulcherrima(100); | Metschnikowia | 13 | Saprotroph | \|Undefined Saprotroph\| | Yeast | NULL | Probable | As nectar-tap_saprotroph (Põlme et al. 2020); Decay sugar-rich substrates (Põlme et al. 2020) | Põlme S, et al. 2020. Fungal Diversity 105:1-16 ((https://doi.org/10.1007/s13225-020-00466-2)) |
| Otu13593 | 1 | k__Fungi(100);p__Ascomycota(100);c__Saccharomycetes(100);o__Saccharomycetales(100);f__Metschnikowiaceae(100);g__Metschnikowia(100);s__Metschnikowia_pulcherrima(100); | Metschnikowia | 13 | Saprotroph | \|Undefined Saprotroph\| | Yeast | NULL | Probable | As nectar-tap_saprotroph (Põlme et al. 2020); Decay sugar-rich substrates (Põlme et al. 2020) | Põlme S, et al. 2020. Fungal Diversity 105:1-16 ((https://doi.org/10.1007/s13225-020-00466-2)) |
| Otu13594 | 1 | k__Fungi(100);p__Ascomycota(100);c__Saccharomycetes(100);o__Saccharomycetales(100);f__Metschnikowiaceae(100);g__Metschnikowia(100);s__Metschnikowia_pulcherrima(100); | Metschnikowia | 13 | Saprotroph | \|Undefined Saprotroph\| | Yeast | NULL | Probable | As nectar-tap_saprotroph (Põlme et al. 2020); Decay sugar-rich substrates (Põlme et al. 2020) | Põlme S, et al. 2020. Fungal Diversity 105:1-16 ((https://doi.org/10.1007/s13225-020-00466-2)) |
| Otu13595 | 1 | k__Fungi(100);p__Ascomycota(100);c__Saccharomycetes(100);o__Saccharomycetales(100);f__Metschnikowiaceae(100);g__Metschnikowia(100);g__Metschnikowia_unclassified(100); | Metschnikowia | 13 | Saprotroph | \|Undefined Saprotroph\| | Yeast | NULL | Probable | As nectar-tap_saprotroph (Põlme et al. 2020); Decay sugar-rich substrates (Põlme et al. 2020) | Põlme S, et al. 2020. Fungal Diversity 105:1-16 ((https://doi.org/10.1007/s13225-020-00466-2)) |
| Otu13597 | 1 | k__Fungi(100);p__Ascomycota(100);c__Saccharomycetes(100);o__Saccharomycetales(100);f__Metschnikowiaceae(100);g__Metschnikowia(100);s__Metschnikowia_pulcherrima(100); | Metschnikowia | 13 | Saprotroph | \|Undefined Saprotroph\| | Yeast | NULL | Probable | As nectar-tap_saprotroph (Põlme et al. 2020); Decay sugar-rich substrates (Põlme et al. 2020) | Põlme S, et al. 2020. Fungal Diversity 105:1-16 ((https://doi.org/10.1007/s13225-020-00466-2)) |
| Otu13598 | 1 | k__Fungi(100);p__Ascomycota(100);c__Saccharomycetes(100);o__Saccharomycetales(100);f__Metschnikowiaceae(100);g__Metschnikowia(100);s__Metschnikowia_pulcherrima(100); | Metschnikowia | 13 | Saprotroph | \|Undefined Saprotroph\| | Yeast | NULL | Probable | As nectar-tap_saprotroph (Põlme et al. 2020); Decay sugar-rich substrates (Põlme et al. 2020) | Põlme S, et al. 2020. Fungal Diversity 105:1-16 ((https://doi.org/10.1007/s13225-020-00466-2)) |
| Otu13600 | 1 | k__Fungi(100);p__Ascomycota(100);c__Saccharomycetes(100);o__Saccharomycetales(100);f__Metschnikowiaceae(100);g__Metschnikowia(100);s__Metschnikowia_pulcherrima(100); | Metschnikowia | 13 | Saprotroph | \|Undefined Saprotroph\| | Yeast | NULL | Probable | As nectar-tap_saprotroph (Põlme et al. 2020); Decay sugar-rich substrates (Põlme et al. 2020) | Põlme S, et al. 2020. Fungal Diversity 105:1-16 ((https://doi.org/10.1007/s13225-020-00466-2)) |
| Otu13601 | 1 | k__Fungi(100);p__Ascomycota(100);c__Saccharomycetes(100);o__Saccharomycetales(100);f__Metschnikowiaceae(100);g__Metschnikowia(100);s__Metschnikowia_pulcherrima(100); | Metschnikowia | 13 | Saprotroph | \|Undefined Saprotroph\| | Yeast | NULL | Probable | As nectar-tap_saprotroph (Põlme et al. 2020); Decay sugar-rich substrates (Põlme et al. 2020) | Põlme S, et al. 2020. Fungal Diversity 105:1-16 ((https://doi.org/10.1007/s13225-020-00466-2)) |
| Otu13602 | 1 | k__Fungi(100);p__Ascomycota(100);c__Saccharomycetes(100);o__Saccharomycetales(100);f__Metschnikowiaceae(100);g__Metschnikowia(100);s__Metschnikowia_pulcherrima(100); | Metschnikowia | 13 | Saprotroph | \|Undefined Saprotroph\| | Yeast | NULL | Probable | As nectar-tap_saprotroph (Põlme et al. 2020); Decay sugar-rich substrates (Põlme et al. 2020) | Põlme S, et al. 2020. Fungal Diversity 105:1-16 ((https://doi.org/10.1007/s13225-020-00466-2)) |
| Otu13603 | 1 | k__Fungi(100);p__Ascomycota(100);c__Saccharomycetes(100);o__Saccharomycetales(100);f__Metschnikowiaceae(100);g__Metschnikowia(100);s__Metschnikowia_pulcherrima(100); | Metschnikowia | 13 | Saprotroph | \|Undefined Saprotroph\| | Yeast | NULL | Probable | As nectar-tap_saprotroph (Põlme et al. 2020); Decay sugar-rich substrates (Põlme et al. 2020) | Põlme S, et al. 2020. Fungal Diversity 105:1-16 ((https://doi.org/10.1007/s13225-020-00466-2)) |
| Otu13604 | 1 | k__Fungi(100);p__Ascomycota(100);c__Saccharomycetes(100);o__Saccharomycetales(100);f__Metschnikowiaceae(100);g__Metschnikowia(100);s__Metschnikowia_pulcherrima(100); | Metschnikowia | 13 | Saprotroph | \|Undefined Saprotroph\| | Yeast | NULL | Probable | As nectar-tap_saprotroph (Põlme et al. 2020); Decay sugar-rich substrates (Põlme et al. 2020) | Põlme S, et al. 2020. Fungal Diversity 105:1-16 ((https://doi.org/10.1007/s13225-020-00466-2)) |
| Otu13606 | 1 | k__Fungi(100);p__Ascomycota(100);c__Saccharomycetes(100);o__Saccharomycetales(100);f__Metschnikowiaceae(100);g__Metschnikowia(100);s__Metschnikowia_pulcherrima(100); | Metschnikowia | 13 | Saprotroph | \|Undefined Saprotroph\| | Yeast | NULL | Probable | As nectar-tap_saprotroph (Põlme et al. 2020); Decay sugar-rich substrates (Põlme et al. 2020) | Põlme S, et al. 2020. Fungal Diversity 105:1-16 ((https://doi.org/10.1007/s13225-020-00466-2)) |
| Otu13607 | 1 | k__Fungi(100);p__Ascomycota(100);c__Saccharomycetes(100);o__Saccharomycetales(100);f__Metschnikowiaceae(100);g__Metschnikowia(100);g__Metschnikowia_unclassified(100); | Metschnikowia | 13 | Saprotroph | \|Undefined Saprotroph\| | Yeast | NULL | Probable | As nectar-tap_saprotroph (Põlme et al. 2020); Decay sugar-rich substrates (Põlme et al. 2020) | Põlme S, et al. 2020. Fungal Diversity 105:1-16 ((https://doi.org/10.1007/s13225-020-00466-2)) |
| Otu13610 | 1 | k__Fungi(100);p__Ascomycota(100);c__Saccharomycetes(100);o__Saccharomycetales(100);f__Metschnikowiaceae(100);g__Metschnikowia(100);s__Metschnikowia_pulcherrima(100); | Metschnikowia | 13 | Saprotroph | \|Undefined Saprotroph\| | Yeast | NULL | Probable | As nectar-tap_saprotroph (Põlme et al. 2020); Decay sugar-rich substrates (Põlme et al. 2020) | Põlme S, et al. 2020. Fungal Diversity 105:1-16 ((https://doi.org/10.1007/s13225-020-00466-2)) |
| Otu13616 | 1 | k__Fungi(100);p__Ascomycota(100);c__Saccharomycetes(100);o__Saccharomycetales(100);f__Metschnikowiaceae(100);g__Metschnikowia(100);s__Metschnikowia_pulcherrima(100); | Metschnikowia | 13 | Saprotroph | \|Undefined Saprotroph\| | Yeast | NULL | Probable | As nectar-tap_saprotroph (Põlme et al. 2020); Decay sugar-rich substrates (Põlme et al. 2020) | Põlme S, et al. 2020. Fungal Diversity 105:1-16 ((https://doi.org/10.1007/s13225-020-00466-2)) |
| Otu13617 | 1 | k__Fungi(100);p__Ascomycota(100);c__Saccharomycetes(100);o__Saccharomycetales(100);f__Metschnikowiaceae(100);g__Metschnikowia(100);s__Metschnikowia_pulcherrima(100); | Metschnikowia | 13 | Saprotroph | \|Undefined Saprotroph\| | Yeast | NULL | Probable | As nectar-tap_saprotroph (Põlme et al. 2020); Decay sugar-rich substrates (Põlme et al. 2020) | Põlme S, et al. 2020. Fungal Diversity 105:1-16 ((https://doi.org/10.1007/s13225-020-00466-2)) |
| Otu13618 | 1 | k__Fungi(100);p__Ascomycota(100);c__Saccharomycetes(100);o__Saccharomycetales(100);f__Metschnikowiaceae(100);g__Metschnikowia(100);s__Metschnikowia_pulcherrima(100); | Metschnikowia | 13 | Saprotroph | \|Undefined Saprotroph\| | Yeast | NULL | Probable | As nectar-tap_saprotroph (Põlme et al. 2020); Decay sugar-rich substrates (Põlme et al. 2020) | Põlme S, et al. 2020. Fungal Diversity 105:1-16 ((https://doi.org/10.1007/s13225-020-00466-2)) |
| Otu13619 | 1 | k__Fungi(100);p__Ascomycota(100);c__Saccharomycetes(100);o__Saccharomycetales(100);f__Metschnikowiaceae(100);g__Metschnikowia(100);g__Metschnikowia_unclassified(100); | Metschnikowia | 13 | Saprotroph | \|Undefined Saprotroph\| | Yeast | NULL | Probable | As nectar-tap_saprotroph (Põlme et al. 2020); Decay sugar-rich substrates (Põlme et al. 2020) | Põlme S, et al. 2020. Fungal Diversity 105:1-16 ((https://doi.org/10.1007/s13225-020-00466-2)) |
| Otu13621 | 1 | k__Fungi(100);p__Ascomycota(100);c__Saccharomycetes(100);o__Saccharomycetales(100);f__Metschnikowiaceae(100);g__Metschnikowia(100);s__Metschnikowia_pulcherrima(100); | Metschnikowia | 13 | Saprotroph | \|Undefined Saprotroph\| | Yeast | NULL | Probable | As nectar-tap_saprotroph (Põlme et al. 2020); Decay sugar-rich substrates (Põlme et al. 2020) | Põlme S, et al. 2020. Fungal Diversity 105:1-16 ((https://doi.org/10.1007/s13225-020-00466-2)) |
| Otu13624 | 1 | k__Fungi(100);p__Ascomycota(100);c__Saccharomycetes(100);o__Saccharomycetales(100);f__Metschnikowiaceae(100);g__Metschnikowia(100);s__Metschnikowia_pulcherrima(100); | Metschnikowia | 13 | Saprotroph | \|Undefined Saprotroph\| | Yeast | NULL | Probable | As nectar-tap_saprotroph (Põlme et al. 2020); Decay sugar-rich substrates (Põlme et al. 2020) | Põlme S, et al. 2020. Fungal Diversity 105:1-16 ((https://doi.org/10.1007/s13225-020-00466-2)) |
| Otu13625 | 1 | k__Fungi(100);p__Ascomycota(100);c__Saccharomycetes(100);o__Saccharomycetales(100);f__Metschnikowiaceae(100);g__Metschnikowia(100);s__Metschnikowia_pulcherrima(100); | Metschnikowia | 13 | Saprotroph | \|Undefined Saprotroph\| | Yeast | NULL | Probable | As nectar-tap_saprotroph (Põlme et al. 2020); Decay sugar-rich substrates (Põlme et al. 2020) | Põlme S, et al. 2020. Fungal Diversity 105:1-16 ((https://doi.org/10.1007/s13225-020-00466-2)) |
| Otu13626 | 1 | k__Fungi(100);p__Ascomycota(100);c__Saccharomycetes(100);o__Saccharomycetales(100);f__Metschnikowiaceae(100);g__Metschnikowia(100);g__Metschnikowia_unclassified(100); | Metschnikowia | 13 | Saprotroph | \|Undefined Saprotroph\| | Yeast | NULL | Probable | As nectar-tap_saprotroph (Põlme et al. 2020); Decay sugar-rich substrates (Põlme et al. 2020) | Põlme S, et al. 2020. Fungal Diversity 105:1-16 ((https://doi.org/10.1007/s13225-020-00466-2)) |
| Otu13628 | 1 | k__Fungi(100);p__Ascomycota(100);c__Saccharomycetes(100);o__Saccharomycetales(100);f__Metschnikowiaceae(100);g__Metschnikowia(100);s__Metschnikowia_pulcherrima(100); | Metschnikowia | 13 | Saprotroph | \|Undefined Saprotroph\| | Yeast | NULL | Probable | As nectar-tap_saprotroph (Põlme et al. 2020); Decay sugar-rich substrates (Põlme et al. 2020) | Põlme S, et al. 2020. Fungal Diversity 105:1-16 ((https://doi.org/10.1007/s13225-020-00466-2)) |
| Otu13630 | 1 | k__Fungi(100);p__Ascomycota(100);c__Saccharomycetes(100);o__Saccharomycetales(100);f__Metschnikowiaceae(100);g__Metschnikowia(100);s__Metschnikowia_pulcherrima(100); | Metschnikowia | 13 | Saprotroph | \|Undefined Saprotroph\| | Yeast | NULL | Probable | As nectar-tap_saprotroph (Põlme et al. 2020); Decay sugar-rich substrates (Põlme et al. 2020) | Põlme S, et al. 2020. Fungal Diversity 105:1-16 ((https://doi.org/10.1007/s13225-020-00466-2)) |
| Otu13631 | 1 | k__Fungi(100);p__Ascomycota(100);c__Saccharomycetes(100);o__Saccharomycetales(100);f__Saccharomycetales_family_Incertae_sedis(100);g__Candida(100);g__Candida_unclassified(100); | Candida | 13 | Pathotroph-Saprotroph-Symbiotroph | Animal Pathogen-Endophyte-Endosymbiont-Epiphyte-\|Undefined Saprotroph\| | Dimorphic Yeast-Yeast | NULL | Probable | As nectar-tap_saprotroph (Põlme et al. 2020); Decay sugar-rich substrates (Põlme et al. 2020); This asexual genus is polyphyletic and contain species that function very differently depend on what they are. Caution should be taken when interpreting guilds for Candida species. | Manolakaki D, et al. 2010. Virulence. 1: 367-375 ((https://doi.org/10.4161/viru.1.5.12796)); Mekbib SB, et al. 2011. Tropical Plant Pathology 36:233-240 ((https://doi.org/10.1590/S1982-56762011000400004)); Põlme S, et al. 2020. Fungal Diversity 105:1-16 ((https://doi.org/10.1007/s13225-020-00466-2)) |
| Otu13637 | 1 | k__Fungi(100);p__Ascomycota(100);c__Saccharomycetes(100);o__Saccharomycetales(100);f__Metschnikowiaceae(100);g__Metschnikowia(100);g__Metschnikowia_unclassified(100); | Metschnikowia | 13 | Saprotroph | \|Undefined Saprotroph\| | Yeast | NULL | Probable | As nectar-tap_saprotroph (Põlme et al. 2020); Decay sugar-rich substrates (Põlme et al. 2020) | Põlme S, et al. 2020. Fungal Diversity 105:1-16 ((https://doi.org/10.1007/s13225-020-00466-2)) |
| Otu13638 | 1 | k__Fungi(100);p__Ascomycota(100);c__Saccharomycetes(100);o__Saccharomycetales(100);f__Metschnikowiaceae(100);g__Metschnikowia(100);g__Metschnikowia_unclassified(100); | Metschnikowia | 13 | Saprotroph | \|Undefined Saprotroph\| | Yeast | NULL | Probable | As nectar-tap_saprotroph (Põlme et al. 2020); Decay sugar-rich substrates (Põlme et al. 2020) | Põlme S, et al. 2020. Fungal Diversity 105:1-16 ((https://doi.org/10.1007/s13225-020-00466-2)) |
| Otu13645 | 1 | k__Fungi(100);p__Ascomycota(100);c__Saccharomycetes(100);o__Saccharomycetales(100);f__Saccharomycetales_family_Incertae_sedis(100);f__Saccharomycetales_family_Incertae_sedis_unclassified(100);f__Saccharomycetales_family_Incertae_sedis_unclassified(100); | Saccharomycetales | 7 | Saprotroph | Undefined Saprotroph | Yeast | NULL | Possible | NULL | Sterkenburg E, et al. 2015. New Phytologist 207:1145-1158 ((https://doi.org/10.1111/nph.13426)) |
| Otu13646 | 1 | k__Fungi(100);p__Ascomycota(100);c__Saccharomycetes(100);o__Saccharomycetales(100);f__Metschnikowiaceae(100);g__Metschnikowia(100);g__Metschnikowia_unclassified(100); | Metschnikowia | 13 | Saprotroph | \|Undefined Saprotroph\| | Yeast | NULL | Probable | As nectar-tap_saprotroph (Põlme et al. 2020); Decay sugar-rich substrates (Põlme et al. 2020) | Põlme S, et al. 2020. Fungal Diversity 105:1-16 ((https://doi.org/10.1007/s13225-020-00466-2)) |
| Otu13648 | 1 | k__Fungi(100);p__Ascomycota(100);c__Saccharomycetes(100);o__Saccharomycetales(100);f__Metschnikowiaceae(100);g__Metschnikowia(100);g__Metschnikowia_unclassified(100); | Metschnikowia | 13 | Saprotroph | \|Undefined Saprotroph\| | Yeast | NULL | Probable | As nectar-tap_saprotroph (Põlme et al. 2020); Decay sugar-rich substrates (Põlme et al. 2020) | Põlme S, et al. 2020. Fungal Diversity 105:1-16 ((https://doi.org/10.1007/s13225-020-00466-2)) |
| Otu13649 | 1 | k__Fungi(100);p__Ascomycota(100);c__Saccharomycetes(100);o__Saccharomycetales(100);f__Metschnikowiaceae(100);g__Metschnikowia(100);g__Metschnikowia_unclassified(100); | Metschnikowia | 13 | Saprotroph | \|Undefined Saprotroph\| | Yeast | NULL | Probable | As nectar-tap_saprotroph (Põlme et al. 2020); Decay sugar-rich substrates (Põlme et al. 2020) | Põlme S, et al. 2020. Fungal Diversity 105:1-16 ((https://doi.org/10.1007/s13225-020-00466-2)) |
| Otu13650 | 1 | k__Fungi(100);p__Ascomycota(100);c__Saccharomycetes(100);o__Saccharomycetales(100);f__Metschnikowiaceae(100);g__Metschnikowia(100);s__Metschnikowia_pulcherrima(100); | Metschnikowia | 13 | Saprotroph | \|Undefined Saprotroph\| | Yeast | NULL | Probable | As nectar-tap_saprotroph (Põlme et al. 2020); Decay sugar-rich substrates (Põlme et al. 2020) | Põlme S, et al. 2020. Fungal Diversity 105:1-16 ((https://doi.org/10.1007/s13225-020-00466-2)) |
| Otu13651 | 1 | k__Fungi(100);p__Ascomycota(100);c__Saccharomycetes(100);o__Saccharomycetales(100);f__Metschnikowiaceae(100);g__Metschnikowia(100);s__Metschnikowia_pulcherrima(100); | Metschnikowia | 13 | Saprotroph | \|Undefined Saprotroph\| | Yeast | NULL | Probable | As nectar-tap_saprotroph (Põlme et al. 2020); Decay sugar-rich substrates (Põlme et al. 2020) | Põlme S, et al. 2020. Fungal Diversity 105:1-16 ((https://doi.org/10.1007/s13225-020-00466-2)) |
| Otu13652 | 1 | k__Fungi(100);p__Ascomycota(100);c__Saccharomycetes(100);o__Saccharomycetales(100);f__Metschnikowiaceae(100);g__Metschnikowia(100);g__Metschnikowia_unclassified(100); | Metschnikowia | 13 | Saprotroph | \|Undefined Saprotroph\| | Yeast | NULL | Probable | As nectar-tap_saprotroph (Põlme et al. 2020); Decay sugar-rich substrates (Põlme et al. 2020) | Põlme S, et al. 2020. Fungal Diversity 105:1-16 ((https://doi.org/10.1007/s13225-020-00466-2)) |
| Otu13660 | 1 | k__Fungi(100);p__Ascomycota(100);c__Saccharomycetes(100);o__Saccharomycetales(100);f__Metschnikowiaceae(100);g__Metschnikowia(100);s__Metschnikowia_pulcherrima(100); | Metschnikowia | 13 | Saprotroph | \|Undefined Saprotroph\| | Yeast | NULL | Probable | As nectar-tap_saprotroph (Põlme et al. 2020); Decay sugar-rich substrates (Põlme et al. 2020) | Põlme S, et al. 2020. Fungal Diversity 105:1-16 ((https://doi.org/10.1007/s13225-020-00466-2)) |
| Otu13661 | 1 | k__Fungi(100);p__Ascomycota(100);c__Saccharomycetes(100);o__Saccharomycetales(100);f__Metschnikowiaceae(100);g__Metschnikowia(100);s__Metschnikowia_pulcherrima(100); | Metschnikowia | 13 | Saprotroph | \|Undefined Saprotroph\| | Yeast | NULL | Probable | As nectar-tap_saprotroph (Põlme et al. 2020); Decay sugar-rich substrates (Põlme et al. 2020) | Põlme S, et al. 2020. Fungal Diversity 105:1-16 ((https://doi.org/10.1007/s13225-020-00466-2)) |
| Otu13665 | 1 | k__Fungi(100);p__Ascomycota(100);c__Saccharomycetes(100);o__Saccharomycetales(100);f__Metschnikowiaceae(100);g__Metschnikowia(100);s__Metschnikowia_pulcherrima(100); | Metschnikowia | 13 | Saprotroph | \|Undefined Saprotroph\| | Yeast | NULL | Probable | As nectar-tap_saprotroph (Põlme et al. 2020); Decay sugar-rich substrates (Põlme et al. 2020) | Põlme S, et al. 2020. Fungal Diversity 105:1-16 ((https://doi.org/10.1007/s13225-020-00466-2)) |
| Otu13666 | 1 | k__Fungi(100);p__Ascomycota(100);c__Saccharomycetes(100);o__Saccharomycetales(100);f__Metschnikowiaceae(100);g__Metschnikowia(100);s__Metschnikowia_pulcherrima(100); | Metschnikowia | 13 | Saprotroph | \|Undefined Saprotroph\| | Yeast | NULL | Probable | As nectar-tap_saprotroph (Põlme et al. 2020); Decay sugar-rich substrates (Põlme et al. 2020) | Põlme S, et al. 2020. Fungal Diversity 105:1-16 ((https://doi.org/10.1007/s13225-020-00466-2)) |
| Otu13667 | 1 | k__Fungi(100);p__Ascomycota(100);c__Saccharomycetes(100);o__Saccharomycetales(100);f__Metschnikowiaceae(100);g__Metschnikowia(100);s__Metschnikowia_pulcherrima(100); | Metschnikowia | 13 | Saprotroph | \|Undefined Saprotroph\| | Yeast | NULL | Probable | As nectar-tap_saprotroph (Põlme et al. 2020); Decay sugar-rich substrates (Põlme et al. 2020) | Põlme S, et al. 2020. Fungal Diversity 105:1-16 ((https://doi.org/10.1007/s13225-020-00466-2)) |
| Otu13668 | 1 | k__Fungi(100);p__Ascomycota(100);c__Saccharomycetes(100);o__Saccharomycetales(100);f__Metschnikowiaceae(100);g__Metschnikowia(100);s__Metschnikowia_pulcherrima(100); | Metschnikowia | 13 | Saprotroph | \|Undefined Saprotroph\| | Yeast | NULL | Probable | As nectar-tap_saprotroph (Põlme et al. 2020); Decay sugar-rich substrates (Põlme et al. 2020) | Põlme S, et al. 2020. Fungal Diversity 105:1-16 ((https://doi.org/10.1007/s13225-020-00466-2)) |
| Otu13670 | 1 | k__Fungi(100);p__Ascomycota(100);c__Saccharomycetes(100);o__Saccharomycetales(100);f__Metschnikowiaceae(100);g__Metschnikowia(100);s__Metschnikowia_pulcherrima(100); | Metschnikowia | 13 | Saprotroph | \|Undefined Saprotroph\| | Yeast | NULL | Probable | As nectar-tap_saprotroph (Põlme et al. 2020); Decay sugar-rich substrates (Põlme et al. 2020) | Põlme S, et al. 2020. Fungal Diversity 105:1-16 ((https://doi.org/10.1007/s13225-020-00466-2)) |
| Otu13671 | 1 | k__Fungi(100);p__Ascomycota(100);c__Saccharomycetes(100);o__Saccharomycetales(100);f__Metschnikowiaceae(100);g__Metschnikowia(100);s__Metschnikowia_pulcherrima(100); | Metschnikowia | 13 | Saprotroph | \|Undefined Saprotroph\| | Yeast | NULL | Probable | As nectar-tap_saprotroph (Põlme et al. 2020); Decay sugar-rich substrates (Põlme et al. 2020) | Põlme S, et al. 2020. Fungal Diversity 105:1-16 ((https://doi.org/10.1007/s13225-020-00466-2)) |
| Otu13672 | 1 | k__Fungi(100);p__Ascomycota(100);c__Saccharomycetes(100);o__Saccharomycetales(100);f__Metschnikowiaceae(100);g__Metschnikowia(100);s__Metschnikowia_pulcherrima(100); | Metschnikowia | 13 | Saprotroph | \|Undefined Saprotroph\| | Yeast | NULL | Probable | As nectar-tap_saprotroph (Põlme et al. 2020); Decay sugar-rich substrates (Põlme et al. 2020) | Põlme S, et al. 2020. Fungal Diversity 105:1-16 ((https://doi.org/10.1007/s13225-020-00466-2)) |
| Otu13673 | 1 | k__Fungi(100);p__Ascomycota(100);c__Saccharomycetes(100);o__Saccharomycetales(100);f__Metschnikowiaceae(100);g__Metschnikowia(100);g__Metschnikowia_unclassified(100); | Metschnikowia | 13 | Saprotroph | \|Undefined Saprotroph\| | Yeast | NULL | Probable | As nectar-tap_saprotroph (Põlme et al. 2020); Decay sugar-rich substrates (Põlme et al. 2020) | Põlme S, et al. 2020. Fungal Diversity 105:1-16 ((https://doi.org/10.1007/s13225-020-00466-2)) |
| Otu13674 | 1 | k__Fungi(100);p__Ascomycota(100);c__Saccharomycetes(100);o__Saccharomycetales(100);f__Metschnikowiaceae(100);g__Metschnikowia(100);s__Metschnikowia_pulcherrima(100); | Metschnikowia | 13 | Saprotroph | \|Undefined Saprotroph\| | Yeast | NULL | Probable | As nectar-tap_saprotroph (Põlme et al. 2020); Decay sugar-rich substrates (Põlme et al. 2020) | Põlme S, et al. 2020. Fungal Diversity 105:1-16 ((https://doi.org/10.1007/s13225-020-00466-2)) |
| Otu13676 | 1 | k__Fungi(100);p__Ascomycota(100);c__Saccharomycetes(100);o__Saccharomycetales(100);f__Metschnikowiaceae(100);g__Metschnikowia(100);s__Metschnikowia_pulcherrima(100); | Metschnikowia | 13 | Saprotroph | \|Undefined Saprotroph\| | Yeast | NULL | Probable | As nectar-tap_saprotroph (Põlme et al. 2020); Decay sugar-rich substrates (Põlme et al. 2020) | Põlme S, et al. 2020. Fungal Diversity 105:1-16 ((https://doi.org/10.1007/s13225-020-00466-2)) |
| Otu13677 | 1 | k__Fungi(100);p__Ascomycota(100);c__Saccharomycetes(100);o__Saccharomycetales(100);o__Saccharomycetales_unclassified(100);o__Saccharomycetales_unclassified(100);o__Saccharomycetales_unclassified(100); | Saccharomycetales | 7 | Saprotroph | Undefined Saprotroph | Yeast | NULL | Possible | NULL | Sterkenburg E, et al. 2015. New Phytologist 207:1145-1158 ((https://doi.org/10.1111/nph.13426)) |
| Otu13681 | 1 | k__Fungi(100);p__Ascomycota(100);c__Saccharomycetes(100);o__Saccharomycetales(100);f__Metschnikowiaceae(100);g__Metschnikowia(100);s__Metschnikowia_pulcherrima(100); | Metschnikowia | 13 | Saprotroph | \|Undefined Saprotroph\| | Yeast | NULL | Probable | As nectar-tap_saprotroph (Põlme et al. 2020); Decay sugar-rich substrates (Põlme et al. 2020) | Põlme S, et al. 2020. Fungal Diversity 105:1-16 ((https://doi.org/10.1007/s13225-020-00466-2)) |
| Otu13682 | 1 | k__Fungi(100);p__Ascomycota(100);c__Saccharomycetes(100);o__Saccharomycetales(100);f__Saccharomycetales_family_Incertae_sedis(100);g__Candida(100);s__Candida_cretensis(100); | Candida | 13 | Pathotroph-Saprotroph-Symbiotroph | Animal Pathogen-Endophyte-Endosymbiont-Epiphyte-\|Undefined Saprotroph\| | Dimorphic Yeast-Yeast | NULL | Probable | As nectar-tap_saprotroph (Põlme et al. 2020); Decay sugar-rich substrates (Põlme et al. 2020); This asexual genus is polyphyletic and contain species that function very differently depend on what they are. Caution should be taken when interpreting guilds for Candida species. | Manolakaki D, et al. 2010. Virulence. 1: 367-375 ((https://doi.org/10.4161/viru.1.5.12796)); Mekbib SB, et al. 2011. Tropical Plant Pathology 36:233-240 ((https://doi.org/10.1590/S1982-56762011000400004)); Põlme S, et al. 2020. Fungal Diversity 105:1-16 ((https://doi.org/10.1007/s13225-020-00466-2)) |
| Otu13683 | 1 | k__Fungi(100);p__Ascomycota(100);c__Saccharomycetes(100);o__Saccharomycetales(100);f__Metschnikowiaceae(100);g__Metschnikowia(100);s__Metschnikowia_pulcherrima(100); | Metschnikowia | 13 | Saprotroph | \|Undefined Saprotroph\| | Yeast | NULL | Probable | As nectar-tap_saprotroph (Põlme et al. 2020); Decay sugar-rich substrates (Põlme et al. 2020) | Põlme S, et al. 2020. Fungal Diversity 105:1-16 ((https://doi.org/10.1007/s13225-020-00466-2)) |
| Otu13684 | 1 | k__Fungi(100);p__Ascomycota(100);c__Saccharomycetes(100);o__Saccharomycetales(100);f__Metschnikowiaceae(100);g__Metschnikowia(100);g__Metschnikowia_unclassified(100); | Metschnikowia | 13 | Saprotroph | \|Undefined Saprotroph\| | Yeast | NULL | Probable | As nectar-tap_saprotroph (Põlme et al. 2020); Decay sugar-rich substrates (Põlme et al. 2020) | Põlme S, et al. 2020. Fungal Diversity 105:1-16 ((https://doi.org/10.1007/s13225-020-00466-2)) |
| Otu13685 | 1 | k__Fungi(100);p__Ascomycota(100);c__Saccharomycetes(100);o__Saccharomycetales(100);f__Saccharomycetales_family_Incertae_sedis(100);g__Candida(100);g__Candida_unclassified(100); | Candida | 13 | Pathotroph-Saprotroph-Symbiotroph | Animal Pathogen-Endophyte-Endosymbiont-Epiphyte-\|Undefined Saprotroph\| | Dimorphic Yeast-Yeast | NULL | Probable | As nectar-tap_saprotroph (Põlme et al. 2020); Decay sugar-rich substrates (Põlme et al. 2020); This asexual genus is polyphyletic and contain species that function very differently depend on what they are. Caution should be taken when interpreting guilds for Candida species. | Manolakaki D, et al. 2010. Virulence. 1: 367-375 ((https://doi.org/10.4161/viru.1.5.12796)); Mekbib SB, et al. 2011. Tropical Plant Pathology 36:233-240 ((https://doi.org/10.1590/S1982-56762011000400004)); Põlme S, et al. 2020. Fungal Diversity 105:1-16 ((https://doi.org/10.1007/s13225-020-00466-2)) |
| Otu13688 | 1 | k__Fungi(100);p__Ascomycota(100);c__Saccharomycetes(100);o__Saccharomycetales(100);f__Saccharomycetales_family_Incertae_sedis(100);g__Candida(100);g__Candida_unclassified(100); | Candida | 13 | Pathotroph-Saprotroph-Symbiotroph | Animal Pathogen-Endophyte-Endosymbiont-Epiphyte-\|Undefined Saprotroph\| | Dimorphic Yeast-Yeast | NULL | Probable | As nectar-tap_saprotroph (Põlme et al. 2020); Decay sugar-rich substrates (Põlme et al. 2020); This asexual genus is polyphyletic and contain species that function very differently depend on what they are. Caution should be taken when interpreting guilds for Candida species. | Manolakaki D, et al. 2010. Virulence. 1: 367-375 ((https://doi.org/10.4161/viru.1.5.12796)); Mekbib SB, et al. 2011. Tropical Plant Pathology 36:233-240 ((https://doi.org/10.1590/S1982-56762011000400004)); Põlme S, et al. 2020. Fungal Diversity 105:1-16 ((https://doi.org/10.1007/s13225-020-00466-2)) |
| Otu13689 | 1 | k__Fungi(100);p__Ascomycota(100);c__Saccharomycetes(100);o__Saccharomycetales(100);f__Metschnikowiaceae(100);g__Metschnikowia(100);s__Metschnikowia_pulcherrima(100); | Metschnikowia | 13 | Saprotroph | \|Undefined Saprotroph\| | Yeast | NULL | Probable | As nectar-tap_saprotroph (Põlme et al. 2020); Decay sugar-rich substrates (Põlme et al. 2020) | Põlme S, et al. 2020. Fungal Diversity 105:1-16 ((https://doi.org/10.1007/s13225-020-00466-2)) |
| Otu13691 | 1 | k__Fungi(100);p__Ascomycota(100);c__Saccharomycetes(100);o__Saccharomycetales(100);f__Metschnikowiaceae(100);g__Metschnikowia(100);g__Metschnikowia_unclassified(100); | Metschnikowia | 13 | Saprotroph | \|Undefined Saprotroph\| | Yeast | NULL | Probable | As nectar-tap_saprotroph (Põlme et al. 2020); Decay sugar-rich substrates (Põlme et al. 2020) | Põlme S, et al. 2020. Fungal Diversity 105:1-16 ((https://doi.org/10.1007/s13225-020-00466-2)) |
| Otu13693 | 1 | k__Fungi(100);p__Ascomycota(100);c__Saccharomycetes(100);o__Saccharomycetales(100);f__Metschnikowiaceae(100);g__Metschnikowia(100);s__Metschnikowia_pulcherrima(100); | Metschnikowia | 13 | Saprotroph | \|Undefined Saprotroph\| | Yeast | NULL | Probable | As nectar-tap_saprotroph (Põlme et al. 2020); Decay sugar-rich substrates (Põlme et al. 2020) | Põlme S, et al. 2020. Fungal Diversity 105:1-16 ((https://doi.org/10.1007/s13225-020-00466-2)) |
| Otu13695 | 1 | k__Fungi(100);p__Ascomycota(100);c__Saccharomycetes(100);o__Saccharomycetales(100);f__Metschnikowiaceae(100);g__Metschnikowia(100);s__Metschnikowia_pulcherrima(100); | Metschnikowia | 13 | Saprotroph | \|Undefined Saprotroph\| | Yeast | NULL | Probable | As nectar-tap_saprotroph (Põlme et al. 2020); Decay sugar-rich substrates (Põlme et al. 2020) | Põlme S, et al. 2020. Fungal Diversity 105:1-16 ((https://doi.org/10.1007/s13225-020-00466-2)) |
| Otu13696 | 1 | k__Fungi(100);p__Ascomycota(100);c__Saccharomycetes(100);o__Saccharomycetales(100);f__Metschnikowiaceae(100);g__Metschnikowia(100);s__Metschnikowia_pulcherrima(100); | Metschnikowia | 13 | Saprotroph | \|Undefined Saprotroph\| | Yeast | NULL | Probable | As nectar-tap_saprotroph (Põlme et al. 2020); Decay sugar-rich substrates (Põlme et al. 2020) | Põlme S, et al. 2020. Fungal Diversity 105:1-16 ((https://doi.org/10.1007/s13225-020-00466-2)) |
| Otu13697 | 1 | k__Fungi(100);p__Ascomycota(100);c__Saccharomycetes(100);o__Saccharomycetales(100);f__Metschnikowiaceae(100);g__Metschnikowia(100);s__Metschnikowia_pulcherrima(100); | Metschnikowia | 13 | Saprotroph | \|Undefined Saprotroph\| | Yeast | NULL | Probable | As nectar-tap_saprotroph (Põlme et al. 2020); Decay sugar-rich substrates (Põlme et al. 2020) | Põlme S, et al. 2020. Fungal Diversity 105:1-16 ((https://doi.org/10.1007/s13225-020-00466-2)) |
| Otu13698 | 1 | k__Fungi(100);p__Ascomycota(100);c__Saccharomycetes(100);o__Saccharomycetales(100);f__Metschnikowiaceae(100);g__Metschnikowia(100);s__Metschnikowia_pulcherrima(100); | Metschnikowia | 13 | Saprotroph | \|Undefined Saprotroph\| | Yeast | NULL | Probable | As nectar-tap_saprotroph (Põlme et al. 2020); Decay sugar-rich substrates (Põlme et al. 2020) | Põlme S, et al. 2020. Fungal Diversity 105:1-16 ((https://doi.org/10.1007/s13225-020-00466-2)) |
| Otu13699 | 1 | k__Fungi(100);p__Ascomycota(100);c__Saccharomycetes(100);o__Saccharomycetales(100);f__Saccharomycetales_family_Incertae_sedis(100);g__Candida(100);s__Candida_cretensis(100); | Candida | 13 | Pathotroph-Saprotroph-Symbiotroph | Animal Pathogen-Endophyte-Endosymbiont-Epiphyte-\|Undefined Saprotroph\| | Dimorphic Yeast-Yeast | NULL | Probable | As nectar-tap_saprotroph (Põlme et al. 2020); Decay sugar-rich substrates (Põlme et al. 2020); This asexual genus is polyphyletic and contain species that function very differently depend on what they are. Caution should be taken when interpreting guilds for Candida species. | Manolakaki D, et al. 2010. Virulence. 1: 367-375 ((https://doi.org/10.4161/viru.1.5.12796)); Mekbib SB, et al. 2011. Tropical Plant Pathology 36:233-240 ((https://doi.org/10.1590/S1982-56762011000400004)); Põlme S, et al. 2020. Fungal Diversity 105:1-16 ((https://doi.org/10.1007/s13225-020-00466-2)) |
| Otu13701 | 1 | k__Fungi(100);p__Ascomycota(100);c__Saccharomycetes(100);o__Saccharomycetales(100);f__Metschnikowiaceae(100);g__Metschnikowia(100);s__Metschnikowia_pulcherrima(100); | Metschnikowia | 13 | Saprotroph | \|Undefined Saprotroph\| | Yeast | NULL | Probable | As nectar-tap_saprotroph (Põlme et al. 2020); Decay sugar-rich substrates (Põlme et al. 2020) | Põlme S, et al. 2020. Fungal Diversity 105:1-16 ((https://doi.org/10.1007/s13225-020-00466-2)) |
| Otu13702 | 1 | k__Fungi(100);p__Ascomycota(100);c__Saccharomycetes(100);o__Saccharomycetales(100);f__Metschnikowiaceae(100);g__Metschnikowia(100);s__Metschnikowia_pulcherrima(100); | Metschnikowia | 13 | Saprotroph | \|Undefined Saprotroph\| | Yeast | NULL | Probable | As nectar-tap_saprotroph (Põlme et al. 2020); Decay sugar-rich substrates (Põlme et al. 2020) | Põlme S, et al. 2020. Fungal Diversity 105:1-16 ((https://doi.org/10.1007/s13225-020-00466-2)) |
| Otu13704 | 1 | k__Fungi(100);p__Ascomycota(100);c__Saccharomycetes(100);o__Saccharomycetales(100);f__Metschnikowiaceae(100);g__Metschnikowia(100);s__Metschnikowia_pulcherrima(100); | Metschnikowia | 13 | Saprotroph | \|Undefined Saprotroph\| | Yeast | NULL | Probable | As nectar-tap_saprotroph (Põlme et al. 2020); Decay sugar-rich substrates (Põlme et al. 2020) | Põlme S, et al. 2020. Fungal Diversity 105:1-16 ((https://doi.org/10.1007/s13225-020-00466-2)) |
| Otu13705 | 1 | k__Fungi(100);p__Ascomycota(100);c__Saccharomycetes(100);o__Saccharomycetales(100);f__Metschnikowiaceae(100);g__Metschnikowia(100);s__Metschnikowia_pulcherrima(100); | Metschnikowia | 13 | Saprotroph | \|Undefined Saprotroph\| | Yeast | NULL | Probable | As nectar-tap_saprotroph (Põlme et al. 2020); Decay sugar-rich substrates (Põlme et al. 2020) | Põlme S, et al. 2020. Fungal Diversity 105:1-16 ((https://doi.org/10.1007/s13225-020-00466-2)) |
| Otu13706 | 1 | k__Fungi(100);p__Ascomycota(100);c__Saccharomycetes(100);o__Saccharomycetales(100);f__Metschnikowiaceae(100);g__Metschnikowia(100);g__Metschnikowia_unclassified(100); | Metschnikowia | 13 | Saprotroph | \|Undefined Saprotroph\| | Yeast | NULL | Probable | As nectar-tap_saprotroph (Põlme et al. 2020); Decay sugar-rich substrates (Põlme et al. 2020) | Põlme S, et al. 2020. Fungal Diversity 105:1-16 ((https://doi.org/10.1007/s13225-020-00466-2)) |
| Otu13707 | 1 | k__Fungi(100);p__Ascomycota(100);c__Saccharomycetes(100);o__Saccharomycetales(100);f__Saccharomycetales_family_Incertae_sedis(100);g__Candida(100);g__Candida_unclassified(100); | Candida | 13 | Pathotroph-Saprotroph-Symbiotroph | Animal Pathogen-Endophyte-Endosymbiont-Epiphyte-\|Undefined Saprotroph\| | Dimorphic Yeast-Yeast | NULL | Probable | As nectar-tap_saprotroph (Põlme et al. 2020); Decay sugar-rich substrates (Põlme et al. 2020); This asexual genus is polyphyletic and contain species that function very differently depend on what they are. Caution should be taken when interpreting guilds for Candida species. | Manolakaki D, et al. 2010. Virulence. 1: 367-375 ((https://doi.org/10.4161/viru.1.5.12796)); Mekbib SB, et al. 2011. Tropical Plant Pathology 36:233-240 ((https://doi.org/10.1590/S1982-56762011000400004)); Põlme S, et al. 2020. Fungal Diversity 105:1-16 ((https://doi.org/10.1007/s13225-020-00466-2)) |
| Otu13708 | 1 | k__Fungi(100);p__Ascomycota(100);c__Saccharomycetes(100);o__Saccharomycetales(100);f__Metschnikowiaceae(100);g__Metschnikowia(100);s__Metschnikowia_pulcherrima(100); | Metschnikowia | 13 | Saprotroph | \|Undefined Saprotroph\| | Yeast | NULL | Probable | As nectar-tap_saprotroph (Põlme et al. 2020); Decay sugar-rich substrates (Põlme et al. 2020) | Põlme S, et al. 2020. Fungal Diversity 105:1-16 ((https://doi.org/10.1007/s13225-020-00466-2)) |
| Otu13710 | 1 | k__Fungi(100);p__Ascomycota(100);c__Saccharomycetes(100);o__Saccharomycetales(100);f__Metschnikowiaceae(100);g__Metschnikowia(100);g__Metschnikowia_unclassified(100); | Metschnikowia | 13 | Saprotroph | \|Undefined Saprotroph\| | Yeast | NULL | Probable | As nectar-tap_saprotroph (Põlme et al. 2020); Decay sugar-rich substrates (Põlme et al. 2020) | Põlme S, et al. 2020. Fungal Diversity 105:1-16 ((https://doi.org/10.1007/s13225-020-00466-2)) |
| Otu13711 | 1 | k__Fungi(100);p__Ascomycota(100);c__Saccharomycetes(100);o__Saccharomycetales(100);f__Metschnikowiaceae(100);g__Metschnikowia(100);g__Metschnikowia_unclassified(100); | Metschnikowia | 13 | Saprotroph | \|Undefined Saprotroph\| | Yeast | NULL | Probable | As nectar-tap_saprotroph (Põlme et al. 2020); Decay sugar-rich substrates (Põlme et al. 2020) | Põlme S, et al. 2020. Fungal Diversity 105:1-16 ((https://doi.org/10.1007/s13225-020-00466-2)) |
| Otu13712 | 1 | k__Fungi(100);p__Ascomycota(100);c__Saccharomycetes(100);o__Saccharomycetales(100);f__Metschnikowiaceae(100);g__Metschnikowia(100);s__Metschnikowia_pulcherrima(100); | Metschnikowia | 13 | Saprotroph | \|Undefined Saprotroph\| | Yeast | NULL | Probable | As nectar-tap_saprotroph (Põlme et al. 2020); Decay sugar-rich substrates (Põlme et al. 2020) | Põlme S, et al. 2020. Fungal Diversity 105:1-16 ((https://doi.org/10.1007/s13225-020-00466-2)) |
| Otu13713 | 1 | k__Fungi(100);p__Ascomycota(100);c__Saccharomycetes(100);o__Saccharomycetales(100);f__Metschnikowiaceae(100);g__Metschnikowia(100);g__Metschnikowia_unclassified(100); | Metschnikowia | 13 | Saprotroph | \|Undefined Saprotroph\| | Yeast | NULL | Probable | As nectar-tap_saprotroph (Põlme et al. 2020); Decay sugar-rich substrates (Põlme et al. 2020) | Põlme S, et al. 2020. Fungal Diversity 105:1-16 ((https://doi.org/10.1007/s13225-020-00466-2)) |
| Otu13714 | 1 | k__Fungi(100);p__Ascomycota(100);c__Saccharomycetes(100);o__Saccharomycetales(100);f__Metschnikowiaceae(100);g__Metschnikowia(100);s__Metschnikowia_pulcherrima(100); | Metschnikowia | 13 | Saprotroph | \|Undefined Saprotroph\| | Yeast | NULL | Probable | As nectar-tap_saprotroph (Põlme et al. 2020); Decay sugar-rich substrates (Põlme et al. 2020) | Põlme S, et al. 2020. Fungal Diversity 105:1-16 ((https://doi.org/10.1007/s13225-020-00466-2)) |
| Otu13715 | 1 | k__Fungi(100);p__Ascomycota(100);c__Saccharomycetes(100);o__Saccharomycetales(100);f__Metschnikowiaceae(100);g__Metschnikowia(100);s__Metschnikowia_pulcherrima(100); | Metschnikowia | 13 | Saprotroph | \|Undefined Saprotroph\| | Yeast | NULL | Probable | As nectar-tap_saprotroph (Põlme et al. 2020); Decay sugar-rich substrates (Põlme et al. 2020) | Põlme S, et al. 2020. Fungal Diversity 105:1-16 ((https://doi.org/10.1007/s13225-020-00466-2)) |
| Otu13716 | 1 | k__Fungi(100);p__Ascomycota(100);c__Saccharomycetes(100);o__Saccharomycetales(100);f__Metschnikowiaceae(100);g__Metschnikowia(100);s__Metschnikowia_pulcherrima(100); | Metschnikowia | 13 | Saprotroph | \|Undefined Saprotroph\| | Yeast | NULL | Probable | As nectar-tap_saprotroph (Põlme et al. 2020); Decay sugar-rich substrates (Põlme et al. 2020) | Põlme S, et al. 2020. Fungal Diversity 105:1-16 ((https://doi.org/10.1007/s13225-020-00466-2)) |
| Otu13717 | 1 | k__Fungi(100);p__Ascomycota(100);c__Saccharomycetes(100);o__Saccharomycetales(100);f__Metschnikowiaceae(100);g__Metschnikowia(100);s__Metschnikowia_pulcherrima(100); | Metschnikowia | 13 | Saprotroph | \|Undefined Saprotroph\| | Yeast | NULL | Probable | As nectar-tap_saprotroph (Põlme et al. 2020); Decay sugar-rich substrates (Põlme et al. 2020) | Põlme S, et al. 2020. Fungal Diversity 105:1-16 ((https://doi.org/10.1007/s13225-020-00466-2)) |
| Otu13719 | 1 | k__Fungi(100);p__Ascomycota(100);c__Saccharomycetes(100);o__Saccharomycetales(100);f__Saccharomycetales_family_Incertae_sedis(100);g__Candida(100);g__Candida_unclassified(100); | Candida | 13 | Pathotroph-Saprotroph-Symbiotroph | Animal Pathogen-Endophyte-Endosymbiont-Epiphyte-\|Undefined Saprotroph\| | Dimorphic Yeast-Yeast | NULL | Probable | As nectar-tap_saprotroph (Põlme et al. 2020); Decay sugar-rich substrates (Põlme et al. 2020); This asexual genus is polyphyletic and contain species that function very differently depend on what they are. Caution should be taken when interpreting guilds for Candida species. | Manolakaki D, et al. 2010. Virulence. 1: 367-375 ((https://doi.org/10.4161/viru.1.5.12796)); Mekbib SB, et al. 2011. Tropical Plant Pathology 36:233-240 ((https://doi.org/10.1590/S1982-56762011000400004)); Põlme S, et al. 2020. Fungal Diversity 105:1-16 ((https://doi.org/10.1007/s13225-020-00466-2)) |
| Otu13721 | 1 | k__Fungi(100);p__Ascomycota(100);c__Saccharomycetes(100);o__Saccharomycetales(100);f__Metschnikowiaceae(100);g__Metschnikowia(100);g__Metschnikowia_unclassified(100); | Metschnikowia | 13 | Saprotroph | \|Undefined Saprotroph\| | Yeast | NULL | Probable | As nectar-tap_saprotroph (Põlme et al. 2020); Decay sugar-rich substrates (Põlme et al. 2020) | Põlme S, et al. 2020. Fungal Diversity 105:1-16 ((https://doi.org/10.1007/s13225-020-00466-2)) |
| Otu13722 | 1 | k__Fungi(100);p__Ascomycota(100);c__Saccharomycetes(100);o__Saccharomycetales(100);f__Metschnikowiaceae(100);g__Metschnikowia(100);s__Metschnikowia_pulcherrima(100); | Metschnikowia | 13 | Saprotroph | \|Undefined Saprotroph\| | Yeast | NULL | Probable | As nectar-tap_saprotroph (Põlme et al. 2020); Decay sugar-rich substrates (Põlme et al. 2020) | Põlme S, et al. 2020. Fungal Diversity 105:1-16 ((https://doi.org/10.1007/s13225-020-00466-2)) |
| Otu13724 | 1 | k__Fungi(100);p__Ascomycota(100);c__Saccharomycetes(100);o__Saccharomycetales(100);f__Metschnikowiaceae(100);g__Metschnikowia(100);g__Metschnikowia_unclassified(100); | Metschnikowia | 13 | Saprotroph | \|Undefined Saprotroph\| | Yeast | NULL | Probable | As nectar-tap_saprotroph (Põlme et al. 2020); Decay sugar-rich substrates (Põlme et al. 2020) | Põlme S, et al. 2020. Fungal Diversity 105:1-16 ((https://doi.org/10.1007/s13225-020-00466-2)) |
| Otu13726 | 1 | k__Fungi(100);p__Ascomycota(100);c__Saccharomycetes(100);o__Saccharomycetales(100);f__Metschnikowiaceae(100);g__Metschnikowia(100);s__Metschnikowia_pulcherrima(100); | Metschnikowia | 13 | Saprotroph | \|Undefined Saprotroph\| | Yeast | NULL | Probable | As nectar-tap_saprotroph (Põlme et al. 2020); Decay sugar-rich substrates (Põlme et al. 2020) | Põlme S, et al. 2020. Fungal Diversity 105:1-16 ((https://doi.org/10.1007/s13225-020-00466-2)) |
| Otu13727 | 1 | k__Fungi(100);p__Ascomycota(100);c__Saccharomycetes(100);o__Saccharomycetales(100);f__Metschnikowiaceae(100);g__Metschnikowia(100);s__Metschnikowia_pulcherrima(100); | Metschnikowia | 13 | Saprotroph | \|Undefined Saprotroph\| | Yeast | NULL | Probable | As nectar-tap_saprotroph (Põlme et al. 2020); Decay sugar-rich substrates (Põlme et al. 2020) | Põlme S, et al. 2020. Fungal Diversity 105:1-16 ((https://doi.org/10.1007/s13225-020-00466-2)) |
| Otu13728 | 1 | k__Fungi(100);p__Ascomycota(100);c__Saccharomycetes(100);o__Saccharomycetales(100);f__Saccharomycetales_family_Incertae_sedis(100);g__Candida(100);g__Candida_unclassified(100); | Candida | 13 | Pathotroph-Saprotroph-Symbiotroph | Animal Pathogen-Endophyte-Endosymbiont-Epiphyte-\|Undefined Saprotroph\| | Dimorphic Yeast-Yeast | NULL | Probable | As nectar-tap_saprotroph (Põlme et al. 2020); Decay sugar-rich substrates (Põlme et al. 2020); This asexual genus is polyphyletic and contain species that function very differently depend on what they are. Caution should be taken when interpreting guilds for Candida species. | Manolakaki D, et al. 2010. Virulence. 1: 367-375 ((https://doi.org/10.4161/viru.1.5.12796)); Mekbib SB, et al. 2011. Tropical Plant Pathology 36:233-240 ((https://doi.org/10.1590/S1982-56762011000400004)); Põlme S, et al. 2020. Fungal Diversity 105:1-16 ((https://doi.org/10.1007/s13225-020-00466-2)) |
| Otu13731 | 1 | k__Fungi(100);p__Ascomycota(100);c__Saccharomycetes(100);o__Saccharomycetales(100);f__Metschnikowiaceae(100);g__Metschnikowia(100);s__Metschnikowia_pulcherrima(100); | Metschnikowia | 13 | Saprotroph | \|Undefined Saprotroph\| | Yeast | NULL | Probable | As nectar-tap_saprotroph (Põlme et al. 2020); Decay sugar-rich substrates (Põlme et al. 2020) | Põlme S, et al. 2020. Fungal Diversity 105:1-16 ((https://doi.org/10.1007/s13225-020-00466-2)) |
| Otu13734 | 1 | k__Fungi(100);p__Ascomycota(100);c__Saccharomycetes(100);o__Saccharomycetales(100);f__Metschnikowiaceae(100);g__Metschnikowia(100);g__Metschnikowia_unclassified(100); | Metschnikowia | 13 | Saprotroph | \|Undefined Saprotroph\| | Yeast | NULL | Probable | As nectar-tap_saprotroph (Põlme et al. 2020); Decay sugar-rich substrates (Põlme et al. 2020) | Põlme S, et al. 2020. Fungal Diversity 105:1-16 ((https://doi.org/10.1007/s13225-020-00466-2)) |
| Otu13735 | 1 | k__Fungi(100);p__Ascomycota(100);c__Saccharomycetes(100);o__Saccharomycetales(100);f__Metschnikowiaceae(100);g__Metschnikowia(100);s__Metschnikowia_pulcherrima(100); | Metschnikowia | 13 | Saprotroph | \|Undefined Saprotroph\| | Yeast | NULL | Probable | As nectar-tap_saprotroph (Põlme et al. 2020); Decay sugar-rich substrates (Põlme et al. 2020) | Põlme S, et al. 2020. Fungal Diversity 105:1-16 ((https://doi.org/10.1007/s13225-020-00466-2)) |
| Otu13737 | 1 | k__Fungi(100);p__Ascomycota(100);c__Saccharomycetes(100);o__Saccharomycetales(100);f__Metschnikowiaceae(100);g__Metschnikowia(100);g__Metschnikowia_unclassified(100); | Metschnikowia | 13 | Saprotroph | \|Undefined Saprotroph\| | Yeast | NULL | Probable | As nectar-tap_saprotroph (Põlme et al. 2020); Decay sugar-rich substrates (Põlme et al. 2020) | Põlme S, et al. 2020. Fungal Diversity 105:1-16 ((https://doi.org/10.1007/s13225-020-00466-2)) |
| Otu13738 | 1 | k__Fungi(100);p__Ascomycota(100);c__Saccharomycetes(100);o__Saccharomycetales(100);f__Metschnikowiaceae(100);g__Metschnikowia(100);s__Metschnikowia_pulcherrima(100); | Metschnikowia | 13 | Saprotroph | \|Undefined Saprotroph\| | Yeast | NULL | Probable | As nectar-tap_saprotroph (Põlme et al. 2020); Decay sugar-rich substrates (Põlme et al. 2020) | Põlme S, et al. 2020. Fungal Diversity 105:1-16 ((https://doi.org/10.1007/s13225-020-00466-2)) |
| Otu13739 | 1 | k__Fungi(100);p__Ascomycota(100);c__Saccharomycetes(100);o__Saccharomycetales(100);f__Metschnikowiaceae(100);g__Metschnikowia(100);s__Metschnikowia_pulcherrima(100); | Metschnikowia | 13 | Saprotroph | \|Undefined Saprotroph\| | Yeast | NULL | Probable | As nectar-tap_saprotroph (Põlme et al. 2020); Decay sugar-rich substrates (Põlme et al. 2020) | Põlme S, et al. 2020. Fungal Diversity 105:1-16 ((https://doi.org/10.1007/s13225-020-00466-2)) |
| Otu13740 | 1 | k__Fungi(100);p__Ascomycota(100);c__Saccharomycetes(100);o__Saccharomycetales(100);f__Metschnikowiaceae(100);g__Metschnikowia(100);s__Metschnikowia_pulcherrima(100); | Metschnikowia | 13 | Saprotroph | \|Undefined Saprotroph\| | Yeast | NULL | Probable | As nectar-tap_saprotroph (Põlme et al. 2020); Decay sugar-rich substrates (Põlme et al. 2020) | Põlme S, et al. 2020. Fungal Diversity 105:1-16 ((https://doi.org/10.1007/s13225-020-00466-2)) |
| Otu13741 | 1 | k__Fungi(100);p__Ascomycota(100);c__Saccharomycetes(100);o__Saccharomycetales(100);f__Metschnikowiaceae(100);g__Metschnikowia(100);g__Metschnikowia_unclassified(100); | Metschnikowia | 13 | Saprotroph | \|Undefined Saprotroph\| | Yeast | NULL | Probable | As nectar-tap_saprotroph (Põlme et al. 2020); Decay sugar-rich substrates (Põlme et al. 2020) | Põlme S, et al. 2020. Fungal Diversity 105:1-16 ((https://doi.org/10.1007/s13225-020-00466-2)) |
| Otu13743 | 1 | k__Fungi(100);p__Ascomycota(100);c__Saccharomycetes(100);o__Saccharomycetales(100);f__Metschnikowiaceae(100);g__Metschnikowia(100);s__Metschnikowia_pulcherrima(100); | Metschnikowia | 13 | Saprotroph | \|Undefined Saprotroph\| | Yeast | NULL | Probable | As nectar-tap_saprotroph (Põlme et al. 2020); Decay sugar-rich substrates (Põlme et al. 2020) | Põlme S, et al. 2020. Fungal Diversity 105:1-16 ((https://doi.org/10.1007/s13225-020-00466-2)) |
| Otu13744 | 1 | k__Fungi(100);p__Ascomycota(100);c__Saccharomycetes(100);o__Saccharomycetales(100);f__Metschnikowiaceae(100);g__Metschnikowia(100);s__Metschnikowia_pulcherrima(100); | Metschnikowia | 13 | Saprotroph | \|Undefined Saprotroph\| | Yeast | NULL | Probable | As nectar-tap_saprotroph (Põlme et al. 2020); Decay sugar-rich substrates (Põlme et al. 2020) | Põlme S, et al. 2020. Fungal Diversity 105:1-16 ((https://doi.org/10.1007/s13225-020-00466-2)) |
| Otu13745 | 1 | k__Fungi(100);p__Ascomycota(100);c__Saccharomycetes(100);o__Saccharomycetales(100);f__Metschnikowiaceae(100);g__Metschnikowia(100);s__Metschnikowia_pulcherrima(100); | Metschnikowia | 13 | Saprotroph | \|Undefined Saprotroph\| | Yeast | NULL | Probable | As nectar-tap_saprotroph (Põlme et al. 2020); Decay sugar-rich substrates (Põlme et al. 2020) | Põlme S, et al. 2020. Fungal Diversity 105:1-16 ((https://doi.org/10.1007/s13225-020-00466-2)) |
| Otu13746 | 1 | k__Fungi(100);p__Ascomycota(100);c__Saccharomycetes(100);o__Saccharomycetales(100);f__Metschnikowiaceae(100);g__Metschnikowia(100);s__Metschnikowia_pulcherrima(100); | Metschnikowia | 13 | Saprotroph | \|Undefined Saprotroph\| | Yeast | NULL | Probable | As nectar-tap_saprotroph (Põlme et al. 2020); Decay sugar-rich substrates (Põlme et al. 2020) | Põlme S, et al. 2020. Fungal Diversity 105:1-16 ((https://doi.org/10.1007/s13225-020-00466-2)) |
| Otu13748 | 1 | k__Fungi(100);p__Ascomycota(100);c__Saccharomycetes(100);o__Saccharomycetales(100);f__Metschnikowiaceae(100);g__Metschnikowia(100);s__Metschnikowia_pulcherrima(100); | Metschnikowia | 13 | Saprotroph | \|Undefined Saprotroph\| | Yeast | NULL | Probable | As nectar-tap_saprotroph (Põlme et al. 2020); Decay sugar-rich substrates (Põlme et al. 2020) | Põlme S, et al. 2020. Fungal Diversity 105:1-16 ((https://doi.org/10.1007/s13225-020-00466-2)) |
| Otu13752 | 1 | k__Fungi(100);p__Ascomycota(100);c__Saccharomycetes(100);o__Saccharomycetales(100);f__Metschnikowiaceae(100);g__Metschnikowia(100);s__Metschnikowia_pulcherrima(100); | Metschnikowia | 13 | Saprotroph | \|Undefined Saprotroph\| | Yeast | NULL | Probable | As nectar-tap_saprotroph (Põlme et al. 2020); Decay sugar-rich substrates (Põlme et al. 2020) | Põlme S, et al. 2020. Fungal Diversity 105:1-16 ((https://doi.org/10.1007/s13225-020-00466-2)) |
| Otu13753 | 1 | k__Fungi(100);p__Ascomycota(100);c__Saccharomycetes(100);o__Saccharomycetales(100);f__Metschnikowiaceae(100);g__Metschnikowia(100);g__Metschnikowia_unclassified(100); | Metschnikowia | 13 | Saprotroph | \|Undefined Saprotroph\| | Yeast | NULL | Probable | As nectar-tap_saprotroph (Põlme et al. 2020); Decay sugar-rich substrates (Põlme et al. 2020) | Põlme S, et al. 2020. Fungal Diversity 105:1-16 ((https://doi.org/10.1007/s13225-020-00466-2)) |
| Otu13755 | 1 | k__Fungi(100);p__Ascomycota(100);c__Saccharomycetes(100);o__Saccharomycetales(100);f__Metschnikowiaceae(100);g__Metschnikowia(100);g__Metschnikowia_unclassified(100); | Metschnikowia | 13 | Saprotroph | \|Undefined Saprotroph\| | Yeast | NULL | Probable | As nectar-tap_saprotroph (Põlme et al. 2020); Decay sugar-rich substrates (Põlme et al. 2020) | Põlme S, et al. 2020. Fungal Diversity 105:1-16 ((https://doi.org/10.1007/s13225-020-00466-2)) |
| Otu13756 | 1 | k__Fungi(100);p__Ascomycota(100);c__Saccharomycetes(100);o__Saccharomycetales(100);f__Saccharomycetales_family_Incertae_sedis(100);g__Candida(100);g__Candida_unclassified(100); | Candida | 13 | Pathotroph-Saprotroph-Symbiotroph | Animal Pathogen-Endophyte-Endosymbiont-Epiphyte-\|Undefined Saprotroph\| | Dimorphic Yeast-Yeast | NULL | Probable | As nectar-tap_saprotroph (Põlme et al. 2020); Decay sugar-rich substrates (Põlme et al. 2020); This asexual genus is polyphyletic and contain species that function very differently depend on what they are. Caution should be taken when interpreting guilds for Candida species. | Manolakaki D, et al. 2010. Virulence. 1: 367-375 ((https://doi.org/10.4161/viru.1.5.12796)); Mekbib SB, et al. 2011. Tropical Plant Pathology 36:233-240 ((https://doi.org/10.1590/S1982-56762011000400004)); Põlme S, et al. 2020. Fungal Diversity 105:1-16 ((https://doi.org/10.1007/s13225-020-00466-2)) |
| Otu13757 | 1 | k__Fungi(100);p__Ascomycota(100);c__Saccharomycetes(100);o__Saccharomycetales(100);f__Metschnikowiaceae(100);g__Metschnikowia(100);g__Metschnikowia_unclassified(100); | Metschnikowia | 13 | Saprotroph | \|Undefined Saprotroph\| | Yeast | NULL | Probable | As nectar-tap_saprotroph (Põlme et al. 2020); Decay sugar-rich substrates (Põlme et al. 2020) | Põlme S, et al. 2020. Fungal Diversity 105:1-16 ((https://doi.org/10.1007/s13225-020-00466-2)) |
| Otu13760 | 1 | k__Fungi(100);p__Ascomycota(100);c__Saccharomycetes(100);o__Saccharomycetales(100);f__Metschnikowiaceae(100);g__Metschnikowia(100);s__Metschnikowia_pulcherrima(100); | Metschnikowia | 13 | Saprotroph | \|Undefined Saprotroph\| | Yeast | NULL | Probable | As nectar-tap_saprotroph (Põlme et al. 2020); Decay sugar-rich substrates (Põlme et al. 2020) | Põlme S, et al. 2020. Fungal Diversity 105:1-16 ((https://doi.org/10.1007/s13225-020-00466-2)) |
| Otu13761 | 1 | k__Fungi(100);p__Ascomycota(100);c__Saccharomycetes(100);o__Saccharomycetales(100);f__Metschnikowiaceae(100);g__Metschnikowia(100);s__Metschnikowia_pulcherrima(100); | Metschnikowia | 13 | Saprotroph | \|Undefined Saprotroph\| | Yeast | NULL | Probable | As nectar-tap_saprotroph (Põlme et al. 2020); Decay sugar-rich substrates (Põlme et al. 2020) | Põlme S, et al. 2020. Fungal Diversity 105:1-16 ((https://doi.org/10.1007/s13225-020-00466-2)) |
| Otu13762 | 1 | k__Fungi(100);p__Ascomycota(100);c__Saccharomycetes(100);o__Saccharomycetales(100);f__Metschnikowiaceae(100);g__Metschnikowia(100);s__Metschnikowia_pulcherrima(100); | Metschnikowia | 13 | Saprotroph | \|Undefined Saprotroph\| | Yeast | NULL | Probable | As nectar-tap_saprotroph (Põlme et al. 2020); Decay sugar-rich substrates (Põlme et al. 2020) | Põlme S, et al. 2020. Fungal Diversity 105:1-16 ((https://doi.org/10.1007/s13225-020-00466-2)) |
| Otu13764 | 1 | k__Fungi(100);p__Ascomycota(100);c__Saccharomycetes(100);o__Saccharomycetales(100);f__Metschnikowiaceae(100);g__Metschnikowia(100);s__Metschnikowia_pulcherrima(100); | Metschnikowia | 13 | Saprotroph | \|Undefined Saprotroph\| | Yeast | NULL | Probable | As nectar-tap_saprotroph (Põlme et al. 2020); Decay sugar-rich substrates (Põlme et al. 2020) | Põlme S, et al. 2020. Fungal Diversity 105:1-16 ((https://doi.org/10.1007/s13225-020-00466-2)) |
| Otu13766 | 1 | k__Fungi(100);p__Ascomycota(100);c__Saccharomycetes(100);o__Saccharomycetales(100);o__Saccharomycetales_unclassified(100);o__Saccharomycetales_unclassified(100);o__Saccharomycetales_unclassified(100); | Saccharomycetales | 7 | Saprotroph | Undefined Saprotroph | Yeast | NULL | Possible | NULL | Sterkenburg E, et al. 2015. New Phytologist 207:1145-1158 ((https://doi.org/10.1111/nph.13426)) |
| Otu13767 | 1 | k__Fungi(100);p__Ascomycota(100);c__Saccharomycetes(100);o__Saccharomycetales(100);f__Metschnikowiaceae(100);g__Metschnikowia(100);s__Metschnikowia_pulcherrima(100); | Metschnikowia | 13 | Saprotroph | \|Undefined Saprotroph\| | Yeast | NULL | Probable | As nectar-tap_saprotroph (Põlme et al. 2020); Decay sugar-rich substrates (Põlme et al. 2020) | Põlme S, et al. 2020. Fungal Diversity 105:1-16 ((https://doi.org/10.1007/s13225-020-00466-2)) |
| Otu13768 | 1 | k__Fungi(100);p__Ascomycota(100);c__Saccharomycetes(100);o__Saccharomycetales(100);f__Metschnikowiaceae(100);g__Metschnikowia(100);s__Metschnikowia_pulcherrima(100); | Metschnikowia | 13 | Saprotroph | \|Undefined Saprotroph\| | Yeast | NULL | Probable | As nectar-tap_saprotroph (Põlme et al. 2020); Decay sugar-rich substrates (Põlme et al. 2020) | Põlme S, et al. 2020. Fungal Diversity 105:1-16 ((https://doi.org/10.1007/s13225-020-00466-2)) |
| Otu13769 | 1 | k__Fungi(100);p__Ascomycota(100);c__Saccharomycetes(100);o__Saccharomycetales(100);f__Metschnikowiaceae(100);g__Metschnikowia(100);g__Metschnikowia_unclassified(100); | Metschnikowia | 13 | Saprotroph | \|Undefined Saprotroph\| | Yeast | NULL | Probable | As nectar-tap_saprotroph (Põlme et al. 2020); Decay sugar-rich substrates (Põlme et al. 2020) | Põlme S, et al. 2020. Fungal Diversity 105:1-16 ((https://doi.org/10.1007/s13225-020-00466-2)) |
| Otu13770 | 1 | k__Fungi(100);p__Ascomycota(100);c__Saccharomycetes(100);o__Saccharomycetales(100);f__Saccharomycetales_family_Incertae_sedis(100);g__Candida(100);g__Candida_unclassified(100); | Candida | 13 | Pathotroph-Saprotroph-Symbiotroph | Animal Pathogen-Endophyte-Endosymbiont-Epiphyte-\|Undefined Saprotroph\| | Dimorphic Yeast-Yeast | NULL | Probable | As nectar-tap_saprotroph (Põlme et al. 2020); Decay sugar-rich substrates (Põlme et al. 2020); This asexual genus is polyphyletic and contain species that function very differently depend on what they are. Caution should be taken when interpreting guilds for Candida species. | Manolakaki D, et al. 2010. Virulence. 1: 367-375 ((https://doi.org/10.4161/viru.1.5.12796)); Mekbib SB, et al. 2011. Tropical Plant Pathology 36:233-240 ((https://doi.org/10.1590/S1982-56762011000400004)); Põlme S, et al. 2020. Fungal Diversity 105:1-16 ((https://doi.org/10.1007/s13225-020-00466-2)) |
| Otu13771 | 1 | k__Fungi(100);p__Ascomycota(100);c__Saccharomycetes(100);o__Saccharomycetales(100);f__Metschnikowiaceae(100);g__Metschnikowia(100);s__Metschnikowia_pulcherrima(100); | Metschnikowia | 13 | Saprotroph | \|Undefined Saprotroph\| | Yeast | NULL | Probable | As nectar-tap_saprotroph (Põlme et al. 2020); Decay sugar-rich substrates (Põlme et al. 2020) | Põlme S, et al. 2020. Fungal Diversity 105:1-16 ((https://doi.org/10.1007/s13225-020-00466-2)) |
| Otu13773 | 1 | k__Fungi(100);p__Ascomycota(100);c__Saccharomycetes(100);o__Saccharomycetales(100);f__Saccharomycetales_family_Incertae_sedis(100);g__Candida(100);g__Candida_unclassified(100); | Candida | 13 | Pathotroph-Saprotroph-Symbiotroph | Animal Pathogen-Endophyte-Endosymbiont-Epiphyte-\|Undefined Saprotroph\| | Dimorphic Yeast-Yeast | NULL | Probable | As nectar-tap_saprotroph (Põlme et al. 2020); Decay sugar-rich substrates (Põlme et al. 2020); This asexual genus is polyphyletic and contain species that function very differently depend on what they are. Caution should be taken when interpreting guilds for Candida species. | Manolakaki D, et al. 2010. Virulence. 1: 367-375 ((https://doi.org/10.4161/viru.1.5.12796)); Mekbib SB, et al. 2011. Tropical Plant Pathology 36:233-240 ((https://doi.org/10.1590/S1982-56762011000400004)); Põlme S, et al. 2020. Fungal Diversity 105:1-16 ((https://doi.org/10.1007/s13225-020-00466-2)) |
| Otu13774 | 1 | k__Fungi(100);p__Ascomycota(100);c__Saccharomycetes(100);o__Saccharomycetales(100);o__Saccharomycetales_unclassified(100);o__Saccharomycetales_unclassified(100);o__Saccharomycetales_unclassified(100); | Saccharomycetales | 7 | Saprotroph | Undefined Saprotroph | Yeast | NULL | Possible | NULL | Sterkenburg E, et al. 2015. New Phytologist 207:1145-1158 ((https://doi.org/10.1111/nph.13426)) |
| Otu13775 | 1 | k__Fungi(100);p__Ascomycota(100);c__Saccharomycetes(100);o__Saccharomycetales(100);f__Metschnikowiaceae(100);g__Metschnikowia(100);s__Metschnikowia_pulcherrima(100); | Metschnikowia | 13 | Saprotroph | \|Undefined Saprotroph\| | Yeast | NULL | Probable | As nectar-tap_saprotroph (Põlme et al. 2020); Decay sugar-rich substrates (Põlme et al. 2020) | Põlme S, et al. 2020. Fungal Diversity 105:1-16 ((https://doi.org/10.1007/s13225-020-00466-2)) |
| Otu13776 | 1 | k__Fungi(100);p__Ascomycota(100);c__Saccharomycetes(100);o__Saccharomycetales(100);f__Metschnikowiaceae(100);g__Metschnikowia(100);g__Metschnikowia_unclassified(100); | Metschnikowia | 13 | Saprotroph | \|Undefined Saprotroph\| | Yeast | NULL | Probable | As nectar-tap_saprotroph (Põlme et al. 2020); Decay sugar-rich substrates (Põlme et al. 2020) | Põlme S, et al. 2020. Fungal Diversity 105:1-16 ((https://doi.org/10.1007/s13225-020-00466-2)) |
| Otu13777 | 1 | k__Fungi(100);p__Ascomycota(100);c__Saccharomycetes(100);o__Saccharomycetales(100);f__Metschnikowiaceae(100);g__Metschnikowia(100);s__Metschnikowia_pulcherrima(100); | Metschnikowia | 13 | Saprotroph | \|Undefined Saprotroph\| | Yeast | NULL | Probable | As nectar-tap_saprotroph (Põlme et al. 2020); Decay sugar-rich substrates (Põlme et al. 2020) | Põlme S, et al. 2020. Fungal Diversity 105:1-16 ((https://doi.org/10.1007/s13225-020-00466-2)) |
| Otu13779 | 1 | k__Fungi(100);p__Ascomycota(100);c__Saccharomycetes(100);o__Saccharomycetales(100);f__Metschnikowiaceae(100);g__Metschnikowia(100);s__Metschnikowia_pulcherrima(100); | Metschnikowia | 13 | Saprotroph | \|Undefined Saprotroph\| | Yeast | NULL | Probable | As nectar-tap_saprotroph (Põlme et al. 2020); Decay sugar-rich substrates (Põlme et al. 2020) | Põlme S, et al. 2020. Fungal Diversity 105:1-16 ((https://doi.org/10.1007/s13225-020-00466-2)) |
| Otu13780 | 1 | k__Fungi(100);p__Ascomycota(100);c__Saccharomycetes(100);o__Saccharomycetales(100);f__Metschnikowiaceae(100);g__Metschnikowia(100);s__Metschnikowia_pulcherrima(100); | Metschnikowia | 13 | Saprotroph | \|Undefined Saprotroph\| | Yeast | NULL | Probable | As nectar-tap_saprotroph (Põlme et al. 2020); Decay sugar-rich substrates (Põlme et al. 2020) | Põlme S, et al. 2020. Fungal Diversity 105:1-16 ((https://doi.org/10.1007/s13225-020-00466-2)) |
| Otu13781 | 1 | k__Fungi(100);p__Ascomycota(100);c__Saccharomycetes(100);o__Saccharomycetales(100);f__Metschnikowiaceae(100);g__Metschnikowia(100);g__Metschnikowia_unclassified(100); | Metschnikowia | 13 | Saprotroph | \|Undefined Saprotroph\| | Yeast | NULL | Probable | As nectar-tap_saprotroph (Põlme et al. 2020); Decay sugar-rich substrates (Põlme et al. 2020) | Põlme S, et al. 2020. Fungal Diversity 105:1-16 ((https://doi.org/10.1007/s13225-020-00466-2)) |
| Otu13782 | 1 | k__Fungi(100);p__Ascomycota(100);c__Saccharomycetes(100);o__Saccharomycetales(100);f__Metschnikowiaceae(100);g__Metschnikowia(100);s__Metschnikowia_pulcherrima(100); | Metschnikowia | 13 | Saprotroph | \|Undefined Saprotroph\| | Yeast | NULL | Probable | As nectar-tap_saprotroph (Põlme et al. 2020); Decay sugar-rich substrates (Põlme et al. 2020) | Põlme S, et al. 2020. Fungal Diversity 105:1-16 ((https://doi.org/10.1007/s13225-020-00466-2)) |
| Otu13783 | 1 | k__Fungi(100);p__Ascomycota(100);c__Saccharomycetes(100);o__Saccharomycetales(100);f__Metschnikowiaceae(100);g__Metschnikowia(100);g__Metschnikowia_unclassified(100); | Metschnikowia | 13 | Saprotroph | \|Undefined Saprotroph\| | Yeast | NULL | Probable | As nectar-tap_saprotroph (Põlme et al. 2020); Decay sugar-rich substrates (Põlme et al. 2020) | Põlme S, et al. 2020. Fungal Diversity 105:1-16 ((https://doi.org/10.1007/s13225-020-00466-2)) |
| Otu13784 | 1 | k__Fungi(100);p__Ascomycota(100);c__Saccharomycetes(100);o__Saccharomycetales(100);f__Metschnikowiaceae(100);g__Metschnikowia(100);s__Metschnikowia_pulcherrima(100); | Metschnikowia | 13 | Saprotroph | \|Undefined Saprotroph\| | Yeast | NULL | Probable | As nectar-tap_saprotroph (Põlme et al. 2020); Decay sugar-rich substrates (Põlme et al. 2020) | Põlme S, et al. 2020. Fungal Diversity 105:1-16 ((https://doi.org/10.1007/s13225-020-00466-2)) |
| Otu13785 | 1 | k__Fungi(100);p__Ascomycota(100);c__Saccharomycetes(100);o__Saccharomycetales(100);f__Saccharomycetales_family_Incertae_sedis(100);g__Candida(100);g__Candida_unclassified(100); | Candida | 13 | Pathotroph-Saprotroph-Symbiotroph | Animal Pathogen-Endophyte-Endosymbiont-Epiphyte-\|Undefined Saprotroph\| | Dimorphic Yeast-Yeast | NULL | Probable | As nectar-tap_saprotroph (Põlme et al. 2020); Decay sugar-rich substrates (Põlme et al. 2020); This asexual genus is polyphyletic and contain species that function very differently depend on what they are. Caution should be taken when interpreting guilds for Candida species. | Manolakaki D, et al. 2010. Virulence. 1: 367-375 ((https://doi.org/10.4161/viru.1.5.12796)); Mekbib SB, et al. 2011. Tropical Plant Pathology 36:233-240 ((https://doi.org/10.1590/S1982-56762011000400004)); Põlme S, et al. 2020. Fungal Diversity 105:1-16 ((https://doi.org/10.1007/s13225-020-00466-2)) |
| Otu13786 | 1 | k__Fungi(100);p__Ascomycota(100);c__Saccharomycetes(100);o__Saccharomycetales(100);f__Metschnikowiaceae(100);g__Metschnikowia(100);s__Metschnikowia_pulcherrima(100); | Metschnikowia | 13 | Saprotroph | \|Undefined Saprotroph\| | Yeast | NULL | Probable | As nectar-tap_saprotroph (Põlme et al. 2020); Decay sugar-rich substrates (Põlme et al. 2020) | Põlme S, et al. 2020. Fungal Diversity 105:1-16 ((https://doi.org/10.1007/s13225-020-00466-2)) |
| Otu13787 | 1 | k__Fungi(100);p__Ascomycota(100);c__Saccharomycetes(100);o__Saccharomycetales(100);f__Metschnikowiaceae(100);g__Metschnikowia(100);g__Metschnikowia_unclassified(100); | Metschnikowia | 13 | Saprotroph | \|Undefined Saprotroph\| | Yeast | NULL | Probable | As nectar-tap_saprotroph (Põlme et al. 2020); Decay sugar-rich substrates (Põlme et al. 2020) | Põlme S, et al. 2020. Fungal Diversity 105:1-16 ((https://doi.org/10.1007/s13225-020-00466-2)) |
| Otu13788 | 1 | k__Fungi(100);p__Ascomycota(100);c__Saccharomycetes(100);o__Saccharomycetales(100);f__Saccharomycetaceae(100);g__Kazachstania(100);s__Kazachstania_intestinalis(100); | Kazachstania | 13 | Saprotroph | \|Undefined Saprotroph\| | Yeast | NULL | Probable | As nectar-tap_saprotroph (Põlme et al. 2020); Decay sugar-rich substrates (Põlme et al. 2020) | Põlme S, et al. 2020. Fungal Diversity 105:1-16 ((https://doi.org/10.1007/s13225-020-00466-2)) |
| Otu13789 | 1 | k__Fungi(100);p__Ascomycota(100);c__Saccharomycetes(100);o__Saccharomycetales(100);f__Metschnikowiaceae(100);g__Metschnikowia(100);s__Metschnikowia_pulcherrima(100); | Metschnikowia | 13 | Saprotroph | \|Undefined Saprotroph\| | Yeast | NULL | Probable | As nectar-tap_saprotroph (Põlme et al. 2020); Decay sugar-rich substrates (Põlme et al. 2020) | Põlme S, et al. 2020. Fungal Diversity 105:1-16 ((https://doi.org/10.1007/s13225-020-00466-2)) |
| Otu13790 | 1 | k__Fungi(100);p__Ascomycota(100);c__Saccharomycetes(100);o__Saccharomycetales(100);f__Metschnikowiaceae(100);g__Metschnikowia(100);g__Metschnikowia_unclassified(100); | Metschnikowia | 13 | Saprotroph | \|Undefined Saprotroph\| | Yeast | NULL | Probable | As nectar-tap_saprotroph (Põlme et al. 2020); Decay sugar-rich substrates (Põlme et al. 2020) | Põlme S, et al. 2020. Fungal Diversity 105:1-16 ((https://doi.org/10.1007/s13225-020-00466-2)) |
| Otu13791 | 1 | k__Fungi(100);p__Ascomycota(100);c__Saccharomycetes(100);o__Saccharomycetales(100);f__Metschnikowiaceae(100);g__Metschnikowia(100);s__Metschnikowia_pulcherrima(100); | Metschnikowia | 13 | Saprotroph | \|Undefined Saprotroph\| | Yeast | NULL | Probable | As nectar-tap_saprotroph (Põlme et al. 2020); Decay sugar-rich substrates (Põlme et al. 2020) | Põlme S, et al. 2020. Fungal Diversity 105:1-16 ((https://doi.org/10.1007/s13225-020-00466-2)) |
| Otu13793 | 1 | k__Fungi(100);p__Ascomycota(100);c__Saccharomycetes(100);o__Saccharomycetales(100);f__Metschnikowiaceae(100);g__Metschnikowia(100);s__Metschnikowia_pulcherrima(100); | Metschnikowia | 13 | Saprotroph | \|Undefined Saprotroph\| | Yeast | NULL | Probable | As nectar-tap_saprotroph (Põlme et al. 2020); Decay sugar-rich substrates (Põlme et al. 2020) | Põlme S, et al. 2020. Fungal Diversity 105:1-16 ((https://doi.org/10.1007/s13225-020-00466-2)) |
| Otu13794 | 1 | k__Fungi(100);p__Ascomycota(100);c__Saccharomycetes(100);o__Saccharomycetales(100);f__Metschnikowiaceae(100);g__Metschnikowia(100);s__Metschnikowia_pulcherrima(100); | Metschnikowia | 13 | Saprotroph | \|Undefined Saprotroph\| | Yeast | NULL | Probable | As nectar-tap_saprotroph (Põlme et al. 2020); Decay sugar-rich substrates (Põlme et al. 2020) | Põlme S, et al. 2020. Fungal Diversity 105:1-16 ((https://doi.org/10.1007/s13225-020-00466-2)) |
| Otu13795 | 1 | k__Fungi(100);p__Ascomycota(100);c__Saccharomycetes(100);o__Saccharomycetales(100);f__Metschnikowiaceae(100);g__Metschnikowia(100);s__Metschnikowia_pulcherrima(100); | Metschnikowia | 13 | Saprotroph | \|Undefined Saprotroph\| | Yeast | NULL | Probable | As nectar-tap_saprotroph (Põlme et al. 2020); Decay sugar-rich substrates (Põlme et al. 2020) | Põlme S, et al. 2020. Fungal Diversity 105:1-16 ((https://doi.org/10.1007/s13225-020-00466-2)) |
| Otu13796 | 1 | k__Fungi(100);p__Ascomycota(100);c__Saccharomycetes(100);o__Saccharomycetales(100);f__Metschnikowiaceae(100);g__Metschnikowia(100);g__Metschnikowia_unclassified(100); | Metschnikowia | 13 | Saprotroph | \|Undefined Saprotroph\| | Yeast | NULL | Probable | As nectar-tap_saprotroph (Põlme et al. 2020); Decay sugar-rich substrates (Põlme et al. 2020) | Põlme S, et al. 2020. Fungal Diversity 105:1-16 ((https://doi.org/10.1007/s13225-020-00466-2)) |
| Otu13797 | 1 | k__Fungi(100);p__Ascomycota(100);c__Saccharomycetes(100);o__Saccharomycetales(100);f__Metschnikowiaceae(100);g__Metschnikowia(100);s__Metschnikowia_pulcherrima(100); | Metschnikowia | 13 | Saprotroph | \|Undefined Saprotroph\| | Yeast | NULL | Probable | As nectar-tap_saprotroph (Põlme et al. 2020); Decay sugar-rich substrates (Põlme et al. 2020) | Põlme S, et al. 2020. Fungal Diversity 105:1-16 ((https://doi.org/10.1007/s13225-020-00466-2)) |
| Otu13798 | 1 | k__Fungi(100);p__Ascomycota(100);c__Saccharomycetes(100);o__Saccharomycetales(100);f__Metschnikowiaceae(100);g__Metschnikowia(100);g__Metschnikowia_unclassified(100); | Metschnikowia | 13 | Saprotroph | \|Undefined Saprotroph\| | Yeast | NULL | Probable | As nectar-tap_saprotroph (Põlme et al. 2020); Decay sugar-rich substrates (Põlme et al. 2020) | Põlme S, et al. 2020. Fungal Diversity 105:1-16 ((https://doi.org/10.1007/s13225-020-00466-2)) |
| Otu13799 | 1 | k__Fungi(100);p__Ascomycota(100);c__Saccharomycetes(100);o__Saccharomycetales(100);f__Metschnikowiaceae(100);g__Metschnikowia(100);s__Metschnikowia_pulcherrima(100); | Metschnikowia | 13 | Saprotroph | \|Undefined Saprotroph\| | Yeast | NULL | Probable | As nectar-tap_saprotroph (Põlme et al. 2020); Decay sugar-rich substrates (Põlme et al. 2020) | Põlme S, et al. 2020. Fungal Diversity 105:1-16 ((https://doi.org/10.1007/s13225-020-00466-2)) |
| Otu13800 | 1 | k__Fungi(100);p__Ascomycota(100);c__Saccharomycetes(100);o__Saccharomycetales(100);f__Metschnikowiaceae(100);g__Metschnikowia(100);s__Metschnikowia_pulcherrima(100); | Metschnikowia | 13 | Saprotroph | \|Undefined Saprotroph\| | Yeast | NULL | Probable | As nectar-tap_saprotroph (Põlme et al. 2020); Decay sugar-rich substrates (Põlme et al. 2020) | Põlme S, et al. 2020. Fungal Diversity 105:1-16 ((https://doi.org/10.1007/s13225-020-00466-2)) |
| Otu13801 | 1 | k__Fungi(100);p__Ascomycota(100);c__Saccharomycetes(100);o__Saccharomycetales(100);f__Metschnikowiaceae(100);g__Metschnikowia(100);g__Metschnikowia_unclassified(100); | Metschnikowia | 13 | Saprotroph | \|Undefined Saprotroph\| | Yeast | NULL | Probable | As nectar-tap_saprotroph (Põlme et al. 2020); Decay sugar-rich substrates (Põlme et al. 2020) | Põlme S, et al. 2020. Fungal Diversity 105:1-16 ((https://doi.org/10.1007/s13225-020-00466-2)) |
| Otu13802 | 1 | k__Fungi(100);p__Ascomycota(100);c__Saccharomycetes(100);o__Saccharomycetales(100);f__Metschnikowiaceae(100);g__Metschnikowia(100);g__Metschnikowia_unclassified(100); | Metschnikowia | 13 | Saprotroph | \|Undefined Saprotroph\| | Yeast | NULL | Probable | As nectar-tap_saprotroph (Põlme et al. 2020); Decay sugar-rich substrates (Põlme et al. 2020) | Põlme S, et al. 2020. Fungal Diversity 105:1-16 ((https://doi.org/10.1007/s13225-020-00466-2)) |
| Otu13803 | 1 | k__Fungi(100);p__Ascomycota(100);c__Saccharomycetes(100);o__Saccharomycetales(100);f__Metschnikowiaceae(100);g__Metschnikowia(100);s__Metschnikowia_pulcherrima(100); | Metschnikowia | 13 | Saprotroph | \|Undefined Saprotroph\| | Yeast | NULL | Probable | As nectar-tap_saprotroph (Põlme et al. 2020); Decay sugar-rich substrates (Põlme et al. 2020) | Põlme S, et al. 2020. Fungal Diversity 105:1-16 ((https://doi.org/10.1007/s13225-020-00466-2)) |
| Otu13804 | 1 | k__Fungi(100);p__Ascomycota(100);c__Saccharomycetes(100);o__Saccharomycetales(100);f__Metschnikowiaceae(100);g__Metschnikowia(100);g__Metschnikowia_unclassified(100); | Metschnikowia | 13 | Saprotroph | \|Undefined Saprotroph\| | Yeast | NULL | Probable | As nectar-tap_saprotroph (Põlme et al. 2020); Decay sugar-rich substrates (Põlme et al. 2020) | Põlme S, et al. 2020. Fungal Diversity 105:1-16 ((https://doi.org/10.1007/s13225-020-00466-2)) |
| Otu13805 | 1 | k__Fungi(100);p__Ascomycota(100);c__Saccharomycetes(100);o__Saccharomycetales(100);f__Metschnikowiaceae(100);g__Metschnikowia(100);g__Metschnikowia_unclassified(100); | Metschnikowia | 13 | Saprotroph | \|Undefined Saprotroph\| | Yeast | NULL | Probable | As nectar-tap_saprotroph (Põlme et al. 2020); Decay sugar-rich substrates (Põlme et al. 2020) | Põlme S, et al. 2020. Fungal Diversity 105:1-16 ((https://doi.org/10.1007/s13225-020-00466-2)) |
| Otu13808 | 1 | k__Fungi(100);p__Ascomycota(100);c__Saccharomycetes(100);o__Saccharomycetales(100);f__Metschnikowiaceae(100);g__Metschnikowia(100);s__Metschnikowia_pulcherrima(100); | Metschnikowia | 13 | Saprotroph | \|Undefined Saprotroph\| | Yeast | NULL | Probable | As nectar-tap_saprotroph (Põlme et al. 2020); Decay sugar-rich substrates (Põlme et al. 2020) | Põlme S, et al. 2020. Fungal Diversity 105:1-16 ((https://doi.org/10.1007/s13225-020-00466-2)) |
| Otu13809 | 1 | k__Fungi(100);p__Ascomycota(100);c__Saccharomycetes(100);o__Saccharomycetales(100);f__Metschnikowiaceae(100);g__Metschnikowia(100);s__Metschnikowia_pulcherrima(100); | Metschnikowia | 13 | Saprotroph | \|Undefined Saprotroph\| | Yeast | NULL | Probable | As nectar-tap_saprotroph (Põlme et al. 2020); Decay sugar-rich substrates (Põlme et al. 2020) | Põlme S, et al. 2020. Fungal Diversity 105:1-16 ((https://doi.org/10.1007/s13225-020-00466-2)) |
| Otu13810 | 1 | k__Fungi(100);p__Ascomycota(100);c__Saccharomycetes(100);o__Saccharomycetales(100);f__Metschnikowiaceae(100);g__Metschnikowia(100);s__Metschnikowia_pulcherrima(100); | Metschnikowia | 13 | Saprotroph | \|Undefined Saprotroph\| | Yeast | NULL | Probable | As nectar-tap_saprotroph (Põlme et al. 2020); Decay sugar-rich substrates (Põlme et al. 2020) | Põlme S, et al. 2020. Fungal Diversity 105:1-16 ((https://doi.org/10.1007/s13225-020-00466-2)) |
| Otu13811 | 1 | k__Fungi(100);p__Ascomycota(100);c__Saccharomycetes(100);o__Saccharomycetales(100);f__Metschnikowiaceae(100);g__Metschnikowia(100);g__Metschnikowia_unclassified(100); | Metschnikowia | 13 | Saprotroph | \|Undefined Saprotroph\| | Yeast | NULL | Probable | As nectar-tap_saprotroph (Põlme et al. 2020); Decay sugar-rich substrates (Põlme et al. 2020) | Põlme S, et al. 2020. Fungal Diversity 105:1-16 ((https://doi.org/10.1007/s13225-020-00466-2)) |
| Otu13812 | 1 | k__Fungi(100);p__Ascomycota(100);c__Saccharomycetes(100);o__Saccharomycetales(100);f__Metschnikowiaceae(100);g__Metschnikowia(100);s__Metschnikowia_pulcherrima(100); | Metschnikowia | 13 | Saprotroph | \|Undefined Saprotroph\| | Yeast | NULL | Probable | As nectar-tap_saprotroph (Põlme et al. 2020); Decay sugar-rich substrates (Põlme et al. 2020) | Põlme S, et al. 2020. Fungal Diversity 105:1-16 ((https://doi.org/10.1007/s13225-020-00466-2)) |
| Otu13813 | 1 | k__Fungi(100);p__Ascomycota(100);c__Saccharomycetes(100);o__Saccharomycetales(100);f__Metschnikowiaceae(100);g__Metschnikowia(100);s__Metschnikowia_pulcherrima(100); | Metschnikowia | 13 | Saprotroph | \|Undefined Saprotroph\| | Yeast | NULL | Probable | As nectar-tap_saprotroph (Põlme et al. 2020); Decay sugar-rich substrates (Põlme et al. 2020) | Põlme S, et al. 2020. Fungal Diversity 105:1-16 ((https://doi.org/10.1007/s13225-020-00466-2)) |
| Otu13814 | 1 | k__Fungi(100);p__Ascomycota(100);c__Saccharomycetes(100);o__Saccharomycetales(100);f__Metschnikowiaceae(100);g__Metschnikowia(100);s__Metschnikowia_pulcherrima(100); | Metschnikowia | 13 | Saprotroph | \|Undefined Saprotroph\| | Yeast | NULL | Probable | As nectar-tap_saprotroph (Põlme et al. 2020); Decay sugar-rich substrates (Põlme et al. 2020) | Põlme S, et al. 2020. Fungal Diversity 105:1-16 ((https://doi.org/10.1007/s13225-020-00466-2)) |
| Otu13815 | 1 | k__Fungi(100);p__Ascomycota(100);c__Saccharomycetes(100);o__Saccharomycetales(100);f__Metschnikowiaceae(100);g__Metschnikowia(100);g__Metschnikowia_unclassified(100); | Metschnikowia | 13 | Saprotroph | \|Undefined Saprotroph\| | Yeast | NULL | Probable | As nectar-tap_saprotroph (Põlme et al. 2020); Decay sugar-rich substrates (Põlme et al. 2020) | Põlme S, et al. 2020. Fungal Diversity 105:1-16 ((https://doi.org/10.1007/s13225-020-00466-2)) |
| Otu13816 | 1 | k__Fungi(100);p__Ascomycota(100);c__Saccharomycetes(100);o__Saccharomycetales(100);f__Metschnikowiaceae(100);g__Metschnikowia(100);g__Metschnikowia_unclassified(100); | Metschnikowia | 13 | Saprotroph | \|Undefined Saprotroph\| | Yeast | NULL | Probable | As nectar-tap_saprotroph (Põlme et al. 2020); Decay sugar-rich substrates (Põlme et al. 2020) | Põlme S, et al. 2020. Fungal Diversity 105:1-16 ((https://doi.org/10.1007/s13225-020-00466-2)) |
| Otu13817 | 1 | k__Fungi(100);p__Ascomycota(100);c__Saccharomycetes(100);o__Saccharomycetales(100);f__Metschnikowiaceae(100);g__Metschnikowia(100);g__Metschnikowia_unclassified(100); | Metschnikowia | 13 | Saprotroph | \|Undefined Saprotroph\| | Yeast | NULL | Probable | As nectar-tap_saprotroph (Põlme et al. 2020); Decay sugar-rich substrates (Põlme et al. 2020) | Põlme S, et al. 2020. Fungal Diversity 105:1-16 ((https://doi.org/10.1007/s13225-020-00466-2)) |
| Otu13818 | 1 | k__Fungi(100);p__Ascomycota(100);c__Saccharomycetes(100);o__Saccharomycetales(100);f__Metschnikowiaceae(100);g__Metschnikowia(100);g__Metschnikowia_unclassified(100); | Metschnikowia | 13 | Saprotroph | \|Undefined Saprotroph\| | Yeast | NULL | Probable | As nectar-tap_saprotroph (Põlme et al. 2020); Decay sugar-rich substrates (Põlme et al. 2020) | Põlme S, et al. 2020. Fungal Diversity 105:1-16 ((https://doi.org/10.1007/s13225-020-00466-2)) |
| Otu13819 | 1 | k__Fungi(100);p__Ascomycota(100);c__Saccharomycetes(100);o__Saccharomycetales(100);f__Saccharomycetales_family_Incertae_sedis(100);g__Candida(100);g__Candida_unclassified(100); | Candida | 13 | Pathotroph-Saprotroph-Symbiotroph | Animal Pathogen-Endophyte-Endosymbiont-Epiphyte-\|Undefined Saprotroph\| | Dimorphic Yeast-Yeast | NULL | Probable | As nectar-tap_saprotroph (Põlme et al. 2020); Decay sugar-rich substrates (Põlme et al. 2020); This asexual genus is polyphyletic and contain species that function very differently depend on what they are. Caution should be taken when interpreting guilds for Candida species. | Manolakaki D, et al. 2010. Virulence. 1: 367-375 ((https://doi.org/10.4161/viru.1.5.12796)); Mekbib SB, et al. 2011. Tropical Plant Pathology 36:233-240 ((https://doi.org/10.1590/S1982-56762011000400004)); Põlme S, et al. 2020. Fungal Diversity 105:1-16 ((https://doi.org/10.1007/s13225-020-00466-2)) |
| Otu13820 | 1 | k__Fungi(100);p__Ascomycota(100);c__Saccharomycetes(100);o__Saccharomycetales(100);f__Metschnikowiaceae(100);g__Metschnikowia(100);s__Metschnikowia_pulcherrima(100); | Metschnikowia | 13 | Saprotroph | \|Undefined Saprotroph\| | Yeast | NULL | Probable | As nectar-tap_saprotroph (Põlme et al. 2020); Decay sugar-rich substrates (Põlme et al. 2020) | Põlme S, et al. 2020. Fungal Diversity 105:1-16 ((https://doi.org/10.1007/s13225-020-00466-2)) |
| Otu13821 | 1 | k__Fungi(100);p__Ascomycota(100);c__Saccharomycetes(100);o__Saccharomycetales(100);f__Metschnikowiaceae(100);g__Metschnikowia(100);s__Metschnikowia_pulcherrima(100); | Metschnikowia | 13 | Saprotroph | \|Undefined Saprotroph\| | Yeast | NULL | Probable | As nectar-tap_saprotroph (Põlme et al. 2020); Decay sugar-rich substrates (Põlme et al. 2020) | Põlme S, et al. 2020. Fungal Diversity 105:1-16 ((https://doi.org/10.1007/s13225-020-00466-2)) |
| Otu13822 | 1 | k__Fungi(100);p__Ascomycota(100);c__Saccharomycetes(100);o__Saccharomycetales(100);f__Metschnikowiaceae(100);g__Metschnikowia(100);s__Metschnikowia_pulcherrima(100); | Metschnikowia | 13 | Saprotroph | \|Undefined Saprotroph\| | Yeast | NULL | Probable | As nectar-tap_saprotroph (Põlme et al. 2020); Decay sugar-rich substrates (Põlme et al. 2020) | Põlme S, et al. 2020. Fungal Diversity 105:1-16 ((https://doi.org/10.1007/s13225-020-00466-2)) |
| Otu13823 | 1 | k__Fungi(100);p__Ascomycota(100);c__Saccharomycetes(100);o__Saccharomycetales(100);f__Metschnikowiaceae(100);g__Metschnikowia(100);s__Metschnikowia_pulcherrima(100); | Metschnikowia | 13 | Saprotroph | \|Undefined Saprotroph\| | Yeast | NULL | Probable | As nectar-tap_saprotroph (Põlme et al. 2020); Decay sugar-rich substrates (Põlme et al. 2020) | Põlme S, et al. 2020. Fungal Diversity 105:1-16 ((https://doi.org/10.1007/s13225-020-00466-2)) |
| Otu13824 | 1 | k__Fungi(100);p__Ascomycota(100);c__Saccharomycetes(100);o__Saccharomycetales(100);f__Metschnikowiaceae(100);g__Metschnikowia(100);s__Metschnikowia_pulcherrima(100); | Metschnikowia | 13 | Saprotroph | \|Undefined Saprotroph\| | Yeast | NULL | Probable | As nectar-tap_saprotroph (Põlme et al. 2020); Decay sugar-rich substrates (Põlme et al. 2020) | Põlme S, et al. 2020. Fungal Diversity 105:1-16 ((https://doi.org/10.1007/s13225-020-00466-2)) |
| Otu13825 | 1 | k__Fungi(100);p__Ascomycota(100);c__Saccharomycetes(100);o__Saccharomycetales(100);f__Metschnikowiaceae(100);g__Metschnikowia(100);g__Metschnikowia_unclassified(100); | Metschnikowia | 13 | Saprotroph | \|Undefined Saprotroph\| | Yeast | NULL | Probable | As nectar-tap_saprotroph (Põlme et al. 2020); Decay sugar-rich substrates (Põlme et al. 2020) | Põlme S, et al. 2020. Fungal Diversity 105:1-16 ((https://doi.org/10.1007/s13225-020-00466-2)) |
| Otu13827 | 1 | k__Fungi(100);p__Ascomycota(100);c__Saccharomycetes(100);o__Saccharomycetales(100);f__Metschnikowiaceae(100);g__Metschnikowia(100);s__Metschnikowia_pulcherrima(100); | Metschnikowia | 13 | Saprotroph | \|Undefined Saprotroph\| | Yeast | NULL | Probable | As nectar-tap_saprotroph (Põlme et al. 2020); Decay sugar-rich substrates (Põlme et al. 2020) | Põlme S, et al. 2020. Fungal Diversity 105:1-16 ((https://doi.org/10.1007/s13225-020-00466-2)) |
| Otu13828 | 1 | k__Fungi(100);p__Ascomycota(100);c__Saccharomycetes(100);o__Saccharomycetales(100);f__Metschnikowiaceae(100);g__Metschnikowia(100);s__Metschnikowia_pulcherrima(100); | Metschnikowia | 13 | Saprotroph | \|Undefined Saprotroph\| | Yeast | NULL | Probable | As nectar-tap_saprotroph (Põlme et al. 2020); Decay sugar-rich substrates (Põlme et al. 2020) | Põlme S, et al. 2020. Fungal Diversity 105:1-16 ((https://doi.org/10.1007/s13225-020-00466-2)) |
| Otu13829 | 1 | k__Fungi(100);p__Ascomycota(100);c__Saccharomycetes(100);o__Saccharomycetales(100);f__Metschnikowiaceae(100);g__Metschnikowia(100);s__Metschnikowia_pulcherrima(100); | Metschnikowia | 13 | Saprotroph | \|Undefined Saprotroph\| | Yeast | NULL | Probable | As nectar-tap_saprotroph (Põlme et al. 2020); Decay sugar-rich substrates (Põlme et al. 2020) | Põlme S, et al. 2020. Fungal Diversity 105:1-16 ((https://doi.org/10.1007/s13225-020-00466-2)) |
| Otu13830 | 1 | k__Fungi(100);p__Ascomycota(100);c__Saccharomycetes(100);o__Saccharomycetales(100);f__Metschnikowiaceae(100);g__Metschnikowia(100);s__Metschnikowia_pulcherrima(100); | Metschnikowia | 13 | Saprotroph | \|Undefined Saprotroph\| | Yeast | NULL | Probable | As nectar-tap_saprotroph (Põlme et al. 2020); Decay sugar-rich substrates (Põlme et al. 2020) | Põlme S, et al. 2020. Fungal Diversity 105:1-16 ((https://doi.org/10.1007/s13225-020-00466-2)) |
| Otu13832 | 1 | k__Fungi(100);p__Ascomycota(100);c__Saccharomycetes(100);o__Saccharomycetales(100);o__Saccharomycetales_unclassified(100);o__Saccharomycetales_unclassified(100);o__Saccharomycetales_unclassified(100); | Saccharomycetales | 7 | Saprotroph | Undefined Saprotroph | Yeast | NULL | Possible | NULL | Sterkenburg E, et al. 2015. New Phytologist 207:1145-1158 ((https://doi.org/10.1111/nph.13426)) |
| Otu13833 | 1 | k__Fungi(100);p__Ascomycota(100);c__Saccharomycetes(100);o__Saccharomycetales(100);f__Metschnikowiaceae(100);g__Metschnikowia(100);g__Metschnikowia_unclassified(100); | Metschnikowia | 13 | Saprotroph | \|Undefined Saprotroph\| | Yeast | NULL | Probable | As nectar-tap_saprotroph (Põlme et al. 2020); Decay sugar-rich substrates (Põlme et al. 2020) | Põlme S, et al. 2020. Fungal Diversity 105:1-16 ((https://doi.org/10.1007/s13225-020-00466-2)) |
| Otu13834 | 1 | k__Fungi(100);p__Ascomycota(100);c__Saccharomycetes(100);o__Saccharomycetales(100);f__Metschnikowiaceae(100);g__Metschnikowia(100);s__Metschnikowia_pulcherrima(100); | Metschnikowia | 13 | Saprotroph | \|Undefined Saprotroph\| | Yeast | NULL | Probable | As nectar-tap_saprotroph (Põlme et al. 2020); Decay sugar-rich substrates (Põlme et al. 2020) | Põlme S, et al. 2020. Fungal Diversity 105:1-16 ((https://doi.org/10.1007/s13225-020-00466-2)) |
| Otu13835 | 1 | k__Fungi(100);p__Ascomycota(100);c__Saccharomycetes(100);o__Saccharomycetales(100);f__Metschnikowiaceae(100);g__Metschnikowia(100);g__Metschnikowia_unclassified(100); | Metschnikowia | 13 | Saprotroph | \|Undefined Saprotroph\| | Yeast | NULL | Probable | As nectar-tap_saprotroph (Põlme et al. 2020); Decay sugar-rich substrates (Põlme et al. 2020) | Põlme S, et al. 2020. Fungal Diversity 105:1-16 ((https://doi.org/10.1007/s13225-020-00466-2)) |
| Otu13836 | 1 | k__Fungi(100);p__Ascomycota(100);c__Saccharomycetes(100);o__Saccharomycetales(100);f__Metschnikowiaceae(100);g__Metschnikowia(100);g__Metschnikowia_unclassified(100); | Metschnikowia | 13 | Saprotroph | \|Undefined Saprotroph\| | Yeast | NULL | Probable | As nectar-tap_saprotroph (Põlme et al. 2020); Decay sugar-rich substrates (Põlme et al. 2020) | Põlme S, et al. 2020. Fungal Diversity 105:1-16 ((https://doi.org/10.1007/s13225-020-00466-2)) |
| Otu13837 | 1 | k__Fungi(100);p__Ascomycota(100);c__Saccharomycetes(100);o__Saccharomycetales(100);f__Metschnikowiaceae(100);g__Metschnikowia(100);s__Metschnikowia_pulcherrima(100); | Metschnikowia | 13 | Saprotroph | \|Undefined Saprotroph\| | Yeast | NULL | Probable | As nectar-tap_saprotroph (Põlme et al. 2020); Decay sugar-rich substrates (Põlme et al. 2020) | Põlme S, et al. 2020. Fungal Diversity 105:1-16 ((https://doi.org/10.1007/s13225-020-00466-2)) |
| Otu13838 | 1 | k__Fungi(100);p__Ascomycota(100);c__Saccharomycetes(100);o__Saccharomycetales(100);f__Metschnikowiaceae(100);g__Metschnikowia(100);s__Metschnikowia_pulcherrima(100); | Metschnikowia | 13 | Saprotroph | \|Undefined Saprotroph\| | Yeast | NULL | Probable | As nectar-tap_saprotroph (Põlme et al. 2020); Decay sugar-rich substrates (Põlme et al. 2020) | Põlme S, et al. 2020. Fungal Diversity 105:1-16 ((https://doi.org/10.1007/s13225-020-00466-2)) |
| Otu13840 | 1 | k__Fungi(100);p__Ascomycota(100);c__Saccharomycetes(100);o__Saccharomycetales(100);f__Metschnikowiaceae(100);g__Metschnikowia(100);g__Metschnikowia_unclassified(100); | Metschnikowia | 13 | Saprotroph | \|Undefined Saprotroph\| | Yeast | NULL | Probable | As nectar-tap_saprotroph (Põlme et al. 2020); Decay sugar-rich substrates (Põlme et al. 2020) | Põlme S, et al. 2020. Fungal Diversity 105:1-16 ((https://doi.org/10.1007/s13225-020-00466-2)) |
| Otu13842 | 1 | k__Fungi(100);p__Ascomycota(100);c__Saccharomycetes(100);o__Saccharomycetales(100);f__Metschnikowiaceae(100);g__Metschnikowia(100);s__Metschnikowia_pulcherrima(100); | Metschnikowia | 13 | Saprotroph | \|Undefined Saprotroph\| | Yeast | NULL | Probable | As nectar-tap_saprotroph (Põlme et al. 2020); Decay sugar-rich substrates (Põlme et al. 2020) | Põlme S, et al. 2020. Fungal Diversity 105:1-16 ((https://doi.org/10.1007/s13225-020-00466-2)) |
| Otu13843 | 1 | k__Fungi(100);p__Ascomycota(100);c__Saccharomycetes(100);o__Saccharomycetales(100);f__Metschnikowiaceae(100);g__Metschnikowia(100);s__Metschnikowia_pulcherrima(100); | Metschnikowia | 13 | Saprotroph | \|Undefined Saprotroph\| | Yeast | NULL | Probable | As nectar-tap_saprotroph (Põlme et al. 2020); Decay sugar-rich substrates (Põlme et al. 2020) | Põlme S, et al. 2020. Fungal Diversity 105:1-16 ((https://doi.org/10.1007/s13225-020-00466-2)) |
| Otu13844 | 1 | k__Fungi(100);p__Ascomycota(100);c__Saccharomycetes(100);o__Saccharomycetales(100);f__Saccharomycetales_family_Incertae_sedis(100);g__Candida(100);g__Candida_unclassified(100); | Candida | 13 | Pathotroph-Saprotroph-Symbiotroph | Animal Pathogen-Endophyte-Endosymbiont-Epiphyte-\|Undefined Saprotroph\| | Dimorphic Yeast-Yeast | NULL | Probable | As nectar-tap_saprotroph (Põlme et al. 2020); Decay sugar-rich substrates (Põlme et al. 2020); This asexual genus is polyphyletic and contain species that function very differently depend on what they are. Caution should be taken when interpreting guilds for Candida species. | Manolakaki D, et al. 2010. Virulence. 1: 367-375 ((https://doi.org/10.4161/viru.1.5.12796)); Mekbib SB, et al. 2011. Tropical Plant Pathology 36:233-240 ((https://doi.org/10.1590/S1982-56762011000400004)); Põlme S, et al. 2020. Fungal Diversity 105:1-16 ((https://doi.org/10.1007/s13225-020-00466-2)) |
| Otu13846 | 1 | k__Fungi(100);p__Ascomycota(100);c__Saccharomycetes(100);o__Saccharomycetales(100);f__Metschnikowiaceae(100);g__Metschnikowia(100);s__Metschnikowia_pulcherrima(100); | Metschnikowia | 13 | Saprotroph | \|Undefined Saprotroph\| | Yeast | NULL | Probable | As nectar-tap_saprotroph (Põlme et al. 2020); Decay sugar-rich substrates (Põlme et al. 2020) | Põlme S, et al. 2020. Fungal Diversity 105:1-16 ((https://doi.org/10.1007/s13225-020-00466-2)) |
| Otu13847 | 1 | k__Fungi(100);p__Ascomycota(100);c__Saccharomycetes(100);o__Saccharomycetales(100);f__Metschnikowiaceae(100);g__Metschnikowia(100);s__Metschnikowia_pulcherrima(100); | Metschnikowia | 13 | Saprotroph | \|Undefined Saprotroph\| | Yeast | NULL | Probable | As nectar-tap_saprotroph (Põlme et al. 2020); Decay sugar-rich substrates (Põlme et al. 2020) | Põlme S, et al. 2020. Fungal Diversity 105:1-16 ((https://doi.org/10.1007/s13225-020-00466-2)) |
| Otu13849 | 1 | k__Fungi(100);p__Ascomycota(100);c__Saccharomycetes(100);o__Saccharomycetales(100);f__Metschnikowiaceae(100);g__Metschnikowia(100);g__Metschnikowia_unclassified(100); | Metschnikowia | 13 | Saprotroph | \|Undefined Saprotroph\| | Yeast | NULL | Probable | As nectar-tap_saprotroph (Põlme et al. 2020); Decay sugar-rich substrates (Põlme et al. 2020) | Põlme S, et al. 2020. Fungal Diversity 105:1-16 ((https://doi.org/10.1007/s13225-020-00466-2)) |
| Otu13851 | 1 | k__Fungi(100);p__Ascomycota(100);c__Saccharomycetes(100);o__Saccharomycetales(100);f__Metschnikowiaceae(100);g__Metschnikowia(100);g__Metschnikowia_unclassified(100); | Metschnikowia | 13 | Saprotroph | \|Undefined Saprotroph\| | Yeast | NULL | Probable | As nectar-tap_saprotroph (Põlme et al. 2020); Decay sugar-rich substrates (Põlme et al. 2020) | Põlme S, et al. 2020. Fungal Diversity 105:1-16 ((https://doi.org/10.1007/s13225-020-00466-2)) |
| Otu13852 | 1 | k__Fungi(100);p__Ascomycota(100);c__Saccharomycetes(100);o__Saccharomycetales(100);f__Metschnikowiaceae(100);g__Metschnikowia(100);s__Metschnikowia_pulcherrima(100); | Metschnikowia | 13 | Saprotroph | \|Undefined Saprotroph\| | Yeast | NULL | Probable | As nectar-tap_saprotroph (Põlme et al. 2020); Decay sugar-rich substrates (Põlme et al. 2020) | Põlme S, et al. 2020. Fungal Diversity 105:1-16 ((https://doi.org/10.1007/s13225-020-00466-2)) |
| Otu13854 | 1 | k__Fungi(100);p__Ascomycota(100);c__Saccharomycetes(100);o__Saccharomycetales(100);f__Metschnikowiaceae(100);g__Metschnikowia(100);s__Metschnikowia_pulcherrima(100); | Metschnikowia | 13 | Saprotroph | \|Undefined Saprotroph\| | Yeast | NULL | Probable | As nectar-tap_saprotroph (Põlme et al. 2020); Decay sugar-rich substrates (Põlme et al. 2020) | Põlme S, et al. 2020. Fungal Diversity 105:1-16 ((https://doi.org/10.1007/s13225-020-00466-2)) |
| Otu13855 | 1 | k__Fungi(100);p__Ascomycota(100);c__Saccharomycetes(100);o__Saccharomycetales(100);f__Metschnikowiaceae(100);g__Metschnikowia(100);s__Metschnikowia_pulcherrima(100); | Metschnikowia | 13 | Saprotroph | \|Undefined Saprotroph\| | Yeast | NULL | Probable | As nectar-tap_saprotroph (Põlme et al. 2020); Decay sugar-rich substrates (Põlme et al. 2020) | Põlme S, et al. 2020. Fungal Diversity 105:1-16 ((https://doi.org/10.1007/s13225-020-00466-2)) |
| Otu13859 | 1 | k__Fungi(100);p__Ascomycota(100);c__Saccharomycetes(100);o__Saccharomycetales(100);f__Metschnikowiaceae(100);g__Metschnikowia(100);g__Metschnikowia_unclassified(100); | Metschnikowia | 13 | Saprotroph | \|Undefined Saprotroph\| | Yeast | NULL | Probable | As nectar-tap_saprotroph (Põlme et al. 2020); Decay sugar-rich substrates (Põlme et al. 2020) | Põlme S, et al. 2020. Fungal Diversity 105:1-16 ((https://doi.org/10.1007/s13225-020-00466-2)) |
| Otu13860 | 1 | k__Fungi(100);p__Ascomycota(100);c__Saccharomycetes(100);o__Saccharomycetales(100);f__Metschnikowiaceae(100);g__Metschnikowia(100);s__Metschnikowia_pulcherrima(100); | Metschnikowia | 13 | Saprotroph | \|Undefined Saprotroph\| | Yeast | NULL | Probable | As nectar-tap_saprotroph (Põlme et al. 2020); Decay sugar-rich substrates (Põlme et al. 2020) | Põlme S, et al. 2020. Fungal Diversity 105:1-16 ((https://doi.org/10.1007/s13225-020-00466-2)) |
| Otu13861 | 1 | k__Fungi(100);p__Ascomycota(100);c__Saccharomycetes(100);o__Saccharomycetales(100);f__Metschnikowiaceae(100);g__Metschnikowia(100);s__Metschnikowia_pulcherrima(100); | Metschnikowia | 13 | Saprotroph | \|Undefined Saprotroph\| | Yeast | NULL | Probable | As nectar-tap_saprotroph (Põlme et al. 2020); Decay sugar-rich substrates (Põlme et al. 2020) | Põlme S, et al. 2020. Fungal Diversity 105:1-16 ((https://doi.org/10.1007/s13225-020-00466-2)) |
| Otu13862 | 1 | k__Fungi(100);p__Ascomycota(100);c__Saccharomycetes(100);o__Saccharomycetales(100);f__Metschnikowiaceae(100);g__Metschnikowia(100);s__Metschnikowia_pulcherrima(100); | Metschnikowia | 13 | Saprotroph | \|Undefined Saprotroph\| | Yeast | NULL | Probable | As nectar-tap_saprotroph (Põlme et al. 2020); Decay sugar-rich substrates (Põlme et al. 2020) | Põlme S, et al. 2020. Fungal Diversity 105:1-16 ((https://doi.org/10.1007/s13225-020-00466-2)) |
| Otu13863 | 1 | k__Fungi(100);p__Ascomycota(100);c__Saccharomycetes(100);o__Saccharomycetales(100);f__Metschnikowiaceae(100);g__Metschnikowia(100);s__Metschnikowia_pulcherrima(100); | Metschnikowia | 13 | Saprotroph | \|Undefined Saprotroph\| | Yeast | NULL | Probable | As nectar-tap_saprotroph (Põlme et al. 2020); Decay sugar-rich substrates (Põlme et al. 2020) | Põlme S, et al. 2020. Fungal Diversity 105:1-16 ((https://doi.org/10.1007/s13225-020-00466-2)) |
| Otu13864 | 1 | k__Fungi(100);p__Ascomycota(100);c__Saccharomycetes(100);o__Saccharomycetales(100);f__Metschnikowiaceae(100);g__Metschnikowia(100);g__Metschnikowia_unclassified(100); | Metschnikowia | 13 | Saprotroph | \|Undefined Saprotroph\| | Yeast | NULL | Probable | As nectar-tap_saprotroph (Põlme et al. 2020); Decay sugar-rich substrates (Põlme et al. 2020) | Põlme S, et al. 2020. Fungal Diversity 105:1-16 ((https://doi.org/10.1007/s13225-020-00466-2)) |
| Otu13865 | 1 | k__Fungi(100);p__Ascomycota(100);c__Saccharomycetes(100);o__Saccharomycetales(100);f__Metschnikowiaceae(100);g__Metschnikowia(100);g__Metschnikowia_unclassified(100); | Metschnikowia | 13 | Saprotroph | \|Undefined Saprotroph\| | Yeast | NULL | Probable | As nectar-tap_saprotroph (Põlme et al. 2020); Decay sugar-rich substrates (Põlme et al. 2020) | Põlme S, et al. 2020. Fungal Diversity 105:1-16 ((https://doi.org/10.1007/s13225-020-00466-2)) |
| Otu13866 | 1 | k__Fungi(100);p__Ascomycota(100);c__Saccharomycetes(100);o__Saccharomycetales(100);f__Metschnikowiaceae(100);g__Metschnikowia(100);s__Metschnikowia_pulcherrima(100); | Metschnikowia | 13 | Saprotroph | \|Undefined Saprotroph\| | Yeast | NULL | Probable | As nectar-tap_saprotroph (Põlme et al. 2020); Decay sugar-rich substrates (Põlme et al. 2020) | Põlme S, et al. 2020. Fungal Diversity 105:1-16 ((https://doi.org/10.1007/s13225-020-00466-2)) |
| Otu13867 | 1 | k__Fungi(100);p__Ascomycota(100);c__Saccharomycetes(100);o__Saccharomycetales(100);f__Metschnikowiaceae(100);g__Metschnikowia(100);g__Metschnikowia_unclassified(100); | Metschnikowia | 13 | Saprotroph | \|Undefined Saprotroph\| | Yeast | NULL | Probable | As nectar-tap_saprotroph (Põlme et al. 2020); Decay sugar-rich substrates (Põlme et al. 2020) | Põlme S, et al. 2020. Fungal Diversity 105:1-16 ((https://doi.org/10.1007/s13225-020-00466-2)) |
| Otu13868 | 1 | k__Fungi(100);p__Ascomycota(100);c__Saccharomycetes(100);o__Saccharomycetales(100);f__Metschnikowiaceae(100);g__Metschnikowia(100);g__Metschnikowia_unclassified(100); | Metschnikowia | 13 | Saprotroph | \|Undefined Saprotroph\| | Yeast | NULL | Probable | As nectar-tap_saprotroph (Põlme et al. 2020); Decay sugar-rich substrates (Põlme et al. 2020) | Põlme S, et al. 2020. Fungal Diversity 105:1-16 ((https://doi.org/10.1007/s13225-020-00466-2)) |
| Otu13869 | 1 | k__Fungi(100);p__Ascomycota(100);c__Saccharomycetes(100);o__Saccharomycetales(100);f__Metschnikowiaceae(100);g__Metschnikowia(100);s__Metschnikowia_pulcherrima(100); | Metschnikowia | 13 | Saprotroph | \|Undefined Saprotroph\| | Yeast | NULL | Probable | As nectar-tap_saprotroph (Põlme et al. 2020); Decay sugar-rich substrates (Põlme et al. 2020) | Põlme S, et al. 2020. Fungal Diversity 105:1-16 ((https://doi.org/10.1007/s13225-020-00466-2)) |
| Otu13871 | 1 | k__Fungi(100);p__Ascomycota(100);c__Saccharomycetes(100);o__Saccharomycetales(100);f__Metschnikowiaceae(100);g__Metschnikowia(100);g__Metschnikowia_unclassified(100); | Metschnikowia | 13 | Saprotroph | \|Undefined Saprotroph\| | Yeast | NULL | Probable | As nectar-tap_saprotroph (Põlme et al. 2020); Decay sugar-rich substrates (Põlme et al. 2020) | Põlme S, et al. 2020. Fungal Diversity 105:1-16 ((https://doi.org/10.1007/s13225-020-00466-2)) |
| Otu13872 | 1 | k__Fungi(100);p__Ascomycota(100);c__Saccharomycetes(100);o__Saccharomycetales(100);f__Metschnikowiaceae(100);g__Metschnikowia(100);s__Metschnikowia_pulcherrima(100); | Metschnikowia | 13 | Saprotroph | \|Undefined Saprotroph\| | Yeast | NULL | Probable | As nectar-tap_saprotroph (Põlme et al. 2020); Decay sugar-rich substrates (Põlme et al. 2020) | Põlme S, et al. 2020. Fungal Diversity 105:1-16 ((https://doi.org/10.1007/s13225-020-00466-2)) |
| Otu13873 | 1 | k__Fungi(100);p__Ascomycota(100);c__Saccharomycetes(100);o__Saccharomycetales(100);o__Saccharomycetales_unclassified(100);o__Saccharomycetales_unclassified(100);o__Saccharomycetales_unclassified(100); | Saccharomycetales | 7 | Saprotroph | Undefined Saprotroph | Yeast | NULL | Possible | NULL | Sterkenburg E, et al. 2015. New Phytologist 207:1145-1158 ((https://doi.org/10.1111/nph.13426)) |
| Otu13874 | 1 | k__Fungi(100);p__Ascomycota(100);c__Saccharomycetes(100);o__Saccharomycetales(100);f__Metschnikowiaceae(100);g__Metschnikowia(100);s__Metschnikowia_pulcherrima(100); | Metschnikowia | 13 | Saprotroph | \|Undefined Saprotroph\| | Yeast | NULL | Probable | As nectar-tap_saprotroph (Põlme et al. 2020); Decay sugar-rich substrates (Põlme et al. 2020) | Põlme S, et al. 2020. Fungal Diversity 105:1-16 ((https://doi.org/10.1007/s13225-020-00466-2)) |

| | | | | | | | | | | | |
|---|---|---|---|---|---|---|---|---|---|---|---|
| Otu13875 | 1 | k__Fungi(100);p__Ascomycota(100);c__Saccharomycetes(100);o__Saccharomycetales(100);f__Metschnikowiaceae(100);g__Metschnikowia(100);g__Metschnikowia_unclassified(100); | Metschnikowia | 13 | Saprotroph | \|Undefined Saprotroph\| | Yeast | NULL | Probable | As nectar-tap_saprotroph (Põlme et al. 2020); Decay sugar-rich substrates (Põlme et al. 2020) | Põlme S, et al. 2020. Fungal Diversity 105:1-16 ((https://doi.org/10.1007/s13225-020-00466-2)) |
| Otu13876 | 1 | k__Fungi(100);p__Ascomycota(100);c__Saccharomycetes(100);o__Saccharomycetales(100);f__Metschnikowiaceae(100);g__Metschnikowia(100);s__Metschnikowia_pulcherrima(100); | Metschnikowia | 13 | Saprotroph | \|Undefined Saprotroph\| | Yeast | NULL | Probable | As nectar-tap_saprotroph (Põlme et al. 2020); Decay sugar-rich substrates (Põlme et al. 2020) | Põlme S, et al. 2020. Fungal Diversity 105:1-16 ((https://doi.org/10.1007/s13225-020-00466-2)) |
| Otu13878 | 1 | k__Fungi(100);p__Ascomycota(100);c__Saccharomycetes(100);o__Saccharomycetales(100);f__Metschnikowiaceae(100);g__Metschnikowia(100);s__Metschnikowia_pulcherrima(100); | Metschnikowia | 13 | Saprotroph | \|Undefined Saprotroph\| | Yeast | NULL | Probable | As nectar-tap_saprotroph (Põlme et al. 2020); Decay sugar-rich substrates (Põlme et al. 2020) | Põlme S, et al. 2020. Fungal Diversity 105:1-16 ((https://doi.org/10.1007/s13225-020-00466-2)) |
| Otu13879 | 1 | k__Fungi(100);p__Ascomycota(100);c__Saccharomycetes(100);o__Saccharomycetales(100);f__Metschnikowiaceae(100);g__Metschnikowia(100);s__Metschnikowia_pulcherrima(100); | Metschnikowia | 13 | Saprotroph | \|Undefined Saprotroph\| | Yeast | NULL | Probable | As nectar-tap_saprotroph (Põlme et al. 2020); Decay sugar-rich substrates (Põlme et al. 2020) | Põlme S, et al. 2020. Fungal Diversity 105:1-16 ((https://doi.org/10.1007/s13225-020-00466-2)) |
| Otu13880 | 1 | k__Fungi(100);p__Ascomycota(100);c__Saccharomycetes(100);o__Saccharomycetales(100);f__Metschnikowiaceae(100);g__Metschnikowia(100);g__Metschnikowia_unclassified(100); | Metschnikowia | 13 | Saprotroph | \|Undefined Saprotroph\| | Yeast | NULL | Probable | As nectar-tap_saprotroph (Põlme et al. 2020); Decay sugar-rich substrates (Põlme et al. 2020) | Põlme S, et al. 2020. Fungal Diversity 105:1-16 ((https://doi.org/10.1007/s13225-020-00466-2)) |
| Otu13881 | 1 | k__Fungi(100);p__Ascomycota(100);c__Saccharomycetes(100);o__Saccharomycetales(100);f__Metschnikowiaceae(100);g__Metschnikowia(100);s__Metschnikowia_pulcherrima(100); | Metschnikowia | 13 | Saprotroph | \|Undefined Saprotroph\| | Yeast | NULL | Probable | As nectar-tap_saprotroph (Põlme et al. 2020); Decay sugar-rich substrates (Põlme et al. 2020) | Põlme S, et al. 2020. Fungal Diversity 105:1-16 ((https://doi.org/10.1007/s13225-020-00466-2)) |
| Otu13882 | 1 | k__Fungi(100);p__Ascomycota(100);c__Saccharomycetes(100);o__Saccharomycetales(100);f__Metschnikowiaceae(100);g__Metschnikowia(100);s__Metschnikowia_pulcherrima(100); | Metschnikowia | 13 | Saprotroph | \|Undefined Saprotroph\| | Yeast | NULL | Probable | As nectar-tap_saprotroph (Põlme et al. 2020); Decay sugar-rich substrates (Põlme et al. 2020) | Põlme S, et al. 2020. Fungal Diversity 105:1-16 ((https://doi.org/10.1007/s13225-020-00466-2)) |
| Otu13883 | 1 | k__Fungi(100);p__Ascomycota(100);c__Saccharomycetes(100);o__Saccharomycetales(100);f__Metschnikowiaceae(100);g__Metschnikowia(100);s__Metschnikowia_pulcherrima(100); | Metschnikowia | 13 | Saprotroph | \|Undefined Saprotroph\| | Yeast | NULL | Probable | As nectar-tap_saprotroph (Põlme et al. 2020); Decay sugar-rich substrates (Põlme et al. 2020) | Põlme S, et al. 2020. Fungal Diversity 105:1-16 ((https://doi.org/10.1007/s13225-020-00466-2)) |
| Otu13884 | 1 | k__Fungi(100);p__Ascomycota(100);c__Saccharomycetes(100);o__Saccharomycetales(100);f__Metschnikowiaceae(100);g__Metschnikowia(100);s__Metschnikowia_pulcherrima(100); | Metschnikowia | 13 | Saprotroph | \|Undefined Saprotroph\| | Yeast | NULL | Probable | As nectar-tap_saprotroph (Põlme et al. 2020); Decay sugar-rich substrates (Põlme et al. 2020) | Põlme S, et al. 2020. Fungal Diversity 105:1-16 ((https://doi.org/10.1007/s13225-020-00466-2)) |
| Otu13885 | 1 | k__Fungi(100);p__Ascomycota(100);c__Saccharomycetes(100);o__Saccharomycetales(100);f__Metschnikowiaceae(100);g__Metschnikowia(100);s__Metschnikowia_pulcherrima(100); | Metschnikowia | 13 | Saprotroph | \|Undefined Saprotroph\| | Yeast | NULL | Probable | As nectar-tap_saprotroph (Põlme et al. 2020); Decay sugar-rich substrates (Põlme et al. 2020) | Põlme S, et al. 2020. Fungal Diversity 105:1-16 ((https://doi.org/10.1007/s13225-020-00466-2)) |
| Otu13886 | 1 | k__Fungi(100);p__Ascomycota(100);c__Saccharomycetes(100);o__Saccharomycetales(100);f__Metschnikowiaceae(100);g__Metschnikowia(100);s__Metschnikowia_pulcherrima(100); | Metschnikowia | 13 | Saprotroph | \|Undefined Saprotroph\| | Yeast | NULL | Probable | As nectar-tap_saprotroph (Põlme et al. 2020); Decay sugar-rich substrates (Põlme et al. 2020) | Põlme S, et al. 2020. Fungal Diversity 105:1-16 ((https://doi.org/10.1007/s13225-020-00466-2)) |
| Otu13887 | 1 | k__Fungi(100);p__Ascomycota(100);c__Saccharomycetes(100);o__Saccharomycetales(100);f__Metschnikowiaceae(100);g__Metschnikowia(100);s__Metschnikowia_pulcherrima(100); | Metschnikowia | 13 | Saprotroph | \|Undefined Saprotroph\| | Yeast | NULL | Probable | As nectar-tap_saprotroph (Põlme et al. 2020); Decay sugar-rich substrates (Põlme et al. 2020) | Põlme S, et al. 2020. Fungal Diversity 105:1-16 ((https://doi.org/10.1007/s13225-020-00466-2)) |
| Otu13888 | 1 | k__Fungi(100);p__Ascomycota(100);c__Saccharomycetes(100);o__Saccharomycetales(100);f__Metschnikowiaceae(100);g__Metschnikowia(100);s__Metschnikowia_pulcherrima(100); | Metschnikowia | 13 | Saprotroph | \|Undefined Saprotroph\| | Yeast | NULL | Probable | As nectar-tap_saprotroph (Põlme et al. 2020); Decay sugar-rich substrates (Põlme et al. 2020) | Põlme S, et al. 2020. Fungal Diversity 105:1-16 ((https://doi.org/10.1007/s13225-020-00466-2)) |
| Otu13890 | 1 | k__Fungi(100);p__Ascomycota(100);c__Saccharomycetes(100);o__Saccharomycetales(100);f__Metschnikowiaceae(100);g__Metschnikowia(100);s__Metschnikowia_pulcherrima(100); | Metschnikowia | 13 | Saprotroph | \|Undefined Saprotroph\| | Yeast | NULL | Probable | As nectar-tap_saprotroph (Põlme et al. 2020); Decay sugar-rich substrates (Põlme et al. 2020) | Põlme S, et al. 2020. Fungal Diversity 105:1-16 ((https://doi.org/10.1007/s13225-020-00466-2)) |
| Otu13891 | 1 | k__Fungi(100);p__Ascomycota(100);c__Saccharomycetes(100);o__Saccharomycetales(100);f__Metschnikowiaceae(100);g__Metschnikowia(100);g__Metschnikowia_unclassified(100); | Metschnikowia | 13 | Saprotroph | \|Undefined Saprotroph\| | Yeast | NULL | Probable | As nectar-tap_saprotroph (Põlme et al. 2020); Decay sugar-rich substrates (Põlme et al. 2020) | Põlme S, et al. 2020. Fungal Diversity 105:1-16 ((https://doi.org/10.1007/s13225-020-00466-2)) |
| Otu13892 | 1 | k__Fungi(100);p__Ascomycota(100);c__Saccharomycetes(100);o__Saccharomycetales(100);f__Metschnikowiaceae(100);g__Metschnikowia(100);g__Metschnikowia_unclassified(100); | Metschnikowia | 13 | Saprotroph | \|Undefined Saprotroph\| | Yeast | NULL | Probable | As nectar-tap_saprotroph (Põlme et al. 2020); Decay sugar-rich substrates (Põlme et al. 2020) | Põlme S, et al. 2020. Fungal Diversity 105:1-16 ((https://doi.org/10.1007/s13225-020-00466-2)) |
| Otu13893 | 1 | k__Fungi(100);p__Ascomycota(100);c__Saccharomycetes(100);o__Saccharomycetales(100);f__Metschnikowiaceae(100);g__Metschnikowia(100);s__Metschnikowia_pulcherrima(100); | Metschnikowia | 13 | Saprotroph | \|Undefined Saprotroph\| | Yeast | NULL | Probable | As nectar-tap_saprotroph (Põlme et al. 2020); Decay sugar-rich substrates (Põlme et al. 2020) | Põlme S, et al. 2020. Fungal Diversity 105:1-16 ((https://doi.org/10.1007/s13225-020-00466-2)) |
| Otu13895 | 1 | k__Fungi(100);p__Ascomycota(100);c__Saccharomycetes(100);o__Saccharomycetales(100);f__Metschnikowiaceae(100);g__Metschnikowia(100);s__Metschnikowia_pulcherrima(100); | Metschnikowia | 13 | Saprotroph | \|Undefined Saprotroph\| | Yeast | NULL | Probable | As nectar-tap_saprotroph (Põlme et al. 2020); Decay sugar-rich substrates (Põlme et al. 2020) | Põlme S, et al. 2020. Fungal Diversity 105:1-16 ((https://doi.org/10.1007/s13225-020-00466-2)) |
| Otu13896 | 1 | k__Fungi(100);p__Ascomycota(100);c__Saccharomycetes(100);o__Saccharomycetales(100);f__Metschnikowiaceae(100);g__Metschnikowia(100);s__Metschnikowia_pulcherrima(100); | Metschnikowia | 13 | Saprotroph | \|Undefined Saprotroph\| | Yeast | NULL | Probable | As nectar-tap_saprotroph (Põlme et al. 2020); Decay sugar-rich substrates (Põlme et al. 2020) | Põlme S, et al. 2020. Fungal Diversity 105:1-16 ((https://doi.org/10.1007/s13225-020-00466-2)) |
| Otu13899 | 1 | k__Fungi(100);p__Ascomycota(100);c__Saccharomycetes(100);o__Saccharomycetales(100);f__Metschnikowiaceae(100);g__Metschnikowia(100);g__Metschnikowia_unclassified(100); | Metschnikowia | 13 | Saprotroph | \|Undefined Saprotroph\| | Yeast | NULL | Probable | As nectar-tap_saprotroph (Põlme et al. 2020); Decay sugar-rich substrates (Põlme et al. 2020) | Põlme S, et al. 2020. Fungal Diversity 105:1-16 ((https://doi.org/10.1007/s13225-020-00466-2)) |
| Otu13901 | 1 | k__Fungi(100);p__Ascomycota(100);c__Saccharomycetes(100);o__Saccharomycetales(100);f__Metschnikowiaceae(100);g__Metschnikowia(100);s__Metschnikowia_pulcherrima(100); | Metschnikowia | 13 | Saprotroph | \|Undefined Saprotroph\| | Yeast | NULL | Probable | As nectar-tap_saprotroph (Põlme et al. 2020); Decay sugar-rich substrates (Põlme et al. 2020) | Põlme S, et al. 2020. Fungal Diversity 105:1-16 ((https://doi.org/10.1007/s13225-020-00466-2)) |
| Otu13902 | 1 | k__Fungi(100);p__Ascomycota(100);c__Saccharomycetes(100);o__Saccharomycetales(100);f__Metschnikowiaceae(100);g__Metschnikowia(100);s__Metschnikowia_pulcherrima(100); | Metschnikowia | 13 | Saprotroph | \|Undefined Saprotroph\| | Yeast | NULL | Probable | As nectar-tap_saprotroph (Põlme et al. 2020); Decay sugar-rich substrates (Põlme et al. 2020) | Põlme S, et al. 2020. Fungal Diversity 105:1-16 ((https://doi.org/10.1007/s13225-020-00466-2)) |
| Otu13904 | 1 | k__Fungi(100);p__Ascomycota(100);c__Saccharomycetes(100);o__Saccharomycetales(100);f__Metschnikowiaceae(100);g__Metschnikowia(100);s__Metschnikowia_pulcherrima(100); | Metschnikowia | 13 | Saprotroph | \|Undefined Saprotroph\| | Yeast | NULL | Probable | As nectar-tap_saprotroph (Põlme et al. 2020); Decay sugar-rich substrates (Põlme et al. 2020) | Põlme S, et al. 2020. Fungal Diversity 105:1-16 ((https://doi.org/10.1007/s13225-020-00466-2)) |
| Otu13905 | 1 | k__Fungi(100);p__Ascomycota(100);c__Saccharomycetes(100);o__Saccharomycetales(100);f__Metschnikowiaceae(100);g__Metschnikowia(100);g__Metschnikowia_unclassified(100); | Metschnikowia | 13 | Saprotroph | \|Undefined Saprotroph\| | Yeast | NULL | Probable | As nectar-tap_saprotroph (Põlme et al. 2020); Decay sugar-rich substrates (Põlme et al. 2020) | Põlme S, et al. 2020. Fungal Diversity 105:1-16 ((https://doi.org/10.1007/s13225-020-00466-2)) |
| Otu13906 | 1 | k__Fungi(100);p__Ascomycota(100);c__Saccharomycetes(100);o__Saccharomycetales(100);f__Metschnikowiaceae(100);g__Metschnikowia(100);s__Metschnikowia_pulcherrima(100); | Metschnikowia | 13 | Saprotroph | \|Undefined Saprotroph\| | Yeast | NULL | Probable | As nectar-tap_saprotroph (Põlme et al. 2020); Decay sugar-rich substrates (Põlme et al. 2020) | Põlme S, et al. 2020. Fungal Diversity 105:1-16 ((https://doi.org/10.1007/s13225-020-00466-2)) |
| Otu13907 | 1 | k__Fungi(100);p__Ascomycota(100);c__Saccharomycetes(100);o__Saccharomycetales(100);f__Metschnikowiaceae(100);g__Metschnikowia(100);g__Metschnikowia_unclassified(100); | Metschnikowia | 13 | Saprotroph | \|Undefined Saprotroph\| | Yeast | NULL | Probable | As nectar-tap_saprotroph (Põlme et al. 2020); Decay sugar-rich substrates (Põlme et al. 2020) | Põlme S, et al. 2020. Fungal Diversity 105:1-16 ((https://doi.org/10.1007/s13225-020-00466-2)) |
| Otu13908 | 1 | k__Fungi(100);p__Ascomycota(100);c__Saccharomycetes(100);o__Saccharomycetales(100);f__Metschnikowiaceae(100);g__Metschnikowia(100);s__Metschnikowia_pulcherrima(100); | Metschnikowia | 13 | Saprotroph | \|Undefined Saprotroph\| | Yeast | NULL | Probable | As nectar-tap_saprotroph (Põlme et al. 2020); Decay sugar-rich substrates (Põlme et al. 2020) | Põlme S, et al. 2020. Fungal Diversity 105:1-16 ((https://doi.org/10.1007/s13225-020-00466-2)) |
| Otu13909 | 1 | k__Fungi(100);p__Ascomycota(100);c__Saccharomycetes(100);o__Saccharomycetales(100);f__Metschnikowiaceae(100);g__Metschnikowia(100);g__Metschnikowia_unclassified(100); | Metschnikowia | 13 | Saprotroph | \|Undefined Saprotroph\| | Yeast | NULL | Probable | As nectar-tap_saprotroph (Põlme et al. 2020); Decay sugar-rich substrates (Põlme et al. 2020) | Põlme S, et al. 2020. Fungal Diversity 105:1-16 ((https://doi.org/10.1007/s13225-020-00466-2)) |
| Otu13910 | 1 | k__Fungi(100);p__Ascomycota(100);c__Saccharomycetes(100);o__Saccharomycetales(100);f__Metschnikowiaceae(100);g__Metschnikowia(100);s__Metschnikowia_pulcherrima(100); | Metschnikowia | 13 | Saprotroph | \|Undefined Saprotroph\| | Yeast | NULL | Probable | As nectar-tap_saprotroph (Põlme et al. 2020); Decay sugar-rich substrates (Põlme et al. 2020) | Põlme S, et al. 2020. Fungal Diversity 105:1-16 ((https://doi.org/10.1007/s13225-020-00466-2)) |
| Otu13911 | 1 | k__Fungi(100);p__Ascomycota(100);c__Saccharomycetes(100);o__Saccharomycetales(100);f__Metschnikowiaceae(100);g__Metschnikowia(100);g__Metschnikowia_unclassified(100); | Metschnikowia | 13 | Saprotroph | \|Undefined Saprotroph\| | Yeast | NULL | Probable | As nectar-tap_saprotroph (Põlme et al. 2020); Decay sugar-rich substrates (Põlme et al. 2020) | Põlme S, et al. 2020. Fungal Diversity 105:1-16 ((https://doi.org/10.1007/s13225-020-00466-2)) |
| Otu13912 | 1 | k__Fungi(100);p__Ascomycota(100);c__Saccharomycetes(100);o__Saccharomycetales(100);f__Metschnikowiaceae(100);g__Metschnikowia(100);g__Metschnikowia_unclassified(100); | Metschnikowia | 13 | Saprotroph | \|Undefined Saprotroph\| | Yeast | NULL | Probable | As nectar-tap_saprotroph (Põlme et al. 2020); Decay sugar-rich substrates (Põlme et al. 2020) | Põlme S, et al. 2020. Fungal Diversity 105:1-16 ((https://doi.org/10.1007/s13225-020-00466-2)) |
| Otu13913 | 1 | k__Fungi(100);p__Ascomycota(100);c__Saccharomycetes(100);o__Saccharomycetales(100);f__Saccharomycetales_family_Incertae_sedis(100);g__Candida(100);s__Candida_cretensis(100); | Candida | 13 | Pathotroph-Saprotroph-Symbiotroph | Animal Pathogen-Endophyte-Endosymbiont-Epiphyte-\|Undefined Saprotroph\| | Dimorphic Yeast-Yeast | NULL | Probable | As nectar-tap_saprotroph (Põlme et al. 2020); Decay sugar-rich substrates (Põlme et al. 2020); This asexual genus is polyphyletic and contain species that function very differently depend on what they are. Caution should be taken when interpreting guilds for Candida species. | Manolakaki D, et al. 2010. Virulence. 1: 367-375 ((https://doi.org/10.4161/viru.1.5.12796)); Mekbib SB, et al. 2011. Tropical Plant Pathology 36:233-240 ((https://doi.org/10.1590/S1982-56762011000400004)); Põlme S, et al. 2020. Fungal Diversity 105:1-16 ((https://doi.org/10.1007/s13225-020-00466-2)) |
| Otu13915 | 1 | k__Fungi(100);p__Ascomycota(100);c__Saccharomycetes(100);o__Saccharomycetales(100);f__Metschnikowiaceae(100);g__Metschnikowia(100);s__Metschnikowia_pulcherrima(100); | Metschnikowia | 13 | Saprotroph | \|Undefined Saprotroph\| | Yeast | NULL | Probable | As nectar-tap_saprotroph (Põlme et al. 2020); Decay sugar-rich substrates (Põlme et al. 2020) | Põlme S, et al. 2020. Fungal Diversity 105:1-16 ((https://doi.org/10.1007/s13225-020-00466-2)) |
| Otu13916 | 1 | k__Fungi(100);p__Ascomycota(100);c__Saccharomycetes(100);o__Saccharomycetales(100);f__Metschnikowiaceae(100);g__Metschnikowia(100);s__Metschnikowia_pulcherrima(100); | Metschnikowia | 13 | Saprotroph | \|Undefined Saprotroph\| | Yeast | NULL | Probable | As nectar-tap_saprotroph (Põlme et al. 2020); Decay sugar-rich substrates (Põlme et al. 2020) | Põlme S, et al. 2020. Fungal Diversity 105:1-16 ((https://doi.org/10.1007/s13225-020-00466-2)) |
| Otu13918 | 1 | k__Fungi(100);p__Ascomycota(100);c__Saccharomycetes(100);o__Saccharomycetales(100);f__Metschnikowiaceae(100);g__Metschnikowia(100);s__Metschnikowia_pulcherrima(100); | Metschnikowia | 13 | Saprotroph | \|Undefined Saprotroph\| | Yeast | NULL | Probable | As nectar-tap_saprotroph (Põlme et al. 2020); Decay sugar-rich substrates (Põlme et al. 2020) | Põlme S, et al. 2020. Fungal Diversity 105:1-16 ((https://doi.org/10.1007/s13225-020-00466-2)) |
| Otu13919 | 1 | k__Fungi(100);p__Ascomycota(100);c__Saccharomycetes(100);o__Saccharomycetales(100);f__Metschnikowiaceae(100);g__Metschnikowia(100);s__Metschnikowia_pulcherrima(100); | Metschnikowia | 13 | Saprotroph | \|Undefined Saprotroph\| | Yeast | NULL | Probable | As nectar-tap_saprotroph (Põlme et al. 2020); Decay sugar-rich substrates (Põlme et al. 2020) | Põlme S, et al. 2020. Fungal Diversity 105:1-16 ((https://doi.org/10.1007/s13225-020-00466-2)) |
| Otu13920 | 1 | k__Fungi(100);p__Ascomycota(100);c__Saccharomycetes(100);o__Saccharomycetales(100);f__Metschnikowiaceae(100);g__Metschnikowia(100);s__Metschnikowia_pulcherrima(100); | Metschnikowia | 13 | Saprotroph | \|Undefined Saprotroph\| | Yeast | NULL | Probable | As nectar-tap_saprotroph (Põlme et al. 2020); Decay sugar-rich substrates (Põlme et al. 2020) | Põlme S, et al. 2020. Fungal Diversity 105:1-16 ((https://doi.org/10.1007/s13225-020-00466-2)) |
| Otu13921 | 1 | k__Fungi(100);p__Ascomycota(100);c__Saccharomycetes(100);o__Saccharomycetales(100);f__Metschnikowiaceae(100);g__Metschnikowia(100);s__Metschnikowia_pulcherrima(100); | Metschnikowia | 13 | Saprotroph | \|Undefined Saprotroph\| | Yeast | NULL | Probable | As nectar-tap_saprotroph (Põlme et al. 2020); Decay sugar-rich substrates (Põlme et al. 2020) | Põlme S, et al. 2020. Fungal Diversity 105:1-16 ((https://doi.org/10.1007/s13225-020-00466-2)) |
| Otu13925 | 1 | k__Fungi(100);p__Ascomycota(100);c__Saccharomycetes(100);o__Saccharomycetales(100);f__Metschnikowiaceae(100);g__Metschnikowia(100);s__Metschnikowia_pulcherrima(100); | Metschnikowia | 13 | Saprotroph | \|Undefined Saprotroph\| | Yeast | NULL | Probable | As nectar-tap_saprotroph (Põlme et al. 2020); Decay sugar-rich substrates (Põlme et al. 2020) | Põlme S, et al. 2020. Fungal Diversity 105:1-16 ((https://doi.org/10.1007/s13225-020-00466-2)) |
| Otu13926 | 1 | k__Fungi(100);p__Ascomycota(100);c__Saccharomycetes(100);o__Saccharomycetales(100);f__Metschnikowiaceae(100);g__Metschnikowia(100);s__Metschnikowia_pulcherrima(100); | Metschnikowia | 13 | Saprotroph | \|Undefined Saprotroph\| | Yeast | NULL | Probable | As nectar-tap_saprotroph (Põlme et al. 2020); Decay sugar-rich substrates (Põlme et al. 2020) | Põlme S, et al. 2020. Fungal Diversity 105:1-16 ((https://doi.org/10.1007/s13225-020-00466-2)) |
| Otu13927 | 1 | k__Fungi(100);p__Ascomycota(100);c__Saccharomycetes(100);o__Saccharomycetales(100);f__Metschnikowiaceae(100);g__Metschnikowia(100);s__Metschnikowia_pulcherrima(100); | Metschnikowia | 13 | Saprotroph | \|Undefined Saprotroph\| | Yeast | NULL | Probable | As nectar-tap_saprotroph (Põlme et al. 2020); Decay sugar-rich substrates (Põlme et al. 2020) | Põlme S, et al. 2020. Fungal Diversity 105:1-16 ((https://doi.org/10.1007/s13225-020-00466-2)) |
| Otu13930 | 1 | k__Fungi(100);p__Ascomycota(100);c__Saccharomycetes(100);o__Saccharomycetales(100);f__Metschnikowiaceae(100);g__Metschnikowia(100);s__Metschnikowia_pulcherrima(100); | Metschnikowia | 13 | Saprotroph | \|Undefined Saprotroph\| | Yeast | NULL | Probable | As nectar-tap_saprotroph (Põlme et al. 2020); Decay sugar-rich substrates (Põlme et al. 2020) | Põlme S, et al. 2020. Fungal Diversity 105:1-16 ((https://doi.org/10.1007/s13225-020-00466-2)) |
| Otu13931 | 1 | k__Fungi(100);p__Ascomycota(100);c__Saccharomycetes(100);o__Saccharomycetales(100);f__Metschnikowiaceae(100);g__Metschnikowia(100);s__Metschnikowia_pulcherrima(100); | Metschnikowia | 13 | Saprotroph | \|Undefined Saprotroph\| | Yeast | NULL | Probable | As nectar-tap_saprotroph (Põlme et al. 2020); Decay sugar-rich substrates (Põlme et al. 2020) | Põlme S, et al. 2020. Fungal Diversity 105:1-16 ((https://doi.org/10.1007/s13225-020-00466-2)) |
| Otu13933 | 1 | k__Fungi(100);p__Ascomycota(100);c__Saccharomycetes(100);o__Saccharomycetales(100);f__Metschnikowiaceae(100);g__Metschnikowia(100);s__Metschnikowia_pulcherrima(100); | Metschnikowia | 13 | Saprotroph | \|Undefined Saprotroph\| | Yeast | NULL | Probable | As nectar-tap_saprotroph (Põlme et al. 2020); Decay sugar-rich substrates (Põlme et al. 2020) | Põlme S, et al. 2020. Fungal Diversity 105:1-16 ((https://doi.org/10.1007/s13225-020-00466-2)) |
| Otu13934 | 1 | k__Fungi(100);p__Ascomycota(100);c__Saccharomycetes(100);o__Saccharomycetales(100);f__Metschnikowiaceae(100);g__Metschnikowia(100);g__Metschnikowia_unclassified(100); | Metschnikowia | 13 | Saprotroph | \|Undefined Saprotroph\| | Yeast | NULL | Probable | As nectar-tap_saprotroph (Põlme et al. 2020); Decay sugar-rich substrates (Põlme et al. 2020) | Põlme S, et al. 2020. Fungal Diversity 105:1-16 ((https://doi.org/10.1007/s13225-020-00466-2)) |
| Otu13935 | 1 | k__Fungi(100);p__Ascomycota(100);c__Saccharomycetes(100);o__Saccharomycetales(100);f__Metschnikowiaceae(100);g__Metschnikowia(100);s__Metschnikowia_pulcherrima(100); | Metschnikowia | 13 | Saprotroph | \|Undefined Saprotroph\| | Yeast | NULL | Probable | As nectar-tap_saprotroph (Põlme et al. 2020); Decay sugar-rich substrates (Põlme et al. 2020) | Põlme S, et al. 2020. Fungal Diversity 105:1-16 ((https://doi.org/10.1007/s13225-020-00466-2)) |
| Otu13936 | 1 | k__Fungi(100);p__Ascomycota(100);c__Saccharomycetes(100);o__Saccharomycetales(100);f__Metschnikowiaceae(100);g__Metschnikowia(100);s__Metschnikowia_pulcherrima(100); | Metschnikowia | 13 | Saprotroph | \|Undefined Saprotroph\| | Yeast | NULL | Probable | As nectar-tap_saprotroph (Põlme et al. 2020); Decay sugar-rich substrates (Põlme et al. 2020) | Põlme S, et al. 2020. Fungal Diversity 105:1-16 ((https://doi.org/10.1007/s13225-020-00466-2)) |
| Otu13937 | 1 | k__Fungi(100);p__Ascomycota(100);c__Saccharomycetes(100);o__Saccharomycetales(100);f__Metschnikowiaceae(100);g__Metschnikowia(100);s__Metschnikowia_pulcherrima(100); | Metschnikowia | 13 | Saprotroph | \|Undefined Saprotroph\| | Yeast | NULL | Probable | As nectar-tap_saprotroph (Põlme et al. 2020); Decay sugar-rich substrates (Põlme et al. 2020) | Põlme S, et al. 2020. Fungal Diversity 105:1-16 ((https://doi.org/10.1007/s13225-020-00466-2)) |
| Otu13938 | 1 | k__Fungi(100);p__Ascomycota(100);c__Saccharomycetes(100);o__Saccharomycetales(100);f__Metschnikowiaceae(100);g__Metschnikowia(100);s__Metschnikowia_pulcherrima(100); | Metschnikowia | 13 | Saprotroph | \|Undefined Saprotroph\| | Yeast | NULL | Probable | As nectar-tap_saprotroph (Põlme et al. 2020); Decay sugar-rich substrates (Põlme et al. 2020) | Põlme S, et al. 2020. Fungal Diversity 105:1-16 ((https://doi.org/10.1007/s13225-020-00466-2)) |
| Otu13939 | 1 | k__Fungi(100);p__Ascomycota(100);c__Saccharomycetes(100);o__Saccharomycetales(100);f__Metschnikowiaceae(100);g__Metschnikowia(100);s__Metschnikowia_pulcherrima(100); | Metschnikowia | 13 | Saprotroph | \|Undefined Saprotroph\| | Yeast | NULL | Probable | As nectar-tap_saprotroph (Põlme et al. 2020); Decay sugar-rich substrates (Põlme et al. 2020) | Põlme S, et al. 2020. Fungal Diversity 105:1-16 ((https://doi.org/10.1007/s13225-020-00466-2)) |
| Otu13940 | 1 | k__Fungi(100);p__Ascomycota(100);c__Saccharomycetes(100);o__Saccharomycetales(100);f__Metschnikowiaceae(100);g__Metschnikowia(100);s__Metschnikowia_pulcherrima(100); | Metschnikowia | 13 | Saprotroph | \|Undefined Saprotroph\| | Yeast | NULL | Probable | As nectar-tap_saprotroph (Põlme et al. 2020); Decay sugar-rich substrates (Põlme et al. 2020) | Põlme S, et al. 2020. Fungal Diversity 105:1-16 ((https://doi.org/10.1007/s13225-020-00466-2)) |
| Otu13941 | 1 | k__Fungi(100);p__Ascomycota(100);c__Saccharomycetes(100);o__Saccharomycetales(100);f__Metschnikowiaceae(100);g__Metschnikowia(100);s__Metschnikowia_pulcherrima(100); | Metschnikowia | 13 | Saprotroph | \|Undefined Saprotroph\| | Yeast | NULL | Probable | As nectar-tap_saprotroph (Põlme et al. 2020); Decay sugar-rich substrates (Põlme et al. 2020) | Põlme S, et al. 2020. Fungal Diversity 105:1-16 ((https://doi.org/10.1007/s13225-020-00466-2)) |
| Otu13942 | 1 | k__Fungi(100);p__Ascomycota(100);c__Saccharomycetes(100);o__Saccharomycetales(100);f__Metschnikowiaceae(100);g__Metschnikowia(100);g__Metschnikowia_unclassified(100); | Metschnikowia | 13 | Saprotroph | \|Undefined Saprotroph\| | Yeast | NULL | Probable | As nectar-tap_saprotroph (Põlme et al. 2020); Decay sugar-rich substrates (Põlme et al. 2020) | Põlme S, et al. 2020. Fungal Diversity 105:1-16 ((https://doi.org/10.1007/s13225-020-00466-2)) |
| Otu13944 | 1 | k__Fungi(100);p__Ascomycota(100);c__Saccharomycetes(100);o__Saccharomycetales(100);f__Metschnikowiaceae(100);g__Metschnikowia(100);g__Metschnikowia_unclassified(100); | Metschnikowia | 13 | Saprotroph | \|Undefined Saprotroph\| | Yeast | NULL | Probable | As nectar-tap_saprotroph (Põlme et al. 2020); Decay sugar-rich substrates (Põlme et al. 2020) | Põlme S, et al. 2020. Fungal Diversity 105:1-16 ((https://doi.org/10.1007/s13225-020-00466-2)) |
| Otu13945 | 1 | k__Fungi(100);p__Ascomycota(100);c__Saccharomycetes(100);o__Saccharomycetales(100);f__Metschnikowiaceae(100);g__Metschnikowia(100);s__Metschnikowia_pulcherrima(100); | Metschnikowia | 13 | Saprotroph | \|Undefined Saprotroph\| | Yeast | NULL | Probable | As nectar-tap_saprotroph (Põlme et al. 2020); Decay sugar-rich substrates (Põlme et al. 2020) | Põlme S, et al. 2020. Fungal Diversity 105:1-16 ((https://doi.org/10.1007/s13225-020-00466-2)) |
| Otu13947 | 1 | k__Fungi(100);p__Ascomycota(100);c__Saccharomycetes(100);o__Saccharomycetales(100);f__Metschnikowiaceae(100);g__Metschnikowia(100);s__Metschnikowia_pulcherrima(100); | Metschnikowia | 13 | Saprotroph | \|Undefined Saprotroph\| | Yeast | NULL | Probable | As nectar-tap_saprotroph (Põlme et al. 2020); Decay sugar-rich substrates (Põlme et al. 2020) | Põlme S, et al. 2020. Fungal Diversity 105:1-16 ((https://doi.org/10.1007/s13225-020-00466-2)) |
| Otu13948 | 1 | k__Fungi(100);p__Ascomycota(100);c__Saccharomycetes(100);o__Saccharomycetales(100);f__Metschnikowiaceae(100);g__Metschnikowia(100);g__Metschnikowia_unclassified(100); | Metschnikowia | 13 | Saprotroph | \|Undefined Saprotroph\| | Yeast | NULL | Probable | As nectar-tap_saprotroph (Põlme et al. 2020); Decay sugar-rich substrates (Põlme et al. 2020) | Põlme S, et al. 2020. Fungal Diversity 105:1-16 ((https://doi.org/10.1007/s13225-020-00466-2)) |
| Otu13949 | 1 | k__Fungi(100);p__Ascomycota(100);c__Saccharomycetes(100);o__Saccharomycetales(100);f__Metschnikowiaceae(100);g__Metschnikowia(100);s__Metschnikowia_pulcherrima(100); | Metschnikowia | 13 | Saprotroph | \|Undefined Saprotroph\| | Yeast | NULL | Probable | As nectar-tap_saprotroph (Põlme et al. 2020); Decay sugar-rich substrates (Põlme et al. 2020) | Põlme S, et al. 2020. Fungal Diversity 105:1-16 ((https://doi.org/10.1007/s13225-020-00466-2)) |
| Otu13950 | 1 | k__Fungi(100);p__Ascomycota(100);c__Saccharomycetes(100);o__Saccharomycetales(100);f__Metschnikowiaceae(100);g__Metschnikowia(100);s__Metschnikowia_pulcherrima(100); | Metschnikowia | 13 | Saprotroph | \|Undefined Saprotroph\| | Yeast | NULL | Probable | As nectar-tap_saprotroph (Põlme et al. 2020); Decay sugar-rich substrates (Põlme et al. 2020) | Põlme S, et al. 2020. Fungal Diversity 105:1-16 ((https://doi.org/10.1007/s13225-020-00466-2)) |
| Otu13956 | 1 | k__Fungi(100);p__Ascomycota(100);c__Saccharomycetes(100);o__Saccharomycetales(100);f__Metschnikowiaceae(100);g__Metschnikowia(100);g__Metschnikowia_unclassified(100); | Metschnikowia | 13 | Saprotroph | \|Undefined Saprotroph\| | Yeast | NULL | Probable | As nectar-tap_saprotroph (Põlme et al. 2020); Decay sugar-rich substrates (Põlme et al. 2020) | Põlme S, et al. 2020. Fungal Diversity 105:1-16 ((https://doi.org/10.1007/s13225-020-00466-2)) |
| Otu13957 | 1 | k__Fungi(100);p__Ascomycota(100);c__Saccharomycetes(100);o__Saccharomycetales(100);f__Metschnikowiaceae(100);g__Metschnikowia(100);s__Metschnikowia_pulcherrima(100); | Metschnikowia | 13 | Saprotroph | \|Undefined Saprotroph\| | Yeast | NULL | Probable | As nectar-tap_saprotroph (Põlme et al. 2020); Decay sugar-rich substrates (Põlme et al. 2020) | Põlme S, et al. 2020. Fungal Diversity 105:1-16 ((https://doi.org/10.1007/s13225-020-00466-2)) |
| Otu13959 | 1 | k__Fungi(100);p__Ascomycota(100);c__Saccharomycetes(100);o__Saccharomycetales(100);f__Metschnikowiaceae(100);g__Metschnikowia(100);s__Metschnikowia_pulcherrima(100); | Metschnikowia | 13 | Saprotroph | \|Undefined Saprotroph\| | Yeast | NULL | Probable | As nectar-tap_saprotroph (Põlme et al. 2020); Decay sugar-rich substrates (Põlme et al. 2020) | Põlme S, et al. 2020. Fungal Diversity 105:1-16 ((https://doi.org/10.1007/s13225-020-00466-2)) |
| Otu13965 | 1 | k__Fungi(100);p__Ascomycota(100);c__Saccharomycetes(100);o__Saccharomycetales(100);f__Metschnikowiaceae(100);g__Metschnikowia(100);s__Metschnikowia_pulcherrima(100); | Metschnikowia | 13 | Saprotroph | \|Undefined Saprotroph\| | Yeast | NULL | Probable | As nectar-tap_saprotroph (Põlme et al. 2020); Decay sugar-rich substrates (Põlme et al. 2020) | Põlme S, et al. 2020. Fungal Diversity 105:1-16 ((https://doi.org/10.1007/s13225-020-00466-2)) |
| Otu13967 | 1 | k__Fungi(100);p__Ascomycota(100);c__Saccharomycetes(100);o__Saccharomycetales(100);o__Saccharomycetales_unclassified(100);o__Saccharomycetales_unclassified(100);o__Saccharomycetales_unclassified(100); | Saccharomycetales | 7 | Saprotroph | Undefined Saprotroph | Yeast | NULL | Possible | NULL | Sterkenburg E, et al. 2015. New Phytologist 207:1145-1158 ((https://doi.org/10.1111/nph.13426)) |
| Otu13970 | 1 | k__Fungi(100);p__Ascomycota(100);c__Saccharomycetes(100);o__Saccharomycetales(100);f__Metschnikowiaceae(100);g__Metschnikowia(100);s__Metschnikowia_pulcherrima(100); | Metschnikowia | 13 | Saprotroph | \|Undefined Saprotroph\| | Yeast | NULL | Probable | As nectar-tap_saprotroph (Põlme et al. 2020); Decay sugar-rich substrates (Põlme et al. 2020) | Põlme S, et al. 2020. Fungal Diversity 105:1-16 ((https://doi.org/10.1007/s13225-020-00466-2)) |
| Otu13974 | 1 | k__Fungi(100);p__Ascomycota(100);c__Saccharomycetes(100);o__Saccharomycetales(100);f__Saccharomycetales_family_Incertae_sedis(100);f__Saccharomycetales_family_Incertae_sedis_unclassified(100);f__Saccharomycetales_family_Incertae_sedis_unclassified(100); | Saccharomycetales | 7 | Saprotroph | Undefined Saprotroph | Yeast | NULL | Possible | NULL | Sterkenburg E, et al. 2015. New Phytologist 207:1145-1158 ((https://doi.org/10.1111/nph.13426)) |
| Otu13976 | 1 | k__Fungi(100);p__Ascomycota(100);c__Saccharomycetes(100);o__Saccharomycetales(100);f__Metschnikowiaceae(100);g__Metschnikowia(100);s__Metschnikowia_pulcherrima(100); | Metschnikowia | 13 | Saprotroph | \|Undefined Saprotroph\| | Yeast | NULL | Probable | As nectar-tap_saprotroph (Põlme et al. 2020); Decay sugar-rich substrates (Põlme et al. 2020) | Põlme S, et al. 2020. Fungal Diversity 105:1-16 ((https://doi.org/10.1007/s13225-020-00466-2)) |
| Otu13978 | 1 | k__Fungi(100);p__Ascomycota(100);c__Saccharomycetes(100);o__Saccharomycetales(100);f__Metschnikowiaceae(100);g__Metschnikowia(100);s__Metschnikowia_pulcherrima(100); | Metschnikowia | 13 | Saprotroph | \|Undefined Saprotroph\| | Yeast | NULL | Probable | As nectar-tap_saprotroph (Põlme et al. 2020); Decay sugar-rich substrates (Põlme et al. 2020) | Põlme S, et al. 2020. Fungal Diversity 105:1-16 ((https://doi.org/10.1007/s13225-020-00466-2)) |
| Otu13979 | 1 | k__Fungi(100);p__Ascomycota(100);c__Saccharomycetes(100);o__Saccharomycetales(100);f__Saccharomycetales_family_Incertae_sedis(100);g__Candida(100);g__Candida_unclassified(100); | Candida | 13 | Pathotroph-Saprotroph-Symbiotroph | Animal Pathogen-Endophyte-Endosymbiont-Epiphyte-\|Undefined Saprotroph\| | Dimorphic Yeast-Yeast | NULL | Probable | As nectar-tap_saprotroph (Põlme et al. 2020); Decay sugar-rich substrates (Põlme et al. 2020); This asexual genus is polyphyletic and contain species that function very differently depend on what they are. Caution should be taken when interpreting guilds for Candida species. | Manolakaki D, et al. 2010. Virulence. 1: 367-375 ((https://doi.org/10.4161/viru.1.5.12796)); Mekbib SB, et al. 2011. Tropical Plant Pathology 36:233-240 ((https://doi.org/10.1590/S1982-56762011000400004)); Põlme S, et al. 2020. Fungal Diversity 105:1-16 ((https://doi.org/10.1007/s13225-020-00466-2)) |
| Otu13980 | 1 | k__Fungi(100);p__Ascomycota(100);c__Saccharomycetes(100);o__Saccharomycetales(100);o__Saccharomycetales_unclassified(100);o__Saccharomycetales_unclassified(100);o__Saccharomycetales_unclassified(100); | Saccharomycetales | 7 | Saprotroph | Undefined Saprotroph | Yeast | NULL | Possible | NULL | Sterkenburg E, et al. 2015. New Phytologist 207:1145-1158 ((https://doi.org/10.1111/nph.13426)) |
| Otu13982 | 1 | k__Fungi(100);p__Ascomycota(100);c__Saccharomycetes(100);o__Saccharomycetales(100);f__Metschnikowiaceae(100);g__Metschnikowia(100);g__Metschnikowia_unclassified(100); | Metschnikowia | 13 | Saprotroph | \|Undefined Saprotroph\| | Yeast | NULL | Probable | As nectar-tap_saprotroph (Põlme et al. 2020); Decay sugar-rich substrates (Põlme et al. 2020) | Põlme S, et al. 2020. Fungal Diversity 105:1-16 ((https://doi.org/10.1007/s13225-020-00466-2)) |
| Otu13985 | 1 | k__Fungi(100);p__Ascomycota(100);c__Saccharomycetes(100);o__Saccharomycetales(100);f__Metschnikowiaceae(100);g__Metschnikowia(100);s__Metschnikowia_pulcherrima(100); | Metschnikowia | 13 | Saprotroph | \|Undefined Saprotroph\| | Yeast | NULL | Probable | As nectar-tap_saprotroph (Põlme et al. 2020); Decay sugar-rich substrates (Põlme et al. 2020) | Põlme S, et al. 2020. Fungal Diversity 105:1-16 ((https://doi.org/10.1007/s13225-020-00466-2)) |
| Otu13986 | 1 | k__Fungi(100);p__Ascomycota(100);c__Saccharomycetes(100);o__Saccharomycetales(100);f__Metschnikowiaceae(100);g__Metschnikowia(100);s__Metschnikowia_pulcherrima(100); | Metschnikowia | 13 | Saprotroph | \|Undefined Saprotroph\| | Yeast | NULL | Probable | As nectar-tap_saprotroph (Põlme et al. 2020); Decay sugar-rich substrates (Põlme et al. 2020) | Põlme S, et al. 2020. Fungal Diversity 105:1-16 ((https://doi.org/10.1007/s13225-020-00466-2)) |
| Otu13987 | 1 | k__Fungi(100);p__Ascomycota(100);c__Saccharomycetes(100);o__Saccharomycetales(100);f__Saccharomycetales_family_Incertae_sedis(100);g__Candida(100);s__Candida_cretensis(100); | Candida | 13 | Pathotroph-Saprotroph-Symbiotroph | Animal Pathogen-Endophyte-Endosymbiont-Epiphyte-\|Undefined Saprotroph\| | Dimorphic Yeast-Yeast | NULL | Probable | As nectar-tap_saprotroph (Põlme et al. 2020); Decay sugar-rich substrates (Põlme et al. 2020); This asexual genus is polyphyletic and contain species that function very differently depend on what they are. Caution should be taken when interpreting guilds for Candida species. | Manolakaki D, et al. 2010. Virulence. 1: 367-375 ((https://doi.org/10.4161/viru.1.5.12796)); Mekbib SB, et al. 2011. Tropical Plant Pathology 36:233-240 ((https://doi.org/10.1590/S1982-56762011000400004)); Põlme S, et al. 2020. Fungal Diversity 105:1-16 ((https://doi.org/10.1007/s13225-020-00466-2)) |
| Otu13988 | 1 | k__Fungi(100);p__Ascomycota(100);c__Saccharomycetes(100);o__Saccharomycetales(100);f__Metschnikowiaceae(100);g__Metschnikowia(100);s__Metschnikowia_pulcherrima(100); | Metschnikowia | 13 | Saprotroph | \|Undefined Saprotroph\| | Yeast | NULL | Probable | As nectar-tap_saprotroph (Põlme et al. 2020); Decay sugar-rich substrates (Põlme et al. 2020) | Põlme S, et al. 2020. Fungal Diversity 105:1-16 ((https://doi.org/10.1007/s13225-020-00466-2)) |
| Otu13991 | 1 | k__Fungi(100);p__Ascomycota(100);c__Saccharomycetes(100);o__Saccharomycetales(100);f__Metschnikowiaceae(100);g__Metschnikowia(100);s__Metschnikowia_pulcherrima(100); | Metschnikowia | 13 | Saprotroph | \|Undefined Saprotroph\| | Yeast | NULL | Probable | As nectar-tap_saprotroph (Põlme et al. 2020); Decay sugar-rich substrates (Põlme et al. 2020) | Põlme S, et al. 2020. Fungal Diversity 105:1-16 ((https://doi.org/10.1007/s13225-020-00466-2)) |
| Otu13996 | 1 | k__Fungi(100);p__Ascomycota(100);c__Saccharomycetes(100);o__Saccharomycetales(100);f__Metschnikowiaceae(100);g__Metschnikowia(100);s__Metschnikowia_pulcherrima(100); | Metschnikowia | 13 | Saprotroph | \|Undefined Saprotroph\| | Yeast | NULL | Probable | As nectar-tap_saprotroph (Põlme et al. 2020); Decay sugar-rich substrates (Põlme et al. 2020) | Põlme S, et al. 2020. Fungal Diversity 105:1-16 ((https://doi.org/10.1007/s13225-020-00466-2)) |
| Otu13997 | 1 | k__Fungi(100);p__Ascomycota(100);c__Saccharomycetes(100);o__Saccharomycetales(100);f__Metschnikowiaceae(100);g__Metschnikowia(100);s__Metschnikowia_pulcherrima(100); | Metschnikowia | 13 | Saprotroph | \|Undefined Saprotroph\| | Yeast | NULL | Probable | As nectar-tap_saprotroph (Põlme et al. 2020); Decay sugar-rich substrates (Põlme et al. 2020) | Põlme S, et al. 2020. Fungal Diversity 105:1-16 ((https://doi.org/10.1007/s13225-020-00466-2)) |
| Otu13998 | 1 | k__Fungi(100);p__Ascomycota(100);c__Saccharomycetes(100);o__Saccharomycetales(100);f__Metschnikowiaceae(100);g__Metschnikowia(100);s__Metschnikowia_pulcherrima(100); | Metschnikowia | 13 | Saprotroph | \|Undefined Saprotroph\| | Yeast | NULL | Probable | As nectar-tap_saprotroph (Põlme et al. 2020); Decay sugar-rich substrates (Põlme et al. 2020) | Põlme S, et al. 2020. Fungal Diversity 105:1-16 ((https://doi.org/10.1007/s13225-020-00466-2)) |
| Otu13999 | 1 | k__Fungi(100);p__Ascomycota(100);c__Saccharomycetes(100);o__Saccharomycetales(100);f__Metschnikowiaceae(100);g__Metschnikowia(100);s__Metschnikowia_pulcherrima(100); | Metschnikowia | 13 | Saprotroph | \|Undefined Saprotroph\| | Yeast | NULL | Probable | As nectar-tap_saprotroph (Põlme et al. 2020); Decay sugar-rich substrates (Põlme et al. 2020) | Põlme S, et al. 2020. Fungal Diversity 105:1-16 ((https://doi.org/10.1007/s13225-020-00466-2)) |
| Otu14002 | 1 | k__Fungi(100);p__Ascomycota(100);c__Saccharomycetes(100);o__Saccharomycetales(100);f__Metschnikowiaceae(100);g__Metschnikowia(100);g__Metschnikowia_unclassified(100); | Metschnikowia | 13 | Saprotroph | \|Undefined Saprotroph\| | Yeast | NULL | Probable | As nectar-tap_saprotroph (Põlme et al. 2020); Decay sugar-rich substrates (Põlme et al. 2020) | Põlme S, et al. 2020. Fungal Diversity 105:1-16 ((https://doi.org/10.1007/s13225-020-00466-2)) |
| Otu14003 | 1 | k__Fungi(100);p__Ascomycota(100);c__Saccharomycetes(100);o__Saccharomycetales(100);f__Metschnikowiaceae(100);g__Metschnikowia(100);s__Metschnikowia_pulcherrima(100); | Metschnikowia | 13 | Saprotroph | \|Undefined Saprotroph\| | Yeast | NULL | Probable | As nectar-tap_saprotroph (Põlme et al. 2020); Decay sugar-rich substrates (Põlme et al. 2020) | Põlme S, et al. 2020. Fungal Diversity 105:1-16 ((https://doi.org/10.1007/s13225-020-00466-2)) |
| Otu14004 | 1 | k__Fungi(100);p__Ascomycota(100);c__Saccharomycetes(100);o__Saccharomycetales(100);f__Metschnikowiaceae(100);g__Metschnikowia(100);g__Metschnikowia_unclassified(100); | Metschnikowia | 13 | Saprotroph | \|Undefined Saprotroph\| | Yeast | NULL | Probable | As nectar-tap_saprotroph (Põlme et al. 2020); Decay sugar-rich substrates (Põlme et al. 2020) | Põlme S, et al. 2020. Fungal Diversity 105:1-16 ((https://doi.org/10.1007/s13225-020-00466-2)) |
| Otu14005 | 1 | k__Fungi(100);p__Ascomycota(100);c__Saccharomycetes(100);o__Saccharomycetales(100);f__Metschnikowiaceae(100);g__Metschnikowia(100);s__Metschnikowia_pulcherrima(100); | Metschnikowia | 13 | Saprotroph | \|Undefined Saprotroph\| | Yeast | NULL | Probable | As nectar-tap_saprotroph (Põlme et al. 2020); Decay sugar-rich substrates (Põlme et al. 2020) | Põlme S, et al. 2020. Fungal Diversity 105:1-16 ((https://doi.org/10.1007/s13225-020-00466-2)) |
| Otu14006 | 1 | k__Fungi(100);p__Ascomycota(100);c__Saccharomycetes(100);o__Saccharomycetales(100);f__Metschnikowiaceae(100);g__Metschnikowia(100);s__Metschnikowia_pulcherrima(100); | Metschnikowia | 13 | Saprotroph | \|Undefined Saprotroph\| | Yeast | NULL | Probable | As nectar-tap_saprotroph (Põlme et al. 2020); Decay sugar-rich substrates (Põlme et al. 2020) | Põlme S, et al. 2020. Fungal Diversity 105:1-16 ((https://doi.org/10.1007/s13225-020-00466-2)) |
| Otu14007 | 1 | k__Fungi(100);p__Ascomycota(100);c__Saccharomycetes(100);o__Saccharomycetales(100);f__Metschnikowiaceae(100);g__Metschnikowia(100);s__Metschnikowia_pulcherrima(100); | Metschnikowia | 13 | Saprotroph | \|Undefined Saprotroph\| | Yeast | NULL | Probable | As nectar-tap_saprotroph (Põlme et al. 2020); Decay sugar-rich substrates (Põlme et al. 2020) | Põlme S, et al. 2020. Fungal Diversity 105:1-16 ((https://doi.org/10.1007/s13225-020-00466-2)) |
| Otu14008 | 1 | k__Fungi(100);p__Ascomycota(100);c__Saccharomycetes(100);o__Saccharomycetales(100);f__Metschnikowiaceae(100);g__Metschnikowia(100);s__Metschnikowia_pulcherrima(100); | Metschnikowia | 13 | Saprotroph | \|Undefined Saprotroph\| | Yeast | NULL | Probable | As nectar-tap_saprotroph (Põlme et al. 2020); Decay sugar-rich substrates (Põlme et al. 2020) | Põlme S, et al. 2020. Fungal Diversity 105:1-16 ((https://doi.org/10.1007/s13225-020-00466-2)) |
| Otu14009 | 1 | k__Fungi(100);p__Ascomycota(100);c__Saccharomycetes(100);o__Saccharomycetales(100);f__Metschnikowiaceae(100);g__Metschnikowia(100);s__Metschnikowia_pulcherrima(100); | Metschnikowia | 13 | Saprotroph | \|Undefined Saprotroph\| | Yeast | NULL | Probable | As nectar-tap_saprotroph (Põlme et al. 2020); Decay sugar-rich substrates (Põlme et al. 2020) | Põlme S, et al. 2020. Fungal Diversity 105:1-16 ((https://doi.org/10.1007/s13225-020-00466-2)) |
| Otu14010 | 1 | k__Fungi(100);p__Ascomycota(100);c__Saccharomycetes(100);o__Saccharomycetales(100);f__Metschnikowiaceae(100);g__Metschnikowia(100);s__Metschnikowia_pulcherrima(100); | Metschnikowia | 13 | Saprotroph | \|Undefined Saprotroph\| | Yeast | NULL | Probable | As nectar-tap_saprotroph (Põlme et al. 2020); Decay sugar-rich substrates (Põlme et al. 2020) | Põlme S, et al. 2020. Fungal Diversity 105:1-16 ((https://doi.org/10.1007/s13225-020-00466-2)) |
| Otu14012 | 1 | k__Fungi(100);p__Ascomycota(100);c__Saccharomycetes(100);o__Saccharomycetales(100);f__Metschnikowiaceae(100);g__Metschnikowia(100);s__Metschnikowia_pulcherrima(100); | Metschnikowia | 13 | Saprotroph | \|Undefined Saprotroph\| | Yeast | NULL | Probable | As nectar-tap_saprotroph (Põlme et al. 2020); Decay sugar-rich substrates (Põlme et al. 2020) | Põlme S, et al. 2020. Fungal Diversity 105:1-16 ((https://doi.org/10.1007/s13225-020-00466-2)) |
| Otu14013 | 1 | k__Fungi(100);p__Ascomycota(100);c__Saccharomycetes(100);o__Saccharomycetales(100);f__Saccharomycetales_family_Incertae_sedis(100);g__Candida(100);g__Candida_unclassified(100); | Candida | 13 | Pathotroph-Saprotroph-Symbiotroph | Animal Pathogen-Endophyte-Endosymbiont-Epiphyte-\|Undefined Saprotroph\| | Dimorphic Yeast-Yeast | NULL | Probable | As nectar-tap_saprotroph (Põlme et al. 2020); Decay sugar-rich substrates (Põlme et al. 2020); This asexual genus is polyphyletic and contain species that function very differently depend on what they are. Caution should be taken when interpreting guilds for Candida species. | Manolakaki D, et al. 2010. Virulence. 1: 367-375 ((https://doi.org/10.4161/viru.1.5.12796)); Mekbib SB, et al. 2011. Tropical Plant Pathology 36:233-240 ((https://doi.org/10.1590/S1982-56762011000400004)); Põlme S, et al. 2020. Fungal Diversity 105:1-16 ((https://doi.org/10.1007/s13225-020-00466-2)) |
| Otu14014 | 1 | k__Fungi(100);p__Ascomycota(100);c__Saccharomycetes(100);o__Saccharomycetales(100);f__Metschnikowiaceae(100);g__Metschnikowia(100);g__Metschnikowia_unclassified(100); | Metschnikowia | 13 | Saprotroph | \|Undefined Saprotroph\| | Yeast | NULL | Probable | As nectar-tap_saprotroph (Põlme et al. 2020); Decay sugar-rich substrates (Põlme et al. 2020) | Põlme S, et al. 2020. Fungal Diversity 105:1-16 ((https://doi.org/10.1007/s13225-020-00466-2)) |
| Otu14015 | 1 | k__Fungi(100);p__Ascomycota(100);c__Saccharomycetes(100);o__Saccharomycetales(100);f__Metschnikowiaceae(100);g__Metschnikowia(100);s__Metschnikowia_pulcherrima(100); | Metschnikowia | 13 | Saprotroph | \|Undefined Saprotroph\| | Yeast | NULL | Probable | As nectar-tap_saprotroph (Põlme et al. 2020); Decay sugar-rich substrates (Põlme et al. 2020) | Põlme S, et al. 2020. Fungal Diversity 105:1-16 ((https://doi.org/10.1007/s13225-020-00466-2)) |
| Otu14017 | 1 | k__Fungi(100);p__Ascomycota(100);c__Saccharomycetes(100);o__Saccharomycetales(100);f__Metschnikowiaceae(100);g__Metschnikowia(100);s__Metschnikowia_pulcherrima(100); | Metschnikowia | 13 | Saprotroph | \|Undefined Saprotroph\| | Yeast | NULL | Probable | As nectar-tap_saprotroph (Põlme et al. 2020); Decay sugar-rich substrates (Põlme et al. 2020) | Põlme S, et al. 2020. Fungal Diversity 105:1-16 ((https://doi.org/10.1007/s13225-020-00466-2)) |
| Otu14019 | 1 | k__Fungi(100);p__Ascomycota(100);c__Saccharomycetes(100);o__Saccharomycetales(100);f__Metschnikowiaceae(100);g__Metschnikowia(100);g__Metschnikowia_unclassified(100); | Metschnikowia | 13 | Saprotroph | \|Undefined Saprotroph\| | Yeast | NULL | Probable | As nectar-tap_saprotroph (Põlme et al. 2020); Decay sugar-rich substrates (Põlme et al. 2020) | Põlme S, et al. 2020. Fungal Diversity 105:1-16 ((https://doi.org/10.1007/s13225-020-00466-2)) |
| Otu14021 | 1 | k__Fungi(100);p__Ascomycota(100);c__Saccharomycetes(100);o__Saccharomycetales(100);f__Metschnikowiaceae(100);g__Metschnikowia(100);g__Metschnikowia_unclassified(100); | Metschnikowia | 13 | Saprotroph | \|Undefined Saprotroph\| | Yeast | NULL | Probable | As nectar-tap_saprotroph (Põlme et al. 2020); Decay sugar-rich substrates (Põlme et al. 2020) | Põlme S, et al. 2020. Fungal Diversity 105:1-16 ((https://doi.org/10.1007/s13225-020-00466-2)) |
| Otu14022 | 1 | k__Fungi(100);p__Ascomycota(100);c__Saccharomycetes(100);o__Saccharomycetales(100);f__Metschnikowiaceae(100);g__Metschnikowia(100);g__Metschnikowia_unclassified(100); | Metschnikowia | 13 | Saprotroph | \|Undefined Saprotroph\| | Yeast | NULL | Probable | As nectar-tap_saprotroph (Põlme et al. 2020); Decay sugar-rich substrates (Põlme et al. 2020) | Põlme S, et al. 2020. Fungal Diversity 105:1-16 ((https://doi.org/10.1007/s13225-020-00466-2)) |
| Otu14023 | 1 | k__Fungi(100);p__Ascomycota(100);c__Saccharomycetes(100);o__Saccharomycetales(100);f__Metschnikowiaceae(100);g__Metschnikowia(100);s__Metschnikowia_pulcherrima(100); | Metschnikowia | 13 | Saprotroph | \|Undefined Saprotroph\| | Yeast | NULL | Probable | As nectar-tap_saprotroph (Põlme et al. 2020); Decay sugar-rich substrates (Põlme et al. 2020) | Põlme S, et al. 2020. Fungal Diversity 105:1-16 ((https://doi.org/10.1007/s13225-020-00466-2)) |
| Otu14024 | 1 | k__Fungi(100);p__Ascomycota(100);c__Saccharomycetes(100);o__Saccharomycetales(100);f__Metschnikowiaceae(100);g__Metschnikowia(100);s__Metschnikowia_pulcherrima(100); | Metschnikowia | 13 | Saprotroph | \|Undefined Saprotroph\| | Yeast | NULL | Probable | As nectar-tap_saprotroph (Põlme et al. 2020); Decay sugar-rich substrates (Põlme et al. 2020) | Põlme S, et al. 2020. Fungal Diversity 105:1-16 ((https://doi.org/10.1007/s13225-020-00466-2)) |
| Otu14025 | 1 | k__Fungi(100);p__Ascomycota(100);c__Saccharomycetes(100);o__Saccharomycetales(100);f__Metschnikowiaceae(100);g__Metschnikowia(100);s__Metschnikowia_pulcherrima(100); | Metschnikowia | 13 | Saprotroph | \|Undefined Saprotroph\| | Yeast | NULL | Probable | As nectar-tap_saprotroph (Põlme et al. 2020); Decay sugar-rich substrates (Põlme et al. 2020) | Põlme S, et al. 2020. Fungal Diversity 105:1-16 ((https://doi.org/10.1007/s13225-020-00466-2)) |
| Otu14026 | 1 | k__Fungi(100);p__Ascomycota(100);c__Saccharomycetes(100);o__Saccharomycetales(100);f__Metschnikowiaceae(100);g__Metschnikowia(100);g__Metschnikowia_unclassified(100); | Metschnikowia | 13 | Saprotroph | \|Undefined Saprotroph\| | Yeast | NULL | Probable | As nectar-tap_saprotroph (Põlme et al. 2020); Decay sugar-rich substrates (Põlme et al. 2020) | Põlme S, et al. 2020. Fungal Diversity 105:1-16 ((https://doi.org/10.1007/s13225-020-00466-2)) |
| Otu14027 | 1 | k__Fungi(100);p__Ascomycota(100);c__Saccharomycetes(100);o__Saccharomycetales(100);f__Metschnikowiaceae(100);g__Metschnikowia(100);g__Metschnikowia_unclassified(100); | Metschnikowia | 13 | Saprotroph | \|Undefined Saprotroph\| | Yeast | NULL | Probable | As nectar-tap_saprotroph (Põlme et al. 2020); Decay sugar-rich substrates (Põlme et al. 2020) | Põlme S, et al. 2020. Fungal Diversity 105:1-16 ((https://doi.org/10.1007/s13225-020-00466-2)) |
| Otu14028 | 1 | k__Fungi(100);p__Ascomycota(100);c__Saccharomycetes(100);o__Saccharomycetales(100);f__Metschnikowiaceae(100);g__Metschnikowia(100);s__Metschnikowia_pulcherrima(100); | Metschnikowia | 13 | Saprotroph | \|Undefined Saprotroph\| | Yeast | NULL | Probable | As nectar-tap_saprotroph (Põlme et al. 2020); Decay sugar-rich substrates (Põlme et al. 2020) | Põlme S, et al. 2020. Fungal Diversity 105:1-16 ((https://doi.org/10.1007/s13225-020-00466-2)) |
| Otu14029 | 1 | k__Fungi(100);p__Ascomycota(100);c__Saccharomycetes(100);o__Saccharomycetales(100);f__Metschnikowiaceae(100);g__Metschnikowia(100);g__Metschnikowia_unclassified(100); | Metschnikowia | 13 | Saprotroph | \|Undefined Saprotroph\| | Yeast | NULL | Probable | As nectar-tap_saprotroph (Põlme et al. 2020); Decay sugar-rich substrates (Põlme et al. 2020) | Põlme S, et al. 2020. Fungal Diversity 105:1-16 ((https://doi.org/10.1007/s13225-020-00466-2)) |
| Otu14031 | 1 | k__Fungi(100);p__Ascomycota(100);c__Saccharomycetes(100);o__Saccharomycetales(100);f__Metschnikowiaceae(100);g__Metschnikowia(100);g__Metschnikowia_unclassified(100); | Metschnikowia | 13 | Saprotroph | \|Undefined Saprotroph\| | Yeast | NULL | Probable | As nectar-tap_saprotroph (Põlme et al. 2020); Decay sugar-rich substrates (Põlme et al. 2020) | Põlme S, et al. 2020. Fungal Diversity 105:1-16 ((https://doi.org/10.1007/s13225-020-00466-2)) |
| Otu14032 | 1 | k__Fungi(100);p__Ascomycota(100);c__Saccharomycetes(100);o__Saccharomycetales(100);f__Metschnikowiaceae(100);g__Metschnikowia(100);s__Metschnikowia_pulcherrima(100); | Metschnikowia | 13 | Saprotroph | \|Undefined Saprotroph\| | Yeast | NULL | Probable | As nectar-tap_saprotroph (Põlme et al. 2020); Decay sugar-rich substrates (Põlme et al. 2020) | Põlme S, et al. 2020. Fungal Diversity 105:1-16 ((https://doi.org/10.1007/s13225-020-00466-2)) |
| Otu14033 | 1 | k__Fungi(100);p__Ascomycota(100);c__Saccharomycetes(100);o__Saccharomycetales(100);f__Metschnikowiaceae(100);g__Metschnikowia(100);s__Metschnikowia_pulcherrima(100); | Metschnikowia | 13 | Saprotroph | \|Undefined Saprotroph\| | Yeast | NULL | Probable | As nectar-tap_saprotroph (Põlme et al. 2020); Decay sugar-rich substrates (Põlme et al. 2020) | Põlme S, et al. 2020. Fungal Diversity 105:1-16 ((https://doi.org/10.1007/s13225-020-00466-2)) |
| Otu14035 | 1 | k__Fungi(100);p__Ascomycota(100);c__Saccharomycetes(100);o__Saccharomycetales(100);f__Saccharomycetales_family_Incertae_sedis(100);g__Candida(100);g__Candida_unclassified(100); | Candida | 13 | Pathotroph-Saprotroph-Symbiotroph | Animal Pathogen-Endophyte-Endosymbiont-Epiphyte-\|Undefined Saprotroph\| | Dimorphic Yeast-Yeast | NULL | Probable | As nectar-tap_saprotroph (Põlme et al. 2020); Decay sugar-rich substrates (Põlme et al. 2020); This asexual genus is polyphyletic and contain species that function very differently depend on what they are. Caution should be taken when interpreting guilds for Candida species. | Manolakaki D, et al. 2010. Virulence. 1: 367-375 ((https://doi.org/10.4161/viru.1.5.12796)); Mekbib SB, et al. 2011. Tropical Plant Pathology 36:233-240 ((https://doi.org/10.1590/S1982-56762011000400004)); Põlme S, et al. 2020. Fungal Diversity 105:1-16 ((https://doi.org/10.1007/s13225-020-00466-2)) |
| Otu14036 | 1 | k__Fungi(100);p__Ascomycota(100);c__Saccharomycetes(100);o__Saccharomycetales(100);f__Metschnikowiaceae(100);g__Metschnikowia(100);s__Metschnikowia_pulcherrima(100); | Metschnikowia | 13 | Saprotroph | \|Undefined Saprotroph\| | Yeast | NULL | Probable | As nectar-tap_saprotroph (Põlme et al. 2020); Decay sugar-rich substrates (Põlme et al. 2020) | Põlme S, et al. 2020. Fungal Diversity 105:1-16 ((https://doi.org/10.1007/s13225-020-00466-2)) |
| Otu14037 | 1 | k__Fungi(100);p__Ascomycota(100);c__Saccharomycetes(100);o__Saccharomycetales(100);f__Metschnikowiaceae(100);g__Metschnikowia(100);g__Metschnikowia_unclassified(100); | Metschnikowia | 13 | Saprotroph | \|Undefined Saprotroph\| | Yeast | NULL | Probable | As nectar-tap_saprotroph (Põlme et al. 2020); Decay sugar-rich substrates (Põlme et al. 2020) | Põlme S, et al. 2020. Fungal Diversity 105:1-16 ((https://doi.org/10.1007/s13225-020-00466-2)) |
| Otu14038 | 1 | k__Fungi(100);p__Ascomycota(100);c__Saccharomycetes(100);o__Saccharomycetales(100);f__Metschnikowiaceae(100);g__Metschnikowia(100);s__Metschnikowia_pulcherrima(100); | Metschnikowia | 13 | Saprotroph | \|Undefined Saprotroph\| | Yeast | NULL | Probable | As nectar-tap_saprotroph (Põlme et al. 2020); Decay sugar-rich substrates (Põlme et al. 2020) | Põlme S, et al. 2020. Fungal Diversity 105:1-16 ((https://doi.org/10.1007/s13225-020-00466-2)) |
| Otu14039 | 1 | k__Fungi(100);p__Ascomycota(100);c__Saccharomycetes(100);o__Saccharomycetales(100);f__Metschnikowiaceae(100);g__Metschnikowia(100);g__Metschnikowia_unclassified(100); | Metschnikowia | 13 | Saprotroph | \|Undefined Saprotroph\| | Yeast | NULL | Probable | As nectar-tap_saprotroph (Põlme et al. 2020); Decay sugar-rich substrates (Põlme et al. 2020) | Põlme S, et al. 2020. Fungal Diversity 105:1-16 ((https://doi.org/10.1007/s13225-020-00466-2)) |
| Otu14040 | 1 | k__Fungi(100);p__Ascomycota(100);c__Saccharomycetes(100);o__Saccharomycetales(100);f__Metschnikowiaceae(100);g__Metschnikowia(100);g__Metschnikowia_unclassified(100); | Metschnikowia | 13 | Saprotroph | \|Undefined Saprotroph\| | Yeast | NULL | Probable | As nectar-tap_saprotroph (Põlme et al. 2020); Decay sugar-rich substrates (Põlme et al. 2020) | Põlme S, et al. 2020. Fungal Diversity 105:1-16 ((https://doi.org/10.1007/s13225-020-00466-2)) |
| Otu14041 | 1 | k__Fungi(100);p__Ascomycota(100);c__Saccharomycetes(100);o__Saccharomycetales(100);f__Metschnikowiaceae(100);g__Metschnikowia(100);s__Metschnikowia_pulcherrima(100); | Metschnikowia | 13 | Saprotroph | \|Undefined Saprotroph\| | Yeast | NULL | Probable | As nectar-tap_saprotroph (Põlme et al. 2020); Decay sugar-rich substrates (Põlme et al. 2020) | Põlme S, et al. 2020. Fungal Diversity 105:1-16 ((https://doi.org/10.1007/s13225-020-00466-2)) |
| Otu14042 | 1 | k__Fungi(100);p__Ascomycota(100);c__Saccharomycetes(100);o__Saccharomycetales(100);f__Metschnikowiaceae(100);g__Metschnikowia(100);s__Metschnikowia_pulcherrima(100); | Metschnikowia | 13 | Saprotroph | \|Undefined Saprotroph\| | Yeast | NULL | Probable | As nectar-tap_saprotroph (Põlme et al. 2020); Decay sugar-rich substrates (Põlme et al. 2020) | Põlme S, et al. 2020. Fungal Diversity 105:1-16 ((https://doi.org/10.1007/s13225-020-00466-2)) |
| Otu14043 | 1 | k__Fungi(100);p__Ascomycota(100);c__Saccharomycetes(100);o__Saccharomycetales(100);f__Metschnikowiaceae(100);g__Metschnikowia(100);s__Metschnikowia_pulcherrima(100); | Metschnikowia | 13 | Saprotroph | \|Undefined Saprotroph\| | Yeast | NULL | Probable | As nectar-tap_saprotroph (Põlme et al. 2020); Decay sugar-rich substrates (Põlme et al. 2020) | Põlme S, et al. 2020. Fungal Diversity 105:1-16 ((https://doi.org/10.1007/s13225-020-00466-2)) |
| Otu14044 | 1 | k__Fungi(100);p__Ascomycota(100);c__Saccharomycetes(100);o__Saccharomycetales(100);f__Metschnikowiaceae(100);g__Metschnikowia(100);g__Metschnikowia_unclassified(100); | Metschnikowia | 13 | Saprotroph | \|Undefined Saprotroph\| | Yeast | NULL | Probable | As nectar-tap_saprotroph (Põlme et al. 2020); Decay sugar-rich substrates (Põlme et al. 2020) | Põlme S, et al. 2020. Fungal Diversity 105:1-16 ((https://doi.org/10.1007/s13225-020-00466-2)) |
| Otu14045 | 1 | k__Fungi(100);p__Ascomycota(100);c__Saccharomycetes(100);o__Saccharomycetales(100);f__Metschnikowiaceae(100);g__Metschnikowia(100);s__Metschnikowia_pulcherrima(100); | Metschnikowia | 13 | Saprotroph | \|Undefined Saprotroph\| | Yeast | NULL | Probable | As nectar-tap_saprotroph (Põlme et al. 2020); Decay sugar-rich substrates (Põlme et al. 2020) | Põlme S, et al. 2020. Fungal Diversity 105:1-16 ((https://doi.org/10.1007/s13225-020-00466-2)) |
| Otu14049 | 1 | k__Fungi(100);p__Ascomycota(100);c__Saccharomycetes(100);o__Saccharomycetales(100);f__Metschnikowiaceae(100);g__Metschnikowia(100);s__Metschnikowia_pulcherrima(100); | Metschnikowia | 13 | Saprotroph | \|Undefined Saprotroph\| | Yeast | NULL | Probable | As nectar-tap_saprotroph (Põlme et al. 2020); Decay sugar-rich substrates (Põlme et al. 2020) | Põlme S, et al. 2020. Fungal Diversity 105:1-16 ((https://doi.org/10.1007/s13225-020-00466-2)) |
| Otu14050 | 1 | k__Fungi(100);p__Ascomycota(100);c__Saccharomycetes(100);o__Saccharomycetales(100);f__Metschnikowiaceae(100);g__Metschnikowia(100);s__Metschnikowia_pulcherrima(100); | Metschnikowia | 13 | Saprotroph | \|Undefined Saprotroph\| | Yeast | NULL | Probable | As nectar-tap_saprotroph (Põlme et al. 2020); Decay sugar-rich substrates (Põlme et al. 2020) | Põlme S, et al. 2020. Fungal Diversity 105:1-16 ((https://doi.org/10.1007/s13225-020-00466-2)) |
| Otu14051 | 1 | k__Fungi(100);p__Ascomycota(100);c__Saccharomycetes(100);o__Saccharomycetales(100);f__Metschnikowiaceae(100);g__Metschnikowia(100);s__Metschnikowia_pulcherrima(100); | Metschnikowia | 13 | Saprotroph | \|Undefined Saprotroph\| | Yeast | NULL | Probable | As nectar-tap_saprotroph (Põlme et al. 2020); Decay sugar-rich substrates (Põlme et al. 2020) | Põlme S, et al. 2020. Fungal Diversity 105:1-16 ((https://doi.org/10.1007/s13225-020-00466-2)) |
| Otu14052 | 1 | k__Fungi(100);p__Ascomycota(100);c__Saccharomycetes(100);o__Saccharomycetales(100);f__Metschnikowiaceae(100);g__Metschnikowia(100);s__Metschnikowia_pulcherrima(100); | Metschnikowia | 13 | Saprotroph | \|Undefined Saprotroph\| | Yeast | NULL | Probable | As nectar-tap_saprotroph (Põlme et al. 2020); Decay sugar-rich substrates (Põlme et al. 2020) | Põlme S, et al. 2020. Fungal Diversity 105:1-16 ((https://doi.org/10.1007/s13225-020-00466-2)) |
| Otu14053 | 1 | k__Fungi(100);p__Ascomycota(100);c__Saccharomycetes(100);o__Saccharomycetales(100);f__Metschnikowiaceae(100);g__Metschnikowia(100);g__Metschnikowia_unclassified(100); | Metschnikowia | 13 | Saprotroph | \|Undefined Saprotroph\| | Yeast | NULL | Probable | As nectar-tap_saprotroph (Põlme et al. 2020); Decay sugar-rich substrates (Põlme et al. 2020) | Põlme S, et al. 2020. Fungal Diversity 105:1-16 ((https://doi.org/10.1007/s13225-020-00466-2)) |
| Otu14054 | 1 | k__Fungi(100);p__Ascomycota(100);c__Saccharomycetes(100);o__Saccharomycetales(100);f__Metschnikowiaceae(100);g__Metschnikowia(100);s__Metschnikowia_pulcherrima(100); | Metschnikowia | 13 | Saprotroph | \|Undefined Saprotroph\| | Yeast | NULL | Probable | As nectar-tap_saprotroph (Põlme et al. 2020); Decay sugar-rich substrates (Põlme et al. 2020) | Põlme S, et al. 2020. Fungal Diversity 105:1-16 ((https://doi.org/10.1007/s13225-020-00466-2)) |
| Otu14055 | 1 | k__Fungi(100);p__Ascomycota(100);c__Saccharomycetes(100);o__Saccharomycetales(100);f__Metschnikowiaceae(100);g__Metschnikowia(100);s__Metschnikowia_pulcherrima(100); | Metschnikowia | 13 | Saprotroph | \|Undefined Saprotroph\| | Yeast | NULL | Probable | As nectar-tap_saprotroph (Põlme et al. 2020); Decay sugar-rich substrates (Põlme et al. 2020) | Põlme S, et al. 2020. Fungal Diversity 105:1-16 ((https://doi.org/10.1007/s13225-020-00466-2)) |
| Otu14056 | 1 | k__Fungi(100);p__Ascomycota(100);c__Saccharomycetes(100);o__Saccharomycetales(100);f__Metschnikowiaceae(100);g__Metschnikowia(100);s__Metschnikowia_pulcherrima(100); | Metschnikowia | 13 | Saprotroph | \|Undefined Saprotroph\| | Yeast | NULL | Probable | As nectar-tap_saprotroph (Põlme et al. 2020); Decay sugar-rich substrates (Põlme et al. 2020) | Põlme S, et al. 2020. Fungal Diversity 105:1-16 ((https://doi.org/10.1007/s13225-020-00466-2)) |
| Otu14057 | 1 | k__Fungi(100);p__Ascomycota(100);c__Saccharomycetes(100);o__Saccharomycetales(100);f__Metschnikowiaceae(100);g__Metschnikowia(100);g__Metschnikowia_unclassified(100); | Metschnikowia | 13 | Saprotroph | \|Undefined Saprotroph\| | Yeast | NULL | Probable | As nectar-tap_saprotroph (Põlme et al. 2020); Decay sugar-rich substrates (Põlme et al. 2020) | Põlme S, et al. 2020. Fungal Diversity 105:1-16 ((https://doi.org/10.1007/s13225-020-00466-2)) |
| Otu14058 | 1 | k__Fungi(100);p__Ascomycota(100);c__Saccharomycetes(100);o__Saccharomycetales(100);f__Metschnikowiaceae(100);g__Metschnikowia(100);g__Metschnikowia_unclassified(100); | Metschnikowia | 13 | Saprotroph | \|Undefined Saprotroph\| | Yeast | NULL | Probable | As nectar-tap_saprotroph (Põlme et al. 2020); Decay sugar-rich substrates (Põlme et al. 2020) | Põlme S, et al. 2020. Fungal Diversity 105:1-16 ((https://doi.org/10.1007/s13225-020-00466-2)) |
| Otu14059 | 1 | k__Fungi(100);p__Ascomycota(100);c__Saccharomycetes(100);o__Saccharomycetales(100);f__Metschnikowiaceae(100);g__Metschnikowia(100);s__Metschnikowia_pulcherrima(100); | Metschnikowia | 13 | Saprotroph | \|Undefined Saprotroph\| | Yeast | NULL | Probable | As nectar-tap_saprotroph (Põlme et al. 2020); Decay sugar-rich substrates (Põlme et al. 2020) | Põlme S, et al. 2020. Fungal Diversity 105:1-16 ((https://doi.org/10.1007/s13225-020-00466-2)) |
| Otu14060 | 1 | k__Fungi(100);p__Ascomycota(100);c__Saccharomycetes(100);o__Saccharomycetales(100);f__Metschnikowiaceae(100);g__Metschnikowia(100);g__Metschnikowia_unclassified(100); | Metschnikowia | 13 | Saprotroph | \|Undefined Saprotroph\| | Yeast | NULL | Probable | As nectar-tap_saprotroph (Põlme et al. 2020); Decay sugar-rich substrates (Põlme et al. 2020) | Põlme S, et al. 2020. Fungal Diversity 105:1-16 ((https://doi.org/10.1007/s13225-020-00466-2)) |
| Otu14062 | 1 | k__Fungi(100);p__Basidiomycota(100);c__Microbotryomycetes(100);o__Sporidiobolales(100);f__Sporidiobolales_family_Incertae_sedis(100);g__Sporobolomyces(100);s__Sporobolomyces_lactosus(100); | Sporobolomyces | 13 | Pathotroph-Saprotroph | \|Fungal Parasite\|-Plant Saprotroph | Yeast | NULL | Probable | As litter_saprotroph (Põlme et al. 2020); As mycoparasite (Põlme et al. 2020); Decay fungal material (Põlme et al. 2020) | [illegible] |
| Otu14065 | 1 | k__Fungi(100);p__Ascomycota(100);c__Saccharomycetes(100);o__Saccharomycetales(100);f__Metschnikowiaceae(100);g__Metschnikowia(100);s__Metschnikowia_pulcherrima(100); | Metschnikowia | 13 | Saprotroph | \|Undefined Saprotroph\| | Yeast | NULL | Probable | As nectar-tap_saprotroph (Põlme et al. 2020); Decay sugar-rich substrates (Põlme et al. 2020) | Põlme S, et al. 2020. Fungal Diversity 105:1-16 ((https://doi.org/10.1007/s13225-020-00466-2)) |
| Otu14066 | 1 | k__Fungi(100);p__Ascomycota(100);c__Saccharomycetes(100);o__Saccharomycetales(100);f__Metschnikowiaceae(100);g__Metschnikowia(100);s__Metschnikowia_pulcherrima(100); | Metschnikowia | 13 | Saprotroph | \|Undefined Saprotroph\| | Yeast | NULL | Probable | As nectar-tap_saprotroph (Põlme et al. 2020); Decay sugar-rich substrates (Põlme et al. 2020) | Põlme S, et al. 2020. Fungal Diversity 105:1-16 ((https://doi.org/10.1007/s13225-020-00466-2)) |
| Otu14067 | 1 | k__Fungi(100);p__Ascomycota(100);c__Saccharomycetes(100);o__Saccharomycetales(100);f__Metschnikowiaceae(100);g__Metschnikowia(100);s__Metschnikowia_pulcherrima(100); | Metschnikowia | 13 | Saprotroph | \|Undefined Saprotroph\| | Yeast | NULL | Probable | As nectar-tap_saprotroph (Põlme et al. 2020); Decay sugar-rich substrates (Põlme et al. 2020) | Põlme S, et al. 2020. Fungal Diversity 105:1-16 ((https://doi.org/10.1007/s13225-020-00466-2)) |
| Otu14068 | 1 | k__Fungi(100);p__Ascomycota(100);c__Saccharomycetes(100);o__Saccharomycetales(100);f__Metschnikowiaceae(100);g__Metschnikowia(100);g__Metschnikowia_unclassified(100); | Metschnikowia | 13 | Saprotroph | \|Undefined Saprotroph\| | Yeast | NULL | Probable | As nectar-tap_saprotroph (Põlme et al. 2020); Decay sugar-rich substrates (Põlme et al. 2020) | Põlme S, et al. 2020. Fungal Diversity 105:1-16 ((https://doi.org/10.1007/s13225-020-00466-2)) |
| Otu14070 | 1 | k__Fungi(100);p__Ascomycota(100);c__Saccharomycetes(100);o__Saccharomycetales(100);f__Metschnikowiaceae(100);g__Metschnikowia(100);s__Metschnikowia_pulcherrima(100); | Metschnikowia | 13 | Saprotroph | \|Undefined Saprotroph\| | Yeast | NULL | Probable | As nectar-tap_saprotroph (Põlme et al. 2020); Decay sugar-rich substrates (Põlme et al. 2020) | Põlme S, et al. 2020. Fungal Diversity 105:1-16 ((https://doi.org/10.1007/s13225-020-00466-2)) |
| Otu14071 | 1 | k__Fungi(100);p__Ascomycota(100);c__Saccharomycetes(100);o__Saccharomycetales(100);f__Metschnikowiaceae(100);g__Metschnikowia(100);s__Metschnikowia_pulcherrima(100); | Metschnikowia | 13 | Saprotroph | \|Undefined Saprotroph\| | Yeast | NULL | Probable | As nectar-tap_saprotroph (Põlme et al. 2020); Decay sugar-rich substrates (Põlme et al. 2020) | Põlme S, et al. 2020. Fungal Diversity 105:1-16 ((https://doi.org/10.1007/s13225-020-00466-2)) |
| Otu14072 | 1 | k__Fungi(100);p__Ascomycota(100);c__Saccharomycetes(100);o__Saccharomycetales(100);f__Metschnikowiaceae(100);g__Metschnikowia(100);s__Metschnikowia_pulcherrima(100); | Metschnikowia | 13 | Saprotroph | \|Undefined Saprotroph\| | Yeast | NULL | Probable | As nectar-tap_saprotroph (Põlme et al. 2020); Decay sugar-rich substrates (Põlme et al. 2020) | Põlme S, et al. 2020. Fungal Diversity 105:1-16 ((https://doi.org/10.1007/s13225-020-00466-2)) |
| Otu14073 | 1 | k__Fungi(100);p__Ascomycota(100);c__Saccharomycetes(100);o__Saccharomycetales(100);f__Saccharomycetales_family_Incertae_sedis(100);g__Candida(100);g__Candida_unclassified(100); | Candida | 13 | Pathotroph-Saprotroph-Symbiotroph | Animal Pathogen-Endophyte-Endosymbiont-Epiphyte-\|Undefined Saprotroph\| | Dimorphic Yeast-Yeast | NULL | Probable | As nectar-tap_saprotroph (Põlme et al. 2020); Decay sugar-rich substrates (Põlme et al. 2020); This asexual genus is polyphyletic and contain species that function very differently depend on what they are. Caution should be taken when interpreting guilds for Candida species. | Manolakaki D, et al. 2010. Virulence. 1: 367-375 ((https://doi.org/10.4161/viru.1.5.12796)); Mekbib SB, et al. 2011. Tropical Plant Pathology 36:233-240 ((https://doi.org/10.1590/S1982-56762011000400004)); Põlme S, et al. 2020. Fungal Diversity 105:1-16 ((https://doi.org/10.1007/s13225-020-00466-2)) |
| Otu14074 | 1 | k__Fungi(100);p__Ascomycota(100);c__Saccharomycetes(100);o__Saccharomycetales(100);f__Metschnikowiaceae(100);g__Metschnikowia(100);s__Metschnikowia_pulcherrima(100); | Metschnikowia | 13 | Saprotroph | \|Undefined Saprotroph\| | Yeast | NULL | Probable | As nectar-tap_saprotroph (Põlme et al. 2020); Decay sugar-rich substrates (Põlme et al. 2020) | Põlme S, et al. 2020. Fungal Diversity 105:1-16 ((https://doi.org/10.1007/s13225-020-00466-2)) |
| Otu14077 | 1 | k__Fungi(100);p__Ascomycota(100);c__Saccharomycetes(100);o__Saccharomycetales(100);f__Metschnikowiaceae(100);g__Metschnikowia(100);s__Metschnikowia_pulcherrima(100); | Metschnikowia | 13 | Saprotroph | \|Undefined Saprotroph\| | Yeast | NULL | Probable | As nectar-tap_saprotroph (Põlme et al. 2020); Decay sugar-rich substrates (Põlme et al. 2020) | Põlme S, et al. 2020. Fungal Diversity 105:1-16 ((https://doi.org/10.1007/s13225-020-00466-2)) |
| Otu14079 | 1 | k__Fungi(100);p__Ascomycota(100);c__Saccharomycetes(100);o__Saccharomycetales(100);f__Metschnikowiaceae(100);g__Metschnikowia(100);s__Metschnikowia_pulcherrima(100); | Metschnikowia | 13 | Saprotroph | \|Undefined Saprotroph\| | Yeast | NULL | Probable | As nectar-tap_saprotroph (Põlme et al. 2020); Decay sugar-rich substrates (Põlme et al. 2020) | Põlme S, et al. 2020. Fungal Diversity 105:1-16 ((https://doi.org/10.1007/s13225-020-00466-2)) |
| Otu14081 | 1 | k__Fungi(100);p__Ascomycota(100);c__Saccharomycetes(100);o__Saccharomycetales(100);f__Metschnikowiaceae(100);g__Metschnikowia(100);s__Metschnikowia_pulcherrima(100); | Metschnikowia | 13 | Saprotroph | \|Undefined Saprotroph\| | Yeast | NULL | Probable | As nectar-tap_saprotroph (Põlme et al. 2020); Decay sugar-rich substrates (Põlme et al. 2020) | Põlme S, et al. 2020. Fungal Diversity 105:1-16 ((https://doi.org/10.1007/s13225-020-00466-2)) |
| Otu14082 | 1 | k__Fungi(100);p__Ascomycota(100);c__Saccharomycetes(100);o__Saccharomycetales(100);f__Metschnikowiaceae(100);g__Metschnikowia(100);g__Metschnikowia_unclassified(100); | Metschnikowia | 13 | Saprotroph | \|Undefined Saprotroph\| | Yeast | NULL | Probable | As nectar-tap_saprotroph (Põlme et al. 2020); Decay sugar-rich substrates (Põlme et al. 2020) | Põlme S, et al. 2020. Fungal Diversity 105:1-16 ((https://doi.org/10.1007/s13225-020-00466-2)) |
| Otu14084 | 1 | k__Fungi(100);p__Ascomycota(100);c__Saccharomycetes(100);o__Saccharomycetales(100);f__Metschnikowiaceae(100);g__Metschnikowia(100);s__Metschnikowia_pulcherrima(100); | Metschnikowia | 13 | Saprotroph | \|Undefined Saprotroph\| | Yeast | NULL | Probable | As nectar-tap_saprotroph (Põlme et al. 2020); Decay sugar-rich substrates (Põlme et al. 2020) | Põlme S, et al. 2020. Fungal Diversity 105:1-16 ((https://doi.org/10.1007/s13225-020-00466-2)) |
| Otu14085 | 1 | k__Fungi(100);p__Ascomycota(100);c__Saccharomycetes(100);o__Saccharomycetales(100);f__Metschnikowiaceae(100);g__Metschnikowia(100);s__Metschnikowia_pulcherrima(100); | Metschnikowia | 13 | Saprotroph | \|Undefined Saprotroph\| | Yeast | NULL | Probable | As nectar-tap_saprotroph (Põlme et al. 2020); Decay sugar-rich substrates (Põlme et al. 2020) | Põlme S, et al. 2020. Fungal Diversity 105:1-16 ((https://doi.org/10.1007/s13225-020-00466-2)) |
| Otu14086 | 1 | k__Fungi(100);p__Ascomycota(100);c__Saccharomycetes(100);o__Saccharomycetales(100);f__Metschnikowiaceae(100);g__Metschnikowia(100);s__Metschnikowia_pulcherrima(100); | Metschnikowia | 13 | Saprotroph | \|Undefined Saprotroph\| | Yeast | NULL | Probable | As nectar-tap_saprotroph (Põlme et al. 2020); Decay sugar-rich substrates (Põlme et al. 2020) | Põlme S, et al. 2020. Fungal Diversity 105:1-16 ((https://doi.org/10.1007/s13225-020-00466-2)) |
| Otu14087 | 1 | k__Fungi(100);p__Ascomycota(100);c__Saccharomycetes(100);o__Saccharomycetales(100);f__Metschnikowiaceae(100);g__Metschnikowia(100);s__Metschnikowia_pulcherrima(100); | Metschnikowia | 13 | Saprotroph | \|Undefined Saprotroph\| | Yeast | NULL | Probable | As nectar-tap_saprotroph (Põlme et al. 2020); Decay sugar-rich substrates (Põlme et al. 2020) | Põlme S, et al. 2020. Fungal Diversity 105:1-16 ((https://doi.org/10.1007/s13225-020-00466-2)) |
| Otu14088 | 1 | k__Fungi(100);p__Ascomycota(100);c__Saccharomycetes(100);o__Saccharomycetales(100);f__Metschnikowiaceae(100);g__Metschnikowia(100);s__Metschnikowia_pulcherrima(100); | Metschnikowia | 13 | Saprotroph | \|Undefined Saprotroph\| | Yeast | NULL | Probable | As nectar-tap_saprotroph (Põlme et al. 2020); Decay sugar-rich substrates (Põlme et al. 2020) | Põlme S, et al. 2020. Fungal Diversity 105:1-16 ((https://doi.org/10.1007/s13225-020-00466-2)) |
| Otu14090 | 1 | k__Fungi(100);p__Ascomycota(100);c__Saccharomycetes(100);o__Saccharomycetales(100);f__Metschnikowiaceae(100);g__Metschnikowia(100);s__Metschnikowia_pulcherrima(100); | Metschnikowia | 13 | Saprotroph | \|Undefined Saprotroph\| | Yeast | NULL | Probable | As nectar-tap_saprotroph (Põlme et al. 2020); Decay sugar-rich substrates (Põlme et al. 2020) | Põlme S, et al. 2020. Fungal Diversity 105:1-16 ((https://doi.org/10.1007/s13225-020-00466-2)) |
| Otu14091 | 1 | k__Fungi(100);p__Ascomycota(100);c__Saccharomycetes(100);o__Saccharomycetales(100);f__Metschnikowiaceae(100);g__Metschnikowia(100);s__Metschnikowia_pulcherrima(100); | Metschnikowia | 13 | Saprotroph | \|Undefined Saprotroph\| | Yeast | NULL | Probable | As nectar-tap_saprotroph (Põlme et al. 2020); Decay sugar-rich substrates (Põlme et al. 2020) | Põlme S, et al. 2020. Fungal Diversity 105:1-16 ((https://doi.org/10.1007/s13225-020-00466-2)) |
| Otu14092 | 1 | k__Fungi(100);p__Ascomycota(100);c__Saccharomycetes(100);o__Saccharomycetales(100);f__Metschnikowiaceae(100);g__Metschnikowia(100);g__Metschnikowia_unclassified(100); | Metschnikowia | 13 | Saprotroph | \|Undefined Saprotroph\| | Yeast | NULL | Probable | As nectar-tap_saprotroph (Põlme et al. 2020); Decay sugar-rich substrates (Põlme et al. 2020) | Põlme S, et al. 2020. Fungal Diversity 105:1-16 ((https://doi.org/10.1007/s13225-020-00466-2)) |
| Otu14093 | 1 | k__Fungi(100);p__Ascomycota(100);c__Saccharomycetes(100);o__Saccharomycetales(100);f__Metschnikowiaceae(100);g__Metschnikowia(100);g__Metschnikowia_unclassified(100); | Metschnikowia | 13 | Saprotroph | \|Undefined Saprotroph\| | Yeast | NULL | Probable | As nectar-tap_saprotroph (Põlme et al. 2020); Decay sugar-rich substrates (Põlme et al. 2020) | Põlme S, et al. 2020. Fungal Diversity 105:1-16 ((https://doi.org/10.1007/s13225-020-00466-2)) |
| Otu14095 | 1 | k__Fungi(100);p__Ascomycota(100);c__Saccharomycetes(100);o__Saccharomycetales(100);f__Metschnikowiaceae(100);g__Metschnikowia(100);s__Metschnikowia_pulcherrima(100); | Metschnikowia | 13 | Saprotroph | \|Undefined Saprotroph\| | Yeast | NULL | Probable | As nectar-tap_saprotroph (Põlme et al. 2020); Decay sugar-rich substrates (Põlme et al. 2020) | Põlme S, et al. 2020. Fungal Diversity 105:1-16 ((https://doi.org/10.1007/s13225-020-00466-2)) |
| Otu14097 | 1 | k__Fungi(100);p__Ascomycota(100);c__Saccharomycetes(100);o__Saccharomycetales(100);f__Metschnikowiaceae(100);g__Metschnikowia(100);s__Metschnikowia_pulcherrima(100); | Metschnikowia | 13 | Saprotroph | \|Undefined Saprotroph\| | Yeast | NULL | Probable | As nectar-tap_saprotroph (Põlme et al. 2020); Decay sugar-rich substrates (Põlme et al. 2020) | Põlme S, et al. 2020. Fungal Diversity 105:1-16 ((https://doi.org/10.1007/s13225-020-00466-2)) |
| Otu14098 | 1 | k__Fungi(100);p__Ascomycota(100);c__Saccharomycetes(100);o__Saccharomycetales(100);f__Metschnikowiaceae(100);g__Metschnikowia(100);s__Metschnikowia_pulcherrima(100); | Metschnikowia | 13 | Saprotroph | \|Undefined Saprotroph\| | Yeast | NULL | Probable | As nectar-tap_saprotroph (Põlme et al. 2020); Decay sugar-rich substrates (Põlme et al. 2020) | Põlme S, et al. 2020. Fungal Diversity 105:1-16 ((https://doi.org/10.1007/s13225-020-00466-2)) |
| Otu14099 | 1 | k__Fungi(100);p__Ascomycota(100);c__Saccharomycetes(100);o__Saccharomycetales(100);f__Saccharomycetales_family_Incertae_sedis(100);f__Saccharomycetales_family_Incertae_sedis_unclassified(100);f__Saccharomycetales_family_Incertae_sedis_unclassified(100); | Saccharomycetales | 7 | Saprotroph | Undefined Saprotroph | Yeast | NULL | Possible | NULL | Sterkenburg E, et al. 2015. New Phytologist 207:1145-1158 ((https://doi.org/10.1111/nph.13426)) |
| Otu14101 | 1 | k__Fungi(100);p__Ascomycota(100);c__Saccharomycetes(100);o__Saccharomycetales(100);f__Metschnikowiaceae(100);g__Metschnikowia(100);s__Metschnikowia_pulcherrima(100); | Metschnikowia | 13 | Saprotroph | \|Undefined Saprotroph\| | Yeast | NULL | Probable | As nectar-tap_saprotroph (Põlme et al. 2020); Decay sugar-rich substrates (Põlme et al. 2020) | Põlme S, et al. 2020. Fungal Diversity 105:1-16 ((https://doi.org/10.1007/s13225-020-00466-2)) |
| Otu14102 | 1 | k__Fungi(100);p__Ascomycota(100);c__Saccharomycetes(100);o__Saccharomycetales(100);f__Metschnikowiaceae(100);g__Metschnikowia(100);s__Metschnikowia_pulcherrima(100); | Metschnikowia | 13 | Saprotroph | \|Undefined Saprotroph\| | Yeast | NULL | Probable | As nectar-tap_saprotroph (Põlme et al. 2020); Decay sugar-rich substrates (Põlme et al. 2020) | Põlme S, et al. 2020. Fungal Diversity 105:1-16 ((https://doi.org/10.1007/s13225-020-00466-2)) |
| Otu14106 | 1 | k__Fungi(100);p__Ascomycota(100);c__Saccharomycetes(100);o__Saccharomycetales(100);f__Metschnikowiaceae(100);g__Metschnikowia(100);s__Metschnikowia_pulcherrima(100); | Metschnikowia | 13 | Saprotroph | \|Undefined Saprotroph\| | Yeast | NULL | Probable | As nectar-tap_saprotroph (Põlme et al. 2020); Decay sugar-rich substrates (Põlme et al. 2020) | Põlme S, et al. 2020. Fungal Diversity 105:1-16 ((https://doi.org/10.1007/s13225-020-00466-2)) |
| Otu14107 | 1 | k__Fungi(100);p__Ascomycota(100);c__Saccharomycetes(100);o__Saccharomycetales(100);f__Metschnikowiaceae(100);g__Metschnikowia(100);s__Metschnikowia_pulcherrima(100); | Metschnikowia | 13 | Saprotroph | \|Undefined Saprotroph\| | Yeast | NULL | Probable | As nectar-tap_saprotroph (Põlme et al. 2020); Decay sugar-rich substrates (Põlme et al. 2020) | Põlme S, et al. 2020. Fungal Diversity 105:1-16 ((https://doi.org/10.1007/s13225-020-00466-2)) |
| Otu14116 | 1 | k__Fungi(100);p__Ascomycota(100);c__Saccharomycetes(100);o__Saccharomycetales(100);f__Metschnikowiaceae(100);g__Metschnikowia(100);g__Metschnikowia_unclassified(100); | Metschnikowia | 13 | Saprotroph | \|Undefined Saprotroph\| | Yeast | NULL | Probable | As nectar-tap_saprotroph (Põlme et al. 2020); Decay sugar-rich substrates (Põlme et al. 2020) | Põlme S, et al. 2020. Fungal Diversity 105:1-16 ((https://doi.org/10.1007/s13225-020-00466-2)) |
| Otu14117 | 1 | k__Fungi(100);p__Ascomycota(100);c__Saccharomycetes(100);o__Saccharomycetales(100);f__Metschnikowiaceae(100);g__Metschnikowia(100);s__Metschnikowia_pulcherrima(100); | Metschnikowia | 13 | Saprotroph | \|Undefined Saprotroph\| | Yeast | NULL | Probable | As nectar-tap_saprotroph (Põlme et al. 2020); Decay sugar-rich substrates (Põlme et al. 2020) | Põlme S, et al. 2020. Fungal Diversity 105:1-16 ((https://doi.org/10.1007/s13225-020-00466-2)) |
| Otu14122 | 1 | k__Fungi(100);p__Ascomycota(100);c__Saccharomycetes(100);o__Saccharomycetales(100);f__Metschnikowiaceae(100);g__Metschnikowia(100);g__Metschnikowia_unclassified(100); | Metschnikowia | 13 | Saprotroph | \|Undefined Saprotroph\| | Yeast | NULL | Probable | As nectar-tap_saprotroph (Põlme et al. 2020); Decay sugar-rich substrates (Põlme et al. 2020) | Põlme S, et al. 2020. Fungal Diversity 105:1-16 ((https://doi.org/10.1007/s13225-020-00466-2)) |
| Otu14125 | 1 | k__Fungi(100);p__Ascomycota(100);c__Saccharomycetes(100);o__Saccharomycetales(100);f__Metschnikowiaceae(100);g__Metschnikowia(100);s__Metschnikowia_pulcherrima(100); | Metschnikowia | 13 | Saprotroph | \|Undefined Saprotroph\| | Yeast | NULL | Probable | As nectar-tap_saprotroph (Põlme et al. 2020); Decay sugar-rich substrates (Põlme et al. 2020) | Põlme S, et al. 2020. Fungal Diversity 105:1-16 ((https://doi.org/10.1007/s13225-020-00466-2)) |
| Otu14127 | 1 | k__Fungi(100);p__Ascomycota(100);c__Saccharomycetes(100);o__Saccharomycetales(100);f__Metschnikowiaceae(100);g__Metschnikowia(100);g__Metschnikowia_unclassified(100); | Metschnikowia | 13 | Saprotroph | \|Undefined Saprotroph\| | Yeast | NULL | Probable | As nectar-tap_saprotroph (Põlme et al. 2020); Decay sugar-rich substrates (Põlme et al. 2020) | Põlme S, et al. 2020. Fungal Diversity 105:1-16 ((https://doi.org/10.1007/s13225-020-00466-2)) |
| Otu14129 | 1 | k__Fungi(100);p__Ascomycota(100);c__Saccharomycetes(100);o__Saccharomycetales(100);f__Metschnikowiaceae(100);g__Metschnikowia(100);s__Metschnikowia_pulcherrima(100); | Metschnikowia | 13 | Saprotroph | \|Undefined Saprotroph\| | Yeast | NULL | Probable | As nectar-tap_saprotroph (Põlme et al. 2020); Decay sugar-rich substrates (Põlme et al. 2020) | Põlme S, et al. 2020. Fungal Diversity 105:1-16 ((https://doi.org/10.1007/s13225-020-00466-2)) |
| Otu14135 | 1 | k__Fungi(100);p__Ascomycota(100);c__Saccharomycetes(100);o__Saccharomycetales(100);f__Metschnikowiaceae(100);g__Metschnikowia(100);s__Metschnikowia_pulcherrima(100); | Metschnikowia | 13 | Saprotroph | \|Undefined Saprotroph\| | Yeast | NULL | Probable | As nectar-tap_saprotroph (Põlme et al. 2020); Decay sugar-rich substrates (Põlme et al. 2020) | Põlme S, et al. 2020. Fungal Diversity 105:1-16 ((https://doi.org/10.1007/s13225-020-00466-2)) |
| Otu14137 | 1 | k__Fungi(100);p__Ascomycota(100);c__Saccharomycetes(100);o__Saccharomycetales(100);f__Metschnikowiaceae(100);g__Metschnikowia(100);s__Metschnikowia_pulcherrima(100); | Metschnikowia | 13 | Saprotroph | \|Undefined Saprotroph\| | Yeast | NULL | Probable | As nectar-tap_saprotroph (Põlme et al. 2020); Decay sugar-rich substrates (Põlme et al. 2020) | Põlme S, et al. 2020. Fungal Diversity 105:1-16 ((https://doi.org/10.1007/s13225-020-00466-2)) |
| Otu14138 | 1 | k__Fungi(100);p__Ascomycota(100);c__Saccharomycetes(100);o__Saccharomycetales(100);f__Metschnikowiaceae(100);g__Metschnikowia(100);g__Metschnikowia_unclassified(100); | Metschnikowia | 13 | Saprotroph | \|Undefined Saprotroph\| | Yeast | NULL | Probable | As nectar-tap_saprotroph (Põlme et al. 2020); Decay sugar-rich substrates (Põlme et al. 2020) | Põlme S, et al. 2020. Fungal Diversity 105:1-16 ((https://doi.org/10.1007/s13225-020-00466-2)) |
| Otu14139 | 1 | k__Fungi(100);p__Ascomycota(100);c__Saccharomycetes(100);o__Saccharomycetales(100);f__Metschnikowiaceae(100);g__Metschnikowia(100);s__Metschnikowia_pulcherrima(100); | Metschnikowia | 13 | Saprotroph | \|Undefined Saprotroph\| | Yeast | NULL | Probable | As nectar-tap_saprotroph (Põlme et al. 2020); Decay sugar-rich substrates (Põlme et al. 2020) | Põlme S, et al. 2020. Fungal Diversity 105:1-16 ((https://doi.org/10.1007/s13225-020-00466-2)) |
| Otu14140 | 1 | k__Fungi(100);p__Ascomycota(100);c__Saccharomycetes(100);o__Saccharomycetales(100);f__Metschnikowiaceae(100);g__Metschnikowia(100);s__Metschnikowia_pulcherrima(100); | Metschnikowia | 13 | Saprotroph | \|Undefined Saprotroph\| | Yeast | NULL | Probable | As nectar-tap_saprotroph (Põlme et al. 2020); Decay sugar-rich substrates (Põlme et al. 2020) | Põlme S, et al. 2020. Fungal Diversity 105:1-16 ((https://doi.org/10.1007/s13225-020-00466-2)) |
| Otu14141 | 1 | k__Fungi(100);p__Ascomycota(100);c__Saccharomycetes(100);o__Saccharomycetales(100);f__Metschnikowiaceae(100);g__Metschnikowia(100);s__Metschnikowia_pulcherrima(100); | Metschnikowia | 13 | Saprotroph | \|Undefined Saprotroph\| | Yeast | NULL | Probable | As nectar-tap_saprotroph (Põlme et al. 2020); Decay sugar-rich substrates (Põlme et al. 2020) | Põlme S, et al. 2020. Fungal Diversity 105:1-16 ((https://doi.org/10.1007/s13225-020-00466-2)) |
| Otu14146 | 1 | k__Fungi(100);p__Ascomycota(100);c__Saccharomycetes(100);o__Saccharomycetales(100);f__Metschnikowiaceae(100);g__Metschnikowia(100);s__Metschnikowia_pulcherrima(100); | Metschnikowia | 13 | Saprotroph | \|Undefined Saprotroph\| | Yeast | NULL | Probable | As nectar-tap_saprotroph (Põlme et al. 2020); Decay sugar-rich substrates (Põlme et al. 2020) | Põlme S, et al. 2020. Fungal Diversity 105:1-16 ((https://doi.org/10.1007/s13225-020-00466-2)) |
| Otu14147 | 1 | k__Fungi(100);p__Ascomycota(100);c__Saccharomycetes(100);o__Saccharomycetales(100);f__Metschnikowiaceae(100);g__Metschnikowia(100);s__Metschnikowia_pulcherrima(100); | Metschnikowia | 13 | Saprotroph | \|Undefined Saprotroph\| | Yeast | NULL | Probable | As nectar-tap_saprotroph (Põlme et al. 2020); Decay sugar-rich substrates (Põlme et al. 2020) | Põlme S, et al. 2020. Fungal Diversity 105:1-16 ((https://doi.org/10.1007/s13225-020-00466-2)) |
| Otu14150 | 1 | k__Fungi(100);p__Ascomycota(100);c__Saccharomycetes(100);o__Saccharomycetales(100);f__Metschnikowiaceae(100);g__Metschnikowia(100);s__Metschnikowia_pulcherrima(100); | Metschnikowia | 13 | Saprotroph | \|Undefined Saprotroph\| | Yeast | NULL | Probable | As nectar-tap_saprotroph (Põlme et al. 2020); Decay sugar-rich substrates (Põlme et al. 2020) | Põlme S, et al. 2020. Fungal Diversity 105:1-16 ((https://doi.org/10.1007/s13225-020-00466-2)) |
| Otu14151 | 1 | k__Fungi(100);p__Ascomycota(100);c__Saccharomycetes(100);o__Saccharomycetales(100);f__Metschnikowiaceae(100);g__Metschnikowia(100);g__Metschnikowia_unclassified(100); | Metschnikowia | 13 | Saprotroph | \|Undefined Saprotroph\| | Yeast | NULL | Probable | As nectar-tap_saprotroph (Põlme et al. 2020); Decay sugar-rich substrates (Põlme et al. 2020) | Põlme S, et al. 2020. Fungal Diversity 105:1-16 ((https://doi.org/10.1007/s13225-020-00466-2)) |
| Otu14153 | 1 | k__Fungi(100);p__Ascomycota(100);c__Saccharomycetes(100);o__Saccharomycetales(100);f__Metschnikowiaceae(100);g__Metschnikowia(100);s__Metschnikowia_pulcherrima(100); | Metschnikowia | 13 | Saprotroph | \|Undefined Saprotroph\| | Yeast | NULL | Probable | As nectar-tap_saprotroph (Põlme et al. 2020); Decay sugar-rich substrates (Põlme et al. 2020) | Põlme S, et al. 2020. Fungal Diversity 105:1-16 ((https://doi.org/10.1007/s13225-020-00466-2)) |
| Otu14155 | 1 | k__Fungi(100);p__Ascomycota(100);c__Saccharomycetes(100);o__Saccharomycetales(100);f__Metschnikowiaceae(100);g__Metschnikowia(100);g__Metschnikowia_unclassified(100); | Metschnikowia | 13 | Saprotroph | \|Undefined Saprotroph\| | Yeast | NULL | Probable | As nectar-tap_saprotroph (Põlme et al. 2020); Decay sugar-rich substrates (Põlme et al. 2020) | Põlme S, et al. 2020. Fungal Diversity 105:1-16 ((https://doi.org/10.1007/s13225-020-00466-2)) |
| Otu14159 | 1 | k__Fungi(100);p__Ascomycota(100);c__Saccharomycetes(100);o__Saccharomycetales(100);f__Metschnikowiaceae(100);g__Metschnikowia(100);s__Metschnikowia_pulcherrima(100); | Metschnikowia | 13 | Saprotroph | \|Undefined Saprotroph\| | Yeast | NULL | Probable | As nectar-tap_saprotroph (Põlme et al. 2020); Decay sugar-rich substrates (Põlme et al. 2020) | Põlme S, et al. 2020. Fungal Diversity 105:1-16 ((https://doi.org/10.1007/s13225-020-00466-2)) |
| Otu14161 | 1 | k__Fungi(100);p__Ascomycota(100);c__Saccharomycetes(100);o__Saccharomycetales(100);f__Metschnikowiaceae(100);g__Metschnikowia(100);s__Metschnikowia_pulcherrima(100); | Metschnikowia | 13 | Saprotroph | \|Undefined Saprotroph\| | Yeast | NULL | Probable | As nectar-tap_saprotroph (Põlme et al. 2020); Decay sugar-rich substrates (Põlme et al. 2020) | Põlme S, et al. 2020. Fungal Diversity 105:1-16 ((https://doi.org/10.1007/s13225-020-00466-2)) |
| Otu14162 | 1 | k__Fungi(100);p__Ascomycota(100);c__Saccharomycetes(100);o__Saccharomycetales(100);f__Saccharomycetales_family_Incertae_sedis(100);g__Candida(100);g__Candida_unclassified(100); | Candida | 13 | Pathotroph-Saprotroph-Symbiotroph | Animal Pathogen-Endophyte-Endosymbiont-Epiphyte-\|Undefined Saprotroph\| | Dimorphic Yeast-Yeast | NULL | Probable | As nectar-tap_saprotroph (Põlme et al. 2020); Decay sugar-rich substrates (Põlme et al. 2020); This asexual genus is polyphyletic and contain species that function very differently depend on what they are. Caution should be taken when interpreting guilds for Candida species. | Manolakaki D, et al. 2010. Virulence. 1: 367-375 ((https://doi.org/10.4161/viru.1.5.12796)); Mekbib SB, et al. 2011. Tropical Plant Pathology 36:233-240 ((https://doi.org/10.1590/S1982-56762011000400004)); Põlme S, et al. 2020. Fungal Diversity 105:1-16 ((https://doi.org/10.1007/s13225-020-00466-2)) |
| Otu14163 | 1 | k__Fungi(100);p__Ascomycota(100);c__Saccharomycetes(100);o__Saccharomycetales(100);f__Metschnikowiaceae(100);g__Metschnikowia(100);s__Metschnikowia_pulcherrima(100); | Metschnikowia | 13 | Saprotroph | \|Undefined Saprotroph\| | Yeast | NULL | Probable | As nectar-tap_saprotroph (Põlme et al. 2020); Decay sugar-rich substrates (Põlme et al. 2020) | Põlme S, et al. 2020. Fungal Diversity 105:1-16 ((https://doi.org/10.1007/s13225-020-00466-2)) |
| Otu14164 | 1 | k__Fungi(100);p__Ascomycota(100);c__Saccharomycetes(100);o__Saccharomycetales(100);f__Metschnikowiaceae(100);g__Metschnikowia(100);s__Metschnikowia_pulcherrima(100); | Metschnikowia | 13 | Saprotroph | \|Undefined Saprotroph\| | Yeast | NULL | Probable | As nectar-tap_saprotroph (Põlme et al. 2020); Decay sugar-rich substrates (Põlme et al. 2020) | Põlme S, et al. 2020. Fungal Diversity 105:1-16 ((https://doi.org/10.1007/s13225-020-00466-2)) |
| Otu14165 | 1 | k__Fungi(100);p__Ascomycota(100);c__Saccharomycetes(100);o__Saccharomycetales(100);f__Metschnikowiaceae(100);g__Metschnikowia(100);s__Metschnikowia_pulcherrima(100); | Metschnikowia | 13 | Saprotroph | \|Undefined Saprotroph\| | Yeast | NULL | Probable | As nectar-tap_saprotroph (Põlme et al. 2020); Decay sugar-rich substrates (Põlme et al. 2020) | Põlme S, et al. 2020. Fungal Diversity 105:1-16 ((https://doi.org/10.1007/s13225-020-00466-2)) |
| Otu14166 | 1 | k__Fungi(100);p__Ascomycota(100);c__Saccharomycetes(100);o__Saccharomycetales(100);f__Metschnikowiaceae(100);g__Metschnikowia(100);s__Metschnikowia_pulcherrima(100); | Metschnikowia | 13 | Saprotroph | \|Undefined Saprotroph\| | Yeast | NULL | Probable | As nectar-tap_saprotroph (Põlme et al. 2020); Decay sugar-rich substrates (Põlme et al. 2020) | Põlme S, et al. 2020. Fungal Diversity 105:1-16 ((https://doi.org/10.1007/s13225-020-00466-2)) |
| Otu14167 | 1 | k__Fungi(100);p__Ascomycota(100);c__Saccharomycetes(100);o__Saccharomycetales(100);f__Metschnikowiaceae(100);g__Metschnikowia(100);s__Metschnikowia_pulcherrima(100); | Metschnikowia | 13 | Saprotroph | \|Undefined Saprotroph\| | Yeast | NULL | Probable | As nectar-tap_saprotroph (Põlme et al. 2020); Decay sugar-rich substrates (Põlme et al. 2020) | Põlme S, et al. 2020. Fungal Diversity 105:1-16 ((https://doi.org/10.1007/s13225-020-00466-2)) |
| Otu14169 | 1 | k__Fungi(100);p__Ascomycota(100);c__Saccharomycetes(100);o__Saccharomycetales(100);o__Saccharomycetales_unclassified(100);o__Saccharomycetales_unclassified(100);o__Saccharomycetales_unclassified(100); | Saccharomycetales | 7 | Saprotroph | Undefined Saprotroph | Yeast | NULL | Possible | NULL | Sterkenburg E, et al. 2015. New Phytologist 207:1145-1158 ((https://doi.org/10.1111/nph.13426)) |
| Otu14170 | 1 | k__Fungi(100);p__Ascomycota(100);c__Saccharomycetes(100);o__Saccharomycetales(100);f__Metschnikowiaceae(100);g__Metschnikowia(100);s__Metschnikowia_pulcherrima(100); | Metschnikowia | 13 | Saprotroph | \|Undefined Saprotroph\| | Yeast | NULL | Probable | As nectar-tap_saprotroph (Põlme et al. 2020); Decay sugar-rich substrates (Põlme et al. 2020) | Põlme S, et al. 2020. Fungal Diversity 105:1-16 ((https://doi.org/10.1007/s13225-020-00466-2)) |
| Otu14174 | 1 | k__Fungi(100);p__Ascomycota(100);c__Saccharomycetes(100);o__Saccharomycetales(100);f__Metschnikowiaceae(100);g__Metschnikowia(100);s__Metschnikowia_pulcherrima(100); | Metschnikowia | 13 | Saprotroph | \|Undefined Saprotroph\| | Yeast | NULL | Probable | As nectar-tap_saprotroph (Põlme et al. 2020); Decay sugar-rich substrates (Põlme et al. 2020) | Põlme S, et al. 2020. Fungal Diversity 105:1-16 ((https://doi.org/10.1007/s13225-020-00466-2)) |
| Otu14175 | 1 | k__Fungi(100);p__Ascomycota(100);c__Saccharomycetes(100);o__Saccharomycetales(100);f__Metschnikowiaceae(100);g__Metschnikowia(100);s__Metschnikowia_pulcherrima(100); | Metschnikowia | 13 | Saprotroph | \|Undefined Saprotroph\| | Yeast | NULL | Probable | As nectar-tap_saprotroph (Põlme et al. 2020); Decay sugar-rich substrates (Põlme et al. 2020) | Põlme S, et al. 2020. Fungal Diversity 105:1-16 ((https://doi.org/10.1007/s13225-020-00466-2)) |
| Otu14180 | 1 | k__Fungi(100);p__Ascomycota(100);c__Saccharomycetes(100);o__Saccharomycetales(100);f__Metschnikowiaceae(100);g__Metschnikowia(100);s__Metschnikowia_pulcherrima(100); | Metschnikowia | 13 | Saprotroph | \|Undefined Saprotroph\| | Yeast | NULL | Probable | As nectar-tap_saprotroph (Põlme et al. 2020); Decay sugar-rich substrates (Põlme et al. 2020) | Põlme S, et al. 2020. Fungal Diversity 105:1-16 ((https://doi.org/10.1007/s13225-020-00466-2)) |
| Otu14181 | 1 | k__Fungi(100);p__Ascomycota(100);c__Saccharomycetes(100);o__Saccharomycetales(100);f__Metschnikowiaceae(100);g__Metschnikowia(100);s__Metschnikowia_pulcherrima(100); | Metschnikowia | 13 | Saprotroph | \|Undefined Saprotroph\| | Yeast | NULL | Probable | As nectar-tap_saprotroph (Põlme et al. 2020); Decay sugar-rich substrates (Põlme et al. 2020) | Põlme S, et al. 2020. Fungal Diversity 105:1-16 ((https://doi.org/10.1007/s13225-020-00466-2)) |
| Otu14183 | 1 | k__Fungi(100);p__Ascomycota(100);c__Saccharomycetes(100);o__Saccharomycetales(100);f__Metschnikowiaceae(100);g__Metschnikowia(100);s__Metschnikowia_pulcherrima(100); | Metschnikowia | 13 | Saprotroph | \|Undefined Saprotroph\| | Yeast | NULL | Probable | As nectar-tap_saprotroph (Põlme et al. 2020); Decay sugar-rich substrates (Põlme et al. 2020) | Põlme S, et al. 2020. Fungal Diversity 105:1-16 ((https://doi.org/10.1007/s13225-020-00466-2)) |
| Otu14184 | 1 | k__Fungi(100);p__Ascomycota(100);c__Saccharomycetes(100);o__Saccharomycetales(100);f__Metschnikowiaceae(100);g__Metschnikowia(100);s__Metschnikowia_pulcherrima(100); | Metschnikowia | 13 | Saprotroph | \|Undefined Saprotroph\| | Yeast | NULL | Probable | As nectar-tap_saprotroph (Põlme et al. 2020); Decay sugar-rich substrates (Põlme et al. 2020) | Põlme S, et al. 2020. Fungal Diversity 105:1-16 ((https://doi.org/10.1007/s13225-020-00466-2)) |
| Otu14191 | 1 | k__Fungi(100);p__Ascomycota(100);c__Saccharomycetes(100);o__Saccharomycetales(100);f__Metschnikowiaceae(100);g__Metschnikowia(100);g__Metschnikowia_unclassified(100); | Metschnikowia | 13 | Saprotroph | \|Undefined Saprotroph\| | Yeast | NULL | Probable | As nectar-tap_saprotroph (Põlme et al. 2020); Decay sugar-rich substrates (Põlme et al. 2020) | Põlme S, et al. 2020. Fungal Diversity 105:1-16 ((https://doi.org/10.1007/s13225-020-00466-2)) |
| Otu14195 | 1 | k__Fungi(100);p__Ascomycota(100);c__Saccharomycetes(100);o__Saccharomycetales(100);f__Metschnikowiaceae(100);g__Metschnikowia(100);s__Metschnikowia_pulcherrima(100); | Metschnikowia | 13 | Saprotroph | \|Undefined Saprotroph\| | Yeast | NULL | Probable | As nectar-tap_saprotroph (Põlme et al. 2020); Decay sugar-rich substrates (Põlme et al. 2020) | Põlme S, et al. 2020. Fungal Diversity 105:1-16 ((https://doi.org/10.1007/s13225-020-00466-2)) |
| Otu14196 | 1 | k__Fungi(100);p__Ascomycota(100);c__Saccharomycetes(100);o__Saccharomycetales(100);f__Metschnikowiaceae(100);g__Metschnikowia(100);s__Metschnikowia_pulcherrima(100); | Metschnikowia | 13 | Saprotroph | \|Undefined Saprotroph\| | Yeast | NULL | Probable | As nectar-tap_saprotroph (Põlme et al. 2020); Decay sugar-rich substrates (Põlme et al. 2020) | Põlme S, et al. 2020. Fungal Diversity 105:1-16 ((https://doi.org/10.1007/s13225-020-00466-2)) |
| Otu14198 | 1 | k__Fungi(100);p__Ascomycota(100);c__Saccharomycetes(100);o__Saccharomycetales(100);f__Metschnikowiaceae(100);g__Metschnikowia(100);g__Metschnikowia_unclassified(100); | Metschnikowia | 13 | Saprotroph | \|Undefined Saprotroph\| | Yeast | NULL | Probable | As nectar-tap_saprotroph (Põlme et al. 2020); Decay sugar-rich substrates (Põlme et al. 2020) | Põlme S, et al. 2020. Fungal Diversity 105:1-16 ((https://doi.org/10.1007/s13225-020-00466-2)) |
| Otu14199 | 1 | k__Fungi(100);p__Ascomycota(100);c__Saccharomycetes(100);o__Saccharomycetales(100);f__Metschnikowiaceae(100);g__Metschnikowia(100);s__Metschnikowia_pulcherrima(100); | Metschnikowia | 13 | Saprotroph | \|Undefined Saprotroph\| | Yeast | NULL | Probable | As nectar-tap_saprotroph (Põlme et al. 2020); Decay sugar-rich substrates (Põlme et al. 2020) | Põlme S, et al. 2020. Fungal Diversity 105:1-16 ((https://doi.org/10.1007/s13225-020-00466-2)) |
| Otu14201 | 1 | k__Fungi(100);p__Ascomycota(100);c__Saccharomycetes(100);o__Saccharomycetales(100);f__Metschnikowiaceae(100);g__Metschnikowia(100);s__Metschnikowia_pulcherrima(100); | Metschnikowia | 13 | Saprotroph | \|Undefined Saprotroph\| | Yeast | NULL | Probable | As nectar-tap_saprotroph (Põlme et al. 2020); Decay sugar-rich substrates (Põlme et al. 2020) | Põlme S, et al. 2020. Fungal Diversity 105:1-16 ((https://doi.org/10.1007/s13225-020-00466-2)) |
| Otu14202 | 1 | k__Fungi(100);p__Ascomycota(100);c__Saccharomycetes(100);o__Saccharomycetales(100);f__Metschnikowiaceae(100);g__Metschnikowia(100);s__Metschnikowia_pulcherrima(100); | Metschnikowia | 13 | Saprotroph | \|Undefined Saprotroph\| | Yeast | NULL | Probable | As nectar-tap_saprotroph (Põlme et al. 2020); Decay sugar-rich substrates (Põlme et al. 2020) | Põlme S, et al. 2020. Fungal Diversity 105:1-16 ((https://doi.org/10.1007/s13225-020-00466-2)) |
| Otu14204 | 1 | k__Fungi(100);p__Ascomycota(100);c__Saccharomycetes(100);o__Saccharomycetales(100);f__Metschnikowiaceae(100);g__Metschnikowia(100);s__Metschnikowia_pulcherrima(100); | Metschnikowia | 13 | Saprotroph | \|Undefined Saprotroph\| | Yeast | NULL | Probable | As nectar-tap_saprotroph (Põlme et al. 2020); Decay sugar-rich substrates (Põlme et al. 2020) | Põlme S, et al. 2020. Fungal Diversity 105:1-16 ((https://doi.org/10.1007/s13225-020-00466-2)) |
| Otu14205 | 1 | k__Fungi(100);p__Ascomycota(100);c__Saccharomycetes(100);o__Saccharomycetales(100);f__Metschnikowiaceae(100);g__Metschnikowia(100);s__Metschnikowia_pulcherrima(100); | Metschnikowia | 13 | Saprotroph | \|Undefined Saprotroph\| | Yeast | NULL | Probable | As nectar-tap_saprotroph (Põlme et al. 2020); Decay sugar-rich substrates (Põlme et al. 2020) | Põlme S, et al. 2020. Fungal Diversity 105:1-16 ((https://doi.org/10.1007/s13225-020-00466-2)) |
| Otu14206 | 1 | k__Fungi(100);p__Ascomycota(100);c__Saccharomycetes(100);o__Saccharomycetales(100);f__Metschnikowiaceae(100);g__Metschnikowia(100);s__Metschnikowia_pulcherrima(100); | Metschnikowia | 13 | Saprotroph | \|Undefined Saprotroph\| | Yeast | NULL | Probable | As nectar-tap_saprotroph (Põlme et al. 2020); Decay sugar-rich substrates (Põlme et al. 2020) | Põlme S, et al. 2020. Fungal Diversity 105:1-16 ((https://doi.org/10.1007/s13225-020-00466-2)) |

| | | | | | | | | | | | |
|---|---|---|---|---|---|---|---|---|---|---|---|
| Otu14208 | 1 | k__Fungi(100);p__Basidiomycota(100);c__Agaricomycetes(100);o__Polyporales(100);f__Meruliaceae(100);g__Steccherinum(100);s__Steccherinum_sp(100); | Steccherinum | 13 | Saprotroph | Undefined Saprotroph-\|Wood Saprotroph\| | Corticioid | White Rot | Probable | As white_rot (Põlme et al. 2020); Decay wood (Põlme et al. 2020) | Tedersoo L, et al. 2014. Science 346:e1256688 ((https://doi.org/10.1126/science.1256688)); Põlme S, et al. 2020. Fungal Diversity 105:1-16 ((https://doi.org/10.1007/s13225-020-00466-2)) |
| Otu14209 | 1 | k__Fungi(100);p__Basidiomycota(100);c__Agaricomycetes(100);o__Atheliales(100);f__Atheliaceae(100);g__Tylospora(100);s__Tylospora_sp(100); | Tylospora | 13 | Symbiotroph | \|Ectomycorrhizal\| | Corticioid | NULL | Highly Probable | NULL | Rinaldi AC, et al. 2008. Fungal Diversity 33:1-45 ((http://www.fungaldiversity.org/fdp/sfdp/33-1.pdf)); Tedersoo L, et al. 2010. Mycorrhiza 20:217-263 ((https://doi.org/10.1007/s00572-009-0274-x)); Põlme S, et al. 2020. Fungal Diversity 105:1-16 ((https://doi.org/10.1007/s13225-020-00466-2)); http://mycorrhizas.info/ecmf.html ((http://mycorrhizas.info/ecmf.html)) |
| Otu14211 | 1 | k__Fungi(100);p__Ascomycota(100);c__Saccharomycetes(100);o__Saccharomycetales(100);f__Metschnikowiaceae(100);g__Metschnikowia(100);s__Metschnikowia_pulcherrima(100); | Metschnikowia | 13 | Saprotroph | \|Undefined Saprotroph\| | Yeast | NULL | Probable | As nectar-tap_saprotroph (Põlme et al. 2020); Decay sugar-rich substrates (Põlme et al. 2020) | Põlme S, et al. 2020. Fungal Diversity 105:1-16 ((https://doi.org/10.1007/s13225-020-00466-2)) |
| Otu14212 | 1 | k__Fungi(100);p__Ascomycota(100);c__Saccharomycetes(100);o__Saccharomycetales(100);f__Metschnikowiaceae(100);g__Metschnikowia(100);s__Metschnikowia_pulcherrima(100); | Metschnikowia | 13 | Saprotroph | \|Undefined Saprotroph\| | Yeast | NULL | Probable | As nectar-tap_saprotroph (Põlme et al. 2020); Decay sugar-rich substrates (Põlme et al. 2020) | Põlme S, et al. 2020. Fungal Diversity 105:1-16 ((https://doi.org/10.1007/s13225-020-00466-2)) |
| Otu14214 | 1 | k__Fungi(100);p__Ascomycota(100);c__Saccharomycetes(100);o__Saccharomycetales(100);f__Metschnikowiaceae(100);g__Metschnikowia(100);s__Metschnikowia_pulcherrima(100); | Metschnikowia | 13 | Saprotroph | \|Undefined Saprotroph\| | Yeast | NULL | Probable | As nectar-tap_saprotroph (Põlme et al. 2020); Decay sugar-rich substrates (Põlme et al. 2020) | Põlme S, et al. 2020. Fungal Diversity 105:1-16 ((https://doi.org/10.1007/s13225-020-00466-2)) |
| Otu14221 | 1 | k__Fungi(100);p__Ascomycota(100);c__Saccharomycetes(100);o__Saccharomycetales(100);f__Metschnikowiaceae(100);g__Metschnikowia(100);s__Metschnikowia_pulcherrima(100); | Metschnikowia | 13 | Saprotroph | \|Undefined Saprotroph\| | Yeast | NULL | Probable | As nectar-tap_saprotroph (Põlme et al. 2020); Decay sugar-rich substrates (Põlme et al. 2020) | Põlme S, et al. 2020. Fungal Diversity 105:1-16 ((https://doi.org/10.1007/s13225-020-00466-2)) |
| Otu14223 | 1 | k__Fungi(100);p__Ascomycota(100);c__Saccharomycetes(100);o__Saccharomycetales(100);f__Metschnikowiaceae(100);g__Metschnikowia(100);s__Metschnikowia_pulcherrima(100); | Metschnikowia | 13 | Saprotroph | \|Undefined Saprotroph\| | Yeast | NULL | Probable | As nectar-tap_saprotroph (Põlme et al. 2020); Decay sugar-rich substrates (Põlme et al. 2020) | Põlme S, et al. 2020. Fungal Diversity 105:1-16 ((https://doi.org/10.1007/s13225-020-00466-2)) |
| Otu14225 | 1 | k__Fungi(100);p__Ascomycota(100);c__Saccharomycetes(100);o__Saccharomycetales(100);f__Metschnikowiaceae(100);g__Metschnikowia(100);s__Metschnikowia_pulcherrima(100); | Metschnikowia | 13 | Saprotroph | \|Undefined Saprotroph\| | Yeast | NULL | Probable | As nectar-tap_saprotroph (Põlme et al. 2020); Decay sugar-rich substrates (Põlme et al. 2020) | Põlme S, et al. 2020. Fungal Diversity 105:1-16 ((https://doi.org/10.1007/s13225-020-00466-2)) |
| Otu14226 | 1 | k__Fungi(100);p__Ascomycota(100);c__Saccharomycetes(100);o__Saccharomycetales(100);f__Metschnikowiaceae(100);g__Metschnikowia(100);s__Metschnikowia_pulcherrima(100); | Metschnikowia | 13 | Saprotroph | \|Undefined Saprotroph\| | Yeast | NULL | Probable | As nectar-tap_saprotroph (Põlme et al. 2020); Decay sugar-rich substrates (Põlme et al. 2020) | Põlme S, et al. 2020. Fungal Diversity 105:1-16 ((https://doi.org/10.1007/s13225-020-00466-2)) |
| Otu14232 | 1 | k__Fungi(100);p__Ascomycota(100);c__Saccharomycetes(100);o__Saccharomycetales(100);f__Metschnikowiaceae(100);g__Metschnikowia(100);s__Metschnikowia_pulcherrima(100); | Metschnikowia | 13 | Saprotroph | \|Undefined Saprotroph\| | Yeast | NULL | Probable | As nectar-tap_saprotroph (Põlme et al. 2020); Decay sugar-rich substrates (Põlme et al. 2020) | Põlme S, et al. 2020. Fungal Diversity 105:1-16 ((https://doi.org/10.1007/s13225-020-00466-2)) |
| Otu14233 | 1 | k__Fungi(100);p__Ascomycota(100);c__Saccharomycetes(100);o__Saccharomycetales(100);f__Metschnikowiaceae(100);g__Metschnikowia(100);s__Metschnikowia_pulcherrima(100); | Metschnikowia | 13 | Saprotroph | \|Undefined Saprotroph\| | Yeast | NULL | Probable | As nectar-tap_saprotroph (Põlme et al. 2020); Decay sugar-rich substrates (Põlme et al. 2020) | Põlme S, et al. 2020. Fungal Diversity 105:1-16 ((https://doi.org/10.1007/s13225-020-00466-2)) |
| Otu14234 | 1 | k__Fungi(100);p__Ascomycota(100);c__Saccharomycetes(100);o__Saccharomycetales(100);f__Metschnikowiaceae(100);g__Metschnikowia(100);s__Metschnikowia_pulcherrima(100); | Metschnikowia | 13 | Saprotroph | \|Undefined Saprotroph\| | Yeast | NULL | Probable | As nectar-tap_saprotroph (Põlme et al. 2020); Decay sugar-rich substrates (Põlme et al. 2020) | Põlme S, et al. 2020. Fungal Diversity 105:1-16 ((https://doi.org/10.1007/s13225-020-00466-2)) |
| Otu14235 | 1 | k__Fungi(100);p__Ascomycota(100);c__Saccharomycetes(100);o__Saccharomycetales(100);f__Metschnikowiaceae(100);g__Metschnikowia(100);s__Metschnikowia_pulcherrima(100); | Metschnikowia | 13 | Saprotroph | \|Undefined Saprotroph\| | Yeast | NULL | Probable | As nectar-tap_saprotroph (Põlme et al. 2020); Decay sugar-rich substrates (Põlme et al. 2020) | Põlme S, et al. 2020. Fungal Diversity 105:1-16 ((https://doi.org/10.1007/s13225-020-00466-2)) |
| Otu14236 | 1 | k__Fungi(100);p__Ascomycota(100);c__Saccharomycetes(100);o__Saccharomycetales(100);f__Metschnikowiaceae(100);g__Metschnikowia(100);s__Metschnikowia_pulcherrima(100); | Metschnikowia | 13 | Saprotroph | \|Undefined Saprotroph\| | Yeast | NULL | Probable | As nectar-tap_saprotroph (Põlme et al. 2020); Decay sugar-rich substrates (Põlme et al. 2020) | Põlme S, et al. 2020. Fungal Diversity 105:1-16 ((https://doi.org/10.1007/s13225-020-00466-2)) |
| Otu14239 | 1 | k__Fungi(100);p__Ascomycota(100);c__Saccharomycetes(100);o__Saccharomycetales(100);f__Metschnikowiaceae(100);g__Metschnikowia(100);g__Metschnikowia_unclassified(100); | Metschnikowia | 13 | Saprotroph | \|Undefined Saprotroph\| | Yeast | NULL | Probable | As nectar-tap_saprotroph (Põlme et al. 2020); Decay sugar-rich substrates (Põlme et al. 2020) | Põlme S, et al. 2020. Fungal Diversity 105:1-16 ((https://doi.org/10.1007/s13225-020-00466-2)) |
| Otu14240 | 1 | k__Fungi(100);p__Ascomycota(100);c__Saccharomycetes(100);o__Saccharomycetales(100);f__Metschnikowiaceae(100);g__Metschnikowia(100);s__Metschnikowia_pulcherrima(100); | Metschnikowia | 13 | Saprotroph | \|Undefined Saprotroph\| | Yeast | NULL | Probable | As nectar-tap_saprotroph (Põlme et al. 2020); Decay sugar-rich substrates (Põlme et al. 2020) | Põlme S, et al. 2020. Fungal Diversity 105:1-16 ((https://doi.org/10.1007/s13225-020-00466-2)) |
| Otu14243 | 1 | k__Fungi(100);p__Ascomycota(100);c__Saccharomycetes(100);o__Saccharomycetales(100);f__Saccharomycetales_family_Incertae_sedis(100);g__Candida(100);g__Candida_unclassified(100); | Candida | 13 | Pathotroph-Saprotroph-Symbiotroph | Animal Pathogen-Endophyte-Endosymbiont-Epiphyte-\|Undefined Saprotroph\| | Dimorphic Yeast-Yeast | NULL | Probable | As nectar-tap_saprotroph (Põlme et al. 2020); Decay sugar-rich substrates (Põlme et al. 2020); This asexual genus is polyphyletic and contain species that function very differently depend on what they are. Caution should be taken when interpreting guilds for Candida species. | Manolakaki D, et al. 2010. Virulence. 1: 367-375 ((https://doi.org/10.4161/viru.1.5.12796)); Mekbib SB, et al. 2011. Tropical Plant Pathology 36:233-240 ((https://doi.org/10.1590/S1982-56762011000400004)); Põlme S, et al. 2020. Fungal Diversity 105:1-16 ((https://doi.org/10.1007/s13225-020-00466-2)) |
| Otu14245 | 1 | k__Fungi(100);p__Ascomycota(100);c__Saccharomycetes(100);o__Saccharomycetales(100);f__Metschnikowiaceae(100);g__Metschnikowia(100);s__Metschnikowia_pulcherrima(100); | Metschnikowia | 13 | Saprotroph | \|Undefined Saprotroph\| | Yeast | NULL | Probable | As nectar-tap_saprotroph (Põlme et al. 2020); Decay sugar-rich substrates (Põlme et al. 2020) | Põlme S, et al. 2020. Fungal Diversity 105:1-16 ((https://doi.org/10.1007/s13225-020-00466-2)) |
| Otu14247 | 1 | k__Fungi(100);p__Ascomycota(100);c__Saccharomycetes(100);o__Saccharomycetales(100);f__Metschnikowiaceae(100);g__Metschnikowia(100);s__Metschnikowia_pulcherrima(100); | Metschnikowia | 13 | Saprotroph | \|Undefined Saprotroph\| | Yeast | NULL | Probable | As nectar-tap_saprotroph (Põlme et al. 2020); Decay sugar-rich substrates (Põlme et al. 2020) | Põlme S, et al. 2020. Fungal Diversity 105:1-16 ((https://doi.org/10.1007/s13225-020-00466-2)) |
| Otu14248 | 1 | k__Fungi(100);p__Ascomycota(100);c__Saccharomycetes(100);o__Saccharomycetales(100);f__Metschnikowiaceae(100);g__Metschnikowia(100);s__Metschnikowia_pulcherrima(100); | Metschnikowia | 13 | Saprotroph | \|Undefined Saprotroph\| | Yeast | NULL | Probable | As nectar-tap_saprotroph (Põlme et al. 2020); Decay sugar-rich substrates (Põlme et al. 2020) | Põlme S, et al. 2020. Fungal Diversity 105:1-16 ((https://doi.org/10.1007/s13225-020-00466-2)) |
| Otu14249 | 1 | k__Fungi(100);p__Ascomycota(100);c__Saccharomycetes(100);o__Saccharomycetales(100);f__Metschnikowiaceae(100);g__Metschnikowia(100);s__Metschnikowia_pulcherrima(100); | Metschnikowia | 13 | Saprotroph | \|Undefined Saprotroph\| | Yeast | NULL | Probable | As nectar-tap_saprotroph (Põlme et al. 2020); Decay sugar-rich substrates (Põlme et al. 2020) | Põlme S, et al. 2020. Fungal Diversity 105:1-16 ((https://doi.org/10.1007/s13225-020-00466-2)) |
| Otu14250 | 1 | k__Fungi(100);p__Ascomycota(100);c__Saccharomycetes(100);o__Saccharomycetales(100);f__Metschnikowiaceae(100);g__Metschnikowia(100);g__Metschnikowia_unclassified(100); | Metschnikowia | 13 | Saprotroph | \|Undefined Saprotroph\| | Yeast | NULL | Probable | As nectar-tap_saprotroph (Põlme et al. 2020); Decay sugar-rich substrates (Põlme et al. 2020) | Põlme S, et al. 2020. Fungal Diversity 105:1-16 ((https://doi.org/10.1007/s13225-020-00466-2)) |
| Otu14251 | 1 | k__Fungi(100);p__Ascomycota(100);c__Saccharomycetes(100);o__Saccharomycetales(100);f__Metschnikowiaceae(100);g__Metschnikowia(100);g__Metschnikowia_unclassified(100); | Metschnikowia | 13 | Saprotroph | \|Undefined Saprotroph\| | Yeast | NULL | Probable | As nectar-tap_saprotroph (Põlme et al. 2020); Decay sugar-rich substrates (Põlme et al. 2020) | Põlme S, et al. 2020. Fungal Diversity 105:1-16 ((https://doi.org/10.1007/s13225-020-00466-2)) |
| Otu14255 | 1 | k__Fungi(100);p__Ascomycota(100);c__Saccharomycetes(100);o__Saccharomycetales(100);f__Metschnikowiaceae(100);g__Metschnikowia(100);s__Metschnikowia_pulcherrima(100); | Metschnikowia | 13 | Saprotroph | \|Undefined Saprotroph\| | Yeast | NULL | Probable | As nectar-tap_saprotroph (Põlme et al. 2020); Decay sugar-rich substrates (Põlme et al. 2020) | Põlme S, et al. 2020. Fungal Diversity 105:1-16 ((https://doi.org/10.1007/s13225-020-00466-2)) |
| Otu14256 | 1 | k__Fungi(100);p__Ascomycota(100);c__Saccharomycetes(100);o__Saccharomycetales(100);f__Metschnikowiaceae(100);g__Metschnikowia(100);s__Metschnikowia_pulcherrima(100); | Metschnikowia | 13 | Saprotroph | \|Undefined Saprotroph\| | Yeast | NULL | Probable | As nectar-tap_saprotroph (Põlme et al. 2020); Decay sugar-rich substrates (Põlme et al. 2020) | Põlme S, et al. 2020. Fungal Diversity 105:1-16 ((https://doi.org/10.1007/s13225-020-00466-2)) |
| Otu14257 | 1 | k__Fungi(100);p__Ascomycota(100);c__Saccharomycetes(100);o__Saccharomycetales(100);f__Metschnikowiaceae(100);g__Metschnikowia(100);s__Metschnikowia_pulcherrima(100); | Metschnikowia | 13 | Saprotroph | \|Undefined Saprotroph\| | Yeast | NULL | Probable | As nectar-tap_saprotroph (Põlme et al. 2020); Decay sugar-rich substrates (Põlme et al. 2020) | Põlme S, et al. 2020. Fungal Diversity 105:1-16 ((https://doi.org/10.1007/s13225-020-00466-2)) |
| Otu14258 | 1 | k__Fungi(100);p__Ascomycota(100);c__Saccharomycetes(100);o__Saccharomycetales(100);f__Metschnikowiaceae(100);g__Metschnikowia(100);s__Metschnikowia_pulcherrima(100); | Metschnikowia | 13 | Saprotroph | \|Undefined Saprotroph\| | Yeast | NULL | Probable | As nectar-tap_saprotroph (Põlme et al. 2020); Decay sugar-rich substrates (Põlme et al. 2020) | Põlme S, et al. 2020. Fungal Diversity 105:1-16 ((https://doi.org/10.1007/s13225-020-00466-2)) |
| Otu14259 | 1 | k__Fungi(100);p__Ascomycota(100);c__Saccharomycetes(100);o__Saccharomycetales(100);f__Metschnikowiaceae(100);g__Metschnikowia(100);s__Metschnikowia_pulcherrima(100); | Metschnikowia | 13 | Saprotroph | \|Undefined Saprotroph\| | Yeast | NULL | Probable | As nectar-tap_saprotroph (Põlme et al. 2020); Decay sugar-rich substrates (Põlme et al. 2020) | Põlme S, et al. 2020. Fungal Diversity 105:1-16 ((https://doi.org/10.1007/s13225-020-00466-2)) |
| Otu14260 | 1 | k__Fungi(100);p__Ascomycota(100);c__Saccharomycetes(100);o__Saccharomycetales(100);f__Metschnikowiaceae(100);g__Metschnikowia(100);s__Metschnikowia_pulcherrima(100); | Metschnikowia | 13 | Saprotroph | \|Undefined Saprotroph\| | Yeast | NULL | Probable | As nectar-tap_saprotroph (Põlme et al. 2020); Decay sugar-rich substrates (Põlme et al. 2020) | Põlme S, et al. 2020. Fungal Diversity 105:1-16 ((https://doi.org/10.1007/s13225-020-00466-2)) |
| Otu14261 | 1 | k__Fungi(100);p__Ascomycota(100);c__Saccharomycetes(100);o__Saccharomycetales(100);f__Metschnikowiaceae(100);g__Metschnikowia(100);s__Metschnikowia_pulcherrima(100); | Metschnikowia | 13 | Saprotroph | \|Undefined Saprotroph\| | Yeast | NULL | Probable | As nectar-tap_saprotroph (Põlme et al. 2020); Decay sugar-rich substrates (Põlme et al. 2020) | Põlme S, et al. 2020. Fungal Diversity 105:1-16 ((https://doi.org/10.1007/s13225-020-00466-2)) |
| Otu14263 | 1 | k__Fungi(100);p__Ascomycota(100);c__Saccharomycetes(100);o__Saccharomycetales(100);f__Metschnikowiaceae(100);g__Metschnikowia(100);s__Metschnikowia_pulcherrima(100); | Metschnikowia | 13 | Saprotroph | \|Undefined Saprotroph\| | Yeast | NULL | Probable | As nectar-tap_saprotroph (Põlme et al. 2020); Decay sugar-rich substrates (Põlme et al. 2020) | Põlme S, et al. 2020. Fungal Diversity 105:1-16 ((https://doi.org/10.1007/s13225-020-00466-2)) |
| Otu14264 | 1 | k__Fungi(100);p__Ascomycota(100);c__Saccharomycetes(100);o__Saccharomycetales(100);f__Metschnikowiaceae(100);g__Metschnikowia(100);s__Metschnikowia_pulcherrima(100); | Metschnikowia | 13 | Saprotroph | \|Undefined Saprotroph\| | Yeast | NULL | Probable | As nectar-tap_saprotroph (Põlme et al. 2020); Decay sugar-rich substrates (Põlme et al. 2020) | Põlme S, et al. 2020. Fungal Diversity 105:1-16 ((https://doi.org/10.1007/s13225-020-00466-2)) |
| Otu14265 | 1 | k__Fungi(100);p__Ascomycota(100);c__Saccharomycetes(100);o__Saccharomycetales(100);f__Metschnikowiaceae(100);g__Metschnikowia(100);g__Metschnikowia_unclassified(100); | Metschnikowia | 13 | Saprotroph | \|Undefined Saprotroph\| | Yeast | NULL | Probable | As nectar-tap_saprotroph (Põlme et al. 2020); Decay sugar-rich substrates (Põlme et al. 2020) | Põlme S, et al. 2020. Fungal Diversity 105:1-16 ((https://doi.org/10.1007/s13225-020-00466-2)) |
| Otu14267 | 1 | k__Fungi(100);p__Ascomycota(100);c__Saccharomycetes(100);o__Saccharomycetales(100);f__Metschnikowiaceae(100);g__Metschnikowia(100);g__Metschnikowia_unclassified(100); | Metschnikowia | 13 | Saprotroph | \|Undefined Saprotroph\| | Yeast | NULL | Probable | As nectar-tap_saprotroph (Põlme et al. 2020); Decay sugar-rich substrates (Põlme et al. 2020) | Põlme S, et al. 2020. Fungal Diversity 105:1-16 ((https://doi.org/10.1007/s13225-020-00466-2)) |
| Otu14270 | 1 | k__Fungi(100);p__Ascomycota(100);c__Saccharomycetes(100);o__Saccharomycetales(100);f__Metschnikowiaceae(100);g__Metschnikowia(100);s__Metschnikowia_pulcherrima(100); | Metschnikowia | 13 | Saprotroph | \|Undefined Saprotroph\| | Yeast | NULL | Probable | As nectar-tap_saprotroph (Põlme et al. 2020); Decay sugar-rich substrates (Põlme et al. 2020) | Põlme S, et al. 2020. Fungal Diversity 105:1-16 ((https://doi.org/10.1007/s13225-020-00466-2)) |
| Otu14271 | 1 | k__Fungi(100);p__Ascomycota(100);c__Saccharomycetes(100);o__Saccharomycetales(100);f__Metschnikowiaceae(100);g__Metschnikowia(100);s__Metschnikowia_pulcherrima(100); | Metschnikowia | 13 | Saprotroph | \|Undefined Saprotroph\| | Yeast | NULL | Probable | As nectar-tap_saprotroph (Põlme et al. 2020); Decay sugar-rich substrates (Põlme et al. 2020) | Põlme S, et al. 2020. Fungal Diversity 105:1-16 ((https://doi.org/10.1007/s13225-020-00466-2)) |
| Otu14273 | 1 | k__Fungi(100);p__Ascomycota(100);c__Saccharomycetes(100);o__Saccharomycetales(100);f__Metschnikowiaceae(100);g__Metschnikowia(100);g__Metschnikowia_unclassified(100); | Metschnikowia | 13 | Saprotroph | \|Undefined Saprotroph\| | Yeast | NULL | Probable | As nectar-tap_saprotroph (Põlme et al. 2020); Decay sugar-rich substrates (Põlme et al. 2020) | Põlme S, et al. 2020. Fungal Diversity 105:1-16 ((https://doi.org/10.1007/s13225-020-00466-2)) |
| Otu14274 | 1 | k__Fungi(100);p__Ascomycota(100);c__Saccharomycetes(100);o__Saccharomycetales(100);f__Metschnikowiaceae(100);g__Metschnikowia(100);s__Metschnikowia_pulcherrima(100); | Metschnikowia | 13 | Saprotroph | \|Undefined Saprotroph\| | Yeast | NULL | Probable | As nectar-tap_saprotroph (Põlme et al. 2020); Decay sugar-rich substrates (Põlme et al. 2020) | Põlme S, et al. 2020. Fungal Diversity 105:1-16 ((https://doi.org/10.1007/s13225-020-00466-2)) |
| Otu14276 | 1 | k__Fungi(100);p__Ascomycota(100);c__Saccharomycetes(100);o__Saccharomycetales(100);f__Saccharomycetales_family_Incertae_sedis(100);g__Candida(100);g__Candida_unclassified(100); | Candida | 13 | Pathotroph-Saprotroph-Symbiotroph | Animal Pathogen-Endophyte-Endosymbiont-Epiphyte-\|Undefined Saprotroph\| | Dimorphic Yeast-Yeast | NULL | Probable | As nectar-tap_saprotroph (Põlme et al. 2020); Decay sugar-rich substrates (Põlme et al. 2020); This asexual genus is polyphyletic and contain species that function very differently depend on what they are. Caution should be taken when interpreting guilds for Candida species. | Manolakaki D, et al. 2010. Virulence. 1: 367-375 ((https://doi.org/10.4161/viru.1.5.12796)); Mekbib SB, et al. 2011. Tropical Plant Pathology 36:233-240 ((https://doi.org/10.1590/S1982-56762011000400004)); Põlme S, et al. 2020. Fungal Diversity 105:1-16 ((https://doi.org/10.1007/s13225-020-00466-2)) |
| Otu14279 | 1 | k__Fungi(100);p__Ascomycota(100);c__Saccharomycetes(100);o__Saccharomycetales(100);f__Metschnikowiaceae(100);g__Metschnikowia(100);g__Metschnikowia_unclassified(100); | Metschnikowia | 13 | Saprotroph | \|Undefined Saprotroph\| | Yeast | NULL | Probable | As nectar-tap_saprotroph (Põlme et al. 2020); Decay sugar-rich substrates (Põlme et al. 2020) | Põlme S, et al. 2020. Fungal Diversity 105:1-16 ((https://doi.org/10.1007/s13225-020-00466-2)) |
| Otu14281 | 1 | k__Fungi(100);p__Ascomycota(100);c__Saccharomycetes(100);o__Saccharomycetales(100);f__Metschnikowiaceae(100);g__Metschnikowia(100);s__Metschnikowia_pulcherrima(100); | Metschnikowia | 13 | Saprotroph | \|Undefined Saprotroph\| | Yeast | NULL | Probable | As nectar-tap_saprotroph (Põlme et al. 2020); Decay sugar-rich substrates (Põlme et al. 2020) | Põlme S, et al. 2020. Fungal Diversity 105:1-16 ((https://doi.org/10.1007/s13225-020-00466-2)) |
| Otu14282 | 1 | k__Fungi(100);p__Ascomycota(100);c__Saccharomycetes(100);o__Saccharomycetales(100);f__Metschnikowiaceae(100);g__Metschnikowia(100);s__Metschnikowia_pulcherrima(100); | Metschnikowia | 13 | Saprotroph | \|Undefined Saprotroph\| | Yeast | NULL | Probable | As nectar-tap_saprotroph (Põlme et al. 2020); Decay sugar-rich substrates (Põlme et al. 2020) | Põlme S, et al. 2020. Fungal Diversity 105:1-16 ((https://doi.org/10.1007/s13225-020-00466-2)) |
| Otu14284 | 1 | k__Fungi(100);p__Ascomycota(100);c__Saccharomycetes(100);o__Saccharomycetales(100);f__Metschnikowiaceae(100);g__Metschnikowia(100);s__Metschnikowia_pulcherrima(100); | Metschnikowia | 13 | Saprotroph | \|Undefined Saprotroph\| | Yeast | NULL | Probable | As nectar-tap_saprotroph (Põlme et al. 2020); Decay sugar-rich substrates (Põlme et al. 2020) | Põlme S, et al. 2020. Fungal Diversity 105:1-16 ((https://doi.org/10.1007/s13225-020-00466-2)) |
| Otu14286 | 1 | k__Fungi(100);p__Ascomycota(100);c__Saccharomycetes(100);o__Saccharomycetales(100);f__Metschnikowiaceae(100);g__Metschnikowia(100);s__Metschnikowia_pulcherrima(100); | Metschnikowia | 13 | Saprotroph | \|Undefined Saprotroph\| | Yeast | NULL | Probable | As nectar-tap_saprotroph (Põlme et al. 2020); Decay sugar-rich substrates (Põlme et al. 2020) | Põlme S, et al. 2020. Fungal Diversity 105:1-16 ((https://doi.org/10.1007/s13225-020-00466-2)) |
| Otu14287 | 1 | k__Fungi(100);p__Ascomycota(100);c__Saccharomycetes(100);o__Saccharomycetales(100);f__Metschnikowiaceae(100);g__Metschnikowia(100);s__Metschnikowia_pulcherrima(100); | Metschnikowia | 13 | Saprotroph | \|Undefined Saprotroph\| | Yeast | NULL | Probable | As nectar-tap_saprotroph (Põlme et al. 2020); Decay sugar-rich substrates (Põlme et al. 2020) | Põlme S, et al. 2020. Fungal Diversity 105:1-16 ((https://doi.org/10.1007/s13225-020-00466-2)) |
| Otu14288 | 1 | k__Fungi(100);p__Ascomycota(100);c__Saccharomycetes(100);o__Saccharomycetales(100);f__Metschnikowiaceae(100);g__Metschnikowia(100);g__Metschnikowia_unclassified(100); | Metschnikowia | 13 | Saprotroph | \|Undefined Saprotroph\| | Yeast | NULL | Probable | As nectar-tap_saprotroph (Põlme et al. 2020); Decay sugar-rich substrates (Põlme et al. 2020) | Põlme S, et al. 2020. Fungal Diversity 105:1-16 ((https://doi.org/10.1007/s13225-020-00466-2)) |
| Otu14289 | 1 | k__Fungi(100);p__Ascomycota(100);c__Saccharomycetes(100);o__Saccharomycetales(100);f__Metschnikowiaceae(100);g__Metschnikowia(100);s__Metschnikowia_pulcherrima(100); | Metschnikowia | 13 | Saprotroph | \|Undefined Saprotroph\| | Yeast | NULL | Probable | As nectar-tap_saprotroph (Põlme et al. 2020); Decay sugar-rich substrates (Põlme et al. 2020) | Põlme S, et al. 2020. Fungal Diversity 105:1-16 ((https://doi.org/10.1007/s13225-020-00466-2)) |
| Otu14290 | 1 | k__Fungi(100);p__Ascomycota(100);c__Saccharomycetes(100);o__Saccharomycetales(100);f__Metschnikowiaceae(100);g__Metschnikowia(100);g__Metschnikowia_unclassified(100); | Metschnikowia | 13 | Saprotroph | \|Undefined Saprotroph\| | Yeast | NULL | Probable | As nectar-tap_saprotroph (Põlme et al. 2020); Decay sugar-rich substrates (Põlme et al. 2020) | Põlme S, et al. 2020. Fungal Diversity 105:1-16 ((https://doi.org/10.1007/s13225-020-00466-2)) |
| Otu14291 | 1 | k__Fungi(100);p__Ascomycota(100);c__Saccharomycetes(100);o__Saccharomycetales(100);f__Metschnikowiaceae(100);g__Metschnikowia(100);s__Metschnikowia_pulcherrima(100); | Metschnikowia | 13 | Saprotroph | \|Undefined Saprotroph\| | Yeast | NULL | Probable | As nectar-tap_saprotroph (Põlme et al. 2020); Decay sugar-rich substrates (Põlme et al. 2020) | Põlme S, et al. 2020. Fungal Diversity 105:1-16 ((https://doi.org/10.1007/s13225-020-00466-2)) |
| Otu14293 | 1 | k__Fungi(100);p__Ascomycota(100);c__Saccharomycetes(100);o__Saccharomycetales(100);f__Metschnikowiaceae(100);g__Metschnikowia(100);s__Metschnikowia_pulcherrima(100); | Metschnikowia | 13 | Saprotroph | \|Undefined Saprotroph\| | Yeast | NULL | Probable | As nectar-tap_saprotroph (Põlme et al. 2020); Decay sugar-rich substrates (Põlme et al. 2020) | Põlme S, et al. 2020. Fungal Diversity 105:1-16 ((https://doi.org/10.1007/s13225-020-00466-2)) |
| Otu14294 | 1 | k__Fungi(100);p__Ascomycota(100);c__Saccharomycetes(100);o__Saccharomycetales(100);f__Metschnikowiaceae(100);g__Metschnikowia(100);g__Metschnikowia_unclassified(100); | Metschnikowia | 13 | Saprotroph | \|Undefined Saprotroph\| | Yeast | NULL | Probable | As nectar-tap_saprotroph (Põlme et al. 2020); Decay sugar-rich substrates (Põlme et al. 2020) | Põlme S, et al. 2020. Fungal Diversity 105:1-16 ((https://doi.org/10.1007/s13225-020-00466-2)) |
| Otu14295 | 1 | k__Fungi(100);p__Ascomycota(100);c__Saccharomycetes(100);o__Saccharomycetales(100);f__Metschnikowiaceae(100);g__Metschnikowia(100);s__Metschnikowia_pulcherrima(100); | Metschnikowia | 13 | Saprotroph | \|Undefined Saprotroph\| | Yeast | NULL | Probable | As nectar-tap_saprotroph (Põlme et al. 2020); Decay sugar-rich substrates (Põlme et al. 2020) | Põlme S, et al. 2020. Fungal Diversity 105:1-16 ((https://doi.org/10.1007/s13225-020-00466-2)) |
| Otu14296 | 1 | k__Fungi(100);p__Ascomycota(100);c__Saccharomycetes(100);o__Saccharomycetales(100);f__Metschnikowiaceae(100);g__Metschnikowia(100);g__Metschnikowia_unclassified(100); | Metschnikowia | 13 | Saprotroph | \|Undefined Saprotroph\| | Yeast | NULL | Probable | As nectar-tap_saprotroph (Põlme et al. 2020); Decay sugar-rich substrates (Põlme et al. 2020) | Põlme S, et al. 2020. Fungal Diversity 105:1-16 ((https://doi.org/10.1007/s13225-020-00466-2)) |
| Otu14297 | 1 | k__Fungi(100);p__Ascomycota(100);c__Saccharomycetes(100);o__Saccharomycetales(100);f__Metschnikowiaceae(100);g__Metschnikowia(100);s__Metschnikowia_pulcherrima(100); | Metschnikowia | 13 | Saprotroph | \|Undefined Saprotroph\| | Yeast | NULL | Probable | As nectar-tap_saprotroph (Põlme et al. 2020); Decay sugar-rich substrates (Põlme et al. 2020) | Põlme S, et al. 2020. Fungal Diversity 105:1-16 ((https://doi.org/10.1007/s13225-020-00466-2)) |
| Otu14298 | 1 | k__Fungi(100);p__Ascomycota(100);c__Saccharomycetes(100);o__Saccharomycetales(100);f__Metschnikowiaceae(100);g__Metschnikowia(100);g__Metschnikowia_unclassified(100); | Metschnikowia | 13 | Saprotroph | \|Undefined Saprotroph\| | Yeast | NULL | Probable | As nectar-tap_saprotroph (Põlme et al. 2020); Decay sugar-rich substrates (Põlme et al. 2020) | Põlme S, et al. 2020. Fungal Diversity 105:1-16 ((https://doi.org/10.1007/s13225-020-00466-2)) |
| Otu14299 | 1 | k__Fungi(100);p__Ascomycota(100);c__Saccharomycetes(100);o__Saccharomycetales(100);f__Metschnikowiaceae(100);g__Metschnikowia(100);s__Metschnikowia_pulcherrima(100); | Metschnikowia | 13 | Saprotroph | \|Undefined Saprotroph\| | Yeast | NULL | Probable | As nectar-tap_saprotroph (Põlme et al. 2020); Decay sugar-rich substrates (Põlme et al. 2020) | Põlme S, et al. 2020. Fungal Diversity 105:1-16 ((https://doi.org/10.1007/s13225-020-00466-2)) |
| Otu14300 | 1 | k__Fungi(100);p__Ascomycota(100);c__Saccharomycetes(100);o__Saccharomycetales(100);f__Metschnikowiaceae(100);g__Metschnikowia(100);g__Metschnikowia_unclassified(100); | Metschnikowia | 13 | Saprotroph | \|Undefined Saprotroph\| | Yeast | NULL | Probable | As nectar-tap_saprotroph (Põlme et al. 2020); Decay sugar-rich substrates (Põlme et al. 2020) | Põlme S, et al. 2020. Fungal Diversity 105:1-16 ((https://doi.org/10.1007/s13225-020-00466-2)) |
| Otu14301 | 1 | k__Fungi(100);p__Ascomycota(100);c__Saccharomycetes(100);o__Saccharomycetales(100);f__Metschnikowiaceae(100);g__Metschnikowia(100);g__Metschnikowia_unclassified(100); | Metschnikowia | 13 | Saprotroph | \|Undefined Saprotroph\| | Yeast | NULL | Probable | As nectar-tap_saprotroph (Põlme et al. 2020); Decay sugar-rich substrates (Põlme et al. 2020) | Põlme S, et al. 2020. Fungal Diversity 105:1-16 ((https://doi.org/10.1007/s13225-020-00466-2)) |
| Otu14302 | 1 | k__Fungi(100);p__Ascomycota(100);c__Saccharomycetes(100);o__Saccharomycetales(100);f__Metschnikowiaceae(100);g__Metschnikowia(100);s__Metschnikowia_pulcherrima(100); | Metschnikowia | 13 | Saprotroph | \|Undefined Saprotroph\| | Yeast | NULL | Probable | As nectar-tap_saprotroph (Põlme et al. 2020); Decay sugar-rich substrates (Põlme et al. 2020) | Põlme S, et al. 2020. Fungal Diversity 105:1-16 ((https://doi.org/10.1007/s13225-020-00466-2)) |
| Otu14303 | 1 | k__Fungi(100);p__Ascomycota(100);c__Saccharomycetes(100);o__Saccharomycetales(100);f__Metschnikowiaceae(100);g__Metschnikowia(100);s__Metschnikowia_pulcherrima(100); | Metschnikowia | 13 | Saprotroph | \|Undefined Saprotroph\| | Yeast | NULL | Probable | As nectar-tap_saprotroph (Põlme et al. 2020); Decay sugar-rich substrates (Põlme et al. 2020) | Põlme S, et al. 2020. Fungal Diversity 105:1-16 ((https://doi.org/10.1007/s13225-020-00466-2)) |
| Otu14304 | 1 | k__Fungi(100);p__Ascomycota(100);c__Saccharomycetes(100);o__Saccharomycetales(100);f__Metschnikowiaceae(100);g__Metschnikowia(100);s__Metschnikowia_pulcherrima(100); | Metschnikowia | 13 | Saprotroph | \|Undefined Saprotroph\| | Yeast | NULL | Probable | As nectar-tap_saprotroph (Põlme et al. 2020); Decay sugar-rich substrates (Põlme et al. 2020) | Põlme S, et al. 2020. Fungal Diversity 105:1-16 ((https://doi.org/10.1007/s13225-020-00466-2)) |
| Otu14305 | 1 | k__Fungi(100);p__Ascomycota(100);c__Saccharomycetes(100);o__Saccharomycetales(100);f__Metschnikowiaceae(100);g__Metschnikowia(100);s__Metschnikowia_pulcherrima(100); | Metschnikowia | 13 | Saprotroph | \|Undefined Saprotroph\| | Yeast | NULL | Probable | As nectar-tap_saprotroph (Põlme et al. 2020); Decay sugar-rich substrates (Põlme et al. 2020) | Põlme S, et al. 2020. Fungal Diversity 105:1-16 ((https://doi.org/10.1007/s13225-020-00466-2)) |
| Otu14307 | 1 | k__Fungi(100);p__Ascomycota(100);c__Saccharomycetes(100);o__Saccharomycetales(100);f__Saccharomycetales_family_Incertae_sedis(100);g__Candida(100);g__Candida_unclassified(100); | Candida | 13 | Pathotroph-Saprotroph-Symbiotroph | Animal Pathogen-Endophyte-Endosymbiont-Epiphyte-\|Undefined Saprotroph\| | Dimorphic Yeast-Yeast | NULL | Probable | As nectar-tap_saprotroph (Põlme et al. 2020); Decay sugar-rich substrates (Põlme et al. 2020); This asexual genus is polyphyletic and contain species that function very differently depend on what they are. Caution should be taken when interpreting guilds for Candida species. | Manolakaki D, et al. 2010. Virulence. 1: 367-375 ((https://doi.org/10.4161/viru.1.5.12796)); Mekbib SB, et al. 2011. Tropical Plant Pathology 36:233-240 ((https://doi.org/10.1590/S1982-56762011000400004)); Põlme S, et al. 2020. Fungal Diversity 105:1-16 ((https://doi.org/10.1007/s13225-020-00466-2)) |
| Otu14309 | 1 | k__Fungi(100);p__Ascomycota(100);c__Saccharomycetes(100);o__Saccharomycetales(100);f__Metschnikowiaceae(100);g__Metschnikowia(100);s__Metschnikowia_pulcherrima(100); | Metschnikowia | 13 | Saprotroph | \|Undefined Saprotroph\| | Yeast | NULL | Probable | As nectar-tap_saprotroph (Põlme et al. 2020); Decay sugar-rich substrates (Põlme et al. 2020) | Põlme S, et al. 2020. Fungal Diversity 105:1-16 ((https://doi.org/10.1007/s13225-020-00466-2)) |
| Otu14317 | 1 | k__Fungi(100);p__Ascomycota(100);c__Saccharomycetes(100);o__Saccharomycetales(100);f__Metschnikowiaceae(100);g__Metschnikowia(100);s__Metschnikowia_pulcherrima(100); | Metschnikowia | 13 | Saprotroph | \|Undefined Saprotroph\| | Yeast | NULL | Probable | As nectar-tap_saprotroph (Põlme et al. 2020); Decay sugar-rich substrates (Põlme et al. 2020) | Põlme S, et al. 2020. Fungal Diversity 105:1-16 ((https://doi.org/10.1007/s13225-020-00466-2)) |
| Otu14321 | 1 | k__Fungi(100);p__Ascomycota(100);c__Saccharomycetes(100);o__Saccharomycetales(100);f__Metschnikowiaceae(100);g__Metschnikowia(100);g__Metschnikowia_unclassified(100); | Metschnikowia | 13 | Saprotroph | \|Undefined Saprotroph\| | Yeast | NULL | Probable | As nectar-tap_saprotroph (Põlme et al. 2020); Decay sugar-rich substrates (Põlme et al. 2020) | Põlme S, et al. 2020. Fungal Diversity 105:1-16 ((https://doi.org/10.1007/s13225-020-00466-2)) |
| Otu14322 | 1 | k__Fungi(100);p__Ascomycota(100);c__Saccharomycetes(100);o__Saccharomycetales(100);f__Metschnikowiaceae(100);g__Metschnikowia(100);s__Metschnikowia_pulcherrima(100); | Metschnikowia | 13 | Saprotroph | \|Undefined Saprotroph\| | Yeast | NULL | Probable | As nectar-tap_saprotroph (Põlme et al. 2020); Decay sugar-rich substrates (Põlme et al. 2020) | Põlme S, et al. 2020. Fungal Diversity 105:1-16 ((https://doi.org/10.1007/s13225-020-00466-2)) |
| Otu14323 | 1 | k__Fungi(100);p__Ascomycota(100);c__Saccharomycetes(100);o__Saccharomycetales(100);f__Metschnikowiaceae(100);g__Metschnikowia(100);s__Metschnikowia_pulcherrima(100); | Metschnikowia | 13 | Saprotroph | \|Undefined Saprotroph\| | Yeast | NULL | Probable | As nectar-tap_saprotroph (Põlme et al. 2020); Decay sugar-rich substrates (Põlme et al. 2020) | Põlme S, et al. 2020. Fungal Diversity 105:1-16 ((https://doi.org/10.1007/s13225-020-00466-2)) |
| Otu14330 | 1 | k__Fungi(100);p__Ascomycota(100);c__Saccharomycetes(100);o__Saccharomycetales(100);f__Metschnikowiaceae(100);g__Metschnikowia(100);s__Metschnikowia_pulcherrima(100); | Metschnikowia | 13 | Saprotroph | \|Undefined Saprotroph\| | Yeast | NULL | Probable | As nectar-tap_saprotroph (Põlme et al. 2020); Decay sugar-rich substrates (Põlme et al. 2020) | Põlme S, et al. 2020. Fungal Diversity 105:1-16 ((https://doi.org/10.1007/s13225-020-00466-2)) |
| Otu14331 | 1 | k__Fungi(100);p__Ascomycota(100);c__Saccharomycetes(100);o__Saccharomycetales(100);f__Metschnikowiaceae(100);g__Metschnikowia(100);s__Metschnikowia_pulcherrima(100); | Metschnikowia | 13 | Saprotroph | \|Undefined Saprotroph\| | Yeast | NULL | Probable | As nectar-tap_saprotroph (Põlme et al. 2020); Decay sugar-rich substrates (Põlme et al. 2020) | Põlme S, et al. 2020. Fungal Diversity 105:1-16 ((https://doi.org/10.1007/s13225-020-00466-2)) |
| Otu14333 | 1 | k__Fungi(100);p__Ascomycota(100);c__Saccharomycetes(100);o__Saccharomycetales(100);f__Metschnikowiaceae(100);g__Metschnikowia(100);s__Metschnikowia_pulcherrima(100); | Metschnikowia | 13 | Saprotroph | \|Undefined Saprotroph\| | Yeast | NULL | Probable | As nectar-tap_saprotroph (Põlme et al. 2020); Decay sugar-rich substrates (Põlme et al. 2020) | Põlme S, et al. 2020. Fungal Diversity 105:1-16 ((https://doi.org/10.1007/s13225-020-00466-2)) |
| Otu14335 | 1 | k__Fungi(100);p__Ascomycota(100);c__Saccharomycetes(100);o__Saccharomycetales(100);f__Metschnikowiaceae(100);g__Metschnikowia(100);s__Metschnikowia_pulcherrima(100); | Metschnikowia | 13 | Saprotroph | \|Undefined Saprotroph\| | Yeast | NULL | Probable | As nectar-tap_saprotroph (Põlme et al. 2020); Decay sugar-rich substrates (Põlme et al. 2020) | Põlme S, et al. 2020. Fungal Diversity 105:1-16 ((https://doi.org/10.1007/s13225-020-00466-2)) |
| Otu14337 | 1 | k__Fungi(100);p__Ascomycota(100);c__Saccharomycetes(100);o__Saccharomycetales(100);f__Metschnikowiaceae(100);g__Metschnikowia(100);s__Metschnikowia_pulcherrima(100); | Metschnikowia | 13 | Saprotroph | \|Undefined Saprotroph\| | Yeast | NULL | Probable | As nectar-tap_saprotroph (Põlme et al. 2020); Decay sugar-rich substrates (Põlme et al. 2020) | Põlme S, et al. 2020. Fungal Diversity 105:1-16 ((https://doi.org/10.1007/s13225-020-00466-2)) |
| Otu14338 | 1 | k__Fungi(100);p__Ascomycota(100);c__Saccharomycetes(100);o__Saccharomycetales(100);f__Metschnikowiaceae(100);g__Metschnikowia(100);s__Metschnikowia_pulcherrima(100); | Metschnikowia | 13 | Saprotroph | \|Undefined Saprotroph\| | Yeast | NULL | Probable | As nectar-tap_saprotroph (Põlme et al. 2020); Decay sugar-rich substrates (Põlme et al. 2020) | Põlme S, et al. 2020. Fungal Diversity 105:1-16 ((https://doi.org/10.1007/s13225-020-00466-2)) |
| Otu14339 | 1 | k__Fungi(100);p__Ascomycota(100);c__Saccharomycetes(100);o__Saccharomycetales(100);f__Metschnikowiaceae(100);g__Metschnikowia(100);s__Metschnikowia_pulcherrima(100); | Metschnikowia | 13 | Saprotroph | \|Undefined Saprotroph\| | Yeast | NULL | Probable | As nectar-tap_saprotroph (Põlme et al. 2020); Decay sugar-rich substrates (Põlme et al. 2020) | Põlme S, et al. 2020. Fungal Diversity 105:1-16 ((https://doi.org/10.1007/s13225-020-00466-2)) |
| Otu14340 | 1 | k__Fungi(100);p__Ascomycota(100);c__Saccharomycetes(100);o__Saccharomycetales(100);f__Metschnikowiaceae(100);g__Metschnikowia(100);s__Metschnikowia_pulcherrima(100); | Metschnikowia | 13 | Saprotroph | \|Undefined Saprotroph\| | Yeast | NULL | Probable | As nectar-tap_saprotroph (Põlme et al. 2020); Decay sugar-rich substrates (Põlme et al. 2020) | Põlme S, et al. 2020. Fungal Diversity 105:1-16 ((https://doi.org/10.1007/s13225-020-00466-2)) |
| Otu14341 | 1 | k__Fungi(100);p__Ascomycota(100);c__Saccharomycetes(100);o__Saccharomycetales(100);f__Metschnikowiaceae(100);g__Metschnikowia(100);g__Metschnikowia_unclassified(100); | Metschnikowia | 13 | Saprotroph | \|Undefined Saprotroph\| | Yeast | NULL | Probable | As nectar-tap_saprotroph (Põlme et al. 2020); Decay sugar-rich substrates (Põlme et al. 2020) | Põlme S, et al. 2020. Fungal Diversity 105:1-16 ((https://doi.org/10.1007/s13225-020-00466-2)) |
| Otu14342 | 1 | k__Fungi(100);p__Ascomycota(100);c__Saccharomycetes(100);o__Saccharomycetales(100);f__Metschnikowiaceae(100);g__Metschnikowia(100);g__Metschnikowia_unclassified(100); | Metschnikowia | 13 | Saprotroph | \|Undefined Saprotroph\| | Yeast | NULL | Probable | As nectar-tap_saprotroph (Põlme et al. 2020); Decay sugar-rich substrates (Põlme et al. 2020) | Põlme S, et al. 2020. Fungal Diversity 105:1-16 ((https://doi.org/10.1007/s13225-020-00466-2)) |
| Otu14343 | 1 | k__Fungi(100);p__Ascomycota(100);c__Saccharomycetes(100);o__Saccharomycetales(100);f__Metschnikowiaceae(100);g__Metschnikowia(100);s__Metschnikowia_pulcherrima(100); | Metschnikowia | 13 | Saprotroph | \|Undefined Saprotroph\| | Yeast | NULL | Probable | As nectar-tap_saprotroph (Põlme et al. 2020); Decay sugar-rich substrates (Põlme et al. 2020) | Põlme S, et al. 2020. Fungal Diversity 105:1-16 ((https://doi.org/10.1007/s13225-020-00466-2)) |
| Otu14344 | 1 | k__Fungi(100);p__Ascomycota(100);c__Saccharomycetes(100);o__Saccharomycetales(100);f__Metschnikowiaceae(100);g__Metschnikowia(100);g__Metschnikowia_unclassified(100); | Metschnikowia | 13 | Saprotroph | \|Undefined Saprotroph\| | Yeast | NULL | Probable | As nectar-tap_saprotroph (Põlme et al. 2020); Decay sugar-rich substrates (Põlme et al. 2020) | Põlme S, et al. 2020. Fungal Diversity 105:1-16 ((https://doi.org/10.1007/s13225-020-00466-2)) |
| Otu14345 | 1 | k__Fungi(100);p__Ascomycota(100);c__Saccharomycetes(100);o__Saccharomycetales(100);f__Metschnikowiaceae(100);g__Metschnikowia(100);s__Metschnikowia_pulcherrima(100); | Metschnikowia | 13 | Saprotroph | \|Undefined Saprotroph\| | Yeast | NULL | Probable | As nectar-tap_saprotroph (Põlme et al. 2020); Decay sugar-rich substrates (Põlme et al. 2020) | Põlme S, et al. 2020. Fungal Diversity 105:1-16 ((https://doi.org/10.1007/s13225-020-00466-2)) |
| Otu14346 | 1 | k__Fungi(100);p__Ascomycota(100);c__Saccharomycetes(100);o__Saccharomycetales(100);f__Metschnikowiaceae(100);g__Metschnikowia(100);s__Metschnikowia_pulcherrima(100); | Metschnikowia | 13 | Saprotroph | \|Undefined Saprotroph\| | Yeast | NULL | Probable | As nectar-tap_saprotroph (Põlme et al. 2020); Decay sugar-rich substrates (Põlme et al. 2020) | Põlme S, et al. 2020. Fungal Diversity 105:1-16 ((https://doi.org/10.1007/s13225-020-00466-2)) |
| Otu14347 | 1 | k__Fungi(100);p__Ascomycota(100);c__Saccharomycetes(100);o__Saccharomycetales(100);f__Metschnikowiaceae(100);g__Metschnikowia(100);g__Metschnikowia_unclassified(100); | Metschnikowia | 13 | Saprotroph | \|Undefined Saprotroph\| | Yeast | NULL | Probable | As nectar-tap_saprotroph (Põlme et al. 2020); Decay sugar-rich substrates (Põlme et al. 2020) | Põlme S, et al. 2020. Fungal Diversity 105:1-16 ((https://doi.org/10.1007/s13225-020-00466-2)) |
| Otu14348 | 1 | k__Fungi(100);p__Ascomycota(100);c__Saccharomycetes(100);o__Saccharomycetales(100);f__Metschnikowiaceae(100);g__Metschnikowia(100);s__Metschnikowia_pulcherrima(100); | Metschnikowia | 13 | Saprotroph | \|Undefined Saprotroph\| | Yeast | NULL | Probable | As nectar-tap_saprotroph (Põlme et al. 2020); Decay sugar-rich substrates (Põlme et al. 2020) | Põlme S, et al. 2020. Fungal Diversity 105:1-16 ((https://doi.org/10.1007/s13225-020-00466-2)) |
| Otu14350 | 1 | k__Fungi(100);p__Ascomycota(100);c__Saccharomycetes(100);o__Saccharomycetales(100);f__Metschnikowiaceae(100);g__Metschnikowia(100);s__Metschnikowia_pulcherrima(100); | Metschnikowia | 13 | Saprotroph | \|Undefined Saprotroph\| | Yeast | NULL | Probable | As nectar-tap_saprotroph (Põlme et al. 2020); Decay sugar-rich substrates (Põlme et al. 2020) | Põlme S, et al. 2020. Fungal Diversity 105:1-16 ((https://doi.org/10.1007/s13225-020-00466-2)) |
| Otu14351 | 1 | k__Fungi(100);p__Ascomycota(100);c__Saccharomycetes(100);o__Saccharomycetales(100);f__Metschnikowiaceae(100);g__Metschnikowia(100);s__Metschnikowia_pulcherrima(100); | Metschnikowia | 13 | Saprotroph | \|Undefined Saprotroph\| | Yeast | NULL | Probable | As nectar-tap_saprotroph (Põlme et al. 2020); Decay sugar-rich substrates (Põlme et al. 2020) | Põlme S, et al. 2020. Fungal Diversity 105:1-16 ((https://doi.org/10.1007/s13225-020-00466-2)) |
| Otu14352 | 1 | k__Fungi(100);p__Ascomycota(100);c__Saccharomycetes(100);o__Saccharomycetales(100);f__Metschnikowiaceae(100);g__Metschnikowia(100);s__Metschnikowia_pulcherrima(100); | Metschnikowia | 13 | Saprotroph | \|Undefined Saprotroph\| | Yeast | NULL | Probable | As nectar-tap_saprotroph (Põlme et al. 2020); Decay sugar-rich substrates (Põlme et al. 2020) | Põlme S, et al. 2020. Fungal Diversity 105:1-16 ((https://doi.org/10.1007/s13225-020-00466-2)) |
| Otu14353 | 1 | k__Fungi(100);p__Ascomycota(100);c__Saccharomycetes(100);o__Saccharomycetales(100);f__Metschnikowiaceae(100);g__Metschnikowia(100);s__Metschnikowia_pulcherrima(100); | Metschnikowia | 13 | Saprotroph | \|Undefined Saprotroph\| | Yeast | NULL | Probable | As nectar-tap_saprotroph (Põlme et al. 2020); Decay sugar-rich substrates (Põlme et al. 2020) | Põlme S, et al. 2020. Fungal Diversity 105:1-16 ((https://doi.org/10.1007/s13225-020-00466-2)) |
| Otu14355 | 1 | k__Fungi(100);p__Ascomycota(100);c__Saccharomycetes(100);o__Saccharomycetales(100);f__Metschnikowiaceae(100);g__Metschnikowia(100);g__Metschnikowia_unclassified(100); | Metschnikowia | 13 | Saprotroph | \|Undefined Saprotroph\| | Yeast | NULL | Probable | As nectar-tap_saprotroph (Põlme et al. 2020); Decay sugar-rich substrates (Põlme et al. 2020) | Põlme S, et al. 2020. Fungal Diversity 105:1-16 ((https://doi.org/10.1007/s13225-020-00466-2)) |
| Otu14356 | 1 | k__Fungi(100);p__Ascomycota(100);c__Saccharomycetes(100);o__Saccharomycetales(100);f__Metschnikowiaceae(100);g__Metschnikowia(100);s__Metschnikowia_pulcherrima(100); | Metschnikowia | 13 | Saprotroph | \|Undefined Saprotroph\| | Yeast | NULL | Probable | As nectar-tap_saprotroph (Põlme et al. 2020); Decay sugar-rich substrates (Põlme et al. 2020) | Põlme S, et al. 2020. Fungal Diversity 105:1-16 ((https://doi.org/10.1007/s13225-020-00466-2)) |
| Otu14358 | 1 | k__Fungi(100);p__Ascomycota(100);c__Saccharomycetes(100);o__Saccharomycetales(100);f__Metschnikowiaceae(100);g__Metschnikowia(100);s__Metschnikowia_pulcherrima(100); | Metschnikowia | 13 | Saprotroph | \|Undefined Saprotroph\| | Yeast | NULL | Probable | As nectar-tap_saprotroph (Põlme et al. 2020); Decay sugar-rich substrates (Põlme et al. 2020) | Põlme S, et al. 2020. Fungal Diversity 105:1-16 ((https://doi.org/10.1007/s13225-020-00466-2)) |
| Otu14359 | 1 | k__Fungi(100);p__Ascomycota(100);c__Saccharomycetes(100);o__Saccharomycetales(100);f__Metschnikowiaceae(100);g__Metschnikowia(100);s__Metschnikowia_pulcherrima(100); | Metschnikowia | 13 | Saprotroph | \|Undefined Saprotroph\| | Yeast | NULL | Probable | As nectar-tap_saprotroph (Põlme et al. 2020); Decay sugar-rich substrates (Põlme et al. 2020) | Põlme S, et al. 2020. Fungal Diversity 105:1-16 ((https://doi.org/10.1007/s13225-020-00466-2)) |
| Otu14360 | 1 | k__Fungi(100);p__Ascomycota(100);c__Saccharomycetes(100);o__Saccharomycetales(100);f__Metschnikowiaceae(100);g__Metschnikowia(100);s__Metschnikowia_pulcherrima(100); | Metschnikowia | 13 | Saprotroph | \|Undefined Saprotroph\| | Yeast | NULL | Probable | As nectar-tap_saprotroph (Põlme et al. 2020); Decay sugar-rich substrates (Põlme et al. 2020) | Põlme S, et al. 2020. Fungal Diversity 105:1-16 ((https://doi.org/10.1007/s13225-020-00466-2)) |
| Otu14362 | 1 | k__Fungi(100);p__Ascomycota(100);c__Saccharomycetes(100);o__Saccharomycetales(100);f__Metschnikowiaceae(100);g__Metschnikowia(100);s__Metschnikowia_pulcherrima(100); | Metschnikowia | 13 | Saprotroph | \|Undefined Saprotroph\| | Yeast | NULL | Probable | As nectar-tap_saprotroph (Põlme et al. 2020); Decay sugar-rich substrates (Põlme et al. 2020) | Põlme S, et al. 2020. Fungal Diversity 105:1-16 ((https://doi.org/10.1007/s13225-020-00466-2)) |
| Otu14363 | 1 | k__Fungi(100);p__Ascomycota(100);c__Saccharomycetes(100);o__Saccharomycetales(100);f__Metschnikowiaceae(100);g__Metschnikowia(100);g__Metschnikowia_unclassified(100); | Metschnikowia | 13 | Saprotroph | \|Undefined Saprotroph\| | Yeast | NULL | Probable | As nectar-tap_saprotroph (Põlme et al. 2020); Decay sugar-rich substrates (Põlme et al. 2020) | Põlme S, et al. 2020. Fungal Diversity 105:1-16 ((https://doi.org/10.1007/s13225-020-00466-2)) |
| Otu14364 | 1 | k__Fungi(100);p__Ascomycota(100);c__Saccharomycetes(100);o__Saccharomycetales(100);f__Metschnikowiaceae(100);g__Metschnikowia(100);s__Metschnikowia_pulcherrima(100); | Metschnikowia | 13 | Saprotroph | \|Undefined Saprotroph\| | Yeast | NULL | Probable | As nectar-tap_saprotroph (Põlme et al. 2020); Decay sugar-rich substrates (Põlme et al. 2020) | Põlme S, et al. 2020. Fungal Diversity 105:1-16 ((https://doi.org/10.1007/s13225-020-00466-2)) |
| Otu14365 | 1 | k__Fungi(100);p__Ascomycota(100);c__Saccharomycetes(100);o__Saccharomycetales(100);f__Metschnikowiaceae(100);g__Metschnikowia(100);s__Metschnikowia_pulcherrima(100); | Metschnikowia | 13 | Saprotroph | \|Undefined Saprotroph\| | Yeast | NULL | Probable | As nectar-tap_saprotroph (Põlme et al. 2020); Decay sugar-rich substrates (Põlme et al. 2020) | Põlme S, et al. 2020. Fungal Diversity 105:1-16 ((https://doi.org/10.1007/s13225-020-00466-2)) |
| Otu14366 | 1 | k__Fungi(100);p__Ascomycota(100);c__Saccharomycetes(100);o__Saccharomycetales(100);f__Metschnikowiaceae(100);g__Metschnikowia(100);g__Metschnikowia_unclassified(100); | Metschnikowia | 13 | Saprotroph | \|Undefined Saprotroph\| | Yeast | NULL | Probable | As nectar-tap_saprotroph (Põlme et al. 2020); Decay sugar-rich substrates (Põlme et al. 2020) | Põlme S, et al. 2020. Fungal Diversity 105:1-16 ((https://doi.org/10.1007/s13225-020-00466-2)) |
| Otu14368 | 1 | k__Fungi(100);p__Ascomycota(100);c__Saccharomycetes(100);o__Saccharomycetales(100);f__Metschnikowiaceae(100);g__Metschnikowia(100);s__Metschnikowia_pulcherrima(100); | Metschnikowia | 13 | Saprotroph | \|Undefined Saprotroph\| | Yeast | NULL | Probable | As nectar-tap_saprotroph (Põlme et al. 2020); Decay sugar-rich substrates (Põlme et al. 2020) | Põlme S, et al. 2020. Fungal Diversity 105:1-16 ((https://doi.org/10.1007/s13225-020-00466-2)) |
| Otu14369 | 1 | k__Fungi(100);p__Ascomycota(100);c__Saccharomycetes(100);o__Saccharomycetales(100);f__Metschnikowiaceae(100);g__Metschnikowia(100);s__Metschnikowia_pulcherrima(100); | Metschnikowia | 13 | Saprotroph | \|Undefined Saprotroph\| | Yeast | NULL | Probable | As nectar-tap_saprotroph (Põlme et al. 2020); Decay sugar-rich substrates (Põlme et al. 2020) | Põlme S, et al. 2020. Fungal Diversity 105:1-16 ((https://doi.org/10.1007/s13225-020-00466-2)) |
| Otu14370 | 1 | k__Fungi(100);p__Ascomycota(100);c__Saccharomycetes(100);o__Saccharomycetales(100);f__Metschnikowiaceae(100);g__Metschnikowia(100);s__Metschnikowia_pulcherrima(100); | Metschnikowia | 13 | Saprotroph | \|Undefined Saprotroph\| | Yeast | NULL | Probable | As nectar-tap_saprotroph (Põlme et al. 2020); Decay sugar-rich substrates (Põlme et al. 2020) | Põlme S, et al. 2020. Fungal Diversity 105:1-16 ((https://doi.org/10.1007/s13225-020-00466-2)) |
| Otu14371 | 1 | k__Fungi(100);p__Ascomycota(100);c__Saccharomycetes(100);o__Saccharomycetales(100);f__Metschnikowiaceae(100);g__Metschnikowia(100);s__Metschnikowia_pulcherrima(100); | Metschnikowia | 13 | Saprotroph | \|Undefined Saprotroph\| | Yeast | NULL | Probable | As nectar-tap_saprotroph (Põlme et al. 2020); Decay sugar-rich substrates (Põlme et al. 2020) | Põlme S, et al. 2020. Fungal Diversity 105:1-16 ((https://doi.org/10.1007/s13225-020-00466-2)) |
| Otu14372 | 1 | k__Fungi(100);p__Ascomycota(100);c__Saccharomycetes(100);o__Saccharomycetales(100);f__Metschnikowiaceae(100);g__Metschnikowia(100);g__Metschnikowia_unclassified(100); | Metschnikowia | 13 | Saprotroph | \|Undefined Saprotroph\| | Yeast | NULL | Probable | As nectar-tap_saprotroph (Põlme et al. 2020); Decay sugar-rich substrates (Põlme et al. 2020) | Põlme S, et al. 2020. Fungal Diversity 105:1-16 ((https://doi.org/10.1007/s13225-020-00466-2)) |
| Otu14373 | 1 | k__Fungi(100);p__Ascomycota(100);c__Saccharomycetes(100);o__Saccharomycetales(100);f__Metschnikowiaceae(100);g__Metschnikowia(100);s__Metschnikowia_pulcherrima(100); | Metschnikowia | 13 | Saprotroph | \|Undefined Saprotroph\| | Yeast | NULL | Probable | As nectar-tap_saprotroph (Põlme et al. 2020); Decay sugar-rich substrates (Põlme et al. 2020) | Põlme S, et al. 2020. Fungal Diversity 105:1-16 ((https://doi.org/10.1007/s13225-020-00466-2)) |
| Otu14375 | 1 | k__Fungi(100);p__Ascomycota(100);c__Saccharomycetes(100);o__Saccharomycetales(100);f__Metschnikowiaceae(100);g__Metschnikowia(100);s__Metschnikowia_pulcherrima(100); | Metschnikowia | 13 | Saprotroph | \|Undefined Saprotroph\| | Yeast | NULL | Probable | As nectar-tap_saprotroph (Põlme et al. 2020); Decay sugar-rich substrates (Põlme et al. 2020) | Põlme S, et al. 2020. Fungal Diversity 105:1-16 ((https://doi.org/10.1007/s13225-020-00466-2)) |
| Otu14377 | 1 | k__Fungi(100);p__Ascomycota(100);c__Saccharomycetes(100);o__Saccharomycetales(100);f__Metschnikowiaceae(100);g__Metschnikowia(100);s__Metschnikowia_pulcherrima(100); | Metschnikowia | 13 | Saprotroph | \|Undefined Saprotroph\| | Yeast | NULL | Probable | As nectar-tap_saprotroph (Põlme et al. 2020); Decay sugar-rich substrates (Põlme et al. 2020) | Põlme S, et al. 2020. Fungal Diversity 105:1-16 ((https://doi.org/10.1007/s13225-020-00466-2)) |
| Otu14379 | 1 | k__Fungi(100);p__Ascomycota(100);c__Saccharomycetes(100);o__Saccharomycetales(100);f__Metschnikowiaceae(100);g__Metschnikowia(100);s__Metschnikowia_pulcherrima(100); | Metschnikowia | 13 | Saprotroph | \|Undefined Saprotroph\| | Yeast | NULL | Probable | As nectar-tap_saprotroph (Põlme et al. 2020); Decay sugar-rich substrates (Põlme et al. 2020) | Põlme S, et al. 2020. Fungal Diversity 105:1-16 ((https://doi.org/10.1007/s13225-020-00466-2)) |
| Otu14382 | 1 | k__Fungi(100);p__Ascomycota(100);c__Saccharomycetes(100);o__Saccharomycetales(100);f__Metschnikowiaceae(100);g__Metschnikowia(100);s__Metschnikowia_pulcherrima(100); | Metschnikowia | 13 | Saprotroph | \|Undefined Saprotroph\| | Yeast | NULL | Probable | As nectar-tap_saprotroph (Põlme et al. 2020); Decay sugar-rich substrates (Põlme et al. 2020) | Põlme S, et al. 2020. Fungal Diversity 105:1-16 ((https://doi.org/10.1007/s13225-020-00466-2)) |
| Otu14384 | 1 | k__Fungi(100);p__Ascomycota(100);c__Saccharomycetes(100);o__Saccharomycetales(100);f__Metschnikowiaceae(100);g__Metschnikowia(100);s__Metschnikowia_pulcherrima(100); | Metschnikowia | 13 | Saprotroph | \|Undefined Saprotroph\| | Yeast | NULL | Probable | As nectar-tap_saprotroph (Põlme et al. 2020); Decay sugar-rich substrates (Põlme et al. 2020) | Põlme S, et al. 2020. Fungal Diversity 105:1-16 ((https://doi.org/10.1007/s13225-020-00466-2)) |
| Otu14385 | 1 | k__Fungi(100);p__Ascomycota(100);c__Saccharomycetes(100);o__Saccharomycetales(100);f__Metschnikowiaceae(100);g__Metschnikowia(100);s__Metschnikowia_pulcherrima(100); | Metschnikowia | 13 | Saprotroph | \|Undefined Saprotroph\| | Yeast | NULL | Probable | As nectar-tap_saprotroph (Põlme et al. 2020); Decay sugar-rich substrates (Põlme et al. 2020) | Põlme S, et al. 2020. Fungal Diversity 105:1-16 ((https://doi.org/10.1007/s13225-020-00466-2)) |
| Otu14386 | 1 | k__Fungi(100);p__Ascomycota(100);c__Saccharomycetes(100);o__Saccharomycetales(100);f__Metschnikowiaceae(100);g__Metschnikowia(100);s__Metschnikowia_pulcherrima(100); | Metschnikowia | 13 | Saprotroph | \|Undefined Saprotroph\| | Yeast | NULL | Probable | As nectar-tap_saprotroph (Põlme et al. 2020); Decay sugar-rich substrates (Põlme et al. 2020) | Põlme S, et al. 2020. Fungal Diversity 105:1-16 ((https://doi.org/10.1007/s13225-020-00466-2)) |
| Otu14387 | 1 | k__Fungi(100);p__Ascomycota(100);c__Saccharomycetes(100);o__Saccharomycetales(100);f__Metschnikowiaceae(100);g__Metschnikowia(100);s__Metschnikowia_pulcherrima(100); | Metschnikowia | 13 | Saprotroph | \|Undefined Saprotroph\| | Yeast | NULL | Probable | As nectar-tap_saprotroph (Põlme et al. 2020); Decay sugar-rich substrates (Põlme et al. 2020) | Põlme S, et al. 2020. Fungal Diversity 105:1-16 ((https://doi.org/10.1007/s13225-020-00466-2)) |
| Otu14388 | 1 | k__Fungi(100);p__Ascomycota(100);c__Saccharomycetes(100);o__Saccharomycetales(100);f__Metschnikowiaceae(100);g__Metschnikowia(100);s__Metschnikowia_pulcherrima(100); | Metschnikowia | 13 | Saprotroph | \|Undefined Saprotroph\| | Yeast | NULL | Probable | As nectar-tap_saprotroph (Põlme et al. 2020); Decay sugar-rich substrates (Põlme et al. 2020) | Põlme S, et al. 2020. Fungal Diversity 105:1-16 ((https://doi.org/10.1007/s13225-020-00466-2)) |
| Otu14389 | 1 | k__Fungi(100);p__Ascomycota(100);c__Saccharomycetes(100);o__Saccharomycetales(100);f__Metschnikowiaceae(100);g__Metschnikowia(100);s__Metschnikowia_pulcherrima(100); | Metschnikowia | 13 | Saprotroph | \|Undefined Saprotroph\| | Yeast | NULL | Probable | As nectar-tap_saprotroph (Põlme et al. 2020); Decay sugar-rich substrates (Põlme et al. 2020) | Põlme S, et al. 2020. Fungal Diversity 105:1-16 ((https://doi.org/10.1007/s13225-020-00466-2)) |
| Otu14390 | 1 | k__Fungi(100);p__Ascomycota(100);c__Saccharomycetes(100);o__Saccharomycetales(100);f__Metschnikowiaceae(100);g__Metschnikowia(100);s__Metschnikowia_pulcherrima(100); | Metschnikowia | 13 | Saprotroph | \|Undefined Saprotroph\| | Yeast | NULL | Probable | As nectar-tap_saprotroph (Põlme et al. 2020); Decay sugar-rich substrates (Põlme et al. 2020) | Põlme S, et al. 2020. Fungal Diversity 105:1-16 ((https://doi.org/10.1007/s13225-020-00466-2)) |
| Otu14391 | 1 | k__Fungi(100);p__Ascomycota(100);c__Saccharomycetes(100);o__Saccharomycetales(100);f__Metschnikowiaceae(100);g__Metschnikowia(100);s__Metschnikowia_pulcherrima(100); | Metschnikowia | 13 | Saprotroph | \|Undefined Saprotroph\| | Yeast | NULL | Probable | As nectar-tap_saprotroph (Põlme et al. 2020); Decay sugar-rich substrates (Põlme et al. 2020) | Põlme S, et al. 2020. Fungal Diversity 105:1-16 ((https://doi.org/10.1007/s13225-020-00466-2)) |
| Otu14393 | 1 | k__Fungi(100);p__Ascomycota(100);c__Saccharomycetes(100);o__Saccharomycetales(100);f__Metschnikowiaceae(100);g__Metschnikowia(100);s__Metschnikowia_pulcherrima(100); | Metschnikowia | 13 | Saprotroph | \|Undefined Saprotroph\| | Yeast | NULL | Probable | As nectar-tap_saprotroph (Põlme et al. 2020); Decay sugar-rich substrates (Põlme et al. 2020) | Põlme S, et al. 2020. Fungal Diversity 105:1-16 ((https://doi.org/10.1007/s13225-020-00466-2)) |
| Otu14398 | 1 | k__Fungi(100);p__Ascomycota(100);c__Saccharomycetes(100);o__Saccharomycetales(100);f__Metschnikowiaceae(100);g__Metschnikowia(100);g__Metschnikowia_unclassified(100); | Metschnikowia | 13 | Saprotroph | \|Undefined Saprotroph\| | Yeast | NULL | Probable | As nectar-tap_saprotroph (Põlme et al. 2020); Decay sugar-rich substrates (Põlme et al. 2020) | Põlme S, et al. 2020. Fungal Diversity 105:1-16 ((https://doi.org/10.1007/s13225-020-00466-2)) |
| Otu14399 | 1 | k__Fungi(100);p__Ascomycota(100);c__Saccharomycetes(100);o__Saccharomycetales(100);f__Saccharomycetales_family_Incertae_sedis(100);g__Candida(100);g__Candida_unclassified(100); | Candida | 13 | Pathotroph-Saprotroph-Symbiotroph | Animal Pathogen-Endophyte-Endosymbiont-Epiphyte-\|Undefined Saprotroph\| | Dimorphic Yeast-Yeast | NULL | Probable | As nectar-tap_saprotroph (Põlme et al. 2020); Decay sugar-rich substrates (Põlme et al. 2020); This asexual genus is polyphyletic and contain species that function very differently depend on what they are. Caution should be taken when interpreting guilds for Candida species. | Manolakaki D, et al. 2010. Virulence. 1: 367-375 ((https://doi.org/10.4161/viru.1.5.12796)); Mekbib SB, et al. 2011. Tropical Plant Pathology 36:233-240 ((https://doi.org/10.1590/S1982-56762011000400004)); Põlme S, et al. 2020. Fungal Diversity 105:1-16 ((https://doi.org/10.1007/s13225-020-00466-2)) |
| Otu14400 | 1 | k__Fungi(100);p__Ascomycota(100);c__Saccharomycetes(100);o__Saccharomycetales(100);f__Metschnikowiaceae(100);g__Metschnikowia(100);s__Metschnikowia_pulcherrima(100); | Metschnikowia | 13 | Saprotroph | \|Undefined Saprotroph\| | Yeast | NULL | Probable | As nectar-tap_saprotroph (Põlme et al. 2020); Decay sugar-rich substrates (Põlme et al. 2020) | Põlme S, et al. 2020. Fungal Diversity 105:1-16 ((https://doi.org/10.1007/s13225-020-00466-2)) |
| Otu14401 | 1 | k__Fungi(100);p__Ascomycota(100);c__Saccharomycetes(100);o__Saccharomycetales(100);f__Metschnikowiaceae(100);g__Metschnikowia(100);s__Metschnikowia_pulcherrima(100); | Metschnikowia | 13 | Saprotroph | \|Undefined Saprotroph\| | Yeast | NULL | Probable | As nectar-tap_saprotroph (Põlme et al. 2020); Decay sugar-rich substrates (Põlme et al. 2020) | Põlme S, et al. 2020. Fungal Diversity 105:1-16 ((https://doi.org/10.1007/s13225-020-00466-2)) |
| Otu14402 | 1 | k__Fungi(100);p__Ascomycota(100);c__Saccharomycetes(100);o__Saccharomycetales(100);f__Metschnikowiaceae(100);g__Metschnikowia(100);s__Metschnikowia_pulcherrima(100); | Metschnikowia | 13 | Saprotroph | \|Undefined Saprotroph\| | Yeast | NULL | Probable | As nectar-tap_saprotroph (Põlme et al. 2020); Decay sugar-rich substrates (Põlme et al. 2020) | Põlme S, et al. 2020. Fungal Diversity 105:1-16 ((https://doi.org/10.1007/s13225-020-00466-2)) |
| Otu14404 | 1 | k__Fungi(100);p__Ascomycota(100);c__Saccharomycetes(100);o__Saccharomycetales(100);f__Metschnikowiaceae(100);g__Metschnikowia(100);g__Metschnikowia_unclassified(100); | Metschnikowia | 13 | Saprotroph | \|Undefined Saprotroph\| | Yeast | NULL | Probable | As nectar-tap_saprotroph (Põlme et al. 2020); Decay sugar-rich substrates (Põlme et al. 2020) | Põlme S, et al. 2020. Fungal Diversity 105:1-16 ((https://doi.org/10.1007/s13225-020-00466-2)) |
| Otu14405 | 1 | k__Fungi(100);p__Ascomycota(100);c__Saccharomycetes(100);o__Saccharomycetales(100);f__Metschnikowiaceae(100);g__Metschnikowia(100);g__Metschnikowia_unclassified(100); | Metschnikowia | 13 | Saprotroph | \|Undefined Saprotroph\| | Yeast | NULL | Probable | As nectar-tap_saprotroph (Põlme et al. 2020); Decay sugar-rich substrates (Põlme et al. 2020) | Põlme S, et al. 2020. Fungal Diversity 105:1-16 ((https://doi.org/10.1007/s13225-020-00466-2)) |
| Otu14406 | 1 | k__Fungi(100);p__Ascomycota(100);c__Saccharomycetes(100);o__Saccharomycetales(100);f__Metschnikowiaceae(100);g__Metschnikowia(100);g__Metschnikowia_unclassified(100); | Metschnikowia | 13 | Saprotroph | \|Undefined Saprotroph\| | Yeast | NULL | Probable | As nectar-tap_saprotroph (Põlme et al. 2020); Decay sugar-rich substrates (Põlme et al. 2020) | Põlme S, et al. 2020. Fungal Diversity 105:1-16 ((https://doi.org/10.1007/s13225-020-00466-2)) |
| Otu14408 | 1 | k__Fungi(100);p__Ascomycota(100);c__Saccharomycetes(100);o__Saccharomycetales(100);f__Metschnikowiaceae(100);g__Metschnikowia(100);g__Metschnikowia_unclassified(100); | Metschnikowia | 13 | Saprotroph | \|Undefined Saprotroph\| | Yeast | NULL | Probable | As nectar-tap_saprotroph (Põlme et al. 2020); Decay sugar-rich substrates (Põlme et al. 2020) | Põlme S, et al. 2020. Fungal Diversity 105:1-16 ((https://doi.org/10.1007/s13225-020-00466-2)) |
| Otu14409 | 1 | k__Fungi(100);p__Ascomycota(100);c__Saccharomycetes(100);o__Saccharomycetales(100);f__Metschnikowiaceae(100);g__Metschnikowia(100);s__Metschnikowia_pulcherrima(100); | Metschnikowia | 13 | Saprotroph | \|Undefined Saprotroph\| | Yeast | NULL | Probable | As nectar-tap_saprotroph (Põlme et al. 2020); Decay sugar-rich substrates (Põlme et al. 2020) | Põlme S, et al. 2020. Fungal Diversity 105:1-16 ((https://doi.org/10.1007/s13225-020-00466-2)) |
| Otu14410 | 1 | k__Fungi(100);p__Ascomycota(100);c__Saccharomycetes(100);o__Saccharomycetales(100);f__Metschnikowiaceae(100);g__Metschnikowia(100);s__Metschnikowia_pulcherrima(100); | Metschnikowia | 13 | Saprotroph | \|Undefined Saprotroph\| | Yeast | NULL | Probable | As nectar-tap_saprotroph (Põlme et al. 2020); Decay sugar-rich substrates (Põlme et al. 2020) | Põlme S, et al. 2020. Fungal Diversity 105:1-16 ((https://doi.org/10.1007/s13225-020-00466-2)) |
| Otu14411 | 1 | k__Fungi(100);p__Ascomycota(100);c__Saccharomycetes(100);o__Saccharomycetales(100);f__Metschnikowiaceae(100);g__Metschnikowia(100);s__Metschnikowia_pulcherrima(100); | Metschnikowia | 13 | Saprotroph | \|Undefined Saprotroph\| | Yeast | NULL | Probable | As nectar-tap_saprotroph (Põlme et al. 2020); Decay sugar-rich substrates (Põlme et al. 2020) | Põlme S, et al. 2020. Fungal Diversity 105:1-16 ((https://doi.org/10.1007/s13225-020-00466-2)) |
| Otu14412 | 1 | k__Fungi(100);p__Ascomycota(100);c__Saccharomycetes(100);o__Saccharomycetales(100);f__Metschnikowiaceae(100);g__Metschnikowia(100);s__Metschnikowia_pulcherrima(100); | Metschnikowia | 13 | Saprotroph | \|Undefined Saprotroph\| | Yeast | NULL | Probable | As nectar-tap_saprotroph (Põlme et al. 2020); Decay sugar-rich substrates (Põlme et al. 2020) | Põlme S, et al. 2020. Fungal Diversity 105:1-16 ((https://doi.org/10.1007/s13225-020-00466-2)) |
| Otu14414 | 1 | k__Fungi(100);p__Ascomycota(100);c__Saccharomycetes(100);o__Saccharomycetales(100);f__Metschnikowiaceae(100);g__Metschnikowia(100);s__Metschnikowia_pulcherrima(100); | Metschnikowia | 13 | Saprotroph | \|Undefined Saprotroph\| | Yeast | NULL | Probable | As nectar-tap_saprotroph (Põlme et al. 2020); Decay sugar-rich substrates (Põlme et al. 2020) | Põlme S, et al. 2020. Fungal Diversity 105:1-16 ((https://doi.org/10.1007/s13225-020-00466-2)) |
| Otu14415 | 1 | k__Fungi(100);p__Ascomycota(100);c__Saccharomycetes(100);o__Saccharomycetales(100);f__Metschnikowiaceae(100);g__Metschnikowia(100);s__Metschnikowia_pulcherrima(100); | Metschnikowia | 13 | Saprotroph | \|Undefined Saprotroph\| | Yeast | NULL | Probable | As nectar-tap_saprotroph (Põlme et al. 2020); Decay sugar-rich substrates (Põlme et al. 2020) | Põlme S, et al. 2020. Fungal Diversity 105:1-16 ((https://doi.org/10.1007/s13225-020-00466-2)) |
| Otu14416 | 1 | k__Fungi(100);p__Ascomycota(100);c__Saccharomycetes(100);o__Saccharomycetales(100);f__Metschnikowiaceae(100);g__Metschnikowia(100);s__Metschnikowia_pulcherrima(100); | Metschnikowia | 13 | Saprotroph | \|Undefined Saprotroph\| | Yeast | NULL | Probable | As nectar-tap_saprotroph (Põlme et al. 2020); Decay sugar-rich substrates (Põlme et al. 2020) | Põlme S, et al. 2020. Fungal Diversity 105:1-16 ((https://doi.org/10.1007/s13225-020-00466-2)) |
| Otu14417 | 1 | k__Fungi(100);p__Ascomycota(100);c__Saccharomycetes(100);o__Saccharomycetales(100);f__Metschnikowiaceae(100);g__Metschnikowia(100);g__Metschnikowia_unclassified(100); | Metschnikowia | 13 | Saprotroph | \|Undefined Saprotroph\| | Yeast | NULL | Probable | As nectar-tap_saprotroph (Põlme et al. 2020); Decay sugar-rich substrates (Põlme et al. 2020) | Põlme S, et al. 2020. Fungal Diversity 105:1-16 ((https://doi.org/10.1007/s13225-020-00466-2)) |
| Otu14418 | 1 | k__Fungi(100);p__Ascomycota(100);c__Saccharomycetes(100);o__Saccharomycetales(100);f__Metschnikowiaceae(100);g__Metschnikowia(100);s__Metschnikowia_pulcherrima(100); | Metschnikowia | 13 | Saprotroph | \|Undefined Saprotroph\| | Yeast | NULL | Probable | As nectar-tap_saprotroph (Põlme et al. 2020); Decay sugar-rich substrates (Põlme et al. 2020) | Põlme S, et al. 2020. Fungal Diversity 105:1-16 ((https://doi.org/10.1007/s13225-020-00466-2)) |
| Otu14420 | 1 | k__Fungi(100);p__Ascomycota(100);c__Saccharomycetes(100);o__Saccharomycetales(100);f__Metschnikowiaceae(100);g__Metschnikowia(100);s__Metschnikowia_pulcherrima(100); | Metschnikowia | 13 | Saprotroph | \|Undefined Saprotroph\| | Yeast | NULL | Probable | As nectar-tap_saprotroph (Põlme et al. 2020); Decay sugar-rich substrates (Põlme et al. 2020) | Põlme S, et al. 2020. Fungal Diversity 105:1-16 ((https://doi.org/10.1007/s13225-020-00466-2)) |
| Otu14421 | 1 | k__Fungi(100);p__Ascomycota(100);c__Saccharomycetes(100);o__Saccharomycetales(100);f__Metschnikowiaceae(100);g__Metschnikowia(100);s__Metschnikowia_pulcherrima(100); | Metschnikowia | 13 | Saprotroph | \|Undefined Saprotroph\| | Yeast | NULL | Probable | As nectar-tap_saprotroph (Põlme et al. 2020); Decay sugar-rich substrates (Põlme et al. 2020) | Põlme S, et al. 2020. Fungal Diversity 105:1-16 ((https://doi.org/10.1007/s13225-020-00466-2)) |
| Otu14422 | 1 | k__Fungi(100);p__Ascomycota(100);c__Saccharomycetes(100);o__Saccharomycetales(100);f__Metschnikowiaceae(100);g__Metschnikowia(100);g__Metschnikowia_unclassified(100); | Metschnikowia | 13 | Saprotroph | \|Undefined Saprotroph\| | Yeast | NULL | Probable | As nectar-tap_saprotroph (Põlme et al. 2020); Decay sugar-rich substrates (Põlme et al. 2020) | Põlme S, et al. 2020. Fungal Diversity 105:1-16 ((https://doi.org/10.1007/s13225-020-00466-2)) |
| Otu14423 | 1 | k__Fungi(100);p__Ascomycota(100);c__Saccharomycetes(100);o__Saccharomycetales(100);f__Metschnikowiaceae(100);g__Metschnikowia(100);s__Metschnikowia_pulcherrima(100); | Metschnikowia | 13 | Saprotroph | \|Undefined Saprotroph\| | Yeast | NULL | Probable | As nectar-tap_saprotroph (Põlme et al. 2020); Decay sugar-rich substrates (Põlme et al. 2020) | Põlme S, et al. 2020. Fungal Diversity 105:1-16 ((https://doi.org/10.1007/s13225-020-00466-2)) |
| Otu14424 | 1 | k__Fungi(100);p__Ascomycota(100);c__Saccharomycetes(100);o__Saccharomycetales(100);f__Metschnikowiaceae(100);g__Metschnikowia(100);s__Metschnikowia_pulcherrima(100); | Metschnikowia | 13 | Saprotroph | \|Undefined Saprotroph\| | Yeast | NULL | Probable | As nectar-tap_saprotroph (Põlme et al. 2020); Decay sugar-rich substrates (Põlme et al. 2020) | Põlme S, et al. 2020. Fungal Diversity 105:1-16 ((https://doi.org/10.1007/s13225-020-00466-2)) |
| Otu14425 | 1 | k__Fungi(100);p__Ascomycota(100);c__Saccharomycetes(100);o__Saccharomycetales(100);f__Metschnikowiaceae(100);g__Metschnikowia(100);s__Metschnikowia_pulcherrima(100); | Metschnikowia | 13 | Saprotroph | \|Undefined Saprotroph\| | Yeast | NULL | Probable | As nectar-tap_saprotroph (Põlme et al. 2020); Decay sugar-rich substrates (Põlme et al. 2020) | Põlme S, et al. 2020. Fungal Diversity 105:1-16 ((https://doi.org/10.1007/s13225-020-00466-2)) |
| Otu14426 | 1 | k__Fungi(100);p__Ascomycota(100);c__Saccharomycetes(100);o__Saccharomycetales(100);f__Metschnikowiaceae(100);g__Metschnikowia(100);s__Metschnikowia_pulcherrima(100); | Metschnikowia | 13 | Saprotroph | \|Undefined Saprotroph\| | Yeast | NULL | Probable | As nectar-tap_saprotroph (Põlme et al. 2020); Decay sugar-rich substrates (Põlme et al. 2020) | Põlme S, et al. 2020. Fungal Diversity 105:1-16 ((https://doi.org/10.1007/s13225-020-00466-2)) |
| Otu14427 | 1 | k__Fungi(100);p__Ascomycota(100);c__Saccharomycetes(100);o__Saccharomycetales(100);f__Metschnikowiaceae(100);g__Metschnikowia(100);g__Metschnikowia_unclassified(100); | Metschnikowia | 13 | Saprotroph | \|Undefined Saprotroph\| | Yeast | NULL | Probable | As nectar-tap_saprotroph (Põlme et al. 2020); Decay sugar-rich substrates (Põlme et al. 2020) | Põlme S, et al. 2020. Fungal Diversity 105:1-16 ((https://doi.org/10.1007/s13225-020-00466-2)) |
| Otu14431 | 1 | k__Fungi(100);p__Ascomycota(100);c__Saccharomycetes(100);o__Saccharomycetales(100);f__Metschnikowiaceae(100);g__Metschnikowia(100);s__Metschnikowia_pulcherrima(100); | Metschnikowia | 13 | Saprotroph | \|Undefined Saprotroph\| | Yeast | NULL | Probable | As nectar-tap_saprotroph (Põlme et al. 2020); Decay sugar-rich substrates (Põlme et al. 2020) | Põlme S, et al. 2020. Fungal Diversity 105:1-16 ((https://doi.org/10.1007/s13225-020-00466-2)) |
| Otu14438 | 1 | k__Fungi(100);p__Ascomycota(100);c__Saccharomycetes(100);o__Saccharomycetales(100);f__Metschnikowiaceae(100);g__Metschnikowia(100);g__Metschnikowia_unclassified(100); | Metschnikowia | 13 | Saprotroph | \|Undefined Saprotroph\| | Yeast | NULL | Probable | As nectar-tap_saprotroph (Põlme et al. 2020); Decay sugar-rich substrates (Põlme et al. 2020) | Põlme S, et al. 2020. Fungal Diversity 105:1-16 ((https://doi.org/10.1007/s13225-020-00466-2)) |
| Otu14440 | 1 | k__Fungi(100);p__Ascomycota(100);c__Saccharomycetes(100);o__Saccharomycetales(100);f__Metschnikowiaceae(100);g__Metschnikowia(100);s__Metschnikowia_pulcherrima(100); | Metschnikowia | 13 | Saprotroph | \|Undefined Saprotroph\| | Yeast | NULL | Probable | As nectar-tap_saprotroph (Põlme et al. 2020); Decay sugar-rich substrates (Põlme et al. 2020) | Põlme S, et al. 2020. Fungal Diversity 105:1-16 ((https://doi.org/10.1007/s13225-020-00466-2)) |
| Otu14443 | 1 | k__Fungi(100);p__Ascomycota(100);c__Saccharomycetes(100);o__Saccharomycetales(100);f__Metschnikowiaceae(100);g__Metschnikowia(100);s__Metschnikowia_pulcherrima(100); | Metschnikowia | 13 | Saprotroph | \|Undefined Saprotroph\| | Yeast | NULL | Probable | As nectar-tap_saprotroph (Põlme et al. 2020); Decay sugar-rich substrates (Põlme et al. 2020) | Põlme S, et al. 2020. Fungal Diversity 105:1-16 ((https://doi.org/10.1007/s13225-020-00466-2)) |
| Otu14444 | 1 | k__Fungi(100);p__Ascomycota(100);c__Saccharomycetes(100);o__Saccharomycetales(100);f__Metschnikowiaceae(100);g__Metschnikowia(100);s__Metschnikowia_pulcherrima(100); | Metschnikowia | 13 | Saprotroph | \|Undefined Saprotroph\| | Yeast | NULL | Probable | As nectar-tap_saprotroph (Põlme et al. 2020); Decay sugar-rich substrates (Põlme et al. 2020) | Põlme S, et al. 2020. Fungal Diversity 105:1-16 ((https://doi.org/10.1007/s13225-020-00466-2)) |
| Otu14446 | 1 | k__Fungi(100);p__Ascomycota(100);c__Saccharomycetes(100);o__Saccharomycetales(100);f__Metschnikowiaceae(100);g__Metschnikowia(100);s__Metschnikowia_pulcherrima(100); | Metschnikowia | 13 | Saprotroph | \|Undefined Saprotroph\| | Yeast | NULL | Probable | As nectar-tap_saprotroph (Põlme et al. 2020); Decay sugar-rich substrates (Põlme et al. 2020) | Põlme S, et al. 2020. Fungal Diversity 105:1-16 ((https://doi.org/10.1007/s13225-020-00466-2)) |
| Otu14447 | 1 | k__Fungi(100);p__Ascomycota(100);c__Saccharomycetes(100);o__Saccharomycetales(100);f__Metschnikowiaceae(100);g__Metschnikowia(100);s__Metschnikowia_pulcherrima(100); | Metschnikowia | 13 | Saprotroph | \|Undefined Saprotroph\| | Yeast | NULL | Probable | As nectar-tap_saprotroph (Põlme et al. 2020); Decay sugar-rich substrates (Põlme et al. 2020) | Põlme S, et al. 2020. Fungal Diversity 105:1-16 ((https://doi.org/10.1007/s13225-020-00466-2)) |
| Otu14448 | 1 | k__Fungi(100);p__Ascomycota(100);c__Saccharomycetes(100);o__Saccharomycetales(100);f__Metschnikowiaceae(100);g__Metschnikowia(100);s__Metschnikowia_pulcherrima(100); | Metschnikowia | 13 | Saprotroph | \|Undefined Saprotroph\| | Yeast | NULL | Probable | As nectar-tap_saprotroph (Põlme et al. 2020); Decay sugar-rich substrates (Põlme et al. 2020) | Põlme S, et al. 2020. Fungal Diversity 105:1-16 ((https://doi.org/10.1007/s13225-020-00466-2)) |
| Otu14452 | 1 | k__Fungi(100);p__Ascomycota(100);c__Saccharomycetes(100);o__Saccharomycetales(100);f__Metschnikowiaceae(100);g__Metschnikowia(100);s__Metschnikowia_pulcherrima(100); | Metschnikowia | 13 | Saprotroph | \|Undefined Saprotroph\| | Yeast | NULL | Probable | As nectar-tap_saprotroph (Põlme et al. 2020); Decay sugar-rich substrates (Põlme et al. 2020) | Põlme S, et al. 2020. Fungal Diversity 105:1-16 ((https://doi.org/10.1007/s13225-020-00466-2)) |
| Otu14453 | 1 | k__Fungi(100);p__Ascomycota(100);c__Saccharomycetes(100);o__Saccharomycetales(100);f__Metschnikowiaceae(100);g__Metschnikowia(100);g__Metschnikowia_unclassified(100); | Metschnikowia | 13 | Saprotroph | \|Undefined Saprotroph\| | Yeast | NULL | Probable | As nectar-tap_saprotroph (Põlme et al. 2020); Decay sugar-rich substrates (Põlme et al. 2020) | Põlme S, et al. 2020. Fungal Diversity 105:1-16 ((https://doi.org/10.1007/s13225-020-00466-2)) |
| Otu14454 | 1 | k__Fungi(100);p__Ascomycota(100);c__Saccharomycetes(100);o__Saccharomycetales(100);f__Metschnikowiaceae(100);g__Metschnikowia(100);s__Metschnikowia_pulcherrima(100); | Metschnikowia | 13 | Saprotroph | \|Undefined Saprotroph\| | Yeast | NULL | Probable | As nectar-tap_saprotroph (Põlme et al. 2020); Decay sugar-rich substrates (Põlme et al. 2020) | Põlme S, et al. 2020. Fungal Diversity 105:1-16 ((https://doi.org/10.1007/s13225-020-00466-2)) |
| Otu14455 | 1 | k__Fungi(100);p__Ascomycota(100);c__Saccharomycetes(100);o__Saccharomycetales(100);f__Metschnikowiaceae(100);g__Metschnikowia(100);s__Metschnikowia_pulcherrima(100); | Metschnikowia | 13 | Saprotroph | \|Undefined Saprotroph\| | Yeast | NULL | Probable | As nectar-tap_saprotroph (Põlme et al. 2020); Decay sugar-rich substrates (Põlme et al. 2020) | Põlme S, et al. 2020. Fungal Diversity 105:1-16 ((https://doi.org/10.1007/s13225-020-00466-2)) |
| Otu14456 | 1 | k__Fungi(100);p__Ascomycota(100);c__Saccharomycetes(100);o__Saccharomycetales(100);f__Metschnikowiaceae(100);g__Metschnikowia(100);s__Metschnikowia_pulcherrima(100); | Metschnikowia | 13 | Saprotroph | \|Undefined Saprotroph\| | Yeast | NULL | Probable | As nectar-tap_saprotroph (Põlme et al. 2020); Decay sugar-rich substrates (Põlme et al. 2020) | Põlme S, et al. 2020. Fungal Diversity 105:1-16 ((https://doi.org/10.1007/s13225-020-00466-2)) |
| Otu14460 | 1 | k__Fungi(100);p__Ascomycota(100);c__Saccharomycetes(100);o__Saccharomycetales(100);f__Metschnikowiaceae(100);g__Metschnikowia(100);s__Metschnikowia_pulcherrima(100); | Metschnikowia | 13 | Saprotroph | \|Undefined Saprotroph\| | Yeast | NULL | Probable | As nectar-tap_saprotroph (Põlme et al. 2020); Decay sugar-rich substrates (Põlme et al. 2020) | Põlme S, et al. 2020. Fungal Diversity 105:1-16 ((https://doi.org/10.1007/s13225-020-00466-2)) |
| Otu14464 | 1 | k__Fungi(100);p__Ascomycota(100);c__Saccharomycetes(100);o__Saccharomycetales(100);f__Metschnikowiaceae(100);g__Metschnikowia(100);s__Metschnikowia_pulcherrima(100); | Metschnikowia | 13 | Saprotroph | \|Undefined Saprotroph\| | Yeast | NULL | Probable | As nectar-tap_saprotroph (Põlme et al. 2020); Decay sugar-rich substrates (Põlme et al. 2020) | Põlme S, et al. 2020. Fungal Diversity 105:1-16 ((https://doi.org/10.1007/s13225-020-00466-2)) |
| Otu14465 | 1 | k__Fungi(100);p__Ascomycota(100);c__Saccharomycetes(100);o__Saccharomycetales(100);f__Metschnikowiaceae(100);g__Metschnikowia(100);g__Metschnikowia_unclassified(100); | Metschnikowia | 13 | Saprotroph | \|Undefined Saprotroph\| | Yeast | NULL | Probable | As nectar-tap_saprotroph (Põlme et al. 2020); Decay sugar-rich substrates (Põlme et al. 2020) | Põlme S, et al. 2020. Fungal Diversity 105:1-16 ((https://doi.org/10.1007/s13225-020-00466-2)) |
| Otu14467 | 1 | k__Fungi(100);p__Ascomycota(100);c__Saccharomycetes(100);o__Saccharomycetales(100);f__Metschnikowiaceae(100);g__Metschnikowia(100);s__Metschnikowia_pulcherrima(100); | Metschnikowia | 13 | Saprotroph | \|Undefined Saprotroph\| | Yeast | NULL | Probable | As nectar-tap_saprotroph (Põlme et al. 2020); Decay sugar-rich substrates (Põlme et al. 2020) | Põlme S, et al. 2020. Fungal Diversity 105:1-16 ((https://doi.org/10.1007/s13225-020-00466-2)) |
| Otu14471 | 1 | k__Fungi(100);p__Ascomycota(100);c__Saccharomycetes(100);o__Saccharomycetales(100);f__Metschnikowiaceae(100);g__Metschnikowia(100);g__Metschnikowia_unclassified(100); | Metschnikowia | 13 | Saprotroph | \|Undefined Saprotroph\| | Yeast | NULL | Probable | As nectar-tap_saprotroph (Põlme et al. 2020); Decay sugar-rich substrates (Põlme et al. 2020) | Põlme S, et al. 2020. Fungal Diversity 105:1-16 ((https://doi.org/10.1007/s13225-020-00466-2)) |
| Otu14472 | 1 | k__Fungi(100);p__Ascomycota(100);c__Saccharomycetes(100);o__Saccharomycetales(100);f__Metschnikowiaceae(100);g__Metschnikowia(100);s__Metschnikowia_pulcherrima(100); | Metschnikowia | 13 | Saprotroph | \|Undefined Saprotroph\| | Yeast | NULL | Probable | As nectar-tap_saprotroph (Põlme et al. 2020); Decay sugar-rich substrates (Põlme et al. 2020) | Põlme S, et al. 2020. Fungal Diversity 105:1-16 ((https://doi.org/10.1007/s13225-020-00466-2)) |
| Otu14473 | 1 | k__Fungi(100);p__Ascomycota(100);c__Saccharomycetes(100);o__Saccharomycetales(100);f__Metschnikowiaceae(100);g__Metschnikowia(100);s__Metschnikowia_pulcherrima(100); | Metschnikowia | 13 | Saprotroph | \|Undefined Saprotroph\| | Yeast | NULL | Probable | As nectar-tap_saprotroph (Põlme et al. 2020); Decay sugar-rich substrates (Põlme et al. 2020) | Põlme S, et al. 2020. Fungal Diversity 105:1-16 ((https://doi.org/10.1007/s13225-020-00466-2)) |
| Otu14474 | 1 | k__Fungi(100);p__Basidiomycota(100);c__Wallemiomycetes(100);o__Wallemiales(100);f__Wallemiaceae(100);g__Wallemia(100);s__Wallemia_sebi(100); | Wallemia | 13 | Saprotroph | \|Undefined Saprotroph\| | Microfungus | NULL | Probable | Decay sugar-rich substrates (Põlme et al. 2020) | Tedersoo L, et al. 2014. Science 346:e1256688 ((https://doi.org/10.1126/science.1256688)); Põlme S, et al. 2020. Fungal Diversity 105:1-16 ((https://doi.org/10.1007/s13225-020-00466-2)) |
| Otu14475 | 1 | k__Fungi(100);p__Ascomycota(100);c__Saccharomycetes(100);o__Saccharomycetales(100);f__Metschnikowiaceae(100);g__Metschnikowia(100);s__Metschnikowia_pulcherrima(100); | Metschnikowia | 13 | Saprotroph | \|Undefined Saprotroph\| | Yeast | NULL | Probable | As nectar-tap_saprotroph (Põlme et al. 2020); Decay sugar-rich substrates (Põlme et al. 2020) | Põlme S, et al. 2020. Fungal Diversity 105:1-16 ((https://doi.org/10.1007/s13225-020-00466-2)) |
| Otu14476 | 1 | k__Fungi(100);p__Ascomycota(100);c__Saccharomycetes(100);o__Saccharomycetales(100);f__Metschnikowiaceae(100);g__Metschnikowia(100);s__Metschnikowia_pulcherrima(100); | Metschnikowia | 13 | Saprotroph | \|Undefined Saprotroph\| | Yeast | NULL | Probable | As nectar-tap_saprotroph (Põlme et al. 2020); Decay sugar-rich substrates (Põlme et al. 2020) | Põlme S, et al. 2020. Fungal Diversity 105:1-16 ((https://doi.org/10.1007/s13225-020-00466-2)) |
| Otu14482 | 1 | k__Fungi(100);p__Ascomycota(100);c__Saccharomycetes(100);o__Saccharomycetales(100);f__Metschnikowiaceae(100);g__Metschnikowia(100);s__Metschnikowia_pulcherrima(100); | Metschnikowia | 13 | Saprotroph | \|Undefined Saprotroph\| | Yeast | NULL | Probable | As nectar-tap_saprotroph (Põlme et al. 2020); Decay sugar-rich substrates (Põlme et al. 2020) | Põlme S, et al. 2020. Fungal Diversity 105:1-16 ((https://doi.org/10.1007/s13225-020-00466-2)) |
| Otu14483 | 1 | k__Fungi(100);p__Ascomycota(100);c__Saccharomycetes(100);o__Saccharomycetales(100);f__Metschnikowiaceae(100);g__Metschnikowia(100);g__Metschnikowia_unclassified(100); | Metschnikowia | 13 | Saprotroph | \|Undefined Saprotroph\| | Yeast | NULL | Probable | As nectar-tap_saprotroph (Põlme et al. 2020); Decay sugar-rich substrates (Põlme et al. 2020) | Põlme S, et al. 2020. Fungal Diversity 105:1-16 ((https://doi.org/10.1007/s13225-020-00466-2)) |
| Otu14484 | 1 | k__Fungi(100);p__Ascomycota(100);c__Saccharomycetes(100);o__Saccharomycetales(100);f__Metschnikowiaceae(100);g__Metschnikowia(100);s__Metschnikowia_pulcherrima(100); | Metschnikowia | 13 | Saprotroph | \|Undefined Saprotroph\| | Yeast | NULL | Probable | As nectar-tap_saprotroph (Põlme et al. 2020); Decay sugar-rich substrates (Põlme et al. 2020) | Põlme S, et al. 2020. Fungal Diversity 105:1-16 ((https://doi.org/10.1007/s13225-020-00466-2)) |
| Otu14485 | 1 | k__Fungi(100);p__Ascomycota(100);c__Saccharomycetes(100);o__Saccharomycetales(100);f__Metschnikowiaceae(100);g__Metschnikowia(100);g__Metschnikowia_unclassified(100); | Metschnikowia | 13 | Saprotroph | \|Undefined Saprotroph\| | Yeast | NULL | Probable | As nectar-tap_saprotroph (Põlme et al. 2020); Decay sugar-rich substrates (Põlme et al. 2020) | Põlme S, et al. 2020. Fungal Diversity 105:1-16 ((https://doi.org/10.1007/s13225-020-00466-2)) |
| Otu14486 | 1 | k__Fungi(100);p__Ascomycota(100);c__Saccharomycetes(100);o__Saccharomycetales(100);f__Metschnikowiaceae(100);g__Metschnikowia(100);s__Metschnikowia_pulcherrima(100); | Metschnikowia | 13 | Saprotroph | \|Undefined Saprotroph\| | Yeast | NULL | Probable | As nectar-tap_saprotroph (Põlme et al. 2020); Decay sugar-rich substrates (Põlme et al. 2020) | Põlme S, et al. 2020. Fungal Diversity 105:1-16 ((https://doi.org/10.1007/s13225-020-00466-2)) |
| Otu14487 | 1 | k__Fungi(100);p__Ascomycota(100);c__Saccharomycetes(100);o__Saccharomycetales(100);f__Metschnikowiaceae(100);g__Metschnikowia(100);s__Metschnikowia_pulcherrima(100); | Metschnikowia | 13 | Saprotroph | \|Undefined Saprotroph\| | Yeast | NULL | Probable | As nectar-tap_saprotroph (Põlme et al. 2020); Decay sugar-rich substrates (Põlme et al. 2020) | Põlme S, et al. 2020. Fungal Diversity 105:1-16 ((https://doi.org/10.1007/s13225-020-00466-2)) |
| Otu14488 | 1 | k__Fungi(100);p__Ascomycota(100);c__Saccharomycetes(100);o__Saccharomycetales(100);f__Metschnikowiaceae(100);g__Metschnikowia(100);g__Metschnikowia_unclassified(100); | Metschnikowia | 13 | Saprotroph | \|Undefined Saprotroph\| | Yeast | NULL | Probable | As nectar-tap_saprotroph (Põlme et al. 2020); Decay sugar-rich substrates (Põlme et al. 2020) | Põlme S, et al. 2020. Fungal Diversity 105:1-16 ((https://doi.org/10.1007/s13225-020-00466-2)) |
| Otu14489 | 1 | k__Fungi(100);p__Ascomycota(100);c__Saccharomycetes(100);o__Saccharomycetales(100);f__Metschnikowiaceae(100);g__Metschnikowia(100);s__Metschnikowia_pulcherrima(100); | Metschnikowia | 13 | Saprotroph | \|Undefined Saprotroph\| | Yeast | NULL | Probable | As nectar-tap_saprotroph (Põlme et al. 2020); Decay sugar-rich substrates (Põlme et al. 2020) | Põlme S, et al. 2020. Fungal Diversity 105:1-16 ((https://doi.org/10.1007/s13225-020-00466-2)) |
| Otu14491 | 1 | k__Fungi(100);p__Ascomycota(100);c__Saccharomycetes(100);o__Saccharomycetales(100);f__Metschnikowiaceae(100);g__Metschnikowia(100);g__Metschnikowia_unclassified(100); | Metschnikowia | 13 | Saprotroph | \|Undefined Saprotroph\| | Yeast | NULL | Probable | As nectar-tap_saprotroph (Põlme et al. 2020); Decay sugar-rich substrates (Põlme et al. 2020) | Põlme S, et al. 2020. Fungal Diversity 105:1-16 ((https://doi.org/10.1007/s13225-020-00466-2)) |
| Otu14492 | 1 | k__Fungi(100);p__Ascomycota(100);c__Saccharomycetes(100);o__Saccharomycetales(100);f__Metschnikowiaceae(100);g__Metschnikowia(100);g__Metschnikowia_unclassified(100); | Metschnikowia | 13 | Saprotroph | \|Undefined Saprotroph\| | Yeast | NULL | Probable | As nectar-tap_saprotroph (Põlme et al. 2020); Decay sugar-rich substrates (Põlme et al. 2020) | Põlme S, et al. 2020. Fungal Diversity 105:1-16 ((https://doi.org/10.1007/s13225-020-00466-2)) |
| Otu14493 | 1 | k__Fungi(100);p__Ascomycota(100);c__Saccharomycetes(100);o__Saccharomycetales(100);f__Metschnikowiaceae(100);g__Metschnikowia(100);s__Metschnikowia_pulcherrima(100); | Metschnikowia | 13 | Saprotroph | \|Undefined Saprotroph\| | Yeast | NULL | Probable | As nectar-tap_saprotroph (Põlme et al. 2020); Decay sugar-rich substrates (Põlme et al. 2020) | Põlme S, et al. 2020. Fungal Diversity 105:1-16 ((https://doi.org/10.1007/s13225-020-00466-2)) |
| Otu14495 | 1 | k__Fungi(100);p__Ascomycota(100);c__Saccharomycetes(100);o__Saccharomycetales(100);f__Metschnikowiaceae(100);g__Metschnikowia(100);s__Metschnikowia_pulcherrima(100); | Metschnikowia | 13 | Saprotroph | \|Undefined Saprotroph\| | Yeast | NULL | Probable | As nectar-tap_saprotroph (Põlme et al. 2020); Decay sugar-rich substrates (Põlme et al. 2020) | Põlme S, et al. 2020. Fungal Diversity 105:1-16 ((https://doi.org/10.1007/s13225-020-00466-2)) |
| Otu14496 | 1 | k__Fungi(100);p__Ascomycota(100);c__Saccharomycetes(100);o__Saccharomycetales(100);f__Metschnikowiaceae(100);g__Metschnikowia(100);s__Metschnikowia_pulcherrima(100); | Metschnikowia | 13 | Saprotroph | \|Undefined Saprotroph\| | Yeast | NULL | Probable | As nectar-tap_saprotroph (Põlme et al. 2020); Decay sugar-rich substrates (Põlme et al. 2020) | Põlme S, et al. 2020. Fungal Diversity 105:1-16 ((https://doi.org/10.1007/s13225-020-00466-2)) |
| Otu14497 | 1 | k__Fungi(100);p__Ascomycota(100);c__Saccharomycetes(100);o__Saccharomycetales(100);f__Metschnikowiaceae(100);g__Metschnikowia(100);s__Metschnikowia_pulcherrima(100); | Metschnikowia | 13 | Saprotroph | \|Undefined Saprotroph\| | Yeast | NULL | Probable | As nectar-tap_saprotroph (Põlme et al. 2020); Decay sugar-rich substrates (Põlme et al. 2020) | Põlme S, et al. 2020. Fungal Diversity 105:1-16 ((https://doi.org/10.1007/s13225-020-00466-2)) |
| Otu14499 | 1 | k__Fungi(100);p__Ascomycota(100);c__Saccharomycetes(100);o__Saccharomycetales(100);f__Metschnikowiaceae(100);g__Metschnikowia(100);g__Metschnikowia_unclassified(100); | Metschnikowia | 13 | Saprotroph | \|Undefined Saprotroph\| | Yeast | NULL | Probable | As nectar-tap_saprotroph (Põlme et al. 2020); Decay sugar-rich substrates (Põlme et al. 2020) | Põlme S, et al. 2020. Fungal Diversity 105:1-16 ((https://doi.org/10.1007/s13225-020-00466-2)) |
| Otu14500 | 1 | k__Fungi(100);p__Ascomycota(100);c__Saccharomycetes(100);o__Saccharomycetales(100);f__Metschnikowiaceae(100);g__Metschnikowia(100);s__Metschnikowia_pulcherrima(100); | Metschnikowia | 13 | Saprotroph | \|Undefined Saprotroph\| | Yeast | NULL | Probable | As nectar-tap_saprotroph (Põlme et al. 2020); Decay sugar-rich substrates (Põlme et al. 2020) | Põlme S, et al. 2020. Fungal Diversity 105:1-16 ((https://doi.org/10.1007/s13225-020-00466-2)) |
| Otu14501 | 1 | k__Fungi(100);p__Ascomycota(100);c__Saccharomycetes(100);o__Saccharomycetales(100);f__Metschnikowiaceae(100);g__Metschnikowia(100);s__Metschnikowia_pulcherrima(100); | Metschnikowia | 13 | Saprotroph | \|Undefined Saprotroph\| | Yeast | NULL | Probable | As nectar-tap_saprotroph (Põlme et al. 2020); Decay sugar-rich substrates (Põlme et al. 2020) | Põlme S, et al. 2020. Fungal Diversity 105:1-16 ((https://doi.org/10.1007/s13225-020-00466-2)) |
| Otu14502 | 1 | k__Fungi(100);p__Ascomycota(100);c__Saccharomycetes(100);o__Saccharomycetales(100);f__Metschnikowiaceae(100);g__Metschnikowia(100);g__Metschnikowia_unclassified(100); | Metschnikowia | 13 | Saprotroph | \|Undefined Saprotroph\| | Yeast | NULL | Probable | As nectar-tap_saprotroph (Põlme et al. 2020); Decay sugar-rich substrates (Põlme et al. 2020) | Põlme S, et al. 2020. Fungal Diversity 105:1-16 ((https://doi.org/10.1007/s13225-020-00466-2)) |
| Otu14503 | 1 | k__Fungi(100);p__Ascomycota(100);c__Saccharomycetes(100);o__Saccharomycetales(100);f__Metschnikowiaceae(100);g__Metschnikowia(100);s__Metschnikowia_pulcherrima(100); | Metschnikowia | 13 | Saprotroph | \|Undefined Saprotroph\| | Yeast | NULL | Probable | As nectar-tap_saprotroph (Põlme et al. 2020); Decay sugar-rich substrates (Põlme et al. 2020) | Põlme S, et al. 2020. Fungal Diversity 105:1-16 ((https://doi.org/10.1007/s13225-020-00466-2)) |
| Otu14504 | 1 | k__Fungi(100);p__Ascomycota(100);c__Saccharomycetes(100);o__Saccharomycetales(100);f__Metschnikowiaceae(100);g__Metschnikowia(100);s__Metschnikowia_pulcherrima(100); | Metschnikowia | 13 | Saprotroph | \|Undefined Saprotroph\| | Yeast | NULL | Probable | As nectar-tap_saprotroph (Põlme et al. 2020); Decay sugar-rich substrates (Põlme et al. 2020) | Põlme S, et al. 2020. Fungal Diversity 105:1-16 ((https://doi.org/10.1007/s13225-020-00466-2)) |
| Otu14505 | 1 | k__Fungi(100);p__Ascomycota(100);c__Saccharomycetes(100);o__Saccharomycetales(100);f__Metschnikowiaceae(100);g__Metschnikowia(100);s__Metschnikowia_pulcherrima(100); | Metschnikowia | 13 | Saprotroph | \|Undefined Saprotroph\| | Yeast | NULL | Probable | As nectar-tap_saprotroph (Põlme et al. 2020); Decay sugar-rich substrates (Põlme et al. 2020) | Põlme S, et al. 2020. Fungal Diversity 105:1-16 ((https://doi.org/10.1007/s13225-020-00466-2)) |
| Otu14506 | 1 | k__Fungi(100);p__Ascomycota(100);c__Saccharomycetes(100);o__Saccharomycetales(100);f__Metschnikowiaceae(100);g__Metschnikowia(100);s__Metschnikowia_pulcherrima(100); | Metschnikowia | 13 | Saprotroph | \|Undefined Saprotroph\| | Yeast | NULL | Probable | As nectar-tap_saprotroph (Põlme et al. 2020); Decay sugar-rich substrates (Põlme et al. 2020) | Põlme S, et al. 2020. Fungal Diversity 105:1-16 ((https://doi.org/10.1007/s13225-020-00466-2)) |
| Otu14507 | 1 | k__Fungi(100);p__Ascomycota(100);c__Saccharomycetes(100);o__Saccharomycetales(100);f__Metschnikowiaceae(100);g__Metschnikowia(100);s__Metschnikowia_pulcherrima(100); | Metschnikowia | 13 | Saprotroph | \|Undefined Saprotroph\| | Yeast | NULL | Probable | As nectar-tap_saprotroph (Põlme et al. 2020); Decay sugar-rich substrates (Põlme et al. 2020) | Põlme S, et al. 2020. Fungal Diversity 105:1-16 ((https://doi.org/10.1007/s13225-020-00466-2)) |
| Otu14508 | 1 | k__Fungi(100);p__Ascomycota(100);c__Saccharomycetes(100);o__Saccharomycetales(100);f__Metschnikowiaceae(100);g__Metschnikowia(100);g__Metschnikowia_unclassified(100); | Metschnikowia | 13 | Saprotroph | \|Undefined Saprotroph\| | Yeast | NULL | Probable | As nectar-tap_saprotroph (Põlme et al. 2020); Decay sugar-rich substrates (Põlme et al. 2020) | Põlme S, et al. 2020. Fungal Diversity 105:1-16 ((https://doi.org/10.1007/s13225-020-00466-2)) |
| Otu14509 | 1 | k__Fungi(100);p__Ascomycota(100);c__Saccharomycetes(100);o__Saccharomycetales(100);f__Metschnikowiaceae(100);g__Metschnikowia(100);s__Metschnikowia_pulcherrima(100); | Metschnikowia | 13 | Saprotroph | \|Undefined Saprotroph\| | Yeast | NULL | Probable | As nectar-tap_saprotroph (Põlme et al. 2020); Decay sugar-rich substrates (Põlme et al. 2020) | Põlme S, et al. 2020. Fungal Diversity 105:1-16 ((https://doi.org/10.1007/s13225-020-00466-2)) |
| Otu14511 | 1 | k__Fungi(100);p__Ascomycota(100);c__Saccharomycetes(100);o__Saccharomycetales(100);f__Saccharomycetaceae(100);g__Lachancea(100);s__Lachancea_thermotolerans(100); | Lachancea | 13 | Pathotroph-Saprotroph | Animal Parasite-Animal Pathogen-\|Undefined Saprotroph\| | Yeast | NULL | Possible | As nectar-tap_saprotroph (Põlme et al. 2020); Decay animal material, sugar-rich substrates (Põlme et al. 2020); Likely opportunistic human pathogen (Irinyi et al. 2015; see L. fermentati) | Irinyi L, et al. 2015. Medical Mycology 53:313-337 ((https://doi.org/10.1093/mmy/myv008)); Põlme S, et al. 2020. Fungal Diversity 105:1-16 ((https://doi.org/10.1007/s13225-020-00466-2)) |
| Otu14512 | 1 | k__Fungi(100);p__Zygomycota(100);c__Zygomycota_class_Incertae_sedis(100);o__Mucorales(100);f__Umbelopsidaceae(100);g__Umbelopsis(100);s__Umbelopsis_angularis(100); | Umbelopsis | 13 | Saprotroph | \|Undefined Saprotroph\| | Microfungus | NULL | Probable | As soil_saprotroph (Põlme et al. 2020); Decay roots, soil (Põlme et al. 2020) | Tedersoo L, et al. 2014. Science 346:e1256688 ((https://doi.org/10.1126/science.1256688)); Põlme S, et al. 2020. Fungal Diversity 105:1-16 ((https://doi.org/10.1007/s13225-020-00466-2)) |
| Otu14513 | 1 | k__Fungi(100);p__Ascomycota(100);c__Eurotiomycetes(100);o__Chaetothyriales(100);f__Herpotrichiellaceae(100);f__Herpotrichiellaceae_unclassified(100);f__Herpotrichiellaceae_unclassified(100); | Herpotrichiellaceae | 9 | Pathotroph-Saprotroph | Animal Pathogen-Fungal Parasite-Undefined Saprotroph | Facultative Yeast-Microfungus | NULL | Probable | Mostly saprobic on plants or other fungi, isolated from soil etc., and from medical sources (Cannon & Kirk 2007) | Cannon PF, Kirk PM. 2007. Fungal Families of the World. CAB International, Cambridge (ISBN: 978-0851998275) |
| Otu14514 | 1 | k__Fungi(100);p__Ascomycota(100);c__Saccharomycetes(100);o__Saccharomycetales(100);f__Metschnikowiaceae(100);g__Metschnikowia(100);s__Metschnikowia_pulcherrima(100); | Metschnikowia | 13 | Saprotroph | \|Undefined Saprotroph\| | Yeast | NULL | Probable | As nectar-tap_saprotroph (Põlme et al. 2020); Decay sugar-rich substrates (Põlme et al. 2020) | Põlme S, et al. 2020. Fungal Diversity 105:1-16 ((https://doi.org/10.1007/s13225-020-00466-2)) |
| Otu14515 | 1 | k__Fungi(100);p__Ascomycota(100);c__Saccharomycetes(100);o__Saccharomycetales(100);f__Metschnikowiaceae(100);g__Metschnikowia(100);g__Metschnikowia_unclassified(100); | Metschnikowia | 13 | Saprotroph | \|Undefined Saprotroph\| | Yeast | NULL | Probable | As nectar-tap_saprotroph (Põlme et al. 2020); Decay sugar-rich substrates (Põlme et al. 2020) | Põlme S, et al. 2020. Fungal Diversity 105:1-16 ((https://doi.org/10.1007/s13225-020-00466-2)) |
| Otu14516 | 1 | k__Fungi(100);p__Ascomycota(100);c__Saccharomycetes(100);o__Saccharomycetales(100);f__Metschnikowiaceae(100);g__Metschnikowia(100);s__Metschnikowia_pulcherrima(100); | Metschnikowia | 13 | Saprotroph | \|Undefined Saprotroph\| | Yeast | NULL | Probable | As nectar-tap_saprotroph (Põlme et al. 2020); Decay sugar-rich substrates (Põlme et al. 2020) | Põlme S, et al. 2020. Fungal Diversity 105:1-16 ((https://doi.org/10.1007/s13225-020-00466-2)) |
| Otu14517 | 1 | k__Fungi(100);p__Ascomycota(100);c__Saccharomycetes(100);o__Saccharomycetales(100);f__Metschnikowiaceae(100);g__Metschnikowia(100);g__Metschnikowia_unclassified(100); | Metschnikowia | 13 | Saprotroph | \|Undefined Saprotroph\| | Yeast | NULL | Probable | As nectar-tap_saprotroph (Põlme et al. 2020); Decay sugar-rich substrates (Põlme et al. 2020) | Põlme S, et al. 2020. Fungal Diversity 105:1-16 ((https://doi.org/10.1007/s13225-020-00466-2)) |
| Otu14518 | 1 | k__Fungi(100);p__Ascomycota(100);c__Saccharomycetes(100);o__Saccharomycetales(100);f__Metschnikowiaceae(100);g__Metschnikowia(100);s__Metschnikowia_pulcherrima(100); | Metschnikowia | 13 | Saprotroph | \|Undefined Saprotroph\| | Yeast | NULL | Probable | As nectar-tap_saprotroph (Põlme et al. 2020); Decay sugar-rich substrates (Põlme et al. 2020) | Põlme S, et al. 2020. Fungal Diversity 105:1-16 ((https://doi.org/10.1007/s13225-020-00466-2)) |
| Otu14519 | 1 | k__Fungi(100);p__Ascomycota(100);c__Saccharomycetes(100);o__Saccharomycetales(100);f__Metschnikowiaceae(100);g__Metschnikowia(100);s__Metschnikowia_pulcherrima(100); | Metschnikowia | 13 | Saprotroph | \|Undefined Saprotroph\| | Yeast | NULL | Probable | As nectar-tap_saprotroph (Põlme et al. 2020); Decay sugar-rich substrates (Põlme et al. 2020) | Põlme S, et al. 2020. Fungal Diversity 105:1-16 ((https://doi.org/10.1007/s13225-020-00466-2)) |
| Otu14520 | 1 | k__Fungi(100);p__Ascomycota(100);c__Saccharomycetes(100);o__Saccharomycetales(100);f__Metschnikowiaceae(100);g__Metschnikowia(100);s__Metschnikowia_pulcherrima(100); | Metschnikowia | 13 | Saprotroph | \|Undefined Saprotroph\| | Yeast | NULL | Probable | As nectar-tap_saprotroph (Põlme et al. 2020); Decay sugar-rich substrates (Põlme et al. 2020) | Põlme S, et al. 2020. Fungal Diversity 105:1-16 ((https://doi.org/10.1007/s13225-020-00466-2)) |
| Otu14521 | 1 | k__Fungi(100);p__Ascomycota(100);c__Saccharomycetes(100);o__Saccharomycetales(100);f__Metschnikowiaceae(100);g__Metschnikowia(100);s__Metschnikowia_pulcherrima(100); | Metschnikowia | 13 | Saprotroph | \|Undefined Saprotroph\| | Yeast | NULL | Probable | As nectar-tap_saprotroph (Põlme et al. 2020); Decay sugar-rich substrates (Põlme et al. 2020) | Põlme S, et al. 2020. Fungal Diversity 105:1-16 ((https://doi.org/10.1007/s13225-020-00466-2)) |
| Otu14522 | 1 | k__Fungi(100);p__Ascomycota(100);c__Saccharomycetes(100);o__Saccharomycetales(100);f__Saccharomycetales_family_Incertae_sedis(100);g__Candida(100);g__Candida_unclassified(100); | Candida | 13 | Pathotroph-Saprotroph-Symbiotroph | Animal Pathogen-Endophyte-Endosymbiont-Epiphyte-\|Undefined Saprotroph\| | Dimorphic Yeast-Yeast | NULL | Probable | As nectar-tap_saprotroph (Põlme et al. 2020); Decay sugar-rich substrates (Põlme et al. 2020); This asexual genus is polyphyletic and contain species that function very differently depend on what they are. Caution should be taken when interpreting guilds for Candida species. | Manolakaki D, et al. 2010. Virulence. 1: 367-375 ((https://doi.org/10.4161/viru.1.5.12796)); Mekbib SB, et al. 2011. Tropical Plant Pathology 36:233-240 ((https://doi.org/10.1590/S1982-56762011000400004)); Põlme S, et al. 2020. Fungal Diversity 105:1-16 ((https://doi.org/10.1007/s13225-020-00466-2)) |
| Otu14525 | 1 | k__Fungi(100);p__Ascomycota(100);c__Saccharomycetes(100);o__Saccharomycetales(100);f__Metschnikowiaceae(100);g__Metschnikowia(100);s__Metschnikowia_pulcherrima(100); | Metschnikowia | 13 | Saprotroph | \|Undefined Saprotroph\| | Yeast | NULL | Probable | As nectar-tap_saprotroph (Põlme et al. 2020); Decay sugar-rich substrates (Põlme et al. 2020) | Põlme S, et al. 2020. Fungal Diversity 105:1-16 ((https://doi.org/10.1007/s13225-020-00466-2)) |
| Otu14526 | 1 | k__Fungi(100);p__Ascomycota(100);c__Saccharomycetes(100);o__Saccharomycetales(100);f__Metschnikowiaceae(100);g__Metschnikowia(100);s__Metschnikowia_pulcherrima(100); | Metschnikowia | 13 | Saprotroph | \|Undefined Saprotroph\| | Yeast | NULL | Probable | As nectar-tap_saprotroph (Põlme et al. 2020); Decay sugar-rich substrates (Põlme et al. 2020) | Põlme S, et al. 2020. Fungal Diversity 105:1-16 ((https://doi.org/10.1007/s13225-020-00466-2)) |

| | | | | | | | | | | | |
|---|---|---|---|---|---|---|---|---|---|---|---|
| Otu14528 | 1 | k__Fungi(100);p__Ascomycota(100);c__Saccharomycetes(100);o__Saccharomycetales(100);f__Metschnikowiaceae(100);g__Metschnikowia(100);s__Metschnikowia_pulcherrima(100); | Metschnikowia | 13 | Saprotroph | \|Undefined Saprotroph\| | Yeast | NULL | Probable | As nectar-tap_saprotroph (Põlme et al. 2020); Decay sugar-rich substrates (Põlme et al. 2020) | Põlme S, et al. 2020. Fungal Diversity 105:1-16 ((https://doi.org/10.1007/s13225-020-00466-2)) |
| Otu14530 | 1 | k__Fungi(100);p__Ascomycota(100);c__Saccharomycetes(100);o__Saccharomycetales(100);f__Metschnikowiaceae(100);g__Metschnikowia(100);s__Metschnikowia_pulcherrima(100); | Metschnikowia | 13 | Saprotroph | \|Undefined Saprotroph\| | Yeast | NULL | Probable | As nectar-tap_saprotroph (Põlme et al. 2020); Decay sugar-rich substrates (Põlme et al. 2020) | Põlme S, et al. 2020. Fungal Diversity 105:1-16 ((https://doi.org/10.1007/s13225-020-00466-2)) |
| Otu14531 | 1 | k__Fungi(100);p__Ascomycota(100);c__Saccharomycetes(100);o__Saccharomycetales(100);f__Metschnikowiaceae(100);g__Metschnikowia(100);s__Metschnikowia_pulcherrima(100); | Metschnikowia | 13 | Saprotroph | \|Undefined Saprotroph\| | Yeast | NULL | Probable | As nectar-tap_saprotroph (Põlme et al. 2020); Decay sugar-rich substrates (Põlme et al. 2020) | Põlme S, et al. 2020. Fungal Diversity 105:1-16 ((https://doi.org/10.1007/s13225-020-00466-2)) |
| Otu14532 | 1 | k__Fungi(100);p__Ascomycota(100);c__Saccharomycetes(100);o__Saccharomycetales(100);f__Metschnikowiaceae(100);g__Metschnikowia(100);s__Metschnikowia_pulcherrima(100); | Metschnikowia | 13 | Saprotroph | \|Undefined Saprotroph\| | Yeast | NULL | Probable | As nectar-tap_saprotroph (Põlme et al. 2020); Decay sugar-rich substrates (Põlme et al. 2020) | Põlme S, et al. 2020. Fungal Diversity 105:1-16 ((https://doi.org/10.1007/s13225-020-00466-2)) |
| Otu14533 | 1 | k__Fungi(100);p__Ascomycota(100);c__Saccharomycetes(100);o__Saccharomycetales(100);f__Metschnikowiaceae(100);g__Metschnikowia(100);g__Metschnikowia_unclassified(100); | Metschnikowia | 13 | Saprotroph | \|Undefined Saprotroph\| | Yeast | NULL | Probable | As nectar-tap_saprotroph (Põlme et al. 2020); Decay sugar-rich substrates (Põlme et al. 2020) | Põlme S, et al. 2020. Fungal Diversity 105:1-16 ((https://doi.org/10.1007/s13225-020-00466-2)) |
| Otu14534 | 1 | k__Fungi(100);p__Ascomycota(100);c__Saccharomycetes(100);o__Saccharomycetales(100);f__Metschnikowiaceae(100);g__Metschnikowia(100);s__Metschnikowia_pulcherrima(100); | Metschnikowia | 13 | Saprotroph | \|Undefined Saprotroph\| | Yeast | NULL | Probable | As nectar-tap_saprotroph (Põlme et al. 2020); Decay sugar-rich substrates (Põlme et al. 2020) | Põlme S, et al. 2020. Fungal Diversity 105:1-16 ((https://doi.org/10.1007/s13225-020-00466-2)) |
| Otu14536 | 1 | k__Fungi(100);p__Ascomycota(100);c__Saccharomycetes(100);o__Saccharomycetales(100);f__Metschnikowiaceae(100);g__Metschnikowia(100);s__Metschnikowia_pulcherrima(100); | Metschnikowia | 13 | Saprotroph | \|Undefined Saprotroph\| | Yeast | NULL | Probable | As nectar-tap_saprotroph (Põlme et al. 2020); Decay sugar-rich substrates (Põlme et al. 2020) | Põlme S, et al. 2020. Fungal Diversity 105:1-16 ((https://doi.org/10.1007/s13225-020-00466-2)) |
| Otu14537 | 1 | k__Fungi(100);p__Ascomycota(100);c__Saccharomycetes(100);o__Saccharomycetales(100);f__Metschnikowiaceae(100);g__Metschnikowia(100);g__Metschnikowia_unclassified(100); | Metschnikowia | 13 | Saprotroph | \|Undefined Saprotroph\| | Yeast | NULL | Probable | As nectar-tap_saprotroph (Põlme et al. 2020); Decay sugar-rich substrates (Põlme et al. 2020) | Põlme S, et al. 2020. Fungal Diversity 105:1-16 ((https://doi.org/10.1007/s13225-020-00466-2)) |
| Otu14538 | 1 | k__Fungi(100);p__Ascomycota(100);c__Saccharomycetes(100);o__Saccharomycetales(100);f__Metschnikowiaceae(100);g__Metschnikowia(100);s__Metschnikowia_pulcherrima(100); | Metschnikowia | 13 | Saprotroph | \|Undefined Saprotroph\| | Yeast | NULL | Probable | As nectar-tap_saprotroph (Põlme et al. 2020); Decay sugar-rich substrates (Põlme et al. 2020) | Põlme S, et al. 2020. Fungal Diversity 105:1-16 ((https://doi.org/10.1007/s13225-020-00466-2)) |
| Otu14540 | 1 | k__Fungi(100);p__Ascomycota(100);c__Saccharomycetes(100);o__Saccharomycetales(100);f__Metschnikowiaceae(100);g__Metschnikowia(100);s__Metschnikowia_pulcherrima(100); | Metschnikowia | 13 | Saprotroph | \|Undefined Saprotroph\| | Yeast | NULL | Probable | As nectar-tap_saprotroph (Põlme et al. 2020); Decay sugar-rich substrates (Põlme et al. 2020) | Põlme S, et al. 2020. Fungal Diversity 105:1-16 ((https://doi.org/10.1007/s13225-020-00466-2)) |
| Otu14541 | 1 | k__Fungi(100);p__Ascomycota(100);c__Saccharomycetes(100);o__Saccharomycetales(100);f__Metschnikowiaceae(100);g__Metschnikowia(100);s__Metschnikowia_pulcherrima(100); | Metschnikowia | 13 | Saprotroph | \|Undefined Saprotroph\| | Yeast | NULL | Probable | As nectar-tap_saprotroph (Põlme et al. 2020); Decay sugar-rich substrates (Põlme et al. 2020) | Põlme S, et al. 2020. Fungal Diversity 105:1-16 ((https://doi.org/10.1007/s13225-020-00466-2)) |
| Otu14543 | 1 | k__Fungi(100);p__Ascomycota(100);c__Saccharomycetes(100);o__Saccharomycetales(100);f__Metschnikowiaceae(100);g__Metschnikowia(100);s__Metschnikowia_pulcherrima(100); | Metschnikowia | 13 | Saprotroph | \|Undefined Saprotroph\| | Yeast | NULL | Probable | As nectar-tap_saprotroph (Põlme et al. 2020); Decay sugar-rich substrates (Põlme et al. 2020) | Põlme S, et al. 2020. Fungal Diversity 105:1-16 ((https://doi.org/10.1007/s13225-020-00466-2)) |
| Otu14546 | 1 | k__Fungi(100);p__Ascomycota(100);c__Saccharomycetes(100);o__Saccharomycetales(100);f__Metschnikowiaceae(100);g__Metschnikowia(100);s__Metschnikowia_pulcherrima(100); | Metschnikowia | 13 | Saprotroph | \|Undefined Saprotroph\| | Yeast | NULL | Probable | As nectar-tap_saprotroph (Põlme et al. 2020); Decay sugar-rich substrates (Põlme et al. 2020) | Põlme S, et al. 2020. Fungal Diversity 105:1-16 ((https://doi.org/10.1007/s13225-020-00466-2)) |
| Otu14550 | 1 | k__Fungi(100);p__Ascomycota(100);c__Saccharomycetes(100);o__Saccharomycetales(100);f__Metschnikowiaceae(100);g__Metschnikowia(100);s__Metschnikowia_pulcherrima(100); | Metschnikowia | 13 | Saprotroph | \|Undefined Saprotroph\| | Yeast | NULL | Probable | As nectar-tap_saprotroph (Põlme et al. 2020); Decay sugar-rich substrates (Põlme et al. 2020) | Põlme S, et al. 2020. Fungal Diversity 105:1-16 ((https://doi.org/10.1007/s13225-020-00466-2)) |
| Otu14555 | 1 | k__Fungi(100);p__Ascomycota(100);c__Saccharomycetes(100);o__Saccharomycetales(100);f__Metschnikowiaceae(100);g__Metschnikowia(100);s__Metschnikowia_pulcherrima(100); | Metschnikowia | 13 | Saprotroph | \|Undefined Saprotroph\| | Yeast | NULL | Probable | As nectar-tap_saprotroph (Põlme et al. 2020); Decay sugar-rich substrates (Põlme et al. 2020) | Põlme S, et al. 2020. Fungal Diversity 105:1-16 ((https://doi.org/10.1007/s13225-020-00466-2)) |
| Otu14557 | 1 | k__Fungi(100);p__Ascomycota(100);c__Saccharomycetes(100);o__Saccharomycetales(100);f__Metschnikowiaceae(100);g__Metschnikowia(100);s__Metschnikowia_pulcherrima(100); | Metschnikowia | 13 | Saprotroph | \|Undefined Saprotroph\| | Yeast | NULL | Probable | As nectar-tap_saprotroph (Põlme et al. 2020); Decay sugar-rich substrates (Põlme et al. 2020) | Põlme S, et al. 2020. Fungal Diversity 105:1-16 ((https://doi.org/10.1007/s13225-020-00466-2)) |
| Otu14558 | 1 | k__Fungi(100);p__Ascomycota(100);c__Saccharomycetes(100);o__Saccharomycetales(100);f__Saccharomycetales_family_Incertae_sedis(100);g__Candida(100);g__Candida_unclassified(100); | Candida | 13 | Pathotroph-Saprotroph-Symbiotroph | Animal Pathogen-Endophyte-Endosymbiont-Epiphyte-\|Undefined Saprotroph\| | Dimorphic Yeast-Yeast | NULL | Probable | As nectar-tap_saprotroph (Põlme et al. 2020); Decay sugar-rich substrates (Põlme et al. 2020); This asexual genus is polyphyletic and contain species that function very differently depend on what they are. Caution should be taken when interpreting guilds for Candida species. | Manolakaki D, et al. 2010. Virulence. 1: 367-375 ((https://doi.org/10.4161/viru.1.5.12796)); Mekbib SB, et al. 2011. Tropical Plant Pathology 36:233-240 ((https://doi.org/10.1590/S1982-56762011000400004)); Põlme S, et al. 2020. Fungal Diversity 105:1-16 ((https://doi.org/10.1007/s13225-020-00466-2)) |
| Otu14560 | 1 | k__Fungi(100);p__Ascomycota(100);c__Saccharomycetes(100);o__Saccharomycetales(100);f__Metschnikowiaceae(100);g__Metschnikowia(100);g__Metschnikowia_unclassified(100); | Metschnikowia | 13 | Saprotroph | \|Undefined Saprotroph\| | Yeast | NULL | Probable | As nectar-tap_saprotroph (Põlme et al. 2020); Decay sugar-rich substrates (Põlme et al. 2020) | Põlme S, et al. 2020. Fungal Diversity 105:1-16 ((https://doi.org/10.1007/s13225-020-00466-2)) |
| Otu14561 | 1 | k__Fungi(100);p__Ascomycota(100);c__Saccharomycetes(100);o__Saccharomycetales(100);f__Metschnikowiaceae(100);g__Metschnikowia(100);s__Metschnikowia_pulcherrima(100); | Metschnikowia | 13 | Saprotroph | \|Undefined Saprotroph\| | Yeast | NULL | Probable | As nectar-tap_saprotroph (Põlme et al. 2020); Decay sugar-rich substrates (Põlme et al. 2020) | Põlme S, et al. 2020. Fungal Diversity 105:1-16 ((https://doi.org/10.1007/s13225-020-00466-2)) |
| Otu14562 | 1 | k__Fungi(100);p__Ascomycota(100);c__Saccharomycetes(100);o__Saccharomycetales(100);f__Metschnikowiaceae(100);g__Metschnikowia(100);s__Metschnikowia_pulcherrima(100); | Metschnikowia | 13 | Saprotroph | \|Undefined Saprotroph\| | Yeast | NULL | Probable | As nectar-tap_saprotroph (Põlme et al. 2020); Decay sugar-rich substrates (Põlme et al. 2020) | Põlme S, et al. 2020. Fungal Diversity 105:1-16 ((https://doi.org/10.1007/s13225-020-00466-2)) |
| Otu14563 | 1 | k__Fungi(100);p__Ascomycota(100);c__Saccharomycetes(100);o__Saccharomycetales(100);f__Metschnikowiaceae(100);g__Metschnikowia(100);g__Metschnikowia_unclassified(100); | Metschnikowia | 13 | Saprotroph | \|Undefined Saprotroph\| | Yeast | NULL | Probable | As nectar-tap_saprotroph (Põlme et al. 2020); Decay sugar-rich substrates (Põlme et al. 2020) | Põlme S, et al. 2020. Fungal Diversity 105:1-16 ((https://doi.org/10.1007/s13225-020-00466-2)) |
| Otu14564 | 1 | k__Fungi(100);p__Ascomycota(100);c__Saccharomycetes(100);o__Saccharomycetales(100);f__Metschnikowiaceae(100);g__Metschnikowia(100);s__Metschnikowia_pulcherrima(100); | Metschnikowia | 13 | Saprotroph | \|Undefined Saprotroph\| | Yeast | NULL | Probable | As nectar-tap_saprotroph (Põlme et al. 2020); Decay sugar-rich substrates (Põlme et al. 2020) | Põlme S, et al. 2020. Fungal Diversity 105:1-16 ((https://doi.org/10.1007/s13225-020-00466-2)) |
| Otu14565 | 1 | k__Fungi(100);p__Ascomycota(100);c__Saccharomycetes(100);o__Saccharomycetales(100);f__Metschnikowiaceae(100);g__Metschnikowia(100);g__Metschnikowia_unclassified(100); | Metschnikowia | 13 | Saprotroph | \|Undefined Saprotroph\| | Yeast | NULL | Probable | As nectar-tap_saprotroph (Põlme et al. 2020); Decay sugar-rich substrates (Põlme et al. 2020) | Põlme S, et al. 2020. Fungal Diversity 105:1-16 ((https://doi.org/10.1007/s13225-020-00466-2)) |
| Otu14566 | 1 | k__Fungi(100);p__Ascomycota(100);c__Saccharomycetes(100);o__Saccharomycetales(100);f__Metschnikowiaceae(100);g__Metschnikowia(100);s__Metschnikowia_pulcherrima(100); | Metschnikowia | 13 | Saprotroph | \|Undefined Saprotroph\| | Yeast | NULL | Probable | As nectar-tap_saprotroph (Põlme et al. 2020); Decay sugar-rich substrates (Põlme et al. 2020) | Põlme S, et al. 2020. Fungal Diversity 105:1-16 ((https://doi.org/10.1007/s13225-020-00466-2)) |
| Otu14567 | 1 | k__Fungi(100);p__Ascomycota(100);c__Saccharomycetes(100);o__Saccharomycetales(100);f__Metschnikowiaceae(100);g__Metschnikowia(100);s__Metschnikowia_pulcherrima(100); | Metschnikowia | 13 | Saprotroph | \|Undefined Saprotroph\| | Yeast | NULL | Probable | As nectar-tap_saprotroph (Põlme et al. 2020); Decay sugar-rich substrates (Põlme et al. 2020) | Põlme S, et al. 2020. Fungal Diversity 105:1-16 ((https://doi.org/10.1007/s13225-020-00466-2)) |
| Otu14568 | 1 | k__Fungi(100);p__Ascomycota(100);c__Saccharomycetes(100);o__Saccharomycetales(100);f__Metschnikowiaceae(100);g__Metschnikowia(100);s__Metschnikowia_pulcherrima(100); | Metschnikowia | 13 | Saprotroph | \|Undefined Saprotroph\| | Yeast | NULL | Probable | As nectar-tap_saprotroph (Põlme et al. 2020); Decay sugar-rich substrates (Põlme et al. 2020) | Põlme S, et al. 2020. Fungal Diversity 105:1-16 ((https://doi.org/10.1007/s13225-020-00466-2)) |
| Otu14569 | 1 | k__Fungi(100);p__Ascomycota(100);c__Saccharomycetes(100);o__Saccharomycetales(100);f__Metschnikowiaceae(100);g__Metschnikowia(100);s__Metschnikowia_pulcherrima(100); | Metschnikowia | 13 | Saprotroph | \|Undefined Saprotroph\| | Yeast | NULL | Probable | As nectar-tap_saprotroph (Põlme et al. 2020); Decay sugar-rich substrates (Põlme et al. 2020) | Põlme S, et al. 2020. Fungal Diversity 105:1-16 ((https://doi.org/10.1007/s13225-020-00466-2)) |
| Otu14570 | 1 | k__Fungi(100);p__Ascomycota(100);c__Saccharomycetes(100);o__Saccharomycetales(100);f__Metschnikowiaceae(100);g__Metschnikowia(100);s__Metschnikowia_pulcherrima(100); | Metschnikowia | 13 | Saprotroph | \|Undefined Saprotroph\| | Yeast | NULL | Probable | As nectar-tap_saprotroph (Põlme et al. 2020); Decay sugar-rich substrates (Põlme et al. 2020) | Põlme S, et al. 2020. Fungal Diversity 105:1-16 ((https://doi.org/10.1007/s13225-020-00466-2)) |
| Otu14572 | 1 | k__Fungi(100);p__Ascomycota(100);c__Saccharomycetes(100);o__Saccharomycetales(100);f__Metschnikowiaceae(100);g__Metschnikowia(100);s__Metschnikowia_pulcherrima(100); | Metschnikowia | 13 | Saprotroph | \|Undefined Saprotroph\| | Yeast | NULL | Probable | As nectar-tap_saprotroph (Põlme et al. 2020); Decay sugar-rich substrates (Põlme et al. 2020) | Põlme S, et al. 2020. Fungal Diversity 105:1-16 ((https://doi.org/10.1007/s13225-020-00466-2)) |
| Otu14573 | 1 | k__Fungi(100);p__Ascomycota(100);c__Saccharomycetes(100);o__Saccharomycetales(100);f__Metschnikowiaceae(100);g__Metschnikowia(100);s__Metschnikowia_pulcherrima(100); | Metschnikowia | 13 | Saprotroph | \|Undefined Saprotroph\| | Yeast | NULL | Probable | As nectar-tap_saprotroph (Põlme et al. 2020); Decay sugar-rich substrates (Põlme et al. 2020) | Põlme S, et al. 2020. Fungal Diversity 105:1-16 ((https://doi.org/10.1007/s13225-020-00466-2)) |
| Otu14574 | 1 | k__Fungi(100);p__Ascomycota(100);c__Saccharomycetes(100);o__Saccharomycetales(100);f__Metschnikowiaceae(100);g__Metschnikowia(100);s__Metschnikowia_pulcherrima(100); | Metschnikowia | 13 | Saprotroph | \|Undefined Saprotroph\| | Yeast | NULL | Probable | As nectar-tap_saprotroph (Põlme et al. 2020); Decay sugar-rich substrates (Põlme et al. 2020) | Põlme S, et al. 2020. Fungal Diversity 105:1-16 ((https://doi.org/10.1007/s13225-020-00466-2)) |
| Otu14575 | 1 | k__Fungi(100);p__Ascomycota(100);c__Saccharomycetes(100);o__Saccharomycetales(100);f__Metschnikowiaceae(100);g__Metschnikowia(100);s__Metschnikowia_pulcherrima(100); | Metschnikowia | 13 | Saprotroph | \|Undefined Saprotroph\| | Yeast | NULL | Probable | As nectar-tap_saprotroph (Põlme et al. 2020); Decay sugar-rich substrates (Põlme et al. 2020) | Põlme S, et al. 2020. Fungal Diversity 105:1-16 ((https://doi.org/10.1007/s13225-020-00466-2)) |
| Otu14576 | 1 | k__Fungi(100);p__Ascomycota(100);c__Saccharomycetes(100);o__Saccharomycetales(100);f__Metschnikowiaceae(100);g__Metschnikowia(100);g__Metschnikowia_unclassified(100); | Metschnikowia | 13 | Saprotroph | \|Undefined Saprotroph\| | Yeast | NULL | Probable | As nectar-tap_saprotroph (Põlme et al. 2020); Decay sugar-rich substrates (Põlme et al. 2020) | Põlme S, et al. 2020. Fungal Diversity 105:1-16 ((https://doi.org/10.1007/s13225-020-00466-2)) |
| Otu14577 | 1 | k__Fungi(100);p__Ascomycota(100);c__Saccharomycetes(100);o__Saccharomycetales(100);f__Metschnikowiaceae(100);g__Metschnikowia(100);s__Metschnikowia_pulcherrima(100); | Metschnikowia | 13 | Saprotroph | \|Undefined Saprotroph\| | Yeast | NULL | Probable | As nectar-tap_saprotroph (Põlme et al. 2020); Decay sugar-rich substrates (Põlme et al. 2020) | Põlme S, et al. 2020. Fungal Diversity 105:1-16 ((https://doi.org/10.1007/s13225-020-00466-2)) |
| Otu14578 | 1 | k__Fungi(100);p__Ascomycota(100);c__Saccharomycetes(100);o__Saccharomycetales(100);f__Metschnikowiaceae(100);g__Metschnikowia(100);s__Metschnikowia_pulcherrima(100); | Metschnikowia | 13 | Saprotroph | \|Undefined Saprotroph\| | Yeast | NULL | Probable | As nectar-tap_saprotroph (Põlme et al. 2020); Decay sugar-rich substrates (Põlme et al. 2020) | Põlme S, et al. 2020. Fungal Diversity 105:1-16 ((https://doi.org/10.1007/s13225-020-00466-2)) |
| Otu14582 | 1 | k__Fungi(100);p__Ascomycota(100);c__Saccharomycetes(100);o__Saccharomycetales(100);f__Metschnikowiaceae(100);g__Metschnikowia(100);s__Metschnikowia_pulcherrima(100); | Metschnikowia | 13 | Saprotroph | \|Undefined Saprotroph\| | Yeast | NULL | Probable | As nectar-tap_saprotroph (Põlme et al. 2020); Decay sugar-rich substrates (Põlme et al. 2020) | Põlme S, et al. 2020. Fungal Diversity 105:1-16 ((https://doi.org/10.1007/s13225-020-00466-2)) |
| Otu14583 | 1 | k__Fungi(100);p__Ascomycota(100);c__Saccharomycetes(100);o__Saccharomycetales(100);f__Metschnikowiaceae(100);g__Metschnikowia(100);s__Metschnikowia_pulcherrima(100); | Metschnikowia | 13 | Saprotroph | \|Undefined Saprotroph\| | Yeast | NULL | Probable | As nectar-tap_saprotroph (Põlme et al. 2020); Decay sugar-rich substrates (Põlme et al. 2020) | Põlme S, et al. 2020. Fungal Diversity 105:1-16 ((https://doi.org/10.1007/s13225-020-00466-2)) |
| Otu14585 | 1 | k__Fungi(100);p__Ascomycota(100);c__Saccharomycetes(100);o__Saccharomycetales(100);f__Metschnikowiaceae(100);g__Metschnikowia(100);s__Metschnikowia_pulcherrima(100); | Metschnikowia | 13 | Saprotroph | \|Undefined Saprotroph\| | Yeast | NULL | Probable | As nectar-tap_saprotroph (Põlme et al. 2020); Decay sugar-rich substrates (Põlme et al. 2020) | Põlme S, et al. 2020. Fungal Diversity 105:1-16 ((https://doi.org/10.1007/s13225-020-00466-2)) |
| Otu14586 | 1 | k__Fungi(100);p__Ascomycota(100);c__Saccharomycetes(100);o__Saccharomycetales(100);f__Metschnikowiaceae(100);g__Metschnikowia(100);s__Metschnikowia_pulcherrima(100); | Metschnikowia | 13 | Saprotroph | \|Undefined Saprotroph\| | Yeast | NULL | Probable | As nectar-tap_saprotroph (Põlme et al. 2020); Decay sugar-rich substrates (Põlme et al. 2020) | Põlme S, et al. 2020. Fungal Diversity 105:1-16 ((https://doi.org/10.1007/s13225-020-00466-2)) |
| Otu14587 | 1 | k__Fungi(100);p__Ascomycota(100);c__Saccharomycetes(100);o__Saccharomycetales(100);f__Metschnikowiaceae(100);g__Metschnikowia(100);s__Metschnikowia_pulcherrima(100); | Metschnikowia | 13 | Saprotroph | \|Undefined Saprotroph\| | Yeast | NULL | Probable | As nectar-tap_saprotroph (Põlme et al. 2020); Decay sugar-rich substrates (Põlme et al. 2020) | Põlme S, et al. 2020. Fungal Diversity 105:1-16 ((https://doi.org/10.1007/s13225-020-00466-2)) |
| Otu14588 | 1 | k__Fungi(100);p__Ascomycota(100);c__Saccharomycetes(100);o__Saccharomycetales(100);f__Metschnikowiaceae(100);g__Metschnikowia(100);g__Metschnikowia_unclassified(100); | Metschnikowia | 13 | Saprotroph | \|Undefined Saprotroph\| | Yeast | NULL | Probable | As nectar-tap_saprotroph (Põlme et al. 2020); Decay sugar-rich substrates (Põlme et al. 2020) | Põlme S, et al. 2020. Fungal Diversity 105:1-16 ((https://doi.org/10.1007/s13225-020-00466-2)) |
| Otu14590 | 1 | k__Fungi(100);p__Ascomycota(100);c__Saccharomycetes(100);o__Saccharomycetales(100);f__Metschnikowiaceae(100);g__Metschnikowia(100);s__Metschnikowia_pulcherrima(100); | Metschnikowia | 13 | Saprotroph | \|Undefined Saprotroph\| | Yeast | NULL | Probable | As nectar-tap_saprotroph (Põlme et al. 2020); Decay sugar-rich substrates (Põlme et al. 2020) | Põlme S, et al. 2020. Fungal Diversity 105:1-16 ((https://doi.org/10.1007/s13225-020-00466-2)) |
| Otu14591 | 1 | k__Fungi(100);p__Ascomycota(100);c__Saccharomycetes(100);o__Saccharomycetales(100);f__Metschnikowiaceae(100);g__Metschnikowia(100);s__Metschnikowia_pulcherrima(100); | Metschnikowia | 13 | Saprotroph | \|Undefined Saprotroph\| | Yeast | NULL | Probable | As nectar-tap_saprotroph (Põlme et al. 2020); Decay sugar-rich substrates (Põlme et al. 2020) | Põlme S, et al. 2020. Fungal Diversity 105:1-16 ((https://doi.org/10.1007/s13225-020-00466-2)) |
| Otu14592 | 1 | k__Fungi(100);p__Ascomycota(100);c__Saccharomycetes(100);o__Saccharomycetales(100);f__Metschnikowiaceae(100);g__Metschnikowia(100);g__Metschnikowia_unclassified(100); | Metschnikowia | 13 | Saprotroph | \|Undefined Saprotroph\| | Yeast | NULL | Probable | As nectar-tap_saprotroph (Põlme et al. 2020); Decay sugar-rich substrates (Põlme et al. 2020) | Põlme S, et al. 2020. Fungal Diversity 105:1-16 ((https://doi.org/10.1007/s13225-020-00466-2)) |
| Otu14593 | 1 | k__Fungi(100);p__Ascomycota(100);c__Saccharomycetes(100);o__Saccharomycetales(100);f__Metschnikowiaceae(100);g__Metschnikowia(100);s__Metschnikowia_pulcherrima(100); | Metschnikowia | 13 | Saprotroph | \|Undefined Saprotroph\| | Yeast | NULL | Probable | As nectar-tap_saprotroph (Põlme et al. 2020); Decay sugar-rich substrates (Põlme et al. 2020) | Põlme S, et al. 2020. Fungal Diversity 105:1-16 ((https://doi.org/10.1007/s13225-020-00466-2)) |
| Otu14595 | 1 | k__Fungi(100);p__Ascomycota(100);c__Saccharomycetes(100);o__Saccharomycetales(100);f__Metschnikowiaceae(100);g__Metschnikowia(100);s__Metschnikowia_pulcherrima(100); | Metschnikowia | 13 | Saprotroph | \|Undefined Saprotroph\| | Yeast | NULL | Probable | As nectar-tap_saprotroph (Põlme et al. 2020); Decay sugar-rich substrates (Põlme et al. 2020) | Põlme S, et al. 2020. Fungal Diversity 105:1-16 ((https://doi.org/10.1007/s13225-020-00466-2)) |
| Otu14597 | 1 | k__Fungi(100);p__Ascomycota(100);c__Saccharomycetes(100);o__Saccharomycetales(100);f__Metschnikowiaceae(100);g__Metschnikowia(100);s__Metschnikowia_pulcherrima(100); | Metschnikowia | 13 | Saprotroph | \|Undefined Saprotroph\| | Yeast | NULL | Probable | As nectar-tap_saprotroph (Põlme et al. 2020); Decay sugar-rich substrates (Põlme et al. 2020) | Põlme S, et al. 2020. Fungal Diversity 105:1-16 ((https://doi.org/10.1007/s13225-020-00466-2)) |
| Otu14598 | 1 | k__Fungi(100);p__Ascomycota(100);c__Saccharomycetes(100);o__Saccharomycetales(100);f__Metschnikowiaceae(100);g__Metschnikowia(100);s__Metschnikowia_pulcherrima(100); | Metschnikowia | 13 | Saprotroph | \|Undefined Saprotroph\| | Yeast | NULL | Probable | As nectar-tap_saprotroph (Põlme et al. 2020); Decay sugar-rich substrates (Põlme et al. 2020) | Põlme S, et al. 2020. Fungal Diversity 105:1-16 ((https://doi.org/10.1007/s13225-020-00466-2)) |
| Otu14599 | 1 | k__Fungi(100);p__Ascomycota(100);c__Saccharomycetes(100);o__Saccharomycetales(100);f__Metschnikowiaceae(100);g__Metschnikowia(100);s__Metschnikowia_pulcherrima(100); | Metschnikowia | 13 | Saprotroph | \|Undefined Saprotroph\| | Yeast | NULL | Probable | As nectar-tap_saprotroph (Põlme et al. 2020); Decay sugar-rich substrates (Põlme et al. 2020) | Põlme S, et al. 2020. Fungal Diversity 105:1-16 ((https://doi.org/10.1007/s13225-020-00466-2)) |
| Otu14600 | 1 | k__Fungi(100);p__Ascomycota(100);c__Saccharomycetes(100);o__Saccharomycetales(100);f__Metschnikowiaceae(100);g__Metschnikowia(100);g__Metschnikowia_unclassified(100); | Metschnikowia | 13 | Saprotroph | \|Undefined Saprotroph\| | Yeast | NULL | Probable | As nectar-tap_saprotroph (Põlme et al. 2020); Decay sugar-rich substrates (Põlme et al. 2020) | Põlme S, et al. 2020. Fungal Diversity 105:1-16 ((https://doi.org/10.1007/s13225-020-00466-2)) |
| Otu14603 | 1 | k__Fungi(100);p__Ascomycota(100);c__Saccharomycetes(100);o__Saccharomycetales(100);f__Metschnikowiaceae(100);g__Metschnikowia(100);s__Metschnikowia_pulcherrima(100); | Metschnikowia | 13 | Saprotroph | \|Undefined Saprotroph\| | Yeast | NULL | Probable | As nectar-tap_saprotroph (Põlme et al. 2020); Decay sugar-rich substrates (Põlme et al. 2020) | Põlme S, et al. 2020. Fungal Diversity 105:1-16 ((https://doi.org/10.1007/s13225-020-00466-2)) |
| Otu14604 | 1 | k__Fungi(100);p__Ascomycota(100);c__Saccharomycetes(100);o__Saccharomycetales(100);f__Metschnikowiaceae(100);g__Metschnikowia(100);s__Metschnikowia_pulcherrima(100); | Metschnikowia | 13 | Saprotroph | \|Undefined Saprotroph\| | Yeast | NULL | Probable | As nectar-tap_saprotroph (Põlme et al. 2020); Decay sugar-rich substrates (Põlme et al. 2020) | Põlme S, et al. 2020. Fungal Diversity 105:1-16 ((https://doi.org/10.1007/s13225-020-00466-2)) |
| Otu14605 | 1 | k__Fungi(100);p__Ascomycota(100);c__Saccharomycetes(100);o__Saccharomycetales(100);f__Metschnikowiaceae(100);g__Metschnikowia(100);g__Metschnikowia_unclassified(100); | Metschnikowia | 13 | Saprotroph | \|Undefined Saprotroph\| | Yeast | NULL | Probable | As nectar-tap_saprotroph (Põlme et al. 2020); Decay sugar-rich substrates (Põlme et al. 2020) | Põlme S, et al. 2020. Fungal Diversity 105:1-16 ((https://doi.org/10.1007/s13225-020-00466-2)) |
| Otu14606 | 1 | k__Fungi(100);p__Ascomycota(100);c__Saccharomycetes(100);o__Saccharomycetales(100);f__Metschnikowiaceae(100);g__Metschnikowia(100);s__Metschnikowia_pulcherrima(100); | Metschnikowia | 13 | Saprotroph | \|Undefined Saprotroph\| | Yeast | NULL | Probable | As nectar-tap_saprotroph (Põlme et al. 2020); Decay sugar-rich substrates (Põlme et al. 2020) | Põlme S, et al. 2020. Fungal Diversity 105:1-16 ((https://doi.org/10.1007/s13225-020-00466-2)) |
| Otu14607 | 1 | k__Fungi(100);p__Ascomycota(100);c__Saccharomycetes(100);o__Saccharomycetales(100);f__Metschnikowiaceae(100);g__Metschnikowia(100);g__Metschnikowia_unclassified(100); | Metschnikowia | 13 | Saprotroph | \|Undefined Saprotroph\| | Yeast | NULL | Probable | As nectar-tap_saprotroph (Põlme et al. 2020); Decay sugar-rich substrates (Põlme et al. 2020) | Põlme S, et al. 2020. Fungal Diversity 105:1-16 ((https://doi.org/10.1007/s13225-020-00466-2)) |
| Otu14610 | 1 | k__Fungi(100);p__Ascomycota(100);c__Saccharomycetes(100);o__Saccharomycetales(100);f__Metschnikowiaceae(100);g__Metschnikowia(100);s__Metschnikowia_pulcherrima(100); | Metschnikowia | 13 | Saprotroph | \|Undefined Saprotroph\| | Yeast | NULL | Probable | As nectar-tap_saprotroph (Põlme et al. 2020); Decay sugar-rich substrates (Põlme et al. 2020) | Põlme S, et al. 2020. Fungal Diversity 105:1-16 ((https://doi.org/10.1007/s13225-020-00466-2)) |
| Otu14611 | 1 | k__Fungi(100);p__Ascomycota(100);c__Saccharomycetes(100);o__Saccharomycetales(100);f__Metschnikowiaceae(100);g__Metschnikowia(100);g__Metschnikowia_unclassified(100); | Metschnikowia | 13 | Saprotroph | \|Undefined Saprotroph\| | Yeast | NULL | Probable | As nectar-tap_saprotroph (Põlme et al. 2020); Decay sugar-rich substrates (Põlme et al. 2020) | Põlme S, et al. 2020. Fungal Diversity 105:1-16 ((https://doi.org/10.1007/s13225-020-00466-2)) |
| Otu14612 | 1 | k__Fungi(100);p__Ascomycota(100);c__Saccharomycetes(100);o__Saccharomycetales(100);f__Metschnikowiaceae(100);g__Metschnikowia(100);s__Metschnikowia_pulcherrima(100); | Metschnikowia | 13 | Saprotroph | \|Undefined Saprotroph\| | Yeast | NULL | Probable | As nectar-tap_saprotroph (Põlme et al. 2020); Decay sugar-rich substrates (Põlme et al. 2020) | Põlme S, et al. 2020. Fungal Diversity 105:1-16 ((https://doi.org/10.1007/s13225-020-00466-2)) |
| Otu14613 | 1 | k__Fungi(100);p__Ascomycota(100);c__Saccharomycetes(100);o__Saccharomycetales(100);f__Metschnikowiaceae(100);g__Metschnikowia(100);s__Metschnikowia_pulcherrima(100); | Metschnikowia | 13 | Saprotroph | \|Undefined Saprotroph\| | Yeast | NULL | Probable | As nectar-tap_saprotroph (Põlme et al. 2020); Decay sugar-rich substrates (Põlme et al. 2020) | Põlme S, et al. 2020. Fungal Diversity 105:1-16 ((https://doi.org/10.1007/s13225-020-00466-2)) |
| Otu14614 | 1 | k__Fungi(100);p__Ascomycota(100);c__Saccharomycetes(100);o__Saccharomycetales(100);f__Metschnikowiaceae(100);g__Metschnikowia(100);s__Metschnikowia_pulcherrima(100); | Metschnikowia | 13 | Saprotroph | \|Undefined Saprotroph\| | Yeast | NULL | Probable | As nectar-tap_saprotroph (Põlme et al. 2020); Decay sugar-rich substrates (Põlme et al. 2020) | Põlme S, et al. 2020. Fungal Diversity 105:1-16 ((https://doi.org/10.1007/s13225-020-00466-2)) |
| Otu14615 | 1 | k__Fungi(100);p__Ascomycota(100);c__Saccharomycetes(100);o__Saccharomycetales(100);f__Metschnikowiaceae(100);g__Metschnikowia(100);s__Metschnikowia_pulcherrima(100); | Metschnikowia | 13 | Saprotroph | \|Undefined Saprotroph\| | Yeast | NULL | Probable | As nectar-tap_saprotroph (Põlme et al. 2020); Decay sugar-rich substrates (Põlme et al. 2020) | Põlme S, et al. 2020. Fungal Diversity 105:1-16 ((https://doi.org/10.1007/s13225-020-00466-2)) |
| Otu14616 | 1 | k__Fungi(100);p__Ascomycota(100);c__Saccharomycetes(100);o__Saccharomycetales(100);f__Metschnikowiaceae(100);g__Metschnikowia(100);s__Metschnikowia_pulcherrima(100); | Metschnikowia | 13 | Saprotroph | \|Undefined Saprotroph\| | Yeast | NULL | Probable | As nectar-tap_saprotroph (Põlme et al. 2020); Decay sugar-rich substrates (Põlme et al. 2020) | Põlme S, et al. 2020. Fungal Diversity 105:1-16 ((https://doi.org/10.1007/s13225-020-00466-2)) |
| Otu14618 | 1 | k__Fungi(100);p__Ascomycota(100);c__Saccharomycetes(100);o__Saccharomycetales(100);f__Metschnikowiaceae(100);g__Metschnikowia(100);s__Metschnikowia_pulcherrima(100); | Metschnikowia | 13 | Saprotroph | \|Undefined Saprotroph\| | Yeast | NULL | Probable | As nectar-tap_saprotroph (Põlme et al. 2020); Decay sugar-rich substrates (Põlme et al. 2020) | Põlme S, et al. 2020. Fungal Diversity 105:1-16 ((https://doi.org/10.1007/s13225-020-00466-2)) |
| Otu14619 | 1 | k__Fungi(100);p__Ascomycota(100);c__Saccharomycetes(100);o__Saccharomycetales(100);f__Metschnikowiaceae(100);g__Metschnikowia(100);s__Metschnikowia_pulcherrima(100); | Metschnikowia | 13 | Saprotroph | \|Undefined Saprotroph\| | Yeast | NULL | Probable | As nectar-tap_saprotroph (Põlme et al. 2020); Decay sugar-rich substrates (Põlme et al. 2020) | Põlme S, et al. 2020. Fungal Diversity 105:1-16 ((https://doi.org/10.1007/s13225-020-00466-2)) |
| Otu14621 | 1 | k__Fungi(100);p__Ascomycota(100);c__Saccharomycetes(100);o__Saccharomycetales(100);f__Metschnikowiaceae(100);g__Metschnikowia(100);g__Metschnikowia_unclassified(100); | Metschnikowia | 13 | Saprotroph | \|Undefined Saprotroph\| | Yeast | NULL | Probable | As nectar-tap_saprotroph (Põlme et al. 2020); Decay sugar-rich substrates (Põlme et al. 2020) | Põlme S, et al. 2020. Fungal Diversity 105:1-16 ((https://doi.org/10.1007/s13225-020-00466-2)) |
| Otu14622 | 1 | k__Fungi(100);p__Ascomycota(100);c__Saccharomycetes(100);o__Saccharomycetales(100);f__Metschnikowiaceae(100);g__Metschnikowia(100);s__Metschnikowia_pulcherrima(100); | Metschnikowia | 13 | Saprotroph | \|Undefined Saprotroph\| | Yeast | NULL | Probable | As nectar-tap_saprotroph (Põlme et al. 2020); Decay sugar-rich substrates (Põlme et al. 2020) | Põlme S, et al. 2020. Fungal Diversity 105:1-16 ((https://doi.org/10.1007/s13225-020-00466-2)) |
| Otu14623 | 1 | k__Fungi(100);p__Ascomycota(100);c__Saccharomycetes(100);o__Saccharomycetales(100);f__Saccharomycetales_family_Incertae_sedis(100);g__Candida(100);g__Candida_unclassified(100); | Candida | 13 | Pathotroph-Saprotroph-Symbiotroph | Animal Pathogen-Endophyte-Endosymbiont-Epiphyte-\|Undefined Saprotroph\| | Dimorphic Yeast-Yeast | NULL | Probable | As nectar-tap_saprotroph (Põlme et al. 2020); Decay sugar-rich substrates (Põlme et al. 2020); This asexual genus is polyphyletic and contain species that function very differently depend on what they are. Caution should be taken when interpreting guilds for Candida species. | Manolakaki D, et al. 2010. Virulence. 1: 367-375 ((https://doi.org/10.4161/viru.1.5.12796)); Mekbib SB, et al. 2011. Tropical Plant Pathology 36:233-240 ((https://doi.org/10.1590/S1982-56762011000400004)); Põlme S, et al. 2020. Fungal Diversity 105:1-16 ((https://doi.org/10.1007/s13225-020-00466-2)) |
| Otu14624 | 1 | k__Fungi(100);p__Ascomycota(100);c__Saccharomycetes(100);o__Saccharomycetales(100);f__Metschnikowiaceae(100);g__Metschnikowia(100);s__Metschnikowia_pulcherrima(100); | Metschnikowia | 13 | Saprotroph | \|Undefined Saprotroph\| | Yeast | NULL | Probable | As nectar-tap_saprotroph (Põlme et al. 2020); Decay sugar-rich substrates (Põlme et al. 2020) | Põlme S, et al. 2020. Fungal Diversity 105:1-16 ((https://doi.org/10.1007/s13225-020-00466-2)) |
| Otu14625 | 1 | k__Fungi(100);p__Ascomycota(100);c__Saccharomycetes(100);o__Saccharomycetales(100);f__Metschnikowiaceae(100);g__Metschnikowia(100);g__Metschnikowia_unclassified(100); | Metschnikowia | 13 | Saprotroph | \|Undefined Saprotroph\| | Yeast | NULL | Probable | As nectar-tap_saprotroph (Põlme et al. 2020); Decay sugar-rich substrates (Põlme et al. 2020) | Põlme S, et al. 2020. Fungal Diversity 105:1-16 ((https://doi.org/10.1007/s13225-020-00466-2)) |
| Otu14626 | 1 | k__Fungi(100);p__Ascomycota(100);c__Saccharomycetes(100);o__Saccharomycetales(100);f__Saccharomycetales_family_Incertae_sedis(100);g__Candida(100);g__Candida_unclassified(100); | Candida | 13 | Pathotroph-Saprotroph-Symbiotroph | Animal Pathogen-Endophyte-Endosymbiont-Epiphyte-\|Undefined Saprotroph\| | Dimorphic Yeast-Yeast | NULL | Probable | As nectar-tap_saprotroph (Põlme et al. 2020); Decay sugar-rich substrates (Põlme et al. 2020); This asexual genus is polyphyletic and contain species that function very differently depend on what they are. Caution should be taken when interpreting guilds for Candida species. | Manolakaki D, et al. 2010. Virulence. 1: 367-375 ((https://doi.org/10.4161/viru.1.5.12796)); Mekbib SB, et al. 2011. Tropical Plant Pathology 36:233-240 ((https://doi.org/10.1590/S1982-56762011000400004)); Põlme S, et al. 2020. Fungal Diversity 105:1-16 ((https://doi.org/10.1007/s13225-020-00466-2)) |
| Otu14627 | 1 | k__Fungi(100);p__Ascomycota(100);c__Saccharomycetes(100);o__Saccharomycetales(100);f__Metschnikowiaceae(100);g__Metschnikowia(100);g__Metschnikowia_unclassified(100); | Metschnikowia | 13 | Saprotroph | \|Undefined Saprotroph\| | Yeast | NULL | Probable | As nectar-tap_saprotroph (Põlme et al. 2020); Decay sugar-rich substrates (Põlme et al. 2020) | Põlme S, et al. 2020. Fungal Diversity 105:1-16 ((https://doi.org/10.1007/s13225-020-00466-2)) |
| Otu14628 | 1 | k__Fungi(100);p__Ascomycota(100);c__Saccharomycetes(100);o__Saccharomycetales(100);f__Metschnikowiaceae(100);g__Metschnikowia(100);s__Metschnikowia_pulcherrima(100); | Metschnikowia | 13 | Saprotroph | \|Undefined Saprotroph\| | Yeast | NULL | Probable | As nectar-tap_saprotroph (Põlme et al. 2020); Decay sugar-rich substrates (Põlme et al. 2020) | Põlme S, et al. 2020. Fungal Diversity 105:1-16 ((https://doi.org/10.1007/s13225-020-00466-2)) |
| Otu14629 | 1 | k__Fungi(100);p__Ascomycota(100);c__Saccharomycetes(100);o__Saccharomycetales(100);f__Metschnikowiaceae(100);g__Metschnikowia(100);s__Metschnikowia_pulcherrima(100); | Metschnikowia | 13 | Saprotroph | \|Undefined Saprotroph\| | Yeast | NULL | Probable | As nectar-tap_saprotroph (Põlme et al. 2020); Decay sugar-rich substrates (Põlme et al. 2020) | Põlme S, et al. 2020. Fungal Diversity 105:1-16 ((https://doi.org/10.1007/s13225-020-00466-2)) |
| Otu14630 | 1 | k__Fungi(100);p__Ascomycota(100);c__Saccharomycetes(100);o__Saccharomycetales(100);f__Metschnikowiaceae(100);g__Metschnikowia(100);s__Metschnikowia_pulcherrima(100); | Metschnikowia | 13 | Saprotroph | \|Undefined Saprotroph\| | Yeast | NULL | Probable | As nectar-tap_saprotroph (Põlme et al. 2020); Decay sugar-rich substrates (Põlme et al. 2020) | Põlme S, et al. 2020. Fungal Diversity 105:1-16 ((https://doi.org/10.1007/s13225-020-00466-2)) |
| Otu14631 | 1 | k__Fungi(100);p__Ascomycota(100);c__Saccharomycetes(100);o__Saccharomycetales(100);f__Metschnikowiaceae(100);g__Metschnikowia(100);s__Metschnikowia_pulcherrima(100); | Metschnikowia | 13 | Saprotroph | \|Undefined Saprotroph\| | Yeast | NULL | Probable | As nectar-tap_saprotroph (Põlme et al. 2020); Decay sugar-rich substrates (Põlme et al. 2020) | Põlme S, et al. 2020. Fungal Diversity 105:1-16 ((https://doi.org/10.1007/s13225-020-00466-2)) |
| Otu14632 | 1 | k__Fungi(100);p__Ascomycota(100);c__Saccharomycetes(100);o__Saccharomycetales(100);f__Metschnikowiaceae(100);g__Metschnikowia(100);s__Metschnikowia_pulcherrima(100); | Metschnikowia | 13 | Saprotroph | \|Undefined Saprotroph\| | Yeast | NULL | Probable | As nectar-tap_saprotroph (Põlme et al. 2020); Decay sugar-rich substrates (Põlme et al. 2020) | Põlme S, et al. 2020. Fungal Diversity 105:1-16 ((https://doi.org/10.1007/s13225-020-00466-2)) |
| Otu14636 | 1 | k__Fungi(100);p__Ascomycota(100);c__Saccharomycetes(100);o__Saccharomycetales(100);f__Metschnikowiaceae(100);g__Metschnikowia(100);g__Metschnikowia_unclassified(100); | Metschnikowia | 13 | Saprotroph | \|Undefined Saprotroph\| | Yeast | NULL | Probable | As nectar-tap_saprotroph (Põlme et al. 2020); Decay sugar-rich substrates (Põlme et al. 2020) | Põlme S, et al. 2020. Fungal Diversity 105:1-16 ((https://doi.org/10.1007/s13225-020-00466-2)) |
| Otu14637 | 1 | k__Fungi(100);p__Ascomycota(100);c__Saccharomycetes(100);o__Saccharomycetales(100);f__Metschnikowiaceae(100);g__Metschnikowia(100);s__Metschnikowia_pulcherrima(100); | Metschnikowia | 13 | Saprotroph | \|Undefined Saprotroph\| | Yeast | NULL | Probable | As nectar-tap_saprotroph (Põlme et al. 2020); Decay sugar-rich substrates (Põlme et al. 2020) | Põlme S, et al. 2020. Fungal Diversity 105:1-16 ((https://doi.org/10.1007/s13225-020-00466-2)) |
| Otu14638 | 1 | k__Fungi(100);p__Ascomycota(100);c__Saccharomycetes(100);o__Saccharomycetales(100);f__Metschnikowiaceae(100);g__Metschnikowia(100);g__Metschnikowia_unclassified(100); | Metschnikowia | 13 | Saprotroph | \|Undefined Saprotroph\| | Yeast | NULL | Probable | As nectar-tap_saprotroph (Põlme et al. 2020); Decay sugar-rich substrates (Põlme et al. 2020) | Põlme S, et al. 2020. Fungal Diversity 105:1-16 ((https://doi.org/10.1007/s13225-020-00466-2)) |
| Otu14639 | 1 | k__Fungi(100);p__Ascomycota(100);c__Saccharomycetes(100);o__Saccharomycetales(100);f__Metschnikowiaceae(100);g__Metschnikowia(100);s__Metschnikowia_pulcherrima(100); | Metschnikowia | 13 | Saprotroph | \|Undefined Saprotroph\| | Yeast | NULL | Probable | As nectar-tap_saprotroph (Põlme et al. 2020); Decay sugar-rich substrates (Põlme et al. 2020) | Põlme S, et al. 2020. Fungal Diversity 105:1-16 ((https://doi.org/10.1007/s13225-020-00466-2)) |
| Otu14640 | 1 | k__Fungi(100);p__Ascomycota(100);c__Saccharomycetes(100);o__Saccharomycetales(100);f__Metschnikowiaceae(100);g__Metschnikowia(100);s__Metschnikowia_pulcherrima(100); | Metschnikowia | 13 | Saprotroph | \|Undefined Saprotroph\| | Yeast | NULL | Probable | As nectar-tap_saprotroph (Põlme et al. 2020); Decay sugar-rich substrates (Põlme et al. 2020) | Põlme S, et al. 2020. Fungal Diversity 105:1-16 ((https://doi.org/10.1007/s13225-020-00466-2)) |
| Otu14641 | 1 | k__Fungi(100);p__Ascomycota(100);c__Saccharomycetes(100);o__Saccharomycetales(100);f__Metschnikowiaceae(100);g__Metschnikowia(100);s__Metschnikowia_pulcherrima(100); | Metschnikowia | 13 | Saprotroph | \|Undefined Saprotroph\| | Yeast | NULL | Probable | As nectar-tap_saprotroph (Põlme et al. 2020); Decay sugar-rich substrates (Põlme et al. 2020) | Põlme S, et al. 2020. Fungal Diversity 105:1-16 ((https://doi.org/10.1007/s13225-020-00466-2)) |
| Otu14642 | 1 | k__Fungi(100);p__Ascomycota(100);c__Saccharomycetes(100);o__Saccharomycetales(100);f__Metschnikowiaceae(100);g__Metschnikowia(100);s__Metschnikowia_pulcherrima(100); | Metschnikowia | 13 | Saprotroph | \|Undefined Saprotroph\| | Yeast | NULL | Probable | As nectar-tap_saprotroph (Põlme et al. 2020); Decay sugar-rich substrates (Põlme et al. 2020) | Põlme S, et al. 2020. Fungal Diversity 105:1-16 ((https://doi.org/10.1007/s13225-020-00466-2)) |
| Otu14643 | 1 | k__Fungi(100);p__Ascomycota(100);c__Saccharomycetes(100);o__Saccharomycetales(100);f__Metschnikowiaceae(100);g__Metschnikowia(100);s__Metschnikowia_pulcherrima(100); | Metschnikowia | 13 | Saprotroph | \|Undefined Saprotroph\| | Yeast | NULL | Probable | As nectar-tap_saprotroph (Põlme et al. 2020); Decay sugar-rich substrates (Põlme et al. 2020) | Põlme S, et al. 2020. Fungal Diversity 105:1-16 ((https://doi.org/10.1007/s13225-020-00466-2)) |
| Otu14644 | 1 | k__Fungi(100);p__Ascomycota(100);c__Saccharomycetes(100);o__Saccharomycetales(100);f__Metschnikowiaceae(100);g__Metschnikowia(100);s__Metschnikowia_pulcherrima(100); | Metschnikowia | 13 | Saprotroph | \|Undefined Saprotroph\| | Yeast | NULL | Probable | As nectar-tap_saprotroph (Põlme et al. 2020); Decay sugar-rich substrates (Põlme et al. 2020) | Põlme S, et al. 2020. Fungal Diversity 105:1-16 ((https://doi.org/10.1007/s13225-020-00466-2)) |
| Otu14645 | 1 | k__Fungi(100);p__Ascomycota(100);c__Saccharomycetes(100);o__Saccharomycetales(100);f__Metschnikowiaceae(100);g__Metschnikowia(100);g__Metschnikowia_unclassified(100); | Metschnikowia | 13 | Saprotroph | \|Undefined Saprotroph\| | Yeast | NULL | Probable | As nectar-tap_saprotroph (Põlme et al. 2020); Decay sugar-rich substrates (Põlme et al. 2020) | Põlme S, et al. 2020. Fungal Diversity 105:1-16 ((https://doi.org/10.1007/s13225-020-00466-2)) |
| Otu14646 | 1 | k__Fungi(100);p__Ascomycota(100);c__Saccharomycetes(100);o__Saccharomycetales(100);f__Metschnikowiaceae(100);g__Metschnikowia(100);s__Metschnikowia_pulcherrima(100); | Metschnikowia | 13 | Saprotroph | \|Undefined Saprotroph\| | Yeast | NULL | Probable | As nectar-tap_saprotroph (Põlme et al. 2020); Decay sugar-rich substrates (Põlme et al. 2020) | Põlme S, et al. 2020. Fungal Diversity 105:1-16 ((https://doi.org/10.1007/s13225-020-00466-2)) |
| Otu14647 | 1 | k__Fungi(100);p__Ascomycota(100);c__Saccharomycetes(100);o__Saccharomycetales(100);f__Metschnikowiaceae(100);g__Metschnikowia(100);s__Metschnikowia_pulcherrima(100); | Metschnikowia | 13 | Saprotroph | \|Undefined Saprotroph\| | Yeast | NULL | Probable | As nectar-tap_saprotroph (Põlme et al. 2020); Decay sugar-rich substrates (Põlme et al. 2020) | Põlme S, et al. 2020. Fungal Diversity 105:1-16 ((https://doi.org/10.1007/s13225-020-00466-2)) |
| Otu14649 | 1 | k__Fungi(100);p__Ascomycota(100);c__Saccharomycetes(100);o__Saccharomycetales(100);f__Metschnikowiaceae(100);g__Metschnikowia(100);s__Metschnikowia_pulcherrima(100); | Metschnikowia | 13 | Saprotroph | \|Undefined Saprotroph\| | Yeast | NULL | Probable | As nectar-tap_saprotroph (Põlme et al. 2020); Decay sugar-rich substrates (Põlme et al. 2020) | Põlme S, et al. 2020. Fungal Diversity 105:1-16 ((https://doi.org/10.1007/s13225-020-00466-2)) |
| Otu14650 | 1 | k__Fungi(100);p__Ascomycota(100);c__Saccharomycetes(100);o__Saccharomycetales(100);f__Metschnikowiaceae(100);g__Metschnikowia(100);g__Metschnikowia_unclassified(100); | Metschnikowia | 13 | Saprotroph | \|Undefined Saprotroph\| | Yeast | NULL | Probable | As nectar-tap_saprotroph (Põlme et al. 2020); Decay sugar-rich substrates (Põlme et al. 2020) | Põlme S, et al. 2020. Fungal Diversity 105:1-16 ((https://doi.org/10.1007/s13225-020-00466-2)) |
| Otu14651 | 1 | k__Fungi(100);p__Ascomycota(100);c__Saccharomycetes(100);o__Saccharomycetales(100);f__Metschnikowiaceae(100);g__Metschnikowia(100);s__Metschnikowia_pulcherrima(100); | Metschnikowia | 13 | Saprotroph | \|Undefined Saprotroph\| | Yeast | NULL | Probable | As nectar-tap_saprotroph (Põlme et al. 2020); Decay sugar-rich substrates (Põlme et al. 2020) | Põlme S, et al. 2020. Fungal Diversity 105:1-16 ((https://doi.org/10.1007/s13225-020-00466-2)) |
| Otu14652 | 1 | k__Fungi(100);p__Ascomycota(100);c__Saccharomycetes(100);o__Saccharomycetales(100);f__Metschnikowiaceae(100);g__Metschnikowia(100);s__Metschnikowia_pulcherrima(100); | Metschnikowia | 13 | Saprotroph | \|Undefined Saprotroph\| | Yeast | NULL | Probable | As nectar-tap_saprotroph (Põlme et al. 2020); Decay sugar-rich substrates (Põlme et al. 2020) | Põlme S, et al. 2020. Fungal Diversity 105:1-16 ((https://doi.org/10.1007/s13225-020-00466-2)) |
| Otu14653 | 1 | k__Fungi(100);p__Ascomycota(100);c__Saccharomycetes(100);o__Saccharomycetales(100);f__Metschnikowiaceae(100);g__Metschnikowia(100);s__Metschnikowia_pulcherrima(100); | Metschnikowia | 13 | Saprotroph | \|Undefined Saprotroph\| | Yeast | NULL | Probable | As nectar-tap_saprotroph (Põlme et al. 2020); Decay sugar-rich substrates (Põlme et al. 2020) | Põlme S, et al. 2020. Fungal Diversity 105:1-16 ((https://doi.org/10.1007/s13225-020-00466-2)) |
| Otu14654 | 1 | k__Fungi(100);p__Ascomycota(100);c__Saccharomycetes(100);o__Saccharomycetales(100);f__Metschnikowiaceae(100);g__Metschnikowia(100);s__Metschnikowia_pulcherrima(100); | Metschnikowia | 13 | Saprotroph | \|Undefined Saprotroph\| | Yeast | NULL | Probable | As nectar-tap_saprotroph (Põlme et al. 2020); Decay sugar-rich substrates (Põlme et al. 2020) | Põlme S, et al. 2020. Fungal Diversity 105:1-16 ((https://doi.org/10.1007/s13225-020-00466-2)) |
| Otu14655 | 1 | k__Fungi(100);p__Ascomycota(100);c__Saccharomycetes(100);o__Saccharomycetales(100);f__Metschnikowiaceae(100);g__Metschnikowia(100);g__Metschnikowia_unclassified(100); | Metschnikowia | 13 | Saprotroph | \|Undefined Saprotroph\| | Yeast | NULL | Probable | As nectar-tap_saprotroph (Põlme et al. 2020); Decay sugar-rich substrates (Põlme et al. 2020) | Põlme S, et al. 2020. Fungal Diversity 105:1-16 ((https://doi.org/10.1007/s13225-020-00466-2)) |
| Otu14656 | 1 | k__Fungi(100);p__Ascomycota(100);c__Saccharomycetes(100);o__Saccharomycetales(100);f__Metschnikowiaceae(100);g__Metschnikowia(100);s__Metschnikowia_pulcherrima(100); | Metschnikowia | 13 | Saprotroph | \|Undefined Saprotroph\| | Yeast | NULL | Probable | As nectar-tap_saprotroph (Põlme et al. 2020); Decay sugar-rich substrates (Põlme et al. 2020) | Põlme S, et al. 2020. Fungal Diversity 105:1-16 ((https://doi.org/10.1007/s13225-020-00466-2)) |
| Otu14657 | 1 | k__Fungi(100);p__Ascomycota(100);c__Saccharomycetes(100);o__Saccharomycetales(100);f__Metschnikowiaceae(100);g__Metschnikowia(100);s__Metschnikowia_pulcherrima(100); | Metschnikowia | 13 | Saprotroph | \|Undefined Saprotroph\| | Yeast | NULL | Probable | As nectar-tap_saprotroph (Põlme et al. 2020); Decay sugar-rich substrates (Põlme et al. 2020) | Põlme S, et al. 2020. Fungal Diversity 105:1-16 ((https://doi.org/10.1007/s13225-020-00466-2)) |
| Otu14658 | 1 | k__Fungi(100);p__Ascomycota(100);c__Saccharomycetes(100);o__Saccharomycetales(100);f__Metschnikowiaceae(100);g__Metschnikowia(100);s__Metschnikowia_pulcherrima(100); | Metschnikowia | 13 | Saprotroph | \|Undefined Saprotroph\| | Yeast | NULL | Probable | As nectar-tap_saprotroph (Põlme et al. 2020); Decay sugar-rich substrates (Põlme et al. 2020) | Põlme S, et al. 2020. Fungal Diversity 105:1-16 ((https://doi.org/10.1007/s13225-020-00466-2)) |
| Otu14659 | 1 | k__Fungi(100);p__Ascomycota(100);c__Saccharomycetes(100);o__Saccharomycetales(100);f__Metschnikowiaceae(100);g__Metschnikowia(100);g__Metschnikowia_unclassified(100); | Metschnikowia | 13 | Saprotroph | \|Undefined Saprotroph\| | Yeast | NULL | Probable | As nectar-tap_saprotroph (Põlme et al. 2020); Decay sugar-rich substrates (Põlme et al. 2020) | Põlme S, et al. 2020. Fungal Diversity 105:1-16 ((https://doi.org/10.1007/s13225-020-00466-2)) |
| Otu14660 | 1 | k__Fungi(100);p__Ascomycota(100);c__Saccharomycetes(100);o__Saccharomycetales(100);f__Metschnikowiaceae(100);g__Metschnikowia(100);s__Metschnikowia_pulcherrima(100); | Metschnikowia | 13 | Saprotroph | \|Undefined Saprotroph\| | Yeast | NULL | Probable | As nectar-tap_saprotroph (Põlme et al. 2020); Decay sugar-rich substrates (Põlme et al. 2020) | Põlme S, et al. 2020. Fungal Diversity 105:1-16 ((https://doi.org/10.1007/s13225-020-00466-2)) |
| Otu14663 | 1 | k__Fungi(100);p__Ascomycota(100);c__Saccharomycetes(100);o__Saccharomycetales(100);f__Metschnikowiaceae(100);g__Metschnikowia(100);s__Metschnikowia_pulcherrima(100); | Metschnikowia | 13 | Saprotroph | \|Undefined Saprotroph\| | Yeast | NULL | Probable | As nectar-tap_saprotroph (Põlme et al. 2020); Decay sugar-rich substrates (Põlme et al. 2020) | Põlme S, et al. 2020. Fungal Diversity 105:1-16 ((https://doi.org/10.1007/s13225-020-00466-2)) |
| Otu14664 | 1 | k__Fungi(100);p__Ascomycota(100);c__Saccharomycetes(100);o__Saccharomycetales(100);f__Metschnikowiaceae(100);g__Metschnikowia(100);g__Metschnikowia_unclassified(100); | Metschnikowia | 13 | Saprotroph | \|Undefined Saprotroph\| | Yeast | NULL | Probable | As nectar-tap_saprotroph (Põlme et al. 2020); Decay sugar-rich substrates (Põlme et al. 2020) | Põlme S, et al. 2020. Fungal Diversity 105:1-16 ((https://doi.org/10.1007/s13225-020-00466-2)) |
| Otu14665 | 1 | k__Fungi(100);p__Ascomycota(100);c__Saccharomycetes(100);o__Saccharomycetales(100);f__Metschnikowiaceae(100);g__Metschnikowia(100);s__Metschnikowia_pulcherrima(100); | Metschnikowia | 13 | Saprotroph | \|Undefined Saprotroph\| | Yeast | NULL | Probable | As nectar-tap_saprotroph (Põlme et al. 2020); Decay sugar-rich substrates (Põlme et al. 2020) | Põlme S, et al. 2020. Fungal Diversity 105:1-16 ((https://doi.org/10.1007/s13225-020-00466-2)) |
| Otu14666 | 1 | k__Fungi(100);p__Ascomycota(100);c__Saccharomycetes(100);o__Saccharomycetales(100);f__Metschnikowiaceae(100);g__Metschnikowia(100);s__Metschnikowia_pulcherrima(100); | Metschnikowia | 13 | Saprotroph | \|Undefined Saprotroph\| | Yeast | NULL | Probable | As nectar-tap_saprotroph (Põlme et al. 2020); Decay sugar-rich substrates (Põlme et al. 2020) | Põlme S, et al. 2020. Fungal Diversity 105:1-16 ((https://doi.org/10.1007/s13225-020-00466-2)) |
| Otu14668 | 1 | k__Fungi(100);p__Ascomycota(100);c__Saccharomycetes(100);o__Saccharomycetales(100);f__Metschnikowiaceae(100);g__Metschnikowia(100);s__Metschnikowia_pulcherrima(100); | Metschnikowia | 13 | Saprotroph | \|Undefined Saprotroph\| | Yeast | NULL | Probable | As nectar-tap_saprotroph (Põlme et al. 2020); Decay sugar-rich substrates (Põlme et al. 2020) | Põlme S, et al. 2020. Fungal Diversity 105:1-16 ((https://doi.org/10.1007/s13225-020-00466-2)) |
| Otu14669 | 1 | k__Fungi(100);p__Ascomycota(100);c__Saccharomycetes(100);o__Saccharomycetales(100);f__Metschnikowiaceae(100);g__Metschnikowia(100);s__Metschnikowia_pulcherrima(100); | Metschnikowia | 13 | Saprotroph | \|Undefined Saprotroph\| | Yeast | NULL | Probable | As nectar-tap_saprotroph (Põlme et al. 2020); Decay sugar-rich substrates (Põlme et al. 2020) | Põlme S, et al. 2020. Fungal Diversity 105:1-16 ((https://doi.org/10.1007/s13225-020-00466-2)) |
| Otu14670 | 1 | k__Fungi(100);p__Ascomycota(100);c__Saccharomycetes(100);o__Saccharomycetales(100);f__Metschnikowiaceae(100);g__Metschnikowia(100);s__Metschnikowia_pulcherrima(100); | Metschnikowia | 13 | Saprotroph | \|Undefined Saprotroph\| | Yeast | NULL | Probable | As nectar-tap_saprotroph (Põlme et al. 2020); Decay sugar-rich substrates (Põlme et al. 2020) | Põlme S, et al. 2020. Fungal Diversity 105:1-16 ((https://doi.org/10.1007/s13225-020-00466-2)) |
| Otu14671 | 1 | k__Fungi(100);p__Ascomycota(100);c__Saccharomycetes(100);o__Saccharomycetales(100);f__Metschnikowiaceae(100);g__Metschnikowia(100);s__Metschnikowia_pulcherrima(100); | Metschnikowia | 13 | Saprotroph | \|Undefined Saprotroph\| | Yeast | NULL | Probable | As nectar-tap_saprotroph (Põlme et al. 2020); Decay sugar-rich substrates (Põlme et al. 2020) | Põlme S, et al. 2020. Fungal Diversity 105:1-16 ((https://doi.org/10.1007/s13225-020-00466-2)) |
| Otu14672 | 1 | k__Fungi(100);p__Ascomycota(100);c__Saccharomycetes(100);o__Saccharomycetales(100);f__Metschnikowiaceae(100);g__Metschnikowia(100);s__Metschnikowia_pulcherrima(100); | Metschnikowia | 13 | Saprotroph | \|Undefined Saprotroph\| | Yeast | NULL | Probable | As nectar-tap_saprotroph (Põlme et al. 2020); Decay sugar-rich substrates (Põlme et al. 2020) | Põlme S, et al. 2020. Fungal Diversity 105:1-16 ((https://doi.org/10.1007/s13225-020-00466-2)) |
| Otu14673 | 1 | k__Fungi(100);p__Ascomycota(100);c__Saccharomycetes(100);o__Saccharomycetales(100);f__Metschnikowiaceae(100);g__Metschnikowia(100);g__Metschnikowia_unclassified(100); | Metschnikowia | 13 | Saprotroph | \|Undefined Saprotroph\| | Yeast | NULL | Probable | As nectar-tap_saprotroph (Põlme et al. 2020); Decay sugar-rich substrates (Põlme et al. 2020) | Põlme S, et al. 2020. Fungal Diversity 105:1-16 ((https://doi.org/10.1007/s13225-020-00466-2)) |
| Otu14674 | 1 | k__Fungi(100);p__Ascomycota(100);c__Saccharomycetes(100);o__Saccharomycetales(100);f__Metschnikowiaceae(100);g__Metschnikowia(100);s__Metschnikowia_pulcherrima(100); | Metschnikowia | 13 | Saprotroph | \|Undefined Saprotroph\| | Yeast | NULL | Probable | As nectar-tap_saprotroph (Põlme et al. 2020); Decay sugar-rich substrates (Põlme et al. 2020) | Põlme S, et al. 2020. Fungal Diversity 105:1-16 ((https://doi.org/10.1007/s13225-020-00466-2)) |
| Otu14675 | 1 | k__Fungi(100);p__Ascomycota(100);c__Saccharomycetes(100);o__Saccharomycetales(100);f__Metschnikowiaceae(100);g__Metschnikowia(100);s__Metschnikowia_pulcherrima(100); | Metschnikowia | 13 | Saprotroph | \|Undefined Saprotroph\| | Yeast | NULL | Probable | As nectar-tap_saprotroph (Põlme et al. 2020); Decay sugar-rich substrates (Põlme et al. 2020) | Põlme S, et al. 2020. Fungal Diversity 105:1-16 ((https://doi.org/10.1007/s13225-020-00466-2)) |
| Otu14676 | 1 | k__Fungi(100);p__Ascomycota(100);c__Saccharomycetes(100);o__Saccharomycetales(100);f__Metschnikowiaceae(100);g__Metschnikowia(100);g__Metschnikowia_unclassified(100); | Metschnikowia | 13 | Saprotroph | \|Undefined Saprotroph\| | Yeast | NULL | Probable | As nectar-tap_saprotroph (Põlme et al. 2020); Decay sugar-rich substrates (Põlme et al. 2020) | Põlme S, et al. 2020. Fungal Diversity 105:1-16 ((https://doi.org/10.1007/s13225-020-00466-2)) |
| Otu14677 | 1 | k__Fungi(100);p__Ascomycota(100);c__Saccharomycetes(100);o__Saccharomycetales(100);f__Metschnikowiaceae(100);g__Metschnikowia(100);g__Metschnikowia_unclassified(100); | Metschnikowia | 13 | Saprotroph | \|Undefined Saprotroph\| | Yeast | NULL | Probable | As nectar-tap_saprotroph (Põlme et al. 2020); Decay sugar-rich substrates (Põlme et al. 2020) | Põlme S, et al. 2020. Fungal Diversity 105:1-16 ((https://doi.org/10.1007/s13225-020-00466-2)) |
| Otu14678 | 1 | k__Fungi(100);p__Ascomycota(100);c__Saccharomycetes(100);o__Saccharomycetales(100);f__Metschnikowiaceae(100);g__Metschnikowia(100);s__Metschnikowia_pulcherrima(100); | Metschnikowia | 13 | Saprotroph | \|Undefined Saprotroph\| | Yeast | NULL | Probable | As nectar-tap_saprotroph (Põlme et al. 2020); Decay sugar-rich substrates (Põlme et al. 2020) | Põlme S, et al. 2020. Fungal Diversity 105:1-16 ((https://doi.org/10.1007/s13225-020-00466-2)) |
| Otu14679 | 1 | k__Fungi(100);p__Ascomycota(100);c__Saccharomycetes(100);o__Saccharomycetales(100);f__Metschnikowiaceae(100);g__Metschnikowia(100);s__Metschnikowia_pulcherrima(100); | Metschnikowia | 13 | Saprotroph | \|Undefined Saprotroph\| | Yeast | NULL | Probable | As nectar-tap_saprotroph (Põlme et al. 2020); Decay sugar-rich substrates (Põlme et al. 2020) | Põlme S, et al. 2020. Fungal Diversity 105:1-16 ((https://doi.org/10.1007/s13225-020-00466-2)) |
| Otu14682 | 1 | k__Fungi(100);p__Ascomycota(100);c__Saccharomycetes(100);o__Saccharomycetales(100);f__Saccharomycetales_family_Incertae_sedis(100);g__Candida(100);g__Candida_unclassified(100); | Candida | 13 | Pathotroph-Saprotroph-Symbiotroph | Animal Pathogen-Endophyte-Endosymbiont-Epiphyte-\|Undefined Saprotroph\| | Dimorphic Yeast-Yeast | NULL | Probable | As nectar-tap_saprotroph (Põlme et al. 2020); Decay sugar-rich substrates (Põlme et al. 2020); This asexual genus is polyphyletic and contain species that function very differently depend on what they are. Caution should be taken when interpreting guilds for Candida species. | Manolakaki D, et al. 2010. Virulence. 1: 367-375 ((https://doi.org/10.4161/viru.1.5.12796)); Mekbib SB, et al. 2011. Tropical Plant Pathology 36:233-240 ((https://doi.org/10.1590/S1982-56762011000400004)); Põlme S, et al. 2020. Fungal Diversity 105:1-16 ((https://doi.org/10.1007/s13225-020-00466-2)) |
| Otu14684 | 1 | k__Fungi(100);p__Ascomycota(100);c__Saccharomycetes(100);o__Saccharomycetales(100);f__Metschnikowiaceae(100);g__Metschnikowia(100);s__Metschnikowia_pulcherrima(100); | Metschnikowia | 13 | Saprotroph | \|Undefined Saprotroph\| | Yeast | NULL | Probable | As nectar-tap_saprotroph (Põlme et al. 2020); Decay sugar-rich substrates (Põlme et al. 2020) | Põlme S, et al. 2020. Fungal Diversity 105:1-16 ((https://doi.org/10.1007/s13225-020-00466-2)) |
| Otu14686 | 1 | k__Fungi(100);p__Ascomycota(100);c__Saccharomycetes(100);o__Saccharomycetales(100);o__Saccharomycetales_unclassified(100);o__Saccharomycetales_unclassified(100);o__Saccharomycetales_unclassified(100); | Saccharomycetales | 7 | Saprotroph | Undefined Saprotroph | Yeast | NULL | Possible | NULL | Sterkenburg E, et al. 2015. New Phytologist 207:1145-1158 ((https://doi.org/10.1111/nph.13426)) |
| Otu14687 | 1 | k__Fungi(100);p__Ascomycota(100);c__Saccharomycetes(100);o__Saccharomycetales(100);f__Metschnikowiaceae(100);g__Metschnikowia(100);s__Metschnikowia_pulcherrima(100); | Metschnikowia | 13 | Saprotroph | \|Undefined Saprotroph\| | Yeast | NULL | Probable | As nectar-tap_saprotroph (Põlme et al. 2020); Decay sugar-rich substrates (Põlme et al. 2020) | Põlme S, et al. 2020. Fungal Diversity 105:1-16 ((https://doi.org/10.1007/s13225-020-00466-2)) |
| Otu14688 | 1 | k__Fungi(100);p__Ascomycota(100);c__Saccharomycetes(100);o__Saccharomycetales(100);f__Metschnikowiaceae(100);g__Metschnikowia(100);s__Metschnikowia_pulcherrima(100); | Metschnikowia | 13 | Saprotroph | \|Undefined Saprotroph\| | Yeast | NULL | Probable | As nectar-tap_saprotroph (Põlme et al. 2020); Decay sugar-rich substrates (Põlme et al. 2020) | Põlme S, et al. 2020. Fungal Diversity 105:1-16 ((https://doi.org/10.1007/s13225-020-00466-2)) |
| Otu14689 | 1 | k__Fungi(100);p__Ascomycota(100);c__Saccharomycetes(100);o__Saccharomycetales(100);f__Metschnikowiaceae(100);g__Metschnikowia(100);s__Metschnikowia_pulcherrima(100); | Metschnikowia | 13 | Saprotroph | \|Undefined Saprotroph\| | Yeast | NULL | Probable | As nectar-tap_saprotroph (Põlme et al. 2020); Decay sugar-rich substrates (Põlme et al. 2020) | Põlme S, et al. 2020. Fungal Diversity 105:1-16 ((https://doi.org/10.1007/s13225-020-00466-2)) |
| Otu14691 | 1 | k__Fungi(100);p__Ascomycota(100);c__Saccharomycetes(100);o__Saccharomycetales(100);f__Metschnikowiaceae(100);g__Metschnikowia(100);g__Metschnikowia_unclassified(100); | Metschnikowia | 13 | Saprotroph | \|Undefined Saprotroph\| | Yeast | NULL | Probable | As nectar-tap_saprotroph (Põlme et al. 2020); Decay sugar-rich substrates (Põlme et al. 2020) | Põlme S, et al. 2020. Fungal Diversity 105:1-16 ((https://doi.org/10.1007/s13225-020-00466-2)) |
| Otu14693 | 1 | k__Fungi(100);p__Ascomycota(100);c__Saccharomycetes(100);o__Saccharomycetales(100);f__Metschnikowiaceae(100);g__Metschnikowia(100);g__Metschnikowia_unclassified(100); | Metschnikowia | 13 | Saprotroph | \|Undefined Saprotroph\| | Yeast | NULL | Probable | As nectar-tap_saprotroph (Põlme et al. 2020); Decay sugar-rich substrates (Põlme et al. 2020) | Põlme S, et al. 2020. Fungal Diversity 105:1-16 ((https://doi.org/10.1007/s13225-020-00466-2)) |
| Otu14694 | 1 | k__Fungi(100);p__Ascomycota(100);c__Saccharomycetes(100);o__Saccharomycetales(100);f__Metschnikowiaceae(100);g__Metschnikowia(100);s__Metschnikowia_pulcherrima(100); | Metschnikowia | 13 | Saprotroph | \|Undefined Saprotroph\| | Yeast | NULL | Probable | As nectar-tap_saprotroph (Põlme et al. 2020); Decay sugar-rich substrates (Põlme et al. 2020) | Põlme S, et al. 2020. Fungal Diversity 105:1-16 ((https://doi.org/10.1007/s13225-020-00466-2)) |
| Otu14695 | 1 | k__Fungi(100);p__Ascomycota(100);c__Saccharomycetes(100);o__Saccharomycetales(100);f__Metschnikowiaceae(100);g__Metschnikowia(100);s__Metschnikowia_pulcherrima(100); | Metschnikowia | 13 | Saprotroph | \|Undefined Saprotroph\| | Yeast | NULL | Probable | As nectar-tap_saprotroph (Põlme et al. 2020); Decay sugar-rich substrates (Põlme et al. 2020) | Põlme S, et al. 2020. Fungal Diversity 105:1-16 ((https://doi.org/10.1007/s13225-020-00466-2)) |
| Otu14698 | 1 | k__Fungi(100);p__Ascomycota(100);c__Saccharomycetes(100);o__Saccharomycetales(100);f__Metschnikowiaceae(100);g__Metschnikowia(100);g__Metschnikowia_unclassified(100); | Metschnikowia | 13 | Saprotroph | \|Undefined Saprotroph\| | Yeast | NULL | Probable | As nectar-tap_saprotroph (Põlme et al. 2020); Decay sugar-rich substrates (Põlme et al. 2020) | Põlme S, et al. 2020. Fungal Diversity 105:1-16 ((https://doi.org/10.1007/s13225-020-00466-2)) |
| Otu14701 | 1 | k__Fungi(100);p__Ascomycota(100);c__Saccharomycetes(100);o__Saccharomycetales(100);f__Metschnikowiaceae(100);g__Metschnikowia(100);s__Metschnikowia_pulcherrima(100); | Metschnikowia | 13 | Saprotroph | \|Undefined Saprotroph\| | Yeast | NULL | Probable | As nectar-tap_saprotroph (Põlme et al. 2020); Decay sugar-rich substrates (Põlme et al. 2020) | Põlme S, et al. 2020. Fungal Diversity 105:1-16 ((https://doi.org/10.1007/s13225-020-00466-2)) |
| Otu14702 | 1 | k__Fungi(100);p__Ascomycota(100);c__Saccharomycetes(100);o__Saccharomycetales(100);f__Metschnikowiaceae(100);g__Metschnikowia(100);s__Metschnikowia_pulcherrima(100); | Metschnikowia | 13 | Saprotroph | \|Undefined Saprotroph\| | Yeast | NULL | Probable | As nectar-tap_saprotroph (Põlme et al. 2020); Decay sugar-rich substrates (Põlme et al. 2020) | Põlme S, et al. 2020. Fungal Diversity 105:1-16 ((https://doi.org/10.1007/s13225-020-00466-2)) |
| Otu14704 | 1 | k__Fungi(100);p__Ascomycota(100);c__Saccharomycetes(100);o__Saccharomycetales(100);f__Metschnikowiaceae(100);g__Metschnikowia(100);s__Metschnikowia_pulcherrima(100); | Metschnikowia | 13 | Saprotroph | \|Undefined Saprotroph\| | Yeast | NULL | Probable | As nectar-tap_saprotroph (Põlme et al. 2020); Decay sugar-rich substrates (Põlme et al. 2020) | Põlme S, et al. 2020. Fungal Diversity 105:1-16 ((https://doi.org/10.1007/s13225-020-00466-2)) |
| Otu14707 | 1 | k__Fungi(100);p__Ascomycota(100);c__Saccharomycetes(100);o__Saccharomycetales(100);f__Metschnikowiaceae(100);g__Metschnikowia(100);s__Metschnikowia_pulcherrima(100); | Metschnikowia | 13 | Saprotroph | \|Undefined Saprotroph\| | Yeast | NULL | Probable | As nectar-tap_saprotroph (Põlme et al. 2020); Decay sugar-rich substrates (Põlme et al. 2020) | Põlme S, et al. 2020. Fungal Diversity 105:1-16 ((https://doi.org/10.1007/s13225-020-00466-2)) |
| Otu14708 | 1 | k__Fungi(100);p__Ascomycota(100);c__Saccharomycetes(100);o__Saccharomycetales(100);f__Saccharomycetales_family_Incertae_sedis(100);f__Saccharomycetales_family_Incertae_sedis_unclassified(100);f__Saccharomycetales_family_Incertae_sedis_unclassified(100); | Saccharomycetales | 7 | Saprotroph | Undefined Saprotroph | Yeast | NULL | Possible | NULL | Sterkenburg E, et al. 2015. New Phytologist 207:1145-1158 ((https://doi.org/10.1111/nph.13426)) |
| Otu14713 | 1 | k__Fungi(100);p__Ascomycota(100);c__Sordariomycetes(100);o__Hypocreales(100);f__Nectriaceae(100);g__Gibberella(100);s__Gibberella_intricans(100); | Gibberella | 13 | Pathotroph-Saprotroph-Symbiotroph | Endophyte-\|Plant Pathogen\|-Plant Saprotroph-Undefined Saprotroph | Microfungus | Soft Rot | Probable | As foliar_endophyte (Põlme et al. 2020); As litter_saprotroph (Põlme et al. 2020); Causing damping-off disease (Põlme et al. 2020); Decay leaf, fruit, seed, roots, soil, animal material (Põlme et al. 2020) | Tedersoo L, et al. 2014. Science 346:e1256688 ((https://doi.org/10.1126/science.1256688)); Põlme S, et al. 2020. Fungal Diversity 105:1-16 ((https://doi.org/10.1007/s13225-020-00466-2)) |
| Otu14714 | 1 | k__Fungi(100);p__Ascomycota(100);c__Saccharomycetes(100);o__Saccharomycetales(100);f__Metschnikowiaceae(100);g__Metschnikowia(100);s__Metschnikowia_pulcherrima(100); | Metschnikowia | 13 | Saprotroph | \|Undefined Saprotroph\| | Yeast | NULL | Probable | As nectar-tap_saprotroph (Põlme et al. 2020); Decay sugar-rich substrates (Põlme et al. 2020) | Põlme S, et al. 2020. Fungal Diversity 105:1-16 ((https://doi.org/10.1007/s13225-020-00466-2)) |
| Otu14715 | 1 | k__Fungi(100);p__Ascomycota(100);c__Saccharomycetes(100);o__Saccharomycetales(100);f__Metschnikowiaceae(100);g__Metschnikowia(100);s__Metschnikowia_pulcherrima(100); | Metschnikowia | 13 | Saprotroph | \|Undefined Saprotroph\| | Yeast | NULL | Probable | As nectar-tap_saprotroph (Põlme et al. 2020); Decay sugar-rich substrates (Põlme et al. 2020) | Põlme S, et al. 2020. Fungal Diversity 105:1-16 ((https://doi.org/10.1007/s13225-020-00466-2)) |
| Otu14716 | 1 | k__Fungi(100);p__Ascomycota(100);c__Saccharomycetes(100);o__Saccharomycetales(100);f__Metschnikowiaceae(100);g__Metschnikowia(100);g__Metschnikowia_unclassified(100); | Metschnikowia | 13 | Saprotroph | \|Undefined Saprotroph\| | Yeast | NULL | Probable | As nectar-tap_saprotroph (Põlme et al. 2020); Decay sugar-rich substrates (Põlme et al. 2020) | Põlme S, et al. 2020. Fungal Diversity 105:1-16 ((https://doi.org/10.1007/s13225-020-00466-2)) |
| Otu14717 | 1 | k__Fungi(100);p__Ascomycota(100);c__Saccharomycetes(100);o__Saccharomycetales(100);f__Metschnikowiaceae(100);g__Metschnikowia(100);s__Metschnikowia_pulcherrima(100); | Metschnikowia | 13 | Saprotroph | \|Undefined Saprotroph\| | Yeast | NULL | Probable | As nectar-tap_saprotroph (Põlme et al. 2020); Decay sugar-rich substrates (Põlme et al. 2020) | Põlme S, et al. 2020. Fungal Diversity 105:1-16 ((https://doi.org/10.1007/s13225-020-00466-2)) |
| Otu14719 | 1 | k__Fungi(100);p__Ascomycota(100);c__Saccharomycetes(100);o__Saccharomycetales(100);f__Metschnikowiaceae(100);g__Metschnikowia(100);s__Metschnikowia_pulcherrima(100); | Metschnikowia | 13 | Saprotroph | \|Undefined Saprotroph\| | Yeast | NULL | Probable | As nectar-tap_saprotroph (Põlme et al. 2020); Decay sugar-rich substrates (Põlme et al. 2020) | Põlme S, et al. 2020. Fungal Diversity 105:1-16 ((https://doi.org/10.1007/s13225-020-00466-2)) |
| Otu14720 | 1 | k__Fungi(100);p__Ascomycota(100);c__Saccharomycetes(100);o__Saccharomycetales(100);f__Metschnikowiaceae(100);g__Metschnikowia(100);g__Metschnikowia_unclassified(100); | Metschnikowia | 13 | Saprotroph | \|Undefined Saprotroph\| | Yeast | NULL | Probable | As nectar-tap_saprotroph (Põlme et al. 2020); Decay sugar-rich substrates (Põlme et al. 2020) | Põlme S, et al. 2020. Fungal Diversity 105:1-16 ((https://doi.org/10.1007/s13225-020-00466-2)) |
| Otu14721 | 1 | k__Fungi(100);p__Ascomycota(100);c__Saccharomycetes(100);o__Saccharomycetales(100);f__Metschnikowiaceae(100);g__Metschnikowia(100);s__Metschnikowia_pulcherrima(100); | Metschnikowia | 13 | Saprotroph | \|Undefined Saprotroph\| | Yeast | NULL | Probable | As nectar-tap_saprotroph (Põlme et al. 2020); Decay sugar-rich substrates (Põlme et al. 2020) | Põlme S, et al. 2020. Fungal Diversity 105:1-16 ((https://doi.org/10.1007/s13225-020-00466-2)) |
| Otu14722 | 1 | k__Fungi(100);p__Ascomycota(100);c__Saccharomycetes(100);o__Saccharomycetales(100);f__Metschnikowiaceae(100);g__Metschnikowia(100);s__Metschnikowia_pulcherrima(100); | Metschnikowia | 13 | Saprotroph | \|Undefined Saprotroph\| | Yeast | NULL | Probable | As nectar-tap_saprotroph (Põlme et al. 2020); Decay sugar-rich substrates (Põlme et al. 2020) | Põlme S, et al. 2020. Fungal Diversity 105:1-16 ((https://doi.org/10.1007/s13225-020-00466-2)) |
| Otu14723 | 1 | k__Fungi(100);p__Ascomycota(100);c__Saccharomycetes(100);o__Saccharomycetales(100);f__Metschnikowiaceae(100);g__Metschnikowia(100);s__Metschnikowia_pulcherrima(100); | Metschnikowia | 13 | Saprotroph | \|Undefined Saprotroph\| | Yeast | NULL | Probable | As nectar-tap_saprotroph (Põlme et al. 2020); Decay sugar-rich substrates (Põlme et al. 2020) | Põlme S, et al. 2020. Fungal Diversity 105:1-16 ((https://doi.org/10.1007/s13225-020-00466-2)) |
| Otu14725 | 1 | k__Fungi(100);p__Ascomycota(100);c__Saccharomycetes(100);o__Saccharomycetales(100);f__Metschnikowiaceae(100);g__Metschnikowia(100);g__Metschnikowia_unclassified(100); | Metschnikowia | 13 | Saprotroph | \|Undefined Saprotroph\| | Yeast | NULL | Probable | As nectar-tap_saprotroph (Põlme et al. 2020); Decay sugar-rich substrates (Põlme et al. 2020) | Põlme S, et al. 2020. Fungal Diversity 105:1-16 ((https://doi.org/10.1007/s13225-020-00466-2)) |
| Otu14726 | 1 | k__Fungi(100);p__Ascomycota(100);c__Saccharomycetes(100);o__Saccharomycetales(100);f__Metschnikowiaceae(100);g__Metschnikowia(100);s__Metschnikowia_pulcherrima(100); | Metschnikowia | 13 | Saprotroph | \|Undefined Saprotroph\| | Yeast | NULL | Probable | As nectar-tap_saprotroph (Põlme et al. 2020); Decay sugar-rich substrates (Põlme et al. 2020) | Põlme S, et al. 2020. Fungal Diversity 105:1-16 ((https://doi.org/10.1007/s13225-020-00466-2)) |
| Otu14727 | 1 | k__Fungi(100);p__Ascomycota(100);c__Saccharomycetes(100);o__Saccharomycetales(100);f__Metschnikowiaceae(100);g__Metschnikowia(100);s__Metschnikowia_pulcherrima(100); | Metschnikowia | 13 | Saprotroph | \|Undefined Saprotroph\| | Yeast | NULL | Probable | As nectar-tap_saprotroph (Põlme et al. 2020); Decay sugar-rich substrates (Põlme et al. 2020) | Põlme S, et al. 2020. Fungal Diversity 105:1-16 ((https://doi.org/10.1007/s13225-020-00466-2)) |
| Otu14728 | 1 | k__Fungi(100);p__Ascomycota(100);c__Saccharomycetes(100);o__Saccharomycetales(100);f__Metschnikowiaceae(100);g__Metschnikowia(100);s__Metschnikowia_pulcherrima(100); | Metschnikowia | 13 | Saprotroph | \|Undefined Saprotroph\| | Yeast | NULL | Probable | As nectar-tap_saprotroph (Põlme et al. 2020); Decay sugar-rich substrates (Põlme et al. 2020) | Põlme S, et al. 2020. Fungal Diversity 105:1-16 ((https://doi.org/10.1007/s13225-020-00466-2)) |
| Otu14734 | 1 | k__Fungi(100);p__Ascomycota(100);c__Saccharomycetes(100);o__Saccharomycetales(100);f__Metschnikowiaceae(100);g__Metschnikowia(100);g__Metschnikowia_unclassified(100); | Metschnikowia | 13 | Saprotroph | \|Undefined Saprotroph\| | Yeast | NULL | Probable | As nectar-tap_saprotroph (Põlme et al. 2020); Decay sugar-rich substrates (Põlme et al. 2020) | Põlme S, et al. 2020. Fungal Diversity 105:1-16 ((https://doi.org/10.1007/s13225-020-00466-2)) |
| Otu14735 | 1 | k__Fungi(100);p__Ascomycota(100);c__Saccharomycetes(100);o__Saccharomycetales(100);f__Metschnikowiaceae(100);g__Metschnikowia(100);s__Metschnikowia_pulcherrima(100); | Metschnikowia | 13 | Saprotroph | \|Undefined Saprotroph\| | Yeast | NULL | Probable | As nectar-tap_saprotroph (Põlme et al. 2020); Decay sugar-rich substrates (Põlme et al. 2020) | Põlme S, et al. 2020. Fungal Diversity 105:1-16 ((https://doi.org/10.1007/s13225-020-00466-2)) |
| Otu14737 | 1 | k__Fungi(100);p__Ascomycota(100);c__Saccharomycetes(100);o__Saccharomycetales(100);f__Metschnikowiaceae(100);g__Metschnikowia(100);s__Metschnikowia_pulcherrima(100); | Metschnikowia | 13 | Saprotroph | \|Undefined Saprotroph\| | Yeast | NULL | Probable | As nectar-tap_saprotroph (Põlme et al. 2020); Decay sugar-rich substrates (Põlme et al. 2020) | Põlme S, et al. 2020. Fungal Diversity 105:1-16 ((https://doi.org/10.1007/s13225-020-00466-2)) |
| Otu14740 | 1 | k__Fungi(100);p__Ascomycota(100);c__Saccharomycetes(100);o__Saccharomycetales(100);f__Metschnikowiaceae(100);g__Metschnikowia(100);g__Metschnikowia_unclassified(100); | Metschnikowia | 13 | Saprotroph | \|Undefined Saprotroph\| | Yeast | NULL | Probable | As nectar-tap_saprotroph (Põlme et al. 2020); Decay sugar-rich substrates (Põlme et al. 2020) | Põlme S, et al. 2020. Fungal Diversity 105:1-16 ((https://doi.org/10.1007/s13225-020-00466-2)) |
| Otu14741 | 1 | k__Fungi(100);p__Ascomycota(100);c__Saccharomycetes(100);o__Saccharomycetales(100);f__Metschnikowiaceae(100);g__Metschnikowia(100);s__Metschnikowia_pulcherrima(100); | Metschnikowia | 13 | Saprotroph | \|Undefined Saprotroph\| | Yeast | NULL | Probable | As nectar-tap_saprotroph (Põlme et al. 2020); Decay sugar-rich substrates (Põlme et al. 2020) | Põlme S, et al. 2020. Fungal Diversity 105:1-16 ((https://doi.org/10.1007/s13225-020-00466-2)) |
| Otu14743 | 1 | k__Fungi(100);p__Ascomycota(100);c__Sordariomycetes(100);o__Hypocreales(100);f__Nectriaceae(100);f__Nectriaceae_unclassified(100);f__Nectriaceae_unclassified(100); | Nectriaceae | 9 | Pathotroph-Saprotroph-Symbiotroph | Animal Pathogen-Endophyte-Fungal Parasite-Lichen Parasite-Plant Pathogen-Wood Saprotroph | Microfungus | NULL | Possible | Fusarium species are amongst the economically significant plant pathogens (Cannon & Kirk 2007) | Cannon PF, Kirk PM. 2007. Fungal Families of the World. CAB International, Cambridge (ISBN: 978-0851998275); Hirooka Y, et al. 2012. Studies in Mycology 711-210 ((https://doi.org/10.3114/sim0001)); Tedersoo L, et al. 2014. Science 346:e1256688 ((https://doi.org/10.1126/science.1256688)); The American Phytopathological Society Common Names of Plant Diseases ((http://www.apsnet.org/publications/commonnames/Pages/default.aspx)); https://www.cdc.gov ((https://www.cdc.gov)) |
| Otu14745 | 1 | k__Fungi(100);p__Ascomycota(100);c__Saccharomycetes(100);o__Saccharomycetales(100);f__Metschnikowiaceae(100);g__Metschnikowia(100);g__Metschnikowia_unclassified(100); | Metschnikowia | 13 | Saprotroph | \|Undefined Saprotroph\| | Yeast | NULL | Probable | As nectar-tap_saprotroph (Põlme et al. 2020); Decay sugar-rich substrates (Põlme et al. 2020) | Põlme S, et al. 2020. Fungal Diversity 105:1-16 ((https://doi.org/10.1007/s13225-020-00466-2)) |
| Otu14746 | 1 | k__Fungi(100);p__Ascomycota(100);c__Saccharomycetes(100);o__Saccharomycetales(100);f__Metschnikowiaceae(100);g__Metschnikowia(100);s__Metschnikowia_pulcherrima(100); | Metschnikowia | 13 | Saprotroph | \|Undefined Saprotroph\| | Yeast | NULL | Probable | As nectar-tap_saprotroph (Põlme et al. 2020); Decay sugar-rich substrates (Põlme et al. 2020) | Põlme S, et al. 2020. Fungal Diversity 105:1-16 ((https://doi.org/10.1007/s13225-020-00466-2)) |
| Otu14747 | 1 | k__Fungi(100);p__Ascomycota(100);c__Saccharomycetes(100);o__Saccharomycetales(100);f__Saccharomycetales_family_Incertae_sedis(100);g__Candida(100);g__Candida_unclassified(100); | Candida | 13 | Pathotroph-Saprotroph-Symbiotroph | Animal Pathogen-Endophyte-Endosymbiont-Epiphyte-\|Undefined Saprotroph\| | Dimorphic Yeast-Yeast | NULL | Probable | As nectar-tap_saprotroph (Põlme et al. 2020); Decay sugar-rich substrates (Põlme et al. 2020); This asexual genus is polyphyletic and contain species that function very differently depend on what they are. Caution should be taken when interpreting guilds for Candida species. | Manolakaki D, et al. 2010. Virulence. 1: 367-375 ((https://doi.org/10.4161/viru.1.5.12796)); Mekbib SB, et al. 2011. Tropical Plant Pathology 36:233-240 ((https://doi.org/10.1590/S1982-56762011000400004)); Põlme S, et al. 2020. Fungal Diversity 105:1-16 ((https://doi.org/10.1007/s13225-020-00466-2)) |
| Otu14748 | 1 | k__Fungi(100);p__Ascomycota(100);c__Saccharomycetes(100);o__Saccharomycetales(100);f__Metschnikowiaceae(100);g__Metschnikowia(100);s__Metschnikowia_pulcherrima(100); | Metschnikowia | 13 | Saprotroph | \|Undefined Saprotroph\| | Yeast | NULL | Probable | As nectar-tap_saprotroph (Põlme et al. 2020); Decay sugar-rich substrates (Põlme et al. 2020) | Põlme S, et al. 2020. Fungal Diversity 105:1-16 ((https://doi.org/10.1007/s13225-020-00466-2)) |
| Otu14752 | 1 | k__Fungi(100);p__Ascomycota(100);c__Saccharomycetes(100);o__Saccharomycetales(100);f__Metschnikowiaceae(100);g__Metschnikowia(100);g__Metschnikowia_unclassified(100); | Metschnikowia | 13 | Saprotroph | \|Undefined Saprotroph\| | Yeast | NULL | Probable | As nectar-tap_saprotroph (Põlme et al. 2020); Decay sugar-rich substrates (Põlme et al. 2020) | Põlme S, et al. 2020. Fungal Diversity 105:1-16 ((https://doi.org/10.1007/s13225-020-00466-2)) |
| Otu14753 | 1 | k__Fungi(100);p__Ascomycota(100);c__Saccharomycetes(100);o__Saccharomycetales(100);f__Metschnikowiaceae(100);g__Metschnikowia(100);s__Metschnikowia_pulcherrima(100); | Metschnikowia | 13 | Saprotroph | \|Undefined Saprotroph\| | Yeast | NULL | Probable | As nectar-tap_saprotroph (Põlme et al. 2020); Decay sugar-rich substrates (Põlme et al. 2020) | Põlme S, et al. 2020. Fungal Diversity 105:1-16 ((https://doi.org/10.1007/s13225-020-00466-2)) |
| Otu14755 | 1 | k__Fungi(100);p__Ascomycota(100);c__Saccharomycetes(100);o__Saccharomycetales(100);f__Metschnikowiaceae(100);g__Metschnikowia(100);g__Metschnikowia_unclassified(100); | Metschnikowia | 13 | Saprotroph | \|Undefined Saprotroph\| | Yeast | NULL | Probable | As nectar-tap_saprotroph (Põlme et al. 2020); Decay sugar-rich substrates (Põlme et al. 2020) | Põlme S, et al. 2020. Fungal Diversity 105:1-16 ((https://doi.org/10.1007/s13225-020-00466-2)) |
| Otu14756 | 1 | k__Fungi(100);p__Ascomycota(100);c__Saccharomycetes(100);o__Saccharomycetales(100);f__Metschnikowiaceae(100);g__Metschnikowia(100);g__Metschnikowia_unclassified(100); | Metschnikowia | 13 | Saprotroph | \|Undefined Saprotroph\| | Yeast | NULL | Probable | As nectar-tap_saprotroph (Põlme et al. 2020); Decay sugar-rich substrates (Põlme et al. 2020) | Põlme S, et al. 2020. Fungal Diversity 105:1-16 ((https://doi.org/10.1007/s13225-020-00466-2)) |
| Otu14759 | 1 | k__Fungi(100);p__Ascomycota(100);c__Saccharomycetes(100);o__Saccharomycetales(100);f__Saccharomycetales_family_Incertae_sedis(100);g__Candida(100);g__Candida_unclassified(100); | Candida | 13 | Pathotroph-Saprotroph-Symbiotroph | Animal Pathogen-Endophyte-Endosymbiont-Epiphyte-\|Undefined Saprotroph\| | Dimorphic Yeast-Yeast | NULL | Probable | As nectar-tap_saprotroph (Põlme et al. 2020); Decay sugar-rich substrates (Põlme et al. 2020); This asexual genus is polyphyletic and contain species that function very differently depend on what they are. Caution should be taken when interpreting guilds for Candida species. | Manolakaki D, et al. 2010. Virulence. 1: 367-375 ((https://doi.org/10.4161/viru.1.5.12796)); Mekbib SB, et al. 2011. Tropical Plant Pathology 36:233-240 ((https://doi.org/10.1590/S1982-56762011000400004)); Põlme S, et al. 2020. Fungal Diversity 105:1-16 ((https://doi.org/10.1007/s13225-020-00466-2)) |
| Otu14764 | 1 | k__Fungi(100);p__Ascomycota(100);c__Saccharomycetes(100);o__Saccharomycetales(100);f__Metschnikowiaceae(100);g__Metschnikowia(100);s__Metschnikowia_pulcherrima(100); | Metschnikowia | 13 | Saprotroph | \|Undefined Saprotroph\| | Yeast | NULL | Probable | As nectar-tap_saprotroph (Põlme et al. 2020); Decay sugar-rich substrates (Põlme et al. 2020) | Põlme S, et al. 2020. Fungal Diversity 105:1-16 ((https://doi.org/10.1007/s13225-020-00466-2)) |
| Otu14765 | 1 | k__Fungi(100);p__Ascomycota(100);c__Saccharomycetes(100);o__Saccharomycetales(100);f__Metschnikowiaceae(100);g__Metschnikowia(100);g__Metschnikowia_unclassified(100); | Metschnikowia | 13 | Saprotroph | \|Undefined Saprotroph\| | Yeast | NULL | Probable | As nectar-tap_saprotroph (Põlme et al. 2020); Decay sugar-rich substrates (Põlme et al. 2020) | Põlme S, et al. 2020. Fungal Diversity 105:1-16 ((https://doi.org/10.1007/s13225-020-00466-2)) |
| Otu14768 | 1 | k__Fungi(100);p__Ascomycota(100);c__Saccharomycetes(100);o__Saccharomycetales(100);f__Metschnikowiaceae(100);g__Metschnikowia(100);s__Metschnikowia_pulcherrima(100); | Metschnikowia | 13 | Saprotroph | \|Undefined Saprotroph\| | Yeast | NULL | Probable | As nectar-tap_saprotroph (Põlme et al. 2020); Decay sugar-rich substrates (Põlme et al. 2020) | Põlme S, et al. 2020. Fungal Diversity 105:1-16 ((https://doi.org/10.1007/s13225-020-00466-2)) |
| Otu14769 | 1 | k__Fungi(100);p__Ascomycota(100);c__Saccharomycetes(100);o__Saccharomycetales(100);f__Metschnikowiaceae(100);g__Metschnikowia(100);g__Metschnikowia_unclassified(100); | Metschnikowia | 13 | Saprotroph | \|Undefined Saprotroph\| | Yeast | NULL | Probable | As nectar-tap_saprotroph (Põlme et al. 2020); Decay sugar-rich substrates (Põlme et al. 2020) | Põlme S, et al. 2020. Fungal Diversity 105:1-16 ((https://doi.org/10.1007/s13225-020-00466-2)) |
| Otu14770 | 1 | k__Fungi(100);p__Ascomycota(100);c__Saccharomycetes(100);o__Saccharomycetales(100);f__Metschnikowiaceae(100);g__Metschnikowia(100);s__Metschnikowia_pulcherrima(100); | Metschnikowia | 13 | Saprotroph | \|Undefined Saprotroph\| | Yeast | NULL | Probable | As nectar-tap_saprotroph (Põlme et al. 2020); Decay sugar-rich substrates (Põlme et al. 2020) | Põlme S, et al. 2020. Fungal Diversity 105:1-16 ((https://doi.org/10.1007/s13225-020-00466-2)) |
| Otu14773 | 1 | k__Fungi(100);p__Ascomycota(100);c__Saccharomycetes(100);o__Saccharomycetales(100);f__Metschnikowiaceae(100);g__Metschnikowia(100);s__Metschnikowia_pulcherrima(100); | Metschnikowia | 13 | Saprotroph | \|Undefined Saprotroph\| | Yeast | NULL | Probable | As nectar-tap_saprotroph (Põlme et al. 2020); Decay sugar-rich substrates (Põlme et al. 2020) | Põlme S, et al. 2020. Fungal Diversity 105:1-16 ((https://doi.org/10.1007/s13225-020-00466-2)) |
| Otu14774 | 1 | k__Fungi(100);p__Ascomycota(100);c__Saccharomycetes(100);o__Saccharomycetales(100);f__Metschnikowiaceae(100);g__Metschnikowia(100);g__Metschnikowia_unclassified(100); | Metschnikowia | 13 | Saprotroph | \|Undefined Saprotroph\| | Yeast | NULL | Probable | As nectar-tap_saprotroph (Põlme et al. 2020); Decay sugar-rich substrates (Põlme et al. 2020) | Põlme S, et al. 2020. Fungal Diversity 105:1-16 ((https://doi.org/10.1007/s13225-020-00466-2)) |
| Otu14775 | 1 | k__Fungi(100);p__Ascomycota(100);c__Saccharomycetes(100);o__Saccharomycetales(100);f__Metschnikowiaceae(100);g__Metschnikowia(100);s__Metschnikowia_pulcherrima(100); | Metschnikowia | 13 | Saprotroph | \|Undefined Saprotroph\| | Yeast | NULL | Probable | As nectar-tap_saprotroph (Põlme et al. 2020); Decay sugar-rich substrates (Põlme et al. 2020) | Põlme S, et al. 2020. Fungal Diversity 105:1-16 ((https://doi.org/10.1007/s13225-020-00466-2)) |
| Otu14776 | 1 | k__Fungi(100);p__Ascomycota(100);c__Saccharomycetes(100);o__Saccharomycetales(100);f__Metschnikowiaceae(100);g__Metschnikowia(100);g__Metschnikowia_unclassified(100); | Metschnikowia | 13 | Saprotroph | \|Undefined Saprotroph\| | Yeast | NULL | Probable | As nectar-tap_saprotroph (Põlme et al. 2020); Decay sugar-rich substrates (Põlme et al. 2020) | Põlme S, et al. 2020. Fungal Diversity 105:1-16 ((https://doi.org/10.1007/s13225-020-00466-2)) |
| Otu14777 | 1 | k__Fungi(100);p__Ascomycota(100);c__Saccharomycetes(100);o__Saccharomycetales(100);f__Metschnikowiaceae(100);g__Metschnikowia(100);s__Metschnikowia_pulcherrima(100); | Metschnikowia | 13 | Saprotroph | \|Undefined Saprotroph\| | Yeast | NULL | Probable | As nectar-tap_saprotroph (Põlme et al. 2020); Decay sugar-rich substrates (Põlme et al. 2020) | Põlme S, et al. 2020. Fungal Diversity 105:1-16 ((https://doi.org/10.1007/s13225-020-00466-2)) |
| Otu14780 | 1 | k__Fungi(100);p__Ascomycota(100);c__Saccharomycetes(100);o__Saccharomycetales(100);f__Metschnikowiaceae(100);g__Metschnikowia(100);g__Metschnikowia_unclassified(100); | Metschnikowia | 13 | Saprotroph | \|Undefined Saprotroph\| | Yeast | NULL | Probable | As nectar-tap_saprotroph (Põlme et al. 2020); Decay sugar-rich substrates (Põlme et al. 2020) | Põlme S, et al. 2020. Fungal Diversity 105:1-16 ((https://doi.org/10.1007/s13225-020-00466-2)) |
| Otu14781 | 1 | k__Fungi(100);p__Ascomycota(100);c__Saccharomycetes(100);o__Saccharomycetales(100);f__Metschnikowiaceae(100);g__Metschnikowia(100);s__Metschnikowia_pulcherrima(100); | Metschnikowia | 13 | Saprotroph | \|Undefined Saprotroph\| | Yeast | NULL | Probable | As nectar-tap_saprotroph (Põlme et al. 2020); Decay sugar-rich substrates (Põlme et al. 2020) | Põlme S, et al. 2020. Fungal Diversity 105:1-16 ((https://doi.org/10.1007/s13225-020-00466-2)) |
| Otu14783 | 1 | k__Fungi(100);p__Ascomycota(100);c__Saccharomycetes(100);o__Saccharomycetales(100);f__Metschnikowiaceae(100);g__Metschnikowia(100);s__Metschnikowia_pulcherrima(100); | Metschnikowia | 13 | Saprotroph | \|Undefined Saprotroph\| | Yeast | NULL | Probable | As nectar-tap_saprotroph (Põlme et al. 2020); Decay sugar-rich substrates (Põlme et al. 2020) | Põlme S, et al. 2020. Fungal Diversity 105:1-16 ((https://doi.org/10.1007/s13225-020-00466-2)) |
| Otu14784 | 1 | k__Fungi(100);p__Ascomycota(100);c__Saccharomycetes(100);o__Saccharomycetales(100);f__Metschnikowiaceae(100);g__Metschnikowia(100);s__Metschnikowia_pulcherrima(100); | Metschnikowia | 13 | Saprotroph | \|Undefined Saprotroph\| | Yeast | NULL | Probable | As nectar-tap_saprotroph (Põlme et al. 2020); Decay sugar-rich substrates (Põlme et al. 2020) | Põlme S, et al. 2020. Fungal Diversity 105:1-16 ((https://doi.org/10.1007/s13225-020-00466-2)) |
| Otu14786 | 1 | k__Fungi(100);p__Ascomycota(100);c__Saccharomycetes(100);o__Saccharomycetales(100);f__Metschnikowiaceae(100);g__Metschnikowia(100);g__Metschnikowia_unclassified(100); | Metschnikowia | 13 | Saprotroph | \|Undefined Saprotroph\| | Yeast | NULL | Probable | As nectar-tap_saprotroph (Põlme et al. 2020); Decay sugar-rich substrates (Põlme et al. 2020) | Põlme S, et al. 2020. Fungal Diversity 105:1-16 ((https://doi.org/10.1007/s13225-020-00466-2)) |
| Otu14787 | 1 | k__Fungi(100);p__Ascomycota(100);c__Saccharomycetes(100);o__Saccharomycetales(100);f__Metschnikowiaceae(100);g__Metschnikowia(100);s__Metschnikowia_pulcherrima(100); | Metschnikowia | 13 | Saprotroph | \|Undefined Saprotroph\| | Yeast | NULL | Probable | As nectar-tap_saprotroph (Põlme et al. 2020); Decay sugar-rich substrates (Põlme et al. 2020) | Põlme S, et al. 2020. Fungal Diversity 105:1-16 ((https://doi.org/10.1007/s13225-020-00466-2)) |
| Otu14788 | 1 | k__Fungi(100);p__Ascomycota(100);c__Saccharomycetes(100);o__Saccharomycetales(100);f__Metschnikowiaceae(100);g__Metschnikowia(100);s__Metschnikowia_pulcherrima(100); | Metschnikowia | 13 | Saprotroph | \|Undefined Saprotroph\| | Yeast | NULL | Probable | As nectar-tap_saprotroph (Põlme et al. 2020); Decay sugar-rich substrates (Põlme et al. 2020) | Põlme S, et al. 2020. Fungal Diversity 105:1-16 ((https://doi.org/10.1007/s13225-020-00466-2)) |
| Otu14789 | 1 | k__Fungi(100);p__Ascomycota(100);c__Saccharomycetes(100);o__Saccharomycetales(100);f__Metschnikowiaceae(100);g__Metschnikowia(100);g__Metschnikowia_unclassified(100); | Metschnikowia | 13 | Saprotroph | \|Undefined Saprotroph\| | Yeast | NULL | Probable | As nectar-tap_saprotroph (Põlme et al. 2020); Decay sugar-rich substrates (Põlme et al. 2020) | Põlme S, et al. 2020. Fungal Diversity 105:1-16 ((https://doi.org/10.1007/s13225-020-00466-2)) |
| Otu14790 | 1 | k__Fungi(100);p__Ascomycota(100);c__Saccharomycetes(100);o__Saccharomycetales(100);f__Metschnikowiaceae(100);g__Metschnikowia(100);s__Metschnikowia_pulcherrima(100); | Metschnikowia | 13 | Saprotroph | \|Undefined Saprotroph\| | Yeast | NULL | Probable | As nectar-tap_saprotroph (Põlme et al. 2020); Decay sugar-rich substrates (Põlme et al. 2020) | Põlme S, et al. 2020. Fungal Diversity 105:1-16 ((https://doi.org/10.1007/s13225-020-00466-2)) |
| Otu14794 | 1 | k__Fungi(100);p__Ascomycota(100);c__Saccharomycetes(100);o__Saccharomycetales(100);f__Metschnikowiaceae(100);g__Metschnikowia(100);s__Metschnikowia_pulcherrima(100); | Metschnikowia | 13 | Saprotroph | \|Undefined Saprotroph\| | Yeast | NULL | Probable | As nectar-tap_saprotroph (Põlme et al. 2020); Decay sugar-rich substrates (Põlme et al. 2020) | Põlme S, et al. 2020. Fungal Diversity 105:1-16 ((https://doi.org/10.1007/s13225-020-00466-2)) |
| Otu14795 | 1 | k__Fungi(100);p__Ascomycota(100);c__Saccharomycetes(100);o__Saccharomycetales(100);f__Metschnikowiaceae(100);g__Metschnikowia(100);s__Metschnikowia_pulcherrima(100); | Metschnikowia | 13 | Saprotroph | \|Undefined Saprotroph\| | Yeast | NULL | Probable | As nectar-tap_saprotroph (Põlme et al. 2020); Decay sugar-rich substrates (Põlme et al. 2020) | Põlme S, et al. 2020. Fungal Diversity 105:1-16 ((https://doi.org/10.1007/s13225-020-00466-2)) |
| Otu14799 | 1 | k__Fungi(100);p__Ascomycota(100);c__Saccharomycetes(100);o__Saccharomycetales(100);f__Metschnikowiaceae(100);g__Metschnikowia(100);s__Metschnikowia_pulcherrima(100); | Metschnikowia | 13 | Saprotroph | \|Undefined Saprotroph\| | Yeast | NULL | Probable | As nectar-tap_saprotroph (Põlme et al. 2020); Decay sugar-rich substrates (Põlme et al. 2020) | Põlme S, et al. 2020. Fungal Diversity 105:1-16 ((https://doi.org/10.1007/s13225-020-00466-2)) |
| Otu14800 | 1 | k__Fungi(100);p__Ascomycota(100);c__Saccharomycetes(100);o__Saccharomycetales(100);f__Metschnikowiaceae(100);g__Metschnikowia(100);s__Metschnikowia_pulcherrima(100); | Metschnikowia | 13 | Saprotroph | \|Undefined Saprotroph\| | Yeast | NULL | Probable | As nectar-tap_saprotroph (Põlme et al. 2020); Decay sugar-rich substrates (Põlme et al. 2020) | Põlme S, et al. 2020. Fungal Diversity 105:1-16 ((https://doi.org/10.1007/s13225-020-00466-2)) |
| Otu14801 | 1 | k__Fungi(100);p__Ascomycota(100);c__Saccharomycetes(100);o__Saccharomycetales(100);f__Metschnikowiaceae(100);g__Metschnikowia(100);s__Metschnikowia_pulcherrima(100); | Metschnikowia | 13 | Saprotroph | \|Undefined Saprotroph\| | Yeast | NULL | Probable | As nectar-tap_saprotroph (Põlme et al. 2020); Decay sugar-rich substrates (Põlme et al. 2020) | Põlme S, et al. 2020. Fungal Diversity 105:1-16 ((https://doi.org/10.1007/s13225-020-00466-2)) |
| Otu14802 | 1 | k__Fungi(100);p__Ascomycota(100);c__Saccharomycetes(100);o__Saccharomycetales(100);f__Saccharomycetales_family_Incertae_sedis(100);g__Candida(100);g__Candida_unclassified(100); | Candida | 13 | Pathotroph-Saprotroph-Symbiotroph | Animal Pathogen-Endophyte-Endosymbiont-Epiphyte-\|Undefined Saprotroph\| | Dimorphic Yeast-Yeast | NULL | Probable | As nectar-tap_saprotroph (Põlme et al. 2020); Decay sugar-rich substrates (Põlme et al. 2020); This asexual genus is polyphyletic and contain species that function very differently depend on what they are. Caution should be taken when interpreting guilds for Candida species. | Manolakaki D, et al. 2010. Virulence. 1: 367-375 ((https://doi.org/10.4161/viru.1.5.12796)); Mekbib SB, et al. 2011. Tropical Plant Pathology 36:233-240 ((https://doi.org/10.1590/S1982-56762011000400004)); Põlme S, et al. 2020. Fungal Diversity 105:1-16 ((https://doi.org/10.1007/s13225-020-00466-2)) |
| Otu14803 | 1 | k__Fungi(100);p__Ascomycota(100);c__Saccharomycetes(100);o__Saccharomycetales(100);f__Metschnikowiaceae(100);g__Metschnikowia(100);g__Metschnikowia_unclassified(100); | Metschnikowia | 13 | Saprotroph | \|Undefined Saprotroph\| | Yeast | NULL | Probable | As nectar-tap_saprotroph (Põlme et al. 2020); Decay sugar-rich substrates (Põlme et al. 2020) | Põlme S, et al. 2020. Fungal Diversity 105:1-16 ((https://doi.org/10.1007/s13225-020-00466-2)) |
| Otu14807 | 1 | k__Fungi(100);p__Ascomycota(100);c__Saccharomycetes(100);o__Saccharomycetales(100);f__Metschnikowiaceae(100);g__Metschnikowia(100);s__Metschnikowia_pulcherrima(100); | Metschnikowia | 13 | Saprotroph | \|Undefined Saprotroph\| | Yeast | NULL | Probable | As nectar-tap_saprotroph (Põlme et al. 2020); Decay sugar-rich substrates (Põlme et al. 2020) | Põlme S, et al. 2020. Fungal Diversity 105:1-16 ((https://doi.org/10.1007/s13225-020-00466-2)) |
| Otu14812 | 1 | k__Fungi(100);p__Ascomycota(100);c__Saccharomycetes(100);o__Saccharomycetales(100);f__Metschnikowiaceae(100);g__Metschnikowia(100);s__Metschnikowia_pulcherrima(100); | Metschnikowia | 13 | Saprotroph | \|Undefined Saprotroph\| | Yeast | NULL | Probable | As nectar-tap_saprotroph (Põlme et al. 2020); Decay sugar-rich substrates (Põlme et al. 2020) | Põlme S, et al. 2020. Fungal Diversity 105:1-16 ((https://doi.org/10.1007/s13225-020-00466-2)) |
| Otu14813 | 1 | k__Fungi(100);p__Ascomycota(100);c__Saccharomycetes(100);o__Saccharomycetales(100);f__Saccharomycetales_family_Incertae_sedis(100);g__Candida(100);g__Candida_unclassified(100); | Candida | 13 | Pathotroph-Saprotroph-Symbiotroph | Animal Pathogen-Endophyte-Endosymbiont-Epiphyte-\|Undefined Saprotroph\| | Dimorphic Yeast-Yeast | NULL | Probable | As nectar-tap_saprotroph (Põlme et al. 2020); Decay sugar-rich substrates (Põlme et al. 2020); This asexual genus is polyphyletic and contain species that function very differently depend on what they are. Caution should be taken when interpreting guilds for Candida species. | Manolakaki D, et al. 2010. Virulence. 1: 367-375 ((https://doi.org/10.4161/viru.1.5.12796)); Mekbib SB, et al. 2011. Tropical Plant Pathology 36:233-240 ((https://doi.org/10.1590/S1982-56762011000400004)); Põlme S, et al. 2020. Fungal Diversity 105:1-16 ((https://doi.org/10.1007/s13225-020-00466-2)) |
| Otu14814 | 1 | k__Fungi(100);p__Ascomycota(100);c__Saccharomycetes(100);o__Saccharomycetales(100);f__Metschnikowiaceae(100);g__Metschnikowia(100);s__Metschnikowia_pulcherrima(100); | Metschnikowia | 13 | Saprotroph | \|Undefined Saprotroph\| | Yeast | NULL | Probable | As nectar-tap_saprotroph (Põlme et al. 2020); Decay sugar-rich substrates (Põlme et al. 2020) | Põlme S, et al. 2020. Fungal Diversity 105:1-16 ((https://doi.org/10.1007/s13225-020-00466-2)) |
| Otu14815 | 1 | k__Fungi(100);p__Ascomycota(100);c__Saccharomycetes(100);o__Saccharomycetales(100);f__Metschnikowiaceae(100);g__Metschnikowia(100);s__Metschnikowia_pulcherrima(100); | Metschnikowia | 13 | Saprotroph | \|Undefined Saprotroph\| | Yeast | NULL | Probable | As nectar-tap_saprotroph (Põlme et al. 2020); Decay sugar-rich substrates (Põlme et al. 2020) | Põlme S, et al. 2020. Fungal Diversity 105:1-16 ((https://doi.org/10.1007/s13225-020-00466-2)) |
| Otu14820 | 1 | k__Fungi(100);p__Ascomycota(100);c__Saccharomycetes(100);o__Saccharomycetales(100);f__Saccharomycetales_family_Incertae_sedis(100);f__Saccharomycetales_family_Incertae_sedis_unclassified(100);f__Saccharomycetales_family_Incertae_sedis_unclassified(100); | Saccharomycetales | 7 | Saprotroph | Undefined Saprotroph | Yeast | NULL | Possible | NULL | Sterkenburg E, et al. 2015. New Phytologist 207:1145-1158 ((https://doi.org/10.1111/nph.13426)) |
| Otu14823 | 1 | k__Fungi(100);p__Ascomycota(100);c__Saccharomycetes(100);o__Saccharomycetales(100);f__Metschnikowiaceae(100);g__Metschnikowia(100);g__Metschnikowia_unclassified(100); | Metschnikowia | 13 | Saprotroph | \|Undefined Saprotroph\| | Yeast | NULL | Probable | As nectar-tap_saprotroph (Põlme et al. 2020); Decay sugar-rich substrates (Põlme et al. 2020) | Põlme S, et al. 2020. Fungal Diversity 105:1-16 ((https://doi.org/10.1007/s13225-020-00466-2)) |
| Otu14825 | 1 | k__Fungi(100);p__Ascomycota(100);c__Saccharomycetes(100);o__Saccharomycetales(100);f__Metschnikowiaceae(100);g__Metschnikowia(100);g__Metschnikowia_unclassified(100); | Metschnikowia | 13 | Saprotroph | \|Undefined Saprotroph\| | Yeast | NULL | Probable | As nectar-tap_saprotroph (Põlme et al. 2020); Decay sugar-rich substrates (Põlme et al. 2020) | Põlme S, et al. 2020. Fungal Diversity 105:1-16 ((https://doi.org/10.1007/s13225-020-00466-2)) |
| Otu14826 | 1 | k__Fungi(100);p__Ascomycota(100);c__Saccharomycetes(100);o__Saccharomycetales(100);f__Metschnikowiaceae(100);g__Metschnikowia(100);g__Metschnikowia_unclassified(100); | Metschnikowia | 13 | Saprotroph | \|Undefined Saprotroph\| | Yeast | NULL | Probable | As nectar-tap_saprotroph (Põlme et al. 2020); Decay sugar-rich substrates (Põlme et al. 2020) | Põlme S, et al. 2020. Fungal Diversity 105:1-16 ((https://doi.org/10.1007/s13225-020-00466-2)) |
| Otu14827 | 1 | k__Fungi(100);p__Ascomycota(100);c__Saccharomycetes(100);o__Saccharomycetales(100);f__Metschnikowiaceae(100);g__Metschnikowia(100);s__Metschnikowia_pulcherrima(100); | Metschnikowia | 13 | Saprotroph | \|Undefined Saprotroph\| | Yeast | NULL | Probable | As nectar-tap_saprotroph (Põlme et al. 2020); Decay sugar-rich substrates (Põlme et al. 2020) | Põlme S, et al. 2020. Fungal Diversity 105:1-16 ((https://doi.org/10.1007/s13225-020-00466-2)) |
| Otu14828 | 1 | k__Fungi(100);p__Ascomycota(100);c__Saccharomycetes(100);o__Saccharomycetales(100);f__Metschnikowiaceae(100);g__Metschnikowia(100);g__Metschnikowia_unclassified(100); | Metschnikowia | 13 | Saprotroph | \|Undefined Saprotroph\| | Yeast | NULL | Probable | As nectar-tap_saprotroph (Põlme et al. 2020); Decay sugar-rich substrates (Põlme et al. 2020) | Põlme S, et al. 2020. Fungal Diversity 105:1-16 ((https://doi.org/10.1007/s13225-020-00466-2)) |
| Otu14829 | 1 | k__Fungi(100);p__Ascomycota(100);c__Saccharomycetes(100);o__Saccharomycetales(100);f__Metschnikowiaceae(100);g__Metschnikowia(100);g__Metschnikowia_unclassified(100); | Metschnikowia | 13 | Saprotroph | \|Undefined Saprotroph\| | Yeast | NULL | Probable | As nectar-tap_saprotroph (Põlme et al. 2020); Decay sugar-rich substrates (Põlme et al. 2020) | Põlme S, et al. 2020. Fungal Diversity 105:1-16 ((https://doi.org/10.1007/s13225-020-00466-2)) |
| Otu14830 | 1 | k__Fungi(100);p__Ascomycota(100);c__Saccharomycetes(100);o__Saccharomycetales(100);f__Metschnikowiaceae(100);g__Metschnikowia(100);s__Metschnikowia_pulcherrima(100); | Metschnikowia | 13 | Saprotroph | \|Undefined Saprotroph\| | Yeast | NULL | Probable | As nectar-tap_saprotroph (Põlme et al. 2020); Decay sugar-rich substrates (Põlme et al. 2020) | Põlme S, et al. 2020. Fungal Diversity 105:1-16 ((https://doi.org/10.1007/s13225-020-00466-2)) |
| Otu14831 | 1 | k__Fungi(100);p__Ascomycota(100);c__Saccharomycetes(100);o__Saccharomycetales(100);f__Metschnikowiaceae(100);g__Metschnikowia(100);s__Metschnikowia_pulcherrima(100); | Metschnikowia | 13 | Saprotroph | \|Undefined Saprotroph\| | Yeast | NULL | Probable | As nectar-tap_saprotroph (Põlme et al. 2020); Decay sugar-rich substrates (Põlme et al. 2020) | Põlme S, et al. 2020. Fungal Diversity 105:1-16 ((https://doi.org/10.1007/s13225-020-00466-2)) |
| Otu14832 | 1 | k__Fungi(100);p__Ascomycota(100);c__Saccharomycetes(100);o__Saccharomycetales(100);f__Metschnikowiaceae(100);g__Metschnikowia(100);s__Metschnikowia_pulcherrima(100); | Metschnikowia | 13 | Saprotroph | \|Undefined Saprotroph\| | Yeast | NULL | Probable | As nectar-tap_saprotroph (Põlme et al. 2020); Decay sugar-rich substrates (Põlme et al. 2020) | Põlme S, et al. 2020. Fungal Diversity 105:1-16 ((https://doi.org/10.1007/s13225-020-00466-2)) |
| Otu14833 | 1 | k__Fungi(100);p__Ascomycota(100);c__Saccharomycetes(100);o__Saccharomycetales(100);f__Metschnikowiaceae(100);g__Metschnikowia(100);s__Metschnikowia_pulcherrima(100); | Metschnikowia | 13 | Saprotroph | \|Undefined Saprotroph\| | Yeast | NULL | Probable | As nectar-tap_saprotroph (Põlme et al. 2020); Decay sugar-rich substrates (Põlme et al. 2020) | Põlme S, et al. 2020. Fungal Diversity 105:1-16 ((https://doi.org/10.1007/s13225-020-00466-2)) |

| | | | | | | | | | | | |
|---|---|---|---|---|---|---|---|---|---|---|---|
| Otu14834 | 1 | k__Fungi(100);p__Ascomycota(100);c__Saccharomycetes(100);o__Saccharomycetales(100);f__Metschnikowiaceae(100);g__Metschnikowia(100);s__Metschnikowia_pulcherrima(100); | Metschnikowia | 13 | Saprotroph | \|Undefined Saprotroph\| | Yeast | NULL | Probable | As nectar-tap_saprotroph (Põlme et al. 2020); Decay sugar-rich substrates (Põlme et al. 2020) | Põlme S, et al. 2020. Fungal Diversity 105:1-16 ((https://doi.org/10.1007/s13225-020-00466-2)) |
| Otu14837 | 1 | k__Fungi(100);p__Ascomycota(100);c__Saccharomycetes(100);o__Saccharomycetales(100);f__Metschnikowiaceae(100);g__Metschnikowia(100);s__Metschnikowia_pulcherrima(100); | Metschnikowia | 13 | Saprotroph | \|Undefined Saprotroph\| | Yeast | NULL | Probable | As nectar-tap_saprotroph (Põlme et al. 2020); Decay sugar-rich substrates (Põlme et al. 2020) | Põlme S, et al. 2020. Fungal Diversity 105:1-16 ((https://doi.org/10.1007/s13225-020-00466-2)) |
| Otu14839 | 1 | k__Fungi(100);p__Ascomycota(100);c__Saccharomycetes(100);o__Saccharomycetales(100);f__Metschnikowiaceae(100);g__Metschnikowia(100);g__Metschnikowia_unclassified(100); | Metschnikowia | 13 | Saprotroph | \|Undefined Saprotroph\| | Yeast | NULL | Probable | As nectar-tap_saprotroph (Põlme et al. 2020); Decay sugar-rich substrates (Põlme et al. 2020) | Põlme S, et al. 2020. Fungal Diversity 105:1-16 ((https://doi.org/10.1007/s13225-020-00466-2)) |
| Otu14841 | 1 | k__Fungi(100);p__Ascomycota(100);c__Saccharomycetes(100);o__Saccharomycetales(100);f__Metschnikowiaceae(100);g__Metschnikowia(100);g__Metschnikowia_unclassified(100); | Metschnikowia | 13 | Saprotroph | \|Undefined Saprotroph\| | Yeast | NULL | Probable | As nectar-tap_saprotroph (Põlme et al. 2020); Decay sugar-rich substrates (Põlme et al. 2020) | Põlme S, et al. 2020. Fungal Diversity 105:1-16 ((https://doi.org/10.1007/s13225-020-00466-2)) |
| Otu14843 | 1 | k__Fungi(100);p__Ascomycota(100);c__Saccharomycetes(100);o__Saccharomycetales(100);f__Metschnikowiaceae(100);g__Metschnikowia(100);g__Metschnikowia_unclassified(100); | Metschnikowia | 13 | Saprotroph | \|Undefined Saprotroph\| | Yeast | NULL | Probable | As nectar-tap_saprotroph (Põlme et al. 2020); Decay sugar-rich substrates (Põlme et al. 2020) | Põlme S, et al. 2020. Fungal Diversity 105:1-16 ((https://doi.org/10.1007/s13225-020-00466-2)) |
| Otu14846 | 1 | k__Fungi(100);p__Ascomycota(100);c__Saccharomycetes(100);o__Saccharomycetales(100);f__Metschnikowiaceae(100);g__Metschnikowia(100);g__Metschnikowia_unclassified(100); | Metschnikowia | 13 | Saprotroph | \|Undefined Saprotroph\| | Yeast | NULL | Probable | As nectar-tap_saprotroph (Põlme et al. 2020); Decay sugar-rich substrates (Põlme et al. 2020) | Põlme S, et al. 2020. Fungal Diversity 105:1-16 ((https://doi.org/10.1007/s13225-020-00466-2)) |
| Otu14848 | 1 | k__Fungi(100);p__Ascomycota(100);c__Saccharomycetes(100);o__Saccharomycetales(100);f__Metschnikowiaceae(100);g__Metschnikowia(100);s__Metschnikowia_pulcherrima(100); | Metschnikowia | 13 | Saprotroph | \|Undefined Saprotroph\| | Yeast | NULL | Probable | As nectar-tap_saprotroph (Põlme et al. 2020); Decay sugar-rich substrates (Põlme et al. 2020) | Põlme S, et al. 2020. Fungal Diversity 105:1-16 ((https://doi.org/10.1007/s13225-020-00466-2)) |
| Otu14849 | 1 | k__Fungi(100);p__Ascomycota(100);c__Saccharomycetes(100);o__Saccharomycetales(100);f__Metschnikowiaceae(100);g__Metschnikowia(100);s__Metschnikowia_pulcherrima(100); | Metschnikowia | 13 | Saprotroph | \|Undefined Saprotroph\| | Yeast | NULL | Probable | As nectar-tap_saprotroph (Põlme et al. 2020); Decay sugar-rich substrates (Põlme et al. 2020) | Põlme S, et al. 2020. Fungal Diversity 105:1-16 ((https://doi.org/10.1007/s13225-020-00466-2)) |
| Otu14850 | 1 | k__Fungi(100);p__Ascomycota(100);c__Saccharomycetes(100);o__Saccharomycetales(100);f__Metschnikowiaceae(100);g__Metschnikowia(100);g__Metschnikowia_unclassified(100); | Metschnikowia | 13 | Saprotroph | \|Undefined Saprotroph\| | Yeast | NULL | Probable | As nectar-tap_saprotroph (Põlme et al. 2020); Decay sugar-rich substrates (Põlme et al. 2020) | Põlme S, et al. 2020. Fungal Diversity 105:1-16 ((https://doi.org/10.1007/s13225-020-00466-2)) |
| Otu14851 | 1 | k__Fungi(100);p__Ascomycota(100);c__Saccharomycetes(100);o__Saccharomycetales(100);f__Saccharomycetales_family_Incertae_sedis(100);g__Candida(100);g__Candida_unclassified(100); | Candida | 13 | Pathotroph-Saprotroph-Symbiotroph | Animal Pathogen-Endophyte-Endosymbiont-Epiphyte-\|Undefined Saprotroph\| | Dimorphic Yeast-Yeast | NULL | Probable | As nectar-tap_saprotroph (Põlme et al. 2020); Decay sugar-rich substrates (Põlme et al. 2020); This asexual genus is polyphyletic and contain species that function very differently depend on what they are. Caution should be taken when interpreting guilds for Candida species. | Manolakaki D, et al. 2010. Virulence. 1: 367-375 ((https://doi.org/10.4161/viru.1.5.12796)); Mekbib SB, et al. 2011. Tropical Plant Pathology 36:233-240 ((https://doi.org/10.1590/S1982-56762011000400004)); Põlme S, et al. 2020. Fungal Diversity 105:1-16 ((https://doi.org/10.1007/s13225-020-00466-2)) |
| Otu14852 | 1 | k__Fungi(100);p__Ascomycota(100);c__Saccharomycetes(100);o__Saccharomycetales(100);f__Metschnikowiaceae(100);g__Metschnikowia(100);s__Metschnikowia_pulcherrima(100); | Metschnikowia | 13 | Saprotroph | \|Undefined Saprotroph\| | Yeast | NULL | Probable | As nectar-tap_saprotroph (Põlme et al. 2020); Decay sugar-rich substrates (Põlme et al. 2020) | Põlme S, et al. 2020. Fungal Diversity 105:1-16 ((https://doi.org/10.1007/s13225-020-00466-2)) |
| Otu14854 | 1 | k__Fungi(100);p__Ascomycota(100);c__Saccharomycetes(100);o__Saccharomycetales(100);f__Metschnikowiaceae(100);g__Metschnikowia(100);g__Metschnikowia_unclassified(100); | Metschnikowia | 13 | Saprotroph | \|Undefined Saprotroph\| | Yeast | NULL | Probable | As nectar-tap_saprotroph (Põlme et al. 2020); Decay sugar-rich substrates (Põlme et al. 2020) | Põlme S, et al. 2020. Fungal Diversity 105:1-16 ((https://doi.org/10.1007/s13225-020-00466-2)) |
| Otu14857 | 1 | k__Fungi(100);p__Ascomycota(100);c__Saccharomycetes(100);o__Saccharomycetales(100);f__Metschnikowiaceae(100);g__Metschnikowia(100);g__Metschnikowia_unclassified(100); | Metschnikowia | 13 | Saprotroph | \|Undefined Saprotroph\| | Yeast | NULL | Probable | As nectar-tap_saprotroph (Põlme et al. 2020); Decay sugar-rich substrates (Põlme et al. 2020) | Põlme S, et al. 2020. Fungal Diversity 105:1-16 ((https://doi.org/10.1007/s13225-020-00466-2)) |
| Otu14858 | 1 | k__Fungi(100);p__Ascomycota(100);c__Saccharomycetes(100);o__Saccharomycetales(100);f__Metschnikowiaceae(100);g__Metschnikowia(100);g__Metschnikowia_unclassified(100); | Metschnikowia | 13 | Saprotroph | \|Undefined Saprotroph\| | Yeast | NULL | Probable | As nectar-tap_saprotroph (Põlme et al. 2020); Decay sugar-rich substrates (Põlme et al. 2020) | Põlme S, et al. 2020. Fungal Diversity 105:1-16 ((https://doi.org/10.1007/s13225-020-00466-2)) |
| Otu14859 | 1 | k__Fungi(100);p__Ascomycota(100);c__Saccharomycetes(100);o__Saccharomycetales(100);f__Metschnikowiaceae(100);g__Metschnikowia(100);s__Metschnikowia_pulcherrima(100); | Metschnikowia | 13 | Saprotroph | \|Undefined Saprotroph\| | Yeast | NULL | Probable | As nectar-tap_saprotroph (Põlme et al. 2020); Decay sugar-rich substrates (Põlme et al. 2020) | Põlme S, et al. 2020. Fungal Diversity 105:1-16 ((https://doi.org/10.1007/s13225-020-00466-2)) |
| Otu14860 | 1 | k__Fungi(100);p__Ascomycota(100);c__Saccharomycetes(100);o__Saccharomycetales(100);f__Metschnikowiaceae(100);g__Metschnikowia(100);s__Metschnikowia_pulcherrima(100); | Metschnikowia | 13 | Saprotroph | \|Undefined Saprotroph\| | Yeast | NULL | Probable | As nectar-tap_saprotroph (Põlme et al. 2020); Decay sugar-rich substrates (Põlme et al. 2020) | Põlme S, et al. 2020. Fungal Diversity 105:1-16 ((https://doi.org/10.1007/s13225-020-00466-2)) |
| Otu14861 | 1 | k__Fungi(100);p__Ascomycota(100);c__Saccharomycetes(100);o__Saccharomycetales(100);f__Metschnikowiaceae(100);g__Metschnikowia(100);s__Metschnikowia_pulcherrima(100); | Metschnikowia | 13 | Saprotroph | \|Undefined Saprotroph\| | Yeast | NULL | Probable | As nectar-tap_saprotroph (Põlme et al. 2020); Decay sugar-rich substrates (Põlme et al. 2020) | Põlme S, et al. 2020. Fungal Diversity 105:1-16 ((https://doi.org/10.1007/s13225-020-00466-2)) |
| Otu14862 | 1 | k__Fungi(100);p__Ascomycota(100);c__Saccharomycetes(100);o__Saccharomycetales(100);f__Metschnikowiaceae(100);g__Metschnikowia(100);g__Metschnikowia_unclassified(100); | Metschnikowia | 13 | Saprotroph | \|Undefined Saprotroph\| | Yeast | NULL | Probable | As nectar-tap_saprotroph (Põlme et al. 2020); Decay sugar-rich substrates (Põlme et al. 2020) | Põlme S, et al. 2020. Fungal Diversity 105:1-16 ((https://doi.org/10.1007/s13225-020-00466-2)) |
| Otu14863 | 1 | k__Fungi(100);p__Ascomycota(100);c__Saccharomycetes(100);o__Saccharomycetales(100);f__Metschnikowiaceae(100);g__Metschnikowia(100);s__Metschnikowia_pulcherrima(100); | Metschnikowia | 13 | Saprotroph | \|Undefined Saprotroph\| | Yeast | NULL | Probable | As nectar-tap_saprotroph (Põlme et al. 2020); Decay sugar-rich substrates (Põlme et al. 2020) | Põlme S, et al. 2020. Fungal Diversity 105:1-16 ((https://doi.org/10.1007/s13225-020-00466-2)) |
| Otu14864 | 1 | k__Fungi(100);p__Ascomycota(100);c__Saccharomycetes(100);o__Saccharomycetales(100);f__Metschnikowiaceae(100);g__Metschnikowia(100);s__Metschnikowia_pulcherrima(100); | Metschnikowia | 13 | Saprotroph | \|Undefined Saprotroph\| | Yeast | NULL | Probable | As nectar-tap_saprotroph (Põlme et al. 2020); Decay sugar-rich substrates (Põlme et al. 2020) | Põlme S, et al. 2020. Fungal Diversity 105:1-16 ((https://doi.org/10.1007/s13225-020-00466-2)) |
| Otu14866 | 1 | k__Fungi(100);p__Ascomycota(100);c__Saccharomycetes(100);o__Saccharomycetales(100);f__Metschnikowiaceae(100);g__Metschnikowia(100);g__Metschnikowia_unclassified(100); | Metschnikowia | 13 | Saprotroph | \|Undefined Saprotroph\| | Yeast | NULL | Probable | As nectar-tap_saprotroph (Põlme et al. 2020); Decay sugar-rich substrates (Põlme et al. 2020) | Põlme S, et al. 2020. Fungal Diversity 105:1-16 ((https://doi.org/10.1007/s13225-020-00466-2)) |
| Otu14867 | 1 | k__Fungi(100);p__Ascomycota(100);c__Saccharomycetes(100);o__Saccharomycetales(100);f__Metschnikowiaceae(100);g__Metschnikowia(100);s__Metschnikowia_pulcherrima(100); | Metschnikowia | 13 | Saprotroph | \|Undefined Saprotroph\| | Yeast | NULL | Probable | As nectar-tap_saprotroph (Põlme et al. 2020); Decay sugar-rich substrates (Põlme et al. 2020) | Põlme S, et al. 2020. Fungal Diversity 105:1-16 ((https://doi.org/10.1007/s13225-020-00466-2)) |
| Otu14868 | 1 | k__Fungi(100);p__Ascomycota(100);c__Saccharomycetes(100);o__Saccharomycetales(100);f__Metschnikowiaceae(100);g__Metschnikowia(100);s__Metschnikowia_pulcherrima(100); | Metschnikowia | 13 | Saprotroph | \|Undefined Saprotroph\| | Yeast | NULL | Probable | As nectar-tap_saprotroph (Põlme et al. 2020); Decay sugar-rich substrates (Põlme et al. 2020) | Põlme S, et al. 2020. Fungal Diversity 105:1-16 ((https://doi.org/10.1007/s13225-020-00466-2)) |
| Otu14869 | 1 | k__Fungi(100);p__Ascomycota(100);c__Saccharomycetes(100);o__Saccharomycetales(100);f__Metschnikowiaceae(100);g__Metschnikowia(100);s__Metschnikowia_pulcherrima(100); | Metschnikowia | 13 | Saprotroph | \|Undefined Saprotroph\| | Yeast | NULL | Probable | As nectar-tap_saprotroph (Põlme et al. 2020); Decay sugar-rich substrates (Põlme et al. 2020) | Põlme S, et al. 2020. Fungal Diversity 105:1-16 ((https://doi.org/10.1007/s13225-020-00466-2)) |
| Otu14870 | 1 | k__Fungi(100);p__Ascomycota(100);c__Saccharomycetes(100);o__Saccharomycetales(100);f__Metschnikowiaceae(100);g__Metschnikowia(100);s__Metschnikowia_pulcherrima(100); | Metschnikowia | 13 | Saprotroph | \|Undefined Saprotroph\| | Yeast | NULL | Probable | As nectar-tap_saprotroph (Põlme et al. 2020); Decay sugar-rich substrates (Põlme et al. 2020) | Põlme S, et al. 2020. Fungal Diversity 105:1-16 ((https://doi.org/10.1007/s13225-020-00466-2)) |
| Otu14871 | 1 | k__Fungi(100);p__Ascomycota(100);c__Saccharomycetes(100);o__Saccharomycetales(100);f__Metschnikowiaceae(100);g__Metschnikowia(100);s__Metschnikowia_pulcherrima(100); | Metschnikowia | 13 | Saprotroph | \|Undefined Saprotroph\| | Yeast | NULL | Probable | As nectar-tap_saprotroph (Põlme et al. 2020); Decay sugar-rich substrates (Põlme et al. 2020) | Põlme S, et al. 2020. Fungal Diversity 105:1-16 ((https://doi.org/10.1007/s13225-020-00466-2)) |
| Otu14872 | 1 | k__Fungi(100);p__Ascomycota(100);c__Saccharomycetes(100);o__Saccharomycetales(100);f__Metschnikowiaceae(100);g__Metschnikowia(100);s__Metschnikowia_pulcherrima(100); | Metschnikowia | 13 | Saprotroph | \|Undefined Saprotroph\| | Yeast | NULL | Probable | As nectar-tap_saprotroph (Põlme et al. 2020); Decay sugar-rich substrates (Põlme et al. 2020) | Põlme S, et al. 2020. Fungal Diversity 105:1-16 ((https://doi.org/10.1007/s13225-020-00466-2)) |
| Otu14874 | 1 | k__Fungi(100);p__Ascomycota(100);c__Saccharomycetes(100);o__Saccharomycetales(100);f__Metschnikowiaceae(100);g__Metschnikowia(100);g__Metschnikowia_unclassified(100); | Metschnikowia | 13 | Saprotroph | \|Undefined Saprotroph\| | Yeast | NULL | Probable | As nectar-tap_saprotroph (Põlme et al. 2020); Decay sugar-rich substrates (Põlme et al. 2020) | Põlme S, et al. 2020. Fungal Diversity 105:1-16 ((https://doi.org/10.1007/s13225-020-00466-2)) |
| Otu14875 | 1 | k__Fungi(100);p__Ascomycota(100);c__Saccharomycetes(100);o__Saccharomycetales(100);f__Metschnikowiaceae(100);g__Metschnikowia(100);g__Metschnikowia_unclassified(100); | Metschnikowia | 13 | Saprotroph | \|Undefined Saprotroph\| | Yeast | NULL | Probable | As nectar-tap_saprotroph (Põlme et al. 2020); Decay sugar-rich substrates (Põlme et al. 2020) | Põlme S, et al. 2020. Fungal Diversity 105:1-16 ((https://doi.org/10.1007/s13225-020-00466-2)) |
| Otu14877 | 1 | k__Fungi(100);p__Ascomycota(100);c__Saccharomycetes(100);o__Saccharomycetales(100);f__Metschnikowiaceae(100);g__Metschnikowia(100);s__Metschnikowia_pulcherrima(100); | Metschnikowia | 13 | Saprotroph | \|Undefined Saprotroph\| | Yeast | NULL | Probable | As nectar-tap_saprotroph (Põlme et al. 2020); Decay sugar-rich substrates (Põlme et al. 2020) | Põlme S, et al. 2020. Fungal Diversity 105:1-16 ((https://doi.org/10.1007/s13225-020-00466-2)) |
| Otu14879 | 1 | k__Fungi(100);p__Ascomycota(100);c__Saccharomycetes(100);o__Saccharomycetales(100);f__Metschnikowiaceae(100);g__Metschnikowia(100);s__Metschnikowia_pulcherrima(100); | Metschnikowia | 13 | Saprotroph | \|Undefined Saprotroph\| | Yeast | NULL | Probable | As nectar-tap_saprotroph (Põlme et al. 2020); Decay sugar-rich substrates (Põlme et al. 2020) | Põlme S, et al. 2020. Fungal Diversity 105:1-16 ((https://doi.org/10.1007/s13225-020-00466-2)) |
| Otu14881 | 1 | k__Fungi(100);p__Ascomycota(100);c__Saccharomycetes(100);o__Saccharomycetales(100);f__Metschnikowiaceae(100);g__Metschnikowia(100);g__Metschnikowia_unclassified(100); | Metschnikowia | 13 | Saprotroph | \|Undefined Saprotroph\| | Yeast | NULL | Probable | As nectar-tap_saprotroph (Põlme et al. 2020); Decay sugar-rich substrates (Põlme et al. 2020) | Põlme S, et al. 2020. Fungal Diversity 105:1-16 ((https://doi.org/10.1007/s13225-020-00466-2)) |
| Otu14882 | 1 | k__Fungi(100);p__Ascomycota(100);c__Saccharomycetes(100);o__Saccharomycetales(100);f__Metschnikowiaceae(100);g__Metschnikowia(100);s__Metschnikowia_pulcherrima(100); | Metschnikowia | 13 | Saprotroph | \|Undefined Saprotroph\| | Yeast | NULL | Probable | As nectar-tap_saprotroph (Põlme et al. 2020); Decay sugar-rich substrates (Põlme et al. 2020) | Põlme S, et al. 2020. Fungal Diversity 105:1-16 ((https://doi.org/10.1007/s13225-020-00466-2)) |
| Otu14883 | 1 | k__Fungi(100);p__Ascomycota(100);c__Saccharomycetes(100);o__Saccharomycetales(100);f__Metschnikowiaceae(100);g__Metschnikowia(100);g__Metschnikowia_unclassified(100); | Metschnikowia | 13 | Saprotroph | \|Undefined Saprotroph\| | Yeast | NULL | Probable | As nectar-tap_saprotroph (Põlme et al. 2020); Decay sugar-rich substrates (Põlme et al. 2020) | Põlme S, et al. 2020. Fungal Diversity 105:1-16 ((https://doi.org/10.1007/s13225-020-00466-2)) |
| Otu14884 | 1 | k__Fungi(100);p__Ascomycota(100);c__Saccharomycetes(100);o__Saccharomycetales(100);f__Metschnikowiaceae(100);g__Metschnikowia(100);s__Metschnikowia_pulcherrima(100); | Metschnikowia | 13 | Saprotroph | \|Undefined Saprotroph\| | Yeast | NULL | Probable | As nectar-tap_saprotroph (Põlme et al. 2020); Decay sugar-rich substrates (Põlme et al. 2020) | Põlme S, et al. 2020. Fungal Diversity 105:1-16 ((https://doi.org/10.1007/s13225-020-00466-2)) |
| Otu14886 | 1 | k__Fungi(100);p__Ascomycota(100);c__Saccharomycetes(100);o__Saccharomycetales(100);f__Metschnikowiaceae(100);g__Metschnikowia(100);g__Metschnikowia_unclassified(100); | Metschnikowia | 13 | Saprotroph | \|Undefined Saprotroph\| | Yeast | NULL | Probable | As nectar-tap_saprotroph (Põlme et al. 2020); Decay sugar-rich substrates (Põlme et al. 2020) | Põlme S, et al. 2020. Fungal Diversity 105:1-16 ((https://doi.org/10.1007/s13225-020-00466-2)) |
| Otu14888 | 1 | k__Fungi(100);p__Ascomycota(100);c__Saccharomycetes(100);o__Saccharomycetales(100);f__Metschnikowiaceae(100);g__Metschnikowia(100);s__Metschnikowia_pulcherrima(100); | Metschnikowia | 13 | Saprotroph | \|Undefined Saprotroph\| | Yeast | NULL | Probable | As nectar-tap_saprotroph (Põlme et al. 2020); Decay sugar-rich substrates (Põlme et al. 2020) | Põlme S, et al. 2020. Fungal Diversity 105:1-16 ((https://doi.org/10.1007/s13225-020-00466-2)) |
| Otu14889 | 1 | k__Fungi(100);p__Ascomycota(100);c__Saccharomycetes(100);o__Saccharomycetales(100);f__Metschnikowiaceae(100);g__Metschnikowia(100);s__Metschnikowia_pulcherrima(100); | Metschnikowia | 13 | Saprotroph | \|Undefined Saprotroph\| | Yeast | NULL | Probable | As nectar-tap_saprotroph (Põlme et al. 2020); Decay sugar-rich substrates (Põlme et al. 2020) | Põlme S, et al. 2020. Fungal Diversity 105:1-16 ((https://doi.org/10.1007/s13225-020-00466-2)) |
| Otu14890 | 1 | k__Fungi(100);p__Ascomycota(100);c__Saccharomycetes(100);o__Saccharomycetales(100);f__Metschnikowiaceae(100);g__Metschnikowia(100);s__Metschnikowia_pulcherrima(100); | Metschnikowia | 13 | Saprotroph | \|Undefined Saprotroph\| | Yeast | NULL | Probable | As nectar-tap_saprotroph (Põlme et al. 2020); Decay sugar-rich substrates (Põlme et al. 2020) | Põlme S, et al. 2020. Fungal Diversity 105:1-16 ((https://doi.org/10.1007/s13225-020-00466-2)) |
| Otu14891 | 1 | k__Fungi(100);p__Ascomycota(100);c__Saccharomycetes(100);o__Saccharomycetales(100);f__Metschnikowiaceae(100);g__Metschnikowia(100);s__Metschnikowia_pulcherrima(100); | Metschnikowia | 13 | Saprotroph | \|Undefined Saprotroph\| | Yeast | NULL | Probable | As nectar-tap_saprotroph (Põlme et al. 2020); Decay sugar-rich substrates (Põlme et al. 2020) | Põlme S, et al. 2020. Fungal Diversity 105:1-16 ((https://doi.org/10.1007/s13225-020-00466-2)) |
| Otu14892 | 1 | k__Fungi(100);p__Ascomycota(100);c__Saccharomycetes(100);o__Saccharomycetales(100);f__Metschnikowiaceae(100);g__Metschnikowia(100);s__Metschnikowia_pulcherrima(100); | Metschnikowia | 13 | Saprotroph | \|Undefined Saprotroph\| | Yeast | NULL | Probable | As nectar-tap_saprotroph (Põlme et al. 2020); Decay sugar-rich substrates (Põlme et al. 2020) | Põlme S, et al. 2020. Fungal Diversity 105:1-16 ((https://doi.org/10.1007/s13225-020-00466-2)) |
| Otu14894 | 1 | k__Fungi(100);p__Ascomycota(100);c__Saccharomycetes(100);o__Saccharomycetales(100);f__Saccharomycetales_family_Incertae_sedis(100);g__Candida(100);g__Candida_unclassified(100); | Candida | 13 | Pathotroph-Saprotroph-Symbiotroph | Animal Pathogen-Endophyte-Endosymbiont-Epiphyte-\|Undefined Saprotroph\| | Dimorphic Yeast-Yeast | NULL | Probable | As nectar-tap_saprotroph (Põlme et al. 2020); Decay sugar-rich substrates (Põlme et al. 2020); This asexual genus is polyphyletic and contain species that function very differently depend on what they are. Caution should be taken when interpreting guilds for Candida species. | Manolakaki D, et al. 2010. Virulence. 1: 367-375 ((https://doi.org/10.4161/viru.1.5.12796)); Mekbib SB, et al. 2011. Tropical Plant Pathology 36:233-240 ((https://doi.org/10.1590/S1982-56762011000400004)); Põlme S, et al. 2020. Fungal Diversity 105:1-16 ((https://doi.org/10.1007/s13225-020-00466-2)) |
| Otu14896 | 1 | k__Fungi(100);p__Ascomycota(100);c__Saccharomycetes(100);o__Saccharomycetales(100);f__Metschnikowiaceae(100);g__Metschnikowia(100);s__Metschnikowia_pulcherrima(100); | Metschnikowia | 13 | Saprotroph | \|Undefined Saprotroph\| | Yeast | NULL | Probable | As nectar-tap_saprotroph (Põlme et al. 2020); Decay sugar-rich substrates (Põlme et al. 2020) | Põlme S, et al. 2020. Fungal Diversity 105:1-16 ((https://doi.org/10.1007/s13225-020-00466-2)) |
| Otu14898 | 1 | k__Fungi(100);p__Ascomycota(100);c__Saccharomycetes(100);o__Saccharomycetales(100);f__Metschnikowiaceae(100);g__Metschnikowia(100);s__Metschnikowia_pulcherrima(100); | Metschnikowia | 13 | Saprotroph | \|Undefined Saprotroph\| | Yeast | NULL | Probable | As nectar-tap_saprotroph (Põlme et al. 2020); Decay sugar-rich substrates (Põlme et al. 2020) | Põlme S, et al. 2020. Fungal Diversity 105:1-16 ((https://doi.org/10.1007/s13225-020-00466-2)) |
| Otu14899 | 1 | k__Fungi(100);p__Ascomycota(100);c__Saccharomycetes(100);o__Saccharomycetales(100);f__Saccharomycetales_family_Incertae_sedis(100);g__Candida(100);g__Candida_unclassified(100); | Candida | 13 | Pathotroph-Saprotroph-Symbiotroph | Animal Pathogen-Endophyte-Endosymbiont-Epiphyte-\|Undefined Saprotroph\| | Dimorphic Yeast-Yeast | NULL | Probable | As nectar-tap_saprotroph (Põlme et al. 2020); Decay sugar-rich substrates (Põlme et al. 2020); This asexual genus is polyphyletic and contain species that function very differently depend on what they are. Caution should be taken when interpreting guilds for Candida species. | Manolakaki D, et al. 2010. Virulence. 1: 367-375 ((https://doi.org/10.4161/viru.1.5.12796)); Mekbib SB, et al. 2011. Tropical Plant Pathology 36:233-240 ((https://doi.org/10.1590/S1982-56762011000400004)); Põlme S, et al. 2020. Fungal Diversity 105:1-16 ((https://doi.org/10.1007/s13225-020-00466-2)) |
| Otu14900 | 1 | k__Fungi(100);p__Ascomycota(100);c__Saccharomycetes(100);o__Saccharomycetales(100);f__Saccharomycetales_family_Incertae_sedis(100);f__Saccharomycetales_family_Incertae_sedis_unclassified(100);f__Saccharomycetales_family_Incertae_sedis_unclassified(100); | Saccharomycetales | 7 | Saprotroph | Undefined Saprotroph | Yeast | NULL | Possible | NULL | Sterkenburg E, et al. 2015. New Phytologist 207:1145-1158 ((https://doi.org/10.1111/nph.13426)) |
| Otu14901 | 1 | k__Fungi(100);p__Ascomycota(100);c__Saccharomycetes(100);o__Saccharomycetales(100);f__Metschnikowiaceae(100);g__Metschnikowia(100);s__Metschnikowia_pulcherrima(100); | Metschnikowia | 13 | Saprotroph | \|Undefined Saprotroph\| | Yeast | NULL | Probable | As nectar-tap_saprotroph (Põlme et al. 2020); Decay sugar-rich substrates (Põlme et al. 2020) | Põlme S, et al. 2020. Fungal Diversity 105:1-16 ((https://doi.org/10.1007/s13225-020-00466-2)) |
| Otu14902 | 1 | k__Fungi(100);p__Ascomycota(100);c__Saccharomycetes(100);o__Saccharomycetales(100);f__Metschnikowiaceae(100);g__Metschnikowia(100);s__Metschnikowia_pulcherrima(100); | Metschnikowia | 13 | Saprotroph | \|Undefined Saprotroph\| | Yeast | NULL | Probable | As nectar-tap_saprotroph (Põlme et al. 2020); Decay sugar-rich substrates (Põlme et al. 2020) | Põlme S, et al. 2020. Fungal Diversity 105:1-16 ((https://doi.org/10.1007/s13225-020-00466-2)) |
| Otu14903 | 1 | k__Fungi(100);p__Ascomycota(100);c__Saccharomycetes(100);o__Saccharomycetales(100);f__Metschnikowiaceae(100);g__Metschnikowia(100);s__Metschnikowia_pulcherrima(100); | Metschnikowia | 13 | Saprotroph | \|Undefined Saprotroph\| | Yeast | NULL | Probable | As nectar-tap_saprotroph (Põlme et al. 2020); Decay sugar-rich substrates (Põlme et al. 2020) | Põlme S, et al. 2020. Fungal Diversity 105:1-16 ((https://doi.org/10.1007/s13225-020-00466-2)) |
| Otu14904 | 1 | k__Fungi(100);p__Ascomycota(100);c__Saccharomycetes(100);o__Saccharomycetales(100);f__Metschnikowiaceae(100);g__Metschnikowia(100);g__Metschnikowia_unclassified(100); | Metschnikowia | 13 | Saprotroph | \|Undefined Saprotroph\| | Yeast | NULL | Probable | As nectar-tap_saprotroph (Põlme et al. 2020); Decay sugar-rich substrates (Põlme et al. 2020) | Põlme S, et al. 2020. Fungal Diversity 105:1-16 ((https://doi.org/10.1007/s13225-020-00466-2)) |
| Otu14905 | 1 | k__Fungi(100);p__Ascomycota(100);c__Saccharomycetes(100);o__Saccharomycetales(100);f__Metschnikowiaceae(100);g__Metschnikowia(100);s__Metschnikowia_pulcherrima(100); | Metschnikowia | 13 | Saprotroph | \|Undefined Saprotroph\| | Yeast | NULL | Probable | As nectar-tap_saprotroph (Põlme et al. 2020); Decay sugar-rich substrates (Põlme et al. 2020) | Põlme S, et al. 2020. Fungal Diversity 105:1-16 ((https://doi.org/10.1007/s13225-020-00466-2)) |
| Otu14906 | 1 | k__Fungi(100);p__Ascomycota(100);c__Saccharomycetes(100);o__Saccharomycetales(100);f__Metschnikowiaceae(100);g__Metschnikowia(100);g__Metschnikowia_unclassified(100); | Metschnikowia | 13 | Saprotroph | \|Undefined Saprotroph\| | Yeast | NULL | Probable | As nectar-tap_saprotroph (Põlme et al. 2020); Decay sugar-rich substrates (Põlme et al. 2020) | Põlme S, et al. 2020. Fungal Diversity 105:1-16 ((https://doi.org/10.1007/s13225-020-00466-2)) |
| Otu14907 | 1 | k__Fungi(100);p__Ascomycota(100);c__Saccharomycetes(100);o__Saccharomycetales(100);f__Metschnikowiaceae(100);g__Metschnikowia(100);s__Metschnikowia_pulcherrima(100); | Metschnikowia | 13 | Saprotroph | \|Undefined Saprotroph\| | Yeast | NULL | Probable | As nectar-tap_saprotroph (Põlme et al. 2020); Decay sugar-rich substrates (Põlme et al. 2020) | Põlme S, et al. 2020. Fungal Diversity 105:1-16 ((https://doi.org/10.1007/s13225-020-00466-2)) |
| Otu14908 | 1 | k__Fungi(100);p__Ascomycota(100);c__Saccharomycetes(100);o__Saccharomycetales(100);f__Metschnikowiaceae(100);g__Metschnikowia(100);s__Metschnikowia_pulcherrima(100); | Metschnikowia | 13 | Saprotroph | \|Undefined Saprotroph\| | Yeast | NULL | Probable | As nectar-tap_saprotroph (Põlme et al. 2020); Decay sugar-rich substrates (Põlme et al. 2020) | Põlme S, et al. 2020. Fungal Diversity 105:1-16 ((https://doi.org/10.1007/s13225-020-00466-2)) |
| Otu14911 | 1 | k__Fungi(100);p__Ascomycota(100);c__Saccharomycetes(100);o__Saccharomycetales(100);f__Metschnikowiaceae(100);g__Metschnikowia(100);s__Metschnikowia_pulcherrima(100); | Metschnikowia | 13 | Saprotroph | \|Undefined Saprotroph\| | Yeast | NULL | Probable | As nectar-tap_saprotroph (Põlme et al. 2020); Decay sugar-rich substrates (Põlme et al. 2020) | Põlme S, et al. 2020. Fungal Diversity 105:1-16 ((https://doi.org/10.1007/s13225-020-00466-2)) |
| Otu14912 | 1 | k__Fungi(100);p__Ascomycota(100);c__Saccharomycetes(100);o__Saccharomycetales(100);f__Metschnikowiaceae(100);g__Metschnikowia(100);s__Metschnikowia_pulcherrima(100); | Metschnikowia | 13 | Saprotroph | \|Undefined Saprotroph\| | Yeast | NULL | Probable | As nectar-tap_saprotroph (Põlme et al. 2020); Decay sugar-rich substrates (Põlme et al. 2020) | Põlme S, et al. 2020. Fungal Diversity 105:1-16 ((https://doi.org/10.1007/s13225-020-00466-2)) |
| Otu14913 | 1 | k__Fungi(100);p__Ascomycota(100);c__Saccharomycetes(100);o__Saccharomycetales(100);f__Metschnikowiaceae(100);g__Metschnikowia(100);s__Metschnikowia_pulcherrima(100); | Metschnikowia | 13 | Saprotroph | \|Undefined Saprotroph\| | Yeast | NULL | Probable | As nectar-tap_saprotroph (Põlme et al. 2020); Decay sugar-rich substrates (Põlme et al. 2020) | Põlme S, et al. 2020. Fungal Diversity 105:1-16 ((https://doi.org/10.1007/s13225-020-00466-2)) |
| Otu14918 | 1 | k__Fungi(100);p__Ascomycota(100);c__Saccharomycetes(100);o__Saccharomycetales(100);f__Saccharomycetales_family_Incertae_sedis(100);f__Saccharomycetales_family_Incertae_sedis_unclassified(100);f__Saccharomycetales_family_Incertae_sedis_unclassified(100); | Saccharomycetales | 7 | Saprotroph | Undefined Saprotroph | Yeast | NULL | Possible | NULL | Sterkenburg E, et al. 2015. New Phytologist 207:1145-1158 ((https://doi.org/10.1111/nph.13426)) |
| Otu14919 | 1 | k__Fungi(100);p__Ascomycota(100);c__Saccharomycetes(100);o__Saccharomycetales(100);f__Metschnikowiaceae(100);g__Metschnikowia(100);s__Metschnikowia_pulcherrima(100); | Metschnikowia | 13 | Saprotroph | \|Undefined Saprotroph\| | Yeast | NULL | Probable | As nectar-tap_saprotroph (Põlme et al. 2020); Decay sugar-rich substrates (Põlme et al. 2020) | Põlme S, et al. 2020. Fungal Diversity 105:1-16 ((https://doi.org/10.1007/s13225-020-00466-2)) |
| Otu14923 | 1 | k__Fungi(100);p__Ascomycota(100);c__Saccharomycetes(100);o__Saccharomycetales(100);f__Metschnikowiaceae(100);g__Metschnikowia(100);s__Metschnikowia_pulcherrima(100); | Metschnikowia | 13 | Saprotroph | \|Undefined Saprotroph\| | Yeast | NULL | Probable | As nectar-tap_saprotroph (Põlme et al. 2020); Decay sugar-rich substrates (Põlme et al. 2020) | Põlme S, et al. 2020. Fungal Diversity 105:1-16 ((https://doi.org/10.1007/s13225-020-00466-2)) |
| Otu14924 | 1 | k__Fungi(100);p__Ascomycota(100);c__Saccharomycetes(100);o__Saccharomycetales(100);f__Metschnikowiaceae(100);g__Metschnikowia(100);s__Metschnikowia_pulcherrima(100); | Metschnikowia | 13 | Saprotroph | \|Undefined Saprotroph\| | Yeast | NULL | Probable | As nectar-tap_saprotroph (Põlme et al. 2020); Decay sugar-rich substrates (Põlme et al. 2020) | Põlme S, et al. 2020. Fungal Diversity 105:1-16 ((https://doi.org/10.1007/s13225-020-00466-2)) |
| Otu14925 | 1 | k__Fungi(100);p__Ascomycota(100);c__Saccharomycetes(100);o__Saccharomycetales(100);f__Metschnikowiaceae(100);g__Metschnikowia(100);s__Metschnikowia_pulcherrima(100); | Metschnikowia | 13 | Saprotroph | \|Undefined Saprotroph\| | Yeast | NULL | Probable | As nectar-tap_saprotroph (Põlme et al. 2020); Decay sugar-rich substrates (Põlme et al. 2020) | Põlme S, et al. 2020. Fungal Diversity 105:1-16 ((https://doi.org/10.1007/s13225-020-00466-2)) |
| Otu14927 | 1 | k__Fungi(100);p__Ascomycota(100);c__Saccharomycetes(100);o__Saccharomycetales(100);f__Metschnikowiaceae(100);g__Metschnikowia(100);s__Metschnikowia_pulcherrima(100); | Metschnikowia | 13 | Saprotroph | \|Undefined Saprotroph\| | Yeast | NULL | Probable | As nectar-tap_saprotroph (Põlme et al. 2020); Decay sugar-rich substrates (Põlme et al. 2020) | Põlme S, et al. 2020. Fungal Diversity 105:1-16 ((https://doi.org/10.1007/s13225-020-00466-2)) |
| Otu14929 | 1 | k__Fungi(100);p__Ascomycota(100);c__Saccharomycetes(100);o__Saccharomycetales(100);f__Metschnikowiaceae(100);g__Metschnikowia(100);s__Metschnikowia_pulcherrima(100); | Metschnikowia | 13 | Saprotroph | \|Undefined Saprotroph\| | Yeast | NULL | Probable | As nectar-tap_saprotroph (Põlme et al. 2020); Decay sugar-rich substrates (Põlme et al. 2020) | Põlme S, et al. 2020. Fungal Diversity 105:1-16 ((https://doi.org/10.1007/s13225-020-00466-2)) |
| Otu14930 | 1 | k__Fungi(100);p__Ascomycota(100);c__Saccharomycetes(100);o__Saccharomycetales(100);f__Metschnikowiaceae(100);g__Metschnikowia(100);g__Metschnikowia_unclassified(100); | Metschnikowia | 13 | Saprotroph | \|Undefined Saprotroph\| | Yeast | NULL | Probable | As nectar-tap_saprotroph (Põlme et al. 2020); Decay sugar-rich substrates (Põlme et al. 2020) | Põlme S, et al. 2020. Fungal Diversity 105:1-16 ((https://doi.org/10.1007/s13225-020-00466-2)) |
| Otu14933 | 1 | k__Fungi(100);p__Ascomycota(100);c__Saccharomycetes(100);o__Saccharomycetales(100);f__Metschnikowiaceae(100);g__Metschnikowia(100);s__Metschnikowia_pulcherrima(100); | Metschnikowia | 13 | Saprotroph | \|Undefined Saprotroph\| | Yeast | NULL | Probable | As nectar-tap_saprotroph (Põlme et al. 2020); Decay sugar-rich substrates (Põlme et al. 2020) | Põlme S, et al. 2020. Fungal Diversity 105:1-16 ((https://doi.org/10.1007/s13225-020-00466-2)) |
| Otu14935 | 1 | k__Fungi(100);p__Ascomycota(100);c__Saccharomycetes(100);o__Saccharomycetales(100);f__Metschnikowiaceae(100);g__Metschnikowia(100);g__Metschnikowia_unclassified(100); | Metschnikowia | 13 | Saprotroph | \|Undefined Saprotroph\| | Yeast | NULL | Probable | As nectar-tap_saprotroph (Põlme et al. 2020); Decay sugar-rich substrates (Põlme et al. 2020) | Põlme S, et al. 2020. Fungal Diversity 105:1-16 ((https://doi.org/10.1007/s13225-020-00466-2)) |
| Otu14937 | 1 | k__Fungi(100);p__Ascomycota(100);c__Saccharomycetes(100);o__Saccharomycetales(100);f__Metschnikowiaceae(100);g__Metschnikowia(100);s__Metschnikowia_pulcherrima(100); | Metschnikowia | 13 | Saprotroph | \|Undefined Saprotroph\| | Yeast | NULL | Probable | As nectar-tap_saprotroph (Põlme et al. 2020); Decay sugar-rich substrates (Põlme et al. 2020) | Põlme S, et al. 2020. Fungal Diversity 105:1-16 ((https://doi.org/10.1007/s13225-020-00466-2)) |
| Otu14938 | 1 | k__Fungi(100);p__Ascomycota(100);c__Saccharomycetes(100);o__Saccharomycetales(100);f__Metschnikowiaceae(100);g__Metschnikowia(100);s__Metschnikowia_pulcherrima(100); | Metschnikowia | 13 | Saprotroph | \|Undefined Saprotroph\| | Yeast | NULL | Probable | As nectar-tap_saprotroph (Põlme et al. 2020); Decay sugar-rich substrates (Põlme et al. 2020) | Põlme S, et al. 2020. Fungal Diversity 105:1-16 ((https://doi.org/10.1007/s13225-020-00466-2)) |
| Otu14939 | 1 | k__Fungi(100);p__Ascomycota(100);c__Saccharomycetes(100);o__Saccharomycetales(100);f__Metschnikowiaceae(100);g__Metschnikowia(100);s__Metschnikowia_pulcherrima(100); | Metschnikowia | 13 | Saprotroph | \|Undefined Saprotroph\| | Yeast | NULL | Probable | As nectar-tap_saprotroph (Põlme et al. 2020); Decay sugar-rich substrates (Põlme et al. 2020) | Põlme S, et al. 2020. Fungal Diversity 105:1-16 ((https://doi.org/10.1007/s13225-020-00466-2)) |
| Otu14940 | 1 | k__Fungi(100);p__Ascomycota(100);c__Saccharomycetes(100);o__Saccharomycetales(100);f__Metschnikowiaceae(100);g__Metschnikowia(100);s__Metschnikowia_pulcherrima(100); | Metschnikowia | 13 | Saprotroph | \|Undefined Saprotroph\| | Yeast | NULL | Probable | As nectar-tap_saprotroph (Põlme et al. 2020); Decay sugar-rich substrates (Põlme et al. 2020) | Põlme S, et al. 2020. Fungal Diversity 105:1-16 ((https://doi.org/10.1007/s13225-020-00466-2)) |
| Otu14943 | 1 | k__Fungi(100);p__Ascomycota(100);c__Saccharomycetes(100);o__Saccharomycetales(100);f__Metschnikowiaceae(100);g__Metschnikowia(100);s__Metschnikowia_pulcherrima(100); | Metschnikowia | 13 | Saprotroph | \|Undefined Saprotroph\| | Yeast | NULL | Probable | As nectar-tap_saprotroph (Põlme et al. 2020); Decay sugar-rich substrates (Põlme et al. 2020) | Põlme S, et al. 2020. Fungal Diversity 105:1-16 ((https://doi.org/10.1007/s13225-020-00466-2)) |
| Otu14946 | 1 | k__Fungi(100);p__Ascomycota(100);c__Saccharomycetes(100);o__Saccharomycetales(100);f__Metschnikowiaceae(100);g__Metschnikowia(100);s__Metschnikowia_pulcherrima(100); | Metschnikowia | 13 | Saprotroph | \|Undefined Saprotroph\| | Yeast | NULL | Probable | As nectar-tap_saprotroph (Põlme et al. 2020); Decay sugar-rich substrates (Põlme et al. 2020) | Põlme S, et al. 2020. Fungal Diversity 105:1-16 ((https://doi.org/10.1007/s13225-020-00466-2)) |
| Otu14948 | 1 | k__Fungi(100);p__Ascomycota(100);c__Saccharomycetes(100);o__Saccharomycetales(100);f__Metschnikowiaceae(100);g__Metschnikowia(100);g__Metschnikowia_unclassified(100); | Metschnikowia | 13 | Saprotroph | \|Undefined Saprotroph\| | Yeast | NULL | Probable | As nectar-tap_saprotroph (Põlme et al. 2020); Decay sugar-rich substrates (Põlme et al. 2020) | Põlme S, et al. 2020. Fungal Diversity 105:1-16 ((https://doi.org/10.1007/s13225-020-00466-2)) |
| Otu14949 | 1 | k__Fungi(100);p__Ascomycota(100);c__Saccharomycetes(100);o__Saccharomycetales(100);f__Metschnikowiaceae(100);g__Metschnikowia(100);g__Metschnikowia_unclassified(100); | Metschnikowia | 13 | Saprotroph | \|Undefined Saprotroph\| | Yeast | NULL | Probable | As nectar-tap_saprotroph (Põlme et al. 2020); Decay sugar-rich substrates (Põlme et al. 2020) | Põlme S, et al. 2020. Fungal Diversity 105:1-16 ((https://doi.org/10.1007/s13225-020-00466-2)) |
| Otu14951 | 1 | k__Fungi(100);p__Ascomycota(100);c__Saccharomycetes(100);o__Saccharomycetales(100);f__Saccharomycetales_family_Incertae_sedis(100);f__Saccharomycetales_family_Incertae_sedis_unclassified(100);f__Saccharomycetales_family_Incertae_sedis_unclassified(100); | Saccharomycetales | 7 | Saprotroph | Undefined Saprotroph | Yeast | NULL | Possible | NULL | Sterkenburg E, et al. 2015. New Phytologist 207:1145-1158 ((https://doi.org/10.1111/nph.13426)) |
| Otu14952 | 1 | k__Fungi(100);p__Ascomycota(100);c__Saccharomycetes(100);o__Saccharomycetales(100);f__Metschnikowiaceae(100);g__Metschnikowia(100);s__Metschnikowia_pulcherrima(100); | Metschnikowia | 13 | Saprotroph | \|Undefined Saprotroph\| | Yeast | NULL | Probable | As nectar-tap_saprotroph (Põlme et al. 2020); Decay sugar-rich substrates (Põlme et al. 2020) | Põlme S, et al. 2020. Fungal Diversity 105:1-16 ((https://doi.org/10.1007/s13225-020-00466-2)) |
| Otu14953 | 1 | k__Fungi(100);p__Ascomycota(100);c__Saccharomycetes(100);o__Saccharomycetales(100);f__Saccharomycetales_family_Incertae_sedis(100);f__Saccharomycetales_family_Incertae_sedis_unclassified(100);f__Saccharomycetales_family_Incertae_sedis_unclassified(100); | Saccharomycetales | 7 | Saprotroph | Undefined Saprotroph | Yeast | NULL | Possible | NULL | Sterkenburg E, et al. 2015. New Phytologist 207:1145-1158 ((https://doi.org/10.1111/nph.13426)) |
| Otu14957 | 1 | k__Fungi(100);p__Ascomycota(100);c__Saccharomycetes(100);o__Saccharomycetales(100);f__Metschnikowiaceae(100);g__Metschnikowia(100);s__Metschnikowia_pulcherrima(100); | Metschnikowia | 13 | Saprotroph | \|Undefined Saprotroph\| | Yeast | NULL | Probable | As nectar-tap_saprotroph (Põlme et al. 2020); Decay sugar-rich substrates (Põlme et al. 2020) | Põlme S, et al. 2020. Fungal Diversity 105:1-16 ((https://doi.org/10.1007/s13225-020-00466-2)) |
| Otu14959 | 1 | k__Fungi(100);p__Ascomycota(100);c__Saccharomycetes(100);o__Saccharomycetales(100);f__Metschnikowiaceae(100);g__Metschnikowia(100);s__Metschnikowia_pulcherrima(100); | Metschnikowia | 13 | Saprotroph | \|Undefined Saprotroph\| | Yeast | NULL | Probable | As nectar-tap_saprotroph (Põlme et al. 2020); Decay sugar-rich substrates (Põlme et al. 2020) | Põlme S, et al. 2020. Fungal Diversity 105:1-16 ((https://doi.org/10.1007/s13225-020-00466-2)) |
| Otu14960 | 1 | k__Fungi(100);p__Ascomycota(100);c__Saccharomycetes(100);o__Saccharomycetales(100);f__Metschnikowiaceae(100);g__Metschnikowia(100);g__Metschnikowia_unclassified(100); | Metschnikowia | 13 | Saprotroph | \|Undefined Saprotroph\| | Yeast | NULL | Probable | As nectar-tap_saprotroph (Põlme et al. 2020); Decay sugar-rich substrates (Põlme et al. 2020) | Põlme S, et al. 2020. Fungal Diversity 105:1-16 ((https://doi.org/10.1007/s13225-020-00466-2)) |
| Otu14964 | 1 | k__Fungi(100);p__Ascomycota(100);c__Saccharomycetes(100);o__Saccharomycetales(100);f__Saccharomycetales_family_Incertae_sedis(100);g__Candida(100);g__Candida_unclassified(100); | Candida | 13 | Pathotroph-Saprotroph-Symbiotroph | Animal Pathogen-Endophyte-Endosymbiont-Epiphyte-\|Undefined Saprotroph\| | Dimorphic Yeast-Yeast | NULL | Probable | As nectar-tap_saprotroph (Põlme et al. 2020); Decay sugar-rich substrates (Põlme et al. 2020); This asexual genus is polyphyletic and contain species that function very differently depend on what they are. Caution should be taken when interpreting guilds for Candida species. | Manolakaki D, et al. 2010. Virulence. 1: 367-375 ((https://doi.org/10.4161/viru.1.5.12796)); Mekbib SB, et al. 2011. Tropical Plant Pathology 36:233-240 ((https://doi.org/10.1590/S1982-56762011000400004)); Põlme S, et al. 2020. Fungal Diversity 105:1-16 ((https://doi.org/10.1007/s13225-020-00466-2)) |
| Otu14965 | 1 | k__Fungi(100);p__Ascomycota(100);c__Saccharomycetes(100);o__Saccharomycetales(100);f__Metschnikowiaceae(100);g__Metschnikowia(100);s__Metschnikowia_pulcherrima(100); | Metschnikowia | 13 | Saprotroph | \|Undefined Saprotroph\| | Yeast | NULL | Probable | As nectar-tap_saprotroph (Põlme et al. 2020); Decay sugar-rich substrates (Põlme et al. 2020) | Põlme S, et al. 2020. Fungal Diversity 105:1-16 ((https://doi.org/10.1007/s13225-020-00466-2)) |
| Otu14967 | 1 | k__Fungi(100);p__Ascomycota(100);c__Saccharomycetes(100);o__Saccharomycetales(100);f__Metschnikowiaceae(100);g__Metschnikowia(100);g__Metschnikowia_unclassified(100); | Metschnikowia | 13 | Saprotroph | \|Undefined Saprotroph\| | Yeast | NULL | Probable | As nectar-tap_saprotroph (Põlme et al. 2020); Decay sugar-rich substrates (Põlme et al. 2020) | Põlme S, et al. 2020. Fungal Diversity 105:1-16 ((https://doi.org/10.1007/s13225-020-00466-2)) |
| Otu14968 | 1 | k__Fungi(100);p__Ascomycota(100);c__Saccharomycetes(100);o__Saccharomycetales(100);o__Saccharomycetales_unclassified(100);o__Saccharomycetales_unclassified(100);o__Saccharomycetales_unclassified(100); | Saccharomycetales | 7 | Saprotroph | Undefined Saprotroph | Yeast | NULL | Possible | NULL | Sterkenburg E, et al. 2015. New Phytologist 207:1145-1158 ((https://doi.org/10.1111/nph.13426)) |
| Otu14969 | 1 | k__Fungi(100);p__Ascomycota(100);c__Saccharomycetes(100);o__Saccharomycetales(100);f__Saccharomycetales_family_Incertae_sedis(100);g__Candida(100);g__Candida_unclassified(100); | Candida | 13 | Pathotroph-Saprotroph-Symbiotroph | Animal Pathogen-Endophyte-Endosymbiont-Epiphyte-\|Undefined Saprotroph\| | Dimorphic Yeast-Yeast | NULL | Probable | As nectar-tap_saprotroph (Põlme et al. 2020); Decay sugar-rich substrates (Põlme et al. 2020); This asexual genus is polyphyletic and contain species that function very differently depend on what they are. Caution should be taken when interpreting guilds for Candida species. | Manolakaki D, et al. 2010. Virulence. 1: 367-375 ((https://doi.org/10.4161/viru.1.5.12796)); Mekbib SB, et al. 2011. Tropical Plant Pathology 36:233-240 ((https://doi.org/10.1590/S1982-56762011000400004)); Põlme S, et al. 2020. Fungal Diversity 105:1-16 ((https://doi.org/10.1007/s13225-020-00466-2)) |
| Otu14970 | 1 | k__Fungi(100);p__Ascomycota(100);c__Saccharomycetes(100);o__Saccharomycetales(100);f__Metschnikowiaceae(100);g__Metschnikowia(100);s__Metschnikowia_pulcherrima(100); | Metschnikowia | 13 | Saprotroph | \|Undefined Saprotroph\| | Yeast | NULL | Probable | As nectar-tap_saprotroph (Põlme et al. 2020); Decay sugar-rich substrates (Põlme et al. 2020) | Põlme S, et al. 2020. Fungal Diversity 105:1-16 ((https://doi.org/10.1007/s13225-020-00466-2)) |
| Otu14971 | 1 | k__Fungi(100);p__Ascomycota(100);c__Saccharomycetes(100);o__Saccharomycetales(100);f__Metschnikowiaceae(100);g__Metschnikowia(100);s__Metschnikowia_pulcherrima(100); | Metschnikowia | 13 | Saprotroph | \|Undefined Saprotroph\| | Yeast | NULL | Probable | As nectar-tap_saprotroph (Põlme et al. 2020); Decay sugar-rich substrates (Põlme et al. 2020) | Põlme S, et al. 2020. Fungal Diversity 105:1-16 ((https://doi.org/10.1007/s13225-020-00466-2)) |
| Otu14973 | 1 | k__Fungi(100);p__Ascomycota(100);c__Saccharomycetes(100);o__Saccharomycetales(100);f__Saccharomycetales_family_Incertae_sedis(100);f__Saccharomycetales_family_Incertae_sedis_unclassified(100);f__Saccharomycetales_family_Incertae_sedis_unclassified(100); | Saccharomycetales | 7 | Saprotroph | Undefined Saprotroph | Yeast | NULL | Possible | NULL | Sterkenburg E, et al. 2015. New Phytologist 207:1145-1158 ((https://doi.org/10.1111/nph.13426)) |
| Otu14974 | 1 | k__Fungi(100);p__Ascomycota(100);c__Saccharomycetes(100);o__Saccharomycetales(100);f__Metschnikowiaceae(100);g__Metschnikowia(100);s__Metschnikowia_pulcherrima(100); | Metschnikowia | 13 | Saprotroph | \|Undefined Saprotroph\| | Yeast | NULL | Probable | As nectar-tap_saprotroph (Põlme et al. 2020); Decay sugar-rich substrates (Põlme et al. 2020) | Põlme S, et al. 2020. Fungal Diversity 105:1-16 ((https://doi.org/10.1007/s13225-020-00466-2)) |
| Otu14975 | 1 | k__Fungi(100);p__Ascomycota(100);c__Saccharomycetes(100);o__Saccharomycetales(100);f__Metschnikowiaceae(100);g__Metschnikowia(100);s__Metschnikowia_pulcherrima(100); | Metschnikowia | 13 | Saprotroph | \|Undefined Saprotroph\| | Yeast | NULL | Probable | As nectar-tap_saprotroph (Põlme et al. 2020); Decay sugar-rich substrates (Põlme et al. 2020) | Põlme S, et al. 2020. Fungal Diversity 105:1-16 ((https://doi.org/10.1007/s13225-020-00466-2)) |
| Otu14976 | 1 | k__Fungi(100);p__Ascomycota(100);c__Saccharomycetes(100);o__Saccharomycetales(100);f__Metschnikowiaceae(100);g__Metschnikowia(100);s__Metschnikowia_pulcherrima(100); | Metschnikowia | 13 | Saprotroph | \|Undefined Saprotroph\| | Yeast | NULL | Probable | As nectar-tap_saprotroph (Põlme et al. 2020); Decay sugar-rich substrates (Põlme et al. 2020) | Põlme S, et al. 2020. Fungal Diversity 105:1-16 ((https://doi.org/10.1007/s13225-020-00466-2)) |
| Otu14977 | 1 | k__Fungi(100);p__Ascomycota(100);c__Saccharomycetes(100);o__Saccharomycetales(100);f__Metschnikowiaceae(100);g__Metschnikowia(100);s__Metschnikowia_pulcherrima(100); | Metschnikowia | 13 | Saprotroph | \|Undefined Saprotroph\| | Yeast | NULL | Probable | As nectar-tap_saprotroph (Põlme et al. 2020); Decay sugar-rich substrates (Põlme et al. 2020) | Põlme S, et al. 2020. Fungal Diversity 105:1-16 ((https://doi.org/10.1007/s13225-020-00466-2)) |
| Otu14979 | 1 | k__Fungi(100);p__Ascomycota(100);c__Saccharomycetes(100);o__Saccharomycetales(100);f__Metschnikowiaceae(100);g__Metschnikowia(100);s__Metschnikowia_pulcherrima(100); | Metschnikowia | 13 | Saprotroph | \|Undefined Saprotroph\| | Yeast | NULL | Probable | As nectar-tap_saprotroph (Põlme et al. 2020); Decay sugar-rich substrates (Põlme et al. 2020) | Põlme S, et al. 2020. Fungal Diversity 105:1-16 ((https://doi.org/10.1007/s13225-020-00466-2)) |
| Otu14980 | 1 | k__Fungi(100);p__Ascomycota(100);c__Saccharomycetes(100);o__Saccharomycetales(100);f__Metschnikowiaceae(100);g__Metschnikowia(100);s__Metschnikowia_pulcherrima(100); | Metschnikowia | 13 | Saprotroph | \|Undefined Saprotroph\| | Yeast | NULL | Probable | As nectar-tap_saprotroph (Põlme et al. 2020); Decay sugar-rich substrates (Põlme et al. 2020) | Põlme S, et al. 2020. Fungal Diversity 105:1-16 ((https://doi.org/10.1007/s13225-020-00466-2)) |
| Otu14982 | 1 | k__Fungi(100);p__Ascomycota(100);c__Saccharomycetes(100);o__Saccharomycetales(100);f__Metschnikowiaceae(100);g__Metschnikowia(100);s__Metschnikowia_pulcherrima(100); | Metschnikowia | 13 | Saprotroph | \|Undefined Saprotroph\| | Yeast | NULL | Probable | As nectar-tap_saprotroph (Põlme et al. 2020); Decay sugar-rich substrates (Põlme et al. 2020) | Põlme S, et al. 2020. Fungal Diversity 105:1-16 ((https://doi.org/10.1007/s13225-020-00466-2)) |
| Otu14984 | 1 | k__Fungi(100);p__Ascomycota(100);c__Saccharomycetes(100);o__Saccharomycetales(100);f__Metschnikowiaceae(100);g__Metschnikowia(100);g__Metschnikowia_unclassified(100); | Metschnikowia | 13 | Saprotroph | \|Undefined Saprotroph\| | Yeast | NULL | Probable | As nectar-tap_saprotroph (Põlme et al. 2020); Decay sugar-rich substrates (Põlme et al. 2020) | Põlme S, et al. 2020. Fungal Diversity 105:1-16 ((https://doi.org/10.1007/s13225-020-00466-2)) |
| Otu14985 | 1 | k__Fungi(100);p__Ascomycota(100);c__Saccharomycetes(100);o__Saccharomycetales(100);f__Metschnikowiaceae(100);g__Metschnikowia(100);s__Metschnikowia_pulcherrima(100); | Metschnikowia | 13 | Saprotroph | \|Undefined Saprotroph\| | Yeast | NULL | Probable | As nectar-tap_saprotroph (Põlme et al. 2020); Decay sugar-rich substrates (Põlme et al. 2020) | Põlme S, et al. 2020. Fungal Diversity 105:1-16 ((https://doi.org/10.1007/s13225-020-00466-2)) |
| Otu14986 | 1 | k__Fungi(100);p__Ascomycota(100);c__Saccharomycetes(100);o__Saccharomycetales(100);f__Metschnikowiaceae(100);g__Metschnikowia(100);g__Metschnikowia_unclassified(100); | Metschnikowia | 13 | Saprotroph | \|Undefined Saprotroph\| | Yeast | NULL | Probable | As nectar-tap_saprotroph (Põlme et al. 2020); Decay sugar-rich substrates (Põlme et al. 2020) | Põlme S, et al. 2020. Fungal Diversity 105:1-16 ((https://doi.org/10.1007/s13225-020-00466-2)) |
| Otu14987 | 1 | k__Fungi(100);p__Ascomycota(100);c__Saccharomycetes(100);o__Saccharomycetales(100);o__Saccharomycetales_unclassified(100);o__Saccharomycetales_unclassified(100);o__Saccharomycetales_unclassified(100); | Saccharomycetales | 7 | Saprotroph | Undefined Saprotroph | Yeast | NULL | Possible | NULL | Sterkenburg E, et al. 2015. New Phytologist 207:1145-1158 ((https://doi.org/10.1111/nph.13426)) |
| Otu14989 | 1 | k__Fungi(100);p__Ascomycota(100);c__Saccharomycetes(100);o__Saccharomycetales(100);f__Metschnikowiaceae(100);g__Metschnikowia(100);s__Metschnikowia_pulcherrima(100); | Metschnikowia | 13 | Saprotroph | \|Undefined Saprotroph\| | Yeast | NULL | Probable | As nectar-tap_saprotroph (Põlme et al. 2020); Decay sugar-rich substrates (Põlme et al. 2020) | Põlme S, et al. 2020. Fungal Diversity 105:1-16 ((https://doi.org/10.1007/s13225-020-00466-2)) |
| Otu14993 | 1 | k__Fungi(100);p__Ascomycota(100);c__Saccharomycetes(100);o__Saccharomycetales(100);f__Saccharomycetales_family_Incertae_sedis(100);g__Candida(100);g__Candida_unclassified(100); | Candida | 13 | Pathotroph-Saprotroph-Symbiotroph | Animal Pathogen-Endophyte-Endosymbiont-Epiphyte-\|Undefined Saprotroph\| | Dimorphic Yeast-Yeast | NULL | Probable | As nectar-tap_saprotroph (Põlme et al. 2020); Decay sugar-rich substrates (Põlme et al. 2020); This asexual genus is polyphyletic and contain species that function very differently depend on what they are. Caution should be taken when interpreting guilds for Candida species. | Manolakaki D, et al. 2010. Virulence. 1: 367-375 ((https://doi.org/10.4161/viru.1.5.12796)); Mekbib SB, et al. 2011. Tropical Plant Pathology 36:233-240 ((https://doi.org/10.1590/S1982-56762011000400004)); Põlme S, et al. 2020. Fungal Diversity 105:1-16 ((https://doi.org/10.1007/s13225-020-00466-2)) |
| Otu14995 | 1 | k__Fungi(100);p__Ascomycota(100);c__Saccharomycetes(100);o__Saccharomycetales(100);f__Metschnikowiaceae(100);g__Metschnikowia(100);g__Metschnikowia_unclassified(100); | Metschnikowia | 13 | Saprotroph | \|Undefined Saprotroph\| | Yeast | NULL | Probable | As nectar-tap_saprotroph (Põlme et al. 2020); Decay sugar-rich substrates (Põlme et al. 2020) | Põlme S, et al. 2020. Fungal Diversity 105:1-16 ((https://doi.org/10.1007/s13225-020-00466-2)) |
| Otu14997 | 1 | k__Fungi(100);p__Ascomycota(100);c__Saccharomycetes(100);o__Saccharomycetales(100);f__Saccharomycetales_family_Incertae_sedis(100);g__Candida(100);g__Candida_unclassified(100); | Candida | 13 | Pathotroph-Saprotroph-Symbiotroph | Animal Pathogen-Endophyte-Endosymbiont-Epiphyte-\|Undefined Saprotroph\| | Dimorphic Yeast-Yeast | NULL | Probable | As nectar-tap_saprotroph (Põlme et al. 2020); Decay sugar-rich substrates (Põlme et al. 2020); This asexual genus is polyphyletic and contain species that function very differently depend on what they are. Caution should be taken when interpreting guilds for Candida species. | Manolakaki D, et al. 2010. Virulence. 1: 367-375 ((https://doi.org/10.4161/viru.1.5.12796)); Mekbib SB, et al. 2011. Tropical Plant Pathology 36:233-240 ((https://doi.org/10.1590/S1982-56762011000400004)); Põlme S, et al. 2020. Fungal Diversity 105:1-16 ((https://doi.org/10.1007/s13225-020-00466-2)) |
| Otu14999 | 1 | k__Fungi(100);p__Ascomycota(100);c__Saccharomycetes(100);o__Saccharomycetales(100);f__Metschnikowiaceae(100);g__Metschnikowia(100);s__Metschnikowia_pulcherrima(100); | Metschnikowia | 13 | Saprotroph | \|Undefined Saprotroph\| | Yeast | NULL | Probable | As nectar-tap_saprotroph (Põlme et al. 2020); Decay sugar-rich substrates (Põlme et al. 2020) | Põlme S, et al. 2020. Fungal Diversity 105:1-16 ((https://doi.org/10.1007/s13225-020-00466-2)) |
| Otu15002 | 1 | k__Fungi(100);p__Ascomycota(100);c__Saccharomycetes(100);o__Saccharomycetales(100);f__Metschnikowiaceae(100);g__Metschnikowia(100);s__Metschnikowia_pulcherrima(100); | Metschnikowia | 13 | Saprotroph | \|Undefined Saprotroph\| | Yeast | NULL | Probable | As nectar-tap_saprotroph (Põlme et al. 2020); Decay sugar-rich substrates (Põlme et al. 2020) | Põlme S, et al. 2020. Fungal Diversity 105:1-16 ((https://doi.org/10.1007/s13225-020-00466-2)) |
| Otu15005 | 1 | k__Fungi(100);p__Ascomycota(100);c__Leotiomycetes(100);o__Helotiales(100);f__Helotiales_family_Incertae_sedis(100);g__Infundichalara(100);s__Infundichalara_sp(100); | Infundichalara | 13 | Saprotroph | \|Plant Saprotroph\|-Undefined Saprotroph | Microfungus | NULL | Probable | As litter_saprotroph (Põlme et al. 2020); Decay leaf, fruit, seed (Põlme et al. 2020); Host - Pinus (Põlme et al. 2020) | Tedersoo L, et al. 2014. Science 346:e1256688 ((https://doi.org/10.1126/science.1256688)); Põlme S, et al. 2020. Fungal Diversity 105:1-16 ((https://doi.org/10.1007/s13225-020-00466-2)) |
| Otu15007 | 1 | k__Fungi(100);p__Ascomycota(100);c__Saccharomycetes(100);o__Saccharomycetales(100);f__Metschnikowiaceae(100);g__Metschnikowia(100);s__Metschnikowia_pulcherrima(100); | Metschnikowia | 13 | Saprotroph | \|Undefined Saprotroph\| | Yeast | NULL | Probable | As nectar-tap_saprotroph (Põlme et al. 2020); Decay sugar-rich substrates (Põlme et al. 2020) | Põlme S, et al. 2020. Fungal Diversity 105:1-16 ((https://doi.org/10.1007/s13225-020-00466-2)) |
| Otu15012 | 1 | k__Fungi(100);p__Ascomycota(100);c__Saccharomycetes(100);o__Saccharomycetales(100);f__Metschnikowiaceae(100);g__Metschnikowia(100);s__Metschnikowia_pulcherrima(100); | Metschnikowia | 13 | Saprotroph | \|Undefined Saprotroph\| | Yeast | NULL | Probable | As nectar-tap_saprotroph (Põlme et al. 2020); Decay sugar-rich substrates (Põlme et al. 2020) | Põlme S, et al. 2020. Fungal Diversity 105:1-16 ((https://doi.org/10.1007/s13225-020-00466-2)) |
| Otu15013 | 1 | k__Fungi(100);p__Ascomycota(100);c__Saccharomycetes(100);o__Saccharomycetales(100);f__Metschnikowiaceae(100);g__Metschnikowia(100);g__Metschnikowia_unclassified(100); | Metschnikowia | 13 | Saprotroph | \|Undefined Saprotroph\| | Yeast | NULL | Probable | As nectar-tap_saprotroph (Põlme et al. 2020); Decay sugar-rich substrates (Põlme et al. 2020) | Põlme S, et al. 2020. Fungal Diversity 105:1-16 ((https://doi.org/10.1007/s13225-020-00466-2)) |
| Otu15014 | 1 | k__Fungi(100);p__Ascomycota(100);c__Saccharomycetes(100);o__Saccharomycetales(100);f__Metschnikowiaceae(100);g__Metschnikowia(100);g__Metschnikowia_unclassified(100); | Metschnikowia | 13 | Saprotroph | \|Undefined Saprotroph\| | Yeast | NULL | Probable | As nectar-tap_saprotroph (Põlme et al. 2020); Decay sugar-rich substrates (Põlme et al. 2020) | Põlme S, et al. 2020. Fungal Diversity 105:1-16 ((https://doi.org/10.1007/s13225-020-00466-2)) |
| Otu15016 | 1 | k__Fungi(100);p__Ascomycota(100);c__Saccharomycetes(100);o__Saccharomycetales(100);f__Metschnikowiaceae(100);g__Metschnikowia(100);s__Metschnikowia_pulcherrima(100); | Metschnikowia | 13 | Saprotroph | \|Undefined Saprotroph\| | Yeast | NULL | Probable | As nectar-tap_saprotroph (Põlme et al. 2020); Decay sugar-rich substrates (Põlme et al. 2020) | Põlme S, et al. 2020. Fungal Diversity 105:1-16 ((https://doi.org/10.1007/s13225-020-00466-2)) |
| Otu15020 | 1 | k__Fungi(100);p__Ascomycota(100);c__Saccharomycetes(100);o__Saccharomycetales(100);f__Metschnikowiaceae(100);g__Metschnikowia(100);s__Metschnikowia_pulcherrima(100); | Metschnikowia | 13 | Saprotroph | \|Undefined Saprotroph\| | Yeast | NULL | Probable | As nectar-tap_saprotroph (Põlme et al. 2020); Decay sugar-rich substrates (Põlme et al. 2020) | Põlme S, et al. 2020. Fungal Diversity 105:1-16 ((https://doi.org/10.1007/s13225-020-00466-2)) |
| Otu15021 | 1 | k__Fungi(100);p__Ascomycota(100);c__Saccharomycetes(100);o__Saccharomycetales(100);f__Metschnikowiaceae(100);g__Metschnikowia(100);g__Metschnikowia_unclassified(100); | Metschnikowia | 13 | Saprotroph | \|Undefined Saprotroph\| | Yeast | NULL | Probable | As nectar-tap_saprotroph (Põlme et al. 2020); Decay sugar-rich substrates (Põlme et al. 2020) | Põlme S, et al. 2020. Fungal Diversity 105:1-16 ((https://doi.org/10.1007/s13225-020-00466-2)) |
| Otu15025 | 1 | k__Fungi(100);p__Ascomycota(100);c__Saccharomycetes(100);o__Saccharomycetales(100);f__Metschnikowiaceae(100);g__Metschnikowia(100);s__Metschnikowia_pulcherrima(100); | Metschnikowia | 13 | Saprotroph | \|Undefined Saprotroph\| | Yeast | NULL | Probable | As nectar-tap_saprotroph (Põlme et al. 2020); Decay sugar-rich substrates (Põlme et al. 2020) | Põlme S, et al. 2020. Fungal Diversity 105:1-16 ((https://doi.org/10.1007/s13225-020-00466-2)) |
| Otu15026 | 1 | k__Fungi(100);p__Ascomycota(100);c__Saccharomycetes(100);o__Saccharomycetales(100);f__Metschnikowiaceae(100);g__Metschnikowia(100);s__Metschnikowia_pulcherrima(100); | Metschnikowia | 13 | Saprotroph | \|Undefined Saprotroph\| | Yeast | NULL | Probable | As nectar-tap_saprotroph (Põlme et al. 2020); Decay sugar-rich substrates (Põlme et al. 2020) | Põlme S, et al. 2020. Fungal Diversity 105:1-16 ((https://doi.org/10.1007/s13225-020-00466-2)) |
| Otu15027 | 1 | k__Fungi(100);p__Ascomycota(100);c__Saccharomycetes(100);o__Saccharomycetales(100);f__Metschnikowiaceae(100);g__Metschnikowia(100);s__Metschnikowia_pulcherrima(100); | Metschnikowia | 13 | Saprotroph | \|Undefined Saprotroph\| | Yeast | NULL | Probable | As nectar-tap_saprotroph (Põlme et al. 2020); Decay sugar-rich substrates (Põlme et al. 2020) | Põlme S, et al. 2020. Fungal Diversity 105:1-16 ((https://doi.org/10.1007/s13225-020-00466-2)) |
| Otu15028 | 1 | k__Fungi(100);p__Ascomycota(100);c__Saccharomycetes(100);o__Saccharomycetales(100);f__Metschnikowiaceae(100);g__Metschnikowia(100);s__Metschnikowia_pulcherrima(100); | Metschnikowia | 13 | Saprotroph | \|Undefined Saprotroph\| | Yeast | NULL | Probable | As nectar-tap_saprotroph (Põlme et al. 2020); Decay sugar-rich substrates (Põlme et al. 2020) | Põlme S, et al. 2020. Fungal Diversity 105:1-16 ((https://doi.org/10.1007/s13225-020-00466-2)) |
| Otu15031 | 1 | k__Fungi(100);p__Ascomycota(100);c__Saccharomycetes(100);o__Saccharomycetales(100);f__Metschnikowiaceae(100);g__Metschnikowia(100);s__Metschnikowia_pulcherrima(100); | Metschnikowia | 13 | Saprotroph | \|Undefined Saprotroph\| | Yeast | NULL | Probable | As nectar-tap_saprotroph (Põlme et al. 2020); Decay sugar-rich substrates (Põlme et al. 2020) | Põlme S, et al. 2020. Fungal Diversity 105:1-16 ((https://doi.org/10.1007/s13225-020-00466-2)) |
| Otu15032 | 1 | k__Fungi(100);p__Ascomycota(100);c__Saccharomycetes(100);o__Saccharomycetales(100);f__Metschnikowiaceae(100);g__Metschnikowia(100);s__Metschnikowia_pulcherrima(100); | Metschnikowia | 13 | Saprotroph | \|Undefined Saprotroph\| | Yeast | NULL | Probable | As nectar-tap_saprotroph (Põlme et al. 2020); Decay sugar-rich substrates (Põlme et al. 2020) | Põlme S, et al. 2020. Fungal Diversity 105:1-16 ((https://doi.org/10.1007/s13225-020-00466-2)) |
| Otu15033 | 1 | k__Fungi(100);p__Ascomycota(100);c__Saccharomycetes(100);o__Saccharomycetales(100);f__Metschnikowiaceae(100);g__Metschnikowia(100);g__Metschnikowia_unclassified(100); | Metschnikowia | 13 | Saprotroph | \|Undefined Saprotroph\| | Yeast | NULL | Probable | As nectar-tap_saprotroph (Põlme et al. 2020); Decay sugar-rich substrates (Põlme et al. 2020) | Põlme S, et al. 2020. Fungal Diversity 105:1-16 ((https://doi.org/10.1007/s13225-020-00466-2)) |
| Otu15034 | 1 | k__Fungi(100);p__Ascomycota(100);c__Saccharomycetes(100);o__Saccharomycetales(100);o__Saccharomycetales_unclassified(100);o__Saccharomycetales_unclassified(100);o__Saccharomycetales_unclassified(100); | Saccharomycetales | 7 | Saprotroph | Undefined Saprotroph | Yeast | NULL | Possible | NULL | Sterkenburg E, et al. 2015. New Phytologist 207:1145-1158 ((https://doi.org/10.1111/nph.13426)) |
| Otu15035 | 1 | k__Fungi(100);p__Ascomycota(100);c__Saccharomycetes(100);o__Saccharomycetales(100);f__Metschnikowiaceae(100);g__Metschnikowia(100);g__Metschnikowia_unclassified(100); | Metschnikowia | 13 | Saprotroph | \|Undefined Saprotroph\| | Yeast | NULL | Probable | As nectar-tap_saprotroph (Põlme et al. 2020); Decay sugar-rich substrates (Põlme et al. 2020) | Põlme S, et al. 2020. Fungal Diversity 105:1-16 ((https://doi.org/10.1007/s13225-020-00466-2)) |
| Otu15036 | 1 | k__Fungi(100);p__Ascomycota(100);c__Saccharomycetes(100);o__Saccharomycetales(100);f__Metschnikowiaceae(100);g__Metschnikowia(100);s__Metschnikowia_pulcherrima(100); | Metschnikowia | 13 | Saprotroph | \|Undefined Saprotroph\| | Yeast | NULL | Probable | As nectar-tap_saprotroph (Põlme et al. 2020); Decay sugar-rich substrates (Põlme et al. 2020) | Põlme S, et al. 2020. Fungal Diversity 105:1-16 ((https://doi.org/10.1007/s13225-020-00466-2)) |
| Otu15037 | 1 | k__Fungi(100);p__Ascomycota(100);c__Saccharomycetes(100);o__Saccharomycetales(100);f__Saccharomycetales_family_Incertae_sedis(100);g__Candida(100);g__Candida_unclassified(100); | Candida | 13 | Pathotroph-Saprotroph-Symbiotroph | Animal Pathogen-Endophyte-Endosymbiont-Epiphyte-\|Undefined Saprotroph\| | Dimorphic Yeast-Yeast | NULL | Probable | As nectar-tap_saprotroph (Põlme et al. 2020); Decay sugar-rich substrates (Põlme et al. 2020); This asexual genus is polyphyletic and contain species that function very differently depend on what they are. Caution should be taken when interpreting guilds for Candida species. | Manolakaki D, et al. 2010. Virulence. 1: 367-375 ((https://doi.org/10.4161/viru.1.5.12796)); Mekbib SB, et al. 2011. Tropical Plant Pathology 36:233-240 ((https://doi.org/10.1590/S1982-56762011000400004)); Põlme S, et al. 2020. Fungal Diversity 105:1-16 ((https://doi.org/10.1007/s13225-020-00466-2)) |
| Otu15038 | 1 | k__Fungi(100);p__Ascomycota(100);c__Saccharomycetes(100);o__Saccharomycetales(100);f__Metschnikowiaceae(100);g__Metschnikowia(100);s__Metschnikowia_pulcherrima(100); | Metschnikowia | 13 | Saprotroph | \|Undefined Saprotroph\| | Yeast | NULL | Probable | As nectar-tap_saprotroph (Põlme et al. 2020); Decay sugar-rich substrates (Põlme et al. 2020) | Põlme S, et al. 2020. Fungal Diversity 105:1-16 ((https://doi.org/10.1007/s13225-020-00466-2)) |
| Otu15039 | 1 | k__Fungi(100);p__Ascomycota(100);c__Saccharomycetes(100);o__Saccharomycetales(100);f__Metschnikowiaceae(100);g__Metschnikowia(100);s__Metschnikowia_pulcherrima(100); | Metschnikowia | 13 | Saprotroph | \|Undefined Saprotroph\| | Yeast | NULL | Probable | As nectar-tap_saprotroph (Põlme et al. 2020); Decay sugar-rich substrates (Põlme et al. 2020) | Põlme S, et al. 2020. Fungal Diversity 105:1-16 ((https://doi.org/10.1007/s13225-020-00466-2)) |
| Otu15040 | 1 | k__Fungi(100);p__Ascomycota(100);c__Saccharomycetes(100);o__Saccharomycetales(100);f__Metschnikowiaceae(100);g__Metschnikowia(100);s__Metschnikowia_pulcherrima(100); | Metschnikowia | 13 | Saprotroph | \|Undefined Saprotroph\| | Yeast | NULL | Probable | As nectar-tap_saprotroph (Põlme et al. 2020); Decay sugar-rich substrates (Põlme et al. 2020) | Põlme S, et al. 2020. Fungal Diversity 105:1-16 ((https://doi.org/10.1007/s13225-020-00466-2)) |
| Otu15041 | 1 | k__Fungi(100);p__Ascomycota(100);c__Saccharomycetes(100);o__Saccharomycetales(100);f__Metschnikowiaceae(100);g__Metschnikowia(100);s__Metschnikowia_pulcherrima(100); | Metschnikowia | 13 | Saprotroph | \|Undefined Saprotroph\| | Yeast | NULL | Probable | As nectar-tap_saprotroph (Põlme et al. 2020); Decay sugar-rich substrates (Põlme et al. 2020) | Põlme S, et al. 2020. Fungal Diversity 105:1-16 ((https://doi.org/10.1007/s13225-020-00466-2)) |
| Otu15042 | 1 | k__Fungi(100);p__Ascomycota(100);c__Saccharomycetes(100);o__Saccharomycetales(100);f__Metschnikowiaceae(100);g__Metschnikowia(100);g__Metschnikowia_unclassified(100); | Metschnikowia | 13 | Saprotroph | \|Undefined Saprotroph\| | Yeast | NULL | Probable | As nectar-tap_saprotroph (Põlme et al. 2020); Decay sugar-rich substrates (Põlme et al. 2020) | Põlme S, et al. 2020. Fungal Diversity 105:1-16 ((https://doi.org/10.1007/s13225-020-00466-2)) |
| Otu15043 | 1 | k__Fungi(100);p__Ascomycota(100);c__Saccharomycetes(100);o__Saccharomycetales(100);f__Metschnikowiaceae(100);g__Metschnikowia(100);s__Metschnikowia_pulcherrima(100); | Metschnikowia | 13 | Saprotroph | \|Undefined Saprotroph\| | Yeast | NULL | Probable | As nectar-tap_saprotroph (Põlme et al. 2020); Decay sugar-rich substrates (Põlme et al. 2020) | Põlme S, et al. 2020. Fungal Diversity 105:1-16 ((https://doi.org/10.1007/s13225-020-00466-2)) |
| Otu15046 | 1 | k__Fungi(100);p__Ascomycota(100);c__Saccharomycetes(100);o__Saccharomycetales(100);f__Metschnikowiaceae(100);g__Metschnikowia(100);s__Metschnikowia_pulcherrima(100); | Metschnikowia | 13 | Saprotroph | \|Undefined Saprotroph\| | Yeast | NULL | Probable | As nectar-tap_saprotroph (Põlme et al. 2020); Decay sugar-rich substrates (Põlme et al. 2020) | Põlme S, et al. 2020. Fungal Diversity 105:1-16 ((https://doi.org/10.1007/s13225-020-00466-2)) |
| Otu15048 | 1 | k__Fungi(100);p__Ascomycota(100);c__Saccharomycetes(100);o__Saccharomycetales(100);f__Metschnikowiaceae(100);g__Metschnikowia(100);g__Metschnikowia_unclassified(100); | Metschnikowia | 13 | Saprotroph | \|Undefined Saprotroph\| | Yeast | NULL | Probable | As nectar-tap_saprotroph (Põlme et al. 2020); Decay sugar-rich substrates (Põlme et al. 2020) | Põlme S, et al. 2020. Fungal Diversity 105:1-16 ((https://doi.org/10.1007/s13225-020-00466-2)) |
| Otu15049 | 1 | k__Fungi(100);p__Ascomycota(100);c__Saccharomycetes(100);o__Saccharomycetales(100);f__Metschnikowiaceae(100);g__Metschnikowia(100);s__Metschnikowia_pulcherrima(100); | Metschnikowia | 13 | Saprotroph | \|Undefined Saprotroph\| | Yeast | NULL | Probable | As nectar-tap_saprotroph (Põlme et al. 2020); Decay sugar-rich substrates (Põlme et al. 2020) | Põlme S, et al. 2020. Fungal Diversity 105:1-16 ((https://doi.org/10.1007/s13225-020-00466-2)) |
| Otu15050 | 1 | k__Fungi(100);p__Ascomycota(100);c__Saccharomycetes(100);o__Saccharomycetales(100);f__Metschnikowiaceae(100);g__Metschnikowia(100);s__Metschnikowia_pulcherrima(100); | Metschnikowia | 13 | Saprotroph | \|Undefined Saprotroph\| | Yeast | NULL | Probable | As nectar-tap_saprotroph (Põlme et al. 2020); Decay sugar-rich substrates (Põlme et al. 2020) | Põlme S, et al. 2020. Fungal Diversity 105:1-16 ((https://doi.org/10.1007/s13225-020-00466-2)) |
| Otu15051 | 1 | k__Fungi(100);p__Ascomycota(100);c__Saccharomycetes(100);o__Saccharomycetales(100);f__Metschnikowiaceae(100);g__Metschnikowia(100);s__Metschnikowia_pulcherrima(100); | Metschnikowia | 13 | Saprotroph | \|Undefined Saprotroph\| | Yeast | NULL | Probable | As nectar-tap_saprotroph (Põlme et al. 2020); Decay sugar-rich substrates (Põlme et al. 2020) | Põlme S, et al. 2020. Fungal Diversity 105:1-16 ((https://doi.org/10.1007/s13225-020-00466-2)) |
| Otu15052 | 1 | k__Fungi(100);p__Ascomycota(100);c__Saccharomycetes(100);o__Saccharomycetales(100);f__Metschnikowiaceae(100);g__Metschnikowia(100);g__Metschnikowia_unclassified(100); | Metschnikowia | 13 | Saprotroph | \|Undefined Saprotroph\| | Yeast | NULL | Probable | As nectar-tap_saprotroph (Põlme et al. 2020); Decay sugar-rich substrates (Põlme et al. 2020) | Põlme S, et al. 2020. Fungal Diversity 105:1-16 ((https://doi.org/10.1007/s13225-020-00466-2)) |
| Otu15053 | 1 | k__Fungi(100);p__Ascomycota(100);c__Saccharomycetes(100);o__Saccharomycetales(100);f__Metschnikowiaceae(100);g__Metschnikowia(100);s__Metschnikowia_pulcherrima(100); | Metschnikowia | 13 | Saprotroph | \|Undefined Saprotroph\| | Yeast | NULL | Probable | As nectar-tap_saprotroph (Põlme et al. 2020); Decay sugar-rich substrates (Põlme et al. 2020) | Põlme S, et al. 2020. Fungal Diversity 105:1-16 ((https://doi.org/10.1007/s13225-020-00466-2)) |
| Otu15054 | 1 | k__Fungi(100);p__Ascomycota(100);c__Saccharomycetes(100);o__Saccharomycetales(100);f__Metschnikowiaceae(100);g__Metschnikowia(100);s__Metschnikowia_pulcherrima(100); | Metschnikowia | 13 | Saprotroph | \|Undefined Saprotroph\| | Yeast | NULL | Probable | As nectar-tap_saprotroph (Põlme et al. 2020); Decay sugar-rich substrates (Põlme et al. 2020) | Põlme S, et al. 2020. Fungal Diversity 105:1-16 ((https://doi.org/10.1007/s13225-020-00466-2)) |
| Otu15055 | 1 | k__Fungi(100);p__Ascomycota(100);c__Saccharomycetes(100);o__Saccharomycetales(100);f__Metschnikowiaceae(100);g__Metschnikowia(100);s__Metschnikowia_pulcherrima(100); | Metschnikowia | 13 | Saprotroph | \|Undefined Saprotroph\| | Yeast | NULL | Probable | As nectar-tap_saprotroph (Põlme et al. 2020); Decay sugar-rich substrates (Põlme et al. 2020) | Põlme S, et al. 2020. Fungal Diversity 105:1-16 ((https://doi.org/10.1007/s13225-020-00466-2)) |
| Otu15057 | 1 | k__Fungi(100);p__Ascomycota(100);c__Saccharomycetes(100);o__Saccharomycetales(100);f__Metschnikowiaceae(100);g__Metschnikowia(100);s__Metschnikowia_pulcherrima(100); | Metschnikowia | 13 | Saprotroph | \|Undefined Saprotroph\| | Yeast | NULL | Probable | As nectar-tap_saprotroph (Põlme et al. 2020); Decay sugar-rich substrates (Põlme et al. 2020) | Põlme S, et al. 2020. Fungal Diversity 105:1-16 ((https://doi.org/10.1007/s13225-020-00466-2)) |
| Otu15058 | 1 | k__Fungi(100);p__Ascomycota(100);c__Saccharomycetes(100);o__Saccharomycetales(100);f__Metschnikowiaceae(100);g__Metschnikowia(100);s__Metschnikowia_pulcherrima(100); | Metschnikowia | 13 | Saprotroph | \|Undefined Saprotroph\| | Yeast | NULL | Probable | As nectar-tap_saprotroph (Põlme et al. 2020); Decay sugar-rich substrates (Põlme et al. 2020) | Põlme S, et al. 2020. Fungal Diversity 105:1-16 ((https://doi.org/10.1007/s13225-020-00466-2)) |
| Otu15061 | 1 | k__Fungi(100);p__Ascomycota(100);c__Saccharomycetes(100);o__Saccharomycetales(100);f__Metschnikowiaceae(100);g__Metschnikowia(100);s__Metschnikowia_pulcherrima(100); | Metschnikowia | 13 | Saprotroph | \|Undefined Saprotroph\| | Yeast | NULL | Probable | As nectar-tap_saprotroph (Põlme et al. 2020); Decay sugar-rich substrates (Põlme et al. 2020) | Põlme S, et al. 2020. Fungal Diversity 105:1-16 ((https://doi.org/10.1007/s13225-020-00466-2)) |
| Otu15062 | 1 | k__Fungi(100);p__Ascomycota(100);c__Saccharomycetes(100);o__Saccharomycetales(100);f__Metschnikowiaceae(100);g__Metschnikowia(100);s__Metschnikowia_pulcherrima(100); | Metschnikowia | 13 | Saprotroph | \|Undefined Saprotroph\| | Yeast | NULL | Probable | As nectar-tap_saprotroph (Põlme et al. 2020); Decay sugar-rich substrates (Põlme et al. 2020) | Põlme S, et al. 2020. Fungal Diversity 105:1-16 ((https://doi.org/10.1007/s13225-020-00466-2)) |
| Otu15063 | 1 | k__Fungi(100);p__Ascomycota(100);c__Saccharomycetes(100);o__Saccharomycetales(100);f__Metschnikowiaceae(100);g__Metschnikowia(100);g__Metschnikowia_unclassified(100); | Metschnikowia | 13 | Saprotroph | \|Undefined Saprotroph\| | Yeast | NULL | Probable | As nectar-tap_saprotroph (Põlme et al. 2020); Decay sugar-rich substrates (Põlme et al. 2020) | Põlme S, et al. 2020. Fungal Diversity 105:1-16 ((https://doi.org/10.1007/s13225-020-00466-2)) |
| Otu15065 | 1 | k__Fungi(100);p__Ascomycota(100);c__Saccharomycetes(100);o__Saccharomycetales(100);f__Metschnikowiaceae(100);g__Metschnikowia(100);s__Metschnikowia_pulcherrima(100); | Metschnikowia | 13 | Saprotroph | \|Undefined Saprotroph\| | Yeast | NULL | Probable | As nectar-tap_saprotroph (Põlme et al. 2020); Decay sugar-rich substrates (Põlme et al. 2020) | Põlme S, et al. 2020. Fungal Diversity 105:1-16 ((https://doi.org/10.1007/s13225-020-00466-2)) |
| Otu15066 | 1 | k__Fungi(100);p__Ascomycota(100);c__Saccharomycetes(100);o__Saccharomycetales(100);f__Metschnikowiaceae(100);g__Metschnikowia(100);s__Metschnikowia_pulcherrima(100); | Metschnikowia | 13 | Saprotroph | \|Undefined Saprotroph\| | Yeast | NULL | Probable | As nectar-tap_saprotroph (Põlme et al. 2020); Decay sugar-rich substrates (Põlme et al. 2020) | Põlme S, et al. 2020. Fungal Diversity 105:1-16 ((https://doi.org/10.1007/s13225-020-00466-2)) |
| Otu15068 | 1 | k__Fungi(100);p__Ascomycota(100);c__Saccharomycetes(100);o__Saccharomycetales(100);f__Metschnikowiaceae(100);g__Metschnikowia(100);g__Metschnikowia_unclassified(100); | Metschnikowia | 13 | Saprotroph | \|Undefined Saprotroph\| | Yeast | NULL | Probable | As nectar-tap_saprotroph (Põlme et al. 2020); Decay sugar-rich substrates (Põlme et al. 2020) | Põlme S, et al. 2020. Fungal Diversity 105:1-16 ((https://doi.org/10.1007/s13225-020-00466-2)) |
| Otu15069 | 1 | k__Fungi(100);p__Ascomycota(100);c__Saccharomycetes(100);o__Saccharomycetales(100);f__Saccharomycetales_family_Incertae_sedis(100);g__Candida(100);g__Candida_unclassified(100); | Candida | 13 | Pathotroph-Saprotroph-Symbiotroph | Animal Pathogen-Endophyte-Endosymbiont-Epiphyte-\|Undefined Saprotroph\| | Dimorphic Yeast-Yeast | NULL | Probable | As nectar-tap_saprotroph (Põlme et al. 2020); Decay sugar-rich substrates (Põlme et al. 2020); This asexual genus is polyphyletic and contain species that function very differently depend on what they are. Caution should be taken when interpreting guilds for Candida species. | Manolakaki D, et al. 2010. Virulence. 1: 367-375 ((https://doi.org/10.4161/viru.1.5.12796)); Mekbib SB, et al. 2011. Tropical Plant Pathology 36:233-240 ((https://doi.org/10.1590/S1982-56762011000400004)); Põlme S, et al. 2020. Fungal Diversity 105:1-16 ((https://doi.org/10.1007/s13225-020-00466-2)) |
| Otu15070 | 1 | k__Fungi(100);p__Ascomycota(100);c__Saccharomycetes(100);o__Saccharomycetales(100);f__Metschnikowiaceae(100);g__Metschnikowia(100);s__Metschnikowia_pulcherrima(100); | Metschnikowia | 13 | Saprotroph | \|Undefined Saprotroph\| | Yeast | NULL | Probable | As nectar-tap_saprotroph (Põlme et al. 2020); Decay sugar-rich substrates (Põlme et al. 2020) | Põlme S, et al. 2020. Fungal Diversity 105:1-16 ((https://doi.org/10.1007/s13225-020-00466-2)) |
| Otu15076 | 1 | k__Fungi(100);p__Ascomycota(100);c__Saccharomycetes(100);o__Saccharomycetales(100);f__Metschnikowiaceae(100);g__Metschnikowia(100);s__Metschnikowia_pulcherrima(100); | Metschnikowia | 13 | Saprotroph | \|Undefined Saprotroph\| | Yeast | NULL | Probable | As nectar-tap_saprotroph (Põlme et al. 2020); Decay sugar-rich substrates (Põlme et al. 2020) | Põlme S, et al. 2020. Fungal Diversity 105:1-16 ((https://doi.org/10.1007/s13225-020-00466-2)) |
| Otu15077 | 1 | k__Fungi(100);p__Ascomycota(100);c__Saccharomycetes(100);o__Saccharomycetales(100);f__Metschnikowiaceae(100);g__Metschnikowia(100);g__Metschnikowia_unclassified(100); | Metschnikowia | 13 | Saprotroph | \|Undefined Saprotroph\| | Yeast | NULL | Probable | As nectar-tap_saprotroph (Põlme et al. 2020); Decay sugar-rich substrates (Põlme et al. 2020) | Põlme S, et al. 2020. Fungal Diversity 105:1-16 ((https://doi.org/10.1007/s13225-020-00466-2)) |
| Otu15078 | 1 | k__Fungi(100);p__Ascomycota(100);c__Saccharomycetes(100);o__Saccharomycetales(100);f__Metschnikowiaceae(100);g__Metschnikowia(100);s__Metschnikowia_pulcherrima(100); | Metschnikowia | 13 | Saprotroph | \|Undefined Saprotroph\| | Yeast | NULL | Probable | As nectar-tap_saprotroph (Põlme et al. 2020); Decay sugar-rich substrates (Põlme et al. 2020) | Põlme S, et al. 2020. Fungal Diversity 105:1-16 ((https://doi.org/10.1007/s13225-020-00466-2)) |
| Otu15080 | 1 | k__Fungi(100);p__Ascomycota(100);c__Saccharomycetes(100);o__Saccharomycetales(100);f__Metschnikowiaceae(100);g__Metschnikowia(100);g__Metschnikowia_unclassified(100); | Metschnikowia | 13 | Saprotroph | \|Undefined Saprotroph\| | Yeast | NULL | Probable | As nectar-tap_saprotroph (Põlme et al. 2020); Decay sugar-rich substrates (Põlme et al. 2020) | Põlme S, et al. 2020. Fungal Diversity 105:1-16 ((https://doi.org/10.1007/s13225-020-00466-2)) |
| Otu15081 | 1 | k__Fungi(100);p__Ascomycota(100);c__Saccharomycetes(100);o__Saccharomycetales(100);f__Metschnikowiaceae(100);g__Metschnikowia(100);s__Metschnikowia_pulcherrima(100); | Metschnikowia | 13 | Saprotroph | \|Undefined Saprotroph\| | Yeast | NULL | Probable | As nectar-tap_saprotroph (Põlme et al. 2020); Decay sugar-rich substrates (Põlme et al. 2020) | Põlme S, et al. 2020. Fungal Diversity 105:1-16 ((https://doi.org/10.1007/s13225-020-00466-2)) |
| Otu15083 | 1 | k__Fungi(100);p__Ascomycota(100);c__Saccharomycetes(100);o__Saccharomycetales(100);f__Metschnikowiaceae(100);g__Metschnikowia(100);s__Metschnikowia_pulcherrima(100); | Metschnikowia | 13 | Saprotroph | \|Undefined Saprotroph\| | Yeast | NULL | Probable | As nectar-tap_saprotroph (Põlme et al. 2020); Decay sugar-rich substrates (Põlme et al. 2020) | Põlme S, et al. 2020. Fungal Diversity 105:1-16 ((https://doi.org/10.1007/s13225-020-00466-2)) |
| Otu15084 | 1 | k__Fungi(100);p__Ascomycota(100);c__Saccharomycetes(100);o__Saccharomycetales(100);f__Metschnikowiaceae(100);g__Metschnikowia(100);s__Metschnikowia_pulcherrima(100); | Metschnikowia | 13 | Saprotroph | \|Undefined Saprotroph\| | Yeast | NULL | Probable | As nectar-tap_saprotroph (Põlme et al. 2020); Decay sugar-rich substrates (Põlme et al. 2020) | Põlme S, et al. 2020. Fungal Diversity 105:1-16 ((https://doi.org/10.1007/s13225-020-00466-2)) |
| Otu15086 | 1 | k__Fungi(100);p__Ascomycota(100);c__Saccharomycetes(100);o__Saccharomycetales(100);f__Metschnikowiaceae(100);g__Metschnikowia(100);s__Metschnikowia_pulcherrima(100); | Metschnikowia | 13 | Saprotroph | \|Undefined Saprotroph\| | Yeast | NULL | Probable | As nectar-tap_saprotroph (Põlme et al. 2020); Decay sugar-rich substrates (Põlme et al. 2020) | Põlme S, et al. 2020. Fungal Diversity 105:1-16 ((https://doi.org/10.1007/s13225-020-00466-2)) |
| Otu15087 | 1 | k__Fungi(100);p__Ascomycota(100);c__Saccharomycetes(100);o__Saccharomycetales(100);f__Metschnikowiaceae(100);g__Metschnikowia(100);s__Metschnikowia_pulcherrima(100); | Metschnikowia | 13 | Saprotroph | \|Undefined Saprotroph\| | Yeast | NULL | Probable | As nectar-tap_saprotroph (Põlme et al. 2020); Decay sugar-rich substrates (Põlme et al. 2020) | Põlme S, et al. 2020. Fungal Diversity 105:1-16 ((https://doi.org/10.1007/s13225-020-00466-2)) |
| Otu15088 | 1 | k__Fungi(100);p__Ascomycota(100);c__Saccharomycetes(100);o__Saccharomycetales(100);f__Metschnikowiaceae(100);g__Metschnikowia(100);s__Metschnikowia_pulcherrima(100); | Metschnikowia | 13 | Saprotroph | \|Undefined Saprotroph\| | Yeast | NULL | Probable | As nectar-tap_saprotroph (Põlme et al. 2020); Decay sugar-rich substrates (Põlme et al. 2020) | Põlme S, et al. 2020. Fungal Diversity 105:1-16 ((https://doi.org/10.1007/s13225-020-00466-2)) |
| Otu15089 | 1 | k__Fungi(100);p__Ascomycota(100);c__Saccharomycetes(100);o__Saccharomycetales(100);f__Metschnikowiaceae(100);g__Metschnikowia(100);g__Metschnikowia_unclassified(100); | Metschnikowia | 13 | Saprotroph | \|Undefined Saprotroph\| | Yeast | NULL | Probable | As nectar-tap_saprotroph (Põlme et al. 2020); Decay sugar-rich substrates (Põlme et al. 2020) | Põlme S, et al. 2020. Fungal Diversity 105:1-16 ((https://doi.org/10.1007/s13225-020-00466-2)) |
| Otu15091 | 1 | k__Fungi(100);p__Ascomycota(100);c__Saccharomycetes(100);o__Saccharomycetales(100);f__Metschnikowiaceae(100);g__Metschnikowia(100);g__Metschnikowia_unclassified(100); | Metschnikowia | 13 | Saprotroph | \|Undefined Saprotroph\| | Yeast | NULL | Probable | As nectar-tap_saprotroph (Põlme et al. 2020); Decay sugar-rich substrates (Põlme et al. 2020) | Põlme S, et al. 2020. Fungal Diversity 105:1-16 ((https://doi.org/10.1007/s13225-020-00466-2)) |
| Otu15092 | 1 | k__Fungi(100);p__Ascomycota(100);c__Saccharomycetes(100);o__Saccharomycetales(100);f__Metschnikowiaceae(100);g__Metschnikowia(100);s__Metschnikowia_pulcherrima(100); | Metschnikowia | 13 | Saprotroph | \|Undefined Saprotroph\| | Yeast | NULL | Probable | As nectar-tap_saprotroph (Põlme et al. 2020); Decay sugar-rich substrates (Põlme et al. 2020) | Põlme S, et al. 2020. Fungal Diversity 105:1-16 ((https://doi.org/10.1007/s13225-020-00466-2)) |
| Otu15093 | 1 | k__Fungi(100);p__Ascomycota(100);c__Saccharomycetes(100);o__Saccharomycetales(100);f__Metschnikowiaceae(100);g__Metschnikowia(100);s__Metschnikowia_pulcherrima(100); | Metschnikowia | 13 | Saprotroph | \|Undefined Saprotroph\| | Yeast | NULL | Probable | As nectar-tap_saprotroph (Põlme et al. 2020); Decay sugar-rich substrates (Põlme et al. 2020) | Põlme S, et al. 2020. Fungal Diversity 105:1-16 ((https://doi.org/10.1007/s13225-020-00466-2)) |
| Otu15094 | 1 | k__Fungi(100);p__Ascomycota(100);c__Saccharomycetes(100);o__Saccharomycetales(100);f__Metschnikowiaceae(100);g__Metschnikowia(100);s__Metschnikowia_pulcherrima(100); | Metschnikowia | 13 | Saprotroph | \|Undefined Saprotroph\| | Yeast | NULL | Probable | As nectar-tap_saprotroph (Põlme et al. 2020); Decay sugar-rich substrates (Põlme et al. 2020) | Põlme S, et al. 2020. Fungal Diversity 105:1-16 ((https://doi.org/10.1007/s13225-020-00466-2)) |
| Otu15095 | 1 | k__Fungi(100);p__Ascomycota(100);c__Saccharomycetes(100);o__Saccharomycetales(100);f__Metschnikowiaceae(100);g__Metschnikowia(100);g__Metschnikowia_unclassified(100); | Metschnikowia | 13 | Saprotroph | \|Undefined Saprotroph\| | Yeast | NULL | Probable | As nectar-tap_saprotroph (Põlme et al. 2020); Decay sugar-rich substrates (Põlme et al. 2020) | Põlme S, et al. 2020. Fungal Diversity 105:1-16 ((https://doi.org/10.1007/s13225-020-00466-2)) |
| Otu15096 | 1 | k__Fungi(100);p__Ascomycota(100);c__Saccharomycetes(100);o__Saccharomycetales(100);f__Metschnikowiaceae(100);g__Metschnikowia(100);s__Metschnikowia_pulcherrima(100); | Metschnikowia | 13 | Saprotroph | \|Undefined Saprotroph\| | Yeast | NULL | Probable | As nectar-tap_saprotroph (Põlme et al. 2020); Decay sugar-rich substrates (Põlme et al. 2020) | Põlme S, et al. 2020. Fungal Diversity 105:1-16 ((https://doi.org/10.1007/s13225-020-00466-2)) |
| Otu15097 | 1 | k__Fungi(100);p__Ascomycota(100);c__Saccharomycetes(100);o__Saccharomycetales(100);f__Metschnikowiaceae(100);g__Metschnikowia(100);s__Metschnikowia_pulcherrima(100); | Metschnikowia | 13 | Saprotroph | \|Undefined Saprotroph\| | Yeast | NULL | Probable | As nectar-tap_saprotroph (Põlme et al. 2020); Decay sugar-rich substrates (Põlme et al. 2020) | Põlme S, et al. 2020. Fungal Diversity 105:1-16 ((https://doi.org/10.1007/s13225-020-00466-2)) |
| Otu15098 | 1 | k__Fungi(100);p__Ascomycota(100);c__Saccharomycetes(100);o__Saccharomycetales(100);f__Metschnikowiaceae(100);g__Metschnikowia(100);s__Metschnikowia_pulcherrima(100); | Metschnikowia | 13 | Saprotroph | \|Undefined Saprotroph\| | Yeast | NULL | Probable | As nectar-tap_saprotroph (Põlme et al. 2020); Decay sugar-rich substrates (Põlme et al. 2020) | Põlme S, et al. 2020. Fungal Diversity 105:1-16 ((https://doi.org/10.1007/s13225-020-00466-2)) |
| Otu15099 | 1 | k__Fungi(100);p__Ascomycota(100);c__Saccharomycetes(100);o__Saccharomycetales(100);f__Metschnikowiaceae(100);g__Metschnikowia(100);s__Metschnikowia_pulcherrima(100); | Metschnikowia | 13 | Saprotroph | \|Undefined Saprotroph\| | Yeast | NULL | Probable | As nectar-tap_saprotroph (Põlme et al. 2020); Decay sugar-rich substrates (Põlme et al. 2020) | Põlme S, et al. 2020. Fungal Diversity 105:1-16 ((https://doi.org/10.1007/s13225-020-00466-2)) |
| Otu15100 | 1 | k__Fungi(100);p__Ascomycota(100);c__Saccharomycetes(100);o__Saccharomycetales(100);f__Metschnikowiaceae(100);g__Metschnikowia(100);s__Metschnikowia_pulcherrima(100); | Metschnikowia | 13 | Saprotroph | \|Undefined Saprotroph\| | Yeast | NULL | Probable | As nectar-tap_saprotroph (Põlme et al. 2020); Decay sugar-rich substrates (Põlme et al. 2020) | Põlme S, et al. 2020. Fungal Diversity 105:1-16 ((https://doi.org/10.1007/s13225-020-00466-2)) |
| Otu15101 | 1 | k__Fungi(100);p__Ascomycota(100);c__Saccharomycetes(100);o__Saccharomycetales(100);f__Metschnikowiaceae(100);g__Metschnikowia(100);s__Metschnikowia_pulcherrima(100); | Metschnikowia | 13 | Saprotroph | \|Undefined Saprotroph\| | Yeast | NULL | Probable | As nectar-tap_saprotroph (Põlme et al. 2020); Decay sugar-rich substrates (Põlme et al. 2020) | Põlme S, et al. 2020. Fungal Diversity 105:1-16 ((https://doi.org/10.1007/s13225-020-00466-2)) |
| Otu15102 | 1 | k__Fungi(100);p__Ascomycota(100);c__Saccharomycetes(100);o__Saccharomycetales(100);f__Metschnikowiaceae(100);g__Metschnikowia(100);s__Metschnikowia_pulcherrima(100); | Metschnikowia | 13 | Saprotroph | \|Undefined Saprotroph\| | Yeast | NULL | Probable | As nectar-tap_saprotroph (Põlme et al. 2020); Decay sugar-rich substrates (Põlme et al. 2020) | Põlme S, et al. 2020. Fungal Diversity 105:1-16 ((https://doi.org/10.1007/s13225-020-00466-2)) |
| Otu15103 | 1 | k__Fungi(100);p__Ascomycota(100);c__Saccharomycetes(100);o__Saccharomycetales(100);f__Metschnikowiaceae(100);g__Metschnikowia(100);s__Metschnikowia_pulcherrima(100); | Metschnikowia | 13 | Saprotroph | \|Undefined Saprotroph\| | Yeast | NULL | Probable | As nectar-tap_saprotroph (Põlme et al. 2020); Decay sugar-rich substrates (Põlme et al. 2020) | Põlme S, et al. 2020. Fungal Diversity 105:1-16 ((https://doi.org/10.1007/s13225-020-00466-2)) |
| Otu15104 | 1 | k__Fungi(100);p__Ascomycota(100);c__Saccharomycetes(100);o__Saccharomycetales(100);f__Metschnikowiaceae(100);g__Metschnikowia(100);s__Metschnikowia_pulcherrima(100); | Metschnikowia | 13 | Saprotroph | \|Undefined Saprotroph\| | Yeast | NULL | Probable | As nectar-tap_saprotroph (Põlme et al. 2020); Decay sugar-rich substrates (Põlme et al. 2020) | Põlme S, et al. 2020. Fungal Diversity 105:1-16 ((https://doi.org/10.1007/s13225-020-00466-2)) |
| Otu15105 | 1 | k__Fungi(100);p__Ascomycota(100);c__Saccharomycetes(100);o__Saccharomycetales(100);f__Metschnikowiaceae(100);g__Metschnikowia(100);s__Metschnikowia_pulcherrima(100); | Metschnikowia | 13 | Saprotroph | \|Undefined Saprotroph\| | Yeast | NULL | Probable | As nectar-tap_saprotroph (Põlme et al. 2020); Decay sugar-rich substrates (Põlme et al. 2020) | Põlme S, et al. 2020. Fungal Diversity 105:1-16 ((https://doi.org/10.1007/s13225-020-00466-2)) |
| Otu15106 | 1 | k__Fungi(100);p__Ascomycota(100);c__Saccharomycetes(100);o__Saccharomycetales(100);f__Metschnikowiaceae(100);g__Metschnikowia(100);s__Metschnikowia_pulcherrima(100); | Metschnikowia | 13 | Saprotroph | \|Undefined Saprotroph\| | Yeast | NULL | Probable | As nectar-tap_saprotroph (Põlme et al. 2020); Decay sugar-rich substrates (Põlme et al. 2020) | Põlme S, et al. 2020. Fungal Diversity 105:1-16 ((https://doi.org/10.1007/s13225-020-00466-2)) |
| Otu15107 | 1 | k__Fungi(100);p__Ascomycota(100);c__Saccharomycetes(100);o__Saccharomycetales(100);f__Metschnikowiaceae(100);g__Metschnikowia(100);s__Metschnikowia_pulcherrima(100); | Metschnikowia | 13 | Saprotroph | \|Undefined Saprotroph\| | Yeast | NULL | Probable | As nectar-tap_saprotroph (Põlme et al. 2020); Decay sugar-rich substrates (Põlme et al. 2020) | Põlme S, et al. 2020. Fungal Diversity 105:1-16 ((https://doi.org/10.1007/s13225-020-00466-2)) |
| Otu15109 | 1 | k__Fungi(100);p__Ascomycota(100);c__Saccharomycetes(100);o__Saccharomycetales(100);f__Metschnikowiaceae(100);g__Metschnikowia(100);s__Metschnikowia_pulcherrima(100); | Metschnikowia | 13 | Saprotroph | \|Undefined Saprotroph\| | Yeast | NULL | Probable | As nectar-tap_saprotroph (Põlme et al. 2020); Decay sugar-rich substrates (Põlme et al. 2020) | Põlme S, et al. 2020. Fungal Diversity 105:1-16 ((https://doi.org/10.1007/s13225-020-00466-2)) |
| Otu15110 | 1 | k__Fungi(100);p__Ascomycota(100);c__Saccharomycetes(100);o__Saccharomycetales(100);f__Metschnikowiaceae(100);g__Metschnikowia(100);s__Metschnikowia_pulcherrima(100); | Metschnikowia | 13 | Saprotroph | \|Undefined Saprotroph\| | Yeast | NULL | Probable | As nectar-tap_saprotroph (Põlme et al. 2020); Decay sugar-rich substrates (Põlme et al. 2020) | Põlme S, et al. 2020. Fungal Diversity 105:1-16 ((https://doi.org/10.1007/s13225-020-00466-2)) |
| Otu15111 | 1 | k__Fungi(100);p__Ascomycota(100);c__Saccharomycetes(100);o__Saccharomycetales(100);f__Metschnikowiaceae(100);g__Metschnikowia(100);s__Metschnikowia_pulcherrima(100); | Metschnikowia | 13 | Saprotroph | \|Undefined Saprotroph\| | Yeast | NULL | Probable | As nectar-tap_saprotroph (Põlme et al. 2020); Decay sugar-rich substrates (Põlme et al. 2020) | Põlme S, et al. 2020. Fungal Diversity 105:1-16 ((https://doi.org/10.1007/s13225-020-00466-2)) |
| Otu15112 | 1 | k__Fungi(100);p__Ascomycota(100);c__Saccharomycetes(100);o__Saccharomycetales(100);f__Metschnikowiaceae(100);g__Metschnikowia(100);s__Metschnikowia_pulcherrima(100); | Metschnikowia | 13 | Saprotroph | \|Undefined Saprotroph\| | Yeast | NULL | Probable | As nectar-tap_saprotroph (Põlme et al. 2020); Decay sugar-rich substrates (Põlme et al. 2020) | Põlme S, et al. 2020. Fungal Diversity 105:1-16 ((https://doi.org/10.1007/s13225-020-00466-2)) |
| Otu15114 | 1 | k__Fungi(100);p__Ascomycota(100);c__Saccharomycetes(100);o__Saccharomycetales(100);f__Metschnikowiaceae(100);g__Metschnikowia(100);s__Metschnikowia_pulcherrima(100); | Metschnikowia | 13 | Saprotroph | \|Undefined Saprotroph\| | Yeast | NULL | Probable | As nectar-tap_saprotroph (Põlme et al. 2020); Decay sugar-rich substrates (Põlme et al. 2020) | Põlme S, et al. 2020. Fungal Diversity 105:1-16 ((https://doi.org/10.1007/s13225-020-00466-2)) |
| Otu15116 | 1 | k__Fungi(100);p__Ascomycota(100);c__Saccharomycetes(100);o__Saccharomycetales(100);f__Metschnikowiaceae(100);g__Metschnikowia(100);g__Metschnikowia_unclassified(100); | Metschnikowia | 13 | Saprotroph | \|Undefined Saprotroph\| | Yeast | NULL | Probable | As nectar-tap_saprotroph (Põlme et al. 2020); Decay sugar-rich substrates (Põlme et al. 2020) | Põlme S, et al. 2020. Fungal Diversity 105:1-16 ((https://doi.org/10.1007/s13225-020-00466-2)) |
| Otu15117 | 1 | k__Fungi(100);p__Ascomycota(100);c__Saccharomycetes(100);o__Saccharomycetales(100);f__Metschnikowiaceae(100);g__Metschnikowia(100);s__Metschnikowia_pulcherrima(100); | Metschnikowia | 13 | Saprotroph | \|Undefined Saprotroph\| | Yeast | NULL | Probable | As nectar-tap_saprotroph (Põlme et al. 2020); Decay sugar-rich substrates (Põlme et al. 2020) | Põlme S, et al. 2020. Fungal Diversity 105:1-16 ((https://doi.org/10.1007/s13225-020-00466-2)) |
| Otu15118 | 1 | k__Fungi(100);p__Ascomycota(100);c__Saccharomycetes(100);o__Saccharomycetales(100);f__Metschnikowiaceae(100);g__Metschnikowia(100);g__Metschnikowia_unclassified(100); | Metschnikowia | 13 | Saprotroph | \|Undefined Saprotroph\| | Yeast | NULL | Probable | As nectar-tap_saprotroph (Põlme et al. 2020); Decay sugar-rich substrates (Põlme et al. 2020) | Põlme S, et al. 2020. Fungal Diversity 105:1-16 ((https://doi.org/10.1007/s13225-020-00466-2)) |
| Otu15120 | 1 | k__Fungi(100);p__Ascomycota(100);c__Saccharomycetes(100);o__Saccharomycetales(100);f__Metschnikowiaceae(100);g__Metschnikowia(100);s__Metschnikowia_pulcherrima(100); | Metschnikowia | 13 | Saprotroph | \|Undefined Saprotroph\| | Yeast | NULL | Probable | As nectar-tap_saprotroph (Põlme et al. 2020); Decay sugar-rich substrates (Põlme et al. 2020) | Põlme S, et al. 2020. Fungal Diversity 105:1-16 ((https://doi.org/10.1007/s13225-020-00466-2)) |
| Otu15121 | 1 | k__Fungi(100);p__Ascomycota(100);c__Saccharomycetes(100);o__Saccharomycetales(100);f__Metschnikowiaceae(100);g__Metschnikowia(100);s__Metschnikowia_pulcherrima(100); | Metschnikowia | 13 | Saprotroph | \|Undefined Saprotroph\| | Yeast | NULL | Probable | As nectar-tap_saprotroph (Põlme et al. 2020); Decay sugar-rich substrates (Põlme et al. 2020) | Põlme S, et al. 2020. Fungal Diversity 105:1-16 ((https://doi.org/10.1007/s13225-020-00466-2)) |
| Otu15122 | 1 | k__Fungi(100);p__Ascomycota(100);c__Saccharomycetes(100);o__Saccharomycetales(100);f__Metschnikowiaceae(100);g__Metschnikowia(100);s__Metschnikowia_pulcherrima(100); | Metschnikowia | 13 | Saprotroph | \|Undefined Saprotroph\| | Yeast | NULL | Probable | As nectar-tap_saprotroph (Põlme et al. 2020); Decay sugar-rich substrates (Põlme et al. 2020) | Põlme S, et al. 2020. Fungal Diversity 105:1-16 ((https://doi.org/10.1007/s13225-020-00466-2)) |
| Otu15123 | 1 | k__Fungi(100);p__Ascomycota(100);c__Saccharomycetes(100);o__Saccharomycetales(100);f__Metschnikowiaceae(100);g__Metschnikowia(100);s__Metschnikowia_pulcherrima(100); | Metschnikowia | 13 | Saprotroph | \|Undefined Saprotroph\| | Yeast | NULL | Probable | As nectar-tap_saprotroph (Põlme et al. 2020); Decay sugar-rich substrates (Põlme et al. 2020) | Põlme S, et al. 2020. Fungal Diversity 105:1-16 ((https://doi.org/10.1007/s13225-020-00466-2)) |
| Otu15124 | 1 | k__Fungi(100);p__Ascomycota(100);c__Saccharomycetes(100);o__Saccharomycetales(100);f__Metschnikowiaceae(100);g__Metschnikowia(100);g__Metschnikowia_unclassified(100); | Metschnikowia | 13 | Saprotroph | \|Undefined Saprotroph\| | Yeast | NULL | Probable | As nectar-tap_saprotroph (Põlme et al. 2020); Decay sugar-rich substrates (Põlme et al. 2020) | Põlme S, et al. 2020. Fungal Diversity 105:1-16 ((https://doi.org/10.1007/s13225-020-00466-2)) |
| Otu15126 | 1 | k__Fungi(100);p__Ascomycota(100);c__Saccharomycetes(100);o__Saccharomycetales(100);f__Metschnikowiaceae(100);g__Metschnikowia(100);g__Metschnikowia_unclassified(100); | Metschnikowia | 13 | Saprotroph | \|Undefined Saprotroph\| | Yeast | NULL | Probable | As nectar-tap_saprotroph (Põlme et al. 2020); Decay sugar-rich substrates (Põlme et al. 2020) | Põlme S, et al. 2020. Fungal Diversity 105:1-16 ((https://doi.org/10.1007/s13225-020-00466-2)) |
| Otu15127 | 1 | k__Fungi(100);p__Ascomycota(100);c__Saccharomycetes(100);o__Saccharomycetales(100);f__Metschnikowiaceae(100);g__Metschnikowia(100);s__Metschnikowia_pulcherrima(100); | Metschnikowia | 13 | Saprotroph | \|Undefined Saprotroph\| | Yeast | NULL | Probable | As nectar-tap_saprotroph (Põlme et al. 2020); Decay sugar-rich substrates (Põlme et al. 2020) | Põlme S, et al. 2020. Fungal Diversity 105:1-16 ((https://doi.org/10.1007/s13225-020-00466-2)) |
| Otu15128 | 1 | k__Fungi(100);p__Ascomycota(100);c__Saccharomycetes(100);o__Saccharomycetales(100);f__Metschnikowiaceae(100);g__Metschnikowia(100);g__Metschnikowia_unclassified(100); | Metschnikowia | 13 | Saprotroph | \|Undefined Saprotroph\| | Yeast | NULL | Probable | As nectar-tap_saprotroph (Põlme et al. 2020); Decay sugar-rich substrates (Põlme et al. 2020) | Põlme S, et al. 2020. Fungal Diversity 105:1-16 ((https://doi.org/10.1007/s13225-020-00466-2)) |
| Otu15129 | 1 | k__Fungi(100);p__Ascomycota(100);c__Saccharomycetes(100);o__Saccharomycetales(100);f__Metschnikowiaceae(100);g__Metschnikowia(100);s__Metschnikowia_pulcherrima(100); | Metschnikowia | 13 | Saprotroph | \|Undefined Saprotroph\| | Yeast | NULL | Probable | As nectar-tap_saprotroph (Põlme et al. 2020); Decay sugar-rich substrates (Põlme et al. 2020) | Põlme S, et al. 2020. Fungal Diversity 105:1-16 ((https://doi.org/10.1007/s13225-020-00466-2)) |
| Otu15131 | 1 | k__Fungi(100);p__Ascomycota(100);c__Saccharomycetes(100);o__Saccharomycetales(100);f__Metschnikowiaceae(100);g__Metschnikowia(100);s__Metschnikowia_pulcherrima(100); | Metschnikowia | 13 | Saprotroph | \|Undefined Saprotroph\| | Yeast | NULL | Probable | As nectar-tap_saprotroph (Põlme et al. 2020); Decay sugar-rich substrates (Põlme et al. 2020) | Põlme S, et al. 2020. Fungal Diversity 105:1-16 ((https://doi.org/10.1007/s13225-020-00466-2)) |
| Otu15132 | 1 | k__Fungi(100);p__Ascomycota(100);c__Saccharomycetes(100);o__Saccharomycetales(100);f__Metschnikowiaceae(100);g__Metschnikowia(100);s__Metschnikowia_pulcherrima(100); | Metschnikowia | 13 | Saprotroph | \|Undefined Saprotroph\| | Yeast | NULL | Probable | As nectar-tap_saprotroph (Põlme et al. 2020); Decay sugar-rich substrates (Põlme et al. 2020) | Põlme S, et al. 2020. Fungal Diversity 105:1-16 ((https://doi.org/10.1007/s13225-020-00466-2)) |
| Otu15133 | 1 | k__Fungi(100);p__Ascomycota(100);c__Saccharomycetes(100);o__Saccharomycetales(100);f__Metschnikowiaceae(100);g__Metschnikowia(100);s__Metschnikowia_pulcherrima(100); | Metschnikowia | 13 | Saprotroph | \|Undefined Saprotroph\| | Yeast | NULL | Probable | As nectar-tap_saprotroph (Põlme et al. 2020); Decay sugar-rich substrates (Põlme et al. 2020) | Põlme S, et al. 2020. Fungal Diversity 105:1-16 ((https://doi.org/10.1007/s13225-020-00466-2)) |
| Otu15134 | 1 | k__Fungi(100);p__Ascomycota(100);c__Saccharomycetes(100);o__Saccharomycetales(100);f__Metschnikowiaceae(100);g__Metschnikowia(100);s__Metschnikowia_pulcherrima(100); | Metschnikowia | 13 | Saprotroph | \|Undefined Saprotroph\| | Yeast | NULL | Probable | As nectar-tap_saprotroph (Põlme et al. 2020); Decay sugar-rich substrates (Põlme et al. 2020) | Põlme S, et al. 2020. Fungal Diversity 105:1-16 ((https://doi.org/10.1007/s13225-020-00466-2)) |
| Otu15135 | 1 | k__Fungi(100);p__Ascomycota(100);c__Saccharomycetes(100);o__Saccharomycetales(100);f__Saccharomycetales_family_Incertae_sedis(100);f__Saccharomycetales_family_Incertae_sedis_unclassified(100);f__Saccharomycetales_family_Incertae_sedis_unclassified(100); | Saccharomycetales | 7 | Saprotroph | Undefined Saprotroph | Yeast | NULL | Possible | NULL | Sterkenburg E, et al. 2015. New Phytologist 207:1145-1158 ((https://doi.org/10.1111/nph.13426)) |
| Otu15136 | 1 | k__Fungi(100);p__Ascomycota(100);c__Saccharomycetes(100);o__Saccharomycetales(100);f__Metschnikowiaceae(100);g__Metschnikowia(100);s__Metschnikowia_pulcherrima(100); | Metschnikowia | 13 | Saprotroph | \|Undefined Saprotroph\| | Yeast | NULL | Probable | As nectar-tap_saprotroph (Põlme et al. 2020); Decay sugar-rich substrates (Põlme et al. 2020) | Põlme S, et al. 2020. Fungal Diversity 105:1-16 ((https://doi.org/10.1007/s13225-020-00466-2)) |
| Otu15137 | 1 | k__Fungi(100);p__Ascomycota(100);c__Saccharomycetes(100);o__Saccharomycetales(100);f__Metschnikowiaceae(100);g__Metschnikowia(100);s__Metschnikowia_pulcherrima(100); | Metschnikowia | 13 | Saprotroph | \|Undefined Saprotroph\| | Yeast | NULL | Probable | As nectar-tap_saprotroph (Põlme et al. 2020); Decay sugar-rich substrates (Põlme et al. 2020) | Põlme S, et al. 2020. Fungal Diversity 105:1-16 ((https://doi.org/10.1007/s13225-020-00466-2)) |
| Otu15138 | 1 | k__Fungi(100);p__Ascomycota(100);c__Saccharomycetes(100);o__Saccharomycetales(100);f__Metschnikowiaceae(100);g__Metschnikowia(100);s__Metschnikowia_pulcherrima(100); | Metschnikowia | 13 | Saprotroph | \|Undefined Saprotroph\| | Yeast | NULL | Probable | As nectar-tap_saprotroph (Põlme et al. 2020); Decay sugar-rich substrates (Põlme et al. 2020) | Põlme S, et al. 2020. Fungal Diversity 105:1-16 ((https://doi.org/10.1007/s13225-020-00466-2)) |

| | | | | | | | | | | | |
|---|---|---|---|---|---|---|---|---|---|---|---|
| Otu15139 | 1 | k__Fungi(100);p__Ascomycota(100);c__Saccharomycetes(100);o__Saccharomycetales(100);f__Metschnikowiaceae(100);g__Metschnikowia(100);g__Metschnikowia_unclassified(100); | Metschnikowia | 13 | Saprotroph | \|Undefined Saprotroph\| | Yeast | NULL | Probable | As nectar-tap_saprotroph (Põlme et al. 2020); Decay sugar-rich substrates (Põlme et al. 2020) | Põlme S, et al. 2020. Fungal Diversity 105:1-16 ((https://doi.org/10.1007/s13225-020-00466-2)) |
| Otu15140 | 1 | k__Fungi(100);p__Ascomycota(100);c__Saccharomycetes(100);o__Saccharomycetales(100);f__Metschnikowiaceae(100);g__Metschnikowia(100);s__Metschnikowia_pulcherrima(100); | Metschnikowia | 13 | Saprotroph | \|Undefined Saprotroph\| | Yeast | NULL | Probable | As nectar-tap_saprotroph (Põlme et al. 2020); Decay sugar-rich substrates (Põlme et al. 2020) | Põlme S, et al. 2020. Fungal Diversity 105:1-16 ((https://doi.org/10.1007/s13225-020-00466-2)) |
| Otu15141 | 1 | k__Fungi(100);p__Ascomycota(100);c__Saccharomycetes(100);o__Saccharomycetales(100);f__Metschnikowiaceae(100);g__Metschnikowia(100);g__Metschnikowia_unclassified(100); | Metschnikowia | 13 | Saprotroph | \|Undefined Saprotroph\| | Yeast | NULL | Probable | As nectar-tap_saprotroph (Põlme et al. 2020); Decay sugar-rich substrates (Põlme et al. 2020) | Põlme S, et al. 2020. Fungal Diversity 105:1-16 ((https://doi.org/10.1007/s13225-020-00466-2)) |
| Otu15142 | 1 | k__Fungi(100);p__Ascomycota(100);c__Saccharomycetes(100);o__Saccharomycetales(100);f__Saccharomycetales_family_Incertae_sedis(100);g__Candida(100);g__Candida_unclassified(100); | Candida | 13 | Pathotroph-Saprotroph-Symbiotroph | Animal Pathogen-Endophyte-Endosymbiont-Epiphyte-\|Undefined Saprotroph\| | Dimorphic Yeast-Yeast | NULL | Probable | As nectar-tap_saprotroph (Põlme et al. 2020); Decay sugar-rich substrates (Põlme et al. 2020); This asexual genus is polyphyletic and contain species that function very differently depend on what they are. Caution should be taken when interpreting guilds for Candida species. | Manolakaki D, et al. 2010. Virulence. 1: 367-375 ((https://doi.org/10.4161/viru.1.5.12796)); Mekbib SB, et al. 2011. Tropical Plant Pathology 36:233-240 ((https://doi.org/10.1590/S1982-56762011000400004)); Põlme S, et al. 2020. Fungal Diversity 105:1-16 ((https://doi.org/10.1007/s13225-020-00466-2)) |
| Otu15143 | 1 | k__Fungi(100);p__Ascomycota(100);c__Saccharomycetes(100);o__Saccharomycetales(100);f__Metschnikowiaceae(100);g__Metschnikowia(100);s__Metschnikowia_pulcherrima(100); | Metschnikowia | 13 | Saprotroph | \|Undefined Saprotroph\| | Yeast | NULL | Probable | As nectar-tap_saprotroph (Põlme et al. 2020); Decay sugar-rich substrates (Põlme et al. 2020) | Põlme S, et al. 2020. Fungal Diversity 105:1-16 ((https://doi.org/10.1007/s13225-020-00466-2)) |
| Otu15144 | 1 | k__Fungi(100);p__Ascomycota(100);c__Saccharomycetes(100);o__Saccharomycetales(100);f__Metschnikowiaceae(100);g__Metschnikowia(100);s__Metschnikowia_pulcherrima(100); | Metschnikowia | 13 | Saprotroph | \|Undefined Saprotroph\| | Yeast | NULL | Probable | As nectar-tap_saprotroph (Põlme et al. 2020); Decay sugar-rich substrates (Põlme et al. 2020) | Põlme S, et al. 2020. Fungal Diversity 105:1-16 ((https://doi.org/10.1007/s13225-020-00466-2)) |
| Otu15147 | 1 | k__Fungi(100);p__Ascomycota(100);c__Saccharomycetes(100);o__Saccharomycetales(100);f__Metschnikowiaceae(100);g__Metschnikowia(100);s__Metschnikowia_pulcherrima(100); | Metschnikowia | 13 | Saprotroph | \|Undefined Saprotroph\| | Yeast | NULL | Probable | As nectar-tap_saprotroph (Põlme et al. 2020); Decay sugar-rich substrates (Põlme et al. 2020) | Põlme S, et al. 2020. Fungal Diversity 105:1-16 ((https://doi.org/10.1007/s13225-020-00466-2)) |
| Otu15148 | 1 | k__Fungi(100);p__Ascomycota(100);c__Saccharomycetes(100);o__Saccharomycetales(100);f__Metschnikowiaceae(100);g__Metschnikowia(100);s__Metschnikowia_pulcherrima(100); | Metschnikowia | 13 | Saprotroph | \|Undefined Saprotroph\| | Yeast | NULL | Probable | As nectar-tap_saprotroph (Põlme et al. 2020); Decay sugar-rich substrates (Põlme et al. 2020) | Põlme S, et al. 2020. Fungal Diversity 105:1-16 ((https://doi.org/10.1007/s13225-020-00466-2)) |
| Otu15149 | 1 | k__Fungi(100);p__Ascomycota(100);c__Saccharomycetes(100);o__Saccharomycetales(100);f__Metschnikowiaceae(100);g__Metschnikowia(100);s__Metschnikowia_pulcherrima(100); | Metschnikowia | 13 | Saprotroph | \|Undefined Saprotroph\| | Yeast | NULL | Probable | As nectar-tap_saprotroph (Põlme et al. 2020); Decay sugar-rich substrates (Põlme et al. 2020) | Põlme S, et al. 2020. Fungal Diversity 105:1-16 ((https://doi.org/10.1007/s13225-020-00466-2)) |
| Otu15150 | 1 | k__Fungi(100);p__Ascomycota(100);c__Saccharomycetes(100);o__Saccharomycetales(100);f__Metschnikowiaceae(100);g__Metschnikowia(100);s__Metschnikowia_pulcherrima(100); | Metschnikowia | 13 | Saprotroph | \|Undefined Saprotroph\| | Yeast | NULL | Probable | As nectar-tap_saprotroph (Põlme et al. 2020); Decay sugar-rich substrates (Põlme et al. 2020) | Põlme S, et al. 2020. Fungal Diversity 105:1-16 ((https://doi.org/10.1007/s13225-020-00466-2)) |
| Otu15152 | 1 | k__Fungi(100);p__Ascomycota(100);c__Saccharomycetes(100);o__Saccharomycetales(100);f__Metschnikowiaceae(100);g__Metschnikowia(100);s__Metschnikowia_pulcherrima(100); | Metschnikowia | 13 | Saprotroph | \|Undefined Saprotroph\| | Yeast | NULL | Probable | As nectar-tap_saprotroph (Põlme et al. 2020); Decay sugar-rich substrates (Põlme et al. 2020) | Põlme S, et al. 2020. Fungal Diversity 105:1-16 ((https://doi.org/10.1007/s13225-020-00466-2)) |
| Otu15153 | 1 | k__Fungi(100);p__Ascomycota(100);c__Saccharomycetes(100);o__Saccharomycetales(100);f__Metschnikowiaceae(100);g__Metschnikowia(100);s__Metschnikowia_pulcherrima(100); | Metschnikowia | 13 | Saprotroph | \|Undefined Saprotroph\| | Yeast | NULL | Probable | As nectar-tap_saprotroph (Põlme et al. 2020); Decay sugar-rich substrates (Põlme et al. 2020) | Põlme S, et al. 2020. Fungal Diversity 105:1-16 ((https://doi.org/10.1007/s13225-020-00466-2)) |
| Otu15154 | 1 | k__Fungi(100);p__Ascomycota(100);c__Saccharomycetes(100);o__Saccharomycetales(100);f__Metschnikowiaceae(100);g__Metschnikowia(100);s__Metschnikowia_pulcherrima(100); | Metschnikowia | 13 | Saprotroph | \|Undefined Saprotroph\| | Yeast | NULL | Probable | As nectar-tap_saprotroph (Põlme et al. 2020); Decay sugar-rich substrates (Põlme et al. 2020) | Põlme S, et al. 2020. Fungal Diversity 105:1-16 ((https://doi.org/10.1007/s13225-020-00466-2)) |
| Otu15156 | 1 | k__Fungi(100);p__Ascomycota(100);c__Saccharomycetes(100);o__Saccharomycetales(100);f__Saccharomycetales_family_Incertae_sedis(100);g__Candida(100);g__Candida_unclassified(100); | Candida | 13 | Pathotroph-Saprotroph-Symbiotroph | Animal Pathogen-Endophyte-Endosymbiont-Epiphyte-\|Undefined Saprotroph\| | Dimorphic Yeast-Yeast | NULL | Probable | As nectar-tap_saprotroph (Põlme et al. 2020); Decay sugar-rich substrates (Põlme et al. 2020); This asexual genus is polyphyletic and contain species that function very differently depend on what they are. Caution should be taken when interpreting guilds for Candida species. | Manolakaki D, et al. 2010. Virulence. 1: 367-375 ((https://doi.org/10.4161/viru.1.5.12796)); Mekbib SB, et al. 2011. Tropical Plant Pathology 36:233-240 ((https://doi.org/10.1590/S1982-56762011000400004)); Põlme S, et al. 2020. Fungal Diversity 105:1-16 ((https://doi.org/10.1007/s13225-020-00466-2)) |
| Otu15157 | 1 | k__Fungi(100);p__Ascomycota(100);c__Saccharomycetes(100);o__Saccharomycetales(100);f__Metschnikowiaceae(100);g__Metschnikowia(100);s__Metschnikowia_pulcherrima(100); | Metschnikowia | 13 | Saprotroph | \|Undefined Saprotroph\| | Yeast | NULL | Probable | As nectar-tap_saprotroph (Põlme et al. 2020); Decay sugar-rich substrates (Põlme et al. 2020) | Põlme S, et al. 2020. Fungal Diversity 105:1-16 ((https://doi.org/10.1007/s13225-020-00466-2)) |
| Otu15158 | 1 | k__Fungi(100);p__Ascomycota(100);c__Saccharomycetes(100);o__Saccharomycetales(100);f__Metschnikowiaceae(100);g__Metschnikowia(100);s__Metschnikowia_pulcherrima(100); | Metschnikowia | 13 | Saprotroph | \|Undefined Saprotroph\| | Yeast | NULL | Probable | As nectar-tap_saprotroph (Põlme et al. 2020); Decay sugar-rich substrates (Põlme et al. 2020) | Põlme S, et al. 2020. Fungal Diversity 105:1-16 ((https://doi.org/10.1007/s13225-020-00466-2)) |
| Otu15159 | 1 | k__Fungi(100);p__Ascomycota(100);c__Saccharomycetes(100);o__Saccharomycetales(100);f__Metschnikowiaceae(100);g__Metschnikowia(100);s__Metschnikowia_pulcherrima(100); | Metschnikowia | 13 | Saprotroph | \|Undefined Saprotroph\| | Yeast | NULL | Probable | As nectar-tap_saprotroph (Põlme et al. 2020); Decay sugar-rich substrates (Põlme et al. 2020) | Põlme S, et al. 2020. Fungal Diversity 105:1-16 ((https://doi.org/10.1007/s13225-020-00466-2)) |
| Otu15160 | 1 | k__Fungi(100);p__Ascomycota(100);c__Saccharomycetes(100);o__Saccharomycetales(100);f__Metschnikowiaceae(100);g__Metschnikowia(100);s__Metschnikowia_pulcherrima(100); | Metschnikowia | 13 | Saprotroph | \|Undefined Saprotroph\| | Yeast | NULL | Probable | As nectar-tap_saprotroph (Põlme et al. 2020); Decay sugar-rich substrates (Põlme et al. 2020) | Põlme S, et al. 2020. Fungal Diversity 105:1-16 ((https://doi.org/10.1007/s13225-020-00466-2)) |
| Otu15161 | 1 | k__Fungi(100);p__Ascomycota(100);c__Saccharomycetes(100);o__Saccharomycetales(100);f__Metschnikowiaceae(100);g__Metschnikowia(100);s__Metschnikowia_pulcherrima(100); | Metschnikowia | 13 | Saprotroph | \|Undefined Saprotroph\| | Yeast | NULL | Probable | As nectar-tap_saprotroph (Põlme et al. 2020); Decay sugar-rich substrates (Põlme et al. 2020) | Põlme S, et al. 2020. Fungal Diversity 105:1-16 ((https://doi.org/10.1007/s13225-020-00466-2)) |
| Otu15162 | 1 | k__Fungi(100);p__Ascomycota(100);c__Saccharomycetes(100);o__Saccharomycetales(100);f__Metschnikowiaceae(100);g__Metschnikowia(100);s__Metschnikowia_pulcherrima(100); | Metschnikowia | 13 | Saprotroph | \|Undefined Saprotroph\| | Yeast | NULL | Probable | As nectar-tap_saprotroph (Põlme et al. 2020); Decay sugar-rich substrates (Põlme et al. 2020) | Põlme S, et al. 2020. Fungal Diversity 105:1-16 ((https://doi.org/10.1007/s13225-020-00466-2)) |
| Otu15164 | 1 | k__Fungi(100);p__Ascomycota(100);c__Saccharomycetes(100);o__Saccharomycetales(100);f__Metschnikowiaceae(100);g__Metschnikowia(100);g__Metschnikowia_unclassified(100); | Metschnikowia | 13 | Saprotroph | \|Undefined Saprotroph\| | Yeast | NULL | Probable | As nectar-tap_saprotroph (Põlme et al. 2020); Decay sugar-rich substrates (Põlme et al. 2020) | Põlme S, et al. 2020. Fungal Diversity 105:1-16 ((https://doi.org/10.1007/s13225-020-00466-2)) |
| Otu15165 | 1 | k__Fungi(100);p__Ascomycota(100);c__Saccharomycetes(100);o__Saccharomycetales(100);f__Metschnikowiaceae(100);g__Metschnikowia(100);s__Metschnikowia_pulcherrima(100); | Metschnikowia | 13 | Saprotroph | \|Undefined Saprotroph\| | Yeast | NULL | Probable | As nectar-tap_saprotroph (Põlme et al. 2020); Decay sugar-rich substrates (Põlme et al. 2020) | Põlme S, et al. 2020. Fungal Diversity 105:1-16 ((https://doi.org/10.1007/s13225-020-00466-2)) |
| Otu15166 | 1 | k__Fungi(100);p__Ascomycota(100);c__Saccharomycetes(100);o__Saccharomycetales(100);f__Saccharomycetales_family_Incertae_sedis(100);g__Candida(100);g__Candida_unclassified(100); | Candida | 13 | Pathotroph-Saprotroph-Symbiotroph | Animal Pathogen-Endophyte-Endosymbiont-Epiphyte-\|Undefined Saprotroph\| | Dimorphic Yeast-Yeast | NULL | Probable | As nectar-tap_saprotroph (Põlme et al. 2020); Decay sugar-rich substrates (Põlme et al. 2020); This asexual genus is polyphyletic and contain species that function very differently depend on what they are. Caution should be taken when interpreting guilds for Candida species. | Manolakaki D, et al. 2010. Virulence. 1: 367-375 ((https://doi.org/10.4161/viru.1.5.12796)); Mekbib SB, et al. 2011. Tropical Plant Pathology 36:233-240 ((https://doi.org/10.1590/S1982-56762011000400004)); Põlme S, et al. 2020. Fungal Diversity 105:1-16 ((https://doi.org/10.1007/s13225-020-00466-2)) |
| Otu15167 | 1 | k__Fungi(100);p__Ascomycota(100);c__Saccharomycetes(100);o__Saccharomycetales(100);f__Metschnikowiaceae(100);g__Metschnikowia(100);s__Metschnikowia_pulcherrima(100); | Metschnikowia | 13 | Saprotroph | \|Undefined Saprotroph\| | Yeast | NULL | Probable | As nectar-tap_saprotroph (Põlme et al. 2020); Decay sugar-rich substrates (Põlme et al. 2020) | Põlme S, et al. 2020. Fungal Diversity 105:1-16 ((https://doi.org/10.1007/s13225-020-00466-2)) |
| Otu15168 | 1 | k__Fungi(100);p__Ascomycota(100);c__Saccharomycetes(100);o__Saccharomycetales(100);f__Metschnikowiaceae(100);g__Metschnikowia(100);s__Metschnikowia_pulcherrima(100); | Metschnikowia | 13 | Saprotroph | \|Undefined Saprotroph\| | Yeast | NULL | Probable | As nectar-tap_saprotroph (Põlme et al. 2020); Decay sugar-rich substrates (Põlme et al. 2020) | Põlme S, et al. 2020. Fungal Diversity 105:1-16 ((https://doi.org/10.1007/s13225-020-00466-2)) |
| Otu15169 | 1 | k__Fungi(100);p__Ascomycota(100);c__Saccharomycetes(100);o__Saccharomycetales(100);f__Metschnikowiaceae(100);g__Metschnikowia(100);s__Metschnikowia_pulcherrima(100); | Metschnikowia | 13 | Saprotroph | \|Undefined Saprotroph\| | Yeast | NULL | Probable | As nectar-tap_saprotroph (Põlme et al. 2020); Decay sugar-rich substrates (Põlme et al. 2020) | Põlme S, et al. 2020. Fungal Diversity 105:1-16 ((https://doi.org/10.1007/s13225-020-00466-2)) |
| Otu15170 | 1 | k__Fungi(100);p__Ascomycota(100);c__Saccharomycetes(100);o__Saccharomycetales(100);f__Metschnikowiaceae(100);g__Metschnikowia(100);s__Metschnikowia_pulcherrima(100); | Metschnikowia | 13 | Saprotroph | \|Undefined Saprotroph\| | Yeast | NULL | Probable | As nectar-tap_saprotroph (Põlme et al. 2020); Decay sugar-rich substrates (Põlme et al. 2020) | Põlme S, et al. 2020. Fungal Diversity 105:1-16 ((https://doi.org/10.1007/s13225-020-00466-2)) |
| Otu15171 | 1 | k__Fungi(100);p__Ascomycota(100);c__Saccharomycetes(100);o__Saccharomycetales(100);f__Metschnikowiaceae(100);g__Metschnikowia(100);s__Metschnikowia_pulcherrima(100); | Metschnikowia | 13 | Saprotroph | \|Undefined Saprotroph\| | Yeast | NULL | Probable | As nectar-tap_saprotroph (Põlme et al. 2020); Decay sugar-rich substrates (Põlme et al. 2020) | Põlme S, et al. 2020. Fungal Diversity 105:1-16 ((https://doi.org/10.1007/s13225-020-00466-2)) |
| Otu15172 | 1 | k__Fungi(100);p__Ascomycota(100);c__Saccharomycetes(100);o__Saccharomycetales(100);f__Metschnikowiaceae(100);g__Metschnikowia(100);s__Metschnikowia_pulcherrima(100); | Metschnikowia | 13 | Saprotroph | \|Undefined Saprotroph\| | Yeast | NULL | Probable | As nectar-tap_saprotroph (Põlme et al. 2020); Decay sugar-rich substrates (Põlme et al. 2020) | Põlme S, et al. 2020. Fungal Diversity 105:1-16 ((https://doi.org/10.1007/s13225-020-00466-2)) |
| Otu15173 | 1 | k__Fungi(100);p__Ascomycota(100);c__Saccharomycetes(100);o__Saccharomycetales(100);f__Metschnikowiaceae(100);g__Metschnikowia(100);s__Metschnikowia_pulcherrima(100); | Metschnikowia | 13 | Saprotroph | \|Undefined Saprotroph\| | Yeast | NULL | Probable | As nectar-tap_saprotroph (Põlme et al. 2020); Decay sugar-rich substrates (Põlme et al. 2020) | Põlme S, et al. 2020. Fungal Diversity 105:1-16 ((https://doi.org/10.1007/s13225-020-00466-2)) |
| Otu15174 | 1 | k__Fungi(100);p__Ascomycota(100);c__Saccharomycetes(100);o__Saccharomycetales(100);f__Metschnikowiaceae(100);g__Metschnikowia(100);s__Metschnikowia_pulcherrima(100); | Metschnikowia | 13 | Saprotroph | \|Undefined Saprotroph\| | Yeast | NULL | Probable | As nectar-tap_saprotroph (Põlme et al. 2020); Decay sugar-rich substrates (Põlme et al. 2020) | Põlme S, et al. 2020. Fungal Diversity 105:1-16 ((https://doi.org/10.1007/s13225-020-00466-2)) |
| Otu15175 | 1 | k__Fungi(100);p__Ascomycota(100);c__Saccharomycetes(100);o__Saccharomycetales(100);f__Metschnikowiaceae(100);g__Metschnikowia(100);g__Metschnikowia_unclassified(100); | Metschnikowia | 13 | Saprotroph | \|Undefined Saprotroph\| | Yeast | NULL | Probable | As nectar-tap_saprotroph (Põlme et al. 2020); Decay sugar-rich substrates (Põlme et al. 2020) | Põlme S, et al. 2020. Fungal Diversity 105:1-16 ((https://doi.org/10.1007/s13225-020-00466-2)) |
| Otu15176 | 1 | k__Fungi(100);p__Ascomycota(100);c__Saccharomycetes(100);o__Saccharomycetales(100);f__Metschnikowiaceae(100);g__Metschnikowia(100);s__Metschnikowia_pulcherrima(100); | Metschnikowia | 13 | Saprotroph | \|Undefined Saprotroph\| | Yeast | NULL | Probable | As nectar-tap_saprotroph (Põlme et al. 2020); Decay sugar-rich substrates (Põlme et al. 2020) | Põlme S, et al. 2020. Fungal Diversity 105:1-16 ((https://doi.org/10.1007/s13225-020-00466-2)) |
| Otu15177 | 1 | k__Fungi(100);p__Ascomycota(100);c__Saccharomycetes(100);o__Saccharomycetales(100);f__Metschnikowiaceae(100);g__Metschnikowia(100);s__Metschnikowia_pulcherrima(100); | Metschnikowia | 13 | Saprotroph | \|Undefined Saprotroph\| | Yeast | NULL | Probable | As nectar-tap_saprotroph (Põlme et al. 2020); Decay sugar-rich substrates (Põlme et al. 2020) | Põlme S, et al. 2020. Fungal Diversity 105:1-16 ((https://doi.org/10.1007/s13225-020-00466-2)) |
| Otu15178 | 1 | k__Fungi(100);p__Ascomycota(100);c__Saccharomycetes(100);o__Saccharomycetales(100);f__Metschnikowiaceae(100);g__Metschnikowia(100);s__Metschnikowia_pulcherrima(100); | Metschnikowia | 13 | Saprotroph | \|Undefined Saprotroph\| | Yeast | NULL | Probable | As nectar-tap_saprotroph (Põlme et al. 2020); Decay sugar-rich substrates (Põlme et al. 2020) | Põlme S, et al. 2020. Fungal Diversity 105:1-16 ((https://doi.org/10.1007/s13225-020-00466-2)) |
| Otu15180 | 1 | k__Fungi(100);p__Ascomycota(100);c__Saccharomycetes(100);o__Saccharomycetales(100);f__Metschnikowiaceae(100);g__Metschnikowia(100);g__Metschnikowia_unclassified(100); | Metschnikowia | 13 | Saprotroph | \|Undefined Saprotroph\| | Yeast | NULL | Probable | As nectar-tap_saprotroph (Põlme et al. 2020); Decay sugar-rich substrates (Põlme et al. 2020) | Põlme S, et al. 2020. Fungal Diversity 105:1-16 ((https://doi.org/10.1007/s13225-020-00466-2)) |
| Otu15181 | 1 | k__Fungi(100);p__Ascomycota(100);c__Saccharomycetes(100);o__Saccharomycetales(100);f__Metschnikowiaceae(100);g__Metschnikowia(100);g__Metschnikowia_unclassified(100); | Metschnikowia | 13 | Saprotroph | \|Undefined Saprotroph\| | Yeast | NULL | Probable | As nectar-tap_saprotroph (Põlme et al. 2020); Decay sugar-rich substrates (Põlme et al. 2020) | Põlme S, et al. 2020. Fungal Diversity 105:1-16 ((https://doi.org/10.1007/s13225-020-00466-2)) |
| Otu15182 | 1 | k__Fungi(100);p__Ascomycota(100);c__Saccharomycetes(100);o__Saccharomycetales(100);f__Metschnikowiaceae(100);g__Metschnikowia(100);s__Metschnikowia_pulcherrima(100); | Metschnikowia | 13 | Saprotroph | \|Undefined Saprotroph\| | Yeast | NULL | Probable | As nectar-tap_saprotroph (Põlme et al. 2020); Decay sugar-rich substrates (Põlme et al. 2020) | Põlme S, et al. 2020. Fungal Diversity 105:1-16 ((https://doi.org/10.1007/s13225-020-00466-2)) |
| Otu15183 | 1 | k__Fungi(100);p__Ascomycota(100);c__Saccharomycetes(100);o__Saccharomycetales(100);f__Metschnikowiaceae(100);g__Metschnikowia(100);s__Metschnikowia_pulcherrima(100); | Metschnikowia | 13 | Saprotroph | \|Undefined Saprotroph\| | Yeast | NULL | Probable | As nectar-tap_saprotroph (Põlme et al. 2020); Decay sugar-rich substrates (Põlme et al. 2020) | Põlme S, et al. 2020. Fungal Diversity 105:1-16 ((https://doi.org/10.1007/s13225-020-00466-2)) |
| Otu15184 | 1 | k__Fungi(100);p__Ascomycota(100);c__Saccharomycetes(100);o__Saccharomycetales(100);f__Metschnikowiaceae(100);g__Metschnikowia(100);s__Metschnikowia_pulcherrima(100); | Metschnikowia | 13 | Saprotroph | \|Undefined Saprotroph\| | Yeast | NULL | Probable | As nectar-tap_saprotroph (Põlme et al. 2020); Decay sugar-rich substrates (Põlme et al. 2020) | Põlme S, et al. 2020. Fungal Diversity 105:1-16 ((https://doi.org/10.1007/s13225-020-00466-2)) |
| Otu15185 | 1 | k__Fungi(100);p__Ascomycota(100);c__Saccharomycetes(100);o__Saccharomycetales(100);f__Metschnikowiaceae(100);g__Metschnikowia(100);g__Metschnikowia_unclassified(100); | Metschnikowia | 13 | Saprotroph | \|Undefined Saprotroph\| | Yeast | NULL | Probable | As nectar-tap_saprotroph (Põlme et al. 2020); Decay sugar-rich substrates (Põlme et al. 2020) | Põlme S, et al. 2020. Fungal Diversity 105:1-16 ((https://doi.org/10.1007/s13225-020-00466-2)) |
| Otu15186 | 1 | k__Fungi(100);p__Ascomycota(100);c__Saccharomycetes(100);o__Saccharomycetales(100);f__Metschnikowiaceae(100);g__Metschnikowia(100);s__Metschnikowia_pulcherrima(100); | Metschnikowia | 13 | Saprotroph | \|Undefined Saprotroph\| | Yeast | NULL | Probable | As nectar-tap_saprotroph (Põlme et al. 2020); Decay sugar-rich substrates (Põlme et al. 2020) | Põlme S, et al. 2020. Fungal Diversity 105:1-16 ((https://doi.org/10.1007/s13225-020-00466-2)) |
| Otu15187 | 1 | k__Fungi(100);p__Ascomycota(100);c__Saccharomycetes(100);o__Saccharomycetales(100);f__Metschnikowiaceae(100);g__Metschnikowia(100);s__Metschnikowia_pulcherrima(100); | Metschnikowia | 13 | Saprotroph | \|Undefined Saprotroph\| | Yeast | NULL | Probable | As nectar-tap_saprotroph (Põlme et al. 2020); Decay sugar-rich substrates (Põlme et al. 2020) | Põlme S, et al. 2020. Fungal Diversity 105:1-16 ((https://doi.org/10.1007/s13225-020-00466-2)) |
| Otu15188 | 1 | k__Fungi(100);p__Ascomycota(100);c__Saccharomycetes(100);o__Saccharomycetales(100);f__Metschnikowiaceae(100);g__Metschnikowia(100);s__Metschnikowia_pulcherrima(100); | Metschnikowia | 13 | Saprotroph | \|Undefined Saprotroph\| | Yeast | NULL | Probable | As nectar-tap_saprotroph (Põlme et al. 2020); Decay sugar-rich substrates (Põlme et al. 2020) | Põlme S, et al. 2020. Fungal Diversity 105:1-16 ((https://doi.org/10.1007/s13225-020-00466-2)) |
| Otu15189 | 1 | k__Fungi(100);p__Ascomycota(100);c__Saccharomycetes(100);o__Saccharomycetales(100);f__Metschnikowiaceae(100);g__Metschnikowia(100);g__Metschnikowia_unclassified(100); | Metschnikowia | 13 | Saprotroph | \|Undefined Saprotroph\| | Yeast | NULL | Probable | As nectar-tap_saprotroph (Põlme et al. 2020); Decay sugar-rich substrates (Põlme et al. 2020) | Põlme S, et al. 2020. Fungal Diversity 105:1-16 ((https://doi.org/10.1007/s13225-020-00466-2)) |
| Otu15190 | 1 | k__Fungi(100);p__Ascomycota(100);c__Saccharomycetes(100);o__Saccharomycetales(100);f__Metschnikowiaceae(100);g__Metschnikowia(100);g__Metschnikowia_unclassified(100); | Metschnikowia | 13 | Saprotroph | \|Undefined Saprotroph\| | Yeast | NULL | Probable | As nectar-tap_saprotroph (Põlme et al. 2020); Decay sugar-rich substrates (Põlme et al. 2020) | Põlme S, et al. 2020. Fungal Diversity 105:1-16 ((https://doi.org/10.1007/s13225-020-00466-2)) |
| Otu15191 | 1 | k__Fungi(100);p__Basidiomycota(100);c__Microbotryomycetes(100);o__Sporidiobolales(100);f__Sporidiobolales_family_Incertae_sedis(100);g__Sporobolomyces(100);s__Sporobolomyces_lactosus(100); | Sporobolomyces | 13 | Pathotroph-Saprotroph | \|Fungal Parasite\|-Plant Saprotroph | Yeast | NULL | Probable | As litter_saprotroph (Põlme et al. 2020); As mycoparasite (Põlme et al. 2020); Decay fungal material (Põlme et al. 2020) | Tedersoo L, et al. 2014. Science 346:e1256688 ((https://doi.org/10.1126/science.1256688)); Purahong W, et al. 2016. Molecular Ecology 25:4059-4074 ((https://doi.org/10.1111/mec.13739)); Põlme S, et al. 2020. Fungal Diversity 105:1-16 ((https://doi.org/10.1007/s13225-020-00466-2)) |
| Otu15192 | 1 | k__Fungi(100);p__Ascomycota(100);c__Saccharomycetes(100);o__Saccharomycetales(100);f__Metschnikowiaceae(100);g__Metschnikowia(100);s__Metschnikowia_pulcherrima(100); | Metschnikowia | 13 | Saprotroph | \|Undefined Saprotroph\| | Yeast | NULL | Probable | As nectar-tap_saprotroph (Põlme et al. 2020); Decay sugar-rich substrates (Põlme et al. 2020) | Põlme S, et al. 2020. Fungal Diversity 105:1-16 ((https://doi.org/10.1007/s13225-020-00466-2)) |
| Otu15193 | 1 | k__Fungi(100);p__Ascomycota(100);c__Saccharomycetes(100);o__Saccharomycetales(100);f__Metschnikowiaceae(100);g__Metschnikowia(100);s__Metschnikowia_pulcherrima(100); | Metschnikowia | 13 | Saprotroph | \|Undefined Saprotroph\| | Yeast | NULL | Probable | As nectar-tap_saprotroph (Põlme et al. 2020); Decay sugar-rich substrates (Põlme et al. 2020) | Põlme S, et al. 2020. Fungal Diversity 105:1-16 ((https://doi.org/10.1007/s13225-020-00466-2)) |
| Otu15194 | 1 | k__Fungi(100);p__Ascomycota(100);c__Saccharomycetes(100);o__Saccharomycetales(100);f__Metschnikowiaceae(100);g__Metschnikowia(100);s__Metschnikowia_pulcherrima(100); | Metschnikowia | 13 | Saprotroph | \|Undefined Saprotroph\| | Yeast | NULL | Probable | As nectar-tap_saprotroph (Põlme et al. 2020); Decay sugar-rich substrates (Põlme et al. 2020) | Põlme S, et al. 2020. Fungal Diversity 105:1-16 ((https://doi.org/10.1007/s13225-020-00466-2)) |
| Otu15195 | 1 | k__Fungi(100);p__Ascomycota(100);c__Saccharomycetes(100);o__Saccharomycetales(100);f__Metschnikowiaceae(100);g__Metschnikowia(100);s__Metschnikowia_pulcherrima(100); | Metschnikowia | 13 | Saprotroph | \|Undefined Saprotroph\| | Yeast | NULL | Probable | As nectar-tap_saprotroph (Põlme et al. 2020); Decay sugar-rich substrates (Põlme et al. 2020) | Põlme S, et al. 2020. Fungal Diversity 105:1-16 ((https://doi.org/10.1007/s13225-020-00466-2)) |
| Otu15196 | 1 | k__Fungi(100);p__Ascomycota(100);c__Saccharomycetes(100);o__Saccharomycetales(100);f__Metschnikowiaceae(100);g__Metschnikowia(100);s__Metschnikowia_pulcherrima(100); | Metschnikowia | 13 | Saprotroph | \|Undefined Saprotroph\| | Yeast | NULL | Probable | As nectar-tap_saprotroph (Põlme et al. 2020); Decay sugar-rich substrates (Põlme et al. 2020) | Põlme S, et al. 2020. Fungal Diversity 105:1-16 ((https://doi.org/10.1007/s13225-020-00466-2)) |
| Otu15197 | 1 | k__Fungi(100);p__Ascomycota(100);c__Saccharomycetes(100);o__Saccharomycetales(100);f__Metschnikowiaceae(100);g__Metschnikowia(100);s__Metschnikowia_pulcherrima(100); | Metschnikowia | 13 | Saprotroph | \|Undefined Saprotroph\| | Yeast | NULL | Probable | As nectar-tap_saprotroph (Põlme et al. 2020); Decay sugar-rich substrates (Põlme et al. 2020) | Põlme S, et al. 2020. Fungal Diversity 105:1-16 ((https://doi.org/10.1007/s13225-020-00466-2)) |
| Otu15198 | 1 | k__Fungi(100);p__Ascomycota(100);c__Saccharomycetes(100);o__Saccharomycetales(100);f__Saccharomycetales_family_Incertae_sedis(100);g__Candida(100);s__Candida_stellata(100); | Candida | 13 | Pathotroph-Saprotroph-Symbiotroph | Animal Pathogen-Endophyte-Endosymbiont-Epiphyte-\|Undefined Saprotroph\| | Dimorphic Yeast-Yeast | NULL | Probable | As nectar-tap_saprotroph (Põlme et al. 2020); Decay sugar-rich substrates (Põlme et al. 2020); This asexual genus is polyphyletic and contain species that function very differently depend on what they are. Caution should be taken when interpreting guilds for Candida species. | Manolakaki D, et al. 2010. Virulence. 1: 367-375 ((https://doi.org/10.4161/viru.1.5.12796)); Mekbib SB, et al. 2011. Tropical Plant Pathology 36:233-240 ((https://doi.org/10.1590/S1982-56762011000400004)); Põlme S, et al. 2020. Fungal Diversity 105:1-16 ((https://doi.org/10.1007/s13225-020-00466-2)) |
| Otu15199 | 1 | k__Fungi(100);p__Ascomycota(100);c__Saccharomycetes(100);o__Saccharomycetales(100);f__Metschnikowiaceae(100);g__Metschnikowia(100);g__Metschnikowia_unclassified(100); | Metschnikowia | 13 | Saprotroph | \|Undefined Saprotroph\| | Yeast | NULL | Probable | As nectar-tap_saprotroph (Põlme et al. 2020); Decay sugar-rich substrates (Põlme et al. 2020) | Põlme S, et al. 2020. Fungal Diversity 105:1-16 ((https://doi.org/10.1007/s13225-020-00466-2)) |
| Otu15200 | 1 | k__Fungi(100);p__Ascomycota(100);c__Saccharomycetes(100);o__Saccharomycetales(100);f__Metschnikowiaceae(100);g__Metschnikowia(100);s__Metschnikowia_pulcherrima(100); | Metschnikowia | 13 | Saprotroph | \|Undefined Saprotroph\| | Yeast | NULL | Probable | As nectar-tap_saprotroph (Põlme et al. 2020); Decay sugar-rich substrates (Põlme et al. 2020) | Põlme S, et al. 2020. Fungal Diversity 105:1-16 ((https://doi.org/10.1007/s13225-020-00466-2)) |
| Otu15201 | 1 | k__Fungi(100);p__Ascomycota(100);c__Saccharomycetes(100);o__Saccharomycetales(100);f__Metschnikowiaceae(100);g__Metschnikowia(100);s__Metschnikowia_pulcherrima(100); | Metschnikowia | 13 | Saprotroph | \|Undefined Saprotroph\| | Yeast | NULL | Probable | As nectar-tap_saprotroph (Põlme et al. 2020); Decay sugar-rich substrates (Põlme et al. 2020) | Põlme S, et al. 2020. Fungal Diversity 105:1-16 ((https://doi.org/10.1007/s13225-020-00466-2)) |
| Otu15202 | 1 | k__Fungi(100);p__Ascomycota(100);c__Saccharomycetes(100);o__Saccharomycetales(100);f__Metschnikowiaceae(100);g__Metschnikowia(100);s__Metschnikowia_pulcherrima(100); | Metschnikowia | 13 | Saprotroph | \|Undefined Saprotroph\| | Yeast | NULL | Probable | As nectar-tap_saprotroph (Põlme et al. 2020); Decay sugar-rich substrates (Põlme et al. 2020) | Põlme S, et al. 2020. Fungal Diversity 105:1-16 ((https://doi.org/10.1007/s13225-020-00466-2)) |
| Otu15203 | 1 | k__Fungi(100);p__Ascomycota(100);c__Saccharomycetes(100);o__Saccharomycetales(100);f__Metschnikowiaceae(100);g__Metschnikowia(100);g__Metschnikowia_unclassified(100); | Metschnikowia | 13 | Saprotroph | \|Undefined Saprotroph\| | Yeast | NULL | Probable | As nectar-tap_saprotroph (Põlme et al. 2020); Decay sugar-rich substrates (Põlme et al. 2020) | Põlme S, et al. 2020. Fungal Diversity 105:1-16 ((https://doi.org/10.1007/s13225-020-00466-2)) |
| Otu15204 | 1 | k__Fungi(100);p__Ascomycota(100);c__Saccharomycetes(100);o__Saccharomycetales(100);f__Metschnikowiaceae(100);g__Metschnikowia(100);s__Metschnikowia_pulcherrima(100); | Metschnikowia | 13 | Saprotroph | \|Undefined Saprotroph\| | Yeast | NULL | Probable | As nectar-tap_saprotroph (Põlme et al. 2020); Decay sugar-rich substrates (Põlme et al. 2020) | Põlme S, et al. 2020. Fungal Diversity 105:1-16 ((https://doi.org/10.1007/s13225-020-00466-2)) |
| Otu15205 | 1 | k__Fungi(100);p__Ascomycota(100);c__Saccharomycetes(100);o__Saccharomycetales(100);f__Metschnikowiaceae(100);g__Metschnikowia(100);g__Metschnikowia_unclassified(100); | Metschnikowia | 13 | Saprotroph | \|Undefined Saprotroph\| | Yeast | NULL | Probable | As nectar-tap_saprotroph (Põlme et al. 2020); Decay sugar-rich substrates (Põlme et al. 2020) | Põlme S, et al. 2020. Fungal Diversity 105:1-16 ((https://doi.org/10.1007/s13225-020-00466-2)) |
| Otu15206 | 1 | k__Fungi(100);p__Ascomycota(100);c__Saccharomycetes(100);o__Saccharomycetales(100);f__Metschnikowiaceae(100);g__Metschnikowia(100);s__Metschnikowia_pulcherrima(100); | Metschnikowia | 13 | Saprotroph | \|Undefined Saprotroph\| | Yeast | NULL | Probable | As nectar-tap_saprotroph (Põlme et al. 2020); Decay sugar-rich substrates (Põlme et al. 2020) | Põlme S, et al. 2020. Fungal Diversity 105:1-16 ((https://doi.org/10.1007/s13225-020-00466-2)) |
| Otu15207 | 1 | k__Fungi(100);p__Ascomycota(100);c__Saccharomycetes(100);o__Saccharomycetales(100);f__Metschnikowiaceae(100);g__Metschnikowia(100);s__Metschnikowia_pulcherrima(100); | Metschnikowia | 13 | Saprotroph | \|Undefined Saprotroph\| | Yeast | NULL | Probable | As nectar-tap_saprotroph (Põlme et al. 2020); Decay sugar-rich substrates (Põlme et al. 2020) | Põlme S, et al. 2020. Fungal Diversity 105:1-16 ((https://doi.org/10.1007/s13225-020-00466-2)) |
| Otu15208 | 1 | k__Fungi(100);p__Ascomycota(100);c__Saccharomycetes(100);o__Saccharomycetales(100);f__Metschnikowiaceae(100);g__Metschnikowia(100);s__Metschnikowia_pulcherrima(100); | Metschnikowia | 13 | Saprotroph | \|Undefined Saprotroph\| | Yeast | NULL | Probable | As nectar-tap_saprotroph (Põlme et al. 2020); Decay sugar-rich substrates (Põlme et al. 2020) | Põlme S, et al. 2020. Fungal Diversity 105:1-16 ((https://doi.org/10.1007/s13225-020-00466-2)) |
| Otu15210 | 1 | k__Fungi(100);p__Ascomycota(100);c__Saccharomycetes(100);o__Saccharomycetales(100);f__Metschnikowiaceae(100);g__Metschnikowia(100);g__Metschnikowia_unclassified(100); | Metschnikowia | 13 | Saprotroph | \|Undefined Saprotroph\| | Yeast | NULL | Probable | As nectar-tap_saprotroph (Põlme et al. 2020); Decay sugar-rich substrates (Põlme et al. 2020) | Põlme S, et al. 2020. Fungal Diversity 105:1-16 ((https://doi.org/10.1007/s13225-020-00466-2)) |
| Otu15211 | 1 | k__Fungi(100);p__Ascomycota(100);c__Saccharomycetes(100);o__Saccharomycetales(100);f__Metschnikowiaceae(100);g__Metschnikowia(100);s__Metschnikowia_pulcherrima(100); | Metschnikowia | 13 | Saprotroph | \|Undefined Saprotroph\| | Yeast | NULL | Probable | As nectar-tap_saprotroph (Põlme et al. 2020); Decay sugar-rich substrates (Põlme et al. 2020) | Põlme S, et al. 2020. Fungal Diversity 105:1-16 ((https://doi.org/10.1007/s13225-020-00466-2)) |
| Otu15212 | 1 | k__Fungi(100);p__Ascomycota(100);c__Saccharomycetes(100);o__Saccharomycetales(100);f__Metschnikowiaceae(100);g__Metschnikowia(100);g__Metschnikowia_unclassified(100); | Metschnikowia | 13 | Saprotroph | \|Undefined Saprotroph\| | Yeast | NULL | Probable | As nectar-tap_saprotroph (Põlme et al. 2020); Decay sugar-rich substrates (Põlme et al. 2020) | Põlme S, et al. 2020. Fungal Diversity 105:1-16 ((https://doi.org/10.1007/s13225-020-00466-2)) |
| Otu15213 | 1 | k__Fungi(100);p__Ascomycota(100);c__Saccharomycetes(100);o__Saccharomycetales(100);f__Metschnikowiaceae(100);g__Metschnikowia(100);s__Metschnikowia_pulcherrima(100); | Metschnikowia | 13 | Saprotroph | \|Undefined Saprotroph\| | Yeast | NULL | Probable | As nectar-tap_saprotroph (Põlme et al. 2020); Decay sugar-rich substrates (Põlme et al. 2020) | Põlme S, et al. 2020. Fungal Diversity 105:1-16 ((https://doi.org/10.1007/s13225-020-00466-2)) |
| Otu15215 | 1 | k__Fungi(100);p__Ascomycota(100);c__Saccharomycetes(100);o__Saccharomycetales(100);f__Metschnikowiaceae(100);g__Metschnikowia(100);g__Metschnikowia_unclassified(100); | Metschnikowia | 13 | Saprotroph | \|Undefined Saprotroph\| | Yeast | NULL | Probable | As nectar-tap_saprotroph (Põlme et al. 2020); Decay sugar-rich substrates (Põlme et al. 2020) | Põlme S, et al. 2020. Fungal Diversity 105:1-16 ((https://doi.org/10.1007/s13225-020-00466-2)) |
| Otu15216 | 1 | k__Fungi(100);p__Ascomycota(100);c__Saccharomycetes(100);o__Saccharomycetales(100);f__Metschnikowiaceae(100);g__Metschnikowia(100);s__Metschnikowia_pulcherrima(100); | Metschnikowia | 13 | Saprotroph | \|Undefined Saprotroph\| | Yeast | NULL | Probable | As nectar-tap_saprotroph (Põlme et al. 2020); Decay sugar-rich substrates (Põlme et al. 2020) | Põlme S, et al. 2020. Fungal Diversity 105:1-16 ((https://doi.org/10.1007/s13225-020-00466-2)) |
| Otu15217 | 1 | k__Fungi(100);p__Ascomycota(100);c__Saccharomycetes(100);o__Saccharomycetales(100);f__Metschnikowiaceae(100);g__Metschnikowia(100);s__Metschnikowia_pulcherrima(100); | Metschnikowia | 13 | Saprotroph | \|Undefined Saprotroph\| | Yeast | NULL | Probable | As nectar-tap_saprotroph (Põlme et al. 2020); Decay sugar-rich substrates (Põlme et al. 2020) | Põlme S, et al. 2020. Fungal Diversity 105:1-16 ((https://doi.org/10.1007/s13225-020-00466-2)) |
| Otu15218 | 1 | k__Fungi(100);p__Ascomycota(100);c__Saccharomycetes(100);o__Saccharomycetales(100);f__Metschnikowiaceae(100);g__Metschnikowia(100);s__Metschnikowia_pulcherrima(100); | Metschnikowia | 13 | Saprotroph | \|Undefined Saprotroph\| | Yeast | NULL | Probable | As nectar-tap_saprotroph (Põlme et al. 2020); Decay sugar-rich substrates (Põlme et al. 2020) | Põlme S, et al. 2020. Fungal Diversity 105:1-16 ((https://doi.org/10.1007/s13225-020-00466-2)) |
| Otu15219 | 1 | k__Fungi(100);p__Ascomycota(100);c__Saccharomycetes(100);o__Saccharomycetales(100);f__Metschnikowiaceae(100);g__Metschnikowia(100);s__Metschnikowia_pulcherrima(100); | Metschnikowia | 13 | Saprotroph | \|Undefined Saprotroph\| | Yeast | NULL | Probable | As nectar-tap_saprotroph (Põlme et al. 2020); Decay sugar-rich substrates (Põlme et al. 2020) | Põlme S, et al. 2020. Fungal Diversity 105:1-16 ((https://doi.org/10.1007/s13225-020-00466-2)) |
| Otu15220 | 1 | k__Fungi(100);p__Ascomycota(100);c__Saccharomycetes(100);o__Saccharomycetales(100);f__Metschnikowiaceae(100);g__Metschnikowia(100);s__Metschnikowia_pulcherrima(100); | Metschnikowia | 13 | Saprotroph | \|Undefined Saprotroph\| | Yeast | NULL | Probable | As nectar-tap_saprotroph (Põlme et al. 2020); Decay sugar-rich substrates (Põlme et al. 2020) | Põlme S, et al. 2020. Fungal Diversity 105:1-16 ((https://doi.org/10.1007/s13225-020-00466-2)) |
| Otu15221 | 1 | k__Fungi(100);p__Ascomycota(100);c__Saccharomycetes(100);o__Saccharomycetales(100);f__Metschnikowiaceae(100);g__Metschnikowia(100);g__Metschnikowia_unclassified(100); | Metschnikowia | 13 | Saprotroph | \|Undefined Saprotroph\| | Yeast | NULL | Probable | As nectar-tap_saprotroph (Põlme et al. 2020); Decay sugar-rich substrates (Põlme et al. 2020) | Põlme S, et al. 2020. Fungal Diversity 105:1-16 ((https://doi.org/10.1007/s13225-020-00466-2)) |
| Otu15223 | 1 | k__Fungi(100);p__Ascomycota(100);c__Saccharomycetes(100);o__Saccharomycetales(100);f__Metschnikowiaceae(100);g__Metschnikowia(100);s__Metschnikowia_pulcherrima(100); | Metschnikowia | 13 | Saprotroph | \|Undefined Saprotroph\| | Yeast | NULL | Probable | As nectar-tap_saprotroph (Põlme et al. 2020); Decay sugar-rich substrates (Põlme et al. 2020) | Põlme S, et al. 2020. Fungal Diversity 105:1-16 ((https://doi.org/10.1007/s13225-020-00466-2)) |
| Otu15224 | 1 | k__Fungi(100);p__Ascomycota(100);c__Saccharomycetes(100);o__Saccharomycetales(100);f__Metschnikowiaceae(100);g__Metschnikowia(100);s__Metschnikowia_pulcherrima(100); | Metschnikowia | 13 | Saprotroph | \|Undefined Saprotroph\| | Yeast | NULL | Probable | As nectar-tap_saprotroph (Põlme et al. 2020); Decay sugar-rich substrates (Põlme et al. 2020) | Põlme S, et al. 2020. Fungal Diversity 105:1-16 ((https://doi.org/10.1007/s13225-020-00466-2)) |
| Otu15225 | 1 | k__Fungi(100);p__Ascomycota(100);c__Saccharomycetes(100);o__Saccharomycetales(100);f__Metschnikowiaceae(100);g__Metschnikowia(100);s__Metschnikowia_pulcherrima(100); | Metschnikowia | 13 | Saprotroph | \|Undefined Saprotroph\| | Yeast | NULL | Probable | As nectar-tap_saprotroph (Põlme et al. 2020); Decay sugar-rich substrates (Põlme et al. 2020) | Põlme S, et al. 2020. Fungal Diversity 105:1-16 ((https://doi.org/10.1007/s13225-020-00466-2)) |
| Otu15226 | 1 | k__Fungi(100);p__Ascomycota(100);c__Saccharomycetes(100);o__Saccharomycetales(100);f__Metschnikowiaceae(100);g__Metschnikowia(100);s__Metschnikowia_pulcherrima(100); | Metschnikowia | 13 | Saprotroph | \|Undefined Saprotroph\| | Yeast | NULL | Probable | As nectar-tap_saprotroph (Põlme et al. 2020); Decay sugar-rich substrates (Põlme et al. 2020) | Põlme S, et al. 2020. Fungal Diversity 105:1-16 ((https://doi.org/10.1007/s13225-020-00466-2)) |
| Otu15227 | 1 | k__Fungi(100);p__Ascomycota(100);c__Saccharomycetes(100);o__Saccharomycetales(100);f__Metschnikowiaceae(100);g__Metschnikowia(100);s__Metschnikowia_pulcherrima(100); | Metschnikowia | 13 | Saprotroph | \|Undefined Saprotroph\| | Yeast | NULL | Probable | As nectar-tap_saprotroph (Põlme et al. 2020); Decay sugar-rich substrates (Põlme et al. 2020) | Põlme S, et al. 2020. Fungal Diversity 105:1-16 ((https://doi.org/10.1007/s13225-020-00466-2)) |
| Otu15228 | 1 | k__Fungi(100);p__Ascomycota(100);c__Saccharomycetes(100);o__Saccharomycetales(100);f__Metschnikowiaceae(100);g__Metschnikowia(100);s__Metschnikowia_pulcherrima(100); | Metschnikowia | 13 | Saprotroph | \|Undefined Saprotroph\| | Yeast | NULL | Probable | As nectar-tap_saprotroph (Põlme et al. 2020); Decay sugar-rich substrates (Põlme et al. 2020) | Põlme S, et al. 2020. Fungal Diversity 105:1-16 ((https://doi.org/10.1007/s13225-020-00466-2)) |
| Otu15229 | 1 | k__Fungi(100);p__Ascomycota(100);c__Saccharomycetes(100);o__Saccharomycetales(100);f__Metschnikowiaceae(100);g__Metschnikowia(100);s__Metschnikowia_pulcherrima(100); | Metschnikowia | 13 | Saprotroph | \|Undefined Saprotroph\| | Yeast | NULL | Probable | As nectar-tap_saprotroph (Põlme et al. 2020); Decay sugar-rich substrates (Põlme et al. 2020) | Põlme S, et al. 2020. Fungal Diversity 105:1-16 ((https://doi.org/10.1007/s13225-020-00466-2)) |
| Otu15230 | 1 | k__Fungi(100);p__Ascomycota(100);c__Saccharomycetes(100);o__Saccharomycetales(100);f__Metschnikowiaceae(100);g__Metschnikowia(100);s__Metschnikowia_pulcherrima(100); | Metschnikowia | 13 | Saprotroph | \|Undefined Saprotroph\| | Yeast | NULL | Probable | As nectar-tap_saprotroph (Põlme et al. 2020); Decay sugar-rich substrates (Põlme et al. 2020) | Põlme S, et al. 2020. Fungal Diversity 105:1-16 ((https://doi.org/10.1007/s13225-020-00466-2)) |
| Otu15231 | 1 | k__Fungi(100);p__Ascomycota(100);c__Saccharomycetes(100);o__Saccharomycetales(100);f__Metschnikowiaceae(100);g__Metschnikowia(100);s__Metschnikowia_pulcherrima(100); | Metschnikowia | 13 | Saprotroph | \|Undefined Saprotroph\| | Yeast | NULL | Probable | As nectar-tap_saprotroph (Põlme et al. 2020); Decay sugar-rich substrates (Põlme et al. 2020) | Põlme S, et al. 2020. Fungal Diversity 105:1-16 ((https://doi.org/10.1007/s13225-020-00466-2)) |
| Otu15232 | 1 | k__Fungi(100);p__Ascomycota(100);c__Saccharomycetes(100);o__Saccharomycetales(100);f__Metschnikowiaceae(100);g__Metschnikowia(100);s__Metschnikowia_pulcherrima(100); | Metschnikowia | 13 | Saprotroph | \|Undefined Saprotroph\| | Yeast | NULL | Probable | As nectar-tap_saprotroph (Põlme et al. 2020); Decay sugar-rich substrates (Põlme et al. 2020) | Põlme S, et al. 2020. Fungal Diversity 105:1-16 ((https://doi.org/10.1007/s13225-020-00466-2)) |
| Otu15233 | 1 | k__Fungi(100);p__Ascomycota(100);c__Saccharomycetes(100);o__Saccharomycetales(100);f__Metschnikowiaceae(100);g__Metschnikowia(100);s__Metschnikowia_pulcherrima(100); | Metschnikowia | 13 | Saprotroph | \|Undefined Saprotroph\| | Yeast | NULL | Probable | As nectar-tap_saprotroph (Põlme et al. 2020); Decay sugar-rich substrates (Põlme et al. 2020) | Põlme S, et al. 2020. Fungal Diversity 105:1-16 ((https://doi.org/10.1007/s13225-020-00466-2)) |
| Otu15234 | 1 | k__Fungi(100);p__Ascomycota(100);c__Saccharomycetes(100);o__Saccharomycetales(100);f__Metschnikowiaceae(100);g__Metschnikowia(100);s__Metschnikowia_pulcherrima(100); | Metschnikowia | 13 | Saprotroph | \|Undefined Saprotroph\| | Yeast | NULL | Probable | As nectar-tap_saprotroph (Põlme et al. 2020); Decay sugar-rich substrates (Põlme et al. 2020) | Põlme S, et al. 2020. Fungal Diversity 105:1-16 ((https://doi.org/10.1007/s13225-020-00466-2)) |
| Otu15235 | 1 | k__Fungi(100);p__Ascomycota(100);c__Saccharomycetes(100);o__Saccharomycetales(100);f__Metschnikowiaceae(100);g__Metschnikowia(100);s__Metschnikowia_pulcherrima(100); | Metschnikowia | 13 | Saprotroph | \|Undefined Saprotroph\| | Yeast | NULL | Probable | As nectar-tap_saprotroph (Põlme et al. 2020); Decay sugar-rich substrates (Põlme et al. 2020) | Põlme S, et al. 2020. Fungal Diversity 105:1-16 ((https://doi.org/10.1007/s13225-020-00466-2)) |
| Otu15236 | 1 | k__Fungi(100);p__Ascomycota(100);c__Saccharomycetes(100);o__Saccharomycetales(100);f__Metschnikowiaceae(100);g__Metschnikowia(100);g__Metschnikowia_unclassified(100); | Metschnikowia | 13 | Saprotroph | \|Undefined Saprotroph\| | Yeast | NULL | Probable | As nectar-tap_saprotroph (Põlme et al. 2020); Decay sugar-rich substrates (Põlme et al. 2020) | Põlme S, et al. 2020. Fungal Diversity 105:1-16 ((https://doi.org/10.1007/s13225-020-00466-2)) |
| Otu15237 | 1 | k__Fungi(100);p__Ascomycota(100);c__Saccharomycetes(100);o__Saccharomycetales(100);f__Metschnikowiaceae(100);g__Metschnikowia(100);s__Metschnikowia_pulcherrima(100); | Metschnikowia | 13 | Saprotroph | \|Undefined Saprotroph\| | Yeast | NULL | Probable | As nectar-tap_saprotroph (Põlme et al. 2020); Decay sugar-rich substrates (Põlme et al. 2020) | Põlme S, et al. 2020. Fungal Diversity 105:1-16 ((https://doi.org/10.1007/s13225-020-00466-2)) |
| Otu15238 | 1 | k__Fungi(100);p__Ascomycota(100);c__Saccharomycetes(100);o__Saccharomycetales(100);f__Metschnikowiaceae(100);g__Metschnikowia(100);s__Metschnikowia_pulcherrima(100); | Metschnikowia | 13 | Saprotroph | \|Undefined Saprotroph\| | Yeast | NULL | Probable | As nectar-tap_saprotroph (Põlme et al. 2020); Decay sugar-rich substrates (Põlme et al. 2020) | Põlme S, et al. 2020. Fungal Diversity 105:1-16 ((https://doi.org/10.1007/s13225-020-00466-2)) |
| Otu15239 | 1 | k__Fungi(100);p__Ascomycota(100);c__Saccharomycetes(100);o__Saccharomycetales(100);f__Metschnikowiaceae(100);g__Metschnikowia(100);s__Metschnikowia_pulcherrima(100); | Metschnikowia | 13 | Saprotroph | \|Undefined Saprotroph\| | Yeast | NULL | Probable | As nectar-tap_saprotroph (Põlme et al. 2020); Decay sugar-rich substrates (Põlme et al. 2020) | Põlme S, et al. 2020. Fungal Diversity 105:1-16 ((https://doi.org/10.1007/s13225-020-00466-2)) |
| Otu15240 | 1 | k__Fungi(100);p__Ascomycota(100);c__Saccharomycetes(100);o__Saccharomycetales(100);f__Metschnikowiaceae(100);g__Metschnikowia(100);g__Metschnikowia_unclassified(100); | Metschnikowia | 13 | Saprotroph | \|Undefined Saprotroph\| | Yeast | NULL | Probable | As nectar-tap_saprotroph (Põlme et al. 2020); Decay sugar-rich substrates (Põlme et al. 2020) | Põlme S, et al. 2020. Fungal Diversity 105:1-16 ((https://doi.org/10.1007/s13225-020-00466-2)) |
| Otu15241 | 1 | k__Fungi(100);p__Ascomycota(100);c__Saccharomycetes(100);o__Saccharomycetales(100);f__Metschnikowiaceae(100);g__Metschnikowia(100);s__Metschnikowia_pulcherrima(100); | Metschnikowia | 13 | Saprotroph | \|Undefined Saprotroph\| | Yeast | NULL | Probable | As nectar-tap_saprotroph (Põlme et al. 2020); Decay sugar-rich substrates (Põlme et al. 2020) | Põlme S, et al. 2020. Fungal Diversity 105:1-16 ((https://doi.org/10.1007/s13225-020-00466-2)) |
| Otu15242 | 1 | k__Fungi(100);p__Ascomycota(100);c__Saccharomycetes(100);o__Saccharomycetales(100);f__Metschnikowiaceae(100);g__Metschnikowia(100);s__Metschnikowia_pulcherrima(100); | Metschnikowia | 13 | Saprotroph | \|Undefined Saprotroph\| | Yeast | NULL | Probable | As nectar-tap_saprotroph (Põlme et al. 2020); Decay sugar-rich substrates (Põlme et al. 2020) | Põlme S, et al. 2020. Fungal Diversity 105:1-16 ((https://doi.org/10.1007/s13225-020-00466-2)) |
| Otu15245 | 1 | k__Fungi(100);p__Ascomycota(100);c__Saccharomycetes(100);o__Saccharomycetales(100);f__Metschnikowiaceae(100);g__Metschnikowia(100);g__Metschnikowia_unclassified(100); | Metschnikowia | 13 | Saprotroph | \|Undefined Saprotroph\| | Yeast | NULL | Probable | As nectar-tap_saprotroph (Põlme et al. 2020); Decay sugar-rich substrates (Põlme et al. 2020) | Põlme S, et al. 2020. Fungal Diversity 105:1-16 ((https://doi.org/10.1007/s13225-020-00466-2)) |
| Otu15246 | 1 | k__Fungi(100);p__Ascomycota(100);c__Saccharomycetes(100);o__Saccharomycetales(100);f__Metschnikowiaceae(100);g__Metschnikowia(100);s__Metschnikowia_pulcherrima(100); | Metschnikowia | 13 | Saprotroph | \|Undefined Saprotroph\| | Yeast | NULL | Probable | As nectar-tap_saprotroph (Põlme et al. 2020); Decay sugar-rich substrates (Põlme et al. 2020) | Põlme S, et al. 2020. Fungal Diversity 105:1-16 ((https://doi.org/10.1007/s13225-020-00466-2)) |
| Otu15247 | 1 | k__Fungi(100);p__Ascomycota(100);c__Saccharomycetes(100);o__Saccharomycetales(100);f__Metschnikowiaceae(100);g__Metschnikowia(100);s__Metschnikowia_pulcherrima(100); | Metschnikowia | 13 | Saprotroph | \|Undefined Saprotroph\| | Yeast | NULL | Probable | As nectar-tap_saprotroph (Põlme et al. 2020); Decay sugar-rich substrates (Põlme et al. 2020) | Põlme S, et al. 2020. Fungal Diversity 105:1-16 ((https://doi.org/10.1007/s13225-020-00466-2)) |
| Otu15248 | 1 | k__Fungi(100);p__Ascomycota(100);c__Saccharomycetes(100);o__Saccharomycetales(100);f__Metschnikowiaceae(100);g__Metschnikowia(100);s__Metschnikowia_pulcherrima(100); | Metschnikowia | 13 | Saprotroph | \|Undefined Saprotroph\| | Yeast | NULL | Probable | As nectar-tap_saprotroph (Põlme et al. 2020); Decay sugar-rich substrates (Põlme et al. 2020) | Põlme S, et al. 2020. Fungal Diversity 105:1-16 ((https://doi.org/10.1007/s13225-020-00466-2)) |
| Otu15249 | 1 | k__Fungi(100);p__Ascomycota(100);c__Saccharomycetes(100);o__Saccharomycetales(100);f__Metschnikowiaceae(100);g__Metschnikowia(100);s__Metschnikowia_pulcherrima(100); | Metschnikowia | 13 | Saprotroph | \|Undefined Saprotroph\| | Yeast | NULL | Probable | As nectar-tap_saprotroph (Põlme et al. 2020); Decay sugar-rich substrates (Põlme et al. 2020) | Põlme S, et al. 2020. Fungal Diversity 105:1-16 ((https://doi.org/10.1007/s13225-020-00466-2)) |
| Otu15250 | 1 | k__Fungi(100);p__Ascomycota(100);c__Saccharomycetes(100);o__Saccharomycetales(100);f__Metschnikowiaceae(100);g__Metschnikowia(100);s__Metschnikowia_pulcherrima(100); | Metschnikowia | 13 | Saprotroph | \|Undefined Saprotroph\| | Yeast | NULL | Probable | As nectar-tap_saprotroph (Põlme et al. 2020); Decay sugar-rich substrates (Põlme et al. 2020) | Põlme S, et al. 2020. Fungal Diversity 105:1-16 ((https://doi.org/10.1007/s13225-020-00466-2)) |
| Otu15252 | 1 | k__Fungi(100);p__Ascomycota(100);c__Saccharomycetes(100);o__Saccharomycetales(100);f__Metschnikowiaceae(100);g__Metschnikowia(100);s__Metschnikowia_pulcherrima(100); | Metschnikowia | 13 | Saprotroph | \|Undefined Saprotroph\| | Yeast | NULL | Probable | As nectar-tap_saprotroph (Põlme et al. 2020); Decay sugar-rich substrates (Põlme et al. 2020) | Põlme S, et al. 2020. Fungal Diversity 105:1-16 ((https://doi.org/10.1007/s13225-020-00466-2)) |
| Otu15254 | 1 | k__Fungi(100);p__Ascomycota(100);c__Saccharomycetes(100);o__Saccharomycetales(100);f__Metschnikowiaceae(100);g__Metschnikowia(100);s__Metschnikowia_pulcherrima(100); | Metschnikowia | 13 | Saprotroph | \|Undefined Saprotroph\| | Yeast | NULL | Probable | As nectar-tap_saprotroph (Põlme et al. 2020); Decay sugar-rich substrates (Põlme et al. 2020) | Põlme S, et al. 2020. Fungal Diversity 105:1-16 ((https://doi.org/10.1007/s13225-020-00466-2)) |
| Otu15255 | 1 | k__Fungi(100);p__Ascomycota(100);c__Saccharomycetes(100);o__Saccharomycetales(100);f__Metschnikowiaceae(100);g__Metschnikowia(100);g__Metschnikowia_unclassified(100); | Metschnikowia | 13 | Saprotroph | \|Undefined Saprotroph\| | Yeast | NULL | Probable | As nectar-tap_saprotroph (Põlme et al. 2020); Decay sugar-rich substrates (Põlme et al. 2020) | Põlme S, et al. 2020. Fungal Diversity 105:1-16 ((https://doi.org/10.1007/s13225-020-00466-2)) |
| Otu15257 | 1 | k__Fungi(100);p__Ascomycota(100);c__Saccharomycetes(100);o__Saccharomycetales(100);f__Metschnikowiaceae(100);g__Metschnikowia(100);g__Metschnikowia_unclassified(100); | Metschnikowia | 13 | Saprotroph | \|Undefined Saprotroph\| | Yeast | NULL | Probable | As nectar-tap_saprotroph (Põlme et al. 2020); Decay sugar-rich substrates (Põlme et al. 2020) | Põlme S, et al. 2020. Fungal Diversity 105:1-16 ((https://doi.org/10.1007/s13225-020-00466-2)) |
| Otu15258 | 1 | k__Fungi(100);p__Ascomycota(100);c__Saccharomycetes(100);o__Saccharomycetales(100);f__Metschnikowiaceae(100);g__Metschnikowia(100);s__Metschnikowia_pulcherrima(100); | Metschnikowia | 13 | Saprotroph | \|Undefined Saprotroph\| | Yeast | NULL | Probable | As nectar-tap_saprotroph (Põlme et al. 2020); Decay sugar-rich substrates (Põlme et al. 2020) | Põlme S, et al. 2020. Fungal Diversity 105:1-16 ((https://doi.org/10.1007/s13225-020-00466-2)) |
| Otu15259 | 1 | k__Fungi(100);p__Ascomycota(100);c__Saccharomycetes(100);o__Saccharomycetales(100);f__Metschnikowiaceae(100);g__Metschnikowia(100);s__Metschnikowia_pulcherrima(100); | Metschnikowia | 13 | Saprotroph | \|Undefined Saprotroph\| | Yeast | NULL | Probable | As nectar-tap_saprotroph (Põlme et al. 2020); Decay sugar-rich substrates (Põlme et al. 2020) | Põlme S, et al. 2020. Fungal Diversity 105:1-16 ((https://doi.org/10.1007/s13225-020-00466-2)) |
| Otu15260 | 1 | k__Fungi(100);p__Ascomycota(100);c__Saccharomycetes(100);o__Saccharomycetales(100);f__Metschnikowiaceae(100);g__Metschnikowia(100);g__Metschnikowia_unclassified(100); | Metschnikowia | 13 | Saprotroph | \|Undefined Saprotroph\| | Yeast | NULL | Probable | As nectar-tap_saprotroph (Põlme et al. 2020); Decay sugar-rich substrates (Põlme et al. 2020) | Põlme S, et al. 2020. Fungal Diversity 105:1-16 ((https://doi.org/10.1007/s13225-020-00466-2)) |
| Otu15262 | 1 | k__Fungi(100);p__Ascomycota(100);c__Saccharomycetes(100);o__Saccharomycetales(100);f__Metschnikowiaceae(100);g__Metschnikowia(100);s__Metschnikowia_pulcherrima(100); | Metschnikowia | 13 | Saprotroph | \|Undefined Saprotroph\| | Yeast | NULL | Probable | As nectar-tap_saprotroph (Põlme et al. 2020); Decay sugar-rich substrates (Põlme et al. 2020) | Põlme S, et al. 2020. Fungal Diversity 105:1-16 ((https://doi.org/10.1007/s13225-020-00466-2)) |
| Otu15264 | 1 | k__Fungi(100);p__Ascomycota(100);c__Saccharomycetes(100);o__Saccharomycetales(100);f__Metschnikowiaceae(100);g__Metschnikowia(100);s__Metschnikowia_pulcherrima(100); | Metschnikowia | 13 | Saprotroph | \|Undefined Saprotroph\| | Yeast | NULL | Probable | As nectar-tap_saprotroph (Põlme et al. 2020); Decay sugar-rich substrates (Põlme et al. 2020) | Põlme S, et al. 2020. Fungal Diversity 105:1-16 ((https://doi.org/10.1007/s13225-020-00466-2)) |
| Otu15265 | 1 | k__Fungi(100);p__Ascomycota(100);c__Saccharomycetes(100);o__Saccharomycetales(100);f__Metschnikowiaceae(100);g__Metschnikowia(100);s__Metschnikowia_pulcherrima(100); | Metschnikowia | 13 | Saprotroph | \|Undefined Saprotroph\| | Yeast | NULL | Probable | As nectar-tap_saprotroph (Põlme et al. 2020); Decay sugar-rich substrates (Põlme et al. 2020) | Põlme S, et al. 2020. Fungal Diversity 105:1-16 ((https://doi.org/10.1007/s13225-020-00466-2)) |
| Otu15266 | 1 | k__Fungi(100);p__Ascomycota(100);c__Saccharomycetes(100);o__Saccharomycetales(100);f__Metschnikowiaceae(100);g__Metschnikowia(100);s__Metschnikowia_pulcherrima(100); | Metschnikowia | 13 | Saprotroph | \|Undefined Saprotroph\| | Yeast | NULL | Probable | As nectar-tap_saprotroph (Põlme et al. 2020); Decay sugar-rich substrates (Põlme et al. 2020) | Põlme S, et al. 2020. Fungal Diversity 105:1-16 ((https://doi.org/10.1007/s13225-020-00466-2)) |
| Otu15267 | 1 | k__Fungi(100);p__Ascomycota(100);c__Saccharomycetes(100);o__Saccharomycetales(100);f__Metschnikowiaceae(100);g__Metschnikowia(100);g__Metschnikowia_unclassified(100); | Metschnikowia | 13 | Saprotroph | \|Undefined Saprotroph\| | Yeast | NULL | Probable | As nectar-tap_saprotroph (Põlme et al. 2020); Decay sugar-rich substrates (Põlme et al. 2020) | Põlme S, et al. 2020. Fungal Diversity 105:1-16 ((https://doi.org/10.1007/s13225-020-00466-2)) |
| Otu15268 | 1 | k__Fungi(100);p__Ascomycota(100);c__Saccharomycetes(100);o__Saccharomycetales(100);f__Metschnikowiaceae(100);g__Metschnikowia(100);s__Metschnikowia_pulcherrima(100); | Metschnikowia | 13 | Saprotroph | \|Undefined Saprotroph\| | Yeast | NULL | Probable | As nectar-tap_saprotroph (Põlme et al. 2020); Decay sugar-rich substrates (Põlme et al. 2020) | Põlme S, et al. 2020. Fungal Diversity 105:1-16 ((https://doi.org/10.1007/s13225-020-00466-2)) |
| Otu15269 | 1 | k__Fungi(100);p__Ascomycota(100);c__Saccharomycetes(100);o__Saccharomycetales(100);f__Metschnikowiaceae(100);g__Metschnikowia(100);s__Metschnikowia_pulcherrima(100); | Metschnikowia | 13 | Saprotroph | \|Undefined Saprotroph\| | Yeast | NULL | Probable | As nectar-tap_saprotroph (Põlme et al. 2020); Decay sugar-rich substrates (Põlme et al. 2020) | Põlme S, et al. 2020. Fungal Diversity 105:1-16 ((https://doi.org/10.1007/s13225-020-00466-2)) |
| Otu15270 | 1 | k__Fungi(100);p__Ascomycota(100);c__Saccharomycetes(100);o__Saccharomycetales(100);f__Metschnikowiaceae(100);g__Metschnikowia(100);g__Metschnikowia_unclassified(100); | Metschnikowia | 13 | Saprotroph | \|Undefined Saprotroph\| | Yeast | NULL | Probable | As nectar-tap_saprotroph (Põlme et al. 2020); Decay sugar-rich substrates (Põlme et al. 2020) | Põlme S, et al. 2020. Fungal Diversity 105:1-16 ((https://doi.org/10.1007/s13225-020-00466-2)) |
| Otu15271 | 1 | k__Fungi(100);p__Ascomycota(100);c__Saccharomycetes(100);o__Saccharomycetales(100);f__Metschnikowiaceae(100);g__Metschnikowia(100);g__Metschnikowia_unclassified(100); | Metschnikowia | 13 | Saprotroph | \|Undefined Saprotroph\| | Yeast | NULL | Probable | As nectar-tap_saprotroph (Põlme et al. 2020); Decay sugar-rich substrates (Põlme et al. 2020) | Põlme S, et al. 2020. Fungal Diversity 105:1-16 ((https://doi.org/10.1007/s13225-020-00466-2)) |
| Otu15272 | 1 | k__Fungi(100);p__Ascomycota(100);c__Saccharomycetes(100);o__Saccharomycetales(100);f__Metschnikowiaceae(100);g__Metschnikowia(100);g__Metschnikowia_unclassified(100); | Metschnikowia | 13 | Saprotroph | \|Undefined Saprotroph\| | Yeast | NULL | Probable | As nectar-tap_saprotroph (Põlme et al. 2020); Decay sugar-rich substrates (Põlme et al. 2020) | Põlme S, et al. 2020. Fungal Diversity 105:1-16 ((https://doi.org/10.1007/s13225-020-00466-2)) |
| Otu15273 | 1 | k__Fungi(100);p__Ascomycota(100);c__Saccharomycetes(100);o__Saccharomycetales(100);f__Metschnikowiaceae(100);g__Metschnikowia(100);g__Metschnikowia_unclassified(100); | Metschnikowia | 13 | Saprotroph | \|Undefined Saprotroph\| | Yeast | NULL | Probable | As nectar-tap_saprotroph (Põlme et al. 2020); Decay sugar-rich substrates (Põlme et al. 2020) | Põlme S, et al. 2020. Fungal Diversity 105:1-16 ((https://doi.org/10.1007/s13225-020-00466-2)) |
| Otu15274 | 1 | k__Fungi(100);p__Ascomycota(100);c__Saccharomycetes(100);o__Saccharomycetales(100);f__Metschnikowiaceae(100);g__Metschnikowia(100);s__Metschnikowia_pulcherrima(100); | Metschnikowia | 13 | Saprotroph | \|Undefined Saprotroph\| | Yeast | NULL | Probable | As nectar-tap_saprotroph (Põlme et al. 2020); Decay sugar-rich substrates (Põlme et al. 2020) | Põlme S, et al. 2020. Fungal Diversity 105:1-16 ((https://doi.org/10.1007/s13225-020-00466-2)) |
| Otu15275 | 1 | k__Fungi(100);p__Ascomycota(100);c__Saccharomycetes(100);o__Saccharomycetales(100);f__Metschnikowiaceae(100);g__Metschnikowia(100);s__Metschnikowia_pulcherrima(100); | Metschnikowia | 13 | Saprotroph | \|Undefined Saprotroph\| | Yeast | NULL | Probable | As nectar-tap_saprotroph (Põlme et al. 2020); Decay sugar-rich substrates (Põlme et al. 2020) | Põlme S, et al. 2020. Fungal Diversity 105:1-16 ((https://doi.org/10.1007/s13225-020-00466-2)) |
| Otu15276 | 1 | k__Fungi(100);p__Ascomycota(100);c__Saccharomycetes(100);o__Saccharomycetales(100);f__Metschnikowiaceae(100);g__Metschnikowia(100);g__Metschnikowia_unclassified(100); | Metschnikowia | 13 | Saprotroph | \|Undefined Saprotroph\| | Yeast | NULL | Probable | As nectar-tap_saprotroph (Põlme et al. 2020); Decay sugar-rich substrates (Põlme et al. 2020) | Põlme S, et al. 2020. Fungal Diversity 105:1-16 ((https://doi.org/10.1007/s13225-020-00466-2)) |
| Otu15277 | 1 | k__Fungi(100);p__Ascomycota(100);c__Saccharomycetes(100);o__Saccharomycetales(100);f__Metschnikowiaceae(100);g__Metschnikowia(100);s__Metschnikowia_pulcherrima(100); | Metschnikowia | 13 | Saprotroph | \|Undefined Saprotroph\| | Yeast | NULL | Probable | As nectar-tap_saprotroph (Põlme et al. 2020); Decay sugar-rich substrates (Põlme et al. 2020) | Põlme S, et al. 2020. Fungal Diversity 105:1-16 ((https://doi.org/10.1007/s13225-020-00466-2)) |
| Otu15278 | 1 | k__Fungi(100);p__Ascomycota(100);c__Saccharomycetes(100);o__Saccharomycetales(100);f__Metschnikowiaceae(100);g__Metschnikowia(100);s__Metschnikowia_pulcherrima(100); | Metschnikowia | 13 | Saprotroph | \|Undefined Saprotroph\| | Yeast | NULL | Probable | As nectar-tap_saprotroph (Põlme et al. 2020); Decay sugar-rich substrates (Põlme et al. 2020) | Põlme S, et al. 2020. Fungal Diversity 105:1-16 ((https://doi.org/10.1007/s13225-020-00466-2)) |
| Otu15279 | 1 | k__Fungi(100);p__Ascomycota(100);c__Saccharomycetes(100);o__Saccharomycetales(100);f__Metschnikowiaceae(100);g__Metschnikowia(100);s__Metschnikowia_pulcherrima(100); | Metschnikowia | 13 | Saprotroph | \|Undefined Saprotroph\| | Yeast | NULL | Probable | As nectar-tap_saprotroph (Põlme et al. 2020); Decay sugar-rich substrates (Põlme et al. 2020) | Põlme S, et al. 2020. Fungal Diversity 105:1-16 ((https://doi.org/10.1007/s13225-020-00466-2)) |
| Otu15280 | 1 | k__Fungi(100);p__Ascomycota(100);c__Saccharomycetes(100);o__Saccharomycetales(100);f__Metschnikowiaceae(100);g__Metschnikowia(100);s__Metschnikowia_pulcherrima(100); | Metschnikowia | 13 | Saprotroph | \|Undefined Saprotroph\| | Yeast | NULL | Probable | As nectar-tap_saprotroph (Põlme et al. 2020); Decay sugar-rich substrates (Põlme et al. 2020) | Põlme S, et al. 2020. Fungal Diversity 105:1-16 ((https://doi.org/10.1007/s13225-020-00466-2)) |
| Otu15281 | 1 | k__Fungi(100);p__Ascomycota(100);c__Saccharomycetes(100);o__Saccharomycetales(100);f__Metschnikowiaceae(100);g__Metschnikowia(100);g__Metschnikowia_unclassified(100); | Metschnikowia | 13 | Saprotroph | \|Undefined Saprotroph\| | Yeast | NULL | Probable | As nectar-tap_saprotroph (Põlme et al. 2020); Decay sugar-rich substrates (Põlme et al. 2020) | Põlme S, et al. 2020. Fungal Diversity 105:1-16 ((https://doi.org/10.1007/s13225-020-00466-2)) |
| Otu15282 | 1 | k__Fungi(100);p__Ascomycota(100);c__Saccharomycetes(100);o__Saccharomycetales(100);f__Metschnikowiaceae(100);g__Metschnikowia(100);s__Metschnikowia_pulcherrima(100); | Metschnikowia | 13 | Saprotroph | \|Undefined Saprotroph\| | Yeast | NULL | Probable | As nectar-tap_saprotroph (Põlme et al. 2020); Decay sugar-rich substrates (Põlme et al. 2020) | Põlme S, et al. 2020. Fungal Diversity 105:1-16 ((https://doi.org/10.1007/s13225-020-00466-2)) |
| Otu15284 | 1 | k__Fungi(100);p__Ascomycota(100);c__Saccharomycetes(100);o__Saccharomycetales(100);f__Metschnikowiaceae(100);g__Metschnikowia(100);s__Metschnikowia_pulcherrima(100); | Metschnikowia | 13 | Saprotroph | \|Undefined Saprotroph\| | Yeast | NULL | Probable | As nectar-tap_saprotroph (Põlme et al. 2020); Decay sugar-rich substrates (Põlme et al. 2020) | Põlme S, et al. 2020. Fungal Diversity 105:1-16 ((https://doi.org/10.1007/s13225-020-00466-2)) |
| Otu15285 | 1 | k__Fungi(100);p__Ascomycota(100);c__Saccharomycetes(100);o__Saccharomycetales(100);f__Metschnikowiaceae(100);g__Metschnikowia(100);g__Metschnikowia_unclassified(100); | Metschnikowia | 13 | Saprotroph | \|Undefined Saprotroph\| | Yeast | NULL | Probable | As nectar-tap_saprotroph (Põlme et al. 2020); Decay sugar-rich substrates (Põlme et al. 2020) | Põlme S, et al. 2020. Fungal Diversity 105:1-16 ((https://doi.org/10.1007/s13225-020-00466-2)) |
| Otu15286 | 1 | k__Fungi(100);p__Ascomycota(100);c__Saccharomycetes(100);o__Saccharomycetales(100);f__Metschnikowiaceae(100);g__Metschnikowia(100);s__Metschnikowia_pulcherrima(100); | Metschnikowia | 13 | Saprotroph | \|Undefined Saprotroph\| | Yeast | NULL | Probable | As nectar-tap_saprotroph (Põlme et al. 2020); Decay sugar-rich substrates (Põlme et al. 2020) | Põlme S, et al. 2020. Fungal Diversity 105:1-16 ((https://doi.org/10.1007/s13225-020-00466-2)) |
| Otu15287 | 1 | k__Fungi(100);p__Ascomycota(100);c__Saccharomycetes(100);o__Saccharomycetales(100);f__Metschnikowiaceae(100);g__Metschnikowia(100);s__Metschnikowia_pulcherrima(100); | Metschnikowia | 13 | Saprotroph | \|Undefined Saprotroph\| | Yeast | NULL | Probable | As nectar-tap_saprotroph (Põlme et al. 2020); Decay sugar-rich substrates (Põlme et al. 2020) | Põlme S, et al. 2020. Fungal Diversity 105:1-16 ((https://doi.org/10.1007/s13225-020-00466-2)) |
| Otu15288 | 1 | k__Fungi(100);p__Ascomycota(100);c__Saccharomycetes(100);o__Saccharomycetales(100);f__Metschnikowiaceae(100);g__Metschnikowia(100);s__Metschnikowia_pulcherrima(100); | Metschnikowia | 13 | Saprotroph | \|Undefined Saprotroph\| | Yeast | NULL | Probable | As nectar-tap_saprotroph (Põlme et al. 2020); Decay sugar-rich substrates (Põlme et al. 2020) | Põlme S, et al. 2020. Fungal Diversity 105:1-16 ((https://doi.org/10.1007/s13225-020-00466-2)) |
| Otu15289 | 1 | k__Fungi(100);p__Ascomycota(100);c__Saccharomycetes(100);o__Saccharomycetales(100);f__Metschnikowiaceae(100);g__Metschnikowia(100);s__Metschnikowia_pulcherrima(100); | Metschnikowia | 13 | Saprotroph | \|Undefined Saprotroph\| | Yeast | NULL | Probable | As nectar-tap_saprotroph (Põlme et al. 2020); Decay sugar-rich substrates (Põlme et al. 2020) | Põlme S, et al. 2020. Fungal Diversity 105:1-16 ((https://doi.org/10.1007/s13225-020-00466-2)) |
| Otu15290 | 1 | k__Fungi(100);p__Ascomycota(100);c__Saccharomycetes(100);o__Saccharomycetales(100);f__Metschnikowiaceae(100);g__Metschnikowia(100);s__Metschnikowia_pulcherrima(100); | Metschnikowia | 13 | Saprotroph | \|Undefined Saprotroph\| | Yeast | NULL | Probable | As nectar-tap_saprotroph (Põlme et al. 2020); Decay sugar-rich substrates (Põlme et al. 2020) | Põlme S, et al. 2020. Fungal Diversity 105:1-16 ((https://doi.org/10.1007/s13225-020-00466-2)) |
| Otu15291 | 1 | k__Fungi(100);p__Ascomycota(100);c__Saccharomycetes(100);o__Saccharomycetales(100);f__Metschnikowiaceae(100);g__Metschnikowia(100);s__Metschnikowia_pulcherrima(100); | Metschnikowia | 13 | Saprotroph | \|Undefined Saprotroph\| | Yeast | NULL | Probable | As nectar-tap_saprotroph (Põlme et al. 2020); Decay sugar-rich substrates (Põlme et al. 2020) | Põlme S, et al. 2020. Fungal Diversity 105:1-16 ((https://doi.org/10.1007/s13225-020-00466-2)) |
| Otu15292 | 1 | k__Fungi(100);p__Ascomycota(100);c__Saccharomycetes(100);o__Saccharomycetales(100);f__Saccharomycetales_family_Incertae_sedis(100);g__Candida(100);g__Candida_unclassified(100); | Candida | 13 | Pathotroph-Saprotroph-Symbiotroph | Animal Pathogen-Endophyte-Endosymbiont-Epiphyte-\|Undefined Saprotroph\| | Dimorphic Yeast-Yeast | NULL | Probable | As nectar-tap_saprotroph (Põlme et al. 2020); Decay sugar-rich substrates (Põlme et al. 2020); This asexual genus is polyphyletic and contain species that function very differently depend on what they are. Caution should be taken when interpreting guilds for Candida species. | Manolakaki D, et al. 2010. Virulence. 1: 367-375 ((https://doi.org/10.4161/viru.1.5.12796)); Mekbib SB, et al. 2011. Tropical Plant Pathology 36:233-240 ((https://doi.org/10.1590/S1982-56762011000400004)); Põlme S, et al. 2020. Fungal Diversity 105:1-16 ((https://doi.org/10.1007/s13225-020-00466-2)) |
| Otu15294 | 1 | k__Fungi(100);p__Ascomycota(100);c__Saccharomycetes(100);o__Saccharomycetales(100);f__Metschnikowiaceae(100);g__Metschnikowia(100);s__Metschnikowia_pulcherrima(100); | Metschnikowia | 13 | Saprotroph | \|Undefined Saprotroph\| | Yeast | NULL | Probable | As nectar-tap_saprotroph (Põlme et al. 2020); Decay sugar-rich substrates (Põlme et al. 2020) | Põlme S, et al. 2020. Fungal Diversity 105:1-16 ((https://doi.org/10.1007/s13225-020-00466-2)) |
| Otu15295 | 1 | k__Fungi(100);p__Ascomycota(100);c__Saccharomycetes(100);o__Saccharomycetales(100);f__Metschnikowiaceae(100);g__Metschnikowia(100);s__Metschnikowia_pulcherrima(100); | Metschnikowia | 13 | Saprotroph | \|Undefined Saprotroph\| | Yeast | NULL | Probable | As nectar-tap_saprotroph (Põlme et al. 2020); Decay sugar-rich substrates (Põlme et al. 2020) | Põlme S, et al. 2020. Fungal Diversity 105:1-16 ((https://doi.org/10.1007/s13225-020-00466-2)) |
| Otu15296 | 1 | k__Fungi(100);p__Ascomycota(100);c__Saccharomycetes(100);o__Saccharomycetales(100);f__Metschnikowiaceae(100);g__Metschnikowia(100);g__Metschnikowia_unclassified(100); | Metschnikowia | 13 | Saprotroph | \|Undefined Saprotroph\| | Yeast | NULL | Probable | As nectar-tap_saprotroph (Põlme et al. 2020); Decay sugar-rich substrates (Põlme et al. 2020) | Põlme S, et al. 2020. Fungal Diversity 105:1-16 ((https://doi.org/10.1007/s13225-020-00466-2)) |
| Otu15297 | 1 | k__Fungi(100);p__Ascomycota(100);c__Saccharomycetes(100);o__Saccharomycetales(100);f__Metschnikowiaceae(100);g__Metschnikowia(100);s__Metschnikowia_pulcherrima(100); | Metschnikowia | 13 | Saprotroph | \|Undefined Saprotroph\| | Yeast | NULL | Probable | As nectar-tap_saprotroph (Põlme et al. 2020); Decay sugar-rich substrates (Põlme et al. 2020) | Põlme S, et al. 2020. Fungal Diversity 105:1-16 ((https://doi.org/10.1007/s13225-020-00466-2)) |
| Otu15298 | 1 | k__Fungi(100);p__Ascomycota(100);c__Saccharomycetes(100);o__Saccharomycetales(100);f__Metschnikowiaceae(100);g__Metschnikowia(100);s__Metschnikowia_pulcherrima(100); | Metschnikowia | 13 | Saprotroph | \|Undefined Saprotroph\| | Yeast | NULL | Probable | As nectar-tap_saprotroph (Põlme et al. 2020); Decay sugar-rich substrates (Põlme et al. 2020) | Põlme S, et al. 2020. Fungal Diversity 105:1-16 ((https://doi.org/10.1007/s13225-020-00466-2)) |
| Otu15299 | 1 | k__Fungi(100);p__Ascomycota(100);c__Saccharomycetes(100);o__Saccharomycetales(100);f__Saccharomycetales_family_Incertae_sedis(100);g__Candida(100);g__Candida_unclassified(100); | Candida | 13 | Pathotroph-Saprotroph-Symbiotroph | Animal Pathogen-Endophyte-Endosymbiont-Epiphyte-\|Undefined Saprotroph\| | Dimorphic Yeast-Yeast | NULL | Probable | As nectar-tap_saprotroph (Põlme et al. 2020); Decay sugar-rich substrates (Põlme et al. 2020); This asexual genus is polyphyletic and contain species that function very differently depend on what they are. Caution should be taken when interpreting guilds for Candida species. | Manolakaki D, et al. 2010. Virulence. 1: 367-375 ((https://doi.org/10.4161/viru.1.5.12796)); Mekbib SB, et al. 2011. Tropical Plant Pathology 36:233-240 ((https://doi.org/10.1590/S1982-56762011000400004)); Põlme S, et al. 2020. Fungal Diversity 105:1-16 ((https://doi.org/10.1007/s13225-020-00466-2)) |
| Otu15300 | 1 | k__Fungi(100);p__Ascomycota(100);c__Saccharomycetes(100);o__Saccharomycetales(100);f__Metschnikowiaceae(100);g__Metschnikowia(100);s__Metschnikowia_pulcherrima(100); | Metschnikowia | 13 | Saprotroph | \|Undefined Saprotroph\| | Yeast | NULL | Probable | As nectar-tap_saprotroph (Põlme et al. 2020); Decay sugar-rich substrates (Põlme et al. 2020) | Põlme S, et al. 2020. Fungal Diversity 105:1-16 ((https://doi.org/10.1007/s13225-020-00466-2)) |
| Otu15301 | 1 | k__Fungi(100);p__Ascomycota(100);c__Saccharomycetes(100);o__Saccharomycetales(100);f__Saccharomycetales_family_Incertae_sedis(100);g__Candida(100);g__Candida_unclassified(100); | Candida | 13 | Pathotroph-Saprotroph-Symbiotroph | Animal Pathogen-Endophyte-Endosymbiont-Epiphyte-\|Undefined Saprotroph\| | Dimorphic Yeast-Yeast | NULL | Probable | As nectar-tap_saprotroph (Põlme et al. 2020); Decay sugar-rich substrates (Põlme et al. 2020); This asexual genus is polyphyletic and contain species that function very differently depend on what they are. Caution should be taken when interpreting guilds for Candida species. | Manolakaki D, et al. 2010. Virulence. 1: 367-375 ((https://doi.org/10.4161/viru.1.5.12796)); Mekbib SB, et al. 2011. Tropical Plant Pathology 36:233-240 ((https://doi.org/10.1590/S1982-56762011000400004)); Põlme S, et al. 2020. Fungal Diversity 105:1-16 ((https://doi.org/10.1007/s13225-020-00466-2)) |
| Otu15302 | 1 | k__Fungi(100);p__Ascomycota(100);c__Saccharomycetes(100);o__Saccharomycetales(100);f__Metschnikowiaceae(100);g__Metschnikowia(100);s__Metschnikowia_pulcherrima(100); | Metschnikowia | 13 | Saprotroph | \|Undefined Saprotroph\| | Yeast | NULL | Probable | As nectar-tap_saprotroph (Põlme et al. 2020); Decay sugar-rich substrates (Põlme et al. 2020) | Põlme S, et al. 2020. Fungal Diversity 105:1-16 ((https://doi.org/10.1007/s13225-020-00466-2)) |
| Otu15303 | 1 | k__Fungi(100);p__Ascomycota(100);c__Saccharomycetes(100);o__Saccharomycetales(100);f__Metschnikowiaceae(100);g__Metschnikowia(100);g__Metschnikowia_unclassified(100); | Metschnikowia | 13 | Saprotroph | \|Undefined Saprotroph\| | Yeast | NULL | Probable | As nectar-tap_saprotroph (Põlme et al. 2020); Decay sugar-rich substrates (Põlme et al. 2020) | Põlme S, et al. 2020. Fungal Diversity 105:1-16 ((https://doi.org/10.1007/s13225-020-00466-2)) |
| Otu15304 | 1 | k__Fungi(100);p__Ascomycota(100);c__Saccharomycetes(100);o__Saccharomycetales(100);f__Metschnikowiaceae(100);g__Metschnikowia(100);g__Metschnikowia_unclassified(100); | Metschnikowia | 13 | Saprotroph | \|Undefined Saprotroph\| | Yeast | NULL | Probable | As nectar-tap_saprotroph (Põlme et al. 2020); Decay sugar-rich substrates (Põlme et al. 2020) | Põlme S, et al. 2020. Fungal Diversity 105:1-16 ((https://doi.org/10.1007/s13225-020-00466-2)) |
| Otu15305 | 1 | k__Fungi(100);p__Ascomycota(100);c__Saccharomycetes(100);o__Saccharomycetales(100);f__Metschnikowiaceae(100);g__Metschnikowia(100);s__Metschnikowia_pulcherrima(100); | Metschnikowia | 13 | Saprotroph | \|Undefined Saprotroph\| | Yeast | NULL | Probable | As nectar-tap_saprotroph (Põlme et al. 2020); Decay sugar-rich substrates (Põlme et al. 2020) | Põlme S, et al. 2020. Fungal Diversity 105:1-16 ((https://doi.org/10.1007/s13225-020-00466-2)) |
| Otu15307 | 1 | k__Fungi(100);p__Ascomycota(100);c__Saccharomycetes(100);o__Saccharomycetales(100);f__Metschnikowiaceae(100);g__Metschnikowia(100);s__Metschnikowia_pulcherrima(100); | Metschnikowia | 13 | Saprotroph | \|Undefined Saprotroph\| | Yeast | NULL | Probable | As nectar-tap_saprotroph (Põlme et al. 2020); Decay sugar-rich substrates (Põlme et al. 2020) | Põlme S, et al. 2020. Fungal Diversity 105:1-16 ((https://doi.org/10.1007/s13225-020-00466-2)) |
| Otu15308 | 1 | k__Fungi(100);p__Ascomycota(100);c__Saccharomycetes(100);o__Saccharomycetales(100);f__Metschnikowiaceae(100);g__Metschnikowia(100);s__Metschnikowia_pulcherrima(100); | Metschnikowia | 13 | Saprotroph | \|Undefined Saprotroph\| | Yeast | NULL | Probable | As nectar-tap_saprotroph (Põlme et al. 2020); Decay sugar-rich substrates (Põlme et al. 2020) | Põlme S, et al. 2020. Fungal Diversity 105:1-16 ((https://doi.org/10.1007/s13225-020-00466-2)) |
| Otu15309 | 1 | k__Fungi(100);p__Ascomycota(100);c__Saccharomycetes(100);o__Saccharomycetales(100);f__Metschnikowiaceae(100);g__Metschnikowia(100);s__Metschnikowia_pulcherrima(100); | Metschnikowia | 13 | Saprotroph | \|Undefined Saprotroph\| | Yeast | NULL | Probable | As nectar-tap_saprotroph (Põlme et al. 2020); Decay sugar-rich substrates (Põlme et al. 2020) | Põlme S, et al. 2020. Fungal Diversity 105:1-16 ((https://doi.org/10.1007/s13225-020-00466-2)) |
| Otu15310 | 1 | k__Fungi(100);p__Ascomycota(100);c__Saccharomycetes(100);o__Saccharomycetales(100);f__Metschnikowiaceae(100);g__Metschnikowia(100);s__Metschnikowia_pulcherrima(100); | Metschnikowia | 13 | Saprotroph | \|Undefined Saprotroph\| | Yeast | NULL | Probable | As nectar-tap_saprotroph (Põlme et al. 2020); Decay sugar-rich substrates (Põlme et al. 2020) | Põlme S, et al. 2020. Fungal Diversity 105:1-16 ((https://doi.org/10.1007/s13225-020-00466-2)) |
| Otu15313 | 1 | k__Fungi(100);p__Ascomycota(100);c__Saccharomycetes(100);o__Saccharomycetales(100);f__Metschnikowiaceae(100);g__Metschnikowia(100);s__Metschnikowia_pulcherrima(100); | Metschnikowia | 13 | Saprotroph | \|Undefined Saprotroph\| | Yeast | NULL | Probable | As nectar-tap_saprotroph (Põlme et al. 2020); Decay sugar-rich substrates (Põlme et al. 2020) | Põlme S, et al. 2020. Fungal Diversity 105:1-16 ((https://doi.org/10.1007/s13225-020-00466-2)) |
| Otu15314 | 1 | k__Fungi(100);p__Ascomycota(100);c__Saccharomycetes(100);o__Saccharomycetales(100);f__Metschnikowiaceae(100);g__Metschnikowia(100);s__Metschnikowia_pulcherrima(100); | Metschnikowia | 13 | Saprotroph | \|Undefined Saprotroph\| | Yeast | NULL | Probable | As nectar-tap_saprotroph (Põlme et al. 2020); Decay sugar-rich substrates (Põlme et al. 2020) | Põlme S, et al. 2020. Fungal Diversity 105:1-16 ((https://doi.org/10.1007/s13225-020-00466-2)) |
| Otu15315 | 1 | k__Fungi(100);p__Ascomycota(100);c__Saccharomycetes(100);o__Saccharomycetales(100);f__Saccharomycetales_family_Incertae_sedis(100);g__Candida(100);s__Candida_cretensis(100); | Candida | 13 | Pathotroph-Saprotroph-Symbiotroph | Animal Pathogen-Endophyte-Endosymbiont-Epiphyte-\|Undefined Saprotroph\| | Dimorphic Yeast-Yeast | NULL | Probable | As nectar-tap_saprotroph (Põlme et al. 2020); Decay sugar-rich substrates (Põlme et al. 2020); This asexual genus is polyphyletic and contain species that function very differently depend on what they are. Caution should be taken when interpreting guilds for Candida species. | Manolakaki D, et al. 2010. Virulence. 1: 367-375 ((https://doi.org/10.4161/viru.1.5.12796)); Mekbib SB, et al. 2011. Tropical Plant Pathology 36:233-240 ((https://doi.org/10.1590/S1982-56762011000400004)); Põlme S, et al. 2020. Fungal Diversity 105:1-16 ((https://doi.org/10.1007/s13225-020-00466-2)) |
| Otu15317 | 1 | k__Fungi(100);p__Ascomycota(100);c__Saccharomycetes(100);o__Saccharomycetales(100);f__Metschnikowiaceae(100);g__Metschnikowia(100);s__Metschnikowia_pulcherrima(100); | Metschnikowia | 13 | Saprotroph | \|Undefined Saprotroph\| | Yeast | NULL | Probable | As nectar-tap_saprotroph (Põlme et al. 2020); Decay sugar-rich substrates (Põlme et al. 2020) | Põlme S, et al. 2020. Fungal Diversity 105:1-16 ((https://doi.org/10.1007/s13225-020-00466-2)) |
| Otu15320 | 1 | k__Fungi(100);p__Ascomycota(100);c__Saccharomycetes(100);o__Saccharomycetales(100);f__Metschnikowiaceae(100);g__Metschnikowia(100);s__Metschnikowia_pulcherrima(100); | Metschnikowia | 13 | Saprotroph | \|Undefined Saprotroph\| | Yeast | NULL | Probable | As nectar-tap_saprotroph (Põlme et al. 2020); Decay sugar-rich substrates (Põlme et al. 2020) | Põlme S, et al. 2020. Fungal Diversity 105:1-16 ((https://doi.org/10.1007/s13225-020-00466-2)) |
| Otu15321 | 1 | k__Fungi(100);p__Ascomycota(100);c__Saccharomycetes(100);o__Saccharomycetales(100);f__Metschnikowiaceae(100);g__Metschnikowia(100);s__Metschnikowia_pulcherrima(100); | Metschnikowia | 13 | Saprotroph | \|Undefined Saprotroph\| | Yeast | NULL | Probable | As nectar-tap_saprotroph (Põlme et al. 2020); Decay sugar-rich substrates (Põlme et al. 2020) | Põlme S, et al. 2020. Fungal Diversity 105:1-16 ((https://doi.org/10.1007/s13225-020-00466-2)) |
| Otu15322 | 1 | k__Fungi(100);p__Ascomycota(100);c__Saccharomycetes(100);o__Saccharomycetales(100);f__Saccharomycetales_family_Incertae_sedis(100);g__Candida(100);s__Candida_cretensis(100); | Candida | 13 | Pathotroph-Saprotroph-Symbiotroph | Animal Pathogen-Endophyte-Endosymbiont-Epiphyte-\|Undefined Saprotroph\| | Dimorphic Yeast-Yeast | NULL | Probable | As nectar-tap_saprotroph (Põlme et al. 2020); Decay sugar-rich substrates (Põlme et al. 2020); This asexual genus is polyphyletic and contain species that function very differently depend on what they are. Caution should be taken when interpreting guilds for Candida species. | Manolakaki D, et al. 2010. Virulence. 1: 367-375 ((https://doi.org/10.4161/viru.1.5.12796)); Mekbib SB, et al. 2011. Tropical Plant Pathology 36:233-240 ((https://doi.org/10.1590/S1982-56762011000400004)); Põlme S, et al. 2020. Fungal Diversity 105:1-16 ((https://doi.org/10.1007/s13225-020-00466-2)) |
| Otu15323 | 1 | k__Fungi(100);p__Ascomycota(100);c__Saccharomycetes(100);o__Saccharomycetales(100);f__Metschnikowiaceae(100);g__Metschnikowia(100);s__Metschnikowia_pulcherrima(100); | Metschnikowia | 13 | Saprotroph | \|Undefined Saprotroph\| | Yeast | NULL | Probable | As nectar-tap_saprotroph (Põlme et al. 2020); Decay sugar-rich substrates (Põlme et al. 2020) | Põlme S, et al. 2020. Fungal Diversity 105:1-16 ((https://doi.org/10.1007/s13225-020-00466-2)) |
| Otu15324 | 1 | k__Fungi(100);p__Ascomycota(100);c__Saccharomycetes(100);o__Saccharomycetales(100);f__Metschnikowiaceae(100);g__Metschnikowia(100);s__Metschnikowia_pulcherrima(100); | Metschnikowia | 13 | Saprotroph | \|Undefined Saprotroph\| | Yeast | NULL | Probable | As nectar-tap_saprotroph (Põlme et al. 2020); Decay sugar-rich substrates (Põlme et al. 2020) | Põlme S, et al. 2020. Fungal Diversity 105:1-16 ((https://doi.org/10.1007/s13225-020-00466-2)) |
| Otu15325 | 1 | k__Fungi(100);p__Ascomycota(100);c__Saccharomycetes(100);o__Saccharomycetales(100);f__Metschnikowiaceae(100);g__Metschnikowia(100);s__Metschnikowia_pulcherrima(100); | Metschnikowia | 13 | Saprotroph | \|Undefined Saprotroph\| | Yeast | NULL | Probable | As nectar-tap_saprotroph (Põlme et al. 2020); Decay sugar-rich substrates (Põlme et al. 2020) | Põlme S, et al. 2020. Fungal Diversity 105:1-16 ((https://doi.org/10.1007/s13225-020-00466-2)) |
| Otu15327 | 1 | k__Fungi(100);p__Ascomycota(100);c__Saccharomycetes(100);o__Saccharomycetales(100);f__Saccharomycetales_family_Incertae_sedis(100);f__Saccharomycetales_family_Incertae_sedis_unclassified(100);f__Saccharomycetales_family_Incertae_sedis_unclassified(100); | Saccharomycetales | 7 | Saprotroph | Undefined Saprotroph | Yeast | NULL | Possible | NULL | Sterkenburg E, et al. 2015. New Phytologist 207:1145-1158 ((https://doi.org/10.1111/nph.13426)) |
| Otu15330 | 1 | k__Fungi(100);p__Ascomycota(100);c__Saccharomycetes(100);o__Saccharomycetales(100);f__Metschnikowiaceae(100);g__Metschnikowia(100);s__Metschnikowia_pulcherrima(100); | Metschnikowia | 13 | Saprotroph | \|Undefined Saprotroph\| | Yeast | NULL | Probable | As nectar-tap_saprotroph (Põlme et al. 2020); Decay sugar-rich substrates (Põlme et al. 2020) | Põlme S, et al. 2020. Fungal Diversity 105:1-16 ((https://doi.org/10.1007/s13225-020-00466-2)) |
| Otu15334 | 1 | k__Fungi(100);p__Ascomycota(100);c__Saccharomycetes(100);o__Saccharomycetales(100);f__Metschnikowiaceae(100);g__Metschnikowia(100);s__Metschnikowia_pulcherrima(100); | Metschnikowia | 13 | Saprotroph | \|Undefined Saprotroph\| | Yeast | NULL | Probable | As nectar-tap_saprotroph (Põlme et al. 2020); Decay sugar-rich substrates (Põlme et al. 2020) | Põlme S, et al. 2020. Fungal Diversity 105:1-16 ((https://doi.org/10.1007/s13225-020-00466-2)) |
| Otu15336 | 1 | k__Fungi(100);p__Ascomycota(100);c__Saccharomycetes(100);o__Saccharomycetales(100);f__Metschnikowiaceae(100);g__Metschnikowia(100);s__Metschnikowia_pulcherrima(100); | Metschnikowia | 13 | Saprotroph | \|Undefined Saprotroph\| | Yeast | NULL | Probable | As nectar-tap_saprotroph (Põlme et al. 2020); Decay sugar-rich substrates (Põlme et al. 2020) | Põlme S, et al. 2020. Fungal Diversity 105:1-16 ((https://doi.org/10.1007/s13225-020-00466-2)) |
| Otu15337 | 1 | k__Fungi(100);p__Ascomycota(100);c__Saccharomycetes(100);o__Saccharomycetales(100);f__Metschnikowiaceae(100);g__Metschnikowia(100);s__Metschnikowia_pulcherrima(100); | Metschnikowia | 13 | Saprotroph | \|Undefined Saprotroph\| | Yeast | NULL | Probable | As nectar-tap_saprotroph (Põlme et al. 2020); Decay sugar-rich substrates (Põlme et al. 2020) | Põlme S, et al. 2020. Fungal Diversity 105:1-16 ((https://doi.org/10.1007/s13225-020-00466-2)) |
| Otu15338 | 1 | k__Fungi(100);p__Ascomycota(100);c__Saccharomycetes(100);o__Saccharomycetales(100);f__Metschnikowiaceae(100);g__Metschnikowia(100);s__Metschnikowia_pulcherrima(100); | Metschnikowia | 13 | Saprotroph | \|Undefined Saprotroph\| | Yeast | NULL | Probable | As nectar-tap_saprotroph (Põlme et al. 2020); Decay sugar-rich substrates (Põlme et al. 2020) | Põlme S, et al. 2020. Fungal Diversity 105:1-16 ((https://doi.org/10.1007/s13225-020-00466-2)) |
| Otu15340 | 1 | k__Fungi(100);p__Ascomycota(100);c__Saccharomycetes(100);o__Saccharomycetales(100);f__Metschnikowiaceae(100);g__Metschnikowia(100);s__Metschnikowia_pulcherrima(100); | Metschnikowia | 13 | Saprotroph | \|Undefined Saprotroph\| | Yeast | NULL | Probable | As nectar-tap_saprotroph (Põlme et al. 2020); Decay sugar-rich substrates (Põlme et al. 2020) | Põlme S, et al. 2020. Fungal Diversity 105:1-16 ((https://doi.org/10.1007/s13225-020-00466-2)) |
| Otu15341 | 1 | k__Fungi(100);p__Ascomycota(100);c__Saccharomycetes(100);o__Saccharomycetales(100);f__Metschnikowiaceae(100);g__Metschnikowia(100);s__Metschnikowia_pulcherrima(100); | Metschnikowia | 13 | Saprotroph | \|Undefined Saprotroph\| | Yeast | NULL | Probable | As nectar-tap_saprotroph (Põlme et al. 2020); Decay sugar-rich substrates (Põlme et al. 2020) | Põlme S, et al. 2020. Fungal Diversity 105:1-16 ((https://doi.org/10.1007/s13225-020-00466-2)) |
| Otu15342 | 1 | k__Fungi(100);p__Ascomycota(100);c__Saccharomycetes(100);o__Saccharomycetales(100);f__Metschnikowiaceae(100);g__Metschnikowia(100);s__Metschnikowia_pulcherrima(100); | Metschnikowia | 13 | Saprotroph | \|Undefined Saprotroph\| | Yeast | NULL | Probable | As nectar-tap_saprotroph (Põlme et al. 2020); Decay sugar-rich substrates (Põlme et al. 2020) | Põlme S, et al. 2020. Fungal Diversity 105:1-16 ((https://doi.org/10.1007/s13225-020-00466-2)) |
| Otu15343 | 1 | k__Fungi(100);p__Ascomycota(100);c__Saccharomycetes(100);o__Saccharomycetales(100);f__Metschnikowiaceae(100);g__Metschnikowia(100);s__Metschnikowia_pulcherrima(100); | Metschnikowia | 13 | Saprotroph | \|Undefined Saprotroph\| | Yeast | NULL | Probable | As nectar-tap_saprotroph (Põlme et al. 2020); Decay sugar-rich substrates (Põlme et al. 2020) | Põlme S, et al. 2020. Fungal Diversity 105:1-16 ((https://doi.org/10.1007/s13225-020-00466-2)) |
| Otu15344 | 1 | k__Fungi(100);p__Ascomycota(100);c__Saccharomycetes(100);o__Saccharomycetales(100);f__Metschnikowiaceae(100);g__Metschnikowia(100);s__Metschnikowia_pulcherrima(100); | Metschnikowia | 13 | Saprotroph | \|Undefined Saprotroph\| | Yeast | NULL | Probable | As nectar-tap_saprotroph (Põlme et al. 2020); Decay sugar-rich substrates (Põlme et al. 2020) | Põlme S, et al. 2020. Fungal Diversity 105:1-16 ((https://doi.org/10.1007/s13225-020-00466-2)) |
| Otu15345 | 1 | k__Fungi(100);p__Ascomycota(100);c__Saccharomycetes(100);o__Saccharomycetales(100);f__Metschnikowiaceae(100);g__Metschnikowia(100);s__Metschnikowia_pulcherrima(100); | Metschnikowia | 13 | Saprotroph | \|Undefined Saprotroph\| | Yeast | NULL | Probable | As nectar-tap_saprotroph (Põlme et al. 2020); Decay sugar-rich substrates (Põlme et al. 2020) | Põlme S, et al. 2020. Fungal Diversity 105:1-16 ((https://doi.org/10.1007/s13225-020-00466-2)) |
| Otu15347 | 1 | k__Fungi(100);p__Ascomycota(100);c__Saccharomycetes(100);o__Saccharomycetales(100);f__Metschnikowiaceae(100);g__Metschnikowia(100);g__Metschnikowia_unclassified(100); | Metschnikowia | 13 | Saprotroph | \|Undefined Saprotroph\| | Yeast | NULL | Probable | As nectar-tap_saprotroph (Põlme et al. 2020); Decay sugar-rich substrates (Põlme et al. 2020) | Põlme S, et al. 2020. Fungal Diversity 105:1-16 ((https://doi.org/10.1007/s13225-020-00466-2)) |
| Otu15348 | 1 | k__Fungi(100);p__Ascomycota(100);c__Saccharomycetes(100);o__Saccharomycetales(100);f__Metschnikowiaceae(100);g__Metschnikowia(100);s__Metschnikowia_pulcherrima(100); | Metschnikowia | 13 | Saprotroph | \|Undefined Saprotroph\| | Yeast | NULL | Probable | As nectar-tap_saprotroph (Põlme et al. 2020); Decay sugar-rich substrates (Põlme et al. 2020) | Põlme S, et al. 2020. Fungal Diversity 105:1-16 ((https://doi.org/10.1007/s13225-020-00466-2)) |
| Otu15349 | 1 | k__Fungi(100);p__Ascomycota(100);c__Saccharomycetes(100);o__Saccharomycetales(100);f__Saccharomycetales_family_Incertae_sedis(100);g__Candida(100);g__Candida_unclassified(100); | Candida | 13 | Pathotroph-Saprotroph-Symbiotroph | Animal Pathogen-Endophyte-Endosymbiont-Epiphyte-\|Undefined Saprotroph\| | Dimorphic Yeast-Yeast | NULL | Probable | As nectar-tap_saprotroph (Põlme et al. 2020); Decay sugar-rich substrates (Põlme et al. 2020); This asexual genus is polyphyletic and contain species that function very differently depend on what they are. Caution should be taken when interpreting guilds for Candida species. | Manolakaki D, et al. 2010. Virulence. 1: 367-375 ((https://doi.org/10.4161/viru.1.5.12796)); Mekbib SB, et al. 2011. Tropical Plant Pathology 36:233-240 ((https://doi.org/10.1590/S1982-56762011000400004)); Põlme S, et al. 2020. Fungal Diversity 105:1-16 ((https://doi.org/10.1007/s13225-020-00466-2)) |
| Otu15351 | 1 | k__Fungi(100);p__Ascomycota(100);c__Saccharomycetes(100);o__Saccharomycetales(100);f__Metschnikowiaceae(100);g__Metschnikowia(100);s__Metschnikowia_pulcherrima(100); | Metschnikowia | 13 | Saprotroph | \|Undefined Saprotroph\| | Yeast | NULL | Probable | As nectar-tap_saprotroph (Põlme et al. 2020); Decay sugar-rich substrates (Põlme et al. 2020) | Põlme S, et al. 2020. Fungal Diversity 105:1-16 ((https://doi.org/10.1007/s13225-020-00466-2)) |
| Otu15352 | 1 | k__Fungi(100);p__Ascomycota(100);c__Saccharomycetes(100);o__Saccharomycetales(100);f__Metschnikowiaceae(100);g__Metschnikowia(100);s__Metschnikowia_pulcherrima(100); | Metschnikowia | 13 | Saprotroph | \|Undefined Saprotroph\| | Yeast | NULL | Probable | As nectar-tap_saprotroph (Põlme et al. 2020); Decay sugar-rich substrates (Põlme et al. 2020) | Põlme S, et al. 2020. Fungal Diversity 105:1-16 ((https://doi.org/10.1007/s13225-020-00466-2)) |
| Otu15353 | 1 | k__Fungi(100);p__Ascomycota(100);c__Saccharomycetes(100);o__Saccharomycetales(100);f__Metschnikowiaceae(100);g__Metschnikowia(100);s__Metschnikowia_pulcherrima(100); | Metschnikowia | 13 | Saprotroph | \|Undefined Saprotroph\| | Yeast | NULL | Probable | As nectar-tap_saprotroph (Põlme et al. 2020); Decay sugar-rich substrates (Põlme et al. 2020) | Põlme S, et al. 2020. Fungal Diversity 105:1-16 ((https://doi.org/10.1007/s13225-020-00466-2)) |
| Otu15355 | 1 | k__Fungi(100);p__Ascomycota(100);c__Saccharomycetes(100);o__Saccharomycetales(100);f__Metschnikowiaceae(100);g__Metschnikowia(100);s__Metschnikowia_pulcherrima(100); | Metschnikowia | 13 | Saprotroph | \|Undefined Saprotroph\| | Yeast | NULL | Probable | As nectar-tap_saprotroph (Põlme et al. 2020); Decay sugar-rich substrates (Põlme et al. 2020) | Põlme S, et al. 2020. Fungal Diversity 105:1-16 ((https://doi.org/10.1007/s13225-020-00466-2)) |
| Otu15356 | 1 | k__Fungi(100);p__Ascomycota(100);c__Saccharomycetes(100);o__Saccharomycetales(100);f__Metschnikowiaceae(100);g__Metschnikowia(100);s__Metschnikowia_pulcherrima(100); | Metschnikowia | 13 | Saprotroph | \|Undefined Saprotroph\| | Yeast | NULL | Probable | As nectar-tap_saprotroph (Põlme et al. 2020); Decay sugar-rich substrates (Põlme et al. 2020) | Põlme S, et al. 2020. Fungal Diversity 105:1-16 ((https://doi.org/10.1007/s13225-020-00466-2)) |
| Otu15359 | 1 | k__Fungi(100);p__Ascomycota(100);c__Saccharomycetes(100);o__Saccharomycetales(100);f__Metschnikowiaceae(100);g__Metschnikowia(100);g__Metschnikowia_unclassified(100); | Metschnikowia | 13 | Saprotroph | \|Undefined Saprotroph\| | Yeast | NULL | Probable | As nectar-tap_saprotroph (Põlme et al. 2020); Decay sugar-rich substrates (Põlme et al. 2020) | Põlme S, et al. 2020. Fungal Diversity 105:1-16 ((https://doi.org/10.1007/s13225-020-00466-2)) |
| Otu15360 | 1 | k__Fungi(100);p__Ascomycota(100);c__Saccharomycetes(100);o__Saccharomycetales(100);f__Metschnikowiaceae(100);g__Metschnikowia(100);s__Metschnikowia_pulcherrima(100); | Metschnikowia | 13 | Saprotroph | \|Undefined Saprotroph\| | Yeast | NULL | Probable | As nectar-tap_saprotroph (Põlme et al. 2020); Decay sugar-rich substrates (Põlme et al. 2020) | Põlme S, et al. 2020. Fungal Diversity 105:1-16 ((https://doi.org/10.1007/s13225-020-00466-2)) |
| Otu15361 | 1 | k__Fungi(100);p__Ascomycota(100);c__Saccharomycetes(100);o__Saccharomycetales(100);f__Metschnikowiaceae(100);g__Metschnikowia(100);s__Metschnikowia_pulcherrima(100); | Metschnikowia | 13 | Saprotroph | \|Undefined Saprotroph\| | Yeast | NULL | Probable | As nectar-tap_saprotroph (Põlme et al. 2020); Decay sugar-rich substrates (Põlme et al. 2020) | Põlme S, et al. 2020. Fungal Diversity 105:1-16 ((https://doi.org/10.1007/s13225-020-00466-2)) |
| Otu15362 | 1 | k__Fungi(100);p__Ascomycota(100);c__Saccharomycetes(100);o__Saccharomycetales(100);f__Metschnikowiaceae(100);g__Metschnikowia(100);s__Metschnikowia_pulcherrima(100); | Metschnikowia | 13 | Saprotroph | \|Undefined Saprotroph\| | Yeast | NULL | Probable | As nectar-tap_saprotroph (Põlme et al. 2020); Decay sugar-rich substrates (Põlme et al. 2020) | Põlme S, et al. 2020. Fungal Diversity 105:1-16 ((https://doi.org/10.1007/s13225-020-00466-2)) |
| Otu15364 | 1 | k__Fungi(100);p__Ascomycota(100);c__Saccharomycetes(100);o__Saccharomycetales(100);f__Metschnikowiaceae(100);g__Metschnikowia(100);g__Metschnikowia_unclassified(100); | Metschnikowia | 13 | Saprotroph | \|Undefined Saprotroph\| | Yeast | NULL | Probable | As nectar-tap_saprotroph (Põlme et al. 2020); Decay sugar-rich substrates (Põlme et al. 2020) | Põlme S, et al. 2020. Fungal Diversity 105:1-16 ((https://doi.org/10.1007/s13225-020-00466-2)) |
| Otu15366 | 1 | k__Fungi(100);p__Ascomycota(100);c__Saccharomycetes(100);o__Saccharomycetales(100);f__Metschnikowiaceae(100);g__Metschnikowia(100);s__Metschnikowia_pulcherrima(100); | Metschnikowia | 13 | Saprotroph | \|Undefined Saprotroph\| | Yeast | NULL | Probable | As nectar-tap_saprotroph (Põlme et al. 2020); Decay sugar-rich substrates (Põlme et al. 2020) | Põlme S, et al. 2020. Fungal Diversity 105:1-16 ((https://doi.org/10.1007/s13225-020-00466-2)) |
| Otu15367 | 1 | k__Fungi(100);p__Ascomycota(100);c__Saccharomycetes(100);o__Saccharomycetales(100);f__Metschnikowiaceae(100);g__Metschnikowia(100);s__Metschnikowia_pulcherrima(100); | Metschnikowia | 13 | Saprotroph | \|Undefined Saprotroph\| | Yeast | NULL | Probable | As nectar-tap_saprotroph (Põlme et al. 2020); Decay sugar-rich substrates (Põlme et al. 2020) | Põlme S, et al. 2020. Fungal Diversity 105:1-16 ((https://doi.org/10.1007/s13225-020-00466-2)) |
| Otu15368 | 1 | k__Fungi(100);p__Ascomycota(100);c__Saccharomycetes(100);o__Saccharomycetales(100);f__Metschnikowiaceae(100);g__Metschnikowia(100);s__Metschnikowia_pulcherrima(100); | Metschnikowia | 13 | Saprotroph | \|Undefined Saprotroph\| | Yeast | NULL | Probable | As nectar-tap_saprotroph (Põlme et al. 2020); Decay sugar-rich substrates (Põlme et al. 2020) | Põlme S, et al. 2020. Fungal Diversity 105:1-16 ((https://doi.org/10.1007/s13225-020-00466-2)) |
| Otu15369 | 1 | k__Fungi(100);p__Ascomycota(100);c__Saccharomycetes(100);o__Saccharomycetales(100);f__Metschnikowiaceae(100);g__Metschnikowia(100);g__Metschnikowia_unclassified(100); | Metschnikowia | 13 | Saprotroph | \|Undefined Saprotroph\| | Yeast | NULL | Probable | As nectar-tap_saprotroph (Põlme et al. 2020); Decay sugar-rich substrates (Põlme et al. 2020) | Põlme S, et al. 2020. Fungal Diversity 105:1-16 ((https://doi.org/10.1007/s13225-020-00466-2)) |
| Otu15370 | 1 | k__Fungi(100);p__Ascomycota(100);c__Saccharomycetes(100);o__Saccharomycetales(100);f__Metschnikowiaceae(100);g__Metschnikowia(100);s__Metschnikowia_pulcherrima(100); | Metschnikowia | 13 | Saprotroph | \|Undefined Saprotroph\| | Yeast | NULL | Probable | As nectar-tap_saprotroph (Põlme et al. 2020); Decay sugar-rich substrates (Põlme et al. 2020) | Põlme S, et al. 2020. Fungal Diversity 105:1-16 ((https://doi.org/10.1007/s13225-020-00466-2)) |
| Otu15373 | 1 | k__Fungi(100);p__Ascomycota(100);c__Saccharomycetes(100);o__Saccharomycetales(100);f__Metschnikowiaceae(100);g__Metschnikowia(100);s__Metschnikowia_pulcherrima(100); | Metschnikowia | 13 | Saprotroph | \|Undefined Saprotroph\| | Yeast | NULL | Probable | As nectar-tap_saprotroph (Põlme et al. 2020); Decay sugar-rich substrates (Põlme et al. 2020) | Põlme S, et al. 2020. Fungal Diversity 105:1-16 ((https://doi.org/10.1007/s13225-020-00466-2)) |
| Otu15374 | 1 | k__Fungi(100);p__Ascomycota(100);c__Saccharomycetes(100);o__Saccharomycetales(100);f__Metschnikowiaceae(100);g__Metschnikowia(100);s__Metschnikowia_pulcherrima(100); | Metschnikowia | 13 | Saprotroph | \|Undefined Saprotroph\| | Yeast | NULL | Probable | As nectar-tap_saprotroph (Põlme et al. 2020); Decay sugar-rich substrates (Põlme et al. 2020) | Põlme S, et al. 2020. Fungal Diversity 105:1-16 ((https://doi.org/10.1007/s13225-020-00466-2)) |
| Otu15375 | 1 | k__Fungi(100);p__Ascomycota(100);c__Saccharomycetes(100);o__Saccharomycetales(100);f__Metschnikowiaceae(100);g__Metschnikowia(100);s__Metschnikowia_pulcherrima(100); | Metschnikowia | 13 | Saprotroph | \|Undefined Saprotroph\| | Yeast | NULL | Probable | As nectar-tap_saprotroph (Põlme et al. 2020); Decay sugar-rich substrates (Põlme et al. 2020) | Põlme S, et al. 2020. Fungal Diversity 105:1-16 ((https://doi.org/10.1007/s13225-020-00466-2)) |
| Otu15377 | 1 | k__Fungi(100);p__Ascomycota(100);c__Saccharomycetes(100);o__Saccharomycetales(100);f__Metschnikowiaceae(100);g__Metschnikowia(100);s__Metschnikowia_pulcherrima(100); | Metschnikowia | 13 | Saprotroph | \|Undefined Saprotroph\| | Yeast | NULL | Probable | As nectar-tap_saprotroph (Põlme et al. 2020); Decay sugar-rich substrates (Põlme et al. 2020) | Põlme S, et al. 2020. Fungal Diversity 105:1-16 ((https://doi.org/10.1007/s13225-020-00466-2)) |
| Otu15381 | 1 | k__Fungi(100);p__Ascomycota(100);c__Saccharomycetes(100);o__Saccharomycetales(100);f__Metschnikowiaceae(100);g__Metschnikowia(100);s__Metschnikowia_pulcherrima(100); | Metschnikowia | 13 | Saprotroph | \|Undefined Saprotroph\| | Yeast | NULL | Probable | As nectar-tap_saprotroph (Põlme et al. 2020); Decay sugar-rich substrates (Põlme et al. 2020) | Põlme S, et al. 2020. Fungal Diversity 105:1-16 ((https://doi.org/10.1007/s13225-020-00466-2)) |
| Otu15382 | 1 | k__Fungi(100);p__Ascomycota(100);c__Saccharomycetes(100);o__Saccharomycetales(100);f__Metschnikowiaceae(100);g__Metschnikowia(100);g__Metschnikowia_unclassified(100); | Metschnikowia | 13 | Saprotroph | \|Undefined Saprotroph\| | Yeast | NULL | Probable | As nectar-tap_saprotroph (Põlme et al. 2020); Decay sugar-rich substrates (Põlme et al. 2020) | Põlme S, et al. 2020. Fungal Diversity 105:1-16 ((https://doi.org/10.1007/s13225-020-00466-2)) |
| Otu15383 | 1 | k__Fungi(100);p__Ascomycota(100);c__Saccharomycetes(100);o__Saccharomycetales(100);f__Metschnikowiaceae(100);g__Metschnikowia(100);s__Metschnikowia_pulcherrima(100); | Metschnikowia | 13 | Saprotroph | \|Undefined Saprotroph\| | Yeast | NULL | Probable | As nectar-tap_saprotroph (Põlme et al. 2020); Decay sugar-rich substrates (Põlme et al. 2020) | Põlme S, et al. 2020. Fungal Diversity 105:1-16 ((https://doi.org/10.1007/s13225-020-00466-2)) |
| Otu15384 | 1 | k__Fungi(100);p__Ascomycota(100);c__Saccharomycetes(100);o__Saccharomycetales(100);f__Metschnikowiaceae(100);g__Metschnikowia(100);s__Metschnikowia_pulcherrima(100); | Metschnikowia | 13 | Saprotroph | \|Undefined Saprotroph\| | Yeast | NULL | Probable | As nectar-tap_saprotroph (Põlme et al. 2020); Decay sugar-rich substrates (Põlme et al. 2020) | Põlme S, et al. 2020. Fungal Diversity 105:1-16 ((https://doi.org/10.1007/s13225-020-00466-2)) |

| | | | | | | | | | | | |
|---|---|---|---|---|---|---|---|---|---|---|---|
| Otu15388 | 1 | k__Fungi(100);p__Ascomycota(100);c__Saccharomycetes(100);o__Saccharomycetales(100);f__Metschnikowiaceae(100);g__Metschnikowia(100);s__Metschnikowia_pulcherrima(100); | Metschnikowia | 13 | Saprotroph | \|Undefined Saprotroph\| | Yeast | NULL | Probable | As nectar-tap_saprotroph (Põlme et al. 2020); Decay sugar-rich substrates (Põlme et al. 2020) | Põlme S, et al. 2020. Fungal Diversity 105:1-16 ((https://doi.org/10.1007/s13225-020-00466-2)) |
| Otu15389 | 1 | k__Fungi(100);p__Ascomycota(100);c__Saccharomycetes(100);o__Saccharomycetales(100);f__Metschnikowiaceae(100);g__Metschnikowia(100);s__Metschnikowia_pulcherrima(100); | Metschnikowia | 13 | Saprotroph | \|Undefined Saprotroph\| | Yeast | NULL | Probable | As nectar-tap_saprotroph (Põlme et al. 2020); Decay sugar-rich substrates (Põlme et al. 2020) | Põlme S, et al. 2020. Fungal Diversity 105:1-16 ((https://doi.org/10.1007/s13225-020-00466-2)) |
| Otu15390 | 1 | k__Fungi(100);p__Ascomycota(100);c__Saccharomycetes(100);o__Saccharomycetales(100);f__Metschnikowiaceae(100);g__Metschnikowia(100);g__Metschnikowia_unclassified(100); | Metschnikowia | 13 | Saprotroph | \|Undefined Saprotroph\| | Yeast | NULL | Probable | As nectar-tap_saprotroph (Põlme et al. 2020); Decay sugar-rich substrates (Põlme et al. 2020) | Põlme S, et al. 2020. Fungal Diversity 105:1-16 ((https://doi.org/10.1007/s13225-020-00466-2)) |
| Otu15391 | 1 | k__Fungi(100);p__Ascomycota(100);c__Saccharomycetes(100);o__Saccharomycetales(100);f__Metschnikowiaceae(100);g__Metschnikowia(100);g__Metschnikowia_unclassified(100); | Metschnikowia | 13 | Saprotroph | \|Undefined Saprotroph\| | Yeast | NULL | Probable | As nectar-tap_saprotroph (Põlme et al. 2020); Decay sugar-rich substrates (Põlme et al. 2020) | Põlme S, et al. 2020. Fungal Diversity 105:1-16 ((https://doi.org/10.1007/s13225-020-00466-2)) |
| Otu15392 | 1 | k__Fungi(100);p__Ascomycota(100);c__Saccharomycetes(100);o__Saccharomycetales(100);f__Metschnikowiaceae(100);g__Metschnikowia(100);s__Metschnikowia_pulcherrima(100); | Metschnikowia | 13 | Saprotroph | \|Undefined Saprotroph\| | Yeast | NULL | Probable | As nectar-tap_saprotroph (Põlme et al. 2020); Decay sugar-rich substrates (Põlme et al. 2020) | Põlme S, et al. 2020. Fungal Diversity 105:1-16 ((https://doi.org/10.1007/s13225-020-00466-2)) |
| Otu15394 | 1 | k__Fungi(100);p__Ascomycota(100);c__Saccharomycetes(100);o__Saccharomycetales(100);f__Metschnikowiaceae(100);g__Metschnikowia(100);s__Metschnikowia_pulcherrima(100); | Metschnikowia | 13 | Saprotroph | \|Undefined Saprotroph\| | Yeast | NULL | Probable | As nectar-tap_saprotroph (Põlme et al. 2020); Decay sugar-rich substrates (Põlme et al. 2020) | Põlme S, et al. 2020. Fungal Diversity 105:1-16 ((https://doi.org/10.1007/s13225-020-00466-2)) |
| Otu15395 | 1 | k__Fungi(100);p__Ascomycota(100);c__Saccharomycetes(100);o__Saccharomycetales(100);f__Metschnikowiaceae(100);g__Metschnikowia(100);s__Metschnikowia_pulcherrima(100); | Metschnikowia | 13 | Saprotroph | \|Undefined Saprotroph\| | Yeast | NULL | Probable | As nectar-tap_saprotroph (Põlme et al. 2020); Decay sugar-rich substrates (Põlme et al. 2020) | Põlme S, et al. 2020. Fungal Diversity 105:1-16 ((https://doi.org/10.1007/s13225-020-00466-2)) |
| Otu15396 | 1 | k__Fungi(100);p__Ascomycota(100);c__Saccharomycetes(100);o__Saccharomycetales(100);f__Metschnikowiaceae(100);g__Metschnikowia(100);g__Metschnikowia_unclassified(100); | Metschnikowia | 13 | Saprotroph | \|Undefined Saprotroph\| | Yeast | NULL | Probable | As nectar-tap_saprotroph (Põlme et al. 2020); Decay sugar-rich substrates (Põlme et al. 2020) | Põlme S, et al. 2020. Fungal Diversity 105:1-16 ((https://doi.org/10.1007/s13225-020-00466-2)) |
| Otu15398 | 1 | k__Fungi(100);p__Ascomycota(100);c__Saccharomycetes(100);o__Saccharomycetales(100);f__Metschnikowiaceae(100);g__Metschnikowia(100);s__Metschnikowia_pulcherrima(100); | Metschnikowia | 13 | Saprotroph | \|Undefined Saprotroph\| | Yeast | NULL | Probable | As nectar-tap_saprotroph (Põlme et al. 2020); Decay sugar-rich substrates (Põlme et al. 2020) | Põlme S, et al. 2020. Fungal Diversity 105:1-16 ((https://doi.org/10.1007/s13225-020-00466-2)) |
| Otu15399 | 1 | k__Fungi(100);p__Ascomycota(100);c__Saccharomycetes(100);o__Saccharomycetales(100);f__Metschnikowiaceae(100);g__Metschnikowia(100);g__Metschnikowia_unclassified(100); | Metschnikowia | 13 | Saprotroph | \|Undefined Saprotroph\| | Yeast | NULL | Probable | As nectar-tap_saprotroph (Põlme et al. 2020); Decay sugar-rich substrates (Põlme et al. 2020) | Põlme S, et al. 2020. Fungal Diversity 105:1-16 ((https://doi.org/10.1007/s13225-020-00466-2)) |
| Otu15400 | 1 | k__Fungi(100);p__Ascomycota(100);c__Saccharomycetes(100);o__Saccharomycetales(100);f__Metschnikowiaceae(100);g__Metschnikowia(100);g__Metschnikowia_unclassified(100); | Metschnikowia | 13 | Saprotroph | \|Undefined Saprotroph\| | Yeast | NULL | Probable | As nectar-tap_saprotroph (Põlme et al. 2020); Decay sugar-rich substrates (Põlme et al. 2020) | Põlme S, et al. 2020. Fungal Diversity 105:1-16 ((https://doi.org/10.1007/s13225-020-00466-2)) |
| Otu15401 | 1 | k__Fungi(100);p__Ascomycota(100);c__Saccharomycetes(100);o__Saccharomycetales(100);f__Metschnikowiaceae(100);g__Metschnikowia(100);s__Metschnikowia_pulcherrima(100); | Metschnikowia | 13 | Saprotroph | \|Undefined Saprotroph\| | Yeast | NULL | Probable | As nectar-tap_saprotroph (Põlme et al. 2020); Decay sugar-rich substrates (Põlme et al. 2020) | Põlme S, et al. 2020. Fungal Diversity 105:1-16 ((https://doi.org/10.1007/s13225-020-00466-2)) |
| Otu15402 | 1 | k__Fungi(100);p__Ascomycota(100);c__Saccharomycetes(100);o__Saccharomycetales(100);f__Metschnikowiaceae(100);g__Metschnikowia(100);g__Metschnikowia_unclassified(100); | Metschnikowia | 13 | Saprotroph | \|Undefined Saprotroph\| | Yeast | NULL | Probable | As nectar-tap_saprotroph (Põlme et al. 2020); Decay sugar-rich substrates (Põlme et al. 2020) | Põlme S, et al. 2020. Fungal Diversity 105:1-16 ((https://doi.org/10.1007/s13225-020-00466-2)) |
| Otu15403 | 1 | k__Fungi(100);p__Ascomycota(100);c__Saccharomycetes(100);o__Saccharomycetales(100);f__Metschnikowiaceae(100);g__Metschnikowia(100);s__Metschnikowia_pulcherrima(100); | Metschnikowia | 13 | Saprotroph | \|Undefined Saprotroph\| | Yeast | NULL | Probable | As nectar-tap_saprotroph (Põlme et al. 2020); Decay sugar-rich substrates (Põlme et al. 2020) | Põlme S, et al. 2020. Fungal Diversity 105:1-16 ((https://doi.org/10.1007/s13225-020-00466-2)) |
| Otu15404 | 1 | k__Fungi(100);p__Ascomycota(100);c__Saccharomycetes(100);o__Saccharomycetales(100);f__Metschnikowiaceae(100);g__Metschnikowia(100);s__Metschnikowia_pulcherrima(100); | Metschnikowia | 13 | Saprotroph | \|Undefined Saprotroph\| | Yeast | NULL | Probable | As nectar-tap_saprotroph (Põlme et al. 2020); Decay sugar-rich substrates (Põlme et al. 2020) | Põlme S, et al. 2020. Fungal Diversity 105:1-16 ((https://doi.org/10.1007/s13225-020-00466-2)) |
| Otu15405 | 1 | k__Fungi(100);p__Ascomycota(100);c__Saccharomycetes(100);o__Saccharomycetales(100);f__Metschnikowiaceae(100);g__Metschnikowia(100);s__Metschnikowia_pulcherrima(100); | Metschnikowia | 13 | Saprotroph | \|Undefined Saprotroph\| | Yeast | NULL | Probable | As nectar-tap_saprotroph (Põlme et al. 2020); Decay sugar-rich substrates (Põlme et al. 2020) | Põlme S, et al. 2020. Fungal Diversity 105:1-16 ((https://doi.org/10.1007/s13225-020-00466-2)) |
| Otu15406 | 1 | k__Fungi(100);p__Ascomycota(100);c__Saccharomycetes(100);o__Saccharomycetales(100);f__Metschnikowiaceae(100);g__Metschnikowia(100);s__Metschnikowia_pulcherrima(100); | Metschnikowia | 13 | Saprotroph | \|Undefined Saprotroph\| | Yeast | NULL | Probable | As nectar-tap_saprotroph (Põlme et al. 2020); Decay sugar-rich substrates (Põlme et al. 2020) | Põlme S, et al. 2020. Fungal Diversity 105:1-16 ((https://doi.org/10.1007/s13225-020-00466-2)) |
| Otu15409 | 1 | k__Fungi(100);p__Ascomycota(100);c__Saccharomycetes(100);o__Saccharomycetales(100);f__Metschnikowiaceae(100);g__Metschnikowia(100);s__Metschnikowia_pulcherrima(100); | Metschnikowia | 13 | Saprotroph | \|Undefined Saprotroph\| | Yeast | NULL | Probable | As nectar-tap_saprotroph (Põlme et al. 2020); Decay sugar-rich substrates (Põlme et al. 2020) | Põlme S, et al. 2020. Fungal Diversity 105:1-16 ((https://doi.org/10.1007/s13225-020-00466-2)) |
| Otu15410 | 1 | k__Fungi(100);p__Ascomycota(100);c__Saccharomycetes(100);o__Saccharomycetales(100);f__Metschnikowiaceae(100);g__Metschnikowia(100);s__Metschnikowia_pulcherrima(100); | Metschnikowia | 13 | Saprotroph | \|Undefined Saprotroph\| | Yeast | NULL | Probable | As nectar-tap_saprotroph (Põlme et al. 2020); Decay sugar-rich substrates (Põlme et al. 2020) | Põlme S, et al. 2020. Fungal Diversity 105:1-16 ((https://doi.org/10.1007/s13225-020-00466-2)) |
| Otu15411 | 1 | k__Fungi(100);p__Ascomycota(100);c__Saccharomycetes(100);o__Saccharomycetales(100);f__Metschnikowiaceae(100);g__Metschnikowia(100);g__Metschnikowia_unclassified(100); | Metschnikowia | 13 | Saprotroph | \|Undefined Saprotroph\| | Yeast | NULL | Probable | As nectar-tap_saprotroph (Põlme et al. 2020); Decay sugar-rich substrates (Põlme et al. 2020) | Põlme S, et al. 2020. Fungal Diversity 105:1-16 ((https://doi.org/10.1007/s13225-020-00466-2)) |
| Otu15412 | 1 | k__Fungi(100);p__Ascomycota(100);c__Saccharomycetes(100);o__Saccharomycetales(100);f__Metschnikowiaceae(100);g__Metschnikowia(100);g__Metschnikowia_unclassified(100); | Metschnikowia | 13 | Saprotroph | \|Undefined Saprotroph\| | Yeast | NULL | Probable | As nectar-tap_saprotroph (Põlme et al. 2020); Decay sugar-rich substrates (Põlme et al. 2020) | Põlme S, et al. 2020. Fungal Diversity 105:1-16 ((https://doi.org/10.1007/s13225-020-00466-2)) |
| Otu15413 | 1 | k__Fungi(100);p__Ascomycota(100);c__Saccharomycetes(100);o__Saccharomycetales(100);f__Metschnikowiaceae(100);g__Metschnikowia(100);s__Metschnikowia_pulcherrima(100); | Metschnikowia | 13 | Saprotroph | \|Undefined Saprotroph\| | Yeast | NULL | Probable | As nectar-tap_saprotroph (Põlme et al. 2020); Decay sugar-rich substrates (Põlme et al. 2020) | Põlme S, et al. 2020. Fungal Diversity 105:1-16 ((https://doi.org/10.1007/s13225-020-00466-2)) |
| Otu15414 | 1 | k__Fungi(100);p__Ascomycota(100);c__Saccharomycetes(100);o__Saccharomycetales(100);f__Metschnikowiaceae(100);g__Metschnikowia(100);s__Metschnikowia_pulcherrima(100); | Metschnikowia | 13 | Saprotroph | \|Undefined Saprotroph\| | Yeast | NULL | Probable | As nectar-tap_saprotroph (Põlme et al. 2020); Decay sugar-rich substrates (Põlme et al. 2020) | Põlme S, et al. 2020. Fungal Diversity 105:1-16 ((https://doi.org/10.1007/s13225-020-00466-2)) |
| Otu15415 | 1 | k__Fungi(100);p__Ascomycota(100);c__Saccharomycetes(100);o__Saccharomycetales(100);f__Metschnikowiaceae(100);g__Metschnikowia(100);g__Metschnikowia_unclassified(100); | Metschnikowia | 13 | Saprotroph | \|Undefined Saprotroph\| | Yeast | NULL | Probable | As nectar-tap_saprotroph (Põlme et al. 2020); Decay sugar-rich substrates (Põlme et al. 2020) | Põlme S, et al. 2020. Fungal Diversity 105:1-16 ((https://doi.org/10.1007/s13225-020-00466-2)) |
| Otu15416 | 1 | k__Fungi(100);p__Ascomycota(100);c__Saccharomycetes(100);o__Saccharomycetales(100);f__Metschnikowiaceae(100);g__Metschnikowia(100);g__Metschnikowia_unclassified(100); | Metschnikowia | 13 | Saprotroph | \|Undefined Saprotroph\| | Yeast | NULL | Probable | As nectar-tap_saprotroph (Põlme et al. 2020); Decay sugar-rich substrates (Põlme et al. 2020) | Põlme S, et al. 2020. Fungal Diversity 105:1-16 ((https://doi.org/10.1007/s13225-020-00466-2)) |
| Otu15417 | 1 | k__Fungi(100);p__Ascomycota(100);c__Saccharomycetes(100);o__Saccharomycetales(100);o__Saccharomycetales_unclassified(100);o__Saccharomycetales_unclassified(100);o__Saccharomycetales_unclassified(100); | Saccharomycetales | 7 | Saprotroph | Undefined Saprotroph | Yeast | NULL | Possible | NULL | Sterkenburg E, et al. 2015. New Phytologist 207:1145-1158 ((https://doi.org/10.1111/nph.13426)) |
| Otu15418 | 1 | k__Fungi(100);p__Ascomycota(100);c__Saccharomycetes(100);o__Saccharomycetales(100);f__Metschnikowiaceae(100);g__Metschnikowia(100);g__Metschnikowia_unclassified(100); | Metschnikowia | 13 | Saprotroph | \|Undefined Saprotroph\| | Yeast | NULL | Probable | As nectar-tap_saprotroph (Põlme et al. 2020); Decay sugar-rich substrates (Põlme et al. 2020) | Põlme S, et al. 2020. Fungal Diversity 105:1-16 ((https://doi.org/10.1007/s13225-020-00466-2)) |
| Otu15419 | 1 | k__Fungi(100);p__Ascomycota(100);c__Saccharomycetes(100);o__Saccharomycetales(100);f__Metschnikowiaceae(100);g__Metschnikowia(100);g__Metschnikowia_unclassified(100); | Metschnikowia | 13 | Saprotroph | \|Undefined Saprotroph\| | Yeast | NULL | Probable | As nectar-tap_saprotroph (Põlme et al. 2020); Decay sugar-rich substrates (Põlme et al. 2020) | Põlme S, et al. 2020. Fungal Diversity 105:1-16 ((https://doi.org/10.1007/s13225-020-00466-2)) |
| Otu15420 | 1 | k__Fungi(100);p__Ascomycota(100);c__Saccharomycetes(100);o__Saccharomycetales(100);f__Metschnikowiaceae(100);g__Metschnikowia(100);g__Metschnikowia_unclassified(100); | Metschnikowia | 13 | Saprotroph | \|Undefined Saprotroph\| | Yeast | NULL | Probable | As nectar-tap_saprotroph (Põlme et al. 2020); Decay sugar-rich substrates (Põlme et al. 2020) | Põlme S, et al. 2020. Fungal Diversity 105:1-16 ((https://doi.org/10.1007/s13225-020-00466-2)) |
| Otu15421 | 1 | k__Fungi(100);p__Ascomycota(100);c__Saccharomycetes(100);o__Saccharomycetales(100);f__Metschnikowiaceae(100);g__Metschnikowia(100);g__Metschnikowia_unclassified(100); | Metschnikowia | 13 | Saprotroph | \|Undefined Saprotroph\| | Yeast | NULL | Probable | As nectar-tap_saprotroph (Põlme et al. 2020); Decay sugar-rich substrates (Põlme et al. 2020) | Põlme S, et al. 2020. Fungal Diversity 105:1-16 ((https://doi.org/10.1007/s13225-020-00466-2)) |
| Otu15423 | 1 | k__Fungi(100);p__Ascomycota(100);c__Saccharomycetes(100);o__Saccharomycetales(100);f__Metschnikowiaceae(100);g__Metschnikowia(100);s__Metschnikowia_pulcherrima(100); | Metschnikowia | 13 | Saprotroph | \|Undefined Saprotroph\| | Yeast | NULL | Probable | As nectar-tap_saprotroph (Põlme et al. 2020); Decay sugar-rich substrates (Põlme et al. 2020) | Põlme S, et al. 2020. Fungal Diversity 105:1-16 ((https://doi.org/10.1007/s13225-020-00466-2)) |
| Otu15424 | 1 | k__Fungi(100);p__Ascomycota(100);c__Saccharomycetes(100);o__Saccharomycetales(100);f__Metschnikowiaceae(100);g__Metschnikowia(100);g__Metschnikowia_unclassified(100); | Metschnikowia | 13 | Saprotroph | \|Undefined Saprotroph\| | Yeast | NULL | Probable | As nectar-tap_saprotroph (Põlme et al. 2020); Decay sugar-rich substrates (Põlme et al. 2020) | Põlme S, et al. 2020. Fungal Diversity 105:1-16 ((https://doi.org/10.1007/s13225-020-00466-2)) |
| Otu15425 | 1 | k__Fungi(100);p__Ascomycota(100);c__Saccharomycetes(100);o__Saccharomycetales(100);f__Metschnikowiaceae(100);g__Metschnikowia(100);s__Metschnikowia_pulcherrima(100); | Metschnikowia | 13 | Saprotroph | \|Undefined Saprotroph\| | Yeast | NULL | Probable | As nectar-tap_saprotroph (Põlme et al. 2020); Decay sugar-rich substrates (Põlme et al. 2020) | Põlme S, et al. 2020. Fungal Diversity 105:1-16 ((https://doi.org/10.1007/s13225-020-00466-2)) |
| Otu15429 | 1 | k__Fungi(100);p__Ascomycota(100);c__Saccharomycetes(100);o__Saccharomycetales(100);f__Metschnikowiaceae(100);g__Metschnikowia(100);g__Metschnikowia_unclassified(100); | Metschnikowia | 13 | Saprotroph | \|Undefined Saprotroph\| | Yeast | NULL | Probable | As nectar-tap_saprotroph (Põlme et al. 2020); Decay sugar-rich substrates (Põlme et al. 2020) | Põlme S, et al. 2020. Fungal Diversity 105:1-16 ((https://doi.org/10.1007/s13225-020-00466-2)) |
| Otu15430 | 1 | k__Fungi(100);p__Ascomycota(100);c__Saccharomycetes(100);o__Saccharomycetales(100);f__Metschnikowiaceae(100);g__Metschnikowia(100);g__Metschnikowia_unclassified(100); | Metschnikowia | 13 | Saprotroph | \|Undefined Saprotroph\| | Yeast | NULL | Probable | As nectar-tap_saprotroph (Põlme et al. 2020); Decay sugar-rich substrates (Põlme et al. 2020) | Põlme S, et al. 2020. Fungal Diversity 105:1-16 ((https://doi.org/10.1007/s13225-020-00466-2)) |
| Otu15431 | 1 | k__Fungi(100);p__Ascomycota(100);c__Saccharomycetes(100);o__Saccharomycetales(100);f__Metschnikowiaceae(100);g__Metschnikowia(100);g__Metschnikowia_unclassified(100); | Metschnikowia | 13 | Saprotroph | \|Undefined Saprotroph\| | Yeast | NULL | Probable | As nectar-tap_saprotroph (Põlme et al. 2020); Decay sugar-rich substrates (Põlme et al. 2020) | Põlme S, et al. 2020. Fungal Diversity 105:1-16 ((https://doi.org/10.1007/s13225-020-00466-2)) |
| Otu15433 | 1 | k__Fungi(100);p__Ascomycota(100);c__Saccharomycetes(100);o__Saccharomycetales(100);f__Metschnikowiaceae(100);g__Metschnikowia(100);s__Metschnikowia_pulcherrima(100); | Metschnikowia | 13 | Saprotroph | \|Undefined Saprotroph\| | Yeast | NULL | Probable | As nectar-tap_saprotroph (Põlme et al. 2020); Decay sugar-rich substrates (Põlme et al. 2020) | Põlme S, et al. 2020. Fungal Diversity 105:1-16 ((https://doi.org/10.1007/s13225-020-00466-2)) |
| Otu15435 | 1 | k__Fungi(100);p__Ascomycota(100);c__Saccharomycetes(100);o__Saccharomycetales(100);f__Metschnikowiaceae(100);g__Metschnikowia(100);s__Metschnikowia_pulcherrima(100); | Metschnikowia | 13 | Saprotroph | \|Undefined Saprotroph\| | Yeast | NULL | Probable | As nectar-tap_saprotroph (Põlme et al. 2020); Decay sugar-rich substrates (Põlme et al. 2020) | Põlme S, et al. 2020. Fungal Diversity 105:1-16 ((https://doi.org/10.1007/s13225-020-00466-2)) |
| Otu15437 | 1 | k__Fungi(100);p__Ascomycota(100);c__Saccharomycetes(100);o__Saccharomycetales(100);f__Metschnikowiaceae(100);g__Metschnikowia(100);s__Metschnikowia_pulcherrima(100); | Metschnikowia | 13 | Saprotroph | \|Undefined Saprotroph\| | Yeast | NULL | Probable | As nectar-tap_saprotroph (Põlme et al. 2020); Decay sugar-rich substrates (Põlme et al. 2020) | Põlme S, et al. 2020. Fungal Diversity 105:1-16 ((https://doi.org/10.1007/s13225-020-00466-2)) |
| Otu15438 | 1 | k__Fungi(100);p__Ascomycota(100);c__Saccharomycetes(100);o__Saccharomycetales(100);f__Metschnikowiaceae(100);g__Metschnikowia(100);g__Metschnikowia_unclassified(100); | Metschnikowia | 13 | Saprotroph | \|Undefined Saprotroph\| | Yeast | NULL | Probable | As nectar-tap_saprotroph (Põlme et al. 2020); Decay sugar-rich substrates (Põlme et al. 2020) | Põlme S, et al. 2020. Fungal Diversity 105:1-16 ((https://doi.org/10.1007/s13225-020-00466-2)) |
| Otu15439 | 1 | k__Fungi(100);p__Ascomycota(100);c__Saccharomycetes(100);o__Saccharomycetales(100);f__Metschnikowiaceae(100);g__Metschnikowia(100);s__Metschnikowia_pulcherrima(100); | Metschnikowia | 13 | Saprotroph | \|Undefined Saprotroph\| | Yeast | NULL | Probable | As nectar-tap_saprotroph (Põlme et al. 2020); Decay sugar-rich substrates (Põlme et al. 2020) | Põlme S, et al. 2020. Fungal Diversity 105:1-16 ((https://doi.org/10.1007/s13225-020-00466-2)) |
| Otu15442 | 1 | k__Fungi(100);p__Ascomycota(100);c__Saccharomycetes(100);o__Saccharomycetales(100);f__Metschnikowiaceae(100);g__Metschnikowia(100);g__Metschnikowia_unclassified(100); | Metschnikowia | 13 | Saprotroph | \|Undefined Saprotroph\| | Yeast | NULL | Probable | As nectar-tap_saprotroph (Põlme et al. 2020); Decay sugar-rich substrates (Põlme et al. 2020) | Põlme S, et al. 2020. Fungal Diversity 105:1-16 ((https://doi.org/10.1007/s13225-020-00466-2)) |
| Otu15443 | 1 | k__Fungi(100);p__Ascomycota(100);c__Saccharomycetes(100);o__Saccharomycetales(100);f__Metschnikowiaceae(100);g__Metschnikowia(100);g__Metschnikowia_unclassified(100); | Metschnikowia | 13 | Saprotroph | \|Undefined Saprotroph\| | Yeast | NULL | Probable | As nectar-tap_saprotroph (Põlme et al. 2020); Decay sugar-rich substrates (Põlme et al. 2020) | Põlme S, et al. 2020. Fungal Diversity 105:1-16 ((https://doi.org/10.1007/s13225-020-00466-2)) |
| Otu15444 | 1 | k__Fungi(100);p__Ascomycota(100);c__Saccharomycetes(100);o__Saccharomycetales(100);f__Metschnikowiaceae(100);g__Metschnikowia(100);s__Metschnikowia_pulcherrima(100); | Metschnikowia | 13 | Saprotroph | \|Undefined Saprotroph\| | Yeast | NULL | Probable | As nectar-tap_saprotroph (Põlme et al. 2020); Decay sugar-rich substrates (Põlme et al. 2020) | Põlme S, et al. 2020. Fungal Diversity 105:1-16 ((https://doi.org/10.1007/s13225-020-00466-2)) |
| Otu15445 | 1 | k__Fungi(100);p__Ascomycota(100);c__Saccharomycetes(100);o__Saccharomycetales(100);f__Metschnikowiaceae(100);g__Metschnikowia(100);s__Metschnikowia_pulcherrima(100); | Metschnikowia | 13 | Saprotroph | \|Undefined Saprotroph\| | Yeast | NULL | Probable | As nectar-tap_saprotroph (Põlme et al. 2020); Decay sugar-rich substrates (Põlme et al. 2020) | Põlme S, et al. 2020. Fungal Diversity 105:1-16 ((https://doi.org/10.1007/s13225-020-00466-2)) |
| Otu15446 | 1 | k__Fungi(100);p__Ascomycota(100);c__Saccharomycetes(100);o__Saccharomycetales(100);f__Metschnikowiaceae(100);g__Metschnikowia(100);s__Metschnikowia_pulcherrima(100); | Metschnikowia | 13 | Saprotroph | \|Undefined Saprotroph\| | Yeast | NULL | Probable | As nectar-tap_saprotroph (Põlme et al. 2020); Decay sugar-rich substrates (Põlme et al. 2020) | Põlme S, et al. 2020. Fungal Diversity 105:1-16 ((https://doi.org/10.1007/s13225-020-00466-2)) |
| Otu15447 | 1 | k__Fungi(100);p__Ascomycota(100);c__Saccharomycetes(100);o__Saccharomycetales(100);f__Metschnikowiaceae(100);g__Metschnikowia(100);s__Metschnikowia_pulcherrima(100); | Metschnikowia | 13 | Saprotroph | \|Undefined Saprotroph\| | Yeast | NULL | Probable | As nectar-tap_saprotroph (Põlme et al. 2020); Decay sugar-rich substrates (Põlme et al. 2020) | Põlme S, et al. 2020. Fungal Diversity 105:1-16 ((https://doi.org/10.1007/s13225-020-00466-2)) |
| Otu15448 | 1 | k__Fungi(100);p__Ascomycota(100);c__Saccharomycetes(100);o__Saccharomycetales(100);f__Metschnikowiaceae(100);g__Metschnikowia(100);s__Metschnikowia_pulcherrima(100); | Metschnikowia | 13 | Saprotroph | \|Undefined Saprotroph\| | Yeast | NULL | Probable | As nectar-tap_saprotroph (Põlme et al. 2020); Decay sugar-rich substrates (Põlme et al. 2020) | Põlme S, et al. 2020. Fungal Diversity 105:1-16 ((https://doi.org/10.1007/s13225-020-00466-2)) |
| Otu15449 | 1 | k__Fungi(100);p__Ascomycota(100);c__Saccharomycetes(100);o__Saccharomycetales(100);f__Metschnikowiaceae(100);g__Metschnikowia(100);g__Metschnikowia_unclassified(100); | Metschnikowia | 13 | Saprotroph | \|Undefined Saprotroph\| | Yeast | NULL | Probable | As nectar-tap_saprotroph (Põlme et al. 2020); Decay sugar-rich substrates (Põlme et al. 2020) | Põlme S, et al. 2020. Fungal Diversity 105:1-16 ((https://doi.org/10.1007/s13225-020-00466-2)) |
| Otu15450 | 1 | k__Fungi(100);p__Ascomycota(100);c__Saccharomycetes(100);o__Saccharomycetales(100);o__Saccharomycetales_unclassified(100);o__Saccharomycetales_unclassified(100);o__Saccharomycetales_unclassified(100); | Saccharomycetales | 7 | Saprotroph | Undefined Saprotroph | Yeast | NULL | Possible | NULL | Sterkenburg E, et al. 2015. New Phytologist 207:1145-1158 ((https://doi.org/10.1111/nph.13426)) |
| Otu15451 | 1 | k__Fungi(100);p__Ascomycota(100);c__Saccharomycetes(100);o__Saccharomycetales(100);f__Metschnikowiaceae(100);g__Metschnikowia(100);s__Metschnikowia_pulcherrima(100); | Metschnikowia | 13 | Saprotroph | \|Undefined Saprotroph\| | Yeast | NULL | Probable | As nectar-tap_saprotroph (Põlme et al. 2020); Decay sugar-rich substrates (Põlme et al. 2020) | Põlme S, et al. 2020. Fungal Diversity 105:1-16 ((https://doi.org/10.1007/s13225-020-00466-2)) |
| Otu15452 | 1 | k__Fungi(100);p__Ascomycota(100);c__Saccharomycetes(100);o__Saccharomycetales(100);f__Metschnikowiaceae(100);g__Metschnikowia(100);s__Metschnikowia_pulcherrima(100); | Metschnikowia | 13 | Saprotroph | \|Undefined Saprotroph\| | Yeast | NULL | Probable | As nectar-tap_saprotroph (Põlme et al. 2020); Decay sugar-rich substrates (Põlme et al. 2020) | Põlme S, et al. 2020. Fungal Diversity 105:1-16 ((https://doi.org/10.1007/s13225-020-00466-2)) |
| Otu15453 | 1 | k__Fungi(100);p__Ascomycota(100);c__Saccharomycetes(100);o__Saccharomycetales(100);f__Metschnikowiaceae(100);g__Metschnikowia(100);s__Metschnikowia_pulcherrima(100); | Metschnikowia | 13 | Saprotroph | \|Undefined Saprotroph\| | Yeast | NULL | Probable | As nectar-tap_saprotroph (Põlme et al. 2020); Decay sugar-rich substrates (Põlme et al. 2020) | Põlme S, et al. 2020. Fungal Diversity 105:1-16 ((https://doi.org/10.1007/s13225-020-00466-2)) |
| Otu15454 | 1 | k__Fungi(100);p__Ascomycota(100);c__Saccharomycetes(100);o__Saccharomycetales(100);f__Metschnikowiaceae(100);g__Metschnikowia(100);s__Metschnikowia_pulcherrima(100); | Metschnikowia | 13 | Saprotroph | \|Undefined Saprotroph\| | Yeast | NULL | Probable | As nectar-tap_saprotroph (Põlme et al. 2020); Decay sugar-rich substrates (Põlme et al. 2020) | Põlme S, et al. 2020. Fungal Diversity 105:1-16 ((https://doi.org/10.1007/s13225-020-00466-2)) |
| Otu15456 | 1 | k__Fungi(100);p__Ascomycota(100);c__Saccharomycetes(100);o__Saccharomycetales(100);f__Metschnikowiaceae(100);g__Metschnikowia(100);s__Metschnikowia_pulcherrima(100); | Metschnikowia | 13 | Saprotroph | \|Undefined Saprotroph\| | Yeast | NULL | Probable | As nectar-tap_saprotroph (Põlme et al. 2020); Decay sugar-rich substrates (Põlme et al. 2020) | Põlme S, et al. 2020. Fungal Diversity 105:1-16 ((https://doi.org/10.1007/s13225-020-00466-2)) |
| Otu15457 | 1 | k__Fungi(100);p__Ascomycota(100);c__Saccharomycetes(100);o__Saccharomycetales(100);f__Metschnikowiaceae(100);g__Metschnikowia(100);s__Metschnikowia_pulcherrima(100); | Metschnikowia | 13 | Saprotroph | \|Undefined Saprotroph\| | Yeast | NULL | Probable | As nectar-tap_saprotroph (Põlme et al. 2020); Decay sugar-rich substrates (Põlme et al. 2020) | Põlme S, et al. 2020. Fungal Diversity 105:1-16 ((https://doi.org/10.1007/s13225-020-00466-2)) |
| Otu15458 | 1 | k__Fungi(100);p__Ascomycota(100);c__Saccharomycetes(100);o__Saccharomycetales(100);f__Metschnikowiaceae(100);g__Metschnikowia(100);s__Metschnikowia_pulcherrima(100); | Metschnikowia | 13 | Saprotroph | \|Undefined Saprotroph\| | Yeast | NULL | Probable | As nectar-tap_saprotroph (Põlme et al. 2020); Decay sugar-rich substrates (Põlme et al. 2020) | Põlme S, et al. 2020. Fungal Diversity 105:1-16 ((https://doi.org/10.1007/s13225-020-00466-2)) |
| Otu15460 | 1 | k__Fungi(100);p__Ascomycota(100);c__Saccharomycetes(100);o__Saccharomycetales(100);f__Metschnikowiaceae(100);g__Metschnikowia(100);s__Metschnikowia_pulcherrima(100); | Metschnikowia | 13 | Saprotroph | \|Undefined Saprotroph\| | Yeast | NULL | Probable | As nectar-tap_saprotroph (Põlme et al. 2020); Decay sugar-rich substrates (Põlme et al. 2020) | Põlme S, et al. 2020. Fungal Diversity 105:1-16 ((https://doi.org/10.1007/s13225-020-00466-2)) |
| Otu15461 | 1 | k__Fungi(100);p__Ascomycota(100);c__Saccharomycetes(100);o__Saccharomycetales(100);f__Metschnikowiaceae(100);g__Metschnikowia(100);s__Metschnikowia_pulcherrima(100); | Metschnikowia | 13 | Saprotroph | \|Undefined Saprotroph\| | Yeast | NULL | Probable | As nectar-tap_saprotroph (Põlme et al. 2020); Decay sugar-rich substrates (Põlme et al. 2020) | Põlme S, et al. 2020. Fungal Diversity 105:1-16 ((https://doi.org/10.1007/s13225-020-00466-2)) |
| Otu15462 | 1 | k__Fungi(100);p__Ascomycota(100);c__Saccharomycetes(100);o__Saccharomycetales(100);f__Metschnikowiaceae(100);g__Metschnikowia(100);s__Metschnikowia_pulcherrima(100); | Metschnikowia | 13 | Saprotroph | \|Undefined Saprotroph\| | Yeast | NULL | Probable | As nectar-tap_saprotroph (Põlme et al. 2020); Decay sugar-rich substrates (Põlme et al. 2020) | Põlme S, et al. 2020. Fungal Diversity 105:1-16 ((https://doi.org/10.1007/s13225-020-00466-2)) |
| Otu15463 | 1 | k__Fungi(100);p__Ascomycota(100);c__Saccharomycetes(100);o__Saccharomycetales(100);f__Metschnikowiaceae(100);g__Metschnikowia(100);s__Metschnikowia_pulcherrima(100); | Metschnikowia | 13 | Saprotroph | \|Undefined Saprotroph\| | Yeast | NULL | Probable | As nectar-tap_saprotroph (Põlme et al. 2020); Decay sugar-rich substrates (Põlme et al. 2020) | Põlme S, et al. 2020. Fungal Diversity 105:1-16 ((https://doi.org/10.1007/s13225-020-00466-2)) |
| Otu15465 | 1 | k__Fungi(100);p__Ascomycota(100);c__Saccharomycetes(100);o__Saccharomycetales(100);f__Metschnikowiaceae(100);g__Metschnikowia(100);s__Metschnikowia_pulcherrima(100); | Metschnikowia | 13 | Saprotroph | \|Undefined Saprotroph\| | Yeast | NULL | Probable | As nectar-tap_saprotroph (Põlme et al. 2020); Decay sugar-rich substrates (Põlme et al. 2020) | Põlme S, et al. 2020. Fungal Diversity 105:1-16 ((https://doi.org/10.1007/s13225-020-00466-2)) |
| Otu15466 | 1 | k__Fungi(100);p__Ascomycota(100);c__Saccharomycetes(100);o__Saccharomycetales(100);f__Saccharomycetales_family_Incertae_sedis(100);g__Candida(100);g__Candida_unclassified(100); | Candida | 13 | Pathotroph-Saprotroph-Symbiotroph | Animal Pathogen-Endophyte-Endosymbiont-Epiphyte-\|Undefined Saprotroph\| | Dimorphic Yeast-Yeast | NULL | Probable | As nectar-tap_saprotroph (Põlme et al. 2020); Decay sugar-rich substrates (Põlme et al. 2020); This asexual genus is polyphyletic and contain species that function very differently depend on what they are. Caution should be taken when interpreting guilds for Candida species. | Manolakaki D, et al. 2010. Virulence. 1: 367-375 ((https://doi.org/10.4161/viru.1.5.12796)); Mekbib SB, et al. 2011. Tropical Plant Pathology 36:233-240 ((https://doi.org/10.1590/S1982-56762011000400004)); Põlme S, et al. 2020. Fungal Diversity 105:1-16 ((https://doi.org/10.1007/s13225-020-00466-2)) |
| Otu15468 | 1 | k__Fungi(100);p__Ascomycota(100);c__Saccharomycetes(100);o__Saccharomycetales(100);f__Metschnikowiaceae(100);g__Metschnikowia(100);g__Metschnikowia_unclassified(100); | Metschnikowia | 13 | Saprotroph | \|Undefined Saprotroph\| | Yeast | NULL | Probable | As nectar-tap_saprotroph (Põlme et al. 2020); Decay sugar-rich substrates (Põlme et al. 2020) | Põlme S, et al. 2020. Fungal Diversity 105:1-16 ((https://doi.org/10.1007/s13225-020-00466-2)) |
| Otu15469 | 1 | k__Fungi(100);p__Ascomycota(100);c__Saccharomycetes(100);o__Saccharomycetales(100);f__Metschnikowiaceae(100);g__Metschnikowia(100);s__Metschnikowia_pulcherrima(100); | Metschnikowia | 13 | Saprotroph | \|Undefined Saprotroph\| | Yeast | NULL | Probable | As nectar-tap_saprotroph (Põlme et al. 2020); Decay sugar-rich substrates (Põlme et al. 2020) | Põlme S, et al. 2020. Fungal Diversity 105:1-16 ((https://doi.org/10.1007/s13225-020-00466-2)) |
| Otu15470 | 1 | k__Fungi(100);p__Ascomycota(100);c__Saccharomycetes(100);o__Saccharomycetales(100);f__Metschnikowiaceae(100);g__Metschnikowia(100);s__Metschnikowia_pulcherrima(100); | Metschnikowia | 13 | Saprotroph | \|Undefined Saprotroph\| | Yeast | NULL | Probable | As nectar-tap_saprotroph (Põlme et al. 2020); Decay sugar-rich substrates (Põlme et al. 2020) | Põlme S, et al. 2020. Fungal Diversity 105:1-16 ((https://doi.org/10.1007/s13225-020-00466-2)) |
| Otu15471 | 1 | k__Fungi(100);p__Ascomycota(100);c__Saccharomycetes(100);o__Saccharomycetales(100);f__Metschnikowiaceae(100);g__Metschnikowia(100);s__Metschnikowia_pulcherrima(100); | Metschnikowia | 13 | Saprotroph | \|Undefined Saprotroph\| | Yeast | NULL | Probable | As nectar-tap_saprotroph (Põlme et al. 2020); Decay sugar-rich substrates (Põlme et al. 2020) | Põlme S, et al. 2020. Fungal Diversity 105:1-16 ((https://doi.org/10.1007/s13225-020-00466-2)) |
| Otu15472 | 1 | k__Fungi(100);p__Ascomycota(100);c__Saccharomycetes(100);o__Saccharomycetales(100);f__Metschnikowiaceae(100);g__Metschnikowia(100);g__Metschnikowia_unclassified(100); | Metschnikowia | 13 | Saprotroph | \|Undefined Saprotroph\| | Yeast | NULL | Probable | As nectar-tap_saprotroph (Põlme et al. 2020); Decay sugar-rich substrates (Põlme et al. 2020) | Põlme S, et al. 2020. Fungal Diversity 105:1-16 ((https://doi.org/10.1007/s13225-020-00466-2)) |
| Otu15473 | 1 | k__Fungi(100);p__Ascomycota(100);c__Saccharomycetes(100);o__Saccharomycetales(100);f__Metschnikowiaceae(100);g__Metschnikowia(100);g__Metschnikowia_unclassified(100); | Metschnikowia | 13 | Saprotroph | \|Undefined Saprotroph\| | Yeast | NULL | Probable | As nectar-tap_saprotroph (Põlme et al. 2020); Decay sugar-rich substrates (Põlme et al. 2020) | Põlme S, et al. 2020. Fungal Diversity 105:1-16 ((https://doi.org/10.1007/s13225-020-00466-2)) |
| Otu15474 | 1 | k__Fungi(100);p__Ascomycota(100);c__Saccharomycetes(100);o__Saccharomycetales(100);f__Metschnikowiaceae(100);g__Metschnikowia(100);s__Metschnikowia_pulcherrima(100); | Metschnikowia | 13 | Saprotroph | \|Undefined Saprotroph\| | Yeast | NULL | Probable | As nectar-tap_saprotroph (Põlme et al. 2020); Decay sugar-rich substrates (Põlme et al. 2020) | Põlme S, et al. 2020. Fungal Diversity 105:1-16 ((https://doi.org/10.1007/s13225-020-00466-2)) |
| Otu15475 | 1 | k__Fungi(100);p__Ascomycota(100);c__Saccharomycetes(100);o__Saccharomycetales(100);f__Metschnikowiaceae(100);g__Metschnikowia(100);s__Metschnikowia_pulcherrima(100); | Metschnikowia | 13 | Saprotroph | \|Undefined Saprotroph\| | Yeast | NULL | Probable | As nectar-tap_saprotroph (Põlme et al. 2020); Decay sugar-rich substrates (Põlme et al. 2020) | Põlme S, et al. 2020. Fungal Diversity 105:1-16 ((https://doi.org/10.1007/s13225-020-00466-2)) |
| Otu15476 | 1 | k__Fungi(100);p__Ascomycota(100);c__Saccharomycetes(100);o__Saccharomycetales(100);f__Metschnikowiaceae(100);g__Metschnikowia(100);s__Metschnikowia_pulcherrima(100); | Metschnikowia | 13 | Saprotroph | \|Undefined Saprotroph\| | Yeast | NULL | Probable | As nectar-tap_saprotroph (Põlme et al. 2020); Decay sugar-rich substrates (Põlme et al. 2020) | Põlme S, et al. 2020. Fungal Diversity 105:1-16 ((https://doi.org/10.1007/s13225-020-00466-2)) |
| Otu15477 | 1 | k__Fungi(100);p__Ascomycota(100);c__Saccharomycetes(100);o__Saccharomycetales(100);f__Metschnikowiaceae(100);g__Metschnikowia(100);s__Metschnikowia_pulcherrima(100); | Metschnikowia | 13 | Saprotroph | \|Undefined Saprotroph\| | Yeast | NULL | Probable | As nectar-tap_saprotroph (Põlme et al. 2020); Decay sugar-rich substrates (Põlme et al. 2020) | Põlme S, et al. 2020. Fungal Diversity 105:1-16 ((https://doi.org/10.1007/s13225-020-00466-2)) |
| Otu15478 | 1 | k__Fungi(100);p__Ascomycota(100);c__Saccharomycetes(100);o__Saccharomycetales(100);f__Metschnikowiaceae(100);g__Metschnikowia(100);s__Metschnikowia_pulcherrima(100); | Metschnikowia | 13 | Saprotroph | \|Undefined Saprotroph\| | Yeast | NULL | Probable | As nectar-tap_saprotroph (Põlme et al. 2020); Decay sugar-rich substrates (Põlme et al. 2020) | Põlme S, et al. 2020. Fungal Diversity 105:1-16 ((https://doi.org/10.1007/s13225-020-00466-2)) |
| Otu15479 | 1 | k__Fungi(100);p__Ascomycota(100);c__Saccharomycetes(100);o__Saccharomycetales(100);f__Metschnikowiaceae(100);g__Metschnikowia(100);g__Metschnikowia_unclassified(100); | Metschnikowia | 13 | Saprotroph | \|Undefined Saprotroph\| | Yeast | NULL | Probable | As nectar-tap_saprotroph (Põlme et al. 2020); Decay sugar-rich substrates (Põlme et al. 2020) | Põlme S, et al. 2020. Fungal Diversity 105:1-16 ((https://doi.org/10.1007/s13225-020-00466-2)) |
| Otu15480 | 1 | k__Fungi(100);p__Ascomycota(100);c__Saccharomycetes(100);o__Saccharomycetales(100);f__Metschnikowiaceae(100);g__Metschnikowia(100);s__Metschnikowia_pulcherrima(100); | Metschnikowia | 13 | Saprotroph | \|Undefined Saprotroph\| | Yeast | NULL | Probable | As nectar-tap_saprotroph (Põlme et al. 2020); Decay sugar-rich substrates (Põlme et al. 2020) | Põlme S, et al. 2020. Fungal Diversity 105:1-16 ((https://doi.org/10.1007/s13225-020-00466-2)) |
| Otu15482 | 1 | k__Fungi(100);p__Ascomycota(100);c__Saccharomycetes(100);o__Saccharomycetales(100);f__Metschnikowiaceae(100);g__Metschnikowia(100);s__Metschnikowia_pulcherrima(100); | Metschnikowia | 13 | Saprotroph | \|Undefined Saprotroph\| | Yeast | NULL | Probable | As nectar-tap_saprotroph (Põlme et al. 2020); Decay sugar-rich substrates (Põlme et al. 2020) | Põlme S, et al. 2020. Fungal Diversity 105:1-16 ((https://doi.org/10.1007/s13225-020-00466-2)) |
| Otu15483 | 1 | k__Fungi(100);p__Ascomycota(100);c__Saccharomycetes(100);o__Saccharomycetales(100);f__Metschnikowiaceae(100);g__Metschnikowia(100);g__Metschnikowia_unclassified(100); | Metschnikowia | 13 | Saprotroph | \|Undefined Saprotroph\| | Yeast | NULL | Probable | As nectar-tap_saprotroph (Põlme et al. 2020); Decay sugar-rich substrates (Põlme et al. 2020) | Põlme S, et al. 2020. Fungal Diversity 105:1-16 ((https://doi.org/10.1007/s13225-020-00466-2)) |
| Otu15484 | 1 | k__Fungi(100);p__Ascomycota(100);c__Saccharomycetes(100);o__Saccharomycetales(100);f__Metschnikowiaceae(100);g__Metschnikowia(100);s__Metschnikowia_pulcherrima(100); | Metschnikowia | 13 | Saprotroph | \|Undefined Saprotroph\| | Yeast | NULL | Probable | As nectar-tap_saprotroph (Põlme et al. 2020); Decay sugar-rich substrates (Põlme et al. 2020) | Põlme S, et al. 2020. Fungal Diversity 105:1-16 ((https://doi.org/10.1007/s13225-020-00466-2)) |
| Otu15485 | 1 | k__Fungi(100);p__Ascomycota(100);c__Saccharomycetes(100);o__Saccharomycetales(100);f__Metschnikowiaceae(100);g__Metschnikowia(100);g__Metschnikowia_unclassified(100); | Metschnikowia | 13 | Saprotroph | \|Undefined Saprotroph\| | Yeast | NULL | Probable | As nectar-tap_saprotroph (Põlme et al. 2020); Decay sugar-rich substrates (Põlme et al. 2020) | Põlme S, et al. 2020. Fungal Diversity 105:1-16 ((https://doi.org/10.1007/s13225-020-00466-2)) |
| Otu15486 | 1 | k__Fungi(100);p__Ascomycota(100);c__Saccharomycetes(100);o__Saccharomycetales(100);f__Metschnikowiaceae(100);g__Metschnikowia(100);s__Metschnikowia_pulcherrima(100); | Metschnikowia | 13 | Saprotroph | \|Undefined Saprotroph\| | Yeast | NULL | Probable | As nectar-tap_saprotroph (Põlme et al. 2020); Decay sugar-rich substrates (Põlme et al. 2020) | Põlme S, et al. 2020. Fungal Diversity 105:1-16 ((https://doi.org/10.1007/s13225-020-00466-2)) |
| Otu15487 | 1 | k__Fungi(100);p__Ascomycota(100);c__Saccharomycetes(100);o__Saccharomycetales(100);f__Metschnikowiaceae(100);g__Metschnikowia(100);s__Metschnikowia_pulcherrima(100); | Metschnikowia | 13 | Saprotroph | \|Undefined Saprotroph\| | Yeast | NULL | Probable | As nectar-tap_saprotroph (Põlme et al. 2020); Decay sugar-rich substrates (Põlme et al. 2020) | Põlme S, et al. 2020. Fungal Diversity 105:1-16 ((https://doi.org/10.1007/s13225-020-00466-2)) |
| Otu15488 | 1 | k__Fungi(100);p__Ascomycota(100);c__Saccharomycetes(100);o__Saccharomycetales(100);f__Metschnikowiaceae(100);g__Metschnikowia(100);g__Metschnikowia_unclassified(100); | Metschnikowia | 13 | Saprotroph | \|Undefined Saprotroph\| | Yeast | NULL | Probable | As nectar-tap_saprotroph (Põlme et al. 2020); Decay sugar-rich substrates (Põlme et al. 2020) | Põlme S, et al. 2020. Fungal Diversity 105:1-16 ((https://doi.org/10.1007/s13225-020-00466-2)) |
| Otu15490 | 1 | k__Fungi(100);p__Ascomycota(100);c__Saccharomycetes(100);o__Saccharomycetales(100);f__Metschnikowiaceae(100);g__Metschnikowia(100);s__Metschnikowia_pulcherrima(100); | Metschnikowia | 13 | Saprotroph | \|Undefined Saprotroph\| | Yeast | NULL | Probable | As nectar-tap_saprotroph (Põlme et al. 2020); Decay sugar-rich substrates (Põlme et al. 2020) | Põlme S, et al. 2020. Fungal Diversity 105:1-16 ((https://doi.org/10.1007/s13225-020-00466-2)) |
| Otu15491 | 1 | k__Fungi(100);p__Ascomycota(100);c__Saccharomycetes(100);o__Saccharomycetales(100);f__Metschnikowiaceae(100);g__Metschnikowia(100);s__Metschnikowia_pulcherrima(100); | Metschnikowia | 13 | Saprotroph | \|Undefined Saprotroph\| | Yeast | NULL | Probable | As nectar-tap_saprotroph (Põlme et al. 2020); Decay sugar-rich substrates (Põlme et al. 2020) | Põlme S, et al. 2020. Fungal Diversity 105:1-16 ((https://doi.org/10.1007/s13225-020-00466-2)) |
| Otu15492 | 1 | k__Fungi(100);p__Ascomycota(100);c__Saccharomycetes(100);o__Saccharomycetales(100);f__Metschnikowiaceae(100);g__Metschnikowia(100);s__Metschnikowia_pulcherrima(100); | Metschnikowia | 13 | Saprotroph | \|Undefined Saprotroph\| | Yeast | NULL | Probable | As nectar-tap_saprotroph (Põlme et al. 2020); Decay sugar-rich substrates (Põlme et al. 2020) | Põlme S, et al. 2020. Fungal Diversity 105:1-16 ((https://doi.org/10.1007/s13225-020-00466-2)) |
| Otu15493 | 1 | k__Fungi(100);p__Ascomycota(100);c__Saccharomycetes(100);o__Saccharomycetales(100);f__Metschnikowiaceae(100);g__Metschnikowia(100);s__Metschnikowia_pulcherrima(100); | Metschnikowia | 13 | Saprotroph | \|Undefined Saprotroph\| | Yeast | NULL | Probable | As nectar-tap_saprotroph (Põlme et al. 2020); Decay sugar-rich substrates (Põlme et al. 2020) | Põlme S, et al. 2020. Fungal Diversity 105:1-16 ((https://doi.org/10.1007/s13225-020-00466-2)) |
| Otu15494 | 1 | k__Fungi(100);p__Ascomycota(100);c__Saccharomycetes(100);o__Saccharomycetales(100);f__Metschnikowiaceae(100);g__Metschnikowia(100);g__Metschnikowia_unclassified(100); | Metschnikowia | 13 | Saprotroph | \|Undefined Saprotroph\| | Yeast | NULL | Probable | As nectar-tap_saprotroph (Põlme et al. 2020); Decay sugar-rich substrates (Põlme et al. 2020) | Põlme S, et al. 2020. Fungal Diversity 105:1-16 ((https://doi.org/10.1007/s13225-020-00466-2)) |
| Otu15495 | 1 | k__Fungi(100);p__Ascomycota(100);c__Saccharomycetes(100);o__Saccharomycetales(100);f__Metschnikowiaceae(100);g__Metschnikowia(100);g__Metschnikowia_unclassified(100); | Metschnikowia | 13 | Saprotroph | \|Undefined Saprotroph\| | Yeast | NULL | Probable | As nectar-tap_saprotroph (Põlme et al. 2020); Decay sugar-rich substrates (Põlme et al. 2020) | Põlme S, et al. 2020. Fungal Diversity 105:1-16 ((https://doi.org/10.1007/s13225-020-00466-2)) |
| Otu15496 | 1 | k__Fungi(100);p__Ascomycota(100);c__Saccharomycetes(100);o__Saccharomycetales(100);f__Metschnikowiaceae(100);g__Metschnikowia(100);g__Metschnikowia_unclassified(100); | Metschnikowia | 13 | Saprotroph | \|Undefined Saprotroph\| | Yeast | NULL | Probable | As nectar-tap_saprotroph (Põlme et al. 2020); Decay sugar-rich substrates (Põlme et al. 2020) | Põlme S, et al. 2020. Fungal Diversity 105:1-16 ((https://doi.org/10.1007/s13225-020-00466-2)) |
| Otu15497 | 1 | k__Fungi(100);p__Ascomycota(100);c__Saccharomycetes(100);o__Saccharomycetales(100);f__Metschnikowiaceae(100);g__Metschnikowia(100);s__Metschnikowia_pulcherrima(100); | Metschnikowia | 13 | Saprotroph | \|Undefined Saprotroph\| | Yeast | NULL | Probable | As nectar-tap_saprotroph (Põlme et al. 2020); Decay sugar-rich substrates (Põlme et al. 2020) | Põlme S, et al. 2020. Fungal Diversity 105:1-16 ((https://doi.org/10.1007/s13225-020-00466-2)) |
| Otu15498 | 1 | k__Fungi(100);p__Ascomycota(100);c__Saccharomycetes(100);o__Saccharomycetales(100);f__Metschnikowiaceae(100);g__Metschnikowia(100);s__Metschnikowia_pulcherrima(100); | Metschnikowia | 13 | Saprotroph | \|Undefined Saprotroph\| | Yeast | NULL | Probable | As nectar-tap_saprotroph (Põlme et al. 2020); Decay sugar-rich substrates (Põlme et al. 2020) | Põlme S, et al. 2020. Fungal Diversity 105:1-16 ((https://doi.org/10.1007/s13225-020-00466-2)) |
| Otu15499 | 1 | k__Fungi(100);p__Ascomycota(100);c__Saccharomycetes(100);o__Saccharomycetales(100);f__Metschnikowiaceae(100);g__Metschnikowia(100);s__Metschnikowia_pulcherrima(100); | Metschnikowia | 13 | Saprotroph | \|Undefined Saprotroph\| | Yeast | NULL | Probable | As nectar-tap_saprotroph (Põlme et al. 2020); Decay sugar-rich substrates (Põlme et al. 2020) | Põlme S, et al. 2020. Fungal Diversity 105:1-16 ((https://doi.org/10.1007/s13225-020-00466-2)) |
| Otu15500 | 1 | k__Fungi(100);p__Ascomycota(100);c__Saccharomycetes(100);o__Saccharomycetales(100);f__Metschnikowiaceae(100);g__Metschnikowia(100);g__Metschnikowia_unclassified(100); | Metschnikowia | 13 | Saprotroph | \|Undefined Saprotroph\| | Yeast | NULL | Probable | As nectar-tap_saprotroph (Põlme et al. 2020); Decay sugar-rich substrates (Põlme et al. 2020) | Põlme S, et al. 2020. Fungal Diversity 105:1-16 ((https://doi.org/10.1007/s13225-020-00466-2)) |
| Otu15502 | 1 | k__Fungi(100);p__Ascomycota(100);c__Saccharomycetes(100);o__Saccharomycetales(100);f__Metschnikowiaceae(100);g__Metschnikowia(100);g__Metschnikowia_unclassified(100); | Metschnikowia | 13 | Saprotroph | \|Undefined Saprotroph\| | Yeast | NULL | Probable | As nectar-tap_saprotroph (Põlme et al. 2020); Decay sugar-rich substrates (Põlme et al. 2020) | Põlme S, et al. 2020. Fungal Diversity 105:1-16 ((https://doi.org/10.1007/s13225-020-00466-2)) |
| Otu15503 | 1 | k__Fungi(100);p__Ascomycota(100);c__Saccharomycetes(100);o__Saccharomycetales(100);f__Metschnikowiaceae(100);g__Metschnikowia(100);s__Metschnikowia_pulcherrima(100); | Metschnikowia | 13 | Saprotroph | \|Undefined Saprotroph\| | Yeast | NULL | Probable | As nectar-tap_saprotroph (Põlme et al. 2020); Decay sugar-rich substrates (Põlme et al. 2020) | Põlme S, et al. 2020. Fungal Diversity 105:1-16 ((https://doi.org/10.1007/s13225-020-00466-2)) |
| Otu15504 | 1 | k__Fungi(100);p__Ascomycota(100);c__Saccharomycetes(100);o__Saccharomycetales(100);f__Metschnikowiaceae(100);g__Metschnikowia(100);s__Metschnikowia_pulcherrima(100); | Metschnikowia | 13 | Saprotroph | \|Undefined Saprotroph\| | Yeast | NULL | Probable | As nectar-tap_saprotroph (Põlme et al. 2020); Decay sugar-rich substrates (Põlme et al. 2020) | Põlme S, et al. 2020. Fungal Diversity 105:1-16 ((https://doi.org/10.1007/s13225-020-00466-2)) |
| Otu15506 | 1 | k__Fungi(100);p__Ascomycota(100);c__Saccharomycetes(100);o__Saccharomycetales(100);f__Metschnikowiaceae(100);g__Metschnikowia(100);s__Metschnikowia_pulcherrima(100); | Metschnikowia | 13 | Saprotroph | \|Undefined Saprotroph\| | Yeast | NULL | Probable | As nectar-tap_saprotroph (Põlme et al. 2020); Decay sugar-rich substrates (Põlme et al. 2020) | Põlme S, et al. 2020. Fungal Diversity 105:1-16 ((https://doi.org/10.1007/s13225-020-00466-2)) |
| Otu15507 | 1 | k__Fungi(100);p__Ascomycota(100);c__Saccharomycetes(100);o__Saccharomycetales(100);f__Metschnikowiaceae(100);g__Metschnikowia(100);s__Metschnikowia_pulcherrima(100); | Metschnikowia | 13 | Saprotroph | \|Undefined Saprotroph\| | Yeast | NULL | Probable | As nectar-tap_saprotroph (Põlme et al. 2020); Decay sugar-rich substrates (Põlme et al. 2020) | Põlme S, et al. 2020. Fungal Diversity 105:1-16 ((https://doi.org/10.1007/s13225-020-00466-2)) |
| Otu15508 | 1 | k__Fungi(100);p__Ascomycota(100);c__Saccharomycetes(100);o__Saccharomycetales(100);f__Metschnikowiaceae(100);g__Metschnikowia(100);s__Metschnikowia_pulcherrima(100); | Metschnikowia | 13 | Saprotroph | \|Undefined Saprotroph\| | Yeast | NULL | Probable | As nectar-tap_saprotroph (Põlme et al. 2020); Decay sugar-rich substrates (Põlme et al. 2020) | Põlme S, et al. 2020. Fungal Diversity 105:1-16 ((https://doi.org/10.1007/s13225-020-00466-2)) |
| Otu15510 | 1 | k__Fungi(100);p__Ascomycota(100);c__Saccharomycetes(100);o__Saccharomycetales(100);f__Metschnikowiaceae(100);g__Metschnikowia(100);s__Metschnikowia_pulcherrima(100); | Metschnikowia | 13 | Saprotroph | \|Undefined Saprotroph\| | Yeast | NULL | Probable | As nectar-tap_saprotroph (Põlme et al. 2020); Decay sugar-rich substrates (Põlme et al. 2020) | Põlme S, et al. 2020. Fungal Diversity 105:1-16 ((https://doi.org/10.1007/s13225-020-00466-2)) |
| Otu15511 | 1 | k__Fungi(100);p__Ascomycota(100);c__Saccharomycetes(100);o__Saccharomycetales(100);f__Metschnikowiaceae(100);g__Metschnikowia(100);s__Metschnikowia_pulcherrima(100); | Metschnikowia | 13 | Saprotroph | \|Undefined Saprotroph\| | Yeast | NULL | Probable | As nectar-tap_saprotroph (Põlme et al. 2020); Decay sugar-rich substrates (Põlme et al. 2020) | Põlme S, et al. 2020. Fungal Diversity 105:1-16 ((https://doi.org/10.1007/s13225-020-00466-2)) |
| Otu15513 | 1 | k__Fungi(100);p__Ascomycota(100);c__Saccharomycetes(100);o__Saccharomycetales(100);f__Metschnikowiaceae(100);g__Metschnikowia(100);s__Metschnikowia_pulcherrima(100); | Metschnikowia | 13 | Saprotroph | \|Undefined Saprotroph\| | Yeast | NULL | Probable | As nectar-tap_saprotroph (Põlme et al. 2020); Decay sugar-rich substrates (Põlme et al. 2020) | Põlme S, et al. 2020. Fungal Diversity 105:1-16 ((https://doi.org/10.1007/s13225-020-00466-2)) |
| Otu15514 | 1 | k__Fungi(100);p__Ascomycota(100);c__Saccharomycetes(100);o__Saccharomycetales(100);f__Metschnikowiaceae(100);g__Metschnikowia(100);s__Metschnikowia_pulcherrima(100); | Metschnikowia | 13 | Saprotroph | \|Undefined Saprotroph\| | Yeast | NULL | Probable | As nectar-tap_saprotroph (Põlme et al. 2020); Decay sugar-rich substrates (Põlme et al. 2020) | Põlme S, et al. 2020. Fungal Diversity 105:1-16 ((https://doi.org/10.1007/s13225-020-00466-2)) |
| Otu15515 | 1 | k__Fungi(100);p__Ascomycota(100);c__Saccharomycetes(100);o__Saccharomycetales(100);f__Metschnikowiaceae(100);g__Metschnikowia(100);s__Metschnikowia_pulcherrima(100); | Metschnikowia | 13 | Saprotroph | \|Undefined Saprotroph\| | Yeast | NULL | Probable | As nectar-tap_saprotroph (Põlme et al. 2020); Decay sugar-rich substrates (Põlme et al. 2020) | Põlme S, et al. 2020. Fungal Diversity 105:1-16 ((https://doi.org/10.1007/s13225-020-00466-2)) |
| Otu15516 | 1 | k__Fungi(100);p__Ascomycota(100);c__Saccharomycetes(100);o__Saccharomycetales(100);f__Metschnikowiaceae(100);g__Metschnikowia(100);s__Metschnikowia_pulcherrima(100); | Metschnikowia | 13 | Saprotroph | \|Undefined Saprotroph\| | Yeast | NULL | Probable | As nectar-tap_saprotroph (Põlme et al. 2020); Decay sugar-rich substrates (Põlme et al. 2020) | Põlme S, et al. 2020. Fungal Diversity 105:1-16 ((https://doi.org/10.1007/s13225-020-00466-2)) |
| Otu15518 | 1 | k__Fungi(100);p__Ascomycota(100);c__Saccharomycetes(100);o__Saccharomycetales(100);f__Metschnikowiaceae(100);g__Metschnikowia(100);s__Metschnikowia_pulcherrima(100); | Metschnikowia | 13 | Saprotroph | \|Undefined Saprotroph\| | Yeast | NULL | Probable | As nectar-tap_saprotroph (Põlme et al. 2020); Decay sugar-rich substrates (Põlme et al. 2020) | Põlme S, et al. 2020. Fungal Diversity 105:1-16 ((https://doi.org/10.1007/s13225-020-00466-2)) |
| Otu15519 | 1 | k__Fungi(100);p__Ascomycota(100);c__Saccharomycetes(100);o__Saccharomycetales(100);f__Metschnikowiaceae(100);g__Metschnikowia(100);g__Metschnikowia_unclassified(100); | Metschnikowia | 13 | Saprotroph | \|Undefined Saprotroph\| | Yeast | NULL | Probable | As nectar-tap_saprotroph (Põlme et al. 2020); Decay sugar-rich substrates (Põlme et al. 2020) | Põlme S, et al. 2020. Fungal Diversity 105:1-16 ((https://doi.org/10.1007/s13225-020-00466-2)) |
| Otu15520 | 1 | k__Fungi(100);p__Ascomycota(100);c__Saccharomycetes(100);o__Saccharomycetales(100);f__Metschnikowiaceae(100);g__Metschnikowia(100);s__Metschnikowia_pulcherrima(100); | Metschnikowia | 13 | Saprotroph | \|Undefined Saprotroph\| | Yeast | NULL | Probable | As nectar-tap_saprotroph (Põlme et al. 2020); Decay sugar-rich substrates (Põlme et al. 2020) | Põlme S, et al. 2020. Fungal Diversity 105:1-16 ((https://doi.org/10.1007/s13225-020-00466-2)) |
| Otu15521 | 1 | k__Fungi(100);p__Ascomycota(100);c__Saccharomycetes(100);o__Saccharomycetales(100);f__Metschnikowiaceae(100);g__Metschnikowia(100);s__Metschnikowia_pulcherrima(100); | Metschnikowia | 13 | Saprotroph | \|Undefined Saprotroph\| | Yeast | NULL | Probable | As nectar-tap_saprotroph (Põlme et al. 2020); Decay sugar-rich substrates (Põlme et al. 2020) | Põlme S, et al. 2020. Fungal Diversity 105:1-16 ((https://doi.org/10.1007/s13225-020-00466-2)) |
| Otu15523 | 1 | k__Fungi(100);p__Ascomycota(100);c__Saccharomycetes(100);o__Saccharomycetales(100);f__Metschnikowiaceae(100);g__Metschnikowia(100);g__Metschnikowia_unclassified(100); | Metschnikowia | 13 | Saprotroph | \|Undefined Saprotroph\| | Yeast | NULL | Probable | As nectar-tap_saprotroph (Põlme et al. 2020); Decay sugar-rich substrates (Põlme et al. 2020) | Põlme S, et al. 2020. Fungal Diversity 105:1-16 ((https://doi.org/10.1007/s13225-020-00466-2)) |
| Otu15524 | 1 | k__Fungi(100);p__Ascomycota(100);c__Saccharomycetes(100);o__Saccharomycetales(100);f__Metschnikowiaceae(100);g__Metschnikowia(100);s__Metschnikowia_pulcherrima(100); | Metschnikowia | 13 | Saprotroph | \|Undefined Saprotroph\| | Yeast | NULL | Probable | As nectar-tap_saprotroph (Põlme et al. 2020); Decay sugar-rich substrates (Põlme et al. 2020) | Põlme S, et al. 2020. Fungal Diversity 105:1-16 ((https://doi.org/10.1007/s13225-020-00466-2)) |
| Otu15525 | 1 | k__Fungi(100);p__Ascomycota(100);c__Saccharomycetes(100);o__Saccharomycetales(100);f__Metschnikowiaceae(100);g__Metschnikowia(100);s__Metschnikowia_pulcherrima(100); | Metschnikowia | 13 | Saprotroph | \|Undefined Saprotroph\| | Yeast | NULL | Probable | As nectar-tap_saprotroph (Põlme et al. 2020); Decay sugar-rich substrates (Põlme et al. 2020) | Põlme S, et al. 2020. Fungal Diversity 105:1-16 ((https://doi.org/10.1007/s13225-020-00466-2)) |
| Otu15526 | 1 | k__Fungi(100);p__Ascomycota(100);c__Saccharomycetes(100);o__Saccharomycetales(100);f__Metschnikowiaceae(100);g__Metschnikowia(100);s__Metschnikowia_pulcherrima(100); | Metschnikowia | 13 | Saprotroph | \|Undefined Saprotroph\| | Yeast | NULL | Probable | As nectar-tap_saprotroph (Põlme et al. 2020); Decay sugar-rich substrates (Põlme et al. 2020) | Põlme S, et al. 2020. Fungal Diversity 105:1-16 ((https://doi.org/10.1007/s13225-020-00466-2)) |
| Otu15527 | 1 | k__Fungi(100);p__Ascomycota(100);c__Saccharomycetes(100);o__Saccharomycetales(100);f__Metschnikowiaceae(100);g__Metschnikowia(100);s__Metschnikowia_pulcherrima(100); | Metschnikowia | 13 | Saprotroph | \|Undefined Saprotroph\| | Yeast | NULL | Probable | As nectar-tap_saprotroph (Põlme et al. 2020); Decay sugar-rich substrates (Põlme et al. 2020) | Põlme S, et al. 2020. Fungal Diversity 105:1-16 ((https://doi.org/10.1007/s13225-020-00466-2)) |
| Otu15528 | 1 | k__Fungi(100);p__Ascomycota(100);c__Saccharomycetes(100);o__Saccharomycetales(100);f__Metschnikowiaceae(100);g__Metschnikowia(100);s__Metschnikowia_pulcherrima(100); | Metschnikowia | 13 | Saprotroph | \|Undefined Saprotroph\| | Yeast | NULL | Probable | As nectar-tap_saprotroph (Põlme et al. 2020); Decay sugar-rich substrates (Põlme et al. 2020) | Põlme S, et al. 2020. Fungal Diversity 105:1-16 ((https://doi.org/10.1007/s13225-020-00466-2)) |
| Otu15529 | 1 | k__Fungi(100);p__Ascomycota(100);c__Saccharomycetes(100);o__Saccharomycetales(100);f__Metschnikowiaceae(100);g__Metschnikowia(100);s__Metschnikowia_pulcherrima(100); | Metschnikowia | 13 | Saprotroph | \|Undefined Saprotroph\| | Yeast | NULL | Probable | As nectar-tap_saprotroph (Põlme et al. 2020); Decay sugar-rich substrates (Põlme et al. 2020) | Põlme S, et al. 2020. Fungal Diversity 105:1-16 ((https://doi.org/10.1007/s13225-020-00466-2)) |
| Otu15530 | 1 | k__Fungi(100);p__Ascomycota(100);c__Saccharomycetes(100);o__Saccharomycetales(100);f__Metschnikowiaceae(100);g__Metschnikowia(100);s__Metschnikowia_pulcherrima(100); | Metschnikowia | 13 | Saprotroph | \|Undefined Saprotroph\| | Yeast | NULL | Probable | As nectar-tap_saprotroph (Põlme et al. 2020); Decay sugar-rich substrates (Põlme et al. 2020) | Põlme S, et al. 2020. Fungal Diversity 105:1-16 ((https://doi.org/10.1007/s13225-020-00466-2)) |
| Otu15532 | 1 | k__Fungi(100);p__Ascomycota(100);c__Saccharomycetes(100);o__Saccharomycetales(100);f__Metschnikowiaceae(100);g__Metschnikowia(100);s__Metschnikowia_pulcherrima(100); | Metschnikowia | 13 | Saprotroph | \|Undefined Saprotroph\| | Yeast | NULL | Probable | As nectar-tap_saprotroph (Põlme et al. 2020); Decay sugar-rich substrates (Põlme et al. 2020) | Põlme S, et al. 2020. Fungal Diversity 105:1-16 ((https://doi.org/10.1007/s13225-020-00466-2)) |
| Otu15533 | 1 | k__Fungi(100);p__Ascomycota(100);c__Saccharomycetes(100);o__Saccharomycetales(100);f__Metschnikowiaceae(100);g__Metschnikowia(100);g__Metschnikowia_unclassified(100); | Metschnikowia | 13 | Saprotroph | \|Undefined Saprotroph\| | Yeast | NULL | Probable | As nectar-tap_saprotroph (Põlme et al. 2020); Decay sugar-rich substrates (Põlme et al. 2020) | Põlme S, et al. 2020. Fungal Diversity 105:1-16 ((https://doi.org/10.1007/s13225-020-00466-2)) |
| Otu15535 | 1 | k__Fungi(100);p__Ascomycota(100);c__Saccharomycetes(100);o__Saccharomycetales(100);f__Metschnikowiaceae(100);g__Metschnikowia(100);s__Metschnikowia_pulcherrima(100); | Metschnikowia | 13 | Saprotroph | \|Undefined Saprotroph\| | Yeast | NULL | Probable | As nectar-tap_saprotroph (Põlme et al. 2020); Decay sugar-rich substrates (Põlme et al. 2020) | Põlme S, et al. 2020. Fungal Diversity 105:1-16 ((https://doi.org/10.1007/s13225-020-00466-2)) |
| Otu15536 | 1 | k__Fungi(100);p__Ascomycota(100);c__Saccharomycetes(100);o__Saccharomycetales(100);f__Metschnikowiaceae(100);g__Metschnikowia(100);s__Metschnikowia_pulcherrima(100); | Metschnikowia | 13 | Saprotroph | \|Undefined Saprotroph\| | Yeast | NULL | Probable | As nectar-tap_saprotroph (Põlme et al. 2020); Decay sugar-rich substrates (Põlme et al. 2020) | Põlme S, et al. 2020. Fungal Diversity 105:1-16 ((https://doi.org/10.1007/s13225-020-00466-2)) |
| Otu15537 | 1 | k__Fungi(100);p__Ascomycota(100);c__Saccharomycetes(100);o__Saccharomycetales(100);f__Metschnikowiaceae(100);g__Metschnikowia(100);g__Metschnikowia_unclassified(100); | Metschnikowia | 13 | Saprotroph | \|Undefined Saprotroph\| | Yeast | NULL | Probable | As nectar-tap_saprotroph (Põlme et al. 2020); Decay sugar-rich substrates (Põlme et al. 2020) | Põlme S, et al. 2020. Fungal Diversity 105:1-16 ((https://doi.org/10.1007/s13225-020-00466-2)) |
| Otu15538 | 1 | k__Fungi(100);p__Ascomycota(100);c__Saccharomycetes(100);o__Saccharomycetales(100);f__Metschnikowiaceae(100);g__Metschnikowia(100);s__Metschnikowia_pulcherrima(100); | Metschnikowia | 13 | Saprotroph | \|Undefined Saprotroph\| | Yeast | NULL | Probable | As nectar-tap_saprotroph (Põlme et al. 2020); Decay sugar-rich substrates (Põlme et al. 2020) | Põlme S, et al. 2020. Fungal Diversity 105:1-16 ((https://doi.org/10.1007/s13225-020-00466-2)) |
| Otu15539 | 1 | k__Fungi(100);p__Ascomycota(100);c__Saccharomycetes(100);o__Saccharomycetales(100);f__Metschnikowiaceae(100);g__Metschnikowia(100);s__Metschnikowia_pulcherrima(100); | Metschnikowia | 13 | Saprotroph | \|Undefined Saprotroph\| | Yeast | NULL | Probable | As nectar-tap_saprotroph (Põlme et al. 2020); Decay sugar-rich substrates (Põlme et al. 2020) | Põlme S, et al. 2020. Fungal Diversity 105:1-16 ((https://doi.org/10.1007/s13225-020-00466-2)) |
| Otu15540 | 1 | k__Fungi(100);p__Ascomycota(100);c__Saccharomycetes(100);o__Saccharomycetales(100);f__Metschnikowiaceae(100);g__Metschnikowia(100);g__Metschnikowia_unclassified(100); | Metschnikowia | 13 | Saprotroph | \|Undefined Saprotroph\| | Yeast | NULL | Probable | As nectar-tap_saprotroph (Põlme et al. 2020); Decay sugar-rich substrates (Põlme et al. 2020) | Põlme S, et al. 2020. Fungal Diversity 105:1-16 ((https://doi.org/10.1007/s13225-020-00466-2)) |
| Otu15541 | 1 | k__Fungi(100);p__Ascomycota(100);c__Saccharomycetes(100);o__Saccharomycetales(100);f__Metschnikowiaceae(100);g__Metschnikowia(100);s__Metschnikowia_pulcherrima(100); | Metschnikowia | 13 | Saprotroph | \|Undefined Saprotroph\| | Yeast | NULL | Probable | As nectar-tap_saprotroph (Põlme et al. 2020); Decay sugar-rich substrates (Põlme et al. 2020) | Põlme S, et al. 2020. Fungal Diversity 105:1-16 ((https://doi.org/10.1007/s13225-020-00466-2)) |
| Otu15542 | 1 | k__Fungi(100);p__Ascomycota(100);c__Saccharomycetes(100);o__Saccharomycetales(100);f__Metschnikowiaceae(100);g__Metschnikowia(100);s__Metschnikowia_pulcherrima(100); | Metschnikowia | 13 | Saprotroph | \|Undefined Saprotroph\| | Yeast | NULL | Probable | As nectar-tap_saprotroph (Põlme et al. 2020); Decay sugar-rich substrates (Põlme et al. 2020) | Põlme S, et al. 2020. Fungal Diversity 105:1-16 ((https://doi.org/10.1007/s13225-020-00466-2)) |
| Otu15543 | 1 | k__Fungi(100);p__Ascomycota(100);c__Saccharomycetes(100);o__Saccharomycetales(100);f__Metschnikowiaceae(100);g__Metschnikowia(100);s__Metschnikowia_pulcherrima(100); | Metschnikowia | 13 | Saprotroph | \|Undefined Saprotroph\| | Yeast | NULL | Probable | As nectar-tap_saprotroph (Põlme et al. 2020); Decay sugar-rich substrates (Põlme et al. 2020) | Põlme S, et al. 2020. Fungal Diversity 105:1-16 ((https://doi.org/10.1007/s13225-020-00466-2)) |
| Otu15544 | 1 | k__Fungi(100);p__Ascomycota(100);c__Saccharomycetes(100);o__Saccharomycetales(100);f__Metschnikowiaceae(100);g__Metschnikowia(100);g__Metschnikowia_unclassified(100); | Metschnikowia | 13 | Saprotroph | \|Undefined Saprotroph\| | Yeast | NULL | Probable | As nectar-tap_saprotroph (Põlme et al. 2020); Decay sugar-rich substrates (Põlme et al. 2020) | Põlme S, et al. 2020. Fungal Diversity 105:1-16 ((https://doi.org/10.1007/s13225-020-00466-2)) |
| Otu15545 | 1 | k__Fungi(100);p__Ascomycota(100);c__Saccharomycetes(100);o__Saccharomycetales(100);f__Metschnikowiaceae(100);g__Metschnikowia(100);s__Metschnikowia_pulcherrima(100); | Metschnikowia | 13 | Saprotroph | \|Undefined Saprotroph\| | Yeast | NULL | Probable | As nectar-tap_saprotroph (Põlme et al. 2020); Decay sugar-rich substrates (Põlme et al. 2020) | Põlme S, et al. 2020. Fungal Diversity 105:1-16 ((https://doi.org/10.1007/s13225-020-00466-2)) |
| Otu15546 | 1 | k__Fungi(100);p__Ascomycota(100);c__Saccharomycetes(100);o__Saccharomycetales(100);f__Metschnikowiaceae(100);g__Metschnikowia(100);s__Metschnikowia_pulcherrima(100); | Metschnikowia | 13 | Saprotroph | \|Undefined Saprotroph\| | Yeast | NULL | Probable | As nectar-tap_saprotroph (Põlme et al. 2020); Decay sugar-rich substrates (Põlme et al. 2020) | Põlme S, et al. 2020. Fungal Diversity 105:1-16 ((https://doi.org/10.1007/s13225-020-00466-2)) |
| Otu15547 | 1 | k__Fungi(100);p__Ascomycota(100);c__Saccharomycetes(100);o__Saccharomycetales(100);f__Metschnikowiaceae(100);g__Metschnikowia(100);s__Metschnikowia_pulcherrima(100); | Metschnikowia | 13 | Saprotroph | \|Undefined Saprotroph\| | Yeast | NULL | Probable | As nectar-tap_saprotroph (Põlme et al. 2020); Decay sugar-rich substrates (Põlme et al. 2020) | Põlme S, et al. 2020. Fungal Diversity 105:1-16 ((https://doi.org/10.1007/s13225-020-00466-2)) |
| Otu15548 | 1 | k__Fungi(100);p__Ascomycota(100);c__Saccharomycetes(100);o__Saccharomycetales(100);f__Metschnikowiaceae(100);g__Metschnikowia(100);g__Metschnikowia_unclassified(100); | Metschnikowia | 13 | Saprotroph | \|Undefined Saprotroph\| | Yeast | NULL | Probable | As nectar-tap_saprotroph (Põlme et al. 2020); Decay sugar-rich substrates (Põlme et al. 2020) | Põlme S, et al. 2020. Fungal Diversity 105:1-16 ((https://doi.org/10.1007/s13225-020-00466-2)) |
| Otu15549 | 1 | k__Fungi(100);p__Ascomycota(100);c__Saccharomycetes(100);o__Saccharomycetales(100);f__Metschnikowiaceae(100);g__Metschnikowia(100);s__Metschnikowia_pulcherrima(100); | Metschnikowia | 13 | Saprotroph | \|Undefined Saprotroph\| | Yeast | NULL | Probable | As nectar-tap_saprotroph (Põlme et al. 2020); Decay sugar-rich substrates (Põlme et al. 2020) | Põlme S, et al. 2020. Fungal Diversity 105:1-16 ((https://doi.org/10.1007/s13225-020-00466-2)) |
| Otu15550 | 1 | k__Fungi(100);p__Ascomycota(100);c__Saccharomycetes(100);o__Saccharomycetales(100);f__Metschnikowiaceae(100);g__Metschnikowia(100);s__Metschnikowia_pulcherrima(100); | Metschnikowia | 13 | Saprotroph | \|Undefined Saprotroph\| | Yeast | NULL | Probable | As nectar-tap_saprotroph (Põlme et al. 2020); Decay sugar-rich substrates (Põlme et al. 2020) | Põlme S, et al. 2020. Fungal Diversity 105:1-16 ((https://doi.org/10.1007/s13225-020-00466-2)) |
| Otu15551 | 1 | k__Fungi(100);p__Ascomycota(100);c__Saccharomycetes(100);o__Saccharomycetales(100);f__Metschnikowiaceae(100);g__Metschnikowia(100);g__Metschnikowia_unclassified(100); | Metschnikowia | 13 | Saprotroph | \|Undefined Saprotroph\| | Yeast | NULL | Probable | As nectar-tap_saprotroph (Põlme et al. 2020); Decay sugar-rich substrates (Põlme et al. 2020) | Põlme S, et al. 2020. Fungal Diversity 105:1-16 ((https://doi.org/10.1007/s13225-020-00466-2)) |
| Otu15552 | 1 | k__Fungi(100);p__Ascomycota(100);c__Saccharomycetes(100);o__Saccharomycetales(100);f__Metschnikowiaceae(100);g__Metschnikowia(100);s__Metschnikowia_pulcherrima(100); | Metschnikowia | 13 | Saprotroph | \|Undefined Saprotroph\| | Yeast | NULL | Probable | As nectar-tap_saprotroph (Põlme et al. 2020); Decay sugar-rich substrates (Põlme et al. 2020) | Põlme S, et al. 2020. Fungal Diversity 105:1-16 ((https://doi.org/10.1007/s13225-020-00466-2)) |
| Otu15553 | 1 | k__Fungi(100);p__Ascomycota(100);c__Saccharomycetes(100);o__Saccharomycetales(100);f__Metschnikowiaceae(100);g__Metschnikowia(100);g__Metschnikowia_unclassified(100); | Metschnikowia | 13 | Saprotroph | \|Undefined Saprotroph\| | Yeast | NULL | Probable | As nectar-tap_saprotroph (Põlme et al. 2020); Decay sugar-rich substrates (Põlme et al. 2020) | Põlme S, et al. 2020. Fungal Diversity 105:1-16 ((https://doi.org/10.1007/s13225-020-00466-2)) |
| Otu15554 | 1 | k__Fungi(100);p__Ascomycota(100);c__Saccharomycetes(100);o__Saccharomycetales(100);f__Metschnikowiaceae(100);g__Metschnikowia(100);s__Metschnikowia_pulcherrima(100); | Metschnikowia | 13 | Saprotroph | \|Undefined Saprotroph\| | Yeast | NULL | Probable | As nectar-tap_saprotroph (Põlme et al. 2020); Decay sugar-rich substrates (Põlme et al. 2020) | Põlme S, et al. 2020. Fungal Diversity 105:1-16 ((https://doi.org/10.1007/s13225-020-00466-2)) |
| Otu15555 | 1 | k__Fungi(100);p__Ascomycota(100);c__Saccharomycetes(100);o__Saccharomycetales(100);f__Metschnikowiaceae(100);g__Metschnikowia(100);s__Metschnikowia_pulcherrima(100); | Metschnikowia | 13 | Saprotroph | \|Undefined Saprotroph\| | Yeast | NULL | Probable | As nectar-tap_saprotroph (Põlme et al. 2020); Decay sugar-rich substrates (Põlme et al. 2020) | Põlme S, et al. 2020. Fungal Diversity 105:1-16 ((https://doi.org/10.1007/s13225-020-00466-2)) |
| Otu15556 | 1 | k__Fungi(100);p__Ascomycota(100);c__Saccharomycetes(100);o__Saccharomycetales(100);f__Metschnikowiaceae(100);g__Metschnikowia(100);s__Metschnikowia_pulcherrima(100); | Metschnikowia | 13 | Saprotroph | \|Undefined Saprotroph\| | Yeast | NULL | Probable | As nectar-tap_saprotroph (Põlme et al. 2020); Decay sugar-rich substrates (Põlme et al. 2020) | Põlme S, et al. 2020. Fungal Diversity 105:1-16 ((https://doi.org/10.1007/s13225-020-00466-2)) |
| Otu15559 | 1 | k__Fungi(100);p__Ascomycota(100);c__Saccharomycetes(100);o__Saccharomycetales(100);f__Metschnikowiaceae(100);g__Metschnikowia(100);s__Metschnikowia_pulcherrima(100); | Metschnikowia | 13 | Saprotroph | \|Undefined Saprotroph\| | Yeast | NULL | Probable | As nectar-tap_saprotroph (Põlme et al. 2020); Decay sugar-rich substrates (Põlme et al. 2020) | Põlme S, et al. 2020. Fungal Diversity 105:1-16 ((https://doi.org/10.1007/s13225-020-00466-2)) |
| Otu15560 | 1 | k__Fungi(100);p__Ascomycota(100);c__Saccharomycetes(100);o__Saccharomycetales(100);f__Metschnikowiaceae(100);g__Metschnikowia(100);s__Metschnikowia_pulcherrima(100); | Metschnikowia | 13 | Saprotroph | \|Undefined Saprotroph\| | Yeast | NULL | Probable | As nectar-tap_saprotroph (Põlme et al. 2020); Decay sugar-rich substrates (Põlme et al. 2020) | Põlme S, et al. 2020. Fungal Diversity 105:1-16 ((https://doi.org/10.1007/s13225-020-00466-2)) |
| Otu15561 | 1 | k__Fungi(100);p__Ascomycota(100);c__Saccharomycetes(100);o__Saccharomycetales(100);f__Metschnikowiaceae(100);g__Metschnikowia(100);s__Metschnikowia_pulcherrima(100); | Metschnikowia | 13 | Saprotroph | \|Undefined Saprotroph\| | Yeast | NULL | Probable | As nectar-tap_saprotroph (Põlme et al. 2020); Decay sugar-rich substrates (Põlme et al. 2020) | Põlme S, et al. 2020. Fungal Diversity 105:1-16 ((https://doi.org/10.1007/s13225-020-00466-2)) |
| Otu15562 | 1 | k__Fungi(100);p__Ascomycota(100);c__Saccharomycetes(100);o__Saccharomycetales(100);f__Metschnikowiaceae(100);g__Metschnikowia(100);g__Metschnikowia_unclassified(100); | Metschnikowia | 13 | Saprotroph | \|Undefined Saprotroph\| | Yeast | NULL | Probable | As nectar-tap_saprotroph (Põlme et al. 2020); Decay sugar-rich substrates (Põlme et al. 2020) | Põlme S, et al. 2020. Fungal Diversity 105:1-16 ((https://doi.org/10.1007/s13225-020-00466-2)) |
| Otu15563 | 1 | k__Fungi(100);p__Ascomycota(100);c__Saccharomycetes(100);o__Saccharomycetales(100);f__Metschnikowiaceae(100);g__Metschnikowia(100);s__Metschnikowia_pulcherrima(100); | Metschnikowia | 13 | Saprotroph | \|Undefined Saprotroph\| | Yeast | NULL | Probable | As nectar-tap_saprotroph (Põlme et al. 2020); Decay sugar-rich substrates (Põlme et al. 2020) | Põlme S, et al. 2020. Fungal Diversity 105:1-16 ((https://doi.org/10.1007/s13225-020-00466-2)) |
| Otu15564 | 1 | k__Fungi(100);p__Ascomycota(100);c__Saccharomycetes(100);o__Saccharomycetales(100);f__Metschnikowiaceae(100);g__Metschnikowia(100);s__Metschnikowia_pulcherrima(100); | Metschnikowia | 13 | Saprotroph | \|Undefined Saprotroph\| | Yeast | NULL | Probable | As nectar-tap_saprotroph (Põlme et al. 2020); Decay sugar-rich substrates (Põlme et al. 2020) | Põlme S, et al. 2020. Fungal Diversity 105:1-16 ((https://doi.org/10.1007/s13225-020-00466-2)) |
| Otu15567 | 1 | k__Fungi(100);p__Ascomycota(100);c__Saccharomycetes(100);o__Saccharomycetales(100);f__Metschnikowiaceae(100);g__Metschnikowia(100);g__Metschnikowia_unclassified(100); | Metschnikowia | 13 | Saprotroph | \|Undefined Saprotroph\| | Yeast | NULL | Probable | As nectar-tap_saprotroph (Põlme et al. 2020); Decay sugar-rich substrates (Põlme et al. 2020) | Põlme S, et al. 2020. Fungal Diversity 105:1-16 ((https://doi.org/10.1007/s13225-020-00466-2)) |
| Otu15568 | 1 | k__Fungi(100);p__Ascomycota(100);c__Saccharomycetes(100);o__Saccharomycetales(100);f__Metschnikowiaceae(100);g__Metschnikowia(100);s__Metschnikowia_pulcherrima(100); | Metschnikowia | 13 | Saprotroph | \|Undefined Saprotroph\| | Yeast | NULL | Probable | As nectar-tap_saprotroph (Põlme et al. 2020); Decay sugar-rich substrates (Põlme et al. 2020) | Põlme S, et al. 2020. Fungal Diversity 105:1-16 ((https://doi.org/10.1007/s13225-020-00466-2)) |
| Otu15569 | 1 | k__Fungi(100);p__Ascomycota(100);c__Saccharomycetes(100);o__Saccharomycetales(100);f__Metschnikowiaceae(100);g__Metschnikowia(100);g__Metschnikowia_unclassified(100); | Metschnikowia | 13 | Saprotroph | \|Undefined Saprotroph\| | Yeast | NULL | Probable | As nectar-tap_saprotroph (Põlme et al. 2020); Decay sugar-rich substrates (Põlme et al. 2020) | Põlme S, et al. 2020. Fungal Diversity 105:1-16 ((https://doi.org/10.1007/s13225-020-00466-2)) |
| Otu15570 | 1 | k__Fungi(100);p__Ascomycota(100);c__Saccharomycetes(100);o__Saccharomycetales(100);f__Metschnikowiaceae(100);g__Metschnikowia(100);g__Metschnikowia_unclassified(100); | Metschnikowia | 13 | Saprotroph | \|Undefined Saprotroph\| | Yeast | NULL | Probable | As nectar-tap_saprotroph (Põlme et al. 2020); Decay sugar-rich substrates (Põlme et al. 2020) | Põlme S, et al. 2020. Fungal Diversity 105:1-16 ((https://doi.org/10.1007/s13225-020-00466-2)) |
| Otu15572 | 1 | k__Fungi(100);p__Ascomycota(100);c__Saccharomycetes(100);o__Saccharomycetales(100);f__Metschnikowiaceae(100);g__Metschnikowia(100);s__Metschnikowia_pulcherrima(100); | Metschnikowia | 13 | Saprotroph | \|Undefined Saprotroph\| | Yeast | NULL | Probable | As nectar-tap_saprotroph (Põlme et al. 2020); Decay sugar-rich substrates (Põlme et al. 2020) | Põlme S, et al. 2020. Fungal Diversity 105:1-16 ((https://doi.org/10.1007/s13225-020-00466-2)) |
| Otu15574 | 1 | k__Fungi(100);p__Ascomycota(100);c__Saccharomycetes(100);o__Saccharomycetales(100);f__Metschnikowiaceae(100);g__Metschnikowia(100);g__Metschnikowia_unclassified(100); | Metschnikowia | 13 | Saprotroph | \|Undefined Saprotroph\| | Yeast | NULL | Probable | As nectar-tap_saprotroph (Põlme et al. 2020); Decay sugar-rich substrates (Põlme et al. 2020) | Põlme S, et al. 2020. Fungal Diversity 105:1-16 ((https://doi.org/10.1007/s13225-020-00466-2)) |
| Otu15576 | 1 | k__Fungi(100);p__Ascomycota(100);c__Saccharomycetes(100);o__Saccharomycetales(100);f__Metschnikowiaceae(100);g__Metschnikowia(100);s__Metschnikowia_pulcherrima(100); | Metschnikowia | 13 | Saprotroph | \|Undefined Saprotroph\| | Yeast | NULL | Probable | As nectar-tap_saprotroph (Põlme et al. 2020); Decay sugar-rich substrates (Põlme et al. 2020) | Põlme S, et al. 2020. Fungal Diversity 105:1-16 ((https://doi.org/10.1007/s13225-020-00466-2)) |
| Otu15577 | 1 | k__Fungi(100);p__Ascomycota(100);c__Saccharomycetes(100);o__Saccharomycetales(100);f__Metschnikowiaceae(100);g__Metschnikowia(100);g__Metschnikowia_unclassified(100); | Metschnikowia | 13 | Saprotroph | \|Undefined Saprotroph\| | Yeast | NULL | Probable | As nectar-tap_saprotroph (Põlme et al. 2020); Decay sugar-rich substrates (Põlme et al. 2020) | Põlme S, et al. 2020. Fungal Diversity 105:1-16 ((https://doi.org/10.1007/s13225-020-00466-2)) |
| Otu15578 | 1 | k__Fungi(100);p__Ascomycota(100);c__Saccharomycetes(100);o__Saccharomycetales(100);f__Metschnikowiaceae(100);g__Metschnikowia(100);s__Metschnikowia_pulcherrima(100); | Metschnikowia | 13 | Saprotroph | \|Undefined Saprotroph\| | Yeast | NULL | Probable | As nectar-tap_saprotroph (Põlme et al. 2020); Decay sugar-rich substrates (Põlme et al. 2020) | Põlme S, et al. 2020. Fungal Diversity 105:1-16 ((https://doi.org/10.1007/s13225-020-00466-2)) |
| Otu15579 | 1 | k__Fungi(100);p__Ascomycota(100);c__Saccharomycetes(100);o__Saccharomycetales(100);f__Metschnikowiaceae(100);g__Metschnikowia(100);s__Metschnikowia_pulcherrima(100); | Metschnikowia | 13 | Saprotroph | \|Undefined Saprotroph\| | Yeast | NULL | Probable | As nectar-tap_saprotroph (Põlme et al. 2020); Decay sugar-rich substrates (Põlme et al. 2020) | Põlme S, et al. 2020. Fungal Diversity 105:1-16 ((https://doi.org/10.1007/s13225-020-00466-2)) |
| Otu15580 | 1 | k__Fungi(100);p__Ascomycota(100);c__Saccharomycetes(100);o__Saccharomycetales(100);f__Metschnikowiaceae(100);g__Metschnikowia(100);s__Metschnikowia_pulcherrima(100); | Metschnikowia | 13 | Saprotroph | \|Undefined Saprotroph\| | Yeast | NULL | Probable | As nectar-tap_saprotroph (Põlme et al. 2020); Decay sugar-rich substrates (Põlme et al. 2020) | Põlme S, et al. 2020. Fungal Diversity 105:1-16 ((https://doi.org/10.1007/s13225-020-00466-2)) |
| Otu15581 | 1 | k__Fungi(100);p__Ascomycota(100);c__Saccharomycetes(100);o__Saccharomycetales(100);f__Metschnikowiaceae(100);g__Metschnikowia(100);s__Metschnikowia_pulcherrima(100); | Metschnikowia | 13 | Saprotroph | \|Undefined Saprotroph\| | Yeast | NULL | Probable | As nectar-tap_saprotroph (Põlme et al. 2020); Decay sugar-rich substrates (Põlme et al. 2020) | Põlme S, et al. 2020. Fungal Diversity 105:1-16 ((https://doi.org/10.1007/s13225-020-00466-2)) |
| Otu15582 | 1 | k__Fungi(100);p__Ascomycota(100);c__Saccharomycetes(100);o__Saccharomycetales(100);f__Metschnikowiaceae(100);g__Metschnikowia(100);s__Metschnikowia_pulcherrima(100); | Metschnikowia | 13 | Saprotroph | \|Undefined Saprotroph\| | Yeast | NULL | Probable | As nectar-tap_saprotroph (Põlme et al. 2020); Decay sugar-rich substrates (Põlme et al. 2020) | Põlme S, et al. 2020. Fungal Diversity 105:1-16 ((https://doi.org/10.1007/s13225-020-00466-2)) |
| Otu15583 | 1 | k__Fungi(100);p__Ascomycota(100);c__Saccharomycetes(100);o__Saccharomycetales(100);f__Metschnikowiaceae(100);g__Metschnikowia(100);s__Metschnikowia_pulcherrima(100); | Metschnikowia | 13 | Saprotroph | \|Undefined Saprotroph\| | Yeast | NULL | Probable | As nectar-tap_saprotroph (Põlme et al. 2020); Decay sugar-rich substrates (Põlme et al. 2020) | Põlme S, et al. 2020. Fungal Diversity 105:1-16 ((https://doi.org/10.1007/s13225-020-00466-2)) |
| Otu15584 | 1 | k__Fungi(100);p__Ascomycota(100);c__Saccharomycetes(100);o__Saccharomycetales(100);f__Metschnikowiaceae(100);g__Metschnikowia(100);s__Metschnikowia_pulcherrima(100); | Metschnikowia | 13 | Saprotroph | \|Undefined Saprotroph\| | Yeast | NULL | Probable | As nectar-tap_saprotroph (Põlme et al. 2020); Decay sugar-rich substrates (Põlme et al. 2020) | Põlme S, et al. 2020. Fungal Diversity 105:1-16 ((https://doi.org/10.1007/s13225-020-00466-2)) |
| Otu15585 | 1 | k__Fungi(100);p__Ascomycota(100);c__Saccharomycetes(100);o__Saccharomycetales(100);f__Metschnikowiaceae(100);g__Metschnikowia(100);s__Metschnikowia_pulcherrima(100); | Metschnikowia | 13 | Saprotroph | \|Undefined Saprotroph\| | Yeast | NULL | Probable | As nectar-tap_saprotroph (Põlme et al. 2020); Decay sugar-rich substrates (Põlme et al. 2020) | Põlme S, et al. 2020. Fungal Diversity 105:1-16 ((https://doi.org/10.1007/s13225-020-00466-2)) |
| Otu15586 | 1 | k__Fungi(100);p__Ascomycota(100);c__Saccharomycetes(100);o__Saccharomycetales(100);f__Metschnikowiaceae(100);g__Metschnikowia(100);s__Metschnikowia_pulcherrima(100); | Metschnikowia | 13 | Saprotroph | \|Undefined Saprotroph\| | Yeast | NULL | Probable | As nectar-tap_saprotroph (Põlme et al. 2020); Decay sugar-rich substrates (Põlme et al. 2020) | Põlme S, et al. 2020. Fungal Diversity 105:1-16 ((https://doi.org/10.1007/s13225-020-00466-2)) |
| Otu15587 | 1 | k__Fungi(100);p__Ascomycota(100);c__Saccharomycetes(100);o__Saccharomycetales(100);f__Metschnikowiaceae(100);g__Metschnikowia(100);g__Metschnikowia_unclassified(100); | Metschnikowia | 13 | Saprotroph | \|Undefined Saprotroph\| | Yeast | NULL | Probable | As nectar-tap_saprotroph (Põlme et al. 2020); Decay sugar-rich substrates (Põlme et al. 2020) | Põlme S, et al. 2020. Fungal Diversity 105:1-16 ((https://doi.org/10.1007/s13225-020-00466-2)) |
| Otu15588 | 1 | k__Fungi(100);p__Ascomycota(100);c__Saccharomycetes(100);o__Saccharomycetales(100);f__Metschnikowiaceae(100);g__Metschnikowia(100);s__Metschnikowia_pulcherrima(100); | Metschnikowia | 13 | Saprotroph | \|Undefined Saprotroph\| | Yeast | NULL | Probable | As nectar-tap_saprotroph (Põlme et al. 2020); Decay sugar-rich substrates (Põlme et al. 2020) | Põlme S, et al. 2020. Fungal Diversity 105:1-16 ((https://doi.org/10.1007/s13225-020-00466-2)) |
| Otu15589 | 1 | k__Fungi(100);p__Ascomycota(100);c__Saccharomycetes(100);o__Saccharomycetales(100);f__Metschnikowiaceae(100);g__Metschnikowia(100);g__Metschnikowia_unclassified(100); | Metschnikowia | 13 | Saprotroph | \|Undefined Saprotroph\| | Yeast | NULL | Probable | As nectar-tap_saprotroph (Põlme et al. 2020); Decay sugar-rich substrates (Põlme et al. 2020) | Põlme S, et al. 2020. Fungal Diversity 105:1-16 ((https://doi.org/10.1007/s13225-020-00466-2)) |
| Otu15590 | 1 | k__Fungi(100);p__Ascomycota(100);c__Saccharomycetes(100);o__Saccharomycetales(100);f__Metschnikowiaceae(100);g__Metschnikowia(100);s__Metschnikowia_pulcherrima(100); | Metschnikowia | 13 | Saprotroph | \|Undefined Saprotroph\| | Yeast | NULL | Probable | As nectar-tap_saprotroph (Põlme et al. 2020); Decay sugar-rich substrates (Põlme et al. 2020) | Põlme S, et al. 2020. Fungal Diversity 105:1-16 ((https://doi.org/10.1007/s13225-020-00466-2)) |
| Otu15591 | 1 | k__Fungi(100);p__Ascomycota(100);c__Saccharomycetes(100);o__Saccharomycetales(100);f__Metschnikowiaceae(100);g__Metschnikowia(100);g__Metschnikowia_unclassified(100); | Metschnikowia | 13 | Saprotroph | \|Undefined Saprotroph\| | Yeast | NULL | Probable | As nectar-tap_saprotroph (Põlme et al. 2020); Decay sugar-rich substrates (Põlme et al. 2020) | Põlme S, et al. 2020. Fungal Diversity 105:1-16 ((https://doi.org/10.1007/s13225-020-00466-2)) |
| Otu15593 | 1 | k__Fungi(100);p__Ascomycota(100);c__Saccharomycetes(100);o__Saccharomycetales(100);f__Metschnikowiaceae(100);g__Metschnikowia(100);g__Metschnikowia_unclassified(100); | Metschnikowia | 13 | Saprotroph | \|Undefined Saprotroph\| | Yeast | NULL | Probable | As nectar-tap_saprotroph (Põlme et al. 2020); Decay sugar-rich substrates (Põlme et al. 2020) | Põlme S, et al. 2020. Fungal Diversity 105:1-16 ((https://doi.org/10.1007/s13225-020-00466-2)) |
| Otu15594 | 1 | k__Fungi(100);p__Ascomycota(100);c__Saccharomycetes(100);o__Saccharomycetales(100);f__Metschnikowiaceae(100);g__Metschnikowia(100);s__Metschnikowia_pulcherrima(100); | Metschnikowia | 13 | Saprotroph | \|Undefined Saprotroph\| | Yeast | NULL | Probable | As nectar-tap_saprotroph (Põlme et al. 2020); Decay sugar-rich substrates (Põlme et al. 2020) | Põlme S, et al. 2020. Fungal Diversity 105:1-16 ((https://doi.org/10.1007/s13225-020-00466-2)) |
| Otu15595 | 1 | k__Fungi(100);p__Ascomycota(100);c__Saccharomycetes(100);o__Saccharomycetales(100);f__Metschnikowiaceae(100);g__Metschnikowia(100);s__Metschnikowia_pulcherrima(100); | Metschnikowia | 13 | Saprotroph | \|Undefined Saprotroph\| | Yeast | NULL | Probable | As nectar-tap_saprotroph (Põlme et al. 2020); Decay sugar-rich substrates (Põlme et al. 2020) | Põlme S, et al. 2020. Fungal Diversity 105:1-16 ((https://doi.org/10.1007/s13225-020-00466-2)) |
| Otu15596 | 1 | k__Fungi(100);p__Ascomycota(100);c__Saccharomycetes(100);o__Saccharomycetales(100);f__Metschnikowiaceae(100);g__Metschnikowia(100);s__Metschnikowia_pulcherrima(100); | Metschnikowia | 13 | Saprotroph | \|Undefined Saprotroph\| | Yeast | NULL | Probable | As nectar-tap_saprotroph (Põlme et al. 2020); Decay sugar-rich substrates (Põlme et al. 2020) | Põlme S, et al. 2020. Fungal Diversity 105:1-16 ((https://doi.org/10.1007/s13225-020-00466-2)) |
| Otu15597 | 1 | k__Fungi(100);p__Ascomycota(100);c__Saccharomycetes(100);o__Saccharomycetales(100);f__Metschnikowiaceae(100);g__Metschnikowia(100);s__Metschnikowia_pulcherrima(100); | Metschnikowia | 13 | Saprotroph | \|Undefined Saprotroph\| | Yeast | NULL | Probable | As nectar-tap_saprotroph (Põlme et al. 2020); Decay sugar-rich substrates (Põlme et al. 2020) | Põlme S, et al. 2020. Fungal Diversity 105:1-16 ((https://doi.org/10.1007/s13225-020-00466-2)) |
| Otu15599 | 1 | k__Fungi(100);p__Ascomycota(100);c__Saccharomycetes(100);o__Saccharomycetales(100);f__Metschnikowiaceae(100);g__Metschnikowia(100);g__Metschnikowia_unclassified(100); | Metschnikowia | 13 | Saprotroph | \|Undefined Saprotroph\| | Yeast | NULL | Probable | As nectar-tap_saprotroph (Põlme et al. 2020); Decay sugar-rich substrates (Põlme et al. 2020) | Põlme S, et al. 2020. Fungal Diversity 105:1-16 ((https://doi.org/10.1007/s13225-020-00466-2)) |
| Otu15600 | 1 | k__Fungi(100);p__Ascomycota(100);c__Saccharomycetes(100);o__Saccharomycetales(100);f__Metschnikowiaceae(100);g__Metschnikowia(100);g__Metschnikowia_unclassified(100); | Metschnikowia | 13 | Saprotroph | \|Undefined Saprotroph\| | Yeast | NULL | Probable | As nectar-tap_saprotroph (Põlme et al. 2020); Decay sugar-rich substrates (Põlme et al. 2020) | Põlme S, et al. 2020. Fungal Diversity 105:1-16 ((https://doi.org/10.1007/s13225-020-00466-2)) |
| Otu15601 | 1 | k__Fungi(100);p__Ascomycota(100);c__Saccharomycetes(100);o__Saccharomycetales(100);f__Saccharomycetales_family_Incertae_sedis(100);g__Candida(100);g__Candida_unclassified(100); | Candida | 13 | Pathotroph-Saprotroph-Symbiotroph | Animal Pathogen-Endophyte-Endosymbiont-Epiphyte-\|Undefined Saprotroph\| | Dimorphic Yeast-Yeast | NULL | Probable | As nectar-tap_saprotroph (Põlme et al. 2020); Decay sugar-rich substrates (Põlme et al. 2020); This asexual genus is polyphyletic and contain species that function very differently depend on what they are. Caution should be taken when interpreting guilds for Candida species. | Manolakaki D, et al. 2010. Virulence. 1: 367-375 ((https://doi.org/10.4161/viru.1.5.12796)); Mekbib SB, et al. 2011. Tropical Plant Pathology 36:233-240 ((https://doi.org/10.1590/S1982-56762011000400004)); Põlme S, et al. 2020. Fungal Diversity 105:1-16 ((https://doi.org/10.1007/s13225-020-00466-2)) |
| Otu15602 | 1 | k__Fungi(100);p__Ascomycota(100);c__Saccharomycetes(100);o__Saccharomycetales(100);f__Metschnikowiaceae(100);g__Metschnikowia(100);s__Metschnikowia_pulcherrima(100); | Metschnikowia | 13 | Saprotroph | \|Undefined Saprotroph\| | Yeast | NULL | Probable | As nectar-tap_saprotroph (Põlme et al. 2020); Decay sugar-rich substrates (Põlme et al. 2020) | Põlme S, et al. 2020. Fungal Diversity 105:1-16 ((https://doi.org/10.1007/s13225-020-00466-2)) |
| Otu15603 | 1 | k__Fungi(100);p__Ascomycota(100);c__Saccharomycetes(100);o__Saccharomycetales(100);f__Metschnikowiaceae(100);g__Metschnikowia(100);g__Metschnikowia_unclassified(100); | Metschnikowia | 13 | Saprotroph | \|Undefined Saprotroph\| | Yeast | NULL | Probable | As nectar-tap_saprotroph (Põlme et al. 2020); Decay sugar-rich substrates (Põlme et al. 2020) | Põlme S, et al. 2020. Fungal Diversity 105:1-16 ((https://doi.org/10.1007/s13225-020-00466-2)) |
| Otu15604 | 1 | k__Fungi(100);p__Ascomycota(100);c__Saccharomycetes(100);o__Saccharomycetales(100);f__Metschnikowiaceae(100);g__Metschnikowia(100);s__Metschnikowia_pulcherrima(100); | Metschnikowia | 13 | Saprotroph | \|Undefined Saprotroph\| | Yeast | NULL | Probable | As nectar-tap_saprotroph (Põlme et al. 2020); Decay sugar-rich substrates (Põlme et al. 2020) | Põlme S, et al. 2020. Fungal Diversity 105:1-16 ((https://doi.org/10.1007/s13225-020-00466-2)) |
| Otu15606 | 1 | k__Fungi(100);p__Ascomycota(100);c__Saccharomycetes(100);o__Saccharomycetales(100);f__Metschnikowiaceae(100);g__Metschnikowia(100);g__Metschnikowia_unclassified(100); | Metschnikowia | 13 | Saprotroph | \|Undefined Saprotroph\| | Yeast | NULL | Probable | As nectar-tap_saprotroph (Põlme et al. 2020); Decay sugar-rich substrates (Põlme et al. 2020) | Põlme S, et al. 2020. Fungal Diversity 105:1-16 ((https://doi.org/10.1007/s13225-020-00466-2)) |
| Otu15607 | 1 | k__Fungi(100);p__Ascomycota(100);c__Saccharomycetes(100);o__Saccharomycetales(100);f__Metschnikowiaceae(100);g__Metschnikowia(100);g__Metschnikowia_unclassified(100); | Metschnikowia | 13 | Saprotroph | \|Undefined Saprotroph\| | Yeast | NULL | Probable | As nectar-tap_saprotroph (Põlme et al. 2020); Decay sugar-rich substrates (Põlme et al. 2020) | Põlme S, et al. 2020. Fungal Diversity 105:1-16 ((https://doi.org/10.1007/s13225-020-00466-2)) |
| Otu15608 | 1 | k__Fungi(100);p__Ascomycota(100);c__Saccharomycetes(100);o__Saccharomycetales(100);f__Metschnikowiaceae(100);g__Metschnikowia(100);s__Metschnikowia_pulcherrima(100); | Metschnikowia | 13 | Saprotroph | \|Undefined Saprotroph\| | Yeast | NULL | Probable | As nectar-tap_saprotroph (Põlme et al. 2020); Decay sugar-rich substrates (Põlme et al. 2020) | Põlme S, et al. 2020. Fungal Diversity 105:1-16 ((https://doi.org/10.1007/s13225-020-00466-2)) |
| Otu15609 | 1 | k__Fungi(100);p__Ascomycota(100);c__Saccharomycetes(100);o__Saccharomycetales(100);f__Metschnikowiaceae(100);g__Metschnikowia(100);s__Metschnikowia_pulcherrima(100); | Metschnikowia | 13 | Saprotroph | \|Undefined Saprotroph\| | Yeast | NULL | Probable | As nectar-tap_saprotroph (Põlme et al. 2020); Decay sugar-rich substrates (Põlme et al. 2020) | Põlme S, et al. 2020. Fungal Diversity 105:1-16 ((https://doi.org/10.1007/s13225-020-00466-2)) |
| Otu15610 | 1 | k__Fungi(100);p__Ascomycota(100);c__Saccharomycetes(100);o__Saccharomycetales(100);f__Metschnikowiaceae(100);g__Metschnikowia(100);g__Metschnikowia_unclassified(100); | Metschnikowia | 13 | Saprotroph | \|Undefined Saprotroph\| | Yeast | NULL | Probable | As nectar-tap_saprotroph (Põlme et al. 2020); Decay sugar-rich substrates (Põlme et al. 2020) | Põlme S, et al. 2020. Fungal Diversity 105:1-16 ((https://doi.org/10.1007/s13225-020-00466-2)) |
| Otu15612 | 1 | k__Fungi(100);p__Ascomycota(100);c__Saccharomycetes(100);o__Saccharomycetales(100);f__Metschnikowiaceae(100);g__Metschnikowia(100);s__Metschnikowia_pulcherrima(100); | Metschnikowia | 13 | Saprotroph | \|Undefined Saprotroph\| | Yeast | NULL | Probable | As nectar-tap_saprotroph (Põlme et al. 2020); Decay sugar-rich substrates (Põlme et al. 2020) | Põlme S, et al. 2020. Fungal Diversity 105:1-16 ((https://doi.org/10.1007/s13225-020-00466-2)) |
| Otu15613 | 1 | k__Fungi(100);p__Ascomycota(100);c__Saccharomycetes(100);o__Saccharomycetales(100);f__Metschnikowiaceae(100);g__Metschnikowia(100);g__Metschnikowia_unclassified(100); | Metschnikowia | 13 | Saprotroph | \|Undefined Saprotroph\| | Yeast | NULL | Probable | As nectar-tap_saprotroph (Põlme et al. 2020); Decay sugar-rich substrates (Põlme et al. 2020) | Põlme S, et al. 2020. Fungal Diversity 105:1-16 ((https://doi.org/10.1007/s13225-020-00466-2)) |
| Otu15614 | 1 | k__Fungi(100);p__Ascomycota(100);c__Saccharomycetes(100);o__Saccharomycetales(100);f__Metschnikowiaceae(100);g__Metschnikowia(100);s__Metschnikowia_pulcherrima(100); | Metschnikowia | 13 | Saprotroph | \|Undefined Saprotroph\| | Yeast | NULL | Probable | As nectar-tap_saprotroph (Põlme et al. 2020); Decay sugar-rich substrates (Põlme et al. 2020) | Põlme S, et al. 2020. Fungal Diversity 105:1-16 ((https://doi.org/10.1007/s13225-020-00466-2)) |
| Otu15615 | 1 | k__Fungi(100);p__Ascomycota(100);c__Saccharomycetes(100);o__Saccharomycetales(100);f__Metschnikowiaceae(100);g__Metschnikowia(100);s__Metschnikowia_pulcherrima(100); | Metschnikowia | 13 | Saprotroph | \|Undefined Saprotroph\| | Yeast | NULL | Probable | As nectar-tap_saprotroph (Põlme et al. 2020); Decay sugar-rich substrates (Põlme et al. 2020) | Põlme S, et al. 2020. Fungal Diversity 105:1-16 ((https://doi.org/10.1007/s13225-020-00466-2)) |
| Otu15616 | 1 | k__Fungi(100);p__Ascomycota(100);c__Saccharomycetes(100);o__Saccharomycetales(100);f__Metschnikowiaceae(100);g__Metschnikowia(100);g__Metschnikowia_unclassified(100); | Metschnikowia | 13 | Saprotroph | \|Undefined Saprotroph\| | Yeast | NULL | Probable | As nectar-tap_saprotroph (Põlme et al. 2020); Decay sugar-rich substrates (Põlme et al. 2020) | Põlme S, et al. 2020. Fungal Diversity 105:1-16 ((https://doi.org/10.1007/s13225-020-00466-2)) |
| Otu15618 | 1 | k__Fungi(100);p__Ascomycota(100);c__Saccharomycetes(100);o__Saccharomycetales(100);f__Metschnikowiaceae(100);g__Metschnikowia(100);g__Metschnikowia_unclassified(100); | Metschnikowia | 13 | Saprotroph | \|Undefined Saprotroph\| | Yeast | NULL | Probable | As nectar-tap_saprotroph (Põlme et al. 2020); Decay sugar-rich substrates (Põlme et al. 2020) | Põlme S, et al. 2020. Fungal Diversity 105:1-16 ((https://doi.org/10.1007/s13225-020-00466-2)) |
| Otu15619 | 1 | k__Fungi(100);p__Ascomycota(100);c__Saccharomycetes(100);o__Saccharomycetales(100);f__Metschnikowiaceae(100);g__Metschnikowia(100);s__Metschnikowia_pulcherrima(100); | Metschnikowia | 13 | Saprotroph | \|Undefined Saprotroph\| | Yeast | NULL | Probable | As nectar-tap_saprotroph (Põlme et al. 2020); Decay sugar-rich substrates (Põlme et al. 2020) | Põlme S, et al. 2020. Fungal Diversity 105:1-16 ((https://doi.org/10.1007/s13225-020-00466-2)) |
| Otu15621 | 1 | k__Fungi(100);p__Ascomycota(100);c__Saccharomycetes(100);o__Saccharomycetales(100);f__Metschnikowiaceae(100);g__Metschnikowia(100);s__Metschnikowia_pulcherrima(100); | Metschnikowia | 13 | Saprotroph | \|Undefined Saprotroph\| | Yeast | NULL | Probable | As nectar-tap_saprotroph (Põlme et al. 2020); Decay sugar-rich substrates (Põlme et al. 2020) | Põlme S, et al. 2020. Fungal Diversity 105:1-16 ((https://doi.org/10.1007/s13225-020-00466-2)) |
| Otu15622 | 1 | k__Fungi(100);p__Ascomycota(100);c__Saccharomycetes(100);o__Saccharomycetales(100);f__Metschnikowiaceae(100);g__Metschnikowia(100);s__Metschnikowia_pulcherrima(100); | Metschnikowia | 13 | Saprotroph | \|Undefined Saprotroph\| | Yeast | NULL | Probable | As nectar-tap_saprotroph (Põlme et al. 2020); Decay sugar-rich substrates (Põlme et al. 2020) | Põlme S, et al. 2020. Fungal Diversity 105:1-16 ((https://doi.org/10.1007/s13225-020-00466-2)) |
| Otu15623 | 1 | k__Fungi(100);p__Ascomycota(100);c__Saccharomycetes(100);o__Saccharomycetales(100);f__Metschnikowiaceae(100);g__Metschnikowia(100);g__Metschnikowia_unclassified(100); | Metschnikowia | 13 | Saprotroph | \|Undefined Saprotroph\| | Yeast | NULL | Probable | As nectar-tap_saprotroph (Põlme et al. 2020); Decay sugar-rich substrates (Põlme et al. 2020) | Põlme S, et al. 2020. Fungal Diversity 105:1-16 ((https://doi.org/10.1007/s13225-020-00466-2)) |
| Otu15624 | 1 | k__Fungi(100);p__Ascomycota(100);c__Saccharomycetes(100);o__Saccharomycetales(100);f__Metschnikowiaceae(100);g__Metschnikowia(100);s__Metschnikowia_pulcherrima(100); | Metschnikowia | 13 | Saprotroph | \|Undefined Saprotroph\| | Yeast | NULL | Probable | As nectar-tap_saprotroph (Põlme et al. 2020); Decay sugar-rich substrates (Põlme et al. 2020) | Põlme S, et al. 2020. Fungal Diversity 105:1-16 ((https://doi.org/10.1007/s13225-020-00466-2)) |
| Otu15625 | 1 | k__Fungi(100);p__Ascomycota(100);c__Saccharomycetes(100);o__Saccharomycetales(100);f__Metschnikowiaceae(100);g__Metschnikowia(100);g__Metschnikowia_unclassified(100); | Metschnikowia | 13 | Saprotroph | \|Undefined Saprotroph\| | Yeast | NULL | Probable | As nectar-tap_saprotroph (Põlme et al. 2020); Decay sugar-rich substrates (Põlme et al. 2020) | Põlme S, et al. 2020. Fungal Diversity 105:1-16 ((https://doi.org/10.1007/s13225-020-00466-2)) |
| Otu15626 | 1 | k__Fungi(100);p__Ascomycota(100);c__Saccharomycetes(100);o__Saccharomycetales(100);f__Metschnikowiaceae(100);g__Metschnikowia(100);s__Metschnikowia_pulcherrima(100); | Metschnikowia | 13 | Saprotroph | \|Undefined Saprotroph\| | Yeast | NULL | Probable | As nectar-tap_saprotroph (Põlme et al. 2020); Decay sugar-rich substrates (Põlme et al. 2020) | Põlme S, et al. 2020. Fungal Diversity 105:1-16 ((https://doi.org/10.1007/s13225-020-00466-2)) |
| Otu15627 | 1 | k__Fungi(100);p__Ascomycota(100);c__Saccharomycetes(100);o__Saccharomycetales(100);f__Metschnikowiaceae(100);g__Metschnikowia(100);s__Metschnikowia_pulcherrima(100); | Metschnikowia | 13 | Saprotroph | \|Undefined Saprotroph\| | Yeast | NULL | Probable | As nectar-tap_saprotroph (Põlme et al. 2020); Decay sugar-rich substrates (Põlme et al. 2020) | Põlme S, et al. 2020. Fungal Diversity 105:1-16 ((https://doi.org/10.1007/s13225-020-00466-2)) |
| Otu15628 | 1 | k__Fungi(100);p__Ascomycota(100);c__Saccharomycetes(100);o__Saccharomycetales(100);f__Metschnikowiaceae(100);g__Metschnikowia(100);s__Metschnikowia_pulcherrima(100); | Metschnikowia | 13 | Saprotroph | \|Undefined Saprotroph\| | Yeast | NULL | Probable | As nectar-tap_saprotroph (Põlme et al. 2020); Decay sugar-rich substrates (Põlme et al. 2020) | Põlme S, et al. 2020. Fungal Diversity 105:1-16 ((https://doi.org/10.1007/s13225-020-00466-2)) |
| Otu15629 | 1 | k__Fungi(100);p__Ascomycota(100);c__Saccharomycetes(100);o__Saccharomycetales(100);f__Metschnikowiaceae(100);g__Metschnikowia(100);s__Metschnikowia_pulcherrima(100); | Metschnikowia | 13 | Saprotroph | \|Undefined Saprotroph\| | Yeast | NULL | Probable | As nectar-tap_saprotroph (Põlme et al. 2020); Decay sugar-rich substrates (Põlme et al. 2020) | Põlme S, et al. 2020. Fungal Diversity 105:1-16 ((https://doi.org/10.1007/s13225-020-00466-2)) |
| Otu15630 | 1 | k__Fungi(100);p__Ascomycota(100);c__Saccharomycetes(100);o__Saccharomycetales(100);f__Metschnikowiaceae(100);g__Metschnikowia(100);s__Metschnikowia_pulcherrima(100); | Metschnikowia | 13 | Saprotroph | \|Undefined Saprotroph\| | Yeast | NULL | Probable | As nectar-tap_saprotroph (Põlme et al. 2020); Decay sugar-rich substrates (Põlme et al. 2020) | Põlme S, et al. 2020. Fungal Diversity 105:1-16 ((https://doi.org/10.1007/s13225-020-00466-2)) |
| Otu15631 | 1 | k__Fungi(100);p__Ascomycota(100);c__Saccharomycetes(100);o__Saccharomycetales(100);f__Metschnikowiaceae(100);g__Metschnikowia(100);s__Metschnikowia_pulcherrima(100); | Metschnikowia | 13 | Saprotroph | \|Undefined Saprotroph\| | Yeast | NULL | Probable | As nectar-tap_saprotroph (Põlme et al. 2020); Decay sugar-rich substrates (Põlme et al. 2020) | Põlme S, et al. 2020. Fungal Diversity 105:1-16 ((https://doi.org/10.1007/s13225-020-00466-2)) |
| Otu15632 | 1 | k__Fungi(100);p__Ascomycota(100);c__Saccharomycetes(100);o__Saccharomycetales(100);f__Metschnikowiaceae(100);g__Metschnikowia(100);s__Metschnikowia_pulcherrima(100); | Metschnikowia | 13 | Saprotroph | \|Undefined Saprotroph\| | Yeast | NULL | Probable | As nectar-tap_saprotroph (Põlme et al. 2020); Decay sugar-rich substrates (Põlme et al. 2020) | Põlme S, et al. 2020. Fungal Diversity 105:1-16 ((https://doi.org/10.1007/s13225-020-00466-2)) |
| Otu15633 | 1 | k__Fungi(100);p__Ascomycota(100);c__Saccharomycetes(100);o__Saccharomycetales(100);f__Metschnikowiaceae(100);g__Metschnikowia(100);g__Metschnikowia_unclassified(100); | Metschnikowia | 13 | Saprotroph | \|Undefined Saprotroph\| | Yeast | NULL | Probable | As nectar-tap_saprotroph (Põlme et al. 2020); Decay sugar-rich substrates (Põlme et al. 2020) | Põlme S, et al. 2020. Fungal Diversity 105:1-16 ((https://doi.org/10.1007/s13225-020-00466-2)) |
| Otu15634 | 1 | k__Fungi(100);p__Ascomycota(100);c__Saccharomycetes(100);o__Saccharomycetales(100);f__Metschnikowiaceae(100);g__Metschnikowia(100);g__Metschnikowia_unclassified(100); | Metschnikowia | 13 | Saprotroph | \|Undefined Saprotroph\| | Yeast | NULL | Probable | As nectar-tap_saprotroph (Põlme et al. 2020); Decay sugar-rich substrates (Põlme et al. 2020) | Põlme S, et al. 2020. Fungal Diversity 105:1-16 ((https://doi.org/10.1007/s13225-020-00466-2)) |
| Otu15635 | 1 | k__Fungi(100);p__Ascomycota(100);c__Saccharomycetes(100);o__Saccharomycetales(100);f__Metschnikowiaceae(100);g__Metschnikowia(100);s__Metschnikowia_pulcherrima(100); | Metschnikowia | 13 | Saprotroph | \|Undefined Saprotroph\| | Yeast | NULL | Probable | As nectar-tap_saprotroph (Põlme et al. 2020); Decay sugar-rich substrates (Põlme et al. 2020) | Põlme S, et al. 2020. Fungal Diversity 105:1-16 ((https://doi.org/10.1007/s13225-020-00466-2)) |
| Otu15636 | 1 | k__Fungi(100);p__Ascomycota(100);c__Saccharomycetes(100);o__Saccharomycetales(100);f__Metschnikowiaceae(100);g__Metschnikowia(100);s__Metschnikowia_pulcherrima(100); | Metschnikowia | 13 | Saprotroph | \|Undefined Saprotroph\| | Yeast | NULL | Probable | As nectar-tap_saprotroph (Põlme et al. 2020); Decay sugar-rich substrates (Põlme et al. 2020) | Põlme S, et al. 2020. Fungal Diversity 105:1-16 ((https://doi.org/10.1007/s13225-020-00466-2)) |
| Otu15637 | 1 | k__Fungi(100);p__Ascomycota(100);c__Saccharomycetes(100);o__Saccharomycetales(100);f__Saccharomycetales_family_Incertae_sedis(100);g__Candida(100);g__Candida_unclassified(100); | Candida | 13 | Pathotroph-Saprotroph-Symbiotroph | Animal Pathogen-Endophyte-Endosymbiont-Epiphyte-\|Undefined Saprotroph\| | Dimorphic Yeast-Yeast | NULL | Probable | As nectar-tap_saprotroph (Põlme et al. 2020); Decay sugar-rich substrates (Põlme et al. 2020); This asexual genus is polyphyletic and contain species that function very differently depend on what they are. Caution should be taken when interpreting guilds for Candida species. | Manolakaki D, et al. 2010. Virulence. 1: 367-375 ((https://doi.org/10.4161/viru.1.5.12796)); Mekbib SB, et al. 2011. Tropical Plant Pathology 36:233-240 ((https://doi.org/10.1590/S1982-56762011000400004)); Põlme S, et al. 2020. Fungal Diversity 105:1-16 ((https://doi.org/10.1007/s13225-020-00466-2)) |
| Otu15638 | 1 | k__Fungi(100);p__Ascomycota(100);c__Saccharomycetes(100);o__Saccharomycetales(100);f__Metschnikowiaceae(100);g__Metschnikowia(100);g__Metschnikowia_unclassified(100); | Metschnikowia | 13 | Saprotroph | \|Undefined Saprotroph\| | Yeast | NULL | Probable | As nectar-tap_saprotroph (Põlme et al. 2020); Decay sugar-rich substrates (Põlme et al. 2020) | Põlme S, et al. 2020. Fungal Diversity 105:1-16 ((https://doi.org/10.1007/s13225-020-00466-2)) |
| Otu15643 | 1 | k__Fungi(100);p__Ascomycota(100);c__Saccharomycetes(100);o__Saccharomycetales(100);f__Metschnikowiaceae(100);g__Metschnikowia(100);s__Metschnikowia_pulcherrima(100); | Metschnikowia | 13 | Saprotroph | \|Undefined Saprotroph\| | Yeast | NULL | Probable | As nectar-tap_saprotroph (Põlme et al. 2020); Decay sugar-rich substrates (Põlme et al. 2020) | Põlme S, et al. 2020. Fungal Diversity 105:1-16 ((https://doi.org/10.1007/s13225-020-00466-2)) |

| | | | | | | | | | | | |
|---|---|---|---|---|---|---|---|---|---|---|---|
| Otu15644 | 1 | k__Fungi(100);p__Ascomycota(100);c__Saccharomycetes(100);o__Saccharomycetales(100);f__Metschnikowiaceae(100);g__Metschnikowia(100);g__Metschnikowia_unclassified(100); | Metschnikowia | 13 | Saprotroph | \|Undefined Saprotroph\| | Yeast | NULL | Probable | As nectar-tap_saprotroph (Põlme et al. 2020); Decay sugar-rich substrates (Põlme et al. 2020) | Põlme S, et al. 2020. Fungal Diversity 105:1-16 ((https://doi.org/10.1007/s13225-020-00466-2)) |
| Otu15646 | 1 | k__Fungi(100);p__Ascomycota(100);c__Saccharomycetes(100);o__Saccharomycetales(100);f__Metschnikowiaceae(100);g__Metschnikowia(100);s__Metschnikowia_pulcherrima(100); | Metschnikowia | 13 | Saprotroph | \|Undefined Saprotroph\| | Yeast | NULL | Probable | As nectar-tap_saprotroph (Põlme et al. 2020); Decay sugar-rich substrates (Põlme et al. 2020) | Põlme S, et al. 2020. Fungal Diversity 105:1-16 ((https://doi.org/10.1007/s13225-020-00466-2)) |
| Otu15648 | 1 | k__Fungi(100);p__Ascomycota(100);c__Saccharomycetes(100);o__Saccharomycetales(100);f__Metschnikowiaceae(100);g__Metschnikowia(100);s__Metschnikowia_pulcherrima(100); | Metschnikowia | 13 | Saprotroph | \|Undefined Saprotroph\| | Yeast | NULL | Probable | As nectar-tap_saprotroph (Põlme et al. 2020); Decay sugar-rich substrates (Põlme et al. 2020) | Põlme S, et al. 2020. Fungal Diversity 105:1-16 ((https://doi.org/10.1007/s13225-020-00466-2)) |
| Otu15649 | 1 | k__Fungi(100);p__Ascomycota(100);c__Saccharomycetes(100);o__Saccharomycetales(100);f__Metschnikowiaceae(100);g__Metschnikowia(100);s__Metschnikowia_pulcherrima(100); | Metschnikowia | 13 | Saprotroph | \|Undefined Saprotroph\| | Yeast | NULL | Probable | As nectar-tap_saprotroph (Põlme et al. 2020); Decay sugar-rich substrates (Põlme et al. 2020) | Põlme S, et al. 2020. Fungal Diversity 105:1-16 ((https://doi.org/10.1007/s13225-020-00466-2)) |
| Otu15652 | 1 | k__Fungi(100);p__Ascomycota(100);c__Saccharomycetes(100);o__Saccharomycetales(100);f__Metschnikowiaceae(100);g__Metschnikowia(100);s__Metschnikowia_pulcherrima(100); | Metschnikowia | 13 | Saprotroph | \|Undefined Saprotroph\| | Yeast | NULL | Probable | As nectar-tap_saprotroph (Põlme et al. 2020); Decay sugar-rich substrates (Põlme et al. 2020) | Põlme S, et al. 2020. Fungal Diversity 105:1-16 ((https://doi.org/10.1007/s13225-020-00466-2)) |
| Otu15656 | 1 | k__Fungi(100);p__Ascomycota(100);c__Saccharomycetes(100);o__Saccharomycetales(100);f__Metschnikowiaceae(100);g__Metschnikowia(100);g__Metschnikowia_unclassified(100); | Metschnikowia | 13 | Saprotroph | \|Undefined Saprotroph\| | Yeast | NULL | Probable | As nectar-tap_saprotroph (Põlme et al. 2020); Decay sugar-rich substrates (Põlme et al. 2020) | Põlme S, et al. 2020. Fungal Diversity 105:1-16 ((https://doi.org/10.1007/s13225-020-00466-2)) |
| Otu15657 | 1 | k__Fungi(100);p__Ascomycota(100);c__Saccharomycetes(100);o__Saccharomycetales(100);f__Metschnikowiaceae(100);g__Metschnikowia(100);g__Metschnikowia_unclassified(100); | Metschnikowia | 13 | Saprotroph | \|Undefined Saprotroph\| | Yeast | NULL | Probable | As nectar-tap_saprotroph (Põlme et al. 2020); Decay sugar-rich substrates (Põlme et al. 2020) | Põlme S, et al. 2020. Fungal Diversity 105:1-16 ((https://doi.org/10.1007/s13225-020-00466-2)) |
| Otu15660 | 1 | k__Fungi(100);p__Ascomycota(100);c__Saccharomycetes(100);o__Saccharomycetales(100);o__Saccharomycetales_unclassified(100);o__Saccharomycetales_unclassified(100);o__Saccharomycetales_unclassified(100); | Saccharomycetales | 7 | Saprotroph | Undefined Saprotroph | Yeast | NULL | Possible | NULL | Sterkenburg E, et al. 2015. New Phytologist 207:1145-1158 ((https://doi.org/10.1111/nph.13426)) |
| Otu15663 | 1 | k__Fungi(100);p__Ascomycota(100);c__Saccharomycetes(100);o__Saccharomycetales(100);f__Metschnikowiaceae(100);g__Metschnikowia(100);g__Metschnikowia_unclassified(100); | Metschnikowia | 13 | Saprotroph | \|Undefined Saprotroph\| | Yeast | NULL | Probable | As nectar-tap_saprotroph (Põlme et al. 2020); Decay sugar-rich substrates (Põlme et al. 2020) | Põlme S, et al. 2020. Fungal Diversity 105:1-16 ((https://doi.org/10.1007/s13225-020-00466-2)) |
| Otu15664 | 1 | k__Fungi(100);p__Ascomycota(100);c__Saccharomycetes(100);o__Saccharomycetales(100);f__Metschnikowiaceae(100);g__Metschnikowia(100);s__Metschnikowia_pulcherrima(100); | Metschnikowia | 13 | Saprotroph | \|Undefined Saprotroph\| | Yeast | NULL | Probable | As nectar-tap_saprotroph (Põlme et al. 2020); Decay sugar-rich substrates (Põlme et al. 2020) | Põlme S, et al. 2020. Fungal Diversity 105:1-16 ((https://doi.org/10.1007/s13225-020-00466-2)) |
| Otu15666 | 1 | k__Fungi(100);p__Ascomycota(100);c__Saccharomycetes(100);o__Saccharomycetales(100);f__Metschnikowiaceae(100);g__Metschnikowia(100);g__Metschnikowia_unclassified(100); | Metschnikowia | 13 | Saprotroph | \|Undefined Saprotroph\| | Yeast | NULL | Probable | As nectar-tap_saprotroph (Põlme et al. 2020); Decay sugar-rich substrates (Põlme et al. 2020) | Põlme S, et al. 2020. Fungal Diversity 105:1-16 ((https://doi.org/10.1007/s13225-020-00466-2)) |
| Otu15667 | 1 | k__Fungi(100);p__Ascomycota(100);c__Saccharomycetes(100);o__Saccharomycetales(100);f__Metschnikowiaceae(100);g__Metschnikowia(100);s__Metschnikowia_pulcherrima(100); | Metschnikowia | 13 | Saprotroph | \|Undefined Saprotroph\| | Yeast | NULL | Probable | As nectar-tap_saprotroph (Põlme et al. 2020); Decay sugar-rich substrates (Põlme et al. 2020) | Põlme S, et al. 2020. Fungal Diversity 105:1-16 ((https://doi.org/10.1007/s13225-020-00466-2)) |
| Otu15668 | 1 | k__Fungi(100);p__Ascomycota(100);c__Saccharomycetes(100);o__Saccharomycetales(100);f__Metschnikowiaceae(100);g__Metschnikowia(100);s__Metschnikowia_pulcherrima(100); | Metschnikowia | 13 | Saprotroph | \|Undefined Saprotroph\| | Yeast | NULL | Probable | As nectar-tap_saprotroph (Põlme et al. 2020); Decay sugar-rich substrates (Põlme et al. 2020) | Põlme S, et al. 2020. Fungal Diversity 105:1-16 ((https://doi.org/10.1007/s13225-020-00466-2)) |
| Otu15669 | 1 | k__Fungi(100);p__Ascomycota(100);c__Saccharomycetes(100);o__Saccharomycetales(100);f__Metschnikowiaceae(100);g__Metschnikowia(100);s__Metschnikowia_pulcherrima(100); | Metschnikowia | 13 | Saprotroph | \|Undefined Saprotroph\| | Yeast | NULL | Probable | As nectar-tap_saprotroph (Põlme et al. 2020); Decay sugar-rich substrates (Põlme et al. 2020) | Põlme S, et al. 2020. Fungal Diversity 105:1-16 ((https://doi.org/10.1007/s13225-020-00466-2)) |
| Otu15670 | 1 | k__Fungi(100);p__Ascomycota(100);c__Saccharomycetes(100);o__Saccharomycetales(100);f__Metschnikowiaceae(100);g__Metschnikowia(100);g__Metschnikowia_unclassified(100); | Metschnikowia | 13 | Saprotroph | \|Undefined Saprotroph\| | Yeast | NULL | Probable | As nectar-tap_saprotroph (Põlme et al. 2020); Decay sugar-rich substrates (Põlme et al. 2020) | Põlme S, et al. 2020. Fungal Diversity 105:1-16 ((https://doi.org/10.1007/s13225-020-00466-2)) |
| Otu15671 | 1 | k__Fungi(100);p__Ascomycota(100);c__Saccharomycetes(100);o__Saccharomycetales(100);f__Metschnikowiaceae(100);g__Metschnikowia(100);g__Metschnikowia_unclassified(100); | Metschnikowia | 13 | Saprotroph | \|Undefined Saprotroph\| | Yeast | NULL | Probable | As nectar-tap_saprotroph (Põlme et al. 2020); Decay sugar-rich substrates (Põlme et al. 2020) | Põlme S, et al. 2020. Fungal Diversity 105:1-16 ((https://doi.org/10.1007/s13225-020-00466-2)) |
| Otu15672 | 1 | k__Fungi(100);p__Ascomycota(100);c__Saccharomycetes(100);o__Saccharomycetales(100);f__Metschnikowiaceae(100);g__Metschnikowia(100);s__Metschnikowia_pulcherrima(100); | Metschnikowia | 13 | Saprotroph | \|Undefined Saprotroph\| | Yeast | NULL | Probable | As nectar-tap_saprotroph (Põlme et al. 2020); Decay sugar-rich substrates (Põlme et al. 2020) | Põlme S, et al. 2020. Fungal Diversity 105:1-16 ((https://doi.org/10.1007/s13225-020-00466-2)) |
| Otu15673 | 1 | k__Fungi(100);p__Ascomycota(100);c__Saccharomycetes(100);o__Saccharomycetales(100);f__Metschnikowiaceae(100);g__Metschnikowia(100);s__Metschnikowia_pulcherrima(100); | Metschnikowia | 13 | Saprotroph | \|Undefined Saprotroph\| | Yeast | NULL | Probable | As nectar-tap_saprotroph (Põlme et al. 2020); Decay sugar-rich substrates (Põlme et al. 2020) | Põlme S, et al. 2020. Fungal Diversity 105:1-16 ((https://doi.org/10.1007/s13225-020-00466-2)) |
| Otu15674 | 1 | k__Fungi(100);p__Ascomycota(100);c__Saccharomycetes(100);o__Saccharomycetales(100);f__Metschnikowiaceae(100);g__Metschnikowia(100);s__Metschnikowia_pulcherrima(100); | Metschnikowia | 13 | Saprotroph | \|Undefined Saprotroph\| | Yeast | NULL | Probable | As nectar-tap_saprotroph (Põlme et al. 2020); Decay sugar-rich substrates (Põlme et al. 2020) | Põlme S, et al. 2020. Fungal Diversity 105:1-16 ((https://doi.org/10.1007/s13225-020-00466-2)) |
| Otu15677 | 1 | k__Fungi(100);p__Ascomycota(100);c__Saccharomycetes(100);o__Saccharomycetales(100);f__Metschnikowiaceae(100);g__Metschnikowia(100);g__Metschnikowia_unclassified(100); | Metschnikowia | 13 | Saprotroph | \|Undefined Saprotroph\| | Yeast | NULL | Probable | As nectar-tap_saprotroph (Põlme et al. 2020); Decay sugar-rich substrates (Põlme et al. 2020) | Põlme S, et al. 2020. Fungal Diversity 105:1-16 ((https://doi.org/10.1007/s13225-020-00466-2)) |
| Otu15679 | 1 | k__Fungi(100);p__Ascomycota(100);c__Saccharomycetes(100);o__Saccharomycetales(100);f__Metschnikowiaceae(100);g__Metschnikowia(100);g__Metschnikowia_unclassified(100); | Metschnikowia | 13 | Saprotroph | \|Undefined Saprotroph\| | Yeast | NULL | Probable | As nectar-tap_saprotroph (Põlme et al. 2020); Decay sugar-rich substrates (Põlme et al. 2020) | Põlme S, et al. 2020. Fungal Diversity 105:1-16 ((https://doi.org/10.1007/s13225-020-00466-2)) |
| Otu15680 | 1 | k__Fungi(100);p__Ascomycota(100);c__Saccharomycetes(100);o__Saccharomycetales(100);f__Metschnikowiaceae(100);g__Metschnikowia(100);g__Metschnikowia_unclassified(100); | Metschnikowia | 13 | Saprotroph | \|Undefined Saprotroph\| | Yeast | NULL | Probable | As nectar-tap_saprotroph (Põlme et al. 2020); Decay sugar-rich substrates (Põlme et al. 2020) | Põlme S, et al. 2020. Fungal Diversity 105:1-16 ((https://doi.org/10.1007/s13225-020-00466-2)) |
| Otu15681 | 1 | k__Fungi(100);p__Ascomycota(100);c__Saccharomycetes(100);o__Saccharomycetales(100);f__Metschnikowiaceae(100);g__Metschnikowia(100);s__Metschnikowia_pulcherrima(100); | Metschnikowia | 13 | Saprotroph | \|Undefined Saprotroph\| | Yeast | NULL | Probable | As nectar-tap_saprotroph (Põlme et al. 2020); Decay sugar-rich substrates (Põlme et al. 2020) | Põlme S, et al. 2020. Fungal Diversity 105:1-16 ((https://doi.org/10.1007/s13225-020-00466-2)) |
| Otu15682 | 1 | k__Fungi(100);p__Ascomycota(100);c__Saccharomycetes(100);o__Saccharomycetales(100);f__Metschnikowiaceae(100);g__Metschnikowia(100);g__Metschnikowia_unclassified(100); | Metschnikowia | 13 | Saprotroph | \|Undefined Saprotroph\| | Yeast | NULL | Probable | As nectar-tap_saprotroph (Põlme et al. 2020); Decay sugar-rich substrates (Põlme et al. 2020) | Põlme S, et al. 2020. Fungal Diversity 105:1-16 ((https://doi.org/10.1007/s13225-020-00466-2)) |
| Otu15683 | 1 | k__Fungi(100);p__Ascomycota(100);c__Saccharomycetes(100);o__Saccharomycetales(100);f__Metschnikowiaceae(100);g__Metschnikowia(100);s__Metschnikowia_pulcherrima(100); | Metschnikowia | 13 | Saprotroph | \|Undefined Saprotroph\| | Yeast | NULL | Probable | As nectar-tap_saprotroph (Põlme et al. 2020); Decay sugar-rich substrates (Põlme et al. 2020) | Põlme S, et al. 2020. Fungal Diversity 105:1-16 ((https://doi.org/10.1007/s13225-020-00466-2)) |
| Otu15686 | 1 | k__Fungi(100);p__Ascomycota(100);c__Saccharomycetes(100);o__Saccharomycetales(100);f__Metschnikowiaceae(100);g__Metschnikowia(100);g__Metschnikowia_unclassified(100); | Metschnikowia | 13 | Saprotroph | \|Undefined Saprotroph\| | Yeast | NULL | Probable | As nectar-tap_saprotroph (Põlme et al. 2020); Decay sugar-rich substrates (Põlme et al. 2020) | Põlme S, et al. 2020. Fungal Diversity 105:1-16 ((https://doi.org/10.1007/s13225-020-00466-2)) |
| Otu15687 | 1 | k__Fungi(100);p__Ascomycota(100);c__Saccharomycetes(100);o__Saccharomycetales(100);f__Metschnikowiaceae(100);g__Metschnikowia(100);s__Metschnikowia_pulcherrima(100); | Metschnikowia | 13 | Saprotroph | \|Undefined Saprotroph\| | Yeast | NULL | Probable | As nectar-tap_saprotroph (Põlme et al. 2020); Decay sugar-rich substrates (Põlme et al. 2020) | Põlme S, et al. 2020. Fungal Diversity 105:1-16 ((https://doi.org/10.1007/s13225-020-00466-2)) |
| Otu15691 | 1 | k__Fungi(100);p__Ascomycota(100);c__Saccharomycetes(100);o__Saccharomycetales(100);f__Metschnikowiaceae(100);g__Metschnikowia(100);s__Metschnikowia_pulcherrima(100); | Metschnikowia | 13 | Saprotroph | \|Undefined Saprotroph\| | Yeast | NULL | Probable | As nectar-tap_saprotroph (Põlme et al. 2020); Decay sugar-rich substrates (Põlme et al. 2020) | Põlme S, et al. 2020. Fungal Diversity 105:1-16 ((https://doi.org/10.1007/s13225-020-00466-2)) |
| Otu15692 | 1 | k__Fungi(100);p__Ascomycota(100);c__Saccharomycetes(100);o__Saccharomycetales(100);f__Metschnikowiaceae(100);g__Metschnikowia(100);s__Metschnikowia_pulcherrima(100); | Metschnikowia | 13 | Saprotroph | \|Undefined Saprotroph\| | Yeast | NULL | Probable | As nectar-tap_saprotroph (Põlme et al. 2020); Decay sugar-rich substrates (Põlme et al. 2020) | Põlme S, et al. 2020. Fungal Diversity 105:1-16 ((https://doi.org/10.1007/s13225-020-00466-2)) |
| Otu15693 | 1 | k__Fungi(100);p__Ascomycota(100);c__Saccharomycetes(100);o__Saccharomycetales(100);f__Metschnikowiaceae(100);g__Metschnikowia(100);s__Metschnikowia_pulcherrima(100); | Metschnikowia | 13 | Saprotroph | \|Undefined Saprotroph\| | Yeast | NULL | Probable | As nectar-tap_saprotroph (Põlme et al. 2020); Decay sugar-rich substrates (Põlme et al. 2020) | Põlme S, et al. 2020. Fungal Diversity 105:1-16 ((https://doi.org/10.1007/s13225-020-00466-2)) |
| Otu15694 | 1 | k__Fungi(100);p__Ascomycota(100);c__Saccharomycetes(100);o__Saccharomycetales(100);f__Metschnikowiaceae(100);g__Metschnikowia(100);g__Metschnikowia_unclassified(100); | Metschnikowia | 13 | Saprotroph | \|Undefined Saprotroph\| | Yeast | NULL | Probable | As nectar-tap_saprotroph (Põlme et al. 2020); Decay sugar-rich substrates (Põlme et al. 2020) | Põlme S, et al. 2020. Fungal Diversity 105:1-16 ((https://doi.org/10.1007/s13225-020-00466-2)) |
| Otu15695 | 1 | k__Fungi(100);p__Ascomycota(100);c__Saccharomycetes(100);o__Saccharomycetales(100);f__Metschnikowiaceae(100);g__Metschnikowia(100);g__Metschnikowia_unclassified(100); | Metschnikowia | 13 | Saprotroph | \|Undefined Saprotroph\| | Yeast | NULL | Probable | As nectar-tap_saprotroph (Põlme et al. 2020); Decay sugar-rich substrates (Põlme et al. 2020) | Põlme S, et al. 2020. Fungal Diversity 105:1-16 ((https://doi.org/10.1007/s13225-020-00466-2)) |
| Otu15697 | 1 | k__Fungi(100);p__Ascomycota(100);c__Saccharomycetes(100);o__Saccharomycetales(100);f__Metschnikowiaceae(100);g__Metschnikowia(100);s__Metschnikowia_pulcherrima(100); | Metschnikowia | 13 | Saprotroph | \|Undefined Saprotroph\| | Yeast | NULL | Probable | As nectar-tap_saprotroph (Põlme et al. 2020); Decay sugar-rich substrates (Põlme et al. 2020) | Põlme S, et al. 2020. Fungal Diversity 105:1-16 ((https://doi.org/10.1007/s13225-020-00466-2)) |
| Otu15698 | 1 | k__Fungi(100);p__Ascomycota(100);c__Saccharomycetes(100);o__Saccharomycetales(100);f__Saccharomycetales_family_Incertae_sedis(100);g__Candida(100);g__Candida_unclassified(100); | Candida | 13 | Pathotroph-Saprotroph-Symbiotroph | Animal Pathogen-Endophyte-Endosymbiont-Epiphyte-\|Undefined Saprotroph\| | Dimorphic Yeast-Yeast | NULL | Probable | As nectar-tap_saprotroph (Põlme et al. 2020); Decay sugar-rich substrates (Põlme et al. 2020); This asexual genus is polyphyletic and contain species that function very differently depend on what they are. Caution should be taken when interpreting guilds for Candida species. | Manolakaki D, et al. 2010. Virulence. 1: 367-375 ((https://doi.org/10.4161/viru.1.5.12796)); Mekbib SB, et al. 2011. Tropical Plant Pathology 36:233-240 ((https://doi.org/10.1590/S1982-56762011000400004)); Põlme S, et al. 2020. Fungal Diversity 105:1-16 ((https://doi.org/10.1007/s13225-020-00466-2)) |
| Otu15699 | 1 | k__Fungi(100);p__Ascomycota(100);c__Saccharomycetes(100);o__Saccharomycetales(100);f__Metschnikowiaceae(100);g__Metschnikowia(100);s__Metschnikowia_pulcherrima(100); | Metschnikowia | 13 | Saprotroph | \|Undefined Saprotroph\| | Yeast | NULL | Probable | As nectar-tap_saprotroph (Põlme et al. 2020); Decay sugar-rich substrates (Põlme et al. 2020) | Põlme S, et al. 2020. Fungal Diversity 105:1-16 ((https://doi.org/10.1007/s13225-020-00466-2)) |
| Otu15700 | 1 | k__Fungi(100);p__Ascomycota(100);c__Saccharomycetes(100);o__Saccharomycetales(100);f__Metschnikowiaceae(100);g__Metschnikowia(100);s__Metschnikowia_pulcherrima(100); | Metschnikowia | 13 | Saprotroph | \|Undefined Saprotroph\| | Yeast | NULL | Probable | As nectar-tap_saprotroph (Põlme et al. 2020); Decay sugar-rich substrates (Põlme et al. 2020) | Põlme S, et al. 2020. Fungal Diversity 105:1-16 ((https://doi.org/10.1007/s13225-020-00466-2)) |
| Otu15701 | 1 | k__Fungi(100);p__Ascomycota(100);c__Saccharomycetes(100);o__Saccharomycetales(100);f__Metschnikowiaceae(100);g__Metschnikowia(100);s__Metschnikowia_pulcherrima(100); | Metschnikowia | 13 | Saprotroph | \|Undefined Saprotroph\| | Yeast | NULL | Probable | As nectar-tap_saprotroph (Põlme et al. 2020); Decay sugar-rich substrates (Põlme et al. 2020) | Põlme S, et al. 2020. Fungal Diversity 105:1-16 ((https://doi.org/10.1007/s13225-020-00466-2)) |
| Otu15702 | 1 | k__Fungi(100);p__Ascomycota(100);c__Saccharomycetes(100);o__Saccharomycetales(100);f__Metschnikowiaceae(100);g__Metschnikowia(100);s__Metschnikowia_pulcherrima(100); | Metschnikowia | 13 | Saprotroph | \|Undefined Saprotroph\| | Yeast | NULL | Probable | As nectar-tap_saprotroph (Põlme et al. 2020); Decay sugar-rich substrates (Põlme et al. 2020) | Põlme S, et al. 2020. Fungal Diversity 105:1-16 ((https://doi.org/10.1007/s13225-020-00466-2)) |
| Otu15703 | 1 | k__Fungi(100);p__Ascomycota(100);c__Saccharomycetes(100);o__Saccharomycetales(100);f__Metschnikowiaceae(100);g__Metschnikowia(100);s__Metschnikowia_pulcherrima(100); | Metschnikowia | 13 | Saprotroph | \|Undefined Saprotroph\| | Yeast | NULL | Probable | As nectar-tap_saprotroph (Põlme et al. 2020); Decay sugar-rich substrates (Põlme et al. 2020) | Põlme S, et al. 2020. Fungal Diversity 105:1-16 ((https://doi.org/10.1007/s13225-020-00466-2)) |
| Otu15706 | 1 | k__Fungi(100);p__Ascomycota(100);c__Saccharomycetes(100);o__Saccharomycetales(100);f__Metschnikowiaceae(100);g__Metschnikowia(100);s__Metschnikowia_pulcherrima(100); | Metschnikowia | 13 | Saprotroph | \|Undefined Saprotroph\| | Yeast | NULL | Probable | As nectar-tap_saprotroph (Põlme et al. 2020); Decay sugar-rich substrates (Põlme et al. 2020) | Põlme S, et al. 2020. Fungal Diversity 105:1-16 ((https://doi.org/10.1007/s13225-020-00466-2)) |
| Otu15707 | 1 | k__Fungi(100);p__Ascomycota(100);c__Saccharomycetes(100);o__Saccharomycetales(100);f__Metschnikowiaceae(100);g__Metschnikowia(100);s__Metschnikowia_pulcherrima(100); | Metschnikowia | 13 | Saprotroph | \|Undefined Saprotroph\| | Yeast | NULL | Probable | As nectar-tap_saprotroph (Põlme et al. 2020); Decay sugar-rich substrates (Põlme et al. 2020) | Põlme S, et al. 2020. Fungal Diversity 105:1-16 ((https://doi.org/10.1007/s13225-020-00466-2)) |
| Otu15708 | 1 | k__Fungi(100);p__Ascomycota(100);c__Saccharomycetes(100);o__Saccharomycetales(100);f__Metschnikowiaceae(100);g__Metschnikowia(100);s__Metschnikowia_pulcherrima(100); | Metschnikowia | 13 | Saprotroph | \|Undefined Saprotroph\| | Yeast | NULL | Probable | As nectar-tap_saprotroph (Põlme et al. 2020); Decay sugar-rich substrates (Põlme et al. 2020) | Põlme S, et al. 2020. Fungal Diversity 105:1-16 ((https://doi.org/10.1007/s13225-020-00466-2)) |
| Otu15709 | 1 | k__Fungi(100);p__Ascomycota(100);c__Saccharomycetes(100);o__Saccharomycetales(100);f__Metschnikowiaceae(100);g__Metschnikowia(100);s__Metschnikowia_pulcherrima(100); | Metschnikowia | 13 | Saprotroph | \|Undefined Saprotroph\| | Yeast | NULL | Probable | As nectar-tap_saprotroph (Põlme et al. 2020); Decay sugar-rich substrates (Põlme et al. 2020) | Põlme S, et al. 2020. Fungal Diversity 105:1-16 ((https://doi.org/10.1007/s13225-020-00466-2)) |
| Otu15711 | 1 | k__Fungi(100);p__Ascomycota(100);c__Saccharomycetes(100);o__Saccharomycetales(100);f__Metschnikowiaceae(100);g__Metschnikowia(100);s__Metschnikowia_pulcherrima(100); | Metschnikowia | 13 | Saprotroph | \|Undefined Saprotroph\| | Yeast | NULL | Probable | As nectar-tap_saprotroph (Põlme et al. 2020); Decay sugar-rich substrates (Põlme et al. 2020) | Põlme S, et al. 2020. Fungal Diversity 105:1-16 ((https://doi.org/10.1007/s13225-020-00466-2)) |
| Otu15713 | 1 | k__Fungi(100);p__Ascomycota(100);c__Saccharomycetes(100);o__Saccharomycetales(100);f__Metschnikowiaceae(100);g__Metschnikowia(100);s__Metschnikowia_pulcherrima(100); | Metschnikowia | 13 | Saprotroph | \|Undefined Saprotroph\| | Yeast | NULL | Probable | As nectar-tap_saprotroph (Põlme et al. 2020); Decay sugar-rich substrates (Põlme et al. 2020) | Põlme S, et al. 2020. Fungal Diversity 105:1-16 ((https://doi.org/10.1007/s13225-020-00466-2)) |
| Otu15716 | 1 | k__Fungi(100);p__Ascomycota(100);c__Saccharomycetes(100);o__Saccharomycetales(100);f__Metschnikowiaceae(100);g__Metschnikowia(100);s__Metschnikowia_pulcherrima(100); | Metschnikowia | 13 | Saprotroph | \|Undefined Saprotroph\| | Yeast | NULL | Probable | As nectar-tap_saprotroph (Põlme et al. 2020); Decay sugar-rich substrates (Põlme et al. 2020) | Põlme S, et al. 2020. Fungal Diversity 105:1-16 ((https://doi.org/10.1007/s13225-020-00466-2)) |
| Otu15719 | 1 | k__Fungi(100);p__Ascomycota(100);c__Saccharomycetes(100);o__Saccharomycetales(100);f__Metschnikowiaceae(100);g__Metschnikowia(100);g__Metschnikowia_unclassified(100); | Metschnikowia | 13 | Saprotroph | \|Undefined Saprotroph\| | Yeast | NULL | Probable | As nectar-tap_saprotroph (Põlme et al. 2020); Decay sugar-rich substrates (Põlme et al. 2020) | Põlme S, et al. 2020. Fungal Diversity 105:1-16 ((https://doi.org/10.1007/s13225-020-00466-2)) |
| Otu15721 | 1 | k__Fungi(100);p__Ascomycota(100);c__Saccharomycetes(100);o__Saccharomycetales(100);f__Saccharomycetales_family_Incertae_sedis(100);g__Candida(100);g__Candida_unclassified(100); | Candida | 13 | Pathotroph-Saprotroph-Symbiotroph | Animal Pathogen-Endophyte-Endosymbiont-Epiphyte-\|Undefined Saprotroph\| | Dimorphic Yeast-Yeast | NULL | Probable | As nectar-tap_saprotroph (Põlme et al. 2020); Decay sugar-rich substrates (Põlme et al. 2020); This asexual genus is polyphyletic and contain species that function very differently depend on what they are. Caution should be taken when interpreting guilds for Candida species. | Manolakaki D, et al. 2010. Virulence. 1: 367-375 ((https://doi.org/10.4161/viru.1.5.12796)); Mekbib SB, et al. 2011. Tropical Plant Pathology 36:233-240 ((https://doi.org/10.1590/S1982-56762011000400004)); Põlme S, et al. 2020. Fungal Diversity 105:1-16 ((https://doi.org/10.1007/s13225-020-00466-2)) |
| Otu15722 | 1 | k__Fungi(100);p__Ascomycota(100);c__Saccharomycetes(100);o__Saccharomycetales(100);o__Saccharomycetales_unclassified(100);o__Saccharomycetales_unclassified(100);o__Saccharomycetales_unclassified(100); | Saccharomycetales | 7 | Saprotroph | Undefined Saprotroph | Yeast | NULL | Possible | NULL | Sterkenburg E, et al. 2015. New Phytologist 207:1145-1158 ((https://doi.org/10.1111/nph.13426)) |
| Otu15723 | 1 | k__Fungi(100);p__Ascomycota(100);c__Saccharomycetes(100);o__Saccharomycetales(100);f__Metschnikowiaceae(100);g__Metschnikowia(100);s__Metschnikowia_pulcherrima(100); | Metschnikowia | 13 | Saprotroph | \|Undefined Saprotroph\| | Yeast | NULL | Probable | As nectar-tap_saprotroph (Põlme et al. 2020); Decay sugar-rich substrates (Põlme et al. 2020) | Põlme S, et al. 2020. Fungal Diversity 105:1-16 ((https://doi.org/10.1007/s13225-020-00466-2)) |
| Otu15726 | 1 | k__Fungi(100);p__Ascomycota(100);c__Saccharomycetes(100);o__Saccharomycetales(100);f__Metschnikowiaceae(100);g__Metschnikowia(100);s__Metschnikowia_pulcherrima(100); | Metschnikowia | 13 | Saprotroph | \|Undefined Saprotroph\| | Yeast | NULL | Probable | As nectar-tap_saprotroph (Põlme et al. 2020); Decay sugar-rich substrates (Põlme et al. 2020) | Põlme S, et al. 2020. Fungal Diversity 105:1-16 ((https://doi.org/10.1007/s13225-020-00466-2)) |
| Otu15728 | 1 | k__Fungi(100);p__Ascomycota(100);c__Saccharomycetes(100);o__Saccharomycetales(100);f__Metschnikowiaceae(100);g__Metschnikowia(100);g__Metschnikowia_unclassified(100); | Metschnikowia | 13 | Saprotroph | \|Undefined Saprotroph\| | Yeast | NULL | Probable | As nectar-tap_saprotroph (Põlme et al. 2020); Decay sugar-rich substrates (Põlme et al. 2020) | Põlme S, et al. 2020. Fungal Diversity 105:1-16 ((https://doi.org/10.1007/s13225-020-00466-2)) |
| Otu15730 | 1 | k__Fungi(100);p__Ascomycota(100);c__Saccharomycetes(100);o__Saccharomycetales(100);f__Metschnikowiaceae(100);g__Metschnikowia(100);g__Metschnikowia_unclassified(100); | Metschnikowia | 13 | Saprotroph | \|Undefined Saprotroph\| | Yeast | NULL | Probable | As nectar-tap_saprotroph (Põlme et al. 2020); Decay sugar-rich substrates (Põlme et al. 2020) | Põlme S, et al. 2020. Fungal Diversity 105:1-16 ((https://doi.org/10.1007/s13225-020-00466-2)) |
| Otu15731 | 1 | k__Fungi(100);p__Ascomycota(100);c__Saccharomycetes(100);o__Saccharomycetales(100);f__Metschnikowiaceae(100);g__Metschnikowia(100);s__Metschnikowia_pulcherrima(100); | Metschnikowia | 13 | Saprotroph | \|Undefined Saprotroph\| | Yeast | NULL | Probable | As nectar-tap_saprotroph (Põlme et al. 2020); Decay sugar-rich substrates (Põlme et al. 2020) | Põlme S, et al. 2020. Fungal Diversity 105:1-16 ((https://doi.org/10.1007/s13225-020-00466-2)) |
| Otu15732 | 1 | k__Fungi(100);p__Ascomycota(100);c__Saccharomycetes(100);o__Saccharomycetales(100);f__Metschnikowiaceae(100);g__Metschnikowia(100);g__Metschnikowia_unclassified(100); | Metschnikowia | 13 | Saprotroph | \|Undefined Saprotroph\| | Yeast | NULL | Probable | As nectar-tap_saprotroph (Põlme et al. 2020); Decay sugar-rich substrates (Põlme et al. 2020) | Põlme S, et al. 2020. Fungal Diversity 105:1-16 ((https://doi.org/10.1007/s13225-020-00466-2)) |
| Otu15733 | 1 | k__Fungi(100);p__Ascomycota(100);c__Saccharomycetes(100);o__Saccharomycetales(100);f__Metschnikowiaceae(100);g__Metschnikowia(100);s__Metschnikowia_pulcherrima(100); | Metschnikowia | 13 | Saprotroph | \|Undefined Saprotroph\| | Yeast | NULL | Probable | As nectar-tap_saprotroph (Põlme et al. 2020); Decay sugar-rich substrates (Põlme et al. 2020) | Põlme S, et al. 2020. Fungal Diversity 105:1-16 ((https://doi.org/10.1007/s13225-020-00466-2)) |
| Otu15734 | 1 | k__Fungi(100);p__Ascomycota(100);c__Saccharomycetes(100);o__Saccharomycetales(100);f__Metschnikowiaceae(100);g__Metschnikowia(100);s__Metschnikowia_pulcherrima(100); | Metschnikowia | 13 | Saprotroph | \|Undefined Saprotroph\| | Yeast | NULL | Probable | As nectar-tap_saprotroph (Põlme et al. 2020); Decay sugar-rich substrates (Põlme et al. 2020) | Põlme S, et al. 2020. Fungal Diversity 105:1-16 ((https://doi.org/10.1007/s13225-020-00466-2)) |
| Otu15735 | 1 | k__Fungi(100);p__Ascomycota(100);c__Saccharomycetes(100);o__Saccharomycetales(100);f__Metschnikowiaceae(100);g__Metschnikowia(100);s__Metschnikowia_pulcherrima(100); | Metschnikowia | 13 | Saprotroph | \|Undefined Saprotroph\| | Yeast | NULL | Probable | As nectar-tap_saprotroph (Põlme et al. 2020); Decay sugar-rich substrates (Põlme et al. 2020) | Põlme S, et al. 2020. Fungal Diversity 105:1-16 ((https://doi.org/10.1007/s13225-020-00466-2)) |
| Otu15736 | 1 | k__Fungi(100);p__Ascomycota(100);c__Saccharomycetes(100);o__Saccharomycetales(100);f__Metschnikowiaceae(100);g__Metschnikowia(100);g__Metschnikowia_unclassified(100); | Metschnikowia | 13 | Saprotroph | \|Undefined Saprotroph\| | Yeast | NULL | Probable | As nectar-tap_saprotroph (Põlme et al. 2020); Decay sugar-rich substrates (Põlme et al. 2020) | Põlme S, et al. 2020. Fungal Diversity 105:1-16 ((https://doi.org/10.1007/s13225-020-00466-2)) |
| Otu15737 | 1 | k__Fungi(100);p__Ascomycota(100);c__Saccharomycetes(100);o__Saccharomycetales(100);f__Metschnikowiaceae(100);g__Metschnikowia(100);s__Metschnikowia_pulcherrima(100); | Metschnikowia | 13 | Saprotroph | \|Undefined Saprotroph\| | Yeast | NULL | Probable | As nectar-tap_saprotroph (Põlme et al. 2020); Decay sugar-rich substrates (Põlme et al. 2020) | Põlme S, et al. 2020. Fungal Diversity 105:1-16 ((https://doi.org/10.1007/s13225-020-00466-2)) |
| Otu15738 | 1 | k__Fungi(100);p__Ascomycota(100);c__Saccharomycetes(100);o__Saccharomycetales(100);f__Metschnikowiaceae(100);g__Metschnikowia(100);g__Metschnikowia_unclassified(100); | Metschnikowia | 13 | Saprotroph | \|Undefined Saprotroph\| | Yeast | NULL | Probable | As nectar-tap_saprotroph (Põlme et al. 2020); Decay sugar-rich substrates (Põlme et al. 2020) | Põlme S, et al. 2020. Fungal Diversity 105:1-16 ((https://doi.org/10.1007/s13225-020-00466-2)) |
| Otu15739 | 1 | k__Fungi(100);p__Ascomycota(100);c__Saccharomycetes(100);o__Saccharomycetales(100);f__Metschnikowiaceae(100);g__Metschnikowia(100);s__Metschnikowia_pulcherrima(100); | Metschnikowia | 13 | Saprotroph | \|Undefined Saprotroph\| | Yeast | NULL | Probable | As nectar-tap_saprotroph (Põlme et al. 2020); Decay sugar-rich substrates (Põlme et al. 2020) | Põlme S, et al. 2020. Fungal Diversity 105:1-16 ((https://doi.org/10.1007/s13225-020-00466-2)) |
| Otu15740 | 1 | k__Fungi(100);p__Ascomycota(100);c__Saccharomycetes(100);o__Saccharomycetales(100);f__Metschnikowiaceae(100);g__Metschnikowia(100);s__Metschnikowia_pulcherrima(100); | Metschnikowia | 13 | Saprotroph | \|Undefined Saprotroph\| | Yeast | NULL | Probable | As nectar-tap_saprotroph (Põlme et al. 2020); Decay sugar-rich substrates (Põlme et al. 2020) | Põlme S, et al. 2020. Fungal Diversity 105:1-16 ((https://doi.org/10.1007/s13225-020-00466-2)) |
| Otu15741 | 1 | k__Fungi(100);p__Ascomycota(100);c__Saccharomycetes(100);o__Saccharomycetales(100);f__Metschnikowiaceae(100);g__Metschnikowia(100);g__Metschnikowia_unclassified(100); | Metschnikowia | 13 | Saprotroph | \|Undefined Saprotroph\| | Yeast | NULL | Probable | As nectar-tap_saprotroph (Põlme et al. 2020); Decay sugar-rich substrates (Põlme et al. 2020) | Põlme S, et al. 2020. Fungal Diversity 105:1-16 ((https://doi.org/10.1007/s13225-020-00466-2)) |
| Otu15743 | 1 | k__Fungi(100);p__Ascomycota(100);c__Saccharomycetes(100);o__Saccharomycetales(100);f__Metschnikowiaceae(100);g__Metschnikowia(100);s__Metschnikowia_pulcherrima(100); | Metschnikowia | 13 | Saprotroph | \|Undefined Saprotroph\| | Yeast | NULL | Probable | As nectar-tap_saprotroph (Põlme et al. 2020); Decay sugar-rich substrates (Põlme et al. 2020) | Põlme S, et al. 2020. Fungal Diversity 105:1-16 ((https://doi.org/10.1007/s13225-020-00466-2)) |
| Otu15745 | 1 | k__Fungi(100);p__Ascomycota(100);c__Saccharomycetes(100);o__Saccharomycetales(100);f__Metschnikowiaceae(100);g__Metschnikowia(100);s__Metschnikowia_pulcherrima(100); | Metschnikowia | 13 | Saprotroph | \|Undefined Saprotroph\| | Yeast | NULL | Probable | As nectar-tap_saprotroph (Põlme et al. 2020); Decay sugar-rich substrates (Põlme et al. 2020) | Põlme S, et al. 2020. Fungal Diversity 105:1-16 ((https://doi.org/10.1007/s13225-020-00466-2)) |
| Otu15747 | 1 | k__Fungi(100);p__Ascomycota(100);c__Saccharomycetes(100);o__Saccharomycetales(100);f__Metschnikowiaceae(100);g__Metschnikowia(100);g__Metschnikowia_unclassified(100); | Metschnikowia | 13 | Saprotroph | \|Undefined Saprotroph\| | Yeast | NULL | Probable | As nectar-tap_saprotroph (Põlme et al. 2020); Decay sugar-rich substrates (Põlme et al. 2020) | Põlme S, et al. 2020. Fungal Diversity 105:1-16 ((https://doi.org/10.1007/s13225-020-00466-2)) |
| Otu15748 | 1 | k__Fungi(100);p__Ascomycota(100);c__Saccharomycetes(100);o__Saccharomycetales(100);f__Metschnikowiaceae(100);g__Metschnikowia(100);g__Metschnikowia_unclassified(100); | Metschnikowia | 13 | Saprotroph | \|Undefined Saprotroph\| | Yeast | NULL | Probable | As nectar-tap_saprotroph (Põlme et al. 2020); Decay sugar-rich substrates (Põlme et al. 2020) | Põlme S, et al. 2020. Fungal Diversity 105:1-16 ((https://doi.org/10.1007/s13225-020-00466-2)) |
| Otu15749 | 1 | k__Fungi(100);p__Ascomycota(100);c__Saccharomycetes(100);o__Saccharomycetales(100);f__Metschnikowiaceae(100);g__Metschnikowia(100);s__Metschnikowia_pulcherrima(100); | Metschnikowia | 13 | Saprotroph | \|Undefined Saprotroph\| | Yeast | NULL | Probable | As nectar-tap_saprotroph (Põlme et al. 2020); Decay sugar-rich substrates (Põlme et al. 2020) | Põlme S, et al. 2020. Fungal Diversity 105:1-16 ((https://doi.org/10.1007/s13225-020-00466-2)) |
| Otu15750 | 1 | k__Fungi(100);p__Ascomycota(100);c__Saccharomycetes(100);o__Saccharomycetales(100);f__Metschnikowiaceae(100);g__Metschnikowia(100);s__Metschnikowia_pulcherrima(100); | Metschnikowia | 13 | Saprotroph | \|Undefined Saprotroph\| | Yeast | NULL | Probable | As nectar-tap_saprotroph (Põlme et al. 2020); Decay sugar-rich substrates (Põlme et al. 2020) | Põlme S, et al. 2020. Fungal Diversity 105:1-16 ((https://doi.org/10.1007/s13225-020-00466-2)) |
| Otu15751 | 1 | k__Fungi(100);p__Ascomycota(100);c__Saccharomycetes(100);o__Saccharomycetales(100);f__Metschnikowiaceae(100);g__Metschnikowia(100);g__Metschnikowia_unclassified(100); | Metschnikowia | 13 | Saprotroph | \|Undefined Saprotroph\| | Yeast | NULL | Probable | As nectar-tap_saprotroph (Põlme et al. 2020); Decay sugar-rich substrates (Põlme et al. 2020) | Põlme S, et al. 2020. Fungal Diversity 105:1-16 ((https://doi.org/10.1007/s13225-020-00466-2)) |
| Otu15753 | 1 | k__Fungi(100);p__Ascomycota(100);c__Saccharomycetes(100);o__Saccharomycetales(100);f__Metschnikowiaceae(100);g__Metschnikowia(100);s__Metschnikowia_pulcherrima(100); | Metschnikowia | 13 | Saprotroph | \|Undefined Saprotroph\| | Yeast | NULL | Probable | As nectar-tap_saprotroph (Põlme et al. 2020); Decay sugar-rich substrates (Põlme et al. 2020) | Põlme S, et al. 2020. Fungal Diversity 105:1-16 ((https://doi.org/10.1007/s13225-020-00466-2)) |
| Otu15754 | 1 | k__Fungi(100);p__Ascomycota(100);c__Saccharomycetes(100);o__Saccharomycetales(100);f__Metschnikowiaceae(100);g__Metschnikowia(100);s__Metschnikowia_pulcherrima(100); | Metschnikowia | 13 | Saprotroph | \|Undefined Saprotroph\| | Yeast | NULL | Probable | As nectar-tap_saprotroph (Põlme et al. 2020); Decay sugar-rich substrates (Põlme et al. 2020) | Põlme S, et al. 2020. Fungal Diversity 105:1-16 ((https://doi.org/10.1007/s13225-020-00466-2)) |
| Otu15755 | 1 | k__Fungi(100);p__Ascomycota(100);c__Saccharomycetes(100);o__Saccharomycetales(100);f__Metschnikowiaceae(100);g__Metschnikowia(100);s__Metschnikowia_pulcherrima(100); | Metschnikowia | 13 | Saprotroph | \|Undefined Saprotroph\| | Yeast | NULL | Probable | As nectar-tap_saprotroph (Põlme et al. 2020); Decay sugar-rich substrates (Põlme et al. 2020) | Põlme S, et al. 2020. Fungal Diversity 105:1-16 ((https://doi.org/10.1007/s13225-020-00466-2)) |
| Otu15756 | 1 | k__Fungi(100);p__Ascomycota(100);c__Saccharomycetes(100);o__Saccharomycetales(100);f__Metschnikowiaceae(100);g__Metschnikowia(100);s__Metschnikowia_pulcherrima(100); | Metschnikowia | 13 | Saprotroph | \|Undefined Saprotroph\| | Yeast | NULL | Probable | As nectar-tap_saprotroph (Põlme et al. 2020); Decay sugar-rich substrates (Põlme et al. 2020) | Põlme S, et al. 2020. Fungal Diversity 105:1-16 ((https://doi.org/10.1007/s13225-020-00466-2)) |
| Otu15757 | 1 | k__Fungi(100);p__Ascomycota(100);c__Saccharomycetes(100);o__Saccharomycetales(100);f__Metschnikowiaceae(100);g__Metschnikowia(100);g__Metschnikowia_unclassified(100); | Metschnikowia | 13 | Saprotroph | \|Undefined Saprotroph\| | Yeast | NULL | Probable | As nectar-tap_saprotroph (Põlme et al. 2020); Decay sugar-rich substrates (Põlme et al. 2020) | Põlme S, et al. 2020. Fungal Diversity 105:1-16 ((https://doi.org/10.1007/s13225-020-00466-2)) |
| Otu15758 | 1 | k__Fungi(100);p__Ascomycota(100);c__Saccharomycetes(100);o__Saccharomycetales(100);f__Metschnikowiaceae(100);g__Metschnikowia(100);s__Metschnikowia_pulcherrima(100); | Metschnikowia | 13 | Saprotroph | \|Undefined Saprotroph\| | Yeast | NULL | Probable | As nectar-tap_saprotroph (Põlme et al. 2020); Decay sugar-rich substrates (Põlme et al. 2020) | Põlme S, et al. 2020. Fungal Diversity 105:1-16 ((https://doi.org/10.1007/s13225-020-00466-2)) |
| Otu15760 | 1 | k__Fungi(100);p__Ascomycota(100);c__Saccharomycetes(100);o__Saccharomycetales(100);f__Metschnikowiaceae(100);g__Metschnikowia(100);g__Metschnikowia_unclassified(100); | Metschnikowia | 13 | Saprotroph | \|Undefined Saprotroph\| | Yeast | NULL | Probable | As nectar-tap_saprotroph (Põlme et al. 2020); Decay sugar-rich substrates (Põlme et al. 2020) | Põlme S, et al. 2020. Fungal Diversity 105:1-16 ((https://doi.org/10.1007/s13225-020-00466-2)) |
| Otu15761 | 1 | k__Fungi(100);p__Ascomycota(100);c__Saccharomycetes(100);o__Saccharomycetales(100);f__Metschnikowiaceae(100);g__Metschnikowia(100);s__Metschnikowia_pulcherrima(100); | Metschnikowia | 13 | Saprotroph | \|Undefined Saprotroph\| | Yeast | NULL | Probable | As nectar-tap_saprotroph (Põlme et al. 2020); Decay sugar-rich substrates (Põlme et al. 2020) | Põlme S, et al. 2020. Fungal Diversity 105:1-16 ((https://doi.org/10.1007/s13225-020-00466-2)) |
| Otu15763 | 1 | k__Fungi(100);p__Ascomycota(100);c__Saccharomycetes(100);o__Saccharomycetales(100);f__Metschnikowiaceae(100);g__Metschnikowia(100);s__Metschnikowia_pulcherrima(100); | Metschnikowia | 13 | Saprotroph | \|Undefined Saprotroph\| | Yeast | NULL | Probable | As nectar-tap_saprotroph (Põlme et al. 2020); Decay sugar-rich substrates (Põlme et al. 2020) | Põlme S, et al. 2020. Fungal Diversity 105:1-16 ((https://doi.org/10.1007/s13225-020-00466-2)) |
| Otu15764 | 1 | k__Fungi(100);p__Ascomycota(100);c__Saccharomycetes(100);o__Saccharomycetales(100);f__Metschnikowiaceae(100);g__Metschnikowia(100);s__Metschnikowia_pulcherrima(100); | Metschnikowia | 13 | Saprotroph | \|Undefined Saprotroph\| | Yeast | NULL | Probable | As nectar-tap_saprotroph (Põlme et al. 2020); Decay sugar-rich substrates (Põlme et al. 2020) | Põlme S, et al. 2020. Fungal Diversity 105:1-16 ((https://doi.org/10.1007/s13225-020-00466-2)) |
| Otu15765 | 1 | k__Fungi(100);p__Ascomycota(100);c__Saccharomycetes(100);o__Saccharomycetales(100);f__Metschnikowiaceae(100);g__Metschnikowia(100);s__Metschnikowia_pulcherrima(100); | Metschnikowia | 13 | Saprotroph | \|Undefined Saprotroph\| | Yeast | NULL | Probable | As nectar-tap_saprotroph (Põlme et al. 2020); Decay sugar-rich substrates (Põlme et al. 2020) | Põlme S, et al. 2020. Fungal Diversity 105:1-16 ((https://doi.org/10.1007/s13225-020-00466-2)) |
| Otu15766 | 1 | k__Fungi(100);p__Ascomycota(100);c__Saccharomycetes(100);o__Saccharomycetales(100);f__Metschnikowiaceae(100);g__Metschnikowia(100);s__Metschnikowia_pulcherrima(100); | Metschnikowia | 13 | Saprotroph | \|Undefined Saprotroph\| | Yeast | NULL | Probable | As nectar-tap_saprotroph (Põlme et al. 2020); Decay sugar-rich substrates (Põlme et al. 2020) | Põlme S, et al. 2020. Fungal Diversity 105:1-16 ((https://doi.org/10.1007/s13225-020-00466-2)) |
| Otu15768 | 1 | k__Fungi(100);p__Ascomycota(100);c__Saccharomycetes(100);o__Saccharomycetales(100);f__Metschnikowiaceae(100);g__Metschnikowia(100);s__Metschnikowia_pulcherrima(100); | Metschnikowia | 13 | Saprotroph | \|Undefined Saprotroph\| | Yeast | NULL | Probable | As nectar-tap_saprotroph (Põlme et al. 2020); Decay sugar-rich substrates (Põlme et al. 2020) | Põlme S, et al. 2020. Fungal Diversity 105:1-16 ((https://doi.org/10.1007/s13225-020-00466-2)) |
| Otu15770 | 1 | k__Fungi(100);p__Ascomycota(100);c__Saccharomycetes(100);o__Saccharomycetales(100);f__Metschnikowiaceae(100);g__Metschnikowia(100);g__Metschnikowia_unclassified(100); | Metschnikowia | 13 | Saprotroph | \|Undefined Saprotroph\| | Yeast | NULL | Probable | As nectar-tap_saprotroph (Põlme et al. 2020); Decay sugar-rich substrates (Põlme et al. 2020) | Põlme S, et al. 2020. Fungal Diversity 105:1-16 ((https://doi.org/10.1007/s13225-020-00466-2)) |
| Otu15771 | 1 | k__Fungi(100);p__Ascomycota(100);c__Saccharomycetes(100);o__Saccharomycetales(100);f__Metschnikowiaceae(100);g__Metschnikowia(100);s__Metschnikowia_pulcherrima(100); | Metschnikowia | 13 | Saprotroph | \|Undefined Saprotroph\| | Yeast | NULL | Probable | As nectar-tap_saprotroph (Põlme et al. 2020); Decay sugar-rich substrates (Põlme et al. 2020) | Põlme S, et al. 2020. Fungal Diversity 105:1-16 ((https://doi.org/10.1007/s13225-020-00466-2)) |
| Otu15772 | 1 | k__Fungi(100);p__Ascomycota(100);c__Saccharomycetes(100);o__Saccharomycetales(100);f__Metschnikowiaceae(100);g__Metschnikowia(100);s__Metschnikowia_pulcherrima(100); | Metschnikowia | 13 | Saprotroph | \|Undefined Saprotroph\| | Yeast | NULL | Probable | As nectar-tap_saprotroph (Põlme et al. 2020); Decay sugar-rich substrates (Põlme et al. 2020) | Põlme S, et al. 2020. Fungal Diversity 105:1-16 ((https://doi.org/10.1007/s13225-020-00466-2)) |
| Otu15773 | 1 | k__Fungi(100);p__Ascomycota(100);c__Saccharomycetes(100);o__Saccharomycetales(100);f__Metschnikowiaceae(100);g__Metschnikowia(100);s__Metschnikowia_pulcherrima(100); | Metschnikowia | 13 | Saprotroph | \|Undefined Saprotroph\| | Yeast | NULL | Probable | As nectar-tap_saprotroph (Põlme et al. 2020); Decay sugar-rich substrates (Põlme et al. 2020) | Põlme S, et al. 2020. Fungal Diversity 105:1-16 ((https://doi.org/10.1007/s13225-020-00466-2)) |
| Otu15774 | 1 | k__Fungi(100);p__Ascomycota(100);c__Saccharomycetes(100);o__Saccharomycetales(100);f__Metschnikowiaceae(100);g__Metschnikowia(100);s__Metschnikowia_pulcherrima(100); | Metschnikowia | 13 | Saprotroph | \|Undefined Saprotroph\| | Yeast | NULL | Probable | As nectar-tap_saprotroph (Põlme et al. 2020); Decay sugar-rich substrates (Põlme et al. 2020) | Põlme S, et al. 2020. Fungal Diversity 105:1-16 ((https://doi.org/10.1007/s13225-020-00466-2)) |
| Otu15775 | 1 | k__Fungi(100);p__Ascomycota(100);c__Saccharomycetes(100);o__Saccharomycetales(100);f__Metschnikowiaceae(100);g__Metschnikowia(100);s__Metschnikowia_pulcherrima(100); | Metschnikowia | 13 | Saprotroph | \|Undefined Saprotroph\| | Yeast | NULL | Probable | As nectar-tap_saprotroph (Põlme et al. 2020); Decay sugar-rich substrates (Põlme et al. 2020) | Põlme S, et al. 2020. Fungal Diversity 105:1-16 ((https://doi.org/10.1007/s13225-020-00466-2)) |
| Otu15776 | 1 | k__Fungi(100);p__Ascomycota(100);c__Saccharomycetes(100);o__Saccharomycetales(100);f__Metschnikowiaceae(100);g__Metschnikowia(100);g__Metschnikowia_unclassified(100); | Metschnikowia | 13 | Saprotroph | \|Undefined Saprotroph\| | Yeast | NULL | Probable | As nectar-tap_saprotroph (Põlme et al. 2020); Decay sugar-rich substrates (Põlme et al. 2020) | Põlme S, et al. 2020. Fungal Diversity 105:1-16 ((https://doi.org/10.1007/s13225-020-00466-2)) |
| Otu15777 | 1 | k__Fungi(100);p__Ascomycota(100);c__Saccharomycetes(100);o__Saccharomycetales(100);f__Metschnikowiaceae(100);g__Metschnikowia(100);s__Metschnikowia_pulcherrima(100); | Metschnikowia | 13 | Saprotroph | \|Undefined Saprotroph\| | Yeast | NULL | Probable | As nectar-tap_saprotroph (Põlme et al. 2020); Decay sugar-rich substrates (Põlme et al. 2020) | Põlme S, et al. 2020. Fungal Diversity 105:1-16 ((https://doi.org/10.1007/s13225-020-00466-2)) |
| Otu15778 | 1 | k__Fungi(100);p__Ascomycota(100);c__Saccharomycetes(100);o__Saccharomycetales(100);f__Metschnikowiaceae(100);g__Metschnikowia(100);g__Metschnikowia_unclassified(100); | Metschnikowia | 13 | Saprotroph | \|Undefined Saprotroph\| | Yeast | NULL | Probable | As nectar-tap_saprotroph (Põlme et al. 2020); Decay sugar-rich substrates (Põlme et al. 2020) | Põlme S, et al. 2020. Fungal Diversity 105:1-16 ((https://doi.org/10.1007/s13225-020-00466-2)) |
| Otu15779 | 1 | k__Fungi(100);p__Ascomycota(100);c__Saccharomycetes(100);o__Saccharomycetales(100);f__Metschnikowiaceae(100);g__Metschnikowia(100);s__Metschnikowia_pulcherrima(100); | Metschnikowia | 13 | Saprotroph | \|Undefined Saprotroph\| | Yeast | NULL | Probable | As nectar-tap_saprotroph (Põlme et al. 2020); Decay sugar-rich substrates (Põlme et al. 2020) | Põlme S, et al. 2020. Fungal Diversity 105:1-16 ((https://doi.org/10.1007/s13225-020-00466-2)) |
| Otu15780 | 1 | k__Fungi(100);p__Ascomycota(100);c__Saccharomycetes(100);o__Saccharomycetales(100);f__Metschnikowiaceae(100);g__Metschnikowia(100);g__Metschnikowia_unclassified(100); | Metschnikowia | 13 | Saprotroph | \|Undefined Saprotroph\| | Yeast | NULL | Probable | As nectar-tap_saprotroph (Põlme et al. 2020); Decay sugar-rich substrates (Põlme et al. 2020) | Põlme S, et al. 2020. Fungal Diversity 105:1-16 ((https://doi.org/10.1007/s13225-020-00466-2)) |
| Otu15781 | 1 | k__Fungi(100);p__Ascomycota(100);c__Saccharomycetes(100);o__Saccharomycetales(100);f__Metschnikowiaceae(100);g__Metschnikowia(100);s__Metschnikowia_pulcherrima(100); | Metschnikowia | 13 | Saprotroph | \|Undefined Saprotroph\| | Yeast | NULL | Probable | As nectar-tap_saprotroph (Põlme et al. 2020); Decay sugar-rich substrates (Põlme et al. 2020) | Põlme S, et al. 2020. Fungal Diversity 105:1-16 ((https://doi.org/10.1007/s13225-020-00466-2)) |
| Otu15782 | 1 | k__Fungi(100);p__Ascomycota(100);c__Saccharomycetes(100);o__Saccharomycetales(100);f__Metschnikowiaceae(100);g__Metschnikowia(100);s__Metschnikowia_pulcherrima(100); | Metschnikowia | 13 | Saprotroph | \|Undefined Saprotroph\| | Yeast | NULL | Probable | As nectar-tap_saprotroph (Põlme et al. 2020); Decay sugar-rich substrates (Põlme et al. 2020) | Põlme S, et al. 2020. Fungal Diversity 105:1-16 ((https://doi.org/10.1007/s13225-020-00466-2)) |
| Otu15783 | 1 | k__Fungi(100);p__Ascomycota(100);c__Saccharomycetes(100);o__Saccharomycetales(100);f__Metschnikowiaceae(100);g__Metschnikowia(100);g__Metschnikowia_unclassified(100); | Metschnikowia | 13 | Saprotroph | \|Undefined Saprotroph\| | Yeast | NULL | Probable | As nectar-tap_saprotroph (Põlme et al. 2020); Decay sugar-rich substrates (Põlme et al. 2020) | Põlme S, et al. 2020. Fungal Diversity 105:1-16 ((https://doi.org/10.1007/s13225-020-00466-2)) |
| Otu15784 | 1 | k__Fungi(100);p__Ascomycota(100);c__Saccharomycetes(100);o__Saccharomycetales(100);f__Metschnikowiaceae(100);g__Metschnikowia(100);s__Metschnikowia_pulcherrima(100); | Metschnikowia | 13 | Saprotroph | \|Undefined Saprotroph\| | Yeast | NULL | Probable | As nectar-tap_saprotroph (Põlme et al. 2020); Decay sugar-rich substrates (Põlme et al. 2020) | Põlme S, et al. 2020. Fungal Diversity 105:1-16 ((https://doi.org/10.1007/s13225-020-00466-2)) |
| Otu15785 | 1 | k__Fungi(100);p__Ascomycota(100);c__Saccharomycetes(100);o__Saccharomycetales(100);f__Metschnikowiaceae(100);g__Metschnikowia(100);s__Metschnikowia_pulcherrima(100); | Metschnikowia | 13 | Saprotroph | \|Undefined Saprotroph\| | Yeast | NULL | Probable | As nectar-tap_saprotroph (Põlme et al. 2020); Decay sugar-rich substrates (Põlme et al. 2020) | Põlme S, et al. 2020. Fungal Diversity 105:1-16 ((https://doi.org/10.1007/s13225-020-00466-2)) |
| Otu15786 | 1 | k__Fungi(100);p__Ascomycota(100);c__Saccharomycetes(100);o__Saccharomycetales(100);f__Metschnikowiaceae(100);g__Metschnikowia(100);s__Metschnikowia_pulcherrima(100); | Metschnikowia | 13 | Saprotroph | \|Undefined Saprotroph\| | Yeast | NULL | Probable | As nectar-tap_saprotroph (Põlme et al. 2020); Decay sugar-rich substrates (Põlme et al. 2020) | Põlme S, et al. 2020. Fungal Diversity 105:1-16 ((https://doi.org/10.1007/s13225-020-00466-2)) |
| Otu15788 | 1 | k__Fungi(100);p__Ascomycota(100);c__Saccharomycetes(100);o__Saccharomycetales(100);f__Metschnikowiaceae(100);g__Metschnikowia(100);s__Metschnikowia_pulcherrima(100); | Metschnikowia | 13 | Saprotroph | \|Undefined Saprotroph\| | Yeast | NULL | Probable | As nectar-tap_saprotroph (Põlme et al. 2020); Decay sugar-rich substrates (Põlme et al. 2020) | Põlme S, et al. 2020. Fungal Diversity 105:1-16 ((https://doi.org/10.1007/s13225-020-00466-2)) |
| Otu15789 | 1 | k__Fungi(100);p__Ascomycota(100);c__Saccharomycetes(100);o__Saccharomycetales(100);f__Metschnikowiaceae(100);g__Metschnikowia(100);s__Metschnikowia_pulcherrima(100); | Metschnikowia | 13 | Saprotroph | \|Undefined Saprotroph\| | Yeast | NULL | Probable | As nectar-tap_saprotroph (Põlme et al. 2020); Decay sugar-rich substrates (Põlme et al. 2020) | Põlme S, et al. 2020. Fungal Diversity 105:1-16 ((https://doi.org/10.1007/s13225-020-00466-2)) |
| Otu15790 | 1 | k__Fungi(100);p__Ascomycota(100);c__Saccharomycetes(100);o__Saccharomycetales(100);f__Metschnikowiaceae(100);g__Metschnikowia(100);s__Metschnikowia_pulcherrima(100); | Metschnikowia | 13 | Saprotroph | \|Undefined Saprotroph\| | Yeast | NULL | Probable | As nectar-tap_saprotroph (Põlme et al. 2020); Decay sugar-rich substrates (Põlme et al. 2020) | Põlme S, et al. 2020. Fungal Diversity 105:1-16 ((https://doi.org/10.1007/s13225-020-00466-2)) |
| Otu15791 | 1 | k__Fungi(100);p__Ascomycota(100);c__Saccharomycetes(100);o__Saccharomycetales(100);f__Metschnikowiaceae(100);g__Metschnikowia(100);s__Metschnikowia_pulcherrima(100); | Metschnikowia | 13 | Saprotroph | \|Undefined Saprotroph\| | Yeast | NULL | Probable | As nectar-tap_saprotroph (Põlme et al. 2020); Decay sugar-rich substrates (Põlme et al. 2020) | Põlme S, et al. 2020. Fungal Diversity 105:1-16 ((https://doi.org/10.1007/s13225-020-00466-2)) |
| Otu15792 | 1 | k__Fungi(100);p__Ascomycota(100);c__Saccharomycetes(100);o__Saccharomycetales(100);f__Metschnikowiaceae(100);g__Metschnikowia(100);s__Metschnikowia_pulcherrima(100); | Metschnikowia | 13 | Saprotroph | \|Undefined Saprotroph\| | Yeast | NULL | Probable | As nectar-tap_saprotroph (Põlme et al. 2020); Decay sugar-rich substrates (Põlme et al. 2020) | Põlme S, et al. 2020. Fungal Diversity 105:1-16 ((https://doi.org/10.1007/s13225-020-00466-2)) |
| Otu15793 | 1 | k__Fungi(100);p__Ascomycota(100);c__Saccharomycetes(100);o__Saccharomycetales(100);f__Metschnikowiaceae(100);g__Metschnikowia(100);s__Metschnikowia_pulcherrima(100); | Metschnikowia | 13 | Saprotroph | \|Undefined Saprotroph\| | Yeast | NULL | Probable | As nectar-tap_saprotroph (Põlme et al. 2020); Decay sugar-rich substrates (Põlme et al. 2020) | Põlme S, et al. 2020. Fungal Diversity 105:1-16 ((https://doi.org/10.1007/s13225-020-00466-2)) |
| Otu15794 | 1 | k__Fungi(100);p__Ascomycota(100);c__Saccharomycetes(100);o__Saccharomycetales(100);f__Metschnikowiaceae(100);g__Metschnikowia(100);s__Metschnikowia_pulcherrima(100); | Metschnikowia | 13 | Saprotroph | \|Undefined Saprotroph\| | Yeast | NULL | Probable | As nectar-tap_saprotroph (Põlme et al. 2020); Decay sugar-rich substrates (Põlme et al. 2020) | Põlme S, et al. 2020. Fungal Diversity 105:1-16 ((https://doi.org/10.1007/s13225-020-00466-2)) |
| Otu15795 | 1 | k__Fungi(100);p__Ascomycota(100);c__Saccharomycetes(100);o__Saccharomycetales(100);f__Metschnikowiaceae(100);g__Metschnikowia(100);s__Metschnikowia_pulcherrima(100); | Metschnikowia | 13 | Saprotroph | \|Undefined Saprotroph\| | Yeast | NULL | Probable | As nectar-tap_saprotroph (Põlme et al. 2020); Decay sugar-rich substrates (Põlme et al. 2020) | Põlme S, et al. 2020. Fungal Diversity 105:1-16 ((https://doi.org/10.1007/s13225-020-00466-2)) |
| Otu15796 | 1 | k__Fungi(100);p__Ascomycota(100);c__Saccharomycetes(100);o__Saccharomycetales(100);f__Metschnikowiaceae(100);g__Metschnikowia(100);s__Metschnikowia_pulcherrima(100); | Metschnikowia | 13 | Saprotroph | \|Undefined Saprotroph\| | Yeast | NULL | Probable | As nectar-tap_saprotroph (Põlme et al. 2020); Decay sugar-rich substrates (Põlme et al. 2020) | Põlme S, et al. 2020. Fungal Diversity 105:1-16 ((https://doi.org/10.1007/s13225-020-00466-2)) |
| Otu15797 | 1 | k__Fungi(100);p__Ascomycota(100);c__Saccharomycetes(100);o__Saccharomycetales(100);f__Metschnikowiaceae(100);g__Metschnikowia(100);s__Metschnikowia_pulcherrima(100); | Metschnikowia | 13 | Saprotroph | \|Undefined Saprotroph\| | Yeast | NULL | Probable | As nectar-tap_saprotroph (Põlme et al. 2020); Decay sugar-rich substrates (Põlme et al. 2020) | Põlme S, et al. 2020. Fungal Diversity 105:1-16 ((https://doi.org/10.1007/s13225-020-00466-2)) |
| Otu15799 | 1 | k__Fungi(100);p__Ascomycota(100);c__Saccharomycetes(100);o__Saccharomycetales(100);f__Metschnikowiaceae(100);g__Metschnikowia(100);g__Metschnikowia_unclassified(100); | Metschnikowia | 13 | Saprotroph | \|Undefined Saprotroph\| | Yeast | NULL | Probable | As nectar-tap_saprotroph (Põlme et al. 2020); Decay sugar-rich substrates (Põlme et al. 2020) | Põlme S, et al. 2020. Fungal Diversity 105:1-16 ((https://doi.org/10.1007/s13225-020-00466-2)) |
| Otu15801 | 1 | k__Fungi(100);p__Ascomycota(100);c__Saccharomycetes(100);o__Saccharomycetales(100);f__Metschnikowiaceae(100);g__Metschnikowia(100);s__Metschnikowia_pulcherrima(100); | Metschnikowia | 13 | Saprotroph | \|Undefined Saprotroph\| | Yeast | NULL | Probable | As nectar-tap_saprotroph (Põlme et al. 2020); Decay sugar-rich substrates (Põlme et al. 2020) | Põlme S, et al. 2020. Fungal Diversity 105:1-16 ((https://doi.org/10.1007/s13225-020-00466-2)) |
| Otu15802 | 1 | k__Fungi(100);p__Ascomycota(100);c__Saccharomycetes(100);o__Saccharomycetales(100);f__Metschnikowiaceae(100);g__Metschnikowia(100);s__Metschnikowia_pulcherrima(100); | Metschnikowia | 13 | Saprotroph | \|Undefined Saprotroph\| | Yeast | NULL | Probable | As nectar-tap_saprotroph (Põlme et al. 2020); Decay sugar-rich substrates (Põlme et al. 2020) | Põlme S, et al. 2020. Fungal Diversity 105:1-16 ((https://doi.org/10.1007/s13225-020-00466-2)) |
| Otu15803 | 1 | k__Fungi(100);p__Ascomycota(100);c__Saccharomycetes(100);o__Saccharomycetales(100);f__Metschnikowiaceae(100);g__Metschnikowia(100);s__Metschnikowia_pulcherrima(100); | Metschnikowia | 13 | Saprotroph | \|Undefined Saprotroph\| | Yeast | NULL | Probable | As nectar-tap_saprotroph (Põlme et al. 2020); Decay sugar-rich substrates (Põlme et al. 2020) | Põlme S, et al. 2020. Fungal Diversity 105:1-16 ((https://doi.org/10.1007/s13225-020-00466-2)) |
| Otu15804 | 1 | k__Fungi(100);p__Ascomycota(100);c__Saccharomycetes(100);o__Saccharomycetales(100);f__Metschnikowiaceae(100);g__Metschnikowia(100);g__Metschnikowia_unclassified(100); | Metschnikowia | 13 | Saprotroph | \|Undefined Saprotroph\| | Yeast | NULL | Probable | As nectar-tap_saprotroph (Põlme et al. 2020); Decay sugar-rich substrates (Põlme et al. 2020) | Põlme S, et al. 2020. Fungal Diversity 105:1-16 ((https://doi.org/10.1007/s13225-020-00466-2)) |
| Otu15806 | 1 | k__Fungi(100);p__Ascomycota(100);c__Saccharomycetes(100);o__Saccharomycetales(100);f__Metschnikowiaceae(100);g__Metschnikowia(100);g__Metschnikowia_unclassified(100); | Metschnikowia | 13 | Saprotroph | \|Undefined Saprotroph\| | Yeast | NULL | Probable | As nectar-tap_saprotroph (Põlme et al. 2020); Decay sugar-rich substrates (Põlme et al. 2020) | Põlme S, et al. 2020. Fungal Diversity 105:1-16 ((https://doi.org/10.1007/s13225-020-00466-2)) |
| Otu15807 | 1 | k__Fungi(100);p__Ascomycota(100);c__Saccharomycetes(100);o__Saccharomycetales(100);f__Metschnikowiaceae(100);g__Metschnikowia(100);s__Metschnikowia_pulcherrima(100); | Metschnikowia | 13 | Saprotroph | \|Undefined Saprotroph\| | Yeast | NULL | Probable | As nectar-tap_saprotroph (Põlme et al. 2020); Decay sugar-rich substrates (Põlme et al. 2020) | Põlme S, et al. 2020. Fungal Diversity 105:1-16 ((https://doi.org/10.1007/s13225-020-00466-2)) |
| Otu15808 | 1 | k__Fungi(100);p__Ascomycota(100);c__Saccharomycetes(100);o__Saccharomycetales(100);f__Metschnikowiaceae(100);g__Metschnikowia(100);g__Metschnikowia_unclassified(100); | Metschnikowia | 13 | Saprotroph | \|Undefined Saprotroph\| | Yeast | NULL | Probable | As nectar-tap_saprotroph (Põlme et al. 2020); Decay sugar-rich substrates (Põlme et al. 2020) | Põlme S, et al. 2020. Fungal Diversity 105:1-16 ((https://doi.org/10.1007/s13225-020-00466-2)) |
| Otu15809 | 1 | k__Fungi(100);p__Ascomycota(100);c__Saccharomycetes(100);o__Saccharomycetales(100);f__Metschnikowiaceae(100);g__Metschnikowia(100);s__Metschnikowia_pulcherrima(100); | Metschnikowia | 13 | Saprotroph | \|Undefined Saprotroph\| | Yeast | NULL | Probable | As nectar-tap_saprotroph (Põlme et al. 2020); Decay sugar-rich substrates (Põlme et al. 2020) | Põlme S, et al. 2020. Fungal Diversity 105:1-16 ((https://doi.org/10.1007/s13225-020-00466-2)) |
| Otu15810 | 1 | k__Fungi(100);p__Ascomycota(100);c__Saccharomycetes(100);o__Saccharomycetales(100);f__Metschnikowiaceae(100);g__Metschnikowia(100);g__Metschnikowia_unclassified(100); | Metschnikowia | 13 | Saprotroph | \|Undefined Saprotroph\| | Yeast | NULL | Probable | As nectar-tap_saprotroph (Põlme et al. 2020); Decay sugar-rich substrates (Põlme et al. 2020) | Põlme S, et al. 2020. Fungal Diversity 105:1-16 ((https://doi.org/10.1007/s13225-020-00466-2)) |
| Otu15811 | 1 | k__Fungi(100);p__Ascomycota(100);c__Saccharomycetes(100);o__Saccharomycetales(100);f__Metschnikowiaceae(100);g__Metschnikowia(100);s__Metschnikowia_pulcherrima(100); | Metschnikowia | 13 | Saprotroph | \|Undefined Saprotroph\| | Yeast | NULL | Probable | As nectar-tap_saprotroph (Põlme et al. 2020); Decay sugar-rich substrates (Põlme et al. 2020) | Põlme S, et al. 2020. Fungal Diversity 105:1-16 ((https://doi.org/10.1007/s13225-020-00466-2)) |
| Otu15812 | 1 | k__Fungi(100);p__Ascomycota(100);c__Saccharomycetes(100);o__Saccharomycetales(100);f__Metschnikowiaceae(100);g__Metschnikowia(100);s__Metschnikowia_pulcherrima(100); | Metschnikowia | 13 | Saprotroph | \|Undefined Saprotroph\| | Yeast | NULL | Probable | As nectar-tap_saprotroph (Põlme et al. 2020); Decay sugar-rich substrates (Põlme et al. 2020) | Põlme S, et al. 2020. Fungal Diversity 105:1-16 ((https://doi.org/10.1007/s13225-020-00466-2)) |
| Otu15813 | 1 | k__Fungi(100);p__Ascomycota(100);c__Saccharomycetes(100);o__Saccharomycetales(100);f__Metschnikowiaceae(100);g__Metschnikowia(100);g__Metschnikowia_unclassified(100); | Metschnikowia | 13 | Saprotroph | \|Undefined Saprotroph\| | Yeast | NULL | Probable | As nectar-tap_saprotroph (Põlme et al. 2020); Decay sugar-rich substrates (Põlme et al. 2020) | Põlme S, et al. 2020. Fungal Diversity 105:1-16 ((https://doi.org/10.1007/s13225-020-00466-2)) |
| Otu15814 | 1 | k__Fungi(100);p__Ascomycota(100);c__Saccharomycetes(100);o__Saccharomycetales(100);f__Metschnikowiaceae(100);g__Metschnikowia(100);s__Metschnikowia_pulcherrima(100); | Metschnikowia | 13 | Saprotroph | \|Undefined Saprotroph\| | Yeast | NULL | Probable | As nectar-tap_saprotroph (Põlme et al. 2020); Decay sugar-rich substrates (Põlme et al. 2020) | Põlme S, et al. 2020. Fungal Diversity 105:1-16 ((https://doi.org/10.1007/s13225-020-00466-2)) |
| Otu15815 | 1 | k__Fungi(100);p__Ascomycota(100);c__Saccharomycetes(100);o__Saccharomycetales(100);f__Metschnikowiaceae(100);g__Metschnikowia(100);g__Metschnikowia_unclassified(100); | Metschnikowia | 13 | Saprotroph | \|Undefined Saprotroph\| | Yeast | NULL | Probable | As nectar-tap_saprotroph (Põlme et al. 2020); Decay sugar-rich substrates (Põlme et al. 2020) | Põlme S, et al. 2020. Fungal Diversity 105:1-16 ((https://doi.org/10.1007/s13225-020-00466-2)) |
| Otu15816 | 1 | k__Fungi(100);p__Ascomycota(100);c__Saccharomycetes(100);o__Saccharomycetales(100);f__Metschnikowiaceae(100);g__Metschnikowia(100);s__Metschnikowia_pulcherrima(100); | Metschnikowia | 13 | Saprotroph | \|Undefined Saprotroph\| | Yeast | NULL | Probable | As nectar-tap_saprotroph (Põlme et al. 2020); Decay sugar-rich substrates (Põlme et al. 2020) | Põlme S, et al. 2020. Fungal Diversity 105:1-16 ((https://doi.org/10.1007/s13225-020-00466-2)) |
| Otu15817 | 1 | k__Fungi(100);p__Ascomycota(100);c__Dothideomycetes(100);o__Pleosporales(100);f__Sporormiaceae(100);g__Sporormiella(100);s__Sporormiella_sp(100); | Sporormiella | 13 | Saprotroph | Dung Saprotroph-Endophyte-Lichen Parasite-\|Plant Saprotroph\| | Microfungus | NULL | Probable | As litter_saprotroph (Põlme et al. 2020); Decay leaf, fruit, seed (Põlme et al. 2020) | Bell A. 1983. Dung Fungi: An Illustrated Guide to Coprophilous Fungi in New Zealand. Victoria University Press, Wellington (ISBN: 978-0864730015); Costa IPMW, et al. 2012. Mycotaxon 119:e26 ((http://mycotaxon.com/resources/checklists/costa_v119_checklist.pdf)); Bills GF, et al. 2013. Current Opinion in Microbiology 16:549-565 ((https://doi.org/10.1016/j.mib.2013.08.001)); Lawrey JD, Diederich P. 2016. http://www.lichenicolous.net (accessed 24 May 2017) ((http://www.lichenicolous.net/)); Põlme S, et al. 2020. Fungal Diversity 105:1-16 ((https://doi.org/10.1007/s13225-020-00466-2)) |
| Otu15818 | 1 | k__Fungi(100);p__Ascomycota(100);c__Saccharomycetes(100);o__Saccharomycetales(100);f__Metschnikowiaceae(100);g__Metschnikowia(100);s__Metschnikowia_pulcherrima(100); | Metschnikowia | 13 | Saprotroph | \|Undefined Saprotroph\| | Yeast | NULL | Probable | As nectar-tap_saprotroph (Põlme et al. 2020); Decay sugar-rich substrates (Põlme et al. 2020) | Põlme S, et al. 2020. Fungal Diversity 105:1-16 ((https://doi.org/10.1007/s13225-020-00466-2)) |
| Otu15819 | 1 | k__Fungi(100);p__Ascomycota(100);c__Saccharomycetes(100);o__Saccharomycetales(100);f__Metschnikowiaceae(100);g__Metschnikowia(100);s__Metschnikowia_pulcherrima(100); | Metschnikowia | 13 | Saprotroph | \|Undefined Saprotroph\| | Yeast | NULL | Probable | As nectar-tap_saprotroph (Põlme et al. 2020); Decay sugar-rich substrates (Põlme et al. 2020) | Põlme S, et al. 2020. Fungal Diversity 105:1-16 ((https://doi.org/10.1007/s13225-020-00466-2)) |
| Otu15820 | 1 | k__Fungi(100);p__Ascomycota(100);c__Saccharomycetes(100);o__Saccharomycetales(100);f__Metschnikowiaceae(100);g__Metschnikowia(100);s__Metschnikowia_pulcherrima(100); | Metschnikowia | 13 | Saprotroph | \|Undefined Saprotroph\| | Yeast | NULL | Probable | As nectar-tap_saprotroph (Põlme et al. 2020); Decay sugar-rich substrates (Põlme et al. 2020) | Põlme S, et al. 2020. Fungal Diversity 105:1-16 ((https://doi.org/10.1007/s13225-020-00466-2)) |
| Otu15821 | 1 | k__Fungi(100);p__Ascomycota(100);c__Saccharomycetes(100);o__Saccharomycetales(100);f__Metschnikowiaceae(100);g__Metschnikowia(100);s__Metschnikowia_pulcherrima(100); | Metschnikowia | 13 | Saprotroph | \|Undefined Saprotroph\| | Yeast | NULL | Probable | As nectar-tap_saprotroph (Põlme et al. 2020); Decay sugar-rich substrates (Põlme et al. 2020) | Põlme S, et al. 2020. Fungal Diversity 105:1-16 ((https://doi.org/10.1007/s13225-020-00466-2)) |
| Otu15822 | 1 | k__Fungi(100);p__Ascomycota(100);c__Saccharomycetes(100);o__Saccharomycetales(100);f__Metschnikowiaceae(100);g__Metschnikowia(100);s__Metschnikowia_pulcherrima(100); | Metschnikowia | 13 | Saprotroph | \|Undefined Saprotroph\| | Yeast | NULL | Probable | As nectar-tap_saprotroph (Põlme et al. 2020); Decay sugar-rich substrates (Põlme et al. 2020) | Põlme S, et al. 2020. Fungal Diversity 105:1-16 ((https://doi.org/10.1007/s13225-020-00466-2)) |
| Otu15825 | 1 | k__Fungi(100);p__Ascomycota(100);c__Saccharomycetes(100);o__Saccharomycetales(100);f__Metschnikowiaceae(100);g__Metschnikowia(100);g__Metschnikowia_unclassified(100); | Metschnikowia | 13 | Saprotroph | \|Undefined Saprotroph\| | Yeast | NULL | Probable | As nectar-tap_saprotroph (Põlme et al. 2020); Decay sugar-rich substrates (Põlme et al. 2020) | Põlme S, et al. 2020. Fungal Diversity 105:1-16 ((https://doi.org/10.1007/s13225-020-00466-2)) |
| Otu15826 | 1 | k__Fungi(100);p__Ascomycota(100);c__Saccharomycetes(100);o__Saccharomycetales(100);f__Metschnikowiaceae(100);g__Metschnikowia(100);s__Metschnikowia_pulcherrima(100); | Metschnikowia | 13 | Saprotroph | \|Undefined Saprotroph\| | Yeast | NULL | Probable | As nectar-tap_saprotroph (Põlme et al. 2020); Decay sugar-rich substrates (Põlme et al. 2020) | Põlme S, et al. 2020. Fungal Diversity 105:1-16 ((https://doi.org/10.1007/s13225-020-00466-2)) |
| Otu15827 | 1 | k__Fungi(100);p__Ascomycota(100);c__Saccharomycetes(100);o__Saccharomycetales(100);f__Metschnikowiaceae(100);g__Metschnikowia(100);s__Metschnikowia_pulcherrima(100); | Metschnikowia | 13 | Saprotroph | \|Undefined Saprotroph\| | Yeast | NULL | Probable | As nectar-tap_saprotroph (Põlme et al. 2020); Decay sugar-rich substrates (Põlme et al. 2020) | Põlme S, et al. 2020. Fungal Diversity 105:1-16 ((https://doi.org/10.1007/s13225-020-00466-2)) |
| Otu15828 | 1 | k__Fungi(100);p__Ascomycota(100);c__Saccharomycetes(100);o__Saccharomycetales(100);f__Metschnikowiaceae(100);g__Metschnikowia(100);g__Metschnikowia_unclassified(100); | Metschnikowia | 13 | Saprotroph | \|Undefined Saprotroph\| | Yeast | NULL | Probable | As nectar-tap_saprotroph (Põlme et al. 2020); Decay sugar-rich substrates (Põlme et al. 2020) | Põlme S, et al. 2020. Fungal Diversity 105:1-16 ((https://doi.org/10.1007/s13225-020-00466-2)) |
| Otu15829 | 1 | k__Fungi(100);p__Ascomycota(100);c__Saccharomycetes(100);o__Saccharomycetales(100);f__Metschnikowiaceae(100);g__Metschnikowia(100);s__Metschnikowia_pulcherrima(100); | Metschnikowia | 13 | Saprotroph | \|Undefined Saprotroph\| | Yeast | NULL | Probable | As nectar-tap_saprotroph (Põlme et al. 2020); Decay sugar-rich substrates (Põlme et al. 2020) | Põlme S, et al. 2020. Fungal Diversity 105:1-16 ((https://doi.org/10.1007/s13225-020-00466-2)) |
| Otu15830 | 1 | k__Fungi(100);p__Ascomycota(100);c__Saccharomycetes(100);o__Saccharomycetales(100);f__Metschnikowiaceae(100);g__Metschnikowia(100);s__Metschnikowia_pulcherrima(100); | Metschnikowia | 13 | Saprotroph | \|Undefined Saprotroph\| | Yeast | NULL | Probable | As nectar-tap_saprotroph (Põlme et al. 2020); Decay sugar-rich substrates (Põlme et al. 2020) | Põlme S, et al. 2020. Fungal Diversity 105:1-16 ((https://doi.org/10.1007/s13225-020-00466-2)) |
| Otu15831 | 1 | k__Fungi(100);p__Ascomycota(100);c__Saccharomycetes(100);o__Saccharomycetales(100);f__Metschnikowiaceae(100);g__Metschnikowia(100);s__Metschnikowia_pulcherrima(100); | Metschnikowia | 13 | Saprotroph | \|Undefined Saprotroph\| | Yeast | NULL | Probable | As nectar-tap_saprotroph (Põlme et al. 2020); Decay sugar-rich substrates (Põlme et al. 2020) | Põlme S, et al. 2020. Fungal Diversity 105:1-16 ((https://doi.org/10.1007/s13225-020-00466-2)) |
| Otu15833 | 1 | k__Fungi(100);p__Ascomycota(100);c__Saccharomycetes(100);o__Saccharomycetales(100);f__Metschnikowiaceae(100);g__Metschnikowia(100);g__Metschnikowia_unclassified(100); | Metschnikowia | 13 | Saprotroph | \|Undefined Saprotroph\| | Yeast | NULL | Probable | As nectar-tap_saprotroph (Põlme et al. 2020); Decay sugar-rich substrates (Põlme et al. 2020) | Põlme S, et al. 2020. Fungal Diversity 105:1-16 ((https://doi.org/10.1007/s13225-020-00466-2)) |
| Otu15834 | 1 | k__Fungi(100);p__Ascomycota(100);c__Saccharomycetes(100);o__Saccharomycetales(100);f__Metschnikowiaceae(100);g__Metschnikowia(100);g__Metschnikowia_unclassified(100); | Metschnikowia | 13 | Saprotroph | \|Undefined Saprotroph\| | Yeast | NULL | Probable | As nectar-tap_saprotroph (Põlme et al. 2020); Decay sugar-rich substrates (Põlme et al. 2020) | Põlme S, et al. 2020. Fungal Diversity 105:1-16 ((https://doi.org/10.1007/s13225-020-00466-2)) |
| Otu15836 | 1 | k__Fungi(100);p__Ascomycota(100);c__Saccharomycetes(100);o__Saccharomycetales(100);f__Metschnikowiaceae(100);g__Metschnikowia(100);s__Metschnikowia_pulcherrima(100); | Metschnikowia | 13 | Saprotroph | \|Undefined Saprotroph\| | Yeast | NULL | Probable | As nectar-tap_saprotroph (Põlme et al. 2020); Decay sugar-rich substrates (Põlme et al. 2020) | Põlme S, et al. 2020. Fungal Diversity 105:1-16 ((https://doi.org/10.1007/s13225-020-00466-2)) |
| Otu15838 | 1 | k__Fungi(100);p__Ascomycota(100);c__Saccharomycetes(100);o__Saccharomycetales(100);f__Metschnikowiaceae(100);g__Metschnikowia(100);g__Metschnikowia_unclassified(100); | Metschnikowia | 13 | Saprotroph | \|Undefined Saprotroph\| | Yeast | NULL | Probable | As nectar-tap_saprotroph (Põlme et al. 2020); Decay sugar-rich substrates (Põlme et al. 2020) | Põlme S, et al. 2020. Fungal Diversity 105:1-16 ((https://doi.org/10.1007/s13225-020-00466-2)) |
| Otu15839 | 1 | k__Fungi(100);p__Ascomycota(100);c__Saccharomycetes(100);o__Saccharomycetales(100);f__Metschnikowiaceae(100);g__Metschnikowia(100);s__Metschnikowia_pulcherrima(100); | Metschnikowia | 13 | Saprotroph | \|Undefined Saprotroph\| | Yeast | NULL | Probable | As nectar-tap_saprotroph (Põlme et al. 2020); Decay sugar-rich substrates (Põlme et al. 2020) | Põlme S, et al. 2020. Fungal Diversity 105:1-16 ((https://doi.org/10.1007/s13225-020-00466-2)) |
| Otu15840 | 1 | k__Fungi(100);p__Ascomycota(100);c__Saccharomycetes(100);o__Saccharomycetales(100);f__Metschnikowiaceae(100);g__Metschnikowia(100);s__Metschnikowia_pulcherrima(100); | Metschnikowia | 13 | Saprotroph | \|Undefined Saprotroph\| | Yeast | NULL | Probable | As nectar-tap_saprotroph (Põlme et al. 2020); Decay sugar-rich substrates (Põlme et al. 2020) | Põlme S, et al. 2020. Fungal Diversity 105:1-16 ((https://doi.org/10.1007/s13225-020-00466-2)) |
| Otu15841 | 1 | k__Fungi(100);p__Ascomycota(100);c__Saccharomycetes(100);o__Saccharomycetales(100);f__Metschnikowiaceae(100);g__Metschnikowia(100);s__Metschnikowia_pulcherrima(100); | Metschnikowia | 13 | Saprotroph | \|Undefined Saprotroph\| | Yeast | NULL | Probable | As nectar-tap_saprotroph (Põlme et al. 2020); Decay sugar-rich substrates (Põlme et al. 2020) | Põlme S, et al. 2020. Fungal Diversity 105:1-16 ((https://doi.org/10.1007/s13225-020-00466-2)) |
| Otu15842 | 1 | k__Fungi(100);p__Ascomycota(100);c__Saccharomycetes(100);o__Saccharomycetales(100);f__Metschnikowiaceae(100);g__Metschnikowia(100);g__Metschnikowia_unclassified(100); | Metschnikowia | 13 | Saprotroph | \|Undefined Saprotroph\| | Yeast | NULL | Probable | As nectar-tap_saprotroph (Põlme et al. 2020); Decay sugar-rich substrates (Põlme et al. 2020) | Põlme S, et al. 2020. Fungal Diversity 105:1-16 ((https://doi.org/10.1007/s13225-020-00466-2)) |
| Otu15843 | 1 | k__Fungi(100);p__Ascomycota(100);c__Saccharomycetes(100);o__Saccharomycetales(100);f__Metschnikowiaceae(100);g__Metschnikowia(100);s__Metschnikowia_pulcherrima(100); | Metschnikowia | 13 | Saprotroph | \|Undefined Saprotroph\| | Yeast | NULL | Probable | As nectar-tap_saprotroph (Põlme et al. 2020); Decay sugar-rich substrates (Põlme et al. 2020) | Põlme S, et al. 2020. Fungal Diversity 105:1-16 ((https://doi.org/10.1007/s13225-020-00466-2)) |
| Otu15844 | 1 | k__Fungi(100);p__Ascomycota(100);c__Saccharomycetes(100);o__Saccharomycetales(100);f__Metschnikowiaceae(100);g__Metschnikowia(100);s__Metschnikowia_pulcherrima(100); | Metschnikowia | 13 | Saprotroph | \|Undefined Saprotroph\| | Yeast | NULL | Probable | As nectar-tap_saprotroph (Põlme et al. 2020); Decay sugar-rich substrates (Põlme et al. 2020) | Põlme S, et al. 2020. Fungal Diversity 105:1-16 ((https://doi.org/10.1007/s13225-020-00466-2)) |
| Otu15845 | 1 | k__Fungi(100);p__Ascomycota(100);c__Saccharomycetes(100);o__Saccharomycetales(100);f__Metschnikowiaceae(100);g__Metschnikowia(100);g__Metschnikowia_unclassified(100); | Metschnikowia | 13 | Saprotroph | \|Undefined Saprotroph\| | Yeast | NULL | Probable | As nectar-tap_saprotroph (Põlme et al. 2020); Decay sugar-rich substrates (Põlme et al. 2020) | Põlme S, et al. 2020. Fungal Diversity 105:1-16 ((https://doi.org/10.1007/s13225-020-00466-2)) |
| Otu15846 | 1 | k__Fungi(100);p__Ascomycota(100);c__Saccharomycetes(100);o__Saccharomycetales(100);f__Metschnikowiaceae(100);g__Metschnikowia(100);s__Metschnikowia_pulcherrima(100); | Metschnikowia | 13 | Saprotroph | \|Undefined Saprotroph\| | Yeast | NULL | Probable | As nectar-tap_saprotroph (Põlme et al. 2020); Decay sugar-rich substrates (Põlme et al. 2020) | Põlme S, et al. 2020. Fungal Diversity 105:1-16 ((https://doi.org/10.1007/s13225-020-00466-2)) |
| Otu15848 | 1 | k__Fungi(100);p__Ascomycota(100);c__Saccharomycetes(100);o__Saccharomycetales(100);f__Metschnikowiaceae(100);g__Metschnikowia(100);s__Metschnikowia_pulcherrima(100); | Metschnikowia | 13 | Saprotroph | \|Undefined Saprotroph\| | Yeast | NULL | Probable | As nectar-tap_saprotroph (Põlme et al. 2020); Decay sugar-rich substrates (Põlme et al. 2020) | Põlme S, et al. 2020. Fungal Diversity 105:1-16 ((https://doi.org/10.1007/s13225-020-00466-2)) |
| Otu15849 | 1 | k__Fungi(100);p__Ascomycota(100);c__Saccharomycetes(100);o__Saccharomycetales(100);f__Metschnikowiaceae(100);g__Metschnikowia(100);g__Metschnikowia_unclassified(100); | Metschnikowia | 13 | Saprotroph | \|Undefined Saprotroph\| | Yeast | NULL | Probable | As nectar-tap_saprotroph (Põlme et al. 2020); Decay sugar-rich substrates (Põlme et al. 2020) | Põlme S, et al. 2020. Fungal Diversity 105:1-16 ((https://doi.org/10.1007/s13225-020-00466-2)) |
| Otu15850 | 1 | k__Fungi(100);p__Ascomycota(100);c__Saccharomycetes(100);o__Saccharomycetales(100);f__Metschnikowiaceae(100);g__Metschnikowia(100);s__Metschnikowia_pulcherrima(100); | Metschnikowia | 13 | Saprotroph | \|Undefined Saprotroph\| | Yeast | NULL | Probable | As nectar-tap_saprotroph (Põlme et al. 2020); Decay sugar-rich substrates (Põlme et al. 2020) | Põlme S, et al. 2020. Fungal Diversity 105:1-16 ((https://doi.org/10.1007/s13225-020-00466-2)) |
| Otu15851 | 1 | k__Fungi(100);p__Ascomycota(100);c__Saccharomycetes(100);o__Saccharomycetales(100);f__Metschnikowiaceae(100);g__Metschnikowia(100);s__Metschnikowia_pulcherrima(100); | Metschnikowia | 13 | Saprotroph | \|Undefined Saprotroph\| | Yeast | NULL | Probable | As nectar-tap_saprotroph (Põlme et al. 2020); Decay sugar-rich substrates (Põlme et al. 2020) | Põlme S, et al. 2020. Fungal Diversity 105:1-16 ((https://doi.org/10.1007/s13225-020-00466-2)) |
| Otu15852 | 1 | k__Fungi(100);p__Ascomycota(100);c__Saccharomycetes(100);o__Saccharomycetales(100);f__Metschnikowiaceae(100);g__Metschnikowia(100);s__Metschnikowia_pulcherrima(100); | Metschnikowia | 13 | Saprotroph | \|Undefined Saprotroph\| | Yeast | NULL | Probable | As nectar-tap_saprotroph (Põlme et al. 2020); Decay sugar-rich substrates (Põlme et al. 2020) | Põlme S, et al. 2020. Fungal Diversity 105:1-16 ((https://doi.org/10.1007/s13225-020-00466-2)) |
| Otu15853 | 1 | k__Fungi(100);p__Ascomycota(100);c__Saccharomycetes(100);o__Saccharomycetales(100);f__Metschnikowiaceae(100);g__Metschnikowia(100);g__Metschnikowia_unclassified(100); | Metschnikowia | 13 | Saprotroph | \|Undefined Saprotroph\| | Yeast | NULL | Probable | As nectar-tap_saprotroph (Põlme et al. 2020); Decay sugar-rich substrates (Põlme et al. 2020) | Põlme S, et al. 2020. Fungal Diversity 105:1-16 ((https://doi.org/10.1007/s13225-020-00466-2)) |
| Otu15854 | 1 | k__Fungi(100);p__Ascomycota(100);c__Saccharomycetes(100);o__Saccharomycetales(100);f__Metschnikowiaceae(100);g__Metschnikowia(100);s__Metschnikowia_pulcherrima(100); | Metschnikowia | 13 | Saprotroph | \|Undefined Saprotroph\| | Yeast | NULL | Probable | As nectar-tap_saprotroph (Põlme et al. 2020); Decay sugar-rich substrates (Põlme et al. 2020) | Põlme S, et al. 2020. Fungal Diversity 105:1-16 ((https://doi.org/10.1007/s13225-020-00466-2)) |
| Otu15855 | 1 | k__Fungi(100);p__Ascomycota(100);c__Saccharomycetes(100);o__Saccharomycetales(100);f__Metschnikowiaceae(100);g__Metschnikowia(100);s__Metschnikowia_pulcherrima(100); | Metschnikowia | 13 | Saprotroph | \|Undefined Saprotroph\| | Yeast | NULL | Probable | As nectar-tap_saprotroph (Põlme et al. 2020); Decay sugar-rich substrates (Põlme et al. 2020) | Põlme S, et al. 2020. Fungal Diversity 105:1-16 ((https://doi.org/10.1007/s13225-020-00466-2)) |
| Otu15856 | 1 | k__Fungi(100);p__Ascomycota(100);c__Saccharomycetes(100);o__Saccharomycetales(100);f__Metschnikowiaceae(100);g__Metschnikowia(100);s__Metschnikowia_pulcherrima(100); | Metschnikowia | 13 | Saprotroph | \|Undefined Saprotroph\| | Yeast | NULL | Probable | As nectar-tap_saprotroph (Põlme et al. 2020); Decay sugar-rich substrates (Põlme et al. 2020) | Põlme S, et al. 2020. Fungal Diversity 105:1-16 ((https://doi.org/10.1007/s13225-020-00466-2)) |
| Otu15857 | 1 | k__Fungi(100);p__Ascomycota(100);c__Saccharomycetes(100);o__Saccharomycetales(100);f__Metschnikowiaceae(100);g__Metschnikowia(100);s__Metschnikowia_pulcherrima(100); | Metschnikowia | 13 | Saprotroph | \|Undefined Saprotroph\| | Yeast | NULL | Probable | As nectar-tap_saprotroph (Põlme et al. 2020); Decay sugar-rich substrates (Põlme et al. 2020) | Põlme S, et al. 2020. Fungal Diversity 105:1-16 ((https://doi.org/10.1007/s13225-020-00466-2)) |
| Otu15858 | 1 | k__Fungi(100);p__Ascomycota(100);c__Saccharomycetes(100);o__Saccharomycetales(100);f__Metschnikowiaceae(100);g__Metschnikowia(100);g__Metschnikowia_unclassified(100); | Metschnikowia | 13 | Saprotroph | \|Undefined Saprotroph\| | Yeast | NULL | Probable | As nectar-tap_saprotroph (Põlme et al. 2020); Decay sugar-rich substrates (Põlme et al. 2020) | Põlme S, et al. 2020. Fungal Diversity 105:1-16 ((https://doi.org/10.1007/s13225-020-00466-2)) |
| Otu15859 | 1 | k__Fungi(100);p__Ascomycota(100);c__Saccharomycetes(100);o__Saccharomycetales(100);f__Metschnikowiaceae(100);g__Metschnikowia(100);s__Metschnikowia_pulcherrima(100); | Metschnikowia | 13 | Saprotroph | \|Undefined Saprotroph\| | Yeast | NULL | Probable | As nectar-tap_saprotroph (Põlme et al. 2020); Decay sugar-rich substrates (Põlme et al. 2020) | Põlme S, et al. 2020. Fungal Diversity 105:1-16 ((https://doi.org/10.1007/s13225-020-00466-2)) |
| Otu15860 | 1 | k__Fungi(100);p__Ascomycota(100);c__Saccharomycetes(100);o__Saccharomycetales(100);f__Metschnikowiaceae(100);g__Metschnikowia(100);s__Metschnikowia_pulcherrima(100); | Metschnikowia | 13 | Saprotroph | \|Undefined Saprotroph\| | Yeast | NULL | Probable | As nectar-tap_saprotroph (Põlme et al. 2020); Decay sugar-rich substrates (Põlme et al. 2020) | Põlme S, et al. 2020. Fungal Diversity 105:1-16 ((https://doi.org/10.1007/s13225-020-00466-2)) |
| Otu15861 | 1 | k__Fungi(100);p__Ascomycota(100);c__Saccharomycetes(100);o__Saccharomycetales(100);f__Metschnikowiaceae(100);g__Metschnikowia(100);s__Metschnikowia_pulcherrima(100); | Metschnikowia | 13 | Saprotroph | \|Undefined Saprotroph\| | Yeast | NULL | Probable | As nectar-tap_saprotroph (Põlme et al. 2020); Decay sugar-rich substrates (Põlme et al. 2020) | Põlme S, et al. 2020. Fungal Diversity 105:1-16 ((https://doi.org/10.1007/s13225-020-00466-2)) |
| Otu15862 | 1 | k__Fungi(100);p__Ascomycota(100);c__Saccharomycetes(100);o__Saccharomycetales(100);f__Metschnikowiaceae(100);g__Metschnikowia(100);s__Metschnikowia_pulcherrima(100); | Metschnikowia | 13 | Saprotroph | \|Undefined Saprotroph\| | Yeast | NULL | Probable | As nectar-tap_saprotroph (Põlme et al. 2020); Decay sugar-rich substrates (Põlme et al. 2020) | Põlme S, et al. 2020. Fungal Diversity 105:1-16 ((https://doi.org/10.1007/s13225-020-00466-2)) |
| Otu15863 | 1 | k__Fungi(100);p__Ascomycota(100);c__Saccharomycetes(100);o__Saccharomycetales(100);f__Metschnikowiaceae(100);g__Metschnikowia(100);s__Metschnikowia_pulcherrima(100); | Metschnikowia | 13 | Saprotroph | \|Undefined Saprotroph\| | Yeast | NULL | Probable | As nectar-tap_saprotroph (Põlme et al. 2020); Decay sugar-rich substrates (Põlme et al. 2020) | Põlme S, et al. 2020. Fungal Diversity 105:1-16 ((https://doi.org/10.1007/s13225-020-00466-2)) |
| Otu15864 | 1 | k__Fungi(100);p__Ascomycota(100);c__Saccharomycetes(100);o__Saccharomycetales(100);f__Metschnikowiaceae(100);g__Metschnikowia(100);g__Metschnikowia_unclassified(100); | Metschnikowia | 13 | Saprotroph | \|Undefined Saprotroph\| | Yeast | NULL | Probable | As nectar-tap_saprotroph (Põlme et al. 2020); Decay sugar-rich substrates (Põlme et al. 2020) | Põlme S, et al. 2020. Fungal Diversity 105:1-16 ((https://doi.org/10.1007/s13225-020-00466-2)) |
| Otu15865 | 1 | k__Fungi(100);p__Ascomycota(100);c__Saccharomycetes(100);o__Saccharomycetales(100);f__Metschnikowiaceae(100);g__Metschnikowia(100);s__Metschnikowia_pulcherrima(100); | Metschnikowia | 13 | Saprotroph | \|Undefined Saprotroph\| | Yeast | NULL | Probable | As nectar-tap_saprotroph (Põlme et al. 2020); Decay sugar-rich substrates (Põlme et al. 2020) | Põlme S, et al. 2020. Fungal Diversity 105:1-16 ((https://doi.org/10.1007/s13225-020-00466-2)) |
| Otu15866 | 1 | k__Fungi(100);p__Ascomycota(100);c__Saccharomycetes(100);o__Saccharomycetales(100);f__Metschnikowiaceae(100);g__Metschnikowia(100);s__Metschnikowia_pulcherrima(100); | Metschnikowia | 13 | Saprotroph | \|Undefined Saprotroph\| | Yeast | NULL | Probable | As nectar-tap_saprotroph (Põlme et al. 2020); Decay sugar-rich substrates (Põlme et al. 2020) | Põlme S, et al. 2020. Fungal Diversity 105:1-16 ((https://doi.org/10.1007/s13225-020-00466-2)) |
| Otu15867 | 1 | k__Fungi(100);p__Ascomycota(100);c__Saccharomycetes(100);o__Saccharomycetales(100);f__Metschnikowiaceae(100);g__Metschnikowia(100);s__Metschnikowia_pulcherrima(100); | Metschnikowia | 13 | Saprotroph | \|Undefined Saprotroph\| | Yeast | NULL | Probable | As nectar-tap_saprotroph (Põlme et al. 2020); Decay sugar-rich substrates (Põlme et al. 2020) | Põlme S, et al. 2020. Fungal Diversity 105:1-16 ((https://doi.org/10.1007/s13225-020-00466-2)) |
| Otu15868 | 1 | k__Fungi(100);p__Ascomycota(100);c__Saccharomycetes(100);o__Saccharomycetales(100);f__Metschnikowiaceae(100);g__Metschnikowia(100);s__Metschnikowia_pulcherrima(100); | Metschnikowia | 13 | Saprotroph | \|Undefined Saprotroph\| | Yeast | NULL | Probable | As nectar-tap_saprotroph (Põlme et al. 2020); Decay sugar-rich substrates (Põlme et al. 2020) | Põlme S, et al. 2020. Fungal Diversity 105:1-16 ((https://doi.org/10.1007/s13225-020-00466-2)) |
| Otu15869 | 1 | k__Fungi(100);p__Ascomycota(100);c__Saccharomycetes(100);o__Saccharomycetales(100);f__Metschnikowiaceae(100);g__Metschnikowia(100);s__Metschnikowia_pulcherrima(100); | Metschnikowia | 13 | Saprotroph | \|Undefined Saprotroph\| | Yeast | NULL | Probable | As nectar-tap_saprotroph (Põlme et al. 2020); Decay sugar-rich substrates (Põlme et al. 2020) | Põlme S, et al. 2020. Fungal Diversity 105:1-16 ((https://doi.org/10.1007/s13225-020-00466-2)) |
| Otu15870 | 1 | k__Fungi(100);p__Ascomycota(100);c__Saccharomycetes(100);o__Saccharomycetales(100);f__Metschnikowiaceae(100);g__Metschnikowia(100);s__Metschnikowia_pulcherrima(100); | Metschnikowia | 13 | Saprotroph | \|Undefined Saprotroph\| | Yeast | NULL | Probable | As nectar-tap_saprotroph (Põlme et al. 2020); Decay sugar-rich substrates (Põlme et al. 2020) | Põlme S, et al. 2020. Fungal Diversity 105:1-16 ((https://doi.org/10.1007/s13225-020-00466-2)) |
| Otu15871 | 1 | k__Fungi(100);p__Ascomycota(100);c__Saccharomycetes(100);o__Saccharomycetales(100);f__Metschnikowiaceae(100);g__Metschnikowia(100);s__Metschnikowia_pulcherrima(100); | Metschnikowia | 13 | Saprotroph | \|Undefined Saprotroph\| | Yeast | NULL | Probable | As nectar-tap_saprotroph (Põlme et al. 2020); Decay sugar-rich substrates (Põlme et al. 2020) | Põlme S, et al. 2020. Fungal Diversity 105:1-16 ((https://doi.org/10.1007/s13225-020-00466-2)) |
| Otu15872 | 1 | k__Fungi(100);p__Ascomycota(100);c__Saccharomycetes(100);o__Saccharomycetales(100);f__Metschnikowiaceae(100);g__Metschnikowia(100);s__Metschnikowia_pulcherrima(100); | Metschnikowia | 13 | Saprotroph | \|Undefined Saprotroph\| | Yeast | NULL | Probable | As nectar-tap_saprotroph (Põlme et al. 2020); Decay sugar-rich substrates (Põlme et al. 2020) | Põlme S, et al. 2020. Fungal Diversity 105:1-16 ((https://doi.org/10.1007/s13225-020-00466-2)) |
| Otu15873 | 1 | k__Fungi(100);p__Ascomycota(100);c__Saccharomycetes(100);o__Saccharomycetales(100);f__Metschnikowiaceae(100);g__Metschnikowia(100);s__Metschnikowia_pulcherrima(100); | Metschnikowia | 13 | Saprotroph | \|Undefined Saprotroph\| | Yeast | NULL | Probable | As nectar-tap_saprotroph (Põlme et al. 2020); Decay sugar-rich substrates (Põlme et al. 2020) | Põlme S, et al. 2020. Fungal Diversity 105:1-16 ((https://doi.org/10.1007/s13225-020-00466-2)) |
| Otu15874 | 1 | k__Fungi(100);p__Ascomycota(100);c__Saccharomycetes(100);o__Saccharomycetales(100);f__Metschnikowiaceae(100);g__Metschnikowia(100);s__Metschnikowia_pulcherrima(100); | Metschnikowia | 13 | Saprotroph | \|Undefined Saprotroph\| | Yeast | NULL | Probable | As nectar-tap_saprotroph (Põlme et al. 2020); Decay sugar-rich substrates (Põlme et al. 2020) | Põlme S, et al. 2020. Fungal Diversity 105:1-16 ((https://doi.org/10.1007/s13225-020-00466-2)) |
| Otu15875 | 1 | k__Fungi(100);p__Ascomycota(100);c__Saccharomycetes(100);o__Saccharomycetales(100);f__Metschnikowiaceae(100);g__Metschnikowia(100);s__Metschnikowia_pulcherrima(100); | Metschnikowia | 13 | Saprotroph | \|Undefined Saprotroph\| | Yeast | NULL | Probable | As nectar-tap_saprotroph (Põlme et al. 2020); Decay sugar-rich substrates (Põlme et al. 2020) | Põlme S, et al. 2020. Fungal Diversity 105:1-16 ((https://doi.org/10.1007/s13225-020-00466-2)) |
| Otu15876 | 1 | k__Fungi(100);p__Ascomycota(100);c__Saccharomycetes(100);o__Saccharomycetales(100);f__Metschnikowiaceae(100);g__Metschnikowia(100);s__Metschnikowia_pulcherrima(100); | Metschnikowia | 13 | Saprotroph | \|Undefined Saprotroph\| | Yeast | NULL | Probable | As nectar-tap_saprotroph (Põlme et al. 2020); Decay sugar-rich substrates (Põlme et al. 2020) | Põlme S, et al. 2020. Fungal Diversity 105:1-16 ((https://doi.org/10.1007/s13225-020-00466-2)) |
| Otu15877 | 1 | k__Fungi(100);p__Ascomycota(100);c__Saccharomycetes(100);o__Saccharomycetales(100);f__Metschnikowiaceae(100);g__Metschnikowia(100);g__Metschnikowia_unclassified(100); | Metschnikowia | 13 | Saprotroph | \|Undefined Saprotroph\| | Yeast | NULL | Probable | As nectar-tap_saprotroph (Põlme et al. 2020); Decay sugar-rich substrates (Põlme et al. 2020) | Põlme S, et al. 2020. Fungal Diversity 105:1-16 ((https://doi.org/10.1007/s13225-020-00466-2)) |
| Otu15878 | 1 | k__Fungi(100);p__Ascomycota(100);c__Saccharomycetes(100);o__Saccharomycetales(100);f__Metschnikowiaceae(100);g__Metschnikowia(100);s__Metschnikowia_pulcherrima(100); | Metschnikowia | 13 | Saprotroph | \|Undefined Saprotroph\| | Yeast | NULL | Probable | As nectar-tap_saprotroph (Põlme et al. 2020); Decay sugar-rich substrates (Põlme et al. 2020) | Põlme S, et al. 2020. Fungal Diversity 105:1-16 ((https://doi.org/10.1007/s13225-020-00466-2)) |
| Otu15879 | 1 | k__Fungi(100);p__Ascomycota(100);c__Saccharomycetes(100);o__Saccharomycetales(100);f__Metschnikowiaceae(100);g__Metschnikowia(100);s__Metschnikowia_pulcherrima(100); | Metschnikowia | 13 | Saprotroph | \|Undefined Saprotroph\| | Yeast | NULL | Probable | As nectar-tap_saprotroph (Põlme et al. 2020); Decay sugar-rich substrates (Põlme et al. 2020) | Põlme S, et al. 2020. Fungal Diversity 105:1-16 ((https://doi.org/10.1007/s13225-020-00466-2)) |
| Otu15880 | 1 | k__Fungi(100);p__Ascomycota(100);c__Saccharomycetes(100);o__Saccharomycetales(100);f__Metschnikowiaceae(100);g__Metschnikowia(100);s__Metschnikowia_pulcherrima(100); | Metschnikowia | 13 | Saprotroph | \|Undefined Saprotroph\| | Yeast | NULL | Probable | As nectar-tap_saprotroph (Põlme et al. 2020); Decay sugar-rich substrates (Põlme et al. 2020) | Põlme S, et al. 2020. Fungal Diversity 105:1-16 ((https://doi.org/10.1007/s13225-020-00466-2)) |
| Otu15881 | 1 | k__Fungi(100);p__Ascomycota(100);c__Saccharomycetes(100);o__Saccharomycetales(100);f__Metschnikowiaceae(100);g__Metschnikowia(100);s__Metschnikowia_pulcherrima(100); | Metschnikowia | 13 | Saprotroph | \|Undefined Saprotroph\| | Yeast | NULL | Probable | As nectar-tap_saprotroph (Põlme et al. 2020); Decay sugar-rich substrates (Põlme et al. 2020) | Põlme S, et al. 2020. Fungal Diversity 105:1-16 ((https://doi.org/10.1007/s13225-020-00466-2)) |
| Otu15882 | 1 | k__Fungi(100);p__Ascomycota(100);c__Saccharomycetes(100);o__Saccharomycetales(100);f__Metschnikowiaceae(100);g__Metschnikowia(100);s__Metschnikowia_pulcherrima(100); | Metschnikowia | 13 | Saprotroph | \|Undefined Saprotroph\| | Yeast | NULL | Probable | As nectar-tap_saprotroph (Põlme et al. 2020); Decay sugar-rich substrates (Põlme et al. 2020) | Põlme S, et al. 2020. Fungal Diversity 105:1-16 ((https://doi.org/10.1007/s13225-020-00466-2)) |
| Otu15883 | 1 | k__Fungi(100);p__Ascomycota(100);c__Saccharomycetes(100);o__Saccharomycetales(100);f__Metschnikowiaceae(100);g__Metschnikowia(100);g__Metschnikowia_unclassified(100); | Metschnikowia | 13 | Saprotroph | \|Undefined Saprotroph\| | Yeast | NULL | Probable | As nectar-tap_saprotroph (Põlme et al. 2020); Decay sugar-rich substrates (Põlme et al. 2020) | Põlme S, et al. 2020. Fungal Diversity 105:1-16 ((https://doi.org/10.1007/s13225-020-00466-2)) |
| Otu15884 | 1 | k__Fungi(100);p__Ascomycota(100);c__Saccharomycetes(100);o__Saccharomycetales(100);f__Metschnikowiaceae(100);g__Metschnikowia(100);s__Metschnikowia_pulcherrima(100); | Metschnikowia | 13 | Saprotroph | \|Undefined Saprotroph\| | Yeast | NULL | Probable | As nectar-tap_saprotroph (Põlme et al. 2020); Decay sugar-rich substrates (Põlme et al. 2020) | Põlme S, et al. 2020. Fungal Diversity 105:1-16 ((https://doi.org/10.1007/s13225-020-00466-2)) |
| Otu15885 | 1 | k__Fungi(100);p__Ascomycota(100);c__Saccharomycetes(100);o__Saccharomycetales(100);f__Metschnikowiaceae(100);g__Metschnikowia(100);s__Metschnikowia_pulcherrima(100); | Metschnikowia | 13 | Saprotroph | \|Undefined Saprotroph\| | Yeast | NULL | Probable | As nectar-tap_saprotroph (Põlme et al. 2020); Decay sugar-rich substrates (Põlme et al. 2020) | Põlme S, et al. 2020. Fungal Diversity 105:1-16 ((https://doi.org/10.1007/s13225-020-00466-2)) |
| Otu15886 | 1 | k__Fungi(100);p__Ascomycota(100);c__Saccharomycetes(100);o__Saccharomycetales(100);f__Metschnikowiaceae(100);g__Metschnikowia(100);s__Metschnikowia_pulcherrima(100); | Metschnikowia | 13 | Saprotroph | \|Undefined Saprotroph\| | Yeast | NULL | Probable | As nectar-tap_saprotroph (Põlme et al. 2020); Decay sugar-rich substrates (Põlme et al. 2020) | Põlme S, et al. 2020. Fungal Diversity 105:1-16 ((https://doi.org/10.1007/s13225-020-00466-2)) |
| Otu15887 | 1 | k__Fungi(100);p__Ascomycota(100);c__Saccharomycetes(100);o__Saccharomycetales(100);f__Metschnikowiaceae(100);g__Metschnikowia(100);s__Metschnikowia_pulcherrima(100); | Metschnikowia | 13 | Saprotroph | \|Undefined Saprotroph\| | Yeast | NULL | Probable | As nectar-tap_saprotroph (Põlme et al. 2020); Decay sugar-rich substrates (Põlme et al. 2020) | Põlme S, et al. 2020. Fungal Diversity 105:1-16 ((https://doi.org/10.1007/s13225-020-00466-2)) |
| Otu15888 | 1 | k__Fungi(100);p__Ascomycota(100);c__Saccharomycetes(100);o__Saccharomycetales(100);f__Metschnikowiaceae(100);g__Metschnikowia(100);s__Metschnikowia_pulcherrima(100); | Metschnikowia | 13 | Saprotroph | \|Undefined Saprotroph\| | Yeast | NULL | Probable | As nectar-tap_saprotroph (Põlme et al. 2020); Decay sugar-rich substrates (Põlme et al. 2020) | Põlme S, et al. 2020. Fungal Diversity 105:1-16 ((https://doi.org/10.1007/s13225-020-00466-2)) |
| Otu15889 | 1 | k__Fungi(100);p__Ascomycota(100);c__Saccharomycetes(100);o__Saccharomycetales(100);f__Metschnikowiaceae(100);g__Metschnikowia(100);g__Metschnikowia_unclassified(100); | Metschnikowia | 13 | Saprotroph | \|Undefined Saprotroph\| | Yeast | NULL | Probable | As nectar-tap_saprotroph (Põlme et al. 2020); Decay sugar-rich substrates (Põlme et al. 2020) | Põlme S, et al. 2020. Fungal Diversity 105:1-16 ((https://doi.org/10.1007/s13225-020-00466-2)) |
| Otu15890 | 1 | k__Fungi(100);p__Ascomycota(100);c__Saccharomycetes(100);o__Saccharomycetales(100);f__Metschnikowiaceae(100);g__Metschnikowia(100);s__Metschnikowia_pulcherrima(100); | Metschnikowia | 13 | Saprotroph | \|Undefined Saprotroph\| | Yeast | NULL | Probable | As nectar-tap_saprotroph (Põlme et al. 2020); Decay sugar-rich substrates (Põlme et al. 2020) | Põlme S, et al. 2020. Fungal Diversity 105:1-16 ((https://doi.org/10.1007/s13225-020-00466-2)) |
| Otu15891 | 1 | k__Fungi(100);p__Ascomycota(100);c__Saccharomycetes(100);o__Saccharomycetales(100);f__Metschnikowiaceae(100);g__Metschnikowia(100);s__Metschnikowia_pulcherrima(100); | Metschnikowia | 13 | Saprotroph | \|Undefined Saprotroph\| | Yeast | NULL | Probable | As nectar-tap_saprotroph (Põlme et al. 2020); Decay sugar-rich substrates (Põlme et al. 2020) | Põlme S, et al. 2020. Fungal Diversity 105:1-16 ((https://doi.org/10.1007/s13225-020-00466-2)) |
| Otu15892 | 1 | k__Fungi(100);p__Ascomycota(100);c__Saccharomycetes(100);o__Saccharomycetales(100);f__Metschnikowiaceae(100);g__Metschnikowia(100);g__Metschnikowia_unclassified(100); | Metschnikowia | 13 | Saprotroph | \|Undefined Saprotroph\| | Yeast | NULL | Probable | As nectar-tap_saprotroph (Põlme et al. 2020); Decay sugar-rich substrates (Põlme et al. 2020) | Põlme S, et al. 2020. Fungal Diversity 105:1-16 ((https://doi.org/10.1007/s13225-020-00466-2)) |
| Otu15893 | 1 | k__Fungi(100);p__Ascomycota(100);c__Saccharomycetes(100);o__Saccharomycetales(100);f__Metschnikowiaceae(100);g__Metschnikowia(100);s__Metschnikowia_pulcherrima(100); | Metschnikowia | 13 | Saprotroph | \|Undefined Saprotroph\| | Yeast | NULL | Probable | As nectar-tap_saprotroph (Põlme et al. 2020); Decay sugar-rich substrates (Põlme et al. 2020) | Põlme S, et al. 2020. Fungal Diversity 105:1-16 ((https://doi.org/10.1007/s13225-020-00466-2)) |
| Otu15894 | 1 | k__Fungi(100);p__Ascomycota(100);c__Saccharomycetes(100);o__Saccharomycetales(100);f__Metschnikowiaceae(100);g__Metschnikowia(100);g__Metschnikowia_unclassified(100); | Metschnikowia | 13 | Saprotroph | \|Undefined Saprotroph\| | Yeast | NULL | Probable | As nectar-tap_saprotroph (Põlme et al. 2020); Decay sugar-rich substrates (Põlme et al. 2020) | Põlme S, et al. 2020. Fungal Diversity 105:1-16 ((https://doi.org/10.1007/s13225-020-00466-2)) |
| Otu15895 | 1 | k__Fungi(100);p__Ascomycota(100);c__Saccharomycetes(100);o__Saccharomycetales(100);f__Metschnikowiaceae(100);g__Metschnikowia(100);s__Metschnikowia_pulcherrima(100); | Metschnikowia | 13 | Saprotroph | \|Undefined Saprotroph\| | Yeast | NULL | Probable | As nectar-tap_saprotroph (Põlme et al. 2020); Decay sugar-rich substrates (Põlme et al. 2020) | Põlme S, et al. 2020. Fungal Diversity 105:1-16 ((https://doi.org/10.1007/s13225-020-00466-2)) |
| Otu15897 | 1 | k__Fungi(100);p__Ascomycota(100);c__Saccharomycetes(100);o__Saccharomycetales(100);f__Metschnikowiaceae(100);g__Metschnikowia(100);s__Metschnikowia_pulcherrima(100); | Metschnikowia | 13 | Saprotroph | \|Undefined Saprotroph\| | Yeast | NULL | Probable | As nectar-tap_saprotroph (Põlme et al. 2020); Decay sugar-rich substrates (Põlme et al. 2020) | Põlme S, et al. 2020. Fungal Diversity 105:1-16 ((https://doi.org/10.1007/s13225-020-00466-2)) |
| Otu15898 | 1 | k__Fungi(100);p__Ascomycota(100);c__Saccharomycetes(100);o__Saccharomycetales(100);f__Metschnikowiaceae(100);g__Metschnikowia(100);g__Metschnikowia_unclassified(100); | Metschnikowia | 13 | Saprotroph | \|Undefined Saprotroph\| | Yeast | NULL | Probable | As nectar-tap_saprotroph (Põlme et al. 2020); Decay sugar-rich substrates (Põlme et al. 2020) | Põlme S, et al. 2020. Fungal Diversity 105:1-16 ((https://doi.org/10.1007/s13225-020-00466-2)) |
| Otu15899 | 1 | k__Fungi(100);p__Ascomycota(100);c__Saccharomycetes(100);o__Saccharomycetales(100);f__Metschnikowiaceae(100);g__Metschnikowia(100);g__Metschnikowia_unclassified(100); | Metschnikowia | 13 | Saprotroph | \|Undefined Saprotroph\| | Yeast | NULL | Probable | As nectar-tap_saprotroph (Põlme et al. 2020); Decay sugar-rich substrates (Põlme et al. 2020) | Põlme S, et al. 2020. Fungal Diversity 105:1-16 ((https://doi.org/10.1007/s13225-020-00466-2)) |
| Otu15900 | 1 | k__Fungi(100);p__Ascomycota(100);c__Saccharomycetes(100);o__Saccharomycetales(100);f__Metschnikowiaceae(100);g__Metschnikowia(100);g__Metschnikowia_unclassified(100); | Metschnikowia | 13 | Saprotroph | \|Undefined Saprotroph\| | Yeast | NULL | Probable | As nectar-tap_saprotroph (Põlme et al. 2020); Decay sugar-rich substrates (Põlme et al. 2020) | Põlme S, et al. 2020. Fungal Diversity 105:1-16 ((https://doi.org/10.1007/s13225-020-00466-2)) |
| Otu15901 | 1 | k__Fungi(100);p__Ascomycota(100);c__Saccharomycetes(100);o__Saccharomycetales(100);f__Metschnikowiaceae(100);g__Metschnikowia(100);s__Metschnikowia_pulcherrima(100); | Metschnikowia | 13 | Saprotroph | \|Undefined Saprotroph\| | Yeast | NULL | Probable | As nectar-tap_saprotroph (Põlme et al. 2020); Decay sugar-rich substrates (Põlme et al. 2020) | Põlme S, et al. 2020. Fungal Diversity 105:1-16 ((https://doi.org/10.1007/s13225-020-00466-2)) |
| Otu15902 | 1 | k__Fungi(100);p__Ascomycota(100);c__Saccharomycetes(100);o__Saccharomycetales(100);f__Metschnikowiaceae(100);g__Metschnikowia(100);s__Metschnikowia_pulcherrima(100); | Metschnikowia | 13 | Saprotroph | \|Undefined Saprotroph\| | Yeast | NULL | Probable | As nectar-tap_saprotroph (Põlme et al. 2020); Decay sugar-rich substrates (Põlme et al. 2020) | Põlme S, et al. 2020. Fungal Diversity 105:1-16 ((https://doi.org/10.1007/s13225-020-00466-2)) |
| Otu15903 | 1 | k__Fungi(100);p__Ascomycota(100);c__Saccharomycetes(100);o__Saccharomycetales(100);f__Metschnikowiaceae(100);g__Metschnikowia(100);s__Metschnikowia_pulcherrima(100); | Metschnikowia | 13 | Saprotroph | \|Undefined Saprotroph\| | Yeast | NULL | Probable | As nectar-tap_saprotroph (Põlme et al. 2020); Decay sugar-rich substrates (Põlme et al. 2020) | Põlme S, et al. 2020. Fungal Diversity 105:1-16 ((https://doi.org/10.1007/s13225-020-00466-2)) |
| Otu15904 | 1 | k__Fungi(100);p__Ascomycota(100);c__Saccharomycetes(100);o__Saccharomycetales(100);f__Metschnikowiaceae(100);g__Metschnikowia(100);s__Metschnikowia_pulcherrima(100); | Metschnikowia | 13 | Saprotroph | \|Undefined Saprotroph\| | Yeast | NULL | Probable | As nectar-tap_saprotroph (Põlme et al. 2020); Decay sugar-rich substrates (Põlme et al. 2020) | Põlme S, et al. 2020. Fungal Diversity 105:1-16 ((https://doi.org/10.1007/s13225-020-00466-2)) |
| Otu15905 | 1 | k__Fungi(100);p__Ascomycota(100);c__Saccharomycetes(100);o__Saccharomycetales(100);f__Metschnikowiaceae(100);g__Metschnikowia(100);s__Metschnikowia_pulcherrima(100); | Metschnikowia | 13 | Saprotroph | \|Undefined Saprotroph\| | Yeast | NULL | Probable | As nectar-tap_saprotroph (Põlme et al. 2020); Decay sugar-rich substrates (Põlme et al. 2020) | Põlme S, et al. 2020. Fungal Diversity 105:1-16 ((https://doi.org/10.1007/s13225-020-00466-2)) |
| Otu15906 | 1 | k__Fungi(100);p__Ascomycota(100);c__Saccharomycetes(100);o__Saccharomycetales(100);f__Metschnikowiaceae(100);g__Metschnikowia(100);s__Metschnikowia_pulcherrima(100); | Metschnikowia | 13 | Saprotroph | \|Undefined Saprotroph\| | Yeast | NULL | Probable | As nectar-tap_saprotroph (Põlme et al. 2020); Decay sugar-rich substrates (Põlme et al. 2020) | Põlme S, et al. 2020. Fungal Diversity 105:1-16 ((https://doi.org/10.1007/s13225-020-00466-2)) |
| Otu15909 | 1 | k__Fungi(100);p__Ascomycota(100);c__Saccharomycetes(100);o__Saccharomycetales(100);f__Metschnikowiaceae(100);g__Metschnikowia(100);s__Metschnikowia_pulcherrima(100); | Metschnikowia | 13 | Saprotroph | \|Undefined Saprotroph\| | Yeast | NULL | Probable | As nectar-tap_saprotroph (Põlme et al. 2020); Decay sugar-rich substrates (Põlme et al. 2020) | Põlme S, et al. 2020. Fungal Diversity 105:1-16 ((https://doi.org/10.1007/s13225-020-00466-2)) |

| | | | | | | | | | | | |
|---|---|---|---|---|---|---|---|---|---|---|---|
| Otu15910 | 1 | k__Fungi(100);p__Ascomycota(100);c__Saccharomycetes(100);o__Saccharomycetales(100);f__Metschnikowiaceae(100);g__Metschnikowia(100);s__Metschnikowia_pulcherrima(100); | Metschnikowia | 13 | Saprotroph | \|Undefined Saprotroph\| | Yeast | NULL | Probable | As nectar-tap_saprotroph (Põlme et al. 2020); Decay sugar-rich substrates (Põlme et al. 2020) | Põlme S, et al. 2020. Fungal Diversity 105:1-16 ((https://doi.org/10.1007/s13225-020-00466-2)) |
| Otu15911 | 1 | k__Fungi(100);p__Ascomycota(100);c__Saccharomycetes(100);o__Saccharomycetales(100);f__Metschnikowiaceae(100);g__Metschnikowia(100);s__Metschnikowia_pulcherrima(100); | Metschnikowia | 13 | Saprotroph | \|Undefined Saprotroph\| | Yeast | NULL | Probable | As nectar-tap_saprotroph (Põlme et al. 2020); Decay sugar-rich substrates (Põlme et al. 2020) | Põlme S, et al. 2020. Fungal Diversity 105:1-16 ((https://doi.org/10.1007/s13225-020-00466-2)) |
| Otu15912 | 1 | k__Fungi(100);p__Ascomycota(100);c__Saccharomycetes(100);o__Saccharomycetales(100);f__Metschnikowiaceae(100);g__Metschnikowia(100);g__Metschnikowia_unclassified(100); | Metschnikowia | 13 | Saprotroph | \|Undefined Saprotroph\| | Yeast | NULL | Probable | As nectar-tap_saprotroph (Põlme et al. 2020); Decay sugar-rich substrates (Põlme et al. 2020) | Põlme S, et al. 2020. Fungal Diversity 105:1-16 ((https://doi.org/10.1007/s13225-020-00466-2)) |
| Otu15913 | 1 | k__Fungi(100);p__Ascomycota(100);c__Saccharomycetes(100);o__Saccharomycetales(100);f__Metschnikowiaceae(100);g__Metschnikowia(100);g__Metschnikowia_unclassified(100); | Metschnikowia | 13 | Saprotroph | \|Undefined Saprotroph\| | Yeast | NULL | Probable | As nectar-tap_saprotroph (Põlme et al. 2020); Decay sugar-rich substrates (Põlme et al. 2020) | Põlme S, et al. 2020. Fungal Diversity 105:1-16 ((https://doi.org/10.1007/s13225-020-00466-2)) |
| Otu15914 | 1 | k__Fungi(100);p__Ascomycota(100);c__Saccharomycetes(100);o__Saccharomycetales(100);f__Metschnikowiaceae(100);g__Metschnikowia(100);g__Metschnikowia_unclassified(100); | Metschnikowia | 13 | Saprotroph | \|Undefined Saprotroph\| | Yeast | NULL | Probable | As nectar-tap_saprotroph (Põlme et al. 2020); Decay sugar-rich substrates (Põlme et al. 2020) | Põlme S, et al. 2020. Fungal Diversity 105:1-16 ((https://doi.org/10.1007/s13225-020-00466-2)) |
| Otu15915 | 1 | k__Fungi(100);p__Ascomycota(100);c__Saccharomycetes(100);o__Saccharomycetales(100);f__Saccharomycetales_family_Incertae_sedis(100);g__Candida(100);s__Candida_cretensis(100); | Candida | 13 | Pathotroph-Saprotroph-Symbiotroph | Animal Pathogen-Endophyte-Endosymbiont-Epiphyte-\|Undefined Saprotroph\| | Dimorphic Yeast-Yeast | NULL | Probable | As nectar-tap_saprotroph (Põlme et al. 2020); Decay sugar-rich substrates (Põlme et al. 2020); This asexual genus is polyphyletic and contain species that function very differently depend on what they are. Caution should be taken when interpreting guilds for Candida species. | Manolakaki D, et al. 2010. Virulence. 1: 367-375 ((https://doi.org/10.4161/viru.1.5.12796)); Mekbib SB, et al. 2011. Tropical Plant Pathology 36:233-240 ((https://doi.org/10.1590/S1982-56762011000400004)); Põlme S, et al. 2020. Fungal Diversity 105:1-16 ((https://doi.org/10.1007/s13225-020-00466-2)) |
| Otu15916 | 1 | k__Fungi(100);p__Ascomycota(100);c__Saccharomycetes(100);o__Saccharomycetales(100);f__Metschnikowiaceae(100);g__Metschnikowia(100);s__Metschnikowia_pulcherrima(100); | Metschnikowia | 13 | Saprotroph | \|Undefined Saprotroph\| | Yeast | NULL | Probable | As nectar-tap_saprotroph (Põlme et al. 2020); Decay sugar-rich substrates (Põlme et al. 2020) | Põlme S, et al. 2020. Fungal Diversity 105:1-16 ((https://doi.org/10.1007/s13225-020-00466-2)) |
| Otu15917 | 1 | k__Fungi(100);p__Ascomycota(100);c__Saccharomycetes(100);o__Saccharomycetales(100);f__Metschnikowiaceae(100);g__Metschnikowia(100);s__Metschnikowia_pulcherrima(100); | Metschnikowia | 13 | Saprotroph | \|Undefined Saprotroph\| | Yeast | NULL | Probable | As nectar-tap_saprotroph (Põlme et al. 2020); Decay sugar-rich substrates (Põlme et al. 2020) | Põlme S, et al. 2020. Fungal Diversity 105:1-16 ((https://doi.org/10.1007/s13225-020-00466-2)) |
| Otu15918 | 1 | k__Fungi(100);p__Ascomycota(100);c__Saccharomycetes(100);o__Saccharomycetales(100);f__Metschnikowiaceae(100);g__Metschnikowia(100);s__Metschnikowia_pulcherrima(100); | Metschnikowia | 13 | Saprotroph | \|Undefined Saprotroph\| | Yeast | NULL | Probable | As nectar-tap_saprotroph (Põlme et al. 2020); Decay sugar-rich substrates (Põlme et al. 2020) | Põlme S, et al. 2020. Fungal Diversity 105:1-16 ((https://doi.org/10.1007/s13225-020-00466-2)) |
| Otu15919 | 1 | k__Fungi(100);p__Ascomycota(100);c__Saccharomycetes(100);o__Saccharomycetales(100);f__Metschnikowiaceae(100);g__Metschnikowia(100);g__Metschnikowia_unclassified(100); | Metschnikowia | 13 | Saprotroph | \|Undefined Saprotroph\| | Yeast | NULL | Probable | As nectar-tap_saprotroph (Põlme et al. 2020); Decay sugar-rich substrates (Põlme et al. 2020) | Põlme S, et al. 2020. Fungal Diversity 105:1-16 ((https://doi.org/10.1007/s13225-020-00466-2)) |
| Otu15920 | 1 | k__Fungi(100);p__Ascomycota(100);c__Saccharomycetes(100);o__Saccharomycetales(100);f__Metschnikowiaceae(100);g__Metschnikowia(100);g__Metschnikowia_unclassified(100); | Metschnikowia | 13 | Saprotroph | \|Undefined Saprotroph\| | Yeast | NULL | Probable | As nectar-tap_saprotroph (Põlme et al. 2020); Decay sugar-rich substrates (Põlme et al. 2020) | Põlme S, et al. 2020. Fungal Diversity 105:1-16 ((https://doi.org/10.1007/s13225-020-00466-2)) |
| Otu15921 | 1 | k__Fungi(100);p__Ascomycota(100);c__Saccharomycetes(100);o__Saccharomycetales(100);f__Metschnikowiaceae(100);g__Metschnikowia(100);s__Metschnikowia_pulcherrima(100); | Metschnikowia | 13 | Saprotroph | \|Undefined Saprotroph\| | Yeast | NULL | Probable | As nectar-tap_saprotroph (Põlme et al. 2020); Decay sugar-rich substrates (Põlme et al. 2020) | Põlme S, et al. 2020. Fungal Diversity 105:1-16 ((https://doi.org/10.1007/s13225-020-00466-2)) |
| Otu15922 | 1 | k__Fungi(100);p__Ascomycota(100);c__Saccharomycetes(100);o__Saccharomycetales(100);f__Metschnikowiaceae(100);g__Metschnikowia(100);g__Metschnikowia_unclassified(100); | Metschnikowia | 13 | Saprotroph | \|Undefined Saprotroph\| | Yeast | NULL | Probable | As nectar-tap_saprotroph (Põlme et al. 2020); Decay sugar-rich substrates (Põlme et al. 2020) | Põlme S, et al. 2020. Fungal Diversity 105:1-16 ((https://doi.org/10.1007/s13225-020-00466-2)) |
| Otu15923 | 1 | k__Fungi(100);p__Ascomycota(100);c__Saccharomycetes(100);o__Saccharomycetales(100);f__Metschnikowiaceae(100);g__Metschnikowia(100);s__Metschnikowia_pulcherrima(100); | Metschnikowia | 13 | Saprotroph | \|Undefined Saprotroph\| | Yeast | NULL | Probable | As nectar-tap_saprotroph (Põlme et al. 2020); Decay sugar-rich substrates (Põlme et al. 2020) | Põlme S, et al. 2020. Fungal Diversity 105:1-16 ((https://doi.org/10.1007/s13225-020-00466-2)) |
| Otu15924 | 1 | k__Fungi(100);p__Ascomycota(100);c__Saccharomycetes(100);o__Saccharomycetales(100);f__Metschnikowiaceae(100);g__Metschnikowia(100);s__Metschnikowia_pulcherrima(100); | Metschnikowia | 13 | Saprotroph | \|Undefined Saprotroph\| | Yeast | NULL | Probable | As nectar-tap_saprotroph (Põlme et al. 2020); Decay sugar-rich substrates (Põlme et al. 2020) | Põlme S, et al. 2020. Fungal Diversity 105:1-16 ((https://doi.org/10.1007/s13225-020-00466-2)) |
| Otu15925 | 1 | k__Fungi(100);p__Ascomycota(100);c__Saccharomycetes(100);o__Saccharomycetales(100);f__Metschnikowiaceae(100);g__Metschnikowia(100);g__Metschnikowia_unclassified(100); | Metschnikowia | 13 | Saprotroph | \|Undefined Saprotroph\| | Yeast | NULL | Probable | As nectar-tap_saprotroph (Põlme et al. 2020); Decay sugar-rich substrates (Põlme et al. 2020) | Põlme S, et al. 2020. Fungal Diversity 105:1-16 ((https://doi.org/10.1007/s13225-020-00466-2)) |
| Otu15927 | 1 | k__Fungi(100);p__Ascomycota(100);c__Saccharomycetes(100);o__Saccharomycetales(100);f__Metschnikowiaceae(100);g__Metschnikowia(100);s__Metschnikowia_pulcherrima(100); | Metschnikowia | 13 | Saprotroph | \|Undefined Saprotroph\| | Yeast | NULL | Probable | As nectar-tap_saprotroph (Põlme et al. 2020); Decay sugar-rich substrates (Põlme et al. 2020) | Põlme S, et al. 2020. Fungal Diversity 105:1-16 ((https://doi.org/10.1007/s13225-020-00466-2)) |
| Otu15930 | 1 | k__Fungi(100);p__Ascomycota(100);c__Saccharomycetes(100);o__Saccharomycetales(100);f__Metschnikowiaceae(100);g__Metschnikowia(100);s__Metschnikowia_pulcherrima(100); | Metschnikowia | 13 | Saprotroph | \|Undefined Saprotroph\| | Yeast | NULL | Probable | As nectar-tap_saprotroph (Põlme et al. 2020); Decay sugar-rich substrates (Põlme et al. 2020) | Põlme S, et al. 2020. Fungal Diversity 105:1-16 ((https://doi.org/10.1007/s13225-020-00466-2)) |
| Otu15932 | 1 | k__Fungi(100);p__Ascomycota(100);c__Saccharomycetes(100);o__Saccharomycetales(100);f__Metschnikowiaceae(100);g__Metschnikowia(100);s__Metschnikowia_pulcherrima(100); | Metschnikowia | 13 | Saprotroph | \|Undefined Saprotroph\| | Yeast | NULL | Probable | As nectar-tap_saprotroph (Põlme et al. 2020); Decay sugar-rich substrates (Põlme et al. 2020) | Põlme S, et al. 2020. Fungal Diversity 105:1-16 ((https://doi.org/10.1007/s13225-020-00466-2)) |
| Otu15933 | 1 | k__Fungi(100);p__Ascomycota(100);c__Saccharomycetes(100);o__Saccharomycetales(100);f__Metschnikowiaceae(100);g__Metschnikowia(100);s__Metschnikowia_pulcherrima(100); | Metschnikowia | 13 | Saprotroph | \|Undefined Saprotroph\| | Yeast | NULL | Probable | As nectar-tap_saprotroph (Põlme et al. 2020); Decay sugar-rich substrates (Põlme et al. 2020) | Põlme S, et al. 2020. Fungal Diversity 105:1-16 ((https://doi.org/10.1007/s13225-020-00466-2)) |
| Otu15934 | 1 | k__Fungi(100);p__Ascomycota(100);c__Saccharomycetes(100);o__Saccharomycetales(100);f__Metschnikowiaceae(100);g__Metschnikowia(100);g__Metschnikowia_unclassified(100); | Metschnikowia | 13 | Saprotroph | \|Undefined Saprotroph\| | Yeast | NULL | Probable | As nectar-tap_saprotroph (Põlme et al. 2020); Decay sugar-rich substrates (Põlme et al. 2020) | Põlme S, et al. 2020. Fungal Diversity 105:1-16 ((https://doi.org/10.1007/s13225-020-00466-2)) |
| Otu15935 | 1 | k__Fungi(100);p__Ascomycota(100);c__Saccharomycetes(100);o__Saccharomycetales(100);f__Metschnikowiaceae(100);g__Metschnikowia(100);s__Metschnikowia_pulcherrima(100); | Metschnikowia | 13 | Saprotroph | \|Undefined Saprotroph\| | Yeast | NULL | Probable | As nectar-tap_saprotroph (Põlme et al. 2020); Decay sugar-rich substrates (Põlme et al. 2020) | Põlme S, et al. 2020. Fungal Diversity 105:1-16 ((https://doi.org/10.1007/s13225-020-00466-2)) |
| Otu15937 | 1 | k__Fungi(100);p__Ascomycota(100);c__Saccharomycetes(100);o__Saccharomycetales(100);f__Metschnikowiaceae(100);g__Metschnikowia(100);s__Metschnikowia_pulcherrima(100); | Metschnikowia | 13 | Saprotroph | \|Undefined Saprotroph\| | Yeast | NULL | Probable | As nectar-tap_saprotroph (Põlme et al. 2020); Decay sugar-rich substrates (Põlme et al. 2020) | Põlme S, et al. 2020. Fungal Diversity 105:1-16 ((https://doi.org/10.1007/s13225-020-00466-2)) |
| Otu15938 | 1 | k__Fungi(100);p__Ascomycota(100);c__Saccharomycetes(100);o__Saccharomycetales(100);f__Metschnikowiaceae(100);g__Metschnikowia(100);s__Metschnikowia_pulcherrima(100); | Metschnikowia | 13 | Saprotroph | \|Undefined Saprotroph\| | Yeast | NULL | Probable | As nectar-tap_saprotroph (Põlme et al. 2020); Decay sugar-rich substrates (Põlme et al. 2020) | Põlme S, et al. 2020. Fungal Diversity 105:1-16 ((https://doi.org/10.1007/s13225-020-00466-2)) |
| Otu15940 | 1 | k__Fungi(100);p__Ascomycota(100);c__Saccharomycetes(100);o__Saccharomycetales(100);f__Metschnikowiaceae(100);g__Metschnikowia(100);s__Metschnikowia_pulcherrima(100); | Metschnikowia | 13 | Saprotroph | \|Undefined Saprotroph\| | Yeast | NULL | Probable | As nectar-tap_saprotroph (Põlme et al. 2020); Decay sugar-rich substrates (Põlme et al. 2020) | Põlme S, et al. 2020. Fungal Diversity 105:1-16 ((https://doi.org/10.1007/s13225-020-00466-2)) |
| Otu15941 | 1 | k__Fungi(100);p__Ascomycota(100);c__Saccharomycetes(100);o__Saccharomycetales(100);f__Metschnikowiaceae(100);g__Metschnikowia(100);s__Metschnikowia_pulcherrima(100); | Metschnikowia | 13 | Saprotroph | \|Undefined Saprotroph\| | Yeast | NULL | Probable | As nectar-tap_saprotroph (Põlme et al. 2020); Decay sugar-rich substrates (Põlme et al. 2020) | Põlme S, et al. 2020. Fungal Diversity 105:1-16 ((https://doi.org/10.1007/s13225-020-00466-2)) |
| Otu15942 | 1 | k__Fungi(100);p__Ascomycota(100);c__Saccharomycetes(100);o__Saccharomycetales(100);f__Metschnikowiaceae(100);g__Metschnikowia(100);s__Metschnikowia_pulcherrima(100); | Metschnikowia | 13 | Saprotroph | \|Undefined Saprotroph\| | Yeast | NULL | Probable | As nectar-tap_saprotroph (Põlme et al. 2020); Decay sugar-rich substrates (Põlme et al. 2020) | Põlme S, et al. 2020. Fungal Diversity 105:1-16 ((https://doi.org/10.1007/s13225-020-00466-2)) |
| Otu15943 | 1 | k__Fungi(100);p__Ascomycota(100);c__Saccharomycetes(100);o__Saccharomycetales(100);f__Metschnikowiaceae(100);g__Metschnikowia(100);g__Metschnikowia_unclassified(100); | Metschnikowia | 13 | Saprotroph | \|Undefined Saprotroph\| | Yeast | NULL | Probable | As nectar-tap_saprotroph (Põlme et al. 2020); Decay sugar-rich substrates (Põlme et al. 2020) | Põlme S, et al. 2020. Fungal Diversity 105:1-16 ((https://doi.org/10.1007/s13225-020-00466-2)) |
| Otu15944 | 1 | k__Fungi(100);p__Ascomycota(100);c__Saccharomycetes(100);o__Saccharomycetales(100);f__Metschnikowiaceae(100);g__Metschnikowia(100);s__Metschnikowia_pulcherrima(100); | Metschnikowia | 13 | Saprotroph | \|Undefined Saprotroph\| | Yeast | NULL | Probable | As nectar-tap_saprotroph (Põlme et al. 2020); Decay sugar-rich substrates (Põlme et al. 2020) | Põlme S, et al. 2020. Fungal Diversity 105:1-16 ((https://doi.org/10.1007/s13225-020-00466-2)) |
| Otu15945 | 1 | k__Fungi(100);p__Ascomycota(100);c__Saccharomycetes(100);o__Saccharomycetales(100);f__Metschnikowiaceae(100);g__Metschnikowia(100);s__Metschnikowia_pulcherrima(100); | Metschnikowia | 13 | Saprotroph | \|Undefined Saprotroph\| | Yeast | NULL | Probable | As nectar-tap_saprotroph (Põlme et al. 2020); Decay sugar-rich substrates (Põlme et al. 2020) | Põlme S, et al. 2020. Fungal Diversity 105:1-16 ((https://doi.org/10.1007/s13225-020-00466-2)) |
| Otu15946 | 1 | k__Fungi(100);p__Ascomycota(100);c__Saccharomycetes(100);o__Saccharomycetales(100);f__Metschnikowiaceae(100);g__Metschnikowia(100);g__Metschnikowia_unclassified(100); | Metschnikowia | 13 | Saprotroph | \|Undefined Saprotroph\| | Yeast | NULL | Probable | As nectar-tap_saprotroph (Põlme et al. 2020); Decay sugar-rich substrates (Põlme et al. 2020) | Põlme S, et al. 2020. Fungal Diversity 105:1-16 ((https://doi.org/10.1007/s13225-020-00466-2)) |
| Otu15947 | 1 | k__Fungi(100);p__Ascomycota(100);c__Saccharomycetes(100);o__Saccharomycetales(100);f__Metschnikowiaceae(100);g__Metschnikowia(100);g__Metschnikowia_unclassified(100); | Metschnikowia | 13 | Saprotroph | \|Undefined Saprotroph\| | Yeast | NULL | Probable | As nectar-tap_saprotroph (Põlme et al. 2020); Decay sugar-rich substrates (Põlme et al. 2020) | Põlme S, et al. 2020. Fungal Diversity 105:1-16 ((https://doi.org/10.1007/s13225-020-00466-2)) |
| Otu15948 | 1 | k__Fungi(100);p__Ascomycota(100);c__Saccharomycetes(100);o__Saccharomycetales(100);f__Metschnikowiaceae(100);g__Metschnikowia(100);g__Metschnikowia_unclassified(100); | Metschnikowia | 13 | Saprotroph | \|Undefined Saprotroph\| | Yeast | NULL | Probable | As nectar-tap_saprotroph (Põlme et al. 2020); Decay sugar-rich substrates (Põlme et al. 2020) | Põlme S, et al. 2020. Fungal Diversity 105:1-16 ((https://doi.org/10.1007/s13225-020-00466-2)) |
| Otu15949 | 1 | k__Fungi(100);p__Ascomycota(100);c__Saccharomycetes(100);o__Saccharomycetales(100);f__Metschnikowiaceae(100);g__Metschnikowia(100);s__Metschnikowia_pulcherrima(100); | Metschnikowia | 13 | Saprotroph | \|Undefined Saprotroph\| | Yeast | NULL | Probable | As nectar-tap_saprotroph (Põlme et al. 2020); Decay sugar-rich substrates (Põlme et al. 2020) | Põlme S, et al. 2020. Fungal Diversity 105:1-16 ((https://doi.org/10.1007/s13225-020-00466-2)) |
| Otu15950 | 1 | k__Fungi(100);p__Ascomycota(100);c__Saccharomycetes(100);o__Saccharomycetales(100);f__Metschnikowiaceae(100);g__Metschnikowia(100);s__Metschnikowia_pulcherrima(100); | Metschnikowia | 13 | Saprotroph | \|Undefined Saprotroph\| | Yeast | NULL | Probable | As nectar-tap_saprotroph (Põlme et al. 2020); Decay sugar-rich substrates (Põlme et al. 2020) | Põlme S, et al. 2020. Fungal Diversity 105:1-16 ((https://doi.org/10.1007/s13225-020-00466-2)) |
| Otu15951 | 1 | k__Fungi(100);p__Ascomycota(100);c__Saccharomycetes(100);o__Saccharomycetales(100);f__Metschnikowiaceae(100);g__Metschnikowia(100);s__Metschnikowia_pulcherrima(100); | Metschnikowia | 13 | Saprotroph | \|Undefined Saprotroph\| | Yeast | NULL | Probable | As nectar-tap_saprotroph (Põlme et al. 2020); Decay sugar-rich substrates (Põlme et al. 2020) | Põlme S, et al. 2020. Fungal Diversity 105:1-16 ((https://doi.org/10.1007/s13225-020-00466-2)) |
| Otu15952 | 1 | k__Fungi(100);p__Ascomycota(100);c__Saccharomycetes(100);o__Saccharomycetales(100);f__Metschnikowiaceae(100);g__Metschnikowia(100);g__Metschnikowia_unclassified(100); | Metschnikowia | 13 | Saprotroph | \|Undefined Saprotroph\| | Yeast | NULL | Probable | As nectar-tap_saprotroph (Põlme et al. 2020); Decay sugar-rich substrates (Põlme et al. 2020) | Põlme S, et al. 2020. Fungal Diversity 105:1-16 ((https://doi.org/10.1007/s13225-020-00466-2)) |
| Otu15953 | 1 | k__Fungi(100);p__Ascomycota(100);c__Saccharomycetes(100);o__Saccharomycetales(100);f__Metschnikowiaceae(100);g__Metschnikowia(100);s__Metschnikowia_pulcherrima(100); | Metschnikowia | 13 | Saprotroph | \|Undefined Saprotroph\| | Yeast | NULL | Probable | As nectar-tap_saprotroph (Põlme et al. 2020); Decay sugar-rich substrates (Põlme et al. 2020) | Põlme S, et al. 2020. Fungal Diversity 105:1-16 ((https://doi.org/10.1007/s13225-020-00466-2)) |
| Otu15954 | 1 | k__Fungi(100);p__Ascomycota(100);c__Saccharomycetes(100);o__Saccharomycetales(100);f__Metschnikowiaceae(100);g__Metschnikowia(100);s__Metschnikowia_pulcherrima(100); | Metschnikowia | 13 | Saprotroph | \|Undefined Saprotroph\| | Yeast | NULL | Probable | As nectar-tap_saprotroph (Põlme et al. 2020); Decay sugar-rich substrates (Põlme et al. 2020) | Põlme S, et al. 2020. Fungal Diversity 105:1-16 ((https://doi.org/10.1007/s13225-020-00466-2)) |
| Otu15955 | 1 | k__Fungi(100);p__Ascomycota(100);c__Saccharomycetes(100);o__Saccharomycetales(100);f__Metschnikowiaceae(100);g__Metschnikowia(100);s__Metschnikowia_pulcherrima(100); | Metschnikowia | 13 | Saprotroph | \|Undefined Saprotroph\| | Yeast | NULL | Probable | As nectar-tap_saprotroph (Põlme et al. 2020); Decay sugar-rich substrates (Põlme et al. 2020) | Põlme S, et al. 2020. Fungal Diversity 105:1-16 ((https://doi.org/10.1007/s13225-020-00466-2)) |
| Otu15956 | 1 | k__Fungi(100);p__Ascomycota(100);c__Saccharomycetes(100);o__Saccharomycetales(100);f__Metschnikowiaceae(100);g__Metschnikowia(100);s__Metschnikowia_pulcherrima(100); | Metschnikowia | 13 | Saprotroph | \|Undefined Saprotroph\| | Yeast | NULL | Probable | As nectar-tap_saprotroph (Põlme et al. 2020); Decay sugar-rich substrates (Põlme et al. 2020) | Põlme S, et al. 2020. Fungal Diversity 105:1-16 ((https://doi.org/10.1007/s13225-020-00466-2)) |
| Otu15957 | 1 | k__Fungi(100);p__Ascomycota(100);c__Saccharomycetes(100);o__Saccharomycetales(100);f__Metschnikowiaceae(100);g__Metschnikowia(100);s__Metschnikowia_pulcherrima(100); | Metschnikowia | 13 | Saprotroph | \|Undefined Saprotroph\| | Yeast | NULL | Probable | As nectar-tap_saprotroph (Põlme et al. 2020); Decay sugar-rich substrates (Põlme et al. 2020) | Põlme S, et al. 2020. Fungal Diversity 105:1-16 ((https://doi.org/10.1007/s13225-020-00466-2)) |
| Otu15958 | 1 | k__Fungi(100);p__Ascomycota(100);c__Saccharomycetes(100);o__Saccharomycetales(100);f__Metschnikowiaceae(100);g__Metschnikowia(100);s__Metschnikowia_pulcherrima(100); | Metschnikowia | 13 | Saprotroph | \|Undefined Saprotroph\| | Yeast | NULL | Probable | As nectar-tap_saprotroph (Põlme et al. 2020); Decay sugar-rich substrates (Põlme et al. 2020) | Põlme S, et al. 2020. Fungal Diversity 105:1-16 ((https://doi.org/10.1007/s13225-020-00466-2)) |
| Otu15959 | 1 | k__Fungi(100);p__Ascomycota(100);c__Saccharomycetes(100);o__Saccharomycetales(100);f__Metschnikowiaceae(100);g__Metschnikowia(100);s__Metschnikowia_pulcherrima(100); | Metschnikowia | 13 | Saprotroph | \|Undefined Saprotroph\| | Yeast | NULL | Probable | As nectar-tap_saprotroph (Põlme et al. 2020); Decay sugar-rich substrates (Põlme et al. 2020) | Põlme S, et al. 2020. Fungal Diversity 105:1-16 ((https://doi.org/10.1007/s13225-020-00466-2)) |
| Otu15960 | 1 | k__Fungi(100);p__Ascomycota(100);c__Saccharomycetes(100);o__Saccharomycetales(100);f__Metschnikowiaceae(100);g__Metschnikowia(100);s__Metschnikowia_pulcherrima(100); | Metschnikowia | 13 | Saprotroph | \|Undefined Saprotroph\| | Yeast | NULL | Probable | As nectar-tap_saprotroph (Põlme et al. 2020); Decay sugar-rich substrates (Põlme et al. 2020) | Põlme S, et al. 2020. Fungal Diversity 105:1-16 ((https://doi.org/10.1007/s13225-020-00466-2)) |
| Otu15961 | 1 | k__Fungi(100);p__Ascomycota(100);c__Saccharomycetes(100);o__Saccharomycetales(100);f__Metschnikowiaceae(100);g__Metschnikowia(100);g__Metschnikowia_unclassified(100); | Metschnikowia | 13 | Saprotroph | \|Undefined Saprotroph\| | Yeast | NULL | Probable | As nectar-tap_saprotroph (Põlme et al. 2020); Decay sugar-rich substrates (Põlme et al. 2020) | Põlme S, et al. 2020. Fungal Diversity 105:1-16 ((https://doi.org/10.1007/s13225-020-00466-2)) |
| Otu15962 | 1 | k__Fungi(100);p__Ascomycota(100);c__Saccharomycetes(100);o__Saccharomycetales(100);f__Metschnikowiaceae(100);g__Metschnikowia(100);g__Metschnikowia_unclassified(100); | Metschnikowia | 13 | Saprotroph | \|Undefined Saprotroph\| | Yeast | NULL | Probable | As nectar-tap_saprotroph (Põlme et al. 2020); Decay sugar-rich substrates (Põlme et al. 2020) | Põlme S, et al. 2020. Fungal Diversity 105:1-16 ((https://doi.org/10.1007/s13225-020-00466-2)) |
| Otu15964 | 1 | k__Fungi(100);p__Ascomycota(100);c__Saccharomycetes(100);o__Saccharomycetales(100);f__Metschnikowiaceae(100);g__Metschnikowia(100);g__Metschnikowia_unclassified(100); | Metschnikowia | 13 | Saprotroph | \|Undefined Saprotroph\| | Yeast | NULL | Probable | As nectar-tap_saprotroph (Põlme et al. 2020); Decay sugar-rich substrates (Põlme et al. 2020) | Põlme S, et al. 2020. Fungal Diversity 105:1-16 ((https://doi.org/10.1007/s13225-020-00466-2)) |
| Otu15965 | 1 | k__Fungi(100);p__Ascomycota(100);c__Saccharomycetes(100);o__Saccharomycetales(100);f__Metschnikowiaceae(100);g__Metschnikowia(100);s__Metschnikowia_pulcherrima(100); | Metschnikowia | 13 | Saprotroph | \|Undefined Saprotroph\| | Yeast | NULL | Probable | As nectar-tap_saprotroph (Põlme et al. 2020); Decay sugar-rich substrates (Põlme et al. 2020) | Põlme S, et al. 2020. Fungal Diversity 105:1-16 ((https://doi.org/10.1007/s13225-020-00466-2)) |
| Otu15966 | 1 | k__Fungi(100);p__Ascomycota(100);c__Saccharomycetes(100);o__Saccharomycetales(100);f__Metschnikowiaceae(100);g__Metschnikowia(100);g__Metschnikowia_unclassified(100); | Metschnikowia | 13 | Saprotroph | \|Undefined Saprotroph\| | Yeast | NULL | Probable | As nectar-tap_saprotroph (Põlme et al. 2020); Decay sugar-rich substrates (Põlme et al. 2020) | Põlme S, et al. 2020. Fungal Diversity 105:1-16 ((https://doi.org/10.1007/s13225-020-00466-2)) |
| Otu15967 | 1 | k__Fungi(100);p__Ascomycota(100);c__Saccharomycetes(100);o__Saccharomycetales(100);f__Metschnikowiaceae(100);g__Metschnikowia(100);s__Metschnikowia_pulcherrima(100); | Metschnikowia | 13 | Saprotroph | \|Undefined Saprotroph\| | Yeast | NULL | Probable | As nectar-tap_saprotroph (Põlme et al. 2020); Decay sugar-rich substrates (Põlme et al. 2020) | Põlme S, et al. 2020. Fungal Diversity 105:1-16 ((https://doi.org/10.1007/s13225-020-00466-2)) |
| Otu15968 | 1 | k__Fungi(100);p__Ascomycota(100);c__Saccharomycetes(100);o__Saccharomycetales(100);f__Metschnikowiaceae(100);g__Metschnikowia(100);s__Metschnikowia_pulcherrima(100); | Metschnikowia | 13 | Saprotroph | \|Undefined Saprotroph\| | Yeast | NULL | Probable | As nectar-tap_saprotroph (Põlme et al. 2020); Decay sugar-rich substrates (Põlme et al. 2020) | Põlme S, et al. 2020. Fungal Diversity 105:1-16 ((https://doi.org/10.1007/s13225-020-00466-2)) |
| Otu15969 | 1 | k__Fungi(100);p__Ascomycota(100);c__Saccharomycetes(100);o__Saccharomycetales(100);f__Metschnikowiaceae(100);g__Metschnikowia(100);g__Metschnikowia_unclassified(100); | Metschnikowia | 13 | Saprotroph | \|Undefined Saprotroph\| | Yeast | NULL | Probable | As nectar-tap_saprotroph (Põlme et al. 2020); Decay sugar-rich substrates (Põlme et al. 2020) | Põlme S, et al. 2020. Fungal Diversity 105:1-16 ((https://doi.org/10.1007/s13225-020-00466-2)) |
| Otu15970 | 1 | k__Fungi(100);p__Ascomycota(100);c__Saccharomycetes(100);o__Saccharomycetales(100);f__Metschnikowiaceae(100);g__Metschnikowia(100);s__Metschnikowia_pulcherrima(100); | Metschnikowia | 13 | Saprotroph | \|Undefined Saprotroph\| | Yeast | NULL | Probable | As nectar-tap_saprotroph (Põlme et al. 2020); Decay sugar-rich substrates (Põlme et al. 2020) | Põlme S, et al. 2020. Fungal Diversity 105:1-16 ((https://doi.org/10.1007/s13225-020-00466-2)) |
| Otu15971 | 1 | k__Fungi(100);p__Ascomycota(100);c__Saccharomycetes(100);o__Saccharomycetales(100);f__Metschnikowiaceae(100);g__Metschnikowia(100);s__Metschnikowia_pulcherrima(100); | Metschnikowia | 13 | Saprotroph | \|Undefined Saprotroph\| | Yeast | NULL | Probable | As nectar-tap_saprotroph (Põlme et al. 2020); Decay sugar-rich substrates (Põlme et al. 2020) | Põlme S, et al. 2020. Fungal Diversity 105:1-16 ((https://doi.org/10.1007/s13225-020-00466-2)) |
| Otu15972 | 1 | k__Fungi(100);p__Ascomycota(100);c__Saccharomycetes(100);o__Saccharomycetales(100);f__Metschnikowiaceae(100);g__Metschnikowia(100);g__Metschnikowia_unclassified(100); | Metschnikowia | 13 | Saprotroph | \|Undefined Saprotroph\| | Yeast | NULL | Probable | As nectar-tap_saprotroph (Põlme et al. 2020); Decay sugar-rich substrates (Põlme et al. 2020) | Põlme S, et al. 2020. Fungal Diversity 105:1-16 ((https://doi.org/10.1007/s13225-020-00466-2)) |
| Otu15973 | 1 | k__Fungi(100);p__Ascomycota(100);c__Saccharomycetes(100);o__Saccharomycetales(100);f__Metschnikowiaceae(100);g__Metschnikowia(100);s__Metschnikowia_pulcherrima(100); | Metschnikowia | 13 | Saprotroph | \|Undefined Saprotroph\| | Yeast | NULL | Probable | As nectar-tap_saprotroph (Põlme et al. 2020); Decay sugar-rich substrates (Põlme et al. 2020) | Põlme S, et al. 2020. Fungal Diversity 105:1-16 ((https://doi.org/10.1007/s13225-020-00466-2)) |
| Otu15974 | 1 | k__Fungi(100);p__Ascomycota(100);c__Saccharomycetes(100);o__Saccharomycetales(100);f__Metschnikowiaceae(100);g__Metschnikowia(100);g__Metschnikowia_unclassified(100); | Metschnikowia | 13 | Saprotroph | \|Undefined Saprotroph\| | Yeast | NULL | Probable | As nectar-tap_saprotroph (Põlme et al. 2020); Decay sugar-rich substrates (Põlme et al. 2020) | Põlme S, et al. 2020. Fungal Diversity 105:1-16 ((https://doi.org/10.1007/s13225-020-00466-2)) |
| Otu15975 | 1 | k__Fungi(100);p__Ascomycota(100);c__Saccharomycetes(100);o__Saccharomycetales(100);f__Metschnikowiaceae(100);g__Metschnikowia(100);s__Metschnikowia_pulcherrima(100); | Metschnikowia | 13 | Saprotroph | \|Undefined Saprotroph\| | Yeast | NULL | Probable | As nectar-tap_saprotroph (Põlme et al. 2020); Decay sugar-rich substrates (Põlme et al. 2020) | Põlme S, et al. 2020. Fungal Diversity 105:1-16 ((https://doi.org/10.1007/s13225-020-00466-2)) |
| Otu15976 | 1 | k__Fungi(100);p__Ascomycota(100);c__Saccharomycetes(100);o__Saccharomycetales(100);f__Metschnikowiaceae(100);g__Metschnikowia(100);g__Metschnikowia_unclassified(100); | Metschnikowia | 13 | Saprotroph | \|Undefined Saprotroph\| | Yeast | NULL | Probable | As nectar-tap_saprotroph (Põlme et al. 2020); Decay sugar-rich substrates (Põlme et al. 2020) | Põlme S, et al. 2020. Fungal Diversity 105:1-16 ((https://doi.org/10.1007/s13225-020-00466-2)) |
| Otu15977 | 1 | k__Fungi(100);p__Ascomycota(100);c__Saccharomycetes(100);o__Saccharomycetales(100);f__Metschnikowiaceae(100);g__Metschnikowia(100);s__Metschnikowia_pulcherrima(100); | Metschnikowia | 13 | Saprotroph | \|Undefined Saprotroph\| | Yeast | NULL | Probable | As nectar-tap_saprotroph (Põlme et al. 2020); Decay sugar-rich substrates (Põlme et al. 2020) | Põlme S, et al. 2020. Fungal Diversity 105:1-16 ((https://doi.org/10.1007/s13225-020-00466-2)) |
| Otu15978 | 1 | k__Fungi(100);p__Ascomycota(100);c__Saccharomycetes(100);o__Saccharomycetales(100);f__Metschnikowiaceae(100);g__Metschnikowia(100);s__Metschnikowia_pulcherrima(100); | Metschnikowia | 13 | Saprotroph | \|Undefined Saprotroph\| | Yeast | NULL | Probable | As nectar-tap_saprotroph (Põlme et al. 2020); Decay sugar-rich substrates (Põlme et al. 2020) | Põlme S, et al. 2020. Fungal Diversity 105:1-16 ((https://doi.org/10.1007/s13225-020-00466-2)) |
| Otu15979 | 1 | k__Fungi(100);p__Ascomycota(100);c__Saccharomycetes(100);o__Saccharomycetales(100);f__Metschnikowiaceae(100);g__Metschnikowia(100);s__Metschnikowia_pulcherrima(100); | Metschnikowia | 13 | Saprotroph | \|Undefined Saprotroph\| | Yeast | NULL | Probable | As nectar-tap_saprotroph (Põlme et al. 2020); Decay sugar-rich substrates (Põlme et al. 2020) | Põlme S, et al. 2020. Fungal Diversity 105:1-16 ((https://doi.org/10.1007/s13225-020-00466-2)) |
| Otu15981 | 1 | k__Fungi(100);p__Ascomycota(100);c__Saccharomycetes(100);o__Saccharomycetales(100);f__Metschnikowiaceae(100);g__Metschnikowia(100);g__Metschnikowia_unclassified(100); | Metschnikowia | 13 | Saprotroph | \|Undefined Saprotroph\| | Yeast | NULL | Probable | As nectar-tap_saprotroph (Põlme et al. 2020); Decay sugar-rich substrates (Põlme et al. 2020) | Põlme S, et al. 2020. Fungal Diversity 105:1-16 ((https://doi.org/10.1007/s13225-020-00466-2)) |
| Otu15982 | 1 | k__Fungi(100);p__Ascomycota(100);c__Saccharomycetes(100);o__Saccharomycetales(100);f__Metschnikowiaceae(100);g__Metschnikowia(100);s__Metschnikowia_pulcherrima(100); | Metschnikowia | 13 | Saprotroph | \|Undefined Saprotroph\| | Yeast | NULL | Probable | As nectar-tap_saprotroph (Põlme et al. 2020); Decay sugar-rich substrates (Põlme et al. 2020) | Põlme S, et al. 2020. Fungal Diversity 105:1-16 ((https://doi.org/10.1007/s13225-020-00466-2)) |
| Otu15983 | 1 | k__Fungi(100);p__Ascomycota(100);c__Saccharomycetes(100);o__Saccharomycetales(100);f__Metschnikowiaceae(100);g__Metschnikowia(100);s__Metschnikowia_pulcherrima(100); | Metschnikowia | 13 | Saprotroph | \|Undefined Saprotroph\| | Yeast | NULL | Probable | As nectar-tap_saprotroph (Põlme et al. 2020); Decay sugar-rich substrates (Põlme et al. 2020) | Põlme S, et al. 2020. Fungal Diversity 105:1-16 ((https://doi.org/10.1007/s13225-020-00466-2)) |
| Otu15985 | 1 | k__Fungi(100);p__Ascomycota(100);c__Saccharomycetes(100);o__Saccharomycetales(100);f__Metschnikowiaceae(100);g__Metschnikowia(100);g__Metschnikowia_unclassified(100); | Metschnikowia | 13 | Saprotroph | \|Undefined Saprotroph\| | Yeast | NULL | Probable | As nectar-tap_saprotroph (Põlme et al. 2020); Decay sugar-rich substrates (Põlme et al. 2020) | Põlme S, et al. 2020. Fungal Diversity 105:1-16 ((https://doi.org/10.1007/s13225-020-00466-2)) |
| Otu15987 | 1 | k__Fungi(100);p__Ascomycota(100);c__Saccharomycetes(100);o__Saccharomycetales(100);f__Metschnikowiaceae(100);g__Metschnikowia(100);s__Metschnikowia_pulcherrima(100); | Metschnikowia | 13 | Saprotroph | \|Undefined Saprotroph\| | Yeast | NULL | Probable | As nectar-tap_saprotroph (Põlme et al. 2020); Decay sugar-rich substrates (Põlme et al. 2020) | Põlme S, et al. 2020. Fungal Diversity 105:1-16 ((https://doi.org/10.1007/s13225-020-00466-2)) |
| Otu15988 | 1 | k__Fungi(100);p__Ascomycota(100);c__Saccharomycetes(100);o__Saccharomycetales(100);f__Metschnikowiaceae(100);g__Metschnikowia(100);s__Metschnikowia_pulcherrima(100); | Metschnikowia | 13 | Saprotroph | \|Undefined Saprotroph\| | Yeast | NULL | Probable | As nectar-tap_saprotroph (Põlme et al. 2020); Decay sugar-rich substrates (Põlme et al. 2020) | Põlme S, et al. 2020. Fungal Diversity 105:1-16 ((https://doi.org/10.1007/s13225-020-00466-2)) |
| Otu15991 | 1 | k__Fungi(100);p__Ascomycota(100);c__Saccharomycetes(100);o__Saccharomycetales(100);f__Metschnikowiaceae(100);g__Metschnikowia(100);s__Metschnikowia_pulcherrima(100); | Metschnikowia | 13 | Saprotroph | \|Undefined Saprotroph\| | Yeast | NULL | Probable | As nectar-tap_saprotroph (Põlme et al. 2020); Decay sugar-rich substrates (Põlme et al. 2020) | Põlme S, et al. 2020. Fungal Diversity 105:1-16 ((https://doi.org/10.1007/s13225-020-00466-2)) |
| Otu15992 | 1 | k__Fungi(100);p__Ascomycota(100);c__Saccharomycetes(100);o__Saccharomycetales(100);f__Metschnikowiaceae(100);g__Metschnikowia(100);g__Metschnikowia_unclassified(100); | Metschnikowia | 13 | Saprotroph | \|Undefined Saprotroph\| | Yeast | NULL | Probable | As nectar-tap_saprotroph (Põlme et al. 2020); Decay sugar-rich substrates (Põlme et al. 2020) | Põlme S, et al. 2020. Fungal Diversity 105:1-16 ((https://doi.org/10.1007/s13225-020-00466-2)) |
| Otu15993 | 1 | k__Fungi(100);p__Ascomycota(100);c__Saccharomycetes(100);o__Saccharomycetales(100);o__Saccharomycetales_unclassified(100);o__Saccharomycetales_unclassified(100);o__Saccharomycetales_unclassified(100); | Saccharomycetales | 7 | Saprotroph | Undefined Saprotroph | Yeast | NULL | Possible | NULL | Sterkenburg E, et al. 2015. New Phytologist 207:1145-1158 ((https://doi.org/10.1111/nph.13426)) |
| Otu15994 | 1 | k__Fungi(100);p__Ascomycota(100);c__Saccharomycetes(100);o__Saccharomycetales(100);f__Metschnikowiaceae(100);g__Metschnikowia(100);s__Metschnikowia_pulcherrima(100); | Metschnikowia | 13 | Saprotroph | \|Undefined Saprotroph\| | Yeast | NULL | Probable | As nectar-tap_saprotroph (Põlme et al. 2020); Decay sugar-rich substrates (Põlme et al. 2020) | Põlme S, et al. 2020. Fungal Diversity 105:1-16 ((https://doi.org/10.1007/s13225-020-00466-2)) |
| Otu15995 | 1 | k__Fungi(100);p__Ascomycota(100);c__Saccharomycetes(100);o__Saccharomycetales(100);f__Metschnikowiaceae(100);g__Metschnikowia(100);g__Metschnikowia_unclassified(100); | Metschnikowia | 13 | Saprotroph | \|Undefined Saprotroph\| | Yeast | NULL | Probable | As nectar-tap_saprotroph (Põlme et al. 2020); Decay sugar-rich substrates (Põlme et al. 2020) | Põlme S, et al. 2020. Fungal Diversity 105:1-16 ((https://doi.org/10.1007/s13225-020-00466-2)) |
| Otu15996 | 1 | k__Fungi(100);p__Ascomycota(100);c__Saccharomycetes(100);o__Saccharomycetales(100);f__Metschnikowiaceae(100);g__Metschnikowia(100);g__Metschnikowia_unclassified(100); | Metschnikowia | 13 | Saprotroph | \|Undefined Saprotroph\| | Yeast | NULL | Probable | As nectar-tap_saprotroph (Põlme et al. 2020); Decay sugar-rich substrates (Põlme et al. 2020) | Põlme S, et al. 2020. Fungal Diversity 105:1-16 ((https://doi.org/10.1007/s13225-020-00466-2)) |
| Otu15997 | 1 | k__Fungi(100);p__Ascomycota(100);c__Saccharomycetes(100);o__Saccharomycetales(100);f__Metschnikowiaceae(100);g__Metschnikowia(100);g__Metschnikowia_unclassified(100); | Metschnikowia | 13 | Saprotroph | \|Undefined Saprotroph\| | Yeast | NULL | Probable | As nectar-tap_saprotroph (Põlme et al. 2020); Decay sugar-rich substrates (Põlme et al. 2020) | Põlme S, et al. 2020. Fungal Diversity 105:1-16 ((https://doi.org/10.1007/s13225-020-00466-2)) |
| Otu15999 | 1 | k__Fungi(100);p__Ascomycota(100);c__Saccharomycetes(100);o__Saccharomycetales(100);f__Metschnikowiaceae(100);g__Metschnikowia(100);g__Metschnikowia_unclassified(100); | Metschnikowia | 13 | Saprotroph | \|Undefined Saprotroph\| | Yeast | NULL | Probable | As nectar-tap_saprotroph (Põlme et al. 2020); Decay sugar-rich substrates (Põlme et al. 2020) | Põlme S, et al. 2020. Fungal Diversity 105:1-16 ((https://doi.org/10.1007/s13225-020-00466-2)) |
| Otu16000 | 1 | k__Fungi(100);p__Ascomycota(100);c__Saccharomycetes(100);o__Saccharomycetales(100);f__Metschnikowiaceae(100);g__Metschnikowia(100);g__Metschnikowia_unclassified(100); | Metschnikowia | 13 | Saprotroph | \|Undefined Saprotroph\| | Yeast | NULL | Probable | As nectar-tap_saprotroph (Põlme et al. 2020); Decay sugar-rich substrates (Põlme et al. 2020) | Põlme S, et al. 2020. Fungal Diversity 105:1-16 ((https://doi.org/10.1007/s13225-020-00466-2)) |
| Otu16001 | 1 | k__Fungi(100);p__Ascomycota(100);c__Saccharomycetes(100);o__Saccharomycetales(100);f__Metschnikowiaceae(100);g__Metschnikowia(100);g__Metschnikowia_unclassified(100); | Metschnikowia | 13 | Saprotroph | \|Undefined Saprotroph\| | Yeast | NULL | Probable | As nectar-tap_saprotroph (Põlme et al. 2020); Decay sugar-rich substrates (Põlme et al. 2020) | Põlme S, et al. 2020. Fungal Diversity 105:1-16 ((https://doi.org/10.1007/s13225-020-00466-2)) |
| Otu16002 | 1 | k__Fungi(100);p__Ascomycota(100);c__Saccharomycetes(100);o__Saccharomycetales(100);f__Metschnikowiaceae(100);g__Metschnikowia(100);s__Metschnikowia_pulcherrima(100); | Metschnikowia | 13 | Saprotroph | \|Undefined Saprotroph\| | Yeast | NULL | Probable | As nectar-tap_saprotroph (Põlme et al. 2020); Decay sugar-rich substrates (Põlme et al. 2020) | Põlme S, et al. 2020. Fungal Diversity 105:1-16 ((https://doi.org/10.1007/s13225-020-00466-2)) |
| Otu16003 | 1 | k__Fungi(100);p__Ascomycota(100);c__Saccharomycetes(100);o__Saccharomycetales(100);f__Metschnikowiaceae(100);g__Metschnikowia(100);s__Metschnikowia_pulcherrima(100); | Metschnikowia | 13 | Saprotroph | \|Undefined Saprotroph\| | Yeast | NULL | Probable | As nectar-tap_saprotroph (Põlme et al. 2020); Decay sugar-rich substrates (Põlme et al. 2020) | Põlme S, et al. 2020. Fungal Diversity 105:1-16 ((https://doi.org/10.1007/s13225-020-00466-2)) |
| Otu16007 | 1 | k__Fungi(100);p__Ascomycota(100);c__Saccharomycetes(100);o__Saccharomycetales(100);f__Metschnikowiaceae(100);g__Metschnikowia(100);s__Metschnikowia_pulcherrima(100); | Metschnikowia | 13 | Saprotroph | \|Undefined Saprotroph\| | Yeast | NULL | Probable | As nectar-tap_saprotroph (Põlme et al. 2020); Decay sugar-rich substrates (Põlme et al. 2020) | Põlme S, et al. 2020. Fungal Diversity 105:1-16 ((https://doi.org/10.1007/s13225-020-00466-2)) |
| Otu16008 | 1 | k__Fungi(100);p__Ascomycota(100);c__Saccharomycetes(100);o__Saccharomycetales(100);f__Metschnikowiaceae(100);g__Metschnikowia(100);s__Metschnikowia_pulcherrima(100); | Metschnikowia | 13 | Saprotroph | \|Undefined Saprotroph\| | Yeast | NULL | Probable | As nectar-tap_saprotroph (Põlme et al. 2020); Decay sugar-rich substrates (Põlme et al. 2020) | Põlme S, et al. 2020. Fungal Diversity 105:1-16 ((https://doi.org/10.1007/s13225-020-00466-2)) |
| Otu16013 | 1 | k__Fungi(100);p__Ascomycota(100);c__Saccharomycetes(100);o__Saccharomycetales(100);f__Metschnikowiaceae(100);g__Metschnikowia(100);g__Metschnikowia_unclassified(100); | Metschnikowia | 13 | Saprotroph | \|Undefined Saprotroph\| | Yeast | NULL | Probable | As nectar-tap_saprotroph (Põlme et al. 2020); Decay sugar-rich substrates (Põlme et al. 2020) | Põlme S, et al. 2020. Fungal Diversity 105:1-16 ((https://doi.org/10.1007/s13225-020-00466-2)) |
| Otu16017 | 1 | k__Fungi(100);p__Ascomycota(100);c__Saccharomycetes(100);o__Saccharomycetales(100);f__Metschnikowiaceae(100);g__Metschnikowia(100);s__Metschnikowia_pulcherrima(100); | Metschnikowia | 13 | Saprotroph | \|Undefined Saprotroph\| | Yeast | NULL | Probable | As nectar-tap_saprotroph (Põlme et al. 2020); Decay sugar-rich substrates (Põlme et al. 2020) | Põlme S, et al. 2020. Fungal Diversity 105:1-16 ((https://doi.org/10.1007/s13225-020-00466-2)) |
| Otu16018 | 1 | k__Fungi(100);p__Ascomycota(100);c__Saccharomycetes(100);o__Saccharomycetales(100);f__Metschnikowiaceae(100);g__Metschnikowia(100);s__Metschnikowia_pulcherrima(100); | Metschnikowia | 13 | Saprotroph | \|Undefined Saprotroph\| | Yeast | NULL | Probable | As nectar-tap_saprotroph (Põlme et al. 2020); Decay sugar-rich substrates (Põlme et al. 2020) | Põlme S, et al. 2020. Fungal Diversity 105:1-16 ((https://doi.org/10.1007/s13225-020-00466-2)) |
| Otu16020 | 1 | k__Fungi(100);p__Ascomycota(100);c__Saccharomycetes(100);o__Saccharomycetales(100);f__Metschnikowiaceae(100);g__Metschnikowia(100);s__Metschnikowia_pulcherrima(100); | Metschnikowia | 13 | Saprotroph | \|Undefined Saprotroph\| | Yeast | NULL | Probable | As nectar-tap_saprotroph (Põlme et al. 2020); Decay sugar-rich substrates (Põlme et al. 2020) | Põlme S, et al. 2020. Fungal Diversity 105:1-16 ((https://doi.org/10.1007/s13225-020-00466-2)) |
| Otu16022 | 1 | k__Fungi(100);p__Ascomycota(100);c__Saccharomycetes(100);o__Saccharomycetales(100);f__Metschnikowiaceae(100);g__Metschnikowia(100);g__Metschnikowia_unclassified(100); | Metschnikowia | 13 | Saprotroph | \|Undefined Saprotroph\| | Yeast | NULL | Probable | As nectar-tap_saprotroph (Põlme et al. 2020); Decay sugar-rich substrates (Põlme et al. 2020) | Põlme S, et al. 2020. Fungal Diversity 105:1-16 ((https://doi.org/10.1007/s13225-020-00466-2)) |
| Otu16023 | 1 | k__Fungi(100);p__Ascomycota(100);c__Saccharomycetes(100);o__Saccharomycetales(100);f__Metschnikowiaceae(100);g__Metschnikowia(100);s__Metschnikowia_pulcherrima(100); | Metschnikowia | 13 | Saprotroph | \|Undefined Saprotroph\| | Yeast | NULL | Probable | As nectar-tap_saprotroph (Põlme et al. 2020); Decay sugar-rich substrates (Põlme et al. 2020) | Põlme S, et al. 2020. Fungal Diversity 105:1-16 ((https://doi.org/10.1007/s13225-020-00466-2)) |
| Otu16025 | 1 | k__Fungi(100);p__Ascomycota(100);c__Saccharomycetes(100);o__Saccharomycetales(100);f__Metschnikowiaceae(100);g__Metschnikowia(100);g__Metschnikowia_unclassified(100); | Metschnikowia | 13 | Saprotroph | \|Undefined Saprotroph\| | Yeast | NULL | Probable | As nectar-tap_saprotroph (Põlme et al. 2020); Decay sugar-rich substrates (Põlme et al. 2020) | Põlme S, et al. 2020. Fungal Diversity 105:1-16 ((https://doi.org/10.1007/s13225-020-00466-2)) |
| Otu16026 | 1 | k__Fungi(100);p__Ascomycota(100);c__Saccharomycetes(100);o__Saccharomycetales(100);f__Metschnikowiaceae(100);g__Metschnikowia(100);g__Metschnikowia_unclassified(100); | Metschnikowia | 13 | Saprotroph | \|Undefined Saprotroph\| | Yeast | NULL | Probable | As nectar-tap_saprotroph (Põlme et al. 2020); Decay sugar-rich substrates (Põlme et al. 2020) | Põlme S, et al. 2020. Fungal Diversity 105:1-16 ((https://doi.org/10.1007/s13225-020-00466-2)) |
| Otu16028 | 1 | k__Fungi(100);p__Ascomycota(100);c__Saccharomycetes(100);o__Saccharomycetales(100);f__Metschnikowiaceae(100);g__Metschnikowia(100);s__Metschnikowia_pulcherrima(100); | Metschnikowia | 13 | Saprotroph | \|Undefined Saprotroph\| | Yeast | NULL | Probable | As nectar-tap_saprotroph (Põlme et al. 2020); Decay sugar-rich substrates (Põlme et al. 2020) | Põlme S, et al. 2020. Fungal Diversity 105:1-16 ((https://doi.org/10.1007/s13225-020-00466-2)) |
| Otu16033 | 1 | k__Fungi(100);p__Ascomycota(100);c__Saccharomycetes(100);o__Saccharomycetales(100);f__Metschnikowiaceae(100);g__Metschnikowia(100);s__Metschnikowia_pulcherrima(100); | Metschnikowia | 13 | Saprotroph | \|Undefined Saprotroph\| | Yeast | NULL | Probable | As nectar-tap_saprotroph (Põlme et al. 2020); Decay sugar-rich substrates (Põlme et al. 2020) | Põlme S, et al. 2020. Fungal Diversity 105:1-16 ((https://doi.org/10.1007/s13225-020-00466-2)) |
| Otu16034 | 1 | k__Fungi(100);p__Ascomycota(100);c__Saccharomycetes(100);o__Saccharomycetales(100);f__Metschnikowiaceae(100);g__Metschnikowia(100);s__Metschnikowia_pulcherrima(100); | Metschnikowia | 13 | Saprotroph | \|Undefined Saprotroph\| | Yeast | NULL | Probable | As nectar-tap_saprotroph (Põlme et al. 2020); Decay sugar-rich substrates (Põlme et al. 2020) | Põlme S, et al. 2020. Fungal Diversity 105:1-16 ((https://doi.org/10.1007/s13225-020-00466-2)) |
| Otu16035 | 1 | k__Fungi(100);p__Ascomycota(100);c__Saccharomycetes(100);o__Saccharomycetales(100);f__Metschnikowiaceae(100);g__Metschnikowia(100);s__Metschnikowia_pulcherrima(100); | Metschnikowia | 13 | Saprotroph | \|Undefined Saprotroph\| | Yeast | NULL | Probable | As nectar-tap_saprotroph (Põlme et al. 2020); Decay sugar-rich substrates (Põlme et al. 2020) | Põlme S, et al. 2020. Fungal Diversity 105:1-16 ((https://doi.org/10.1007/s13225-020-00466-2)) |
| Otu16036 | 1 | k__Fungi(100);p__Ascomycota(100);c__Saccharomycetes(100);o__Saccharomycetales(100);f__Saccharomycetales_family_Incertae_sedis(100);g__Candida(100);g__Candida_unclassified(100); | Candida | 13 | Pathotroph-Saprotroph-Symbiotroph | Animal Pathogen-Endophyte-Endosymbiont-Epiphyte-\|Undefined Saprotroph\| | Dimorphic Yeast-Yeast | NULL | Probable | As nectar-tap_saprotroph (Põlme et al. 2020); Decay sugar-rich substrates (Põlme et al. 2020); This asexual genus is polyphyletic and contain species that function very differently depend on what they are. Caution should be taken when interpreting guilds for Candida species. | Manolakaki D, et al. 2010. Virulence. 1: 367-375 ((https://doi.org/10.4161/viru.1.5.12796)); Mekbib SB, et al. 2011. Tropical Plant Pathology 36:233-240 ((https://doi.org/10.1590/S1982-56762011000400004)); Põlme S, et al. 2020. Fungal Diversity 105:1-16 ((https://doi.org/10.1007/s13225-020-00466-2)) |
| Otu16038 | 1 | k__Fungi(100);p__Ascomycota(100);c__Saccharomycetes(100);o__Saccharomycetales(100);f__Metschnikowiaceae(100);g__Metschnikowia(100);g__Metschnikowia_unclassified(100); | Metschnikowia | 13 | Saprotroph | \|Undefined Saprotroph\| | Yeast | NULL | Probable | As nectar-tap_saprotroph (Põlme et al. 2020); Decay sugar-rich substrates (Põlme et al. 2020) | Põlme S, et al. 2020. Fungal Diversity 105:1-16 ((https://doi.org/10.1007/s13225-020-00466-2)) |
| Otu16039 | 1 | k__Fungi(100);p__Ascomycota(100);c__Saccharomycetes(100);o__Saccharomycetales(100);f__Saccharomycetales_family_Incertae_sedis(100);g__Candida(100);s__Candida_cretensis(100); | Candida | 13 | Pathotroph-Saprotroph-Symbiotroph | Animal Pathogen-Endophyte-Endosymbiont-Epiphyte-\|Undefined Saprotroph\| | Dimorphic Yeast-Yeast | NULL | Probable | As nectar-tap_saprotroph (Põlme et al. 2020); Decay sugar-rich substrates (Põlme et al. 2020); This asexual genus is polyphyletic and contain species that function very differently depend on what they are. Caution should be taken when interpreting guilds for Candida species. | Manolakaki D, et al. 2010. Virulence. 1: 367-375 ((https://doi.org/10.4161/viru.1.5.12796)); Mekbib SB, et al. 2011. Tropical Plant Pathology 36:233-240 ((https://doi.org/10.1590/S1982-56762011000400004)); Põlme S, et al. 2020. Fungal Diversity 105:1-16 ((https://doi.org/10.1007/s13225-020-00466-2)) |
| Otu16040 | 1 | k__Fungi(100);p__Ascomycota(100);c__Saccharomycetes(100);o__Saccharomycetales(100);f__Metschnikowiaceae(100);g__Metschnikowia(100);s__Metschnikowia_pulcherrima(100); | Metschnikowia | 13 | Saprotroph | \|Undefined Saprotroph\| | Yeast | NULL | Probable | As nectar-tap_saprotroph (Põlme et al. 2020); Decay sugar-rich substrates (Põlme et al. 2020) | Põlme S, et al. 2020. Fungal Diversity 105:1-16 ((https://doi.org/10.1007/s13225-020-00466-2)) |
| Otu16042 | 1 | k__Fungi(100);p__Ascomycota(100);c__Saccharomycetes(100);o__Saccharomycetales(100);f__Metschnikowiaceae(100);g__Metschnikowia(100);s__Metschnikowia_pulcherrima(100); | Metschnikowia | 13 | Saprotroph | \|Undefined Saprotroph\| | Yeast | NULL | Probable | As nectar-tap_saprotroph (Põlme et al. 2020); Decay sugar-rich substrates (Põlme et al. 2020) | Põlme S, et al. 2020. Fungal Diversity 105:1-16 ((https://doi.org/10.1007/s13225-020-00466-2)) |
| Otu16043 | 1 | k__Fungi(100);p__Ascomycota(100);c__Saccharomycetes(100);o__Saccharomycetales(100);f__Metschnikowiaceae(100);g__Metschnikowia(100);s__Metschnikowia_pulcherrima(100); | Metschnikowia | 13 | Saprotroph | \|Undefined Saprotroph\| | Yeast | NULL | Probable | As nectar-tap_saprotroph (Põlme et al. 2020); Decay sugar-rich substrates (Põlme et al. 2020) | Põlme S, et al. 2020. Fungal Diversity 105:1-16 ((https://doi.org/10.1007/s13225-020-00466-2)) |
| Otu16044 | 1 | k__Fungi(100);p__Ascomycota(100);c__Saccharomycetes(100);o__Saccharomycetales(100);f__Metschnikowiaceae(100);g__Metschnikowia(100);g__Metschnikowia_unclassified(100); | Metschnikowia | 13 | Saprotroph | \|Undefined Saprotroph\| | Yeast | NULL | Probable | As nectar-tap_saprotroph (Põlme et al. 2020); Decay sugar-rich substrates (Põlme et al. 2020) | Põlme S, et al. 2020. Fungal Diversity 105:1-16 ((https://doi.org/10.1007/s13225-020-00466-2)) |
| Otu16047 | 1 | k__Fungi(100);p__Ascomycota(100);c__Saccharomycetes(100);o__Saccharomycetales(100);f__Saccharomycetales_family_Incertae_sedis(100);g__Candida(100);g__Candida_unclassified(100); | Candida | 13 | Pathotroph-Saprotroph-Symbiotroph | Animal Pathogen-Endophyte-Endosymbiont-Epiphyte-\|Undefined Saprotroph\| | Dimorphic Yeast-Yeast | NULL | Probable | As nectar-tap_saprotroph (Põlme et al. 2020); Decay sugar-rich substrates (Põlme et al. 2020); This asexual genus is polyphyletic and contain species that function very differently depend on what they are. Caution should be taken when interpreting guilds for Candida species. | Manolakaki D, et al. 2010. Virulence. 1: 367-375 ((https://doi.org/10.4161/viru.1.5.12796)); Mekbib SB, et al. 2011. Tropical Plant Pathology 36:233-240 ((https://doi.org/10.1590/S1982-56762011000400004)); Põlme S, et al. 2020. Fungal Diversity 105:1-16 ((https://doi.org/10.1007/s13225-020-00466-2)) |
| Otu16049 | 1 | k__Fungi(100);p__Ascomycota(100);c__Saccharomycetes(100);o__Saccharomycetales(100);f__Metschnikowiaceae(100);g__Metschnikowia(100);s__Metschnikowia_pulcherrima(100); | Metschnikowia | 13 | Saprotroph | \|Undefined Saprotroph\| | Yeast | NULL | Probable | As nectar-tap_saprotroph (Põlme et al. 2020); Decay sugar-rich substrates (Põlme et al. 2020) | Põlme S, et al. 2020. Fungal Diversity 105:1-16 ((https://doi.org/10.1007/s13225-020-00466-2)) |
| Otu16050 | 1 | k__Fungi(100);p__Ascomycota(100);c__Saccharomycetes(100);o__Saccharomycetales(100);f__Metschnikowiaceae(100);g__Metschnikowia(100);s__Metschnikowia_pulcherrima(100); | Metschnikowia | 13 | Saprotroph | \|Undefined Saprotroph\| | Yeast | NULL | Probable | As nectar-tap_saprotroph (Põlme et al. 2020); Decay sugar-rich substrates (Põlme et al. 2020) | Põlme S, et al. 2020. Fungal Diversity 105:1-16 ((https://doi.org/10.1007/s13225-020-00466-2)) |
| Otu16051 | 1 | k__Fungi(100);p__Ascomycota(100);c__Saccharomycetes(100);o__Saccharomycetales(100);f__Metschnikowiaceae(100);g__Metschnikowia(100);s__Metschnikowia_pulcherrima(100); | Metschnikowia | 13 | Saprotroph | \|Undefined Saprotroph\| | Yeast | NULL | Probable | As nectar-tap_saprotroph (Põlme et al. 2020); Decay sugar-rich substrates (Põlme et al. 2020) | Põlme S, et al. 2020. Fungal Diversity 105:1-16 ((https://doi.org/10.1007/s13225-020-00466-2)) |
| Otu16053 | 1 | k__Fungi(100);p__Ascomycota(100);c__Saccharomycetes(100);o__Saccharomycetales(100);f__Saccharomycetales_family_Incertae_sedis(100);g__Candida(100);g__Candida_unclassified(100); | Candida | 13 | Pathotroph-Saprotroph-Symbiotroph | Animal Pathogen-Endophyte-Endosymbiont-Epiphyte-\|Undefined Saprotroph\| | Dimorphic Yeast-Yeast | NULL | Probable | As nectar-tap_saprotroph (Põlme et al. 2020); Decay sugar-rich substrates (Põlme et al. 2020); This asexual genus is polyphyletic and contain species that function very differently depend on what they are. Caution should be taken when interpreting guilds for Candida species. | Manolakaki D, et al. 2010. Virulence. 1: 367-375 ((https://doi.org/10.4161/viru.1.5.12796)); Mekbib SB, et al. 2011. Tropical Plant Pathology 36:233-240 ((https://doi.org/10.1590/S1982-56762011000400004)); Põlme S, et al. 2020. Fungal Diversity 105:1-16 ((https://doi.org/10.1007/s13225-020-00466-2)) |
| Otu16054 | 1 | k__Fungi(100);p__Ascomycota(100);c__Saccharomycetes(100);o__Saccharomycetales(100);f__Metschnikowiaceae(100);g__Metschnikowia(100);s__Metschnikowia_pulcherrima(100); | Metschnikowia | 13 | Saprotroph | \|Undefined Saprotroph\| | Yeast | NULL | Probable | As nectar-tap_saprotroph (Põlme et al. 2020); Decay sugar-rich substrates (Põlme et al. 2020) | Põlme S, et al. 2020. Fungal Diversity 105:1-16 ((https://doi.org/10.1007/s13225-020-00466-2)) |
| Otu16055 | 1 | k__Fungi(100);p__Ascomycota(100);c__Saccharomycetes(100);o__Saccharomycetales(100);f__Metschnikowiaceae(100);g__Metschnikowia(100);s__Metschnikowia_pulcherrima(100); | Metschnikowia | 13 | Saprotroph | \|Undefined Saprotroph\| | Yeast | NULL | Probable | As nectar-tap_saprotroph (Põlme et al. 2020); Decay sugar-rich substrates (Põlme et al. 2020) | Põlme S, et al. 2020. Fungal Diversity 105:1-16 ((https://doi.org/10.1007/s13225-020-00466-2)) |
| Otu16058 | 1 | k__Fungi(100);p__Ascomycota(100);c__Saccharomycetes(100);o__Saccharomycetales(100);f__Metschnikowiaceae(100);g__Metschnikowia(100);g__Metschnikowia_unclassified(100); | Metschnikowia | 13 | Saprotroph | \|Undefined Saprotroph\| | Yeast | NULL | Probable | As nectar-tap_saprotroph (Põlme et al. 2020); Decay sugar-rich substrates (Põlme et al. 2020) | Põlme S, et al. 2020. Fungal Diversity 105:1-16 ((https://doi.org/10.1007/s13225-020-00466-2)) |
| Otu16059 | 1 | k__Fungi(100);p__Ascomycota(100);c__Saccharomycetes(100);o__Saccharomycetales(100);f__Saccharomycetales_family_Incertae_sedis(100);g__Candida(100);g__Candida_unclassified(100); | Candida | 13 | Pathotroph-Saprotroph-Symbiotroph | Animal Pathogen-Endophyte-Endosymbiont-Epiphyte-\|Undefined Saprotroph\| | Dimorphic Yeast-Yeast | NULL | Probable | As nectar-tap_saprotroph (Põlme et al. 2020); Decay sugar-rich substrates (Põlme et al. 2020); This asexual genus is polyphyletic and contain species that function very differently depend on what they are. Caution should be taken when interpreting guilds for Candida species. | Manolakaki D, et al. 2010. Virulence. 1: 367-375 ((https://doi.org/10.4161/viru.1.5.12796)); Mekbib SB, et al. 2011. Tropical Plant Pathology 36:233-240 ((https://doi.org/10.1590/S1982-56762011000400004)); Põlme S, et al. 2020. Fungal Diversity 105:1-16 ((https://doi.org/10.1007/s13225-020-00466-2)) |
| Otu16061 | 1 | k__Fungi(100);p__Ascomycota(100);c__Saccharomycetes(100);o__Saccharomycetales(100);f__Metschnikowiaceae(100);g__Metschnikowia(100);g__Metschnikowia_unclassified(100); | Metschnikowia | 13 | Saprotroph | \|Undefined Saprotroph\| | Yeast | NULL | Probable | As nectar-tap_saprotroph (Põlme et al. 2020); Decay sugar-rich substrates (Põlme et al. 2020) | Põlme S, et al. 2020. Fungal Diversity 105:1-16 ((https://doi.org/10.1007/s13225-020-00466-2)) |
| Otu16062 | 1 | k__Fungi(100);p__Ascomycota(100);c__Saccharomycetes(100);o__Saccharomycetales(100);f__Metschnikowiaceae(100);g__Metschnikowia(100);s__Metschnikowia_pulcherrima(100); | Metschnikowia | 13 | Saprotroph | \|Undefined Saprotroph\| | Yeast | NULL | Probable | As nectar-tap_saprotroph (Põlme et al. 2020); Decay sugar-rich substrates (Põlme et al. 2020) | Põlme S, et al. 2020. Fungal Diversity 105:1-16 ((https://doi.org/10.1007/s13225-020-00466-2)) |
| Otu16064 | 1 | k__Fungi(100);p__Ascomycota(100);c__Saccharomycetes(100);o__Saccharomycetales(100);f__Metschnikowiaceae(100);g__Metschnikowia(100);s__Metschnikowia_pulcherrima(100); | Metschnikowia | 13 | Saprotroph | \|Undefined Saprotroph\| | Yeast | NULL | Probable | As nectar-tap_saprotroph (Põlme et al. 2020); Decay sugar-rich substrates (Põlme et al. 2020) | Põlme S, et al. 2020. Fungal Diversity 105:1-16 ((https://doi.org/10.1007/s13225-020-00466-2)) |
| Otu16065 | 1 | k__Fungi(100);p__Ascomycota(100);c__Saccharomycetes(100);o__Saccharomycetales(100);f__Metschnikowiaceae(100);g__Metschnikowia(100);s__Metschnikowia_pulcherrima(100); | Metschnikowia | 13 | Saprotroph | \|Undefined Saprotroph\| | Yeast | NULL | Probable | As nectar-tap_saprotroph (Põlme et al. 2020); Decay sugar-rich substrates (Põlme et al. 2020) | Põlme S, et al. 2020. Fungal Diversity 105:1-16 ((https://doi.org/10.1007/s13225-020-00466-2)) |
| Otu16066 | 1 | k__Fungi(100);p__Ascomycota(100);c__Saccharomycetes(100);o__Saccharomycetales(100);f__Metschnikowiaceae(100);g__Metschnikowia(100);s__Metschnikowia_pulcherrima(100); | Metschnikowia | 13 | Saprotroph | \|Undefined Saprotroph\| | Yeast | NULL | Probable | As nectar-tap_saprotroph (Põlme et al. 2020); Decay sugar-rich substrates (Põlme et al. 2020) | Põlme S, et al. 2020. Fungal Diversity 105:1-16 ((https://doi.org/10.1007/s13225-020-00466-2)) |
| Otu16070 | 1 | k__Fungi(100);p__Ascomycota(100);c__Saccharomycetes(100);o__Saccharomycetales(100);f__Metschnikowiaceae(100);g__Metschnikowia(100);g__Metschnikowia_unclassified(100); | Metschnikowia | 13 | Saprotroph | \|Undefined Saprotroph\| | Yeast | NULL | Probable | As nectar-tap_saprotroph (Põlme et al. 2020); Decay sugar-rich substrates (Põlme et al. 2020) | Põlme S, et al. 2020. Fungal Diversity 105:1-16 ((https://doi.org/10.1007/s13225-020-00466-2)) |
| Otu16071 | 1 | k__Fungi(100);p__Ascomycota(100);c__Saccharomycetes(100);o__Saccharomycetales(100);f__Metschnikowiaceae(100);g__Metschnikowia(100);g__Metschnikowia_unclassified(100); | Metschnikowia | 13 | Saprotroph | \|Undefined Saprotroph\| | Yeast | NULL | Probable | As nectar-tap_saprotroph (Põlme et al. 2020); Decay sugar-rich substrates (Põlme et al. 2020) | Põlme S, et al. 2020. Fungal Diversity 105:1-16 ((https://doi.org/10.1007/s13225-020-00466-2)) |
| Otu16074 | 1 | k__Fungi(100);p__Ascomycota(100);c__Saccharomycetes(100);o__Saccharomycetales(100);f__Metschnikowiaceae(100);g__Metschnikowia(100);s__Metschnikowia_pulcherrima(100); | Metschnikowia | 13 | Saprotroph | \|Undefined Saprotroph\| | Yeast | NULL | Probable | As nectar-tap_saprotroph (Põlme et al. 2020); Decay sugar-rich substrates (Põlme et al. 2020) | Põlme S, et al. 2020. Fungal Diversity 105:1-16 ((https://doi.org/10.1007/s13225-020-00466-2)) |
| Otu16076 | 1 | k__Fungi(100);p__Ascomycota(100);c__Saccharomycetes(100);o__Saccharomycetales(100);f__Metschnikowiaceae(100);g__Metschnikowia(100);g__Metschnikowia_unclassified(100); | Metschnikowia | 13 | Saprotroph | \|Undefined Saprotroph\| | Yeast | NULL | Probable | As nectar-tap_saprotroph (Põlme et al. 2020); Decay sugar-rich substrates (Põlme et al. 2020) | Põlme S, et al. 2020. Fungal Diversity 105:1-16 ((https://doi.org/10.1007/s13225-020-00466-2)) |
| Otu16078 | 1 | k__Fungi(100);p__Ascomycota(100);c__Saccharomycetes(100);o__Saccharomycetales(100);f__Metschnikowiaceae(100);g__Metschnikowia(100);g__Metschnikowia_unclassified(100); | Metschnikowia | 13 | Saprotroph | \|Undefined Saprotroph\| | Yeast | NULL | Probable | As nectar-tap_saprotroph (Põlme et al. 2020); Decay sugar-rich substrates (Põlme et al. 2020) | Põlme S, et al. 2020. Fungal Diversity 105:1-16 ((https://doi.org/10.1007/s13225-020-00466-2)) |
| Otu16079 | 1 | k__Fungi(100);p__Ascomycota(100);c__Ascomycota_class_Incertae_sedis(100);o__Ascomycota_order_Incertae_sedis(100);f__Ascomycota_family_Incertae_sedis(100);g__Pyrigemmula(100);s__Pyrigemmula_aurantiaca(100); | Pyrigemmula | 13 | Saprotroph | \|Undefined Saprotroph\|-Wood Saprotroph | Microfungus | NULL | Probable | As nectar-tap_saprotroph (Põlme et al. 2020); Decay wood, sugar-rich substrates (Põlme et al. 2020) | Põlme S, et al. 2020. Fungal Diversity 105:1-16 ((https://doi.org/10.1007/s13225-020-00466-2)) |
| Otu16081 | 1 | k__Fungi(100);p__Ascomycota(100);c__Saccharomycetes(100);o__Saccharomycetales(100);f__Metschnikowiaceae(100);g__Metschnikowia(100);s__Metschnikowia_pulcherrima(100); | Metschnikowia | 13 | Saprotroph | \|Undefined Saprotroph\| | Yeast | NULL | Probable | As nectar-tap_saprotroph (Põlme et al. 2020); Decay sugar-rich substrates (Põlme et al. 2020) | Põlme S, et al. 2020. Fungal Diversity 105:1-16 ((https://doi.org/10.1007/s13225-020-00466-2)) |
| Otu16082 | 1 | k__Fungi(100);p__Ascomycota(100);c__Saccharomycetes(100);o__Saccharomycetales(100);f__Metschnikowiaceae(100);g__Metschnikowia(100);g__Metschnikowia_unclassified(100); | Metschnikowia | 13 | Saprotroph | \|Undefined Saprotroph\| | Yeast | NULL | Probable | As nectar-tap_saprotroph (Põlme et al. 2020); Decay sugar-rich substrates (Põlme et al. 2020) | Põlme S, et al. 2020. Fungal Diversity 105:1-16 ((https://doi.org/10.1007/s13225-020-00466-2)) |
| Otu16083 | 1 | k__Fungi(100);p__Ascomycota(100);c__Saccharomycetes(100);o__Saccharomycetales(100);f__Metschnikowiaceae(100);g__Metschnikowia(100);s__Metschnikowia_pulcherrima(100); | Metschnikowia | 13 | Saprotroph | \|Undefined Saprotroph\| | Yeast | NULL | Probable | As nectar-tap_saprotroph (Põlme et al. 2020); Decay sugar-rich substrates (Põlme et al. 2020) | Põlme S, et al. 2020. Fungal Diversity 105:1-16 ((https://doi.org/10.1007/s13225-020-00466-2)) |
| Otu16084 | 1 | k__Fungi(100);p__Ascomycota(100);c__Saccharomycetes(100);o__Saccharomycetales(100);f__Metschnikowiaceae(100);g__Metschnikowia(100);s__Metschnikowia_pulcherrima(100); | Metschnikowia | 13 | Saprotroph | \|Undefined Saprotroph\| | Yeast | NULL | Probable | As nectar-tap_saprotroph (Põlme et al. 2020); Decay sugar-rich substrates (Põlme et al. 2020) | Põlme S, et al. 2020. Fungal Diversity 105:1-16 ((https://doi.org/10.1007/s13225-020-00466-2)) |
| Otu16086 | 1 | k__Fungi(100);p__Ascomycota(100);c__Saccharomycetes(100);o__Saccharomycetales(100);f__Metschnikowiaceae(100);g__Metschnikowia(100);g__Metschnikowia_unclassified(100); | Metschnikowia | 13 | Saprotroph | \|Undefined Saprotroph\| | Yeast | NULL | Probable | As nectar-tap_saprotroph (Põlme et al. 2020); Decay sugar-rich substrates (Põlme et al. 2020) | Põlme S, et al. 2020. Fungal Diversity 105:1-16 ((https://doi.org/10.1007/s13225-020-00466-2)) |
| Otu16089 | 1 | k__Fungi(100);p__Ascomycota(100);c__Saccharomycetes(100);o__Saccharomycetales(100);f__Metschnikowiaceae(100);g__Metschnikowia(100);g__Metschnikowia_unclassified(100); | Metschnikowia | 13 | Saprotroph | \|Undefined Saprotroph\| | Yeast | NULL | Probable | As nectar-tap_saprotroph (Põlme et al. 2020); Decay sugar-rich substrates (Põlme et al. 2020) | Põlme S, et al. 2020. Fungal Diversity 105:1-16 ((https://doi.org/10.1007/s13225-020-00466-2)) |
| Otu16093 | 1 | k__Fungi(100);p__Ascomycota(100);c__Saccharomycetes(100);o__Saccharomycetales(100);f__Metschnikowiaceae(100);g__Metschnikowia(100);s__Metschnikowia_pulcherrima(100); | Metschnikowia | 13 | Saprotroph | \|Undefined Saprotroph\| | Yeast | NULL | Probable | As nectar-tap_saprotroph (Põlme et al. 2020); Decay sugar-rich substrates (Põlme et al. 2020) | Põlme S, et al. 2020. Fungal Diversity 105:1-16 ((https://doi.org/10.1007/s13225-020-00466-2)) |
| Otu16096 | 1 | k__Fungi(100);p__Ascomycota(100);c__Saccharomycetes(100);o__Saccharomycetales(100);f__Metschnikowiaceae(100);g__Metschnikowia(100);s__Metschnikowia_pulcherrima(100); | Metschnikowia | 13 | Saprotroph | \|Undefined Saprotroph\| | Yeast | NULL | Probable | As nectar-tap_saprotroph (Põlme et al. 2020); Decay sugar-rich substrates (Põlme et al. 2020) | Põlme S, et al. 2020. Fungal Diversity 105:1-16 ((https://doi.org/10.1007/s13225-020-00466-2)) |
| Otu16099 | 1 | k__Fungi(100);p__Ascomycota(100);c__Saccharomycetes(100);o__Saccharomycetales(100);f__Metschnikowiaceae(100);g__Metschnikowia(100);s__Metschnikowia_pulcherrima(100); | Metschnikowia | 13 | Saprotroph | \|Undefined Saprotroph\| | Yeast | NULL | Probable | As nectar-tap_saprotroph (Põlme et al. 2020); Decay sugar-rich substrates (Põlme et al. 2020) | Põlme S, et al. 2020. Fungal Diversity 105:1-16 ((https://doi.org/10.1007/s13225-020-00466-2)) |
| Otu16102 | 1 | k__Fungi(100);p__Ascomycota(100);c__Saccharomycetes(100);o__Saccharomycetales(100);f__Metschnikowiaceae(100);g__Metschnikowia(100);s__Metschnikowia_pulcherrima(100); | Metschnikowia | 13 | Saprotroph | \|Undefined Saprotroph\| | Yeast | NULL | Probable | As nectar-tap_saprotroph (Põlme et al. 2020); Decay sugar-rich substrates (Põlme et al. 2020) | Põlme S, et al. 2020. Fungal Diversity 105:1-16 ((https://doi.org/10.1007/s13225-020-00466-2)) |
| Otu16103 | 1 | k__Fungi(100);p__Ascomycota(100);c__Saccharomycetes(100);o__Saccharomycetales(100);f__Metschnikowiaceae(100);g__Metschnikowia(100);s__Metschnikowia_pulcherrima(100); | Metschnikowia | 13 | Saprotroph | \|Undefined Saprotroph\| | Yeast | NULL | Probable | As nectar-tap_saprotroph (Põlme et al. 2020); Decay sugar-rich substrates (Põlme et al. 2020) | Põlme S, et al. 2020. Fungal Diversity 105:1-16 ((https://doi.org/10.1007/s13225-020-00466-2)) |
| Otu16108 | 1 | k__Fungi(100);p__Ascomycota(100);c__Saccharomycetes(100);o__Saccharomycetales(100);f__Metschnikowiaceae(100);g__Metschnikowia(100);g__Metschnikowia_unclassified(100); | Metschnikowia | 13 | Saprotroph | \|Undefined Saprotroph\| | Yeast | NULL | Probable | As nectar-tap_saprotroph (Põlme et al. 2020); Decay sugar-rich substrates (Põlme et al. 2020) | Põlme S, et al. 2020. Fungal Diversity 105:1-16 ((https://doi.org/10.1007/s13225-020-00466-2)) |
| Otu16109 | 1 | k__Fungi(100);p__Ascomycota(100);c__Saccharomycetes(100);o__Saccharomycetales(100);f__Metschnikowiaceae(100);g__Metschnikowia(100);s__Metschnikowia_pulcherrima(100); | Metschnikowia | 13 | Saprotroph | \|Undefined Saprotroph\| | Yeast | NULL | Probable | As nectar-tap_saprotroph (Põlme et al. 2020); Decay sugar-rich substrates (Põlme et al. 2020) | Põlme S, et al. 2020. Fungal Diversity 105:1-16 ((https://doi.org/10.1007/s13225-020-00466-2)) |
| Otu16110 | 1 | k__Fungi(100);p__Ascomycota(100);c__Saccharomycetes(100);o__Saccharomycetales(100);f__Metschnikowiaceae(100);g__Metschnikowia(100);s__Metschnikowia_pulcherrima(100); | Metschnikowia | 13 | Saprotroph | \|Undefined Saprotroph\| | Yeast | NULL | Probable | As nectar-tap_saprotroph (Põlme et al. 2020); Decay sugar-rich substrates (Põlme et al. 2020) | Põlme S, et al. 2020. Fungal Diversity 105:1-16 ((https://doi.org/10.1007/s13225-020-00466-2)) |
| Otu16111 | 1 | k__Fungi(100);p__Ascomycota(100);c__Saccharomycetes(100);o__Saccharomycetales(100);f__Metschnikowiaceae(100);g__Metschnikowia(100);s__Metschnikowia_pulcherrima(100); | Metschnikowia | 13 | Saprotroph | \|Undefined Saprotroph\| | Yeast | NULL | Probable | As nectar-tap_saprotroph (Põlme et al. 2020); Decay sugar-rich substrates (Põlme et al. 2020) | Põlme S, et al. 2020. Fungal Diversity 105:1-16 ((https://doi.org/10.1007/s13225-020-00466-2)) |
| Otu16112 | 1 | k__Fungi(100);p__Ascomycota(100);c__Saccharomycetes(100);o__Saccharomycetales(100);f__Metschnikowiaceae(100);g__Metschnikowia(100);g__Metschnikowia_unclassified(100); | Metschnikowia | 13 | Saprotroph | \|Undefined Saprotroph\| | Yeast | NULL | Probable | As nectar-tap_saprotroph (Põlme et al. 2020); Decay sugar-rich substrates (Põlme et al. 2020) | Põlme S, et al. 2020. Fungal Diversity 105:1-16 ((https://doi.org/10.1007/s13225-020-00466-2)) |
| Otu16113 | 1 | k__Fungi(100);p__Ascomycota(100);c__Saccharomycetes(100);o__Saccharomycetales(100);f__Metschnikowiaceae(100);g__Metschnikowia(100);g__Metschnikowia_unclassified(100); | Metschnikowia | 13 | Saprotroph | \|Undefined Saprotroph\| | Yeast | NULL | Probable | As nectar-tap_saprotroph (Põlme et al. 2020); Decay sugar-rich substrates (Põlme et al. 2020) | Põlme S, et al. 2020. Fungal Diversity 105:1-16 ((https://doi.org/10.1007/s13225-020-00466-2)) |
| Otu16114 | 1 | k__Fungi(100);p__Ascomycota(100);c__Saccharomycetes(100);o__Saccharomycetales(100);f__Metschnikowiaceae(100);g__Metschnikowia(100);s__Metschnikowia_pulcherrima(100); | Metschnikowia | 13 | Saprotroph | \|Undefined Saprotroph\| | Yeast | NULL | Probable | As nectar-tap_saprotroph (Põlme et al. 2020); Decay sugar-rich substrates (Põlme et al. 2020) | Põlme S, et al. 2020. Fungal Diversity 105:1-16 ((https://doi.org/10.1007/s13225-020-00466-2)) |
| Otu16115 | 1 | k__Fungi(100);p__Ascomycota(100);c__Saccharomycetes(100);o__Saccharomycetales(100);f__Metschnikowiaceae(100);g__Metschnikowia(100);g__Metschnikowia_unclassified(100); | Metschnikowia | 13 | Saprotroph | \|Undefined Saprotroph\| | Yeast | NULL | Probable | As nectar-tap_saprotroph (Põlme et al. 2020); Decay sugar-rich substrates (Põlme et al. 2020) | Põlme S, et al. 2020. Fungal Diversity 105:1-16 ((https://doi.org/10.1007/s13225-020-00466-2)) |
| Otu16116 | 1 | k__Fungi(100);p__Ascomycota(100);c__Saccharomycetes(100);o__Saccharomycetales(100);f__Metschnikowiaceae(100);g__Metschnikowia(100);s__Metschnikowia_pulcherrima(100); | Metschnikowia | 13 | Saprotroph | \|Undefined Saprotroph\| | Yeast | NULL | Probable | As nectar-tap_saprotroph (Põlme et al. 2020); Decay sugar-rich substrates (Põlme et al. 2020) | Põlme S, et al. 2020. Fungal Diversity 105:1-16 ((https://doi.org/10.1007/s13225-020-00466-2)) |
| Otu16117 | 1 | k__Fungi(100);p__Ascomycota(100);c__Saccharomycetes(100);o__Saccharomycetales(100);f__Metschnikowiaceae(100);g__Metschnikowia(100);s__Metschnikowia_pulcherrima(100); | Metschnikowia | 13 | Saprotroph | \|Undefined Saprotroph\| | Yeast | NULL | Probable | As nectar-tap_saprotroph (Põlme et al. 2020); Decay sugar-rich substrates (Põlme et al. 2020) | Põlme S, et al. 2020. Fungal Diversity 105:1-16 ((https://doi.org/10.1007/s13225-020-00466-2)) |
| Otu16120 | 1 | k__Fungi(100);p__Ascomycota(100);c__Saccharomycetes(100);o__Saccharomycetales(100);f__Saccharomycetales_family_Incertae_sedis(100);g__Candida(100);g__Candida_unclassified(100); | Candida | 13 | Pathotroph-Saprotroph-Symbiotroph | Animal Pathogen-Endophyte-Endosymbiont-Epiphyte-\|Undefined Saprotroph\| | Dimorphic Yeast-Yeast | NULL | Probable | As nectar-tap_saprotroph (Põlme et al. 2020); Decay sugar-rich substrates (Põlme et al. 2020); This asexual genus is polyphyletic and contain species that function very differently depend on what they are. Caution should be taken when interpreting guilds for Candida species. | Manolakaki D, et al. 2010. Virulence. 1: 367-375 ((https://doi.org/10.4161/viru.1.5.12796)); Mekbib SB, et al. 2011. Tropical Plant Pathology 36:233-240 ((https://doi.org/10.1590/S1982-56762011000400004)); Põlme S, et al. 2020. Fungal Diversity 105:1-16 ((https://doi.org/10.1007/s13225-020-00466-2)) |
| Otu16122 | 1 | k__Fungi(100);p__Ascomycota(100);c__Saccharomycetes(100);o__Saccharomycetales(100);f__Metschnikowiaceae(100);g__Metschnikowia(100);s__Metschnikowia_pulcherrima(100); | Metschnikowia | 13 | Saprotroph | \|Undefined Saprotroph\| | Yeast | NULL | Probable | As nectar-tap_saprotroph (Põlme et al. 2020); Decay sugar-rich substrates (Põlme et al. 2020) | Põlme S, et al. 2020. Fungal Diversity 105:1-16 ((https://doi.org/10.1007/s13225-020-00466-2)) |
| Otu16123 | 1 | k__Fungi(100);p__Ascomycota(100);c__Saccharomycetes(100);o__Saccharomycetales(100);f__Metschnikowiaceae(100);g__Metschnikowia(100);g__Metschnikowia_unclassified(100); | Metschnikowia | 13 | Saprotroph | \|Undefined Saprotroph\| | Yeast | NULL | Probable | As nectar-tap_saprotroph (Põlme et al. 2020); Decay sugar-rich substrates (Põlme et al. 2020) | Põlme S, et al. 2020. Fungal Diversity 105:1-16 ((https://doi.org/10.1007/s13225-020-00466-2)) |
| Otu16128 | 1 | k__Fungi(100);p__Ascomycota(100);c__Saccharomycetes(100);o__Saccharomycetales(100);f__Metschnikowiaceae(100);g__Metschnikowia(100);g__Metschnikowia_unclassified(100); | Metschnikowia | 13 | Saprotroph | \|Undefined Saprotroph\| | Yeast | NULL | Probable | As nectar-tap_saprotroph (Põlme et al. 2020); Decay sugar-rich substrates (Põlme et al. 2020) | Põlme S, et al. 2020. Fungal Diversity 105:1-16 ((https://doi.org/10.1007/s13225-020-00466-2)) |
| Otu16131 | 1 | k__Fungi(100);p__Ascomycota(100);c__Saccharomycetes(100);o__Saccharomycetales(100);f__Metschnikowiaceae(100);g__Metschnikowia(100);s__Metschnikowia_pulcherrima(100); | Metschnikowia | 13 | Saprotroph | \|Undefined Saprotroph\| | Yeast | NULL | Probable | As nectar-tap_saprotroph (Põlme et al. 2020); Decay sugar-rich substrates (Põlme et al. 2020) | Põlme S, et al. 2020. Fungal Diversity 105:1-16 ((https://doi.org/10.1007/s13225-020-00466-2)) |
| Otu16132 | 1 | k__Fungi(100);p__Ascomycota(100);c__Saccharomycetes(100);o__Saccharomycetales(100);f__Metschnikowiaceae(100);g__Metschnikowia(100);s__Metschnikowia_pulcherrima(100); | Metschnikowia | 13 | Saprotroph | \|Undefined Saprotroph\| | Yeast | NULL | Probable | As nectar-tap_saprotroph (Põlme et al. 2020); Decay sugar-rich substrates (Põlme et al. 2020) | Põlme S, et al. 2020. Fungal Diversity 105:1-16 ((https://doi.org/10.1007/s13225-020-00466-2)) |
| Otu16133 | 1 | k__Fungi(100);p__Ascomycota(100);c__Saccharomycetes(100);o__Saccharomycetales(100);f__Metschnikowiaceae(100);g__Metschnikowia(100);g__Metschnikowia_unclassified(100); | Metschnikowia | 13 | Saprotroph | \|Undefined Saprotroph\| | Yeast | NULL | Probable | As nectar-tap_saprotroph (Põlme et al. 2020); Decay sugar-rich substrates (Põlme et al. 2020) | Põlme S, et al. 2020. Fungal Diversity 105:1-16 ((https://doi.org/10.1007/s13225-020-00466-2)) |
| Otu16134 | 1 | k__Fungi(100);p__Ascomycota(100);c__Saccharomycetes(100);o__Saccharomycetales(100);f__Metschnikowiaceae(100);g__Metschnikowia(100);s__Metschnikowia_pulcherrima(100); | Metschnikowia | 13 | Saprotroph | \|Undefined Saprotroph\| | Yeast | NULL | Probable | As nectar-tap_saprotroph (Põlme et al. 2020); Decay sugar-rich substrates (Põlme et al. 2020) | Põlme S, et al. 2020. Fungal Diversity 105:1-16 ((https://doi.org/10.1007/s13225-020-00466-2)) |
| Otu16136 | 1 | k__Fungi(100);p__Ascomycota(100);c__Saccharomycetes(100);o__Saccharomycetales(100);o__Saccharomycetales_unclassified(100);o__Saccharomycetales_unclassified(100);o__Saccharomycetales_unclassified(100); | Saccharomycetales | 7 | Saprotroph | Undefined Saprotroph | Yeast | NULL | Possible | NULL | Sterkenburg E, et al. 2015. New Phytologist 207:1145-1158 ((https://doi.org/10.1111/nph.13426)) |
| Otu16137 | 1 | k__Fungi(100);p__Ascomycota(100);c__Saccharomycetes(100);o__Saccharomycetales(100);f__Metschnikowiaceae(100);g__Metschnikowia(100);g__Metschnikowia_unclassified(100); | Metschnikowia | 13 | Saprotroph | \|Undefined Saprotroph\| | Yeast | NULL | Probable | As nectar-tap_saprotroph (Põlme et al. 2020); Decay sugar-rich substrates (Põlme et al. 2020) | Põlme S, et al. 2020. Fungal Diversity 105:1-16 ((https://doi.org/10.1007/s13225-020-00466-2)) |
| Otu16139 | 1 | k__Fungi(100);p__Ascomycota(100);c__Saccharomycetes(100);o__Saccharomycetales(100);f__Metschnikowiaceae(100);g__Metschnikowia(100);g__Metschnikowia_unclassified(100); | Metschnikowia | 13 | Saprotroph | \|Undefined Saprotroph\| | Yeast | NULL | Probable | As nectar-tap_saprotroph (Põlme et al. 2020); Decay sugar-rich substrates (Põlme et al. 2020) | Põlme S, et al. 2020. Fungal Diversity 105:1-16 ((https://doi.org/10.1007/s13225-020-00466-2)) |
| Otu16141 | 1 | k__Fungi(100);p__Ascomycota(100);c__Saccharomycetes(100);o__Saccharomycetales(100);f__Metschnikowiaceae(100);g__Metschnikowia(100);s__Metschnikowia_pulcherrima(100); | Metschnikowia | 13 | Saprotroph | \|Undefined Saprotroph\| | Yeast | NULL | Probable | As nectar-tap_saprotroph (Põlme et al. 2020); Decay sugar-rich substrates (Põlme et al. 2020) | Põlme S, et al. 2020. Fungal Diversity 105:1-16 ((https://doi.org/10.1007/s13225-020-00466-2)) |
| Otu16142 | 1 | k__Fungi(100);p__Ascomycota(100);c__Saccharomycetes(100);o__Saccharomycetales(100);f__Metschnikowiaceae(100);g__Metschnikowia(100);s__Metschnikowia_pulcherrima(100); | Metschnikowia | 13 | Saprotroph | \|Undefined Saprotroph\| | Yeast | NULL | Probable | As nectar-tap_saprotroph (Põlme et al. 2020); Decay sugar-rich substrates (Põlme et al. 2020) | Põlme S, et al. 2020. Fungal Diversity 105:1-16 ((https://doi.org/10.1007/s13225-020-00466-2)) |
| Otu16143 | 1 | k__Fungi(100);p__Ascomycota(100);c__Saccharomycetes(100);o__Saccharomycetales(100);f__Metschnikowiaceae(100);g__Metschnikowia(100);g__Metschnikowia_unclassified(100); | Metschnikowia | 13 | Saprotroph | \|Undefined Saprotroph\| | Yeast | NULL | Probable | As nectar-tap_saprotroph (Põlme et al. 2020); Decay sugar-rich substrates (Põlme et al. 2020) | Põlme S, et al. 2020. Fungal Diversity 105:1-16 ((https://doi.org/10.1007/s13225-020-00466-2)) |
| Otu16144 | 1 | k__Fungi(100);p__Ascomycota(100);c__Saccharomycetes(100);o__Saccharomycetales(100);f__Metschnikowiaceae(100);g__Metschnikowia(100);s__Metschnikowia_pulcherrima(100); | Metschnikowia | 13 | Saprotroph | \|Undefined Saprotroph\| | Yeast | NULL | Probable | As nectar-tap_saprotroph (Põlme et al. 2020); Decay sugar-rich substrates (Põlme et al. 2020) | Põlme S, et al. 2020. Fungal Diversity 105:1-16 ((https://doi.org/10.1007/s13225-020-00466-2)) |
| Otu16145 | 1 | k__Fungi(100);p__Ascomycota(100);c__Saccharomycetes(100);o__Saccharomycetales(100);f__Metschnikowiaceae(100);g__Metschnikowia(100);s__Metschnikowia_pulcherrima(100); | Metschnikowia | 13 | Saprotroph | \|Undefined Saprotroph\| | Yeast | NULL | Probable | As nectar-tap_saprotroph (Põlme et al. 2020); Decay sugar-rich substrates (Põlme et al. 2020) | Põlme S, et al. 2020. Fungal Diversity 105:1-16 ((https://doi.org/10.1007/s13225-020-00466-2)) |
| Otu16147 | 1 | k__Fungi(100);p__Ascomycota(100);c__Saccharomycetes(100);o__Saccharomycetales(100);f__Metschnikowiaceae(100);g__Metschnikowia(100);s__Metschnikowia_pulcherrima(100); | Metschnikowia | 13 | Saprotroph | \|Undefined Saprotroph\| | Yeast | NULL | Probable | As nectar-tap_saprotroph (Põlme et al. 2020); Decay sugar-rich substrates (Põlme et al. 2020) | Põlme S, et al. 2020. Fungal Diversity 105:1-16 ((https://doi.org/10.1007/s13225-020-00466-2)) |
| Otu16148 | 1 | k__Fungi(100);p__Ascomycota(100);c__Saccharomycetes(100);o__Saccharomycetales(100);f__Metschnikowiaceae(100);g__Metschnikowia(100);g__Metschnikowia_unclassified(100); | Metschnikowia | 13 | Saprotroph | \|Undefined Saprotroph\| | Yeast | NULL | Probable | As nectar-tap_saprotroph (Põlme et al. 2020); Decay sugar-rich substrates (Põlme et al. 2020) | Põlme S, et al. 2020. Fungal Diversity 105:1-16 ((https://doi.org/10.1007/s13225-020-00466-2)) |
| Otu16150 | 1 | k__Fungi(100);p__Ascomycota(100);c__Saccharomycetes(100);o__Saccharomycetales(100);f__Metschnikowiaceae(100);g__Metschnikowia(100);s__Metschnikowia_pulcherrima(100); | Metschnikowia | 13 | Saprotroph | \|Undefined Saprotroph\| | Yeast | NULL | Probable | As nectar-tap_saprotroph (Põlme et al. 2020); Decay sugar-rich substrates (Põlme et al. 2020) | Põlme S, et al. 2020. Fungal Diversity 105:1-16 ((https://doi.org/10.1007/s13225-020-00466-2)) |
| Otu16151 | 1 | k__Fungi(100);p__Ascomycota(100);c__Saccharomycetes(100);o__Saccharomycetales(100);f__Metschnikowiaceae(100);g__Metschnikowia(100);g__Metschnikowia_unclassified(100); | Metschnikowia | 13 | Saprotroph | \|Undefined Saprotroph\| | Yeast | NULL | Probable | As nectar-tap_saprotroph (Põlme et al. 2020); Decay sugar-rich substrates (Põlme et al. 2020) | Põlme S, et al. 2020. Fungal Diversity 105:1-16 ((https://doi.org/10.1007/s13225-020-00466-2)) |
| Otu16153 | 1 | k__Fungi(100);p__Ascomycota(100);c__Saccharomycetes(100);o__Saccharomycetales(100);f__Metschnikowiaceae(100);g__Metschnikowia(100);s__Metschnikowia_pulcherrima(100); | Metschnikowia | 13 | Saprotroph | \|Undefined Saprotroph\| | Yeast | NULL | Probable | As nectar-tap_saprotroph (Põlme et al. 2020); Decay sugar-rich substrates (Põlme et al. 2020) | Põlme S, et al. 2020. Fungal Diversity 105:1-16 ((https://doi.org/10.1007/s13225-020-00466-2)) |
| Otu16154 | 1 | k__Fungi(100);p__Ascomycota(100);c__Saccharomycetes(100);o__Saccharomycetales(100);f__Metschnikowiaceae(100);g__Metschnikowia(100);s__Metschnikowia_pulcherrima(100); | Metschnikowia | 13 | Saprotroph | \|Undefined Saprotroph\| | Yeast | NULL | Probable | As nectar-tap_saprotroph (Põlme et al. 2020); Decay sugar-rich substrates (Põlme et al. 2020) | Põlme S, et al. 2020. Fungal Diversity 105:1-16 ((https://doi.org/10.1007/s13225-020-00466-2)) |
| Otu16155 | 1 | k__Fungi(100);p__Ascomycota(100);c__Saccharomycetes(100);o__Saccharomycetales(100);f__Metschnikowiaceae(100);g__Metschnikowia(100);s__Metschnikowia_pulcherrima(100); | Metschnikowia | 13 | Saprotroph | \|Undefined Saprotroph\| | Yeast | NULL | Probable | As nectar-tap_saprotroph (Põlme et al. 2020); Decay sugar-rich substrates (Põlme et al. 2020) | Põlme S, et al. 2020. Fungal Diversity 105:1-16 ((https://doi.org/10.1007/s13225-020-00466-2)) |
| Otu16156 | 1 | k__Fungi(100);p__Ascomycota(100);c__Saccharomycetes(100);o__Saccharomycetales(100);f__Metschnikowiaceae(100);g__Metschnikowia(100);g__Metschnikowia_unclassified(100); | Metschnikowia | 13 | Saprotroph | \|Undefined Saprotroph\| | Yeast | NULL | Probable | As nectar-tap_saprotroph (Põlme et al. 2020); Decay sugar-rich substrates (Põlme et al. 2020) | Põlme S, et al. 2020. Fungal Diversity 105:1-16 ((https://doi.org/10.1007/s13225-020-00466-2)) |
| Otu16157 | 1 | k__Fungi(100);p__Ascomycota(100);c__Saccharomycetes(100);o__Saccharomycetales(100);f__Metschnikowiaceae(100);g__Metschnikowia(100);s__Metschnikowia_pulcherrima(100); | Metschnikowia | 13 | Saprotroph | \|Undefined Saprotroph\| | Yeast | NULL | Probable | As nectar-tap_saprotroph (Põlme et al. 2020); Decay sugar-rich substrates (Põlme et al. 2020) | Põlme S, et al. 2020. Fungal Diversity 105:1-16 ((https://doi.org/10.1007/s13225-020-00466-2)) |
| Otu16158 | 1 | k__Fungi(100);p__Ascomycota(100);c__Saccharomycetes(100);o__Saccharomycetales(100);f__Metschnikowiaceae(100);g__Metschnikowia(100);s__Metschnikowia_pulcherrima(100); | Metschnikowia | 13 | Saprotroph | \|Undefined Saprotroph\| | Yeast | NULL | Probable | As nectar-tap_saprotroph (Põlme et al. 2020); Decay sugar-rich substrates (Põlme et al. 2020) | Põlme S, et al. 2020. Fungal Diversity 105:1-16 ((https://doi.org/10.1007/s13225-020-00466-2)) |
| Otu16159 | 1 | k__Fungi(100);p__Ascomycota(100);c__Saccharomycetes(100);o__Saccharomycetales(100);f__Metschnikowiaceae(100);g__Metschnikowia(100);g__Metschnikowia_unclassified(100); | Metschnikowia | 13 | Saprotroph | \|Undefined Saprotroph\| | Yeast | NULL | Probable | As nectar-tap_saprotroph (Põlme et al. 2020); Decay sugar-rich substrates (Põlme et al. 2020) | Põlme S, et al. 2020. Fungal Diversity 105:1-16 ((https://doi.org/10.1007/s13225-020-00466-2)) |
| Otu16162 | 1 | k__Fungi(100);p__Ascomycota(100);c__Saccharomycetes(100);o__Saccharomycetales(100);f__Metschnikowiaceae(100);g__Metschnikowia(100);g__Metschnikowia_unclassified(100); | Metschnikowia | 13 | Saprotroph | \|Undefined Saprotroph\| | Yeast | NULL | Probable | As nectar-tap_saprotroph (Põlme et al. 2020); Decay sugar-rich substrates (Põlme et al. 2020) | Põlme S, et al. 2020. Fungal Diversity 105:1-16 ((https://doi.org/10.1007/s13225-020-00466-2)) |
| Otu16165 | 1 | k__Fungi(100);p__Ascomycota(100);c__Saccharomycetes(100);o__Saccharomycetales(100);f__Metschnikowiaceae(100);g__Metschnikowia(100);s__Metschnikowia_pulcherrima(100); | Metschnikowia | 13 | Saprotroph | \|Undefined Saprotroph\| | Yeast | NULL | Probable | As nectar-tap_saprotroph (Põlme et al. 2020); Decay sugar-rich substrates (Põlme et al. 2020) | Põlme S, et al. 2020. Fungal Diversity 105:1-16 ((https://doi.org/10.1007/s13225-020-00466-2)) |
| Otu16167 | 1 | k__Fungi(100);p__Ascomycota(100);c__Saccharomycetes(100);o__Saccharomycetales(100);f__Metschnikowiaceae(100);g__Metschnikowia(100);s__Metschnikowia_pulcherrima(100); | Metschnikowia | 13 | Saprotroph | \|Undefined Saprotroph\| | Yeast | NULL | Probable | As nectar-tap_saprotroph (Põlme et al. 2020); Decay sugar-rich substrates (Põlme et al. 2020) | Põlme S, et al. 2020. Fungal Diversity 105:1-16 ((https://doi.org/10.1007/s13225-020-00466-2)) |
| Otu16168 | 1 | k__Fungi(100);p__Ascomycota(100);c__Saccharomycetes(100);o__Saccharomycetales(100);f__Metschnikowiaceae(100);g__Metschnikowia(100);s__Metschnikowia_pulcherrima(100); | Metschnikowia | 13 | Saprotroph | \|Undefined Saprotroph\| | Yeast | NULL | Probable | As nectar-tap_saprotroph (Põlme et al. 2020); Decay sugar-rich substrates (Põlme et al. 2020) | Põlme S, et al. 2020. Fungal Diversity 105:1-16 ((https://doi.org/10.1007/s13225-020-00466-2)) |
| Otu16170 | 1 | k__Fungi(100);p__Ascomycota(100);c__Saccharomycetes(100);o__Saccharomycetales(100);f__Metschnikowiaceae(100);g__Metschnikowia(100);s__Metschnikowia_pulcherrima(100); | Metschnikowia | 13 | Saprotroph | \|Undefined Saprotroph\| | Yeast | NULL | Probable | As nectar-tap_saprotroph (Põlme et al. 2020); Decay sugar-rich substrates (Põlme et al. 2020) | Põlme S, et al. 2020. Fungal Diversity 105:1-16 ((https://doi.org/10.1007/s13225-020-00466-2)) |
| Otu16171 | 1 | k__Fungi(100);p__Ascomycota(100);c__Saccharomycetes(100);o__Saccharomycetales(100);f__Metschnikowiaceae(100);g__Metschnikowia(100);s__Metschnikowia_pulcherrima(100); | Metschnikowia | 13 | Saprotroph | \|Undefined Saprotroph\| | Yeast | NULL | Probable | As nectar-tap_saprotroph (Põlme et al. 2020); Decay sugar-rich substrates (Põlme et al. 2020) | Põlme S, et al. 2020. Fungal Diversity 105:1-16 ((https://doi.org/10.1007/s13225-020-00466-2)) |
| Otu16172 | 1 | k__Fungi(100);p__Ascomycota(100);c__Saccharomycetes(100);o__Saccharomycetales(100);f__Metschnikowiaceae(100);g__Metschnikowia(100);g__Metschnikowia_unclassified(100); | Metschnikowia | 13 | Saprotroph | \|Undefined Saprotroph\| | Yeast | NULL | Probable | As nectar-tap_saprotroph (Põlme et al. 2020); Decay sugar-rich substrates (Põlme et al. 2020) | Põlme S, et al. 2020. Fungal Diversity 105:1-16 ((https://doi.org/10.1007/s13225-020-00466-2)) |
| Otu16174 | 1 | k__Fungi(100);p__Ascomycota(100);c__Saccharomycetes(100);o__Saccharomycetales(100);f__Metschnikowiaceae(100);g__Metschnikowia(100);s__Metschnikowia_pulcherrima(100); | Metschnikowia | 13 | Saprotroph | \|Undefined Saprotroph\| | Yeast | NULL | Probable | As nectar-tap_saprotroph (Põlme et al. 2020); Decay sugar-rich substrates (Põlme et al. 2020) | Põlme S, et al. 2020. Fungal Diversity 105:1-16 ((https://doi.org/10.1007/s13225-020-00466-2)) |
| Otu16175 | 1 | k__Fungi(100);p__Ascomycota(100);c__Saccharomycetes(100);o__Saccharomycetales(100);f__Metschnikowiaceae(100);g__Metschnikowia(100);s__Metschnikowia_pulcherrima(100); | Metschnikowia | 13 | Saprotroph | \|Undefined Saprotroph\| | Yeast | NULL | Probable | As nectar-tap_saprotroph (Põlme et al. 2020); Decay sugar-rich substrates (Põlme et al. 2020) | Põlme S, et al. 2020. Fungal Diversity 105:1-16 ((https://doi.org/10.1007/s13225-020-00466-2)) |
| Otu16178 | 1 | k__Fungi(100);p__Ascomycota(100);c__Saccharomycetes(100);o__Saccharomycetales(100);f__Metschnikowiaceae(100);g__Metschnikowia(100);g__Metschnikowia_unclassified(100); | Metschnikowia | 13 | Saprotroph | \|Undefined Saprotroph\| | Yeast | NULL | Probable | As nectar-tap_saprotroph (Põlme et al. 2020); Decay sugar-rich substrates (Põlme et al. 2020) | Põlme S, et al. 2020. Fungal Diversity 105:1-16 ((https://doi.org/10.1007/s13225-020-00466-2)) |
| Otu16179 | 1 | k__Fungi(100);p__Ascomycota(100);c__Sordariomycetes(100);o__Hypocreales(100);f__Ophiocordycipitaceae(100);g__Ophiocordyceps(100);s__Ophiocordyceps_heteropoda_var_langyashanensis(100); | Ophiocordyceps | 13 | Pathotroph-Saprotroph-Symbiotroph | Endophyte-\|Insect Pathogen\|-Undefined Saprotroph | Microfungus | NULL | Probable | As animal_decomposer (Põlme et al. 2020); Contains about 140 species that grow on insects (Kirk et al. 2008); Decay animal material (Põlme et al. 2020) | Kirk PM et al. 2008. Dictionary of the Fungi. Tenth Edition. CAB International, Wallingford (ISBN: 978-0851998268); Costa IPMW, et al. 2012. Mycotaxon 119:e26 ((http://mycotaxon.com/resources/checklists/costa_v119_checklist.pdf)); Tedersoo L, et al. 2014. Science 346: e1256688 ((https://doi.org/10.1126/science.1256688)); Põlme S, et al. 2020. Fungal Diversity 105:1-16 ((https://doi.org/10.1007/s13225-020-00466-2)) |
| Otu16180 | 1 | k__Fungi(100);p__Ascomycota(100);c__Saccharomycetes(100);o__Saccharomycetales(100);f__Metschnikowiaceae(100);g__Metschnikowia(100);s__Metschnikowia_pulcherrima(100); | Metschnikowia | 13 | Saprotroph | \|Undefined Saprotroph\| | Yeast | NULL | Probable | As nectar-tap_saprotroph (Põlme et al. 2020); Decay sugar-rich substrates (Põlme et al. 2020) | Põlme S, et al. 2020. Fungal Diversity 105:1-16 ((https://doi.org/10.1007/s13225-020-00466-2)) |
| Otu16182 | 1 | k__Fungi(100);p__Ascomycota(100);c__Saccharomycetes(100);o__Saccharomycetales(100);f__Metschnikowiaceae(100);g__Metschnikowia(100);s__Metschnikowia_pulcherrima(100); | Metschnikowia | 13 | Saprotroph | \|Undefined Saprotroph\| | Yeast | NULL | Probable | As nectar-tap_saprotroph (Põlme et al. 2020); Decay sugar-rich substrates (Põlme et al. 2020) | Põlme S, et al. 2020. Fungal Diversity 105:1-16 ((https://doi.org/10.1007/s13225-020-00466-2)) |
| Otu16183 | 1 | k__Fungi(100);p__Ascomycota(100);c__Saccharomycetes(100);o__Saccharomycetales(100);f__Metschnikowiaceae(100);g__Metschnikowia(100);s__Metschnikowia_pulcherrima(100); | Metschnikowia | 13 | Saprotroph | \|Undefined Saprotroph\| | Yeast | NULL | Probable | As nectar-tap_saprotroph (Põlme et al. 2020); Decay sugar-rich substrates (Põlme et al. 2020) | Põlme S, et al. 2020. Fungal Diversity 105:1-16 ((https://doi.org/10.1007/s13225-020-00466-2)) |
| Otu16184 | 1 | k__Fungi(100);p__Ascomycota(100);c__Saccharomycetes(100);o__Saccharomycetales(100);f__Metschnikowiaceae(100);g__Metschnikowia(100);s__Metschnikowia_pulcherrima(100); | Metschnikowia | 13 | Saprotroph | \|Undefined Saprotroph\| | Yeast | NULL | Probable | As nectar-tap_saprotroph (Põlme et al. 2020); Decay sugar-rich substrates (Põlme et al. 2020) | Põlme S, et al. 2020. Fungal Diversity 105:1-16 ((https://doi.org/10.1007/s13225-020-00466-2)) |
| Otu16186 | 1 | k__Fungi(100);p__Ascomycota(100);c__Saccharomycetes(100);o__Saccharomycetales(100);f__Metschnikowiaceae(100);g__Metschnikowia(100);s__Metschnikowia_pulcherrima(100); | Metschnikowia | 13 | Saprotroph | \|Undefined Saprotroph\| | Yeast | NULL | Probable | As nectar-tap_saprotroph (Põlme et al. 2020); Decay sugar-rich substrates (Põlme et al. 2020) | Põlme S, et al. 2020. Fungal Diversity 105:1-16 ((https://doi.org/10.1007/s13225-020-00466-2)) |
| Otu16187 | 1 | k__Fungi(100);p__Ascomycota(100);c__Saccharomycetes(100);o__Saccharomycetales(100);f__Metschnikowiaceae(100);g__Metschnikowia(100);s__Metschnikowia_pulcherrima(100); | Metschnikowia | 13 | Saprotroph | \|Undefined Saprotroph\| | Yeast | NULL | Probable | As nectar-tap_saprotroph (Põlme et al. 2020); Decay sugar-rich substrates (Põlme et al. 2020) | Põlme S, et al. 2020. Fungal Diversity 105:1-16 ((https://doi.org/10.1007/s13225-020-00466-2)) |
| Otu16194 | 1 | k__Fungi(100);p__Ascomycota(100);c__Saccharomycetes(100);o__Saccharomycetales(100);f__Metschnikowiaceae(100);g__Metschnikowia(100);s__Metschnikowia_pulcherrima(100); | Metschnikowia | 13 | Saprotroph | \|Undefined Saprotroph\| | Yeast | NULL | Probable | As nectar-tap_saprotroph (Põlme et al. 2020); Decay sugar-rich substrates (Põlme et al. 2020) | Põlme S, et al. 2020. Fungal Diversity 105:1-16 ((https://doi.org/10.1007/s13225-020-00466-2)) |
| Otu16198 | 1 | k__Fungi(100);p__Ascomycota(100);c__Saccharomycetes(100);o__Saccharomycetales(100);f__Metschnikowiaceae(100);g__Metschnikowia(100);s__Metschnikowia_pulcherrima(100); | Metschnikowia | 13 | Saprotroph | \|Undefined Saprotroph\| | Yeast | NULL | Probable | As nectar-tap_saprotroph (Põlme et al. 2020); Decay sugar-rich substrates (Põlme et al. 2020) | Põlme S, et al. 2020. Fungal Diversity 105:1-16 ((https://doi.org/10.1007/s13225-020-00466-2)) |
| Otu16200 | 1 | k__Fungi(100);p__Ascomycota(100);c__Saccharomycetes(100);o__Saccharomycetales(100);f__Metschnikowiaceae(100);g__Metschnikowia(100);s__Metschnikowia_pulcherrima(100); | Metschnikowia | 13 | Saprotroph | \|Undefined Saprotroph\| | Yeast | NULL | Probable | As nectar-tap_saprotroph (Põlme et al. 2020); Decay sugar-rich substrates (Põlme et al. 2020) | Põlme S, et al. 2020. Fungal Diversity 105:1-16 ((https://doi.org/10.1007/s13225-020-00466-2)) |
| Otu16204 | 1 | k__Fungi(100);p__Ascomycota(100);c__Saccharomycetes(100);o__Saccharomycetales(100);f__Metschnikowiaceae(100);g__Metschnikowia(100);s__Metschnikowia_pulcherrima(100); | Metschnikowia | 13 | Saprotroph | \|Undefined Saprotroph\| | Yeast | NULL | Probable | As nectar-tap_saprotroph (Põlme et al. 2020); Decay sugar-rich substrates (Põlme et al. 2020) | Põlme S, et al. 2020. Fungal Diversity 105:1-16 ((https://doi.org/10.1007/s13225-020-00466-2)) |
| Otu16207 | 1 | k__Fungi(100);p__Ascomycota(100);c__Saccharomycetes(100);o__Saccharomycetales(100);f__Metschnikowiaceae(100);g__Metschnikowia(100);s__Metschnikowia_pulcherrima(100); | Metschnikowia | 13 | Saprotroph | \|Undefined Saprotroph\| | Yeast | NULL | Probable | As nectar-tap_saprotroph (Põlme et al. 2020); Decay sugar-rich substrates (Põlme et al. 2020) | Põlme S, et al. 2020. Fungal Diversity 105:1-16 ((https://doi.org/10.1007/s13225-020-00466-2)) |
| Otu16209 | 1 | k__Fungi(100);p__Ascomycota(100);c__Saccharomycetes(100);o__Saccharomycetales(100);f__Metschnikowiaceae(100);g__Metschnikowia(100);s__Metschnikowia_pulcherrima(100); | Metschnikowia | 13 | Saprotroph | \|Undefined Saprotroph\| | Yeast | NULL | Probable | As nectar-tap_saprotroph (Põlme et al. 2020); Decay sugar-rich substrates (Põlme et al. 2020) | Põlme S, et al. 2020. Fungal Diversity 105:1-16 ((https://doi.org/10.1007/s13225-020-00466-2)) |
| Otu16210 | 1 | k__Fungi(100);p__Ascomycota(100);c__Saccharomycetes(100);o__Saccharomycetales(100);f__Metschnikowiaceae(100);g__Metschnikowia(100);s__Metschnikowia_pulcherrima(100); | Metschnikowia | 13 | Saprotroph | \|Undefined Saprotroph\| | Yeast | NULL | Probable | As nectar-tap_saprotroph (Põlme et al. 2020); Decay sugar-rich substrates (Põlme et al. 2020) | Põlme S, et al. 2020. Fungal Diversity 105:1-16 ((https://doi.org/10.1007/s13225-020-00466-2)) |
| Otu16211 | 1 | k__Fungi(100);p__Ascomycota(100);c__Saccharomycetes(100);o__Saccharomycetales(100);f__Metschnikowiaceae(100);g__Metschnikowia(100);g__Metschnikowia_unclassified(100); | Metschnikowia | 13 | Saprotroph | \|Undefined Saprotroph\| | Yeast | NULL | Probable | As nectar-tap_saprotroph (Põlme et al. 2020); Decay sugar-rich substrates (Põlme et al. 2020) | Põlme S, et al. 2020. Fungal Diversity 105:1-16 ((https://doi.org/10.1007/s13225-020-00466-2)) |
| Otu16212 | 1 | k__Fungi(100);p__Ascomycota(100);c__Saccharomycetes(100);o__Saccharomycetales(100);f__Metschnikowiaceae(100);g__Metschnikowia(100);s__Metschnikowia_pulcherrima(100); | Metschnikowia | 13 | Saprotroph | \|Undefined Saprotroph\| | Yeast | NULL | Probable | As nectar-tap_saprotroph (Põlme et al. 2020); Decay sugar-rich substrates (Põlme et al. 2020) | Põlme S, et al. 2020. Fungal Diversity 105:1-16 ((https://doi.org/10.1007/s13225-020-00466-2)) |
| Otu16213 | 1 | k__Fungi(100);p__Ascomycota(100);c__Saccharomycetes(100);o__Saccharomycetales(100);f__Metschnikowiaceae(100);g__Metschnikowia(100);g__Metschnikowia_unclassified(100); | Metschnikowia | 13 | Saprotroph | \|Undefined Saprotroph\| | Yeast | NULL | Probable | As nectar-tap_saprotroph (Põlme et al. 2020); Decay sugar-rich substrates (Põlme et al. 2020) | Põlme S, et al. 2020. Fungal Diversity 105:1-16 ((https://doi.org/10.1007/s13225-020-00466-2)) |
| Otu16214 | 1 | k__Fungi(100);p__Ascomycota(100);c__Saccharomycetes(100);o__Saccharomycetales(100);f__Metschnikowiaceae(100);g__Metschnikowia(100);s__Metschnikowia_pulcherrima(100); | Metschnikowia | 13 | Saprotroph | \|Undefined Saprotroph\| | Yeast | NULL | Probable | As nectar-tap_saprotroph (Põlme et al. 2020); Decay sugar-rich substrates (Põlme et al. 2020) | Põlme S, et al. 2020. Fungal Diversity 105:1-16 ((https://doi.org/10.1007/s13225-020-00466-2)) |
| Otu16215 | 1 | k__Fungi(100);p__Ascomycota(100);c__Saccharomycetes(100);o__Saccharomycetales(100);f__Metschnikowiaceae(100);g__Metschnikowia(100);g__Metschnikowia_unclassified(100); | Metschnikowia | 13 | Saprotroph | \|Undefined Saprotroph\| | Yeast | NULL | Probable | As nectar-tap_saprotroph (Põlme et al. 2020); Decay sugar-rich substrates (Põlme et al. 2020) | Põlme S, et al. 2020. Fungal Diversity 105:1-16 ((https://doi.org/10.1007/s13225-020-00466-2)) |
| Otu16216 | 1 | k__Fungi(100);p__Ascomycota(100);c__Saccharomycetes(100);o__Saccharomycetales(100);f__Metschnikowiaceae(100);g__Metschnikowia(100);s__Metschnikowia_pulcherrima(100); | Metschnikowia | 13 | Saprotroph | \|Undefined Saprotroph\| | Yeast | NULL | Probable | As nectar-tap_saprotroph (Põlme et al. 2020); Decay sugar-rich substrates (Põlme et al. 2020) | Põlme S, et al. 2020. Fungal Diversity 105:1-16 ((https://doi.org/10.1007/s13225-020-00466-2)) |
| Otu16217 | 1 | k__Fungi(100);p__Ascomycota(100);c__Saccharomycetes(100);o__Saccharomycetales(100);f__Metschnikowiaceae(100);g__Metschnikowia(100);s__Metschnikowia_pulcherrima(100); | Metschnikowia | 13 | Saprotroph | \|Undefined Saprotroph\| | Yeast | NULL | Probable | As nectar-tap_saprotroph (Põlme et al. 2020); Decay sugar-rich substrates (Põlme et al. 2020) | Põlme S, et al. 2020. Fungal Diversity 105:1-16 ((https://doi.org/10.1007/s13225-020-00466-2)) |
| Otu16218 | 1 | k__Fungi(100);p__Ascomycota(100);c__Saccharomycetes(100);o__Saccharomycetales(100);f__Metschnikowiaceae(100);g__Metschnikowia(100);g__Metschnikowia_unclassified(100); | Metschnikowia | 13 | Saprotroph | \|Undefined Saprotroph\| | Yeast | NULL | Probable | As nectar-tap_saprotroph (Põlme et al. 2020); Decay sugar-rich substrates (Põlme et al. 2020) | Põlme S, et al. 2020. Fungal Diversity 105:1-16 ((https://doi.org/10.1007/s13225-020-00466-2)) |
| Otu16220 | 1 | k__Fungi(100);p__Ascomycota(100);c__Saccharomycetes(100);o__Saccharomycetales(100);f__Metschnikowiaceae(100);g__Metschnikowia(100);s__Metschnikowia_pulcherrima(100); | Metschnikowia | 13 | Saprotroph | \|Undefined Saprotroph\| | Yeast | NULL | Probable | As nectar-tap_saprotroph (Põlme et al. 2020); Decay sugar-rich substrates (Põlme et al. 2020) | Põlme S, et al. 2020. Fungal Diversity 105:1-16 ((https://doi.org/10.1007/s13225-020-00466-2)) |
| Otu16221 | 1 | k__Fungi(100);p__Ascomycota(100);c__Saccharomycetes(100);o__Saccharomycetales(100);f__Metschnikowiaceae(100);g__Metschnikowia(100);g__Metschnikowia_unclassified(100); | Metschnikowia | 13 | Saprotroph | \|Undefined Saprotroph\| | Yeast | NULL | Probable | As nectar-tap_saprotroph (Põlme et al. 2020); Decay sugar-rich substrates (Põlme et al. 2020) | Põlme S, et al. 2020. Fungal Diversity 105:1-16 ((https://doi.org/10.1007/s13225-020-00466-2)) |
| Otu16222 | 1 | k__Fungi(100);p__Ascomycota(100);c__Saccharomycetes(100);o__Saccharomycetales(100);f__Metschnikowiaceae(100);g__Metschnikowia(100);g__Metschnikowia_unclassified(100); | Metschnikowia | 13 | Saprotroph | \|Undefined Saprotroph\| | Yeast | NULL | Probable | As nectar-tap_saprotroph (Põlme et al. 2020); Decay sugar-rich substrates (Põlme et al. 2020) | Põlme S, et al. 2020. Fungal Diversity 105:1-16 ((https://doi.org/10.1007/s13225-020-00466-2)) |

| | | | | | | | | | | | |
|---|---|---|---|---|---|---|---|---|---|---|---|
| Otu16223 | 1 | k__Fungi(100);p__Ascomycota(100);c__Saccharomycetes(100);o__Saccharomycetales(100);f__Saccharomycetaceae(100);g__Kazachstania(100);s__Kazachstania_intestinalis(100); | Kazachstania | 13 | Saprotroph | \|Undefined Saprotroph\| | Yeast | NULL | Probable | As nectar-tap_saprotroph (Põlme et al. 2020); Decay sugar-rich substrates (Põlme et al. 2020) | Põlme S, et al. 2020. Fungal Diversity 105:1-16 ((https://doi.org/10.1007/s13225-020-00466-2)) |
| Otu16224 | 1 | k__Fungi(100);p__Ascomycota(100);c__Saccharomycetes(100);o__Saccharomycetales(100);f__Metschnikowiaceae(100);g__Metschnikowia(100);g__Metschnikowia_unclassified(100); | Metschnikowia | 13 | Saprotroph | \|Undefined Saprotroph\| | Yeast | NULL | Probable | As nectar-tap_saprotroph (Põlme et al. 2020); Decay sugar-rich substrates (Põlme et al. 2020) | Põlme S, et al. 2020. Fungal Diversity 105:1-16 ((https://doi.org/10.1007/s13225-020-00466-2)) |
| Otu16225 | 1 | k__Fungi(100);p__Ascomycota(100);c__Saccharomycetes(100);o__Saccharomycetales(100);f__Metschnikowiaceae(100);g__Metschnikowia(100);s__Metschnikowia_pulcherrima(100); | Metschnikowia | 13 | Saprotroph | \|Undefined Saprotroph\| | Yeast | NULL | Probable | As nectar-tap_saprotroph (Põlme et al. 2020); Decay sugar-rich substrates (Põlme et al. 2020) | Põlme S, et al. 2020. Fungal Diversity 105:1-16 ((https://doi.org/10.1007/s13225-020-00466-2)) |
| Otu16226 | 1 | k__Fungi(100);p__Ascomycota(100);c__Saccharomycetes(100);o__Saccharomycetales(100);f__Metschnikowiaceae(100);g__Metschnikowia(100);s__Metschnikowia_pulcherrima(100); | Metschnikowia | 13 | Saprotroph | \|Undefined Saprotroph\| | Yeast | NULL | Probable | As nectar-tap_saprotroph (Põlme et al. 2020); Decay sugar-rich substrates (Põlme et al. 2020) | Põlme S, et al. 2020. Fungal Diversity 105:1-16 ((https://doi.org/10.1007/s13225-020-00466-2)) |
| Otu16227 | 1 | k__Fungi(100);p__Ascomycota(100);c__Saccharomycetes(100);o__Saccharomycetales(100);f__Metschnikowiaceae(100);g__Metschnikowia(100);s__Metschnikowia_pulcherrima(100); | Metschnikowia | 13 | Saprotroph | \|Undefined Saprotroph\| | Yeast | NULL | Probable | As nectar-tap_saprotroph (Põlme et al. 2020); Decay sugar-rich substrates (Põlme et al. 2020) | Põlme S, et al. 2020. Fungal Diversity 105:1-16 ((https://doi.org/10.1007/s13225-020-00466-2)) |
| Otu16228 | 1 | k__Fungi(100);p__Ascomycota(100);c__Saccharomycetes(100);o__Saccharomycetales(100);f__Metschnikowiaceae(100);g__Metschnikowia(100);s__Metschnikowia_pulcherrima(100); | Metschnikowia | 13 | Saprotroph | \|Undefined Saprotroph\| | Yeast | NULL | Probable | As nectar-tap_saprotroph (Põlme et al. 2020); Decay sugar-rich substrates (Põlme et al. 2020) | Põlme S, et al. 2020. Fungal Diversity 105:1-16 ((https://doi.org/10.1007/s13225-020-00466-2)) |
| Otu16229 | 1 | k__Fungi(100);p__Ascomycota(100);c__Saccharomycetes(100);o__Saccharomycetales(100);f__Metschnikowiaceae(100);g__Metschnikowia(100);g__Metschnikowia_unclassified(100); | Metschnikowia | 13 | Saprotroph | \|Undefined Saprotroph\| | Yeast | NULL | Probable | As nectar-tap_saprotroph (Põlme et al. 2020); Decay sugar-rich substrates (Põlme et al. 2020) | Põlme S, et al. 2020. Fungal Diversity 105:1-16 ((https://doi.org/10.1007/s13225-020-00466-2)) |
| Otu16230 | 1 | k__Fungi(100);p__Ascomycota(100);c__Saccharomycetes(100);o__Saccharomycetales(100);f__Metschnikowiaceae(100);g__Metschnikowia(100);g__Metschnikowia_unclassified(100); | Metschnikowia | 13 | Saprotroph | \|Undefined Saprotroph\| | Yeast | NULL | Probable | As nectar-tap_saprotroph (Põlme et al. 2020); Decay sugar-rich substrates (Põlme et al. 2020) | Põlme S, et al. 2020. Fungal Diversity 105:1-16 ((https://doi.org/10.1007/s13225-020-00466-2)) |
| Otu16231 | 1 | k__Fungi(100);p__Ascomycota(100);c__Saccharomycetes(100);o__Saccharomycetales(100);f__Metschnikowiaceae(100);g__Metschnikowia(100);s__Metschnikowia_pulcherrima(100); | Metschnikowia | 13 | Saprotroph | \|Undefined Saprotroph\| | Yeast | NULL | Probable | As nectar-tap_saprotroph (Põlme et al. 2020); Decay sugar-rich substrates (Põlme et al. 2020) | Põlme S, et al. 2020. Fungal Diversity 105:1-16 ((https://doi.org/10.1007/s13225-020-00466-2)) |
| Otu16232 | 1 | k__Fungi(100);p__Ascomycota(100);c__Saccharomycetes(100);o__Saccharomycetales(100);f__Metschnikowiaceae(100);g__Metschnikowia(100);g__Metschnikowia_unclassified(100); | Metschnikowia | 13 | Saprotroph | \|Undefined Saprotroph\| | Yeast | NULL | Probable | As nectar-tap_saprotroph (Põlme et al. 2020); Decay sugar-rich substrates (Põlme et al. 2020) | Põlme S, et al. 2020. Fungal Diversity 105:1-16 ((https://doi.org/10.1007/s13225-020-00466-2)) |
| Otu16233 | 1 | k__Fungi(100);p__Ascomycota(100);c__Saccharomycetes(100);o__Saccharomycetales(100);f__Metschnikowiaceae(100);g__Metschnikowia(100);s__Metschnikowia_pulcherrima(100); | Metschnikowia | 13 | Saprotroph | \|Undefined Saprotroph\| | Yeast | NULL | Probable | As nectar-tap_saprotroph (Põlme et al. 2020); Decay sugar-rich substrates (Põlme et al. 2020) | Põlme S, et al. 2020. Fungal Diversity 105:1-16 ((https://doi.org/10.1007/s13225-020-00466-2)) |
| Otu16234 | 1 | k__Fungi(100);p__Ascomycota(100);c__Saccharomycetes(100);o__Saccharomycetales(100);f__Metschnikowiaceae(100);g__Metschnikowia(100);s__Metschnikowia_pulcherrima(100); | Metschnikowia | 13 | Saprotroph | \|Undefined Saprotroph\| | Yeast | NULL | Probable | As nectar-tap_saprotroph (Põlme et al. 2020); Decay sugar-rich substrates (Põlme et al. 2020) | Põlme S, et al. 2020. Fungal Diversity 105:1-16 ((https://doi.org/10.1007/s13225-020-00466-2)) |
| Otu16236 | 1 | k__Fungi(100);p__Ascomycota(100);c__Saccharomycetes(100);o__Saccharomycetales(100);f__Metschnikowiaceae(100);g__Metschnikowia(100);s__Metschnikowia_pulcherrima(100); | Metschnikowia | 13 | Saprotroph | \|Undefined Saprotroph\| | Yeast | NULL | Probable | As nectar-tap_saprotroph (Põlme et al. 2020); Decay sugar-rich substrates (Põlme et al. 2020) | Põlme S, et al. 2020. Fungal Diversity 105:1-16 ((https://doi.org/10.1007/s13225-020-00466-2)) |
| Otu16237 | 1 | k__Fungi(100);p__Ascomycota(100);c__Saccharomycetes(100);o__Saccharomycetales(100);f__Metschnikowiaceae(100);g__Metschnikowia(100);s__Metschnikowia_pulcherrima(100); | Metschnikowia | 13 | Saprotroph | \|Undefined Saprotroph\| | Yeast | NULL | Probable | As nectar-tap_saprotroph (Põlme et al. 2020); Decay sugar-rich substrates (Põlme et al. 2020) | Põlme S, et al. 2020. Fungal Diversity 105:1-16 ((https://doi.org/10.1007/s13225-020-00466-2)) |
| Otu16242 | 1 | k__Fungi(100);p__Ascomycota(100);c__Saccharomycetes(100);o__Saccharomycetales(100);f__Metschnikowiaceae(100);g__Metschnikowia(100);s__Metschnikowia_pulcherrima(100); | Metschnikowia | 13 | Saprotroph | \|Undefined Saprotroph\| | Yeast | NULL | Probable | As nectar-tap_saprotroph (Põlme et al. 2020); Decay sugar-rich substrates (Põlme et al. 2020) | Põlme S, et al. 2020. Fungal Diversity 105:1-16 ((https://doi.org/10.1007/s13225-020-00466-2)) |
| Otu16243 | 1 | k__Fungi(100);p__Ascomycota(100);c__Saccharomycetes(100);o__Saccharomycetales(100);f__Metschnikowiaceae(100);g__Metschnikowia(100);s__Metschnikowia_pulcherrima(100); | Metschnikowia | 13 | Saprotroph | \|Undefined Saprotroph\| | Yeast | NULL | Probable | As nectar-tap_saprotroph (Põlme et al. 2020); Decay sugar-rich substrates (Põlme et al. 2020) | Põlme S, et al. 2020. Fungal Diversity 105:1-16 ((https://doi.org/10.1007/s13225-020-00466-2)) |
| Otu16244 | 1 | k__Fungi(100);p__Ascomycota(100);c__Saccharomycetes(100);o__Saccharomycetales(100);f__Metschnikowiaceae(100);g__Metschnikowia(100);s__Metschnikowia_pulcherrima(100); | Metschnikowia | 13 | Saprotroph | \|Undefined Saprotroph\| | Yeast | NULL | Probable | As nectar-tap_saprotroph (Põlme et al. 2020); Decay sugar-rich substrates (Põlme et al. 2020) | Põlme S, et al. 2020. Fungal Diversity 105:1-16 ((https://doi.org/10.1007/s13225-020-00466-2)) |
| Otu16245 | 1 | k__Fungi(100);p__Ascomycota(100);c__Saccharomycetes(100);o__Saccharomycetales(100);f__Metschnikowiaceae(100);g__Metschnikowia(100);s__Metschnikowia_pulcherrima(100); | Metschnikowia | 13 | Saprotroph | \|Undefined Saprotroph\| | Yeast | NULL | Probable | As nectar-tap_saprotroph (Põlme et al. 2020); Decay sugar-rich substrates (Põlme et al. 2020) | Põlme S, et al. 2020. Fungal Diversity 105:1-16 ((https://doi.org/10.1007/s13225-020-00466-2)) |
| Otu16246 | 1 | k__Fungi(100);p__Ascomycota(100);c__Saccharomycetes(100);o__Saccharomycetales(100);f__Metschnikowiaceae(100);g__Metschnikowia(100);s__Metschnikowia_pulcherrima(100); | Metschnikowia | 13 | Saprotroph | \|Undefined Saprotroph\| | Yeast | NULL | Probable | As nectar-tap_saprotroph (Põlme et al. 2020); Decay sugar-rich substrates (Põlme et al. 2020) | Põlme S, et al. 2020. Fungal Diversity 105:1-16 ((https://doi.org/10.1007/s13225-020-00466-2)) |
| Otu16247 | 1 | k__Fungi(100);p__Ascomycota(100);c__Saccharomycetes(100);o__Saccharomycetales(100);f__Metschnikowiaceae(100);g__Metschnikowia(100);s__Metschnikowia_pulcherrima(100); | Metschnikowia | 13 | Saprotroph | \|Undefined Saprotroph\| | Yeast | NULL | Probable | As nectar-tap_saprotroph (Põlme et al. 2020); Decay sugar-rich substrates (Põlme et al. 2020) | Põlme S, et al. 2020. Fungal Diversity 105:1-16 ((https://doi.org/10.1007/s13225-020-00466-2)) |
| Otu16249 | 1 | k__Fungi(100);p__Ascomycota(100);c__Saccharomycetes(100);o__Saccharomycetales(100);f__Metschnikowiaceae(100);g__Metschnikowia(100);g__Metschnikowia_unclassified(100); | Metschnikowia | 13 | Saprotroph | \|Undefined Saprotroph\| | Yeast | NULL | Probable | As nectar-tap_saprotroph (Põlme et al. 2020); Decay sugar-rich substrates (Põlme et al. 2020) | Põlme S, et al. 2020. Fungal Diversity 105:1-16 ((https://doi.org/10.1007/s13225-020-00466-2)) |
| Otu16250 | 1 | k__Fungi(100);p__Ascomycota(100);c__Saccharomycetes(100);o__Saccharomycetales(100);f__Metschnikowiaceae(100);g__Metschnikowia(100);s__Metschnikowia_pulcherrima(100); | Metschnikowia | 13 | Saprotroph | \|Undefined Saprotroph\| | Yeast | NULL | Probable | As nectar-tap_saprotroph (Põlme et al. 2020); Decay sugar-rich substrates (Põlme et al. 2020) | Põlme S, et al. 2020. Fungal Diversity 105:1-16 ((https://doi.org/10.1007/s13225-020-00466-2)) |
| Otu16251 | 1 | k__Fungi(100);p__Ascomycota(100);c__Saccharomycetes(100);o__Saccharomycetales(100);f__Metschnikowiaceae(100);g__Metschnikowia(100);s__Metschnikowia_pulcherrima(100); | Metschnikowia | 13 | Saprotroph | \|Undefined Saprotroph\| | Yeast | NULL | Probable | As nectar-tap_saprotroph (Põlme et al. 2020); Decay sugar-rich substrates (Põlme et al. 2020) | Põlme S, et al. 2020. Fungal Diversity 105:1-16 ((https://doi.org/10.1007/s13225-020-00466-2)) |
| Otu16252 | 1 | k__Fungi(100);p__Ascomycota(100);c__Saccharomycetes(100);o__Saccharomycetales(100);f__Metschnikowiaceae(100);g__Metschnikowia(100);s__Metschnikowia_pulcherrima(100); | Metschnikowia | 13 | Saprotroph | \|Undefined Saprotroph\| | Yeast | NULL | Probable | As nectar-tap_saprotroph (Põlme et al. 2020); Decay sugar-rich substrates (Põlme et al. 2020) | Põlme S, et al. 2020. Fungal Diversity 105:1-16 ((https://doi.org/10.1007/s13225-020-00466-2)) |
| Otu16254 | 1 | k__Fungi(100);p__Ascomycota(100);c__Saccharomycetes(100);o__Saccharomycetales(100);f__Metschnikowiaceae(100);g__Metschnikowia(100);s__Metschnikowia_pulcherrima(100); | Metschnikowia | 13 | Saprotroph | \|Undefined Saprotroph\| | Yeast | NULL | Probable | As nectar-tap_saprotroph (Põlme et al. 2020); Decay sugar-rich substrates (Põlme et al. 2020) | Põlme S, et al. 2020. Fungal Diversity 105:1-16 ((https://doi.org/10.1007/s13225-020-00466-2)) |
| Otu16255 | 1 | k__Fungi(100);p__Ascomycota(100);c__Saccharomycetes(100);o__Saccharomycetales(100);f__Metschnikowiaceae(100);g__Metschnikowia(100);s__Metschnikowia_pulcherrima(100); | Metschnikowia | 13 | Saprotroph | \|Undefined Saprotroph\| | Yeast | NULL | Probable | As nectar-tap_saprotroph (Põlme et al. 2020); Decay sugar-rich substrates (Põlme et al. 2020) | Põlme S, et al. 2020. Fungal Diversity 105:1-16 ((https://doi.org/10.1007/s13225-020-00466-2)) |
| Otu16257 | 1 | k__Fungi(100);p__Ascomycota(100);c__Saccharomycetes(100);o__Saccharomycetales(100);f__Metschnikowiaceae(100);g__Metschnikowia(100);s__Metschnikowia_pulcherrima(100); | Metschnikowia | 13 | Saprotroph | \|Undefined Saprotroph\| | Yeast | NULL | Probable | As nectar-tap_saprotroph (Põlme et al. 2020); Decay sugar-rich substrates (Põlme et al. 2020) | Põlme S, et al. 2020. Fungal Diversity 105:1-16 ((https://doi.org/10.1007/s13225-020-00466-2)) |
| Otu16262 | 1 | k__Fungi(100);p__Ascomycota(100);c__Saccharomycetes(100);o__Saccharomycetales(100);f__Metschnikowiaceae(100);g__Metschnikowia(100);s__Metschnikowia_pulcherrima(100); | Metschnikowia | 13 | Saprotroph | \|Undefined Saprotroph\| | Yeast | NULL | Probable | As nectar-tap_saprotroph (Põlme et al. 2020); Decay sugar-rich substrates (Põlme et al. 2020) | Põlme S, et al. 2020. Fungal Diversity 105:1-16 ((https://doi.org/10.1007/s13225-020-00466-2)) |
| Otu16266 | 1 | k__Fungi(100);p__Ascomycota(100);c__Saccharomycetes(100);o__Saccharomycetales(100);f__Metschnikowiaceae(100);g__Metschnikowia(100);g__Metschnikowia_unclassified(100); | Metschnikowia | 13 | Saprotroph | \|Undefined Saprotroph\| | Yeast | NULL | Probable | As nectar-tap_saprotroph (Põlme et al. 2020); Decay sugar-rich substrates (Põlme et al. 2020) | Põlme S, et al. 2020. Fungal Diversity 105:1-16 ((https://doi.org/10.1007/s13225-020-00466-2)) |
| Otu16267 | 1 | k__Fungi(100);p__Ascomycota(100);c__Saccharomycetes(100);o__Saccharomycetales(100);f__Metschnikowiaceae(100);g__Metschnikowia(100);g__Metschnikowia_unclassified(100); | Metschnikowia | 13 | Saprotroph | \|Undefined Saprotroph\| | Yeast | NULL | Probable | As nectar-tap_saprotroph (Põlme et al. 2020); Decay sugar-rich substrates (Põlme et al. 2020) | Põlme S, et al. 2020. Fungal Diversity 105:1-16 ((https://doi.org/10.1007/s13225-020-00466-2)) |
| Otu16268 | 1 | k__Fungi(100);p__Ascomycota(100);c__Saccharomycetes(100);o__Saccharomycetales(100);f__Metschnikowiaceae(100);g__Metschnikowia(100);g__Metschnikowia_unclassified(100); | Metschnikowia | 13 | Saprotroph | \|Undefined Saprotroph\| | Yeast | NULL | Probable | As nectar-tap_saprotroph (Põlme et al. 2020); Decay sugar-rich substrates (Põlme et al. 2020) | Põlme S, et al. 2020. Fungal Diversity 105:1-16 ((https://doi.org/10.1007/s13225-020-00466-2)) |
| Otu16269 | 1 | k__Fungi(100);p__Ascomycota(100);c__Saccharomycetes(100);o__Saccharomycetales(100);f__Metschnikowiaceae(100);g__Metschnikowia(100);g__Metschnikowia_unclassified(100); | Metschnikowia | 13 | Saprotroph | \|Undefined Saprotroph\| | Yeast | NULL | Probable | As nectar-tap_saprotroph (Põlme et al. 2020); Decay sugar-rich substrates (Põlme et al. 2020) | Põlme S, et al. 2020. Fungal Diversity 105:1-16 ((https://doi.org/10.1007/s13225-020-00466-2)) |
| Otu16271 | 1 | k__Fungi(100);p__Ascomycota(100);c__Saccharomycetes(100);o__Saccharomycetales(100);f__Metschnikowiaceae(100);g__Metschnikowia(100);s__Metschnikowia_pulcherrima(100); | Metschnikowia | 13 | Saprotroph | \|Undefined Saprotroph\| | Yeast | NULL | Probable | As nectar-tap_saprotroph (Põlme et al. 2020); Decay sugar-rich substrates (Põlme et al. 2020) | Põlme S, et al. 2020. Fungal Diversity 105:1-16 ((https://doi.org/10.1007/s13225-020-00466-2)) |
| Otu16272 | 1 | k__Fungi(100);p__Ascomycota(100);c__Saccharomycetes(100);o__Saccharomycetales(100);f__Metschnikowiaceae(100);g__Metschnikowia(100);g__Metschnikowia_unclassified(100); | Metschnikowia | 13 | Saprotroph | \|Undefined Saprotroph\| | Yeast | NULL | Probable | As nectar-tap_saprotroph (Põlme et al. 2020); Decay sugar-rich substrates (Põlme et al. 2020) | Põlme S, et al. 2020. Fungal Diversity 105:1-16 ((https://doi.org/10.1007/s13225-020-00466-2)) |
| Otu16275 | 1 | k__Fungi(100);p__Ascomycota(100);c__Saccharomycetes(100);o__Saccharomycetales(100);f__Metschnikowiaceae(100);g__Metschnikowia(100);s__Metschnikowia_pulcherrima(100); | Metschnikowia | 13 | Saprotroph | \|Undefined Saprotroph\| | Yeast | NULL | Probable | As nectar-tap_saprotroph (Põlme et al. 2020); Decay sugar-rich substrates (Põlme et al. 2020) | Põlme S, et al. 2020. Fungal Diversity 105:1-16 ((https://doi.org/10.1007/s13225-020-00466-2)) |
| Otu16276 | 1 | k__Fungi(100);p__Ascomycota(100);c__Saccharomycetes(100);o__Saccharomycetales(100);f__Metschnikowiaceae(100);g__Metschnikowia(100);s__Metschnikowia_pulcherrima(100); | Metschnikowia | 13 | Saprotroph | \|Undefined Saprotroph\| | Yeast | NULL | Probable | As nectar-tap_saprotroph (Põlme et al. 2020); Decay sugar-rich substrates (Põlme et al. 2020) | Põlme S, et al. 2020. Fungal Diversity 105:1-16 ((https://doi.org/10.1007/s13225-020-00466-2)) |
| Otu16277 | 1 | k__Fungi(100);p__Ascomycota(100);c__Saccharomycetes(100);o__Saccharomycetales(100);f__Metschnikowiaceae(100);g__Metschnikowia(100);s__Metschnikowia_pulcherrima(100); | Metschnikowia | 13 | Saprotroph | \|Undefined Saprotroph\| | Yeast | NULL | Probable | As nectar-tap_saprotroph (Põlme et al. 2020); Decay sugar-rich substrates (Põlme et al. 2020) | Põlme S, et al. 2020. Fungal Diversity 105:1-16 ((https://doi.org/10.1007/s13225-020-00466-2)) |
| Otu16278 | 1 | k__Fungi(100);p__Ascomycota(100);c__Leotiomycetes(100);o__Helotiales(100);f__Vibrisseaceae(100);g__Vibrissea(100);g__Vibrissea_unclassified(100); | Vibrissea | 13 | Saprotroph | \|Plant Saprotroph\|-Undefined Saprotroph | Microfungus | NULL | Probable | As litter_saprotroph (Põlme et al. 2020); Decay leaf, fruit, seed (Põlme et al. 2020) | Tedersoo L, et al. 2014. Science 346:e1256688 ((https://doi.org/10.1126/science.1256688)); Põlme S, et al. 2020. Fungal Diversity 105:1-16 ((https://doi.org/10.1007/s13225-020-00466-2)) |
| Otu16280 | 1 | k__Fungi(100);p__Ascomycota(100);c__Saccharomycetes(100);o__Saccharomycetales(100);f__Metschnikowiaceae(100);g__Metschnikowia(100);s__Metschnikowia_pulcherrima(100); | Metschnikowia | 13 | Saprotroph | \|Undefined Saprotroph\| | Yeast | NULL | Probable | As nectar-tap_saprotroph (Põlme et al. 2020); Decay sugar-rich substrates (Põlme et al. 2020) | Põlme S, et al. 2020. Fungal Diversity 105:1-16 ((https://doi.org/10.1007/s13225-020-00466-2)) |
| Otu16281 | 1 | k__Fungi(100);p__Ascomycota(100);c__Saccharomycetes(100);o__Saccharomycetales(100);f__Metschnikowiaceae(100);g__Metschnikowia(100);s__Metschnikowia_pulcherrima(100); | Metschnikowia | 13 | Saprotroph | \|Undefined Saprotroph\| | Yeast | NULL | Probable | As nectar-tap_saprotroph (Põlme et al. 2020); Decay sugar-rich substrates (Põlme et al. 2020) | Põlme S, et al. 2020. Fungal Diversity 105:1-16 ((https://doi.org/10.1007/s13225-020-00466-2)) |
| Otu16282 | 1 | k__Fungi(100);p__Ascomycota(100);c__Saccharomycetes(100);o__Saccharomycetales(100);f__Metschnikowiaceae(100);g__Metschnikowia(100);g__Metschnikowia_unclassified(100); | Metschnikowia | 13 | Saprotroph | \|Undefined Saprotroph\| | Yeast | NULL | Probable | As nectar-tap_saprotroph (Põlme et al. 2020); Decay sugar-rich substrates (Põlme et al. 2020) | Põlme S, et al. 2020. Fungal Diversity 105:1-16 ((https://doi.org/10.1007/s13225-020-00466-2)) |
| Otu16283 | 1 | k__Fungi(100);p__Ascomycota(100);c__Saccharomycetes(100);o__Saccharomycetales(100);f__Metschnikowiaceae(100);g__Metschnikowia(100);g__Metschnikowia_unclassified(100); | Metschnikowia | 13 | Saprotroph | \|Undefined Saprotroph\| | Yeast | NULL | Probable | As nectar-tap_saprotroph (Põlme et al. 2020); Decay sugar-rich substrates (Põlme et al. 2020) | Põlme S, et al. 2020. Fungal Diversity 105:1-16 ((https://doi.org/10.1007/s13225-020-00466-2)) |
| Otu16284 | 1 | k__Fungi(100);p__Ascomycota(100);c__Saccharomycetes(100);o__Saccharomycetales(100);f__Metschnikowiaceae(100);g__Metschnikowia(100);g__Metschnikowia_unclassified(100); | Metschnikowia | 13 | Saprotroph | \|Undefined Saprotroph\| | Yeast | NULL | Probable | As nectar-tap_saprotroph (Põlme et al. 2020); Decay sugar-rich substrates (Põlme et al. 2020) | Põlme S, et al. 2020. Fungal Diversity 105:1-16 ((https://doi.org/10.1007/s13225-020-00466-2)) |
| Otu16285 | 1 | k__Fungi(100);p__Ascomycota(100);c__Saccharomycetes(100);o__Saccharomycetales(100);f__Metschnikowiaceae(100);g__Metschnikowia(100);s__Metschnikowia_pulcherrima(100); | Metschnikowia | 13 | Saprotroph | \|Undefined Saprotroph\| | Yeast | NULL | Probable | As nectar-tap_saprotroph (Põlme et al. 2020); Decay sugar-rich substrates (Põlme et al. 2020) | Põlme S, et al. 2020. Fungal Diversity 105:1-16 ((https://doi.org/10.1007/s13225-020-00466-2)) |
| Otu16286 | 1 | k__Fungi(100);p__Ascomycota(100);c__Saccharomycetes(100);o__Saccharomycetales(100);f__Metschnikowiaceae(100);g__Metschnikowia(100);s__Metschnikowia_pulcherrima(100); | Metschnikowia | 13 | Saprotroph | \|Undefined Saprotroph\| | Yeast | NULL | Probable | As nectar-tap_saprotroph (Põlme et al. 2020); Decay sugar-rich substrates (Põlme et al. 2020) | Põlme S, et al. 2020. Fungal Diversity 105:1-16 ((https://doi.org/10.1007/s13225-020-00466-2)) |
| Otu16287 | 1 | k__Fungi(100);p__Ascomycota(100);c__Saccharomycetes(100);o__Saccharomycetales(100);f__Metschnikowiaceae(100);g__Metschnikowia(100);s__Metschnikowia_pulcherrima(100); | Metschnikowia | 13 | Saprotroph | \|Undefined Saprotroph\| | Yeast | NULL | Probable | As nectar-tap_saprotroph (Põlme et al. 2020); Decay sugar-rich substrates (Põlme et al. 2020) | Põlme S, et al. 2020. Fungal Diversity 105:1-16 ((https://doi.org/10.1007/s13225-020-00466-2)) |
| Otu16288 | 1 | k__Fungi(100);p__Ascomycota(100);c__Saccharomycetes(100);o__Saccharomycetales(100);f__Metschnikowiaceae(100);g__Metschnikowia(100);g__Metschnikowia_unclassified(100); | Metschnikowia | 13 | Saprotroph | \|Undefined Saprotroph\| | Yeast | NULL | Probable | As nectar-tap_saprotroph (Põlme et al. 2020); Decay sugar-rich substrates (Põlme et al. 2020) | Põlme S, et al. 2020. Fungal Diversity 105:1-16 ((https://doi.org/10.1007/s13225-020-00466-2)) |
| Otu16289 | 1 | k__Fungi(100);p__Ascomycota(100);c__Saccharomycetes(100);o__Saccharomycetales(100);f__Metschnikowiaceae(100);g__Metschnikowia(100);s__Metschnikowia_pulcherrima(100); | Metschnikowia | 13 | Saprotroph | \|Undefined Saprotroph\| | Yeast | NULL | Probable | As nectar-tap_saprotroph (Põlme et al. 2020); Decay sugar-rich substrates (Põlme et al. 2020) | Põlme S, et al. 2020. Fungal Diversity 105:1-16 ((https://doi.org/10.1007/s13225-020-00466-2)) |
| Otu16290 | 1 | k__Fungi(100);p__Ascomycota(100);c__Saccharomycetes(100);o__Saccharomycetales(100);f__Metschnikowiaceae(100);g__Metschnikowia(100);s__Metschnikowia_pulcherrima(100); | Metschnikowia | 13 | Saprotroph | \|Undefined Saprotroph\| | Yeast | NULL | Probable | As nectar-tap_saprotroph (Põlme et al. 2020); Decay sugar-rich substrates (Põlme et al. 2020) | Põlme S, et al. 2020. Fungal Diversity 105:1-16 ((https://doi.org/10.1007/s13225-020-00466-2)) |
| Otu16292 | 1 | k__Fungi(100);p__Ascomycota(100);c__Saccharomycetes(100);o__Saccharomycetales(100);f__Metschnikowiaceae(100);g__Metschnikowia(100);s__Metschnikowia_pulcherrima(100); | Metschnikowia | 13 | Saprotroph | \|Undefined Saprotroph\| | Yeast | NULL | Probable | As nectar-tap_saprotroph (Põlme et al. 2020); Decay sugar-rich substrates (Põlme et al. 2020) | Põlme S, et al. 2020. Fungal Diversity 105:1-16 ((https://doi.org/10.1007/s13225-020-00466-2)) |
| Otu16294 | 1 | k__Fungi(100);p__Ascomycota(100);c__Saccharomycetes(100);o__Saccharomycetales(100);f__Metschnikowiaceae(100);g__Metschnikowia(100);s__Metschnikowia_pulcherrima(100); | Metschnikowia | 13 | Saprotroph | \|Undefined Saprotroph\| | Yeast | NULL | Probable | As nectar-tap_saprotroph (Põlme et al. 2020); Decay sugar-rich substrates (Põlme et al. 2020) | Põlme S, et al. 2020. Fungal Diversity 105:1-16 ((https://doi.org/10.1007/s13225-020-00466-2)) |
| Otu16295 | 1 | k__Fungi(100);p__Ascomycota(100);c__Saccharomycetes(100);o__Saccharomycetales(100);f__Metschnikowiaceae(100);g__Metschnikowia(100);s__Metschnikowia_pulcherrima(100); | Metschnikowia | 13 | Saprotroph | \|Undefined Saprotroph\| | Yeast | NULL | Probable | As nectar-tap_saprotroph (Põlme et al. 2020); Decay sugar-rich substrates (Põlme et al. 2020) | Põlme S, et al. 2020. Fungal Diversity 105:1-16 ((https://doi.org/10.1007/s13225-020-00466-2)) |
| Otu16296 | 1 | k__Fungi(100);p__Ascomycota(100);c__Saccharomycetes(100);o__Saccharomycetales(100);f__Metschnikowiaceae(100);g__Metschnikowia(100);g__Metschnikowia_unclassified(100); | Metschnikowia | 13 | Saprotroph | \|Undefined Saprotroph\| | Yeast | NULL | Probable | As nectar-tap_saprotroph (Põlme et al. 2020); Decay sugar-rich substrates (Põlme et al. 2020) | Põlme S, et al. 2020. Fungal Diversity 105:1-16 ((https://doi.org/10.1007/s13225-020-00466-2)) |
| Otu16298 | 1 | k__Fungi(100);p__Ascomycota(100);c__Saccharomycetes(100);o__Saccharomycetales(100);f__Metschnikowiaceae(100);g__Metschnikowia(100);g__Metschnikowia_unclassified(100); | Metschnikowia | 13 | Saprotroph | \|Undefined Saprotroph\| | Yeast | NULL | Probable | As nectar-tap_saprotroph (Põlme et al. 2020); Decay sugar-rich substrates (Põlme et al. 2020) | Põlme S, et al. 2020. Fungal Diversity 105:1-16 ((https://doi.org/10.1007/s13225-020-00466-2)) |
| Otu16299 | 1 | k__Fungi(100);p__Ascomycota(100);c__Saccharomycetes(100);o__Saccharomycetales(100);f__Metschnikowiaceae(100);g__Metschnikowia(100);s__Metschnikowia_pulcherrima(100); | Metschnikowia | 13 | Saprotroph | \|Undefined Saprotroph\| | Yeast | NULL | Probable | As nectar-tap_saprotroph (Põlme et al. 2020); Decay sugar-rich substrates (Põlme et al. 2020) | Põlme S, et al. 2020. Fungal Diversity 105:1-16 ((https://doi.org/10.1007/s13225-020-00466-2)) |
| Otu16300 | 1 | k__Fungi(100);p__Ascomycota(100);c__Saccharomycetes(100);o__Saccharomycetales(100);f__Metschnikowiaceae(100);g__Metschnikowia(100);s__Metschnikowia_pulcherrima(100); | Metschnikowia | 13 | Saprotroph | \|Undefined Saprotroph\| | Yeast | NULL | Probable | As nectar-tap_saprotroph (Põlme et al. 2020); Decay sugar-rich substrates (Põlme et al. 2020) | Põlme S, et al. 2020. Fungal Diversity 105:1-16 ((https://doi.org/10.1007/s13225-020-00466-2)) |
| Otu16301 | 1 | k__Fungi(100);p__Ascomycota(100);c__Saccharomycetes(100);o__Saccharomycetales(100);f__Metschnikowiaceae(100);g__Metschnikowia(100);g__Metschnikowia_unclassified(100); | Metschnikowia | 13 | Saprotroph | \|Undefined Saprotroph\| | Yeast | NULL | Probable | As nectar-tap_saprotroph (Põlme et al. 2020); Decay sugar-rich substrates (Põlme et al. 2020) | Põlme S, et al. 2020. Fungal Diversity 105:1-16 ((https://doi.org/10.1007/s13225-020-00466-2)) |
| Otu16302 | 1 | k__Fungi(100);p__Ascomycota(100);c__Saccharomycetes(100);o__Saccharomycetales(100);f__Metschnikowiaceae(100);g__Metschnikowia(100);s__Metschnikowia_pulcherrima(100); | Metschnikowia | 13 | Saprotroph | \|Undefined Saprotroph\| | Yeast | NULL | Probable | As nectar-tap_saprotroph (Põlme et al. 2020); Decay sugar-rich substrates (Põlme et al. 2020) | Põlme S, et al. 2020. Fungal Diversity 105:1-16 ((https://doi.org/10.1007/s13225-020-00466-2)) |
| Otu16304 | 1 | k__Fungi(100);p__Ascomycota(100);c__Saccharomycetes(100);o__Saccharomycetales(100);f__Metschnikowiaceae(100);g__Metschnikowia(100);s__Metschnikowia_pulcherrima(100); | Metschnikowia | 13 | Saprotroph | \|Undefined Saprotroph\| | Yeast | NULL | Probable | As nectar-tap_saprotroph (Põlme et al. 2020); Decay sugar-rich substrates (Põlme et al. 2020) | Põlme S, et al. 2020. Fungal Diversity 105:1-16 ((https://doi.org/10.1007/s13225-020-00466-2)) |
| Otu16305 | 1 | k__Fungi(100);p__Ascomycota(100);c__Saccharomycetes(100);o__Saccharomycetales(100);f__Metschnikowiaceae(100);g__Metschnikowia(100);g__Metschnikowia_unclassified(100); | Metschnikowia | 13 | Saprotroph | \|Undefined Saprotroph\| | Yeast | NULL | Probable | As nectar-tap_saprotroph (Põlme et al. 2020); Decay sugar-rich substrates (Põlme et al. 2020) | Põlme S, et al. 2020. Fungal Diversity 105:1-16 ((https://doi.org/10.1007/s13225-020-00466-2)) |
| Otu16306 | 1 | k__Fungi(100);p__Ascomycota(100);c__Saccharomycetes(100);o__Saccharomycetales(100);f__Metschnikowiaceae(100);g__Metschnikowia(100);s__Metschnikowia_pulcherrima(100); | Metschnikowia | 13 | Saprotroph | \|Undefined Saprotroph\| | Yeast | NULL | Probable | As nectar-tap_saprotroph (Põlme et al. 2020); Decay sugar-rich substrates (Põlme et al. 2020) | Põlme S, et al. 2020. Fungal Diversity 105:1-16 ((https://doi.org/10.1007/s13225-020-00466-2)) |
| Otu16307 | 1 | k__Fungi(100);p__Ascomycota(100);c__Saccharomycetes(100);o__Saccharomycetales(100);f__Metschnikowiaceae(100);g__Metschnikowia(100);g__Metschnikowia_unclassified(100); | Metschnikowia | 13 | Saprotroph | \|Undefined Saprotroph\| | Yeast | NULL | Probable | As nectar-tap_saprotroph (Põlme et al. 2020); Decay sugar-rich substrates (Põlme et al. 2020) | Põlme S, et al. 2020. Fungal Diversity 105:1-16 ((https://doi.org/10.1007/s13225-020-00466-2)) |
| Otu16308 | 1 | k__Fungi(100);p__Ascomycota(100);c__Saccharomycetes(100);o__Saccharomycetales(100);f__Metschnikowiaceae(100);g__Metschnikowia(100);s__Metschnikowia_pulcherrima(100); | Metschnikowia | 13 | Saprotroph | \|Undefined Saprotroph\| | Yeast | NULL | Probable | As nectar-tap_saprotroph (Põlme et al. 2020); Decay sugar-rich substrates (Põlme et al. 2020) | Põlme S, et al. 2020. Fungal Diversity 105:1-16 ((https://doi.org/10.1007/s13225-020-00466-2)) |
| Otu16309 | 1 | k__Fungi(100);p__Ascomycota(100);c__Saccharomycetes(100);o__Saccharomycetales(100);f__Metschnikowiaceae(100);g__Metschnikowia(100);s__Metschnikowia_pulcherrima(100); | Metschnikowia | 13 | Saprotroph | \|Undefined Saprotroph\| | Yeast | NULL | Probable | As nectar-tap_saprotroph (Põlme et al. 2020); Decay sugar-rich substrates (Põlme et al. 2020) | Põlme S, et al. 2020. Fungal Diversity 105:1-16 ((https://doi.org/10.1007/s13225-020-00466-2)) |
| Otu16311 | 1 | k__Fungi(100);p__Ascomycota(100);c__Saccharomycetes(100);o__Saccharomycetales(100);f__Metschnikowiaceae(100);g__Metschnikowia(100);s__Metschnikowia_pulcherrima(100); | Metschnikowia | 13 | Saprotroph | \|Undefined Saprotroph\| | Yeast | NULL | Probable | As nectar-tap_saprotroph (Põlme et al. 2020); Decay sugar-rich substrates (Põlme et al. 2020) | Põlme S, et al. 2020. Fungal Diversity 105:1-16 ((https://doi.org/10.1007/s13225-020-00466-2)) |
| Otu16312 | 1 | k__Fungi(100);p__Ascomycota(100);c__Saccharomycetes(100);o__Saccharomycetales(100);f__Metschnikowiaceae(100);g__Metschnikowia(100);g__Metschnikowia_unclassified(100); | Metschnikowia | 13 | Saprotroph | \|Undefined Saprotroph\| | Yeast | NULL | Probable | As nectar-tap_saprotroph (Põlme et al. 2020); Decay sugar-rich substrates (Põlme et al. 2020) | Põlme S, et al. 2020. Fungal Diversity 105:1-16 ((https://doi.org/10.1007/s13225-020-00466-2)) |
| Otu16313 | 1 | k__Fungi(100);p__Ascomycota(100);c__Saccharomycetes(100);o__Saccharomycetales(100);f__Metschnikowiaceae(100);g__Metschnikowia(100);g__Metschnikowia_unclassified(100); | Metschnikowia | 13 | Saprotroph | \|Undefined Saprotroph\| | Yeast | NULL | Probable | As nectar-tap_saprotroph (Põlme et al. 2020); Decay sugar-rich substrates (Põlme et al. 2020) | Põlme S, et al. 2020. Fungal Diversity 105:1-16 ((https://doi.org/10.1007/s13225-020-00466-2)) |
| Otu16314 | 1 | k__Fungi(100);p__Ascomycota(100);c__Saccharomycetes(100);o__Saccharomycetales(100);f__Metschnikowiaceae(100);g__Metschnikowia(100);s__Metschnikowia_pulcherrima(100); | Metschnikowia | 13 | Saprotroph | \|Undefined Saprotroph\| | Yeast | NULL | Probable | As nectar-tap_saprotroph (Põlme et al. 2020); Decay sugar-rich substrates (Põlme et al. 2020) | Põlme S, et al. 2020. Fungal Diversity 105:1-16 ((https://doi.org/10.1007/s13225-020-00466-2)) |
| Otu16315 | 1 | k__Fungi(100);p__Ascomycota(100);c__Saccharomycetes(100);o__Saccharomycetales(100);f__Metschnikowiaceae(100);g__Metschnikowia(100);s__Metschnikowia_pulcherrima(100); | Metschnikowia | 13 | Saprotroph | \|Undefined Saprotroph\| | Yeast | NULL | Probable | As nectar-tap_saprotroph (Põlme et al. 2020); Decay sugar-rich substrates (Põlme et al. 2020) | Põlme S, et al. 2020. Fungal Diversity 105:1-16 ((https://doi.org/10.1007/s13225-020-00466-2)) |
| Otu16316 | 1 | k__Fungi(100);p__Ascomycota(100);c__Saccharomycetes(100);o__Saccharomycetales(100);f__Metschnikowiaceae(100);g__Metschnikowia(100);s__Metschnikowia_pulcherrima(100); | Metschnikowia | 13 | Saprotroph | \|Undefined Saprotroph\| | Yeast | NULL | Probable | As nectar-tap_saprotroph (Põlme et al. 2020); Decay sugar-rich substrates (Põlme et al. 2020) | Põlme S, et al. 2020. Fungal Diversity 105:1-16 ((https://doi.org/10.1007/s13225-020-00466-2)) |
| Otu16317 | 1 | k__Fungi(100);p__Ascomycota(100);c__Saccharomycetes(100);o__Saccharomycetales(100);f__Metschnikowiaceae(100);g__Metschnikowia(100);g__Metschnikowia_unclassified(100); | Metschnikowia | 13 | Saprotroph | \|Undefined Saprotroph\| | Yeast | NULL | Probable | As nectar-tap_saprotroph (Põlme et al. 2020); Decay sugar-rich substrates (Põlme et al. 2020) | Põlme S, et al. 2020. Fungal Diversity 105:1-16 ((https://doi.org/10.1007/s13225-020-00466-2)) |
| Otu16318 | 1 | k__Fungi(100);p__Ascomycota(100);c__Saccharomycetes(100);o__Saccharomycetales(100);f__Metschnikowiaceae(100);g__Metschnikowia(100);s__Metschnikowia_pulcherrima(100); | Metschnikowia | 13 | Saprotroph | \|Undefined Saprotroph\| | Yeast | NULL | Probable | As nectar-tap_saprotroph (Põlme et al. 2020); Decay sugar-rich substrates (Põlme et al. 2020) | Põlme S, et al. 2020. Fungal Diversity 105:1-16 ((https://doi.org/10.1007/s13225-020-00466-2)) |
| Otu16319 | 1 | k__Fungi(100);p__Ascomycota(100);c__Saccharomycetes(100);o__Saccharomycetales(100);f__Metschnikowiaceae(100);g__Metschnikowia(100);s__Metschnikowia_pulcherrima(100); | Metschnikowia | 13 | Saprotroph | \|Undefined Saprotroph\| | Yeast | NULL | Probable | As nectar-tap_saprotroph (Põlme et al. 2020); Decay sugar-rich substrates (Põlme et al. 2020) | Põlme S, et al. 2020. Fungal Diversity 105:1-16 ((https://doi.org/10.1007/s13225-020-00466-2)) |
| Otu16320 | 1 | k__Fungi(100);p__Ascomycota(100);c__Saccharomycetes(100);o__Saccharomycetales(100);f__Metschnikowiaceae(100);g__Metschnikowia(100);s__Metschnikowia_pulcherrima(100); | Metschnikowia | 13 | Saprotroph | \|Undefined Saprotroph\| | Yeast | NULL | Probable | As nectar-tap_saprotroph (Põlme et al. 2020); Decay sugar-rich substrates (Põlme et al. 2020) | Põlme S, et al. 2020. Fungal Diversity 105:1-16 ((https://doi.org/10.1007/s13225-020-00466-2)) |
| Otu16323 | 1 | k__Fungi(100);p__Ascomycota(100);c__Saccharomycetes(100);o__Saccharomycetales(100);f__Metschnikowiaceae(100);g__Metschnikowia(100);s__Metschnikowia_pulcherrima(100); | Metschnikowia | 13 | Saprotroph | \|Undefined Saprotroph\| | Yeast | NULL | Probable | As nectar-tap_saprotroph (Põlme et al. 2020); Decay sugar-rich substrates (Põlme et al. 2020) | Põlme S, et al. 2020. Fungal Diversity 105:1-16 ((https://doi.org/10.1007/s13225-020-00466-2)) |
| Otu16325 | 1 | k__Fungi(100);p__Ascomycota(100);c__Saccharomycetes(100);o__Saccharomycetales(100);f__Metschnikowiaceae(100);g__Metschnikowia(100);s__Metschnikowia_pulcherrima(100); | Metschnikowia | 13 | Saprotroph | \|Undefined Saprotroph\| | Yeast | NULL | Probable | As nectar-tap_saprotroph (Põlme et al. 2020); Decay sugar-rich substrates (Põlme et al. 2020) | Põlme S, et al. 2020. Fungal Diversity 105:1-16 ((https://doi.org/10.1007/s13225-020-00466-2)) |
| Otu16326 | 1 | k__Fungi(100);p__Ascomycota(100);c__Saccharomycetes(100);o__Saccharomycetales(100);f__Saccharomycetales_family_Incertae_sedis(100);g__Candida(100);s__Candida_cretensis(100); | Candida | 13 | Pathotroph-Saprotroph-Symbiotroph | Animal Pathogen-Endophyte-Endosymbiont-Epiphyte-\|Undefined Saprotroph\| | Dimorphic Yeast-Yeast | NULL | Probable | As nectar-tap_saprotroph (Põlme et al. 2020); Decay sugar-rich substrates (Põlme et al. 2020); This asexual genus is polyphyletic and contain species that function very differently depend on what they are. Caution should be taken when interpreting guilds for Candida species. | Manolakaki D, et al. 2010. Virulence. 1: 367-375 ((https://doi.org/10.4161/viru.1.5.12796)); Mekbib SB, et al. 2011. Tropical Plant Pathology 36:233-240 ((https://doi.org/10.1590/S1982-56762011000400004)); Põlme S, et al. 2020. Fungal Diversity 105:1-16 ((https://doi.org/10.1007/s13225-020-00466-2)) |
| Otu16327 | 1 | k__Fungi(100);p__Ascomycota(100);c__Saccharomycetes(100);o__Saccharomycetales(100);f__Metschnikowiaceae(100);g__Metschnikowia(100);s__Metschnikowia_pulcherrima(100); | Metschnikowia | 13 | Saprotroph | \|Undefined Saprotroph\| | Yeast | NULL | Probable | As nectar-tap_saprotroph (Põlme et al. 2020); Decay sugar-rich substrates (Põlme et al. 2020) | Põlme S, et al. 2020. Fungal Diversity 105:1-16 ((https://doi.org/10.1007/s13225-020-00466-2)) |
| Otu16328 | 1 | k__Fungi(100);p__Ascomycota(100);c__Saccharomycetes(100);o__Saccharomycetales(100);f__Metschnikowiaceae(100);g__Metschnikowia(100);g__Metschnikowia_unclassified(100); | Metschnikowia | 13 | Saprotroph | \|Undefined Saprotroph\| | Yeast | NULL | Probable | As nectar-tap_saprotroph (Põlme et al. 2020); Decay sugar-rich substrates (Põlme et al. 2020) | Põlme S, et al. 2020. Fungal Diversity 105:1-16 ((https://doi.org/10.1007/s13225-020-00466-2)) |
| Otu16329 | 1 | k__Fungi(100);p__Ascomycota(100);c__Saccharomycetes(100);o__Saccharomycetales(100);f__Metschnikowiaceae(100);g__Metschnikowia(100);s__Metschnikowia_pulcherrima(100); | Metschnikowia | 13 | Saprotroph | \|Undefined Saprotroph\| | Yeast | NULL | Probable | As nectar-tap_saprotroph (Põlme et al. 2020); Decay sugar-rich substrates (Põlme et al. 2020) | Põlme S, et al. 2020. Fungal Diversity 105:1-16 ((https://doi.org/10.1007/s13225-020-00466-2)) |
| Otu16332 | 1 | k__Fungi(100);p__Ascomycota(100);c__Saccharomycetes(100);o__Saccharomycetales(100);f__Metschnikowiaceae(100);g__Metschnikowia(100);g__Metschnikowia_unclassified(100); | Metschnikowia | 13 | Saprotroph | \|Undefined Saprotroph\| | Yeast | NULL | Probable | As nectar-tap_saprotroph (Põlme et al. 2020); Decay sugar-rich substrates (Põlme et al. 2020) | Põlme S, et al. 2020. Fungal Diversity 105:1-16 ((https://doi.org/10.1007/s13225-020-00466-2)) |
| Otu16335 | 1 | k__Fungi(100);p__Ascomycota(100);c__Saccharomycetes(100);o__Saccharomycetales(100);f__Metschnikowiaceae(100);g__Metschnikowia(100);s__Metschnikowia_pulcherrima(100); | Metschnikowia | 13 | Saprotroph | \|Undefined Saprotroph\| | Yeast | NULL | Probable | As nectar-tap_saprotroph (Põlme et al. 2020); Decay sugar-rich substrates (Põlme et al. 2020) | Põlme S, et al. 2020. Fungal Diversity 105:1-16 ((https://doi.org/10.1007/s13225-020-00466-2)) |
| Otu16340 | 1 | k__Fungi(100);p__Ascomycota(100);c__Saccharomycetes(100);o__Saccharomycetales(100);f__Metschnikowiaceae(100);g__Metschnikowia(100);s__Metschnikowia_pulcherrima(100); | Metschnikowia | 13 | Saprotroph | \|Undefined Saprotroph\| | Yeast | NULL | Probable | As nectar-tap_saprotroph (Põlme et al. 2020); Decay sugar-rich substrates (Põlme et al. 2020) | Põlme S, et al. 2020. Fungal Diversity 105:1-16 ((https://doi.org/10.1007/s13225-020-00466-2)) |
| Otu16341 | 1 | k__Fungi(100);p__Ascomycota(100);c__Saccharomycetes(100);o__Saccharomycetales(100);f__Metschnikowiaceae(100);g__Metschnikowia(100);s__Metschnikowia_pulcherrima(100); | Metschnikowia | 13 | Saprotroph | \|Undefined Saprotroph\| | Yeast | NULL | Probable | As nectar-tap_saprotroph (Põlme et al. 2020); Decay sugar-rich substrates (Põlme et al. 2020) | Põlme S, et al. 2020. Fungal Diversity 105:1-16 ((https://doi.org/10.1007/s13225-020-00466-2)) |
| Otu16342 | 1 | k__Fungi(100);p__Ascomycota(100);c__Saccharomycetes(100);o__Saccharomycetales(100);f__Metschnikowiaceae(100);g__Metschnikowia(100);s__Metschnikowia_pulcherrima(100); | Metschnikowia | 13 | Saprotroph | \|Undefined Saprotroph\| | Yeast | NULL | Probable | As nectar-tap_saprotroph (Põlme et al. 2020); Decay sugar-rich substrates (Põlme et al. 2020) | Põlme S, et al. 2020. Fungal Diversity 105:1-16 ((https://doi.org/10.1007/s13225-020-00466-2)) |
| Otu16344 | 1 | k__Fungi(100);p__Ascomycota(100);c__Saccharomycetes(100);o__Saccharomycetales(100);f__Metschnikowiaceae(100);g__Metschnikowia(100);s__Metschnikowia_pulcherrima(100); | Metschnikowia | 13 | Saprotroph | \|Undefined Saprotroph\| | Yeast | NULL | Probable | As nectar-tap_saprotroph (Põlme et al. 2020); Decay sugar-rich substrates (Põlme et al. 2020) | Põlme S, et al. 2020. Fungal Diversity 105:1-16 ((https://doi.org/10.1007/s13225-020-00466-2)) |
| Otu16345 | 1 | k__Fungi(100);p__Ascomycota(100);c__Saccharomycetes(100);o__Saccharomycetales(100);f__Metschnikowiaceae(100);g__Metschnikowia(100);s__Metschnikowia_pulcherrima(100); | Metschnikowia | 13 | Saprotroph | \|Undefined Saprotroph\| | Yeast | NULL | Probable | As nectar-tap_saprotroph (Põlme et al. 2020); Decay sugar-rich substrates (Põlme et al. 2020) | Põlme S, et al. 2020. Fungal Diversity 105:1-16 ((https://doi.org/10.1007/s13225-020-00466-2)) |
| Otu16346 | 1 | k__Fungi(100);p__Ascomycota(100);c__Saccharomycetes(100);o__Saccharomycetales(100);f__Metschnikowiaceae(100);g__Metschnikowia(100);g__Metschnikowia_unclassified(100); | Metschnikowia | 13 | Saprotroph | \|Undefined Saprotroph\| | Yeast | NULL | Probable | As nectar-tap_saprotroph (Põlme et al. 2020); Decay sugar-rich substrates (Põlme et al. 2020) | Põlme S, et al. 2020. Fungal Diversity 105:1-16 ((https://doi.org/10.1007/s13225-020-00466-2)) |
| Otu16347 | 1 | k__Fungi(100);p__Ascomycota(100);c__Saccharomycetes(100);o__Saccharomycetales(100);f__Metschnikowiaceae(100);g__Metschnikowia(100);g__Metschnikowia_unclassified(100); | Metschnikowia | 13 | Saprotroph | \|Undefined Saprotroph\| | Yeast | NULL | Probable | As nectar-tap_saprotroph (Põlme et al. 2020); Decay sugar-rich substrates (Põlme et al. 2020) | Põlme S, et al. 2020. Fungal Diversity 105:1-16 ((https://doi.org/10.1007/s13225-020-00466-2)) |
| Otu16349 | 1 | k__Fungi(100);p__Ascomycota(100);c__Saccharomycetes(100);o__Saccharomycetales(100);f__Metschnikowiaceae(100);g__Metschnikowia(100);g__Metschnikowia_unclassified(100); | Metschnikowia | 13 | Saprotroph | \|Undefined Saprotroph\| | Yeast | NULL | Probable | As nectar-tap_saprotroph (Põlme et al. 2020); Decay sugar-rich substrates (Põlme et al. 2020) | Põlme S, et al. 2020. Fungal Diversity 105:1-16 ((https://doi.org/10.1007/s13225-020-00466-2)) |
| Otu16350 | 1 | k__Fungi(100);p__Ascomycota(100);c__Saccharomycetes(100);o__Saccharomycetales(100);f__Saccharomycetales_family_Incertae_sedis(100);g__Candida(100);g__Candida_unclassified(100); | Candida | 13 | Pathotroph-Saprotroph-Symbiotroph | Animal Pathogen-Endophyte-Endosymbiont-Epiphyte-\|Undefined Saprotroph\| | Dimorphic Yeast-Yeast | NULL | Probable | As nectar-tap_saprotroph (Põlme et al. 2020); Decay sugar-rich substrates (Põlme et al. 2020); This asexual genus is polyphyletic and contain species that function very differently depend on what they are. Caution should be taken when interpreting guilds for Candida species. | Manolakaki D, et al. 2010. Virulence. 1: 367-375 ((https://doi.org/10.4161/viru.1.5.12796)); Mekbib SB, et al. 2011. Tropical Plant Pathology 36:233-240 ((https://doi.org/10.1590/S1982-56762011000400004)); Põlme S, et al. 2020. Fungal Diversity 105:1-16 ((https://doi.org/10.1007/s13225-020-00466-2)) |
| Otu16352 | 1 | k__Fungi(100);p__Ascomycota(100);c__Saccharomycetes(100);o__Saccharomycetales(100);f__Metschnikowiaceae(100);g__Metschnikowia(100);s__Metschnikowia_pulcherrima(100); | Metschnikowia | 13 | Saprotroph | \|Undefined Saprotroph\| | Yeast | NULL | Probable | As nectar-tap_saprotroph (Põlme et al. 2020); Decay sugar-rich substrates (Põlme et al. 2020) | Põlme S, et al. 2020. Fungal Diversity 105:1-16 ((https://doi.org/10.1007/s13225-020-00466-2)) |
| Otu16353 | 1 | k__Fungi(100);p__Ascomycota(100);c__Saccharomycetes(100);o__Saccharomycetales(100);f__Metschnikowiaceae(100);g__Metschnikowia(100);g__Metschnikowia_unclassified(100); | Metschnikowia | 13 | Saprotroph | \|Undefined Saprotroph\| | Yeast | NULL | Probable | As nectar-tap_saprotroph (Põlme et al. 2020); Decay sugar-rich substrates (Põlme et al. 2020) | Põlme S, et al. 2020. Fungal Diversity 105:1-16 ((https://doi.org/10.1007/s13225-020-00466-2)) |
| Otu16354 | 1 | k__Fungi(100);p__Ascomycota(100);c__Saccharomycetes(100);o__Saccharomycetales(100);f__Metschnikowiaceae(100);g__Metschnikowia(100);s__Metschnikowia_pulcherrima(100); | Metschnikowia | 13 | Saprotroph | \|Undefined Saprotroph\| | Yeast | NULL | Probable | As nectar-tap_saprotroph (Põlme et al. 2020); Decay sugar-rich substrates (Põlme et al. 2020) | Põlme S, et al. 2020. Fungal Diversity 105:1-16 ((https://doi.org/10.1007/s13225-020-00466-2)) |
| Otu16355 | 1 | k__Fungi(100);p__Ascomycota(100);c__Saccharomycetes(100);o__Saccharomycetales(100);f__Metschnikowiaceae(100);g__Metschnikowia(100);s__Metschnikowia_pulcherrima(100); | Metschnikowia | 13 | Saprotroph | \|Undefined Saprotroph\| | Yeast | NULL | Probable | As nectar-tap_saprotroph (Põlme et al. 2020); Decay sugar-rich substrates (Põlme et al. 2020) | Põlme S, et al. 2020. Fungal Diversity 105:1-16 ((https://doi.org/10.1007/s13225-020-00466-2)) |
| Otu16356 | 1 | k__Fungi(100);p__Ascomycota(100);c__Saccharomycetes(100);o__Saccharomycetales(100);f__Metschnikowiaceae(100);g__Metschnikowia(100);s__Metschnikowia_pulcherrima(100); | Metschnikowia | 13 | Saprotroph | \|Undefined Saprotroph\| | Yeast | NULL | Probable | As nectar-tap_saprotroph (Põlme et al. 2020); Decay sugar-rich substrates (Põlme et al. 2020) | Põlme S, et al. 2020. Fungal Diversity 105:1-16 ((https://doi.org/10.1007/s13225-020-00466-2)) |
| Otu16357 | 1 | k__Fungi(100);p__Ascomycota(100);c__Saccharomycetes(100);o__Saccharomycetales(100);f__Saccharomycetales_family_Incertae_sedis(100);g__Candida(100);s__Candida_cretensis(100); | Candida | 13 | Pathotroph-Saprotroph-Symbiotroph | Animal Pathogen-Endophyte-Endosymbiont-Epiphyte-\|Undefined Saprotroph\| | Dimorphic Yeast-Yeast | NULL | Probable | As nectar-tap_saprotroph (Põlme et al. 2020); Decay sugar-rich substrates (Põlme et al. 2020); This asexual genus is polyphyletic and contain species that function very differently depend on what they are. Caution should be taken when interpreting guilds for Candida species. | Manolakaki D, et al. 2010. Virulence. 1: 367-375 ((https://doi.org/10.4161/viru.1.5.12796)); Mekbib SB, et al. 2011. Tropical Plant Pathology 36:233-240 ((https://doi.org/10.1590/S1982-56762011000400004)); Põlme S, et al. 2020. Fungal Diversity 105:1-16 ((https://doi.org/10.1007/s13225-020-00466-2)) |
| Otu16359 | 1 | k__Fungi(100);p__Ascomycota(100);c__Saccharomycetes(100);o__Saccharomycetales(100);f__Metschnikowiaceae(100);g__Metschnikowia(100);g__Metschnikowia_unclassified(100); | Metschnikowia | 13 | Saprotroph | \|Undefined Saprotroph\| | Yeast | NULL | Probable | As nectar-tap_saprotroph (Põlme et al. 2020); Decay sugar-rich substrates (Põlme et al. 2020) | Põlme S, et al. 2020. Fungal Diversity 105:1-16 ((https://doi.org/10.1007/s13225-020-00466-2)) |
| Otu16360 | 1 | k__Fungi(100);p__Ascomycota(100);c__Saccharomycetes(100);o__Saccharomycetales(100);f__Metschnikowiaceae(100);g__Metschnikowia(100);g__Metschnikowia_unclassified(100); | Metschnikowia | 13 | Saprotroph | \|Undefined Saprotroph\| | Yeast | NULL | Probable | As nectar-tap_saprotroph (Põlme et al. 2020); Decay sugar-rich substrates (Põlme et al. 2020) | Põlme S, et al. 2020. Fungal Diversity 105:1-16 ((https://doi.org/10.1007/s13225-020-00466-2)) |
| Otu16362 | 1 | k__Fungi(100);p__Ascomycota(100);c__Saccharomycetes(100);o__Saccharomycetales(100);f__Metschnikowiaceae(100);g__Metschnikowia(100);s__Metschnikowia_pulcherrima(100); | Metschnikowia | 13 | Saprotroph | \|Undefined Saprotroph\| | Yeast | NULL | Probable | As nectar-tap_saprotroph (Põlme et al. 2020); Decay sugar-rich substrates (Põlme et al. 2020) | Põlme S, et al. 2020. Fungal Diversity 105:1-16 ((https://doi.org/10.1007/s13225-020-00466-2)) |
| Otu16364 | 1 | k__Fungi(100);p__Ascomycota(100);c__Saccharomycetes(100);o__Saccharomycetales(100);f__Metschnikowiaceae(100);g__Metschnikowia(100);s__Metschnikowia_pulcherrima(100); | Metschnikowia | 13 | Saprotroph | \|Undefined Saprotroph\| | Yeast | NULL | Probable | As nectar-tap_saprotroph (Põlme et al. 2020); Decay sugar-rich substrates (Põlme et al. 2020) | Põlme S, et al. 2020. Fungal Diversity 105:1-16 ((https://doi.org/10.1007/s13225-020-00466-2)) |
| Otu16365 | 1 | k__Fungi(100);p__Ascomycota(100);c__Saccharomycetes(100);o__Saccharomycetales(100);f__Metschnikowiaceae(100);g__Metschnikowia(100);s__Metschnikowia_pulcherrima(100); | Metschnikowia | 13 | Saprotroph | \|Undefined Saprotroph\| | Yeast | NULL | Probable | As nectar-tap_saprotroph (Põlme et al. 2020); Decay sugar-rich substrates (Põlme et al. 2020) | Põlme S, et al. 2020. Fungal Diversity 105:1-16 ((https://doi.org/10.1007/s13225-020-00466-2)) |
| Otu16366 | 1 | k__Fungi(100);p__Ascomycota(100);c__Saccharomycetes(100);o__Saccharomycetales(100);f__Metschnikowiaceae(100);g__Metschnikowia(100);g__Metschnikowia_unclassified(100); | Metschnikowia | 13 | Saprotroph | \|Undefined Saprotroph\| | Yeast | NULL | Probable | As nectar-tap_saprotroph (Põlme et al. 2020); Decay sugar-rich substrates (Põlme et al. 2020) | Põlme S, et al. 2020. Fungal Diversity 105:1-16 ((https://doi.org/10.1007/s13225-020-00466-2)) |
| Otu16367 | 1 | k__Fungi(100);p__Ascomycota(100);c__Saccharomycetes(100);o__Saccharomycetales(100);f__Metschnikowiaceae(100);g__Metschnikowia(100);s__Metschnikowia_pulcherrima(100); | Metschnikowia | 13 | Saprotroph | \|Undefined Saprotroph\| | Yeast | NULL | Probable | As nectar-tap_saprotroph (Põlme et al. 2020); Decay sugar-rich substrates (Põlme et al. 2020) | Põlme S, et al. 2020. Fungal Diversity 105:1-16 ((https://doi.org/10.1007/s13225-020-00466-2)) |
| Otu16368 | 1 | k__Fungi(100);p__Ascomycota(100);c__Saccharomycetes(100);o__Saccharomycetales(100);f__Metschnikowiaceae(100);g__Metschnikowia(100);s__Metschnikowia_pulcherrima(100); | Metschnikowia | 13 | Saprotroph | \|Undefined Saprotroph\| | Yeast | NULL | Probable | As nectar-tap_saprotroph (Põlme et al. 2020); Decay sugar-rich substrates (Põlme et al. 2020) | Põlme S, et al. 2020. Fungal Diversity 105:1-16 ((https://doi.org/10.1007/s13225-020-00466-2)) |
| Otu16369 | 1 | k__Fungi(100);p__Ascomycota(100);c__Saccharomycetes(100);o__Saccharomycetales(100);f__Metschnikowiaceae(100);g__Metschnikowia(100);s__Metschnikowia_pulcherrima(100); | Metschnikowia | 13 | Saprotroph | \|Undefined Saprotroph\| | Yeast | NULL | Probable | As nectar-tap_saprotroph (Põlme et al. 2020); Decay sugar-rich substrates (Põlme et al. 2020) | Põlme S, et al. 2020. Fungal Diversity 105:1-16 ((https://doi.org/10.1007/s13225-020-00466-2)) |
| Otu16370 | 1 | k__Fungi(100);p__Ascomycota(100);c__Saccharomycetes(100);o__Saccharomycetales(100);f__Metschnikowiaceae(100);g__Metschnikowia(100);s__Metschnikowia_pulcherrima(100); | Metschnikowia | 13 | Saprotroph | \|Undefined Saprotroph\| | Yeast | NULL | Probable | As nectar-tap_saprotroph (Põlme et al. 2020); Decay sugar-rich substrates (Põlme et al. 2020) | Põlme S, et al. 2020. Fungal Diversity 105:1-16 ((https://doi.org/10.1007/s13225-020-00466-2)) |
| Otu16371 | 1 | k__Fungi(100);p__Ascomycota(100);c__Saccharomycetes(100);o__Saccharomycetales(100);f__Metschnikowiaceae(100);g__Metschnikowia(100);s__Metschnikowia_pulcherrima(100); | Metschnikowia | 13 | Saprotroph | \|Undefined Saprotroph\| | Yeast | NULL | Probable | As nectar-tap_saprotroph (Põlme et al. 2020); Decay sugar-rich substrates (Põlme et al. 2020) | Põlme S, et al. 2020. Fungal Diversity 105:1-16 ((https://doi.org/10.1007/s13225-020-00466-2)) |
| Otu16372 | 1 | k__Fungi(100);p__Ascomycota(100);c__Saccharomycetes(100);o__Saccharomycetales(100);f__Metschnikowiaceae(100);g__Metschnikowia(100);g__Metschnikowia_unclassified(100); | Metschnikowia | 13 | Saprotroph | \|Undefined Saprotroph\| | Yeast | NULL | Probable | As nectar-tap_saprotroph (Põlme et al. 2020); Decay sugar-rich substrates (Põlme et al. 2020) | Põlme S, et al. 2020. Fungal Diversity 105:1-16 ((https://doi.org/10.1007/s13225-020-00466-2)) |
| Otu16374 | 1 | k__Fungi(100);p__Ascomycota(100);c__Saccharomycetes(100);o__Saccharomycetales(100);f__Metschnikowiaceae(100);g__Metschnikowia(100);s__Metschnikowia_pulcherrima(100); | Metschnikowia | 13 | Saprotroph | \|Undefined Saprotroph\| | Yeast | NULL | Probable | As nectar-tap_saprotroph (Põlme et al. 2020); Decay sugar-rich substrates (Põlme et al. 2020) | Põlme S, et al. 2020. Fungal Diversity 105:1-16 ((https://doi.org/10.1007/s13225-020-00466-2)) |
| Otu16375 | 1 | k__Fungi(100);p__Ascomycota(100);c__Saccharomycetes(100);o__Saccharomycetales(100);f__Metschnikowiaceae(100);g__Metschnikowia(100);s__Metschnikowia_pulcherrima(100); | Metschnikowia | 13 | Saprotroph | \|Undefined Saprotroph\| | Yeast | NULL | Probable | As nectar-tap_saprotroph (Põlme et al. 2020); Decay sugar-rich substrates (Põlme et al. 2020) | Põlme S, et al. 2020. Fungal Diversity 105:1-16 ((https://doi.org/10.1007/s13225-020-00466-2)) |
| Otu16376 | 1 | k__Fungi(100);p__Ascomycota(100);c__Saccharomycetes(100);o__Saccharomycetales(100);f__Metschnikowiaceae(100);g__Metschnikowia(100);s__Metschnikowia_pulcherrima(100); | Metschnikowia | 13 | Saprotroph | \|Undefined Saprotroph\| | Yeast | NULL | Probable | As nectar-tap_saprotroph (Põlme et al. 2020); Decay sugar-rich substrates (Põlme et al. 2020) | Põlme S, et al. 2020. Fungal Diversity 105:1-16 ((https://doi.org/10.1007/s13225-020-00466-2)) |
| Otu16377 | 1 | k__Fungi(100);p__Ascomycota(100);c__Saccharomycetes(100);o__Saccharomycetales(100);f__Metschnikowiaceae(100);g__Metschnikowia(100);g__Metschnikowia_unclassified(100); | Metschnikowia | 13 | Saprotroph | \|Undefined Saprotroph\| | Yeast | NULL | Probable | As nectar-tap_saprotroph (Põlme et al. 2020); Decay sugar-rich substrates (Põlme et al. 2020) | Põlme S, et al. 2020. Fungal Diversity 105:1-16 ((https://doi.org/10.1007/s13225-020-00466-2)) |
| Otu16378 | 1 | k__Fungi(100);p__Ascomycota(100);c__Saccharomycetes(100);o__Saccharomycetales(100);f__Metschnikowiaceae(100);g__Metschnikowia(100);g__Metschnikowia_unclassified(100); | Metschnikowia | 13 | Saprotroph | \|Undefined Saprotroph\| | Yeast | NULL | Probable | As nectar-tap_saprotroph (Põlme et al. 2020); Decay sugar-rich substrates (Põlme et al. 2020) | Põlme S, et al. 2020. Fungal Diversity 105:1-16 ((https://doi.org/10.1007/s13225-020-00466-2)) |
| Otu16379 | 1 | k__Fungi(100);p__Ascomycota(100);c__Saccharomycetes(100);o__Saccharomycetales(100);f__Metschnikowiaceae(100);g__Metschnikowia(100);g__Metschnikowia_unclassified(100); | Metschnikowia | 13 | Saprotroph | \|Undefined Saprotroph\| | Yeast | NULL | Probable | As nectar-tap_saprotroph (Põlme et al. 2020); Decay sugar-rich substrates (Põlme et al. 2020) | Põlme S, et al. 2020. Fungal Diversity 105:1-16 ((https://doi.org/10.1007/s13225-020-00466-2)) |
| Otu16380 | 1 | k__Fungi(100);p__Ascomycota(100);c__Saccharomycetes(100);o__Saccharomycetales(100);f__Metschnikowiaceae(100);g__Metschnikowia(100);g__Metschnikowia_unclassified(100); | Metschnikowia | 13 | Saprotroph | \|Undefined Saprotroph\| | Yeast | NULL | Probable | As nectar-tap_saprotroph (Põlme et al. 2020); Decay sugar-rich substrates (Põlme et al. 2020) | Põlme S, et al. 2020. Fungal Diversity 105:1-16 ((https://doi.org/10.1007/s13225-020-00466-2)) |
| Otu16382 | 1 | k__Fungi(100);p__Ascomycota(100);c__Saccharomycetes(100);o__Saccharomycetales(100);f__Metschnikowiaceae(100);g__Metschnikowia(100);s__Metschnikowia_pulcherrima(100); | Metschnikowia | 13 | Saprotroph | \|Undefined Saprotroph\| | Yeast | NULL | Probable | As nectar-tap_saprotroph (Põlme et al. 2020); Decay sugar-rich substrates (Põlme et al. 2020) | Põlme S, et al. 2020. Fungal Diversity 105:1-16 ((https://doi.org/10.1007/s13225-020-00466-2)) |
| Otu16383 | 1 | k__Fungi(100);p__Ascomycota(100);c__Saccharomycetes(100);o__Saccharomycetales(100);f__Metschnikowiaceae(100);g__Metschnikowia(100);s__Metschnikowia_pulcherrima(100); | Metschnikowia | 13 | Saprotroph | \|Undefined Saprotroph\| | Yeast | NULL | Probable | As nectar-tap_saprotroph (Põlme et al. 2020); Decay sugar-rich substrates (Põlme et al. 2020) | Põlme S, et al. 2020. Fungal Diversity 105:1-16 ((https://doi.org/10.1007/s13225-020-00466-2)) |
| Otu16384 | 1 | k__Fungi(100);p__Ascomycota(100);c__Saccharomycetes(100);o__Saccharomycetales(100);f__Metschnikowiaceae(100);g__Metschnikowia(100);s__Metschnikowia_pulcherrima(100); | Metschnikowia | 13 | Saprotroph | \|Undefined Saprotroph\| | Yeast | NULL | Probable | As nectar-tap_saprotroph (Põlme et al. 2020); Decay sugar-rich substrates (Põlme et al. 2020) | Põlme S, et al. 2020. Fungal Diversity 105:1-16 ((https://doi.org/10.1007/s13225-020-00466-2)) |
| Otu16385 | 1 | k__Fungi(100);p__Ascomycota(100);c__Saccharomycetes(100);o__Saccharomycetales(100);f__Metschnikowiaceae(100);g__Metschnikowia(100);s__Metschnikowia_pulcherrima(100); | Metschnikowia | 13 | Saprotroph | \|Undefined Saprotroph\| | Yeast | NULL | Probable | As nectar-tap_saprotroph (Põlme et al. 2020); Decay sugar-rich substrates (Põlme et al. 2020) | Põlme S, et al. 2020. Fungal Diversity 105:1-16 ((https://doi.org/10.1007/s13225-020-00466-2)) |
| Otu16386 | 1 | k__Fungi(100);p__Ascomycota(100);c__Saccharomycetes(100);o__Saccharomycetales(100);f__Metschnikowiaceae(100);g__Metschnikowia(100);s__Metschnikowia_pulcherrima(100); | Metschnikowia | 13 | Saprotroph | \|Undefined Saprotroph\| | Yeast | NULL | Probable | As nectar-tap_saprotroph (Põlme et al. 2020); Decay sugar-rich substrates (Põlme et al. 2020) | Põlme S, et al. 2020. Fungal Diversity 105:1-16 ((https://doi.org/10.1007/s13225-020-00466-2)) |
| Otu16387 | 1 | k__Fungi(100);p__Ascomycota(100);c__Saccharomycetes(100);o__Saccharomycetales(100);f__Metschnikowiaceae(100);g__Metschnikowia(100);s__Metschnikowia_pulcherrima(100); | Metschnikowia | 13 | Saprotroph | \|Undefined Saprotroph\| | Yeast | NULL | Probable | As nectar-tap_saprotroph (Põlme et al. 2020); Decay sugar-rich substrates (Põlme et al. 2020) | Põlme S, et al. 2020. Fungal Diversity 105:1-16 ((https://doi.org/10.1007/s13225-020-00466-2)) |
| Otu16389 | 1 | k__Fungi(100);p__Ascomycota(100);c__Saccharomycetes(100);o__Saccharomycetales(100);f__Metschnikowiaceae(100);g__Metschnikowia(100);s__Metschnikowia_pulcherrima(100); | Metschnikowia | 13 | Saprotroph | \|Undefined Saprotroph\| | Yeast | NULL | Probable | As nectar-tap_saprotroph (Põlme et al. 2020); Decay sugar-rich substrates (Põlme et al. 2020) | Põlme S, et al. 2020. Fungal Diversity 105:1-16 ((https://doi.org/10.1007/s13225-020-00466-2)) |
| Otu16391 | 1 | k__Fungi(100);p__Ascomycota(100);c__Saccharomycetes(100);o__Saccharomycetales(100);f__Metschnikowiaceae(100);g__Metschnikowia(100);g__Metschnikowia_unclassified(100); | Metschnikowia | 13 | Saprotroph | \|Undefined Saprotroph\| | Yeast | NULL | Probable | As nectar-tap_saprotroph (Põlme et al. 2020); Decay sugar-rich substrates (Põlme et al. 2020) | Põlme S, et al. 2020. Fungal Diversity 105:1-16 ((https://doi.org/10.1007/s13225-020-00466-2)) |
| Otu16392 | 1 | k__Fungi(100);p__Ascomycota(100);c__Saccharomycetes(100);o__Saccharomycetales(100);f__Metschnikowiaceae(100);g__Metschnikowia(100);s__Metschnikowia_pulcherrima(100); | Metschnikowia | 13 | Saprotroph | \|Undefined Saprotroph\| | Yeast | NULL | Probable | As nectar-tap_saprotroph (Põlme et al. 2020); Decay sugar-rich substrates (Põlme et al. 2020) | Põlme S, et al. 2020. Fungal Diversity 105:1-16 ((https://doi.org/10.1007/s13225-020-00466-2)) |
| Otu16393 | 1 | k__Fungi(100);p__Ascomycota(100);c__Saccharomycetes(100);o__Saccharomycetales(100);f__Metschnikowiaceae(100);g__Metschnikowia(100);s__Metschnikowia_pulcherrima(100); | Metschnikowia | 13 | Saprotroph | \|Undefined Saprotroph\| | Yeast | NULL | Probable | As nectar-tap_saprotroph (Põlme et al. 2020); Decay sugar-rich substrates (Põlme et al. 2020) | Põlme S, et al. 2020. Fungal Diversity 105:1-16 ((https://doi.org/10.1007/s13225-020-00466-2)) |
| Otu16395 | 1 | k__Fungi(100);p__Ascomycota(100);c__Saccharomycetes(100);o__Saccharomycetales(100);f__Metschnikowiaceae(100);g__Metschnikowia(100);g__Metschnikowia_unclassified(100); | Metschnikowia | 13 | Saprotroph | \|Undefined Saprotroph\| | Yeast | NULL | Probable | As nectar-tap_saprotroph (Põlme et al. 2020); Decay sugar-rich substrates (Põlme et al. 2020) | Põlme S, et al. 2020. Fungal Diversity 105:1-16 ((https://doi.org/10.1007/s13225-020-00466-2)) |
| Otu16396 | 1 | k__Fungi(100);p__Ascomycota(100);c__Saccharomycetes(100);o__Saccharomycetales(100);f__Metschnikowiaceae(100);g__Metschnikowia(100);s__Metschnikowia_pulcherrima(100); | Metschnikowia | 13 | Saprotroph | \|Undefined Saprotroph\| | Yeast | NULL | Probable | As nectar-tap_saprotroph (Põlme et al. 2020); Decay sugar-rich substrates (Põlme et al. 2020) | Põlme S, et al. 2020. Fungal Diversity 105:1-16 ((https://doi.org/10.1007/s13225-020-00466-2)) |
| Otu16397 | 1 | k__Fungi(100);p__Ascomycota(100);c__Saccharomycetes(100);o__Saccharomycetales(100);f__Metschnikowiaceae(100);g__Metschnikowia(100);s__Metschnikowia_pulcherrima(100); | Metschnikowia | 13 | Saprotroph | \|Undefined Saprotroph\| | Yeast | NULL | Probable | As nectar-tap_saprotroph (Põlme et al. 2020); Decay sugar-rich substrates (Põlme et al. 2020) | Põlme S, et al. 2020. Fungal Diversity 105:1-16 ((https://doi.org/10.1007/s13225-020-00466-2)) |
| Otu16398 | 1 | k__Fungi(100);p__Ascomycota(100);c__Saccharomycetes(100);o__Saccharomycetales(100);f__Metschnikowiaceae(100);g__Metschnikowia(100);s__Metschnikowia_pulcherrima(100); | Metschnikowia | 13 | Saprotroph | \|Undefined Saprotroph\| | Yeast | NULL | Probable | As nectar-tap_saprotroph (Põlme et al. 2020); Decay sugar-rich substrates (Põlme et al. 2020) | Põlme S, et al. 2020. Fungal Diversity 105:1-16 ((https://doi.org/10.1007/s13225-020-00466-2)) |
| Otu16399 | 1 | k__Fungi(100);p__Ascomycota(100);c__Saccharomycetes(100);o__Saccharomycetales(100);f__Metschnikowiaceae(100);g__Metschnikowia(100);s__Metschnikowia_pulcherrima(100); | Metschnikowia | 13 | Saprotroph | \|Undefined Saprotroph\| | Yeast | NULL | Probable | As nectar-tap_saprotroph (Põlme et al. 2020); Decay sugar-rich substrates (Põlme et al. 2020) | Põlme S, et al. 2020. Fungal Diversity 105:1-16 ((https://doi.org/10.1007/s13225-020-00466-2)) |
| Otu16400 | 1 | k__Fungi(100);p__Ascomycota(100);c__Saccharomycetes(100);o__Saccharomycetales(100);f__Metschnikowiaceae(100);g__Metschnikowia(100);g__Metschnikowia_unclassified(100); | Metschnikowia | 13 | Saprotroph | \|Undefined Saprotroph\| | Yeast | NULL | Probable | As nectar-tap_saprotroph (Põlme et al. 2020); Decay sugar-rich substrates (Põlme et al. 2020) | Põlme S, et al. 2020. Fungal Diversity 105:1-16 ((https://doi.org/10.1007/s13225-020-00466-2)) |
| Otu16401 | 1 | k__Fungi(100);p__Ascomycota(100);c__Saccharomycetes(100);o__Saccharomycetales(100);f__Metschnikowiaceae(100);g__Metschnikowia(100);g__Metschnikowia_unclassified(100); | Metschnikowia | 13 | Saprotroph | \|Undefined Saprotroph\| | Yeast | NULL | Probable | As nectar-tap_saprotroph (Põlme et al. 2020); Decay sugar-rich substrates (Põlme et al. 2020) | Põlme S, et al. 2020. Fungal Diversity 105:1-16 ((https://doi.org/10.1007/s13225-020-00466-2)) |
| Otu16402 | 1 | k__Fungi(100);p__Ascomycota(100);c__Saccharomycetes(100);o__Saccharomycetales(100);f__Metschnikowiaceae(100);g__Metschnikowia(100);s__Metschnikowia_pulcherrima(100); | Metschnikowia | 13 | Saprotroph | \|Undefined Saprotroph\| | Yeast | NULL | Probable | As nectar-tap_saprotroph (Põlme et al. 2020); Decay sugar-rich substrates (Põlme et al. 2020) | Põlme S, et al. 2020. Fungal Diversity 105:1-16 ((https://doi.org/10.1007/s13225-020-00466-2)) |
| Otu16403 | 1 | k__Fungi(100);p__Ascomycota(100);c__Saccharomycetes(100);o__Saccharomycetales(100);f__Metschnikowiaceae(100);g__Metschnikowia(100);g__Metschnikowia_unclassified(100); | Metschnikowia | 13 | Saprotroph | \|Undefined Saprotroph\| | Yeast | NULL | Probable | As nectar-tap_saprotroph (Põlme et al. 2020); Decay sugar-rich substrates (Põlme et al. 2020) | Põlme S, et al. 2020. Fungal Diversity 105:1-16 ((https://doi.org/10.1007/s13225-020-00466-2)) |
| Otu16404 | 1 | k__Fungi(100);p__Ascomycota(100);c__Saccharomycetes(100);o__Saccharomycetales(100);f__Metschnikowiaceae(100);g__Metschnikowia(100);s__Metschnikowia_pulcherrima(100); | Metschnikowia | 13 | Saprotroph | \|Undefined Saprotroph\| | Yeast | NULL | Probable | As nectar-tap_saprotroph (Põlme et al. 2020); Decay sugar-rich substrates (Põlme et al. 2020) | Põlme S, et al. 2020. Fungal Diversity 105:1-16 ((https://doi.org/10.1007/s13225-020-00466-2)) |
| Otu16405 | 1 | k__Fungi(100);p__Ascomycota(100);c__Saccharomycetes(100);o__Saccharomycetales(100);f__Metschnikowiaceae(100);g__Metschnikowia(100);s__Metschnikowia_pulcherrima(100); | Metschnikowia | 13 | Saprotroph | \|Undefined Saprotroph\| | Yeast | NULL | Probable | As nectar-tap_saprotroph (Põlme et al. 2020); Decay sugar-rich substrates (Põlme et al. 2020) | Põlme S, et al. 2020. Fungal Diversity 105:1-16 ((https://doi.org/10.1007/s13225-020-00466-2)) |
| Otu16406 | 1 | k__Fungi(100);p__Ascomycota(100);c__Saccharomycetes(100);o__Saccharomycetales(100);f__Metschnikowiaceae(100);g__Metschnikowia(100);g__Metschnikowia_unclassified(100); | Metschnikowia | 13 | Saprotroph | \|Undefined Saprotroph\| | Yeast | NULL | Probable | As nectar-tap_saprotroph (Põlme et al. 2020); Decay sugar-rich substrates (Põlme et al. 2020) | Põlme S, et al. 2020. Fungal Diversity 105:1-16 ((https://doi.org/10.1007/s13225-020-00466-2)) |
| Otu16408 | 1 | k__Fungi(100);p__Ascomycota(100);c__Saccharomycetes(100);o__Saccharomycetales(100);f__Metschnikowiaceae(100);g__Metschnikowia(100);s__Metschnikowia_pulcherrima(100); | Metschnikowia | 13 | Saprotroph | \|Undefined Saprotroph\| | Yeast | NULL | Probable | As nectar-tap_saprotroph (Põlme et al. 2020); Decay sugar-rich substrates (Põlme et al. 2020) | Põlme S, et al. 2020. Fungal Diversity 105:1-16 ((https://doi.org/10.1007/s13225-020-00466-2)) |
| Otu16409 | 1 | k__Fungi(100);p__Ascomycota(100);c__Saccharomycetes(100);o__Saccharomycetales(100);f__Metschnikowiaceae(100);g__Metschnikowia(100);s__Metschnikowia_pulcherrima(100); | Metschnikowia | 13 | Saprotroph | \|Undefined Saprotroph\| | Yeast | NULL | Probable | As nectar-tap_saprotroph (Põlme et al. 2020); Decay sugar-rich substrates (Põlme et al. 2020) | Põlme S, et al. 2020. Fungal Diversity 105:1-16 ((https://doi.org/10.1007/s13225-020-00466-2)) |
| Otu16411 | 1 | k__Fungi(100);p__Ascomycota(100);c__Saccharomycetes(100);o__Saccharomycetales(100);f__Metschnikowiaceae(100);g__Metschnikowia(100);g__Metschnikowia_unclassified(100); | Metschnikowia | 13 | Saprotroph | \|Undefined Saprotroph\| | Yeast | NULL | Probable | As nectar-tap_saprotroph (Põlme et al. 2020); Decay sugar-rich substrates (Põlme et al. 2020) | Põlme S, et al. 2020. Fungal Diversity 105:1-16 ((https://doi.org/10.1007/s13225-020-00466-2)) |
| Otu16412 | 1 | k__Fungi(100);p__Ascomycota(100);c__Saccharomycetes(100);o__Saccharomycetales(100);f__Metschnikowiaceae(100);g__Metschnikowia(100);s__Metschnikowia_pulcherrima(100); | Metschnikowia | 13 | Saprotroph | \|Undefined Saprotroph\| | Yeast | NULL | Probable | As nectar-tap_saprotroph (Põlme et al. 2020); Decay sugar-rich substrates (Põlme et al. 2020) | Põlme S, et al. 2020. Fungal Diversity 105:1-16 ((https://doi.org/10.1007/s13225-020-00466-2)) |
| Otu16414 | 1 | k__Fungi(100);p__Ascomycota(100);c__Saccharomycetes(100);o__Saccharomycetales(100);f__Metschnikowiaceae(100);g__Metschnikowia(100);s__Metschnikowia_pulcherrima(100); | Metschnikowia | 13 | Saprotroph | \|Undefined Saprotroph\| | Yeast | NULL | Probable | As nectar-tap_saprotroph (Põlme et al. 2020); Decay sugar-rich substrates (Põlme et al. 2020) | Põlme S, et al. 2020. Fungal Diversity 105:1-16 ((https://doi.org/10.1007/s13225-020-00466-2)) |
| Otu16416 | 1 | k__Fungi(100);p__Ascomycota(100);c__Saccharomycetes(100);o__Saccharomycetales(100);f__Metschnikowiaceae(100);g__Metschnikowia(100);g__Metschnikowia_unclassified(100); | Metschnikowia | 13 | Saprotroph | \|Undefined Saprotroph\| | Yeast | NULL | Probable | As nectar-tap_saprotroph (Põlme et al. 2020); Decay sugar-rich substrates (Põlme et al. 2020) | Põlme S, et al. 2020. Fungal Diversity 105:1-16 ((https://doi.org/10.1007/s13225-020-00466-2)) |
| Otu16417 | 1 | k__Fungi(100);p__Ascomycota(100);c__Saccharomycetes(100);o__Saccharomycetales(100);f__Metschnikowiaceae(100);g__Metschnikowia(100);g__Metschnikowia_unclassified(100); | Metschnikowia | 13 | Saprotroph | \|Undefined Saprotroph\| | Yeast | NULL | Probable | As nectar-tap_saprotroph (Põlme et al. 2020); Decay sugar-rich substrates (Põlme et al. 2020) | Põlme S, et al. 2020. Fungal Diversity 105:1-16 ((https://doi.org/10.1007/s13225-020-00466-2)) |
| Otu16418 | 1 | k__Fungi(100);p__Ascomycota(100);c__Saccharomycetes(100);o__Saccharomycetales(100);f__Metschnikowiaceae(100);g__Metschnikowia(100);s__Metschnikowia_pulcherrima(100); | Metschnikowia | 13 | Saprotroph | \|Undefined Saprotroph\| | Yeast | NULL | Probable | As nectar-tap_saprotroph (Põlme et al. 2020); Decay sugar-rich substrates (Põlme et al. 2020) | Põlme S, et al. 2020. Fungal Diversity 105:1-16 ((https://doi.org/10.1007/s13225-020-00466-2)) |
| Otu16419 | 1 | k__Fungi(100);p__Ascomycota(100);c__Saccharomycetes(100);o__Saccharomycetales(100);f__Metschnikowiaceae(100);g__Metschnikowia(100);s__Metschnikowia_pulcherrima(100); | Metschnikowia | 13 | Saprotroph | \|Undefined Saprotroph\| | Yeast | NULL | Probable | As nectar-tap_saprotroph (Põlme et al. 2020); Decay sugar-rich substrates (Põlme et al. 2020) | Põlme S, et al. 2020. Fungal Diversity 105:1-16 ((https://doi.org/10.1007/s13225-020-00466-2)) |
| Otu16420 | 1 | k__Fungi(100);p__Ascomycota(100);c__Saccharomycetes(100);o__Saccharomycetales(100);f__Metschnikowiaceae(100);g__Metschnikowia(100);s__Metschnikowia_pulcherrima(100); | Metschnikowia | 13 | Saprotroph | \|Undefined Saprotroph\| | Yeast | NULL | Probable | As nectar-tap_saprotroph (Põlme et al. 2020); Decay sugar-rich substrates (Põlme et al. 2020) | Põlme S, et al. 2020. Fungal Diversity 105:1-16 ((https://doi.org/10.1007/s13225-020-00466-2)) |
| Otu16421 | 1 | k__Fungi(100);p__Ascomycota(100);c__Saccharomycetes(100);o__Saccharomycetales(100);f__Metschnikowiaceae(100);g__Metschnikowia(100);s__Metschnikowia_pulcherrima(100); | Metschnikowia | 13 | Saprotroph | \|Undefined Saprotroph\| | Yeast | NULL | Probable | As nectar-tap_saprotroph (Põlme et al. 2020); Decay sugar-rich substrates (Põlme et al. 2020) | Põlme S, et al. 2020. Fungal Diversity 105:1-16 ((https://doi.org/10.1007/s13225-020-00466-2)) |
| Otu16422 | 1 | k__Fungi(100);p__Ascomycota(100);c__Saccharomycetes(100);o__Saccharomycetales(100);f__Metschnikowiaceae(100);g__Metschnikowia(100);g__Metschnikowia_unclassified(100); | Metschnikowia | 13 | Saprotroph | \|Undefined Saprotroph\| | Yeast | NULL | Probable | As nectar-tap_saprotroph (Põlme et al. 2020); Decay sugar-rich substrates (Põlme et al. 2020) | Põlme S, et al. 2020. Fungal Diversity 105:1-16 ((https://doi.org/10.1007/s13225-020-00466-2)) |
| Otu16423 | 1 | k__Fungi(100);p__Ascomycota(100);c__Saccharomycetes(100);o__Saccharomycetales(100);f__Metschnikowiaceae(100);g__Metschnikowia(100);s__Metschnikowia_pulcherrima(100); | Metschnikowia | 13 | Saprotroph | \|Undefined Saprotroph\| | Yeast | NULL | Probable | As nectar-tap_saprotroph (Põlme et al. 2020); Decay sugar-rich substrates (Põlme et al. 2020) | Põlme S, et al. 2020. Fungal Diversity 105:1-16 ((https://doi.org/10.1007/s13225-020-00466-2)) |
| Otu16424 | 1 | k__Fungi(100);p__Ascomycota(100);c__Saccharomycetes(100);o__Saccharomycetales(100);f__Metschnikowiaceae(100);g__Metschnikowia(100);s__Metschnikowia_pulcherrima(100); | Metschnikowia | 13 | Saprotroph | \|Undefined Saprotroph\| | Yeast | NULL | Probable | As nectar-tap_saprotroph (Põlme et al. 2020); Decay sugar-rich substrates (Põlme et al. 2020) | Põlme S, et al. 2020. Fungal Diversity 105:1-16 ((https://doi.org/10.1007/s13225-020-00466-2)) |
| Otu16425 | 1 | k__Fungi(100);p__Ascomycota(100);c__Saccharomycetes(100);o__Saccharomycetales(100);f__Metschnikowiaceae(100);g__Metschnikowia(100);g__Metschnikowia_unclassified(100); | Metschnikowia | 13 | Saprotroph | \|Undefined Saprotroph\| | Yeast | NULL | Probable | As nectar-tap_saprotroph (Põlme et al. 2020); Decay sugar-rich substrates (Põlme et al. 2020) | Põlme S, et al. 2020. Fungal Diversity 105:1-16 ((https://doi.org/10.1007/s13225-020-00466-2)) |
| Otu16426 | 1 | k__Fungi(100);p__Ascomycota(100);c__Saccharomycetes(100);o__Saccharomycetales(100);f__Metschnikowiaceae(100);g__Metschnikowia(100);s__Metschnikowia_pulcherrima(100); | Metschnikowia | 13 | Saprotroph | \|Undefined Saprotroph\| | Yeast | NULL | Probable | As nectar-tap_saprotroph (Põlme et al. 2020); Decay sugar-rich substrates (Põlme et al. 2020) | Põlme S, et al. 2020. Fungal Diversity 105:1-16 ((https://doi.org/10.1007/s13225-020-00466-2)) |
| Otu16428 | 1 | k__Fungi(100);p__Ascomycota(100);c__Saccharomycetes(100);o__Saccharomycetales(100);f__Metschnikowiaceae(100);g__Metschnikowia(100);s__Metschnikowia_pulcherrima(100); | Metschnikowia | 13 | Saprotroph | \|Undefined Saprotroph\| | Yeast | NULL | Probable | As nectar-tap_saprotroph (Põlme et al. 2020); Decay sugar-rich substrates (Põlme et al. 2020) | Põlme S, et al. 2020. Fungal Diversity 105:1-16 ((https://doi.org/10.1007/s13225-020-00466-2)) |
| Otu16429 | 1 | k__Fungi(100);p__Ascomycota(100);c__Saccharomycetes(100);o__Saccharomycetales(100);f__Metschnikowiaceae(100);g__Metschnikowia(100);s__Metschnikowia_pulcherrima(100); | Metschnikowia | 13 | Saprotroph | \|Undefined Saprotroph\| | Yeast | NULL | Probable | As nectar-tap_saprotroph (Põlme et al. 2020); Decay sugar-rich substrates (Põlme et al. 2020) | Põlme S, et al. 2020. Fungal Diversity 105:1-16 ((https://doi.org/10.1007/s13225-020-00466-2)) |
| Otu16433 | 1 | k__Fungi(100);p__Ascomycota(100);c__Saccharomycetes(100);o__Saccharomycetales(100);f__Metschnikowiaceae(100);g__Metschnikowia(100);s__Metschnikowia_pulcherrima(100); | Metschnikowia | 13 | Saprotroph | \|Undefined Saprotroph\| | Yeast | NULL | Probable | As nectar-tap_saprotroph (Põlme et al. 2020); Decay sugar-rich substrates (Põlme et al. 2020) | Põlme S, et al. 2020. Fungal Diversity 105:1-16 ((https://doi.org/10.1007/s13225-020-00466-2)) |
| Otu16435 | 1 | k__Fungi(100);p__Ascomycota(100);c__Saccharomycetes(100);o__Saccharomycetales(100);f__Metschnikowiaceae(100);g__Metschnikowia(100);s__Metschnikowia_pulcherrima(100); | Metschnikowia | 13 | Saprotroph | \|Undefined Saprotroph\| | Yeast | NULL | Probable | As nectar-tap_saprotroph (Põlme et al. 2020); Decay sugar-rich substrates (Põlme et al. 2020) | Põlme S, et al. 2020. Fungal Diversity 105:1-16 ((https://doi.org/10.1007/s13225-020-00466-2)) |
| Otu16436 | 1 | k__Fungi(100);p__Ascomycota(100);c__Saccharomycetes(100);o__Saccharomycetales(100);f__Metschnikowiaceae(100);g__Metschnikowia(100);s__Metschnikowia_pulcherrima(100); | Metschnikowia | 13 | Saprotroph | \|Undefined Saprotroph\| | Yeast | NULL | Probable | As nectar-tap_saprotroph (Põlme et al. 2020); Decay sugar-rich substrates (Põlme et al. 2020) | Põlme S, et al. 2020. Fungal Diversity 105:1-16 ((https://doi.org/10.1007/s13225-020-00466-2)) |
| Otu16437 | 1 | k__Fungi(100);p__Ascomycota(100);c__Saccharomycetes(100);o__Saccharomycetales(100);f__Metschnikowiaceae(100);g__Metschnikowia(100);s__Metschnikowia_pulcherrima(100); | Metschnikowia | 13 | Saprotroph | \|Undefined Saprotroph\| | Yeast | NULL | Probable | As nectar-tap_saprotroph (Põlme et al. 2020); Decay sugar-rich substrates (Põlme et al. 2020) | Põlme S, et al. 2020. Fungal Diversity 105:1-16 ((https://doi.org/10.1007/s13225-020-00466-2)) |
| Otu16438 | 1 | k__Fungi(100);p__Ascomycota(100);c__Saccharomycetes(100);o__Saccharomycetales(100);f__Metschnikowiaceae(100);g__Metschnikowia(100);s__Metschnikowia_pulcherrima(100); | Metschnikowia | 13 | Saprotroph | \|Undefined Saprotroph\| | Yeast | NULL | Probable | As nectar-tap_saprotroph (Põlme et al. 2020); Decay sugar-rich substrates (Põlme et al. 2020) | Põlme S, et al. 2020. Fungal Diversity 105:1-16 ((https://doi.org/10.1007/s13225-020-00466-2)) |
| Otu16439 | 1 | k__Fungi(100);p__Ascomycota(100);c__Leotiomycetes(100);o__Helotiales(100);f__Helotiaceae(100);f__Helotiaceae_unclassified(100);f__Helotiaceae_unclassified(100); | Helotiaceae | 9 | Saprotroph-Symbiotroph | Ectomycorrhizal-Fungal Parasite-Plant Pathogen-Wood Saprotroph | Microfungus | NULL | Probable | Usually saprobic on herbaceous or woody tissues, some species fungicolous. A few species are known to form mycorrhizae (Cannon & Kirk 2007) | Cannon PF, Kirk PM. 2007. Fungal Families of the World. CAB International, Cambridge (ISBN: 978-0851998275) |
| Otu16440 | 1 | k__Fungi(100);p__Ascomycota(100);c__Saccharomycetes(100);o__Saccharomycetales(100);f__Metschnikowiaceae(100);g__Metschnikowia(100);s__Metschnikowia_pulcherrima(100); | Metschnikowia | 13 | Saprotroph | \|Undefined Saprotroph\| | Yeast | NULL | Probable | As nectar-tap_saprotroph (Põlme et al. 2020); Decay sugar-rich substrates (Põlme et al. 2020) | Põlme S, et al. 2020. Fungal Diversity 105:1-16 ((https://doi.org/10.1007/s13225-020-00466-2)) |
| Otu16442 | 1 | k__Fungi(100);p__Ascomycota(100);c__Saccharomycetes(100);o__Saccharomycetales(100);f__Metschnikowiaceae(100);g__Metschnikowia(100);s__Metschnikowia_pulcherrima(100); | Metschnikowia | 13 | Saprotroph | \|Undefined Saprotroph\| | Yeast | NULL | Probable | As nectar-tap_saprotroph (Põlme et al. 2020); Decay sugar-rich substrates (Põlme et al. 2020) | Põlme S, et al. 2020. Fungal Diversity 105:1-16 ((https://doi.org/10.1007/s13225-020-00466-2)) |
| Otu16443 | 1 | k__Fungi(100);p__Ascomycota(100);c__Dothideomycetes(100);o__Capnodiales(100);f__Mycosphaerellaceae(100);g__Stenella(100);s__Stenella_queenslandica(100); | Stenella | 13 | Pathotroph-Saprotroph | Plant Pathogen-\|Plant Saprotroph\|-Undefined Saprotroph | Microfungus | NULL | Probable | As litter_saprotroph (Põlme et al. 2020); Decay leaf, fruit, seed (Põlme et al. 2020) | Tedersoo L, et al. 2014. Science 346:e1256688 ((https://doi.org/10.1126/science.1256688)); Põlme S, et al. 2020. Fungal Diversity 105:1-16 ((https://doi.org/10.1007/s13225-020-00466-2)) |
| Otu16444 | 1 | k__Fungi(100);p__Ascomycota(100);c__Saccharomycetes(100);o__Saccharomycetales(100);f__Metschnikowiaceae(100);g__Metschnikowia(100);s__Metschnikowia_pulcherrima(100); | Metschnikowia | 13 | Saprotroph | \|Undefined Saprotroph\| | Yeast | NULL | Probable | As nectar-tap_saprotroph (Põlme et al. 2020); Decay sugar-rich substrates (Põlme et al. 2020) | Põlme S, et al. 2020. Fungal Diversity 105:1-16 ((https://doi.org/10.1007/s13225-020-00466-2)) |
| Otu16445 | 1 | k__Fungi(100);p__Ascomycota(100);c__Saccharomycetes(100);o__Saccharomycetales(100);f__Metschnikowiaceae(100);g__Metschnikowia(100);s__Metschnikowia_pulcherrima(100); | Metschnikowia | 13 | Saprotroph | \|Undefined Saprotroph\| | Yeast | NULL | Probable | As nectar-tap_saprotroph (Põlme et al. 2020); Decay sugar-rich substrates (Põlme et al. 2020) | Põlme S, et al. 2020. Fungal Diversity 105:1-16 ((https://doi.org/10.1007/s13225-020-00466-2)) |
| Otu16446 | 1 | k__Fungi(100);p__Ascomycota(100);c__Saccharomycetes(100);o__Saccharomycetales(100);f__Metschnikowiaceae(100);g__Metschnikowia(100);g__Metschnikowia_unclassified(100); | Metschnikowia | 13 | Saprotroph | \|Undefined Saprotroph\| | Yeast | NULL | Probable | As nectar-tap_saprotroph (Põlme et al. 2020); Decay sugar-rich substrates (Põlme et al. 2020) | Põlme S, et al. 2020. Fungal Diversity 105:1-16 ((https://doi.org/10.1007/s13225-020-00466-2)) |
| Otu16447 | 1 | k__Fungi(100);p__Ascomycota(100);c__Saccharomycetes(100);o__Saccharomycetales(100);f__Metschnikowiaceae(100);g__Metschnikowia(100);s__Metschnikowia_pulcherrima(100); | Metschnikowia | 13 | Saprotroph | \|Undefined Saprotroph\| | Yeast | NULL | Probable | As nectar-tap_saprotroph (Põlme et al. 2020); Decay sugar-rich substrates (Põlme et al. 2020) | Põlme S, et al. 2020. Fungal Diversity 105:1-16 ((https://doi.org/10.1007/s13225-020-00466-2)) |
| Otu16448 | 1 | k__Fungi(100);p__Ascomycota(100);c__Saccharomycetes(100);o__Saccharomycetales(100);f__Metschnikowiaceae(100);g__Metschnikowia(100);s__Metschnikowia_pulcherrima(100); | Metschnikowia | 13 | Saprotroph | \|Undefined Saprotroph\| | Yeast | NULL | Probable | As nectar-tap_saprotroph (Põlme et al. 2020); Decay sugar-rich substrates (Põlme et al. 2020) | Põlme S, et al. 2020. Fungal Diversity 105:1-16 ((https://doi.org/10.1007/s13225-020-00466-2)) |
| Otu16449 | 1 | k__Fungi(100);p__Ascomycota(100);c__Saccharomycetes(100);o__Saccharomycetales(100);f__Metschnikowiaceae(100);g__Metschnikowia(100);s__Metschnikowia_pulcherrima(100); | Metschnikowia | 13 | Saprotroph | \|Undefined Saprotroph\| | Yeast | NULL | Probable | As nectar-tap_saprotroph (Põlme et al. 2020); Decay sugar-rich substrates (Põlme et al. 2020) | Põlme S, et al. 2020. Fungal Diversity 105:1-16 ((https://doi.org/10.1007/s13225-020-00466-2)) |
| Otu16450 | 1 | k__Fungi(100);p__Ascomycota(100);c__Saccharomycetes(100);o__Saccharomycetales(100);f__Metschnikowiaceae(100);g__Metschnikowia(100);s__Metschnikowia_pulcherrima(100); | Metschnikowia | 13 | Saprotroph | \|Undefined Saprotroph\| | Yeast | NULL | Probable | As nectar-tap_saprotroph (Põlme et al. 2020); Decay sugar-rich substrates (Põlme et al. 2020) | Põlme S, et al. 2020. Fungal Diversity 105:1-16 ((https://doi.org/10.1007/s13225-020-00466-2)) |
| Otu16451 | 1 | k__Fungi(100);p__Ascomycota(100);c__Saccharomycetes(100);o__Saccharomycetales(100);f__Metschnikowiaceae(100);g__Metschnikowia(100);s__Metschnikowia_pulcherrima(100); | Metschnikowia | 13 | Saprotroph | \|Undefined Saprotroph\| | Yeast | NULL | Probable | As nectar-tap_saprotroph (Põlme et al. 2020); Decay sugar-rich substrates (Põlme et al. 2020) | Põlme S, et al. 2020. Fungal Diversity 105:1-16 ((https://doi.org/10.1007/s13225-020-00466-2)) |
| Otu16452 | 1 | k__Fungi(100);p__Ascomycota(100);c__Saccharomycetes(100);o__Saccharomycetales(100);f__Metschnikowiaceae(100);g__Metschnikowia(100);s__Metschnikowia_pulcherrima(100); | Metschnikowia | 13 | Saprotroph | \|Undefined Saprotroph\| | Yeast | NULL | Probable | As nectar-tap_saprotroph (Põlme et al. 2020); Decay sugar-rich substrates (Põlme et al. 2020) | Põlme S, et al. 2020. Fungal Diversity 105:1-16 ((https://doi.org/10.1007/s13225-020-00466-2)) |
| Otu16453 | 1 | k__Fungi(100);p__Ascomycota(100);c__Saccharomycetes(100);o__Saccharomycetales(100);f__Metschnikowiaceae(100);g__Metschnikowia(100);s__Metschnikowia_pulcherrima(100); | Metschnikowia | 13 | Saprotroph | \|Undefined Saprotroph\| | Yeast | NULL | Probable | As nectar-tap_saprotroph (Põlme et al. 2020); Decay sugar-rich substrates (Põlme et al. 2020) | Põlme S, et al. 2020. Fungal Diversity 105:1-16 ((https://doi.org/10.1007/s13225-020-00466-2)) |
| Otu16454 | 1 | k__Fungi(100);p__Ascomycota(100);c__Saccharomycetes(100);o__Saccharomycetales(100);f__Metschnikowiaceae(100);g__Metschnikowia(100);g__Metschnikowia_unclassified(100); | Metschnikowia | 13 | Saprotroph | \|Undefined Saprotroph\| | Yeast | NULL | Probable | As nectar-tap_saprotroph (Põlme et al. 2020); Decay sugar-rich substrates (Põlme et al. 2020) | Põlme S, et al. 2020. Fungal Diversity 105:1-16 ((https://doi.org/10.1007/s13225-020-00466-2)) |
| Otu16455 | 1 | k__Fungi(100);p__Ascomycota(100);c__Saccharomycetes(100);o__Saccharomycetales(100);f__Metschnikowiaceae(100);g__Metschnikowia(100);s__Metschnikowia_pulcherrima(100); | Metschnikowia | 13 | Saprotroph | \|Undefined Saprotroph\| | Yeast | NULL | Probable | As nectar-tap_saprotroph (Põlme et al. 2020); Decay sugar-rich substrates (Põlme et al. 2020) | Põlme S, et al. 2020. Fungal Diversity 105:1-16 ((https://doi.org/10.1007/s13225-020-00466-2)) |
| Otu16457 | 1 | k__Fungi(100);p__Ascomycota(100);c__Saccharomycetes(100);o__Saccharomycetales(100);f__Metschnikowiaceae(100);g__Metschnikowia(100);s__Metschnikowia_pulcherrima(100); | Metschnikowia | 13 | Saprotroph | \|Undefined Saprotroph\| | Yeast | NULL | Probable | As nectar-tap_saprotroph (Põlme et al. 2020); Decay sugar-rich substrates (Põlme et al. 2020) | Põlme S, et al. 2020. Fungal Diversity 105:1-16 ((https://doi.org/10.1007/s13225-020-00466-2)) |
| Otu16458 | 1 | k__Fungi(100);p__Ascomycota(100);c__Saccharomycetes(100);o__Saccharomycetales(100);f__Metschnikowiaceae(100);g__Metschnikowia(100);s__Metschnikowia_pulcherrima(100); | Metschnikowia | 13 | Saprotroph | \|Undefined Saprotroph\| | Yeast | NULL | Probable | As nectar-tap_saprotroph (Põlme et al. 2020); Decay sugar-rich substrates (Põlme et al. 2020) | Põlme S, et al. 2020. Fungal Diversity 105:1-16 ((https://doi.org/10.1007/s13225-020-00466-2)) |
| Otu16459 | 1 | k__Fungi(100);p__Ascomycota(100);c__Saccharomycetes(100);o__Saccharomycetales(100);f__Metschnikowiaceae(100);g__Metschnikowia(100);s__Metschnikowia_pulcherrima(100); | Metschnikowia | 13 | Saprotroph | \|Undefined Saprotroph\| | Yeast | NULL | Probable | As nectar-tap_saprotroph (Põlme et al. 2020); Decay sugar-rich substrates (Põlme et al. 2020) | Põlme S, et al. 2020. Fungal Diversity 105:1-16 ((https://doi.org/10.1007/s13225-020-00466-2)) |
| Otu16460 | 1 | k__Fungi(100);p__Ascomycota(100);c__Saccharomycetes(100);o__Saccharomycetales(100);f__Metschnikowiaceae(100);g__Metschnikowia(100);s__Metschnikowia_pulcherrima(100); | Metschnikowia | 13 | Saprotroph | \|Undefined Saprotroph\| | Yeast | NULL | Probable | As nectar-tap_saprotroph (Põlme et al. 2020); Decay sugar-rich substrates (Põlme et al. 2020) | Põlme S, et al. 2020. Fungal Diversity 105:1-16 ((https://doi.org/10.1007/s13225-020-00466-2)) |
| Otu16461 | 1 | k__Fungi(100);p__Ascomycota(100);c__Saccharomycetes(100);o__Saccharomycetales(100);f__Metschnikowiaceae(100);g__Metschnikowia(100);g__Metschnikowia_unclassified(100); | Metschnikowia | 13 | Saprotroph | \|Undefined Saprotroph\| | Yeast | NULL | Probable | As nectar-tap_saprotroph (Põlme et al. 2020); Decay sugar-rich substrates (Põlme et al. 2020) | Põlme S, et al. 2020. Fungal Diversity 105:1-16 ((https://doi.org/10.1007/s13225-020-00466-2)) |
| Otu16462 | 1 | k__Fungi(100);p__Ascomycota(100);c__Saccharomycetes(100);o__Saccharomycetales(100);f__Metschnikowiaceae(100);g__Metschnikowia(100);g__Metschnikowia_unclassified(100); | Metschnikowia | 13 | Saprotroph | \|Undefined Saprotroph\| | Yeast | NULL | Probable | As nectar-tap_saprotroph (Põlme et al. 2020); Decay sugar-rich substrates (Põlme et al. 2020) | Põlme S, et al. 2020. Fungal Diversity 105:1-16 ((https://doi.org/10.1007/s13225-020-00466-2)) |
| Otu16463 | 1 | k__Fungi(100);p__Ascomycota(100);c__Saccharomycetes(100);o__Saccharomycetales(100);f__Metschnikowiaceae(100);g__Metschnikowia(100);s__Metschnikowia_pulcherrima(100); | Metschnikowia | 13 | Saprotroph | \|Undefined Saprotroph\| | Yeast | NULL | Probable | As nectar-tap_saprotroph (Põlme et al. 2020); Decay sugar-rich substrates (Põlme et al. 2020) | Põlme S, et al. 2020. Fungal Diversity 105:1-16 ((https://doi.org/10.1007/s13225-020-00466-2)) |
| Otu16464 | 1 | k__Fungi(100);p__Ascomycota(100);c__Saccharomycetes(100);o__Saccharomycetales(100);f__Metschnikowiaceae(100);g__Metschnikowia(100);g__Metschnikowia_unclassified(100); | Metschnikowia | 13 | Saprotroph | \|Undefined Saprotroph\| | Yeast | NULL | Probable | As nectar-tap_saprotroph (Põlme et al. 2020); Decay sugar-rich substrates (Põlme et al. 2020) | Põlme S, et al. 2020. Fungal Diversity 105:1-16 ((https://doi.org/10.1007/s13225-020-00466-2)) |
| Otu16465 | 1 | k__Fungi(100);p__Ascomycota(100);c__Saccharomycetes(100);o__Saccharomycetales(100);f__Metschnikowiaceae(100);g__Metschnikowia(100);g__Metschnikowia_unclassified(100); | Metschnikowia | 13 | Saprotroph | \|Undefined Saprotroph\| | Yeast | NULL | Probable | As nectar-tap_saprotroph (Põlme et al. 2020); Decay sugar-rich substrates (Põlme et al. 2020) | Põlme S, et al. 2020. Fungal Diversity 105:1-16 ((https://doi.org/10.1007/s13225-020-00466-2)) |
| Otu16466 | 1 | k__Fungi(100);p__Ascomycota(100);c__Saccharomycetes(100);o__Saccharomycetales(100);f__Metschnikowiaceae(100);g__Metschnikowia(100);g__Metschnikowia_unclassified(100); | Metschnikowia | 13 | Saprotroph | \|Undefined Saprotroph\| | Yeast | NULL | Probable | As nectar-tap_saprotroph (Põlme et al. 2020); Decay sugar-rich substrates (Põlme et al. 2020) | Põlme S, et al. 2020. Fungal Diversity 105:1-16 ((https://doi.org/10.1007/s13225-020-00466-2)) |
| Otu16467 | 1 | k__Fungi(100);p__Ascomycota(100);c__Saccharomycetes(100);o__Saccharomycetales(100);f__Metschnikowiaceae(100);g__Metschnikowia(100);g__Metschnikowia_unclassified(100); | Metschnikowia | 13 | Saprotroph | \|Undefined Saprotroph\| | Yeast | NULL | Probable | As nectar-tap_saprotroph (Põlme et al. 2020); Decay sugar-rich substrates (Põlme et al. 2020) | Põlme S, et al. 2020. Fungal Diversity 105:1-16 ((https://doi.org/10.1007/s13225-020-00466-2)) |
| Otu16468 | 1 | k__Fungi(100);p__Ascomycota(100);c__Saccharomycetes(100);o__Saccharomycetales(100);f__Metschnikowiaceae(100);g__Metschnikowia(100);g__Metschnikowia_unclassified(100); | Metschnikowia | 13 | Saprotroph | \|Undefined Saprotroph\| | Yeast | NULL | Probable | As nectar-tap_saprotroph (Põlme et al. 2020); Decay sugar-rich substrates (Põlme et al. 2020) | Põlme S, et al. 2020. Fungal Diversity 105:1-16 ((https://doi.org/10.1007/s13225-020-00466-2)) |
| Otu16469 | 1 | k__Fungi(100);p__Ascomycota(100);c__Saccharomycetes(100);o__Saccharomycetales(100);f__Metschnikowiaceae(100);g__Metschnikowia(100);g__Metschnikowia_unclassified(100); | Metschnikowia | 13 | Saprotroph | \|Undefined Saprotroph\| | Yeast | NULL | Probable | As nectar-tap_saprotroph (Põlme et al. 2020); Decay sugar-rich substrates (Põlme et al. 2020) | Põlme S, et al. 2020. Fungal Diversity 105:1-16 ((https://doi.org/10.1007/s13225-020-00466-2)) |
| Otu16470 | 1 | k__Fungi(100);p__Ascomycota(100);c__Saccharomycetes(100);o__Saccharomycetales(100);f__Metschnikowiaceae(100);g__Metschnikowia(100);s__Metschnikowia_pulcherrima(100); | Metschnikowia | 13 | Saprotroph | \|Undefined Saprotroph\| | Yeast | NULL | Probable | As nectar-tap_saprotroph (Põlme et al. 2020); Decay sugar-rich substrates (Põlme et al. 2020) | Põlme S, et al. 2020. Fungal Diversity 105:1-16 ((https://doi.org/10.1007/s13225-020-00466-2)) |
| Otu16471 | 1 | k__Fungi(100);p__Ascomycota(100);c__Saccharomycetes(100);o__Saccharomycetales(100);f__Metschnikowiaceae(100);g__Metschnikowia(100);s__Metschnikowia_pulcherrima(100); | Metschnikowia | 13 | Saprotroph | \|Undefined Saprotroph\| | Yeast | NULL | Probable | As nectar-tap_saprotroph (Põlme et al. 2020); Decay sugar-rich substrates (Põlme et al. 2020) | Põlme S, et al. 2020. Fungal Diversity 105:1-16 ((https://doi.org/10.1007/s13225-020-00466-2)) |
| Otu16472 | 1 | k__Fungi(100);p__Ascomycota(100);c__Saccharomycetes(100);o__Saccharomycetales(100);f__Metschnikowiaceae(100);g__Metschnikowia(100);s__Metschnikowia_pulcherrima(100); | Metschnikowia | 13 | Saprotroph | \|Undefined Saprotroph\| | Yeast | NULL | Probable | As nectar-tap_saprotroph (Põlme et al. 2020); Decay sugar-rich substrates (Põlme et al. 2020) | Põlme S, et al. 2020. Fungal Diversity 105:1-16 ((https://doi.org/10.1007/s13225-020-00466-2)) |
| Otu16473 | 1 | k__Fungi(100);p__Ascomycota(100);c__Saccharomycetes(100);o__Saccharomycetales(100);f__Metschnikowiaceae(100);g__Metschnikowia(100);g__Metschnikowia_unclassified(100); | Metschnikowia | 13 | Saprotroph | \|Undefined Saprotroph\| | Yeast | NULL | Probable | As nectar-tap_saprotroph (Põlme et al. 2020); Decay sugar-rich substrates (Põlme et al. 2020) | Põlme S, et al. 2020. Fungal Diversity 105:1-16 ((https://doi.org/10.1007/s13225-020-00466-2)) |
| Otu16475 | 1 | k__Fungi(100);p__Ascomycota(100);c__Saccharomycetes(100);o__Saccharomycetales(100);f__Metschnikowiaceae(100);g__Metschnikowia(100);s__Metschnikowia_pulcherrima(100); | Metschnikowia | 13 | Saprotroph | \|Undefined Saprotroph\| | Yeast | NULL | Probable | As nectar-tap_saprotroph (Põlme et al. 2020); Decay sugar-rich substrates (Põlme et al. 2020) | Põlme S, et al. 2020. Fungal Diversity 105:1-16 ((https://doi.org/10.1007/s13225-020-00466-2)) |
| Otu16476 | 1 | k__Fungi(100);p__Ascomycota(100);c__Saccharomycetes(100);o__Saccharomycetales(100);f__Metschnikowiaceae(100);g__Metschnikowia(100);g__Metschnikowia_unclassified(100); | Metschnikowia | 13 | Saprotroph | \|Undefined Saprotroph\| | Yeast | NULL | Probable | As nectar-tap_saprotroph (Põlme et al. 2020); Decay sugar-rich substrates (Põlme et al. 2020) | Põlme S, et al. 2020. Fungal Diversity 105:1-16 ((https://doi.org/10.1007/s13225-020-00466-2)) |
| Otu16477 | 1 | k__Fungi(100);p__Ascomycota(100);c__Saccharomycetes(100);o__Saccharomycetales(100);f__Metschnikowiaceae(100);g__Metschnikowia(100);g__Metschnikowia_unclassified(100); | Metschnikowia | 13 | Saprotroph | \|Undefined Saprotroph\| | Yeast | NULL | Probable | As nectar-tap_saprotroph (Põlme et al. 2020); Decay sugar-rich substrates (Põlme et al. 2020) | Põlme S, et al. 2020. Fungal Diversity 105:1-16 ((https://doi.org/10.1007/s13225-020-00466-2)) |
| Otu16481 | 1 | k__Fungi(100);p__Ascomycota(100);c__Saccharomycetes(100);o__Saccharomycetales(100);f__Metschnikowiaceae(100);g__Metschnikowia(100);s__Metschnikowia_pulcherrima(100); | Metschnikowia | 13 | Saprotroph | \|Undefined Saprotroph\| | Yeast | NULL | Probable | As nectar-tap_saprotroph (Põlme et al. 2020); Decay sugar-rich substrates (Põlme et al. 2020) | Põlme S, et al. 2020. Fungal Diversity 105:1-16 ((https://doi.org/10.1007/s13225-020-00466-2)) |
| Otu16482 | 1 | k__Fungi(100);p__Ascomycota(100);c__Saccharomycetes(100);o__Saccharomycetales(100);f__Metschnikowiaceae(100);g__Metschnikowia(100);s__Metschnikowia_pulcherrima(100); | Metschnikowia | 13 | Saprotroph | \|Undefined Saprotroph\| | Yeast | NULL | Probable | As nectar-tap_saprotroph (Põlme et al. 2020); Decay sugar-rich substrates (Põlme et al. 2020) | Põlme S, et al. 2020. Fungal Diversity 105:1-16 ((https://doi.org/10.1007/s13225-020-00466-2)) |
| Otu16483 | 1 | k__Fungi(100);p__Ascomycota(100);c__Saccharomycetes(100);o__Saccharomycetales(100);f__Metschnikowiaceae(100);g__Metschnikowia(100);g__Metschnikowia_unclassified(100); | Metschnikowia | 13 | Saprotroph | \|Undefined Saprotroph\| | Yeast | NULL | Probable | As nectar-tap_saprotroph (Põlme et al. 2020); Decay sugar-rich substrates (Põlme et al. 2020) | Põlme S, et al. 2020. Fungal Diversity 105:1-16 ((https://doi.org/10.1007/s13225-020-00466-2)) |
| Otu16485 | 1 | k__Fungi(100);p__Ascomycota(100);c__Saccharomycetes(100);o__Saccharomycetales(100);f__Metschnikowiaceae(100);g__Metschnikowia(100);g__Metschnikowia_unclassified(100); | Metschnikowia | 13 | Saprotroph | \|Undefined Saprotroph\| | Yeast | NULL | Probable | As nectar-tap_saprotroph (Põlme et al. 2020); Decay sugar-rich substrates (Põlme et al. 2020) | Põlme S, et al. 2020. Fungal Diversity 105:1-16 ((https://doi.org/10.1007/s13225-020-00466-2)) |
| Otu16487 | 1 | k__Fungi(100);p__Ascomycota(100);c__Saccharomycetes(100);o__Saccharomycetales(100);f__Metschnikowiaceae(100);g__Metschnikowia(100);s__Metschnikowia_pulcherrima(100); | Metschnikowia | 13 | Saprotroph | \|Undefined Saprotroph\| | Yeast | NULL | Probable | As nectar-tap_saprotroph (Põlme et al. 2020); Decay sugar-rich substrates (Põlme et al. 2020) | Põlme S, et al. 2020. Fungal Diversity 105:1-16 ((https://doi.org/10.1007/s13225-020-00466-2)) |
| Otu16488 | 1 | k__Fungi(100);p__Ascomycota(100);c__Saccharomycetes(100);o__Saccharomycetales(100);f__Metschnikowiaceae(100);g__Metschnikowia(100);g__Metschnikowia_unclassified(100); | Metschnikowia | 13 | Saprotroph | \|Undefined Saprotroph\| | Yeast | NULL | Probable | As nectar-tap_saprotroph (Põlme et al. 2020); Decay sugar-rich substrates (Põlme et al. 2020) | Põlme S, et al. 2020. Fungal Diversity 105:1-16 ((https://doi.org/10.1007/s13225-020-00466-2)) |
| Otu16489 | 1 | k__Fungi(100);p__Ascomycota(100);c__Saccharomycetes(100);o__Saccharomycetales(100);f__Metschnikowiaceae(100);g__Metschnikowia(100);s__Metschnikowia_pulcherrima(100); | Metschnikowia | 13 | Saprotroph | \|Undefined Saprotroph\| | Yeast | NULL | Probable | As nectar-tap_saprotroph (Põlme et al. 2020); Decay sugar-rich substrates (Põlme et al. 2020) | Põlme S, et al. 2020. Fungal Diversity 105:1-16 ((https://doi.org/10.1007/s13225-020-00466-2)) |
| Otu16490 | 1 | k__Fungi(100);p__Ascomycota(100);c__Saccharomycetes(100);o__Saccharomycetales(100);f__Metschnikowiaceae(100);g__Metschnikowia(100);s__Metschnikowia_pulcherrima(100); | Metschnikowia | 13 | Saprotroph | \|Undefined Saprotroph\| | Yeast | NULL | Probable | As nectar-tap_saprotroph (Põlme et al. 2020); Decay sugar-rich substrates (Põlme et al. 2020) | Põlme S, et al. 2020. Fungal Diversity 105:1-16 ((https://doi.org/10.1007/s13225-020-00466-2)) |
| Otu16491 | 1 | k__Fungi(100);p__Ascomycota(100);c__Saccharomycetes(100);o__Saccharomycetales(100);f__Metschnikowiaceae(100);g__Metschnikowia(100);s__Metschnikowia_pulcherrima(100); | Metschnikowia | 13 | Saprotroph | \|Undefined Saprotroph\| | Yeast | NULL | Probable | As nectar-tap_saprotroph (Põlme et al. 2020); Decay sugar-rich substrates (Põlme et al. 2020) | Põlme S, et al. 2020. Fungal Diversity 105:1-16 ((https://doi.org/10.1007/s13225-020-00466-2)) |
| Otu16493 | 1 | k__Fungi(100);p__Ascomycota(100);c__Saccharomycetes(100);o__Saccharomycetales(100);f__Metschnikowiaceae(100);g__Metschnikowia(100);s__Metschnikowia_pulcherrima(100); | Metschnikowia | 13 | Saprotroph | \|Undefined Saprotroph\| | Yeast | NULL | Probable | As nectar-tap_saprotroph (Põlme et al. 2020); Decay sugar-rich substrates (Põlme et al. 2020) | Põlme S, et al. 2020. Fungal Diversity 105:1-16 ((https://doi.org/10.1007/s13225-020-00466-2)) |
| Otu16494 | 1 | k__Fungi(100);p__Ascomycota(100);c__Saccharomycetes(100);o__Saccharomycetales(100);f__Metschnikowiaceae(100);g__Metschnikowia(100);s__Metschnikowia_pulcherrima(100); | Metschnikowia | 13 | Saprotroph | \|Undefined Saprotroph\| | Yeast | NULL | Probable | As nectar-tap_saprotroph (Põlme et al. 2020); Decay sugar-rich substrates (Põlme et al. 2020) | Põlme S, et al. 2020. Fungal Diversity 105:1-16 ((https://doi.org/10.1007/s13225-020-00466-2)) |
| Otu16495 | 1 | k__Fungi(100);p__Ascomycota(100);c__Saccharomycetes(100);o__Saccharomycetales(100);f__Metschnikowiaceae(100);g__Metschnikowia(100);s__Metschnikowia_pulcherrima(100); | Metschnikowia | 13 | Saprotroph | \|Undefined Saprotroph\| | Yeast | NULL | Probable | As nectar-tap_saprotroph (Põlme et al. 2020); Decay sugar-rich substrates (Põlme et al. 2020) | Põlme S, et al. 2020. Fungal Diversity 105:1-16 ((https://doi.org/10.1007/s13225-020-00466-2)) |
| Otu16496 | 1 | k__Fungi(100);p__Ascomycota(100);c__Saccharomycetes(100);o__Saccharomycetales(100);f__Metschnikowiaceae(100);g__Metschnikowia(100);g__Metschnikowia_unclassified(100); | Metschnikowia | 13 | Saprotroph | \|Undefined Saprotroph\| | Yeast | NULL | Probable | As nectar-tap_saprotroph (Põlme et al. 2020); Decay sugar-rich substrates (Põlme et al. 2020) | Põlme S, et al. 2020. Fungal Diversity 105:1-16 ((https://doi.org/10.1007/s13225-020-00466-2)) |
| Otu16497 | 1 | k__Fungi(100);p__Ascomycota(100);c__Saccharomycetes(100);o__Saccharomycetales(100);f__Metschnikowiaceae(100);g__Metschnikowia(100);g__Metschnikowia_unclassified(100); | Metschnikowia | 13 | Saprotroph | \|Undefined Saprotroph\| | Yeast | NULL | Probable | As nectar-tap_saprotroph (Põlme et al. 2020); Decay sugar-rich substrates (Põlme et al. 2020) | Põlme S, et al. 2020. Fungal Diversity 105:1-16 ((https://doi.org/10.1007/s13225-020-00466-2)) |

| | | | | | | | | | | | |
|---|---|---|---|---|---|---|---|---|---|---|---|
| Otu16498 | 1 | k__Fungi(100);p__Ascomycota(100);c__Saccharomycetes(100);o__Saccharomycetales(100);f__Metschnikowiaceae(100);g__Metschnikowia(100);s__Metschnikowia_pulcherrima(100); | Metschnikowia | 13 | Saprotroph | \|Undefined Saprotroph\| | Yeast | NULL | Probable | As nectar-tap_saprotroph (Põlme et al. 2020); Decay sugar-rich substrates (Põlme et al. 2020) | Põlme S, et al. 2020. Fungal Diversity 105:1-16 ((https://doi.org/10.1007/s13225-020-00466-2)) |
| Otu16499 | 1 | k__Fungi(100);p__Ascomycota(100);c__Saccharomycetes(100);o__Saccharomycetales(100);f__Metschnikowiaceae(100);g__Metschnikowia(100);s__Metschnikowia_pulcherrima(100); | Metschnikowia | 13 | Saprotroph | \|Undefined Saprotroph\| | Yeast | NULL | Probable | As nectar-tap_saprotroph (Põlme et al. 2020); Decay sugar-rich substrates (Põlme et al. 2020) | Põlme S, et al. 2020. Fungal Diversity 105:1-16 ((https://doi.org/10.1007/s13225-020-00466-2)) |
| Otu16501 | 1 | k__Fungi(100);p__Ascomycota(100);c__Saccharomycetes(100);o__Saccharomycetales(100);f__Metschnikowiaceae(100);g__Metschnikowia(100);s__Metschnikowia_pulcherrima(100); | Metschnikowia | 13 | Saprotroph | \|Undefined Saprotroph\| | Yeast | NULL | Probable | As nectar-tap_saprotroph (Põlme et al. 2020); Decay sugar-rich substrates (Põlme et al. 2020) | Põlme S, et al. 2020. Fungal Diversity 105:1-16 ((https://doi.org/10.1007/s13225-020-00466-2)) |
| Otu16502 | 1 | k__Fungi(100);p__Ascomycota(100);c__Saccharomycetes(100);o__Saccharomycetales(100);f__Metschnikowiaceae(100);g__Metschnikowia(100);s__Metschnikowia_pulcherrima(100); | Metschnikowia | 13 | Saprotroph | \|Undefined Saprotroph\| | Yeast | NULL | Probable | As nectar-tap_saprotroph (Põlme et al. 2020); Decay sugar-rich substrates (Põlme et al. 2020) | Põlme S, et al. 2020. Fungal Diversity 105:1-16 ((https://doi.org/10.1007/s13225-020-00466-2)) |
| Otu16503 | 1 | k__Fungi(100);p__Ascomycota(100);c__Saccharomycetes(100);o__Saccharomycetales(100);f__Metschnikowiaceae(100);g__Metschnikowia(100);s__Metschnikowia_pulcherrima(100); | Metschnikowia | 13 | Saprotroph | \|Undefined Saprotroph\| | Yeast | NULL | Probable | As nectar-tap_saprotroph (Põlme et al. 2020); Decay sugar-rich substrates (Põlme et al. 2020) | Põlme S, et al. 2020. Fungal Diversity 105:1-16 ((https://doi.org/10.1007/s13225-020-00466-2)) |
| Otu16505 | 1 | k__Fungi(100);p__Ascomycota(100);c__Saccharomycetes(100);o__Saccharomycetales(100);f__Saccharomycetales_family_Incertae_sedis(100);g__Candida(100);s__Candida_cretensis(100); | Candida | 13 | Pathotroph-Saprotroph-Symbiotroph | Animal Pathogen-Endophyte-Endosymbiont-Epiphyte-\|Undefined Saprotroph\| | Dimorphic Yeast-Yeast | NULL | Probable | As nectar-tap_saprotroph (Põlme et al. 2020); Decay sugar-rich substrates (Põlme et al. 2020); This asexual genus is polyphyletic and contain species that function very differently depend on what they are. Caution should be taken when interpreting guilds for Candida species. | Manolakaki D, et al. 2010. Virulence. 1: 367-375 ((https://doi.org/10.4161/viru.1.5.12796)); Mekbib SB, et al. 2011. Tropical Plant Pathology 36:233-240 ((https://doi.org/10.1590/S1982-56762011000400004)); Põlme S, et al. 2020. Fungal Diversity 105:1-16 ((https://doi.org/10.1007/s13225-020-00466-2)) |
| Otu16506 | 1 | k__Fungi(100);p__Ascomycota(100);c__Saccharomycetes(100);o__Saccharomycetales(100);f__Metschnikowiaceae(100);g__Metschnikowia(100);s__Metschnikowia_pulcherrima(100); | Metschnikowia | 13 | Saprotroph | \|Undefined Saprotroph\| | Yeast | NULL | Probable | As nectar-tap_saprotroph (Põlme et al. 2020); Decay sugar-rich substrates (Põlme et al. 2020) | Põlme S, et al. 2020. Fungal Diversity 105:1-16 ((https://doi.org/10.1007/s13225-020-00466-2)) |
| Otu16507 | 1 | k__Fungi(100);p__Ascomycota(100);c__Saccharomycetes(100);o__Saccharomycetales(100);f__Metschnikowiaceae(100);g__Metschnikowia(100);s__Metschnikowia_pulcherrima(100); | Metschnikowia | 13 | Saprotroph | \|Undefined Saprotroph\| | Yeast | NULL | Probable | As nectar-tap_saprotroph (Põlme et al. 2020); Decay sugar-rich substrates (Põlme et al. 2020) | Põlme S, et al. 2020. Fungal Diversity 105:1-16 ((https://doi.org/10.1007/s13225-020-00466-2)) |
| Otu16510 | 1 | k__Fungi(100);p__Ascomycota(100);c__Saccharomycetes(100);o__Saccharomycetales(100);f__Metschnikowiaceae(100);g__Metschnikowia(100);s__Metschnikowia_pulcherrima(100); | Metschnikowia | 13 | Saprotroph | \|Undefined Saprotroph\| | Yeast | NULL | Probable | As nectar-tap_saprotroph (Põlme et al. 2020); Decay sugar-rich substrates (Põlme et al. 2020) | Põlme S, et al. 2020. Fungal Diversity 105:1-16 ((https://doi.org/10.1007/s13225-020-00466-2)) |
| Otu16512 | 1 | k__Fungi(100);p__Ascomycota(100);c__Saccharomycetes(100);o__Saccharomycetales(100);f__Metschnikowiaceae(100);g__Metschnikowia(100);s__Metschnikowia_pulcherrima(100); | Metschnikowia | 13 | Saprotroph | \|Undefined Saprotroph\| | Yeast | NULL | Probable | As nectar-tap_saprotroph (Põlme et al. 2020); Decay sugar-rich substrates (Põlme et al. 2020) | Põlme S, et al. 2020. Fungal Diversity 105:1-16 ((https://doi.org/10.1007/s13225-020-00466-2)) |
| Otu16513 | 1 | k__Fungi(100);p__Ascomycota(100);c__Saccharomycetes(100);o__Saccharomycetales(100);f__Metschnikowiaceae(100);g__Metschnikowia(100);s__Metschnikowia_pulcherrima(100); | Metschnikowia | 13 | Saprotroph | \|Undefined Saprotroph\| | Yeast | NULL | Probable | As nectar-tap_saprotroph (Põlme et al. 2020); Decay sugar-rich substrates (Põlme et al. 2020) | Põlme S, et al. 2020. Fungal Diversity 105:1-16 ((https://doi.org/10.1007/s13225-020-00466-2)) |
| Otu16519 | 1 | k__Fungi(100);p__Ascomycota(100);c__Saccharomycetes(100);o__Saccharomycetales(100);f__Metschnikowiaceae(100);g__Metschnikowia(100);s__Metschnikowia_pulcherrima(100); | Metschnikowia | 13 | Saprotroph | \|Undefined Saprotroph\| | Yeast | NULL | Probable | As nectar-tap_saprotroph (Põlme et al. 2020); Decay sugar-rich substrates (Põlme et al. 2020) | Põlme S, et al. 2020. Fungal Diversity 105:1-16 ((https://doi.org/10.1007/s13225-020-00466-2)) |
| Otu16520 | 1 | k__Fungi(100);p__Ascomycota(100);c__Saccharomycetes(100);o__Saccharomycetales(100);f__Metschnikowiaceae(100);g__Metschnikowia(100);s__Metschnikowia_pulcherrima(100); | Metschnikowia | 13 | Saprotroph | \|Undefined Saprotroph\| | Yeast | NULL | Probable | As nectar-tap_saprotroph (Põlme et al. 2020); Decay sugar-rich substrates (Põlme et al. 2020) | Põlme S, et al. 2020. Fungal Diversity 105:1-16 ((https://doi.org/10.1007/s13225-020-00466-2)) |
| Otu16527 | 1 | k__Fungi(100);p__Ascomycota(100);c__Saccharomycetes(100);o__Saccharomycetales(100);f__Metschnikowiaceae(100);g__Metschnikowia(100);s__Metschnikowia_pulcherrima(100); | Metschnikowia | 13 | Saprotroph | \|Undefined Saprotroph\| | Yeast | NULL | Probable | As nectar-tap_saprotroph (Põlme et al. 2020); Decay sugar-rich substrates (Põlme et al. 2020) | Põlme S, et al. 2020. Fungal Diversity 105:1-16 ((https://doi.org/10.1007/s13225-020-00466-2)) |
| Otu16530 | 1 | k__Fungi(100);p__Ascomycota(100);c__Saccharomycetes(100);o__Saccharomycetales(100);f__Metschnikowiaceae(100);g__Metschnikowia(100);s__Metschnikowia_pulcherrima(100); | Metschnikowia | 13 | Saprotroph | \|Undefined Saprotroph\| | Yeast | NULL | Probable | As nectar-tap_saprotroph (Põlme et al. 2020); Decay sugar-rich substrates (Põlme et al. 2020) | Põlme S, et al. 2020. Fungal Diversity 105:1-16 ((https://doi.org/10.1007/s13225-020-00466-2)) |
| Otu16532 | 1 | k__Fungi(100);p__Ascomycota(100);c__Saccharomycetes(100);o__Saccharomycetales(100);f__Metschnikowiaceae(100);g__Metschnikowia(100);s__Metschnikowia_pulcherrima(100); | Metschnikowia | 13 | Saprotroph | \|Undefined Saprotroph\| | Yeast | NULL | Probable | As nectar-tap_saprotroph (Põlme et al. 2020); Decay sugar-rich substrates (Põlme et al. 2020) | Põlme S, et al. 2020. Fungal Diversity 105:1-16 ((https://doi.org/10.1007/s13225-020-00466-2)) |
| Otu16533 | 1 | k__Fungi(100);p__Ascomycota(100);c__Saccharomycetes(100);o__Saccharomycetales(100);f__Metschnikowiaceae(100);g__Metschnikowia(100);s__Metschnikowia_pulcherrima(100); | Metschnikowia | 13 | Saprotroph | \|Undefined Saprotroph\| | Yeast | NULL | Probable | As nectar-tap_saprotroph (Põlme et al. 2020); Decay sugar-rich substrates (Põlme et al. 2020) | Põlme S, et al. 2020. Fungal Diversity 105:1-16 ((https://doi.org/10.1007/s13225-020-00466-2)) |
| Otu16534 | 1 | k__Fungi(100);p__Ascomycota(100);c__Saccharomycetes(100);o__Saccharomycetales(100);f__Metschnikowiaceae(100);g__Metschnikowia(100);g__Metschnikowia_unclassified(100); | Metschnikowia | 13 | Saprotroph | \|Undefined Saprotroph\| | Yeast | NULL | Probable | As nectar-tap_saprotroph (Põlme et al. 2020); Decay sugar-rich substrates (Põlme et al. 2020) | Põlme S, et al. 2020. Fungal Diversity 105:1-16 ((https://doi.org/10.1007/s13225-020-00466-2)) |
| Otu16535 | 1 | k__Fungi(100);p__Ascomycota(100);c__Saccharomycetes(100);o__Saccharomycetales(100);f__Metschnikowiaceae(100);g__Metschnikowia(100);s__Metschnikowia_pulcherrima(100); | Metschnikowia | 13 | Saprotroph | \|Undefined Saprotroph\| | Yeast | NULL | Probable | As nectar-tap_saprotroph (Põlme et al. 2020); Decay sugar-rich substrates (Põlme et al. 2020) | Põlme S, et al. 2020. Fungal Diversity 105:1-16 ((https://doi.org/10.1007/s13225-020-00466-2)) |
| Otu16536 | 1 | k__Fungi(100);p__Ascomycota(100);c__Saccharomycetes(100);o__Saccharomycetales(100);f__Metschnikowiaceae(100);g__Metschnikowia(100);s__Metschnikowia_pulcherrima(100); | Metschnikowia | 13 | Saprotroph | \|Undefined Saprotroph\| | Yeast | NULL | Probable | As nectar-tap_saprotroph (Põlme et al. 2020); Decay sugar-rich substrates (Põlme et al. 2020) | Põlme S, et al. 2020. Fungal Diversity 105:1-16 ((https://doi.org/10.1007/s13225-020-00466-2)) |
| Otu16539 | 1 | k__Fungi(100);p__Ascomycota(100);c__Saccharomycetes(100);o__Saccharomycetales(100);f__Metschnikowiaceae(100);g__Metschnikowia(100);g__Metschnikowia_unclassified(100); | Metschnikowia | 13 | Saprotroph | \|Undefined Saprotroph\| | Yeast | NULL | Probable | As nectar-tap_saprotroph (Põlme et al. 2020); Decay sugar-rich substrates (Põlme et al. 2020) | Põlme S, et al. 2020. Fungal Diversity 105:1-16 ((https://doi.org/10.1007/s13225-020-00466-2)) |
| Otu16542 | 1 | k__Fungi(100);p__Ascomycota(100);c__Saccharomycetes(100);o__Saccharomycetales(100);f__Metschnikowiaceae(100);g__Metschnikowia(100);s__Metschnikowia_pulcherrima(100); | Metschnikowia | 13 | Saprotroph | \|Undefined Saprotroph\| | Yeast | NULL | Probable | As nectar-tap_saprotroph (Põlme et al. 2020); Decay sugar-rich substrates (Põlme et al. 2020) | Põlme S, et al. 2020. Fungal Diversity 105:1-16 ((https://doi.org/10.1007/s13225-020-00466-2)) |
| Otu16545 | 1 | k__Fungi(100);p__Ascomycota(100);c__Saccharomycetes(100);o__Saccharomycetales(100);f__Metschnikowiaceae(100);g__Metschnikowia(100);g__Metschnikowia_unclassified(100); | Metschnikowia | 13 | Saprotroph | \|Undefined Saprotroph\| | Yeast | NULL | Probable | As nectar-tap_saprotroph (Põlme et al. 2020); Decay sugar-rich substrates (Põlme et al. 2020) | Põlme S, et al. 2020. Fungal Diversity 105:1-16 ((https://doi.org/10.1007/s13225-020-00466-2)) |
| Otu16548 | 1 | k__Fungi(100);p__Ascomycota(100);c__Saccharomycetes(100);o__Saccharomycetales(100);f__Metschnikowiaceae(100);g__Metschnikowia(100);g__Metschnikowia_unclassified(100); | Metschnikowia | 13 | Saprotroph | \|Undefined Saprotroph\| | Yeast | NULL | Probable | As nectar-tap_saprotroph (Põlme et al. 2020); Decay sugar-rich substrates (Põlme et al. 2020) | Põlme S, et al. 2020. Fungal Diversity 105:1-16 ((https://doi.org/10.1007/s13225-020-00466-2)) |
| Otu16549 | 1 | k__Fungi(100);p__Ascomycota(100);c__Saccharomycetes(100);o__Saccharomycetales(100);f__Metschnikowiaceae(100);g__Metschnikowia(100);s__Metschnikowia_pulcherrima(100); | Metschnikowia | 13 | Saprotroph | \|Undefined Saprotroph\| | Yeast | NULL | Probable | As nectar-tap_saprotroph (Põlme et al. 2020); Decay sugar-rich substrates (Põlme et al. 2020) | Põlme S, et al. 2020. Fungal Diversity 105:1-16 ((https://doi.org/10.1007/s13225-020-00466-2)) |
| Otu16550 | 1 | k__Fungi(100);p__Ascomycota(100);c__Saccharomycetes(100);o__Saccharomycetales(100);f__Metschnikowiaceae(100);g__Metschnikowia(100);s__Metschnikowia_pulcherrima(100); | Metschnikowia | 13 | Saprotroph | \|Undefined Saprotroph\| | Yeast | NULL | Probable | As nectar-tap_saprotroph (Põlme et al. 2020); Decay sugar-rich substrates (Põlme et al. 2020) | Põlme S, et al. 2020. Fungal Diversity 105:1-16 ((https://doi.org/10.1007/s13225-020-00466-2)) |
| Otu16551 | 1 | k__Fungi(100);p__Ascomycota(100);c__Saccharomycetes(100);o__Saccharomycetales(100);f__Metschnikowiaceae(100);g__Metschnikowia(100);s__Metschnikowia_pulcherrima(100); | Metschnikowia | 13 | Saprotroph | \|Undefined Saprotroph\| | Yeast | NULL | Probable | As nectar-tap_saprotroph (Põlme et al. 2020); Decay sugar-rich substrates (Põlme et al. 2020) | Põlme S, et al. 2020. Fungal Diversity 105:1-16 ((https://doi.org/10.1007/s13225-020-00466-2)) |
| Otu16552 | 1 | k__Fungi(100);p__Ascomycota(100);c__Saccharomycetes(100);o__Saccharomycetales(100);f__Metschnikowiaceae(100);g__Metschnikowia(100);s__Metschnikowia_pulcherrima(100); | Metschnikowia | 13 | Saprotroph | \|Undefined Saprotroph\| | Yeast | NULL | Probable | As nectar-tap_saprotroph (Põlme et al. 2020); Decay sugar-rich substrates (Põlme et al. 2020) | Põlme S, et al. 2020. Fungal Diversity 105:1-16 ((https://doi.org/10.1007/s13225-020-00466-2)) |
| Otu16553 | 1 | k__Fungi(100);p__Ascomycota(100);c__Saccharomycetes(100);o__Saccharomycetales(100);f__Metschnikowiaceae(100);g__Metschnikowia(100);s__Metschnikowia_pulcherrima(100); | Metschnikowia | 13 | Saprotroph | \|Undefined Saprotroph\| | Yeast | NULL | Probable | As nectar-tap_saprotroph (Põlme et al. 2020); Decay sugar-rich substrates (Põlme et al. 2020) | Põlme S, et al. 2020. Fungal Diversity 105:1-16 ((https://doi.org/10.1007/s13225-020-00466-2)) |
| Otu16554 | 1 | k__Fungi(100);p__Ascomycota(100);c__Saccharomycetes(100);o__Saccharomycetales(100);f__Metschnikowiaceae(100);g__Metschnikowia(100);s__Metschnikowia_pulcherrima(100); | Metschnikowia | 13 | Saprotroph | \|Undefined Saprotroph\| | Yeast | NULL | Probable | As nectar-tap_saprotroph (Põlme et al. 2020); Decay sugar-rich substrates (Põlme et al. 2020) | Põlme S, et al. 2020. Fungal Diversity 105:1-16 ((https://doi.org/10.1007/s13225-020-00466-2)) |
| Otu16555 | 1 | k__Fungi(100);p__Ascomycota(100);c__Saccharomycetes(100);o__Saccharomycetales(100);f__Metschnikowiaceae(100);g__Metschnikowia(100);g__Metschnikowia_unclassified(100); | Metschnikowia | 13 | Saprotroph | \|Undefined Saprotroph\| | Yeast | NULL | Probable | As nectar-tap_saprotroph (Põlme et al. 2020); Decay sugar-rich substrates (Põlme et al. 2020) | Põlme S, et al. 2020. Fungal Diversity 105:1-16 ((https://doi.org/10.1007/s13225-020-00466-2)) |
| Otu16556 | 1 | k__Fungi(100);p__Ascomycota(100);c__Saccharomycetes(100);o__Saccharomycetales(100);f__Metschnikowiaceae(100);g__Metschnikowia(100);s__Metschnikowia_pulcherrima(100); | Metschnikowia | 13 | Saprotroph | \|Undefined Saprotroph\| | Yeast | NULL | Probable | As nectar-tap_saprotroph (Põlme et al. 2020); Decay sugar-rich substrates (Põlme et al. 2020) | Põlme S, et al. 2020. Fungal Diversity 105:1-16 ((https://doi.org/10.1007/s13225-020-00466-2)) |
| Otu16557 | 1 | k__Fungi(100);p__Ascomycota(100);c__Saccharomycetes(100);o__Saccharomycetales(100);f__Metschnikowiaceae(100);g__Metschnikowia(100);s__Metschnikowia_pulcherrima(100); | Metschnikowia | 13 | Saprotroph | \|Undefined Saprotroph\| | Yeast | NULL | Probable | As nectar-tap_saprotroph (Põlme et al. 2020); Decay sugar-rich substrates (Põlme et al. 2020) | Põlme S, et al. 2020. Fungal Diversity 105:1-16 ((https://doi.org/10.1007/s13225-020-00466-2)) |
| Otu16559 | 1 | k__Fungi(100);p__Ascomycota(100);c__Saccharomycetes(100);o__Saccharomycetales(100);f__Metschnikowiaceae(100);g__Metschnikowia(100);g__Metschnikowia_unclassified(100); | Metschnikowia | 13 | Saprotroph | \|Undefined Saprotroph\| | Yeast | NULL | Probable | As nectar-tap_saprotroph (Põlme et al. 2020); Decay sugar-rich substrates (Põlme et al. 2020) | Põlme S, et al. 2020. Fungal Diversity 105:1-16 ((https://doi.org/10.1007/s13225-020-00466-2)) |
| Otu16560 | 1 | k__Fungi(100);p__Ascomycota(100);c__Saccharomycetes(100);o__Saccharomycetales(100);f__Metschnikowiaceae(100);g__Metschnikowia(100);g__Metschnikowia_unclassified(100); | Metschnikowia | 13 | Saprotroph | \|Undefined Saprotroph\| | Yeast | NULL | Probable | As nectar-tap_saprotroph (Põlme et al. 2020); Decay sugar-rich substrates (Põlme et al. 2020) | Põlme S, et al. 2020. Fungal Diversity 105:1-16 ((https://doi.org/10.1007/s13225-020-00466-2)) |
| Otu16562 | 1 | k__Fungi(100);p__Ascomycota(100);c__Saccharomycetes(100);o__Saccharomycetales(100);f__Metschnikowiaceae(100);g__Metschnikowia(100);s__Metschnikowia_pulcherrima(100); | Metschnikowia | 13 | Saprotroph | \|Undefined Saprotroph\| | Yeast | NULL | Probable | As nectar-tap_saprotroph (Põlme et al. 2020); Decay sugar-rich substrates (Põlme et al. 2020) | Põlme S, et al. 2020. Fungal Diversity 105:1-16 ((https://doi.org/10.1007/s13225-020-00466-2)) |
| Otu16563 | 1 | k__Fungi(100);p__Ascomycota(100);c__Saccharomycetes(100);o__Saccharomycetales(100);f__Metschnikowiaceae(100);g__Metschnikowia(100);g__Metschnikowia_unclassified(100); | Metschnikowia | 13 | Saprotroph | \|Undefined Saprotroph\| | Yeast | NULL | Probable | As nectar-tap_saprotroph (Põlme et al. 2020); Decay sugar-rich substrates (Põlme et al. 2020) | Põlme S, et al. 2020. Fungal Diversity 105:1-16 ((https://doi.org/10.1007/s13225-020-00466-2)) |
| Otu16564 | 1 | k__Fungi(100);p__Ascomycota(100);c__Saccharomycetes(100);o__Saccharomycetales(100);f__Metschnikowiaceae(100);g__Metschnikowia(100);g__Metschnikowia_unclassified(100); | Metschnikowia | 13 | Saprotroph | \|Undefined Saprotroph\| | Yeast | NULL | Probable | As nectar-tap_saprotroph (Põlme et al. 2020); Decay sugar-rich substrates (Põlme et al. 2020) | Põlme S, et al. 2020. Fungal Diversity 105:1-16 ((https://doi.org/10.1007/s13225-020-00466-2)) |
| Otu16565 | 1 | k__Fungi(100);p__Ascomycota(100);c__Saccharomycetes(100);o__Saccharomycetales(100);f__Metschnikowiaceae(100);g__Metschnikowia(100);s__Metschnikowia_pulcherrima(100); | Metschnikowia | 13 | Saprotroph | \|Undefined Saprotroph\| | Yeast | NULL | Probable | As nectar-tap_saprotroph (Põlme et al. 2020); Decay sugar-rich substrates (Põlme et al. 2020) | Põlme S, et al. 2020. Fungal Diversity 105:1-16 ((https://doi.org/10.1007/s13225-020-00466-2)) |
| Otu16566 | 1 | k__Fungi(100);p__Ascomycota(100);c__Saccharomycetes(100);o__Saccharomycetales(100);f__Metschnikowiaceae(100);g__Metschnikowia(100);s__Metschnikowia_pulcherrima(100); | Metschnikowia | 13 | Saprotroph | \|Undefined Saprotroph\| | Yeast | NULL | Probable | As nectar-tap_saprotroph (Põlme et al. 2020); Decay sugar-rich substrates (Põlme et al. 2020) | Põlme S, et al. 2020. Fungal Diversity 105:1-16 ((https://doi.org/10.1007/s13225-020-00466-2)) |
| Otu16570 | 1 | k__Fungi(100);p__Ascomycota(100);c__Saccharomycetes(100);o__Saccharomycetales(100);f__Metschnikowiaceae(100);g__Metschnikowia(100);s__Metschnikowia_pulcherrima(100); | Metschnikowia | 13 | Saprotroph | \|Undefined Saprotroph\| | Yeast | NULL | Probable | As nectar-tap_saprotroph (Põlme et al. 2020); Decay sugar-rich substrates (Põlme et al. 2020) | Põlme S, et al. 2020. Fungal Diversity 105:1-16 ((https://doi.org/10.1007/s13225-020-00466-2)) |
| Otu16571 | 1 | k__Fungi(100);p__Ascomycota(100);c__Saccharomycetes(100);o__Saccharomycetales(100);f__Metschnikowiaceae(100);g__Metschnikowia(100);s__Metschnikowia_pulcherrima(100); | Metschnikowia | 13 | Saprotroph | \|Undefined Saprotroph\| | Yeast | NULL | Probable | As nectar-tap_saprotroph (Põlme et al. 2020); Decay sugar-rich substrates (Põlme et al. 2020) | Põlme S, et al. 2020. Fungal Diversity 105:1-16 ((https://doi.org/10.1007/s13225-020-00466-2)) |
| Otu16572 | 1 | k__Fungi(100);p__Ascomycota(100);c__Saccharomycetes(100);o__Saccharomycetales(100);f__Metschnikowiaceae(100);g__Metschnikowia(100);s__Metschnikowia_pulcherrima(100); | Metschnikowia | 13 | Saprotroph | \|Undefined Saprotroph\| | Yeast | NULL | Probable | As nectar-tap_saprotroph (Põlme et al. 2020); Decay sugar-rich substrates (Põlme et al. 2020) | Põlme S, et al. 2020. Fungal Diversity 105:1-16 ((https://doi.org/10.1007/s13225-020-00466-2)) |
| Otu16573 | 1 | k__Fungi(100);p__Ascomycota(100);c__Saccharomycetes(100);o__Saccharomycetales(100);f__Metschnikowiaceae(100);g__Metschnikowia(100);s__Metschnikowia_pulcherrima(100); | Metschnikowia | 13 | Saprotroph | \|Undefined Saprotroph\| | Yeast | NULL | Probable | As nectar-tap_saprotroph (Põlme et al. 2020); Decay sugar-rich substrates (Põlme et al. 2020) | Põlme S, et al. 2020. Fungal Diversity 105:1-16 ((https://doi.org/10.1007/s13225-020-00466-2)) |
| Otu16574 | 1 | k__Fungi(100);p__Ascomycota(100);c__Saccharomycetes(100);o__Saccharomycetales(100);f__Metschnikowiaceae(100);g__Metschnikowia(100);s__Metschnikowia_pulcherrima(100); | Metschnikowia | 13 | Saprotroph | \|Undefined Saprotroph\| | Yeast | NULL | Probable | As nectar-tap_saprotroph (Põlme et al. 2020); Decay sugar-rich substrates (Põlme et al. 2020) | Põlme S, et al. 2020. Fungal Diversity 105:1-16 ((https://doi.org/10.1007/s13225-020-00466-2)) |
| Otu16575 | 1 | k__Fungi(100);p__Ascomycota(100);c__Saccharomycetes(100);o__Saccharomycetales(100);f__Metschnikowiaceae(100);g__Metschnikowia(100);s__Metschnikowia_pulcherrima(100); | Metschnikowia | 13 | Saprotroph | \|Undefined Saprotroph\| | Yeast | NULL | Probable | As nectar-tap_saprotroph (Põlme et al. 2020); Decay sugar-rich substrates (Põlme et al. 2020) | Põlme S, et al. 2020. Fungal Diversity 105:1-16 ((https://doi.org/10.1007/s13225-020-00466-2)) |
| Otu16576 | 1 | k__Fungi(100);p__Ascomycota(100);c__Saccharomycetes(100);o__Saccharomycetales(100);f__Metschnikowiaceae(100);g__Metschnikowia(100);s__Metschnikowia_pulcherrima(100); | Metschnikowia | 13 | Saprotroph | \|Undefined Saprotroph\| | Yeast | NULL | Probable | As nectar-tap_saprotroph (Põlme et al. 2020); Decay sugar-rich substrates (Põlme et al. 2020) | Põlme S, et al. 2020. Fungal Diversity 105:1-16 ((https://doi.org/10.1007/s13225-020-00466-2)) |
| Otu16577 | 1 | k__Fungi(100);p__Ascomycota(100);c__Saccharomycetes(100);o__Saccharomycetales(100);f__Metschnikowiaceae(100);g__Metschnikowia(100);s__Metschnikowia_pulcherrima(100); | Metschnikowia | 13 | Saprotroph | \|Undefined Saprotroph\| | Yeast | NULL | Probable | As nectar-tap_saprotroph (Põlme et al. 2020); Decay sugar-rich substrates (Põlme et al. 2020) | Põlme S, et al. 2020. Fungal Diversity 105:1-16 ((https://doi.org/10.1007/s13225-020-00466-2)) |
| Otu16578 | 1 | k__Fungi(100);p__Ascomycota(100);c__Saccharomycetes(100);o__Saccharomycetales(100);f__Metschnikowiaceae(100);g__Metschnikowia(100);s__Metschnikowia_pulcherrima(100); | Metschnikowia | 13 | Saprotroph | \|Undefined Saprotroph\| | Yeast | NULL | Probable | As nectar-tap_saprotroph (Põlme et al. 2020); Decay sugar-rich substrates (Põlme et al. 2020) | Põlme S, et al. 2020. Fungal Diversity 105:1-16 ((https://doi.org/10.1007/s13225-020-00466-2)) |
| Otu16579 | 1 | k__Fungi(100);p__Ascomycota(100);c__Saccharomycetes(100);o__Saccharomycetales(100);f__Metschnikowiaceae(100);g__Metschnikowia(100);s__Metschnikowia_pulcherrima(100); | Metschnikowia | 13 | Saprotroph | \|Undefined Saprotroph\| | Yeast | NULL | Probable | As nectar-tap_saprotroph (Põlme et al. 2020); Decay sugar-rich substrates (Põlme et al. 2020) | Põlme S, et al. 2020. Fungal Diversity 105:1-16 ((https://doi.org/10.1007/s13225-020-00466-2)) |
| Otu16580 | 1 | k__Fungi(100);p__Ascomycota(100);c__Saccharomycetes(100);o__Saccharomycetales(100);f__Metschnikowiaceae(100);g__Metschnikowia(100);g__Metschnikowia_unclassified(100); | Metschnikowia | 13 | Saprotroph | \|Undefined Saprotroph\| | Yeast | NULL | Probable | As nectar-tap_saprotroph (Põlme et al. 2020); Decay sugar-rich substrates (Põlme et al. 2020) | Põlme S, et al. 2020. Fungal Diversity 105:1-16 ((https://doi.org/10.1007/s13225-020-00466-2)) |
| Otu16581 | 1 | k__Fungi(100);p__Ascomycota(100);c__Saccharomycetes(100);o__Saccharomycetales(100);f__Metschnikowiaceae(100);g__Metschnikowia(100);s__Metschnikowia_pulcherrima(100); | Metschnikowia | 13 | Saprotroph | \|Undefined Saprotroph\| | Yeast | NULL | Probable | As nectar-tap_saprotroph (Põlme et al. 2020); Decay sugar-rich substrates (Põlme et al. 2020) | Põlme S, et al. 2020. Fungal Diversity 105:1-16 ((https://doi.org/10.1007/s13225-020-00466-2)) |
| Otu16582 | 1 | k__Fungi(100);p__Ascomycota(100);c__Saccharomycetes(100);o__Saccharomycetales(100);f__Saccharomycetales_family_Incertae_sedis(100);g__Candida(100);g__Candida_unclassified(100); | Candida | 13 | Pathotroph-Saprotroph-Symbiotroph | Animal Pathogen-Endophyte-Endosymbiont-Epiphyte-\|Undefined Saprotroph\| | Dimorphic Yeast-Yeast | NULL | Probable | As nectar-tap_saprotroph (Põlme et al. 2020); Decay sugar-rich substrates (Põlme et al. 2020); This asexual genus is polyphyletic and contain species that function very differently depend on what they are. Caution should be taken when interpreting guilds for Candida species. | Manolakaki D, et al. 2010. Virulence. 1: 367-375 ((https://doi.org/10.4161/viru.1.5.12796)); Mekbib SB, et al. 2011. Tropical Plant Pathology 36:233-240 ((https://doi.org/10.1590/S1982-56762011000400004)); Põlme S, et al. 2020. Fungal Diversity 105:1-16 ((https://doi.org/10.1007/s13225-020-00466-2)) |
| Otu16585 | 1 | k__Fungi(100);p__Ascomycota(100);c__Saccharomycetes(100);o__Saccharomycetales(100);f__Metschnikowiaceae(100);g__Metschnikowia(100);s__Metschnikowia_pulcherrima(100); | Metschnikowia | 13 | Saprotroph | \|Undefined Saprotroph\| | Yeast | NULL | Probable | As nectar-tap_saprotroph (Põlme et al. 2020); Decay sugar-rich substrates (Põlme et al. 2020) | Põlme S, et al. 2020. Fungal Diversity 105:1-16 ((https://doi.org/10.1007/s13225-020-00466-2)) |
| Otu16586 | 1 | k__Fungi(100);p__Ascomycota(100);c__Saccharomycetes(100);o__Saccharomycetales(100);f__Metschnikowiaceae(100);g__Metschnikowia(100);g__Metschnikowia_unclassified(100); | Metschnikowia | 13 | Saprotroph | \|Undefined Saprotroph\| | Yeast | NULL | Probable | As nectar-tap_saprotroph (Põlme et al. 2020); Decay sugar-rich substrates (Põlme et al. 2020) | Põlme S, et al. 2020. Fungal Diversity 105:1-16 ((https://doi.org/10.1007/s13225-020-00466-2)) |
| Otu16587 | 1 | k__Fungi(100);p__Ascomycota(100);c__Saccharomycetes(100);o__Saccharomycetales(100);f__Metschnikowiaceae(100);g__Metschnikowia(100);s__Metschnikowia_pulcherrima(100); | Metschnikowia | 13 | Saprotroph | \|Undefined Saprotroph\| | Yeast | NULL | Probable | As nectar-tap_saprotroph (Põlme et al. 2020); Decay sugar-rich substrates (Põlme et al. 2020) | Põlme S, et al. 2020. Fungal Diversity 105:1-16 ((https://doi.org/10.1007/s13225-020-00466-2)) |
| Otu16588 | 1 | k__Fungi(100);p__Ascomycota(100);c__Saccharomycetes(100);o__Saccharomycetales(100);f__Metschnikowiaceae(100);g__Metschnikowia(100);s__Metschnikowia_pulcherrima(100); | Metschnikowia | 13 | Saprotroph | \|Undefined Saprotroph\| | Yeast | NULL | Probable | As nectar-tap_saprotroph (Põlme et al. 2020); Decay sugar-rich substrates (Põlme et al. 2020) | Põlme S, et al. 2020. Fungal Diversity 105:1-16 ((https://doi.org/10.1007/s13225-020-00466-2)) |
| Otu16589 | 1 | k__Fungi(100);p__Ascomycota(100);c__Saccharomycetes(100);o__Saccharomycetales(100);f__Metschnikowiaceae(100);g__Metschnikowia(100);s__Metschnikowia_pulcherrima(100); | Metschnikowia | 13 | Saprotroph | \|Undefined Saprotroph\| | Yeast | NULL | Probable | As nectar-tap_saprotroph (Põlme et al. 2020); Decay sugar-rich substrates (Põlme et al. 2020) | Põlme S, et al. 2020. Fungal Diversity 105:1-16 ((https://doi.org/10.1007/s13225-020-00466-2)) |
| Otu16590 | 1 | k__Fungi(100);p__Ascomycota(100);c__Saccharomycetes(100);o__Saccharomycetales(100);f__Metschnikowiaceae(100);g__Metschnikowia(100);s__Metschnikowia_pulcherrima(100); | Metschnikowia | 13 | Saprotroph | \|Undefined Saprotroph\| | Yeast | NULL | Probable | As nectar-tap_saprotroph (Põlme et al. 2020); Decay sugar-rich substrates (Põlme et al. 2020) | Põlme S, et al. 2020. Fungal Diversity 105:1-16 ((https://doi.org/10.1007/s13225-020-00466-2)) |
| Otu16591 | 1 | k__Fungi(100);p__Ascomycota(100);c__Saccharomycetes(100);o__Saccharomycetales(100);f__Saccharomycetales_family_Incertae_sedis(100);g__Candida(100);s__Candida_cretensis(100); | Candida | 13 | Pathotroph-Saprotroph-Symbiotroph | Animal Pathogen-Endophyte-Endosymbiont-Epiphyte-\|Undefined Saprotroph\| | Dimorphic Yeast-Yeast | NULL | Probable | As nectar-tap_saprotroph (Põlme et al. 2020); Decay sugar-rich substrates (Põlme et al. 2020); This asexual genus is polyphyletic and contain species that function very differently depend on what they are. Caution should be taken when interpreting guilds for Candida species. | Manolakaki D, et al. 2010. Virulence. 1: 367-375 ((https://doi.org/10.4161/viru.1.5.12796)); Mekbib SB, et al. 2011. Tropical Plant Pathology 36:233-240 ((https://doi.org/10.1590/S1982-56762011000400004)); Põlme S, et al. 2020. Fungal Diversity 105:1-16 ((https://doi.org/10.1007/s13225-020-00466-2)) |
| Otu16592 | 1 | k__Fungi(100);p__Ascomycota(100);c__Saccharomycetes(100);o__Saccharomycetales(100);f__Metschnikowiaceae(100);g__Metschnikowia(100);s__Metschnikowia_pulcherrima(100); | Metschnikowia | 13 | Saprotroph | \|Undefined Saprotroph\| | Yeast | NULL | Probable | As nectar-tap_saprotroph (Põlme et al. 2020); Decay sugar-rich substrates (Põlme et al. 2020) | Põlme S, et al. 2020. Fungal Diversity 105:1-16 ((https://doi.org/10.1007/s13225-020-00466-2)) |
| Otu16593 | 1 | k__Fungi(100);p__Ascomycota(100);c__Saccharomycetes(100);o__Saccharomycetales(100);f__Metschnikowiaceae(100);g__Metschnikowia(100);s__Metschnikowia_pulcherrima(100); | Metschnikowia | 13 | Saprotroph | \|Undefined Saprotroph\| | Yeast | NULL | Probable | As nectar-tap_saprotroph (Põlme et al. 2020); Decay sugar-rich substrates (Põlme et al. 2020) | Põlme S, et al. 2020. Fungal Diversity 105:1-16 ((https://doi.org/10.1007/s13225-020-00466-2)) |
| Otu16594 | 1 | k__Fungi(100);p__Ascomycota(100);c__Saccharomycetes(100);o__Saccharomycetales(100);f__Metschnikowiaceae(100);g__Metschnikowia(100);s__Metschnikowia_pulcherrima(100); | Metschnikowia | 13 | Saprotroph | \|Undefined Saprotroph\| | Yeast | NULL | Probable | As nectar-tap_saprotroph (Põlme et al. 2020); Decay sugar-rich substrates (Põlme et al. 2020) | Põlme S, et al. 2020. Fungal Diversity 105:1-16 ((https://doi.org/10.1007/s13225-020-00466-2)) |
| Otu16595 | 1 | k__Fungi(100);p__Ascomycota(100);c__Saccharomycetes(100);o__Saccharomycetales(100);f__Metschnikowiaceae(100);g__Metschnikowia(100);s__Metschnikowia_pulcherrima(100); | Metschnikowia | 13 | Saprotroph | \|Undefined Saprotroph\| | Yeast | NULL | Probable | As nectar-tap_saprotroph (Põlme et al. 2020); Decay sugar-rich substrates (Põlme et al. 2020) | Põlme S, et al. 2020. Fungal Diversity 105:1-16 ((https://doi.org/10.1007/s13225-020-00466-2)) |
| Otu16596 | 1 | k__Fungi(100);p__Ascomycota(100);c__Saccharomycetes(100);o__Saccharomycetales(100);f__Metschnikowiaceae(100);g__Metschnikowia(100);s__Metschnikowia_pulcherrima(100); | Metschnikowia | 13 | Saprotroph | \|Undefined Saprotroph\| | Yeast | NULL | Probable | As nectar-tap_saprotroph (Põlme et al. 2020); Decay sugar-rich substrates (Põlme et al. 2020) | Põlme S, et al. 2020. Fungal Diversity 105:1-16 ((https://doi.org/10.1007/s13225-020-00466-2)) |
| Otu16597 | 1 | k__Fungi(100);p__Ascomycota(100);c__Leotiomycetes(100);o__Helotiales(100);f__Helotiaceae(100);g__Hymenoscyphus(100);g__Hymenoscyphus_unclassified(100); | Hymenoscyphus | 13 | Pathotroph-Saprotroph-Symbiotroph | Bryophyte Parasite-Ectomycorrhizal-Ericoid Mycorrhizal-Plant Pathogen-\|Plant Saprotroph\|-Undefined Saprotroph | Microfungus | NULL | Possible | As litter_saprotroph (Põlme et al. 2020); Dark Septate Microfungus; Syn. Pezoloma among other (see e.g., Pezoloma ericae); Decay leaf, fruit, seed (Põlme et al. 2020) | Baral HO, Krieglsteiner L. 2006. Acta Mycologica 41:11-20 ((https://doi.org/10.5586/am.2006.003)) (as a bryophyte parasite); Huhtinen S, et al. 2010. Nova Hedwigia 90:413-431 ((https://doi.org/10.1127/0029-5035/2010/0090-0413)); Bent E, et al. 2011. Applied and Environmental Microbiology 77:3351-3359 ((https://doi.org/10.1128/AEM.02575-10)) (as ectomycorrhizae); Tedersoo L, et al. 2014. Science 346:1256688 ((https://doi.org/10.1126/science.1256688)) (as saprotroph); Põlme S, et al. 2020. Fungal Diversity 105:1-16 ((https://doi.org/10.1007/s13225-020-00466-2)) |
| Otu16598 | 1 | k__Fungi(100);p__Ascomycota(100);c__Saccharomycetes(100);o__Saccharomycetales(100);f__Metschnikowiaceae(100);g__Metschnikowia(100);s__Metschnikowia_pulcherrima(100); | Metschnikowia | 13 | Saprotroph | \|Undefined Saprotroph\| | Yeast | NULL | Probable | As nectar-tap_saprotroph (Põlme et al. 2020); Decay sugar-rich substrates (Põlme et al. 2020) | Põlme S, et al. 2020. Fungal Diversity 105:1-16 ((https://doi.org/10.1007/s13225-020-00466-2)) |
| Otu16599 | 1 | k__Fungi(100);p__Ascomycota(100);c__Saccharomycetes(100);o__Saccharomycetales(100);f__Metschnikowiaceae(100);g__Metschnikowia(100);s__Metschnikowia_pulcherrima(100); | Metschnikowia | 13 | Saprotroph | \|Undefined Saprotroph\| | Yeast | NULL | Probable | As nectar-tap_saprotroph (Põlme et al. 2020); Decay sugar-rich substrates (Põlme et al. 2020) | Põlme S, et al. 2020. Fungal Diversity 105:1-16 ((https://doi.org/10.1007/s13225-020-00466-2)) |
| Otu16600 | 1 | k__Fungi(100);p__Ascomycota(100);c__Saccharomycetes(100);o__Saccharomycetales(100);f__Metschnikowiaceae(100);g__Metschnikowia(100);s__Metschnikowia_pulcherrima(100); | Metschnikowia | 13 | Saprotroph | \|Undefined Saprotroph\| | Yeast | NULL | Probable | As nectar-tap_saprotroph (Põlme et al. 2020); Decay sugar-rich substrates (Põlme et al. 2020) | Põlme S, et al. 2020. Fungal Diversity 105:1-16 ((https://doi.org/10.1007/s13225-020-00466-2)) |
| Otu16601 | 1 | k__Fungi(100);p__Ascomycota(100);c__Saccharomycetes(100);o__Saccharomycetales(100);f__Metschnikowiaceae(100);g__Metschnikowia(100);s__Metschnikowia_pulcherrima(100); | Metschnikowia | 13 | Saprotroph | \|Undefined Saprotroph\| | Yeast | NULL | Probable | As nectar-tap_saprotroph (Põlme et al. 2020); Decay sugar-rich substrates (Põlme et al. 2020) | Põlme S, et al. 2020. Fungal Diversity 105:1-16 ((https://doi.org/10.1007/s13225-020-00466-2)) |
| Otu16602 | 1 | k__Fungi(100);p__Ascomycota(100);c__Saccharomycetes(100);o__Saccharomycetales(100);f__Metschnikowiaceae(100);g__Metschnikowia(100);g__Metschnikowia_unclassified(100); | Metschnikowia | 13 | Saprotroph | \|Undefined Saprotroph\| | Yeast | NULL | Probable | As nectar-tap_saprotroph (Põlme et al. 2020); Decay sugar-rich substrates (Põlme et al. 2020) | Põlme S, et al. 2020. Fungal Diversity 105:1-16 ((https://doi.org/10.1007/s13225-020-00466-2)) |
| Otu16603 | 1 | k__Fungi(100);p__Ascomycota(100);c__Saccharomycetes(100);o__Saccharomycetales(100);f__Metschnikowiaceae(100);g__Metschnikowia(100);s__Metschnikowia_pulcherrima(100); | Metschnikowia | 13 | Saprotroph | \|Undefined Saprotroph\| | Yeast | NULL | Probable | As nectar-tap_saprotroph (Põlme et al. 2020); Decay sugar-rich substrates (Põlme et al. 2020) | Põlme S, et al. 2020. Fungal Diversity 105:1-16 ((https://doi.org/10.1007/s13225-020-00466-2)) |
| Otu16604 | 1 | k__Fungi(100);p__Ascomycota(100);c__Saccharomycetes(100);o__Saccharomycetales(100);f__Metschnikowiaceae(100);g__Metschnikowia(100);g__Metschnikowia_unclassified(100); | Metschnikowia | 13 | Saprotroph | \|Undefined Saprotroph\| | Yeast | NULL | Probable | As nectar-tap_saprotroph (Põlme et al. 2020); Decay sugar-rich substrates (Põlme et al. 2020) | Põlme S, et al. 2020. Fungal Diversity 105:1-16 ((https://doi.org/10.1007/s13225-020-00466-2)) |
| Otu16605 | 1 | k__Fungi(100);p__Ascomycota(100);c__Saccharomycetes(100);o__Saccharomycetales(100);f__Metschnikowiaceae(100);g__Metschnikowia(100);s__Metschnikowia_pulcherrima(100); | Metschnikowia | 13 | Saprotroph | \|Undefined Saprotroph\| | Yeast | NULL | Probable | As nectar-tap_saprotroph (Põlme et al. 2020); Decay sugar-rich substrates (Põlme et al. 2020) | Põlme S, et al. 2020. Fungal Diversity 105:1-16 ((https://doi.org/10.1007/s13225-020-00466-2)) |
| Otu16606 | 1 | k__Fungi(100);p__Ascomycota(100);c__Saccharomycetes(100);o__Saccharomycetales(100);f__Saccharomycetales_family_Incertae_sedis(100);g__Candida(100);s__Candida_cretensis(100); | Candida | 13 | Pathotroph-Saprotroph-Symbiotroph | Animal Pathogen-Endophyte-Endosymbiont-Epiphyte-\|Undefined Saprotroph\| | Dimorphic Yeast-Yeast | NULL | Probable | As nectar-tap_saprotroph (Põlme et al. 2020); Decay sugar-rich substrates (Põlme et al. 2020); This asexual genus is polyphyletic and contain species that function very differently depend on what they are. Caution should be taken when interpreting guilds for Candida species. | Manolakaki D, et al. 2010. Virulence. 1: 367-375 ((https://doi.org/10.4161/viru.1.5.12796)); Mekbib SB, et al. 2011. Tropical Plant Pathology 36:233-240 ((https://doi.org/10.1590/S1982-56762011000400004)); Põlme S, et al. 2020. Fungal Diversity 105:1-16 ((https://doi.org/10.1007/s13225-020-00466-2)) |
| Otu16607 | 1 | k__Fungi(100);p__Ascomycota(100);c__Saccharomycetes(100);o__Saccharomycetales(100);f__Metschnikowiaceae(100);g__Metschnikowia(100);s__Metschnikowia_sp(100); | Metschnikowia | 13 | Saprotroph | \|Undefined Saprotroph\| | Yeast | NULL | Probable | As nectar-tap_saprotroph (Põlme et al. 2020); Decay sugar-rich substrates (Põlme et al. 2020) | Põlme S, et al. 2020. Fungal Diversity 105:1-16 ((https://doi.org/10.1007/s13225-020-00466-2)) |
| Otu16608 | 1 | k__Fungi(100);p__Ascomycota(100);c__Saccharomycetes(100);o__Saccharomycetales(100);f__Metschnikowiaceae(100);g__Metschnikowia(100);g__Metschnikowia_unclassified(100); | Metschnikowia | 13 | Saprotroph | \|Undefined Saprotroph\| | Yeast | NULL | Probable | As nectar-tap_saprotroph (Põlme et al. 2020); Decay sugar-rich substrates (Põlme et al. 2020) | Põlme S, et al. 2020. Fungal Diversity 105:1-16 ((https://doi.org/10.1007/s13225-020-00466-2)) |
| Otu16609 | 1 | k__Fungi(100);p__Ascomycota(100);c__Saccharomycetes(100);o__Saccharomycetales(100);f__Metschnikowiaceae(100);g__Metschnikowia(100);s__Metschnikowia_pulcherrima(100); | Metschnikowia | 13 | Saprotroph | \|Undefined Saprotroph\| | Yeast | NULL | Probable | As nectar-tap_saprotroph (Põlme et al. 2020); Decay sugar-rich substrates (Põlme et al. 2020) | Põlme S, et al. 2020. Fungal Diversity 105:1-16 ((https://doi.org/10.1007/s13225-020-00466-2)) |
| Otu16610 | 1 | k__Fungi(100);p__Ascomycota(100);c__Saccharomycetes(100);o__Saccharomycetales(100);f__Metschnikowiaceae(100);g__Metschnikowia(100);s__Metschnikowia_pulcherrima(100); | Metschnikowia | 13 | Saprotroph | \|Undefined Saprotroph\| | Yeast | NULL | Probable | As nectar-tap_saprotroph (Põlme et al. 2020); Decay sugar-rich substrates (Põlme et al. 2020) | Põlme S, et al. 2020. Fungal Diversity 105:1-16 ((https://doi.org/10.1007/s13225-020-00466-2)) |
| Otu16611 | 1 | k__Fungi(100);p__Ascomycota(100);c__Saccharomycetes(100);o__Saccharomycetales(100);f__Metschnikowiaceae(100);g__Metschnikowia(100);s__Metschnikowia_pulcherrima(100); | Metschnikowia | 13 | Saprotroph | \|Undefined Saprotroph\| | Yeast | NULL | Probable | As nectar-tap_saprotroph (Põlme et al. 2020); Decay sugar-rich substrates (Põlme et al. 2020) | Põlme S, et al. 2020. Fungal Diversity 105:1-16 ((https://doi.org/10.1007/s13225-020-00466-2)) |
| Otu16612 | 1 | k__Fungi(100);p__Ascomycota(100);c__Saccharomycetes(100);o__Saccharomycetales(100);f__Metschnikowiaceae(100);g__Metschnikowia(100);s__Metschnikowia_pulcherrima(100); | Metschnikowia | 13 | Saprotroph | \|Undefined Saprotroph\| | Yeast | NULL | Probable | As nectar-tap_saprotroph (Põlme et al. 2020); Decay sugar-rich substrates (Põlme et al. 2020) | Põlme S, et al. 2020. Fungal Diversity 105:1-16 ((https://doi.org/10.1007/s13225-020-00466-2)) |
| Otu16615 | 1 | k__Fungi(100);p__Ascomycota(100);c__Saccharomycetes(100);o__Saccharomycetales(100);f__Metschnikowiaceae(100);g__Metschnikowia(100);s__Metschnikowia_pulcherrima(100); | Metschnikowia | 13 | Saprotroph | \|Undefined Saprotroph\| | Yeast | NULL | Probable | As nectar-tap_saprotroph (Põlme et al. 2020); Decay sugar-rich substrates (Põlme et al. 2020) | Põlme S, et al. 2020. Fungal Diversity 105:1-16 ((https://doi.org/10.1007/s13225-020-00466-2)) |
| Otu16616 | 1 | k__Fungi(100);p__Ascomycota(100);c__Saccharomycetes(100);o__Saccharomycetales(100);f__Metschnikowiaceae(100);g__Metschnikowia(100);s__Metschnikowia_pulcherrima(100); | Metschnikowia | 13 | Saprotroph | \|Undefined Saprotroph\| | Yeast | NULL | Probable | As nectar-tap_saprotroph (Põlme et al. 2020); Decay sugar-rich substrates (Põlme et al. 2020) | Põlme S, et al. 2020. Fungal Diversity 105:1-16 ((https://doi.org/10.1007/s13225-020-00466-2)) |
| Otu16617 | 1 | k__Fungi(100);p__Ascomycota(100);c__Saccharomycetes(100);o__Saccharomycetales(100);f__Metschnikowiaceae(100);g__Metschnikowia(100);s__Metschnikowia_pulcherrima(100); | Metschnikowia | 13 | Saprotroph | \|Undefined Saprotroph\| | Yeast | NULL | Probable | As nectar-tap_saprotroph (Põlme et al. 2020); Decay sugar-rich substrates (Põlme et al. 2020) | Põlme S, et al. 2020. Fungal Diversity 105:1-16 ((https://doi.org/10.1007/s13225-020-00466-2)) |
| Otu16618 | 1 | k__Fungi(100);p__Ascomycota(100);c__Saccharomycetes(100);o__Saccharomycetales(100);f__Metschnikowiaceae(100);g__Metschnikowia(100);g__Metschnikowia_unclassified(100); | Metschnikowia | 13 | Saprotroph | \|Undefined Saprotroph\| | Yeast | NULL | Probable | As nectar-tap_saprotroph (Põlme et al. 2020); Decay sugar-rich substrates (Põlme et al. 2020) | Põlme S, et al. 2020. Fungal Diversity 105:1-16 ((https://doi.org/10.1007/s13225-020-00466-2)) |
| Otu16619 | 1 | k__Fungi(100);p__Ascomycota(100);c__Saccharomycetes(100);o__Saccharomycetales(100);f__Metschnikowiaceae(100);g__Metschnikowia(100);g__Metschnikowia_unclassified(100); | Metschnikowia | 13 | Saprotroph | \|Undefined Saprotroph\| | Yeast | NULL | Probable | As nectar-tap_saprotroph (Põlme et al. 2020); Decay sugar-rich substrates (Põlme et al. 2020) | Põlme S, et al. 2020. Fungal Diversity 105:1-16 ((https://doi.org/10.1007/s13225-020-00466-2)) |
| Otu16622 | 1 | k__Fungi(100);p__Ascomycota(100);c__Saccharomycetes(100);o__Saccharomycetales(100);f__Metschnikowiaceae(100);g__Metschnikowia(100);s__Metschnikowia_pulcherrima(100); | Metschnikowia | 13 | Saprotroph | \|Undefined Saprotroph\| | Yeast | NULL | Probable | As nectar-tap_saprotroph (Põlme et al. 2020); Decay sugar-rich substrates (Põlme et al. 2020) | Põlme S, et al. 2020. Fungal Diversity 105:1-16 ((https://doi.org/10.1007/s13225-020-00466-2)) |
| Otu16623 | 1 | k__Fungi(100);p__Ascomycota(100);c__Saccharomycetes(100);o__Saccharomycetales(100);f__Metschnikowiaceae(100);g__Metschnikowia(100);s__Metschnikowia_pulcherrima(100); | Metschnikowia | 13 | Saprotroph | \|Undefined Saprotroph\| | Yeast | NULL | Probable | As nectar-tap_saprotroph (Põlme et al. 2020); Decay sugar-rich substrates (Põlme et al. 2020) | Põlme S, et al. 2020. Fungal Diversity 105:1-16 ((https://doi.org/10.1007/s13225-020-00466-2)) |
| Otu16624 | 1 | k__Fungi(100);p__Ascomycota(100);c__Saccharomycetes(100);o__Saccharomycetales(100);f__Metschnikowiaceae(100);g__Metschnikowia(100);s__Metschnikowia_pulcherrima(100); | Metschnikowia | 13 | Saprotroph | \|Undefined Saprotroph\| | Yeast | NULL | Probable | As nectar-tap_saprotroph (Põlme et al. 2020); Decay sugar-rich substrates (Põlme et al. 2020) | Põlme S, et al. 2020. Fungal Diversity 105:1-16 ((https://doi.org/10.1007/s13225-020-00466-2)) |
| Otu16625 | 1 | k__Fungi(100);p__Ascomycota(100);c__Saccharomycetes(100);o__Saccharomycetales(100);f__Metschnikowiaceae(100);g__Metschnikowia(100);s__Metschnikowia_pulcherrima(100); | Metschnikowia | 13 | Saprotroph | \|Undefined Saprotroph\| | Yeast | NULL | Probable | As nectar-tap_saprotroph (Põlme et al. 2020); Decay sugar-rich substrates (Põlme et al. 2020) | Põlme S, et al. 2020. Fungal Diversity 105:1-16 ((https://doi.org/10.1007/s13225-020-00466-2)) |
| Otu16626 | 1 | k__Fungi(100);p__Ascomycota(100);c__Saccharomycetes(100);o__Saccharomycetales(100);f__Metschnikowiaceae(100);g__Metschnikowia(100);s__Metschnikowia_pulcherrima(100); | Metschnikowia | 13 | Saprotroph | \|Undefined Saprotroph\| | Yeast | NULL | Probable | As nectar-tap_saprotroph (Põlme et al. 2020); Decay sugar-rich substrates (Põlme et al. 2020) | Põlme S, et al. 2020. Fungal Diversity 105:1-16 ((https://doi.org/10.1007/s13225-020-00466-2)) |
| Otu16627 | 1 | k__Fungi(100);p__Ascomycota(100);c__Saccharomycetes(100);o__Saccharomycetales(100);f__Metschnikowiaceae(100);g__Metschnikowia(100);s__Metschnikowia_pulcherrima(100); | Metschnikowia | 13 | Saprotroph | \|Undefined Saprotroph\| | Yeast | NULL | Probable | As nectar-tap_saprotroph (Põlme et al. 2020); Decay sugar-rich substrates (Põlme et al. 2020) | Põlme S, et al. 2020. Fungal Diversity 105:1-16 ((https://doi.org/10.1007/s13225-020-00466-2)) |
| Otu16628 | 1 | k__Fungi(100);p__Ascomycota(100);c__Saccharomycetes(100);o__Saccharomycetales(100);f__Metschnikowiaceae(100);g__Metschnikowia(100);s__Metschnikowia_pulcherrima(100); | Metschnikowia | 13 | Saprotroph | \|Undefined Saprotroph\| | Yeast | NULL | Probable | As nectar-tap_saprotroph (Põlme et al. 2020); Decay sugar-rich substrates (Põlme et al. 2020) | Põlme S, et al. 2020. Fungal Diversity 105:1-16 ((https://doi.org/10.1007/s13225-020-00466-2)) |
| Otu16629 | 1 | k__Fungi(100);p__Ascomycota(100);c__Saccharomycetes(100);o__Saccharomycetales(100);f__Metschnikowiaceae(100);g__Metschnikowia(100);s__Metschnikowia_pulcherrima(100); | Metschnikowia | 13 | Saprotroph | \|Undefined Saprotroph\| | Yeast | NULL | Probable | As nectar-tap_saprotroph (Põlme et al. 2020); Decay sugar-rich substrates (Põlme et al. 2020) | Põlme S, et al. 2020. Fungal Diversity 105:1-16 ((https://doi.org/10.1007/s13225-020-00466-2)) |
| Otu16630 | 1 | k__Fungi(100);p__Ascomycota(100);c__Saccharomycetes(100);o__Saccharomycetales(100);f__Metschnikowiaceae(100);g__Metschnikowia(100);s__Metschnikowia_pulcherrima(100); | Metschnikowia | 13 | Saprotroph | \|Undefined Saprotroph\| | Yeast | NULL | Probable | As nectar-tap_saprotroph (Põlme et al. 2020); Decay sugar-rich substrates (Põlme et al. 2020) | Põlme S, et al. 2020. Fungal Diversity 105:1-16 ((https://doi.org/10.1007/s13225-020-00466-2)) |
| Otu16631 | 1 | k__Fungi(100);p__Ascomycota(100);c__Saccharomycetes(100);o__Saccharomycetales(100);f__Metschnikowiaceae(100);g__Metschnikowia(100);g__Metschnikowia_unclassified(100); | Metschnikowia | 13 | Saprotroph | \|Undefined Saprotroph\| | Yeast | NULL | Probable | As nectar-tap_saprotroph (Põlme et al. 2020); Decay sugar-rich substrates (Põlme et al. 2020) | Põlme S, et al. 2020. Fungal Diversity 105:1-16 ((https://doi.org/10.1007/s13225-020-00466-2)) |
| Otu16632 | 1 | k__Fungi(100);p__Ascomycota(100);c__Saccharomycetes(100);o__Saccharomycetales(100);f__Metschnikowiaceae(100);g__Metschnikowia(100);s__Metschnikowia_pulcherrima(100); | Metschnikowia | 13 | Saprotroph | \|Undefined Saprotroph\| | Yeast | NULL | Probable | As nectar-tap_saprotroph (Põlme et al. 2020); Decay sugar-rich substrates (Põlme et al. 2020) | Põlme S, et al. 2020. Fungal Diversity 105:1-16 ((https://doi.org/10.1007/s13225-020-00466-2)) |
| Otu16633 | 1 | k__Fungi(100);p__Ascomycota(100);c__Saccharomycetes(100);o__Saccharomycetales(100);f__Metschnikowiaceae(100);g__Metschnikowia(100);g__Metschnikowia_unclassified(100); | Metschnikowia | 13 | Saprotroph | \|Undefined Saprotroph\| | Yeast | NULL | Probable | As nectar-tap_saprotroph (Põlme et al. 2020); Decay sugar-rich substrates (Põlme et al. 2020) | Põlme S, et al. 2020. Fungal Diversity 105:1-16 ((https://doi.org/10.1007/s13225-020-00466-2)) |
| Otu16634 | 1 | k__Fungi(100);p__Ascomycota(100);c__Saccharomycetes(100);o__Saccharomycetales(100);f__Metschnikowiaceae(100);g__Metschnikowia(100);s__Metschnikowia_pulcherrima(100); | Metschnikowia | 13 | Saprotroph | \|Undefined Saprotroph\| | Yeast | NULL | Probable | As nectar-tap_saprotroph (Põlme et al. 2020); Decay sugar-rich substrates (Põlme et al. 2020) | Põlme S, et al. 2020. Fungal Diversity 105:1-16 ((https://doi.org/10.1007/s13225-020-00466-2)) |
| Otu16635 | 1 | k__Fungi(100);p__Ascomycota(100);c__Saccharomycetes(100);o__Saccharomycetales(100);f__Metschnikowiaceae(100);g__Metschnikowia(100);s__Metschnikowia_pulcherrima(100); | Metschnikowia | 13 | Saprotroph | \|Undefined Saprotroph\| | Yeast | NULL | Probable | As nectar-tap_saprotroph (Põlme et al. 2020); Decay sugar-rich substrates (Põlme et al. 2020) | Põlme S, et al. 2020. Fungal Diversity 105:1-16 ((https://doi.org/10.1007/s13225-020-00466-2)) |
| Otu16636 | 1 | k__Fungi(100);p__Ascomycota(100);c__Saccharomycetes(100);o__Saccharomycetales(100);f__Metschnikowiaceae(100);g__Metschnikowia(100);s__Metschnikowia_pulcherrima(100); | Metschnikowia | 13 | Saprotroph | \|Undefined Saprotroph\| | Yeast | NULL | Probable | As nectar-tap_saprotroph (Põlme et al. 2020); Decay sugar-rich substrates (Põlme et al. 2020) | Põlme S, et al. 2020. Fungal Diversity 105:1-16 ((https://doi.org/10.1007/s13225-020-00466-2)) |
| Otu16637 | 1 | k__Fungi(100);p__Ascomycota(100);c__Saccharomycetes(100);o__Saccharomycetales(100);f__Metschnikowiaceae(100);g__Metschnikowia(100);s__Metschnikowia_pulcherrima(100); | Metschnikowia | 13 | Saprotroph | \|Undefined Saprotroph\| | Yeast | NULL | Probable | As nectar-tap_saprotroph (Põlme et al. 2020); Decay sugar-rich substrates (Põlme et al. 2020) | Põlme S, et al. 2020. Fungal Diversity 105:1-16 ((https://doi.org/10.1007/s13225-020-00466-2)) |
| Otu16638 | 1 | k__Fungi(100);p__Ascomycota(100);c__Saccharomycetes(100);o__Saccharomycetales(100);f__Metschnikowiaceae(100);g__Metschnikowia(100);s__Metschnikowia_pulcherrima(100); | Metschnikowia | 13 | Saprotroph | \|Undefined Saprotroph\| | Yeast | NULL | Probable | As nectar-tap_saprotroph (Põlme et al. 2020); Decay sugar-rich substrates (Põlme et al. 2020) | Põlme S, et al. 2020. Fungal Diversity 105:1-16 ((https://doi.org/10.1007/s13225-020-00466-2)) |
| Otu16639 | 1 | k__Fungi(100);p__Ascomycota(100);c__Saccharomycetes(100);o__Saccharomycetales(100);f__Metschnikowiaceae(100);g__Metschnikowia(100);s__Metschnikowia_pulcherrima(100); | Metschnikowia | 13 | Saprotroph | \|Undefined Saprotroph\| | Yeast | NULL | Probable | As nectar-tap_saprotroph (Põlme et al. 2020); Decay sugar-rich substrates (Põlme et al. 2020) | Põlme S, et al. 2020. Fungal Diversity 105:1-16 ((https://doi.org/10.1007/s13225-020-00466-2)) |
| Otu16641 | 1 | k__Fungi(100);p__Ascomycota(100);c__Saccharomycetes(100);o__Saccharomycetales(100);f__Metschnikowiaceae(100);g__Metschnikowia(100);s__Metschnikowia_pulcherrima(100); | Metschnikowia | 13 | Saprotroph | \|Undefined Saprotroph\| | Yeast | NULL | Probable | As nectar-tap_saprotroph (Põlme et al. 2020); Decay sugar-rich substrates (Põlme et al. 2020) | Põlme S, et al. 2020. Fungal Diversity 105:1-16 ((https://doi.org/10.1007/s13225-020-00466-2)) |
| Otu16642 | 1 | k__Fungi(100);p__Ascomycota(100);c__Saccharomycetes(100);o__Saccharomycetales(100);f__Metschnikowiaceae(100);g__Metschnikowia(100);g__Metschnikowia_unclassified(100); | Metschnikowia | 13 | Saprotroph | \|Undefined Saprotroph\| | Yeast | NULL | Probable | As nectar-tap_saprotroph (Põlme et al. 2020); Decay sugar-rich substrates (Põlme et al. 2020) | Põlme S, et al. 2020. Fungal Diversity 105:1-16 ((https://doi.org/10.1007/s13225-020-00466-2)) |
| Otu16643 | 1 | k__Fungi(100);p__Basidiomycota(100);c__Microbotryomycetes(100);o__Sporidiobolales(100);f__Sporidiobolales_family_Incertae_sedis(100);g__Sporobolomyces(100);s__Sporobolomyces_lactosus(100); | Sporobolomyces | 13 | Pathotroph-Saprotroph | \|Fungal Parasite\|-Plant Saprotroph | Yeast | NULL | Probable | As litter_saprotroph (Põlme et al. 2020); As mycoparasite (Põlme et al. 2020); Decay fungal material (Põlme et al. 2020) | Tedersoo L, et al. 2014. Science 346:e1256688 ((https://doi.org/10.1126/science.1256688)); Purahong W, et al. 2016. Molecular Ecology 25:4059-4074 ((https://doi.org/10.1111/mec.13739)); Põlme S, et al. 2020. Fungal Diversity 105:1-16 ((https://doi.org/10.1007/s13225-020-00466-2)) |
| Otu16644 | 1 | k__Fungi(100);p__Ascomycota(100);c__Saccharomycetes(100);o__Saccharomycetales(100);f__Metschnikowiaceae(100);g__Metschnikowia(100);s__Metschnikowia_pulcherrima(100); | Metschnikowia | 13 | Saprotroph | \|Undefined Saprotroph\| | Yeast | NULL | Probable | As nectar-tap_saprotroph (Põlme et al. 2020); Decay sugar-rich substrates (Põlme et al. 2020) | Põlme S, et al. 2020. Fungal Diversity 105:1-16 ((https://doi.org/10.1007/s13225-020-00466-2)) |
| Otu16645 | 1 | k__Fungi(100);p__Ascomycota(100);c__Saccharomycetes(100);o__Saccharomycetales(100);f__Metschnikowiaceae(100);g__Metschnikowia(100);s__Metschnikowia_pulcherrima(100); | Metschnikowia | 13 | Saprotroph | \|Undefined Saprotroph\| | Yeast | NULL | Probable | As nectar-tap_saprotroph (Põlme et al. 2020); Decay sugar-rich substrates (Põlme et al. 2020) | Põlme S, et al. 2020. Fungal Diversity 105:1-16 ((https://doi.org/10.1007/s13225-020-00466-2)) |
| Otu16646 | 1 | k__Fungi(100);p__Ascomycota(100);c__Saccharomycetes(100);o__Saccharomycetales(100);f__Metschnikowiaceae(100);g__Metschnikowia(100);g__Metschnikowia_unclassified(100); | Metschnikowia | 13 | Saprotroph | \|Undefined Saprotroph\| | Yeast | NULL | Probable | As nectar-tap_saprotroph (Põlme et al. 2020); Decay sugar-rich substrates (Põlme et al. 2020) | Põlme S, et al. 2020. Fungal Diversity 105:1-16 ((https://doi.org/10.1007/s13225-020-00466-2)) |
| Otu16647 | 1 | k__Fungi(100);p__Ascomycota(100);c__Saccharomycetes(100);o__Saccharomycetales(100);f__Metschnikowiaceae(100);g__Metschnikowia(100);s__Metschnikowia_pulcherrima(100); | Metschnikowia | 13 | Saprotroph | \|Undefined Saprotroph\| | Yeast | NULL | Probable | As nectar-tap_saprotroph (Põlme et al. 2020); Decay sugar-rich substrates (Põlme et al. 2020) | Põlme S, et al. 2020. Fungal Diversity 105:1-16 ((https://doi.org/10.1007/s13225-020-00466-2)) |
| Otu16648 | 1 | k__Fungi(100);p__Ascomycota(100);c__Saccharomycetes(100);o__Saccharomycetales(100);f__Metschnikowiaceae(100);g__Metschnikowia(100);s__Metschnikowia_pulcherrima(100); | Metschnikowia | 13 | Saprotroph | \|Undefined Saprotroph\| | Yeast | NULL | Probable | As nectar-tap_saprotroph (Põlme et al. 2020); Decay sugar-rich substrates (Põlme et al. 2020) | Põlme S, et al. 2020. Fungal Diversity 105:1-16 ((https://doi.org/10.1007/s13225-020-00466-2)) |
| Otu16649 | 1 | k__Fungi(100);p__Ascomycota(100);c__Saccharomycetes(100);o__Saccharomycetales(100);f__Metschnikowiaceae(100);g__Metschnikowia(100);s__Metschnikowia_pulcherrima(100); | Metschnikowia | 13 | Saprotroph | \|Undefined Saprotroph\| | Yeast | NULL | Probable | As nectar-tap_saprotroph (Põlme et al. 2020); Decay sugar-rich substrates (Põlme et al. 2020) | Põlme S, et al. 2020. Fungal Diversity 105:1-16 ((https://doi.org/10.1007/s13225-020-00466-2)) |
| Otu16650 | 1 | k__Fungi(100);p__Ascomycota(100);c__Saccharomycetes(100);o__Saccharomycetales(100);f__Metschnikowiaceae(100);g__Metschnikowia(100);s__Metschnikowia_pulcherrima(100); | Metschnikowia | 13 | Saprotroph | \|Undefined Saprotroph\| | Yeast | NULL | Probable | As nectar-tap_saprotroph (Põlme et al. 2020); Decay sugar-rich substrates (Põlme et al. 2020) | Põlme S, et al. 2020. Fungal Diversity 105:1-16 ((https://doi.org/10.1007/s13225-020-00466-2)) |
| Otu16651 | 1 | k__Fungi(100);p__Ascomycota(100);c__Saccharomycetes(100);o__Saccharomycetales(100);f__Metschnikowiaceae(100);g__Metschnikowia(100);s__Metschnikowia_pulcherrima(100); | Metschnikowia | 13 | Saprotroph | \|Undefined Saprotroph\| | Yeast | NULL | Probable | As nectar-tap_saprotroph (Põlme et al. 2020); Decay sugar-rich substrates (Põlme et al. 2020) | Põlme S, et al. 2020. Fungal Diversity 105:1-16 ((https://doi.org/10.1007/s13225-020-00466-2)) |
| Otu16652 | 1 | k__Fungi(100);p__Ascomycota(100);c__Saccharomycetes(100);o__Saccharomycetales(100);f__Metschnikowiaceae(100);g__Metschnikowia(100);s__Metschnikowia_pulcherrima(100); | Metschnikowia | 13 | Saprotroph | \|Undefined Saprotroph\| | Yeast | NULL | Probable | As nectar-tap_saprotroph (Põlme et al. 2020); Decay sugar-rich substrates (Põlme et al. 2020) | Põlme S, et al. 2020. Fungal Diversity 105:1-16 ((https://doi.org/10.1007/s13225-020-00466-2)) |
| Otu16653 | 1 | k__Fungi(100);p__Ascomycota(100);c__Saccharomycetes(100);o__Saccharomycetales(100);f__Metschnikowiaceae(100);g__Metschnikowia(100);s__Metschnikowia_pulcherrima(100); | Metschnikowia | 13 | Saprotroph | \|Undefined Saprotroph\| | Yeast | NULL | Probable | As nectar-tap_saprotroph (Põlme et al. 2020); Decay sugar-rich substrates (Põlme et al. 2020) | Põlme S, et al. 2020. Fungal Diversity 105:1-16 ((https://doi.org/10.1007/s13225-020-00466-2)) |
| Otu16654 | 1 | k__Fungi(100);p__Ascomycota(100);c__Saccharomycetes(100);o__Saccharomycetales(100);f__Metschnikowiaceae(100);g__Metschnikowia(100);s__Metschnikowia_pulcherrima(100); | Metschnikowia | 13 | Saprotroph | \|Undefined Saprotroph\| | Yeast | NULL | Probable | As nectar-tap_saprotroph (Põlme et al. 2020); Decay sugar-rich substrates (Põlme et al. 2020) | Põlme S, et al. 2020. Fungal Diversity 105:1-16 ((https://doi.org/10.1007/s13225-020-00466-2)) |
| Otu16655 | 1 | k__Fungi(100);p__Ascomycota(100);c__Saccharomycetes(100);o__Saccharomycetales(100);f__Metschnikowiaceae(100);g__Metschnikowia(100);g__Metschnikowia_unclassified(100); | Metschnikowia | 13 | Saprotroph | \|Undefined Saprotroph\| | Yeast | NULL | Probable | As nectar-tap_saprotroph (Põlme et al. 2020); Decay sugar-rich substrates (Põlme et al. 2020) | Põlme S, et al. 2020. Fungal Diversity 105:1-16 ((https://doi.org/10.1007/s13225-020-00466-2)) |
| Otu16656 | 1 | k__Fungi(100);p__Ascomycota(100);c__Saccharomycetes(100);o__Saccharomycetales(100);f__Metschnikowiaceae(100);g__Metschnikowia(100);s__Metschnikowia_pulcherrima(100); | Metschnikowia | 13 | Saprotroph | \|Undefined Saprotroph\| | Yeast | NULL | Probable | As nectar-tap_saprotroph (Põlme et al. 2020); Decay sugar-rich substrates (Põlme et al. 2020) | Põlme S, et al. 2020. Fungal Diversity 105:1-16 ((https://doi.org/10.1007/s13225-020-00466-2)) |
| Otu16657 | 1 | k__Fungi(100);p__Ascomycota(100);c__Saccharomycetes(100);o__Saccharomycetales(100);f__Metschnikowiaceae(100);g__Metschnikowia(100);s__Metschnikowia_pulcherrima(100); | Metschnikowia | 13 | Saprotroph | \|Undefined Saprotroph\| | Yeast | NULL | Probable | As nectar-tap_saprotroph (Põlme et al. 2020); Decay sugar-rich substrates (Põlme et al. 2020) | Põlme S, et al. 2020. Fungal Diversity 105:1-16 ((https://doi.org/10.1007/s13225-020-00466-2)) |
| Otu16658 | 1 | k__Fungi(100);p__Ascomycota(100);c__Saccharomycetes(100);o__Saccharomycetales(100);f__Saccharomycetales_family_Incertae_sedis(100);g__Candida(100);g__Candida_unclassified(100); | Candida | 13 | Pathotroph-Saprotroph-Symbiotroph | Animal Pathogen-Endophyte-Endosymbiont-Epiphyte-\|Undefined Saprotroph\| | Dimorphic Yeast-Yeast | NULL | Probable | As nectar-tap_saprotroph (Põlme et al. 2020); Decay sugar-rich substrates (Põlme et al. 2020); This asexual genus is polyphyletic and contain species that function very differently depend on what they are. Caution should be taken when interpreting guilds for Candida species. | Manolakaki D, et al. 2010. Virulence. 1: 367-375 ((https://doi.org/10.4161/viru.1.5.12796)); Mekbib SB, et al. 2011. Tropical Plant Pathology 36:233-240 ((https://doi.org/10.1590/S1982-56762011000400004)); Põlme S, et al. 2020. Fungal Diversity 105:1-16 ((https://doi.org/10.1007/s13225-020-00466-2)) |
| Otu16659 | 1 | k__Fungi(100);p__Ascomycota(100);c__Saccharomycetes(100);o__Saccharomycetales(100);f__Metschnikowiaceae(100);g__Metschnikowia(100);s__Metschnikowia_pulcherrima(100); | Metschnikowia | 13 | Saprotroph | \|Undefined Saprotroph\| | Yeast | NULL | Probable | As nectar-tap_saprotroph (Põlme et al. 2020); Decay sugar-rich substrates (Põlme et al. 2020) | Põlme S, et al. 2020. Fungal Diversity 105:1-16 ((https://doi.org/10.1007/s13225-020-00466-2)) |
| Otu16660 | 1 | k__Fungi(100);p__Ascomycota(100);c__Saccharomycetes(100);o__Saccharomycetales(100);f__Metschnikowiaceae(100);g__Metschnikowia(100);s__Metschnikowia_pulcherrima(100); | Metschnikowia | 13 | Saprotroph | \|Undefined Saprotroph\| | Yeast | NULL | Probable | As nectar-tap_saprotroph (Põlme et al. 2020); Decay sugar-rich substrates (Põlme et al. 2020) | Põlme S, et al. 2020. Fungal Diversity 105:1-16 ((https://doi.org/10.1007/s13225-020-00466-2)) |
| Otu16661 | 1 | k__Fungi(100);p__Ascomycota(100);c__Saccharomycetes(100);o__Saccharomycetales(100);f__Metschnikowiaceae(100);g__Metschnikowia(100);s__Metschnikowia_pulcherrima(100); | Metschnikowia | 13 | Saprotroph | \|Undefined Saprotroph\| | Yeast | NULL | Probable | As nectar-tap_saprotroph (Põlme et al. 2020); Decay sugar-rich substrates (Põlme et al. 2020) | Põlme S, et al. 2020. Fungal Diversity 105:1-16 ((https://doi.org/10.1007/s13225-020-00466-2)) |
| Otu16662 | 1 | k__Fungi(100);p__Ascomycota(100);c__Saccharomycetes(100);o__Saccharomycetales(100);f__Metschnikowiaceae(100);g__Metschnikowia(100);g__Metschnikowia_unclassified(100); | Metschnikowia | 13 | Saprotroph | \|Undefined Saprotroph\| | Yeast | NULL | Probable | As nectar-tap_saprotroph (Põlme et al. 2020); Decay sugar-rich substrates (Põlme et al. 2020) | Põlme S, et al. 2020. Fungal Diversity 105:1-16 ((https://doi.org/10.1007/s13225-020-00466-2)) |
| Otu16663 | 1 | k__Fungi(100);p__Ascomycota(100);c__Saccharomycetes(100);o__Saccharomycetales(100);f__Metschnikowiaceae(100);g__Metschnikowia(100);s__Metschnikowia_pulcherrima(100); | Metschnikowia | 13 | Saprotroph | \|Undefined Saprotroph\| | Yeast | NULL | Probable | As nectar-tap_saprotroph (Põlme et al. 2020); Decay sugar-rich substrates (Põlme et al. 2020) | Põlme S, et al. 2020. Fungal Diversity 105:1-16 ((https://doi.org/10.1007/s13225-020-00466-2)) |
| Otu16664 | 1 | k__Fungi(100);p__Ascomycota(100);c__Saccharomycetes(100);o__Saccharomycetales(100);f__Metschnikowiaceae(100);g__Metschnikowia(100);g__Metschnikowia_unclassified(100); | Metschnikowia | 13 | Saprotroph | \|Undefined Saprotroph\| | Yeast | NULL | Probable | As nectar-tap_saprotroph (Põlme et al. 2020); Decay sugar-rich substrates (Põlme et al. 2020) | Põlme S, et al. 2020. Fungal Diversity 105:1-16 ((https://doi.org/10.1007/s13225-020-00466-2)) |
| Otu16665 | 1 | k__Fungi(100);p__Ascomycota(100);c__Saccharomycetes(100);o__Saccharomycetales(100);f__Metschnikowiaceae(100);g__Metschnikowia(100);s__Metschnikowia_pulcherrima(100); | Metschnikowia | 13 | Saprotroph | \|Undefined Saprotroph\| | Yeast | NULL | Probable | As nectar-tap_saprotroph (Põlme et al. 2020); Decay sugar-rich substrates (Põlme et al. 2020) | Põlme S, et al. 2020. Fungal Diversity 105:1-16 ((https://doi.org/10.1007/s13225-020-00466-2)) |
| Otu16667 | 1 | k__Fungi(100);p__Ascomycota(100);c__Saccharomycetes(100);o__Saccharomycetales(100);f__Metschnikowiaceae(100);g__Metschnikowia(100);g__Metschnikowia_unclassified(100); | Metschnikowia | 13 | Saprotroph | \|Undefined Saprotroph\| | Yeast | NULL | Probable | As nectar-tap_saprotroph (Põlme et al. 2020); Decay sugar-rich substrates (Põlme et al. 2020) | Põlme S, et al. 2020. Fungal Diversity 105:1-16 ((https://doi.org/10.1007/s13225-020-00466-2)) |
| Otu16668 | 1 | k__Fungi(100);p__Ascomycota(100);c__Saccharomycetes(100);o__Saccharomycetales(100);f__Metschnikowiaceae(100);g__Metschnikowia(100);s__Metschnikowia_pulcherrima(100); | Metschnikowia | 13 | Saprotroph | \|Undefined Saprotroph\| | Yeast | NULL | Probable | As nectar-tap_saprotroph (Põlme et al. 2020); Decay sugar-rich substrates (Põlme et al. 2020) | Põlme S, et al. 2020. Fungal Diversity 105:1-16 ((https://doi.org/10.1007/s13225-020-00466-2)) |
| Otu16669 | 1 | k__Fungi(100);p__Ascomycota(100);c__Saccharomycetes(100);o__Saccharomycetales(100);f__Metschnikowiaceae(100);g__Metschnikowia(100);s__Metschnikowia_pulcherrima(100); | Metschnikowia | 13 | Saprotroph | \|Undefined Saprotroph\| | Yeast | NULL | Probable | As nectar-tap_saprotroph (Põlme et al. 2020); Decay sugar-rich substrates (Põlme et al. 2020) | Põlme S, et al. 2020. Fungal Diversity 105:1-16 ((https://doi.org/10.1007/s13225-020-00466-2)) |
| Otu16670 | 1 | k__Fungi(100);p__Ascomycota(100);c__Saccharomycetes(100);o__Saccharomycetales(100);f__Metschnikowiaceae(100);g__Metschnikowia(100);g__Metschnikowia_unclassified(100); | Metschnikowia | 13 | Saprotroph | \|Undefined Saprotroph\| | Yeast | NULL | Probable | As nectar-tap_saprotroph (Põlme et al. 2020); Decay sugar-rich substrates (Põlme et al. 2020) | Põlme S, et al. 2020. Fungal Diversity 105:1-16 ((https://doi.org/10.1007/s13225-020-00466-2)) |
| Otu16671 | 1 | k__Fungi(100);p__Ascomycota(100);c__Saccharomycetes(100);o__Saccharomycetales(100);f__Metschnikowiaceae(100);g__Metschnikowia(100);s__Metschnikowia_pulcherrima(100); | Metschnikowia | 13 | Saprotroph | \|Undefined Saprotroph\| | Yeast | NULL | Probable | As nectar-tap_saprotroph (Põlme et al. 2020); Decay sugar-rich substrates (Põlme et al. 2020) | Põlme S, et al. 2020. Fungal Diversity 105:1-16 ((https://doi.org/10.1007/s13225-020-00466-2)) |
| Otu16672 | 1 | k__Fungi(100);p__Ascomycota(100);c__Saccharomycetes(100);o__Saccharomycetales(100);f__Metschnikowiaceae(100);g__Metschnikowia(100);g__Metschnikowia_unclassified(100); | Metschnikowia | 13 | Saprotroph | \|Undefined Saprotroph\| | Yeast | NULL | Probable | As nectar-tap_saprotroph (Põlme et al. 2020); Decay sugar-rich substrates (Põlme et al. 2020) | Põlme S, et al. 2020. Fungal Diversity 105:1-16 ((https://doi.org/10.1007/s13225-020-00466-2)) |
| Otu16673 | 1 | k__Fungi(100);p__Ascomycota(100);c__Saccharomycetes(100);o__Saccharomycetales(100);f__Metschnikowiaceae(100);g__Metschnikowia(100);s__Metschnikowia_pulcherrima(100); | Metschnikowia | 13 | Saprotroph | \|Undefined Saprotroph\| | Yeast | NULL | Probable | As nectar-tap_saprotroph (Põlme et al. 2020); Decay sugar-rich substrates (Põlme et al. 2020) | Põlme S, et al. 2020. Fungal Diversity 105:1-16 ((https://doi.org/10.1007/s13225-020-00466-2)) |
| Otu16674 | 1 | k__Fungi(100);p__Basidiomycota(100);c__Microbotryomycetes(100);o__Sporidiobolales(100);f__Sporidiobolales_family_Incertae_sedis(100);g__Sporobolomyces(100);s__Sporobolomyces_lactosus(100); | Sporobolomyces | 13 | Pathotroph-Saprotroph | \|Fungal Parasite\|-Plant Saprotroph | Yeast | NULL | Probable | As litter_saprotroph (Põlme et al. 2020); As mycoparasite (Põlme et al. 2020); Decay fungal material (Põlme et al. 2020) | Tedersoo L, et al. 2014. Science 346:e1256688 ((https://doi.org/10.1126/science.1256688)); Purahong W, et al. 2016. Molecular Ecology 25:4059-4074 ((https://doi.org/10.1111/mec.13739)); Põlme S, et al. 2020. Fungal Diversity 105:1-16 ((https://doi.org/10.1007/s13225-020-00466-2)) |
| Otu16675 | 1 | k__Fungi(100);p__Ascomycota(100);c__Saccharomycetes(100);o__Saccharomycetales(100);f__Metschnikowiaceae(100);g__Metschnikowia(100);s__Metschnikowia_pulcherrima(100); | Metschnikowia | 13 | Saprotroph | \|Undefined Saprotroph\| | Yeast | NULL | Probable | As nectar-tap_saprotroph (Põlme et al. 2020); Decay sugar-rich substrates (Põlme et al. 2020) | Põlme S, et al. 2020. Fungal Diversity 105:1-16 ((https://doi.org/10.1007/s13225-020-00466-2)) |
| Otu16676 | 1 | k__Fungi(100);p__Ascomycota(100);c__Saccharomycetes(100);o__Saccharomycetales(100);f__Metschnikowiaceae(100);g__Metschnikowia(100);g__Metschnikowia_unclassified(100); | Metschnikowia | 13 | Saprotroph | \|Undefined Saprotroph\| | Yeast | NULL | Probable | As nectar-tap_saprotroph (Põlme et al. 2020); Decay sugar-rich substrates (Põlme et al. 2020) | Põlme S, et al. 2020. Fungal Diversity 105:1-16 ((https://doi.org/10.1007/s13225-020-00466-2)) |
| Otu16677 | 1 | k__Fungi(100);p__Ascomycota(100);c__Saccharomycetes(100);o__Saccharomycetales(100);f__Metschnikowiaceae(100);g__Metschnikowia(100);s__Metschnikowia_pulcherrima(100); | Metschnikowia | 13 | Saprotroph | \|Undefined Saprotroph\| | Yeast | NULL | Probable | As nectar-tap_saprotroph (Põlme et al. 2020); Decay sugar-rich substrates (Põlme et al. 2020) | Põlme S, et al. 2020. Fungal Diversity 105:1-16 ((https://doi.org/10.1007/s13225-020-00466-2)) |
| Otu16678 | 1 | k__Fungi(100);p__Ascomycota(100);c__Saccharomycetes(100);o__Saccharomycetales(100);f__Metschnikowiaceae(100);g__Metschnikowia(100);s__Metschnikowia_pulcherrima(100); | Metschnikowia | 13 | Saprotroph | \|Undefined Saprotroph\| | Yeast | NULL | Probable | As nectar-tap_saprotroph (Põlme et al. 2020); Decay sugar-rich substrates (Põlme et al. 2020) | Põlme S, et al. 2020. Fungal Diversity 105:1-16 ((https://doi.org/10.1007/s13225-020-00466-2)) |
| Otu16679 | 1 | k__Fungi(100);p__Ascomycota(100);c__Saccharomycetes(100);o__Saccharomycetales(100);f__Metschnikowiaceae(100);g__Metschnikowia(100);s__Metschnikowia_pulcherrima(100); | Metschnikowia | 13 | Saprotroph | \|Undefined Saprotroph\| | Yeast | NULL | Probable | As nectar-tap_saprotroph (Põlme et al. 2020); Decay sugar-rich substrates (Põlme et al. 2020) | Põlme S, et al. 2020. Fungal Diversity 105:1-16 ((https://doi.org/10.1007/s13225-020-00466-2)) |
| Otu16680 | 1 | k__Fungi(100);p__Ascomycota(100);c__Saccharomycetes(100);o__Saccharomycetales(100);f__Metschnikowiaceae(100);g__Metschnikowia(100);g__Metschnikowia_unclassified(100); | Metschnikowia | 13 | Saprotroph | \|Undefined Saprotroph\| | Yeast | NULL | Probable | As nectar-tap_saprotroph (Põlme et al. 2020); Decay sugar-rich substrates (Põlme et al. 2020) | Põlme S, et al. 2020. Fungal Diversity 105:1-16 ((https://doi.org/10.1007/s13225-020-00466-2)) |
| Otu16681 | 1 | k__Fungi(100);p__Ascomycota(100);c__Saccharomycetes(100);o__Saccharomycetales(100);f__Metschnikowiaceae(100);g__Metschnikowia(100);s__Metschnikowia_pulcherrima(100); | Metschnikowia | 13 | Saprotroph | \|Undefined Saprotroph\| | Yeast | NULL | Probable | As nectar-tap_saprotroph (Põlme et al. 2020); Decay sugar-rich substrates (Põlme et al. 2020) | Põlme S, et al. 2020. Fungal Diversity 105:1-16 ((https://doi.org/10.1007/s13225-020-00466-2)) |
| Otu16682 | 1 | k__Fungi(100);p__Ascomycota(100);c__Saccharomycetes(100);o__Saccharomycetales(100);f__Metschnikowiaceae(100);g__Metschnikowia(100);s__Metschnikowia_pulcherrima(100); | Metschnikowia | 13 | Saprotroph | \|Undefined Saprotroph\| | Yeast | NULL | Probable | As nectar-tap_saprotroph (Põlme et al. 2020); Decay sugar-rich substrates (Põlme et al. 2020) | Põlme S, et al. 2020. Fungal Diversity 105:1-16 ((https://doi.org/10.1007/s13225-020-00466-2)) |
| Otu16683 | 1 | k__Fungi(100);p__Ascomycota(100);c__Saccharomycetes(100);o__Saccharomycetales(100);f__Metschnikowiaceae(100);g__Metschnikowia(100);g__Metschnikowia_unclassified(100); | Metschnikowia | 13 | Saprotroph | \|Undefined Saprotroph\| | Yeast | NULL | Probable | As nectar-tap_saprotroph (Põlme et al. 2020); Decay sugar-rich substrates (Põlme et al. 2020) | Põlme S, et al. 2020. Fungal Diversity 105:1-16 ((https://doi.org/10.1007/s13225-020-00466-2)) |
| Otu16684 | 1 | k__Fungi(100);p__Ascomycota(100);c__Saccharomycetes(100);o__Saccharomycetales(100);f__Metschnikowiaceae(100);g__Metschnikowia(100);g__Metschnikowia_unclassified(100); | Metschnikowia | 13 | Saprotroph | \|Undefined Saprotroph\| | Yeast | NULL | Probable | As nectar-tap_saprotroph (Põlme et al. 2020); Decay sugar-rich substrates (Põlme et al. 2020) | Põlme S, et al. 2020. Fungal Diversity 105:1-16 ((https://doi.org/10.1007/s13225-020-00466-2)) |
| Otu16685 | 1 | k__Fungi(100);p__Ascomycota(100);c__Saccharomycetes(100);o__Saccharomycetales(100);f__Metschnikowiaceae(100);g__Metschnikowia(100);s__Metschnikowia_pulcherrima(100); | Metschnikowia | 13 | Saprotroph | \|Undefined Saprotroph\| | Yeast | NULL | Probable | As nectar-tap_saprotroph (Põlme et al. 2020); Decay sugar-rich substrates (Põlme et al. 2020) | Põlme S, et al. 2020. Fungal Diversity 105:1-16 ((https://doi.org/10.1007/s13225-020-00466-2)) |
| Otu16686 | 1 | k__Fungi(100);p__Ascomycota(100);c__Saccharomycetes(100);o__Saccharomycetales(100);f__Metschnikowiaceae(100);g__Metschnikowia(100);s__Metschnikowia_pulcherrima(100); | Metschnikowia | 13 | Saprotroph | \|Undefined Saprotroph\| | Yeast | NULL | Probable | As nectar-tap_saprotroph (Põlme et al. 2020); Decay sugar-rich substrates (Põlme et al. 2020) | Põlme S, et al. 2020. Fungal Diversity 105:1-16 ((https://doi.org/10.1007/s13225-020-00466-2)) |
| Otu16687 | 1 | k__Fungi(100);p__Ascomycota(100);c__Saccharomycetes(100);o__Saccharomycetales(100);f__Metschnikowiaceae(100);g__Metschnikowia(100);s__Metschnikowia_pulcherrima(100); | Metschnikowia | 13 | Saprotroph | \|Undefined Saprotroph\| | Yeast | NULL | Probable | As nectar-tap_saprotroph (Põlme et al. 2020); Decay sugar-rich substrates (Põlme et al. 2020) | Põlme S, et al. 2020. Fungal Diversity 105:1-16 ((https://doi.org/10.1007/s13225-020-00466-2)) |
| Otu16688 | 1 | k__Fungi(100);p__Ascomycota(100);c__Saccharomycetes(100);o__Saccharomycetales(100);f__Metschnikowiaceae(100);g__Metschnikowia(100);s__Metschnikowia_pulcherrima(100); | Metschnikowia | 13 | Saprotroph | \|Undefined Saprotroph\| | Yeast | NULL | Probable | As nectar-tap_saprotroph (Põlme et al. 2020); Decay sugar-rich substrates (Põlme et al. 2020) | Põlme S, et al. 2020. Fungal Diversity 105:1-16 ((https://doi.org/10.1007/s13225-020-00466-2)) |
| Otu16689 | 1 | k__Fungi(100);p__Ascomycota(100);c__Saccharomycetes(100);o__Saccharomycetales(100);f__Metschnikowiaceae(100);g__Metschnikowia(100);g__Metschnikowia_unclassified(100); | Metschnikowia | 13 | Saprotroph | \|Undefined Saprotroph\| | Yeast | NULL | Probable | As nectar-tap_saprotroph (Põlme et al. 2020); Decay sugar-rich substrates (Põlme et al. 2020) | Põlme S, et al. 2020. Fungal Diversity 105:1-16 ((https://doi.org/10.1007/s13225-020-00466-2)) |
| Otu16690 | 1 | k__Fungi(100);p__Ascomycota(100);c__Saccharomycetes(100);o__Saccharomycetales(100);f__Metschnikowiaceae(100);g__Metschnikowia(100);s__Metschnikowia_pulcherrima(100); | Metschnikowia | 13 | Saprotroph | \|Undefined Saprotroph\| | Yeast | NULL | Probable | As nectar-tap_saprotroph (Põlme et al. 2020); Decay sugar-rich substrates (Põlme et al. 2020) | Põlme S, et al. 2020. Fungal Diversity 105:1-16 ((https://doi.org/10.1007/s13225-020-00466-2)) |
| Otu16691 | 1 | k__Fungi(100);p__Ascomycota(100);c__Saccharomycetes(100);o__Saccharomycetales(100);f__Metschnikowiaceae(100);g__Metschnikowia(100);g__Metschnikowia_unclassified(100); | Metschnikowia | 13 | Saprotroph | \|Undefined Saprotroph\| | Yeast | NULL | Probable | As nectar-tap_saprotroph (Põlme et al. 2020); Decay sugar-rich substrates (Põlme et al. 2020) | Põlme S, et al. 2020. Fungal Diversity 105:1-16 ((https://doi.org/10.1007/s13225-020-00466-2)) |
| Otu16692 | 1 | k__Fungi(100);p__Ascomycota(100);c__Saccharomycetes(100);o__Saccharomycetales(100);f__Metschnikowiaceae(100);g__Metschnikowia(100);s__Metschnikowia_pulcherrima(100); | Metschnikowia | 13 | Saprotroph | \|Undefined Saprotroph\| | Yeast | NULL | Probable | As nectar-tap_saprotroph (Põlme et al. 2020); Decay sugar-rich substrates (Põlme et al. 2020) | Põlme S, et al. 2020. Fungal Diversity 105:1-16 ((https://doi.org/10.1007/s13225-020-00466-2)) |
| Otu16693 | 1 | k__Fungi(100);p__Ascomycota(100);c__Saccharomycetes(100);o__Saccharomycetales(100);f__Metschnikowiaceae(100);g__Metschnikowia(100);s__Metschnikowia_pulcherrima(100); | Metschnikowia | 13 | Saprotroph | \|Undefined Saprotroph\| | Yeast | NULL | Probable | As nectar-tap_saprotroph (Põlme et al. 2020); Decay sugar-rich substrates (Põlme et al. 2020) | Põlme S, et al. 2020. Fungal Diversity 105:1-16 ((https://doi.org/10.1007/s13225-020-00466-2)) |
| Otu16694 | 1 | k__Fungi(100);p__Ascomycota(100);c__Saccharomycetes(100);o__Saccharomycetales(100);f__Metschnikowiaceae(100);g__Metschnikowia(100);s__Metschnikowia_pulcherrima(100); | Metschnikowia | 13 | Saprotroph | \|Undefined Saprotroph\| | Yeast | NULL | Probable | As nectar-tap_saprotroph (Põlme et al. 2020); Decay sugar-rich substrates (Põlme et al. 2020) | Põlme S, et al. 2020. Fungal Diversity 105:1-16 ((https://doi.org/10.1007/s13225-020-00466-2)) |
| Otu16695 | 1 | k__Fungi(100);p__Ascomycota(100);c__Saccharomycetes(100);o__Saccharomycetales(100);f__Metschnikowiaceae(100);g__Metschnikowia(100);s__Metschnikowia_pulcherrima(100); | Metschnikowia | 13 | Saprotroph | \|Undefined Saprotroph\| | Yeast | NULL | Probable | As nectar-tap_saprotroph (Põlme et al. 2020); Decay sugar-rich substrates (Põlme et al. 2020) | Põlme S, et al. 2020. Fungal Diversity 105:1-16 ((https://doi.org/10.1007/s13225-020-00466-2)) |
| Otu16696 | 1 | k__Fungi(100);p__Ascomycota(100);c__Saccharomycetes(100);o__Saccharomycetales(100);f__Metschnikowiaceae(100);g__Metschnikowia(100);s__Metschnikowia_pulcherrima(100); | Metschnikowia | 13 | Saprotroph | \|Undefined Saprotroph\| | Yeast | NULL | Probable | As nectar-tap_saprotroph (Põlme et al. 2020); Decay sugar-rich substrates (Põlme et al. 2020) | Põlme S, et al. 2020. Fungal Diversity 105:1-16 ((https://doi.org/10.1007/s13225-020-00466-2)) |
| Otu16697 | 1 | k__Fungi(100);p__Ascomycota(100);c__Saccharomycetes(100);o__Saccharomycetales(100);f__Metschnikowiaceae(100);g__Metschnikowia(100);g__Metschnikowia_unclassified(100); | Metschnikowia | 13 | Saprotroph | \|Undefined Saprotroph\| | Yeast | NULL | Probable | As nectar-tap_saprotroph (Põlme et al. 2020); Decay sugar-rich substrates (Põlme et al. 2020) | Põlme S, et al. 2020. Fungal Diversity 105:1-16 ((https://doi.org/10.1007/s13225-020-00466-2)) |
| Otu16698 | 1 | k__Fungi(100);p__Ascomycota(100);c__Saccharomycetes(100);o__Saccharomycetales(100);f__Metschnikowiaceae(100);g__Metschnikowia(100);s__Metschnikowia_pulcherrima(100); | Metschnikowia | 13 | Saprotroph | \|Undefined Saprotroph\| | Yeast | NULL | Probable | As nectar-tap_saprotroph (Põlme et al. 2020); Decay sugar-rich substrates (Põlme et al. 2020) | Põlme S, et al. 2020. Fungal Diversity 105:1-16 ((https://doi.org/10.1007/s13225-020-00466-2)) |
| Otu16699 | 1 | k__Fungi(100);p__Ascomycota(100);c__Saccharomycetes(100);o__Saccharomycetales(100);f__Metschnikowiaceae(100);g__Metschnikowia(100);s__Metschnikowia_pulcherrima(100); | Metschnikowia | 13 | Saprotroph | \|Undefined Saprotroph\| | Yeast | NULL | Probable | As nectar-tap_saprotroph (Põlme et al. 2020); Decay sugar-rich substrates (Põlme et al. 2020) | Põlme S, et al. 2020. Fungal Diversity 105:1-16 ((https://doi.org/10.1007/s13225-020-00466-2)) |
| Otu16700 | 1 | k__Fungi(100);p__Ascomycota(100);c__Saccharomycetes(100);o__Saccharomycetales(100);f__Metschnikowiaceae(100);g__Metschnikowia(100);s__Metschnikowia_pulcherrima(100); | Metschnikowia | 13 | Saprotroph | \|Undefined Saprotroph\| | Yeast | NULL | Probable | As nectar-tap_saprotroph (Põlme et al. 2020); Decay sugar-rich substrates (Põlme et al. 2020) | Põlme S, et al. 2020. Fungal Diversity 105:1-16 ((https://doi.org/10.1007/s13225-020-00466-2)) |
| Otu16701 | 1 | k__Fungi(100);p__Ascomycota(100);c__Saccharomycetes(100);o__Saccharomycetales(100);f__Metschnikowiaceae(100);g__Metschnikowia(100);s__Metschnikowia_pulcherrima(100); | Metschnikowia | 13 | Saprotroph | \|Undefined Saprotroph\| | Yeast | NULL | Probable | As nectar-tap_saprotroph (Põlme et al. 2020); Decay sugar-rich substrates (Põlme et al. 2020) | Põlme S, et al. 2020. Fungal Diversity 105:1-16 ((https://doi.org/10.1007/s13225-020-00466-2)) |
| Otu16702 | 1 | k__Fungi(100);p__Ascomycota(100);c__Saccharomycetes(100);o__Saccharomycetales(100);f__Metschnikowiaceae(100);g__Metschnikowia(100);s__Metschnikowia_pulcherrima(100); | Metschnikowia | 13 | Saprotroph | \|Undefined Saprotroph\| | Yeast | NULL | Probable | As nectar-tap_saprotroph (Põlme et al. 2020); Decay sugar-rich substrates (Põlme et al. 2020) | Põlme S, et al. 2020. Fungal Diversity 105:1-16 ((https://doi.org/10.1007/s13225-020-00466-2)) |
| Otu16703 | 1 | k__Fungi(100);p__Ascomycota(100);c__Saccharomycetes(100);o__Saccharomycetales(100);f__Metschnikowiaceae(100);g__Metschnikowia(100);g__Metschnikowia_unclassified(100); | Metschnikowia | 13 | Saprotroph | \|Undefined Saprotroph\| | Yeast | NULL | Probable | As nectar-tap_saprotroph (Põlme et al. 2020); Decay sugar-rich substrates (Põlme et al. 2020) | Põlme S, et al. 2020. Fungal Diversity 105:1-16 ((https://doi.org/10.1007/s13225-020-00466-2)) |
| Otu16704 | 1 | k__Fungi(100);p__Ascomycota(100);c__Saccharomycetes(100);o__Saccharomycetales(100);f__Metschnikowiaceae(100);g__Metschnikowia(100);s__Metschnikowia_pulcherrima(100); | Metschnikowia | 13 | Saprotroph | \|Undefined Saprotroph\| | Yeast | NULL | Probable | As nectar-tap_saprotroph (Põlme et al. 2020); Decay sugar-rich substrates (Põlme et al. 2020) | Põlme S, et al. 2020. Fungal Diversity 105:1-16 ((https://doi.org/10.1007/s13225-020-00466-2)) |
| Otu16705 | 1 | k__Fungi(100);p__Ascomycota(100);c__Saccharomycetes(100);o__Saccharomycetales(100);f__Metschnikowiaceae(100);g__Metschnikowia(100);s__Metschnikowia_pulcherrima(100); | Metschnikowia | 13 | Saprotroph | \|Undefined Saprotroph\| | Yeast | NULL | Probable | As nectar-tap_saprotroph (Põlme et al. 2020); Decay sugar-rich substrates (Põlme et al. 2020) | Põlme S, et al. 2020. Fungal Diversity 105:1-16 ((https://doi.org/10.1007/s13225-020-00466-2)) |
| Otu16706 | 1 | k__Fungi(100);p__Ascomycota(100);c__Saccharomycetes(100);o__Saccharomycetales(100);f__Metschnikowiaceae(100);g__Metschnikowia(100);g__Metschnikowia_unclassified(100); | Metschnikowia | 13 | Saprotroph | \|Undefined Saprotroph\| | Yeast | NULL | Probable | As nectar-tap_saprotroph (Põlme et al. 2020); Decay sugar-rich substrates (Põlme et al. 2020) | Põlme S, et al. 2020. Fungal Diversity 105:1-16 ((https://doi.org/10.1007/s13225-020-00466-2)) |
| Otu16707 | 1 | k__Fungi(100);p__Ascomycota(100);c__Saccharomycetes(100);o__Saccharomycetales(100);f__Metschnikowiaceae(100);g__Metschnikowia(100);s__Metschnikowia_pulcherrima(100); | Metschnikowia | 13 | Saprotroph | \|Undefined Saprotroph\| | Yeast | NULL | Probable | As nectar-tap_saprotroph (Põlme et al. 2020); Decay sugar-rich substrates (Põlme et al. 2020) | Põlme S, et al. 2020. Fungal Diversity 105:1-16 ((https://doi.org/10.1007/s13225-020-00466-2)) |
| Otu16708 | 1 | k__Fungi(100);p__Ascomycota(100);c__Saccharomycetes(100);o__Saccharomycetales(100);f__Metschnikowiaceae(100);g__Metschnikowia(100);s__Metschnikowia_pulcherrima(100); | Metschnikowia | 13 | Saprotroph | \|Undefined Saprotroph\| | Yeast | NULL | Probable | As nectar-tap_saprotroph (Põlme et al. 2020); Decay sugar-rich substrates (Põlme et al. 2020) | Põlme S, et al. 2020. Fungal Diversity 105:1-16 ((https://doi.org/10.1007/s13225-020-00466-2)) |
| Otu16711 | 1 | k__Fungi(100);p__Ascomycota(100);c__Saccharomycetes(100);o__Saccharomycetales(100);f__Metschnikowiaceae(100);g__Metschnikowia(100);s__Metschnikowia_pulcherrima(100); | Metschnikowia | 13 | Saprotroph | \|Undefined Saprotroph\| | Yeast | NULL | Probable | As nectar-tap_saprotroph (Põlme et al. 2020); Decay sugar-rich substrates (Põlme et al. 2020) | Põlme S, et al. 2020. Fungal Diversity 105:1-16 ((https://doi.org/10.1007/s13225-020-00466-2)) |
| Otu16714 | 1 | k__Fungi(100);p__Ascomycota(100);c__Saccharomycetes(100);o__Saccharomycetales(100);f__Metschnikowiaceae(100);g__Metschnikowia(100);s__Metschnikowia_pulcherrima(100); | Metschnikowia | 13 | Saprotroph | \|Undefined Saprotroph\| | Yeast | NULL | Probable | As nectar-tap_saprotroph (Põlme et al. 2020); Decay sugar-rich substrates (Põlme et al. 2020) | Põlme S, et al. 2020. Fungal Diversity 105:1-16 ((https://doi.org/10.1007/s13225-020-00466-2)) |
| Otu16715 | 1 | k__Fungi(100);p__Ascomycota(100);c__Saccharomycetes(100);o__Saccharomycetales(100);f__Metschnikowiaceae(100);g__Metschnikowia(100);s__Metschnikowia_pulcherrima(100); | Metschnikowia | 13 | Saprotroph | \|Undefined Saprotroph\| | Yeast | NULL | Probable | As nectar-tap_saprotroph (Põlme et al. 2020); Decay sugar-rich substrates (Põlme et al. 2020) | Põlme S, et al. 2020. Fungal Diversity 105:1-16 ((https://doi.org/10.1007/s13225-020-00466-2)) |
| Otu16716 | 1 | k__Fungi(100);p__Ascomycota(100);c__Saccharomycetes(100);o__Saccharomycetales(100);f__Metschnikowiaceae(100);g__Metschnikowia(100);g__Metschnikowia_unclassified(100); | Metschnikowia | 13 | Saprotroph | \|Undefined Saprotroph\| | Yeast | NULL | Probable | As nectar-tap_saprotroph (Põlme et al. 2020); Decay sugar-rich substrates (Põlme et al. 2020) | Põlme S, et al. 2020. Fungal Diversity 105:1-16 ((https://doi.org/10.1007/s13225-020-00466-2)) |
| Otu16717 | 1 | k__Fungi(100);p__Ascomycota(100);c__Saccharomycetes(100);o__Saccharomycetales(100);f__Metschnikowiaceae(100);g__Metschnikowia(100);g__Metschnikowia_unclassified(100); | Metschnikowia | 13 | Saprotroph | \|Undefined Saprotroph\| | Yeast | NULL | Probable | As nectar-tap_saprotroph (Põlme et al. 2020); Decay sugar-rich substrates (Põlme et al. 2020) | Põlme S, et al. 2020. Fungal Diversity 105:1-16 ((https://doi.org/10.1007/s13225-020-00466-2)) |
| Otu16718 | 1 | k__Fungi(100);p__Ascomycota(100);c__Saccharomycetes(100);o__Saccharomycetales(100);f__Metschnikowiaceae(100);g__Metschnikowia(100);g__Metschnikowia_unclassified(100); | Metschnikowia | 13 | Saprotroph | \|Undefined Saprotroph\| | Yeast | NULL | Probable | As nectar-tap_saprotroph (Põlme et al. 2020); Decay sugar-rich substrates (Põlme et al. 2020) | Põlme S, et al. 2020. Fungal Diversity 105:1-16 ((https://doi.org/10.1007/s13225-020-00466-2)) |
| Otu16719 | 1 | k__Fungi(100);p__Ascomycota(100);c__Saccharomycetes(100);o__Saccharomycetales(100);f__Metschnikowiaceae(100);g__Metschnikowia(100);s__Metschnikowia_pulcherrima(100); | Metschnikowia | 13 | Saprotroph | \|Undefined Saprotroph\| | Yeast | NULL | Probable | As nectar-tap_saprotroph (Põlme et al. 2020); Decay sugar-rich substrates (Põlme et al. 2020) | Põlme S, et al. 2020. Fungal Diversity 105:1-16 ((https://doi.org/10.1007/s13225-020-00466-2)) |
| Otu16720 | 1 | k__Fungi(100);p__Ascomycota(100);c__Saccharomycetes(100);o__Saccharomycetales(100);f__Metschnikowiaceae(100);g__Metschnikowia(100);s__Metschnikowia_pulcherrima(100); | Metschnikowia | 13 | Saprotroph | \|Undefined Saprotroph\| | Yeast | NULL | Probable | As nectar-tap_saprotroph (Põlme et al. 2020); Decay sugar-rich substrates (Põlme et al. 2020) | Põlme S, et al. 2020. Fungal Diversity 105:1-16 ((https://doi.org/10.1007/s13225-020-00466-2)) |
| Otu16721 | 1 | k__Fungi(100);p__Ascomycota(100);c__Saccharomycetes(100);o__Saccharomycetales(100);f__Metschnikowiaceae(100);g__Metschnikowia(100);s__Metschnikowia_pulcherrima(100); | Metschnikowia | 13 | Saprotroph | \|Undefined Saprotroph\| | Yeast | NULL | Probable | As nectar-tap_saprotroph (Põlme et al. 2020); Decay sugar-rich substrates (Põlme et al. 2020) | Põlme S, et al. 2020. Fungal Diversity 105:1-16 ((https://doi.org/10.1007/s13225-020-00466-2)) |
| Otu16722 | 1 | k__Fungi(100);p__Ascomycota(100);c__Saccharomycetes(100);o__Saccharomycetales(100);f__Metschnikowiaceae(100);g__Metschnikowia(100);g__Metschnikowia_unclassified(100); | Metschnikowia | 13 | Saprotroph | \|Undefined Saprotroph\| | Yeast | NULL | Probable | As nectar-tap_saprotroph (Põlme et al. 2020); Decay sugar-rich substrates (Põlme et al. 2020) | Põlme S, et al. 2020. Fungal Diversity 105:1-16 ((https://doi.org/10.1007/s13225-020-00466-2)) |
| Otu16723 | 1 | k__Fungi(100);p__Ascomycota(100);c__Saccharomycetes(100);o__Saccharomycetales(100);f__Metschnikowiaceae(100);g__Metschnikowia(100);s__Metschnikowia_pulcherrima(100); | Metschnikowia | 13 | Saprotroph | \|Undefined Saprotroph\| | Yeast | NULL | Probable | As nectar-tap_saprotroph (Põlme et al. 2020); Decay sugar-rich substrates (Põlme et al. 2020) | Põlme S, et al. 2020. Fungal Diversity 105:1-16 ((https://doi.org/10.1007/s13225-020-00466-2)) |
| Otu16724 | 1 | k__Fungi(100);p__Ascomycota(100);c__Saccharomycetes(100);o__Saccharomycetales(100);f__Metschnikowiaceae(100);g__Metschnikowia(100);s__Metschnikowia_pulcherrima(100); | Metschnikowia | 13 | Saprotroph | \|Undefined Saprotroph\| | Yeast | NULL | Probable | As nectar-tap_saprotroph (Põlme et al. 2020); Decay sugar-rich substrates (Põlme et al. 2020) | Põlme S, et al. 2020. Fungal Diversity 105:1-16 ((https://doi.org/10.1007/s13225-020-00466-2)) |
| Otu16725 | 1 | k__Fungi(100);p__Ascomycota(100);c__Saccharomycetes(100);o__Saccharomycetales(100);f__Metschnikowiaceae(100);g__Metschnikowia(100);s__Metschnikowia_pulcherrima(100); | Metschnikowia | 13 | Saprotroph | \|Undefined Saprotroph\| | Yeast | NULL | Probable | As nectar-tap_saprotroph (Põlme et al. 2020); Decay sugar-rich substrates (Põlme et al. 2020) | Põlme S, et al. 2020. Fungal Diversity 105:1-16 ((https://doi.org/10.1007/s13225-020-00466-2)) |
| Otu16726 | 1 | k__Fungi(100);p__Ascomycota(100);c__Saccharomycetes(100);o__Saccharomycetales(100);f__Metschnikowiaceae(100);g__Metschnikowia(100);g__Metschnikowia_unclassified(100); | Metschnikowia | 13 | Saprotroph | \|Undefined Saprotroph\| | Yeast | NULL | Probable | As nectar-tap_saprotroph (Põlme et al. 2020); Decay sugar-rich substrates (Põlme et al. 2020) | Põlme S, et al. 2020. Fungal Diversity 105:1-16 ((https://doi.org/10.1007/s13225-020-00466-2)) |
| Otu16727 | 1 | k__Fungi(100);p__Ascomycota(100);c__Saccharomycetes(100);o__Saccharomycetales(100);f__Metschnikowiaceae(100);g__Metschnikowia(100);s__Metschnikowia_pulcherrima(100); | Metschnikowia | 13 | Saprotroph | \|Undefined Saprotroph\| | Yeast | NULL | Probable | As nectar-tap_saprotroph (Põlme et al. 2020); Decay sugar-rich substrates (Põlme et al. 2020) | Põlme S, et al. 2020. Fungal Diversity 105:1-16 ((https://doi.org/10.1007/s13225-020-00466-2)) |
| Otu16728 | 1 | k__Fungi(100);p__Ascomycota(100);c__Saccharomycetes(100);o__Saccharomycetales(100);f__Metschnikowiaceae(100);g__Metschnikowia(100);s__Metschnikowia_pulcherrima(100); | Metschnikowia | 13 | Saprotroph | \|Undefined Saprotroph\| | Yeast | NULL | Probable | As nectar-tap_saprotroph (Põlme et al. 2020); Decay sugar-rich substrates (Põlme et al. 2020) | Põlme S, et al. 2020. Fungal Diversity 105:1-16 ((https://doi.org/10.1007/s13225-020-00466-2)) |
| Otu16729 | 1 | k__Fungi(100);p__Ascomycota(100);c__Saccharomycetes(100);o__Saccharomycetales(100);f__Metschnikowiaceae(100);g__Metschnikowia(100);g__Metschnikowia_unclassified(100); | Metschnikowia | 13 | Saprotroph | \|Undefined Saprotroph\| | Yeast | NULL | Probable | As nectar-tap_saprotroph (Põlme et al. 2020); Decay sugar-rich substrates (Põlme et al. 2020) | Põlme S, et al. 2020. Fungal Diversity 105:1-16 ((https://doi.org/10.1007/s13225-020-00466-2)) |
| Otu16730 | 1 | k__Fungi(100);p__Ascomycota(100);c__Saccharomycetes(100);o__Saccharomycetales(100);f__Metschnikowiaceae(100);g__Metschnikowia(100);s__Metschnikowia_pulcherrima(100); | Metschnikowia | 13 | Saprotroph | \|Undefined Saprotroph\| | Yeast | NULL | Probable | As nectar-tap_saprotroph (Põlme et al. 2020); Decay sugar-rich substrates (Põlme et al. 2020) | Põlme S, et al. 2020. Fungal Diversity 105:1-16 ((https://doi.org/10.1007/s13225-020-00466-2)) |
| Otu16731 | 1 | k__Fungi(100);p__Ascomycota(100);c__Saccharomycetes(100);o__Saccharomycetales(100);f__Metschnikowiaceae(100);g__Metschnikowia(100);g__Metschnikowia_unclassified(100); | Metschnikowia | 13 | Saprotroph | \|Undefined Saprotroph\| | Yeast | NULL | Probable | As nectar-tap_saprotroph (Põlme et al. 2020); Decay sugar-rich substrates (Põlme et al. 2020) | Põlme S, et al. 2020. Fungal Diversity 105:1-16 ((https://doi.org/10.1007/s13225-020-00466-2)) |
| Otu16732 | 1 | k__Fungi(100);p__Ascomycota(100);c__Saccharomycetes(100);o__Saccharomycetales(100);f__Metschnikowiaceae(100);g__Metschnikowia(100);s__Metschnikowia_pulcherrima(100); | Metschnikowia | 13 | Saprotroph | \|Undefined Saprotroph\| | Yeast | NULL | Probable | As nectar-tap_saprotroph (Põlme et al. 2020); Decay sugar-rich substrates (Põlme et al. 2020) | Põlme S, et al. 2020. Fungal Diversity 105:1-16 ((https://doi.org/10.1007/s13225-020-00466-2)) |
| Otu16733 | 1 | k__Fungi(100);p__Ascomycota(100);c__Saccharomycetes(100);o__Saccharomycetales(100);f__Metschnikowiaceae(100);g__Metschnikowia(100);s__Metschnikowia_pulcherrima(100); | Metschnikowia | 13 | Saprotroph | \|Undefined Saprotroph\| | Yeast | NULL | Probable | As nectar-tap_saprotroph (Põlme et al. 2020); Decay sugar-rich substrates (Põlme et al. 2020) | Põlme S, et al. 2020. Fungal Diversity 105:1-16 ((https://doi.org/10.1007/s13225-020-00466-2)) |
| Otu16736 | 1 | k__Fungi(100);p__Ascomycota(100);c__Saccharomycetes(100);o__Saccharomycetales(100);f__Metschnikowiaceae(100);g__Metschnikowia(100);s__Metschnikowia_pulcherrima(100); | Metschnikowia | 13 | Saprotroph | \|Undefined Saprotroph\| | Yeast | NULL | Probable | As nectar-tap_saprotroph (Põlme et al. 2020); Decay sugar-rich substrates (Põlme et al. 2020) | Põlme S, et al. 2020. Fungal Diversity 105:1-16 ((https://doi.org/10.1007/s13225-020-00466-2)) |
| Otu16737 | 1 | k__Fungi(100);p__Ascomycota(100);c__Saccharomycetes(100);o__Saccharomycetales(100);f__Metschnikowiaceae(100);g__Metschnikowia(100);s__Metschnikowia_pulcherrima(100); | Metschnikowia | 13 | Saprotroph | \|Undefined Saprotroph\| | Yeast | NULL | Probable | As nectar-tap_saprotroph (Põlme et al. 2020); Decay sugar-rich substrates (Põlme et al. 2020) | Põlme S, et al. 2020. Fungal Diversity 105:1-16 ((https://doi.org/10.1007/s13225-020-00466-2)) |
| Otu16738 | 1 | k__Fungi(100);p__Ascomycota(100);c__Saccharomycetes(100);o__Saccharomycetales(100);f__Metschnikowiaceae(100);g__Metschnikowia(100);g__Metschnikowia_unclassified(100); | Metschnikowia | 13 | Saprotroph | \|Undefined Saprotroph\| | Yeast | NULL | Probable | As nectar-tap_saprotroph (Põlme et al. 2020); Decay sugar-rich substrates (Põlme et al. 2020) | Põlme S, et al. 2020. Fungal Diversity 105:1-16 ((https://doi.org/10.1007/s13225-020-00466-2)) |
| Otu16739 | 1 | k__Fungi(100);p__Ascomycota(100);c__Saccharomycetes(100);o__Saccharomycetales(100);f__Metschnikowiaceae(100);g__Metschnikowia(100);s__Metschnikowia_pulcherrima(100); | Metschnikowia | 13 | Saprotroph | \|Undefined Saprotroph\| | Yeast | NULL | Probable | As nectar-tap_saprotroph (Põlme et al. 2020); Decay sugar-rich substrates (Põlme et al. 2020) | Põlme S, et al. 2020. Fungal Diversity 105:1-16 ((https://doi.org/10.1007/s13225-020-00466-2)) |
| Otu16740 | 1 | k__Fungi(100);p__Ascomycota(100);c__Saccharomycetes(100);o__Saccharomycetales(100);f__Metschnikowiaceae(100);g__Metschnikowia(100);s__Metschnikowia_pulcherrima(100); | Metschnikowia | 13 | Saprotroph | \|Undefined Saprotroph\| | Yeast | NULL | Probable | As nectar-tap_saprotroph (Põlme et al. 2020); Decay sugar-rich substrates (Põlme et al. 2020) | Põlme S, et al. 2020. Fungal Diversity 105:1-16 ((https://doi.org/10.1007/s13225-020-00466-2)) |
| Otu16742 | 1 | k__Fungi(100);p__Ascomycota(100);c__Saccharomycetes(100);o__Saccharomycetales(100);f__Metschnikowiaceae(100);g__Metschnikowia(100);s__Metschnikowia_pulcherrima(100); | Metschnikowia | 13 | Saprotroph | \|Undefined Saprotroph\| | Yeast | NULL | Probable | As nectar-tap_saprotroph (Põlme et al. 2020); Decay sugar-rich substrates (Põlme et al. 2020) | Põlme S, et al. 2020. Fungal Diversity 105:1-16 ((https://doi.org/10.1007/s13225-020-00466-2)) |
| Otu16753 | 1 | k__Fungi(100);p__Ascomycota(100);c__Saccharomycetes(100);o__Saccharomycetales(100);f__Metschnikowiaceae(100);g__Metschnikowia(100);g__Metschnikowia_unclassified(100); | Metschnikowia | 13 | Saprotroph | \|Undefined Saprotroph\| | Yeast | NULL | Probable | As nectar-tap_saprotroph (Põlme et al. 2020); Decay sugar-rich substrates (Põlme et al. 2020) | Põlme S, et al. 2020. Fungal Diversity 105:1-16 ((https://doi.org/10.1007/s13225-020-00466-2)) |
| Otu16754 | 1 | k__Fungi(100);p__Ascomycota(100);c__Saccharomycetes(100);o__Saccharomycetales(100);f__Metschnikowiaceae(100);g__Metschnikowia(100);g__Metschnikowia_unclassified(100); | Metschnikowia | 13 | Saprotroph | \|Undefined Saprotroph\| | Yeast | NULL | Probable | As nectar-tap_saprotroph (Põlme et al. 2020); Decay sugar-rich substrates (Põlme et al. 2020) | Põlme S, et al. 2020. Fungal Diversity 105:1-16 ((https://doi.org/10.1007/s13225-020-00466-2)) |
| Otu16756 | 1 | k__Fungi(100);p__Ascomycota(100);c__Saccharomycetes(100);o__Saccharomycetales(100);f__Metschnikowiaceae(100);g__Metschnikowia(100);s__Metschnikowia_pulcherrima(100); | Metschnikowia | 13 | Saprotroph | \|Undefined Saprotroph\| | Yeast | NULL | Probable | As nectar-tap_saprotroph (Põlme et al. 2020); Decay sugar-rich substrates (Põlme et al. 2020) | Põlme S, et al. 2020. Fungal Diversity 105:1-16 ((https://doi.org/10.1007/s13225-020-00466-2)) |
| Otu16758 | 1 | k__Fungi(100);p__Ascomycota(100);c__Saccharomycetes(100);o__Saccharomycetales(100);f__Metschnikowiaceae(100);g__Metschnikowia(100);s__Metschnikowia_pulcherrima(100); | Metschnikowia | 13 | Saprotroph | \|Undefined Saprotroph\| | Yeast | NULL | Probable | As nectar-tap_saprotroph (Põlme et al. 2020); Decay sugar-rich substrates (Põlme et al. 2020) | Põlme S, et al. 2020. Fungal Diversity 105:1-16 ((https://doi.org/10.1007/s13225-020-00466-2)) |
| Otu16760 | 1 | k__Fungi(100);p__Ascomycota(100);c__Saccharomycetes(100);o__Saccharomycetales(100);f__Metschnikowiaceae(100);g__Metschnikowia(100);s__Metschnikowia_pulcherrima(100); | Metschnikowia | 13 | Saprotroph | \|Undefined Saprotroph\| | Yeast | NULL | Probable | As nectar-tap_saprotroph (Põlme et al. 2020); Decay sugar-rich substrates (Põlme et al. 2020) | Põlme S, et al. 2020. Fungal Diversity 105:1-16 ((https://doi.org/10.1007/s13225-020-00466-2)) |
| Otu16761 | 1 | k__Fungi(100);p__Ascomycota(100);c__Saccharomycetes(100);o__Saccharomycetales(100);f__Metschnikowiaceae(100);g__Metschnikowia(100);g__Metschnikowia_unclassified(100); | Metschnikowia | 13 | Saprotroph | \|Undefined Saprotroph\| | Yeast | NULL | Probable | As nectar-tap_saprotroph (Põlme et al. 2020); Decay sugar-rich substrates (Põlme et al. 2020) | Põlme S, et al. 2020. Fungal Diversity 105:1-16 ((https://doi.org/10.1007/s13225-020-00466-2)) |

| | | | | | | | | | | | |
|---|---|---|---|---|---|---|---|---|---|---|---|
| Otu16762 | 1 | k__Fungi(100);p__Ascomycota(100);c__Saccharomycetes(100);o__Saccharomycetales(100);f__Metschnikowiaceae(100);g__Metschnikowia(100);g__Metschnikowia_unclassified(100); | Metschnikowia | 13 | Saprotroph | \|Undefined Saprotroph\| | Yeast | NULL | Probable | As nectar-tap_saprotroph (Põlme et al. 2020); Decay sugar-rich substrates (Põlme et al. 2020) | Põlme S, et al. 2020. Fungal Diversity 105:1-16 ((https://doi.org/10.1007/s13225-020-00466-2)) |
| Otu16763 | 1 | k__Fungi(100);p__Ascomycota(100);c__Saccharomycetes(100);o__Saccharomycetales(100);f__Metschnikowiaceae(100);g__Metschnikowia(100);s__Metschnikowia_pulcherrima(100); | Metschnikowia | 13 | Saprotroph | \|Undefined Saprotroph\| | Yeast | NULL | Probable | As nectar-tap_saprotroph (Põlme et al. 2020); Decay sugar-rich substrates (Põlme et al. 2020) | Põlme S, et al. 2020. Fungal Diversity 105:1-16 ((https://doi.org/10.1007/s13225-020-00466-2)) |
| Otu16764 | 1 | k__Fungi(100);p__Ascomycota(100);c__Saccharomycetes(100);o__Saccharomycetales(100);f__Metschnikowiaceae(100);g__Metschnikowia(100);s__Metschnikowia_pulcherrima(100); | Metschnikowia | 13 | Saprotroph | \|Undefined Saprotroph\| | Yeast | NULL | Probable | As nectar-tap_saprotroph (Põlme et al. 2020); Decay sugar-rich substrates (Põlme et al. 2020) | Põlme S, et al. 2020. Fungal Diversity 105:1-16 ((https://doi.org/10.1007/s13225-020-00466-2)) |
| Otu16765 | 1 | k__Fungi(100);p__Ascomycota(100);c__Saccharomycetes(100);o__Saccharomycetales(100);f__Metschnikowiaceae(100);g__Metschnikowia(100);g__Metschnikowia_unclassified(100); | Metschnikowia | 13 | Saprotroph | \|Undefined Saprotroph\| | Yeast | NULL | Probable | As nectar-tap_saprotroph (Põlme et al. 2020); Decay sugar-rich substrates (Põlme et al. 2020) | Põlme S, et al. 2020. Fungal Diversity 105:1-16 ((https://doi.org/10.1007/s13225-020-00466-2)) |
| Otu16766 | 1 | k__Fungi(100);p__Ascomycota(100);c__Saccharomycetes(100);o__Saccharomycetales(100);f__Metschnikowiaceae(100);g__Metschnikowia(100);s__Metschnikowia_pulcherrima(100); | Metschnikowia | 13 | Saprotroph | \|Undefined Saprotroph\| | Yeast | NULL | Probable | As nectar-tap_saprotroph (Põlme et al. 2020); Decay sugar-rich substrates (Põlme et al. 2020) | Põlme S, et al. 2020. Fungal Diversity 105:1-16 ((https://doi.org/10.1007/s13225-020-00466-2)) |
| Otu16767 | 1 | k__Fungi(100);p__Ascomycota(100);c__Saccharomycetes(100);o__Saccharomycetales(100);f__Metschnikowiaceae(100);g__Metschnikowia(100);g__Metschnikowia_unclassified(100); | Metschnikowia | 13 | Saprotroph | \|Undefined Saprotroph\| | Yeast | NULL | Probable | As nectar-tap_saprotroph (Põlme et al. 2020); Decay sugar-rich substrates (Põlme et al. 2020) | Põlme S, et al. 2020. Fungal Diversity 105:1-16 ((https://doi.org/10.1007/s13225-020-00466-2)) |
| Otu16768 | 1 | k__Fungi(100);p__Ascomycota(100);c__Saccharomycetes(100);o__Saccharomycetales(100);f__Metschnikowiaceae(100);g__Metschnikowia(100);g__Metschnikowia_unclassified(100); | Metschnikowia | 13 | Saprotroph | \|Undefined Saprotroph\| | Yeast | NULL | Probable | As nectar-tap_saprotroph (Põlme et al. 2020); Decay sugar-rich substrates (Põlme et al. 2020) | Põlme S, et al. 2020. Fungal Diversity 105:1-16 ((https://doi.org/10.1007/s13225-020-00466-2)) |
| Otu16769 | 1 | k__Fungi(100);p__Ascomycota(100);c__Saccharomycetes(100);o__Saccharomycetales(100);f__Metschnikowiaceae(100);g__Metschnikowia(100);g__Metschnikowia_unclassified(100); | Metschnikowia | 13 | Saprotroph | \|Undefined Saprotroph\| | Yeast | NULL | Probable | As nectar-tap_saprotroph (Põlme et al. 2020); Decay sugar-rich substrates (Põlme et al. 2020) | Põlme S, et al. 2020. Fungal Diversity 105:1-16 ((https://doi.org/10.1007/s13225-020-00466-2)) |
| Otu16770 | 1 | k__Fungi(100);p__Ascomycota(100);c__Saccharomycetes(100);o__Saccharomycetales(100);f__Metschnikowiaceae(100);g__Metschnikowia(100);s__Metschnikowia_pulcherrima(100); | Metschnikowia | 13 | Saprotroph | \|Undefined Saprotroph\| | Yeast | NULL | Probable | As nectar-tap_saprotroph (Põlme et al. 2020); Decay sugar-rich substrates (Põlme et al. 2020) | Põlme S, et al. 2020. Fungal Diversity 105:1-16 ((https://doi.org/10.1007/s13225-020-00466-2)) |
| Otu16771 | 1 | k__Fungi(100);p__Ascomycota(100);c__Saccharomycetes(100);o__Saccharomycetales(100);f__Metschnikowiaceae(100);g__Metschnikowia(100);s__Metschnikowia_pulcherrima(100); | Metschnikowia | 13 | Saprotroph | \|Undefined Saprotroph\| | Yeast | NULL | Probable | As nectar-tap_saprotroph (Põlme et al. 2020); Decay sugar-rich substrates (Põlme et al. 2020) | Põlme S, et al. 2020. Fungal Diversity 105:1-16 ((https://doi.org/10.1007/s13225-020-00466-2)) |
| Otu16772 | 1 | k__Fungi(100);p__Ascomycota(100);c__Saccharomycetes(100);o__Saccharomycetales(100);f__Metschnikowiaceae(100);g__Metschnikowia(100);s__Metschnikowia_pulcherrima(100); | Metschnikowia | 13 | Saprotroph | \|Undefined Saprotroph\| | Yeast | NULL | Probable | As nectar-tap_saprotroph (Põlme et al. 2020); Decay sugar-rich substrates (Põlme et al. 2020) | Põlme S, et al. 2020. Fungal Diversity 105:1-16 ((https://doi.org/10.1007/s13225-020-00466-2)) |
| Otu16774 | 1 | k__Fungi(100);p__Ascomycota(100);c__Saccharomycetes(100);o__Saccharomycetales(100);f__Metschnikowiaceae(100);g__Metschnikowia(100);s__Metschnikowia_pulcherrima(100); | Metschnikowia | 13 | Saprotroph | \|Undefined Saprotroph\| | Yeast | NULL | Probable | As nectar-tap_saprotroph (Põlme et al. 2020); Decay sugar-rich substrates (Põlme et al. 2020) | Põlme S, et al. 2020. Fungal Diversity 105:1-16 ((https://doi.org/10.1007/s13225-020-00466-2)) |
| Otu16775 | 1 | k__Fungi(100);p__Ascomycota(100);c__Saccharomycetes(100);o__Saccharomycetales(100);f__Metschnikowiaceae(100);g__Metschnikowia(100);s__Metschnikowia_pulcherrima(100); | Metschnikowia | 13 | Saprotroph | \|Undefined Saprotroph\| | Yeast | NULL | Probable | As nectar-tap_saprotroph (Põlme et al. 2020); Decay sugar-rich substrates (Põlme et al. 2020) | Põlme S, et al. 2020. Fungal Diversity 105:1-16 ((https://doi.org/10.1007/s13225-020-00466-2)) |
| Otu16776 | 1 | k__Fungi(100);p__Ascomycota(100);c__Saccharomycetes(100);o__Saccharomycetales(100);f__Metschnikowiaceae(100);g__Metschnikowia(100);s__Metschnikowia_pulcherrima(100); | Metschnikowia | 13 | Saprotroph | \|Undefined Saprotroph\| | Yeast | NULL | Probable | As nectar-tap_saprotroph (Põlme et al. 2020); Decay sugar-rich substrates (Põlme et al. 2020) | Põlme S, et al. 2020. Fungal Diversity 105:1-16 ((https://doi.org/10.1007/s13225-020-00466-2)) |
| Otu16778 | 1 | k__Fungi(100);p__Ascomycota(100);c__Saccharomycetes(100);o__Saccharomycetales(100);f__Metschnikowiaceae(100);g__Metschnikowia(100);g__Metschnikowia_unclassified(100); | Metschnikowia | 13 | Saprotroph | \|Undefined Saprotroph\| | Yeast | NULL | Probable | As nectar-tap_saprotroph (Põlme et al. 2020); Decay sugar-rich substrates (Põlme et al. 2020) | Põlme S, et al. 2020. Fungal Diversity 105:1-16 ((https://doi.org/10.1007/s13225-020-00466-2)) |
| Otu16779 | 1 | k__Fungi(100);p__Ascomycota(100);c__Saccharomycetes(100);o__Saccharomycetales(100);f__Metschnikowiaceae(100);g__Metschnikowia(100);s__Metschnikowia_pulcherrima(100); | Metschnikowia | 13 | Saprotroph | \|Undefined Saprotroph\| | Yeast | NULL | Probable | As nectar-tap_saprotroph (Põlme et al. 2020); Decay sugar-rich substrates (Põlme et al. 2020) | Põlme S, et al. 2020. Fungal Diversity 105:1-16 ((https://doi.org/10.1007/s13225-020-00466-2)) |
| Otu16780 | 1 | k__Fungi(100);p__Ascomycota(100);c__Saccharomycetes(100);o__Saccharomycetales(100);f__Metschnikowiaceae(100);g__Metschnikowia(100);s__Metschnikowia_pulcherrima(100); | Metschnikowia | 13 | Saprotroph | \|Undefined Saprotroph\| | Yeast | NULL | Probable | As nectar-tap_saprotroph (Põlme et al. 2020); Decay sugar-rich substrates (Põlme et al. 2020) | Põlme S, et al. 2020. Fungal Diversity 105:1-16 ((https://doi.org/10.1007/s13225-020-00466-2)) |
| Otu16781 | 1 | k__Fungi(100);p__Ascomycota(100);c__Saccharomycetes(100);o__Saccharomycetales(100);f__Metschnikowiaceae(100);g__Metschnikowia(100);s__Metschnikowia_pulcherrima(100); | Metschnikowia | 13 | Saprotroph | \|Undefined Saprotroph\| | Yeast | NULL | Probable | As nectar-tap_saprotroph (Põlme et al. 2020); Decay sugar-rich substrates (Põlme et al. 2020) | Põlme S, et al. 2020. Fungal Diversity 105:1-16 ((https://doi.org/10.1007/s13225-020-00466-2)) |
| Otu16782 | 1 | k__Fungi(100);p__Ascomycota(100);c__Saccharomycetes(100);o__Saccharomycetales(100);f__Metschnikowiaceae(100);g__Metschnikowia(100);s__Metschnikowia_pulcherrima(100); | Metschnikowia | 13 | Saprotroph | \|Undefined Saprotroph\| | Yeast | NULL | Probable | As nectar-tap_saprotroph (Põlme et al. 2020); Decay sugar-rich substrates (Põlme et al. 2020) | Põlme S, et al. 2020. Fungal Diversity 105:1-16 ((https://doi.org/10.1007/s13225-020-00466-2)) |
| Otu16784 | 1 | k__Fungi(100);p__Ascomycota(100);c__Saccharomycetes(100);o__Saccharomycetales(100);f__Metschnikowiaceae(100);g__Metschnikowia(100);s__Metschnikowia_pulcherrima(100); | Metschnikowia | 13 | Saprotroph | \|Undefined Saprotroph\| | Yeast | NULL | Probable | As nectar-tap_saprotroph (Põlme et al. 2020); Decay sugar-rich substrates (Põlme et al. 2020) | Põlme S, et al. 2020. Fungal Diversity 105:1-16 ((https://doi.org/10.1007/s13225-020-00466-2)) |
| Otu16785 | 1 | k__Fungi(100);p__Ascomycota(100);c__Saccharomycetes(100);o__Saccharomycetales(100);f__Metschnikowiaceae(100);g__Metschnikowia(100);s__Metschnikowia_pulcherrima(100); | Metschnikowia | 13 | Saprotroph | \|Undefined Saprotroph\| | Yeast | NULL | Probable | As nectar-tap_saprotroph (Põlme et al. 2020); Decay sugar-rich substrates (Põlme et al. 2020) | Põlme S, et al. 2020. Fungal Diversity 105:1-16 ((https://doi.org/10.1007/s13225-020-00466-2)) |
| Otu16786 | 1 | k__Fungi(100);p__Ascomycota(100);c__Saccharomycetes(100);o__Saccharomycetales(100);f__Metschnikowiaceae(100);g__Metschnikowia(100);g__Metschnikowia_unclassified(100); | Metschnikowia | 13 | Saprotroph | \|Undefined Saprotroph\| | Yeast | NULL | Probable | As nectar-tap_saprotroph (Põlme et al. 2020); Decay sugar-rich substrates (Põlme et al. 2020) | Põlme S, et al. 2020. Fungal Diversity 105:1-16 ((https://doi.org/10.1007/s13225-020-00466-2)) |
| Otu16787 | 1 | k__Fungi(100);p__Ascomycota(100);c__Saccharomycetes(100);o__Saccharomycetales(100);f__Metschnikowiaceae(100);g__Metschnikowia(100);s__Metschnikowia_pulcherrima(100); | Metschnikowia | 13 | Saprotroph | \|Undefined Saprotroph\| | Yeast | NULL | Probable | As nectar-tap_saprotroph (Põlme et al. 2020); Decay sugar-rich substrates (Põlme et al. 2020) | Põlme S, et al. 2020. Fungal Diversity 105:1-16 ((https://doi.org/10.1007/s13225-020-00466-2)) |
| Otu16789 | 1 | k__Fungi(100);p__Ascomycota(100);c__Saccharomycetes(100);o__Saccharomycetales(100);f__Metschnikowiaceae(100);g__Metschnikowia(100);s__Metschnikowia_pulcherrima(100); | Metschnikowia | 13 | Saprotroph | \|Undefined Saprotroph\| | Yeast | NULL | Probable | As nectar-tap_saprotroph (Põlme et al. 2020); Decay sugar-rich substrates (Põlme et al. 2020) | Põlme S, et al. 2020. Fungal Diversity 105:1-16 ((https://doi.org/10.1007/s13225-020-00466-2)) |
| Otu16790 | 1 | k__Fungi(100);p__Ascomycota(100);c__Saccharomycetes(100);o__Saccharomycetales(100);f__Saccharomycetales_family_Incertae_sedis(100);g__Candida(100);g__Candida_unclassified(100); | Candida | 13 | Pathotroph-Saprotroph-Symbiotroph | Animal Pathogen-Endophyte-Endosymbiont-Epiphyte-\|Undefined Saprotroph\| | Dimorphic Yeast-Yeast | NULL | Probable | As nectar-tap_saprotroph (Põlme et al. 2020); Decay sugar-rich substrates (Põlme et al. 2020); This asexual genus is polyphyletic and contain species that function very differently depend on what they are. Caution should be taken when interpreting guilds for Candida species. | Manolakaki D, et al. 2010. Virulence. 1: 367-375 ((https://doi.org/10.4161/viru.1.5.12796)); Mekbib SB, et al. 2011. Tropical Plant Pathology 36:233-240 ((https://doi.org/10.1590/S1982-56762011000400004)); Põlme S, et al. 2020. Fungal Diversity 105:1-16 ((https://doi.org/10.1007/s13225-020-00466-2)) |
| Otu16791 | 1 | k__Fungi(100);p__Ascomycota(100);c__Saccharomycetes(100);o__Saccharomycetales(100);f__Saccharomycetales_family_Incertae_sedis(100);f__Saccharomycetales_family_Incertae_sedis_unclassified(100);f__Saccharomycetales_family_Incertae_sedis_unclassified(100); | Saccharomycetales | 7 | Saprotroph | Undefined Saprotroph | Yeast | NULL | Possible | NULL | Sterkenburg E, et al. 2015. New Phytologist 207:1145-1158 ((https://doi.org/10.1111/nph.13426)) |
| Otu16799 | 1 | k__Fungi(100);p__Ascomycota(100);c__Saccharomycetes(100);o__Saccharomycetales(100);f__Metschnikowiaceae(100);g__Metschnikowia(100);s__Metschnikowia_pulcherrima(100); | Metschnikowia | 13 | Saprotroph | \|Undefined Saprotroph\| | Yeast | NULL | Probable | As nectar-tap_saprotroph (Põlme et al. 2020); Decay sugar-rich substrates (Põlme et al. 2020) | Põlme S, et al. 2020. Fungal Diversity 105:1-16 ((https://doi.org/10.1007/s13225-020-00466-2)) |
| Otu16800 | 1 | k__Fungi(100);p__Ascomycota(100);c__Saccharomycetes(100);o__Saccharomycetales(100);f__Metschnikowiaceae(100);g__Metschnikowia(100);g__Metschnikowia_unclassified(100); | Metschnikowia | 13 | Saprotroph | \|Undefined Saprotroph\| | Yeast | NULL | Probable | As nectar-tap_saprotroph (Põlme et al. 2020); Decay sugar-rich substrates (Põlme et al. 2020) | Põlme S, et al. 2020. Fungal Diversity 105:1-16 ((https://doi.org/10.1007/s13225-020-00466-2)) |
| Otu16801 | 1 | k__Fungi(100);p__Ascomycota(100);c__Saccharomycetes(100);o__Saccharomycetales(100);f__Metschnikowiaceae(100);g__Metschnikowia(100);s__Metschnikowia_pulcherrima(100); | Metschnikowia | 13 | Saprotroph | \|Undefined Saprotroph\| | Yeast | NULL | Probable | As nectar-tap_saprotroph (Põlme et al. 2020); Decay sugar-rich substrates (Põlme et al. 2020) | Põlme S, et al. 2020. Fungal Diversity 105:1-16 ((https://doi.org/10.1007/s13225-020-00466-2)) |
| Otu16802 | 1 | k__Fungi(100);p__Ascomycota(100);c__Saccharomycetes(100);o__Saccharomycetales(100);f__Metschnikowiaceae(100);g__Metschnikowia(100);s__Metschnikowia_pulcherrima(100); | Metschnikowia | 13 | Saprotroph | \|Undefined Saprotroph\| | Yeast | NULL | Probable | As nectar-tap_saprotroph (Põlme et al. 2020); Decay sugar-rich substrates (Põlme et al. 2020) | Põlme S, et al. 2020. Fungal Diversity 105:1-16 ((https://doi.org/10.1007/s13225-020-00466-2)) |
| Otu16803 | 1 | k__Fungi(100);p__Ascomycota(100);c__Saccharomycetes(100);o__Saccharomycetales(100);f__Metschnikowiaceae(100);g__Metschnikowia(100);s__Metschnikowia_pulcherrima(100); | Metschnikowia | 13 | Saprotroph | \|Undefined Saprotroph\| | Yeast | NULL | Probable | As nectar-tap_saprotroph (Põlme et al. 2020); Decay sugar-rich substrates (Põlme et al. 2020) | Põlme S, et al. 2020. Fungal Diversity 105:1-16 ((https://doi.org/10.1007/s13225-020-00466-2)) |
| Otu16805 | 1 | k__Fungi(100);p__Ascomycota(100);c__Saccharomycetes(100);o__Saccharomycetales(100);f__Metschnikowiaceae(100);g__Metschnikowia(100);g__Metschnikowia_unclassified(100); | Metschnikowia | 13 | Saprotroph | \|Undefined Saprotroph\| | Yeast | NULL | Probable | As nectar-tap_saprotroph (Põlme et al. 2020); Decay sugar-rich substrates (Põlme et al. 2020) | Põlme S, et al. 2020. Fungal Diversity 105:1-16 ((https://doi.org/10.1007/s13225-020-00466-2)) |
| Otu16806 | 1 | k__Fungi(100);p__Ascomycota(100);c__Saccharomycetes(100);o__Saccharomycetales(100);f__Metschnikowiaceae(100);g__Metschnikowia(100);s__Metschnikowia_pulcherrima(100); | Metschnikowia | 13 | Saprotroph | \|Undefined Saprotroph\| | Yeast | NULL | Probable | As nectar-tap_saprotroph (Põlme et al. 2020); Decay sugar-rich substrates (Põlme et al. 2020) | Põlme S, et al. 2020. Fungal Diversity 105:1-16 ((https://doi.org/10.1007/s13225-020-00466-2)) |
| Otu16807 | 1 | k__Fungi(100);p__Ascomycota(100);c__Saccharomycetes(100);o__Saccharomycetales(100);f__Metschnikowiaceae(100);g__Metschnikowia(100);s__Metschnikowia_pulcherrima(100); | Metschnikowia | 13 | Saprotroph | \|Undefined Saprotroph\| | Yeast | NULL | Probable | As nectar-tap_saprotroph (Põlme et al. 2020); Decay sugar-rich substrates (Põlme et al. 2020) | Põlme S, et al. 2020. Fungal Diversity 105:1-16 ((https://doi.org/10.1007/s13225-020-00466-2)) |
| Otu16808 | 1 | k__Fungi(100);p__Ascomycota(100);c__Saccharomycetes(100);o__Saccharomycetales(100);f__Metschnikowiaceae(100);g__Metschnikowia(100);s__Metschnikowia_pulcherrima(100); | Metschnikowia | 13 | Saprotroph | \|Undefined Saprotroph\| | Yeast | NULL | Probable | As nectar-tap_saprotroph (Põlme et al. 2020); Decay sugar-rich substrates (Põlme et al. 2020) | Põlme S, et al. 2020. Fungal Diversity 105:1-16 ((https://doi.org/10.1007/s13225-020-00466-2)) |
| Otu16810 | 1 | k__Fungi(100);p__Ascomycota(100);c__Saccharomycetes(100);o__Saccharomycetales(100);f__Saccharomycetales_family_Incertae_sedis(100);g__Candida(100);s__Candida_cretensis(100); | Candida | 13 | Pathotroph-Saprotroph-Symbiotroph | Animal Pathogen-Endophyte-Endosymbiont-Epiphyte-\|Undefined Saprotroph\| | Dimorphic Yeast-Yeast | NULL | Probable | As nectar-tap_saprotroph (Põlme et al. 2020); Decay sugar-rich substrates (Põlme et al. 2020); This asexual genus is polyphyletic and contain species that function very differently depend on what they are. Caution should be taken when interpreting guilds for Candida species. | Manolakaki D, et al. 2010. Virulence. 1: 367-375 ((https://doi.org/10.4161/viru.1.5.12796)); Mekbib SB, et al. 2011. Tropical Plant Pathology 36:233-240 ((https://doi.org/10.1590/S1982-56762011000400004)); Põlme S, et al. 2020. Fungal Diversity 105:1-16 ((https://doi.org/10.1007/s13225-020-00466-2)) |
| Otu16811 | 1 | k__Fungi(100);p__Ascomycota(100);c__Saccharomycetes(100);o__Saccharomycetales(100);f__Metschnikowiaceae(100);g__Metschnikowia(100);s__Metschnikowia_pulcherrima(100); | Metschnikowia | 13 | Saprotroph | \|Undefined Saprotroph\| | Yeast | NULL | Probable | As nectar-tap_saprotroph (Põlme et al. 2020); Decay sugar-rich substrates (Põlme et al. 2020) | Põlme S, et al. 2020. Fungal Diversity 105:1-16 ((https://doi.org/10.1007/s13225-020-00466-2)) |
| Otu16812 | 1 | k__Fungi(100);p__Ascomycota(100);c__Saccharomycetes(100);o__Saccharomycetales(100);f__Saccharomycetales_family_Incertae_sedis(100);g__Candida(100);g__Candida_unclassified(100); | Candida | 13 | Pathotroph-Saprotroph-Symbiotroph | Animal Pathogen-Endophyte-Endosymbiont-Epiphyte-\|Undefined Saprotroph\| | Dimorphic Yeast-Yeast | NULL | Probable | As nectar-tap_saprotroph (Põlme et al. 2020); Decay sugar-rich substrates (Põlme et al. 2020); This asexual genus is polyphyletic and contain species that function very differently depend on what they are. Caution should be taken when interpreting guilds for Candida species. | Manolakaki D, et al. 2010. Virulence. 1: 367-375 ((https://doi.org/10.4161/viru.1.5.12796)); Mekbib SB, et al. 2011. Tropical Plant Pathology 36:233-240 ((https://doi.org/10.1590/S1982-56762011000400004)); Põlme S, et al. 2020. Fungal Diversity 105:1-16 ((https://doi.org/10.1007/s13225-020-00466-2)) |
| Otu16813 | 1 | k__Fungi(100);p__Ascomycota(100);c__Saccharomycetes(100);o__Saccharomycetales(100);f__Metschnikowiaceae(100);g__Metschnikowia(100);s__Metschnikowia_pulcherrima(100); | Metschnikowia | 13 | Saprotroph | \|Undefined Saprotroph\| | Yeast | NULL | Probable | As nectar-tap_saprotroph (Põlme et al. 2020); Decay sugar-rich substrates (Põlme et al. 2020) | Põlme S, et al. 2020. Fungal Diversity 105:1-16 ((https://doi.org/10.1007/s13225-020-00466-2)) |
| Otu16816 | 1 | k__Fungi(100);p__Ascomycota(100);c__Saccharomycetes(100);o__Saccharomycetales(100);f__Metschnikowiaceae(100);g__Metschnikowia(100);g__Metschnikowia_unclassified(100); | Metschnikowia | 13 | Saprotroph | \|Undefined Saprotroph\| | Yeast | NULL | Probable | As nectar-tap_saprotroph (Põlme et al. 2020); Decay sugar-rich substrates (Põlme et al. 2020) | Põlme S, et al. 2020. Fungal Diversity 105:1-16 ((https://doi.org/10.1007/s13225-020-00466-2)) |
| Otu16817 | 1 | k__Fungi(100);p__Ascomycota(100);c__Saccharomycetes(100);o__Saccharomycetales(100);f__Metschnikowiaceae(100);g__Metschnikowia(100);s__Metschnikowia_pulcherrima(100); | Metschnikowia | 13 | Saprotroph | \|Undefined Saprotroph\| | Yeast | NULL | Probable | As nectar-tap_saprotroph (Põlme et al. 2020); Decay sugar-rich substrates (Põlme et al. 2020) | Põlme S, et al. 2020. Fungal Diversity 105:1-16 ((https://doi.org/10.1007/s13225-020-00466-2)) |
| Otu16818 | 1 | k__Fungi(100);p__Ascomycota(100);c__Saccharomycetes(100);o__Saccharomycetales(100);f__Metschnikowiaceae(100);g__Metschnikowia(100);s__Metschnikowia_pulcherrima(100); | Metschnikowia | 13 | Saprotroph | \|Undefined Saprotroph\| | Yeast | NULL | Probable | As nectar-tap_saprotroph (Põlme et al. 2020); Decay sugar-rich substrates (Põlme et al. 2020) | Põlme S, et al. 2020. Fungal Diversity 105:1-16 ((https://doi.org/10.1007/s13225-020-00466-2)) |
| Otu16820 | 1 | k__Fungi(100);p__Ascomycota(100);c__Saccharomycetes(100);o__Saccharomycetales(100);f__Metschnikowiaceae(100);g__Metschnikowia(100);g__Metschnikowia_unclassified(100); | Metschnikowia | 13 | Saprotroph | \|Undefined Saprotroph\| | Yeast | NULL | Probable | As nectar-tap_saprotroph (Põlme et al. 2020); Decay sugar-rich substrates (Põlme et al. 2020) | Põlme S, et al. 2020. Fungal Diversity 105:1-16 ((https://doi.org/10.1007/s13225-020-00466-2)) |
| Otu16823 | 1 | k__Fungi(100);p__Ascomycota(100);c__Saccharomycetes(100);o__Saccharomycetales(100);f__Metschnikowiaceae(100);g__Metschnikowia(100);s__Metschnikowia_pulcherrima(100); | Metschnikowia | 13 | Saprotroph | \|Undefined Saprotroph\| | Yeast | NULL | Probable | As nectar-tap_saprotroph (Põlme et al. 2020); Decay sugar-rich substrates (Põlme et al. 2020) | Põlme S, et al. 2020. Fungal Diversity 105:1-16 ((https://doi.org/10.1007/s13225-020-00466-2)) |
| Otu16824 | 1 | k__Fungi(100);p__Ascomycota(100);c__Saccharomycetes(100);o__Saccharomycetales(100);f__Saccharomycetales_family_Incertae_sedis(100);g__Candida(100);g__Candida_unclassified(100); | Candida | 13 | Pathotroph-Saprotroph-Symbiotroph | Animal Pathogen-Endophyte-Endosymbiont-Epiphyte-\|Undefined Saprotroph\| | Dimorphic Yeast-Yeast | NULL | Probable | As nectar-tap_saprotroph (Põlme et al. 2020); Decay sugar-rich substrates (Põlme et al. 2020); This asexual genus is polyphyletic and contain species that function very differently depend on what they are. Caution should be taken when interpreting guilds for Candida species. | Manolakaki D, et al. 2010. Virulence. 1: 367-375 ((https://doi.org/10.4161/viru.1.5.12796)); Mekbib SB, et al. 2011. Tropical Plant Pathology 36:233-240 ((https://doi.org/10.1590/S1982-56762011000400004)); Põlme S, et al. 2020. Fungal Diversity 105:1-16 ((https://doi.org/10.1007/s13225-020-00466-2)) |
| Otu16826 | 1 | k__Fungi(100);p__Ascomycota(100);c__Saccharomycetes(100);o__Saccharomycetales(100);f__Metschnikowiaceae(100);g__Metschnikowia(100);s__Metschnikowia_pulcherrima(100); | Metschnikowia | 13 | Saprotroph | \|Undefined Saprotroph\| | Yeast | NULL | Probable | As nectar-tap_saprotroph (Põlme et al. 2020); Decay sugar-rich substrates (Põlme et al. 2020) | Põlme S, et al. 2020. Fungal Diversity 105:1-16 ((https://doi.org/10.1007/s13225-020-00466-2)) |
| Otu16830 | 1 | k__Fungi(100);p__Ascomycota(100);c__Saccharomycetes(100);o__Saccharomycetales(100);f__Saccharomycetales_family_Incertae_sedis(100);f__Saccharomycetales_family_Incertae_sedis_unclassified(100);f__Saccharomycetales_family_Incertae_sedis_unclassified(100); | Saccharomycetales | 7 | Saprotroph | Undefined Saprotroph | Yeast | NULL | Possible | NULL | Sterkenburg E, et al. 2015. New Phytologist 207:1145-1158 ((https://doi.org/10.1111/nph.13426)) |
| Otu16832 | 1 | k__Fungi(100);p__Ascomycota(100);c__Saccharomycetes(100);o__Saccharomycetales(100);f__Metschnikowiaceae(100);g__Metschnikowia(100);s__Metschnikowia_pulcherrima(100); | Metschnikowia | 13 | Saprotroph | \|Undefined Saprotroph\| | Yeast | NULL | Probable | As nectar-tap_saprotroph (Põlme et al. 2020); Decay sugar-rich substrates (Põlme et al. 2020) | Põlme S, et al. 2020. Fungal Diversity 105:1-16 ((https://doi.org/10.1007/s13225-020-00466-2)) |
| Otu16833 | 1 | k__Fungi(100);p__Ascomycota(100);c__Saccharomycetes(100);o__Saccharomycetales(100);f__Metschnikowiaceae(100);g__Metschnikowia(100);s__Metschnikowia_pulcherrima(100); | Metschnikowia | 13 | Saprotroph | \|Undefined Saprotroph\| | Yeast | NULL | Probable | As nectar-tap_saprotroph (Põlme et al. 2020); Decay sugar-rich substrates (Põlme et al. 2020) | Põlme S, et al. 2020. Fungal Diversity 105:1-16 ((https://doi.org/10.1007/s13225-020-00466-2)) |
| Otu16834 | 1 | k__Fungi(100);p__Ascomycota(100);c__Saccharomycetes(100);o__Saccharomycetales(100);f__Metschnikowiaceae(100);g__Metschnikowia(100);s__Metschnikowia_pulcherrima(100); | Metschnikowia | 13 | Saprotroph | \|Undefined Saprotroph\| | Yeast | NULL | Probable | As nectar-tap_saprotroph (Põlme et al. 2020); Decay sugar-rich substrates (Põlme et al. 2020) | Põlme S, et al. 2020. Fungal Diversity 105:1-16 ((https://doi.org/10.1007/s13225-020-00466-2)) |
| Otu16835 | 1 | k__Fungi(100);p__Ascomycota(100);c__Saccharomycetes(100);o__Saccharomycetales(100);f__Metschnikowiaceae(100);g__Metschnikowia(100);g__Metschnikowia_unclassified(100); | Metschnikowia | 13 | Saprotroph | \|Undefined Saprotroph\| | Yeast | NULL | Probable | As nectar-tap_saprotroph (Põlme et al. 2020); Decay sugar-rich substrates (Põlme et al. 2020) | Põlme S, et al. 2020. Fungal Diversity 105:1-16 ((https://doi.org/10.1007/s13225-020-00466-2)) |
| Otu16836 | 1 | k__Fungi(100);p__Ascomycota(100);c__Saccharomycetes(100);o__Saccharomycetales(100);f__Metschnikowiaceae(100);g__Metschnikowia(100);s__Metschnikowia_pulcherrima(100); | Metschnikowia | 13 | Saprotroph | \|Undefined Saprotroph\| | Yeast | NULL | Probable | As nectar-tap_saprotroph (Põlme et al. 2020); Decay sugar-rich substrates (Põlme et al. 2020) | Põlme S, et al. 2020. Fungal Diversity 105:1-16 ((https://doi.org/10.1007/s13225-020-00466-2)) |
| Otu16837 | 1 | k__Fungi(100);p__Ascomycota(100);c__Saccharomycetes(100);o__Saccharomycetales(100);f__Metschnikowiaceae(100);g__Metschnikowia(100);s__Metschnikowia_pulcherrima(100); | Metschnikowia | 13 | Saprotroph | \|Undefined Saprotroph\| | Yeast | NULL | Probable | As nectar-tap_saprotroph (Põlme et al. 2020); Decay sugar-rich substrates (Põlme et al. 2020) | Põlme S, et al. 2020. Fungal Diversity 105:1-16 ((https://doi.org/10.1007/s13225-020-00466-2)) |
| Otu16838 | 1 | k__Fungi(100);p__Ascomycota(100);c__Saccharomycetes(100);o__Saccharomycetales(100);f__Metschnikowiaceae(100);g__Metschnikowia(100);s__Metschnikowia_pulcherrima(100); | Metschnikowia | 13 | Saprotroph | \|Undefined Saprotroph\| | Yeast | NULL | Probable | As nectar-tap_saprotroph (Põlme et al. 2020); Decay sugar-rich substrates (Põlme et al. 2020) | Põlme S, et al. 2020. Fungal Diversity 105:1-16 ((https://doi.org/10.1007/s13225-020-00466-2)) |
| Otu16840 | 1 | k__Fungi(100);p__Ascomycota(100);c__Leotiomycetes(100);o__Helotiales(100);f__Helotiales_family_Incertae_sedis(100);g__Naevala(100);s__Naevala_minutissima(100); | Naevala | 13 | Pathotroph-Symbiotroph | Endophyte-\|Plant Pathogen\| | Microfungus | NULL | Probable | As foliar_endophyte (Põlme et al. 2020); Decay leaf, fruit, seed (Põlme et al. 2020) | Põlme S, et al. 2020. Fungal Diversity 105:1-16 ((https://doi.org/10.1007/s13225-020-00466-2)) |
| Otu16843 | 1 | k__Fungi(100);p__Ascomycota(100);c__Dothideomycetes(100);o__Capnodiales(100);f__Mycosphaerellaceae(100);f__Mycosphaerellaceae_unclassified(100);f__Mycosphaerellaceae_unclassified(100); | Mycosphaerellaceae | 9 | Pathotroph-Saprotroph | Plant Pathogen-Undefined Saprotroph | Microfungus | NULL | Probable | Biotrophic, necrotrophic or saprobic on various plant tissues (Cannon & Kirk 2007) | Cannon PF, Kirk PM. 2007. Fungal Families of the World. CAB International, Cambridge (ISBN: 978-0851998275) |
| Otu16844 | 1 | k__Fungi(100);p__Ascomycota(100);c__Saccharomycetes(100);o__Saccharomycetales(100);f__Metschnikowiaceae(100);g__Metschnikowia(100);s__Metschnikowia_pulcherrima(100); | Metschnikowia | 13 | Saprotroph | \|Undefined Saprotroph\| | Yeast | NULL | Probable | As nectar-tap_saprotroph (Põlme et al. 2020); Decay sugar-rich substrates (Põlme et al. 2020) | Põlme S, et al. 2020. Fungal Diversity 105:1-16 ((https://doi.org/10.1007/s13225-020-00466-2)) |
| Otu16845 | 1 | k__Fungi(100);p__Ascomycota(100);c__Saccharomycetes(100);o__Saccharomycetales(100);f__Metschnikowiaceae(100);g__Metschnikowia(100);g__Metschnikowia_unclassified(100); | Metschnikowia | 13 | Saprotroph | \|Undefined Saprotroph\| | Yeast | NULL | Probable | As nectar-tap_saprotroph (Põlme et al. 2020); Decay sugar-rich substrates (Põlme et al. 2020) | Põlme S, et al. 2020. Fungal Diversity 105:1-16 ((https://doi.org/10.1007/s13225-020-00466-2)) |
| Otu16846 | 1 | k__Fungi(100);p__Ascomycota(100);c__Saccharomycetes(100);o__Saccharomycetales(100);f__Metschnikowiaceae(100);g__Metschnikowia(100);s__Metschnikowia_pulcherrima(100); | Metschnikowia | 13 | Saprotroph | \|Undefined Saprotroph\| | Yeast | NULL | Probable | As nectar-tap_saprotroph (Põlme et al. 2020); Decay sugar-rich substrates (Põlme et al. 2020) | Põlme S, et al. 2020. Fungal Diversity 105:1-16 ((https://doi.org/10.1007/s13225-020-00466-2)) |
| Otu16847 | 1 | k__Fungi(100);p__Ascomycota(100);c__Saccharomycetes(100);o__Saccharomycetales(100);f__Metschnikowiaceae(100);g__Metschnikowia(100);g__Metschnikowia_unclassified(100); | Metschnikowia | 13 | Saprotroph | \|Undefined Saprotroph\| | Yeast | NULL | Probable | As nectar-tap_saprotroph (Põlme et al. 2020); Decay sugar-rich substrates (Põlme et al. 2020) | Põlme S, et al. 2020. Fungal Diversity 105:1-16 ((https://doi.org/10.1007/s13225-020-00466-2)) |
| Otu16848 | 1 | k__Fungi(100);p__Ascomycota(100);c__Saccharomycetes(100);o__Saccharomycetales(100);f__Metschnikowiaceae(100);g__Metschnikowia(100);g__Metschnikowia_unclassified(100); | Metschnikowia | 13 | Saprotroph | \|Undefined Saprotroph\| | Yeast | NULL | Probable | As nectar-tap_saprotroph (Põlme et al. 2020); Decay sugar-rich substrates (Põlme et al. 2020) | Põlme S, et al. 2020. Fungal Diversity 105:1-16 ((https://doi.org/10.1007/s13225-020-00466-2)) |
| Otu16849 | 1 | k__Fungi(100);p__Ascomycota(100);c__Saccharomycetes(100);o__Saccharomycetales(100);f__Metschnikowiaceae(100);g__Metschnikowia(100);s__Metschnikowia_pulcherrima(100); | Metschnikowia | 13 | Saprotroph | \|Undefined Saprotroph\| | Yeast | NULL | Probable | As nectar-tap_saprotroph (Põlme et al. 2020); Decay sugar-rich substrates (Põlme et al. 2020) | Põlme S, et al. 2020. Fungal Diversity 105:1-16 ((https://doi.org/10.1007/s13225-020-00466-2)) |
| Otu16850 | 1 | k__Fungi(100);p__Ascomycota(100);c__Saccharomycetes(100);o__Saccharomycetales(100);f__Metschnikowiaceae(100);g__Metschnikowia(100);s__Metschnikowia_pulcherrima(100); | Metschnikowia | 13 | Saprotroph | \|Undefined Saprotroph\| | Yeast | NULL | Probable | As nectar-tap_saprotroph (Põlme et al. 2020); Decay sugar-rich substrates (Põlme et al. 2020) | Põlme S, et al. 2020. Fungal Diversity 105:1-16 ((https://doi.org/10.1007/s13225-020-00466-2)) |
| Otu16853 | 1 | k__Fungi(100);p__Ascomycota(100);c__Saccharomycetes(100);o__Saccharomycetales(100);f__Metschnikowiaceae(100);g__Metschnikowia(100);s__Metschnikowia_pulcherrima(100); | Metschnikowia | 13 | Saprotroph | \|Undefined Saprotroph\| | Yeast | NULL | Probable | As nectar-tap_saprotroph (Põlme et al. 2020); Decay sugar-rich substrates (Põlme et al. 2020) | Põlme S, et al. 2020. Fungal Diversity 105:1-16 ((https://doi.org/10.1007/s13225-020-00466-2)) |
| Otu16855 | 1 | k__Fungi(100);p__Ascomycota(100);c__Saccharomycetes(100);o__Saccharomycetales(100);f__Metschnikowiaceae(100);g__Metschnikowia(100);s__Metschnikowia_pulcherrima(100); | Metschnikowia | 13 | Saprotroph | \|Undefined Saprotroph\| | Yeast | NULL | Probable | As nectar-tap_saprotroph (Põlme et al. 2020); Decay sugar-rich substrates (Põlme et al. 2020) | Põlme S, et al. 2020. Fungal Diversity 105:1-16 ((https://doi.org/10.1007/s13225-020-00466-2)) |
| Otu16856 | 1 | k__Fungi(100);p__Ascomycota(100);c__Saccharomycetes(100);o__Saccharomycetales(100);f__Metschnikowiaceae(100);g__Metschnikowia(100);s__Metschnikowia_pulcherrima(100); | Metschnikowia | 13 | Saprotroph | \|Undefined Saprotroph\| | Yeast | NULL | Probable | As nectar-tap_saprotroph (Põlme et al. 2020); Decay sugar-rich substrates (Põlme et al. 2020) | Põlme S, et al. 2020. Fungal Diversity 105:1-16 ((https://doi.org/10.1007/s13225-020-00466-2)) |
| Otu16857 | 1 | k__Fungi(100);p__Ascomycota(100);c__Saccharomycetes(100);o__Saccharomycetales(100);f__Metschnikowiaceae(100);g__Metschnikowia(100);s__Metschnikowia_pulcherrima(100); | Metschnikowia | 13 | Saprotroph | \|Undefined Saprotroph\| | Yeast | NULL | Probable | As nectar-tap_saprotroph (Põlme et al. 2020); Decay sugar-rich substrates (Põlme et al. 2020) | Põlme S, et al. 2020. Fungal Diversity 105:1-16 ((https://doi.org/10.1007/s13225-020-00466-2)) |
| Otu16859 | 1 | k__Fungi(100);p__Basidiomycota(100);c__Microbotryomycetes(100);o__Sporidiobolales(100);f__Sporidiobolales_family_Incertae_sedis(100);g__Sporobolomyces(100);s__Sporobolomyces_lactosus(100); | Sporobolomyces | 13 | Pathotroph-Saprotroph | \|Fungal Parasite\|-Plant Saprotroph | Yeast | NULL | Probable | As litter_saprotroph (Põlme et al. 2020); As mycoparasite (Põlme et al. 2020); Decay fungal material (Põlme et al. 2020) | Tedersoo L, et al. 2014. Science 346:e1256688 ((https://doi.org/10.1126/science.1256688)); Purahong W, et al. 2016. Molecular Ecology 25:4059-4074 ((https://doi.org/10.1111/mec.13739)); Põlme S, et al. 2020. Fungal Diversity 105:1-16 ((https://doi.org/10.1007/s13225-020-00466-2)) |
| Otu16860 | 1 | k__Fungi(100);p__Basidiomycota(100);c__Microbotryomycetes(100);o__Sporidiobolales(100);f__Sporidiobolales_family_Incertae_sedis(100);g__Sporobolomyces(100);s__Sporobolomyces_lactosus(100); | Sporobolomyces | 13 | Pathotroph-Saprotroph | \|Fungal Parasite\|-Plant Saprotroph | Yeast | NULL | Probable | As litter_saprotroph (Põlme et al. 2020); As mycoparasite (Põlme et al. 2020); Decay fungal material (Põlme et al. 2020) | Tedersoo L, et al. 2014. Science 346:e1256688 ((https://doi.org/10.1126/science.1256688)); Purahong W, et al. 2016. Molecular Ecology 25:4059-4074 ((https://doi.org/10.1111/mec.13739)); Põlme S, et al. 2020. Fungal Diversity 105:1-16 ((https://doi.org/10.1007/s13225-020-00466-2)) |
| Otu16863 | 1 | k__Fungi(100);p__Ascomycota(100);c__Saccharomycetes(100);o__Saccharomycetales(100);f__Metschnikowiaceae(100);g__Metschnikowia(100);g__Metschnikowia_unclassified(100); | Metschnikowia | 13 | Saprotroph | \|Undefined Saprotroph\| | Yeast | NULL | Probable | As nectar-tap_saprotroph (Põlme et al. 2020); Decay sugar-rich substrates (Põlme et al. 2020) | Põlme S, et al. 2020. Fungal Diversity 105:1-16 ((https://doi.org/10.1007/s13225-020-00466-2)) |
| Otu16864 | 1 | k__Fungi(100);p__Ascomycota(100);c__Saccharomycetes(100);o__Saccharomycetales(100);f__Metschnikowiaceae(100);g__Metschnikowia(100);g__Metschnikowia_unclassified(100); | Metschnikowia | 13 | Saprotroph | \|Undefined Saprotroph\| | Yeast | NULL | Probable | As nectar-tap_saprotroph (Põlme et al. 2020); Decay sugar-rich substrates (Põlme et al. 2020) | Põlme S, et al. 2020. Fungal Diversity 105:1-16 ((https://doi.org/10.1007/s13225-020-00466-2)) |
| Otu16865 | 1 | k__Fungi(100);p__Ascomycota(100);c__Saccharomycetes(100);o__Saccharomycetales(100);f__Metschnikowiaceae(100);g__Metschnikowia(100);g__Metschnikowia_unclassified(100); | Metschnikowia | 13 | Saprotroph | \|Undefined Saprotroph\| | Yeast | NULL | Probable | As nectar-tap_saprotroph (Põlme et al. 2020); Decay sugar-rich substrates (Põlme et al. 2020) | Põlme S, et al. 2020. Fungal Diversity 105:1-16 ((https://doi.org/10.1007/s13225-020-00466-2)) |
| Otu16866 | 1 | k__Fungi(100);p__Ascomycota(100);c__Saccharomycetes(100);o__Saccharomycetales(100);f__Metschnikowiaceae(100);g__Metschnikowia(100);s__Metschnikowia_pulcherrima(100); | Metschnikowia | 13 | Saprotroph | \|Undefined Saprotroph\| | Yeast | NULL | Probable | As nectar-tap_saprotroph (Põlme et al. 2020); Decay sugar-rich substrates (Põlme et al. 2020) | Põlme S, et al. 2020. Fungal Diversity 105:1-16 ((https://doi.org/10.1007/s13225-020-00466-2)) |
| Otu16867 | 1 | k__Fungi(100);p__Ascomycota(100);c__Saccharomycetes(100);o__Saccharomycetales(100);f__Metschnikowiaceae(100);g__Metschnikowia(100);g__Metschnikowia_unclassified(100); | Metschnikowia | 13 | Saprotroph | \|Undefined Saprotroph\| | Yeast | NULL | Probable | As nectar-tap_saprotroph (Põlme et al. 2020); Decay sugar-rich substrates (Põlme et al. 2020) | Põlme S, et al. 2020. Fungal Diversity 105:1-16 ((https://doi.org/10.1007/s13225-020-00466-2)) |
| Otu16868 | 1 | k__Fungi(100);p__Ascomycota(100);c__Saccharomycetes(100);o__Saccharomycetales(100);f__Metschnikowiaceae(100);g__Metschnikowia(100);s__Metschnikowia_pulcherrima(100); | Metschnikowia | 13 | Saprotroph | \|Undefined Saprotroph\| | Yeast | NULL | Probable | As nectar-tap_saprotroph (Põlme et al. 2020); Decay sugar-rich substrates (Põlme et al. 2020) | Põlme S, et al. 2020. Fungal Diversity 105:1-16 ((https://doi.org/10.1007/s13225-020-00466-2)) |
| Otu16870 | 1 | k__Fungi(100);p__Ascomycota(100);c__Saccharomycetes(100);o__Saccharomycetales(100);f__Metschnikowiaceae(100);g__Metschnikowia(100);s__Metschnikowia_pulcherrima(100); | Metschnikowia | 13 | Saprotroph | \|Undefined Saprotroph\| | Yeast | NULL | Probable | As nectar-tap_saprotroph (Põlme et al. 2020); Decay sugar-rich substrates (Põlme et al. 2020) | Põlme S, et al. 2020. Fungal Diversity 105:1-16 ((https://doi.org/10.1007/s13225-020-00466-2)) |
| Otu16871 | 1 | k__Fungi(100);p__Ascomycota(100);c__Saccharomycetes(100);o__Saccharomycetales(100);f__Metschnikowiaceae(100);g__Metschnikowia(100);s__Metschnikowia_pulcherrima(100); | Metschnikowia | 13 | Saprotroph | \|Undefined Saprotroph\| | Yeast | NULL | Probable | As nectar-tap_saprotroph (Põlme et al. 2020); Decay sugar-rich substrates (Põlme et al. 2020) | Põlme S, et al. 2020. Fungal Diversity 105:1-16 ((https://doi.org/10.1007/s13225-020-00466-2)) |
| Otu16872 | 1 | k__Fungi(100);p__Ascomycota(100);c__Saccharomycetes(100);o__Saccharomycetales(100);f__Metschnikowiaceae(100);g__Metschnikowia(100);s__Metschnikowia_pulcherrima(100); | Metschnikowia | 13 | Saprotroph | \|Undefined Saprotroph\| | Yeast | NULL | Probable | As nectar-tap_saprotroph (Põlme et al. 2020); Decay sugar-rich substrates (Põlme et al. 2020) | Põlme S, et al. 2020. Fungal Diversity 105:1-16 ((https://doi.org/10.1007/s13225-020-00466-2)) |
| Otu16873 | 1 | k__Fungi(100);p__Ascomycota(100);c__Saccharomycetes(100);o__Saccharomycetales(100);f__Metschnikowiaceae(100);g__Metschnikowia(100);g__Metschnikowia_unclassified(100); | Metschnikowia | 13 | Saprotroph | \|Undefined Saprotroph\| | Yeast | NULL | Probable | As nectar-tap_saprotroph (Põlme et al. 2020); Decay sugar-rich substrates (Põlme et al. 2020) | Põlme S, et al. 2020. Fungal Diversity 105:1-16 ((https://doi.org/10.1007/s13225-020-00466-2)) |
| Otu16874 | 1 | k__Fungi(100);p__Ascomycota(100);c__Saccharomycetes(100);o__Saccharomycetales(100);f__Saccharomycetales_family_Incertae_sedis(100);g__Candida(100);g__Candida_unclassified(100); | Candida | 13 | Pathotroph-Saprotroph-Symbiotroph | Animal Pathogen-Endophyte-Endosymbiont-Epiphyte-\|Undefined Saprotroph\| | Dimorphic Yeast-Yeast | NULL | Probable | As nectar-tap_saprotroph (Põlme et al. 2020); Decay sugar-rich substrates (Põlme et al. 2020); This asexual genus is polyphyletic and contain species that function very differently depend on what they are. Caution should be taken when interpreting guilds for Candida species. | Manolakaki D, et al. 2010. Virulence. 1: 367-375 ((https://doi.org/10.4161/viru.1.5.12796)); Mekbib SB, et al. 2011. Tropical Plant Pathology 36:233-240 ((https://doi.org/10.1590/S1982-56762011000400004)); Põlme S, et al. 2020. Fungal Diversity 105:1-16 ((https://doi.org/10.1007/s13225-020-00466-2)) |
| Otu16875 | 1 | k__Fungi(100);p__Ascomycota(100);c__Saccharomycetes(100);o__Saccharomycetales(100);f__Saccharomycetales_family_Incertae_sedis(100);f__Saccharomycetales_family_Incertae_sedis_unclassified(100);f__Saccharomycetales_family_Incertae_sedis_unclassified(100); | Saccharomycetales | 7 | Saprotroph | Undefined Saprotroph | Yeast | NULL | Possible | NULL | Sterkenburg E, et al. 2015. New Phytologist 207:1145-1158 ((https://doi.org/10.1111/nph.13426)) |
| Otu16877 | 1 | k__Fungi(100);p__Ascomycota(100);c__Saccharomycetes(100);o__Saccharomycetales(100);f__Metschnikowiaceae(100);g__Metschnikowia(100);s__Metschnikowia_pulcherrima(100); | Metschnikowia | 13 | Saprotroph | \|Undefined Saprotroph\| | Yeast | NULL | Probable | As nectar-tap_saprotroph (Põlme et al. 2020); Decay sugar-rich substrates (Põlme et al. 2020) | Põlme S, et al. 2020. Fungal Diversity 105:1-16 ((https://doi.org/10.1007/s13225-020-00466-2)) |
| Otu16878 | 1 | k__Fungi(100);p__Ascomycota(100);c__Saccharomycetes(100);o__Saccharomycetales(100);f__Metschnikowiaceae(100);g__Metschnikowia(100);g__Metschnikowia_unclassified(100); | Metschnikowia | 13 | Saprotroph | \|Undefined Saprotroph\| | Yeast | NULL | Probable | As nectar-tap_saprotroph (Põlme et al. 2020); Decay sugar-rich substrates (Põlme et al. 2020) | Põlme S, et al. 2020. Fungal Diversity 105:1-16 ((https://doi.org/10.1007/s13225-020-00466-2)) |
| Otu16879 | 1 | k__Fungi(100);p__Ascomycota(100);c__Saccharomycetes(100);o__Saccharomycetales(100);f__Metschnikowiaceae(100);g__Metschnikowia(100);g__Metschnikowia_unclassified(100); | Metschnikowia | 13 | Saprotroph | \|Undefined Saprotroph\| | Yeast | NULL | Probable | As nectar-tap_saprotroph (Põlme et al. 2020); Decay sugar-rich substrates (Põlme et al. 2020) | Põlme S, et al. 2020. Fungal Diversity 105:1-16 ((https://doi.org/10.1007/s13225-020-00466-2)) |
| Otu16882 | 1 | k__Fungi(100);p__Ascomycota(100);c__Saccharomycetes(100);o__Saccharomycetales(100);f__Metschnikowiaceae(100);g__Metschnikowia(100);g__Metschnikowia_unclassified(100); | Metschnikowia | 13 | Saprotroph | \|Undefined Saprotroph\| | Yeast | NULL | Probable | As nectar-tap_saprotroph (Põlme et al. 2020); Decay sugar-rich substrates (Põlme et al. 2020) | Põlme S, et al. 2020. Fungal Diversity 105:1-16 ((https://doi.org/10.1007/s13225-020-00466-2)) |
| Otu16883 | 1 | k__Fungi(100);p__Ascomycota(100);c__Saccharomycetes(100);o__Saccharomycetales(100);f__Metschnikowiaceae(100);g__Metschnikowia(100);s__Metschnikowia_pulcherrima(100); | Metschnikowia | 13 | Saprotroph | \|Undefined Saprotroph\| | Yeast | NULL | Probable | As nectar-tap_saprotroph (Põlme et al. 2020); Decay sugar-rich substrates (Põlme et al. 2020) | Põlme S, et al. 2020. Fungal Diversity 105:1-16 ((https://doi.org/10.1007/s13225-020-00466-2)) |
| Otu16884 | 1 | k__Fungi(100);p__Basidiomycota(100);c__Microbotryomycetes(100);o__Sporidiobolales(100);f__Sporidiobolales_family_Incertae_sedis(100);g__Rhodotorula(100);s__Rhodotorula_nothofagi(100); | Rhodotorula | 13 | Pathotroph-Saprotroph-Symbiotroph | Animal Pathogen-Animal Symbiotroph-Endophyte-Plant Pathogen-\|Undefined Saprotroph\| | Yeast | NULL | Probable | As foliar_endophyte (Põlme et al. 2020); Opportunistic human pathogen | Põlme S, et al. 2020. Fungal Diversity 105:1-16 ((https://doi.org/10.1007/s13225-020-00466-2)); https://www.cdc.gov ((https://www.cdc.gov)) |
| Otu16885 | 1 | k__Fungi(100);p__Ascomycota(100);c__Saccharomycetes(100);o__Saccharomycetales(100);f__Metschnikowiaceae(100);g__Metschnikowia(100);g__Metschnikowia_unclassified(100); | Metschnikowia | 13 | Saprotroph | \|Undefined Saprotroph\| | Yeast | NULL | Probable | As nectar-tap_saprotroph (Põlme et al. 2020); Decay sugar-rich substrates (Põlme et al. 2020) | Põlme S, et al. 2020. Fungal Diversity 105:1-16 ((https://doi.org/10.1007/s13225-020-00466-2)) |
| Otu16886 | 1 | k__Fungi(100);p__Ascomycota(100);c__Saccharomycetes(100);o__Saccharomycetales(100);f__Metschnikowiaceae(100);g__Metschnikowia(100);s__Metschnikowia_pulcherrima(100); | Metschnikowia | 13 | Saprotroph | \|Undefined Saprotroph\| | Yeast | NULL | Probable | As nectar-tap_saprotroph (Põlme et al. 2020); Decay sugar-rich substrates (Põlme et al. 2020) | Põlme S, et al. 2020. Fungal Diversity 105:1-16 ((https://doi.org/10.1007/s13225-020-00466-2)) |
| Otu16887 | 1 | k__Fungi(100);p__Ascomycota(100);c__Saccharomycetes(100);o__Saccharomycetales(100);f__Metschnikowiaceae(100);g__Metschnikowia(100);s__Metschnikowia_pulcherrima(100); | Metschnikowia | 13 | Saprotroph | \|Undefined Saprotroph\| | Yeast | NULL | Probable | As nectar-tap_saprotroph (Põlme et al. 2020); Decay sugar-rich substrates (Põlme et al. 2020) | Põlme S, et al. 2020. Fungal Diversity 105:1-16 ((https://doi.org/10.1007/s13225-020-00466-2)) |
| Otu16888 | 1 | k__Fungi(100);p__Ascomycota(100);c__Saccharomycetes(100);o__Saccharomycetales(100);f__Metschnikowiaceae(100);g__Metschnikowia(100);s__Metschnikowia_pulcherrima(100); | Metschnikowia | 13 | Saprotroph | \|Undefined Saprotroph\| | Yeast | NULL | Probable | As nectar-tap_saprotroph (Põlme et al. 2020); Decay sugar-rich substrates (Põlme et al. 2020) | Põlme S, et al. 2020. Fungal Diversity 105:1-16 ((https://doi.org/10.1007/s13225-020-00466-2)) |
| Otu16889 | 1 | k__Fungi(100);p__Ascomycota(100);c__Dothideomycetes(100);o__Dothideomycetes_order_Incertae_sedis(100);f__Pseudeurotiaceae(100);g__unclassified_Pseudeurotiaceae(100);s__Pseudeurotiaceae_sp(100); | Pseudeurotiaceae | 9 | Saprotroph | Plant Saprotroph-Wood Saprotroph | Microfungus | NULL | Probable | Saprobic on woody tissue and rotting vegetation, often isolated from soil (Cannon & Kirk 2007) | Cannon PF, Kirk PM. 2007. Fungal Families of the World. CAB International, Cambridge (ISBN: 978-0851998275) |
| Otu16890 | 1 | k__Fungi(100);p__Ascomycota(100);c__Saccharomycetes(100);o__Saccharomycetales(100);f__Metschnikowiaceae(100);g__Metschnikowia(100);s__Metschnikowia_pulcherrima(100); | Metschnikowia | 13 | Saprotroph | \|Undefined Saprotroph\| | Yeast | NULL | Probable | As nectar-tap_saprotroph (Põlme et al. 2020); Decay sugar-rich substrates (Põlme et al. 2020) | Põlme S, et al. 2020. Fungal Diversity 105:1-16 ((https://doi.org/10.1007/s13225-020-00466-2)) |
| Otu16891 | 1 | k__Fungi(100);p__Ascomycota(100);c__Saccharomycetes(100);o__Saccharomycetales(100);f__Metschnikowiaceae(100);g__Metschnikowia(100);s__Metschnikowia_pulcherrima(100); | Metschnikowia | 13 | Saprotroph | \|Undefined Saprotroph\| | Yeast | NULL | Probable | As nectar-tap_saprotroph (Põlme et al. 2020); Decay sugar-rich substrates (Põlme et al. 2020) | Põlme S, et al. 2020. Fungal Diversity 105:1-16 ((https://doi.org/10.1007/s13225-020-00466-2)) |
| Otu16892 | 1 | k__Fungi(100);p__Ascomycota(100);c__Saccharomycetes(100);o__Saccharomycetales(100);f__Metschnikowiaceae(100);g__Metschnikowia(100);s__Metschnikowia_pulcherrima(100); | Metschnikowia | 13 | Saprotroph | \|Undefined Saprotroph\| | Yeast | NULL | Probable | As nectar-tap_saprotroph (Põlme et al. 2020); Decay sugar-rich substrates (Põlme et al. 2020) | Põlme S, et al. 2020. Fungal Diversity 105:1-16 ((https://doi.org/10.1007/s13225-020-00466-2)) |
| Otu16893 | 1 | k__Fungi(100);p__Ascomycota(100);c__Saccharomycetes(100);o__Saccharomycetales(100);f__Metschnikowiaceae(100);g__Metschnikowia(100);g__Metschnikowia_unclassified(100); | Metschnikowia | 13 | Saprotroph | \|Undefined Saprotroph\| | Yeast | NULL | Probable | As nectar-tap_saprotroph (Põlme et al. 2020); Decay sugar-rich substrates (Põlme et al. 2020) | Põlme S, et al. 2020. Fungal Diversity 105:1-16 ((https://doi.org/10.1007/s13225-020-00466-2)) |
| Otu16894 | 1 | k__Fungi(100);p__Ascomycota(100);c__Saccharomycetes(100);o__Saccharomycetales(100);f__Metschnikowiaceae(100);g__Metschnikowia(100);g__Metschnikowia_unclassified(100); | Metschnikowia | 13 | Saprotroph | \|Undefined Saprotroph\| | Yeast | NULL | Probable | As nectar-tap_saprotroph (Põlme et al. 2020); Decay sugar-rich substrates (Põlme et al. 2020) | Põlme S, et al. 2020. Fungal Diversity 105:1-16 ((https://doi.org/10.1007/s13225-020-00466-2)) |
| Otu16895 | 1 | k__Fungi(100);p__Ascomycota(100);c__Saccharomycetes(100);o__Saccharomycetales(100);f__Metschnikowiaceae(100);g__Metschnikowia(100);g__Metschnikowia_unclassified(100); | Metschnikowia | 13 | Saprotroph | \|Undefined Saprotroph\| | Yeast | NULL | Probable | As nectar-tap_saprotroph (Põlme et al. 2020); Decay sugar-rich substrates (Põlme et al. 2020) | Põlme S, et al. 2020. Fungal Diversity 105:1-16 ((https://doi.org/10.1007/s13225-020-00466-2)) |
| Otu16896 | 1 | k__Fungi(100);p__Ascomycota(100);c__Saccharomycetes(100);o__Saccharomycetales(100);f__Metschnikowiaceae(100);g__Metschnikowia(100);s__Metschnikowia_pulcherrima(100); | Metschnikowia | 13 | Saprotroph | \|Undefined Saprotroph\| | Yeast | NULL | Probable | As nectar-tap_saprotroph (Põlme et al. 2020); Decay sugar-rich substrates (Põlme et al. 2020) | Põlme S, et al. 2020. Fungal Diversity 105:1-16 ((https://doi.org/10.1007/s13225-020-00466-2)) |
| Otu16897 | 1 | k__Fungi(100);p__Ascomycota(100);c__Saccharomycetes(100);o__Saccharomycetales(100);f__Metschnikowiaceae(100);g__Metschnikowia(100);s__Metschnikowia_pulcherrima(100); | Metschnikowia | 13 | Saprotroph | \|Undefined Saprotroph\| | Yeast | NULL | Probable | As nectar-tap_saprotroph (Põlme et al. 2020); Decay sugar-rich substrates (Põlme et al. 2020) | Põlme S, et al. 2020. Fungal Diversity 105:1-16 ((https://doi.org/10.1007/s13225-020-00466-2)) |
| Otu16898 | 1 | k__Fungi(100);p__Ascomycota(100);c__Saccharomycetes(100);o__Saccharomycetales(100);f__Metschnikowiaceae(100);g__Metschnikowia(100);s__Metschnikowia_pulcherrima(100); | Metschnikowia | 13 | Saprotroph | \|Undefined Saprotroph\| | Yeast | NULL | Probable | As nectar-tap_saprotroph (Põlme et al. 2020); Decay sugar-rich substrates (Põlme et al. 2020) | Põlme S, et al. 2020. Fungal Diversity 105:1-16 ((https://doi.org/10.1007/s13225-020-00466-2)) |
| Otu16899 | 1 | k__Fungi(100);p__Ascomycota(100);c__Saccharomycetes(100);o__Saccharomycetales(100);f__Metschnikowiaceae(100);g__Metschnikowia(100);s__Metschnikowia_pulcherrima(100); | Metschnikowia | 13 | Saprotroph | \|Undefined Saprotroph\| | Yeast | NULL | Probable | As nectar-tap_saprotroph (Põlme et al. 2020); Decay sugar-rich substrates (Põlme et al. 2020) | Põlme S, et al. 2020. Fungal Diversity 105:1-16 ((https://doi.org/10.1007/s13225-020-00466-2)) |
| Otu16901 | 1 | k__Fungi(100);p__Ascomycota(100);c__Saccharomycetes(100);o__Saccharomycetales(100);f__Metschnikowiaceae(100);g__Metschnikowia(100);g__Metschnikowia_unclassified(100); | Metschnikowia | 13 | Saprotroph | \|Undefined Saprotroph\| | Yeast | NULL | Probable | As nectar-tap_saprotroph (Põlme et al. 2020); Decay sugar-rich substrates (Põlme et al. 2020) | Põlme S, et al. 2020. Fungal Diversity 105:1-16 ((https://doi.org/10.1007/s13225-020-00466-2)) |
| Otu16902 | 1 | k__Fungi(100);p__Ascomycota(100);c__Saccharomycetes(100);o__Saccharomycetales(100);f__Metschnikowiaceae(100);g__Metschnikowia(100);g__Metschnikowia_unclassified(100); | Metschnikowia | 13 | Saprotroph | \|Undefined Saprotroph\| | Yeast | NULL | Probable | As nectar-tap_saprotroph (Põlme et al. 2020); Decay sugar-rich substrates (Põlme et al. 2020) | Põlme S, et al. 2020. Fungal Diversity 105:1-16 ((https://doi.org/10.1007/s13225-020-00466-2)) |
| Otu16903 | 1 | k__Fungi(100);p__Ascomycota(100);c__Saccharomycetes(100);o__Saccharomycetales(100);f__Metschnikowiaceae(100);g__Metschnikowia(100);s__Metschnikowia_pulcherrima(100); | Metschnikowia | 13 | Saprotroph | \|Undefined Saprotroph\| | Yeast | NULL | Probable | As nectar-tap_saprotroph (Põlme et al. 2020); Decay sugar-rich substrates (Põlme et al. 2020) | Põlme S, et al. 2020. Fungal Diversity 105:1-16 ((https://doi.org/10.1007/s13225-020-00466-2)) |
| Otu16904 | 1 | k__Fungi(100);p__Ascomycota(100);c__Saccharomycetes(100);o__Saccharomycetales(100);f__Metschnikowiaceae(100);g__Metschnikowia(100);s__Metschnikowia_pulcherrima(100); | Metschnikowia | 13 | Saprotroph | \|Undefined Saprotroph\| | Yeast | NULL | Probable | As nectar-tap_saprotroph (Põlme et al. 2020); Decay sugar-rich substrates (Põlme et al. 2020) | Põlme S, et al. 2020. Fungal Diversity 105:1-16 ((https://doi.org/10.1007/s13225-020-00466-2)) |
| Otu16905 | 1 | k__Fungi(100);p__Ascomycota(100);c__Saccharomycetes(100);o__Saccharomycetales(100);f__Metschnikowiaceae(100);g__Metschnikowia(100);g__Metschnikowia_unclassified(100); | Metschnikowia | 13 | Saprotroph | \|Undefined Saprotroph\| | Yeast | NULL | Probable | As nectar-tap_saprotroph (Põlme et al. 2020); Decay sugar-rich substrates (Põlme et al. 2020) | Põlme S, et al. 2020. Fungal Diversity 105:1-16 ((https://doi.org/10.1007/s13225-020-00466-2)) |
| Otu16906 | 1 | k__Fungi(100);p__Ascomycota(100);c__Saccharomycetes(100);o__Saccharomycetales(100);f__Metschnikowiaceae(100);g__Metschnikowia(100);s__Metschnikowia_pulcherrima(100); | Metschnikowia | 13 | Saprotroph | \|Undefined Saprotroph\| | Yeast | NULL | Probable | As nectar-tap_saprotroph (Põlme et al. 2020); Decay sugar-rich substrates (Põlme et al. 2020) | Põlme S, et al. 2020. Fungal Diversity 105:1-16 ((https://doi.org/10.1007/s13225-020-00466-2)) |
| Otu16907 | 1 | k__Fungi(100);p__Ascomycota(100);c__Saccharomycetes(100);o__Saccharomycetales(100);f__Metschnikowiaceae(100);g__Metschnikowia(100);s__Metschnikowia_pulcherrima(100); | Metschnikowia | 13 | Saprotroph | \|Undefined Saprotroph\| | Yeast | NULL | Probable | As nectar-tap_saprotroph (Põlme et al. 2020); Decay sugar-rich substrates (Põlme et al. 2020) | Põlme S, et al. 2020. Fungal Diversity 105:1-16 ((https://doi.org/10.1007/s13225-020-00466-2)) |
| Otu16908 | 1 | k__Fungi(100);p__Ascomycota(100);c__Saccharomycetes(100);o__Saccharomycetales(100);f__Metschnikowiaceae(100);g__Metschnikowia(100);s__Metschnikowia_pulcherrima(100); | Metschnikowia | 13 | Saprotroph | \|Undefined Saprotroph\| | Yeast | NULL | Probable | As nectar-tap_saprotroph (Põlme et al. 2020); Decay sugar-rich substrates (Põlme et al. 2020) | Põlme S, et al. 2020. Fungal Diversity 105:1-16 ((https://doi.org/10.1007/s13225-020-00466-2)) |
| Otu16909 | 1 | k__Fungi(100);p__Ascomycota(100);c__Saccharomycetes(100);o__Saccharomycetales(100);f__Metschnikowiaceae(100);g__Metschnikowia(100);s__Metschnikowia_pulcherrima(100); | Metschnikowia | 13 | Saprotroph | \|Undefined Saprotroph\| | Yeast | NULL | Probable | As nectar-tap_saprotroph (Põlme et al. 2020); Decay sugar-rich substrates (Põlme et al. 2020) | Põlme S, et al. 2020. Fungal Diversity 105:1-16 ((https://doi.org/10.1007/s13225-020-00466-2)) |
| Otu16910 | 1 | k__Fungi(100);p__Ascomycota(100);c__Saccharomycetes(100);o__Saccharomycetales(100);f__Metschnikowiaceae(100);g__Metschnikowia(100);s__Metschnikowia_pulcherrima(100); | Metschnikowia | 13 | Saprotroph | \|Undefined Saprotroph\| | Yeast | NULL | Probable | As nectar-tap_saprotroph (Põlme et al. 2020); Decay sugar-rich substrates (Põlme et al. 2020) | Põlme S, et al. 2020. Fungal Diversity 105:1-16 ((https://doi.org/10.1007/s13225-020-00466-2)) |
| Otu16911 | 1 | k__Fungi(100);p__Ascomycota(100);c__Saccharomycetes(100);o__Saccharomycetales(100);f__Metschnikowiaceae(100);g__Metschnikowia(100);s__Metschnikowia_pulcherrima(100); | Metschnikowia | 13 | Saprotroph | \|Undefined Saprotroph\| | Yeast | NULL | Probable | As nectar-tap_saprotroph (Põlme et al. 2020); Decay sugar-rich substrates (Põlme et al. 2020) | Põlme S, et al. 2020. Fungal Diversity 105:1-16 ((https://doi.org/10.1007/s13225-020-00466-2)) |
| Otu16912 | 1 | k__Fungi(100);p__Ascomycota(100);c__Saccharomycetes(100);o__Saccharomycetales(100);f__Metschnikowiaceae(100);g__Metschnikowia(100);s__Metschnikowia_pulcherrima(100); | Metschnikowia | 13 | Saprotroph | \|Undefined Saprotroph\| | Yeast | NULL | Probable | As nectar-tap_saprotroph (Põlme et al. 2020); Decay sugar-rich substrates (Põlme et al. 2020) | Põlme S, et al. 2020. Fungal Diversity 105:1-16 ((https://doi.org/10.1007/s13225-020-00466-2)) |
| Otu16913 | 1 | k__Fungi(100);p__Ascomycota(100);c__Saccharomycetes(100);o__Saccharomycetales(100);f__Metschnikowiaceae(100);g__Metschnikowia(100);s__Metschnikowia_pulcherrima(100); | Metschnikowia | 13 | Saprotroph | \|Undefined Saprotroph\| | Yeast | NULL | Probable | As nectar-tap_saprotroph (Põlme et al. 2020); Decay sugar-rich substrates (Põlme et al. 2020) | Põlme S, et al. 2020. Fungal Diversity 105:1-16 ((https://doi.org/10.1007/s13225-020-00466-2)) |
| Otu16915 | 1 | k__Fungi(100);p__Ascomycota(100);c__Saccharomycetes(100);o__Saccharomycetales(100);f__Metschnikowiaceae(100);g__Metschnikowia(100);s__Metschnikowia_pulcherrima(100); | Metschnikowia | 13 | Saprotroph | \|Undefined Saprotroph\| | Yeast | NULL | Probable | As nectar-tap_saprotroph (Põlme et al. 2020); Decay sugar-rich substrates (Põlme et al. 2020) | Põlme S, et al. 2020. Fungal Diversity 105:1-16 ((https://doi.org/10.1007/s13225-020-00466-2)) |
| Otu16916 | 1 | k__Fungi(100);p__Ascomycota(100);c__Saccharomycetes(100);o__Saccharomycetales(100);f__Metschnikowiaceae(100);g__Metschnikowia(100);g__Metschnikowia_unclassified(100); | Metschnikowia | 13 | Saprotroph | \|Undefined Saprotroph\| | Yeast | NULL | Probable | As nectar-tap_saprotroph (Põlme et al. 2020); Decay sugar-rich substrates (Põlme et al. 2020) | Põlme S, et al. 2020. Fungal Diversity 105:1-16 ((https://doi.org/10.1007/s13225-020-00466-2)) |
| Otu16917 | 1 | k__Fungi(100);p__Ascomycota(100);c__Saccharomycetes(100);o__Saccharomycetales(100);f__Metschnikowiaceae(100);g__Metschnikowia(100);s__Metschnikowia_pulcherrima(100); | Metschnikowia | 13 | Saprotroph | \|Undefined Saprotroph\| | Yeast | NULL | Probable | As nectar-tap_saprotroph (Põlme et al. 2020); Decay sugar-rich substrates (Põlme et al. 2020) | Põlme S, et al. 2020. Fungal Diversity 105:1-16 ((https://doi.org/10.1007/s13225-020-00466-2)) |
| Otu16918 | 1 | k__Fungi(100);p__Ascomycota(100);c__Saccharomycetes(100);o__Saccharomycetales(100);f__Metschnikowiaceae(100);g__Metschnikowia(100);s__Metschnikowia_pulcherrima(100); | Metschnikowia | 13 | Saprotroph | \|Undefined Saprotroph\| | Yeast | NULL | Probable | As nectar-tap_saprotroph (Põlme et al. 2020); Decay sugar-rich substrates (Põlme et al. 2020) | Põlme S, et al. 2020. Fungal Diversity 105:1-16 ((https://doi.org/10.1007/s13225-020-00466-2)) |
| Otu16919 | 1 | k__Fungi(100);p__Ascomycota(100);c__Saccharomycetes(100);o__Saccharomycetales(100);f__Metschnikowiaceae(100);g__Metschnikowia(100);g__Metschnikowia_unclassified(100); | Metschnikowia | 13 | Saprotroph | \|Undefined Saprotroph\| | Yeast | NULL | Probable | As nectar-tap_saprotroph (Põlme et al. 2020); Decay sugar-rich substrates (Põlme et al. 2020) | Põlme S, et al. 2020. Fungal Diversity 105:1-16 ((https://doi.org/10.1007/s13225-020-00466-2)) |
| Otu16920 | 1 | k__Fungi(100);p__Ascomycota(100);c__Saccharomycetes(100);o__Saccharomycetales(100);f__Metschnikowiaceae(100);g__Metschnikowia(100);g__Metschnikowia_unclassified(100); | Metschnikowia | 13 | Saprotroph | \|Undefined Saprotroph\| | Yeast | NULL | Probable | As nectar-tap_saprotroph (Põlme et al. 2020); Decay sugar-rich substrates (Põlme et al. 2020) | Põlme S, et al. 2020. Fungal Diversity 105:1-16 ((https://doi.org/10.1007/s13225-020-00466-2)) |
| Otu16921 | 1 | k__Fungi(100);p__Ascomycota(100);c__Saccharomycetes(100);o__Saccharomycetales(100);f__Metschnikowiaceae(100);g__Metschnikowia(100);s__Metschnikowia_pulcherrima(100); | Metschnikowia | 13 | Saprotroph | \|Undefined Saprotroph\| | Yeast | NULL | Probable | As nectar-tap_saprotroph (Põlme et al. 2020); Decay sugar-rich substrates (Põlme et al. 2020) | Põlme S, et al. 2020. Fungal Diversity 105:1-16 ((https://doi.org/10.1007/s13225-020-00466-2)) |
| Otu16922 | 1 | k__Fungi(100);p__Ascomycota(100);c__Saccharomycetes(100);o__Saccharomycetales(100);f__Metschnikowiaceae(100);g__Metschnikowia(100);s__Metschnikowia_pulcherrima(100); | Metschnikowia | 13 | Saprotroph | \|Undefined Saprotroph\| | Yeast | NULL | Probable | As nectar-tap_saprotroph (Põlme et al. 2020); Decay sugar-rich substrates (Põlme et al. 2020) | Põlme S, et al. 2020. Fungal Diversity 105:1-16 ((https://doi.org/10.1007/s13225-020-00466-2)) |
| Otu16923 | 1 | k__Fungi(100);p__Ascomycota(100);c__Saccharomycetes(100);o__Saccharomycetales(100);f__Metschnikowiaceae(100);g__Metschnikowia(100);s__Metschnikowia_pulcherrima(100); | Metschnikowia | 13 | Saprotroph | \|Undefined Saprotroph\| | Yeast | NULL | Probable | As nectar-tap_saprotroph (Põlme et al. 2020); Decay sugar-rich substrates (Põlme et al. 2020) | Põlme S, et al. 2020. Fungal Diversity 105:1-16 ((https://doi.org/10.1007/s13225-020-00466-2)) |
| Otu16924 | 1 | k__Fungi(100);p__Ascomycota(100);c__Saccharomycetes(100);o__Saccharomycetales(100);f__Metschnikowiaceae(100);g__Metschnikowia(100);s__Metschnikowia_pulcherrima(100); | Metschnikowia | 13 | Saprotroph | \|Undefined Saprotroph\| | Yeast | NULL | Probable | As nectar-tap_saprotroph (Põlme et al. 2020); Decay sugar-rich substrates (Põlme et al. 2020) | Põlme S, et al. 2020. Fungal Diversity 105:1-16 ((https://doi.org/10.1007/s13225-020-00466-2)) |
| Otu16925 | 1 | k__Fungi(100);p__Ascomycota(100);c__Saccharomycetes(100);o__Saccharomycetales(100);f__Metschnikowiaceae(100);g__Metschnikowia(100);g__Metschnikowia_unclassified(100); | Metschnikowia | 13 | Saprotroph | \|Undefined Saprotroph\| | Yeast | NULL | Probable | As nectar-tap_saprotroph (Põlme et al. 2020); Decay sugar-rich substrates (Põlme et al. 2020) | Põlme S, et al. 2020. Fungal Diversity 105:1-16 ((https://doi.org/10.1007/s13225-020-00466-2)) |
| Otu16926 | 1 | k__Fungi(100);p__Ascomycota(100);c__Saccharomycetes(100);o__Saccharomycetales(100);f__Metschnikowiaceae(100);g__Metschnikowia(100);s__Metschnikowia_pulcherrima(100); | Metschnikowia | 13 | Saprotroph | \|Undefined Saprotroph\| | Yeast | NULL | Probable | As nectar-tap_saprotroph (Põlme et al. 2020); Decay sugar-rich substrates (Põlme et al. 2020) | Põlme S, et al. 2020. Fungal Diversity 105:1-16 ((https://doi.org/10.1007/s13225-020-00466-2)) |
| Otu16927 | 1 | k__Fungi(100);p__Ascomycota(100);c__Saccharomycetes(100);o__Saccharomycetales(100);f__Saccharomycetales_family_Incertae_sedis(100);g__Candida(100);g__Candida_unclassified(100); | Candida | 13 | Pathotroph-Saprotroph-Symbiotroph | Animal Pathogen-Endophyte-Endosymbiont-Epiphyte-\|Undefined Saprotroph\| | Dimorphic Yeast-Yeast | NULL | Probable | As nectar-tap_saprotroph (Põlme et al. 2020); Decay sugar-rich substrates (Põlme et al. 2020); This asexual genus is polyphyletic and contain species that function very differently depend on what they are. Caution should be taken when interpreting guilds for Candida species. | Manolakaki D, et al. 2010. Virulence. 1: 367-375 ((https://doi.org/10.4161/viru.1.5.12796)); Mekbib SB, et al. 2011. Tropical Plant Pathology 36:233-240 ((https://doi.org/10.1590/S1982-56762011000400004)); Põlme S, et al. 2020. Fungal Diversity 105:1-16 ((https://doi.org/10.1007/s13225-020-00466-2)) |
| Otu16928 | 1 | k__Fungi(100);p__Ascomycota(100);c__Saccharomycetes(100);o__Saccharomycetales(100);f__Metschnikowiaceae(100);g__Metschnikowia(100);s__Metschnikowia_pulcherrima(100); | Metschnikowia | 13 | Saprotroph | \|Undefined Saprotroph\| | Yeast | NULL | Probable | As nectar-tap_saprotroph (Põlme et al. 2020); Decay sugar-rich substrates (Põlme et al. 2020) | Põlme S, et al. 2020. Fungal Diversity 105:1-16 ((https://doi.org/10.1007/s13225-020-00466-2)) |
| Otu16929 | 1 | k__Fungi(100);p__Ascomycota(100);c__Saccharomycetes(100);o__Saccharomycetales(100);f__Metschnikowiaceae(100);g__Metschnikowia(100);s__Metschnikowia_pulcherrima(100); | Metschnikowia | 13 | Saprotroph | \|Undefined Saprotroph\| | Yeast | NULL | Probable | As nectar-tap_saprotroph (Põlme et al. 2020); Decay sugar-rich substrates (Põlme et al. 2020) | Põlme S, et al. 2020. Fungal Diversity 105:1-16 ((https://doi.org/10.1007/s13225-020-00466-2)) |
| Otu16930 | 1 | k__Fungi(100);p__Ascomycota(100);c__Saccharomycetes(100);o__Saccharomycetales(100);f__Metschnikowiaceae(100);g__Metschnikowia(100);s__Metschnikowia_pulcherrima(100); | Metschnikowia | 13 | Saprotroph | \|Undefined Saprotroph\| | Yeast | NULL | Probable | As nectar-tap_saprotroph (Põlme et al. 2020); Decay sugar-rich substrates (Põlme et al. 2020) | Põlme S, et al. 2020. Fungal Diversity 105:1-16 ((https://doi.org/10.1007/s13225-020-00466-2)) |
| Otu16931 | 1 | k__Fungi(100);p__Ascomycota(100);c__Saccharomycetes(100);o__Saccharomycetales(100);f__Metschnikowiaceae(100);g__Metschnikowia(100);g__Metschnikowia_unclassified(100); | Metschnikowia | 13 | Saprotroph | \|Undefined Saprotroph\| | Yeast | NULL | Probable | As nectar-tap_saprotroph (Põlme et al. 2020); Decay sugar-rich substrates (Põlme et al. 2020) | Põlme S, et al. 2020. Fungal Diversity 105:1-16 ((https://doi.org/10.1007/s13225-020-00466-2)) |
| Otu16932 | 1 | k__Fungi(100);p__Ascomycota(100);c__Saccharomycetes(100);o__Saccharomycetales(100);f__Metschnikowiaceae(100);g__Metschnikowia(100);g__Metschnikowia_unclassified(100); | Metschnikowia | 13 | Saprotroph | \|Undefined Saprotroph\| | Yeast | NULL | Probable | As nectar-tap_saprotroph (Põlme et al. 2020); Decay sugar-rich substrates (Põlme et al. 2020) | Põlme S, et al. 2020. Fungal Diversity 105:1-16 ((https://doi.org/10.1007/s13225-020-00466-2)) |
| Otu16933 | 1 | k__Fungi(100);p__Ascomycota(100);c__Saccharomycetes(100);o__Saccharomycetales(100);f__Metschnikowiaceae(100);g__Metschnikowia(100);s__Metschnikowia_pulcherrima(100); | Metschnikowia | 13 | Saprotroph | \|Undefined Saprotroph\| | Yeast | NULL | Probable | As nectar-tap_saprotroph (Põlme et al. 2020); Decay sugar-rich substrates (Põlme et al. 2020) | Põlme S, et al. 2020. Fungal Diversity 105:1-16 ((https://doi.org/10.1007/s13225-020-00466-2)) |
| Otu16934 | 1 | k__Fungi(100);p__Ascomycota(100);c__Saccharomycetes(100);o__Saccharomycetales(100);f__Metschnikowiaceae(100);g__Metschnikowia(100);s__Metschnikowia_pulcherrima(100); | Metschnikowia | 13 | Saprotroph | \|Undefined Saprotroph\| | Yeast | NULL | Probable | As nectar-tap_saprotroph (Põlme et al. 2020); Decay sugar-rich substrates (Põlme et al. 2020) | Põlme S, et al. 2020. Fungal Diversity 105:1-16 ((https://doi.org/10.1007/s13225-020-00466-2)) |
| Otu16937 | 1 | k__Fungi(100);p__Ascomycota(100);c__Saccharomycetes(100);o__Saccharomycetales(100);f__Metschnikowiaceae(100);g__Metschnikowia(100);s__Metschnikowia_pulcherrima(100); | Metschnikowia | 13 | Saprotroph | \|Undefined Saprotroph\| | Yeast | NULL | Probable | As nectar-tap_saprotroph (Põlme et al. 2020); Decay sugar-rich substrates (Põlme et al. 2020) | Põlme S, et al. 2020. Fungal Diversity 105:1-16 ((https://doi.org/10.1007/s13225-020-00466-2)) |
| Otu16939 | 1 | k__Fungi(100);p__Ascomycota(100);c__Saccharomycetes(100);o__Saccharomycetales(100);f__Metschnikowiaceae(100);g__Metschnikowia(100);s__Metschnikowia_pulcherrima(100); | Metschnikowia | 13 | Saprotroph | \|Undefined Saprotroph\| | Yeast | NULL | Probable | As nectar-tap_saprotroph (Põlme et al. 2020); Decay sugar-rich substrates (Põlme et al. 2020) | Põlme S, et al. 2020. Fungal Diversity 105:1-16 ((https://doi.org/10.1007/s13225-020-00466-2)) |
| Otu16940 | 1 | k__Fungi(100);p__Ascomycota(100);c__Saccharomycetes(100);o__Saccharomycetales(100);f__Metschnikowiaceae(100);g__Metschnikowia(100);s__Metschnikowia_pulcherrima(100); | Metschnikowia | 13 | Saprotroph | \|Undefined Saprotroph\| | Yeast | NULL | Probable | As nectar-tap_saprotroph (Põlme et al. 2020); Decay sugar-rich substrates (Põlme et al. 2020) | Põlme S, et al. 2020. Fungal Diversity 105:1-16 ((https://doi.org/10.1007/s13225-020-00466-2)) |
| Otu16941 | 1 | k__Fungi(100);p__Ascomycota(100);c__Saccharomycetes(100);o__Saccharomycetales(100);f__Metschnikowiaceae(100);g__Metschnikowia(100);s__Metschnikowia_pulcherrima(100); | Metschnikowia | 13 | Saprotroph | \|Undefined Saprotroph\| | Yeast | NULL | Probable | As nectar-tap_saprotroph (Põlme et al. 2020); Decay sugar-rich substrates (Põlme et al. 2020) | Põlme S, et al. 2020. Fungal Diversity 105:1-16 ((https://doi.org/10.1007/s13225-020-00466-2)) |
| Otu16942 | 1 | k__Fungi(100);p__Ascomycota(100);c__Saccharomycetes(100);o__Saccharomycetales(100);f__Metschnikowiaceae(100);g__Metschnikowia(100);s__Metschnikowia_pulcherrima(100); | Metschnikowia | 13 | Saprotroph | \|Undefined Saprotroph\| | Yeast | NULL | Probable | As nectar-tap_saprotroph (Põlme et al. 2020); Decay sugar-rich substrates (Põlme et al. 2020) | Põlme S, et al. 2020. Fungal Diversity 105:1-16 ((https://doi.org/10.1007/s13225-020-00466-2)) |
| Otu16946 | 1 | k__Fungi(100);p__Ascomycota(100);c__Saccharomycetes(100);o__Saccharomycetales(100);f__Metschnikowiaceae(100);g__Metschnikowia(100);g__Metschnikowia_unclassified(100); | Metschnikowia | 13 | Saprotroph | \|Undefined Saprotroph\| | Yeast | NULL | Probable | As nectar-tap_saprotroph (Põlme et al. 2020); Decay sugar-rich substrates (Põlme et al. 2020) | Põlme S, et al. 2020. Fungal Diversity 105:1-16 ((https://doi.org/10.1007/s13225-020-00466-2)) |
| Otu16947 | 1 | k__Fungi(100);p__Ascomycota(100);c__Saccharomycetes(100);o__Saccharomycetales(100);f__Metschnikowiaceae(100);g__Metschnikowia(100);s__Metschnikowia_pulcherrima(100); | Metschnikowia | 13 | Saprotroph | \|Undefined Saprotroph\| | Yeast | NULL | Probable | As nectar-tap_saprotroph (Põlme et al. 2020); Decay sugar-rich substrates (Põlme et al. 2020) | Põlme S, et al. 2020. Fungal Diversity 105:1-16 ((https://doi.org/10.1007/s13225-020-00466-2)) |
| Otu16948 | 1 | k__Fungi(100);p__Ascomycota(100);c__Saccharomycetes(100);o__Saccharomycetales(100);f__Metschnikowiaceae(100);g__Metschnikowia(100);g__Metschnikowia_unclassified(100); | Metschnikowia | 13 | Saprotroph | \|Undefined Saprotroph\| | Yeast | NULL | Probable | As nectar-tap_saprotroph (Põlme et al. 2020); Decay sugar-rich substrates (Põlme et al. 2020) | Põlme S, et al. 2020. Fungal Diversity 105:1-16 ((https://doi.org/10.1007/s13225-020-00466-2)) |
| Otu16949 | 1 | k__Fungi(100);p__Ascomycota(100);c__Saccharomycetes(100);o__Saccharomycetales(100);f__Metschnikowiaceae(100);g__Metschnikowia(100);s__Metschnikowia_pulcherrima(100); | Metschnikowia | 13 | Saprotroph | \|Undefined Saprotroph\| | Yeast | NULL | Probable | As nectar-tap_saprotroph (Põlme et al. 2020); Decay sugar-rich substrates (Põlme et al. 2020) | Põlme S, et al. 2020. Fungal Diversity 105:1-16 ((https://doi.org/10.1007/s13225-020-00466-2)) |
| Otu16952 | 1 | k__Fungi(100);p__Ascomycota(100);c__Saccharomycetes(100);o__Saccharomycetales(100);f__Metschnikowiaceae(100);g__Metschnikowia(100);s__Metschnikowia_pulcherrima(100); | Metschnikowia | 13 | Saprotroph | \|Undefined Saprotroph\| | Yeast | NULL | Probable | As nectar-tap_saprotroph (Põlme et al. 2020); Decay sugar-rich substrates (Põlme et al. 2020) | Põlme S, et al. 2020. Fungal Diversity 105:1-16 ((https://doi.org/10.1007/s13225-020-00466-2)) |
| Otu16953 | 1 | k__Fungi(100);p__Ascomycota(100);c__Saccharomycetes(100);o__Saccharomycetales(100);f__Metschnikowiaceae(100);g__Metschnikowia(100);s__Metschnikowia_pulcherrima(100); | Metschnikowia | 13 | Saprotroph | \|Undefined Saprotroph\| | Yeast | NULL | Probable | As nectar-tap_saprotroph (Põlme et al. 2020); Decay sugar-rich substrates (Põlme et al. 2020) | Põlme S, et al. 2020. Fungal Diversity 105:1-16 ((https://doi.org/10.1007/s13225-020-00466-2)) |
| Otu16954 | 1 | k__Fungi(100);p__Ascomycota(100);c__Saccharomycetes(100);o__Saccharomycetales(100);f__Metschnikowiaceae(100);g__Metschnikowia(100);g__Metschnikowia_unclassified(100); | Metschnikowia | 13 | Saprotroph | \|Undefined Saprotroph\| | Yeast | NULL | Probable | As nectar-tap_saprotroph (Põlme et al. 2020); Decay sugar-rich substrates (Põlme et al. 2020) | Põlme S, et al. 2020. Fungal Diversity 105:1-16 ((https://doi.org/10.1007/s13225-020-00466-2)) |
| Otu16957 | 1 | k__Fungi(100);p__Ascomycota(100);c__Saccharomycetes(100);o__Saccharomycetales(100);f__Metschnikowiaceae(100);g__Metschnikowia(100);s__Metschnikowia_pulcherrima(100); | Metschnikowia | 13 | Saprotroph | \|Undefined Saprotroph\| | Yeast | NULL | Probable | As nectar-tap_saprotroph (Põlme et al. 2020); Decay sugar-rich substrates (Põlme et al. 2020) | Põlme S, et al. 2020. Fungal Diversity 105:1-16 ((https://doi.org/10.1007/s13225-020-00466-2)) |
| Otu16958 | 1 | k__Fungi(100);p__Ascomycota(100);c__Saccharomycetes(100);o__Saccharomycetales(100);f__Metschnikowiaceae(100);g__Metschnikowia(100);s__Metschnikowia_pulcherrima(100); | Metschnikowia | 13 | Saprotroph | \|Undefined Saprotroph\| | Yeast | NULL | Probable | As nectar-tap_saprotroph (Põlme et al. 2020); Decay sugar-rich substrates (Põlme et al. 2020) | Põlme S, et al. 2020. Fungal Diversity 105:1-16 ((https://doi.org/10.1007/s13225-020-00466-2)) |
| Otu16959 | 1 | k__Fungi(100);p__Ascomycota(100);c__Saccharomycetes(100);o__Saccharomycetales(100);f__Metschnikowiaceae(100);g__Metschnikowia(100);s__Metschnikowia_pulcherrima(100); | Metschnikowia | 13 | Saprotroph | \|Undefined Saprotroph\| | Yeast | NULL | Probable | As nectar-tap_saprotroph (Põlme et al. 2020); Decay sugar-rich substrates (Põlme et al. 2020) | Põlme S, et al. 2020. Fungal Diversity 105:1-16 ((https://doi.org/10.1007/s13225-020-00466-2)) |
| Otu16960 | 1 | k__Fungi(100);p__Ascomycota(100);c__Saccharomycetes(100);o__Saccharomycetales(100);f__Metschnikowiaceae(100);g__Metschnikowia(100);s__Metschnikowia_pulcherrima(100); | Metschnikowia | 13 | Saprotroph | \|Undefined Saprotroph\| | Yeast | NULL | Probable | As nectar-tap_saprotroph (Põlme et al. 2020); Decay sugar-rich substrates (Põlme et al. 2020) | Põlme S, et al. 2020. Fungal Diversity 105:1-16 ((https://doi.org/10.1007/s13225-020-00466-2)) |
| Otu16961 | 1 | k__Fungi(100);p__Ascomycota(100);c__Saccharomycetes(100);o__Saccharomycetales(100);f__Metschnikowiaceae(100);g__Metschnikowia(100);s__Metschnikowia_pulcherrima(100); | Metschnikowia | 13 | Saprotroph | \|Undefined Saprotroph\| | Yeast | NULL | Probable | As nectar-tap_saprotroph (Põlme et al. 2020); Decay sugar-rich substrates (Põlme et al. 2020) | Põlme S, et al. 2020. Fungal Diversity 105:1-16 ((https://doi.org/10.1007/s13225-020-00466-2)) |
| Otu16962 | 1 | k__Fungi(100);p__Ascomycota(100);c__Saccharomycetes(100);o__Saccharomycetales(100);f__Metschnikowiaceae(100);g__Metschnikowia(100);s__Metschnikowia_pulcherrima(100); | Metschnikowia | 13 | Saprotroph | \|Undefined Saprotroph\| | Yeast | NULL | Probable | As nectar-tap_saprotroph (Põlme et al. 2020); Decay sugar-rich substrates (Põlme et al. 2020) | Põlme S, et al. 2020. Fungal Diversity 105:1-16 ((https://doi.org/10.1007/s13225-020-00466-2)) |
| Otu16963 | 1 | k__Fungi(100);p__Ascomycota(100);c__Saccharomycetes(100);o__Saccharomycetales(100);f__Metschnikowiaceae(100);g__Metschnikowia(100);s__Metschnikowia_pulcherrima(100); | Metschnikowia | 13 | Saprotroph | \|Undefined Saprotroph\| | Yeast | NULL | Probable | As nectar-tap_saprotroph (Põlme et al. 2020); Decay sugar-rich substrates (Põlme et al. 2020) | Põlme S, et al. 2020. Fungal Diversity 105:1-16 ((https://doi.org/10.1007/s13225-020-00466-2)) |
| Otu16964 | 1 | k__Fungi(100);p__Ascomycota(100);c__Saccharomycetes(100);o__Saccharomycetales(100);f__Metschnikowiaceae(100);g__Metschnikowia(100);g__Metschnikowia_unclassified(100); | Metschnikowia | 13 | Saprotroph | \|Undefined Saprotroph\| | Yeast | NULL | Probable | As nectar-tap_saprotroph (Põlme et al. 2020); Decay sugar-rich substrates (Põlme et al. 2020) | Põlme S, et al. 2020. Fungal Diversity 105:1-16 ((https://doi.org/10.1007/s13225-020-00466-2)) |
| Otu16965 | 1 | k__Fungi(100);p__Ascomycota(100);c__Saccharomycetes(100);o__Saccharomycetales(100);f__Metschnikowiaceae(100);g__Metschnikowia(100);g__Metschnikowia_unclassified(100); | Metschnikowia | 13 | Saprotroph | \|Undefined Saprotroph\| | Yeast | NULL | Probable | As nectar-tap_saprotroph (Põlme et al. 2020); Decay sugar-rich substrates (Põlme et al. 2020) | Põlme S, et al. 2020. Fungal Diversity 105:1-16 ((https://doi.org/10.1007/s13225-020-00466-2)) |
| Otu16966 | 1 | k__Fungi(100);p__Ascomycota(100);c__Saccharomycetes(100);o__Saccharomycetales(100);f__Metschnikowiaceae(100);g__Metschnikowia(100);s__Metschnikowia_pulcherrima(100); | Metschnikowia | 13 | Saprotroph | \|Undefined Saprotroph\| | Yeast | NULL | Probable | As nectar-tap_saprotroph (Põlme et al. 2020); Decay sugar-rich substrates (Põlme et al. 2020) | Põlme S, et al. 2020. Fungal Diversity 105:1-16 ((https://doi.org/10.1007/s13225-020-00466-2)) |
| Otu16967 | 1 | k__Fungi(100);p__Ascomycota(100);c__Saccharomycetes(100);o__Saccharomycetales(100);f__Metschnikowiaceae(100);g__Metschnikowia(100);s__Metschnikowia_pulcherrima(100); | Metschnikowia | 13 | Saprotroph | \|Undefined Saprotroph\| | Yeast | NULL | Probable | As nectar-tap_saprotroph (Põlme et al. 2020); Decay sugar-rich substrates (Põlme et al. 2020) | Põlme S, et al. 2020. Fungal Diversity 105:1-16 ((https://doi.org/10.1007/s13225-020-00466-2)) |
| Otu16969 | 1 | k__Fungi(100);p__Ascomycota(100);c__Saccharomycetes(100);o__Saccharomycetales(100);f__Metschnikowiaceae(100);g__Metschnikowia(100);s__Metschnikowia_pulcherrima(100); | Metschnikowia | 13 | Saprotroph | \|Undefined Saprotroph\| | Yeast | NULL | Probable | As nectar-tap_saprotroph (Põlme et al. 2020); Decay sugar-rich substrates (Põlme et al. 2020) | Põlme S, et al. 2020. Fungal Diversity 105:1-16 ((https://doi.org/10.1007/s13225-020-00466-2)) |
| Otu16970 | 1 | k__Fungi(100);p__Ascomycota(100);c__Saccharomycetes(100);o__Saccharomycetales(100);f__Metschnikowiaceae(100);g__Metschnikowia(100);s__Metschnikowia_pulcherrima(100); | Metschnikowia | 13 | Saprotroph | \|Undefined Saprotroph\| | Yeast | NULL | Probable | As nectar-tap_saprotroph (Põlme et al. 2020); Decay sugar-rich substrates (Põlme et al. 2020) | Põlme S, et al. 2020. Fungal Diversity 105:1-16 ((https://doi.org/10.1007/s13225-020-00466-2)) |
| Otu16971 | 1 | k__Fungi(100);p__Ascomycota(100);c__Saccharomycetes(100);o__Saccharomycetales(100);f__Metschnikowiaceae(100);g__Metschnikowia(100);s__Metschnikowia_pulcherrima(100); | Metschnikowia | 13 | Saprotroph | \|Undefined Saprotroph\| | Yeast | NULL | Probable | As nectar-tap_saprotroph (Põlme et al. 2020); Decay sugar-rich substrates (Põlme et al. 2020) | Põlme S, et al. 2020. Fungal Diversity 105:1-16 ((https://doi.org/10.1007/s13225-020-00466-2)) |
| Otu16972 | 1 | k__Fungi(100);p__Ascomycota(100);c__Saccharomycetes(100);o__Saccharomycetales(100);f__Metschnikowiaceae(100);g__Metschnikowia(100);s__Metschnikowia_pulcherrima(100); | Metschnikowia | 13 | Saprotroph | \|Undefined Saprotroph\| | Yeast | NULL | Probable | As nectar-tap_saprotroph (Põlme et al. 2020); Decay sugar-rich substrates (Põlme et al. 2020) | Põlme S, et al. 2020. Fungal Diversity 105:1-16 ((https://doi.org/10.1007/s13225-020-00466-2)) |
| Otu16973 | 1 | k__Fungi(100);p__Ascomycota(100);c__Saccharomycetes(100);o__Saccharomycetales(100);f__Metschnikowiaceae(100);g__Metschnikowia(100);g__Metschnikowia_unclassified(100); | Metschnikowia | 13 | Saprotroph | \|Undefined Saprotroph\| | Yeast | NULL | Probable | As nectar-tap_saprotroph (Põlme et al. 2020); Decay sugar-rich substrates (Põlme et al. 2020) | Põlme S, et al. 2020. Fungal Diversity 105:1-16 ((https://doi.org/10.1007/s13225-020-00466-2)) |
| Otu16974 | 1 | k__Fungi(100);p__Ascomycota(100);c__Saccharomycetes(100);o__Saccharomycetales(100);f__Metschnikowiaceae(100);g__Metschnikowia(100);g__Metschnikowia_unclassified(100); | Metschnikowia | 13 | Saprotroph | \|Undefined Saprotroph\| | Yeast | NULL | Probable | As nectar-tap_saprotroph (Põlme et al. 2020); Decay sugar-rich substrates (Põlme et al. 2020) | Põlme S, et al. 2020. Fungal Diversity 105:1-16 ((https://doi.org/10.1007/s13225-020-00466-2)) |
| Otu16981 | 1 | k__Fungi(100);p__Ascomycota(100);c__Saccharomycetes(100);o__Saccharomycetales(100);f__Metschnikowiaceae(100);g__Metschnikowia(100);s__Metschnikowia_pulcherrima(100); | Metschnikowia | 13 | Saprotroph | \|Undefined Saprotroph\| | Yeast | NULL | Probable | As nectar-tap_saprotroph (Põlme et al. 2020); Decay sugar-rich substrates (Põlme et al. 2020) | Põlme S, et al. 2020. Fungal Diversity 105:1-16 ((https://doi.org/10.1007/s13225-020-00466-2)) |
| Otu16986 | 1 | k__Fungi(100);p__Ascomycota(100);c__Saccharomycetes(100);o__Saccharomycetales(100);f__Metschnikowiaceae(100);g__Metschnikowia(100);s__Metschnikowia_pulcherrima(100); | Metschnikowia | 13 | Saprotroph | \|Undefined Saprotroph\| | Yeast | NULL | Probable | As nectar-tap_saprotroph (Põlme et al. 2020); Decay sugar-rich substrates (Põlme et al. 2020) | Põlme S, et al. 2020. Fungal Diversity 105:1-16 ((https://doi.org/10.1007/s13225-020-00466-2)) |
| Otu16987 | 1 | k__Fungi(100);p__Ascomycota(100);c__Saccharomycetes(100);o__Saccharomycetales(100);f__Metschnikowiaceae(100);g__Metschnikowia(100);g__Metschnikowia_unclassified(100); | Metschnikowia | 13 | Saprotroph | \|Undefined Saprotroph\| | Yeast | NULL | Probable | As nectar-tap_saprotroph (Põlme et al. 2020); Decay sugar-rich substrates (Põlme et al. 2020) | Põlme S, et al. 2020. Fungal Diversity 105:1-16 ((https://doi.org/10.1007/s13225-020-00466-2)) |
| Otu16988 | 1 | k__Fungi(100);p__Ascomycota(100);c__Saccharomycetes(100);o__Saccharomycetales(100);f__Metschnikowiaceae(100);g__Metschnikowia(100);s__Metschnikowia_pulcherrima(100); | Metschnikowia | 13 | Saprotroph | \|Undefined Saprotroph\| | Yeast | NULL | Probable | As nectar-tap_saprotroph (Põlme et al. 2020); Decay sugar-rich substrates (Põlme et al. 2020) | Põlme S, et al. 2020. Fungal Diversity 105:1-16 ((https://doi.org/10.1007/s13225-020-00466-2)) |
| Otu16989 | 1 | k__Fungi(100);p__Ascomycota(100);c__Saccharomycetes(100);o__Saccharomycetales(100);f__Metschnikowiaceae(100);g__Metschnikowia(100);s__Metschnikowia_pulcherrima(100); | Metschnikowia | 13 | Saprotroph | \|Undefined Saprotroph\| | Yeast | NULL | Probable | As nectar-tap_saprotroph (Põlme et al. 2020); Decay sugar-rich substrates (Põlme et al. 2020) | Põlme S, et al. 2020. Fungal Diversity 105:1-16 ((https://doi.org/10.1007/s13225-020-00466-2)) |
| Otu16994 | 1 | k__Fungi(100);p__Ascomycota(100);c__Saccharomycetes(100);o__Saccharomycetales(100);f__Metschnikowiaceae(100);g__Metschnikowia(100);s__Metschnikowia_pulcherrima(100); | Metschnikowia | 13 | Saprotroph | \|Undefined Saprotroph\| | Yeast | NULL | Probable | As nectar-tap_saprotroph (Põlme et al. 2020); Decay sugar-rich substrates (Põlme et al. 2020) | Põlme S, et al. 2020. Fungal Diversity 105:1-16 ((https://doi.org/10.1007/s13225-020-00466-2)) |
| Otu16995 | 1 | k__Fungi(100);p__Ascomycota(100);c__Saccharomycetes(100);o__Saccharomycetales(100);f__Metschnikowiaceae(100);g__Metschnikowia(100);g__Metschnikowia_unclassified(100); | Metschnikowia | 13 | Saprotroph | \|Undefined Saprotroph\| | Yeast | NULL | Probable | As nectar-tap_saprotroph (Põlme et al. 2020); Decay sugar-rich substrates (Põlme et al. 2020) | Põlme S, et al. 2020. Fungal Diversity 105:1-16 ((https://doi.org/10.1007/s13225-020-00466-2)) |
| Otu16996 | 1 | k__Fungi(100);p__Ascomycota(100);c__Saccharomycetes(100);o__Saccharomycetales(100);f__Metschnikowiaceae(100);g__Metschnikowia(100);s__Metschnikowia_pulcherrima(100); | Metschnikowia | 13 | Saprotroph | \|Undefined Saprotroph\| | Yeast | NULL | Probable | As nectar-tap_saprotroph (Põlme et al. 2020); Decay sugar-rich substrates (Põlme et al. 2020) | Põlme S, et al. 2020. Fungal Diversity 105:1-16 ((https://doi.org/10.1007/s13225-020-00466-2)) |
| Otu16997 | 1 | k__Fungi(100);p__Ascomycota(100);c__Saccharomycetes(100);o__Saccharomycetales(100);f__Metschnikowiaceae(100);g__Metschnikowia(100);s__Metschnikowia_pulcherrima(100); | Metschnikowia | 13 | Saprotroph | \|Undefined Saprotroph\| | Yeast | NULL | Probable | As nectar-tap_saprotroph (Põlme et al. 2020); Decay sugar-rich substrates (Põlme et al. 2020) | Põlme S, et al. 2020. Fungal Diversity 105:1-16 ((https://doi.org/10.1007/s13225-020-00466-2)) |
| Otu16998 | 1 | k__Fungi(100);p__Ascomycota(100);c__Saccharomycetes(100);o__Saccharomycetales(100);f__Metschnikowiaceae(100);g__Metschnikowia(100);s__Metschnikowia_pulcherrima(100); | Metschnikowia | 13 | Saprotroph | \|Undefined Saprotroph\| | Yeast | NULL | Probable | As nectar-tap_saprotroph (Põlme et al. 2020); Decay sugar-rich substrates (Põlme et al. 2020) | Põlme S, et al. 2020. Fungal Diversity 105:1-16 ((https://doi.org/10.1007/s13225-020-00466-2)) |
| Otu16999 | 1 | k__Fungi(100);p__Ascomycota(100);c__Saccharomycetes(100);o__Saccharomycetales(100);f__Metschnikowiaceae(100);g__Metschnikowia(100);s__Metschnikowia_pulcherrima(100); | Metschnikowia | 13 | Saprotroph | \|Undefined Saprotroph\| | Yeast | NULL | Probable | As nectar-tap_saprotroph (Põlme et al. 2020); Decay sugar-rich substrates (Põlme et al. 2020) | Põlme S, et al. 2020. Fungal Diversity 105:1-16 ((https://doi.org/10.1007/s13225-020-00466-2)) |
| Otu17000 | 1 | k__Fungi(100);p__Ascomycota(100);c__Saccharomycetes(100);o__Saccharomycetales(100);f__Metschnikowiaceae(100);g__Metschnikowia(100);s__Metschnikowia_pulcherrima(100); | Metschnikowia | 13 | Saprotroph | \|Undefined Saprotroph\| | Yeast | NULL | Probable | As nectar-tap_saprotroph (Põlme et al. 2020); Decay sugar-rich substrates (Põlme et al. 2020) | Põlme S, et al. 2020. Fungal Diversity 105:1-16 ((https://doi.org/10.1007/s13225-020-00466-2)) |
| Otu17001 | 1 | k__Fungi(100);p__Ascomycota(100);c__Saccharomycetes(100);o__Saccharomycetales(100);f__Metschnikowiaceae(100);g__Metschnikowia(100);s__Metschnikowia_pulcherrima(100); | Metschnikowia | 13 | Saprotroph | \|Undefined Saprotroph\| | Yeast | NULL | Probable | As nectar-tap_saprotroph (Põlme et al. 2020); Decay sugar-rich substrates (Põlme et al. 2020) | Põlme S, et al. 2020. Fungal Diversity 105:1-16 ((https://doi.org/10.1007/s13225-020-00466-2)) |
| Otu17002 | 1 | k__Fungi(100);p__Ascomycota(100);c__Saccharomycetes(100);o__Saccharomycetales(100);f__Metschnikowiaceae(100);g__Metschnikowia(100);s__Metschnikowia_pulcherrima(100); | Metschnikowia | 13 | Saprotroph | \|Undefined Saprotroph\| | Yeast | NULL | Probable | As nectar-tap_saprotroph (Põlme et al. 2020); Decay sugar-rich substrates (Põlme et al. 2020) | Põlme S, et al. 2020. Fungal Diversity 105:1-16 ((https://doi.org/10.1007/s13225-020-00466-2)) |
| Otu17004 | 1 | k__Fungi(100);p__Ascomycota(100);c__Saccharomycetes(100);o__Saccharomycetales(100);f__Metschnikowiaceae(100);g__Metschnikowia(100);s__Metschnikowia_pulcherrima(100); | Metschnikowia | 13 | Saprotroph | \|Undefined Saprotroph\| | Yeast | NULL | Probable | As nectar-tap_saprotroph (Põlme et al. 2020); Decay sugar-rich substrates (Põlme et al. 2020) | Põlme S, et al. 2020. Fungal Diversity 105:1-16 ((https://doi.org/10.1007/s13225-020-00466-2)) |
| Otu17005 | 1 | k__Fungi(100);p__Ascomycota(100);c__Saccharomycetes(100);o__Saccharomycetales(100);f__Metschnikowiaceae(100);g__Metschnikowia(100);s__Metschnikowia_pulcherrima(100); | Metschnikowia | 13 | Saprotroph | \|Undefined Saprotroph\| | Yeast | NULL | Probable | As nectar-tap_saprotroph (Põlme et al. 2020); Decay sugar-rich substrates (Põlme et al. 2020) | Põlme S, et al. 2020. Fungal Diversity 105:1-16 ((https://doi.org/10.1007/s13225-020-00466-2)) |
| Otu17006 | 1 | k__Fungi(100);p__Ascomycota(100);c__Saccharomycetes(100);o__Saccharomycetales(100);f__Metschnikowiaceae(100);g__Metschnikowia(100);s__Metschnikowia_pulcherrima(100); | Metschnikowia | 13 | Saprotroph | \|Undefined Saprotroph\| | Yeast | NULL | Probable | As nectar-tap_saprotroph (Põlme et al. 2020); Decay sugar-rich substrates (Põlme et al. 2020) | Põlme S, et al. 2020. Fungal Diversity 105:1-16 ((https://doi.org/10.1007/s13225-020-00466-2)) |
| Otu17007 | 1 | k__Fungi(100);p__Ascomycota(100);c__Saccharomycetes(100);o__Saccharomycetales(100);f__Metschnikowiaceae(100);g__Metschnikowia(100);s__Metschnikowia_pulcherrima(100); | Metschnikowia | 13 | Saprotroph | \|Undefined Saprotroph\| | Yeast | NULL | Probable | As nectar-tap_saprotroph (Põlme et al. 2020); Decay sugar-rich substrates (Põlme et al. 2020) | Põlme S, et al. 2020. Fungal Diversity 105:1-16 ((https://doi.org/10.1007/s13225-020-00466-2)) |
| Otu17008 | 1 | k__Fungi(100);p__Ascomycota(100);c__Saccharomycetes(100);o__Saccharomycetales(100);f__Metschnikowiaceae(100);g__Metschnikowia(100);g__Metschnikowia_unclassified(100); | Metschnikowia | 13 | Saprotroph | \|Undefined Saprotroph\| | Yeast | NULL | Probable | As nectar-tap_saprotroph (Põlme et al. 2020); Decay sugar-rich substrates (Põlme et al. 2020) | Põlme S, et al. 2020. Fungal Diversity 105:1-16 ((https://doi.org/10.1007/s13225-020-00466-2)) |
| Otu17009 | 1 | k__Fungi(100);p__Ascomycota(100);c__Saccharomycetes(100);o__Saccharomycetales(100);f__Metschnikowiaceae(100);g__Metschnikowia(100);g__Metschnikowia_unclassified(100); | Metschnikowia | 13 | Saprotroph | \|Undefined Saprotroph\| | Yeast | NULL | Probable | As nectar-tap_saprotroph (Põlme et al. 2020); Decay sugar-rich substrates (Põlme et al. 2020) | Põlme S, et al. 2020. Fungal Diversity 105:1-16 ((https://doi.org/10.1007/s13225-020-00466-2)) |
| Otu17010 | 1 | k__Fungi(100);p__Ascomycota(100);c__Saccharomycetes(100);o__Saccharomycetales(100);f__Metschnikowiaceae(100);g__Metschnikowia(100);s__Metschnikowia_pulcherrima(100); | Metschnikowia | 13 | Saprotroph | \|Undefined Saprotroph\| | Yeast | NULL | Probable | As nectar-tap_saprotroph (Põlme et al. 2020); Decay sugar-rich substrates (Põlme et al. 2020) | Põlme S, et al. 2020. Fungal Diversity 105:1-16 ((https://doi.org/10.1007/s13225-020-00466-2)) |
| Otu17011 | 1 | k__Fungi(100);p__Ascomycota(100);c__Saccharomycetes(100);o__Saccharomycetales(100);f__Metschnikowiaceae(100);g__Metschnikowia(100);g__Metschnikowia_unclassified(100); | Metschnikowia | 13 | Saprotroph | \|Undefined Saprotroph\| | Yeast | NULL | Probable | As nectar-tap_saprotroph (Põlme et al. 2020); Decay sugar-rich substrates (Põlme et al. 2020) | Põlme S, et al. 2020. Fungal Diversity 105:1-16 ((https://doi.org/10.1007/s13225-020-00466-2)) |
| Otu17012 | 1 | k__Fungi(100);p__Ascomycota(100);c__Saccharomycetes(100);o__Saccharomycetales(100);f__Metschnikowiaceae(100);g__Metschnikowia(100);g__Metschnikowia_unclassified(100); | Metschnikowia | 13 | Saprotroph | \|Undefined Saprotroph\| | Yeast | NULL | Probable | As nectar-tap_saprotroph (Põlme et al. 2020); Decay sugar-rich substrates (Põlme et al. 2020) | Põlme S, et al. 2020. Fungal Diversity 105:1-16 ((https://doi.org/10.1007/s13225-020-00466-2)) |
| Otu17013 | 1 | k__Fungi(100);p__Ascomycota(100);c__Saccharomycetes(100);o__Saccharomycetales(100);f__Saccharomycetales_family_Incertae_sedis(100);f__Saccharomycetales_family_Incertae_sedis_unclassified(100);f__Saccharomycetales_family_Incertae_sedis_unclassified(100); | Saccharomycetales | 7 | Saprotroph | Undefined Saprotroph | Yeast | NULL | Possible | NULL | Sterkenburg E, et al. 2015. New Phytologist 207:1145-1158 ((https://doi.org/10.1111/nph.13426)) |
| Otu17015 | 1 | k__Fungi(100);p__Ascomycota(100);c__Saccharomycetes(100);o__Saccharomycetales(100);f__Metschnikowiaceae(100);g__Metschnikowia(100);g__Metschnikowia_unclassified(100); | Metschnikowia | 13 | Saprotroph | \|Undefined Saprotroph\| | Yeast | NULL | Probable | As nectar-tap_saprotroph (Põlme et al. 2020); Decay sugar-rich substrates (Põlme et al. 2020) | Põlme S, et al. 2020. Fungal Diversity 105:1-16 ((https://doi.org/10.1007/s13225-020-00466-2)) |
| Otu17018 | 1 | k__Fungi(100);p__Ascomycota(100);c__Saccharomycetes(100);o__Saccharomycetales(100);f__Metschnikowiaceae(100);g__Metschnikowia(100);s__Metschnikowia_pulcherrima(100); | Metschnikowia | 13 | Saprotroph | \|Undefined Saprotroph\| | Yeast | NULL | Probable | As nectar-tap_saprotroph (Põlme et al. 2020); Decay sugar-rich substrates (Põlme et al. 2020) | Põlme S, et al. 2020. Fungal Diversity 105:1-16 ((https://doi.org/10.1007/s13225-020-00466-2)) |
| Otu17020 | 1 | k__Fungi(100);p__Ascomycota(100);c__Saccharomycetes(100);o__Saccharomycetales(100);f__Metschnikowiaceae(100);g__Metschnikowia(100);g__Metschnikowia_unclassified(100); | Metschnikowia | 13 | Saprotroph | \|Undefined Saprotroph\| | Yeast | NULL | Probable | As nectar-tap_saprotroph (Põlme et al. 2020); Decay sugar-rich substrates (Põlme et al. 2020) | Põlme S, et al. 2020. Fungal Diversity 105:1-16 ((https://doi.org/10.1007/s13225-020-00466-2)) |
| Otu17023 | 1 | k__Fungi(100);p__Ascomycota(100);c__Saccharomycetes(100);o__Saccharomycetales(100);f__Metschnikowiaceae(100);g__Metschnikowia(100);s__Metschnikowia_pulcherrima(100); | Metschnikowia | 13 | Saprotroph | \|Undefined Saprotroph\| | Yeast | NULL | Probable | As nectar-tap_saprotroph (Põlme et al. 2020); Decay sugar-rich substrates (Põlme et al. 2020) | Põlme S, et al. 2020. Fungal Diversity 105:1-16 ((https://doi.org/10.1007/s13225-020-00466-2)) |
| Otu17024 | 1 | k__Fungi(100);p__Ascomycota(100);c__Saccharomycetes(100);o__Saccharomycetales(100);f__Metschnikowiaceae(100);g__Metschnikowia(100);s__Metschnikowia_pulcherrima(100); | Metschnikowia | 13 | Saprotroph | \|Undefined Saprotroph\| | Yeast | NULL | Probable | As nectar-tap_saprotroph (Põlme et al. 2020); Decay sugar-rich substrates (Põlme et al. 2020) | Põlme S, et al. 2020. Fungal Diversity 105:1-16 ((https://doi.org/10.1007/s13225-020-00466-2)) |
| Otu17025 | 1 | k__Fungi(100);p__Ascomycota(100);c__Saccharomycetes(100);o__Saccharomycetales(100);f__Metschnikowiaceae(100);g__Metschnikowia(100);g__Metschnikowia_unclassified(100); | Metschnikowia | 13 | Saprotroph | \|Undefined Saprotroph\| | Yeast | NULL | Probable | As nectar-tap_saprotroph (Põlme et al. 2020); Decay sugar-rich substrates (Põlme et al. 2020) | Põlme S, et al. 2020. Fungal Diversity 105:1-16 ((https://doi.org/10.1007/s13225-020-00466-2)) |
| Otu17026 | 1 | k__Fungi(100);p__Ascomycota(100);c__Saccharomycetes(100);o__Saccharomycetales(100);f__Metschnikowiaceae(100);g__Metschnikowia(100);g__Metschnikowia_unclassified(100); | Metschnikowia | 13 | Saprotroph | \|Undefined Saprotroph\| | Yeast | NULL | Probable | As nectar-tap_saprotroph (Põlme et al. 2020); Decay sugar-rich substrates (Põlme et al. 2020) | Põlme S, et al. 2020. Fungal Diversity 105:1-16 ((https://doi.org/10.1007/s13225-020-00466-2)) |
| Otu17028 | 1 | k__Fungi(100);p__Ascomycota(100);c__Saccharomycetes(100);o__Saccharomycetales(100);f__Metschnikowiaceae(100);g__Metschnikowia(100);s__Metschnikowia_pulcherrima(100); | Metschnikowia | 13 | Saprotroph | \|Undefined Saprotroph\| | Yeast | NULL | Probable | As nectar-tap_saprotroph (Põlme et al. 2020); Decay sugar-rich substrates (Põlme et al. 2020) | Põlme S, et al. 2020. Fungal Diversity 105:1-16 ((https://doi.org/10.1007/s13225-020-00466-2)) |
| Otu17029 | 1 | k__Fungi(100);p__Ascomycota(100);c__Saccharomycetes(100);o__Saccharomycetales(100);f__Metschnikowiaceae(100);g__Metschnikowia(100);g__Metschnikowia_unclassified(100); | Metschnikowia | 13 | Saprotroph | \|Undefined Saprotroph\| | Yeast | NULL | Probable | As nectar-tap_saprotroph (Põlme et al. 2020); Decay sugar-rich substrates (Põlme et al. 2020) | Põlme S, et al. 2020. Fungal Diversity 105:1-16 ((https://doi.org/10.1007/s13225-020-00466-2)) |
| Otu17030 | 1 | k__Fungi(100);p__Ascomycota(100);c__Saccharomycetes(100);o__Saccharomycetales(100);f__Metschnikowiaceae(100);g__Metschnikowia(100);s__Metschnikowia_pulcherrima(100); | Metschnikowia | 13 | Saprotroph | \|Undefined Saprotroph\| | Yeast | NULL | Probable | As nectar-tap_saprotroph (Põlme et al. 2020); Decay sugar-rich substrates (Põlme et al. 2020) | Põlme S, et al. 2020. Fungal Diversity 105:1-16 ((https://doi.org/10.1007/s13225-020-00466-2)) |
| Otu17032 | 1 | k__Fungi(100);p__Ascomycota(100);c__Saccharomycetes(100);o__Saccharomycetales(100);f__Metschnikowiaceae(100);g__Metschnikowia(100);g__Metschnikowia_unclassified(100); | Metschnikowia | 13 | Saprotroph | \|Undefined Saprotroph\| | Yeast | NULL | Probable | As nectar-tap_saprotroph (Põlme et al. 2020); Decay sugar-rich substrates (Põlme et al. 2020) | Põlme S, et al. 2020. Fungal Diversity 105:1-16 ((https://doi.org/10.1007/s13225-020-00466-2)) |
| Otu17034 | 1 | k__Fungi(100);p__Ascomycota(100);c__Saccharomycetes(100);o__Saccharomycetales(100);f__Metschnikowiaceae(100);g__Metschnikowia(100);s__Metschnikowia_pulcherrima(100); | Metschnikowia | 13 | Saprotroph | \|Undefined Saprotroph\| | Yeast | NULL | Probable | As nectar-tap_saprotroph (Põlme et al. 2020); Decay sugar-rich substrates (Põlme et al. 2020) | Põlme S, et al. 2020. Fungal Diversity 105:1-16 ((https://doi.org/10.1007/s13225-020-00466-2)) |
| Otu17035 | 1 | k__Fungi(100);p__Ascomycota(100);c__Saccharomycetes(100);o__Saccharomycetales(100);f__Metschnikowiaceae(100);g__Metschnikowia(100);s__Metschnikowia_pulcherrima(100); | Metschnikowia | 13 | Saprotroph | \|Undefined Saprotroph\| | Yeast | NULL | Probable | As nectar-tap_saprotroph (Põlme et al. 2020); Decay sugar-rich substrates (Põlme et al. 2020) | Põlme S, et al. 2020. Fungal Diversity 105:1-16 ((https://doi.org/10.1007/s13225-020-00466-2)) |
| Otu17036 | 1 | k__Fungi(100);p__Ascomycota(100);c__Saccharomycetes(100);o__Saccharomycetales(100);f__Metschnikowiaceae(100);g__Metschnikowia(100);s__Metschnikowia_pulcherrima(100); | Metschnikowia | 13 | Saprotroph | \|Undefined Saprotroph\| | Yeast | NULL | Probable | As nectar-tap_saprotroph (Põlme et al. 2020); Decay sugar-rich substrates (Põlme et al. 2020) | Põlme S, et al. 2020. Fungal Diversity 105:1-16 ((https://doi.org/10.1007/s13225-020-00466-2)) |
| Otu17037 | 1 | k__Fungi(100);p__Ascomycota(100);c__Saccharomycetes(100);o__Saccharomycetales(100);f__Metschnikowiaceae(100);g__Metschnikowia(100);s__Metschnikowia_pulcherrima(100); | Metschnikowia | 13 | Saprotroph | \|Undefined Saprotroph\| | Yeast | NULL | Probable | As nectar-tap_saprotroph (Põlme et al. 2020); Decay sugar-rich substrates (Põlme et al. 2020) | Põlme S, et al. 2020. Fungal Diversity 105:1-16 ((https://doi.org/10.1007/s13225-020-00466-2)) |
| Otu17038 | 1 | k__Fungi(100);p__Ascomycota(100);c__Saccharomycetes(100);o__Saccharomycetales(100);f__Metschnikowiaceae(100);g__Metschnikowia(100);g__Metschnikowia_unclassified(100); | Metschnikowia | 13 | Saprotroph | \|Undefined Saprotroph\| | Yeast | NULL | Probable | As nectar-tap_saprotroph (Põlme et al. 2020); Decay sugar-rich substrates (Põlme et al. 2020) | Põlme S, et al. 2020. Fungal Diversity 105:1-16 ((https://doi.org/10.1007/s13225-020-00466-2)) |
| Otu17039 | 1 | k__Fungi(100);p__Ascomycota(100);c__Saccharomycetes(100);o__Saccharomycetales(100);f__Metschnikowiaceae(100);g__Metschnikowia(100);s__Metschnikowia_pulcherrima(100); | Metschnikowia | 13 | Saprotroph | \|Undefined Saprotroph\| | Yeast | NULL | Probable | As nectar-tap_saprotroph (Põlme et al. 2020); Decay sugar-rich substrates (Põlme et al. 2020) | Põlme S, et al. 2020. Fungal Diversity 105:1-16 ((https://doi.org/10.1007/s13225-020-00466-2)) |

| | | | | | | | | | | | |
|---|---|---|---|---|---|---|---|---|---|---|---|
| Otu17040 | 1 | k__Fungi(100);p__Ascomycota(100);c__Saccharomycetes(100);o__Saccharomycetales(100);f__Metschnikowiaceae(100);g__Metschnikowia(100);s__Metschnikowia_pulcherrima(100); | Metschnikowia | 13 | Saprotroph | \|Undefined Saprotroph\| | Yeast | NULL | Probable | As nectar-tap_saprotroph (Põlme et al. 2020); Decay sugar-rich substrates (Põlme et al. 2020) | Põlme S, et al. 2020. Fungal Diversity 105:1-16 ((https://doi.org/10.1007/s13225-020-00466-2)) |
| Otu17041 | 1 | k__Fungi(100);p__Ascomycota(100);c__Saccharomycetes(100);o__Saccharomycetales(100);f__Metschnikowiaceae(100);g__Metschnikowia(100);s__Metschnikowia_pulcherrima(100); | Metschnikowia | 13 | Saprotroph | \|Undefined Saprotroph\| | Yeast | NULL | Probable | As nectar-tap_saprotroph (Põlme et al. 2020); Decay sugar-rich substrates (Põlme et al. 2020) | Põlme S, et al. 2020. Fungal Diversity 105:1-16 ((https://doi.org/10.1007/s13225-020-00466-2)) |
| Otu17042 | 1 | k__Fungi(100);p__Ascomycota(100);c__Saccharomycetes(100);o__Saccharomycetales(100);f__Metschnikowiaceae(100);g__Metschnikowia(100);s__Metschnikowia_pulcherrima(100); | Metschnikowia | 13 | Saprotroph | \|Undefined Saprotroph\| | Yeast | NULL | Probable | As nectar-tap_saprotroph (Põlme et al. 2020); Decay sugar-rich substrates (Põlme et al. 2020) | Põlme S, et al. 2020. Fungal Diversity 105:1-16 ((https://doi.org/10.1007/s13225-020-00466-2)) |
| Otu17043 | 1 | k__Fungi(100);p__Ascomycota(100);c__Saccharomycetes(100);o__Saccharomycetales(100);f__Metschnikowiaceae(100);g__Metschnikowia(100);g__Metschnikowia_unclassified(100); | Metschnikowia | 13 | Saprotroph | \|Undefined Saprotroph\| | Yeast | NULL | Probable | As nectar-tap_saprotroph (Põlme et al. 2020); Decay sugar-rich substrates (Põlme et al. 2020) | Põlme S, et al. 2020. Fungal Diversity 105:1-16 ((https://doi.org/10.1007/s13225-020-00466-2)) |
| Otu17044 | 1 | k__Fungi(100);p__Ascomycota(100);c__Saccharomycetes(100);o__Saccharomycetales(100);f__Metschnikowiaceae(100);g__Metschnikowia(100);s__Metschnikowia_pulcherrima(100); | Metschnikowia | 13 | Saprotroph | \|Undefined Saprotroph\| | Yeast | NULL | Probable | As nectar-tap_saprotroph (Põlme et al. 2020); Decay sugar-rich substrates (Põlme et al. 2020) | Põlme S, et al. 2020. Fungal Diversity 105:1-16 ((https://doi.org/10.1007/s13225-020-00466-2)) |
| Otu17045 | 1 | k__Fungi(100);p__Ascomycota(100);c__Saccharomycetes(100);o__Saccharomycetales(100);f__Metschnikowiaceae(100);g__Metschnikowia(100);s__Metschnikowia_pulcherrima(100); | Metschnikowia | 13 | Saprotroph | \|Undefined Saprotroph\| | Yeast | NULL | Probable | As nectar-tap_saprotroph (Põlme et al. 2020); Decay sugar-rich substrates (Põlme et al. 2020) | Põlme S, et al. 2020. Fungal Diversity 105:1-16 ((https://doi.org/10.1007/s13225-020-00466-2)) |
| Otu17046 | 1 | k__Fungi(100);p__Ascomycota(100);c__Saccharomycetes(100);o__Saccharomycetales(100);f__Metschnikowiaceae(100);g__Metschnikowia(100);s__Metschnikowia_pulcherrima(100); | Metschnikowia | 13 | Saprotroph | \|Undefined Saprotroph\| | Yeast | NULL | Probable | As nectar-tap_saprotroph (Põlme et al. 2020); Decay sugar-rich substrates (Põlme et al. 2020) | Põlme S, et al. 2020. Fungal Diversity 105:1-16 ((https://doi.org/10.1007/s13225-020-00466-2)) |
| Otu17047 | 1 | k__Fungi(100);p__Ascomycota(100);c__Saccharomycetes(100);o__Saccharomycetales(100);f__Metschnikowiaceae(100);g__Metschnikowia(100);g__Metschnikowia_unclassified(100); | Metschnikowia | 13 | Saprotroph | \|Undefined Saprotroph\| | Yeast | NULL | Probable | As nectar-tap_saprotroph (Põlme et al. 2020); Decay sugar-rich substrates (Põlme et al. 2020) | Põlme S, et al. 2020. Fungal Diversity 105:1-16 ((https://doi.org/10.1007/s13225-020-00466-2)) |
| Otu17048 | 1 | k__Fungi(100);p__Ascomycota(100);c__Saccharomycetes(100);o__Saccharomycetales(100);f__Metschnikowiaceae(100);g__Metschnikowia(100);g__Metschnikowia_unclassified(100); | Metschnikowia | 13 | Saprotroph | \|Undefined Saprotroph\| | Yeast | NULL | Probable | As nectar-tap_saprotroph (Põlme et al. 2020); Decay sugar-rich substrates (Põlme et al. 2020) | Põlme S, et al. 2020. Fungal Diversity 105:1-16 ((https://doi.org/10.1007/s13225-020-00466-2)) |
| Otu17049 | 1 | k__Fungi(100);p__Ascomycota(100);c__Saccharomycetes(100);o__Saccharomycetales(100);f__Metschnikowiaceae(100);g__Metschnikowia(100);s__Metschnikowia_pulcherrima(100); | Metschnikowia | 13 | Saprotroph | \|Undefined Saprotroph\| | Yeast | NULL | Probable | As nectar-tap_saprotroph (Põlme et al. 2020); Decay sugar-rich substrates (Põlme et al. 2020) | Põlme S, et al. 2020. Fungal Diversity 105:1-16 ((https://doi.org/10.1007/s13225-020-00466-2)) |
| Otu17050 | 1 | k__Fungi(100);p__Ascomycota(100);c__Saccharomycetes(100);o__Saccharomycetales(100);f__Metschnikowiaceae(100);g__Metschnikowia(100);g__Metschnikowia_unclassified(100); | Metschnikowia | 13 | Saprotroph | \|Undefined Saprotroph\| | Yeast | NULL | Probable | As nectar-tap_saprotroph (Põlme et al. 2020); Decay sugar-rich substrates (Põlme et al. 2020) | Põlme S, et al. 2020. Fungal Diversity 105:1-16 ((https://doi.org/10.1007/s13225-020-00466-2)) |
| Otu17051 | 1 | k__Fungi(100);p__Ascomycota(100);c__Saccharomycetes(100);o__Saccharomycetales(100);f__Metschnikowiaceae(100);g__Metschnikowia(100);s__Metschnikowia_pulcherrima(100); | Metschnikowia | 13 | Saprotroph | \|Undefined Saprotroph\| | Yeast | NULL | Probable | As nectar-tap_saprotroph (Põlme et al. 2020); Decay sugar-rich substrates (Põlme et al. 2020) | Põlme S, et al. 2020. Fungal Diversity 105:1-16 ((https://doi.org/10.1007/s13225-020-00466-2)) |
| Otu17052 | 1 | k__Fungi(100);p__Ascomycota(100);c__Saccharomycetes(100);o__Saccharomycetales(100);f__Metschnikowiaceae(100);g__Metschnikowia(100);g__Metschnikowia_unclassified(100); | Metschnikowia | 13 | Saprotroph | \|Undefined Saprotroph\| | Yeast | NULL | Probable | As nectar-tap_saprotroph (Põlme et al. 2020); Decay sugar-rich substrates (Põlme et al. 2020) | Põlme S, et al. 2020. Fungal Diversity 105:1-16 ((https://doi.org/10.1007/s13225-020-00466-2)) |
| Otu17053 | 1 | k__Fungi(100);p__Ascomycota(100);c__Saccharomycetes(100);o__Saccharomycetales(100);f__Metschnikowiaceae(100);g__Metschnikowia(100);g__Metschnikowia_unclassified(100); | Metschnikowia | 13 | Saprotroph | \|Undefined Saprotroph\| | Yeast | NULL | Probable | As nectar-tap_saprotroph (Põlme et al. 2020); Decay sugar-rich substrates (Põlme et al. 2020) | Põlme S, et al. 2020. Fungal Diversity 105:1-16 ((https://doi.org/10.1007/s13225-020-00466-2)) |
| Otu17054 | 1 | k__Fungi(100);p__Ascomycota(100);c__Saccharomycetes(100);o__Saccharomycetales(100);f__Metschnikowiaceae(100);g__Metschnikowia(100);g__Metschnikowia_unclassified(100); | Metschnikowia | 13 | Saprotroph | \|Undefined Saprotroph\| | Yeast | NULL | Probable | As nectar-tap_saprotroph (Põlme et al. 2020); Decay sugar-rich substrates (Põlme et al. 2020) | Põlme S, et al. 2020. Fungal Diversity 105:1-16 ((https://doi.org/10.1007/s13225-020-00466-2)) |
| Otu17055 | 1 | k__Fungi(100);p__Ascomycota(100);c__Saccharomycetes(100);o__Saccharomycetales(100);f__Metschnikowiaceae(100);g__Metschnikowia(100);g__Metschnikowia_unclassified(100); | Metschnikowia | 13 | Saprotroph | \|Undefined Saprotroph\| | Yeast | NULL | Probable | As nectar-tap_saprotroph (Põlme et al. 2020); Decay sugar-rich substrates (Põlme et al. 2020) | Põlme S, et al. 2020. Fungal Diversity 105:1-16 ((https://doi.org/10.1007/s13225-020-00466-2)) |
| Otu17056 | 1 | k__Fungi(100);p__Ascomycota(100);c__Saccharomycetes(100);o__Saccharomycetales(100);f__Metschnikowiaceae(100);g__Metschnikowia(100);s__Metschnikowia_pulcherrima(100); | Metschnikowia | 13 | Saprotroph | \|Undefined Saprotroph\| | Yeast | NULL | Probable | As nectar-tap_saprotroph (Põlme et al. 2020); Decay sugar-rich substrates (Põlme et al. 2020) | Põlme S, et al. 2020. Fungal Diversity 105:1-16 ((https://doi.org/10.1007/s13225-020-00466-2)) |
| Otu17057 | 1 | k__Fungi(100);p__Ascomycota(100);c__Saccharomycetes(100);o__Saccharomycetales(100);f__Metschnikowiaceae(100);g__Metschnikowia(100);s__Metschnikowia_pulcherrima(100); | Metschnikowia | 13 | Saprotroph | \|Undefined Saprotroph\| | Yeast | NULL | Probable | As nectar-tap_saprotroph (Põlme et al. 2020); Decay sugar-rich substrates (Põlme et al. 2020) | Põlme S, et al. 2020. Fungal Diversity 105:1-16 ((https://doi.org/10.1007/s13225-020-00466-2)) |
| Otu17059 | 1 | k__Fungi(100);p__Ascomycota(100);c__Saccharomycetes(100);o__Saccharomycetales(100);f__Metschnikowiaceae(100);g__Metschnikowia(100);s__Metschnikowia_pulcherrima(100); | Metschnikowia | 13 | Saprotroph | \|Undefined Saprotroph\| | Yeast | NULL | Probable | As nectar-tap_saprotroph (Põlme et al. 2020); Decay sugar-rich substrates (Põlme et al. 2020) | Põlme S, et al. 2020. Fungal Diversity 105:1-16 ((https://doi.org/10.1007/s13225-020-00466-2)) |
| Otu17060 | 1 | k__Fungi(100);p__Ascomycota(100);c__Saccharomycetes(100);o__Saccharomycetales(100);f__Metschnikowiaceae(100);g__Metschnikowia(100);s__Metschnikowia_pulcherrima(100); | Metschnikowia | 13 | Saprotroph | \|Undefined Saprotroph\| | Yeast | NULL | Probable | As nectar-tap_saprotroph (Põlme et al. 2020); Decay sugar-rich substrates (Põlme et al. 2020) | Põlme S, et al. 2020. Fungal Diversity 105:1-16 ((https://doi.org/10.1007/s13225-020-00466-2)) |
| Otu17061 | 1 | k__Fungi(100);p__Ascomycota(100);c__Saccharomycetes(100);o__Saccharomycetales(100);f__Metschnikowiaceae(100);g__Metschnikowia(100);s__Metschnikowia_pulcherrima(100); | Metschnikowia | 13 | Saprotroph | \|Undefined Saprotroph\| | Yeast | NULL | Probable | As nectar-tap_saprotroph (Põlme et al. 2020); Decay sugar-rich substrates (Põlme et al. 2020) | Põlme S, et al. 2020. Fungal Diversity 105:1-16 ((https://doi.org/10.1007/s13225-020-00466-2)) |
| Otu17062 | 1 | k__Fungi(100);p__Ascomycota(100);c__Saccharomycetes(100);o__Saccharomycetales(100);f__Metschnikowiaceae(100);g__Metschnikowia(100);s__Metschnikowia_pulcherrima(100); | Metschnikowia | 13 | Saprotroph | \|Undefined Saprotroph\| | Yeast | NULL | Probable | As nectar-tap_saprotroph (Põlme et al. 2020); Decay sugar-rich substrates (Põlme et al. 2020) | Põlme S, et al. 2020. Fungal Diversity 105:1-16 ((https://doi.org/10.1007/s13225-020-00466-2)) |
| Otu17063 | 1 | k__Fungi(100);p__Ascomycota(100);c__Saccharomycetes(100);o__Saccharomycetales(100);f__Metschnikowiaceae(100);g__Metschnikowia(100);s__Metschnikowia_pulcherrima(100); | Metschnikowia | 13 | Saprotroph | \|Undefined Saprotroph\| | Yeast | NULL | Probable | As nectar-tap_saprotroph (Põlme et al. 2020); Decay sugar-rich substrates (Põlme et al. 2020) | Põlme S, et al. 2020. Fungal Diversity 105:1-16 ((https://doi.org/10.1007/s13225-020-00466-2)) |
| Otu17064 | 1 | k__Fungi(100);p__Ascomycota(100);c__Saccharomycetes(100);o__Saccharomycetales(100);f__Metschnikowiaceae(100);g__Metschnikowia(100);s__Metschnikowia_pulcherrima(100); | Metschnikowia | 13 | Saprotroph | \|Undefined Saprotroph\| | Yeast | NULL | Probable | As nectar-tap_saprotroph (Põlme et al. 2020); Decay sugar-rich substrates (Põlme et al. 2020) | Põlme S, et al. 2020. Fungal Diversity 105:1-16 ((https://doi.org/10.1007/s13225-020-00466-2)) |
| Otu17066 | 1 | k__Fungi(100);p__Ascomycota(100);c__Saccharomycetes(100);o__Saccharomycetales(100);f__Metschnikowiaceae(100);g__Metschnikowia(100);g__Metschnikowia_unclassified(100); | Metschnikowia | 13 | Saprotroph | \|Undefined Saprotroph\| | Yeast | NULL | Probable | As nectar-tap_saprotroph (Põlme et al. 2020); Decay sugar-rich substrates (Põlme et al. 2020) | Põlme S, et al. 2020. Fungal Diversity 105:1-16 ((https://doi.org/10.1007/s13225-020-00466-2)) |
| Otu17067 | 1 | k__Fungi(100);p__Ascomycota(100);c__Saccharomycetes(100);o__Saccharomycetales(100);f__Metschnikowiaceae(100);g__Metschnikowia(100);g__Metschnikowia_unclassified(100); | Metschnikowia | 13 | Saprotroph | \|Undefined Saprotroph\| | Yeast | NULL | Probable | As nectar-tap_saprotroph (Põlme et al. 2020); Decay sugar-rich substrates (Põlme et al. 2020) | Põlme S, et al. 2020. Fungal Diversity 105:1-16 ((https://doi.org/10.1007/s13225-020-00466-2)) |
| Otu17068 | 1 | k__Fungi(100);p__Ascomycota(100);c__Saccharomycetes(100);o__Saccharomycetales(100);f__Metschnikowiaceae(100);g__Metschnikowia(100);s__Metschnikowia_pulcherrima(100); | Metschnikowia | 13 | Saprotroph | \|Undefined Saprotroph\| | Yeast | NULL | Probable | As nectar-tap_saprotroph (Põlme et al. 2020); Decay sugar-rich substrates (Põlme et al. 2020) | Põlme S, et al. 2020. Fungal Diversity 105:1-16 ((https://doi.org/10.1007/s13225-020-00466-2)) |
| Otu17069 | 1 | k__Fungi(100);p__Ascomycota(100);c__Saccharomycetes(100);o__Saccharomycetales(100);f__Metschnikowiaceae(100);g__Metschnikowia(100);s__Metschnikowia_pulcherrima(100); | Metschnikowia | 13 | Saprotroph | \|Undefined Saprotroph\| | Yeast | NULL | Probable | As nectar-tap_saprotroph (Põlme et al. 2020); Decay sugar-rich substrates (Põlme et al. 2020) | Põlme S, et al. 2020. Fungal Diversity 105:1-16 ((https://doi.org/10.1007/s13225-020-00466-2)) |
| Otu17070 | 1 | k__Fungi(100);p__Ascomycota(100);c__Saccharomycetes(100);o__Saccharomycetales(100);f__Metschnikowiaceae(100);g__Metschnikowia(100);g__Metschnikowia_unclassified(100); | Metschnikowia | 13 | Saprotroph | \|Undefined Saprotroph\| | Yeast | NULL | Probable | As nectar-tap_saprotroph (Põlme et al. 2020); Decay sugar-rich substrates (Põlme et al. 2020) | Põlme S, et al. 2020. Fungal Diversity 105:1-16 ((https://doi.org/10.1007/s13225-020-00466-2)) |
| Otu17071 | 1 | k__Fungi(100);p__Ascomycota(100);c__Saccharomycetes(100);o__Saccharomycetales(100);f__Metschnikowiaceae(100);g__Metschnikowia(100);s__Metschnikowia_pulcherrima(100); | Metschnikowia | 13 | Saprotroph | \|Undefined Saprotroph\| | Yeast | NULL | Probable | As nectar-tap_saprotroph (Põlme et al. 2020); Decay sugar-rich substrates (Põlme et al. 2020) | Põlme S, et al. 2020. Fungal Diversity 105:1-16 ((https://doi.org/10.1007/s13225-020-00466-2)) |
| Otu17073 | 1 | k__Fungi(100);p__Ascomycota(100);c__Saccharomycetes(100);o__Saccharomycetales(100);f__Metschnikowiaceae(100);g__Metschnikowia(100);s__Metschnikowia_pulcherrima(100); | Metschnikowia | 13 | Saprotroph | \|Undefined Saprotroph\| | Yeast | NULL | Probable | As nectar-tap_saprotroph (Põlme et al. 2020); Decay sugar-rich substrates (Põlme et al. 2020) | Põlme S, et al. 2020. Fungal Diversity 105:1-16 ((https://doi.org/10.1007/s13225-020-00466-2)) |
| Otu17074 | 1 | k__Fungi(100);p__Ascomycota(100);c__Saccharomycetes(100);o__Saccharomycetales(100);f__Metschnikowiaceae(100);g__Metschnikowia(100);g__Metschnikowia_unclassified(100); | Metschnikowia | 13 | Saprotroph | \|Undefined Saprotroph\| | Yeast | NULL | Probable | As nectar-tap_saprotroph (Põlme et al. 2020); Decay sugar-rich substrates (Põlme et al. 2020) | Põlme S, et al. 2020. Fungal Diversity 105:1-16 ((https://doi.org/10.1007/s13225-020-00466-2)) |
| Otu17075 | 1 | k__Fungi(100);p__Ascomycota(100);c__Saccharomycetes(100);o__Saccharomycetales(100);f__Metschnikowiaceae(100);g__Metschnikowia(100);g__Metschnikowia_unclassified(100); | Metschnikowia | 13 | Saprotroph | \|Undefined Saprotroph\| | Yeast | NULL | Probable | As nectar-tap_saprotroph (Põlme et al. 2020); Decay sugar-rich substrates (Põlme et al. 2020) | Põlme S, et al. 2020. Fungal Diversity 105:1-16 ((https://doi.org/10.1007/s13225-020-00466-2)) |
| Otu17076 | 1 | k__Fungi(100);p__Ascomycota(100);c__Saccharomycetes(100);o__Saccharomycetales(100);f__Metschnikowiaceae(100);g__Metschnikowia(100);s__Metschnikowia_pulcherrima(100); | Metschnikowia | 13 | Saprotroph | \|Undefined Saprotroph\| | Yeast | NULL | Probable | As nectar-tap_saprotroph (Põlme et al. 2020); Decay sugar-rich substrates (Põlme et al. 2020) | Põlme S, et al. 2020. Fungal Diversity 105:1-16 ((https://doi.org/10.1007/s13225-020-00466-2)) |
| Otu17077 | 1 | k__Fungi(100);p__Ascomycota(100);c__Saccharomycetes(100);o__Saccharomycetales(100);f__Metschnikowiaceae(100);g__Metschnikowia(100);s__Metschnikowia_pulcherrima(100); | Metschnikowia | 13 | Saprotroph | \|Undefined Saprotroph\| | Yeast | NULL | Probable | As nectar-tap_saprotroph (Põlme et al. 2020); Decay sugar-rich substrates (Põlme et al. 2020) | Põlme S, et al. 2020. Fungal Diversity 105:1-16 ((https://doi.org/10.1007/s13225-020-00466-2)) |
| Otu17078 | 1 | k__Fungi(100);p__Ascomycota(100);c__Saccharomycetes(100);o__Saccharomycetales(100);f__Metschnikowiaceae(100);g__Metschnikowia(100);s__Metschnikowia_pulcherrima(100); | Metschnikowia | 13 | Saprotroph | \|Undefined Saprotroph\| | Yeast | NULL | Probable | As nectar-tap_saprotroph (Põlme et al. 2020); Decay sugar-rich substrates (Põlme et al. 2020) | Põlme S, et al. 2020. Fungal Diversity 105:1-16 ((https://doi.org/10.1007/s13225-020-00466-2)) |
| Otu17079 | 1 | k__Fungi(100);p__Ascomycota(100);c__Saccharomycetes(100);o__Saccharomycetales(100);f__Metschnikowiaceae(100);g__Metschnikowia(100);s__Metschnikowia_pulcherrima(100); | Metschnikowia | 13 | Saprotroph | \|Undefined Saprotroph\| | Yeast | NULL | Probable | As nectar-tap_saprotroph (Põlme et al. 2020); Decay sugar-rich substrates (Põlme et al. 2020) | Põlme S, et al. 2020. Fungal Diversity 105:1-16 ((https://doi.org/10.1007/s13225-020-00466-2)) |
| Otu17082 | 1 | k__Fungi(100);p__Ascomycota(100);c__Saccharomycetes(100);o__Saccharomycetales(100);f__Metschnikowiaceae(100);g__Metschnikowia(100);s__Metschnikowia_pulcherrima(100); | Metschnikowia | 13 | Saprotroph | \|Undefined Saprotroph\| | Yeast | NULL | Probable | As nectar-tap_saprotroph (Põlme et al. 2020); Decay sugar-rich substrates (Põlme et al. 2020) | Põlme S, et al. 2020. Fungal Diversity 105:1-16 ((https://doi.org/10.1007/s13225-020-00466-2)) |
| Otu17085 | 1 | k__Fungi(100);p__Ascomycota(100);c__Saccharomycetes(100);o__Saccharomycetales(100);f__Metschnikowiaceae(100);g__Metschnikowia(100);g__Metschnikowia_unclassified(100); | Metschnikowia | 13 | Saprotroph | \|Undefined Saprotroph\| | Yeast | NULL | Probable | As nectar-tap_saprotroph (Põlme et al. 2020); Decay sugar-rich substrates (Põlme et al. 2020) | Põlme S, et al. 2020. Fungal Diversity 105:1-16 ((https://doi.org/10.1007/s13225-020-00466-2)) |
| Otu17086 | 1 | k__Fungi(100);p__Ascomycota(100);c__Saccharomycetes(100);o__Saccharomycetales(100);f__Metschnikowiaceae(100);g__Metschnikowia(100);g__Metschnikowia_unclassified(100); | Metschnikowia | 13 | Saprotroph | \|Undefined Saprotroph\| | Yeast | NULL | Probable | As nectar-tap_saprotroph (Põlme et al. 2020); Decay sugar-rich substrates (Põlme et al. 2020) | Põlme S, et al. 2020. Fungal Diversity 105:1-16 ((https://doi.org/10.1007/s13225-020-00466-2)) |
| Otu17087 | 1 | k__Fungi(100);p__Ascomycota(100);c__Saccharomycetes(100);o__Saccharomycetales(100);f__Metschnikowiaceae(100);g__Metschnikowia(100);s__Metschnikowia_pulcherrima(100); | Metschnikowia | 13 | Saprotroph | \|Undefined Saprotroph\| | Yeast | NULL | Probable | As nectar-tap_saprotroph (Põlme et al. 2020); Decay sugar-rich substrates (Põlme et al. 2020) | Põlme S, et al. 2020. Fungal Diversity 105:1-16 ((https://doi.org/10.1007/s13225-020-00466-2)) |
| Otu17088 | 1 | k__Fungi(100);p__Ascomycota(100);c__Saccharomycetes(100);o__Saccharomycetales(100);f__Metschnikowiaceae(100);g__Metschnikowia(100);g__Metschnikowia_unclassified(100); | Metschnikowia | 13 | Saprotroph | \|Undefined Saprotroph\| | Yeast | NULL | Probable | As nectar-tap_saprotroph (Põlme et al. 2020); Decay sugar-rich substrates (Põlme et al. 2020) | Põlme S, et al. 2020. Fungal Diversity 105:1-16 ((https://doi.org/10.1007/s13225-020-00466-2)) |
| Otu17090 | 1 | k__Fungi(100);p__Ascomycota(100);c__Saccharomycetes(100);o__Saccharomycetales(100);f__Metschnikowiaceae(100);g__Metschnikowia(100);g__Metschnikowia_unclassified(100); | Metschnikowia | 13 | Saprotroph | \|Undefined Saprotroph\| | Yeast | NULL | Probable | As nectar-tap_saprotroph (Põlme et al. 2020); Decay sugar-rich substrates (Põlme et al. 2020) | Põlme S, et al. 2020. Fungal Diversity 105:1-16 ((https://doi.org/10.1007/s13225-020-00466-2)) |
| Otu17091 | 1 | k__Fungi(100);p__Ascomycota(100);c__Saccharomycetes(100);o__Saccharomycetales(100);f__Metschnikowiaceae(100);g__Metschnikowia(100);s__Metschnikowia_pulcherrima(100); | Metschnikowia | 13 | Saprotroph | \|Undefined Saprotroph\| | Yeast | NULL | Probable | As nectar-tap_saprotroph (Põlme et al. 2020); Decay sugar-rich substrates (Põlme et al. 2020) | Põlme S, et al. 2020. Fungal Diversity 105:1-16 ((https://doi.org/10.1007/s13225-020-00466-2)) |
| Otu17092 | 1 | k__Fungi(100);p__Ascomycota(100);c__Saccharomycetes(100);o__Saccharomycetales(100);f__Metschnikowiaceae(100);g__Metschnikowia(100);s__Metschnikowia_pulcherrima(100); | Metschnikowia | 13 | Saprotroph | \|Undefined Saprotroph\| | Yeast | NULL | Probable | As nectar-tap_saprotroph (Põlme et al. 2020); Decay sugar-rich substrates (Põlme et al. 2020) | Põlme S, et al. 2020. Fungal Diversity 105:1-16 ((https://doi.org/10.1007/s13225-020-00466-2)) |
| Otu17093 | 1 | k__Fungi(100);p__Ascomycota(100);c__Saccharomycetes(100);o__Saccharomycetales(100);f__Metschnikowiaceae(100);g__Metschnikowia(100);s__Metschnikowia_pulcherrima(100); | Metschnikowia | 13 | Saprotroph | \|Undefined Saprotroph\| | Yeast | NULL | Probable | As nectar-tap_saprotroph (Põlme et al. 2020); Decay sugar-rich substrates (Põlme et al. 2020) | Põlme S, et al. 2020. Fungal Diversity 105:1-16 ((https://doi.org/10.1007/s13225-020-00466-2)) |
| Otu17094 | 1 | k__Fungi(100);p__Ascomycota(100);c__Saccharomycetes(100);o__Saccharomycetales(100);f__Metschnikowiaceae(100);g__Metschnikowia(100);s__Metschnikowia_pulcherrima(100); | Metschnikowia | 13 | Saprotroph | \|Undefined Saprotroph\| | Yeast | NULL | Probable | As nectar-tap_saprotroph (Põlme et al. 2020); Decay sugar-rich substrates (Põlme et al. 2020) | Põlme S, et al. 2020. Fungal Diversity 105:1-16 ((https://doi.org/10.1007/s13225-020-00466-2)) |
| Otu17096 | 1 | k__Fungi(100);p__Ascomycota(100);c__Saccharomycetes(100);o__Saccharomycetales(100);f__Metschnikowiaceae(100);g__Metschnikowia(100);s__Metschnikowia_pulcherrima(100); | Metschnikowia | 13 | Saprotroph | \|Undefined Saprotroph\| | Yeast | NULL | Probable | As nectar-tap_saprotroph (Põlme et al. 2020); Decay sugar-rich substrates (Põlme et al. 2020) | Põlme S, et al. 2020. Fungal Diversity 105:1-16 ((https://doi.org/10.1007/s13225-020-00466-2)) |
| Otu17097 | 1 | k__Fungi(100);p__Ascomycota(100);c__Saccharomycetes(100);o__Saccharomycetales(100);f__Metschnikowiaceae(100);g__Metschnikowia(100);s__Metschnikowia_pulcherrima(100); | Metschnikowia | 13 | Saprotroph | \|Undefined Saprotroph\| | Yeast | NULL | Probable | As nectar-tap_saprotroph (Põlme et al. 2020); Decay sugar-rich substrates (Põlme et al. 2020) | Põlme S, et al. 2020. Fungal Diversity 105:1-16 ((https://doi.org/10.1007/s13225-020-00466-2)) |
| Otu17098 | 1 | k__Fungi(100);p__Ascomycota(100);c__Saccharomycetes(100);o__Saccharomycetales(100);o__Saccharomycetales_unclassified(100);o__Saccharomycetales_unclassified(100);o__Saccharomycetales_unclassified(100); | Saccharomycetales | 7 | Saprotroph | Undefined Saprotroph | Yeast | NULL | Possible | NULL | Sterkenburg E, et al. 2015. New Phytologist 207:1145-1158 ((https://doi.org/10.1111/nph.13426)) |
| Otu17101 | 1 | k__Fungi(100);p__Ascomycota(100);c__Saccharomycetes(100);o__Saccharomycetales(100);f__Saccharomycetales_family_Incertae_sedis(100);g__Candida(100);g__Candida_unclassified(100); | Candida | 13 | Pathotroph-Saprotroph-Symbiotroph | Animal Pathogen-Endophyte-Endosymbiont-Epiphyte-\|Undefined Saprotroph\| | Dimorphic Yeast-Yeast | NULL | Probable | As nectar-tap_saprotroph (Põlme et al. 2020); Decay sugar-rich substrates (Põlme et al. 2020); This asexual genus is polyphyletic and contain species that function very differently depend on what they are. Caution should be taken when interpreting guilds for Candida species. | Manolakaki D, et al. 2010. Virulence. 1: 367-375 ((https://doi.org/10.4161/viru.1.5.12796)); Mekbib SB, et al. 2011. Tropical Plant Pathology 36:233-240 ((https://doi.org/10.1590/S1982-56762011000400004)); Põlme S, et al. 2020. Fungal Diversity 105:1-16 ((https://doi.org/10.1007/s13225-020-00466-2)) |
| Otu17104 | 1 | k__Fungi(100);p__Ascomycota(100);c__Saccharomycetes(100);o__Saccharomycetales(100);f__Metschnikowiaceae(100);g__Metschnikowia(100);g__Metschnikowia_unclassified(100); | Metschnikowia | 13 | Saprotroph | \|Undefined Saprotroph\| | Yeast | NULL | Probable | As nectar-tap_saprotroph (Põlme et al. 2020); Decay sugar-rich substrates (Põlme et al. 2020) | Põlme S, et al. 2020. Fungal Diversity 105:1-16 ((https://doi.org/10.1007/s13225-020-00466-2)) |
| Otu17107 | 1 | k__Fungi(100);p__Ascomycota(100);c__Saccharomycetes(100);o__Saccharomycetales(100);f__Saccharomycetales_family_Incertae_sedis(100);g__Candida(100);s__Candida_cretensis(100); | Candida | 13 | Pathotroph-Saprotroph-Symbiotroph | Animal Pathogen-Endophyte-Endosymbiont-Epiphyte-\|Undefined Saprotroph\| | Dimorphic Yeast-Yeast | NULL | Probable | As nectar-tap_saprotroph (Põlme et al. 2020); Decay sugar-rich substrates (Põlme et al. 2020); This asexual genus is polyphyletic and contain species that function very differently depend on what they are. Caution should be taken when interpreting guilds for Candida species. | Manolakaki D, et al. 2010. Virulence. 1: 367-375 ((https://doi.org/10.4161/viru.1.5.12796)); Mekbib SB, et al. 2011. Tropical Plant Pathology 36:233-240 ((https://doi.org/10.1590/S1982-56762011000400004)); Põlme S, et al. 2020. Fungal Diversity 105:1-16 ((https://doi.org/10.1007/s13225-020-00466-2)) |
| Otu17108 | 1 | k__Fungi(100);p__Ascomycota(100);c__Saccharomycetes(100);o__Saccharomycetales(100);f__Saccharomycetales_family_Incertae_sedis(100);g__Candida(100);s__Candida_cretensis(100); | Candida | 13 | Pathotroph-Saprotroph-Symbiotroph | Animal Pathogen-Endophyte-Endosymbiont-Epiphyte-\|Undefined Saprotroph\| | Dimorphic Yeast-Yeast | NULL | Probable | As nectar-tap_saprotroph (Põlme et al. 2020); Decay sugar-rich substrates (Põlme et al. 2020); This asexual genus is polyphyletic and contain species that function very differently depend on what they are. Caution should be taken when interpreting guilds for Candida species. | Manolakaki D, et al. 2010. Virulence. 1: 367-375 ((https://doi.org/10.4161/viru.1.5.12796)); Mekbib SB, et al. 2011. Tropical Plant Pathology 36:233-240 ((https://doi.org/10.1590/S1982-56762011000400004)); Põlme S, et al. 2020. Fungal Diversity 105:1-16 ((https://doi.org/10.1007/s13225-020-00466-2)) |
| Otu17109 | 1 | k__Fungi(100);p__Ascomycota(100);c__Saccharomycetes(100);o__Saccharomycetales(100);f__Saccharomycetales_family_Incertae_sedis(100);f__Saccharomycetales_family_Incertae_sedis_unclassified(100);f__Saccharomycetales_family_Incertae_sedis_unclassified(100); | Saccharomycetales | 7 | Saprotroph | Undefined Saprotroph | Yeast | NULL | Possible | NULL | Sterkenburg E, et al. 2015. New Phytologist 207:1145-1158 ((https://doi.org/10.1111/nph.13426)) |
| Otu17112 | 1 | k__Fungi(100);p__Ascomycota(100);c__Saccharomycetes(100);o__Saccharomycetales(100);f__Metschnikowiaceae(100);g__Metschnikowia(100);g__Metschnikowia_unclassified(100); | Metschnikowia | 13 | Saprotroph | \|Undefined Saprotroph\| | Yeast | NULL | Probable | As nectar-tap_saprotroph (Põlme et al. 2020); Decay sugar-rich substrates (Põlme et al. 2020) | Põlme S, et al. 2020. Fungal Diversity 105:1-16 ((https://doi.org/10.1007/s13225-020-00466-2)) |
| Otu17113 | 1 | k__Fungi(100);p__Ascomycota(100);c__Saccharomycetes(100);o__Saccharomycetales(100);f__Metschnikowiaceae(100);g__Metschnikowia(100);s__Metschnikowia_pulcherrima(100); | Metschnikowia | 13 | Saprotroph | \|Undefined Saprotroph\| | Yeast | NULL | Probable | As nectar-tap_saprotroph (Põlme et al. 2020); Decay sugar-rich substrates (Põlme et al. 2020) | Põlme S, et al. 2020. Fungal Diversity 105:1-16 ((https://doi.org/10.1007/s13225-020-00466-2)) |
| Otu17114 | 1 | k__Fungi(100);p__Ascomycota(100);c__Saccharomycetes(100);o__Saccharomycetales(100);f__Metschnikowiaceae(100);g__Metschnikowia(100);s__Metschnikowia_pulcherrima(100); | Metschnikowia | 13 | Saprotroph | \|Undefined Saprotroph\| | Yeast | NULL | Probable | As nectar-tap_saprotroph (Põlme et al. 2020); Decay sugar-rich substrates (Põlme et al. 2020) | Põlme S, et al. 2020. Fungal Diversity 105:1-16 ((https://doi.org/10.1007/s13225-020-00466-2)) |
| Otu17115 | 1 | k__Fungi(100);p__Ascomycota(100);c__Saccharomycetes(100);o__Saccharomycetales(100);f__Metschnikowiaceae(100);g__Metschnikowia(100);s__Metschnikowia_pulcherrima(100); | Metschnikowia | 13 | Saprotroph | \|Undefined Saprotroph\| | Yeast | NULL | Probable | As nectar-tap_saprotroph (Põlme et al. 2020); Decay sugar-rich substrates (Põlme et al. 2020) | Põlme S, et al. 2020. Fungal Diversity 105:1-16 ((https://doi.org/10.1007/s13225-020-00466-2)) |
| Otu17121 | 1 | k__Fungi(100);p__Ascomycota(100);c__Saccharomycetes(100);o__Saccharomycetales(100);f__Metschnikowiaceae(100);g__Metschnikowia(100);s__Metschnikowia_pulcherrima(100); | Metschnikowia | 13 | Saprotroph | \|Undefined Saprotroph\| | Yeast | NULL | Probable | As nectar-tap_saprotroph (Põlme et al. 2020); Decay sugar-rich substrates (Põlme et al. 2020) | Põlme S, et al. 2020. Fungal Diversity 105:1-16 ((https://doi.org/10.1007/s13225-020-00466-2)) |
| Otu17122 | 1 | k__Fungi(100);p__Ascomycota(100);c__Saccharomycetes(100);o__Saccharomycetales(100);f__Metschnikowiaceae(100);g__Metschnikowia(100);s__Metschnikowia_pulcherrima(100); | Metschnikowia | 13 | Saprotroph | \|Undefined Saprotroph\| | Yeast | NULL | Probable | As nectar-tap_saprotroph (Põlme et al. 2020); Decay sugar-rich substrates (Põlme et al. 2020) | Põlme S, et al. 2020. Fungal Diversity 105:1-16 ((https://doi.org/10.1007/s13225-020-00466-2)) |
| Otu17123 | 1 | k__Fungi(100);p__Ascomycota(100);c__Saccharomycetes(100);o__Saccharomycetales(100);f__Metschnikowiaceae(100);g__Metschnikowia(100);s__Metschnikowia_pulcherrima(100); | Metschnikowia | 13 | Saprotroph | \|Undefined Saprotroph\| | Yeast | NULL | Probable | As nectar-tap_saprotroph (Põlme et al. 2020); Decay sugar-rich substrates (Põlme et al. 2020) | Põlme S, et al. 2020. Fungal Diversity 105:1-16 ((https://doi.org/10.1007/s13225-020-00466-2)) |
| Otu17124 | 1 | k__Fungi(100);p__Ascomycota(100);c__Saccharomycetes(100);o__Saccharomycetales(100);f__Metschnikowiaceae(100);g__Metschnikowia(100);s__Metschnikowia_pulcherrima(100); | Metschnikowia | 13 | Saprotroph | \|Undefined Saprotroph\| | Yeast | NULL | Probable | As nectar-tap_saprotroph (Põlme et al. 2020); Decay sugar-rich substrates (Põlme et al. 2020) | Põlme S, et al. 2020. Fungal Diversity 105:1-16 ((https://doi.org/10.1007/s13225-020-00466-2)) |
| Otu17125 | 1 | k__Fungi(100);p__Ascomycota(100);c__Saccharomycetes(100);o__Saccharomycetales(100);f__Metschnikowiaceae(100);g__Metschnikowia(100);g__Metschnikowia_unclassified(100); | Metschnikowia | 13 | Saprotroph | \|Undefined Saprotroph\| | Yeast | NULL | Probable | As nectar-tap_saprotroph (Põlme et al. 2020); Decay sugar-rich substrates (Põlme et al. 2020) | Põlme S, et al. 2020. Fungal Diversity 105:1-16 ((https://doi.org/10.1007/s13225-020-00466-2)) |
| Otu17126 | 1 | k__Fungi(100);p__Ascomycota(100);c__Saccharomycetes(100);o__Saccharomycetales(100);f__Metschnikowiaceae(100);g__Metschnikowia(100);s__Metschnikowia_pulcherrima(100); | Metschnikowia | 13 | Saprotroph | \|Undefined Saprotroph\| | Yeast | NULL | Probable | As nectar-tap_saprotroph (Põlme et al. 2020); Decay sugar-rich substrates (Põlme et al. 2020) | Põlme S, et al. 2020. Fungal Diversity 105:1-16 ((https://doi.org/10.1007/s13225-020-00466-2)) |
| Otu17127 | 1 | k__Fungi(100);p__Ascomycota(100);c__Saccharomycetes(100);o__Saccharomycetales(100);f__Metschnikowiaceae(100);g__Metschnikowia(100);s__Metschnikowia_pulcherrima(100); | Metschnikowia | 13 | Saprotroph | \|Undefined Saprotroph\| | Yeast | NULL | Probable | As nectar-tap_saprotroph (Põlme et al. 2020); Decay sugar-rich substrates (Põlme et al. 2020) | Põlme S, et al. 2020. Fungal Diversity 105:1-16 ((https://doi.org/10.1007/s13225-020-00466-2)) |
| Otu17128 | 1 | k__Fungi(100);p__Ascomycota(100);c__Saccharomycetes(100);o__Saccharomycetales(100);f__Metschnikowiaceae(100);g__Metschnikowia(100);s__Metschnikowia_pulcherrima(100); | Metschnikowia | 13 | Saprotroph | \|Undefined Saprotroph\| | Yeast | NULL | Probable | As nectar-tap_saprotroph (Põlme et al. 2020); Decay sugar-rich substrates (Põlme et al. 2020) | Põlme S, et al. 2020. Fungal Diversity 105:1-16 ((https://doi.org/10.1007/s13225-020-00466-2)) |
| Otu17129 | 1 | k__Fungi(100);p__Ascomycota(100);c__Saccharomycetes(100);o__Saccharomycetales(100);f__Metschnikowiaceae(100);g__Metschnikowia(100);s__Metschnikowia_pulcherrima(100); | Metschnikowia | 13 | Saprotroph | \|Undefined Saprotroph\| | Yeast | NULL | Probable | As nectar-tap_saprotroph (Põlme et al. 2020); Decay sugar-rich substrates (Põlme et al. 2020) | Põlme S, et al. 2020. Fungal Diversity 105:1-16 ((https://doi.org/10.1007/s13225-020-00466-2)) |
| Otu17130 | 1 | k__Fungi(100);p__Ascomycota(100);c__Saccharomycetes(100);o__Saccharomycetales(100);f__Saccharomycetales_family_Incertae_sedis(100);f__Saccharomycetales_family_Incertae_sedis_unclassified(100);f__Saccharomycetales_family_Incertae_sedis_unclassified(100); | Saccharomycetales | 7 | Saprotroph | Undefined Saprotroph | Yeast | NULL | Possible | NULL | Sterkenburg E, et al. 2015. New Phytologist 207:1145-1158 ((https://doi.org/10.1111/nph.13426)) |
| Otu17132 | 1 | k__Fungi(100);p__Ascomycota(100);c__Saccharomycetes(100);o__Saccharomycetales(100);f__Metschnikowiaceae(100);g__Metschnikowia(100);s__Metschnikowia_pulcherrima(100); | Metschnikowia | 13 | Saprotroph | \|Undefined Saprotroph\| | Yeast | NULL | Probable | As nectar-tap_saprotroph (Põlme et al. 2020); Decay sugar-rich substrates (Põlme et al. 2020) | Põlme S, et al. 2020. Fungal Diversity 105:1-16 ((https://doi.org/10.1007/s13225-020-00466-2)) |
| Otu17133 | 1 | k__Fungi(100);p__Ascomycota(100);c__Saccharomycetes(100);o__Saccharomycetales(100);f__Metschnikowiaceae(100);g__Metschnikowia(100);s__Metschnikowia_pulcherrima(100); | Metschnikowia | 13 | Saprotroph | \|Undefined Saprotroph\| | Yeast | NULL | Probable | As nectar-tap_saprotroph (Põlme et al. 2020); Decay sugar-rich substrates (Põlme et al. 2020) | Põlme S, et al. 2020. Fungal Diversity 105:1-16 ((https://doi.org/10.1007/s13225-020-00466-2)) |
| Otu17135 | 1 | k__Fungi(100);p__Ascomycota(100);c__Saccharomycetes(100);o__Saccharomycetales(100);f__Metschnikowiaceae(100);g__Metschnikowia(100);s__Metschnikowia_pulcherrima(100); | Metschnikowia | 13 | Saprotroph | \|Undefined Saprotroph\| | Yeast | NULL | Probable | As nectar-tap_saprotroph (Põlme et al. 2020); Decay sugar-rich substrates (Põlme et al. 2020) | Põlme S, et al. 2020. Fungal Diversity 105:1-16 ((https://doi.org/10.1007/s13225-020-00466-2)) |
| Otu17137 | 1 | k__Fungi(100);p__Ascomycota(100);c__Saccharomycetes(100);o__Saccharomycetales(100);f__Metschnikowiaceae(100);g__Metschnikowia(100);s__Metschnikowia_pulcherrima(100); | Metschnikowia | 13 | Saprotroph | \|Undefined Saprotroph\| | Yeast | NULL | Probable | As nectar-tap_saprotroph (Põlme et al. 2020); Decay sugar-rich substrates (Põlme et al. 2020) | Põlme S, et al. 2020. Fungal Diversity 105:1-16 ((https://doi.org/10.1007/s13225-020-00466-2)) |
| Otu17138 | 1 | k__Fungi(100);p__Ascomycota(100);c__Saccharomycetes(100);o__Saccharomycetales(100);f__Saccharomycetales_family_Incertae_sedis(100);g__Candida(100);s__Candida_cretensis(100); | Candida | 13 | Pathotroph-Saprotroph-Symbiotroph | Animal Pathogen-Endophyte-Endosymbiont-Epiphyte-\|Undefined Saprotroph\| | Dimorphic Yeast-Yeast | NULL | Probable | As nectar-tap_saprotroph (Põlme et al. 2020); Decay sugar-rich substrates (Põlme et al. 2020); This asexual genus is polyphyletic and contain species that function very differently depend on what they are. Caution should be taken when interpreting guilds for Candida species. | Manolakaki D, et al. 2010. Virulence. 1: 367-375 ((https://doi.org/10.4161/viru.1.5.12796)); Mekbib SB, et al. 2011. Tropical Plant Pathology 36:233-240 ((https://doi.org/10.1590/S1982-56762011000400004)); Põlme S, et al. 2020. Fungal Diversity 105:1-16 ((https://doi.org/10.1007/s13225-020-00466-2)) |
| Otu17143 | 1 | k__Fungi(100);p__Ascomycota(100);c__Saccharomycetes(100);o__Saccharomycetales(100);f__Saccharomycetales_family_Incertae_sedis(100);g__Candida(100);g__Candida_unclassified(100); | Candida | 13 | Pathotroph-Saprotroph-Symbiotroph | Animal Pathogen-Endophyte-Endosymbiont-Epiphyte-\|Undefined Saprotroph\| | Dimorphic Yeast-Yeast | NULL | Probable | As nectar-tap_saprotroph (Põlme et al. 2020); Decay sugar-rich substrates (Põlme et al. 2020); This asexual genus is polyphyletic and contain species that function very differently depend on what they are. Caution should be taken when interpreting guilds for Candida species. | Manolakaki D, et al. 2010. Virulence. 1: 367-375 ((https://doi.org/10.4161/viru.1.5.12796)); Mekbib SB, et al. 2011. Tropical Plant Pathology 36:233-240 ((https://doi.org/10.1590/S1982-56762011000400004)); Põlme S, et al. 2020. Fungal Diversity 105:1-16 ((https://doi.org/10.1007/s13225-020-00466-2)) |
| Otu17145 | 1 | k__Fungi(100);p__Ascomycota(100);c__Saccharomycetes(100);o__Saccharomycetales(100);f__Metschnikowiaceae(100);g__Metschnikowia(100);s__Metschnikowia_pulcherrima(100); | Metschnikowia | 13 | Saprotroph | \|Undefined Saprotroph\| | Yeast | NULL | Probable | As nectar-tap_saprotroph (Põlme et al. 2020); Decay sugar-rich substrates (Põlme et al. 2020) | Põlme S, et al. 2020. Fungal Diversity 105:1-16 ((https://doi.org/10.1007/s13225-020-00466-2)) |
| Otu17148 | 1 | k__Fungi(100);p__Ascomycota(100);c__Saccharomycetes(100);o__Saccharomycetales(100);f__Saccharomycetales_family_Incertae_sedis(100);g__Candida(100);g__Candida_unclassified(100); | Candida | 13 | Pathotroph-Saprotroph-Symbiotroph | Animal Pathogen-Endophyte-Endosymbiont-Epiphyte-\|Undefined Saprotroph\| | Dimorphic Yeast-Yeast | NULL | Probable | As nectar-tap_saprotroph (Põlme et al. 2020); Decay sugar-rich substrates (Põlme et al. 2020); This asexual genus is polyphyletic and contain species that function very differently depend on what they are. Caution should be taken when interpreting guilds for Candida species. | Manolakaki D, et al. 2010. Virulence. 1: 367-375 ((https://doi.org/10.4161/viru.1.5.12796)); Mekbib SB, et al. 2011. Tropical Plant Pathology 36:233-240 ((https://doi.org/10.1590/S1982-56762011000400004)); Põlme S, et al. 2020. Fungal Diversity 105:1-16 ((https://doi.org/10.1007/s13225-020-00466-2)) |
| Otu17150 | 1 | k__Fungi(100);p__Ascomycota(100);c__Saccharomycetes(100);o__Saccharomycetales(100);f__Metschnikowiaceae(100);g__Metschnikowia(100);s__Metschnikowia_pulcherrima(100); | Metschnikowia | 13 | Saprotroph | \|Undefined Saprotroph\| | Yeast | NULL | Probable | As nectar-tap_saprotroph (Põlme et al. 2020); Decay sugar-rich substrates (Põlme et al. 2020) | Põlme S, et al. 2020. Fungal Diversity 105:1-16 ((https://doi.org/10.1007/s13225-020-00466-2)) |
| Otu17151 | 1 | k__Fungi(100);p__Ascomycota(100);c__Saccharomycetes(100);o__Saccharomycetales(100);f__Saccharomycetales_family_Incertae_sedis(100);g__Candida(100);g__Candida_unclassified(100); | Candida | 13 | Pathotroph-Saprotroph-Symbiotroph | Animal Pathogen-Endophyte-Endosymbiont-Epiphyte-\|Undefined Saprotroph\| | Dimorphic Yeast-Yeast | NULL | Probable | As nectar-tap_saprotroph (Põlme et al. 2020); Decay sugar-rich substrates (Põlme et al. 2020); This asexual genus is polyphyletic and contain species that function very differently depend on what they are. Caution should be taken when interpreting guilds for Candida species. | Manolakaki D, et al. 2010. Virulence. 1: 367-375 ((https://doi.org/10.4161/viru.1.5.12796)); Mekbib SB, et al. 2011. Tropical Plant Pathology 36:233-240 ((https://doi.org/10.1590/S1982-56762011000400004)); Põlme S, et al. 2020. Fungal Diversity 105:1-16 ((https://doi.org/10.1007/s13225-020-00466-2)) |
| Otu17152 | 1 | k__Fungi(100);p__Ascomycota(100);c__Saccharomycetes(100);o__Saccharomycetales(100);f__Metschnikowiaceae(100);g__Metschnikowia(100);s__Metschnikowia_pulcherrima(100); | Metschnikowia | 13 | Saprotroph | \|Undefined Saprotroph\| | Yeast | NULL | Probable | As nectar-tap_saprotroph (Põlme et al. 2020); Decay sugar-rich substrates (Põlme et al. 2020) | Põlme S, et al. 2020. Fungal Diversity 105:1-16 ((https://doi.org/10.1007/s13225-020-00466-2)) |
| Otu17153 | 1 | k__Fungi(100);p__Ascomycota(100);c__Saccharomycetes(100);o__Saccharomycetales(100);f__Metschnikowiaceae(100);g__Metschnikowia(100);s__Metschnikowia_pulcherrima(100); | Metschnikowia | 13 | Saprotroph | \|Undefined Saprotroph\| | Yeast | NULL | Probable | As nectar-tap_saprotroph (Põlme et al. 2020); Decay sugar-rich substrates (Põlme et al. 2020) | Põlme S, et al. 2020. Fungal Diversity 105:1-16 ((https://doi.org/10.1007/s13225-020-00466-2)) |
| Otu17154 | 1 | k__Fungi(100);p__Ascomycota(100);c__Saccharomycetes(100);o__Saccharomycetales(100);f__Metschnikowiaceae(100);g__Metschnikowia(100);s__Metschnikowia_pulcherrima(100); | Metschnikowia | 13 | Saprotroph | \|Undefined Saprotroph\| | Yeast | NULL | Probable | As nectar-tap_saprotroph (Põlme et al. 2020); Decay sugar-rich substrates (Põlme et al. 2020) | Põlme S, et al. 2020. Fungal Diversity 105:1-16 ((https://doi.org/10.1007/s13225-020-00466-2)) |
| Otu17155 | 1 | k__Fungi(100);p__Ascomycota(100);c__Saccharomycetes(100);o__Saccharomycetales(100);f__Metschnikowiaceae(100);g__Metschnikowia(100);s__Metschnikowia_pulcherrima(100); | Metschnikowia | 13 | Saprotroph | \|Undefined Saprotroph\| | Yeast | NULL | Probable | As nectar-tap_saprotroph (Põlme et al. 2020); Decay sugar-rich substrates (Põlme et al. 2020) | Põlme S, et al. 2020. Fungal Diversity 105:1-16 ((https://doi.org/10.1007/s13225-020-00466-2)) |
| Otu17156 | 1 | k__Fungi(100);p__Ascomycota(100);c__Saccharomycetes(100);o__Saccharomycetales(100);f__Metschnikowiaceae(100);g__Metschnikowia(100);g__Metschnikowia_unclassified(100); | Metschnikowia | 13 | Saprotroph | \|Undefined Saprotroph\| | Yeast | NULL | Probable | As nectar-tap_saprotroph (Põlme et al. 2020); Decay sugar-rich substrates (Põlme et al. 2020) | Põlme S, et al. 2020. Fungal Diversity 105:1-16 ((https://doi.org/10.1007/s13225-020-00466-2)) |
| Otu17157 | 1 | k__Fungi(100);p__Ascomycota(100);c__Saccharomycetes(100);o__Saccharomycetales(100);f__Metschnikowiaceae(100);g__Metschnikowia(100);g__Metschnikowia_unclassified(100); | Metschnikowia | 13 | Saprotroph | \|Undefined Saprotroph\| | Yeast | NULL | Probable | As nectar-tap_saprotroph (Põlme et al. 2020); Decay sugar-rich substrates (Põlme et al. 2020) | Põlme S, et al. 2020. Fungal Diversity 105:1-16 ((https://doi.org/10.1007/s13225-020-00466-2)) |
| Otu17159 | 1 | k__Fungi(100);p__Ascomycota(100);c__Saccharomycetes(100);o__Saccharomycetales(100);f__Saccharomycetales_family_Incertae_sedis(100);g__Candida(100);s__Candida_cretensis(100); | Candida | 13 | Pathotroph-Saprotroph-Symbiotroph | Animal Pathogen-Endophyte-Endosymbiont-Epiphyte-\|Undefined Saprotroph\| | Dimorphic Yeast-Yeast | NULL | Probable | As nectar-tap_saprotroph (Põlme et al. 2020); Decay sugar-rich substrates (Põlme et al. 2020); This asexual genus is polyphyletic and contain species that function very differently depend on what they are. Caution should be taken when interpreting guilds for Candida species. | Manolakaki D, et al. 2010. Virulence. 1: 367-375 ((https://doi.org/10.4161/viru.1.5.12796)); Mekbib SB, et al. 2011. Tropical Plant Pathology 36:233-240 ((https://doi.org/10.1590/S1982-56762011000400004)); Põlme S, et al. 2020. Fungal Diversity 105:1-16 ((https://doi.org/10.1007/s13225-020-00466-2)) |
| Otu17160 | 1 | k__Fungi(100);p__Ascomycota(100);c__Saccharomycetes(100);o__Saccharomycetales(100);f__Saccharomycetaceae(100);g__Kazachstania(100);s__Kazachstania_intestinalis(100); | Kazachstania | 13 | Saprotroph | \|Undefined Saprotroph\| | Yeast | NULL | Probable | As nectar-tap_saprotroph (Põlme et al. 2020); Decay sugar-rich substrates (Põlme et al. 2020) | Põlme S, et al. 2020. Fungal Diversity 105:1-16 ((https://doi.org/10.1007/s13225-020-00466-2)) |
| Otu17161 | 1 | k__Fungi(100);p__Ascomycota(100);c__Saccharomycetes(100);o__Saccharomycetales(100);f__Metschnikowiaceae(100);g__Metschnikowia(100);s__Metschnikowia_pulcherrima(100); | Metschnikowia | 13 | Saprotroph | \|Undefined Saprotroph\| | Yeast | NULL | Probable | As nectar-tap_saprotroph (Põlme et al. 2020); Decay sugar-rich substrates (Põlme et al. 2020) | Põlme S, et al. 2020. Fungal Diversity 105:1-16 ((https://doi.org/10.1007/s13225-020-00466-2)) |
| Otu17162 | 1 | k__Fungi(100);p__Ascomycota(100);c__Saccharomycetes(100);o__Saccharomycetales(100);f__Metschnikowiaceae(100);g__Metschnikowia(100);g__Metschnikowia_unclassified(100); | Metschnikowia | 13 | Saprotroph | \|Undefined Saprotroph\| | Yeast | NULL | Probable | As nectar-tap_saprotroph (Põlme et al. 2020); Decay sugar-rich substrates (Põlme et al. 2020) | Põlme S, et al. 2020. Fungal Diversity 105:1-16 ((https://doi.org/10.1007/s13225-020-00466-2)) |
| Otu17163 | 1 | k__Fungi(100);p__Ascomycota(100);c__Saccharomycetes(100);o__Saccharomycetales(100);f__Metschnikowiaceae(100);g__Metschnikowia(100);s__Metschnikowia_pulcherrima(100); | Metschnikowia | 13 | Saprotroph | \|Undefined Saprotroph\| | Yeast | NULL | Probable | As nectar-tap_saprotroph (Põlme et al. 2020); Decay sugar-rich substrates (Põlme et al. 2020) | Põlme S, et al. 2020. Fungal Diversity 105:1-16 ((https://doi.org/10.1007/s13225-020-00466-2)) |
| Otu17164 | 1 | k__Fungi(100);p__Ascomycota(100);c__Saccharomycetes(100);o__Saccharomycetales(100);f__Saccharomycetales_family_Incertae_sedis(100);g__Candida(100);g__Candida_unclassified(100); | Candida | 13 | Pathotroph-Saprotroph-Symbiotroph | Animal Pathogen-Endophyte-Endosymbiont-Epiphyte-\|Undefined Saprotroph\| | Dimorphic Yeast-Yeast | NULL | Probable | As nectar-tap_saprotroph (Põlme et al. 2020); Decay sugar-rich substrates (Põlme et al. 2020); This asexual genus is polyphyletic and contain species that function very differently depend on what they are. Caution should be taken when interpreting guilds for Candida species. | Manolakaki D, et al. 2010. Virulence. 1: 367-375 ((https://doi.org/10.4161/viru.1.5.12796)); Mekbib SB, et al. 2011. Tropical Plant Pathology 36:233-240 ((https://doi.org/10.1590/S1982-56762011000400004)); Põlme S, et al. 2020. Fungal Diversity 105:1-16 ((https://doi.org/10.1007/s13225-020-00466-2)) |
| Otu17166 | 1 | k__Fungi(100);p__Ascomycota(100);c__Saccharomycetes(100);o__Saccharomycetales(100);f__Metschnikowiaceae(100);g__Metschnikowia(100);s__Metschnikowia_pulcherrima(100); | Metschnikowia | 13 | Saprotroph | \|Undefined Saprotroph\| | Yeast | NULL | Probable | As nectar-tap_saprotroph (Põlme et al. 2020); Decay sugar-rich substrates (Põlme et al. 2020) | Põlme S, et al. 2020. Fungal Diversity 105:1-16 ((https://doi.org/10.1007/s13225-020-00466-2)) |
| Otu17168 | 1 | k__Fungi(100);p__Ascomycota(100);c__Saccharomycetes(100);o__Saccharomycetales(100);f__Metschnikowiaceae(100);g__Metschnikowia(100);s__Metschnikowia_pulcherrima(100); | Metschnikowia | 13 | Saprotroph | \|Undefined Saprotroph\| | Yeast | NULL | Probable | As nectar-tap_saprotroph (Põlme et al. 2020); Decay sugar-rich substrates (Põlme et al. 2020) | Põlme S, et al. 2020. Fungal Diversity 105:1-16 ((https://doi.org/10.1007/s13225-020-00466-2)) |
| Otu17169 | 1 | k__Fungi(100);p__Ascomycota(100);c__Saccharomycetes(100);o__Saccharomycetales(100);f__Metschnikowiaceae(100);g__Metschnikowia(100);s__Metschnikowia_pulcherrima(100); | Metschnikowia | 13 | Saprotroph | \|Undefined Saprotroph\| | Yeast | NULL | Probable | As nectar-tap_saprotroph (Põlme et al. 2020); Decay sugar-rich substrates (Põlme et al. 2020) | Põlme S, et al. 2020. Fungal Diversity 105:1-16 ((https://doi.org/10.1007/s13225-020-00466-2)) |
| Otu17170 | 1 | k__Fungi(100);p__Ascomycota(100);c__Saccharomycetes(100);o__Saccharomycetales(100);f__Metschnikowiaceae(100);g__Metschnikowia(100);g__Metschnikowia_unclassified(100); | Metschnikowia | 13 | Saprotroph | \|Undefined Saprotroph\| | Yeast | NULL | Probable | As nectar-tap_saprotroph (Põlme et al. 2020); Decay sugar-rich substrates (Põlme et al. 2020) | Põlme S, et al. 2020. Fungal Diversity 105:1-16 ((https://doi.org/10.1007/s13225-020-00466-2)) |
| Otu17171 | 1 | k__Fungi(100);p__Ascomycota(100);c__Saccharomycetes(100);o__Saccharomycetales(100);f__Metschnikowiaceae(100);g__Metschnikowia(100);s__Metschnikowia_pulcherrima(100); | Metschnikowia | 13 | Saprotroph | \|Undefined Saprotroph\| | Yeast | NULL | Probable | As nectar-tap_saprotroph (Põlme et al. 2020); Decay sugar-rich substrates (Põlme et al. 2020) | Põlme S, et al. 2020. Fungal Diversity 105:1-16 ((https://doi.org/10.1007/s13225-020-00466-2)) |
| Otu17173 | 1 | k__Fungi(100);p__Ascomycota(100);c__Saccharomycetes(100);o__Saccharomycetales(100);f__Metschnikowiaceae(100);g__Metschnikowia(100);g__Metschnikowia_unclassified(100); | Metschnikowia | 13 | Saprotroph | \|Undefined Saprotroph\| | Yeast | NULL | Probable | As nectar-tap_saprotroph (Põlme et al. 2020); Decay sugar-rich substrates (Põlme et al. 2020) | Põlme S, et al. 2020. Fungal Diversity 105:1-16 ((https://doi.org/10.1007/s13225-020-00466-2)) |
| Otu17175 | 1 | k__Fungi(100);p__Ascomycota(100);c__Saccharomycetes(100);o__Saccharomycetales(100);f__Saccharomycetales_family_Incertae_sedis(100);g__Candida(100);s__Candida_cretensis(100); | Candida | 13 | Pathotroph-Saprotroph-Symbiotroph | Animal Pathogen-Endophyte-Endosymbiont-Epiphyte-\|Undefined Saprotroph\| | Dimorphic Yeast-Yeast | NULL | Probable | As nectar-tap_saprotroph (Põlme et al. 2020); Decay sugar-rich substrates (Põlme et al. 2020); This asexual genus is polyphyletic and contain species that function very differently depend on what they are. Caution should be taken when interpreting guilds for Candida species. | Manolakaki D, et al. 2010. Virulence. 1: 367-375 ((https://doi.org/10.4161/viru.1.5.12796)); Mekbib SB, et al. 2011. Tropical Plant Pathology 36:233-240 ((https://doi.org/10.1590/S1982-56762011000400004)); Põlme S, et al. 2020. Fungal Diversity 105:1-16 ((https://doi.org/10.1007/s13225-020-00466-2)) |
| Otu17176 | 1 | k__Fungi(100);p__Ascomycota(100);c__Saccharomycetes(100);o__Saccharomycetales(100);f__Metschnikowiaceae(100);g__Metschnikowia(100);s__Metschnikowia_pulcherrima(100); | Metschnikowia | 13 | Saprotroph | \|Undefined Saprotroph\| | Yeast | NULL | Probable | As nectar-tap_saprotroph (Põlme et al. 2020); Decay sugar-rich substrates (Põlme et al. 2020) | Põlme S, et al. 2020. Fungal Diversity 105:1-16 ((https://doi.org/10.1007/s13225-020-00466-2)) |
| Otu17177 | 1 | k__Fungi(100);p__Ascomycota(100);c__Saccharomycetes(100);o__Saccharomycetales(100);f__Metschnikowiaceae(100);g__Metschnikowia(100);s__Metschnikowia_pulcherrima(100); | Metschnikowia | 13 | Saprotroph | \|Undefined Saprotroph\| | Yeast | NULL | Probable | As nectar-tap_saprotroph (Põlme et al. 2020); Decay sugar-rich substrates (Põlme et al. 2020) | Põlme S, et al. 2020. Fungal Diversity 105:1-16 ((https://doi.org/10.1007/s13225-020-00466-2)) |
| Otu17180 | 1 | k__Fungi(100);p__Ascomycota(100);c__Saccharomycetes(100);o__Saccharomycetales(100);f__Metschnikowiaceae(100);g__Metschnikowia(100);s__Metschnikowia_pulcherrima(100); | Metschnikowia | 13 | Saprotroph | \|Undefined Saprotroph\| | Yeast | NULL | Probable | As nectar-tap_saprotroph (Põlme et al. 2020); Decay sugar-rich substrates (Põlme et al. 2020) | Põlme S, et al. 2020. Fungal Diversity 105:1-16 ((https://doi.org/10.1007/s13225-020-00466-2)) |
| Otu17181 | 1 | k__Fungi(100);p__Ascomycota(100);c__Saccharomycetes(100);o__Saccharomycetales(100);f__Metschnikowiaceae(100);g__Metschnikowia(100);s__Metschnikowia_pulcherrima(100); | Metschnikowia | 13 | Saprotroph | \|Undefined Saprotroph\| | Yeast | NULL | Probable | As nectar-tap_saprotroph (Põlme et al. 2020); Decay sugar-rich substrates (Põlme et al. 2020) | Põlme S, et al. 2020. Fungal Diversity 105:1-16 ((https://doi.org/10.1007/s13225-020-00466-2)) |
| Otu17182 | 1 | k__Fungi(100);p__Ascomycota(100);c__Saccharomycetes(100);o__Saccharomycetales(100);f__Metschnikowiaceae(100);g__Metschnikowia(100);s__Metschnikowia_pulcherrima(100); | Metschnikowia | 13 | Saprotroph | \|Undefined Saprotroph\| | Yeast | NULL | Probable | As nectar-tap_saprotroph (Põlme et al. 2020); Decay sugar-rich substrates (Põlme et al. 2020) | Põlme S, et al. 2020. Fungal Diversity 105:1-16 ((https://doi.org/10.1007/s13225-020-00466-2)) |
| Otu17184 | 1 | k__Fungi(100);p__Ascomycota(100);c__Saccharomycetes(100);o__Saccharomycetales(100);f__Saccharomycetales_family_Incertae_sedis(100);g__Candida(100);g__Candida_unclassified(100); | Candida | 13 | Pathotroph-Saprotroph-Symbiotroph | Animal Pathogen-Endophyte-Endosymbiont-Epiphyte-\|Undefined Saprotroph\| | Dimorphic Yeast-Yeast | NULL | Probable | As nectar-tap_saprotroph (Põlme et al. 2020); Decay sugar-rich substrates (Põlme et al. 2020); This asexual genus is polyphyletic and contain species that function very differently depend on what they are. Caution should be taken when interpreting guilds for Candida species. | Manolakaki D, et al. 2010. Virulence. 1: 367-375 ((https://doi.org/10.4161/viru.1.5.12796)); Mekbib SB, et al. 2011. Tropical Plant Pathology 36:233-240 ((https://doi.org/10.1590/S1982-56762011000400004)); Põlme S, et al. 2020. Fungal Diversity 105:1-16 ((https://doi.org/10.1007/s13225-020-00466-2)) |
| Otu17187 | 1 | k__Fungi(100);p__Ascomycota(100);c__Saccharomycetes(100);o__Saccharomycetales(100);f__Saccharomycetales_family_Incertae_sedis(100);g__Candida(100);g__Candida_unclassified(100); | Candida | 13 | Pathotroph-Saprotroph-Symbiotroph | Animal Pathogen-Endophyte-Endosymbiont-Epiphyte-\|Undefined Saprotroph\| | Dimorphic Yeast-Yeast | NULL | Probable | As nectar-tap_saprotroph (Põlme et al. 2020); Decay sugar-rich substrates (Põlme et al. 2020); This asexual genus is polyphyletic and contain species that function very differently depend on what they are. Caution should be taken when interpreting guilds for Candida species. | Manolakaki D, et al. 2010. Virulence. 1: 367-375 ((https://doi.org/10.4161/viru.1.5.12796)); Mekbib SB, et al. 2011. Tropical Plant Pathology 36:233-240 ((https://doi.org/10.1590/S1982-56762011000400004)); Põlme S, et al. 2020. Fungal Diversity 105:1-16 ((https://doi.org/10.1007/s13225-020-00466-2)) |
| Otu17188 | 1 | k__Fungi(100);p__Ascomycota(100);c__Saccharomycetes(100);o__Saccharomycetales(100);f__Metschnikowiaceae(100);g__Metschnikowia(100);s__Metschnikowia_pulcherrima(100); | Metschnikowia | 13 | Saprotroph | \|Undefined Saprotroph\| | Yeast | NULL | Probable | As nectar-tap_saprotroph (Põlme et al. 2020); Decay sugar-rich substrates (Põlme et al. 2020) | Põlme S, et al. 2020. Fungal Diversity 105:1-16 ((https://doi.org/10.1007/s13225-020-00466-2)) |
| Otu17190 | 1 | k__Fungi(100);p__Ascomycota(100);c__Saccharomycetes(100);o__Saccharomycetales(100);f__Metschnikowiaceae(100);g__Metschnikowia(100);s__Metschnikowia_pulcherrima(100); | Metschnikowia | 13 | Saprotroph | \|Undefined Saprotroph\| | Yeast | NULL | Probable | As nectar-tap_saprotroph (Põlme et al. 2020); Decay sugar-rich substrates (Põlme et al. 2020) | Põlme S, et al. 2020. Fungal Diversity 105:1-16 ((https://doi.org/10.1007/s13225-020-00466-2)) |
| Otu17191 | 1 | k__Fungi(100);p__Ascomycota(100);c__Saccharomycetes(100);o__Saccharomycetales(100);f__Metschnikowiaceae(100);g__Metschnikowia(100);s__Metschnikowia_pulcherrima(100); | Metschnikowia | 13 | Saprotroph | \|Undefined Saprotroph\| | Yeast | NULL | Probable | As nectar-tap_saprotroph (Põlme et al. 2020); Decay sugar-rich substrates (Põlme et al. 2020) | Põlme S, et al. 2020. Fungal Diversity 105:1-16 ((https://doi.org/10.1007/s13225-020-00466-2)) |
| Otu17192 | 1 | k__Fungi(100);p__Ascomycota(100);c__Saccharomycetes(100);o__Saccharomycetales(100);f__Metschnikowiaceae(100);g__Metschnikowia(100);s__Metschnikowia_pulcherrima(100); | Metschnikowia | 13 | Saprotroph | \|Undefined Saprotroph\| | Yeast | NULL | Probable | As nectar-tap_saprotroph (Põlme et al. 2020); Decay sugar-rich substrates (Põlme et al. 2020) | Põlme S, et al. 2020. Fungal Diversity 105:1-16 ((https://doi.org/10.1007/s13225-020-00466-2)) |
| Otu17193 | 1 | k__Fungi(100);p__Ascomycota(100);c__Saccharomycetes(100);o__Saccharomycetales(100);f__Metschnikowiaceae(100);g__Metschnikowia(100);s__Metschnikowia_pulcherrima(100); | Metschnikowia | 13 | Saprotroph | \|Undefined Saprotroph\| | Yeast | NULL | Probable | As nectar-tap_saprotroph (Põlme et al. 2020); Decay sugar-rich substrates (Põlme et al. 2020) | Põlme S, et al. 2020. Fungal Diversity 105:1-16 ((https://doi.org/10.1007/s13225-020-00466-2)) |
| Otu17194 | 1 | k__Fungi(100);p__Ascomycota(100);c__Saccharomycetes(100);o__Saccharomycetales(100);f__Metschnikowiaceae(100);g__Metschnikowia(100);g__Metschnikowia_unclassified(100); | Metschnikowia | 13 | Saprotroph | \|Undefined Saprotroph\| | Yeast | NULL | Probable | As nectar-tap_saprotroph (Põlme et al. 2020); Decay sugar-rich substrates (Põlme et al. 2020) | Põlme S, et al. 2020. Fungal Diversity 105:1-16 ((https://doi.org/10.1007/s13225-020-00466-2)) |
| Otu17195 | 1 | k__Fungi(100);p__Ascomycota(100);c__Saccharomycetes(100);o__Saccharomycetales(100);f__Metschnikowiaceae(100);g__Metschnikowia(100);g__Metschnikowia_unclassified(100); | Metschnikowia | 13 | Saprotroph | \|Undefined Saprotroph\| | Yeast | NULL | Probable | As nectar-tap_saprotroph (Põlme et al. 2020); Decay sugar-rich substrates (Põlme et al. 2020) | Põlme S, et al. 2020. Fungal Diversity 105:1-16 ((https://doi.org/10.1007/s13225-020-00466-2)) |
| Otu17196 | 1 | k__Fungi(100);p__Ascomycota(100);c__Saccharomycetes(100);o__Saccharomycetales(100);f__Metschnikowiaceae(100);g__Metschnikowia(100);s__Metschnikowia_pulcherrima(100); | Metschnikowia | 13 | Saprotroph | \|Undefined Saprotroph\| | Yeast | NULL | Probable | As nectar-tap_saprotroph (Põlme et al. 2020); Decay sugar-rich substrates (Põlme et al. 2020) | Põlme S, et al. 2020. Fungal Diversity 105:1-16 ((https://doi.org/10.1007/s13225-020-00466-2)) |
| Otu17197 | 1 | k__Fungi(100);p__Ascomycota(100);c__Saccharomycetes(100);o__Saccharomycetales(100);f__Metschnikowiaceae(100);g__Metschnikowia(100);s__Metschnikowia_pulcherrima(100); | Metschnikowia | 13 | Saprotroph | \|Undefined Saprotroph\| | Yeast | NULL | Probable | As nectar-tap_saprotroph (Põlme et al. 2020); Decay sugar-rich substrates (Põlme et al. 2020) | Põlme S, et al. 2020. Fungal Diversity 105:1-16 ((https://doi.org/10.1007/s13225-020-00466-2)) |
| Otu17198 | 1 | k__Fungi(100);p__Ascomycota(100);c__Saccharomycetes(100);o__Saccharomycetales(100);f__Metschnikowiaceae(100);g__Metschnikowia(100);s__Metschnikowia_pulcherrima(100); | Metschnikowia | 13 | Saprotroph | \|Undefined Saprotroph\| | Yeast | NULL | Probable | As nectar-tap_saprotroph (Põlme et al. 2020); Decay sugar-rich substrates (Põlme et al. 2020) | Põlme S, et al. 2020. Fungal Diversity 105:1-16 ((https://doi.org/10.1007/s13225-020-00466-2)) |
| Otu17199 | 1 | k__Fungi(100);p__Ascomycota(100);c__Saccharomycetes(100);o__Saccharomycetales(100);f__Metschnikowiaceae(100);g__Metschnikowia(100);s__Metschnikowia_pulcherrima(100); | Metschnikowia | 13 | Saprotroph | \|Undefined Saprotroph\| | Yeast | NULL | Probable | As nectar-tap_saprotroph (Põlme et al. 2020); Decay sugar-rich substrates (Põlme et al. 2020) | Põlme S, et al. 2020. Fungal Diversity 105:1-16 ((https://doi.org/10.1007/s13225-020-00466-2)) |
| Otu17200 | 1 | k__Fungi(100);p__Ascomycota(100);c__Leotiomycetes(100);o__Helotiales(100);f__Rutstroemiaceae(100);g__Lambertella(100);s__Lambertella_corni_maris(100); | Lambertella | 13 | Pathotroph-Saprotroph | Plant Pathogen-\|Plant Saprotroph\|-Undefined Saprotroph | Microfungus | NULL | Probable | As litter_saprotroph (Põlme et al. 2020); Decay leaf, fruit, seed (Põlme et al. 2020) | Tedersoo L, et al. 2014. Science 346:e1256688 ((https://doi.org/10.1126/science.1256688)); Põlme S, et al. 2020. Fungal Diversity 105:1-16 ((https://doi.org/10.1007/s13225-020-00466-2)) |
| Otu17201 | 1 | k__Fungi(100);p__Ascomycota(100);c__Saccharomycetes(100);o__Saccharomycetales(100);f__Metschnikowiaceae(100);g__Metschnikowia(100);s__Metschnikowia_pulcherrima(100); | Metschnikowia | 13 | Saprotroph | \|Undefined Saprotroph\| | Yeast | NULL | Probable | As nectar-tap_saprotroph (Põlme et al. 2020); Decay sugar-rich substrates (Põlme et al. 2020) | Põlme S, et al. 2020. Fungal Diversity 105:1-16 ((https://doi.org/10.1007/s13225-020-00466-2)) |
| Otu17202 | 1 | k__Fungi(100);p__Ascomycota(100);c__Saccharomycetes(100);o__Saccharomycetales(100);f__Metschnikowiaceae(100);g__Metschnikowia(100);s__Metschnikowia_pulcherrima(100); | Metschnikowia | 13 | Saprotroph | \|Undefined Saprotroph\| | Yeast | NULL | Probable | As nectar-tap_saprotroph (Põlme et al. 2020); Decay sugar-rich substrates (Põlme et al. 2020) | Põlme S, et al. 2020. Fungal Diversity 105:1-16 ((https://doi.org/10.1007/s13225-020-00466-2)) |
| Otu17203 | 1 | k__Fungi(100);p__Ascomycota(100);c__Saccharomycetes(100);o__Saccharomycetales(100);f__Metschnikowiaceae(100);g__Metschnikowia(100);g__Metschnikowia_unclassified(100); | Metschnikowia | 13 | Saprotroph | \|Undefined Saprotroph\| | Yeast | NULL | Probable | As nectar-tap_saprotroph (Põlme et al. 2020); Decay sugar-rich substrates (Põlme et al. 2020) | Põlme S, et al. 2020. Fungal Diversity 105:1-16 ((https://doi.org/10.1007/s13225-020-00466-2)) |
| Otu17204 | 1 | k__Fungi(100);p__Ascomycota(100);c__Saccharomycetes(100);o__Saccharomycetales(100);f__Metschnikowiaceae(100);g__Metschnikowia(100);s__Metschnikowia_pulcherrima(100); | Metschnikowia | 13 | Saprotroph | \|Undefined Saprotroph\| | Yeast | NULL | Probable | As nectar-tap_saprotroph (Põlme et al. 2020); Decay sugar-rich substrates (Põlme et al. 2020) | Põlme S, et al. 2020. Fungal Diversity 105:1-16 ((https://doi.org/10.1007/s13225-020-00466-2)) |
| Otu17205 | 1 | k__Fungi(100);p__Ascomycota(100);c__Saccharomycetes(100);o__Saccharomycetales(100);f__Metschnikowiaceae(100);g__Metschnikowia(100);g__Metschnikowia_unclassified(100); | Metschnikowia | 13 | Saprotroph | \|Undefined Saprotroph\| | Yeast | NULL | Probable | As nectar-tap_saprotroph (Põlme et al. 2020); Decay sugar-rich substrates (Põlme et al. 2020) | Põlme S, et al. 2020. Fungal Diversity 105:1-16 ((https://doi.org/10.1007/s13225-020-00466-2)) |
| Otu17206 | 1 | k__Fungi(100);p__Ascomycota(100);c__Saccharomycetes(100);o__Saccharomycetales(100);f__Metschnikowiaceae(100);g__Metschnikowia(100);g__Metschnikowia_unclassified(100); | Metschnikowia | 13 | Saprotroph | \|Undefined Saprotroph\| | Yeast | NULL | Probable | As nectar-tap_saprotroph (Põlme et al. 2020); Decay sugar-rich substrates (Põlme et al. 2020) | Põlme S, et al. 2020. Fungal Diversity 105:1-16 ((https://doi.org/10.1007/s13225-020-00466-2)) |
| Otu17207 | 1 | k__Fungi(100);p__Ascomycota(100);c__Saccharomycetes(100);o__Saccharomycetales(100);f__Metschnikowiaceae(100);g__Metschnikowia(100);g__Metschnikowia_unclassified(100); | Metschnikowia | 13 | Saprotroph | \|Undefined Saprotroph\| | Yeast | NULL | Probable | As nectar-tap_saprotroph (Põlme et al. 2020); Decay sugar-rich substrates (Põlme et al. 2020) | Põlme S, et al. 2020. Fungal Diversity 105:1-16 ((https://doi.org/10.1007/s13225-020-00466-2)) |
| Otu17208 | 1 | k__Fungi(100);p__Ascomycota(100);c__Saccharomycetes(100);o__Saccharomycetales(100);f__Metschnikowiaceae(100);g__Metschnikowia(100);s__Metschnikowia_pulcherrima(100); | Metschnikowia | 13 | Saprotroph | \|Undefined Saprotroph\| | Yeast | NULL | Probable | As nectar-tap_saprotroph (Põlme et al. 2020); Decay sugar-rich substrates (Põlme et al. 2020) | Põlme S, et al. 2020. Fungal Diversity 105:1-16 ((https://doi.org/10.1007/s13225-020-00466-2)) |
| Otu17209 | 1 | k__Fungi(100);p__Ascomycota(100);c__Saccharomycetes(100);o__Saccharomycetales(100);f__Metschnikowiaceae(100);g__Metschnikowia(100);g__Metschnikowia_unclassified(100); | Metschnikowia | 13 | Saprotroph | \|Undefined Saprotroph\| | Yeast | NULL | Probable | As nectar-tap_saprotroph (Põlme et al. 2020); Decay sugar-rich substrates (Põlme et al. 2020) | Põlme S, et al. 2020. Fungal Diversity 105:1-16 ((https://doi.org/10.1007/s13225-020-00466-2)) |
| Otu17210 | 1 | k__Fungi(100);p__Ascomycota(100);c__Saccharomycetes(100);o__Saccharomycetales(100);f__Metschnikowiaceae(100);g__Metschnikowia(100);g__Metschnikowia_unclassified(100); | Metschnikowia | 13 | Saprotroph | \|Undefined Saprotroph\| | Yeast | NULL | Probable | As nectar-tap_saprotroph (Põlme et al. 2020); Decay sugar-rich substrates (Põlme et al. 2020) | Põlme S, et al. 2020. Fungal Diversity 105:1-16 ((https://doi.org/10.1007/s13225-020-00466-2)) |
| Otu17211 | 1 | k__Fungi(100);p__Ascomycota(100);c__Saccharomycetes(100);o__Saccharomycetales(100);f__Metschnikowiaceae(100);g__Metschnikowia(100);g__Metschnikowia_unclassified(100); | Metschnikowia | 13 | Saprotroph | \|Undefined Saprotroph\| | Yeast | NULL | Probable | As nectar-tap_saprotroph (Põlme et al. 2020); Decay sugar-rich substrates (Põlme et al. 2020) | Põlme S, et al. 2020. Fungal Diversity 105:1-16 ((https://doi.org/10.1007/s13225-020-00466-2)) |
| Otu17212 | 1 | k__Fungi(100);p__Ascomycota(100);c__Saccharomycetes(100);o__Saccharomycetales(100);f__Metschnikowiaceae(100);g__Metschnikowia(100);g__Metschnikowia_unclassified(100); | Metschnikowia | 13 | Saprotroph | \|Undefined Saprotroph\| | Yeast | NULL | Probable | As nectar-tap_saprotroph (Põlme et al. 2020); Decay sugar-rich substrates (Põlme et al. 2020) | Põlme S, et al. 2020. Fungal Diversity 105:1-16 ((https://doi.org/10.1007/s13225-020-00466-2)) |
| Otu17213 | 1 | k__Fungi(100);p__Ascomycota(100);c__Saccharomycetes(100);o__Saccharomycetales(100);f__Saccharomycetales_family_Incertae_sedis(100);f__Saccharomycetales_family_Incertae_sedis_unclassified(100);f__Saccharomycetales_family_Incertae_sedis_unclassified(100); | Saccharomycetales | 7 | Saprotroph | Undefined Saprotroph | Yeast | NULL | Possible | NULL | Sterkenburg E, et al. 2015. New Phytologist 207:1145-1158 ((https://doi.org/10.1111/nph.13426)) |
| Otu17214 | 1 | k__Fungi(100);p__Ascomycota(100);c__Saccharomycetes(100);o__Saccharomycetales(100);f__Metschnikowiaceae(100);g__Metschnikowia(100);s__Metschnikowia_pulcherrima(100); | Metschnikowia | 13 | Saprotroph | \|Undefined Saprotroph\| | Yeast | NULL | Probable | As nectar-tap_saprotroph (Põlme et al. 2020); Decay sugar-rich substrates (Põlme et al. 2020) | Põlme S, et al. 2020. Fungal Diversity 105:1-16 ((https://doi.org/10.1007/s13225-020-00466-2)) |
| Otu17215 | 1 | k__Fungi(100);p__Ascomycota(100);c__Saccharomycetes(100);o__Saccharomycetales(100);f__Metschnikowiaceae(100);g__Metschnikowia(100);s__Metschnikowia_pulcherrima(100); | Metschnikowia | 13 | Saprotroph | \|Undefined Saprotroph\| | Yeast | NULL | Probable | As nectar-tap_saprotroph (Põlme et al. 2020); Decay sugar-rich substrates (Põlme et al. 2020) | Põlme S, et al. 2020. Fungal Diversity 105:1-16 ((https://doi.org/10.1007/s13225-020-00466-2)) |
| Otu17216 | 1 | k__Fungi(100);p__Ascomycota(100);c__Saccharomycetes(100);o__Saccharomycetales(100);f__Metschnikowiaceae(100);g__Metschnikowia(100);s__Metschnikowia_pulcherrima(100); | Metschnikowia | 13 | Saprotroph | \|Undefined Saprotroph\| | Yeast | NULL | Probable | As nectar-tap_saprotroph (Põlme et al. 2020); Decay sugar-rich substrates (Põlme et al. 2020) | Põlme S, et al. 2020. Fungal Diversity 105:1-16 ((https://doi.org/10.1007/s13225-020-00466-2)) |
| Otu17217 | 1 | k__Fungi(100);p__Ascomycota(100);c__Saccharomycetes(100);o__Saccharomycetales(100);f__Metschnikowiaceae(100);g__Metschnikowia(100);s__Metschnikowia_pulcherrima(100); | Metschnikowia | 13 | Saprotroph | \|Undefined Saprotroph\| | Yeast | NULL | Probable | As nectar-tap_saprotroph (Põlme et al. 2020); Decay sugar-rich substrates (Põlme et al. 2020) | Põlme S, et al. 2020. Fungal Diversity 105:1-16 ((https://doi.org/10.1007/s13225-020-00466-2)) |
| Otu17218 | 1 | k__Fungi(100);p__Ascomycota(100);c__Saccharomycetes(100);o__Saccharomycetales(100);f__Metschnikowiaceae(100);g__Metschnikowia(100);s__Metschnikowia_pulcherrima(100); | Metschnikowia | 13 | Saprotroph | \|Undefined Saprotroph\| | Yeast | NULL | Probable | As nectar-tap_saprotroph (Põlme et al. 2020); Decay sugar-rich substrates (Põlme et al. 2020) | Põlme S, et al. 2020. Fungal Diversity 105:1-16 ((https://doi.org/10.1007/s13225-020-00466-2)) |
| Otu17219 | 1 | k__Fungi(100);p__Ascomycota(100);c__Saccharomycetes(100);o__Saccharomycetales(100);f__Metschnikowiaceae(100);g__Metschnikowia(100);g__Metschnikowia_unclassified(100); | Metschnikowia | 13 | Saprotroph | \|Undefined Saprotroph\| | Yeast | NULL | Probable | As nectar-tap_saprotroph (Põlme et al. 2020); Decay sugar-rich substrates (Põlme et al. 2020) | Põlme S, et al. 2020. Fungal Diversity 105:1-16 ((https://doi.org/10.1007/s13225-020-00466-2)) |
| Otu17220 | 1 | k__Fungi(100);p__Ascomycota(100);c__Saccharomycetes(100);o__Saccharomycetales(100);f__Metschnikowiaceae(100);g__Metschnikowia(100);g__Metschnikowia_unclassified(100); | Metschnikowia | 13 | Saprotroph | \|Undefined Saprotroph\| | Yeast | NULL | Probable | As nectar-tap_saprotroph (Põlme et al. 2020); Decay sugar-rich substrates (Põlme et al. 2020) | Põlme S, et al. 2020. Fungal Diversity 105:1-16 ((https://doi.org/10.1007/s13225-020-00466-2)) |
| Otu17221 | 1 | k__Fungi(100);p__Ascomycota(100);c__Saccharomycetes(100);o__Saccharomycetales(100);f__Metschnikowiaceae(100);g__Metschnikowia(100);s__Metschnikowia_pulcherrima(100); | Metschnikowia | 13 | Saprotroph | \|Undefined Saprotroph\| | Yeast | NULL | Probable | As nectar-tap_saprotroph (Põlme et al. 2020); Decay sugar-rich substrates (Põlme et al. 2020) | Põlme S, et al. 2020. Fungal Diversity 105:1-16 ((https://doi.org/10.1007/s13225-020-00466-2)) |
| Otu17222 | 1 | k__Fungi(100);p__Ascomycota(100);c__Saccharomycetes(100);o__Saccharomycetales(100);f__Metschnikowiaceae(100);g__Metschnikowia(100);g__Metschnikowia_unclassified(100); | Metschnikowia | 13 | Saprotroph | \|Undefined Saprotroph\| | Yeast | NULL | Probable | As nectar-tap_saprotroph (Põlme et al. 2020); Decay sugar-rich substrates (Põlme et al. 2020) | Põlme S, et al. 2020. Fungal Diversity 105:1-16 ((https://doi.org/10.1007/s13225-020-00466-2)) |
| Otu17223 | 1 | k__Fungi(100);p__Ascomycota(100);c__Saccharomycetes(100);o__Saccharomycetales(100);f__Metschnikowiaceae(100);g__Metschnikowia(100);s__Metschnikowia_pulcherrima(100); | Metschnikowia | 13 | Saprotroph | \|Undefined Saprotroph\| | Yeast | NULL | Probable | As nectar-tap_saprotroph (Põlme et al. 2020); Decay sugar-rich substrates (Põlme et al. 2020) | Põlme S, et al. 2020. Fungal Diversity 105:1-16 ((https://doi.org/10.1007/s13225-020-00466-2)) |
| Otu17224 | 1 | k__Fungi(100);p__Ascomycota(100);c__Saccharomycetes(100);o__Saccharomycetales(100);f__Metschnikowiaceae(100);g__Metschnikowia(100);g__Metschnikowia_unclassified(100); | Metschnikowia | 13 | Saprotroph | \|Undefined Saprotroph\| | Yeast | NULL | Probable | As nectar-tap_saprotroph (Põlme et al. 2020); Decay sugar-rich substrates (Põlme et al. 2020) | Põlme S, et al. 2020. Fungal Diversity 105:1-16 ((https://doi.org/10.1007/s13225-020-00466-2)) |
| Otu17225 | 1 | k__Fungi(100);p__Ascomycota(100);c__Saccharomycetes(100);o__Saccharomycetales(100);f__Metschnikowiaceae(100);g__Metschnikowia(100);s__Metschnikowia_pulcherrima(100); | Metschnikowia | 13 | Saprotroph | \|Undefined Saprotroph\| | Yeast | NULL | Probable | As nectar-tap_saprotroph (Põlme et al. 2020); Decay sugar-rich substrates (Põlme et al. 2020) | Põlme S, et al. 2020. Fungal Diversity 105:1-16 ((https://doi.org/10.1007/s13225-020-00466-2)) |
| Otu17227 | 1 | k__Fungi(100);p__Ascomycota(100);c__Saccharomycetes(100);o__Saccharomycetales(100);f__Metschnikowiaceae(100);g__Metschnikowia(100);s__Metschnikowia_pulcherrima(100); | Metschnikowia | 13 | Saprotroph | \|Undefined Saprotroph\| | Yeast | NULL | Probable | As nectar-tap_saprotroph (Põlme et al. 2020); Decay sugar-rich substrates (Põlme et al. 2020) | Põlme S, et al. 2020. Fungal Diversity 105:1-16 ((https://doi.org/10.1007/s13225-020-00466-2)) |
| Otu17228 | 1 | k__Fungi(100);p__Ascomycota(100);c__Saccharomycetes(100);o__Saccharomycetales(100);f__Metschnikowiaceae(100);g__Metschnikowia(100);g__Metschnikowia_unclassified(100); | Metschnikowia | 13 | Saprotroph | \|Undefined Saprotroph\| | Yeast | NULL | Probable | As nectar-tap_saprotroph (Põlme et al. 2020); Decay sugar-rich substrates (Põlme et al. 2020) | Põlme S, et al. 2020. Fungal Diversity 105:1-16 ((https://doi.org/10.1007/s13225-020-00466-2)) |
| Otu17229 | 1 | k__Fungi(100);p__Ascomycota(100);c__Saccharomycetes(100);o__Saccharomycetales(100);f__Metschnikowiaceae(100);g__Metschnikowia(100);s__Metschnikowia_pulcherrima(100); | Metschnikowia | 13 | Saprotroph | \|Undefined Saprotroph\| | Yeast | NULL | Probable | As nectar-tap_saprotroph (Põlme et al. 2020); Decay sugar-rich substrates (Põlme et al. 2020) | Põlme S, et al. 2020. Fungal Diversity 105:1-16 ((https://doi.org/10.1007/s13225-020-00466-2)) |
| Otu17230 | 1 | k__Fungi(100);p__Ascomycota(100);c__Saccharomycetes(100);o__Saccharomycetales(100);f__Metschnikowiaceae(100);g__Metschnikowia(100);s__Metschnikowia_pulcherrima(100); | Metschnikowia | 13 | Saprotroph | \|Undefined Saprotroph\| | Yeast | NULL | Probable | As nectar-tap_saprotroph (Põlme et al. 2020); Decay sugar-rich substrates (Põlme et al. 2020) | Põlme S, et al. 2020. Fungal Diversity 105:1-16 ((https://doi.org/10.1007/s13225-020-00466-2)) |
| Otu17231 | 1 | k__Fungi(100);p__Ascomycota(100);c__Saccharomycetes(100);o__Saccharomycetales(100);f__Metschnikowiaceae(100);g__Metschnikowia(100);s__Metschnikowia_pulcherrima(100); | Metschnikowia | 13 | Saprotroph | \|Undefined Saprotroph\| | Yeast | NULL | Probable | As nectar-tap_saprotroph (Põlme et al. 2020); Decay sugar-rich substrates (Põlme et al. 2020) | Põlme S, et al. 2020. Fungal Diversity 105:1-16 ((https://doi.org/10.1007/s13225-020-00466-2)) |
| Otu17232 | 1 | k__Fungi(100);p__Ascomycota(100);c__Saccharomycetes(100);o__Saccharomycetales(100);f__Metschnikowiaceae(100);g__Metschnikowia(100);s__Metschnikowia_pulcherrima(100); | Metschnikowia | 13 | Saprotroph | \|Undefined Saprotroph\| | Yeast | NULL | Probable | As nectar-tap_saprotroph (Põlme et al. 2020); Decay sugar-rich substrates (Põlme et al. 2020) | Põlme S, et al. 2020. Fungal Diversity 105:1-16 ((https://doi.org/10.1007/s13225-020-00466-2)) |
| Otu17233 | 1 | k__Fungi(100);p__Ascomycota(100);c__Saccharomycetes(100);o__Saccharomycetales(100);f__Metschnikowiaceae(100);g__Metschnikowia(100);s__Metschnikowia_pulcherrima(100); | Metschnikowia | 13 | Saprotroph | \|Undefined Saprotroph\| | Yeast | NULL | Probable | As nectar-tap_saprotroph (Põlme et al. 2020); Decay sugar-rich substrates (Põlme et al. 2020) | Põlme S, et al. 2020. Fungal Diversity 105:1-16 ((https://doi.org/10.1007/s13225-020-00466-2)) |
| Otu17234 | 1 | k__Fungi(100);p__Ascomycota(100);c__Saccharomycetes(100);o__Saccharomycetales(100);f__Metschnikowiaceae(100);g__Metschnikowia(100);s__Metschnikowia_pulcherrima(100); | Metschnikowia | 13 | Saprotroph | \|Undefined Saprotroph\| | Yeast | NULL | Probable | As nectar-tap_saprotroph (Põlme et al. 2020); Decay sugar-rich substrates (Põlme et al. 2020) | Põlme S, et al. 2020. Fungal Diversity 105:1-16 ((https://doi.org/10.1007/s13225-020-00466-2)) |
| Otu17235 | 1 | k__Fungi(100);p__Ascomycota(100);c__Saccharomycetes(100);o__Saccharomycetales(100);f__Metschnikowiaceae(100);g__Metschnikowia(100);s__Metschnikowia_pulcherrima(100); | Metschnikowia | 13 | Saprotroph | \|Undefined Saprotroph\| | Yeast | NULL | Probable | As nectar-tap_saprotroph (Põlme et al. 2020); Decay sugar-rich substrates (Põlme et al. 2020) | Põlme S, et al. 2020. Fungal Diversity 105:1-16 ((https://doi.org/10.1007/s13225-020-00466-2)) |
| Otu17238 | 1 | k__Fungi(100);p__Ascomycota(100);c__Saccharomycetes(100);o__Saccharomycetales(100);f__Metschnikowiaceae(100);g__Metschnikowia(100);s__Metschnikowia_pulcherrima(100); | Metschnikowia | 13 | Saprotroph | \|Undefined Saprotroph\| | Yeast | NULL | Probable | As nectar-tap_saprotroph (Põlme et al. 2020); Decay sugar-rich substrates (Põlme et al. 2020) | Põlme S, et al. 2020. Fungal Diversity 105:1-16 ((https://doi.org/10.1007/s13225-020-00466-2)) |
| Otu17239 | 1 | k__Fungi(100);p__Ascomycota(100);c__Saccharomycetes(100);o__Saccharomycetales(100);f__Metschnikowiaceae(100);g__Metschnikowia(100);s__Metschnikowia_pulcherrima(100); | Metschnikowia | 13 | Saprotroph | \|Undefined Saprotroph\| | Yeast | NULL | Probable | As nectar-tap_saprotroph (Põlme et al. 2020); Decay sugar-rich substrates (Põlme et al. 2020) | Põlme S, et al. 2020. Fungal Diversity 105:1-16 ((https://doi.org/10.1007/s13225-020-00466-2)) |
| Otu17240 | 1 | k__Fungi(100);p__Ascomycota(100);c__Saccharomycetes(100);o__Saccharomycetales(100);f__Metschnikowiaceae(100);g__Metschnikowia(100);s__Metschnikowia_pulcherrima(100); | Metschnikowia | 13 | Saprotroph | \|Undefined Saprotroph\| | Yeast | NULL | Probable | As nectar-tap_saprotroph (Põlme et al. 2020); Decay sugar-rich substrates (Põlme et al. 2020) | Põlme S, et al. 2020. Fungal Diversity 105:1-16 ((https://doi.org/10.1007/s13225-020-00466-2)) |
| Otu17241 | 1 | k__Fungi(100);p__Ascomycota(100);c__Saccharomycetes(100);o__Saccharomycetales(100);f__Metschnikowiaceae(100);g__Metschnikowia(100);g__Metschnikowia_unclassified(100); | Metschnikowia | 13 | Saprotroph | \|Undefined Saprotroph\| | Yeast | NULL | Probable | As nectar-tap_saprotroph (Põlme et al. 2020); Decay sugar-rich substrates (Põlme et al. 2020) | Põlme S, et al. 2020. Fungal Diversity 105:1-16 ((https://doi.org/10.1007/s13225-020-00466-2)) |
| Otu17242 | 1 | k__Fungi(100);p__Ascomycota(100);c__Saccharomycetes(100);o__Saccharomycetales(100);f__Metschnikowiaceae(100);g__Metschnikowia(100);s__Metschnikowia_pulcherrima(100); | Metschnikowia | 13 | Saprotroph | \|Undefined Saprotroph\| | Yeast | NULL | Probable | As nectar-tap_saprotroph (Põlme et al. 2020); Decay sugar-rich substrates (Põlme et al. 2020) | Põlme S, et al. 2020. Fungal Diversity 105:1-16 ((https://doi.org/10.1007/s13225-020-00466-2)) |
| Otu17243 | 1 | k__Fungi(100);p__Ascomycota(100);c__Saccharomycetes(100);o__Saccharomycetales(100);f__Metschnikowiaceae(100);g__Metschnikowia(100);s__Metschnikowia_pulcherrima(100); | Metschnikowia | 13 | Saprotroph | \|Undefined Saprotroph\| | Yeast | NULL | Probable | As nectar-tap_saprotroph (Põlme et al. 2020); Decay sugar-rich substrates (Põlme et al. 2020) | Põlme S, et al. 2020. Fungal Diversity 105:1-16 ((https://doi.org/10.1007/s13225-020-00466-2)) |
| Otu17244 | 1 | k__Fungi(100);p__Ascomycota(100);c__Saccharomycetes(100);o__Saccharomycetales(100);f__Metschnikowiaceae(100);g__Metschnikowia(100);s__Metschnikowia_pulcherrima(100); | Metschnikowia | 13 | Saprotroph | \|Undefined Saprotroph\| | Yeast | NULL | Probable | As nectar-tap_saprotroph (Põlme et al. 2020); Decay sugar-rich substrates (Põlme et al. 2020) | Põlme S, et al. 2020. Fungal Diversity 105:1-16 ((https://doi.org/10.1007/s13225-020-00466-2)) |
| Otu17245 | 1 | k__Fungi(100);p__Ascomycota(100);c__Saccharomycetes(100);o__Saccharomycetales(100);f__Metschnikowiaceae(100);g__Metschnikowia(100);s__Metschnikowia_pulcherrima(100); | Metschnikowia | 13 | Saprotroph | \|Undefined Saprotroph\| | Yeast | NULL | Probable | As nectar-tap_saprotroph (Põlme et al. 2020); Decay sugar-rich substrates (Põlme et al. 2020) | Põlme S, et al. 2020. Fungal Diversity 105:1-16 ((https://doi.org/10.1007/s13225-020-00466-2)) |
| Otu17247 | 1 | k__Fungi(100);p__Ascomycota(100);c__Saccharomycetes(100);o__Saccharomycetales(100);f__Metschnikowiaceae(100);g__Metschnikowia(100);g__Metschnikowia_unclassified(100); | Metschnikowia | 13 | Saprotroph | \|Undefined Saprotroph\| | Yeast | NULL | Probable | As nectar-tap_saprotroph (Põlme et al. 2020); Decay sugar-rich substrates (Põlme et al. 2020) | Põlme S, et al. 2020. Fungal Diversity 105:1-16 ((https://doi.org/10.1007/s13225-020-00466-2)) |
| Otu17248 | 1 | k__Fungi(100);p__Ascomycota(100);c__Saccharomycetes(100);o__Saccharomycetales(100);f__Metschnikowiaceae(100);g__Metschnikowia(100);s__Metschnikowia_pulcherrima(100); | Metschnikowia | 13 | Saprotroph | \|Undefined Saprotroph\| | Yeast | NULL | Probable | As nectar-tap_saprotroph (Põlme et al. 2020); Decay sugar-rich substrates (Põlme et al. 2020) | Põlme S, et al. 2020. Fungal Diversity 105:1-16 ((https://doi.org/10.1007/s13225-020-00466-2)) |
| Otu17249 | 1 | k__Fungi(100);p__Ascomycota(100);c__Saccharomycetes(100);o__Saccharomycetales(100);f__Metschnikowiaceae(100);g__Metschnikowia(100);s__Metschnikowia_pulcherrima(100); | Metschnikowia | 13 | Saprotroph | \|Undefined Saprotroph\| | Yeast | NULL | Probable | As nectar-tap_saprotroph (Põlme et al. 2020); Decay sugar-rich substrates (Põlme et al. 2020) | Põlme S, et al. 2020. Fungal Diversity 105:1-16 ((https://doi.org/10.1007/s13225-020-00466-2)) |
| Otu17250 | 1 | k__Fungi(100);p__Ascomycota(100);c__Saccharomycetes(100);o__Saccharomycetales(100);f__Metschnikowiaceae(100);g__Metschnikowia(100);g__Metschnikowia_unclassified(100); | Metschnikowia | 13 | Saprotroph | \|Undefined Saprotroph\| | Yeast | NULL | Probable | As nectar-tap_saprotroph (Põlme et al. 2020); Decay sugar-rich substrates (Põlme et al. 2020) | Põlme S, et al. 2020. Fungal Diversity 105:1-16 ((https://doi.org/10.1007/s13225-020-00466-2)) |
| Otu17251 | 1 | k__Fungi(100);p__Ascomycota(100);c__Saccharomycetes(100);o__Saccharomycetales(100);f__Metschnikowiaceae(100);g__Metschnikowia(100);s__Metschnikowia_pulcherrima(100); | Metschnikowia | 13 | Saprotroph | \|Undefined Saprotroph\| | Yeast | NULL | Probable | As nectar-tap_saprotroph (Põlme et al. 2020); Decay sugar-rich substrates (Põlme et al. 2020) | Põlme S, et al. 2020. Fungal Diversity 105:1-16 ((https://doi.org/10.1007/s13225-020-00466-2)) |
| Otu17252 | 1 | k__Fungi(100);p__Ascomycota(100);c__Saccharomycetes(100);o__Saccharomycetales(100);f__Metschnikowiaceae(100);g__Metschnikowia(100);s__Metschnikowia_pulcherrima(100); | Metschnikowia | 13 | Saprotroph | \|Undefined Saprotroph\| | Yeast | NULL | Probable | As nectar-tap_saprotroph (Põlme et al. 2020); Decay sugar-rich substrates (Põlme et al. 2020) | Põlme S, et al. 2020. Fungal Diversity 105:1-16 ((https://doi.org/10.1007/s13225-020-00466-2)) |
| Otu17253 | 1 | k__Fungi(100);p__Ascomycota(100);c__Saccharomycetes(100);o__Saccharomycetales(100);f__Saccharomycetales_family_Incertae_sedis(100);g__Candida(100);g__Candida_unclassified(100); | Candida | 13 | Pathotroph-Saprotroph-Symbiotroph | Animal Pathogen-Endophyte-Endosymbiont-Epiphyte-\|Undefined Saprotroph\| | Dimorphic Yeast-Yeast | NULL | Probable | As nectar-tap_saprotroph (Põlme et al. 2020); Decay sugar-rich substrates (Põlme et al. 2020); This asexual genus is polyphyletic and contain species that function very differently depend on what they are. Caution should be taken when interpreting guilds for Candida species. | Manolakaki D, et al. 2010. Virulence. 1: 367-375 ((https://doi.org/10.4161/viru.1.5.12796)); Mekbib SB, et al. 2011. Tropical Plant Pathology 36:233-240 ((https://doi.org/10.1590/S1982-56762011000400004)); Põlme S, et al. 2020. Fungal Diversity 105:1-16 ((https://doi.org/10.1007/s13225-020-00466-2)) |
| Otu17254 | 1 | k__Fungi(100);p__Ascomycota(100);c__Saccharomycetes(100);o__Saccharomycetales(100);f__Metschnikowiaceae(100);g__Metschnikowia(100);g__Metschnikowia_unclassified(100); | Metschnikowia | 13 | Saprotroph | \|Undefined Saprotroph\| | Yeast | NULL | Probable | As nectar-tap_saprotroph (Põlme et al. 2020); Decay sugar-rich substrates (Põlme et al. 2020) | Põlme S, et al. 2020. Fungal Diversity 105:1-16 ((https://doi.org/10.1007/s13225-020-00466-2)) |
| Otu17255 | 1 | k__Fungi(100);p__Ascomycota(100);c__Saccharomycetes(100);o__Saccharomycetales(100);f__Metschnikowiaceae(100);g__Metschnikowia(100);s__Metschnikowia_pulcherrima(100); | Metschnikowia | 13 | Saprotroph | \|Undefined Saprotroph\| | Yeast | NULL | Probable | As nectar-tap_saprotroph (Põlme et al. 2020); Decay sugar-rich substrates (Põlme et al. 2020) | Põlme S, et al. 2020. Fungal Diversity 105:1-16 ((https://doi.org/10.1007/s13225-020-00466-2)) |
| Otu17256 | 1 | k__Fungi(100);p__Ascomycota(100);c__Saccharomycetes(100);o__Saccharomycetales(100);f__Saccharomycetales_family_Incertae_sedis(100);f__Saccharomycetales_family_Incertae_sedis_unclassified(100);f__Saccharomycetales_family_Incertae_sedis_unclassified(100); | Saccharomycetales | 7 | Saprotroph | Undefined Saprotroph | Yeast | NULL | Possible | NULL | Sterkenburg E, et al. 2015. New Phytologist 207:1145-1158 ((https://doi.org/10.1111/nph.13426)) |
| Otu17257 | 1 | k__Fungi(100);p__Ascomycota(100);c__Saccharomycetes(100);o__Saccharomycetales(100);f__Saccharomycetales_family_Incertae_sedis(100);g__Candida(100);g__Candida_unclassified(100); | Candida | 13 | Pathotroph-Saprotroph-Symbiotroph | Animal Pathogen-Endophyte-Endosymbiont-Epiphyte-\|Undefined Saprotroph\| | Dimorphic Yeast-Yeast | NULL | Probable | As nectar-tap_saprotroph (Põlme et al. 2020); Decay sugar-rich substrates (Põlme et al. 2020); This asexual genus is polyphyletic and contain species that function very differently depend on what they are. Caution should be taken when interpreting guilds for Candida species. | Manolakaki D, et al. 2010. Virulence. 1: 367-375 ((https://doi.org/10.4161/viru.1.5.12796)); Mekbib SB, et al. 2011. Tropical Plant Pathology 36:233-240 ((https://doi.org/10.1590/S1982-56762011000400004)); Põlme S, et al. 2020. Fungal Diversity 105:1-16 ((https://doi.org/10.1007/s13225-020-00466-2)) |
| Otu17260 | 1 | k__Fungi(100);p__Ascomycota(100);c__Saccharomycetes(100);o__Saccharomycetales(100);f__Metschnikowiaceae(100);g__Metschnikowia(100);s__Metschnikowia_pulcherrima(100); | Metschnikowia | 13 | Saprotroph | \|Undefined Saprotroph\| | Yeast | NULL | Probable | As nectar-tap_saprotroph (Põlme et al. 2020); Decay sugar-rich substrates (Põlme et al. 2020) | Põlme S, et al. 2020. Fungal Diversity 105:1-16 ((https://doi.org/10.1007/s13225-020-00466-2)) |
| Otu17263 | 1 | k__Fungi(100);p__Ascomycota(100);c__Saccharomycetes(100);o__Saccharomycetales(100);f__Metschnikowiaceae(100);g__Metschnikowia(100);s__Metschnikowia_pulcherrima(100); | Metschnikowia | 13 | Saprotroph | \|Undefined Saprotroph\| | Yeast | NULL | Probable | As nectar-tap_saprotroph (Põlme et al. 2020); Decay sugar-rich substrates (Põlme et al. 2020) | Põlme S, et al. 2020. Fungal Diversity 105:1-16 ((https://doi.org/10.1007/s13225-020-00466-2)) |
| Otu17264 | 1 | k__Fungi(100);p__Ascomycota(100);c__Saccharomycetes(100);o__Saccharomycetales(100);f__Metschnikowiaceae(100);g__Metschnikowia(100);s__Metschnikowia_pulcherrima(100); | Metschnikowia | 13 | Saprotroph | \|Undefined Saprotroph\| | Yeast | NULL | Probable | As nectar-tap_saprotroph (Põlme et al. 2020); Decay sugar-rich substrates (Põlme et al. 2020) | Põlme S, et al. 2020. Fungal Diversity 105:1-16 ((https://doi.org/10.1007/s13225-020-00466-2)) |
| Otu17265 | 1 | k__Fungi(100);p__Ascomycota(100);c__Saccharomycetes(100);o__Saccharomycetales(100);f__Metschnikowiaceae(100);g__Metschnikowia(100);g__Metschnikowia_unclassified(100); | Metschnikowia | 13 | Saprotroph | \|Undefined Saprotroph\| | Yeast | NULL | Probable | As nectar-tap_saprotroph (Põlme et al. 2020); Decay sugar-rich substrates (Põlme et al. 2020) | Põlme S, et al. 2020. Fungal Diversity 105:1-16 ((https://doi.org/10.1007/s13225-020-00466-2)) |
| Otu17266 | 1 | k__Fungi(100);p__Ascomycota(100);c__Saccharomycetes(100);o__Saccharomycetales(100);f__Metschnikowiaceae(100);g__Metschnikowia(100);g__Metschnikowia_unclassified(100); | Metschnikowia | 13 | Saprotroph | \|Undefined Saprotroph\| | Yeast | NULL | Probable | As nectar-tap_saprotroph (Põlme et al. 2020); Decay sugar-rich substrates (Põlme et al. 2020) | Põlme S, et al. 2020. Fungal Diversity 105:1-16 ((https://doi.org/10.1007/s13225-020-00466-2)) |
| Otu17267 | 1 | k__Fungi(100);p__Ascomycota(100);c__Saccharomycetes(100);o__Saccharomycetales(100);f__Metschnikowiaceae(100);g__Metschnikowia(100);s__Metschnikowia_pulcherrima(100); | Metschnikowia | 13 | Saprotroph | \|Undefined Saprotroph\| | Yeast | NULL | Probable | As nectar-tap_saprotroph (Põlme et al. 2020); Decay sugar-rich substrates (Põlme et al. 2020) | Põlme S, et al. 2020. Fungal Diversity 105:1-16 ((https://doi.org/10.1007/s13225-020-00466-2)) |
| Otu17268 | 1 | k__Fungi(100);p__Basidiomycota(100);c__Tremellomycetes(100);o__Filobasidiales(100);f__Filobasidiaceae(100);g__Cryptococcus(100);s__Cryptococcus_heimaeyensis(100); | Cryptococcus | 13 | Pathotroph-Saprotroph-Symbiotroph | Animal Pathogen-Endophyte-Epiphyte-Undefined Saprotroph | Dimorphic Yeast-Yeast | NULL | Possible | As foliar_endophyte (Põlme et al. 2020); Decay leaf, fruit, seed, soil, dung, animal material (Põlme et al. 2020) | Andrews JH, et al. 1983. Phytopathology 73:228-234 ((https://doi.org/10.1094/Phyto-73-228)); Kurtzman CP, et al. (eds.) 2011. The Yeasts, a Taxonomic Study. Fifth Edition. Vols 1-3. Elsevier, San Diego; ISBN: 9780444521491; Põlme S, et al. 2020. Fungal Diversity 105:1-16 ((https://doi.org/10.1007/s13225-020-00466-2)) |
| Otu17270 | 1 | k__Fungi(100);p__Ascomycota(100);c__Saccharomycetes(100);o__Saccharomycetales(100);f__Metschnikowiaceae(100);g__Metschnikowia(100);g__Metschnikowia_unclassified(100); | Metschnikowia | 13 | Saprotroph | \|Undefined Saprotroph\| | Yeast | NULL | Probable | As nectar-tap_saprotroph (Põlme et al. 2020); Decay sugar-rich substrates (Põlme et al. 2020) | Põlme S, et al. 2020. Fungal Diversity 105:1-16 ((https://doi.org/10.1007/s13225-020-00466-2)) |
| Otu17271 | 1 | k__Fungi(100);p__Ascomycota(100);c__Saccharomycetes(100);o__Saccharomycetales(100);f__Metschnikowiaceae(100);g__Metschnikowia(100);s__Metschnikowia_pulcherrima(100); | Metschnikowia | 13 | Saprotroph | \|Undefined Saprotroph\| | Yeast | NULL | Probable | As nectar-tap_saprotroph (Põlme et al. 2020); Decay sugar-rich substrates (Põlme et al. 2020) | Põlme S, et al. 2020. Fungal Diversity 105:1-16 ((https://doi.org/10.1007/s13225-020-00466-2)) |
| Otu17272 | 1 | k__Fungi(100);p__Ascomycota(100);c__Saccharomycetes(100);o__Saccharomycetales(100);f__Metschnikowiaceae(100);g__Metschnikowia(100);s__Metschnikowia_pulcherrima(100); | Metschnikowia | 13 | Saprotroph | \|Undefined Saprotroph\| | Yeast | NULL | Probable | As nectar-tap_saprotroph (Põlme et al. 2020); Decay sugar-rich substrates (Põlme et al. 2020) | Põlme S, et al. 2020. Fungal Diversity 105:1-16 ((https://doi.org/10.1007/s13225-020-00466-2)) |
| Otu17273 | 1 | k__Fungi(100);p__Ascomycota(100);c__Saccharomycetes(100);o__Saccharomycetales(100);f__Metschnikowiaceae(100);g__Metschnikowia(100);g__Metschnikowia_unclassified(100); | Metschnikowia | 13 | Saprotroph | \|Undefined Saprotroph\| | Yeast | NULL | Probable | As nectar-tap_saprotroph (Põlme et al. 2020); Decay sugar-rich substrates (Põlme et al. 2020) | Põlme S, et al. 2020. Fungal Diversity 105:1-16 ((https://doi.org/10.1007/s13225-020-00466-2)) |
| Otu17274 | 1 | k__Fungi(100);p__Ascomycota(100);c__Saccharomycetes(100);o__Saccharomycetales(100);f__Metschnikowiaceae(100);g__Metschnikowia(100);s__Metschnikowia_pulcherrima(100); | Metschnikowia | 13 | Saprotroph | \|Undefined Saprotroph\| | Yeast | NULL | Probable | As nectar-tap_saprotroph (Põlme et al. 2020); Decay sugar-rich substrates (Põlme et al. 2020) | Põlme S, et al. 2020. Fungal Diversity 105:1-16 ((https://doi.org/10.1007/s13225-020-00466-2)) |
| Otu17275 | 1 | k__Fungi(100);p__Ascomycota(100);c__Saccharomycetes(100);o__Saccharomycetales(100);f__Metschnikowiaceae(100);g__Metschnikowia(100);g__Metschnikowia_unclassified(100); | Metschnikowia | 13 | Saprotroph | \|Undefined Saprotroph\| | Yeast | NULL | Probable | As nectar-tap_saprotroph (Põlme et al. 2020); Decay sugar-rich substrates (Põlme et al. 2020) | Põlme S, et al. 2020. Fungal Diversity 105:1-16 ((https://doi.org/10.1007/s13225-020-00466-2)) |
| Otu17276 | 1 | k__Fungi(100);p__Ascomycota(100);c__Saccharomycetes(100);o__Saccharomycetales(100);f__Metschnikowiaceae(100);g__Metschnikowia(100);g__Metschnikowia_unclassified(100); | Metschnikowia | 13 | Saprotroph | \|Undefined Saprotroph\| | Yeast | NULL | Probable | As nectar-tap_saprotroph (Põlme et al. 2020); Decay sugar-rich substrates (Põlme et al. 2020) | Põlme S, et al. 2020. Fungal Diversity 105:1-16 ((https://doi.org/10.1007/s13225-020-00466-2)) |
| Otu17277 | 1 | k__Fungi(100);p__Ascomycota(100);c__Saccharomycetes(100);o__Saccharomycetales(100);f__Metschnikowiaceae(100);g__Metschnikowia(100);s__Metschnikowia_pulcherrima(100); | Metschnikowia | 13 | Saprotroph | \|Undefined Saprotroph\| | Yeast | NULL | Probable | As nectar-tap_saprotroph (Põlme et al. 2020); Decay sugar-rich substrates (Põlme et al. 2020) | Põlme S, et al. 2020. Fungal Diversity 105:1-16 ((https://doi.org/10.1007/s13225-020-00466-2)) |
| Otu17278 | 1 | k__Fungi(100);p__Ascomycota(100);c__Saccharomycetes(100);o__Saccharomycetales(100);f__Metschnikowiaceae(100);g__Metschnikowia(100);s__Metschnikowia_pulcherrima(100); | Metschnikowia | 13 | Saprotroph | \|Undefined Saprotroph\| | Yeast | NULL | Probable | As nectar-tap_saprotroph (Põlme et al. 2020); Decay sugar-rich substrates (Põlme et al. 2020) | Põlme S, et al. 2020. Fungal Diversity 105:1-16 ((https://doi.org/10.1007/s13225-020-00466-2)) |
| Otu17279 | 1 | k__Fungi(100);p__Ascomycota(100);c__Saccharomycetes(100);o__Saccharomycetales(100);f__Metschnikowiaceae(100);g__Metschnikowia(100);s__Metschnikowia_pulcherrima(100); | Metschnikowia | 13 | Saprotroph | \|Undefined Saprotroph\| | Yeast | NULL | Probable | As nectar-tap_saprotroph (Põlme et al. 2020); Decay sugar-rich substrates (Põlme et al. 2020) | Põlme S, et al. 2020. Fungal Diversity 105:1-16 ((https://doi.org/10.1007/s13225-020-00466-2)) |
| Otu17281 | 1 | k__Fungi(100);p__Ascomycota(100);c__Saccharomycetes(100);o__Saccharomycetales(100);f__Metschnikowiaceae(100);g__Metschnikowia(100);s__Metschnikowia_pulcherrima(100); | Metschnikowia | 13 | Saprotroph | \|Undefined Saprotroph\| | Yeast | NULL | Probable | As nectar-tap_saprotroph (Põlme et al. 2020); Decay sugar-rich substrates (Põlme et al. 2020) | Põlme S, et al. 2020. Fungal Diversity 105:1-16 ((https://doi.org/10.1007/s13225-020-00466-2)) |
| Otu17284 | 1 | k__Fungi(100);p__Ascomycota(100);c__Saccharomycetes(100);o__Saccharomycetales(100);o__Saccharomycetales_unclassified(100);o__Saccharomycetales_unclassified(100);o__Saccharomycetales_unclassified(100); | Saccharomycetales | 7 | Saprotroph | Undefined Saprotroph | Yeast | NULL | Possible | NULL | Sterkenburg E, et al. 2015. New Phytologist 207:1145-1158 ((https://doi.org/10.1111/nph.13426)) |
| Otu17285 | 1 | k__Fungi(100);p__Ascomycota(100);c__Saccharomycetes(100);o__Saccharomycetales(100);f__Metschnikowiaceae(100);g__Metschnikowia(100);g__Metschnikowia_unclassified(100); | Metschnikowia | 13 | Saprotroph | \|Undefined Saprotroph\| | Yeast | NULL | Probable | As nectar-tap_saprotroph (Põlme et al. 2020); Decay sugar-rich substrates (Põlme et al. 2020) | Põlme S, et al. 2020. Fungal Diversity 105:1-16 ((https://doi.org/10.1007/s13225-020-00466-2)) |
| Otu17295 | 1 | k__Fungi(100);p__Ascomycota(100);c__Saccharomycetes(100);o__Saccharomycetales(100);f__Metschnikowiaceae(100);g__Metschnikowia(100);g__Metschnikowia_unclassified(100); | Metschnikowia | 13 | Saprotroph | \|Undefined Saprotroph\| | Yeast | NULL | Probable | As nectar-tap_saprotroph (Põlme et al. 2020); Decay sugar-rich substrates (Põlme et al. 2020) | Põlme S, et al. 2020. Fungal Diversity 105:1-16 ((https://doi.org/10.1007/s13225-020-00466-2)) |
| Otu17298 | 1 | k__Fungi(100);p__Ascomycota(100);c__Saccharomycetes(100);o__Saccharomycetales(100);f__Metschnikowiaceae(100);g__Metschnikowia(100);g__Metschnikowia_unclassified(100); | Metschnikowia | 13 | Saprotroph | \|Undefined Saprotroph\| | Yeast | NULL | Probable | As nectar-tap_saprotroph (Põlme et al. 2020); Decay sugar-rich substrates (Põlme et al. 2020) | Põlme S, et al. 2020. Fungal Diversity 105:1-16 ((https://doi.org/10.1007/s13225-020-00466-2)) |
| Otu17299 | 1 | k__Fungi(100);p__Ascomycota(100);c__Saccharomycetes(100);o__Saccharomycetales(100);f__Metschnikowiaceae(100);g__Metschnikowia(100);s__Metschnikowia_pulcherrima(100); | Metschnikowia | 13 | Saprotroph | \|Undefined Saprotroph\| | Yeast | NULL | Probable | As nectar-tap_saprotroph (Põlme et al. 2020); Decay sugar-rich substrates (Põlme et al. 2020) | Põlme S, et al. 2020. Fungal Diversity 105:1-16 ((https://doi.org/10.1007/s13225-020-00466-2)) |
| Otu17300 | 1 | k__Fungi(100);p__Ascomycota(100);c__Saccharomycetes(100);o__Saccharomycetales(100);f__Metschnikowiaceae(100);g__Metschnikowia(100);s__Metschnikowia_pulcherrima(100); | Metschnikowia | 13 | Saprotroph | \|Undefined Saprotroph\| | Yeast | NULL | Probable | As nectar-tap_saprotroph (Põlme et al. 2020); Decay sugar-rich substrates (Põlme et al. 2020) | Põlme S, et al. 2020. Fungal Diversity 105:1-16 ((https://doi.org/10.1007/s13225-020-00466-2)) |
| Otu17301 | 1 | k__Fungi(100);p__Ascomycota(100);c__Saccharomycetes(100);o__Saccharomycetales(100);f__Metschnikowiaceae(100);g__Metschnikowia(100);g__Metschnikowia_unclassified(100); | Metschnikowia | 13 | Saprotroph | \|Undefined Saprotroph\| | Yeast | NULL | Probable | As nectar-tap_saprotroph (Põlme et al. 2020); Decay sugar-rich substrates (Põlme et al. 2020) | Põlme S, et al. 2020. Fungal Diversity 105:1-16 ((https://doi.org/10.1007/s13225-020-00466-2)) |

| | | | | | | | | | | | |
|---|---|---|---|---|---|---|---|---|---|---|---|
| Otu17302 | 1 | k__Fungi(100);p__Ascomycota(100);c__Saccharomycetes(100);o__Saccharomycetales(100);f__Metschnikowiaceae(100);g__Metschnikowia(100);g__Metschnikowia_unclassified(100); | Metschnikowia | 13 | Saprotroph | \|Undefined Saprotroph\| | Yeast | NULL | Probable | As nectar-tap_saprotroph (Põlme et al. 2020); Decay sugar-rich substrates (Põlme et al. 2020) | Põlme S, et al. 2020. Fungal Diversity 105:1-16 ((https://doi.org/10.1007/s13225-020-00466-2)) |
| Otu17305 | 1 | k__Fungi(100);p__Ascomycota(100);c__Saccharomycetes(100);o__Saccharomycetales(100);f__Metschnikowiaceae(100);g__Metschnikowia(100);s__Metschnikowia_pulcherrima(100); | Metschnikowia | 13 | Saprotroph | \|Undefined Saprotroph\| | Yeast | NULL | Probable | As nectar-tap_saprotroph (Põlme et al. 2020); Decay sugar-rich substrates (Põlme et al. 2020) | Põlme S, et al. 2020. Fungal Diversity 105:1-16 ((https://doi.org/10.1007/s13225-020-00466-2)) |
| Otu17306 | 1 | k__Fungi(100);p__Ascomycota(100);c__Saccharomycetes(100);o__Saccharomycetales(100);f__Metschnikowiaceae(100);g__Metschnikowia(100);s__Metschnikowia_pulcherrima(100); | Metschnikowia | 13 | Saprotroph | \|Undefined Saprotroph\| | Yeast | NULL | Probable | As nectar-tap_saprotroph (Põlme et al. 2020); Decay sugar-rich substrates (Põlme et al. 2020) | Põlme S, et al. 2020. Fungal Diversity 105:1-16 ((https://doi.org/10.1007/s13225-020-00466-2)) |
| Otu17313 | 1 | k__Fungi(100);p__Ascomycota(100);c__Saccharomycetes(100);o__Saccharomycetales(100);f__Metschnikowiaceae(100);g__Metschnikowia(100);g__Metschnikowia_unclassified(100); | Metschnikowia | 13 | Saprotroph | \|Undefined Saprotroph\| | Yeast | NULL | Probable | As nectar-tap_saprotroph (Põlme et al. 2020); Decay sugar-rich substrates (Põlme et al. 2020) | Põlme S, et al. 2020. Fungal Diversity 105:1-16 ((https://doi.org/10.1007/s13225-020-00466-2)) |
| Otu17314 | 1 | k__Fungi(100);p__Ascomycota(100);c__Saccharomycetes(100);o__Saccharomycetales(100);f__Metschnikowiaceae(100);g__Metschnikowia(100);s__Metschnikowia_pulcherrima(100); | Metschnikowia | 13 | Saprotroph | \|Undefined Saprotroph\| | Yeast | NULL | Probable | As nectar-tap_saprotroph (Põlme et al. 2020); Decay sugar-rich substrates (Põlme et al. 2020) | Põlme S, et al. 2020. Fungal Diversity 105:1-16 ((https://doi.org/10.1007/s13225-020-00466-2)) |
| Otu17315 | 1 | k__Fungi(100);p__Ascomycota(100);c__Saccharomycetes(100);o__Saccharomycetales(100);f__Metschnikowiaceae(100);g__Metschnikowia(100);g__Metschnikowia_unclassified(100); | Metschnikowia | 13 | Saprotroph | \|Undefined Saprotroph\| | Yeast | NULL | Probable | As nectar-tap_saprotroph (Põlme et al. 2020); Decay sugar-rich substrates (Põlme et al. 2020) | Põlme S, et al. 2020. Fungal Diversity 105:1-16 ((https://doi.org/10.1007/s13225-020-00466-2)) |
| Otu17316 | 1 | k__Fungi(100);p__Ascomycota(100);c__Saccharomycetes(100);o__Saccharomycetales(100);f__Metschnikowiaceae(100);g__Metschnikowia(100);g__Metschnikowia_unclassified(100); | Metschnikowia | 13 | Saprotroph | \|Undefined Saprotroph\| | Yeast | NULL | Probable | As nectar-tap_saprotroph (Põlme et al. 2020); Decay sugar-rich substrates (Põlme et al. 2020) | Põlme S, et al. 2020. Fungal Diversity 105:1-16 ((https://doi.org/10.1007/s13225-020-00466-2)) |
| Otu17317 | 1 | k__Fungi(100);p__Ascomycota(100);c__Saccharomycetes(100);o__Saccharomycetales(100);f__Metschnikowiaceae(100);g__Metschnikowia(100);s__Metschnikowia_pulcherrima(100); | Metschnikowia | 13 | Saprotroph | \|Undefined Saprotroph\| | Yeast | NULL | Probable | As nectar-tap_saprotroph (Põlme et al. 2020); Decay sugar-rich substrates (Põlme et al. 2020) | Põlme S, et al. 2020. Fungal Diversity 105:1-16 ((https://doi.org/10.1007/s13225-020-00466-2)) |
| Otu17318 | 1 | k__Fungi(100);p__Ascomycota(100);c__Saccharomycetes(100);o__Saccharomycetales(100);f__Metschnikowiaceae(100);g__Metschnikowia(100);s__Metschnikowia_pulcherrima(100); | Metschnikowia | 13 | Saprotroph | \|Undefined Saprotroph\| | Yeast | NULL | Probable | As nectar-tap_saprotroph (Põlme et al. 2020); Decay sugar-rich substrates (Põlme et al. 2020) | Põlme S, et al. 2020. Fungal Diversity 105:1-16 ((https://doi.org/10.1007/s13225-020-00466-2)) |
| Otu17320 | 1 | k__Fungi(100);p__Ascomycota(100);c__Saccharomycetes(100);o__Saccharomycetales(100);f__Metschnikowiaceae(100);g__Metschnikowia(100);s__Metschnikowia_pulcherrima(100); | Metschnikowia | 13 | Saprotroph | \|Undefined Saprotroph\| | Yeast | NULL | Probable | As nectar-tap_saprotroph (Põlme et al. 2020); Decay sugar-rich substrates (Põlme et al. 2020) | Põlme S, et al. 2020. Fungal Diversity 105:1-16 ((https://doi.org/10.1007/s13225-020-00466-2)) |
| Otu17321 | 1 | k__Fungi(100);p__Ascomycota(100);c__Saccharomycetes(100);o__Saccharomycetales(100);f__Metschnikowiaceae(100);g__Metschnikowia(100);s__Metschnikowia_pulcherrima(100); | Metschnikowia | 13 | Saprotroph | \|Undefined Saprotroph\| | Yeast | NULL | Probable | As nectar-tap_saprotroph (Põlme et al. 2020); Decay sugar-rich substrates (Põlme et al. 2020) | Põlme S, et al. 2020. Fungal Diversity 105:1-16 ((https://doi.org/10.1007/s13225-020-00466-2)) |
| Otu17323 | 1 | k__Fungi(100);p__Ascomycota(100);c__Saccharomycetes(100);o__Saccharomycetales(100);f__Metschnikowiaceae(100);g__Metschnikowia(100);s__Metschnikowia_pulcherrima(100); | Metschnikowia | 13 | Saprotroph | \|Undefined Saprotroph\| | Yeast | NULL | Probable | As nectar-tap_saprotroph (Põlme et al. 2020); Decay sugar-rich substrates (Põlme et al. 2020) | Põlme S, et al. 2020. Fungal Diversity 105:1-16 ((https://doi.org/10.1007/s13225-020-00466-2)) |
| Otu17324 | 1 | k__Fungi(100);p__Basidiomycota(100);c__Agaricomycetes(100);o__Boletales(100);f__Suillaceae(100);g__Suillus(100);s__Suillus_placidus(100); | Suillus | 13 | Symbiotroph | \|Ectomycorrhizal\| | Agaricoid-Boletoid | NULL | Highly Probable | Host - Pinaceae (Põlme et al. 2020) | Rinaldi AC, et al. 2008. Fungal Diversity 33:1-45 ((http://www.fungaldiversity.org/fdp/sfdp/33-1.pdf)); Tedersoo L, et al. 2010. Mycorrhiza 20:217-263 ((https://doi.org/10.1007/s00572-009-0274-x)); Põlme S, et al. 2020. Fungal Diversity 105:1-16 ((https://doi.org/10.1007/s13225-020-00466-2)); http://mycorrhizas.info/ecmf.html ((http://mycorrhizas.info/ecmf.html)) |
| Otu17325 | 1 | k__Fungi(100);p__Ascomycota(100);c__Saccharomycetes(100);o__Saccharomycetales(100);f__Metschnikowiaceae(100);g__Metschnikowia(100);g__Metschnikowia_unclassified(100); | Metschnikowia | 13 | Saprotroph | \|Undefined Saprotroph\| | Yeast | NULL | Probable | As nectar-tap_saprotroph (Põlme et al. 2020); Decay sugar-rich substrates (Põlme et al. 2020) | Põlme S, et al. 2020. Fungal Diversity 105:1-16 ((https://doi.org/10.1007/s13225-020-00466-2)) |
| Otu17326 | 1 | k__Fungi(100);p__Ascomycota(100);c__Saccharomycetes(100);o__Saccharomycetales(100);f__Metschnikowiaceae(100);g__Metschnikowia(100);s__Metschnikowia_pulcherrima(100); | Metschnikowia | 13 | Saprotroph | \|Undefined Saprotroph\| | Yeast | NULL | Probable | As nectar-tap_saprotroph (Põlme et al. 2020); Decay sugar-rich substrates (Põlme et al. 2020) | Põlme S, et al. 2020. Fungal Diversity 105:1-16 ((https://doi.org/10.1007/s13225-020-00466-2)) |
| Otu17327 | 1 | k__Fungi(100);p__Ascomycota(100);c__Saccharomycetes(100);o__Saccharomycetales(100);f__Metschnikowiaceae(100);g__Metschnikowia(100);s__Metschnikowia_pulcherrima(100); | Metschnikowia | 13 | Saprotroph | \|Undefined Saprotroph\| | Yeast | NULL | Probable | As nectar-tap_saprotroph (Põlme et al. 2020); Decay sugar-rich substrates (Põlme et al. 2020) | Põlme S, et al. 2020. Fungal Diversity 105:1-16 ((https://doi.org/10.1007/s13225-020-00466-2)) |
| Otu17328 | 1 | k__Fungi(100);p__Ascomycota(100);c__Saccharomycetes(100);o__Saccharomycetales(100);f__Metschnikowiaceae(100);g__Metschnikowia(100);s__Metschnikowia_pulcherrima(100); | Metschnikowia | 13 | Saprotroph | \|Undefined Saprotroph\| | Yeast | NULL | Probable | As nectar-tap_saprotroph (Põlme et al. 2020); Decay sugar-rich substrates (Põlme et al. 2020) | Põlme S, et al. 2020. Fungal Diversity 105:1-16 ((https://doi.org/10.1007/s13225-020-00466-2)) |
| Otu17330 | 1 | k__Fungi(100);p__Ascomycota(100);c__Saccharomycetes(100);o__Saccharomycetales(100);f__Metschnikowiaceae(100);g__Metschnikowia(100);g__Metschnikowia_unclassified(100); | Metschnikowia | 13 | Saprotroph | \|Undefined Saprotroph\| | Yeast | NULL | Probable | As nectar-tap_saprotroph (Põlme et al. 2020); Decay sugar-rich substrates (Põlme et al. 2020) | Põlme S, et al. 2020. Fungal Diversity 105:1-16 ((https://doi.org/10.1007/s13225-020-00466-2)) |
| Otu17334 | 1 | k__Fungi(100);p__Zygomycota(100);c__Zygomycota_class_Incertae_sedis(100);o__Mortierellales(100);f__Mortierellaceae(100);g__unclassified_Mortierellaceae(100);s__Mortierellaceae_sp(100); | Mortierellaceae | 9 | Saprotroph-Symbiotroph | Endophyte-Litter Saprotroph-Soil Saprotroph-Undefined Saprotroph | Microfungus | NULL | Possible | Saprobic in soil and thus frequently isolated from soil and material contaminated with soil such as hay or rotten silage and spoil tips (Cannon & Kirk 2007) | Cannon PF, Kirk PM. 2007. Fungal Families of the World. CAB International, Cambridge (ISBN: 978-0851998275); Purahong W, et al. 2016. Molecular Ecology 25:4059-4074 ((https://doi.org/10.1111/mec.13739)) |
| Otu17335 | 1 | k__Fungi(100);p__Ascomycota(100);c__Saccharomycetes(100);o__Saccharomycetales(100);o__Saccharomycetales_unclassified(100);o__Saccharomycetales_unclassified(100);o__Saccharomycetales_unclassified(100); | Saccharomycetales | 7 | Saprotroph | Undefined Saprotroph | Yeast | NULL | Possible | NULL | Sterkenburg E, et al. 2015. New Phytologist 207:1145-1158 ((https://doi.org/10.1111/nph.13426)) |
| Otu17344 | 1 | k__Fungi(100);p__Ascomycota(100);c__Saccharomycetes(100);o__Saccharomycetales(100);f__Metschnikowiaceae(100);g__Metschnikowia(100);s__Metschnikowia_pulcherrima(100); | Metschnikowia | 13 | Saprotroph | \|Undefined Saprotroph\| | Yeast | NULL | Probable | As nectar-tap_saprotroph (Põlme et al. 2020); Decay sugar-rich substrates (Põlme et al. 2020) | Põlme S, et al. 2020. Fungal Diversity 105:1-16 ((https://doi.org/10.1007/s13225-020-00466-2)) |
| Otu17345 | 1 | k__Fungi(100);p__Ascomycota(100);c__Saccharomycetes(100);o__Saccharomycetales(100);f__Metschnikowiaceae(100);g__Metschnikowia(100);s__Metschnikowia_pulcherrima(100); | Metschnikowia | 13 | Saprotroph | \|Undefined Saprotroph\| | Yeast | NULL | Probable | As nectar-tap_saprotroph (Põlme et al. 2020); Decay sugar-rich substrates (Põlme et al. 2020) | Põlme S, et al. 2020. Fungal Diversity 105:1-16 ((https://doi.org/10.1007/s13225-020-00466-2)) |
| Otu17352 | 1 | k__Fungi(100);p__Ascomycota(100);c__Saccharomycetes(100);o__Saccharomycetales(100);f__Metschnikowiaceae(100);g__Metschnikowia(100);s__Metschnikowia_pulcherrima(100); | Metschnikowia | 13 | Saprotroph | \|Undefined Saprotroph\| | Yeast | NULL | Probable | As nectar-tap_saprotroph (Põlme et al. 2020); Decay sugar-rich substrates (Põlme et al. 2020) | Põlme S, et al. 2020. Fungal Diversity 105:1-16 ((https://doi.org/10.1007/s13225-020-00466-2)) |
| Otu17355 | 1 | k__Fungi(100);p__Ascomycota(100);c__Saccharomycetes(100);o__Saccharomycetales(100);f__Metschnikowiaceae(100);g__Metschnikowia(100);s__Metschnikowia_pulcherrima(100); | Metschnikowia | 13 | Saprotroph | \|Undefined Saprotroph\| | Yeast | NULL | Probable | As nectar-tap_saprotroph (Põlme et al. 2020); Decay sugar-rich substrates (Põlme et al. 2020) | Põlme S, et al. 2020. Fungal Diversity 105:1-16 ((https://doi.org/10.1007/s13225-020-00466-2)) |
| Otu17356 | 1 | k__Fungi(100);p__Ascomycota(100);c__Saccharomycetes(100);o__Saccharomycetales(100);f__Metschnikowiaceae(100);g__Metschnikowia(100);s__Metschnikowia_pulcherrima(100); | Metschnikowia | 13 | Saprotroph | \|Undefined Saprotroph\| | Yeast | NULL | Probable | As nectar-tap_saprotroph (Põlme et al. 2020); Decay sugar-rich substrates (Põlme et al. 2020) | Põlme S, et al. 2020. Fungal Diversity 105:1-16 ((https://doi.org/10.1007/s13225-020-00466-2)) |
| Otu17357 | 1 | k__Fungi(100);p__Ascomycota(100);c__Saccharomycetes(100);o__Saccharomycetales(100);f__Metschnikowiaceae(100);g__Metschnikowia(100);s__Metschnikowia_pulcherrima(100); | Metschnikowia | 13 | Saprotroph | \|Undefined Saprotroph\| | Yeast | NULL | Probable | As nectar-tap_saprotroph (Põlme et al. 2020); Decay sugar-rich substrates (Põlme et al. 2020) | Põlme S, et al. 2020. Fungal Diversity 105:1-16 ((https://doi.org/10.1007/s13225-020-00466-2)) |
| Otu17362 | 1 | k__Fungi(100);p__Ascomycota(100);c__Saccharomycetes(100);o__Saccharomycetales(100);f__Metschnikowiaceae(100);g__Metschnikowia(100);g__Metschnikowia_unclassified(100); | Metschnikowia | 13 | Saprotroph | \|Undefined Saprotroph\| | Yeast | NULL | Probable | As nectar-tap_saprotroph (Põlme et al. 2020); Decay sugar-rich substrates (Põlme et al. 2020) | Põlme S, et al. 2020. Fungal Diversity 105:1-16 ((https://doi.org/10.1007/s13225-020-00466-2)) |
| Otu17366 | 1 | k__Fungi(100);p__Ascomycota(100);c__Saccharomycetes(100);o__Saccharomycetales(100);f__Metschnikowiaceae(100);g__Metschnikowia(100);s__Metschnikowia_pulcherrima(100); | Metschnikowia | 13 | Saprotroph | \|Undefined Saprotroph\| | Yeast | NULL | Probable | As nectar-tap_saprotroph (Põlme et al. 2020); Decay sugar-rich substrates (Põlme et al. 2020) | Põlme S, et al. 2020. Fungal Diversity 105:1-16 ((https://doi.org/10.1007/s13225-020-00466-2)) |
| Otu17371 | 1 | k__Fungi(100);p__Ascomycota(100);c__Saccharomycetes(100);o__Saccharomycetales(100);f__Metschnikowiaceae(100);g__Metschnikowia(100);s__Metschnikowia_pulcherrima(100); | Metschnikowia | 13 | Saprotroph | \|Undefined Saprotroph\| | Yeast | NULL | Probable | As nectar-tap_saprotroph (Põlme et al. 2020); Decay sugar-rich substrates (Põlme et al. 2020) | Põlme S, et al. 2020. Fungal Diversity 105:1-16 ((https://doi.org/10.1007/s13225-020-00466-2)) |
| Otu17373 | 1 | k__Fungi(100);p__Ascomycota(100);c__Saccharomycetes(100);o__Saccharomycetales(100);f__Saccharomycetaceae(100);g__Kazachstania(100);s__Kazachstania_intestinalis(100); | Kazachstania | 13 | Saprotroph | \|Undefined Saprotroph\| | Yeast | NULL | Probable | As nectar-tap_saprotroph (Põlme et al. 2020); Decay sugar-rich substrates (Põlme et al. 2020) | Põlme S, et al. 2020. Fungal Diversity 105:1-16 ((https://doi.org/10.1007/s13225-020-00466-2)) |
| Otu17376 | 1 | k__Fungi(100);p__Ascomycota(100);c__Saccharomycetes(100);o__Saccharomycetales(100);f__Metschnikowiaceae(100);g__Metschnikowia(100);s__Metschnikowia_pulcherrima(100); | Metschnikowia | 13 | Saprotroph | \|Undefined Saprotroph\| | Yeast | NULL | Probable | As nectar-tap_saprotroph (Põlme et al. 2020); Decay sugar-rich substrates (Põlme et al. 2020) | Põlme S, et al. 2020. Fungal Diversity 105:1-16 ((https://doi.org/10.1007/s13225-020-00466-2)) |
| Otu17381 | 1 | k__Fungi(100);p__Ascomycota(100);c__Saccharomycetes(100);o__Saccharomycetales(100);f__Metschnikowiaceae(100);g__Metschnikowia(100);s__Metschnikowia_pulcherrima(100); | Metschnikowia | 13 | Saprotroph | \|Undefined Saprotroph\| | Yeast | NULL | Probable | As nectar-tap_saprotroph (Põlme et al. 2020); Decay sugar-rich substrates (Põlme et al. 2020) | Põlme S, et al. 2020. Fungal Diversity 105:1-16 ((https://doi.org/10.1007/s13225-020-00466-2)) |
| Otu17382 | 1 | k__Fungi(100);p__Ascomycota(100);c__Saccharomycetes(100);o__Saccharomycetales(100);f__Metschnikowiaceae(100);g__Metschnikowia(100);s__Metschnikowia_pulcherrima(100); | Metschnikowia | 13 | Saprotroph | \|Undefined Saprotroph\| | Yeast | NULL | Probable | As nectar-tap_saprotroph (Põlme et al. 2020); Decay sugar-rich substrates (Põlme et al. 2020) | Põlme S, et al. 2020. Fungal Diversity 105:1-16 ((https://doi.org/10.1007/s13225-020-00466-2)) |
| Otu17383 | 1 | k__Fungi(100);p__Ascomycota(100);c__Saccharomycetes(100);o__Saccharomycetales(100);f__Metschnikowiaceae(100);g__Metschnikowia(100);s__Metschnikowia_pulcherrima(100); | Metschnikowia | 13 | Saprotroph | \|Undefined Saprotroph\| | Yeast | NULL | Probable | As nectar-tap_saprotroph (Põlme et al. 2020); Decay sugar-rich substrates (Põlme et al. 2020) | Põlme S, et al. 2020. Fungal Diversity 105:1-16 ((https://doi.org/10.1007/s13225-020-00466-2)) |
| Otu17384 | 1 | k__Fungi(100);p__Ascomycota(100);c__Saccharomycetes(100);o__Saccharomycetales(100);f__Metschnikowiaceae(100);g__Metschnikowia(100);s__Metschnikowia_pulcherrima(100); | Metschnikowia | 13 | Saprotroph | \|Undefined Saprotroph\| | Yeast | NULL | Probable | As nectar-tap_saprotroph (Põlme et al. 2020); Decay sugar-rich substrates (Põlme et al. 2020) | Põlme S, et al. 2020. Fungal Diversity 105:1-16 ((https://doi.org/10.1007/s13225-020-00466-2)) |
| Otu17385 | 1 | k__Fungi(100);p__Ascomycota(100);c__Saccharomycetes(100);o__Saccharomycetales(100);f__Metschnikowiaceae(100);g__Metschnikowia(100);g__Metschnikowia_unclassified(100); | Metschnikowia | 13 | Saprotroph | \|Undefined Saprotroph\| | Yeast | NULL | Probable | As nectar-tap_saprotroph (Põlme et al. 2020); Decay sugar-rich substrates (Põlme et al. 2020) | Põlme S, et al. 2020. Fungal Diversity 105:1-16 ((https://doi.org/10.1007/s13225-020-00466-2)) |
| Otu17387 | 1 | k__Fungi(100);p__Ascomycota(100);c__Saccharomycetes(100);o__Saccharomycetales(100);f__Metschnikowiaceae(100);g__Metschnikowia(100);g__Metschnikowia_unclassified(100); | Metschnikowia | 13 | Saprotroph | \|Undefined Saprotroph\| | Yeast | NULL | Probable | As nectar-tap_saprotroph (Põlme et al. 2020); Decay sugar-rich substrates (Põlme et al. 2020) | Põlme S, et al. 2020. Fungal Diversity 105:1-16 ((https://doi.org/10.1007/s13225-020-00466-2)) |
| Otu17388 | 1 | k__Fungi(100);p__Ascomycota(100);c__Leotiomycetes(100);o__Helotiales(100);f__Dermateaceae(100);g__Mollisia(100);g__Mollisia_unclassified(100); | Mollisia | 13 | Pathotroph-Saprotroph-Symbiotroph | Endophyte-\|Plant Pathogen\|-Plant Saprotroph | Microfungus-Pezizoid | Soft Rot | Probable | As foliar_endophyte (Põlme et al. 2020); As litter_saprotroph (Põlme et al. 2020); Dark Septate Endophyte, known from plant roots.; Decay leaf, fruit, seed (Põlme et al. 2020) | James TY, et al. 2006. Nature 443:818-822 ((https://doi.org/10.1038/nature05110)); Tedersoo L, et al. 2014. Science 346:1256688 ((https://doi.org/10.1126/science.1256688)) (as Plant Pathogen); Põlme S, et al. 2020. Fungal Diversity 105:1-16 ((https://doi.org/10.1007/s13225-020-00466-2)) |
| Otu17389 | 1 | k__Fungi(100);p__Ascomycota(100);c__Saccharomycetes(100);o__Saccharomycetales(100);f__Metschnikowiaceae(100);g__Metschnikowia(100);g__Metschnikowia_unclassified(100); | Metschnikowia | 13 | Saprotroph | \|Undefined Saprotroph\| | Yeast | NULL | Probable | As nectar-tap_saprotroph (Põlme et al. 2020); Decay sugar-rich substrates (Põlme et al. 2020) | Põlme S, et al. 2020. Fungal Diversity 105:1-16 ((https://doi.org/10.1007/s13225-020-00466-2)) |
| Otu17390 | 1 | k__Fungi(100);p__Ascomycota(100);c__Saccharomycetes(100);o__Saccharomycetales(100);f__Metschnikowiaceae(100);g__Metschnikowia(100);s__Metschnikowia_pulcherrima(100); | Metschnikowia | 13 | Saprotroph | \|Undefined Saprotroph\| | Yeast | NULL | Probable | As nectar-tap_saprotroph (Põlme et al. 2020); Decay sugar-rich substrates (Põlme et al. 2020) | Põlme S, et al. 2020. Fungal Diversity 105:1-16 ((https://doi.org/10.1007/s13225-020-00466-2)) |
| Otu17392 | 1 | k__Fungi(100);p__Ascomycota(100);c__Saccharomycetes(100);o__Saccharomycetales(100);f__Metschnikowiaceae(100);g__Metschnikowia(100);s__Metschnikowia_pulcherrima(100); | Metschnikowia | 13 | Saprotroph | \|Undefined Saprotroph\| | Yeast | NULL | Probable | As nectar-tap_saprotroph (Põlme et al. 2020); Decay sugar-rich substrates (Põlme et al. 2020) | Põlme S, et al. 2020. Fungal Diversity 105:1-16 ((https://doi.org/10.1007/s13225-020-00466-2)) |
| Otu17393 | 1 | k__Fungi(100);p__Ascomycota(100);c__Saccharomycetes(100);o__Saccharomycetales(100);f__Metschnikowiaceae(100);g__Metschnikowia(100);s__Metschnikowia_pulcherrima(100); | Metschnikowia | 13 | Saprotroph | \|Undefined Saprotroph\| | Yeast | NULL | Probable | As nectar-tap_saprotroph (Põlme et al. 2020); Decay sugar-rich substrates (Põlme et al. 2020) | Põlme S, et al. 2020. Fungal Diversity 105:1-16 ((https://doi.org/10.1007/s13225-020-00466-2)) |
| Otu17394 | 1 | k__Fungi(100);p__Ascomycota(100);c__Saccharomycetes(100);o__Saccharomycetales(100);f__Metschnikowiaceae(100);g__Metschnikowia(100);s__Metschnikowia_pulcherrima(100); | Metschnikowia | 13 | Saprotroph | \|Undefined Saprotroph\| | Yeast | NULL | Probable | As nectar-tap_saprotroph (Põlme et al. 2020); Decay sugar-rich substrates (Põlme et al. 2020) | Põlme S, et al. 2020. Fungal Diversity 105:1-16 ((https://doi.org/10.1007/s13225-020-00466-2)) |
| Otu17395 | 1 | k__Fungi(100);p__Ascomycota(100);c__Saccharomycetes(100);o__Saccharomycetales(100);f__Metschnikowiaceae(100);g__Metschnikowia(100);s__Metschnikowia_pulcherrima(100); | Metschnikowia | 13 | Saprotroph | \|Undefined Saprotroph\| | Yeast | NULL | Probable | As nectar-tap_saprotroph (Põlme et al. 2020); Decay sugar-rich substrates (Põlme et al. 2020) | Põlme S, et al. 2020. Fungal Diversity 105:1-16 ((https://doi.org/10.1007/s13225-020-00466-2)) |
| Otu17396 | 1 | k__Fungi(100);p__Ascomycota(100);c__Saccharomycetes(100);o__Saccharomycetales(100);f__Metschnikowiaceae(100);g__Metschnikowia(100);g__Metschnikowia_unclassified(100); | Metschnikowia | 13 | Saprotroph | \|Undefined Saprotroph\| | Yeast | NULL | Probable | As nectar-tap_saprotroph (Põlme et al. 2020); Decay sugar-rich substrates (Põlme et al. 2020) | Põlme S, et al. 2020. Fungal Diversity 105:1-16 ((https://doi.org/10.1007/s13225-020-00466-2)) |
| Otu17397 | 1 | k__Fungi(100);p__Ascomycota(100);c__Saccharomycetes(100);o__Saccharomycetales(100);f__Metschnikowiaceae(100);g__Metschnikowia(100);s__Metschnikowia_pulcherrima(100); | Metschnikowia | 13 | Saprotroph | \|Undefined Saprotroph\| | Yeast | NULL | Probable | As nectar-tap_saprotroph (Põlme et al. 2020); Decay sugar-rich substrates (Põlme et al. 2020) | Põlme S, et al. 2020. Fungal Diversity 105:1-16 ((https://doi.org/10.1007/s13225-020-00466-2)) |
| Otu17398 | 1 | k__Fungi(100);p__Ascomycota(100);c__Saccharomycetes(100);o__Saccharomycetales(100);f__Metschnikowiaceae(100);g__Metschnikowia(100);s__Metschnikowia_pulcherrima(100); | Metschnikowia | 13 | Saprotroph | \|Undefined Saprotroph\| | Yeast | NULL | Probable | As nectar-tap_saprotroph (Põlme et al. 2020); Decay sugar-rich substrates (Põlme et al. 2020) | Põlme S, et al. 2020. Fungal Diversity 105:1-16 ((https://doi.org/10.1007/s13225-020-00466-2)) |
| Otu17399 | 1 | k__Fungi(100);p__Ascomycota(100);c__Saccharomycetes(100);o__Saccharomycetales(100);f__Metschnikowiaceae(100);g__Metschnikowia(100);s__Metschnikowia_pulcherrima(100); | Metschnikowia | 13 | Saprotroph | \|Undefined Saprotroph\| | Yeast | NULL | Probable | As nectar-tap_saprotroph (Põlme et al. 2020); Decay sugar-rich substrates (Põlme et al. 2020) | Põlme S, et al. 2020. Fungal Diversity 105:1-16 ((https://doi.org/10.1007/s13225-020-00466-2)) |
| Otu17400 | 1 | k__Fungi(100);p__Ascomycota(100);c__Saccharomycetes(100);o__Saccharomycetales(100);f__Metschnikowiaceae(100);g__Metschnikowia(100);g__Metschnikowia_unclassified(100); | Metschnikowia | 13 | Saprotroph | \|Undefined Saprotroph\| | Yeast | NULL | Probable | As nectar-tap_saprotroph (Põlme et al. 2020); Decay sugar-rich substrates (Põlme et al. 2020) | Põlme S, et al. 2020. Fungal Diversity 105:1-16 ((https://doi.org/10.1007/s13225-020-00466-2)) |
| Otu17401 | 1 | k__Fungi(100);p__Ascomycota(100);c__Saccharomycetes(100);o__Saccharomycetales(100);f__Metschnikowiaceae(100);g__Metschnikowia(100);s__Metschnikowia_pulcherrima(100); | Metschnikowia | 13 | Saprotroph | \|Undefined Saprotroph\| | Yeast | NULL | Probable | As nectar-tap_saprotroph (Põlme et al. 2020); Decay sugar-rich substrates (Põlme et al. 2020) | Põlme S, et al. 2020. Fungal Diversity 105:1-16 ((https://doi.org/10.1007/s13225-020-00466-2)) |
| Otu17402 | 1 | k__Fungi(100);p__Ascomycota(100);c__Saccharomycetes(100);o__Saccharomycetales(100);f__Metschnikowiaceae(100);g__Metschnikowia(100);s__Metschnikowia_pulcherrima(100); | Metschnikowia | 13 | Saprotroph | \|Undefined Saprotroph\| | Yeast | NULL | Probable | As nectar-tap_saprotroph (Põlme et al. 2020); Decay sugar-rich substrates (Põlme et al. 2020) | Põlme S, et al. 2020. Fungal Diversity 105:1-16 ((https://doi.org/10.1007/s13225-020-00466-2)) |
| Otu17404 | 1 | k__Fungi(100);p__Ascomycota(100);c__Saccharomycetes(100);o__Saccharomycetales(100);f__Metschnikowiaceae(100);g__Metschnikowia(100);g__Metschnikowia_unclassified(100); | Metschnikowia | 13 | Saprotroph | \|Undefined Saprotroph\| | Yeast | NULL | Probable | As nectar-tap_saprotroph (Põlme et al. 2020); Decay sugar-rich substrates (Põlme et al. 2020) | Põlme S, et al. 2020. Fungal Diversity 105:1-16 ((https://doi.org/10.1007/s13225-020-00466-2)) |
| Otu17407 | 1 | k__Fungi(100);p__Ascomycota(100);c__Saccharomycetes(100);o__Saccharomycetales(100);f__Metschnikowiaceae(100);g__Metschnikowia(100);g__Metschnikowia_unclassified(100); | Metschnikowia | 13 | Saprotroph | \|Undefined Saprotroph\| | Yeast | NULL | Probable | As nectar-tap_saprotroph (Põlme et al. 2020); Decay sugar-rich substrates (Põlme et al. 2020) | Põlme S, et al. 2020. Fungal Diversity 105:1-16 ((https://doi.org/10.1007/s13225-020-00466-2)) |
| Otu17408 | 1 | k__Fungi(100);p__Ascomycota(100);c__Saccharomycetes(100);o__Saccharomycetales(100);f__Metschnikowiaceae(100);g__Metschnikowia(100);g__Metschnikowia_unclassified(100); | Metschnikowia | 13 | Saprotroph | \|Undefined Saprotroph\| | Yeast | NULL | Probable | As nectar-tap_saprotroph (Põlme et al. 2020); Decay sugar-rich substrates (Põlme et al. 2020) | Põlme S, et al. 2020. Fungal Diversity 105:1-16 ((https://doi.org/10.1007/s13225-020-00466-2)) |
| Otu17409 | 1 | k__Fungi(100);p__Ascomycota(100);c__Saccharomycetes(100);o__Saccharomycetales(100);f__Metschnikowiaceae(100);g__Metschnikowia(100);s__Metschnikowia_pulcherrima(100); | Metschnikowia | 13 | Saprotroph | \|Undefined Saprotroph\| | Yeast | NULL | Probable | As nectar-tap_saprotroph (Põlme et al. 2020); Decay sugar-rich substrates (Põlme et al. 2020) | Põlme S, et al. 2020. Fungal Diversity 105:1-16 ((https://doi.org/10.1007/s13225-020-00466-2)) |
| Otu17417 | 1 | k__Fungi(100);p__Ascomycota(100);c__Saccharomycetes(100);o__Saccharomycetales(100);f__Saccharomycetales_family_Incertae_sedis(100);g__Candida(100);g__Candida_unclassified(100); | Candida | 13 | Pathotroph-Saprotroph-Symbiotroph | Animal Pathogen-Endophyte-Endosymbiont-Epiphyte-\|Undefined Saprotroph\| | Dimorphic Yeast-Yeast | NULL | Probable | As nectar-tap_saprotroph (Põlme et al. 2020); Decay sugar-rich substrates (Põlme et al. 2020); This asexual genus is polyphyletic and contain species that function very differently depend on what they are. Caution should be taken when interpreting guilds for Candida species. | Manolakaki D, et al. 2010. Virulence. 1: 367-375 ((https://doi.org/10.4161/viru.1.5.12796)); Mekbib SB, et al. 2011. Tropical Plant Pathology 36:233-240 ((https://doi.org/10.1590/S1982-56762011000400004)); Põlme S, et al. 2020. Fungal Diversity 105:1-16 ((https://doi.org/10.1007/s13225-020-00466-2)) |
| Otu17431 | 1 | k__Fungi(100);p__Ascomycota(100);c__Saccharomycetes(100);o__Saccharomycetales(100);f__Metschnikowiaceae(100);g__Metschnikowia(100);g__Metschnikowia_unclassified(100); | Metschnikowia | 13 | Saprotroph | \|Undefined Saprotroph\| | Yeast | NULL | Probable | As nectar-tap_saprotroph (Põlme et al. 2020); Decay sugar-rich substrates (Põlme et al. 2020) | Põlme S, et al. 2020. Fungal Diversity 105:1-16 ((https://doi.org/10.1007/s13225-020-00466-2)) |
| Otu17433 | 1 | k__Fungi(100);p__Ascomycota(100);c__Saccharomycetes(100);o__Saccharomycetales(100);f__Metschnikowiaceae(100);g__Metschnikowia(100);s__Metschnikowia_pulcherrima(100); | Metschnikowia | 13 | Saprotroph | \|Undefined Saprotroph\| | Yeast | NULL | Probable | As nectar-tap_saprotroph (Põlme et al. 2020); Decay sugar-rich substrates (Põlme et al. 2020) | Põlme S, et al. 2020. Fungal Diversity 105:1-16 ((https://doi.org/10.1007/s13225-020-00466-2)) |
| Otu17434 | 1 | k__Fungi(100);p__Ascomycota(100);c__Saccharomycetes(100);o__Saccharomycetales(100);f__Metschnikowiaceae(100);g__Metschnikowia(100);s__Metschnikowia_pulcherrima(100); | Metschnikowia | 13 | Saprotroph | \|Undefined Saprotroph\| | Yeast | NULL | Probable | As nectar-tap_saprotroph (Põlme et al. 2020); Decay sugar-rich substrates (Põlme et al. 2020) | Põlme S, et al. 2020. Fungal Diversity 105:1-16 ((https://doi.org/10.1007/s13225-020-00466-2)) |
| Otu17435 | 1 | k__Fungi(100);p__Ascomycota(100);c__Saccharomycetes(100);o__Saccharomycetales(100);f__Metschnikowiaceae(100);g__Metschnikowia(100);g__Metschnikowia_unclassified(100); | Metschnikowia | 13 | Saprotroph | \|Undefined Saprotroph\| | Yeast | NULL | Probable | As nectar-tap_saprotroph (Põlme et al. 2020); Decay sugar-rich substrates (Põlme et al. 2020) | Põlme S, et al. 2020. Fungal Diversity 105:1-16 ((https://doi.org/10.1007/s13225-020-00466-2)) |
| Otu17437 | 1 | k__Fungi(100);p__Ascomycota(100);c__Saccharomycetes(100);o__Saccharomycetales(100);f__Metschnikowiaceae(100);g__Metschnikowia(100);s__Metschnikowia_pulcherrima(100); | Metschnikowia | 13 | Saprotroph | \|Undefined Saprotroph\| | Yeast | NULL | Probable | As nectar-tap_saprotroph (Põlme et al. 2020); Decay sugar-rich substrates (Põlme et al. 2020) | Põlme S, et al. 2020. Fungal Diversity 105:1-16 ((https://doi.org/10.1007/s13225-020-00466-2)) |
| Otu17439 | 1 | k__Fungi(100);p__Ascomycota(100);c__Saccharomycetes(100);o__Saccharomycetales(100);f__Metschnikowiaceae(100);g__Metschnikowia(100);s__Metschnikowia_pulcherrima(100); | Metschnikowia | 13 | Saprotroph | \|Undefined Saprotroph\| | Yeast | NULL | Probable | As nectar-tap_saprotroph (Põlme et al. 2020); Decay sugar-rich substrates (Põlme et al. 2020) | Põlme S, et al. 2020. Fungal Diversity 105:1-16 ((https://doi.org/10.1007/s13225-020-00466-2)) |
| Otu17440 | 1 | k__Fungi(100);p__Ascomycota(100);c__Saccharomycetes(100);o__Saccharomycetales(100);f__Metschnikowiaceae(100);g__Metschnikowia(100);s__Metschnikowia_pulcherrima(100); | Metschnikowia | 13 | Saprotroph | \|Undefined Saprotroph\| | Yeast | NULL | Probable | As nectar-tap_saprotroph (Põlme et al. 2020); Decay sugar-rich substrates (Põlme et al. 2020) | Põlme S, et al. 2020. Fungal Diversity 105:1-16 ((https://doi.org/10.1007/s13225-020-00466-2)) |
| Otu17442 | 1 | k__Fungi(100);p__Ascomycota(100);c__Saccharomycetes(100);o__Saccharomycetales(100);f__Metschnikowiaceae(100);g__Metschnikowia(100);s__Metschnikowia_pulcherrima(100); | Metschnikowia | 13 | Saprotroph | \|Undefined Saprotroph\| | Yeast | NULL | Probable | As nectar-tap_saprotroph (Põlme et al. 2020); Decay sugar-rich substrates (Põlme et al. 2020) | Põlme S, et al. 2020. Fungal Diversity 105:1-16 ((https://doi.org/10.1007/s13225-020-00466-2)) |
| Otu17443 | 1 | k__Fungi(100);p__Basidiomycota(100);c__Tremellomycetes(100);o__Tremellales(100);f__Tremellales_family_Incertae_sedis(100);g__Dioszegia(100);s__Dioszegia_changbaiensis(100); | Dioszegia | 13 | Saprotroph | \|Plant Saprotroph\|-Undefined Saprotroph | Yeast | NULL | Probable | As litter_saprotroph (Põlme et al. 2020); Decay leaf, fruit, seed (Põlme et al. 2020); The Dioszegia subclade sensu Inacio (Inacio et al. 20015) is characterized by orange; yellow or orange to deep orange colonies. Isolated from plant leaves and soil. | Põlme S, et al. 2020. Fungal Diversity 105:1-16 ((https://doi.org/10.1007/s13225-020-00466-2)); Takashima M, et al. 2011 International Journal of Systematic and Evolutionary Microbiology 61:1736-1739 ((https://doi.org/10.1099/ijs.0.025254-0)) |
| Otu17445 | 1 | k__Fungi(100);p__Ascomycota(100);c__Saccharomycetes(100);o__Saccharomycetales(100);f__Metschnikowiaceae(100);g__Metschnikowia(100);s__Metschnikowia_pulcherrima(100); | Metschnikowia | 13 | Saprotroph | \|Undefined Saprotroph\| | Yeast | NULL | Probable | As nectar-tap_saprotroph (Põlme et al. 2020); Decay sugar-rich substrates (Põlme et al. 2020) | Põlme S, et al. 2020. Fungal Diversity 105:1-16 ((https://doi.org/10.1007/s13225-020-00466-2)) |
| Otu17446 | 1 | k__Fungi(100);p__Ascomycota(100);c__Saccharomycetes(100);o__Saccharomycetales(100);f__Metschnikowiaceae(100);g__Metschnikowia(100);s__Metschnikowia_pulcherrima(100); | Metschnikowia | 13 | Saprotroph | \|Undefined Saprotroph\| | Yeast | NULL | Probable | As nectar-tap_saprotroph (Põlme et al. 2020); Decay sugar-rich substrates (Põlme et al. 2020) | Põlme S, et al. 2020. Fungal Diversity 105:1-16 ((https://doi.org/10.1007/s13225-020-00466-2)) |
| Otu17449 | 1 | k__Fungi(100);p__Ascomycota(100);c__Saccharomycetes(100);o__Saccharomycetales(100);f__Metschnikowiaceae(100);g__Metschnikowia(100);s__Metschnikowia_pulcherrima(100); | Metschnikowia | 13 | Saprotroph | \|Undefined Saprotroph\| | Yeast | NULL | Probable | As nectar-tap_saprotroph (Põlme et al. 2020); Decay sugar-rich substrates (Põlme et al. 2020) | Põlme S, et al. 2020. Fungal Diversity 105:1-16 ((https://doi.org/10.1007/s13225-020-00466-2)) |
| Otu17450 | 1 | k__Fungi(100);p__Ascomycota(100);c__Saccharomycetes(100);o__Saccharomycetales(100);f__Metschnikowiaceae(100);g__Metschnikowia(100);s__Metschnikowia_pulcherrima(100); | Metschnikowia | 13 | Saprotroph | \|Undefined Saprotroph\| | Yeast | NULL | Probable | As nectar-tap_saprotroph (Põlme et al. 2020); Decay sugar-rich substrates (Põlme et al. 2020) | Põlme S, et al. 2020. Fungal Diversity 105:1-16 ((https://doi.org/10.1007/s13225-020-00466-2)) |
| Otu17457 | 1 | k__Fungi(100);p__Basidiomycota(100);c__Tremellomycetes(100);o__Tremellales(100);f__Tremellales_family_Incertae_sedis(100);g__Dioszegia(100);s__Dioszegia_changbaiensis(100); | Dioszegia | 13 | Saprotroph | \|Plant Saprotroph\|-Undefined Saprotroph | Yeast | NULL | Probable | As litter_saprotroph (Põlme et al. 2020); Decay leaf, fruit, seed (Põlme et al. 2020); The Dioszegia subclade sensu Inacio (Inacio et al. 20015) is characterized by orange; yellow or orange to deep orange colonies. Isolated from plant leaves and soil. | Põlme S, et al. 2020. Fungal Diversity 105:1-16 ((https://doi.org/10.1007/s13225-020-00466-2)); Takashima M, et al. 2011 International Journal of Systematic and Evolutionary Microbiology 61:1736-1739 ((https://doi.org/10.1099/ijs.0.025254-0)) |
| Otu17463 | 1 | k__Fungi(100);p__Ascomycota(100);c__Saccharomycetes(100);o__Saccharomycetales(100);f__Metschnikowiaceae(100);g__Metschnikowia(100);s__Metschnikowia_pulcherrima(100); | Metschnikowia | 13 | Saprotroph | \|Undefined Saprotroph\| | Yeast | NULL | Probable | As nectar-tap_saprotroph (Põlme et al. 2020); Decay sugar-rich substrates (Põlme et al. 2020) | Põlme S, et al. 2020. Fungal Diversity 105:1-16 ((https://doi.org/10.1007/s13225-020-00466-2)) |
| Otu17464 | 1 | k__Fungi(100);p__Ascomycota(100);c__Saccharomycetes(100);o__Saccharomycetales(100);f__Metschnikowiaceae(100);g__Metschnikowia(100);s__Metschnikowia_pulcherrima(100); | Metschnikowia | 13 | Saprotroph | \|Undefined Saprotroph\| | Yeast | NULL | Probable | As nectar-tap_saprotroph (Põlme et al. 2020); Decay sugar-rich substrates (Põlme et al. 2020) | Põlme S, et al. 2020. Fungal Diversity 105:1-16 ((https://doi.org/10.1007/s13225-020-00466-2)) |
| Otu17465 | 1 | k__Fungi(100);p__Ascomycota(100);c__Saccharomycetes(100);o__Saccharomycetales(100);f__Metschnikowiaceae(100);g__Metschnikowia(100);s__Metschnikowia_pulcherrima(100); | Metschnikowia | 13 | Saprotroph | \|Undefined Saprotroph\| | Yeast | NULL | Probable | As nectar-tap_saprotroph (Põlme et al. 2020); Decay sugar-rich substrates (Põlme et al. 2020) | Põlme S, et al. 2020. Fungal Diversity 105:1-16 ((https://doi.org/10.1007/s13225-020-00466-2)) |
| Otu17466 | 1 | k__Fungi(100);p__Ascomycota(100);c__Saccharomycetes(100);o__Saccharomycetales(100);f__Metschnikowiaceae(100);g__Metschnikowia(100);g__Metschnikowia_unclassified(100); | Metschnikowia | 13 | Saprotroph | \|Undefined Saprotroph\| | Yeast | NULL | Probable | As nectar-tap_saprotroph (Põlme et al. 2020); Decay sugar-rich substrates (Põlme et al. 2020) | Põlme S, et al. 2020. Fungal Diversity 105:1-16 ((https://doi.org/10.1007/s13225-020-00466-2)) |
| Otu17467 | 1 | k__Fungi(100);p__Ascomycota(100);c__Saccharomycetes(100);o__Saccharomycetales(100);f__Metschnikowiaceae(100);g__Metschnikowia(100);s__Metschnikowia_pulcherrima(100); | Metschnikowia | 13 | Saprotroph | \|Undefined Saprotroph\| | Yeast | NULL | Probable | As nectar-tap_saprotroph (Põlme et al. 2020); Decay sugar-rich substrates (Põlme et al. 2020) | Põlme S, et al. 2020. Fungal Diversity 105:1-16 ((https://doi.org/10.1007/s13225-020-00466-2)) |
| Otu17474 | 1 | k__Fungi(100);p__Ascomycota(100);c__Saccharomycetes(100);o__Saccharomycetales(100);f__Metschnikowiaceae(100);g__Metschnikowia(100);s__Metschnikowia_pulcherrima(100); | Metschnikowia | 13 | Saprotroph | \|Undefined Saprotroph\| | Yeast | NULL | Probable | As nectar-tap_saprotroph (Põlme et al. 2020); Decay sugar-rich substrates (Põlme et al. 2020) | Põlme S, et al. 2020. Fungal Diversity 105:1-16 ((https://doi.org/10.1007/s13225-020-00466-2)) |
| Otu17475 | 1 | k__Fungi(100);p__Ascomycota(100);c__Saccharomycetes(100);o__Saccharomycetales(100);f__Metschnikowiaceae(100);g__Metschnikowia(100);g__Metschnikowia_unclassified(100); | Metschnikowia | 13 | Saprotroph | \|Undefined Saprotroph\| | Yeast | NULL | Probable | As nectar-tap_saprotroph (Põlme et al. 2020); Decay sugar-rich substrates (Põlme et al. 2020) | Põlme S, et al. 2020. Fungal Diversity 105:1-16 ((https://doi.org/10.1007/s13225-020-00466-2)) |
| Otu17476 | 1 | k__Fungi(100);p__Ascomycota(100);c__Saccharomycetes(100);o__Saccharomycetales(100);f__Metschnikowiaceae(100);g__Metschnikowia(100);s__Metschnikowia_pulcherrima(100); | Metschnikowia | 13 | Saprotroph | \|Undefined Saprotroph\| | Yeast | NULL | Probable | As nectar-tap_saprotroph (Põlme et al. 2020); Decay sugar-rich substrates (Põlme et al. 2020) | Põlme S, et al. 2020. Fungal Diversity 105:1-16 ((https://doi.org/10.1007/s13225-020-00466-2)) |
| Otu17488 | 1 | k__Fungi(100);p__Ascomycota(100);c__Saccharomycetes(100);o__Saccharomycetales(100);f__Metschnikowiaceae(100);g__Metschnikowia(100);s__Metschnikowia_pulcherrima(100); | Metschnikowia | 13 | Saprotroph | \|Undefined Saprotroph\| | Yeast | NULL | Probable | As nectar-tap_saprotroph (Põlme et al. 2020); Decay sugar-rich substrates (Põlme et al. 2020) | Põlme S, et al. 2020. Fungal Diversity 105:1-16 ((https://doi.org/10.1007/s13225-020-00466-2)) |
| Otu17489 | 1 | k__Fungi(100);p__Ascomycota(100);c__Saccharomycetes(100);o__Saccharomycetales(100);f__Metschnikowiaceae(100);g__Metschnikowia(100);s__Metschnikowia_pulcherrima(100); | Metschnikowia | 13 | Saprotroph | \|Undefined Saprotroph\| | Yeast | NULL | Probable | As nectar-tap_saprotroph (Põlme et al. 2020); Decay sugar-rich substrates (Põlme et al. 2020) | Põlme S, et al. 2020. Fungal Diversity 105:1-16 ((https://doi.org/10.1007/s13225-020-00466-2)) |
| Otu17493 | 1 | k__Fungi(100);p__Ascomycota(100);c__Saccharomycetes(100);o__Saccharomycetales(100);f__Metschnikowiaceae(100);g__Metschnikowia(100);s__Metschnikowia_pulcherrima(100); | Metschnikowia | 13 | Saprotroph | \|Undefined Saprotroph\| | Yeast | NULL | Probable | As nectar-tap_saprotroph (Põlme et al. 2020); Decay sugar-rich substrates (Põlme et al. 2020) | Põlme S, et al. 2020. Fungal Diversity 105:1-16 ((https://doi.org/10.1007/s13225-020-00466-2)) |
| Otu17495 | 1 | k__Fungi(100);p__Ascomycota(100);c__Saccharomycetes(100);o__Saccharomycetales(100);f__Metschnikowiaceae(100);g__Metschnikowia(100);s__Metschnikowia_pulcherrima(100); | Metschnikowia | 13 | Saprotroph | \|Undefined Saprotroph\| | Yeast | NULL | Probable | As nectar-tap_saprotroph (Põlme et al. 2020); Decay sugar-rich substrates (Põlme et al. 2020) | Põlme S, et al. 2020. Fungal Diversity 105:1-16 ((https://doi.org/10.1007/s13225-020-00466-2)) |
| Otu17496 | 1 | k__Fungi(100);p__Ascomycota(100);c__Saccharomycetes(100);o__Saccharomycetales(100);f__Metschnikowiaceae(100);g__Metschnikowia(100);g__Metschnikowia_unclassified(100); | Metschnikowia | 13 | Saprotroph | \|Undefined Saprotroph\| | Yeast | NULL | Probable | As nectar-tap_saprotroph (Põlme et al. 2020); Decay sugar-rich substrates (Põlme et al. 2020) | Põlme S, et al. 2020. Fungal Diversity 105:1-16 ((https://doi.org/10.1007/s13225-020-00466-2)) |
| Otu17497 | 1 | k__Fungi(100);p__Ascomycota(100);c__Saccharomycetes(100);o__Saccharomycetales(100);f__Metschnikowiaceae(100);g__Metschnikowia(100);s__Metschnikowia_pulcherrima(100); | Metschnikowia | 13 | Saprotroph | \|Undefined Saprotroph\| | Yeast | NULL | Probable | As nectar-tap_saprotroph (Põlme et al. 2020); Decay sugar-rich substrates (Põlme et al. 2020) | Põlme S, et al. 2020. Fungal Diversity 105:1-16 ((https://doi.org/10.1007/s13225-020-00466-2)) |
| Otu17498 | 1 | k__Fungi(100);p__Ascomycota(100);c__Saccharomycetes(100);o__Saccharomycetales(100);f__Metschnikowiaceae(100);g__Metschnikowia(100);s__Metschnikowia_pulcherrima(100); | Metschnikowia | 13 | Saprotroph | \|Undefined Saprotroph\| | Yeast | NULL | Probable | As nectar-tap_saprotroph (Põlme et al. 2020); Decay sugar-rich substrates (Põlme et al. 2020) | Põlme S, et al. 2020. Fungal Diversity 105:1-16 ((https://doi.org/10.1007/s13225-020-00466-2)) |
| Otu17500 | 1 | k__Fungi(100);p__Ascomycota(100);c__Saccharomycetes(100);o__Saccharomycetales(100);f__Metschnikowiaceae(100);g__Metschnikowia(100);s__Metschnikowia_pulcherrima(100); | Metschnikowia | 13 | Saprotroph | \|Undefined Saprotroph\| | Yeast | NULL | Probable | As nectar-tap_saprotroph (Põlme et al. 2020); Decay sugar-rich substrates (Põlme et al. 2020) | Põlme S, et al. 2020. Fungal Diversity 105:1-16 ((https://doi.org/10.1007/s13225-020-00466-2)) |
| Otu17504 | 1 | k__Fungi(100);p__Ascomycota(100);c__Saccharomycetes(100);o__Saccharomycetales(100);f__Metschnikowiaceae(100);g__Metschnikowia(100);g__Metschnikowia_unclassified(100); | Metschnikowia | 13 | Saprotroph | \|Undefined Saprotroph\| | Yeast | NULL | Probable | As nectar-tap_saprotroph (Põlme et al. 2020); Decay sugar-rich substrates (Põlme et al. 2020) | Põlme S, et al. 2020. Fungal Diversity 105:1-16 ((https://doi.org/10.1007/s13225-020-00466-2)) |
| Otu17507 | 1 | k__Fungi(100);p__Ascomycota(100);c__Saccharomycetes(100);o__Saccharomycetales(100);f__Metschnikowiaceae(100);g__Metschnikowia(100);s__Metschnikowia_pulcherrima(100); | Metschnikowia | 13 | Saprotroph | \|Undefined Saprotroph\| | Yeast | NULL | Probable | As nectar-tap_saprotroph (Põlme et al. 2020); Decay sugar-rich substrates (Põlme et al. 2020) | Põlme S, et al. 2020. Fungal Diversity 105:1-16 ((https://doi.org/10.1007/s13225-020-00466-2)) |
| Otu17509 | 1 | k__Fungi(100);p__Ascomycota(100);c__Saccharomycetes(100);o__Saccharomycetales(100);f__Metschnikowiaceae(100);g__Metschnikowia(100);s__Metschnikowia_pulcherrima(100); | Metschnikowia | 13 | Saprotroph | \|Undefined Saprotroph\| | Yeast | NULL | Probable | As nectar-tap_saprotroph (Põlme et al. 2020); Decay sugar-rich substrates (Põlme et al. 2020) | Põlme S, et al. 2020. Fungal Diversity 105:1-16 ((https://doi.org/10.1007/s13225-020-00466-2)) |
| Otu17517 | 1 | k__Fungi(100);p__Ascomycota(100);c__Saccharomycetes(100);o__Saccharomycetales(100);f__Metschnikowiaceae(100);g__Metschnikowia(100);s__Metschnikowia_pulcherrima(100); | Metschnikowia | 13 | Saprotroph | \|Undefined Saprotroph\| | Yeast | NULL | Probable | As nectar-tap_saprotroph (Põlme et al. 2020); Decay sugar-rich substrates (Põlme et al. 2020) | Põlme S, et al. 2020. Fungal Diversity 105:1-16 ((https://doi.org/10.1007/s13225-020-00466-2)) |
| Otu17518 | 1 | k__Fungi(100);p__Ascomycota(100);c__Saccharomycetes(100);o__Saccharomycetales(100);f__Metschnikowiaceae(100);g__Metschnikowia(100);s__Metschnikowia_pulcherrima(100); | Metschnikowia | 13 | Saprotroph | \|Undefined Saprotroph\| | Yeast | NULL | Probable | As nectar-tap_saprotroph (Põlme et al. 2020); Decay sugar-rich substrates (Põlme et al. 2020) | Põlme S, et al. 2020. Fungal Diversity 105:1-16 ((https://doi.org/10.1007/s13225-020-00466-2)) |
| Otu17522 | 1 | k__Fungi(100);p__Ascomycota(100);c__Saccharomycetes(100);o__Saccharomycetales(100);f__Metschnikowiaceae(100);g__Metschnikowia(100);s__Metschnikowia_pulcherrima(100); | Metschnikowia | 13 | Saprotroph | \|Undefined Saprotroph\| | Yeast | NULL | Probable | As nectar-tap_saprotroph (Põlme et al. 2020); Decay sugar-rich substrates (Põlme et al. 2020) | Põlme S, et al. 2020. Fungal Diversity 105:1-16 ((https://doi.org/10.1007/s13225-020-00466-2)) |
| Otu17539 | 1 | k__Fungi(100);p__Ascomycota(100);c__Saccharomycetes(100);o__Saccharomycetales(100);f__Metschnikowiaceae(100);g__Metschnikowia(100);s__Metschnikowia_pulcherrima(100); | Metschnikowia | 13 | Saprotroph | \|Undefined Saprotroph\| | Yeast | NULL | Probable | As nectar-tap_saprotroph (Põlme et al. 2020); Decay sugar-rich substrates (Põlme et al. 2020) | Põlme S, et al. 2020. Fungal Diversity 105:1-16 ((https://doi.org/10.1007/s13225-020-00466-2)) |
| Otu17541 | 1 | k__Fungi(100);p__Ascomycota(100);c__Saccharomycetes(100);o__Saccharomycetales(100);f__Metschnikowiaceae(100);g__Metschnikowia(100);s__Metschnikowia_pulcherrima(100); | Metschnikowia | 13 | Saprotroph | \|Undefined Saprotroph\| | Yeast | NULL | Probable | As nectar-tap_saprotroph (Põlme et al. 2020); Decay sugar-rich substrates (Põlme et al. 2020) | Põlme S, et al. 2020. Fungal Diversity 105:1-16 ((https://doi.org/10.1007/s13225-020-00466-2)) |
| Otu17542 | 1 | k__Fungi(100);p__Ascomycota(100);c__Saccharomycetes(100);o__Saccharomycetales(100);f__Metschnikowiaceae(100);g__Metschnikowia(100);s__Metschnikowia_pulcherrima(100); | Metschnikowia | 13 | Saprotroph | \|Undefined Saprotroph\| | Yeast | NULL | Probable | As nectar-tap_saprotroph (Põlme et al. 2020); Decay sugar-rich substrates (Põlme et al. 2020) | Põlme S, et al. 2020. Fungal Diversity 105:1-16 ((https://doi.org/10.1007/s13225-020-00466-2)) |
| Otu17543 | 1 | k__Fungi(100);p__Ascomycota(100);c__Saccharomycetes(100);o__Saccharomycetales(100);f__Metschnikowiaceae(100);g__Metschnikowia(100);s__Metschnikowia_pulcherrima(100); | Metschnikowia | 13 | Saprotroph | \|Undefined Saprotroph\| | Yeast | NULL | Probable | As nectar-tap_saprotroph (Põlme et al. 2020); Decay sugar-rich substrates (Põlme et al. 2020) | Põlme S, et al. 2020. Fungal Diversity 105:1-16 ((https://doi.org/10.1007/s13225-020-00466-2)) |
| Otu17544 | 1 | k__Fungi(100);p__Ascomycota(100);c__Saccharomycetes(100);o__Saccharomycetales(100);f__Metschnikowiaceae(100);g__Metschnikowia(100);s__Metschnikowia_pulcherrima(100); | Metschnikowia | 13 | Saprotroph | \|Undefined Saprotroph\| | Yeast | NULL | Probable | As nectar-tap_saprotroph (Põlme et al. 2020); Decay sugar-rich substrates (Põlme et al. 2020) | Põlme S, et al. 2020. Fungal Diversity 105:1-16 ((https://doi.org/10.1007/s13225-020-00466-2)) |
| Otu17545 | 1 | k__Fungi(100);p__Ascomycota(100);c__Saccharomycetes(100);o__Saccharomycetales(100);f__Metschnikowiaceae(100);g__Metschnikowia(100);s__Metschnikowia_pulcherrima(100); | Metschnikowia | 13 | Saprotroph | \|Undefined Saprotroph\| | Yeast | NULL | Probable | As nectar-tap_saprotroph (Põlme et al. 2020); Decay sugar-rich substrates (Põlme et al. 2020) | Põlme S, et al. 2020. Fungal Diversity 105:1-16 ((https://doi.org/10.1007/s13225-020-00466-2)) |
| Otu17546 | 1 | k__Fungi(100);p__Ascomycota(100);c__Saccharomycetes(100);o__Saccharomycetales(100);f__Metschnikowiaceae(100);g__Metschnikowia(100);s__Metschnikowia_pulcherrima(100); | Metschnikowia | 13 | Saprotroph | \|Undefined Saprotroph\| | Yeast | NULL | Probable | As nectar-tap_saprotroph (Põlme et al. 2020); Decay sugar-rich substrates (Põlme et al. 2020) | Põlme S, et al. 2020. Fungal Diversity 105:1-16 ((https://doi.org/10.1007/s13225-020-00466-2)) |
| Otu17547 | 1 | k__Fungi(100);p__Ascomycota(100);c__Saccharomycetes(100);o__Saccharomycetales(100);f__Metschnikowiaceae(100);g__Metschnikowia(100);s__Metschnikowia_pulcherrima(100); | Metschnikowia | 13 | Saprotroph | \|Undefined Saprotroph\| | Yeast | NULL | Probable | As nectar-tap_saprotroph (Põlme et al. 2020); Decay sugar-rich substrates (Põlme et al. 2020) | Põlme S, et al. 2020. Fungal Diversity 105:1-16 ((https://doi.org/10.1007/s13225-020-00466-2)) |
| Otu17548 | 1 | k__Fungi(100);p__Ascomycota(100);c__Saccharomycetes(100);o__Saccharomycetales(100);f__Metschnikowiaceae(100);g__Metschnikowia(100);s__Metschnikowia_pulcherrima(100); | Metschnikowia | 13 | Saprotroph | \|Undefined Saprotroph\| | Yeast | NULL | Probable | As nectar-tap_saprotroph (Põlme et al. 2020); Decay sugar-rich substrates (Põlme et al. 2020) | Põlme S, et al. 2020. Fungal Diversity 105:1-16 ((https://doi.org/10.1007/s13225-020-00466-2)) |
| Otu17549 | 1 | k__Fungi(100);p__Ascomycota(100);c__Saccharomycetes(100);o__Saccharomycetales(100);f__Metschnikowiaceae(100);g__Metschnikowia(100);s__Metschnikowia_pulcherrima(100); | Metschnikowia | 13 | Saprotroph | \|Undefined Saprotroph\| | Yeast | NULL | Probable | As nectar-tap_saprotroph (Põlme et al. 2020); Decay sugar-rich substrates (Põlme et al. 2020) | Põlme S, et al. 2020. Fungal Diversity 105:1-16 ((https://doi.org/10.1007/s13225-020-00466-2)) |
| Otu17550 | 1 | k__Fungi(100);p__Ascomycota(100);c__Saccharomycetes(100);o__Saccharomycetales(100);f__Metschnikowiaceae(100);g__Metschnikowia(100);s__Metschnikowia_pulcherrima(100); | Metschnikowia | 13 | Saprotroph | \|Undefined Saprotroph\| | Yeast | NULL | Probable | As nectar-tap_saprotroph (Põlme et al. 2020); Decay sugar-rich substrates (Põlme et al. 2020) | Põlme S, et al. 2020. Fungal Diversity 105:1-16 ((https://doi.org/10.1007/s13225-020-00466-2)) |
| Otu17551 | 1 | k__Fungi(100);p__Ascomycota(100);c__Saccharomycetes(100);o__Saccharomycetales(100);f__Metschnikowiaceae(100);g__Metschnikowia(100);g__Metschnikowia_unclassified(100); | Metschnikowia | 13 | Saprotroph | \|Undefined Saprotroph\| | Yeast | NULL | Probable | As nectar-tap_saprotroph (Põlme et al. 2020); Decay sugar-rich substrates (Põlme et al. 2020) | Põlme S, et al. 2020. Fungal Diversity 105:1-16 ((https://doi.org/10.1007/s13225-020-00466-2)) |
| Otu17566 | 1 | k__Fungi(100);p__Ascomycota(100);c__Saccharomycetes(100);o__Saccharomycetales(100);f__Metschnikowiaceae(100);g__Metschnikowia(100);g__Metschnikowia_unclassified(100); | Metschnikowia | 13 | Saprotroph | \|Undefined Saprotroph\| | Yeast | NULL | Probable | As nectar-tap_saprotroph (Põlme et al. 2020); Decay sugar-rich substrates (Põlme et al. 2020) | Põlme S, et al. 2020. Fungal Diversity 105:1-16 ((https://doi.org/10.1007/s13225-020-00466-2)) |
| Otu17571 | 1 | k__Fungi(100);p__Ascomycota(100);c__Saccharomycetes(100);o__Saccharomycetales(100);f__Metschnikowiaceae(100);g__Metschnikowia(100);s__Metschnikowia_pulcherrima(100); | Metschnikowia | 13 | Saprotroph | \|Undefined Saprotroph\| | Yeast | NULL | Probable | As nectar-tap_saprotroph (Põlme et al. 2020); Decay sugar-rich substrates (Põlme et al. 2020) | Põlme S, et al. 2020. Fungal Diversity 105:1-16 ((https://doi.org/10.1007/s13225-020-00466-2)) |
| Otu17580 | 1 | k__Fungi(100);p__Ascomycota(100);c__Saccharomycetes(100);o__Saccharomycetales(100);f__Metschnikowiaceae(100);g__Metschnikowia(100);s__Metschnikowia_pulcherrima(100); | Metschnikowia | 13 | Saprotroph | \|Undefined Saprotroph\| | Yeast | NULL | Probable | As nectar-tap_saprotroph (Põlme et al. 2020); Decay sugar-rich substrates (Põlme et al. 2020) | Põlme S, et al. 2020. Fungal Diversity 105:1-16 ((https://doi.org/10.1007/s13225-020-00466-2)) |
| Otu17581 | 1 | k__Fungi(100);p__Ascomycota(100);c__Saccharomycetes(100);o__Saccharomycetales(100);f__Metschnikowiaceae(100);g__Metschnikowia(100);s__Metschnikowia_pulcherrima(100); | Metschnikowia | 13 | Saprotroph | \|Undefined Saprotroph\| | Yeast | NULL | Probable | As nectar-tap_saprotroph (Põlme et al. 2020); Decay sugar-rich substrates (Põlme et al. 2020) | Põlme S, et al. 2020. Fungal Diversity 105:1-16 ((https://doi.org/10.1007/s13225-020-00466-2)) |
| Otu17582 | 1 | k__Fungi(100);p__Ascomycota(100);c__Saccharomycetes(100);o__Saccharomycetales(100);f__Metschnikowiaceae(100);g__Metschnikowia(100);s__Metschnikowia_pulcherrima(100); | Metschnikowia | 13 | Saprotroph | \|Undefined Saprotroph\| | Yeast | NULL | Probable | As nectar-tap_saprotroph (Põlme et al. 2020); Decay sugar-rich substrates (Põlme et al. 2020) | Põlme S, et al. 2020. Fungal Diversity 105:1-16 ((https://doi.org/10.1007/s13225-020-00466-2)) |
| Otu17583 | 1 | k__Fungi(100);p__Ascomycota(100);c__Saccharomycetes(100);o__Saccharomycetales(100);f__Metschnikowiaceae(100);g__Metschnikowia(100);s__Metschnikowia_pulcherrima(100); | Metschnikowia | 13 | Saprotroph | \|Undefined Saprotroph\| | Yeast | NULL | Probable | As nectar-tap_saprotroph (Põlme et al. 2020); Decay sugar-rich substrates (Põlme et al. 2020) | Põlme S, et al. 2020. Fungal Diversity 105:1-16 ((https://doi.org/10.1007/s13225-020-00466-2)) |
| Otu17584 | 1 | k__Fungi(100);p__Ascomycota(100);c__Saccharomycetes(100);o__Saccharomycetales(100);f__Metschnikowiaceae(100);g__Metschnikowia(100);s__Metschnikowia_pulcherrima(100); | Metschnikowia | 13 | Saprotroph | \|Undefined Saprotroph\| | Yeast | NULL | Probable | As nectar-tap_saprotroph (Põlme et al. 2020); Decay sugar-rich substrates (Põlme et al. 2020) | Põlme S, et al. 2020. Fungal Diversity 105:1-16 ((https://doi.org/10.1007/s13225-020-00466-2)) |
| Otu17585 | 1 | k__Fungi(100);p__Ascomycota(100);c__Saccharomycetes(100);o__Saccharomycetales(100);f__Metschnikowiaceae(100);g__Metschnikowia(100);s__Metschnikowia_pulcherrima(100); | Metschnikowia | 13 | Saprotroph | \|Undefined Saprotroph\| | Yeast | NULL | Probable | As nectar-tap_saprotroph (Põlme et al. 2020); Decay sugar-rich substrates (Põlme et al. 2020) | Põlme S, et al. 2020. Fungal Diversity 105:1-16 ((https://doi.org/10.1007/s13225-020-00466-2)) |
| Otu17586 | 1 | k__Fungi(100);p__Ascomycota(100);c__Saccharomycetes(100);o__Saccharomycetales(100);f__Metschnikowiaceae(100);g__Metschnikowia(100);s__Metschnikowia_pulcherrima(100); | Metschnikowia | 13 | Saprotroph | \|Undefined Saprotroph\| | Yeast | NULL | Probable | As nectar-tap_saprotroph (Põlme et al. 2020); Decay sugar-rich substrates (Põlme et al. 2020) | Põlme S, et al. 2020. Fungal Diversity 105:1-16 ((https://doi.org/10.1007/s13225-020-00466-2)) |
| Otu17587 | 1 | k__Fungi(100);p__Ascomycota(100);c__Saccharomycetes(100);o__Saccharomycetales(100);f__Metschnikowiaceae(100);g__Metschnikowia(100);g__Metschnikowia_unclassified(100); | Metschnikowia | 13 | Saprotroph | \|Undefined Saprotroph\| | Yeast | NULL | Probable | As nectar-tap_saprotroph (Põlme et al. 2020); Decay sugar-rich substrates (Põlme et al. 2020) | Põlme S, et al. 2020. Fungal Diversity 105:1-16 ((https://doi.org/10.1007/s13225-020-00466-2)) |
| Otu17588 | 1 | k__Fungi(100);p__Ascomycota(100);c__Saccharomycetes(100);o__Saccharomycetales(100);f__Metschnikowiaceae(100);g__Metschnikowia(100);g__Metschnikowia_unclassified(100); | Metschnikowia | 13 | Saprotroph | \|Undefined Saprotroph\| | Yeast | NULL | Probable | As nectar-tap_saprotroph (Põlme et al. 2020); Decay sugar-rich substrates (Põlme et al. 2020) | Põlme S, et al. 2020. Fungal Diversity 105:1-16 ((https://doi.org/10.1007/s13225-020-00466-2)) |
| Otu17589 | 1 | k__Fungi(100);p__Ascomycota(100);c__Saccharomycetes(100);o__Saccharomycetales(100);f__Metschnikowiaceae(100);g__Metschnikowia(100);s__Metschnikowia_pulcherrima(100); | Metschnikowia | 13 | Saprotroph | \|Undefined Saprotroph\| | Yeast | NULL | Probable | As nectar-tap_saprotroph (Põlme et al. 2020); Decay sugar-rich substrates (Põlme et al. 2020) | Põlme S, et al. 2020. Fungal Diversity 105:1-16 ((https://doi.org/10.1007/s13225-020-00466-2)) |
| Otu17590 | 1 | k__Fungi(100);p__Ascomycota(100);c__Saccharomycetes(100);o__Saccharomycetales(100);f__Metschnikowiaceae(100);g__Metschnikowia(100);s__Metschnikowia_pulcherrima(100); | Metschnikowia | 13 | Saprotroph | \|Undefined Saprotroph\| | Yeast | NULL | Probable | As nectar-tap_saprotroph (Põlme et al. 2020); Decay sugar-rich substrates (Põlme et al. 2020) | Põlme S, et al. 2020. Fungal Diversity 105:1-16 ((https://doi.org/10.1007/s13225-020-00466-2)) |
| Otu17591 | 1 | k__Fungi(100);p__Ascomycota(100);c__Saccharomycetes(100);o__Saccharomycetales(100);f__Metschnikowiaceae(100);g__Metschnikowia(100);s__Metschnikowia_pulcherrima(100); | Metschnikowia | 13 | Saprotroph | \|Undefined Saprotroph\| | Yeast | NULL | Probable | As nectar-tap_saprotroph (Põlme et al. 2020); Decay sugar-rich substrates (Põlme et al. 2020) | Põlme S, et al. 2020. Fungal Diversity 105:1-16 ((https://doi.org/10.1007/s13225-020-00466-2)) |
| Otu17595 | 1 | k__Fungi(100);p__Ascomycota(100);c__Saccharomycetes(100);o__Saccharomycetales(100);f__Metschnikowiaceae(100);g__Metschnikowia(100);g__Metschnikowia_unclassified(100); | Metschnikowia | 13 | Saprotroph | \|Undefined Saprotroph\| | Yeast | NULL | Probable | As nectar-tap_saprotroph (Põlme et al. 2020); Decay sugar-rich substrates (Põlme et al. 2020) | Põlme S, et al. 2020. Fungal Diversity 105:1-16 ((https://doi.org/10.1007/s13225-020-00466-2)) |
| Otu17602 | 1 | k__Fungi(100);p__Ascomycota(100);c__Saccharomycetes(100);o__Saccharomycetales(100);f__Metschnikowiaceae(100);g__Metschnikowia(100);s__Metschnikowia_pulcherrima(100); | Metschnikowia | 13 | Saprotroph | \|Undefined Saprotroph\| | Yeast | NULL | Probable | As nectar-tap_saprotroph (Põlme et al. 2020); Decay sugar-rich substrates (Põlme et al. 2020) | Põlme S, et al. 2020. Fungal Diversity 105:1-16 ((https://doi.org/10.1007/s13225-020-00466-2)) |
| Otu17604 | 1 | k__Fungi(100);p__Ascomycota(100);c__Saccharomycetes(100);o__Saccharomycetales(100);f__Metschnikowiaceae(100);g__Metschnikowia(100);s__Metschnikowia_pulcherrima(100); | Metschnikowia | 13 | Saprotroph | \|Undefined Saprotroph\| | Yeast | NULL | Probable | As nectar-tap_saprotroph (Põlme et al. 2020); Decay sugar-rich substrates (Põlme et al. 2020) | Põlme S, et al. 2020. Fungal Diversity 105:1-16 ((https://doi.org/10.1007/s13225-020-00466-2)) |
| Otu17605 | 1 | k__Fungi(100);p__Ascomycota(100);c__Saccharomycetes(100);o__Saccharomycetales(100);f__Metschnikowiaceae(100);g__Metschnikowia(100);s__Metschnikowia_pulcherrima(100); | Metschnikowia | 13 | Saprotroph | \|Undefined Saprotroph\| | Yeast | NULL | Probable | As nectar-tap_saprotroph (Põlme et al. 2020); Decay sugar-rich substrates (Põlme et al. 2020) | Põlme S, et al. 2020. Fungal Diversity 105:1-16 ((https://doi.org/10.1007/s13225-020-00466-2)) |
| Otu17610 | 1 | k__Fungi(100);p__Ascomycota(100);c__Saccharomycetes(100);o__Saccharomycetales(100);f__Metschnikowiaceae(100);g__Metschnikowia(100);s__Metschnikowia_pulcherrima(100); | Metschnikowia | 13 | Saprotroph | \|Undefined Saprotroph\| | Yeast | NULL | Probable | As nectar-tap_saprotroph (Põlme et al. 2020); Decay sugar-rich substrates (Põlme et al. 2020) | Põlme S, et al. 2020. Fungal Diversity 105:1-16 ((https://doi.org/10.1007/s13225-020-00466-2)) |
| Otu17616 | 1 | k__Fungi(100);p__Ascomycota(100);c__Saccharomycetes(100);o__Saccharomycetales(100);f__Metschnikowiaceae(100);g__Metschnikowia(100);s__Metschnikowia_pulcherrima(100); | Metschnikowia | 13 | Saprotroph | \|Undefined Saprotroph\| | Yeast | NULL | Probable | As nectar-tap_saprotroph (Põlme et al. 2020); Decay sugar-rich substrates (Põlme et al. 2020) | Põlme S, et al. 2020. Fungal Diversity 105:1-16 ((https://doi.org/10.1007/s13225-020-00466-2)) |
| Otu17618 | 1 | k__Fungi(100);p__Ascomycota(100);c__Saccharomycetes(100);o__Saccharomycetales(100);f__Metschnikowiaceae(100);g__Metschnikowia(100);s__Metschnikowia_pulcherrima(100); | Metschnikowia | 13 | Saprotroph | \|Undefined Saprotroph\| | Yeast | NULL | Probable | As nectar-tap_saprotroph (Põlme et al. 2020); Decay sugar-rich substrates (Põlme et al. 2020) | Põlme S, et al. 2020. Fungal Diversity 105:1-16 ((https://doi.org/10.1007/s13225-020-00466-2)) |
| Otu17619 | 1 | k__Fungi(100);p__Ascomycota(100);c__Saccharomycetes(100);o__Saccharomycetales(100);f__Metschnikowiaceae(100);g__Metschnikowia(100);s__Metschnikowia_pulcherrima(100); | Metschnikowia | 13 | Saprotroph | \|Undefined Saprotroph\| | Yeast | NULL | Probable | As nectar-tap_saprotroph (Põlme et al. 2020); Decay sugar-rich substrates (Põlme et al. 2020) | Põlme S, et al. 2020. Fungal Diversity 105:1-16 ((https://doi.org/10.1007/s13225-020-00466-2)) |
| Otu17620 | 1 | k__Fungi(100);p__Ascomycota(100);c__Saccharomycetes(100);o__Saccharomycetales(100);f__Metschnikowiaceae(100);g__Metschnikowia(100);s__Metschnikowia_pulcherrima(100); | Metschnikowia | 13 | Saprotroph | \|Undefined Saprotroph\| | Yeast | NULL | Probable | As nectar-tap_saprotroph (Põlme et al. 2020); Decay sugar-rich substrates (Põlme et al. 2020) | Põlme S, et al. 2020. Fungal Diversity 105:1-16 ((https://doi.org/10.1007/s13225-020-00466-2)) |
| Otu17621 | 1 | k__Fungi(100);p__Ascomycota(100);c__Saccharomycetes(100);o__Saccharomycetales(100);f__Metschnikowiaceae(100);g__Metschnikowia(100);s__Metschnikowia_pulcherrima(100); | Metschnikowia | 13 | Saprotroph | \|Undefined Saprotroph\| | Yeast | NULL | Probable | As nectar-tap_saprotroph (Põlme et al. 2020); Decay sugar-rich substrates (Põlme et al. 2020) | Põlme S, et al. 2020. Fungal Diversity 105:1-16 ((https://doi.org/10.1007/s13225-020-00466-2)) |
| Otu17622 | 1 | k__Fungi(100);p__Ascomycota(100);c__Saccharomycetes(100);o__Saccharomycetales(100);f__Metschnikowiaceae(100);g__Metschnikowia(100);s__Metschnikowia_pulcherrima(100); | Metschnikowia | 13 | Saprotroph | \|Undefined Saprotroph\| | Yeast | NULL | Probable | As nectar-tap_saprotroph (Põlme et al. 2020); Decay sugar-rich substrates (Põlme et al. 2020) | Põlme S, et al. 2020. Fungal Diversity 105:1-16 ((https://doi.org/10.1007/s13225-020-00466-2)) |
| Otu17623 | 1 | k__Fungi(100);p__Ascomycota(100);c__Saccharomycetes(100);o__Saccharomycetales(100);f__Metschnikowiaceae(100);g__Metschnikowia(100);s__Metschnikowia_pulcherrima(100); | Metschnikowia | 13 | Saprotroph | \|Undefined Saprotroph\| | Yeast | NULL | Probable | As nectar-tap_saprotroph (Põlme et al. 2020); Decay sugar-rich substrates (Põlme et al. 2020) | Põlme S, et al. 2020. Fungal Diversity 105:1-16 ((https://doi.org/10.1007/s13225-020-00466-2)) |
| Otu17624 | 1 | k__Fungi(100);p__Ascomycota(100);c__Saccharomycetes(100);o__Saccharomycetales(100);f__Metschnikowiaceae(100);g__Metschnikowia(100);s__Metschnikowia_pulcherrima(100); | Metschnikowia | 13 | Saprotroph | \|Undefined Saprotroph\| | Yeast | NULL | Probable | As nectar-tap_saprotroph (Põlme et al. 2020); Decay sugar-rich substrates (Põlme et al. 2020) | Põlme S, et al. 2020. Fungal Diversity 105:1-16 ((https://doi.org/10.1007/s13225-020-00466-2)) |
| Otu17625 | 1 | k__Fungi(100);p__Ascomycota(100);c__Saccharomycetes(100);o__Saccharomycetales(100);f__Metschnikowiaceae(100);g__Metschnikowia(100);g__Metschnikowia_unclassified(100); | Metschnikowia | 13 | Saprotroph | \|Undefined Saprotroph\| | Yeast | NULL | Probable | As nectar-tap_saprotroph (Põlme et al. 2020); Decay sugar-rich substrates (Põlme et al. 2020) | Põlme S, et al. 2020. Fungal Diversity 105:1-16 ((https://doi.org/10.1007/s13225-020-00466-2)) |
| Otu17626 | 1 | k__Fungi(100);p__Ascomycota(100);c__Saccharomycetes(100);o__Saccharomycetales(100);f__Metschnikowiaceae(100);g__Metschnikowia(100);g__Metschnikowia_unclassified(100); | Metschnikowia | 13 | Saprotroph | \|Undefined Saprotroph\| | Yeast | NULL | Probable | As nectar-tap_saprotroph (Põlme et al. 2020); Decay sugar-rich substrates (Põlme et al. 2020) | Põlme S, et al. 2020. Fungal Diversity 105:1-16 ((https://doi.org/10.1007/s13225-020-00466-2)) |
| Otu17627 | 1 | k__Fungi(100);p__Ascomycota(100);c__Saccharomycetes(100);o__Saccharomycetales(100);f__Metschnikowiaceae(100);g__Metschnikowia(100);g__Metschnikowia_unclassified(100); | Metschnikowia | 13 | Saprotroph | \|Undefined Saprotroph\| | Yeast | NULL | Probable | As nectar-tap_saprotroph (Põlme et al. 2020); Decay sugar-rich substrates (Põlme et al. 2020) | Põlme S, et al. 2020. Fungal Diversity 105:1-16 ((https://doi.org/10.1007/s13225-020-00466-2)) |
| Otu17628 | 1 | k__Fungi(100);p__Ascomycota(100);c__Saccharomycetes(100);o__Saccharomycetales(100);f__Metschnikowiaceae(100);g__Metschnikowia(100);g__Metschnikowia_unclassified(100); | Metschnikowia | 13 | Saprotroph | \|Undefined Saprotroph\| | Yeast | NULL | Probable | As nectar-tap_saprotroph (Põlme et al. 2020); Decay sugar-rich substrates (Põlme et al. 2020) | Põlme S, et al. 2020. Fungal Diversity 105:1-16 ((https://doi.org/10.1007/s13225-020-00466-2)) |
| Otu17629 | 1 | k__Fungi(100);p__Ascomycota(100);c__Saccharomycetes(100);o__Saccharomycetales(100);f__Metschnikowiaceae(100);g__Metschnikowia(100);g__Metschnikowia_unclassified(100); | Metschnikowia | 13 | Saprotroph | \|Undefined Saprotroph\| | Yeast | NULL | Probable | As nectar-tap_saprotroph (Põlme et al. 2020); Decay sugar-rich substrates (Põlme et al. 2020) | Põlme S, et al. 2020. Fungal Diversity 105:1-16 ((https://doi.org/10.1007/s13225-020-00466-2)) |
| Otu17630 | 1 | k__Fungi(100);p__Ascomycota(100);c__Saccharomycetes(100);o__Saccharomycetales(100);f__Metschnikowiaceae(100);g__Metschnikowia(100);s__Metschnikowia_pulcherrima(100); | Metschnikowia | 13 | Saprotroph | \|Undefined Saprotroph\| | Yeast | NULL | Probable | As nectar-tap_saprotroph (Põlme et al. 2020); Decay sugar-rich substrates (Põlme et al. 2020) | Põlme S, et al. 2020. Fungal Diversity 105:1-16 ((https://doi.org/10.1007/s13225-020-00466-2)) |
| Otu17631 | 1 | k__Fungi(100);p__Ascomycota(100);c__Saccharomycetes(100);o__Saccharomycetales(100);f__Metschnikowiaceae(100);g__Metschnikowia(100);s__Metschnikowia_pulcherrima(100); | Metschnikowia | 13 | Saprotroph | \|Undefined Saprotroph\| | Yeast | NULL | Probable | As nectar-tap_saprotroph (Põlme et al. 2020); Decay sugar-rich substrates (Põlme et al. 2020) | Põlme S, et al. 2020. Fungal Diversity 105:1-16 ((https://doi.org/10.1007/s13225-020-00466-2)) |
| Otu17632 | 1 | k__Fungi(100);p__Ascomycota(100);c__Saccharomycetes(100);o__Saccharomycetales(100);f__Metschnikowiaceae(100);g__Metschnikowia(100);s__Metschnikowia_pulcherrima(100); | Metschnikowia | 13 | Saprotroph | \|Undefined Saprotroph\| | Yeast | NULL | Probable | As nectar-tap_saprotroph (Põlme et al. 2020); Decay sugar-rich substrates (Põlme et al. 2020) | Põlme S, et al. 2020. Fungal Diversity 105:1-16 ((https://doi.org/10.1007/s13225-020-00466-2)) |
| Otu17633 | 1 | k__Fungi(100);p__Ascomycota(100);c__Saccharomycetes(100);o__Saccharomycetales(100);f__Metschnikowiaceae(100);g__Metschnikowia(100);s__Metschnikowia_pulcherrima(100); | Metschnikowia | 13 | Saprotroph | \|Undefined Saprotroph\| | Yeast | NULL | Probable | As nectar-tap_saprotroph (Põlme et al. 2020); Decay sugar-rich substrates (Põlme et al. 2020) | Põlme S, et al. 2020. Fungal Diversity 105:1-16 ((https://doi.org/10.1007/s13225-020-00466-2)) |
| Otu17634 | 1 | k__Fungi(100);p__Ascomycota(100);c__Saccharomycetes(100);o__Saccharomycetales(100);f__Metschnikowiaceae(100);g__Metschnikowia(100);s__Metschnikowia_pulcherrima(100); | Metschnikowia | 13 | Saprotroph | \|Undefined Saprotroph\| | Yeast | NULL | Probable | As nectar-tap_saprotroph (Põlme et al. 2020); Decay sugar-rich substrates (Põlme et al. 2020) | Põlme S, et al. 2020. Fungal Diversity 105:1-16 ((https://doi.org/10.1007/s13225-020-00466-2)) |
| Otu17635 | 1 | k__Fungi(100);p__Ascomycota(100);c__Saccharomycetes(100);o__Saccharomycetales(100);f__Metschnikowiaceae(100);g__Metschnikowia(100);s__Metschnikowia_pulcherrima(100); | Metschnikowia | 13 | Saprotroph | \|Undefined Saprotroph\| | Yeast | NULL | Probable | As nectar-tap_saprotroph (Põlme et al. 2020); Decay sugar-rich substrates (Põlme et al. 2020) | Põlme S, et al. 2020. Fungal Diversity 105:1-16 ((https://doi.org/10.1007/s13225-020-00466-2)) |
| Otu17636 | 1 | k__Fungi(100);p__Ascomycota(100);c__Saccharomycetes(100);o__Saccharomycetales(100);f__Metschnikowiaceae(100);g__Metschnikowia(100);s__Metschnikowia_pulcherrima(100); | Metschnikowia | 13 | Saprotroph | \|Undefined Saprotroph\| | Yeast | NULL | Probable | As nectar-tap_saprotroph (Põlme et al. 2020); Decay sugar-rich substrates (Põlme et al. 2020) | Põlme S, et al. 2020. Fungal Diversity 105:1-16 ((https://doi.org/10.1007/s13225-020-00466-2)) |
| Otu17637 | 1 | k__Fungi(100);p__Ascomycota(100);c__Saccharomycetes(100);o__Saccharomycetales(100);f__Metschnikowiaceae(100);g__Metschnikowia(100);s__Metschnikowia_pulcherrima(100); | Metschnikowia | 13 | Saprotroph | \|Undefined Saprotroph\| | Yeast | NULL | Probable | As nectar-tap_saprotroph (Põlme et al. 2020); Decay sugar-rich substrates (Põlme et al. 2020) | Põlme S, et al. 2020. Fungal Diversity 105:1-16 ((https://doi.org/10.1007/s13225-020-00466-2)) |
| Otu17638 | 1 | k__Fungi(100);p__Ascomycota(100);c__Saccharomycetes(100);o__Saccharomycetales(100);f__Metschnikowiaceae(100);g__Metschnikowia(100);s__Metschnikowia_pulcherrima(100); | Metschnikowia | 13 | Saprotroph | \|Undefined Saprotroph\| | Yeast | NULL | Probable | As nectar-tap_saprotroph (Põlme et al. 2020); Decay sugar-rich substrates (Põlme et al. 2020) | Põlme S, et al. 2020. Fungal Diversity 105:1-16 ((https://doi.org/10.1007/s13225-020-00466-2)) |
| Otu17640 | 1 | k__Fungi(100);p__Ascomycota(100);c__Saccharomycetes(100);o__Saccharomycetales(100);f__Metschnikowiaceae(100);g__Metschnikowia(100);g__Metschnikowia_unclassified(100); | Metschnikowia | 13 | Saprotroph | \|Undefined Saprotroph\| | Yeast | NULL | Probable | As nectar-tap_saprotroph (Põlme et al. 2020); Decay sugar-rich substrates (Põlme et al. 2020) | Põlme S, et al. 2020. Fungal Diversity 105:1-16 ((https://doi.org/10.1007/s13225-020-00466-2)) |
| Otu17642 | 1 | k__Fungi(100);p__Ascomycota(100);c__Saccharomycetes(100);o__Saccharomycetales(100);f__Metschnikowiaceae(100);g__Metschnikowia(100);g__Metschnikowia_unclassified(100); | Metschnikowia | 13 | Saprotroph | \|Undefined Saprotroph\| | Yeast | NULL | Probable | As nectar-tap_saprotroph (Põlme et al. 2020); Decay sugar-rich substrates (Põlme et al. 2020) | Põlme S, et al. 2020. Fungal Diversity 105:1-16 ((https://doi.org/10.1007/s13225-020-00466-2)) |
| Otu17643 | 1 | k__Fungi(100);p__Ascomycota(100);c__Saccharomycetes(100);o__Saccharomycetales(100);f__Metschnikowiaceae(100);g__Metschnikowia(100);g__Metschnikowia_unclassified(100); | Metschnikowia | 13 | Saprotroph | \|Undefined Saprotroph\| | Yeast | NULL | Probable | As nectar-tap_saprotroph (Põlme et al. 2020); Decay sugar-rich substrates (Põlme et al. 2020) | Põlme S, et al. 2020. Fungal Diversity 105:1-16 ((https://doi.org/10.1007/s13225-020-00466-2)) |
| Otu17644 | 1 | k__Fungi(100);p__Ascomycota(100);c__Saccharomycetes(100);o__Saccharomycetales(100);f__Saccharomycetales_family_Incertae_sedis(100);g__Candida(100);g__Candida_unclassified(100); | Candida | 13 | Pathotroph-Saprotroph-Symbiotroph | Animal Pathogen-Endophyte-Endosymbiont-Epiphyte-\|Undefined Saprotroph\| | Dimorphic Yeast-Yeast | NULL | Probable | As nectar-tap_saprotroph (Põlme et al. 2020); Decay sugar-rich substrates (Põlme et al. 2020); This asexual genus is polyphyletic and contain species that function very differently depend on what they are. Caution should be taken when interpreting guilds for Candida species. | Manolakaki D, et al. 2010. Virulence. 1: 367-375 ((https://doi.org/10.4161/viru.1.5.12796)); Mekbib SB, et al. 2011. Tropical Plant Pathology 36:233-240 ((https://doi.org/10.1590/S1982-56762011000400004)); Põlme S, et al. 2020. Fungal Diversity 105:1-16 ((https://doi.org/10.1007/s13225-020-00466-2)) |
| Otu17646 | 1 | k__Fungi(100);p__Ascomycota(100);c__Saccharomycetes(100);o__Saccharomycetales(100);f__Metschnikowiaceae(100);g__Metschnikowia(100);g__Metschnikowia_unclassified(100); | Metschnikowia | 13 | Saprotroph | \|Undefined Saprotroph\| | Yeast | NULL | Probable | As nectar-tap_saprotroph (Põlme et al. 2020); Decay sugar-rich substrates (Põlme et al. 2020) | Põlme S, et al. 2020. Fungal Diversity 105:1-16 ((https://doi.org/10.1007/s13225-020-00466-2)) |
| Otu17647 | 1 | k__Fungi(100);p__Ascomycota(100);c__Saccharomycetes(100);o__Saccharomycetales(100);f__Metschnikowiaceae(100);g__Metschnikowia(100);g__Metschnikowia_unclassified(100); | Metschnikowia | 13 | Saprotroph | \|Undefined Saprotroph\| | Yeast | NULL | Probable | As nectar-tap_saprotroph (Põlme et al. 2020); Decay sugar-rich substrates (Põlme et al. 2020) | Põlme S, et al. 2020. Fungal Diversity 105:1-16 ((https://doi.org/10.1007/s13225-020-00466-2)) |
| Otu17652 | 1 | k__Fungi(100);p__Ascomycota(100);c__Saccharomycetes(100);o__Saccharomycetales(100);f__Metschnikowiaceae(100);g__Metschnikowia(100);s__Metschnikowia_pulcherrima(100); | Metschnikowia | 13 | Saprotroph | \|Undefined Saprotroph\| | Yeast | NULL | Probable | As nectar-tap_saprotroph (Põlme et al. 2020); Decay sugar-rich substrates (Põlme et al. 2020) | Põlme S, et al. 2020. Fungal Diversity 105:1-16 ((https://doi.org/10.1007/s13225-020-00466-2)) |
| Otu17653 | 1 | k__Fungi(100);p__Ascomycota(100);c__Saccharomycetes(100);o__Saccharomycetales(100);f__Metschnikowiaceae(100);g__Metschnikowia(100);s__Metschnikowia_pulcherrima(100); | Metschnikowia | 13 | Saprotroph | \|Undefined Saprotroph\| | Yeast | NULL | Probable | As nectar-tap_saprotroph (Põlme et al. 2020); Decay sugar-rich substrates (Põlme et al. 2020) | Põlme S, et al. 2020. Fungal Diversity 105:1-16 ((https://doi.org/10.1007/s13225-020-00466-2)) |
| Otu17654 | 1 | k__Fungi(100);p__Ascomycota(100);c__Saccharomycetes(100);o__Saccharomycetales(100);f__Metschnikowiaceae(100);g__Metschnikowia(100);g__Metschnikowia_unclassified(100); | Metschnikowia | 13 | Saprotroph | \|Undefined Saprotroph\| | Yeast | NULL | Probable | As nectar-tap_saprotroph (Põlme et al. 2020); Decay sugar-rich substrates (Põlme et al. 2020) | Põlme S, et al. 2020. Fungal Diversity 105:1-16 ((https://doi.org/10.1007/s13225-020-00466-2)) |
| Otu17655 | 1 | k__Fungi(100);p__Ascomycota(100);c__Saccharomycetes(100);o__Saccharomycetales(100);f__Metschnikowiaceae(100);g__Metschnikowia(100);s__Metschnikowia_pulcherrima(100); | Metschnikowia | 13 | Saprotroph | \|Undefined Saprotroph\| | Yeast | NULL | Probable | As nectar-tap_saprotroph (Põlme et al. 2020); Decay sugar-rich substrates (Põlme et al. 2020) | Põlme S, et al. 2020. Fungal Diversity 105:1-16 ((https://doi.org/10.1007/s13225-020-00466-2)) |
| Otu17657 | 1 | k__Fungi(100);p__Ascomycota(100);c__Saccharomycetes(100);o__Saccharomycetales(100);f__Metschnikowiaceae(100);g__Metschnikowia(100);s__Metschnikowia_pulcherrima(100); | Metschnikowia | 13 | Saprotroph | \|Undefined Saprotroph\| | Yeast | NULL | Probable | As nectar-tap_saprotroph (Põlme et al. 2020); Decay sugar-rich substrates (Põlme et al. 2020) | Põlme S, et al. 2020. Fungal Diversity 105:1-16 ((https://doi.org/10.1007/s13225-020-00466-2)) |
| Otu17658 | 1 | k__Fungi(100);p__Ascomycota(100);c__Saccharomycetes(100);o__Saccharomycetales(100);f__Metschnikowiaceae(100);g__Metschnikowia(100);g__Metschnikowia_unclassified(100); | Metschnikowia | 13 | Saprotroph | \|Undefined Saprotroph\| | Yeast | NULL | Probable | As nectar-tap_saprotroph (Põlme et al. 2020); Decay sugar-rich substrates (Põlme et al. 2020) | Põlme S, et al. 2020. Fungal Diversity 105:1-16 ((https://doi.org/10.1007/s13225-020-00466-2)) |
| Otu17659 | 1 | k__Fungi(100);p__Ascomycota(100);c__Saccharomycetes(100);o__Saccharomycetales(100);f__Metschnikowiaceae(100);g__Metschnikowia(100);s__Metschnikowia_pulcherrima(100); | Metschnikowia | 13 | Saprotroph | \|Undefined Saprotroph\| | Yeast | NULL | Probable | As nectar-tap_saprotroph (Põlme et al. 2020); Decay sugar-rich substrates (Põlme et al. 2020) | Põlme S, et al. 2020. Fungal Diversity 105:1-16 ((https://doi.org/10.1007/s13225-020-00466-2)) |
| Otu17660 | 1 | k__Fungi(100);p__Ascomycota(100);c__Saccharomycetes(100);o__Saccharomycetales(100);f__Metschnikowiaceae(100);g__Metschnikowia(100);s__Metschnikowia_pulcherrima(100); | Metschnikowia | 13 | Saprotroph | \|Undefined Saprotroph\| | Yeast | NULL | Probable | As nectar-tap_saprotroph (Põlme et al. 2020); Decay sugar-rich substrates (Põlme et al. 2020) | Põlme S, et al. 2020. Fungal Diversity 105:1-16 ((https://doi.org/10.1007/s13225-020-00466-2)) |
| Otu17661 | 1 | k__Fungi(100);p__Ascomycota(100);c__Saccharomycetes(100);o__Saccharomycetales(100);f__Metschnikowiaceae(100);g__Metschnikowia(100);g__Metschnikowia_unclassified(100); | Metschnikowia | 13 | Saprotroph | \|Undefined Saprotroph\| | Yeast | NULL | Probable | As nectar-tap_saprotroph (Põlme et al. 2020); Decay sugar-rich substrates (Põlme et al. 2020) | Põlme S, et al. 2020. Fungal Diversity 105:1-16 ((https://doi.org/10.1007/s13225-020-00466-2)) |
| Otu17662 | 1 | k__Fungi(100);p__Ascomycota(100);c__Saccharomycetes(100);o__Saccharomycetales(100);f__Metschnikowiaceae(100);g__Metschnikowia(100);g__Metschnikowia_unclassified(100); | Metschnikowia | 13 | Saprotroph | \|Undefined Saprotroph\| | Yeast | NULL | Probable | As nectar-tap_saprotroph (Põlme et al. 2020); Decay sugar-rich substrates (Põlme et al. 2020) | Põlme S, et al. 2020. Fungal Diversity 105:1-16 ((https://doi.org/10.1007/s13225-020-00466-2)) |
| Otu17663 | 1 | k__Fungi(100);p__Ascomycota(100);c__Saccharomycetes(100);o__Saccharomycetales(100);f__Metschnikowiaceae(100);g__Metschnikowia(100);s__Metschnikowia_pulcherrima(100); | Metschnikowia | 13 | Saprotroph | \|Undefined Saprotroph\| | Yeast | NULL | Probable | As nectar-tap_saprotroph (Põlme et al. 2020); Decay sugar-rich substrates (Põlme et al. 2020) | Põlme S, et al. 2020. Fungal Diversity 105:1-16 ((https://doi.org/10.1007/s13225-020-00466-2)) |
| Otu17664 | 1 | k__Fungi(100);p__Ascomycota(100);c__Saccharomycetes(100);o__Saccharomycetales(100);f__Metschnikowiaceae(100);g__Metschnikowia(100);s__Metschnikowia_pulcherrima(100); | Metschnikowia | 13 | Saprotroph | \|Undefined Saprotroph\| | Yeast | NULL | Probable | As nectar-tap_saprotroph (Põlme et al. 2020); Decay sugar-rich substrates (Põlme et al. 2020) | Põlme S, et al. 2020. Fungal Diversity 105:1-16 ((https://doi.org/10.1007/s13225-020-00466-2)) |
| Otu17665 | 1 | k__Fungi(100);p__Ascomycota(100);c__Saccharomycetes(100);o__Saccharomycetales(100);f__Metschnikowiaceae(100);g__Metschnikowia(100);s__Metschnikowia_pulcherrima(100); | Metschnikowia | 13 | Saprotroph | \|Undefined Saprotroph\| | Yeast | NULL | Probable | As nectar-tap_saprotroph (Põlme et al. 2020); Decay sugar-rich substrates (Põlme et al. 2020) | Põlme S, et al. 2020. Fungal Diversity 105:1-16 ((https://doi.org/10.1007/s13225-020-00466-2)) |
| Otu17666 | 1 | k__Fungi(100);p__Ascomycota(100);c__Saccharomycetes(100);o__Saccharomycetales(100);f__Metschnikowiaceae(100);g__Metschnikowia(100);s__Metschnikowia_pulcherrima(100); | Metschnikowia | 13 | Saprotroph | \|Undefined Saprotroph\| | Yeast | NULL | Probable | As nectar-tap_saprotroph (Põlme et al. 2020); Decay sugar-rich substrates (Põlme et al. 2020) | Põlme S, et al. 2020. Fungal Diversity 105:1-16 ((https://doi.org/10.1007/s13225-020-00466-2)) |
| Otu17667 | 1 | k__Fungi(100);p__Ascomycota(100);c__Saccharomycetes(100);o__Saccharomycetales(100);f__Metschnikowiaceae(100);g__Metschnikowia(100);g__Metschnikowia_unclassified(100); | Metschnikowia | 13 | Saprotroph | \|Undefined Saprotroph\| | Yeast | NULL | Probable | As nectar-tap_saprotroph (Põlme et al. 2020); Decay sugar-rich substrates (Põlme et al. 2020) | Põlme S, et al. 2020. Fungal Diversity 105:1-16 ((https://doi.org/10.1007/s13225-020-00466-2)) |
| Otu17668 | 1 | k__Fungi(100);p__Ascomycota(100);c__Saccharomycetes(100);o__Saccharomycetales(100);f__Metschnikowiaceae(100);g__Metschnikowia(100);s__Metschnikowia_pulcherrima(100); | Metschnikowia | 13 | Saprotroph | \|Undefined Saprotroph\| | Yeast | NULL | Probable | As nectar-tap_saprotroph (Põlme et al. 2020); Decay sugar-rich substrates (Põlme et al. 2020) | Põlme S, et al. 2020. Fungal Diversity 105:1-16 ((https://doi.org/10.1007/s13225-020-00466-2)) |
| Otu17669 | 1 | k__Fungi(100);p__Ascomycota(100);c__Saccharomycetes(100);o__Saccharomycetales(100);f__Metschnikowiaceae(100);g__Metschnikowia(100);g__Metschnikowia_unclassified(100); | Metschnikowia | 13 | Saprotroph | \|Undefined Saprotroph\| | Yeast | NULL | Probable | As nectar-tap_saprotroph (Põlme et al. 2020); Decay sugar-rich substrates (Põlme et al. 2020) | Põlme S, et al. 2020. Fungal Diversity 105:1-16 ((https://doi.org/10.1007/s13225-020-00466-2)) |
| Otu17670 | 1 | k__Fungi(100);p__Ascomycota(100);c__Saccharomycetes(100);o__Saccharomycetales(100);f__Metschnikowiaceae(100);g__Metschnikowia(100);s__Metschnikowia_pulcherrima(100); | Metschnikowia | 13 | Saprotroph | \|Undefined Saprotroph\| | Yeast | NULL | Probable | As nectar-tap_saprotroph (Põlme et al. 2020); Decay sugar-rich substrates (Põlme et al. 2020) | Põlme S, et al. 2020. Fungal Diversity 105:1-16 ((https://doi.org/10.1007/s13225-020-00466-2)) |
| Otu17673 | 1 | k__Fungi(100);p__Ascomycota(100);c__Saccharomycetes(100);o__Saccharomycetales(100);f__Metschnikowiaceae(100);g__Metschnikowia(100);s__Metschnikowia_pulcherrima(100); | Metschnikowia | 13 | Saprotroph | \|Undefined Saprotroph\| | Yeast | NULL | Probable | As nectar-tap_saprotroph (Põlme et al. 2020); Decay sugar-rich substrates (Põlme et al. 2020) | Põlme S, et al. 2020. Fungal Diversity 105:1-16 ((https://doi.org/10.1007/s13225-020-00466-2)) |
| Otu17674 | 1 | k__Fungi(100);p__Ascomycota(100);c__Saccharomycetes(100);o__Saccharomycetales(100);o__Saccharomycetales_unclassified(100);o__Saccharomycetales_unclassified(100);o__Saccharomycetales_unclassified(100); | Saccharomycetales | 7 | Saprotroph | Undefined Saprotroph | Yeast | NULL | Possible | NULL | Sterkenburg E, et al. 2015. New Phytologist 207:1145-1158 ((https://doi.org/10.1111/nph.13426)) |
| Otu17675 | 1 | k__Fungi(100);p__Ascomycota(100);c__Saccharomycetes(100);o__Saccharomycetales(100);f__Metschnikowiaceae(100);g__Metschnikowia(100);s__Metschnikowia_pulcherrima(100); | Metschnikowia | 13 | Saprotroph | \|Undefined Saprotroph\| | Yeast | NULL | Probable | As nectar-tap_saprotroph (Põlme et al. 2020); Decay sugar-rich substrates (Põlme et al. 2020) | Põlme S, et al. 2020. Fungal Diversity 105:1-16 ((https://doi.org/10.1007/s13225-020-00466-2)) |
| Otu17676 | 1 | k__Fungi(100);p__Ascomycota(100);c__Saccharomycetes(100);o__Saccharomycetales(100);f__Metschnikowiaceae(100);g__Metschnikowia(100);s__Metschnikowia_pulcherrima(100); | Metschnikowia | 13 | Saprotroph | \|Undefined Saprotroph\| | Yeast | NULL | Probable | As nectar-tap_saprotroph (Põlme et al. 2020); Decay sugar-rich substrates (Põlme et al. 2020) | Põlme S, et al. 2020. Fungal Diversity 105:1-16 ((https://doi.org/10.1007/s13225-020-00466-2)) |
| Otu17677 | 1 | k__Fungi(100);p__Ascomycota(100);c__Saccharomycetes(100);o__Saccharomycetales(100);f__Metschnikowiaceae(100);g__Metschnikowia(100);s__Metschnikowia_pulcherrima(100); | Metschnikowia | 13 | Saprotroph | \|Undefined Saprotroph\| | Yeast | NULL | Probable | As nectar-tap_saprotroph (Põlme et al. 2020); Decay sugar-rich substrates (Põlme et al. 2020) | Põlme S, et al. 2020. Fungal Diversity 105:1-16 ((https://doi.org/10.1007/s13225-020-00466-2)) |
| Otu17678 | 1 | k__Fungi(100);p__Ascomycota(100);c__Saccharomycetes(100);o__Saccharomycetales(100);f__Metschnikowiaceae(100);g__Metschnikowia(100);s__Metschnikowia_pulcherrima(100); | Metschnikowia | 13 | Saprotroph | \|Undefined Saprotroph\| | Yeast | NULL | Probable | As nectar-tap_saprotroph (Põlme et al. 2020); Decay sugar-rich substrates (Põlme et al. 2020) | Põlme S, et al. 2020. Fungal Diversity 105:1-16 ((https://doi.org/10.1007/s13225-020-00466-2)) |
| Otu17679 | 1 | k__Fungi(100);p__Ascomycota(100);c__Saccharomycetes(100);o__Saccharomycetales(100);f__Metschnikowiaceae(100);g__Metschnikowia(100);s__Metschnikowia_pulcherrima(100); | Metschnikowia | 13 | Saprotroph | \|Undefined Saprotroph\| | Yeast | NULL | Probable | As nectar-tap_saprotroph (Põlme et al. 2020); Decay sugar-rich substrates (Põlme et al. 2020) | Põlme S, et al. 2020. Fungal Diversity 105:1-16 ((https://doi.org/10.1007/s13225-020-00466-2)) |
| Otu17680 | 1 | k__Fungi(100);p__Ascomycota(100);c__Saccharomycetes(100);o__Saccharomycetales(100);f__Metschnikowiaceae(100);g__Metschnikowia(100);g__Metschnikowia_unclassified(100); | Metschnikowia | 13 | Saprotroph | \|Undefined Saprotroph\| | Yeast | NULL | Probable | As nectar-tap_saprotroph (Põlme et al. 2020); Decay sugar-rich substrates (Põlme et al. 2020) | Põlme S, et al. 2020. Fungal Diversity 105:1-16 ((https://doi.org/10.1007/s13225-020-00466-2)) |
| Otu17681 | 1 | k__Fungi(100);p__Ascomycota(100);c__Saccharomycetes(100);o__Saccharomycetales(100);f__Metschnikowiaceae(100);g__Metschnikowia(100);s__Metschnikowia_pulcherrima(100); | Metschnikowia | 13 | Saprotroph | \|Undefined Saprotroph\| | Yeast | NULL | Probable | As nectar-tap_saprotroph (Põlme et al. 2020); Decay sugar-rich substrates (Põlme et al. 2020) | Põlme S, et al. 2020. Fungal Diversity 105:1-16 ((https://doi.org/10.1007/s13225-020-00466-2)) |
| Otu17682 | 1 | k__Fungi(100);p__Ascomycota(100);c__Saccharomycetes(100);o__Saccharomycetales(100);f__Metschnikowiaceae(100);g__Metschnikowia(100);g__Metschnikowia_unclassified(100); | Metschnikowia | 13 | Saprotroph | \|Undefined Saprotroph\| | Yeast | NULL | Probable | As nectar-tap_saprotroph (Põlme et al. 2020); Decay sugar-rich substrates (Põlme et al. 2020) | Põlme S, et al. 2020. Fungal Diversity 105:1-16 ((https://doi.org/10.1007/s13225-020-00466-2)) |
| Otu17683 | 1 | k__Fungi(100);p__Ascomycota(100);c__Saccharomycetes(100);o__Saccharomycetales(100);f__Metschnikowiaceae(100);g__Metschnikowia(100);g__Metschnikowia_unclassified(100); | Metschnikowia | 13 | Saprotroph | \|Undefined Saprotroph\| | Yeast | NULL | Probable | As nectar-tap_saprotroph (Põlme et al. 2020); Decay sugar-rich substrates (Põlme et al. 2020) | Põlme S, et al. 2020. Fungal Diversity 105:1-16 ((https://doi.org/10.1007/s13225-020-00466-2)) |
| Otu17684 | 1 | k__Fungi(100);p__Ascomycota(100);c__Saccharomycetes(100);o__Saccharomycetales(100);f__Metschnikowiaceae(100);g__Metschnikowia(100);s__Metschnikowia_pulcherrima(100); | Metschnikowia | 13 | Saprotroph | \|Undefined Saprotroph\| | Yeast | NULL | Probable | As nectar-tap_saprotroph (Põlme et al. 2020); Decay sugar-rich substrates (Põlme et al. 2020) | Põlme S, et al. 2020. Fungal Diversity 105:1-16 ((https://doi.org/10.1007/s13225-020-00466-2)) |
| Otu17685 | 1 | k__Fungi(100);p__Ascomycota(100);c__Saccharomycetes(100);o__Saccharomycetales(100);f__Saccharomycetales_family_Incertae_sedis(100);g__Candida(100);g__Candida_unclassified(100); | Candida | 13 | Pathotroph-Saprotroph-Symbiotroph | Animal Pathogen-Endophyte-Endosymbiont-Epiphyte-\|Undefined Saprotroph\| | Dimorphic Yeast-Yeast | NULL | Probable | As nectar-tap_saprotroph (Põlme et al. 2020); Decay sugar-rich substrates (Põlme et al. 2020); This asexual genus is polyphyletic and contain species that function very differently depend on what they are. Caution should be taken when interpreting guilds for Candida species. | Manolakaki D, et al. 2010. Virulence. 1: 367-375 ((https://doi.org/10.4161/viru.1.5.12796)); Mekbib SB, et al. 2011. Tropical Plant Pathology 36:233-240 ((https://doi.org/10.1590/S1982-56762011000400004)); Põlme S, et al. 2020. Fungal Diversity 105:1-16 ((https://doi.org/10.1007/s13225-020-00466-2)) |
| Otu17686 | 1 | k__Fungi(100);p__Ascomycota(100);c__Saccharomycetes(100);o__Saccharomycetales(100);f__Metschnikowiaceae(100);g__Metschnikowia(100);s__Metschnikowia_pulcherrima(100); | Metschnikowia | 13 | Saprotroph | \|Undefined Saprotroph\| | Yeast | NULL | Probable | As nectar-tap_saprotroph (Põlme et al. 2020); Decay sugar-rich substrates (Põlme et al. 2020) | Põlme S, et al. 2020. Fungal Diversity 105:1-16 ((https://doi.org/10.1007/s13225-020-00466-2)) |
| Otu17687 | 1 | k__Fungi(100);p__Ascomycota(100);c__Saccharomycetes(100);o__Saccharomycetales(100);f__Metschnikowiaceae(100);g__Metschnikowia(100);s__Metschnikowia_pulcherrima(100); | Metschnikowia | 13 | Saprotroph | \|Undefined Saprotroph\| | Yeast | NULL | Probable | As nectar-tap_saprotroph (Põlme et al. 2020); Decay sugar-rich substrates (Põlme et al. 2020) | Põlme S, et al. 2020. Fungal Diversity 105:1-16 ((https://doi.org/10.1007/s13225-020-00466-2)) |
| Otu17688 | 1 | k__Fungi(100);p__Ascomycota(100);c__Saccharomycetes(100);o__Saccharomycetales(100);f__Metschnikowiaceae(100);g__Metschnikowia(100);s__Metschnikowia_pulcherrima(100); | Metschnikowia | 13 | Saprotroph | \|Undefined Saprotroph\| | Yeast | NULL | Probable | As nectar-tap_saprotroph (Põlme et al. 2020); Decay sugar-rich substrates (Põlme et al. 2020) | Põlme S, et al. 2020. Fungal Diversity 105:1-16 ((https://doi.org/10.1007/s13225-020-00466-2)) |
| Otu17689 | 1 | k__Fungi(100);p__Ascomycota(100);c__Saccharomycetes(100);o__Saccharomycetales(100);f__Metschnikowiaceae(100);g__Metschnikowia(100);g__Metschnikowia_unclassified(100); | Metschnikowia | 13 | Saprotroph | \|Undefined Saprotroph\| | Yeast | NULL | Probable | As nectar-tap_saprotroph (Põlme et al. 2020); Decay sugar-rich substrates (Põlme et al. 2020) | Põlme S, et al. 2020. Fungal Diversity 105:1-16 ((https://doi.org/10.1007/s13225-020-00466-2)) |
| Otu17690 | 1 | k__Fungi(100);p__Ascomycota(100);c__Saccharomycetes(100);o__Saccharomycetales(100);f__Metschnikowiaceae(100);g__Metschnikowia(100);s__Metschnikowia_pulcherrima(100); | Metschnikowia | 13 | Saprotroph | \|Undefined Saprotroph\| | Yeast | NULL | Probable | As nectar-tap_saprotroph (Põlme et al. 2020); Decay sugar-rich substrates (Põlme et al. 2020) | Põlme S, et al. 2020. Fungal Diversity 105:1-16 ((https://doi.org/10.1007/s13225-020-00466-2)) |
| Otu17691 | 1 | k__Fungi(100);p__Ascomycota(100);c__Saccharomycetes(100);o__Saccharomycetales(100);f__Metschnikowiaceae(100);g__Metschnikowia(100);s__Metschnikowia_pulcherrima(100); | Metschnikowia | 13 | Saprotroph | \|Undefined Saprotroph\| | Yeast | NULL | Probable | As nectar-tap_saprotroph (Põlme et al. 2020); Decay sugar-rich substrates (Põlme et al. 2020) | Põlme S, et al. 2020. Fungal Diversity 105:1-16 ((https://doi.org/10.1007/s13225-020-00466-2)) |
| Otu17692 | 1 | k__Fungi(100);p__Ascomycota(100);c__Saccharomycetes(100);o__Saccharomycetales(100);f__Metschnikowiaceae(100);g__Metschnikowia(100);s__Metschnikowia_pulcherrima(100); | Metschnikowia | 13 | Saprotroph | \|Undefined Saprotroph\| | Yeast | NULL | Probable | As nectar-tap_saprotroph (Põlme et al. 2020); Decay sugar-rich substrates (Põlme et al. 2020) | Põlme S, et al. 2020. Fungal Diversity 105:1-16 ((https://doi.org/10.1007/s13225-020-00466-2)) |
| Otu17693 | 1 | k__Fungi(100);p__Ascomycota(100);c__Saccharomycetes(100);o__Saccharomycetales(100);f__Metschnikowiaceae(100);g__Metschnikowia(100);s__Metschnikowia_pulcherrima(100); | Metschnikowia | 13 | Saprotroph | \|Undefined Saprotroph\| | Yeast | NULL | Probable | As nectar-tap_saprotroph (Põlme et al. 2020); Decay sugar-rich substrates (Põlme et al. 2020) | Põlme S, et al. 2020. Fungal Diversity 105:1-16 ((https://doi.org/10.1007/s13225-020-00466-2)) |
| Otu17694 | 1 | k__Fungi(100);p__Ascomycota(100);c__Saccharomycetes(100);o__Saccharomycetales(100);f__Metschnikowiaceae(100);g__Metschnikowia(100);g__Metschnikowia_unclassified(100); | Metschnikowia | 13 | Saprotroph | \|Undefined Saprotroph\| | Yeast | NULL | Probable | As nectar-tap_saprotroph (Põlme et al. 2020); Decay sugar-rich substrates (Põlme et al. 2020) | Põlme S, et al. 2020. Fungal Diversity 105:1-16 ((https://doi.org/10.1007/s13225-020-00466-2)) |
| Otu17695 | 1 | k__Fungi(100);p__Ascomycota(100);c__Saccharomycetes(100);o__Saccharomycetales(100);f__Metschnikowiaceae(100);g__Metschnikowia(100);s__Metschnikowia_pulcherrima(100); | Metschnikowia | 13 | Saprotroph | \|Undefined Saprotroph\| | Yeast | NULL | Probable | As nectar-tap_saprotroph (Põlme et al. 2020); Decay sugar-rich substrates (Põlme et al. 2020) | Põlme S, et al. 2020. Fungal Diversity 105:1-16 ((https://doi.org/10.1007/s13225-020-00466-2)) |
| Otu17696 | 1 | k__Fungi(100);p__Ascomycota(100);c__Saccharomycetes(100);o__Saccharomycetales(100);f__Metschnikowiaceae(100);g__Metschnikowia(100);s__Metschnikowia_pulcherrima(100); | Metschnikowia | 13 | Saprotroph | \|Undefined Saprotroph\| | Yeast | NULL | Probable | As nectar-tap_saprotroph (Põlme et al. 2020); Decay sugar-rich substrates (Põlme et al. 2020) | Põlme S, et al. 2020. Fungal Diversity 105:1-16 ((https://doi.org/10.1007/s13225-020-00466-2)) |
| Otu17697 | 1 | k__Fungi(100);p__Ascomycota(100);c__Saccharomycetes(100);o__Saccharomycetales(100);f__Metschnikowiaceae(100);g__Metschnikowia(100);s__Metschnikowia_pulcherrima(100); | Metschnikowia | 13 | Saprotroph | \|Undefined Saprotroph\| | Yeast | NULL | Probable | As nectar-tap_saprotroph (Põlme et al. 2020); Decay sugar-rich substrates (Põlme et al. 2020) | Põlme S, et al. 2020. Fungal Diversity 105:1-16 ((https://doi.org/10.1007/s13225-020-00466-2)) |
| Otu17698 | 1 | k__Fungi(100);p__Ascomycota(100);c__Saccharomycetes(100);o__Saccharomycetales(100);f__Metschnikowiaceae(100);g__Metschnikowia(100);g__Metschnikowia_unclassified(100); | Metschnikowia | 13 | Saprotroph | \|Undefined Saprotroph\| | Yeast | NULL | Probable | As nectar-tap_saprotroph (Põlme et al. 2020); Decay sugar-rich substrates (Põlme et al. 2020) | Põlme S, et al. 2020. Fungal Diversity 105:1-16 ((https://doi.org/10.1007/s13225-020-00466-2)) |
| Otu17699 | 1 | k__Fungi(100);p__Ascomycota(100);c__Saccharomycetes(100);o__Saccharomycetales(100);f__Metschnikowiaceae(100);g__Metschnikowia(100);g__Metschnikowia_unclassified(100); | Metschnikowia | 13 | Saprotroph | \|Undefined Saprotroph\| | Yeast | NULL | Probable | As nectar-tap_saprotroph (Põlme et al. 2020); Decay sugar-rich substrates (Põlme et al. 2020) | Põlme S, et al. 2020. Fungal Diversity 105:1-16 ((https://doi.org/10.1007/s13225-020-00466-2)) |
| Otu17700 | 1 | k__Fungi(100);p__Ascomycota(100);c__Saccharomycetes(100);o__Saccharomycetales(100);f__Metschnikowiaceae(100);g__Metschnikowia(100);g__Metschnikowia_unclassified(100); | Metschnikowia | 13 | Saprotroph | \|Undefined Saprotroph\| | Yeast | NULL | Probable | As nectar-tap_saprotroph (Põlme et al. 2020); Decay sugar-rich substrates (Põlme et al. 2020) | Põlme S, et al. 2020. Fungal Diversity 105:1-16 ((https://doi.org/10.1007/s13225-020-00466-2)) |
| Otu17701 | 1 | k__Fungi(100);p__Ascomycota(100);c__Saccharomycetes(100);o__Saccharomycetales(100);f__Metschnikowiaceae(100);g__Metschnikowia(100);g__Metschnikowia_unclassified(100); | Metschnikowia | 13 | Saprotroph | \|Undefined Saprotroph\| | Yeast | NULL | Probable | As nectar-tap_saprotroph (Põlme et al. 2020); Decay sugar-rich substrates (Põlme et al. 2020) | Põlme S, et al. 2020. Fungal Diversity 105:1-16 ((https://doi.org/10.1007/s13225-020-00466-2)) |
| Otu17702 | 1 | k__Fungi(100);p__Ascomycota(100);c__Saccharomycetes(100);o__Saccharomycetales(100);f__Metschnikowiaceae(100);g__Metschnikowia(100);s__Metschnikowia_pulcherrima(100); | Metschnikowia | 13 | Saprotroph | \|Undefined Saprotroph\| | Yeast | NULL | Probable | As nectar-tap_saprotroph (Põlme et al. 2020); Decay sugar-rich substrates (Põlme et al. 2020) | Põlme S, et al. 2020. Fungal Diversity 105:1-16 ((https://doi.org/10.1007/s13225-020-00466-2)) |
| Otu17703 | 1 | k__Fungi(100);p__Ascomycota(100);c__Saccharomycetes(100);o__Saccharomycetales(100);f__Metschnikowiaceae(100);g__Metschnikowia(100);s__Metschnikowia_pulcherrima(100); | Metschnikowia | 13 | Saprotroph | \|Undefined Saprotroph\| | Yeast | NULL | Probable | As nectar-tap_saprotroph (Põlme et al. 2020); Decay sugar-rich substrates (Põlme et al. 2020) | Põlme S, et al. 2020. Fungal Diversity 105:1-16 ((https://doi.org/10.1007/s13225-020-00466-2)) |
| Otu17704 | 1 | k__Fungi(100);p__Ascomycota(100);c__Saccharomycetes(100);o__Saccharomycetales(100);f__Metschnikowiaceae(100);g__Metschnikowia(100);s__Metschnikowia_pulcherrima(100); | Metschnikowia | 13 | Saprotroph | \|Undefined Saprotroph\| | Yeast | NULL | Probable | As nectar-tap_saprotroph (Põlme et al. 2020); Decay sugar-rich substrates (Põlme et al. 2020) | Põlme S, et al. 2020. Fungal Diversity 105:1-16 ((https://doi.org/10.1007/s13225-020-00466-2)) |
| Otu17705 | 1 | k__Fungi(100);p__Ascomycota(100);c__Saccharomycetes(100);o__Saccharomycetales(100);f__Metschnikowiaceae(100);g__Metschnikowia(100);s__Metschnikowia_pulcherrima(100); | Metschnikowia | 13 | Saprotroph | \|Undefined Saprotroph\| | Yeast | NULL | Probable | As nectar-tap_saprotroph (Põlme et al. 2020); Decay sugar-rich substrates (Põlme et al. 2020) | Põlme S, et al. 2020. Fungal Diversity 105:1-16 ((https://doi.org/10.1007/s13225-020-00466-2)) |
| Otu17706 | 1 | k__Fungi(100);p__Ascomycota(100);c__Saccharomycetes(100);o__Saccharomycetales(100);f__Metschnikowiaceae(100);g__Metschnikowia(100);s__Metschnikowia_pulcherrima(100); | Metschnikowia | 13 | Saprotroph | \|Undefined Saprotroph\| | Yeast | NULL | Probable | As nectar-tap_saprotroph (Põlme et al. 2020); Decay sugar-rich substrates (Põlme et al. 2020) | Põlme S, et al. 2020. Fungal Diversity 105:1-16 ((https://doi.org/10.1007/s13225-020-00466-2)) |
| Otu17707 | 1 | k__Fungi(100);p__Ascomycota(100);c__Saccharomycetes(100);o__Saccharomycetales(100);f__Metschnikowiaceae(100);g__Metschnikowia(100);s__Metschnikowia_pulcherrima(100); | Metschnikowia | 13 | Saprotroph | \|Undefined Saprotroph\| | Yeast | NULL | Probable | As nectar-tap_saprotroph (Põlme et al. 2020); Decay sugar-rich substrates (Põlme et al. 2020) | Põlme S, et al. 2020. Fungal Diversity 105:1-16 ((https://doi.org/10.1007/s13225-020-00466-2)) |
| Otu17708 | 1 | k__Fungi(100);p__Ascomycota(100);c__Saccharomycetes(100);o__Saccharomycetales(100);f__Metschnikowiaceae(100);g__Metschnikowia(100);s__Metschnikowia_pulcherrima(100); | Metschnikowia | 13 | Saprotroph | \|Undefined Saprotroph\| | Yeast | NULL | Probable | As nectar-tap_saprotroph (Põlme et al. 2020); Decay sugar-rich substrates (Põlme et al. 2020) | Põlme S, et al. 2020. Fungal Diversity 105:1-16 ((https://doi.org/10.1007/s13225-020-00466-2)) |
| Otu17709 | 1 | k__Fungi(100);p__Ascomycota(100);c__Saccharomycetes(100);o__Saccharomycetales(100);f__Metschnikowiaceae(100);g__Metschnikowia(100);s__Metschnikowia_pulcherrima(100); | Metschnikowia | 13 | Saprotroph | \|Undefined Saprotroph\| | Yeast | NULL | Probable | As nectar-tap_saprotroph (Põlme et al. 2020); Decay sugar-rich substrates (Põlme et al. 2020) | Põlme S, et al. 2020. Fungal Diversity 105:1-16 ((https://doi.org/10.1007/s13225-020-00466-2)) |
| Otu17711 | 1 | k__Fungi(100);p__Ascomycota(100);c__Saccharomycetes(100);o__Saccharomycetales(100);o__Saccharomycetales_unclassified(100);o__Saccharomycetales_unclassified(100);o__Saccharomycetales_unclassified(100); | Saccharomycetales | 7 | Saprotroph | Undefined Saprotroph | Yeast | NULL | Possible | NULL | Sterkenburg E, et al. 2015. New Phytologist 207:1145-1158 ((https://doi.org/10.1111/nph.13426)) |

| | | | | | | | | | | | |
|---|---|---|---|---|---|---|---|---|---|---|---|
| Otu17712 | 1 | k__Fungi(100);p__Ascomycota(100);c__Saccharomycetes(100);o__Saccharomycetales(100);f__Metschnikowiaceae(100);g__Metschnikowia(100);g__Metschnikowia_unclassified(100); | Metschnikowia | 13 | Saprotroph | \|Undefined Saprotroph\| | Yeast | NULL | Probable | As nectar-tap_saprotroph (Põlme et al. 2020); Decay sugar-rich substrates (Põlme et al. 2020) | Põlme S, et al. 2020. Fungal Diversity 105:1-16 ((https://doi.org/10.1007/s13225-020-00466-2)) |
| Otu17713 | 1 | k__Fungi(100);p__Ascomycota(100);c__Saccharomycetes(100);o__Saccharomycetales(100);f__Metschnikowiaceae(100);g__Metschnikowia(100);s__Metschnikowia_pulcherrima(100); | Metschnikowia | 13 | Saprotroph | \|Undefined Saprotroph\| | Yeast | NULL | Probable | As nectar-tap_saprotroph (Põlme et al. 2020); Decay sugar-rich substrates (Põlme et al. 2020) | Põlme S, et al. 2020. Fungal Diversity 105:1-16 ((https://doi.org/10.1007/s13225-020-00466-2)) |
| Otu17714 | 1 | k__Fungi(100);p__Ascomycota(100);c__Saccharomycetes(100);o__Saccharomycetales(100);f__Metschnikowiaceae(100);g__Metschnikowia(100);s__Metschnikowia_pulcherrima(100); | Metschnikowia | 13 | Saprotroph | \|Undefined Saprotroph\| | Yeast | NULL | Probable | As nectar-tap_saprotroph (Põlme et al. 2020); Decay sugar-rich substrates (Põlme et al. 2020) | Põlme S, et al. 2020. Fungal Diversity 105:1-16 ((https://doi.org/10.1007/s13225-020-00466-2)) |
| Otu17715 | 1 | k__Fungi(100);p__Ascomycota(100);c__Saccharomycetes(100);o__Saccharomycetales(100);f__Metschnikowiaceae(100);g__Metschnikowia(100);g__Metschnikowia_unclassified(100); | Metschnikowia | 13 | Saprotroph | \|Undefined Saprotroph\| | Yeast | NULL | Probable | As nectar-tap_saprotroph (Põlme et al. 2020); Decay sugar-rich substrates (Põlme et al. 2020) | Põlme S, et al. 2020. Fungal Diversity 105:1-16 ((https://doi.org/10.1007/s13225-020-00466-2)) |
| Otu17716 | 1 | k__Fungi(100);p__Ascomycota(100);c__Saccharomycetes(100);o__Saccharomycetales(100);f__Metschnikowiaceae(100);g__Metschnikowia(100);s__Metschnikowia_pulcherrima(100); | Metschnikowia | 13 | Saprotroph | \|Undefined Saprotroph\| | Yeast | NULL | Probable | As nectar-tap_saprotroph (Põlme et al. 2020); Decay sugar-rich substrates (Põlme et al. 2020) | Põlme S, et al. 2020. Fungal Diversity 105:1-16 ((https://doi.org/10.1007/s13225-020-00466-2)) |
| Otu17717 | 1 | k__Fungi(100);p__Ascomycota(100);c__Saccharomycetes(100);o__Saccharomycetales(100);f__Metschnikowiaceae(100);g__Metschnikowia(100);s__Metschnikowia_pulcherrima(100); | Metschnikowia | 13 | Saprotroph | \|Undefined Saprotroph\| | Yeast | NULL | Probable | As nectar-tap_saprotroph (Põlme et al. 2020); Decay sugar-rich substrates (Põlme et al. 2020) | Põlme S, et al. 2020. Fungal Diversity 105:1-16 ((https://doi.org/10.1007/s13225-020-00466-2)) |
| Otu17718 | 1 | k__Fungi(100);p__Ascomycota(100);c__Saccharomycetes(100);o__Saccharomycetales(100);f__Metschnikowiaceae(100);g__Metschnikowia(100);g__Metschnikowia_unclassified(100); | Metschnikowia | 13 | Saprotroph | \|Undefined Saprotroph\| | Yeast | NULL | Probable | As nectar-tap_saprotroph (Põlme et al. 2020); Decay sugar-rich substrates (Põlme et al. 2020) | Põlme S, et al. 2020. Fungal Diversity 105:1-16 ((https://doi.org/10.1007/s13225-020-00466-2)) |
| Otu17719 | 1 | k__Fungi(100);p__Ascomycota(100);c__Saccharomycetes(100);o__Saccharomycetales(100);f__Metschnikowiaceae(100);g__Metschnikowia(100);s__Metschnikowia_pulcherrima(100); | Metschnikowia | 13 | Saprotroph | \|Undefined Saprotroph\| | Yeast | NULL | Probable | As nectar-tap_saprotroph (Põlme et al. 2020); Decay sugar-rich substrates (Põlme et al. 2020) | Põlme S, et al. 2020. Fungal Diversity 105:1-16 ((https://doi.org/10.1007/s13225-020-00466-2)) |
| Otu17720 | 1 | k__Fungi(100);p__Ascomycota(100);c__Saccharomycetes(100);o__Saccharomycetales(100);f__Metschnikowiaceae(100);g__Metschnikowia(100);s__Metschnikowia_pulcherrima(100); | Metschnikowia | 13 | Saprotroph | \|Undefined Saprotroph\| | Yeast | NULL | Probable | As nectar-tap_saprotroph (Põlme et al. 2020); Decay sugar-rich substrates (Põlme et al. 2020) | Põlme S, et al. 2020. Fungal Diversity 105:1-16 ((https://doi.org/10.1007/s13225-020-00466-2)) |
| Otu17721 | 1 | k__Fungi(100);p__Ascomycota(100);c__Saccharomycetes(100);o__Saccharomycetales(100);f__Metschnikowiaceae(100);g__Metschnikowia(100);s__Metschnikowia_pulcherrima(100); | Metschnikowia | 13 | Saprotroph | \|Undefined Saprotroph\| | Yeast | NULL | Probable | As nectar-tap_saprotroph (Põlme et al. 2020); Decay sugar-rich substrates (Põlme et al. 2020) | Põlme S, et al. 2020. Fungal Diversity 105:1-16 ((https://doi.org/10.1007/s13225-020-00466-2)) |
| Otu17722 | 1 | k__Fungi(100);p__Ascomycota(100);c__Saccharomycetes(100);o__Saccharomycetales(100);f__Metschnikowiaceae(100);g__Metschnikowia(100);s__Metschnikowia_pulcherrima(100); | Metschnikowia | 13 | Saprotroph | \|Undefined Saprotroph\| | Yeast | NULL | Probable | As nectar-tap_saprotroph (Põlme et al. 2020); Decay sugar-rich substrates (Põlme et al. 2020) | Põlme S, et al. 2020. Fungal Diversity 105:1-16 ((https://doi.org/10.1007/s13225-020-00466-2)) |
| Otu17723 | 1 | k__Fungi(100);p__Ascomycota(100);c__Saccharomycetes(100);o__Saccharomycetales(100);f__Metschnikowiaceae(100);g__Metschnikowia(100);s__Metschnikowia_pulcherrima(100); | Metschnikowia | 13 | Saprotroph | \|Undefined Saprotroph\| | Yeast | NULL | Probable | As nectar-tap_saprotroph (Põlme et al. 2020); Decay sugar-rich substrates (Põlme et al. 2020) | Põlme S, et al. 2020. Fungal Diversity 105:1-16 ((https://doi.org/10.1007/s13225-020-00466-2)) |
| Otu17724 | 1 | k__Fungi(100);p__Ascomycota(100);c__Saccharomycetes(100);o__Saccharomycetales(100);f__Metschnikowiaceae(100);g__Metschnikowia(100);g__Metschnikowia_unclassified(100); | Metschnikowia | 13 | Saprotroph | \|Undefined Saprotroph\| | Yeast | NULL | Probable | As nectar-tap_saprotroph (Põlme et al. 2020); Decay sugar-rich substrates (Põlme et al. 2020) | Põlme S, et al. 2020. Fungal Diversity 105:1-16 ((https://doi.org/10.1007/s13225-020-00466-2)) |
| Otu17726 | 1 | k__Fungi(100);p__Ascomycota(100);c__Saccharomycetes(100);o__Saccharomycetales(100);f__Metschnikowiaceae(100);g__Metschnikowia(100);g__Metschnikowia_unclassified(100); | Metschnikowia | 13 | Saprotroph | \|Undefined Saprotroph\| | Yeast | NULL | Probable | As nectar-tap_saprotroph (Põlme et al. 2020); Decay sugar-rich substrates (Põlme et al. 2020) | Põlme S, et al. 2020. Fungal Diversity 105:1-16 ((https://doi.org/10.1007/s13225-020-00466-2)) |
| Otu17727 | 1 | k__Fungi(100);p__Ascomycota(100);c__Saccharomycetes(100);o__Saccharomycetales(100);f__Metschnikowiaceae(100);g__Metschnikowia(100);s__Metschnikowia_pulcherrima(100); | Metschnikowia | 13 | Saprotroph | \|Undefined Saprotroph\| | Yeast | NULL | Probable | As nectar-tap_saprotroph (Põlme et al. 2020); Decay sugar-rich substrates (Põlme et al. 2020) | Põlme S, et al. 2020. Fungal Diversity 105:1-16 ((https://doi.org/10.1007/s13225-020-00466-2)) |
| Otu17728 | 1 | k__Fungi(100);p__Ascomycota(100);c__Saccharomycetes(100);o__Saccharomycetales(100);f__Metschnikowiaceae(100);g__Metschnikowia(100);s__Metschnikowia_pulcherrima(100); | Metschnikowia | 13 | Saprotroph | \|Undefined Saprotroph\| | Yeast | NULL | Probable | As nectar-tap_saprotroph (Põlme et al. 2020); Decay sugar-rich substrates (Põlme et al. 2020) | Põlme S, et al. 2020. Fungal Diversity 105:1-16 ((https://doi.org/10.1007/s13225-020-00466-2)) |
| Otu17729 | 1 | k__Fungi(100);p__Ascomycota(100);c__Saccharomycetes(100);o__Saccharomycetales(100);f__Metschnikowiaceae(100);g__Metschnikowia(100);s__Metschnikowia_pulcherrima(100); | Metschnikowia | 13 | Saprotroph | \|Undefined Saprotroph\| | Yeast | NULL | Probable | As nectar-tap_saprotroph (Põlme et al. 2020); Decay sugar-rich substrates (Põlme et al. 2020) | Põlme S, et al. 2020. Fungal Diversity 105:1-16 ((https://doi.org/10.1007/s13225-020-00466-2)) |
| Otu17730 | 1 | k__Fungi(100);p__Ascomycota(100);c__Saccharomycetes(100);o__Saccharomycetales(100);f__Metschnikowiaceae(100);g__Metschnikowia(100);g__Metschnikowia_unclassified(100); | Metschnikowia | 13 | Saprotroph | \|Undefined Saprotroph\| | Yeast | NULL | Probable | As nectar-tap_saprotroph (Põlme et al. 2020); Decay sugar-rich substrates (Põlme et al. 2020) | Põlme S, et al. 2020. Fungal Diversity 105:1-16 ((https://doi.org/10.1007/s13225-020-00466-2)) |
| Otu17731 | 1 | k__Fungi(100);p__Ascomycota(100);c__Saccharomycetes(100);o__Saccharomycetales(100);f__Metschnikowiaceae(100);g__Metschnikowia(100);s__Metschnikowia_pulcherrima(100); | Metschnikowia | 13 | Saprotroph | \|Undefined Saprotroph\| | Yeast | NULL | Probable | As nectar-tap_saprotroph (Põlme et al. 2020); Decay sugar-rich substrates (Põlme et al. 2020) | Põlme S, et al. 2020. Fungal Diversity 105:1-16 ((https://doi.org/10.1007/s13225-020-00466-2)) |
| Otu17732 | 1 | k__Fungi(100);p__Ascomycota(100);c__Saccharomycetes(100);o__Saccharomycetales(100);f__Metschnikowiaceae(100);g__Metschnikowia(100);g__Metschnikowia_unclassified(100); | Metschnikowia | 13 | Saprotroph | \|Undefined Saprotroph\| | Yeast | NULL | Probable | As nectar-tap_saprotroph (Põlme et al. 2020); Decay sugar-rich substrates (Põlme et al. 2020) | Põlme S, et al. 2020. Fungal Diversity 105:1-16 ((https://doi.org/10.1007/s13225-020-00466-2)) |
| Otu17733 | 1 | k__Fungi(100);p__Ascomycota(100);c__Saccharomycetes(100);o__Saccharomycetales(100);f__Metschnikowiaceae(100);g__Metschnikowia(100);g__Metschnikowia_unclassified(100); | Metschnikowia | 13 | Saprotroph | \|Undefined Saprotroph\| | Yeast | NULL | Probable | As nectar-tap_saprotroph (Põlme et al. 2020); Decay sugar-rich substrates (Põlme et al. 2020) | Põlme S, et al. 2020. Fungal Diversity 105:1-16 ((https://doi.org/10.1007/s13225-020-00466-2)) |
| Otu17734 | 1 | k__Fungi(100);p__Ascomycota(100);c__Saccharomycetes(100);o__Saccharomycetales(100);f__Metschnikowiaceae(100);g__Metschnikowia(100);s__Metschnikowia_pulcherrima(100); | Metschnikowia | 13 | Saprotroph | \|Undefined Saprotroph\| | Yeast | NULL | Probable | As nectar-tap_saprotroph (Põlme et al. 2020); Decay sugar-rich substrates (Põlme et al. 2020) | Põlme S, et al. 2020. Fungal Diversity 105:1-16 ((https://doi.org/10.1007/s13225-020-00466-2)) |
| Otu17735 | 1 | k__Fungi(100);p__Ascomycota(100);c__Saccharomycetes(100);o__Saccharomycetales(100);f__Metschnikowiaceae(100);g__Metschnikowia(100);s__Metschnikowia_pulcherrima(100); | Metschnikowia | 13 | Saprotroph | \|Undefined Saprotroph\| | Yeast | NULL | Probable | As nectar-tap_saprotroph (Põlme et al. 2020); Decay sugar-rich substrates (Põlme et al. 2020) | Põlme S, et al. 2020. Fungal Diversity 105:1-16 ((https://doi.org/10.1007/s13225-020-00466-2)) |
| Otu17736 | 1 | k__Fungi(100);p__Ascomycota(100);c__Saccharomycetes(100);o__Saccharomycetales(100);f__Metschnikowiaceae(100);g__Metschnikowia(100);s__Metschnikowia_pulcherrima(100); | Metschnikowia | 13 | Saprotroph | \|Undefined Saprotroph\| | Yeast | NULL | Probable | As nectar-tap_saprotroph (Põlme et al. 2020); Decay sugar-rich substrates (Põlme et al. 2020) | Põlme S, et al. 2020. Fungal Diversity 105:1-16 ((https://doi.org/10.1007/s13225-020-00466-2)) |
| Otu17737 | 1 | k__Fungi(100);p__Ascomycota(100);c__Saccharomycetes(100);o__Saccharomycetales(100);f__Metschnikowiaceae(100);g__Metschnikowia(100);s__Metschnikowia_pulcherrima(100); | Metschnikowia | 13 | Saprotroph | \|Undefined Saprotroph\| | Yeast | NULL | Probable | As nectar-tap_saprotroph (Põlme et al. 2020); Decay sugar-rich substrates (Põlme et al. 2020) | Põlme S, et al. 2020. Fungal Diversity 105:1-16 ((https://doi.org/10.1007/s13225-020-00466-2)) |
| Otu17738 | 1 | k__Fungi(100);p__Ascomycota(100);c__Saccharomycetes(100);o__Saccharomycetales(100);f__Metschnikowiaceae(100);g__Metschnikowia(100);s__Metschnikowia_pulcherrima(100); | Metschnikowia | 13 | Saprotroph | \|Undefined Saprotroph\| | Yeast | NULL | Probable | As nectar-tap_saprotroph (Põlme et al. 2020); Decay sugar-rich substrates (Põlme et al. 2020) | Põlme S, et al. 2020. Fungal Diversity 105:1-16 ((https://doi.org/10.1007/s13225-020-00466-2)) |
| Otu17739 | 1 | k__Fungi(100);p__Ascomycota(100);c__Saccharomycetes(100);o__Saccharomycetales(100);f__Metschnikowiaceae(100);g__Metschnikowia(100);s__Metschnikowia_pulcherrima(100); | Metschnikowia | 13 | Saprotroph | \|Undefined Saprotroph\| | Yeast | NULL | Probable | As nectar-tap_saprotroph (Põlme et al. 2020); Decay sugar-rich substrates (Põlme et al. 2020) | Põlme S, et al. 2020. Fungal Diversity 105:1-16 ((https://doi.org/10.1007/s13225-020-00466-2)) |
| Otu17740 | 1 | k__Fungi(100);p__Ascomycota(100);c__Saccharomycetes(100);o__Saccharomycetales(100);f__Metschnikowiaceae(100);g__Metschnikowia(100);g__Metschnikowia_unclassified(100); | Metschnikowia | 13 | Saprotroph | \|Undefined Saprotroph\| | Yeast | NULL | Probable | As nectar-tap_saprotroph (Põlme et al. 2020); Decay sugar-rich substrates (Põlme et al. 2020) | Põlme S, et al. 2020. Fungal Diversity 105:1-16 ((https://doi.org/10.1007/s13225-020-00466-2)) |
| Otu17741 | 1 | k__Fungi(100);p__Ascomycota(100);c__Saccharomycetes(100);o__Saccharomycetales(100);f__Metschnikowiaceae(100);g__Metschnikowia(100);s__Metschnikowia_pulcherrima(100); | Metschnikowia | 13 | Saprotroph | \|Undefined Saprotroph\| | Yeast | NULL | Probable | As nectar-tap_saprotroph (Põlme et al. 2020); Decay sugar-rich substrates (Põlme et al. 2020) | Põlme S, et al. 2020. Fungal Diversity 105:1-16 ((https://doi.org/10.1007/s13225-020-00466-2)) |
| Otu17742 | 1 | k__Fungi(100);p__Zygomycota(100);c__Zygomycota_class_Incertae_sedis(100);o__Mucorales(100);f__Mucoraceae(100);g__Mucor(100);g__Mucor_unclassified(100); | Mucor | 13 | Saprotroph | \|Undefined Saprotroph\| | Microfungus | NULL | Probable | As soil_saprotroph (Põlme et al. 2020); Decay soil (Põlme et al. 2020) | Tedersoo L, et al. 2014. Science 346:e1256688 ((https://doi.org/10.1126/science.1256688)); Põlme S, et al. 2020. Fungal Diversity 105:1-16 ((https://doi.org/10.1007/s13225-020-00466-2)) |
| Otu17743 | 1 | k__Fungi(100);p__Ascomycota(100);c__Saccharomycetes(100);o__Saccharomycetales(100);f__Metschnikowiaceae(100);g__Metschnikowia(100);s__Metschnikowia_pulcherrima(100); | Metschnikowia | 13 | Saprotroph | \|Undefined Saprotroph\| | Yeast | NULL | Probable | As nectar-tap_saprotroph (Põlme et al. 2020); Decay sugar-rich substrates (Põlme et al. 2020) | Põlme S, et al. 2020. Fungal Diversity 105:1-16 ((https://doi.org/10.1007/s13225-020-00466-2)) |
| Otu17744 | 1 | k__Fungi(100);p__Ascomycota(100);c__Saccharomycetes(100);o__Saccharomycetales(100);f__Metschnikowiaceae(100);g__Metschnikowia(100);s__Metschnikowia_pulcherrima(100); | Metschnikowia | 13 | Saprotroph | \|Undefined Saprotroph\| | Yeast | NULL | Probable | As nectar-tap_saprotroph (Põlme et al. 2020); Decay sugar-rich substrates (Põlme et al. 2020) | Põlme S, et al. 2020. Fungal Diversity 105:1-16 ((https://doi.org/10.1007/s13225-020-00466-2)) |
| Otu17745 | 1 | k__Fungi(100);p__Ascomycota(100);c__Saccharomycetes(100);o__Saccharomycetales(100);f__Metschnikowiaceae(100);g__Metschnikowia(100);s__Metschnikowia_pulcherrima(100); | Metschnikowia | 13 | Saprotroph | \|Undefined Saprotroph\| | Yeast | NULL | Probable | As nectar-tap_saprotroph (Põlme et al. 2020); Decay sugar-rich substrates (Põlme et al. 2020) | Põlme S, et al. 2020. Fungal Diversity 105:1-16 ((https://doi.org/10.1007/s13225-020-00466-2)) |
| Otu17746 | 1 | k__Fungi(100);p__Ascomycota(100);c__Saccharomycetes(100);o__Saccharomycetales(100);f__Metschnikowiaceae(100);g__Metschnikowia(100);g__Metschnikowia_unclassified(100); | Metschnikowia | 13 | Saprotroph | \|Undefined Saprotroph\| | Yeast | NULL | Probable | As nectar-tap_saprotroph (Põlme et al. 2020); Decay sugar-rich substrates (Põlme et al. 2020) | Põlme S, et al. 2020. Fungal Diversity 105:1-16 ((https://doi.org/10.1007/s13225-020-00466-2)) |
| Otu17747 | 1 | k__Fungi(100);p__Ascomycota(100);c__Saccharomycetes(100);o__Saccharomycetales(100);f__Metschnikowiaceae(100);g__Metschnikowia(100);s__Metschnikowia_pulcherrima(100); | Metschnikowia | 13 | Saprotroph | \|Undefined Saprotroph\| | Yeast | NULL | Probable | As nectar-tap_saprotroph (Põlme et al. 2020); Decay sugar-rich substrates (Põlme et al. 2020) | Põlme S, et al. 2020. Fungal Diversity 105:1-16 ((https://doi.org/10.1007/s13225-020-00466-2)) |
| Otu17748 | 1 | k__Fungi(100);p__Ascomycota(100);c__Saccharomycetes(100);o__Saccharomycetales(100);f__Metschnikowiaceae(100);g__Metschnikowia(100);g__Metschnikowia_unclassified(100); | Metschnikowia | 13 | Saprotroph | \|Undefined Saprotroph\| | Yeast | NULL | Probable | As nectar-tap_saprotroph (Põlme et al. 2020); Decay sugar-rich substrates (Põlme et al. 2020) | Põlme S, et al. 2020. Fungal Diversity 105:1-16 ((https://doi.org/10.1007/s13225-020-00466-2)) |
| Otu17749 | 1 | k__Fungi(100);p__Ascomycota(100);c__Saccharomycetes(100);o__Saccharomycetales(100);f__Metschnikowiaceae(100);g__Metschnikowia(100);s__Metschnikowia_pulcherrima(100); | Metschnikowia | 13 | Saprotroph | \|Undefined Saprotroph\| | Yeast | NULL | Probable | As nectar-tap_saprotroph (Põlme et al. 2020); Decay sugar-rich substrates (Põlme et al. 2020) | Põlme S, et al. 2020. Fungal Diversity 105:1-16 ((https://doi.org/10.1007/s13225-020-00466-2)) |
| Otu17750 | 1 | k__Fungi(100);p__Ascomycota(100);c__Saccharomycetes(100);o__Saccharomycetales(100);f__Metschnikowiaceae(100);g__Metschnikowia(100);g__Metschnikowia_unclassified(100); | Metschnikowia | 13 | Saprotroph | \|Undefined Saprotroph\| | Yeast | NULL | Probable | As nectar-tap_saprotroph (Põlme et al. 2020); Decay sugar-rich substrates (Põlme et al. 2020) | Põlme S, et al. 2020. Fungal Diversity 105:1-16 ((https://doi.org/10.1007/s13225-020-00466-2)) |
| Otu17751 | 1 | k__Fungi(100);p__Ascomycota(100);c__Saccharomycetes(100);o__Saccharomycetales(100);f__Metschnikowiaceae(100);g__Metschnikowia(100);s__Metschnikowia_pulcherrima(100); | Metschnikowia | 13 | Saprotroph | \|Undefined Saprotroph\| | Yeast | NULL | Probable | As nectar-tap_saprotroph (Põlme et al. 2020); Decay sugar-rich substrates (Põlme et al. 2020) | Põlme S, et al. 2020. Fungal Diversity 105:1-16 ((https://doi.org/10.1007/s13225-020-00466-2)) |
| Otu17752 | 1 | k__Fungi(100);p__Ascomycota(100);c__Saccharomycetes(100);o__Saccharomycetales(100);f__Metschnikowiaceae(100);g__Metschnikowia(100);s__Metschnikowia_pulcherrima(100); | Metschnikowia | 13 | Saprotroph | \|Undefined Saprotroph\| | Yeast | NULL | Probable | As nectar-tap_saprotroph (Põlme et al. 2020); Decay sugar-rich substrates (Põlme et al. 2020) | Põlme S, et al. 2020. Fungal Diversity 105:1-16 ((https://doi.org/10.1007/s13225-020-00466-2)) |
| Otu17753 | 1 | k__Fungi(100);p__Ascomycota(100);c__Saccharomycetes(100);o__Saccharomycetales(100);f__Metschnikowiaceae(100);g__Metschnikowia(100);s__Metschnikowia_pulcherrima(100); | Metschnikowia | 13 | Saprotroph | \|Undefined Saprotroph\| | Yeast | NULL | Probable | As nectar-tap_saprotroph (Põlme et al. 2020); Decay sugar-rich substrates (Põlme et al. 2020) | Põlme S, et al. 2020. Fungal Diversity 105:1-16 ((https://doi.org/10.1007/s13225-020-00466-2)) |
| Otu17754 | 1 | k__Fungi(100);p__Ascomycota(100);c__Saccharomycetes(100);o__Saccharomycetales(100);f__Metschnikowiaceae(100);g__Metschnikowia(100);s__Metschnikowia_pulcherrima(100); | Metschnikowia | 13 | Saprotroph | \|Undefined Saprotroph\| | Yeast | NULL | Probable | As nectar-tap_saprotroph (Põlme et al. 2020); Decay sugar-rich substrates (Põlme et al. 2020) | Põlme S, et al. 2020. Fungal Diversity 105:1-16 ((https://doi.org/10.1007/s13225-020-00466-2)) |
| Otu17755 | 1 | k__Fungi(100);p__Ascomycota(100);c__Saccharomycetes(100);o__Saccharomycetales(100);f__Metschnikowiaceae(100);g__Metschnikowia(100);s__Metschnikowia_pulcherrima(100); | Metschnikowia | 13 | Saprotroph | \|Undefined Saprotroph\| | Yeast | NULL | Probable | As nectar-tap_saprotroph (Põlme et al. 2020); Decay sugar-rich substrates (Põlme et al. 2020) | Põlme S, et al. 2020. Fungal Diversity 105:1-16 ((https://doi.org/10.1007/s13225-020-00466-2)) |
| Otu17756 | 1 | k__Fungi(100);p__Ascomycota(100);c__Saccharomycetes(100);o__Saccharomycetales(100);f__Metschnikowiaceae(100);g__Metschnikowia(100);s__Metschnikowia_pulcherrima(100); | Metschnikowia | 13 | Saprotroph | \|Undefined Saprotroph\| | Yeast | NULL | Probable | As nectar-tap_saprotroph (Põlme et al. 2020); Decay sugar-rich substrates (Põlme et al. 2020) | Põlme S, et al. 2020. Fungal Diversity 105:1-16 ((https://doi.org/10.1007/s13225-020-00466-2)) |
| Otu17757 | 1 | k__Fungi(100);p__Ascomycota(100);c__Saccharomycetes(100);o__Saccharomycetales(100);f__Metschnikowiaceae(100);g__Metschnikowia(100);s__Metschnikowia_pulcherrima(100); | Metschnikowia | 13 | Saprotroph | \|Undefined Saprotroph\| | Yeast | NULL | Probable | As nectar-tap_saprotroph (Põlme et al. 2020); Decay sugar-rich substrates (Põlme et al. 2020) | Põlme S, et al. 2020. Fungal Diversity 105:1-16 ((https://doi.org/10.1007/s13225-020-00466-2)) |
| Otu17759 | 1 | k__Fungi(100);p__Ascomycota(100);c__Saccharomycetes(100);o__Saccharomycetales(100);f__Metschnikowiaceae(100);g__Metschnikowia(100);s__Metschnikowia_pulcherrima(100); | Metschnikowia | 13 | Saprotroph | \|Undefined Saprotroph\| | Yeast | NULL | Probable | As nectar-tap_saprotroph (Põlme et al. 2020); Decay sugar-rich substrates (Põlme et al. 2020) | Põlme S, et al. 2020. Fungal Diversity 105:1-16 ((https://doi.org/10.1007/s13225-020-00466-2)) |
| Otu17760 | 1 | k__Fungi(100);p__Ascomycota(100);c__Saccharomycetes(100);o__Saccharomycetales(100);f__Metschnikowiaceae(100);g__Metschnikowia(100);s__Metschnikowia_pulcherrima(100); | Metschnikowia | 13 | Saprotroph | \|Undefined Saprotroph\| | Yeast | NULL | Probable | As nectar-tap_saprotroph (Põlme et al. 2020); Decay sugar-rich substrates (Põlme et al. 2020) | Põlme S, et al. 2020. Fungal Diversity 105:1-16 ((https://doi.org/10.1007/s13225-020-00466-2)) |
| Otu17761 | 1 | k__Fungi(100);p__Ascomycota(100);c__Saccharomycetes(100);o__Saccharomycetales(100);f__Metschnikowiaceae(100);g__Metschnikowia(100);s__Metschnikowia_pulcherrima(100); | Metschnikowia | 13 | Saprotroph | \|Undefined Saprotroph\| | Yeast | NULL | Probable | As nectar-tap_saprotroph (Põlme et al. 2020); Decay sugar-rich substrates (Põlme et al. 2020) | Põlme S, et al. 2020. Fungal Diversity 105:1-16 ((https://doi.org/10.1007/s13225-020-00466-2)) |
| Otu17762 | 1 | k__Fungi(100);p__Ascomycota(100);c__Saccharomycetes(100);o__Saccharomycetales(100);f__Metschnikowiaceae(100);g__Metschnikowia(100);s__Metschnikowia_pulcherrima(100); | Metschnikowia | 13 | Saprotroph | \|Undefined Saprotroph\| | Yeast | NULL | Probable | As nectar-tap_saprotroph (Põlme et al. 2020); Decay sugar-rich substrates (Põlme et al. 2020) | Põlme S, et al. 2020. Fungal Diversity 105:1-16 ((https://doi.org/10.1007/s13225-020-00466-2)) |
| Otu17763 | 1 | k__Fungi(100);p__Ascomycota(100);c__Saccharomycetes(100);o__Saccharomycetales(100);f__Metschnikowiaceae(100);g__Metschnikowia(100);s__Metschnikowia_pulcherrima(100); | Metschnikowia | 13 | Saprotroph | \|Undefined Saprotroph\| | Yeast | NULL | Probable | As nectar-tap_saprotroph (Põlme et al. 2020); Decay sugar-rich substrates (Põlme et al. 2020) | Põlme S, et al. 2020. Fungal Diversity 105:1-16 ((https://doi.org/10.1007/s13225-020-00466-2)) |
| Otu17764 | 1 | k__Fungi(100);p__Ascomycota(100);c__Saccharomycetes(100);o__Saccharomycetales(100);f__Metschnikowiaceae(100);g__Metschnikowia(100);s__Metschnikowia_pulcherrima(100); | Metschnikowia | 13 | Saprotroph | \|Undefined Saprotroph\| | Yeast | NULL | Probable | As nectar-tap_saprotroph (Põlme et al. 2020); Decay sugar-rich substrates (Põlme et al. 2020) | Põlme S, et al. 2020. Fungal Diversity 105:1-16 ((https://doi.org/10.1007/s13225-020-00466-2)) |
| Otu17765 | 1 | k__Fungi(100);p__Ascomycota(100);c__Saccharomycetes(100);o__Saccharomycetales(100);f__Metschnikowiaceae(100);g__Metschnikowia(100);s__Metschnikowia_pulcherrima(100); | Metschnikowia | 13 | Saprotroph | \|Undefined Saprotroph\| | Yeast | NULL | Probable | As nectar-tap_saprotroph (Põlme et al. 2020); Decay sugar-rich substrates (Põlme et al. 2020) | Põlme S, et al. 2020. Fungal Diversity 105:1-16 ((https://doi.org/10.1007/s13225-020-00466-2)) |
| Otu17766 | 1 | k__Fungi(100);p__Ascomycota(100);c__Saccharomycetes(100);o__Saccharomycetales(100);f__Metschnikowiaceae(100);g__Metschnikowia(100);s__Metschnikowia_pulcherrima(100); | Metschnikowia | 13 | Saprotroph | \|Undefined Saprotroph\| | Yeast | NULL | Probable | As nectar-tap_saprotroph (Põlme et al. 2020); Decay sugar-rich substrates (Põlme et al. 2020) | Põlme S, et al. 2020. Fungal Diversity 105:1-16 ((https://doi.org/10.1007/s13225-020-00466-2)) |
| Otu17767 | 1 | k__Fungi(100);p__Ascomycota(100);c__Saccharomycetes(100);o__Saccharomycetales(100);f__Metschnikowiaceae(100);g__Metschnikowia(100);s__Metschnikowia_pulcherrima(100); | Metschnikowia | 13 | Saprotroph | \|Undefined Saprotroph\| | Yeast | NULL | Probable | As nectar-tap_saprotroph (Põlme et al. 2020); Decay sugar-rich substrates (Põlme et al. 2020) | Põlme S, et al. 2020. Fungal Diversity 105:1-16 ((https://doi.org/10.1007/s13225-020-00466-2)) |
| Otu17768 | 1 | k__Fungi(100);p__Ascomycota(100);c__Saccharomycetes(100);o__Saccharomycetales(100);f__Metschnikowiaceae(100);g__Metschnikowia(100);s__Metschnikowia_pulcherrima(100); | Metschnikowia | 13 | Saprotroph | \|Undefined Saprotroph\| | Yeast | NULL | Probable | As nectar-tap_saprotroph (Põlme et al. 2020); Decay sugar-rich substrates (Põlme et al. 2020) | Põlme S, et al. 2020. Fungal Diversity 105:1-16 ((https://doi.org/10.1007/s13225-020-00466-2)) |
| Otu17769 | 1 | k__Fungi(100);p__Ascomycota(100);c__Saccharomycetes(100);o__Saccharomycetales(100);f__Metschnikowiaceae(100);g__Metschnikowia(100);s__Metschnikowia_pulcherrima(100); | Metschnikowia | 13 | Saprotroph | \|Undefined Saprotroph\| | Yeast | NULL | Probable | As nectar-tap_saprotroph (Põlme et al. 2020); Decay sugar-rich substrates (Põlme et al. 2020) | Põlme S, et al. 2020. Fungal Diversity 105:1-16 ((https://doi.org/10.1007/s13225-020-00466-2)) |
| Otu17770 | 1 | k__Fungi(100);p__Ascomycota(100);c__Saccharomycetes(100);o__Saccharomycetales(100);f__Metschnikowiaceae(100);g__Metschnikowia(100);s__Metschnikowia_pulcherrima(100); | Metschnikowia | 13 | Saprotroph | \|Undefined Saprotroph\| | Yeast | NULL | Probable | As nectar-tap_saprotroph (Põlme et al. 2020); Decay sugar-rich substrates (Põlme et al. 2020) | Põlme S, et al. 2020. Fungal Diversity 105:1-16 ((https://doi.org/10.1007/s13225-020-00466-2)) |
| Otu17771 | 1 | k__Fungi(100);p__Ascomycota(100);c__Saccharomycetes(100);o__Saccharomycetales(100);f__Metschnikowiaceae(100);g__Metschnikowia(100);s__Metschnikowia_pulcherrima(100); | Metschnikowia | 13 | Saprotroph | \|Undefined Saprotroph\| | Yeast | NULL | Probable | As nectar-tap_saprotroph (Põlme et al. 2020); Decay sugar-rich substrates (Põlme et al. 2020) | Põlme S, et al. 2020. Fungal Diversity 105:1-16 ((https://doi.org/10.1007/s13225-020-00466-2)) |
| Otu17772 | 1 | k__Fungi(100);p__Ascomycota(100);c__Saccharomycetes(100);o__Saccharomycetales(100);f__Metschnikowiaceae(100);g__Metschnikowia(100);s__Metschnikowia_pulcherrima(100); | Metschnikowia | 13 | Saprotroph | \|Undefined Saprotroph\| | Yeast | NULL | Probable | As nectar-tap_saprotroph (Põlme et al. 2020); Decay sugar-rich substrates (Põlme et al. 2020) | Põlme S, et al. 2020. Fungal Diversity 105:1-16 ((https://doi.org/10.1007/s13225-020-00466-2)) |
| Otu17773 | 1 | k__Fungi(100);p__Ascomycota(100);c__Saccharomycetes(100);o__Saccharomycetales(100);f__Metschnikowiaceae(100);g__Metschnikowia(100);s__Metschnikowia_pulcherrima(100); | Metschnikowia | 13 | Saprotroph | \|Undefined Saprotroph\| | Yeast | NULL | Probable | As nectar-tap_saprotroph (Põlme et al. 2020); Decay sugar-rich substrates (Põlme et al. 2020) | Põlme S, et al. 2020. Fungal Diversity 105:1-16 ((https://doi.org/10.1007/s13225-020-00466-2)) |
| Otu17774 | 1 | k__Fungi(100);p__Basidiomycota(100);c__Agaricomycetes(100);o__Hymenochaetales(100);f__Schizoporaceae(100);g__Hyphodontia(100);s__Hyphodontia_sp_DLL2011_1(100); | Hyphodontia | 13 | Saprotroph | Undefined Saprotroph-\|Wood Saprotroph\| | Corticioid | White Rot | Probable | As white_rot (Põlme et al. 2020); Decay wood (Põlme et al. 2020) | Tedersoo L, et al. 2014. Science 346:e1256688 ((https://doi.org/10.1126/science.1256688)); Põlme S, et al. 2020. Fungal Diversity 105:1-16 ((https://doi.org/10.1007/s13225-020-00466-2)) |
| Otu17775 | 1 | k__Fungi(100);p__Ascomycota(100);c__Saccharomycetes(100);o__Saccharomycetales(100);f__Metschnikowiaceae(100);g__Metschnikowia(100);s__Metschnikowia_pulcherrima(100); | Metschnikowia | 13 | Saprotroph | \|Undefined Saprotroph\| | Yeast | NULL | Probable | As nectar-tap_saprotroph (Põlme et al. 2020); Decay sugar-rich substrates (Põlme et al. 2020) | Põlme S, et al. 2020. Fungal Diversity 105:1-16 ((https://doi.org/10.1007/s13225-020-00466-2)) |
| Otu17776 | 1 | k__Fungi(100);p__Ascomycota(100);c__Saccharomycetes(100);o__Saccharomycetales(100);f__Metschnikowiaceae(100);g__Metschnikowia(100);s__Metschnikowia_pulcherrima(100); | Metschnikowia | 13 | Saprotroph | \|Undefined Saprotroph\| | Yeast | NULL | Probable | As nectar-tap_saprotroph (Põlme et al. 2020); Decay sugar-rich substrates (Põlme et al. 2020) | Põlme S, et al. 2020. Fungal Diversity 105:1-16 ((https://doi.org/10.1007/s13225-020-00466-2)) |
| Otu17777 | 1 | k__Fungi(100);p__Ascomycota(100);c__Saccharomycetes(100);o__Saccharomycetales(100);f__Metschnikowiaceae(100);g__Metschnikowia(100);s__Metschnikowia_pulcherrima(100); | Metschnikowia | 13 | Saprotroph | \|Undefined Saprotroph\| | Yeast | NULL | Probable | As nectar-tap_saprotroph (Põlme et al. 2020); Decay sugar-rich substrates (Põlme et al. 2020) | Põlme S, et al. 2020. Fungal Diversity 105:1-16 ((https://doi.org/10.1007/s13225-020-00466-2)) |
| Otu17778 | 1 | k__Fungi(100);p__Ascomycota(100);c__Saccharomycetes(100);o__Saccharomycetales(100);f__Metschnikowiaceae(100);g__Metschnikowia(100);g__Metschnikowia_unclassified(100); | Metschnikowia | 13 | Saprotroph | \|Undefined Saprotroph\| | Yeast | NULL | Probable | As nectar-tap_saprotroph (Põlme et al. 2020); Decay sugar-rich substrates (Põlme et al. 2020) | Põlme S, et al. 2020. Fungal Diversity 105:1-16 ((https://doi.org/10.1007/s13225-020-00466-2)) |
| Otu17779 | 1 | k__Fungi(100);p__Ascomycota(100);c__Saccharomycetes(100);o__Saccharomycetales(100);f__Metschnikowiaceae(100);g__Metschnikowia(100);g__Metschnikowia_unclassified(100); | Metschnikowia | 13 | Saprotroph | \|Undefined Saprotroph\| | Yeast | NULL | Probable | As nectar-tap_saprotroph (Põlme et al. 2020); Decay sugar-rich substrates (Põlme et al. 2020) | Põlme S, et al. 2020. Fungal Diversity 105:1-16 ((https://doi.org/10.1007/s13225-020-00466-2)) |
| Otu17780 | 1 | k__Fungi(100);p__Ascomycota(100);c__Saccharomycetes(100);o__Saccharomycetales(100);f__Metschnikowiaceae(100);g__Metschnikowia(100);s__Metschnikowia_pulcherrima(100); | Metschnikowia | 13 | Saprotroph | \|Undefined Saprotroph\| | Yeast | NULL | Probable | As nectar-tap_saprotroph (Põlme et al. 2020); Decay sugar-rich substrates (Põlme et al. 2020) | Põlme S, et al. 2020. Fungal Diversity 105:1-16 ((https://doi.org/10.1007/s13225-020-00466-2)) |
| Otu17782 | 1 | k__Fungi(100);p__Ascomycota(100);c__Saccharomycetes(100);o__Saccharomycetales(100);f__Metschnikowiaceae(100);g__Metschnikowia(100);s__Metschnikowia_pulcherrima(100); | Metschnikowia | 13 | Saprotroph | \|Undefined Saprotroph\| | Yeast | NULL | Probable | As nectar-tap_saprotroph (Põlme et al. 2020); Decay sugar-rich substrates (Põlme et al. 2020) | Põlme S, et al. 2020. Fungal Diversity 105:1-16 ((https://doi.org/10.1007/s13225-020-00466-2)) |
| Otu17785 | 1 | k__Fungi(100);p__Zygomycota(100);c__Zygomycota_class_Incertae_sedis(100);o__Mortierellales(100);f__Mortierellaceae(100);g__unclassified_Mortierellaceae(100);s__Mortierellaceae_sp(100); | Mortierellaceae | 9 | Saprotroph-Symbiotroph | Endophyte-Litter Saprotroph-Soil Saprotroph-Undefined Saprotroph | Microfungus | NULL | Possible | Saprobic in soil and thus frequently isolated from soil and material contaminated with soil such as hay or rotten silage and spoil tips (Cannon & Kirk 2007) | Cannon PF, Kirk PM. 2007. Fungal Families of the World. CAB International, Cambridge (ISBN: 978-0851998275); Purahong W, et al. 2016. Molecular Ecology 25:4059-4074 ((https://doi.org/10.1111/mec.13739)) |
| Otu17786 | 1 | k__Fungi(100);p__Basidiomycota(100);c__Microbotryomycetes(100);o__Sporidiobolales(100);f__Sporidiobolales_family_Incertae_sedis(100);g__Rhodotorula(100);s__Rhodotorula_sp_JCM_11351(100); | Rhodotorula | 13 | Pathotroph-Saprotroph-Symbiotroph | Animal Pathogen-Animal Symbiotroph-Endophyte-Plant Pathogen-\|Undefined Saprotroph\| | Yeast | NULL | Probable | As foliar_endophyte (Põlme et al. 2020); Opportunistic human pathogen | Põlme S, et al. 2020. Fungal Diversity 105:1-16 ((https://doi.org/10.1007/s13225-020-00466-2)); https://www.cdc.gov ((https://www.cdc.gov)) |
| Otu17788 | 1 | k__Fungi(100);p__Ascomycota(100);c__Eurotiomycetes(100);o__Chaetothyriales(100);f__Herpotrichiellaceae(100);f__Herpotrichiellaceae_unclassified(100);f__Herpotrichiellaceae_unclassified(100); | Herpotrichiellaceae | 9 | Pathotroph-Saprotroph | Animal Pathogen-Fungal Parasite-Undefined Saprotroph | Facultative Yeast-Microfungus | NULL | Probable | Mostly saprobic on plants or other fungi, isolated from soil etc., and from medical sources (Cannon & Kirk 2007) | Cannon PF, Kirk PM. 2007. Fungal Families of the World. CAB International, Cambridge (ISBN: 978-0851998275) |
| Otu17789 | 1 | k__Fungi(100);p__Ascomycota(100);c__Saccharomycetes(100);o__Saccharomycetales(100);f__Saccharomycetaceae(100);g__Kazachstania(100);s__Kazachstania_intestinalis(100); | Kazachstania | 13 | Saprotroph | \|Undefined Saprotroph\| | Yeast | NULL | Probable | As nectar-tap_saprotroph (Põlme et al. 2020); Decay sugar-rich substrates (Põlme et al. 2020) | Põlme S, et al. 2020. Fungal Diversity 105:1-16 ((https://doi.org/10.1007/s13225-020-00466-2)) |
| Otu17790 | 1 | k__Fungi(100);p__Ascomycota(100);c__Saccharomycetes(100);o__Saccharomycetales(100);f__Metschnikowiaceae(100);g__Metschnikowia(100);s__Metschnikowia_pulcherrima(100); | Metschnikowia | 13 | Saprotroph | \|Undefined Saprotroph\| | Yeast | NULL | Probable | As nectar-tap_saprotroph (Põlme et al. 2020); Decay sugar-rich substrates (Põlme et al. 2020) | Põlme S, et al. 2020. Fungal Diversity 105:1-16 ((https://doi.org/10.1007/s13225-020-00466-2)) |
| Otu17791 | 1 | k__Fungi(100);p__Ascomycota(100);c__Saccharomycetes(100);o__Saccharomycetales(100);f__Metschnikowiaceae(100);g__Metschnikowia(100);s__Metschnikowia_pulcherrima(100); | Metschnikowia | 13 | Saprotroph | \|Undefined Saprotroph\| | Yeast | NULL | Probable | As nectar-tap_saprotroph (Põlme et al. 2020); Decay sugar-rich substrates (Põlme et al. 2020) | Põlme S, et al. 2020. Fungal Diversity 105:1-16 ((https://doi.org/10.1007/s13225-020-00466-2)) |
| Otu17792 | 1 | k__Fungi(100);p__Ascomycota(100);c__Saccharomycetes(100);o__Saccharomycetales(100);f__Metschnikowiaceae(100);g__Metschnikowia(100);s__Metschnikowia_pulcherrima(100); | Metschnikowia | 13 | Saprotroph | \|Undefined Saprotroph\| | Yeast | NULL | Probable | As nectar-tap_saprotroph (Põlme et al. 2020); Decay sugar-rich substrates (Põlme et al. 2020) | Põlme S, et al. 2020. Fungal Diversity 105:1-16 ((https://doi.org/10.1007/s13225-020-00466-2)) |
| Otu17793 | 1 | k__Fungi(100);p__Ascomycota(100);c__Saccharomycetes(100);o__Saccharomycetales(100);f__Metschnikowiaceae(100);g__Metschnikowia(100);s__Metschnikowia_pulcherrima(100); | Metschnikowia | 13 | Saprotroph | \|Undefined Saprotroph\| | Yeast | NULL | Probable | As nectar-tap_saprotroph (Põlme et al. 2020); Decay sugar-rich substrates (Põlme et al. 2020) | Põlme S, et al. 2020. Fungal Diversity 105:1-16 ((https://doi.org/10.1007/s13225-020-00466-2)) |
| Otu17794 | 1 | k__Fungi(100);p__Ascomycota(100);c__Saccharomycetes(100);o__Saccharomycetales(100);f__Metschnikowiaceae(100);g__Metschnikowia(100);g__Metschnikowia_unclassified(100); | Metschnikowia | 13 | Saprotroph | \|Undefined Saprotroph\| | Yeast | NULL | Probable | As nectar-tap_saprotroph (Põlme et al. 2020); Decay sugar-rich substrates (Põlme et al. 2020) | Põlme S, et al. 2020. Fungal Diversity 105:1-16 ((https://doi.org/10.1007/s13225-020-00466-2)) |
| Otu17795 | 1 | k__Fungi(100);p__Ascomycota(100);c__Saccharomycetes(100);o__Saccharomycetales(100);f__Metschnikowiaceae(100);g__Metschnikowia(100);g__Metschnikowia_unclassified(100); | Metschnikowia | 13 | Saprotroph | \|Undefined Saprotroph\| | Yeast | NULL | Probable | As nectar-tap_saprotroph (Põlme et al. 2020); Decay sugar-rich substrates (Põlme et al. 2020) | Põlme S, et al. 2020. Fungal Diversity 105:1-16 ((https://doi.org/10.1007/s13225-020-00466-2)) |
| Otu17799 | 1 | k__Fungi(100);p__Ascomycota(100);c__Saccharomycetes(100);o__Saccharomycetales(100);f__Saccharomycetales_family_Incertae_sedis(100);f__Saccharomycetales_family_Incertae_sedis_unclassified(100);f__Saccharomycetales_family_Incertae_sedis_unclassified(100); | Saccharomycetales | 7 | Saprotroph | Undefined Saprotroph | Yeast | NULL | Possible | NULL | Sterkenburg E, et al. 2015. New Phytologist 207:1145-1158 ((https://doi.org/10.1111/nph.13426)) |
| Otu17800 | 1 | k__Fungi(100);p__Ascomycota(100);c__Saccharomycetes(100);o__Saccharomycetales(100);f__Metschnikowiaceae(100);g__Metschnikowia(100);g__Metschnikowia_unclassified(100); | Metschnikowia | 13 | Saprotroph | \|Undefined Saprotroph\| | Yeast | NULL | Probable | As nectar-tap_saprotroph (Põlme et al. 2020); Decay sugar-rich substrates (Põlme et al. 2020) | Põlme S, et al. 2020. Fungal Diversity 105:1-16 ((https://doi.org/10.1007/s13225-020-00466-2)) |
| Otu17801 | 1 | k__Fungi(100);p__Ascomycota(100);c__Saccharomycetes(100);o__Saccharomycetales(100);f__Metschnikowiaceae(100);g__Metschnikowia(100);g__Metschnikowia_unclassified(100); | Metschnikowia | 13 | Saprotroph | \|Undefined Saprotroph\| | Yeast | NULL | Probable | As nectar-tap_saprotroph (Põlme et al. 2020); Decay sugar-rich substrates (Põlme et al. 2020) | Põlme S, et al. 2020. Fungal Diversity 105:1-16 ((https://doi.org/10.1007/s13225-020-00466-2)) |
| Otu17803 | 1 | k__Fungi(100);p__Ascomycota(100);c__Saccharomycetes(100);o__Saccharomycetales(100);f__Metschnikowiaceae(100);g__Metschnikowia(100);s__Metschnikowia_pulcherrima(100); | Metschnikowia | 13 | Saprotroph | \|Undefined Saprotroph\| | Yeast | NULL | Probable | As nectar-tap_saprotroph (Põlme et al. 2020); Decay sugar-rich substrates (Põlme et al. 2020) | Põlme S, et al. 2020. Fungal Diversity 105:1-16 ((https://doi.org/10.1007/s13225-020-00466-2)) |
| Otu17804 | 1 | k__Fungi(100);p__Ascomycota(100);c__Saccharomycetes(100);o__Saccharomycetales(100);f__Metschnikowiaceae(100);g__Metschnikowia(100);s__Metschnikowia_pulcherrima(100); | Metschnikowia | 13 | Saprotroph | \|Undefined Saprotroph\| | Yeast | NULL | Probable | As nectar-tap_saprotroph (Põlme et al. 2020); Decay sugar-rich substrates (Põlme et al. 2020) | Põlme S, et al. 2020. Fungal Diversity 105:1-16 ((https://doi.org/10.1007/s13225-020-00466-2)) |
| Otu17805 | 1 | k__Fungi(100);p__Ascomycota(100);c__Saccharomycetes(100);o__Saccharomycetales(100);f__Metschnikowiaceae(100);g__Metschnikowia(100);g__Metschnikowia_unclassified(100); | Metschnikowia | 13 | Saprotroph | \|Undefined Saprotroph\| | Yeast | NULL | Probable | As nectar-tap_saprotroph (Põlme et al. 2020); Decay sugar-rich substrates (Põlme et al. 2020) | Põlme S, et al. 2020. Fungal Diversity 105:1-16 ((https://doi.org/10.1007/s13225-020-00466-2)) |
| Otu17806 | 1 | k__Fungi(100);p__Ascomycota(100);c__Saccharomycetes(100);o__Saccharomycetales(100);f__Metschnikowiaceae(100);g__Metschnikowia(100);s__Metschnikowia_pulcherrima(100); | Metschnikowia | 13 | Saprotroph | \|Undefined Saprotroph\| | Yeast | NULL | Probable | As nectar-tap_saprotroph (Põlme et al. 2020); Decay sugar-rich substrates (Põlme et al. 2020) | Põlme S, et al. 2020. Fungal Diversity 105:1-16 ((https://doi.org/10.1007/s13225-020-00466-2)) |
| Otu17807 | 1 | k__Fungi(100);p__Ascomycota(100);c__Saccharomycetes(100);o__Saccharomycetales(100);f__Metschnikowiaceae(100);g__Metschnikowia(100);s__Metschnikowia_pulcherrima(100); | Metschnikowia | 13 | Saprotroph | \|Undefined Saprotroph\| | Yeast | NULL | Probable | As nectar-tap_saprotroph (Põlme et al. 2020); Decay sugar-rich substrates (Põlme et al. 2020) | Põlme S, et al. 2020. Fungal Diversity 105:1-16 ((https://doi.org/10.1007/s13225-020-00466-2)) |
| Otu17808 | 1 | k__Fungi(100);p__Ascomycota(100);c__Saccharomycetes(100);o__Saccharomycetales(100);f__Metschnikowiaceae(100);g__Metschnikowia(100);g__Metschnikowia_unclassified(100); | Metschnikowia | 13 | Saprotroph | \|Undefined Saprotroph\| | Yeast | NULL | Probable | As nectar-tap_saprotroph (Põlme et al. 2020); Decay sugar-rich substrates (Põlme et al. 2020) | Põlme S, et al. 2020. Fungal Diversity 105:1-16 ((https://doi.org/10.1007/s13225-020-00466-2)) |
| Otu17809 | 1 | k__Fungi(100);p__Ascomycota(100);c__Saccharomycetes(100);o__Saccharomycetales(100);f__Metschnikowiaceae(100);g__Metschnikowia(100);s__Metschnikowia_pulcherrima(100); | Metschnikowia | 13 | Saprotroph | \|Undefined Saprotroph\| | Yeast | NULL | Probable | As nectar-tap_saprotroph (Põlme et al. 2020); Decay sugar-rich substrates (Põlme et al. 2020) | Põlme S, et al. 2020. Fungal Diversity 105:1-16 ((https://doi.org/10.1007/s13225-020-00466-2)) |
| Otu17810 | 1 | k__Fungi(100);p__Ascomycota(100);c__Saccharomycetes(100);o__Saccharomycetales(100);f__Metschnikowiaceae(100);g__Metschnikowia(100);s__Metschnikowia_pulcherrima(100); | Metschnikowia | 13 | Saprotroph | \|Undefined Saprotroph\| | Yeast | NULL | Probable | As nectar-tap_saprotroph (Põlme et al. 2020); Decay sugar-rich substrates (Põlme et al. 2020) | Põlme S, et al. 2020. Fungal Diversity 105:1-16 ((https://doi.org/10.1007/s13225-020-00466-2)) |
| Otu17811 | 1 | k__Fungi(100);p__Ascomycota(100);c__Saccharomycetes(100);o__Saccharomycetales(100);f__Metschnikowiaceae(100);g__Metschnikowia(100);s__Metschnikowia_pulcherrima(100); | Metschnikowia | 13 | Saprotroph | \|Undefined Saprotroph\| | Yeast | NULL | Probable | As nectar-tap_saprotroph (Põlme et al. 2020); Decay sugar-rich substrates (Põlme et al. 2020) | Põlme S, et al. 2020. Fungal Diversity 105:1-16 ((https://doi.org/10.1007/s13225-020-00466-2)) |
| Otu17812 | 1 | k__Fungi(100);p__Ascomycota(100);c__Saccharomycetes(100);o__Saccharomycetales(100);f__Metschnikowiaceae(100);g__Metschnikowia(100);g__Metschnikowia_unclassified(100); | Metschnikowia | 13 | Saprotroph | \|Undefined Saprotroph\| | Yeast | NULL | Probable | As nectar-tap_saprotroph (Põlme et al. 2020); Decay sugar-rich substrates (Põlme et al. 2020) | Põlme S, et al. 2020. Fungal Diversity 105:1-16 ((https://doi.org/10.1007/s13225-020-00466-2)) |
| Otu17813 | 1 | k__Fungi(100);p__Ascomycota(100);c__Saccharomycetes(100);o__Saccharomycetales(100);f__Metschnikowiaceae(100);g__Metschnikowia(100);g__Metschnikowia_unclassified(100); | Metschnikowia | 13 | Saprotroph | \|Undefined Saprotroph\| | Yeast | NULL | Probable | As nectar-tap_saprotroph (Põlme et al. 2020); Decay sugar-rich substrates (Põlme et al. 2020) | Põlme S, et al. 2020. Fungal Diversity 105:1-16 ((https://doi.org/10.1007/s13225-020-00466-2)) |
| Otu17814 | 1 | k__Fungi(100);p__Ascomycota(100);c__Saccharomycetes(100);o__Saccharomycetales(100);f__Metschnikowiaceae(100);g__Metschnikowia(100);g__Metschnikowia_unclassified(100); | Metschnikowia | 13 | Saprotroph | \|Undefined Saprotroph\| | Yeast | NULL | Probable | As nectar-tap_saprotroph (Põlme et al. 2020); Decay sugar-rich substrates (Põlme et al. 2020) | Põlme S, et al. 2020. Fungal Diversity 105:1-16 ((https://doi.org/10.1007/s13225-020-00466-2)) |
| Otu17816 | 1 | k__Fungi(100);p__Ascomycota(100);c__Saccharomycetes(100);o__Saccharomycetales(100);f__Metschnikowiaceae(100);g__Metschnikowia(100);s__Metschnikowia_pulcherrima(100); | Metschnikowia | 13 | Saprotroph | \|Undefined Saprotroph\| | Yeast | NULL | Probable | As nectar-tap_saprotroph (Põlme et al. 2020); Decay sugar-rich substrates (Põlme et al. 2020) | Põlme S, et al. 2020. Fungal Diversity 105:1-16 ((https://doi.org/10.1007/s13225-020-00466-2)) |
| Otu17818 | 1 | k__Fungi(100);p__Ascomycota(100);c__Saccharomycetes(100);o__Saccharomycetales(100);f__Metschnikowiaceae(100);g__Metschnikowia(100);g__Metschnikowia_unclassified(100); | Metschnikowia | 13 | Saprotroph | \|Undefined Saprotroph\| | Yeast | NULL | Probable | As nectar-tap_saprotroph (Põlme et al. 2020); Decay sugar-rich substrates (Põlme et al. 2020) | Põlme S, et al. 2020. Fungal Diversity 105:1-16 ((https://doi.org/10.1007/s13225-020-00466-2)) |
| Otu17822 | 1 | k__Fungi(100);p__Ascomycota(100);c__Saccharomycetes(100);o__Saccharomycetales(100);f__Metschnikowiaceae(100);g__Metschnikowia(100);g__Metschnikowia_unclassified(100); | Metschnikowia | 13 | Saprotroph | \|Undefined Saprotroph\| | Yeast | NULL | Probable | As nectar-tap_saprotroph (Põlme et al. 2020); Decay sugar-rich substrates (Põlme et al. 2020) | Põlme S, et al. 2020. Fungal Diversity 105:1-16 ((https://doi.org/10.1007/s13225-020-00466-2)) |
| Otu17823 | 1 | k__Fungi(100);p__Ascomycota(100);c__Saccharomycetes(100);o__Saccharomycetales(100);f__Metschnikowiaceae(100);g__Metschnikowia(100);g__Metschnikowia_unclassified(100); | Metschnikowia | 13 | Saprotroph | \|Undefined Saprotroph\| | Yeast | NULL | Probable | As nectar-tap_saprotroph (Põlme et al. 2020); Decay sugar-rich substrates (Põlme et al. 2020) | Põlme S, et al. 2020. Fungal Diversity 105:1-16 ((https://doi.org/10.1007/s13225-020-00466-2)) |
| Otu17824 | 1 | k__Fungi(100);p__Ascomycota(100);c__Saccharomycetes(100);o__Saccharomycetales(100);f__Metschnikowiaceae(100);g__Metschnikowia(100);s__Metschnikowia_pulcherrima(100); | Metschnikowia | 13 | Saprotroph | \|Undefined Saprotroph\| | Yeast | NULL | Probable | As nectar-tap_saprotroph (Põlme et al. 2020); Decay sugar-rich substrates (Põlme et al. 2020) | Põlme S, et al. 2020. Fungal Diversity 105:1-16 ((https://doi.org/10.1007/s13225-020-00466-2)) |
| Otu17826 | 1 | k__Fungi(100);p__Ascomycota(100);c__Saccharomycetes(100);o__Saccharomycetales(100);f__Metschnikowiaceae(100);g__Metschnikowia(100);s__Metschnikowia_pulcherrima(100); | Metschnikowia | 13 | Saprotroph | \|Undefined Saprotroph\| | Yeast | NULL | Probable | As nectar-tap_saprotroph (Põlme et al. 2020); Decay sugar-rich substrates (Põlme et al. 2020) | Põlme S, et al. 2020. Fungal Diversity 105:1-16 ((https://doi.org/10.1007/s13225-020-00466-2)) |
| Otu17827 | 1 | k__Fungi(100);p__Ascomycota(100);c__Saccharomycetes(100);o__Saccharomycetales(100);f__Metschnikowiaceae(100);g__Metschnikowia(100);s__Metschnikowia_pulcherrima(100); | Metschnikowia | 13 | Saprotroph | \|Undefined Saprotroph\| | Yeast | NULL | Probable | As nectar-tap_saprotroph (Põlme et al. 2020); Decay sugar-rich substrates (Põlme et al. 2020) | Põlme S, et al. 2020. Fungal Diversity 105:1-16 ((https://doi.org/10.1007/s13225-020-00466-2)) |
| Otu17828 | 1 | k__Fungi(100);p__Ascomycota(100);c__Saccharomycetes(100);o__Saccharomycetales(100);f__Metschnikowiaceae(100);g__Metschnikowia(100);s__Metschnikowia_pulcherrima(100); | Metschnikowia | 13 | Saprotroph | \|Undefined Saprotroph\| | Yeast | NULL | Probable | As nectar-tap_saprotroph (Põlme et al. 2020); Decay sugar-rich substrates (Põlme et al. 2020) | Põlme S, et al. 2020. Fungal Diversity 105:1-16 ((https://doi.org/10.1007/s13225-020-00466-2)) |
| Otu17829 | 1 | k__Fungi(100);p__Ascomycota(100);c__Saccharomycetes(100);o__Saccharomycetales(100);f__Metschnikowiaceae(100);g__Metschnikowia(100);g__Metschnikowia_unclassified(100); | Metschnikowia | 13 | Saprotroph | \|Undefined Saprotroph\| | Yeast | NULL | Probable | As nectar-tap_saprotroph (Põlme et al. 2020); Decay sugar-rich substrates (Põlme et al. 2020) | Põlme S, et al. 2020. Fungal Diversity 105:1-16 ((https://doi.org/10.1007/s13225-020-00466-2)) |
| Otu17830 | 1 | k__Fungi(100);p__Ascomycota(100);c__Saccharomycetes(100);o__Saccharomycetales(100);f__Metschnikowiaceae(100);g__Metschnikowia(100);g__Metschnikowia_unclassified(100); | Metschnikowia | 13 | Saprotroph | \|Undefined Saprotroph\| | Yeast | NULL | Probable | As nectar-tap_saprotroph (Põlme et al. 2020); Decay sugar-rich substrates (Põlme et al. 2020) | Põlme S, et al. 2020. Fungal Diversity 105:1-16 ((https://doi.org/10.1007/s13225-020-00466-2)) |
| Otu17831 | 1 | k__Fungi(100);p__Ascomycota(100);c__Saccharomycetes(100);o__Saccharomycetales(100);f__Metschnikowiaceae(100);g__Metschnikowia(100);g__Metschnikowia_unclassified(100); | Metschnikowia | 13 | Saprotroph | \|Undefined Saprotroph\| | Yeast | NULL | Probable | As nectar-tap_saprotroph (Põlme et al. 2020); Decay sugar-rich substrates (Põlme et al. 2020) | Põlme S, et al. 2020. Fungal Diversity 105:1-16 ((https://doi.org/10.1007/s13225-020-00466-2)) |
| Otu17832 | 1 | k__Fungi(100);p__Ascomycota(100);c__Saccharomycetes(100);o__Saccharomycetales(100);f__Metschnikowiaceae(100);g__Metschnikowia(100);g__Metschnikowia_unclassified(100); | Metschnikowia | 13 | Saprotroph | \|Undefined Saprotroph\| | Yeast | NULL | Probable | As nectar-tap_saprotroph (Põlme et al. 2020); Decay sugar-rich substrates (Põlme et al. 2020) | Põlme S, et al. 2020. Fungal Diversity 105:1-16 ((https://doi.org/10.1007/s13225-020-00466-2)) |
| Otu17833 | 1 | k__Fungi(100);p__Ascomycota(100);c__Saccharomycetes(100);o__Saccharomycetales(100);f__Metschnikowiaceae(100);g__Metschnikowia(100);g__Metschnikowia_unclassified(100); | Metschnikowia | 13 | Saprotroph | \|Undefined Saprotroph\| | Yeast | NULL | Probable | As nectar-tap_saprotroph (Põlme et al. 2020); Decay sugar-rich substrates (Põlme et al. 2020) | Põlme S, et al. 2020. Fungal Diversity 105:1-16 ((https://doi.org/10.1007/s13225-020-00466-2)) |
| Otu17834 | 1 | k__Fungi(100);p__Ascomycota(100);c__Saccharomycetes(100);o__Saccharomycetales(100);f__Metschnikowiaceae(100);g__Metschnikowia(100);s__Metschnikowia_pulcherrima(100); | Metschnikowia | 13 | Saprotroph | \|Undefined Saprotroph\| | Yeast | NULL | Probable | As nectar-tap_saprotroph (Põlme et al. 2020); Decay sugar-rich substrates (Põlme et al. 2020) | Põlme S, et al. 2020. Fungal Diversity 105:1-16 ((https://doi.org/10.1007/s13225-020-00466-2)) |
| Otu17835 | 1 | k__Fungi(100);p__Ascomycota(100);c__Saccharomycetes(100);o__Saccharomycetales(100);f__Metschnikowiaceae(100);g__Metschnikowia(100);s__Metschnikowia_pulcherrima(100); | Metschnikowia | 13 | Saprotroph | \|Undefined Saprotroph\| | Yeast | NULL | Probable | As nectar-tap_saprotroph (Põlme et al. 2020); Decay sugar-rich substrates (Põlme et al. 2020) | Põlme S, et al. 2020. Fungal Diversity 105:1-16 ((https://doi.org/10.1007/s13225-020-00466-2)) |
| Otu17836 | 1 | k__Fungi(100);p__Ascomycota(100);c__Saccharomycetes(100);o__Saccharomycetales(100);f__Metschnikowiaceae(100);g__Metschnikowia(100);g__Metschnikowia_unclassified(100); | Metschnikowia | 13 | Saprotroph | \|Undefined Saprotroph\| | Yeast | NULL | Probable | As nectar-tap_saprotroph (Põlme et al. 2020); Decay sugar-rich substrates (Põlme et al. 2020) | Põlme S, et al. 2020. Fungal Diversity 105:1-16 ((https://doi.org/10.1007/s13225-020-00466-2)) |
| Otu17837 | 1 | k__Fungi(100);p__Ascomycota(100);c__Saccharomycetes(100);o__Saccharomycetales(100);f__Metschnikowiaceae(100);g__Metschnikowia(100);s__Metschnikowia_pulcherrima(100); | Metschnikowia | 13 | Saprotroph | \|Undefined Saprotroph\| | Yeast | NULL | Probable | As nectar-tap_saprotroph (Põlme et al. 2020); Decay sugar-rich substrates (Põlme et al. 2020) | Põlme S, et al. 2020. Fungal Diversity 105:1-16 ((https://doi.org/10.1007/s13225-020-00466-2)) |
| Otu17838 | 1 | k__Fungi(100);p__Ascomycota(100);c__Saccharomycetes(100);o__Saccharomycetales(100);f__Metschnikowiaceae(100);g__Metschnikowia(100);s__Metschnikowia_pulcherrima(100); | Metschnikowia | 13 | Saprotroph | \|Undefined Saprotroph\| | Yeast | NULL | Probable | As nectar-tap_saprotroph (Põlme et al. 2020); Decay sugar-rich substrates (Põlme et al. 2020) | Põlme S, et al. 2020. Fungal Diversity 105:1-16 ((https://doi.org/10.1007/s13225-020-00466-2)) |
| Otu17839 | 1 | k__Fungi(100);p__Ascomycota(100);c__Saccharomycetes(100);o__Saccharomycetales(100);f__Metschnikowiaceae(100);g__Metschnikowia(100);s__Metschnikowia_pulcherrima(100); | Metschnikowia | 13 | Saprotroph | \|Undefined Saprotroph\| | Yeast | NULL | Probable | As nectar-tap_saprotroph (Põlme et al. 2020); Decay sugar-rich substrates (Põlme et al. 2020) | Põlme S, et al. 2020. Fungal Diversity 105:1-16 ((https://doi.org/10.1007/s13225-020-00466-2)) |
| Otu17840 | 1 | k__Fungi(100);p__Ascomycota(100);c__Saccharomycetes(100);o__Saccharomycetales(100);f__Metschnikowiaceae(100);g__Metschnikowia(100);s__Metschnikowia_pulcherrima(100); | Metschnikowia | 13 | Saprotroph | \|Undefined Saprotroph\| | Yeast | NULL | Probable | As nectar-tap_saprotroph (Põlme et al. 2020); Decay sugar-rich substrates (Põlme et al. 2020) | Põlme S, et al. 2020. Fungal Diversity 105:1-16 ((https://doi.org/10.1007/s13225-020-00466-2)) |
| Otu17841 | 1 | k__Fungi(100);p__Ascomycota(100);c__Saccharomycetes(100);o__Saccharomycetales(100);f__Metschnikowiaceae(100);g__Metschnikowia(100);s__Metschnikowia_pulcherrima(100); | Metschnikowia | 13 | Saprotroph | \|Undefined Saprotroph\| | Yeast | NULL | Probable | As nectar-tap_saprotroph (Põlme et al. 2020); Decay sugar-rich substrates (Põlme et al. 2020) | Põlme S, et al. 2020. Fungal Diversity 105:1-16 ((https://doi.org/10.1007/s13225-020-00466-2)) |
| Otu17842 | 1 | k__Fungi(100);p__Ascomycota(100);c__Saccharomycetes(100);o__Saccharomycetales(100);f__Metschnikowiaceae(100);g__Metschnikowia(100);s__Metschnikowia_pulcherrima(100); | Metschnikowia | 13 | Saprotroph | \|Undefined Saprotroph\| | Yeast | NULL | Probable | As nectar-tap_saprotroph (Põlme et al. 2020); Decay sugar-rich substrates (Põlme et al. 2020) | Põlme S, et al. 2020. Fungal Diversity 105:1-16 ((https://doi.org/10.1007/s13225-020-00466-2)) |
| Otu17843 | 1 | k__Fungi(100);p__Ascomycota(100);c__Saccharomycetes(100);o__Saccharomycetales(100);f__Metschnikowiaceae(100);g__Metschnikowia(100);g__Metschnikowia_unclassified(100); | Metschnikowia | 13 | Saprotroph | \|Undefined Saprotroph\| | Yeast | NULL | Probable | As nectar-tap_saprotroph (Põlme et al. 2020); Decay sugar-rich substrates (Põlme et al. 2020) | Põlme S, et al. 2020. Fungal Diversity 105:1-16 ((https://doi.org/10.1007/s13225-020-00466-2)) |
| Otu17844 | 1 | k__Fungi(100);p__Ascomycota(100);c__Saccharomycetes(100);o__Saccharomycetales(100);f__Metschnikowiaceae(100);g__Metschnikowia(100);g__Metschnikowia_unclassified(100); | Metschnikowia | 13 | Saprotroph | \|Undefined Saprotroph\| | Yeast | NULL | Probable | As nectar-tap_saprotroph (Põlme et al. 2020); Decay sugar-rich substrates (Põlme et al. 2020) | Põlme S, et al. 2020. Fungal Diversity 105:1-16 ((https://doi.org/10.1007/s13225-020-00466-2)) |
| Otu17845 | 1 | k__Fungi(100);p__Ascomycota(100);c__Saccharomycetes(100);o__Saccharomycetales(100);f__Metschnikowiaceae(100);g__Metschnikowia(100);s__Metschnikowia_pulcherrima(100); | Metschnikowia | 13 | Saprotroph | \|Undefined Saprotroph\| | Yeast | NULL | Probable | As nectar-tap_saprotroph (Põlme et al. 2020); Decay sugar-rich substrates (Põlme et al. 2020) | Põlme S, et al. 2020. Fungal Diversity 105:1-16 ((https://doi.org/10.1007/s13225-020-00466-2)) |
| Otu17846 | 1 | k__Fungi(100);p__Ascomycota(100);c__Saccharomycetes(100);o__Saccharomycetales(100);f__Metschnikowiaceae(100);g__Metschnikowia(100);s__Metschnikowia_pulcherrima(100); | Metschnikowia | 13 | Saprotroph | \|Undefined Saprotroph\| | Yeast | NULL | Probable | As nectar-tap_saprotroph (Põlme et al. 2020); Decay sugar-rich substrates (Põlme et al. 2020) | Põlme S, et al. 2020. Fungal Diversity 105:1-16 ((https://doi.org/10.1007/s13225-020-00466-2)) |
| Otu17847 | 1 | k__Fungi(100);p__Ascomycota(100);c__Saccharomycetes(100);o__Saccharomycetales(100);f__Metschnikowiaceae(100);g__Metschnikowia(100);g__Metschnikowia_unclassified(100); | Metschnikowia | 13 | Saprotroph | \|Undefined Saprotroph\| | Yeast | NULL | Probable | As nectar-tap_saprotroph (Põlme et al. 2020); Decay sugar-rich substrates (Põlme et al. 2020) | Põlme S, et al. 2020. Fungal Diversity 105:1-16 ((https://doi.org/10.1007/s13225-020-00466-2)) |
| Otu17848 | 1 | k__Fungi(100);p__Ascomycota(100);c__Saccharomycetes(100);o__Saccharomycetales(100);f__Metschnikowiaceae(100);g__Metschnikowia(100);g__Metschnikowia_unclassified(100); | Metschnikowia | 13 | Saprotroph | \|Undefined Saprotroph\| | Yeast | NULL | Probable | As nectar-tap_saprotroph (Põlme et al. 2020); Decay sugar-rich substrates (Põlme et al. 2020) | Põlme S, et al. 2020. Fungal Diversity 105:1-16 ((https://doi.org/10.1007/s13225-020-00466-2)) |
| Otu17849 | 1 | k__Fungi(100);p__Ascomycota(100);c__Saccharomycetes(100);o__Saccharomycetales(100);f__Metschnikowiaceae(100);g__Metschnikowia(100);s__Metschnikowia_pulcherrima(100); | Metschnikowia | 13 | Saprotroph | \|Undefined Saprotroph\| | Yeast | NULL | Probable | As nectar-tap_saprotroph (Põlme et al. 2020); Decay sugar-rich substrates (Põlme et al. 2020) | Põlme S, et al. 2020. Fungal Diversity 105:1-16 ((https://doi.org/10.1007/s13225-020-00466-2)) |
| Otu17850 | 1 | k__Fungi(100);p__Ascomycota(100);c__Saccharomycetes(100);o__Saccharomycetales(100);f__Metschnikowiaceae(100);g__Metschnikowia(100);g__Metschnikowia_unclassified(100); | Metschnikowia | 13 | Saprotroph | \|Undefined Saprotroph\| | Yeast | NULL | Probable | As nectar-tap_saprotroph (Põlme et al. 2020); Decay sugar-rich substrates (Põlme et al. 2020) | Põlme S, et al. 2020. Fungal Diversity 105:1-16 ((https://doi.org/10.1007/s13225-020-00466-2)) |
| Otu17851 | 1 | k__Fungi(100);p__Ascomycota(100);c__Saccharomycetes(100);o__Saccharomycetales(100);f__Metschnikowiaceae(100);g__Metschnikowia(100);s__Metschnikowia_pulcherrima(100); | Metschnikowia | 13 | Saprotroph | \|Undefined Saprotroph\| | Yeast | NULL | Probable | As nectar-tap_saprotroph (Põlme et al. 2020); Decay sugar-rich substrates (Põlme et al. 2020) | Põlme S, et al. 2020. Fungal Diversity 105:1-16 ((https://doi.org/10.1007/s13225-020-00466-2)) |
| Otu17852 | 1 | k__Fungi(100);p__Ascomycota(100);c__Saccharomycetes(100);o__Saccharomycetales(100);f__Metschnikowiaceae(100);g__Metschnikowia(100);s__Metschnikowia_pulcherrima(100); | Metschnikowia | 13 | Saprotroph | \|Undefined Saprotroph\| | Yeast | NULL | Probable | As nectar-tap_saprotroph (Põlme et al. 2020); Decay sugar-rich substrates (Põlme et al. 2020) | Põlme S, et al. 2020. Fungal Diversity 105:1-16 ((https://doi.org/10.1007/s13225-020-00466-2)) |
| Otu17853 | 1 | k__Fungi(100);p__Ascomycota(100);c__Saccharomycetes(100);o__Saccharomycetales(100);f__Metschnikowiaceae(100);g__Metschnikowia(100);g__Metschnikowia_unclassified(100); | Metschnikowia | 13 | Saprotroph | \|Undefined Saprotroph\| | Yeast | NULL | Probable | As nectar-tap_saprotroph (Põlme et al. 2020); Decay sugar-rich substrates (Põlme et al. 2020) | Põlme S, et al. 2020. Fungal Diversity 105:1-16 ((https://doi.org/10.1007/s13225-020-00466-2)) |
| Otu17854 | 1 | k__Fungi(100);p__Ascomycota(100);c__Saccharomycetes(100);o__Saccharomycetales(100);f__Metschnikowiaceae(100);g__Metschnikowia(100);g__Metschnikowia_unclassified(100); | Metschnikowia | 13 | Saprotroph | \|Undefined Saprotroph\| | Yeast | NULL | Probable | As nectar-tap_saprotroph (Põlme et al. 2020); Decay sugar-rich substrates (Põlme et al. 2020) | Põlme S, et al. 2020. Fungal Diversity 105:1-16 ((https://doi.org/10.1007/s13225-020-00466-2)) |
| Otu17855 | 1 | k__Fungi(100);p__Ascomycota(100);c__Saccharomycetes(100);o__Saccharomycetales(100);f__Metschnikowiaceae(100);g__Metschnikowia(100);s__Metschnikowia_pulcherrima(100); | Metschnikowia | 13 | Saprotroph | \|Undefined Saprotroph\| | Yeast | NULL | Probable | As nectar-tap_saprotroph (Põlme et al. 2020); Decay sugar-rich substrates (Põlme et al. 2020) | Põlme S, et al. 2020. Fungal Diversity 105:1-16 ((https://doi.org/10.1007/s13225-020-00466-2)) |
| Otu17856 | 1 | k__Fungi(100);p__Ascomycota(100);c__Saccharomycetes(100);o__Saccharomycetales(100);f__Metschnikowiaceae(100);g__Metschnikowia(100);g__Metschnikowia_unclassified(100); | Metschnikowia | 13 | Saprotroph | \|Undefined Saprotroph\| | Yeast | NULL | Probable | As nectar-tap_saprotroph (Põlme et al. 2020); Decay sugar-rich substrates (Põlme et al. 2020) | Põlme S, et al. 2020. Fungal Diversity 105:1-16 ((https://doi.org/10.1007/s13225-020-00466-2)) |
| Otu17860 | 1 | k__Fungi(100);p__Ascomycota(100);c__Saccharomycetes(100);o__Saccharomycetales(100);f__Metschnikowiaceae(100);g__Metschnikowia(100);s__Metschnikowia_pulcherrima(100); | Metschnikowia | 13 | Saprotroph | \|Undefined Saprotroph\| | Yeast | NULL | Probable | As nectar-tap_saprotroph (Põlme et al. 2020); Decay sugar-rich substrates (Põlme et al. 2020) | Põlme S, et al. 2020. Fungal Diversity 105:1-16 ((https://doi.org/10.1007/s13225-020-00466-2)) |
| Otu17861 | 1 | k__Fungi(100);p__Ascomycota(100);c__Saccharomycetes(100);o__Saccharomycetales(100);f__Metschnikowiaceae(100);g__Metschnikowia(100);s__Metschnikowia_pulcherrima(100); | Metschnikowia | 13 | Saprotroph | \|Undefined Saprotroph\| | Yeast | NULL | Probable | As nectar-tap_saprotroph (Põlme et al. 2020); Decay sugar-rich substrates (Põlme et al. 2020) | Põlme S, et al. 2020. Fungal Diversity 105:1-16 ((https://doi.org/10.1007/s13225-020-00466-2)) |
| Otu17862 | 1 | k__Fungi(100);p__Ascomycota(100);c__Saccharomycetes(100);o__Saccharomycetales(100);f__Metschnikowiaceae(100);g__Metschnikowia(100);s__Metschnikowia_pulcherrima(100); | Metschnikowia | 13 | Saprotroph | \|Undefined Saprotroph\| | Yeast | NULL | Probable | As nectar-tap_saprotroph (Põlme et al. 2020); Decay sugar-rich substrates (Põlme et al. 2020) | Põlme S, et al. 2020. Fungal Diversity 105:1-16 ((https://doi.org/10.1007/s13225-020-00466-2)) |
| Otu17864 | 1 | k__Fungi(100);p__Ascomycota(100);c__Saccharomycetes(100);o__Saccharomycetales(100);f__Metschnikowiaceae(100);g__Metschnikowia(100);g__Metschnikowia_unclassified(100); | Metschnikowia | 13 | Saprotroph | \|Undefined Saprotroph\| | Yeast | NULL | Probable | As nectar-tap_saprotroph (Põlme et al. 2020); Decay sugar-rich substrates (Põlme et al. 2020) | Põlme S, et al. 2020. Fungal Diversity 105:1-16 ((https://doi.org/10.1007/s13225-020-00466-2)) |
| Otu17865 | 1 | k__Fungi(100);p__Ascomycota(100);c__Saccharomycetes(100);o__Saccharomycetales(100);f__Metschnikowiaceae(100);g__Metschnikowia(100);s__Metschnikowia_pulcherrima(100); | Metschnikowia | 13 | Saprotroph | \|Undefined Saprotroph\| | Yeast | NULL | Probable | As nectar-tap_saprotroph (Põlme et al. 2020); Decay sugar-rich substrates (Põlme et al. 2020) | Põlme S, et al. 2020. Fungal Diversity 105:1-16 ((https://doi.org/10.1007/s13225-020-00466-2)) |
| Otu17866 | 1 | k__Fungi(100);p__Ascomycota(100);c__Saccharomycetes(100);o__Saccharomycetales(100);f__Metschnikowiaceae(100);g__Metschnikowia(100);s__Metschnikowia_pulcherrima(100); | Metschnikowia | 13 | Saprotroph | \|Undefined Saprotroph\| | Yeast | NULL | Probable | As nectar-tap_saprotroph (Põlme et al. 2020); Decay sugar-rich substrates (Põlme et al. 2020) | Põlme S, et al. 2020. Fungal Diversity 105:1-16 ((https://doi.org/10.1007/s13225-020-00466-2)) |
| Otu17867 | 1 | k__Fungi(100);p__Ascomycota(100);c__Saccharomycetes(100);o__Saccharomycetales(100);f__Metschnikowiaceae(100);g__Metschnikowia(100);s__Metschnikowia_pulcherrima(100); | Metschnikowia | 13 | Saprotroph | \|Undefined Saprotroph\| | Yeast | NULL | Probable | As nectar-tap_saprotroph (Põlme et al. 2020); Decay sugar-rich substrates (Põlme et al. 2020) | Põlme S, et al. 2020. Fungal Diversity 105:1-16 ((https://doi.org/10.1007/s13225-020-00466-2)) |
| Otu17868 | 1 | k__Fungi(100);p__Ascomycota(100);c__Saccharomycetes(100);o__Saccharomycetales(100);f__Metschnikowiaceae(100);g__Metschnikowia(100);s__Metschnikowia_pulcherrima(100); | Metschnikowia | 13 | Saprotroph | \|Undefined Saprotroph\| | Yeast | NULL | Probable | As nectar-tap_saprotroph (Põlme et al. 2020); Decay sugar-rich substrates (Põlme et al. 2020) | Põlme S, et al. 2020. Fungal Diversity 105:1-16 ((https://doi.org/10.1007/s13225-020-00466-2)) |
| Otu17869 | 1 | k__Fungi(100);p__Ascomycota(100);c__Saccharomycetes(100);o__Saccharomycetales(100);f__Metschnikowiaceae(100);g__Metschnikowia(100);s__Metschnikowia_pulcherrima(100); | Metschnikowia | 13 | Saprotroph | \|Undefined Saprotroph\| | Yeast | NULL | Probable | As nectar-tap_saprotroph (Põlme et al. 2020); Decay sugar-rich substrates (Põlme et al. 2020) | Põlme S, et al. 2020. Fungal Diversity 105:1-16 ((https://doi.org/10.1007/s13225-020-00466-2)) |
| Otu17870 | 1 | k__Fungi(100);p__Ascomycota(100);c__Saccharomycetes(100);o__Saccharomycetales(100);f__Metschnikowiaceae(100);g__Metschnikowia(100);s__Metschnikowia_pulcherrima(100); | Metschnikowia | 13 | Saprotroph | \|Undefined Saprotroph\| | Yeast | NULL | Probable | As nectar-tap_saprotroph (Põlme et al. 2020); Decay sugar-rich substrates (Põlme et al. 2020) | Põlme S, et al. 2020. Fungal Diversity 105:1-16 ((https://doi.org/10.1007/s13225-020-00466-2)) |
| Otu17871 | 1 | k__Fungi(100);p__Ascomycota(100);c__Saccharomycetes(100);o__Saccharomycetales(100);f__Metschnikowiaceae(100);g__Metschnikowia(100);g__Metschnikowia_unclassified(100); | Metschnikowia | 13 | Saprotroph | \|Undefined Saprotroph\| | Yeast | NULL | Probable | As nectar-tap_saprotroph (Põlme et al. 2020); Decay sugar-rich substrates (Põlme et al. 2020) | Põlme S, et al. 2020. Fungal Diversity 105:1-16 ((https://doi.org/10.1007/s13225-020-00466-2)) |
| Otu17872 | 1 | k__Fungi(100);p__Ascomycota(100);c__Saccharomycetes(100);o__Saccharomycetales(100);f__Metschnikowiaceae(100);g__Metschnikowia(100);s__Metschnikowia_pulcherrima(100); | Metschnikowia | 13 | Saprotroph | \|Undefined Saprotroph\| | Yeast | NULL | Probable | As nectar-tap_saprotroph (Põlme et al. 2020); Decay sugar-rich substrates (Põlme et al. 2020) | Põlme S, et al. 2020. Fungal Diversity 105:1-16 ((https://doi.org/10.1007/s13225-020-00466-2)) |
| Otu17873 | 1 | k__Fungi(100);p__Ascomycota(100);c__Saccharomycetes(100);o__Saccharomycetales(100);f__Metschnikowiaceae(100);g__Metschnikowia(100);s__Metschnikowia_pulcherrima(100); | Metschnikowia | 13 | Saprotroph | \|Undefined Saprotroph\| | Yeast | NULL | Probable | As nectar-tap_saprotroph (Põlme et al. 2020); Decay sugar-rich substrates (Põlme et al. 2020) | Põlme S, et al. 2020. Fungal Diversity 105:1-16 ((https://doi.org/10.1007/s13225-020-00466-2)) |
| Otu17874 | 1 | k__Fungi(100);p__Ascomycota(100);c__Saccharomycetes(100);o__Saccharomycetales(100);f__Metschnikowiaceae(100);g__Metschnikowia(100);s__Metschnikowia_pulcherrima(100); | Metschnikowia | 13 | Saprotroph | \|Undefined Saprotroph\| | Yeast | NULL | Probable | As nectar-tap_saprotroph (Põlme et al. 2020); Decay sugar-rich substrates (Põlme et al. 2020) | Põlme S, et al. 2020. Fungal Diversity 105:1-16 ((https://doi.org/10.1007/s13225-020-00466-2)) |
| Otu17875 | 1 | k__Fungi(100);p__Ascomycota(100);c__Saccharomycetes(100);o__Saccharomycetales(100);f__Metschnikowiaceae(100);g__Metschnikowia(100);s__Metschnikowia_pulcherrima(100); | Metschnikowia | 13 | Saprotroph | \|Undefined Saprotroph\| | Yeast | NULL | Probable | As nectar-tap_saprotroph (Põlme et al. 2020); Decay sugar-rich substrates (Põlme et al. 2020) | Põlme S, et al. 2020. Fungal Diversity 105:1-16 ((https://doi.org/10.1007/s13225-020-00466-2)) |
| Otu17876 | 1 | k__Fungi(100);p__Ascomycota(100);c__Saccharomycetes(100);o__Saccharomycetales(100);f__Metschnikowiaceae(100);g__Metschnikowia(100);s__Metschnikowia_pulcherrima(100); | Metschnikowia | 13 | Saprotroph | \|Undefined Saprotroph\| | Yeast | NULL | Probable | As nectar-tap_saprotroph (Põlme et al. 2020); Decay sugar-rich substrates (Põlme et al. 2020) | Põlme S, et al. 2020. Fungal Diversity 105:1-16 ((https://doi.org/10.1007/s13225-020-00466-2)) |
| Otu17877 | 1 | k__Fungi(100);p__Ascomycota(100);c__Saccharomycetes(100);o__Saccharomycetales(100);f__Metschnikowiaceae(100);g__Metschnikowia(100);g__Metschnikowia_unclassified(100); | Metschnikowia | 13 | Saprotroph | \|Undefined Saprotroph\| | Yeast | NULL | Probable | As nectar-tap_saprotroph (Põlme et al. 2020); Decay sugar-rich substrates (Põlme et al. 2020) | Põlme S, et al. 2020. Fungal Diversity 105:1-16 ((https://doi.org/10.1007/s13225-020-00466-2)) |
| Otu17878 | 1 | k__Fungi(100);p__Ascomycota(100);c__Saccharomycetes(100);o__Saccharomycetales(100);f__Metschnikowiaceae(100);g__Metschnikowia(100);g__Metschnikowia_unclassified(100); | Metschnikowia | 13 | Saprotroph | \|Undefined Saprotroph\| | Yeast | NULL | Probable | As nectar-tap_saprotroph (Põlme et al. 2020); Decay sugar-rich substrates (Põlme et al. 2020) | Põlme S, et al. 2020. Fungal Diversity 105:1-16 ((https://doi.org/10.1007/s13225-020-00466-2)) |
| Otu17879 | 1 | k__Fungi(100);p__Ascomycota(100);c__Saccharomycetes(100);o__Saccharomycetales(100);f__Metschnikowiaceae(100);g__Metschnikowia(100);s__Metschnikowia_pulcherrima(100); | Metschnikowia | 13 | Saprotroph | \|Undefined Saprotroph\| | Yeast | NULL | Probable | As nectar-tap_saprotroph (Põlme et al. 2020); Decay sugar-rich substrates (Põlme et al. 2020) | Põlme S, et al. 2020. Fungal Diversity 105:1-16 ((https://doi.org/10.1007/s13225-020-00466-2)) |
| Otu17880 | 1 | k__Fungi(100);p__Ascomycota(100);c__Saccharomycetes(100);o__Saccharomycetales(100);f__Metschnikowiaceae(100);g__Metschnikowia(100);s__Metschnikowia_pulcherrima(100); | Metschnikowia | 13 | Saprotroph | \|Undefined Saprotroph\| | Yeast | NULL | Probable | As nectar-tap_saprotroph (Põlme et al. 2020); Decay sugar-rich substrates (Põlme et al. 2020) | Põlme S, et al. 2020. Fungal Diversity 105:1-16 ((https://doi.org/10.1007/s13225-020-00466-2)) |
| Otu17881 | 1 | k__Fungi(100);p__Ascomycota(100);c__Saccharomycetes(100);o__Saccharomycetales(100);f__Metschnikowiaceae(100);g__Metschnikowia(100);g__Metschnikowia_unclassified(100); | Metschnikowia | 13 | Saprotroph | \|Undefined Saprotroph\| | Yeast | NULL | Probable | As nectar-tap_saprotroph (Põlme et al. 2020); Decay sugar-rich substrates (Põlme et al. 2020) | Põlme S, et al. 2020. Fungal Diversity 105:1-16 ((https://doi.org/10.1007/s13225-020-00466-2)) |
| Otu17882 | 1 | k__Fungi(100);p__Ascomycota(100);c__Saccharomycetes(100);o__Saccharomycetales(100);f__Metschnikowiaceae(100);g__Metschnikowia(100);s__Metschnikowia_pulcherrima(100); | Metschnikowia | 13 | Saprotroph | \|Undefined Saprotroph\| | Yeast | NULL | Probable | As nectar-tap_saprotroph (Põlme et al. 2020); Decay sugar-rich substrates (Põlme et al. 2020) | Põlme S, et al. 2020. Fungal Diversity 105:1-16 ((https://doi.org/10.1007/s13225-020-00466-2)) |
| Otu17883 | 1 | k__Fungi(100);p__Ascomycota(100);c__Saccharomycetes(100);o__Saccharomycetales(100);f__Metschnikowiaceae(100);g__Metschnikowia(100);s__Metschnikowia_pulcherrima(100); | Metschnikowia | 13 | Saprotroph | \|Undefined Saprotroph\| | Yeast | NULL | Probable | As nectar-tap_saprotroph (Põlme et al. 2020); Decay sugar-rich substrates (Põlme et al. 2020) | Põlme S, et al. 2020. Fungal Diversity 105:1-16 ((https://doi.org/10.1007/s13225-020-00466-2)) |
| Otu17884 | 1 | k__Fungi(100);p__Ascomycota(100);c__Saccharomycetes(100);o__Saccharomycetales(100);f__Metschnikowiaceae(100);g__Metschnikowia(100);s__Metschnikowia_pulcherrima(100); | Metschnikowia | 13 | Saprotroph | \|Undefined Saprotroph\| | Yeast | NULL | Probable | As nectar-tap_saprotroph (Põlme et al. 2020); Decay sugar-rich substrates (Põlme et al. 2020) | Põlme S, et al. 2020. Fungal Diversity 105:1-16 ((https://doi.org/10.1007/s13225-020-00466-2)) |
| Otu17885 | 1 | k__Fungi(100);p__Ascomycota(100);c__Saccharomycetes(100);o__Saccharomycetales(100);f__Metschnikowiaceae(100);g__Metschnikowia(100);s__Metschnikowia_pulcherrima(100); | Metschnikowia | 13 | Saprotroph | \|Undefined Saprotroph\| | Yeast | NULL | Probable | As nectar-tap_saprotroph (Põlme et al. 2020); Decay sugar-rich substrates (Põlme et al. 2020) | Põlme S, et al. 2020. Fungal Diversity 105:1-16 ((https://doi.org/10.1007/s13225-020-00466-2)) |
| Otu17886 | 1 | k__Fungi(100);p__Ascomycota(100);c__Saccharomycetes(100);o__Saccharomycetales(100);f__Metschnikowiaceae(100);g__Metschnikowia(100);s__Metschnikowia_pulcherrima(100); | Metschnikowia | 13 | Saprotroph | \|Undefined Saprotroph\| | Yeast | NULL | Probable | As nectar-tap_saprotroph (Põlme et al. 2020); Decay sugar-rich substrates (Põlme et al. 2020) | Põlme S, et al. 2020. Fungal Diversity 105:1-16 ((https://doi.org/10.1007/s13225-020-00466-2)) |
| Otu17887 | 1 | k__Fungi(100);p__Ascomycota(100);c__Saccharomycetes(100);o__Saccharomycetales(100);f__Metschnikowiaceae(100);g__Metschnikowia(100);s__Metschnikowia_pulcherrima(100); | Metschnikowia | 13 | Saprotroph | \|Undefined Saprotroph\| | Yeast | NULL | Probable | As nectar-tap_saprotroph (Põlme et al. 2020); Decay sugar-rich substrates (Põlme et al. 2020) | Põlme S, et al. 2020. Fungal Diversity 105:1-16 ((https://doi.org/10.1007/s13225-020-00466-2)) |
| Otu17888 | 1 | k__Fungi(100);p__Ascomycota(100);c__Saccharomycetes(100);o__Saccharomycetales(100);f__Metschnikowiaceae(100);g__Metschnikowia(100);s__Metschnikowia_pulcherrima(100); | Metschnikowia | 13 | Saprotroph | \|Undefined Saprotroph\| | Yeast | NULL | Probable | As nectar-tap_saprotroph (Põlme et al. 2020); Decay sugar-rich substrates (Põlme et al. 2020) | Põlme S, et al. 2020. Fungal Diversity 105:1-16 ((https://doi.org/10.1007/s13225-020-00466-2)) |
| Otu17889 | 1 | k__Fungi(100);p__Ascomycota(100);c__Saccharomycetes(100);o__Saccharomycetales(100);f__Metschnikowiaceae(100);g__Metschnikowia(100);g__Metschnikowia_unclassified(100); | Metschnikowia | 13 | Saprotroph | \|Undefined Saprotroph\| | Yeast | NULL | Probable | As nectar-tap_saprotroph (Põlme et al. 2020); Decay sugar-rich substrates (Põlme et al. 2020) | Põlme S, et al. 2020. Fungal Diversity 105:1-16 ((https://doi.org/10.1007/s13225-020-00466-2)) |
| Otu17890 | 1 | k__Fungi(100);p__Ascomycota(100);c__Saccharomycetes(100);o__Saccharomycetales(100);f__Metschnikowiaceae(100);g__Metschnikowia(100);s__Metschnikowia_pulcherrima(100); | Metschnikowia | 13 | Saprotroph | \|Undefined Saprotroph\| | Yeast | NULL | Probable | As nectar-tap_saprotroph (Põlme et al. 2020); Decay sugar-rich substrates (Põlme et al. 2020) | Põlme S, et al. 2020. Fungal Diversity 105:1-16 ((https://doi.org/10.1007/s13225-020-00466-2)) |
| Otu17891 | 1 | k__Fungi(100);p__Ascomycota(100);c__Saccharomycetes(100);o__Saccharomycetales(100);f__Metschnikowiaceae(100);g__Metschnikowia(100);s__Metschnikowia_pulcherrima(100); | Metschnikowia | 13 | Saprotroph | \|Undefined Saprotroph\| | Yeast | NULL | Probable | As nectar-tap_saprotroph (Põlme et al. 2020); Decay sugar-rich substrates (Põlme et al. 2020) | Põlme S, et al. 2020. Fungal Diversity 105:1-16 ((https://doi.org/10.1007/s13225-020-00466-2)) |
| Otu17892 | 1 | k__Fungi(100);p__Ascomycota(100);c__Saccharomycetes(100);o__Saccharomycetales(100);f__Metschnikowiaceae(100);g__Metschnikowia(100);s__Metschnikowia_pulcherrima(100); | Metschnikowia | 13 | Saprotroph | \|Undefined Saprotroph\| | Yeast | NULL | Probable | As nectar-tap_saprotroph (Põlme et al. 2020); Decay sugar-rich substrates (Põlme et al. 2020) | Põlme S, et al. 2020. Fungal Diversity 105:1-16 ((https://doi.org/10.1007/s13225-020-00466-2)) |
| Otu17893 | 1 | k__Fungi(100);p__Ascomycota(100);c__Saccharomycetes(100);o__Saccharomycetales(100);f__Metschnikowiaceae(100);g__Metschnikowia(100);s__Metschnikowia_pulcherrima(100); | Metschnikowia | 13 | Saprotroph | \|Undefined Saprotroph\| | Yeast | NULL | Probable | As nectar-tap_saprotroph (Põlme et al. 2020); Decay sugar-rich substrates (Põlme et al. 2020) | Põlme S, et al. 2020. Fungal Diversity 105:1-16 ((https://doi.org/10.1007/s13225-020-00466-2)) |
| Otu17894 | 1 | k__Fungi(100);p__Ascomycota(100);c__Saccharomycetes(100);o__Saccharomycetales(100);f__Metschnikowiaceae(100);g__Metschnikowia(100);g__Metschnikowia_unclassified(100); | Metschnikowia | 13 | Saprotroph | \|Undefined Saprotroph\| | Yeast | NULL | Probable | As nectar-tap_saprotroph (Põlme et al. 2020); Decay sugar-rich substrates (Põlme et al. 2020) | Põlme S, et al. 2020. Fungal Diversity 105:1-16 ((https://doi.org/10.1007/s13225-020-00466-2)) |
| Otu17895 | 1 | k__Fungi(100);p__Ascomycota(100);c__Saccharomycetes(100);o__Saccharomycetales(100);f__Metschnikowiaceae(100);g__Metschnikowia(100);s__Metschnikowia_pulcherrima(100); | Metschnikowia | 13 | Saprotroph | \|Undefined Saprotroph\| | Yeast | NULL | Probable | As nectar-tap_saprotroph (Põlme et al. 2020); Decay sugar-rich substrates (Põlme et al. 2020) | Põlme S, et al. 2020. Fungal Diversity 105:1-16 ((https://doi.org/10.1007/s13225-020-00466-2)) |
| Otu17896 | 1 | k__Fungi(100);p__Ascomycota(100);c__Saccharomycetes(100);o__Saccharomycetales(100);f__Metschnikowiaceae(100);g__Metschnikowia(100);g__Metschnikowia_unclassified(100); | Metschnikowia | 13 | Saprotroph | \|Undefined Saprotroph\| | Yeast | NULL | Probable | As nectar-tap_saprotroph (Põlme et al. 2020); Decay sugar-rich substrates (Põlme et al. 2020) | Põlme S, et al. 2020. Fungal Diversity 105:1-16 ((https://doi.org/10.1007/s13225-020-00466-2)) |
| Otu17897 | 1 | k__Fungi(100);p__Ascomycota(100);c__Saccharomycetes(100);o__Saccharomycetales(100);f__Metschnikowiaceae(100);g__Metschnikowia(100);s__Metschnikowia_pulcherrima(100); | Metschnikowia | 13 | Saprotroph | \|Undefined Saprotroph\| | Yeast | NULL | Probable | As nectar-tap_saprotroph (Põlme et al. 2020); Decay sugar-rich substrates (Põlme et al. 2020) | Põlme S, et al. 2020. Fungal Diversity 105:1-16 ((https://doi.org/10.1007/s13225-020-00466-2)) |
| Otu17898 | 1 | k__Fungi(100);p__Ascomycota(100);c__Saccharomycetes(100);o__Saccharomycetales(100);f__Metschnikowiaceae(100);g__Metschnikowia(100);s__Metschnikowia_pulcherrima(100); | Metschnikowia | 13 | Saprotroph | \|Undefined Saprotroph\| | Yeast | NULL | Probable | As nectar-tap_saprotroph (Põlme et al. 2020); Decay sugar-rich substrates (Põlme et al. 2020) | Põlme S, et al. 2020. Fungal Diversity 105:1-16 ((https://doi.org/10.1007/s13225-020-00466-2)) |
| Otu17899 | 1 | k__Fungi(100);p__Ascomycota(100);c__Saccharomycetes(100);o__Saccharomycetales(100);f__Metschnikowiaceae(100);g__Metschnikowia(100);s__Metschnikowia_pulcherrima(100); | Metschnikowia | 13 | Saprotroph | \|Undefined Saprotroph\| | Yeast | NULL | Probable | As nectar-tap_saprotroph (Põlme et al. 2020); Decay sugar-rich substrates (Põlme et al. 2020) | Põlme S, et al. 2020. Fungal Diversity 105:1-16 ((https://doi.org/10.1007/s13225-020-00466-2)) |
| Otu17900 | 1 | k__Fungi(100);p__Ascomycota(100);c__Saccharomycetes(100);o__Saccharomycetales(100);f__Metschnikowiaceae(100);g__Metschnikowia(100);s__Metschnikowia_pulcherrima(100); | Metschnikowia | 13 | Saprotroph | \|Undefined Saprotroph\| | Yeast | NULL | Probable | As nectar-tap_saprotroph (Põlme et al. 2020); Decay sugar-rich substrates (Põlme et al. 2020) | Põlme S, et al. 2020. Fungal Diversity 105:1-16 ((https://doi.org/10.1007/s13225-020-00466-2)) |
| Otu17901 | 1 | k__Fungi(100);p__Ascomycota(100);c__Saccharomycetes(100);o__Saccharomycetales(100);f__Metschnikowiaceae(100);g__Metschnikowia(100);s__Metschnikowia_pulcherrima(100); | Metschnikowia | 13 | Saprotroph | \|Undefined Saprotroph\| | Yeast | NULL | Probable | As nectar-tap_saprotroph (Põlme et al. 2020); Decay sugar-rich substrates (Põlme et al. 2020) | Põlme S, et al. 2020. Fungal Diversity 105:1-16 ((https://doi.org/10.1007/s13225-020-00466-2)) |
| Otu17902 | 1 | k__Fungi(100);p__Ascomycota(100);c__Saccharomycetes(100);o__Saccharomycetales(100);f__Metschnikowiaceae(100);g__Metschnikowia(100);s__Metschnikowia_pulcherrima(100); | Metschnikowia | 13 | Saprotroph | \|Undefined Saprotroph\| | Yeast | NULL | Probable | As nectar-tap_saprotroph (Põlme et al. 2020); Decay sugar-rich substrates (Põlme et al. 2020) | Põlme S, et al. 2020. Fungal Diversity 105:1-16 ((https://doi.org/10.1007/s13225-020-00466-2)) |
| Otu17903 | 1 | k__Fungi(100);p__Ascomycota(100);c__Saccharomycetes(100);o__Saccharomycetales(100);f__Metschnikowiaceae(100);g__Metschnikowia(100);s__Metschnikowia_pulcherrima(100); | Metschnikowia | 13 | Saprotroph | \|Undefined Saprotroph\| | Yeast | NULL | Probable | As nectar-tap_saprotroph (Põlme et al. 2020); Decay sugar-rich substrates (Põlme et al. 2020) | Põlme S, et al. 2020. Fungal Diversity 105:1-16 ((https://doi.org/10.1007/s13225-020-00466-2)) |
| Otu17904 | 1 | k__Fungi(100);p__Ascomycota(100);c__Saccharomycetes(100);o__Saccharomycetales(100);f__Metschnikowiaceae(100);g__Metschnikowia(100);g__Metschnikowia_unclassified(100); | Metschnikowia | 13 | Saprotroph | \|Undefined Saprotroph\| | Yeast | NULL | Probable | As nectar-tap_saprotroph (Põlme et al. 2020); Decay sugar-rich substrates (Põlme et al. 2020) | Põlme S, et al. 2020. Fungal Diversity 105:1-16 ((https://doi.org/10.1007/s13225-020-00466-2)) |
| Otu17905 | 1 | k__Fungi(100);p__Ascomycota(100);c__Saccharomycetes(100);o__Saccharomycetales(100);f__Metschnikowiaceae(100);g__Metschnikowia(100);s__Metschnikowia_pulcherrima(100); | Metschnikowia | 13 | Saprotroph | \|Undefined Saprotroph\| | Yeast | NULL | Probable | As nectar-tap_saprotroph (Põlme et al. 2020); Decay sugar-rich substrates (Põlme et al. 2020) | Põlme S, et al. 2020. Fungal Diversity 105:1-16 ((https://doi.org/10.1007/s13225-020-00466-2)) |
| Otu17906 | 1 | k__Fungi(100);p__Ascomycota(100);c__Saccharomycetes(100);o__Saccharomycetales(100);f__Metschnikowiaceae(100);g__Metschnikowia(100);s__Metschnikowia_pulcherrima(100); | Metschnikowia | 13 | Saprotroph | \|Undefined Saprotroph\| | Yeast | NULL | Probable | As nectar-tap_saprotroph (Põlme et al. 2020); Decay sugar-rich substrates (Põlme et al. 2020) | Põlme S, et al. 2020. Fungal Diversity 105:1-16 ((https://doi.org/10.1007/s13225-020-00466-2)) |
| Otu17907 | 1 | k__Fungi(100);p__Ascomycota(100);c__Saccharomycetes(100);o__Saccharomycetales(100);f__Metschnikowiaceae(100);g__Metschnikowia(100);s__Metschnikowia_pulcherrima(100); | Metschnikowia | 13 | Saprotroph | \|Undefined Saprotroph\| | Yeast | NULL | Probable | As nectar-tap_saprotroph (Põlme et al. 2020); Decay sugar-rich substrates (Põlme et al. 2020) | Põlme S, et al. 2020. Fungal Diversity 105:1-16 ((https://doi.org/10.1007/s13225-020-00466-2)) |
| Otu17908 | 1 | k__Fungi(100);p__Ascomycota(100);c__Saccharomycetes(100);o__Saccharomycetales(100);f__Metschnikowiaceae(100);g__Metschnikowia(100);s__Metschnikowia_pulcherrima(100); | Metschnikowia | 13 | Saprotroph | \|Undefined Saprotroph\| | Yeast | NULL | Probable | As nectar-tap_saprotroph (Põlme et al. 2020); Decay sugar-rich substrates (Põlme et al. 2020) | Põlme S, et al. 2020. Fungal Diversity 105:1-16 ((https://doi.org/10.1007/s13225-020-00466-2)) |
| Otu17909 | 1 | k__Fungi(100);p__Ascomycota(100);c__Saccharomycetes(100);o__Saccharomycetales(100);f__Metschnikowiaceae(100);g__Metschnikowia(100);g__Metschnikowia_unclassified(100); | Metschnikowia | 13 | Saprotroph | \|Undefined Saprotroph\| | Yeast | NULL | Probable | As nectar-tap_saprotroph (Põlme et al. 2020); Decay sugar-rich substrates (Põlme et al. 2020) | Põlme S, et al. 2020. Fungal Diversity 105:1-16 ((https://doi.org/10.1007/s13225-020-00466-2)) |
| Otu17910 | 1 | k__Fungi(100);p__Ascomycota(100);c__Saccharomycetes(100);o__Saccharomycetales(100);f__Metschnikowiaceae(100);g__Metschnikowia(100);s__Metschnikowia_pulcherrima(100); | Metschnikowia | 13 | Saprotroph | \|Undefined Saprotroph\| | Yeast | NULL | Probable | As nectar-tap_saprotroph (Põlme et al. 2020); Decay sugar-rich substrates (Põlme et al. 2020) | Põlme S, et al. 2020. Fungal Diversity 105:1-16 ((https://doi.org/10.1007/s13225-020-00466-2)) |
| Otu17911 | 1 | k__Fungi(100);p__Ascomycota(100);c__Saccharomycetes(100);o__Saccharomycetales(100);f__Metschnikowiaceae(100);g__Metschnikowia(100);g__Metschnikowia_unclassified(100); | Metschnikowia | 13 | Saprotroph | \|Undefined Saprotroph\| | Yeast | NULL | Probable | As nectar-tap_saprotroph (Põlme et al. 2020); Decay sugar-rich substrates (Põlme et al. 2020) | Põlme S, et al. 2020. Fungal Diversity 105:1-16 ((https://doi.org/10.1007/s13225-020-00466-2)) |
| Otu17912 | 1 | k__Fungi(100);p__Ascomycota(100);c__Saccharomycetes(100);o__Saccharomycetales(100);f__Metschnikowiaceae(100);g__Metschnikowia(100);s__Metschnikowia_pulcherrima(100); | Metschnikowia | 13 | Saprotroph | \|Undefined Saprotroph\| | Yeast | NULL | Probable | As nectar-tap_saprotroph (Põlme et al. 2020); Decay sugar-rich substrates (Põlme et al. 2020) | Põlme S, et al. 2020. Fungal Diversity 105:1-16 ((https://doi.org/10.1007/s13225-020-00466-2)) |
| Otu17913 | 1 | k__Fungi(100);p__Ascomycota(100);c__Saccharomycetes(100);o__Saccharomycetales(100);f__Metschnikowiaceae(100);g__Metschnikowia(100);s__Metschnikowia_pulcherrima(100); | Metschnikowia | 13 | Saprotroph | \|Undefined Saprotroph\| | Yeast | NULL | Probable | As nectar-tap_saprotroph (Põlme et al. 2020); Decay sugar-rich substrates (Põlme et al. 2020) | Põlme S, et al. 2020. Fungal Diversity 105:1-16 ((https://doi.org/10.1007/s13225-020-00466-2)) |
| Otu17914 | 1 | k__Fungi(100);p__Ascomycota(100);c__Saccharomycetes(100);o__Saccharomycetales(100);f__Metschnikowiaceae(100);g__Metschnikowia(100);s__Metschnikowia_pulcherrima(100); | Metschnikowia | 13 | Saprotroph | \|Undefined Saprotroph\| | Yeast | NULL | Probable | As nectar-tap_saprotroph (Põlme et al. 2020); Decay sugar-rich substrates (Põlme et al. 2020) | Põlme S, et al. 2020. Fungal Diversity 105:1-16 ((https://doi.org/10.1007/s13225-020-00466-2)) |
| Otu17915 | 1 | k__Fungi(100);p__Ascomycota(100);c__Saccharomycetes(100);o__Saccharomycetales(100);f__Metschnikowiaceae(100);g__Metschnikowia(100);g__Metschnikowia_unclassified(100); | Metschnikowia | 13 | Saprotroph | \|Undefined Saprotroph\| | Yeast | NULL | Probable | As nectar-tap_saprotroph (Põlme et al. 2020); Decay sugar-rich substrates (Põlme et al. 2020) | Põlme S, et al. 2020. Fungal Diversity 105:1-16 ((https://doi.org/10.1007/s13225-020-00466-2)) |
| Otu17916 | 1 | k__Fungi(100);p__Ascomycota(100);c__Saccharomycetes(100);o__Saccharomycetales(100);f__Metschnikowiaceae(100);g__Metschnikowia(100);g__Metschnikowia_unclassified(100); | Metschnikowia | 13 | Saprotroph | \|Undefined Saprotroph\| | Yeast | NULL | Probable | As nectar-tap_saprotroph (Põlme et al. 2020); Decay sugar-rich substrates (Põlme et al. 2020) | Põlme S, et al. 2020. Fungal Diversity 105:1-16 ((https://doi.org/10.1007/s13225-020-00466-2)) |
| Otu17917 | 1 | k__Fungi(100);p__Ascomycota(100);c__Saccharomycetes(100);o__Saccharomycetales(100);f__Metschnikowiaceae(100);g__Metschnikowia(100);s__Metschnikowia_pulcherrima(100); | Metschnikowia | 13 | Saprotroph | \|Undefined Saprotroph\| | Yeast | NULL | Probable | As nectar-tap_saprotroph (Põlme et al. 2020); Decay sugar-rich substrates (Põlme et al. 2020) | Põlme S, et al. 2020. Fungal Diversity 105:1-16 ((https://doi.org/10.1007/s13225-020-00466-2)) |
| Otu17918 | 1 | k__Fungi(100);p__Ascomycota(100);c__Saccharomycetes(100);o__Saccharomycetales(100);f__Metschnikowiaceae(100);g__Metschnikowia(100);s__Metschnikowia_pulcherrima(100); | Metschnikowia | 13 | Saprotroph | \|Undefined Saprotroph\| | Yeast | NULL | Probable | As nectar-tap_saprotroph (Põlme et al. 2020); Decay sugar-rich substrates (Põlme et al. 2020) | Põlme S, et al. 2020. Fungal Diversity 105:1-16 ((https://doi.org/10.1007/s13225-020-00466-2)) |
| Otu17919 | 1 | k__Fungi(100);p__Ascomycota(100);c__Saccharomycetes(100);o__Saccharomycetales(100);f__Metschnikowiaceae(100);g__Metschnikowia(100);g__Metschnikowia_unclassified(100); | Metschnikowia | 13 | Saprotroph | \|Undefined Saprotroph\| | Yeast | NULL | Probable | As nectar-tap_saprotroph (Põlme et al. 2020); Decay sugar-rich substrates (Põlme et al. 2020) | Põlme S, et al. 2020. Fungal Diversity 105:1-16 ((https://doi.org/10.1007/s13225-020-00466-2)) |
| Otu17920 | 1 | k__Fungi(100);p__Ascomycota(100);c__Saccharomycetes(100);o__Saccharomycetales(100);f__Metschnikowiaceae(100);g__Metschnikowia(100);s__Metschnikowia_pulcherrima(100); | Metschnikowia | 13 | Saprotroph | \|Undefined Saprotroph\| | Yeast | NULL | Probable | As nectar-tap_saprotroph (Põlme et al. 2020); Decay sugar-rich substrates (Põlme et al. 2020) | Põlme S, et al. 2020. Fungal Diversity 105:1-16 ((https://doi.org/10.1007/s13225-020-00466-2)) |
| Otu17921 | 1 | k__Fungi(100);p__Ascomycota(100);c__Saccharomycetes(100);o__Saccharomycetales(100);f__Metschnikowiaceae(100);g__Metschnikowia(100);s__Metschnikowia_pulcherrima(100); | Metschnikowia | 13 | Saprotroph | \|Undefined Saprotroph\| | Yeast | NULL | Probable | As nectar-tap_saprotroph (Põlme et al. 2020); Decay sugar-rich substrates (Põlme et al. 2020) | Põlme S, et al. 2020. Fungal Diversity 105:1-16 ((https://doi.org/10.1007/s13225-020-00466-2)) |
| Otu17922 | 1 | k__Fungi(100);p__Ascomycota(100);c__Saccharomycetes(100);o__Saccharomycetales(100);f__Metschnikowiaceae(100);g__Metschnikowia(100);g__Metschnikowia_unclassified(100); | Metschnikowia | 13 | Saprotroph | \|Undefined Saprotroph\| | Yeast | NULL | Probable | As nectar-tap_saprotroph (Põlme et al. 2020); Decay sugar-rich substrates (Põlme et al. 2020) | Põlme S, et al. 2020. Fungal Diversity 105:1-16 ((https://doi.org/10.1007/s13225-020-00466-2)) |
| Otu17923 | 1 | k__Fungi(100);p__Ascomycota(100);c__Saccharomycetes(100);o__Saccharomycetales(100);f__Metschnikowiaceae(100);g__Metschnikowia(100);s__Metschnikowia_pulcherrima(100); | Metschnikowia | 13 | Saprotroph | \|Undefined Saprotroph\| | Yeast | NULL | Probable | As nectar-tap_saprotroph (Põlme et al. 2020); Decay sugar-rich substrates (Põlme et al. 2020) | Põlme S, et al. 2020. Fungal Diversity 105:1-16 ((https://doi.org/10.1007/s13225-020-00466-2)) |
| Otu17924 | 1 | k__Fungi(100);p__Ascomycota(100);c__Saccharomycetes(100);o__Saccharomycetales(100);f__Metschnikowiaceae(100);g__Metschnikowia(100);s__Metschnikowia_pulcherrima(100); | Metschnikowia | 13 | Saprotroph | \|Undefined Saprotroph\| | Yeast | NULL | Probable | As nectar-tap_saprotroph (Põlme et al. 2020); Decay sugar-rich substrates (Põlme et al. 2020) | Põlme S, et al. 2020. Fungal Diversity 105:1-16 ((https://doi.org/10.1007/s13225-020-00466-2)) |
| Otu17925 | 1 | k__Fungi(100);p__Ascomycota(100);c__Saccharomycetes(100);o__Saccharomycetales(100);f__Metschnikowiaceae(100);g__Metschnikowia(100);s__Metschnikowia_pulcherrima(100); | Metschnikowia | 13 | Saprotroph | \|Undefined Saprotroph\| | Yeast | NULL | Probable | As nectar-tap_saprotroph (Põlme et al. 2020); Decay sugar-rich substrates (Põlme et al. 2020) | Põlme S, et al. 2020. Fungal Diversity 105:1-16 ((https://doi.org/10.1007/s13225-020-00466-2)) |
| Otu17926 | 1 | k__Fungi(100);p__Ascomycota(100);c__Saccharomycetes(100);o__Saccharomycetales(100);f__Metschnikowiaceae(100);g__Metschnikowia(100);s__Metschnikowia_pulcherrima(100); | Metschnikowia | 13 | Saprotroph | \|Undefined Saprotroph\| | Yeast | NULL | Probable | As nectar-tap_saprotroph (Põlme et al. 2020); Decay sugar-rich substrates (Põlme et al. 2020) | Põlme S, et al. 2020. Fungal Diversity 105:1-16 ((https://doi.org/10.1007/s13225-020-00466-2)) |
| Otu17927 | 1 | k__Fungi(100);p__Ascomycota(100);c__Saccharomycetes(100);o__Saccharomycetales(100);f__Metschnikowiaceae(100);g__Metschnikowia(100);s__Metschnikowia_pulcherrima(100); | Metschnikowia | 13 | Saprotroph | \|Undefined Saprotroph\| | Yeast | NULL | Probable | As nectar-tap_saprotroph (Põlme et al. 2020); Decay sugar-rich substrates (Põlme et al. 2020) | Põlme S, et al. 2020. Fungal Diversity 105:1-16 ((https://doi.org/10.1007/s13225-020-00466-2)) |
| Otu17928 | 1 | k__Fungi(100);p__Ascomycota(100);c__Saccharomycetes(100);o__Saccharomycetales(100);f__Metschnikowiaceae(100);g__Metschnikowia(100);s__Metschnikowia_pulcherrima(100); | Metschnikowia | 13 | Saprotroph | \|Undefined Saprotroph\| | Yeast | NULL | Probable | As nectar-tap_saprotroph (Põlme et al. 2020); Decay sugar-rich substrates (Põlme et al. 2020) | Põlme S, et al. 2020. Fungal Diversity 105:1-16 ((https://doi.org/10.1007/s13225-020-00466-2)) |
| Otu17929 | 1 | k__Fungi(100);p__Ascomycota(100);c__Saccharomycetes(100);o__Saccharomycetales(100);f__Metschnikowiaceae(100);g__Metschnikowia(100);s__Metschnikowia_pulcherrima(100); | Metschnikowia | 13 | Saprotroph | \|Undefined Saprotroph\| | Yeast | NULL | Probable | As nectar-tap_saprotroph (Põlme et al. 2020); Decay sugar-rich substrates (Põlme et al. 2020) | Põlme S, et al. 2020. Fungal Diversity 105:1-16 ((https://doi.org/10.1007/s13225-020-00466-2)) |
| Otu17930 | 1 | k__Fungi(100);p__Ascomycota(100);c__Saccharomycetes(100);o__Saccharomycetales(100);f__Metschnikowiaceae(100);g__Metschnikowia(100);g__Metschnikowia_unclassified(100); | Metschnikowia | 13 | Saprotroph | \|Undefined Saprotroph\| | Yeast | NULL | Probable | As nectar-tap_saprotroph (Põlme et al. 2020); Decay sugar-rich substrates (Põlme et al. 2020) | Põlme S, et al. 2020. Fungal Diversity 105:1-16 ((https://doi.org/10.1007/s13225-020-00466-2)) |
| Otu17931 | 1 | k__Fungi(100);p__Ascomycota(100);c__Saccharomycetes(100);o__Saccharomycetales(100);f__Metschnikowiaceae(100);g__Metschnikowia(100);g__Metschnikowia_unclassified(100); | Metschnikowia | 13 | Saprotroph | \|Undefined Saprotroph\| | Yeast | NULL | Probable | As nectar-tap_saprotroph (Põlme et al. 2020); Decay sugar-rich substrates (Põlme et al. 2020) | Põlme S, et al. 2020. Fungal Diversity 105:1-16 ((https://doi.org/10.1007/s13225-020-00466-2)) |
| Otu17932 | 1 | k__Fungi(100);p__Ascomycota(100);c__Saccharomycetes(100);o__Saccharomycetales(100);f__Metschnikowiaceae(100);g__Metschnikowia(100);s__Metschnikowia_pulcherrima(100); | Metschnikowia | 13 | Saprotroph | \|Undefined Saprotroph\| | Yeast | NULL | Probable | As nectar-tap_saprotroph (Põlme et al. 2020); Decay sugar-rich substrates (Põlme et al. 2020) | Põlme S, et al. 2020. Fungal Diversity 105:1-16 ((https://doi.org/10.1007/s13225-020-00466-2)) |
| Otu17933 | 1 | k__Fungi(100);p__Ascomycota(100);c__Saccharomycetes(100);o__Saccharomycetales(100);f__Metschnikowiaceae(100);g__Metschnikowia(100);g__Metschnikowia_unclassified(100); | Metschnikowia | 13 | Saprotroph | \|Undefined Saprotroph\| | Yeast | NULL | Probable | As nectar-tap_saprotroph (Põlme et al. 2020); Decay sugar-rich substrates (Põlme et al. 2020) | Põlme S, et al. 2020. Fungal Diversity 105:1-16 ((https://doi.org/10.1007/s13225-020-00466-2)) |
| Otu17934 | 1 | k__Fungi(100);p__Ascomycota(100);c__Saccharomycetes(100);o__Saccharomycetales(100);f__Metschnikowiaceae(100);g__Metschnikowia(100);g__Metschnikowia_unclassified(100); | Metschnikowia | 13 | Saprotroph | \|Undefined Saprotroph\| | Yeast | NULL | Probable | As nectar-tap_saprotroph (Põlme et al. 2020); Decay sugar-rich substrates (Põlme et al. 2020) | Põlme S, et al. 2020. Fungal Diversity 105:1-16 ((https://doi.org/10.1007/s13225-020-00466-2)) |
| Otu17935 | 1 | k__Fungi(100);p__Ascomycota(100);c__Saccharomycetes(100);o__Saccharomycetales(100);f__Metschnikowiaceae(100);g__Metschnikowia(100);s__Metschnikowia_pulcherrima(100); | Metschnikowia | 13 | Saprotroph | \|Undefined Saprotroph\| | Yeast | NULL | Probable | As nectar-tap_saprotroph (Põlme et al. 2020); Decay sugar-rich substrates (Põlme et al. 2020) | Põlme S, et al. 2020. Fungal Diversity 105:1-16 ((https://doi.org/10.1007/s13225-020-00466-2)) |
| Otu17936 | 1 | k__Fungi(100);p__Ascomycota(100);c__Saccharomycetes(100);o__Saccharomycetales(100);f__Metschnikowiaceae(100);g__Metschnikowia(100);s__Metschnikowia_pulcherrima(100); | Metschnikowia | 13 | Saprotroph | \|Undefined Saprotroph\| | Yeast | NULL | Probable | As nectar-tap_saprotroph (Põlme et al. 2020); Decay sugar-rich substrates (Põlme et al. 2020) | Põlme S, et al. 2020. Fungal Diversity 105:1-16 ((https://doi.org/10.1007/s13225-020-00466-2)) |
| Otu17937 | 1 | k__Fungi(100);p__Ascomycota(100);c__Saccharomycetes(100);o__Saccharomycetales(100);f__Metschnikowiaceae(100);g__Metschnikowia(100);s__Metschnikowia_pulcherrima(100); | Metschnikowia | 13 | Saprotroph | \|Undefined Saprotroph\| | Yeast | NULL | Probable | As nectar-tap_saprotroph (Põlme et al. 2020); Decay sugar-rich substrates (Põlme et al. 2020) | Põlme S, et al. 2020. Fungal Diversity 105:1-16 ((https://doi.org/10.1007/s13225-020-00466-2)) |
| Otu17938 | 1 | k__Fungi(100);p__Ascomycota(100);c__Saccharomycetes(100);o__Saccharomycetales(100);f__Metschnikowiaceae(100);g__Metschnikowia(100);s__Metschnikowia_pulcherrima(100); | Metschnikowia | 13 | Saprotroph | \|Undefined Saprotroph\| | Yeast | NULL | Probable | As nectar-tap_saprotroph (Põlme et al. 2020); Decay sugar-rich substrates (Põlme et al. 2020) | Põlme S, et al. 2020. Fungal Diversity 105:1-16 ((https://doi.org/10.1007/s13225-020-00466-2)) |
| Otu17939 | 1 | k__Fungi(100);p__Ascomycota(100);c__Saccharomycetes(100);o__Saccharomycetales(100);f__Metschnikowiaceae(100);g__Metschnikowia(100);s__Metschnikowia_pulcherrima(100); | Metschnikowia | 13 | Saprotroph | \|Undefined Saprotroph\| | Yeast | NULL | Probable | As nectar-tap_saprotroph (Põlme et al. 2020); Decay sugar-rich substrates (Põlme et al. 2020) | Põlme S, et al. 2020. Fungal Diversity 105:1-16 ((https://doi.org/10.1007/s13225-020-00466-2)) |
| Otu17940 | 1 | k__Fungi(100);p__Ascomycota(100);c__Saccharomycetes(100);o__Saccharomycetales(100);f__Saccharomycetales_family_Incertae_sedis(100);g__Candida(100);g__Candida_unclassified(100); | Candida | 13 | Pathotroph-Saprotroph-Symbiotroph | Animal Pathogen-Endophyte-Endosymbiont-Epiphyte-\|Undefined Saprotroph\| | Dimorphic Yeast-Yeast | NULL | Probable | As nectar-tap_saprotroph (Põlme et al. 2020); Decay sugar-rich substrates (Põlme et al. 2020); This asexual genus is polyphyletic and contain species that function very differently depend on what they are. Caution should be taken when interpreting guilds for Candida species. | Manolakaki D, et al. 2010. Virulence. 1: 367-375 ((https://doi.org/10.4161/viru.1.5.12796)); Mekibib SB, et al. 2011. Tropical Plant Pathology 36:233-240 ((https://doi.org/10.1590/S1982-56762011000400004)); Põlme S, et al. 2020. Fungal Diversity 105:1-16 ((https://doi.org/10.1007/s13225-020-00466-2)) |
| Otu17941 | 1 | k__Fungi(100);p__Ascomycota(100);c__Saccharomycetes(100);o__Saccharomycetales(100);f__Metschnikowiaceae(100);g__Metschnikowia(100);s__Metschnikowia_pulcherrima(100); | Metschnikowia | 13 | Saprotroph | \|Undefined Saprotroph\| | Yeast | NULL | Probable | As nectar-tap_saprotroph (Põlme et al. 2020); Decay sugar-rich substrates (Põlme et al. 2020) | Põlme S, et al. 2020. Fungal Diversity 105:1-16 ((https://doi.org/10.1007/s13225-020-00466-2)) |
| Otu17942 | 1 | k__Fungi(100);p__Ascomycota(100);c__Saccharomycetes(100);o__Saccharomycetales(100);f__Metschnikowiaceae(100);g__Metschnikowia(100);s__Metschnikowia_pulcherrima(100); | Metschnikowia | 13 | Saprotroph | \|Undefined Saprotroph\| | Yeast | NULL | Probable | As nectar-tap_saprotroph (Põlme et al. 2020); Decay sugar-rich substrates (Põlme et al. 2020) | Põlme S, et al. 2020. Fungal Diversity 105:1-16 ((https://doi.org/10.1007/s13225-020-00466-2)) |
| Otu17943 | 1 | k__Fungi(100);p__Ascomycota(100);c__Saccharomycetes(100);o__Saccharomycetales(100);f__Metschnikowiaceae(100);g__Metschnikowia(100);s__Metschnikowia_pulcherrima(100); | Metschnikowia | 13 | Saprotroph | \|Undefined Saprotroph\| | Yeast | NULL | Probable | As nectar-tap_saprotroph (Põlme et al. 2020); Decay sugar-rich substrates (Põlme et al. 2020) | Põlme S, et al. 2020. Fungal Diversity 105:1-16 ((https://doi.org/10.1007/s13225-020-00466-2)) |

| | | | | | | | | | | | |
|---|---|---|---|---|---|---|---|---|---|---|---|
| Otu17944 | 1 | k__Fungi(100);p__Ascomycota(100);c__Saccharomycetes(100);o__Saccharomycetales(100);f__Metschnikowiaceae(100);g__Metschnikowia(100);s__Metschnikowia_pulcherrima(100); | Metschnikowia | 13 | Saprotroph | \|Undefined Saprotroph\| | Yeast | NULL | Probable | As nectar-tap_saprotroph (Põlme et al. 2020); Decay sugar-rich substrates (Põlme et al. 2020) | Põlme S, et al. 2020. Fungal Diversity 105:1-16 ((https://doi.org/10.1007/s13225-020-00466-2)) |
| Otu17945 | 1 | k__Fungi(100);p__Ascomycota(100);c__Saccharomycetes(100);o__Saccharomycetales(100);f__Metschnikowiaceae(100);g__Metschnikowia(100);s__Metschnikowia_pulcherrima(100); | Metschnikowia | 13 | Saprotroph | \|Undefined Saprotroph\| | Yeast | NULL | Probable | As nectar-tap_saprotroph (Põlme et al. 2020); Decay sugar-rich substrates (Põlme et al. 2020) | Põlme S, et al. 2020. Fungal Diversity 105:1-16 ((https://doi.org/10.1007/s13225-020-00466-2)) |
| Otu17946 | 1 | k__Fungi(100);p__Ascomycota(100);c__Saccharomycetes(100);o__Saccharomycetales(100);f__Metschnikowiaceae(100);g__Metschnikowia(100);s__Metschnikowia_pulcherrima(100); | Metschnikowia | 13 | Saprotroph | \|Undefined Saprotroph\| | Yeast | NULL | Probable | As nectar-tap_saprotroph (Põlme et al. 2020); Decay sugar-rich substrates (Põlme et al. 2020) | Põlme S, et al. 2020. Fungal Diversity 105:1-16 ((https://doi.org/10.1007/s13225-020-00466-2)) |
| Otu17947 | 1 | k__Fungi(100);p__Ascomycota(100);c__Saccharomycetes(100);o__Saccharomycetales(100);f__Metschnikowiaceae(100);g__Metschnikowia(100);g__Metschnikowia_unclassified(100); | Metschnikowia | 13 | Saprotroph | \|Undefined Saprotroph\| | Yeast | NULL | Probable | As nectar-tap_saprotroph (Põlme et al. 2020); Decay sugar-rich substrates (Põlme et al. 2020) | Põlme S, et al. 2020. Fungal Diversity 105:1-16 ((https://doi.org/10.1007/s13225-020-00466-2)) |
| Otu17948 | 1 | k__Fungi(100);p__Ascomycota(100);c__Saccharomycetes(100);o__Saccharomycetales(100);f__Metschnikowiaceae(100);g__Metschnikowia(100);g__Metschnikowia_unclassified(100); | Metschnikowia | 13 | Saprotroph | \|Undefined Saprotroph\| | Yeast | NULL | Probable | As nectar-tap_saprotroph (Põlme et al. 2020); Decay sugar-rich substrates (Põlme et al. 2020) | Põlme S, et al. 2020. Fungal Diversity 105:1-16 ((https://doi.org/10.1007/s13225-020-00466-2)) |
| Otu17949 | 1 | k__Fungi(100);p__Ascomycota(100);c__Saccharomycetes(100);o__Saccharomycetales(100);f__Metschnikowiaceae(100);g__Metschnikowia(100);s__Metschnikowia_pulcherrima(100); | Metschnikowia | 13 | Saprotroph | \|Undefined Saprotroph\| | Yeast | NULL | Probable | As nectar-tap_saprotroph (Põlme et al. 2020); Decay sugar-rich substrates (Põlme et al. 2020) | Põlme S, et al. 2020. Fungal Diversity 105:1-16 ((https://doi.org/10.1007/s13225-020-00466-2)) |
| Otu17950 | 1 | k__Fungi(100);p__Ascomycota(100);c__Saccharomycetes(100);o__Saccharomycetales(100);f__Saccharomycetales_family_Incertae_sedis(100);g__Candida(100);g__Candida_unclassified(100); | Candida | 13 | Pathotroph-Saprotroph-Symbiotroph | Animal Pathogen-Endophyte-Endosymbiont-Epiphyte-\|Undefined Saprotroph\| | Dimorphic Yeast-Yeast | NULL | Probable | As nectar-tap_saprotroph (Põlme et al. 2020); Decay sugar-rich substrates (Põlme et al. 2020); This asexual genus is polyphyletic and contain species that function very differently depend on what they are. Caution should be taken when interpreting guilds for Candida species. | Manolakaki D, et al. 2010. Virulence. 1: 367-375 ((https://doi.org/10.4161/viru.1.5.12796)); Mekbib SB, et al. 2011. Tropical Plant Pathology 36:233-240 ((https://doi.org/10.1590/S1982-56762011000400004)); Põlme S, et al. 2020. Fungal Diversity 105:1-16 ((https://doi.org/10.1007/s13225-020-00466-2)) |
| Otu17951 | 1 | k__Fungi(100);p__Ascomycota(100);c__Saccharomycetes(100);o__Saccharomycetales(100);f__Metschnikowiaceae(100);g__Metschnikowia(100);s__Metschnikowia_pulcherrima(100); | Metschnikowia | 13 | Saprotroph | \|Undefined Saprotroph\| | Yeast | NULL | Probable | As nectar-tap_saprotroph (Põlme et al. 2020); Decay sugar-rich substrates (Põlme et al. 2020) | Põlme S, et al. 2020. Fungal Diversity 105:1-16 ((https://doi.org/10.1007/s13225-020-00466-2)) |
| Otu17952 | 1 | k__Fungi(100);p__Ascomycota(100);c__Saccharomycetes(100);o__Saccharomycetales(100);f__Metschnikowiaceae(100);g__Metschnikowia(100);s__Metschnikowia_pulcherrima(100); | Metschnikowia | 13 | Saprotroph | \|Undefined Saprotroph\| | Yeast | NULL | Probable | As nectar-tap_saprotroph (Põlme et al. 2020); Decay sugar-rich substrates (Põlme et al. 2020) | Põlme S, et al. 2020. Fungal Diversity 105:1-16 ((https://doi.org/10.1007/s13225-020-00466-2)) |
| Otu17953 | 1 | k__Fungi(100);p__Ascomycota(100);c__Saccharomycetes(100);o__Saccharomycetales(100);f__Metschnikowiaceae(100);g__Metschnikowia(100);s__Metschnikowia_pulcherrima(100); | Metschnikowia | 13 | Saprotroph | \|Undefined Saprotroph\| | Yeast | NULL | Probable | As nectar-tap_saprotroph (Põlme et al. 2020); Decay sugar-rich substrates (Põlme et al. 2020) | Põlme S, et al. 2020. Fungal Diversity 105:1-16 ((https://doi.org/10.1007/s13225-020-00466-2)) |
| Otu17954 | 1 | k__Fungi(100);p__Ascomycota(100);c__Saccharomycetes(100);o__Saccharomycetales(100);f__Metschnikowiaceae(100);g__Metschnikowia(100);s__Metschnikowia_pulcherrima(100); | Metschnikowia | 13 | Saprotroph | \|Undefined Saprotroph\| | Yeast | NULL | Probable | As nectar-tap_saprotroph (Põlme et al. 2020); Decay sugar-rich substrates (Põlme et al. 2020) | Põlme S, et al. 2020. Fungal Diversity 105:1-16 ((https://doi.org/10.1007/s13225-020-00466-2)) |
| Otu17955 | 1 | k__Fungi(100);p__Ascomycota(100);c__Saccharomycetes(100);o__Saccharomycetales(100);f__Metschnikowiaceae(100);g__Metschnikowia(100);s__Metschnikowia_pulcherrima(100); | Metschnikowia | 13 | Saprotroph | \|Undefined Saprotroph\| | Yeast | NULL | Probable | As nectar-tap_saprotroph (Põlme et al. 2020); Decay sugar-rich substrates (Põlme et al. 2020) | Põlme S, et al. 2020. Fungal Diversity 105:1-16 ((https://doi.org/10.1007/s13225-020-00466-2)) |
| Otu17956 | 1 | k__Fungi(100);p__Ascomycota(100);c__Saccharomycetes(100);o__Saccharomycetales(100);f__Metschnikowiaceae(100);g__Metschnikowia(100);s__Metschnikowia_pulcherrima(100); | Metschnikowia | 13 | Saprotroph | \|Undefined Saprotroph\| | Yeast | NULL | Probable | As nectar-tap_saprotroph (Põlme et al. 2020); Decay sugar-rich substrates (Põlme et al. 2020) | Põlme S, et al. 2020. Fungal Diversity 105:1-16 ((https://doi.org/10.1007/s13225-020-00466-2)) |
| Otu17957 | 1 | k__Fungi(100);p__Ascomycota(100);c__Saccharomycetes(100);o__Saccharomycetales(100);f__Metschnikowiaceae(100);g__Metschnikowia(100);g__Metschnikowia_unclassified(100); | Metschnikowia | 13 | Saprotroph | \|Undefined Saprotroph\| | Yeast | NULL | Probable | As nectar-tap_saprotroph (Põlme et al. 2020); Decay sugar-rich substrates (Põlme et al. 2020) | Põlme S, et al. 2020. Fungal Diversity 105:1-16 ((https://doi.org/10.1007/s13225-020-00466-2)) |
| Otu17958 | 1 | k__Fungi(100);p__Ascomycota(100);c__Saccharomycetes(100);o__Saccharomycetales(100);f__Metschnikowiaceae(100);g__Metschnikowia(100);s__Metschnikowia_pulcherrima(100); | Metschnikowia | 13 | Saprotroph | \|Undefined Saprotroph\| | Yeast | NULL | Probable | As nectar-tap_saprotroph (Põlme et al. 2020); Decay sugar-rich substrates (Põlme et al. 2020) | Põlme S, et al. 2020. Fungal Diversity 105:1-16 ((https://doi.org/10.1007/s13225-020-00466-2)) |
| Otu17959 | 1 | k__Fungi(100);p__Ascomycota(100);c__Saccharomycetes(100);o__Saccharomycetales(100);f__Metschnikowiaceae(100);g__Metschnikowia(100);g__Metschnikowia_unclassified(100); | Metschnikowia | 13 | Saprotroph | \|Undefined Saprotroph\| | Yeast | NULL | Probable | As nectar-tap_saprotroph (Põlme et al. 2020); Decay sugar-rich substrates (Põlme et al. 2020) | Põlme S, et al. 2020. Fungal Diversity 105:1-16 ((https://doi.org/10.1007/s13225-020-00466-2)) |
| Otu17960 | 1 | k__Fungi(100);p__Ascomycota(100);c__Saccharomycetes(100);o__Saccharomycetales(100);f__Metschnikowiaceae(100);g__Metschnikowia(100);s__Metschnikowia_pulcherrima(100); | Metschnikowia | 13 | Saprotroph | \|Undefined Saprotroph\| | Yeast | NULL | Probable | As nectar-tap_saprotroph (Põlme et al. 2020); Decay sugar-rich substrates (Põlme et al. 2020) | Põlme S, et al. 2020. Fungal Diversity 105:1-16 ((https://doi.org/10.1007/s13225-020-00466-2)) |
| Otu17961 | 1 | k__Fungi(100);p__Ascomycota(100);c__Saccharomycetes(100);o__Saccharomycetales(100);f__Metschnikowiaceae(100);g__Metschnikowia(100);s__Metschnikowia_pulcherrima(100); | Metschnikowia | 13 | Saprotroph | \|Undefined Saprotroph\| | Yeast | NULL | Probable | As nectar-tap_saprotroph (Põlme et al. 2020); Decay sugar-rich substrates (Põlme et al. 2020) | Põlme S, et al. 2020. Fungal Diversity 105:1-16 ((https://doi.org/10.1007/s13225-020-00466-2)) |
| Otu17962 | 1 | k__Fungi(100);p__Ascomycota(100);c__Saccharomycetes(100);o__Saccharomycetales(100);f__Metschnikowiaceae(100);g__Metschnikowia(100);s__Metschnikowia_pulcherrima(100); | Metschnikowia | 13 | Saprotroph | \|Undefined Saprotroph\| | Yeast | NULL | Probable | As nectar-tap_saprotroph (Põlme et al. 2020); Decay sugar-rich substrates (Põlme et al. 2020) | Põlme S, et al. 2020. Fungal Diversity 105:1-16 ((https://doi.org/10.1007/s13225-020-00466-2)) |
| Otu17963 | 1 | k__Fungi(100);p__Ascomycota(100);c__Saccharomycetes(100);o__Saccharomycetales(100);f__Metschnikowiaceae(100);g__Metschnikowia(100);g__Metschnikowia_unclassified(100); | Metschnikowia | 13 | Saprotroph | \|Undefined Saprotroph\| | Yeast | NULL | Probable | As nectar-tap_saprotroph (Põlme et al. 2020); Decay sugar-rich substrates (Põlme et al. 2020) | Põlme S, et al. 2020. Fungal Diversity 105:1-16 ((https://doi.org/10.1007/s13225-020-00466-2)) |
| Otu17965 | 1 | k__Fungi(100);p__Ascomycota(100);c__Saccharomycetes(100);o__Saccharomycetales(100);f__Metschnikowiaceae(100);g__Metschnikowia(100);s__Metschnikowia_pulcherrima(100); | Metschnikowia | 13 | Saprotroph | \|Undefined Saprotroph\| | Yeast | NULL | Probable | As nectar-tap_saprotroph (Põlme et al. 2020); Decay sugar-rich substrates (Põlme et al. 2020) | Põlme S, et al. 2020. Fungal Diversity 105:1-16 ((https://doi.org/10.1007/s13225-020-00466-2)) |
| Otu17966 | 1 | k__Fungi(100);p__Ascomycota(100);c__Saccharomycetes(100);o__Saccharomycetales(100);f__Metschnikowiaceae(100);g__Metschnikowia(100);s__Metschnikowia_pulcherrima(100); | Metschnikowia | 13 | Saprotroph | \|Undefined Saprotroph\| | Yeast | NULL | Probable | As nectar-tap_saprotroph (Põlme et al. 2020); Decay sugar-rich substrates (Põlme et al. 2020) | Põlme S, et al. 2020. Fungal Diversity 105:1-16 ((https://doi.org/10.1007/s13225-020-00466-2)) |
| Otu17967 | 1 | k__Fungi(100);p__Ascomycota(100);c__Saccharomycetes(100);o__Saccharomycetales(100);f__Metschnikowiaceae(100);g__Metschnikowia(100);s__Metschnikowia_pulcherrima(100); | Metschnikowia | 13 | Saprotroph | \|Undefined Saprotroph\| | Yeast | NULL | Probable | As nectar-tap_saprotroph (Põlme et al. 2020); Decay sugar-rich substrates (Põlme et al. 2020) | Põlme S, et al. 2020. Fungal Diversity 105:1-16 ((https://doi.org/10.1007/s13225-020-00466-2)) |
| Otu17968 | 1 | k__Fungi(100);p__Ascomycota(100);c__Saccharomycetes(100);o__Saccharomycetales(100);f__Metschnikowiaceae(100);g__Metschnikowia(100);g__Metschnikowia_unclassified(100); | Metschnikowia | 13 | Saprotroph | \|Undefined Saprotroph\| | Yeast | NULL | Probable | As nectar-tap_saprotroph (Põlme et al. 2020); Decay sugar-rich substrates (Põlme et al. 2020) | Põlme S, et al. 2020. Fungal Diversity 105:1-16 ((https://doi.org/10.1007/s13225-020-00466-2)) |
| Otu17969 | 1 | k__Fungi(100);p__Ascomycota(100);c__Saccharomycetes(100);o__Saccharomycetales(100);f__Metschnikowiaceae(100);g__Metschnikowia(100);s__Metschnikowia_pulcherrima(100); | Metschnikowia | 13 | Saprotroph | \|Undefined Saprotroph\| | Yeast | NULL | Probable | As nectar-tap_saprotroph (Põlme et al. 2020); Decay sugar-rich substrates (Põlme et al. 2020) | Põlme S, et al. 2020. Fungal Diversity 105:1-16 ((https://doi.org/10.1007/s13225-020-00466-2)) |
| Otu17970 | 1 | k__Fungi(100);p__Ascomycota(100);c__Saccharomycetes(100);o__Saccharomycetales(100);f__Metschnikowiaceae(100);g__Metschnikowia(100);s__Metschnikowia_pulcherrima(100); | Metschnikowia | 13 | Saprotroph | \|Undefined Saprotroph\| | Yeast | NULL | Probable | As nectar-tap_saprotroph (Põlme et al. 2020); Decay sugar-rich substrates (Põlme et al. 2020) | Põlme S, et al. 2020. Fungal Diversity 105:1-16 ((https://doi.org/10.1007/s13225-020-00466-2)) |
| Otu17971 | 1 | k__Fungi(100);p__Ascomycota(100);c__Saccharomycetes(100);o__Saccharomycetales(100);f__Metschnikowiaceae(100);g__Metschnikowia(100);s__Metschnikowia_pulcherrima(100); | Metschnikowia | 13 | Saprotroph | \|Undefined Saprotroph\| | Yeast | NULL | Probable | As nectar-tap_saprotroph (Põlme et al. 2020); Decay sugar-rich substrates (Põlme et al. 2020) | Põlme S, et al. 2020. Fungal Diversity 105:1-16 ((https://doi.org/10.1007/s13225-020-00466-2)) |
| Otu17972 | 1 | k__Fungi(100);p__Ascomycota(100);c__Saccharomycetes(100);o__Saccharomycetales(100);f__Metschnikowiaceae(100);g__Metschnikowia(100);g__Metschnikowia_unclassified(100); | Metschnikowia | 13 | Saprotroph | \|Undefined Saprotroph\| | Yeast | NULL | Probable | As nectar-tap_saprotroph (Põlme et al. 2020); Decay sugar-rich substrates (Põlme et al. 2020) | Põlme S, et al. 2020. Fungal Diversity 105:1-16 ((https://doi.org/10.1007/s13225-020-00466-2)) |
| Otu17973 | 1 | k__Fungi(100);p__Ascomycota(100);c__Saccharomycetes(100);o__Saccharomycetales(100);f__Metschnikowiaceae(100);g__Metschnikowia(100);s__Metschnikowia_pulcherrima(100); | Metschnikowia | 13 | Saprotroph | \|Undefined Saprotroph\| | Yeast | NULL | Probable | As nectar-tap_saprotroph (Põlme et al. 2020); Decay sugar-rich substrates (Põlme et al. 2020) | Põlme S, et al. 2020. Fungal Diversity 105:1-16 ((https://doi.org/10.1007/s13225-020-00466-2)) |
| Otu17974 | 1 | k__Fungi(100);p__Ascomycota(100);c__Saccharomycetes(100);o__Saccharomycetales(100);f__Metschnikowiaceae(100);g__Metschnikowia(100);s__Metschnikowia_pulcherrima(100); | Metschnikowia | 13 | Saprotroph | \|Undefined Saprotroph\| | Yeast | NULL | Probable | As nectar-tap_saprotroph (Põlme et al. 2020); Decay sugar-rich substrates (Põlme et al. 2020) | Põlme S, et al. 2020. Fungal Diversity 105:1-16 ((https://doi.org/10.1007/s13225-020-00466-2)) |
| Otu17975 | 1 | k__Fungi(100);p__Ascomycota(100);c__Saccharomycetes(100);o__Saccharomycetales(100);f__Metschnikowiaceae(100);g__Metschnikowia(100);s__Metschnikowia_pulcherrima(100); | Metschnikowia | 13 | Saprotroph | \|Undefined Saprotroph\| | Yeast | NULL | Probable | As nectar-tap_saprotroph (Põlme et al. 2020); Decay sugar-rich substrates (Põlme et al. 2020) | Põlme S, et al. 2020. Fungal Diversity 105:1-16 ((https://doi.org/10.1007/s13225-020-00466-2)) |
| Otu17976 | 1 | k__Fungi(100);p__Ascomycota(100);c__Saccharomycetes(100);o__Saccharomycetales(100);f__Metschnikowiaceae(100);g__Metschnikowia(100);s__Metschnikowia_pulcherrima(100); | Metschnikowia | 13 | Saprotroph | \|Undefined Saprotroph\| | Yeast | NULL | Probable | As nectar-tap_saprotroph (Põlme et al. 2020); Decay sugar-rich substrates (Põlme et al. 2020) | Põlme S, et al. 2020. Fungal Diversity 105:1-16 ((https://doi.org/10.1007/s13225-020-00466-2)) |
| Otu17977 | 1 | k__Fungi(100);p__Ascomycota(100);c__Saccharomycetes(100);o__Saccharomycetales(100);f__Metschnikowiaceae(100);g__Metschnikowia(100);g__Metschnikowia_unclassified(100); | Metschnikowia | 13 | Saprotroph | \|Undefined Saprotroph\| | Yeast | NULL | Probable | As nectar-tap_saprotroph (Põlme et al. 2020); Decay sugar-rich substrates (Põlme et al. 2020) | Põlme S, et al. 2020. Fungal Diversity 105:1-16 ((https://doi.org/10.1007/s13225-020-00466-2)) |
| Otu17978 | 1 | k__Fungi(100);p__Ascomycota(100);c__Saccharomycetes(100);o__Saccharomycetales(100);f__Metschnikowiaceae(100);g__Metschnikowia(100);s__Metschnikowia_pulcherrima(100); | Metschnikowia | 13 | Saprotroph | \|Undefined Saprotroph\| | Yeast | NULL | Probable | As nectar-tap_saprotroph (Põlme et al. 2020); Decay sugar-rich substrates (Põlme et al. 2020) | Põlme S, et al. 2020. Fungal Diversity 105:1-16 ((https://doi.org/10.1007/s13225-020-00466-2)) |
| Otu17979 | 1 | k__Fungi(100);p__Ascomycota(100);c__Saccharomycetes(100);o__Saccharomycetales(100);f__Metschnikowiaceae(100);g__Metschnikowia(100);s__Metschnikowia_pulcherrima(100); | Metschnikowia | 13 | Saprotroph | \|Undefined Saprotroph\| | Yeast | NULL | Probable | As nectar-tap_saprotroph (Põlme et al. 2020); Decay sugar-rich substrates (Põlme et al. 2020) | Põlme S, et al. 2020. Fungal Diversity 105:1-16 ((https://doi.org/10.1007/s13225-020-00466-2)) |
| Otu17980 | 1 | k__Fungi(100);p__Ascomycota(100);c__Saccharomycetes(100);o__Saccharomycetales(100);f__Metschnikowiaceae(100);g__Metschnikowia(100);s__Metschnikowia_pulcherrima(100); | Metschnikowia | 13 | Saprotroph | \|Undefined Saprotroph\| | Yeast | NULL | Probable | As nectar-tap_saprotroph (Põlme et al. 2020); Decay sugar-rich substrates (Põlme et al. 2020) | Põlme S, et al. 2020. Fungal Diversity 105:1-16 ((https://doi.org/10.1007/s13225-020-00466-2)) |
| Otu17981 | 1 | k__Fungi(100);p__Ascomycota(100);c__Saccharomycetes(100);o__Saccharomycetales(100);f__Metschnikowiaceae(100);g__Metschnikowia(100);s__Metschnikowia_pulcherrima(100); | Metschnikowia | 13 | Saprotroph | \|Undefined Saprotroph\| | Yeast | NULL | Probable | As nectar-tap_saprotroph (Põlme et al. 2020); Decay sugar-rich substrates (Põlme et al. 2020) | Põlme S, et al. 2020. Fungal Diversity 105:1-16 ((https://doi.org/10.1007/s13225-020-00466-2)) |
| Otu17982 | 1 | k__Fungi(100);p__Ascomycota(100);c__Saccharomycetes(100);o__Saccharomycetales(100);f__Metschnikowiaceae(100);g__Metschnikowia(100);s__Metschnikowia_pulcherrima(100); | Metschnikowia | 13 | Saprotroph | \|Undefined Saprotroph\| | Yeast | NULL | Probable | As nectar-tap_saprotroph (Põlme et al. 2020); Decay sugar-rich substrates (Põlme et al. 2020) | Põlme S, et al. 2020. Fungal Diversity 105:1-16 ((https://doi.org/10.1007/s13225-020-00466-2)) |
| Otu17983 | 1 | k__Fungi(100);p__Ascomycota(100);c__Saccharomycetes(100);o__Saccharomycetales(100);f__Metschnikowiaceae(100);g__Metschnikowia(100);s__Metschnikowia_pulcherrima(100); | Metschnikowia | 13 | Saprotroph | \|Undefined Saprotroph\| | Yeast | NULL | Probable | As nectar-tap_saprotroph (Põlme et al. 2020); Decay sugar-rich substrates (Põlme et al. 2020) | Põlme S, et al. 2020. Fungal Diversity 105:1-16 ((https://doi.org/10.1007/s13225-020-00466-2)) |
| Otu17984 | 1 | k__Fungi(100);p__Ascomycota(100);c__Saccharomycetes(100);o__Saccharomycetales(100);f__Metschnikowiaceae(100);g__Metschnikowia(100);s__Metschnikowia_pulcherrima(100); | Metschnikowia | 13 | Saprotroph | \|Undefined Saprotroph\| | Yeast | NULL | Probable | As nectar-tap_saprotroph (Põlme et al. 2020); Decay sugar-rich substrates (Põlme et al. 2020) | Põlme S, et al. 2020. Fungal Diversity 105:1-16 ((https://doi.org/10.1007/s13225-020-00466-2)) |
| Otu17985 | 1 | k__Fungi(100);p__Ascomycota(100);c__Saccharomycetes(100);o__Saccharomycetales(100);f__Metschnikowiaceae(100);g__Metschnikowia(100);s__Metschnikowia_pulcherrima(100); | Metschnikowia | 13 | Saprotroph | \|Undefined Saprotroph\| | Yeast | NULL | Probable | As nectar-tap_saprotroph (Põlme et al. 2020); Decay sugar-rich substrates (Põlme et al. 2020) | Põlme S, et al. 2020. Fungal Diversity 105:1-16 ((https://doi.org/10.1007/s13225-020-00466-2)) |
| Otu17986 | 1 | k__Fungi(100);p__Ascomycota(100);c__Saccharomycetes(100);o__Saccharomycetales(100);f__Metschnikowiaceae(100);g__Metschnikowia(100);s__Metschnikowia_pulcherrima(100); | Metschnikowia | 13 | Saprotroph | \|Undefined Saprotroph\| | Yeast | NULL | Probable | As nectar-tap_saprotroph (Põlme et al. 2020); Decay sugar-rich substrates (Põlme et al. 2020) | Põlme S, et al. 2020. Fungal Diversity 105:1-16 ((https://doi.org/10.1007/s13225-020-00466-2)) |
| Otu17987 | 1 | k__Fungi(100);p__Ascomycota(100);c__Saccharomycetes(100);o__Saccharomycetales(100);f__Metschnikowiaceae(100);g__Metschnikowia(100);s__Metschnikowia_pulcherrima(100); | Metschnikowia | 13 | Saprotroph | \|Undefined Saprotroph\| | Yeast | NULL | Probable | As nectar-tap_saprotroph (Põlme et al. 2020); Decay sugar-rich substrates (Põlme et al. 2020) | Põlme S, et al. 2020. Fungal Diversity 105:1-16 ((https://doi.org/10.1007/s13225-020-00466-2)) |
| Otu17988 | 1 | k__Fungi(100);p__Ascomycota(100);c__Saccharomycetes(100);o__Saccharomycetales(100);f__Metschnikowiaceae(100);g__Metschnikowia(100);g__Metschnikowia_unclassified(100); | Metschnikowia | 13 | Saprotroph | \|Undefined Saprotroph\| | Yeast | NULL | Probable | As nectar-tap_saprotroph (Põlme et al. 2020); Decay sugar-rich substrates (Põlme et al. 2020) | Põlme S, et al. 2020. Fungal Diversity 105:1-16 ((https://doi.org/10.1007/s13225-020-00466-2)) |
| Otu17989 | 1 | k__Fungi(100);p__Basidiomycota(100);c__Tremellomycetes(100);o__Tremellales(100);f__Tremellales_family_Incertae_sedis(100);g__Dioszegia(100);s__Dioszegia_changbaiensis(100); | Dioszegia | 13 | Saprotroph | \|Plant Saprotroph\|-Undefined Saprotroph | Yeast | NULL | Probable | As litter_saprotroph (Põlme et al. 2020); Decay leaf, fruit, seed (Põlme et al. 2020); The Dioszegia subclade sensu Inacio (Inacio et al. 20015) is characterized by orange; yellow or orange to deep orange colonies. Isolated from plant leaves and soil. | Põlme S, et al. 2020. Fungal Diversity 105:1-16 ((https://doi.org/10.1007/s13225-020-00466-2)); Takashima M, et al. 2011 International Journal of Systematic and Evolutionary Microbiology 61:1736-1739 ((https://doi.org/10.1099/ijs.0.025254-0)) |
| Otu17990 | 1 | k__Fungi(100);p__Ascomycota(100);c__Saccharomycetes(100);o__Saccharomycetales(100);f__Metschnikowiaceae(100);g__Metschnikowia(100);s__Metschnikowia_pulcherrima(100); | Metschnikowia | 13 | Saprotroph | \|Undefined Saprotroph\| | Yeast | NULL | Probable | As nectar-tap_saprotroph (Põlme et al. 2020); Decay sugar-rich substrates (Põlme et al. 2020) | Põlme S, et al. 2020. Fungal Diversity 105:1-16 ((https://doi.org/10.1007/s13225-020-00466-2)) |
| Otu17991 | 1 | k__Fungi(100);p__Ascomycota(100);c__Saccharomycetes(100);o__Saccharomycetales(100);f__Metschnikowiaceae(100);g__Metschnikowia(100);s__Metschnikowia_pulcherrima(100); | Metschnikowia | 13 | Saprotroph | \|Undefined Saprotroph\| | Yeast | NULL | Probable | As nectar-tap_saprotroph (Põlme et al. 2020); Decay sugar-rich substrates (Põlme et al. 2020) | Põlme S, et al. 2020. Fungal Diversity 105:1-16 ((https://doi.org/10.1007/s13225-020-00466-2)) |
| Otu17992 | 1 | k__Fungi(100);p__Ascomycota(100);c__Saccharomycetes(100);o__Saccharomycetales(100);f__Metschnikowiaceae(100);g__Metschnikowia(100);s__Metschnikowia_pulcherrima(100); | Metschnikowia | 13 | Saprotroph | \|Undefined Saprotroph\| | Yeast | NULL | Probable | As nectar-tap_saprotroph (Põlme et al. 2020); Decay sugar-rich substrates (Põlme et al. 2020) | Põlme S, et al. 2020. Fungal Diversity 105:1-16 ((https://doi.org/10.1007/s13225-020-00466-2)) |
| Otu17993 | 1 | k__Fungi(100);p__Ascomycota(100);c__Saccharomycetes(100);o__Saccharomycetales(100);f__Metschnikowiaceae(100);g__Metschnikowia(100);g__Metschnikowia_unclassified(100); | Metschnikowia | 13 | Saprotroph | \|Undefined Saprotroph\| | Yeast | NULL | Probable | As nectar-tap_saprotroph (Põlme et al. 2020); Decay sugar-rich substrates (Põlme et al. 2020) | Põlme S, et al. 2020. Fungal Diversity 105:1-16 ((https://doi.org/10.1007/s13225-020-00466-2)) |
| Otu17995 | 1 | k__Fungi(100);p__Ascomycota(100);c__Saccharomycetes(100);o__Saccharomycetales(100);f__Metschnikowiaceae(100);g__Metschnikowia(100);s__Metschnikowia_pulcherrima(100); | Metschnikowia | 13 | Saprotroph | \|Undefined Saprotroph\| | Yeast | NULL | Probable | As nectar-tap_saprotroph (Põlme et al. 2020); Decay sugar-rich substrates (Põlme et al. 2020) | Põlme S, et al. 2020. Fungal Diversity 105:1-16 ((https://doi.org/10.1007/s13225-020-00466-2)) |
| Otu17996 | 1 | k__Fungi(100);p__Ascomycota(100);c__Saccharomycetes(100);o__Saccharomycetales(100);f__Metschnikowiaceae(100);g__Metschnikowia(100);s__Metschnikowia_pulcherrima(100); | Metschnikowia | 13 | Saprotroph | \|Undefined Saprotroph\| | Yeast | NULL | Probable | As nectar-tap_saprotroph (Põlme et al. 2020); Decay sugar-rich substrates (Põlme et al. 2020) | Põlme S, et al. 2020. Fungal Diversity 105:1-16 ((https://doi.org/10.1007/s13225-020-00466-2)) |
| Otu17997 | 1 | k__Fungi(100);p__Ascomycota(100);c__Saccharomycetes(100);o__Saccharomycetales(100);f__Metschnikowiaceae(100);g__Metschnikowia(100);g__Metschnikowia_unclassified(100); | Metschnikowia | 13 | Saprotroph | \|Undefined Saprotroph\| | Yeast | NULL | Probable | As nectar-tap_saprotroph (Põlme et al. 2020); Decay sugar-rich substrates (Põlme et al. 2020) | Põlme S, et al. 2020. Fungal Diversity 105:1-16 ((https://doi.org/10.1007/s13225-020-00466-2)) |
| Otu17998 | 1 | k__Fungi(100);p__Ascomycota(100);c__Saccharomycetes(100);o__Saccharomycetales(100);f__Metschnikowiaceae(100);g__Metschnikowia(100);s__Metschnikowia_pulcherrima(100); | Metschnikowia | 13 | Saprotroph | \|Undefined Saprotroph\| | Yeast | NULL | Probable | As nectar-tap_saprotroph (Põlme et al. 2020); Decay sugar-rich substrates (Põlme et al. 2020) | Põlme S, et al. 2020. Fungal Diversity 105:1-16 ((https://doi.org/10.1007/s13225-020-00466-2)) |
| Otu18002 | 1 | k__Fungi(100);p__Ascomycota(100);c__Saccharomycetes(100);o__Saccharomycetales(100);f__Metschnikowiaceae(100);g__Metschnikowia(100);s__Metschnikowia_pulcherrima(100); | Metschnikowia | 13 | Saprotroph | \|Undefined Saprotroph\| | Yeast | NULL | Probable | As nectar-tap_saprotroph (Põlme et al. 2020); Decay sugar-rich substrates (Põlme et al. 2020) | Põlme S, et al. 2020. Fungal Diversity 105:1-16 ((https://doi.org/10.1007/s13225-020-00466-2)) |
| Otu18003 | 1 | k__Fungi(100);p__Ascomycota(100);c__Saccharomycetes(100);o__Saccharomycetales(100);f__Metschnikowiaceae(100);g__Metschnikowia(100);s__Metschnikowia_pulcherrima(100); | Metschnikowia | 13 | Saprotroph | \|Undefined Saprotroph\| | Yeast | NULL | Probable | As nectar-tap_saprotroph (Põlme et al. 2020); Decay sugar-rich substrates (Põlme et al. 2020) | Põlme S, et al. 2020. Fungal Diversity 105:1-16 ((https://doi.org/10.1007/s13225-020-00466-2)) |
| Otu18005 | 1 | k__Fungi(100);p__Ascomycota(100);c__Saccharomycetes(100);o__Saccharomycetales(100);f__Metschnikowiaceae(100);g__Metschnikowia(100);g__Metschnikowia_unclassified(100); | Metschnikowia | 13 | Saprotroph | \|Undefined Saprotroph\| | Yeast | NULL | Probable | As nectar-tap_saprotroph (Põlme et al. 2020); Decay sugar-rich substrates (Põlme et al. 2020) | Põlme S, et al. 2020. Fungal Diversity 105:1-16 ((https://doi.org/10.1007/s13225-020-00466-2)) |
| Otu18008 | 1 | k__Fungi(100);p__Ascomycota(100);c__Saccharomycetes(100);o__Saccharomycetales(100);f__Metschnikowiaceae(100);g__Metschnikowia(100);g__Metschnikowia_unclassified(100); | Metschnikowia | 13 | Saprotroph | \|Undefined Saprotroph\| | Yeast | NULL | Probable | As nectar-tap_saprotroph (Põlme et al. 2020); Decay sugar-rich substrates (Põlme et al. 2020) | Põlme S, et al. 2020. Fungal Diversity 105:1-16 ((https://doi.org/10.1007/s13225-020-00466-2)) |
| Otu18010 | 1 | k__Fungi(100);p__Ascomycota(100);c__Saccharomycetes(100);o__Saccharomycetales(100);f__Metschnikowiaceae(100);g__Metschnikowia(100);g__Metschnikowia_unclassified(100); | Metschnikowia | 13 | Saprotroph | \|Undefined Saprotroph\| | Yeast | NULL | Probable | As nectar-tap_saprotroph (Põlme et al. 2020); Decay sugar-rich substrates (Põlme et al. 2020) | Põlme S, et al. 2020. Fungal Diversity 105:1-16 ((https://doi.org/10.1007/s13225-020-00466-2)) |
| Otu18011 | 1 | k__Fungi(100);p__Ascomycota(100);c__Saccharomycetes(100);o__Saccharomycetales(100);f__Metschnikowiaceae(100);g__Metschnikowia(100);s__Metschnikowia_pulcherrima(100); | Metschnikowia | 13 | Saprotroph | \|Undefined Saprotroph\| | Yeast | NULL | Probable | As nectar-tap_saprotroph (Põlme et al. 2020); Decay sugar-rich substrates (Põlme et al. 2020) | Põlme S, et al. 2020. Fungal Diversity 105:1-16 ((https://doi.org/10.1007/s13225-020-00466-2)) |
| Otu18012 | 1 | k__Fungi(100);p__Ascomycota(100);c__Saccharomycetes(100);o__Saccharomycetales(100);f__Metschnikowiaceae(100);g__Metschnikowia(100);g__Metschnikowia_unclassified(100); | Metschnikowia | 13 | Saprotroph | \|Undefined Saprotroph\| | Yeast | NULL | Probable | As nectar-tap_saprotroph (Põlme et al. 2020); Decay sugar-rich substrates (Põlme et al. 2020) | Põlme S, et al. 2020. Fungal Diversity 105:1-16 ((https://doi.org/10.1007/s13225-020-00466-2)) |
| Otu18013 | 1 | k__Fungi(100);p__Ascomycota(100);c__Saccharomycetes(100);o__Saccharomycetales(100);f__Metschnikowiaceae(100);g__Metschnikowia(100);s__Metschnikowia_pulcherrima(100); | Metschnikowia | 13 | Saprotroph | \|Undefined Saprotroph\| | Yeast | NULL | Probable | As nectar-tap_saprotroph (Põlme et al. 2020); Decay sugar-rich substrates (Põlme et al. 2020) | Põlme S, et al. 2020. Fungal Diversity 105:1-16 ((https://doi.org/10.1007/s13225-020-00466-2)) |
| Otu18014 | 1 | k__Fungi(100);p__Ascomycota(100);c__Saccharomycetes(100);o__Saccharomycetales(100);f__Metschnikowiaceae(100);g__Metschnikowia(100);s__Metschnikowia_pulcherrima(100); | Metschnikowia | 13 | Saprotroph | \|Undefined Saprotroph\| | Yeast | NULL | Probable | As nectar-tap_saprotroph (Põlme et al. 2020); Decay sugar-rich substrates (Põlme et al. 2020) | Põlme S, et al. 2020. Fungal Diversity 105:1-16 ((https://doi.org/10.1007/s13225-020-00466-2)) |
| Otu18015 | 1 | k__Fungi(100);p__Ascomycota(100);c__Saccharomycetes(100);o__Saccharomycetales(100);f__Metschnikowiaceae(100);g__Metschnikowia(100);g__Metschnikowia_unclassified(100); | Metschnikowia | 13 | Saprotroph | \|Undefined Saprotroph\| | Yeast | NULL | Probable | As nectar-tap_saprotroph (Põlme et al. 2020); Decay sugar-rich substrates (Põlme et al. 2020) | Põlme S, et al. 2020. Fungal Diversity 105:1-16 ((https://doi.org/10.1007/s13225-020-00466-2)) |
| Otu18016 | 1 | k__Fungi(100);p__Ascomycota(100);c__Saccharomycetes(100);o__Saccharomycetales(100);f__Metschnikowiaceae(100);g__Metschnikowia(100);g__Metschnikowia_unclassified(100); | Metschnikowia | 13 | Saprotroph | \|Undefined Saprotroph\| | Yeast | NULL | Probable | As nectar-tap_saprotroph (Põlme et al. 2020); Decay sugar-rich substrates (Põlme et al. 2020) | Põlme S, et al. 2020. Fungal Diversity 105:1-16 ((https://doi.org/10.1007/s13225-020-00466-2)) |
| Otu18017 | 1 | k__Fungi(100);p__Ascomycota(100);c__Saccharomycetes(100);o__Saccharomycetales(100);f__Metschnikowiaceae(100);g__Metschnikowia(100);s__Metschnikowia_pulcherrima(100); | Metschnikowia | 13 | Saprotroph | \|Undefined Saprotroph\| | Yeast | NULL | Probable | As nectar-tap_saprotroph (Põlme et al. 2020); Decay sugar-rich substrates (Põlme et al. 2020) | Põlme S, et al. 2020. Fungal Diversity 105:1-16 ((https://doi.org/10.1007/s13225-020-00466-2)) |
| Otu18018 | 1 | k__Fungi(100);p__Ascomycota(100);c__Saccharomycetes(100);o__Saccharomycetales(100);f__Metschnikowiaceae(100);g__Metschnikowia(100);g__Metschnikowia_unclassified(100); | Metschnikowia | 13 | Saprotroph | \|Undefined Saprotroph\| | Yeast | NULL | Probable | As nectar-tap_saprotroph (Põlme et al. 2020); Decay sugar-rich substrates (Põlme et al. 2020) | Põlme S, et al. 2020. Fungal Diversity 105:1-16 ((https://doi.org/10.1007/s13225-020-00466-2)) |
| Otu18019 | 1 | k__Fungi(100);p__Ascomycota(100);c__Saccharomycetes(100);o__Saccharomycetales(100);f__Metschnikowiaceae(100);g__Metschnikowia(100);g__Metschnikowia_unclassified(100); | Metschnikowia | 13 | Saprotroph | \|Undefined Saprotroph\| | Yeast | NULL | Probable | As nectar-tap_saprotroph (Põlme et al. 2020); Decay sugar-rich substrates (Põlme et al. 2020) | Põlme S, et al. 2020. Fungal Diversity 105:1-16 ((https://doi.org/10.1007/s13225-020-00466-2)) |
| Otu18020 | 1 | k__Fungi(100);p__Ascomycota(100);c__Saccharomycetes(100);o__Saccharomycetales(100);f__Metschnikowiaceae(100);g__Metschnikowia(100);s__Metschnikowia_pulcherrima(100); | Metschnikowia | 13 | Saprotroph | \|Undefined Saprotroph\| | Yeast | NULL | Probable | As nectar-tap_saprotroph (Põlme et al. 2020); Decay sugar-rich substrates (Põlme et al. 2020) | Põlme S, et al. 2020. Fungal Diversity 105:1-16 ((https://doi.org/10.1007/s13225-020-00466-2)) |
| Otu18021 | 1 | k__Fungi(100);p__Ascomycota(100);c__Saccharomycetes(100);o__Saccharomycetales(100);f__Metschnikowiaceae(100);g__Metschnikowia(100);g__Metschnikowia_unclassified(100); | Metschnikowia | 13 | Saprotroph | \|Undefined Saprotroph\| | Yeast | NULL | Probable | As nectar-tap_saprotroph (Põlme et al. 2020); Decay sugar-rich substrates (Põlme et al. 2020) | Põlme S, et al. 2020. Fungal Diversity 105:1-16 ((https://doi.org/10.1007/s13225-020-00466-2)) |
| Otu18022 | 1 | k__Fungi(100);p__Ascomycota(100);c__Saccharomycetes(100);o__Saccharomycetales(100);f__Metschnikowiaceae(100);g__Metschnikowia(100);g__Metschnikowia_unclassified(100); | Metschnikowia | 13 | Saprotroph | \|Undefined Saprotroph\| | Yeast | NULL | Probable | As nectar-tap_saprotroph (Põlme et al. 2020); Decay sugar-rich substrates (Põlme et al. 2020) | Põlme S, et al. 2020. Fungal Diversity 105:1-16 ((https://doi.org/10.1007/s13225-020-00466-2)) |
| Otu18023 | 1 | k__Fungi(100);p__Ascomycota(100);c__Saccharomycetes(100);o__Saccharomycetales(100);f__Metschnikowiaceae(100);g__Metschnikowia(100);g__Metschnikowia_unclassified(100); | Metschnikowia | 13 | Saprotroph | \|Undefined Saprotroph\| | Yeast | NULL | Probable | As nectar-tap_saprotroph (Põlme et al. 2020); Decay sugar-rich substrates (Põlme et al. 2020) | Põlme S, et al. 2020. Fungal Diversity 105:1-16 ((https://doi.org/10.1007/s13225-020-00466-2)) |
| Otu18024 | 1 | k__Fungi(100);p__Ascomycota(100);c__Saccharomycetes(100);o__Saccharomycetales(100);f__Metschnikowiaceae(100);g__Metschnikowia(100);g__Metschnikowia_unclassified(100); | Metschnikowia | 13 | Saprotroph | \|Undefined Saprotroph\| | Yeast | NULL | Probable | As nectar-tap_saprotroph (Põlme et al. 2020); Decay sugar-rich substrates (Põlme et al. 2020) | Põlme S, et al. 2020. Fungal Diversity 105:1-16 ((https://doi.org/10.1007/s13225-020-00466-2)) |
| Otu18025 | 1 | k__Fungi(100);p__Ascomycota(100);c__Saccharomycetes(100);o__Saccharomycetales(100);f__Metschnikowiaceae(100);g__Metschnikowia(100);s__Metschnikowia_pulcherrima(100); | Metschnikowia | 13 | Saprotroph | \|Undefined Saprotroph\| | Yeast | NULL | Probable | As nectar-tap_saprotroph (Põlme et al. 2020); Decay sugar-rich substrates (Põlme et al. 2020) | Põlme S, et al. 2020. Fungal Diversity 105:1-16 ((https://doi.org/10.1007/s13225-020-00466-2)) |
| Otu18026 | 1 | k__Fungi(100);p__Ascomycota(100);c__Saccharomycetes(100);o__Saccharomycetales(100);f__Metschnikowiaceae(100);g__Metschnikowia(100);s__Metschnikowia_pulcherrima(100); | Metschnikowia | 13 | Saprotroph | \|Undefined Saprotroph\| | Yeast | NULL | Probable | As nectar-tap_saprotroph (Põlme et al. 2020); Decay sugar-rich substrates (Põlme et al. 2020) | Põlme S, et al. 2020. Fungal Diversity 105:1-16 ((https://doi.org/10.1007/s13225-020-00466-2)) |
| Otu18027 | 1 | k__Fungi(100);p__Ascomycota(100);c__Saccharomycetes(100);o__Saccharomycetales(100);f__Metschnikowiaceae(100);g__Metschnikowia(100);s__Metschnikowia_pulcherrima(100); | Metschnikowia | 13 | Saprotroph | \|Undefined Saprotroph\| | Yeast | NULL | Probable | As nectar-tap_saprotroph (Põlme et al. 2020); Decay sugar-rich substrates (Põlme et al. 2020) | Põlme S, et al. 2020. Fungal Diversity 105:1-16 ((https://doi.org/10.1007/s13225-020-00466-2)) |
| Otu18028 | 1 | k__Fungi(100);p__Ascomycota(100);c__Saccharomycetes(100);o__Saccharomycetales(100);f__Metschnikowiaceae(100);g__Metschnikowia(100);s__Metschnikowia_pulcherrima(100); | Metschnikowia | 13 | Saprotroph | \|Undefined Saprotroph\| | Yeast | NULL | Probable | As nectar-tap_saprotroph (Põlme et al. 2020); Decay sugar-rich substrates (Põlme et al. 2020) | Põlme S, et al. 2020. Fungal Diversity 105:1-16 ((https://doi.org/10.1007/s13225-020-00466-2)) |
| Otu18029 | 1 | k__Fungi(100);p__Ascomycota(100);c__Saccharomycetes(100);o__Saccharomycetales(100);f__Metschnikowiaceae(100);g__Metschnikowia(100);s__Metschnikowia_pulcherrima(100); | Metschnikowia | 13 | Saprotroph | \|Undefined Saprotroph\| | Yeast | NULL | Probable | As nectar-tap_saprotroph (Põlme et al. 2020); Decay sugar-rich substrates (Põlme et al. 2020) | Põlme S, et al. 2020. Fungal Diversity 105:1-16 ((https://doi.org/10.1007/s13225-020-00466-2)) |
| Otu18030 | 1 | k__Fungi(100);p__Ascomycota(100);c__Saccharomycetes(100);o__Saccharomycetales(100);f__Metschnikowiaceae(100);g__Metschnikowia(100);s__Metschnikowia_pulcherrima(100); | Metschnikowia | 13 | Saprotroph | \|Undefined Saprotroph\| | Yeast | NULL | Probable | As nectar-tap_saprotroph (Põlme et al. 2020); Decay sugar-rich substrates (Põlme et al. 2020) | Põlme S, et al. 2020. Fungal Diversity 105:1-16 ((https://doi.org/10.1007/s13225-020-00466-2)) |
| Otu18031 | 1 | k__Fungi(100);p__Ascomycota(100);c__Saccharomycetes(100);o__Saccharomycetales(100);f__Metschnikowiaceae(100);g__Metschnikowia(100);s__Metschnikowia_pulcherrima(100); | Metschnikowia | 13 | Saprotroph | \|Undefined Saprotroph\| | Yeast | NULL | Probable | As nectar-tap_saprotroph (Põlme et al. 2020); Decay sugar-rich substrates (Põlme et al. 2020) | Põlme S, et al. 2020. Fungal Diversity 105:1-16 ((https://doi.org/10.1007/s13225-020-00466-2)) |
| Otu18032 | 1 | k__Fungi(100);p__Ascomycota(100);c__Saccharomycetes(100);o__Saccharomycetales(100);f__Metschnikowiaceae(100);g__Metschnikowia(100);s__Metschnikowia_pulcherrima(100); | Metschnikowia | 13 | Saprotroph | \|Undefined Saprotroph\| | Yeast | NULL | Probable | As nectar-tap_saprotroph (Põlme et al. 2020); Decay sugar-rich substrates (Põlme et al. 2020) | Põlme S, et al. 2020. Fungal Diversity 105:1-16 ((https://doi.org/10.1007/s13225-020-00466-2)) |
| Otu18033 | 1 | k__Fungi(100);p__Ascomycota(100);c__Saccharomycetes(100);o__Saccharomycetales(100);f__Metschnikowiaceae(100);g__Metschnikowia(100);s__Metschnikowia_pulcherrima(100); | Metschnikowia | 13 | Saprotroph | \|Undefined Saprotroph\| | Yeast | NULL | Probable | As nectar-tap_saprotroph (Põlme et al. 2020); Decay sugar-rich substrates (Põlme et al. 2020) | Põlme S, et al. 2020. Fungal Diversity 105:1-16 ((https://doi.org/10.1007/s13225-020-00466-2)) |
| Otu18034 | 1 | k__Fungi(100);p__Ascomycota(100);c__Saccharomycetes(100);o__Saccharomycetales(100);f__Metschnikowiaceae(100);g__Metschnikowia(100);s__Metschnikowia_pulcherrima(100); | Metschnikowia | 13 | Saprotroph | \|Undefined Saprotroph\| | Yeast | NULL | Probable | As nectar-tap_saprotroph (Põlme et al. 2020); Decay sugar-rich substrates (Põlme et al. 2020) | Põlme S, et al. 2020. Fungal Diversity 105:1-16 ((https://doi.org/10.1007/s13225-020-00466-2)) |
| Otu18035 | 1 | k__Fungi(100);p__Ascomycota(100);c__Saccharomycetes(100);o__Saccharomycetales(100);f__Metschnikowiaceae(100);g__Metschnikowia(100);s__Metschnikowia_pulcherrima(100); | Metschnikowia | 13 | Saprotroph | \|Undefined Saprotroph\| | Yeast | NULL | Probable | As nectar-tap_saprotroph (Põlme et al. 2020); Decay sugar-rich substrates (Põlme et al. 2020) | Põlme S, et al. 2020. Fungal Diversity 105:1-16 ((https://doi.org/10.1007/s13225-020-00466-2)) |
| Otu18036 | 1 | k__Fungi(100);p__Ascomycota(100);c__Saccharomycetes(100);o__Saccharomycetales(100);f__Metschnikowiaceae(100);g__Metschnikowia(100);g__Metschnikowia_unclassified(100); | Metschnikowia | 13 | Saprotroph | \|Undefined Saprotroph\| | Yeast | NULL | Probable | As nectar-tap_saprotroph (Põlme et al. 2020); Decay sugar-rich substrates (Põlme et al. 2020) | Põlme S, et al. 2020. Fungal Diversity 105:1-16 ((https://doi.org/10.1007/s13225-020-00466-2)) |
| Otu18037 | 1 | k__Fungi(100);p__Ascomycota(100);c__Saccharomycetes(100);o__Saccharomycetales(100);f__Metschnikowiaceae(100);g__Metschnikowia(100);s__Metschnikowia_pulcherrima(100); | Metschnikowia | 13 | Saprotroph | \|Undefined Saprotroph\| | Yeast | NULL | Probable | As nectar-tap_saprotroph (Põlme et al. 2020); Decay sugar-rich substrates (Põlme et al. 2020) | Põlme S, et al. 2020. Fungal Diversity 105:1-16 ((https://doi.org/10.1007/s13225-020-00466-2)) |
| Otu18038 | 1 | k__Fungi(100);p__Ascomycota(100);c__Saccharomycetes(100);o__Saccharomycetales(100);f__Metschnikowiaceae(100);g__Metschnikowia(100);s__Metschnikowia_pulcherrima(100); | Metschnikowia | 13 | Saprotroph | \|Undefined Saprotroph\| | Yeast | NULL | Probable | As nectar-tap_saprotroph (Põlme et al. 2020); Decay sugar-rich substrates (Põlme et al. 2020) | Põlme S, et al. 2020. Fungal Diversity 105:1-16 ((https://doi.org/10.1007/s13225-020-00466-2)) |
| Otu18039 | 1 | k__Fungi(100);p__Ascomycota(100);c__Saccharomycetes(100);o__Saccharomycetales(100);f__Metschnikowiaceae(100);g__Metschnikowia(100);g__Metschnikowia_unclassified(100); | Metschnikowia | 13 | Saprotroph | \|Undefined Saprotroph\| | Yeast | NULL | Probable | As nectar-tap_saprotroph (Põlme et al. 2020); Decay sugar-rich substrates (Põlme et al. 2020) | Põlme S, et al. 2020. Fungal Diversity 105:1-16 ((https://doi.org/10.1007/s13225-020-00466-2)) |
| Otu18040 | 1 | k__Fungi(100);p__Ascomycota(100);c__Saccharomycetes(100);o__Saccharomycetales(100);f__Metschnikowiaceae(100);g__Metschnikowia(100);s__Metschnikowia_pulcherrima(100); | Metschnikowia | 13 | Saprotroph | \|Undefined Saprotroph\| | Yeast | NULL | Probable | As nectar-tap_saprotroph (Põlme et al. 2020); Decay sugar-rich substrates (Põlme et al. 2020) | Põlme S, et al. 2020. Fungal Diversity 105:1-16 ((https://doi.org/10.1007/s13225-020-00466-2)) |
| Otu18041 | 1 | k__Fungi(100);p__Ascomycota(100);c__Saccharomycetes(100);o__Saccharomycetales(100);f__Metschnikowiaceae(100);g__Metschnikowia(100);s__Metschnikowia_pulcherrima(100); | Metschnikowia | 13 | Saprotroph | \|Undefined Saprotroph\| | Yeast | NULL | Probable | As nectar-tap_saprotroph (Põlme et al. 2020); Decay sugar-rich substrates (Põlme et al. 2020) | Põlme S, et al. 2020. Fungal Diversity 105:1-16 ((https://doi.org/10.1007/s13225-020-00466-2)) |
| Otu18042 | 1 | k__Fungi(100);p__Ascomycota(100);c__Saccharomycetes(100);o__Saccharomycetales(100);f__Metschnikowiaceae(100);g__Metschnikowia(100);s__Metschnikowia_pulcherrima(100); | Metschnikowia | 13 | Saprotroph | \|Undefined Saprotroph\| | Yeast | NULL | Probable | As nectar-tap_saprotroph (Põlme et al. 2020); Decay sugar-rich substrates (Põlme et al. 2020) | Põlme S, et al. 2020. Fungal Diversity 105:1-16 ((https://doi.org/10.1007/s13225-020-00466-2)) |
| Otu18044 | 1 | k__Fungi(100);p__Ascomycota(100);c__Saccharomycetes(100);o__Saccharomycetales(100);f__Saccharomycetales_family_Incertae_sedis(100);f__Saccharomycetales_family_Incertae_sedis_unclassified(100);f__Saccharomycetales_family_Incertae_sedis_unclassified(100); | Saccharomycetales | 7 | Saprotroph | Undefined Saprotroph | Yeast | NULL | Possible | NULL | Sterkenburg E, et al. 2015. New Phytologist 207:1145-1158 ((https://doi.org/10.1111/nph.13426)) |
| Otu18045 | 1 | k__Fungi(100);p__Ascomycota(100);c__Saccharomycetes(100);o__Saccharomycetales(100);f__Metschnikowiaceae(100);g__Metschnikowia(100);s__Metschnikowia_pulcherrima(100); | Metschnikowia | 13 | Saprotroph | \|Undefined Saprotroph\| | Yeast | NULL | Probable | As nectar-tap_saprotroph (Põlme et al. 2020); Decay sugar-rich substrates (Põlme et al. 2020) | Põlme S, et al. 2020. Fungal Diversity 105:1-16 ((https://doi.org/10.1007/s13225-020-00466-2)) |
| Otu18046 | 1 | k__Fungi(100);p__Ascomycota(100);c__Saccharomycetes(100);o__Saccharomycetales(100);f__Metschnikowiaceae(100);g__Metschnikowia(100);s__Metschnikowia_pulcherrima(100); | Metschnikowia | 13 | Saprotroph | \|Undefined Saprotroph\| | Yeast | NULL | Probable | As nectar-tap_saprotroph (Põlme et al. 2020); Decay sugar-rich substrates (Põlme et al. 2020) | Põlme S, et al. 2020. Fungal Diversity 105:1-16 ((https://doi.org/10.1007/s13225-020-00466-2)) |
| Otu18047 | 1 | k__Fungi(100);p__Ascomycota(100);c__Saccharomycetes(100);o__Saccharomycetales(100);f__Metschnikowiaceae(100);g__Metschnikowia(100);g__Metschnikowia_unclassified(100); | Metschnikowia | 13 | Saprotroph | \|Undefined Saprotroph\| | Yeast | NULL | Probable | As nectar-tap_saprotroph (Põlme et al. 2020); Decay sugar-rich substrates (Põlme et al. 2020) | Põlme S, et al. 2020. Fungal Diversity 105:1-16 ((https://doi.org/10.1007/s13225-020-00466-2)) |
| Otu18049 | 1 | k__Fungi(100);p__Ascomycota(100);c__Saccharomycetes(100);o__Saccharomycetales(100);f__Metschnikowiaceae(100);g__Metschnikowia(100);s__Metschnikowia_pulcherrima(100); | Metschnikowia | 13 | Saprotroph | \|Undefined Saprotroph\| | Yeast | NULL | Probable | As nectar-tap_saprotroph (Põlme et al. 2020); Decay sugar-rich substrates (Põlme et al. 2020) | Põlme S, et al. 2020. Fungal Diversity 105:1-16 ((https://doi.org/10.1007/s13225-020-00466-2)) |
| Otu18050 | 1 | k__Fungi(100);p__Ascomycota(100);c__Saccharomycetes(100);o__Saccharomycetales(100);f__Metschnikowiaceae(100);g__Metschnikowia(100);s__Metschnikowia_pulcherrima(100); | Metschnikowia | 13 | Saprotroph | \|Undefined Saprotroph\| | Yeast | NULL | Probable | As nectar-tap_saprotroph (Põlme et al. 2020); Decay sugar-rich substrates (Põlme et al. 2020) | Põlme S, et al. 2020. Fungal Diversity 105:1-16 ((https://doi.org/10.1007/s13225-020-00466-2)) |
| Otu18053 | 1 | k__Fungi(100);p__Ascomycota(100);c__Saccharomycetes(100);o__Saccharomycetales(100);f__Metschnikowiaceae(100);g__Metschnikowia(100);s__Metschnikowia_pulcherrima(100); | Metschnikowia | 13 | Saprotroph | \|Undefined Saprotroph\| | Yeast | NULL | Probable | As nectar-tap_saprotroph (Põlme et al. 2020); Decay sugar-rich substrates (Põlme et al. 2020) | Põlme S, et al. 2020. Fungal Diversity 105:1-16 ((https://doi.org/10.1007/s13225-020-00466-2)) |
| Otu18057 | 1 | k__Fungi(100);p__Ascomycota(100);c__Saccharomycetes(100);o__Saccharomycetales(100);f__Metschnikowiaceae(100);g__Metschnikowia(100);g__Metschnikowia_unclassified(100); | Metschnikowia | 13 | Saprotroph | \|Undefined Saprotroph\| | Yeast | NULL | Probable | As nectar-tap_saprotroph (Põlme et al. 2020); Decay sugar-rich substrates (Põlme et al. 2020) | Põlme S, et al. 2020. Fungal Diversity 105:1-16 ((https://doi.org/10.1007/s13225-020-00466-2)) |
| Otu18058 | 1 | k__Fungi(100);p__Ascomycota(100);c__Saccharomycetes(100);o__Saccharomycetales(100);f__Metschnikowiaceae(100);g__Metschnikowia(100);s__Metschnikowia_pulcherrima(100); | Metschnikowia | 13 | Saprotroph | \|Undefined Saprotroph\| | Yeast | NULL | Probable | As nectar-tap_saprotroph (Põlme et al. 2020); Decay sugar-rich substrates (Põlme et al. 2020) | Põlme S, et al. 2020. Fungal Diversity 105:1-16 ((https://doi.org/10.1007/s13225-020-00466-2)) |
| Otu18059 | 1 | k__Fungi(100);p__Ascomycota(100);c__Saccharomycetes(100);o__Saccharomycetales(100);f__Metschnikowiaceae(100);g__Metschnikowia(100);s__Metschnikowia_pulcherrima(100); | Metschnikowia | 13 | Saprotroph | \|Undefined Saprotroph\| | Yeast | NULL | Probable | As nectar-tap_saprotroph (Põlme et al. 2020); Decay sugar-rich substrates (Põlme et al. 2020) | Põlme S, et al. 2020. Fungal Diversity 105:1-16 ((https://doi.org/10.1007/s13225-020-00466-2)) |
| Otu18060 | 1 | k__Fungi(100);p__Ascomycota(100);c__Saccharomycetes(100);o__Saccharomycetales(100);f__Metschnikowiaceae(100);g__Metschnikowia(100);s__Metschnikowia_pulcherrima(100); | Metschnikowia | 13 | Saprotroph | \|Undefined Saprotroph\| | Yeast | NULL | Probable | As nectar-tap_saprotroph (Põlme et al. 2020); Decay sugar-rich substrates (Põlme et al. 2020) | Põlme S, et al. 2020. Fungal Diversity 105:1-16 ((https://doi.org/10.1007/s13225-020-00466-2)) |
| Otu18061 | 1 | k__Fungi(100);p__Ascomycota(100);c__Saccharomycetes(100);o__Saccharomycetales(100);f__Metschnikowiaceae(100);g__Metschnikowia(100);g__Metschnikowia_unclassified(100); | Metschnikowia | 13 | Saprotroph | \|Undefined Saprotroph\| | Yeast | NULL | Probable | As nectar-tap_saprotroph (Põlme et al. 2020); Decay sugar-rich substrates (Põlme et al. 2020) | Põlme S, et al. 2020. Fungal Diversity 105:1-16 ((https://doi.org/10.1007/s13225-020-00466-2)) |
| Otu18062 | 1 | k__Fungi(100);p__Ascomycota(100);c__Saccharomycetes(100);o__Saccharomycetales(100);f__Metschnikowiaceae(100);g__Metschnikowia(100);s__Metschnikowia_pulcherrima(100); | Metschnikowia | 13 | Saprotroph | \|Undefined Saprotroph\| | Yeast | NULL | Probable | As nectar-tap_saprotroph (Põlme et al. 2020); Decay sugar-rich substrates (Põlme et al. 2020) | Põlme S, et al. 2020. Fungal Diversity 105:1-16 ((https://doi.org/10.1007/s13225-020-00466-2)) |
| Otu18063 | 1 | k__Fungi(100);p__Ascomycota(100);c__Saccharomycetes(100);o__Saccharomycetales(100);f__Metschnikowiaceae(100);g__Metschnikowia(100);g__Metschnikowia_unclassified(100); | Metschnikowia | 13 | Saprotroph | \|Undefined Saprotroph\| | Yeast | NULL | Probable | As nectar-tap_saprotroph (Põlme et al. 2020); Decay sugar-rich substrates (Põlme et al. 2020) | Põlme S, et al. 2020. Fungal Diversity 105:1-16 ((https://doi.org/10.1007/s13225-020-00466-2)) |
| Otu18064 | 1 | k__Fungi(100);p__Ascomycota(100);c__Saccharomycetes(100);o__Saccharomycetales(100);f__Metschnikowiaceae(100);g__Metschnikowia(100);g__Metschnikowia_unclassified(100); | Metschnikowia | 13 | Saprotroph | \|Undefined Saprotroph\| | Yeast | NULL | Probable | As nectar-tap_saprotroph (Põlme et al. 2020); Decay sugar-rich substrates (Põlme et al. 2020) | Põlme S, et al. 2020. Fungal Diversity 105:1-16 ((https://doi.org/10.1007/s13225-020-00466-2)) |
| Otu18065 | 1 | k__Fungi(100);p__Ascomycota(100);c__Saccharomycetes(100);o__Saccharomycetales(100);f__Metschnikowiaceae(100);g__Metschnikowia(100);s__Metschnikowia_pulcherrima(100); | Metschnikowia | 13 | Saprotroph | \|Undefined Saprotroph\| | Yeast | NULL | Probable | As nectar-tap_saprotroph (Põlme et al. 2020); Decay sugar-rich substrates (Põlme et al. 2020) | Põlme S, et al. 2020. Fungal Diversity 105:1-16 ((https://doi.org/10.1007/s13225-020-00466-2)) |
| Otu18066 | 1 | k__Fungi(100);p__Ascomycota(100);c__Saccharomycetes(100);o__Saccharomycetales(100);f__Metschnikowiaceae(100);g__Metschnikowia(100);s__Metschnikowia_pulcherrima(100); | Metschnikowia | 13 | Saprotroph | \|Undefined Saprotroph\| | Yeast | NULL | Probable | As nectar-tap_saprotroph (Põlme et al. 2020); Decay sugar-rich substrates (Põlme et al. 2020) | Põlme S, et al. 2020. Fungal Diversity 105:1-16 ((https://doi.org/10.1007/s13225-020-00466-2)) |
| Otu18067 | 1 | k__Fungi(100);p__Ascomycota(100);c__Saccharomycetes(100);o__Saccharomycetales(100);f__Metschnikowiaceae(100);g__Metschnikowia(100);g__Metschnikowia_unclassified(100); | Metschnikowia | 13 | Saprotroph | \|Undefined Saprotroph\| | Yeast | NULL | Probable | As nectar-tap_saprotroph (Põlme et al. 2020); Decay sugar-rich substrates (Põlme et al. 2020) | Põlme S, et al. 2020. Fungal Diversity 105:1-16 ((https://doi.org/10.1007/s13225-020-00466-2)) |
| Otu18068 | 1 | k__Fungi(100);p__Ascomycota(100);c__Saccharomycetes(100);o__Saccharomycetales(100);f__Metschnikowiaceae(100);g__Metschnikowia(100);s__Metschnikowia_pulcherrima(100); | Metschnikowia | 13 | Saprotroph | \|Undefined Saprotroph\| | Yeast | NULL | Probable | As nectar-tap_saprotroph (Põlme et al. 2020); Decay sugar-rich substrates (Põlme et al. 2020) | Põlme S, et al. 2020. Fungal Diversity 105:1-16 ((https://doi.org/10.1007/s13225-020-00466-2)) |
| Otu18069 | 1 | k__Fungi(100);p__Ascomycota(100);c__Saccharomycetes(100);o__Saccharomycetales(100);f__Metschnikowiaceae(100);g__Metschnikowia(100);s__Metschnikowia_pulcherrima(100); | Metschnikowia | 13 | Saprotroph | \|Undefined Saprotroph\| | Yeast | NULL | Probable | As nectar-tap_saprotroph (Põlme et al. 2020); Decay sugar-rich substrates (Põlme et al. 2020) | Põlme S, et al. 2020. Fungal Diversity 105:1-16 ((https://doi.org/10.1007/s13225-020-00466-2)) |
| Otu18070 | 1 | k__Fungi(100);p__Ascomycota(100);c__Saccharomycetes(100);o__Saccharomycetales(100);f__Metschnikowiaceae(100);g__Metschnikowia(100);s__Metschnikowia_pulcherrima(100); | Metschnikowia | 13 | Saprotroph | \|Undefined Saprotroph\| | Yeast | NULL | Probable | As nectar-tap_saprotroph (Põlme et al. 2020); Decay sugar-rich substrates (Põlme et al. 2020) | Põlme S, et al. 2020. Fungal Diversity 105:1-16 ((https://doi.org/10.1007/s13225-020-00466-2)) |
| Otu18071 | 1 | k__Fungi(100);p__Ascomycota(100);c__Saccharomycetes(100);o__Saccharomycetales(100);f__Metschnikowiaceae(100);g__Metschnikowia(100);s__Metschnikowia_pulcherrima(100); | Metschnikowia | 13 | Saprotroph | \|Undefined Saprotroph\| | Yeast | NULL | Probable | As nectar-tap_saprotroph (Põlme et al. 2020); Decay sugar-rich substrates (Põlme et al. 2020) | Põlme S, et al. 2020. Fungal Diversity 105:1-16 ((https://doi.org/10.1007/s13225-020-00466-2)) |
| Otu18072 | 1 | k__Fungi(100);p__Ascomycota(100);c__Saccharomycetes(100);o__Saccharomycetales(100);f__Metschnikowiaceae(100);g__Metschnikowia(100);s__Metschnikowia_pulcherrima(100); | Metschnikowia | 13 | Saprotroph | \|Undefined Saprotroph\| | Yeast | NULL | Probable | As nectar-tap_saprotroph (Põlme et al. 2020); Decay sugar-rich substrates (Põlme et al. 2020) | Põlme S, et al. 2020. Fungal Diversity 105:1-16 ((https://doi.org/10.1007/s13225-020-00466-2)) |
| Otu18073 | 1 | k__Fungi(100);p__Ascomycota(100);c__Saccharomycetes(100);o__Saccharomycetales(100);f__Metschnikowiaceae(100);g__Metschnikowia(100);s__Metschnikowia_pulcherrima(100); | Metschnikowia | 13 | Saprotroph | \|Undefined Saprotroph\| | Yeast | NULL | Probable | As nectar-tap_saprotroph (Põlme et al. 2020); Decay sugar-rich substrates (Põlme et al. 2020) | Põlme S, et al. 2020. Fungal Diversity 105:1-16 ((https://doi.org/10.1007/s13225-020-00466-2)) |
| Otu18074 | 1 | k__Fungi(100);p__Ascomycota(100);c__Saccharomycetes(100);o__Saccharomycetales(100);f__Metschnikowiaceae(100);g__Metschnikowia(100);s__Metschnikowia_pulcherrima(100); | Metschnikowia | 13 | Saprotroph | \|Undefined Saprotroph\| | Yeast | NULL | Probable | As nectar-tap_saprotroph (Põlme et al. 2020); Decay sugar-rich substrates (Põlme et al. 2020) | Põlme S, et al. 2020. Fungal Diversity 105:1-16 ((https://doi.org/10.1007/s13225-020-00466-2)) |
| Otu18075 | 1 | k__Fungi(100);p__Ascomycota(100);c__Saccharomycetes(100);o__Saccharomycetales(100);f__Metschnikowiaceae(100);g__Metschnikowia(100);s__Metschnikowia_pulcherrima(100); | Metschnikowia | 13 | Saprotroph | \|Undefined Saprotroph\| | Yeast | NULL | Probable | As nectar-tap_saprotroph (Põlme et al. 2020); Decay sugar-rich substrates (Põlme et al. 2020) | Põlme S, et al. 2020. Fungal Diversity 105:1-16 ((https://doi.org/10.1007/s13225-020-00466-2)) |
| Otu18076 | 1 | k__Fungi(100);p__Ascomycota(100);c__Saccharomycetes(100);o__Saccharomycetales(100);f__Metschnikowiaceae(100);g__Metschnikowia(100);g__Metschnikowia_unclassified(100); | Metschnikowia | 13 | Saprotroph | \|Undefined Saprotroph\| | Yeast | NULL | Probable | As nectar-tap_saprotroph (Põlme et al. 2020); Decay sugar-rich substrates (Põlme et al. 2020) | Põlme S, et al. 2020. Fungal Diversity 105:1-16 ((https://doi.org/10.1007/s13225-020-00466-2)) |
| Otu18080 | 1 | k__Fungi(100);p__Ascomycota(100);c__Saccharomycetes(100);o__Saccharomycetales(100);f__Metschnikowiaceae(100);g__Metschnikowia(100);g__Metschnikowia_unclassified(100); | Metschnikowia | 13 | Saprotroph | \|Undefined Saprotroph\| | Yeast | NULL | Probable | As nectar-tap_saprotroph (Põlme et al. 2020); Decay sugar-rich substrates (Põlme et al. 2020) | Põlme S, et al. 2020. Fungal Diversity 105:1-16 ((https://doi.org/10.1007/s13225-020-00466-2)) |
| Otu18082 | 1 | k__Fungi(100);p__Ascomycota(100);c__Saccharomycetes(100);o__Saccharomycetales(100);f__Metschnikowiaceae(100);g__Metschnikowia(100);g__Metschnikowia_unclassified(100); | Metschnikowia | 13 | Saprotroph | \|Undefined Saprotroph\| | Yeast | NULL | Probable | As nectar-tap_saprotroph (Põlme et al. 2020); Decay sugar-rich substrates (Põlme et al. 2020) | Põlme S, et al. 2020. Fungal Diversity 105:1-16 ((https://doi.org/10.1007/s13225-020-00466-2)) |
| Otu18083 | 1 | k__Fungi(100);p__Ascomycota(100);c__Saccharomycetes(100);o__Saccharomycetales(100);f__Metschnikowiaceae(100);g__Metschnikowia(100);g__Metschnikowia_unclassified(100); | Metschnikowia | 13 | Saprotroph | \|Undefined Saprotroph\| | Yeast | NULL | Probable | As nectar-tap_saprotroph (Põlme et al. 2020); Decay sugar-rich substrates (Põlme et al. 2020) | Põlme S, et al. 2020. Fungal Diversity 105:1-16 ((https://doi.org/10.1007/s13225-020-00466-2)) |
| Otu18084 | 1 | k__Fungi(100);p__Ascomycota(100);c__Saccharomycetes(100);o__Saccharomycetales(100);f__Metschnikowiaceae(100);g__Metschnikowia(100);s__Metschnikowia_pulcherrima(100); | Metschnikowia | 13 | Saprotroph | \|Undefined Saprotroph\| | Yeast | NULL | Probable | As nectar-tap_saprotroph (Põlme et al. 2020); Decay sugar-rich substrates (Põlme et al. 2020) | Põlme S, et al. 2020. Fungal Diversity 105:1-16 ((https://doi.org/10.1007/s13225-020-00466-2)) |
| Otu18085 | 1 | k__Fungi(100);p__Ascomycota(100);c__Saccharomycetes(100);o__Saccharomycetales(100);f__Metschnikowiaceae(100);g__Metschnikowia(100);g__Metschnikowia_unclassified(100); | Metschnikowia | 13 | Saprotroph | \|Undefined Saprotroph\| | Yeast | NULL | Probable | As nectar-tap_saprotroph (Põlme et al. 2020); Decay sugar-rich substrates (Põlme et al. 2020) | Põlme S, et al. 2020. Fungal Diversity 105:1-16 ((https://doi.org/10.1007/s13225-020-00466-2)) |
| Otu18086 | 1 | k__Fungi(100);p__Ascomycota(100);c__Saccharomycetes(100);o__Saccharomycetales(100);f__Metschnikowiaceae(100);g__Metschnikowia(100);s__Metschnikowia_pulcherrima(100); | Metschnikowia | 13 | Saprotroph | \|Undefined Saprotroph\| | Yeast | NULL | Probable | As nectar-tap_saprotroph (Põlme et al. 2020); Decay sugar-rich substrates (Põlme et al. 2020) | Põlme S, et al. 2020. Fungal Diversity 105:1-16 ((https://doi.org/10.1007/s13225-020-00466-2)) |
| Otu18087 | 1 | k__Fungi(100);p__Ascomycota(100);c__Saccharomycetes(100);o__Saccharomycetales(100);f__Metschnikowiaceae(100);g__Metschnikowia(100);s__Metschnikowia_pulcherrima(100); | Metschnikowia | 13 | Saprotroph | \|Undefined Saprotroph\| | Yeast | NULL | Probable | As nectar-tap_saprotroph (Põlme et al. 2020); Decay sugar-rich substrates (Põlme et al. 2020) | Põlme S, et al. 2020. Fungal Diversity 105:1-16 ((https://doi.org/10.1007/s13225-020-00466-2)) |
| Otu18088 | 1 | k__Fungi(100);p__Ascomycota(100);c__Saccharomycetes(100);o__Saccharomycetales(100);f__Metschnikowiaceae(100);g__Metschnikowia(100);s__Metschnikowia_pulcherrima(100); | Metschnikowia | 13 | Saprotroph | \|Undefined Saprotroph\| | Yeast | NULL | Probable | As nectar-tap_saprotroph (Põlme et al. 2020); Decay sugar-rich substrates (Põlme et al. 2020) | Põlme S, et al. 2020. Fungal Diversity 105:1-16 ((https://doi.org/10.1007/s13225-020-00466-2)) |
| Otu18089 | 1 | k__Fungi(100);p__Ascomycota(100);c__Saccharomycetes(100);o__Saccharomycetales(100);f__Metschnikowiaceae(100);g__Metschnikowia(100);s__Metschnikowia_pulcherrima(100); | Metschnikowia | 13 | Saprotroph | \|Undefined Saprotroph\| | Yeast | NULL | Probable | As nectar-tap_saprotroph (Põlme et al. 2020); Decay sugar-rich substrates (Põlme et al. 2020) | Põlme S, et al. 2020. Fungal Diversity 105:1-16 ((https://doi.org/10.1007/s13225-020-00466-2)) |
| Otu18090 | 1 | k__Fungi(100);p__Ascomycota(100);c__Saccharomycetes(100);o__Saccharomycetales(100);f__Metschnikowiaceae(100);g__Metschnikowia(100);s__Metschnikowia_pulcherrima(100); | Metschnikowia | 13 | Saprotroph | \|Undefined Saprotroph\| | Yeast | NULL | Probable | As nectar-tap_saprotroph (Põlme et al. 2020); Decay sugar-rich substrates (Põlme et al. 2020) | Põlme S, et al. 2020. Fungal Diversity 105:1-16 ((https://doi.org/10.1007/s13225-020-00466-2)) |
| Otu18091 | 1 | k__Fungi(100);p__Ascomycota(100);c__Saccharomycetes(100);o__Saccharomycetales(100);f__Metschnikowiaceae(100);g__Metschnikowia(100);s__Metschnikowia_pulcherrima(100); | Metschnikowia | 13 | Saprotroph | \|Undefined Saprotroph\| | Yeast | NULL | Probable | As nectar-tap_saprotroph (Põlme et al. 2020); Decay sugar-rich substrates (Põlme et al. 2020) | Põlme S, et al. 2020. Fungal Diversity 105:1-16 ((https://doi.org/10.1007/s13225-020-00466-2)) |
| Otu18092 | 1 | k__Fungi(100);p__Ascomycota(100);c__Saccharomycetes(100);o__Saccharomycetales(100);f__Metschnikowiaceae(100);g__Metschnikowia(100);s__Metschnikowia_pulcherrima(100); | Metschnikowia | 13 | Saprotroph | \|Undefined Saprotroph\| | Yeast | NULL | Probable | As nectar-tap_saprotroph (Põlme et al. 2020); Decay sugar-rich substrates (Põlme et al. 2020) | Põlme S, et al. 2020. Fungal Diversity 105:1-16 ((https://doi.org/10.1007/s13225-020-00466-2)) |
| Otu18093 | 1 | k__Fungi(100);p__Ascomycota(100);c__Saccharomycetes(100);o__Saccharomycetales(100);f__Metschnikowiaceae(100);g__Metschnikowia(100);s__Metschnikowia_pulcherrima(100); | Metschnikowia | 13 | Saprotroph | \|Undefined Saprotroph\| | Yeast | NULL | Probable | As nectar-tap_saprotroph (Põlme et al. 2020); Decay sugar-rich substrates (Põlme et al. 2020) | Põlme S, et al. 2020. Fungal Diversity 105:1-16 ((https://doi.org/10.1007/s13225-020-00466-2)) |
| Otu18094 | 1 | k__Fungi(100);p__Ascomycota(100);c__Saccharomycetes(100);o__Saccharomycetales(100);f__Metschnikowiaceae(100);g__Metschnikowia(100);g__Metschnikowia_unclassified(100); | Metschnikowia | 13 | Saprotroph | \|Undefined Saprotroph\| | Yeast | NULL | Probable | As nectar-tap_saprotroph (Põlme et al. 2020); Decay sugar-rich substrates (Põlme et al. 2020) | Põlme S, et al. 2020. Fungal Diversity 105:1-16 ((https://doi.org/10.1007/s13225-020-00466-2)) |
| Otu18095 | 1 | k__Fungi(100);p__Ascomycota(100);c__Saccharomycetes(100);o__Saccharomycetales(100);f__Metschnikowiaceae(100);g__Metschnikowia(100);g__Metschnikowia_unclassified(100); | Metschnikowia | 13 | Saprotroph | \|Undefined Saprotroph\| | Yeast | NULL | Probable | As nectar-tap_saprotroph (Põlme et al. 2020); Decay sugar-rich substrates (Põlme et al. 2020) | Põlme S, et al. 2020. Fungal Diversity 105:1-16 ((https://doi.org/10.1007/s13225-020-00466-2)) |
| Otu18096 | 1 | k__Fungi(100);p__Ascomycota(100);c__Saccharomycetes(100);o__Saccharomycetales(100);f__Metschnikowiaceae(100);g__Metschnikowia(100);s__Metschnikowia_pulcherrima(100); | Metschnikowia | 13 | Saprotroph | \|Undefined Saprotroph\| | Yeast | NULL | Probable | As nectar-tap_saprotroph (Põlme et al. 2020); Decay sugar-rich substrates (Põlme et al. 2020) | Põlme S, et al. 2020. Fungal Diversity 105:1-16 ((https://doi.org/10.1007/s13225-020-00466-2)) |
| Otu18097 | 1 | k__Fungi(100);p__Ascomycota(100);c__Saccharomycetes(100);o__Saccharomycetales(100);f__Metschnikowiaceae(100);g__Metschnikowia(100);s__Metschnikowia_pulcherrima(100); | Metschnikowia | 13 | Saprotroph | \|Undefined Saprotroph\| | Yeast | NULL | Probable | As nectar-tap_saprotroph (Põlme et al. 2020); Decay sugar-rich substrates (Põlme et al. 2020) | Põlme S, et al. 2020. Fungal Diversity 105:1-16 ((https://doi.org/10.1007/s13225-020-00466-2)) |
| Otu18098 | 1 | k__Fungi(100);p__Ascomycota(100);c__Saccharomycetes(100);o__Saccharomycetales(100);f__Metschnikowiaceae(100);g__Metschnikowia(100);g__Metschnikowia_unclassified(100); | Metschnikowia | 13 | Saprotroph | \|Undefined Saprotroph\| | Yeast | NULL | Probable | As nectar-tap_saprotroph (Põlme et al. 2020); Decay sugar-rich substrates (Põlme et al. 2020) | Põlme S, et al. 2020. Fungal Diversity 105:1-16 ((https://doi.org/10.1007/s13225-020-00466-2)) |
| Otu18099 | 1 | k__Fungi(100);p__Ascomycota(100);c__Saccharomycetes(100);o__Saccharomycetales(100);f__Metschnikowiaceae(100);g__Metschnikowia(100);s__Metschnikowia_pulcherrima(100); | Metschnikowia | 13 | Saprotroph | \|Undefined Saprotroph\| | Yeast | NULL | Probable | As nectar-tap_saprotroph (Põlme et al. 2020); Decay sugar-rich substrates (Põlme et al. 2020) | Põlme S, et al. 2020. Fungal Diversity 105:1-16 ((https://doi.org/10.1007/s13225-020-00466-2)) |
| Otu18100 | 1 | k__Fungi(100);p__Ascomycota(100);c__Saccharomycetes(100);o__Saccharomycetales(100);f__Metschnikowiaceae(100);g__Metschnikowia(100);s__Metschnikowia_pulcherrima(100); | Metschnikowia | 13 | Saprotroph | \|Undefined Saprotroph\| | Yeast | NULL | Probable | As nectar-tap_saprotroph (Põlme et al. 2020); Decay sugar-rich substrates (Põlme et al. 2020) | Põlme S, et al. 2020. Fungal Diversity 105:1-16 ((https://doi.org/10.1007/s13225-020-00466-2)) |
| Otu18101 | 1 | k__Fungi(100);p__Ascomycota(100);c__Saccharomycetes(100);o__Saccharomycetales(100);f__Metschnikowiaceae(100);g__Metschnikowia(100);g__Metschnikowia_unclassified(100); | Metschnikowia | 13 | Saprotroph | \|Undefined Saprotroph\| | Yeast | NULL | Probable | As nectar-tap_saprotroph (Põlme et al. 2020); Decay sugar-rich substrates (Põlme et al. 2020) | Põlme S, et al. 2020. Fungal Diversity 105:1-16 ((https://doi.org/10.1007/s13225-020-00466-2)) |
| Otu18102 | 1 | k__Fungi(100);p__Ascomycota(100);c__Saccharomycetes(100);o__Saccharomycetales(100);f__Metschnikowiaceae(100);g__Metschnikowia(100);s__Metschnikowia_pulcherrima(100); | Metschnikowia | 13 | Saprotroph | \|Undefined Saprotroph\| | Yeast | NULL | Probable | As nectar-tap_saprotroph (Põlme et al. 2020); Decay sugar-rich substrates (Põlme et al. 2020) | Põlme S, et al. 2020. Fungal Diversity 105:1-16 ((https://doi.org/10.1007/s13225-020-00466-2)) |
| Otu18103 | 1 | k__Fungi(100);p__Ascomycota(100);c__Saccharomycetes(100);o__Saccharomycetales(100);f__Metschnikowiaceae(100);g__Metschnikowia(100);g__Metschnikowia_unclassified(100); | Metschnikowia | 13 | Saprotroph | \|Undefined Saprotroph\| | Yeast | NULL | Probable | As nectar-tap_saprotroph (Põlme et al. 2020); Decay sugar-rich substrates (Põlme et al. 2020) | Põlme S, et al. 2020. Fungal Diversity 105:1-16 ((https://doi.org/10.1007/s13225-020-00466-2)) |
| Otu18104 | 1 | k__Fungi(100);p__Ascomycota(100);c__Saccharomycetes(100);o__Saccharomycetales(100);f__Metschnikowiaceae(100);g__Metschnikowia(100);s__Metschnikowia_pulcherrima(100); | Metschnikowia | 13 | Saprotroph | \|Undefined Saprotroph\| | Yeast | NULL | Probable | As nectar-tap_saprotroph (Põlme et al. 2020); Decay sugar-rich substrates (Põlme et al. 2020) | Põlme S, et al. 2020. Fungal Diversity 105:1-16 ((https://doi.org/10.1007/s13225-020-00466-2)) |
| Otu18105 | 1 | k__Fungi(100);p__Ascomycota(100);c__Saccharomycetes(100);o__Saccharomycetales(100);f__Metschnikowiaceae(100);g__Metschnikowia(100);g__Metschnikowia_unclassified(100); | Metschnikowia | 13 | Saprotroph | \|Undefined Saprotroph\| | Yeast | NULL | Probable | As nectar-tap_saprotroph (Põlme et al. 2020); Decay sugar-rich substrates (Põlme et al. 2020) | Põlme S, et al. 2020. Fungal Diversity 105:1-16 ((https://doi.org/10.1007/s13225-020-00466-2)) |
| Otu18106 | 1 | k__Fungi(100);p__Ascomycota(100);c__Saccharomycetes(100);o__Saccharomycetales(100);f__Metschnikowiaceae(100);g__Metschnikowia(100);s__Metschnikowia_pulcherrima(100); | Metschnikowia | 13 | Saprotroph | \|Undefined Saprotroph\| | Yeast | NULL | Probable | As nectar-tap_saprotroph (Põlme et al. 2020); Decay sugar-rich substrates (Põlme et al. 2020) | Põlme S, et al. 2020. Fungal Diversity 105:1-16 ((https://doi.org/10.1007/s13225-020-00466-2)) |
| Otu18107 | 1 | k__Fungi(100);p__Ascomycota(100);c__Saccharomycetes(100);o__Saccharomycetales(100);f__Metschnikowiaceae(100);g__Metschnikowia(100);s__Metschnikowia_pulcherrima(100); | Metschnikowia | 13 | Saprotroph | \|Undefined Saprotroph\| | Yeast | NULL | Probable | As nectar-tap_saprotroph (Põlme et al. 2020); Decay sugar-rich substrates (Põlme et al. 2020) | Põlme S, et al. 2020. Fungal Diversity 105:1-16 ((https://doi.org/10.1007/s13225-020-00466-2)) |
| Otu18108 | 1 | k__Fungi(100);p__Ascomycota(100);c__Saccharomycetes(100);o__Saccharomycetales(100);f__Metschnikowiaceae(100);g__Metschnikowia(100);s__Metschnikowia_pulcherrima(100); | Metschnikowia | 13 | Saprotroph | \|Undefined Saprotroph\| | Yeast | NULL | Probable | As nectar-tap_saprotroph (Põlme et al. 2020); Decay sugar-rich substrates (Põlme et al. 2020) | Põlme S, et al. 2020. Fungal Diversity 105:1-16 ((https://doi.org/10.1007/s13225-020-00466-2)) |
| Otu18109 | 1 | k__Fungi(100);p__Ascomycota(100);c__Saccharomycetes(100);o__Saccharomycetales(100);f__Metschnikowiaceae(100);g__Metschnikowia(100);g__Metschnikowia_unclassified(100); | Metschnikowia | 13 | Saprotroph | \|Undefined Saprotroph\| | Yeast | NULL | Probable | As nectar-tap_saprotroph (Põlme et al. 2020); Decay sugar-rich substrates (Põlme et al. 2020) | Põlme S, et al. 2020. Fungal Diversity 105:1-16 ((https://doi.org/10.1007/s13225-020-00466-2)) |
| Otu18110 | 1 | k__Fungi(100);p__Ascomycota(100);c__Saccharomycetes(100);o__Saccharomycetales(100);f__Metschnikowiaceae(100);g__Metschnikowia(100);s__Metschnikowia_pulcherrima(100); | Metschnikowia | 13 | Saprotroph | \|Undefined Saprotroph\| | Yeast | NULL | Probable | As nectar-tap_saprotroph (Põlme et al. 2020); Decay sugar-rich substrates (Põlme et al. 2020) | Põlme S, et al. 2020. Fungal Diversity 105:1-16 ((https://doi.org/10.1007/s13225-020-00466-2)) |
| Otu18111 | 1 | k__Fungi(100);p__Ascomycota(100);c__Saccharomycetes(100);o__Saccharomycetales(100);f__Metschnikowiaceae(100);g__Metschnikowia(100);s__Metschnikowia_pulcherrima(100); | Metschnikowia | 13 | Saprotroph | \|Undefined Saprotroph\| | Yeast | NULL | Probable | As nectar-tap_saprotroph (Põlme et al. 2020); Decay sugar-rich substrates (Põlme et al. 2020) | Põlme S, et al. 2020. Fungal Diversity 105:1-16 ((https://doi.org/10.1007/s13225-020-00466-2)) |
| Otu18112 | 1 | k__Fungi(100);p__Ascomycota(100);c__Sordariomycetes(100);o__Hypocreales(100);f__Nectriaceae(100);g__Fusarium(100);g__Fusarium_unclassified(100); | Fusarium | 13 | Pathotroph-Saprotroph-Symbiotroph | Animal Pathogen-Endophyte-Lichen Parasite-\|Plant Pathogen\|-Plant Saprotroph-Undefined Saprotroph-Wood Saprotroph | Microfungus | Soft Rot | Possible | As foliar_endophyte (Põlme et al. 2020); As litter_saprotroph (Põlme et al. 2020); Causing damping-off disease (Põlme et al. 2020); Decay leaf, fruit, seed, roots, soil, animal material (Põlme et al. 2020); Likely opportunistic human pathogen (Irinyi et al. 2015) | Duncan CG, Eslyn WE. 1966. Mycologia 58:642-664 ((https://doi.org/10.2307/3757045)); Seehann G, et al. 1975. List of Fungi in Soft-Rot Tissue Tests. Institute fur Holzbiologie und Holzschutz, Bundesforschungsanstalt fur Forst - und Holzwirtschaft, Hamburg - Lohbrugge, Germany; Costa IPMW, et al. 2012. Mycotaxon 119:e26 ((http://mycotaxon.com/resources/checklists/costa_v119_checklist.pdf)); Tedersoo L, et al. 2014. Science 346:e1256688 ((https://doi.org/10.1126/science.1256688)); Irinyi L, et al. 2015. Medical Mycology 53:313-337 ((https://doi.org/10.1093/mmy/myv008)); Schroers, et al. 2016. Mycologia 108:806-819 ((https://doi.org/10.3852/15-255)); Põlme S, et al. 2020. Fungal Diversity 105:1-16 ((https://doi.org/10.1007/s13225-020-00466-2)); The American Phytopathological Society Common Names of Plant Diseases ((http://www.apsnet.org/publications/commonnames/Pages/default.aspx)); Lawrey JD, Diederich P. 2016. http://www.lichenicolous.net (accessed 24 May 2017) ((http://www.lichenicolous.net)) |
| Otu18113 | 1 | k__Fungi(100);p__Ascomycota(100);c__Saccharomycetes(100);o__Saccharomycetales(100);f__Metschnikowiaceae(100);g__Metschnikowia(100);g__Metschnikowia_unclassified(100); | Metschnikowia | 13 | Saprotroph | \|Undefined Saprotroph\| | Yeast | NULL | Probable | As nectar-tap_saprotroph (Põlme et al. 2020); Decay sugar-rich substrates (Põlme et al. 2020) | Põlme S, et al. 2020. Fungal Diversity 105:1-16 ((https://doi.org/10.1007/s13225-020-00466-2)) |
| Otu18114 | 1 | k__Fungi(100);p__Ascomycota(100);c__Saccharomycetes(100);o__Saccharomycetales(100);f__Metschnikowiaceae(100);g__Metschnikowia(100);g__Metschnikowia_unclassified(100); | Metschnikowia | 13 | Saprotroph | \|Undefined Saprotroph\| | Yeast | NULL | Probable | As nectar-tap_saprotroph (Põlme et al. 2020); Decay sugar-rich substrates (Põlme et al. 2020) | Põlme S, et al. 2020. Fungal Diversity 105:1-16 ((https://doi.org/10.1007/s13225-020-00466-2)) |
| Otu18115 | 1 | k__Fungi(100);p__Ascomycota(100);c__Saccharomycetes(100);o__Saccharomycetales(100);f__Metschnikowiaceae(100);g__Metschnikowia(100);s__Metschnikowia_pulcherrima(100); | Metschnikowia | 13 | Saprotroph | \|Undefined Saprotroph\| | Yeast | NULL | Probable | As nectar-tap_saprotroph (Põlme et al. 2020); Decay sugar-rich substrates (Põlme et al. 2020) | Põlme S, et al. 2020. Fungal Diversity 105:1-16 ((https://doi.org/10.1007/s13225-020-00466-2)) |
| Otu18117 | 1 | k__Fungi(100);p__Ascomycota(100);c__Saccharomycetes(100);o__Saccharomycetales(100);f__Saccharomycetales_family_Incertae_sedis(100);g__Candida(100);s__Candida_cretensis(100); | Candida | 13 | Pathotroph-Saprotroph-Symbiotroph | Animal Pathogen-Endophyte-Endosymbiont-Epiphyte-\|Undefined Saprotroph\| | Dimorphic Yeast-Yeast | NULL | Probable | As nectar-tap_saprotroph (Põlme et al. 2020); Decay sugar-rich substrates (Põlme et al. 2020); This asexual genus is polyphyletic and contain species that function very differently depend on what they are. Caution should be taken when interpreting guilds for Candida species. | Manolakaki D, et al. 2010. Virulence. 1: 367-375 ((https://doi.org/10.4161/viru.1.5.12796)); Mekbib SB, et al. 2011. Tropical Plant Pathology 36:233-240 ((https://doi.org/10.1590/S1982-56762011000400004)); Põlme S, et al. 2020. Fungal Diversity 105:1-16 ((https://doi.org/10.1007/s13225-020-00466-2)) |
| Otu18118 | 1 | k__Fungi(100);p__Ascomycota(100);c__Saccharomycetes(100);o__Saccharomycetales(100);f__Saccharomycetales_family_Incertae_sedis(100);g__Candida(100);g__Candida_unclassified(100); | Candida | 13 | Pathotroph-Saprotroph-Symbiotroph | Animal Pathogen-Endophyte-Endosymbiont-Epiphyte-\|Undefined Saprotroph\| | Dimorphic Yeast-Yeast | NULL | Probable | As nectar-tap_saprotroph (Põlme et al. 2020); Decay sugar-rich substrates (Põlme et al. 2020); This asexual genus is polyphyletic and contain species that function very differently depend on what they are. Caution should be taken when interpreting guilds for Candida species. | Manolakaki D, et al. 2010. Virulence. 1: 367-375 ((https://doi.org/10.4161/viru.1.5.12796)); Mekbib SB, et al. 2011. Tropical Plant Pathology 36:233-240 ((https://doi.org/10.1590/S1982-56762011000400004)); Põlme S, et al. 2020. Fungal Diversity 105:1-16 ((https://doi.org/10.1007/s13225-020-00466-2)) |
| Otu18119 | 1 | k__Fungi(100);p__Ascomycota(100);c__Saccharomycetes(100);o__Saccharomycetales(100);f__Metschnikowiaceae(100);g__Metschnikowia(100);s__Metschnikowia_pulcherrima(100); | Metschnikowia | 13 | Saprotroph | \|Undefined Saprotroph\| | Yeast | NULL | Probable | As nectar-tap_saprotroph (Põlme et al. 2020); Decay sugar-rich substrates (Põlme et al. 2020) | Põlme S, et al. 2020. Fungal Diversity 105:1-16 ((https://doi.org/10.1007/s13225-020-00466-2)) |
| Otu18120 | 1 | k__Fungi(100);p__Ascomycota(100);c__Saccharomycetes(100);o__Saccharomycetales(100);f__Metschnikowiaceae(100);g__Metschnikowia(100);g__Metschnikowia_unclassified(100); | Metschnikowia | 13 | Saprotroph | \|Undefined Saprotroph\| | Yeast | NULL | Probable | As nectar-tap_saprotroph (Põlme et al. 2020); Decay sugar-rich substrates (Põlme et al. 2020) | Põlme S, et al. 2020. Fungal Diversity 105:1-16 ((https://doi.org/10.1007/s13225-020-00466-2)) |
| Otu18121 | 1 | k__Fungi(100);p__Ascomycota(100);c__Saccharomycetes(100);o__Saccharomycetales(100);f__Saccharomycetales_family_Incertae_sedis(100);g__Candida(100);g__Candida_unclassified(100); | Candida | 13 | Pathotroph-Saprotroph-Symbiotroph | Animal Pathogen-Endophyte-Endosymbiont-Epiphyte-\|Undefined Saprotroph\| | Dimorphic Yeast-Yeast | NULL | Probable | As nectar-tap_saprotroph (Põlme et al. 2020); Decay sugar-rich substrates (Põlme et al. 2020); This asexual genus is polyphyletic and contain species that function very differently depend on what they are. Caution should be taken when interpreting guilds for Candida species. | Manolakaki D, et al. 2010. Virulence. 1: 367-375 ((https://doi.org/10.4161/viru.1.5.12796)); Mekbib SB, et al. 2011. Tropical Plant Pathology 36:233-240 ((https://doi.org/10.1590/S1982-56762011000400004)); Põlme S, et al. 2020. Fungal Diversity 105:1-16 ((https://doi.org/10.1007/s13225-020-00466-2)) |
| Otu18122 | 1 | k__Fungi(100);p__Ascomycota(100);c__Saccharomycetes(100);o__Saccharomycetales(100);f__Metschnikowiaceae(100);g__Metschnikowia(100);s__Metschnikowia_pulcherrima(100); | Metschnikowia | 13 | Saprotroph | \|Undefined Saprotroph\| | Yeast | NULL | Probable | As nectar-tap_saprotroph (Põlme et al. 2020); Decay sugar-rich substrates (Põlme et al. 2020) | Põlme S, et al. 2020. Fungal Diversity 105:1-16 ((https://doi.org/10.1007/s13225-020-00466-2)) |
| Otu18123 | 1 | k__Fungi(100);p__Ascomycota(100);c__Saccharomycetes(100);o__Saccharomycetales(100);f__Saccharomycetales_family_Incertae_sedis(100);g__Candida(100);g__Candida_unclassified(100); | Candida | 13 | Pathotroph-Saprotroph-Symbiotroph | Animal Pathogen-Endophyte-Endosymbiont-Epiphyte-\|Undefined Saprotroph\| | Dimorphic Yeast-Yeast | NULL | Probable | As nectar-tap_saprotroph (Põlme et al. 2020); Decay sugar-rich substrates (Põlme et al. 2020); This asexual genus is polyphyletic and contain species that function very differently depend on what they are. Caution should be taken when interpreting guilds for Candida species. | Manolakaki D, et al. 2010. Virulence. 1: 367-375 ((https://doi.org/10.4161/viru.1.5.12796)); Mekbib SB, et al. 2011. Tropical Plant Pathology 36:233-240 ((https://doi.org/10.1590/S1982-56762011000400004)); Põlme S, et al. 2020. Fungal Diversity 105:1-16 ((https://doi.org/10.1007/s13225-020-00466-2)) |
| Otu18124 | 1 | k__Fungi(100);p__Ascomycota(100);c__Saccharomycetes(100);o__Saccharomycetales(100);f__Metschnikowiaceae(100);g__Metschnikowia(100);s__Metschnikowia_pulcherrima(100); | Metschnikowia | 13 | Saprotroph | \|Undefined Saprotroph\| | Yeast | NULL | Probable | As nectar-tap_saprotroph (Põlme et al. 2020); Decay sugar-rich substrates (Põlme et al. 2020) | Põlme S, et al. 2020. Fungal Diversity 105:1-16 ((https://doi.org/10.1007/s13225-020-00466-2)) |
| Otu18126 | 1 | k__Fungi(100);p__Ascomycota(100);c__Saccharomycetes(100);o__Saccharomycetales(100);f__Metschnikowiaceae(100);g__Metschnikowia(100);s__Metschnikowia_pulcherrima(100); | Metschnikowia | 13 | Saprotroph | \|Undefined Saprotroph\| | Yeast | NULL | Probable | As nectar-tap_saprotroph (Põlme et al. 2020); Decay sugar-rich substrates (Põlme et al. 2020) | Põlme S, et al. 2020. Fungal Diversity 105:1-16 ((https://doi.org/10.1007/s13225-020-00466-2)) |
| Otu18127 | 1 | k__Fungi(100);p__Ascomycota(100);c__Saccharomycetes(100);o__Saccharomycetales(100);f__Metschnikowiaceae(100);g__Metschnikowia(100);s__Metschnikowia_pulcherrima(100); | Metschnikowia | 13 | Saprotroph | \|Undefined Saprotroph\| | Yeast | NULL | Probable | As nectar-tap_saprotroph (Põlme et al. 2020); Decay sugar-rich substrates (Põlme et al. 2020) | Põlme S, et al. 2020. Fungal Diversity 105:1-16 ((https://doi.org/10.1007/s13225-020-00466-2)) |
| Otu18128 | 1 | k__Fungi(100);p__Ascomycota(100);c__Saccharomycetes(100);o__Saccharomycetales(100);f__Metschnikowiaceae(100);g__Metschnikowia(100);s__Metschnikowia_pulcherrima(100); | Metschnikowia | 13 | Saprotroph | \|Undefined Saprotroph\| | Yeast | NULL | Probable | As nectar-tap_saprotroph (Põlme et al. 2020); Decay sugar-rich substrates (Põlme et al. 2020) | Põlme S, et al. 2020. Fungal Diversity 105:1-16 ((https://doi.org/10.1007/s13225-020-00466-2)) |
| Otu18129 | 1 | k__Fungi(100);p__Ascomycota(100);c__Saccharomycetes(100);o__Saccharomycetales(100);f__Metschnikowiaceae(100);g__Metschnikowia(100);s__Metschnikowia_pulcherrima(100); | Metschnikowia | 13 | Saprotroph | \|Undefined Saprotroph\| | Yeast | NULL | Probable | As nectar-tap_saprotroph (Põlme et al. 2020); Decay sugar-rich substrates (Põlme et al. 2020) | Põlme S, et al. 2020. Fungal Diversity 105:1-16 ((https://doi.org/10.1007/s13225-020-00466-2)) |
| Otu18130 | 1 | k__Fungi(100);p__Ascomycota(100);c__Saccharomycetes(100);o__Saccharomycetales(100);f__Metschnikowiaceae(100);g__Metschnikowia(100);g__Metschnikowia_unclassified(100); | Metschnikowia | 13 | Saprotroph | \|Undefined Saprotroph\| | Yeast | NULL | Probable | As nectar-tap_saprotroph (Põlme et al. 2020); Decay sugar-rich substrates (Põlme et al. 2020) | Põlme S, et al. 2020. Fungal Diversity 105:1-16 ((https://doi.org/10.1007/s13225-020-00466-2)) |
| Otu18131 | 1 | k__Fungi(100);p__Ascomycota(100);c__Saccharomycetes(100);o__Saccharomycetales(100);f__Metschnikowiaceae(100);g__Metschnikowia(100);g__Metschnikowia_unclassified(100); | Metschnikowia | 13 | Saprotroph | \|Undefined Saprotroph\| | Yeast | NULL | Probable | As nectar-tap_saprotroph (Põlme et al. 2020); Decay sugar-rich substrates (Põlme et al. 2020) | Põlme S, et al. 2020. Fungal Diversity 105:1-16 ((https://doi.org/10.1007/s13225-020-00466-2)) |
| Otu18132 | 1 | k__Fungi(100);p__Ascomycota(100);c__Saccharomycetes(100);o__Saccharomycetales(100);f__Metschnikowiaceae(100);g__Metschnikowia(100);s__Metschnikowia_pulcherrima(100); | Metschnikowia | 13 | Saprotroph | \|Undefined Saprotroph\| | Yeast | NULL | Probable | As nectar-tap_saprotroph (Põlme et al. 2020); Decay sugar-rich substrates (Põlme et al. 2020) | Põlme S, et al. 2020. Fungal Diversity 105:1-16 ((https://doi.org/10.1007/s13225-020-00466-2)) |
| Otu18133 | 1 | k__Fungi(100);p__Ascomycota(100);c__Saccharomycetes(100);o__Saccharomycetales(100);f__Metschnikowiaceae(100);g__Metschnikowia(100);g__Metschnikowia_unclassified(100); | Metschnikowia | 13 | Saprotroph | \|Undefined Saprotroph\| | Yeast | NULL | Probable | As nectar-tap_saprotroph (Põlme et al. 2020); Decay sugar-rich substrates (Põlme et al. 2020) | Põlme S, et al. 2020. Fungal Diversity 105:1-16 ((https://doi.org/10.1007/s13225-020-00466-2)) |
| Otu18134 | 1 | k__Fungi(100);p__Ascomycota(100);c__Saccharomycetes(100);o__Saccharomycetales(100);f__Metschnikowiaceae(100);g__Metschnikowia(100);g__Metschnikowia_unclassified(100); | Metschnikowia | 13 | Saprotroph | \|Undefined Saprotroph\| | Yeast | NULL | Probable | As nectar-tap_saprotroph (Põlme et al. 2020); Decay sugar-rich substrates (Põlme et al. 2020) | Põlme S, et al. 2020. Fungal Diversity 105:1-16 ((https://doi.org/10.1007/s13225-020-00466-2)) |
| Otu18135 | 1 | k__Fungi(100);p__Ascomycota(100);c__Saccharomycetes(100);o__Saccharomycetales(100);f__Metschnikowiaceae(100);g__Metschnikowia(100);s__Metschnikowia_pulcherrima(100); | Metschnikowia | 13 | Saprotroph | \|Undefined Saprotroph\| | Yeast | NULL | Probable | As nectar-tap_saprotroph (Põlme et al. 2020); Decay sugar-rich substrates (Põlme et al. 2020) | Põlme S, et al. 2020. Fungal Diversity 105:1-16 ((https://doi.org/10.1007/s13225-020-00466-2)) |
| Otu18136 | 1 | k__Fungi(100);p__Ascomycota(100);c__Saccharomycetes(100);o__Saccharomycetales(100);f__Metschnikowiaceae(100);g__Metschnikowia(100);g__Metschnikowia_unclassified(100); | Metschnikowia | 13 | Saprotroph | \|Undefined Saprotroph\| | Yeast | NULL | Probable | As nectar-tap_saprotroph (Põlme et al. 2020); Decay sugar-rich substrates (Põlme et al. 2020) | Põlme S, et al. 2020. Fungal Diversity 105:1-16 ((https://doi.org/10.1007/s13225-020-00466-2)) |
| Otu18138 | 1 | k__Fungi(100);p__Ascomycota(100);c__Saccharomycetes(100);o__Saccharomycetales(100);f__Metschnikowiaceae(100);g__Metschnikowia(100);g__Metschnikowia_unclassified(100); | Metschnikowia | 13 | Saprotroph | \|Undefined Saprotroph\| | Yeast | NULL | Probable | As nectar-tap_saprotroph (Põlme et al. 2020); Decay sugar-rich substrates (Põlme et al. 2020) | Põlme S, et al. 2020. Fungal Diversity 105:1-16 ((https://doi.org/10.1007/s13225-020-00466-2)) |
| Otu18139 | 1 | k__Fungi(100);p__Ascomycota(100);c__Saccharomycetes(100);o__Saccharomycetales(100);f__Metschnikowiaceae(100);g__Metschnikowia(100);s__Metschnikowia_pulcherrima(100); | Metschnikowia | 13 | Saprotroph | \|Undefined Saprotroph\| | Yeast | NULL | Probable | As nectar-tap_saprotroph (Põlme et al. 2020); Decay sugar-rich substrates (Põlme et al. 2020) | Põlme S, et al. 2020. Fungal Diversity 105:1-16 ((https://doi.org/10.1007/s13225-020-00466-2)) |
| Otu18140 | 1 | k__Fungi(100);p__Ascomycota(100);c__Saccharomycetes(100);o__Saccharomycetales(100);f__Metschnikowiaceae(100);g__Metschnikowia(100);g__Metschnikowia_unclassified(100); | Metschnikowia | 13 | Saprotroph | \|Undefined Saprotroph\| | Yeast | NULL | Probable | As nectar-tap_saprotroph (Põlme et al. 2020); Decay sugar-rich substrates (Põlme et al. 2020) | Põlme S, et al. 2020. Fungal Diversity 105:1-16 ((https://doi.org/10.1007/s13225-020-00466-2)) |
| Otu18141 | 1 | k__Fungi(100);p__Ascomycota(100);c__Saccharomycetes(100);o__Saccharomycetales(100);f__Metschnikowiaceae(100);g__Metschnikowia(100);s__Metschnikowia_pulcherrima(100); | Metschnikowia | 13 | Saprotroph | \|Undefined Saprotroph\| | Yeast | NULL | Probable | As nectar-tap_saprotroph (Põlme et al. 2020); Decay sugar-rich substrates (Põlme et al. 2020) | Põlme S, et al. 2020. Fungal Diversity 105:1-16 ((https://doi.org/10.1007/s13225-020-00466-2)) |
| Otu18142 | 1 | k__Fungi(100);p__Ascomycota(100);c__Saccharomycetes(100);o__Saccharomycetales(100);f__Metschnikowiaceae(100);g__Metschnikowia(100);g__Metschnikowia_unclassified(100); | Metschnikowia | 13 | Saprotroph | \|Undefined Saprotroph\| | Yeast | NULL | Probable | As nectar-tap_saprotroph (Põlme et al. 2020); Decay sugar-rich substrates (Põlme et al. 2020) | Põlme S, et al. 2020. Fungal Diversity 105:1-16 ((https://doi.org/10.1007/s13225-020-00466-2)) |
| Otu18143 | 1 | k__Fungi(100);p__Ascomycota(100);c__Saccharomycetes(100);o__Saccharomycetales(100);f__Metschnikowiaceae(100);g__Metschnikowia(100);s__Metschnikowia_pulcherrima(100); | Metschnikowia | 13 | Saprotroph | \|Undefined Saprotroph\| | Yeast | NULL | Probable | As nectar-tap_saprotroph (Põlme et al. 2020); Decay sugar-rich substrates (Põlme et al. 2020) | Põlme S, et al. 2020. Fungal Diversity 105:1-16 ((https://doi.org/10.1007/s13225-020-00466-2)) |
| Otu18145 | 1 | k__Fungi(100);p__Ascomycota(100);c__Saccharomycetes(100);o__Saccharomycetales(100);f__Metschnikowiaceae(100);g__Metschnikowia(100);s__Metschnikowia_pulcherrima(100); | Metschnikowia | 13 | Saprotroph | \|Undefined Saprotroph\| | Yeast | NULL | Probable | As nectar-tap_saprotroph (Põlme et al. 2020); Decay sugar-rich substrates (Põlme et al. 2020) | Põlme S, et al. 2020. Fungal Diversity 105:1-16 ((https://doi.org/10.1007/s13225-020-00466-2)) |
| Otu18147 | 1 | k__Fungi(100);p__Ascomycota(100);c__Saccharomycetes(100);o__Saccharomycetales(100);f__Metschnikowiaceae(100);g__Metschnikowia(100);s__Metschnikowia_pulcherrima(100); | Metschnikowia | 13 | Saprotroph | \|Undefined Saprotroph\| | Yeast | NULL | Probable | As nectar-tap_saprotroph (Põlme et al. 2020); Decay sugar-rich substrates (Põlme et al. 2020) | Põlme S, et al. 2020. Fungal Diversity 105:1-16 ((https://doi.org/10.1007/s13225-020-00466-2)) |
| Otu18148 | 1 | k__Fungi(100);p__Ascomycota(100);c__Saccharomycetes(100);o__Saccharomycetales(100);f__Metschnikowiaceae(100);g__Metschnikowia(100);s__Metschnikowia_pulcherrima(100); | Metschnikowia | 13 | Saprotroph | \|Undefined Saprotroph\| | Yeast | NULL | Probable | As nectar-tap_saprotroph (Põlme et al. 2020); Decay sugar-rich substrates (Põlme et al. 2020) | Põlme S, et al. 2020. Fungal Diversity 105:1-16 ((https://doi.org/10.1007/s13225-020-00466-2)) |
| Otu18150 | 1 | k__Fungi(100);p__Ascomycota(100);c__Saccharomycetes(100);o__Saccharomycetales(100);f__Metschnikowiaceae(100);g__Metschnikowia(100);s__Metschnikowia_pulcherrima(100); | Metschnikowia | 13 | Saprotroph | \|Undefined Saprotroph\| | Yeast | NULL | Probable | As nectar-tap_saprotroph (Põlme et al. 2020); Decay sugar-rich substrates (Põlme et al. 2020) | Põlme S, et al. 2020. Fungal Diversity 105:1-16 ((https://doi.org/10.1007/s13225-020-00466-2)) |
| Otu18153 | 1 | k__Fungi(100);p__Ascomycota(100);c__Saccharomycetes(100);o__Saccharomycetales(100);f__Saccharomycetales_family_Incertae_sedis(100);g__Candida(100);g__Candida_unclassified(100); | Candida | 13 | Pathotroph-Saprotroph-Symbiotroph | Animal Pathogen-Endophyte-Endosymbiont-Epiphyte-\|Undefined Saprotroph\| | Dimorphic Yeast-Yeast | NULL | Probable | As nectar-tap_saprotroph (Põlme et al. 2020); Decay sugar-rich substrates (Põlme et al. 2020); This asexual genus is polyphyletic and contain species that function very differently depend on what they are. Caution should be taken when interpreting guilds for Candida species. | Manolakaki D, et al. 2010. Virulence. 1: 367-375 ((https://doi.org/10.4161/viru.1.5.12796)); Mekbib SB, et al. 2011. Tropical Plant Pathology 36:233-240 ((https://doi.org/10.1590/S1982-56762011000400004)); Põlme S, et al. 2020. Fungal Diversity 105:1-16 ((https://doi.org/10.1007/s13225-020-00466-2)) |
| Otu18155 | 1 | k__Fungi(100);p__Ascomycota(100);c__Saccharomycetes(100);o__Saccharomycetales(100);f__Metschnikowiaceae(100);g__Metschnikowia(100);s__Metschnikowia_pulcherrima(100); | Metschnikowia | 13 | Saprotroph | \|Undefined Saprotroph\| | Yeast | NULL | Probable | As nectar-tap_saprotroph (Põlme et al. 2020); Decay sugar-rich substrates (Põlme et al. 2020) | Põlme S, et al. 2020. Fungal Diversity 105:1-16 ((https://doi.org/10.1007/s13225-020-00466-2)) |
| Otu18163 | 1 | k__Fungi(100);p__Ascomycota(100);c__Saccharomycetes(100);o__Saccharomycetales(100);f__Metschnikowiaceae(100);g__Metschnikowia(100);s__Metschnikowia_pulcherrima(100); | Metschnikowia | 13 | Saprotroph | \|Undefined Saprotroph\| | Yeast | NULL | Probable | As nectar-tap_saprotroph (Põlme et al. 2020); Decay sugar-rich substrates (Põlme et al. 2020) | Põlme S, et al. 2020. Fungal Diversity 105:1-16 ((https://doi.org/10.1007/s13225-020-00466-2)) |
| Otu18164 | 1 | k__Fungi(100);p__Ascomycota(100);c__Saccharomycetes(100);o__Saccharomycetales(100);f__Metschnikowiaceae(100);g__Metschnikowia(100);s__Metschnikowia_pulcherrima(100); | Metschnikowia | 13 | Saprotroph | \|Undefined Saprotroph\| | Yeast | NULL | Probable | As nectar-tap_saprotroph (Põlme et al. 2020); Decay sugar-rich substrates (Põlme et al. 2020) | Põlme S, et al. 2020. Fungal Diversity 105:1-16 ((https://doi.org/10.1007/s13225-020-00466-2)) |
| Otu18165 | 1 | k__Fungi(100);p__Ascomycota(100);c__Saccharomycetes(100);o__Saccharomycetales(100);f__Metschnikowiaceae(100);g__Metschnikowia(100);s__Metschnikowia_pulcherrima(100); | Metschnikowia | 13 | Saprotroph | \|Undefined Saprotroph\| | Yeast | NULL | Probable | As nectar-tap_saprotroph (Põlme et al. 2020); Decay sugar-rich substrates (Põlme et al. 2020) | Põlme S, et al. 2020. Fungal Diversity 105:1-16 ((https://doi.org/10.1007/s13225-020-00466-2)) |
| Otu18166 | 1 | k__Fungi(100);p__Ascomycota(100);c__Saccharomycetes(100);o__Saccharomycetales(100);f__Metschnikowiaceae(100);g__Metschnikowia(100);s__Metschnikowia_pulcherrima(100); | Metschnikowia | 13 | Saprotroph | \|Undefined Saprotroph\| | Yeast | NULL | Probable | As nectar-tap_saprotroph (Põlme et al. 2020); Decay sugar-rich substrates (Põlme et al. 2020) | Põlme S, et al. 2020. Fungal Diversity 105:1-16 ((https://doi.org/10.1007/s13225-020-00466-2)) |
| Otu18167 | 1 | k__Fungi(100);p__Ascomycota(100);c__Saccharomycetes(100);o__Saccharomycetales(100);f__Metschnikowiaceae(100);g__Metschnikowia(100);s__Metschnikowia_pulcherrima(100); | Metschnikowia | 13 | Saprotroph | \|Undefined Saprotroph\| | Yeast | NULL | Probable | As nectar-tap_saprotroph (Põlme et al. 2020); Decay sugar-rich substrates (Põlme et al. 2020) | Põlme S, et al. 2020. Fungal Diversity 105:1-16 ((https://doi.org/10.1007/s13225-020-00466-2)) |
| Otu18168 | 1 | k__Fungi(100);p__Ascomycota(100);c__Saccharomycetes(100);o__Saccharomycetales(100);f__Metschnikowiaceae(100);g__Metschnikowia(100);s__Metschnikowia_pulcherrima(100); | Metschnikowia | 13 | Saprotroph | \|Undefined Saprotroph\| | Yeast | NULL | Probable | As nectar-tap_saprotroph (Põlme et al. 2020); Decay sugar-rich substrates (Põlme et al. 2020) | Põlme S, et al. 2020. Fungal Diversity 105:1-16 ((https://doi.org/10.1007/s13225-020-00466-2)) |
| Otu18169 | 1 | k__Fungi(100);p__Ascomycota(100);c__Saccharomycetes(100);o__Saccharomycetales(100);f__Metschnikowiaceae(100);g__Metschnikowia(100);s__Metschnikowia_pulcherrima(100); | Metschnikowia | 13 | Saprotroph | \|Undefined Saprotroph\| | Yeast | NULL | Probable | As nectar-tap_saprotroph (Põlme et al. 2020); Decay sugar-rich substrates (Põlme et al. 2020) | Põlme S, et al. 2020. Fungal Diversity 105:1-16 ((https://doi.org/10.1007/s13225-020-00466-2)) |
| Otu18170 | 1 | k__Fungi(100);p__Ascomycota(100);c__Saccharomycetes(100);o__Saccharomycetales(100);f__Metschnikowiaceae(100);g__Metschnikowia(100);g__Metschnikowia_unclassified(100); | Metschnikowia | 13 | Saprotroph | \|Undefined Saprotroph\| | Yeast | NULL | Probable | As nectar-tap_saprotroph (Põlme et al. 2020); Decay sugar-rich substrates (Põlme et al. 2020) | Põlme S, et al. 2020. Fungal Diversity 105:1-16 ((https://doi.org/10.1007/s13225-020-00466-2)) |
| Otu18171 | 1 | k__Fungi(100);p__Ascomycota(100);c__Saccharomycetes(100);o__Saccharomycetales(100);f__Metschnikowiaceae(100);g__Metschnikowia(100);s__Metschnikowia_pulcherrima(100); | Metschnikowia | 13 | Saprotroph | \|Undefined Saprotroph\| | Yeast | NULL | Probable | As nectar-tap_saprotroph (Põlme et al. 2020); Decay sugar-rich substrates (Põlme et al. 2020) | Põlme S, et al. 2020. Fungal Diversity 105:1-16 ((https://doi.org/10.1007/s13225-020-00466-2)) |
| Otu18172 | 1 | k__Fungi(100);p__Ascomycota(100);c__Saccharomycetes(100);o__Saccharomycetales(100);f__Saccharomycetales_family_Incertae_sedis(100);f__Saccharomycetales_family_Incertae_sedis_unclassified(100);f__Saccharomycetales_family_Incertae_sedis_unclassified(100); | Saccharomycetales | 7 | Saprotroph | Undefined Saprotroph | Yeast | NULL | Possible | NULL | Sterkenburg E, et al. 2015. New Phytologist 207:1145-1158 ((https://doi.org/10.1111/nph.13426)) |
| Otu18175 | 1 | k__Fungi(100);p__Ascomycota(100);c__Saccharomycetes(100);o__Saccharomycetales(100);f__Metschnikowiaceae(100);g__Metschnikowia(100);s__Metschnikowia_pulcherrima(100); | Metschnikowia | 13 | Saprotroph | \|Undefined Saprotroph\| | Yeast | NULL | Probable | As nectar-tap_saprotroph (Põlme et al. 2020); Decay sugar-rich substrates (Põlme et al. 2020) | Põlme S, et al. 2020. Fungal Diversity 105:1-16 ((https://doi.org/10.1007/s13225-020-00466-2)) |
| Otu18176 | 1 | k__Fungi(100);p__Ascomycota(100);c__Saccharomycetes(100);o__Saccharomycetales(100);f__Metschnikowiaceae(100);g__Metschnikowia(100);s__Metschnikowia_pulcherrima(100); | Metschnikowia | 13 | Saprotroph | \|Undefined Saprotroph\| | Yeast | NULL | Probable | As nectar-tap_saprotroph (Põlme et al. 2020); Decay sugar-rich substrates (Põlme et al. 2020) | Põlme S, et al. 2020. Fungal Diversity 105:1-16 ((https://doi.org/10.1007/s13225-020-00466-2)) |
| Otu18177 | 1 | k__Fungi(100);p__Ascomycota(100);c__Saccharomycetes(100);o__Saccharomycetales(100);f__Metschnikowiaceae(100);g__Metschnikowia(100);s__Metschnikowia_pulcherrima(100); | Metschnikowia | 13 | Saprotroph | \|Undefined Saprotroph\| | Yeast | NULL | Probable | As nectar-tap_saprotroph (Põlme et al. 2020); Decay sugar-rich substrates (Põlme et al. 2020) | Põlme S, et al. 2020. Fungal Diversity 105:1-16 ((https://doi.org/10.1007/s13225-020-00466-2)) |
| Otu18178 | 1 | k__Fungi(100);p__Ascomycota(100);c__Saccharomycetes(100);o__Saccharomycetales(100);f__Metschnikowiaceae(100);g__Metschnikowia(100);s__Metschnikowia_pulcherrima(100); | Metschnikowia | 13 | Saprotroph | \|Undefined Saprotroph\| | Yeast | NULL | Probable | As nectar-tap_saprotroph (Põlme et al. 2020); Decay sugar-rich substrates (Põlme et al. 2020) | Põlme S, et al. 2020. Fungal Diversity 105:1-16 ((https://doi.org/10.1007/s13225-020-00466-2)) |
| Otu18179 | 1 | k__Fungi(100);p__Ascomycota(100);c__Dothideomycetes(100);o__Capnodiales(100);f__Mycosphaerellaceae(100);f__Mycosphaerellaceae_unclassified(100);f__Mycosphaerellaceae_unclassified(100); | Mycosphaerellaceae | 9 | Pathotroph-Saprotroph | Plant Pathogen-Undefined Saprotroph | Microfungus | NULL | Probable | Biotrophic, necrotrophic or saprobic on various plant tissues (Cannon & Kirk 2007) | Cannon PF, Kirk PM. 2007. Fungal Families of the World. CAB International, Cambridge (ISBN: 978-0851998275) |
| Otu18180 | 1 | k__Fungi(100);p__Ascomycota(100);c__Saccharomycetes(100);o__Saccharomycetales(100);f__Metschnikowiaceae(100);g__Metschnikowia(100);s__Metschnikowia_pulcherrima(100); | Metschnikowia | 13 | Saprotroph | \|Undefined Saprotroph\| | Yeast | NULL | Probable | As nectar-tap_saprotroph (Põlme et al. 2020); Decay sugar-rich substrates (Põlme et al. 2020) | Põlme S, et al. 2020. Fungal Diversity 105:1-16 ((https://doi.org/10.1007/s13225-020-00466-2)) |
| Otu18181 | 1 | k__Fungi(100);p__Ascomycota(100);c__Saccharomycetes(100);o__Saccharomycetales(100);f__Metschnikowiaceae(100);g__Metschnikowia(100);g__Metschnikowia_unclassified(100); | Metschnikowia | 13 | Saprotroph | \|Undefined Saprotroph\| | Yeast | NULL | Probable | As nectar-tap_saprotroph (Põlme et al. 2020); Decay sugar-rich substrates (Põlme et al. 2020) | Põlme S, et al. 2020. Fungal Diversity 105:1-16 ((https://doi.org/10.1007/s13225-020-00466-2)) |
| Otu18182 | 1 | k__Fungi(100);p__Ascomycota(100);c__Saccharomycetes(100);o__Saccharomycetales(100);f__Metschnikowiaceae(100);g__Metschnikowia(100);s__Metschnikowia_pulcherrima(100); | Metschnikowia | 13 | Saprotroph | \|Undefined Saprotroph\| | Yeast | NULL | Probable | As nectar-tap_saprotroph (Põlme et al. 2020); Decay sugar-rich substrates (Põlme et al. 2020) | Põlme S, et al. 2020. Fungal Diversity 105:1-16 ((https://doi.org/10.1007/s13225-020-00466-2)) |
| Otu18183 | 1 | k__Fungi(100);p__Ascomycota(100);c__Saccharomycetes(100);o__Saccharomycetales(100);f__Metschnikowiaceae(100);g__Metschnikowia(100);s__Metschnikowia_pulcherrima(100); | Metschnikowia | 13 | Saprotroph | \|Undefined Saprotroph\| | Yeast | NULL | Probable | As nectar-tap_saprotroph (Põlme et al. 2020); Decay sugar-rich substrates (Põlme et al. 2020) | Põlme S, et al. 2020. Fungal Diversity 105:1-16 ((https://doi.org/10.1007/s13225-020-00466-2)) |

| | | | | | | | | | | | |
|---|---|---|---|---|---|---|---|---|---|---|---|
| Otu18185 | 1 | k__Fungi(100);p__Ascomycota(100);c__Saccharomycetes(100);o__Saccharomycetales(100);f__Metschnikowiaceae(100);g__Metschnikowia(100);g__Metschnikowia_unclassified(100); | Metschnikowia | 13 | Saprotroph | \|Undefined Saprotroph\| | Yeast | NULL | Probable | As nectar-tap_saprotroph (Põlme et al. 2020); Decay sugar-rich substrates (Põlme et al. 2020) | Põlme S, et al. 2020. Fungal Diversity 105:1-16 ((https://doi.org/10.1007/s13225-020-00466-2)) |
| Otu18186 | 1 | k__Fungi(100);p__Ascomycota(100);c__Saccharomycetes(100);o__Saccharomycetales(100);f__Metschnikowiaceae(100);g__Metschnikowia(100);s__Metschnikowia_pulcherrima(100); | Metschnikowia | 13 | Saprotroph | \|Undefined Saprotroph\| | Yeast | NULL | Probable | As nectar-tap_saprotroph (Põlme et al. 2020); Decay sugar-rich substrates (Põlme et al. 2020) | Põlme S, et al. 2020. Fungal Diversity 105:1-16 ((https://doi.org/10.1007/s13225-020-00466-2)) |
| Otu18187 | 1 | k__Fungi(100);p__Ascomycota(100);c__Saccharomycetes(100);o__Saccharomycetales(100);f__Metschnikowiaceae(100);g__Metschnikowia(100);g__Metschnikowia_unclassified(100); | Metschnikowia | 13 | Saprotroph | \|Undefined Saprotroph\| | Yeast | NULL | Probable | As nectar-tap_saprotroph (Põlme et al. 2020); Decay sugar-rich substrates (Põlme et al. 2020) | Põlme S, et al. 2020. Fungal Diversity 105:1-16 ((https://doi.org/10.1007/s13225-020-00466-2)) |
| Otu18188 | 1 | k__Fungi(100);p__Ascomycota(100);c__Saccharomycetes(100);o__Saccharomycetales(100);f__Metschnikowiaceae(100);g__Metschnikowia(100);s__Metschnikowia_pulcherrima(100); | Metschnikowia | 13 | Saprotroph | \|Undefined Saprotroph\| | Yeast | NULL | Probable | As nectar-tap_saprotroph (Põlme et al. 2020); Decay sugar-rich substrates (Põlme et al. 2020) | Põlme S, et al. 2020. Fungal Diversity 105:1-16 ((https://doi.org/10.1007/s13225-020-00466-2)) |
| Otu18189 | 1 | k__Fungi(100);p__Ascomycota(100);c__Saccharomycetes(100);o__Saccharomycetales(100);f__Metschnikowiaceae(100);g__Metschnikowia(100);s__Metschnikowia_pulcherrima(100); | Metschnikowia | 13 | Saprotroph | \|Undefined Saprotroph\| | Yeast | NULL | Probable | As nectar-tap_saprotroph (Põlme et al. 2020); Decay sugar-rich substrates (Põlme et al. 2020) | Põlme S, et al. 2020. Fungal Diversity 105:1-16 ((https://doi.org/10.1007/s13225-020-00466-2)) |
| Otu18190 | 1 | k__Fungi(100);p__Ascomycota(100);c__Saccharomycetes(100);o__Saccharomycetales(100);f__Metschnikowiaceae(100);g__Metschnikowia(100);g__Metschnikowia_unclassified(100); | Metschnikowia | 13 | Saprotroph | \|Undefined Saprotroph\| | Yeast | NULL | Probable | As nectar-tap_saprotroph (Põlme et al. 2020); Decay sugar-rich substrates (Põlme et al. 2020) | Põlme S, et al. 2020. Fungal Diversity 105:1-16 ((https://doi.org/10.1007/s13225-020-00466-2)) |
| Otu18191 | 1 | k__Fungi(100);p__Ascomycota(100);c__Saccharomycetes(100);o__Saccharomycetales(100);f__Metschnikowiaceae(100);g__Metschnikowia(100);s__Metschnikowia_pulcherrima(100); | Metschnikowia | 13 | Saprotroph | \|Undefined Saprotroph\| | Yeast | NULL | Probable | As nectar-tap_saprotroph (Põlme et al. 2020); Decay sugar-rich substrates (Põlme et al. 2020) | Põlme S, et al. 2020. Fungal Diversity 105:1-16 ((https://doi.org/10.1007/s13225-020-00466-2)) |
| Otu18192 | 1 | k__Fungi(100);p__Ascomycota(100);c__Saccharomycetes(100);o__Saccharomycetales(100);f__Metschnikowiaceae(100);g__Metschnikowia(100);s__Metschnikowia_pulcherrima(100); | Metschnikowia | 13 | Saprotroph | \|Undefined Saprotroph\| | Yeast | NULL | Probable | As nectar-tap_saprotroph (Põlme et al. 2020); Decay sugar-rich substrates (Põlme et al. 2020) | Põlme S, et al. 2020. Fungal Diversity 105:1-16 ((https://doi.org/10.1007/s13225-020-00466-2)) |
| Otu18193 | 1 | k__Fungi(100);p__Ascomycota(100);c__Saccharomycetes(100);o__Saccharomycetales(100);f__Metschnikowiaceae(100);g__Metschnikowia(100);g__Metschnikowia_unclassified(100); | Metschnikowia | 13 | Saprotroph | \|Undefined Saprotroph\| | Yeast | NULL | Probable | As nectar-tap_saprotroph (Põlme et al. 2020); Decay sugar-rich substrates (Põlme et al. 2020) | Põlme S, et al. 2020. Fungal Diversity 105:1-16 ((https://doi.org/10.1007/s13225-020-00466-2)) |
| Otu18194 | 1 | k__Fungi(100);p__Ascomycota(100);c__Saccharomycetes(100);o__Saccharomycetales(100);f__Metschnikowiaceae(100);g__Metschnikowia(100);s__Metschnikowia_pulcherrima(100); | Metschnikowia | 13 | Saprotroph | \|Undefined Saprotroph\| | Yeast | NULL | Probable | As nectar-tap_saprotroph (Põlme et al. 2020); Decay sugar-rich substrates (Põlme et al. 2020) | Põlme S, et al. 2020. Fungal Diversity 105:1-16 ((https://doi.org/10.1007/s13225-020-00466-2)) |
| Otu18197 | 1 | k__Fungi(100);p__Ascomycota(100);c__Saccharomycetes(100);o__Saccharomycetales(100);f__Metschnikowiaceae(100);g__Metschnikowia(100);s__Metschnikowia_pulcherrima(100); | Metschnikowia | 13 | Saprotroph | \|Undefined Saprotroph\| | Yeast | NULL | Probable | As nectar-tap_saprotroph (Põlme et al. 2020); Decay sugar-rich substrates (Põlme et al. 2020) | Põlme S, et al. 2020. Fungal Diversity 105:1-16 ((https://doi.org/10.1007/s13225-020-00466-2)) |
| Otu18198 | 1 | k__Fungi(100);p__Ascomycota(100);c__Saccharomycetes(100);o__Saccharomycetales(100);f__Metschnikowiaceae(100);g__Metschnikowia(100);s__Metschnikowia_pulcherrima(100); | Metschnikowia | 13 | Saprotroph | \|Undefined Saprotroph\| | Yeast | NULL | Probable | As nectar-tap_saprotroph (Põlme et al. 2020); Decay sugar-rich substrates (Põlme et al. 2020) | Põlme S, et al. 2020. Fungal Diversity 105:1-16 ((https://doi.org/10.1007/s13225-020-00466-2)) |
| Otu18199 | 1 | k__Fungi(100);p__Ascomycota(100);c__Saccharomycetes(100);o__Saccharomycetales(100);f__Metschnikowiaceae(100);g__Metschnikowia(100);g__Metschnikowia_unclassified(100); | Metschnikowia | 13 | Saprotroph | \|Undefined Saprotroph\| | Yeast | NULL | Probable | As nectar-tap_saprotroph (Põlme et al. 2020); Decay sugar-rich substrates (Põlme et al. 2020) | Põlme S, et al. 2020. Fungal Diversity 105:1-16 ((https://doi.org/10.1007/s13225-020-00466-2)) |
| Otu18200 | 1 | k__Fungi(100);p__Ascomycota(100);c__Saccharomycetes(100);o__Saccharomycetales(100);f__Metschnikowiaceae(100);g__Metschnikowia(100);g__Metschnikowia_unclassified(100); | Metschnikowia | 13 | Saprotroph | \|Undefined Saprotroph\| | Yeast | NULL | Probable | As nectar-tap_saprotroph (Põlme et al. 2020); Decay sugar-rich substrates (Põlme et al. 2020) | Põlme S, et al. 2020. Fungal Diversity 105:1-16 ((https://doi.org/10.1007/s13225-020-00466-2)) |
| Otu18201 | 1 | k__Fungi(100);p__Ascomycota(100);c__Saccharomycetes(100);o__Saccharomycetales(100);f__Metschnikowiaceae(100);g__Metschnikowia(100);s__Metschnikowia_pulcherrima(100); | Metschnikowia | 13 | Saprotroph | \|Undefined Saprotroph\| | Yeast | NULL | Probable | As nectar-tap_saprotroph (Põlme et al. 2020); Decay sugar-rich substrates (Põlme et al. 2020) | Põlme S, et al. 2020. Fungal Diversity 105:1-16 ((https://doi.org/10.1007/s13225-020-00466-2)) |
| Otu18202 | 1 | k__Fungi(100);p__Ascomycota(100);c__Saccharomycetes(100);o__Saccharomycetales(100);f__Metschnikowiaceae(100);g__Metschnikowia(100);g__Metschnikowia_unclassified(100); | Metschnikowia | 13 | Saprotroph | \|Undefined Saprotroph\| | Yeast | NULL | Probable | As nectar-tap_saprotroph (Põlme et al. 2020); Decay sugar-rich substrates (Põlme et al. 2020) | Põlme S, et al. 2020. Fungal Diversity 105:1-16 ((https://doi.org/10.1007/s13225-020-00466-2)) |
| Otu18203 | 1 | k__Fungi(100);p__Ascomycota(100);c__Saccharomycetes(100);o__Saccharomycetales(100);f__Metschnikowiaceae(100);g__Metschnikowia(100);g__Metschnikowia_unclassified(100); | Metschnikowia | 13 | Saprotroph | \|Undefined Saprotroph\| | Yeast | NULL | Probable | As nectar-tap_saprotroph (Põlme et al. 2020); Decay sugar-rich substrates (Põlme et al. 2020) | Põlme S, et al. 2020. Fungal Diversity 105:1-16 ((https://doi.org/10.1007/s13225-020-00466-2)) |
| Otu18204 | 1 | k__Fungi(100);p__Ascomycota(100);c__Saccharomycetes(100);o__Saccharomycetales(100);f__Metschnikowiaceae(100);g__Metschnikowia(100);s__Metschnikowia_pulcherrima(100); | Metschnikowia | 13 | Saprotroph | \|Undefined Saprotroph\| | Yeast | NULL | Probable | As nectar-tap_saprotroph (Põlme et al. 2020); Decay sugar-rich substrates (Põlme et al. 2020) | Põlme S, et al. 2020. Fungal Diversity 105:1-16 ((https://doi.org/10.1007/s13225-020-00466-2)) |
| Otu18205 | 1 | k__Fungi(100);p__Ascomycota(100);c__Saccharomycetes(100);o__Saccharomycetales(100);f__Metschnikowiaceae(100);g__Metschnikowia(100);s__Metschnikowia_pulcherrima(100); | Metschnikowia | 13 | Saprotroph | \|Undefined Saprotroph\| | Yeast | NULL | Probable | As nectar-tap_saprotroph (Põlme et al. 2020); Decay sugar-rich substrates (Põlme et al. 2020) | Põlme S, et al. 2020. Fungal Diversity 105:1-16 ((https://doi.org/10.1007/s13225-020-00466-2)) |
| Otu18206 | 1 | k__Fungi(100);p__Ascomycota(100);c__Saccharomycetes(100);o__Saccharomycetales(100);f__Saccharomycetales_family_Incertae_sedis(100);g__Candida(100);g__Candida_unclassified(100); | Candida | 13 | Pathotroph-Saprotroph-Symbiotroph | Animal Pathogen-Endophyte-Endosymbiont-Epiphyte-\|Undefined Saprotroph\| | Dimorphic Yeast-Yeast | NULL | Probable | As nectar-tap_saprotroph (Põlme et al. 2020); Decay sugar-rich substrates (Põlme et al. 2020); This asexual genus is polyphyletic and contain species that function very differently depend on what they are. Caution should be taken when interpreting guilds for Candida species. | Manolakaki D, et al. 2010. Virulence. 1: 367-375 ((https://doi.org/10.4161/viru.1.5.12796)); Mekbib SB, et al. 2011. Tropical Plant Pathology 36:233-240 ((https://doi.org/10.1590/S1982-56762011000400004)); Põlme S, et al. 2020. Fungal Diversity 105:1-16 ((https://doi.org/10.1007/s13225-020-00466-2)) |
| Otu18207 | 1 | k__Fungi(100);p__Ascomycota(100);c__Saccharomycetes(100);o__Saccharomycetales(100);f__Metschnikowiaceae(100);g__Metschnikowia(100);s__Metschnikowia_pulcherrima(100); | Metschnikowia | 13 | Saprotroph | \|Undefined Saprotroph\| | Yeast | NULL | Probable | As nectar-tap_saprotroph (Põlme et al. 2020); Decay sugar-rich substrates (Põlme et al. 2020) | Põlme S, et al. 2020. Fungal Diversity 105:1-16 ((https://doi.org/10.1007/s13225-020-00466-2)) |
| Otu18208 | 1 | k__Fungi(100);p__Ascomycota(100);c__Saccharomycetes(100);o__Saccharomycetales(100);f__Metschnikowiaceae(100);g__Metschnikowia(100);s__Metschnikowia_pulcherrima(100); | Metschnikowia | 13 | Saprotroph | \|Undefined Saprotroph\| | Yeast | NULL | Probable | As nectar-tap_saprotroph (Põlme et al. 2020); Decay sugar-rich substrates (Põlme et al. 2020) | Põlme S, et al. 2020. Fungal Diversity 105:1-16 ((https://doi.org/10.1007/s13225-020-00466-2)) |
| Otu18209 | 1 | k__Fungi(100);p__Ascomycota(100);c__Saccharomycetes(100);o__Saccharomycetales(100);f__Metschnikowiaceae(100);g__Metschnikowia(100);s__Metschnikowia_pulcherrima(100); | Metschnikowia | 13 | Saprotroph | \|Undefined Saprotroph\| | Yeast | NULL | Probable | As nectar-tap_saprotroph (Põlme et al. 2020); Decay sugar-rich substrates (Põlme et al. 2020) | Põlme S, et al. 2020. Fungal Diversity 105:1-16 ((https://doi.org/10.1007/s13225-020-00466-2)) |
| Otu18210 | 1 | k__Fungi(100);p__Ascomycota(100);c__Saccharomycetes(100);o__Saccharomycetales(100);f__Metschnikowiaceae(100);g__Metschnikowia(100);s__Metschnikowia_pulcherrima(100); | Metschnikowia | 13 | Saprotroph | \|Undefined Saprotroph\| | Yeast | NULL | Probable | As nectar-tap_saprotroph (Põlme et al. 2020); Decay sugar-rich substrates (Põlme et al. 2020) | Põlme S, et al. 2020. Fungal Diversity 105:1-16 ((https://doi.org/10.1007/s13225-020-00466-2)) |
| Otu18211 | 1 | k__Fungi(100);p__Ascomycota(100);c__Saccharomycetes(100);o__Saccharomycetales(100);f__Saccharomycetales_family_Incertae_sedis(100);g__Candida(100);g__Candida_unclassified(100); | Candida | 13 | Pathotroph-Saprotroph-Symbiotroph | Animal Pathogen-Endophyte-Endosymbiont-Epiphyte-\|Undefined Saprotroph\| | Dimorphic Yeast-Yeast | NULL | Probable | As nectar-tap_saprotroph (Põlme et al. 2020); Decay sugar-rich substrates (Põlme et al. 2020); This asexual genus is polyphyletic and contain species that function very differently depend on what they are. Caution should be taken when interpreting guilds for Candida species. | Manolakaki D, et al. 2010. Virulence. 1: 367-375 ((https://doi.org/10.4161/viru.1.5.12796)); Mekbib SB, et al. 2011. Tropical Plant Pathology 36:233-240 ((https://doi.org/10.1590/S1982-56762011000400004)); Põlme S, et al. 2020. Fungal Diversity 105:1-16 ((https://doi.org/10.1007/s13225-020-00466-2)) |
| Otu18212 | 1 | k__Fungi(100);p__Ascomycota(100);c__Saccharomycetes(100);o__Saccharomycetales(100);f__Saccharomycetales_family_Incertae_sedis(100);g__Candida(100);g__Candida_unclassified(100); | Candida | 13 | Pathotroph-Saprotroph-Symbiotroph | Animal Pathogen-Endophyte-Endosymbiont-Epiphyte-\|Undefined Saprotroph\| | Dimorphic Yeast-Yeast | NULL | Probable | As nectar-tap_saprotroph (Põlme et al. 2020); Decay sugar-rich substrates (Põlme et al. 2020); This asexual genus is polyphyletic and contain species that function very differently depend on what they are. Caution should be taken when interpreting guilds for Candida species. | Manolakaki D, et al. 2010. Virulence. 1: 367-375 ((https://doi.org/10.4161/viru.1.5.12796)); Mekbib SB, et al. 2011. Tropical Plant Pathology 36:233-240 ((https://doi.org/10.1590/S1982-56762011000400004)); Põlme S, et al. 2020. Fungal Diversity 105:1-16 ((https://doi.org/10.1007/s13225-020-00466-2)) |
| Otu18213 | 1 | k__Fungi(100);p__Ascomycota(100);c__Saccharomycetes(100);o__Saccharomycetales(100);f__Metschnikowiaceae(100);g__Metschnikowia(100);s__Metschnikowia_pulcherrima(100); | Metschnikowia | 13 | Saprotroph | \|Undefined Saprotroph\| | Yeast | NULL | Probable | As nectar-tap_saprotroph (Põlme et al. 2020); Decay sugar-rich substrates (Põlme et al. 2020) | Põlme S, et al. 2020. Fungal Diversity 105:1-16 ((https://doi.org/10.1007/s13225-020-00466-2)) |
| Otu18215 | 1 | k__Fungi(100);p__Ascomycota(100);c__Saccharomycetes(100);o__Saccharomycetales(100);f__Metschnikowiaceae(100);g__Metschnikowia(100);s__Metschnikowia_pulcherrima(100); | Metschnikowia | 13 | Saprotroph | \|Undefined Saprotroph\| | Yeast | NULL | Probable | As nectar-tap_saprotroph (Põlme et al. 2020); Decay sugar-rich substrates (Põlme et al. 2020) | Põlme S, et al. 2020. Fungal Diversity 105:1-16 ((https://doi.org/10.1007/s13225-020-00466-2)) |
| Otu18217 | 1 | k__Fungi(100);p__Ascomycota(100);c__Saccharomycetes(100);o__Saccharomycetales(100);f__Metschnikowiaceae(100);g__Metschnikowia(100);g__Metschnikowia_unclassified(100); | Metschnikowia | 13 | Saprotroph | \|Undefined Saprotroph\| | Yeast | NULL | Probable | As nectar-tap_saprotroph (Põlme et al. 2020); Decay sugar-rich substrates (Põlme et al. 2020) | Põlme S, et al. 2020. Fungal Diversity 105:1-16 ((https://doi.org/10.1007/s13225-020-00466-2)) |
| Otu18218 | 1 | k__Fungi(100);p__Ascomycota(100);c__Saccharomycetes(100);o__Saccharomycetales(100);f__Metschnikowiaceae(100);g__Metschnikowia(100);s__Metschnikowia_pulcherrima(100); | Metschnikowia | 13 | Saprotroph | \|Undefined Saprotroph\| | Yeast | NULL | Probable | As nectar-tap_saprotroph (Põlme et al. 2020); Decay sugar-rich substrates (Põlme et al. 2020) | Põlme S, et al. 2020. Fungal Diversity 105:1-16 ((https://doi.org/10.1007/s13225-020-00466-2)) |
| Otu18219 | 1 | k__Fungi(100);p__Ascomycota(100);c__Saccharomycetes(100);o__Saccharomycetales(100);f__Metschnikowiaceae(100);g__Metschnikowia(100);g__Metschnikowia_unclassified(100); | Metschnikowia | 13 | Saprotroph | \|Undefined Saprotroph\| | Yeast | NULL | Probable | As nectar-tap_saprotroph (Põlme et al. 2020); Decay sugar-rich substrates (Põlme et al. 2020) | Põlme S, et al. 2020. Fungal Diversity 105:1-16 ((https://doi.org/10.1007/s13225-020-00466-2)) |
| Otu18220 | 1 | k__Fungi(100);p__Ascomycota(100);c__Saccharomycetes(100);o__Saccharomycetales(100);f__Metschnikowiaceae(100);g__Metschnikowia(100);s__Metschnikowia_pulcherrima(100); | Metschnikowia | 13 | Saprotroph | \|Undefined Saprotroph\| | Yeast | NULL | Probable | As nectar-tap_saprotroph (Põlme et al. 2020); Decay sugar-rich substrates (Põlme et al. 2020) | Põlme S, et al. 2020. Fungal Diversity 105:1-16 ((https://doi.org/10.1007/s13225-020-00466-2)) |
| Otu18221 | 1 | k__Fungi(100);p__Ascomycota(100);c__Saccharomycetes(100);o__Saccharomycetales(100);f__Metschnikowiaceae(100);g__Metschnikowia(100);g__Metschnikowia_unclassified(100); | Metschnikowia | 13 | Saprotroph | \|Undefined Saprotroph\| | Yeast | NULL | Probable | As nectar-tap_saprotroph (Põlme et al. 2020); Decay sugar-rich substrates (Põlme et al. 2020) | Põlme S, et al. 2020. Fungal Diversity 105:1-16 ((https://doi.org/10.1007/s13225-020-00466-2)) |
| Otu18222 | 1 | k__Fungi(100);p__Ascomycota(100);c__Saccharomycetes(100);o__Saccharomycetales(100);f__Metschnikowiaceae(100);g__Metschnikowia(100);s__Metschnikowia_pulcherrima(100); | Metschnikowia | 13 | Saprotroph | \|Undefined Saprotroph\| | Yeast | NULL | Probable | As nectar-tap_saprotroph (Põlme et al. 2020); Decay sugar-rich substrates (Põlme et al. 2020) | Põlme S, et al. 2020. Fungal Diversity 105:1-16 ((https://doi.org/10.1007/s13225-020-00466-2)) |
| Otu18223 | 1 | k__Fungi(100);p__Ascomycota(100);c__Saccharomycetes(100);o__Saccharomycetales(100);f__Metschnikowiaceae(100);g__Metschnikowia(100);s__Metschnikowia_pulcherrima(100); | Metschnikowia | 13 | Saprotroph | \|Undefined Saprotroph\| | Yeast | NULL | Probable | As nectar-tap_saprotroph (Põlme et al. 2020); Decay sugar-rich substrates (Põlme et al. 2020) | Põlme S, et al. 2020. Fungal Diversity 105:1-16 ((https://doi.org/10.1007/s13225-020-00466-2)) |
| Otu18224 | 1 | k__Fungi(100);p__Ascomycota(100);c__Saccharomycetes(100);o__Saccharomycetales(100);f__Metschnikowiaceae(100);g__Metschnikowia(100);g__Metschnikowia_unclassified(100); | Metschnikowia | 13 | Saprotroph | \|Undefined Saprotroph\| | Yeast | NULL | Probable | As nectar-tap_saprotroph (Põlme et al. 2020); Decay sugar-rich substrates (Põlme et al. 2020) | Põlme S, et al. 2020. Fungal Diversity 105:1-16 ((https://doi.org/10.1007/s13225-020-00466-2)) |
| Otu18225 | 1 | k__Fungi(100);p__Ascomycota(100);c__Saccharomycetes(100);o__Saccharomycetales(100);f__Metschnikowiaceae(100);g__Metschnikowia(100);s__Metschnikowia_pulcherrima(100); | Metschnikowia | 13 | Saprotroph | \|Undefined Saprotroph\| | Yeast | NULL | Probable | As nectar-tap_saprotroph (Põlme et al. 2020); Decay sugar-rich substrates (Põlme et al. 2020) | Põlme S, et al. 2020. Fungal Diversity 105:1-16 ((https://doi.org/10.1007/s13225-020-00466-2)) |
| Otu18226 | 1 | k__Fungi(100);p__Ascomycota(100);c__Saccharomycetes(100);o__Saccharomycetales(100);f__Metschnikowiaceae(100);g__Metschnikowia(100);g__Metschnikowia_unclassified(100); | Metschnikowia | 13 | Saprotroph | \|Undefined Saprotroph\| | Yeast | NULL | Probable | As nectar-tap_saprotroph (Põlme et al. 2020); Decay sugar-rich substrates (Põlme et al. 2020) | Põlme S, et al. 2020. Fungal Diversity 105:1-16 ((https://doi.org/10.1007/s13225-020-00466-2)) |
| Otu18227 | 1 | k__Fungi(100);p__Ascomycota(100);c__Saccharomycetes(100);o__Saccharomycetales(100);f__Metschnikowiaceae(100);g__Metschnikowia(100);s__Metschnikowia_pulcherrima(100); | Metschnikowia | 13 | Saprotroph | \|Undefined Saprotroph\| | Yeast | NULL | Probable | As nectar-tap_saprotroph (Põlme et al. 2020); Decay sugar-rich substrates (Põlme et al. 2020) | Põlme S, et al. 2020. Fungal Diversity 105:1-16 ((https://doi.org/10.1007/s13225-020-00466-2)) |
| Otu18228 | 1 | k__Fungi(100);p__Ascomycota(100);c__Saccharomycetes(100);o__Saccharomycetales(100);f__Metschnikowiaceae(100);g__Metschnikowia(100);s__Metschnikowia_pulcherrima(100); | Metschnikowia | 13 | Saprotroph | \|Undefined Saprotroph\| | Yeast | NULL | Probable | As nectar-tap_saprotroph (Põlme et al. 2020); Decay sugar-rich substrates (Põlme et al. 2020) | Põlme S, et al. 2020. Fungal Diversity 105:1-16 ((https://doi.org/10.1007/s13225-020-00466-2)) |
| Otu18229 | 1 | k__Fungi(100);p__Ascomycota(100);c__Saccharomycetes(100);o__Saccharomycetales(100);f__Metschnikowiaceae(100);g__Metschnikowia(100);s__Metschnikowia_pulcherrima(100); | Metschnikowia | 13 | Saprotroph | \|Undefined Saprotroph\| | Yeast | NULL | Probable | As nectar-tap_saprotroph (Põlme et al. 2020); Decay sugar-rich substrates (Põlme et al. 2020) | Põlme S, et al. 2020. Fungal Diversity 105:1-16 ((https://doi.org/10.1007/s13225-020-00466-2)) |
| Otu18230 | 1 | k__Fungi(100);p__Ascomycota(100);c__Saccharomycetes(100);o__Saccharomycetales(100);f__Metschnikowiaceae(100);g__Metschnikowia(100);s__Metschnikowia_pulcherrima(100); | Metschnikowia | 13 | Saprotroph | \|Undefined Saprotroph\| | Yeast | NULL | Probable | As nectar-tap_saprotroph (Põlme et al. 2020); Decay sugar-rich substrates (Põlme et al. 2020) | Põlme S, et al. 2020. Fungal Diversity 105:1-16 ((https://doi.org/10.1007/s13225-020-00466-2)) |
| Otu18231 | 1 | k__Fungi(100);p__Ascomycota(100);c__Saccharomycetes(100);o__Saccharomycetales(100);f__Metschnikowiaceae(100);g__Metschnikowia(100);s__Metschnikowia_pulcherrima(100); | Metschnikowia | 13 | Saprotroph | \|Undefined Saprotroph\| | Yeast | NULL | Probable | As nectar-tap_saprotroph (Põlme et al. 2020); Decay sugar-rich substrates (Põlme et al. 2020) | Põlme S, et al. 2020. Fungal Diversity 105:1-16 ((https://doi.org/10.1007/s13225-020-00466-2)) |
| Otu18232 | 1 | k__Fungi(100);p__Ascomycota(100);c__Saccharomycetes(100);o__Saccharomycetales(100);f__Saccharomycetales_family_Incertae_sedis(100);g__Candida(100);g__Candida_unclassified(100); | Candida | 13 | Pathotroph-Saprotroph-Symbiotroph | Animal Pathogen-Endophyte-Endosymbiont-Epiphyte-\|Undefined Saprotroph\| | Dimorphic Yeast-Yeast | NULL | Probable | As nectar-tap_saprotroph (Põlme et al. 2020); Decay sugar-rich substrates (Põlme et al. 2020); This asexual genus is polyphyletic and contain species that function very differently depend on what they are. Caution should be taken when interpreting guilds for Candida species. | Manolakaki D, et al. 2010. Virulence. 1: 367-375 ((https://doi.org/10.4161/viru.1.5.12796)); Mekbib SB, et al. 2011. Tropical Plant Pathology 36:233-240 ((https://doi.org/10.1590/S1982-56762011000400004)); Põlme S, et al. 2020. Fungal Diversity 105:1-16 ((https://doi.org/10.1007/s13225-020-00466-2)) |
| Otu18233 | 1 | k__Fungi(100);p__Ascomycota(100);c__Saccharomycetes(100);o__Saccharomycetales(100);f__Metschnikowiaceae(100);g__Metschnikowia(100);s__Metschnikowia_pulcherrima(100); | Metschnikowia | 13 | Saprotroph | \|Undefined Saprotroph\| | Yeast | NULL | Probable | As nectar-tap_saprotroph (Põlme et al. 2020); Decay sugar-rich substrates (Põlme et al. 2020) | Põlme S, et al. 2020. Fungal Diversity 105:1-16 ((https://doi.org/10.1007/s13225-020-00466-2)) |
| Otu18234 | 1 | k__Fungi(100);p__Ascomycota(100);c__Saccharomycetes(100);o__Saccharomycetales(100);f__Metschnikowiaceae(100);g__Metschnikowia(100);s__Metschnikowia_pulcherrima(100); | Metschnikowia | 13 | Saprotroph | \|Undefined Saprotroph\| | Yeast | NULL | Probable | As nectar-tap_saprotroph (Põlme et al. 2020); Decay sugar-rich substrates (Põlme et al. 2020) | Põlme S, et al. 2020. Fungal Diversity 105:1-16 ((https://doi.org/10.1007/s13225-020-00466-2)) |
| Otu18237 | 1 | k__Fungi(100);p__Ascomycota(100);c__Saccharomycetes(100);o__Saccharomycetales(100);f__Metschnikowiaceae(100);g__Metschnikowia(100);g__Metschnikowia_unclassified(100); | Metschnikowia | 13 | Saprotroph | \|Undefined Saprotroph\| | Yeast | NULL | Probable | As nectar-tap_saprotroph (Põlme et al. 2020); Decay sugar-rich substrates (Põlme et al. 2020) | Põlme S, et al. 2020. Fungal Diversity 105:1-16 ((https://doi.org/10.1007/s13225-020-00466-2)) |
| Otu18238 | 1 | k__Fungi(100);p__Ascomycota(100);c__Saccharomycetes(100);o__Saccharomycetales(100);f__Metschnikowiaceae(100);g__Metschnikowia(100);s__Metschnikowia_pulcherrima(100); | Metschnikowia | 13 | Saprotroph | \|Undefined Saprotroph\| | Yeast | NULL | Probable | As nectar-tap_saprotroph (Põlme et al. 2020); Decay sugar-rich substrates (Põlme et al. 2020) | Põlme S, et al. 2020. Fungal Diversity 105:1-16 ((https://doi.org/10.1007/s13225-020-00466-2)) |
| Otu18239 | 1 | k__Fungi(100);p__Ascomycota(100);c__Saccharomycetes(100);o__Saccharomycetales(100);f__Metschnikowiaceae(100);g__Metschnikowia(100);s__Metschnikowia_pulcherrima(100); | Metschnikowia | 13 | Saprotroph | \|Undefined Saprotroph\| | Yeast | NULL | Probable | As nectar-tap_saprotroph (Põlme et al. 2020); Decay sugar-rich substrates (Põlme et al. 2020) | Põlme S, et al. 2020. Fungal Diversity 105:1-16 ((https://doi.org/10.1007/s13225-020-00466-2)) |
| Otu18240 | 1 | k__Fungi(100);p__Ascomycota(100);c__Saccharomycetes(100);o__Saccharomycetales(100);f__Metschnikowiaceae(100);g__Metschnikowia(100);s__Metschnikowia_pulcherrima(100); | Metschnikowia | 13 | Saprotroph | \|Undefined Saprotroph\| | Yeast | NULL | Probable | As nectar-tap_saprotroph (Põlme et al. 2020); Decay sugar-rich substrates (Põlme et al. 2020) | Põlme S, et al. 2020. Fungal Diversity 105:1-16 ((https://doi.org/10.1007/s13225-020-00466-2)) |
| Otu18241 | 1 | k__Fungi(100);p__Ascomycota(100);c__Saccharomycetes(100);o__Saccharomycetales(100);f__Metschnikowiaceae(100);g__Metschnikowia(100);s__Metschnikowia_pulcherrima(100); | Metschnikowia | 13 | Saprotroph | \|Undefined Saprotroph\| | Yeast | NULL | Probable | As nectar-tap_saprotroph (Põlme et al. 2020); Decay sugar-rich substrates (Põlme et al. 2020) | Põlme S, et al. 2020. Fungal Diversity 105:1-16 ((https://doi.org/10.1007/s13225-020-00466-2)) |
| Otu18243 | 1 | k__Fungi(100);p__Ascomycota(100);c__Saccharomycetes(100);o__Saccharomycetales(100);f__Metschnikowiaceae(100);g__Metschnikowia(100);s__Metschnikowia_pulcherrima(100); | Metschnikowia | 13 | Saprotroph | \|Undefined Saprotroph\| | Yeast | NULL | Probable | As nectar-tap_saprotroph (Põlme et al. 2020); Decay sugar-rich substrates (Põlme et al. 2020) | Põlme S, et al. 2020. Fungal Diversity 105:1-16 ((https://doi.org/10.1007/s13225-020-00466-2)) |
| Otu18244 | 1 | k__Fungi(100);p__Ascomycota(100);c__Saccharomycetes(100);o__Saccharomycetales(100);f__Metschnikowiaceae(100);g__Metschnikowia(100);s__Metschnikowia_pulcherrima(100); | Metschnikowia | 13 | Saprotroph | \|Undefined Saprotroph\| | Yeast | NULL | Probable | As nectar-tap_saprotroph (Põlme et al. 2020); Decay sugar-rich substrates (Põlme et al. 2020) | Põlme S, et al. 2020. Fungal Diversity 105:1-16 ((https://doi.org/10.1007/s13225-020-00466-2)) |
| Otu18245 | 1 | k__Fungi(100);p__Ascomycota(100);c__Saccharomycetes(100);o__Saccharomycetales(100);f__Metschnikowiaceae(100);g__Metschnikowia(100);s__Metschnikowia_pulcherrima(100); | Metschnikowia | 13 | Saprotroph | \|Undefined Saprotroph\| | Yeast | NULL | Probable | As nectar-tap_saprotroph (Põlme et al. 2020); Decay sugar-rich substrates (Põlme et al. 2020) | Põlme S, et al. 2020. Fungal Diversity 105:1-16 ((https://doi.org/10.1007/s13225-020-00466-2)) |
| Otu18246 | 1 | k__Fungi(100);p__Ascomycota(100);c__Saccharomycetes(100);o__Saccharomycetales(100);f__Metschnikowiaceae(100);g__Metschnikowia(100);s__Metschnikowia_pulcherrima(100); | Metschnikowia | 13 | Saprotroph | \|Undefined Saprotroph\| | Yeast | NULL | Probable | As nectar-tap_saprotroph (Põlme et al. 2020); Decay sugar-rich substrates (Põlme et al. 2020) | Põlme S, et al. 2020. Fungal Diversity 105:1-16 ((https://doi.org/10.1007/s13225-020-00466-2)) |
| Otu18247 | 1 | k__Fungi(100);p__Ascomycota(100);c__Saccharomycetes(100);o__Saccharomycetales(100);f__Metschnikowiaceae(100);g__Metschnikowia(100);s__Metschnikowia_pulcherrima(100); | Metschnikowia | 13 | Saprotroph | \|Undefined Saprotroph\| | Yeast | NULL | Probable | As nectar-tap_saprotroph (Põlme et al. 2020); Decay sugar-rich substrates (Põlme et al. 2020) | Põlme S, et al. 2020. Fungal Diversity 105:1-16 ((https://doi.org/10.1007/s13225-020-00466-2)) |
| Otu18248 | 1 | k__Fungi(100);p__Ascomycota(100);c__Saccharomycetes(100);o__Saccharomycetales(100);f__Metschnikowiaceae(100);g__Metschnikowia(100);s__Metschnikowia_pulcherrima(100); | Metschnikowia | 13 | Saprotroph | \|Undefined Saprotroph\| | Yeast | NULL | Probable | As nectar-tap_saprotroph (Põlme et al. 2020); Decay sugar-rich substrates (Põlme et al. 2020) | Põlme S, et al. 2020. Fungal Diversity 105:1-16 ((https://doi.org/10.1007/s13225-020-00466-2)) |
| Otu18251 | 1 | k__Fungi(100);p__Ascomycota(100);c__Saccharomycetes(100);o__Saccharomycetales(100);f__Metschnikowiaceae(100);g__Metschnikowia(100);g__Metschnikowia_unclassified(100); | Metschnikowia | 13 | Saprotroph | \|Undefined Saprotroph\| | Yeast | NULL | Probable | As nectar-tap_saprotroph (Põlme et al. 2020); Decay sugar-rich substrates (Põlme et al. 2020) | Põlme S, et al. 2020. Fungal Diversity 105:1-16 ((https://doi.org/10.1007/s13225-020-00466-2)) |
| Otu18252 | 1 | k__Fungi(100);p__Ascomycota(100);c__Saccharomycetes(100);o__Saccharomycetales(100);f__Metschnikowiaceae(100);g__Metschnikowia(100);g__Metschnikowia_unclassified(100); | Metschnikowia | 13 | Saprotroph | \|Undefined Saprotroph\| | Yeast | NULL | Probable | As nectar-tap_saprotroph (Põlme et al. 2020); Decay sugar-rich substrates (Põlme et al. 2020) | Põlme S, et al. 2020. Fungal Diversity 105:1-16 ((https://doi.org/10.1007/s13225-020-00466-2)) |
| Otu18253 | 1 | k__Fungi(100);p__Ascomycota(100);c__Saccharomycetes(100);o__Saccharomycetales(100);f__Metschnikowiaceae(100);g__Metschnikowia(100);s__Metschnikowia_pulcherrima(100); | Metschnikowia | 13 | Saprotroph | \|Undefined Saprotroph\| | Yeast | NULL | Probable | As nectar-tap_saprotroph (Põlme et al. 2020); Decay sugar-rich substrates (Põlme et al. 2020) | Põlme S, et al. 2020. Fungal Diversity 105:1-16 ((https://doi.org/10.1007/s13225-020-00466-2)) |
| Otu18254 | 1 | k__Fungi(100);p__Ascomycota(100);c__Saccharomycetes(100);o__Saccharomycetales(100);f__Saccharomycetales_family_Incertae_sedis(100);g__Candida(100);g__Candida_unclassified(100); | Candida | 13 | Pathotroph-Saprotroph-Symbiotroph | Animal Pathogen-Endophyte-Endosymbiont-Epiphyte-\|Undefined Saprotroph\| | Dimorphic Yeast-Yeast | NULL | Probable | As nectar-tap_saprotroph (Põlme et al. 2020); Decay sugar-rich substrates (Põlme et al. 2020); This asexual genus is polyphyletic and contain species that function very differently depend on what they are. Caution should be taken when interpreting guilds for Candida species. | Manolakaki D, et al. 2010. Virulence. 1: 367-375 ((https://doi.org/10.4161/viru.1.5.12796)); Mekbib SB, et al. 2011. Tropical Plant Pathology 36:233-240 ((https://doi.org/10.1590/S1982-56762011000400004)); Põlme S, et al. 2020. Fungal Diversity 105:1-16 ((https://doi.org/10.1007/s13225-020-00466-2)) |
| Otu18255 | 1 | k__Fungi(100);p__Ascomycota(100);c__Saccharomycetes(100);o__Saccharomycetales(100);f__Metschnikowiaceae(100);g__Metschnikowia(100);s__Metschnikowia_pulcherrima(100); | Metschnikowia | 13 | Saprotroph | \|Undefined Saprotroph\| | Yeast | NULL | Probable | As nectar-tap_saprotroph (Põlme et al. 2020); Decay sugar-rich substrates (Põlme et al. 2020) | Põlme S, et al. 2020. Fungal Diversity 105:1-16 ((https://doi.org/10.1007/s13225-020-00466-2)) |
| Otu18256 | 1 | k__Fungi(100);p__Ascomycota(100);c__Saccharomycetes(100);o__Saccharomycetales(100);f__Metschnikowiaceae(100);g__Metschnikowia(100);s__Metschnikowia_pulcherrima(100); | Metschnikowia | 13 | Saprotroph | \|Undefined Saprotroph\| | Yeast | NULL | Probable | As nectar-tap_saprotroph (Põlme et al. 2020); Decay sugar-rich substrates (Põlme et al. 2020) | Põlme S, et al. 2020. Fungal Diversity 105:1-16 ((https://doi.org/10.1007/s13225-020-00466-2)) |
| Otu18257 | 1 | k__Fungi(100);p__Ascomycota(100);c__Saccharomycetes(100);o__Saccharomycetales(100);f__Metschnikowiaceae(100);g__Metschnikowia(100);s__Metschnikowia_pulcherrima(100); | Metschnikowia | 13 | Saprotroph | \|Undefined Saprotroph\| | Yeast | NULL | Probable | As nectar-tap_saprotroph (Põlme et al. 2020); Decay sugar-rich substrates (Põlme et al. 2020) | Põlme S, et al. 2020. Fungal Diversity 105:1-16 ((https://doi.org/10.1007/s13225-020-00466-2)) |
| Otu18258 | 1 | k__Fungi(100);p__Ascomycota(100);c__Saccharomycetes(100);o__Saccharomycetales(100);f__Metschnikowiaceae(100);g__Metschnikowia(100);s__Metschnikowia_pulcherrima(100); | Metschnikowia | 13 | Saprotroph | \|Undefined Saprotroph\| | Yeast | NULL | Probable | As nectar-tap_saprotroph (Põlme et al. 2020); Decay sugar-rich substrates (Põlme et al. 2020) | Põlme S, et al. 2020. Fungal Diversity 105:1-16 ((https://doi.org/10.1007/s13225-020-00466-2)) |
| Otu18259 | 1 | k__Fungi(100);p__Ascomycota(100);c__Saccharomycetes(100);o__Saccharomycetales(100);f__Metschnikowiaceae(100);g__Metschnikowia(100);g__Metschnikowia_unclassified(100); | Metschnikowia | 13 | Saprotroph | \|Undefined Saprotroph\| | Yeast | NULL | Probable | As nectar-tap_saprotroph (Põlme et al. 2020); Decay sugar-rich substrates (Põlme et al. 2020) | Põlme S, et al. 2020. Fungal Diversity 105:1-16 ((https://doi.org/10.1007/s13225-020-00466-2)) |
| Otu18260 | 1 | k__Fungi(100);p__Ascomycota(100);c__Saccharomycetes(100);o__Saccharomycetales(100);f__Metschnikowiaceae(100);g__Metschnikowia(100);g__Metschnikowia_unclassified(100); | Metschnikowia | 13 | Saprotroph | \|Undefined Saprotroph\| | Yeast | NULL | Probable | As nectar-tap_saprotroph (Põlme et al. 2020); Decay sugar-rich substrates (Põlme et al. 2020) | Põlme S, et al. 2020. Fungal Diversity 105:1-16 ((https://doi.org/10.1007/s13225-020-00466-2)) |
| Otu18261 | 1 | k__Fungi(100);p__Ascomycota(100);c__Saccharomycetes(100);o__Saccharomycetales(100);f__Metschnikowiaceae(100);g__Metschnikowia(100);s__Metschnikowia_pulcherrima(100); | Metschnikowia | 13 | Saprotroph | \|Undefined Saprotroph\| | Yeast | NULL | Probable | As nectar-tap_saprotroph (Põlme et al. 2020); Decay sugar-rich substrates (Põlme et al. 2020) | Põlme S, et al. 2020. Fungal Diversity 105:1-16 ((https://doi.org/10.1007/s13225-020-00466-2)) |
| Otu18262 | 1 | k__Fungi(100);p__Ascomycota(100);c__Saccharomycetes(100);o__Saccharomycetales(100);f__Metschnikowiaceae(100);g__Metschnikowia(100);s__Metschnikowia_pulcherrima(100); | Metschnikowia | 13 | Saprotroph | \|Undefined Saprotroph\| | Yeast | NULL | Probable | As nectar-tap_saprotroph (Põlme et al. 2020); Decay sugar-rich substrates (Põlme et al. 2020) | Põlme S, et al. 2020. Fungal Diversity 105:1-16 ((https://doi.org/10.1007/s13225-020-00466-2)) |
| Otu18263 | 1 | k__Fungi(100);p__Ascomycota(100);c__Saccharomycetes(100);o__Saccharomycetales(100);f__Metschnikowiaceae(100);g__Metschnikowia(100);g__Metschnikowia_unclassified(100); | Metschnikowia | 13 | Saprotroph | \|Undefined Saprotroph\| | Yeast | NULL | Probable | As nectar-tap_saprotroph (Põlme et al. 2020); Decay sugar-rich substrates (Põlme et al. 2020) | Põlme S, et al. 2020. Fungal Diversity 105:1-16 ((https://doi.org/10.1007/s13225-020-00466-2)) |
| Otu18264 | 1 | k__Fungi(100);p__Ascomycota(100);c__Saccharomycetes(100);o__Saccharomycetales(100);f__Metschnikowiaceae(100);g__Metschnikowia(100);s__Metschnikowia_pulcherrima(100); | Metschnikowia | 13 | Saprotroph | \|Undefined Saprotroph\| | Yeast | NULL | Probable | As nectar-tap_saprotroph (Põlme et al. 2020); Decay sugar-rich substrates (Põlme et al. 2020) | Põlme S, et al. 2020. Fungal Diversity 105:1-16 ((https://doi.org/10.1007/s13225-020-00466-2)) |
| Otu18265 | 1 | k__Fungi(100);p__Ascomycota(100);c__Saccharomycetes(100);o__Saccharomycetales(100);f__Metschnikowiaceae(100);g__Metschnikowia(100);s__Metschnikowia_pulcherrima(100); | Metschnikowia | 13 | Saprotroph | \|Undefined Saprotroph\| | Yeast | NULL | Probable | As nectar-tap_saprotroph (Põlme et al. 2020); Decay sugar-rich substrates (Põlme et al. 2020) | Põlme S, et al. 2020. Fungal Diversity 105:1-16 ((https://doi.org/10.1007/s13225-020-00466-2)) |
| Otu18266 | 1 | k__Fungi(100);p__Ascomycota(100);c__Saccharomycetes(100);o__Saccharomycetales(100);f__Metschnikowiaceae(100);g__Metschnikowia(100);s__Metschnikowia_pulcherrima(100); | Metschnikowia | 13 | Saprotroph | \|Undefined Saprotroph\| | Yeast | NULL | Probable | As nectar-tap_saprotroph (Põlme et al. 2020); Decay sugar-rich substrates (Põlme et al. 2020) | Põlme S, et al. 2020. Fungal Diversity 105:1-16 ((https://doi.org/10.1007/s13225-020-00466-2)) |
| Otu18267 | 1 | k__Fungi(100);p__Ascomycota(100);c__Saccharomycetes(100);o__Saccharomycetales(100);f__Metschnikowiaceae(100);g__Metschnikowia(100);s__Metschnikowia_pulcherrima(100); | Metschnikowia | 13 | Saprotroph | \|Undefined Saprotroph\| | Yeast | NULL | Probable | As nectar-tap_saprotroph (Põlme et al. 2020); Decay sugar-rich substrates (Põlme et al. 2020) | Põlme S, et al. 2020. Fungal Diversity 105:1-16 ((https://doi.org/10.1007/s13225-020-00466-2)) |
| Otu18270 | 1 | k__Fungi(100);p__Ascomycota(100);c__Saccharomycetes(100);o__Saccharomycetales(100);f__Metschnikowiaceae(100);g__Metschnikowia(100);g__Metschnikowia_unclassified(100); | Metschnikowia | 13 | Saprotroph | \|Undefined Saprotroph\| | Yeast | NULL | Probable | As nectar-tap_saprotroph (Põlme et al. 2020); Decay sugar-rich substrates (Põlme et al. 2020) | Põlme S, et al. 2020. Fungal Diversity 105:1-16 ((https://doi.org/10.1007/s13225-020-00466-2)) |
| Otu18271 | 1 | k__Fungi(100);p__Ascomycota(100);c__Saccharomycetes(100);o__Saccharomycetales(100);f__Metschnikowiaceae(100);g__Metschnikowia(100);s__Metschnikowia_pulcherrima(100); | Metschnikowia | 13 | Saprotroph | \|Undefined Saprotroph\| | Yeast | NULL | Probable | As nectar-tap_saprotroph (Põlme et al. 2020); Decay sugar-rich substrates (Põlme et al. 2020) | Põlme S, et al. 2020. Fungal Diversity 105:1-16 ((https://doi.org/10.1007/s13225-020-00466-2)) |
| Otu18272 | 1 | k__Fungi(100);p__Ascomycota(100);c__Saccharomycetes(100);o__Saccharomycetales(100);f__Metschnikowiaceae(100);g__Metschnikowia(100);g__Metschnikowia_unclassified(100); | Metschnikowia | 13 | Saprotroph | \|Undefined Saprotroph\| | Yeast | NULL | Probable | As nectar-tap_saprotroph (Põlme et al. 2020); Decay sugar-rich substrates (Põlme et al. 2020) | Põlme S, et al. 2020. Fungal Diversity 105:1-16 ((https://doi.org/10.1007/s13225-020-00466-2)) |
| Otu18274 | 1 | k__Fungi(100);p__Ascomycota(100);c__Saccharomycetes(100);o__Saccharomycetales(100);f__Metschnikowiaceae(100);g__Metschnikowia(100);s__Metschnikowia_pulcherrima(100); | Metschnikowia | 13 | Saprotroph | \|Undefined Saprotroph\| | Yeast | NULL | Probable | As nectar-tap_saprotroph (Põlme et al. 2020); Decay sugar-rich substrates (Põlme et al. 2020) | Põlme S, et al. 2020. Fungal Diversity 105:1-16 ((https://doi.org/10.1007/s13225-020-00466-2)) |
| Otu18276 | 1 | k__Fungi(100);p__Ascomycota(100);c__Saccharomycetes(100);o__Saccharomycetales(100);f__Metschnikowiaceae(100);g__Metschnikowia(100);s__Metschnikowia_pulcherrima(100); | Metschnikowia | 13 | Saprotroph | \|Undefined Saprotroph\| | Yeast | NULL | Probable | As nectar-tap_saprotroph (Põlme et al. 2020); Decay sugar-rich substrates (Põlme et al. 2020) | Põlme S, et al. 2020. Fungal Diversity 105:1-16 ((https://doi.org/10.1007/s13225-020-00466-2)) |
| Otu18280 | 1 | k__Fungi(100);p__Ascomycota(100);c__Saccharomycetes(100);o__Saccharomycetales(100);f__Metschnikowiaceae(100);g__Metschnikowia(100);s__Metschnikowia_pulcherrima(100); | Metschnikowia | 13 | Saprotroph | \|Undefined Saprotroph\| | Yeast | NULL | Probable | As nectar-tap_saprotroph (Põlme et al. 2020); Decay sugar-rich substrates (Põlme et al. 2020) | Põlme S, et al. 2020. Fungal Diversity 105:1-16 ((https://doi.org/10.1007/s13225-020-00466-2)) |
| Otu18281 | 1 | k__Fungi(100);p__Ascomycota(100);c__Saccharomycetes(100);o__Saccharomycetales(100);f__Saccharomycetaceae(100);g__Kazachstania(100);s__Kazachstania_intestinalis(100); | Kazachstania | 13 | Saprotroph | \|Undefined Saprotroph\| | Yeast | NULL | Probable | As nectar-tap_saprotroph (Põlme et al. 2020); Decay sugar-rich substrates (Põlme et al. 2020) | Põlme S, et al. 2020. Fungal Diversity 105:1-16 ((https://doi.org/10.1007/s13225-020-00466-2)) |
| Otu18282 | 1 | k__Fungi(100);p__Ascomycota(100);c__Saccharomycetes(100);o__Saccharomycetales(100);f__Metschnikowiaceae(100);g__Metschnikowia(100);s__Metschnikowia_pulcherrima(100); | Metschnikowia | 13 | Saprotroph | \|Undefined Saprotroph\| | Yeast | NULL | Probable | As nectar-tap_saprotroph (Põlme et al. 2020); Decay sugar-rich substrates (Põlme et al. 2020) | Põlme S, et al. 2020. Fungal Diversity 105:1-16 ((https://doi.org/10.1007/s13225-020-00466-2)) |
| Otu18283 | 1 | k__Fungi(100);p__Ascomycota(100);c__Saccharomycetes(100);o__Saccharomycetales(100);f__Metschnikowiaceae(100);g__Metschnikowia(100);s__Metschnikowia_pulcherrima(100); | Metschnikowia | 13 | Saprotroph | \|Undefined Saprotroph\| | Yeast | NULL | Probable | As nectar-tap_saprotroph (Põlme et al. 2020); Decay sugar-rich substrates (Põlme et al. 2020) | Põlme S, et al. 2020. Fungal Diversity 105:1-16 ((https://doi.org/10.1007/s13225-020-00466-2)) |
| Otu18284 | 1 | k__Fungi(100);p__Ascomycota(100);c__Saccharomycetes(100);o__Saccharomycetales(100);f__Metschnikowiaceae(100);g__Metschnikowia(100);s__Metschnikowia_pulcherrima(100); | Metschnikowia | 13 | Saprotroph | \|Undefined Saprotroph\| | Yeast | NULL | Probable | As nectar-tap_saprotroph (Põlme et al. 2020); Decay sugar-rich substrates (Põlme et al. 2020) | Põlme S, et al. 2020. Fungal Diversity 105:1-16 ((https://doi.org/10.1007/s13225-020-00466-2)) |
| Otu18285 | 1 | k__Fungi(100);p__Ascomycota(100);c__Saccharomycetes(100);o__Saccharomycetales(100);f__Metschnikowiaceae(100);g__Metschnikowia(100);s__Metschnikowia_pulcherrima(100); | Metschnikowia | 13 | Saprotroph | \|Undefined Saprotroph\| | Yeast | NULL | Probable | As nectar-tap_saprotroph (Põlme et al. 2020); Decay sugar-rich substrates (Põlme et al. 2020) | Põlme S, et al. 2020. Fungal Diversity 105:1-16 ((https://doi.org/10.1007/s13225-020-00466-2)) |
| Otu18286 | 1 | k__Fungi(100);p__Ascomycota(100);c__Saccharomycetes(100);o__Saccharomycetales(100);f__Metschnikowiaceae(100);g__Metschnikowia(100);s__Metschnikowia_pulcherrima(100); | Metschnikowia | 13 | Saprotroph | \|Undefined Saprotroph\| | Yeast | NULL | Probable | As nectar-tap_saprotroph (Põlme et al. 2020); Decay sugar-rich substrates (Põlme et al. 2020) | Põlme S, et al. 2020. Fungal Diversity 105:1-16 ((https://doi.org/10.1007/s13225-020-00466-2)) |
| Otu18287 | 1 | k__Fungi(100);p__Ascomycota(100);c__Saccharomycetes(100);o__Saccharomycetales(100);f__Metschnikowiaceae(100);g__Metschnikowia(100);s__Metschnikowia_pulcherrima(100); | Metschnikowia | 13 | Saprotroph | \|Undefined Saprotroph\| | Yeast | NULL | Probable | As nectar-tap_saprotroph (Põlme et al. 2020); Decay sugar-rich substrates (Põlme et al. 2020) | Põlme S, et al. 2020. Fungal Diversity 105:1-16 ((https://doi.org/10.1007/s13225-020-00466-2)) |
| Otu18290 | 1 | k__Fungi(100);p__Ascomycota(100);c__Saccharomycetes(100);o__Saccharomycetales(100);f__Metschnikowiaceae(100);g__Metschnikowia(100);g__Metschnikowia_unclassified(100); | Metschnikowia | 13 | Saprotroph | \|Undefined Saprotroph\| | Yeast | NULL | Probable | As nectar-tap_saprotroph (Põlme et al. 2020); Decay sugar-rich substrates (Põlme et al. 2020) | Põlme S, et al. 2020. Fungal Diversity 105:1-16 ((https://doi.org/10.1007/s13225-020-00466-2)) |
| Otu18291 | 1 | k__Fungi(100);p__Ascomycota(100);c__Saccharomycetes(100);o__Saccharomycetales(100);f__Metschnikowiaceae(100);g__Metschnikowia(100);g__Metschnikowia_unclassified(100); | Metschnikowia | 13 | Saprotroph | \|Undefined Saprotroph\| | Yeast | NULL | Probable | As nectar-tap_saprotroph (Põlme et al. 2020); Decay sugar-rich substrates (Põlme et al. 2020) | Põlme S, et al. 2020. Fungal Diversity 105:1-16 ((https://doi.org/10.1007/s13225-020-00466-2)) |
| Otu18292 | 1 | k__Fungi(100);p__Ascomycota(100);c__Saccharomycetes(100);o__Saccharomycetales(100);f__Metschnikowiaceae(100);g__Metschnikowia(100);s__Metschnikowia_pulcherrima(100); | Metschnikowia | 13 | Saprotroph | \|Undefined Saprotroph\| | Yeast | NULL | Probable | As nectar-tap_saprotroph (Põlme et al. 2020); Decay sugar-rich substrates (Põlme et al. 2020) | Põlme S, et al. 2020. Fungal Diversity 105:1-16 ((https://doi.org/10.1007/s13225-020-00466-2)) |
| Otu18293 | 1 | k__Fungi(100);p__Ascomycota(100);c__Saccharomycetes(100);o__Saccharomycetales(100);f__Metschnikowiaceae(100);g__Metschnikowia(100);s__Metschnikowia_pulcherrima(100); | Metschnikowia | 13 | Saprotroph | \|Undefined Saprotroph\| | Yeast | NULL | Probable | As nectar-tap_saprotroph (Põlme et al. 2020); Decay sugar-rich substrates (Põlme et al. 2020) | Põlme S, et al. 2020. Fungal Diversity 105:1-16 ((https://doi.org/10.1007/s13225-020-00466-2)) |
| Otu18294 | 1 | k__Fungi(100);p__Ascomycota(100);c__Saccharomycetes(100);o__Saccharomycetales(100);f__Metschnikowiaceae(100);g__Metschnikowia(100);s__Metschnikowia_pulcherrima(100); | Metschnikowia | 13 | Saprotroph | \|Undefined Saprotroph\| | Yeast | NULL | Probable | As nectar-tap_saprotroph (Põlme et al. 2020); Decay sugar-rich substrates (Põlme et al. 2020) | Põlme S, et al. 2020. Fungal Diversity 105:1-16 ((https://doi.org/10.1007/s13225-020-00466-2)) |
| Otu18295 | 1 | k__Fungi(100);p__Ascomycota(100);c__Saccharomycetes(100);o__Saccharomycetales(100);f__Metschnikowiaceae(100);g__Metschnikowia(100);g__Metschnikowia_unclassified(100); | Metschnikowia | 13 | Saprotroph | \|Undefined Saprotroph\| | Yeast | NULL | Probable | As nectar-tap_saprotroph (Põlme et al. 2020); Decay sugar-rich substrates (Põlme et al. 2020) | Põlme S, et al. 2020. Fungal Diversity 105:1-16 ((https://doi.org/10.1007/s13225-020-00466-2)) |
| Otu18296 | 1 | k__Fungi(100);p__Ascomycota(100);c__Saccharomycetes(100);o__Saccharomycetales(100);f__Metschnikowiaceae(100);g__Metschnikowia(100);g__Metschnikowia_unclassified(100); | Metschnikowia | 13 | Saprotroph | \|Undefined Saprotroph\| | Yeast | NULL | Probable | As nectar-tap_saprotroph (Põlme et al. 2020); Decay sugar-rich substrates (Põlme et al. 2020) | Põlme S, et al. 2020. Fungal Diversity 105:1-16 ((https://doi.org/10.1007/s13225-020-00466-2)) |
| Otu18297 | 1 | k__Fungi(100);p__Ascomycota(100);c__Saccharomycetes(100);o__Saccharomycetales(100);f__Metschnikowiaceae(100);g__Metschnikowia(100);g__Metschnikowia_unclassified(100); | Metschnikowia | 13 | Saprotroph | \|Undefined Saprotroph\| | Yeast | NULL | Probable | As nectar-tap_saprotroph (Põlme et al. 2020); Decay sugar-rich substrates (Põlme et al. 2020) | Põlme S, et al. 2020. Fungal Diversity 105:1-16 ((https://doi.org/10.1007/s13225-020-00466-2)) |
| Otu18298 | 1 | k__Fungi(100);p__Ascomycota(100);c__Saccharomycetes(100);o__Saccharomycetales(100);f__Metschnikowiaceae(100);g__Metschnikowia(100);s__Metschnikowia_pulcherrima(100); | Metschnikowia | 13 | Saprotroph | \|Undefined Saprotroph\| | Yeast | NULL | Probable | As nectar-tap_saprotroph (Põlme et al. 2020); Decay sugar-rich substrates (Põlme et al. 2020) | Põlme S, et al. 2020. Fungal Diversity 105:1-16 ((https://doi.org/10.1007/s13225-020-00466-2)) |
| Otu18299 | 1 | k__Fungi(100);p__Ascomycota(100);c__Saccharomycetes(100);o__Saccharomycetales(100);f__Metschnikowiaceae(100);g__Metschnikowia(100);s__Metschnikowia_pulcherrima(100); | Metschnikowia | 13 | Saprotroph | \|Undefined Saprotroph\| | Yeast | NULL | Probable | As nectar-tap_saprotroph (Põlme et al. 2020); Decay sugar-rich substrates (Põlme et al. 2020) | Põlme S, et al. 2020. Fungal Diversity 105:1-16 ((https://doi.org/10.1007/s13225-020-00466-2)) |
| Otu18300 | 1 | k__Fungi(100);p__Ascomycota(100);c__Saccharomycetes(100);o__Saccharomycetales(100);f__Metschnikowiaceae(100);g__Metschnikowia(100);s__Metschnikowia_pulcherrima(100); | Metschnikowia | 13 | Saprotroph | \|Undefined Saprotroph\| | Yeast | NULL | Probable | As nectar-tap_saprotroph (Põlme et al. 2020); Decay sugar-rich substrates (Põlme et al. 2020) | Põlme S, et al. 2020. Fungal Diversity 105:1-16 ((https://doi.org/10.1007/s13225-020-00466-2)) |
| Otu18301 | 1 | k__Fungi(100);p__Ascomycota(100);c__Saccharomycetes(100);o__Saccharomycetales(100);f__Metschnikowiaceae(100);g__Metschnikowia(100);s__Metschnikowia_pulcherrima(100); | Metschnikowia | 13 | Saprotroph | \|Undefined Saprotroph\| | Yeast | NULL | Probable | As nectar-tap_saprotroph (Põlme et al. 2020); Decay sugar-rich substrates (Põlme et al. 2020) | Põlme S, et al. 2020. Fungal Diversity 105:1-16 ((https://doi.org/10.1007/s13225-020-00466-2)) |
| Otu18302 | 1 | k__Fungi(100);p__Ascomycota(100);c__Saccharomycetes(100);o__Saccharomycetales(100);f__Metschnikowiaceae(100);g__Metschnikowia(100);s__Metschnikowia_pulcherrima(100); | Metschnikowia | 13 | Saprotroph | \|Undefined Saprotroph\| | Yeast | NULL | Probable | As nectar-tap_saprotroph (Põlme et al. 2020); Decay sugar-rich substrates (Põlme et al. 2020) | Põlme S, et al. 2020. Fungal Diversity 105:1-16 ((https://doi.org/10.1007/s13225-020-00466-2)) |
| Otu18303 | 1 | k__Fungi(100);p__Ascomycota(100);c__Saccharomycetes(100);o__Saccharomycetales(100);f__Metschnikowiaceae(100);g__Metschnikowia(100);s__Metschnikowia_pulcherrima(100); | Metschnikowia | 13 | Saprotroph | \|Undefined Saprotroph\| | Yeast | NULL | Probable | As nectar-tap_saprotroph (Põlme et al. 2020); Decay sugar-rich substrates (Põlme et al. 2020) | Põlme S, et al. 2020. Fungal Diversity 105:1-16 ((https://doi.org/10.1007/s13225-020-00466-2)) |
| Otu18304 | 1 | k__Fungi(100);p__Ascomycota(100);c__Saccharomycetes(100);o__Saccharomycetales(100);f__Metschnikowiaceae(100);g__Metschnikowia(100);s__Metschnikowia_pulcherrima(100); | Metschnikowia | 13 | Saprotroph | \|Undefined Saprotroph\| | Yeast | NULL | Probable | As nectar-tap_saprotroph (Põlme et al. 2020); Decay sugar-rich substrates (Põlme et al. 2020) | Põlme S, et al. 2020. Fungal Diversity 105:1-16 ((https://doi.org/10.1007/s13225-020-00466-2)) |
| Otu18305 | 1 | k__Fungi(100);p__Ascomycota(100);c__Saccharomycetes(100);o__Saccharomycetales(100);f__Metschnikowiaceae(100);g__Metschnikowia(100);s__Metschnikowia_pulcherrima(100); | Metschnikowia | 13 | Saprotroph | \|Undefined Saprotroph\| | Yeast | NULL | Probable | As nectar-tap_saprotroph (Põlme et al. 2020); Decay sugar-rich substrates (Põlme et al. 2020) | Põlme S, et al. 2020. Fungal Diversity 105:1-16 ((https://doi.org/10.1007/s13225-020-00466-2)) |
| Otu18306 | 1 | k__Fungi(100);p__Ascomycota(100);c__Saccharomycetes(100);o__Saccharomycetales(100);f__Metschnikowiaceae(100);g__Metschnikowia(100);g__Metschnikowia_unclassified(100); | Metschnikowia | 13 | Saprotroph | \|Undefined Saprotroph\| | Yeast | NULL | Probable | As nectar-tap_saprotroph (Põlme et al. 2020); Decay sugar-rich substrates (Põlme et al. 2020) | Põlme S, et al. 2020. Fungal Diversity 105:1-16 ((https://doi.org/10.1007/s13225-020-00466-2)) |
| Otu18307 | 1 | k__Fungi(100);p__Ascomycota(100);c__Saccharomycetes(100);o__Saccharomycetales(100);f__Metschnikowiaceae(100);g__Metschnikowia(100);s__Metschnikowia_pulcherrima(100); | Metschnikowia | 13 | Saprotroph | \|Undefined Saprotroph\| | Yeast | NULL | Probable | As nectar-tap_saprotroph (Põlme et al. 2020); Decay sugar-rich substrates (Põlme et al. 2020) | Põlme S, et al. 2020. Fungal Diversity 105:1-16 ((https://doi.org/10.1007/s13225-020-00466-2)) |
| Otu18308 | 1 | k__Fungi(100);p__Ascomycota(100);c__Saccharomycetes(100);o__Saccharomycetales(100);f__Metschnikowiaceae(100);g__Metschnikowia(100);s__Metschnikowia_pulcherrima(100); | Metschnikowia | 13 | Saprotroph | \|Undefined Saprotroph\| | Yeast | NULL | Probable | As nectar-tap_saprotroph (Põlme et al. 2020); Decay sugar-rich substrates (Põlme et al. 2020) | Põlme S, et al. 2020. Fungal Diversity 105:1-16 ((https://doi.org/10.1007/s13225-020-00466-2)) |
| Otu18309 | 1 | k__Fungi(100);p__Ascomycota(100);c__Saccharomycetes(100);o__Saccharomycetales(100);f__Metschnikowiaceae(100);g__Metschnikowia(100);s__Metschnikowia_pulcherrima(100); | Metschnikowia | 13 | Saprotroph | \|Undefined Saprotroph\| | Yeast | NULL | Probable | As nectar-tap_saprotroph (Põlme et al. 2020); Decay sugar-rich substrates (Põlme et al. 2020) | Põlme S, et al. 2020. Fungal Diversity 105:1-16 ((https://doi.org/10.1007/s13225-020-00466-2)) |
| Otu18310 | 1 | k__Fungi(100);p__Ascomycota(100);c__Saccharomycetes(100);o__Saccharomycetales(100);f__Metschnikowiaceae(100);g__Metschnikowia(100);s__Metschnikowia_pulcherrima(100); | Metschnikowia | 13 | Saprotroph | \|Undefined Saprotroph\| | Yeast | NULL | Probable | As nectar-tap_saprotroph (Põlme et al. 2020); Decay sugar-rich substrates (Põlme et al. 2020) | Põlme S, et al. 2020. Fungal Diversity 105:1-16 ((https://doi.org/10.1007/s13225-020-00466-2)) |
| Otu18311 | 1 | k__Fungi(100);p__Ascomycota(100);c__Saccharomycetes(100);o__Saccharomycetales(100);f__Metschnikowiaceae(100);g__Metschnikowia(100);g__Metschnikowia_unclassified(100); | Metschnikowia | 13 | Saprotroph | \|Undefined Saprotroph\| | Yeast | NULL | Probable | As nectar-tap_saprotroph (Põlme et al. 2020); Decay sugar-rich substrates (Põlme et al. 2020) | Põlme S, et al. 2020. Fungal Diversity 105:1-16 ((https://doi.org/10.1007/s13225-020-00466-2)) |
| Otu18312 | 1 | k__Fungi(100);p__Ascomycota(100);c__Saccharomycetes(100);o__Saccharomycetales(100);f__Metschnikowiaceae(100);g__Metschnikowia(100);s__Metschnikowia_pulcherrima(100); | Metschnikowia | 13 | Saprotroph | \|Undefined Saprotroph\| | Yeast | NULL | Probable | As nectar-tap_saprotroph (Põlme et al. 2020); Decay sugar-rich substrates (Põlme et al. 2020) | Põlme S, et al. 2020. Fungal Diversity 105:1-16 ((https://doi.org/10.1007/s13225-020-00466-2)) |
| Otu18313 | 1 | k__Fungi(100);p__Ascomycota(100);c__Saccharomycetes(100);o__Saccharomycetales(100);f__Metschnikowiaceae(100);g__Metschnikowia(100);g__Metschnikowia_unclassified(100); | Metschnikowia | 13 | Saprotroph | \|Undefined Saprotroph\| | Yeast | NULL | Probable | As nectar-tap_saprotroph (Põlme et al. 2020); Decay sugar-rich substrates (Põlme et al. 2020) | Põlme S, et al. 2020. Fungal Diversity 105:1-16 ((https://doi.org/10.1007/s13225-020-00466-2)) |
| Otu18314 | 1 | k__Fungi(100);p__Ascomycota(100);c__Saccharomycetes(100);o__Saccharomycetales(100);f__Metschnikowiaceae(100);g__Metschnikowia(100);s__Metschnikowia_pulcherrima(100); | Metschnikowia | 13 | Saprotroph | \|Undefined Saprotroph\| | Yeast | NULL | Probable | As nectar-tap_saprotroph (Põlme et al. 2020); Decay sugar-rich substrates (Põlme et al. 2020) | Põlme S, et al. 2020. Fungal Diversity 105:1-16 ((https://doi.org/10.1007/s13225-020-00466-2)) |
| Otu18315 | 1 | k__Fungi(100);p__Ascomycota(100);c__Saccharomycetes(100);o__Saccharomycetales(100);f__Metschnikowiaceae(100);g__Metschnikowia(100);s__Metschnikowia_pulcherrima(100); | Metschnikowia | 13 | Saprotroph | \|Undefined Saprotroph\| | Yeast | NULL | Probable | As nectar-tap_saprotroph (Põlme et al. 2020); Decay sugar-rich substrates (Põlme et al. 2020) | Põlme S, et al. 2020. Fungal Diversity 105:1-16 ((https://doi.org/10.1007/s13225-020-00466-2)) |
| Otu18316 | 1 | k__Fungi(100);p__Ascomycota(100);c__Saccharomycetes(100);o__Saccharomycetales(100);f__Metschnikowiaceae(100);g__Metschnikowia(100);s__Metschnikowia_pulcherrima(100); | Metschnikowia | 13 | Saprotroph | \|Undefined Saprotroph\| | Yeast | NULL | Probable | As nectar-tap_saprotroph (Põlme et al. 2020); Decay sugar-rich substrates (Põlme et al. 2020) | Põlme S, et al. 2020. Fungal Diversity 105:1-16 ((https://doi.org/10.1007/s13225-020-00466-2)) |
| Otu18317 | 1 | k__Fungi(100);p__Ascomycota(100);c__Saccharomycetes(100);o__Saccharomycetales(100);f__Metschnikowiaceae(100);g__Metschnikowia(100);s__Metschnikowia_pulcherrima(100); | Metschnikowia | 13 | Saprotroph | \|Undefined Saprotroph\| | Yeast | NULL | Probable | As nectar-tap_saprotroph (Põlme et al. 2020); Decay sugar-rich substrates (Põlme et al. 2020) | Põlme S, et al. 2020. Fungal Diversity 105:1-16 ((https://doi.org/10.1007/s13225-020-00466-2)) |
| Otu18318 | 1 | k__Fungi(100);p__Ascomycota(100);c__Saccharomycetes(100);o__Saccharomycetales(100);f__Metschnikowiaceae(100);g__Metschnikowia(100);s__Metschnikowia_pulcherrima(100); | Metschnikowia | 13 | Saprotroph | \|Undefined Saprotroph\| | Yeast | NULL | Probable | As nectar-tap_saprotroph (Põlme et al. 2020); Decay sugar-rich substrates (Põlme et al. 2020) | Põlme S, et al. 2020. Fungal Diversity 105:1-16 ((https://doi.org/10.1007/s13225-020-00466-2)) |
| Otu18319 | 1 | k__Fungi(100);p__Ascomycota(100);c__Saccharomycetes(100);o__Saccharomycetales(100);f__Metschnikowiaceae(100);g__Metschnikowia(100);g__Metschnikowia_unclassified(100); | Metschnikowia | 13 | Saprotroph | \|Undefined Saprotroph\| | Yeast | NULL | Probable | As nectar-tap_saprotroph (Põlme et al. 2020); Decay sugar-rich substrates (Põlme et al. 2020) | Põlme S, et al. 2020. Fungal Diversity 105:1-16 ((https://doi.org/10.1007/s13225-020-00466-2)) |
| Otu18320 | 1 | k__Fungi(100);p__Ascomycota(100);c__Saccharomycetes(100);o__Saccharomycetales(100);f__Metschnikowiaceae(100);g__Metschnikowia(100);s__Metschnikowia_pulcherrima(100); | Metschnikowia | 13 | Saprotroph | \|Undefined Saprotroph\| | Yeast | NULL | Probable | As nectar-tap_saprotroph (Põlme et al. 2020); Decay sugar-rich substrates (Põlme et al. 2020) | Põlme S, et al. 2020. Fungal Diversity 105:1-16 ((https://doi.org/10.1007/s13225-020-00466-2)) |
| Otu18321 | 1 | k__Fungi(100);p__Ascomycota(100);c__Saccharomycetes(100);o__Saccharomycetales(100);f__Metschnikowiaceae(100);g__Metschnikowia(100);g__Metschnikowia_unclassified(100); | Metschnikowia | 13 | Saprotroph | \|Undefined Saprotroph\| | Yeast | NULL | Probable | As nectar-tap_saprotroph (Põlme et al. 2020); Decay sugar-rich substrates (Põlme et al. 2020) | Põlme S, et al. 2020. Fungal Diversity 105:1-16 ((https://doi.org/10.1007/s13225-020-00466-2)) |
| Otu18322 | 1 | k__Fungi(100);p__Ascomycota(100);c__Saccharomycetes(100);o__Saccharomycetales(100);f__Metschnikowiaceae(100);g__Metschnikowia(100);g__Metschnikowia_unclassified(100); | Metschnikowia | 13 | Saprotroph | \|Undefined Saprotroph\| | Yeast | NULL | Probable | As nectar-tap_saprotroph (Põlme et al. 2020); Decay sugar-rich substrates (Põlme et al. 2020) | Põlme S, et al. 2020. Fungal Diversity 105:1-16 ((https://doi.org/10.1007/s13225-020-00466-2)) |
| Otu18323 | 1 | k__Fungi(100);p__Ascomycota(100);c__Saccharomycetes(100);o__Saccharomycetales(100);f__Metschnikowiaceae(100);g__Metschnikowia(100);s__Metschnikowia_pulcherrima(100); | Metschnikowia | 13 | Saprotroph | \|Undefined Saprotroph\| | Yeast | NULL | Probable | As nectar-tap_saprotroph (Põlme et al. 2020); Decay sugar-rich substrates (Põlme et al. 2020) | Põlme S, et al. 2020. Fungal Diversity 105:1-16 ((https://doi.org/10.1007/s13225-020-00466-2)) |
| Otu18324 | 1 | k__Fungi(100);p__Ascomycota(100);c__Saccharomycetes(100);o__Saccharomycetales(100);f__Metschnikowiaceae(100);g__Metschnikowia(100);s__Metschnikowia_pulcherrima(100); | Metschnikowia | 13 | Saprotroph | \|Undefined Saprotroph\| | Yeast | NULL | Probable | As nectar-tap_saprotroph (Põlme et al. 2020); Decay sugar-rich substrates (Põlme et al. 2020) | Põlme S, et al. 2020. Fungal Diversity 105:1-16 ((https://doi.org/10.1007/s13225-020-00466-2)) |
| Otu18325 | 1 | k__Fungi(100);p__Ascomycota(100);c__Saccharomycetes(100);o__Saccharomycetales(100);f__Metschnikowiaceae(100);g__Metschnikowia(100);s__Metschnikowia_pulcherrima(100); | Metschnikowia | 13 | Saprotroph | \|Undefined Saprotroph\| | Yeast | NULL | Probable | As nectar-tap_saprotroph (Põlme et al. 2020); Decay sugar-rich substrates (Põlme et al. 2020) | Põlme S, et al. 2020. Fungal Diversity 105:1-16 ((https://doi.org/10.1007/s13225-020-00466-2)) |
| Otu18326 | 1 | k__Fungi(100);p__Ascomycota(100);c__Saccharomycetes(100);o__Saccharomycetales(100);f__Metschnikowiaceae(100);g__Metschnikowia(100);g__Metschnikowia_unclassified(100); | Metschnikowia | 13 | Saprotroph | \|Undefined Saprotroph\| | Yeast | NULL | Probable | As nectar-tap_saprotroph (Põlme et al. 2020); Decay sugar-rich substrates (Põlme et al. 2020) | Põlme S, et al. 2020. Fungal Diversity 105:1-16 ((https://doi.org/10.1007/s13225-020-00466-2)) |
| Otu18327 | 1 | k__Fungi(100);p__Ascomycota(100);c__Saccharomycetes(100);o__Saccharomycetales(100);f__Metschnikowiaceae(100);g__Metschnikowia(100);s__Metschnikowia_pulcherrima(100); | Metschnikowia | 13 | Saprotroph | \|Undefined Saprotroph\| | Yeast | NULL | Probable | As nectar-tap_saprotroph (Põlme et al. 2020); Decay sugar-rich substrates (Põlme et al. 2020) | Põlme S, et al. 2020. Fungal Diversity 105:1-16 ((https://doi.org/10.1007/s13225-020-00466-2)) |
| Otu18328 | 1 | k__Fungi(100);p__Ascomycota(100);c__Saccharomycetes(100);o__Saccharomycetales(100);f__Metschnikowiaceae(100);g__Metschnikowia(100);s__Metschnikowia_pulcherrima(100); | Metschnikowia | 13 | Saprotroph | \|Undefined Saprotroph\| | Yeast | NULL | Probable | As nectar-tap_saprotroph (Põlme et al. 2020); Decay sugar-rich substrates (Põlme et al. 2020) | Põlme S, et al. 2020. Fungal Diversity 105:1-16 ((https://doi.org/10.1007/s13225-020-00466-2)) |
| Otu18329 | 1 | k__Fungi(100);p__Ascomycota(100);c__Saccharomycetes(100);o__Saccharomycetales(100);f__Metschnikowiaceae(100);g__Metschnikowia(100);s__Metschnikowia_pulcherrima(100); | Metschnikowia | 13 | Saprotroph | \|Undefined Saprotroph\| | Yeast | NULL | Probable | As nectar-tap_saprotroph (Põlme et al. 2020); Decay sugar-rich substrates (Põlme et al. 2020) | Põlme S, et al. 2020. Fungal Diversity 105:1-16 ((https://doi.org/10.1007/s13225-020-00466-2)) |
| Otu18330 | 1 | k__Fungi(100);p__Ascomycota(100);c__Saccharomycetes(100);o__Saccharomycetales(100);f__Metschnikowiaceae(100);g__Metschnikowia(100);g__Metschnikowia_unclassified(100); | Metschnikowia | 13 | Saprotroph | \|Undefined Saprotroph\| | Yeast | NULL | Probable | As nectar-tap_saprotroph (Põlme et al. 2020); Decay sugar-rich substrates (Põlme et al. 2020) | Põlme S, et al. 2020. Fungal Diversity 105:1-16 ((https://doi.org/10.1007/s13225-020-00466-2)) |
| Otu18331 | 1 | k__Fungi(100);p__Ascomycota(100);c__Saccharomycetes(100);o__Saccharomycetales(100);f__Metschnikowiaceae(100);g__Metschnikowia(100);s__Metschnikowia_pulcherrima(100); | Metschnikowia | 13 | Saprotroph | \|Undefined Saprotroph\| | Yeast | NULL | Probable | As nectar-tap_saprotroph (Põlme et al. 2020); Decay sugar-rich substrates (Põlme et al. 2020) | Põlme S, et al. 2020. Fungal Diversity 105:1-16 ((https://doi.org/10.1007/s13225-020-00466-2)) |
| Otu18333 | 1 | k__Fungi(100);p__Ascomycota(100);c__Saccharomycetes(100);o__Saccharomycetales(100);f__Metschnikowiaceae(100);g__Metschnikowia(100);s__Metschnikowia_pulcherrima(100); | Metschnikowia | 13 | Saprotroph | \|Undefined Saprotroph\| | Yeast | NULL | Probable | As nectar-tap_saprotroph (Põlme et al. 2020); Decay sugar-rich substrates (Põlme et al. 2020) | Põlme S, et al. 2020. Fungal Diversity 105:1-16 ((https://doi.org/10.1007/s13225-020-00466-2)) |
| Otu18334 | 1 | k__Fungi(100);p__Ascomycota(100);c__Saccharomycetes(100);o__Saccharomycetales(100);f__Metschnikowiaceae(100);g__Metschnikowia(100);g__Metschnikowia_unclassified(100); | Metschnikowia | 13 | Saprotroph | \|Undefined Saprotroph\| | Yeast | NULL | Probable | As nectar-tap_saprotroph (Põlme et al. 2020); Decay sugar-rich substrates (Põlme et al. 2020) | Põlme S, et al. 2020. Fungal Diversity 105:1-16 ((https://doi.org/10.1007/s13225-020-00466-2)) |
| Otu18335 | 1 | k__Fungi(100);p__Ascomycota(100);c__Saccharomycetes(100);o__Saccharomycetales(100);f__Metschnikowiaceae(100);g__Metschnikowia(100);g__Metschnikowia_unclassified(100); | Metschnikowia | 13 | Saprotroph | \|Undefined Saprotroph\| | Yeast | NULL | Probable | As nectar-tap_saprotroph (Põlme et al. 2020); Decay sugar-rich substrates (Põlme et al. 2020) | Põlme S, et al. 2020. Fungal Diversity 105:1-16 ((https://doi.org/10.1007/s13225-020-00466-2)) |
| Otu18336 | 1 | k__Fungi(100);p__Ascomycota(100);c__Saccharomycetes(100);o__Saccharomycetales(100);f__Metschnikowiaceae(100);g__Metschnikowia(100);s__Metschnikowia_pulcherrima(100); | Metschnikowia | 13 | Saprotroph | \|Undefined Saprotroph\| | Yeast | NULL | Probable | As nectar-tap_saprotroph (Põlme et al. 2020); Decay sugar-rich substrates (Põlme et al. 2020) | Põlme S, et al. 2020. Fungal Diversity 105:1-16 ((https://doi.org/10.1007/s13225-020-00466-2)) |
| Otu18337 | 1 | k__Fungi(100);p__Ascomycota(100);c__Saccharomycetes(100);o__Saccharomycetales(100);f__Metschnikowiaceae(100);g__Metschnikowia(100);s__Metschnikowia_pulcherrima(100); | Metschnikowia | 13 | Saprotroph | \|Undefined Saprotroph\| | Yeast | NULL | Probable | As nectar-tap_saprotroph (Põlme et al. 2020); Decay sugar-rich substrates (Põlme et al. 2020) | Põlme S, et al. 2020. Fungal Diversity 105:1-16 ((https://doi.org/10.1007/s13225-020-00466-2)) |
| Otu18338 | 1 | k__Fungi(100);p__Ascomycota(100);c__Saccharomycetes(100);o__Saccharomycetales(100);f__Metschnikowiaceae(100);g__Metschnikowia(100);s__Metschnikowia_pulcherrima(100); | Metschnikowia | 13 | Saprotroph | \|Undefined Saprotroph\| | Yeast | NULL | Probable | As nectar-tap_saprotroph (Põlme et al. 2020); Decay sugar-rich substrates (Põlme et al. 2020) | Põlme S, et al. 2020. Fungal Diversity 105:1-16 ((https://doi.org/10.1007/s13225-020-00466-2)) |
| Otu18339 | 1 | k__Fungi(100);p__Ascomycota(100);c__Saccharomycetes(100);o__Saccharomycetales(100);f__Metschnikowiaceae(100);g__Metschnikowia(100);s__Metschnikowia_pulcherrima(100); | Metschnikowia | 13 | Saprotroph | \|Undefined Saprotroph\| | Yeast | NULL | Probable | As nectar-tap_saprotroph (Põlme et al. 2020); Decay sugar-rich substrates (Põlme et al. 2020) | Põlme S, et al. 2020. Fungal Diversity 105:1-16 ((https://doi.org/10.1007/s13225-020-00466-2)) |
| Otu18340 | 1 | k__Fungi(100);p__Ascomycota(100);c__Saccharomycetes(100);o__Saccharomycetales(100);f__Metschnikowiaceae(100);g__Metschnikowia(100);s__Metschnikowia_pulcherrima(100); | Metschnikowia | 13 | Saprotroph | \|Undefined Saprotroph\| | Yeast | NULL | Probable | As nectar-tap_saprotroph (Põlme et al. 2020); Decay sugar-rich substrates (Põlme et al. 2020) | Põlme S, et al. 2020. Fungal Diversity 105:1-16 ((https://doi.org/10.1007/s13225-020-00466-2)) |
| Otu18341 | 1 | k__Fungi(100);p__Ascomycota(100);c__Saccharomycetes(100);o__Saccharomycetales(100);f__Metschnikowiaceae(100);g__Metschnikowia(100);g__Metschnikowia_unclassified(100); | Metschnikowia | 13 | Saprotroph | \|Undefined Saprotroph\| | Yeast | NULL | Probable | As nectar-tap_saprotroph (Põlme et al. 2020); Decay sugar-rich substrates (Põlme et al. 2020) | Põlme S, et al. 2020. Fungal Diversity 105:1-16 ((https://doi.org/10.1007/s13225-020-00466-2)) |
| Otu18342 | 1 | k__Fungi(100);p__Ascomycota(100);c__Saccharomycetes(100);o__Saccharomycetales(100);f__Metschnikowiaceae(100);g__Metschnikowia(100);s__Metschnikowia_pulcherrima(100); | Metschnikowia | 13 | Saprotroph | \|Undefined Saprotroph\| | Yeast | NULL | Probable | As nectar-tap_saprotroph (Põlme et al. 2020); Decay sugar-rich substrates (Põlme et al. 2020) | Põlme S, et al. 2020. Fungal Diversity 105:1-16 ((https://doi.org/10.1007/s13225-020-00466-2)) |
| Otu18343 | 1 | k__Fungi(100);p__Ascomycota(100);c__Saccharomycetes(100);o__Saccharomycetales(100);f__Metschnikowiaceae(100);g__Metschnikowia(100);s__Metschnikowia_pulcherrima(100); | Metschnikowia | 13 | Saprotroph | \|Undefined Saprotroph\| | Yeast | NULL | Probable | As nectar-tap_saprotroph (Põlme et al. 2020); Decay sugar-rich substrates (Põlme et al. 2020) | Põlme S, et al. 2020. Fungal Diversity 105:1-16 ((https://doi.org/10.1007/s13225-020-00466-2)) |
| Otu18345 | 1 | k__Fungi(100);p__Ascomycota(100);c__Saccharomycetes(100);o__Saccharomycetales(100);f__Metschnikowiaceae(100);g__Metschnikowia(100);s__Metschnikowia_pulcherrima(100); | Metschnikowia | 13 | Saprotroph | \|Undefined Saprotroph\| | Yeast | NULL | Probable | As nectar-tap_saprotroph (Põlme et al. 2020); Decay sugar-rich substrates (Põlme et al. 2020) | Põlme S, et al. 2020. Fungal Diversity 105:1-16 ((https://doi.org/10.1007/s13225-020-00466-2)) |
| Otu18346 | 1 | k__Fungi(100);p__Ascomycota(100);c__Saccharomycetes(100);o__Saccharomycetales(100);f__Metschnikowiaceae(100);g__Metschnikowia(100);s__Metschnikowia_pulcherrima(100); | Metschnikowia | 13 | Saprotroph | \|Undefined Saprotroph\| | Yeast | NULL | Probable | As nectar-tap_saprotroph (Põlme et al. 2020); Decay sugar-rich substrates (Põlme et al. 2020) | Põlme S, et al. 2020. Fungal Diversity 105:1-16 ((https://doi.org/10.1007/s13225-020-00466-2)) |
| Otu18347 | 1 | k__Fungi(100);p__Ascomycota(100);c__Saccharomycetes(100);o__Saccharomycetales(100);f__Metschnikowiaceae(100);g__Metschnikowia(100);s__Metschnikowia_pulcherrima(100); | Metschnikowia | 13 | Saprotroph | \|Undefined Saprotroph\| | Yeast | NULL | Probable | As nectar-tap_saprotroph (Põlme et al. 2020); Decay sugar-rich substrates (Põlme et al. 2020) | Põlme S, et al. 2020. Fungal Diversity 105:1-16 ((https://doi.org/10.1007/s13225-020-00466-2)) |
| Otu18348 | 1 | k__Fungi(100);p__Ascomycota(100);c__Saccharomycetes(100);o__Saccharomycetales(100);f__Metschnikowiaceae(100);g__Metschnikowia(100);g__Metschnikowia_unclassified(100); | Metschnikowia | 13 | Saprotroph | \|Undefined Saprotroph\| | Yeast | NULL | Probable | As nectar-tap_saprotroph (Põlme et al. 2020); Decay sugar-rich substrates (Põlme et al. 2020) | Põlme S, et al. 2020. Fungal Diversity 105:1-16 ((https://doi.org/10.1007/s13225-020-00466-2)) |
| Otu18349 | 1 | k__Fungi(100);p__Ascomycota(100);c__Saccharomycetes(100);o__Saccharomycetales(100);f__Metschnikowiaceae(100);g__Metschnikowia(100);s__Metschnikowia_pulcherrima(100); | Metschnikowia | 13 | Saprotroph | \|Undefined Saprotroph\| | Yeast | NULL | Probable | As nectar-tap_saprotroph (Põlme et al. 2020); Decay sugar-rich substrates (Põlme et al. 2020) | Põlme S, et al. 2020. Fungal Diversity 105:1-16 ((https://doi.org/10.1007/s13225-020-00466-2)) |
| Otu18350 | 1 | k__Fungi(100);p__Ascomycota(100);c__Saccharomycetes(100);o__Saccharomycetales(100);f__Metschnikowiaceae(100);g__Metschnikowia(100);g__Metschnikowia_unclassified(100); | Metschnikowia | 13 | Saprotroph | \|Undefined Saprotroph\| | Yeast | NULL | Probable | As nectar-tap_saprotroph (Põlme et al. 2020); Decay sugar-rich substrates (Põlme et al. 2020) | Põlme S, et al. 2020. Fungal Diversity 105:1-16 ((https://doi.org/10.1007/s13225-020-00466-2)) |
| Otu18351 | 1 | k__Fungi(100);p__Ascomycota(100);c__Saccharomycetes(100);o__Saccharomycetales(100);f__Metschnikowiaceae(100);g__Metschnikowia(100);s__Metschnikowia_pulcherrima(100); | Metschnikowia | 13 | Saprotroph | \|Undefined Saprotroph\| | Yeast | NULL | Probable | As nectar-tap_saprotroph (Põlme et al. 2020); Decay sugar-rich substrates (Põlme et al. 2020) | Põlme S, et al. 2020. Fungal Diversity 105:1-16 ((https://doi.org/10.1007/s13225-020-00466-2)) |
| Otu18352 | 1 | k__Fungi(100);p__Ascomycota(100);c__Saccharomycetes(100);o__Saccharomycetales(100);f__Metschnikowiaceae(100);g__Metschnikowia(100);s__Metschnikowia_pulcherrima(100); | Metschnikowia | 13 | Saprotroph | \|Undefined Saprotroph\| | Yeast | NULL | Probable | As nectar-tap_saprotroph (Põlme et al. 2020); Decay sugar-rich substrates (Põlme et al. 2020) | Põlme S, et al. 2020. Fungal Diversity 105:1-16 ((https://doi.org/10.1007/s13225-020-00466-2)) |
| Otu18353 | 1 | k__Fungi(100);p__Ascomycota(100);c__Saccharomycetes(100);o__Saccharomycetales(100);f__Metschnikowiaceae(100);g__Metschnikowia(100);s__Metschnikowia_pulcherrima(100); | Metschnikowia | 13 | Saprotroph | \|Undefined Saprotroph\| | Yeast | NULL | Probable | As nectar-tap_saprotroph (Põlme et al. 2020); Decay sugar-rich substrates (Põlme et al. 2020) | Põlme S, et al. 2020. Fungal Diversity 105:1-16 ((https://doi.org/10.1007/s13225-020-00466-2)) |
| Otu18354 | 1 | k__Fungi(100);p__Ascomycota(100);c__Saccharomycetes(100);o__Saccharomycetales(100);f__Metschnikowiaceae(100);g__Metschnikowia(100);g__Metschnikowia_unclassified(100); | Metschnikowia | 13 | Saprotroph | \|Undefined Saprotroph\| | Yeast | NULL | Probable | As nectar-tap_saprotroph (Põlme et al. 2020); Decay sugar-rich substrates (Põlme et al. 2020) | Põlme S, et al. 2020. Fungal Diversity 105:1-16 ((https://doi.org/10.1007/s13225-020-00466-2)) |
| Otu18355 | 1 | k__Fungi(100);p__Ascomycota(100);c__Saccharomycetes(100);o__Saccharomycetales(100);f__Metschnikowiaceae(100);g__Metschnikowia(100);s__Metschnikowia_pulcherrima(100); | Metschnikowia | 13 | Saprotroph | \|Undefined Saprotroph\| | Yeast | NULL | Probable | As nectar-tap_saprotroph (Põlme et al. 2020); Decay sugar-rich substrates (Põlme et al. 2020) | Põlme S, et al. 2020. Fungal Diversity 105:1-16 ((https://doi.org/10.1007/s13225-020-00466-2)) |
| Otu18356 | 1 | k__Fungi(100);p__Ascomycota(100);c__Saccharomycetes(100);o__Saccharomycetales(100);f__Metschnikowiaceae(100);g__Metschnikowia(100);s__Metschnikowia_pulcherrima(100); | Metschnikowia | 13 | Saprotroph | \|Undefined Saprotroph\| | Yeast | NULL | Probable | As nectar-tap_saprotroph (Põlme et al. 2020); Decay sugar-rich substrates (Põlme et al. 2020) | Põlme S, et al. 2020. Fungal Diversity 105:1-16 ((https://doi.org/10.1007/s13225-020-00466-2)) |
| Otu18357 | 1 | k__Fungi(100);p__Ascomycota(100);c__Saccharomycetes(100);o__Saccharomycetales(100);f__Metschnikowiaceae(100);g__Metschnikowia(100);g__Metschnikowia_unclassified(100); | Metschnikowia | 13 | Saprotroph | \|Undefined Saprotroph\| | Yeast | NULL | Probable | As nectar-tap_saprotroph (Põlme et al. 2020); Decay sugar-rich substrates (Põlme et al. 2020) | Põlme S, et al. 2020. Fungal Diversity 105:1-16 ((https://doi.org/10.1007/s13225-020-00466-2)) |
| Otu18359 | 1 | k__Fungi(100);p__Ascomycota(100);c__Saccharomycetes(100);o__Saccharomycetales(100);f__Metschnikowiaceae(100);g__Metschnikowia(100);s__Metschnikowia_pulcherrima(100); | Metschnikowia | 13 | Saprotroph | \|Undefined Saprotroph\| | Yeast | NULL | Probable | As nectar-tap_saprotroph (Põlme et al. 2020); Decay sugar-rich substrates (Põlme et al. 2020) | Põlme S, et al. 2020. Fungal Diversity 105:1-16 ((https://doi.org/10.1007/s13225-020-00466-2)) |
| Otu18360 | 1 | k__Fungi(100);p__Ascomycota(100);c__Saccharomycetes(100);o__Saccharomycetales(100);f__Metschnikowiaceae(100);g__Metschnikowia(100);g__Metschnikowia_unclassified(100); | Metschnikowia | 13 | Saprotroph | \|Undefined Saprotroph\| | Yeast | NULL | Probable | As nectar-tap_saprotroph (Põlme et al. 2020); Decay sugar-rich substrates (Põlme et al. 2020) | Põlme S, et al. 2020. Fungal Diversity 105:1-16 ((https://doi.org/10.1007/s13225-020-00466-2)) |
| Otu18361 | 1 | k__Fungi(100);p__Ascomycota(100);c__Saccharomycetes(100);o__Saccharomycetales(100);f__Metschnikowiaceae(100);g__Metschnikowia(100);s__Metschnikowia_pulcherrima(100); | Metschnikowia | 13 | Saprotroph | \|Undefined Saprotroph\| | Yeast | NULL | Probable | As nectar-tap_saprotroph (Põlme et al. 2020); Decay sugar-rich substrates (Põlme et al. 2020) | Põlme S, et al. 2020. Fungal Diversity 105:1-16 ((https://doi.org/10.1007/s13225-020-00466-2)) |
| Otu18362 | 1 | k__Fungi(100);p__Ascomycota(100);c__Saccharomycetes(100);o__Saccharomycetales(100);f__Metschnikowiaceae(100);g__Metschnikowia(100);s__Metschnikowia_pulcherrima(100); | Metschnikowia | 13 | Saprotroph | \|Undefined Saprotroph\| | Yeast | NULL | Probable | As nectar-tap_saprotroph (Põlme et al. 2020); Decay sugar-rich substrates (Põlme et al. 2020) | Põlme S, et al. 2020. Fungal Diversity 105:1-16 ((https://doi.org/10.1007/s13225-020-00466-2)) |
| Otu18363 | 1 | k__Fungi(100);p__Ascomycota(100);c__Saccharomycetes(100);o__Saccharomycetales(100);f__Metschnikowiaceae(100);g__Metschnikowia(100);s__Metschnikowia_pulcherrima(100); | Metschnikowia | 13 | Saprotroph | \|Undefined Saprotroph\| | Yeast | NULL | Probable | As nectar-tap_saprotroph (Põlme et al. 2020); Decay sugar-rich substrates (Põlme et al. 2020) | Põlme S, et al. 2020. Fungal Diversity 105:1-16 ((https://doi.org/10.1007/s13225-020-00466-2)) |
| Otu18364 | 1 | k__Fungi(100);p__Ascomycota(100);c__Saccharomycetes(100);o__Saccharomycetales(100);f__Metschnikowiaceae(100);g__Metschnikowia(100);g__Metschnikowia_unclassified(100); | Metschnikowia | 13 | Saprotroph | \|Undefined Saprotroph\| | Yeast | NULL | Probable | As nectar-tap_saprotroph (Põlme et al. 2020); Decay sugar-rich substrates (Põlme et al. 2020) | Põlme S, et al. 2020. Fungal Diversity 105:1-16 ((https://doi.org/10.1007/s13225-020-00466-2)) |
| Otu18365 | 1 | k__Fungi(100);p__Ascomycota(100);c__Saccharomycetes(100);o__Saccharomycetales(100);f__Metschnikowiaceae(100);g__Metschnikowia(100);g__Metschnikowia_unclassified(100); | Metschnikowia | 13 | Saprotroph | \|Undefined Saprotroph\| | Yeast | NULL | Probable | As nectar-tap_saprotroph (Põlme et al. 2020); Decay sugar-rich substrates (Põlme et al. 2020) | Põlme S, et al. 2020. Fungal Diversity 105:1-16 ((https://doi.org/10.1007/s13225-020-00466-2)) |
| Otu18366 | 1 | k__Fungi(100);p__Ascomycota(100);c__Saccharomycetes(100);o__Saccharomycetales(100);f__Metschnikowiaceae(100);g__Metschnikowia(100);s__Metschnikowia_pulcherrima(100); | Metschnikowia | 13 | Saprotroph | \|Undefined Saprotroph\| | Yeast | NULL | Probable | As nectar-tap_saprotroph (Põlme et al. 2020); Decay sugar-rich substrates (Põlme et al. 2020) | Põlme S, et al. 2020. Fungal Diversity 105:1-16 ((https://doi.org/10.1007/s13225-020-00466-2)) |
| Otu18367 | 1 | k__Fungi(100);p__Ascomycota(100);c__Saccharomycetes(100);o__Saccharomycetales(100);f__Metschnikowiaceae(100);g__Metschnikowia(100);s__Metschnikowia_pulcherrima(100); | Metschnikowia | 13 | Saprotroph | \|Undefined Saprotroph\| | Yeast | NULL | Probable | As nectar-tap_saprotroph (Põlme et al. 2020); Decay sugar-rich substrates (Põlme et al. 2020) | Põlme S, et al. 2020. Fungal Diversity 105:1-16 ((https://doi.org/10.1007/s13225-020-00466-2)) |
| Otu18368 | 1 | k__Fungi(100);p__Ascomycota(100);c__Saccharomycetes(100);o__Saccharomycetales(100);f__Metschnikowiaceae(100);g__Metschnikowia(100);s__Metschnikowia_pulcherrima(100); | Metschnikowia | 13 | Saprotroph | \|Undefined Saprotroph\| | Yeast | NULL | Probable | As nectar-tap_saprotroph (Põlme et al. 2020); Decay sugar-rich substrates (Põlme et al. 2020) | Põlme S, et al. 2020. Fungal Diversity 105:1-16 ((https://doi.org/10.1007/s13225-020-00466-2)) |
| Otu18369 | 1 | k__Fungi(100);p__Ascomycota(100);c__Saccharomycetes(100);o__Saccharomycetales(100);f__Metschnikowiaceae(100);g__Metschnikowia(100);s__Metschnikowia_pulcherrima(100); | Metschnikowia | 13 | Saprotroph | \|Undefined Saprotroph\| | Yeast | NULL | Probable | As nectar-tap_saprotroph (Põlme et al. 2020); Decay sugar-rich substrates (Põlme et al. 2020) | Põlme S, et al. 2020. Fungal Diversity 105:1-16 ((https://doi.org/10.1007/s13225-020-00466-2)) |
| Otu18370 | 1 | k__Fungi(100);p__Ascomycota(100);c__Saccharomycetes(100);o__Saccharomycetales(100);f__Metschnikowiaceae(100);g__Metschnikowia(100);g__Metschnikowia_unclassified(100); | Metschnikowia | 13 | Saprotroph | \|Undefined Saprotroph\| | Yeast | NULL | Probable | As nectar-tap_saprotroph (Põlme et al. 2020); Decay sugar-rich substrates (Põlme et al. 2020) | Põlme S, et al. 2020. Fungal Diversity 105:1-16 ((https://doi.org/10.1007/s13225-020-00466-2)) |
| Otu18371 | 1 | k__Fungi(100);p__Ascomycota(100);c__Saccharomycetes(100);o__Saccharomycetales(100);f__Metschnikowiaceae(100);g__Metschnikowia(100);g__Metschnikowia_unclassified(100); | Metschnikowia | 13 | Saprotroph | \|Undefined Saprotroph\| | Yeast | NULL | Probable | As nectar-tap_saprotroph (Põlme et al. 2020); Decay sugar-rich substrates (Põlme et al. 2020) | Põlme S, et al. 2020. Fungal Diversity 105:1-16 ((https://doi.org/10.1007/s13225-020-00466-2)) |
| Otu18372 | 1 | k__Fungi(100);p__Ascomycota(100);c__Saccharomycetes(100);o__Saccharomycetales(100);f__Metschnikowiaceae(100);g__Metschnikowia(100);g__Metschnikowia_unclassified(100); | Metschnikowia | 13 | Saprotroph | \|Undefined Saprotroph\| | Yeast | NULL | Probable | As nectar-tap_saprotroph (Põlme et al. 2020); Decay sugar-rich substrates (Põlme et al. 2020) | Põlme S, et al. 2020. Fungal Diversity 105:1-16 ((https://doi.org/10.1007/s13225-020-00466-2)) |
| Otu18373 | 1 | k__Fungi(100);p__Ascomycota(100);c__Saccharomycetes(100);o__Saccharomycetales(100);f__Metschnikowiaceae(100);g__Metschnikowia(100);s__Metschnikowia_pulcherrima(100); | Metschnikowia | 13 | Saprotroph | \|Undefined Saprotroph\| | Yeast | NULL | Probable | As nectar-tap_saprotroph (Põlme et al. 2020); Decay sugar-rich substrates (Põlme et al. 2020) | Põlme S, et al. 2020. Fungal Diversity 105:1-16 ((https://doi.org/10.1007/s13225-020-00466-2)) |
| Otu18374 | 1 | k__Fungi(100);p__Ascomycota(100);c__Saccharomycetes(100);o__Saccharomycetales(100);f__Metschnikowiaceae(100);g__Metschnikowia(100);g__Metschnikowia_unclassified(100); | Metschnikowia | 13 | Saprotroph | \|Undefined Saprotroph\| | Yeast | NULL | Probable | As nectar-tap_saprotroph (Põlme et al. 2020); Decay sugar-rich substrates (Põlme et al. 2020) | Põlme S, et al. 2020. Fungal Diversity 105:1-16 ((https://doi.org/10.1007/s13225-020-00466-2)) |
| Otu18375 | 1 | k__Fungi(100);p__Ascomycota(100);c__Saccharomycetes(100);o__Saccharomycetales(100);f__Metschnikowiaceae(100);g__Metschnikowia(100);s__Metschnikowia_pulcherrima(100); | Metschnikowia | 13 | Saprotroph | \|Undefined Saprotroph\| | Yeast | NULL | Probable | As nectar-tap_saprotroph (Põlme et al. 2020); Decay sugar-rich substrates (Põlme et al. 2020) | Põlme S, et al. 2020. Fungal Diversity 105:1-16 ((https://doi.org/10.1007/s13225-020-00466-2)) |
| Otu18376 | 1 | k__Fungi(100);p__Ascomycota(100);c__Saccharomycetes(100);o__Saccharomycetales(100);f__Metschnikowiaceae(100);g__Metschnikowia(100);g__Metschnikowia_unclassified(100); | Metschnikowia | 13 | Saprotroph | \|Undefined Saprotroph\| | Yeast | NULL | Probable | As nectar-tap_saprotroph (Põlme et al. 2020); Decay sugar-rich substrates (Põlme et al. 2020) | Põlme S, et al. 2020. Fungal Diversity 105:1-16 ((https://doi.org/10.1007/s13225-020-00466-2)) |
| Otu18377 | 1 | k__Fungi(100);p__Ascomycota(100);c__Saccharomycetes(100);o__Saccharomycetales(100);f__Metschnikowiaceae(100);g__Metschnikowia(100);g__Metschnikowia_unclassified(100); | Metschnikowia | 13 | Saprotroph | \|Undefined Saprotroph\| | Yeast | NULL | Probable | As nectar-tap_saprotroph (Põlme et al. 2020); Decay sugar-rich substrates (Põlme et al. 2020) | Põlme S, et al. 2020. Fungal Diversity 105:1-16 ((https://doi.org/10.1007/s13225-020-00466-2)) |
| Otu18378 | 1 | k__Fungi(100);p__Ascomycota(100);c__Saccharomycetes(100);o__Saccharomycetales(100);f__Metschnikowiaceae(100);g__Metschnikowia(100);s__Metschnikowia_pulcherrima(100); | Metschnikowia | 13 | Saprotroph | \|Undefined Saprotroph\| | Yeast | NULL | Probable | As nectar-tap_saprotroph (Põlme et al. 2020); Decay sugar-rich substrates (Põlme et al. 2020) | Põlme S, et al. 2020. Fungal Diversity 105:1-16 ((https://doi.org/10.1007/s13225-020-00466-2)) |
| Otu18379 | 1 | k__Fungi(100);p__Ascomycota(100);c__Saccharomycetes(100);o__Saccharomycetales(100);f__Metschnikowiaceae(100);g__Metschnikowia(100);s__Metschnikowia_pulcherrima(100); | Metschnikowia | 13 | Saprotroph | \|Undefined Saprotroph\| | Yeast | NULL | Probable | As nectar-tap_saprotroph (Põlme et al. 2020); Decay sugar-rich substrates (Põlme et al. 2020) | Põlme S, et al. 2020. Fungal Diversity 105:1-16 ((https://doi.org/10.1007/s13225-020-00466-2)) |
| Otu18380 | 1 | k__Fungi(100);p__Ascomycota(100);c__Saccharomycetes(100);o__Saccharomycetales(100);f__Metschnikowiaceae(100);g__Metschnikowia(100);g__Metschnikowia_unclassified(100); | Metschnikowia | 13 | Saprotroph | \|Undefined Saprotroph\| | Yeast | NULL | Probable | As nectar-tap_saprotroph (Põlme et al. 2020); Decay sugar-rich substrates (Põlme et al. 2020) | Põlme S, et al. 2020. Fungal Diversity 105:1-16 ((https://doi.org/10.1007/s13225-020-00466-2)) |
| Otu18381 | 1 | k__Fungi(100);p__Ascomycota(100);c__Saccharomycetes(100);o__Saccharomycetales(100);f__Metschnikowiaceae(100);g__Metschnikowia(100);s__Metschnikowia_pulcherrima(100); | Metschnikowia | 13 | Saprotroph | \|Undefined Saprotroph\| | Yeast | NULL | Probable | As nectar-tap_saprotroph (Põlme et al. 2020); Decay sugar-rich substrates (Põlme et al. 2020) | Põlme S, et al. 2020. Fungal Diversity 105:1-16 ((https://doi.org/10.1007/s13225-020-00466-2)) |
| Otu18382 | 1 | k__Fungi(100);p__Ascomycota(100);c__Saccharomycetes(100);o__Saccharomycetales(100);f__Metschnikowiaceae(100);g__Metschnikowia(100);g__Metschnikowia_unclassified(100); | Metschnikowia | 13 | Saprotroph | \|Undefined Saprotroph\| | Yeast | NULL | Probable | As nectar-tap_saprotroph (Põlme et al. 2020); Decay sugar-rich substrates (Põlme et al. 2020) | Põlme S, et al. 2020. Fungal Diversity 105:1-16 ((https://doi.org/10.1007/s13225-020-00466-2)) |
| Otu18383 | 1 | k__Fungi(100);p__Ascomycota(100);c__Saccharomycetes(100);o__Saccharomycetales(100);f__Metschnikowiaceae(100);g__Metschnikowia(100);s__Metschnikowia_pulcherrima(100); | Metschnikowia | 13 | Saprotroph | \|Undefined Saprotroph\| | Yeast | NULL | Probable | As nectar-tap_saprotroph (Põlme et al. 2020); Decay sugar-rich substrates (Põlme et al. 2020) | Põlme S, et al. 2020. Fungal Diversity 105:1-16 ((https://doi.org/10.1007/s13225-020-00466-2)) |
| Otu18384 | 1 | k__Fungi(100);p__Ascomycota(100);c__Saccharomycetes(100);o__Saccharomycetales(100);f__Metschnikowiaceae(100);g__Metschnikowia(100);g__Metschnikowia_unclassified(100); | Metschnikowia | 13 | Saprotroph | \|Undefined Saprotroph\| | Yeast | NULL | Probable | As nectar-tap_saprotroph (Põlme et al. 2020); Decay sugar-rich substrates (Põlme et al. 2020) | Põlme S, et al. 2020. Fungal Diversity 105:1-16 ((https://doi.org/10.1007/s13225-020-00466-2)) |
| Otu18385 | 1 | k__Fungi(100);p__Ascomycota(100);c__Saccharomycetes(100);o__Saccharomycetales(100);f__Metschnikowiaceae(100);g__Metschnikowia(100);g__Metschnikowia_unclassified(100); | Metschnikowia | 13 | Saprotroph | \|Undefined Saprotroph\| | Yeast | NULL | Probable | As nectar-tap_saprotroph (Põlme et al. 2020); Decay sugar-rich substrates (Põlme et al. 2020) | Põlme S, et al. 2020. Fungal Diversity 105:1-16 ((https://doi.org/10.1007/s13225-020-00466-2)) |
| Otu18386 | 1 | k__Fungi(100);p__Ascomycota(100);c__Saccharomycetes(100);o__Saccharomycetales(100);f__Metschnikowiaceae(100);g__Metschnikowia(100);g__Metschnikowia_unclassified(100); | Metschnikowia | 13 | Saprotroph | \|Undefined Saprotroph\| | Yeast | NULL | Probable | As nectar-tap_saprotroph (Põlme et al. 2020); Decay sugar-rich substrates (Põlme et al. 2020) | Põlme S, et al. 2020. Fungal Diversity 105:1-16 ((https://doi.org/10.1007/s13225-020-00466-2)) |
| Otu18387 | 1 | k__Fungi(100);p__Ascomycota(100);c__Saccharomycetes(100);o__Saccharomycetales(100);f__Metschnikowiaceae(100);g__Metschnikowia(100);g__Metschnikowia_unclassified(100); | Metschnikowia | 13 | Saprotroph | \|Undefined Saprotroph\| | Yeast | NULL | Probable | As nectar-tap_saprotroph (Põlme et al. 2020); Decay sugar-rich substrates (Põlme et al. 2020) | Põlme S, et al. 2020. Fungal Diversity 105:1-16 ((https://doi.org/10.1007/s13225-020-00466-2)) |
| Otu18388 | 1 | k__Fungi(100);p__Ascomycota(100);c__Saccharomycetes(100);o__Saccharomycetales(100);f__Metschnikowiaceae(100);g__Metschnikowia(100);s__Metschnikowia_pulcherrima(100); | Metschnikowia | 13 | Saprotroph | \|Undefined Saprotroph\| | Yeast | NULL | Probable | As nectar-tap_saprotroph (Põlme et al. 2020); Decay sugar-rich substrates (Põlme et al. 2020) | Põlme S, et al. 2020. Fungal Diversity 105:1-16 ((https://doi.org/10.1007/s13225-020-00466-2)) |
| Otu18389 | 1 | k__Fungi(100);p__Ascomycota(100);c__Saccharomycetes(100);o__Saccharomycetales(100);f__Metschnikowiaceae(100);g__Metschnikowia(100);s__Metschnikowia_pulcherrima(100); | Metschnikowia | 13 | Saprotroph | \|Undefined Saprotroph\| | Yeast | NULL | Probable | As nectar-tap_saprotroph (Põlme et al. 2020); Decay sugar-rich substrates (Põlme et al. 2020) | Põlme S, et al. 2020. Fungal Diversity 105:1-16 ((https://doi.org/10.1007/s13225-020-00466-2)) |
| Otu18390 | 1 | k__Fungi(100);p__Ascomycota(100);c__Saccharomycetes(100);o__Saccharomycetales(100);f__Metschnikowiaceae(100);g__Metschnikowia(100);s__Metschnikowia_pulcherrima(100); | Metschnikowia | 13 | Saprotroph | \|Undefined Saprotroph\| | Yeast | NULL | Probable | As nectar-tap_saprotroph (Põlme et al. 2020); Decay sugar-rich substrates (Põlme et al. 2020) | Põlme S, et al. 2020. Fungal Diversity 105:1-16 ((https://doi.org/10.1007/s13225-020-00466-2)) |
| Otu18391 | 1 | k__Fungi(100);p__Ascomycota(100);c__Saccharomycetes(100);o__Saccharomycetales(100);f__Metschnikowiaceae(100);g__Metschnikowia(100);s__Metschnikowia_pulcherrima(100); | Metschnikowia | 13 | Saprotroph | \|Undefined Saprotroph\| | Yeast | NULL | Probable | As nectar-tap_saprotroph (Põlme et al. 2020); Decay sugar-rich substrates (Põlme et al. 2020) | Põlme S, et al. 2020. Fungal Diversity 105:1-16 ((https://doi.org/10.1007/s13225-020-00466-2)) |
| Otu18392 | 1 | k__Fungi(100);p__Ascomycota(100);c__Saccharomycetes(100);o__Saccharomycetales(100);f__Metschnikowiaceae(100);g__Metschnikowia(100);s__Metschnikowia_pulcherrima(100); | Metschnikowia | 13 | Saprotroph | \|Undefined Saprotroph\| | Yeast | NULL | Probable | As nectar-tap_saprotroph (Põlme et al. 2020); Decay sugar-rich substrates (Põlme et al. 2020) | Põlme S, et al. 2020. Fungal Diversity 105:1-16 ((https://doi.org/10.1007/s13225-020-00466-2)) |
| Otu18393 | 1 | k__Fungi(100);p__Ascomycota(100);c__Saccharomycetes(100);o__Saccharomycetales(100);f__Metschnikowiaceae(100);g__Metschnikowia(100);s__Metschnikowia_pulcherrima(100); | Metschnikowia | 13 | Saprotroph | \|Undefined Saprotroph\| | Yeast | NULL | Probable | As nectar-tap_saprotroph (Põlme et al. 2020); Decay sugar-rich substrates (Põlme et al. 2020) | Põlme S, et al. 2020. Fungal Diversity 105:1-16 ((https://doi.org/10.1007/s13225-020-00466-2)) |
| Otu18394 | 1 | k__Fungi(100);p__Ascomycota(100);c__Saccharomycetes(100);o__Saccharomycetales(100);f__Metschnikowiaceae(100);g__Metschnikowia(100);s__Metschnikowia_pulcherrima(100); | Metschnikowia | 13 | Saprotroph | \|Undefined Saprotroph\| | Yeast | NULL | Probable | As nectar-tap_saprotroph (Põlme et al. 2020); Decay sugar-rich substrates (Põlme et al. 2020) | Põlme S, et al. 2020. Fungal Diversity 105:1-16 ((https://doi.org/10.1007/s13225-020-00466-2)) |
| Otu18395 | 1 | k__Fungi(100);p__Ascomycota(100);c__Saccharomycetes(100);o__Saccharomycetales(100);f__Metschnikowiaceae(100);g__Metschnikowia(100);s__Metschnikowia_pulcherrima(100); | Metschnikowia | 13 | Saprotroph | \|Undefined Saprotroph\| | Yeast | NULL | Probable | As nectar-tap_saprotroph (Põlme et al. 2020); Decay sugar-rich substrates (Põlme et al. 2020) | Põlme S, et al. 2020. Fungal Diversity 105:1-16 ((https://doi.org/10.1007/s13225-020-00466-2)) |
| Otu18396 | 1 | k__Fungi(100);p__Ascomycota(100);c__Saccharomycetes(100);o__Saccharomycetales(100);f__Metschnikowiaceae(100);g__Metschnikowia(100);s__Metschnikowia_pulcherrima(100); | Metschnikowia | 13 | Saprotroph | \|Undefined Saprotroph\| | Yeast | NULL | Probable | As nectar-tap_saprotroph (Põlme et al. 2020); Decay sugar-rich substrates (Põlme et al. 2020) | Põlme S, et al. 2020. Fungal Diversity 105:1-16 ((https://doi.org/10.1007/s13225-020-00466-2)) |
| Otu18397 | 1 | k__Fungi(100);p__Ascomycota(100);c__Saccharomycetes(100);o__Saccharomycetales(100);f__Metschnikowiaceae(100);g__Metschnikowia(100);s__Metschnikowia_pulcherrima(100); | Metschnikowia | 13 | Saprotroph | \|Undefined Saprotroph\| | Yeast | NULL | Probable | As nectar-tap_saprotroph (Põlme et al. 2020); Decay sugar-rich substrates (Põlme et al. 2020) | Põlme S, et al. 2020. Fungal Diversity 105:1-16 ((https://doi.org/10.1007/s13225-020-00466-2)) |
| Otu18401 | 1 | k__Fungi(100);p__Ascomycota(100);c__Saccharomycetes(100);o__Saccharomycetales(100);f__Metschnikowiaceae(100);g__Metschnikowia(100);s__Metschnikowia_pulcherrima(100); | Metschnikowia | 13 | Saprotroph | \|Undefined Saprotroph\| | Yeast | NULL | Probable | As nectar-tap_saprotroph (Põlme et al. 2020); Decay sugar-rich substrates (Põlme et al. 2020) | Põlme S, et al. 2020. Fungal Diversity 105:1-16 ((https://doi.org/10.1007/s13225-020-00466-2)) |
| Otu18402 | 1 | k__Fungi(100);p__Ascomycota(100);c__Saccharomycetes(100);o__Saccharomycetales(100);f__Metschnikowiaceae(100);g__Metschnikowia(100);g__Metschnikowia_unclassified(100); | Metschnikowia | 13 | Saprotroph | \|Undefined Saprotroph\| | Yeast | NULL | Probable | As nectar-tap_saprotroph (Põlme et al. 2020); Decay sugar-rich substrates (Põlme et al. 2020) | Põlme S, et al. 2020. Fungal Diversity 105:1-16 ((https://doi.org/10.1007/s13225-020-00466-2)) |
| Otu18403 | 1 | k__Fungi(100);p__Ascomycota(100);c__Saccharomycetes(100);o__Saccharomycetales(100);f__Metschnikowiaceae(100);g__Metschnikowia(100);g__Metschnikowia_unclassified(100); | Metschnikowia | 13 | Saprotroph | \|Undefined Saprotroph\| | Yeast | NULL | Probable | As nectar-tap_saprotroph (Põlme et al. 2020); Decay sugar-rich substrates (Põlme et al. 2020) | Põlme S, et al. 2020. Fungal Diversity 105:1-16 ((https://doi.org/10.1007/s13225-020-00466-2)) |
| Otu18404 | 1 | k__Fungi(100);p__Ascomycota(100);c__Saccharomycetes(100);o__Saccharomycetales(100);f__Metschnikowiaceae(100);g__Metschnikowia(100);s__Metschnikowia_pulcherrima(100); | Metschnikowia | 13 | Saprotroph | \|Undefined Saprotroph\| | Yeast | NULL | Probable | As nectar-tap_saprotroph (Põlme et al. 2020); Decay sugar-rich substrates (Põlme et al. 2020) | Põlme S, et al. 2020. Fungal Diversity 105:1-16 ((https://doi.org/10.1007/s13225-020-00466-2)) |
| Otu18405 | 1 | k__Fungi(100);p__Ascomycota(100);c__Saccharomycetes(100);o__Saccharomycetales(100);f__Metschnikowiaceae(100);g__Metschnikowia(100);s__Metschnikowia_pulcherrima(100); | Metschnikowia | 13 | Saprotroph | \|Undefined Saprotroph\| | Yeast | NULL | Probable | As nectar-tap_saprotroph (Põlme et al. 2020); Decay sugar-rich substrates (Põlme et al. 2020) | Põlme S, et al. 2020. Fungal Diversity 105:1-16 ((https://doi.org/10.1007/s13225-020-00466-2)) |
| Otu18406 | 1 | k__Fungi(100);p__Ascomycota(100);c__Saccharomycetes(100);o__Saccharomycetales(100);f__Metschnikowiaceae(100);g__Metschnikowia(100);s__Metschnikowia_pulcherrima(100); | Metschnikowia | 13 | Saprotroph | \|Undefined Saprotroph\| | Yeast | NULL | Probable | As nectar-tap_saprotroph (Põlme et al. 2020); Decay sugar-rich substrates (Põlme et al. 2020) | Põlme S, et al. 2020. Fungal Diversity 105:1-16 ((https://doi.org/10.1007/s13225-020-00466-2)) |
| Otu18407 | 1 | k__Fungi(100);p__Ascomycota(100);c__Saccharomycetes(100);o__Saccharomycetales(100);f__Metschnikowiaceae(100);g__Metschnikowia(100);g__Metschnikowia_unclassified(100); | Metschnikowia | 13 | Saprotroph | \|Undefined Saprotroph\| | Yeast | NULL | Probable | As nectar-tap_saprotroph (Põlme et al. 2020); Decay sugar-rich substrates (Põlme et al. 2020) | Põlme S, et al. 2020. Fungal Diversity 105:1-16 ((https://doi.org/10.1007/s13225-020-00466-2)) |
| Otu18408 | 1 | k__Fungi(100);p__Ascomycota(100);c__Saccharomycetes(100);o__Saccharomycetales(100);f__Metschnikowiaceae(100);g__Metschnikowia(100);s__Metschnikowia_pulcherrima(100); | Metschnikowia | 13 | Saprotroph | \|Undefined Saprotroph\| | Yeast | NULL | Probable | As nectar-tap_saprotroph (Põlme et al. 2020); Decay sugar-rich substrates (Põlme et al. 2020) | Põlme S, et al. 2020. Fungal Diversity 105:1-16 ((https://doi.org/10.1007/s13225-020-00466-2)) |
| Otu18409 | 1 | k__Fungi(100);p__Ascomycota(100);c__Saccharomycetes(100);o__Saccharomycetales(100);f__Metschnikowiaceae(100);g__Metschnikowia(100);s__Metschnikowia_pulcherrima(100); | Metschnikowia | 13 | Saprotroph | \|Undefined Saprotroph\| | Yeast | NULL | Probable | As nectar-tap_saprotroph (Põlme et al. 2020); Decay sugar-rich substrates (Põlme et al. 2020) | Põlme S, et al. 2020. Fungal Diversity 105:1-16 ((https://doi.org/10.1007/s13225-020-00466-2)) |
| Otu18410 | 1 | k__Fungi(100);p__Ascomycota(100);c__Saccharomycetes(100);o__Saccharomycetales(100);f__Metschnikowiaceae(100);g__Metschnikowia(100);s__Metschnikowia_pulcherrima(100); | Metschnikowia | 13 | Saprotroph | \|Undefined Saprotroph\| | Yeast | NULL | Probable | As nectar-tap_saprotroph (Põlme et al. 2020); Decay sugar-rich substrates (Põlme et al. 2020) | Põlme S, et al. 2020. Fungal Diversity 105:1-16 ((https://doi.org/10.1007/s13225-020-00466-2)) |
| Otu18411 | 1 | k__Fungi(100);p__Ascomycota(100);c__Saccharomycetes(100);o__Saccharomycetales(100);f__Metschnikowiaceae(100);g__Metschnikowia(100);s__Metschnikowia_pulcherrima(100); | Metschnikowia | 13 | Saprotroph | \|Undefined Saprotroph\| | Yeast | NULL | Probable | As nectar-tap_saprotroph (Põlme et al. 2020); Decay sugar-rich substrates (Põlme et al. 2020) | Põlme S, et al. 2020. Fungal Diversity 105:1-16 ((https://doi.org/10.1007/s13225-020-00466-2)) |
| Otu18412 | 1 | k__Fungi(100);p__Ascomycota(100);c__Saccharomycetes(100);o__Saccharomycetales(100);f__Metschnikowiaceae(100);g__Metschnikowia(100);g__Metschnikowia_unclassified(100); | Metschnikowia | 13 | Saprotroph | \|Undefined Saprotroph\| | Yeast | NULL | Probable | As nectar-tap_saprotroph (Põlme et al. 2020); Decay sugar-rich substrates (Põlme et al. 2020) | Põlme S, et al. 2020. Fungal Diversity 105:1-16 ((https://doi.org/10.1007/s13225-020-00466-2)) |
| Otu18413 | 1 | k__Fungi(100);p__Ascomycota(100);c__Saccharomycetes(100);o__Saccharomycetales(100);f__Metschnikowiaceae(100);g__Metschnikowia(100);g__Metschnikowia_unclassified(100); | Metschnikowia | 13 | Saprotroph | \|Undefined Saprotroph\| | Yeast | NULL | Probable | As nectar-tap_saprotroph (Põlme et al. 2020); Decay sugar-rich substrates (Põlme et al. 2020) | Põlme S, et al. 2020. Fungal Diversity 105:1-16 ((https://doi.org/10.1007/s13225-020-00466-2)) |
| Otu18414 | 1 | k__Fungi(100);p__Ascomycota(100);c__Saccharomycetes(100);o__Saccharomycetales(100);f__Metschnikowiaceae(100);g__Metschnikowia(100);s__Metschnikowia_pulcherrima(100); | Metschnikowia | 13 | Saprotroph | \|Undefined Saprotroph\| | Yeast | NULL | Probable | As nectar-tap_saprotroph (Põlme et al. 2020); Decay sugar-rich substrates (Põlme et al. 2020) | Põlme S, et al. 2020. Fungal Diversity 105:1-16 ((https://doi.org/10.1007/s13225-020-00466-2)) |
| Otu18415 | 1 | k__Fungi(100);p__Ascomycota(100);c__Saccharomycetes(100);o__Saccharomycetales(100);f__Metschnikowiaceae(100);g__Metschnikowia(100);s__Metschnikowia_pulcherrima(100); | Metschnikowia | 13 | Saprotroph | \|Undefined Saprotroph\| | Yeast | NULL | Probable | As nectar-tap_saprotroph (Põlme et al. 2020); Decay sugar-rich substrates (Põlme et al. 2020) | Põlme S, et al. 2020. Fungal Diversity 105:1-16 ((https://doi.org/10.1007/s13225-020-00466-2)) |
| Otu18416 | 1 | k__Fungi(100);p__Ascomycota(100);c__Saccharomycetes(100);o__Saccharomycetales(100);f__Metschnikowiaceae(100);g__Metschnikowia(100);s__Metschnikowia_pulcherrima(100); | Metschnikowia | 13 | Saprotroph | \|Undefined Saprotroph\| | Yeast | NULL | Probable | As nectar-tap_saprotroph (Põlme et al. 2020); Decay sugar-rich substrates (Põlme et al. 2020) | Põlme S, et al. 2020. Fungal Diversity 105:1-16 ((https://doi.org/10.1007/s13225-020-00466-2)) |
| Otu18417 | 1 | k__Fungi(100);p__Ascomycota(100);c__Saccharomycetes(100);o__Saccharomycetales(100);f__Metschnikowiaceae(100);g__Metschnikowia(100);s__Metschnikowia_pulcherrima(100); | Metschnikowia | 13 | Saprotroph | \|Undefined Saprotroph\| | Yeast | NULL | Probable | As nectar-tap_saprotroph (Põlme et al. 2020); Decay sugar-rich substrates (Põlme et al. 2020) | Põlme S, et al. 2020. Fungal Diversity 105:1-16 ((https://doi.org/10.1007/s13225-020-00466-2)) |

| | | | | | | | | | | | |
|---|---|---|---|---|---|---|---|---|---|---|---|
| Otu18418 | 1 | k__Fungi(100);p__Ascomycota(100);c__Saccharomycetes(100);o__Saccharomycetales(100);f__Metschnikowiaceae(100);g__Metschnikowia(100);s__Metschnikowia_pulcherrima(100); | Metschnikowia | 13 | Saprotroph | Undefined Saprotroph | Yeast | NULL | Probable | As nectar-tap_saprotroph (Põlme et al. 2020); Decay sugar-rich substrates (Põlme et al. 2020) | Põlme S, et al. 2020. Fungal Diversity 105:1-16 ((https://doi.org/10.1007/s13225-020-00466-2)) |
| Otu18419 | 1 | k__Fungi(100);p__Ascomycota(100);c__Saccharomycetes(100);o__Saccharomycetales(100);f__Metschnikowiaceae(100);g__Metschnikowia(100);s__Metschnikowia_pulcherrima(100); | Metschnikowia | 13 | Saprotroph | Undefined Saprotroph | Yeast | NULL | Probable | As nectar-tap_saprotroph (Põlme et al. 2020); Decay sugar-rich substrates (Põlme et al. 2020) | Põlme S, et al. 2020. Fungal Diversity 105:1-16 ((https://doi.org/10.1007/s13225-020-00466-2)) |
| Otu18420 | 1 | k__Fungi(100);p__Ascomycota(100);c__Saccharomycetes(100);o__Saccharomycetales(100);f__Metschnikowiaceae(100);g__Metschnikowia(100);s__Metschnikowia_pulcherrima(100); | Metschnikowia | 13 | Saprotroph | Undefined Saprotroph | Yeast | NULL | Probable | As nectar-tap_saprotroph (Põlme et al. 2020); Decay sugar-rich substrates (Põlme et al. 2020) | Põlme S, et al. 2020. Fungal Diversity 105:1-16 ((https://doi.org/10.1007/s13225-020-00466-2)) |
| Otu18421 | 1 | k__Fungi(100);p__Ascomycota(100);c__Saccharomycetes(100);o__Saccharomycetales(100);f__Metschnikowiaceae(100);g__Metschnikowia(100);s__Metschnikowia_pulcherrima(100); | Metschnikowia | 13 | Saprotroph | Undefined Saprotroph | Yeast | NULL | Probable | As nectar-tap_saprotroph (Põlme et al. 2020); Decay sugar-rich substrates (Põlme et al. 2020) | Põlme S, et al. 2020. Fungal Diversity 105:1-16 ((https://doi.org/10.1007/s13225-020-00466-2)) |
| Otu18422 | 1 | k__Fungi(100);p__Ascomycota(100);c__Saccharomycetes(100);o__Saccharomycetales(100);f__Metschnikowiaceae(100);g__Metschnikowia(100);g__Metschnikowia_unclassified(100); | Metschnikowia | 13 | Saprotroph | Undefined Saprotroph | Yeast | NULL | Probable | As nectar-tap_saprotroph (Põlme et al. 2020); Decay sugar-rich substrates (Põlme et al. 2020) | Põlme S, et al. 2020. Fungal Diversity 105:1-16 ((https://doi.org/10.1007/s13225-020-00466-2)) |
| Otu18424 | 1 | k__Fungi(100);p__Ascomycota(100);c__Saccharomycetes(100);o__Saccharomycetales(100);f__Metschnikowiaceae(100);g__Metschnikowia(100);s__Metschnikowia_pulcherrima(100); | Metschnikowia | 13 | Saprotroph | Undefined Saprotroph | Yeast | NULL | Probable | As nectar-tap_saprotroph (Põlme et al. 2020); Decay sugar-rich substrates (Põlme et al. 2020) | Põlme S, et al. 2020. Fungal Diversity 105:1-16 ((https://doi.org/10.1007/s13225-020-00466-2)) |
| Otu18425 | 1 | k__Fungi(100);p__Ascomycota(100);c__Saccharomycetes(100);o__Saccharomycetales(100);f__Metschnikowiaceae(100);g__Metschnikowia(100);s__Metschnikowia_pulcherrima(100); | Metschnikowia | 13 | Saprotroph | Undefined Saprotroph | Yeast | NULL | Probable | As nectar-tap_saprotroph (Põlme et al. 2020); Decay sugar-rich substrates (Põlme et al. 2020) | Põlme S, et al. 2020. Fungal Diversity 105:1-16 ((https://doi.org/10.1007/s13225-020-00466-2)) |
| Otu18426 | 1 | k__Fungi(100);p__Ascomycota(100);c__Saccharomycetes(100);o__Saccharomycetales(100);f__Metschnikowiaceae(100);g__Metschnikowia(100);s__Metschnikowia_pulcherrima(100); | Metschnikowia | 13 | Saprotroph | Undefined Saprotroph | Yeast | NULL | Probable | As nectar-tap_saprotroph (Põlme et al. 2020); Decay sugar-rich substrates (Põlme et al. 2020) | Põlme S, et al. 2020. Fungal Diversity 105:1-16 ((https://doi.org/10.1007/s13225-020-00466-2)) |
| Otu18427 | 1 | k__Fungi(100);p__Ascomycota(100);c__Saccharomycetes(100);o__Saccharomycetales(100);f__Metschnikowiaceae(100);g__Metschnikowia(100);s__Metschnikowia_pulcherrima(100); | Metschnikowia | 13 | Saprotroph | Undefined Saprotroph | Yeast | NULL | Probable | As nectar-tap_saprotroph (Põlme et al. 2020); Decay sugar-rich substrates (Põlme et al. 2020) | Põlme S, et al. 2020. Fungal Diversity 105:1-16 ((https://doi.org/10.1007/s13225-020-00466-2)) |
| Otu18428 | 1 | k__Fungi(100);p__Ascomycota(100);c__Saccharomycetes(100);o__Saccharomycetales(100);f__Metschnikowiaceae(100);g__Metschnikowia(100);s__Metschnikowia_pulcherrima(100); | Metschnikowia | 13 | Saprotroph | Undefined Saprotroph | Yeast | NULL | Probable | As nectar-tap_saprotroph (Põlme et al. 2020); Decay sugar-rich substrates (Põlme et al. 2020) | Põlme S, et al. 2020. Fungal Diversity 105:1-16 ((https://doi.org/10.1007/s13225-020-00466-2)) |
| Otu18429 | 1 | k__Fungi(100);p__Ascomycota(100);c__Saccharomycetes(100);o__Saccharomycetales(100);f__Metschnikowiaceae(100);g__Metschnikowia(100);s__Metschnikowia_pulcherrima(100); | Metschnikowia | 13 | Saprotroph | Undefined Saprotroph | Yeast | NULL | Probable | As nectar-tap_saprotroph (Põlme et al. 2020); Decay sugar-rich substrates (Põlme et al. 2020) | Põlme S, et al. 2020. Fungal Diversity 105:1-16 ((https://doi.org/10.1007/s13225-020-00466-2)) |
| Otu18430 | 1 | k__Fungi(100);p__Ascomycota(100);c__Saccharomycetes(100);o__Saccharomycetales(100);f__Metschnikowiaceae(100);g__Metschnikowia(100);s__Metschnikowia_pulcherrima(100); | Metschnikowia | 13 | Saprotroph | Undefined Saprotroph | Yeast | NULL | Probable | As nectar-tap_saprotroph (Põlme et al. 2020); Decay sugar-rich substrates (Põlme et al. 2020) | Põlme S, et al. 2020. Fungal Diversity 105:1-16 ((https://doi.org/10.1007/s13225-020-00466-2)) |
| Otu18432 | 1 | k__Fungi(100);p__Ascomycota(100);c__Saccharomycetes(100);o__Saccharomycetales(100);f__Metschnikowiaceae(100);g__Metschnikowia(100);s__Metschnikowia_pulcherrima(100); | Metschnikowia | 13 | Saprotroph | Undefined Saprotroph | Yeast | NULL | Probable | As nectar-tap_saprotroph (Põlme et al. 2020); Decay sugar-rich substrates (Põlme et al. 2020) | Põlme S, et al. 2020. Fungal Diversity 105:1-16 ((https://doi.org/10.1007/s13225-020-00466-2)) |
| Otu18433 | 1 | k__Fungi(100);p__Ascomycota(100);c__Saccharomycetes(100);o__Saccharomycetales(100);f__Metschnikowiaceae(100);g__Metschnikowia(100);s__Metschnikowia_pulcherrima(100); | Metschnikowia | 13 | Saprotroph | Undefined Saprotroph | Yeast | NULL | Probable | As nectar-tap_saprotroph (Põlme et al. 2020); Decay sugar-rich substrates (Põlme et al. 2020) | Põlme S, et al. 2020. Fungal Diversity 105:1-16 ((https://doi.org/10.1007/s13225-020-00466-2)) |
| Otu18434 | 1 | k__Fungi(100);p__Ascomycota(100);c__Saccharomycetes(100);o__Saccharomycetales(100);f__Metschnikowiaceae(100);g__Metschnikowia(100);g__Metschnikowia_unclassified(100); | Metschnikowia | 13 | Saprotroph | Undefined Saprotroph | Yeast | NULL | Probable | As nectar-tap_saprotroph (Põlme et al. 2020); Decay sugar-rich substrates (Põlme et al. 2020) | Põlme S, et al. 2020. Fungal Diversity 105:1-16 ((https://doi.org/10.1007/s13225-020-00466-2)) |
| Otu18436 | 1 | k__Fungi(100);p__Ascomycota(100);c__Saccharomycetes(100);o__Saccharomycetales(100);f__Metschnikowiaceae(100);g__Metschnikowia(100);s__Metschnikowia_pulcherrima(100); | Metschnikowia | 13 | Saprotroph | Undefined Saprotroph | Yeast | NULL | Probable | As nectar-tap_saprotroph (Põlme et al. 2020); Decay sugar-rich substrates (Põlme et al. 2020) | Põlme S, et al. 2020. Fungal Diversity 105:1-16 ((https://doi.org/10.1007/s13225-020-00466-2)) |
| Otu18437 | 1 | k__Fungi(100);p__Ascomycota(100);c__Saccharomycetes(100);o__Saccharomycetales(100);f__Metschnikowiaceae(100);g__Metschnikowia(100);s__Metschnikowia_pulcherrima(100); | Metschnikowia | 13 | Saprotroph | Undefined Saprotroph | Yeast | NULL | Probable | As nectar-tap_saprotroph (Põlme et al. 2020); Decay sugar-rich substrates (Põlme et al. 2020) | Põlme S, et al. 2020. Fungal Diversity 105:1-16 ((https://doi.org/10.1007/s13225-020-00466-2)) |
| Otu18439 | 1 | k__Fungi(100);p__Ascomycota(100);c__Saccharomycetes(100);o__Saccharomycetales(100);f__Metschnikowiaceae(100);g__Metschnikowia(100);s__Metschnikowia_pulcherrima(100); | Metschnikowia | 13 | Saprotroph | Undefined Saprotroph | Yeast | NULL | Probable | As nectar-tap_saprotroph (Põlme et al. 2020); Decay sugar-rich substrates (Põlme et al. 2020) | Põlme S, et al. 2020. Fungal Diversity 105:1-16 ((https://doi.org/10.1007/s13225-020-00466-2)) |
| Otu18440 | 1 | k__Fungi(100);p__Ascomycota(100);c__Saccharomycetes(100);o__Saccharomycetales(100);f__Metschnikowiaceae(100);g__Metschnikowia(100);s__Metschnikowia_pulcherrima(100); | Metschnikowia | 13 | Saprotroph | Undefined Saprotroph | Yeast | NULL | Probable | As nectar-tap_saprotroph (Põlme et al. 2020); Decay sugar-rich substrates (Põlme et al. 2020) | Põlme S, et al. 2020. Fungal Diversity 105:1-16 ((https://doi.org/10.1007/s13225-020-00466-2)) |
| Otu18441 | 1 | k__Fungi(100);p__Ascomycota(100);c__Saccharomycetes(100);o__Saccharomycetales(100);f__Metschnikowiaceae(100);g__Metschnikowia(100);s__Metschnikowia_pulcherrima(100); | Metschnikowia | 13 | Saprotroph | Undefined Saprotroph | Yeast | NULL | Probable | As nectar-tap_saprotroph (Põlme et al. 2020); Decay sugar-rich substrates (Põlme et al. 2020) | Põlme S, et al. 2020. Fungal Diversity 105:1-16 ((https://doi.org/10.1007/s13225-020-00466-2)) |
| Otu18443 | 1 | k__Fungi(100);p__Ascomycota(100);c__Saccharomycetes(100);o__Saccharomycetales(100);f__Metschnikowiaceae(100);g__Metschnikowia(100);s__Metschnikowia_pulcherrima(100); | Metschnikowia | 13 | Saprotroph | Undefined Saprotroph | Yeast | NULL | Probable | As nectar-tap_saprotroph (Põlme et al. 2020); Decay sugar-rich substrates (Põlme et al. 2020) | Põlme S, et al. 2020. Fungal Diversity 105:1-16 ((https://doi.org/10.1007/s13225-020-00466-2)) |
| Otu18444 | 1 | k__Fungi(100);p__Ascomycota(100);c__Saccharomycetes(100);o__Saccharomycetales(100);f__Metschnikowiaceae(100);g__Metschnikowia(100);s__Metschnikowia_pulcherrima(100); | Metschnikowia | 13 | Saprotroph | Undefined Saprotroph | Yeast | NULL | Probable | As nectar-tap_saprotroph (Põlme et al. 2020); Decay sugar-rich substrates (Põlme et al. 2020) | Põlme S, et al. 2020. Fungal Diversity 105:1-16 ((https://doi.org/10.1007/s13225-020-00466-2)) |
| Otu18445 | 1 | k__Fungi(100);p__Ascomycota(100);c__Saccharomycetes(100);o__Saccharomycetales(100);f__Metschnikowiaceae(100);g__Metschnikowia(100);s__Metschnikowia_pulcherrima(100); | Metschnikowia | 13 | Saprotroph | Undefined Saprotroph | Yeast | NULL | Probable | As nectar-tap_saprotroph (Põlme et al. 2020); Decay sugar-rich substrates (Põlme et al. 2020) | Põlme S, et al. 2020. Fungal Diversity 105:1-16 ((https://doi.org/10.1007/s13225-020-00466-2)) |
| Otu18446 | 1 | k__Fungi(100);p__Ascomycota(100);c__Saccharomycetes(100);o__Saccharomycetales(100);f__Saccharomycetales_family_Incertae_sedis(100);g__Candida(100);g__Candida_unclassified(100); | Candida | 13 | Pathotroph-Saprotroph-Symbiotroph | Animal Pathogen-Endophyte-Endosymbiont-Epiphyte-Undefined Saprotroph | Dimorphic Yeast-Yeast | NULL | Probable | As nectar-tap_saprotroph (Põlme et al. 2020); Decay sugar-rich substrates (Põlme et al. 2020); This asexual genus is polyphyletic and contain species that function very differently depend on what they are. Caution should be taken when interpreting guilds for Candida species. | Manolakaki D, et al. 2010. Virulence. 1: 367-375 ((https://doi.org/10.4161/viru.1.5.12796)); Mekbib SB, et al. 2011. Tropical Plant Pathology 36:233-240 ((https://doi.org/10.1590/S1982-56762011000400004)); Põlme S, et al. 2020. Fungal Diversity 105:1-16 ((https://doi.org/10.1007/s13225-020-00466-2)) |
| Otu18450 | 1 | k__Fungi(100);p__Ascomycota(100);c__Saccharomycetes(100);o__Saccharomycetales(100);f__Metschnikowiaceae(100);g__Metschnikowia(100);s__Metschnikowia_pulcherrima(100); | Metschnikowia | 13 | Saprotroph | Undefined Saprotroph | Yeast | NULL | Probable | As nectar-tap_saprotroph (Põlme et al. 2020); Decay sugar-rich substrates (Põlme et al. 2020) | Põlme S, et al. 2020. Fungal Diversity 105:1-16 ((https://doi.org/10.1007/s13225-020-00466-2)) |
| Otu18451 | 1 | k__Fungi(100);p__Ascomycota(100);c__Saccharomycetes(100);o__Saccharomycetales(100);f__Metschnikowiaceae(100);g__Metschnikowia(100);s__Metschnikowia_pulcherrima(100); | Metschnikowia | 13 | Saprotroph | Undefined Saprotroph | Yeast | NULL | Probable | As nectar-tap_saprotroph (Põlme et al. 2020); Decay sugar-rich substrates (Põlme et al. 2020) | Põlme S, et al. 2020. Fungal Diversity 105:1-16 ((https://doi.org/10.1007/s13225-020-00466-2)) |
| Otu18452 | 1 | k__Fungi(100);p__Ascomycota(100);c__Saccharomycetes(100);o__Saccharomycetales(100);f__Metschnikowiaceae(100);g__Metschnikowia(100);s__Metschnikowia_pulcherrima(100); | Metschnikowia | 13 | Saprotroph | Undefined Saprotroph | Yeast | NULL | Probable | As nectar-tap_saprotroph (Põlme et al. 2020); Decay sugar-rich substrates (Põlme et al. 2020) | Põlme S, et al. 2020. Fungal Diversity 105:1-16 ((https://doi.org/10.1007/s13225-020-00466-2)) |
| Otu18453 | 1 | k__Fungi(100);p__Ascomycota(100);c__Saccharomycetes(100);o__Saccharomycetales(100);f__Metschnikowiaceae(100);g__Metschnikowia(100);s__Metschnikowia_pulcherrima(100); | Metschnikowia | 13 | Saprotroph | Undefined Saprotroph | Yeast | NULL | Probable | As nectar-tap_saprotroph (Põlme et al. 2020); Decay sugar-rich substrates (Põlme et al. 2020) | Põlme S, et al. 2020. Fungal Diversity 105:1-16 ((https://doi.org/10.1007/s13225-020-00466-2)) |
| Otu18454 | 1 | k__Fungi(100);p__Ascomycota(100);c__Saccharomycetes(100);o__Saccharomycetales(100);f__Metschnikowiaceae(100);g__Metschnikowia(100);s__Metschnikowia_pulcherrima(100); | Metschnikowia | 13 | Saprotroph | Undefined Saprotroph | Yeast | NULL | Probable | As nectar-tap_saprotroph (Põlme et al. 2020); Decay sugar-rich substrates (Põlme et al. 2020) | Põlme S, et al. 2020. Fungal Diversity 105:1-16 ((https://doi.org/10.1007/s13225-020-00466-2)) |
| Otu18455 | 1 | k__Fungi(100);p__Ascomycota(100);c__Saccharomycetes(100);o__Saccharomycetales(100);f__Metschnikowiaceae(100);g__Metschnikowia(100);s__Metschnikowia_pulcherrima(100); | Metschnikowia | 13 | Saprotroph | Undefined Saprotroph | Yeast | NULL | Probable | As nectar-tap_saprotroph (Põlme et al. 2020); Decay sugar-rich substrates (Põlme et al. 2020) | Põlme S, et al. 2020. Fungal Diversity 105:1-16 ((https://doi.org/10.1007/s13225-020-00466-2)) |
| Otu18456 | 1 | k__Fungi(100);p__Ascomycota(100);c__Saccharomycetes(100);o__Saccharomycetales(100);f__Metschnikowiaceae(100);g__Metschnikowia(100);g__Metschnikowia_unclassified(100); | Metschnikowia | 13 | Saprotroph | Undefined Saprotroph | Yeast | NULL | Probable | As nectar-tap_saprotroph (Põlme et al. 2020); Decay sugar-rich substrates (Põlme et al. 2020) | Põlme S, et al. 2020. Fungal Diversity 105:1-16 ((https://doi.org/10.1007/s13225-020-00466-2)) |
| Otu18457 | 1 | k__Fungi(100);p__Ascomycota(100);c__Saccharomycetes(100);o__Saccharomycetales(100);f__Metschnikowiaceae(100);g__Metschnikowia(100);s__Metschnikowia_pulcherrima(100); | Metschnikowia | 13 | Saprotroph | Undefined Saprotroph | Yeast | NULL | Probable | As nectar-tap_saprotroph (Põlme et al. 2020); Decay sugar-rich substrates (Põlme et al. 2020) | Põlme S, et al. 2020. Fungal Diversity 105:1-16 ((https://doi.org/10.1007/s13225-020-00466-2)) |
| Otu18458 | 1 | k__Fungi(100);p__Ascomycota(100);c__Saccharomycetes(100);o__Saccharomycetales(100);f__Metschnikowiaceae(100);g__Metschnikowia(100);g__Metschnikowia_unclassified(100); | Metschnikowia | 13 | Saprotroph | Undefined Saprotroph | Yeast | NULL | Probable | As nectar-tap_saprotroph (Põlme et al. 2020); Decay sugar-rich substrates (Põlme et al. 2020) | Põlme S, et al. 2020. Fungal Diversity 105:1-16 ((https://doi.org/10.1007/s13225-020-00466-2)) |
| Otu18459 | 1 | k__Fungi(100);p__Ascomycota(100);c__Saccharomycetes(100);o__Saccharomycetales(100);f__Metschnikowiaceae(100);g__Metschnikowia(100);g__Metschnikowia_unclassified(100); | Metschnikowia | 13 | Saprotroph | Undefined Saprotroph | Yeast | NULL | Probable | As nectar-tap_saprotroph (Põlme et al. 2020); Decay sugar-rich substrates (Põlme et al. 2020) | Põlme S, et al. 2020. Fungal Diversity 105:1-16 ((https://doi.org/10.1007/s13225-020-00466-2)) |
| Otu18461 | 1 | k__Fungi(100);p__Ascomycota(100);c__Saccharomycetes(100);o__Saccharomycetales(100);f__Metschnikowiaceae(100);g__Metschnikowia(100);s__Metschnikowia_pulcherrima(100); | Metschnikowia | 13 | Saprotroph | Undefined Saprotroph | Yeast | NULL | Probable | As nectar-tap_saprotroph (Põlme et al. 2020); Decay sugar-rich substrates (Põlme et al. 2020) | Põlme S, et al. 2020. Fungal Diversity 105:1-16 ((https://doi.org/10.1007/s13225-020-00466-2)) |
| Otu18462 | 1 | k__Fungi(100);p__Ascomycota(100);c__Saccharomycetes(100);o__Saccharomycetales(100);f__Metschnikowiaceae(100);g__Metschnikowia(100);s__Metschnikowia_pulcherrima(100); | Metschnikowia | 13 | Saprotroph | Undefined Saprotroph | Yeast | NULL | Probable | As nectar-tap_saprotroph (Põlme et al. 2020); Decay sugar-rich substrates (Põlme et al. 2020) | Põlme S, et al. 2020. Fungal Diversity 105:1-16 ((https://doi.org/10.1007/s13225-020-00466-2)) |
| Otu18463 | 1 | k__Fungi(100);p__Ascomycota(100);c__Saccharomycetes(100);o__Saccharomycetales(100);f__Metschnikowiaceae(100);g__Metschnikowia(100);s__Metschnikowia_pulcherrima(100); | Metschnikowia | 13 | Saprotroph | Undefined Saprotroph | Yeast | NULL | Probable | As nectar-tap_saprotroph (Põlme et al. 2020); Decay sugar-rich substrates (Põlme et al. 2020) | Põlme S, et al. 2020. Fungal Diversity 105:1-16 ((https://doi.org/10.1007/s13225-020-00466-2)) |
| Otu18464 | 1 | k__Fungi(100);p__Ascomycota(100);c__Saccharomycetes(100);o__Saccharomycetales(100);f__Metschnikowiaceae(100);g__Metschnikowia(100);s__Metschnikowia_pulcherrima(100); | Metschnikowia | 13 | Saprotroph | Undefined Saprotroph | Yeast | NULL | Probable | As nectar-tap_saprotroph (Põlme et al. 2020); Decay sugar-rich substrates (Põlme et al. 2020) | Põlme S, et al. 2020. Fungal Diversity 105:1-16 ((https://doi.org/10.1007/s13225-020-00466-2)) |
| Otu18465 | 1 | k__Fungi(100);p__Ascomycota(100);c__Saccharomycetes(100);o__Saccharomycetales(100);f__Metschnikowiaceae(100);g__Metschnikowia(100);s__Metschnikowia_pulcherrima(100); | Metschnikowia | 13 | Saprotroph | Undefined Saprotroph | Yeast | NULL | Probable | As nectar-tap_saprotroph (Põlme et al. 2020); Decay sugar-rich substrates (Põlme et al. 2020) | Põlme S, et al. 2020. Fungal Diversity 105:1-16 ((https://doi.org/10.1007/s13225-020-00466-2)) |
| Otu18466 | 1 | k__Fungi(100);p__Ascomycota(100);c__Saccharomycetes(100);o__Saccharomycetales(100);f__Metschnikowiaceae(100);g__Metschnikowia(100);s__Metschnikowia_pulcherrima(100); | Metschnikowia | 13 | Saprotroph | Undefined Saprotroph | Yeast | NULL | Probable | As nectar-tap_saprotroph (Põlme et al. 2020); Decay sugar-rich substrates (Põlme et al. 2020) | Põlme S, et al. 2020. Fungal Diversity 105:1-16 ((https://doi.org/10.1007/s13225-020-00466-2)) |
| Otu18467 | 1 | k__Fungi(100);p__Ascomycota(100);c__Saccharomycetes(100);o__Saccharomycetales(100);f__Metschnikowiaceae(100);g__Metschnikowia(100);s__Metschnikowia_pulcherrima(100); | Metschnikowia | 13 | Saprotroph | Undefined Saprotroph | Yeast | NULL | Probable | As nectar-tap_saprotroph (Põlme et al. 2020); Decay sugar-rich substrates (Põlme et al. 2020) | Põlme S, et al. 2020. Fungal Diversity 105:1-16 ((https://doi.org/10.1007/s13225-020-00466-2)) |
| Otu18468 | 1 | k__Fungi(100);p__Ascomycota(100);c__Saccharomycetes(100);o__Saccharomycetales(100);f__Metschnikowiaceae(100);g__Metschnikowia(100);s__Metschnikowia_pulcherrima(100); | Metschnikowia | 13 | Saprotroph | Undefined Saprotroph | Yeast | NULL | Probable | As nectar-tap_saprotroph (Põlme et al. 2020); Decay sugar-rich substrates (Põlme et al. 2020) | Põlme S, et al. 2020. Fungal Diversity 105:1-16 ((https://doi.org/10.1007/s13225-020-00466-2)) |
| Otu18469 | 1 | k__Fungi(100);p__Ascomycota(100);c__Saccharomycetes(100);o__Saccharomycetales(100);f__Metschnikowiaceae(100);g__Metschnikowia(100);s__Metschnikowia_pulcherrima(100); | Metschnikowia | 13 | Saprotroph | Undefined Saprotroph | Yeast | NULL | Probable | As nectar-tap_saprotroph (Põlme et al. 2020); Decay sugar-rich substrates (Põlme et al. 2020) | Põlme S, et al. 2020. Fungal Diversity 105:1-16 ((https://doi.org/10.1007/s13225-020-00466-2)) |
| Otu18470 | 1 | k__Fungi(100);p__Ascomycota(100);c__Saccharomycetes(100);o__Saccharomycetales(100);f__Metschnikowiaceae(100);g__Metschnikowia(100);s__Metschnikowia_pulcherrima(100); | Metschnikowia | 13 | Saprotroph | Undefined Saprotroph | Yeast | NULL | Probable | As nectar-tap_saprotroph (Põlme et al. 2020); Decay sugar-rich substrates (Põlme et al. 2020) | Põlme S, et al. 2020. Fungal Diversity 105:1-16 ((https://doi.org/10.1007/s13225-020-00466-2)) |
| Otu18471 | 1 | k__Fungi(100);p__Ascomycota(100);c__Saccharomycetes(100);o__Saccharomycetales(100);f__Metschnikowiaceae(100);g__Metschnikowia(100);s__Metschnikowia_pulcherrima(100); | Metschnikowia | 13 | Saprotroph | Undefined Saprotroph | Yeast | NULL | Probable | As nectar-tap_saprotroph (Põlme et al. 2020); Decay sugar-rich substrates (Põlme et al. 2020) | Põlme S, et al. 2020. Fungal Diversity 105:1-16 ((https://doi.org/10.1007/s13225-020-00466-2)) |
| Otu18472 | 1 | k__Fungi(100);p__Ascomycota(100);c__Saccharomycetes(100);o__Saccharomycetales(100);f__Metschnikowiaceae(100);g__Metschnikowia(100);s__Metschnikowia_pulcherrima(100); | Metschnikowia | 13 | Saprotroph | Undefined Saprotroph | Yeast | NULL | Probable | As nectar-tap_saprotroph (Põlme et al. 2020); Decay sugar-rich substrates (Põlme et al. 2020) | Põlme S, et al. 2020. Fungal Diversity 105:1-16 ((https://doi.org/10.1007/s13225-020-00466-2)) |
| Otu18476 | 1 | k__Fungi(100);p__Ascomycota(100);c__Saccharomycetes(100);o__Saccharomycetales(100);f__Saccharomycetales_family_Incertae_sedis(100);g__Candida(100);g__Candida_unclassified(100); | Candida | 13 | Pathotroph-Saprotroph-Symbiotroph | Animal Pathogen-Endophyte-Endosymbiont-Epiphyte-Undefined Saprotroph | Dimorphic Yeast-Yeast | NULL | Probable | As nectar-tap_saprotroph (Põlme et al. 2020); Decay sugar-rich substrates (Põlme et al. 2020); This asexual genus is polyphyletic and contain species that function very differently depend on what they are. Caution should be taken when interpreting guilds for Candida species. | Manolakaki D, et al. 2010. Virulence. 1: 367-375 ((https://doi.org/10.4161/viru.1.5.12796)); Mekbib SB, et al. 2011. Tropical Plant Pathology 36:233-240 ((https://doi.org/10.1590/S1982-56762011000400004)); Põlme S, et al. 2020. Fungal Diversity 105:1-16 ((https://doi.org/10.1007/s13225-020-00466-2)) |
| Otu18478 | 1 | k__Fungi(100);p__Ascomycota(100);c__Saccharomycetes(100);o__Saccharomycetales(100);o__Saccharomycetales_unclassified(100);o__Saccharomycetales_unclassified(100);o__Saccharomycetales_unclassified(100); | Saccharomycetales | 7 | Saprotroph | Undefined Saprotroph | Yeast | NULL | Possible | NULL | Sterrenburg E, et al. 2015. New Phytologist 207:1145-1158 (https://doi.org/10.1111/nph.13426) |
| Otu18479 | 1 | k__Fungi(100);p__Ascomycota(100);c__Saccharomycetes(100);o__Saccharomycetales(100);f__Metschnikowiaceae(100);g__Metschnikowia(100);g__Metschnikowia_unclassified(100); | Metschnikowia | 13 | Saprotroph | Undefined Saprotroph | Yeast | NULL | Probable | As nectar-tap_saprotroph (Põlme et al. 2020); Decay sugar-rich substrates (Põlme et al. 2020) | Põlme S, et al. 2020. Fungal Diversity 105:1-16 ((https://doi.org/10.1007/s13225-020-00466-2)) |
| Otu18480 | 1 | k__Fungi(100);p__Ascomycota(100);c__Saccharomycetes(100);o__Saccharomycetales(100);f__Metschnikowiaceae(100);g__Metschnikowia(100);s__Metschnikowia_pulcherrima(100); | Metschnikowia | 13 | Saprotroph | Undefined Saprotroph | Yeast | NULL | Probable | As nectar-tap_saprotroph (Põlme et al. 2020); Decay sugar-rich substrates (Põlme et al. 2020) | Põlme S, et al. 2020. Fungal Diversity 105:1-16 ((https://doi.org/10.1007/s13225-020-00466-2)) |
| Otu18482 | 1 | k__Fungi(100);p__Ascomycota(100);c__Saccharomycetes(100);o__Saccharomycetales(100);f__Metschnikowiaceae(100);g__Metschnikowia(100);s__Metschnikowia_pulcherrima(100); | Metschnikowia | 13 | Saprotroph | Undefined Saprotroph | Yeast | NULL | Probable | As nectar-tap_saprotroph (Põlme et al. 2020); Decay sugar-rich substrates (Põlme et al. 2020) | Põlme S, et al. 2020. Fungal Diversity 105:1-16 ((https://doi.org/10.1007/s13225-020-00466-2)) |
| Otu18483 | 1 | k__Fungi(100);p__Ascomycota(100);c__Saccharomycetes(100);o__Saccharomycetales(100);f__Metschnikowiaceae(100);g__Metschnikowia(100);g__Metschnikowia_unclassified(100); | Metschnikowia | 13 | Saprotroph | Undefined Saprotroph | Yeast | NULL | Probable | As nectar-tap_saprotroph (Põlme et al. 2020); Decay sugar-rich substrates (Põlme et al. 2020) | Põlme S, et al. 2020. Fungal Diversity 105:1-16 ((https://doi.org/10.1007/s13225-020-00466-2)) |
| Otu18484 | 1 | k__Fungi(100);p__Ascomycota(100);c__Saccharomycetes(100);o__Saccharomycetales(100);f__Metschnikowiaceae(100);g__Metschnikowia(100);s__Metschnikowia_pulcherrima(100); | Metschnikowia | 13 | Saprotroph | Undefined Saprotroph | Yeast | NULL | Probable | As nectar-tap_saprotroph (Põlme et al. 2020); Decay sugar-rich substrates (Põlme et al. 2020) | Põlme S, et al. 2020. Fungal Diversity 105:1-16 ((https://doi.org/10.1007/s13225-020-00466-2)) |
| Otu18485 | 1 | k__Fungi(100);p__Ascomycota(100);c__Saccharomycetes(100);o__Saccharomycetales(100);f__Metschnikowiaceae(100);g__Metschnikowia(100);s__Metschnikowia_pulcherrima(100); | Metschnikowia | 13 | Saprotroph | Undefined Saprotroph | Yeast | NULL | Probable | As nectar-tap_saprotroph (Põlme et al. 2020); Decay sugar-rich substrates (Põlme et al. 2020) | Põlme S, et al. 2020. Fungal Diversity 105:1-16 ((https://doi.org/10.1007/s13225-020-00466-2)) |
| Otu18486 | 1 | k__Fungi(100);p__Ascomycota(100);c__Saccharomycetes(100);o__Saccharomycetales(100);f__Metschnikowiaceae(100);g__Metschnikowia(100);s__Metschnikowia_pulcherrima(100); | Metschnikowia | 13 | Saprotroph | Undefined Saprotroph | Yeast | NULL | Probable | As nectar-tap_saprotroph (Põlme et al. 2020); Decay sugar-rich substrates (Põlme et al. 2020) | Põlme S, et al. 2020. Fungal Diversity 105:1-16 ((https://doi.org/10.1007/s13225-020-00466-2)) |
| Otu18487 | 1 | k__Fungi(100);p__Ascomycota(100);c__Saccharomycetes(100);o__Saccharomycetales(100);f__Metschnikowiaceae(100);g__Metschnikowia(100);g__Metschnikowia_unclassified(100); | Metschnikowia | 13 | Saprotroph | Undefined Saprotroph | Yeast | NULL | Probable | As nectar-tap_saprotroph (Põlme et al. 2020); Decay sugar-rich substrates (Põlme et al. 2020) | Põlme S, et al. 2020. Fungal Diversity 105:1-16 ((https://doi.org/10.1007/s13225-020-00466-2)) |
| Otu18488 | 1 | k__Fungi(100);p__Ascomycota(100);c__Saccharomycetes(100);o__Saccharomycetales(100);f__Metschnikowiaceae(100);g__Metschnikowia(100);s__Metschnikowia_pulcherrima(100); | Metschnikowia | 13 | Saprotroph | Undefined Saprotroph | Yeast | NULL | Probable | As nectar-tap_saprotroph (Põlme et al. 2020); Decay sugar-rich substrates (Põlme et al. 2020) | Põlme S, et al. 2020. Fungal Diversity 105:1-16 ((https://doi.org/10.1007/s13225-020-00466-2)) |
| Otu18489 | 1 | k__Fungi(100);p__Ascomycota(100);c__Saccharomycetes(100);o__Saccharomycetales(100);f__Metschnikowiaceae(100);g__Metschnikowia(100);s__Metschnikowia_pulcherrima(100); | Metschnikowia | 13 | Saprotroph | Undefined Saprotroph | Yeast | NULL | Probable | As nectar-tap_saprotroph (Põlme et al. 2020); Decay sugar-rich substrates (Põlme et al. 2020) | Põlme S, et al. 2020. Fungal Diversity 105:1-16 ((https://doi.org/10.1007/s13225-020-00466-2)) |
| Otu18490 | 1 | k__Fungi(100);p__Ascomycota(100);c__Saccharomycetes(100);o__Saccharomycetales(100);f__Metschnikowiaceae(100);g__Metschnikowia(100);g__Metschnikowia_unclassified(100); | Metschnikowia | 13 | Saprotroph | Undefined Saprotroph | Yeast | NULL | Probable | As nectar-tap_saprotroph (Põlme et al. 2020); Decay sugar-rich substrates (Põlme et al. 2020) | Põlme S, et al. 2020. Fungal Diversity 105:1-16 ((https://doi.org/10.1007/s13225-020-00466-2)) |
| Otu18492 | 1 | k__Fungi(100);p__Ascomycota(100);c__Saccharomycetes(100);o__Saccharomycetales(100);f__Metschnikowiaceae(100);g__Metschnikowia(100);s__Metschnikowia_pulcherrima(100); | Metschnikowia | 13 | Saprotroph | Undefined Saprotroph | Yeast | NULL | Probable | As nectar-tap_saprotroph (Põlme et al. 2020); Decay sugar-rich substrates (Põlme et al. 2020) | Põlme S, et al. 2020. Fungal Diversity 105:1-16 ((https://doi.org/10.1007/s13225-020-00466-2)) |
| Otu18494 | 1 | k__Fungi(100);p__Ascomycota(100);c__Saccharomycetes(100);o__Saccharomycetales(100);f__Metschnikowiaceae(100);g__Metschnikowia(100);s__Metschnikowia_pulcherrima(100); | Metschnikowia | 13 | Saprotroph | Undefined Saprotroph | Yeast | NULL | Probable | As nectar-tap_saprotroph (Põlme et al. 2020); Decay sugar-rich substrates (Põlme et al. 2020) | Põlme S, et al. 2020. Fungal Diversity 105:1-16 ((https://doi.org/10.1007/s13225-020-00466-2)) |
| Otu18495 | 1 | k__Fungi(100);p__Ascomycota(100);c__Saccharomycetes(100);o__Saccharomycetales(100);f__Metschnikowiaceae(100);g__Metschnikowia(100);s__Metschnikowia_pulcherrima(100); | Metschnikowia | 13 | Saprotroph | Undefined Saprotroph | Yeast | NULL | Probable | As nectar-tap_saprotroph (Põlme et al. 2020); Decay sugar-rich substrates (Põlme et al. 2020) | Põlme S, et al. 2020. Fungal Diversity 105:1-16 ((https://doi.org/10.1007/s13225-020-00466-2)) |
| Otu18496 | 1 | k__Fungi(100);p__Ascomycota(100);c__Saccharomycetes(100);o__Saccharomycetales(100);f__Metschnikowiaceae(100);g__Metschnikowia(100);s__Metschnikowia_pulcherrima(100); | Metschnikowia | 13 | Saprotroph | Undefined Saprotroph | Yeast | NULL | Probable | As nectar-tap_saprotroph (Põlme et al. 2020); Decay sugar-rich substrates (Põlme et al. 2020) | Põlme S, et al. 2020. Fungal Diversity 105:1-16 ((https://doi.org/10.1007/s13225-020-00466-2)) |
| Otu18497 | 1 | k__Fungi(100);p__Ascomycota(100);c__Saccharomycetes(100);o__Saccharomycetales(100);f__Metschnikowiaceae(100);g__Metschnikowia(100);s__Metschnikowia_pulcherrima(100); | Metschnikowia | 13 | Saprotroph | Undefined Saprotroph | Yeast | NULL | Probable | As nectar-tap_saprotroph (Põlme et al. 2020); Decay sugar-rich substrates (Põlme et al. 2020) | Põlme S, et al. 2020. Fungal Diversity 105:1-16 ((https://doi.org/10.1007/s13225-020-00466-2)) |
| Otu18498 | 1 | k__Fungi(100);p__Ascomycota(100);c__Saccharomycetes(100);o__Saccharomycetales(100);f__Metschnikowiaceae(100);g__Metschnikowia(100);s__Metschnikowia_pulcherrima(100); | Metschnikowia | 13 | Saprotroph | Undefined Saprotroph | Yeast | NULL | Probable | As nectar-tap_saprotroph (Põlme et al. 2020); Decay sugar-rich substrates (Põlme et al. 2020) | Põlme S, et al. 2020. Fungal Diversity 105:1-16 ((https://doi.org/10.1007/s13225-020-00466-2)) |
| Otu18499 | 1 | k__Fungi(100);p__Ascomycota(100);c__Saccharomycetes(100);o__Saccharomycetales(100);f__Metschnikowiaceae(100);g__Metschnikowia(100);g__Metschnikowia_unclassified(100); | Metschnikowia | 13 | Saprotroph | Undefined Saprotroph | Yeast | NULL | Probable | As nectar-tap_saprotroph (Põlme et al. 2020); Decay sugar-rich substrates (Põlme et al. 2020) | Põlme S, et al. 2020. Fungal Diversity 105:1-16 ((https://doi.org/10.1007/s13225-020-00466-2)) |
| Otu18500 | 1 | k__Fungi(100);p__Ascomycota(100);c__Saccharomycetes(100);o__Saccharomycetales(100);f__Metschnikowiaceae(100);g__Metschnikowia(100);g__Metschnikowia_unclassified(100); | Metschnikowia | 13 | Saprotroph | Undefined Saprotroph | Yeast | NULL | Probable | As nectar-tap_saprotroph (Põlme et al. 2020); Decay sugar-rich substrates (Põlme et al. 2020) | Põlme S, et al. 2020. Fungal Diversity 105:1-16 ((https://doi.org/10.1007/s13225-020-00466-2)) |
| Otu18501 | 1 | k__Fungi(100);p__Ascomycota(100);c__Saccharomycetes(100);o__Saccharomycetales(100);f__Metschnikowiaceae(100);g__Metschnikowia(100);s__Metschnikowia_pulcherrima(100); | Metschnikowia | 13 | Saprotroph | Undefined Saprotroph | Yeast | NULL | Probable | As nectar-tap_saprotroph (Põlme et al. 2020); Decay sugar-rich substrates (Põlme et al. 2020) | Põlme S, et al. 2020. Fungal Diversity 105:1-16 ((https://doi.org/10.1007/s13225-020-00466-2)) |
| Otu18502 | 1 | k__Fungi(100);p__Ascomycota(100);c__Saccharomycetes(100);o__Saccharomycetales(100);f__Metschnikowiaceae(100);g__Metschnikowia(100);s__Metschnikowia_pulcherrima(100); | Metschnikowia | 13 | Saprotroph | Undefined Saprotroph | Yeast | NULL | Probable | As nectar-tap_saprotroph (Põlme et al. 2020); Decay sugar-rich substrates (Põlme et al. 2020) | Põlme S, et al. 2020. Fungal Diversity 105:1-16 ((https://doi.org/10.1007/s13225-020-00466-2)) |
| Otu18503 | 1 | k__Fungi(100);p__Ascomycota(100);c__Saccharomycetes(100);o__Saccharomycetales(100);f__Saccharomycetales_family_Incertae_sedis(100);g__Candida(100);g__Candida_unclassified(100); | Candida | 13 | Pathotroph-Saprotroph-Symbiotroph | Animal Pathogen-Endophyte-Endosymbiont-Epiphyte-Undefined Saprotroph | Dimorphic Yeast-Yeast | NULL | Probable | As nectar-tap_saprotroph (Põlme et al. 2020); Decay sugar-rich substrates (Põlme et al. 2020); This asexual genus is polyphyletic and contain species that function very differently depend on what they are. Caution should be taken when interpreting guilds for Candida species. | Manolakaki D, et al. 2010. Virulence. 1: 367-375 ((https://doi.org/10.4161/viru.1.5.12796)); Mekbib SB, et al. 2011. Tropical Plant Pathology 36:233-240 ((https://doi.org/10.1590/S1982-56762011000400004)); Põlme S, et al. 2020. Fungal Diversity 105:1-16 ((https://doi.org/10.1007/s13225-020-00466-2)) |
| Otu18505 | 1 | k__Fungi(100);p__Ascomycota(100);c__Saccharomycetes(100);o__Saccharomycetales(100);o__Saccharomycetales_unclassified(100);o__Saccharomycetales_unclassified(100);o__Saccharomycetales_unclassified(100); | Saccharomycetales | 7 | Saprotroph | Undefined Saprotroph | Yeast | NULL | Possible | NULL | Sterrenburg E, et al. 2015. New Phytologist 207:1145-1158 (https://doi.org/10.1111/nph.13426) |
| Otu18506 | 1 | k__Fungi(100);p__Ascomycota(100);c__Saccharomycetes(100);o__Saccharomycetales(100);f__Metschnikowiaceae(100);g__Metschnikowia(100);g__Metschnikowia_unclassified(100); | Metschnikowia | 13 | Saprotroph | Undefined Saprotroph | Yeast | NULL | Probable | As nectar-tap_saprotroph (Põlme et al. 2020); Decay sugar-rich substrates (Põlme et al. 2020) | Põlme S, et al. 2020. Fungal Diversity 105:1-16 ((https://doi.org/10.1007/s13225-020-00466-2)) |
| Otu18507 | 1 | k__Fungi(100);p__Ascomycota(100);c__Saccharomycetes(100);o__Saccharomycetales(100);f__Metschnikowiaceae(100);g__Metschnikowia(100);g__Metschnikowia_unclassified(100); | Metschnikowia | 13 | Saprotroph | Undefined Saprotroph | Yeast | NULL | Probable | As nectar-tap_saprotroph (Põlme et al. 2020); Decay sugar-rich substrates (Põlme et al. 2020) | Põlme S, et al. 2020. Fungal Diversity 105:1-16 ((https://doi.org/10.1007/s13225-020-00466-2)) |
| Otu18508 | 1 | k__Fungi(100);p__Ascomycota(100);c__Saccharomycetes(100);o__Saccharomycetales(100);f__Metschnikowiaceae(100);g__Metschnikowia(100);s__Metschnikowia_pulcherrima(100); | Metschnikowia | 13 | Saprotroph | Undefined Saprotroph | Yeast | NULL | Probable | As nectar-tap_saprotroph (Põlme et al. 2020); Decay sugar-rich substrates (Põlme et al. 2020) | Põlme S, et al. 2020. Fungal Diversity 105:1-16 ((https://doi.org/10.1007/s13225-020-00466-2)) |
| Otu18509 | 1 | k__Fungi(100);p__Ascomycota(100);c__Saccharomycetes(100);o__Saccharomycetales(100);f__Metschnikowiaceae(100);g__Metschnikowia(100);g__Metschnikowia_unclassified(100); | Metschnikowia | 13 | Saprotroph | Undefined Saprotroph | Yeast | NULL | Probable | As nectar-tap_saprotroph (Põlme et al. 2020); Decay sugar-rich substrates (Põlme et al. 2020) | Põlme S, et al. 2020. Fungal Diversity 105:1-16 ((https://doi.org/10.1007/s13225-020-00466-2)) |
| Otu18510 | 1 | k__Fungi(100);p__Ascomycota(100);c__Saccharomycetes(100);o__Saccharomycetales(100);f__Metschnikowiaceae(100);g__Metschnikowia(100);s__Metschnikowia_pulcherrima(100); | Metschnikowia | 13 | Saprotroph | Undefined Saprotroph | Yeast | NULL | Probable | As nectar-tap_saprotroph (Põlme et al. 2020); Decay sugar-rich substrates (Põlme et al. 2020) | Põlme S, et al. 2020. Fungal Diversity 105:1-16 ((https://doi.org/10.1007/s13225-020-00466-2)) |
| Otu18511 | 1 | k__Fungi(100);p__Ascomycota(100);c__Saccharomycetes(100);o__Saccharomycetales(100);f__Metschnikowiaceae(100);g__Metschnikowia(100);g__Metschnikowia_unclassified(100); | Metschnikowia | 13 | Saprotroph | Undefined Saprotroph | Yeast | NULL | Probable | As nectar-tap_saprotroph (Põlme et al. 2020); Decay sugar-rich substrates (Põlme et al. 2020) | Põlme S, et al. 2020. Fungal Diversity 105:1-16 ((https://doi.org/10.1007/s13225-020-00466-2)) |
| Otu18512 | 1 | k__Fungi(100);p__Ascomycota(100);c__Saccharomycetes(100);o__Saccharomycetales(100);f__Metschnikowiaceae(100);g__Metschnikowia(100);s__Metschnikowia_pulcherrima(100); | Metschnikowia | 13 | Saprotroph | Undefined Saprotroph | Yeast | NULL | Probable | As nectar-tap_saprotroph (Põlme et al. 2020); Decay sugar-rich substrates (Põlme et al. 2020) | Põlme S, et al. 2020. Fungal Diversity 105:1-16 ((https://doi.org/10.1007/s13225-020-00466-2)) |
| Otu18513 | 1 | k__Fungi(100);p__Ascomycota(100);c__Saccharomycetes(100);o__Saccharomycetales(100);f__Metschnikowiaceae(100);g__Metschnikowia(100);s__Metschnikowia_pulcherrima(100); | Metschnikowia | 13 | Saprotroph | Undefined Saprotroph | Yeast | NULL | Probable | As nectar-tap_saprotroph (Põlme et al. 2020); Decay sugar-rich substrates (Põlme et al. 2020) | Põlme S, et al. 2020. Fungal Diversity 105:1-16 ((https://doi.org/10.1007/s13225-020-00466-2)) |
| Otu18514 | 1 | k__Fungi(100);p__Ascomycota(100);c__Saccharomycetes(100);o__Saccharomycetales(100);f__Metschnikowiaceae(100);g__Metschnikowia(100);g__Metschnikowia_unclassified(100); | Metschnikowia | 13 | Saprotroph | Undefined Saprotroph | Yeast | NULL | Probable | As nectar-tap_saprotroph (Põlme et al. 2020); Decay sugar-rich substrates (Põlme et al. 2020) | Põlme S, et al. 2020. Fungal Diversity 105:1-16 ((https://doi.org/10.1007/s13225-020-00466-2)) |
| Otu18515 | 1 | k__Fungi(100);p__Ascomycota(100);c__Saccharomycetes(100);o__Saccharomycetales(100);f__Metschnikowiaceae(100);g__Metschnikowia(100);s__Metschnikowia_pulcherrima(100); | Metschnikowia | 13 | Saprotroph | Undefined Saprotroph | Yeast | NULL | Probable | As nectar-tap_saprotroph (Põlme et al. 2020); Decay sugar-rich substrates (Põlme et al. 2020) | Põlme S, et al. 2020. Fungal Diversity 105:1-16 ((https://doi.org/10.1007/s13225-020-00466-2)) |
| Otu18516 | 1 | k__Fungi(100);p__Ascomycota(100);c__Saccharomycetes(100);o__Saccharomycetales(100);f__Metschnikowiaceae(100);g__Metschnikowia(100);g__Metschnikowia_unclassified(100); | Metschnikowia | 13 | Saprotroph | Undefined Saprotroph | Yeast | NULL | Probable | As nectar-tap_saprotroph (Põlme et al. 2020); Decay sugar-rich substrates (Põlme et al. 2020) | Põlme S, et al. 2020. Fungal Diversity 105:1-16 ((https://doi.org/10.1007/s13225-020-00466-2)) |
| Otu18517 | 1 | k__Fungi(100);p__Ascomycota(100);c__Saccharomycetes(100);o__Saccharomycetales(100);f__Metschnikowiaceae(100);g__Metschnikowia(100);g__Metschnikowia_unclassified(100); | Metschnikowia | 13 | Saprotroph | Undefined Saprotroph | Yeast | NULL | Probable | As nectar-tap_saprotroph (Põlme et al. 2020); Decay sugar-rich substrates (Põlme et al. 2020) | Põlme S, et al. 2020. Fungal Diversity 105:1-16 ((https://doi.org/10.1007/s13225-020-00466-2)) |
| Otu18518 | 1 | k__Fungi(100);p__Ascomycota(100);c__Saccharomycetes(100);o__Saccharomycetales(100);f__Metschnikowiaceae(100);g__Metschnikowia(100);g__Metschnikowia_unclassified(100); | Metschnikowia | 13 | Saprotroph | Undefined Saprotroph | Yeast | NULL | Probable | As nectar-tap_saprotroph (Põlme et al. 2020); Decay sugar-rich substrates (Põlme et al. 2020) | Põlme S, et al. 2020. Fungal Diversity 105:1-16 ((https://doi.org/10.1007/s13225-020-00466-2)) |
| Otu18519 | 1 | k__Fungi(100);p__Ascomycota(100);c__Saccharomycetes(100);o__Saccharomycetales(100);f__Metschnikowiaceae(100);g__Metschnikowia(100);s__Metschnikowia_pulcherrima(100); | Metschnikowia | 13 | Saprotroph | Undefined Saprotroph | Yeast | NULL | Probable | As nectar-tap_saprotroph (Põlme et al. 2020); Decay sugar-rich substrates (Põlme et al. 2020) | Põlme S, et al. 2020. Fungal Diversity 105:1-16 ((https://doi.org/10.1007/s13225-020-00466-2)) |
| Otu18520 | 1 | k__Fungi(100);p__Ascomycota(100);c__Saccharomycetes(100);o__Saccharomycetales(100);f__Metschnikowiaceae(100);g__Metschnikowia(100);s__Metschnikowia_pulcherrima(100); | Metschnikowia | 13 | Saprotroph | Undefined Saprotroph | Yeast | NULL | Probable | As nectar-tap_saprotroph (Põlme et al. 2020); Decay sugar-rich substrates (Põlme et al. 2020) | Põlme S, et al. 2020. Fungal Diversity 105:1-16 ((https://doi.org/10.1007/s13225-020-00466-2)) |
| Otu18521 | 1 | k__Fungi(100);p__Ascomycota(100);c__Saccharomycetes(100);o__Saccharomycetales(100);f__Metschnikowiaceae(100);g__Metschnikowia(100);s__Metschnikowia_pulcherrima(100); | Metschnikowia | 13 | Saprotroph | Undefined Saprotroph | Yeast | NULL | Probable | As nectar-tap_saprotroph (Põlme et al. 2020); Decay sugar-rich substrates (Põlme et al. 2020) | Põlme S, et al. 2020. Fungal Diversity 105:1-16 ((https://doi.org/10.1007/s13225-020-00466-2)) |
| Otu18522 | 1 | k__Fungi(100);p__Ascomycota(100);c__Saccharomycetes(100);o__Saccharomycetales(100);f__Metschnikowiaceae(100);g__Metschnikowia(100);s__Metschnikowia_pulcherrima(100); | Metschnikowia | 13 | Saprotroph | Undefined Saprotroph | Yeast | NULL | Probable | As nectar-tap_saprotroph (Põlme et al. 2020); Decay sugar-rich substrates (Põlme et al. 2020) | Põlme S, et al. 2020. Fungal Diversity 105:1-16 ((https://doi.org/10.1007/s13225-020-00466-2)) |
| Otu18523 | 1 | k__Fungi(100);p__Ascomycota(100);c__Saccharomycetes(100);o__Saccharomycetales(100);f__Metschnikowiaceae(100);g__Metschnikowia(100);s__Metschnikowia_pulcherrima(100); | Metschnikowia | 13 | Saprotroph | Undefined Saprotroph | Yeast | NULL | Probable | As nectar-tap_saprotroph (Põlme et al. 2020); Decay sugar-rich substrates (Põlme et al. 2020) | Põlme S, et al. 2020. Fungal Diversity 105:1-16 ((https://doi.org/10.1007/s13225-020-00466-2)) |
| Otu18524 | 1 | k__Fungi(100);p__Ascomycota(100);c__Saccharomycetes(100);o__Saccharomycetales(100);f__Metschnikowiaceae(100);g__Metschnikowia(100);s__Metschnikowia_pulcherrima(100); | Metschnikowia | 13 | Saprotroph | Undefined Saprotroph | Yeast | NULL | Probable | As nectar-tap_saprotroph (Põlme et al. 2020); Decay sugar-rich substrates (Põlme et al. 2020) | Põlme S, et al. 2020. Fungal Diversity 105:1-16 ((https://doi.org/10.1007/s13225-020-00466-2)) |
| Otu18525 | 1 | k__Fungi(100);p__Ascomycota(100);c__Saccharomycetes(100);o__Saccharomycetales(100);f__Metschnikowiaceae(100);g__Metschnikowia(100);g__Metschnikowia_unclassified(100); | Metschnikowia | 13 | Saprotroph | Undefined Saprotroph | Yeast | NULL | Probable | As nectar-tap_saprotroph (Põlme et al. 2020); Decay sugar-rich substrates (Põlme et al. 2020) | Põlme S, et al. 2020. Fungal Diversity 105:1-16 ((https://doi.org/10.1007/s13225-020-00466-2)) |
| Otu18526 | 1 | k__Fungi(100);p__Ascomycota(100);c__Saccharomycetes(100);o__Saccharomycetales(100);f__Metschnikowiaceae(100);g__Metschnikowia(100);g__Metschnikowia_unclassified(100); | Metschnikowia | 13 | Saprotroph | Undefined Saprotroph | Yeast | NULL | Probable | As nectar-tap_saprotroph (Põlme et al. 2020); Decay sugar-rich substrates (Põlme et al. 2020) | Põlme S, et al. 2020. Fungal Diversity 105:1-16 ((https://doi.org/10.1007/s13225-020-00466-2)) |
| Otu18528 | 1 | k__Fungi(100);p__Ascomycota(100);c__Saccharomycetes(100);o__Saccharomycetales(100);f__Metschnikowiaceae(100);g__Metschnikowia(100);s__Metschnikowia_pulcherrima(100); | Metschnikowia | 13 | Saprotroph | Undefined Saprotroph | Yeast | NULL | Probable | As nectar-tap_saprotroph (Põlme et al. 2020); Decay sugar-rich substrates (Põlme et al. 2020) | Põlme S, et al. 2020. Fungal Diversity 105:1-16 ((https://doi.org/10.1007/s13225-020-00466-2)) |
| Otu18529 | 1 | k__Fungi(100);p__Ascomycota(100);c__Saccharomycetes(100);o__Saccharomycetales(100);f__Metschnikowiaceae(100);g__Metschnikowia(100);g__Metschnikowia_unclassified(100); | Metschnikowia | 13 | Saprotroph | Undefined Saprotroph | Yeast | NULL | Probable | As nectar-tap_saprotroph (Põlme et al. 2020); Decay sugar-rich substrates (Põlme et al. 2020) | Põlme S, et al. 2020. Fungal Diversity 105:1-16 ((https://doi.org/10.1007/s13225-020-00466-2)) |
| Otu18530 | 1 | k__Fungi(100);p__Ascomycota(100);c__Saccharomycetes(100);o__Saccharomycetales(100);f__Metschnikowiaceae(100);g__Metschnikowia(100);s__Metschnikowia_pulcherrima(100); | Metschnikowia | 13 | Saprotroph | Undefined Saprotroph | Yeast | NULL | Probable | As nectar-tap_saprotroph (Põlme et al. 2020); Decay sugar-rich substrates (Põlme et al. 2020) | Põlme S, et al. 2020. Fungal Diversity 105:1-16 ((https://doi.org/10.1007/s13225-020-00466-2)) |
| Otu18533 | 1 | k__Fungi(100);p__Ascomycota(100);c__Saccharomycetes(100);o__Saccharomycetales(100);f__Metschnikowiaceae(100);g__Metschnikowia(100);g__Metschnikowia_unclassified(100); | Metschnikowia | 13 | Saprotroph | Undefined Saprotroph | Yeast | NULL | Probable | As nectar-tap_saprotroph (Põlme et al. 2020); Decay sugar-rich substrates (Põlme et al. 2020) | Põlme S, et al. 2020. Fungal Diversity 105:1-16 ((https://doi.org/10.1007/s13225-020-00466-2)) |
| Otu18535 | 1 | k__Fungi(100);p__Ascomycota(100);c__Saccharomycetes(100);o__Saccharomycetales(100);f__Metschnikowiaceae(100);g__Metschnikowia(100);g__Metschnikowia_unclassified(100); | Metschnikowia | 13 | Saprotroph | Undefined Saprotroph | Yeast | NULL | Probable | As nectar-tap_saprotroph (Põlme et al. 2020); Decay sugar-rich substrates (Põlme et al. 2020) | Põlme S, et al. 2020. Fungal Diversity 105:1-16 ((https://doi.org/10.1007/s13225-020-00466-2)) |
| Otu18536 | 1 | k__Fungi(100);p__Ascomycota(100);c__Saccharomycetes(100);o__Saccharomycetales(100);f__Metschnikowiaceae(100);g__Metschnikowia(100);s__Metschnikowia_pulcherrima(100); | Metschnikowia | 13 | Saprotroph | Undefined Saprotroph | Yeast | NULL | Probable | As nectar-tap_saprotroph (Põlme et al. 2020); Decay sugar-rich substrates (Põlme et al. 2020) | Põlme S, et al. 2020. Fungal Diversity 105:1-16 ((https://doi.org/10.1007/s13225-020-00466-2)) |
| Otu18538 | 1 | k__Fungi(100);p__Ascomycota(100);c__Saccharomycetes(100);o__Saccharomycetales(100);f__Saccharomycetales_family_Incertae_sedis(100);g__Candida(100);s__Candida_cretensis(100); | Candida | 13 | Pathotroph-Saprotroph-Symbiotroph | Animal Pathogen-Endophyte-Endosymbiont-Epiphyte-Undefined Saprotroph | Dimorphic Yeast-Yeast | NULL | Probable | As nectar-tap_saprotroph (Põlme et al. 2020); Decay sugar-rich substrates (Põlme et al. 2020); This asexual genus is polyphyletic and contain species that function very differently depend on what they are. Caution should be taken when interpreting guilds for Candida species. | Manolakaki D, et al. 2010. Virulence. 1: 367-375 ((https://doi.org/10.4161/viru.1.5.12796)); Mekbib SB, et al. 2011. Tropical Plant Pathology 36:233-240 ((https://doi.org/10.1590/S1982-56762011000400004)); Põlme S, et al. 2020. Fungal Diversity 105:1-16 ((https://doi.org/10.1007/s13225-020-00466-2)) |
| Otu18540 | 1 | k__Fungi(100);p__Ascomycota(100);c__Saccharomycetes(100);o__Saccharomycetales(100);f__Saccharomycetales_family_Incertae_sedis(100);g__Candida(100);g__Candida_unclassified(100); | Candida | 13 | Pathotroph-Saprotroph-Symbiotroph | Animal Pathogen-Endophyte-Endosymbiont-Epiphyte-Undefined Saprotroph | Dimorphic Yeast-Yeast | NULL | Probable | As nectar-tap_saprotroph (Põlme et al. 2020); Decay sugar-rich substrates (Põlme et al. 2020); This asexual genus is polyphyletic and contain species that function very differently depend on what they are. Caution should be taken when interpreting guilds for Candida species. | Manolakaki D, et al. 2010. Virulence. 1: 367-375 ((https://doi.org/10.4161/viru.1.5.12796)); Mekbib SB, et al. 2011. Tropical Plant Pathology 36:233-240 ((https://doi.org/10.1590/S1982-56762011000400004)); Põlme S, et al. 2020. Fungal Diversity 105:1-16 ((https://doi.org/10.1007/s13225-020-00466-2)) |
| Otu18541 | 1 | k__Fungi(100);p__Ascomycota(100);c__Saccharomycetes(100);o__Saccharomycetales(100);f__Metschnikowiaceae(100);g__Metschnikowia(100);g__Metschnikowia_unclassified(100); | Metschnikowia | 13 | Saprotroph | Undefined Saprotroph | Yeast | NULL | Probable | As nectar-tap_saprotroph (Põlme et al. 2020); Decay sugar-rich substrates (Põlme et al. 2020) | Põlme S, et al. 2020. Fungal Diversity 105:1-16 ((https://doi.org/10.1007/s13225-020-00466-2)) |
| Otu18544 | 1 | k__Fungi(100);p__Ascomycota(100);c__Saccharomycetes(100);o__Saccharomycetales(100);f__Metschnikowiaceae(100);g__Metschnikowia(100);s__Metschnikowia_pulcherrima(100); | Metschnikowia | 13 | Saprotroph | Undefined Saprotroph | Yeast | NULL | Probable | As nectar-tap_saprotroph (Põlme et al. 2020); Decay sugar-rich substrates (Põlme et al. 2020) | Põlme S, et al. 2020. Fungal Diversity 105:1-16 ((https://doi.org/10.1007/s13225-020-00466-2)) |
| Otu18545 | 1 | k__Fungi(100);p__Ascomycota(100);c__Saccharomycetes(100);o__Saccharomycetales(100);f__Metschnikowiaceae(100);g__Metschnikowia(100);s__Metschnikowia_pulcherrima(100); | Metschnikowia | 13 | Saprotroph | Undefined Saprotroph | Yeast | NULL | Probable | As nectar-tap_saprotroph (Põlme et al. 2020); Decay sugar-rich substrates (Põlme et al. 2020) | Põlme S, et al. 2020. Fungal Diversity 105:1-16 ((https://doi.org/10.1007/s13225-020-00466-2)) |
| Otu18546 | 1 | k__Fungi(100);p__Ascomycota(100);c__Saccharomycetes(100);o__Saccharomycetales(100);f__Metschnikowiaceae(100);g__Metschnikowia(100);s__Metschnikowia_pulcherrima(100); | Metschnikowia | 13 | Saprotroph | Undefined Saprotroph | Yeast | NULL | Probable | As nectar-tap_saprotroph (Põlme et al. 2020); Decay sugar-rich substrates (Põlme et al. 2020) | Põlme S, et al. 2020. Fungal Diversity 105:1-16 ((https://doi.org/10.1007/s13225-020-00466-2)) |
| Otu18547 | 1 | k__Fungi(100);p__Ascomycota(100);c__Saccharomycetes(100);o__Saccharomycetales(100);f__Metschnikowiaceae(100);g__Metschnikowia(100);s__Metschnikowia_pulcherrima(100); | Metschnikowia | 13 | Saprotroph | Undefined Saprotroph | Yeast | NULL | Probable | As nectar-tap_saprotroph (Põlme et al. 2020); Decay sugar-rich substrates (Põlme et al. 2020) | Põlme S, et al. 2020. Fungal Diversity 105:1-16 ((https://doi.org/10.1007/s13225-020-00466-2)) |
| Otu18549 | 1 | k__Fungi(100);p__Ascomycota(100);c__Saccharomycetes(100);o__Saccharomycetales(100);f__Metschnikowiaceae(100);g__Metschnikowia(100);g__Metschnikowia_unclassified(100); | Metschnikowia | 13 | Saprotroph | Undefined Saprotroph | Yeast | NULL | Probable | As nectar-tap_saprotroph (Põlme et al. 2020); Decay sugar-rich substrates (Põlme et al. 2020) | Põlme S, et al. 2020. Fungal Diversity 105:1-16 ((https://doi.org/10.1007/s13225-020-00466-2)) |
| Otu18550 | 1 | k__Fungi(100);p__Ascomycota(100);c__Saccharomycetes(100);o__Saccharomycetales(100);f__Metschnikowiaceae(100);g__Metschnikowia(100);g__Metschnikowia_unclassified(100); | Metschnikowia | 13 | Saprotroph | Undefined Saprotroph | Yeast | NULL | Probable | As nectar-tap_saprotroph (Põlme et al. 2020); Decay sugar-rich substrates (Põlme et al. 2020) | Põlme S, et al. 2020. Fungal Diversity 105:1-16 ((https://doi.org/10.1007/s13225-020-00466-2)) |
| Otu18551 | 1 | k__Fungi(100);p__Ascomycota(100);c__Saccharomycetes(100);o__Saccharomycetales(100);f__Metschnikowiaceae(100);g__Metschnikowia(100);s__Metschnikowia_pulcherrima(100); | Metschnikowia | 13 | Saprotroph | Undefined Saprotroph | Yeast | NULL | Probable | As nectar-tap_saprotroph (Põlme et al. 2020); Decay sugar-rich substrates (Põlme et al. 2020) | Põlme S, et al. 2020. Fungal Diversity 105:1-16 ((https://doi.org/10.1007/s13225-020-00466-2)) |
| Otu18552 | 1 | k__Fungi(100);p__Ascomycota(100);c__Saccharomycetes(100);o__Saccharomycetales(100);f__Metschnikowiaceae(100);g__Metschnikowia(100);s__Metschnikowia_pulcherrima(100); | Metschnikowia | 13 | Saprotroph | Undefined Saprotroph | Yeast | NULL | Probable | As nectar-tap_saprotroph (Põlme et al. 2020); Decay sugar-rich substrates (Põlme et al. 2020) | Põlme S, et al. 2020. Fungal Diversity 105:1-16 ((https://doi.org/10.1007/s13225-020-00466-2)) |
| Otu18553 | 1 | k__Fungi(100);p__Ascomycota(100);c__Saccharomycetes(100);o__Saccharomycetales(100);f__Metschnikowiaceae(100);g__Metschnikowia(100);s__Metschnikowia_pulcherrima(100); | Metschnikowia | 13 | Saprotroph | Undefined Saprotroph | Yeast | NULL | Probable | As nectar-tap_saprotroph (Põlme et al. 2020); Decay sugar-rich substrates (Põlme et al. 2020) | Põlme S, et al. 2020. Fungal Diversity 105:1-16 ((https://doi.org/10.1007/s13225-020-00466-2)) |
| Otu18554 | 1 | k__Fungi(100);p__Ascomycota(100);c__Saccharomycetes(100);o__Saccharomycetales(100);f__Metschnikowiaceae(100);g__Metschnikowia(100);s__Metschnikowia_pulcherrima(100); | Metschnikowia | 13 | Saprotroph | Undefined Saprotroph | Yeast | NULL | Probable | As nectar-tap_saprotroph (Põlme et al. 2020); Decay sugar-rich substrates (Põlme et al. 2020) | Põlme S, et al. 2020. Fungal Diversity 105:1-16 ((https://doi.org/10.1007/s13225-020-00466-2)) |
| Otu18555 | 1 | k__Fungi(100);p__Ascomycota(100);c__Saccharomycetes(100);o__Saccharomycetales(100);f__Metschnikowiaceae(100);g__Metschnikowia(100);g__Metschnikowia_unclassified(100); | Metschnikowia | 13 | Saprotroph | Undefined Saprotroph | Yeast | NULL | Probable | As nectar-tap_saprotroph (Põlme et al. 2020); Decay sugar-rich substrates (Põlme et al. 2020) | Põlme S, et al. 2020. Fungal Diversity 105:1-16 ((https://doi.org/10.1007/s13225-020-00466-2)) |
| Otu18556 | 1 | k__Fungi(100);p__Ascomycota(100);c__Saccharomycetes(100);o__Saccharomycetales(100);f__Metschnikowiaceae(100);g__Metschnikowia(100);g__Metschnikowia_unclassified(100); | Metschnikowia | 13 | Saprotroph | Undefined Saprotroph | Yeast | NULL | Probable | As nectar-tap_saprotroph (Põlme et al. 2020); Decay sugar-rich substrates (Põlme et al. 2020) | Põlme S, et al. 2020. Fungal Diversity 105:1-16 ((https://doi.org/10.1007/s13225-020-00466-2)) |
| Otu18557 | 1 | k__Fungi(100);p__Ascomycota(100);c__Saccharomycetes(100);o__Saccharomycetales(100);f__Metschnikowiaceae(100);g__Metschnikowia(100);s__Metschnikowia_pulcherrima(100); | Metschnikowia | 13 | Saprotroph | Undefined Saprotroph | Yeast | NULL | Probable | As nectar-tap_saprotroph (Põlme et al. 2020); Decay sugar-rich substrates (Põlme et al. 2020) | Põlme S, et al. 2020. Fungal Diversity 105:1-16 ((https://doi.org/10.1007/s13225-020-00466-2)) |
| Otu18559 | 1 | k__Fungi(100);p__Ascomycota(100);c__Saccharomycetes(100);o__Saccharomycetales(100);f__Metschnikowiaceae(100);g__Metschnikowia(100);g__Metschnikowia_unclassified(100); | Metschnikowia | 13 | Saprotroph | Undefined Saprotroph | Yeast | NULL | Probable | As nectar-tap_saprotroph (Põlme et al. 2020); Decay sugar-rich substrates (Põlme et al. 2020) | Põlme S, et al. 2020. Fungal Diversity 105:1-16 ((https://doi.org/10.1007/s13225-020-00466-2)) |
| Otu18560 | 1 | k__Fungi(100);p__Ascomycota(100);c__Saccharomycetes(100);o__Saccharomycetales(100);f__Metschnikowiaceae(100);g__Metschnikowia(100);s__Metschnikowia_pulcherrima(100); | Metschnikowia | 13 | Saprotroph | Undefined Saprotroph | Yeast | NULL | Probable | As nectar-tap_saprotroph (Põlme et al. 2020); Decay sugar-rich substrates (Põlme et al. 2020) | Põlme S, et al. 2020. Fungal Diversity 105:1-16 ((https://doi.org/10.1007/s13225-020-00466-2)) |
| Otu18561 | 1 | k__Fungi(100);p__Ascomycota(100);c__Saccharomycetes(100);o__Saccharomycetales(100);f__Metschnikowiaceae(100);g__Metschnikowia(100);s__Metschnikowia_pulcherrima(100); | Metschnikowia | 13 | Saprotroph | Undefined Saprotroph | Yeast | NULL | Probable | As nectar-tap_saprotroph (Põlme et al. 2020); Decay sugar-rich substrates (Põlme et al. 2020) | Põlme S, et al. 2020. Fungal Diversity 105:1-16 ((https://doi.org/10.1007/s13225-020-00466-2)) |
| Otu18562 | 1 | k__Fungi(100);p__Ascomycota(100);c__Saccharomycetes(100);o__Saccharomycetales(100);o__Saccharomycetales_unclassified(100);o__Saccharomycetales_unclassified(100);o__Saccharomycetales_unclassified(100); | Saccharomycetales | 7 | Saprotroph | Undefined Saprotroph | Yeast | NULL | Possible | NULL | Sterrenburg E, et al. 2015. New Phytologist 207:1145-1158 (https://doi.org/10.1111/nph.13426) |
| Otu18564 | 1 | k__Fungi(100);p__Ascomycota(100);c__Saccharomycetes(100);o__Saccharomycetales(100);f__Metschnikowiaceae(100);g__Metschnikowia(100);s__Metschnikowia_pulcherrima(100); | Metschnikowia | 13 | Saprotroph | Undefined Saprotroph | Yeast | NULL | Probable | As nectar-tap_saprotroph (Põlme et al. 2020); Decay sugar-rich substrates (Põlme et al. 2020) | Põlme S, et al. 2020. Fungal Diversity 105:1-16 ((https://doi.org/10.1007/s13225-020-00466-2)) |
| Otu18565 | 1 | k__Fungi(100);p__Ascomycota(100);c__Saccharomycetes(100);o__Saccharomycetales(100);f__Metschnikowiaceae(100);g__Metschnikowia(100);s__Metschnikowia_pulcherrima(100); | Metschnikowia | 13 | Saprotroph | Undefined Saprotroph | Yeast | NULL | Probable | As nectar-tap_saprotroph (Põlme et al. 2020); Decay sugar-rich substrates (Põlme et al. 2020) | Põlme S, et al. 2020. Fungal Diversity 105:1-16 ((https://doi.org/10.1007/s13225-020-00466-2)) |
| Otu18566 | 1 | k__Fungi(100);p__Ascomycota(100);c__Saccharomycetes(100);o__Saccharomycetales(100);f__Metschnikowiaceae(100);g__Metschnikowia(100);s__Metschnikowia_pulcherrima(100); | Metschnikowia | 13 | Saprotroph | Undefined Saprotroph | Yeast | NULL | Probable | As nectar-tap_saprotroph (Põlme et al. 2020); Decay sugar-rich substrates (Põlme et al. 2020) | Põlme S, et al. 2020. Fungal Diversity 105:1-16 ((https://doi.org/10.1007/s13225-020-00466-2)) |
| Otu18567 | 1 | k__Fungi(100);p__Ascomycota(100);c__Saccharomycetes(100);o__Saccharomycetales(100);f__Metschnikowiaceae(100);g__Metschnikowia(100);g__Metschnikowia_unclassified(100); | Metschnikowia | 13 | Saprotroph | Undefined Saprotroph | Yeast | NULL | Probable | As nectar-tap_saprotroph (Põlme et al. 2020); Decay sugar-rich substrates (Põlme et al. 2020) | Põlme S, et al. 2020. Fungal Diversity 105:1-16 ((https://doi.org/10.1007/s13225-020-00466-2)) |
| Otu18568 | 1 | k__Fungi(100);p__Ascomycota(100);c__Saccharomycetes(100);o__Saccharomycetales(100);f__Metschnikowiaceae(100);g__Metschnikowia(100);s__Metschnikowia_pulcherrima(100); | Metschnikowia | 13 | Saprotroph | Undefined Saprotroph | Yeast | NULL | Probable | As nectar-tap_saprotroph (Põlme et al. 2020); Decay sugar-rich substrates (Põlme et al. 2020) | Põlme S, et al. 2020. Fungal Diversity 105:1-16 ((https://doi.org/10.1007/s13225-020-00466-2)) |
| Otu18569 | 1 | k__Fungi(100);p__Ascomycota(100);c__Saccharomycetes(100);o__Saccharomycetales(100);f__Metschnikowiaceae(100);g__Metschnikowia(100);s__Metschnikowia_pulcherrima(100); | Metschnikowia | 13 | Saprotroph | Undefined Saprotroph | Yeast | NULL | Probable | As nectar-tap_saprotroph (Põlme et al. 2020); Decay sugar-rich substrates (Põlme et al. 2020) | Põlme S, et al. 2020. Fungal Diversity 105:1-16 ((https://doi.org/10.1007/s13225-020-00466-2)) |
| Otu18570 | 1 | k__Fungi(100);p__Ascomycota(100);c__Saccharomycetes(100);o__Saccharomycetales(100);f__Metschnikowiaceae(100);g__Metschnikowia(100);s__Metschnikowia_pulcherrima(100); | Metschnikowia | 13 | Saprotroph | Undefined Saprotroph | Yeast | NULL | Probable | As nectar-tap_saprotroph (Põlme et al. 2020); Decay sugar-rich substrates (Põlme et al. 2020) | Põlme S, et al. 2020. Fungal Diversity 105:1-16 ((https://doi.org/10.1007/s13225-020-00466-2)) |
| Otu18571 | 1 | k__Fungi(100);p__Ascomycota(100);c__Saccharomycetes(100);o__Saccharomycetales(100);f__Metschnikowiaceae(100);g__Metschnikowia(100);s__Metschnikowia_pulcherrima(100); | Metschnikowia | 13 | Saprotroph | Undefined Saprotroph | Yeast | NULL | Probable | As nectar-tap_saprotroph (Põlme et al. 2020); Decay sugar-rich substrates (Põlme et al. 2020) | Põlme S, et al. 2020. Fungal Diversity 105:1-16 ((https://doi.org/10.1007/s13225-020-00466-2)) |
| Otu18573 | 1 | k__Fungi(100);p__Ascomycota(100);c__Saccharomycetes(100);o__Saccharomycetales(100);f__Metschnikowiaceae(100);g__Metschnikowia(100);s__Metschnikowia_pulcherrima(100); | Metschnikowia | 13 | Saprotroph | Undefined Saprotroph | Yeast | NULL | Probable | As nectar-tap_saprotroph (Põlme et al. 2020); Decay sugar-rich substrates (Põlme et al. 2020) | Põlme S, et al. 2020. Fungal Diversity 105:1-16 ((https://doi.org/10.1007/s13225-020-00466-2)) |
| Otu18574 | 1 | k__Fungi(100);p__Ascomycota(100);c__Saccharomycetes(100);o__Saccharomycetales(100);f__Metschnikowiaceae(100);g__Metschnikowia(100);g__Metschnikowia_unclassified(100); | Metschnikowia | 13 | Saprotroph | Undefined Saprotroph | Yeast | NULL | Probable | As nectar-tap_saprotroph (Põlme et al. 2020); Decay sugar-rich substrates (Põlme et al. 2020) | Põlme S, et al. 2020. Fungal Diversity 105:1-16 ((https://doi.org/10.1007/s13225-020-00466-2)) |
| Otu18575 | 1 | k__Fungi(100);p__Ascomycota(100);c__Saccharomycetes(100);o__Saccharomycetales(100);f__Metschnikowiaceae(100);g__Metschnikowia(100);s__Metschnikowia_pulcherrima(100); | Metschnikowia | 13 | Saprotroph | Undefined Saprotroph | Yeast | NULL | Probable | As nectar-tap_saprotroph (Põlme et al. 2020); Decay sugar-rich substrates (Põlme et al. 2020) | Põlme S, et al. 2020. Fungal Diversity 105:1-16 ((https://doi.org/10.1007/s13225-020-00466-2)) |
| Otu18576 | 1 | k__Fungi(100);p__Ascomycota(100);c__Saccharomycetes(100);o__Saccharomycetales(100);f__Metschnikowiaceae(100);g__Metschnikowia(100);s__Metschnikowia_pulcherrima(100); | Metschnikowia | 13 | Saprotroph | Undefined Saprotroph | Yeast | NULL | Probable | As nectar-tap_saprotroph (Põlme et al. 2020); Decay sugar-rich substrates (Põlme et al. 2020) | Põlme S, et al. 2020. Fungal Diversity 105:1-16 ((https://doi.org/10.1007/s13225-020-00466-2)) |
| Otu18577 | 1 | k__Fungi(100);p__Ascomycota(100);c__Saccharomycetes(100);o__Saccharomycetales(100);f__Metschnikowiaceae(100);g__Metschnikowia(100);s__Metschnikowia_pulcherrima(100); | Metschnikowia | 13 | Saprotroph | Undefined Saprotroph | Yeast | NULL | Probable | As nectar-tap_saprotroph (Põlme et al. 2020); Decay sugar-rich substrates (Põlme et al. 2020) | Põlme S, et al. 2020. Fungal Diversity 105:1-16 ((https://doi.org/10.1007/s13225-020-00466-2)) |
| Otu18578 | 1 | k__Fungi(100);p__Ascomycota(100);c__Saccharomycetes(100);o__Saccharomycetales(100);f__Metschnikowiaceae(100);g__Metschnikowia(100);s__Metschnikowia_pulcherrima(100); | Metschnikowia | 13 | Saprotroph | Undefined Saprotroph | Yeast | NULL | Probable | As nectar-tap_saprotroph (Põlme et al. 2020); Decay sugar-rich substrates (Põlme et al. 2020) | Põlme S, et al. 2020. Fungal Diversity 105:1-16 ((https://doi.org/10.1007/s13225-020-00466-2)) |
| Otu18579 | 1 | k__Fungi(100);p__Ascomycota(100);c__Saccharomycetes(100);o__Saccharomycetales(100);f__Metschnikowiaceae(100);g__Metschnikowia(100);g__Metschnikowia_unclassified(100); | Metschnikowia | 13 | Saprotroph | Undefined Saprotroph | Yeast | NULL | Probable | As nectar-tap_saprotroph (Põlme et al. 2020); Decay sugar-rich substrates (Põlme et al. 2020) | Põlme S, et al. 2020. Fungal Diversity 105:1-16 ((https://doi.org/10.1007/s13225-020-00466-2)) |
| Otu18580 | 1 | k__Fungi(100);p__Ascomycota(100);c__Saccharomycetes(100);o__Saccharomycetales(100);f__Metschnikowiaceae(100);g__Metschnikowia(100);s__Metschnikowia_pulcherrima(100); | Metschnikowia | 13 | Saprotroph | Undefined Saprotroph | Yeast | NULL | Probable | As nectar-tap_saprotroph (Põlme et al. 2020); Decay sugar-rich substrates (Põlme et al. 2020) | Põlme S, et al. 2020. Fungal Diversity 105:1-16 ((https://doi.org/10.1007/s13225-020-00466-2)) |
| Otu18581 | 1 | k__Fungi(100);p__Ascomycota(100);c__Saccharomycetes(100);o__Saccharomycetales(100);f__Metschnikowiaceae(100);g__Metschnikowia(100);s__Metschnikowia_pulcherrima(100); | Metschnikowia | 13 | Saprotroph | Undefined Saprotroph | Yeast | NULL | Probable | As nectar-tap_saprotroph (Põlme et al. 2020); Decay sugar-rich substrates (Põlme et al. 2020) | Põlme S, et al. 2020. Fungal Diversity 105:1-16 ((https://doi.org/10.1007/s13225-020-00466-2)) |
| Otu18582 | 1 | k__Fungi(100);p__Ascomycota(100);c__Saccharomycetes(100);o__Saccharomycetales(100);f__Metschnikowiaceae(100);g__Metschnikowia(100);s__Metschnikowia_pulcherrima(100); | Metschnikowia | 13 | Saprotroph | Undefined Saprotroph | Yeast | NULL | Probable | As nectar-tap_saprotroph (Põlme et al. 2020); Decay sugar-rich substrates (Põlme et al. 2020) | Põlme S, et al. 2020. Fungal Diversity 105:1-16 ((https://doi.org/10.1007/s13225-020-00466-2)) |
| Otu18583 | 1 | k__Fungi(100);p__Ascomycota(100);c__Saccharomycetes(100);o__Saccharomycetales(100);f__Metschnikowiaceae(100);g__Metschnikowia(100);s__Metschnikowia_pulcherrima(100); | Metschnikowia | 13 | Saprotroph | Undefined Saprotroph | Yeast | NULL | Probable | As nectar-tap_saprotroph (Põlme et al. 2020); Decay sugar-rich substrates (Põlme et al. 2020) | Põlme S, et al. 2020. Fungal Diversity 105:1-16 ((https://doi.org/10.1007/s13225-020-00466-2)) |
| Otu18584 | 1 | k__Fungi(100);p__Ascomycota(100);c__Saccharomycetes(100);o__Saccharomycetales(100);f__Metschnikowiaceae(100);g__Metschnikowia(100);s__Metschnikowia_pulcherrima(100); | Metschnikowia | 13 | Saprotroph | Undefined Saprotroph | Yeast | NULL | Probable | As nectar-tap_saprotroph (Põlme et al. 2020); Decay sugar-rich substrates (Põlme et al. 2020) | Põlme S, et al. 2020. Fungal Diversity 105:1-16 ((https://doi.org/10.1007/s13225-020-00466-2)) |
| Otu18585 | 1 | k__Fungi(100);p__Ascomycota(100);c__Saccharomycetes(100);o__Saccharomycetales(100);f__Metschnikowiaceae(100);g__Metschnikowia(100);s__Metschnikowia_pulcherrima(100); | Metschnikowia | 13 | Saprotroph | Undefined Saprotroph | Yeast | NULL | Probable | As nectar-tap_saprotroph (Põlme et al. 2020); Decay sugar-rich substrates (Põlme et al. 2020) | Põlme S, et al. 2020. Fungal Diversity 105:1-16 ((https://doi.org/10.1007/s13225-020-00466-2)) |
| Otu18586 | 1 | k__Fungi(100);p__Ascomycota(100);c__Saccharomycetes(100);o__Saccharomycetales(100);f__Metschnikowiaceae(100);g__Metschnikowia(100);s__Metschnikowia_pulcherrima(100); | Metschnikowia | 13 | Saprotroph | Undefined Saprotroph | Yeast | NULL | Probable | As nectar-tap_saprotroph (Põlme et al. 2020); Decay sugar-rich substrates (Põlme et al. 2020) | Põlme S, et al. 2020. Fungal Diversity 105:1-16 ((https://doi.org/10.1007/s13225-020-00466-2)) |
| Otu18587 | 1 | k__Fungi(100);p__Ascomycota(100);c__Saccharomycetes(100);o__Saccharomycetales(100);f__Metschnikowiaceae(100);g__Metschnikowia(100);g__Metschnikowia_unclassified(100); | Metschnikowia | 13 | Saprotroph | Undefined Saprotroph | Yeast | NULL | Probable | As nectar-tap_saprotroph (Põlme et al. 2020); Decay sugar-rich substrates (Põlme et al. 2020) | Põlme S, et al. 2020. Fungal Diversity 105:1-16 ((https://doi.org/10.1007/s13225-020-00466-2)) |
| Otu18588 | 1 | k__Fungi(100);p__Ascomycota(100);c__Saccharomycetes(100);o__Saccharomycetales(100);f__Metschnikowiaceae(100);g__Metschnikowia(100);g__Metschnikowia_unclassified(100); | Metschnikowia | 13 | Saprotroph | Undefined Saprotroph | Yeast | NULL | Probable | As nectar-tap_saprotroph (Põlme et al. 2020); Decay sugar-rich substrates (Põlme et al. 2020) | Põlme S, et al. 2020. Fungal Diversity 105:1-16 ((https://doi.org/10.1007/s13225-020-00466-2)) |
| Otu18589 | 1 | k__Fungi(100);p__Ascomycota(100);c__Saccharomycetes(100);o__Saccharomycetales(100);f__Metschnikowiaceae(100);g__Metschnikowia(100);s__Metschnikowia_pulcherrima(100); | Metschnikowia | 13 | Saprotroph | Undefined Saprotroph | Yeast | NULL | Probable | As nectar-tap_saprotroph (Põlme et al. 2020); Decay sugar-rich substrates (Põlme et al. 2020) | Põlme S, et al. 2020. Fungal Diversity 105:1-16 ((https://doi.org/10.1007/s13225-020-00466-2)) |
| Otu18590 | 1 | k__Fungi(100);p__Ascomycota(100);c__Saccharomycetes(100);o__Saccharomycetales(100);f__Metschnikowiaceae(100);g__Metschnikowia(100);s__Metschnikowia_pulcherrima(100); | Metschnikowia | 13 | Saprotroph | Undefined Saprotroph | Yeast | NULL | Probable | As nectar-tap_saprotroph (Põlme et al. 2020); Decay sugar-rich substrates (Põlme et al. 2020) | Põlme S, et al. 2020. Fungal Diversity 105:1-16 ((https://doi.org/10.1007/s13225-020-00466-2)) |
| Otu18591 | 1 | k__Fungi(100);p__Ascomycota(100);c__Saccharomycetes(100);o__Saccharomycetales(100);f__Metschnikowiaceae(100);g__Metschnikowia(100);g__Metschnikowia_unclassified(100); | Metschnikowia | 13 | Saprotroph | Undefined Saprotroph | Yeast | NULL | Probable | As nectar-tap_saprotroph (Põlme et al. 2020); Decay sugar-rich substrates (Põlme et al. 2020) | Põlme S, et al. 2020. Fungal Diversity 105:1-16 ((https://doi.org/10.1007/s13225-020-00466-2)) |
| Otu18592 | 1 | k__Fungi(100);p__Ascomycota(100);c__Saccharomycetes(100);o__Saccharomycetales(100);f__Metschnikowiaceae(100);g__Metschnikowia(100);s__Metschnikowia_pulcherrima(100); | Metschnikowia | 13 | Saprotroph | Undefined Saprotroph | Yeast | NULL | Probable | As nectar-tap_saprotroph (Põlme et al. 2020); Decay sugar-rich substrates (Põlme et al. 2020) | Põlme S, et al. 2020. Fungal Diversity 105:1-16 ((https://doi.org/10.1007/s13225-020-00466-2)) |
| Otu18593 | 1 | k__Fungi(100);p__Ascomycota(100);c__Saccharomycetes(100);o__Saccharomycetales(100);f__Metschnikowiaceae(100);g__Metschnikowia(100);s__Metschnikowia_pulcherrima(100); | Metschnikowia | 13 | Saprotroph | Undefined Saprotroph | Yeast | NULL | Probable | As nectar-tap_saprotroph (Põlme et al. 2020); Decay sugar-rich substrates (Põlme et al. 2020) | Põlme S, et al. 2020. Fungal Diversity 105:1-16 ((https://doi.org/10.1007/s13225-020-00466-2)) |
| Otu18594 | 1 | k__Fungi(100);p__Ascomycota(100);c__Saccharomycetes(100);o__Saccharomycetales(100);f__Metschnikowiaceae(100);g__Metschnikowia(100);s__Metschnikowia_pulcherrima(100); | Metschnikowia | 13 | Saprotroph | Undefined Saprotroph | Yeast | NULL | Probable | As nectar-tap_saprotroph (Põlme et al. 2020); Decay sugar-rich substrates (Põlme et al. 2020) | Põlme S, et al. 2020. Fungal Diversity 105:1-16 ((https://doi.org/10.1007/s13225-020-00466-2)) |
| Otu18595 | 1 | k__Fungi(100);p__Ascomycota(100);c__Saccharomycetes(100);o__Saccharomycetales(100);f__Metschnikowiaceae(100);g__Metschnikowia(100);s__Metschnikowia_pulcherrima(100); | Metschnikowia | 13 | Saprotroph | Undefined Saprotroph | Yeast | NULL | Probable | As nectar-tap_saprotroph (Põlme et al. 2020); Decay sugar-rich substrates (Põlme et al. 2020) | Põlme S, et al. 2020. Fungal Diversity 105:1-16 ((https://doi.org/10.1007/s13225-020-00466-2)) |
| Otu18596 | 1 | k__Fungi(100);p__Ascomycota(100);c__Saccharomycetes(100);o__Saccharomycetales(100);f__Metschnikowiaceae(100);g__Metschnikowia(100);s__Metschnikowia_pulcherrima(100); | Metschnikowia | 13 | Saprotroph | Undefined Saprotroph | Yeast | NULL | Probable | As nectar-tap_saprotroph (Põlme et al. 2020); Decay sugar-rich substrates (Põlme et al. 2020) | Põlme S, et al. 2020. Fungal Diversity 105:1-16 ((https://doi.org/10.1007/s13225-020-00466-2)) |
| Otu18597 | 1 | k__Fungi(100);p__Ascomycota(100);c__Saccharomycetes(100);o__Saccharomycetales(100);f__Metschnikowiaceae(100);g__Metschnikowia(100);s__Metschnikowia_pulcherrima(100); | Metschnikowia | 13 | Saprotroph | Undefined Saprotroph | Yeast | NULL | Probable | As nectar-tap_saprotroph (Põlme et al. 2020); Decay sugar-rich substrates (Põlme et al. 2020) | Põlme S, et al. 2020. Fungal Diversity 105:1-16 ((https://doi.org/10.1007/s13225-020-00466-2)) |
| Otu18598 | 1 | k__Fungi(100);p__Ascomycota(100);c__Saccharomycetes(100);o__Saccharomycetales(100);f__Metschnikowiaceae(100);g__Metschnikowia(100);s__Metschnikowia_pulcherrima(100); | Metschnikowia | 13 | Saprotroph | Undefined Saprotroph | Yeast | NULL | Probable | As nectar-tap_saprotroph (Põlme et al. 2020); Decay sugar-rich substrates (Põlme et al. 2020) | Põlme S, et al. 2020. Fungal Diversity 105:1-16 ((https://doi.org/10.1007/s13225-020-00466-2)) |
| Otu18599 | 1 | k__Fungi(100);p__Ascomycota(100);c__Saccharomycetes(100);o__Saccharomycetales(100);f__Metschnikowiaceae(100);g__Metschnikowia(100);s__Metschnikowia_pulcherrima(100); | Metschnikowia | 13 | Saprotroph | Undefined Saprotroph | Yeast | NULL | Probable | As nectar-tap_saprotroph (Põlme et al. 2020); Decay sugar-rich substrates (Põlme et al. 2020) | Põlme S, et al. 2020. Fungal Diversity 105:1-16 ((https://doi.org/10.1007/s13225-020-00466-2)) |
| Otu18600 | 1 | k__Fungi(100);p__Ascomycota(100);c__Saccharomycetes(100);o__Saccharomycetales(100);f__Metschnikowiaceae(100);g__Metschnikowia(100);s__Metschnikowia_pulcherrima(100); | Metschnikowia | 13 | Saprotroph | Undefined Saprotroph | Yeast | NULL | Probable | As nectar-tap_saprotroph (Põlme et al. 2020); Decay sugar-rich substrates (Põlme et al. 2020) | Põlme S, et al. 2020. Fungal Diversity 105:1-16 ((https://doi.org/10.1007/s13225-020-00466-2)) |
| Otu18601 | 1 | k__Fungi(100);p__Ascomycota(100);c__Saccharomycetes(100);o__Saccharomycetales(100);f__Metschnikowiaceae(100);g__Metschnikowia(100);s__Metschnikowia_pulcherrima(100); | Metschnikowia | 13 | Saprotroph | Undefined Saprotroph | Yeast | NULL | Probable | As nectar-tap_saprotroph (Põlme et al. 2020); Decay sugar-rich substrates (Põlme et al. 2020) | Põlme S, et al. 2020. Fungal Diversity 105:1-16 ((https://doi.org/10.1007/s13225-020-00466-2)) |
| Otu18602 | 1 | k__Fungi(100);p__Ascomycota(100);c__Saccharomycetes(100);o__Saccharomycetales(100);f__Metschnikowiaceae(100);g__Metschnikowia(100);s__Metschnikowia_pulcherrima(100); | Metschnikowia | 13 | Saprotroph | Undefined Saprotroph | Yeast | NULL | Probable | As nectar-tap_saprotroph (Põlme et al. 2020); Decay sugar-rich substrates (Põlme et al. 2020) | Põlme S, et al. 2020. Fungal Diversity 105:1-16 ((https://doi.org/10.1007/s13225-020-00466-2)) |
| Otu18603 | 1 | k__Fungi(100);p__Ascomycota(100);c__Saccharomycetes(100);o__Saccharomycetales(100);f__Metschnikowiaceae(100);g__Metschnikowia(100);s__Metschnikowia_pulcherrima(100); | Metschnikowia | 13 | Saprotroph | Undefined Saprotroph | Yeast | NULL | Probable | As nectar-tap_saprotroph (Põlme et al. 2020); Decay sugar-rich substrates (Põlme et al. 2020) | Põlme S, et al. 2020. Fungal Diversity 105:1-16 ((https://doi.org/10.1007/s13225-020-00466-2)) |
| Otu18604 | 1 | k__Fungi(100);p__Ascomycota(100);c__Saccharomycetes(100);o__Saccharomycetales(100);f__Metschnikowiaceae(100);g__Metschnikowia(100);s__Metschnikowia_pulcherrima(100); | Metschnikowia | 13 | Saprotroph | Undefined Saprotroph | Yeast | NULL | Probable | As nectar-tap_saprotroph (Põlme et al. 2020); Decay sugar-rich substrates (Põlme et al. 2020) | Põlme S, et al. 2020. Fungal Diversity 105:1-16 ((https://doi.org/10.1007/s13225-020-00466-2)) |
| Otu18605 | 1 | k__Fungi(100);p__Ascomycota(100);c__Saccharomycetes(100);o__Saccharomycetales(100);f__Metschnikowiaceae(100);g__Metschnikowia(100);s__Metschnikowia_pulcherrima(100); | Metschnikowia | 13 | Saprotroph | Undefined Saprotroph | Yeast | NULL | Probable | As nectar-tap_saprotroph (Põlme et al. 2020); Decay sugar-rich substrates (Põlme et al. 2020) | Põlme S, et al. 2020. Fungal Diversity 105:1-16 ((https://doi.org/10.1007/s13225-020-00466-2)) |
| Otu18606 | 1 | k__Fungi(100);p__Ascomycota(100);c__Saccharomycetes(100);o__Saccharomycetales(100);f__Metschnikowiaceae(100);g__Metschnikowia(100);s__Metschnikowia_pulcherrima(100); | Metschnikowia | 13 | Saprotroph | Undefined Saprotroph | Yeast | NULL | Probable | As nectar-tap_saprotroph (Põlme et al. 2020); Decay sugar-rich substrates (Põlme et al. 2020) | Põlme S, et al. 2020. Fungal Diversity 105:1-16 ((https://doi.org/10.1007/s13225-020-00466-2)) |
| Otu18607 | 1 | k__Fungi(100);p__Ascomycota(100);c__Saccharomycetes(100);o__Saccharomycetales(100);f__Metschnikowiaceae(100);g__Metschnikowia(100);s__Metschnikowia_pulcherrima(100); | Metschnikowia | 13 | Saprotroph | Undefined Saprotroph | Yeast | NULL | Probable | As nectar-tap_saprotroph (Põlme et al. 2020); Decay sugar-rich substrates (Põlme et al. 2020) | Põlme S, et al. 2020. Fungal Diversity 105:1-16 ((https://doi.org/10.1007/s13225-020-00466-2)) |
| Otu18608 | 1 | k__Fungi(100);p__Ascomycota(100);c__Saccharomycetes(100);o__Saccharomycetales(100);f__Metschnikowiaceae(100);g__Metschnikowia(100);g__Metschnikowia_unclassified(100); | Metschnikowia | 13 | Saprotroph | Undefined Saprotroph | Yeast | NULL | Probable | As nectar-tap_saprotroph (Põlme et al. 2020); Decay sugar-rich substrates (Põlme et al. 2020) | Põlme S, et al. 2020. Fungal Diversity 105:1-16 ((https://doi.org/10.1007/s13225-020-00466-2)) |
| Otu18609 | 1 | k__Fungi(100);p__Ascomycota(100);c__Saccharomycetes(100);o__Saccharomycetales(100);f__Metschnikowiaceae(100);g__Metschnikowia(100);g__Metschnikowia_unclassified(100); | Metschnikowia | 13 | Saprotroph | Undefined Saprotroph | Yeast | NULL | Probable | As nectar-tap_saprotroph (Põlme et al. 2020); Decay sugar-rich substrates (Põlme et al. 2020) | Põlme S, et al. 2020. Fungal Diversity 105:1-16 ((https://doi.org/10.1007/s13225-020-00466-2)) |
| Otu18610 | 1 | k__Fungi(100);p__Ascomycota(100);c__Saccharomycetes(100);o__Saccharomycetales(100);f__Metschnikowiaceae(100);g__Metschnikowia(100);s__Metschnikowia_pulcherrima(100); | Metschnikowia | 13 | Saprotroph | Undefined Saprotroph | Yeast | NULL | Probable | As nectar-tap_saprotroph (Põlme et al. 2020); Decay sugar-rich substrates (Põlme et al. 2020) | Põlme S, et al. 2020. Fungal Diversity 105:1-16 ((https://doi.org/10.1007/s13225-020-00466-2)) |
| Otu18611 | 1 | k__Fungi(100);p__Ascomycota(100);c__Saccharomycetes(100);o__Saccharomycetales(100);f__Metschnikowiaceae(100);g__Metschnikowia(100);s__Metschnikowia_pulcherrima(100); | Metschnikowia | 13 | Saprotroph | Undefined Saprotroph | Yeast | NULL | Probable | As nectar-tap_saprotroph (Põlme et al. 2020); Decay sugar-rich substrates (Põlme et al. 2020) | Põlme S, et al. 2020. Fungal Diversity 105:1-16 ((https://doi.org/10.1007/s13225-020-00466-2)) |
| Otu18612 | 1 | k__Fungi(100);p__Ascomycota(100);c__Saccharomycetes(100);o__Saccharomycetales(100);f__Metschnikowiaceae(100);g__Metschnikowia(100);g__Metschnikowia_unclassified(100); | Metschnikowia | 13 | Saprotroph | Undefined Saprotroph | Yeast | NULL | Probable | As nectar-tap_saprotroph (Põlme et al. 2020); Decay sugar-rich substrates (Põlme et al. 2020) | Põlme S, et al. 2020. Fungal Diversity 105:1-16 ((https://doi.org/10.1007/s13225-020-00466-2)) |
| Otu18613 | 1 | k__Fungi(100);p__Ascomycota(100);c__Saccharomycetes(100);o__Saccharomycetales(100);f__Metschnikowiaceae(100);g__Metschnikowia(100);s__Metschnikowia_pulcherrima(100); | Metschnikowia | 13 | Saprotroph | Undefined Saprotroph | Yeast | NULL | Probable | As nectar-tap_saprotroph (Põlme et al. 2020); Decay sugar-rich substrates (Põlme et al. 2020) | Põlme S, et al. 2020. Fungal Diversity 105:1-16 ((https://doi.org/10.1007/s13225-020-00466-2)) |
| Otu18614 | 1 | k__Fungi(100);p__Ascomycota(100);c__Saccharomycetes(100);o__Saccharomycetales(100);f__Metschnikowiaceae(100);g__Metschnikowia(100);s__Metschnikowia_pulcherrima(100); | Metschnikowia | 13 | Saprotroph | Undefined Saprotroph | Yeast | NULL | Probable | As nectar-tap_saprotroph (Põlme et al. 2020); Decay sugar-rich substrates (Põlme et al. 2020) | Põlme S, et al. 2020. Fungal Diversity 105:1-16 ((https://doi.org/10.1007/s13225-020-00466-2)) |
| Otu18615 | 1 | k__Fungi(100);p__Ascomycota(100);c__Saccharomycetes(100);o__Saccharomycetales(100);f__Metschnikowiaceae(100);g__Metschnikowia(100);g__Metschnikowia_unclassified(100); | Metschnikowia | 13 | Saprotroph | Undefined Saprotroph | Yeast | NULL | Probable | As nectar-tap_saprotroph (Põlme et al. 2020); Decay sugar-rich substrates (Põlme et al. 2020) | Põlme S, et al. 2020. Fungal Diversity 105:1-16 ((https://doi.org/10.1007/s13225-020-00466-2)) |
| Otu18616 | 1 | k__Fungi(100);p__Ascomycota(100);c__Saccharomycetes(100);o__Saccharomycetales(100);f__Metschnikowiaceae(100);g__Metschnikowia(100);s__Metschnikowia_pulcherrima(100); | Metschnikowia | 13 | Saprotroph | Undefined Saprotroph | Yeast | NULL | Probable | As nectar-tap_saprotroph (Põlme et al. 2020); Decay sugar-rich substrates (Põlme et al. 2020) | Põlme S, et al. 2020. Fungal Diversity 105:1-16 ((https://doi.org/10.1007/s13225-020-00466-2)) |
| Otu18617 | 1 | k__Fungi(100);p__Ascomycota(100);c__Saccharomycetes(100);o__Saccharomycetales(100);f__Metschnikowiaceae(100);g__Metschnikowia(100);s__Metschnikowia_pulcherrima(100); | Metschnikowia | 13 | Saprotroph | Undefined Saprotroph | Yeast | NULL | Probable | As nectar-tap_saprotroph (Põlme et al. 2020); Decay sugar-rich substrates (Põlme et al. 2020) | Põlme S, et al. 2020. Fungal Diversity 105:1-16 ((https://doi.org/10.1007/s13225-020-00466-2)) |
| Otu18618 | 1 | k__Fungi(100);p__Ascomycota(100);c__Saccharomycetes(100);o__Saccharomycetales(100);f__Metschnikowiaceae(100);g__Metschnikowia(100);g__Metschnikowia_unclassified(100); | Metschnikowia | 13 | Saprotroph | Undefined Saprotroph | Yeast | NULL | Probable | As nectar-tap_saprotroph (Põlme et al. 2020); Decay sugar-rich substrates (Põlme et al. 2020) | Põlme S, et al. 2020. Fungal Diversity 105:1-16 ((https://doi.org/10.1007/s13225-020-00466-2)) |
| Otu18619 | 1 | k__Fungi(100);p__Ascomycota(100);c__Saccharomycetes(100);o__Saccharomycetales(100);f__Metschnikowiaceae(100);g__Metschnikowia(100);s__Metschnikowia_pulcherrima(100); | Metschnikowia | 13 | Saprotroph | Undefined Saprotroph | Yeast | NULL | Probable | As nectar-tap_saprotroph (Põlme et al. 2020); Decay sugar-rich substrates (Põlme et al. 2020) | Põlme S, et al. 2020. Fungal Diversity 105:1-16 ((https://doi.org/10.1007/s13225-020-00466-2)) |
| Otu18620 | 1 | k__Fungi(100);p__Ascomycota(100);c__Saccharomycetes(100);o__Saccharomycetales(100);f__Metschnikowiaceae(100);g__Metschnikowia(100);s__Metschnikowia_pulcherrima(100); | Metschnikowia | 13 | Saprotroph | Undefined Saprotroph | Yeast | NULL | Probable | As nectar-tap_saprotroph (Põlme et al. 2020); Decay sugar-rich substrates (Põlme et al. 2020) | Põlme S, et al. 2020. Fungal Diversity 105:1-16 ((https://doi.org/10.1007/s13225-020-00466-2)) |
| Otu18621 | 1 | k__Fungi(100);p__Ascomycota(100);c__Saccharomycetes(100);o__Saccharomycetales(100);f__Metschnikowiaceae(100);g__Metschnikowia(100);g__Metschnikowia_unclassified(100); | Metschnikowia | 13 | Saprotroph | Undefined Saprotroph | Yeast | NULL | Probable | As nectar-tap_saprotroph (Põlme et al. 2020); Decay sugar-rich substrates (Põlme et al. 2020) | Põlme S, et al. 2020. Fungal Diversity 105:1-16 ((https://doi.org/10.1007/s13225-020-00466-2)) |
| Otu18622 | 1 | k__Fungi(100);p__Ascomycota(100);c__Saccharomycetes(100);o__Saccharomycetales(100);f__Metschnikowiaceae(100);g__Metschnikowia(100);s__Metschnikowia_pulcherrima(100); | Metschnikowia | 13 | Saprotroph | Undefined Saprotroph | Yeast | NULL | Probable | As nectar-tap_saprotroph (Põlme et al. 2020); Decay sugar-rich substrates (Põlme et al. 2020) | Põlme S, et al. 2020. Fungal Diversity 105:1-16 ((https://doi.org/10.1007/s13225-020-00466-2)) |
| Otu18623 | 1 | k__Fungi(100);p__Ascomycota(100);c__Saccharomycetes(100);o__Saccharomycetales(100);f__Metschnikowiaceae(100);g__Metschnikowia(100);s__Metschnikowia_pulcherrima(100); | Metschnikowia | 13 | Saprotroph | Undefined Saprotroph | Yeast | NULL | Probable | As nectar-tap_saprotroph (Põlme et al. 2020); Decay sugar-rich substrates (Põlme et al. 2020) | Põlme S, et al. 2020. Fungal Diversity 105:1-16 ((https://doi.org/10.1007/s13225-020-00466-2)) |
| Otu18624 | 1 | k__Fungi(100);p__Ascomycota(100);c__Saccharomycetes(100);o__Saccharomycetales(100);f__Metschnikowiaceae(100);g__Metschnikowia(100);s__Metschnikowia_pulcherrima(100); | Metschnikowia | 13 | Saprotroph | Undefined Saprotroph | Yeast | NULL | Probable | As nectar-tap_saprotroph (Põlme et al. 2020); Decay sugar-rich substrates (Põlme et al. 2020) | Põlme S, et al. 2020. Fungal Diversity 105:1-16 ((https://doi.org/10.1007/s13225-020-00466-2)) |
| Otu18625 | 1 | k__Fungi(100);p__Ascomycota(100);c__Saccharomycetes(100);o__Saccharomycetales(100);f__Metschnikowiaceae(100);g__Metschnikowia(100);s__Metschnikowia_pulcherrima(100); | Metschnikowia | 13 | Saprotroph | Undefined Saprotroph | Yeast | NULL | Probable | As nectar-tap_saprotroph (Põlme et al. 2020); Decay sugar-rich substrates (Põlme et al. 2020) | Põlme S, et al. 2020. Fungal Diversity 105:1-16 ((https://doi.org/10.1007/s13225-020-00466-2)) |
| Otu18626 | 1 | k__Fungi(100);p__Ascomycota(100);c__Saccharomycetes(100);o__Saccharomycetales(100);f__Metschnikowiaceae(100);g__Metschnikowia(100);g__Metschnikowia_unclassified(100); | Metschnikowia | 13 | Saprotroph | Undefined Saprotroph | Yeast | NULL | Probable | As nectar-tap_saprotroph (Põlme et al. 2020); Decay sugar-rich substrates (Põlme et al. 2020) | Põlme S, et al. 2020. Fungal Diversity 105:1-16 ((https://doi.org/10.1007/s13225-020-00466-2)) |
| Otu18627 | 1 | k__Fungi(100);p__Ascomycota(100);c__Saccharomycetes(100);o__Saccharomycetales(100);f__Metschnikowiaceae(100);g__Metschnikowia(100);s__Metschnikowia_pulcherrima(100); | Metschnikowia | 13 | Saprotroph | Undefined Saprotroph | Yeast | NULL | Probable | As nectar-tap_saprotroph (Põlme et al. 2020); Decay sugar-rich substrates (Põlme et al. 2020) | Põlme S, et al. 2020. Fungal Diversity 105:1-16 ((https://doi.org/10.1007/s13225-020-00466-2)) |
| Otu18628 | 1 | k__Fungi(100);p__Ascomycota(100);c__Saccharomycetes(100);o__Saccharomycetales(100);f__Metschnikowiaceae(100);g__Metschnikowia(100);s__Metschnikowia_pulcherrima(100); | Metschnikowia | 13 | Saprotroph | Undefined Saprotroph | Yeast | NULL | Probable | As nectar-tap_saprotroph (Põlme et al. 2020); Decay sugar-rich substrates (Põlme et al. 2020) | Põlme S, et al. 2020. Fungal Diversity 105:1-16 ((https://doi.org/10.1007/s13225-020-00466-2)) |
| Otu18629 | 1 | k__Fungi(100);p__Ascomycota(100);c__Saccharomycetes(100);o__Saccharomycetales(100);f__Metschnikowiaceae(100);g__Metschnikowia(100);s__Metschnikowia_pulcherrima(100); | Metschnikowia | 13 | Saprotroph | Undefined Saprotroph | Yeast | NULL | Probable | As nectar-tap_saprotroph (Põlme et al. 2020); Decay sugar-rich substrates (Põlme et al. 2020) | Põlme S, et al. 2020. Fungal Diversity 105:1-16 ((https://doi.org/10.1007/s13225-020-00466-2)) |
| Otu18630 | 1 | k__Fungi(100);p__Ascomycota(100);c__Saccharomycetes(100);o__Saccharomycetales(100);f__Metschnikowiaceae(100);g__Metschnikowia(100);g__Metschnikowia_unclassified(100); | Metschnikowia | 13 | Saprotroph | Undefined Saprotroph | Yeast | NULL | Probable | As nectar-tap_saprotroph (Põlme et al. 2020); Decay sugar-rich substrates (Põlme et al. 2020) | Põlme S, et al. 2020. Fungal Diversity 105:1-16 ((https://doi.org/10.1007/s13225-020-00466-2)) |
| Otu18631 | 1 | k__Fungi(100);p__Ascomycota(100);c__Saccharomycetes(100);o__Saccharomycetales(100);f__Metschnikowiaceae(100);g__Metschnikowia(100);g__Metschnikowia_unclassified(100); | Metschnikowia | 13 | Saprotroph | Undefined Saprotroph | Yeast | NULL | Probable | As nectar-tap_saprotroph (Põlme et al. 2020); Decay sugar-rich substrates (Põlme et al. 2020) | Põlme S, et al. 2020. Fungal Diversity 105:1-16 ((https://doi.org/10.1007/s13225-020-00466-2)) |
| Otu18632 | 1 | k__Fungi(100);p__Ascomycota(100);c__Saccharomycetes(100);o__Saccharomycetales(100);f__Metschnikowiaceae(100);g__Metschnikowia(100);g__Metschnikowia_unclassified(100); | Metschnikowia | 13 | Saprotroph | Undefined Saprotroph | Yeast | NULL | Probable | As nectar-tap_saprotroph (Põlme et al. 2020); Decay sugar-rich substrates (Põlme et al. 2020) | Põlme S, et al. 2020. Fungal Diversity 105:1-16 ((https://doi.org/10.1007/s13225-020-00466-2)) |
| Otu18635 | 1 | k__Fungi(100);p__Ascomycota(100);c__Saccharomycetes(100);o__Saccharomycetales(100);f__Metschnikowiaceae(100);g__Metschnikowia(100);s__Metschnikowia_pulcherrima(100); | Metschnikowia | 13 | Saprotroph | Undefined Saprotroph | Yeast | NULL | Probable | As nectar-tap_saprotroph (Põlme et al. 2020); Decay sugar-rich substrates (Põlme et al. 2020) | Põlme S, et al. 2020. Fungal Diversity 105:1-16 ((https://doi.org/10.1007/s13225-020-00466-2)) |
| Otu18636 | 1 | k__Fungi(100);p__Ascomycota(100);c__Saccharomycetes(100);o__Saccharomycetales(100);f__Metschnikowiaceae(100);g__Metschnikowia(100);s__Metschnikowia_pulcherrima(100); | Metschnikowia | 13 | Saprotroph | Undefined Saprotroph | Yeast | NULL | Probable | As nectar-tap_saprotroph (Põlme et al. 2020); Decay sugar-rich substrates (Põlme et al. 2020) | Põlme S, et al. 2020. Fungal Diversity 105:1-16 ((https://doi.org/10.1007/s13225-020-00466-2)) |
| Otu18637 | 1 | k__Fungi(100);p__Ascomycota(100);c__Saccharomycetes(100);o__Saccharomycetales(100);f__Metschnikowiaceae(100);g__Metschnikowia(100);g__Metschnikowia_unclassified(100); | Metschnikowia | 13 | Saprotroph | Undefined Saprotroph | Yeast | NULL | Probable | As nectar-tap_saprotroph (Põlme et al. 2020); Decay sugar-rich substrates (Põlme et al. 2020) | Põlme S, et al. 2020. Fungal Diversity 105:1-16 ((https://doi.org/10.1007/s13225-020-00466-2)) |
| Otu18640 | 1 | k__Fungi(100);p__Ascomycota(100);c__Saccharomycetes(100);o__Saccharomycetales(100);f__Metschnikowiaceae(100);g__Metschnikowia(100);g__Metschnikowia_unclassified(100); | Metschnikowia | 13 | Saprotroph | Undefined Saprotroph | Yeast | NULL | Probable | As nectar-tap_saprotroph (Põlme et al. 2020); Decay sugar-rich substrates (Põlme et al. 2020) | Põlme S, et al. 2020. Fungal Diversity 105:1-16 ((https://doi.org/10.1007/s13225-020-00466-2)) |
| Otu18641 | 1 | k__Fungi(100);p__Ascomycota(100);c__Saccharomycetes(100);o__Saccharomycetales(100);f__Metschnikowiaceae(100);g__Metschnikowia(100);s__Metschnikowia_pulcherrima(100); | Metschnikowia | 13 | Saprotroph | Undefined Saprotroph | Yeast | NULL | Probable | As nectar-tap_saprotroph (Põlme et al. 2020); Decay sugar-rich substrates (Põlme et al. 2020) | Põlme S, et al. 2020. Fungal Diversity 105:1-16 ((https://doi.org/10.1007/s13225-020-00466-2)) |
| Otu18642 | 1 | k__Fungi(100);p__Ascomycota(100);c__Saccharomycetes(100);o__Saccharomycetales(100);f__Metschnikowiaceae(100);g__Metschnikowia(100);s__Metschnikowia_pulcherrima(100); | Metschnikowia | 13 | Saprotroph | Undefined Saprotroph | Yeast | NULL | Probable | As nectar-tap_saprotroph (Põlme et al. 2020); Decay sugar-rich substrates (Põlme et al. 2020) | Põlme S, et al. 2020. Fungal Diversity 105:1-16 ((https://doi.org/10.1007/s13225-020-00466-2)) |
| Otu18643 | 1 | k__Fungi(100);p__Ascomycota(100);c__Saccharomycetes(100);o__Saccharomycetales(100);f__Metschnikowiaceae(100);g__Metschnikowia(100);g__Metschnikowia_unclassified(100); | Metschnikowia | 13 | Saprotroph | Undefined Saprotroph | Yeast | NULL | Probable | As nectar-tap_saprotroph (Põlme et al. 2020); Decay sugar-rich substrates (Põlme et al. 2020) | Põlme S, et al. 2020. Fungal Diversity 105:1-16 ((https://doi.org/10.1007/s13225-020-00466-2)) |
| Otu18644 | 1 | k__Fungi(100);p__Ascomycota(100);c__Saccharomycetes(100);o__Saccharomycetales(100);f__Metschnikowiaceae(100);g__Metschnikowia(100);s__Metschnikowia_pulcherrima(100); | Metschnikowia | 13 | Saprotroph | Undefined Saprotroph | Yeast | NULL | Probable | As nectar-tap_saprotroph (Põlme et al. 2020); Decay sugar-rich substrates (Põlme et al. 2020) | Põlme S, et al. 2020. Fungal Diversity 105:1-16 ((https://doi.org/10.1007/s13225-020-00466-2)) |
| Otu18645 | 1 | k__Fungi(100);p__Ascomycota(100);c__Saccharomycetes(100);o__Saccharomycetales(100);f__Metschnikowiaceae(100);g__Metschnikowia(100);g__Metschnikowia_unclassified(100); | Metschnikowia | 13 | Saprotroph | Undefined Saprotroph | Yeast | NULL | Probable | As nectar-tap_saprotroph (Põlme et al. 2020); Decay sugar-rich substrates (Põlme et al. 2020) | Põlme S, et al. 2020. Fungal Diversity 105:1-16 ((https://doi.org/10.1007/s13225-020-00466-2)) |
| Otu18646 | 1 | k__Fungi(100);p__Ascomycota(100);c__Saccharomycetes(100);o__Saccharomycetales(100);f__Metschnikowiaceae(100);g__Metschnikowia(100);g__Metschnikowia_unclassified(100); | Metschnikowia | 13 | Saprotroph | Undefined Saprotroph | Yeast | NULL | Probable | As nectar-tap_saprotroph (Põlme et al. 2020); Decay sugar-rich substrates (Põlme et al. 2020) | Põlme S, et al. 2020. Fungal Diversity 105:1-16 ((https://doi.org/10.1007/s13225-020-00466-2)) |
| Otu18647 | 1 | k__Fungi(100);p__Ascomycota(100);c__Saccharomycetes(100);o__Saccharomycetales(100);f__Metschnikowiaceae(100);g__Metschnikowia(100);g__Metschnikowia_unclassified(100); | Metschnikowia | 13 | Saprotroph | Undefined Saprotroph | Yeast | NULL | Probable | As nectar-tap_saprotroph (Põlme et al. 2020); Decay sugar-rich substrates (Põlme et al. 2020) | Põlme S, et al. 2020. Fungal Diversity 105:1-16 ((https://doi.org/10.1007/s13225-020-00466-2)) |
| Otu18648 | 1 | k__Fungi(100);p__Ascomycota(100);c__Saccharomycetes(100);o__Saccharomycetales(100);f__Metschnikowiaceae(100);g__Metschnikowia(100);s__Metschnikowia_pulcherrima(100); | Metschnikowia | 13 | Saprotroph | Undefined Saprotroph | Yeast | NULL | Probable | As nectar-tap_saprotroph (Põlme et al. 2020); Decay sugar-rich substrates (Põlme et al. 2020) | Põlme S, et al. 2020. Fungal Diversity 105:1-16 ((https://doi.org/10.1007/s13225-020-00466-2)) |
| Otu18649 | 1 | k__Fungi(100);p__Ascomycota(100);c__Saccharomycetes(100);o__Saccharomycetales(100);f__Metschnikowiaceae(100);g__Metschnikowia(100);s__Metschnikowia_pulcherrima(100); | Metschnikowia | 13 | Saprotroph | Undefined Saprotroph | Yeast | NULL | Probable | As nectar-tap_saprotroph (Põlme et al. 2020); Decay sugar-rich substrates (Põlme et al. 2020) | Põlme S, et al. 2020. Fungal Diversity 105:1-16 ((https://doi.org/10.1007/s13225-020-00466-2)) |
| Otu18650 | 1 | k__Fungi(100);p__Ascomycota(100);c__Saccharomycetes(100);o__Saccharomycetales(100);f__Metschnikowiaceae(100);g__Metschnikowia(100);g__Metschnikowia_unclassified(100); | Metschnikowia | 13 | Saprotroph | Undefined Saprotroph | Yeast | NULL | Probable | As nectar-tap_saprotroph (Põlme et al. 2020); Decay sugar-rich substrates (Põlme et al. 2020) | Põlme S, et al. 2020. Fungal Diversity 105:1-16 ((https://doi.org/10.1007/s13225-020-00466-2)) |
| Otu18651 | 1 | k__Fungi(100);p__Ascomycota(100);c__Saccharomycetes(100);o__Saccharomycetales(100);f__Metschnikowiaceae(100);g__Metschnikowia(100);g__Metschnikowia_unclassified(100); | Metschnikowia | 13 | Saprotroph | Undefined Saprotroph | Yeast | NULL | Probable | As nectar-tap_saprotroph (Põlme et al. 2020); Decay sugar-rich substrates (Põlme et al. 2020) | Põlme S, et al. 2020. Fungal Diversity 105:1-16 ((https://doi.org/10.1007/s13225-020-00466-2)) |
| Otu18652 | 1 | k__Fungi(100);p__Ascomycota(100);c__Saccharomycetes(100);o__Saccharomycetales(100);f__Metschnikowiaceae(100);g__Metschnikowia(100);g__Metschnikowia_unclassified(100); | Metschnikowia | 13 | Saprotroph | Undefined Saprotroph | Yeast | NULL | Probable | As nectar-tap_saprotroph (Põlme et al. 2020); Decay sugar-rich substrates (Põlme et al. 2020) | Põlme S, et al. 2020. Fungal Diversity 105:1-16 ((https://doi.org/10.1007/s13225-020-00466-2)) |
| Otu18653 | 1 | k__Fungi(100);p__Ascomycota(100);c__Saccharomycetes(100);o__Saccharomycetales(100);f__Metschnikowiaceae(100);g__Metschnikowia(100);g__Metschnikowia_unclassified(100); | Metschnikowia | 13 | Saprotroph | Undefined Saprotroph | Yeast | NULL | Probable | As nectar-tap_saprotroph (Põlme et al. 2020); Decay sugar-rich substrates (Põlme et al. 2020) | Põlme S, et al. 2020. Fungal Diversity 105:1-16 ((https://doi.org/10.1007/s13225-020-00466-2)) |
| Otu18654 | 1 | k__Fungi(100);p__Ascomycota(100);c__Saccharomycetes(100);o__Saccharomycetales(100);f__Metschnikowiaceae(100);g__Metschnikowia(100);s__Metschnikowia_pulcherrima(100); | Metschnikowia | 13 | Saprotroph | Undefined Saprotroph | Yeast | NULL | Probable | As nectar-tap_saprotroph (Põlme et al. 2020); Decay sugar-rich substrates (Põlme et al. 2020) | Põlme S, et al. 2020. Fungal Diversity 105:1-16 ((https://doi.org/10.1007/s13225-020-00466-2)) |
| Otu18655 | 1 | k__Fungi(100);p__Ascomycota(100);c__Saccharomycetes(100);o__Saccharomycetales(100);f__Metschnikowiaceae(100);g__Metschnikowia(100);g__Metschnikowia_unclassified(100); | Metschnikowia | 13 | Saprotroph | Undefined Saprotroph | Yeast | NULL | Probable | As nectar-tap_saprotroph (Põlme et al. 2020); Decay sugar-rich substrates (Põlme et al. 2020) | Põlme S, et al. 2020. Fungal Diversity 105:1-16 ((https://doi.org/10.1007/s13225-020-00466-2)) |
| Otu18656 | 1 | k__Fungi(100);p__Ascomycota(100);c__Saccharomycetes(100);o__Saccharomycetales(100);f__Metschnikowiaceae(100);g__Metschnikowia(100);s__Metschnikowia_pulcherrima(100); | Metschnikowia | 13 | Saprotroph | Undefined Saprotroph | Yeast | NULL | Probable | As nectar-tap_saprotroph (Põlme et al. 2020); Decay sugar-rich substrates (Põlme et al. 2020) | Põlme S, et al. 2020. Fungal Diversity 105:1-16 ((https://doi.org/10.1007/s13225-020-00466-2)) |
| Otu18657 | 1 | k__Fungi(100);p__Ascomycota(100);c__Saccharomycetes(100);o__Saccharomycetales(100);f__Metschnikowiaceae(100);g__Metschnikowia(100);g__Metschnikowia_unclassified(100); | Metschnikowia | 13 | Saprotroph | Undefined Saprotroph | Yeast | NULL | Probable | As nectar-tap_saprotroph (Põlme et al. 2020); Decay sugar-rich substrates (Põlme et al. 2020) | Põlme S, et al. 2020. Fungal Diversity 105:1-16 ((https://doi.org/10.1007/s13225-020-00466-2)) |
| Otu18658 | 1 | k__Fungi(100);p__Ascomycota(100);c__Saccharomycetes(100);o__Saccharomycetales(100);f__Metschnikowiaceae(100);g__Metschnikowia(100);s__Metschnikowia_pulcherrima(100); | Metschnikowia | 13 | Saprotroph | Undefined Saprotroph | Yeast | NULL | Probable | As nectar-tap_saprotroph (Põlme et al. 2020); Decay sugar-rich substrates (Põlme et al. 2020) | Põlme S, et al. 2020. Fungal Diversity 105:1-16 ((https://doi.org/10.1007/s13225-020-00466-2)) |
| Otu18662 | 1 | k__Fungi(100);p__Ascomycota(100);c__Saccharomycetes(100);o__Saccharomycetales(100);f__Metschnikowiaceae(100);g__Metschnikowia(100);s__Metschnikowia_pulcherrima(100); | Metschnikowia | 13 | Saprotroph | Undefined Saprotroph | Yeast | NULL | Probable | As nectar-tap_saprotroph (Põlme et al. 2020); Decay sugar-rich substrates (Põlme et al. 2020) | Põlme S, et al. 2020. Fungal Diversity 105:1-16 ((https://doi.org/10.1007/s13225-020-00466-2)) |

| | | | | | | | | | | | |
|---|---|---|---|---|---|---|---|---|---|---|---|
| Otu18663 | 1 | k__Fungi(100);p__Ascomycota(100);c__Saccharomycetes(100);o__Saccharomycetales(100);f__Metschnikowiaceae(100);g__Metschnikowia(100);s__Metschnikowia_pulcherrima(100); | Metschnikowia | 13 | Saprotroph | \|Undefined Saprotroph\| | Yeast | NULL | Probable | As nectar-tap_saprotroph (Põlme et al. 2020); Decay sugar-rich substrates (Põlme et al. 2020) | Põlme S, et al. 2020. Fungal Diversity 105:1-16 ((https://doi.org/10.1007/s13225-020-00466-2)) |
| Otu18664 | 1 | k__Fungi(100);p__Ascomycota(100);c__Saccharomycetes(100);o__Saccharomycetales(100);f__Metschnikowiaceae(100);g__Metschnikowia(100);s__Metschnikowia_pulcherrima(100); | Metschnikowia | 13 | Saprotroph | \|Undefined Saprotroph\| | Yeast | NULL | Probable | As nectar-tap_saprotroph (Põlme et al. 2020); Decay sugar-rich substrates (Põlme et al. 2020) | Põlme S, et al. 2020. Fungal Diversity 105:1-16 ((https://doi.org/10.1007/s13225-020-00466-2)) |
| Otu18665 | 1 | k__Fungi(100);p__Ascomycota(100);c__Saccharomycetes(100);o__Saccharomycetales(100);f__Metschnikowiaceae(100);g__Metschnikowia(100);g__Metschnikowia_unclassified(100); | Metschnikowia | 13 | Saprotroph | \|Undefined Saprotroph\| | Yeast | NULL | Probable | As nectar-tap_saprotroph (Põlme et al. 2020); Decay sugar-rich substrates (Põlme et al. 2020) | Põlme S, et al. 2020. Fungal Diversity 105:1-16 ((https://doi.org/10.1007/s13225-020-00466-2)) |
| Otu18666 | 1 | k__Fungi(100);p__Ascomycota(100);c__Saccharomycetes(100);o__Saccharomycetales(100);f__Metschnikowiaceae(100);g__Metschnikowia(100);s__Metschnikowia_pulcherrima(100); | Metschnikowia | 13 | Saprotroph | \|Undefined Saprotroph\| | Yeast | NULL | Probable | As nectar-tap_saprotroph (Põlme et al. 2020); Decay sugar-rich substrates (Põlme et al. 2020) | Põlme S, et al. 2020. Fungal Diversity 105:1-16 ((https://doi.org/10.1007/s13225-020-00466-2)) |
| Otu18667 | 1 | k__Fungi(100);p__Ascomycota(100);c__Saccharomycetes(100);o__Saccharomycetales(100);f__Metschnikowiaceae(100);g__Metschnikowia(100);s__Metschnikowia_pulcherrima(100); | Metschnikowia | 13 | Saprotroph | \|Undefined Saprotroph\| | Yeast | NULL | Probable | As nectar-tap_saprotroph (Põlme et al. 2020); Decay sugar-rich substrates (Põlme et al. 2020) | Põlme S, et al. 2020. Fungal Diversity 105:1-16 ((https://doi.org/10.1007/s13225-020-00466-2)) |
| Otu18668 | 1 | k__Fungi(100);p__Ascomycota(100);c__Saccharomycetes(100);o__Saccharomycetales(100);f__Metschnikowiaceae(100);g__Metschnikowia(100);s__Metschnikowia_pulcherrima(100); | Metschnikowia | 13 | Saprotroph | \|Undefined Saprotroph\| | Yeast | NULL | Probable | As nectar-tap_saprotroph (Põlme et al. 2020); Decay sugar-rich substrates (Põlme et al. 2020) | Põlme S, et al. 2020. Fungal Diversity 105:1-16 ((https://doi.org/10.1007/s13225-020-00466-2)) |
| Otu18669 | 1 | k__Fungi(100);p__Ascomycota(100);c__Saccharomycetes(100);o__Saccharomycetales(100);f__Metschnikowiaceae(100);g__Metschnikowia(100);s__Metschnikowia_pulcherrima(100); | Metschnikowia | 13 | Saprotroph | \|Undefined Saprotroph\| | Yeast | NULL | Probable | As nectar-tap_saprotroph (Põlme et al. 2020); Decay sugar-rich substrates (Põlme et al. 2020) | Põlme S, et al. 2020. Fungal Diversity 105:1-16 ((https://doi.org/10.1007/s13225-020-00466-2)) |
| Otu18674 | 1 | k__Fungi(100);p__Ascomycota(100);c__Saccharomycetes(100);o__Saccharomycetales(100);f__Metschnikowiaceae(100);g__Metschnikowia(100);s__Metschnikowia_pulcherrima(100); | Metschnikowia | 13 | Saprotroph | \|Undefined Saprotroph\| | Yeast | NULL | Probable | As nectar-tap_saprotroph (Põlme et al. 2020); Decay sugar-rich substrates (Põlme et al. 2020) | Põlme S, et al. 2020. Fungal Diversity 105:1-16 ((https://doi.org/10.1007/s13225-020-00466-2)) |
| Otu18675 | 1 | k__Fungi(100);p__Ascomycota(100);c__Saccharomycetes(100);o__Saccharomycetales(100);f__Metschnikowiaceae(100);g__Metschnikowia(100);g__Metschnikowia_unclassified(100); | Metschnikowia | 13 | Saprotroph | \|Undefined Saprotroph\| | Yeast | NULL | Probable | As nectar-tap_saprotroph (Põlme et al. 2020); Decay sugar-rich substrates (Põlme et al. 2020) | Põlme S, et al. 2020. Fungal Diversity 105:1-16 ((https://doi.org/10.1007/s13225-020-00466-2)) |
| Otu18676 | 1 | k__Fungi(100);p__Ascomycota(100);c__Saccharomycetes(100);o__Saccharomycetales(100);f__Metschnikowiaceae(100);g__Metschnikowia(100);s__Metschnikowia_pulcherrima(100); | Metschnikowia | 13 | Saprotroph | \|Undefined Saprotroph\| | Yeast | NULL | Probable | As nectar-tap_saprotroph (Põlme et al. 2020); Decay sugar-rich substrates (Põlme et al. 2020) | Põlme S, et al. 2020. Fungal Diversity 105:1-16 ((https://doi.org/10.1007/s13225-020-00466-2)) |
| Otu18682 | 1 | k__Fungi(100);p__Ascomycota(100);c__Saccharomycetes(100);o__Saccharomycetales(100);f__Saccharomycetaceae(100);g__Kazachstania(100);s__Kazachstania_intestinalis(100); | Kazachstania | 13 | Saprotroph | \|Undefined Saprotroph\| | Yeast | NULL | Probable | As nectar-tap_saprotroph (Põlme et al. 2020); Decay sugar-rich substrates (Põlme et al. 2020) | Põlme S, et al. 2020. Fungal Diversity 105:1-16 ((https://doi.org/10.1007/s13225-020-00466-2)) |
| Otu18683 | 1 | k__Fungi(100);p__Ascomycota(100);c__Saccharomycetes(100);o__Saccharomycetales(100);f__Metschnikowiaceae(100);g__Metschnikowia(100);g__Metschnikowia_unclassified(100); | Metschnikowia | 13 | Saprotroph | \|Undefined Saprotroph\| | Yeast | NULL | Probable | As nectar-tap_saprotroph (Põlme et al. 2020); Decay sugar-rich substrates (Põlme et al. 2020) | Põlme S, et al. 2020. Fungal Diversity 105:1-16 ((https://doi.org/10.1007/s13225-020-00466-2)) |
| Otu18684 | 1 | k__Fungi(100);p__Ascomycota(100);c__Saccharomycetes(100);o__Saccharomycetales(100);f__Metschnikowiaceae(100);g__Metschnikowia(100);g__Metschnikowia_unclassified(100); | Metschnikowia | 13 | Saprotroph | \|Undefined Saprotroph\| | Yeast | NULL | Probable | As nectar-tap_saprotroph (Põlme et al. 2020); Decay sugar-rich substrates (Põlme et al. 2020) | Põlme S, et al. 2020. Fungal Diversity 105:1-16 ((https://doi.org/10.1007/s13225-020-00466-2)) |
| Otu18685 | 1 | k__Fungi(100);p__Ascomycota(100);c__Saccharomycetes(100);o__Saccharomycetales(100);f__Metschnikowiaceae(100);g__Metschnikowia(100);s__Metschnikowia_pulcherrima(100); | Metschnikowia | 13 | Saprotroph | \|Undefined Saprotroph\| | Yeast | NULL | Probable | As nectar-tap_saprotroph (Põlme et al. 2020); Decay sugar-rich substrates (Põlme et al. 2020) | Põlme S, et al. 2020. Fungal Diversity 105:1-16 ((https://doi.org/10.1007/s13225-020-00466-2)) |
| Otu18686 | 1 | k__Fungi(100);p__Ascomycota(100);c__Saccharomycetes(100);o__Saccharomycetales(100);f__Metschnikowiaceae(100);g__Metschnikowia(100);g__Metschnikowia_unclassified(100); | Metschnikowia | 13 | Saprotroph | \|Undefined Saprotroph\| | Yeast | NULL | Probable | As nectar-tap_saprotroph (Põlme et al. 2020); Decay sugar-rich substrates (Põlme et al. 2020) | Põlme S, et al. 2020. Fungal Diversity 105:1-16 ((https://doi.org/10.1007/s13225-020-00466-2)) |
| Otu18687 | 1 | k__Fungi(100);p__Ascomycota(100);c__Saccharomycetes(100);o__Saccharomycetales(100);f__Metschnikowiaceae(100);g__Metschnikowia(100);g__Metschnikowia_unclassified(100); | Metschnikowia | 13 | Saprotroph | \|Undefined Saprotroph\| | Yeast | NULL | Probable | As nectar-tap_saprotroph (Põlme et al. 2020); Decay sugar-rich substrates (Põlme et al. 2020) | Põlme S, et al. 2020. Fungal Diversity 105:1-16 ((https://doi.org/10.1007/s13225-020-00466-2)) |
| Otu18688 | 1 | k__Fungi(100);p__Ascomycota(100);c__Saccharomycetes(100);o__Saccharomycetales(100);f__Metschnikowiaceae(100);g__Metschnikowia(100);s__Metschnikowia_pulcherrima(100); | Metschnikowia | 13 | Saprotroph | \|Undefined Saprotroph\| | Yeast | NULL | Probable | As nectar-tap_saprotroph (Põlme et al. 2020); Decay sugar-rich substrates (Põlme et al. 2020) | Põlme S, et al. 2020. Fungal Diversity 105:1-16 ((https://doi.org/10.1007/s13225-020-00466-2)) |
| Otu18689 | 1 | k__Fungi(100);p__Ascomycota(100);c__Saccharomycetes(100);o__Saccharomycetales(100);f__Metschnikowiaceae(100);g__Metschnikowia(100);s__Metschnikowia_pulcherrima(100); | Metschnikowia | 13 | Saprotroph | \|Undefined Saprotroph\| | Yeast | NULL | Probable | As nectar-tap_saprotroph (Põlme et al. 2020); Decay sugar-rich substrates (Põlme et al. 2020) | Põlme S, et al. 2020. Fungal Diversity 105:1-16 ((https://doi.org/10.1007/s13225-020-00466-2)) |
| Otu18690 | 1 | k__Fungi(100);p__Ascomycota(100);c__Saccharomycetes(100);o__Saccharomycetales(100);f__Metschnikowiaceae(100);g__Metschnikowia(100);s__Metschnikowia_pulcherrima(100); | Metschnikowia | 13 | Saprotroph | \|Undefined Saprotroph\| | Yeast | NULL | Probable | As nectar-tap_saprotroph (Põlme et al. 2020); Decay sugar-rich substrates (Põlme et al. 2020) | Põlme S, et al. 2020. Fungal Diversity 105:1-16 ((https://doi.org/10.1007/s13225-020-00466-2)) |
| Otu18691 | 1 | k__Fungi(100);p__Ascomycota(100);c__Saccharomycetes(100);o__Saccharomycetales(100);f__Metschnikowiaceae(100);g__Metschnikowia(100);s__Metschnikowia_pulcherrima(100); | Metschnikowia | 13 | Saprotroph | \|Undefined Saprotroph\| | Yeast | NULL | Probable | As nectar-tap_saprotroph (Põlme et al. 2020); Decay sugar-rich substrates (Põlme et al. 2020) | Põlme S, et al. 2020. Fungal Diversity 105:1-16 ((https://doi.org/10.1007/s13225-020-00466-2)) |
| Otu18692 | 1 | k__Fungi(100);p__Ascomycota(100);c__Saccharomycetes(100);o__Saccharomycetales(100);f__Metschnikowiaceae(100);g__Metschnikowia(100);s__Metschnikowia_pulcherrima(100); | Metschnikowia | 13 | Saprotroph | \|Undefined Saprotroph\| | Yeast | NULL | Probable | As nectar-tap_saprotroph (Põlme et al. 2020); Decay sugar-rich substrates (Põlme et al. 2020) | Põlme S, et al. 2020. Fungal Diversity 105:1-16 ((https://doi.org/10.1007/s13225-020-00466-2)) |
| Otu18693 | 1 | k__Fungi(100);p__Ascomycota(100);c__Saccharomycetes(100);o__Saccharomycetales(100);f__Metschnikowiaceae(100);g__Metschnikowia(100);g__Metschnikowia_unclassified(100); | Metschnikowia | 13 | Saprotroph | \|Undefined Saprotroph\| | Yeast | NULL | Probable | As nectar-tap_saprotroph (Põlme et al. 2020); Decay sugar-rich substrates (Põlme et al. 2020) | Põlme S, et al. 2020. Fungal Diversity 105:1-16 ((https://doi.org/10.1007/s13225-020-00466-2)) |
| Otu18694 | 1 | k__Fungi(100);p__Ascomycota(100);c__Saccharomycetes(100);o__Saccharomycetales(100);f__Metschnikowiaceae(100);g__Metschnikowia(100);s__Metschnikowia_pulcherrima(100); | Metschnikowia | 13 | Saprotroph | \|Undefined Saprotroph\| | Yeast | NULL | Probable | As nectar-tap_saprotroph (Põlme et al. 2020); Decay sugar-rich substrates (Põlme et al. 2020) | Põlme S, et al. 2020. Fungal Diversity 105:1-16 ((https://doi.org/10.1007/s13225-020-00466-2)) |
| Otu18695 | 1 | k__Fungi(100);p__Ascomycota(100);c__Saccharomycetes(100);o__Saccharomycetales(100);f__Metschnikowiaceae(100);g__Metschnikowia(100);s__Metschnikowia_pulcherrima(100); | Metschnikowia | 13 | Saprotroph | \|Undefined Saprotroph\| | Yeast | NULL | Probable | As nectar-tap_saprotroph (Põlme et al. 2020); Decay sugar-rich substrates (Põlme et al. 2020) | Põlme S, et al. 2020. Fungal Diversity 105:1-16 ((https://doi.org/10.1007/s13225-020-00466-2)) |
| Otu18700 | 1 | k__Fungi(100);p__Ascomycota(100);c__Saccharomycetes(100);o__Saccharomycetales(100);f__Metschnikowiaceae(100);g__Metschnikowia(100);s__Metschnikowia_pulcherrima(100); | Metschnikowia | 13 | Saprotroph | \|Undefined Saprotroph\| | Yeast | NULL | Probable | As nectar-tap_saprotroph (Põlme et al. 2020); Decay sugar-rich substrates (Põlme et al. 2020) | Põlme S, et al. 2020. Fungal Diversity 105:1-16 ((https://doi.org/10.1007/s13225-020-00466-2)) |
| Otu18702 | 1 | k__Fungi(100);p__Ascomycota(100);c__Saccharomycetes(100);o__Saccharomycetales(100);f__Metschnikowiaceae(100);g__Metschnikowia(100);s__Metschnikowia_pulcherrima(100); | Metschnikowia | 13 | Saprotroph | \|Undefined Saprotroph\| | Yeast | NULL | Probable | As nectar-tap_saprotroph (Põlme et al. 2020); Decay sugar-rich substrates (Põlme et al. 2020) | Põlme S, et al. 2020. Fungal Diversity 105:1-16 ((https://doi.org/10.1007/s13225-020-00466-2)) |
| Otu18703 | 1 | k__Fungi(100);p__Ascomycota(100);c__Saccharomycetes(100);o__Saccharomycetales(100);f__Metschnikowiaceae(100);g__Metschnikowia(100);g__Metschnikowia_unclassified(100); | Metschnikowia | 13 | Saprotroph | \|Undefined Saprotroph\| | Yeast | NULL | Probable | As nectar-tap_saprotroph (Põlme et al. 2020); Decay sugar-rich substrates (Põlme et al. 2020) | Põlme S, et al. 2020. Fungal Diversity 105:1-16 ((https://doi.org/10.1007/s13225-020-00466-2)) |
| Otu18705 | 1 | k__Fungi(100);p__Ascomycota(100);c__Saccharomycetes(100);o__Saccharomycetales(100);f__Metschnikowiaceae(100);g__Metschnikowia(100);g__Metschnikowia_unclassified(100); | Metschnikowia | 13 | Saprotroph | \|Undefined Saprotroph\| | Yeast | NULL | Probable | As nectar-tap_saprotroph (Põlme et al. 2020); Decay sugar-rich substrates (Põlme et al. 2020) | Põlme S, et al. 2020. Fungal Diversity 105:1-16 ((https://doi.org/10.1007/s13225-020-00466-2)) |
| Otu18706 | 1 | k__Fungi(100);p__Ascomycota(100);c__Saccharomycetes(100);o__Saccharomycetales(100);f__Metschnikowiaceae(100);g__Metschnikowia(100);s__Metschnikowia_pulcherrima(100); | Metschnikowia | 13 | Saprotroph | \|Undefined Saprotroph\| | Yeast | NULL | Probable | As nectar-tap_saprotroph (Põlme et al. 2020); Decay sugar-rich substrates (Põlme et al. 2020) | Põlme S, et al. 2020. Fungal Diversity 105:1-16 ((https://doi.org/10.1007/s13225-020-00466-2)) |
| Otu18707 | 1 | k__Fungi(100);p__Ascomycota(100);c__Saccharomycetes(100);o__Saccharomycetales(100);f__Metschnikowiaceae(100);g__Metschnikowia(100);s__Metschnikowia_pulcherrima(100); | Metschnikowia | 13 | Saprotroph | \|Undefined Saprotroph\| | Yeast | NULL | Probable | As nectar-tap_saprotroph (Põlme et al. 2020); Decay sugar-rich substrates (Põlme et al. 2020) | Põlme S, et al. 2020. Fungal Diversity 105:1-16 ((https://doi.org/10.1007/s13225-020-00466-2)) |
| Otu18708 | 1 | k__Fungi(100);p__Ascomycota(100);c__Saccharomycetes(100);o__Saccharomycetales(100);f__Metschnikowiaceae(100);g__Metschnikowia(100);s__Metschnikowia_pulcherrima(100); | Metschnikowia | 13 | Saprotroph | \|Undefined Saprotroph\| | Yeast | NULL | Probable | As nectar-tap_saprotroph (Põlme et al. 2020); Decay sugar-rich substrates (Põlme et al. 2020) | Põlme S, et al. 2020. Fungal Diversity 105:1-16 ((https://doi.org/10.1007/s13225-020-00466-2)) |
| Otu18709 | 1 | k__Fungi(100);p__Ascomycota(100);c__Saccharomycetes(100);o__Saccharomycetales(100);f__Metschnikowiaceae(100);g__Metschnikowia(100);s__Metschnikowia_pulcherrima(100); | Metschnikowia | 13 | Saprotroph | \|Undefined Saprotroph\| | Yeast | NULL | Probable | As nectar-tap_saprotroph (Põlme et al. 2020); Decay sugar-rich substrates (Põlme et al. 2020) | Põlme S, et al. 2020. Fungal Diversity 105:1-16 ((https://doi.org/10.1007/s13225-020-00466-2)) |
| Otu18710 | 1 | k__Fungi(100);p__Ascomycota(100);c__Saccharomycetes(100);o__Saccharomycetales(100);f__Metschnikowiaceae(100);g__Metschnikowia(100);s__Metschnikowia_pulcherrima(100); | Metschnikowia | 13 | Saprotroph | \|Undefined Saprotroph\| | Yeast | NULL | Probable | As nectar-tap_saprotroph (Põlme et al. 2020); Decay sugar-rich substrates (Põlme et al. 2020) | Põlme S, et al. 2020. Fungal Diversity 105:1-16 ((https://doi.org/10.1007/s13225-020-00466-2)) |
| Otu18711 | 1 | k__Fungi(100);p__Ascomycota(100);c__Saccharomycetes(100);o__Saccharomycetales(100);f__Metschnikowiaceae(100);g__Metschnikowia(100);g__Metschnikowia_unclassified(100); | Metschnikowia | 13 | Saprotroph | \|Undefined Saprotroph\| | Yeast | NULL | Probable | As nectar-tap_saprotroph (Põlme et al. 2020); Decay sugar-rich substrates (Põlme et al. 2020) | Põlme S, et al. 2020. Fungal Diversity 105:1-16 ((https://doi.org/10.1007/s13225-020-00466-2)) |
| Otu18714 | 1 | k__Fungi(100);p__Ascomycota(100);c__Saccharomycetes(100);o__Saccharomycetales(100);f__Metschnikowiaceae(100);g__Metschnikowia(100);s__Metschnikowia_pulcherrima(100); | Metschnikowia | 13 | Saprotroph | \|Undefined Saprotroph\| | Yeast | NULL | Probable | As nectar-tap_saprotroph (Põlme et al. 2020); Decay sugar-rich substrates (Põlme et al. 2020) | Põlme S, et al. 2020. Fungal Diversity 105:1-16 ((https://doi.org/10.1007/s13225-020-00466-2)) |
| Otu18715 | 1 | k__Fungi(100);p__Ascomycota(100);c__Saccharomycetes(100);o__Saccharomycetales(100);f__Metschnikowiaceae(100);g__Metschnikowia(100);s__Metschnikowia_pulcherrima(100); | Metschnikowia | 13 | Saprotroph | \|Undefined Saprotroph\| | Yeast | NULL | Probable | As nectar-tap_saprotroph (Põlme et al. 2020); Decay sugar-rich substrates (Põlme et al. 2020) | Põlme S, et al. 2020. Fungal Diversity 105:1-16 ((https://doi.org/10.1007/s13225-020-00466-2)) |
| Otu18716 | 1 | k__Fungi(100);p__Ascomycota(100);c__Saccharomycetes(100);o__Saccharomycetales(100);f__Metschnikowiaceae(100);g__Metschnikowia(100);s__Metschnikowia_pulcherrima(100); | Metschnikowia | 13 | Saprotroph | \|Undefined Saprotroph\| | Yeast | NULL | Probable | As nectar-tap_saprotroph (Põlme et al. 2020); Decay sugar-rich substrates (Põlme et al. 2020) | Põlme S, et al. 2020. Fungal Diversity 105:1-16 ((https://doi.org/10.1007/s13225-020-00466-2)) |
| Otu18718 | 1 | k__Fungi(100);p__Ascomycota(100);c__Saccharomycetes(100);o__Saccharomycetales(100);f__Metschnikowiaceae(100);g__Metschnikowia(100);s__Metschnikowia_pulcherrima(100); | Metschnikowia | 13 | Saprotroph | \|Undefined Saprotroph\| | Yeast | NULL | Probable | As nectar-tap_saprotroph (Põlme et al. 2020); Decay sugar-rich substrates (Põlme et al. 2020) | Põlme S, et al. 2020. Fungal Diversity 105:1-16 ((https://doi.org/10.1007/s13225-020-00466-2)) |
| Otu18719 | 1 | k__Fungi(100);p__Ascomycota(100);c__Saccharomycetes(100);o__Saccharomycetales(100);f__Metschnikowiaceae(100);g__Metschnikowia(100);s__Metschnikowia_pulcherrima(100); | Metschnikowia | 13 | Saprotroph | \|Undefined Saprotroph\| | Yeast | NULL | Probable | As nectar-tap_saprotroph (Põlme et al. 2020); Decay sugar-rich substrates (Põlme et al. 2020) | Põlme S, et al. 2020. Fungal Diversity 105:1-16 ((https://doi.org/10.1007/s13225-020-00466-2)) |
| Otu18720 | 1 | k__Fungi(100);p__Ascomycota(100);c__Saccharomycetes(100);o__Saccharomycetales(100);f__Metschnikowiaceae(100);g__Metschnikowia(100);g__Metschnikowia_unclassified(100); | Metschnikowia | 13 | Saprotroph | \|Undefined Saprotroph\| | Yeast | NULL | Probable | As nectar-tap_saprotroph (Põlme et al. 2020); Decay sugar-rich substrates (Põlme et al. 2020) | Põlme S, et al. 2020. Fungal Diversity 105:1-16 ((https://doi.org/10.1007/s13225-020-00466-2)) |
| Otu18721 | 1 | k__Fungi(100);p__Ascomycota(100);c__Saccharomycetes(100);o__Saccharomycetales(100);f__Metschnikowiaceae(100);g__Metschnikowia(100);s__Metschnikowia_pulcherrima(100); | Metschnikowia | 13 | Saprotroph | \|Undefined Saprotroph\| | Yeast | NULL | Probable | As nectar-tap_saprotroph (Põlme et al. 2020); Decay sugar-rich substrates (Põlme et al. 2020) | Põlme S, et al. 2020. Fungal Diversity 105:1-16 ((https://doi.org/10.1007/s13225-020-00466-2)) |
| Otu18722 | 1 | k__Fungi(100);p__Ascomycota(100);c__Saccharomycetes(100);o__Saccharomycetales(100);f__Metschnikowiaceae(100);g__Metschnikowia(100);s__Metschnikowia_pulcherrima(100); | Metschnikowia | 13 | Saprotroph | \|Undefined Saprotroph\| | Yeast | NULL | Probable | As nectar-tap_saprotroph (Põlme et al. 2020); Decay sugar-rich substrates (Põlme et al. 2020) | Põlme S, et al. 2020. Fungal Diversity 105:1-16 ((https://doi.org/10.1007/s13225-020-00466-2)) |
| Otu18723 | 1 | k__Fungi(100);p__Ascomycota(100);c__Saccharomycetes(100);o__Saccharomycetales(100);f__Metschnikowiaceae(100);g__Metschnikowia(100);s__Metschnikowia_pulcherrima(100); | Metschnikowia | 13 | Saprotroph | \|Undefined Saprotroph\| | Yeast | NULL | Probable | As nectar-tap_saprotroph (Põlme et al. 2020); Decay sugar-rich substrates (Põlme et al. 2020) | Põlme S, et al. 2020. Fungal Diversity 105:1-16 ((https://doi.org/10.1007/s13225-020-00466-2)) |
| Otu18724 | 1 | k__Fungi(100);p__Ascomycota(100);c__Saccharomycetes(100);o__Saccharomycetales(100);f__Metschnikowiaceae(100);g__Metschnikowia(100);s__Metschnikowia_pulcherrima(100); | Metschnikowia | 13 | Saprotroph | \|Undefined Saprotroph\| | Yeast | NULL | Probable | As nectar-tap_saprotroph (Põlme et al. 2020); Decay sugar-rich substrates (Põlme et al. 2020) | Põlme S, et al. 2020. Fungal Diversity 105:1-16 ((https://doi.org/10.1007/s13225-020-00466-2)) |
| Otu18725 | 1 | k__Fungi(100);p__Ascomycota(100);c__Saccharomycetes(100);o__Saccharomycetales(100);f__Metschnikowiaceae(100);g__Metschnikowia(100);s__Metschnikowia_pulcherrima(100); | Metschnikowia | 13 | Saprotroph | \|Undefined Saprotroph\| | Yeast | NULL | Probable | As nectar-tap_saprotroph (Põlme et al. 2020); Decay sugar-rich substrates (Põlme et al. 2020) | Põlme S, et al. 2020. Fungal Diversity 105:1-16 ((https://doi.org/10.1007/s13225-020-00466-2)) |
| Otu18726 | 1 | k__Fungi(100);p__Ascomycota(100);c__Saccharomycetes(100);o__Saccharomycetales(100);f__Metschnikowiaceae(100);g__Metschnikowia(100);s__Metschnikowia_pulcherrima(100); | Metschnikowia | 13 | Saprotroph | \|Undefined Saprotroph\| | Yeast | NULL | Probable | As nectar-tap_saprotroph (Põlme et al. 2020); Decay sugar-rich substrates (Põlme et al. 2020) | Põlme S, et al. 2020. Fungal Diversity 105:1-16 ((https://doi.org/10.1007/s13225-020-00466-2)) |
| Otu18727 | 1 | k__Fungi(100);p__Ascomycota(100);c__Saccharomycetes(100);o__Saccharomycetales(100);f__Metschnikowiaceae(100);g__Metschnikowia(100);s__Metschnikowia_pulcherrima(100); | Metschnikowia | 13 | Saprotroph | \|Undefined Saprotroph\| | Yeast | NULL | Probable | As nectar-tap_saprotroph (Põlme et al. 2020); Decay sugar-rich substrates (Põlme et al. 2020) | Põlme S, et al. 2020. Fungal Diversity 105:1-16 ((https://doi.org/10.1007/s13225-020-00466-2)) |
| Otu18728 | 1 | k__Fungi(100);p__Ascomycota(100);c__Saccharomycetes(100);o__Saccharomycetales(100);f__Metschnikowiaceae(100);g__Metschnikowia(100);s__Metschnikowia_pulcherrima(100); | Metschnikowia | 13 | Saprotroph | \|Undefined Saprotroph\| | Yeast | NULL | Probable | As nectar-tap_saprotroph (Põlme et al. 2020); Decay sugar-rich substrates (Põlme et al. 2020) | Põlme S, et al. 2020. Fungal Diversity 105:1-16 ((https://doi.org/10.1007/s13225-020-00466-2)) |
| Otu18729 | 1 | k__Fungi(100);p__Ascomycota(100);c__Saccharomycetes(100);o__Saccharomycetales(100);f__Metschnikowiaceae(100);g__Metschnikowia(100);s__Metschnikowia_pulcherrima(100); | Metschnikowia | 13 | Saprotroph | \|Undefined Saprotroph\| | Yeast | NULL | Probable | As nectar-tap_saprotroph (Põlme et al. 2020); Decay sugar-rich substrates (Põlme et al. 2020) | Põlme S, et al. 2020. Fungal Diversity 105:1-16 ((https://doi.org/10.1007/s13225-020-00466-2)) |
| Otu18730 | 1 | k__Fungi(100);p__Ascomycota(100);c__Saccharomycetes(100);o__Saccharomycetales(100);f__Metschnikowiaceae(100);g__Metschnikowia(100);g__Metschnikowia_unclassified(100); | Metschnikowia | 13 | Saprotroph | \|Undefined Saprotroph\| | Yeast | NULL | Probable | As nectar-tap_saprotroph (Põlme et al. 2020); Decay sugar-rich substrates (Põlme et al. 2020) | Põlme S, et al. 2020. Fungal Diversity 105:1-16 ((https://doi.org/10.1007/s13225-020-00466-2)) |
| Otu18731 | 1 | k__Fungi(100);p__Ascomycota(100);c__Saccharomycetes(100);o__Saccharomycetales(100);f__Metschnikowiaceae(100);g__Metschnikowia(100);s__Metschnikowia_pulcherrima(100); | Metschnikowia | 13 | Saprotroph | \|Undefined Saprotroph\| | Yeast | NULL | Probable | As nectar-tap_saprotroph (Põlme et al. 2020); Decay sugar-rich substrates (Põlme et al. 2020) | Põlme S, et al. 2020. Fungal Diversity 105:1-16 ((https://doi.org/10.1007/s13225-020-00466-2)) |
| Otu18732 | 1 | k__Fungi(100);p__Ascomycota(100);c__Saccharomycetes(100);o__Saccharomycetales(100);f__Saccharomycetales_family_Incertae_sedis(100);g__Candida(100);g__Candida_unclassified(100); | Candida | 13 | Pathotroph-Saprotroph-Symbiotroph | Animal Pathogen-Endophyte-Endosymbiont-Epiphyte-\|Undefined Saprotroph\| | Dimorphic Yeast-Yeast | NULL | Probable | As nectar-tap_saprotroph (Põlme et al. 2020); Decay sugar-rich substrates (Põlme et al. 2020); This asexual genus is polyphyletic and contain species that function very differently depend on what they are. Caution should be taken when interpreting guilds for Candida species. | Manolakaki D, et al. 2010. Virulence. 1: 367-375 ((https://doi.org/10.4161/viru.1.5.12796)); Mekbib SB, et al. 2011. Tropical Plant Pathology 36:233-240 ((https://doi.org/10.1590/S1982-56762011000400004)); Põlme S, et al. 2020. Fungal Diversity 105:1-16 ((https://doi.org/10.1007/s13225-020-00466-2)) |
| Otu18734 | 1 | k__Fungi(100);p__Ascomycota(100);c__Saccharomycetes(100);o__Saccharomycetales(100);f__Metschnikowiaceae(100);g__Metschnikowia(100);g__Metschnikowia_unclassified(100); | Metschnikowia | 13 | Saprotroph | \|Undefined Saprotroph\| | Yeast | NULL | Probable | As nectar-tap_saprotroph (Põlme et al. 2020); Decay sugar-rich substrates (Põlme et al. 2020) | Põlme S, et al. 2020. Fungal Diversity 105:1-16 ((https://doi.org/10.1007/s13225-020-00466-2)) |
| Otu18735 | 1 | k__Fungi(100);p__Ascomycota(100);c__Saccharomycetes(100);o__Saccharomycetales(100);f__Metschnikowiaceae(100);g__Metschnikowia(100);s__Metschnikowia_pulcherrima(100); | Metschnikowia | 13 | Saprotroph | \|Undefined Saprotroph\| | Yeast | NULL | Probable | As nectar-tap_saprotroph (Põlme et al. 2020); Decay sugar-rich substrates (Põlme et al. 2020) | Põlme S, et al. 2020. Fungal Diversity 105:1-16 ((https://doi.org/10.1007/s13225-020-00466-2)) |
| Otu18736 | 1 | k__Fungi(100);p__Ascomycota(100);c__Saccharomycetes(100);o__Saccharomycetales(100);f__Metschnikowiaceae(100);g__Metschnikowia(100);s__Metschnikowia_pulcherrima(100); | Metschnikowia | 13 | Saprotroph | \|Undefined Saprotroph\| | Yeast | NULL | Probable | As nectar-tap_saprotroph (Põlme et al. 2020); Decay sugar-rich substrates (Põlme et al. 2020) | Põlme S, et al. 2020. Fungal Diversity 105:1-16 ((https://doi.org/10.1007/s13225-020-00466-2)) |
| Otu18737 | 1 | k__Fungi(100);p__Ascomycota(100);c__Saccharomycetes(100);o__Saccharomycetales(100);f__Metschnikowiaceae(100);g__Metschnikowia(100);s__Metschnikowia_pulcherrima(100); | Metschnikowia | 13 | Saprotroph | \|Undefined Saprotroph\| | Yeast | NULL | Probable | As nectar-tap_saprotroph (Põlme et al. 2020); Decay sugar-rich substrates (Põlme et al. 2020) | Põlme S, et al. 2020. Fungal Diversity 105:1-16 ((https://doi.org/10.1007/s13225-020-00466-2)) |
| Otu18738 | 1 | k__Fungi(100);p__Ascomycota(100);c__Saccharomycetes(100);o__Saccharomycetales(100);f__Metschnikowiaceae(100);g__Metschnikowia(100);s__Metschnikowia_pulcherrima(100); | Metschnikowia | 13 | Saprotroph | \|Undefined Saprotroph\| | Yeast | NULL | Probable | As nectar-tap_saprotroph (Põlme et al. 2020); Decay sugar-rich substrates (Põlme et al. 2020) | Põlme S, et al. 2020. Fungal Diversity 105:1-16 ((https://doi.org/10.1007/s13225-020-00466-2)) |
| Otu18739 | 1 | k__Fungi(100);p__Ascomycota(100);c__Saccharomycetes(100);o__Saccharomycetales(100);f__Metschnikowiaceae(100);g__Metschnikowia(100);s__Metschnikowia_pulcherrima(100); | Metschnikowia | 13 | Saprotroph | \|Undefined Saprotroph\| | Yeast | NULL | Probable | As nectar-tap_saprotroph (Põlme et al. 2020); Decay sugar-rich substrates (Põlme et al. 2020) | Põlme S, et al. 2020. Fungal Diversity 105:1-16 ((https://doi.org/10.1007/s13225-020-00466-2)) |
| Otu18740 | 1 | k__Fungi(100);p__Ascomycota(100);c__Saccharomycetes(100);o__Saccharomycetales(100);f__Metschnikowiaceae(100);g__Metschnikowia(100);s__Metschnikowia_pulcherrima(100); | Metschnikowia | 13 | Saprotroph | \|Undefined Saprotroph\| | Yeast | NULL | Probable | As nectar-tap_saprotroph (Põlme et al. 2020); Decay sugar-rich substrates (Põlme et al. 2020) | Põlme S, et al. 2020. Fungal Diversity 105:1-16 ((https://doi.org/10.1007/s13225-020-00466-2)) |
| Otu18741 | 1 | k__Fungi(100);p__Ascomycota(100);c__Saccharomycetes(100);o__Saccharomycetales(100);f__Metschnikowiaceae(100);g__Metschnikowia(100);s__Metschnikowia_pulcherrima(100); | Metschnikowia | 13 | Saprotroph | \|Undefined Saprotroph\| | Yeast | NULL | Probable | As nectar-tap_saprotroph (Põlme et al. 2020); Decay sugar-rich substrates (Põlme et al. 2020) | Põlme S, et al. 2020. Fungal Diversity 105:1-16 ((https://doi.org/10.1007/s13225-020-00466-2)) |
| Otu18743 | 1 | k__Fungi(100);p__Ascomycota(100);c__Saccharomycetes(100);o__Saccharomycetales(100);f__Metschnikowiaceae(100);g__Metschnikowia(100);s__Metschnikowia_pulcherrima(100); | Metschnikowia | 13 | Saprotroph | \|Undefined Saprotroph\| | Yeast | NULL | Probable | As nectar-tap_saprotroph (Põlme et al. 2020); Decay sugar-rich substrates (Põlme et al. 2020) | Põlme S, et al. 2020. Fungal Diversity 105:1-16 ((https://doi.org/10.1007/s13225-020-00466-2)) |
| Otu18744 | 1 | k__Fungi(100);p__Ascomycota(100);c__Saccharomycetes(100);o__Saccharomycetales(100);f__Saccharomycetales_family_Incertae_sedis(100);g__Candida(100);g__Candida_unclassified(100); | Candida | 13 | Pathotroph-Saprotroph-Symbiotroph | Animal Pathogen-Endophyte-Endosymbiont-Epiphyte-\|Undefined Saprotroph\| | Dimorphic Yeast-Yeast | NULL | Probable | As nectar-tap_saprotroph (Põlme et al. 2020); Decay sugar-rich substrates (Põlme et al. 2020); This asexual genus is polyphyletic and contain species that function very differently depend on what they are. Caution should be taken when interpreting guilds for Candida species. | Manolakaki D, et al. 2010. Virulence. 1: 367-375 ((https://doi.org/10.4161/viru.1.5.12796)); Mekbib SB, et al. 2011. Tropical Plant Pathology 36:233-240 ((https://doi.org/10.1590/S1982-56762011000400004)); Põlme S, et al. 2020. Fungal Diversity 105:1-16 ((https://doi.org/10.1007/s13225-020-00466-2)) |
| Otu18748 | 1 | k__Fungi(100);p__Ascomycota(100);c__Saccharomycetes(100);o__Saccharomycetales(100);f__Metschnikowiaceae(100);g__Metschnikowia(100);s__Metschnikowia_pulcherrima(100); | Metschnikowia | 13 | Saprotroph | \|Undefined Saprotroph\| | Yeast | NULL | Probable | As nectar-tap_saprotroph (Põlme et al. 2020); Decay sugar-rich substrates (Põlme et al. 2020) | Põlme S, et al. 2020. Fungal Diversity 105:1-16 ((https://doi.org/10.1007/s13225-020-00466-2)) |
| Otu18749 | 1 | k__Fungi(100);p__Ascomycota(100);c__Saccharomycetes(100);o__Saccharomycetales(100);f__Metschnikowiaceae(100);g__Metschnikowia(100);s__Metschnikowia_pulcherrima(100); | Metschnikowia | 13 | Saprotroph | \|Undefined Saprotroph\| | Yeast | NULL | Probable | As nectar-tap_saprotroph (Põlme et al. 2020); Decay sugar-rich substrates (Põlme et al. 2020) | Põlme S, et al. 2020. Fungal Diversity 105:1-16 ((https://doi.org/10.1007/s13225-020-00466-2)) |
| Otu18750 | 1 | k__Fungi(100);p__Ascomycota(100);c__Saccharomycetes(100);o__Saccharomycetales(100);f__Metschnikowiaceae(100);g__Metschnikowia(100);s__Metschnikowia_pulcherrima(100); | Metschnikowia | 13 | Saprotroph | \|Undefined Saprotroph\| | Yeast | NULL | Probable | As nectar-tap_saprotroph (Põlme et al. 2020); Decay sugar-rich substrates (Põlme et al. 2020) | Põlme S, et al. 2020. Fungal Diversity 105:1-16 ((https://doi.org/10.1007/s13225-020-00466-2)) |
| Otu18751 | 1 | k__Fungi(100);p__Ascomycota(100);c__Saccharomycetes(100);o__Saccharomycetales(100);f__Metschnikowiaceae(100);g__Metschnikowia(100);g__Metschnikowia_unclassified(100); | Metschnikowia | 13 | Saprotroph | \|Undefined Saprotroph\| | Yeast | NULL | Probable | As nectar-tap_saprotroph (Põlme et al. 2020); Decay sugar-rich substrates (Põlme et al. 2020) | Põlme S, et al. 2020. Fungal Diversity 105:1-16 ((https://doi.org/10.1007/s13225-020-00466-2)) |
| Otu18752 | 1 | k__Fungi(100);p__Ascomycota(100);c__Saccharomycetes(100);o__Saccharomycetales(100);f__Metschnikowiaceae(100);g__Metschnikowia(100);s__Metschnikowia_pulcherrima(100); | Metschnikowia | 13 | Saprotroph | \|Undefined Saprotroph\| | Yeast | NULL | Probable | As nectar-tap_saprotroph (Põlme et al. 2020); Decay sugar-rich substrates (Põlme et al. 2020) | Põlme S, et al. 2020. Fungal Diversity 105:1-16 ((https://doi.org/10.1007/s13225-020-00466-2)) |
| Otu18753 | 1 | k__Fungi(100);p__Ascomycota(100);c__Saccharomycetes(100);o__Saccharomycetales(100);f__Metschnikowiaceae(100);g__Metschnikowia(100);s__Metschnikowia_pulcherrima(100); | Metschnikowia | 13 | Saprotroph | \|Undefined Saprotroph\| | Yeast | NULL | Probable | As nectar-tap_saprotroph (Põlme et al. 2020); Decay sugar-rich substrates (Põlme et al. 2020) | Põlme S, et al. 2020. Fungal Diversity 105:1-16 ((https://doi.org/10.1007/s13225-020-00466-2)) |
| Otu18754 | 1 | k__Fungi(100);p__Ascomycota(100);c__Saccharomycetes(100);o__Saccharomycetales(100);f__Metschnikowiaceae(100);g__Metschnikowia(100);s__Metschnikowia_pulcherrima(100); | Metschnikowia | 13 | Saprotroph | \|Undefined Saprotroph\| | Yeast | NULL | Probable | As nectar-tap_saprotroph (Põlme et al. 2020); Decay sugar-rich substrates (Põlme et al. 2020) | Põlme S, et al. 2020. Fungal Diversity 105:1-16 ((https://doi.org/10.1007/s13225-020-00466-2)) |
| Otu18756 | 1 | k__Fungi(100);p__Ascomycota(100);c__Saccharomycetes(100);o__Saccharomycetales(100);f__Metschnikowiaceae(100);g__Metschnikowia(100);s__Metschnikowia_pulcherrima(100); | Metschnikowia | 13 | Saprotroph | \|Undefined Saprotroph\| | Yeast | NULL | Probable | As nectar-tap_saprotroph (Põlme et al. 2020); Decay sugar-rich substrates (Põlme et al. 2020) | Põlme S, et al. 2020. Fungal Diversity 105:1-16 ((https://doi.org/10.1007/s13225-020-00466-2)) |
| Otu18757 | 1 | k__Fungi(100);p__Ascomycota(100);c__Saccharomycetes(100);o__Saccharomycetales(100);f__Metschnikowiaceae(100);g__Metschnikowia(100);g__Metschnikowia_unclassified(100); | Metschnikowia | 13 | Saprotroph | \|Undefined Saprotroph\| | Yeast | NULL | Probable | As nectar-tap_saprotroph (Põlme et al. 2020); Decay sugar-rich substrates (Põlme et al. 2020) | Põlme S, et al. 2020. Fungal Diversity 105:1-16 ((https://doi.org/10.1007/s13225-020-00466-2)) |
| Otu18758 | 1 | k__Fungi(100);p__Ascomycota(100);c__Saccharomycetes(100);o__Saccharomycetales(100);f__Metschnikowiaceae(100);g__Metschnikowia(100);g__Metschnikowia_unclassified(100); | Metschnikowia | 13 | Saprotroph | \|Undefined Saprotroph\| | Yeast | NULL | Probable | As nectar-tap_saprotroph (Põlme et al. 2020); Decay sugar-rich substrates (Põlme et al. 2020) | Põlme S, et al. 2020. Fungal Diversity 105:1-16 ((https://doi.org/10.1007/s13225-020-00466-2)) |
| Otu18759 | 1 | k__Fungi(100);p__Ascomycota(100);c__Saccharomycetes(100);o__Saccharomycetales(100);f__Metschnikowiaceae(100);g__Metschnikowia(100);g__Metschnikowia_unclassified(100); | Metschnikowia | 13 | Saprotroph | \|Undefined Saprotroph\| | Yeast | NULL | Probable | As nectar-tap_saprotroph (Põlme et al. 2020); Decay sugar-rich substrates (Põlme et al. 2020) | Põlme S, et al. 2020. Fungal Diversity 105:1-16 ((https://doi.org/10.1007/s13225-020-00466-2)) |
| Otu18760 | 1 | k__Fungi(100);p__Ascomycota(100);c__Saccharomycetes(100);o__Saccharomycetales(100);f__Metschnikowiaceae(100);g__Metschnikowia(100);s__Metschnikowia_pulcherrima(100); | Metschnikowia | 13 | Saprotroph | \|Undefined Saprotroph\| | Yeast | NULL | Probable | As nectar-tap_saprotroph (Põlme et al. 2020); Decay sugar-rich substrates (Põlme et al. 2020) | Põlme S, et al. 2020. Fungal Diversity 105:1-16 ((https://doi.org/10.1007/s13225-020-00466-2)) |
| Otu18761 | 1 | k__Fungi(100);p__Ascomycota(100);c__Saccharomycetes(100);o__Saccharomycetales(100);f__Metschnikowiaceae(100);g__Metschnikowia(100);s__Metschnikowia_pulcherrima(100); | Metschnikowia | 13 | Saprotroph | \|Undefined Saprotroph\| | Yeast | NULL | Probable | As nectar-tap_saprotroph (Põlme et al. 2020); Decay sugar-rich substrates (Põlme et al. 2020) | Põlme S, et al. 2020. Fungal Diversity 105:1-16 ((https://doi.org/10.1007/s13225-020-00466-2)) |
| Otu18762 | 1 | k__Fungi(100);p__Ascomycota(100);c__Saccharomycetes(100);o__Saccharomycetales(100);f__Metschnikowiaceae(100);g__Metschnikowia(100);g__Metschnikowia_unclassified(100); | Metschnikowia | 13 | Saprotroph | \|Undefined Saprotroph\| | Yeast | NULL | Probable | As nectar-tap_saprotroph (Põlme et al. 2020); Decay sugar-rich substrates (Põlme et al. 2020) | Põlme S, et al. 2020. Fungal Diversity 105:1-16 ((https://doi.org/10.1007/s13225-020-00466-2)) |
| Otu18763 | 1 | k__Fungi(100);p__Ascomycota(100);c__Saccharomycetes(100);o__Saccharomycetales(100);f__Metschnikowiaceae(100);g__Metschnikowia(100);s__Metschnikowia_pulcherrima(100); | Metschnikowia | 13 | Saprotroph | \|Undefined Saprotroph\| | Yeast | NULL | Probable | As nectar-tap_saprotroph (Põlme et al. 2020); Decay sugar-rich substrates (Põlme et al. 2020) | Põlme S, et al. 2020. Fungal Diversity 105:1-16 ((https://doi.org/10.1007/s13225-020-00466-2)) |
| Otu18764 | 1 | k__Fungi(100);p__Ascomycota(100);c__Saccharomycetes(100);o__Saccharomycetales(100);f__Metschnikowiaceae(100);g__Metschnikowia(100);s__Metschnikowia_pulcherrima(100); | Metschnikowia | 13 | Saprotroph | \|Undefined Saprotroph\| | Yeast | NULL | Probable | As nectar-tap_saprotroph (Põlme et al. 2020); Decay sugar-rich substrates (Põlme et al. 2020) | Põlme S, et al. 2020. Fungal Diversity 105:1-16 ((https://doi.org/10.1007/s13225-020-00466-2)) |
| Otu18765 | 1 | k__Fungi(100);p__Ascomycota(100);c__Saccharomycetes(100);o__Saccharomycetales(100);f__Metschnikowiaceae(100);g__Metschnikowia(100);s__Metschnikowia_pulcherrima(100); | Metschnikowia | 13 | Saprotroph | \|Undefined Saprotroph\| | Yeast | NULL | Probable | As nectar-tap_saprotroph (Põlme et al. 2020); Decay sugar-rich substrates (Põlme et al. 2020) | Põlme S, et al. 2020. Fungal Diversity 105:1-16 ((https://doi.org/10.1007/s13225-020-00466-2)) |
| Otu18766 | 1 | k__Fungi(100);p__Ascomycota(100);c__Saccharomycetes(100);o__Saccharomycetales(100);f__Metschnikowiaceae(100);g__Metschnikowia(100);s__Metschnikowia_pulcherrima(100); | Metschnikowia | 13 | Saprotroph | \|Undefined Saprotroph\| | Yeast | NULL | Probable | As nectar-tap_saprotroph (Põlme et al. 2020); Decay sugar-rich substrates (Põlme et al. 2020) | Põlme S, et al. 2020. Fungal Diversity 105:1-16 ((https://doi.org/10.1007/s13225-020-00466-2)) |
| Otu18767 | 1 | k__Fungi(100);p__Ascomycota(100);c__Saccharomycetes(100);o__Saccharomycetales(100);f__Metschnikowiaceae(100);g__Metschnikowia(100);s__Metschnikowia_pulcherrima(100); | Metschnikowia | 13 | Saprotroph | \|Undefined Saprotroph\| | Yeast | NULL | Probable | As nectar-tap_saprotroph (Põlme et al. 2020); Decay sugar-rich substrates (Põlme et al. 2020) | Põlme S, et al. 2020. Fungal Diversity 105:1-16 ((https://doi.org/10.1007/s13225-020-00466-2)) |
| Otu18768 | 1 | k__Fungi(100);p__Ascomycota(100);c__Saccharomycetes(100);o__Saccharomycetales(100);f__Metschnikowiaceae(100);g__Metschnikowia(100);s__Metschnikowia_pulcherrima(100); | Metschnikowia | 13 | Saprotroph | \|Undefined Saprotroph\| | Yeast | NULL | Probable | As nectar-tap_saprotroph (Põlme et al. 2020); Decay sugar-rich substrates (Põlme et al. 2020) | Põlme S, et al. 2020. Fungal Diversity 105:1-16 ((https://doi.org/10.1007/s13225-020-00466-2)) |
| Otu18769 | 1 | k__Fungi(100);p__Ascomycota(100);c__Saccharomycetes(100);o__Saccharomycetales(100);f__Metschnikowiaceae(100);g__Metschnikowia(100);s__Metschnikowia_pulcherrima(100); | Metschnikowia | 13 | Saprotroph | \|Undefined Saprotroph\| | Yeast | NULL | Probable | As nectar-tap_saprotroph (Põlme et al. 2020); Decay sugar-rich substrates (Põlme et al. 2020) | Põlme S, et al. 2020. Fungal Diversity 105:1-16 ((https://doi.org/10.1007/s13225-020-00466-2)) |
| Otu18770 | 1 | k__Fungi(100);p__Ascomycota(100);c__Saccharomycetes(100);o__Saccharomycetales(100);f__Metschnikowiaceae(100);g__Metschnikowia(100);g__Metschnikowia_unclassified(100); | Metschnikowia | 13 | Saprotroph | \|Undefined Saprotroph\| | Yeast | NULL | Probable | As nectar-tap_saprotroph (Põlme et al. 2020); Decay sugar-rich substrates (Põlme et al. 2020) | Põlme S, et al. 2020. Fungal Diversity 105:1-16 ((https://doi.org/10.1007/s13225-020-00466-2)) |
| Otu18771 | 1 | k__Fungi(100);p__Ascomycota(100);c__Saccharomycetes(100);o__Saccharomycetales(100);f__Metschnikowiaceae(100);g__Metschnikowia(100);s__Metschnikowia_pulcherrima(100); | Metschnikowia | 13 | Saprotroph | \|Undefined Saprotroph\| | Yeast | NULL | Probable | As nectar-tap_saprotroph (Põlme et al. 2020); Decay sugar-rich substrates (Põlme et al. 2020) | Põlme S, et al. 2020. Fungal Diversity 105:1-16 ((https://doi.org/10.1007/s13225-020-00466-2)) |
| Otu18772 | 1 | k__Fungi(100);p__Ascomycota(100);c__Saccharomycetes(100);o__Saccharomycetales(100);f__Metschnikowiaceae(100);g__Metschnikowia(100);s__Metschnikowia_pulcherrima(100); | Metschnikowia | 13 | Saprotroph | \|Undefined Saprotroph\| | Yeast | NULL | Probable | As nectar-tap_saprotroph (Põlme et al. 2020); Decay sugar-rich substrates (Põlme et al. 2020) | Põlme S, et al. 2020. Fungal Diversity 105:1-16 ((https://doi.org/10.1007/s13225-020-00466-2)) |
| Otu18773 | 1 | k__Fungi(100);p__Ascomycota(100);c__Saccharomycetes(100);o__Saccharomycetales(100);f__Metschnikowiaceae(100);g__Metschnikowia(100);g__Metschnikowia_unclassified(100); | Metschnikowia | 13 | Saprotroph | \|Undefined Saprotroph\| | Yeast | NULL | Probable | As nectar-tap_saprotroph (Põlme et al. 2020); Decay sugar-rich substrates (Põlme et al. 2020) | Põlme S, et al. 2020. Fungal Diversity 105:1-16 ((https://doi.org/10.1007/s13225-020-00466-2)) |
| Otu18774 | 1 | k__Fungi(100);p__Ascomycota(100);c__Saccharomycetes(100);o__Saccharomycetales(100);f__Metschnikowiaceae(100);g__Metschnikowia(100);s__Metschnikowia_pulcherrima(100); | Metschnikowia | 13 | Saprotroph | \|Undefined Saprotroph\| | Yeast | NULL | Probable | As nectar-tap_saprotroph (Põlme et al. 2020); Decay sugar-rich substrates (Põlme et al. 2020) | Põlme S, et al. 2020. Fungal Diversity 105:1-16 ((https://doi.org/10.1007/s13225-020-00466-2)) |
| Otu18775 | 1 | k__Fungi(100);p__Ascomycota(100);c__Saccharomycetes(100);o__Saccharomycetales(100);f__Metschnikowiaceae(100);g__Metschnikowia(100);s__Metschnikowia_pulcherrima(100); | Metschnikowia | 13 | Saprotroph | \|Undefined Saprotroph\| | Yeast | NULL | Probable | As nectar-tap_saprotroph (Põlme et al. 2020); Decay sugar-rich substrates (Põlme et al. 2020) | Põlme S, et al. 2020. Fungal Diversity 105:1-16 ((https://doi.org/10.1007/s13225-020-00466-2)) |
| Otu18776 | 1 | k__Fungi(100);p__Ascomycota(100);c__Saccharomycetes(100);o__Saccharomycetales(100);f__Metschnikowiaceae(100);g__Metschnikowia(100);s__Metschnikowia_pulcherrima(100); | Metschnikowia | 13 | Saprotroph | \|Undefined Saprotroph\| | Yeast | NULL | Probable | As nectar-tap_saprotroph (Põlme et al. 2020); Decay sugar-rich substrates (Põlme et al. 2020) | Põlme S, et al. 2020. Fungal Diversity 105:1-16 ((https://doi.org/10.1007/s13225-020-00466-2)) |
| Otu18777 | 1 | k__Fungi(100);p__Ascomycota(100);c__Saccharomycetes(100);o__Saccharomycetales(100);f__Metschnikowiaceae(100);g__Metschnikowia(100);g__Metschnikowia_unclassified(100); | Metschnikowia | 13 | Saprotroph | \|Undefined Saprotroph\| | Yeast | NULL | Probable | As nectar-tap_saprotroph (Põlme et al. 2020); Decay sugar-rich substrates (Põlme et al. 2020) | Põlme S, et al. 2020. Fungal Diversity 105:1-16 ((https://doi.org/10.1007/s13225-020-00466-2)) |
| Otu18778 | 1 | k__Fungi(100);p__Ascomycota(100);c__Saccharomycetes(100);o__Saccharomycetales(100);f__Metschnikowiaceae(100);g__Metschnikowia(100);s__Metschnikowia_pulcherrima(100); | Metschnikowia | 13 | Saprotroph | \|Undefined Saprotroph\| | Yeast | NULL | Probable | As nectar-tap_saprotroph (Põlme et al. 2020); Decay sugar-rich substrates (Põlme et al. 2020) | Põlme S, et al. 2020. Fungal Diversity 105:1-16 ((https://doi.org/10.1007/s13225-020-00466-2)) |
| Otu18779 | 1 | k__Fungi(100);p__Ascomycota(100);c__Saccharomycetes(100);o__Saccharomycetales(100);f__Metschnikowiaceae(100);g__Metschnikowia(100);s__Metschnikowia_pulcherrima(100); | Metschnikowia | 13 | Saprotroph | \|Undefined Saprotroph\| | Yeast | NULL | Probable | As nectar-tap_saprotroph (Põlme et al. 2020); Decay sugar-rich substrates (Põlme et al. 2020) | Põlme S, et al. 2020. Fungal Diversity 105:1-16 ((https://doi.org/10.1007/s13225-020-00466-2)) |
| Otu18780 | 1 | k__Fungi(100);p__Ascomycota(100);c__Saccharomycetes(100);o__Saccharomycetales(100);f__Metschnikowiaceae(100);g__Metschnikowia(100);s__Metschnikowia_pulcherrima(100); | Metschnikowia | 13 | Saprotroph | \|Undefined Saprotroph\| | Yeast | NULL | Probable | As nectar-tap_saprotroph (Põlme et al. 2020); Decay sugar-rich substrates (Põlme et al. 2020) | Põlme S, et al. 2020. Fungal Diversity 105:1-16 ((https://doi.org/10.1007/s13225-020-00466-2)) |
| Otu18781 | 1 | k__Fungi(100);p__Ascomycota(100);c__Saccharomycetes(100);o__Saccharomycetales(100);f__Metschnikowiaceae(100);g__Metschnikowia(100);g__Metschnikowia_unclassified(100); | Metschnikowia | 13 | Saprotroph | \|Undefined Saprotroph\| | Yeast | NULL | Probable | As nectar-tap_saprotroph (Põlme et al. 2020); Decay sugar-rich substrates (Põlme et al. 2020) | Põlme S, et al. 2020. Fungal Diversity 105:1-16 ((https://doi.org/10.1007/s13225-020-00466-2)) |
| Otu18782 | 1 | k__Fungi(100);p__Ascomycota(100);c__Saccharomycetes(100);o__Saccharomycetales(100);f__Metschnikowiaceae(100);g__Metschnikowia(100);s__Metschnikowia_pulcherrima(100); | Metschnikowia | 13 | Saprotroph | \|Undefined Saprotroph\| | Yeast | NULL | Probable | As nectar-tap_saprotroph (Põlme et al. 2020); Decay sugar-rich substrates (Põlme et al. 2020) | Põlme S, et al. 2020. Fungal Diversity 105:1-16 ((https://doi.org/10.1007/s13225-020-00466-2)) |
| Otu18783 | 1 | k__Fungi(100);p__Ascomycota(100);c__Saccharomycetes(100);o__Saccharomycetales(100);f__Metschnikowiaceae(100);g__Metschnikowia(100);g__Metschnikowia_unclassified(100); | Metschnikowia | 13 | Saprotroph | \|Undefined Saprotroph\| | Yeast | NULL | Probable | As nectar-tap_saprotroph (Põlme et al. 2020); Decay sugar-rich substrates (Põlme et al. 2020) | Põlme S, et al. 2020. Fungal Diversity 105:1-16 ((https://doi.org/10.1007/s13225-020-00466-2)) |
| Otu18784 | 1 | k__Fungi(100);p__Ascomycota(100);c__Saccharomycetes(100);o__Saccharomycetales(100);f__Metschnikowiaceae(100);g__Metschnikowia(100);g__Metschnikowia_unclassified(100); | Metschnikowia | 13 | Saprotroph | \|Undefined Saprotroph\| | Yeast | NULL | Probable | As nectar-tap_saprotroph (Põlme et al. 2020); Decay sugar-rich substrates (Põlme et al. 2020) | Põlme S, et al. 2020. Fungal Diversity 105:1-16 ((https://doi.org/10.1007/s13225-020-00466-2)) |
| Otu18785 | 1 | k__Fungi(100);p__Ascomycota(100);c__Saccharomycetes(100);o__Saccharomycetales(100);f__Metschnikowiaceae(100);g__Metschnikowia(100);g__Metschnikowia_unclassified(100); | Metschnikowia | 13 | Saprotroph | \|Undefined Saprotroph\| | Yeast | NULL | Probable | As nectar-tap_saprotroph (Põlme et al. 2020); Decay sugar-rich substrates (Põlme et al. 2020) | Põlme S, et al. 2020. Fungal Diversity 105:1-16 ((https://doi.org/10.1007/s13225-020-00466-2)) |
| Otu18786 | 1 | k__Fungi(100);p__Ascomycota(100);c__Saccharomycetes(100);o__Saccharomycetales(100);f__Metschnikowiaceae(100);g__Metschnikowia(100);g__Metschnikowia_unclassified(100); | Metschnikowia | 13 | Saprotroph | \|Undefined Saprotroph\| | Yeast | NULL | Probable | As nectar-tap_saprotroph (Põlme et al. 2020); Decay sugar-rich substrates (Põlme et al. 2020) | Põlme S, et al. 2020. Fungal Diversity 105:1-16 ((https://doi.org/10.1007/s13225-020-00466-2)) |
| Otu18787 | 1 | k__Fungi(100);p__Ascomycota(100);c__Saccharomycetes(100);o__Saccharomycetales(100);f__Metschnikowiaceae(100);g__Metschnikowia(100);s__Metschnikowia_pulcherrima(100); | Metschnikowia | 13 | Saprotroph | \|Undefined Saprotroph\| | Yeast | NULL | Probable | As nectar-tap_saprotroph (Põlme et al. 2020); Decay sugar-rich substrates (Põlme et al. 2020) | Põlme S, et al. 2020. Fungal Diversity 105:1-16 ((https://doi.org/10.1007/s13225-020-00466-2)) |
| Otu18788 | 1 | k__Fungi(100);p__Ascomycota(100);c__Saccharomycetes(100);o__Saccharomycetales(100);f__Metschnikowiaceae(100);g__Metschnikowia(100);s__Metschnikowia_pulcherrima(100); | Metschnikowia | 13 | Saprotroph | \|Undefined Saprotroph\| | Yeast | NULL | Probable | As nectar-tap_saprotroph (Põlme et al. 2020); Decay sugar-rich substrates (Põlme et al. 2020) | Põlme S, et al. 2020. Fungal Diversity 105:1-16 ((https://doi.org/10.1007/s13225-020-00466-2)) |
| Otu18790 | 1 | k__Fungi(100);p__Ascomycota(100);c__Saccharomycetes(100);o__Saccharomycetales(100);f__Metschnikowiaceae(100);g__Metschnikowia(100);g__Metschnikowia_unclassified(100); | Metschnikowia | 13 | Saprotroph | \|Undefined Saprotroph\| | Yeast | NULL | Probable | As nectar-tap_saprotroph (Põlme et al. 2020); Decay sugar-rich substrates (Põlme et al. 2020) | Põlme S, et al. 2020. Fungal Diversity 105:1-16 ((https://doi.org/10.1007/s13225-020-00466-2)) |
| Otu18791 | 1 | k__Fungi(100);p__Ascomycota(100);c__Saccharomycetes(100);o__Saccharomycetales(100);f__Metschnikowiaceae(100);g__Metschnikowia(100);s__Metschnikowia_pulcherrima(100); | Metschnikowia | 13 | Saprotroph | \|Undefined Saprotroph\| | Yeast | NULL | Probable | As nectar-tap_saprotroph (Põlme et al. 2020); Decay sugar-rich substrates (Põlme et al. 2020) | Põlme S, et al. 2020. Fungal Diversity 105:1-16 ((https://doi.org/10.1007/s13225-020-00466-2)) |
| Otu18793 | 1 | k__Fungi(100);p__Ascomycota(100);c__Saccharomycetes(100);o__Saccharomycetales(100);f__Metschnikowiaceae(100);g__Metschnikowia(100);g__Metschnikowia_unclassified(100); | Metschnikowia | 13 | Saprotroph | \|Undefined Saprotroph\| | Yeast | NULL | Probable | As nectar-tap_saprotroph (Põlme et al. 2020); Decay sugar-rich substrates (Põlme et al. 2020) | Põlme S, et al. 2020. Fungal Diversity 105:1-16 ((https://doi.org/10.1007/s13225-020-00466-2)) |
| Otu18794 | 1 | k__Fungi(100);p__Ascomycota(100);c__Saccharomycetes(100);o__Saccharomycetales(100);f__Metschnikowiaceae(100);g__Metschnikowia(100);g__Metschnikowia_unclassified(100); | Metschnikowia | 13 | Saprotroph | \|Undefined Saprotroph\| | Yeast | NULL | Probable | As nectar-tap_saprotroph (Põlme et al. 2020); Decay sugar-rich substrates (Põlme et al. 2020) | Põlme S, et al. 2020. Fungal Diversity 105:1-16 ((https://doi.org/10.1007/s13225-020-00466-2)) |
| Otu18795 | 1 | k__Fungi(100);p__Ascomycota(100);c__Saccharomycetes(100);o__Saccharomycetales(100);f__Metschnikowiaceae(100);g__Metschnikowia(100);s__Metschnikowia_pulcherrima(100); | Metschnikowia | 13 | Saprotroph | \|Undefined Saprotroph\| | Yeast | NULL | Probable | As nectar-tap_saprotroph (Põlme et al. 2020); Decay sugar-rich substrates (Põlme et al. 2020) | Põlme S, et al. 2020. Fungal Diversity 105:1-16 ((https://doi.org/10.1007/s13225-020-00466-2)) |
| Otu18796 | 1 | k__Fungi(100);p__Ascomycota(100);c__Saccharomycetes(100);o__Saccharomycetales(100);f__Metschnikowiaceae(100);g__Metschnikowia(100);s__Metschnikowia_pulcherrima(100); | Metschnikowia | 13 | Saprotroph | \|Undefined Saprotroph\| | Yeast | NULL | Probable | As nectar-tap_saprotroph (Põlme et al. 2020); Decay sugar-rich substrates (Põlme et al. 2020) | Põlme S, et al. 2020. Fungal Diversity 105:1-16 ((https://doi.org/10.1007/s13225-020-00466-2)) |
| Otu18797 | 1 | k__Fungi(100);p__Ascomycota(100);c__Saccharomycetes(100);o__Saccharomycetales(100);f__Metschnikowiaceae(100);g__Metschnikowia(100);g__Metschnikowia_unclassified(100); | Metschnikowia | 13 | Saprotroph | \|Undefined Saprotroph\| | Yeast | NULL | Probable | As nectar-tap_saprotroph (Põlme et al. 2020); Decay sugar-rich substrates (Põlme et al. 2020) | Põlme S, et al. 2020. Fungal Diversity 105:1-16 ((https://doi.org/10.1007/s13225-020-00466-2)) |
| Otu18798 | 1 | k__Fungi(100);p__Ascomycota(100);c__Saccharomycetes(100);o__Saccharomycetales(100);f__Metschnikowiaceae(100);g__Metschnikowia(100);g__Metschnikowia_unclassified(100); | Metschnikowia | 13 | Saprotroph | \|Undefined Saprotroph\| | Yeast | NULL | Probable | As nectar-tap_saprotroph (Põlme et al. 2020); Decay sugar-rich substrates (Põlme et al. 2020) | Põlme S, et al. 2020. Fungal Diversity 105:1-16 ((https://doi.org/10.1007/s13225-020-00466-2)) |
| Otu18799 | 1 | k__Fungi(100);p__Ascomycota(100);c__Saccharomycetes(100);o__Saccharomycetales(100);f__Metschnikowiaceae(100);g__Metschnikowia(100);g__Metschnikowia_unclassified(100); | Metschnikowia | 13 | Saprotroph | \|Undefined Saprotroph\| | Yeast | NULL | Probable | As nectar-tap_saprotroph (Põlme et al. 2020); Decay sugar-rich substrates (Põlme et al. 2020) | Põlme S, et al. 2020. Fungal Diversity 105:1-16 ((https://doi.org/10.1007/s13225-020-00466-2)) |
| Otu18800 | 1 | k__Fungi(100);p__Ascomycota(100);c__Saccharomycetes(100);o__Saccharomycetales(100);f__Metschnikowiaceae(100);g__Metschnikowia(100);s__Metschnikowia_pulcherrima(100); | Metschnikowia | 13 | Saprotroph | \|Undefined Saprotroph\| | Yeast | NULL | Probable | As nectar-tap_saprotroph (Põlme et al. 2020); Decay sugar-rich substrates (Põlme et al. 2020) | Põlme S, et al. 2020. Fungal Diversity 105:1-16 ((https://doi.org/10.1007/s13225-020-00466-2)) |
| Otu18801 | 1 | k__Fungi(100);p__Ascomycota(100);c__Saccharomycetes(100);o__Saccharomycetales(100);f__Metschnikowiaceae(100);g__Metschnikowia(100);s__Metschnikowia_pulcherrima(100); | Metschnikowia | 13 | Saprotroph | \|Undefined Saprotroph\| | Yeast | NULL | Probable | As nectar-tap_saprotroph (Põlme et al. 2020); Decay sugar-rich substrates (Põlme et al. 2020) | Põlme S, et al. 2020. Fungal Diversity 105:1-16 ((https://doi.org/10.1007/s13225-020-00466-2)) |
| Otu18802 | 1 | k__Fungi(100);p__Ascomycota(100);c__Saccharomycetes(100);o__Saccharomycetales(100);f__Metschnikowiaceae(100);g__Metschnikowia(100);s__Metschnikowia_pulcherrima(100); | Metschnikowia | 13 | Saprotroph | \|Undefined Saprotroph\| | Yeast | NULL | Probable | As nectar-tap_saprotroph (Põlme et al. 2020); Decay sugar-rich substrates (Põlme et al. 2020) | Põlme S, et al. 2020. Fungal Diversity 105:1-16 ((https://doi.org/10.1007/s13225-020-00466-2)) |
| Otu18803 | 1 | k__Fungi(100);p__Ascomycota(100);c__Saccharomycetes(100);o__Saccharomycetales(100);f__Metschnikowiaceae(100);g__Metschnikowia(100);s__Metschnikowia_pulcherrima(100); | Metschnikowia | 13 | Saprotroph | \|Undefined Saprotroph\| | Yeast | NULL | Probable | As nectar-tap_saprotroph (Põlme et al. 2020); Decay sugar-rich substrates (Põlme et al. 2020) | Põlme S, et al. 2020. Fungal Diversity 105:1-16 ((https://doi.org/10.1007/s13225-020-00466-2)) |
| Otu18804 | 1 | k__Fungi(100);p__Ascomycota(100);c__Saccharomycetes(100);o__Saccharomycetales(100);f__Metschnikowiaceae(100);g__Metschnikowia(100);s__Metschnikowia_pulcherrima(100); | Metschnikowia | 13 | Saprotroph | \|Undefined Saprotroph\| | Yeast | NULL | Probable | As nectar-tap_saprotroph (Põlme et al. 2020); Decay sugar-rich substrates (Põlme et al. 2020) | Põlme S, et al. 2020. Fungal Diversity 105:1-16 ((https://doi.org/10.1007/s13225-020-00466-2)) |
| Otu18805 | 1 | k__Fungi(100);p__Ascomycota(100);c__Saccharomycetes(100);o__Saccharomycetales(100);f__Metschnikowiaceae(100);g__Metschnikowia(100);s__Metschnikowia_pulcherrima(100); | Metschnikowia | 13 | Saprotroph | \|Undefined Saprotroph\| | Yeast | NULL | Probable | As nectar-tap_saprotroph (Põlme et al. 2020); Decay sugar-rich substrates (Põlme et al. 2020) | Põlme S, et al. 2020. Fungal Diversity 105:1-16 ((https://doi.org/10.1007/s13225-020-00466-2)) |
| Otu18806 | 1 | k__Fungi(100);p__Ascomycota(100);c__Saccharomycetes(100);o__Saccharomycetales(100);f__Metschnikowiaceae(100);g__Metschnikowia(100);g__Metschnikowia_unclassified(100); | Metschnikowia | 13 | Saprotroph | \|Undefined Saprotroph\| | Yeast | NULL | Probable | As nectar-tap_saprotroph (Põlme et al. 2020); Decay sugar-rich substrates (Põlme et al. 2020) | Põlme S, et al. 2020. Fungal Diversity 105:1-16 ((https://doi.org/10.1007/s13225-020-00466-2)) |
| Otu18807 | 1 | k__Fungi(100);p__Ascomycota(100);c__Saccharomycetes(100);o__Saccharomycetales(100);f__Metschnikowiaceae(100);g__Metschnikowia(100);s__Metschnikowia_pulcherrima(100); | Metschnikowia | 13 | Saprotroph | \|Undefined Saprotroph\| | Yeast | NULL | Probable | As nectar-tap_saprotroph (Põlme et al. 2020); Decay sugar-rich substrates (Põlme et al. 2020) | Põlme S, et al. 2020. Fungal Diversity 105:1-16 ((https://doi.org/10.1007/s13225-020-00466-2)) |
| Otu18808 | 1 | k__Fungi(100);p__Ascomycota(100);c__Saccharomycetes(100);o__Saccharomycetales(100);f__Metschnikowiaceae(100);g__Metschnikowia(100);s__Metschnikowia_pulcherrima(100); | Metschnikowia | 13 | Saprotroph | \|Undefined Saprotroph\| | Yeast | NULL | Probable | As nectar-tap_saprotroph (Põlme et al. 2020); Decay sugar-rich substrates (Põlme et al. 2020) | Põlme S, et al. 2020. Fungal Diversity 105:1-16 ((https://doi.org/10.1007/s13225-020-00466-2)) |
| Otu18809 | 1 | k__Fungi(100);p__Ascomycota(100);c__Saccharomycetes(100);o__Saccharomycetales(100);f__Metschnikowiaceae(100);g__Metschnikowia(100);s__Metschnikowia_pulcherrima(100); | Metschnikowia | 13 | Saprotroph | \|Undefined Saprotroph\| | Yeast | NULL | Probable | As nectar-tap_saprotroph (Põlme et al. 2020); Decay sugar-rich substrates (Põlme et al. 2020) | Põlme S, et al. 2020. Fungal Diversity 105:1-16 ((https://doi.org/10.1007/s13225-020-00466-2)) |
| Otu18811 | 1 | k__Fungi(100);p__Ascomycota(100);c__Saccharomycetes(100);o__Saccharomycetales(100);f__Metschnikowiaceae(100);g__Metschnikowia(100);s__Metschnikowia_pulcherrima(100); | Metschnikowia | 13 | Saprotroph | \|Undefined Saprotroph\| | Yeast | NULL | Probable | As nectar-tap_saprotroph (Põlme et al. 2020); Decay sugar-rich substrates (Põlme et al. 2020) | Põlme S, et al. 2020. Fungal Diversity 105:1-16 ((https://doi.org/10.1007/s13225-020-00466-2)) |
| Otu18812 | 1 | k__Fungi(100);p__Ascomycota(100);c__Saccharomycetes(100);o__Saccharomycetales(100);f__Metschnikowiaceae(100);g__Metschnikowia(100);g__Metschnikowia_unclassified(100); | Metschnikowia | 13 | Saprotroph | \|Undefined Saprotroph\| | Yeast | NULL | Probable | As nectar-tap_saprotroph (Põlme et al. 2020); Decay sugar-rich substrates (Põlme et al. 2020) | Põlme S, et al. 2020. Fungal Diversity 105:1-16 ((https://doi.org/10.1007/s13225-020-00466-2)) |
| Otu18813 | 1 | k__Fungi(100);p__Ascomycota(100);c__Saccharomycetes(100);o__Saccharomycetales(100);f__Metschnikowiaceae(100);g__Metschnikowia(100);s__Metschnikowia_pulcherrima(100); | Metschnikowia | 13 | Saprotroph | \|Undefined Saprotroph\| | Yeast | NULL | Probable | As nectar-tap_saprotroph (Põlme et al. 2020); Decay sugar-rich substrates (Põlme et al. 2020) | Põlme S, et al. 2020. Fungal Diversity 105:1-16 ((https://doi.org/10.1007/s13225-020-00466-2)) |
| Otu18814 | 1 | k__Fungi(100);p__Ascomycota(100);c__Saccharomycetes(100);o__Saccharomycetales(100);f__Metschnikowiaceae(100);g__Metschnikowia(100);s__Metschnikowia_pulcherrima(100); | Metschnikowia | 13 | Saprotroph | \|Undefined Saprotroph\| | Yeast | NULL | Probable | As nectar-tap_saprotroph (Põlme et al. 2020); Decay sugar-rich substrates (Põlme et al. 2020) | Põlme S, et al. 2020. Fungal Diversity 105:1-16 ((https://doi.org/10.1007/s13225-020-00466-2)) |
| Otu18815 | 1 | k__Fungi(100);p__Ascomycota(100);c__Saccharomycetes(100);o__Saccharomycetales(100);f__Metschnikowiaceae(100);g__Metschnikowia(100);s__Metschnikowia_pulcherrima(100); | Metschnikowia | 13 | Saprotroph | \|Undefined Saprotroph\| | Yeast | NULL | Probable | As nectar-tap_saprotroph (Põlme et al. 2020); Decay sugar-rich substrates (Põlme et al. 2020) | Põlme S, et al. 2020. Fungal Diversity 105:1-16 ((https://doi.org/10.1007/s13225-020-00466-2)) |
| Otu18816 | 1 | k__Fungi(100);p__Ascomycota(100);c__Saccharomycetes(100);o__Saccharomycetales(100);f__Metschnikowiaceae(100);g__Metschnikowia(100);s__Metschnikowia_pulcherrima(100); | Metschnikowia | 13 | Saprotroph | \|Undefined Saprotroph\| | Yeast | NULL | Probable | As nectar-tap_saprotroph (Põlme et al. 2020); Decay sugar-rich substrates (Põlme et al. 2020) | Põlme S, et al. 2020. Fungal Diversity 105:1-16 ((https://doi.org/10.1007/s13225-020-00466-2)) |
| Otu18817 | 1 | k__Fungi(100);p__Ascomycota(100);c__Saccharomycetes(100);o__Saccharomycetales(100);f__Metschnikowiaceae(100);g__Metschnikowia(100);s__Metschnikowia_pulcherrima(100); | Metschnikowia | 13 | Saprotroph | \|Undefined Saprotroph\| | Yeast | NULL | Probable | As nectar-tap_saprotroph (Põlme et al. 2020); Decay sugar-rich substrates (Põlme et al. 2020) | Põlme S, et al. 2020. Fungal Diversity 105:1-16 ((https://doi.org/10.1007/s13225-020-00466-2)) |
| Otu18818 | 1 | k__Fungi(100);p__Ascomycota(100);c__Saccharomycetes(100);o__Saccharomycetales(100);f__Metschnikowiaceae(100);g__Metschnikowia(100);s__Metschnikowia_pulcherrima(100); | Metschnikowia | 13 | Saprotroph | \|Undefined Saprotroph\| | Yeast | NULL | Probable | As nectar-tap_saprotroph (Põlme et al. 2020); Decay sugar-rich substrates (Põlme et al. 2020) | Põlme S, et al. 2020. Fungal Diversity 105:1-16 ((https://doi.org/10.1007/s13225-020-00466-2)) |
| Otu18819 | 1 | k__Fungi(100);p__Ascomycota(100);c__Saccharomycetes(100);o__Saccharomycetales(100);f__Metschnikowiaceae(100);g__Metschnikowia(100);s__Metschnikowia_pulcherrima(100); | Metschnikowia | 13 | Saprotroph | \|Undefined Saprotroph\| | Yeast | NULL | Probable | As nectar-tap_saprotroph (Põlme et al. 2020); Decay sugar-rich substrates (Põlme et al. 2020) | Põlme S, et al. 2020. Fungal Diversity 105:1-16 ((https://doi.org/10.1007/s13225-020-00466-2)) |
| Otu18820 | 1 | k__Fungi(100);p__Ascomycota(100);c__Saccharomycetes(100);o__Saccharomycetales(100);f__Metschnikowiaceae(100);g__Metschnikowia(100);s__Metschnikowia_pulcherrima(100); | Metschnikowia | 13 | Saprotroph | \|Undefined Saprotroph\| | Yeast | NULL | Probable | As nectar-tap_saprotroph (Põlme et al. 2020); Decay sugar-rich substrates (Põlme et al. 2020) | Põlme S, et al. 2020. Fungal Diversity 105:1-16 ((https://doi.org/10.1007/s13225-020-00466-2)) |
| Otu18821 | 1 | k__Fungi(100);p__Ascomycota(100);c__Saccharomycetes(100);o__Saccharomycetales(100);f__Metschnikowiaceae(100);g__Metschnikowia(100);g__Metschnikowia_unclassified(100); | Metschnikowia | 13 | Saprotroph | \|Undefined Saprotroph\| | Yeast | NULL | Probable | As nectar-tap_saprotroph (Põlme et al. 2020); Decay sugar-rich substrates (Põlme et al. 2020) | Põlme S, et al. 2020. Fungal Diversity 105:1-16 ((https://doi.org/10.1007/s13225-020-00466-2)) |
| Otu18822 | 1 | k__Fungi(100);p__Ascomycota(100);c__Saccharomycetes(100);o__Saccharomycetales(100);f__Metschnikowiaceae(100);g__Metschnikowia(100);s__Metschnikowia_pulcherrima(100); | Metschnikowia | 13 | Saprotroph | \|Undefined Saprotroph\| | Yeast | NULL | Probable | As nectar-tap_saprotroph (Põlme et al. 2020); Decay sugar-rich substrates (Põlme et al. 2020) | Põlme S, et al. 2020. Fungal Diversity 105:1-16 ((https://doi.org/10.1007/s13225-020-00466-2)) |
| Otu18823 | 1 | k__Fungi(100);p__Ascomycota(100);c__Saccharomycetes(100);o__Saccharomycetales(100);f__Metschnikowiaceae(100);g__Metschnikowia(100);g__Metschnikowia_unclassified(100); | Metschnikowia | 13 | Saprotroph | \|Undefined Saprotroph\| | Yeast | NULL | Probable | As nectar-tap_saprotroph (Põlme et al. 2020); Decay sugar-rich substrates (Põlme et al. 2020) | Põlme S, et al. 2020. Fungal Diversity 105:1-16 ((https://doi.org/10.1007/s13225-020-00466-2)) |
| Otu18824 | 1 | k__Fungi(100);p__Ascomycota(100);c__Saccharomycetes(100);o__Saccharomycetales(100);f__Metschnikowiaceae(100);g__Metschnikowia(100);s__Metschnikowia_pulcherrima(100); | Metschnikowia | 13 | Saprotroph | \|Undefined Saprotroph\| | Yeast | NULL | Probable | As nectar-tap_saprotroph (Põlme et al. 2020); Decay sugar-rich substrates (Põlme et al. 2020) | Põlme S, et al. 2020. Fungal Diversity 105:1-16 ((https://doi.org/10.1007/s13225-020-00466-2)) |
| Otu18825 | 1 | k__Fungi(100);p__Ascomycota(100);c__Saccharomycetes(100);o__Saccharomycetales(100);f__Metschnikowiaceae(100);g__Metschnikowia(100);s__Metschnikowia_pulcherrima(100); | Metschnikowia | 13 | Saprotroph | \|Undefined Saprotroph\| | Yeast | NULL | Probable | As nectar-tap_saprotroph (Põlme et al. 2020); Decay sugar-rich substrates (Põlme et al. 2020) | Põlme S, et al. 2020. Fungal Diversity 105:1-16 ((https://doi.org/10.1007/s13225-020-00466-2)) |
| Otu18826 | 1 | k__Fungi(100);p__Ascomycota(100);c__Saccharomycetes(100);o__Saccharomycetales(100);f__Metschnikowiaceae(100);g__Metschnikowia(100);s__Metschnikowia_pulcherrima(100); | Metschnikowia | 13 | Saprotroph | \|Undefined Saprotroph\| | Yeast | NULL | Probable | As nectar-tap_saprotroph (Põlme et al. 2020); Decay sugar-rich substrates (Põlme et al. 2020) | Põlme S, et al. 2020. Fungal Diversity 105:1-16 ((https://doi.org/10.1007/s13225-020-00466-2)) |
| Otu18827 | 1 | k__Fungi(100);p__Ascomycota(100);c__Saccharomycetes(100);o__Saccharomycetales(100);f__Metschnikowiaceae(100);g__Metschnikowia(100);g__Metschnikowia_unclassified(100); | Metschnikowia | 13 | Saprotroph | \|Undefined Saprotroph\| | Yeast | NULL | Probable | As nectar-tap_saprotroph (Põlme et al. 2020); Decay sugar-rich substrates (Põlme et al. 2020) | Põlme S, et al. 2020. Fungal Diversity 105:1-16 ((https://doi.org/10.1007/s13225-020-00466-2)) |
| Otu18828 | 1 | k__Fungi(100);p__Ascomycota(100);c__Saccharomycetes(100);o__Saccharomycetales(100);f__Metschnikowiaceae(100);g__Metschnikowia(100);g__Metschnikowia_unclassified(100); | Metschnikowia | 13 | Saprotroph | \|Undefined Saprotroph\| | Yeast | NULL | Probable | As nectar-tap_saprotroph (Põlme et al. 2020); Decay sugar-rich substrates (Põlme et al. 2020) | Põlme S, et al. 2020. Fungal Diversity 105:1-16 ((https://doi.org/10.1007/s13225-020-00466-2)) |
| Otu18829 | 1 | k__Fungi(100);p__Ascomycota(100);c__Saccharomycetes(100);o__Saccharomycetales(100);f__Metschnikowiaceae(100);g__Metschnikowia(100);s__Metschnikowia_pulcherrima(100); | Metschnikowia | 13 | Saprotroph | \|Undefined Saprotroph\| | Yeast | NULL | Probable | As nectar-tap_saprotroph (Põlme et al. 2020); Decay sugar-rich substrates (Põlme et al. 2020) | Põlme S, et al. 2020. Fungal Diversity 105:1-16 ((https://doi.org/10.1007/s13225-020-00466-2)) |
| Otu18830 | 1 | k__Fungi(100);p__Ascomycota(100);c__Saccharomycetes(100);o__Saccharomycetales(100);f__Metschnikowiaceae(100);g__Metschnikowia(100);g__Metschnikowia_unclassified(100); | Metschnikowia | 13 | Saprotroph | \|Undefined Saprotroph\| | Yeast | NULL | Probable | As nectar-tap_saprotroph (Põlme et al. 2020); Decay sugar-rich substrates (Põlme et al. 2020) | Põlme S, et al. 2020. Fungal Diversity 105:1-16 ((https://doi.org/10.1007/s13225-020-00466-2)) |
| Otu18831 | 1 | k__Fungi(100);p__Ascomycota(100);c__Saccharomycetes(100);o__Saccharomycetales(100);f__Metschnikowiaceae(100);g__Metschnikowia(100);s__Metschnikowia_pulcherrima(100); | Metschnikowia | 13 | Saprotroph | \|Undefined Saprotroph\| | Yeast | NULL | Probable | As nectar-tap_saprotroph (Põlme et al. 2020); Decay sugar-rich substrates (Põlme et al. 2020) | Põlme S, et al. 2020. Fungal Diversity 105:1-16 ((https://doi.org/10.1007/s13225-020-00466-2)) |
| Otu18832 | 1 | k__Fungi(100);p__Ascomycota(100);c__Saccharomycetes(100);o__Saccharomycetales(100);f__Metschnikowiaceae(100);g__Metschnikowia(100);s__Metschnikowia_pulcherrima(100); | Metschnikowia | 13 | Saprotroph | \|Undefined Saprotroph\| | Yeast | NULL | Probable | As nectar-tap_saprotroph (Põlme et al. 2020); Decay sugar-rich substrates (Põlme et al. 2020) | Põlme S, et al. 2020. Fungal Diversity 105:1-16 ((https://doi.org/10.1007/s13225-020-00466-2)) |
| Otu18833 | 1 | k__Fungi(100);p__Ascomycota(100);c__Saccharomycetes(100);o__Saccharomycetales(100);f__Metschnikowiaceae(100);g__Metschnikowia(100);s__Metschnikowia_pulcherrima(100); | Metschnikowia | 13 | Saprotroph | \|Undefined Saprotroph\| | Yeast | NULL | Probable | As nectar-tap_saprotroph (Põlme et al. 2020); Decay sugar-rich substrates (Põlme et al. 2020) | Põlme S, et al. 2020. Fungal Diversity 105:1-16 ((https://doi.org/10.1007/s13225-020-00466-2)) |
| Otu18834 | 1 | k__Fungi(100);p__Ascomycota(100);c__Saccharomycetes(100);o__Saccharomycetales(100);f__Metschnikowiaceae(100);g__Metschnikowia(100);s__Metschnikowia_pulcherrima(100); | Metschnikowia | 13 | Saprotroph | \|Undefined Saprotroph\| | Yeast | NULL | Probable | As nectar-tap_saprotroph (Põlme et al. 2020); Decay sugar-rich substrates (Põlme et al. 2020) | Põlme S, et al. 2020. Fungal Diversity 105:1-16 ((https://doi.org/10.1007/s13225-020-00466-2)) |
| Otu18835 | 1 | k__Fungi(100);p__Ascomycota(100);c__Saccharomycetes(100);o__Saccharomycetales(100);f__Metschnikowiaceae(100);g__Metschnikowia(100);g__Metschnikowia_unclassified(100); | Metschnikowia | 13 | Saprotroph | \|Undefined Saprotroph\| | Yeast | NULL | Probable | As nectar-tap_saprotroph (Põlme et al. 2020); Decay sugar-rich substrates (Põlme et al. 2020) | Põlme S, et al. 2020. Fungal Diversity 105:1-16 ((https://doi.org/10.1007/s13225-020-00466-2)) |
| Otu18838 | 1 | k__Fungi(100);p__Ascomycota(100);c__Saccharomycetes(100);o__Saccharomycetales(100);f__Metschnikowiaceae(100);g__Metschnikowia(100);s__Metschnikowia_pulcherrima(100); | Metschnikowia | 13 | Saprotroph | \|Undefined Saprotroph\| | Yeast | NULL | Probable | As nectar-tap_saprotroph (Põlme et al. 2020); Decay sugar-rich substrates (Põlme et al. 2020) | Põlme S, et al. 2020. Fungal Diversity 105:1-16 ((https://doi.org/10.1007/s13225-020-00466-2)) |
| Otu18839 | 1 | k__Fungi(100);p__Ascomycota(100);c__Saccharomycetes(100);o__Saccharomycetales(100);f__Metschnikowiaceae(100);g__Metschnikowia(100);s__Metschnikowia_pulcherrima(100); | Metschnikowia | 13 | Saprotroph | \|Undefined Saprotroph\| | Yeast | NULL | Probable | As nectar-tap_saprotroph (Põlme et al. 2020); Decay sugar-rich substrates (Põlme et al. 2020) | Põlme S, et al. 2020. Fungal Diversity 105:1-16 ((https://doi.org/10.1007/s13225-020-00466-2)) |
| Otu18840 | 1 | k__Fungi(100);p__Ascomycota(100);c__Saccharomycetes(100);o__Saccharomycetales(100);f__Metschnikowiaceae(100);g__Metschnikowia(100);s__Metschnikowia_pulcherrima(100); | Metschnikowia | 13 | Saprotroph | \|Undefined Saprotroph\| | Yeast | NULL | Probable | As nectar-tap_saprotroph (Põlme et al. 2020); Decay sugar-rich substrates (Põlme et al. 2020) | Põlme S, et al. 2020. Fungal Diversity 105:1-16 ((https://doi.org/10.1007/s13225-020-00466-2)) |
| Otu18841 | 1 | k__Fungi(100);p__Ascomycota(100);c__Saccharomycetes(100);o__Saccharomycetales(100);f__Metschnikowiaceae(100);g__Metschnikowia(100);s__Metschnikowia_pulcherrima(100); | Metschnikowia | 13 | Saprotroph | \|Undefined Saprotroph\| | Yeast | NULL | Probable | As nectar-tap_saprotroph (Põlme et al. 2020); Decay sugar-rich substrates (Põlme et al. 2020) | Põlme S, et al. 2020. Fungal Diversity 105:1-16 ((https://doi.org/10.1007/s13225-020-00466-2)) |
| Otu18842 | 1 | k__Fungi(100);p__Ascomycota(100);c__Saccharomycetes(100);o__Saccharomycetales(100);f__Metschnikowiaceae(100);g__Metschnikowia(100);s__Metschnikowia_pulcherrima(100); | Metschnikowia | 13 | Saprotroph | \|Undefined Saprotroph\| | Yeast | NULL | Probable | As nectar-tap_saprotroph (Põlme et al. 2020); Decay sugar-rich substrates (Põlme et al. 2020) | Põlme S, et al. 2020. Fungal Diversity 105:1-16 ((https://doi.org/10.1007/s13225-020-00466-2)) |
| Otu18843 | 1 | k__Fungi(100);p__Ascomycota(100);c__Saccharomycetes(100);o__Saccharomycetales(100);f__Metschnikowiaceae(100);g__Metschnikowia(100);g__Metschnikowia_unclassified(100); | Metschnikowia | 13 | Saprotroph | \|Undefined Saprotroph\| | Yeast | NULL | Probable | As nectar-tap_saprotroph (Põlme et al. 2020); Decay sugar-rich substrates (Põlme et al. 2020) | Põlme S, et al. 2020. Fungal Diversity 105:1-16 ((https://doi.org/10.1007/s13225-020-00466-2)) |
| Otu18844 | 1 | k__Fungi(100);p__Ascomycota(100);c__Saccharomycetes(100);o__Saccharomycetales(100);f__Metschnikowiaceae(100);g__Metschnikowia(100);s__Metschnikowia_pulcherrima(100); | Metschnikowia | 13 | Saprotroph | \|Undefined Saprotroph\| | Yeast | NULL | Probable | As nectar-tap_saprotroph (Põlme et al. 2020); Decay sugar-rich substrates (Põlme et al. 2020) | Põlme S, et al. 2020. Fungal Diversity 105:1-16 ((https://doi.org/10.1007/s13225-020-00466-2)) |
| Otu18845 | 1 | k__Fungi(100);p__Ascomycota(100);c__Saccharomycetes(100);o__Saccharomycetales(100);f__Metschnikowiaceae(100);g__Metschnikowia(100);s__Metschnikowia_pulcherrima(100); | Metschnikowia | 13 | Saprotroph | \|Undefined Saprotroph\| | Yeast | NULL | Probable | As nectar-tap_saprotroph (Põlme et al. 2020); Decay sugar-rich substrates (Põlme et al. 2020) | Põlme S, et al. 2020. Fungal Diversity 105:1-16 ((https://doi.org/10.1007/s13225-020-00466-2)) |
| Otu18846 | 1 | k__Fungi(100);p__Ascomycota(100);c__Saccharomycetes(100);o__Saccharomycetales(100);f__Metschnikowiaceae(100);g__Metschnikowia(100);s__Metschnikowia_pulcherrima(100); | Metschnikowia | 13 | Saprotroph | \|Undefined Saprotroph\| | Yeast | NULL | Probable | As nectar-tap_saprotroph (Põlme et al. 2020); Decay sugar-rich substrates (Põlme et al. 2020) | Põlme S, et al. 2020. Fungal Diversity 105:1-16 ((https://doi.org/10.1007/s13225-020-00466-2)) |
| Otu18847 | 1 | k__Fungi(100);p__Ascomycota(100);c__Saccharomycetes(100);o__Saccharomycetales(100);f__Metschnikowiaceae(100);g__Metschnikowia(100);s__Metschnikowia_pulcherrima(100); | Metschnikowia | 13 | Saprotroph | \|Undefined Saprotroph\| | Yeast | NULL | Probable | As nectar-tap_saprotroph (Põlme et al. 2020); Decay sugar-rich substrates (Põlme et al. 2020) | Põlme S, et al. 2020. Fungal Diversity 105:1-16 ((https://doi.org/10.1007/s13225-020-00466-2)) |
| Otu18848 | 1 | k__Fungi(100);p__Ascomycota(100);c__Saccharomycetes(100);o__Saccharomycetales(100);f__Metschnikowiaceae(100);g__Metschnikowia(100);s__Metschnikowia_pulcherrima(100); | Metschnikowia | 13 | Saprotroph | \|Undefined Saprotroph\| | Yeast | NULL | Probable | As nectar-tap_saprotroph (Põlme et al. 2020); Decay sugar-rich substrates (Põlme et al. 2020) | Põlme S, et al. 2020. Fungal Diversity 105:1-16 ((https://doi.org/10.1007/s13225-020-00466-2)) |
| Otu18849 | 1 | k__Fungi(100);p__Ascomycota(100);c__Saccharomycetes(100);o__Saccharomycetales(100);f__Metschnikowiaceae(100);g__Metschnikowia(100);g__Metschnikowia_unclassified(100); | Metschnikowia | 13 | Saprotroph | \|Undefined Saprotroph\| | Yeast | NULL | Probable | As nectar-tap_saprotroph (Põlme et al. 2020); Decay sugar-rich substrates (Põlme et al. 2020) | Põlme S, et al. 2020. Fungal Diversity 105:1-16 ((https://doi.org/10.1007/s13225-020-00466-2)) |
| Otu18850 | 1 | k__Fungi(100);p__Ascomycota(100);c__Saccharomycetes(100);o__Saccharomycetales(100);f__Metschnikowiaceae(100);g__Metschnikowia(100);s__Metschnikowia_pulcherrima(100); | Metschnikowia | 13 | Saprotroph | \|Undefined Saprotroph\| | Yeast | NULL | Probable | As nectar-tap_saprotroph (Põlme et al. 2020); Decay sugar-rich substrates (Põlme et al. 2020) | Põlme S, et al. 2020. Fungal Diversity 105:1-16 ((https://doi.org/10.1007/s13225-020-00466-2)) |
| Otu18851 | 1 | k__Fungi(100);p__Ascomycota(100);c__Saccharomycetes(100);o__Saccharomycetales(100);f__Metschnikowiaceae(100);g__Metschnikowia(100);g__Metschnikowia_unclassified(100); | Metschnikowia | 13 | Saprotroph | \|Undefined Saprotroph\| | Yeast | NULL | Probable | As nectar-tap_saprotroph (Põlme et al. 2020); Decay sugar-rich substrates (Põlme et al. 2020) | Põlme S, et al. 2020. Fungal Diversity 105:1-16 ((https://doi.org/10.1007/s13225-020-00466-2)) |
| Otu18852 | 1 | k__Fungi(100);p__Ascomycota(100);c__Saccharomycetes(100);o__Saccharomycetales(100);f__Metschnikowiaceae(100);g__Metschnikowia(100);s__Metschnikowia_pulcherrima(100); | Metschnikowia | 13 | Saprotroph | \|Undefined Saprotroph\| | Yeast | NULL | Probable | As nectar-tap_saprotroph (Põlme et al. 2020); Decay sugar-rich substrates (Põlme et al. 2020) | Põlme S, et al. 2020. Fungal Diversity 105:1-16 ((https://doi.org/10.1007/s13225-020-00466-2)) |
| Otu18853 | 1 | k__Fungi(100);p__Ascomycota(100);c__Saccharomycetes(100);o__Saccharomycetales(100);f__Metschnikowiaceae(100);g__Metschnikowia(100);s__Metschnikowia_pulcherrima(100); | Metschnikowia | 13 | Saprotroph | \|Undefined Saprotroph\| | Yeast | NULL | Probable | As nectar-tap_saprotroph (Põlme et al. 2020); Decay sugar-rich substrates (Põlme et al. 2020) | Põlme S, et al. 2020. Fungal Diversity 105:1-16 ((https://doi.org/10.1007/s13225-020-00466-2)) |
| Otu18854 | 1 | k__Fungi(100);p__Ascomycota(100);c__Saccharomycetes(100);o__Saccharomycetales(100);f__Metschnikowiaceae(100);g__Metschnikowia(100);s__Metschnikowia_pulcherrima(100); | Metschnikowia | 13 | Saprotroph | \|Undefined Saprotroph\| | Yeast | NULL | Probable | As nectar-tap_saprotroph (Põlme et al. 2020); Decay sugar-rich substrates (Põlme et al. 2020) | Põlme S, et al. 2020. Fungal Diversity 105:1-16 ((https://doi.org/10.1007/s13225-020-00466-2)) |
| Otu18855 | 1 | k__Fungi(100);p__Ascomycota(100);c__Saccharomycetes(100);o__Saccharomycetales(100);f__Metschnikowiaceae(100);g__Metschnikowia(100);g__Metschnikowia_unclassified(100); | Metschnikowia | 13 | Saprotroph | \|Undefined Saprotroph\| | Yeast | NULL | Probable | As nectar-tap_saprotroph (Põlme et al. 2020); Decay sugar-rich substrates (Põlme et al. 2020) | Põlme S, et al. 2020. Fungal Diversity 105:1-16 ((https://doi.org/10.1007/s13225-020-00466-2)) |
| Otu18856 | 1 | k__Fungi(100);p__Ascomycota(100);c__Saccharomycetes(100);o__Saccharomycetales(100);f__Metschnikowiaceae(100);g__Metschnikowia(100);s__Metschnikowia_pulcherrima(100); | Metschnikowia | 13 | Saprotroph | \|Undefined Saprotroph\| | Yeast | NULL | Probable | As nectar-tap_saprotroph (Põlme et al. 2020); Decay sugar-rich substrates (Põlme et al. 2020) | Põlme S, et al. 2020. Fungal Diversity 105:1-16 ((https://doi.org/10.1007/s13225-020-00466-2)) |
| Otu18857 | 1 | k__Fungi(100);p__Ascomycota(100);c__Saccharomycetes(100);o__Saccharomycetales(100);f__Metschnikowiaceae(100);g__Metschnikowia(100);s__Metschnikowia_pulcherrima(100); | Metschnikowia | 13 | Saprotroph | \|Undefined Saprotroph\| | Yeast | NULL | Probable | As nectar-tap_saprotroph (Põlme et al. 2020); Decay sugar-rich substrates (Põlme et al. 2020) | Põlme S, et al. 2020. Fungal Diversity 105:1-16 ((https://doi.org/10.1007/s13225-020-00466-2)) |
| Otu18858 | 1 | k__Fungi(100);p__Ascomycota(100);c__Saccharomycetes(100);o__Saccharomycetales(100);f__Metschnikowiaceae(100);g__Metschnikowia(100);s__Metschnikowia_pulcherrima(100); | Metschnikowia | 13 | Saprotroph | \|Undefined Saprotroph\| | Yeast | NULL | Probable | As nectar-tap_saprotroph (Põlme et al. 2020); Decay sugar-rich substrates (Põlme et al. 2020) | Põlme S, et al. 2020. Fungal Diversity 105:1-16 ((https://doi.org/10.1007/s13225-020-00466-2)) |
| Otu18859 | 1 | k__Fungi(100);p__Ascomycota(100);c__Saccharomycetes(100);o__Saccharomycetales(100);f__Metschnikowiaceae(100);g__Metschnikowia(100);s__Metschnikowia_pulcherrima(100); | Metschnikowia | 13 | Saprotroph | \|Undefined Saprotroph\| | Yeast | NULL | Probable | As nectar-tap_saprotroph (Põlme et al. 2020); Decay sugar-rich substrates (Põlme et al. 2020) | Põlme S, et al. 2020. Fungal Diversity 105:1-16 ((https://doi.org/10.1007/s13225-020-00466-2)) |
| Otu18860 | 1 | k__Fungi(100);p__Ascomycota(100);c__Saccharomycetes(100);o__Saccharomycetales(100);f__Metschnikowiaceae(100);g__Metschnikowia(100);s__Metschnikowia_pulcherrima(100); | Metschnikowia | 13 | Saprotroph | \|Undefined Saprotroph\| | Yeast | NULL | Probable | As nectar-tap_saprotroph (Põlme et al. 2020); Decay sugar-rich substrates (Põlme et al. 2020) | Põlme S, et al. 2020. Fungal Diversity 105:1-16 ((https://doi.org/10.1007/s13225-020-00466-2)) |
| Otu18861 | 1 | k__Fungi(100);p__Ascomycota(100);c__Saccharomycetes(100);o__Saccharomycetales(100);f__Metschnikowiaceae(100);g__Metschnikowia(100);s__Metschnikowia_pulcherrima(100); | Metschnikowia | 13 | Saprotroph | \|Undefined Saprotroph\| | Yeast | NULL | Probable | As nectar-tap_saprotroph (Põlme et al. 2020); Decay sugar-rich substrates (Põlme et al. 2020) | Põlme S, et al. 2020. Fungal Diversity 105:1-16 ((https://doi.org/10.1007/s13225-020-00466-2)) |
| Otu18862 | 1 | k__Fungi(100);p__Ascomycota(100);c__Saccharomycetes(100);o__Saccharomycetales(100);f__Metschnikowiaceae(100);g__Metschnikowia(100);g__Metschnikowia_unclassified(100); | Metschnikowia | 13 | Saprotroph | \|Undefined Saprotroph\| | Yeast | NULL | Probable | As nectar-tap_saprotroph (Põlme et al. 2020); Decay sugar-rich substrates (Põlme et al. 2020) | Põlme S, et al. 2020. Fungal Diversity 105:1-16 ((https://doi.org/10.1007/s13225-020-00466-2)) |
| Otu18863 | 1 | k__Fungi(100);p__Ascomycota(100);c__Saccharomycetes(100);o__Saccharomycetales(100);f__Metschnikowiaceae(100);g__Metschnikowia(100);s__Metschnikowia_pulcherrima(100); | Metschnikowia | 13 | Saprotroph | \|Undefined Saprotroph\| | Yeast | NULL | Probable | As nectar-tap_saprotroph (Põlme et al. 2020); Decay sugar-rich substrates (Põlme et al. 2020) | Põlme S, et al. 2020. Fungal Diversity 105:1-16 ((https://doi.org/10.1007/s13225-020-00466-2)) |
| Otu18864 | 1 | k__Fungi(100);p__Ascomycota(100);c__Saccharomycetes(100);o__Saccharomycetales(100);f__Metschnikowiaceae(100);g__Metschnikowia(100);g__Metschnikowia_unclassified(100); | Metschnikowia | 13 | Saprotroph | \|Undefined Saprotroph\| | Yeast | NULL | Probable | As nectar-tap_saprotroph (Põlme et al. 2020); Decay sugar-rich substrates (Põlme et al. 2020) | Põlme S, et al. 2020. Fungal Diversity 105:1-16 ((https://doi.org/10.1007/s13225-020-00466-2)) |
| Otu18865 | 1 | k__Fungi(100);p__Ascomycota(100);c__Saccharomycetes(100);o__Saccharomycetales(100);f__Metschnikowiaceae(100);g__Metschnikowia(100);s__Metschnikowia_pulcherrima(100); | Metschnikowia | 13 | Saprotroph | \|Undefined Saprotroph\| | Yeast | NULL | Probable | As nectar-tap_saprotroph (Põlme et al. 2020); Decay sugar-rich substrates (Põlme et al. 2020) | Põlme S, et al. 2020. Fungal Diversity 105:1-16 ((https://doi.org/10.1007/s13225-020-00466-2)) |
| Otu18866 | 1 | k__Fungi(100);p__Ascomycota(100);c__Saccharomycetes(100);o__Saccharomycetales(100);f__Metschnikowiaceae(100);g__Metschnikowia(100);s__Metschnikowia_pulcherrima(100); | Metschnikowia | 13 | Saprotroph | \|Undefined Saprotroph\| | Yeast | NULL | Probable | As nectar-tap_saprotroph (Põlme et al. 2020); Decay sugar-rich substrates (Põlme et al. 2020) | Põlme S, et al. 2020. Fungal Diversity 105:1-16 ((https://doi.org/10.1007/s13225-020-00466-2)) |
| Otu18867 | 1 | k__Fungi(100);p__Ascomycota(100);c__Saccharomycetes(100);o__Saccharomycetales(100);f__Metschnikowiaceae(100);g__Metschnikowia(100);s__Metschnikowia_pulcherrima(100); | Metschnikowia | 13 | Saprotroph | \|Undefined Saprotroph\| | Yeast | NULL | Probable | As nectar-tap_saprotroph (Põlme et al. 2020); Decay sugar-rich substrates (Põlme et al. 2020) | Põlme S, et al. 2020. Fungal Diversity 105:1-16 ((https://doi.org/10.1007/s13225-020-00466-2)) |
| Otu18868 | 1 | k__Fungi(100);p__Ascomycota(100);c__Saccharomycetes(100);o__Saccharomycetales(100);f__Metschnikowiaceae(100);g__Metschnikowia(100);g__Metschnikowia_unclassified(100); | Metschnikowia | 13 | Saprotroph | \|Undefined Saprotroph\| | Yeast | NULL | Probable | As nectar-tap_saprotroph (Põlme et al. 2020); Decay sugar-rich substrates (Põlme et al. 2020) | Põlme S, et al. 2020. Fungal Diversity 105:1-16 ((https://doi.org/10.1007/s13225-020-00466-2)) |
| Otu18869 | 1 | k__Fungi(100);p__Ascomycota(100);c__Saccharomycetes(100);o__Saccharomycetales(100);f__Metschnikowiaceae(100);g__Metschnikowia(100);g__Metschnikowia_unclassified(100); | Metschnikowia | 13 | Saprotroph | \|Undefined Saprotroph\| | Yeast | NULL | Probable | As nectar-tap_saprotroph (Põlme et al. 2020); Decay sugar-rich substrates (Põlme et al. 2020) | Põlme S, et al. 2020. Fungal Diversity 105:1-16 ((https://doi.org/10.1007/s13225-020-00466-2)) |
| Otu18870 | 1 | k__Fungi(100);p__Ascomycota(100);c__Saccharomycetes(100);o__Saccharomycetales(100);f__Metschnikowiaceae(100);g__Metschnikowia(100);s__Metschnikowia_pulcherrima(100); | Metschnikowia | 13 | Saprotroph | \|Undefined Saprotroph\| | Yeast | NULL | Probable | As nectar-tap_saprotroph (Põlme et al. 2020); Decay sugar-rich substrates (Põlme et al. 2020) | Põlme S, et al. 2020. Fungal Diversity 105:1-16 ((https://doi.org/10.1007/s13225-020-00466-2)) |
| Otu18871 | 1 | k__Fungi(100);p__Ascomycota(100);c__Saccharomycetes(100);o__Saccharomycetales(100);f__Metschnikowiaceae(100);g__Metschnikowia(100);s__Metschnikowia_pulcherrima(100); | Metschnikowia | 13 | Saprotroph | \|Undefined Saprotroph\| | Yeast | NULL | Probable | As nectar-tap_saprotroph (Põlme et al. 2020); Decay sugar-rich substrates (Põlme et al. 2020) | Põlme S, et al. 2020. Fungal Diversity 105:1-16 ((https://doi.org/10.1007/s13225-020-00466-2)) |
| Otu18872 | 1 | k__Fungi(100);p__Ascomycota(100);c__Saccharomycetes(100);o__Saccharomycetales(100);f__Metschnikowiaceae(100);g__Metschnikowia(100);s__Metschnikowia_pulcherrima(100); | Metschnikowia | 13 | Saprotroph | \|Undefined Saprotroph\| | Yeast | NULL | Probable | As nectar-tap_saprotroph (Põlme et al. 2020); Decay sugar-rich substrates (Põlme et al. 2020) | Põlme S, et al. 2020. Fungal Diversity 105:1-16 ((https://doi.org/10.1007/s13225-020-00466-2)) |
| Otu18873 | 1 | k__Fungi(100);p__Ascomycota(100);c__Saccharomycetes(100);o__Saccharomycetales(100);f__Metschnikowiaceae(100);g__Metschnikowia(100);s__Metschnikowia_pulcherrima(100); | Metschnikowia | 13 | Saprotroph | \|Undefined Saprotroph\| | Yeast | NULL | Probable | As nectar-tap_saprotroph (Põlme et al. 2020); Decay sugar-rich substrates (Põlme et al. 2020) | Põlme S, et al. 2020. Fungal Diversity 105:1-16 ((https://doi.org/10.1007/s13225-020-00466-2)) |
| Otu18874 | 1 | k__Fungi(100);p__Ascomycota(100);c__Saccharomycetes(100);o__Saccharomycetales(100);f__Metschnikowiaceae(100);g__Metschnikowia(100);s__Metschnikowia_pulcherrima(100); | Metschnikowia | 13 | Saprotroph | \|Undefined Saprotroph\| | Yeast | NULL | Probable | As nectar-tap_saprotroph (Põlme et al. 2020); Decay sugar-rich substrates (Põlme et al. 2020) | Põlme S, et al. 2020. Fungal Diversity 105:1-16 ((https://doi.org/10.1007/s13225-020-00466-2)) |
| Otu18875 | 1 | k__Fungi(100);p__Ascomycota(100);c__Saccharomycetes(100);o__Saccharomycetales(100);f__Metschnikowiaceae(100);g__Metschnikowia(100);s__Metschnikowia_pulcherrima(100); | Metschnikowia | 13 | Saprotroph | \|Undefined Saprotroph\| | Yeast | NULL | Probable | As nectar-tap_saprotroph (Põlme et al. 2020); Decay sugar-rich substrates (Põlme et al. 2020) | Põlme S, et al. 2020. Fungal Diversity 105:1-16 ((https://doi.org/10.1007/s13225-020-00466-2)) |
| Otu18876 | 1 | k__Fungi(100);p__Ascomycota(100);c__Saccharomycetes(100);o__Saccharomycetales(100);f__Metschnikowiaceae(100);g__Metschnikowia(100);s__Metschnikowia_pulcherrima(100); | Metschnikowia | 13 | Saprotroph | \|Undefined Saprotroph\| | Yeast | NULL | Probable | As nectar-tap_saprotroph (Põlme et al. 2020); Decay sugar-rich substrates (Põlme et al. 2020) | Põlme S, et al. 2020. Fungal Diversity 105:1-16 ((https://doi.org/10.1007/s13225-020-00466-2)) |
| Otu18877 | 1 | k__Fungi(100);p__Ascomycota(100);c__Saccharomycetes(100);o__Saccharomycetales(100);f__Metschnikowiaceae(100);g__Metschnikowia(100);s__Metschnikowia_pulcherrima(100); | Metschnikowia | 13 | Saprotroph | \|Undefined Saprotroph\| | Yeast | NULL | Probable | As nectar-tap_saprotroph (Põlme et al. 2020); Decay sugar-rich substrates (Põlme et al. 2020) | Põlme S, et al. 2020. Fungal Diversity 105:1-16 ((https://doi.org/10.1007/s13225-020-00466-2)) |
| Otu18878 | 1 | k__Fungi(100);p__Ascomycota(100);c__Saccharomycetes(100);o__Saccharomycetales(100);f__Metschnikowiaceae(100);g__Metschnikowia(100);s__Metschnikowia_pulcherrima(100); | Metschnikowia | 13 | Saprotroph | \|Undefined Saprotroph\| | Yeast | NULL | Probable | As nectar-tap_saprotroph (Põlme et al. 2020); Decay sugar-rich substrates (Põlme et al. 2020) | Põlme S, et al. 2020. Fungal Diversity 105:1-16 ((https://doi.org/10.1007/s13225-020-00466-2)) |
| Otu18879 | 1 | k__Fungi(100);p__Ascomycota(100);c__Saccharomycetes(100);o__Saccharomycetales(100);f__Metschnikowiaceae(100);g__Metschnikowia(100);s__Metschnikowia_pulcherrima(100); | Metschnikowia | 13 | Saprotroph | \|Undefined Saprotroph\| | Yeast | NULL | Probable | As nectar-tap_saprotroph (Põlme et al. 2020); Decay sugar-rich substrates (Põlme et al. 2020) | Põlme S, et al. 2020. Fungal Diversity 105:1-16 ((https://doi.org/10.1007/s13225-020-00466-2)) |
| Otu18880 | 1 | k__Fungi(100);p__Ascomycota(100);c__Saccharomycetes(100);o__Saccharomycetales(100);f__Metschnikowiaceae(100);g__Metschnikowia(100);g__Metschnikowia_unclassified(100); | Metschnikowia | 13 | Saprotroph | \|Undefined Saprotroph\| | Yeast | NULL | Probable | As nectar-tap_saprotroph (Põlme et al. 2020); Decay sugar-rich substrates (Põlme et al. 2020) | Põlme S, et al. 2020. Fungal Diversity 105:1-16 ((https://doi.org/10.1007/s13225-020-00466-2)) |
| Otu18881 | 1 | k__Fungi(100);p__Ascomycota(100);c__Saccharomycetes(100);o__Saccharomycetales(100);f__Metschnikowiaceae(100);g__Metschnikowia(100);s__Metschnikowia_pulcherrima(100); | Metschnikowia | 13 | Saprotroph | \|Undefined Saprotroph\| | Yeast | NULL | Probable | As nectar-tap_saprotroph (Põlme et al. 2020); Decay sugar-rich substrates (Põlme et al. 2020) | Põlme S, et al. 2020. Fungal Diversity 105:1-16 ((https://doi.org/10.1007/s13225-020-00466-2)) |
| Otu18882 | 1 | k__Fungi(100);p__Ascomycota(100);c__Saccharomycetes(100);o__Saccharomycetales(100);f__Metschnikowiaceae(100);g__Metschnikowia(100);g__Metschnikowia_unclassified(100); | Metschnikowia | 13 | Saprotroph | \|Undefined Saprotroph\| | Yeast | NULL | Probable | As nectar-tap_saprotroph (Põlme et al. 2020); Decay sugar-rich substrates (Põlme et al. 2020) | Põlme S, et al. 2020. Fungal Diversity 105:1-16 ((https://doi.org/10.1007/s13225-020-00466-2)) |
| Otu18883 | 1 | k__Fungi(100);p__Ascomycota(100);c__Saccharomycetes(100);o__Saccharomycetales(100);f__Metschnikowiaceae(100);g__Metschnikowia(100);s__Metschnikowia_pulcherrima(100); | Metschnikowia | 13 | Saprotroph | \|Undefined Saprotroph\| | Yeast | NULL | Probable | As nectar-tap_saprotroph (Põlme et al. 2020); Decay sugar-rich substrates (Põlme et al. 2020) | Põlme S, et al. 2020. Fungal Diversity 105:1-16 ((https://doi.org/10.1007/s13225-020-00466-2)) |
| Otu18884 | 1 | k__Fungi(100);p__Ascomycota(100);c__Saccharomycetes(100);o__Saccharomycetales(100);f__Metschnikowiaceae(100);g__Metschnikowia(100);g__Metschnikowia_unclassified(100); | Metschnikowia | 13 | Saprotroph | \|Undefined Saprotroph\| | Yeast | NULL | Probable | As nectar-tap_saprotroph (Põlme et al. 2020); Decay sugar-rich substrates (Põlme et al. 2020) | Põlme S, et al. 2020. Fungal Diversity 105:1-16 ((https://doi.org/10.1007/s13225-020-00466-2)) |
| Otu18885 | 1 | k__Fungi(100);p__Ascomycota(100);c__Saccharomycetes(100);o__Saccharomycetales(100);f__Metschnikowiaceae(100);g__Metschnikowia(100);s__Metschnikowia_pulcherrima(100); | Metschnikowia | 13 | Saprotroph | \|Undefined Saprotroph\| | Yeast | NULL | Probable | As nectar-tap_saprotroph (Põlme et al. 2020); Decay sugar-rich substrates (Põlme et al. 2020) | Põlme S, et al. 2020. Fungal Diversity 105:1-16 ((https://doi.org/10.1007/s13225-020-00466-2)) |
| Otu18886 | 1 | k__Fungi(100);p__Ascomycota(100);c__Saccharomycetes(100);o__Saccharomycetales(100);f__Metschnikowiaceae(100);g__Metschnikowia(100);s__Metschnikowia_pulcherrima(100); | Metschnikowia | 13 | Saprotroph | \|Undefined Saprotroph\| | Yeast | NULL | Probable | As nectar-tap_saprotroph (Põlme et al. 2020); Decay sugar-rich substrates (Põlme et al. 2020) | Põlme S, et al. 2020. Fungal Diversity 105:1-16 ((https://doi.org/10.1007/s13225-020-00466-2)) |
| Otu18889 | 1 | k__Fungi(100);p__Ascomycota(100);c__Saccharomycetes(100);o__Saccharomycetales(100);f__Metschnikowiaceae(100);g__Metschnikowia(100);g__Metschnikowia_unclassified(100); | Metschnikowia | 13 | Saprotroph | \|Undefined Saprotroph\| | Yeast | NULL | Probable | As nectar-tap_saprotroph (Põlme et al. 2020); Decay sugar-rich substrates (Põlme et al. 2020) | Põlme S, et al. 2020. Fungal Diversity 105:1-16 ((https://doi.org/10.1007/s13225-020-00466-2)) |
| Otu18890 | 1 | k__Fungi(100);p__Ascomycota(100);c__Saccharomycetes(100);o__Saccharomycetales(100);f__Metschnikowiaceae(100);g__Metschnikowia(100);s__Metschnikowia_pulcherrima(100); | Metschnikowia | 13 | Saprotroph | \|Undefined Saprotroph\| | Yeast | NULL | Probable | As nectar-tap_saprotroph (Põlme et al. 2020); Decay sugar-rich substrates (Põlme et al. 2020) | Põlme S, et al. 2020. Fungal Diversity 105:1-16 ((https://doi.org/10.1007/s13225-020-00466-2)) |
| Otu18891 | 1 | k__Fungi(100);p__Ascomycota(100);c__Saccharomycetes(100);o__Saccharomycetales(100);f__Metschnikowiaceae(100);g__Metschnikowia(100);s__Metschnikowia_pulcherrima(100); | Metschnikowia | 13 | Saprotroph | \|Undefined Saprotroph\| | Yeast | NULL | Probable | As nectar-tap_saprotroph (Põlme et al. 2020); Decay sugar-rich substrates (Põlme et al. 2020) | Põlme S, et al. 2020. Fungal Diversity 105:1-16 ((https://doi.org/10.1007/s13225-020-00466-2)) |
| Otu18892 | 1 | k__Fungi(100);p__Ascomycota(100);c__Saccharomycetes(100);o__Saccharomycetales(100);f__Metschnikowiaceae(100);g__Metschnikowia(100);s__Metschnikowia_pulcherrima(100); | Metschnikowia | 13 | Saprotroph | \|Undefined Saprotroph\| | Yeast | NULL | Probable | As nectar-tap_saprotroph (Põlme et al. 2020); Decay sugar-rich substrates (Põlme et al. 2020) | Põlme S, et al. 2020. Fungal Diversity 105:1-16 ((https://doi.org/10.1007/s13225-020-00466-2)) |
| Otu18893 | 1 | k__Fungi(100);p__Ascomycota(100);c__Saccharomycetes(100);o__Saccharomycetales(100);f__Metschnikowiaceae(100);g__Metschnikowia(100);g__Metschnikowia_unclassified(100); | Metschnikowia | 13 | Saprotroph | \|Undefined Saprotroph\| | Yeast | NULL | Probable | As nectar-tap_saprotroph (Põlme et al. 2020); Decay sugar-rich substrates (Põlme et al. 2020) | Põlme S, et al. 2020. Fungal Diversity 105:1-16 ((https://doi.org/10.1007/s13225-020-00466-2)) |
| Otu18894 | 1 | k__Fungi(100);p__Ascomycota(100);c__Saccharomycetes(100);o__Saccharomycetales(100);f__Metschnikowiaceae(100);g__Metschnikowia(100);s__Metschnikowia_pulcherrima(100); | Metschnikowia | 13 | Saprotroph | \|Undefined Saprotroph\| | Yeast | NULL | Probable | As nectar-tap_saprotroph (Põlme et al. 2020); Decay sugar-rich substrates (Põlme et al. 2020) | Põlme S, et al. 2020. Fungal Diversity 105:1-16 ((https://doi.org/10.1007/s13225-020-00466-2)) |
| Otu18895 | 1 | k__Fungi(100);p__Ascomycota(100);c__Saccharomycetes(100);o__Saccharomycetales(100);f__Metschnikowiaceae(100);g__Metschnikowia(100);g__Metschnikowia_unclassified(100); | Metschnikowia | 13 | Saprotroph | \|Undefined Saprotroph\| | Yeast | NULL | Probable | As nectar-tap_saprotroph (Põlme et al. 2020); Decay sugar-rich substrates (Põlme et al. 2020) | Põlme S, et al. 2020. Fungal Diversity 105:1-16 ((https://doi.org/10.1007/s13225-020-00466-2)) |
| Otu18896 | 1 | k__Fungi(100);p__Ascomycota(100);c__Saccharomycetes(100);o__Saccharomycetales(100);f__Metschnikowiaceae(100);g__Metschnikowia(100);g__Metschnikowia_unclassified(100); | Metschnikowia | 13 | Saprotroph | \|Undefined Saprotroph\| | Yeast | NULL | Probable | As nectar-tap_saprotroph (Põlme et al. 2020); Decay sugar-rich substrates (Põlme et al. 2020) | Põlme S, et al. 2020. Fungal Diversity 105:1-16 ((https://doi.org/10.1007/s13225-020-00466-2)) |
| Otu18897 | 1 | k__Fungi(100);p__Ascomycota(100);c__Saccharomycetes(100);o__Saccharomycetales(100);f__Metschnikowiaceae(100);g__Metschnikowia(100);s__Metschnikowia_pulcherrima(100); | Metschnikowia | 13 | Saprotroph | \|Undefined Saprotroph\| | Yeast | NULL | Probable | As nectar-tap_saprotroph (Põlme et al. 2020); Decay sugar-rich substrates (Põlme et al. 2020) | Põlme S, et al. 2020. Fungal Diversity 105:1-16 ((https://doi.org/10.1007/s13225-020-00466-2)) |
| Otu18898 | 1 | k__Fungi(100);p__Ascomycota(100);c__Saccharomycetes(100);o__Saccharomycetales(100);f__Metschnikowiaceae(100);g__Metschnikowia(100);s__Metschnikowia_pulcherrima(100); | Metschnikowia | 13 | Saprotroph | \|Undefined Saprotroph\| | Yeast | NULL | Probable | As nectar-tap_saprotroph (Põlme et al. 2020); Decay sugar-rich substrates (Põlme et al. 2020) | Põlme S, et al. 2020. Fungal Diversity 105:1-16 ((https://doi.org/10.1007/s13225-020-00466-2)) |
| Otu18899 | 1 | k__Fungi(100);p__Ascomycota(100);c__Saccharomycetes(100);o__Saccharomycetales(100);f__Metschnikowiaceae(100);g__Metschnikowia(100);s__Metschnikowia_pulcherrima(100); | Metschnikowia | 13 | Saprotroph | \|Undefined Saprotroph\| | Yeast | NULL | Probable | As nectar-tap_saprotroph (Põlme et al. 2020); Decay sugar-rich substrates (Põlme et al. 2020) | Põlme S, et al. 2020. Fungal Diversity 105:1-16 ((https://doi.org/10.1007/s13225-020-00466-2)) |
| Otu18900 | 1 | k__Fungi(100);p__Ascomycota(100);c__Saccharomycetes(100);o__Saccharomycetales(100);f__Metschnikowiaceae(100);g__Metschnikowia(100);s__Metschnikowia_pulcherrima(100); | Metschnikowia | 13 | Saprotroph | \|Undefined Saprotroph\| | Yeast | NULL | Probable | As nectar-tap_saprotroph (Põlme et al. 2020); Decay sugar-rich substrates (Põlme et al. 2020) | Põlme S, et al. 2020. Fungal Diversity 105:1-16 ((https://doi.org/10.1007/s13225-020-00466-2)) |
| Otu18901 | 1 | k__Fungi(100);p__Ascomycota(100);c__Saccharomycetes(100);o__Saccharomycetales(100);f__Metschnikowiaceae(100);g__Metschnikowia(100);s__Metschnikowia_pulcherrima(100); | Metschnikowia | 13 | Saprotroph | \|Undefined Saprotroph\| | Yeast | NULL | Probable | As nectar-tap_saprotroph (Põlme et al. 2020); Decay sugar-rich substrates (Põlme et al. 2020) | Põlme S, et al. 2020. Fungal Diversity 105:1-16 ((https://doi.org/10.1007/s13225-020-00466-2)) |
| Otu18902 | 1 | k__Fungi(100);p__Ascomycota(100);c__Saccharomycetes(100);o__Saccharomycetales(100);f__Metschnikowiaceae(100);g__Metschnikowia(100);s__Metschnikowia_pulcherrima(100); | Metschnikowia | 13 | Saprotroph | \|Undefined Saprotroph\| | Yeast | NULL | Probable | As nectar-tap_saprotroph (Põlme et al. 2020); Decay sugar-rich substrates (Põlme et al. 2020) | Põlme S, et al. 2020. Fungal Diversity 105:1-16 ((https://doi.org/10.1007/s13225-020-00466-2)) |
| Otu18903 | 1 | k__Fungi(100);p__Ascomycota(100);c__Saccharomycetes(100);o__Saccharomycetales(100);f__Metschnikowiaceae(100);g__Metschnikowia(100);s__Metschnikowia_pulcherrima(100); | Metschnikowia | 13 | Saprotroph | \|Undefined Saprotroph\| | Yeast | NULL | Probable | As nectar-tap_saprotroph (Põlme et al. 2020); Decay sugar-rich substrates (Põlme et al. 2020) | Põlme S, et al. 2020. Fungal Diversity 105:1-16 ((https://doi.org/10.1007/s13225-020-00466-2)) |
| Otu18904 | 1 | k__Fungi(100);p__Ascomycota(100);c__Saccharomycetes(100);o__Saccharomycetales(100);f__Metschnikowiaceae(100);g__Metschnikowia(100);s__Metschnikowia_pulcherrima(100); | Metschnikowia | 13 | Saprotroph | \|Undefined Saprotroph\| | Yeast | NULL | Probable | As nectar-tap_saprotroph (Põlme et al. 2020); Decay sugar-rich substrates (Põlme et al. 2020) | Põlme S, et al. 2020. Fungal Diversity 105:1-16 ((https://doi.org/10.1007/s13225-020-00466-2)) |
| Otu18905 | 1 | k__Fungi(100);p__Ascomycota(100);c__Saccharomycetes(100);o__Saccharomycetales(100);f__Metschnikowiaceae(100);g__Metschnikowia(100);s__Metschnikowia_pulcherrima(100); | Metschnikowia | 13 | Saprotroph | \|Undefined Saprotroph\| | Yeast | NULL | Probable | As nectar-tap_saprotroph (Põlme et al. 2020); Decay sugar-rich substrates (Põlme et al. 2020) | Põlme S, et al. 2020. Fungal Diversity 105:1-16 ((https://doi.org/10.1007/s13225-020-00466-2)) |
| Otu18906 | 1 | k__Fungi(100);p__Ascomycota(100);c__Saccharomycetes(100);o__Saccharomycetales(100);f__Metschnikowiaceae(100);g__Metschnikowia(100);g__Metschnikowia_unclassified(100); | Metschnikowia | 13 | Saprotroph | \|Undefined Saprotroph\| | Yeast | NULL | Probable | As nectar-tap_saprotroph (Põlme et al. 2020); Decay sugar-rich substrates (Põlme et al. 2020) | Põlme S, et al. 2020. Fungal Diversity 105:1-16 ((https://doi.org/10.1007/s13225-020-00466-2)) |
| Otu18907 | 1 | k__Fungi(100);p__Ascomycota(100);c__Saccharomycetes(100);o__Saccharomycetales(100);f__Metschnikowiaceae(100);g__Metschnikowia(100);s__Metschnikowia_pulcherrima(100); | Metschnikowia | 13 | Saprotroph | \|Undefined Saprotroph\| | Yeast | NULL | Probable | As nectar-tap_saprotroph (Põlme et al. 2020); Decay sugar-rich substrates (Põlme et al. 2020) | Põlme S, et al. 2020. Fungal Diversity 105:1-16 ((https://doi.org/10.1007/s13225-020-00466-2)) |

| | | | | | | | | | | | |
|---|---|---|---|---|---|---|---|---|---|---|---|
| Otu18908 | 1 | k__Fungi(100);p__Ascomycota(100);c__Saccharomycetes(100);o__Saccharomycetales(100);f__Metschnikowiaceae(100);g__Metschnikowia(100);g__Metschnikowia_unclassified(100); | Metschnikowia | 13 | Saprotroph | \|Undefined Saprotroph\| | Yeast | NULL | Probable | As nectar-tap_saprotroph (Põlme et al. 2020); Decay sugar-rich substrates (Põlme et al. 2020) | Põlme S, et al. 2020. Fungal Diversity 105:1-16 ((https://doi.org/10.1007/s13225-020-00466-2)) |
| Otu18909 | 1 | k__Fungi(100);p__Ascomycota(100);c__Saccharomycetes(100);o__Saccharomycetales(100);f__Metschnikowiaceae(100);g__Metschnikowia(100);s__Metschnikowia_pulcherrima(100); | Metschnikowia | 13 | Saprotroph | \|Undefined Saprotroph\| | Yeast | NULL | Probable | As nectar-tap_saprotroph (Põlme et al. 2020); Decay sugar-rich substrates (Põlme et al. 2020) | Põlme S, et al. 2020. Fungal Diversity 105:1-16 ((https://doi.org/10.1007/s13225-020-00466-2)) |
| Otu18910 | 1 | k__Fungi(100);p__Ascomycota(100);c__Saccharomycetes(100);o__Saccharomycetales(100);f__Metschnikowiaceae(100);g__Metschnikowia(100);g__Metschnikowia_unclassified(100); | Metschnikowia | 13 | Saprotroph | \|Undefined Saprotroph\| | Yeast | NULL | Probable | As nectar-tap_saprotroph (Põlme et al. 2020); Decay sugar-rich substrates (Põlme et al. 2020) | Põlme S, et al. 2020. Fungal Diversity 105:1-16 ((https://doi.org/10.1007/s13225-020-00466-2)) |
| Otu18911 | 1 | k__Fungi(100);p__Ascomycota(100);c__Saccharomycetes(100);o__Saccharomycetales(100);f__Metschnikowiaceae(100);g__Metschnikowia(100);s__Metschnikowia_pulcherrima(100); | Metschnikowia | 13 | Saprotroph | \|Undefined Saprotroph\| | Yeast | NULL | Probable | As nectar-tap_saprotroph (Põlme et al. 2020); Decay sugar-rich substrates (Põlme et al. 2020) | Põlme S, et al. 2020. Fungal Diversity 105:1-16 ((https://doi.org/10.1007/s13225-020-00466-2)) |
| Otu18913 | 1 | k__Fungi(100);p__Ascomycota(100);c__Saccharomycetes(100);o__Saccharomycetales(100);f__Metschnikowiaceae(100);g__Metschnikowia(100);g__Metschnikowia_unclassified(100); | Metschnikowia | 13 | Saprotroph | \|Undefined Saprotroph\| | Yeast | NULL | Probable | As nectar-tap_saprotroph (Põlme et al. 2020); Decay sugar-rich substrates (Põlme et al. 2020) | Põlme S, et al. 2020. Fungal Diversity 105:1-16 ((https://doi.org/10.1007/s13225-020-00466-2)) |
| Otu18914 | 1 | k__Fungi(100);p__Ascomycota(100);c__Saccharomycetes(100);o__Saccharomycetales(100);f__Metschnikowiaceae(100);g__Metschnikowia(100);s__Metschnikowia_pulcherrima(100); | Metschnikowia | 13 | Saprotroph | \|Undefined Saprotroph\| | Yeast | NULL | Probable | As nectar-tap_saprotroph (Põlme et al. 2020); Decay sugar-rich substrates (Põlme et al. 2020) | Põlme S, et al. 2020. Fungal Diversity 105:1-16 ((https://doi.org/10.1007/s13225-020-00466-2)) |
| Otu18915 | 1 | k__Fungi(100);p__Ascomycota(100);c__Saccharomycetes(100);o__Saccharomycetales(100);f__Metschnikowiaceae(100);g__Metschnikowia(100);s__Metschnikowia_pulcherrima(100); | Metschnikowia | 13 | Saprotroph | \|Undefined Saprotroph\| | Yeast | NULL | Probable | As nectar-tap_saprotroph (Põlme et al. 2020); Decay sugar-rich substrates (Põlme et al. 2020) | Põlme S, et al. 2020. Fungal Diversity 105:1-16 ((https://doi.org/10.1007/s13225-020-00466-2)) |
| Otu18916 | 1 | k__Fungi(100);p__Ascomycota(100);c__Saccharomycetes(100);o__Saccharomycetales(100);f__Metschnikowiaceae(100);g__Metschnikowia(100);s__Metschnikowia_pulcherrima(100); | Metschnikowia | 13 | Saprotroph | \|Undefined Saprotroph\| | Yeast | NULL | Probable | As nectar-tap_saprotroph (Põlme et al. 2020); Decay sugar-rich substrates (Põlme et al. 2020) | Põlme S, et al. 2020. Fungal Diversity 105:1-16 ((https://doi.org/10.1007/s13225-020-00466-2)) |
| Otu18917 | 1 | k__Fungi(100);p__Ascomycota(100);c__Saccharomycetes(100);o__Saccharomycetales(100);f__Metschnikowiaceae(100);g__Metschnikowia(100);g__Metschnikowia_unclassified(100); | Metschnikowia | 13 | Saprotroph | \|Undefined Saprotroph\| | Yeast | NULL | Probable | As nectar-tap_saprotroph (Põlme et al. 2020); Decay sugar-rich substrates (Põlme et al. 2020) | Põlme S, et al. 2020. Fungal Diversity 105:1-16 ((https://doi.org/10.1007/s13225-020-00466-2)) |
| Otu18918 | 1 | k__Fungi(100);p__Ascomycota(100);c__Saccharomycetes(100);o__Saccharomycetales(100);f__Metschnikowiaceae(100);g__Metschnikowia(100);g__Metschnikowia_unclassified(100); | Metschnikowia | 13 | Saprotroph | \|Undefined Saprotroph\| | Yeast | NULL | Probable | As nectar-tap_saprotroph (Põlme et al. 2020); Decay sugar-rich substrates (Põlme et al. 2020) | Põlme S, et al. 2020. Fungal Diversity 105:1-16 ((https://doi.org/10.1007/s13225-020-00466-2)) |
| Otu18919 | 1 | k__Fungi(100);p__Ascomycota(100);c__Saccharomycetes(100);o__Saccharomycetales(100);f__Metschnikowiaceae(100);g__Metschnikowia(100);s__Metschnikowia_pulcherrima(100); | Metschnikowia | 13 | Saprotroph | \|Undefined Saprotroph\| | Yeast | NULL | Probable | As nectar-tap_saprotroph (Põlme et al. 2020); Decay sugar-rich substrates (Põlme et al. 2020) | Põlme S, et al. 2020. Fungal Diversity 105:1-16 ((https://doi.org/10.1007/s13225-020-00466-2)) |
| Otu18920 | 1 | k__Fungi(100);p__Ascomycota(100);c__Saccharomycetes(100);o__Saccharomycetales(100);f__Metschnikowiaceae(100);g__Metschnikowia(100);s__Metschnikowia_pulcherrima(100); | Metschnikowia | 13 | Saprotroph | \|Undefined Saprotroph\| | Yeast | NULL | Probable | As nectar-tap_saprotroph (Põlme et al. 2020); Decay sugar-rich substrates (Põlme et al. 2020) | Põlme S, et al. 2020. Fungal Diversity 105:1-16 ((https://doi.org/10.1007/s13225-020-00466-2)) |
| Otu18921 | 1 | k__Fungi(100);p__Ascomycota(100);c__Saccharomycetes(100);o__Saccharomycetales(100);f__Metschnikowiaceae(100);g__Metschnikowia(100);s__Metschnikowia_pulcherrima(100); | Metschnikowia | 13 | Saprotroph | \|Undefined Saprotroph\| | Yeast | NULL | Probable | As nectar-tap_saprotroph (Põlme et al. 2020); Decay sugar-rich substrates (Põlme et al. 2020) | Põlme S, et al. 2020. Fungal Diversity 105:1-16 ((https://doi.org/10.1007/s13225-020-00466-2)) |
| Otu18922 | 1 | k__Fungi(100);p__Ascomycota(100);c__Saccharomycetes(100);o__Saccharomycetales(100);f__Metschnikowiaceae(100);g__Metschnikowia(100);s__Metschnikowia_pulcherrima(100); | Metschnikowia | 13 | Saprotroph | \|Undefined Saprotroph\| | Yeast | NULL | Probable | As nectar-tap_saprotroph (Põlme et al. 2020); Decay sugar-rich substrates (Põlme et al. 2020) | Põlme S, et al. 2020. Fungal Diversity 105:1-16 ((https://doi.org/10.1007/s13225-020-00466-2)) |
| Otu18923 | 1 | k__Fungi(100);p__Ascomycota(100);c__Saccharomycetes(100);o__Saccharomycetales(100);f__Metschnikowiaceae(100);g__Metschnikowia(100);g__Metschnikowia_unclassified(100); | Metschnikowia | 13 | Saprotroph | \|Undefined Saprotroph\| | Yeast | NULL | Probable | As nectar-tap_saprotroph (Põlme et al. 2020); Decay sugar-rich substrates (Põlme et al. 2020) | Põlme S, et al. 2020. Fungal Diversity 105:1-16 ((https://doi.org/10.1007/s13225-020-00466-2)) |
| Otu18924 | 1 | k__Fungi(100);p__Ascomycota(100);c__Saccharomycetes(100);o__Saccharomycetales(100);f__Metschnikowiaceae(100);g__Metschnikowia(100);s__Metschnikowia_pulcherrima(100); | Metschnikowia | 13 | Saprotroph | \|Undefined Saprotroph\| | Yeast | NULL | Probable | As nectar-tap_saprotroph (Põlme et al. 2020); Decay sugar-rich substrates (Põlme et al. 2020) | Põlme S, et al. 2020. Fungal Diversity 105:1-16 ((https://doi.org/10.1007/s13225-020-00466-2)) |
| Otu18925 | 1 | k__Fungi(100);p__Ascomycota(100);c__Saccharomycetes(100);o__Saccharomycetales(100);f__Metschnikowiaceae(100);g__Metschnikowia(100);g__Metschnikowia_unclassified(100); | Metschnikowia | 13 | Saprotroph | \|Undefined Saprotroph\| | Yeast | NULL | Probable | As nectar-tap_saprotroph (Põlme et al. 2020); Decay sugar-rich substrates (Põlme et al. 2020) | Põlme S, et al. 2020. Fungal Diversity 105:1-16 ((https://doi.org/10.1007/s13225-020-00466-2)) |
| Otu18926 | 1 | k__Fungi(100);p__Ascomycota(100);c__Saccharomycetes(100);o__Saccharomycetales(100);f__Metschnikowiaceae(100);g__Metschnikowia(100);s__Metschnikowia_pulcherrima(100); | Metschnikowia | 13 | Saprotroph | \|Undefined Saprotroph\| | Yeast | NULL | Probable | As nectar-tap_saprotroph (Põlme et al. 2020); Decay sugar-rich substrates (Põlme et al. 2020) | Põlme S, et al. 2020. Fungal Diversity 105:1-16 ((https://doi.org/10.1007/s13225-020-00466-2)) |
| Otu18927 | 1 | k__Fungi(100);p__Ascomycota(100);c__Saccharomycetes(100);o__Saccharomycetales(100);f__Metschnikowiaceae(100);g__Metschnikowia(100);g__Metschnikowia_unclassified(100); | Metschnikowia | 13 | Saprotroph | \|Undefined Saprotroph\| | Yeast | NULL | Probable | As nectar-tap_saprotroph (Põlme et al. 2020); Decay sugar-rich substrates (Põlme et al. 2020) | Põlme S, et al. 2020. Fungal Diversity 105:1-16 ((https://doi.org/10.1007/s13225-020-00466-2)) |
| Otu18928 | 1 | k__Fungi(100);p__Ascomycota(100);c__Saccharomycetes(100);o__Saccharomycetales(100);f__Metschnikowiaceae(100);g__Metschnikowia(100);s__Metschnikowia_pulcherrima(100); | Metschnikowia | 13 | Saprotroph | \|Undefined Saprotroph\| | Yeast | NULL | Probable | As nectar-tap_saprotroph (Põlme et al. 2020); Decay sugar-rich substrates (Põlme et al. 2020) | Põlme S, et al. 2020. Fungal Diversity 105:1-16 ((https://doi.org/10.1007/s13225-020-00466-2)) |
| Otu18929 | 1 | k__Fungi(100);p__Ascomycota(100);c__Saccharomycetes(100);o__Saccharomycetales(100);f__Metschnikowiaceae(100);g__Metschnikowia(100);g__Metschnikowia_unclassified(100); | Metschnikowia | 13 | Saprotroph | \|Undefined Saprotroph\| | Yeast | NULL | Probable | As nectar-tap_saprotroph (Põlme et al. 2020); Decay sugar-rich substrates (Põlme et al. 2020) | Põlme S, et al. 2020. Fungal Diversity 105:1-16 ((https://doi.org/10.1007/s13225-020-00466-2)) |
| Otu18930 | 1 | k__Fungi(100);p__Ascomycota(100);c__Saccharomycetes(100);o__Saccharomycetales(100);f__Metschnikowiaceae(100);g__Metschnikowia(100);s__Metschnikowia_pulcherrima(100); | Metschnikowia | 13 | Saprotroph | \|Undefined Saprotroph\| | Yeast | NULL | Probable | As nectar-tap_saprotroph (Põlme et al. 2020); Decay sugar-rich substrates (Põlme et al. 2020) | Põlme S, et al. 2020. Fungal Diversity 105:1-16 ((https://doi.org/10.1007/s13225-020-00466-2)) |
| Otu18931 | 1 | k__Fungi(100);p__Ascomycota(100);c__Saccharomycetes(100);o__Saccharomycetales(100);f__Metschnikowiaceae(100);g__Metschnikowia(100);s__Metschnikowia_pulcherrima(100); | Metschnikowia | 13 | Saprotroph | \|Undefined Saprotroph\| | Yeast | NULL | Probable | As nectar-tap_saprotroph (Põlme et al. 2020); Decay sugar-rich substrates (Põlme et al. 2020) | Põlme S, et al. 2020. Fungal Diversity 105:1-16 ((https://doi.org/10.1007/s13225-020-00466-2)) |
| Otu18934 | 1 | k__Fungi(100);p__Ascomycota(100);c__Saccharomycetes(100);o__Saccharomycetales(100);f__Metschnikowiaceae(100);g__Metschnikowia(100);s__Metschnikowia_pulcherrima(100); | Metschnikowia | 13 | Saprotroph | \|Undefined Saprotroph\| | Yeast | NULL | Probable | As nectar-tap_saprotroph (Põlme et al. 2020); Decay sugar-rich substrates (Põlme et al. 2020) | Põlme S, et al. 2020. Fungal Diversity 105:1-16 ((https://doi.org/10.1007/s13225-020-00466-2)) |
| Otu18935 | 1 | k__Fungi(100);p__Ascomycota(100);c__Saccharomycetes(100);o__Saccharomycetales(100);f__Metschnikowiaceae(100);g__Metschnikowia(100);g__Metschnikowia_unclassified(100); | Metschnikowia | 13 | Saprotroph | \|Undefined Saprotroph\| | Yeast | NULL | Probable | As nectar-tap_saprotroph (Põlme et al. 2020); Decay sugar-rich substrates (Põlme et al. 2020) | Põlme S, et al. 2020. Fungal Diversity 105:1-16 ((https://doi.org/10.1007/s13225-020-00466-2)) |
| Otu18936 | 1 | k__Fungi(100);p__Ascomycota(100);c__Saccharomycetes(100);o__Saccharomycetales(100);f__Metschnikowiaceae(100);g__Metschnikowia(100);g__Metschnikowia_unclassified(100); | Metschnikowia | 13 | Saprotroph | \|Undefined Saprotroph\| | Yeast | NULL | Probable | As nectar-tap_saprotroph (Põlme et al. 2020); Decay sugar-rich substrates (Põlme et al. 2020) | Põlme S, et al. 2020. Fungal Diversity 105:1-16 ((https://doi.org/10.1007/s13225-020-00466-2)) |
| Otu18937 | 1 | k__Fungi(100);p__Ascomycota(100);c__Saccharomycetes(100);o__Saccharomycetales(100);f__Metschnikowiaceae(100);g__Metschnikowia(100);s__Metschnikowia_pulcherrima(100); | Metschnikowia | 13 | Saprotroph | \|Undefined Saprotroph\| | Yeast | NULL | Probable | As nectar-tap_saprotroph (Põlme et al. 2020); Decay sugar-rich substrates (Põlme et al. 2020) | Põlme S, et al. 2020. Fungal Diversity 105:1-16 ((https://doi.org/10.1007/s13225-020-00466-2)) |
| Otu18938 | 1 | k__Fungi(100);p__Ascomycota(100);c__Saccharomycetes(100);o__Saccharomycetales(100);f__Metschnikowiaceae(100);g__Metschnikowia(100);s__Metschnikowia_pulcherrima(100); | Metschnikowia | 13 | Saprotroph | \|Undefined Saprotroph\| | Yeast | NULL | Probable | As nectar-tap_saprotroph (Põlme et al. 2020); Decay sugar-rich substrates (Põlme et al. 2020) | Põlme S, et al. 2020. Fungal Diversity 105:1-16 ((https://doi.org/10.1007/s13225-020-00466-2)) |
| Otu18939 | 1 | k__Fungi(100);p__Ascomycota(100);c__Saccharomycetes(100);o__Saccharomycetales(100);f__Metschnikowiaceae(100);g__Metschnikowia(100);g__Metschnikowia_unclassified(100); | Metschnikowia | 13 | Saprotroph | \|Undefined Saprotroph\| | Yeast | NULL | Probable | As nectar-tap_saprotroph (Põlme et al. 2020); Decay sugar-rich substrates (Põlme et al. 2020) | Põlme S, et al. 2020. Fungal Diversity 105:1-16 ((https://doi.org/10.1007/s13225-020-00466-2)) |
| Otu18940 | 1 | k__Fungi(100);p__Ascomycota(100);c__Saccharomycetes(100);o__Saccharomycetales(100);f__Metschnikowiaceae(100);g__Metschnikowia(100);s__Metschnikowia_pulcherrima(100); | Metschnikowia | 13 | Saprotroph | \|Undefined Saprotroph\| | Yeast | NULL | Probable | As nectar-tap_saprotroph (Põlme et al. 2020); Decay sugar-rich substrates (Põlme et al. 2020) | Põlme S, et al. 2020. Fungal Diversity 105:1-16 ((https://doi.org/10.1007/s13225-020-00466-2)) |
| Otu18941 | 1 | k__Fungi(100);p__Ascomycota(100);c__Saccharomycetes(100);o__Saccharomycetales(100);f__Metschnikowiaceae(100);g__Metschnikowia(100);s__Metschnikowia_pulcherrima(100); | Metschnikowia | 13 | Saprotroph | \|Undefined Saprotroph\| | Yeast | NULL | Probable | As nectar-tap_saprotroph (Põlme et al. 2020); Decay sugar-rich substrates (Põlme et al. 2020) | Põlme S, et al. 2020. Fungal Diversity 105:1-16 ((https://doi.org/10.1007/s13225-020-00466-2)) |
| Otu18942 | 1 | k__Fungi(100);p__Ascomycota(100);c__Saccharomycetes(100);o__Saccharomycetales(100);f__Metschnikowiaceae(100);g__Metschnikowia(100);s__Metschnikowia_pulcherrima(100); | Metschnikowia | 13 | Saprotroph | \|Undefined Saprotroph\| | Yeast | NULL | Probable | As nectar-tap_saprotroph (Põlme et al. 2020); Decay sugar-rich substrates (Põlme et al. 2020) | Põlme S, et al. 2020. Fungal Diversity 105:1-16 ((https://doi.org/10.1007/s13225-020-00466-2)) |
| Otu18943 | 1 | k__Fungi(100);p__Ascomycota(100);c__Saccharomycetes(100);o__Saccharomycetales(100);f__Metschnikowiaceae(100);g__Metschnikowia(100);g__Metschnikowia_unclassified(100); | Metschnikowia | 13 | Saprotroph | \|Undefined Saprotroph\| | Yeast | NULL | Probable | As nectar-tap_saprotroph (Põlme et al. 2020); Decay sugar-rich substrates (Põlme et al. 2020) | Põlme S, et al. 2020. Fungal Diversity 105:1-16 ((https://doi.org/10.1007/s13225-020-00466-2)) |
| Otu18944 | 1 | k__Fungi(100);p__Ascomycota(100);c__Saccharomycetes(100);o__Saccharomycetales(100);f__Metschnikowiaceae(100);g__Metschnikowia(100);s__Metschnikowia_pulcherrima(100); | Metschnikowia | 13 | Saprotroph | \|Undefined Saprotroph\| | Yeast | NULL | Probable | As nectar-tap_saprotroph (Põlme et al. 2020); Decay sugar-rich substrates (Põlme et al. 2020) | Põlme S, et al. 2020. Fungal Diversity 105:1-16 ((https://doi.org/10.1007/s13225-020-00466-2)) |
| Otu18945 | 1 | k__Fungi(100);p__Ascomycota(100);c__Saccharomycetes(100);o__Saccharomycetales(100);f__Metschnikowiaceae(100);g__Metschnikowia(100);g__Metschnikowia_unclassified(100); | Metschnikowia | 13 | Saprotroph | \|Undefined Saprotroph\| | Yeast | NULL | Probable | As nectar-tap_saprotroph (Põlme et al. 2020); Decay sugar-rich substrates (Põlme et al. 2020) | Põlme S, et al. 2020. Fungal Diversity 105:1-16 ((https://doi.org/10.1007/s13225-020-00466-2)) |
| Otu18948 | 1 | k__Fungi(100);p__Ascomycota(100);c__Saccharomycetes(100);o__Saccharomycetales(100);f__Metschnikowiaceae(100);g__Metschnikowia(100);g__Metschnikowia_unclassified(100); | Metschnikowia | 13 | Saprotroph | \|Undefined Saprotroph\| | Yeast | NULL | Probable | As nectar-tap_saprotroph (Põlme et al. 2020); Decay sugar-rich substrates (Põlme et al. 2020) | Põlme S, et al. 2020. Fungal Diversity 105:1-16 ((https://doi.org/10.1007/s13225-020-00466-2)) |
| Otu18949 | 1 | k__Fungi(100);p__Ascomycota(100);c__Saccharomycetes(100);o__Saccharomycetales(100);f__Metschnikowiaceae(100);g__Metschnikowia(100);s__Metschnikowia_pulcherrima(100); | Metschnikowia | 13 | Saprotroph | \|Undefined Saprotroph\| | Yeast | NULL | Probable | As nectar-tap_saprotroph (Põlme et al. 2020); Decay sugar-rich substrates (Põlme et al. 2020) | Põlme S, et al. 2020. Fungal Diversity 105:1-16 ((https://doi.org/10.1007/s13225-020-00466-2)) |
| Otu18950 | 1 | k__Fungi(100);p__Ascomycota(100);c__Saccharomycetes(100);o__Saccharomycetales(100);f__Metschnikowiaceae(100);g__Metschnikowia(100);s__Metschnikowia_pulcherrima(100); | Metschnikowia | 13 | Saprotroph | \|Undefined Saprotroph\| | Yeast | NULL | Probable | As nectar-tap_saprotroph (Põlme et al. 2020); Decay sugar-rich substrates (Põlme et al. 2020) | Põlme S, et al. 2020. Fungal Diversity 105:1-16 ((https://doi.org/10.1007/s13225-020-00466-2)) |
| Otu18951 | 1 | k__Fungi(100);p__Ascomycota(100);c__Saccharomycetes(100);o__Saccharomycetales(100);f__Metschnikowiaceae(100);g__Metschnikowia(100);s__Metschnikowia_pulcherrima(100); | Metschnikowia | 13 | Saprotroph | \|Undefined Saprotroph\| | Yeast | NULL | Probable | As nectar-tap_saprotroph (Põlme et al. 2020); Decay sugar-rich substrates (Põlme et al. 2020) | Põlme S, et al. 2020. Fungal Diversity 105:1-16 ((https://doi.org/10.1007/s13225-020-00466-2)) |
| Otu18952 | 1 | k__Fungi(100);p__Ascomycota(100);c__Saccharomycetes(100);o__Saccharomycetales(100);f__Metschnikowiaceae(100);g__Metschnikowia(100);s__Metschnikowia_pulcherrima(100); | Metschnikowia | 13 | Saprotroph | \|Undefined Saprotroph\| | Yeast | NULL | Probable | As nectar-tap_saprotroph (Põlme et al. 2020); Decay sugar-rich substrates (Põlme et al. 2020) | Põlme S, et al. 2020. Fungal Diversity 105:1-16 ((https://doi.org/10.1007/s13225-020-00466-2)) |
| Otu18953 | 1 | k__Fungi(100);p__Ascomycota(100);c__Saccharomycetes(100);o__Saccharomycetales(100);f__Metschnikowiaceae(100);g__Metschnikowia(100);s__Metschnikowia_pulcherrima(100); | Metschnikowia | 13 | Saprotroph | \|Undefined Saprotroph\| | Yeast | NULL | Probable | As nectar-tap_saprotroph (Põlme et al. 2020); Decay sugar-rich substrates (Põlme et al. 2020) | Põlme S, et al. 2020. Fungal Diversity 105:1-16 ((https://doi.org/10.1007/s13225-020-00466-2)) |
| Otu18955 | 1 | k__Fungi(100);p__Ascomycota(100);c__Saccharomycetes(100);o__Saccharomycetales(100);f__Metschnikowiaceae(100);g__Metschnikowia(100);g__Metschnikowia_unclassified(100); | Metschnikowia | 13 | Saprotroph | \|Undefined Saprotroph\| | Yeast | NULL | Probable | As nectar-tap_saprotroph (Põlme et al. 2020); Decay sugar-rich substrates (Põlme et al. 2020) | Põlme S, et al. 2020. Fungal Diversity 105:1-16 ((https://doi.org/10.1007/s13225-020-00466-2)) |
| Otu18956 | 1 | k__Fungi(100);p__Ascomycota(100);c__Saccharomycetes(100);o__Saccharomycetales(100);f__Metschnikowiaceae(100);g__Metschnikowia(100);s__Metschnikowia_pulcherrima(100); | Metschnikowia | 13 | Saprotroph | \|Undefined Saprotroph\| | Yeast | NULL | Probable | As nectar-tap_saprotroph (Põlme et al. 2020); Decay sugar-rich substrates (Põlme et al. 2020) | Põlme S, et al. 2020. Fungal Diversity 105:1-16 ((https://doi.org/10.1007/s13225-020-00466-2)) |
| Otu18957 | 1 | k__Fungi(100);p__Ascomycota(100);c__Saccharomycetes(100);o__Saccharomycetales(100);f__Metschnikowiaceae(100);g__Metschnikowia(100);s__Metschnikowia_pulcherrima(100); | Metschnikowia | 13 | Saprotroph | \|Undefined Saprotroph\| | Yeast | NULL | Probable | As nectar-tap_saprotroph (Põlme et al. 2020); Decay sugar-rich substrates (Põlme et al. 2020) | Põlme S, et al. 2020. Fungal Diversity 105:1-16 ((https://doi.org/10.1007/s13225-020-00466-2)) |
| Otu18958 | 1 | k__Fungi(100);p__Ascomycota(100);c__Saccharomycetes(100);o__Saccharomycetales(100);f__Metschnikowiaceae(100);g__Metschnikowia(100);s__Metschnikowia_pulcherrima(100); | Metschnikowia | 13 | Saprotroph | \|Undefined Saprotroph\| | Yeast | NULL | Probable | As nectar-tap_saprotroph (Põlme et al. 2020); Decay sugar-rich substrates (Põlme et al. 2020) | Põlme S, et al. 2020. Fungal Diversity 105:1-16 ((https://doi.org/10.1007/s13225-020-00466-2)) |
| Otu18959 | 1 | k__Fungi(100);p__Ascomycota(100);c__Saccharomycetes(100);o__Saccharomycetales(100);f__Metschnikowiaceae(100);g__Metschnikowia(100);s__Metschnikowia_pulcherrima(100); | Metschnikowia | 13 | Saprotroph | \|Undefined Saprotroph\| | Yeast | NULL | Probable | As nectar-tap_saprotroph (Põlme et al. 2020); Decay sugar-rich substrates (Põlme et al. 2020) | Põlme S, et al. 2020. Fungal Diversity 105:1-16 ((https://doi.org/10.1007/s13225-020-00466-2)) |
| Otu18960 | 1 | k__Fungi(100);p__Ascomycota(100);c__Saccharomycetes(100);o__Saccharomycetales(100);f__Metschnikowiaceae(100);g__Metschnikowia(100);s__Metschnikowia_pulcherrima(100); | Metschnikowia | 13 | Saprotroph | \|Undefined Saprotroph\| | Yeast | NULL | Probable | As nectar-tap_saprotroph (Põlme et al. 2020); Decay sugar-rich substrates (Põlme et al. 2020) | Põlme S, et al. 2020. Fungal Diversity 105:1-16 ((https://doi.org/10.1007/s13225-020-00466-2)) |
| Otu18961 | 1 | k__Fungi(100);p__Ascomycota(100);c__Saccharomycetes(100);o__Saccharomycetales(100);f__Metschnikowiaceae(100);g__Metschnikowia(100);s__Metschnikowia_pulcherrima(100); | Metschnikowia | 13 | Saprotroph | \|Undefined Saprotroph\| | Yeast | NULL | Probable | As nectar-tap_saprotroph (Põlme et al. 2020); Decay sugar-rich substrates (Põlme et al. 2020) | Põlme S, et al. 2020. Fungal Diversity 105:1-16 ((https://doi.org/10.1007/s13225-020-00466-2)) |
| Otu18962 | 1 | k__Fungi(100);p__Ascomycota(100);c__Saccharomycetes(100);o__Saccharomycetales(100);f__Metschnikowiaceae(100);g__Metschnikowia(100);s__Metschnikowia_pulcherrima(100); | Metschnikowia | 13 | Saprotroph | \|Undefined Saprotroph\| | Yeast | NULL | Probable | As nectar-tap_saprotroph (Põlme et al. 2020); Decay sugar-rich substrates (Põlme et al. 2020) | Põlme S, et al. 2020. Fungal Diversity 105:1-16 ((https://doi.org/10.1007/s13225-020-00466-2)) |
| Otu18963 | 1 | k__Fungi(100);p__Ascomycota(100);c__Saccharomycetes(100);o__Saccharomycetales(100);f__Metschnikowiaceae(100);g__Metschnikowia(100);s__Metschnikowia_pulcherrima(100); | Metschnikowia | 13 | Saprotroph | \|Undefined Saprotroph\| | Yeast | NULL | Probable | As nectar-tap_saprotroph (Põlme et al. 2020); Decay sugar-rich substrates (Põlme et al. 2020) | Põlme S, et al. 2020. Fungal Diversity 105:1-16 ((https://doi.org/10.1007/s13225-020-00466-2)) |
| Otu18964 | 1 | k__Fungi(100);p__Ascomycota(100);c__Saccharomycetes(100);o__Saccharomycetales(100);f__Metschnikowiaceae(100);g__Metschnikowia(100);s__Metschnikowia_pulcherrima(100); | Metschnikowia | 13 | Saprotroph | \|Undefined Saprotroph\| | Yeast | NULL | Probable | As nectar-tap_saprotroph (Põlme et al. 2020); Decay sugar-rich substrates (Põlme et al. 2020) | Põlme S, et al. 2020. Fungal Diversity 105:1-16 ((https://doi.org/10.1007/s13225-020-00466-2)) |
| Otu18965 | 1 | k__Fungi(100);p__Ascomycota(100);c__Saccharomycetes(100);o__Saccharomycetales(100);f__Metschnikowiaceae(100);g__Metschnikowia(100);g__Metschnikowia_unclassified(100); | Metschnikowia | 13 | Saprotroph | \|Undefined Saprotroph\| | Yeast | NULL | Probable | As nectar-tap_saprotroph (Põlme et al. 2020); Decay sugar-rich substrates (Põlme et al. 2020) | Põlme S, et al. 2020. Fungal Diversity 105:1-16 ((https://doi.org/10.1007/s13225-020-00466-2)) |
| Otu18966 | 1 | k__Fungi(100);p__Ascomycota(100);c__Saccharomycetes(100);o__Saccharomycetales(100);f__Metschnikowiaceae(100);g__Metschnikowia(100);s__Metschnikowia_pulcherrima(100); | Metschnikowia | 13 | Saprotroph | \|Undefined Saprotroph\| | Yeast | NULL | Probable | As nectar-tap_saprotroph (Põlme et al. 2020); Decay sugar-rich substrates (Põlme et al. 2020) | Põlme S, et al. 2020. Fungal Diversity 105:1-16 ((https://doi.org/10.1007/s13225-020-00466-2)) |
| Otu18967 | 1 | k__Fungi(100);p__Ascomycota(100);c__Saccharomycetes(100);o__Saccharomycetales(100);f__Metschnikowiaceae(100);g__Metschnikowia(100);s__Metschnikowia_pulcherrima(100); | Metschnikowia | 13 | Saprotroph | \|Undefined Saprotroph\| | Yeast | NULL | Probable | As nectar-tap_saprotroph (Põlme et al. 2020); Decay sugar-rich substrates (Põlme et al. 2020) | Põlme S, et al. 2020. Fungal Diversity 105:1-16 ((https://doi.org/10.1007/s13225-020-00466-2)) |
| Otu18968 | 1 | k__Fungi(100);p__Ascomycota(100);c__Saccharomycetes(100);o__Saccharomycetales(100);f__Metschnikowiaceae(100);g__Metschnikowia(100);s__Metschnikowia_pulcherrima(100); | Metschnikowia | 13 | Saprotroph | \|Undefined Saprotroph\| | Yeast | NULL | Probable | As nectar-tap_saprotroph (Põlme et al. 2020); Decay sugar-rich substrates (Põlme et al. 2020) | Põlme S, et al. 2020. Fungal Diversity 105:1-16 ((https://doi.org/10.1007/s13225-020-00466-2)) |
| Otu18969 | 1 | k__Fungi(100);p__Ascomycota(100);c__Saccharomycetes(100);o__Saccharomycetales(100);f__Metschnikowiaceae(100);g__Metschnikowia(100);s__Metschnikowia_pulcherrima(100); | Metschnikowia | 13 | Saprotroph | \|Undefined Saprotroph\| | Yeast | NULL | Probable | As nectar-tap_saprotroph (Põlme et al. 2020); Decay sugar-rich substrates (Põlme et al. 2020) | Põlme S, et al. 2020. Fungal Diversity 105:1-16 ((https://doi.org/10.1007/s13225-020-00466-2)) |
| Otu18970 | 1 | k__Fungi(100);p__Ascomycota(100);c__Saccharomycetes(100);o__Saccharomycetales(100);f__Metschnikowiaceae(100);g__Metschnikowia(100);s__Metschnikowia_pulcherrima(100); | Metschnikowia | 13 | Saprotroph | \|Undefined Saprotroph\| | Yeast | NULL | Probable | As nectar-tap_saprotroph (Põlme et al. 2020); Decay sugar-rich substrates (Põlme et al. 2020) | Põlme S, et al. 2020. Fungal Diversity 105:1-16 ((https://doi.org/10.1007/s13225-020-00466-2)) |
| Otu18972 | 1 | k__Fungi(100);p__Ascomycota(100);c__Saccharomycetes(100);o__Saccharomycetales(100);f__Metschnikowiaceae(100);g__Metschnikowia(100);g__Metschnikowia_unclassified(100); | Metschnikowia | 13 | Saprotroph | \|Undefined Saprotroph\| | Yeast | NULL | Probable | As nectar-tap_saprotroph (Põlme et al. 2020); Decay sugar-rich substrates (Põlme et al. 2020) | Põlme S, et al. 2020. Fungal Diversity 105:1-16 ((https://doi.org/10.1007/s13225-020-00466-2)) |
| Otu18973 | 1 | k__Fungi(100);p__Ascomycota(100);c__Saccharomycetes(100);o__Saccharomycetales(100);f__Metschnikowiaceae(100);g__Metschnikowia(100);s__Metschnikowia_pulcherrima(100); | Metschnikowia | 13 | Saprotroph | \|Undefined Saprotroph\| | Yeast | NULL | Probable | As nectar-tap_saprotroph (Põlme et al. 2020); Decay sugar-rich substrates (Põlme et al. 2020) | Põlme S, et al. 2020. Fungal Diversity 105:1-16 ((https://doi.org/10.1007/s13225-020-00466-2)) |
| Otu18974 | 1 | k__Fungi(100);p__Ascomycota(100);c__Saccharomycetes(100);o__Saccharomycetales(100);f__Metschnikowiaceae(100);g__Metschnikowia(100);s__Metschnikowia_pulcherrima(100); | Metschnikowia | 13 | Saprotroph | \|Undefined Saprotroph\| | Yeast | NULL | Probable | As nectar-tap_saprotroph (Põlme et al. 2020); Decay sugar-rich substrates (Põlme et al. 2020) | Põlme S, et al. 2020. Fungal Diversity 105:1-16 ((https://doi.org/10.1007/s13225-020-00466-2)) |
| Otu18975 | 1 | k__Fungi(100);p__Ascomycota(100);c__Saccharomycetes(100);o__Saccharomycetales(100);f__Metschnikowiaceae(100);g__Metschnikowia(100);s__Metschnikowia_pulcherrima(100); | Metschnikowia | 13 | Saprotroph | \|Undefined Saprotroph\| | Yeast | NULL | Probable | As nectar-tap_saprotroph (Põlme et al. 2020); Decay sugar-rich substrates (Põlme et al. 2020) | Põlme S, et al. 2020. Fungal Diversity 105:1-16 ((https://doi.org/10.1007/s13225-020-00466-2)) |
| Otu18976 | 1 | k__Fungi(100);p__Ascomycota(100);c__Saccharomycetes(100);o__Saccharomycetales(100);f__Metschnikowiaceae(100);g__Metschnikowia(100);g__Metschnikowia_unclassified(100); | Metschnikowia | 13 | Saprotroph | \|Undefined Saprotroph\| | Yeast | NULL | Probable | As nectar-tap_saprotroph (Põlme et al. 2020); Decay sugar-rich substrates (Põlme et al. 2020) | Põlme S, et al. 2020. Fungal Diversity 105:1-16 ((https://doi.org/10.1007/s13225-020-00466-2)) |
| Otu18977 | 1 | k__Fungi(100);p__Ascomycota(100);c__Saccharomycetes(100);o__Saccharomycetales(100);f__Metschnikowiaceae(100);g__Metschnikowia(100);s__Metschnikowia_pulcherrima(100); | Metschnikowia | 13 | Saprotroph | \|Undefined Saprotroph\| | Yeast | NULL | Probable | As nectar-tap_saprotroph (Põlme et al. 2020); Decay sugar-rich substrates (Põlme et al. 2020) | Põlme S, et al. 2020. Fungal Diversity 105:1-16 ((https://doi.org/10.1007/s13225-020-00466-2)) |
| Otu18978 | 1 | k__Fungi(100);p__Ascomycota(100);c__Saccharomycetes(100);o__Saccharomycetales(100);f__Metschnikowiaceae(100);g__Metschnikowia(100);g__Metschnikowia_unclassified(100); | Metschnikowia | 13 | Saprotroph | \|Undefined Saprotroph\| | Yeast | NULL | Probable | As nectar-tap_saprotroph (Põlme et al. 2020); Decay sugar-rich substrates (Põlme et al. 2020) | Põlme S, et al. 2020. Fungal Diversity 105:1-16 ((https://doi.org/10.1007/s13225-020-00466-2)) |
| Otu18980 | 1 | k__Fungi(100);p__Ascomycota(100);c__Saccharomycetes(100);o__Saccharomycetales(100);f__Metschnikowiaceae(100);g__Metschnikowia(100);s__Metschnikowia_pulcherrima(100); | Metschnikowia | 13 | Saprotroph | \|Undefined Saprotroph\| | Yeast | NULL | Probable | As nectar-tap_saprotroph (Põlme et al. 2020); Decay sugar-rich substrates (Põlme et al. 2020) | Põlme S, et al. 2020. Fungal Diversity 105:1-16 ((https://doi.org/10.1007/s13225-020-00466-2)) |
| Otu18981 | 1 | k__Fungi(100);p__Ascomycota(100);c__Saccharomycetes(100);o__Saccharomycetales(100);f__Metschnikowiaceae(100);g__Metschnikowia(100);s__Metschnikowia_pulcherrima(100); | Metschnikowia | 13 | Saprotroph | \|Undefined Saprotroph\| | Yeast | NULL | Probable | As nectar-tap_saprotroph (Põlme et al. 2020); Decay sugar-rich substrates (Põlme et al. 2020) | Põlme S, et al. 2020. Fungal Diversity 105:1-16 ((https://doi.org/10.1007/s13225-020-00466-2)) |
| Otu18982 | 1 | k__Fungi(100);p__Ascomycota(100);c__Saccharomycetes(100);o__Saccharomycetales(100);f__Metschnikowiaceae(100);g__Metschnikowia(100);s__Metschnikowia_pulcherrima(100); | Metschnikowia | 13 | Saprotroph | \|Undefined Saprotroph\| | Yeast | NULL | Probable | As nectar-tap_saprotroph (Põlme et al. 2020); Decay sugar-rich substrates (Põlme et al. 2020) | Põlme S, et al. 2020. Fungal Diversity 105:1-16 ((https://doi.org/10.1007/s13225-020-00466-2)) |
| Otu18983 | 1 | k__Fungi(100);p__Ascomycota(100);c__Saccharomycetes(100);o__Saccharomycetales(100);f__Metschnikowiaceae(100);g__Metschnikowia(100);s__Metschnikowia_pulcherrima(100); | Metschnikowia | 13 | Saprotroph | \|Undefined Saprotroph\| | Yeast | NULL | Probable | As nectar-tap_saprotroph (Põlme et al. 2020); Decay sugar-rich substrates (Põlme et al. 2020) | Põlme S, et al. 2020. Fungal Diversity 105:1-16 ((https://doi.org/10.1007/s13225-020-00466-2)) |
| Otu18990 | 1 | k__Fungi(100);p__Ascomycota(100);c__Saccharomycetes(100);o__Saccharomycetales(100);f__Saccharomycetales_family_Incertae_sedis(100);g__Candida(100);g__Candida_unclassified(100); | Candida | 13 | Pathotroph-Saprotroph-Symbiotroph | Animal Pathogen-Endophyte-Endosymbiont-Epiphyte-\|Undefined Saprotroph\| | Dimorphic Yeast-Yeast | NULL | Probable | As nectar-tap_saprotroph (Põlme et al. 2020); Decay sugar-rich substrates (Põlme et al. 2020); This asexual genus is polyphyletic and contain species that function very differently depend on what they are. Caution should be taken when interpreting guilds for Candida species. | Manolakaki D, et al. 2010. Virulence. 1: 367-375 ((https://doi.org/10.4161/viru.1.5.12796)); Mekbib SB, et al. 2011. Tropical Plant Pathology 36:233-240 ((https://doi.org/10.1590/S1982-56762011000400004)); Põlme S, et al. 2020. Fungal Diversity 105:1-16 ((https://doi.org/10.1007/s13225-020-00466-2)) |
| Otu18993 | 1 | k__Fungi(100);p__Ascomycota(100);c__Saccharomycetes(100);o__Saccharomycetales(100);f__Metschnikowiaceae(100);g__Metschnikowia(100);s__Metschnikowia_pulcherrima(100); | Metschnikowia | 13 | Saprotroph | \|Undefined Saprotroph\| | Yeast | NULL | Probable | As nectar-tap_saprotroph (Põlme et al. 2020); Decay sugar-rich substrates (Põlme et al. 2020) | Põlme S, et al. 2020. Fungal Diversity 105:1-16 ((https://doi.org/10.1007/s13225-020-00466-2)) |
| Otu18994 | 1 | k__Fungi(100);p__Ascomycota(100);c__Saccharomycetes(100);o__Saccharomycetales(100);f__Metschnikowiaceae(100);g__Metschnikowia(100);s__Metschnikowia_pulcherrima(100); | Metschnikowia | 13 | Saprotroph | \|Undefined Saprotroph\| | Yeast | NULL | Probable | As nectar-tap_saprotroph (Põlme et al. 2020); Decay sugar-rich substrates (Põlme et al. 2020) | Põlme S, et al. 2020. Fungal Diversity 105:1-16 ((https://doi.org/10.1007/s13225-020-00466-2)) |
| Otu18995 | 1 | k__Fungi(100);p__Ascomycota(100);c__Saccharomycetes(100);o__Saccharomycetales(100);f__Metschnikowiaceae(100);g__Metschnikowia(100);g__Metschnikowia_unclassified(100); | Metschnikowia | 13 | Saprotroph | \|Undefined Saprotroph\| | Yeast | NULL | Probable | As nectar-tap_saprotroph (Põlme et al. 2020); Decay sugar-rich substrates (Põlme et al. 2020) | Põlme S, et al. 2020. Fungal Diversity 105:1-16 ((https://doi.org/10.1007/s13225-020-00466-2)) |
| Otu18996 | 1 | k__Fungi(100);p__Ascomycota(100);c__Saccharomycetes(100);o__Saccharomycetales(100);f__Metschnikowiaceae(100);g__Metschnikowia(100);g__Metschnikowia_unclassified(100); | Metschnikowia | 13 | Saprotroph | \|Undefined Saprotroph\| | Yeast | NULL | Probable | As nectar-tap_saprotroph (Põlme et al. 2020); Decay sugar-rich substrates (Põlme et al. 2020) | Põlme S, et al. 2020. Fungal Diversity 105:1-16 ((https://doi.org/10.1007/s13225-020-00466-2)) |
| Otu18997 | 1 | k__Fungi(100);p__Ascomycota(100);c__Saccharomycetes(100);o__Saccharomycetales(100);f__Metschnikowiaceae(100);g__Metschnikowia(100);g__Metschnikowia_unclassified(100); | Metschnikowia | 13 | Saprotroph | \|Undefined Saprotroph\| | Yeast | NULL | Probable | As nectar-tap_saprotroph (Põlme et al. 2020); Decay sugar-rich substrates (Põlme et al. 2020) | Põlme S, et al. 2020. Fungal Diversity 105:1-16 ((https://doi.org/10.1007/s13225-020-00466-2)) |
| Otu18998 | 1 | k__Fungi(100);p__Ascomycota(100);c__Saccharomycetes(100);o__Saccharomycetales(100);f__Metschnikowiaceae(100);g__Metschnikowia(100);g__Metschnikowia_unclassified(100); | Metschnikowia | 13 | Saprotroph | \|Undefined Saprotroph\| | Yeast | NULL | Probable | As nectar-tap_saprotroph (Põlme et al. 2020); Decay sugar-rich substrates (Põlme et al. 2020) | Põlme S, et al. 2020. Fungal Diversity 105:1-16 ((https://doi.org/10.1007/s13225-020-00466-2)) |
| Otu18999 | 1 | k__Fungi(100);p__Ascomycota(100);c__Saccharomycetes(100);o__Saccharomycetales(100);f__Metschnikowiaceae(100);g__Metschnikowia(100);g__Metschnikowia_unclassified(100); | Metschnikowia | 13 | Saprotroph | \|Undefined Saprotroph\| | Yeast | NULL | Probable | As nectar-tap_saprotroph (Põlme et al. 2020); Decay sugar-rich substrates (Põlme et al. 2020) | Põlme S, et al. 2020. Fungal Diversity 105:1-16 ((https://doi.org/10.1007/s13225-020-00466-2)) |
| Otu19000 | 1 | k__Fungi(100);p__Ascomycota(100);c__Saccharomycetes(100);o__Saccharomycetales(100);f__Metschnikowiaceae(100);g__Metschnikowia(100);g__Metschnikowia_unclassified(100); | Metschnikowia | 13 | Saprotroph | \|Undefined Saprotroph\| | Yeast | NULL | Probable | As nectar-tap_saprotroph (Põlme et al. 2020); Decay sugar-rich substrates (Põlme et al. 2020) | Põlme S, et al. 2020. Fungal Diversity 105:1-16 ((https://doi.org/10.1007/s13225-020-00466-2)) |
| Otu19001 | 1 | k__Fungi(100);p__Ascomycota(100);c__Saccharomycetes(100);o__Saccharomycetales(100);f__Metschnikowiaceae(100);g__Metschnikowia(100);s__Metschnikowia_pulcherrima(100); | Metschnikowia | 13 | Saprotroph | \|Undefined Saprotroph\| | Yeast | NULL | Probable | As nectar-tap_saprotroph (Põlme et al. 2020); Decay sugar-rich substrates (Põlme et al. 2020) | Põlme S, et al. 2020. Fungal Diversity 105:1-16 ((https://doi.org/10.1007/s13225-020-00466-2)) |
| Otu19002 | 1 | k__Fungi(100);p__Ascomycota(100);c__Saccharomycetes(100);o__Saccharomycetales(100);f__Metschnikowiaceae(100);g__Metschnikowia(100);s__Metschnikowia_pulcherrima(100); | Metschnikowia | 13 | Saprotroph | \|Undefined Saprotroph\| | Yeast | NULL | Probable | As nectar-tap_saprotroph (Põlme et al. 2020); Decay sugar-rich substrates (Põlme et al. 2020) | Põlme S, et al. 2020. Fungal Diversity 105:1-16 ((https://doi.org/10.1007/s13225-020-00466-2)) |
| Otu19003 | 1 | k__Fungi(100);p__Ascomycota(100);c__Saccharomycetes(100);o__Saccharomycetales(100);f__Metschnikowiaceae(100);g__Metschnikowia(100);s__Metschnikowia_pulcherrima(100); | Metschnikowia | 13 | Saprotroph | \|Undefined Saprotroph\| | Yeast | NULL | Probable | As nectar-tap_saprotroph (Põlme et al. 2020); Decay sugar-rich substrates (Põlme et al. 2020) | Põlme S, et al. 2020. Fungal Diversity 105:1-16 ((https://doi.org/10.1007/s13225-020-00466-2)) |
| Otu19004 | 1 | k__Fungi(100);p__Ascomycota(100);c__Saccharomycetes(100);o__Saccharomycetales(100);f__Metschnikowiaceae(100);g__Metschnikowia(100);s__Metschnikowia_pulcherrima(100); | Metschnikowia | 13 | Saprotroph | \|Undefined Saprotroph\| | Yeast | NULL | Probable | As nectar-tap_saprotroph (Põlme et al. 2020); Decay sugar-rich substrates (Põlme et al. 2020) | Põlme S, et al. 2020. Fungal Diversity 105:1-16 ((https://doi.org/10.1007/s13225-020-00466-2)) |
| Otu19005 | 1 | k__Fungi(100);p__Ascomycota(100);c__Saccharomycetes(100);o__Saccharomycetales(100);f__Metschnikowiaceae(100);g__Metschnikowia(100);s__Metschnikowia_pulcherrima(100); | Metschnikowia | 13 | Saprotroph | \|Undefined Saprotroph\| | Yeast | NULL | Probable | As nectar-tap_saprotroph (Põlme et al. 2020); Decay sugar-rich substrates (Põlme et al. 2020) | Põlme S, et al. 2020. Fungal Diversity 105:1-16 ((https://doi.org/10.1007/s13225-020-00466-2)) |
| Otu19006 | 1 | k__Fungi(100);p__Ascomycota(100);c__Saccharomycetes(100);o__Saccharomycetales(100);f__Metschnikowiaceae(100);g__Metschnikowia(100);s__Metschnikowia_pulcherrima(100); | Metschnikowia | 13 | Saprotroph | \|Undefined Saprotroph\| | Yeast | NULL | Probable | As nectar-tap_saprotroph (Põlme et al. 2020); Decay sugar-rich substrates (Põlme et al. 2020) | Põlme S, et al. 2020. Fungal Diversity 105:1-16 ((https://doi.org/10.1007/s13225-020-00466-2)) |
| Otu19007 | 1 | k__Fungi(100);p__Ascomycota(100);c__Saccharomycetes(100);o__Saccharomycetales(100);f__Metschnikowiaceae(100);g__Metschnikowia(100);g__Metschnikowia_unclassified(100); | Metschnikowia | 13 | Saprotroph | \|Undefined Saprotroph\| | Yeast | NULL | Probable | As nectar-tap_saprotroph (Põlme et al. 2020); Decay sugar-rich substrates (Põlme et al. 2020) | Põlme S, et al. 2020. Fungal Diversity 105:1-16 ((https://doi.org/10.1007/s13225-020-00466-2)) |
| Otu19008 | 1 | k__Fungi(100);p__Ascomycota(100);c__Saccharomycetes(100);o__Saccharomycetales(100);f__Metschnikowiaceae(100);g__Metschnikowia(100);s__Metschnikowia_pulcherrima(100); | Metschnikowia | 13 | Saprotroph | \|Undefined Saprotroph\| | Yeast | NULL | Probable | As nectar-tap_saprotroph (Põlme et al. 2020); Decay sugar-rich substrates (Põlme et al. 2020) | Põlme S, et al. 2020. Fungal Diversity 105:1-16 ((https://doi.org/10.1007/s13225-020-00466-2)) |
| Otu19009 | 1 | k__Fungi(100);p__Ascomycota(100);c__Saccharomycetes(100);o__Saccharomycetales(100);f__Metschnikowiaceae(100);g__Metschnikowia(100);s__Metschnikowia_pulcherrima(100); | Metschnikowia | 13 | Saprotroph | \|Undefined Saprotroph\| | Yeast | NULL | Probable | As nectar-tap_saprotroph (Põlme et al. 2020); Decay sugar-rich substrates (Põlme et al. 2020) | Põlme S, et al. 2020. Fungal Diversity 105:1-16 ((https://doi.org/10.1007/s13225-020-00466-2)) |
| Otu19010 | 1 | k__Fungi(100);p__Ascomycota(100);c__Saccharomycetes(100);o__Saccharomycetales(100);f__Metschnikowiaceae(100);g__Metschnikowia(100);g__Metschnikowia_unclassified(100); | Metschnikowia | 13 | Saprotroph | \|Undefined Saprotroph\| | Yeast | NULL | Probable | As nectar-tap_saprotroph (Põlme et al. 2020); Decay sugar-rich substrates (Põlme et al. 2020) | Põlme S, et al. 2020. Fungal Diversity 105:1-16 ((https://doi.org/10.1007/s13225-020-00466-2)) |
| Otu19011 | 1 | k__Fungi(100);p__Ascomycota(100);c__Saccharomycetes(100);o__Saccharomycetales(100);f__Metschnikowiaceae(100);g__Metschnikowia(100);g__Metschnikowia_unclassified(100); | Metschnikowia | 13 | Saprotroph | \|Undefined Saprotroph\| | Yeast | NULL | Probable | As nectar-tap_saprotroph (Põlme et al. 2020); Decay sugar-rich substrates (Põlme et al. 2020) | Põlme S, et al. 2020. Fungal Diversity 105:1-16 ((https://doi.org/10.1007/s13225-020-00466-2)) |
| Otu19012 | 1 | k__Fungi(100);p__Ascomycota(100);c__Saccharomycetes(100);o__Saccharomycetales(100);f__Metschnikowiaceae(100);g__Metschnikowia(100);s__Metschnikowia_pulcherrima(100); | Metschnikowia | 13 | Saprotroph | \|Undefined Saprotroph\| | Yeast | NULL | Probable | As nectar-tap_saprotroph (Põlme et al. 2020); Decay sugar-rich substrates (Põlme et al. 2020) | Põlme S, et al. 2020. Fungal Diversity 105:1-16 ((https://doi.org/10.1007/s13225-020-00466-2)) |
| Otu19013 | 1 | k__Fungi(100);p__Ascomycota(100);c__Saccharomycetes(100);o__Saccharomycetales(100);f__Metschnikowiaceae(100);g__Metschnikowia(100);s__Metschnikowia_pulcherrima(100); | Metschnikowia | 13 | Saprotroph | \|Undefined Saprotroph\| | Yeast | NULL | Probable | As nectar-tap_saprotroph (Põlme et al. 2020); Decay sugar-rich substrates (Põlme et al. 2020) | Põlme S, et al. 2020. Fungal Diversity 105:1-16 ((https://doi.org/10.1007/s13225-020-00466-2)) |
| Otu19014 | 1 | k__Fungi(100);p__Ascomycota(100);c__Saccharomycetes(100);o__Saccharomycetales(100);f__Metschnikowiaceae(100);g__Metschnikowia(100);s__Metschnikowia_pulcherrima(100); | Metschnikowia | 13 | Saprotroph | \|Undefined Saprotroph\| | Yeast | NULL | Probable | As nectar-tap_saprotroph (Põlme et al. 2020); Decay sugar-rich substrates (Põlme et al. 2020) | Põlme S, et al. 2020. Fungal Diversity 105:1-16 ((https://doi.org/10.1007/s13225-020-00466-2)) |
| Otu19015 | 1 | k__Fungi(100);p__Ascomycota(100);c__Saccharomycetes(100);o__Saccharomycetales(100);f__Metschnikowiaceae(100);g__Metschnikowia(100);s__Metschnikowia_pulcherrima(100); | Metschnikowia | 13 | Saprotroph | \|Undefined Saprotroph\| | Yeast | NULL | Probable | As nectar-tap_saprotroph (Põlme et al. 2020); Decay sugar-rich substrates (Põlme et al. 2020) | Põlme S, et al. 2020. Fungal Diversity 105:1-16 ((https://doi.org/10.1007/s13225-020-00466-2)) |
| Otu19016 | 1 | k__Fungi(100);p__Ascomycota(100);c__Saccharomycetes(100);o__Saccharomycetales(100);f__Metschnikowiaceae(100);g__Metschnikowia(100);s__Metschnikowia_pulcherrima(100); | Metschnikowia | 13 | Saprotroph | \|Undefined Saprotroph\| | Yeast | NULL | Probable | As nectar-tap_saprotroph (Põlme et al. 2020); Decay sugar-rich substrates (Põlme et al. 2020) | Põlme S, et al. 2020. Fungal Diversity 105:1-16 ((https://doi.org/10.1007/s13225-020-00466-2)) |
| Otu19019 | 1 | k__Fungi(100);p__Ascomycota(100);c__Saccharomycetes(100);o__Saccharomycetales(100);f__Metschnikowiaceae(100);g__Metschnikowia(100);s__Metschnikowia_pulcherrima(100); | Metschnikowia | 13 | Saprotroph | \|Undefined Saprotroph\| | Yeast | NULL | Probable | As nectar-tap_saprotroph (Põlme et al. 2020); Decay sugar-rich substrates (Põlme et al. 2020) | Põlme S, et al. 2020. Fungal Diversity 105:1-16 ((https://doi.org/10.1007/s13225-020-00466-2)) |
| Otu19020 | 1 | k__Fungi(100);p__Ascomycota(100);c__Saccharomycetes(100);o__Saccharomycetales(100);f__Metschnikowiaceae(100);g__Metschnikowia(100);s__Metschnikowia_pulcherrima(100); | Metschnikowia | 13 | Saprotroph | \|Undefined Saprotroph\| | Yeast | NULL | Probable | As nectar-tap_saprotroph (Põlme et al. 2020); Decay sugar-rich substrates (Põlme et al. 2020) | Põlme S, et al. 2020. Fungal Diversity 105:1-16 ((https://doi.org/10.1007/s13225-020-00466-2)) |
| Otu19021 | 1 | k__Fungi(100);p__Ascomycota(100);c__Saccharomycetes(100);o__Saccharomycetales(100);f__Metschnikowiaceae(100);g__Metschnikowia(100);s__Metschnikowia_pulcherrima(100); | Metschnikowia | 13 | Saprotroph | \|Undefined Saprotroph\| | Yeast | NULL | Probable | As nectar-tap_saprotroph (Põlme et al. 2020); Decay sugar-rich substrates (Põlme et al. 2020) | Põlme S, et al. 2020. Fungal Diversity 105:1-16 ((https://doi.org/10.1007/s13225-020-00466-2)) |
| Otu19023 | 1 | k__Fungi(100);p__Ascomycota(100);c__Saccharomycetes(100);o__Saccharomycetales(100);f__Metschnikowiaceae(100);g__Metschnikowia(100);s__Metschnikowia_pulcherrima(100); | Metschnikowia | 13 | Saprotroph | \|Undefined Saprotroph\| | Yeast | NULL | Probable | As nectar-tap_saprotroph (Põlme et al. 2020); Decay sugar-rich substrates (Põlme et al. 2020) | Põlme S, et al. 2020. Fungal Diversity 105:1-16 ((https://doi.org/10.1007/s13225-020-00466-2)) |
| Otu19024 | 1 | k__Fungi(100);p__Ascomycota(100);c__Saccharomycetes(100);o__Saccharomycetales(100);f__Metschnikowiaceae(100);g__Metschnikowia(100);g__Metschnikowia_unclassified(100); | Metschnikowia | 13 | Saprotroph | \|Undefined Saprotroph\| | Yeast | NULL | Probable | As nectar-tap_saprotroph (Põlme et al. 2020); Decay sugar-rich substrates (Põlme et al. 2020) | Põlme S, et al. 2020. Fungal Diversity 105:1-16 ((https://doi.org/10.1007/s13225-020-00466-2)) |
| Otu19025 | 1 | k__Fungi(100);p__Ascomycota(100);c__Saccharomycetes(100);o__Saccharomycetales(100);f__Metschnikowiaceae(100);g__Metschnikowia(100);g__Metschnikowia_unclassified(100); | Metschnikowia | 13 | Saprotroph | \|Undefined Saprotroph\| | Yeast | NULL | Probable | As nectar-tap_saprotroph (Põlme et al. 2020); Decay sugar-rich substrates (Põlme et al. 2020) | Põlme S, et al. 2020. Fungal Diversity 105:1-16 ((https://doi.org/10.1007/s13225-020-00466-2)) |
| Otu19026 | 1 | k__Fungi(100);p__Ascomycota(100);c__Saccharomycetes(100);o__Saccharomycetales(100);f__Metschnikowiaceae(100);g__Metschnikowia(100);s__Metschnikowia_pulcherrima(100); | Metschnikowia | 13 | Saprotroph | \|Undefined Saprotroph\| | Yeast | NULL | Probable | As nectar-tap_saprotroph (Põlme et al. 2020); Decay sugar-rich substrates (Põlme et al. 2020) | Põlme S, et al. 2020. Fungal Diversity 105:1-16 ((https://doi.org/10.1007/s13225-020-00466-2)) |
| Otu19027 | 1 | k__Fungi(100);p__Ascomycota(100);c__Saccharomycetes(100);o__Saccharomycetales(100);f__Metschnikowiaceae(100);g__Metschnikowia(100);s__Metschnikowia_pulcherrima(100); | Metschnikowia | 13 | Saprotroph | \|Undefined Saprotroph\| | Yeast | NULL | Probable | As nectar-tap_saprotroph (Põlme et al. 2020); Decay sugar-rich substrates (Põlme et al. 2020) | Põlme S, et al. 2020. Fungal Diversity 105:1-16 ((https://doi.org/10.1007/s13225-020-00466-2)) |
| Otu19028 | 1 | k__Fungi(100);p__Ascomycota(100);c__Saccharomycetes(100);o__Saccharomycetales(100);f__Metschnikowiaceae(100);g__Metschnikowia(100);g__Metschnikowia_unclassified(100); | Metschnikowia | 13 | Saprotroph | \|Undefined Saprotroph\| | Yeast | NULL | Probable | As nectar-tap_saprotroph (Põlme et al. 2020); Decay sugar-rich substrates (Põlme et al. 2020) | Põlme S, et al. 2020. Fungal Diversity 105:1-16 ((https://doi.org/10.1007/s13225-020-00466-2)) |
| Otu19029 | 1 | k__Fungi(100);p__Ascomycota(100);c__Saccharomycetes(100);o__Saccharomycetales(100);f__Metschnikowiaceae(100);g__Metschnikowia(100);g__Metschnikowia_unclassified(100); | Metschnikowia | 13 | Saprotroph | \|Undefined Saprotroph\| | Yeast | NULL | Probable | As nectar-tap_saprotroph (Põlme et al. 2020); Decay sugar-rich substrates (Põlme et al. 2020) | Põlme S, et al. 2020. Fungal Diversity 105:1-16 ((https://doi.org/10.1007/s13225-020-00466-2)) |
| Otu19030 | 1 | k__Fungi(100);p__Ascomycota(100);c__Saccharomycetes(100);o__Saccharomycetales(100);f__Metschnikowiaceae(100);g__Metschnikowia(100);s__Metschnikowia_pulcherrima(100); | Metschnikowia | 13 | Saprotroph | \|Undefined Saprotroph\| | Yeast | NULL | Probable | As nectar-tap_saprotroph (Põlme et al. 2020); Decay sugar-rich substrates (Põlme et al. 2020) | Põlme S, et al. 2020. Fungal Diversity 105:1-16 ((https://doi.org/10.1007/s13225-020-00466-2)) |
| Otu19031 | 1 | k__Fungi(100);p__Ascomycota(100);c__Saccharomycetes(100);o__Saccharomycetales(100);f__Metschnikowiaceae(100);g__Metschnikowia(100);g__Metschnikowia_unclassified(100); | Metschnikowia | 13 | Saprotroph | \|Undefined Saprotroph\| | Yeast | NULL | Probable | As nectar-tap_saprotroph (Põlme et al. 2020); Decay sugar-rich substrates (Põlme et al. 2020) | Põlme S, et al. 2020. Fungal Diversity 105:1-16 ((https://doi.org/10.1007/s13225-020-00466-2)) |
| Otu19032 | 1 | k__Fungi(100);p__Ascomycota(100);c__Saccharomycetes(100);o__Saccharomycetales(100);f__Metschnikowiaceae(100);g__Metschnikowia(100);s__Metschnikowia_pulcherrima(100); | Metschnikowia | 13 | Saprotroph | \|Undefined Saprotroph\| | Yeast | NULL | Probable | As nectar-tap_saprotroph (Põlme et al. 2020); Decay sugar-rich substrates (Põlme et al. 2020) | Põlme S, et al. 2020. Fungal Diversity 105:1-16 ((https://doi.org/10.1007/s13225-020-00466-2)) |
| Otu19033 | 1 | k__Fungi(100);p__Ascomycota(100);c__Saccharomycetes(100);o__Saccharomycetales(100);f__Metschnikowiaceae(100);g__Metschnikowia(100);s__Metschnikowia_pulcherrima(100); | Metschnikowia | 13 | Saprotroph | \|Undefined Saprotroph\| | Yeast | NULL | Probable | As nectar-tap_saprotroph (Põlme et al. 2020); Decay sugar-rich substrates (Põlme et al. 2020) | Põlme S, et al. 2020. Fungal Diversity 105:1-16 ((https://doi.org/10.1007/s13225-020-00466-2)) |
| Otu19034 | 1 | k__Fungi(100);p__Ascomycota(100);c__Saccharomycetes(100);o__Saccharomycetales(100);f__Saccharomycetales_family_Incertae_sedis(100);g__Candida(100);g__Candida_unclassified(100); | Candida | 13 | Pathotroph-Saprotroph-Symbiotroph | Animal Pathogen-Endophyte-Endosymbiont-Epiphyte-\|Undefined Saprotroph\| | Dimorphic Yeast-Yeast | NULL | Probable | As nectar-tap_saprotroph (Põlme et al. 2020); Decay sugar-rich substrates (Põlme et al. 2020); This asexual genus is polyphyletic and contain species that function very differently depend on what they are. Caution should be taken when interpreting guilds for Candida species. | Manolakaki D, et al. 2010. Virulence. 1: 367-375 ((https://doi.org/10.4161/viru.1.5.12796)); Mekbib SB, et al. 2011. Tropical Plant Pathology 36:233-240 ((https://doi.org/10.1590/S1982-56762011000400004)); Põlme S, et al. 2020. Fungal Diversity 105:1-16 ((https://doi.org/10.1007/s13225-020-00466-2)) |
| Otu19040 | 1 | k__Fungi(100);p__Ascomycota(100);c__Saccharomycetes(100);o__Saccharomycetales(100);f__Saccharomycetales_family_Incertae_sedis(100);g__Candida(100);g__Candida_unclassified(100); | Candida | 13 | Pathotroph-Saprotroph-Symbiotroph | Animal Pathogen-Endophyte-Endosymbiont-Epiphyte-\|Undefined Saprotroph\| | Dimorphic Yeast-Yeast | NULL | Probable | As nectar-tap_saprotroph (Põlme et al. 2020); Decay sugar-rich substrates (Põlme et al. 2020); This asexual genus is polyphyletic and contain species that function very differently depend on what they are. Caution should be taken when interpreting guilds for Candida species. | Manolakaki D, et al. 2010. Virulence. 1: 367-375 ((https://doi.org/10.4161/viru.1.5.12796)); Mekbib SB, et al. 2011. Tropical Plant Pathology 36:233-240 ((https://doi.org/10.1590/S1982-56762011000400004)); Põlme S, et al. 2020. Fungal Diversity 105:1-16 ((https://doi.org/10.1007/s13225-020-00466-2)) |
| Otu19042 | 1 | k__Fungi(100);p__Ascomycota(100);c__Saccharomycetes(100);o__Saccharomycetales(100);f__Metschnikowiaceae(100);g__Metschnikowia(100);s__Metschnikowia_pulcherrima(100); | Metschnikowia | 13 | Saprotroph | \|Undefined Saprotroph\| | Yeast | NULL | Probable | As nectar-tap_saprotroph (Põlme et al. 2020); Decay sugar-rich substrates (Põlme et al. 2020) | Põlme S, et al. 2020. Fungal Diversity 105:1-16 ((https://doi.org/10.1007/s13225-020-00466-2)) |
| Otu19044 | 1 | k__Fungi(100);p__Ascomycota(100);c__Saccharomycetes(100);o__Saccharomycetales(100);f__Metschnikowiaceae(100);g__Metschnikowia(100);s__Metschnikowia_pulcherrima(100); | Metschnikowia | 13 | Saprotroph | \|Undefined Saprotroph\| | Yeast | NULL | Probable | As nectar-tap_saprotroph (Põlme et al. 2020); Decay sugar-rich substrates (Põlme et al. 2020) | Põlme S, et al. 2020. Fungal Diversity 105:1-16 ((https://doi.org/10.1007/s13225-020-00466-2)) |
| Otu19045 | 1 | k__Fungi(100);p__Ascomycota(100);c__Saccharomycetes(100);o__Saccharomycetales(100);f__Metschnikowiaceae(100);g__Metschnikowia(100);g__Metschnikowia_unclassified(100); | Metschnikowia | 13 | Saprotroph | \|Undefined Saprotroph\| | Yeast | NULL | Probable | As nectar-tap_saprotroph (Põlme et al. 2020); Decay sugar-rich substrates (Põlme et al. 2020) | Põlme S, et al. 2020. Fungal Diversity 105:1-16 ((https://doi.org/10.1007/s13225-020-00466-2)) |
| Otu19050 | 1 | k__Fungi(100);p__Ascomycota(100);c__Saccharomycetes(100);o__Saccharomycetales(100);f__Metschnikowiaceae(100);g__Metschnikowia(100);s__Metschnikowia_pulcherrima(100); | Metschnikowia | 13 | Saprotroph | \|Undefined Saprotroph\| | Yeast | NULL | Probable | As nectar-tap_saprotroph (Põlme et al. 2020); Decay sugar-rich substrates (Põlme et al. 2020) | Põlme S, et al. 2020. Fungal Diversity 105:1-16 ((https://doi.org/10.1007/s13225-020-00466-2)) |
| Otu19052 | 1 | k__Fungi(100);p__Ascomycota(100);c__Saccharomycetes(100);o__Saccharomycetales(100);f__Metschnikowiaceae(100);g__Metschnikowia(100);g__Metschnikowia_unclassified(100); | Metschnikowia | 13 | Saprotroph | \|Undefined Saprotroph\| | Yeast | NULL | Probable | As nectar-tap_saprotroph (Põlme et al. 2020); Decay sugar-rich substrates (Põlme et al. 2020) | Põlme S, et al. 2020. Fungal Diversity 105:1-16 ((https://doi.org/10.1007/s13225-020-00466-2)) |
| Otu19053 | 1 | k__Fungi(100);p__Ascomycota(100);c__Saccharomycetes(100);o__Saccharomycetales(100);f__Metschnikowiaceae(100);g__Metschnikowia(100);s__Metschnikowia_pulcherrima(100); | Metschnikowia | 13 | Saprotroph | \|Undefined Saprotroph\| | Yeast | NULL | Probable | As nectar-tap_saprotroph (Põlme et al. 2020); Decay sugar-rich substrates (Põlme et al. 2020) | Põlme S, et al. 2020. Fungal Diversity 105:1-16 ((https://doi.org/10.1007/s13225-020-00466-2)) |
| Otu19054 | 1 | k__Fungi(100);p__Ascomycota(100);c__Saccharomycetes(100);o__Saccharomycetales(100);f__Metschnikowiaceae(100);g__Metschnikowia(100);s__Metschnikowia_pulcherrima(100); | Metschnikowia | 13 | Saprotroph | \|Undefined Saprotroph\| | Yeast | NULL | Probable | As nectar-tap_saprotroph (Põlme et al. 2020); Decay sugar-rich substrates (Põlme et al. 2020) | Põlme S, et al. 2020. Fungal Diversity 105:1-16 ((https://doi.org/10.1007/s13225-020-00466-2)) |
| Otu19055 | 1 | k__Fungi(100);p__Ascomycota(100);c__Saccharomycetes(100);o__Saccharomycetales(100);f__Metschnikowiaceae(100);g__Metschnikowia(100);s__Metschnikowia_pulcherrima(100); | Metschnikowia | 13 | Saprotroph | \|Undefined Saprotroph\| | Yeast | NULL | Probable | As nectar-tap_saprotroph (Põlme et al. 2020); Decay sugar-rich substrates (Põlme et al. 2020) | Põlme S, et al. 2020. Fungal Diversity 105:1-16 ((https://doi.org/10.1007/s13225-020-00466-2)) |
| Otu19057 | 1 | k__Fungi(100);p__Ascomycota(100);c__Saccharomycetes(100);o__Saccharomycetales(100);f__Metschnikowiaceae(100);g__Metschnikowia(100);g__Metschnikowia_unclassified(100); | Metschnikowia | 13 | Saprotroph | \|Undefined Saprotroph\| | Yeast | NULL | Probable | As nectar-tap_saprotroph (Põlme et al. 2020); Decay sugar-rich substrates (Põlme et al. 2020) | Põlme S, et al. 2020. Fungal Diversity 105:1-16 ((https://doi.org/10.1007/s13225-020-00466-2)) |
| Otu19058 | 1 | k__Fungi(100);p__Ascomycota(100);c__Saccharomycetes(100);o__Saccharomycetales(100);f__Metschnikowiaceae(100);g__Metschnikowia(100);s__Metschnikowia_pulcherrima(100); | Metschnikowia | 13 | Saprotroph | \|Undefined Saprotroph\| | Yeast | NULL | Probable | As nectar-tap_saprotroph (Põlme et al. 2020); Decay sugar-rich substrates (Põlme et al. 2020) | Põlme S, et al. 2020. Fungal Diversity 105:1-16 ((https://doi.org/10.1007/s13225-020-00466-2)) |
| Otu19059 | 1 | k__Fungi(100);p__Ascomycota(100);c__Saccharomycetes(100);o__Saccharomycetales(100);f__Metschnikowiaceae(100);g__Metschnikowia(100);g__Metschnikowia_unclassified(100); | Metschnikowia | 13 | Saprotroph | \|Undefined Saprotroph\| | Yeast | NULL | Probable | As nectar-tap_saprotroph (Põlme et al. 2020); Decay sugar-rich substrates (Põlme et al. 2020) | Põlme S, et al. 2020. Fungal Diversity 105:1-16 ((https://doi.org/10.1007/s13225-020-00466-2)) |
| Otu19061 | 1 | k__Fungi(100);p__Ascomycota(100);c__Saccharomycetes(100);o__Saccharomycetales(100);f__Saccharomycetales_family_Incertae_sedis(100);g__Candida(100);s__Candida_cretensis(100); | Candida | 13 | Pathotroph-Saprotroph-Symbiotroph | Animal Pathogen-Endophyte-Endosymbiont-Epiphyte-\|Undefined Saprotroph\| | Dimorphic Yeast-Yeast | NULL | Probable | As nectar-tap_saprotroph (Põlme et al. 2020); Decay sugar-rich substrates (Põlme et al. 2020); This asexual genus is polyphyletic and contain species that function very differently depend on what they are. Caution should be taken when interpreting guilds for Candida species. | Manolakaki D, et al. 2010. Virulence. 1: 367-375 ((https://doi.org/10.4161/viru.1.5.12796)); Mekbib SB, et al. 2011. Tropical Plant Pathology 36:233-240 ((https://doi.org/10.1590/S1982-56762011000400004)); Põlme S, et al. 2020. Fungal Diversity 105:1-16 ((https://doi.org/10.1007/s13225-020-00466-2)) |
| Otu19062 | 1 | k__Fungi(100);p__Ascomycota(100);c__Saccharomycetes(100);o__Saccharomycetales(100);f__Saccharomycetales_family_Incertae_sedis(100);f__Saccharomycetales_family_Incertae_sedis_unclassified(100);f__Saccharomycetales_family_Incertae_sedis_unclassified(100); | Saccharomycetales | 7 | Saprotroph | Undefined Saprotroph | Yeast | NULL | Possible | NULL | Sterkenburg E, et al. 2015. New Phytologist 207:1145-1158 ((https://doi.org/10.1111/nph.13426)) |
| Otu19063 | 1 | k__Fungi(100);p__Ascomycota(100);c__Saccharomycetes(100);o__Saccharomycetales(100);f__Metschnikowiaceae(100);g__Metschnikowia(100);g__Metschnikowia_unclassified(100); | Metschnikowia | 13 | Saprotroph | \|Undefined Saprotroph\| | Yeast | NULL | Probable | As nectar-tap_saprotroph (Põlme et al. 2020); Decay sugar-rich substrates (Põlme et al. 2020) | Põlme S, et al. 2020. Fungal Diversity 105:1-16 ((https://doi.org/10.1007/s13225-020-00466-2)) |
| Otu19065 | 1 | k__Fungi(100);p__Ascomycota(100);c__Saccharomycetes(100);o__Saccharomycetales(100);f__Metschnikowiaceae(100);g__Metschnikowia(100);s__Metschnikowia_pulcherrima(100); | Metschnikowia | 13 | Saprotroph | \|Undefined Saprotroph\| | Yeast | NULL | Probable | As nectar-tap_saprotroph (Põlme et al. 2020); Decay sugar-rich substrates (Põlme et al. 2020) | Põlme S, et al. 2020. Fungal Diversity 105:1-16 ((https://doi.org/10.1007/s13225-020-00466-2)) |
| Otu19069 | 1 | k__Fungi(100);p__Ascomycota(100);c__Saccharomycetes(100);o__Saccharomycetales(100);f__Saccharomycetales_family_Incertae_sedis(100);g__Candida(100);g__Candida_unclassified(100); | Candida | 13 | Pathotroph-Saprotroph-Symbiotroph | Animal Pathogen-Endophyte-Endosymbiont-Epiphyte-\|Undefined Saprotroph\| | Dimorphic Yeast-Yeast | NULL | Probable | As nectar-tap_saprotroph (Põlme et al. 2020); Decay sugar-rich substrates (Põlme et al. 2020); This asexual genus is polyphyletic and contain species that function very differently depend on what they are. Caution should be taken when interpreting guilds for Candida species. | Manolakaki D, et al. 2010. Virulence. 1: 367-375 ((https://doi.org/10.4161/viru.1.5.12796)); Mekbib SB, et al. 2011. Tropical Plant Pathology 36:233-240 ((https://doi.org/10.1590/S1982-56762011000400004)); Põlme S, et al. 2020. Fungal Diversity 105:1-16 ((https://doi.org/10.1007/s13225-020-00466-2)) |
| Otu19070 | 1 | k__Fungi(100);p__Ascomycota(100);c__Saccharomycetes(100);o__Saccharomycetales(100);f__Metschnikowiaceae(100);g__Metschnikowia(100);s__Metschnikowia_pulcherrima(100); | Metschnikowia | 13 | Saprotroph | \|Undefined Saprotroph\| | Yeast | NULL | Probable | As nectar-tap_saprotroph (Põlme et al. 2020); Decay sugar-rich substrates (Põlme et al. 2020) | Põlme S, et al. 2020. Fungal Diversity 105:1-16 ((https://doi.org/10.1007/s13225-020-00466-2)) |
| Otu19071 | 1 | k__Fungi(100);p__Ascomycota(100);c__Saccharomycetes(100);o__Saccharomycetales(100);f__Saccharomycetales_family_Incertae_sedis(100);g__Candida(100);g__Candida_unclassified(100); | Candida | 13 | Pathotroph-Saprotroph-Symbiotroph | Animal Pathogen-Endophyte-Endosymbiont-Epiphyte-\|Undefined Saprotroph\| | Dimorphic Yeast-Yeast | NULL | Probable | As nectar-tap_saprotroph (Põlme et al. 2020); Decay sugar-rich substrates (Põlme et al. 2020); This asexual genus is polyphyletic and contain species that function very differently depend on what they are. Caution should be taken when interpreting guilds for Candida species. | Manolakaki D, et al. 2010. Virulence. 1: 367-375 ((https://doi.org/10.4161/viru.1.5.12796)); Mekbib SB, et al. 2011. Tropical Plant Pathology 36:233-240 ((https://doi.org/10.1590/S1982-56762011000400004)); Põlme S, et al. 2020. Fungal Diversity 105:1-16 ((https://doi.org/10.1007/s13225-020-00466-2)) |
| Otu19072 | 1 | k__Fungi(100);p__Ascomycota(100);c__Saccharomycetes(100);o__Saccharomycetales(100);f__Metschnikowiaceae(100);g__Metschnikowia(100);s__Metschnikowia_pulcherrima(100); | Metschnikowia | 13 | Saprotroph | \|Undefined Saprotroph\| | Yeast | NULL | Probable | As nectar-tap_saprotroph (Põlme et al. 2020); Decay sugar-rich substrates (Põlme et al. 2020) | Põlme S, et al. 2020. Fungal Diversity 105:1-16 ((https://doi.org/10.1007/s13225-020-00466-2)) |
| Otu19073 | 1 | k__Fungi(100);p__Ascomycota(100);c__Saccharomycetes(100);o__Saccharomycetales(100);f__Metschnikowiaceae(100);g__Metschnikowia(100);s__Metschnikowia_pulcherrima(100); | Metschnikowia | 13 | Saprotroph | \|Undefined Saprotroph\| | Yeast | NULL | Probable | As nectar-tap_saprotroph (Põlme et al. 2020); Decay sugar-rich substrates (Põlme et al. 2020) | Põlme S, et al. 2020. Fungal Diversity 105:1-16 ((https://doi.org/10.1007/s13225-020-00466-2)) |
| Otu19074 | 1 | k__Fungi(100);p__Ascomycota(100);c__Saccharomycetes(100);o__Saccharomycetales(100);f__Metschnikowiaceae(100);g__Metschnikowia(100);s__Metschnikowia_pulcherrima(100); | Metschnikowia | 13 | Saprotroph | \|Undefined Saprotroph\| | Yeast | NULL | Probable | As nectar-tap_saprotroph (Põlme et al. 2020); Decay sugar-rich substrates (Põlme et al. 2020) | Põlme S, et al. 2020. Fungal Diversity 105:1-16 ((https://doi.org/10.1007/s13225-020-00466-2)) |
| Otu19075 | 1 | k__Fungi(100);p__Ascomycota(100);c__Saccharomycetes(100);o__Saccharomycetales(100);f__Metschnikowiaceae(100);g__Metschnikowia(100);s__Metschnikowia_pulcherrima(100); | Metschnikowia | 13 | Saprotroph | \|Undefined Saprotroph\| | Yeast | NULL | Probable | As nectar-tap_saprotroph (Põlme et al. 2020); Decay sugar-rich substrates (Põlme et al. 2020) | Põlme S, et al. 2020. Fungal Diversity 105:1-16 ((https://doi.org/10.1007/s13225-020-00466-2)) |
| Otu19076 | 1 | k__Fungi(100);p__Ascomycota(100);c__Saccharomycetes(100);o__Saccharomycetales(100);f__Metschnikowiaceae(100);g__Metschnikowia(100);s__Metschnikowia_pulcherrima(100); | Metschnikowia | 13 | Saprotroph | \|Undefined Saprotroph\| | Yeast | NULL | Probable | As nectar-tap_saprotroph (Põlme et al. 2020); Decay sugar-rich substrates (Põlme et al. 2020) | Põlme S, et al. 2020. Fungal Diversity 105:1-16 ((https://doi.org/10.1007/s13225-020-00466-2)) |
| Otu19077 | 1 | k__Fungi(100);p__Ascomycota(100);c__Saccharomycetes(100);o__Saccharomycetales(100);f__Metschnikowiaceae(100);g__Metschnikowia(100);s__Metschnikowia_pulcherrima(100); | Metschnikowia | 13 | Saprotroph | \|Undefined Saprotroph\| | Yeast | NULL | Probable | As nectar-tap_saprotroph (Põlme et al. 2020); Decay sugar-rich substrates (Põlme et al. 2020) | Põlme S, et al. 2020. Fungal Diversity 105:1-16 ((https://doi.org/10.1007/s13225-020-00466-2)) |
| Otu19078 | 1 | k__Fungi(100);p__Ascomycota(100);c__Saccharomycetes(100);o__Saccharomycetales(100);f__Metschnikowiaceae(100);g__Metschnikowia(100);s__Metschnikowia_pulcherrima(100); | Metschnikowia | 13 | Saprotroph | \|Undefined Saprotroph\| | Yeast | NULL | Probable | As nectar-tap_saprotroph (Põlme et al. 2020); Decay sugar-rich substrates (Põlme et al. 2020) | Põlme S, et al. 2020. Fungal Diversity 105:1-16 ((https://doi.org/10.1007/s13225-020-00466-2)) |
| Otu19079 | 1 | k__Fungi(100);p__Ascomycota(100);c__Saccharomycetes(100);o__Saccharomycetales(100);f__Metschnikowiaceae(100);g__Metschnikowia(100);s__Metschnikowia_pulcherrima(100); | Metschnikowia | 13 | Saprotroph | \|Undefined Saprotroph\| | Yeast | NULL | Probable | As nectar-tap_saprotroph (Põlme et al. 2020); Decay sugar-rich substrates (Põlme et al. 2020) | Põlme S, et al. 2020. Fungal Diversity 105:1-16 ((https://doi.org/10.1007/s13225-020-00466-2)) |
| Otu19081 | 1 | k__Fungi(100);p__Ascomycota(100);c__Saccharomycetes(100);o__Saccharomycetales(100);f__Metschnikowiaceae(100);g__Metschnikowia(100);g__Metschnikowia_unclassified(100); | Metschnikowia | 13 | Saprotroph | \|Undefined Saprotroph\| | Yeast | NULL | Probable | As nectar-tap_saprotroph (Põlme et al. 2020); Decay sugar-rich substrates (Põlme et al. 2020) | Põlme S, et al. 2020. Fungal Diversity 105:1-16 ((https://doi.org/10.1007/s13225-020-00466-2)) |
| Otu19082 | 1 | k__Fungi(100);p__Ascomycota(100);c__Saccharomycetes(100);o__Saccharomycetales(100);f__Metschnikowiaceae(100);g__Metschnikowia(100);s__Metschnikowia_pulcherrima(100); | Metschnikowia | 13 | Saprotroph | \|Undefined Saprotroph\| | Yeast | NULL | Probable | As nectar-tap_saprotroph (Põlme et al. 2020); Decay sugar-rich substrates (Põlme et al. 2020) | Põlme S, et al. 2020. Fungal Diversity 105:1-16 ((https://doi.org/10.1007/s13225-020-00466-2)) |
| Otu19083 | 1 | k__Fungi(100);p__Ascomycota(100);c__Saccharomycetes(100);o__Saccharomycetales(100);f__Metschnikowiaceae(100);g__Metschnikowia(100);s__Metschnikowia_pulcherrima(100); | Metschnikowia | 13 | Saprotroph | \|Undefined Saprotroph\| | Yeast | NULL | Probable | As nectar-tap_saprotroph (Põlme et al. 2020); Decay sugar-rich substrates (Põlme et al. 2020) | Põlme S, et al. 2020. Fungal Diversity 105:1-16 ((https://doi.org/10.1007/s13225-020-00466-2)) |
| Otu19084 | 1 | k__Fungi(100);p__Ascomycota(100);c__Saccharomycetes(100);o__Saccharomycetales(100);f__Metschnikowiaceae(100);g__Metschnikowia(100);g__Metschnikowia_unclassified(100); | Metschnikowia | 13 | Saprotroph | \|Undefined Saprotroph\| | Yeast | NULL | Probable | As nectar-tap_saprotroph (Põlme et al. 2020); Decay sugar-rich substrates (Põlme et al. 2020) | Põlme S, et al. 2020. Fungal Diversity 105:1-16 ((https://doi.org/10.1007/s13225-020-00466-2)) |
| Otu19085 | 1 | k__Fungi(100);p__Ascomycota(100);c__Saccharomycetes(100);o__Saccharomycetales(100);f__Metschnikowiaceae(100);g__Metschnikowia(100);s__Metschnikowia_pulcherrima(100); | Metschnikowia | 13 | Saprotroph | \|Undefined Saprotroph\| | Yeast | NULL | Probable | As nectar-tap_saprotroph (Põlme et al. 2020); Decay sugar-rich substrates (Põlme et al. 2020) | Põlme S, et al. 2020. Fungal Diversity 105:1-16 ((https://doi.org/10.1007/s13225-020-00466-2)) |
| Otu19086 | 1 | k__Fungi(100);p__Ascomycota(100);c__Saccharomycetes(100);o__Saccharomycetales(100);f__Metschnikowiaceae(100);g__Metschnikowia(100);s__Metschnikowia_pulcherrima(100); | Metschnikowia | 13 | Saprotroph | \|Undefined Saprotroph\| | Yeast | NULL | Probable | As nectar-tap_saprotroph (Põlme et al. 2020); Decay sugar-rich substrates (Põlme et al. 2020) | Põlme S, et al. 2020. Fungal Diversity 105:1-16 ((https://doi.org/10.1007/s13225-020-00466-2)) |
| Otu19087 | 1 | k__Fungi(100);p__Ascomycota(100);c__Saccharomycetes(100);o__Saccharomycetales(100);f__Metschnikowiaceae(100);g__Metschnikowia(100);s__Metschnikowia_pulcherrima(100); | Metschnikowia | 13 | Saprotroph | \|Undefined Saprotroph\| | Yeast | NULL | Probable | As nectar-tap_saprotroph (Põlme et al. 2020); Decay sugar-rich substrates (Põlme et al. 2020) | Põlme S, et al. 2020. Fungal Diversity 105:1-16 ((https://doi.org/10.1007/s13225-020-00466-2)) |
| Otu19088 | 1 | k__Fungi(100);p__Ascomycota(100);c__Saccharomycetes(100);o__Saccharomycetales(100);f__Metschnikowiaceae(100);g__Metschnikowia(100);g__Metschnikowia_unclassified(100); | Metschnikowia | 13 | Saprotroph | \|Undefined Saprotroph\| | Yeast | NULL | Probable | As nectar-tap_saprotroph (Põlme et al. 2020); Decay sugar-rich substrates (Põlme et al. 2020) | Põlme S, et al. 2020. Fungal Diversity 105:1-16 ((https://doi.org/10.1007/s13225-020-00466-2)) |
| Otu19089 | 1 | k__Fungi(100);p__Ascomycota(100);c__Saccharomycetes(100);o__Saccharomycetales(100);f__Metschnikowiaceae(100);g__Metschnikowia(100);s__Metschnikowia_pulcherrima(100); | Metschnikowia | 13 | Saprotroph | \|Undefined Saprotroph\| | Yeast | NULL | Probable | As nectar-tap_saprotroph (Põlme et al. 2020); Decay sugar-rich substrates (Põlme et al. 2020) | Põlme S, et al. 2020. Fungal Diversity 105:1-16 ((https://doi.org/10.1007/s13225-020-00466-2)) |
| Otu19090 | 1 | k__Fungi(100);p__Ascomycota(100);c__Saccharomycetes(100);o__Saccharomycetales(100);f__Metschnikowiaceae(100);g__Metschnikowia(100);s__Metschnikowia_pulcherrima(100); | Metschnikowia | 13 | Saprotroph | \|Undefined Saprotroph\| | Yeast | NULL | Probable | As nectar-tap_saprotroph (Põlme et al. 2020); Decay sugar-rich substrates (Põlme et al. 2020) | Põlme S, et al. 2020. Fungal Diversity 105:1-16 ((https://doi.org/10.1007/s13225-020-00466-2)) |
| Otu19091 | 1 | k__Fungi(100);p__Ascomycota(100);c__Saccharomycetes(100);o__Saccharomycetales(100);f__Metschnikowiaceae(100);g__Metschnikowia(100);s__Metschnikowia_pulcherrima(100); | Metschnikowia | 13 | Saprotroph | \|Undefined Saprotroph\| | Yeast | NULL | Probable | As nectar-tap_saprotroph (Põlme et al. 2020); Decay sugar-rich substrates (Põlme et al. 2020) | Põlme S, et al. 2020. Fungal Diversity 105:1-16 ((https://doi.org/10.1007/s13225-020-00466-2)) |
| Otu19092 | 1 | k__Fungi(100);p__Ascomycota(100);c__Saccharomycetes(100);o__Saccharomycetales(100);f__Metschnikowiaceae(100);g__Metschnikowia(100);s__Metschnikowia_pulcherrima(100); | Metschnikowia | 13 | Saprotroph | \|Undefined Saprotroph\| | Yeast | NULL | Probable | As nectar-tap_saprotroph (Põlme et al. 2020); Decay sugar-rich substrates (Põlme et al. 2020) | Põlme S, et al. 2020. Fungal Diversity 105:1-16 ((https://doi.org/10.1007/s13225-020-00466-2)) |
| Otu19094 | 1 | k__Fungi(100);p__Ascomycota(100);c__Saccharomycetes(100);o__Saccharomycetales(100);f__Metschnikowiaceae(100);g__Metschnikowia(100);g__Metschnikowia_unclassified(100); | Metschnikowia | 13 | Saprotroph | \|Undefined Saprotroph\| | Yeast | NULL | Probable | As nectar-tap_saprotroph (Põlme et al. 2020); Decay sugar-rich substrates (Põlme et al. 2020) | Põlme S, et al. 2020. Fungal Diversity 105:1-16 ((https://doi.org/10.1007/s13225-020-00466-2)) |
| Otu19095 | 1 | k__Fungi(100);p__Ascomycota(100);c__Saccharomycetes(100);o__Saccharomycetales(100);f__Metschnikowiaceae(100);g__Metschnikowia(100);s__Metschnikowia_pulcherrima(100); | Metschnikowia | 13 | Saprotroph | \|Undefined Saprotroph\| | Yeast | NULL | Probable | As nectar-tap_saprotroph (Põlme et al. 2020); Decay sugar-rich substrates (Põlme et al. 2020) | Põlme S, et al. 2020. Fungal Diversity 105:1-16 ((https://doi.org/10.1007/s13225-020-00466-2)) |
| Otu19096 | 1 | k__Fungi(100);p__Ascomycota(100);c__Saccharomycetes(100);o__Saccharomycetales(100);f__Metschnikowiaceae(100);g__Metschnikowia(100);g__Metschnikowia_unclassified(100); | Metschnikowia | 13 | Saprotroph | \|Undefined Saprotroph\| | Yeast | NULL | Probable | As nectar-tap_saprotroph (Põlme et al. 2020); Decay sugar-rich substrates (Põlme et al. 2020) | Põlme S, et al. 2020. Fungal Diversity 105:1-16 ((https://doi.org/10.1007/s13225-020-00466-2)) |
| Otu19097 | 1 | k__Fungi(100);p__Ascomycota(100);c__Saccharomycetes(100);o__Saccharomycetales(100);f__Metschnikowiaceae(100);g__Metschnikowia(100);g__Metschnikowia_unclassified(100); | Metschnikowia | 13 | Saprotroph | \|Undefined Saprotroph\| | Yeast | NULL | Probable | As nectar-tap_saprotroph (Põlme et al. 2020); Decay sugar-rich substrates (Põlme et al. 2020) | Põlme S, et al. 2020. Fungal Diversity 105:1-16 ((https://doi.org/10.1007/s13225-020-00466-2)) |
| Otu19098 | 1 | k__Fungi(100);p__Ascomycota(100);c__Saccharomycetes(100);o__Saccharomycetales(100);f__Metschnikowiaceae(100);g__Metschnikowia(100);s__Metschnikowia_pulcherrima(100); | Metschnikowia | 13 | Saprotroph | \|Undefined Saprotroph\| | Yeast | NULL | Probable | As nectar-tap_saprotroph (Põlme et al. 2020); Decay sugar-rich substrates (Põlme et al. 2020) | Põlme S, et al. 2020. Fungal Diversity 105:1-16 ((https://doi.org/10.1007/s13225-020-00466-2)) |
| Otu19100 | 1 | k__Fungi(100);p__Ascomycota(100);c__Saccharomycetes(100);o__Saccharomycetales(100);f__Metschnikowiaceae(100);g__Metschnikowia(100);s__Metschnikowia_pulcherrima(100); | Metschnikowia | 13 | Saprotroph | \|Undefined Saprotroph\| | Yeast | NULL | Probable | As nectar-tap_saprotroph (Põlme et al. 2020); Decay sugar-rich substrates (Põlme et al. 2020) | Põlme S, et al. 2020. Fungal Diversity 105:1-16 ((https://doi.org/10.1007/s13225-020-00466-2)) |
| Otu19101 | 1 | k__Fungi(100);p__Ascomycota(100);c__Saccharomycetes(100);o__Saccharomycetales(100);f__Metschnikowiaceae(100);g__Metschnikowia(100);s__Metschnikowia_pulcherrima(100); | Metschnikowia | 13 | Saprotroph | \|Undefined Saprotroph\| | Yeast | NULL | Probable | As nectar-tap_saprotroph (Põlme et al. 2020); Decay sugar-rich substrates (Põlme et al. 2020) | Põlme S, et al. 2020. Fungal Diversity 105:1-16 ((https://doi.org/10.1007/s13225-020-00466-2)) |
| Otu19102 | 1 | k__Fungi(100);p__Ascomycota(100);c__Saccharomycetes(100);o__Saccharomycetales(100);f__Metschnikowiaceae(100);g__Metschnikowia(100);s__Metschnikowia_pulcherrima(100); | Metschnikowia | 13 | Saprotroph | \|Undefined Saprotroph\| | Yeast | NULL | Probable | As nectar-tap_saprotroph (Põlme et al. 2020); Decay sugar-rich substrates (Põlme et al. 2020) | Põlme S, et al. 2020. Fungal Diversity 105:1-16 ((https://doi.org/10.1007/s13225-020-00466-2)) |
| Otu19103 | 1 | k__Fungi(100);p__Ascomycota(100);c__Saccharomycetes(100);o__Saccharomycetales(100);f__Metschnikowiaceae(100);g__Metschnikowia(100);s__Metschnikowia_pulcherrima(100); | Metschnikowia | 13 | Saprotroph | \|Undefined Saprotroph\| | Yeast | NULL | Probable | As nectar-tap_saprotroph (Põlme et al. 2020); Decay sugar-rich substrates (Põlme et al. 2020) | Põlme S, et al. 2020. Fungal Diversity 105:1-16 ((https://doi.org/10.1007/s13225-020-00466-2)) |
| Otu19104 | 1 | k__Fungi(100);p__Ascomycota(100);c__Saccharomycetes(100);o__Saccharomycetales(100);f__Metschnikowiaceae(100);g__Metschnikowia(100);s__Metschnikowia_pulcherrima(100); | Metschnikowia | 13 | Saprotroph | \|Undefined Saprotroph\| | Yeast | NULL | Probable | As nectar-tap_saprotroph (Põlme et al. 2020); Decay sugar-rich substrates (Põlme et al. 2020) | Põlme S, et al. 2020. Fungal Diversity 105:1-16 ((https://doi.org/10.1007/s13225-020-00466-2)) |
| Otu19105 | 1 | k__Fungi(100);p__Ascomycota(100);c__Saccharomycetes(100);o__Saccharomycetales(100);f__Metschnikowiaceae(100);g__Metschnikowia(100);g__Metschnikowia_unclassified(100); | Metschnikowia | 13 | Saprotroph | \|Undefined Saprotroph\| | Yeast | NULL | Probable | As nectar-tap_saprotroph (Põlme et al. 2020); Decay sugar-rich substrates (Põlme et al. 2020) | Põlme S, et al. 2020. Fungal Diversity 105:1-16 ((https://doi.org/10.1007/s13225-020-00466-2)) |
| Otu19106 | 1 | k__Fungi(100);p__Ascomycota(100);c__Saccharomycetes(100);o__Saccharomycetales(100);f__Metschnikowiaceae(100);g__Metschnikowia(100);s__Metschnikowia_pulcherrima(100); | Metschnikowia | 13 | Saprotroph | \|Undefined Saprotroph\| | Yeast | NULL | Probable | As nectar-tap_saprotroph (Põlme et al. 2020); Decay sugar-rich substrates (Põlme et al. 2020) | Põlme S, et al. 2020. Fungal Diversity 105:1-16 ((https://doi.org/10.1007/s13225-020-00466-2)) |
| Otu19107 | 1 | k__Fungi(100);p__Ascomycota(100);c__Saccharomycetes(100);o__Saccharomycetales(100);f__Metschnikowiaceae(100);g__Metschnikowia(100);s__Metschnikowia_pulcherrima(100); | Metschnikowia | 13 | Saprotroph | \|Undefined Saprotroph\| | Yeast | NULL | Probable | As nectar-tap_saprotroph (Põlme et al. 2020); Decay sugar-rich substrates (Põlme et al. 2020) | Põlme S, et al. 2020. Fungal Diversity 105:1-16 ((https://doi.org/10.1007/s13225-020-00466-2)) |
| Otu19108 | 1 | k__Fungi(100);p__Ascomycota(100);c__Saccharomycetes(100);o__Saccharomycetales(100);f__Metschnikowiaceae(100);g__Metschnikowia(100);s__Metschnikowia_pulcherrima(100); | Metschnikowia | 13 | Saprotroph | \|Undefined Saprotroph\| | Yeast | NULL | Probable | As nectar-tap_saprotroph (Põlme et al. 2020); Decay sugar-rich substrates (Põlme et al. 2020) | Põlme S, et al. 2020. Fungal Diversity 105:1-16 ((https://doi.org/10.1007/s13225-020-00466-2)) |
| Otu19109 | 1 | k__Fungi(100);p__Ascomycota(100);c__Saccharomycetes(100);o__Saccharomycetales(100);f__Metschnikowiaceae(100);g__Metschnikowia(100);s__Metschnikowia_pulcherrima(100); | Metschnikowia | 13 | Saprotroph | \|Undefined Saprotroph\| | Yeast | NULL | Probable | As nectar-tap_saprotroph (Põlme et al. 2020); Decay sugar-rich substrates (Põlme et al. 2020) | Põlme S, et al. 2020. Fungal Diversity 105:1-16 ((https://doi.org/10.1007/s13225-020-00466-2)) |
| Otu19110 | 1 | k__Fungi(100);p__Ascomycota(100);c__Saccharomycetes(100);o__Saccharomycetales(100);f__Metschnikowiaceae(100);g__Metschnikowia(100);s__Metschnikowia_pulcherrima(100); | Metschnikowia | 13 | Saprotroph | \|Undefined Saprotroph\| | Yeast | NULL | Probable | As nectar-tap_saprotroph (Põlme et al. 2020); Decay sugar-rich substrates (Põlme et al. 2020) | Põlme S, et al. 2020. Fungal Diversity 105:1-16 ((https://doi.org/10.1007/s13225-020-00466-2)) |
| Otu19112 | 1 | k__Fungi(100);p__Ascomycota(100);c__Saccharomycetes(100);o__Saccharomycetales(100);f__Metschnikowiaceae(100);g__Metschnikowia(100);s__Metschnikowia_pulcherrima(100); | Metschnikowia | 13 | Saprotroph | \|Undefined Saprotroph\| | Yeast | NULL | Probable | As nectar-tap_saprotroph (Põlme et al. 2020); Decay sugar-rich substrates (Põlme et al. 2020) | Põlme S, et al. 2020. Fungal Diversity 105:1-16 ((https://doi.org/10.1007/s13225-020-00466-2)) |
| Otu19113 | 1 | k__Fungi(100);p__Ascomycota(100);c__Saccharomycetes(100);o__Saccharomycetales(100);f__Metschnikowiaceae(100);g__Metschnikowia(100);g__Metschnikowia_unclassified(100); | Metschnikowia | 13 | Saprotroph | \|Undefined Saprotroph\| | Yeast | NULL | Probable | As nectar-tap_saprotroph (Põlme et al. 2020); Decay sugar-rich substrates (Põlme et al. 2020) | Põlme S, et al. 2020. Fungal Diversity 105:1-16 ((https://doi.org/10.1007/s13225-020-00466-2)) |
| Otu19114 | 1 | k__Fungi(100);p__Ascomycota(100);c__Saccharomycetes(100);o__Saccharomycetales(100);f__Metschnikowiaceae(100);g__Metschnikowia(100);s__Metschnikowia_pulcherrima(100); | Metschnikowia | 13 | Saprotroph | \|Undefined Saprotroph\| | Yeast | NULL | Probable | As nectar-tap_saprotroph (Põlme et al. 2020); Decay sugar-rich substrates (Põlme et al. 2020) | Põlme S, et al. 2020. Fungal Diversity 105:1-16 ((https://doi.org/10.1007/s13225-020-00466-2)) |
| Otu19115 | 1 | k__Fungi(100);p__Ascomycota(100);c__Saccharomycetes(100);o__Saccharomycetales(100);f__Metschnikowiaceae(100);g__Metschnikowia(100);s__Metschnikowia_pulcherrima(100); | Metschnikowia | 13 | Saprotroph | \|Undefined Saprotroph\| | Yeast | NULL | Probable | As nectar-tap_saprotroph (Põlme et al. 2020); Decay sugar-rich substrates (Põlme et al. 2020) | Põlme S, et al. 2020. Fungal Diversity 105:1-16 ((https://doi.org/10.1007/s13225-020-00466-2)) |
| Otu19116 | 1 | k__Fungi(100);p__Ascomycota(100);c__Saccharomycetes(100);o__Saccharomycetales(100);f__Metschnikowiaceae(100);g__Metschnikowia(100);s__Metschnikowia_pulcherrima(100); | Metschnikowia | 13 | Saprotroph | \|Undefined Saprotroph\| | Yeast | NULL | Probable | As nectar-tap_saprotroph (Põlme et al. 2020); Decay sugar-rich substrates (Põlme et al. 2020) | Põlme S, et al. 2020. Fungal Diversity 105:1-16 ((https://doi.org/10.1007/s13225-020-00466-2)) |
| Otu19117 | 1 | k__Fungi(100);p__Ascomycota(100);c__Saccharomycetes(100);o__Saccharomycetales(100);f__Metschnikowiaceae(100);g__Metschnikowia(100);s__Metschnikowia_pulcherrima(100); | Metschnikowia | 13 | Saprotroph | \|Undefined Saprotroph\| | Yeast | NULL | Probable | As nectar-tap_saprotroph (Põlme et al. 2020); Decay sugar-rich substrates (Põlme et al. 2020) | Põlme S, et al. 2020. Fungal Diversity 105:1-16 ((https://doi.org/10.1007/s13225-020-00466-2)) |
| Otu19118 | 1 | k__Fungi(100);p__Ascomycota(100);c__Saccharomycetes(100);o__Saccharomycetales(100);f__Metschnikowiaceae(100);g__Metschnikowia(100);s__Metschnikowia_pulcherrima(100); | Metschnikowia | 13 | Saprotroph | \|Undefined Saprotroph\| | Yeast | NULL | Probable | As nectar-tap_saprotroph (Põlme et al. 2020); Decay sugar-rich substrates (Põlme et al. 2020) | Põlme S, et al. 2020. Fungal Diversity 105:1-16 ((https://doi.org/10.1007/s13225-020-00466-2)) |
| Otu19119 | 1 | k__Fungi(100);p__Ascomycota(100);c__Saccharomycetes(100);o__Saccharomycetales(100);f__Metschnikowiaceae(100);g__Metschnikowia(100);s__Metschnikowia_pulcherrima(100); | Metschnikowia | 13 | Saprotroph | \|Undefined Saprotroph\| | Yeast | NULL | Probable | As nectar-tap_saprotroph (Põlme et al. 2020); Decay sugar-rich substrates (Põlme et al. 2020) | Põlme S, et al. 2020. Fungal Diversity 105:1-16 ((https://doi.org/10.1007/s13225-020-00466-2)) |
| Otu19120 | 1 | k__Fungi(100);p__Ascomycota(100);c__Saccharomycetes(100);o__Saccharomycetales(100);f__Metschnikowiaceae(100);g__Metschnikowia(100);g__Metschnikowia_unclassified(100); | Metschnikowia | 13 | Saprotroph | \|Undefined Saprotroph\| | Yeast | NULL | Probable | As nectar-tap_saprotroph (Põlme et al. 2020); Decay sugar-rich substrates (Põlme et al. 2020) | Põlme S, et al. 2020. Fungal Diversity 105:1-16 ((https://doi.org/10.1007/s13225-020-00466-2)) |
| Otu19121 | 1 | k__Fungi(100);p__Ascomycota(100);c__Saccharomycetes(100);o__Saccharomycetales(100);f__Metschnikowiaceae(100);g__Metschnikowia(100);s__Metschnikowia_pulcherrima(100); | Metschnikowia | 13 | Saprotroph | \|Undefined Saprotroph\| | Yeast | NULL | Probable | As nectar-tap_saprotroph (Põlme et al. 2020); Decay sugar-rich substrates (Põlme et al. 2020) | Põlme S, et al. 2020. Fungal Diversity 105:1-16 ((https://doi.org/10.1007/s13225-020-00466-2)) |
| Otu19122 | 1 | k__Fungi(100);p__Ascomycota(100);c__Saccharomycetes(100);o__Saccharomycetales(100);f__Metschnikowiaceae(100);g__Metschnikowia(100);s__Metschnikowia_pulcherrima(100); | Metschnikowia | 13 | Saprotroph | \|Undefined Saprotroph\| | Yeast | NULL | Probable | As nectar-tap_saprotroph (Põlme et al. 2020); Decay sugar-rich substrates (Põlme et al. 2020) | Põlme S, et al. 2020. Fungal Diversity 105:1-16 ((https://doi.org/10.1007/s13225-020-00466-2)) |
| Otu19123 | 1 | k__Fungi(100);p__Ascomycota(100);c__Saccharomycetes(100);o__Saccharomycetales(100);f__Metschnikowiaceae(100);g__Metschnikowia(100);s__Metschnikowia_pulcherrima(100); | Metschnikowia | 13 | Saprotroph | \|Undefined Saprotroph\| | Yeast | NULL | Probable | As nectar-tap_saprotroph (Põlme et al. 2020); Decay sugar-rich substrates (Põlme et al. 2020) | Põlme S, et al. 2020. Fungal Diversity 105:1-16 ((https://doi.org/10.1007/s13225-020-00466-2)) |
| Otu19124 | 1 | k__Fungi(100);p__Ascomycota(100);c__Saccharomycetes(100);o__Saccharomycetales(100);f__Metschnikowiaceae(100);g__Metschnikowia(100);s__Metschnikowia_pulcherrima(100); | Metschnikowia | 13 | Saprotroph | \|Undefined Saprotroph\| | Yeast | NULL | Probable | As nectar-tap_saprotroph (Põlme et al. 2020); Decay sugar-rich substrates (Põlme et al. 2020) | Põlme S, et al. 2020. Fungal Diversity 105:1-16 ((https://doi.org/10.1007/s13225-020-00466-2)) |
| Otu19127 | 1 | k__Fungi(100);p__Ascomycota(100);c__Saccharomycetes(100);o__Saccharomycetales(100);f__Metschnikowiaceae(100);g__Metschnikowia(100);g__Metschnikowia_unclassified(100); | Metschnikowia | 13 | Saprotroph | \|Undefined Saprotroph\| | Yeast | NULL | Probable | As nectar-tap_saprotroph (Põlme et al. 2020); Decay sugar-rich substrates (Põlme et al. 2020) | Põlme S, et al. 2020. Fungal Diversity 105:1-16 ((https://doi.org/10.1007/s13225-020-00466-2)) |
| Otu19128 | 1 | k__Fungi(100);p__Ascomycota(100);c__Saccharomycetes(100);o__Saccharomycetales(100);f__Metschnikowiaceae(100);g__Metschnikowia(100);s__Metschnikowia_pulcherrima(100); | Metschnikowia | 13 | Saprotroph | \|Undefined Saprotroph\| | Yeast | NULL | Probable | As nectar-tap_saprotroph (Põlme et al. 2020); Decay sugar-rich substrates (Põlme et al. 2020) | Põlme S, et al. 2020. Fungal Diversity 105:1-16 ((https://doi.org/10.1007/s13225-020-00466-2)) |
| Otu19129 | 1 | k__Fungi(100);p__Ascomycota(100);c__Saccharomycetes(100);o__Saccharomycetales(100);f__Metschnikowiaceae(100);g__Metschnikowia(100);s__Metschnikowia_pulcherrima(100); | Metschnikowia | 13 | Saprotroph | \|Undefined Saprotroph\| | Yeast | NULL | Probable | As nectar-tap_saprotroph (Põlme et al. 2020); Decay sugar-rich substrates (Põlme et al. 2020) | Põlme S, et al. 2020. Fungal Diversity 105:1-16 ((https://doi.org/10.1007/s13225-020-00466-2)) |
| Otu19130 | 1 | k__Fungi(100);p__Ascomycota(100);c__Saccharomycetes(100);o__Saccharomycetales(100);f__Metschnikowiaceae(100);g__Metschnikowia(100);s__Metschnikowia_pulcherrima(100); | Metschnikowia | 13 | Saprotroph | \|Undefined Saprotroph\| | Yeast | NULL | Probable | As nectar-tap_saprotroph (Põlme et al. 2020); Decay sugar-rich substrates (Põlme et al. 2020) | Põlme S, et al. 2020. Fungal Diversity 105:1-16 ((https://doi.org/10.1007/s13225-020-00466-2)) |
| Otu19131 | 1 | k__Fungi(100);p__Ascomycota(100);c__Saccharomycetes(100);o__Saccharomycetales(100);f__Metschnikowiaceae(100);g__Metschnikowia(100);s__Metschnikowia_pulcherrima(100); | Metschnikowia | 13 | Saprotroph | \|Undefined Saprotroph\| | Yeast | NULL | Probable | As nectar-tap_saprotroph (Põlme et al. 2020); Decay sugar-rich substrates (Põlme et al. 2020) | Põlme S, et al. 2020. Fungal Diversity 105:1-16 ((https://doi.org/10.1007/s13225-020-00466-2)) |
| Otu19134 | 1 | k__Fungi(100);p__Ascomycota(100);c__Saccharomycetes(100);o__Saccharomycetales(100);f__Metschnikowiaceae(100);g__Metschnikowia(100);s__Metschnikowia_pulcherrima(100); | Metschnikowia | 13 | Saprotroph | \|Undefined Saprotroph\| | Yeast | NULL | Probable | As nectar-tap_saprotroph (Põlme et al. 2020); Decay sugar-rich substrates (Põlme et al. 2020) | Põlme S, et al. 2020. Fungal Diversity 105:1-16 ((https://doi.org/10.1007/s13225-020-00466-2)) |
| Otu19135 | 1 | k__Fungi(100);p__Ascomycota(100);c__Saccharomycetes(100);o__Saccharomycetales(100);f__Metschnikowiaceae(100);g__Metschnikowia(100);s__Metschnikowia_pulcherrima(100); | Metschnikowia | 13 | Saprotroph | \|Undefined Saprotroph\| | Yeast | NULL | Probable | As nectar-tap_saprotroph (Põlme et al. 2020); Decay sugar-rich substrates (Põlme et al. 2020) | Põlme S, et al. 2020. Fungal Diversity 105:1-16 ((https://doi.org/10.1007/s13225-020-00466-2)) |
| Otu19136 | 1 | k__Fungi(100);p__Ascomycota(100);c__Saccharomycetes(100);o__Saccharomycetales(100);f__Metschnikowiaceae(100);g__Metschnikowia(100);s__Metschnikowia_pulcherrima(100); | Metschnikowia | 13 | Saprotroph | \|Undefined Saprotroph\| | Yeast | NULL | Probable | As nectar-tap_saprotroph (Põlme et al. 2020); Decay sugar-rich substrates (Põlme et al. 2020) | Põlme S, et al. 2020. Fungal Diversity 105:1-16 ((https://doi.org/10.1007/s13225-020-00466-2)) |
| Otu19138 | 1 | k__Fungi(100);p__Ascomycota(100);c__Saccharomycetes(100);o__Saccharomycetales(100);f__Metschnikowiaceae(100);g__Metschnikowia(100);g__Metschnikowia_unclassified(100); | Metschnikowia | 13 | Saprotroph | \|Undefined Saprotroph\| | Yeast | NULL | Probable | As nectar-tap_saprotroph (Põlme et al. 2020); Decay sugar-rich substrates (Põlme et al. 2020) | Põlme S, et al. 2020. Fungal Diversity 105:1-16 ((https://doi.org/10.1007/s13225-020-00466-2)) |
| Otu19139 | 1 | k__Fungi(100);p__Ascomycota(100);c__Saccharomycetes(100);o__Saccharomycetales(100);f__Metschnikowiaceae(100);g__Metschnikowia(100);s__Metschnikowia_pulcherrima(100); | Metschnikowia | 13 | Saprotroph | \|Undefined Saprotroph\| | Yeast | NULL | Probable | As nectar-tap_saprotroph (Põlme et al. 2020); Decay sugar-rich substrates (Põlme et al. 2020) | Põlme S, et al. 2020. Fungal Diversity 105:1-16 ((https://doi.org/10.1007/s13225-020-00466-2)) |
| Otu19140 | 1 | k__Fungi(100);p__Ascomycota(100);c__Saccharomycetes(100);o__Saccharomycetales(100);f__Metschnikowiaceae(100);g__Metschnikowia(100);s__Metschnikowia_pulcherrima(100); | Metschnikowia | 13 | Saprotroph | \|Undefined Saprotroph\| | Yeast | NULL | Probable | As nectar-tap_saprotroph (Põlme et al. 2020); Decay sugar-rich substrates (Põlme et al. 2020) | Põlme S, et al. 2020. Fungal Diversity 105:1-16 ((https://doi.org/10.1007/s13225-020-00466-2)) |
| Otu19141 | 1 | k__Fungi(100);p__Ascomycota(100);c__Saccharomycetes(100);o__Saccharomycetales(100);f__Metschnikowiaceae(100);g__Metschnikowia(100);s__Metschnikowia_pulcherrima(100); | Metschnikowia | 13 | Saprotroph | \|Undefined Saprotroph\| | Yeast | NULL | Probable | As nectar-tap_saprotroph (Põlme et al. 2020); Decay sugar-rich substrates (Põlme et al. 2020) | Põlme S, et al. 2020. Fungal Diversity 105:1-16 ((https://doi.org/10.1007/s13225-020-00466-2)) |
| Otu19143 | 1 | k__Fungi(100);p__Ascomycota(100);c__Saccharomycetes(100);o__Saccharomycetales(100);f__Metschnikowiaceae(100);g__Metschnikowia(100);g__Metschnikowia_unclassified(100); | Metschnikowia | 13 | Saprotroph | \|Undefined Saprotroph\| | Yeast | NULL | Probable | As nectar-tap_saprotroph (Põlme et al. 2020); Decay sugar-rich substrates (Põlme et al. 2020) | Põlme S, et al. 2020. Fungal Diversity 105:1-16 ((https://doi.org/10.1007/s13225-020-00466-2)) |
| Otu19147 | 1 | k__Fungi(100);p__Ascomycota(100);c__Saccharomycetes(100);o__Saccharomycetales(100);f__Metschnikowiaceae(100);g__Metschnikowia(100);g__Metschnikowia_unclassified(100); | Metschnikowia | 13 | Saprotroph | \|Undefined Saprotroph\| | Yeast | NULL | Probable | As nectar-tap_saprotroph (Põlme et al. 2020); Decay sugar-rich substrates (Põlme et al. 2020) | Põlme S, et al. 2020. Fungal Diversity 105:1-16 ((https://doi.org/10.1007/s13225-020-00466-2)) |
| Otu19149 | 1 | k__Fungi(100);p__Ascomycota(100);c__Saccharomycetes(100);o__Saccharomycetales(100);f__Metschnikowiaceae(100);g__Metschnikowia(100);g__Metschnikowia_unclassified(100); | Metschnikowia | 13 | Saprotroph | \|Undefined Saprotroph\| | Yeast | NULL | Probable | As nectar-tap_saprotroph (Põlme et al. 2020); Decay sugar-rich substrates (Põlme et al. 2020) | Põlme S, et al. 2020. Fungal Diversity 105:1-16 ((https://doi.org/10.1007/s13225-020-00466-2)) |
| Otu19150 | 1 | k__Fungi(100);p__Ascomycota(100);c__Saccharomycetes(100);o__Saccharomycetales(100);f__Metschnikowiaceae(100);g__Metschnikowia(100);g__Metschnikowia_unclassified(100); | Metschnikowia | 13 | Saprotroph | \|Undefined Saprotroph\| | Yeast | NULL | Probable | As nectar-tap_saprotroph (Põlme et al. 2020); Decay sugar-rich substrates (Põlme et al. 2020) | Põlme S, et al. 2020. Fungal Diversity 105:1-16 ((https://doi.org/10.1007/s13225-020-00466-2)) |
| Otu19153 | 1 | k__Fungi(100);p__Ascomycota(100);c__Saccharomycetes(100);o__Saccharomycetales(100);f__Metschnikowiaceae(100);g__Metschnikowia(100);s__Metschnikowia_pulcherrima(100); | Metschnikowia | 13 | Saprotroph | \|Undefined Saprotroph\| | Yeast | NULL | Probable | As nectar-tap_saprotroph (Põlme et al. 2020); Decay sugar-rich substrates (Põlme et al. 2020) | Põlme S, et al. 2020. Fungal Diversity 105:1-16 ((https://doi.org/10.1007/s13225-020-00466-2)) |
| Otu19154 | 1 | k__Fungi(100);p__Ascomycota(100);c__Saccharomycetes(100);o__Saccharomycetales(100);f__Metschnikowiaceae(100);g__Metschnikowia(100);g__Metschnikowia_unclassified(100); | Metschnikowia | 13 | Saprotroph | \|Undefined Saprotroph\| | Yeast | NULL | Probable | As nectar-tap_saprotroph (Põlme et al. 2020); Decay sugar-rich substrates (Põlme et al. 2020) | Põlme S, et al. 2020. Fungal Diversity 105:1-16 ((https://doi.org/10.1007/s13225-020-00466-2)) |
| Otu19160 | 1 | k__Fungi(100);p__Ascomycota(100);c__Saccharomycetes(100);o__Saccharomycetales(100);f__Metschnikowiaceae(100);g__Metschnikowia(100);s__Metschnikowia_pulcherrima(100); | Metschnikowia | 13 | Saprotroph | \|Undefined Saprotroph\| | Yeast | NULL | Probable | As nectar-tap_saprotroph (Põlme et al. 2020); Decay sugar-rich substrates (Põlme et al. 2020) | Põlme S, et al. 2020. Fungal Diversity 105:1-16 ((https://doi.org/10.1007/s13225-020-00466-2)) |
| Otu19161 | 1 | k__Fungi(100);p__Ascomycota(100);c__Saccharomycetes(100);o__Saccharomycetales(100);f__Metschnikowiaceae(100);g__Metschnikowia(100);s__Metschnikowia_pulcherrima(100); | Metschnikowia | 13 | Saprotroph | \|Undefined Saprotroph\| | Yeast | NULL | Probable | As nectar-tap_saprotroph (Põlme et al. 2020); Decay sugar-rich substrates (Põlme et al. 2020) | Põlme S, et al. 2020. Fungal Diversity 105:1-16 ((https://doi.org/10.1007/s13225-020-00466-2)) |
| Otu19162 | 1 | k__Fungi(100);p__Ascomycota(100);c__Saccharomycetes(100);o__Saccharomycetales(100);f__Metschnikowiaceae(100);g__Metschnikowia(100);s__Metschnikowia_pulcherrima(100); | Metschnikowia | 13 | Saprotroph | \|Undefined Saprotroph\| | Yeast | NULL | Probable | As nectar-tap_saprotroph (Põlme et al. 2020); Decay sugar-rich substrates (Põlme et al. 2020) | Põlme S, et al. 2020. Fungal Diversity 105:1-16 ((https://doi.org/10.1007/s13225-020-00466-2)) |
| Otu19163 | 1 | k__Fungi(100);p__Ascomycota(100);c__Saccharomycetes(100);o__Saccharomycetales(100);f__Metschnikowiaceae(100);g__Metschnikowia(100);g__Metschnikowia_unclassified(100); | Metschnikowia | 13 | Saprotroph | \|Undefined Saprotroph\| | Yeast | NULL | Probable | As nectar-tap_saprotroph (Põlme et al. 2020); Decay sugar-rich substrates (Põlme et al. 2020) | Põlme S, et al. 2020. Fungal Diversity 105:1-16 ((https://doi.org/10.1007/s13225-020-00466-2)) |
| Otu19164 | 1 | k__Fungi(100);p__Ascomycota(100);c__Saccharomycetes(100);o__Saccharomycetales(100);f__Metschnikowiaceae(100);g__Metschnikowia(100);s__Metschnikowia_pulcherrima(100); | Metschnikowia | 13 | Saprotroph | \|Undefined Saprotroph\| | Yeast | NULL | Probable | As nectar-tap_saprotroph (Põlme et al. 2020); Decay sugar-rich substrates (Põlme et al. 2020) | Põlme S, et al. 2020. Fungal Diversity 105:1-16 ((https://doi.org/10.1007/s13225-020-00466-2)) |
| Otu19165 | 1 | k__Fungi(100);p__Ascomycota(100);c__Saccharomycetes(100);o__Saccharomycetales(100);f__Metschnikowiaceae(100);g__Metschnikowia(100);s__Metschnikowia_pulcherrima(100); | Metschnikowia | 13 | Saprotroph | \|Undefined Saprotroph\| | Yeast | NULL | Probable | As nectar-tap_saprotroph (Põlme et al. 2020); Decay sugar-rich substrates (Põlme et al. 2020) | Põlme S, et al. 2020. Fungal Diversity 105:1-16 ((https://doi.org/10.1007/s13225-020-00466-2)) |
| Otu19177 | 1 | k__Fungi(100);p__Ascomycota(100);c__Saccharomycetes(100);o__Saccharomycetales(100);f__Metschnikowiaceae(100);g__Metschnikowia(100);g__Metschnikowia_unclassified(100); | Metschnikowia | 13 | Saprotroph | \|Undefined Saprotroph\| | Yeast | NULL | Probable | As nectar-tap_saprotroph (Põlme et al. 2020); Decay sugar-rich substrates (Põlme et al. 2020) | Põlme S, et al. 2020. Fungal Diversity 105:1-16 ((https://doi.org/10.1007/s13225-020-00466-2)) |
| Otu19178 | 1 | k__Fungi(100);p__Ascomycota(100);c__Saccharomycetes(100);o__Saccharomycetales(100);f__Metschnikowiaceae(100);g__Metschnikowia(100);s__Metschnikowia_pulcherrima(100); | Metschnikowia | 13 | Saprotroph | \|Undefined Saprotroph\| | Yeast | NULL | Probable | As nectar-tap_saprotroph (Põlme et al. 2020); Decay sugar-rich substrates (Põlme et al. 2020) | Põlme S, et al. 2020. Fungal Diversity 105:1-16 ((https://doi.org/10.1007/s13225-020-00466-2)) |
| Otu19179 | 1 | k__Fungi(100);p__Ascomycota(100);c__Saccharomycetes(100);o__Saccharomycetales(100);f__Metschnikowiaceae(100);g__Metschnikowia(100);s__Metschnikowia_pulcherrima(100); | Metschnikowia | 13 | Saprotroph | \|Undefined Saprotroph\| | Yeast | NULL | Probable | As nectar-tap_saprotroph (Põlme et al. 2020); Decay sugar-rich substrates (Põlme et al. 2020) | Põlme S, et al. 2020. Fungal Diversity 105:1-16 ((https://doi.org/10.1007/s13225-020-00466-2)) |
| Otu19180 | 1 | k__Fungi(100);p__Ascomycota(100);c__Saccharomycetes(100);o__Saccharomycetales(100);f__Metschnikowiaceae(100);g__Metschnikowia(100);s__Metschnikowia_pulcherrima(100); | Metschnikowia | 13 | Saprotroph | \|Undefined Saprotroph\| | Yeast | NULL | Probable | As nectar-tap_saprotroph (Põlme et al. 2020); Decay sugar-rich substrates (Põlme et al. 2020) | Põlme S, et al. 2020. Fungal Diversity 105:1-16 ((https://doi.org/10.1007/s13225-020-00466-2)) |
| Otu19181 | 1 | k__Fungi(100);p__Ascomycota(100);c__Saccharomycetes(100);o__Saccharomycetales(100);f__Metschnikowiaceae(100);g__Metschnikowia(100);s__Metschnikowia_pulcherrima(100); | Metschnikowia | 13 | Saprotroph | \|Undefined Saprotroph\| | Yeast | NULL | Probable | As nectar-tap_saprotroph (Põlme et al. 2020); Decay sugar-rich substrates (Põlme et al. 2020) | Põlme S, et al. 2020. Fungal Diversity 105:1-16 ((https://doi.org/10.1007/s13225-020-00466-2)) |
| Otu19182 | 1 | k__Fungi(100);p__Ascomycota(100);c__Saccharomycetes(100);o__Saccharomycetales(100);f__Metschnikowiaceae(100);g__Metschnikowia(100);g__Metschnikowia_unclassified(100); | Metschnikowia | 13 | Saprotroph | \|Undefined Saprotroph\| | Yeast | NULL | Probable | As nectar-tap_saprotroph (Põlme et al. 2020); Decay sugar-rich substrates (Põlme et al. 2020) | Põlme S, et al. 2020. Fungal Diversity 105:1-16 ((https://doi.org/10.1007/s13225-020-00466-2)) |
| Otu19183 | 1 | k__Fungi(100);p__Ascomycota(100);c__Saccharomycetes(100);o__Saccharomycetales(100);f__Metschnikowiaceae(100);g__Metschnikowia(100);s__Metschnikowia_pulcherrima(100); | Metschnikowia | 13 | Saprotroph | \|Undefined Saprotroph\| | Yeast | NULL | Probable | As nectar-tap_saprotroph (Põlme et al. 2020); Decay sugar-rich substrates (Põlme et al. 2020) | Põlme S, et al. 2020. Fungal Diversity 105:1-16 ((https://doi.org/10.1007/s13225-020-00466-2)) |
| Otu19184 | 1 | k__Fungi(100);p__Ascomycota(100);c__Saccharomycetes(100);o__Saccharomycetales(100);f__Metschnikowiaceae(100);g__Metschnikowia(100);g__Metschnikowia_unclassified(100); | Metschnikowia | 13 | Saprotroph | \|Undefined Saprotroph\| | Yeast | NULL | Probable | As nectar-tap_saprotroph (Põlme et al. 2020); Decay sugar-rich substrates (Põlme et al. 2020) | Põlme S, et al. 2020. Fungal Diversity 105:1-16 ((https://doi.org/10.1007/s13225-020-00466-2)) |

| | | | | | | | | | | | |
|---|---|---|---|---|---|---|---|---|---|---|---|
| Otu19185 | 1 | k__Fungi(100);p__Ascomycota(100);c__Leotiomycetes(100);o__Helotiales(100);f__Helotiales_family_Incertae_sedis(100);g__Rhexocercosporidium(100);s__Rhexocercosporidium_panacis(100); | Rhexocercosporidium | 13 | Pathotroph-Saprotroph | \|Plant Pathogen-Plant Saprotroph\| | Microfungus | NULL | Probable | As litter_saprotroph (Põlme et al. 2020); As root_pathogen (Põlme et al. 2020); Decay leaf, fruit, seed, roots (Põlme et al. 2020) | Tedersoo L, et al. 2014. Science 346:e1256688 (https://doi.org/10.1126/science.1256688); Põlme S, et al. 2020. Fungal Diversity 105:1-16 (https://doi.org/10.1007/s13225-020-00466-2) |
| Otu19186 | 1 | k__Fungi(100);p__Ascomycota(100);c__Saccharomycetes(100);o__Saccharomycetales(100);f__Metschnikowiaceae(100);g__Metschnikowia(100);g__Metschnikowia_unclassified(100); | Metschnikowia | 13 | Saprotroph | \|Undefined Saprotroph\| | Yeast | NULL | Probable | As nectar-tap_saprotroph (Põlme et al. 2020); Decay sugar-rich substrates (Põlme et al. 2020) | Põlme S, et al. 2020. Fungal Diversity 105:1-16 (https://doi.org/10.1007/s13225-020-00466-2) |
| Otu19187 | 1 | k__Fungi(100);p__Ascomycota(100);c__Saccharomycetes(100);o__Saccharomycetales(100);f__Metschnikowiaceae(100);g__Metschnikowia(100);s__Metschnikowia_pulcherrima(100); | Metschnikowia | 13 | Saprotroph | \|Undefined Saprotroph\| | Yeast | NULL | Probable | As nectar-tap_saprotroph (Põlme et al. 2020); Decay sugar-rich substrates (Põlme et al. 2020) | Põlme S, et al. 2020. Fungal Diversity 105:1-16 (https://doi.org/10.1007/s13225-020-00466-2) |
| Otu19188 | 1 | k__Fungi(100);p__Ascomycota(100);c__Saccharomycetes(100);o__Saccharomycetales(100);f__Metschnikowiaceae(100);g__Metschnikowia(100);s__Metschnikowia_pulcherrima(100); | Metschnikowia | 13 | Saprotroph | \|Undefined Saprotroph\| | Yeast | NULL | Probable | As nectar-tap_saprotroph (Põlme et al. 2020); Decay sugar-rich substrates (Põlme et al. 2020) | Põlme S, et al. 2020. Fungal Diversity 105:1-16 (https://doi.org/10.1007/s13225-020-00466-2) |
| Otu19189 | 1 | k__Fungi(100);p__Ascomycota(100);c__Saccharomycetes(100);o__Saccharomycetales(100);f__Metschnikowiaceae(100);g__Metschnikowia(100);s__Metschnikowia_pulcherrima(100); | Metschnikowia | 13 | Saprotroph | \|Undefined Saprotroph\| | Yeast | NULL | Probable | As nectar-tap_saprotroph (Põlme et al. 2020); Decay sugar-rich substrates (Põlme et al. 2020) | Põlme S, et al. 2020. Fungal Diversity 105:1-16 (https://doi.org/10.1007/s13225-020-00466-2) |
| Otu19190 | 1 | k__Fungi(100);p__Ascomycota(100);c__Saccharomycetes(100);o__Saccharomycetales(100);f__Metschnikowiaceae(100);g__Metschnikowia(100);g__Metschnikowia_unclassified(100); | Metschnikowia | 13 | Saprotroph | \|Undefined Saprotroph\| | Yeast | NULL | Probable | As nectar-tap_saprotroph (Põlme et al. 2020); Decay sugar-rich substrates (Põlme et al. 2020) | Põlme S, et al. 2020. Fungal Diversity 105:1-16 (https://doi.org/10.1007/s13225-020-00466-2) |
| Otu19191 | 1 | k__Fungi(100);p__Ascomycota(100);c__Saccharomycetes(100);o__Saccharomycetales(100);f__Metschnikowiaceae(100);g__Metschnikowia(100);s__Metschnikowia_pulcherrima(100); | Metschnikowia | 13 | Saprotroph | \|Undefined Saprotroph\| | Yeast | NULL | Probable | As nectar-tap_saprotroph (Põlme et al. 2020); Decay sugar-rich substrates (Põlme et al. 2020) | Põlme S, et al. 2020. Fungal Diversity 105:1-16 (https://doi.org/10.1007/s13225-020-00466-2) |
| Otu19192 | 1 | k__Fungi(100);p__Ascomycota(100);c__Saccharomycetes(100);o__Saccharomycetales(100);f__Metschnikowiaceae(100);g__Metschnikowia(100);s__Metschnikowia_pulcherrima(100); | Metschnikowia | 13 | Saprotroph | \|Undefined Saprotroph\| | Yeast | NULL | Probable | As nectar-tap_saprotroph (Põlme et al. 2020); Decay sugar-rich substrates (Põlme et al. 2020) | Põlme S, et al. 2020. Fungal Diversity 105:1-16 (https://doi.org/10.1007/s13225-020-00466-2) |
| Otu19193 | 1 | k__Fungi(100);p__Ascomycota(100);c__Saccharomycetes(100);o__Saccharomycetales(100);f__Metschnikowiaceae(100);g__Metschnikowia(100);s__Metschnikowia_pulcherrima(100); | Metschnikowia | 13 | Saprotroph | \|Undefined Saprotroph\| | Yeast | NULL | Probable | As nectar-tap_saprotroph (Põlme et al. 2020); Decay sugar-rich substrates (Põlme et al. 2020) | Põlme S, et al. 2020. Fungal Diversity 105:1-16 (https://doi.org/10.1007/s13225-020-00466-2) |
| Otu19194 | 1 | k__Fungi(100);p__Ascomycota(100);c__Saccharomycetes(100);o__Saccharomycetales(100);f__Metschnikowiaceae(100);g__Metschnikowia(100);g__Metschnikowia_unclassified(100); | Metschnikowia | 13 | Saprotroph | \|Undefined Saprotroph\| | Yeast | NULL | Probable | As nectar-tap_saprotroph (Põlme et al. 2020); Decay sugar-rich substrates (Põlme et al. 2020) | Põlme S, et al. 2020. Fungal Diversity 105:1-16 (https://doi.org/10.1007/s13225-020-00466-2) |
| Otu19195 | 1 | k__Fungi(100);p__Ascomycota(100);c__Saccharomycetes(100);o__Saccharomycetales(100);f__Metschnikowiaceae(100);g__Metschnikowia(100);s__Metschnikowia_pulcherrima(100); | Metschnikowia | 13 | Saprotroph | \|Undefined Saprotroph\| | Yeast | NULL | Probable | As nectar-tap_saprotroph (Põlme et al. 2020); Decay sugar-rich substrates (Põlme et al. 2020) | Põlme S, et al. 2020. Fungal Diversity 105:1-16 (https://doi.org/10.1007/s13225-020-00466-2) |
| Otu19196 | 1 | k__Fungi(100);p__Ascomycota(100);c__Saccharomycetes(100);o__Saccharomycetales(100);f__Metschnikowiaceae(100);g__Metschnikowia(100);g__Metschnikowia_unclassified(100); | Metschnikowia | 13 | Saprotroph | \|Undefined Saprotroph\| | Yeast | NULL | Probable | As nectar-tap_saprotroph (Põlme et al. 2020); Decay sugar-rich substrates (Põlme et al. 2020) | Põlme S, et al. 2020. Fungal Diversity 105:1-16 (https://doi.org/10.1007/s13225-020-00466-2) |
| Otu19197 | 1 | k__Fungi(100);p__Ascomycota(100);c__Saccharomycetes(100);o__Saccharomycetales(100);f__Metschnikowiaceae(100);g__Metschnikowia(100);s__Metschnikowia_pulcherrima(100); | Metschnikowia | 13 | Saprotroph | \|Undefined Saprotroph\| | Yeast | NULL | Probable | As nectar-tap_saprotroph (Põlme et al. 2020); Decay sugar-rich substrates (Põlme et al. 2020) | Põlme S, et al. 2020. Fungal Diversity 105:1-16 (https://doi.org/10.1007/s13225-020-00466-2) |
| Otu19198 | 1 | k__Fungi(100);p__Ascomycota(100);c__Saccharomycetes(100);o__Saccharomycetales(100);f__Metschnikowiaceae(100);g__Metschnikowia(100);g__Metschnikowia_unclassified(100); | Metschnikowia | 13 | Saprotroph | \|Undefined Saprotroph\| | Yeast | NULL | Probable | As nectar-tap_saprotroph (Põlme et al. 2020); Decay sugar-rich substrates (Põlme et al. 2020) | Põlme S, et al. 2020. Fungal Diversity 105:1-16 (https://doi.org/10.1007/s13225-020-00466-2) |
| Otu19199 | 1 | k__Fungi(100);p__Ascomycota(100);c__Saccharomycetes(100);o__Saccharomycetales(100);f__Metschnikowiaceae(100);g__Metschnikowia(100);s__Metschnikowia_pulcherrima(100); | Metschnikowia | 13 | Saprotroph | \|Undefined Saprotroph\| | Yeast | NULL | Probable | As nectar-tap_saprotroph (Põlme et al. 2020); Decay sugar-rich substrates (Põlme et al. 2020) | Põlme S, et al. 2020. Fungal Diversity 105:1-16 (https://doi.org/10.1007/s13225-020-00466-2) |
| Otu19200 | 1 | k__Fungi(100);p__Ascomycota(100);c__Saccharomycetes(100);o__Saccharomycetales(100);f__Metschnikowiaceae(100);g__Metschnikowia(100);s__Metschnikowia_pulcherrima(100); | Metschnikowia | 13 | Saprotroph | \|Undefined Saprotroph\| | Yeast | NULL | Probable | As nectar-tap_saprotroph (Põlme et al. 2020); Decay sugar-rich substrates (Põlme et al. 2020) | Põlme S, et al. 2020. Fungal Diversity 105:1-16 (https://doi.org/10.1007/s13225-020-00466-2) |
| Otu19201 | 1 | k__Fungi(100);p__Ascomycota(100);c__Saccharomycetes(100);o__Saccharomycetales(100);f__Metschnikowiaceae(100);g__Metschnikowia(100);s__Metschnikowia_pulcherrima(100); | Metschnikowia | 13 | Saprotroph | \|Undefined Saprotroph\| | Yeast | NULL | Probable | As nectar-tap_saprotroph (Põlme et al. 2020); Decay sugar-rich substrates (Põlme et al. 2020) | Põlme S, et al. 2020. Fungal Diversity 105:1-16 (https://doi.org/10.1007/s13225-020-00466-2) |
| Otu19202 | 1 | k__Fungi(100);p__Ascomycota(100);c__Saccharomycetes(100);o__Saccharomycetales(100);f__Metschnikowiaceae(100);g__Metschnikowia(100);s__Metschnikowia_pulcherrima(100); | Metschnikowia | 13 | Saprotroph | \|Undefined Saprotroph\| | Yeast | NULL | Probable | As nectar-tap_saprotroph (Põlme et al. 2020); Decay sugar-rich substrates (Põlme et al. 2020) | Põlme S, et al. 2020. Fungal Diversity 105:1-16 (https://doi.org/10.1007/s13225-020-00466-2) |
| Otu19203 | 1 | k__Fungi(100);p__Ascomycota(100);c__Saccharomycetes(100);o__Saccharomycetales(100);f__Metschnikowiaceae(100);g__Metschnikowia(100);s__Metschnikowia_pulcherrima(100); | Metschnikowia | 13 | Saprotroph | \|Undefined Saprotroph\| | Yeast | NULL | Probable | As nectar-tap_saprotroph (Põlme et al. 2020); Decay sugar-rich substrates (Põlme et al. 2020) | Põlme S, et al. 2020. Fungal Diversity 105:1-16 (https://doi.org/10.1007/s13225-020-00466-2) |
| Otu19204 | 1 | k__Fungi(100);p__Ascomycota(100);c__Saccharomycetes(100);o__Saccharomycetales(100);f__Metschnikowiaceae(100);g__Metschnikowia(100);s__Metschnikowia_pulcherrima(100); | Metschnikowia | 13 | Saprotroph | \|Undefined Saprotroph\| | Yeast | NULL | Probable | As nectar-tap_saprotroph (Põlme et al. 2020); Decay sugar-rich substrates (Põlme et al. 2020) | Põlme S, et al. 2020. Fungal Diversity 105:1-16 (https://doi.org/10.1007/s13225-020-00466-2) |
| Otu19205 | 1 | k__Fungi(100);p__Ascomycota(100);c__Saccharomycetes(100);o__Saccharomycetales(100);f__Metschnikowiaceae(100);g__Metschnikowia(100);s__Metschnikowia_pulcherrima(100); | Metschnikowia | 13 | Saprotroph | \|Undefined Saprotroph\| | Yeast | NULL | Probable | As nectar-tap_saprotroph (Põlme et al. 2020); Decay sugar-rich substrates (Põlme et al. 2020) | Põlme S, et al. 2020. Fungal Diversity 105:1-16 (https://doi.org/10.1007/s13225-020-00466-2) |
| Otu19206 | 1 | k__Fungi(100);p__Ascomycota(100);c__Saccharomycetes(100);o__Saccharomycetales(100);f__Metschnikowiaceae(100);g__Metschnikowia(100);s__Metschnikowia_pulcherrima(100); | Metschnikowia | 13 | Saprotroph | \|Undefined Saprotroph\| | Yeast | NULL | Probable | As nectar-tap_saprotroph (Põlme et al. 2020); Decay sugar-rich substrates (Põlme et al. 2020) | Põlme S, et al. 2020. Fungal Diversity 105:1-16 (https://doi.org/10.1007/s13225-020-00466-2) |
| Otu19207 | 1 | k__Fungi(100);p__Ascomycota(100);c__Saccharomycetes(100);o__Saccharomycetales(100);f__Metschnikowiaceae(100);g__Metschnikowia(100);s__Metschnikowia_pulcherrima(100); | Metschnikowia | 13 | Saprotroph | \|Undefined Saprotroph\| | Yeast | NULL | Probable | As nectar-tap_saprotroph (Põlme et al. 2020); Decay sugar-rich substrates (Põlme et al. 2020) | Põlme S, et al. 2020. Fungal Diversity 105:1-16 (https://doi.org/10.1007/s13225-020-00466-2) |
| Otu19208 | 1 | k__Fungi(100);p__Ascomycota(100);c__Saccharomycetes(100);o__Saccharomycetales(100);f__Metschnikowiaceae(100);g__Metschnikowia(100);s__Metschnikowia_pulcherrima(100); | Metschnikowia | 13 | Saprotroph | \|Undefined Saprotroph\| | Yeast | NULL | Probable | As nectar-tap_saprotroph (Põlme et al. 2020); Decay sugar-rich substrates (Põlme et al. 2020) | Põlme S, et al. 2020. Fungal Diversity 105:1-16 (https://doi.org/10.1007/s13225-020-00466-2) |
| Otu19209 | 1 | k__Fungi(100);p__Ascomycota(100);c__Saccharomycetes(100);o__Saccharomycetales(100);f__Metschnikowiaceae(100);g__Metschnikowia(100);s__Metschnikowia_pulcherrima(100); | Metschnikowia | 13 | Saprotroph | \|Undefined Saprotroph\| | Yeast | NULL | Probable | As nectar-tap_saprotroph (Põlme et al. 2020); Decay sugar-rich substrates (Põlme et al. 2020) | Põlme S, et al. 2020. Fungal Diversity 105:1-16 (https://doi.org/10.1007/s13225-020-00466-2) |
| Otu19210 | 1 | k__Fungi(100);p__Ascomycota(100);c__Saccharomycetes(100);o__Saccharomycetales(100);f__Metschnikowiaceae(100);g__Metschnikowia(100);g__Metschnikowia_unclassified(100); | Metschnikowia | 13 | Saprotroph | \|Undefined Saprotroph\| | Yeast | NULL | Probable | As nectar-tap_saprotroph (Põlme et al. 2020); Decay sugar-rich substrates (Põlme et al. 2020) | Põlme S, et al. 2020. Fungal Diversity 105:1-16 (https://doi.org/10.1007/s13225-020-00466-2) |
| Otu19211 | 1 | k__Fungi(100);p__Ascomycota(100);c__Saccharomycetes(100);o__Saccharomycetales(100);f__Metschnikowiaceae(100);g__Metschnikowia(100);s__Metschnikowia_pulcherrima(100); | Metschnikowia | 13 | Saprotroph | \|Undefined Saprotroph\| | Yeast | NULL | Probable | As nectar-tap_saprotroph (Põlme et al. 2020); Decay sugar-rich substrates (Põlme et al. 2020) | Põlme S, et al. 2020. Fungal Diversity 105:1-16 (https://doi.org/10.1007/s13225-020-00466-2) |
| Otu19212 | 1 | k__Fungi(100);p__Ascomycota(100);c__Saccharomycetes(100);o__Saccharomycetales(100);f__Metschnikowiaceae(100);g__Metschnikowia(100);s__Metschnikowia_pulcherrima(100); | Metschnikowia | 13 | Saprotroph | \|Undefined Saprotroph\| | Yeast | NULL | Probable | As nectar-tap_saprotroph (Põlme et al. 2020); Decay sugar-rich substrates (Põlme et al. 2020) | Põlme S, et al. 2020. Fungal Diversity 105:1-16 (https://doi.org/10.1007/s13225-020-00466-2) |
| Otu19213 | 1 | k__Fungi(100);p__Ascomycota(100);c__Saccharomycetes(100);o__Saccharomycetales(100);f__Metschnikowiaceae(100);g__Metschnikowia(100);s__Metschnikowia_pulcherrima(100); | Metschnikowia | 13 | Saprotroph | \|Undefined Saprotroph\| | Yeast | NULL | Probable | As nectar-tap_saprotroph (Põlme et al. 2020); Decay sugar-rich substrates (Põlme et al. 2020) | Põlme S, et al. 2020. Fungal Diversity 105:1-16 (https://doi.org/10.1007/s13225-020-00466-2) |
| Otu19214 | 1 | k__Fungi(100);p__Ascomycota(100);c__Saccharomycetes(100);o__Saccharomycetales(100);f__Metschnikowiaceae(100);g__Metschnikowia(100);g__Metschnikowia_unclassified(100); | Metschnikowia | 13 | Saprotroph | \|Undefined Saprotroph\| | Yeast | NULL | Probable | As nectar-tap_saprotroph (Põlme et al. 2020); Decay sugar-rich substrates (Põlme et al. 2020) | Põlme S, et al. 2020. Fungal Diversity 105:1-16 (https://doi.org/10.1007/s13225-020-00466-2) |
| Otu19215 | 1 | k__Fungi(100);p__Ascomycota(100);c__Saccharomycetes(100);o__Saccharomycetales(100);f__Metschnikowiaceae(100);g__Metschnikowia(100);s__Metschnikowia_pulcherrima(100); | Metschnikowia | 13 | Saprotroph | \|Undefined Saprotroph\| | Yeast | NULL | Probable | As nectar-tap_saprotroph (Põlme et al. 2020); Decay sugar-rich substrates (Põlme et al. 2020) | Põlme S, et al. 2020. Fungal Diversity 105:1-16 (https://doi.org/10.1007/s13225-020-00466-2) |
| Otu19216 | 1 | k__Fungi(100);p__Ascomycota(100);c__Saccharomycetes(100);o__Saccharomycetales(100);f__Metschnikowiaceae(100);g__Metschnikowia(100);g__Metschnikowia_unclassified(100); | Metschnikowia | 13 | Saprotroph | \|Undefined Saprotroph\| | Yeast | NULL | Probable | As nectar-tap_saprotroph (Põlme et al. 2020); Decay sugar-rich substrates (Põlme et al. 2020) | Põlme S, et al. 2020. Fungal Diversity 105:1-16 (https://doi.org/10.1007/s13225-020-00466-2) |
| Otu19217 | 1 | k__Fungi(100);p__Ascomycota(100);c__Saccharomycetes(100);o__Saccharomycetales(100);f__Metschnikowiaceae(100);g__Metschnikowia(100);s__Metschnikowia_pulcherrima(100); | Metschnikowia | 13 | Saprotroph | \|Undefined Saprotroph\| | Yeast | NULL | Probable | As nectar-tap_saprotroph (Põlme et al. 2020); Decay sugar-rich substrates (Põlme et al. 2020) | Põlme S, et al. 2020. Fungal Diversity 105:1-16 (https://doi.org/10.1007/s13225-020-00466-2) |
| Otu19221 | 1 | k__Fungi(100);p__Ascomycota(100);c__Saccharomycetes(100);o__Saccharomycetales(100);f__Metschnikowiaceae(100);g__Metschnikowia(100);s__Metschnikowia_pulcherrima(100); | Metschnikowia | 13 | Saprotroph | \|Undefined Saprotroph\| | Yeast | NULL | Probable | As nectar-tap_saprotroph (Põlme et al. 2020); Decay sugar-rich substrates (Põlme et al. 2020) | Põlme S, et al. 2020. Fungal Diversity 105:1-16 (https://doi.org/10.1007/s13225-020-00466-2) |
| Otu19222 | 1 | k__Fungi(100);p__Ascomycota(100);c__Saccharomycetes(100);o__Saccharomycetales(100);f__Metschnikowiaceae(100);g__Metschnikowia(100);s__Metschnikowia_pulcherrima(100); | Metschnikowia | 13 | Saprotroph | \|Undefined Saprotroph\| | Yeast | NULL | Probable | As nectar-tap_saprotroph (Põlme et al. 2020); Decay sugar-rich substrates (Põlme et al. 2020) | Põlme S, et al. 2020. Fungal Diversity 105:1-16 (https://doi.org/10.1007/s13225-020-00466-2) |
| Otu19223 | 1 | k__Fungi(100);p__Ascomycota(100);c__Saccharomycetes(100);o__Saccharomycetales(100);f__Metschnikowiaceae(100);g__Metschnikowia(100);g__Metschnikowia_unclassified(100); | Metschnikowia | 13 | Saprotroph | \|Undefined Saprotroph\| | Yeast | NULL | Probable | As nectar-tap_saprotroph (Põlme et al. 2020); Decay sugar-rich substrates (Põlme et al. 2020) | Põlme S, et al. 2020. Fungal Diversity 105:1-16 (https://doi.org/10.1007/s13225-020-00466-2) |
| Otu19224 | 1 | k__Fungi(100);p__Ascomycota(100);c__Saccharomycetes(100);o__Saccharomycetales(100);f__Metschnikowiaceae(100);g__Metschnikowia(100);g__Metschnikowia_unclassified(100); | Metschnikowia | 13 | Saprotroph | \|Undefined Saprotroph\| | Yeast | NULL | Probable | As nectar-tap_saprotroph (Põlme et al. 2020); Decay sugar-rich substrates (Põlme et al. 2020) | Põlme S, et al. 2020. Fungal Diversity 105:1-16 (https://doi.org/10.1007/s13225-020-00466-2) |
| Otu19225 | 1 | k__Fungi(100);p__Ascomycota(100);c__Saccharomycetes(100);o__Saccharomycetales(100);f__Metschnikowiaceae(100);g__Metschnikowia(100);s__Metschnikowia_pulcherrima(100); | Metschnikowia | 13 | Saprotroph | \|Undefined Saprotroph\| | Yeast | NULL | Probable | As nectar-tap_saprotroph (Põlme et al. 2020); Decay sugar-rich substrates (Põlme et al. 2020) | Põlme S, et al. 2020. Fungal Diversity 105:1-16 (https://doi.org/10.1007/s13225-020-00466-2) |
| Otu19226 | 1 | k__Fungi(100);p__Ascomycota(100);c__Saccharomycetes(100);o__Saccharomycetales(100);f__Metschnikowiaceae(100);g__Metschnikowia(100);s__Metschnikowia_pulcherrima(100); | Metschnikowia | 13 | Saprotroph | \|Undefined Saprotroph\| | Yeast | NULL | Probable | As nectar-tap_saprotroph (Põlme et al. 2020); Decay sugar-rich substrates (Põlme et al. 2020) | Põlme S, et al. 2020. Fungal Diversity 105:1-16 (https://doi.org/10.1007/s13225-020-00466-2) |
| Otu19227 | 1 | k__Fungi(100);p__Ascomycota(100);c__Saccharomycetes(100);o__Saccharomycetales(100);f__Metschnikowiaceae(100);g__Metschnikowia(100);s__Metschnikowia_pulcherrima(100); | Metschnikowia | 13 | Saprotroph | \|Undefined Saprotroph\| | Yeast | NULL | Probable | As nectar-tap_saprotroph (Põlme et al. 2020); Decay sugar-rich substrates (Põlme et al. 2020) | Põlme S, et al. 2020. Fungal Diversity 105:1-16 (https://doi.org/10.1007/s13225-020-00466-2) |
| Otu19228 | 1 | k__Fungi(100);p__Ascomycota(100);c__Saccharomycetes(100);o__Saccharomycetales(100);f__Metschnikowiaceae(100);g__Metschnikowia(100);s__Metschnikowia_pulcherrima(100); | Metschnikowia | 13 | Saprotroph | \|Undefined Saprotroph\| | Yeast | NULL | Probable | As nectar-tap_saprotroph (Põlme et al. 2020); Decay sugar-rich substrates (Põlme et al. 2020) | Põlme S, et al. 2020. Fungal Diversity 105:1-16 (https://doi.org/10.1007/s13225-020-00466-2) |
| Otu19229 | 1 | k__Fungi(100);p__Ascomycota(100);c__Saccharomycetes(100);o__Saccharomycetales(100);f__Metschnikowiaceae(100);g__Metschnikowia(100);s__Metschnikowia_pulcherrima(100); | Metschnikowia | 13 | Saprotroph | \|Undefined Saprotroph\| | Yeast | NULL | Probable | As nectar-tap_saprotroph (Põlme et al. 2020); Decay sugar-rich substrates (Põlme et al. 2020) | Põlme S, et al. 2020. Fungal Diversity 105:1-16 (https://doi.org/10.1007/s13225-020-00466-2) |
| Otu19230 | 1 | k__Fungi(100);p__Ascomycota(100);c__Saccharomycetes(100);o__Saccharomycetales(100);f__Metschnikowiaceae(100);g__Metschnikowia(100);s__Metschnikowia_pulcherrima(100); | Metschnikowia | 13 | Saprotroph | \|Undefined Saprotroph\| | Yeast | NULL | Probable | As nectar-tap_saprotroph (Põlme et al. 2020); Decay sugar-rich substrates (Põlme et al. 2020) | Põlme S, et al. 2020. Fungal Diversity 105:1-16 (https://doi.org/10.1007/s13225-020-00466-2) |
| Otu19231 | 1 | k__Fungi(100);p__Ascomycota(100);c__Saccharomycetes(100);o__Saccharomycetales(100);f__Metschnikowiaceae(100);g__Metschnikowia(100);s__Metschnikowia_pulcherrima(100); | Metschnikowia | 13 | Saprotroph | \|Undefined Saprotroph\| | Yeast | NULL | Probable | As nectar-tap_saprotroph (Põlme et al. 2020); Decay sugar-rich substrates (Põlme et al. 2020) | Põlme S, et al. 2020. Fungal Diversity 105:1-16 (https://doi.org/10.1007/s13225-020-00466-2) |
| Otu19232 | 1 | k__Fungi(100);p__Ascomycota(100);c__Saccharomycetes(100);o__Saccharomycetales(100);f__Metschnikowiaceae(100);g__Metschnikowia(100);s__Metschnikowia_pulcherrima(100); | Metschnikowia | 13 | Saprotroph | \|Undefined Saprotroph\| | Yeast | NULL | Probable | As nectar-tap_saprotroph (Põlme et al. 2020); Decay sugar-rich substrates (Põlme et al. 2020) | Põlme S, et al. 2020. Fungal Diversity 105:1-16 (https://doi.org/10.1007/s13225-020-00466-2) |
| Otu19233 | 1 | k__Fungi(100);p__Ascomycota(100);c__Saccharomycetes(100);o__Saccharomycetales(100);f__Metschnikowiaceae(100);g__Metschnikowia(100);s__Metschnikowia_pulcherrima(100); | Metschnikowia | 13 | Saprotroph | \|Undefined Saprotroph\| | Yeast | NULL | Probable | As nectar-tap_saprotroph (Põlme et al. 2020); Decay sugar-rich substrates (Põlme et al. 2020) | Põlme S, et al. 2020. Fungal Diversity 105:1-16 (https://doi.org/10.1007/s13225-020-00466-2) |
| Otu19234 | 1 | k__Fungi(100);p__Ascomycota(100);c__Saccharomycetes(100);o__Saccharomycetales(100);f__Metschnikowiaceae(100);g__Metschnikowia(100);s__Metschnikowia_pulcherrima(100); | Metschnikowia | 13 | Saprotroph | \|Undefined Saprotroph\| | Yeast | NULL | Probable | As nectar-tap_saprotroph (Põlme et al. 2020); Decay sugar-rich substrates (Põlme et al. 2020) | Põlme S, et al. 2020. Fungal Diversity 105:1-16 (https://doi.org/10.1007/s13225-020-00466-2) |
| Otu19235 | 1 | k__Fungi(100);p__Ascomycota(100);c__Saccharomycetes(100);o__Saccharomycetales(100);f__Metschnikowiaceae(100);g__Metschnikowia(100);s__Metschnikowia_pulcherrima(100); | Metschnikowia | 13 | Saprotroph | \|Undefined Saprotroph\| | Yeast | NULL | Probable | As nectar-tap_saprotroph (Põlme et al. 2020); Decay sugar-rich substrates (Põlme et al. 2020) | Põlme S, et al. 2020. Fungal Diversity 105:1-16 (https://doi.org/10.1007/s13225-020-00466-2) |
| Otu19236 | 1 | k__Fungi(100);p__Ascomycota(100);c__Saccharomycetes(100);o__Saccharomycetales(100);f__Metschnikowiaceae(100);g__Metschnikowia(100);s__Metschnikowia_pulcherrima(100); | Metschnikowia | 13 | Saprotroph | \|Undefined Saprotroph\| | Yeast | NULL | Probable | As nectar-tap_saprotroph (Põlme et al. 2020); Decay sugar-rich substrates (Põlme et al. 2020) | Põlme S, et al. 2020. Fungal Diversity 105:1-16 (https://doi.org/10.1007/s13225-020-00466-2) |
| Otu19237 | 1 | k__Fungi(100);p__Ascomycota(100);c__Saccharomycetes(100);o__Saccharomycetales(100);f__Metschnikowiaceae(100);g__Metschnikowia(100);s__Metschnikowia_pulcherrima(100); | Metschnikowia | 13 | Saprotroph | \|Undefined Saprotroph\| | Yeast | NULL | Probable | As nectar-tap_saprotroph (Põlme et al. 2020); Decay sugar-rich substrates (Põlme et al. 2020) | Põlme S, et al. 2020. Fungal Diversity 105:1-16 (https://doi.org/10.1007/s13225-020-00466-2) |
| Otu19238 | 1 | k__Fungi(100);p__Ascomycota(100);c__Saccharomycetes(100);o__Saccharomycetales(100);f__Metschnikowiaceae(100);g__Metschnikowia(100);s__Metschnikowia_pulcherrima(100); | Metschnikowia | 13 | Saprotroph | \|Undefined Saprotroph\| | Yeast | NULL | Probable | As nectar-tap_saprotroph (Põlme et al. 2020); Decay sugar-rich substrates (Põlme et al. 2020) | Põlme S, et al. 2020. Fungal Diversity 105:1-16 (https://doi.org/10.1007/s13225-020-00466-2) |
| Otu19239 | 1 | k__Fungi(100);p__Ascomycota(100);c__Saccharomycetes(100);o__Saccharomycetales(100);f__Metschnikowiaceae(100);g__Metschnikowia(100);g__Metschnikowia_unclassified(100); | Metschnikowia | 13 | Saprotroph | \|Undefined Saprotroph\| | Yeast | NULL | Probable | As nectar-tap_saprotroph (Põlme et al. 2020); Decay sugar-rich substrates (Põlme et al. 2020) | Põlme S, et al. 2020. Fungal Diversity 105:1-16 (https://doi.org/10.1007/s13225-020-00466-2) |
| Otu19240 | 1 | k__Fungi(100);p__Ascomycota(100);c__Saccharomycetes(100);o__Saccharomycetales(100);f__Metschnikowiaceae(100);g__Metschnikowia(100);s__Metschnikowia_pulcherrima(100); | Metschnikowia | 13 | Saprotroph | \|Undefined Saprotroph\| | Yeast | NULL | Probable | As nectar-tap_saprotroph (Põlme et al. 2020); Decay sugar-rich substrates (Põlme et al. 2020) | Põlme S, et al. 2020. Fungal Diversity 105:1-16 (https://doi.org/10.1007/s13225-020-00466-2) |
| Otu19241 | 1 | k__Fungi(100);p__Ascomycota(100);c__Saccharomycetes(100);o__Saccharomycetales(100);f__Metschnikowiaceae(100);g__Metschnikowia(100);s__Metschnikowia_pulcherrima(100); | Metschnikowia | 13 | Saprotroph | \|Undefined Saprotroph\| | Yeast | NULL | Probable | As nectar-tap_saprotroph (Põlme et al. 2020); Decay sugar-rich substrates (Põlme et al. 2020) | Põlme S, et al. 2020. Fungal Diversity 105:1-16 (https://doi.org/10.1007/s13225-020-00466-2) |
| Otu19242 | 1 | k__Fungi(100);p__Ascomycota(100);c__Saccharomycetes(100);o__Saccharomycetales(100);f__Metschnikowiaceae(100);g__Metschnikowia(100);s__Metschnikowia_pulcherrima(100); | Metschnikowia | 13 | Saprotroph | \|Undefined Saprotroph\| | Yeast | NULL | Probable | As nectar-tap_saprotroph (Põlme et al. 2020); Decay sugar-rich substrates (Põlme et al. 2020) | Põlme S, et al. 2020. Fungal Diversity 105:1-16 (https://doi.org/10.1007/s13225-020-00466-2) |
| Otu19243 | 1 | k__Fungi(100);p__Ascomycota(100);c__Saccharomycetes(100);o__Saccharomycetales(100);f__Metschnikowiaceae(100);g__Metschnikowia(100);s__Metschnikowia_pulcherrima(100); | Metschnikowia | 13 | Saprotroph | \|Undefined Saprotroph\| | Yeast | NULL | Probable | As nectar-tap_saprotroph (Põlme et al. 2020); Decay sugar-rich substrates (Põlme et al. 2020) | Põlme S, et al. 2020. Fungal Diversity 105:1-16 (https://doi.org/10.1007/s13225-020-00466-2) |
| Otu19244 | 1 | k__Fungi(100);p__Ascomycota(100);c__Saccharomycetes(100);o__Saccharomycetales(100);f__Metschnikowiaceae(100);g__Metschnikowia(100);s__Metschnikowia_pulcherrima(100); | Metschnikowia | 13 | Saprotroph | \|Undefined Saprotroph\| | Yeast | NULL | Probable | As nectar-tap_saprotroph (Põlme et al. 2020); Decay sugar-rich substrates (Põlme et al. 2020) | Põlme S, et al. 2020. Fungal Diversity 105:1-16 (https://doi.org/10.1007/s13225-020-00466-2) |
| Otu19245 | 1 | k__Fungi(100);p__Ascomycota(100);c__Saccharomycetes(100);o__Saccharomycetales(100);f__Metschnikowiaceae(100);g__Metschnikowia(100);s__Metschnikowia_pulcherrima(100); | Metschnikowia | 13 | Saprotroph | \|Undefined Saprotroph\| | Yeast | NULL | Probable | As nectar-tap_saprotroph (Põlme et al. 2020); Decay sugar-rich substrates (Põlme et al. 2020) | Põlme S, et al. 2020. Fungal Diversity 105:1-16 (https://doi.org/10.1007/s13225-020-00466-2) |
| Otu19247 | 1 | k__Fungi(100);p__Ascomycota(100);c__Saccharomycetes(100);o__Saccharomycetales(100);f__Metschnikowiaceae(100);g__Metschnikowia(100);s__Metschnikowia_pulcherrima(100); | Metschnikowia | 13 | Saprotroph | \|Undefined Saprotroph\| | Yeast | NULL | Probable | As nectar-tap_saprotroph (Põlme et al. 2020); Decay sugar-rich substrates (Põlme et al. 2020) | Põlme S, et al. 2020. Fungal Diversity 105:1-16 (https://doi.org/10.1007/s13225-020-00466-2) |
| Otu19249 | 1 | k__Fungi(100);p__Ascomycota(100);c__Saccharomycetes(100);o__Saccharomycetales(100);f__Metschnikowiaceae(100);g__Metschnikowia(100);s__Metschnikowia_pulcherrima(100); | Metschnikowia | 13 | Saprotroph | \|Undefined Saprotroph\| | Yeast | NULL | Probable | As nectar-tap_saprotroph (Põlme et al. 2020); Decay sugar-rich substrates (Põlme et al. 2020) | Põlme S, et al. 2020. Fungal Diversity 105:1-16 (https://doi.org/10.1007/s13225-020-00466-2) |
| Otu19250 | 1 | k__Fungi(100);p__Ascomycota(100);c__Saccharomycetes(100);o__Saccharomycetales(100);f__Metschnikowiaceae(100);g__Metschnikowia(100);s__Metschnikowia_pulcherrima(100); | Metschnikowia | 13 | Saprotroph | \|Undefined Saprotroph\| | Yeast | NULL | Probable | As nectar-tap_saprotroph (Põlme et al. 2020); Decay sugar-rich substrates (Põlme et al. 2020) | Põlme S, et al. 2020. Fungal Diversity 105:1-16 (https://doi.org/10.1007/s13225-020-00466-2) |
| Otu19254 | 1 | k__Fungi(100);p__Ascomycota(100);c__Saccharomycetes(100);o__Saccharomycetales(100);f__Metschnikowiaceae(100);g__Metschnikowia(100);g__Metschnikowia_unclassified(100); | Metschnikowia | 13 | Saprotroph | \|Undefined Saprotroph\| | Yeast | NULL | Probable | As nectar-tap_saprotroph (Põlme et al. 2020); Decay sugar-rich substrates (Põlme et al. 2020) | Põlme S, et al. 2020. Fungal Diversity 105:1-16 (https://doi.org/10.1007/s13225-020-00466-2) |
| Otu19255 | 1 | k__Fungi(100);p__Ascomycota(100);c__Saccharomycetes(100);o__Saccharomycetales(100);f__Metschnikowiaceae(100);g__Metschnikowia(100);s__Metschnikowia_pulcherrima(100); | Metschnikowia | 13 | Saprotroph | \|Undefined Saprotroph\| | Yeast | NULL | Probable | As nectar-tap_saprotroph (Põlme et al. 2020); Decay sugar-rich substrates (Põlme et al. 2020) | Põlme S, et al. 2020. Fungal Diversity 105:1-16 (https://doi.org/10.1007/s13225-020-00466-2) |
| Otu19256 | 1 | k__Fungi(100);p__Ascomycota(100);c__Saccharomycetes(100);o__Saccharomycetales(100);f__Metschnikowiaceae(100);g__Metschnikowia(100);s__Metschnikowia_pulcherrima(100); | Metschnikowia | 13 | Saprotroph | \|Undefined Saprotroph\| | Yeast | NULL | Probable | As nectar-tap_saprotroph (Põlme et al. 2020); Decay sugar-rich substrates (Põlme et al. 2020) | Põlme S, et al. 2020. Fungal Diversity 105:1-16 (https://doi.org/10.1007/s13225-020-00466-2) |
| Otu19257 | 1 | k__Fungi(100);p__Ascomycota(100);c__Saccharomycetes(100);o__Saccharomycetales(100);f__Metschnikowiaceae(100);g__Metschnikowia(100);s__Metschnikowia_pulcherrima(100); | Metschnikowia | 13 | Saprotroph | \|Undefined Saprotroph\| | Yeast | NULL | Probable | As nectar-tap_saprotroph (Põlme et al. 2020); Decay sugar-rich substrates (Põlme et al. 2020) | Põlme S, et al. 2020. Fungal Diversity 105:1-16 (https://doi.org/10.1007/s13225-020-00466-2) |
| Otu19258 | 1 | k__Fungi(100);p__Ascomycota(100);c__Saccharomycetes(100);o__Saccharomycetales(100);f__Metschnikowiaceae(100);g__Metschnikowia(100);s__Metschnikowia_pulcherrima(100); | Metschnikowia | 13 | Saprotroph | \|Undefined Saprotroph\| | Yeast | NULL | Probable | As nectar-tap_saprotroph (Põlme et al. 2020); Decay sugar-rich substrates (Põlme et al. 2020) | Põlme S, et al. 2020. Fungal Diversity 105:1-16 (https://doi.org/10.1007/s13225-020-00466-2) |
| Otu19259 | 1 | k__Fungi(100);p__Ascomycota(100);c__Saccharomycetes(100);o__Saccharomycetales(100);f__Metschnikowiaceae(100);g__Metschnikowia(100);s__Metschnikowia_pulcherrima(100); | Metschnikowia | 13 | Saprotroph | \|Undefined Saprotroph\| | Yeast | NULL | Probable | As nectar-tap_saprotroph (Põlme et al. 2020); Decay sugar-rich substrates (Põlme et al. 2020) | Põlme S, et al. 2020. Fungal Diversity 105:1-16 (https://doi.org/10.1007/s13225-020-00466-2) |
| Otu19260 | 1 | k__Fungi(100);p__Ascomycota(100);c__Saccharomycetes(100);o__Saccharomycetales(100);f__Metschnikowiaceae(100);g__Metschnikowia(100);s__Metschnikowia_pulcherrima(100); | Metschnikowia | 13 | Saprotroph | \|Undefined Saprotroph\| | Yeast | NULL | Probable | As nectar-tap_saprotroph (Põlme et al. 2020); Decay sugar-rich substrates (Põlme et al. 2020) | Põlme S, et al. 2020. Fungal Diversity 105:1-16 (https://doi.org/10.1007/s13225-020-00466-2) |
| Otu19261 | 1 | k__Fungi(100);p__Ascomycota(100);c__Saccharomycetes(100);o__Saccharomycetales(100);f__Metschnikowiaceae(100);g__Metschnikowia(100);s__Metschnikowia_pulcherrima(100); | Metschnikowia | 13 | Saprotroph | \|Undefined Saprotroph\| | Yeast | NULL | Probable | As nectar-tap_saprotroph (Põlme et al. 2020); Decay sugar-rich substrates (Põlme et al. 2020) | Põlme S, et al. 2020. Fungal Diversity 105:1-16 (https://doi.org/10.1007/s13225-020-00466-2) |
| Otu19262 | 1 | k__Fungi(100);p__Ascomycota(100);c__Saccharomycetes(100);o__Saccharomycetales(100);f__Metschnikowiaceae(100);g__Metschnikowia(100);s__Metschnikowia_pulcherrima(100); | Metschnikowia | 13 | Saprotroph | \|Undefined Saprotroph\| | Yeast | NULL | Probable | As nectar-tap_saprotroph (Põlme et al. 2020); Decay sugar-rich substrates (Põlme et al. 2020) | Põlme S, et al. 2020. Fungal Diversity 105:1-16 (https://doi.org/10.1007/s13225-020-00466-2) |
| Otu19263 | 1 | k__Fungi(100);p__Ascomycota(100);c__Saccharomycetes(100);o__Saccharomycetales(100);f__Metschnikowiaceae(100);g__Metschnikowia(100);g__Metschnikowia_unclassified(100); | Metschnikowia | 13 | Saprotroph | \|Undefined Saprotroph\| | Yeast | NULL | Probable | As nectar-tap_saprotroph (Põlme et al. 2020); Decay sugar-rich substrates (Põlme et al. 2020) | Põlme S, et al. 2020. Fungal Diversity 105:1-16 (https://doi.org/10.1007/s13225-020-00466-2) |
| Otu19264 | 1 | k__Fungi(100);p__Ascomycota(100);c__Saccharomycetes(100);o__Saccharomycetales(100);f__Metschnikowiaceae(100);g__Metschnikowia(100);g__Metschnikowia_unclassified(100); | Metschnikowia | 13 | Saprotroph | \|Undefined Saprotroph\| | Yeast | NULL | Probable | As nectar-tap_saprotroph (Põlme et al. 2020); Decay sugar-rich substrates (Põlme et al. 2020) | Põlme S, et al. 2020. Fungal Diversity 105:1-16 (https://doi.org/10.1007/s13225-020-00466-2) |
| Otu19265 | 1 | k__Fungi(100);p__Ascomycota(100);c__Saccharomycetes(100);o__Saccharomycetales(100);f__Metschnikowiaceae(100);g__Metschnikowia(100);s__Metschnikowia_pulcherrima(100); | Metschnikowia | 13 | Saprotroph | \|Undefined Saprotroph\| | Yeast | NULL | Probable | As nectar-tap_saprotroph (Põlme et al. 2020); Decay sugar-rich substrates (Põlme et al. 2020) | Põlme S, et al. 2020. Fungal Diversity 105:1-16 (https://doi.org/10.1007/s13225-020-00466-2) |
| Otu19266 | 1 | k__Fungi(100);p__Ascomycota(100);c__Saccharomycetes(100);o__Saccharomycetales(100);f__Metschnikowiaceae(100);g__Metschnikowia(100);g__Metschnikowia_unclassified(100); | Metschnikowia | 13 | Saprotroph | \|Undefined Saprotroph\| | Yeast | NULL | Probable | As nectar-tap_saprotroph (Põlme et al. 2020); Decay sugar-rich substrates (Põlme et al. 2020) | Põlme S, et al. 2020. Fungal Diversity 105:1-16 (https://doi.org/10.1007/s13225-020-00466-2) |
| Otu19267 | 1 | k__Fungi(100);p__Ascomycota(100);c__Saccharomycetes(100);o__Saccharomycetales(100);f__Metschnikowiaceae(100);g__Metschnikowia(100);s__Metschnikowia_pulcherrima(100); | Metschnikowia | 13 | Saprotroph | \|Undefined Saprotroph\| | Yeast | NULL | Probable | As nectar-tap_saprotroph (Põlme et al. 2020); Decay sugar-rich substrates (Põlme et al. 2020) | Põlme S, et al. 2020. Fungal Diversity 105:1-16 (https://doi.org/10.1007/s13225-020-00466-2) |
| Otu19268 | 1 | k__Fungi(100);p__Ascomycota(100);c__Saccharomycetes(100);o__Saccharomycetales(100);f__Metschnikowiaceae(100);g__Metschnikowia(100);g__Metschnikowia_unclassified(100); | Metschnikowia | 13 | Saprotroph | \|Undefined Saprotroph\| | Yeast | NULL | Probable | As nectar-tap_saprotroph (Põlme et al. 2020); Decay sugar-rich substrates (Põlme et al. 2020) | Põlme S, et al. 2020. Fungal Diversity 105:1-16 (https://doi.org/10.1007/s13225-020-00466-2) |
| Otu19269 | 1 | k__Fungi(100);p__Ascomycota(100);c__Saccharomycetes(100);o__Saccharomycetales(100);f__Metschnikowiaceae(100);g__Metschnikowia(100);s__Metschnikowia_pulcherrima(100); | Metschnikowia | 13 | Saprotroph | \|Undefined Saprotroph\| | Yeast | NULL | Probable | As nectar-tap_saprotroph (Põlme et al. 2020); Decay sugar-rich substrates (Põlme et al. 2020) | Põlme S, et al. 2020. Fungal Diversity 105:1-16 (https://doi.org/10.1007/s13225-020-00466-2) |
| Otu19270 | 1 | k__Fungi(100);p__Ascomycota(100);c__Saccharomycetes(100);o__Saccharomycetales(100);f__Metschnikowiaceae(100);g__Metschnikowia(100);s__Metschnikowia_pulcherrima(100); | Metschnikowia | 13 | Saprotroph | \|Undefined Saprotroph\| | Yeast | NULL | Probable | As nectar-tap_saprotroph (Põlme et al. 2020); Decay sugar-rich substrates (Põlme et al. 2020) | Põlme S, et al. 2020. Fungal Diversity 105:1-16 (https://doi.org/10.1007/s13225-020-00466-2) |
| Otu19271 | 1 | k__Fungi(100);p__Ascomycota(100);c__Saccharomycetes(100);o__Saccharomycetales(100);f__Metschnikowiaceae(100);g__Metschnikowia(100);s__Metschnikowia_pulcherrima(100); | Metschnikowia | 13 | Saprotroph | \|Undefined Saprotroph\| | Yeast | NULL | Probable | As nectar-tap_saprotroph (Põlme et al. 2020); Decay sugar-rich substrates (Põlme et al. 2020) | Põlme S, et al. 2020. Fungal Diversity 105:1-16 (https://doi.org/10.1007/s13225-020-00466-2) |
| Otu19272 | 1 | k__Fungi(100);p__Ascomycota(100);c__Saccharomycetes(100);o__Saccharomycetales(100);f__Metschnikowiaceae(100);g__Metschnikowia(100);s__Metschnikowia_pulcherrima(100); | Metschnikowia | 13 | Saprotroph | \|Undefined Saprotroph\| | Yeast | NULL | Probable | As nectar-tap_saprotroph (Põlme et al. 2020); Decay sugar-rich substrates (Põlme et al. 2020) | Põlme S, et al. 2020. Fungal Diversity 105:1-16 (https://doi.org/10.1007/s13225-020-00466-2) |
| Otu19273 | 1 | k__Fungi(100);p__Ascomycota(100);c__Saccharomycetes(100);o__Saccharomycetales(100);f__Metschnikowiaceae(100);g__Metschnikowia(100);s__Metschnikowia_pulcherrima(100); | Metschnikowia | 13 | Saprotroph | \|Undefined Saprotroph\| | Yeast | NULL | Probable | As nectar-tap_saprotroph (Põlme et al. 2020); Decay sugar-rich substrates (Põlme et al. 2020) | Põlme S, et al. 2020. Fungal Diversity 105:1-16 (https://doi.org/10.1007/s13225-020-00466-2) |
| Otu19274 | 1 | k__Fungi(100);p__Ascomycota(100);c__Saccharomycetes(100);o__Saccharomycetales(100);f__Metschnikowiaceae(100);g__Metschnikowia(100);s__Metschnikowia_pulcherrima(100); | Metschnikowia | 13 | Saprotroph | \|Undefined Saprotroph\| | Yeast | NULL | Probable | As nectar-tap_saprotroph (Põlme et al. 2020); Decay sugar-rich substrates (Põlme et al. 2020) | Põlme S, et al. 2020. Fungal Diversity 105:1-16 (https://doi.org/10.1007/s13225-020-00466-2) |
| Otu19275 | 1 | k__Fungi(100);p__Ascomycota(100);c__Saccharomycetes(100);o__Saccharomycetales(100);f__Metschnikowiaceae(100);g__Metschnikowia(100);s__Metschnikowia_pulcherrima(100); | Metschnikowia | 13 | Saprotroph | \|Undefined Saprotroph\| | Yeast | NULL | Probable | As nectar-tap_saprotroph (Põlme et al. 2020); Decay sugar-rich substrates (Põlme et al. 2020) | Põlme S, et al. 2020. Fungal Diversity 105:1-16 (https://doi.org/10.1007/s13225-020-00466-2) |
| Otu19276 | 1 | k__Fungi(100);p__Ascomycota(100);c__Saccharomycetes(100);o__Saccharomycetales(100);f__Metschnikowiaceae(100);g__Metschnikowia(100);s__Metschnikowia_pulcherrima(100); | Metschnikowia | 13 | Saprotroph | \|Undefined Saprotroph\| | Yeast | NULL | Probable | As nectar-tap_saprotroph (Põlme et al. 2020); Decay sugar-rich substrates (Põlme et al. 2020) | Põlme S, et al. 2020. Fungal Diversity 105:1-16 (https://doi.org/10.1007/s13225-020-00466-2) |
| Otu19277 | 1 | k__Fungi(100);p__Ascomycota(100);c__Saccharomycetes(100);o__Saccharomycetales(100);f__Metschnikowiaceae(100);g__Metschnikowia(100);s__Metschnikowia_pulcherrima(100); | Metschnikowia | 13 | Saprotroph | \|Undefined Saprotroph\| | Yeast | NULL | Probable | As nectar-tap_saprotroph (Põlme et al. 2020); Decay sugar-rich substrates (Põlme et al. 2020) | Põlme S, et al. 2020. Fungal Diversity 105:1-16 (https://doi.org/10.1007/s13225-020-00466-2) |
| Otu19278 | 1 | k__Fungi(100);p__Ascomycota(100);c__Saccharomycetes(100);o__Saccharomycetales(100);f__Metschnikowiaceae(100);g__Metschnikowia(100);s__Metschnikowia_pulcherrima(100); | Metschnikowia | 13 | Saprotroph | \|Undefined Saprotroph\| | Yeast | NULL | Probable | As nectar-tap_saprotroph (Põlme et al. 2020); Decay sugar-rich substrates (Põlme et al. 2020) | Põlme S, et al. 2020. Fungal Diversity 105:1-16 (https://doi.org/10.1007/s13225-020-00466-2) |
| Otu19279 | 1 | k__Fungi(100);p__Ascomycota(100);c__Saccharomycetes(100);o__Saccharomycetales(100);f__Metschnikowiaceae(100);g__Metschnikowia(100);g__Metschnikowia_unclassified(100); | Metschnikowia | 13 | Saprotroph | \|Undefined Saprotroph\| | Yeast | NULL | Probable | As nectar-tap_saprotroph (Põlme et al. 2020); Decay sugar-rich substrates (Põlme et al. 2020) | Põlme S, et al. 2020. Fungal Diversity 105:1-16 (https://doi.org/10.1007/s13225-020-00466-2) |
| Otu19280 | 1 | k__Fungi(100);p__Ascomycota(100);c__Saccharomycetes(100);o__Saccharomycetales(100);f__Metschnikowiaceae(100);g__Metschnikowia(100);s__Metschnikowia_pulcherrima(100); | Metschnikowia | 13 | Saprotroph | \|Undefined Saprotroph\| | Yeast | NULL | Probable | As nectar-tap_saprotroph (Põlme et al. 2020); Decay sugar-rich substrates (Põlme et al. 2020) | Põlme S, et al. 2020. Fungal Diversity 105:1-16 (https://doi.org/10.1007/s13225-020-00466-2) |
| Otu19281 | 1 | k__Fungi(100);p__Ascomycota(100);c__Saccharomycetes(100);o__Saccharomycetales(100);f__Metschnikowiaceae(100);g__Metschnikowia(100);g__Metschnikowia_unclassified(100); | Metschnikowia | 13 | Saprotroph | \|Undefined Saprotroph\| | Yeast | NULL | Probable | As nectar-tap_saprotroph (Põlme et al. 2020); Decay sugar-rich substrates (Põlme et al. 2020) | Põlme S, et al. 2020. Fungal Diversity 105:1-16 (https://doi.org/10.1007/s13225-020-00466-2) |
| Otu19282 | 1 | k__Fungi(100);p__Ascomycota(100);c__Saccharomycetes(100);o__Saccharomycetales(100);f__Metschnikowiaceae(100);g__Metschnikowia(100);s__Metschnikowia_pulcherrima(100); | Metschnikowia | 13 | Saprotroph | \|Undefined Saprotroph\| | Yeast | NULL | Probable | As nectar-tap_saprotroph (Põlme et al. 2020); Decay sugar-rich substrates (Põlme et al. 2020) | Põlme S, et al. 2020. Fungal Diversity 105:1-16 (https://doi.org/10.1007/s13225-020-00466-2) |
| Otu19283 | 1 | k__Fungi(100);p__Ascomycota(100);c__Saccharomycetes(100);o__Saccharomycetales(100);f__Metschnikowiaceae(100);g__Metschnikowia(100);s__Metschnikowia_pulcherrima(100); | Metschnikowia | 13 | Saprotroph | \|Undefined Saprotroph\| | Yeast | NULL | Probable | As nectar-tap_saprotroph (Põlme et al. 2020); Decay sugar-rich substrates (Põlme et al. 2020) | Põlme S, et al. 2020. Fungal Diversity 105:1-16 (https://doi.org/10.1007/s13225-020-00466-2) |
| Otu19284 | 1 | k__Fungi(100);p__Ascomycota(100);c__Saccharomycetes(100);o__Saccharomycetales(100);f__Metschnikowiaceae(100);g__Metschnikowia(100);s__Metschnikowia_pulcherrima(100); | Metschnikowia | 13 | Saprotroph | \|Undefined Saprotroph\| | Yeast | NULL | Probable | As nectar-tap_saprotroph (Põlme et al. 2020); Decay sugar-rich substrates (Põlme et al. 2020) | Põlme S, et al. 2020. Fungal Diversity 105:1-16 (https://doi.org/10.1007/s13225-020-00466-2) |
| Otu19285 | 1 | k__Fungi(100);p__Ascomycota(100);c__Saccharomycetes(100);o__Saccharomycetales(100);f__Metschnikowiaceae(100);g__Metschnikowia(100);g__Metschnikowia_unclassified(100); | Metschnikowia | 13 | Saprotroph | \|Undefined Saprotroph\| | Yeast | NULL | Probable | As nectar-tap_saprotroph (Põlme et al. 2020); Decay sugar-rich substrates (Põlme et al. 2020) | Põlme S, et al. 2020. Fungal Diversity 105:1-16 (https://doi.org/10.1007/s13225-020-00466-2) |
| Otu19286 | 1 | k__Fungi(100);p__Ascomycota(100);c__Saccharomycetes(100);o__Saccharomycetales(100);f__Metschnikowiaceae(100);g__Metschnikowia(100);s__Metschnikowia_pulcherrima(100); | Metschnikowia | 13 | Saprotroph | \|Undefined Saprotroph\| | Yeast | NULL | Probable | As nectar-tap_saprotroph (Põlme et al. 2020); Decay sugar-rich substrates (Põlme et al. 2020) | Põlme S, et al. 2020. Fungal Diversity 105:1-16 (https://doi.org/10.1007/s13225-020-00466-2) |
| Otu19287 | 1 | k__Fungi(100);p__Ascomycota(100);c__Saccharomycetes(100);o__Saccharomycetales(100);f__Metschnikowiaceae(100);g__Metschnikowia(100);s__Metschnikowia_pulcherrima(100); | Metschnikowia | 13 | Saprotroph | \|Undefined Saprotroph\| | Yeast | NULL | Probable | As nectar-tap_saprotroph (Põlme et al. 2020); Decay sugar-rich substrates (Põlme et al. 2020) | Põlme S, et al. 2020. Fungal Diversity 105:1-16 (https://doi.org/10.1007/s13225-020-00466-2) |
| Otu19292 | 1 | k__Fungi(100);p__Ascomycota(100);c__Saccharomycetes(100);o__Saccharomycetales(100);f__Metschnikowiaceae(100);g__Metschnikowia(100);s__Metschnikowia_pulcherrima(100); | Metschnikowia | 13 | Saprotroph | \|Undefined Saprotroph\| | Yeast | NULL | Probable | As nectar-tap_saprotroph (Põlme et al. 2020); Decay sugar-rich substrates (Põlme et al. 2020) | Põlme S, et al. 2020. Fungal Diversity 105:1-16 (https://doi.org/10.1007/s13225-020-00466-2) |
| Otu19294 | 1 | k__Fungi(100);p__Ascomycota(100);c__Saccharomycetes(100);o__Saccharomycetales(100);f__Metschnikowiaceae(100);g__Metschnikowia(100);s__Metschnikowia_pulcherrima(100); | Metschnikowia | 13 | Saprotroph | \|Undefined Saprotroph\| | Yeast | NULL | Probable | As nectar-tap_saprotroph (Põlme et al. 2020); Decay sugar-rich substrates (Põlme et al. 2020) | Põlme S, et al. 2020. Fungal Diversity 105:1-16 (https://doi.org/10.1007/s13225-020-00466-2) |
| Otu19296 | 1 | k__Fungi(100);p__Ascomycota(100);c__Saccharomycetes(100);o__Saccharomycetales(100);f__Metschnikowiaceae(100);g__Metschnikowia(100);g__Metschnikowia_unclassified(100); | Metschnikowia | 13 | Saprotroph | \|Undefined Saprotroph\| | Yeast | NULL | Probable | As nectar-tap_saprotroph (Põlme et al. 2020); Decay sugar-rich substrates (Põlme et al. 2020) | Põlme S, et al. 2020. Fungal Diversity 105:1-16 (https://doi.org/10.1007/s13225-020-00466-2) |
| Otu19297 | 1 | k__Fungi(100);p__Ascomycota(100);c__Saccharomycetes(100);o__Saccharomycetales(100);f__Metschnikowiaceae(100);g__Metschnikowia(100);g__Metschnikowia_unclassified(100); | Metschnikowia | 13 | Saprotroph | \|Undefined Saprotroph\| | Yeast | NULL | Probable | As nectar-tap_saprotroph (Põlme et al. 2020); Decay sugar-rich substrates (Põlme et al. 2020) | Põlme S, et al. 2020. Fungal Diversity 105:1-16 (https://doi.org/10.1007/s13225-020-00466-2) |
| Otu19298 | 1 | k__Fungi(100);p__Ascomycota(100);c__Saccharomycetes(100);o__Saccharomycetales(100);f__Metschnikowiaceae(100);g__Metschnikowia(100);g__Metschnikowia_unclassified(100); | Metschnikowia | 13 | Saprotroph | \|Undefined Saprotroph\| | Yeast | NULL | Probable | As nectar-tap_saprotroph (Põlme et al. 2020); Decay sugar-rich substrates (Põlme et al. 2020) | Põlme S, et al. 2020. Fungal Diversity 105:1-16 (https://doi.org/10.1007/s13225-020-00466-2) |
| Otu19299 | 1 | k__Fungi(100);p__Ascomycota(100);c__Saccharomycetes(100);o__Saccharomycetales(100);f__Metschnikowiaceae(100);g__Metschnikowia(100);s__Metschnikowia_pulcherrima(100); | Metschnikowia | 13 | Saprotroph | \|Undefined Saprotroph\| | Yeast | NULL | Probable | As nectar-tap_saprotroph (Põlme et al. 2020); Decay sugar-rich substrates (Põlme et al. 2020) | Põlme S, et al. 2020. Fungal Diversity 105:1-16 (https://doi.org/10.1007/s13225-020-00466-2) |
| Otu19300 | 1 | k__Fungi(100);p__Ascomycota(100);c__Saccharomycetes(100);o__Saccharomycetales(100);f__Metschnikowiaceae(100);g__Metschnikowia(100);s__Metschnikowia_pulcherrima(100); | Metschnikowia | 13 | Saprotroph | \|Undefined Saprotroph\| | Yeast | NULL | Probable | As nectar-tap_saprotroph (Põlme et al. 2020); Decay sugar-rich substrates (Põlme et al. 2020) | Põlme S, et al. 2020. Fungal Diversity 105:1-16 (https://doi.org/10.1007/s13225-020-00466-2) |
| Otu19301 | 1 | k__Fungi(100);p__Ascomycota(100);c__Saccharomycetes(100);o__Saccharomycetales(100);f__Metschnikowiaceae(100);g__Metschnikowia(100);s__Metschnikowia_pulcherrima(100); | Metschnikowia | 13 | Saprotroph | \|Undefined Saprotroph\| | Yeast | NULL | Probable | As nectar-tap_saprotroph (Põlme et al. 2020); Decay sugar-rich substrates (Põlme et al. 2020) | Põlme S, et al. 2020. Fungal Diversity 105:1-16 (https://doi.org/10.1007/s13225-020-00466-2) |
| Otu19302 | 1 | k__Fungi(100);p__Ascomycota(100);c__Saccharomycetes(100);o__Saccharomycetales(100);f__Metschnikowiaceae(100);g__Metschnikowia(100);s__Metschnikowia_pulcherrima(100); | Metschnikowia | 13 | Saprotroph | \|Undefined Saprotroph\| | Yeast | NULL | Probable | As nectar-tap_saprotroph (Põlme et al. 2020); Decay sugar-rich substrates (Põlme et al. 2020) | Põlme S, et al. 2020. Fungal Diversity 105:1-16 (https://doi.org/10.1007/s13225-020-00466-2) |
| Otu19303 | 1 | k__Fungi(100);p__Ascomycota(100);c__Saccharomycetes(100);o__Saccharomycetales(100);f__Metschnikowiaceae(100);g__Metschnikowia(100);s__Metschnikowia_pulcherrima(100); | Metschnikowia | 13 | Saprotroph | \|Undefined Saprotroph\| | Yeast | NULL | Probable | As nectar-tap_saprotroph (Põlme et al. 2020); Decay sugar-rich substrates (Põlme et al. 2020) | Põlme S, et al. 2020. Fungal Diversity 105:1-16 (https://doi.org/10.1007/s13225-020-00466-2) |
| Otu19304 | 1 | k__Fungi(100);p__Ascomycota(100);c__Saccharomycetes(100);o__Saccharomycetales(100);f__Metschnikowiaceae(100);g__Metschnikowia(100);s__Metschnikowia_pulcherrima(100); | Metschnikowia | 13 | Saprotroph | \|Undefined Saprotroph\| | Yeast | NULL | Probable | As nectar-tap_saprotroph (Põlme et al. 2020); Decay sugar-rich substrates (Põlme et al. 2020) | Põlme S, et al. 2020. Fungal Diversity 105:1-16 (https://doi.org/10.1007/s13225-020-00466-2) |
| Otu19305 | 1 | k__Fungi(100);p__Ascomycota(100);c__Saccharomycetes(100);o__Saccharomycetales(100);f__Metschnikowiaceae(100);g__Metschnikowia(100);g__Metschnikowia_unclassified(100); | Metschnikowia | 13 | Saprotroph | \|Undefined Saprotroph\| | Yeast | NULL | Probable | As nectar-tap_saprotroph (Põlme et al. 2020); Decay sugar-rich substrates (Põlme et al. 2020) | Põlme S, et al. 2020. Fungal Diversity 105:1-16 (https://doi.org/10.1007/s13225-020-00466-2) |
| Otu19306 | 1 | k__Fungi(100);p__Ascomycota(100);c__Saccharomycetes(100);o__Saccharomycetales(100);f__Metschnikowiaceae(100);g__Metschnikowia(100);s__Metschnikowia_pulcherrima(100); | Metschnikowia | 13 | Saprotroph | \|Undefined Saprotroph\| | Yeast | NULL | Probable | As nectar-tap_saprotroph (Põlme et al. 2020); Decay sugar-rich substrates (Põlme et al. 2020) | Põlme S, et al. 2020. Fungal Diversity 105:1-16 (https://doi.org/10.1007/s13225-020-00466-2) |
| Otu19307 | 1 | k__Fungi(100);p__Ascomycota(100);c__Saccharomycetes(100);o__Saccharomycetales(100);f__Metschnikowiaceae(100);g__Metschnikowia(100);s__Metschnikowia_pulcherrima(100); | Metschnikowia | 13 | Saprotroph | \|Undefined Saprotroph\| | Yeast | NULL | Probable | As nectar-tap_saprotroph (Põlme et al. 2020); Decay sugar-rich substrates (Põlme et al. 2020) | Põlme S, et al. 2020. Fungal Diversity 105:1-16 (https://doi.org/10.1007/s13225-020-00466-2) |
| Otu19308 | 1 | k__Fungi(100);p__Ascomycota(100);c__Saccharomycetes(100);o__Saccharomycetales(100);f__Metschnikowiaceae(100);g__Metschnikowia(100);s__Metschnikowia_pulcherrima(100); | Metschnikowia | 13 | Saprotroph | \|Undefined Saprotroph\| | Yeast | NULL | Probable | As nectar-tap_saprotroph (Põlme et al. 2020); Decay sugar-rich substrates (Põlme et al. 2020) | Põlme S, et al. 2020. Fungal Diversity 105:1-16 (https://doi.org/10.1007/s13225-020-00466-2) |
| Otu19309 | 1 | k__Fungi(100);p__Ascomycota(100);c__Saccharomycetes(100);o__Saccharomycetales(100);f__Metschnikowiaceae(100);g__Metschnikowia(100);s__Metschnikowia_pulcherrima(100); | Metschnikowia | 13 | Saprotroph | \|Undefined Saprotroph\| | Yeast | NULL | Probable | As nectar-tap_saprotroph (Põlme et al. 2020); Decay sugar-rich substrates (Põlme et al. 2020) | Põlme S, et al. 2020. Fungal Diversity 105:1-16 (https://doi.org/10.1007/s13225-020-00466-2) |
| Otu19310 | 1 | k__Fungi(100);p__Ascomycota(100);c__Saccharomycetes(100);o__Saccharomycetales(100);f__Metschnikowiaceae(100);g__Metschnikowia(100);g__Metschnikowia_unclassified(100); | Metschnikowia | 13 | Saprotroph | \|Undefined Saprotroph\| | Yeast | NULL | Probable | As nectar-tap_saprotroph (Põlme et al. 2020); Decay sugar-rich substrates (Põlme et al. 2020) | Põlme S, et al. 2020. Fungal Diversity 105:1-16 (https://doi.org/10.1007/s13225-020-00466-2) |
| Otu19311 | 1 | k__Fungi(100);p__Ascomycota(100);c__Saccharomycetes(100);o__Saccharomycetales(100);f__Metschnikowiaceae(100);g__Metschnikowia(100);s__Metschnikowia_pulcherrima(100); | Metschnikowia | 13 | Saprotroph | \|Undefined Saprotroph\| | Yeast | NULL | Probable | As nectar-tap_saprotroph (Põlme et al. 2020); Decay sugar-rich substrates (Põlme et al. 2020) | Põlme S, et al. 2020. Fungal Diversity 105:1-16 (https://doi.org/10.1007/s13225-020-00466-2) |
| Otu19312 | 1 | k__Fungi(100);p__Ascomycota(100);c__Saccharomycetes(100);o__Saccharomycetales(100);f__Metschnikowiaceae(100);g__Metschnikowia(100);s__Metschnikowia_pulcherrima(100); | Metschnikowia | 13 | Saprotroph | \|Undefined Saprotroph\| | Yeast | NULL | Probable | As nectar-tap_saprotroph (Põlme et al. 2020); Decay sugar-rich substrates (Põlme et al. 2020) | Põlme S, et al. 2020. Fungal Diversity 105:1-16 (https://doi.org/10.1007/s13225-020-00466-2) |
| Otu19313 | 1 | k__Fungi(100);p__Ascomycota(100);c__Saccharomycetes(100);o__Saccharomycetales(100);f__Metschnikowiaceae(100);g__Metschnikowia(100);s__Metschnikowia_pulcherrima(100); | Metschnikowia | 13 | Saprotroph | \|Undefined Saprotroph\| | Yeast | NULL | Probable | As nectar-tap_saprotroph (Põlme et al. 2020); Decay sugar-rich substrates (Põlme et al. 2020) | Põlme S, et al. 2020. Fungal Diversity 105:1-16 (https://doi.org/10.1007/s13225-020-00466-2) |
| Otu19314 | 1 | k__Fungi(100);p__Ascomycota(100);c__Saccharomycetes(100);o__Saccharomycetales(100);f__Metschnikowiaceae(100);g__Metschnikowia(100);g__Metschnikowia_unclassified(100); | Metschnikowia | 13 | Saprotroph | \|Undefined Saprotroph\| | Yeast | NULL | Probable | As nectar-tap_saprotroph (Põlme et al. 2020); Decay sugar-rich substrates (Põlme et al. 2020) | Põlme S, et al. 2020. Fungal Diversity 105:1-16 (https://doi.org/10.1007/s13225-020-00466-2) |
| Otu19315 | 1 | k__Fungi(100);p__Ascomycota(100);c__Saccharomycetes(100);o__Saccharomycetales(100);f__Metschnikowiaceae(100);g__Metschnikowia(100);s__Metschnikowia_pulcherrima(100); | Metschnikowia | 13 | Saprotroph | \|Undefined Saprotroph\| | Yeast | NULL | Probable | As nectar-tap_saprotroph (Põlme et al. 2020); Decay sugar-rich substrates (Põlme et al. 2020) | Põlme S, et al. 2020. Fungal Diversity 105:1-16 (https://doi.org/10.1007/s13225-020-00466-2) |
| Otu19316 | 1 | k__Fungi(100);p__Ascomycota(100);c__Saccharomycetes(100);o__Saccharomycetales(100);f__Metschnikowiaceae(100);g__Metschnikowia(100);s__Metschnikowia_pulcherrima(100); | Metschnikowia | 13 | Saprotroph | \|Undefined Saprotroph\| | Yeast | NULL | Probable | As nectar-tap_saprotroph (Põlme et al. 2020); Decay sugar-rich substrates (Põlme et al. 2020) | Põlme S, et al. 2020. Fungal Diversity 105:1-16 (https://doi.org/10.1007/s13225-020-00466-2) |
| Otu19317 | 1 | k__Fungi(100);p__Ascomycota(100);c__Saccharomycetes(100);o__Saccharomycetales(100);f__Metschnikowiaceae(100);g__Metschnikowia(100);g__Metschnikowia_unclassified(100); | Metschnikowia | 13 | Saprotroph | \|Undefined Saprotroph\| | Yeast | NULL | Probable | As nectar-tap_saprotroph (Põlme et al. 2020); Decay sugar-rich substrates (Põlme et al. 2020) | Põlme S, et al. 2020. Fungal Diversity 105:1-16 (https://doi.org/10.1007/s13225-020-00466-2) |
| Otu19318 | 1 | k__Fungi(100);p__Ascomycota(100);c__Saccharomycetes(100);o__Saccharomycetales(100);f__Metschnikowiaceae(100);g__Metschnikowia(100);s__Metschnikowia_pulcherrima(100); | Metschnikowia | 13 | Saprotroph | \|Undefined Saprotroph\| | Yeast | NULL | Probable | As nectar-tap_saprotroph (Põlme et al. 2020); Decay sugar-rich substrates (Põlme et al. 2020) | Põlme S, et al. 2020. Fungal Diversity 105:1-16 (https://doi.org/10.1007/s13225-020-00466-2) |
| Otu19320 | 1 | k__Fungi(100);p__Ascomycota(100);c__Saccharomycetes(100);o__Saccharomycetales(100);f__Metschnikowiaceae(100);g__Metschnikowia(100);s__Metschnikowia_pulcherrima(100); | Metschnikowia | 13 | Saprotroph | \|Undefined Saprotroph\| | Yeast | NULL | Probable | As nectar-tap_saprotroph (Põlme et al. 2020); Decay sugar-rich substrates (Põlme et al. 2020) | Põlme S, et al. 2020. Fungal Diversity 105:1-16 (https://doi.org/10.1007/s13225-020-00466-2) |
| Otu19321 | 1 | k__Fungi(100);p__Ascomycota(100);c__Saccharomycetes(100);o__Saccharomycetales(100);f__Metschnikowiaceae(100);g__Metschnikowia(100);s__Metschnikowia_pulcherrima(100); | Metschnikowia | 13 | Saprotroph | \|Undefined Saprotroph\| | Yeast | NULL | Probable | As nectar-tap_saprotroph (Põlme et al. 2020); Decay sugar-rich substrates (Põlme et al. 2020) | Põlme S, et al. 2020. Fungal Diversity 105:1-16 (https://doi.org/10.1007/s13225-020-00466-2) |
| Otu19322 | 1 | k__Fungi(100);p__Ascomycota(100);c__Saccharomycetes(100);o__Saccharomycetales(100);f__Metschnikowiaceae(100);g__Metschnikowia(100);g__Metschnikowia_unclassified(100); | Metschnikowia | 13 | Saprotroph | \|Undefined Saprotroph\| | Yeast | NULL | Probable | As nectar-tap_saprotroph (Põlme et al. 2020); Decay sugar-rich substrates (Põlme et al. 2020) | Põlme S, et al. 2020. Fungal Diversity 105:1-16 (https://doi.org/10.1007/s13225-020-00466-2) |
| Otu19323 | 1 | k__Fungi(100);p__Ascomycota(100);c__Saccharomycetes(100);o__Saccharomycetales(100);f__Saccharomycetales_family_Incertae_sedis(100);g__Candida(100);g__Candida_unclassified(100); | Candida | 13 | Pathotroph-Saprotroph-Symbiotroph | Animal Pathogen-Endophyte-Endosymbiont-Epiphyte-\|Undefined Saprotroph\| | Dimorphic Yeast-Yeast | NULL | Probable | As nectar-tap_saprotroph (Põlme et al. 2020); Decay sugar-rich substrates (Põlme et al. 2020); This asexual genus is polyphyletic and contain species that function very differently depend on what they are. Caution should be taken when interpreting guilds for Candida species. | Mandalasi D, et al. 2010. Virulence. 1: 367-375 (https://doi.org/10.4161/viru.1.5.12796); Mekbib SB, et al. 2011. Tropical Plant Pathology 36:233-240 (https://doi.org/10.1590/S1982-56762011000400004); Põlme S, et al. 2020. Fungal Diversity 105:1-16 (https://doi.org/10.1007/s13225-020-00466-2) |
| Otu19324 | 1 | k__Fungi(100);p__Ascomycota(100);c__Saccharomycetes(100);o__Saccharomycetales(100);f__Metschnikowiaceae(100);g__Metschnikowia(100);s__Metschnikowia_pulcherrima(100); | Metschnikowia | 13 | Saprotroph | \|Undefined Saprotroph\| | Yeast | NULL | Probable | As nectar-tap_saprotroph (Põlme et al. 2020); Decay sugar-rich substrates (Põlme et al. 2020) | Põlme S, et al. 2020. Fungal Diversity 105:1-16 (https://doi.org/10.1007/s13225-020-00466-2) |
| Otu19325 | 1 | k__Fungi(100);p__Ascomycota(100);c__Saccharomycetes(100);o__Saccharomycetales(100);f__Metschnikowiaceae(100);g__Metschnikowia(100);s__Metschnikowia_pulcherrima(100); | Metschnikowia | 13 | Saprotroph | \|Undefined Saprotroph\| | Yeast | NULL | Probable | As nectar-tap_saprotroph (Põlme et al. 2020); Decay sugar-rich substrates (Põlme et al. 2020) | Põlme S, et al. 2020. Fungal Diversity 105:1-16 (https://doi.org/10.1007/s13225-020-00466-2) |
| Otu19326 | 1 | k__Fungi(100);p__Ascomycota(100);c__Saccharomycetes(100);o__Saccharomycetales(100);f__Metschnikowiaceae(100);g__Metschnikowia(100);s__Metschnikowia_pulcherrima(100); | Metschnikowia | 13 | Saprotroph | \|Undefined Saprotroph\| | Yeast | NULL | Probable | As nectar-tap_saprotroph (Põlme et al. 2020); Decay sugar-rich substrates (Põlme et al. 2020) | Põlme S, et al. 2020. Fungal Diversity 105:1-16 (https://doi.org/10.1007/s13225-020-00466-2) |
| Otu19327 | 1 | k__Fungi(100);p__Ascomycota(100);c__Saccharomycetes(100);o__Saccharomycetales(100);f__Metschnikowiaceae(100);g__Metschnikowia(100);g__Metschnikowia_unclassified(100); | Metschnikowia | 13 | Saprotroph | \|Undefined Saprotroph\| | Yeast | NULL | Probable | As nectar-tap_saprotroph (Põlme et al. 2020); Decay sugar-rich substrates (Põlme et al. 2020) | Põlme S, et al. 2020. Fungal Diversity 105:1-16 (https://doi.org/10.1007/s13225-020-00466-2) |
| Otu19328 | 1 | k__Fungi(100);p__Ascomycota(100);c__Saccharomycetes(100);o__Saccharomycetales(100);f__Metschnikowiaceae(100);g__Metschnikowia(100);g__Metschnikowia_unclassified(100); | Metschnikowia | 13 | Saprotroph | \|Undefined Saprotroph\| | Yeast | NULL | Probable | As nectar-tap_saprotroph (Põlme et al. 2020); Decay sugar-rich substrates (Põlme et al. 2020) | Põlme S, et al. 2020. Fungal Diversity 105:1-16 (https://doi.org/10.1007/s13225-020-00466-2) |
| Otu19329 | 1 | k__Fungi(100);p__Ascomycota(100);c__Saccharomycetes(100);o__Saccharomycetales(100);f__Metschnikowiaceae(100);g__Metschnikowia(100);s__Metschnikowia_pulcherrima(100); | Metschnikowia | 13 | Saprotroph | \|Undefined Saprotroph\| | Yeast | NULL | Probable | As nectar-tap_saprotroph (Põlme et al. 2020); Decay sugar-rich substrates (Põlme et al. 2020) | Põlme S, et al. 2020. Fungal Diversity 105:1-16 (https://doi.org/10.1007/s13225-020-00466-2) |
| Otu19330 | 1 | k__Fungi(100);p__Ascomycota(100);c__Saccharomycetes(100);o__Saccharomycetales(100);f__Metschnikowiaceae(100);g__Metschnikowia(100);s__Metschnikowia_pulcherrima(100); | Metschnikowia | 13 | Saprotroph | \|Undefined Saprotroph\| | Yeast | NULL | Probable | As nectar-tap_saprotroph (Põlme et al. 2020); Decay sugar-rich substrates (Põlme et al. 2020) | Põlme S, et al. 2020. Fungal Diversity 105:1-16 (https://doi.org/10.1007/s13225-020-00466-2) |
| Otu19331 | 1 | k__Fungi(100);p__Ascomycota(100);c__Saccharomycetes(100);o__Saccharomycetales(100);f__Metschnikowiaceae(100);g__Metschnikowia(100);s__Metschnikowia_pulcherrima(100); | Metschnikowia | 13 | Saprotroph | \|Undefined Saprotroph\| | Yeast | NULL | Probable | As nectar-tap_saprotroph (Põlme et al. 2020); Decay sugar-rich substrates (Põlme et al. 2020) | Põlme S, et al. 2020. Fungal Diversity 105:1-16 (https://doi.org/10.1007/s13225-020-00466-2) |
| Otu19332 | 1 | k__Fungi(100);p__Ascomycota(100);c__Saccharomycetes(100);o__Saccharomycetales(100);f__Metschnikowiaceae(100);g__Metschnikowia(100);s__Metschnikowia_pulcherrima(100); | Metschnikowia | 13 | Saprotroph | \|Undefined Saprotroph\| | Yeast | NULL | Probable | As nectar-tap_saprotroph (Põlme et al. 2020); Decay sugar-rich substrates (Põlme et al. 2020) | Põlme S, et al. 2020. Fungal Diversity 105:1-16 (https://doi.org/10.1007/s13225-020-00466-2) |
| Otu19333 | 1 | k__Fungi(100);p__Ascomycota(100);c__Saccharomycetes(100);o__Saccharomycetales(100);f__Metschnikowiaceae(100);g__Metschnikowia(100);s__Metschnikowia_pulcherrima(100); | Metschnikowia | 13 | Saprotroph | \|Undefined Saprotroph\| | Yeast | NULL | Probable | As nectar-tap_saprotroph (Põlme et al. 2020); Decay sugar-rich substrates (Põlme et al. 2020) | Põlme S, et al. 2020. Fungal Diversity 105:1-16 (https://doi.org/10.1007/s13225-020-00466-2) |
| Otu19336 | 1 | k__Fungi(100);p__Ascomycota(100);c__Saccharomycetes(100);o__Saccharomycetales(100);f__Saccharomycetales_family_Incertae_sedis(100);g__Candida(100);s__Candida_cretensis(100); | Candida | 13 | Pathotroph-Saprotroph-Symbiotroph | Animal Pathogen-Endophyte-Endosymbiont-Epiphyte-\|Undefined Saprotroph\| | Dimorphic Yeast-Yeast | NULL | Probable | As nectar-tap_saprotroph (Põlme et al. 2020); Decay sugar-rich substrates (Põlme et al. 2020); This asexual genus is polyphyletic and contain species that function very differently depend on what they are. Caution should be taken when interpreting guilds for Candida species. | Mandalasi D, et al. 2010. Virulence. 1: 367-375 (https://doi.org/10.4161/viru.1.5.12796); Mekbib SB, et al. 2011. Tropical Plant Pathology 36:233-240 (https://doi.org/10.1590/S1982-56762011000400004); Põlme S, et al. 2020. Fungal Diversity 105:1-16 (https://doi.org/10.1007/s13225-020-00466-2) |
| Otu19337 | 1 | k__Fungi(100);p__Ascomycota(100);c__Saccharomycetes(100);o__Saccharomycetales(100);f__Metschnikowiaceae(100);g__Metschnikowia(100);s__Metschnikowia_pulcherrima(100); | Metschnikowia | 13 | Saprotroph | \|Undefined Saprotroph\| | Yeast | NULL | Probable | As nectar-tap_saprotroph (Põlme et al. 2020); Decay sugar-rich substrates (Põlme et al. 2020) | Põlme S, et al. 2020. Fungal Diversity 105:1-16 (https://doi.org/10.1007/s13225-020-00466-2) |
| Otu19338 | 1 | k__Fungi(100);p__Ascomycota(100);c__Saccharomycetes(100);o__Saccharomycetales(100);f__Metschnikowiaceae(100);g__Metschnikowia(100);s__Metschnikowia_pulcherrima(100); | Metschnikowia | 13 | Saprotroph | \|Undefined Saprotroph\| | Yeast | NULL | Probable | As nectar-tap_saprotroph (Põlme et al. 2020); Decay sugar-rich substrates (Põlme et al. 2020) | Põlme S, et al. 2020. Fungal Diversity 105:1-16 (https://doi.org/10.1007/s13225-020-00466-2) |
| Otu19339 | 1 | k__Fungi(100);p__Ascomycota(100);c__Saccharomycetes(100);o__Saccharomycetales(100);f__Metschnikowiaceae(100);g__Metschnikowia(100);s__Metschnikowia_pulcherrima(100); | Metschnikowia | 13 | Saprotroph | \|Undefined Saprotroph\| | Yeast | NULL | Probable | As nectar-tap_saprotroph (Põlme et al. 2020); Decay sugar-rich substrates (Põlme et al. 2020) | Põlme S, et al. 2020. Fungal Diversity 105:1-16 (https://doi.org/10.1007/s13225-020-00466-2) |
| Otu19340 | 1 | k__Fungi(100);p__Ascomycota(100);c__Saccharomycetes(100);o__Saccharomycetales(100);f__Metschnikowiaceae(100);g__Metschnikowia(100);s__Metschnikowia_pulcherrima(100); | Metschnikowia | 13 | Saprotroph | \|Undefined Saprotroph\| | Yeast | NULL | Probable | As nectar-tap_saprotroph (Põlme et al. 2020); Decay sugar-rich substrates (Põlme et al. 2020) | Põlme S, et al. 2020. Fungal Diversity 105:1-16 (https://doi.org/10.1007/s13225-020-00466-2) |
| Otu19341 | 1 | k__Fungi(100);p__Ascomycota(100);c__Saccharomycetes(100);o__Saccharomycetales(100);f__Metschnikowiaceae(100);g__Metschnikowia(100);s__Metschnikowia_pulcherrima(100); | Metschnikowia | 13 | Saprotroph | \|Undefined Saprotroph\| | Yeast | NULL | Probable | As nectar-tap_saprotroph (Põlme et al. 2020); Decay sugar-rich substrates (Põlme et al. 2020) | Põlme S, et al. 2020. Fungal Diversity 105:1-16 (https://doi.org/10.1007/s13225-020-00466-2) |
| Otu19342 | 1 | k__Fungi(100);p__Ascomycota(100);c__Saccharomycetes(100);o__Saccharomycetales(100);f__Saccharomycetales_family_Incertae_sedis(100);f__Saccharomycetales_family_Incertae_sedis_unclassified(100);f__Saccharomycetales_family_Incertae_sedis_unclassified(100); | Saccharomycetales | 7 | Saprotroph | Undefined Saprotroph | Yeast | NULL | Possible | NULL | [illegible] |
| Otu19343 | 1 | k__Fungi(100);p__Ascomycota(100);c__Saccharomycetes(100);o__Saccharomycetales(100);f__Metschnikowiaceae(100);g__Metschnikowia(100);s__Metschnikowia_pulcherrima(100); | Metschnikowia | 13 | Saprotroph | \|Undefined Saprotroph\| | Yeast | NULL | Probable | As nectar-tap_saprotroph (Põlme et al. 2020); Decay sugar-rich substrates (Põlme et al. 2020) | Põlme S, et al. 2020. Fungal Diversity 105:1-16 (https://doi.org/10.1007/s13225-020-00466-2) |
| Otu19344 | 1 | k__Fungi(100);p__Ascomycota(100);c__Saccharomycetes(100);o__Saccharomycetales(100);f__Metschnikowiaceae(100);g__Metschnikowia(100);s__Metschnikowia_pulcherrima(100); | Metschnikowia | 13 | Saprotroph | \|Undefined Saprotroph\| | Yeast | NULL | Probable | As nectar-tap_saprotroph (Põlme et al. 2020); Decay sugar-rich substrates (Põlme et al. 2020) | Põlme S, et al. 2020. Fungal Diversity 105:1-16 (https://doi.org/10.1007/s13225-020-00466-2) |
| Otu19345 | 1 | k__Fungi(100);p__Ascomycota(100);c__Saccharomycetes(100);o__Saccharomycetales(100);f__Saccharomycetales_family_Incertae_sedis(100);g__Candida(100);g__Candida_unclassified(100); | Candida | 13 | Pathotroph-Saprotroph-Symbiotroph | Animal Pathogen-Endophyte-Endosymbiont-Epiphyte-\|Undefined Saprotroph\| | Dimorphic Yeast-Yeast | NULL | Probable | As nectar-tap_saprotroph (Põlme et al. 2020); Decay sugar-rich substrates (Põlme et al. 2020); This asexual genus is polyphyletic and contain species that function very differently depend on what they are. Caution should be taken when interpreting guilds for Candida species. | Mandalasi D, et al. 2010. Virulence. 1: 367-375 (https://doi.org/10.4161/viru.1.5.12796); Mekbib SB, et al. 2011. Tropical Plant Pathology 36:233-240 (https://doi.org/10.1590/S1982-56762011000400004); Põlme S, et al. 2020. Fungal Diversity 105:1-16 (https://doi.org/10.1007/s13225-020-00466-2) |
| Otu19346 | 1 | k__Fungi(100);p__Ascomycota(100);c__Saccharomycetes(100);o__Saccharomycetales(100);f__Metschnikowiaceae(100);g__Metschnikowia(100);g__Metschnikowia_unclassified(100); | Metschnikowia | 13 | Saprotroph | \|Undefined Saprotroph\| | Yeast | NULL | Probable | As nectar-tap_saprotroph (Põlme et al. 2020); Decay sugar-rich substrates (Põlme et al. 2020) | Põlme S, et al. 2020. Fungal Diversity 105:1-16 (https://doi.org/10.1007/s13225-020-00466-2) |
| Otu19347 | 1 | k__Fungi(100);p__Ascomycota(100);c__Saccharomycetes(100);o__Saccharomycetales(100);f__Metschnikowiaceae(100);g__Metschnikowia(100);s__Metschnikowia_pulcherrima(100); | Metschnikowia | 13 | Saprotroph | \|Undefined Saprotroph\| | Yeast | NULL | Probable | As nectar-tap_saprotroph (Põlme et al. 2020); Decay sugar-rich substrates (Põlme et al. 2020) | Põlme S, et al. 2020. Fungal Diversity 105:1-16 (https://doi.org/10.1007/s13225-020-00466-2) |
| Otu19348 | 1 | k__Fungi(100);p__Ascomycota(100);c__Saccharomycetes(100);o__Saccharomycetales(100);f__Metschnikowiaceae(100);g__Metschnikowia(100);g__Metschnikowia_unclassified(100); | Metschnikowia | 13 | Saprotroph | \|Undefined Saprotroph\| | Yeast | NULL | Probable | As nectar-tap_saprotroph (Põlme et al. 2020); Decay sugar-rich substrates (Põlme et al. 2020) | Põlme S, et al. 2020. Fungal Diversity 105:1-16 (https://doi.org/10.1007/s13225-020-00466-2) |
| Otu19349 | 1 | k__Fungi(100);p__Ascomycota(100);c__Saccharomycetes(100);o__Saccharomycetales(100);f__Metschnikowiaceae(100);g__Metschnikowia(100);s__Metschnikowia_pulcherrima(100); | Metschnikowia | 13 | Saprotroph | \|Undefined Saprotroph\| | Yeast | NULL | Probable | As nectar-tap_saprotroph (Põlme et al. 2020); Decay sugar-rich substrates (Põlme et al. 2020) | Põlme S, et al. 2020. Fungal Diversity 105:1-16 (https://doi.org/10.1007/s13225-020-00466-2) |
| Otu19350 | 1 | k__Fungi(100);p__Ascomycota(100);c__Saccharomycetes(100);o__Saccharomycetales(100);f__Metschnikowiaceae(100);g__Metschnikowia(100);s__Metschnikowia_pulcherrima(100); | Metschnikowia | 13 | Saprotroph | \|Undefined Saprotroph\| | Yeast | NULL | Probable | As nectar-tap_saprotroph (Põlme et al. 2020); Decay sugar-rich substrates (Põlme et al. 2020) | Põlme S, et al. 2020. Fungal Diversity 105:1-16 (https://doi.org/10.1007/s13225-020-00466-2) |
| Otu19351 | 1 | k__Fungi(100);p__Ascomycota(100);c__Saccharomycetes(100);o__Saccharomycetales(100);f__Metschnikowiaceae(100);g__Metschnikowia(100);g__Metschnikowia_unclassified(100); | Metschnikowia | 13 | Saprotroph | \|Undefined Saprotroph\| | Yeast | NULL | Probable | As nectar-tap_saprotroph (Põlme et al. 2020); Decay sugar-rich substrates (Põlme et al. 2020) | Põlme S, et al. 2020. Fungal Diversity 105:1-16 (https://doi.org/10.1007/s13225-020-00466-2) |
| Otu19352 | 1 | k__Fungi(100);p__Ascomycota(100);c__Saccharomycetes(100);o__Saccharomycetales(100);f__Metschnikowiaceae(100);g__Metschnikowia(100);s__Metschnikowia_pulcherrima(100); | Metschnikowia | 13 | Saprotroph | \|Undefined Saprotroph\| | Yeast | NULL | Probable | As nectar-tap_saprotroph (Põlme et al. 2020); Decay sugar-rich substrates (Põlme et al. 2020) | Põlme S, et al. 2020. Fungal Diversity 105:1-16 (https://doi.org/10.1007/s13225-020-00466-2) |
| Otu19354 | 1 | k__Fungi(100);p__Ascomycota(100);c__Saccharomycetes(100);o__Saccharomycetales(100);f__Metschnikowiaceae(100);g__Metschnikowia(100);s__Metschnikowia_pulcherrima(100); | Metschnikowia | 13 | Saprotroph | \|Undefined Saprotroph\| | Yeast | NULL | Probable | As nectar-tap_saprotroph (Põlme et al. 2020); Decay sugar-rich substrates (Põlme et al. 2020) | Põlme S, et al. 2020. Fungal Diversity 105:1-16 (https://doi.org/10.1007/s13225-020-00466-2) |
| Otu19355 | 1 | k__Fungi(100);p__Ascomycota(100);c__Saccharomycetes(100);o__Saccharomycetales(100);f__Metschnikowiaceae(100);g__Metschnikowia(100);s__Metschnikowia_pulcherrima(100); | Metschnikowia | 13 | Saprotroph | \|Undefined Saprotroph\| | Yeast | NULL | Probable | As nectar-tap_saprotroph (Põlme et al. 2020); Decay sugar-rich substrates (Põlme et al. 2020) | Põlme S, et al. 2020. Fungal Diversity 105:1-16 (https://doi.org/10.1007/s13225-020-00466-2) |
| Otu19356 | 1 | k__Fungi(100);p__Ascomycota(100);c__Saccharomycetes(100);o__Saccharomycetales(100);f__Metschnikowiaceae(100);g__Metschnikowia(100);s__Metschnikowia_pulcherrima(100); | Metschnikowia | 13 | Saprotroph | \|Undefined Saprotroph\| | Yeast | NULL | Probable | As nectar-tap_saprotroph (Põlme et al. 2020); Decay sugar-rich substrates (Põlme et al. 2020) | Põlme S, et al. 2020. Fungal Diversity 105:1-16 (https://doi.org/10.1007/s13225-020-00466-2) |
| Otu19357 | 1 | k__Fungi(100);p__Ascomycota(100);c__Saccharomycetes(100);o__Saccharomycetales(100);f__Metschnikowiaceae(100);g__Metschnikowia(100);s__Metschnikowia_pulcherrima(100); | Metschnikowia | 13 | Saprotroph | \|Undefined Saprotroph\| | Yeast | NULL | Probable | As nectar-tap_saprotroph (Põlme et al. 2020); Decay sugar-rich substrates (Põlme et al. 2020) | Põlme S, et al. 2020. Fungal Diversity 105:1-16 (https://doi.org/10.1007/s13225-020-00466-2) |
| Otu19359 | 1 | k__Fungi(100);p__Ascomycota(100);c__Saccharomycetes(100);o__Saccharomycetales(100);f__Metschnikowiaceae(100);g__Metschnikowia(100);s__Metschnikowia_pulcherrima(100); | Metschnikowia | 13 | Saprotroph | \|Undefined Saprotroph\| | Yeast | NULL | Probable | As nectar-tap_saprotroph (Põlme et al. 2020); Decay sugar-rich substrates (Põlme et al. 2020) | Põlme S, et al. 2020. Fungal Diversity 105:1-16 (https://doi.org/10.1007/s13225-020-00466-2) |
| Otu19360 | 1 | k__Fungi(100);p__Ascomycota(100);c__Saccharomycetes(100);o__Saccharomycetales(100);f__Metschnikowiaceae(100);g__Metschnikowia(100);s__Metschnikowia_pulcherrima(100); | Metschnikowia | 13 | Saprotroph | \|Undefined Saprotroph\| | Yeast | NULL | Probable | As nectar-tap_saprotroph (Põlme et al. 2020); Decay sugar-rich substrates (Põlme et al. 2020) | Põlme S, et al. 2020. Fungal Diversity 105:1-16 (https://doi.org/10.1007/s13225-020-00466-2) |
| Otu19361 | 1 | k__Fungi(100);p__Ascomycota(100);c__Saccharomycetes(100);o__Saccharomycetales(100);f__Metschnikowiaceae(100);g__Metschnikowia(100);s__Metschnikowia_pulcherrima(100); | Metschnikowia | 13 | Saprotroph | \|Undefined Saprotroph\| | Yeast | NULL | Probable | As nectar-tap_saprotroph (Põlme et al. 2020); Decay sugar-rich substrates (Põlme et al. 2020) | Põlme S, et al. 2020. Fungal Diversity 105:1-16 (https://doi.org/10.1007/s13225-020-00466-2) |
| Otu19362 | 1 | k__Fungi(100);p__Ascomycota(100);c__Saccharomycetes(100);o__Saccharomycetales(100);f__Metschnikowiaceae(100);g__Metschnikowia(100);s__Metschnikowia_pulcherrima(100); | Metschnikowia | 13 | Saprotroph | \|Undefined Saprotroph\| | Yeast | NULL | Probable | As nectar-tap_saprotroph (Põlme et al. 2020); Decay sugar-rich substrates (Põlme et al. 2020) | Põlme S, et al. 2020. Fungal Diversity 105:1-16 (https://doi.org/10.1007/s13225-020-00466-2) |
| Otu19363 | 1 | k__Fungi(100);p__Ascomycota(100);c__Saccharomycetes(100);o__Saccharomycetales(100);f__Metschnikowiaceae(100);g__Metschnikowia(100);s__Metschnikowia_pulcherrima(100); | Metschnikowia | 13 | Saprotroph | \|Undefined Saprotroph\| | Yeast | NULL | Probable | As nectar-tap_saprotroph (Põlme et al. 2020); Decay sugar-rich substrates (Põlme et al. 2020) | Põlme S, et al. 2020. Fungal Diversity 105:1-16 (https://doi.org/10.1007/s13225-020-00466-2) |
| Otu19364 | 1 | k__Fungi(100);p__Ascomycota(100);c__Saccharomycetes(100);o__Saccharomycetales(100);f__Metschnikowiaceae(100);g__Metschnikowia(100);s__Metschnikowia_pulcherrima(100); | Metschnikowia | 13 | Saprotroph | \|Undefined Saprotroph\| | Yeast | NULL | Probable | As nectar-tap_saprotroph (Põlme et al. 2020); Decay sugar-rich substrates (Põlme et al. 2020) | Põlme S, et al. 2020. Fungal Diversity 105:1-16 (https://doi.org/10.1007/s13225-020-00466-2) |
| Otu19365 | 1 | k__Fungi(100);p__Ascomycota(100);c__Saccharomycetes(100);o__Saccharomycetales(100);f__Metschnikowiaceae(100);g__Metschnikowia(100);g__Metschnikowia_unclassified(100); | Metschnikowia | 13 | Saprotroph | \|Undefined Saprotroph\| | Yeast | NULL | Probable | As nectar-tap_saprotroph (Põlme et al. 2020); Decay sugar-rich substrates (Põlme et al. 2020) | Põlme S, et al. 2020. Fungal Diversity 105:1-16 (https://doi.org/10.1007/s13225-020-00466-2) |
| Otu19366 | 1 | k__Fungi(100);p__Ascomycota(100);c__Saccharomycetes(100);o__Saccharomycetales(100);f__Metschnikowiaceae(100);g__Metschnikowia(100);g__Metschnikowia_unclassified(100); | Metschnikowia | 13 | Saprotroph | \|Undefined Saprotroph\| | Yeast | NULL | Probable | As nectar-tap_saprotroph (Põlme et al. 2020); Decay sugar-rich substrates (Põlme et al. 2020) | Põlme S, et al. 2020. Fungal Diversity 105:1-16 (https://doi.org/10.1007/s13225-020-00466-2) |
| Otu19367 | 1 | k__Fungi(100);p__Ascomycota(100);c__Saccharomycetes(100);o__Saccharomycetales(100);f__Metschnikowiaceae(100);g__Metschnikowia(100);g__Metschnikowia_unclassified(100); | Metschnikowia | 13 | Saprotroph | \|Undefined Saprotroph\| | Yeast | NULL | Probable | As nectar-tap_saprotroph (Põlme et al. 2020); Decay sugar-rich substrates (Põlme et al. 2020) | Põlme S, et al. 2020. Fungal Diversity 105:1-16 (https://doi.org/10.1007/s13225-020-00466-2) |
| Otu19368 | 1 | k__Fungi(100);p__Ascomycota(100);c__Saccharomycetes(100);o__Saccharomycetales(100);f__Metschnikowiaceae(100);g__Metschnikowia(100);s__Metschnikowia_pulcherrima(100); | Metschnikowia | 13 | Saprotroph | \|Undefined Saprotroph\| | Yeast | NULL | Probable | As nectar-tap_saprotroph (Põlme et al. 2020); Decay sugar-rich substrates (Põlme et al. 2020) | Põlme S, et al. 2020. Fungal Diversity 105:1-16 (https://doi.org/10.1007/s13225-020-00466-2) |
| Otu19369 | 1 | k__Fungi(100);p__Ascomycota(100);c__Saccharomycetes(100);o__Saccharomycetales(100);f__Metschnikowiaceae(100);g__Metschnikowia(100);g__Metschnikowia_unclassified(100); | Metschnikowia | 13 | Saprotroph | \|Undefined Saprotroph\| | Yeast | NULL | Probable | As nectar-tap_saprotroph (Põlme et al. 2020); Decay sugar-rich substrates (Põlme et al. 2020) | Põlme S, et al. 2020. Fungal Diversity 105:1-16 (https://doi.org/10.1007/s13225-020-00466-2) |
| Otu19370 | 1 | k__Fungi(100);p__Ascomycota(100);c__Saccharomycetes(100);o__Saccharomycetales(100);f__Metschnikowiaceae(100);g__Metschnikowia(100);s__Metschnikowia_pulcherrima(100); | Metschnikowia | 13 | Saprotroph | \|Undefined Saprotroph\| | Yeast | NULL | Probable | As nectar-tap_saprotroph (Põlme et al. 2020); Decay sugar-rich substrates (Põlme et al. 2020) | Põlme S, et al. 2020. Fungal Diversity 105:1-16 (https://doi.org/10.1007/s13225-020-00466-2) |
| Otu19371 | 1 | k__Fungi(100);p__Ascomycota(100);c__Saccharomycetes(100);o__Saccharomycetales(100);f__Metschnikowiaceae(100);g__Metschnikowia(100);g__Metschnikowia_unclassified(100); | Metschnikowia | 13 | Saprotroph | \|Undefined Saprotroph\| | Yeast | NULL | Probable | As nectar-tap_saprotroph (Põlme et al. 2020); Decay sugar-rich substrates (Põlme et al. 2020) | Põlme S, et al. 2020. Fungal Diversity 105:1-16 (https://doi.org/10.1007/s13225-020-00466-2) |
| Otu19373 | 1 | k__Fungi(100);p__Ascomycota(100);c__Saccharomycetes(100);o__Saccharomycetales(100);f__Metschnikowiaceae(100);g__Metschnikowia(100);s__Metschnikowia_pulcherrima(100); | Metschnikowia | 13 | Saprotroph | \|Undefined Saprotroph\| | Yeast | NULL | Probable | As nectar-tap_saprotroph (Põlme et al. 2020); Decay sugar-rich substrates (Põlme et al. 2020) | Põlme S, et al. 2020. Fungal Diversity 105:1-16 (https://doi.org/10.1007/s13225-020-00466-2) |
| Otu19376 | 1 | k__Fungi(100);p__Ascomycota(100);c__Saccharomycetes(100);o__Saccharomycetales(100);f__Metschnikowiaceae(100);g__Metschnikowia(100);s__Metschnikowia_pulcherrima(100); | Metschnikowia | 13 | Saprotroph | \|Undefined Saprotroph\| | Yeast | NULL | Probable | As nectar-tap_saprotroph (Põlme et al. 2020); Decay sugar-rich substrates (Põlme et al. 2020) | Põlme S, et al. 2020. Fungal Diversity 105:1-16 (https://doi.org/10.1007/s13225-020-00466-2) |
| Otu19378 | 1 | k__Fungi(100);p__Ascomycota(100);c__Saccharomycetes(100);o__Saccharomycetales(100);f__Saccharomycetales_family_Incertae_sedis(100);g__Candida(100);g__Candida_unclassified(100); | Candida | 13 | Pathotroph-Saprotroph-Symbiotroph | Animal Pathogen-Endophyte-Endosymbiont-Epiphyte-\|Undefined Saprotroph\| | Dimorphic Yeast-Yeast | NULL | Probable | As nectar-tap_saprotroph (Põlme et al. 2020); Decay sugar-rich substrates (Põlme et al. 2020); This asexual genus is polyphyletic and contain species that function very differently depend on what they are. Caution should be taken when interpreting guilds for Candida species. | Mandalasi D, et al. 2010. Virulence. 1: 367-375 (https://doi.org/10.4161/viru.1.5.12796); Mekbib SB, et al. 2011. Tropical Plant Pathology 36:233-240 (https://doi.org/10.1590/S1982-56762011000400004); Põlme S, et al. 2020. Fungal Diversity 105:1-16 (https://doi.org/10.1007/s13225-020-00466-2) |
| Otu19381 | 1 | k__Fungi(100);p__Ascomycota(100);c__Saccharomycetes(100);o__Saccharomycetales(100);f__Metschnikowiaceae(100);g__Metschnikowia(100);s__Metschnikowia_pulcherrima(100); | Metschnikowia | 13 | Saprotroph | \|Undefined Saprotroph\| | Yeast | NULL | Probable | As nectar-tap_saprotroph (Põlme et al. 2020); Decay sugar-rich substrates (Põlme et al. 2020) | Põlme S, et al. 2020. Fungal Diversity 105:1-16 (https://doi.org/10.1007/s13225-020-00466-2) |
| Otu19382 | 1 | k__Fungi(100);p__Ascomycota(100);c__Saccharomycetes(100);o__Saccharomycetales(100);f__Metschnikowiaceae(100);g__Metschnikowia(100);s__Metschnikowia_pulcherrima(100); | Metschnikowia | 13 | Saprotroph | \|Undefined Saprotroph\| | Yeast | NULL | Probable | As nectar-tap_saprotroph (Põlme et al. 2020); Decay sugar-rich substrates (Põlme et al. 2020) | Põlme S, et al. 2020. Fungal Diversity 105:1-16 (https://doi.org/10.1007/s13225-020-00466-2) |
| Otu19383 | 1 | k__Fungi(100);p__Ascomycota(100);c__Saccharomycetes(100);o__Saccharomycetales(100);f__Metschnikowiaceae(100);g__Metschnikowia(100);s__Metschnikowia_pulcherrima(100); | Metschnikowia | 13 | Saprotroph | \|Undefined Saprotroph\| | Yeast | NULL | Probable | As nectar-tap_saprotroph (Põlme et al. 2020); Decay sugar-rich substrates (Põlme et al. 2020) | Põlme S, et al. 2020. Fungal Diversity 105:1-16 (https://doi.org/10.1007/s13225-020-00466-2) |
| Otu19384 | 1 | k__Fungi(100);p__Ascomycota(100);c__Saccharomycetes(100);o__Saccharomycetales(100);f__Metschnikowiaceae(100);g__Metschnikowia(100);s__Metschnikowia_pulcherrima(100); | Metschnikowia | 13 | Saprotroph | \|Undefined Saprotroph\| | Yeast | NULL | Probable | As nectar-tap_saprotroph (Põlme et al. 2020); Decay sugar-rich substrates (Põlme et al. 2020) | Põlme S, et al. 2020. Fungal Diversity 105:1-16 (https://doi.org/10.1007/s13225-020-00466-2) |
| Otu19385 | 1 | k__Fungi(100);p__Ascomycota(100);c__Saccharomycetes(100);o__Saccharomycetales(100);f__Metschnikowiaceae(100);g__Metschnikowia(100);s__Metschnikowia_pulcherrima(100); | Metschnikowia | 13 | Saprotroph | \|Undefined Saprotroph\| | Yeast | NULL | Probable | As nectar-tap_saprotroph (Põlme et al. 2020); Decay sugar-rich substrates (Põlme et al. 2020) | Põlme S, et al. 2020. Fungal Diversity 105:1-16 (https://doi.org/10.1007/s13225-020-00466-2) |
| Otu19386 | 1 | k__Fungi(100);p__Ascomycota(100);c__Saccharomycetes(100);o__Saccharomycetales(100);f__Metschnikowiaceae(100);g__Metschnikowia(100);s__Metschnikowia_pulcherrima(100); | Metschnikowia | 13 | Saprotroph | \|Undefined Saprotroph\| | Yeast | NULL | Probable | As nectar-tap_saprotroph (Põlme et al. 2020); Decay sugar-rich substrates (Põlme et al. 2020) | Põlme S, et al. 2020. Fungal Diversity 105:1-16 (https://doi.org/10.1007/s13225-020-00466-2) |
| Otu19387 | 1 | k__Fungi(100);p__Ascomycota(100);c__Saccharomycetes(100);o__Saccharomycetales(100);f__Metschnikowiaceae(100);g__Metschnikowia(100);s__Metschnikowia_pulcherrima(100); | Metschnikowia | 13 | Saprotroph | \|Undefined Saprotroph\| | Yeast | NULL | Probable | As nectar-tap_saprotroph (Põlme et al. 2020); Decay sugar-rich substrates (Põlme et al. 2020) | Põlme S, et al. 2020. Fungal Diversity 105:1-16 (https://doi.org/10.1007/s13225-020-00466-2) |
| Otu19388 | 1 | k__Fungi(100);p__Ascomycota(100);c__Saccharomycetes(100);o__Saccharomycetales(100);f__Metschnikowiaceae(100);g__Metschnikowia(100);s__Metschnikowia_pulcherrima(100); | Metschnikowia | 13 | Saprotroph | \|Undefined Saprotroph\| | Yeast | NULL | Probable | As nectar-tap_saprotroph (Põlme et al. 2020); Decay sugar-rich substrates (Põlme et al. 2020) | Põlme S, et al. 2020. Fungal Diversity 105:1-16 (https://doi.org/10.1007/s13225-020-00466-2) |
| Otu19389 | 1 | k__Fungi(100);p__Ascomycota(100);c__Saccharomycetes(100);o__Saccharomycetales(100);f__Metschnikowiaceae(100);g__Metschnikowia(100);g__Metschnikowia_unclassified(100); | Metschnikowia | 13 | Saprotroph | \|Undefined Saprotroph\| | Yeast | NULL | Probable | As nectar-tap_saprotroph (Põlme et al. 2020); Decay sugar-rich substrates (Põlme et al. 2020) | Põlme S, et al. 2020. Fungal Diversity 105:1-16 (https://doi.org/10.1007/s13225-020-00466-2) |
| Otu19390 | 1 | k__Fungi(100);p__Ascomycota(100);c__Saccharomycetes(100);o__Saccharomycetales(100);f__Metschnikowiaceae(100);g__Metschnikowia(100);s__Metschnikowia_pulcherrima(100); | Metschnikowia | 13 | Saprotroph | \|Undefined Saprotroph\| | Yeast | NULL | Probable | As nectar-tap_saprotroph (Põlme et al. 2020); Decay sugar-rich substrates (Põlme et al. 2020) | Põlme S, et al. 2020. Fungal Diversity 105:1-16 (https://doi.org/10.1007/s13225-020-00466-2) |
| Otu19391 | 1 | k__Fungi(100);p__Ascomycota(100);c__Saccharomycetes(100);o__Saccharomycetales(100);f__Metschnikowiaceae(100);g__Metschnikowia(100);g__Metschnikowia_unclassified(100); | Metschnikowia | 13 | Saprotroph | \|Undefined Saprotroph\| | Yeast | NULL | Probable | As nectar-tap_saprotroph (Põlme et al. 2020); Decay sugar-rich substrates (Põlme et al. 2020) | Põlme S, et al. 2020. Fungal Diversity 105:1-16 (https://doi.org/10.1007/s13225-020-00466-2) |
| Otu19392 | 1 | k__Fungi(100);p__Ascomycota(100);c__Saccharomycetes(100);o__Saccharomycetales(100);f__Metschnikowiaceae(100);g__Metschnikowia(100);s__Metschnikowia_pulcherrima(100); | Metschnikowia | 13 | Saprotroph | \|Undefined Saprotroph\| | Yeast | NULL | Probable | As nectar-tap_saprotroph (Põlme et al. 2020); Decay sugar-rich substrates (Põlme et al. 2020) | Põlme S, et al. 2020. Fungal Diversity 105:1-16 (https://doi.org/10.1007/s13225-020-00466-2) |
| Otu19393 | 1 | k__Fungi(100);p__Ascomycota(100);c__Saccharomycetes(100);o__Saccharomycetales(100);f__Metschnikowiaceae(100);g__Metschnikowia(100);s__Metschnikowia_pulcherrima(100); | Metschnikowia | 13 | Saprotroph | \|Undefined Saprotroph\| | Yeast | NULL | Probable | As nectar-tap_saprotroph (Põlme et al. 2020); Decay sugar-rich substrates (Põlme et al. 2020) | Põlme S, et al. 2020. Fungal Diversity 105:1-16 (https://doi.org/10.1007/s13225-020-00466-2) |
| Otu19394 | 1 | k__Fungi(100);p__Ascomycota(100);c__Saccharomycetes(100);o__Saccharomycetales(100);f__Metschnikowiaceae(100);g__Metschnikowia(100);s__Metschnikowia_pulcherrima(100); | Metschnikowia | 13 | Saprotroph | \|Undefined Saprotroph\| | Yeast | NULL | Probable | As nectar-tap_saprotroph (Põlme et al. 2020); Decay sugar-rich substrates (Põlme et al. 2020) | Põlme S, et al. 2020. Fungal Diversity 105:1-16 (https://doi.org/10.1007/s13225-020-00466-2) |
| Otu19395 | 1 | k__Fungi(100);p__Ascomycota(100);c__Saccharomycetes(100);o__Saccharomycetales(100);f__Metschnikowiaceae(100);g__Metschnikowia(100);s__Metschnikowia_pulcherrima(100); | Metschnikowia | 13 | Saprotroph | \|Undefined Saprotroph\| | Yeast | NULL | Probable | As nectar-tap_saprotroph (Põlme et al. 2020); Decay sugar-rich substrates (Põlme et al. 2020) | Põlme S, et al. 2020. Fungal Diversity 105:1-16 (https://doi.org/10.1007/s13225-020-00466-2) |
| Otu19396 | 1 | k__Fungi(100);p__Ascomycota(100);c__Saccharomycetes(100);o__Saccharomycetales(100);f__Metschnikowiaceae(100);g__Metschnikowia(100);s__Metschnikowia_pulcherrima(100); | Metschnikowia | 13 | Saprotroph | \|Undefined Saprotroph\| | Yeast | NULL | Probable | As nectar-tap_saprotroph (Põlme et al. 2020); Decay sugar-rich substrates (Põlme et al. 2020) | Põlme S, et al. 2020. Fungal Diversity 105:1-16 (https://doi.org/10.1007/s13225-020-00466-2) |
| Otu19397 | 1 | k__Fungi(100);p__Ascomycota(100);c__Saccharomycetes(100);o__Saccharomycetales(100);f__Metschnikowiaceae(100);g__Metschnikowia(100);g__Metschnikowia_unclassified(100); | Metschnikowia | 13 | Saprotroph | \|Undefined Saprotroph\| | Yeast | NULL | Probable | As nectar-tap_saprotroph (Põlme et al. 2020); Decay sugar-rich substrates (Põlme et al. 2020) | Põlme S, et al. 2020. Fungal Diversity 105:1-16 (https://doi.org/10.1007/s13225-020-00466-2) |
| Otu19398 | 1 | k__Fungi(100);p__Ascomycota(100);c__Saccharomycetes(100);o__Saccharomycetales(100);f__Metschnikowiaceae(100);g__Metschnikowia(100);s__Metschnikowia_pulcherrima(100); | Metschnikowia | 13 | Saprotroph | \|Undefined Saprotroph\| | Yeast | NULL | Probable | As nectar-tap_saprotroph (Põlme et al. 2020); Decay sugar-rich substrates (Põlme et al. 2020) | Põlme S, et al. 2020. Fungal Diversity 105:1-16 (https://doi.org/10.1007/s13225-020-00466-2) |
| Otu19399 | 1 | k__Fungi(100);p__Ascomycota(100);c__Saccharomycetes(100);o__Saccharomycetales(100);f__Metschnikowiaceae(100);g__Metschnikowia(100);s__Metschnikowia_pulcherrima(100); | Metschnikowia | 13 | Saprotroph | \|Undefined Saprotroph\| | Yeast | NULL | Probable | As nectar-tap_saprotroph (Põlme et al. 2020); Decay sugar-rich substrates (Põlme et al. 2020) | Põlme S, et al. 2020. Fungal Diversity 105:1-16 (https://doi.org/10.1007/s13225-020-00466-2) |
| Otu19400 | 1 | k__Fungi(100);p__Ascomycota(100);c__Saccharomycetes(100);o__Saccharomycetales(100);f__Metschnikowiaceae(100);g__Metschnikowia(100);g__Metschnikowia_unclassified(100); | Metschnikowia | 13 | Saprotroph | \|Undefined Saprotroph\| | Yeast | NULL | Probable | As nectar-tap_saprotroph (Põlme et al. 2020); Decay sugar-rich substrates (Põlme et al. 2020) | Põlme S, et al. 2020. Fungal Diversity 105:1-16 (https://doi.org/10.1007/s13225-020-00466-2) |
| Otu19401 | 1 | k__Fungi(100);p__Ascomycota(100);c__Saccharomycetes(100);o__Saccharomycetales(100);f__Metschnikowiaceae(100);g__Metschnikowia(100);s__Metschnikowia_pulcherrima(100); | Metschnikowia | 13 | Saprotroph | \|Undefined Saprotroph\| | Yeast | NULL | Probable | As nectar-tap_saprotroph (Põlme et al. 2020); Decay sugar-rich substrates (Põlme et al. 2020) | Põlme S, et al. 2020. Fungal Diversity 105:1-16 (https://doi.org/10.1007/s13225-020-00466-2) |
| Otu19402 | 1 | k__Fungi(100);p__Ascomycota(100);c__Saccharomycetes(100);o__Saccharomycetales(100);f__Metschnikowiaceae(100);g__Metschnikowia(100);s__Metschnikowia_pulcherrima(100); | Metschnikowia | 13 | Saprotroph | \|Undefined Saprotroph\| | Yeast | NULL | Probable | As nectar-tap_saprotroph (Põlme et al. 2020); Decay sugar-rich substrates (Põlme et al. 2020) | Põlme S, et al. 2020. Fungal Diversity 105:1-16 (https://doi.org/10.1007/s13225-020-00466-2) |
| Otu19403 | 1 | k__Fungi(100);p__Ascomycota(100);c__Saccharomycetes(100);o__Saccharomycetales(100);f__Metschnikowiaceae(100);g__Metschnikowia(100);s__Metschnikowia_pulcherrima(100); | Metschnikowia | 13 | Saprotroph | \|Undefined Saprotroph\| | Yeast | NULL | Probable | As nectar-tap_saprotroph (Põlme et al. 2020); Decay sugar-rich substrates (Põlme et al. 2020) | Põlme S, et al. 2020. Fungal Diversity 105:1-16 (https://doi.org/10.1007/s13225-020-00466-2) |
| Otu19404 | 1 | k__Fungi(100);p__Ascomycota(100);c__Saccharomycetes(100);o__Saccharomycetales(100);f__Metschnikowiaceae(100);g__Metschnikowia(100);g__Metschnikowia_unclassified(100); | Metschnikowia | 13 | Saprotroph | \|Undefined Saprotroph\| | Yeast | NULL | Probable | As nectar-tap_saprotroph (Põlme et al. 2020); Decay sugar-rich substrates (Põlme et al. 2020) | Põlme S, et al. 2020. Fungal Diversity 105:1-16 (https://doi.org/10.1007/s13225-020-00466-2) |
| Otu19405 | 1 | k__Fungi(100);p__Ascomycota(100);c__Saccharomycetes(100);o__Saccharomycetales(100);f__Metschnikowiaceae(100);g__Metschnikowia(100);s__Metschnikowia_pulcherrima(100); | Metschnikowia | 13 | Saprotroph | \|Undefined Saprotroph\| | Yeast | NULL | Probable | As nectar-tap_saprotroph (Põlme et al. 2020); Decay sugar-rich substrates (Põlme et al. 2020) | Põlme S, et al. 2020. Fungal Diversity 105:1-16 (https://doi.org/10.1007/s13225-020-00466-2) |
| Otu19406 | 1 | k__Fungi(100);p__Ascomycota(100);c__Saccharomycetes(100);o__Saccharomycetales(100);f__Metschnikowiaceae(100);g__Metschnikowia(100);s__Metschnikowia_pulcherrima(100); | Metschnikowia | 13 | Saprotroph | \|Undefined Saprotroph\| | Yeast | NULL | Probable | As nectar-tap_saprotroph (Põlme et al. 2020); Decay sugar-rich substrates (Põlme et al. 2020) | Põlme S, et al. 2020. Fungal Diversity 105:1-16 (https://doi.org/10.1007/s13225-020-00466-2) |
| Otu19407 | 1 | k__Fungi(100);p__Ascomycota(100);c__Saccharomycetes(100);o__Saccharomycetales(100);f__Metschnikowiaceae(100);g__Metschnikowia(100);s__Metschnikowia_pulcherrima(100); | Metschnikowia | 13 | Saprotroph | \|Undefined Saprotroph\| | Yeast | NULL | Probable | As nectar-tap_saprotroph (Põlme et al. 2020); Decay sugar-rich substrates (Põlme et al. 2020) | Põlme S, et al. 2020. Fungal Diversity 105:1-16 (https://doi.org/10.1007/s13225-020-00466-2) |
| Otu19408 | 1 | k__Fungi(100);p__Ascomycota(100);c__Saccharomycetes(100);o__Saccharomycetales(100);f__Metschnikowiaceae(100);g__Metschnikowia(100);g__Metschnikowia_unclassified(100); | Metschnikowia | 13 | Saprotroph | \|Undefined Saprotroph\| | Yeast | NULL | Probable | As nectar-tap_saprotroph (Põlme et al. 2020); Decay sugar-rich substrates (Põlme et al. 2020) | Põlme S, et al. 2020. Fungal Diversity 105:1-16 (https://doi.org/10.1007/s13225-020-00466-2) |
| Otu19409 | 1 | k__Fungi(100);p__Ascomycota(100);c__Saccharomycetes(100);o__Saccharomycetales(100);f__Metschnikowiaceae(100);g__Metschnikowia(100);g__Metschnikowia_unclassified(100); | Metschnikowia | 13 | Saprotroph | \|Undefined Saprotroph\| | Yeast | NULL | Probable | As nectar-tap_saprotroph (Põlme et al. 2020); Decay sugar-rich substrates (Põlme et al. 2020) | Põlme S, et al. 2020. Fungal Diversity 105:1-16 (https://doi.org/10.1007/s13225-020-00466-2) |
| Otu19410 | 1 | k__Fungi(100);p__Ascomycota(100);c__Saccharomycetes(100);o__Saccharomycetales(100);f__Metschnikowiaceae(100);g__Metschnikowia(100);s__Metschnikowia_pulcherrima(100); | Metschnikowia | 13 | Saprotroph | \|Undefined Saprotroph\| | Yeast | NULL | Probable | As nectar-tap_saprotroph (Põlme et al. 2020); Decay sugar-rich substrates (Põlme et al. 2020) | Põlme S, et al. 2020. Fungal Diversity 105:1-16 (https://doi.org/10.1007/s13225-020-00466-2) |
| Otu19411 | 1 | k__Fungi(100);p__Ascomycota(100);c__Saccharomycetes(100);o__Saccharomycetales(100);f__Metschnikowiaceae(100);g__Metschnikowia(100);s__Metschnikowia_pulcherrima(100); | Metschnikowia | 13 | Saprotroph | \|Undefined Saprotroph\| | Yeast | NULL | Probable | As nectar-tap_saprotroph (Põlme et al. 2020); Decay sugar-rich substrates (Põlme et al. 2020) | Põlme S, et al. 2020. Fungal Diversity 105:1-16 (https://doi.org/10.1007/s13225-020-00466-2) |
| Otu19412 | 1 | k__Fungi(100);p__Ascomycota(100);c__Saccharomycetes(100);o__Saccharomycetales(100);f__Metschnikowiaceae(100);g__Metschnikowia(100);s__Metschnikowia_pulcherrima(100); | Metschnikowia | 13 | Saprotroph | \|Undefined Saprotroph\| | Yeast | NULL | Probable | As nectar-tap_saprotroph (Põlme et al. 2020); Decay sugar-rich substrates (Põlme et al. 2020) | Põlme S, et al. 2020. Fungal Diversity 105:1-16 (https://doi.org/10.1007/s13225-020-00466-2) |
| Otu19413 | 1 | k__Fungi(100);p__Ascomycota(100);c__Saccharomycetes(100);o__Saccharomycetales(100);f__Metschnikowiaceae(100);g__Metschnikowia(100);s__Metschnikowia_pulcherrima(100); | Metschnikowia | 13 | Saprotroph | \|Undefined Saprotroph\| | Yeast | NULL | Probable | As nectar-tap_saprotroph (Põlme et al. 2020); Decay sugar-rich substrates (Põlme et al. 2020) | Põlme S, et al. 2020. Fungal Diversity 105:1-16 (https://doi.org/10.1007/s13225-020-00466-2) |
| Otu19414 | 1 | k__Fungi(100);p__Ascomycota(100);c__Saccharomycetes(100);o__Saccharomycetales(100);f__Metschnikowiaceae(100);g__Metschnikowia(100);s__Metschnikowia_pulcherrima(100); | Metschnikowia | 13 | Saprotroph | \|Undefined Saprotroph\| | Yeast | NULL | Probable | As nectar-tap_saprotroph (Põlme et al. 2020); Decay sugar-rich substrates (Põlme et al. 2020) | Põlme S, et al. 2020. Fungal Diversity 105:1-16 (https://doi.org/10.1007/s13225-020-00466-2) |
| Otu19415 | 1 | k__Fungi(100);p__Ascomycota(100);c__Saccharomycetes(100);o__Saccharomycetales(100);f__Metschnikowiaceae(100);g__Metschnikowia(100);s__Metschnikowia_pulcherrima(100); | Metschnikowia | 13 | Saprotroph | \|Undefined Saprotroph\| | Yeast | NULL | Probable | As nectar-tap_saprotroph (Põlme et al. 2020); Decay sugar-rich substrates (Põlme et al. 2020) | Põlme S, et al. 2020. Fungal Diversity 105:1-16 (https://doi.org/10.1007/s13225-020-00466-2) |
| Otu19416 | 1 | k__Fungi(100);p__Ascomycota(100);c__Saccharomycetes(100);o__Saccharomycetales(100);f__Metschnikowiaceae(100);g__Metschnikowia(100);s__Metschnikowia_pulcherrima(100); | Metschnikowia | 13 | Saprotroph | \|Undefined Saprotroph\| | Yeast | NULL | Probable | As nectar-tap_saprotroph (Põlme et al. 2020); Decay sugar-rich substrates (Põlme et al. 2020) | Põlme S, et al. 2020. Fungal Diversity 105:1-16 (https://doi.org/10.1007/s13225-020-00466-2) |
| Otu19417 | 1 | k__Fungi(100);p__Ascomycota(100);c__Saccharomycetes(100);o__Saccharomycetales(100);f__Metschnikowiaceae(100);g__Metschnikowia(100);s__Metschnikowia_pulcherrima(100); | Metschnikowia | 13 | Saprotroph | \|Undefined Saprotroph\| | Yeast | NULL | Probable | As nectar-tap_saprotroph (Põlme et al. 2020); Decay sugar-rich substrates (Põlme et al. 2020) | Põlme S, et al. 2020. Fungal Diversity 105:1-16 (https://doi.org/10.1007/s13225-020-00466-2) |
| Otu19418 | 1 | k__Fungi(100);p__Ascomycota(100);c__Saccharomycetes(100);o__Saccharomycetales(100);f__Metschnikowiaceae(100);g__Metschnikowia(100);g__Metschnikowia_unclassified(100); | Metschnikowia | 13 | Saprotroph | \|Undefined Saprotroph\| | Yeast | NULL | Probable | As nectar-tap_saprotroph (Põlme et al. 2020); Decay sugar-rich substrates (Põlme et al. 2020) | Põlme S, et al. 2020. Fungal Diversity 105:1-16 (https://doi.org/10.1007/s13225-020-00466-2) |
| Otu19419 | 1 | k__Fungi(100);p__Ascomycota(100);c__Saccharomycetes(100);o__Saccharomycetales(100);f__Metschnikowiaceae(100);g__Metschnikowia(100);g__Metschnikowia_unclassified(100); | Metschnikowia | 13 | Saprotroph | \|Undefined Saprotroph\| | Yeast | NULL | Probable | As nectar-tap_saprotroph (Põlme et al. 2020); Decay sugar-rich substrates (Põlme et al. 2020) | Põlme S, et al. 2020. Fungal Diversity 105:1-16 (https://doi.org/10.1007/s13225-020-00466-2) |

| | | | | | | | | | | | |
|---|---|---|---|---|---|---|---|---|---|---|---|
| Otu19420 | 1 | k__Fungi(100);p__Ascomycota(100);c__Saccharomycetes(100);o__Saccharomycetales(100);f__Metschnikowiaceae(100);g__Metschnikowia(100);g__Metschnikowia_unclassified(100); | Metschnikowia | 13 | Saprotroph | \|Undefined Saprotroph\| | Yeast | NULL | Probable | As nectar-tap_saprotroph (Põlme et al. 2020); Decay sugar-rich substrates (Põlme et al. 2020) | Põlme S, et al. 2020. Fungal Diversity 105:1-16 ((https://doi.org/10.1007/s13225-020-00466-2)) |
| Otu19421 | 1 | k__Fungi(100);p__Ascomycota(100);c__Saccharomycetes(100);o__Saccharomycetales(100);f__Metschnikowiaceae(100);g__Metschnikowia(100);s__Metschnikowia_pulcherrima(100); | Metschnikowia | 13 | Saprotroph | \|Undefined Saprotroph\| | Yeast | NULL | Probable | As nectar-tap_saprotroph (Põlme et al. 2020); Decay sugar-rich substrates (Põlme et al. 2020) | Põlme S, et al. 2020. Fungal Diversity 105:1-16 ((https://doi.org/10.1007/s13225-020-00466-2)) |
| Otu19422 | 1 | k__Fungi(100);p__Ascomycota(100);c__Saccharomycetes(100);o__Saccharomycetales(100);f__Metschnikowiaceae(100);g__Metschnikowia(100);s__Metschnikowia_pulcherrima(100); | Metschnikowia | 13 | Saprotroph | \|Undefined Saprotroph\| | Yeast | NULL | Probable | As nectar-tap_saprotroph (Põlme et al. 2020); Decay sugar-rich substrates (Põlme et al. 2020) | Põlme S, et al. 2020. Fungal Diversity 105:1-16 ((https://doi.org/10.1007/s13225-020-00466-2)) |
| Otu19423 | 1 | k__Fungi(100);p__Ascomycota(100);c__Saccharomycetes(100);o__Saccharomycetales(100);f__Metschnikowiaceae(100);g__Metschnikowia(100);s__Metschnikowia_pulcherrima(100); | Metschnikowia | 13 | Saprotroph | \|Undefined Saprotroph\| | Yeast | NULL | Probable | As nectar-tap_saprotroph (Põlme et al. 2020); Decay sugar-rich substrates (Põlme et al. 2020) | Põlme S, et al. 2020. Fungal Diversity 105:1-16 ((https://doi.org/10.1007/s13225-020-00466-2)) |
| Otu19424 | 1 | k__Fungi(100);p__Ascomycota(100);c__Saccharomycetes(100);o__Saccharomycetales(100);f__Metschnikowiaceae(100);g__Metschnikowia(100);s__Metschnikowia_pulcherrima(100); | Metschnikowia | 13 | Saprotroph | \|Undefined Saprotroph\| | Yeast | NULL | Probable | As nectar-tap_saprotroph (Põlme et al. 2020); Decay sugar-rich substrates (Põlme et al. 2020) | Põlme S, et al. 2020. Fungal Diversity 105:1-16 ((https://doi.org/10.1007/s13225-020-00466-2)) |
| Otu19425 | 1 | k__Fungi(100);p__Ascomycota(100);c__Saccharomycetes(100);o__Saccharomycetales(100);f__Metschnikowiaceae(100);g__Metschnikowia(100);s__Metschnikowia_pulcherrima(100); | Metschnikowia | 13 | Saprotroph | \|Undefined Saprotroph\| | Yeast | NULL | Probable | As nectar-tap_saprotroph (Põlme et al. 2020); Decay sugar-rich substrates (Põlme et al. 2020) | Põlme S, et al. 2020. Fungal Diversity 105:1-16 ((https://doi.org/10.1007/s13225-020-00466-2)) |
| Otu19426 | 1 | k__Fungi(100);p__Ascomycota(100);c__Saccharomycetes(100);o__Saccharomycetales(100);f__Metschnikowiaceae(100);g__Metschnikowia(100);s__Metschnikowia_pulcherrima(100); | Metschnikowia | 13 | Saprotroph | \|Undefined Saprotroph\| | Yeast | NULL | Probable | As nectar-tap_saprotroph (Põlme et al. 2020); Decay sugar-rich substrates (Põlme et al. 2020) | Põlme S, et al. 2020. Fungal Diversity 105:1-16 ((https://doi.org/10.1007/s13225-020-00466-2)) |
| Otu19427 | 1 | k__Fungi(100);p__Ascomycota(100);c__Saccharomycetes(100);o__Saccharomycetales(100);f__Metschnikowiaceae(100);g__Metschnikowia(100);s__Metschnikowia_pulcherrima(100); | Metschnikowia | 13 | Saprotroph | \|Undefined Saprotroph\| | Yeast | NULL | Probable | As nectar-tap_saprotroph (Põlme et al. 2020); Decay sugar-rich substrates (Põlme et al. 2020) | Põlme S, et al. 2020. Fungal Diversity 105:1-16 ((https://doi.org/10.1007/s13225-020-00466-2)) |
| Otu19428 | 1 | k__Fungi(100);p__Ascomycota(100);c__Saccharomycetes(100);o__Saccharomycetales(100);f__Metschnikowiaceae(100);g__Metschnikowia(100);s__Metschnikowia_pulcherrima(100); | Metschnikowia | 13 | Saprotroph | \|Undefined Saprotroph\| | Yeast | NULL | Probable | As nectar-tap_saprotroph (Põlme et al. 2020); Decay sugar-rich substrates (Põlme et al. 2020) | Põlme S, et al. 2020. Fungal Diversity 105:1-16 ((https://doi.org/10.1007/s13225-020-00466-2)) |
| Otu19429 | 1 | k__Fungi(100);p__Ascomycota(100);c__Saccharomycetes(100);o__Saccharomycetales(100);f__Metschnikowiaceae(100);g__Metschnikowia(100);s__Metschnikowia_pulcherrima(100); | Metschnikowia | 13 | Saprotroph | \|Undefined Saprotroph\| | Yeast | NULL | Probable | As nectar-tap_saprotroph (Põlme et al. 2020); Decay sugar-rich substrates (Põlme et al. 2020) | Põlme S, et al. 2020. Fungal Diversity 105:1-16 ((https://doi.org/10.1007/s13225-020-00466-2)) |
| Otu19430 | 1 | k__Fungi(100);p__Ascomycota(100);c__Saccharomycetes(100);o__Saccharomycetales(100);f__Metschnikowiaceae(100);g__Metschnikowia(100);s__Metschnikowia_pulcherrima(100); | Metschnikowia | 13 | Saprotroph | \|Undefined Saprotroph\| | Yeast | NULL | Probable | As nectar-tap_saprotroph (Põlme et al. 2020); Decay sugar-rich substrates (Põlme et al. 2020) | Põlme S, et al. 2020. Fungal Diversity 105:1-16 ((https://doi.org/10.1007/s13225-020-00466-2)) |
| Otu19431 | 1 | k__Fungi(100);p__Ascomycota(100);c__Saccharomycetes(100);o__Saccharomycetales(100);f__Metschnikowiaceae(100);g__Metschnikowia(100);s__Metschnikowia_pulcherrima(100); | Metschnikowia | 13 | Saprotroph | \|Undefined Saprotroph\| | Yeast | NULL | Probable | As nectar-tap_saprotroph (Põlme et al. 2020); Decay sugar-rich substrates (Põlme et al. 2020) | Põlme S, et al. 2020. Fungal Diversity 105:1-16 ((https://doi.org/10.1007/s13225-020-00466-2)) |
| Otu19432 | 1 | k__Fungi(100);p__Ascomycota(100);c__Saccharomycetes(100);o__Saccharomycetales(100);f__Metschnikowiaceae(100);g__Metschnikowia(100);s__Metschnikowia_sp(100); | Metschnikowia | 13 | Saprotroph | \|Undefined Saprotroph\| | Yeast | NULL | Probable | As nectar-tap_saprotroph (Põlme et al. 2020); Decay sugar-rich substrates (Põlme et al. 2020) | Põlme S, et al. 2020. Fungal Diversity 105:1-16 ((https://doi.org/10.1007/s13225-020-00466-2)) |
| Otu19433 | 1 | k__Fungi(100);p__Ascomycota(100);c__Saccharomycetes(100);o__Saccharomycetales(100);f__Metschnikowiaceae(100);g__Metschnikowia(100);g__Metschnikowia_unclassified(100); | Metschnikowia | 13 | Saprotroph | \|Undefined Saprotroph\| | Yeast | NULL | Probable | As nectar-tap_saprotroph (Põlme et al. 2020); Decay sugar-rich substrates (Põlme et al. 2020) | Põlme S, et al. 2020. Fungal Diversity 105:1-16 ((https://doi.org/10.1007/s13225-020-00466-2)) |
| Otu19434 | 1 | k__Fungi(100);p__Ascomycota(100);c__Saccharomycetes(100);o__Saccharomycetales(100);f__Metschnikowiaceae(100);g__Metschnikowia(100);g__Metschnikowia_unclassified(100); | Metschnikowia | 13 | Saprotroph | \|Undefined Saprotroph\| | Yeast | NULL | Probable | As nectar-tap_saprotroph (Põlme et al. 2020); Decay sugar-rich substrates (Põlme et al. 2020) | Põlme S, et al. 2020. Fungal Diversity 105:1-16 ((https://doi.org/10.1007/s13225-020-00466-2)) |
| Otu19435 | 1 | k__Fungi(100);p__Ascomycota(100);c__Saccharomycetes(100);o__Saccharomycetales(100);f__Metschnikowiaceae(100);g__Metschnikowia(100);s__Metschnikowia_pulcherrima(100); | Metschnikowia | 13 | Saprotroph | \|Undefined Saprotroph\| | Yeast | NULL | Probable | As nectar-tap_saprotroph (Põlme et al. 2020); Decay sugar-rich substrates (Põlme et al. 2020) | Põlme S, et al. 2020. Fungal Diversity 105:1-16 ((https://doi.org/10.1007/s13225-020-00466-2)) |
| Otu19436 | 1 | k__Fungi(100);p__Ascomycota(100);c__Saccharomycetes(100);o__Saccharomycetales(100);f__Metschnikowiaceae(100);g__Metschnikowia(100);s__Metschnikowia_pulcherrima(100); | Metschnikowia | 13 | Saprotroph | \|Undefined Saprotroph\| | Yeast | NULL | Probable | As nectar-tap_saprotroph (Põlme et al. 2020); Decay sugar-rich substrates (Põlme et al. 2020) | Põlme S, et al. 2020. Fungal Diversity 105:1-16 ((https://doi.org/10.1007/s13225-020-00466-2)) |
| Otu19437 | 1 | k__Fungi(100);p__Ascomycota(100);c__Saccharomycetes(100);o__Saccharomycetales(100);f__Metschnikowiaceae(100);g__Metschnikowia(100);s__Metschnikowia_pulcherrima(100); | Metschnikowia | 13 | Saprotroph | \|Undefined Saprotroph\| | Yeast | NULL | Probable | As nectar-tap_saprotroph (Põlme et al. 2020); Decay sugar-rich substrates (Põlme et al. 2020) | Põlme S, et al. 2020. Fungal Diversity 105:1-16 ((https://doi.org/10.1007/s13225-020-00466-2)) |
| Otu19438 | 1 | k__Fungi(100);p__Ascomycota(100);c__Saccharomycetes(100);o__Saccharomycetales(100);f__Metschnikowiaceae(100);g__Metschnikowia(100);s__Metschnikowia_pulcherrima(100); | Metschnikowia | 13 | Saprotroph | \|Undefined Saprotroph\| | Yeast | NULL | Probable | As nectar-tap_saprotroph (Põlme et al. 2020); Decay sugar-rich substrates (Põlme et al. 2020) | Põlme S, et al. 2020. Fungal Diversity 105:1-16 ((https://doi.org/10.1007/s13225-020-00466-2)) |
| Otu19439 | 1 | k__Fungi(100);p__Ascomycota(100);c__Saccharomycetes(100);o__Saccharomycetales(100);f__Metschnikowiaceae(100);g__Metschnikowia(100);s__Metschnikowia_pulcherrima(100); | Metschnikowia | 13 | Saprotroph | \|Undefined Saprotroph\| | Yeast | NULL | Probable | As nectar-tap_saprotroph (Põlme et al. 2020); Decay sugar-rich substrates (Põlme et al. 2020) | Põlme S, et al. 2020. Fungal Diversity 105:1-16 ((https://doi.org/10.1007/s13225-020-00466-2)) |
| Otu19440 | 1 | k__Fungi(100);p__Ascomycota(100);c__Saccharomycetes(100);o__Saccharomycetales(100);f__Metschnikowiaceae(100);g__Metschnikowia(100);s__Metschnikowia_pulcherrima(100); | Metschnikowia | 13 | Saprotroph | \|Undefined Saprotroph\| | Yeast | NULL | Probable | As nectar-tap_saprotroph (Põlme et al. 2020); Decay sugar-rich substrates (Põlme et al. 2020) | Põlme S, et al. 2020. Fungal Diversity 105:1-16 ((https://doi.org/10.1007/s13225-020-00466-2)) |
| Otu19441 | 1 | k__Fungi(100);p__Ascomycota(100);c__Saccharomycetes(100);o__Saccharomycetales(100);f__Metschnikowiaceae(100);g__Metschnikowia(100);s__Metschnikowia_pulcherrima(100); | Metschnikowia | 13 | Saprotroph | \|Undefined Saprotroph\| | Yeast | NULL | Probable | As nectar-tap_saprotroph (Põlme et al. 2020); Decay sugar-rich substrates (Põlme et al. 2020) | Põlme S, et al. 2020. Fungal Diversity 105:1-16 ((https://doi.org/10.1007/s13225-020-00466-2)) |
| Otu19442 | 1 | k__Fungi(100);p__Ascomycota(100);c__Saccharomycetes(100);o__Saccharomycetales(100);f__Metschnikowiaceae(100);g__Metschnikowia(100);s__Metschnikowia_pulcherrima(100); | Metschnikowia | 13 | Saprotroph | \|Undefined Saprotroph\| | Yeast | NULL | Probable | As nectar-tap_saprotroph (Põlme et al. 2020); Decay sugar-rich substrates (Põlme et al. 2020) | Põlme S, et al. 2020. Fungal Diversity 105:1-16 ((https://doi.org/10.1007/s13225-020-00466-2)) |
| Otu19443 | 1 | k__Fungi(100);p__Ascomycota(100);c__Saccharomycetes(100);o__Saccharomycetales(100);f__Metschnikowiaceae(100);g__Metschnikowia(100);s__Metschnikowia_pulcherrima(100); | Metschnikowia | 13 | Saprotroph | \|Undefined Saprotroph\| | Yeast | NULL | Probable | As nectar-tap_saprotroph (Põlme et al. 2020); Decay sugar-rich substrates (Põlme et al. 2020) | Põlme S, et al. 2020. Fungal Diversity 105:1-16 ((https://doi.org/10.1007/s13225-020-00466-2)) |
| Otu19444 | 1 | k__Fungi(100);p__Ascomycota(100);c__Saccharomycetes(100);o__Saccharomycetales(100);f__Metschnikowiaceae(100);g__Metschnikowia(100);s__Metschnikowia_pulcherrima(100); | Metschnikowia | 13 | Saprotroph | \|Undefined Saprotroph\| | Yeast | NULL | Probable | As nectar-tap_saprotroph (Põlme et al. 2020); Decay sugar-rich substrates (Põlme et al. 2020) | Põlme S, et al. 2020. Fungal Diversity 105:1-16 ((https://doi.org/10.1007/s13225-020-00466-2)) |
| Otu19445 | 1 | k__Fungi(100);p__Ascomycota(100);c__Saccharomycetes(100);o__Saccharomycetales(100);f__Metschnikowiaceae(100);g__Metschnikowia(100);s__Metschnikowia_pulcherrima(100); | Metschnikowia | 13 | Saprotroph | \|Undefined Saprotroph\| | Yeast | NULL | Probable | As nectar-tap_saprotroph (Põlme et al. 2020); Decay sugar-rich substrates (Põlme et al. 2020) | Põlme S, et al. 2020. Fungal Diversity 105:1-16 ((https://doi.org/10.1007/s13225-020-00466-2)) |
| Otu19446 | 1 | k__Fungi(100);p__Ascomycota(100);c__Saccharomycetes(100);o__Saccharomycetales(100);f__Metschnikowiaceae(100);g__Metschnikowia(100);s__Metschnikowia_pulcherrima(100); | Metschnikowia | 13 | Saprotroph | \|Undefined Saprotroph\| | Yeast | NULL | Probable | As nectar-tap_saprotroph (Põlme et al. 2020); Decay sugar-rich substrates (Põlme et al. 2020) | Põlme S, et al. 2020. Fungal Diversity 105:1-16 ((https://doi.org/10.1007/s13225-020-00466-2)) |
| Otu19447 | 1 | k__Fungi(100);p__Ascomycota(100);c__Saccharomycetes(100);o__Saccharomycetales(100);f__Metschnikowiaceae(100);g__Metschnikowia(100);s__Metschnikowia_pulcherrima(100); | Metschnikowia | 13 | Saprotroph | \|Undefined Saprotroph\| | Yeast | NULL | Probable | As nectar-tap_saprotroph (Põlme et al. 2020); Decay sugar-rich substrates (Põlme et al. 2020) | Põlme S, et al. 2020. Fungal Diversity 105:1-16 ((https://doi.org/10.1007/s13225-020-00466-2)) |
| Otu19448 | 1 | k__Fungi(100);p__Ascomycota(100);c__Saccharomycetes(100);o__Saccharomycetales(100);f__Metschnikowiaceae(100);g__Metschnikowia(100);s__Metschnikowia_pulcherrima(100); | Metschnikowia | 13 | Saprotroph | \|Undefined Saprotroph\| | Yeast | NULL | Probable | As nectar-tap_saprotroph (Põlme et al. 2020); Decay sugar-rich substrates (Põlme et al. 2020) | Põlme S, et al. 2020. Fungal Diversity 105:1-16 ((https://doi.org/10.1007/s13225-020-00466-2)) |
| Otu19449 | 1 | k__Fungi(100);p__Ascomycota(100);c__Saccharomycetes(100);o__Saccharomycetales(100);f__Metschnikowiaceae(100);g__Metschnikowia(100);s__Metschnikowia_pulcherrima(100); | Metschnikowia | 13 | Saprotroph | \|Undefined Saprotroph\| | Yeast | NULL | Probable | As nectar-tap_saprotroph (Põlme et al. 2020); Decay sugar-rich substrates (Põlme et al. 2020) | Põlme S, et al. 2020. Fungal Diversity 105:1-16 ((https://doi.org/10.1007/s13225-020-00466-2)) |
| Otu19450 | 1 | k__Fungi(100);p__Ascomycota(100);c__Saccharomycetes(100);o__Saccharomycetales(100);f__Metschnikowiaceae(100);g__Metschnikowia(100);s__Metschnikowia_pulcherrima(100); | Metschnikowia | 13 | Saprotroph | \|Undefined Saprotroph\| | Yeast | NULL | Probable | As nectar-tap_saprotroph (Põlme et al. 2020); Decay sugar-rich substrates (Põlme et al. 2020) | Põlme S, et al. 2020. Fungal Diversity 105:1-16 ((https://doi.org/10.1007/s13225-020-00466-2)) |
| Otu19451 | 1 | k__Fungi(100);p__Ascomycota(100);c__Saccharomycetes(100);o__Saccharomycetales(100);f__Metschnikowiaceae(100);g__Metschnikowia(100);s__Metschnikowia_pulcherrima(100); | Metschnikowia | 13 | Saprotroph | \|Undefined Saprotroph\| | Yeast | NULL | Probable | As nectar-tap_saprotroph (Põlme et al. 2020); Decay sugar-rich substrates (Põlme et al. 2020) | Põlme S, et al. 2020. Fungal Diversity 105:1-16 ((https://doi.org/10.1007/s13225-020-00466-2)) |
| Otu19452 | 1 | k__Fungi(100);p__Ascomycota(100);c__Saccharomycetes(100);o__Saccharomycetales(100);f__Metschnikowiaceae(100);g__Metschnikowia(100);s__Metschnikowia_pulcherrima(100); | Metschnikowia | 13 | Saprotroph | \|Undefined Saprotroph\| | Yeast | NULL | Probable | As nectar-tap_saprotroph (Põlme et al. 2020); Decay sugar-rich substrates (Põlme et al. 2020) | Põlme S, et al. 2020. Fungal Diversity 105:1-16 ((https://doi.org/10.1007/s13225-020-00466-2)) |
| Otu19453 | 1 | k__Fungi(100);p__Ascomycota(100);c__Saccharomycetes(100);o__Saccharomycetales(100);f__Metschnikowiaceae(100);g__Metschnikowia(100);s__Metschnikowia_pulcherrima(100); | Metschnikowia | 13 | Saprotroph | \|Undefined Saprotroph\| | Yeast | NULL | Probable | As nectar-tap_saprotroph (Põlme et al. 2020); Decay sugar-rich substrates (Põlme et al. 2020) | Põlme S, et al. 2020. Fungal Diversity 105:1-16 ((https://doi.org/10.1007/s13225-020-00466-2)) |
| Otu19454 | 1 | k__Fungi(100);p__Ascomycota(100);c__Saccharomycetes(100);o__Saccharomycetales(100);f__Metschnikowiaceae(100);g__Metschnikowia(100);g__Metschnikowia_unclassified(100); | Metschnikowia | 13 | Saprotroph | \|Undefined Saprotroph\| | Yeast | NULL | Probable | As nectar-tap_saprotroph (Põlme et al. 2020); Decay sugar-rich substrates (Põlme et al. 2020) | Põlme S, et al. 2020. Fungal Diversity 105:1-16 ((https://doi.org/10.1007/s13225-020-00466-2)) |
| Otu19455 | 1 | k__Fungi(100);p__Ascomycota(100);c__Saccharomycetes(100);o__Saccharomycetales(100);f__Metschnikowiaceae(100);g__Metschnikowia(100);s__Metschnikowia_pulcherrima(100); | Metschnikowia | 13 | Saprotroph | \|Undefined Saprotroph\| | Yeast | NULL | Probable | As nectar-tap_saprotroph (Põlme et al. 2020); Decay sugar-rich substrates (Põlme et al. 2020) | Põlme S, et al. 2020. Fungal Diversity 105:1-16 ((https://doi.org/10.1007/s13225-020-00466-2)) |
| Otu19456 | 1 | k__Fungi(100);p__Ascomycota(100);c__Saccharomycetes(100);o__Saccharomycetales(100);f__Metschnikowiaceae(100);g__Metschnikowia(100);s__Metschnikowia_pulcherrima(100); | Metschnikowia | 13 | Saprotroph | \|Undefined Saprotroph\| | Yeast | NULL | Probable | As nectar-tap_saprotroph (Põlme et al. 2020); Decay sugar-rich substrates (Põlme et al. 2020) | Põlme S, et al. 2020. Fungal Diversity 105:1-16 ((https://doi.org/10.1007/s13225-020-00466-2)) |
| Otu19457 | 1 | k__Fungi(100);p__Ascomycota(100);c__Saccharomycetes(100);o__Saccharomycetales(100);f__Metschnikowiaceae(100);g__Metschnikowia(100);g__Metschnikowia_unclassified(100); | Metschnikowia | 13 | Saprotroph | \|Undefined Saprotroph\| | Yeast | NULL | Probable | As nectar-tap_saprotroph (Põlme et al. 2020); Decay sugar-rich substrates (Põlme et al. 2020) | Põlme S, et al. 2020. Fungal Diversity 105:1-16 ((https://doi.org/10.1007/s13225-020-00466-2)) |
| Otu19458 | 1 | k__Fungi(100);p__Ascomycota(100);c__Saccharomycetes(100);o__Saccharomycetales(100);f__Metschnikowiaceae(100);g__Metschnikowia(100);s__Metschnikowia_pulcherrima(100); | Metschnikowia | 13 | Saprotroph | \|Undefined Saprotroph\| | Yeast | NULL | Probable | As nectar-tap_saprotroph (Põlme et al. 2020); Decay sugar-rich substrates (Põlme et al. 2020) | Põlme S, et al. 2020. Fungal Diversity 105:1-16 ((https://doi.org/10.1007/s13225-020-00466-2)) |
| Otu19459 | 1 | k__Fungi(100);p__Ascomycota(100);c__Saccharomycetes(100);o__Saccharomycetales(100);f__Metschnikowiaceae(100);g__Metschnikowia(100);s__Metschnikowia_pulcherrima(100); | Metschnikowia | 13 | Saprotroph | \|Undefined Saprotroph\| | Yeast | NULL | Probable | As nectar-tap_saprotroph (Põlme et al. 2020); Decay sugar-rich substrates (Põlme et al. 2020) | Põlme S, et al. 2020. Fungal Diversity 105:1-16 ((https://doi.org/10.1007/s13225-020-00466-2)) |
| Otu19460 | 1 | k__Fungi(100);p__Ascomycota(100);c__Saccharomycetes(100);o__Saccharomycetales(100);f__Metschnikowiaceae(100);g__Metschnikowia(100);g__Metschnikowia_unclassified(100); | Metschnikowia | 13 | Saprotroph | \|Undefined Saprotroph\| | Yeast | NULL | Probable | As nectar-tap_saprotroph (Põlme et al. 2020); Decay sugar-rich substrates (Põlme et al. 2020) | Põlme S, et al. 2020. Fungal Diversity 105:1-16 ((https://doi.org/10.1007/s13225-020-00466-2)) |
| Otu19461 | 1 | k__Fungi(100);p__Ascomycota(100);c__Saccharomycetes(100);o__Saccharomycetales(100);f__Metschnikowiaceae(100);g__Metschnikowia(100);g__Metschnikowia_unclassified(100); | Metschnikowia | 13 | Saprotroph | \|Undefined Saprotroph\| | Yeast | NULL | Probable | As nectar-tap_saprotroph (Põlme et al. 2020); Decay sugar-rich substrates (Põlme et al. 2020) | Põlme S, et al. 2020. Fungal Diversity 105:1-16 ((https://doi.org/10.1007/s13225-020-00466-2)) |
| Otu19462 | 1 | k__Fungi(100);p__Ascomycota(100);c__Saccharomycetes(100);o__Saccharomycetales(100);f__Metschnikowiaceae(100);g__Metschnikowia(100);g__Metschnikowia_unclassified(100); | Metschnikowia | 13 | Saprotroph | \|Undefined Saprotroph\| | Yeast | NULL | Probable | As nectar-tap_saprotroph (Põlme et al. 2020); Decay sugar-rich substrates (Põlme et al. 2020) | Põlme S, et al. 2020. Fungal Diversity 105:1-16 ((https://doi.org/10.1007/s13225-020-00466-2)) |
| Otu19463 | 1 | k__Fungi(100);p__Ascomycota(100);c__Saccharomycetes(100);o__Saccharomycetales(100);f__Metschnikowiaceae(100);g__Metschnikowia(100);g__Metschnikowia_unclassified(100); | Metschnikowia | 13 | Saprotroph | \|Undefined Saprotroph\| | Yeast | NULL | Probable | As nectar-tap_saprotroph (Põlme et al. 2020); Decay sugar-rich substrates (Põlme et al. 2020) | Põlme S, et al. 2020. Fungal Diversity 105:1-16 ((https://doi.org/10.1007/s13225-020-00466-2)) |
| Otu19464 | 1 | k__Fungi(100);p__Ascomycota(100);c__Saccharomycetes(100);o__Saccharomycetales(100);f__Metschnikowiaceae(100);g__Metschnikowia(100);s__Metschnikowia_pulcherrima(100); | Metschnikowia | 13 | Saprotroph | \|Undefined Saprotroph\| | Yeast | NULL | Probable | As nectar-tap_saprotroph (Põlme et al. 2020); Decay sugar-rich substrates (Põlme et al. 2020) | Põlme S, et al. 2020. Fungal Diversity 105:1-16 ((https://doi.org/10.1007/s13225-020-00466-2)) |
| Otu19465 | 1 | k__Fungi(100);p__Ascomycota(100);c__Saccharomycetes(100);o__Saccharomycetales(100);f__Metschnikowiaceae(100);g__Metschnikowia(100);s__Metschnikowia_pulcherrima(100); | Metschnikowia | 13 | Saprotroph | \|Undefined Saprotroph\| | Yeast | NULL | Probable | As nectar-tap_saprotroph (Põlme et al. 2020); Decay sugar-rich substrates (Põlme et al. 2020) | Põlme S, et al. 2020. Fungal Diversity 105:1-16 ((https://doi.org/10.1007/s13225-020-00466-2)) |
| Otu19466 | 1 | k__Fungi(100);p__Ascomycota(100);c__Saccharomycetes(100);o__Saccharomycetales(100);f__Metschnikowiaceae(100);g__Metschnikowia(100);s__Metschnikowia_pulcherrima(100); | Metschnikowia | 13 | Saprotroph | \|Undefined Saprotroph\| | Yeast | NULL | Probable | As nectar-tap_saprotroph (Põlme et al. 2020); Decay sugar-rich substrates (Põlme et al. 2020) | Põlme S, et al. 2020. Fungal Diversity 105:1-16 ((https://doi.org/10.1007/s13225-020-00466-2)) |
| Otu19467 | 1 | k__Fungi(100);p__Ascomycota(100);c__Saccharomycetes(100);o__Saccharomycetales(100);f__Metschnikowiaceae(100);g__Metschnikowia(100);s__Metschnikowia_pulcherrima(100); | Metschnikowia | 13 | Saprotroph | \|Undefined Saprotroph\| | Yeast | NULL | Probable | As nectar-tap_saprotroph (Põlme et al. 2020); Decay sugar-rich substrates (Põlme et al. 2020) | Põlme S, et al. 2020. Fungal Diversity 105:1-16 ((https://doi.org/10.1007/s13225-020-00466-2)) |
| Otu19468 | 1 | k__Fungi(100);p__Ascomycota(100);c__Saccharomycetes(100);o__Saccharomycetales(100);f__Metschnikowiaceae(100);g__Metschnikowia(100);s__Metschnikowia_pulcherrima(100); | Metschnikowia | 13 | Saprotroph | \|Undefined Saprotroph\| | Yeast | NULL | Probable | As nectar-tap_saprotroph (Põlme et al. 2020); Decay sugar-rich substrates (Põlme et al. 2020) | Põlme S, et al. 2020. Fungal Diversity 105:1-16 ((https://doi.org/10.1007/s13225-020-00466-2)) |
| Otu19469 | 1 | k__Fungi(100);p__Ascomycota(100);c__Saccharomycetes(100);o__Saccharomycetales(100);f__Metschnikowiaceae(100);g__Metschnikowia(100);s__Metschnikowia_pulcherrima(100); | Metschnikowia | 13 | Saprotroph | \|Undefined Saprotroph\| | Yeast | NULL | Probable | As nectar-tap_saprotroph (Põlme et al. 2020); Decay sugar-rich substrates (Põlme et al. 2020) | Põlme S, et al. 2020. Fungal Diversity 105:1-16 ((https://doi.org/10.1007/s13225-020-00466-2)) |
| Otu19470 | 1 | k__Fungi(100);p__Ascomycota(100);c__Saccharomycetes(100);o__Saccharomycetales(100);f__Metschnikowiaceae(100);g__Metschnikowia(100);s__Metschnikowia_pulcherrima(100); | Metschnikowia | 13 | Saprotroph | \|Undefined Saprotroph\| | Yeast | NULL | Probable | As nectar-tap_saprotroph (Põlme et al. 2020); Decay sugar-rich substrates (Põlme et al. 2020) | Põlme S, et al. 2020. Fungal Diversity 105:1-16 ((https://doi.org/10.1007/s13225-020-00466-2)) |
| Otu19471 | 1 | k__Fungi(100);p__Ascomycota(100);c__Saccharomycetes(100);o__Saccharomycetales(100);f__Metschnikowiaceae(100);g__Metschnikowia(100);g__Metschnikowia_unclassified(100); | Metschnikowia | 13 | Saprotroph | \|Undefined Saprotroph\| | Yeast | NULL | Probable | As nectar-tap_saprotroph (Põlme et al. 2020); Decay sugar-rich substrates (Põlme et al. 2020) | Põlme S, et al. 2020. Fungal Diversity 105:1-16 ((https://doi.org/10.1007/s13225-020-00466-2)) |
| Otu19472 | 1 | k__Fungi(100);p__Ascomycota(100);c__Saccharomycetes(100);o__Saccharomycetales(100);f__Metschnikowiaceae(100);g__Metschnikowia(100);s__Metschnikowia_pulcherrima(100); | Metschnikowia | 13 | Saprotroph | \|Undefined Saprotroph\| | Yeast | NULL | Probable | As nectar-tap_saprotroph (Põlme et al. 2020); Decay sugar-rich substrates (Põlme et al. 2020) | Põlme S, et al. 2020. Fungal Diversity 105:1-16 ((https://doi.org/10.1007/s13225-020-00466-2)) |
| Otu19473 | 1 | k__Fungi(100);p__Ascomycota(100);c__Saccharomycetes(100);o__Saccharomycetales(100);f__Metschnikowiaceae(100);g__Metschnikowia(100);s__Metschnikowia_pulcherrima(100); | Metschnikowia | 13 | Saprotroph | \|Undefined Saprotroph\| | Yeast | NULL | Probable | As nectar-tap_saprotroph (Põlme et al. 2020); Decay sugar-rich substrates (Põlme et al. 2020) | Põlme S, et al. 2020. Fungal Diversity 105:1-16 ((https://doi.org/10.1007/s13225-020-00466-2)) |
| Otu19474 | 1 | k__Fungi(100);p__Ascomycota(100);c__Saccharomycetes(100);o__Saccharomycetales(100);f__Metschnikowiaceae(100);g__Metschnikowia(100);s__Metschnikowia_pulcherrima(100); | Metschnikowia | 13 | Saprotroph | \|Undefined Saprotroph\| | Yeast | NULL | Probable | As nectar-tap_saprotroph (Põlme et al. 2020); Decay sugar-rich substrates (Põlme et al. 2020) | Põlme S, et al. 2020. Fungal Diversity 105:1-16 ((https://doi.org/10.1007/s13225-020-00466-2)) |
| Otu19475 | 1 | k__Fungi(100);p__Ascomycota(100);c__Saccharomycetes(100);o__Saccharomycetales(100);f__Metschnikowiaceae(100);g__Metschnikowia(100);s__Metschnikowia_pulcherrima(100); | Metschnikowia | 13 | Saprotroph | \|Undefined Saprotroph\| | Yeast | NULL | Probable | As nectar-tap_saprotroph (Põlme et al. 2020); Decay sugar-rich substrates (Põlme et al. 2020) | Põlme S, et al. 2020. Fungal Diversity 105:1-16 ((https://doi.org/10.1007/s13225-020-00466-2)) |
| Otu19476 | 1 | k__Fungi(100);p__Ascomycota(100);c__Saccharomycetes(100);o__Saccharomycetales(100);f__Metschnikowiaceae(100);g__Metschnikowia(100);g__Metschnikowia_unclassified(100); | Metschnikowia | 13 | Saprotroph | \|Undefined Saprotroph\| | Yeast | NULL | Probable | As nectar-tap_saprotroph (Põlme et al. 2020); Decay sugar-rich substrates (Põlme et al. 2020) | Põlme S, et al. 2020. Fungal Diversity 105:1-16 ((https://doi.org/10.1007/s13225-020-00466-2)) |
| Otu19477 | 1 | k__Fungi(100);p__Ascomycota(100);c__Saccharomycetes(100);o__Saccharomycetales(100);f__Metschnikowiaceae(100);g__Metschnikowia(100);s__Metschnikowia_pulcherrima(100); | Metschnikowia | 13 | Saprotroph | \|Undefined Saprotroph\| | Yeast | NULL | Probable | As nectar-tap_saprotroph (Põlme et al. 2020); Decay sugar-rich substrates (Põlme et al. 2020) | Põlme S, et al. 2020. Fungal Diversity 105:1-16 ((https://doi.org/10.1007/s13225-020-00466-2)) |
| Otu19478 | 1 | k__Fungi(100);p__Ascomycota(100);c__Saccharomycetes(100);o__Saccharomycetales(100);f__Metschnikowiaceae(100);g__Metschnikowia(100);s__Metschnikowia_pulcherrima(100); | Metschnikowia | 13 | Saprotroph | \|Undefined Saprotroph\| | Yeast | NULL | Probable | As nectar-tap_saprotroph (Põlme et al. 2020); Decay sugar-rich substrates (Põlme et al. 2020) | Põlme S, et al. 2020. Fungal Diversity 105:1-16 ((https://doi.org/10.1007/s13225-020-00466-2)) |
| Otu19479 | 1 | k__Fungi(100);p__Ascomycota(100);c__Saccharomycetes(100);o__Saccharomycetales(100);f__Metschnikowiaceae(100);g__Metschnikowia(100);s__Metschnikowia_pulcherrima(100); | Metschnikowia | 13 | Saprotroph | \|Undefined Saprotroph\| | Yeast | NULL | Probable | As nectar-tap_saprotroph (Põlme et al. 2020); Decay sugar-rich substrates (Põlme et al. 2020) | Põlme S, et al. 2020. Fungal Diversity 105:1-16 ((https://doi.org/10.1007/s13225-020-00466-2)) |
| Otu19480 | 1 | k__Fungi(100);p__Ascomycota(100);c__Saccharomycetes(100);o__Saccharomycetales(100);f__Metschnikowiaceae(100);g__Metschnikowia(100);g__Metschnikowia_unclassified(100); | Metschnikowia | 13 | Saprotroph | \|Undefined Saprotroph\| | Yeast | NULL | Probable | As nectar-tap_saprotroph (Põlme et al. 2020); Decay sugar-rich substrates (Põlme et al. 2020) | Põlme S, et al. 2020. Fungal Diversity 105:1-16 ((https://doi.org/10.1007/s13225-020-00466-2)) |
| Otu19481 | 1 | k__Fungi(100);p__Ascomycota(100);c__Saccharomycetes(100);o__Saccharomycetales(100);f__Metschnikowiaceae(100);g__Metschnikowia(100);g__Metschnikowia_unclassified(100); | Metschnikowia | 13 | Saprotroph | \|Undefined Saprotroph\| | Yeast | NULL | Probable | As nectar-tap_saprotroph (Põlme et al. 2020); Decay sugar-rich substrates (Põlme et al. 2020) | Põlme S, et al. 2020. Fungal Diversity 105:1-16 ((https://doi.org/10.1007/s13225-020-00466-2)) |
| Otu19482 | 1 | k__Fungi(100);p__Ascomycota(100);c__Saccharomycetes(100);o__Saccharomycetales(100);f__Metschnikowiaceae(100);g__Metschnikowia(100);g__Metschnikowia_unclassified(100); | Metschnikowia | 13 | Saprotroph | \|Undefined Saprotroph\| | Yeast | NULL | Probable | As nectar-tap_saprotroph (Põlme et al. 2020); Decay sugar-rich substrates (Põlme et al. 2020) | Põlme S, et al. 2020. Fungal Diversity 105:1-16 ((https://doi.org/10.1007/s13225-020-00466-2)) |
| Otu19483 | 1 | k__Fungi(100);p__Zygomycota(100);c__Zygomycota_class_Incertae_sedis(100);o__Mortierellales(100);f__Mortierellaceae(100);g__Mortierella(100);g__Mortierella_unclassified(100); | Mortierella | 13 | Saprotroph-Symbiotroph | Endophyte-Plant Saprotroph-\|Undefined Saprotroph\| | Microfungus | NULL | Probable | As chitinolytic (Põlme et al. 2020); As soil_saprotroph (Põlme et al. 2020); Decay roots, soil (Põlme et al. 2020); Saprobic in soil and thus frequently isolated from soil and material contaminated with soil such as hay or rotten silage and spoil tips (Cannon & Kirk 2007) | Cannon PF, Kirk PM. 2007. Fungal Families of the World. CAB International, Cambridge (ISBN: 978-0851998275); Tedersoo L, et al. 2014. Science 346:e1256688 ((https://doi.org/10.1126/science.1256688)); Purahong W, et al. 2016. Molecular Ecology 25:4059-4074 ((https://doi.org/10.1111/mec.13739)); Põlme S, et al. 2020. Fungal Diversity 105:1-16 ((https://doi.org/10.1007/s13225-020-00466-2)) |
| Otu19484 | 1 | k__Fungi(100);p__Ascomycota(100);c__Saccharomycetes(100);o__Saccharomycetales(100);f__Metschnikowiaceae(100);g__Metschnikowia(100);s__Metschnikowia_pulcherrima(100); | Metschnikowia | 13 | Saprotroph | \|Undefined Saprotroph\| | Yeast | NULL | Probable | As nectar-tap_saprotroph (Põlme et al. 2020); Decay sugar-rich substrates (Põlme et al. 2020) | Põlme S, et al. 2020. Fungal Diversity 105:1-16 ((https://doi.org/10.1007/s13225-020-00466-2)) |
| Otu19485 | 1 | k__Fungi(100);p__Ascomycota(100);c__Saccharomycetes(100);o__Saccharomycetales(100);f__Metschnikowiaceae(100);g__Metschnikowia(100);s__Metschnikowia_pulcherrima(100); | Metschnikowia | 13 | Saprotroph | \|Undefined Saprotroph\| | Yeast | NULL | Probable | As nectar-tap_saprotroph (Põlme et al. 2020); Decay sugar-rich substrates (Põlme et al. 2020) | Põlme S, et al. 2020. Fungal Diversity 105:1-16 ((https://doi.org/10.1007/s13225-020-00466-2)) |
| Otu19486 | 1 | k__Fungi(100);p__Basidiomycota(100);c__Microbotryomycetes(100);o__Sporidiobolales(100);f__Sporidiobolales_family_Incertae_sedis(100);g__Sporobolomyces(100);s__Sporobolomyces_lactosus(100); | Sporobolomyces | 13 | Pathotroph-Saprotroph | \|Fungal Parasite\|-Plant Saprotroph | Yeast | NULL | Probable | As litter_saprotroph (Põlme et al. 2020); As mycoparasite (Põlme et al. 2020); Decay fungal material (Põlme et al. 2020) | Tedersoo L, et al. 2014. Science 346:e1256688 ((https://doi.org/10.1126/science.1256688)); Purahong W, et al. 2016. Molecular Ecology 25:4059-4074 ((https://doi.org/10.1111/mec.13739)); Põlme S, et al. 2020. Fungal Diversity 105:1-16 ((https://doi.org/10.1007/s13225-020-00466-2)) |
| Otu19487 | 1 | k__Fungi(100);p__Ascomycota(100);c__Saccharomycetes(100);o__Saccharomycetales(100);f__Metschnikowiaceae(100);g__Metschnikowia(100);s__Metschnikowia_pulcherrima(100); | Metschnikowia | 13 | Saprotroph | \|Undefined Saprotroph\| | Yeast | NULL | Probable | As nectar-tap_saprotroph (Põlme et al. 2020); Decay sugar-rich substrates (Põlme et al. 2020) | Põlme S, et al. 2020. Fungal Diversity 105:1-16 ((https://doi.org/10.1007/s13225-020-00466-2)) |
| Otu19488 | 1 | k__Fungi(100);p__Ascomycota(100);c__Saccharomycetes(100);o__Saccharomycetales(100);f__Metschnikowiaceae(100);g__Metschnikowia(100);s__Metschnikowia_pulcherrima(100); | Metschnikowia | 13 | Saprotroph | \|Undefined Saprotroph\| | Yeast | NULL | Probable | As nectar-tap_saprotroph (Põlme et al. 2020); Decay sugar-rich substrates (Põlme et al. 2020) | Põlme S, et al. 2020. Fungal Diversity 105:1-16 ((https://doi.org/10.1007/s13225-020-00466-2)) |
| Otu19489 | 1 | k__Fungi(100);p__Ascomycota(100);c__Saccharomycetes(100);o__Saccharomycetales(100);f__Saccharomycetales_family_Incertae_sedis(100);g__Candida(100);s__Candida_cretensis(100); | Candida | 13 | Pathotroph-Saprotroph-Symbiotroph | Animal Pathogen-Endophyte-Endosymbiont-Epiphyte-\|Undefined Saprotroph\| | Dimorphic Yeast-Yeast | NULL | Probable | As nectar-tap_saprotroph (Põlme et al. 2020); Decay sugar-rich substrates (Põlme et al. 2020); This asexual genus is polyphyletic and contain species that function very differently depend on what they are. Caution should be taken when interpreting guilds for Candida species. | Manolakaki D, et al. 2010. Virulence. 1: 367-375 ((https://doi.org/10.4161/viru.1.5.12796)); Mekbib SB, et al. 2011. Tropical Plant Pathology 36:233-240 ((https://doi.org/10.1590/S1982-56762011000400004)); Põlme S, et al. 2020. Fungal Diversity 105:1-16 ((https://doi.org/10.1007/s13225-020-00466-2)) |
| Otu19490 | 1 | k__Fungi(100);p__Ascomycota(100);c__Saccharomycetes(100);o__Saccharomycetales(100);f__Metschnikowiaceae(100);g__Metschnikowia(100);s__Metschnikowia_pulcherrima(100); | Metschnikowia | 13 | Saprotroph | \|Undefined Saprotroph\| | Yeast | NULL | Probable | As nectar-tap_saprotroph (Põlme et al. 2020); Decay sugar-rich substrates (Põlme et al. 2020) | Põlme S, et al. 2020. Fungal Diversity 105:1-16 ((https://doi.org/10.1007/s13225-020-00466-2)) |
| Otu19491 | 1 | k__Fungi(100);p__Ascomycota(100);c__Saccharomycetes(100);o__Saccharomycetales(100);f__Metschnikowiaceae(100);g__Metschnikowia(100);s__Metschnikowia_pulcherrima(100); | Metschnikowia | 13 | Saprotroph | \|Undefined Saprotroph\| | Yeast | NULL | Probable | As nectar-tap_saprotroph (Põlme et al. 2020); Decay sugar-rich substrates (Põlme et al. 2020) | Põlme S, et al. 2020. Fungal Diversity 105:1-16 ((https://doi.org/10.1007/s13225-020-00466-2)) |
| Otu19492 | 1 | k__Fungi(100);p__Ascomycota(100);c__Saccharomycetes(100);o__Saccharomycetales(100);f__Metschnikowiaceae(100);g__Metschnikowia(100);s__Metschnikowia_pulcherrima(100); | Metschnikowia | 13 | Saprotroph | \|Undefined Saprotroph\| | Yeast | NULL | Probable | As nectar-tap_saprotroph (Põlme et al. 2020); Decay sugar-rich substrates (Põlme et al. 2020) | Põlme S, et al. 2020. Fungal Diversity 105:1-16 ((https://doi.org/10.1007/s13225-020-00466-2)) |
| Otu19493 | 1 | k__Fungi(100);p__Ascomycota(100);c__Saccharomycetes(100);o__Saccharomycetales(100);f__Metschnikowiaceae(100);g__Metschnikowia(100);g__Metschnikowia_unclassified(100); | Metschnikowia | 13 | Saprotroph | \|Undefined Saprotroph\| | Yeast | NULL | Probable | As nectar-tap_saprotroph (Põlme et al. 2020); Decay sugar-rich substrates (Põlme et al. 2020) | Põlme S, et al. 2020. Fungal Diversity 105:1-16 ((https://doi.org/10.1007/s13225-020-00466-2)) |
| Otu19494 | 1 | k__Fungi(100);p__Ascomycota(100);c__Saccharomycetes(100);o__Saccharomycetales(100);f__Metschnikowiaceae(100);g__Metschnikowia(100);s__Metschnikowia_pulcherrima(100); | Metschnikowia | 13 | Saprotroph | \|Undefined Saprotroph\| | Yeast | NULL | Probable | As nectar-tap_saprotroph (Põlme et al. 2020); Decay sugar-rich substrates (Põlme et al. 2020) | Põlme S, et al. 2020. Fungal Diversity 105:1-16 ((https://doi.org/10.1007/s13225-020-00466-2)) |
| Otu19495 | 1 | k__Fungi(100);p__Ascomycota(100);c__Saccharomycetes(100);o__Saccharomycetales(100);f__Metschnikowiaceae(100);g__Metschnikowia(100);s__Metschnikowia_pulcherrima(100); | Metschnikowia | 13 | Saprotroph | \|Undefined Saprotroph\| | Yeast | NULL | Probable | As nectar-tap_saprotroph (Põlme et al. 2020); Decay sugar-rich substrates (Põlme et al. 2020) | Põlme S, et al. 2020. Fungal Diversity 105:1-16 ((https://doi.org/10.1007/s13225-020-00466-2)) |
| Otu19496 | 1 | k__Fungi(100);p__Ascomycota(100);c__Saccharomycetes(100);o__Saccharomycetales(100);f__Metschnikowiaceae(100);g__Metschnikowia(100);s__Metschnikowia_pulcherrima(100); | Metschnikowia | 13 | Saprotroph | \|Undefined Saprotroph\| | Yeast | NULL | Probable | As nectar-tap_saprotroph (Põlme et al. 2020); Decay sugar-rich substrates (Põlme et al. 2020) | Põlme S, et al. 2020. Fungal Diversity 105:1-16 ((https://doi.org/10.1007/s13225-020-00466-2)) |
| Otu19497 | 1 | k__Fungi(100);p__Ascomycota(100);c__Saccharomycetes(100);o__Saccharomycetales(100);f__Metschnikowiaceae(100);g__Metschnikowia(100);g__Metschnikowia_unclassified(100); | Metschnikowia | 13 | Saprotroph | \|Undefined Saprotroph\| | Yeast | NULL | Probable | As nectar-tap_saprotroph (Põlme et al. 2020); Decay sugar-rich substrates (Põlme et al. 2020) | Põlme S, et al. 2020. Fungal Diversity 105:1-16 ((https://doi.org/10.1007/s13225-020-00466-2)) |
| Otu19498 | 1 | k__Fungi(100);p__Ascomycota(100);c__Saccharomycetes(100);o__Saccharomycetales(100);f__Metschnikowiaceae(100);g__Metschnikowia(100);g__Metschnikowia_unclassified(100); | Metschnikowia | 13 | Saprotroph | \|Undefined Saprotroph\| | Yeast | NULL | Probable | As nectar-tap_saprotroph (Põlme et al. 2020); Decay sugar-rich substrates (Põlme et al. 2020) | Põlme S, et al. 2020. Fungal Diversity 105:1-16 ((https://doi.org/10.1007/s13225-020-00466-2)) |
| Otu19499 | 1 | k__Fungi(100);p__Ascomycota(100);c__Saccharomycetes(100);o__Saccharomycetales(100);f__Metschnikowiaceae(100);g__Metschnikowia(100);s__Metschnikowia_pulcherrima(100); | Metschnikowia | 13 | Saprotroph | \|Undefined Saprotroph\| | Yeast | NULL | Probable | As nectar-tap_saprotroph (Põlme et al. 2020); Decay sugar-rich substrates (Põlme et al. 2020) | Põlme S, et al. 2020. Fungal Diversity 105:1-16 ((https://doi.org/10.1007/s13225-020-00466-2)) |
| Otu19500 | 1 | k__Fungi(100);p__Ascomycota(100);c__Saccharomycetes(100);o__Saccharomycetales(100);f__Metschnikowiaceae(100);g__Metschnikowia(100);s__Metschnikowia_pulcherrima(100); | Metschnikowia | 13 | Saprotroph | \|Undefined Saprotroph\| | Yeast | NULL | Probable | As nectar-tap_saprotroph (Põlme et al. 2020); Decay sugar-rich substrates (Põlme et al. 2020) | Põlme S, et al. 2020. Fungal Diversity 105:1-16 ((https://doi.org/10.1007/s13225-020-00466-2)) |
| Otu19502 | 1 | k__Fungi(100);p__Ascomycota(100);c__Saccharomycetes(100);o__Saccharomycetales(100);f__Metschnikowiaceae(100);g__Metschnikowia(100);g__Metschnikowia_unclassified(100); | Metschnikowia | 13 | Saprotroph | \|Undefined Saprotroph\| | Yeast | NULL | Probable | As nectar-tap_saprotroph (Põlme et al. 2020); Decay sugar-rich substrates (Põlme et al. 2020) | Põlme S, et al. 2020. Fungal Diversity 105:1-16 ((https://doi.org/10.1007/s13225-020-00466-2)) |
| Otu19503 | 1 | k__Fungi(100);p__Ascomycota(100);c__Saccharomycetes(100);o__Saccharomycetales(100);f__Metschnikowiaceae(100);g__Metschnikowia(100);s__Metschnikowia_pulcherrima(100); | Metschnikowia | 13 | Saprotroph | \|Undefined Saprotroph\| | Yeast | NULL | Probable | As nectar-tap_saprotroph (Põlme et al. 2020); Decay sugar-rich substrates (Põlme et al. 2020) | Põlme S, et al. 2020. Fungal Diversity 105:1-16 ((https://doi.org/10.1007/s13225-020-00466-2)) |
| Otu19504 | 1 | k__Fungi(100);p__Ascomycota(100);c__Saccharomycetes(100);o__Saccharomycetales(100);f__Metschnikowiaceae(100);g__Metschnikowia(100);s__Metschnikowia_pulcherrima(100); | Metschnikowia | 13 | Saprotroph | \|Undefined Saprotroph\| | Yeast | NULL | Probable | As nectar-tap_saprotroph (Põlme et al. 2020); Decay sugar-rich substrates (Põlme et al. 2020) | Põlme S, et al. 2020. Fungal Diversity 105:1-16 ((https://doi.org/10.1007/s13225-020-00466-2)) |
| Otu19505 | 1 | k__Fungi(100);p__Ascomycota(100);c__Saccharomycetes(100);o__Saccharomycetales(100);f__Metschnikowiaceae(100);g__Metschnikowia(100);g__Metschnikowia_unclassified(100); | Metschnikowia | 13 | Saprotroph | \|Undefined Saprotroph\| | Yeast | NULL | Probable | As nectar-tap_saprotroph (Põlme et al. 2020); Decay sugar-rich substrates (Põlme et al. 2020) | Põlme S, et al. 2020. Fungal Diversity 105:1-16 ((https://doi.org/10.1007/s13225-020-00466-2)) |
| Otu19506 | 1 | k__Fungi(100);p__Ascomycota(100);c__Saccharomycetes(100);o__Saccharomycetales(100);f__Metschnikowiaceae(100);g__Metschnikowia(100);s__Metschnikowia_pulcherrima(100); | Metschnikowia | 13 | Saprotroph | \|Undefined Saprotroph\| | Yeast | NULL | Probable | As nectar-tap_saprotroph (Põlme et al. 2020); Decay sugar-rich substrates (Põlme et al. 2020) | Põlme S, et al. 2020. Fungal Diversity 105:1-16 ((https://doi.org/10.1007/s13225-020-00466-2)) |
| Otu19507 | 1 | k__Fungi(100);p__Ascomycota(100);c__Saccharomycetes(100);o__Saccharomycetales(100);f__Metschnikowiaceae(100);g__Metschnikowia(100);s__Metschnikowia_pulcherrima(100); | Metschnikowia | 13 | Saprotroph | \|Undefined Saprotroph\| | Yeast | NULL | Probable | As nectar-tap_saprotroph (Põlme et al. 2020); Decay sugar-rich substrates (Põlme et al. 2020) | Põlme S, et al. 2020. Fungal Diversity 105:1-16 ((https://doi.org/10.1007/s13225-020-00466-2)) |
| Otu19509 | 1 | k__Fungi(100);p__Ascomycota(100);c__Saccharomycetes(100);o__Saccharomycetales(100);f__Metschnikowiaceae(100);g__Metschnikowia(100);g__Metschnikowia_unclassified(100); | Metschnikowia | 13 | Saprotroph | \|Undefined Saprotroph\| | Yeast | NULL | Probable | As nectar-tap_saprotroph (Põlme et al. 2020); Decay sugar-rich substrates (Põlme et al. 2020) | Põlme S, et al. 2020. Fungal Diversity 105:1-16 ((https://doi.org/10.1007/s13225-020-00466-2)) |
| Otu19510 | 1 | k__Fungi(100);p__Ascomycota(100);c__Saccharomycetes(100);o__Saccharomycetales(100);f__Metschnikowiaceae(100);g__Metschnikowia(100);s__Metschnikowia_pulcherrima(100); | Metschnikowia | 13 | Saprotroph | \|Undefined Saprotroph\| | Yeast | NULL | Probable | As nectar-tap_saprotroph (Põlme et al. 2020); Decay sugar-rich substrates (Põlme et al. 2020) | Põlme S, et al. 2020. Fungal Diversity 105:1-16 ((https://doi.org/10.1007/s13225-020-00466-2)) |
| Otu19511 | 1 | k__Fungi(100);p__Ascomycota(100);c__Saccharomycetes(100);o__Saccharomycetales(100);f__Metschnikowiaceae(100);g__Metschnikowia(100);g__Metschnikowia_unclassified(100); | Metschnikowia | 13 | Saprotroph | \|Undefined Saprotroph\| | Yeast | NULL | Probable | As nectar-tap_saprotroph (Põlme et al. 2020); Decay sugar-rich substrates (Põlme et al. 2020) | Põlme S, et al. 2020. Fungal Diversity 105:1-16 ((https://doi.org/10.1007/s13225-020-00466-2)) |
| Otu19512 | 1 | k__Fungi(100);p__Ascomycota(100);c__Saccharomycetes(100);o__Saccharomycetales(100);f__Metschnikowiaceae(100);g__Metschnikowia(100);s__Metschnikowia_pulcherrima(100); | Metschnikowia | 13 | Saprotroph | \|Undefined Saprotroph\| | Yeast | NULL | Probable | As nectar-tap_saprotroph (Põlme et al. 2020); Decay sugar-rich substrates (Põlme et al. 2020) | Põlme S, et al. 2020. Fungal Diversity 105:1-16 ((https://doi.org/10.1007/s13225-020-00466-2)) |
| Otu19513 | 1 | k__Fungi(100);p__Ascomycota(100);c__Saccharomycetes(100);o__Saccharomycetales(100);f__Metschnikowiaceae(100);g__Metschnikowia(100);s__Metschnikowia_pulcherrima(100); | Metschnikowia | 13 | Saprotroph | \|Undefined Saprotroph\| | Yeast | NULL | Probable | As nectar-tap_saprotroph (Põlme et al. 2020); Decay sugar-rich substrates (Põlme et al. 2020) | Põlme S, et al. 2020. Fungal Diversity 105:1-16 ((https://doi.org/10.1007/s13225-020-00466-2)) |
| Otu19514 | 1 | k__Fungi(100);p__Ascomycota(100);c__Saccharomycetes(100);o__Saccharomycetales(100);f__Metschnikowiaceae(100);g__Metschnikowia(100);s__Metschnikowia_pulcherrima(100); | Metschnikowia | 13 | Saprotroph | \|Undefined Saprotroph\| | Yeast | NULL | Probable | As nectar-tap_saprotroph (Põlme et al. 2020); Decay sugar-rich substrates (Põlme et al. 2020) | Põlme S, et al. 2020. Fungal Diversity 105:1-16 ((https://doi.org/10.1007/s13225-020-00466-2)) |
| Otu19515 | 1 | k__Fungi(100);p__Ascomycota(100);c__Saccharomycetes(100);o__Saccharomycetales(100);f__Metschnikowiaceae(100);g__Metschnikowia(100);s__Metschnikowia_pulcherrima(100); | Metschnikowia | 13 | Saprotroph | \|Undefined Saprotroph\| | Yeast | NULL | Probable | As nectar-tap_saprotroph (Põlme et al. 2020); Decay sugar-rich substrates (Põlme et al. 2020) | Põlme S, et al. 2020. Fungal Diversity 105:1-16 ((https://doi.org/10.1007/s13225-020-00466-2)) |
| Otu19516 | 1 | k__Fungi(100);p__Ascomycota(100);c__Saccharomycetes(100);o__Saccharomycetales(100);f__Metschnikowiaceae(100);g__Metschnikowia(100);s__Metschnikowia_pulcherrima(100); | Metschnikowia | 13 | Saprotroph | \|Undefined Saprotroph\| | Yeast | NULL | Probable | As nectar-tap_saprotroph (Põlme et al. 2020); Decay sugar-rich substrates (Põlme et al. 2020) | Põlme S, et al. 2020. Fungal Diversity 105:1-16 ((https://doi.org/10.1007/s13225-020-00466-2)) |
| Otu19517 | 1 | k__Fungi(100);p__Ascomycota(100);c__Saccharomycetes(100);o__Saccharomycetales(100);f__Metschnikowiaceae(100);g__Metschnikowia(100);s__Metschnikowia_pulcherrima(100); | Metschnikowia | 13 | Saprotroph | \|Undefined Saprotroph\| | Yeast | NULL | Probable | As nectar-tap_saprotroph (Põlme et al. 2020); Decay sugar-rich substrates (Põlme et al. 2020) | Põlme S, et al. 2020. Fungal Diversity 105:1-16 ((https://doi.org/10.1007/s13225-020-00466-2)) |
| Otu19518 | 1 | k__Fungi(100);p__Ascomycota(100);c__Saccharomycetes(100);o__Saccharomycetales(100);f__Metschnikowiaceae(100);g__Metschnikowia(100);g__Metschnikowia_unclassified(100); | Metschnikowia | 13 | Saprotroph | \|Undefined Saprotroph\| | Yeast | NULL | Probable | As nectar-tap_saprotroph (Põlme et al. 2020); Decay sugar-rich substrates (Põlme et al. 2020) | Põlme S, et al. 2020. Fungal Diversity 105:1-16 ((https://doi.org/10.1007/s13225-020-00466-2)) |
| Otu19519 | 1 | k__Fungi(100);p__Ascomycota(100);c__Saccharomycetes(100);o__Saccharomycetales(100);f__Metschnikowiaceae(100);g__Metschnikowia(100);g__Metschnikowia_unclassified(100); | Metschnikowia | 13 | Saprotroph | \|Undefined Saprotroph\| | Yeast | NULL | Probable | As nectar-tap_saprotroph (Põlme et al. 2020); Decay sugar-rich substrates (Põlme et al. 2020) | Põlme S, et al. 2020. Fungal Diversity 105:1-16 ((https://doi.org/10.1007/s13225-020-00466-2)) |
| Otu19520 | 1 | k__Fungi(100);p__Ascomycota(100);c__Saccharomycetes(100);o__Saccharomycetales(100);f__Metschnikowiaceae(100);g__Metschnikowia(100);s__Metschnikowia_pulcherrima(100); | Metschnikowia | 13 | Saprotroph | \|Undefined Saprotroph\| | Yeast | NULL | Probable | As nectar-tap_saprotroph (Põlme et al. 2020); Decay sugar-rich substrates (Põlme et al. 2020) | Põlme S, et al. 2020. Fungal Diversity 105:1-16 ((https://doi.org/10.1007/s13225-020-00466-2)) |
| Otu19521 | 1 | k__Fungi(100);p__Ascomycota(100);c__Saccharomycetes(100);o__Saccharomycetales(100);f__Metschnikowiaceae(100);g__Metschnikowia(100);s__Metschnikowia_pulcherrima(100); | Metschnikowia | 13 | Saprotroph | \|Undefined Saprotroph\| | Yeast | NULL | Probable | As nectar-tap_saprotroph (Põlme et al. 2020); Decay sugar-rich substrates (Põlme et al. 2020) | Põlme S, et al. 2020. Fungal Diversity 105:1-16 ((https://doi.org/10.1007/s13225-020-00466-2)) |
| Otu19522 | 1 | k__Fungi(100);p__Ascomycota(100);c__Saccharomycetes(100);o__Saccharomycetales(100);f__Metschnikowiaceae(100);g__Metschnikowia(100);s__Metschnikowia_pulcherrima(100); | Metschnikowia | 13 | Saprotroph | \|Undefined Saprotroph\| | Yeast | NULL | Probable | As nectar-tap_saprotroph (Põlme et al. 2020); Decay sugar-rich substrates (Põlme et al. 2020) | Põlme S, et al. 2020. Fungal Diversity 105:1-16 ((https://doi.org/10.1007/s13225-020-00466-2)) |
| Otu19523 | 1 | k__Fungi(100);p__Ascomycota(100);c__Saccharomycetes(100);o__Saccharomycetales(100);f__Metschnikowiaceae(100);g__Metschnikowia(100);s__Metschnikowia_pulcherrima(100); | Metschnikowia | 13 | Saprotroph | \|Undefined Saprotroph\| | Yeast | NULL | Probable | As nectar-tap_saprotroph (Põlme et al. 2020); Decay sugar-rich substrates (Põlme et al. 2020) | Põlme S, et al. 2020. Fungal Diversity 105:1-16 ((https://doi.org/10.1007/s13225-020-00466-2)) |
| Otu19525 | 1 | k__Fungi(100);p__Ascomycota(100);c__Saccharomycetes(100);o__Saccharomycetales(100);f__Metschnikowiaceae(100);g__Metschnikowia(100);s__Metschnikowia_pulcherrima(100); | Metschnikowia | 13 | Saprotroph | \|Undefined Saprotroph\| | Yeast | NULL | Probable | As nectar-tap_saprotroph (Põlme et al. 2020); Decay sugar-rich substrates (Põlme et al. 2020) | Põlme S, et al. 2020. Fungal Diversity 105:1-16 ((https://doi.org/10.1007/s13225-020-00466-2)) |
| Otu19526 | 1 | k__Fungi(100);p__Ascomycota(100);c__Saccharomycetes(100);o__Saccharomycetales(100);f__Metschnikowiaceae(100);g__Metschnikowia(100);g__Metschnikowia_unclassified(100); | Metschnikowia | 13 | Saprotroph | \|Undefined Saprotroph\| | Yeast | NULL | Probable | As nectar-tap_saprotroph (Põlme et al. 2020); Decay sugar-rich substrates (Põlme et al. 2020) | Põlme S, et al. 2020. Fungal Diversity 105:1-16 ((https://doi.org/10.1007/s13225-020-00466-2)) |
| Otu19527 | 1 | k__Fungi(100);p__Ascomycota(100);c__Saccharomycetes(100);o__Saccharomycetales(100);f__Metschnikowiaceae(100);g__Metschnikowia(100);s__Metschnikowia_pulcherrima(100); | Metschnikowia | 13 | Saprotroph | \|Undefined Saprotroph\| | Yeast | NULL | Probable | As nectar-tap_saprotroph (Põlme et al. 2020); Decay sugar-rich substrates (Põlme et al. 2020) | Põlme S, et al. 2020. Fungal Diversity 105:1-16 ((https://doi.org/10.1007/s13225-020-00466-2)) |
| Otu19528 | 1 | k__Fungi(100);p__Ascomycota(100);c__Saccharomycetes(100);o__Saccharomycetales(100);f__Metschnikowiaceae(100);g__Metschnikowia(100);s__Metschnikowia_pulcherrima(100); | Metschnikowia | 13 | Saprotroph | \|Undefined Saprotroph\| | Yeast | NULL | Probable | As nectar-tap_saprotroph (Põlme et al. 2020); Decay sugar-rich substrates (Põlme et al. 2020) | Põlme S, et al. 2020. Fungal Diversity 105:1-16 ((https://doi.org/10.1007/s13225-020-00466-2)) |
| Otu19529 | 1 | k__Fungi(100);p__Ascomycota(100);c__Saccharomycetes(100);o__Saccharomycetales(100);f__Metschnikowiaceae(100);g__Metschnikowia(100);s__Metschnikowia_pulcherrima(100); | Metschnikowia | 13 | Saprotroph | \|Undefined Saprotroph\| | Yeast | NULL | Probable | As nectar-tap_saprotroph (Põlme et al. 2020); Decay sugar-rich substrates (Põlme et al. 2020) | Põlme S, et al. 2020. Fungal Diversity 105:1-16 ((https://doi.org/10.1007/s13225-020-00466-2)) |
| Otu19530 | 1 | k__Fungi(100);p__Ascomycota(100);c__Saccharomycetes(100);o__Saccharomycetales(100);f__Metschnikowiaceae(100);g__Metschnikowia(100);s__Metschnikowia_pulcherrima(100); | Metschnikowia | 13 | Saprotroph | \|Undefined Saprotroph\| | Yeast | NULL | Probable | As nectar-tap_saprotroph (Põlme et al. 2020); Decay sugar-rich substrates (Põlme et al. 2020) | Põlme S, et al. 2020. Fungal Diversity 105:1-16 ((https://doi.org/10.1007/s13225-020-00466-2)) |
| Otu19531 | 1 | k__Fungi(100);p__Ascomycota(100);c__Saccharomycetes(100);o__Saccharomycetales(100);f__Metschnikowiaceae(100);g__Metschnikowia(100);s__Metschnikowia_pulcherrima(100); | Metschnikowia | 13 | Saprotroph | \|Undefined Saprotroph\| | Yeast | NULL | Probable | As nectar-tap_saprotroph (Põlme et al. 2020); Decay sugar-rich substrates (Põlme et al. 2020) | Põlme S, et al. 2020. Fungal Diversity 105:1-16 ((https://doi.org/10.1007/s13225-020-00466-2)) |
| Otu19532 | 1 | k__Fungi(100);p__Ascomycota(100);c__Saccharomycetes(100);o__Saccharomycetales(100);f__Metschnikowiaceae(100);g__Metschnikowia(100);s__Metschnikowia_pulcherrima(100); | Metschnikowia | 13 | Saprotroph | \|Undefined Saprotroph\| | Yeast | NULL | Probable | As nectar-tap_saprotroph (Põlme et al. 2020); Decay sugar-rich substrates (Põlme et al. 2020) | Põlme S, et al. 2020. Fungal Diversity 105:1-16 ((https://doi.org/10.1007/s13225-020-00466-2)) |
| Otu19533 | 1 | k__Fungi(100);p__Ascomycota(100);c__Saccharomycetes(100);o__Saccharomycetales(100);f__Metschnikowiaceae(100);g__Metschnikowia(100);s__Metschnikowia_pulcherrima(100); | Metschnikowia | 13 | Saprotroph | \|Undefined Saprotroph\| | Yeast | NULL | Probable | As nectar-tap_saprotroph (Põlme et al. 2020); Decay sugar-rich substrates (Põlme et al. 2020) | Põlme S, et al. 2020. Fungal Diversity 105:1-16 ((https://doi.org/10.1007/s13225-020-00466-2)) |
| Otu19534 | 1 | k__Fungi(100);p__Ascomycota(100);c__Saccharomycetes(100);o__Saccharomycetales(100);f__Metschnikowiaceae(100);g__Metschnikowia(100);g__Metschnikowia_unclassified(100); | Metschnikowia | 13 | Saprotroph | \|Undefined Saprotroph\| | Yeast | NULL | Probable | As nectar-tap_saprotroph (Põlme et al. 2020); Decay sugar-rich substrates (Põlme et al. 2020) | Põlme S, et al. 2020. Fungal Diversity 105:1-16 ((https://doi.org/10.1007/s13225-020-00466-2)) |
| Otu19535 | 1 | k__Fungi(100);p__Ascomycota(100);c__Saccharomycetes(100);o__Saccharomycetales(100);f__Metschnikowiaceae(100);g__Metschnikowia(100);s__Metschnikowia_pulcherrima(100); | Metschnikowia | 13 | Saprotroph | \|Undefined Saprotroph\| | Yeast | NULL | Probable | As nectar-tap_saprotroph (Põlme et al. 2020); Decay sugar-rich substrates (Põlme et al. 2020) | Põlme S, et al. 2020. Fungal Diversity 105:1-16 ((https://doi.org/10.1007/s13225-020-00466-2)) |
| Otu19536 | 1 | k__Fungi(100);p__Ascomycota(100);c__Saccharomycetes(100);o__Saccharomycetales(100);f__Metschnikowiaceae(100);g__Metschnikowia(100);s__Metschnikowia_pulcherrima(100); | Metschnikowia | 13 | Saprotroph | \|Undefined Saprotroph\| | Yeast | NULL | Probable | As nectar-tap_saprotroph (Põlme et al. 2020); Decay sugar-rich substrates (Põlme et al. 2020) | Põlme S, et al. 2020. Fungal Diversity 105:1-16 ((https://doi.org/10.1007/s13225-020-00466-2)) |
| Otu19537 | 1 | k__Fungi(100);p__Ascomycota(100);c__Saccharomycetes(100);o__Saccharomycetales(100);f__Metschnikowiaceae(100);g__Metschnikowia(100);s__Metschnikowia_pulcherrima(100); | Metschnikowia | 13 | Saprotroph | \|Undefined Saprotroph\| | Yeast | NULL | Probable | As nectar-tap_saprotroph (Põlme et al. 2020); Decay sugar-rich substrates (Põlme et al. 2020) | Põlme S, et al. 2020. Fungal Diversity 105:1-16 ((https://doi.org/10.1007/s13225-020-00466-2)) |
| Otu19538 | 1 | k__Fungi(100);p__Ascomycota(100);c__Saccharomycetes(100);o__Saccharomycetales(100);f__Saccharomycetales_family_Incertae_sedis(100);g__Candida(100);g__Candida_unclassified(100); | Candida | 13 | Pathotroph-Saprotroph-Symbiotroph | Animal Pathogen-Endophyte-Endosymbiont-Epiphyte-\|Undefined Saprotroph\| | Dimorphic Yeast-Yeast | NULL | Probable | As nectar-tap_saprotroph (Põlme et al. 2020); Decay sugar-rich substrates (Põlme et al. 2020); This asexual genus is polyphyletic and contain species that function very differently depend on what they are. Caution should be taken when interpreting guilds for Candida species. | Manolakaki D, et al. 2010. Virulence. 1: 367-375 ((https://doi.org/10.4161/viru.1.5.12796)); Mekbib SB, et al. 2011. Tropical Plant Pathology 36:233-240 ((https://doi.org/10.1590/S1982-56762011000400004)); Põlme S, et al. 2020. Fungal Diversity 105:1-16 ((https://doi.org/10.1007/s13225-020-00466-2)) |
| Otu19539 | 1 | k__Fungi(100);p__Ascomycota(100);c__Saccharomycetes(100);o__Saccharomycetales(100);f__Saccharomycetales_family_Incertae_sedis(100);g__Candida(100);g__Candida_unclassified(100); | Candida | 13 | Pathotroph-Saprotroph-Symbiotroph | Animal Pathogen-Endophyte-Endosymbiont-Epiphyte-\|Undefined Saprotroph\| | Dimorphic Yeast-Yeast | NULL | Probable | As nectar-tap_saprotroph (Põlme et al. 2020); Decay sugar-rich substrates (Põlme et al. 2020); This asexual genus is polyphyletic and contain species that function very differently depend on what they are. Caution should be taken when interpreting guilds for Candida species. | Manolakaki D, et al. 2010. Virulence. 1: 367-375 ((https://doi.org/10.4161/viru.1.5.12796)); Mekbib SB, et al. 2011. Tropical Plant Pathology 36:233-240 ((https://doi.org/10.1590/S1982-56762011000400004)); Põlme S, et al. 2020. Fungal Diversity 105:1-16 ((https://doi.org/10.1007/s13225-020-00466-2)) |
| Otu19540 | 1 | k__Fungi(100);p__Ascomycota(100);c__Saccharomycetes(100);o__Saccharomycetales(100);f__Metschnikowiaceae(100);g__Metschnikowia(100);s__Metschnikowia_pulcherrima(100); | Metschnikowia | 13 | Saprotroph | \|Undefined Saprotroph\| | Yeast | NULL | Probable | As nectar-tap_saprotroph (Põlme et al. 2020); Decay sugar-rich substrates (Põlme et al. 2020) | Põlme S, et al. 2020. Fungal Diversity 105:1-16 ((https://doi.org/10.1007/s13225-020-00466-2)) |
| Otu19541 | 1 | k__Fungi(100);p__Ascomycota(100);c__Saccharomycetes(100);o__Saccharomycetales(100);f__Metschnikowiaceae(100);g__Metschnikowia(100);s__Metschnikowia_pulcherrima(100); | Metschnikowia | 13 | Saprotroph | \|Undefined Saprotroph\| | Yeast | NULL | Probable | As nectar-tap_saprotroph (Põlme et al. 2020); Decay sugar-rich substrates (Põlme et al. 2020) | Põlme S, et al. 2020. Fungal Diversity 105:1-16 ((https://doi.org/10.1007/s13225-020-00466-2)) |
| Otu19542 | 1 | k__Fungi(100);p__Ascomycota(100);c__Saccharomycetes(100);o__Saccharomycetales(100);f__Metschnikowiaceae(100);g__Metschnikowia(100);g__Metschnikowia_unclassified(100); | Metschnikowia | 13 | Saprotroph | \|Undefined Saprotroph\| | Yeast | NULL | Probable | As nectar-tap_saprotroph (Põlme et al. 2020); Decay sugar-rich substrates (Põlme et al. 2020) | Põlme S, et al. 2020. Fungal Diversity 105:1-16 ((https://doi.org/10.1007/s13225-020-00466-2)) |
| Otu19543 | 1 | k__Fungi(100);p__Ascomycota(100);c__Saccharomycetes(100);o__Saccharomycetales(100);f__Metschnikowiaceae(100);g__Metschnikowia(100);s__Metschnikowia_pulcherrima(100); | Metschnikowia | 13 | Saprotroph | \|Undefined Saprotroph\| | Yeast | NULL | Probable | As nectar-tap_saprotroph (Põlme et al. 2020); Decay sugar-rich substrates (Põlme et al. 2020) | Põlme S, et al. 2020. Fungal Diversity 105:1-16 ((https://doi.org/10.1007/s13225-020-00466-2)) |
| Otu19544 | 1 | k__Fungi(100);p__Ascomycota(100);c__Saccharomycetes(100);o__Saccharomycetales(100);f__Metschnikowiaceae(100);g__Metschnikowia(100);g__Metschnikowia_unclassified(100); | Metschnikowia | 13 | Saprotroph | \|Undefined Saprotroph\| | Yeast | NULL | Probable | As nectar-tap_saprotroph (Põlme et al. 2020); Decay sugar-rich substrates (Põlme et al. 2020) | Põlme S, et al. 2020. Fungal Diversity 105:1-16 ((https://doi.org/10.1007/s13225-020-00466-2)) |
| Otu19545 | 1 | k__Fungi(100);p__Ascomycota(100);c__Saccharomycetes(100);o__Saccharomycetales(100);f__Metschnikowiaceae(100);g__Metschnikowia(100);s__Metschnikowia_pulcherrima(100); | Metschnikowia | 13 | Saprotroph | \|Undefined Saprotroph\| | Yeast | NULL | Probable | As nectar-tap_saprotroph (Põlme et al. 2020); Decay sugar-rich substrates (Põlme et al. 2020) | Põlme S, et al. 2020. Fungal Diversity 105:1-16 ((https://doi.org/10.1007/s13225-020-00466-2)) |
| Otu19546 | 1 | k__Fungi(100);p__Ascomycota(100);c__Saccharomycetes(100);o__Saccharomycetales(100);f__Metschnikowiaceae(100);g__Metschnikowia(100);s__Metschnikowia_pulcherrima(100); | Metschnikowia | 13 | Saprotroph | \|Undefined Saprotroph\| | Yeast | NULL | Probable | As nectar-tap_saprotroph (Põlme et al. 2020); Decay sugar-rich substrates (Põlme et al. 2020) | Põlme S, et al. 2020. Fungal Diversity 105:1-16 ((https://doi.org/10.1007/s13225-020-00466-2)) |
| Otu19547 | 1 | k__Fungi(100);p__Ascomycota(100);c__Saccharomycetes(100);o__Saccharomycetales(100);f__Metschnikowiaceae(100);g__Metschnikowia(100);g__Metschnikowia_unclassified(100); | Metschnikowia | 13 | Saprotroph | \|Undefined Saprotroph\| | Yeast | NULL | Probable | As nectar-tap_saprotroph (Põlme et al. 2020); Decay sugar-rich substrates (Põlme et al. 2020) | Põlme S, et al. 2020. Fungal Diversity 105:1-16 ((https://doi.org/10.1007/s13225-020-00466-2)) |
| Otu19548 | 1 | k__Fungi(100);p__Ascomycota(100);c__Saccharomycetes(100);o__Saccharomycetales(100);f__Metschnikowiaceae(100);g__Metschnikowia(100);s__Metschnikowia_pulcherrima(100); | Metschnikowia | 13 | Saprotroph | \|Undefined Saprotroph\| | Yeast | NULL | Probable | As nectar-tap_saprotroph (Põlme et al. 2020); Decay sugar-rich substrates (Põlme et al. 2020) | Põlme S, et al. 2020. Fungal Diversity 105:1-16 ((https://doi.org/10.1007/s13225-020-00466-2)) |
| Otu19549 | 1 | k__Fungi(100);p__Ascomycota(100);c__Saccharomycetes(100);o__Saccharomycetales(100);f__Metschnikowiaceae(100);g__Metschnikowia(100);s__Metschnikowia_pulcherrima(100); | Metschnikowia | 13 | Saprotroph | \|Undefined Saprotroph\| | Yeast | NULL | Probable | As nectar-tap_saprotroph (Põlme et al. 2020); Decay sugar-rich substrates (Põlme et al. 2020) | Põlme S, et al. 2020. Fungal Diversity 105:1-16 ((https://doi.org/10.1007/s13225-020-00466-2)) |
| Otu19550 | 1 | k__Fungi(100);p__Ascomycota(100);c__Saccharomycetes(100);o__Saccharomycetales(100);f__Metschnikowiaceae(100);g__Metschnikowia(100);s__Metschnikowia_pulcherrima(100); | Metschnikowia | 13 | Saprotroph | \|Undefined Saprotroph\| | Yeast | NULL | Probable | As nectar-tap_saprotroph (Põlme et al. 2020); Decay sugar-rich substrates (Põlme et al. 2020) | Põlme S, et al. 2020. Fungal Diversity 105:1-16 ((https://doi.org/10.1007/s13225-020-00466-2)) |
| Otu19551 | 1 | k__Fungi(100);p__Ascomycota(100);c__Saccharomycetes(100);o__Saccharomycetales(100);f__Metschnikowiaceae(100);g__Metschnikowia(100);s__Metschnikowia_pulcherrima(100); | Metschnikowia | 13 | Saprotroph | \|Undefined Saprotroph\| | Yeast | NULL | Probable | As nectar-tap_saprotroph (Põlme et al. 2020); Decay sugar-rich substrates (Põlme et al. 2020) | Põlme S, et al. 2020. Fungal Diversity 105:1-16 ((https://doi.org/10.1007/s13225-020-00466-2)) |
| Otu19552 | 1 | k__Fungi(100);p__Ascomycota(100);c__Saccharomycetes(100);o__Saccharomycetales(100);f__Metschnikowiaceae(100);g__Metschnikowia(100);s__Metschnikowia_pulcherrima(100); | Metschnikowia | 13 | Saprotroph | \|Undefined Saprotroph\| | Yeast | NULL | Probable | As nectar-tap_saprotroph (Põlme et al. 2020); Decay sugar-rich substrates (Põlme et al. 2020) | Põlme S, et al. 2020. Fungal Diversity 105:1-16 ((https://doi.org/10.1007/s13225-020-00466-2)) |
| Otu19553 | 1 | k__Fungi(100);p__Ascomycota(100);c__Saccharomycetes(100);o__Saccharomycetales(100);f__Metschnikowiaceae(100);g__Metschnikowia(100);s__Metschnikowia_pulcherrima(100); | Metschnikowia | 13 | Saprotroph | \|Undefined Saprotroph\| | Yeast | NULL | Probable | As nectar-tap_saprotroph (Põlme et al. 2020); Decay sugar-rich substrates (Põlme et al. 2020) | Põlme S, et al. 2020. Fungal Diversity 105:1-16 ((https://doi.org/10.1007/s13225-020-00466-2)) |
| Otu19554 | 1 | k__Fungi(100);p__Ascomycota(100);c__Saccharomycetes(100);o__Saccharomycetales(100);f__Metschnikowiaceae(100);g__Metschnikowia(100);s__Metschnikowia_pulcherrima(100); | Metschnikowia | 13 | Saprotroph | \|Undefined Saprotroph\| | Yeast | NULL | Probable | As nectar-tap_saprotroph (Põlme et al. 2020); Decay sugar-rich substrates (Põlme et al. 2020) | Põlme S, et al. 2020. Fungal Diversity 105:1-16 ((https://doi.org/10.1007/s13225-020-00466-2)) |
| Otu19555 | 1 | k__Fungi(100);p__Ascomycota(100);c__Saccharomycetes(100);o__Saccharomycetales(100);f__Metschnikowiaceae(100);g__Metschnikowia(100);s__Metschnikowia_pulcherrima(100); | Metschnikowia | 13 | Saprotroph | \|Undefined Saprotroph\| | Yeast | NULL | Probable | As nectar-tap_saprotroph (Põlme et al. 2020); Decay sugar-rich substrates (Põlme et al. 2020) | Põlme S, et al. 2020. Fungal Diversity 105:1-16 ((https://doi.org/10.1007/s13225-020-00466-2)) |
| Otu19556 | 1 | k__Fungi(100);p__Ascomycota(100);c__Saccharomycetes(100);o__Saccharomycetales(100);f__Metschnikowiaceae(100);g__Metschnikowia(100);s__Metschnikowia_pulcherrima(100); | Metschnikowia | 13 | Saprotroph | \|Undefined Saprotroph\| | Yeast | NULL | Probable | As nectar-tap_saprotroph (Põlme et al. 2020); Decay sugar-rich substrates (Põlme et al. 2020) | Põlme S, et al. 2020. Fungal Diversity 105:1-16 ((https://doi.org/10.1007/s13225-020-00466-2)) |
| Otu19557 | 1 | k__Fungi(100);p__Ascomycota(100);c__Saccharomycetes(100);o__Saccharomycetales(100);f__Metschnikowiaceae(100);g__Metschnikowia(100);g__Metschnikowia_unclassified(100); | Metschnikowia | 13 | Saprotroph | \|Undefined Saprotroph\| | Yeast | NULL | Probable | As nectar-tap_saprotroph (Põlme et al. 2020); Decay sugar-rich substrates (Põlme et al. 2020) | Põlme S, et al. 2020. Fungal Diversity 105:1-16 ((https://doi.org/10.1007/s13225-020-00466-2)) |
| Otu19558 | 1 | k__Fungi(100);p__Ascomycota(100);c__Saccharomycetes(100);o__Saccharomycetales(100);f__Metschnikowiaceae(100);g__Metschnikowia(100);s__Metschnikowia_pulcherrima(100); | Metschnikowia | 13 | Saprotroph | \|Undefined Saprotroph\| | Yeast | NULL | Probable | As nectar-tap_saprotroph (Põlme et al. 2020); Decay sugar-rich substrates (Põlme et al. 2020) | Põlme S, et al. 2020. Fungal Diversity 105:1-16 ((https://doi.org/10.1007/s13225-020-00466-2)) |
| Otu19559 | 1 | k__Fungi(100);p__Ascomycota(100);c__Saccharomycetes(100);o__Saccharomycetales(100);f__Metschnikowiaceae(100);g__Metschnikowia(100);s__Metschnikowia_pulcherrima(100); | Metschnikowia | 13 | Saprotroph | \|Undefined Saprotroph\| | Yeast | NULL | Probable | As nectar-tap_saprotroph (Põlme et al. 2020); Decay sugar-rich substrates (Põlme et al. 2020) | Põlme S, et al. 2020. Fungal Diversity 105:1-16 ((https://doi.org/10.1007/s13225-020-00466-2)) |
| Otu19560 | 1 | k__Fungi(100);p__Ascomycota(100);c__Saccharomycetes(100);o__Saccharomycetales(100);f__Metschnikowiaceae(100);g__Metschnikowia(100);s__Metschnikowia_pulcherrima(100); | Metschnikowia | 13 | Saprotroph | \|Undefined Saprotroph\| | Yeast | NULL | Probable | As nectar-tap_saprotroph (Põlme et al. 2020); Decay sugar-rich substrates (Põlme et al. 2020) | Põlme S, et al. 2020. Fungal Diversity 105:1-16 ((https://doi.org/10.1007/s13225-020-00466-2)) |
| Otu19561 | 1 | k__Fungi(100);p__Ascomycota(100);c__Saccharomycetes(100);o__Saccharomycetales(100);f__Metschnikowiaceae(100);g__Metschnikowia(100);s__Metschnikowia_pulcherrima(100); | Metschnikowia | 13 | Saprotroph | \|Undefined Saprotroph\| | Yeast | NULL | Probable | As nectar-tap_saprotroph (Põlme et al. 2020); Decay sugar-rich substrates (Põlme et al. 2020) | Põlme S, et al. 2020. Fungal Diversity 105:1-16 ((https://doi.org/10.1007/s13225-020-00466-2)) |
| Otu19562 | 1 | k__Fungi(100);p__Ascomycota(100);c__Saccharomycetes(100);o__Saccharomycetales(100);f__Metschnikowiaceae(100);g__Metschnikowia(100);g__Metschnikowia_unclassified(100); | Metschnikowia | 13 | Saprotroph | \|Undefined Saprotroph\| | Yeast | NULL | Probable | As nectar-tap_saprotroph (Põlme et al. 2020); Decay sugar-rich substrates (Põlme et al. 2020) | Põlme S, et al. 2020. Fungal Diversity 105:1-16 ((https://doi.org/10.1007/s13225-020-00466-2)) |
| Otu19563 | 1 | k__Fungi(100);p__Ascomycota(100);c__Saccharomycetes(100);o__Saccharomycetales(100);f__Metschnikowiaceae(100);g__Metschnikowia(100);s__Metschnikowia_pulcherrima(100); | Metschnikowia | 13 | Saprotroph | \|Undefined Saprotroph\| | Yeast | NULL | Probable | As nectar-tap_saprotroph (Põlme et al. 2020); Decay sugar-rich substrates (Põlme et al. 2020) | Põlme S, et al. 2020. Fungal Diversity 105:1-16 ((https://doi.org/10.1007/s13225-020-00466-2)) |
| Otu19564 | 1 | k__Fungi(100);p__Ascomycota(100);c__Saccharomycetes(100);o__Saccharomycetales(100);f__Metschnikowiaceae(100);g__Metschnikowia(100);s__Metschnikowia_pulcherrima(100); | Metschnikowia | 13 | Saprotroph | \|Undefined Saprotroph\| | Yeast | NULL | Probable | As nectar-tap_saprotroph (Põlme et al. 2020); Decay sugar-rich substrates (Põlme et al. 2020) | Põlme S, et al. 2020. Fungal Diversity 105:1-16 ((https://doi.org/10.1007/s13225-020-00466-2)) |
| Otu19565 | 1 | k__Fungi(100);p__Ascomycota(100);c__Saccharomycetes(100);o__Saccharomycetales(100);f__Metschnikowiaceae(100);g__Metschnikowia(100);g__Metschnikowia_unclassified(100); | Metschnikowia | 13 | Saprotroph | \|Undefined Saprotroph\| | Yeast | NULL | Probable | As nectar-tap_saprotroph (Põlme et al. 2020); Decay sugar-rich substrates (Põlme et al. 2020) | Põlme S, et al. 2020. Fungal Diversity 105:1-16 ((https://doi.org/10.1007/s13225-020-00466-2)) |
| Otu19566 | 1 | k__Fungi(100);p__Ascomycota(100);c__Saccharomycetes(100);o__Saccharomycetales(100);f__Metschnikowiaceae(100);g__Metschnikowia(100);s__Metschnikowia_pulcherrima(100); | Metschnikowia | 13 | Saprotroph | \|Undefined Saprotroph\| | Yeast | NULL | Probable | As nectar-tap_saprotroph (Põlme et al. 2020); Decay sugar-rich substrates (Põlme et al. 2020) | Põlme S, et al. 2020. Fungal Diversity 105:1-16 ((https://doi.org/10.1007/s13225-020-00466-2)) |
| Otu19567 | 1 | k__Fungi(100);p__Ascomycota(100);c__Saccharomycetes(100);o__Saccharomycetales(100);f__Metschnikowiaceae(100);g__Metschnikowia(100);s__Metschnikowia_pulcherrima(100); | Metschnikowia | 13 | Saprotroph | \|Undefined Saprotroph\| | Yeast | NULL | Probable | As nectar-tap_saprotroph (Põlme et al. 2020); Decay sugar-rich substrates (Põlme et al. 2020) | Põlme S, et al. 2020. Fungal Diversity 105:1-16 ((https://doi.org/10.1007/s13225-020-00466-2)) |
| Otu19568 | 1 | k__Fungi(100);p__Ascomycota(100);c__Saccharomycetes(100);o__Saccharomycetales(100);f__Metschnikowiaceae(100);g__Metschnikowia(100);g__Metschnikowia_unclassified(100); | Metschnikowia | 13 | Saprotroph | \|Undefined Saprotroph\| | Yeast | NULL | Probable | As nectar-tap_saprotroph (Põlme et al. 2020); Decay sugar-rich substrates (Põlme et al. 2020) | Põlme S, et al. 2020. Fungal Diversity 105:1-16 ((https://doi.org/10.1007/s13225-020-00466-2)) |
| Otu19569 | 1 | k__Fungi(100);p__Ascomycota(100);c__Saccharomycetes(100);o__Saccharomycetales(100);f__Metschnikowiaceae(100);g__Metschnikowia(100);g__Metschnikowia_unclassified(100); | Metschnikowia | 13 | Saprotroph | \|Undefined Saprotroph\| | Yeast | NULL | Probable | As nectar-tap_saprotroph (Põlme et al. 2020); Decay sugar-rich substrates (Põlme et al. 2020) | Põlme S, et al. 2020. Fungal Diversity 105:1-16 ((https://doi.org/10.1007/s13225-020-00466-2)) |
| Otu19570 | 1 | k__Fungi(100);p__Ascomycota(100);c__Saccharomycetes(100);o__Saccharomycetales(100);f__Metschnikowiaceae(100);g__Metschnikowia(100);g__Metschnikowia_unclassified(100); | Metschnikowia | 13 | Saprotroph | \|Undefined Saprotroph\| | Yeast | NULL | Probable | As nectar-tap_saprotroph (Põlme et al. 2020); Decay sugar-rich substrates (Põlme et al. 2020) | Põlme S, et al. 2020. Fungal Diversity 105:1-16 ((https://doi.org/10.1007/s13225-020-00466-2)) |
| Otu19571 | 1 | k__Fungi(100);p__Ascomycota(100);c__Saccharomycetes(100);o__Saccharomycetales(100);f__Metschnikowiaceae(100);g__Metschnikowia(100);s__Metschnikowia_pulcherrima(100); | Metschnikowia | 13 | Saprotroph | \|Undefined Saprotroph\| | Yeast | NULL | Probable | As nectar-tap_saprotroph (Põlme et al. 2020); Decay sugar-rich substrates (Põlme et al. 2020) | Põlme S, et al. 2020. Fungal Diversity 105:1-16 ((https://doi.org/10.1007/s13225-020-00466-2)) |
| Otu19572 | 1 | k__Fungi(100);p__Ascomycota(100);c__Saccharomycetes(100);o__Saccharomycetales(100);f__Metschnikowiaceae(100);g__Metschnikowia(100);s__Metschnikowia_pulcherrima(100); | Metschnikowia | 13 | Saprotroph | \|Undefined Saprotroph\| | Yeast | NULL | Probable | As nectar-tap_saprotroph (Põlme et al. 2020); Decay sugar-rich substrates (Põlme et al. 2020) | Põlme S, et al. 2020. Fungal Diversity 105:1-16 ((https://doi.org/10.1007/s13225-020-00466-2)) |
| Otu19573 | 1 | k__Fungi(100);p__Ascomycota(100);c__Saccharomycetes(100);o__Saccharomycetales(100);f__Metschnikowiaceae(100);g__Metschnikowia(100);s__Metschnikowia_pulcherrima(100); | Metschnikowia | 13 | Saprotroph | \|Undefined Saprotroph\| | Yeast | NULL | Probable | As nectar-tap_saprotroph (Põlme et al. 2020); Decay sugar-rich substrates (Põlme et al. 2020) | Põlme S, et al. 2020. Fungal Diversity 105:1-16 ((https://doi.org/10.1007/s13225-020-00466-2)) |
| Otu19574 | 1 | k__Fungi(100);p__Ascomycota(100);c__Saccharomycetes(100);o__Saccharomycetales(100);f__Metschnikowiaceae(100);g__Metschnikowia(100);s__Metschnikowia_pulcherrima(100); | Metschnikowia | 13 | Saprotroph | \|Undefined Saprotroph\| | Yeast | NULL | Probable | As nectar-tap_saprotroph (Põlme et al. 2020); Decay sugar-rich substrates (Põlme et al. 2020) | Põlme S, et al. 2020. Fungal Diversity 105:1-16 ((https://doi.org/10.1007/s13225-020-00466-2)) |
| Otu19575 | 1 | k__Fungi(100);p__Ascomycota(100);c__Saccharomycetes(100);o__Saccharomycetales(100);f__Metschnikowiaceae(100);g__Metschnikowia(100);s__Metschnikowia_pulcherrima(100); | Metschnikowia | 13 | Saprotroph | \|Undefined Saprotroph\| | Yeast | NULL | Probable | As nectar-tap_saprotroph (Põlme et al. 2020); Decay sugar-rich substrates (Põlme et al. 2020) | Põlme S, et al. 2020. Fungal Diversity 105:1-16 ((https://doi.org/10.1007/s13225-020-00466-2)) |
| Otu19576 | 1 | k__Fungi(100);p__Ascomycota(100);c__Saccharomycetes(100);o__Saccharomycetales(100);f__Metschnikowiaceae(100);g__Metschnikowia(100);s__Metschnikowia_pulcherrima(100); | Metschnikowia | 13 | Saprotroph | \|Undefined Saprotroph\| | Yeast | NULL | Probable | As nectar-tap_saprotroph (Põlme et al. 2020); Decay sugar-rich substrates (Põlme et al. 2020) | Põlme S, et al. 2020. Fungal Diversity 105:1-16 ((https://doi.org/10.1007/s13225-020-00466-2)) |
| Otu19577 | 1 | k__Fungi(100);p__Ascomycota(100);c__Saccharomycetes(100);o__Saccharomycetales(100);f__Metschnikowiaceae(100);g__Metschnikowia(100);s__Metschnikowia_pulcherrima(100); | Metschnikowia | 13 | Saprotroph | \|Undefined Saprotroph\| | Yeast | NULL | Probable | As nectar-tap_saprotroph (Põlme et al. 2020); Decay sugar-rich substrates (Põlme et al. 2020) | Põlme S, et al. 2020. Fungal Diversity 105:1-16 ((https://doi.org/10.1007/s13225-020-00466-2)) |
| Otu19578 | 1 | k__Fungi(100);p__Ascomycota(100);c__Saccharomycetes(100);o__Saccharomycetales(100);f__Metschnikowiaceae(100);g__Metschnikowia(100);g__Metschnikowia_unclassified(100); | Metschnikowia | 13 | Saprotroph | \|Undefined Saprotroph\| | Yeast | NULL | Probable | As nectar-tap_saprotroph (Põlme et al. 2020); Decay sugar-rich substrates (Põlme et al. 2020) | Põlme S, et al. 2020. Fungal Diversity 105:1-16 ((https://doi.org/10.1007/s13225-020-00466-2)) |
| Otu19579 | 1 | k__Fungi(100);p__Ascomycota(100);c__Saccharomycetes(100);o__Saccharomycetales(100);f__Metschnikowiaceae(100);g__Metschnikowia(100);s__Metschnikowia_pulcherrima(100); | Metschnikowia | 13 | Saprotroph | \|Undefined Saprotroph\| | Yeast | NULL | Probable | As nectar-tap_saprotroph (Põlme et al. 2020); Decay sugar-rich substrates (Põlme et al. 2020) | Põlme S, et al. 2020. Fungal Diversity 105:1-16 ((https://doi.org/10.1007/s13225-020-00466-2)) |
| Otu19580 | 1 | k__Fungi(100);p__Ascomycota(100);c__Saccharomycetes(100);o__Saccharomycetales(100);f__Metschnikowiaceae(100);g__Metschnikowia(100);g__Metschnikowia_unclassified(100); | Metschnikowia | 13 | Saprotroph | \|Undefined Saprotroph\| | Yeast | NULL | Probable | As nectar-tap_saprotroph (Põlme et al. 2020); Decay sugar-rich substrates (Põlme et al. 2020) | Põlme S, et al. 2020. Fungal Diversity 105:1-16 ((https://doi.org/10.1007/s13225-020-00466-2)) |
| Otu19581 | 1 | k__Fungi(100);p__Ascomycota(100);c__Saccharomycetes(100);o__Saccharomycetales(100);f__Metschnikowiaceae(100);g__Metschnikowia(100);s__Metschnikowia_pulcherrima(100); | Metschnikowia | 13 | Saprotroph | \|Undefined Saprotroph\| | Yeast | NULL | Probable | As nectar-tap_saprotroph (Põlme et al. 2020); Decay sugar-rich substrates (Põlme et al. 2020) | Põlme S, et al. 2020. Fungal Diversity 105:1-16 ((https://doi.org/10.1007/s13225-020-00466-2)) |
| Otu19582 | 1 | k__Fungi(100);p__Ascomycota(100);c__Saccharomycetes(100);o__Saccharomycetales(100);f__Metschnikowiaceae(100);g__Metschnikowia(100);s__Metschnikowia_pulcherrima(100); | Metschnikowia | 13 | Saprotroph | \|Undefined Saprotroph\| | Yeast | NULL | Probable | As nectar-tap_saprotroph (Põlme et al. 2020); Decay sugar-rich substrates (Põlme et al. 2020) | Põlme S, et al. 2020. Fungal Diversity 105:1-16 ((https://doi.org/10.1007/s13225-020-00466-2)) |
| Otu19583 | 1 | k__Fungi(100);p__Ascomycota(100);c__Saccharomycetes(100);o__Saccharomycetales(100);f__Metschnikowiaceae(100);g__Metschnikowia(100);s__Metschnikowia_pulcherrima(100); | Metschnikowia | 13 | Saprotroph | \|Undefined Saprotroph\| | Yeast | NULL | Probable | As nectar-tap_saprotroph (Põlme et al. 2020); Decay sugar-rich substrates (Põlme et al. 2020) | Põlme S, et al. 2020. Fungal Diversity 105:1-16 ((https://doi.org/10.1007/s13225-020-00466-2)) |
| Otu19584 | 1 | k__Fungi(100);p__Ascomycota(100);c__Saccharomycetes(100);o__Saccharomycetales(100);f__Metschnikowiaceae(100);g__Metschnikowia(100);s__Metschnikowia_pulcherrima(100); | Metschnikowia | 13 | Saprotroph | \|Undefined Saprotroph\| | Yeast | NULL | Probable | As nectar-tap_saprotroph (Põlme et al. 2020); Decay sugar-rich substrates (Põlme et al. 2020) | Põlme S, et al. 2020. Fungal Diversity 105:1-16 ((https://doi.org/10.1007/s13225-020-00466-2)) |
| Otu19585 | 1 | k__Fungi(100);p__Ascomycota(100);c__Saccharomycetes(100);o__Saccharomycetales(100);f__Metschnikowiaceae(100);g__Metschnikowia(100);s__Metschnikowia_pulcherrima(100); | Metschnikowia | 13 | Saprotroph | \|Undefined Saprotroph\| | Yeast | NULL | Probable | As nectar-tap_saprotroph (Põlme et al. 2020); Decay sugar-rich substrates (Põlme et al. 2020) | Põlme S, et al. 2020. Fungal Diversity 105:1-16 ((https://doi.org/10.1007/s13225-020-00466-2)) |
| Otu19586 | 1 | k__Fungi(100);p__Ascomycota(100);c__Saccharomycetes(100);o__Saccharomycetales(100);f__Metschnikowiaceae(100);g__Metschnikowia(100);g__Metschnikowia_unclassified(100); | Metschnikowia | 13 | Saprotroph | \|Undefined Saprotroph\| | Yeast | NULL | Probable | As nectar-tap_saprotroph (Põlme et al. 2020); Decay sugar-rich substrates (Põlme et al. 2020) | Põlme S, et al. 2020. Fungal Diversity 105:1-16 ((https://doi.org/10.1007/s13225-020-00466-2)) |
| Otu19587 | 1 | k__Fungi(100);p__Ascomycota(100);c__Saccharomycetes(100);o__Saccharomycetales(100);f__Metschnikowiaceae(100);g__Metschnikowia(100);g__Metschnikowia_unclassified(100); | Metschnikowia | 13 | Saprotroph | \|Undefined Saprotroph\| | Yeast | NULL | Probable | As nectar-tap_saprotroph (Põlme et al. 2020); Decay sugar-rich substrates (Põlme et al. 2020) | Põlme S, et al. 2020. Fungal Diversity 105:1-16 ((https://doi.org/10.1007/s13225-020-00466-2)) |
| Otu19588 | 1 | k__Fungi(100);p__Ascomycota(100);c__Saccharomycetes(100);o__Saccharomycetales(100);f__Metschnikowiaceae(100);g__Metschnikowia(100);g__Metschnikowia_unclassified(100); | Metschnikowia | 13 | Saprotroph | \|Undefined Saprotroph\| | Yeast | NULL | Probable | As nectar-tap_saprotroph (Põlme et al. 2020); Decay sugar-rich substrates (Põlme et al. 2020) | Põlme S, et al. 2020. Fungal Diversity 105:1-16 ((https://doi.org/10.1007/s13225-020-00466-2)) |
| Otu19589 | 1 | k__Fungi(100);p__Ascomycota(100);c__Saccharomycetes(100);o__Saccharomycetales(100);f__Saccharomycetales_family_Incertae_sedis(100);g__Candida(100);g__Candida_unclassified(100); | Candida | 13 | Pathotroph-Saprotroph-Symbiotroph | Animal Pathogen-Endophyte-Endosymbiont-Epiphyte-\|Undefined Saprotroph\| | Dimorphic Yeast-Yeast | NULL | Probable | As nectar-tap_saprotroph (Põlme et al. 2020); Decay sugar-rich substrates (Põlme et al. 2020); This asexual genus is polyphyletic and contain species that function very differently depend on what they are. Caution should be taken when interpreting guilds for Candida species. | Manolakaki D, et al. 2010. Virulence. 1: 367-375 ((https://doi.org/10.4161/viru.1.5.12796)); Mekbib SB, et al. 2011. Tropical Plant Pathology 36:233-240 ((https://doi.org/10.1590/S1982-56762011000400004)); Põlme S, et al. 2020. Fungal Diversity 105:1-16 ((https://doi.org/10.1007/s13225-020-00466-2)) |
| Otu19590 | 1 | k__Fungi(100);p__Ascomycota(100);c__Saccharomycetes(100);o__Saccharomycetales(100);f__Metschnikowiaceae(100);g__Metschnikowia(100);s__Metschnikowia_pulcherrima(100); | Metschnikowia | 13 | Saprotroph | \|Undefined Saprotroph\| | Yeast | NULL | Probable | As nectar-tap_saprotroph (Põlme et al. 2020); Decay sugar-rich substrates (Põlme et al. 2020) | Põlme S, et al. 2020. Fungal Diversity 105:1-16 ((https://doi.org/10.1007/s13225-020-00466-2)) |
| Otu19591 | 1 | k__Fungi(100);p__Ascomycota(100);c__Saccharomycetes(100);o__Saccharomycetales(100);f__Metschnikowiaceae(100);g__Metschnikowia(100);s__Metschnikowia_pulcherrima(100); | Metschnikowia | 13 | Saprotroph | \|Undefined Saprotroph\| | Yeast | NULL | Probable | As nectar-tap_saprotroph (Põlme et al. 2020); Decay sugar-rich substrates (Põlme et al. 2020) | Põlme S, et al. 2020. Fungal Diversity 105:1-16 ((https://doi.org/10.1007/s13225-020-00466-2)) |
| Otu19592 | 1 | k__Fungi(100);p__Ascomycota(100);c__Saccharomycetes(100);o__Saccharomycetales(100);f__Metschnikowiaceae(100);g__Metschnikowia(100);s__Metschnikowia_pulcherrima(100); | Metschnikowia | 13 | Saprotroph | \|Undefined Saprotroph\| | Yeast | NULL | Probable | As nectar-tap_saprotroph (Põlme et al. 2020); Decay sugar-rich substrates (Põlme et al. 2020) | Põlme S, et al. 2020. Fungal Diversity 105:1-16 ((https://doi.org/10.1007/s13225-020-00466-2)) |
| Otu19593 | 1 | k__Fungi(100);p__Ascomycota(100);c__Saccharomycetes(100);o__Saccharomycetales(100);f__Metschnikowiaceae(100);g__Metschnikowia(100);s__Metschnikowia_pulcherrima(100); | Metschnikowia | 13 | Saprotroph | \|Undefined Saprotroph\| | Yeast | NULL | Probable | As nectar-tap_saprotroph (Põlme et al. 2020); Decay sugar-rich substrates (Põlme et al. 2020) | Põlme S, et al. 2020. Fungal Diversity 105:1-16 ((https://doi.org/10.1007/s13225-020-00466-2)) |
| Otu19594 | 1 | k__Fungi(100);p__Ascomycota(100);c__Saccharomycetes(100);o__Saccharomycetales(100);f__Metschnikowiaceae(100);g__Metschnikowia(100);g__Metschnikowia_unclassified(100); | Metschnikowia | 13 | Saprotroph | \|Undefined Saprotroph\| | Yeast | NULL | Probable | As nectar-tap_saprotroph (Põlme et al. 2020); Decay sugar-rich substrates (Põlme et al. 2020) | Põlme S, et al. 2020. Fungal Diversity 105:1-16 ((https://doi.org/10.1007/s13225-020-00466-2)) |
| Otu19595 | 1 | k__Fungi(100);p__Ascomycota(100);c__Saccharomycetes(100);o__Saccharomycetales(100);f__Metschnikowiaceae(100);g__Metschnikowia(100);g__Metschnikowia_unclassified(100); | Metschnikowia | 13 | Saprotroph | \|Undefined Saprotroph\| | Yeast | NULL | Probable | As nectar-tap_saprotroph (Põlme et al. 2020); Decay sugar-rich substrates (Põlme et al. 2020) | Põlme S, et al. 2020. Fungal Diversity 105:1-16 ((https://doi.org/10.1007/s13225-020-00466-2)) |
| Otu19597 | 1 | k__Fungi(100);p__Ascomycota(100);c__Saccharomycetes(100);o__Saccharomycetales(100);f__Metschnikowiaceae(100);g__Metschnikowia(100);s__Metschnikowia_pulcherrima(100); | Metschnikowia | 13 | Saprotroph | \|Undefined Saprotroph\| | Yeast | NULL | Probable | As nectar-tap_saprotroph (Põlme et al. 2020); Decay sugar-rich substrates (Põlme et al. 2020) | Põlme S, et al. 2020. Fungal Diversity 105:1-16 ((https://doi.org/10.1007/s13225-020-00466-2)) |
| Otu19598 | 1 | k__Fungi(100);p__Ascomycota(100);c__Saccharomycetes(100);o__Saccharomycetales(100);f__Metschnikowiaceae(100);g__Metschnikowia(100);g__Metschnikowia_unclassified(100); | Metschnikowia | 13 | Saprotroph | \|Undefined Saprotroph\| | Yeast | NULL | Probable | As nectar-tap_saprotroph (Põlme et al. 2020); Decay sugar-rich substrates (Põlme et al. 2020) | Põlme S, et al. 2020. Fungal Diversity 105:1-16 ((https://doi.org/10.1007/s13225-020-00466-2)) |
| Otu19599 | 1 | k__Fungi(100);p__Ascomycota(100);c__Saccharomycetes(100);o__Saccharomycetales(100);f__Metschnikowiaceae(100);g__Metschnikowia(100);s__Metschnikowia_pulcherrima(100); | Metschnikowia | 13 | Saprotroph | \|Undefined Saprotroph\| | Yeast | NULL | Probable | As nectar-tap_saprotroph (Põlme et al. 2020); Decay sugar-rich substrates (Põlme et al. 2020) | Põlme S, et al. 2020. Fungal Diversity 105:1-16 ((https://doi.org/10.1007/s13225-020-00466-2)) |
| Otu19600 | 1 | k__Fungi(100);p__Ascomycota(100);c__Saccharomycetes(100);o__Saccharomycetales(100);f__Metschnikowiaceae(100);g__Metschnikowia(100);g__Metschnikowia_unclassified(100); | Metschnikowia | 13 | Saprotroph | \|Undefined Saprotroph\| | Yeast | NULL | Probable | As nectar-tap_saprotroph (Põlme et al. 2020); Decay sugar-rich substrates (Põlme et al. 2020) | Põlme S, et al. 2020. Fungal Diversity 105:1-16 ((https://doi.org/10.1007/s13225-020-00466-2)) |
| Otu19601 | 1 | k__Fungi(100);p__Ascomycota(100);c__Saccharomycetes(100);o__Saccharomycetales(100);f__Metschnikowiaceae(100);g__Metschnikowia(100);g__Metschnikowia_unclassified(100); | Metschnikowia | 13 | Saprotroph | \|Undefined Saprotroph\| | Yeast | NULL | Probable | As nectar-tap_saprotroph (Põlme et al. 2020); Decay sugar-rich substrates (Põlme et al. 2020) | Põlme S, et al. 2020. Fungal Diversity 105:1-16 ((https://doi.org/10.1007/s13225-020-00466-2)) |
| Otu19602 | 1 | k__Fungi(100);p__Ascomycota(100);c__Saccharomycetes(100);o__Saccharomycetales(100);f__Metschnikowiaceae(100);g__Metschnikowia(100);g__Metschnikowia_unclassified(100); | Metschnikowia | 13 | Saprotroph | \|Undefined Saprotroph\| | Yeast | NULL | Probable | As nectar-tap_saprotroph (Põlme et al. 2020); Decay sugar-rich substrates (Põlme et al. 2020) | Põlme S, et al. 2020. Fungal Diversity 105:1-16 ((https://doi.org/10.1007/s13225-020-00466-2)) |
| Otu19603 | 1 | k__Fungi(100);p__Ascomycota(100);c__Saccharomycetes(100);o__Saccharomycetales(100);f__Metschnikowiaceae(100);g__Metschnikowia(100);s__Metschnikowia_pulcherrima(100); | Metschnikowia | 13 | Saprotroph | \|Undefined Saprotroph\| | Yeast | NULL | Probable | As nectar-tap_saprotroph (Põlme et al. 2020); Decay sugar-rich substrates (Põlme et al. 2020) | Põlme S, et al. 2020. Fungal Diversity 105:1-16 ((https://doi.org/10.1007/s13225-020-00466-2)) |
| Otu19604 | 1 | k__Fungi(100);p__Ascomycota(100);c__Saccharomycetes(100);o__Saccharomycetales(100);f__Metschnikowiaceae(100);g__Metschnikowia(100);s__Metschnikowia_pulcherrima(100); | Metschnikowia | 13 | Saprotroph | \|Undefined Saprotroph\| | Yeast | NULL | Probable | As nectar-tap_saprotroph (Põlme et al. 2020); Decay sugar-rich substrates (Põlme et al. 2020) | Põlme S, et al. 2020. Fungal Diversity 105:1-16 ((https://doi.org/10.1007/s13225-020-00466-2)) |
| Otu19605 | 1 | k__Fungi(100);p__Ascomycota(100);c__Saccharomycetes(100);o__Saccharomycetales(100);f__Metschnikowiaceae(100);g__Metschnikowia(100);s__Metschnikowia_pulcherrima(100); | Metschnikowia | 13 | Saprotroph | \|Undefined Saprotroph\| | Yeast | NULL | Probable | As nectar-tap_saprotroph (Põlme et al. 2020); Decay sugar-rich substrates (Põlme et al. 2020) | Põlme S, et al. 2020. Fungal Diversity 105:1-16 ((https://doi.org/10.1007/s13225-020-00466-2)) |
| Otu19606 | 1 | k__Fungi(100);p__Ascomycota(100);c__Saccharomycetes(100);o__Saccharomycetales(100);f__Metschnikowiaceae(100);g__Metschnikowia(100);s__Metschnikowia_pulcherrima(100); | Metschnikowia | 13 | Saprotroph | \|Undefined Saprotroph\| | Yeast | NULL | Probable | As nectar-tap_saprotroph (Põlme et al. 2020); Decay sugar-rich substrates (Põlme et al. 2020) | Põlme S, et al. 2020. Fungal Diversity 105:1-16 ((https://doi.org/10.1007/s13225-020-00466-2)) |
| Otu19607 | 1 | k__Fungi(100);p__Ascomycota(100);c__Saccharomycetes(100);o__Saccharomycetales(100);f__Metschnikowiaceae(100);g__Metschnikowia(100);s__Metschnikowia_pulcherrima(100); | Metschnikowia | 13 | Saprotroph | \|Undefined Saprotroph\| | Yeast | NULL | Probable | As nectar-tap_saprotroph (Põlme et al. 2020); Decay sugar-rich substrates (Põlme et al. 2020) | Põlme S, et al. 2020. Fungal Diversity 105:1-16 ((https://doi.org/10.1007/s13225-020-00466-2)) |
| Otu19608 | 1 | k__Fungi(100);p__Ascomycota(100);c__Saccharomycetes(100);o__Saccharomycetales(100);f__Metschnikowiaceae(100);g__Metschnikowia(100);s__Metschnikowia_pulcherrima(100); | Metschnikowia | 13 | Saprotroph | \|Undefined Saprotroph\| | Yeast | NULL | Probable | As nectar-tap_saprotroph (Põlme et al. 2020); Decay sugar-rich substrates (Põlme et al. 2020) | Põlme S, et al. 2020. Fungal Diversity 105:1-16 ((https://doi.org/10.1007/s13225-020-00466-2)) |
| Otu19609 | 1 | k__Fungi(100);p__Ascomycota(100);c__Saccharomycetes(100);o__Saccharomycetales(100);f__Metschnikowiaceae(100);g__Metschnikowia(100);s__Metschnikowia_pulcherrima(100); | Metschnikowia | 13 | Saprotroph | \|Undefined Saprotroph\| | Yeast | NULL | Probable | As nectar-tap_saprotroph (Põlme et al. 2020); Decay sugar-rich substrates (Põlme et al. 2020) | Põlme S, et al. 2020. Fungal Diversity 105:1-16 ((https://doi.org/10.1007/s13225-020-00466-2)) |
| Otu19610 | 1 | k__Fungi(100);p__Ascomycota(100);c__Saccharomycetes(100);o__Saccharomycetales(100);f__Metschnikowiaceae(100);g__Metschnikowia(100);g__Metschnikowia_unclassified(100); | Metschnikowia | 13 | Saprotroph | \|Undefined Saprotroph\| | Yeast | NULL | Probable | As nectar-tap_saprotroph (Põlme et al. 2020); Decay sugar-rich substrates (Põlme et al. 2020) | Põlme S, et al. 2020. Fungal Diversity 105:1-16 ((https://doi.org/10.1007/s13225-020-00466-2)) |
| Otu19611 | 1 | k__Fungi(100);p__Ascomycota(100);c__Saccharomycetes(100);o__Saccharomycetales(100);f__Metschnikowiaceae(100);g__Metschnikowia(100);g__Metschnikowia_unclassified(100); | Metschnikowia | 13 | Saprotroph | \|Undefined Saprotroph\| | Yeast | NULL | Probable | As nectar-tap_saprotroph (Põlme et al. 2020); Decay sugar-rich substrates (Põlme et al. 2020) | Põlme S, et al. 2020. Fungal Diversity 105:1-16 ((https://doi.org/10.1007/s13225-020-00466-2)) |
| Otu19612 | 1 | k__Fungi(100);p__Ascomycota(100);c__Saccharomycetes(100);o__Saccharomycetales(100);f__Metschnikowiaceae(100);g__Metschnikowia(100);s__Metschnikowia_pulcherrima(100); | Metschnikowia | 13 | Saprotroph | \|Undefined Saprotroph\| | Yeast | NULL | Probable | As nectar-tap_saprotroph (Põlme et al. 2020); Decay sugar-rich substrates (Põlme et al. 2020) | Põlme S, et al. 2020. Fungal Diversity 105:1-16 ((https://doi.org/10.1007/s13225-020-00466-2)) |
| Otu19613 | 1 | k__Fungi(100);p__Ascomycota(100);c__Saccharomycetes(100);o__Saccharomycetales(100);f__Metschnikowiaceae(100);g__Metschnikowia(100);s__Metschnikowia_pulcherrima(100); | Metschnikowia | 13 | Saprotroph | \|Undefined Saprotroph\| | Yeast | NULL | Probable | As nectar-tap_saprotroph (Põlme et al. 2020); Decay sugar-rich substrates (Põlme et al. 2020) | Põlme S, et al. 2020. Fungal Diversity 105:1-16 ((https://doi.org/10.1007/s13225-020-00466-2)) |
| Otu19614 | 1 | k__Fungi(100);p__Ascomycota(100);c__Saccharomycetes(100);o__Saccharomycetales(100);f__Metschnikowiaceae(100);g__Metschnikowia(100);s__Metschnikowia_pulcherrima(100); | Metschnikowia | 13 | Saprotroph | \|Undefined Saprotroph\| | Yeast | NULL | Probable | As nectar-tap_saprotroph (Põlme et al. 2020); Decay sugar-rich substrates (Põlme et al. 2020) | Põlme S, et al. 2020. Fungal Diversity 105:1-16 ((https://doi.org/10.1007/s13225-020-00466-2)) |
| Otu19615 | 1 | k__Fungi(100);p__Ascomycota(100);c__Saccharomycetes(100);o__Saccharomycetales(100);f__Metschnikowiaceae(100);g__Metschnikowia(100);s__Metschnikowia_pulcherrima(100); | Metschnikowia | 13 | Saprotroph | \|Undefined Saprotroph\| | Yeast | NULL | Probable | As nectar-tap_saprotroph (Põlme et al. 2020); Decay sugar-rich substrates (Põlme et al. 2020) | Põlme S, et al. 2020. Fungal Diversity 105:1-16 ((https://doi.org/10.1007/s13225-020-00466-2)) |
| Otu19616 | 1 | k__Fungi(100);p__Ascomycota(100);c__Saccharomycetes(100);o__Saccharomycetales(100);f__Metschnikowiaceae(100);g__Metschnikowia(100);s__Metschnikowia_pulcherrima(100); | Metschnikowia | 13 | Saprotroph | \|Undefined Saprotroph\| | Yeast | NULL | Probable | As nectar-tap_saprotroph (Põlme et al. 2020); Decay sugar-rich substrates (Põlme et al. 2020) | Põlme S, et al. 2020. Fungal Diversity 105:1-16 ((https://doi.org/10.1007/s13225-020-00466-2)) |
| Otu19617 | 1 | k__Fungi(100);p__Ascomycota(100);c__Saccharomycetes(100);o__Saccharomycetales(100);f__Metschnikowiaceae(100);g__Metschnikowia(100);g__Metschnikowia_unclassified(100); | Metschnikowia | 13 | Saprotroph | \|Undefined Saprotroph\| | Yeast | NULL | Probable | As nectar-tap_saprotroph (Põlme et al. 2020); Decay sugar-rich substrates (Põlme et al. 2020) | Põlme S, et al. 2020. Fungal Diversity 105:1-16 ((https://doi.org/10.1007/s13225-020-00466-2)) |
| Otu19619 | 1 | k__Fungi(100);p__Ascomycota(100);c__Saccharomycetes(100);o__Saccharomycetales(100);f__Metschnikowiaceae(100);g__Metschnikowia(100);s__Metschnikowia_pulcherrima(100); | Metschnikowia | 13 | Saprotroph | \|Undefined Saprotroph\| | Yeast | NULL | Probable | As nectar-tap_saprotroph (Põlme et al. 2020); Decay sugar-rich substrates (Põlme et al. 2020) | Põlme S, et al. 2020. Fungal Diversity 105:1-16 ((https://doi.org/10.1007/s13225-020-00466-2)) |
| Otu19620 | 1 | k__Fungi(100);p__Ascomycota(100);c__Saccharomycetes(100);o__Saccharomycetales(100);f__Metschnikowiaceae(100);g__Metschnikowia(100);g__Metschnikowia_unclassified(100); | Metschnikowia | 13 | Saprotroph | \|Undefined Saprotroph\| | Yeast | NULL | Probable | As nectar-tap_saprotroph (Põlme et al. 2020); Decay sugar-rich substrates (Põlme et al. 2020) | Põlme S, et al. 2020. Fungal Diversity 105:1-16 ((https://doi.org/10.1007/s13225-020-00466-2)) |
| Otu19621 | 1 | k__Fungi(100);p__Ascomycota(100);c__Saccharomycetes(100);o__Saccharomycetales(100);f__Metschnikowiaceae(100);g__Metschnikowia(100);s__Metschnikowia_pulcherrima(100); | Metschnikowia | 13 | Saprotroph | \|Undefined Saprotroph\| | Yeast | NULL | Probable | As nectar-tap_saprotroph (Põlme et al. 2020); Decay sugar-rich substrates (Põlme et al. 2020) | Põlme S, et al. 2020. Fungal Diversity 105:1-16 ((https://doi.org/10.1007/s13225-020-00466-2)) |
| Otu19622 | 1 | k__Fungi(100);p__Ascomycota(100);c__Saccharomycetes(100);o__Saccharomycetales(100);f__Metschnikowiaceae(100);g__Metschnikowia(100);s__Metschnikowia_pulcherrima(100); | Metschnikowia | 13 | Saprotroph | \|Undefined Saprotroph\| | Yeast | NULL | Probable | As nectar-tap_saprotroph (Põlme et al. 2020); Decay sugar-rich substrates (Põlme et al. 2020) | Põlme S, et al. 2020. Fungal Diversity 105:1-16 ((https://doi.org/10.1007/s13225-020-00466-2)) |
| Otu19623 | 1 | k__Fungi(100);p__Ascomycota(100);c__Saccharomycetes(100);o__Saccharomycetales(100);f__Metschnikowiaceae(100);g__Metschnikowia(100);s__Metschnikowia_pulcherrima(100); | Metschnikowia | 13 | Saprotroph | \|Undefined Saprotroph\| | Yeast | NULL | Probable | As nectar-tap_saprotroph (Põlme et al. 2020); Decay sugar-rich substrates (Põlme et al. 2020) | Põlme S, et al. 2020. Fungal Diversity 105:1-16 ((https://doi.org/10.1007/s13225-020-00466-2)) |
| Otu19624 | 1 | k__Fungi(100);p__Ascomycota(100);c__Saccharomycetes(100);o__Saccharomycetales(100);f__Metschnikowiaceae(100);g__Metschnikowia(100);s__Metschnikowia_pulcherrima(100); | Metschnikowia | 13 | Saprotroph | \|Undefined Saprotroph\| | Yeast | NULL | Probable | As nectar-tap_saprotroph (Põlme et al. 2020); Decay sugar-rich substrates (Põlme et al. 2020) | Põlme S, et al. 2020. Fungal Diversity 105:1-16 ((https://doi.org/10.1007/s13225-020-00466-2)) |
| Otu19625 | 1 | k__Fungi(100);p__Ascomycota(100);c__Saccharomycetes(100);o__Saccharomycetales(100);f__Metschnikowiaceae(100);g__Metschnikowia(100);s__Metschnikowia_pulcherrima(100); | Metschnikowia | 13 | Saprotroph | \|Undefined Saprotroph\| | Yeast | NULL | Probable | As nectar-tap_saprotroph (Põlme et al. 2020); Decay sugar-rich substrates (Põlme et al. 2020) | Põlme S, et al. 2020. Fungal Diversity 105:1-16 ((https://doi.org/10.1007/s13225-020-00466-2)) |
| Otu19627 | 1 | k__Fungi(100);p__Ascomycota(100);c__Saccharomycetes(100);o__Saccharomycetales(100);f__Metschnikowiaceae(100);g__Metschnikowia(100);g__Metschnikowia_unclassified(100); | Metschnikowia | 13 | Saprotroph | \|Undefined Saprotroph\| | Yeast | NULL | Probable | As nectar-tap_saprotroph (Põlme et al. 2020); Decay sugar-rich substrates (Põlme et al. 2020) | Põlme S, et al. 2020. Fungal Diversity 105:1-16 ((https://doi.org/10.1007/s13225-020-00466-2)) |
| Otu19630 | 1 | k__Fungi(100);p__Ascomycota(100);c__Saccharomycetes(100);o__Saccharomycetales(100);f__Metschnikowiaceae(100);g__Metschnikowia(100);s__Metschnikowia_pulcherrima(100); | Metschnikowia | 13 | Saprotroph | \|Undefined Saprotroph\| | Yeast | NULL | Probable | As nectar-tap_saprotroph (Põlme et al. 2020); Decay sugar-rich substrates (Põlme et al. 2020) | Põlme S, et al. 2020. Fungal Diversity 105:1-16 ((https://doi.org/10.1007/s13225-020-00466-2)) |
| Otu19631 | 1 | k__Fungi(100);p__Ascomycota(100);c__Saccharomycetes(100);o__Saccharomycetales(100);f__Metschnikowiaceae(100);g__Metschnikowia(100);s__Metschnikowia_pulcherrima(100); | Metschnikowia | 13 | Saprotroph | \|Undefined Saprotroph\| | Yeast | NULL | Probable | As nectar-tap_saprotroph (Põlme et al. 2020); Decay sugar-rich substrates (Põlme et al. 2020) | Põlme S, et al. 2020. Fungal Diversity 105:1-16 ((https://doi.org/10.1007/s13225-020-00466-2)) |
| Otu19633 | 1 | k__Fungi(100);p__Ascomycota(100);c__Saccharomycetes(100);o__Saccharomycetales(100);f__Metschnikowiaceae(100);g__Metschnikowia(100);g__Metschnikowia_unclassified(100); | Metschnikowia | 13 | Saprotroph | \|Undefined Saprotroph\| | Yeast | NULL | Probable | As nectar-tap_saprotroph (Põlme et al. 2020); Decay sugar-rich substrates (Põlme et al. 2020) | Põlme S, et al. 2020. Fungal Diversity 105:1-16 ((https://doi.org/10.1007/s13225-020-00466-2)) |
| Otu19634 | 1 | k__Fungi(100);p__Basidiomycota(100);c__Tremellomycetes(100);o__Tremellales(100);f__Tremellales_family_Incertae_sedis(100);g__Hannaella(100);s__Hannaella_zeae(100); | Hannaella | 13 | Pathotroph-Saprotroph | \|Animal Parasite\|-Undefined Saprotroph | Yeast | NULL | Probable | As animal_decomposer (Põlme et al. 2020); Decay animal material (Põlme et al. 2020) | Põlme S, et al. 2020. Fungal Diversity 105:1-16 ((https://doi.org/10.1007/s13225-020-00466-2)) |
| Otu19636 | 1 | k__Fungi(100);p__Ascomycota(100);c__Saccharomycetes(100);o__Saccharomycetales(100);f__Metschnikowiaceae(100);g__Metschnikowia(100);s__Metschnikowia_pulcherrima(100); | Metschnikowia | 13 | Saprotroph | \|Undefined Saprotroph\| | Yeast | NULL | Probable | As nectar-tap_saprotroph (Põlme et al. 2020); Decay sugar-rich substrates (Põlme et al. 2020) | Põlme S, et al. 2020. Fungal Diversity 105:1-16 ((https://doi.org/10.1007/s13225-020-00466-2)) |
| Otu19637 | 1 | k__Fungi(100);p__Ascomycota(100);c__Saccharomycetes(100);o__Saccharomycetales(100);f__Metschnikowiaceae(100);g__Metschnikowia(100);s__Metschnikowia_pulcherrima(100); | Metschnikowia | 13 | Saprotroph | \|Undefined Saprotroph\| | Yeast | NULL | Probable | As nectar-tap_saprotroph (Põlme et al. 2020); Decay sugar-rich substrates (Põlme et al. 2020) | Põlme S, et al. 2020. Fungal Diversity 105:1-16 ((https://doi.org/10.1007/s13225-020-00466-2)) |

| | | | | | | | | | | | |
|---|---|---|---|---|---|---|---|---|---|---|---|
| Otu19638 | 1 | k__Fungi(100);p__Ascomycota(100);c__Saccharomycetes(100);o__Saccharomycetales(100);f__Metschnikowiaceae(100);g__Metschnikowia(100);s__Metschnikowia_pulcherrima(100); | Metschnikowia | 13 | Saprotroph | \|Undefined Saprotroph\| | Yeast | NULL | Probable | As nectar-tap_saprotroph (Põlme et al. 2020); Decay sugar-rich substrates (Põlme et al. 2020) | Põlme S, et al. 2020. Fungal Diversity 105:1-16 ((https://doi.org/10.1007/s13225-020-00466-2)) |
| Otu19639 | 1 | k__Fungi(100);p__Ascomycota(100);c__Saccharomycetes(100);o__Saccharomycetales(100);f__Metschnikowiaceae(100);g__Metschnikowia(100);s__Metschnikowia_pulcherrima(100); | Metschnikowia | 13 | Saprotroph | \|Undefined Saprotroph\| | Yeast | NULL | Probable | As nectar-tap_saprotroph (Põlme et al. 2020); Decay sugar-rich substrates (Põlme et al. 2020) | Põlme S, et al. 2020. Fungal Diversity 105:1-16 ((https://doi.org/10.1007/s13225-020-00466-2)) |
| Otu19640 | 1 | k__Fungi(100);p__Ascomycota(100);c__Saccharomycetes(100);o__Saccharomycetales(100);f__Metschnikowiaceae(100);g__Metschnikowia(100);s__Metschnikowia_pulcherrima(100); | Metschnikowia | 13 | Saprotroph | \|Undefined Saprotroph\| | Yeast | NULL | Probable | As nectar-tap_saprotroph (Põlme et al. 2020); Decay sugar-rich substrates (Põlme et al. 2020) | Põlme S, et al. 2020. Fungal Diversity 105:1-16 ((https://doi.org/10.1007/s13225-020-00466-2)) |
| Otu19641 | 1 | k__Fungi(100);p__Ascomycota(100);c__Saccharomycetes(100);o__Saccharomycetales(100);f__Metschnikowiaceae(100);g__Metschnikowia(100);s__Metschnikowia_pulcherrima(100); | Metschnikowia | 13 | Saprotroph | \|Undefined Saprotroph\| | Yeast | NULL | Probable | As nectar-tap_saprotroph (Põlme et al. 2020); Decay sugar-rich substrates (Põlme et al. 2020) | Põlme S, et al. 2020. Fungal Diversity 105:1-16 ((https://doi.org/10.1007/s13225-020-00466-2)) |
| Otu19642 | 1 | k__Fungi(100);p__Ascomycota(100);c__Saccharomycetes(100);o__Saccharomycetales(100);f__Metschnikowiaceae(100);g__Metschnikowia(100);s__Metschnikowia_pulcherrima(100); | Metschnikowia | 13 | Saprotroph | \|Undefined Saprotroph\| | Yeast | NULL | Probable | As nectar-tap_saprotroph (Põlme et al. 2020); Decay sugar-rich substrates (Põlme et al. 2020) | Põlme S, et al. 2020. Fungal Diversity 105:1-16 ((https://doi.org/10.1007/s13225-020-00466-2)) |
| Otu19643 | 1 | k__Fungi(100);p__Ascomycota(100);c__Saccharomycetes(100);o__Saccharomycetales(100);f__Metschnikowiaceae(100);g__Metschnikowia(100);s__Metschnikowia_pulcherrima(100); | Metschnikowia | 13 | Saprotroph | \|Undefined Saprotroph\| | Yeast | NULL | Probable | As nectar-tap_saprotroph (Põlme et al. 2020); Decay sugar-rich substrates (Põlme et al. 2020) | Põlme S, et al. 2020. Fungal Diversity 105:1-16 ((https://doi.org/10.1007/s13225-020-00466-2)) |
| Otu19644 | 1 | k__Fungi(100);p__Ascomycota(100);c__Saccharomycetes(100);o__Saccharomycetales(100);f__Metschnikowiaceae(100);g__Metschnikowia(100);s__Metschnikowia_pulcherrima(100); | Metschnikowia | 13 | Saprotroph | \|Undefined Saprotroph\| | Yeast | NULL | Probable | As nectar-tap_saprotroph (Põlme et al. 2020); Decay sugar-rich substrates (Põlme et al. 2020) | Põlme S, et al. 2020. Fungal Diversity 105:1-16 ((https://doi.org/10.1007/s13225-020-00466-2)) |
| Otu19645 | 1 | k__Fungi(100);p__Ascomycota(100);c__Saccharomycetes(100);o__Saccharomycetales(100);f__Metschnikowiaceae(100);g__Metschnikowia(100);s__Metschnikowia_pulcherrima(100); | Metschnikowia | 13 | Saprotroph | \|Undefined Saprotroph\| | Yeast | NULL | Probable | As nectar-tap_saprotroph (Põlme et al. 2020); Decay sugar-rich substrates (Põlme et al. 2020) | Põlme S, et al. 2020. Fungal Diversity 105:1-16 ((https://doi.org/10.1007/s13225-020-00466-2)) |
| Otu19646 | 1 | k__Fungi(100);p__Ascomycota(100);c__Saccharomycetes(100);o__Saccharomycetales(100);f__Metschnikowiaceae(100);g__Metschnikowia(100);s__Metschnikowia_pulcherrima(100); | Metschnikowia | 13 | Saprotroph | \|Undefined Saprotroph\| | Yeast | NULL | Probable | As nectar-tap_saprotroph (Põlme et al. 2020); Decay sugar-rich substrates (Põlme et al. 2020) | Põlme S, et al. 2020. Fungal Diversity 105:1-16 ((https://doi.org/10.1007/s13225-020-00466-2)) |
| Otu19647 | 1 | k__Fungi(100);p__Ascomycota(100);c__Saccharomycetes(100);o__Saccharomycetales(100);f__Metschnikowiaceae(100);g__Metschnikowia(100);s__Metschnikowia_pulcherrima(100); | Metschnikowia | 13 | Saprotroph | \|Undefined Saprotroph\| | Yeast | NULL | Probable | As nectar-tap_saprotroph (Põlme et al. 2020); Decay sugar-rich substrates (Põlme et al. 2020) | Põlme S, et al. 2020. Fungal Diversity 105:1-16 ((https://doi.org/10.1007/s13225-020-00466-2)) |
| Otu19648 | 1 | k__Fungi(100);p__Ascomycota(100);c__Saccharomycetes(100);o__Saccharomycetales(100);f__Metschnikowiaceae(100);g__Metschnikowia(100);s__Metschnikowia_pulcherrima(100); | Metschnikowia | 13 | Saprotroph | \|Undefined Saprotroph\| | Yeast | NULL | Probable | As nectar-tap_saprotroph (Põlme et al. 2020); Decay sugar-rich substrates (Põlme et al. 2020) | Põlme S, et al. 2020. Fungal Diversity 105:1-16 ((https://doi.org/10.1007/s13225-020-00466-2)) |
| Otu19649 | 1 | k__Fungi(100);p__Ascomycota(100);c__Saccharomycetes(100);o__Saccharomycetales(100);f__Metschnikowiaceae(100);g__Metschnikowia(100);s__Metschnikowia_pulcherrima(100); | Metschnikowia | 13 | Saprotroph | \|Undefined Saprotroph\| | Yeast | NULL | Probable | As nectar-tap_saprotroph (Põlme et al. 2020); Decay sugar-rich substrates (Põlme et al. 2020) | Põlme S, et al. 2020. Fungal Diversity 105:1-16 ((https://doi.org/10.1007/s13225-020-00466-2)) |
| Otu19650 | 1 | k__Fungi(100);p__Ascomycota(100);c__Saccharomycetes(100);o__Saccharomycetales(100);f__Metschnikowiaceae(100);g__Metschnikowia(100);g__Metschnikowia_unclassified(100); | Metschnikowia | 13 | Saprotroph | \|Undefined Saprotroph\| | Yeast | NULL | Probable | As nectar-tap_saprotroph (Põlme et al. 2020); Decay sugar-rich substrates (Põlme et al. 2020) | Põlme S, et al. 2020. Fungal Diversity 105:1-16 ((https://doi.org/10.1007/s13225-020-00466-2)) |
| Otu19651 | 1 | k__Fungi(100);p__Ascomycota(100);c__Saccharomycetes(100);o__Saccharomycetales(100);f__Metschnikowiaceae(100);g__Metschnikowia(100);s__Metschnikowia_pulcherrima(100); | Metschnikowia | 13 | Saprotroph | \|Undefined Saprotroph\| | Yeast | NULL | Probable | As nectar-tap_saprotroph (Põlme et al. 2020); Decay sugar-rich substrates (Põlme et al. 2020) | Põlme S, et al. 2020. Fungal Diversity 105:1-16 ((https://doi.org/10.1007/s13225-020-00466-2)) |
| Otu19652 | 1 | k__Fungi(100);p__Ascomycota(100);c__Saccharomycetes(100);o__Saccharomycetales(100);f__Metschnikowiaceae(100);g__Metschnikowia(100);s__Metschnikowia_pulcherrima(100); | Metschnikowia | 13 | Saprotroph | \|Undefined Saprotroph\| | Yeast | NULL | Probable | As nectar-tap_saprotroph (Põlme et al. 2020); Decay sugar-rich substrates (Põlme et al. 2020) | Põlme S, et al. 2020. Fungal Diversity 105:1-16 ((https://doi.org/10.1007/s13225-020-00466-2)) |
| Otu19653 | 1 | k__Fungi(100);p__Ascomycota(100);c__Saccharomycetes(100);o__Saccharomycetales(100);f__Metschnikowiaceae(100);g__Metschnikowia(100);s__Metschnikowia_pulcherrima(100); | Metschnikowia | 13 | Saprotroph | \|Undefined Saprotroph\| | Yeast | NULL | Probable | As nectar-tap_saprotroph (Põlme et al. 2020); Decay sugar-rich substrates (Põlme et al. 2020) | Põlme S, et al. 2020. Fungal Diversity 105:1-16 ((https://doi.org/10.1007/s13225-020-00466-2)) |
| Otu19655 | 1 | k__Fungi(100);p__Ascomycota(100);c__Saccharomycetes(100);o__Saccharomycetales(100);f__Metschnikowiaceae(100);g__Metschnikowia(100);g__Metschnikowia_unclassified(100); | Metschnikowia | 13 | Saprotroph | \|Undefined Saprotroph\| | Yeast | NULL | Probable | As nectar-tap_saprotroph (Põlme et al. 2020); Decay sugar-rich substrates (Põlme et al. 2020) | Põlme S, et al. 2020. Fungal Diversity 105:1-16 ((https://doi.org/10.1007/s13225-020-00466-2)) |
| Otu19656 | 1 | k__Fungi(100);p__Ascomycota(100);c__Saccharomycetes(100);o__Saccharomycetales(100);f__Metschnikowiaceae(100);g__Metschnikowia(100);g__Metschnikowia_unclassified(100); | Metschnikowia | 13 | Saprotroph | \|Undefined Saprotroph\| | Yeast | NULL | Probable | As nectar-tap_saprotroph (Põlme et al. 2020); Decay sugar-rich substrates (Põlme et al. 2020) | Põlme S, et al. 2020. Fungal Diversity 105:1-16 ((https://doi.org/10.1007/s13225-020-00466-2)) |
| Otu19657 | 1 | k__Fungi(100);p__Ascomycota(100);c__Saccharomycetes(100);o__Saccharomycetales(100);f__Metschnikowiaceae(100);g__Metschnikowia(100);g__Metschnikowia_unclassified(100); | Metschnikowia | 13 | Saprotroph | \|Undefined Saprotroph\| | Yeast | NULL | Probable | As nectar-tap_saprotroph (Põlme et al. 2020); Decay sugar-rich substrates (Põlme et al. 2020) | Põlme S, et al. 2020. Fungal Diversity 105:1-16 ((https://doi.org/10.1007/s13225-020-00466-2)) |
| Otu19658 | 1 | k__Fungi(100);p__Ascomycota(100);c__Saccharomycetes(100);o__Saccharomycetales(100);f__Metschnikowiaceae(100);g__Metschnikowia(100);s__Metschnikowia_pulcherrima(100); | Metschnikowia | 13 | Saprotroph | \|Undefined Saprotroph\| | Yeast | NULL | Probable | As nectar-tap_saprotroph (Põlme et al. 2020); Decay sugar-rich substrates (Põlme et al. 2020) | Põlme S, et al. 2020. Fungal Diversity 105:1-16 ((https://doi.org/10.1007/s13225-020-00466-2)) |
| Otu19659 | 1 | k__Fungi(100);p__Ascomycota(100);c__Saccharomycetes(100);o__Saccharomycetales(100);f__Metschnikowiaceae(100);g__Metschnikowia(100);g__Metschnikowia_unclassified(100); | Metschnikowia | 13 | Saprotroph | \|Undefined Saprotroph\| | Yeast | NULL | Probable | As nectar-tap_saprotroph (Põlme et al. 2020); Decay sugar-rich substrates (Põlme et al. 2020) | Põlme S, et al. 2020. Fungal Diversity 105:1-16 ((https://doi.org/10.1007/s13225-020-00466-2)) |
| Otu19660 | 1 | k__Fungi(100);p__Ascomycota(100);c__Saccharomycetes(100);o__Saccharomycetales(100);f__Metschnikowiaceae(100);g__Metschnikowia(100);s__Metschnikowia_pulcherrima(100); | Metschnikowia | 13 | Saprotroph | \|Undefined Saprotroph\| | Yeast | NULL | Probable | As nectar-tap_saprotroph (Põlme et al. 2020); Decay sugar-rich substrates (Põlme et al. 2020) | Põlme S, et al. 2020. Fungal Diversity 105:1-16 ((https://doi.org/10.1007/s13225-020-00466-2)) |
| Otu19661 | 1 | k__Fungi(100);p__Ascomycota(100);c__Saccharomycetes(100);o__Saccharomycetales(100);f__Metschnikowiaceae(100);g__Metschnikowia(100);g__Metschnikowia_unclassified(100); | Metschnikowia | 13 | Saprotroph | \|Undefined Saprotroph\| | Yeast | NULL | Probable | As nectar-tap_saprotroph (Põlme et al. 2020); Decay sugar-rich substrates (Põlme et al. 2020) | Põlme S, et al. 2020. Fungal Diversity 105:1-16 ((https://doi.org/10.1007/s13225-020-00466-2)) |
| Otu19662 | 1 | k__Fungi(100);p__Ascomycota(100);c__Saccharomycetes(100);o__Saccharomycetales(100);f__Metschnikowiaceae(100);g__Metschnikowia(100);s__Metschnikowia_pulcherrima(100); | Metschnikowia | 13 | Saprotroph | \|Undefined Saprotroph\| | Yeast | NULL | Probable | As nectar-tap_saprotroph (Põlme et al. 2020); Decay sugar-rich substrates (Põlme et al. 2020) | Põlme S, et al. 2020. Fungal Diversity 105:1-16 ((https://doi.org/10.1007/s13225-020-00466-2)) |
| Otu19663 | 1 | k__Fungi(100);p__Ascomycota(100);c__Saccharomycetes(100);o__Saccharomycetales(100);f__Metschnikowiaceae(100);g__Metschnikowia(100);g__Metschnikowia_unclassified(100); | Metschnikowia | 13 | Saprotroph | \|Undefined Saprotroph\| | Yeast | NULL | Probable | As nectar-tap_saprotroph (Põlme et al. 2020); Decay sugar-rich substrates (Põlme et al. 2020) | Põlme S, et al. 2020. Fungal Diversity 105:1-16 ((https://doi.org/10.1007/s13225-020-00466-2)) |
| Otu19664 | 1 | k__Fungi(100);p__Ascomycota(100);c__Saccharomycetes(100);o__Saccharomycetales(100);f__Metschnikowiaceae(100);g__Metschnikowia(100);s__Metschnikowia_pulcherrima(100); | Metschnikowia | 13 | Saprotroph | \|Undefined Saprotroph\| | Yeast | NULL | Probable | As nectar-tap_saprotroph (Põlme et al. 2020); Decay sugar-rich substrates (Põlme et al. 2020) | Põlme S, et al. 2020. Fungal Diversity 105:1-16 ((https://doi.org/10.1007/s13225-020-00466-2)) |
| Otu19665 | 1 | k__Fungi(100);p__Ascomycota(100);c__Saccharomycetes(100);o__Saccharomycetales(100);f__Metschnikowiaceae(100);g__Metschnikowia(100);g__Metschnikowia_unclassified(100); | Metschnikowia | 13 | Saprotroph | \|Undefined Saprotroph\| | Yeast | NULL | Probable | As nectar-tap_saprotroph (Põlme et al. 2020); Decay sugar-rich substrates (Põlme et al. 2020) | Põlme S, et al. 2020. Fungal Diversity 105:1-16 ((https://doi.org/10.1007/s13225-020-00466-2)) |
| Otu19666 | 1 | k__Fungi(100);p__Ascomycota(100);c__Saccharomycetes(100);o__Saccharomycetales(100);f__Metschnikowiaceae(100);g__Metschnikowia(100);s__Metschnikowia_pulcherrima(100); | Metschnikowia | 13 | Saprotroph | \|Undefined Saprotroph\| | Yeast | NULL | Probable | As nectar-tap_saprotroph (Põlme et al. 2020); Decay sugar-rich substrates (Põlme et al. 2020) | Põlme S, et al. 2020. Fungal Diversity 105:1-16 ((https://doi.org/10.1007/s13225-020-00466-2)) |
| Otu19667 | 1 | k__Fungi(100);p__Ascomycota(100);c__Saccharomycetes(100);o__Saccharomycetales(100);f__Metschnikowiaceae(100);g__Metschnikowia(100);s__Metschnikowia_pulcherrima(100); | Metschnikowia | 13 | Saprotroph | \|Undefined Saprotroph\| | Yeast | NULL | Probable | As nectar-tap_saprotroph (Põlme et al. 2020); Decay sugar-rich substrates (Põlme et al. 2020) | Põlme S, et al. 2020. Fungal Diversity 105:1-16 ((https://doi.org/10.1007/s13225-020-00466-2)) |
| Otu19668 | 1 | k__Fungi(100);p__Ascomycota(100);c__Saccharomycetes(100);o__Saccharomycetales(100);f__Metschnikowiaceae(100);g__Metschnikowia(100);s__Metschnikowia_pulcherrima(100); | Metschnikowia | 13 | Saprotroph | \|Undefined Saprotroph\| | Yeast | NULL | Probable | As nectar-tap_saprotroph (Põlme et al. 2020); Decay sugar-rich substrates (Põlme et al. 2020) | Põlme S, et al. 2020. Fungal Diversity 105:1-16 ((https://doi.org/10.1007/s13225-020-00466-2)) |
| Otu19669 | 1 | k__Fungi(100);p__Ascomycota(100);c__Saccharomycetes(100);o__Saccharomycetales(100);f__Metschnikowiaceae(100);g__Metschnikowia(100);s__Metschnikowia_pulcherrima(100); | Metschnikowia | 13 | Saprotroph | \|Undefined Saprotroph\| | Yeast | NULL | Probable | As nectar-tap_saprotroph (Põlme et al. 2020); Decay sugar-rich substrates (Põlme et al. 2020) | Põlme S, et al. 2020. Fungal Diversity 105:1-16 ((https://doi.org/10.1007/s13225-020-00466-2)) |
| Otu19670 | 1 | k__Fungi(100);p__Ascomycota(100);c__Saccharomycetes(100);o__Saccharomycetales(100);f__Metschnikowiaceae(100);g__Metschnikowia(100);s__Metschnikowia_pulcherrima(100); | Metschnikowia | 13 | Saprotroph | \|Undefined Saprotroph\| | Yeast | NULL | Probable | As nectar-tap_saprotroph (Põlme et al. 2020); Decay sugar-rich substrates (Põlme et al. 2020) | Põlme S, et al. 2020. Fungal Diversity 105:1-16 ((https://doi.org/10.1007/s13225-020-00466-2)) |
| Otu19671 | 1 | k__Fungi(100);p__Ascomycota(100);c__Saccharomycetes(100);o__Saccharomycetales(100);f__Metschnikowiaceae(100);g__Metschnikowia(100);g__Metschnikowia_unclassified(100); | Metschnikowia | 13 | Saprotroph | \|Undefined Saprotroph\| | Yeast | NULL | Probable | As nectar-tap_saprotroph (Põlme et al. 2020); Decay sugar-rich substrates (Põlme et al. 2020) | Põlme S, et al. 2020. Fungal Diversity 105:1-16 ((https://doi.org/10.1007/s13225-020-00466-2)) |
| Otu19672 | 1 | k__Fungi(100);p__Ascomycota(100);c__Saccharomycetes(100);o__Saccharomycetales(100);f__Metschnikowiaceae(100);g__Metschnikowia(100);s__Metschnikowia_pulcherrima(100); | Metschnikowia | 13 | Saprotroph | \|Undefined Saprotroph\| | Yeast | NULL | Probable | As nectar-tap_saprotroph (Põlme et al. 2020); Decay sugar-rich substrates (Põlme et al. 2020) | Põlme S, et al. 2020. Fungal Diversity 105:1-16 ((https://doi.org/10.1007/s13225-020-00466-2)) |
| Otu19673 | 1 | k__Fungi(100);p__Ascomycota(100);c__Saccharomycetes(100);o__Saccharomycetales(100);f__Metschnikowiaceae(100);g__Metschnikowia(100);s__Metschnikowia_pulcherrima(100); | Metschnikowia | 13 | Saprotroph | \|Undefined Saprotroph\| | Yeast | NULL | Probable | As nectar-tap_saprotroph (Põlme et al. 2020); Decay sugar-rich substrates (Põlme et al. 2020) | Põlme S, et al. 2020. Fungal Diversity 105:1-16 ((https://doi.org/10.1007/s13225-020-00466-2)) |
| Otu19674 | 1 | k__Fungi(100);p__Ascomycota(100);c__Saccharomycetes(100);o__Saccharomycetales(100);f__Metschnikowiaceae(100);g__Metschnikowia(100);s__Metschnikowia_pulcherrima(100); | Metschnikowia | 13 | Saprotroph | \|Undefined Saprotroph\| | Yeast | NULL | Probable | As nectar-tap_saprotroph (Põlme et al. 2020); Decay sugar-rich substrates (Põlme et al. 2020) | Põlme S, et al. 2020. Fungal Diversity 105:1-16 ((https://doi.org/10.1007/s13225-020-00466-2)) |
| Otu19675 | 1 | k__Fungi(100);p__Ascomycota(100);c__Saccharomycetes(100);o__Saccharomycetales(100);f__Metschnikowiaceae(100);g__Metschnikowia(100);g__Metschnikowia_unclassified(100); | Metschnikowia | 13 | Saprotroph | \|Undefined Saprotroph\| | Yeast | NULL | Probable | As nectar-tap_saprotroph (Põlme et al. 2020); Decay sugar-rich substrates (Põlme et al. 2020) | Põlme S, et al. 2020. Fungal Diversity 105:1-16 ((https://doi.org/10.1007/s13225-020-00466-2)) |
| Otu19676 | 1 | k__Fungi(100);p__Ascomycota(100);c__Saccharomycetes(100);o__Saccharomycetales(100);f__Metschnikowiaceae(100);g__Metschnikowia(100);g__Metschnikowia_unclassified(100); | Metschnikowia | 13 | Saprotroph | \|Undefined Saprotroph\| | Yeast | NULL | Probable | As nectar-tap_saprotroph (Põlme et al. 2020); Decay sugar-rich substrates (Põlme et al. 2020) | Põlme S, et al. 2020. Fungal Diversity 105:1-16 ((https://doi.org/10.1007/s13225-020-00466-2)) |
| Otu19677 | 1 | k__Fungi(100);p__Ascomycota(100);c__Saccharomycetes(100);o__Saccharomycetales(100);f__Metschnikowiaceae(100);g__Metschnikowia(100);s__Metschnikowia_pulcherrima(100); | Metschnikowia | 13 | Saprotroph | \|Undefined Saprotroph\| | Yeast | NULL | Probable | As nectar-tap_saprotroph (Põlme et al. 2020); Decay sugar-rich substrates (Põlme et al. 2020) | Põlme S, et al. 2020. Fungal Diversity 105:1-16 ((https://doi.org/10.1007/s13225-020-00466-2)) |
| Otu19678 | 1 | k__Fungi(100);p__Ascomycota(100);c__Saccharomycetes(100);o__Saccharomycetales(100);f__Metschnikowiaceae(100);g__Metschnikowia(100);s__Metschnikowia_pulcherrima(100); | Metschnikowia | 13 | Saprotroph | \|Undefined Saprotroph\| | Yeast | NULL | Probable | As nectar-tap_saprotroph (Põlme et al. 2020); Decay sugar-rich substrates (Põlme et al. 2020) | Põlme S, et al. 2020. Fungal Diversity 105:1-16 ((https://doi.org/10.1007/s13225-020-00466-2)) |
| Otu19679 | 1 | k__Fungi(100);p__Ascomycota(100);c__Saccharomycetes(100);o__Saccharomycetales(100);f__Metschnikowiaceae(100);g__Metschnikowia(100);s__Metschnikowia_pulcherrima(100); | Metschnikowia | 13 | Saprotroph | \|Undefined Saprotroph\| | Yeast | NULL | Probable | As nectar-tap_saprotroph (Põlme et al. 2020); Decay sugar-rich substrates (Põlme et al. 2020) | Põlme S, et al. 2020. Fungal Diversity 105:1-16 ((https://doi.org/10.1007/s13225-020-00466-2)) |
| Otu19680 | 1 | k__Fungi(100);p__Ascomycota(100);c__Saccharomycetes(100);o__Saccharomycetales(100);f__Metschnikowiaceae(100);g__Metschnikowia(100);s__Metschnikowia_pulcherrima(100); | Metschnikowia | 13 | Saprotroph | \|Undefined Saprotroph\| | Yeast | NULL | Probable | As nectar-tap_saprotroph (Põlme et al. 2020); Decay sugar-rich substrates (Põlme et al. 2020) | Põlme S, et al. 2020. Fungal Diversity 105:1-16 ((https://doi.org/10.1007/s13225-020-00466-2)) |
| Otu19681 | 1 | k__Fungi(100);p__Ascomycota(100);c__Saccharomycetes(100);o__Saccharomycetales(100);f__Metschnikowiaceae(100);g__Metschnikowia(100);s__Metschnikowia_pulcherrima(100); | Metschnikowia | 13 | Saprotroph | \|Undefined Saprotroph\| | Yeast | NULL | Probable | As nectar-tap_saprotroph (Põlme et al. 2020); Decay sugar-rich substrates (Põlme et al. 2020) | Põlme S, et al. 2020. Fungal Diversity 105:1-16 ((https://doi.org/10.1007/s13225-020-00466-2)) |
| Otu19682 | 1 | k__Fungi(100);p__Ascomycota(100);c__Saccharomycetes(100);o__Saccharomycetales(100);f__Metschnikowiaceae(100);g__Metschnikowia(100);s__Metschnikowia_pulcherrima(100); | Metschnikowia | 13 | Saprotroph | \|Undefined Saprotroph\| | Yeast | NULL | Probable | As nectar-tap_saprotroph (Põlme et al. 2020); Decay sugar-rich substrates (Põlme et al. 2020) | Põlme S, et al. 2020. Fungal Diversity 105:1-16 ((https://doi.org/10.1007/s13225-020-00466-2)) |
| Otu19683 | 1 | k__Fungi(100);p__Ascomycota(100);c__Saccharomycetes(100);o__Saccharomycetales(100);f__Metschnikowiaceae(100);g__Metschnikowia(100);s__Metschnikowia_pulcherrima(100); | Metschnikowia | 13 | Saprotroph | \|Undefined Saprotroph\| | Yeast | NULL | Probable | As nectar-tap_saprotroph (Põlme et al. 2020); Decay sugar-rich substrates (Põlme et al. 2020) | Põlme S, et al. 2020. Fungal Diversity 105:1-16 ((https://doi.org/10.1007/s13225-020-00466-2)) |
| Otu19684 | 1 | k__Fungi(100);p__Ascomycota(100);c__Saccharomycetes(100);o__Saccharomycetales(100);f__Metschnikowiaceae(100);g__Metschnikowia(100);s__Metschnikowia_pulcherrima(100); | Metschnikowia | 13 | Saprotroph | \|Undefined Saprotroph\| | Yeast | NULL | Probable | As nectar-tap_saprotroph (Põlme et al. 2020); Decay sugar-rich substrates (Põlme et al. 2020) | Põlme S, et al. 2020. Fungal Diversity 105:1-16 ((https://doi.org/10.1007/s13225-020-00466-2)) |
| Otu19685 | 1 | k__Fungi(100);p__Ascomycota(100);c__Saccharomycetes(100);o__Saccharomycetales(100);f__Metschnikowiaceae(100);g__Metschnikowia(100);s__Metschnikowia_pulcherrima(100); | Metschnikowia | 13 | Saprotroph | \|Undefined Saprotroph\| | Yeast | NULL | Probable | As nectar-tap_saprotroph (Põlme et al. 2020); Decay sugar-rich substrates (Põlme et al. 2020) | Põlme S, et al. 2020. Fungal Diversity 105:1-16 ((https://doi.org/10.1007/s13225-020-00466-2)) |
| Otu19686 | 1 | k__Fungi(100);p__Ascomycota(100);c__Saccharomycetes(100);o__Saccharomycetales(100);f__Metschnikowiaceae(100);g__Metschnikowia(100);s__Metschnikowia_pulcherrima(100); | Metschnikowia | 13 | Saprotroph | \|Undefined Saprotroph\| | Yeast | NULL | Probable | As nectar-tap_saprotroph (Põlme et al. 2020); Decay sugar-rich substrates (Põlme et al. 2020) | Põlme S, et al. 2020. Fungal Diversity 105:1-16 ((https://doi.org/10.1007/s13225-020-00466-2)) |
| Otu19687 | 1 | k__Fungi(100);p__Ascomycota(100);c__Saccharomycetes(100);o__Saccharomycetales(100);f__Metschnikowiaceae(100);g__Metschnikowia(100);s__Metschnikowia_pulcherrima(100); | Metschnikowia | 13 | Saprotroph | \|Undefined Saprotroph\| | Yeast | NULL | Probable | As nectar-tap_saprotroph (Põlme et al. 2020); Decay sugar-rich substrates (Põlme et al. 2020) | Põlme S, et al. 2020. Fungal Diversity 105:1-16 ((https://doi.org/10.1007/s13225-020-00466-2)) |
| Otu19688 | 1 | k__Fungi(100);p__Ascomycota(100);c__Saccharomycetes(100);o__Saccharomycetales(100);f__Metschnikowiaceae(100);g__Metschnikowia(100);s__Metschnikowia_pulcherrima(100); | Metschnikowia | 13 | Saprotroph | \|Undefined Saprotroph\| | Yeast | NULL | Probable | As nectar-tap_saprotroph (Põlme et al. 2020); Decay sugar-rich substrates (Põlme et al. 2020) | Põlme S, et al. 2020. Fungal Diversity 105:1-16 ((https://doi.org/10.1007/s13225-020-00466-2)) |
| Otu19689 | 1 | k__Fungi(100);p__Ascomycota(100);c__Saccharomycetes(100);o__Saccharomycetales(100);f__Metschnikowiaceae(100);g__Metschnikowia(100);s__Metschnikowia_pulcherrima(100); | Metschnikowia | 13 | Saprotroph | \|Undefined Saprotroph\| | Yeast | NULL | Probable | As nectar-tap_saprotroph (Põlme et al. 2020); Decay sugar-rich substrates (Põlme et al. 2020) | Põlme S, et al. 2020. Fungal Diversity 105:1-16 ((https://doi.org/10.1007/s13225-020-00466-2)) |
| Otu19690 | 1 | k__Fungi(100);p__Ascomycota(100);c__Saccharomycetes(100);o__Saccharomycetales(100);f__Metschnikowiaceae(100);g__Metschnikowia(100);s__Metschnikowia_pulcherrima(100); | Metschnikowia | 13 | Saprotroph | \|Undefined Saprotroph\| | Yeast | NULL | Probable | As nectar-tap_saprotroph (Põlme et al. 2020); Decay sugar-rich substrates (Põlme et al. 2020) | Põlme S, et al. 2020. Fungal Diversity 105:1-16 ((https://doi.org/10.1007/s13225-020-00466-2)) |
| Otu19691 | 1 | k__Fungi(100);p__Ascomycota(100);c__Saccharomycetes(100);o__Saccharomycetales(100);f__Metschnikowiaceae(100);g__Metschnikowia(100);s__Metschnikowia_pulcherrima(100); | Metschnikowia | 13 | Saprotroph | \|Undefined Saprotroph\| | Yeast | NULL | Probable | As nectar-tap_saprotroph (Põlme et al. 2020); Decay sugar-rich substrates (Põlme et al. 2020) | Põlme S, et al. 2020. Fungal Diversity 105:1-16 ((https://doi.org/10.1007/s13225-020-00466-2)) |
| Otu19692 | 1 | k__Fungi(100);p__Ascomycota(100);c__Saccharomycetes(100);o__Saccharomycetales(100);f__Metschnikowiaceae(100);g__Metschnikowia(100);s__Metschnikowia_pulcherrima(100); | Metschnikowia | 13 | Saprotroph | \|Undefined Saprotroph\| | Yeast | NULL | Probable | As nectar-tap_saprotroph (Põlme et al. 2020); Decay sugar-rich substrates (Põlme et al. 2020) | Põlme S, et al. 2020. Fungal Diversity 105:1-16 ((https://doi.org/10.1007/s13225-020-00466-2)) |
| Otu19693 | 1 | k__Fungi(100);p__Ascomycota(100);c__Saccharomycetes(100);o__Saccharomycetales(100);f__Metschnikowiaceae(100);g__Metschnikowia(100);s__Metschnikowia_pulcherrima(100); | Metschnikowia | 13 | Saprotroph | \|Undefined Saprotroph\| | Yeast | NULL | Probable | As nectar-tap_saprotroph (Põlme et al. 2020); Decay sugar-rich substrates (Põlme et al. 2020) | Põlme S, et al. 2020. Fungal Diversity 105:1-16 ((https://doi.org/10.1007/s13225-020-00466-2)) |
| Otu19694 | 1 | k__Fungi(100);p__Ascomycota(100);c__Saccharomycetes(100);o__Saccharomycetales(100);f__Metschnikowiaceae(100);g__Metschnikowia(100);g__Metschnikowia_unclassified(100); | Metschnikowia | 13 | Saprotroph | \|Undefined Saprotroph\| | Yeast | NULL | Probable | As nectar-tap_saprotroph (Põlme et al. 2020); Decay sugar-rich substrates (Põlme et al. 2020) | Põlme S, et al. 2020. Fungal Diversity 105:1-16 ((https://doi.org/10.1007/s13225-020-00466-2)) |
| Otu19695 | 1 | k__Fungi(100);p__Ascomycota(100);c__Saccharomycetes(100);o__Saccharomycetales(100);f__Metschnikowiaceae(100);g__Metschnikowia(100);s__Metschnikowia_pulcherrima(100); | Metschnikowia | 13 | Saprotroph | \|Undefined Saprotroph\| | Yeast | NULL | Probable | As nectar-tap_saprotroph (Põlme et al. 2020); Decay sugar-rich substrates (Põlme et al. 2020) | Põlme S, et al. 2020. Fungal Diversity 105:1-16 ((https://doi.org/10.1007/s13225-020-00466-2)) |
| Otu19696 | 1 | k__Fungi(100);p__Ascomycota(100);c__Saccharomycetes(100);o__Saccharomycetales(100);f__Metschnikowiaceae(100);g__Metschnikowia(100);s__Metschnikowia_pulcherrima(100); | Metschnikowia | 13 | Saprotroph | \|Undefined Saprotroph\| | Yeast | NULL | Probable | As nectar-tap_saprotroph (Põlme et al. 2020); Decay sugar-rich substrates (Põlme et al. 2020) | Põlme S, et al. 2020. Fungal Diversity 105:1-16 ((https://doi.org/10.1007/s13225-020-00466-2)) |
| Otu19697 | 1 | k__Fungi(100);p__Ascomycota(100);c__Saccharomycetes(100);o__Saccharomycetales(100);f__Metschnikowiaceae(100);g__Metschnikowia(100);s__Metschnikowia_pulcherrima(100); | Metschnikowia | 13 | Saprotroph | \|Undefined Saprotroph\| | Yeast | NULL | Probable | As nectar-tap_saprotroph (Põlme et al. 2020); Decay sugar-rich substrates (Põlme et al. 2020) | Põlme S, et al. 2020. Fungal Diversity 105:1-16 ((https://doi.org/10.1007/s13225-020-00466-2)) |
| Otu19698 | 1 | k__Fungi(100);p__Ascomycota(100);c__Saccharomycetes(100);o__Saccharomycetales(100);f__Metschnikowiaceae(100);g__Metschnikowia(100);s__Metschnikowia_pulcherrima(100); | Metschnikowia | 13 | Saprotroph | \|Undefined Saprotroph\| | Yeast | NULL | Probable | As nectar-tap_saprotroph (Põlme et al. 2020); Decay sugar-rich substrates (Põlme et al. 2020) | Põlme S, et al. 2020. Fungal Diversity 105:1-16 ((https://doi.org/10.1007/s13225-020-00466-2)) |
| Otu19699 | 1 | k__Fungi(100);p__Ascomycota(100);c__Saccharomycetes(100);o__Saccharomycetales(100);f__Metschnikowiaceae(100);g__Metschnikowia(100);s__Metschnikowia_pulcherrima(100); | Metschnikowia | 13 | Saprotroph | \|Undefined Saprotroph\| | Yeast | NULL | Probable | As nectar-tap_saprotroph (Põlme et al. 2020); Decay sugar-rich substrates (Põlme et al. 2020) | Põlme S, et al. 2020. Fungal Diversity 105:1-16 ((https://doi.org/10.1007/s13225-020-00466-2)) |
| Otu19700 | 1 | k__Fungi(100);p__Ascomycota(100);c__Saccharomycetes(100);o__Saccharomycetales(100);f__Metschnikowiaceae(100);g__Metschnikowia(100);s__Metschnikowia_pulcherrima(100); | Metschnikowia | 13 | Saprotroph | \|Undefined Saprotroph\| | Yeast | NULL | Probable | As nectar-tap_saprotroph (Põlme et al. 2020); Decay sugar-rich substrates (Põlme et al. 2020) | Põlme S, et al. 2020. Fungal Diversity 105:1-16 ((https://doi.org/10.1007/s13225-020-00466-2)) |
| Otu19701 | 1 | k__Fungi(100);p__Ascomycota(100);c__Saccharomycetes(100);o__Saccharomycetales(100);f__Metschnikowiaceae(100);g__Metschnikowia(100);s__Metschnikowia_pulcherrima(100); | Metschnikowia | 13 | Saprotroph | \|Undefined Saprotroph\| | Yeast | NULL | Probable | As nectar-tap_saprotroph (Põlme et al. 2020); Decay sugar-rich substrates (Põlme et al. 2020) | Põlme S, et al. 2020. Fungal Diversity 105:1-16 ((https://doi.org/10.1007/s13225-020-00466-2)) |
| Otu19702 | 1 | k__Fungi(100);p__Ascomycota(100);c__Saccharomycetes(100);o__Saccharomycetales(100);f__Metschnikowiaceae(100);g__Metschnikowia(100);s__Metschnikowia_pulcherrima(100); | Metschnikowia | 13 | Saprotroph | \|Undefined Saprotroph\| | Yeast | NULL | Probable | As nectar-tap_saprotroph (Põlme et al. 2020); Decay sugar-rich substrates (Põlme et al. 2020) | Põlme S, et al. 2020. Fungal Diversity 105:1-16 ((https://doi.org/10.1007/s13225-020-00466-2)) |
| Otu19703 | 1 | k__Fungi(100);p__Ascomycota(100);c__Saccharomycetes(100);o__Saccharomycetales(100);f__Metschnikowiaceae(100);g__Metschnikowia(100);s__Metschnikowia_pulcherrima(100); | Metschnikowia | 13 | Saprotroph | \|Undefined Saprotroph\| | Yeast | NULL | Probable | As nectar-tap_saprotroph (Põlme et al. 2020); Decay sugar-rich substrates (Põlme et al. 2020) | Põlme S, et al. 2020. Fungal Diversity 105:1-16 ((https://doi.org/10.1007/s13225-020-00466-2)) |
| Otu19704 | 1 | k__Fungi(100);p__Ascomycota(100);c__Saccharomycetes(100);o__Saccharomycetales(100);f__Metschnikowiaceae(100);g__Metschnikowia(100);s__Metschnikowia_pulcherrima(100); | Metschnikowia | 13 | Saprotroph | \|Undefined Saprotroph\| | Yeast | NULL | Probable | As nectar-tap_saprotroph (Põlme et al. 2020); Decay sugar-rich substrates (Põlme et al. 2020) | Põlme S, et al. 2020. Fungal Diversity 105:1-16 ((https://doi.org/10.1007/s13225-020-00466-2)) |
| Otu19705 | 1 | k__Fungi(100);p__Ascomycota(100);c__Saccharomycetes(100);o__Saccharomycetales(100);f__Metschnikowiaceae(100);g__Metschnikowia(100);g__Metschnikowia_unclassified(100); | Metschnikowia | 13 | Saprotroph | \|Undefined Saprotroph\| | Yeast | NULL | Probable | As nectar-tap_saprotroph (Põlme et al. 2020); Decay sugar-rich substrates (Põlme et al. 2020) | Põlme S, et al. 2020. Fungal Diversity 105:1-16 ((https://doi.org/10.1007/s13225-020-00466-2)) |
| Otu19706 | 1 | k__Fungi(100);p__Ascomycota(100);c__Saccharomycetes(100);o__Saccharomycetales(100);f__Metschnikowiaceae(100);g__Metschnikowia(100);s__Metschnikowia_pulcherrima(100); | Metschnikowia | 13 | Saprotroph | \|Undefined Saprotroph\| | Yeast | NULL | Probable | As nectar-tap_saprotroph (Põlme et al. 2020); Decay sugar-rich substrates (Põlme et al. 2020) | Põlme S, et al. 2020. Fungal Diversity 105:1-16 ((https://doi.org/10.1007/s13225-020-00466-2)) |
| Otu19707 | 1 | k__Fungi(100);p__Ascomycota(100);c__Saccharomycetes(100);o__Saccharomycetales(100);f__Metschnikowiaceae(100);g__Metschnikowia(100);s__Metschnikowia_pulcherrima(100); | Metschnikowia | 13 | Saprotroph | \|Undefined Saprotroph\| | Yeast | NULL | Probable | As nectar-tap_saprotroph (Põlme et al. 2020); Decay sugar-rich substrates (Põlme et al. 2020) | Põlme S, et al. 2020. Fungal Diversity 105:1-16 ((https://doi.org/10.1007/s13225-020-00466-2)) |
| Otu19708 | 1 | k__Fungi(100);p__Ascomycota(100);c__Saccharomycetes(100);o__Saccharomycetales(100);f__Metschnikowiaceae(100);g__Metschnikowia(100);g__Metschnikowia_unclassified(100); | Metschnikowia | 13 | Saprotroph | \|Undefined Saprotroph\| | Yeast | NULL | Probable | As nectar-tap_saprotroph (Põlme et al. 2020); Decay sugar-rich substrates (Põlme et al. 2020) | Põlme S, et al. 2020. Fungal Diversity 105:1-16 ((https://doi.org/10.1007/s13225-020-00466-2)) |
| Otu19709 | 1 | k__Fungi(100);p__Ascomycota(100);c__Saccharomycetes(100);o__Saccharomycetales(100);f__Metschnikowiaceae(100);g__Metschnikowia(100);s__Metschnikowia_pulcherrima(100); | Metschnikowia | 13 | Saprotroph | \|Undefined Saprotroph\| | Yeast | NULL | Probable | As nectar-tap_saprotroph (Põlme et al. 2020); Decay sugar-rich substrates (Põlme et al. 2020) | Põlme S, et al. 2020. Fungal Diversity 105:1-16 ((https://doi.org/10.1007/s13225-020-00466-2)) |
| Otu19710 | 1 | k__Fungi(100);p__Ascomycota(100);c__Saccharomycetes(100);o__Saccharomycetales(100);f__Metschnikowiaceae(100);g__Metschnikowia(100);g__Metschnikowia_unclassified(100); | Metschnikowia | 13 | Saprotroph | \|Undefined Saprotroph\| | Yeast | NULL | Probable | As nectar-tap_saprotroph (Põlme et al. 2020); Decay sugar-rich substrates (Põlme et al. 2020) | Põlme S, et al. 2020. Fungal Diversity 105:1-16 ((https://doi.org/10.1007/s13225-020-00466-2)) |
| Otu19711 | 1 | k__Fungi(100);p__Ascomycota(100);c__Saccharomycetes(100);o__Saccharomycetales(100);f__Metschnikowiaceae(100);g__Metschnikowia(100);s__Metschnikowia_pulcherrima(100); | Metschnikowia | 13 | Saprotroph | \|Undefined Saprotroph\| | Yeast | NULL | Probable | As nectar-tap_saprotroph (Põlme et al. 2020); Decay sugar-rich substrates (Põlme et al. 2020) | Põlme S, et al. 2020. Fungal Diversity 105:1-16 ((https://doi.org/10.1007/s13225-020-00466-2)) |
| Otu19712 | 1 | k__Fungi(100);p__Ascomycota(100);c__Saccharomycetes(100);o__Saccharomycetales(100);f__Metschnikowiaceae(100);g__Metschnikowia(100);g__Metschnikowia_unclassified(100); | Metschnikowia | 13 | Saprotroph | \|Undefined Saprotroph\| | Yeast | NULL | Probable | As nectar-tap_saprotroph (Põlme et al. 2020); Decay sugar-rich substrates (Põlme et al. 2020) | Põlme S, et al. 2020. Fungal Diversity 105:1-16 ((https://doi.org/10.1007/s13225-020-00466-2)) |
| Otu19713 | 1 | k__Fungi(100);p__Ascomycota(100);c__Saccharomycetes(100);o__Saccharomycetales(100);f__Metschnikowiaceae(100);g__Metschnikowia(100);s__Metschnikowia_pulcherrima(100); | Metschnikowia | 13 | Saprotroph | \|Undefined Saprotroph\| | Yeast | NULL | Probable | As nectar-tap_saprotroph (Põlme et al. 2020); Decay sugar-rich substrates (Põlme et al. 2020) | Põlme S, et al. 2020. Fungal Diversity 105:1-16 ((https://doi.org/10.1007/s13225-020-00466-2)) |
| Otu19714 | 1 | k__Fungi(100);p__Ascomycota(100);c__Saccharomycetes(100);o__Saccharomycetales(100);f__Metschnikowiaceae(100);g__Metschnikowia(100);s__Metschnikowia_pulcherrima(100); | Metschnikowia | 13 | Saprotroph | \|Undefined Saprotroph\| | Yeast | NULL | Probable | As nectar-tap_saprotroph (Põlme et al. 2020); Decay sugar-rich substrates (Põlme et al. 2020) | Põlme S, et al. 2020. Fungal Diversity 105:1-16 ((https://doi.org/10.1007/s13225-020-00466-2)) |
| Otu19715 | 1 | k__Fungi(100);p__Ascomycota(100);c__Saccharomycetes(100);o__Saccharomycetales(100);f__Metschnikowiaceae(100);g__Metschnikowia(100);g__Metschnikowia_unclassified(100); | Metschnikowia | 13 | Saprotroph | \|Undefined Saprotroph\| | Yeast | NULL | Probable | As nectar-tap_saprotroph (Põlme et al. 2020); Decay sugar-rich substrates (Põlme et al. 2020) | Põlme S, et al. 2020. Fungal Diversity 105:1-16 ((https://doi.org/10.1007/s13225-020-00466-2)) |
| Otu19716 | 1 | k__Fungi(100);p__Ascomycota(100);c__Saccharomycetes(100);o__Saccharomycetales(100);f__Metschnikowiaceae(100);g__Metschnikowia(100);s__Metschnikowia_pulcherrima(100); | Metschnikowia | 13 | Saprotroph | \|Undefined Saprotroph\| | Yeast | NULL | Probable | As nectar-tap_saprotroph (Põlme et al. 2020); Decay sugar-rich substrates (Põlme et al. 2020) | Põlme S, et al. 2020. Fungal Diversity 105:1-16 ((https://doi.org/10.1007/s13225-020-00466-2)) |
| Otu19717 | 1 | k__Fungi(100);p__Ascomycota(100);c__Saccharomycetes(100);o__Saccharomycetales(100);f__Metschnikowiaceae(100);g__Metschnikowia(100);s__Metschnikowia_pulcherrima(100); | Metschnikowia | 13 | Saprotroph | \|Undefined Saprotroph\| | Yeast | NULL | Probable | As nectar-tap_saprotroph (Põlme et al. 2020); Decay sugar-rich substrates (Põlme et al. 2020) | Põlme S, et al. 2020. Fungal Diversity 105:1-16 ((https://doi.org/10.1007/s13225-020-00466-2)) |
| Otu19718 | 1 | k__Fungi(100);p__Ascomycota(100);c__Saccharomycetes(100);o__Saccharomycetales(100);f__Metschnikowiaceae(100);g__Metschnikowia(100);g__Metschnikowia_unclassified(100); | Metschnikowia | 13 | Saprotroph | \|Undefined Saprotroph\| | Yeast | NULL | Probable | As nectar-tap_saprotroph (Põlme et al. 2020); Decay sugar-rich substrates (Põlme et al. 2020) | Põlme S, et al. 2020. Fungal Diversity 105:1-16 ((https://doi.org/10.1007/s13225-020-00466-2)) |
| Otu19719 | 1 | k__Fungi(100);p__Ascomycota(100);c__Saccharomycetes(100);o__Saccharomycetales(100);f__Metschnikowiaceae(100);g__Metschnikowia(100);s__Metschnikowia_pulcherrima(100); | Metschnikowia | 13 | Saprotroph | \|Undefined Saprotroph\| | Yeast | NULL | Probable | As nectar-tap_saprotroph (Põlme et al. 2020); Decay sugar-rich substrates (Põlme et al. 2020) | Põlme S, et al. 2020. Fungal Diversity 105:1-16 ((https://doi.org/10.1007/s13225-020-00466-2)) |
| Otu19720 | 1 | k__Fungi(100);p__Ascomycota(100);c__Saccharomycetes(100);o__Saccharomycetales(100);f__Metschnikowiaceae(100);g__Metschnikowia(100);s__Metschnikowia_pulcherrima(100); | Metschnikowia | 13 | Saprotroph | \|Undefined Saprotroph\| | Yeast | NULL | Probable | As nectar-tap_saprotroph (Põlme et al. 2020); Decay sugar-rich substrates (Põlme et al. 2020) | Põlme S, et al. 2020. Fungal Diversity 105:1-16 ((https://doi.org/10.1007/s13225-020-00466-2)) |
| Otu19721 | 1 | k__Fungi(100);p__Ascomycota(100);c__Saccharomycetes(100);o__Saccharomycetales(100);f__Metschnikowiaceae(100);g__Metschnikowia(100);g__Metschnikowia_unclassified(100); | Metschnikowia | 13 | Saprotroph | \|Undefined Saprotroph\| | Yeast | NULL | Probable | As nectar-tap_saprotroph (Põlme et al. 2020); Decay sugar-rich substrates (Põlme et al. 2020) | Põlme S, et al. 2020. Fungal Diversity 105:1-16 ((https://doi.org/10.1007/s13225-020-00466-2)) |
| Otu19722 | 1 | k__Fungi(100);p__Ascomycota(100);c__Saccharomycetes(100);o__Saccharomycetales(100);f__Metschnikowiaceae(100);g__Metschnikowia(100);s__Metschnikowia_pulcherrima(100); | Metschnikowia | 13 | Saprotroph | \|Undefined Saprotroph\| | Yeast | NULL | Probable | As nectar-tap_saprotroph (Põlme et al. 2020); Decay sugar-rich substrates (Põlme et al. 2020) | Põlme S, et al. 2020. Fungal Diversity 105:1-16 ((https://doi.org/10.1007/s13225-020-00466-2)) |
| Otu19723 | 1 | k__Fungi(100);p__Ascomycota(100);c__Saccharomycetes(100);o__Saccharomycetales(100);f__Metschnikowiaceae(100);g__Metschnikowia(100);s__Metschnikowia_pulcherrima(100); | Metschnikowia | 13 | Saprotroph | \|Undefined Saprotroph\| | Yeast | NULL | Probable | As nectar-tap_saprotroph (Põlme et al. 2020); Decay sugar-rich substrates (Põlme et al. 2020) | Põlme S, et al. 2020. Fungal Diversity 105:1-16 ((https://doi.org/10.1007/s13225-020-00466-2)) |
| Otu19725 | 1 | k__Fungi(100);p__Ascomycota(100);c__Saccharomycetes(100);o__Saccharomycetales(100);f__Metschnikowiaceae(100);g__Metschnikowia(100);s__Metschnikowia_pulcherrima(100); | Metschnikowia | 13 | Saprotroph | \|Undefined Saprotroph\| | Yeast | NULL | Probable | As nectar-tap_saprotroph (Põlme et al. 2020); Decay sugar-rich substrates (Põlme et al. 2020) | Põlme S, et al. 2020. Fungal Diversity 105:1-16 ((https://doi.org/10.1007/s13225-020-00466-2)) |
| Otu19726 | 1 | k__Fungi(100);p__Ascomycota(100);c__Saccharomycetes(100);o__Saccharomycetales(100);f__Metschnikowiaceae(100);g__Metschnikowia(100);s__Metschnikowia_pulcherrima(100); | Metschnikowia | 13 | Saprotroph | \|Undefined Saprotroph\| | Yeast | NULL | Probable | As nectar-tap_saprotroph (Põlme et al. 2020); Decay sugar-rich substrates (Põlme et al. 2020) | Põlme S, et al. 2020. Fungal Diversity 105:1-16 ((https://doi.org/10.1007/s13225-020-00466-2)) |
| Otu19727 | 1 | k__Fungi(100);p__Ascomycota(100);c__Saccharomycetes(100);o__Saccharomycetales(100);f__Metschnikowiaceae(100);g__Metschnikowia(100);s__Metschnikowia_pulcherrima(100); | Metschnikowia | 13 | Saprotroph | \|Undefined Saprotroph\| | Yeast | NULL | Probable | As nectar-tap_saprotroph (Põlme et al. 2020); Decay sugar-rich substrates (Põlme et al. 2020) | Põlme S, et al. 2020. Fungal Diversity 105:1-16 ((https://doi.org/10.1007/s13225-020-00466-2)) |
| Otu19728 | 1 | k__Fungi(100);p__Ascomycota(100);c__Saccharomycetes(100);o__Saccharomycetales(100);f__Metschnikowiaceae(100);g__Metschnikowia(100);s__Metschnikowia_pulcherrima(100); | Metschnikowia | 13 | Saprotroph | \|Undefined Saprotroph\| | Yeast | NULL | Probable | As nectar-tap_saprotroph (Põlme et al. 2020); Decay sugar-rich substrates (Põlme et al. 2020) | Põlme S, et al. 2020. Fungal Diversity 105:1-16 ((https://doi.org/10.1007/s13225-020-00466-2)) |
| Otu19729 | 1 | k__Fungi(100);p__Ascomycota(100);c__Saccharomycetes(100);o__Saccharomycetales(100);f__Metschnikowiaceae(100);g__Metschnikowia(100);s__Metschnikowia_pulcherrima(100); | Metschnikowia | 13 | Saprotroph | \|Undefined Saprotroph\| | Yeast | NULL | Probable | As nectar-tap_saprotroph (Põlme et al. 2020); Decay sugar-rich substrates (Põlme et al. 2020) | Põlme S, et al. 2020. Fungal Diversity 105:1-16 ((https://doi.org/10.1007/s13225-020-00466-2)) |
| Otu19730 | 1 | k__Fungi(100);p__Ascomycota(100);c__Saccharomycetes(100);o__Saccharomycetales(100);f__Metschnikowiaceae(100);g__Metschnikowia(100);g__Metschnikowia_unclassified(100); | Metschnikowia | 13 | Saprotroph | \|Undefined Saprotroph\| | Yeast | NULL | Probable | As nectar-tap_saprotroph (Põlme et al. 2020); Decay sugar-rich substrates (Põlme et al. 2020) | Põlme S, et al. 2020. Fungal Diversity 105:1-16 ((https://doi.org/10.1007/s13225-020-00466-2)) |
| Otu19731 | 1 | k__Fungi(100);p__Ascomycota(100);c__Saccharomycetes(100);o__Saccharomycetales(100);f__Metschnikowiaceae(100);g__Metschnikowia(100);s__Metschnikowia_pulcherrima(100); | Metschnikowia | 13 | Saprotroph | \|Undefined Saprotroph\| | Yeast | NULL | Probable | As nectar-tap_saprotroph (Põlme et al. 2020); Decay sugar-rich substrates (Põlme et al. 2020) | Põlme S, et al. 2020. Fungal Diversity 105:1-16 ((https://doi.org/10.1007/s13225-020-00466-2)) |
| Otu19732 | 1 | k__Fungi(100);p__Ascomycota(100);c__Saccharomycetes(100);o__Saccharomycetales(100);f__Metschnikowiaceae(100);g__Metschnikowia(100);s__Metschnikowia_pulcherrima(100); | Metschnikowia | 13 | Saprotroph | \|Undefined Saprotroph\| | Yeast | NULL | Probable | As nectar-tap_saprotroph (Põlme et al. 2020); Decay sugar-rich substrates (Põlme et al. 2020) | Põlme S, et al. 2020. Fungal Diversity 105:1-16 ((https://doi.org/10.1007/s13225-020-00466-2)) |
| Otu19734 | 1 | k__Fungi(100);p__Ascomycota(100);c__Saccharomycetes(100);o__Saccharomycetales(100);f__Metschnikowiaceae(100);g__Metschnikowia(100);g__Metschnikowia_unclassified(100); | Metschnikowia | 13 | Saprotroph | \|Undefined Saprotroph\| | Yeast | NULL | Probable | As nectar-tap_saprotroph (Põlme et al. 2020); Decay sugar-rich substrates (Põlme et al. 2020) | Põlme S, et al. 2020. Fungal Diversity 105:1-16 ((https://doi.org/10.1007/s13225-020-00466-2)) |
| Otu19735 | 1 | k__Fungi(100);p__Ascomycota(100);c__Saccharomycetes(100);o__Saccharomycetales(100);f__Metschnikowiaceae(100);g__Metschnikowia(100);s__Metschnikowia_pulcherrima(100); | Metschnikowia | 13 | Saprotroph | \|Undefined Saprotroph\| | Yeast | NULL | Probable | As nectar-tap_saprotroph (Põlme et al. 2020); Decay sugar-rich substrates (Põlme et al. 2020) | Põlme S, et al. 2020. Fungal Diversity 105:1-16 ((https://doi.org/10.1007/s13225-020-00466-2)) |
| Otu19736 | 1 | k__Fungi(100);p__Ascomycota(100);c__Saccharomycetes(100);o__Saccharomycetales(100);f__Metschnikowiaceae(100);g__Metschnikowia(100);g__Metschnikowia_unclassified(100); | Metschnikowia | 13 | Saprotroph | \|Undefined Saprotroph\| | Yeast | NULL | Probable | As nectar-tap_saprotroph (Põlme et al. 2020); Decay sugar-rich substrates (Põlme et al. 2020) | Põlme S, et al. 2020. Fungal Diversity 105:1-16 ((https://doi.org/10.1007/s13225-020-00466-2)) |
| Otu19737 | 1 | k__Fungi(100);p__Ascomycota(100);c__Saccharomycetes(100);o__Saccharomycetales(100);f__Metschnikowiaceae(100);g__Metschnikowia(100);s__Metschnikowia_pulcherrima(100); | Metschnikowia | 13 | Saprotroph | \|Undefined Saprotroph\| | Yeast | NULL | Probable | As nectar-tap_saprotroph (Põlme et al. 2020); Decay sugar-rich substrates (Põlme et al. 2020) | Põlme S, et al. 2020. Fungal Diversity 105:1-16 ((https://doi.org/10.1007/s13225-020-00466-2)) |
| Otu19738 | 1 | k__Fungi(100);p__Ascomycota(100);c__Saccharomycetes(100);o__Saccharomycetales(100);f__Metschnikowiaceae(100);g__Metschnikowia(100);s__Metschnikowia_pulcherrima(100); | Metschnikowia | 13 | Saprotroph | \|Undefined Saprotroph\| | Yeast | NULL | Probable | As nectar-tap_saprotroph (Põlme et al. 2020); Decay sugar-rich substrates (Põlme et al. 2020) | Põlme S, et al. 2020. Fungal Diversity 105:1-16 ((https://doi.org/10.1007/s13225-020-00466-2)) |
| Otu19739 | 1 | k__Fungi(100);p__Ascomycota(100);c__Saccharomycetes(100);o__Saccharomycetales(100);f__Metschnikowiaceae(100);g__Metschnikowia(100);s__Metschnikowia_pulcherrima(100); | Metschnikowia | 13 | Saprotroph | \|Undefined Saprotroph\| | Yeast | NULL | Probable | As nectar-tap_saprotroph (Põlme et al. 2020); Decay sugar-rich substrates (Põlme et al. 2020) | Põlme S, et al. 2020. Fungal Diversity 105:1-16 ((https://doi.org/10.1007/s13225-020-00466-2)) |
| Otu19740 | 1 | k__Fungi(100);p__Ascomycota(100);c__Saccharomycetes(100);o__Saccharomycetales(100);f__Metschnikowiaceae(100);g__Metschnikowia(100);s__Metschnikowia_pulcherrima(100); | Metschnikowia | 13 | Saprotroph | \|Undefined Saprotroph\| | Yeast | NULL | Probable | As nectar-tap_saprotroph (Põlme et al. 2020); Decay sugar-rich substrates (Põlme et al. 2020) | Põlme S, et al. 2020. Fungal Diversity 105:1-16 ((https://doi.org/10.1007/s13225-020-00466-2)) |
| Otu19741 | 1 | k__Fungi(100);p__Ascomycota(100);c__Saccharomycetes(100);o__Saccharomycetales(100);f__Metschnikowiaceae(100);g__Metschnikowia(100);g__Metschnikowia_unclassified(100); | Metschnikowia | 13 | Saprotroph | \|Undefined Saprotroph\| | Yeast | NULL | Probable | As nectar-tap_saprotroph (Põlme et al. 2020); Decay sugar-rich substrates (Põlme et al. 2020) | Põlme S, et al. 2020. Fungal Diversity 105:1-16 ((https://doi.org/10.1007/s13225-020-00466-2)) |
| Otu19742 | 1 | k__Fungi(100);p__Ascomycota(100);c__Saccharomycetes(100);o__Saccharomycetales(100);f__Metschnikowiaceae(100);g__Metschnikowia(100);s__Metschnikowia_pulcherrima(100); | Metschnikowia | 13 | Saprotroph | \|Undefined Saprotroph\| | Yeast | NULL | Probable | As nectar-tap_saprotroph (Põlme et al. 2020); Decay sugar-rich substrates (Põlme et al. 2020) | Põlme S, et al. 2020. Fungal Diversity 105:1-16 ((https://doi.org/10.1007/s13225-020-00466-2)) |
| Otu19743 | 1 | k__Fungi(100);p__Ascomycota(100);c__Saccharomycetes(100);o__Saccharomycetales(100);f__Metschnikowiaceae(100);g__Metschnikowia(100);s__Metschnikowia_pulcherrima(100); | Metschnikowia | 13 | Saprotroph | \|Undefined Saprotroph\| | Yeast | NULL | Probable | As nectar-tap_saprotroph (Põlme et al. 2020); Decay sugar-rich substrates (Põlme et al. 2020) | Põlme S, et al. 2020. Fungal Diversity 105:1-16 ((https://doi.org/10.1007/s13225-020-00466-2)) |
| Otu19744 | 1 | k__Fungi(100);p__Ascomycota(100);c__Saccharomycetes(100);o__Saccharomycetales(100);f__Metschnikowiaceae(100);g__Metschnikowia(100);s__Metschnikowia_pulcherrima(100); | Metschnikowia | 13 | Saprotroph | \|Undefined Saprotroph\| | Yeast | NULL | Probable | As nectar-tap_saprotroph (Põlme et al. 2020); Decay sugar-rich substrates (Põlme et al. 2020) | Põlme S, et al. 2020. Fungal Diversity 105:1-16 ((https://doi.org/10.1007/s13225-020-00466-2)) |
| Otu19745 | 1 | k__Fungi(100);p__Ascomycota(100);c__Saccharomycetes(100);o__Saccharomycetales(100);f__Metschnikowiaceae(100);g__Metschnikowia(100);g__Metschnikowia_unclassified(100); | Metschnikowia | 13 | Saprotroph | \|Undefined Saprotroph\| | Yeast | NULL | Probable | As nectar-tap_saprotroph (Põlme et al. 2020); Decay sugar-rich substrates (Põlme et al. 2020) | Põlme S, et al. 2020. Fungal Diversity 105:1-16 ((https://doi.org/10.1007/s13225-020-00466-2)) |
| Otu19746 | 1 | k__Fungi(100);p__Ascomycota(100);c__Saccharomycetes(100);o__Saccharomycetales(100);f__Metschnikowiaceae(100);g__Metschnikowia(100);s__Metschnikowia_pulcherrima(100); | Metschnikowia | 13 | Saprotroph | \|Undefined Saprotroph\| | Yeast | NULL | Probable | As nectar-tap_saprotroph (Põlme et al. 2020); Decay sugar-rich substrates (Põlme et al. 2020) | Põlme S, et al. 2020. Fungal Diversity 105:1-16 ((https://doi.org/10.1007/s13225-020-00466-2)) |
| Otu19749 | 1 | k__Fungi(100);p__Ascomycota(100);c__Saccharomycetes(100);o__Saccharomycetales(100);f__Metschnikowiaceae(100);g__Metschnikowia(100);s__Metschnikowia_pulcherrima(100); | Metschnikowia | 13 | Saprotroph | \|Undefined Saprotroph\| | Yeast | NULL | Probable | As nectar-tap_saprotroph (Põlme et al. 2020); Decay sugar-rich substrates (Põlme et al. 2020) | Põlme S, et al. 2020. Fungal Diversity 105:1-16 ((https://doi.org/10.1007/s13225-020-00466-2)) |
| Otu19750 | 1 | k__Fungi(100);p__Ascomycota(100);c__Saccharomycetes(100);o__Saccharomycetales(100);f__Metschnikowiaceae(100);g__Metschnikowia(100);s__Metschnikowia_pulcherrima(100); | Metschnikowia | 13 | Saprotroph | \|Undefined Saprotroph\| | Yeast | NULL | Probable | As nectar-tap_saprotroph (Põlme et al. 2020); Decay sugar-rich substrates (Põlme et al. 2020) | Põlme S, et al. 2020. Fungal Diversity 105:1-16 ((https://doi.org/10.1007/s13225-020-00466-2)) |
| Otu19751 | 1 | k__Fungi(100);p__Ascomycota(100);c__Saccharomycetes(100);o__Saccharomycetales(100);f__Metschnikowiaceae(100);g__Metschnikowia(100);s__Metschnikowia_pulcherrima(100); | Metschnikowia | 13 | Saprotroph | \|Undefined Saprotroph\| | Yeast | NULL | Probable | As nectar-tap_saprotroph (Põlme et al. 2020); Decay sugar-rich substrates (Põlme et al. 2020) | Põlme S, et al. 2020. Fungal Diversity 105:1-16 ((https://doi.org/10.1007/s13225-020-00466-2)) |
| Otu19752 | 1 | k__Fungi(100);p__Ascomycota(100);c__Saccharomycetes(100);o__Saccharomycetales(100);f__Metschnikowiaceae(100);g__Metschnikowia(100);s__Metschnikowia_pulcherrima(100); | Metschnikowia | 13 | Saprotroph | \|Undefined Saprotroph\| | Yeast | NULL | Probable | As nectar-tap_saprotroph (Põlme et al. 2020); Decay sugar-rich substrates (Põlme et al. 2020) | Põlme S, et al. 2020. Fungal Diversity 105:1-16 ((https://doi.org/10.1007/s13225-020-00466-2)) |
| Otu19753 | 1 | k__Fungi(100);p__Ascomycota(100);c__Saccharomycetes(100);o__Saccharomycetales(100);f__Metschnikowiaceae(100);g__Metschnikowia(100);s__Metschnikowia_pulcherrima(100); | Metschnikowia | 13 | Saprotroph | \|Undefined Saprotroph\| | Yeast | NULL | Probable | As nectar-tap_saprotroph (Põlme et al. 2020); Decay sugar-rich substrates (Põlme et al. 2020) | Põlme S, et al. 2020. Fungal Diversity 105:1-16 ((https://doi.org/10.1007/s13225-020-00466-2)) |
| Otu19754 | 1 | k__Fungi(100);p__Ascomycota(100);c__Saccharomycetes(100);o__Saccharomycetales(100);f__Metschnikowiaceae(100);g__Metschnikowia(100);s__Metschnikowia_pulcherrima(100); | Metschnikowia | 13 | Saprotroph | \|Undefined Saprotroph\| | Yeast | NULL | Probable | As nectar-tap_saprotroph (Põlme et al. 2020); Decay sugar-rich substrates (Põlme et al. 2020) | Põlme S, et al. 2020. Fungal Diversity 105:1-16 ((https://doi.org/10.1007/s13225-020-00466-2)) |
| Otu19755 | 1 | k__Fungi(100);p__Ascomycota(100);c__Saccharomycetes(100);o__Saccharomycetales(100);f__Metschnikowiaceae(100);g__Metschnikowia(100);s__Metschnikowia_pulcherrima(100); | Metschnikowia | 13 | Saprotroph | \|Undefined Saprotroph\| | Yeast | NULL | Probable | As nectar-tap_saprotroph (Põlme et al. 2020); Decay sugar-rich substrates (Põlme et al. 2020) | Põlme S, et al. 2020. Fungal Diversity 105:1-16 ((https://doi.org/10.1007/s13225-020-00466-2)) |
| Otu19756 | 1 | k__Fungi(100);p__Ascomycota(100);c__Saccharomycetes(100);o__Saccharomycetales(100);f__Metschnikowiaceae(100);g__Metschnikowia(100);s__Metschnikowia_pulcherrima(100); | Metschnikowia | 13 | Saprotroph | \|Undefined Saprotroph\| | Yeast | NULL | Probable | As nectar-tap_saprotroph (Põlme et al. 2020); Decay sugar-rich substrates (Põlme et al. 2020) | Põlme S, et al. 2020. Fungal Diversity 105:1-16 ((https://doi.org/10.1007/s13225-020-00466-2)) |
| Otu19757 | 1 | k__Fungi(100);p__Ascomycota(100);c__Saccharomycetes(100);o__Saccharomycetales(100);f__Metschnikowiaceae(100);g__Metschnikowia(100);s__Metschnikowia_pulcherrima(100); | Metschnikowia | 13 | Saprotroph | \|Undefined Saprotroph\| | Yeast | NULL | Probable | As nectar-tap_saprotroph (Põlme et al. 2020); Decay sugar-rich substrates (Põlme et al. 2020) | Põlme S, et al. 2020. Fungal Diversity 105:1-16 ((https://doi.org/10.1007/s13225-020-00466-2)) |
| Otu19759 | 1 | k__Fungi(100);p__Ascomycota(100);c__Saccharomycetes(100);o__Saccharomycetales(100);f__Metschnikowiaceae(100);g__Metschnikowia(100);s__Metschnikowia_pulcherrima(100); | Metschnikowia | 13 | Saprotroph | \|Undefined Saprotroph\| | Yeast | NULL | Probable | As nectar-tap_saprotroph (Põlme et al. 2020); Decay sugar-rich substrates (Põlme et al. 2020) | Põlme S, et al. 2020. Fungal Diversity 105:1-16 ((https://doi.org/10.1007/s13225-020-00466-2)) |
| Otu19760 | 1 | k__Fungi(100);p__Ascomycota(100);c__Saccharomycetes(100);o__Saccharomycetales(100);f__Metschnikowiaceae(100);g__Metschnikowia(100);g__Metschnikowia_unclassified(100); | Metschnikowia | 13 | Saprotroph | \|Undefined Saprotroph\| | Yeast | NULL | Probable | As nectar-tap_saprotroph (Põlme et al. 2020); Decay sugar-rich substrates (Põlme et al. 2020) | Põlme S, et al. 2020. Fungal Diversity 105:1-16 ((https://doi.org/10.1007/s13225-020-00466-2)) |
| Otu19761 | 1 | k__Fungi(100);p__Ascomycota(100);c__Saccharomycetes(100);o__Saccharomycetales(100);f__Metschnikowiaceae(100);g__Metschnikowia(100);g__Metschnikowia_unclassified(100); | Metschnikowia | 13 | Saprotroph | \|Undefined Saprotroph\| | Yeast | NULL | Probable | As nectar-tap_saprotroph (Põlme et al. 2020); Decay sugar-rich substrates (Põlme et al. 2020) | Põlme S, et al. 2020. Fungal Diversity 105:1-16 ((https://doi.org/10.1007/s13225-020-00466-2)) |
| Otu19762 | 1 | k__Fungi(100);p__Ascomycota(100);c__Saccharomycetes(100);o__Saccharomycetales(100);f__Metschnikowiaceae(100);g__Metschnikowia(100);s__Metschnikowia_pulcherrima(100); | Metschnikowia | 13 | Saprotroph | \|Undefined Saprotroph\| | Yeast | NULL | Probable | As nectar-tap_saprotroph (Põlme et al. 2020); Decay sugar-rich substrates (Põlme et al. 2020) | Põlme S, et al. 2020. Fungal Diversity 105:1-16 ((https://doi.org/10.1007/s13225-020-00466-2)) |
| Otu19764 | 1 | k__Fungi(100);p__Ascomycota(100);c__Saccharomycetes(100);o__Saccharomycetales(100);f__Metschnikowiaceae(100);g__Metschnikowia(100);g__Metschnikowia_unclassified(100); | Metschnikowia | 13 | Saprotroph | \|Undefined Saprotroph\| | Yeast | NULL | Probable | As nectar-tap_saprotroph (Põlme et al. 2020); Decay sugar-rich substrates (Põlme et al. 2020) | Põlme S, et al. 2020. Fungal Diversity 105:1-16 ((https://doi.org/10.1007/s13225-020-00466-2)) |
| Otu19766 | 1 | k__Fungi(100);p__Ascomycota(100);c__Saccharomycetes(100);o__Saccharomycetales(100);f__Metschnikowiaceae(100);g__Metschnikowia(100);s__Metschnikowia_pulcherrima(100); | Metschnikowia | 13 | Saprotroph | \|Undefined Saprotroph\| | Yeast | NULL | Probable | As nectar-tap_saprotroph (Põlme et al. 2020); Decay sugar-rich substrates (Põlme et al. 2020) | Põlme S, et al. 2020. Fungal Diversity 105:1-16 ((https://doi.org/10.1007/s13225-020-00466-2)) |
| Otu19767 | 1 | k__Fungi(100);p__Ascomycota(100);c__Saccharomycetes(100);o__Saccharomycetales(100);f__Metschnikowiaceae(100);g__Metschnikowia(100);s__Metschnikowia_pulcherrima(100); | Metschnikowia | 13 | Saprotroph | \|Undefined Saprotroph\| | Yeast | NULL | Probable | As nectar-tap_saprotroph (Põlme et al. 2020); Decay sugar-rich substrates (Põlme et al. 2020) | Põlme S, et al. 2020. Fungal Diversity 105:1-16 ((https://doi.org/10.1007/s13225-020-00466-2)) |
| Otu19768 | 1 | k__Fungi(100);p__Ascomycota(100);c__Saccharomycetes(100);o__Saccharomycetales(100);f__Metschnikowiaceae(100);g__Metschnikowia(100);s__Metschnikowia_pulcherrima(100); | Metschnikowia | 13 | Saprotroph | \|Undefined Saprotroph\| | Yeast | NULL | Probable | As nectar-tap_saprotroph (Põlme et al. 2020); Decay sugar-rich substrates (Põlme et al. 2020) | Põlme S, et al. 2020. Fungal Diversity 105:1-16 ((https://doi.org/10.1007/s13225-020-00466-2)) |
| Otu19769 | 1 | k__Fungi(100);p__Ascomycota(100);c__Saccharomycetes(100);o__Saccharomycetales(100);f__Metschnikowiaceae(100);g__Metschnikowia(100);s__Metschnikowia_pulcherrima(100); | Metschnikowia | 13 | Saprotroph | \|Undefined Saprotroph\| | Yeast | NULL | Probable | As nectar-tap_saprotroph (Põlme et al. 2020); Decay sugar-rich substrates (Põlme et al. 2020) | Põlme S, et al. 2020. Fungal Diversity 105:1-16 ((https://doi.org/10.1007/s13225-020-00466-2)) |
| Otu19770 | 1 | k__Fungi(100);p__Ascomycota(100);c__Saccharomycetes(100);o__Saccharomycetales(100);f__Metschnikowiaceae(100);g__Metschnikowia(100);s__Metschnikowia_pulcherrima(100); | Metschnikowia | 13 | Saprotroph | \|Undefined Saprotroph\| | Yeast | NULL | Probable | As nectar-tap_saprotroph (Põlme et al. 2020); Decay sugar-rich substrates (Põlme et al. 2020) | Põlme S, et al. 2020. Fungal Diversity 105:1-16 ((https://doi.org/10.1007/s13225-020-00466-2)) |
| Otu19771 | 1 | k__Fungi(100);p__Ascomycota(100);c__Saccharomycetes(100);o__Saccharomycetales(100);f__Metschnikowiaceae(100);g__Metschnikowia(100);s__Metschnikowia_pulcherrima(100); | Metschnikowia | 13 | Saprotroph | \|Undefined Saprotroph\| | Yeast | NULL | Probable | As nectar-tap_saprotroph (Põlme et al. 2020); Decay sugar-rich substrates (Põlme et al. 2020) | Põlme S, et al. 2020. Fungal Diversity 105:1-16 ((https://doi.org/10.1007/s13225-020-00466-2)) |
| Otu19772 | 1 | k__Fungi(100);p__Ascomycota(100);c__Saccharomycetes(100);o__Saccharomycetales(100);f__Metschnikowiaceae(100);g__Metschnikowia(100);s__Metschnikowia_pulcherrima(100); | Metschnikowia | 13 | Saprotroph | \|Undefined Saprotroph\| | Yeast | NULL | Probable | As nectar-tap_saprotroph (Põlme et al. 2020); Decay sugar-rich substrates (Põlme et al. 2020) | Põlme S, et al. 2020. Fungal Diversity 105:1-16 ((https://doi.org/10.1007/s13225-020-00466-2)) |
| Otu19773 | 1 | k__Fungi(100);p__Ascomycota(100);c__Saccharomycetes(100);o__Saccharomycetales(100);f__Metschnikowiaceae(100);g__Metschnikowia(100);g__Metschnikowia_unclassified(100); | Metschnikowia | 13 | Saprotroph | \|Undefined Saprotroph\| | Yeast | NULL | Probable | As nectar-tap_saprotroph (Põlme et al. 2020); Decay sugar-rich substrates (Põlme et al. 2020) | Põlme S, et al. 2020. Fungal Diversity 105:1-16 ((https://doi.org/10.1007/s13225-020-00466-2)) |
| Otu19774 | 1 | k__Fungi(100);p__Ascomycota(100);c__Saccharomycetes(100);o__Saccharomycetales(100);f__Metschnikowiaceae(100);g__Metschnikowia(100);s__Metschnikowia_pulcherrima(100); | Metschnikowia | 13 | Saprotroph | \|Undefined Saprotroph\| | Yeast | NULL | Probable | As nectar-tap_saprotroph (Põlme et al. 2020); Decay sugar-rich substrates (Põlme et al. 2020) | Põlme S, et al. 2020. Fungal Diversity 105:1-16 ((https://doi.org/10.1007/s13225-020-00466-2)) |
| Otu19775 | 1 | k__Fungi(100);p__Ascomycota(100);c__Saccharomycetes(100);o__Saccharomycetales(100);f__Metschnikowiaceae(100);g__Metschnikowia(100);s__Metschnikowia_pulcherrima(100); | Metschnikowia | 13 | Saprotroph | \|Undefined Saprotroph\| | Yeast | NULL | Probable | As nectar-tap_saprotroph (Põlme et al. 2020); Decay sugar-rich substrates (Põlme et al. 2020) | Põlme S, et al. 2020. Fungal Diversity 105:1-16 ((https://doi.org/10.1007/s13225-020-00466-2)) |
| Otu19776 | 1 | k__Fungi(100);p__Ascomycota(100);c__Saccharomycetes(100);o__Saccharomycetales(100);f__Metschnikowiaceae(100);g__Metschnikowia(100);g__Metschnikowia_unclassified(100); | Metschnikowia | 13 | Saprotroph | \|Undefined Saprotroph\| | Yeast | NULL | Probable | As nectar-tap_saprotroph (Põlme et al. 2020); Decay sugar-rich substrates (Põlme et al. 2020) | Põlme S, et al. 2020. Fungal Diversity 105:1-16 ((https://doi.org/10.1007/s13225-020-00466-2)) |
| Otu19777 | 1 | k__Fungi(100);p__Ascomycota(100);c__Saccharomycetes(100);o__Saccharomycetales(100);f__Metschnikowiaceae(100);g__Metschnikowia(100);s__Metschnikowia_pulcherrima(100); | Metschnikowia | 13 | Saprotroph | \|Undefined Saprotroph\| | Yeast | NULL | Probable | As nectar-tap_saprotroph (Põlme et al. 2020); Decay sugar-rich substrates (Põlme et al. 2020) | Põlme S, et al. 2020. Fungal Diversity 105:1-16 ((https://doi.org/10.1007/s13225-020-00466-2)) |
| Otu19778 | 1 | k__Fungi(100);p__Ascomycota(100);c__Saccharomycetes(100);o__Saccharomycetales(100);f__Metschnikowiaceae(100);g__Metschnikowia(100);s__Metschnikowia_pulcherrima(100); | Metschnikowia | 13 | Saprotroph | \|Undefined Saprotroph\| | Yeast | NULL | Probable | As nectar-tap_saprotroph (Põlme et al. 2020); Decay sugar-rich substrates (Põlme et al. 2020) | Põlme S, et al. 2020. Fungal Diversity 105:1-16 ((https://doi.org/10.1007/s13225-020-00466-2)) |
| Otu19779 | 1 | k__Fungi(100);p__Ascomycota(100);c__Saccharomycetes(100);o__Saccharomycetales(100);f__Metschnikowiaceae(100);g__Metschnikowia(100);s__Metschnikowia_pulcherrima(100); | Metschnikowia | 13 | Saprotroph | \|Undefined Saprotroph\| | Yeast | NULL | Probable | As nectar-tap_saprotroph (Põlme et al. 2020); Decay sugar-rich substrates (Põlme et al. 2020) | Põlme S, et al. 2020. Fungal Diversity 105:1-16 ((https://doi.org/10.1007/s13225-020-00466-2)) |
| Otu19780 | 1 | k__Fungi(100);p__Ascomycota(100);c__Saccharomycetes(100);o__Saccharomycetales(100);f__Metschnikowiaceae(100);g__Metschnikowia(100);g__Metschnikowia_unclassified(100); | Metschnikowia | 13 | Saprotroph | \|Undefined Saprotroph\| | Yeast | NULL | Probable | As nectar-tap_saprotroph (Põlme et al. 2020); Decay sugar-rich substrates (Põlme et al. 2020) | Põlme S, et al. 2020. Fungal Diversity 105:1-16 ((https://doi.org/10.1007/s13225-020-00466-2)) |
| Otu19781 | 1 | k__Fungi(100);p__Ascomycota(100);c__Saccharomycetes(100);o__Saccharomycetales(100);f__Metschnikowiaceae(100);g__Metschnikowia(100);s__Metschnikowia_pulcherrima(100); | Metschnikowia | 13 | Saprotroph | \|Undefined Saprotroph\| | Yeast | NULL | Probable | As nectar-tap_saprotroph (Põlme et al. 2020); Decay sugar-rich substrates (Põlme et al. 2020) | Põlme S, et al. 2020. Fungal Diversity 105:1-16 ((https://doi.org/10.1007/s13225-020-00466-2)) |
| Otu19782 | 1 | k__Fungi(100);p__Ascomycota(100);c__Saccharomycetes(100);o__Saccharomycetales(100);f__Metschnikowiaceae(100);g__Metschnikowia(100);s__Metschnikowia_pulcherrima(100); | Metschnikowia | 13 | Saprotroph | \|Undefined Saprotroph\| | Yeast | NULL | Probable | As nectar-tap_saprotroph (Põlme et al. 2020); Decay sugar-rich substrates (Põlme et al. 2020) | Põlme S, et al. 2020. Fungal Diversity 105:1-16 ((https://doi.org/10.1007/s13225-020-00466-2)) |
| Otu19783 | 1 | k__Fungi(100);p__Ascomycota(100);c__Saccharomycetes(100);o__Saccharomycetales(100);f__Metschnikowiaceae(100);g__Metschnikowia(100);g__Metschnikowia_unclassified(100); | Metschnikowia | 13 | Saprotroph | \|Undefined Saprotroph\| | Yeast | NULL | Probable | As nectar-tap_saprotroph (Põlme et al. 2020); Decay sugar-rich substrates (Põlme et al. 2020) | Põlme S, et al. 2020. Fungal Diversity 105:1-16 ((https://doi.org/10.1007/s13225-020-00466-2)) |
| Otu19784 | 1 | k__Fungi(100);p__Ascomycota(100);c__Saccharomycetes(100);o__Saccharomycetales(100);f__Metschnikowiaceae(100);g__Metschnikowia(100);s__Metschnikowia_pulcherrima(100); | Metschnikowia | 13 | Saprotroph | \|Undefined Saprotroph\| | Yeast | NULL | Probable | As nectar-tap_saprotroph (Põlme et al. 2020); Decay sugar-rich substrates (Põlme et al. 2020) | Põlme S, et al. 2020. Fungal Diversity 105:1-16 ((https://doi.org/10.1007/s13225-020-00466-2)) |
| Otu19785 | 1 | k__Fungi(100);p__Ascomycota(100);c__Saccharomycetes(100);o__Saccharomycetales(100);f__Metschnikowiaceae(100);g__Metschnikowia(100);s__Metschnikowia_pulcherrima(100); | Metschnikowia | 13 | Saprotroph | \|Undefined Saprotroph\| | Yeast | NULL | Probable | As nectar-tap_saprotroph (Põlme et al. 2020); Decay sugar-rich substrates (Põlme et al. 2020) | Põlme S, et al. 2020. Fungal Diversity 105:1-16 ((https://doi.org/10.1007/s13225-020-00466-2)) |
| Otu19786 | 1 | k__Fungi(100);p__Ascomycota(100);c__Saccharomycetes(100);o__Saccharomycetales(100);f__Metschnikowiaceae(100);g__Metschnikowia(100);g__Metschnikowia_unclassified(100); | Metschnikowia | 13 | Saprotroph | \|Undefined Saprotroph\| | Yeast | NULL | Probable | As nectar-tap_saprotroph (Põlme et al. 2020); Decay sugar-rich substrates (Põlme et al. 2020) | Põlme S, et al. 2020. Fungal Diversity 105:1-16 ((https://doi.org/10.1007/s13225-020-00466-2)) |
| Otu19787 | 1 | k__Fungi(100);p__Ascomycota(100);c__Saccharomycetes(100);o__Saccharomycetales(100);f__Metschnikowiaceae(100);g__Metschnikowia(100);s__Metschnikowia_pulcherrima(100); | Metschnikowia | 13 | Saprotroph | \|Undefined Saprotroph\| | Yeast | NULL | Probable | As nectar-tap_saprotroph (Põlme et al. 2020); Decay sugar-rich substrates (Põlme et al. 2020) | Põlme S, et al. 2020. Fungal Diversity 105:1-16 ((https://doi.org/10.1007/s13225-020-00466-2)) |
| Otu19788 | 1 | k__Fungi(100);p__Ascomycota(100);c__Saccharomycetes(100);o__Saccharomycetales(100);f__Metschnikowiaceae(100);g__Metschnikowia(100);s__Metschnikowia_pulcherrima(100); | Metschnikowia | 13 | Saprotroph | \|Undefined Saprotroph\| | Yeast | NULL | Probable | As nectar-tap_saprotroph (Põlme et al. 2020); Decay sugar-rich substrates (Põlme et al. 2020) | Põlme S, et al. 2020. Fungal Diversity 105:1-16 ((https://doi.org/10.1007/s13225-020-00466-2)) |
| Otu19790 | 1 | k__Fungi(100);p__Ascomycota(100);c__Saccharomycetes(100);o__Saccharomycetales(100);f__Metschnikowiaceae(100);g__Metschnikowia(100);s__Metschnikowia_pulcherrima(100); | Metschnikowia | 13 | Saprotroph | \|Undefined Saprotroph\| | Yeast | NULL | Probable | As nectar-tap_saprotroph (Põlme et al. 2020); Decay sugar-rich substrates (Põlme et al. 2020) | Põlme S, et al. 2020. Fungal Diversity 105:1-16 ((https://doi.org/10.1007/s13225-020-00466-2)) |
| Otu19791 | 1 | k__Fungi(100);p__Ascomycota(100);c__Saccharomycetes(100);o__Saccharomycetales(100);f__Metschnikowiaceae(100);g__Metschnikowia(100);s__Metschnikowia_pulcherrima(100); | Metschnikowia | 13 | Saprotroph | \|Undefined Saprotroph\| | Yeast | NULL | Probable | As nectar-tap_saprotroph (Põlme et al. 2020); Decay sugar-rich substrates (Põlme et al. 2020) | Põlme S, et al. 2020. Fungal Diversity 105:1-16 ((https://doi.org/10.1007/s13225-020-00466-2)) |
| Otu19792 | 1 | k__Fungi(100);p__Ascomycota(100);c__Saccharomycetes(100);o__Saccharomycetales(100);f__Metschnikowiaceae(100);g__Metschnikowia(100);g__Metschnikowia_unclassified(100); | Metschnikowia | 13 | Saprotroph | \|Undefined Saprotroph\| | Yeast | NULL | Probable | As nectar-tap_saprotroph (Põlme et al. 2020); Decay sugar-rich substrates (Põlme et al. 2020) | Põlme S, et al. 2020. Fungal Diversity 105:1-16 ((https://doi.org/10.1007/s13225-020-00466-2)) |
| Otu19793 | 1 | k__Fungi(100);p__Ascomycota(100);c__Saccharomycetes(100);o__Saccharomycetales(100);f__Metschnikowiaceae(100);g__Metschnikowia(100);s__Metschnikowia_pulcherrima(100); | Metschnikowia | 13 | Saprotroph | \|Undefined Saprotroph\| | Yeast | NULL | Probable | As nectar-tap_saprotroph (Põlme et al. 2020); Decay sugar-rich substrates (Põlme et al. 2020) | Põlme S, et al. 2020. Fungal Diversity 105:1-16 ((https://doi.org/10.1007/s13225-020-00466-2)) |
| Otu19794 | 1 | k__Fungi(100);p__Ascomycota(100);c__Saccharomycetes(100);o__Saccharomycetales(100);f__Metschnikowiaceae(100);g__Metschnikowia(100);s__Metschnikowia_pulcherrima(100); | Metschnikowia | 13 | Saprotroph | \|Undefined Saprotroph\| | Yeast | NULL | Probable | As nectar-tap_saprotroph (Põlme et al. 2020); Decay sugar-rich substrates (Põlme et al. 2020) | Põlme S, et al. 2020. Fungal Diversity 105:1-16 ((https://doi.org/10.1007/s13225-020-00466-2)) |
| Otu19795 | 1 | k__Fungi(100);p__Ascomycota(100);c__Saccharomycetes(100);o__Saccharomycetales(100);f__Metschnikowiaceae(100);g__Metschnikowia(100);s__Metschnikowia_pulcherrima(100); | Metschnikowia | 13 | Saprotroph | \|Undefined Saprotroph\| | Yeast | NULL | Probable | As nectar-tap_saprotroph (Põlme et al. 2020); Decay sugar-rich substrates (Põlme et al. 2020) | Põlme S, et al. 2020. Fungal Diversity 105:1-16 ((https://doi.org/10.1007/s13225-020-00466-2)) |
| Otu19796 | 1 | k__Fungi(100);p__Ascomycota(100);c__Saccharomycetes(100);o__Saccharomycetales(100);f__Metschnikowiaceae(100);g__Metschnikowia(100);s__Metschnikowia_pulcherrima(100); | Metschnikowia | 13 | Saprotroph | \|Undefined Saprotroph\| | Yeast | NULL | Probable | As nectar-tap_saprotroph (Põlme et al. 2020); Decay sugar-rich substrates (Põlme et al. 2020) | Põlme S, et al. 2020. Fungal Diversity 105:1-16 ((https://doi.org/10.1007/s13225-020-00466-2)) |
| Otu19797 | 1 | k__Fungi(100);p__Ascomycota(100);c__Saccharomycetes(100);o__Saccharomycetales(100);f__Metschnikowiaceae(100);g__Metschnikowia(100);g__Metschnikowia_unclassified(100); | Metschnikowia | 13 | Saprotroph | \|Undefined Saprotroph\| | Yeast | NULL | Probable | As nectar-tap_saprotroph (Põlme et al. 2020); Decay sugar-rich substrates (Põlme et al. 2020) | Põlme S, et al. 2020. Fungal Diversity 105:1-16 ((https://doi.org/10.1007/s13225-020-00466-2)) |
| Otu19798 | 1 | k__Fungi(100);p__Ascomycota(100);c__Saccharomycetes(100);o__Saccharomycetales(100);f__Metschnikowiaceae(100);g__Metschnikowia(100);s__Metschnikowia_pulcherrima(100); | Metschnikowia | 13 | Saprotroph | \|Undefined Saprotroph\| | Yeast | NULL | Probable | As nectar-tap_saprotroph (Põlme et al. 2020); Decay sugar-rich substrates (Põlme et al. 2020) | Põlme S, et al. 2020. Fungal Diversity 105:1-16 ((https://doi.org/10.1007/s13225-020-00466-2)) |
| Otu19799 | 1 | k__Fungi(100);p__Ascomycota(100);c__Saccharomycetes(100);o__Saccharomycetales(100);f__Metschnikowiaceae(100);g__Metschnikowia(100);s__Metschnikowia_pulcherrima(100); | Metschnikowia | 13 | Saprotroph | \|Undefined Saprotroph\| | Yeast | NULL | Probable | As nectar-tap_saprotroph (Põlme et al. 2020); Decay sugar-rich substrates (Põlme et al. 2020) | Põlme S, et al. 2020. Fungal Diversity 105:1-16 ((https://doi.org/10.1007/s13225-020-00466-2)) |
| Otu19800 | 1 | k__Fungi(100);p__Ascomycota(100);c__Saccharomycetes(100);o__Saccharomycetales(100);f__Metschnikowiaceae(100);g__Metschnikowia(100);g__Metschnikowia_unclassified(100); | Metschnikowia | 13 | Saprotroph | \|Undefined Saprotroph\| | Yeast | NULL | Probable | As nectar-tap_saprotroph (Põlme et al. 2020); Decay sugar-rich substrates (Põlme et al. 2020) | Põlme S, et al. 2020. Fungal Diversity 105:1-16 ((https://doi.org/10.1007/s13225-020-00466-2)) |
| Otu19801 | 1 | k__Fungi(100);p__Ascomycota(100);c__Saccharomycetes(100);o__Saccharomycetales(100);f__Metschnikowiaceae(100);g__Metschnikowia(100);s__Metschnikowia_pulcherrima(100); | Metschnikowia | 13 | Saprotroph | \|Undefined Saprotroph\| | Yeast | NULL | Probable | As nectar-tap_saprotroph (Põlme et al. 2020); Decay sugar-rich substrates (Põlme et al. 2020) | Põlme S, et al. 2020. Fungal Diversity 105:1-16 ((https://doi.org/10.1007/s13225-020-00466-2)) |
| Otu19802 | 1 | k__Fungi(100);p__Ascomycota(100);c__Saccharomycetes(100);o__Saccharomycetales(100);f__Metschnikowiaceae(100);g__Metschnikowia(100);s__Metschnikowia_pulcherrima(100); | Metschnikowia | 13 | Saprotroph | \|Undefined Saprotroph\| | Yeast | NULL | Probable | As nectar-tap_saprotroph (Põlme et al. 2020); Decay sugar-rich substrates (Põlme et al. 2020) | Põlme S, et al. 2020. Fungal Diversity 105:1-16 ((https://doi.org/10.1007/s13225-020-00466-2)) |
| Otu19803 | 1 | k__Fungi(100);p__Ascomycota(100);c__Saccharomycetes(100);o__Saccharomycetales(100);f__Metschnikowiaceae(100);g__Metschnikowia(100);g__Metschnikowia_unclassified(100); | Metschnikowia | 13 | Saprotroph | \|Undefined Saprotroph\| | Yeast | NULL | Probable | As nectar-tap_saprotroph (Põlme et al. 2020); Decay sugar-rich substrates (Põlme et al. 2020) | Põlme S, et al. 2020. Fungal Diversity 105:1-16 ((https://doi.org/10.1007/s13225-020-00466-2)) |
| Otu19804 | 1 | k__Fungi(100);p__Ascomycota(100);c__Saccharomycetes(100);o__Saccharomycetales(100);f__Metschnikowiaceae(100);g__Metschnikowia(100);s__Metschnikowia_pulcherrima(100); | Metschnikowia | 13 | Saprotroph | \|Undefined Saprotroph\| | Yeast | NULL | Probable | As nectar-tap_saprotroph (Põlme et al. 2020); Decay sugar-rich substrates (Põlme et al. 2020) | Põlme S, et al. 2020. Fungal Diversity 105:1-16 ((https://doi.org/10.1007/s13225-020-00466-2)) |
| Otu19805 | 1 | k__Fungi(100);p__Ascomycota(100);c__Saccharomycetes(100);o__Saccharomycetales(100);f__Metschnikowiaceae(100);g__Metschnikowia(100);s__Metschnikowia_pulcherrima(100); | Metschnikowia | 13 | Saprotroph | \|Undefined Saprotroph\| | Yeast | NULL | Probable | As nectar-tap_saprotroph (Põlme et al. 2020); Decay sugar-rich substrates (Põlme et al. 2020) | Põlme S, et al. 2020. Fungal Diversity 105:1-16 ((https://doi.org/10.1007/s13225-020-00466-2)) |
| Otu19806 | 1 | k__Fungi(100);p__Ascomycota(100);c__Saccharomycetes(100);o__Saccharomycetales(100);f__Metschnikowiaceae(100);g__Metschnikowia(100);g__Metschnikowia_unclassified(100); | Metschnikowia | 13 | Saprotroph | \|Undefined Saprotroph\| | Yeast | NULL | Probable | As nectar-tap_saprotroph (Põlme et al. 2020); Decay sugar-rich substrates (Põlme et al. 2020) | Põlme S, et al. 2020. Fungal Diversity 105:1-16 ((https://doi.org/10.1007/s13225-020-00466-2)) |
| Otu19807 | 1 | k__Fungi(100);p__Ascomycota(100);c__Saccharomycetes(100);o__Saccharomycetales(100);f__Metschnikowiaceae(100);g__Metschnikowia(100);s__Metschnikowia_pulcherrima(100); | Metschnikowia | 13 | Saprotroph | \|Undefined Saprotroph\| | Yeast | NULL | Probable | As nectar-tap_saprotroph (Põlme et al. 2020); Decay sugar-rich substrates (Põlme et al. 2020) | Põlme S, et al. 2020. Fungal Diversity 105:1-16 ((https://doi.org/10.1007/s13225-020-00466-2)) |
| Otu19808 | 1 | k__Fungi(100);p__Ascomycota(100);c__Saccharomycetes(100);o__Saccharomycetales(100);f__Metschnikowiaceae(100);g__Metschnikowia(100);s__Metschnikowia_pulcherrima(100); | Metschnikowia | 13 | Saprotroph | \|Undefined Saprotroph\| | Yeast | NULL | Probable | As nectar-tap_saprotroph (Põlme et al. 2020); Decay sugar-rich substrates (Põlme et al. 2020) | Põlme S, et al. 2020. Fungal Diversity 105:1-16 ((https://doi.org/10.1007/s13225-020-00466-2)) |
| Otu19809 | 1 | k__Fungi(100);p__Ascomycota(100);c__Saccharomycetes(100);o__Saccharomycetales(100);f__Metschnikowiaceae(100);g__Metschnikowia(100);g__Metschnikowia_unclassified(100); | Metschnikowia | 13 | Saprotroph | \|Undefined Saprotroph\| | Yeast | NULL | Probable | As nectar-tap_saprotroph (Põlme et al. 2020); Decay sugar-rich substrates (Põlme et al. 2020) | Põlme S, et al. 2020. Fungal Diversity 105:1-16 ((https://doi.org/10.1007/s13225-020-00466-2)) |
| Otu19810 | 1 | k__Fungi(100);p__Ascomycota(100);c__Saccharomycetes(100);o__Saccharomycetales(100);f__Metschnikowiaceae(100);g__Metschnikowia(100);g__Metschnikowia_unclassified(100); | Metschnikowia | 13 | Saprotroph | \|Undefined Saprotroph\| | Yeast | NULL | Probable | As nectar-tap_saprotroph (Põlme et al. 2020); Decay sugar-rich substrates (Põlme et al. 2020) | Põlme S, et al. 2020. Fungal Diversity 105:1-16 ((https://doi.org/10.1007/s13225-020-00466-2)) |
| Otu19811 | 1 | k__Fungi(100);p__Ascomycota(100);c__Saccharomycetes(100);o__Saccharomycetales(100);f__Metschnikowiaceae(100);g__Metschnikowia(100);s__Metschnikowia_pulcherrima(100); | Metschnikowia | 13 | Saprotroph | \|Undefined Saprotroph\| | Yeast | NULL | Probable | As nectar-tap_saprotroph (Põlme et al. 2020); Decay sugar-rich substrates (Põlme et al. 2020) | Põlme S, et al. 2020. Fungal Diversity 105:1-16 ((https://doi.org/10.1007/s13225-020-00466-2)) |
| Otu19812 | 1 | k__Fungi(100);p__Ascomycota(100);c__Saccharomycetes(100);o__Saccharomycetales(100);f__Metschnikowiaceae(100);g__Metschnikowia(100);s__Metschnikowia_pulcherrima(100); | Metschnikowia | 13 | Saprotroph | \|Undefined Saprotroph\| | Yeast | NULL | Probable | As nectar-tap_saprotroph (Põlme et al. 2020); Decay sugar-rich substrates (Põlme et al. 2020) | Põlme S, et al. 2020. Fungal Diversity 105:1-16 ((https://doi.org/10.1007/s13225-020-00466-2)) |
| Otu19813 | 1 | k__Fungi(100);p__Ascomycota(100);c__Saccharomycetes(100);o__Saccharomycetales(100);f__Metschnikowiaceae(100);g__Metschnikowia(100);g__Metschnikowia_unclassified(100); | Metschnikowia | 13 | Saprotroph | \|Undefined Saprotroph\| | Yeast | NULL | Probable | As nectar-tap_saprotroph (Põlme et al. 2020); Decay sugar-rich substrates (Põlme et al. 2020) | Põlme S, et al. 2020. Fungal Diversity 105:1-16 ((https://doi.org/10.1007/s13225-020-00466-2)) |
| Otu19814 | 1 | k__Fungi(100);p__Ascomycota(100);c__Saccharomycetes(100);o__Saccharomycetales(100);f__Metschnikowiaceae(100);g__Metschnikowia(100);s__Metschnikowia_pulcherrima(100); | Metschnikowia | 13 | Saprotroph | \|Undefined Saprotroph\| | Yeast | NULL | Probable | As nectar-tap_saprotroph (Põlme et al. 2020); Decay sugar-rich substrates (Põlme et al. 2020) | Põlme S, et al. 2020. Fungal Diversity 105:1-16 ((https://doi.org/10.1007/s13225-020-00466-2)) |
| Otu19815 | 1 | k__Fungi(100);p__Ascomycota(100);c__Saccharomycetes(100);o__Saccharomycetales(100);f__Metschnikowiaceae(100);g__Metschnikowia(100);s__Metschnikowia_pulcherrima(100); | Metschnikowia | 13 | Saprotroph | \|Undefined Saprotroph\| | Yeast | NULL | Probable | As nectar-tap_saprotroph (Põlme et al. 2020); Decay sugar-rich substrates (Põlme et al. 2020) | Põlme S, et al. 2020. Fungal Diversity 105:1-16 ((https://doi.org/10.1007/s13225-020-00466-2)) |
| Otu19816 | 1 | k__Fungi(100);p__Ascomycota(100);c__Saccharomycetes(100);o__Saccharomycetales(100);f__Metschnikowiaceae(100);g__Metschnikowia(100);s__Metschnikowia_pulcherrima(100); | Metschnikowia | 13 | Saprotroph | \|Undefined Saprotroph\| | Yeast | NULL | Probable | As nectar-tap_saprotroph (Põlme et al. 2020); Decay sugar-rich substrates (Põlme et al. 2020) | Põlme S, et al. 2020. Fungal Diversity 105:1-16 ((https://doi.org/10.1007/s13225-020-00466-2)) |
| Otu19818 | 1 | k__Fungi(100);p__Ascomycota(100);c__Saccharomycetes(100);o__Saccharomycetales(100);f__Metschnikowiaceae(100);g__Metschnikowia(100);s__Metschnikowia_pulcherrima(100); | Metschnikowia | 13 | Saprotroph | \|Undefined Saprotroph\| | Yeast | NULL | Probable | As nectar-tap_saprotroph (Põlme et al. 2020); Decay sugar-rich substrates (Põlme et al. 2020) | Põlme S, et al. 2020. Fungal Diversity 105:1-16 ((https://doi.org/10.1007/s13225-020-00466-2)) |
| Otu19819 | 1 | k__Fungi(100);p__Ascomycota(100);c__Saccharomycetes(100);o__Saccharomycetales(100);f__Metschnikowiaceae(100);g__Metschnikowia(100);s__Metschnikowia_pulcherrima(100); | Metschnikowia | 13 | Saprotroph | \|Undefined Saprotroph\| | Yeast | NULL | Probable | As nectar-tap_saprotroph (Põlme et al. 2020); Decay sugar-rich substrates (Põlme et al. 2020) | Põlme S, et al. 2020. Fungal Diversity 105:1-16 ((https://doi.org/10.1007/s13225-020-00466-2)) |
| Otu19820 | 1 | k__Fungi(100);p__Ascomycota(100);c__Saccharomycetes(100);o__Saccharomycetales(100);f__Metschnikowiaceae(100);g__Metschnikowia(100);s__Metschnikowia_pulcherrima(100); | Metschnikowia | 13 | Saprotroph | \|Undefined Saprotroph\| | Yeast | NULL | Probable | As nectar-tap_saprotroph (Põlme et al. 2020); Decay sugar-rich substrates (Põlme et al. 2020) | Põlme S, et al. 2020. Fungal Diversity 105:1-16 ((https://doi.org/10.1007/s13225-020-00466-2)) |
| Otu19821 | 1 | k__Fungi(100);p__Ascomycota(100);c__Saccharomycetes(100);o__Saccharomycetales(100);f__Metschnikowiaceae(100);g__Metschnikowia(100);s__Metschnikowia_pulcherrima(100); | Metschnikowia | 13 | Saprotroph | \|Undefined Saprotroph\| | Yeast | NULL | Probable | As nectar-tap_saprotroph (Põlme et al. 2020); Decay sugar-rich substrates (Põlme et al. 2020) | Põlme S, et al. 2020. Fungal Diversity 105:1-16 ((https://doi.org/10.1007/s13225-020-00466-2)) |
| Otu19822 | 1 | k__Fungi(100);p__Ascomycota(100);c__Saccharomycetes(100);o__Saccharomycetales(100);f__Metschnikowiaceae(100);g__Metschnikowia(100);s__Metschnikowia_pulcherrima(100); | Metschnikowia | 13 | Saprotroph | \|Undefined Saprotroph\| | Yeast | NULL | Probable | As nectar-tap_saprotroph (Põlme et al. 2020); Decay sugar-rich substrates (Põlme et al. 2020) | Põlme S, et al. 2020. Fungal Diversity 105:1-16 ((https://doi.org/10.1007/s13225-020-00466-2)) |
| Otu19823 | 1 | k__Fungi(100);p__Ascomycota(100);c__Saccharomycetes(100);o__Saccharomycetales(100);f__Metschnikowiaceae(100);g__Metschnikowia(100);s__Metschnikowia_pulcherrima(100); | Metschnikowia | 13 | Saprotroph | \|Undefined Saprotroph\| | Yeast | NULL | Probable | As nectar-tap_saprotroph (Põlme et al. 2020); Decay sugar-rich substrates (Põlme et al. 2020) | Põlme S, et al. 2020. Fungal Diversity 105:1-16 ((https://doi.org/10.1007/s13225-020-00466-2)) |
| Otu19824 | 1 | k__Fungi(100);p__Ascomycota(100);c__Saccharomycetes(100);o__Saccharomycetales(100);f__Saccharomycetales_family_Incertae_sedis(100);g__Candida(100);g__Candida_unclassified(100); | Candida | 13 | Pathotroph-Saprotroph-Symbiotroph | Animal Pathogen-Endophyte-Endosymbiont-Epiphyte-\|Undefined Saprotroph\| | Dimorphic Yeast-Yeast | NULL | Probable | As nectar-tap_saprotroph (Põlme et al. 2020); Decay sugar-rich substrates (Põlme et al. 2020); This asexual genus is polyphyletic and contain species that function very differently depend on what they are. Caution should be taken when interpreting guilds for Candida species. | Manolakaki D, et al. 2010. Virulence. 1: 367-375 ((https://doi.org/10.4161/viru.1.5.12796)); Mekbib SB, et al. 2011. Tropical Plant Pathology 36:233-240 ((https://doi.org/10.1590/S1982-56762011000400004)); Põlme S, et al. 2020. Fungal Diversity 105:1-16 ((https://doi.org/10.1007/s13225-020-00466-2)) |
| Otu19825 | 1 | k__Fungi(100);p__Ascomycota(100);c__Saccharomycetes(100);o__Saccharomycetales(100);f__Metschnikowiaceae(100);g__Metschnikowia(100);s__Metschnikowia_pulcherrima(100); | Metschnikowia | 13 | Saprotroph | \|Undefined Saprotroph\| | Yeast | NULL | Probable | As nectar-tap_saprotroph (Põlme et al. 2020); Decay sugar-rich substrates (Põlme et al. 2020) | Põlme S, et al. 2020. Fungal Diversity 105:1-16 ((https://doi.org/10.1007/s13225-020-00466-2)) |
| Otu19826 | 1 | k__Fungi(100);p__Ascomycota(100);c__Saccharomycetes(100);o__Saccharomycetales(100);f__Metschnikowiaceae(100);g__Metschnikowia(100);g__Metschnikowia_unclassified(100); | Metschnikowia | 13 | Saprotroph | \|Undefined Saprotroph\| | Yeast | NULL | Probable | As nectar-tap_saprotroph (Põlme et al. 2020); Decay sugar-rich substrates (Põlme et al. 2020) | Põlme S, et al. 2020. Fungal Diversity 105:1-16 ((https://doi.org/10.1007/s13225-020-00466-2)) |
| Otu19827 | 1 | k__Fungi(100);p__Ascomycota(100);c__Saccharomycetes(100);o__Saccharomycetales(100);f__Metschnikowiaceae(100);g__Metschnikowia(100);s__Metschnikowia_pulcherrima(100); | Metschnikowia | 13 | Saprotroph | \|Undefined Saprotroph\| | Yeast | NULL | Probable | As nectar-tap_saprotroph (Põlme et al. 2020); Decay sugar-rich substrates (Põlme et al. 2020) | Põlme S, et al. 2020. Fungal Diversity 105:1-16 ((https://doi.org/10.1007/s13225-020-00466-2)) |
| Otu19828 | 1 | k__Fungi(100);p__Ascomycota(100);c__Saccharomycetes(100);o__Saccharomycetales(100);f__Metschnikowiaceae(100);g__Metschnikowia(100);s__Metschnikowia_pulcherrima(100); | Metschnikowia | 13 | Saprotroph | \|Undefined Saprotroph\| | Yeast | NULL | Probable | As nectar-tap_saprotroph (Põlme et al. 2020); Decay sugar-rich substrates (Põlme et al. 2020) | Põlme S, et al. 2020. Fungal Diversity 105:1-16 ((https://doi.org/10.1007/s13225-020-00466-2)) |
| Otu19829 | 1 | k__Fungi(100);p__Ascomycota(100);c__Saccharomycetes(100);o__Saccharomycetales(100);f__Metschnikowiaceae(100);g__Metschnikowia(100);s__Metschnikowia_pulcherrima(100); | Metschnikowia | 13 | Saprotroph | \|Undefined Saprotroph\| | Yeast | NULL | Probable | As nectar-tap_saprotroph (Põlme et al. 2020); Decay sugar-rich substrates (Põlme et al. 2020) | Põlme S, et al. 2020. Fungal Diversity 105:1-16 ((https://doi.org/10.1007/s13225-020-00466-2)) |
| Otu19830 | 1 | k__Fungi(100);p__Ascomycota(100);c__Saccharomycetes(100);o__Saccharomycetales(100);f__Metschnikowiaceae(100);g__Metschnikowia(100);s__Metschnikowia_pulcherrima(100); | Metschnikowia | 13 | Saprotroph | \|Undefined Saprotroph\| | Yeast | NULL | Probable | As nectar-tap_saprotroph (Põlme et al. 2020); Decay sugar-rich substrates (Põlme et al. 2020) | Põlme S, et al. 2020. Fungal Diversity 105:1-16 ((https://doi.org/10.1007/s13225-020-00466-2)) |
| Otu19831 | 1 | k__Fungi(100);p__Ascomycota(100);c__Saccharomycetes(100);o__Saccharomycetales(100);f__Metschnikowiaceae(100);g__Metschnikowia(100);s__Metschnikowia_pulcherrima(100); | Metschnikowia | 13 | Saprotroph | \|Undefined Saprotroph\| | Yeast | NULL | Probable | As nectar-tap_saprotroph (Põlme et al. 2020); Decay sugar-rich substrates (Põlme et al. 2020) | Põlme S, et al. 2020. Fungal Diversity 105:1-16 ((https://doi.org/10.1007/s13225-020-00466-2)) |
| Otu19832 | 1 | k__Fungi(100);p__Ascomycota(100);c__Saccharomycetes(100);o__Saccharomycetales(100);f__Metschnikowiaceae(100);g__Metschnikowia(100);g__Metschnikowia_unclassified(100); | Metschnikowia | 13 | Saprotroph | \|Undefined Saprotroph\| | Yeast | NULL | Probable | As nectar-tap_saprotroph (Põlme et al. 2020); Decay sugar-rich substrates (Põlme et al. 2020) | Põlme S, et al. 2020. Fungal Diversity 105:1-16 ((https://doi.org/10.1007/s13225-020-00466-2)) |
| Otu19833 | 1 | k__Fungi(100);p__Ascomycota(100);c__Saccharomycetes(100);o__Saccharomycetales(100);f__Metschnikowiaceae(100);g__Metschnikowia(100);s__Metschnikowia_pulcherrima(100); | Metschnikowia | 13 | Saprotroph | \|Undefined Saprotroph\| | Yeast | NULL | Probable | As nectar-tap_saprotroph (Põlme et al. 2020); Decay sugar-rich substrates (Põlme et al. 2020) | Põlme S, et al. 2020. Fungal Diversity 105:1-16 ((https://doi.org/10.1007/s13225-020-00466-2)) |
| Otu19834 | 1 | k__Fungi(100);p__Ascomycota(100);c__Saccharomycetes(100);o__Saccharomycetales(100);f__Metschnikowiaceae(100);g__Metschnikowia(100);g__Metschnikowia_unclassified(100); | Metschnikowia | 13 | Saprotroph | \|Undefined Saprotroph\| | Yeast | NULL | Probable | As nectar-tap_saprotroph (Põlme et al. 2020); Decay sugar-rich substrates (Põlme et al. 2020) | Põlme S, et al. 2020. Fungal Diversity 105:1-16 ((https://doi.org/10.1007/s13225-020-00466-2)) |
| Otu19835 | 1 | k__Fungi(100);p__Ascomycota(100);c__Saccharomycetes(100);o__Saccharomycetales(100);f__Metschnikowiaceae(100);g__Metschnikowia(100);s__Metschnikowia_pulcherrima(100); | Metschnikowia | 13 | Saprotroph | \|Undefined Saprotroph\| | Yeast | NULL | Probable | As nectar-tap_saprotroph (Põlme et al. 2020); Decay sugar-rich substrates (Põlme et al. 2020) | Põlme S, et al. 2020. Fungal Diversity 105:1-16 ((https://doi.org/10.1007/s13225-020-00466-2)) |
| Otu19836 | 1 | k__Fungi(100);p__Ascomycota(100);c__Saccharomycetes(100);o__Saccharomycetales(100);f__Metschnikowiaceae(100);g__Metschnikowia(100);g__Metschnikowia_unclassified(100); | Metschnikowia | 13 | Saprotroph | \|Undefined Saprotroph\| | Yeast | NULL | Probable | As nectar-tap_saprotroph (Põlme et al. 2020); Decay sugar-rich substrates (Põlme et al. 2020) | Põlme S, et al. 2020. Fungal Diversity 105:1-16 ((https://doi.org/10.1007/s13225-020-00466-2)) |
| Otu19837 | 1 | k__Fungi(100);p__Ascomycota(100);c__Saccharomycetes(100);o__Saccharomycetales(100);f__Metschnikowiaceae(100);g__Metschnikowia(100);s__Metschnikowia_pulcherrima(100); | Metschnikowia | 13 | Saprotroph | \|Undefined Saprotroph\| | Yeast | NULL | Probable | As nectar-tap_saprotroph (Põlme et al. 2020); Decay sugar-rich substrates (Põlme et al. 2020) | Põlme S, et al. 2020. Fungal Diversity 105:1-16 ((https://doi.org/10.1007/s13225-020-00466-2)) |
| Otu19838 | 1 | k__Fungi(100);p__Ascomycota(100);c__Saccharomycetes(100);o__Saccharomycetales(100);o__Saccharomycetales_unclassified(100);o__Saccharomycetales_unclassified(100);o__Saccharomycetales_unclassified(100); | Saccharomycetales | 7 | Saprotroph | Undefined Saprotroph | Yeast | NULL | Possible | NULL | Sterkenburg E, et al. 2015. New Phytologist 207:1145-1158 ((https://doi.org/10.1111/nph.13426)) |
| Otu19839 | 1 | k__Fungi(100);p__Ascomycota(100);c__Saccharomycetes(100);o__Saccharomycetales(100);f__Metschnikowiaceae(100);g__Metschnikowia(100);g__Metschnikowia_unclassified(100); | Metschnikowia | 13 | Saprotroph | \|Undefined Saprotroph\| | Yeast | NULL | Probable | As nectar-tap_saprotroph (Põlme et al. 2020); Decay sugar-rich substrates (Põlme et al. 2020) | Põlme S, et al. 2020. Fungal Diversity 105:1-16 ((https://doi.org/10.1007/s13225-020-00466-2)) |
| Otu19840 | 1 | k__Fungi(100);p__Ascomycota(100);c__Saccharomycetes(100);o__Saccharomycetales(100);f__Metschnikowiaceae(100);g__Metschnikowia(100);s__Metschnikowia_pulcherrima(100); | Metschnikowia | 13 | Saprotroph | \|Undefined Saprotroph\| | Yeast | NULL | Probable | As nectar-tap_saprotroph (Põlme et al. 2020); Decay sugar-rich substrates (Põlme et al. 2020) | Põlme S, et al. 2020. Fungal Diversity 105:1-16 ((https://doi.org/10.1007/s13225-020-00466-2)) |
| Otu19841 | 1 | k__Fungi(100);p__Ascomycota(100);c__Saccharomycetes(100);o__Saccharomycetales(100);f__Metschnikowiaceae(100);g__Metschnikowia(100);s__Metschnikowia_pulcherrima(100); | Metschnikowia | 13 | Saprotroph | \|Undefined Saprotroph\| | Yeast | NULL | Probable | As nectar-tap_saprotroph (Põlme et al. 2020); Decay sugar-rich substrates (Põlme et al. 2020) | Põlme S, et al. 2020. Fungal Diversity 105:1-16 ((https://doi.org/10.1007/s13225-020-00466-2)) |
| Otu19842 | 1 | k__Fungi(100);p__Ascomycota(100);c__Saccharomycetes(100);o__Saccharomycetales(100);f__Metschnikowiaceae(100);g__Metschnikowia(100);g__Metschnikowia_unclassified(100); | Metschnikowia | 13 | Saprotroph | \|Undefined Saprotroph\| | Yeast | NULL | Probable | As nectar-tap_saprotroph (Põlme et al. 2020); Decay sugar-rich substrates (Põlme et al. 2020) | Põlme S, et al. 2020. Fungal Diversity 105:1-16 ((https://doi.org/10.1007/s13225-020-00466-2)) |
| Otu19843 | 1 | k__Fungi(100);p__Ascomycota(100);c__Saccharomycetes(100);o__Saccharomycetales(100);f__Metschnikowiaceae(100);g__Metschnikowia(100);g__Metschnikowia_unclassified(100); | Metschnikowia | 13 | Saprotroph | \|Undefined Saprotroph\| | Yeast | NULL | Probable | As nectar-tap_saprotroph (Põlme et al. 2020); Decay sugar-rich substrates (Põlme et al. 2020) | Põlme S, et al. 2020. Fungal Diversity 105:1-16 ((https://doi.org/10.1007/s13225-020-00466-2)) |
| Otu19844 | 1 | k__Fungi(100);p__Ascomycota(100);c__Saccharomycetes(100);o__Saccharomycetales(100);f__Metschnikowiaceae(100);g__Metschnikowia(100);s__Metschnikowia_pulcherrima(100); | Metschnikowia | 13 | Saprotroph | \|Undefined Saprotroph\| | Yeast | NULL | Probable | As nectar-tap_saprotroph (Põlme et al. 2020); Decay sugar-rich substrates (Põlme et al. 2020) | Põlme S, et al. 2020. Fungal Diversity 105:1-16 ((https://doi.org/10.1007/s13225-020-00466-2)) |
| Otu19845 | 1 | k__Fungi(100);p__Ascomycota(100);c__Saccharomycetes(100);o__Saccharomycetales(100);f__Metschnikowiaceae(100);g__Metschnikowia(100);g__Metschnikowia_unclassified(100); | Metschnikowia | 13 | Saprotroph | \|Undefined Saprotroph\| | Yeast | NULL | Probable | As nectar-tap_saprotroph (Põlme et al. 2020); Decay sugar-rich substrates (Põlme et al. 2020) | Põlme S, et al. 2020. Fungal Diversity 105:1-16 ((https://doi.org/10.1007/s13225-020-00466-2)) |
| Otu19846 | 1 | k__Fungi(100);p__Ascomycota(100);c__Saccharomycetes(100);o__Saccharomycetales(100);f__Metschnikowiaceae(100);g__Metschnikowia(100);s__Metschnikowia_pulcherrima(100); | Metschnikowia | 13 | Saprotroph | \|Undefined Saprotroph\| | Yeast | NULL | Probable | As nectar-tap_saprotroph (Põlme et al. 2020); Decay sugar-rich substrates (Põlme et al. 2020) | Põlme S, et al. 2020. Fungal Diversity 105:1-16 ((https://doi.org/10.1007/s13225-020-00466-2)) |
| Otu19847 | 1 | k__Fungi(100);p__Ascomycota(100);c__Saccharomycetes(100);o__Saccharomycetales(100);f__Metschnikowiaceae(100);g__Metschnikowia(100);s__Metschnikowia_pulcherrima(100); | Metschnikowia | 13 | Saprotroph | \|Undefined Saprotroph\| | Yeast | NULL | Probable | As nectar-tap_saprotroph (Põlme et al. 2020); Decay sugar-rich substrates (Põlme et al. 2020) | Põlme S, et al. 2020. Fungal Diversity 105:1-16 ((https://doi.org/10.1007/s13225-020-00466-2)) |
| Otu19848 | 1 | k__Fungi(100);p__Ascomycota(100);c__Saccharomycetes(100);o__Saccharomycetales(100);f__Metschnikowiaceae(100);g__Metschnikowia(100);s__Metschnikowia_pulcherrima(100); | Metschnikowia | 13 | Saprotroph | \|Undefined Saprotroph\| | Yeast | NULL | Probable | As nectar-tap_saprotroph (Põlme et al. 2020); Decay sugar-rich substrates (Põlme et al. 2020) | Põlme S, et al. 2020. Fungal Diversity 105:1-16 ((https://doi.org/10.1007/s13225-020-00466-2)) |
| Otu19849 | 1 | k__Fungi(100);p__Ascomycota(100);c__Saccharomycetes(100);o__Saccharomycetales(100);f__Metschnikowiaceae(100);g__Metschnikowia(100);s__Metschnikowia_pulcherrima(100); | Metschnikowia | 13 | Saprotroph | \|Undefined Saprotroph\| | Yeast | NULL | Probable | As nectar-tap_saprotroph (Põlme et al. 2020); Decay sugar-rich substrates (Põlme et al. 2020) | Põlme S, et al. 2020. Fungal Diversity 105:1-16 ((https://doi.org/10.1007/s13225-020-00466-2)) |
| Otu19851 | 1 | k__Fungi(100);p__Ascomycota(100);c__Saccharomycetes(100);o__Saccharomycetales(100);f__Metschnikowiaceae(100);g__Metschnikowia(100);s__Metschnikowia_pulcherrima(100); | Metschnikowia | 13 | Saprotroph | \|Undefined Saprotroph\| | Yeast | NULL | Probable | As nectar-tap_saprotroph (Põlme et al. 2020); Decay sugar-rich substrates (Põlme et al. 2020) | Põlme S, et al. 2020. Fungal Diversity 105:1-16 ((https://doi.org/10.1007/s13225-020-00466-2)) |
| Otu19852 | 1 | k__Fungi(100);p__Ascomycota(100);c__Saccharomycetes(100);o__Saccharomycetales(100);f__Metschnikowiaceae(100);g__Metschnikowia(100);g__Metschnikowia_unclassified(100); | Metschnikowia | 13 | Saprotroph | \|Undefined Saprotroph\| | Yeast | NULL | Probable | As nectar-tap_saprotroph (Põlme et al. 2020); Decay sugar-rich substrates (Põlme et al. 2020) | Põlme S, et al. 2020. Fungal Diversity 105:1-16 ((https://doi.org/10.1007/s13225-020-00466-2)) |
| Otu19853 | 1 | k__Fungi(100);p__Ascomycota(100);c__Saccharomycetes(100);o__Saccharomycetales(100);f__Metschnikowiaceae(100);g__Metschnikowia(100);s__Metschnikowia_pulcherrima(100); | Metschnikowia | 13 | Saprotroph | \|Undefined Saprotroph\| | Yeast | NULL | Probable | As nectar-tap_saprotroph (Põlme et al. 2020); Decay sugar-rich substrates (Põlme et al. 2020) | Põlme S, et al. 2020. Fungal Diversity 105:1-16 ((https://doi.org/10.1007/s13225-020-00466-2)) |
| Otu19854 | 1 | k__Fungi(100);p__Ascomycota(100);c__Saccharomycetes(100);o__Saccharomycetales(100);f__Metschnikowiaceae(100);g__Metschnikowia(100);s__Metschnikowia_pulcherrima(100); | Metschnikowia | 13 | Saprotroph | \|Undefined Saprotroph\| | Yeast | NULL | Probable | As nectar-tap_saprotroph (Põlme et al. 2020); Decay sugar-rich substrates (Põlme et al. 2020) | Põlme S, et al. 2020. Fungal Diversity 105:1-16 ((https://doi.org/10.1007/s13225-020-00466-2)) |
| Otu19855 | 1 | k__Fungi(100);p__Ascomycota(100);c__Saccharomycetes(100);o__Saccharomycetales(100);f__Metschnikowiaceae(100);g__Metschnikowia(100);s__Metschnikowia_pulcherrima(100); | Metschnikowia | 13 | Saprotroph | \|Undefined Saprotroph\| | Yeast | NULL | Probable | As nectar-tap_saprotroph (Põlme et al. 2020); Decay sugar-rich substrates (Põlme et al. 2020) | Põlme S, et al. 2020. Fungal Diversity 105:1-16 ((https://doi.org/10.1007/s13225-020-00466-2)) |
| Otu19856 | 1 | k__Fungi(100);p__Ascomycota(100);c__Saccharomycetes(100);o__Saccharomycetales(100);f__Metschnikowiaceae(100);g__Metschnikowia(100);s__Metschnikowia_pulcherrima(100); | Metschnikowia | 13 | Saprotroph | \|Undefined Saprotroph\| | Yeast | NULL | Probable | As nectar-tap_saprotroph (Põlme et al. 2020); Decay sugar-rich substrates (Põlme et al. 2020) | Põlme S, et al. 2020. Fungal Diversity 105:1-16 ((https://doi.org/10.1007/s13225-020-00466-2)) |
| Otu19857 | 1 | k__Fungi(100);p__Ascomycota(100);c__Saccharomycetes(100);o__Saccharomycetales(100);f__Metschnikowiaceae(100);g__Metschnikowia(100);s__Metschnikowia_pulcherrima(100); | Metschnikowia | 13 | Saprotroph | \|Undefined Saprotroph\| | Yeast | NULL | Probable | As nectar-tap_saprotroph (Põlme et al. 2020); Decay sugar-rich substrates (Põlme et al. 2020) | Põlme S, et al. 2020. Fungal Diversity 105:1-16 ((https://doi.org/10.1007/s13225-020-00466-2)) |
| Otu19859 | 1 | k__Fungi(100);p__Ascomycota(100);c__Saccharomycetes(100);o__Saccharomycetales(100);f__Metschnikowiaceae(100);g__Metschnikowia(100);s__Metschnikowia_pulcherrima(100); | Metschnikowia | 13 | Saprotroph | \|Undefined Saprotroph\| | Yeast | NULL | Probable | As nectar-tap_saprotroph (Põlme et al. 2020); Decay sugar-rich substrates (Põlme et al. 2020) | Põlme S, et al. 2020. Fungal Diversity 105:1-16 ((https://doi.org/10.1007/s13225-020-00466-2)) |
| Otu19860 | 1 | k__Fungi(100);p__Ascomycota(100);c__Saccharomycetes(100);o__Saccharomycetales(100);f__Metschnikowiaceae(100);g__Metschnikowia(100);s__Metschnikowia_pulcherrima(100); | Metschnikowia | 13 | Saprotroph | \|Undefined Saprotroph\| | Yeast | NULL | Probable | As nectar-tap_saprotroph (Põlme et al. 2020); Decay sugar-rich substrates (Põlme et al. 2020) | Põlme S, et al. 2020. Fungal Diversity 105:1-16 ((https://doi.org/10.1007/s13225-020-00466-2)) |
| Otu19861 | 1 | k__Fungi(100);p__Ascomycota(100);c__Saccharomycetes(100);o__Saccharomycetales(100);f__Metschnikowiaceae(100);g__Metschnikowia(100);s__Metschnikowia_pulcherrima(100); | Metschnikowia | 13 | Saprotroph | \|Undefined Saprotroph\| | Yeast | NULL | Probable | As nectar-tap_saprotroph (Põlme et al. 2020); Decay sugar-rich substrates (Põlme et al. 2020) | Põlme S, et al. 2020. Fungal Diversity 105:1-16 ((https://doi.org/10.1007/s13225-020-00466-2)) |
| Otu19862 | 1 | k__Fungi(100);p__Ascomycota(100);c__Saccharomycetes(100);o__Saccharomycetales(100);f__Metschnikowiaceae(100);g__Metschnikowia(100);g__Metschnikowia_unclassified(100); | Metschnikowia | 13 | Saprotroph | \|Undefined Saprotroph\| | Yeast | NULL | Probable | As nectar-tap_saprotroph (Põlme et al. 2020); Decay sugar-rich substrates (Põlme et al. 2020) | Põlme S, et al. 2020. Fungal Diversity 105:1-16 ((https://doi.org/10.1007/s13225-020-00466-2)) |
| Otu19863 | 1 | k__Fungi(100);p__Ascomycota(100);c__Saccharomycetes(100);o__Saccharomycetales(100);f__Metschnikowiaceae(100);g__Metschnikowia(100);g__Metschnikowia_unclassified(100); | Metschnikowia | 13 | Saprotroph | \|Undefined Saprotroph\| | Yeast | NULL | Probable | As nectar-tap_saprotroph (Põlme et al. 2020); Decay sugar-rich substrates (Põlme et al. 2020) | Põlme S, et al. 2020. Fungal Diversity 105:1-16 ((https://doi.org/10.1007/s13225-020-00466-2)) |
| Otu19864 | 1 | k__Fungi(100);p__Ascomycota(100);c__Saccharomycetes(100);o__Saccharomycetales(100);f__Metschnikowiaceae(100);g__Metschnikowia(100);s__Metschnikowia_pulcherrima(100); | Metschnikowia | 13 | Saprotroph | \|Undefined Saprotroph\| | Yeast | NULL | Probable | As nectar-tap_saprotroph (Põlme et al. 2020); Decay sugar-rich substrates (Põlme et al. 2020) | Põlme S, et al. 2020. Fungal Diversity 105:1-16 ((https://doi.org/10.1007/s13225-020-00466-2)) |

| | | | | | | | | | | | |
|---|---|---|---|---|---|---|---|---|---|---|---|
| Otu20150 | 1 | k__Fungi(100);p__Ascomycota(100);c__Leotiomycetes(100);o__Helotiales(100);f__Vibrisseaceae(100);g__Phialocephala(100);g__Phialocephala_unclassified(100); | Phialocephala | 13 | Saprotroph-Symbiotroph | Endophyte-\|Undefined Saprotroph\| | Microfungus | NULL | Highly Probable | As root_endophyte (Põlme et al. 2020); As root_endophyte_dark_septate (Põlme et al. 2020); As soil_saprotroph (Põlme et al. 2020); Dark Septate Endophyte, known from plant roots.; Decay roots, soil (Põlme et al. 2020) | Newsham KK. 2011. New Phytologist 190:783-793 ((https://doi.org/10.1111/j.1469-8137.2010.03611.x)) (as dark septate root endophytes); Põlme S, et al. 2020. Fungal Diversity 105:1-16 ((https://doi.org/10.1007/s13225-020-00466-2)) |
| Otu20155 | 1 | k__Fungi(100);p__Ascomycota(100);c__Saccharomycetes(100);o__Saccharomycetales(100);o__Saccharomycetales_unclassified(100);o__Saccharomycetales_unclassified(100);o__Saccharomycetales_unclassified(100); | Saccharomycetales | 7 | Saprotroph | Undefined Saprotroph | Yeast | NULL | Possible | NULL | Sterkenburg E, et al. 2015. New Phytologist 207:1145-1158 ((https://doi.org/10.1111/nph.13426)) |
| Otu20159 | 1 | k__Fungi(100);p__Ascomycota(100);c__Saccharomycetes(100);o__Saccharomycetales(100);f__Metschnikowiaceae(100);g__Metschnikowia(100);s__Metschnikowia_pulcherrima(100); | Metschnikowia | 13 | Saprotroph | \|Undefined Saprotroph\| | Yeast | NULL | Probable | As nectar-tap_saprotroph (Põlme et al. 2020); Decay sugar-rich substrates (Põlme et al. 2020) | Põlme S, et al. 2020. Fungal Diversity 105:1-16 ((https://doi.org/10.1007/s13225-020-00466-2)) |
| Otu20161 | 1 | k__Fungi(100);p__Ascomycota(100);c__Saccharomycetes(100);o__Saccharomycetales(100);f__Metschnikowiaceae(100);g__Metschnikowia(100);s__Metschnikowia_pulcherrima(100); | Metschnikowia | 13 | Saprotroph | \|Undefined Saprotroph\| | Yeast | NULL | Probable | As nectar-tap_saprotroph (Põlme et al. 2020); Decay sugar-rich substrates (Põlme et al. 2020) | Põlme S, et al. 2020. Fungal Diversity 105:1-16 ((https://doi.org/10.1007/s13225-020-00466-2)) |
| Otu20162 | 1 | k__Fungi(100);p__Ascomycota(100);c__Saccharomycetes(100);o__Saccharomycetales(100);f__Metschnikowiaceae(100);g__Metschnikowia(100);s__Metschnikowia_pulcherrima(100); | Metschnikowia | 13 | Saprotroph | \|Undefined Saprotroph\| | Yeast | NULL | Probable | As nectar-tap_saprotroph (Põlme et al. 2020); Decay sugar-rich substrates (Põlme et al. 2020) | Põlme S, et al. 2020. Fungal Diversity 105:1-16 ((https://doi.org/10.1007/s13225-020-00466-2)) |
| Otu20165 | 1 | k__Fungi(100);p__Ascomycota(100);c__Saccharomycetes(100);o__Saccharomycetales(100);f__Saccharomycetales_family_Incertae_sedis(100);g__Candida(100);g__Candida_unclassified(100); | Candida | 13 | Pathotroph-Saprotroph-Symbiotroph | Animal Pathogen-Endophyte-Endosymbiont-Epiphyte-\|Undefined Saprotroph\| | Dimorphic Yeast-Yeast | NULL | Probable | As nectar-tap_saprotroph (Põlme et al. 2020); Decay sugar-rich substrates (Põlme et al. 2020); This asexual genus is polyphyletic and contain species that function very differently depend on what they are. Caution should be taken when interpreting guilds for Candida species. | Manolakaki D, et al. 2010. Virulence. 1: 367-375 ((https://doi.org/10.4161/viru.1.5.12796)); Mekbib SB, et al. 2011. Tropical Plant Pathology 36:233-240 ((https://doi.org/10.1590/S1982-56762011000400004)); Põlme S, et al. 2020. Fungal Diversity 105:1-16 ((https://doi.org/10.1007/s13225-020-00466-2)) |
| Otu20167 | 1 | k__Fungi(100);p__Ascomycota(100);c__Saccharomycetes(100);o__Saccharomycetales(100);f__Metschnikowiaceae(100);g__Metschnikowia(100);g__Metschnikowia_unclassified(100); | Metschnikowia | 13 | Saprotroph | \|Undefined Saprotroph\| | Yeast | NULL | Probable | As nectar-tap_saprotroph (Põlme et al. 2020); Decay sugar-rich substrates (Põlme et al. 2020) | Põlme S, et al. 2020. Fungal Diversity 105:1-16 ((https://doi.org/10.1007/s13225-020-00466-2)) |
| Otu20168 | 1 | k__Fungi(100);p__Ascomycota(100);c__Saccharomycetes(100);o__Saccharomycetales(100);f__Saccharomycetales_family_Incertae_sedis(100);g__Candida(100);g__Candida_unclassified(100); | Candida | 13 | Pathotroph-Saprotroph-Symbiotroph | Animal Pathogen-Endophyte-Endosymbiont-Epiphyte-\|Undefined Saprotroph\| | Dimorphic Yeast-Yeast | NULL | Probable | As nectar-tap_saprotroph (Põlme et al. 2020); Decay sugar-rich substrates (Põlme et al. 2020); This asexual genus is polyphyletic and contain species that function very differently depend on what they are. Caution should be taken when interpreting guilds for Candida species. | Manolakaki D, et al. 2010. Virulence. 1: 367-375 ((https://doi.org/10.4161/viru.1.5.12796)); Mekbib SB, et al. 2011. Tropical Plant Pathology 36:233-240 ((https://doi.org/10.1590/S1982-56762011000400004)); Põlme S, et al. 2020. Fungal Diversity 105:1-16 ((https://doi.org/10.1007/s13225-020-00466-2)) |
| Otu20169 | 1 | k__Fungi(100);p__Ascomycota(100);c__Saccharomycetes(100);o__Saccharomycetales(100);f__Saccharomycetales_family_Incertae_sedis(100);f__Saccharomycetales_family_Incertae_sedis_unclassified(100);f__Saccharomycetales_family_Incertae_sedis_unclassified(100); | Saccharomycetales | 7 | Saprotroph | Undefined Saprotroph | Yeast | NULL | Possible | NULL | Sterkenburg E, et al. 2015. New Phytologist 207:1145-1158 ((https://doi.org/10.1111/nph.13426)) |
| Otu20170 | 1 | k__Fungi(100);p__Ascomycota(100);c__Saccharomycetes(100);o__Saccharomycetales(100);f__Saccharomycetales_family_Incertae_sedis(100);g__Candida(100);s__Candida_cretensis(100); | Candida | 13 | Pathotroph-Saprotroph-Symbiotroph | Animal Pathogen-Endophyte-Endosymbiont-Epiphyte-\|Undefined Saprotroph\| | Dimorphic Yeast-Yeast | NULL | Probable | As nectar-tap_saprotroph (Põlme et al. 2020); Decay sugar-rich substrates (Põlme et al. 2020); This asexual genus is polyphyletic and contain species that function very differently depend on what they are. Caution should be taken when interpreting guilds for Candida species. | Manolakaki D, et al. 2010. Virulence. 1: 367-375 ((https://doi.org/10.4161/viru.1.5.12796)); Mekbib SB, et al. 2011. Tropical Plant Pathology 36:233-240 ((https://doi.org/10.1590/S1982-56762011000400004)); Põlme S, et al. 2020. Fungal Diversity 105:1-16 ((https://doi.org/10.1007/s13225-020-00466-2)) |
| Otu20176 | 1 | k__Fungi(100);p__Ascomycota(100);c__Saccharomycetes(100);o__Saccharomycetales(100);f__Metschnikowiaceae(100);g__Metschnikowia(100);s__Metschnikowia_pulcherrima(100); | Metschnikowia | 13 | Saprotroph | \|Undefined Saprotroph\| | Yeast | NULL | Probable | As nectar-tap_saprotroph (Põlme et al. 2020); Decay sugar-rich substrates (Põlme et al. 2020) | Põlme S, et al. 2020. Fungal Diversity 105:1-16 ((https://doi.org/10.1007/s13225-020-00466-2)) |
| Otu20177 | 1 | k__Fungi(100);p__Ascomycota(100);c__Saccharomycetes(100);o__Saccharomycetales(100);f__Metschnikowiaceae(100);g__Metschnikowia(100);s__Metschnikowia_pulcherrima(100); | Metschnikowia | 13 | Saprotroph | \|Undefined Saprotroph\| | Yeast | NULL | Probable | As nectar-tap_saprotroph (Põlme et al. 2020); Decay sugar-rich substrates (Põlme et al. 2020) | Põlme S, et al. 2020. Fungal Diversity 105:1-16 ((https://doi.org/10.1007/s13225-020-00466-2)) |
| Otu20180 | 1 | k__Fungi(100);p__Ascomycota(100);c__Saccharomycetes(100);o__Saccharomycetales(100);f__Metschnikowiaceae(100);g__Metschnikowia(100);g__Metschnikowia_unclassified(100); | Metschnikowia | 13 | Saprotroph | \|Undefined Saprotroph\| | Yeast | NULL | Probable | As nectar-tap_saprotroph (Põlme et al. 2020); Decay sugar-rich substrates (Põlme et al. 2020) | Põlme S, et al. 2020. Fungal Diversity 105:1-16 ((https://doi.org/10.1007/s13225-020-00466-2)) |
| Otu20181 | 1 | k__Fungi(100);p__Ascomycota(100);c__Saccharomycetes(100);o__Saccharomycetales(100);f__Metschnikowiaceae(100);g__Metschnikowia(100);s__Metschnikowia_pulcherrima(100); | Metschnikowia | 13 | Saprotroph | \|Undefined Saprotroph\| | Yeast | NULL | Probable | As nectar-tap_saprotroph (Põlme et al. 2020); Decay sugar-rich substrates (Põlme et al. 2020) | Põlme S, et al. 2020. Fungal Diversity 105:1-16 ((https://doi.org/10.1007/s13225-020-00466-2)) |
| Otu20182 | 1 | k__Fungi(100);p__Ascomycota(100);c__Saccharomycetes(100);o__Saccharomycetales(100);f__Metschnikowiaceae(100);g__Metschnikowia(100);g__Metschnikowia_unclassified(100); | Metschnikowia | 13 | Saprotroph | \|Undefined Saprotroph\| | Yeast | NULL | Probable | As nectar-tap_saprotroph (Põlme et al. 2020); Decay sugar-rich substrates (Põlme et al. 2020) | Põlme S, et al. 2020. Fungal Diversity 105:1-16 ((https://doi.org/10.1007/s13225-020-00466-2)) |
| Otu20183 | 1 | k__Fungi(100);p__Ascomycota(100);c__Saccharomycetes(100);o__Saccharomycetales(100);f__Metschnikowiaceae(100);g__Metschnikowia(100);g__Metschnikowia_unclassified(100); | Metschnikowia | 13 | Saprotroph | \|Undefined Saprotroph\| | Yeast | NULL | Probable | As nectar-tap_saprotroph (Põlme et al. 2020); Decay sugar-rich substrates (Põlme et al. 2020) | Põlme S, et al. 2020. Fungal Diversity 105:1-16 ((https://doi.org/10.1007/s13225-020-00466-2)) |
| Otu20184 | 1 | k__Fungi(100);p__Ascomycota(100);c__Saccharomycetes(100);o__Saccharomycetales(100);f__Metschnikowiaceae(100);g__Metschnikowia(100);s__Metschnikowia_pulcherrima(100); | Metschnikowia | 13 | Saprotroph | \|Undefined Saprotroph\| | Yeast | NULL | Probable | As nectar-tap_saprotroph (Põlme et al. 2020); Decay sugar-rich substrates (Põlme et al. 2020) | Põlme S, et al. 2020. Fungal Diversity 105:1-16 ((https://doi.org/10.1007/s13225-020-00466-2)) |
| Otu20186 | 1 | k__Fungi(100);p__Ascomycota(100);c__Saccharomycetes(100);o__Saccharomycetales(100);f__Saccharomycetales_family_Incertae_sedis(100);f__Saccharomycetales_family_Incertae_sedis_unclassified(100);f__Saccharomycetales_family_Incertae_sedis_unclassified(100); | Saccharomycetales | 7 | Saprotroph | Undefined Saprotroph | Yeast | NULL | Possible | NULL | Sterkenburg E, et al. 2015. New Phytologist 207:1145-1158 ((https://doi.org/10.1111/nph.13426)) |
| Otu20187 | 1 | k__Fungi(100);p__Ascomycota(100);c__Saccharomycetes(100);o__Saccharomycetales(100);f__Metschnikowiaceae(100);g__Metschnikowia(100);s__Metschnikowia_pulcherrima(100); | Metschnikowia | 13 | Saprotroph | \|Undefined Saprotroph\| | Yeast | NULL | Probable | As nectar-tap_saprotroph (Põlme et al. 2020); Decay sugar-rich substrates (Põlme et al. 2020) | Põlme S, et al. 2020. Fungal Diversity 105:1-16 ((https://doi.org/10.1007/s13225-020-00466-2)) |
| Otu20189 | 1 | k__Fungi(100);p__Ascomycota(100);c__Saccharomycetes(100);o__Saccharomycetales(100);f__Metschnikowiaceae(100);g__Metschnikowia(100);s__Metschnikowia_pulcherrima(100); | Metschnikowia | 13 | Saprotroph | \|Undefined Saprotroph\| | Yeast | NULL | Probable | As nectar-tap_saprotroph (Põlme et al. 2020); Decay sugar-rich substrates (Põlme et al. 2020) | Põlme S, et al. 2020. Fungal Diversity 105:1-16 ((https://doi.org/10.1007/s13225-020-00466-2)) |
| Otu20190 | 1 | k__Fungi(100);p__Ascomycota(100);c__Saccharomycetes(100);o__Saccharomycetales(100);f__Metschnikowiaceae(100);g__Metschnikowia(100);g__Metschnikowia_unclassified(100); | Metschnikowia | 13 | Saprotroph | \|Undefined Saprotroph\| | Yeast | NULL | Probable | As nectar-tap_saprotroph (Põlme et al. 2020); Decay sugar-rich substrates (Põlme et al. 2020) | Põlme S, et al. 2020. Fungal Diversity 105:1-16 ((https://doi.org/10.1007/s13225-020-00466-2)) |
| Otu20191 | 1 | k__Fungi(100);p__Ascomycota(100);c__Saccharomycetes(100);o__Saccharomycetales(100);f__Metschnikowiaceae(100);g__Metschnikowia(100);s__Metschnikowia_pulcherrima(100); | Metschnikowia | 13 | Saprotroph | \|Undefined Saprotroph\| | Yeast | NULL | Probable | As nectar-tap_saprotroph (Põlme et al. 2020); Decay sugar-rich substrates (Põlme et al. 2020) | Põlme S, et al. 2020. Fungal Diversity 105:1-16 ((https://doi.org/10.1007/s13225-020-00466-2)) |
| Otu20192 | 1 | k__Fungi(100);p__Ascomycota(100);c__Saccharomycetes(100);o__Saccharomycetales(100);f__Metschnikowiaceae(100);g__Metschnikowia(100);s__Metschnikowia_pulcherrima(100); | Metschnikowia | 13 | Saprotroph | \|Undefined Saprotroph\| | Yeast | NULL | Probable | As nectar-tap_saprotroph (Põlme et al. 2020); Decay sugar-rich substrates (Põlme et al. 2020) | Põlme S, et al. 2020. Fungal Diversity 105:1-16 ((https://doi.org/10.1007/s13225-020-00466-2)) |
| Otu20193 | 1 | k__Fungi(100);p__Ascomycota(100);c__Saccharomycetes(100);o__Saccharomycetales(100);f__Metschnikowiaceae(100);g__Metschnikowia(100);g__Metschnikowia_unclassified(100); | Metschnikowia | 13 | Saprotroph | \|Undefined Saprotroph\| | Yeast | NULL | Probable | As nectar-tap_saprotroph (Põlme et al. 2020); Decay sugar-rich substrates (Põlme et al. 2020) | Põlme S, et al. 2020. Fungal Diversity 105:1-16 ((https://doi.org/10.1007/s13225-020-00466-2)) |
| Otu20194 | 1 | k__Fungi(100);p__Ascomycota(100);c__Saccharomycetes(100);o__Saccharomycetales(100);f__Metschnikowiaceae(100);g__Metschnikowia(100);s__Metschnikowia_pulcherrima(100); | Metschnikowia | 13 | Saprotroph | \|Undefined Saprotroph\| | Yeast | NULL | Probable | As nectar-tap_saprotroph (Põlme et al. 2020); Decay sugar-rich substrates (Põlme et al. 2020) | Põlme S, et al. 2020. Fungal Diversity 105:1-16 ((https://doi.org/10.1007/s13225-020-00466-2)) |
| Otu20195 | 1 | k__Fungi(100);p__Ascomycota(100);c__Saccharomycetes(100);o__Saccharomycetales(100);f__Metschnikowiaceae(100);g__Metschnikowia(100);s__Metschnikowia_pulcherrima(100); | Metschnikowia | 13 | Saprotroph | \|Undefined Saprotroph\| | Yeast | NULL | Probable | As nectar-tap_saprotroph (Põlme et al. 2020); Decay sugar-rich substrates (Põlme et al. 2020) | Põlme S, et al. 2020. Fungal Diversity 105:1-16 ((https://doi.org/10.1007/s13225-020-00466-2)) |
| Otu20196 | 1 | k__Fungi(100);p__Ascomycota(100);c__Saccharomycetes(100);o__Saccharomycetales(100);f__Metschnikowiaceae(100);g__Metschnikowia(100);g__Metschnikowia_unclassified(100); | Metschnikowia | 13 | Saprotroph | \|Undefined Saprotroph\| | Yeast | NULL | Probable | As nectar-tap_saprotroph (Põlme et al. 2020); Decay sugar-rich substrates (Põlme et al. 2020) | Põlme S, et al. 2020. Fungal Diversity 105:1-16 ((https://doi.org/10.1007/s13225-020-00466-2)) |
| Otu20197 | 1 | k__Fungi(100);p__Ascomycota(100);c__Saccharomycetes(100);o__Saccharomycetales(100);f__Metschnikowiaceae(100);g__Metschnikowia(100);s__Metschnikowia_pulcherrima(100); | Metschnikowia | 13 | Saprotroph | \|Undefined Saprotroph\| | Yeast | NULL | Probable | As nectar-tap_saprotroph (Põlme et al. 2020); Decay sugar-rich substrates (Põlme et al. 2020) | Põlme S, et al. 2020. Fungal Diversity 105:1-16 ((https://doi.org/10.1007/s13225-020-00466-2)) |
| Otu20198 | 1 | k__Fungi(100);p__Ascomycota(100);c__Saccharomycetes(100);o__Saccharomycetales(100);f__Metschnikowiaceae(100);g__Metschnikowia(100);s__Metschnikowia_pulcherrima(100); | Metschnikowia | 13 | Saprotroph | \|Undefined Saprotroph\| | Yeast | NULL | Probable | As nectar-tap_saprotroph (Põlme et al. 2020); Decay sugar-rich substrates (Põlme et al. 2020) | Põlme S, et al. 2020. Fungal Diversity 105:1-16 ((https://doi.org/10.1007/s13225-020-00466-2)) |
| Otu20199 | 1 | k__Fungi(100);p__Ascomycota(100);c__Saccharomycetes(100);o__Saccharomycetales(100);f__Metschnikowiaceae(100);g__Metschnikowia(100);g__Metschnikowia_unclassified(100); | Metschnikowia | 13 | Saprotroph | \|Undefined Saprotroph\| | Yeast | NULL | Probable | As nectar-tap_saprotroph (Põlme et al. 2020); Decay sugar-rich substrates (Põlme et al. 2020) | Põlme S, et al. 2020. Fungal Diversity 105:1-16 ((https://doi.org/10.1007/s13225-020-00466-2)) |
| Otu20200 | 1 | k__Fungi(100);p__Ascomycota(100);c__Saccharomycetes(100);o__Saccharomycetales(100);f__Metschnikowiaceae(100);g__Metschnikowia(100);s__Metschnikowia_pulcherrima(100); | Metschnikowia | 13 | Saprotroph | \|Undefined Saprotroph\| | Yeast | NULL | Probable | As nectar-tap_saprotroph (Põlme et al. 2020); Decay sugar-rich substrates (Põlme et al. 2020) | Põlme S, et al. 2020. Fungal Diversity 105:1-16 ((https://doi.org/10.1007/s13225-020-00466-2)) |
| Otu20201 | 1 | k__Fungi(100);p__Ascomycota(100);c__Saccharomycetes(100);o__Saccharomycetales(100);f__Metschnikowiaceae(100);g__Metschnikowia(100);s__Metschnikowia_pulcherrima(100); | Metschnikowia | 13 | Saprotroph | \|Undefined Saprotroph\| | Yeast | NULL | Probable | As nectar-tap_saprotroph (Põlme et al. 2020); Decay sugar-rich substrates (Põlme et al. 2020) | Põlme S, et al. 2020. Fungal Diversity 105:1-16 ((https://doi.org/10.1007/s13225-020-00466-2)) |
| Otu20202 | 1 | k__Fungi(100);p__Ascomycota(100);c__Saccharomycetes(100);o__Saccharomycetales(100);f__Metschnikowiaceae(100);g__Metschnikowia(100);s__Metschnikowia_pulcherrima(100); | Metschnikowia | 13 | Saprotroph | \|Undefined Saprotroph\| | Yeast | NULL | Probable | As nectar-tap_saprotroph (Põlme et al. 2020); Decay sugar-rich substrates (Põlme et al. 2020) | Põlme S, et al. 2020. Fungal Diversity 105:1-16 ((https://doi.org/10.1007/s13225-020-00466-2)) |
| Otu20203 | 1 | k__Fungi(100);p__Ascomycota(100);c__Saccharomycetes(100);o__Saccharomycetales(100);f__Metschnikowiaceae(100);g__Metschnikowia(100);s__Metschnikowia_pulcherrima(100); | Metschnikowia | 13 | Saprotroph | \|Undefined Saprotroph\| | Yeast | NULL | Probable | As nectar-tap_saprotroph (Põlme et al. 2020); Decay sugar-rich substrates (Põlme et al. 2020) | Põlme S, et al. 2020. Fungal Diversity 105:1-16 ((https://doi.org/10.1007/s13225-020-00466-2)) |
| Otu20204 | 1 | k__Fungi(100);p__Ascomycota(100);c__Saccharomycetes(100);o__Saccharomycetales(100);f__Metschnikowiaceae(100);g__Metschnikowia(100);s__Metschnikowia_pulcherrima(100); | Metschnikowia | 13 | Saprotroph | \|Undefined Saprotroph\| | Yeast | NULL | Probable | As nectar-tap_saprotroph (Põlme et al. 2020); Decay sugar-rich substrates (Põlme et al. 2020) | Põlme S, et al. 2020. Fungal Diversity 105:1-16 ((https://doi.org/10.1007/s13225-020-00466-2)) |
| Otu20205 | 1 | k__Fungi(100);p__Ascomycota(100);c__Saccharomycetes(100);o__Saccharomycetales(100);f__Metschnikowiaceae(100);g__Metschnikowia(100);s__Metschnikowia_pulcherrima(100); | Metschnikowia | 13 | Saprotroph | \|Undefined Saprotroph\| | Yeast | NULL | Probable | As nectar-tap_saprotroph (Põlme et al. 2020); Decay sugar-rich substrates (Põlme et al. 2020) | Põlme S, et al. 2020. Fungal Diversity 105:1-16 ((https://doi.org/10.1007/s13225-020-00466-2)) |
| Otu20206 | 1 | k__Fungi(100);p__Ascomycota(100);c__Saccharomycetes(100);o__Saccharomycetales(100);f__Metschnikowiaceae(100);g__Metschnikowia(100);g__Metschnikowia_unclassified(100); | Metschnikowia | 13 | Saprotroph | \|Undefined Saprotroph\| | Yeast | NULL | Probable | As nectar-tap_saprotroph (Põlme et al. 2020); Decay sugar-rich substrates (Põlme et al. 2020) | Põlme S, et al. 2020. Fungal Diversity 105:1-16 ((https://doi.org/10.1007/s13225-020-00466-2)) |
| Otu20207 | 1 | k__Fungi(100);p__Ascomycota(100);c__Saccharomycetes(100);o__Saccharomycetales(100);f__Saccharomycetales_family_Incertae_sedis(100);g__Candida(100);g__Candida_unclassified(100); | Candida | 13 | Pathotroph-Saprotroph-Symbiotroph | Animal Pathogen-Endophyte-Endosymbiont-Epiphyte-\|Undefined Saprotroph\| | Dimorphic Yeast-Yeast | NULL | Probable | As nectar-tap_saprotroph (Põlme et al. 2020); Decay sugar-rich substrates (Põlme et al. 2020); This asexual genus is polyphyletic and contain species that function very differently depend on what they are. Caution should be taken when interpreting guilds for Candida species. | Manolakaki D, et al. 2010. Virulence. 1: 367-375 ((https://doi.org/10.4161/viru.1.5.12796)); Mekbib SB, et al. 2011. Tropical Plant Pathology 36:233-240 ((https://doi.org/10.1590/S1982-56762011000400004)); Põlme S, et al. 2020. Fungal Diversity 105:1-16 ((https://doi.org/10.1007/s13225-020-00466-2)) |
| Otu20211 | 1 | k__Fungi(100);p__Ascomycota(100);c__Saccharomycetes(100);o__Saccharomycetales(100);f__Metschnikowiaceae(100);g__Metschnikowia(100);s__Metschnikowia_pulcherrima(100); | Metschnikowia | 13 | Saprotroph | \|Undefined Saprotroph\| | Yeast | NULL | Probable | As nectar-tap_saprotroph (Põlme et al. 2020); Decay sugar-rich substrates (Põlme et al. 2020) | Põlme S, et al. 2020. Fungal Diversity 105:1-16 ((https://doi.org/10.1007/s13225-020-00466-2)) |
| Otu20212 | 1 | k__Fungi(100);p__Ascomycota(100);c__Saccharomycetes(100);o__Saccharomycetales(100);f__Metschnikowiaceae(100);g__Metschnikowia(100);g__Metschnikowia_unclassified(100); | Metschnikowia | 13 | Saprotroph | \|Undefined Saprotroph\| | Yeast | NULL | Probable | As nectar-tap_saprotroph (Põlme et al. 2020); Decay sugar-rich substrates (Põlme et al. 2020) | Põlme S, et al. 2020. Fungal Diversity 105:1-16 ((https://doi.org/10.1007/s13225-020-00466-2)) |
| Otu20213 | 1 | k__Fungi(100);p__Ascomycota(100);c__Saccharomycetes(100);o__Saccharomycetales(100);f__Metschnikowiaceae(100);g__Metschnikowia(100);g__Metschnikowia_unclassified(100); | Metschnikowia | 13 | Saprotroph | \|Undefined Saprotroph\| | Yeast | NULL | Probable | As nectar-tap_saprotroph (Põlme et al. 2020); Decay sugar-rich substrates (Põlme et al. 2020) | Põlme S, et al. 2020. Fungal Diversity 105:1-16 ((https://doi.org/10.1007/s13225-020-00466-2)) |
| Otu20214 | 1 | k__Fungi(100);p__Ascomycota(100);c__Saccharomycetes(100);o__Saccharomycetales(100);f__Metschnikowiaceae(100);g__Metschnikowia(100);s__Metschnikowia_pulcherrima(100); | Metschnikowia | 13 | Saprotroph | \|Undefined Saprotroph\| | Yeast | NULL | Probable | As nectar-tap_saprotroph (Põlme et al. 2020); Decay sugar-rich substrates (Põlme et al. 2020) | Põlme S, et al. 2020. Fungal Diversity 105:1-16 ((https://doi.org/10.1007/s13225-020-00466-2)) |
| Otu20215 | 1 | k__Fungi(100);p__Ascomycota(100);c__Saccharomycetes(100);o__Saccharomycetales(100);f__Metschnikowiaceae(100);g__Metschnikowia(100);s__Metschnikowia_pulcherrima(100); | Metschnikowia | 13 | Saprotroph | \|Undefined Saprotroph\| | Yeast | NULL | Probable | As nectar-tap_saprotroph (Põlme et al. 2020); Decay sugar-rich substrates (Põlme et al. 2020) | Põlme S, et al. 2020. Fungal Diversity 105:1-16 ((https://doi.org/10.1007/s13225-020-00466-2)) |
| Otu20216 | 1 | k__Fungi(100);p__Ascomycota(100);c__Saccharomycetes(100);o__Saccharomycetales(100);f__Metschnikowiaceae(100);g__Metschnikowia(100);g__Metschnikowia_unclassified(100); | Metschnikowia | 13 | Saprotroph | \|Undefined Saprotroph\| | Yeast | NULL | Probable | As nectar-tap_saprotroph (Põlme et al. 2020); Decay sugar-rich substrates (Põlme et al. 2020) | Põlme S, et al. 2020. Fungal Diversity 105:1-16 ((https://doi.org/10.1007/s13225-020-00466-2)) |
| Otu20217 | 1 | k__Fungi(100);p__Ascomycota(100);c__Dothideomycetes(100);o__Pleosporales(100);f__Venturiaceae(100);g__unclassified_Venturiaceae(100);s__Venturiaceae_sp(100); | Venturiaceae | 9 | Pathotroph-Saprotroph-Symbiotroph | Endophyte-Plant Pathogen-Undefined Saprotroph | Microfungus | NULL | Probable | Mostly endophytes and weak parasites of aerial plant parts, including bark and wood, leaf and fruit tissue; rarely biotrophic. Little is known except for the important pathogens (Cannon & Kirk 2007) | Cannon PF, Kirk PM. 2007. Fungal Families of the World. CAB International, Cambridge (ISBN: 978-0851998275) |
| Otu20218 | 1 | k__Fungi(100);p__Ascomycota(100);c__Saccharomycetes(100);o__Saccharomycetales(100);f__Metschnikowiaceae(100);g__Metschnikowia(100);g__Metschnikowia_unclassified(100); | Metschnikowia | 13 | Saprotroph | \|Undefined Saprotroph\| | Yeast | NULL | Probable | As nectar-tap_saprotroph (Põlme et al. 2020); Decay sugar-rich substrates (Põlme et al. 2020) | Põlme S, et al. 2020. Fungal Diversity 105:1-16 ((https://doi.org/10.1007/s13225-020-00466-2)) |
| Otu20219 | 1 | k__Fungi(100);p__Ascomycota(100);c__Saccharomycetes(100);o__Saccharomycetales(100);f__Metschnikowiaceae(100);g__Metschnikowia(100);g__Metschnikowia_unclassified(100); | Metschnikowia | 13 | Saprotroph | \|Undefined Saprotroph\| | Yeast | NULL | Probable | As nectar-tap_saprotroph (Põlme et al. 2020); Decay sugar-rich substrates (Põlme et al. 2020) | Põlme S, et al. 2020. Fungal Diversity 105:1-16 ((https://doi.org/10.1007/s13225-020-00466-2)) |
| Otu20220 | 1 | k__Fungi(100);p__Ascomycota(100);c__Saccharomycetes(100);o__Saccharomycetales(100);f__Metschnikowiaceae(100);g__Metschnikowia(100);g__Metschnikowia_unclassified(100); | Metschnikowia | 13 | Saprotroph | \|Undefined Saprotroph\| | Yeast | NULL | Probable | As nectar-tap_saprotroph (Põlme et al. 2020); Decay sugar-rich substrates (Põlme et al. 2020) | Põlme S, et al. 2020. Fungal Diversity 105:1-16 ((https://doi.org/10.1007/s13225-020-00466-2)) |
| Otu20221 | 1 | k__Fungi(100);p__Ascomycota(100);c__Saccharomycetes(100);o__Saccharomycetales(100);f__Metschnikowiaceae(100);g__Metschnikowia(100);s__Metschnikowia_pulcherrima(100); | Metschnikowia | 13 | Saprotroph | \|Undefined Saprotroph\| | Yeast | NULL | Probable | As nectar-tap_saprotroph (Põlme et al. 2020); Decay sugar-rich substrates (Põlme et al. 2020) | Põlme S, et al. 2020. Fungal Diversity 105:1-16 ((https://doi.org/10.1007/s13225-020-00466-2)) |
| Otu20223 | 1 | k__Fungi(100);p__Ascomycota(100);c__Saccharomycetes(100);o__Saccharomycetales(100);f__Metschnikowiaceae(100);g__Metschnikowia(100);s__Metschnikowia_pulcherrima(100); | Metschnikowia | 13 | Saprotroph | \|Undefined Saprotroph\| | Yeast | NULL | Probable | As nectar-tap_saprotroph (Põlme et al. 2020); Decay sugar-rich substrates (Põlme et al. 2020) | Põlme S, et al. 2020. Fungal Diversity 105:1-16 ((https://doi.org/10.1007/s13225-020-00466-2)) |
| Otu20224 | 1 | k__Fungi(100);p__Ascomycota(100);c__Saccharomycetes(100);o__Saccharomycetales(100);f__Metschnikowiaceae(100);g__Metschnikowia(100);s__Metschnikowia_pulcherrima(100); | Metschnikowia | 13 | Saprotroph | \|Undefined Saprotroph\| | Yeast | NULL | Probable | As nectar-tap_saprotroph (Põlme et al. 2020); Decay sugar-rich substrates (Põlme et al. 2020) | Põlme S, et al. 2020. Fungal Diversity 105:1-16 ((https://doi.org/10.1007/s13225-020-00466-2)) |
| Otu20225 | 1 | k__Fungi(100);p__Ascomycota(100);c__Saccharomycetes(100);o__Saccharomycetales(100);f__Metschnikowiaceae(100);g__Metschnikowia(100);g__Metschnikowia_unclassified(100); | Metschnikowia | 13 | Saprotroph | \|Undefined Saprotroph\| | Yeast | NULL | Probable | As nectar-tap_saprotroph (Põlme et al. 2020); Decay sugar-rich substrates (Põlme et al. 2020) | Põlme S, et al. 2020. Fungal Diversity 105:1-16 ((https://doi.org/10.1007/s13225-020-00466-2)) |
| Otu20226 | 1 | k__Fungi(100);p__Ascomycota(100);c__Saccharomycetes(100);o__Saccharomycetales(100);f__Metschnikowiaceae(100);g__Metschnikowia(100);s__Metschnikowia_pulcherrima(100); | Metschnikowia | 13 | Saprotroph | \|Undefined Saprotroph\| | Yeast | NULL | Probable | As nectar-tap_saprotroph (Põlme et al. 2020); Decay sugar-rich substrates (Põlme et al. 2020) | Põlme S, et al. 2020. Fungal Diversity 105:1-16 ((https://doi.org/10.1007/s13225-020-00466-2)) |
| Otu20227 | 1 | k__Fungi(100);p__Ascomycota(100);c__Saccharomycetes(100);o__Saccharomycetales(100);f__Metschnikowiaceae(100);g__Metschnikowia(100);g__Metschnikowia_unclassified(100); | Metschnikowia | 13 | Saprotroph | \|Undefined Saprotroph\| | Yeast | NULL | Probable | As nectar-tap_saprotroph (Põlme et al. 2020); Decay sugar-rich substrates (Põlme et al. 2020) | Põlme S, et al. 2020. Fungal Diversity 105:1-16 ((https://doi.org/10.1007/s13225-020-00466-2)) |
| Otu20228 | 1 | k__Fungi(100);p__Ascomycota(100);c__Saccharomycetes(100);o__Saccharomycetales(100);f__Metschnikowiaceae(100);g__Metschnikowia(100);s__Metschnikowia_pulcherrima(100); | Metschnikowia | 13 | Saprotroph | \|Undefined Saprotroph\| | Yeast | NULL | Probable | As nectar-tap_saprotroph (Põlme et al. 2020); Decay sugar-rich substrates (Põlme et al. 2020) | Põlme S, et al. 2020. Fungal Diversity 105:1-16 ((https://doi.org/10.1007/s13225-020-00466-2)) |
| Otu20229 | 1 | k__Fungi(100);p__Ascomycota(100);c__Saccharomycetes(100);o__Saccharomycetales(100);f__Metschnikowiaceae(100);g__Metschnikowia(100);s__Metschnikowia_pulcherrima(100); | Metschnikowia | 13 | Saprotroph | \|Undefined Saprotroph\| | Yeast | NULL | Probable | As nectar-tap_saprotroph (Põlme et al. 2020); Decay sugar-rich substrates (Põlme et al. 2020) | Põlme S, et al. 2020. Fungal Diversity 105:1-16 ((https://doi.org/10.1007/s13225-020-00466-2)) |
| Otu20230 | 1 | k__Fungi(100);p__Ascomycota(100);c__Saccharomycetes(100);o__Saccharomycetales(100);f__Metschnikowiaceae(100);g__Metschnikowia(100);s__Metschnikowia_pulcherrima(100); | Metschnikowia | 13 | Saprotroph | \|Undefined Saprotroph\| | Yeast | NULL | Probable | As nectar-tap_saprotroph (Põlme et al. 2020); Decay sugar-rich substrates (Põlme et al. 2020) | Põlme S, et al. 2020. Fungal Diversity 105:1-16 ((https://doi.org/10.1007/s13225-020-00466-2)) |
| Otu20231 | 1 | k__Fungi(100);p__Ascomycota(100);c__Saccharomycetes(100);o__Saccharomycetales(100);f__Metschnikowiaceae(100);g__Metschnikowia(100);s__Metschnikowia_pulcherrima(100); | Metschnikowia | 13 | Saprotroph | \|Undefined Saprotroph\| | Yeast | NULL | Probable | As nectar-tap_saprotroph (Põlme et al. 2020); Decay sugar-rich substrates (Põlme et al. 2020) | Põlme S, et al. 2020. Fungal Diversity 105:1-16 ((https://doi.org/10.1007/s13225-020-00466-2)) |
| Otu20232 | 1 | k__Fungi(100);p__Ascomycota(100);c__Saccharomycetes(100);o__Saccharomycetales(100);f__Metschnikowiaceae(100);g__Metschnikowia(100);s__Metschnikowia_pulcherrima(100); | Metschnikowia | 13 | Saprotroph | \|Undefined Saprotroph\| | Yeast | NULL | Probable | As nectar-tap_saprotroph (Põlme et al. 2020); Decay sugar-rich substrates (Põlme et al. 2020) | Põlme S, et al. 2020. Fungal Diversity 105:1-16 ((https://doi.org/10.1007/s13225-020-00466-2)) |
| Otu20233 | 1 | k__Fungi(100);p__Ascomycota(100);c__Saccharomycetes(100);o__Saccharomycetales(100);f__Metschnikowiaceae(100);g__Metschnikowia(100);s__Metschnikowia_pulcherrima(100); | Metschnikowia | 13 | Saprotroph | \|Undefined Saprotroph\| | Yeast | NULL | Probable | As nectar-tap_saprotroph (Põlme et al. 2020); Decay sugar-rich substrates (Põlme et al. 2020) | Põlme S, et al. 2020. Fungal Diversity 105:1-16 ((https://doi.org/10.1007/s13225-020-00466-2)) |
| Otu20234 | 1 | k__Fungi(100);p__Ascomycota(100);c__Saccharomycetes(100);o__Saccharomycetales(100);f__Metschnikowiaceae(100);g__Metschnikowia(100);s__Metschnikowia_pulcherrima(100); | Metschnikowia | 13 | Saprotroph | \|Undefined Saprotroph\| | Yeast | NULL | Probable | As nectar-tap_saprotroph (Põlme et al. 2020); Decay sugar-rich substrates (Põlme et al. 2020) | Põlme S, et al. 2020. Fungal Diversity 105:1-16 ((https://doi.org/10.1007/s13225-020-00466-2)) |
| Otu20235 | 1 | k__Fungi(100);p__Ascomycota(100);c__Saccharomycetes(100);o__Saccharomycetales(100);f__Metschnikowiaceae(100);g__Metschnikowia(100);g__Metschnikowia_unclassified(100); | Metschnikowia | 13 | Saprotroph | \|Undefined Saprotroph\| | Yeast | NULL | Probable | As nectar-tap_saprotroph (Põlme et al. 2020); Decay sugar-rich substrates (Põlme et al. 2020) | Põlme S, et al. 2020. Fungal Diversity 105:1-16 ((https://doi.org/10.1007/s13225-020-00466-2)) |
| Otu20236 | 1 | k__Fungi(100);p__Ascomycota(100);c__Saccharomycetes(100);o__Saccharomycetales(100);f__Metschnikowiaceae(100);g__Metschnikowia(100);s__Metschnikowia_pulcherrima(100); | Metschnikowia | 13 | Saprotroph | \|Undefined Saprotroph\| | Yeast | NULL | Probable | As nectar-tap_saprotroph (Põlme et al. 2020); Decay sugar-rich substrates (Põlme et al. 2020) | Põlme S, et al. 2020. Fungal Diversity 105:1-16 ((https://doi.org/10.1007/s13225-020-00466-2)) |
| Otu20237 | 1 | k__Fungi(100);p__Ascomycota(100);c__Saccharomycetes(100);o__Saccharomycetales(100);f__Metschnikowiaceae(100);g__Metschnikowia(100);g__Metschnikowia_unclassified(100); | Metschnikowia | 13 | Saprotroph | \|Undefined Saprotroph\| | Yeast | NULL | Probable | As nectar-tap_saprotroph (Põlme et al. 2020); Decay sugar-rich substrates (Põlme et al. 2020) | Põlme S, et al. 2020. Fungal Diversity 105:1-16 ((https://doi.org/10.1007/s13225-020-00466-2)) |
| Otu20238 | 1 | k__Fungi(100);p__Ascomycota(100);c__Saccharomycetes(100);o__Saccharomycetales(100);f__Metschnikowiaceae(100);g__Metschnikowia(100);g__Metschnikowia_unclassified(100); | Metschnikowia | 13 | Saprotroph | \|Undefined Saprotroph\| | Yeast | NULL | Probable | As nectar-tap_saprotroph (Põlme et al. 2020); Decay sugar-rich substrates (Põlme et al. 2020) | Põlme S, et al. 2020. Fungal Diversity 105:1-16 ((https://doi.org/10.1007/s13225-020-00466-2)) |
| Otu20239 | 1 | k__Fungi(100);p__Ascomycota(100);c__Saccharomycetes(100);o__Saccharomycetales(100);f__Metschnikowiaceae(100);g__Metschnikowia(100);g__Metschnikowia_unclassified(100); | Metschnikowia | 13 | Saprotroph | \|Undefined Saprotroph\| | Yeast | NULL | Probable | As nectar-tap_saprotroph (Põlme et al. 2020); Decay sugar-rich substrates (Põlme et al. 2020) | Põlme S, et al. 2020. Fungal Diversity 105:1-16 ((https://doi.org/10.1007/s13225-020-00466-2)) |
| Otu20240 | 1 | k__Fungi(100);p__Ascomycota(100);c__Saccharomycetes(100);o__Saccharomycetales(100);f__Metschnikowiaceae(100);g__Metschnikowia(100);g__Metschnikowia_unclassified(100); | Metschnikowia | 13 | Saprotroph | \|Undefined Saprotroph\| | Yeast | NULL | Probable | As nectar-tap_saprotroph (Põlme et al. 2020); Decay sugar-rich substrates (Põlme et al. 2020) | Põlme S, et al. 2020. Fungal Diversity 105:1-16 ((https://doi.org/10.1007/s13225-020-00466-2)) |
| Otu20241 | 1 | k__Fungi(100);p__Ascomycota(100);c__Saccharomycetes(100);o__Saccharomycetales(100);f__Metschnikowiaceae(100);g__Metschnikowia(100);s__Metschnikowia_pulcherrima(100); | Metschnikowia | 13 | Saprotroph | \|Undefined Saprotroph\| | Yeast | NULL | Probable | As nectar-tap_saprotroph (Põlme et al. 2020); Decay sugar-rich substrates (Põlme et al. 2020) | Põlme S, et al. 2020. Fungal Diversity 105:1-16 ((https://doi.org/10.1007/s13225-020-00466-2)) |
| Otu20242 | 1 | k__Fungi(100);p__Ascomycota(100);c__Saccharomycetes(100);o__Saccharomycetales(100);f__Metschnikowiaceae(100);g__Metschnikowia(100);s__Metschnikowia_pulcherrima(100); | Metschnikowia | 13 | Saprotroph | \|Undefined Saprotroph\| | Yeast | NULL | Probable | As nectar-tap_saprotroph (Põlme et al. 2020); Decay sugar-rich substrates (Põlme et al. 2020) | Põlme S, et al. 2020. Fungal Diversity 105:1-16 ((https://doi.org/10.1007/s13225-020-00466-2)) |
| Otu20243 | 1 | k__Fungi(100);p__Ascomycota(100);c__Saccharomycetes(100);o__Saccharomycetales(100);f__Metschnikowiaceae(100);g__Metschnikowia(100);g__Metschnikowia_unclassified(100); | Metschnikowia | 13 | Saprotroph | \|Undefined Saprotroph\| | Yeast | NULL | Probable | As nectar-tap_saprotroph (Põlme et al. 2020); Decay sugar-rich substrates (Põlme et al. 2020) | Põlme S, et al. 2020. Fungal Diversity 105:1-16 ((https://doi.org/10.1007/s13225-020-00466-2)) |
| Otu20245 | 1 | k__Fungi(100);p__Ascomycota(100);c__Saccharomycetes(100);o__Saccharomycetales(100);f__Metschnikowiaceae(100);g__Metschnikowia(100);s__Metschnikowia_pulcherrima(100); | Metschnikowia | 13 | Saprotroph | \|Undefined Saprotroph\| | Yeast | NULL | Probable | As nectar-tap_saprotroph (Põlme et al. 2020); Decay sugar-rich substrates (Põlme et al. 2020) | Põlme S, et al. 2020. Fungal Diversity 105:1-16 ((https://doi.org/10.1007/s13225-020-00466-2)) |
| Otu20249 | 1 | k__Fungi(100);p__Ascomycota(100);c__Saccharomycetes(100);o__Saccharomycetales(100);f__Metschnikowiaceae(100);g__Metschnikowia(100);g__Metschnikowia_unclassified(100); | Metschnikowia | 13 | Saprotroph | \|Undefined Saprotroph\| | Yeast | NULL | Probable | As nectar-tap_saprotroph (Põlme et al. 2020); Decay sugar-rich substrates (Põlme et al. 2020) | Põlme S, et al. 2020. Fungal Diversity 105:1-16 ((https://doi.org/10.1007/s13225-020-00466-2)) |
| Otu20251 | 1 | k__Fungi(100);p__Ascomycota(100);c__Saccharomycetes(100);o__Saccharomycetales(100);f__Metschnikowiaceae(100);g__Metschnikowia(100);s__Metschnikowia_pulcherrima(100); | Metschnikowia | 13 | Saprotroph | \|Undefined Saprotroph\| | Yeast | NULL | Probable | As nectar-tap_saprotroph (Põlme et al. 2020); Decay sugar-rich substrates (Põlme et al. 2020) | Põlme S, et al. 2020. Fungal Diversity 105:1-16 ((https://doi.org/10.1007/s13225-020-00466-2)) |
| Otu20252 | 1 | k__Fungi(100);p__Ascomycota(100);c__Saccharomycetes(100);o__Saccharomycetales(100);f__Saccharomycetales_family_Incertae_sedis(100);g__Candida(100);g__Candida_unclassified(100); | Candida | 13 | Pathotroph-Saprotroph-Symbiotroph | Animal Pathogen-Endophyte-Endosymbiont-Epiphyte-\|Undefined Saprotroph\| | Dimorphic Yeast-Yeast | NULL | Probable | As nectar-tap_saprotroph (Põlme et al. 2020); Decay sugar-rich substrates (Põlme et al. 2020); This asexual genus is polyphyletic and contain species that function very differently depend on what they are. Caution should be taken when interpreting guilds for Candida species. | Manolakaki D, et al. 2010. Virulence. 1: 367-375 ((https://doi.org/10.4161/viru.1.5.12796)); Mekbib SB, et al. 2011. Tropical Plant Pathology 36:233-240 ((https://doi.org/10.1590/S1982-56762011000400004)); Põlme S, et al. 2020. Fungal Diversity 105:1-16 ((https://doi.org/10.1007/s13225-020-00466-2)) |
| Otu20254 | 1 | k__Fungi(100);p__Ascomycota(100);c__Saccharomycetes(100);o__Saccharomycetales(100);f__Metschnikowiaceae(100);g__Metschnikowia(100);g__Metschnikowia_unclassified(100); | Metschnikowia | 13 | Saprotroph | \|Undefined Saprotroph\| | Yeast | NULL | Probable | As nectar-tap_saprotroph (Põlme et al. 2020); Decay sugar-rich substrates (Põlme et al. 2020) | Põlme S, et al. 2020. Fungal Diversity 105:1-16 ((https://doi.org/10.1007/s13225-020-00466-2)) |
| Otu20255 | 1 | k__Fungi(100);p__Ascomycota(100);c__Saccharomycetes(100);o__Saccharomycetales(100);f__Metschnikowiaceae(100);g__Metschnikowia(100);g__Metschnikowia_unclassified(100); | Metschnikowia | 13 | Saprotroph | \|Undefined Saprotroph\| | Yeast | NULL | Probable | As nectar-tap_saprotroph (Põlme et al. 2020); Decay sugar-rich substrates (Põlme et al. 2020) | Põlme S, et al. 2020. Fungal Diversity 105:1-16 ((https://doi.org/10.1007/s13225-020-00466-2)) |
| Otu20256 | 1 | k__Fungi(100);p__Ascomycota(100);c__Saccharomycetes(100);o__Saccharomycetales(100);f__Metschnikowiaceae(100);g__Metschnikowia(100);g__Metschnikowia_unclassified(100); | Metschnikowia | 13 | Saprotroph | \|Undefined Saprotroph\| | Yeast | NULL | Probable | As nectar-tap_saprotroph (Põlme et al. 2020); Decay sugar-rich substrates (Põlme et al. 2020) | Põlme S, et al. 2020. Fungal Diversity 105:1-16 ((https://doi.org/10.1007/s13225-020-00466-2)) |
| Otu20257 | 1 | k__Fungi(100);p__Ascomycota(100);c__Saccharomycetes(100);o__Saccharomycetales(100);f__Metschnikowiaceae(100);g__Metschnikowia(100);s__Metschnikowia_pulcherrima(100); | Metschnikowia | 13 | Saprotroph | \|Undefined Saprotroph\| | Yeast | NULL | Probable | As nectar-tap_saprotroph (Põlme et al. 2020); Decay sugar-rich substrates (Põlme et al. 2020) | Põlme S, et al. 2020. Fungal Diversity 105:1-16 ((https://doi.org/10.1007/s13225-020-00466-2)) |
| Otu20258 | 1 | k__Fungi(100);p__Ascomycota(100);c__Saccharomycetes(100);o__Saccharomycetales(100);f__Metschnikowiaceae(100);g__Metschnikowia(100);s__Metschnikowia_pulcherrima(100); | Metschnikowia | 13 | Saprotroph | \|Undefined Saprotroph\| | Yeast | NULL | Probable | As nectar-tap_saprotroph (Põlme et al. 2020); Decay sugar-rich substrates (Põlme et al. 2020) | Põlme S, et al. 2020. Fungal Diversity 105:1-16 ((https://doi.org/10.1007/s13225-020-00466-2)) |
| Otu20259 | 1 | k__Fungi(100);p__Ascomycota(100);c__Saccharomycetes(100);o__Saccharomycetales(100);f__Metschnikowiaceae(100);g__Metschnikowia(100);s__Metschnikowia_pulcherrima(100); | Metschnikowia | 13 | Saprotroph | \|Undefined Saprotroph\| | Yeast | NULL | Probable | As nectar-tap_saprotroph (Põlme et al. 2020); Decay sugar-rich substrates (Põlme et al. 2020) | Põlme S, et al. 2020. Fungal Diversity 105:1-16 ((https://doi.org/10.1007/s13225-020-00466-2)) |
| Otu20260 | 1 | k__Fungi(100);p__Ascomycota(100);c__Saccharomycetes(100);o__Saccharomycetales(100);f__Metschnikowiaceae(100);g__Metschnikowia(100);g__Metschnikowia_unclassified(100); | Metschnikowia | 13 | Saprotroph | \|Undefined Saprotroph\| | Yeast | NULL | Probable | As nectar-tap_saprotroph (Põlme et al. 2020); Decay sugar-rich substrates (Põlme et al. 2020) | Põlme S, et al. 2020. Fungal Diversity 105:1-16 ((https://doi.org/10.1007/s13225-020-00466-2)) |
| Otu20261 | 1 | k__Fungi(100);p__Ascomycota(100);c__Saccharomycetes(100);o__Saccharomycetales(100);f__Metschnikowiaceae(100);g__Metschnikowia(100);g__Metschnikowia_unclassified(100); | Metschnikowia | 13 | Saprotroph | \|Undefined Saprotroph\| | Yeast | NULL | Probable | As nectar-tap_saprotroph (Põlme et al. 2020); Decay sugar-rich substrates (Põlme et al. 2020) | Põlme S, et al. 2020. Fungal Diversity 105:1-16 ((https://doi.org/10.1007/s13225-020-00466-2)) |
| Otu20262 | 1 | k__Fungi(100);p__Ascomycota(100);c__Saccharomycetes(100);o__Saccharomycetales(100);f__Metschnikowiaceae(100);g__Metschnikowia(100);g__Metschnikowia_unclassified(100); | Metschnikowia | 13 | Saprotroph | \|Undefined Saprotroph\| | Yeast | NULL | Probable | As nectar-tap_saprotroph (Põlme et al. 2020); Decay sugar-rich substrates (Põlme et al. 2020) | Põlme S, et al. 2020. Fungal Diversity 105:1-16 ((https://doi.org/10.1007/s13225-020-00466-2)) |
| Otu20263 | 1 | k__Fungi(100);p__Ascomycota(100);c__Saccharomycetes(100);o__Saccharomycetales(100);f__Metschnikowiaceae(100);g__Metschnikowia(100);s__Metschnikowia_pulcherrima(100); | Metschnikowia | 13 | Saprotroph | \|Undefined Saprotroph\| | Yeast | NULL | Probable | As nectar-tap_saprotroph (Põlme et al. 2020); Decay sugar-rich substrates (Põlme et al. 2020) | Põlme S, et al. 2020. Fungal Diversity 105:1-16 ((https://doi.org/10.1007/s13225-020-00466-2)) |
| Otu20264 | 1 | k__Fungi(100);p__Ascomycota(100);c__Saccharomycetes(100);o__Saccharomycetales(100);f__Metschnikowiaceae(100);g__Metschnikowia(100);s__Metschnikowia_pulcherrima(100); | Metschnikowia | 13 | Saprotroph | \|Undefined Saprotroph\| | Yeast | NULL | Probable | As nectar-tap_saprotroph (Põlme et al. 2020); Decay sugar-rich substrates (Põlme et al. 2020) | Põlme S, et al. 2020. Fungal Diversity 105:1-16 ((https://doi.org/10.1007/s13225-020-00466-2)) |
| Otu20265 | 1 | k__Fungi(100);p__Ascomycota(100);c__Saccharomycetes(100);o__Saccharomycetales(100);f__Metschnikowiaceae(100);g__Metschnikowia(100);s__Metschnikowia_pulcherrima(100); | Metschnikowia | 13 | Saprotroph | \|Undefined Saprotroph\| | Yeast | NULL | Probable | As nectar-tap_saprotroph (Põlme et al. 2020); Decay sugar-rich substrates (Põlme et al. 2020) | Põlme S, et al. 2020. Fungal Diversity 105:1-16 ((https://doi.org/10.1007/s13225-020-00466-2)) |
| Otu20266 | 1 | k__Fungi(100);p__Ascomycota(100);c__Saccharomycetes(100);o__Saccharomycetales(100);f__Metschnikowiaceae(100);g__Metschnikowia(100);s__Metschnikowia_pulcherrima(100); | Metschnikowia | 13 | Saprotroph | \|Undefined Saprotroph\| | Yeast | NULL | Probable | As nectar-tap_saprotroph (Põlme et al. 2020); Decay sugar-rich substrates (Põlme et al. 2020) | Põlme S, et al. 2020. Fungal Diversity 105:1-16 ((https://doi.org/10.1007/s13225-020-00466-2)) |
| Otu20267 | 1 | k__Fungi(100);p__Ascomycota(100);c__Saccharomycetes(100);o__Saccharomycetales(100);f__Metschnikowiaceae(100);g__Metschnikowia(100);g__Metschnikowia_unclassified(100); | Metschnikowia | 13 | Saprotroph | \|Undefined Saprotroph\| | Yeast | NULL | Probable | As nectar-tap_saprotroph (Põlme et al. 2020); Decay sugar-rich substrates (Põlme et al. 2020) | Põlme S, et al. 2020. Fungal Diversity 105:1-16 ((https://doi.org/10.1007/s13225-020-00466-2)) |
| Otu20268 | 1 | k__Fungi(100);p__Ascomycota(100);c__Saccharomycetes(100);o__Saccharomycetales(100);f__Metschnikowiaceae(100);g__Metschnikowia(100);s__Metschnikowia_pulcherrima(100); | Metschnikowia | 13 | Saprotroph | \|Undefined Saprotroph\| | Yeast | NULL | Probable | As nectar-tap_saprotroph (Põlme et al. 2020); Decay sugar-rich substrates (Põlme et al. 2020) | Põlme S, et al. 2020. Fungal Diversity 105:1-16 ((https://doi.org/10.1007/s13225-020-00466-2)) |
| Otu20269 | 1 | k__Fungi(100);p__Ascomycota(100);c__Saccharomycetes(100);o__Saccharomycetales(100);f__Metschnikowiaceae(100);g__Metschnikowia(100);s__Metschnikowia_pulcherrima(100); | Metschnikowia | 13 | Saprotroph | \|Undefined Saprotroph\| | Yeast | NULL | Probable | As nectar-tap_saprotroph (Põlme et al. 2020); Decay sugar-rich substrates (Põlme et al. 2020) | Põlme S, et al. 2020. Fungal Diversity 105:1-16 ((https://doi.org/10.1007/s13225-020-00466-2)) |
| Otu20270 | 1 | k__Fungi(100);p__Ascomycota(100);c__Saccharomycetes(100);o__Saccharomycetales(100);f__Metschnikowiaceae(100);g__Metschnikowia(100);s__Metschnikowia_pulcherrima(100); | Metschnikowia | 13 | Saprotroph | \|Undefined Saprotroph\| | Yeast | NULL | Probable | As nectar-tap_saprotroph (Põlme et al. 2020); Decay sugar-rich substrates (Põlme et al. 2020) | Põlme S, et al. 2020. Fungal Diversity 105:1-16 ((https://doi.org/10.1007/s13225-020-00466-2)) |
| Otu20271 | 1 | k__Fungi(100);p__Ascomycota(100);c__Saccharomycetes(100);o__Saccharomycetales(100);f__Metschnikowiaceae(100);g__Metschnikowia(100);s__Metschnikowia_pulcherrima(100); | Metschnikowia | 13 | Saprotroph | \|Undefined Saprotroph\| | Yeast | NULL | Probable | As nectar-tap_saprotroph (Põlme et al. 2020); Decay sugar-rich substrates (Põlme et al. 2020) | Põlme S, et al. 2020. Fungal Diversity 105:1-16 ((https://doi.org/10.1007/s13225-020-00466-2)) |
| Otu20272 | 1 | k__Fungi(100);p__Ascomycota(100);c__Saccharomycetes(100);o__Saccharomycetales(100);f__Metschnikowiaceae(100);g__Metschnikowia(100);s__Metschnikowia_pulcherrima(100); | Metschnikowia | 13 | Saprotroph | \|Undefined Saprotroph\| | Yeast | NULL | Probable | As nectar-tap_saprotroph (Põlme et al. 2020); Decay sugar-rich substrates (Põlme et al. 2020) | Põlme S, et al. 2020. Fungal Diversity 105:1-16 ((https://doi.org/10.1007/s13225-020-00466-2)) |
| Otu20273 | 1 | k__Fungi(100);p__Ascomycota(100);c__Saccharomycetes(100);o__Saccharomycetales(100);f__Metschnikowiaceae(100);g__Metschnikowia(100);s__Metschnikowia_pulcherrima(100); | Metschnikowia | 13 | Saprotroph | \|Undefined Saprotroph\| | Yeast | NULL | Probable | As nectar-tap_saprotroph (Põlme et al. 2020); Decay sugar-rich substrates (Põlme et al. 2020) | Põlme S, et al. 2020. Fungal Diversity 105:1-16 ((https://doi.org/10.1007/s13225-020-00466-2)) |
| Otu20274 | 1 | k__Fungi(100);p__Ascomycota(100);c__Saccharomycetes(100);o__Saccharomycetales(100);f__Metschnikowiaceae(100);g__Metschnikowia(100);s__Metschnikowia_pulcherrima(100); | Metschnikowia | 13 | Saprotroph | \|Undefined Saprotroph\| | Yeast | NULL | Probable | As nectar-tap_saprotroph (Põlme et al. 2020); Decay sugar-rich substrates (Põlme et al. 2020) | Põlme S, et al. 2020. Fungal Diversity 105:1-16 ((https://doi.org/10.1007/s13225-020-00466-2)) |
| Otu20275 | 1 | k__Fungi(100);p__Ascomycota(100);c__Saccharomycetes(100);o__Saccharomycetales(100);f__Metschnikowiaceae(100);g__Metschnikowia(100);s__Metschnikowia_pulcherrima(100); | Metschnikowia | 13 | Saprotroph | \|Undefined Saprotroph\| | Yeast | NULL | Probable | As nectar-tap_saprotroph (Põlme et al. 2020); Decay sugar-rich substrates (Põlme et al. 2020) | Põlme S, et al. 2020. Fungal Diversity 105:1-16 ((https://doi.org/10.1007/s13225-020-00466-2)) |
| Otu20276 | 1 | k__Fungi(100);p__Ascomycota(100);c__Saccharomycetes(100);o__Saccharomycetales(100);f__Metschnikowiaceae(100);g__Metschnikowia(100);s__Metschnikowia_pulcherrima(100); | Metschnikowia | 13 | Saprotroph | \|Undefined Saprotroph\| | Yeast | NULL | Probable | As nectar-tap_saprotroph (Põlme et al. 2020); Decay sugar-rich substrates (Põlme et al. 2020) | Põlme S, et al. 2020. Fungal Diversity 105:1-16 ((https://doi.org/10.1007/s13225-020-00466-2)) |
| Otu20277 | 1 | k__Fungi(100);p__Ascomycota(100);c__Saccharomycetes(100);o__Saccharomycetales(100);f__Metschnikowiaceae(100);g__Metschnikowia(100);s__Metschnikowia_pulcherrima(100); | Metschnikowia | 13 | Saprotroph | \|Undefined Saprotroph\| | Yeast | NULL | Probable | As nectar-tap_saprotroph (Põlme et al. 2020); Decay sugar-rich substrates (Põlme et al. 2020) | Põlme S, et al. 2020. Fungal Diversity 105:1-16 ((https://doi.org/10.1007/s13225-020-00466-2)) |
| Otu20278 | 1 | k__Fungi(100);p__Ascomycota(100);c__Saccharomycetes(100);o__Saccharomycetales(100);f__Metschnikowiaceae(100);g__Metschnikowia(100);s__Metschnikowia_pulcherrima(100); | Metschnikowia | 13 | Saprotroph | \|Undefined Saprotroph\| | Yeast | NULL | Probable | As nectar-tap_saprotroph (Põlme et al. 2020); Decay sugar-rich substrates (Põlme et al. 2020) | Põlme S, et al. 2020. Fungal Diversity 105:1-16 ((https://doi.org/10.1007/s13225-020-00466-2)) |
| Otu20279 | 1 | k__Fungi(100);p__Ascomycota(100);c__Saccharomycetes(100);o__Saccharomycetales(100);f__Metschnikowiaceae(100);g__Metschnikowia(100);s__Metschnikowia_pulcherrima(100); | Metschnikowia | 13 | Saprotroph | \|Undefined Saprotroph\| | Yeast | NULL | Probable | As nectar-tap_saprotroph (Põlme et al. 2020); Decay sugar-rich substrates (Põlme et al. 2020) | Põlme S, et al. 2020. Fungal Diversity 105:1-16 ((https://doi.org/10.1007/s13225-020-00466-2)) |
| Otu20280 | 1 | k__Fungi(100);p__Ascomycota(100);c__Saccharomycetes(100);o__Saccharomycetales(100);f__Metschnikowiaceae(100);g__Metschnikowia(100);g__Metschnikowia_unclassified(100); | Metschnikowia | 13 | Saprotroph | \|Undefined Saprotroph\| | Yeast | NULL | Probable | As nectar-tap_saprotroph (Põlme et al. 2020); Decay sugar-rich substrates (Põlme et al. 2020) | Põlme S, et al. 2020. Fungal Diversity 105:1-16 ((https://doi.org/10.1007/s13225-020-00466-2)) |
| Otu20281 | 1 | k__Fungi(100);p__Ascomycota(100);c__Saccharomycetes(100);o__Saccharomycetales(100);f__Metschnikowiaceae(100);g__Metschnikowia(100);s__Metschnikowia_pulcherrima(100); | Metschnikowia | 13 | Saprotroph | \|Undefined Saprotroph\| | Yeast | NULL | Probable | As nectar-tap_saprotroph (Põlme et al. 2020); Decay sugar-rich substrates (Põlme et al. 2020) | Põlme S, et al. 2020. Fungal Diversity 105:1-16 ((https://doi.org/10.1007/s13225-020-00466-2)) |
| Otu20283 | 1 | k__Fungi(100);p__Ascomycota(100);c__Saccharomycetes(100);o__Saccharomycetales(100);f__Metschnikowiaceae(100);g__Metschnikowia(100);s__Metschnikowia_pulcherrima(100); | Metschnikowia | 13 | Saprotroph | \|Undefined Saprotroph\| | Yeast | NULL | Probable | As nectar-tap_saprotroph (Põlme et al. 2020); Decay sugar-rich substrates (Põlme et al. 2020) | Põlme S, et al. 2020. Fungal Diversity 105:1-16 ((https://doi.org/10.1007/s13225-020-00466-2)) |
| Otu20284 | 1 | k__Fungi(100);p__Ascomycota(100);c__Saccharomycetes(100);o__Saccharomycetales(100);f__Metschnikowiaceae(100);g__Metschnikowia(100);g__Metschnikowia_unclassified(100); | Metschnikowia | 13 | Saprotroph | \|Undefined Saprotroph\| | Yeast | NULL | Probable | As nectar-tap_saprotroph (Põlme et al. 2020); Decay sugar-rich substrates (Põlme et al. 2020) | Põlme S, et al. 2020. Fungal Diversity 105:1-16 ((https://doi.org/10.1007/s13225-020-00466-2)) |
| Otu20285 | 1 | k__Fungi(100);p__Ascomycota(100);c__Saccharomycetes(100);o__Saccharomycetales(100);f__Metschnikowiaceae(100);g__Metschnikowia(100);s__Metschnikowia_pulcherrima(100); | Metschnikowia | 13 | Saprotroph | \|Undefined Saprotroph\| | Yeast | NULL | Probable | As nectar-tap_saprotroph (Põlme et al. 2020); Decay sugar-rich substrates (Põlme et al. 2020) | Põlme S, et al. 2020. Fungal Diversity 105:1-16 ((https://doi.org/10.1007/s13225-020-00466-2)) |
| Otu20286 | 1 | k__Fungi(100);p__Ascomycota(100);c__Saccharomycetes(100);o__Saccharomycetales(100);f__Metschnikowiaceae(100);g__Metschnikowia(100);g__Metschnikowia_unclassified(100); | Metschnikowia | 13 | Saprotroph | \|Undefined Saprotroph\| | Yeast | NULL | Probable | As nectar-tap_saprotroph (Põlme et al. 2020); Decay sugar-rich substrates (Põlme et al. 2020) | Põlme S, et al. 2020. Fungal Diversity 105:1-16 ((https://doi.org/10.1007/s13225-020-00466-2)) |
| Otu20288 | 1 | k__Fungi(100);p__Ascomycota(100);c__Saccharomycetes(100);o__Saccharomycetales(100);f__Metschnikowiaceae(100);g__Metschnikowia(100);s__Metschnikowia_pulcherrima(100); | Metschnikowia | 13 | Saprotroph | \|Undefined Saprotroph\| | Yeast | NULL | Probable | As nectar-tap_saprotroph (Põlme et al. 2020); Decay sugar-rich substrates (Põlme et al. 2020) | Põlme S, et al. 2020. Fungal Diversity 105:1-16 ((https://doi.org/10.1007/s13225-020-00466-2)) |
| Otu20289 | 1 | k__Fungi(100);p__Ascomycota(100);c__Saccharomycetes(100);o__Saccharomycetales(100);f__Metschnikowiaceae(100);g__Metschnikowia(100);s__Metschnikowia_pulcherrima(100); | Metschnikowia | 13 | Saprotroph | \|Undefined Saprotroph\| | Yeast | NULL | Probable | As nectar-tap_saprotroph (Põlme et al. 2020); Decay sugar-rich substrates (Põlme et al. 2020) | Põlme S, et al. 2020. Fungal Diversity 105:1-16 ((https://doi.org/10.1007/s13225-020-00466-2)) |
| Otu20291 | 1 | k__Fungi(100);p__Ascomycota(100);c__Saccharomycetes(100);o__Saccharomycetales(100);f__Metschnikowiaceae(100);g__Metschnikowia(100);g__Metschnikowia_unclassified(100); | Metschnikowia | 13 | Saprotroph | \|Undefined Saprotroph\| | Yeast | NULL | Probable | As nectar-tap_saprotroph (Põlme et al. 2020); Decay sugar-rich substrates (Põlme et al. 2020) | Põlme S, et al. 2020. Fungal Diversity 105:1-16 ((https://doi.org/10.1007/s13225-020-00466-2)) |
| Otu20293 | 1 | k__Fungi(100);p__Ascomycota(100);c__Saccharomycetes(100);o__Saccharomycetales(100);f__Metschnikowiaceae(100);g__Metschnikowia(100);s__Metschnikowia_pulcherrima(100); | Metschnikowia | 13 | Saprotroph | \|Undefined Saprotroph\| | Yeast | NULL | Probable | As nectar-tap_saprotroph (Põlme et al. 2020); Decay sugar-rich substrates (Põlme et al. 2020) | Põlme S, et al. 2020. Fungal Diversity 105:1-16 ((https://doi.org/10.1007/s13225-020-00466-2)) |
| Otu20294 | 1 | k__Fungi(100);p__Ascomycota(100);c__Saccharomycetes(100);o__Saccharomycetales(100);f__Metschnikowiaceae(100);g__Metschnikowia(100);g__Metschnikowia_unclassified(100); | Metschnikowia | 13 | Saprotroph | \|Undefined Saprotroph\| | Yeast | NULL | Probable | As nectar-tap_saprotroph (Põlme et al. 2020); Decay sugar-rich substrates (Põlme et al. 2020) | Põlme S, et al. 2020. Fungal Diversity 105:1-16 ((https://doi.org/10.1007/s13225-020-00466-2)) |
| Otu20295 | 1 | k__Fungi(100);p__Ascomycota(100);c__Saccharomycetes(100);o__Saccharomycetales(100);f__Metschnikowiaceae(100);g__Metschnikowia(100);s__Metschnikowia_pulcherrima(100); | Metschnikowia | 13 | Saprotroph | \|Undefined Saprotroph\| | Yeast | NULL | Probable | As nectar-tap_saprotroph (Põlme et al. 2020); Decay sugar-rich substrates (Põlme et al. 2020) | Põlme S, et al. 2020. Fungal Diversity 105:1-16 ((https://doi.org/10.1007/s13225-020-00466-2)) |
| Otu20296 | 1 | k__Fungi(100);p__Ascomycota(100);c__Saccharomycetes(100);o__Saccharomycetales(100);f__Metschnikowiaceae(100);g__Metschnikowia(100);s__Metschnikowia_pulcherrima(100); | Metschnikowia | 13 | Saprotroph | \|Undefined Saprotroph\| | Yeast | NULL | Probable | As nectar-tap_saprotroph (Põlme et al. 2020); Decay sugar-rich substrates (Põlme et al. 2020) | Põlme S, et al. 2020. Fungal Diversity 105:1-16 ((https://doi.org/10.1007/s13225-020-00466-2)) |
| Otu20297 | 1 | k__Fungi(100);p__Ascomycota(100);c__Saccharomycetes(100);o__Saccharomycetales(100);f__Metschnikowiaceae(100);g__Metschnikowia(100);s__Metschnikowia_pulcherrima(100); | Metschnikowia | 13 | Saprotroph | \|Undefined Saprotroph\| | Yeast | NULL | Probable | As nectar-tap_saprotroph (Põlme et al. 2020); Decay sugar-rich substrates (Põlme et al. 2020) | Põlme S, et al. 2020. Fungal Diversity 105:1-16 ((https://doi.org/10.1007/s13225-020-00466-2)) |
| Otu20298 | 1 | k__Fungi(100);p__Ascomycota(100);c__Saccharomycetes(100);o__Saccharomycetales(100);f__Metschnikowiaceae(100);g__Metschnikowia(100);s__Metschnikowia_pulcherrima(100); | Metschnikowia | 13 | Saprotroph | \|Undefined Saprotroph\| | Yeast | NULL | Probable | As nectar-tap_saprotroph (Põlme et al. 2020); Decay sugar-rich substrates (Põlme et al. 2020) | Põlme S, et al. 2020. Fungal Diversity 105:1-16 ((https://doi.org/10.1007/s13225-020-00466-2)) |
| Otu20299 | 1 | k__Fungi(100);p__Ascomycota(100);c__Saccharomycetes(100);o__Saccharomycetales(100);f__Metschnikowiaceae(100);g__Metschnikowia(100);s__Metschnikowia_pulcherrima(100); | Metschnikowia | 13 | Saprotroph | \|Undefined Saprotroph\| | Yeast | NULL | Probable | As nectar-tap_saprotroph (Põlme et al. 2020); Decay sugar-rich substrates (Põlme et al. 2020) | Põlme S, et al. 2020. Fungal Diversity 105:1-16 ((https://doi.org/10.1007/s13225-020-00466-2)) |
| Otu20300 | 1 | k__Fungi(100);p__Ascomycota(100);c__Saccharomycetes(100);o__Saccharomycetales(100);f__Metschnikowiaceae(100);g__Metschnikowia(100);s__Metschnikowia_pulcherrima(100); | Metschnikowia | 13 | Saprotroph | \|Undefined Saprotroph\| | Yeast | NULL | Probable | As nectar-tap_saprotroph (Põlme et al. 2020); Decay sugar-rich substrates (Põlme et al. 2020) | Põlme S, et al. 2020. Fungal Diversity 105:1-16 ((https://doi.org/10.1007/s13225-020-00466-2)) |
| Otu20301 | 1 | k__Fungi(100);p__Ascomycota(100);c__Saccharomycetes(100);o__Saccharomycetales(100);f__Metschnikowiaceae(100);g__Metschnikowia(100);s__Metschnikowia_pulcherrima(100); | Metschnikowia | 13 | Saprotroph | \|Undefined Saprotroph\| | Yeast | NULL | Probable | As nectar-tap_saprotroph (Põlme et al. 2020); Decay sugar-rich substrates (Põlme et al. 2020) | Põlme S, et al. 2020. Fungal Diversity 105:1-16 ((https://doi.org/10.1007/s13225-020-00466-2)) |
| Otu20302 | 1 | k__Fungi(100);p__Ascomycota(100);c__Saccharomycetes(100);o__Saccharomycetales(100);f__Metschnikowiaceae(100);g__Metschnikowia(100);s__Metschnikowia_pulcherrima(100); | Metschnikowia | 13 | Saprotroph | \|Undefined Saprotroph\| | Yeast | NULL | Probable | As nectar-tap_saprotroph (Põlme et al. 2020); Decay sugar-rich substrates (Põlme et al. 2020) | Põlme S, et al. 2020. Fungal Diversity 105:1-16 ((https://doi.org/10.1007/s13225-020-00466-2)) |
| Otu20303 | 1 | k__Fungi(100);p__Ascomycota(100);c__Saccharomycetes(100);o__Saccharomycetales(100);f__Metschnikowiaceae(100);g__Metschnikowia(100);g__Metschnikowia_unclassified(100); | Metschnikowia | 13 | Saprotroph | \|Undefined Saprotroph\| | Yeast | NULL | Probable | As nectar-tap_saprotroph (Põlme et al. 2020); Decay sugar-rich substrates (Põlme et al. 2020) | Põlme S, et al. 2020. Fungal Diversity 105:1-16 ((https://doi.org/10.1007/s13225-020-00466-2)) |
| Otu20304 | 1 | k__Fungi(100);p__Ascomycota(100);c__Saccharomycetes(100);o__Saccharomycetales(100);f__Metschnikowiaceae(100);g__Metschnikowia(100);s__Metschnikowia_pulcherrima(100); | Metschnikowia | 13 | Saprotroph | \|Undefined Saprotroph\| | Yeast | NULL | Probable | As nectar-tap_saprotroph (Põlme et al. 2020); Decay sugar-rich substrates (Põlme et al. 2020) | Põlme S, et al. 2020. Fungal Diversity 105:1-16 ((https://doi.org/10.1007/s13225-020-00466-2)) |
| Otu20305 | 1 | k__Fungi(100);p__Ascomycota(100);c__Saccharomycetes(100);o__Saccharomycetales(100);f__Metschnikowiaceae(100);g__Metschnikowia(100);s__Metschnikowia_pulcherrima(100); | Metschnikowia | 13 | Saprotroph | \|Undefined Saprotroph\| | Yeast | NULL | Probable | As nectar-tap_saprotroph (Põlme et al. 2020); Decay sugar-rich substrates (Põlme et al. 2020) | Põlme S, et al. 2020. Fungal Diversity 105:1-16 ((https://doi.org/10.1007/s13225-020-00466-2)) |
| Otu20306 | 1 | k__Fungi(100);p__Ascomycota(100);c__Saccharomycetes(100);o__Saccharomycetales(100);f__Metschnikowiaceae(100);g__Metschnikowia(100);s__Metschnikowia_pulcherrima(100); | Metschnikowia | 13 | Saprotroph | \|Undefined Saprotroph\| | Yeast | NULL | Probable | As nectar-tap_saprotroph (Põlme et al. 2020); Decay sugar-rich substrates (Põlme et al. 2020) | Põlme S, et al. 2020. Fungal Diversity 105:1-16 ((https://doi.org/10.1007/s13225-020-00466-2)) |
| Otu20307 | 1 | k__Fungi(100);p__Ascomycota(100);c__Saccharomycetes(100);o__Saccharomycetales(100);f__Metschnikowiaceae(100);g__Metschnikowia(100);s__Metschnikowia_pulcherrima(100); | Metschnikowia | 13 | Saprotroph | \|Undefined Saprotroph\| | Yeast | NULL | Probable | As nectar-tap_saprotroph (Põlme et al. 2020); Decay sugar-rich substrates (Põlme et al. 2020) | Põlme S, et al. 2020. Fungal Diversity 105:1-16 ((https://doi.org/10.1007/s13225-020-00466-2)) |
| Otu20308 | 1 | k__Fungi(100);p__Ascomycota(100);c__Saccharomycetes(100);o__Saccharomycetales(100);f__Metschnikowiaceae(100);g__Metschnikowia(100);s__Metschnikowia_pulcherrima(100); | Metschnikowia | 13 | Saprotroph | \|Undefined Saprotroph\| | Yeast | NULL | Probable | As nectar-tap_saprotroph (Põlme et al. 2020); Decay sugar-rich substrates (Põlme et al. 2020) | Põlme S, et al. 2020. Fungal Diversity 105:1-16 ((https://doi.org/10.1007/s13225-020-00466-2)) |
| Otu20309 | 1 | k__Fungi(100);p__Ascomycota(100);c__Saccharomycetes(100);o__Saccharomycetales(100);f__Metschnikowiaceae(100);g__Metschnikowia(100);s__Metschnikowia_pulcherrima(100); | Metschnikowia | 13 | Saprotroph | \|Undefined Saprotroph\| | Yeast | NULL | Probable | As nectar-tap_saprotroph (Põlme et al. 2020); Decay sugar-rich substrates (Põlme et al. 2020) | Põlme S, et al. 2020. Fungal Diversity 105:1-16 ((https://doi.org/10.1007/s13225-020-00466-2)) |
| Otu20310 | 1 | k__Fungi(100);p__Ascomycota(100);c__Saccharomycetes(100);o__Saccharomycetales(100);f__Metschnikowiaceae(100);g__Metschnikowia(100);s__Metschnikowia_pulcherrima(100); | Metschnikowia | 13 | Saprotroph | \|Undefined Saprotroph\| | Yeast | NULL | Probable | As nectar-tap_saprotroph (Põlme et al. 2020); Decay sugar-rich substrates (Põlme et al. 2020) | Põlme S, et al. 2020. Fungal Diversity 105:1-16 ((https://doi.org/10.1007/s13225-020-00466-2)) |
| Otu20311 | 1 | k__Fungi(100);p__Ascomycota(100);c__Saccharomycetes(100);o__Saccharomycetales(100);f__Metschnikowiaceae(100);g__Metschnikowia(100);s__Metschnikowia_pulcherrima(100); | Metschnikowia | 13 | Saprotroph | \|Undefined Saprotroph\| | Yeast | NULL | Probable | As nectar-tap_saprotroph (Põlme et al. 2020); Decay sugar-rich substrates (Põlme et al. 2020) | Põlme S, et al. 2020. Fungal Diversity 105:1-16 ((https://doi.org/10.1007/s13225-020-00466-2)) |
| Otu20312 | 1 | k__Fungi(100);p__Ascomycota(100);c__Saccharomycetes(100);o__Saccharomycetales(100);f__Metschnikowiaceae(100);g__Metschnikowia(100);g__Metschnikowia_unclassified(100); | Metschnikowia | 13 | Saprotroph | \|Undefined Saprotroph\| | Yeast | NULL | Probable | As nectar-tap_saprotroph (Põlme et al. 2020); Decay sugar-rich substrates (Põlme et al. 2020) | Põlme S, et al. 2020. Fungal Diversity 105:1-16 ((https://doi.org/10.1007/s13225-020-00466-2)) |
| Otu20313 | 1 | k__Fungi(100);p__Ascomycota(100);c__Saccharomycetes(100);o__Saccharomycetales(100);f__Metschnikowiaceae(100);g__Metschnikowia(100);g__Metschnikowia_unclassified(100); | Metschnikowia | 13 | Saprotroph | \|Undefined Saprotroph\| | Yeast | NULL | Probable | As nectar-tap_saprotroph (Põlme et al. 2020); Decay sugar-rich substrates (Põlme et al. 2020) | Põlme S, et al. 2020. Fungal Diversity 105:1-16 ((https://doi.org/10.1007/s13225-020-00466-2)) |
| Otu20314 | 1 | k__Fungi(100);p__Ascomycota(100);c__Saccharomycetes(100);o__Saccharomycetales(100);f__Metschnikowiaceae(100);g__Metschnikowia(100);s__Metschnikowia_pulcherrima(100); | Metschnikowia | 13 | Saprotroph | \|Undefined Saprotroph\| | Yeast | NULL | Probable | As nectar-tap_saprotroph (Põlme et al. 2020); Decay sugar-rich substrates (Põlme et al. 2020) | Põlme S, et al. 2020. Fungal Diversity 105:1-16 ((https://doi.org/10.1007/s13225-020-00466-2)) |
| Otu20316 | 1 | k__Fungi(100);p__Ascomycota(100);c__Saccharomycetes(100);o__Saccharomycetales(100);f__Metschnikowiaceae(100);g__Metschnikowia(100);g__Metschnikowia_unclassified(100); | Metschnikowia | 13 | Saprotroph | \|Undefined Saprotroph\| | Yeast | NULL | Probable | As nectar-tap_saprotroph (Põlme et al. 2020); Decay sugar-rich substrates (Põlme et al. 2020) | Põlme S, et al. 2020. Fungal Diversity 105:1-16 ((https://doi.org/10.1007/s13225-020-00466-2)) |
| Otu20317 | 1 | k__Fungi(100);p__Ascomycota(100);c__Saccharomycetes(100);o__Saccharomycetales(100);f__Metschnikowiaceae(100);g__Metschnikowia(100);s__Metschnikowia_pulcherrima(100); | Metschnikowia | 13 | Saprotroph | \|Undefined Saprotroph\| | Yeast | NULL | Probable | As nectar-tap_saprotroph (Põlme et al. 2020); Decay sugar-rich substrates (Põlme et al. 2020) | Põlme S, et al. 2020. Fungal Diversity 105:1-16 ((https://doi.org/10.1007/s13225-020-00466-2)) |
| Otu20318 | 1 | k__Fungi(100);p__Ascomycota(100);c__Saccharomycetes(100);o__Saccharomycetales(100);f__Metschnikowiaceae(100);g__Metschnikowia(100);s__Metschnikowia_pulcherrima(100); | Metschnikowia | 13 | Saprotroph | \|Undefined Saprotroph\| | Yeast | NULL | Probable | As nectar-tap_saprotroph (Põlme et al. 2020); Decay sugar-rich substrates (Põlme et al. 2020) | Põlme S, et al. 2020. Fungal Diversity 105:1-16 ((https://doi.org/10.1007/s13225-020-00466-2)) |
| Otu20319 | 1 | k__Fungi(100);p__Ascomycota(100);c__Saccharomycetes(100);o__Saccharomycetales(100);f__Metschnikowiaceae(100);g__Metschnikowia(100);s__Metschnikowia_pulcherrima(100); | Metschnikowia | 13 | Saprotroph | \|Undefined Saprotroph\| | Yeast | NULL | Probable | As nectar-tap_saprotroph (Põlme et al. 2020); Decay sugar-rich substrates (Põlme et al. 2020) | Põlme S, et al. 2020. Fungal Diversity 105:1-16 ((https://doi.org/10.1007/s13225-020-00466-2)) |
| Otu20321 | 1 | k__Fungi(100);p__Ascomycota(100);c__Saccharomycetes(100);o__Saccharomycetales(100);o__Saccharomycetales_unclassified(100);o__Saccharomycetales_unclassified(100);o__Saccharomycetales_unclassified(100); | Saccharomycetales | 7 | Saprotroph | Undefined Saprotroph | Yeast | NULL | Possible | NULL | Sterkenburg E, et al. 2015. New Phytologist 207:1145-1158 ((https://doi.org/10.1111/nph.13426)) |
| Otu20322 | 1 | k__Fungi(100);p__Ascomycota(100);c__Saccharomycetes(100);o__Saccharomycetales(100);f__Saccharomycetales_family_Incertae_sedis(100);g__Candida(100);s__Candida_cretensis(100); | Candida | 13 | Pathotroph-Saprotroph-Symbiotroph | Animal Pathogen-Endophyte-Endosymbiont-Epiphyte-\|Undefined Saprotroph\| | Dimorphic Yeast-Yeast | NULL | Probable | As nectar-tap_saprotroph (Põlme et al. 2020); Decay sugar-rich substrates (Põlme et al. 2020); This asexual genus is polyphyletic and contain species that function very differently depend on what they are. Caution should be taken when interpreting guilds for Candida species. | Manolakaki D, et al. 2010. Virulence. 1: 367-375 ((https://doi.org/10.4161/viru.1.5.12796)); Mekbib SB, et al. 2011. Tropical Plant Pathology 36:233-240 ((https://doi.org/10.1590/S1982-56762011000400004)); Põlme S, et al. 2020. Fungal Diversity 105:1-16 ((https://doi.org/10.1007/s13225-020-00466-2)) |
| Otu20324 | 1 | k__Fungi(100);p__Ascomycota(100);c__Saccharomycetes(100);o__Saccharomycetales(100);f__Metschnikowiaceae(100);g__Metschnikowia(100);s__Metschnikowia_pulcherrima(100); | Metschnikowia | 13 | Saprotroph | \|Undefined Saprotroph\| | Yeast | NULL | Probable | As nectar-tap_saprotroph (Põlme et al. 2020); Decay sugar-rich substrates (Põlme et al. 2020) | Põlme S, et al. 2020. Fungal Diversity 105:1-16 ((https://doi.org/10.1007/s13225-020-00466-2)) |
| Otu20325 | 1 | k__Fungi(100);p__Ascomycota(100);c__Saccharomycetes(100);o__Saccharomycetales(100);f__Metschnikowiaceae(100);g__Metschnikowia(100);s__Metschnikowia_pulcherrima(100); | Metschnikowia | 13 | Saprotroph | \|Undefined Saprotroph\| | Yeast | NULL | Probable | As nectar-tap_saprotroph (Põlme et al. 2020); Decay sugar-rich substrates (Põlme et al. 2020) | Põlme S, et al. 2020. Fungal Diversity 105:1-16 ((https://doi.org/10.1007/s13225-020-00466-2)) |
| Otu20326 | 1 | k__Fungi(100);p__Ascomycota(100);c__Saccharomycetes(100);o__Saccharomycetales(100);f__Metschnikowiaceae(100);g__Metschnikowia(100);s__Metschnikowia_pulcherrima(100); | Metschnikowia | 13 | Saprotroph | \|Undefined Saprotroph\| | Yeast | NULL | Probable | As nectar-tap_saprotroph (Põlme et al. 2020); Decay sugar-rich substrates (Põlme et al. 2020) | Põlme S, et al. 2020. Fungal Diversity 105:1-16 ((https://doi.org/10.1007/s13225-020-00466-2)) |
| Otu20327 | 1 | k__Fungi(100);p__Ascomycota(100);c__Saccharomycetes(100);o__Saccharomycetales(100);f__Metschnikowiaceae(100);g__Metschnikowia(100);s__Metschnikowia_pulcherrima(100); | Metschnikowia | 13 | Saprotroph | \|Undefined Saprotroph\| | Yeast | NULL | Probable | As nectar-tap_saprotroph (Põlme et al. 2020); Decay sugar-rich substrates (Põlme et al. 2020) | Põlme S, et al. 2020. Fungal Diversity 105:1-16 ((https://doi.org/10.1007/s13225-020-00466-2)) |
| Otu20328 | 1 | k__Fungi(100);p__Ascomycota(100);c__Saccharomycetes(100);o__Saccharomycetales(100);f__Metschnikowiaceae(100);g__Metschnikowia(100);s__Metschnikowia_pulcherrima(100); | Metschnikowia | 13 | Saprotroph | \|Undefined Saprotroph\| | Yeast | NULL | Probable | As nectar-tap_saprotroph (Põlme et al. 2020); Decay sugar-rich substrates (Põlme et al. 2020) | Põlme S, et al. 2020. Fungal Diversity 105:1-16 ((https://doi.org/10.1007/s13225-020-00466-2)) |
| Otu20329 | 1 | k__Fungi(100);p__Ascomycota(100);c__Saccharomycetes(100);o__Saccharomycetales(100);f__Metschnikowiaceae(100);g__Metschnikowia(100);s__Metschnikowia_pulcherrima(100); | Metschnikowia | 13 | Saprotroph | \|Undefined Saprotroph\| | Yeast | NULL | Probable | As nectar-tap_saprotroph (Põlme et al. 2020); Decay sugar-rich substrates (Põlme et al. 2020) | Põlme S, et al. 2020. Fungal Diversity 105:1-16 ((https://doi.org/10.1007/s13225-020-00466-2)) |
| Otu20331 | 1 | k__Fungi(100);p__Basidiomycota(100);c__Tremellomycetes(100);o__Tremellales(100);f__Tremellales_family_Incertae_sedis(100);g__Hannaella(100);s__Hannaella_sp(100); | Hannaella | 13 | Pathotroph-Saprotroph | \|Animal Parasite\|-Undefined Saprotroph | Yeast | NULL | Probable | As animal_decomposer (Põlme et al. 2020); Decay animal material (Põlme et al. 2020) | Põlme S, et al. 2020. Fungal Diversity 105:1-16 ((https://doi.org/10.1007/s13225-020-00466-2)) |
| Otu20338 | 1 | k__Fungi(100);p__Ascomycota(100);c__Saccharomycetes(100);o__Saccharomycetales(100);f__Metschnikowiaceae(100);g__Metschnikowia(100);s__Metschnikowia_pulcherrima(100); | Metschnikowia | 13 | Saprotroph | \|Undefined Saprotroph\| | Yeast | NULL | Probable | As nectar-tap_saprotroph (Põlme et al. 2020); Decay sugar-rich substrates (Põlme et al. 2020) | Põlme S, et al. 2020. Fungal Diversity 105:1-16 ((https://doi.org/10.1007/s13225-020-00466-2)) |
| Otu20342 | 1 | k__Fungi(100);p__Ascomycota(100);c__Saccharomycetes(100);o__Saccharomycetales(100);f__Metschnikowiaceae(100);g__Metschnikowia(100);s__Metschnikowia_pulcherrima(100); | Metschnikowia | 13 | Saprotroph | \|Undefined Saprotroph\| | Yeast | NULL | Probable | As nectar-tap_saprotroph (Põlme et al. 2020); Decay sugar-rich substrates (Põlme et al. 2020) | Põlme S, et al. 2020. Fungal Diversity 105:1-16 ((https://doi.org/10.1007/s13225-020-00466-2)) |
| Otu20344 | 1 | k__Fungi(100);p__Ascomycota(100);c__Saccharomycetes(100);o__Saccharomycetales(100);f__Metschnikowiaceae(100);g__Metschnikowia(100);s__Metschnikowia_pulcherrima(100); | Metschnikowia | 13 | Saprotroph | \|Undefined Saprotroph\| | Yeast | NULL | Probable | As nectar-tap_saprotroph (Põlme et al. 2020); Decay sugar-rich substrates (Põlme et al. 2020) | Põlme S, et al. 2020. Fungal Diversity 105:1-16 ((https://doi.org/10.1007/s13225-020-00466-2)) |
| Otu20346 | 1 | k__Fungi(100);p__Ascomycota(100);c__Saccharomycetes(100);o__Saccharomycetales(100);f__Metschnikowiaceae(100);g__Metschnikowia(100);s__Metschnikowia_pulcherrima(100); | Metschnikowia | 13 | Saprotroph | \|Undefined Saprotroph\| | Yeast | NULL | Probable | As nectar-tap_saprotroph (Põlme et al. 2020); Decay sugar-rich substrates (Põlme et al. 2020) | Põlme S, et al. 2020. Fungal Diversity 105:1-16 ((https://doi.org/10.1007/s13225-020-00466-2)) |
| Otu20347 | 1 | k__Fungi(100);p__Ascomycota(100);c__Saccharomycetes(100);o__Saccharomycetales(100);f__Metschnikowiaceae(100);g__Metschnikowia(100);s__Metschnikowia_pulcherrima(100); | Metschnikowia | 13 | Saprotroph | \|Undefined Saprotroph\| | Yeast | NULL | Probable | As nectar-tap_saprotroph (Põlme et al. 2020); Decay sugar-rich substrates (Põlme et al. 2020) | Põlme S, et al. 2020. Fungal Diversity 105:1-16 ((https://doi.org/10.1007/s13225-020-00466-2)) |
| Otu20352 | 1 | k__Fungi(100);p__Ascomycota(100);c__Saccharomycetes(100);o__Saccharomycetales(100);f__Metschnikowiaceae(100);g__Metschnikowia(100);s__Metschnikowia_pulcherrima(100); | Metschnikowia | 13 | Saprotroph | \|Undefined Saprotroph\| | Yeast | NULL | Probable | As nectar-tap_saprotroph (Põlme et al. 2020); Decay sugar-rich substrates (Põlme et al. 2020) | Põlme S, et al. 2020. Fungal Diversity 105:1-16 ((https://doi.org/10.1007/s13225-020-00466-2)) |
| Otu20353 | 1 | k__Fungi(100);p__Ascomycota(100);c__Saccharomycetes(100);o__Saccharomycetales(100);f__Metschnikowiaceae(100);g__Metschnikowia(100);s__Metschnikowia_pulcherrima(100); | Metschnikowia | 13 | Saprotroph | \|Undefined Saprotroph\| | Yeast | NULL | Probable | As nectar-tap_saprotroph (Põlme et al. 2020); Decay sugar-rich substrates (Põlme et al. 2020) | Põlme S, et al. 2020. Fungal Diversity 105:1-16 ((https://doi.org/10.1007/s13225-020-00466-2)) |
| Otu20355 | 1 | k__Fungi(100);p__Ascomycota(100);c__Saccharomycetes(100);o__Saccharomycetales(100);f__Metschnikowiaceae(100);g__Metschnikowia(100);s__Metschnikowia_pulcherrima(100); | Metschnikowia | 13 | Saprotroph | \|Undefined Saprotroph\| | Yeast | NULL | Probable | As nectar-tap_saprotroph (Põlme et al. 2020); Decay sugar-rich substrates (Põlme et al. 2020) | Põlme S, et al. 2020. Fungal Diversity 105:1-16 ((https://doi.org/10.1007/s13225-020-00466-2)) |
| Otu20356 | 1 | k__Fungi(100);p__Ascomycota(100);c__Saccharomycetes(100);o__Saccharomycetales(100);f__Metschnikowiaceae(100);g__Metschnikowia(100);s__Metschnikowia_pulcherrima(100); | Metschnikowia | 13 | Saprotroph | \|Undefined Saprotroph\| | Yeast | NULL | Probable | As nectar-tap_saprotroph (Põlme et al. 2020); Decay sugar-rich substrates (Põlme et al. 2020) | Põlme S, et al. 2020. Fungal Diversity 105:1-16 ((https://doi.org/10.1007/s13225-020-00466-2)) |
| Otu20359 | 1 | k__Fungi(100);p__Ascomycota(100);c__Saccharomycetes(100);o__Saccharomycetales(100);f__Metschnikowiaceae(100);g__Metschnikowia(100);s__Metschnikowia_pulcherrima(100); | Metschnikowia | 13 | Saprotroph | \|Undefined Saprotroph\| | Yeast | NULL | Probable | As nectar-tap_saprotroph (Põlme et al. 2020); Decay sugar-rich substrates (Põlme et al. 2020) | Põlme S, et al. 2020. Fungal Diversity 105:1-16 ((https://doi.org/10.1007/s13225-020-00466-2)) |
| Otu20360 | 1 | k__Fungi(100);p__Ascomycota(100);c__Saccharomycetes(100);o__Saccharomycetales(100);f__Metschnikowiaceae(100);g__Metschnikowia(100);s__Metschnikowia_pulcherrima(100); | Metschnikowia | 13 | Saprotroph | \|Undefined Saprotroph\| | Yeast | NULL | Probable | As nectar-tap_saprotroph (Põlme et al. 2020); Decay sugar-rich substrates (Põlme et al. 2020) | Põlme S, et al. 2020. Fungal Diversity 105:1-16 ((https://doi.org/10.1007/s13225-020-00466-2)) |
| Otu20361 | 1 | k__Fungi(100);p__Ascomycota(100);c__Saccharomycetes(100);o__Saccharomycetales(100);f__Metschnikowiaceae(100);g__Metschnikowia(100);s__Metschnikowia_pulcherrima(100); | Metschnikowia | 13 | Saprotroph | \|Undefined Saprotroph\| | Yeast | NULL | Probable | As nectar-tap_saprotroph (Põlme et al. 2020); Decay sugar-rich substrates (Põlme et al. 2020) | Põlme S, et al. 2020. Fungal Diversity 105:1-16 ((https://doi.org/10.1007/s13225-020-00466-2)) |
| Otu20362 | 1 | k__Fungi(100);p__Ascomycota(100);c__Saccharomycetes(100);o__Saccharomycetales(100);f__Metschnikowiaceae(100);g__Metschnikowia(100);s__Metschnikowia_pulcherrima(100); | Metschnikowia | 13 | Saprotroph | \|Undefined Saprotroph\| | Yeast | NULL | Probable | As nectar-tap_saprotroph (Põlme et al. 2020); Decay sugar-rich substrates (Põlme et al. 2020) | Põlme S, et al. 2020. Fungal Diversity 105:1-16 ((https://doi.org/10.1007/s13225-020-00466-2)) |
| Otu20363 | 1 | k__Fungi(100);p__Ascomycota(100);c__Saccharomycetes(100);o__Saccharomycetales(100);f__Metschnikowiaceae(100);g__Metschnikowia(100);g__Metschnikowia_unclassified(100); | Metschnikowia | 13 | Saprotroph | \|Undefined Saprotroph\| | Yeast | NULL | Probable | As nectar-tap_saprotroph (Põlme et al. 2020); Decay sugar-rich substrates (Põlme et al. 2020) | Põlme S, et al. 2020. Fungal Diversity 105:1-16 ((https://doi.org/10.1007/s13225-020-00466-2)) |
| Otu20364 | 1 | k__Fungi(100);p__Ascomycota(100);c__Saccharomycetes(100);o__Saccharomycetales(100);f__Metschnikowiaceae(100);g__Metschnikowia(100);s__Metschnikowia_pulcherrima(100); | Metschnikowia | 13 | Saprotroph | \|Undefined Saprotroph\| | Yeast | NULL | Probable | As nectar-tap_saprotroph (Põlme et al. 2020); Decay sugar-rich substrates (Põlme et al. 2020) | Põlme S, et al. 2020. Fungal Diversity 105:1-16 ((https://doi.org/10.1007/s13225-020-00466-2)) |
| Otu20365 | 1 | k__Fungi(100);p__Ascomycota(100);c__Saccharomycetes(100);o__Saccharomycetales(100);f__Metschnikowiaceae(100);g__Metschnikowia(100);s__Metschnikowia_pulcherrima(100); | Metschnikowia | 13 | Saprotroph | \|Undefined Saprotroph\| | Yeast | NULL | Probable | As nectar-tap_saprotroph (Põlme et al. 2020); Decay sugar-rich substrates (Põlme et al. 2020) | Põlme S, et al. 2020. Fungal Diversity 105:1-16 ((https://doi.org/10.1007/s13225-020-00466-2)) |
| Otu20366 | 1 | k__Fungi(100);p__Ascomycota(100);c__Saccharomycetes(100);o__Saccharomycetales(100);f__Metschnikowiaceae(100);g__Metschnikowia(100);s__Metschnikowia_pulcherrima(100); | Metschnikowia | 13 | Saprotroph | \|Undefined Saprotroph\| | Yeast | NULL | Probable | As nectar-tap_saprotroph (Põlme et al. 2020); Decay sugar-rich substrates (Põlme et al. 2020) | Põlme S, et al. 2020. Fungal Diversity 105:1-16 ((https://doi.org/10.1007/s13225-020-00466-2)) |
| Otu20367 | 1 | k__Fungi(100);p__Ascomycota(100);c__Saccharomycetes(100);o__Saccharomycetales(100);f__Metschnikowiaceae(100);g__Metschnikowia(100);s__Metschnikowia_pulcherrima(100); | Metschnikowia | 13 | Saprotroph | \|Undefined Saprotroph\| | Yeast | NULL | Probable | As nectar-tap_saprotroph (Põlme et al. 2020); Decay sugar-rich substrates (Põlme et al. 2020) | Põlme S, et al. 2020. Fungal Diversity 105:1-16 ((https://doi.org/10.1007/s13225-020-00466-2)) |
| Otu20368 | 1 | k__Fungi(100);p__Ascomycota(100);c__Saccharomycetes(100);o__Saccharomycetales(100);f__Metschnikowiaceae(100);g__Metschnikowia(100);s__Metschnikowia_pulcherrima(100); | Metschnikowia | 13 | Saprotroph | \|Undefined Saprotroph\| | Yeast | NULL | Probable | As nectar-tap_saprotroph (Põlme et al. 2020); Decay sugar-rich substrates (Põlme et al. 2020) | Põlme S, et al. 2020. Fungal Diversity 105:1-16 ((https://doi.org/10.1007/s13225-020-00466-2)) |
| Otu20369 | 1 | k__Fungi(100);p__Ascomycota(100);c__Saccharomycetes(100);o__Saccharomycetales(100);f__Saccharomycetales_family_Incertae_sedis(100);g__Candida(100);g__Candida_unclassified(100); | Candida | 13 | Pathotroph-Saprotroph-Symbiotroph | Animal Pathogen-Endophyte-Endosymbiont-Epiphyte-\|Undefined Saprotroph\| | Dimorphic Yeast-Yeast | NULL | Probable | As nectar-tap_saprotroph (Põlme et al. 2020); Decay sugar-rich substrates (Põlme et al. 2020); This asexual genus is polyphyletic and contain species that function very differently depend on what they are. Caution should be taken when interpreting guilds for Candida species. | Manolakaki D, et al. 2010. Virulence. 1: 367-375 ((https://doi.org/10.4161/viru.1.5.12796)); Mekbib SB, et al. 2011. Tropical Plant Pathology 36:233-240 ((https://doi.org/10.1590/S1982-56762011000400004)); Põlme S, et al. 2020. Fungal Diversity 105:1-16 ((https://doi.org/10.1007/s13225-020-00466-2)) |
| Otu20370 | 1 | k__Fungi(100);p__Ascomycota(100);c__Saccharomycetes(100);o__Saccharomycetales(100);f__Metschnikowiaceae(100);g__Metschnikowia(100);s__Metschnikowia_pulcherrima(100); | Metschnikowia | 13 | Saprotroph | \|Undefined Saprotroph\| | Yeast | NULL | Probable | As nectar-tap_saprotroph (Põlme et al. 2020); Decay sugar-rich substrates (Põlme et al. 2020) | Põlme S, et al. 2020. Fungal Diversity 105:1-16 ((https://doi.org/10.1007/s13225-020-00466-2)) |
| Otu20371 | 1 | k__Fungi(100);p__Ascomycota(100);c__Saccharomycetes(100);o__Saccharomycetales(100);f__Metschnikowiaceae(100);g__Metschnikowia(100);s__Metschnikowia_pulcherrima(100); | Metschnikowia | 13 | Saprotroph | \|Undefined Saprotroph\| | Yeast | NULL | Probable | As nectar-tap_saprotroph (Põlme et al. 2020); Decay sugar-rich substrates (Põlme et al. 2020) | Põlme S, et al. 2020. Fungal Diversity 105:1-16 ((https://doi.org/10.1007/s13225-020-00466-2)) |
| Otu20372 | 1 | k__Fungi(100);p__Ascomycota(100);c__Saccharomycetes(100);o__Saccharomycetales(100);f__Metschnikowiaceae(100);g__Metschnikowia(100);g__Metschnikowia_unclassified(100); | Metschnikowia | 13 | Saprotroph | \|Undefined Saprotroph\| | Yeast | NULL | Probable | As nectar-tap_saprotroph (Põlme et al. 2020); Decay sugar-rich substrates (Põlme et al. 2020) | Põlme S, et al. 2020. Fungal Diversity 105:1-16 ((https://doi.org/10.1007/s13225-020-00466-2)) |
| Otu20373 | 1 | k__Fungi(100);p__Ascomycota(100);c__Saccharomycetes(100);o__Saccharomycetales(100);f__Metschnikowiaceae(100);g__Metschnikowia(100);s__Metschnikowia_pulcherrima(100); | Metschnikowia | 13 | Saprotroph | \|Undefined Saprotroph\| | Yeast | NULL | Probable | As nectar-tap_saprotroph (Põlme et al. 2020); Decay sugar-rich substrates (Põlme et al. 2020) | Põlme S, et al. 2020. Fungal Diversity 105:1-16 ((https://doi.org/10.1007/s13225-020-00466-2)) |
| Otu20375 | 1 | k__Fungi(100);p__Ascomycota(100);c__Saccharomycetes(100);o__Saccharomycetales(100);f__Metschnikowiaceae(100);g__Metschnikowia(100);s__Metschnikowia_pulcherrima(100); | Metschnikowia | 13 | Saprotroph | \|Undefined Saprotroph\| | Yeast | NULL | Probable | As nectar-tap_saprotroph (Põlme et al. 2020); Decay sugar-rich substrates (Põlme et al. 2020) | Põlme S, et al. 2020. Fungal Diversity 105:1-16 ((https://doi.org/10.1007/s13225-020-00466-2)) |
| Otu20377 | 1 | k__Fungi(100);p__Ascomycota(100);c__Saccharomycetes(100);o__Saccharomycetales(100);f__Metschnikowiaceae(100);g__Metschnikowia(100);s__Metschnikowia_pulcherrima(100); | Metschnikowia | 13 | Saprotroph | \|Undefined Saprotroph\| | Yeast | NULL | Probable | As nectar-tap_saprotroph (Põlme et al. 2020); Decay sugar-rich substrates (Põlme et al. 2020) | Põlme S, et al. 2020. Fungal Diversity 105:1-16 ((https://doi.org/10.1007/s13225-020-00466-2)) |
| Otu20378 | 1 | k__Fungi(100);p__Ascomycota(100);c__Saccharomycetes(100);o__Saccharomycetales(100);f__Metschnikowiaceae(100);g__Metschnikowia(100);s__Metschnikowia_pulcherrima(100); | Metschnikowia | 13 | Saprotroph | \|Undefined Saprotroph\| | Yeast | NULL | Probable | As nectar-tap_saprotroph (Põlme et al. 2020); Decay sugar-rich substrates (Põlme et al. 2020) | Põlme S, et al. 2020. Fungal Diversity 105:1-16 ((https://doi.org/10.1007/s13225-020-00466-2)) |
| Otu20379 | 1 | k__Fungi(100);p__Ascomycota(100);c__Saccharomycetes(100);o__Saccharomycetales(100);f__Metschnikowiaceae(100);g__Metschnikowia(100);s__Metschnikowia_pulcherrima(100); | Metschnikowia | 13 | Saprotroph | \|Undefined Saprotroph\| | Yeast | NULL | Probable | As nectar-tap_saprotroph (Põlme et al. 2020); Decay sugar-rich substrates (Põlme et al. 2020) | Põlme S, et al. 2020. Fungal Diversity 105:1-16 ((https://doi.org/10.1007/s13225-020-00466-2)) |
| Otu20380 | 1 | k__Fungi(100);p__Ascomycota(100);c__Saccharomycetes(100);o__Saccharomycetales(100);f__Metschnikowiaceae(100);g__Metschnikowia(100);g__Metschnikowia_unclassified(100); | Metschnikowia | 13 | Saprotroph | \|Undefined Saprotroph\| | Yeast | NULL | Probable | As nectar-tap_saprotroph (Põlme et al. 2020); Decay sugar-rich substrates (Põlme et al. 2020) | Põlme S, et al. 2020. Fungal Diversity 105:1-16 ((https://doi.org/10.1007/s13225-020-00466-2)) |
| Otu20382 | 1 | k__Fungi(100);p__Ascomycota(100);c__Saccharomycetes(100);o__Saccharomycetales(100);f__Metschnikowiaceae(100);g__Metschnikowia(100);s__Metschnikowia_pulcherrima(100); | Metschnikowia | 13 | Saprotroph | \|Undefined Saprotroph\| | Yeast | NULL | Probable | As nectar-tap_saprotroph (Põlme et al. 2020); Decay sugar-rich substrates (Põlme et al. 2020) | Põlme S, et al. 2020. Fungal Diversity 105:1-16 ((https://doi.org/10.1007/s13225-020-00466-2)) |
| Otu20383 | 1 | k__Fungi(100);p__Ascomycota(100);c__Saccharomycetes(100);o__Saccharomycetales(100);f__Metschnikowiaceae(100);g__Metschnikowia(100);s__Metschnikowia_pulcherrima(100); | Metschnikowia | 13 | Saprotroph | \|Undefined Saprotroph\| | Yeast | NULL | Probable | As nectar-tap_saprotroph (Põlme et al. 2020); Decay sugar-rich substrates (Põlme et al. 2020) | Põlme S, et al. 2020. Fungal Diversity 105:1-16 ((https://doi.org/10.1007/s13225-020-00466-2)) |
| Otu20384 | 1 | k__Fungi(100);p__Ascomycota(100);c__Saccharomycetes(100);o__Saccharomycetales(100);f__Metschnikowiaceae(100);g__Metschnikowia(100);s__Metschnikowia_pulcherrima(100); | Metschnikowia | 13 | Saprotroph | \|Undefined Saprotroph\| | Yeast | NULL | Probable | As nectar-tap_saprotroph (Põlme et al. 2020); Decay sugar-rich substrates (Põlme et al. 2020) | Põlme S, et al. 2020. Fungal Diversity 105:1-16 ((https://doi.org/10.1007/s13225-020-00466-2)) |
| Otu20385 | 1 | k__Fungi(100);p__Ascomycota(100);c__Saccharomycetes(100);o__Saccharomycetales(100);f__Metschnikowiaceae(100);g__Metschnikowia(100);s__Metschnikowia_pulcherrima(100); | Metschnikowia | 13 | Saprotroph | \|Undefined Saprotroph\| | Yeast | NULL | Probable | As nectar-tap_saprotroph (Põlme et al. 2020); Decay sugar-rich substrates (Põlme et al. 2020) | Põlme S, et al. 2020. Fungal Diversity 105:1-16 ((https://doi.org/10.1007/s13225-020-00466-2)) |
| Otu20391 | 1 | k__Fungi(100);p__Ascomycota(100);c__Saccharomycetes(100);o__Saccharomycetales(100);f__Metschnikowiaceae(100);g__Metschnikowia(100);s__Metschnikowia_pulcherrima(100); | Metschnikowia | 13 | Saprotroph | \|Undefined Saprotroph\| | Yeast | NULL | Probable | As nectar-tap_saprotroph (Põlme et al. 2020); Decay sugar-rich substrates (Põlme et al. 2020) | Põlme S, et al. 2020. Fungal Diversity 105:1-16 ((https://doi.org/10.1007/s13225-020-00466-2)) |
| Otu20396 | 1 | k__Fungi(100);p__Ascomycota(100);c__Saccharomycetes(100);o__Saccharomycetales(100);f__Metschnikowiaceae(100);g__Metschnikowia(100);s__Metschnikowia_pulcherrima(100); | Metschnikowia | 13 | Saprotroph | \|Undefined Saprotroph\| | Yeast | NULL | Probable | As nectar-tap_saprotroph (Põlme et al. 2020); Decay sugar-rich substrates (Põlme et al. 2020) | Põlme S, et al. 2020. Fungal Diversity 105:1-16 ((https://doi.org/10.1007/s13225-020-00466-2)) |
| Otu20397 | 1 | k__Fungi(100);p__Ascomycota(100);c__Saccharomycetes(100);o__Saccharomycetales(100);f__Metschnikowiaceae(100);g__Metschnikowia(100);s__Metschnikowia_pulcherrima(100); | Metschnikowia | 13 | Saprotroph | \|Undefined Saprotroph\| | Yeast | NULL | Probable | As nectar-tap_saprotroph (Põlme et al. 2020); Decay sugar-rich substrates (Põlme et al. 2020) | Põlme S, et al. 2020. Fungal Diversity 105:1-16 ((https://doi.org/10.1007/s13225-020-00466-2)) |
| Otu20398 | 1 | k__Fungi(100);p__Ascomycota(100);c__Saccharomycetes(100);o__Saccharomycetales(100);f__Metschnikowiaceae(100);g__Metschnikowia(100);s__Metschnikowia_pulcherrima(100); | Metschnikowia | 13 | Saprotroph | \|Undefined Saprotroph\| | Yeast | NULL | Probable | As nectar-tap_saprotroph (Põlme et al. 2020); Decay sugar-rich substrates (Põlme et al. 2020) | Põlme S, et al. 2020. Fungal Diversity 105:1-16 ((https://doi.org/10.1007/s13225-020-00466-2)) |
| Otu20400 | 1 | k__Fungi(100);p__Ascomycota(100);c__Saccharomycetes(100);o__Saccharomycetales(100);f__Metschnikowiaceae(100);g__Metschnikowia(100);s__Metschnikowia_pulcherrima(100); | Metschnikowia | 13 | Saprotroph | \|Undefined Saprotroph\| | Yeast | NULL | Probable | As nectar-tap_saprotroph (Põlme et al. 2020); Decay sugar-rich substrates (Põlme et al. 2020) | Põlme S, et al. 2020. Fungal Diversity 105:1-16 ((https://doi.org/10.1007/s13225-020-00466-2)) |
| Otu20401 | 1 | k__Fungi(100);p__Ascomycota(100);c__Saccharomycetes(100);o__Saccharomycetales(100);f__Metschnikowiaceae(100);g__Metschnikowia(100);s__Metschnikowia_pulcherrima(100); | Metschnikowia | 13 | Saprotroph | \|Undefined Saprotroph\| | Yeast | NULL | Probable | As nectar-tap_saprotroph (Põlme et al. 2020); Decay sugar-rich substrates (Põlme et al. 2020) | Põlme S, et al. 2020. Fungal Diversity 105:1-16 ((https://doi.org/10.1007/s13225-020-00466-2)) |
| Otu20402 | 1 | k__Fungi(100);p__Ascomycota(100);c__Saccharomycetes(100);o__Saccharomycetales(100);f__Metschnikowiaceae(100);g__Metschnikowia(100);g__Metschnikowia_unclassified(100); | Metschnikowia | 13 | Saprotroph | \|Undefined Saprotroph\| | Yeast | NULL | Probable | As nectar-tap_saprotroph (Põlme et al. 2020); Decay sugar-rich substrates (Põlme et al. 2020) | Põlme S, et al. 2020. Fungal Diversity 105:1-16 ((https://doi.org/10.1007/s13225-020-00466-2)) |
| Otu20403 | 1 | k__Fungi(100);p__Ascomycota(100);c__Saccharomycetes(100);o__Saccharomycetales(100);f__Metschnikowiaceae(100);g__Metschnikowia(100);s__Metschnikowia_pulcherrima(100); | Metschnikowia | 13 | Saprotroph | \|Undefined Saprotroph\| | Yeast | NULL | Probable | As nectar-tap_saprotroph (Põlme et al. 2020); Decay sugar-rich substrates (Põlme et al. 2020) | Põlme S, et al. 2020. Fungal Diversity 105:1-16 ((https://doi.org/10.1007/s13225-020-00466-2)) |
| Otu20404 | 1 | k__Fungi(100);p__Ascomycota(100);c__Saccharomycetes(100);o__Saccharomycetales(100);f__Metschnikowiaceae(100);g__Metschnikowia(100);s__Metschnikowia_pulcherrima(100); | Metschnikowia | 13 | Saprotroph | \|Undefined Saprotroph\| | Yeast | NULL | Probable | As nectar-tap_saprotroph (Põlme et al. 2020); Decay sugar-rich substrates (Põlme et al. 2020) | Põlme S, et al. 2020. Fungal Diversity 105:1-16 ((https://doi.org/10.1007/s13225-020-00466-2)) |
| Otu20405 | 1 | k__Fungi(100);p__Ascomycota(100);c__Saccharomycetes(100);o__Saccharomycetales(100);f__Metschnikowiaceae(100);g__Metschnikowia(100);g__Metschnikowia_unclassified(100); | Metschnikowia | 13 | Saprotroph | \|Undefined Saprotroph\| | Yeast | NULL | Probable | As nectar-tap_saprotroph (Põlme et al. 2020); Decay sugar-rich substrates (Põlme et al. 2020) | Põlme S, et al. 2020. Fungal Diversity 105:1-16 ((https://doi.org/10.1007/s13225-020-00466-2)) |
| Otu20406 | 1 | k__Fungi(100);p__Ascomycota(100);c__Saccharomycetes(100);o__Saccharomycetales(100);f__Metschnikowiaceae(100);g__Metschnikowia(100);s__Metschnikowia_pulcherrima(100); | Metschnikowia | 13 | Saprotroph | \|Undefined Saprotroph\| | Yeast | NULL | Probable | As nectar-tap_saprotroph (Põlme et al. 2020); Decay sugar-rich substrates (Põlme et al. 2020) | Põlme S, et al. 2020. Fungal Diversity 105:1-16 ((https://doi.org/10.1007/s13225-020-00466-2)) |
| Otu20407 | 1 | k__Fungi(100);p__Ascomycota(100);c__Saccharomycetes(100);o__Saccharomycetales(100);f__Metschnikowiaceae(100);g__Metschnikowia(100);s__Metschnikowia_pulcherrima(100); | Metschnikowia | 13 | Saprotroph | \|Undefined Saprotroph\| | Yeast | NULL | Probable | As nectar-tap_saprotroph (Põlme et al. 2020); Decay sugar-rich substrates (Põlme et al. 2020) | Põlme S, et al. 2020. Fungal Diversity 105:1-16 ((https://doi.org/10.1007/s13225-020-00466-2)) |
| Otu20408 | 1 | k__Fungi(100);p__Ascomycota(100);c__Saccharomycetes(100);o__Saccharomycetales(100);f__Metschnikowiaceae(100);g__Metschnikowia(100);s__Metschnikowia_pulcherrima(100); | Metschnikowia | 13 | Saprotroph | \|Undefined Saprotroph\| | Yeast | NULL | Probable | As nectar-tap_saprotroph (Põlme et al. 2020); Decay sugar-rich substrates (Põlme et al. 2020) | Põlme S, et al. 2020. Fungal Diversity 105:1-16 ((https://doi.org/10.1007/s13225-020-00466-2)) |
| Otu20409 | 1 | k__Fungi(100);p__Ascomycota(100);c__Saccharomycetes(100);o__Saccharomycetales(100);f__Metschnikowiaceae(100);g__Metschnikowia(100);s__Metschnikowia_pulcherrima(100); | Metschnikowia | 13 | Saprotroph | \|Undefined Saprotroph\| | Yeast | NULL | Probable | As nectar-tap_saprotroph (Põlme et al. 2020); Decay sugar-rich substrates (Põlme et al. 2020) | Põlme S, et al. 2020. Fungal Diversity 105:1-16 ((https://doi.org/10.1007/s13225-020-00466-2)) |
| Otu20411 | 1 | k__Fungi(100);p__Ascomycota(100);c__Saccharomycetes(100);o__Saccharomycetales(100);f__Metschnikowiaceae(100);g__Metschnikowia(100);g__Metschnikowia_unclassified(100); | Metschnikowia | 13 | Saprotroph | \|Undefined Saprotroph\| | Yeast | NULL | Probable | As nectar-tap_saprotroph (Põlme et al. 2020); Decay sugar-rich substrates (Põlme et al. 2020) | Põlme S, et al. 2020. Fungal Diversity 105:1-16 ((https://doi.org/10.1007/s13225-020-00466-2)) |
| Otu20412 | 1 | k__Fungi(100);p__Ascomycota(100);c__Saccharomycetes(100);o__Saccharomycetales(100);f__Metschnikowiaceae(100);g__Metschnikowia(100);s__Metschnikowia_pulcherrima(100); | Metschnikowia | 13 | Saprotroph | \|Undefined Saprotroph\| | Yeast | NULL | Probable | As nectar-tap_saprotroph (Põlme et al. 2020); Decay sugar-rich substrates (Põlme et al. 2020) | Põlme S, et al. 2020. Fungal Diversity 105:1-16 ((https://doi.org/10.1007/s13225-020-00466-2)) |
| Otu20413 | 1 | k__Fungi(100);p__Ascomycota(100);c__Saccharomycetes(100);o__Saccharomycetales(100);f__Metschnikowiaceae(100);g__Metschnikowia(100);g__Metschnikowia_unclassified(100); | Metschnikowia | 13 | Saprotroph | \|Undefined Saprotroph\| | Yeast | NULL | Probable | As nectar-tap_saprotroph (Põlme et al. 2020); Decay sugar-rich substrates (Põlme et al. 2020) | Põlme S, et al. 2020. Fungal Diversity 105:1-16 ((https://doi.org/10.1007/s13225-020-00466-2)) |
| Otu20414 | 1 | k__Fungi(100);p__Ascomycota(100);c__Saccharomycetes(100);o__Saccharomycetales(100);f__Metschnikowiaceae(100);g__Metschnikowia(100);s__Metschnikowia_pulcherrima(100); | Metschnikowia | 13 | Saprotroph | \|Undefined Saprotroph\| | Yeast | NULL | Probable | As nectar-tap_saprotroph (Põlme et al. 2020); Decay sugar-rich substrates (Põlme et al. 2020) | Põlme S, et al. 2020. Fungal Diversity 105:1-16 ((https://doi.org/10.1007/s13225-020-00466-2)) |
| Otu20415 | 1 | k__Fungi(100);p__Ascomycota(100);c__Saccharomycetes(100);o__Saccharomycetales(100);f__Metschnikowiaceae(100);g__Metschnikowia(100);s__Metschnikowia_pulcherrima(100); | Metschnikowia | 13 | Saprotroph | \|Undefined Saprotroph\| | Yeast | NULL | Probable | As nectar-tap_saprotroph (Põlme et al. 2020); Decay sugar-rich substrates (Põlme et al. 2020) | Põlme S, et al. 2020. Fungal Diversity 105:1-16 ((https://doi.org/10.1007/s13225-020-00466-2)) |
| Otu20416 | 1 | k__Fungi(100);p__Ascomycota(100);c__Saccharomycetes(100);o__Saccharomycetales(100);f__Metschnikowiaceae(100);g__Metschnikowia(100);s__Metschnikowia_pulcherrima(100); | Metschnikowia | 13 | Saprotroph | \|Undefined Saprotroph\| | Yeast | NULL | Probable | As nectar-tap_saprotroph (Põlme et al. 2020); Decay sugar-rich substrates (Põlme et al. 2020) | Põlme S, et al. 2020. Fungal Diversity 105:1-16 ((https://doi.org/10.1007/s13225-020-00466-2)) |
| Otu20417 | 1 | k__Fungi(100);p__Ascomycota(100);c__Saccharomycetes(100);o__Saccharomycetales(100);f__Metschnikowiaceae(100);g__Metschnikowia(100);s__Metschnikowia_pulcherrima(100); | Metschnikowia | 13 | Saprotroph | \|Undefined Saprotroph\| | Yeast | NULL | Probable | As nectar-tap_saprotroph (Põlme et al. 2020); Decay sugar-rich substrates (Põlme et al. 2020) | Põlme S, et al. 2020. Fungal Diversity 105:1-16 ((https://doi.org/10.1007/s13225-020-00466-2)) |
| Otu20418 | 1 | k__Fungi(100);p__Ascomycota(100);c__Saccharomycetes(100);o__Saccharomycetales(100);f__Metschnikowiaceae(100);g__Metschnikowia(100);s__Metschnikowia_pulcherrima(100); | Metschnikowia | 13 | Saprotroph | \|Undefined Saprotroph\| | Yeast | NULL | Probable | As nectar-tap_saprotroph (Põlme et al. 2020); Decay sugar-rich substrates (Põlme et al. 2020) | Põlme S, et al. 2020. Fungal Diversity 105:1-16 ((https://doi.org/10.1007/s13225-020-00466-2)) |
| Otu20419 | 1 | k__Fungi(100);p__Ascomycota(100);c__Saccharomycetes(100);o__Saccharomycetales(100);f__Metschnikowiaceae(100);g__Metschnikowia(100);s__Metschnikowia_pulcherrima(100); | Metschnikowia | 13 | Saprotroph | \|Undefined Saprotroph\| | Yeast | NULL | Probable | As nectar-tap_saprotroph (Põlme et al. 2020); Decay sugar-rich substrates (Põlme et al. 2020) | Põlme S, et al. 2020. Fungal Diversity 105:1-16 ((https://doi.org/10.1007/s13225-020-00466-2)) |
| Otu20420 | 1 | k__Fungi(100);p__Ascomycota(100);c__Saccharomycetes(100);o__Saccharomycetales(100);f__Metschnikowiaceae(100);g__Metschnikowia(100);s__Metschnikowia_pulcherrima(100); | Metschnikowia | 13 | Saprotroph | \|Undefined Saprotroph\| | Yeast | NULL | Probable | As nectar-tap_saprotroph (Põlme et al. 2020); Decay sugar-rich substrates (Põlme et al. 2020) | Põlme S, et al. 2020. Fungal Diversity 105:1-16 ((https://doi.org/10.1007/s13225-020-00466-2)) |
| Otu20421 | 1 | k__Fungi(100);p__Ascomycota(100);c__Saccharomycetes(100);o__Saccharomycetales(100);f__Metschnikowiaceae(100);g__Metschnikowia(100);s__Metschnikowia_pulcherrima(100); | Metschnikowia | 13 | Saprotroph | \|Undefined Saprotroph\| | Yeast | NULL | Probable | As nectar-tap_saprotroph (Põlme et al. 2020); Decay sugar-rich substrates (Põlme et al. 2020) | Põlme S, et al. 2020. Fungal Diversity 105:1-16 ((https://doi.org/10.1007/s13225-020-00466-2)) |
| Otu20422 | 1 | k__Fungi(100);p__Ascomycota(100);c__Saccharomycetes(100);o__Saccharomycetales(100);f__Metschnikowiaceae(100);g__Metschnikowia(100);g__Metschnikowia_unclassified(100); | Metschnikowia | 13 | Saprotroph | \|Undefined Saprotroph\| | Yeast | NULL | Probable | As nectar-tap_saprotroph (Põlme et al. 2020); Decay sugar-rich substrates (Põlme et al. 2020) | Põlme S, et al. 2020. Fungal Diversity 105:1-16 ((https://doi.org/10.1007/s13225-020-00466-2)) |
| Otu20423 | 1 | k__Fungi(100);p__Ascomycota(100);c__Saccharomycetes(100);o__Saccharomycetales(100);f__Metschnikowiaceae(100);g__Metschnikowia(100);g__Metschnikowia_unclassified(100); | Metschnikowia | 13 | Saprotroph | \|Undefined Saprotroph\| | Yeast | NULL | Probable | As nectar-tap_saprotroph (Põlme et al. 2020); Decay sugar-rich substrates (Põlme et al. 2020) | Põlme S, et al. 2020. Fungal Diversity 105:1-16 ((https://doi.org/10.1007/s13225-020-00466-2)) |
| Otu20424 | 1 | k__Fungi(100);p__Ascomycota(100);c__Saccharomycetes(100);o__Saccharomycetales(100);f__Metschnikowiaceae(100);g__Metschnikowia(100);s__Metschnikowia_pulcherrima(100); | Metschnikowia | 13 | Saprotroph | \|Undefined Saprotroph\| | Yeast | NULL | Probable | As nectar-tap_saprotroph (Põlme et al. 2020); Decay sugar-rich substrates (Põlme et al. 2020) | Põlme S, et al. 2020. Fungal Diversity 105:1-16 ((https://doi.org/10.1007/s13225-020-00466-2)) |
| Otu20425 | 1 | k__Fungi(100);p__Ascomycota(100);c__Saccharomycetes(100);o__Saccharomycetales(100);f__Metschnikowiaceae(100);g__Metschnikowia(100);s__Metschnikowia_pulcherrima(100); | Metschnikowia | 13 | Saprotroph | \|Undefined Saprotroph\| | Yeast | NULL | Probable | As nectar-tap_saprotroph (Põlme et al. 2020); Decay sugar-rich substrates (Põlme et al. 2020) | Põlme S, et al. 2020. Fungal Diversity 105:1-16 ((https://doi.org/10.1007/s13225-020-00466-2)) |

| | | | | | | | | | | | |
|---|---|---|---|---|---|---|---|---|---|---|---|
| Otu20426 | 1 | k__Fungi(100);p__Ascomycota(100);c__Saccharomycetes(100);o__Saccharomycetales(100);f__Metschnikowiaceae(100);g__Metschnikowia(100);s__Metschnikowia_pulcherrima(100); | Metschnikowia | 13 | Saprotroph | \|Undefined Saprotroph\| | Yeast | NULL | Probable | As nectar-tap_saprotroph (Põlme et al. 2020); Decay sugar-rich substrates (Põlme et al. 2020) | Põlme S, et al. 2020. Fungal Diversity 105:1-16 ((https://doi.org/10.1007/s13225-020-00466-2)) |
| Otu20427 | 1 | k__Fungi(100);p__Ascomycota(100);c__Saccharomycetes(100);o__Saccharomycetales(100);f__Metschnikowiaceae(100);g__Metschnikowia(100);s__Metschnikowia_pulcherrima(100); | Metschnikowia | 13 | Saprotroph | \|Undefined Saprotroph\| | Yeast | NULL | Probable | As nectar-tap_saprotroph (Põlme et al. 2020); Decay sugar-rich substrates (Põlme et al. 2020) | Põlme S, et al. 2020. Fungal Diversity 105:1-16 ((https://doi.org/10.1007/s13225-020-00466-2)) |
| Otu20428 | 1 | k__Fungi(100);p__Ascomycota(100);c__Saccharomycetes(100);o__Saccharomycetales(100);f__Metschnikowiaceae(100);g__Metschnikowia(100);s__Metschnikowia_pulcherrima(100); | Metschnikowia | 13 | Saprotroph | \|Undefined Saprotroph\| | Yeast | NULL | Probable | As nectar-tap_saprotroph (Põlme et al. 2020); Decay sugar-rich substrates (Põlme et al. 2020) | Põlme S, et al. 2020. Fungal Diversity 105:1-16 ((https://doi.org/10.1007/s13225-020-00466-2)) |
| Otu20429 | 1 | k__Fungi(100);p__Ascomycota(100);c__Saccharomycetes(100);o__Saccharomycetales(100);f__Metschnikowiaceae(100);g__Metschnikowia(100);s__Metschnikowia_pulcherrima(100); | Metschnikowia | 13 | Saprotroph | \|Undefined Saprotroph\| | Yeast | NULL | Probable | As nectar-tap_saprotroph (Põlme et al. 2020); Decay sugar-rich substrates (Põlme et al. 2020) | Põlme S, et al. 2020. Fungal Diversity 105:1-16 ((https://doi.org/10.1007/s13225-020-00466-2)) |
| Otu20430 | 1 | k__Fungi(100);p__Ascomycota(100);c__Saccharomycetes(100);o__Saccharomycetales(100);f__Metschnikowiaceae(100);g__Metschnikowia(100);s__Metschnikowia_pulcherrima(100); | Metschnikowia | 13 | Saprotroph | \|Undefined Saprotroph\| | Yeast | NULL | Probable | As nectar-tap_saprotroph (Põlme et al. 2020); Decay sugar-rich substrates (Põlme et al. 2020) | Põlme S, et al. 2020. Fungal Diversity 105:1-16 ((https://doi.org/10.1007/s13225-020-00466-2)) |
| Otu20431 | 1 | k__Fungi(100);p__Ascomycota(100);c__Saccharomycetes(100);o__Saccharomycetales(100);f__Metschnikowiaceae(100);g__Metschnikowia(100);s__Metschnikowia_pulcherrima(100); | Metschnikowia | 13 | Saprotroph | \|Undefined Saprotroph\| | Yeast | NULL | Probable | As nectar-tap_saprotroph (Põlme et al. 2020); Decay sugar-rich substrates (Põlme et al. 2020) | Põlme S, et al. 2020. Fungal Diversity 105:1-16 ((https://doi.org/10.1007/s13225-020-00466-2)) |
| Otu20432 | 1 | k__Fungi(100);p__Ascomycota(100);c__Saccharomycetes(100);o__Saccharomycetales(100);f__Metschnikowiaceae(100);g__Metschnikowia(100);g__Metschnikowia_unclassified(100); | Metschnikowia | 13 | Saprotroph | \|Undefined Saprotroph\| | Yeast | NULL | Probable | As nectar-tap_saprotroph (Põlme et al. 2020); Decay sugar-rich substrates (Põlme et al. 2020) | Põlme S, et al. 2020. Fungal Diversity 105:1-16 ((https://doi.org/10.1007/s13225-020-00466-2)) |
| Otu20433 | 1 | k__Fungi(100);p__Ascomycota(100);c__Saccharomycetes(100);o__Saccharomycetales(100);f__Metschnikowiaceae(100);g__Metschnikowia(100);s__Metschnikowia_pulcherrima(100); | Metschnikowia | 13 | Saprotroph | \|Undefined Saprotroph\| | Yeast | NULL | Probable | As nectar-tap_saprotroph (Põlme et al. 2020); Decay sugar-rich substrates (Põlme et al. 2020) | Põlme S, et al. 2020. Fungal Diversity 105:1-16 ((https://doi.org/10.1007/s13225-020-00466-2)) |
| Otu20434 | 1 | k__Fungi(100);p__Ascomycota(100);c__Saccharomycetes(100);o__Saccharomycetales(100);f__Metschnikowiaceae(100);g__Metschnikowia(100);s__Metschnikowia_pulcherrima(100); | Metschnikowia | 13 | Saprotroph | \|Undefined Saprotroph\| | Yeast | NULL | Probable | As nectar-tap_saprotroph (Põlme et al. 2020); Decay sugar-rich substrates (Põlme et al. 2020) | Põlme S, et al. 2020. Fungal Diversity 105:1-16 ((https://doi.org/10.1007/s13225-020-00466-2)) |
| Otu20435 | 1 | k__Fungi(100);p__Ascomycota(100);c__Saccharomycetes(100);o__Saccharomycetales(100);f__Metschnikowiaceae(100);g__Metschnikowia(100);s__Metschnikowia_pulcherrima(100); | Metschnikowia | 13 | Saprotroph | \|Undefined Saprotroph\| | Yeast | NULL | Probable | As nectar-tap_saprotroph (Põlme et al. 2020); Decay sugar-rich substrates (Põlme et al. 2020) | Põlme S, et al. 2020. Fungal Diversity 105:1-16 ((https://doi.org/10.1007/s13225-020-00466-2)) |
| Otu20436 | 1 | k__Fungi(100);p__Ascomycota(100);c__Saccharomycetes(100);o__Saccharomycetales(100);f__Metschnikowiaceae(100);g__Metschnikowia(100);g__Metschnikowia_unclassified(100); | Metschnikowia | 13 | Saprotroph | \|Undefined Saprotroph\| | Yeast | NULL | Probable | As nectar-tap_saprotroph (Põlme et al. 2020); Decay sugar-rich substrates (Põlme et al. 2020) | Põlme S, et al. 2020. Fungal Diversity 105:1-16 ((https://doi.org/10.1007/s13225-020-00466-2)) |
| Otu20437 | 1 | k__Fungi(100);p__Ascomycota(100);c__Saccharomycetes(100);o__Saccharomycetales(100);f__Metschnikowiaceae(100);g__Metschnikowia(100);g__Metschnikowia_unclassified(100); | Metschnikowia | 13 | Saprotroph | \|Undefined Saprotroph\| | Yeast | NULL | Probable | As nectar-tap_saprotroph (Põlme et al. 2020); Decay sugar-rich substrates (Põlme et al. 2020) | Põlme S, et al. 2020. Fungal Diversity 105:1-16 ((https://doi.org/10.1007/s13225-020-00466-2)) |
| Otu20438 | 1 | k__Fungi(100);p__Ascomycota(100);c__Saccharomycetes(100);o__Saccharomycetales(100);f__Metschnikowiaceae(100);g__Metschnikowia(100);s__Metschnikowia_pulcherrima(100); | Metschnikowia | 13 | Saprotroph | \|Undefined Saprotroph\| | Yeast | NULL | Probable | As nectar-tap_saprotroph (Põlme et al. 2020); Decay sugar-rich substrates (Põlme et al. 2020) | Põlme S, et al. 2020. Fungal Diversity 105:1-16 ((https://doi.org/10.1007/s13225-020-00466-2)) |
| Otu20439 | 1 | k__Fungi(100);p__Ascomycota(100);c__Saccharomycetes(100);o__Saccharomycetales(100);f__Metschnikowiaceae(100);g__Metschnikowia(100);g__Metschnikowia_unclassified(100); | Metschnikowia | 13 | Saprotroph | \|Undefined Saprotroph\| | Yeast | NULL | Probable | As nectar-tap_saprotroph (Põlme et al. 2020); Decay sugar-rich substrates (Põlme et al. 2020) | Põlme S, et al. 2020. Fungal Diversity 105:1-16 ((https://doi.org/10.1007/s13225-020-00466-2)) |
| Otu20441 | 1 | k__Fungi(100);p__Ascomycota(100);c__Saccharomycetes(100);o__Saccharomycetales(100);f__Metschnikowiaceae(100);g__Metschnikowia(100);s__Metschnikowia_pulcherrima(100); | Metschnikowia | 13 | Saprotroph | \|Undefined Saprotroph\| | Yeast | NULL | Probable | As nectar-tap_saprotroph (Põlme et al. 2020); Decay sugar-rich substrates (Põlme et al. 2020) | Põlme S, et al. 2020. Fungal Diversity 105:1-16 ((https://doi.org/10.1007/s13225-020-00466-2)) |
| Otu20442 | 1 | k__Fungi(100);p__Ascomycota(100);c__Saccharomycetes(100);o__Saccharomycetales(100);f__Metschnikowiaceae(100);g__Metschnikowia(100);g__Metschnikowia_unclassified(100); | Metschnikowia | 13 | Saprotroph | \|Undefined Saprotroph\| | Yeast | NULL | Probable | As nectar-tap_saprotroph (Põlme et al. 2020); Decay sugar-rich substrates (Põlme et al. 2020) | Põlme S, et al. 2020. Fungal Diversity 105:1-16 ((https://doi.org/10.1007/s13225-020-00466-2)) |
| Otu20443 | 1 | k__Fungi(100);p__Ascomycota(100);c__Saccharomycetes(100);o__Saccharomycetales(100);f__Metschnikowiaceae(100);g__Metschnikowia(100);s__Metschnikowia_pulcherrima(100); | Metschnikowia | 13 | Saprotroph | \|Undefined Saprotroph\| | Yeast | NULL | Probable | As nectar-tap_saprotroph (Põlme et al. 2020); Decay sugar-rich substrates (Põlme et al. 2020) | Põlme S, et al. 2020. Fungal Diversity 105:1-16 ((https://doi.org/10.1007/s13225-020-00466-2)) |
| Otu20444 | 1 | k__Fungi(100);p__Ascomycota(100);c__Saccharomycetes(100);o__Saccharomycetales(100);f__Metschnikowiaceae(100);g__Metschnikowia(100);g__Metschnikowia_unclassified(100); | Metschnikowia | 13 | Saprotroph | \|Undefined Saprotroph\| | Yeast | NULL | Probable | As nectar-tap_saprotroph (Põlme et al. 2020); Decay sugar-rich substrates (Põlme et al. 2020) | Põlme S, et al. 2020. Fungal Diversity 105:1-16 ((https://doi.org/10.1007/s13225-020-00466-2)) |
| Otu20447 | 1 | k__Fungi(100);p__Ascomycota(100);c__Saccharomycetes(100);o__Saccharomycetales(100);f__Metschnikowiaceae(100);g__Metschnikowia(100);s__Metschnikowia_pulcherrima(100); | Metschnikowia | 13 | Saprotroph | \|Undefined Saprotroph\| | Yeast | NULL | Probable | As nectar-tap_saprotroph (Põlme et al. 2020); Decay sugar-rich substrates (Põlme et al. 2020) | Põlme S, et al. 2020. Fungal Diversity 105:1-16 ((https://doi.org/10.1007/s13225-020-00466-2)) |
| Otu20448 | 1 | k__Fungi(100);p__Ascomycota(100);c__Saccharomycetes(100);o__Saccharomycetales(100);f__Metschnikowiaceae(100);g__Metschnikowia(100);s__Metschnikowia_pulcherrima(100); | Metschnikowia | 13 | Saprotroph | \|Undefined Saprotroph\| | Yeast | NULL | Probable | As nectar-tap_saprotroph (Põlme et al. 2020); Decay sugar-rich substrates (Põlme et al. 2020) | Põlme S, et al. 2020. Fungal Diversity 105:1-16 ((https://doi.org/10.1007/s13225-020-00466-2)) |
| Otu20449 | 1 | k__Fungi(100);p__Ascomycota(100);c__Saccharomycetes(100);o__Saccharomycetales(100);f__Metschnikowiaceae(100);g__Metschnikowia(100);s__Metschnikowia_pulcherrima(100); | Metschnikowia | 13 | Saprotroph | \|Undefined Saprotroph\| | Yeast | NULL | Probable | As nectar-tap_saprotroph (Põlme et al. 2020); Decay sugar-rich substrates (Põlme et al. 2020) | Põlme S, et al. 2020. Fungal Diversity 105:1-16 ((https://doi.org/10.1007/s13225-020-00466-2)) |
| Otu20450 | 1 | k__Fungi(100);p__Ascomycota(100);c__Saccharomycetes(100);o__Saccharomycetales(100);f__Metschnikowiaceae(100);g__Metschnikowia(100);s__Metschnikowia_pulcherrima(100); | Metschnikowia | 13 | Saprotroph | \|Undefined Saprotroph\| | Yeast | NULL | Probable | As nectar-tap_saprotroph (Põlme et al. 2020); Decay sugar-rich substrates (Põlme et al. 2020) | Põlme S, et al. 2020. Fungal Diversity 105:1-16 ((https://doi.org/10.1007/s13225-020-00466-2)) |
| Otu20451 | 1 | k__Fungi(100);p__Ascomycota(100);c__Saccharomycetes(100);o__Saccharomycetales(100);f__Metschnikowiaceae(100);g__Metschnikowia(100);s__Metschnikowia_pulcherrima(100); | Metschnikowia | 13 | Saprotroph | \|Undefined Saprotroph\| | Yeast | NULL | Probable | As nectar-tap_saprotroph (Põlme et al. 2020); Decay sugar-rich substrates (Põlme et al. 2020) | Põlme S, et al. 2020. Fungal Diversity 105:1-16 ((https://doi.org/10.1007/s13225-020-00466-2)) |
| Otu20453 | 1 | k__Fungi(100);p__Ascomycota(100);c__Saccharomycetes(100);o__Saccharomycetales(100);f__Metschnikowiaceae(100);g__Metschnikowia(100);s__Metschnikowia_pulcherrima(100); | Metschnikowia | 13 | Saprotroph | \|Undefined Saprotroph\| | Yeast | NULL | Probable | As nectar-tap_saprotroph (Põlme et al. 2020); Decay sugar-rich substrates (Põlme et al. 2020) | Põlme S, et al. 2020. Fungal Diversity 105:1-16 ((https://doi.org/10.1007/s13225-020-00466-2)) |
| Otu20454 | 1 | k__Fungi(100);p__Ascomycota(100);c__Saccharomycetes(100);o__Saccharomycetales(100);f__Metschnikowiaceae(100);g__Metschnikowia(100);g__Metschnikowia_unclassified(100); | Metschnikowia | 13 | Saprotroph | \|Undefined Saprotroph\| | Yeast | NULL | Probable | As nectar-tap_saprotroph (Põlme et al. 2020); Decay sugar-rich substrates (Põlme et al. 2020) | Põlme S, et al. 2020. Fungal Diversity 105:1-16 ((https://doi.org/10.1007/s13225-020-00466-2)) |
| Otu20455 | 1 | k__Fungi(100);p__Ascomycota(100);c__Saccharomycetes(100);o__Saccharomycetales(100);f__Metschnikowiaceae(100);g__Metschnikowia(100);s__Metschnikowia_pulcherrima(100); | Metschnikowia | 13 | Saprotroph | \|Undefined Saprotroph\| | Yeast | NULL | Probable | As nectar-tap_saprotroph (Põlme et al. 2020); Decay sugar-rich substrates (Põlme et al. 2020) | Põlme S, et al. 2020. Fungal Diversity 105:1-16 ((https://doi.org/10.1007/s13225-020-00466-2)) |
| Otu20456 | 1 | k__Fungi(100);p__Ascomycota(100);c__Saccharomycetes(100);o__Saccharomycetales(100);f__Metschnikowiaceae(100);g__Metschnikowia(100);g__Metschnikowia_unclassified(100); | Metschnikowia | 13 | Saprotroph | \|Undefined Saprotroph\| | Yeast | NULL | Probable | As nectar-tap_saprotroph (Põlme et al. 2020); Decay sugar-rich substrates (Põlme et al. 2020) | Põlme S, et al. 2020. Fungal Diversity 105:1-16 ((https://doi.org/10.1007/s13225-020-00466-2)) |
| Otu20457 | 1 | k__Fungi(100);p__Ascomycota(100);c__Saccharomycetes(100);o__Saccharomycetales(100);f__Metschnikowiaceae(100);g__Metschnikowia(100);s__Metschnikowia_pulcherrima(100); | Metschnikowia | 13 | Saprotroph | \|Undefined Saprotroph\| | Yeast | NULL | Probable | As nectar-tap_saprotroph (Põlme et al. 2020); Decay sugar-rich substrates (Põlme et al. 2020) | Põlme S, et al. 2020. Fungal Diversity 105:1-16 ((https://doi.org/10.1007/s13225-020-00466-2)) |
| Otu20458 | 1 | k__Fungi(100);p__Ascomycota(100);c__Saccharomycetes(100);o__Saccharomycetales(100);f__Metschnikowiaceae(100);g__Metschnikowia(100);g__Metschnikowia_unclassified(100); | Metschnikowia | 13 | Saprotroph | \|Undefined Saprotroph\| | Yeast | NULL | Probable | As nectar-tap_saprotroph (Põlme et al. 2020); Decay sugar-rich substrates (Põlme et al. 2020) | Põlme S, et al. 2020. Fungal Diversity 105:1-16 ((https://doi.org/10.1007/s13225-020-00466-2)) |
| Otu20459 | 1 | k__Fungi(100);p__Ascomycota(100);c__Saccharomycetes(100);o__Saccharomycetales(100);f__Metschnikowiaceae(100);g__Metschnikowia(100);s__Metschnikowia_pulcherrima(100); | Metschnikowia | 13 | Saprotroph | \|Undefined Saprotroph\| | Yeast | NULL | Probable | As nectar-tap_saprotroph (Põlme et al. 2020); Decay sugar-rich substrates (Põlme et al. 2020) | Põlme S, et al. 2020. Fungal Diversity 105:1-16 ((https://doi.org/10.1007/s13225-020-00466-2)) |
| Otu20460 | 1 | k__Fungi(100);p__Ascomycota(100);c__Saccharomycetes(100);o__Saccharomycetales(100);f__Metschnikowiaceae(100);g__Metschnikowia(100);s__Metschnikowia_pulcherrima(100); | Metschnikowia | 13 | Saprotroph | \|Undefined Saprotroph\| | Yeast | NULL | Probable | As nectar-tap_saprotroph (Põlme et al. 2020); Decay sugar-rich substrates (Põlme et al. 2020) | Põlme S, et al. 2020. Fungal Diversity 105:1-16 ((https://doi.org/10.1007/s13225-020-00466-2)) |
| Otu20461 | 1 | k__Fungi(100);p__Ascomycota(100);c__Saccharomycetes(100);o__Saccharomycetales(100);f__Metschnikowiaceae(100);g__Metschnikowia(100);s__Metschnikowia_pulcherrima(100); | Metschnikowia | 13 | Saprotroph | \|Undefined Saprotroph\| | Yeast | NULL | Probable | As nectar-tap_saprotroph (Põlme et al. 2020); Decay sugar-rich substrates (Põlme et al. 2020) | Põlme S, et al. 2020. Fungal Diversity 105:1-16 ((https://doi.org/10.1007/s13225-020-00466-2)) |
| Otu20462 | 1 | k__Fungi(100);p__Ascomycota(100);c__Saccharomycetes(100);o__Saccharomycetales(100);f__Metschnikowiaceae(100);g__Metschnikowia(100);g__Metschnikowia_unclassified(100); | Metschnikowia | 13 | Saprotroph | \|Undefined Saprotroph\| | Yeast | NULL | Probable | As nectar-tap_saprotroph (Põlme et al. 2020); Decay sugar-rich substrates (Põlme et al. 2020) | Põlme S, et al. 2020. Fungal Diversity 105:1-16 ((https://doi.org/10.1007/s13225-020-00466-2)) |
| Otu20463 | 1 | k__Fungi(100);p__Ascomycota(100);c__Saccharomycetes(100);o__Saccharomycetales(100);f__Metschnikowiaceae(100);g__Metschnikowia(100);s__Metschnikowia_pulcherrima(100); | Metschnikowia | 13 | Saprotroph | \|Undefined Saprotroph\| | Yeast | NULL | Probable | As nectar-tap_saprotroph (Põlme et al. 2020); Decay sugar-rich substrates (Põlme et al. 2020) | Põlme S, et al. 2020. Fungal Diversity 105:1-16 ((https://doi.org/10.1007/s13225-020-00466-2)) |
| Otu20464 | 1 | k__Fungi(100);p__Ascomycota(100);c__Saccharomycetes(100);o__Saccharomycetales(100);f__Metschnikowiaceae(100);g__Metschnikowia(100);s__Metschnikowia_pulcherrima(100); | Metschnikowia | 13 | Saprotroph | \|Undefined Saprotroph\| | Yeast | NULL | Probable | As nectar-tap_saprotroph (Põlme et al. 2020); Decay sugar-rich substrates (Põlme et al. 2020) | Põlme S, et al. 2020. Fungal Diversity 105:1-16 ((https://doi.org/10.1007/s13225-020-00466-2)) |
| Otu20465 | 1 | k__Fungi(100);p__Ascomycota(100);c__Saccharomycetes(100);o__Saccharomycetales(100);f__Metschnikowiaceae(100);g__Metschnikowia(100);s__Metschnikowia_pulcherrima(100); | Metschnikowia | 13 | Saprotroph | \|Undefined Saprotroph\| | Yeast | NULL | Probable | As nectar-tap_saprotroph (Põlme et al. 2020); Decay sugar-rich substrates (Põlme et al. 2020) | Põlme S, et al. 2020. Fungal Diversity 105:1-16 ((https://doi.org/10.1007/s13225-020-00466-2)) |
| Otu20467 | 1 | k__Fungi(100);p__Ascomycota(100);c__Saccharomycetes(100);o__Saccharomycetales(100);f__Metschnikowiaceae(100);g__Metschnikowia(100);s__Metschnikowia_pulcherrima(100); | Metschnikowia | 13 | Saprotroph | \|Undefined Saprotroph\| | Yeast | NULL | Probable | As nectar-tap_saprotroph (Põlme et al. 2020); Decay sugar-rich substrates (Põlme et al. 2020) | Põlme S, et al. 2020. Fungal Diversity 105:1-16 ((https://doi.org/10.1007/s13225-020-00466-2)) |
| Otu20468 | 1 | k__Fungi(100);p__Ascomycota(100);c__Saccharomycetes(100);o__Saccharomycetales(100);f__Metschnikowiaceae(100);g__Metschnikowia(100);s__Metschnikowia_pulcherrima(100); | Metschnikowia | 13 | Saprotroph | \|Undefined Saprotroph\| | Yeast | NULL | Probable | As nectar-tap_saprotroph (Põlme et al. 2020); Decay sugar-rich substrates (Põlme et al. 2020) | Põlme S, et al. 2020. Fungal Diversity 105:1-16 ((https://doi.org/10.1007/s13225-020-00466-2)) |
| Otu20469 | 1 | k__Fungi(100);p__Ascomycota(100);c__Saccharomycetes(100);o__Saccharomycetales(100);f__Metschnikowiaceae(100);g__Metschnikowia(100);s__Metschnikowia_pulcherrima(100); | Metschnikowia | 13 | Saprotroph | \|Undefined Saprotroph\| | Yeast | NULL | Probable | As nectar-tap_saprotroph (Põlme et al. 2020); Decay sugar-rich substrates (Põlme et al. 2020) | Põlme S, et al. 2020. Fungal Diversity 105:1-16 ((https://doi.org/10.1007/s13225-020-00466-2)) |
| Otu20470 | 1 | k__Fungi(100);p__Ascomycota(100);c__Saccharomycetes(100);o__Saccharomycetales(100);f__Metschnikowiaceae(100);g__Metschnikowia(100);s__Metschnikowia_pulcherrima(100); | Metschnikowia | 13 | Saprotroph | \|Undefined Saprotroph\| | Yeast | NULL | Probable | As nectar-tap_saprotroph (Põlme et al. 2020); Decay sugar-rich substrates (Põlme et al. 2020) | Põlme S, et al. 2020. Fungal Diversity 105:1-16 ((https://doi.org/10.1007/s13225-020-00466-2)) |
| Otu20471 | 1 | k__Fungi(100);p__Ascomycota(100);c__Saccharomycetes(100);o__Saccharomycetales(100);f__Saccharomycetales_family_Incertae_sedis(100);g__Candida(100);g__Candida_unclassified(100); | Candida | 13 | Pathotroph-Saprotroph-Symbiotroph | Animal Pathogen-Endophyte-Endosymbiont-Epiphyte-\|Undefined Saprotroph\| | Dimorphic Yeast-Yeast | NULL | Probable | As nectar-tap_saprotroph (Põlme et al. 2020); Decay sugar-rich substrates (Põlme et al. 2020); This asexual genus is polyphyletic and contain species that function very differently depend on what they are. Caution should be taken when interpreting guilds for Candida species. | Manolakaki D, et al. 2010. Virulence. 1: 367-375 ((https://doi.org/10.4161/viru.1.5.12796)); Mekbib SB, et al. 2011. Tropical Plant Pathology 36:233-240 ((https://doi.org/10.1590/S1982-56762011000400004)); Põlme S, et al. 2020. Fungal Diversity 105:1-16 ((https://doi.org/10.1007/s13225-020-00466-2)) |
| Otu20473 | 1 | k__Fungi(100);p__Ascomycota(100);c__Saccharomycetes(100);o__Saccharomycetales(100);f__Saccharomycetales_family_Incertae_sedis(100);g__Candida(100);g__Candida_unclassified(100); | Candida | 13 | Pathotroph-Saprotroph-Symbiotroph | Animal Pathogen-Endophyte-Endosymbiont-Epiphyte-\|Undefined Saprotroph\| | Dimorphic Yeast-Yeast | NULL | Probable | As nectar-tap_saprotroph (Põlme et al. 2020); Decay sugar-rich substrates (Põlme et al. 2020); This asexual genus is polyphyletic and contain species that function very differently depend on what they are. Caution should be taken when interpreting guilds for Candida species. | Manolakaki D, et al. 2010. Virulence. 1: 367-375 ((https://doi.org/10.4161/viru.1.5.12796)); Mekbib SB, et al. 2011. Tropical Plant Pathology 36:233-240 ((https://doi.org/10.1590/S1982-56762011000400004)); Põlme S, et al. 2020. Fungal Diversity 105:1-16 ((https://doi.org/10.1007/s13225-020-00466-2)) |
| Otu20474 | 1 | k__Fungi(100);p__Ascomycota(100);c__Saccharomycetes(100);o__Saccharomycetales(100);f__Saccharomycetales_family_Incertae_sedis(100);g__Candida(100);g__Candida_unclassified(100); | Candida | 13 | Pathotroph-Saprotroph-Symbiotroph | Animal Pathogen-Endophyte-Endosymbiont-Epiphyte-\|Undefined Saprotroph\| | Dimorphic Yeast-Yeast | NULL | Probable | As nectar-tap_saprotroph (Põlme et al. 2020); Decay sugar-rich substrates (Põlme et al. 2020); This asexual genus is polyphyletic and contain species that function very differently depend on what they are. Caution should be taken when interpreting guilds for Candida species. | Manolakaki D, et al. 2010. Virulence. 1: 367-375 ((https://doi.org/10.4161/viru.1.5.12796)); Mekbib SB, et al. 2011. Tropical Plant Pathology 36:233-240 ((https://doi.org/10.1590/S1982-56762011000400004)); Põlme S, et al. 2020. Fungal Diversity 105:1-16 ((https://doi.org/10.1007/s13225-020-00466-2)) |
| Otu20476 | 1 | k__Fungi(100);p__Ascomycota(100);c__Saccharomycetes(100);o__Saccharomycetales(100);f__Metschnikowiaceae(100);g__Metschnikowia(100);s__Metschnikowia_pulcherrima(100); | Metschnikowia | 13 | Saprotroph | \|Undefined Saprotroph\| | Yeast | NULL | Probable | As nectar-tap_saprotroph (Põlme et al. 2020); Decay sugar-rich substrates (Põlme et al. 2020) | Põlme S, et al. 2020. Fungal Diversity 105:1-16 ((https://doi.org/10.1007/s13225-020-00466-2)) |
| Otu20478 | 1 | k__Fungi(100);p__Ascomycota(100);c__Saccharomycetes(100);o__Saccharomycetales(100);f__Metschnikowiaceae(100);g__Metschnikowia(100);g__Metschnikowia_unclassified(100); | Metschnikowia | 13 | Saprotroph | \|Undefined Saprotroph\| | Yeast | NULL | Probable | As nectar-tap_saprotroph (Põlme et al. 2020); Decay sugar-rich substrates (Põlme et al. 2020) | Põlme S, et al. 2020. Fungal Diversity 105:1-16 ((https://doi.org/10.1007/s13225-020-00466-2)) |
| Otu20483 | 1 | k__Fungi(100);p__Ascomycota(100);c__Saccharomycetes(100);o__Saccharomycetales(100);f__Metschnikowiaceae(100);g__Metschnikowia(100);s__Metschnikowia_pulcherrima(100); | Metschnikowia | 13 | Saprotroph | \|Undefined Saprotroph\| | Yeast | NULL | Probable | As nectar-tap_saprotroph (Põlme et al. 2020); Decay sugar-rich substrates (Põlme et al. 2020) | Põlme S, et al. 2020. Fungal Diversity 105:1-16 ((https://doi.org/10.1007/s13225-020-00466-2)) |
| Otu20489 | 1 | k__Fungi(100);p__Ascomycota(100);c__Leotiomycetes(100);o__Helotiales(100);f__Vibrisseaceae(100);g__Phialocephala(100);g__Phialocephala_unclassified(100); | Phialocephala | 13 | Saprotroph-Symbiotroph | Endophyte-\|Undefined Saprotroph\| | Microfungus | NULL | Highly Probable | As root_endophyte (Põlme et al. 2020); As root_endophyte_dark_septate (Põlme et al. 2020); As soil_saprotroph (Põlme et al. 2020); Dark Septate Endophyte, known from plant roots.; Decay roots, soil (Põlme et al. 2020) | Newsham KK. 2011. New Phytologist 190:783-793 ((https://doi.org/10.1111/j.1469-8137.2010.03611.x)) (as dark septate root endophytes); Põlme S, et al. 2020. Fungal Diversity 105:1-16 ((https://doi.org/10.1007/s13225-020-00466-2)) |
| Otu20492 | 1 | k__Fungi(100);p__Ascomycota(100);c__Saccharomycetes(100);o__Saccharomycetales(100);f__Metschnikowiaceae(100);g__Metschnikowia(100);g__Metschnikowia_unclassified(100); | Metschnikowia | 13 | Saprotroph | \|Undefined Saprotroph\| | Yeast | NULL | Probable | As nectar-tap_saprotroph (Põlme et al. 2020); Decay sugar-rich substrates (Põlme et al. 2020) | Põlme S, et al. 2020. Fungal Diversity 105:1-16 ((https://doi.org/10.1007/s13225-020-00466-2)) |
| Otu20493 | 1 | k__Fungi(100);p__Ascomycota(100);c__Saccharomycetes(100);o__Saccharomycetales(100);f__Metschnikowiaceae(100);g__Metschnikowia(100);s__Metschnikowia_pulcherrima(100); | Metschnikowia | 13 | Saprotroph | \|Undefined Saprotroph\| | Yeast | NULL | Probable | As nectar-tap_saprotroph (Põlme et al. 2020); Decay sugar-rich substrates (Põlme et al. 2020) | Põlme S, et al. 2020. Fungal Diversity 105:1-16 ((https://doi.org/10.1007/s13225-020-00466-2)) |
| Otu20494 | 1 | k__Fungi(100);p__Ascomycota(100);c__Saccharomycetes(100);o__Saccharomycetales(100);f__Metschnikowiaceae(100);g__Metschnikowia(100);s__Metschnikowia_pulcherrima(100); | Metschnikowia | 13 | Saprotroph | \|Undefined Saprotroph\| | Yeast | NULL | Probable | As nectar-tap_saprotroph (Põlme et al. 2020); Decay sugar-rich substrates (Põlme et al. 2020) | Põlme S, et al. 2020. Fungal Diversity 105:1-16 ((https://doi.org/10.1007/s13225-020-00466-2)) |
| Otu20495 | 1 | k__Fungi(100);p__Ascomycota(100);c__Saccharomycetes(100);o__Saccharomycetales(100);f__Metschnikowiaceae(100);g__Metschnikowia(100);s__Metschnikowia_pulcherrima(100); | Metschnikowia | 13 | Saprotroph | \|Undefined Saprotroph\| | Yeast | NULL | Probable | As nectar-tap_saprotroph (Põlme et al. 2020); Decay sugar-rich substrates (Põlme et al. 2020) | Põlme S, et al. 2020. Fungal Diversity 105:1-16 ((https://doi.org/10.1007/s13225-020-00466-2)) |
| Otu20497 | 1 | k__Fungi(100);p__Ascomycota(100);c__Saccharomycetes(100);o__Saccharomycetales(100);f__Metschnikowiaceae(100);g__Metschnikowia(100);s__Metschnikowia_pulcherrima(100); | Metschnikowia | 13 | Saprotroph | \|Undefined Saprotroph\| | Yeast | NULL | Probable | As nectar-tap_saprotroph (Põlme et al. 2020); Decay sugar-rich substrates (Põlme et al. 2020) | Põlme S, et al. 2020. Fungal Diversity 105:1-16 ((https://doi.org/10.1007/s13225-020-00466-2)) |
| Otu20499 | 1 | k__Fungi(100);p__Ascomycota(100);c__Saccharomycetes(100);o__Saccharomycetales(100);f__Metschnikowiaceae(100);g__Metschnikowia(100);g__Metschnikowia_unclassified(100); | Metschnikowia | 13 | Saprotroph | \|Undefined Saprotroph\| | Yeast | NULL | Probable | As nectar-tap_saprotroph (Põlme et al. 2020); Decay sugar-rich substrates (Põlme et al. 2020) | Põlme S, et al. 2020. Fungal Diversity 105:1-16 ((https://doi.org/10.1007/s13225-020-00466-2)) |
| Otu20500 | 1 | k__Fungi(100);p__Ascomycota(100);c__Saccharomycetes(100);o__Saccharomycetales(100);f__Metschnikowiaceae(100);g__Metschnikowia(100);s__Metschnikowia_pulcherrima(100); | Metschnikowia | 13 | Saprotroph | \|Undefined Saprotroph\| | Yeast | NULL | Probable | As nectar-tap_saprotroph (Põlme et al. 2020); Decay sugar-rich substrates (Põlme et al. 2020) | Põlme S, et al. 2020. Fungal Diversity 105:1-16 ((https://doi.org/10.1007/s13225-020-00466-2)) |
| Otu20501 | 1 | k__Fungi(100);p__Ascomycota(100);c__Saccharomycetes(100);o__Saccharomycetales(100);f__Metschnikowiaceae(100);g__Metschnikowia(100);s__Metschnikowia_pulcherrima(100); | Metschnikowia | 13 | Saprotroph | \|Undefined Saprotroph\| | Yeast | NULL | Probable | As nectar-tap_saprotroph (Põlme et al. 2020); Decay sugar-rich substrates (Põlme et al. 2020) | Põlme S, et al. 2020. Fungal Diversity 105:1-16 ((https://doi.org/10.1007/s13225-020-00466-2)) |
| Otu20502 | 1 | k__Fungi(100);p__Ascomycota(100);c__Saccharomycetes(100);o__Saccharomycetales(100);f__Metschnikowiaceae(100);g__Metschnikowia(100);g__Metschnikowia_unclassified(100); | Metschnikowia | 13 | Saprotroph | \|Undefined Saprotroph\| | Yeast | NULL | Probable | As nectar-tap_saprotroph (Põlme et al. 2020); Decay sugar-rich substrates (Põlme et al. 2020) | Põlme S, et al. 2020. Fungal Diversity 105:1-16 ((https://doi.org/10.1007/s13225-020-00466-2)) |
| Otu20503 | 1 | k__Fungi(100);p__Ascomycota(100);c__Saccharomycetes(100);o__Saccharomycetales(100);f__Metschnikowiaceae(100);g__Metschnikowia(100);s__Metschnikowia_pulcherrima(100); | Metschnikowia | 13 | Saprotroph | \|Undefined Saprotroph\| | Yeast | NULL | Probable | As nectar-tap_saprotroph (Põlme et al. 2020); Decay sugar-rich substrates (Põlme et al. 2020) | Põlme S, et al. 2020. Fungal Diversity 105:1-16 ((https://doi.org/10.1007/s13225-020-00466-2)) |
| Otu20504 | 1 | k__Fungi(100);p__Ascomycota(100);c__Saccharomycetes(100);o__Saccharomycetales(100);f__Metschnikowiaceae(100);g__Metschnikowia(100);s__Metschnikowia_pulcherrima(100); | Metschnikowia | 13 | Saprotroph | \|Undefined Saprotroph\| | Yeast | NULL | Probable | As nectar-tap_saprotroph (Põlme et al. 2020); Decay sugar-rich substrates (Põlme et al. 2020) | Põlme S, et al. 2020. Fungal Diversity 105:1-16 ((https://doi.org/10.1007/s13225-020-00466-2)) |
| Otu20505 | 1 | k__Fungi(100);p__Ascomycota(100);c__Saccharomycetes(100);o__Saccharomycetales(100);f__Metschnikowiaceae(100);g__Metschnikowia(100);s__Metschnikowia_pulcherrima(100); | Metschnikowia | 13 | Saprotroph | \|Undefined Saprotroph\| | Yeast | NULL | Probable | As nectar-tap_saprotroph (Põlme et al. 2020); Decay sugar-rich substrates (Põlme et al. 2020) | Põlme S, et al. 2020. Fungal Diversity 105:1-16 ((https://doi.org/10.1007/s13225-020-00466-2)) |
| Otu20506 | 1 | k__Fungi(100);p__Ascomycota(100);c__Saccharomycetes(100);o__Saccharomycetales(100);f__Metschnikowiaceae(100);g__Metschnikowia(100);s__Metschnikowia_pulcherrima(100); | Metschnikowia | 13 | Saprotroph | \|Undefined Saprotroph\| | Yeast | NULL | Probable | As nectar-tap_saprotroph (Põlme et al. 2020); Decay sugar-rich substrates (Põlme et al. 2020) | Põlme S, et al. 2020. Fungal Diversity 105:1-16 ((https://doi.org/10.1007/s13225-020-00466-2)) |
| Otu20508 | 1 | k__Fungi(100);p__Ascomycota(100);c__Saccharomycetes(100);o__Saccharomycetales(100);f__Metschnikowiaceae(100);g__Metschnikowia(100);s__Metschnikowia_pulcherrima(100); | Metschnikowia | 13 | Saprotroph | \|Undefined Saprotroph\| | Yeast | NULL | Probable | As nectar-tap_saprotroph (Põlme et al. 2020); Decay sugar-rich substrates (Põlme et al. 2020) | Põlme S, et al. 2020. Fungal Diversity 105:1-16 ((https://doi.org/10.1007/s13225-020-00466-2)) |
| Otu20509 | 1 | k__Fungi(100);p__Ascomycota(100);c__Saccharomycetes(100);o__Saccharomycetales(100);f__Metschnikowiaceae(100);g__Metschnikowia(100);s__Metschnikowia_pulcherrima(100); | Metschnikowia | 13 | Saprotroph | \|Undefined Saprotroph\| | Yeast | NULL | Probable | As nectar-tap_saprotroph (Põlme et al. 2020); Decay sugar-rich substrates (Põlme et al. 2020) | Põlme S, et al. 2020. Fungal Diversity 105:1-16 ((https://doi.org/10.1007/s13225-020-00466-2)) |
| Otu20510 | 1 | k__Fungi(100);p__Ascomycota(100);c__Saccharomycetes(100);o__Saccharomycetales(100);f__Metschnikowiaceae(100);g__Metschnikowia(100);s__Metschnikowia_pulcherrima(100); | Metschnikowia | 13 | Saprotroph | \|Undefined Saprotroph\| | Yeast | NULL | Probable | As nectar-tap_saprotroph (Põlme et al. 2020); Decay sugar-rich substrates (Põlme et al. 2020) | Põlme S, et al. 2020. Fungal Diversity 105:1-16 ((https://doi.org/10.1007/s13225-020-00466-2)) |
| Otu20511 | 1 | k__Fungi(100);p__Ascomycota(100);c__Saccharomycetes(100);o__Saccharomycetales(100);f__Saccharomycetales_family_Incertae_sedis(100);g__Candida(100);g__Candida_unclassified(100); | Candida | 13 | Pathotroph-Saprotroph-Symbiotroph | Animal Pathogen-Endophyte-Endosymbiont-Epiphyte-\|Undefined Saprotroph\| | Dimorphic Yeast-Yeast | NULL | Probable | As nectar-tap_saprotroph (Põlme et al. 2020); Decay sugar-rich substrates (Põlme et al. 2020); This asexual genus is polyphyletic and contain species that function very differently depend on what they are. Caution should be taken when interpreting guilds for Candida species. | Manolakaki D, et al. 2010. Virulence. 1: 367-375 ((https://doi.org/10.4161/viru.1.5.12796)); Mekbib SB, et al. 2011. Tropical Plant Pathology 36:233-240 ((https://doi.org/10.1590/S1982-56762011000400004)); Põlme S, et al. 2020. Fungal Diversity 105:1-16 ((https://doi.org/10.1007/s13225-020-00466-2)) |
| Otu20512 | 1 | k__Fungi(100);p__Ascomycota(100);c__Saccharomycetes(100);o__Saccharomycetales(100);f__Saccharomycetales_family_Incertae_sedis(100);g__Candida(100);g__Candida_unclassified(100); | Candida | 13 | Pathotroph-Saprotroph-Symbiotroph | Animal Pathogen-Endophyte-Endosymbiont-Epiphyte-\|Undefined Saprotroph\| | Dimorphic Yeast-Yeast | NULL | Probable | As nectar-tap_saprotroph (Põlme et al. 2020); Decay sugar-rich substrates (Põlme et al. 2020); This asexual genus is polyphyletic and contain species that function very differently depend on what they are. Caution should be taken when interpreting guilds for Candida species. | Manolakaki D, et al. 2010. Virulence. 1: 367-375 ((https://doi.org/10.4161/viru.1.5.12796)); Mekbib SB, et al. 2011. Tropical Plant Pathology 36:233-240 ((https://doi.org/10.1590/S1982-56762011000400004)); Põlme S, et al. 2020. Fungal Diversity 105:1-16 ((https://doi.org/10.1007/s13225-020-00466-2)) |
| Otu20513 | 1 | k__Fungi(100);p__Ascomycota(100);c__Saccharomycetes(100);o__Saccharomycetales(100);f__Metschnikowiaceae(100);g__Metschnikowia(100);s__Metschnikowia_pulcherrima(100); | Metschnikowia | 13 | Saprotroph | \|Undefined Saprotroph\| | Yeast | NULL | Probable | As nectar-tap_saprotroph (Põlme et al. 2020); Decay sugar-rich substrates (Põlme et al. 2020) | Põlme S, et al. 2020. Fungal Diversity 105:1-16 ((https://doi.org/10.1007/s13225-020-00466-2)) |
| Otu20514 | 1 | k__Fungi(100);p__Ascomycota(100);c__Saccharomycetes(100);o__Saccharomycetales(100);f__Metschnikowiaceae(100);g__Metschnikowia(100);s__Metschnikowia_pulcherrima(100); | Metschnikowia | 13 | Saprotroph | \|Undefined Saprotroph\| | Yeast | NULL | Probable | As nectar-tap_saprotroph (Põlme et al. 2020); Decay sugar-rich substrates (Põlme et al. 2020) | Põlme S, et al. 2020. Fungal Diversity 105:1-16 ((https://doi.org/10.1007/s13225-020-00466-2)) |
| Otu20515 | 1 | k__Fungi(100);p__Ascomycota(100);c__Saccharomycetes(100);o__Saccharomycetales(100);f__Metschnikowiaceae(100);g__Metschnikowia(100);s__Metschnikowia_pulcherrima(100); | Metschnikowia | 13 | Saprotroph | \|Undefined Saprotroph\| | Yeast | NULL | Probable | As nectar-tap_saprotroph (Põlme et al. 2020); Decay sugar-rich substrates (Põlme et al. 2020) | Põlme S, et al. 2020. Fungal Diversity 105:1-16 ((https://doi.org/10.1007/s13225-020-00466-2)) |
| Otu20516 | 1 | k__Fungi(100);p__Ascomycota(100);c__Saccharomycetes(100);o__Saccharomycetales(100);f__Metschnikowiaceae(100);g__Metschnikowia(100);g__Metschnikowia_unclassified(100); | Metschnikowia | 13 | Saprotroph | \|Undefined Saprotroph\| | Yeast | NULL | Probable | As nectar-tap_saprotroph (Põlme et al. 2020); Decay sugar-rich substrates (Põlme et al. 2020) | Põlme S, et al. 2020. Fungal Diversity 105:1-16 ((https://doi.org/10.1007/s13225-020-00466-2)) |
| Otu20517 | 1 | k__Fungi(100);p__Ascomycota(100);c__Saccharomycetes(100);o__Saccharomycetales(100);f__Metschnikowiaceae(100);g__Metschnikowia(100);s__Metschnikowia_pulcherrima(100); | Metschnikowia | 13 | Saprotroph | \|Undefined Saprotroph\| | Yeast | NULL | Probable | As nectar-tap_saprotroph (Põlme et al. 2020); Decay sugar-rich substrates (Põlme et al. 2020) | Põlme S, et al. 2020. Fungal Diversity 105:1-16 ((https://doi.org/10.1007/s13225-020-00466-2)) |
| Otu20518 | 1 | k__Fungi(100);p__Ascomycota(100);c__Saccharomycetes(100);o__Saccharomycetales(100);f__Metschnikowiaceae(100);g__Metschnikowia(100);s__Metschnikowia_pulcherrima(100); | Metschnikowia | 13 | Saprotroph | \|Undefined Saprotroph\| | Yeast | NULL | Probable | As nectar-tap_saprotroph (Põlme et al. 2020); Decay sugar-rich substrates (Põlme et al. 2020) | Põlme S, et al. 2020. Fungal Diversity 105:1-16 ((https://doi.org/10.1007/s13225-020-00466-2)) |
| Otu20527 | 1 | k__Fungi(100);p__Ascomycota(100);c__Saccharomycetes(100);o__Saccharomycetales(100);f__Metschnikowiaceae(100);g__Metschnikowia(100);g__Metschnikowia_unclassified(100); | Metschnikowia | 13 | Saprotroph | \|Undefined Saprotroph\| | Yeast | NULL | Probable | As nectar-tap_saprotroph (Põlme et al. 2020); Decay sugar-rich substrates (Põlme et al. 2020) | Põlme S, et al. 2020. Fungal Diversity 105:1-16 ((https://doi.org/10.1007/s13225-020-00466-2)) |
| Otu20528 | 1 | k__Fungi(100);p__Ascomycota(100);c__Saccharomycetes(100);o__Saccharomycetales(100);f__Metschnikowiaceae(100);g__Metschnikowia(100);s__Metschnikowia_pulcherrima(100); | Metschnikowia | 13 | Saprotroph | \|Undefined Saprotroph\| | Yeast | NULL | Probable | As nectar-tap_saprotroph (Põlme et al. 2020); Decay sugar-rich substrates (Põlme et al. 2020) | Põlme S, et al. 2020. Fungal Diversity 105:1-16 ((https://doi.org/10.1007/s13225-020-00466-2)) |
| Otu20529 | 1 | k__Fungi(100);p__Ascomycota(100);c__Saccharomycetes(100);o__Saccharomycetales(100);f__Metschnikowiaceae(100);g__Metschnikowia(100);s__Metschnikowia_pulcherrima(100); | Metschnikowia | 13 | Saprotroph | \|Undefined Saprotroph\| | Yeast | NULL | Probable | As nectar-tap_saprotroph (Põlme et al. 2020); Decay sugar-rich substrates (Põlme et al. 2020) | Põlme S, et al. 2020. Fungal Diversity 105:1-16 ((https://doi.org/10.1007/s13225-020-00466-2)) |
| Otu20530 | 1 | k__Fungi(100);p__Ascomycota(100);c__Saccharomycetes(100);o__Saccharomycetales(100);f__Metschnikowiaceae(100);g__Metschnikowia(100);s__Metschnikowia_pulcherrima(100); | Metschnikowia | 13 | Saprotroph | \|Undefined Saprotroph\| | Yeast | NULL | Probable | As nectar-tap_saprotroph (Põlme et al. 2020); Decay sugar-rich substrates (Põlme et al. 2020) | Põlme S, et al. 2020. Fungal Diversity 105:1-16 ((https://doi.org/10.1007/s13225-020-00466-2)) |
| Otu20531 | 1 | k__Fungi(100);p__Ascomycota(100);c__Saccharomycetes(100);o__Saccharomycetales(100);f__Metschnikowiaceae(100);g__Metschnikowia(100);g__Metschnikowia_unclassified(100); | Metschnikowia | 13 | Saprotroph | \|Undefined Saprotroph\| | Yeast | NULL | Probable | As nectar-tap_saprotroph (Põlme et al. 2020); Decay sugar-rich substrates (Põlme et al. 2020) | Põlme S, et al. 2020. Fungal Diversity 105:1-16 ((https://doi.org/10.1007/s13225-020-00466-2)) |
| Otu20532 | 1 | k__Fungi(100);p__Ascomycota(100);c__Saccharomycetes(100);o__Saccharomycetales(100);f__Metschnikowiaceae(100);g__Metschnikowia(100);s__Metschnikowia_pulcherrima(100); | Metschnikowia | 13 | Saprotroph | \|Undefined Saprotroph\| | Yeast | NULL | Probable | As nectar-tap_saprotroph (Põlme et al. 2020); Decay sugar-rich substrates (Põlme et al. 2020) | Põlme S, et al. 2020. Fungal Diversity 105:1-16 ((https://doi.org/10.1007/s13225-020-00466-2)) |
| Otu20533 | 1 | k__Fungi(100);p__Ascomycota(100);c__Saccharomycetes(100);o__Saccharomycetales(100);f__Metschnikowiaceae(100);g__Metschnikowia(100);s__Metschnikowia_pulcherrima(100); | Metschnikowia | 13 | Saprotroph | \|Undefined Saprotroph\| | Yeast | NULL | Probable | As nectar-tap_saprotroph (Põlme et al. 2020); Decay sugar-rich substrates (Põlme et al. 2020) | Põlme S, et al. 2020. Fungal Diversity 105:1-16 ((https://doi.org/10.1007/s13225-020-00466-2)) |
| Otu20534 | 1 | k__Fungi(100);p__Ascomycota(100);c__Saccharomycetes(100);o__Saccharomycetales(100);f__Metschnikowiaceae(100);g__Metschnikowia(100);g__Metschnikowia_unclassified(100); | Metschnikowia | 13 | Saprotroph | \|Undefined Saprotroph\| | Yeast | NULL | Probable | As nectar-tap_saprotroph (Põlme et al. 2020); Decay sugar-rich substrates (Põlme et al. 2020) | Põlme S, et al. 2020. Fungal Diversity 105:1-16 ((https://doi.org/10.1007/s13225-020-00466-2)) |
| Otu20535 | 1 | k__Fungi(100);p__Ascomycota(100);c__Saccharomycetes(100);o__Saccharomycetales(100);f__Metschnikowiaceae(100);g__Metschnikowia(100);s__Metschnikowia_pulcherrima(100); | Metschnikowia | 13 | Saprotroph | \|Undefined Saprotroph\| | Yeast | NULL | Probable | As nectar-tap_saprotroph (Põlme et al. 2020); Decay sugar-rich substrates (Põlme et al. 2020) | Põlme S, et al. 2020. Fungal Diversity 105:1-16 ((https://doi.org/10.1007/s13225-020-00466-2)) |
| Otu20536 | 1 | k__Fungi(100);p__Ascomycota(100);c__Saccharomycetes(100);o__Saccharomycetales(100);f__Metschnikowiaceae(100);g__Metschnikowia(100);g__Metschnikowia_unclassified(100); | Metschnikowia | 13 | Saprotroph | \|Undefined Saprotroph\| | Yeast | NULL | Probable | As nectar-tap_saprotroph (Põlme et al. 2020); Decay sugar-rich substrates (Põlme et al. 2020) | Põlme S, et al. 2020. Fungal Diversity 105:1-16 ((https://doi.org/10.1007/s13225-020-00466-2)) |
| Otu20539 | 1 | k__Fungi(100);p__Ascomycota(100);c__Saccharomycetes(100);o__Saccharomycetales(100);f__Metschnikowiaceae(100);g__Metschnikowia(100);s__Metschnikowia_pulcherrima(100); | Metschnikowia | 13 | Saprotroph | \|Undefined Saprotroph\| | Yeast | NULL | Probable | As nectar-tap_saprotroph (Põlme et al. 2020); Decay sugar-rich substrates (Põlme et al. 2020) | Põlme S, et al. 2020. Fungal Diversity 105:1-16 ((https://doi.org/10.1007/s13225-020-00466-2)) |
| Otu20540 | 1 | k__Fungi(100);p__Ascomycota(100);c__Saccharomycetes(100);o__Saccharomycetales(100);f__Metschnikowiaceae(100);g__Metschnikowia(100);s__Metschnikowia_pulcherrima(100); | Metschnikowia | 13 | Saprotroph | \|Undefined Saprotroph\| | Yeast | NULL | Probable | As nectar-tap_saprotroph (Põlme et al. 2020); Decay sugar-rich substrates (Põlme et al. 2020) | Põlme S, et al. 2020. Fungal Diversity 105:1-16 ((https://doi.org/10.1007/s13225-020-00466-2)) |
| Otu20541 | 1 | k__Fungi(100);p__Ascomycota(100);c__Saccharomycetes(100);o__Saccharomycetales(100);f__Metschnikowiaceae(100);g__Metschnikowia(100);s__Metschnikowia_pulcherrima(100); | Metschnikowia | 13 | Saprotroph | \|Undefined Saprotroph\| | Yeast | NULL | Probable | As nectar-tap_saprotroph (Põlme et al. 2020); Decay sugar-rich substrates (Põlme et al. 2020) | Põlme S, et al. 2020. Fungal Diversity 105:1-16 ((https://doi.org/10.1007/s13225-020-00466-2)) |
| Otu20542 | 1 | k__Fungi(100);p__Ascomycota(100);c__Saccharomycetes(100);o__Saccharomycetales(100);f__Metschnikowiaceae(100);g__Metschnikowia(100);s__Metschnikowia_pulcherrima(100); | Metschnikowia | 13 | Saprotroph | \|Undefined Saprotroph\| | Yeast | NULL | Probable | As nectar-tap_saprotroph (Põlme et al. 2020); Decay sugar-rich substrates (Põlme et al. 2020) | Põlme S, et al. 2020. Fungal Diversity 105:1-16 ((https://doi.org/10.1007/s13225-020-00466-2)) |
| Otu20543 | 1 | k__Fungi(100);p__Ascomycota(100);c__Saccharomycetes(100);o__Saccharomycetales(100);f__Metschnikowiaceae(100);g__Metschnikowia(100);g__Metschnikowia_unclassified(100); | Metschnikowia | 13 | Saprotroph | \|Undefined Saprotroph\| | Yeast | NULL | Probable | As nectar-tap_saprotroph (Põlme et al. 2020); Decay sugar-rich substrates (Põlme et al. 2020) | Põlme S, et al. 2020. Fungal Diversity 105:1-16 ((https://doi.org/10.1007/s13225-020-00466-2)) |
| Otu20544 | 1 | k__Fungi(100);p__Ascomycota(100);c__Saccharomycetes(100);o__Saccharomycetales(100);f__Metschnikowiaceae(100);g__Metschnikowia(100);s__Metschnikowia_pulcherrima(100); | Metschnikowia | 13 | Saprotroph | \|Undefined Saprotroph\| | Yeast | NULL | Probable | As nectar-tap_saprotroph (Põlme et al. 2020); Decay sugar-rich substrates (Põlme et al. 2020) | Põlme S, et al. 2020. Fungal Diversity 105:1-16 ((https://doi.org/10.1007/s13225-020-00466-2)) |
| Otu20545 | 1 | k__Fungi(100);p__Ascomycota(100);c__Saccharomycetes(100);o__Saccharomycetales(100);f__Metschnikowiaceae(100);g__Metschnikowia(100);s__Metschnikowia_pulcherrima(100); | Metschnikowia | 13 | Saprotroph | \|Undefined Saprotroph\| | Yeast | NULL | Probable | As nectar-tap_saprotroph (Põlme et al. 2020); Decay sugar-rich substrates (Põlme et al. 2020) | Põlme S, et al. 2020. Fungal Diversity 105:1-16 ((https://doi.org/10.1007/s13225-020-00466-2)) |
| Otu20546 | 1 | k__Fungi(100);p__Ascomycota(100);c__Saccharomycetes(100);o__Saccharomycetales(100);f__Metschnikowiaceae(100);g__Metschnikowia(100);s__Metschnikowia_pulcherrima(100); | Metschnikowia | 13 | Saprotroph | \|Undefined Saprotroph\| | Yeast | NULL | Probable | As nectar-tap_saprotroph (Põlme et al. 2020); Decay sugar-rich substrates (Põlme et al. 2020) | Põlme S, et al. 2020. Fungal Diversity 105:1-16 ((https://doi.org/10.1007/s13225-020-00466-2)) |
| Otu20547 | 1 | k__Fungi(100);p__Ascomycota(100);c__Saccharomycetes(100);o__Saccharomycetales(100);f__Metschnikowiaceae(100);g__Metschnikowia(100);s__Metschnikowia_pulcherrima(100); | Metschnikowia | 13 | Saprotroph | \|Undefined Saprotroph\| | Yeast | NULL | Probable | As nectar-tap_saprotroph (Põlme et al. 2020); Decay sugar-rich substrates (Põlme et al. 2020) | Põlme S, et al. 2020. Fungal Diversity 105:1-16 ((https://doi.org/10.1007/s13225-020-00466-2)) |
| Otu20548 | 1 | k__Fungi(100);p__Ascomycota(100);c__Saccharomycetes(100);o__Saccharomycetales(100);f__Metschnikowiaceae(100);g__Metschnikowia(100);s__Metschnikowia_pulcherrima(100); | Metschnikowia | 13 | Saprotroph | \|Undefined Saprotroph\| | Yeast | NULL | Probable | As nectar-tap_saprotroph (Põlme et al. 2020); Decay sugar-rich substrates (Põlme et al. 2020) | Põlme S, et al. 2020. Fungal Diversity 105:1-16 ((https://doi.org/10.1007/s13225-020-00466-2)) |
| Otu20549 | 1 | k__Fungi(100);p__Ascomycota(100);c__Saccharomycetes(100);o__Saccharomycetales(100);f__Metschnikowiaceae(100);g__Metschnikowia(100);g__Metschnikowia_unclassified(100); | Metschnikowia | 13 | Saprotroph | \|Undefined Saprotroph\| | Yeast | NULL | Probable | As nectar-tap_saprotroph (Põlme et al. 2020); Decay sugar-rich substrates (Põlme et al. 2020) | Põlme S, et al. 2020. Fungal Diversity 105:1-16 ((https://doi.org/10.1007/s13225-020-00466-2)) |
| Otu20550 | 1 | k__Fungi(100);p__Ascomycota(100);c__Saccharomycetes(100);o__Saccharomycetales(100);f__Metschnikowiaceae(100);g__Metschnikowia(100);g__Metschnikowia_unclassified(100); | Metschnikowia | 13 | Saprotroph | \|Undefined Saprotroph\| | Yeast | NULL | Probable | As nectar-tap_saprotroph (Põlme et al. 2020); Decay sugar-rich substrates (Põlme et al. 2020) | Põlme S, et al. 2020. Fungal Diversity 105:1-16 ((https://doi.org/10.1007/s13225-020-00466-2)) |
| Otu20551 | 1 | k__Fungi(100);p__Ascomycota(100);c__Saccharomycetes(100);o__Saccharomycetales(100);f__Metschnikowiaceae(100);g__Metschnikowia(100);s__Metschnikowia_pulcherrima(100); | Metschnikowia | 13 | Saprotroph | \|Undefined Saprotroph\| | Yeast | NULL | Probable | As nectar-tap_saprotroph (Põlme et al. 2020); Decay sugar-rich substrates (Põlme et al. 2020) | Põlme S, et al. 2020. Fungal Diversity 105:1-16 ((https://doi.org/10.1007/s13225-020-00466-2)) |
| Otu20552 | 1 | k__Fungi(100);p__Ascomycota(100);c__Saccharomycetes(100);o__Saccharomycetales(100);f__Saccharomycetaceae(100);g__Kazachstania(100);s__Kazachstania_intestinalis(100); | Kazachstania | 13 | Saprotroph | \|Undefined Saprotroph\| | Yeast | NULL | Probable | As nectar-tap_saprotroph (Põlme et al. 2020); Decay sugar-rich substrates (Põlme et al. 2020) | Põlme S, et al. 2020. Fungal Diversity 105:1-16 ((https://doi.org/10.1007/s13225-020-00466-2)) |
| Otu20553 | 1 | k__Fungi(100);p__Ascomycota(100);c__Saccharomycetes(100);o__Saccharomycetales(100);f__Metschnikowiaceae(100);g__Metschnikowia(100);s__Metschnikowia_pulcherrima(100); | Metschnikowia | 13 | Saprotroph | \|Undefined Saprotroph\| | Yeast | NULL | Probable | As nectar-tap_saprotroph (Põlme et al. 2020); Decay sugar-rich substrates (Põlme et al. 2020) | Põlme S, et al. 2020. Fungal Diversity 105:1-16 ((https://doi.org/10.1007/s13225-020-00466-2)) |
| Otu20554 | 1 | k__Fungi(100);p__Ascomycota(100);c__Saccharomycetes(100);o__Saccharomycetales(100);f__Metschnikowiaceae(100);g__Metschnikowia(100);s__Metschnikowia_pulcherrima(100); | Metschnikowia | 13 | Saprotroph | \|Undefined Saprotroph\| | Yeast | NULL | Probable | As nectar-tap_saprotroph (Põlme et al. 2020); Decay sugar-rich substrates (Põlme et al. 2020) | Põlme S, et al. 2020. Fungal Diversity 105:1-16 ((https://doi.org/10.1007/s13225-020-00466-2)) |
| Otu20555 | 1 | k__Fungi(100);p__Ascomycota(100);c__Saccharomycetes(100);o__Saccharomycetales(100);f__Metschnikowiaceae(100);g__Metschnikowia(100);g__Metschnikowia_unclassified(100); | Metschnikowia | 13 | Saprotroph | \|Undefined Saprotroph\| | Yeast | NULL | Probable | As nectar-tap_saprotroph (Põlme et al. 2020); Decay sugar-rich substrates (Põlme et al. 2020) | Põlme S, et al. 2020. Fungal Diversity 105:1-16 ((https://doi.org/10.1007/s13225-020-00466-2)) |
| Otu20556 | 1 | k__Fungi(100);p__Ascomycota(100);c__Saccharomycetes(100);o__Saccharomycetales(100);f__Metschnikowiaceae(100);g__Metschnikowia(100);g__Metschnikowia_unclassified(100); | Metschnikowia | 13 | Saprotroph | \|Undefined Saprotroph\| | Yeast | NULL | Probable | As nectar-tap_saprotroph (Põlme et al. 2020); Decay sugar-rich substrates (Põlme et al. 2020) | Põlme S, et al. 2020. Fungal Diversity 105:1-16 ((https://doi.org/10.1007/s13225-020-00466-2)) |
| Otu20557 | 1 | k__Fungi(100);p__Ascomycota(100);c__Saccharomycetes(100);o__Saccharomycetales(100);f__Metschnikowiaceae(100);g__Metschnikowia(100);g__Metschnikowia_unclassified(100); | Metschnikowia | 13 | Saprotroph | \|Undefined Saprotroph\| | Yeast | NULL | Probable | As nectar-tap_saprotroph (Põlme et al. 2020); Decay sugar-rich substrates (Põlme et al. 2020) | Põlme S, et al. 2020. Fungal Diversity 105:1-16 ((https://doi.org/10.1007/s13225-020-00466-2)) |
| Otu20558 | 1 | k__Fungi(100);p__Ascomycota(100);c__Saccharomycetes(100);o__Saccharomycetales(100);f__Metschnikowiaceae(100);g__Metschnikowia(100);s__Metschnikowia_pulcherrima(100); | Metschnikowia | 13 | Saprotroph | \|Undefined Saprotroph\| | Yeast | NULL | Probable | As nectar-tap_saprotroph (Põlme et al. 2020); Decay sugar-rich substrates (Põlme et al. 2020) | Põlme S, et al. 2020. Fungal Diversity 105:1-16 ((https://doi.org/10.1007/s13225-020-00466-2)) |
| Otu20559 | 1 | k__Fungi(100);p__Ascomycota(100);c__Saccharomycetes(100);o__Saccharomycetales(100);f__Metschnikowiaceae(100);g__Metschnikowia(100);s__Metschnikowia_pulcherrima(100); | Metschnikowia | 13 | Saprotroph | \|Undefined Saprotroph\| | Yeast | NULL | Probable | As nectar-tap_saprotroph (Põlme et al. 2020); Decay sugar-rich substrates (Põlme et al. 2020) | Põlme S, et al. 2020. Fungal Diversity 105:1-16 ((https://doi.org/10.1007/s13225-020-00466-2)) |
| Otu20560 | 1 | k__Fungi(100);p__Basidiomycota(100);c__Microbotryomycetes(100);o__Sporidiobolales(100);f__Sporidiobolales_family_Incertae_sedis(100);g__Sporobolomyces(100);s__Sporobolomyces_lactosus(100); | Sporobolomyces | 13 | Pathotroph-Saprotroph | \|Fungal Parasite\|-Plant Saprotroph | Yeast | NULL | Probable | As litter_saprotroph (Põlme et al. 2020); As mycoparasite (Põlme et al. 2020); Decay fungal material (Põlme et al. 2020) | Tedersoo L, et al. 2014. Science 346:e1256688 ((https://doi.org/10.1126/science.1256688)); Purahong W, et al. 2016. Molecular Ecology 25:4059-4074 ((https://doi.org/10.1111/mec.13739)); Põlme S, et al. 2020. Fungal Diversity 105:1-16 ((https://doi.org/10.1007/s13225-020-00466-2)) |
| Otu20561 | 1 | k__Fungi(100);p__Ascomycota(100);c__Saccharomycetes(100);o__Saccharomycetales(100);f__Metschnikowiaceae(100);g__Metschnikowia(100);s__Metschnikowia_pulcherrima(100); | Metschnikowia | 13 | Saprotroph | \|Undefined Saprotroph\| | Yeast | NULL | Probable | As nectar-tap_saprotroph (Põlme et al. 2020); Decay sugar-rich substrates (Põlme et al. 2020) | Põlme S, et al. 2020. Fungal Diversity 105:1-16 ((https://doi.org/10.1007/s13225-020-00466-2)) |
| Otu20562 | 1 | k__Fungi(100);p__Ascomycota(100);c__Saccharomycetes(100);o__Saccharomycetales(100);f__Metschnikowiaceae(100);g__Metschnikowia(100);s__Metschnikowia_pulcherrima(100); | Metschnikowia | 13 | Saprotroph | \|Undefined Saprotroph\| | Yeast | NULL | Probable | As nectar-tap_saprotroph (Põlme et al. 2020); Decay sugar-rich substrates (Põlme et al. 2020) | Põlme S, et al. 2020. Fungal Diversity 105:1-16 ((https://doi.org/10.1007/s13225-020-00466-2)) |
| Otu20563 | 1 | k__Fungi(100);p__Ascomycota(100);c__Saccharomycetes(100);o__Saccharomycetales(100);f__Metschnikowiaceae(100);g__Metschnikowia(100);s__Metschnikowia_pulcherrima(100); | Metschnikowia | 13 | Saprotroph | \|Undefined Saprotroph\| | Yeast | NULL | Probable | As nectar-tap_saprotroph (Põlme et al. 2020); Decay sugar-rich substrates (Põlme et al. 2020) | Põlme S, et al. 2020. Fungal Diversity 105:1-16 ((https://doi.org/10.1007/s13225-020-00466-2)) |
| Otu20564 | 1 | k__Fungi(100);p__Ascomycota(100);c__Saccharomycetes(100);o__Saccharomycetales(100);f__Metschnikowiaceae(100);g__Metschnikowia(100);s__Metschnikowia_pulcherrima(100); | Metschnikowia | 13 | Saprotroph | \|Undefined Saprotroph\| | Yeast | NULL | Probable | As nectar-tap_saprotroph (Põlme et al. 2020); Decay sugar-rich substrates (Põlme et al. 2020) | Põlme S, et al. 2020. Fungal Diversity 105:1-16 ((https://doi.org/10.1007/s13225-020-00466-2)) |
| Otu20565 | 1 | k__Fungi(100);p__Ascomycota(100);c__Saccharomycetes(100);o__Saccharomycetales(100);f__Metschnikowiaceae(100);g__Metschnikowia(100);s__Metschnikowia_pulcherrima(100); | Metschnikowia | 13 | Saprotroph | \|Undefined Saprotroph\| | Yeast | NULL | Probable | As nectar-tap_saprotroph (Põlme et al. 2020); Decay sugar-rich substrates (Põlme et al. 2020) | Põlme S, et al. 2020. Fungal Diversity 105:1-16 ((https://doi.org/10.1007/s13225-020-00466-2)) |
| Otu20566 | 1 | k__Fungi(100);p__Ascomycota(100);c__Saccharomycetes(100);o__Saccharomycetales(100);f__Metschnikowiaceae(100);g__Metschnikowia(100);s__Metschnikowia_pulcherrima(100); | Metschnikowia | 13 | Saprotroph | \|Undefined Saprotroph\| | Yeast | NULL | Probable | As nectar-tap_saprotroph (Põlme et al. 2020); Decay sugar-rich substrates (Põlme et al. 2020) | Põlme S, et al. 2020. Fungal Diversity 105:1-16 ((https://doi.org/10.1007/s13225-020-00466-2)) |
| Otu20567 | 1 | k__Fungi(100);p__Ascomycota(100);c__Saccharomycetes(100);o__Saccharomycetales(100);f__Metschnikowiaceae(100);g__Metschnikowia(100);g__Metschnikowia_unclassified(100); | Metschnikowia | 13 | Saprotroph | \|Undefined Saprotroph\| | Yeast | NULL | Probable | As nectar-tap_saprotroph (Põlme et al. 2020); Decay sugar-rich substrates (Põlme et al. 2020) | Põlme S, et al. 2020. Fungal Diversity 105:1-16 ((https://doi.org/10.1007/s13225-020-00466-2)) |
| Otu20568 | 1 | k__Fungi(100);p__Ascomycota(100);c__Saccharomycetes(100);o__Saccharomycetales(100);f__Metschnikowiaceae(100);g__Metschnikowia(100);s__Metschnikowia_pulcherrima(100); | Metschnikowia | 13 | Saprotroph | \|Undefined Saprotroph\| | Yeast | NULL | Probable | As nectar-tap_saprotroph (Põlme et al. 2020); Decay sugar-rich substrates (Põlme et al. 2020) | Põlme S, et al. 2020. Fungal Diversity 105:1-16 ((https://doi.org/10.1007/s13225-020-00466-2)) |
| Otu20569 | 1 | k__Fungi(100);p__Ascomycota(100);c__Saccharomycetes(100);o__Saccharomycetales(100);f__Metschnikowiaceae(100);g__Metschnikowia(100);s__Metschnikowia_pulcherrima(100); | Metschnikowia | 13 | Saprotroph | \|Undefined Saprotroph\| | Yeast | NULL | Probable | As nectar-tap_saprotroph (Põlme et al. 2020); Decay sugar-rich substrates (Põlme et al. 2020) | Põlme S, et al. 2020. Fungal Diversity 105:1-16 ((https://doi.org/10.1007/s13225-020-00466-2)) |
| Otu20570 | 1 | k__Fungi(100);p__Ascomycota(100);c__Saccharomycetes(100);o__Saccharomycetales(100);f__Metschnikowiaceae(100);g__Metschnikowia(100);s__Metschnikowia_pulcherrima(100); | Metschnikowia | 13 | Saprotroph | \|Undefined Saprotroph\| | Yeast | NULL | Probable | As nectar-tap_saprotroph (Põlme et al. 2020); Decay sugar-rich substrates (Põlme et al. 2020) | Põlme S, et al. 2020. Fungal Diversity 105:1-16 ((https://doi.org/10.1007/s13225-020-00466-2)) |
| Otu20571 | 1 | k__Fungi(100);p__Ascomycota(100);c__Saccharomycetes(100);o__Saccharomycetales(100);f__Metschnikowiaceae(100);g__Metschnikowia(100);s__Metschnikowia_pulcherrima(100); | Metschnikowia | 13 | Saprotroph | \|Undefined Saprotroph\| | Yeast | NULL | Probable | As nectar-tap_saprotroph (Põlme et al. 2020); Decay sugar-rich substrates (Põlme et al. 2020) | Põlme S, et al. 2020. Fungal Diversity 105:1-16 ((https://doi.org/10.1007/s13225-020-00466-2)) |
| Otu20573 | 1 | k__Fungi(100);p__Ascomycota(100);c__Saccharomycetes(100);o__Saccharomycetales(100);f__Metschnikowiaceae(100);g__Metschnikowia(100);g__Metschnikowia_unclassified(100); | Metschnikowia | 13 | Saprotroph | \|Undefined Saprotroph\| | Yeast | NULL | Probable | As nectar-tap_saprotroph (Põlme et al. 2020); Decay sugar-rich substrates (Põlme et al. 2020) | Põlme S, et al. 2020. Fungal Diversity 105:1-16 ((https://doi.org/10.1007/s13225-020-00466-2)) |
| Otu20574 | 1 | k__Fungi(100);p__Ascomycota(100);c__Saccharomycetes(100);o__Saccharomycetales(100);f__Metschnikowiaceae(100);g__Metschnikowia(100);s__Metschnikowia_pulcherrima(100); | Metschnikowia | 13 | Saprotroph | \|Undefined Saprotroph\| | Yeast | NULL | Probable | As nectar-tap_saprotroph (Põlme et al. 2020); Decay sugar-rich substrates (Põlme et al. 2020) | Põlme S, et al. 2020. Fungal Diversity 105:1-16 ((https://doi.org/10.1007/s13225-020-00466-2)) |
| Otu20575 | 1 | k__Fungi(100);p__Ascomycota(100);c__Saccharomycetes(100);o__Saccharomycetales(100);f__Metschnikowiaceae(100);g__Metschnikowia(100);s__Metschnikowia_pulcherrima(100); | Metschnikowia | 13 | Saprotroph | \|Undefined Saprotroph\| | Yeast | NULL | Probable | As nectar-tap_saprotroph (Põlme et al. 2020); Decay sugar-rich substrates (Põlme et al. 2020) | Põlme S, et al. 2020. Fungal Diversity 105:1-16 ((https://doi.org/10.1007/s13225-020-00466-2)) |
| Otu20576 | 1 | k__Fungi(100);p__Ascomycota(100);c__Saccharomycetes(100);o__Saccharomycetales(100);f__Metschnikowiaceae(100);g__Metschnikowia(100);s__Metschnikowia_pulcherrima(100); | Metschnikowia | 13 | Saprotroph | \|Undefined Saprotroph\| | Yeast | NULL | Probable | As nectar-tap_saprotroph (Põlme et al. 2020); Decay sugar-rich substrates (Põlme et al. 2020) | Põlme S, et al. 2020. Fungal Diversity 105:1-16 ((https://doi.org/10.1007/s13225-020-00466-2)) |
| Otu20577 | 1 | k__Fungi(100);p__Ascomycota(100);c__Saccharomycetes(100);o__Saccharomycetales(100);f__Metschnikowiaceae(100);g__Metschnikowia(100);s__Metschnikowia_pulcherrima(100); | Metschnikowia | 13 | Saprotroph | \|Undefined Saprotroph\| | Yeast | NULL | Probable | As nectar-tap_saprotroph (Põlme et al. 2020); Decay sugar-rich substrates (Põlme et al. 2020) | Põlme S, et al. 2020. Fungal Diversity 105:1-16 ((https://doi.org/10.1007/s13225-020-00466-2)) |
| Otu20578 | 1 | k__Fungi(100);p__Ascomycota(100);c__Saccharomycetes(100);o__Saccharomycetales(100);f__Metschnikowiaceae(100);g__Metschnikowia(100);s__Metschnikowia_pulcherrima(100); | Metschnikowia | 13 | Saprotroph | \|Undefined Saprotroph\| | Yeast | NULL | Probable | As nectar-tap_saprotroph (Põlme et al. 2020); Decay sugar-rich substrates (Põlme et al. 2020) | Põlme S, et al. 2020. Fungal Diversity 105:1-16 ((https://doi.org/10.1007/s13225-020-00466-2)) |
| Otu20579 | 1 | k__Fungi(100);p__Ascomycota(100);c__Saccharomycetes(100);o__Saccharomycetales(100);f__Metschnikowiaceae(100);g__Metschnikowia(100);s__Metschnikowia_pulcherrima(100); | Metschnikowia | 13 | Saprotroph | \|Undefined Saprotroph\| | Yeast | NULL | Probable | As nectar-tap_saprotroph (Põlme et al. 2020); Decay sugar-rich substrates (Põlme et al. 2020) | Põlme S, et al. 2020. Fungal Diversity 105:1-16 ((https://doi.org/10.1007/s13225-020-00466-2)) |
| Otu20580 | 1 | k__Fungi(100);p__Ascomycota(100);c__Saccharomycetes(100);o__Saccharomycetales(100);f__Metschnikowiaceae(100);g__Metschnikowia(100);g__Metschnikowia_unclassified(100); | Metschnikowia | 13 | Saprotroph | \|Undefined Saprotroph\| | Yeast | NULL | Probable | As nectar-tap_saprotroph (Põlme et al. 2020); Decay sugar-rich substrates (Põlme et al. 2020) | Põlme S, et al. 2020. Fungal Diversity 105:1-16 ((https://doi.org/10.1007/s13225-020-00466-2)) |
| Otu20582 | 1 | k__Fungi(100);p__Ascomycota(100);c__Saccharomycetes(100);o__Saccharomycetales(100);f__Metschnikowiaceae(100);g__Metschnikowia(100);g__Metschnikowia_unclassified(100); | Metschnikowia | 13 | Saprotroph | \|Undefined Saprotroph\| | Yeast | NULL | Probable | As nectar-tap_saprotroph (Põlme et al. 2020); Decay sugar-rich substrates (Põlme et al. 2020) | Põlme S, et al. 2020. Fungal Diversity 105:1-16 ((https://doi.org/10.1007/s13225-020-00466-2)) |
| Otu20583 | 1 | k__Fungi(100);p__Ascomycota(100);c__Saccharomycetes(100);o__Saccharomycetales(100);f__Metschnikowiaceae(100);g__Metschnikowia(100);g__Metschnikowia_unclassified(100); | Metschnikowia | 13 | Saprotroph | \|Undefined Saprotroph\| | Yeast | NULL | Probable | As nectar-tap_saprotroph (Põlme et al. 2020); Decay sugar-rich substrates (Põlme et al. 2020) | Põlme S, et al. 2020. Fungal Diversity 105:1-16 ((https://doi.org/10.1007/s13225-020-00466-2)) |
| Otu20584 | 1 | k__Fungi(100);p__Ascomycota(100);c__Saccharomycetes(100);o__Saccharomycetales(100);f__Metschnikowiaceae(100);g__Metschnikowia(100);s__Metschnikowia_pulcherrima(100); | Metschnikowia | 13 | Saprotroph | \|Undefined Saprotroph\| | Yeast | NULL | Probable | As nectar-tap_saprotroph (Põlme et al. 2020); Decay sugar-rich substrates (Põlme et al. 2020) | Põlme S, et al. 2020. Fungal Diversity 105:1-16 ((https://doi.org/10.1007/s13225-020-00466-2)) |
| Otu20585 | 1 | k__Fungi(100);p__Ascomycota(100);c__Saccharomycetes(100);o__Saccharomycetales(100);f__Metschnikowiaceae(100);g__Metschnikowia(100);s__Metschnikowia_pulcherrima(100); | Metschnikowia | 13 | Saprotroph | \|Undefined Saprotroph\| | Yeast | NULL | Probable | As nectar-tap_saprotroph (Põlme et al. 2020); Decay sugar-rich substrates (Põlme et al. 2020) | Põlme S, et al. 2020. Fungal Diversity 105:1-16 ((https://doi.org/10.1007/s13225-020-00466-2)) |
| Otu20586 | 1 | k__Fungi(100);p__Ascomycota(100);c__Saccharomycetes(100);o__Saccharomycetales(100);f__Metschnikowiaceae(100);g__Metschnikowia(100);s__Metschnikowia_pulcherrima(100); | Metschnikowia | 13 | Saprotroph | \|Undefined Saprotroph\| | Yeast | NULL | Probable | As nectar-tap_saprotroph (Põlme et al. 2020); Decay sugar-rich substrates (Põlme et al. 2020) | Põlme S, et al. 2020. Fungal Diversity 105:1-16 ((https://doi.org/10.1007/s13225-020-00466-2)) |
| Otu20587 | 1 | k__Fungi(100);p__Ascomycota(100);c__Saccharomycetes(100);o__Saccharomycetales(100);f__Metschnikowiaceae(100);g__Metschnikowia(100);g__Metschnikowia_unclassified(100); | Metschnikowia | 13 | Saprotroph | \|Undefined Saprotroph\| | Yeast | NULL | Probable | As nectar-tap_saprotroph (Põlme et al. 2020); Decay sugar-rich substrates (Põlme et al. 2020) | Põlme S, et al. 2020. Fungal Diversity 105:1-16 ((https://doi.org/10.1007/s13225-020-00466-2)) |
| Otu20589 | 1 | k__Fungi(100);p__Ascomycota(100);c__Saccharomycetes(100);o__Saccharomycetales(100);f__Metschnikowiaceae(100);g__Metschnikowia(100);g__Metschnikowia_unclassified(100); | Metschnikowia | 13 | Saprotroph | \|Undefined Saprotroph\| | Yeast | NULL | Probable | As nectar-tap_saprotroph (Põlme et al. 2020); Decay sugar-rich substrates (Põlme et al. 2020) | Põlme S, et al. 2020. Fungal Diversity 105:1-16 ((https://doi.org/10.1007/s13225-020-00466-2)) |
| Otu20590 | 1 | k__Fungi(100);p__Ascomycota(100);c__Saccharomycetes(100);o__Saccharomycetales(100);f__Metschnikowiaceae(100);g__Metschnikowia(100);g__Metschnikowia_unclassified(100); | Metschnikowia | 13 | Saprotroph | \|Undefined Saprotroph\| | Yeast | NULL | Probable | As nectar-tap_saprotroph (Põlme et al. 2020); Decay sugar-rich substrates (Põlme et al. 2020) | Põlme S, et al. 2020. Fungal Diversity 105:1-16 ((https://doi.org/10.1007/s13225-020-00466-2)) |
| Otu20591 | 1 | k__Fungi(100);p__Ascomycota(100);c__Saccharomycetes(100);o__Saccharomycetales(100);f__Metschnikowiaceae(100);g__Metschnikowia(100);s__Metschnikowia_pulcherrima(100); | Metschnikowia | 13 | Saprotroph | \|Undefined Saprotroph\| | Yeast | NULL | Probable | As nectar-tap_saprotroph (Põlme et al. 2020); Decay sugar-rich substrates (Põlme et al. 2020) | Põlme S, et al. 2020. Fungal Diversity 105:1-16 ((https://doi.org/10.1007/s13225-020-00466-2)) |
| Otu20593 | 1 | k__Fungi(100);p__Ascomycota(100);c__Saccharomycetes(100);o__Saccharomycetales(100);f__Metschnikowiaceae(100);g__Metschnikowia(100);s__Metschnikowia_pulcherrima(100); | Metschnikowia | 13 | Saprotroph | \|Undefined Saprotroph\| | Yeast | NULL | Probable | As nectar-tap_saprotroph (Põlme et al. 2020); Decay sugar-rich substrates (Põlme et al. 2020) | Põlme S, et al. 2020. Fungal Diversity 105:1-16 ((https://doi.org/10.1007/s13225-020-00466-2)) |
| Otu20594 | 1 | k__Fungi(100);p__Ascomycota(100);c__Saccharomycetes(100);o__Saccharomycetales(100);f__Metschnikowiaceae(100);g__Metschnikowia(100);s__Metschnikowia_pulcherrima(100); | Metschnikowia | 13 | Saprotroph | \|Undefined Saprotroph\| | Yeast | NULL | Probable | As nectar-tap_saprotroph (Põlme et al. 2020); Decay sugar-rich substrates (Põlme et al. 2020) | Põlme S, et al. 2020. Fungal Diversity 105:1-16 ((https://doi.org/10.1007/s13225-020-00466-2)) |
| Otu20595 | 1 | k__Fungi(100);p__Ascomycota(100);c__Saccharomycetes(100);o__Saccharomycetales(100);f__Metschnikowiaceae(100);g__Metschnikowia(100);s__Metschnikowia_pulcherrima(100); | Metschnikowia | 13 | Saprotroph | \|Undefined Saprotroph\| | Yeast | NULL | Probable | As nectar-tap_saprotroph (Põlme et al. 2020); Decay sugar-rich substrates (Põlme et al. 2020) | Põlme S, et al. 2020. Fungal Diversity 105:1-16 ((https://doi.org/10.1007/s13225-020-00466-2)) |
| Otu20596 | 1 | k__Fungi(100);p__Ascomycota(100);c__Saccharomycetes(100);o__Saccharomycetales(100);f__Metschnikowiaceae(100);g__Metschnikowia(100);s__Metschnikowia_pulcherrima(100); | Metschnikowia | 13 | Saprotroph | \|Undefined Saprotroph\| | Yeast | NULL | Probable | As nectar-tap_saprotroph (Põlme et al. 2020); Decay sugar-rich substrates (Põlme et al. 2020) | Põlme S, et al. 2020. Fungal Diversity 105:1-16 ((https://doi.org/10.1007/s13225-020-00466-2)) |
| Otu20597 | 1 | k__Fungi(100);p__Ascomycota(100);c__Saccharomycetes(100);o__Saccharomycetales(100);f__Metschnikowiaceae(100);g__Metschnikowia(100);s__Metschnikowia_pulcherrima(100); | Metschnikowia | 13 | Saprotroph | \|Undefined Saprotroph\| | Yeast | NULL | Probable | As nectar-tap_saprotroph (Põlme et al. 2020); Decay sugar-rich substrates (Põlme et al. 2020) | Põlme S, et al. 2020. Fungal Diversity 105:1-16 ((https://doi.org/10.1007/s13225-020-00466-2)) |
| Otu20599 | 1 | k__Fungi(100);p__Ascomycota(100);c__Saccharomycetes(100);o__Saccharomycetales(100);f__Metschnikowiaceae(100);g__Metschnikowia(100);s__Metschnikowia_pulcherrima(100); | Metschnikowia | 13 | Saprotroph | \|Undefined Saprotroph\| | Yeast | NULL | Probable | As nectar-tap_saprotroph (Põlme et al. 2020); Decay sugar-rich substrates (Põlme et al. 2020) | Põlme S, et al. 2020. Fungal Diversity 105:1-16 ((https://doi.org/10.1007/s13225-020-00466-2)) |
| Otu20600 | 1 | k__Fungi(100);p__Ascomycota(100);c__Saccharomycetes(100);o__Saccharomycetales(100);f__Metschnikowiaceae(100);g__Metschnikowia(100);g__Metschnikowia_unclassified(100); | Metschnikowia | 13 | Saprotroph | \|Undefined Saprotroph\| | Yeast | NULL | Probable | As nectar-tap_saprotroph (Põlme et al. 2020); Decay sugar-rich substrates (Põlme et al. 2020) | Põlme S, et al. 2020. Fungal Diversity 105:1-16 ((https://doi.org/10.1007/s13225-020-00466-2)) |
| Otu20602 | 1 | k__Fungi(100);p__Ascomycota(100);c__Saccharomycetes(100);o__Saccharomycetales(100);f__Metschnikowiaceae(100);g__Metschnikowia(100);s__Metschnikowia_pulcherrima(100); | Metschnikowia | 13 | Saprotroph | \|Undefined Saprotroph\| | Yeast | NULL | Probable | As nectar-tap_saprotroph (Põlme et al. 2020); Decay sugar-rich substrates (Põlme et al. 2020) | Põlme S, et al. 2020. Fungal Diversity 105:1-16 ((https://doi.org/10.1007/s13225-020-00466-2)) |
| Otu20603 | 1 | k__Fungi(100);p__Ascomycota(100);c__Saccharomycetes(100);o__Saccharomycetales(100);f__Metschnikowiaceae(100);g__Metschnikowia(100);g__Metschnikowia_unclassified(100); | Metschnikowia | 13 | Saprotroph | \|Undefined Saprotroph\| | Yeast | NULL | Probable | As nectar-tap_saprotroph (Põlme et al. 2020); Decay sugar-rich substrates (Põlme et al. 2020) | Põlme S, et al. 2020. Fungal Diversity 105:1-16 ((https://doi.org/10.1007/s13225-020-00466-2)) |
| Otu20604 | 1 | k__Fungi(100);p__Ascomycota(100);c__Saccharomycetes(100);o__Saccharomycetales(100);f__Metschnikowiaceae(100);g__Metschnikowia(100);s__Metschnikowia_pulcherrima(100); | Metschnikowia | 13 | Saprotroph | \|Undefined Saprotroph\| | Yeast | NULL | Probable | As nectar-tap_saprotroph (Põlme et al. 2020); Decay sugar-rich substrates (Põlme et al. 2020) | Põlme S, et al. 2020. Fungal Diversity 105:1-16 ((https://doi.org/10.1007/s13225-020-00466-2)) |
| Otu20605 | 1 | k__Fungi(100);p__Ascomycota(100);c__Saccharomycetes(100);o__Saccharomycetales(100);f__Metschnikowiaceae(100);g__Metschnikowia(100);s__Metschnikowia_pulcherrima(100); | Metschnikowia | 13 | Saprotroph | \|Undefined Saprotroph\| | Yeast | NULL | Probable | As nectar-tap_saprotroph (Põlme et al. 2020); Decay sugar-rich substrates (Põlme et al. 2020) | Põlme S, et al. 2020. Fungal Diversity 105:1-16 ((https://doi.org/10.1007/s13225-020-00466-2)) |
| Otu20606 | 1 | k__Fungi(100);p__Ascomycota(100);c__Saccharomycetes(100);o__Saccharomycetales(100);f__Metschnikowiaceae(100);g__Metschnikowia(100);s__Metschnikowia_pulcherrima(100); | Metschnikowia | 13 | Saprotroph | \|Undefined Saprotroph\| | Yeast | NULL | Probable | As nectar-tap_saprotroph (Põlme et al. 2020); Decay sugar-rich substrates (Põlme et al. 2020) | Põlme S, et al. 2020. Fungal Diversity 105:1-16 ((https://doi.org/10.1007/s13225-020-00466-2)) |
| Otu20607 | 1 | k__Fungi(100);p__Ascomycota(100);c__Saccharomycetes(100);o__Saccharomycetales(100);f__Metschnikowiaceae(100);g__Metschnikowia(100);g__Metschnikowia_unclassified(100); | Metschnikowia | 13 | Saprotroph | \|Undefined Saprotroph\| | Yeast | NULL | Probable | As nectar-tap_saprotroph (Põlme et al. 2020); Decay sugar-rich substrates (Põlme et al. 2020) | Põlme S, et al. 2020. Fungal Diversity 105:1-16 ((https://doi.org/10.1007/s13225-020-00466-2)) |
| Otu20608 | 1 | k__Fungi(100);p__Ascomycota(100);c__Saccharomycetes(100);o__Saccharomycetales(100);f__Metschnikowiaceae(100);g__Metschnikowia(100);s__Metschnikowia_pulcherrima(100); | Metschnikowia | 13 | Saprotroph | \|Undefined Saprotroph\| | Yeast | NULL | Probable | As nectar-tap_saprotroph (Põlme et al. 2020); Decay sugar-rich substrates (Põlme et al. 2020) | Põlme S, et al. 2020. Fungal Diversity 105:1-16 ((https://doi.org/10.1007/s13225-020-00466-2)) |
| Otu20609 | 1 | k__Fungi(100);p__Ascomycota(100);c__Saccharomycetes(100);o__Saccharomycetales(100);f__Metschnikowiaceae(100);g__Metschnikowia(100);g__Metschnikowia_unclassified(100); | Metschnikowia | 13 | Saprotroph | \|Undefined Saprotroph\| | Yeast | NULL | Probable | As nectar-tap_saprotroph (Põlme et al. 2020); Decay sugar-rich substrates (Põlme et al. 2020) | Põlme S, et al. 2020. Fungal Diversity 105:1-16 ((https://doi.org/10.1007/s13225-020-00466-2)) |
| Otu20611 | 1 | k__Fungi(100);p__Ascomycota(100);c__Saccharomycetes(100);o__Saccharomycetales(100);f__Metschnikowiaceae(100);g__Metschnikowia(100);g__Metschnikowia_unclassified(100); | Metschnikowia | 13 | Saprotroph | \|Undefined Saprotroph\| | Yeast | NULL | Probable | As nectar-tap_saprotroph (Põlme et al. 2020); Decay sugar-rich substrates (Põlme et al. 2020) | Põlme S, et al. 2020. Fungal Diversity 105:1-16 ((https://doi.org/10.1007/s13225-020-00466-2)) |
| Otu20612 | 1 | k__Fungi(100);p__Ascomycota(100);c__Saccharomycetes(100);o__Saccharomycetales(100);f__Metschnikowiaceae(100);g__Metschnikowia(100);s__Metschnikowia_pulcherrima(100); | Metschnikowia | 13 | Saprotroph | \|Undefined Saprotroph\| | Yeast | NULL | Probable | As nectar-tap_saprotroph (Põlme et al. 2020); Decay sugar-rich substrates (Põlme et al. 2020) | Põlme S, et al. 2020. Fungal Diversity 105:1-16 ((https://doi.org/10.1007/s13225-020-00466-2)) |
| Otu20613 | 1 | k__Fungi(100);p__Ascomycota(100);c__Saccharomycetes(100);o__Saccharomycetales(100);f__Metschnikowiaceae(100);g__Metschnikowia(100);s__Metschnikowia_pulcherrima(100); | Metschnikowia | 13 | Saprotroph | \|Undefined Saprotroph\| | Yeast | NULL | Probable | As nectar-tap_saprotroph (Põlme et al. 2020); Decay sugar-rich substrates (Põlme et al. 2020) | Põlme S, et al. 2020. Fungal Diversity 105:1-16 ((https://doi.org/10.1007/s13225-020-00466-2)) |
| Otu20614 | 1 | k__Fungi(100);p__Ascomycota(100);c__Saccharomycetes(100);o__Saccharomycetales(100);f__Metschnikowiaceae(100);g__Metschnikowia(100);g__Metschnikowia_unclassified(100); | Metschnikowia | 13 | Saprotroph | \|Undefined Saprotroph\| | Yeast | NULL | Probable | As nectar-tap_saprotroph (Põlme et al. 2020); Decay sugar-rich substrates (Põlme et al. 2020) | Põlme S, et al. 2020. Fungal Diversity 105:1-16 ((https://doi.org/10.1007/s13225-020-00466-2)) |
| Otu20615 | 1 | k__Fungi(100);p__Ascomycota(100);c__Saccharomycetes(100);o__Saccharomycetales(100);f__Metschnikowiaceae(100);g__Metschnikowia(100);s__Metschnikowia_pulcherrima(100); | Metschnikowia | 13 | Saprotroph | \|Undefined Saprotroph\| | Yeast | NULL | Probable | As nectar-tap_saprotroph (Põlme et al. 2020); Decay sugar-rich substrates (Põlme et al. 2020) | Põlme S, et al. 2020. Fungal Diversity 105:1-16 ((https://doi.org/10.1007/s13225-020-00466-2)) |
| Otu20616 | 1 | k__Fungi(100);p__Ascomycota(100);c__Saccharomycetes(100);o__Saccharomycetales(100);f__Metschnikowiaceae(100);g__Metschnikowia(100);s__Metschnikowia_pulcherrima(100); | Metschnikowia | 13 | Saprotroph | \|Undefined Saprotroph\| | Yeast | NULL | Probable | As nectar-tap_saprotroph (Põlme et al. 2020); Decay sugar-rich substrates (Põlme et al. 2020) | Põlme S, et al. 2020. Fungal Diversity 105:1-16 ((https://doi.org/10.1007/s13225-020-00466-2)) |
| Otu20617 | 1 | k__Fungi(100);p__Ascomycota(100);c__Saccharomycetes(100);o__Saccharomycetales(100);f__Metschnikowiaceae(100);g__Metschnikowia(100);s__Metschnikowia_pulcherrima(100); | Metschnikowia | 13 | Saprotroph | \|Undefined Saprotroph\| | Yeast | NULL | Probable | As nectar-tap_saprotroph (Põlme et al. 2020); Decay sugar-rich substrates (Põlme et al. 2020) | Põlme S, et al. 2020. Fungal Diversity 105:1-16 ((https://doi.org/10.1007/s13225-020-00466-2)) |
| Otu20618 | 1 | k__Fungi(100);p__Ascomycota(100);c__Saccharomycetes(100);o__Saccharomycetales(100);f__Metschnikowiaceae(100);g__Metschnikowia(100);s__Metschnikowia_pulcherrima(100); | Metschnikowia | 13 | Saprotroph | \|Undefined Saprotroph\| | Yeast | NULL | Probable | As nectar-tap_saprotroph (Põlme et al. 2020); Decay sugar-rich substrates (Põlme et al. 2020) | Põlme S, et al. 2020. Fungal Diversity 105:1-16 ((https://doi.org/10.1007/s13225-020-00466-2)) |
| Otu20619 | 1 | k__Fungi(100);p__Ascomycota(100);c__Saccharomycetes(100);o__Saccharomycetales(100);f__Metschnikowiaceae(100);g__Metschnikowia(100);s__Metschnikowia_pulcherrima(100); | Metschnikowia | 13 | Saprotroph | \|Undefined Saprotroph\| | Yeast | NULL | Probable | As nectar-tap_saprotroph (Põlme et al. 2020); Decay sugar-rich substrates (Põlme et al. 2020) | Põlme S, et al. 2020. Fungal Diversity 105:1-16 ((https://doi.org/10.1007/s13225-020-00466-2)) |
| Otu20620 | 1 | k__Fungi(100);p__Ascomycota(100);c__Saccharomycetes(100);o__Saccharomycetales(100);f__Metschnikowiaceae(100);g__Metschnikowia(100);s__Metschnikowia_pulcherrima(100); | Metschnikowia | 13 | Saprotroph | \|Undefined Saprotroph\| | Yeast | NULL | Probable | As nectar-tap_saprotroph (Põlme et al. 2020); Decay sugar-rich substrates (Põlme et al. 2020) | Põlme S, et al. 2020. Fungal Diversity 105:1-16 ((https://doi.org/10.1007/s13225-020-00466-2)) |
| Otu20621 | 1 | k__Fungi(100);p__Ascomycota(100);c__Saccharomycetes(100);o__Saccharomycetales(100);f__Metschnikowiaceae(100);g__Metschnikowia(100);s__Metschnikowia_pulcherrima(100); | Metschnikowia | 13 | Saprotroph | \|Undefined Saprotroph\| | Yeast | NULL | Probable | As nectar-tap_saprotroph (Põlme et al. 2020); Decay sugar-rich substrates (Põlme et al. 2020) | Põlme S, et al. 2020. Fungal Diversity 105:1-16 ((https://doi.org/10.1007/s13225-020-00466-2)) |
| Otu20622 | 1 | k__Fungi(100);p__Ascomycota(100);c__Saccharomycetes(100);o__Saccharomycetales(100);f__Metschnikowiaceae(100);g__Metschnikowia(100);g__Metschnikowia_unclassified(100); | Metschnikowia | 13 | Saprotroph | \|Undefined Saprotroph\| | Yeast | NULL | Probable | As nectar-tap_saprotroph (Põlme et al. 2020); Decay sugar-rich substrates (Põlme et al. 2020) | Põlme S, et al. 2020. Fungal Diversity 105:1-16 ((https://doi.org/10.1007/s13225-020-00466-2)) |
| Otu20623 | 1 | k__Fungi(100);p__Ascomycota(100);c__Saccharomycetes(100);o__Saccharomycetales(100);f__Metschnikowiaceae(100);g__Metschnikowia(100);s__Metschnikowia_pulcherrima(100); | Metschnikowia | 13 | Saprotroph | \|Undefined Saprotroph\| | Yeast | NULL | Probable | As nectar-tap_saprotroph (Põlme et al. 2020); Decay sugar-rich substrates (Põlme et al. 2020) | Põlme S, et al. 2020. Fungal Diversity 105:1-16 ((https://doi.org/10.1007/s13225-020-00466-2)) |
| Otu20624 | 1 | k__Fungi(100);p__Ascomycota(100);c__Saccharomycetes(100);o__Saccharomycetales(100);f__Metschnikowiaceae(100);g__Metschnikowia(100);g__Metschnikowia_unclassified(100); | Metschnikowia | 13 | Saprotroph | \|Undefined Saprotroph\| | Yeast | NULL | Probable | As nectar-tap_saprotroph (Põlme et al. 2020); Decay sugar-rich substrates (Põlme et al. 2020) | Põlme S, et al. 2020. Fungal Diversity 105:1-16 ((https://doi.org/10.1007/s13225-020-00466-2)) |
| Otu20625 | 1 | k__Fungi(100);p__Ascomycota(100);c__Saccharomycetes(100);o__Saccharomycetales(100);f__Metschnikowiaceae(100);g__Metschnikowia(100);s__Metschnikowia_pulcherrima(100); | Metschnikowia | 13 | Saprotroph | \|Undefined Saprotroph\| | Yeast | NULL | Probable | As nectar-tap_saprotroph (Põlme et al. 2020); Decay sugar-rich substrates (Põlme et al. 2020) | Põlme S, et al. 2020. Fungal Diversity 105:1-16 ((https://doi.org/10.1007/s13225-020-00466-2)) |
| Otu20627 | 1 | k__Fungi(100);p__Ascomycota(100);c__Saccharomycetes(100);o__Saccharomycetales(100);f__Metschnikowiaceae(100);g__Metschnikowia(100);s__Metschnikowia_pulcherrima(100); | Metschnikowia | 13 | Saprotroph | \|Undefined Saprotroph\| | Yeast | NULL | Probable | As nectar-tap_saprotroph (Põlme et al. 2020); Decay sugar-rich substrates (Põlme et al. 2020) | Põlme S, et al. 2020. Fungal Diversity 105:1-16 ((https://doi.org/10.1007/s13225-020-00466-2)) |
| Otu20629 | 1 | k__Fungi(100);p__Ascomycota(100);c__Saccharomycetes(100);o__Saccharomycetales(100);f__Metschnikowiaceae(100);g__Metschnikowia(100);s__Metschnikowia_pulcherrima(100); | Metschnikowia | 13 | Saprotroph | \|Undefined Saprotroph\| | Yeast | NULL | Probable | As nectar-tap_saprotroph (Põlme et al. 2020); Decay sugar-rich substrates (Põlme et al. 2020) | Põlme S, et al. 2020. Fungal Diversity 105:1-16 ((https://doi.org/10.1007/s13225-020-00466-2)) |
| Otu20630 | 1 | k__Fungi(100);p__Ascomycota(100);c__Saccharomycetes(100);o__Saccharomycetales(100);f__Metschnikowiaceae(100);g__Metschnikowia(100);g__Metschnikowia_unclassified(100); | Metschnikowia | 13 | Saprotroph | \|Undefined Saprotroph\| | Yeast | NULL | Probable | As nectar-tap_saprotroph (Põlme et al. 2020); Decay sugar-rich substrates (Põlme et al. 2020) | Põlme S, et al. 2020. Fungal Diversity 105:1-16 ((https://doi.org/10.1007/s13225-020-00466-2)) |
| Otu20631 | 1 | k__Fungi(100);p__Ascomycota(100);c__Saccharomycetes(100);o__Saccharomycetales(100);f__Metschnikowiaceae(100);g__Metschnikowia(100);s__Metschnikowia_pulcherrima(100); | Metschnikowia | 13 | Saprotroph | \|Undefined Saprotroph\| | Yeast | NULL | Probable | As nectar-tap_saprotroph (Põlme et al. 2020); Decay sugar-rich substrates (Põlme et al. 2020) | Põlme S, et al. 2020. Fungal Diversity 105:1-16 ((https://doi.org/10.1007/s13225-020-00466-2)) |
| Otu20632 | 1 | k__Fungi(100);p__Ascomycota(100);c__Saccharomycetes(100);o__Saccharomycetales(100);f__Metschnikowiaceae(100);g__Metschnikowia(100);s__Metschnikowia_pulcherrima(100); | Metschnikowia | 13 | Saprotroph | \|Undefined Saprotroph\| | Yeast | NULL | Probable | As nectar-tap_saprotroph (Põlme et al. 2020); Decay sugar-rich substrates (Põlme et al. 2020) | Põlme S, et al. 2020. Fungal Diversity 105:1-16 ((https://doi.org/10.1007/s13225-020-00466-2)) |
| Otu20633 | 1 | k__Fungi(100);p__Ascomycota(100);c__Saccharomycetes(100);o__Saccharomycetales(100);f__Metschnikowiaceae(100);g__Metschnikowia(100);s__Metschnikowia_pulcherrima(100); | Metschnikowia | 13 | Saprotroph | \|Undefined Saprotroph\| | Yeast | NULL | Probable | As nectar-tap_saprotroph (Põlme et al. 2020); Decay sugar-rich substrates (Põlme et al. 2020) | Põlme S, et al. 2020. Fungal Diversity 105:1-16 ((https://doi.org/10.1007/s13225-020-00466-2)) |
| Otu20634 | 1 | k__Fungi(100);p__Ascomycota(100);c__Saccharomycetes(100);o__Saccharomycetales(100);f__Metschnikowiaceae(100);g__Metschnikowia(100);s__Metschnikowia_pulcherrima(100); | Metschnikowia | 13 | Saprotroph | \|Undefined Saprotroph\| | Yeast | NULL | Probable | As nectar-tap_saprotroph (Põlme et al. 2020); Decay sugar-rich substrates (Põlme et al. 2020) | Põlme S, et al. 2020. Fungal Diversity 105:1-16 ((https://doi.org/10.1007/s13225-020-00466-2)) |
| Otu20635 | 1 | k__Fungi(100);p__Ascomycota(100);c__Saccharomycetes(100);o__Saccharomycetales(100);f__Metschnikowiaceae(100);g__Metschnikowia(100);s__Metschnikowia_pulcherrima(100); | Metschnikowia | 13 | Saprotroph | \|Undefined Saprotroph\| | Yeast | NULL | Probable | As nectar-tap_saprotroph (Põlme et al. 2020); Decay sugar-rich substrates (Põlme et al. 2020) | Põlme S, et al. 2020. Fungal Diversity 105:1-16 ((https://doi.org/10.1007/s13225-020-00466-2)) |
| Otu20636 | 1 | k__Fungi(100);p__Ascomycota(100);c__Saccharomycetes(100);o__Saccharomycetales(100);f__Metschnikowiaceae(100);g__Metschnikowia(100);s__Metschnikowia_pulcherrima(100); | Metschnikowia | 13 | Saprotroph | \|Undefined Saprotroph\| | Yeast | NULL | Probable | As nectar-tap_saprotroph (Põlme et al. 2020); Decay sugar-rich substrates (Põlme et al. 2020) | Põlme S, et al. 2020. Fungal Diversity 105:1-16 ((https://doi.org/10.1007/s13225-020-00466-2)) |
| Otu20637 | 1 | k__Fungi(100);p__Ascomycota(100);c__Saccharomycetes(100);o__Saccharomycetales(100);f__Metschnikowiaceae(100);g__Metschnikowia(100);s__Metschnikowia_pulcherrima(100); | Metschnikowia | 13 | Saprotroph | \|Undefined Saprotroph\| | Yeast | NULL | Probable | As nectar-tap_saprotroph (Põlme et al. 2020); Decay sugar-rich substrates (Põlme et al. 2020) | Põlme S, et al. 2020. Fungal Diversity 105:1-16 ((https://doi.org/10.1007/s13225-020-00466-2)) |
| Otu20638 | 1 | k__Fungi(100);p__Ascomycota(100);c__Saccharomycetes(100);o__Saccharomycetales(100);f__Metschnikowiaceae(100);g__Metschnikowia(100);s__Metschnikowia_pulcherrima(100); | Metschnikowia | 13 | Saprotroph | \|Undefined Saprotroph\| | Yeast | NULL | Probable | As nectar-tap_saprotroph (Põlme et al. 2020); Decay sugar-rich substrates (Põlme et al. 2020) | Põlme S, et al. 2020. Fungal Diversity 105:1-16 ((https://doi.org/10.1007/s13225-020-00466-2)) |
| Otu20641 | 1 | k__Fungi(100);p__Ascomycota(100);c__Saccharomycetes(100);o__Saccharomycetales(100);f__Metschnikowiaceae(100);g__Metschnikowia(100);s__Metschnikowia_pulcherrima(100); | Metschnikowia | 13 | Saprotroph | \|Undefined Saprotroph\| | Yeast | NULL | Probable | As nectar-tap_saprotroph (Põlme et al. 2020); Decay sugar-rich substrates (Põlme et al. 2020) | Põlme S, et al. 2020. Fungal Diversity 105:1-16 ((https://doi.org/10.1007/s13225-020-00466-2)) |
| Otu20642 | 1 | k__Fungi(100);p__Ascomycota(100);c__Saccharomycetes(100);o__Saccharomycetales(100);f__Metschnikowiaceae(100);g__Metschnikowia(100);s__Metschnikowia_pulcherrima(100); | Metschnikowia | 13 | Saprotroph | \|Undefined Saprotroph\| | Yeast | NULL | Probable | As nectar-tap_saprotroph (Põlme et al. 2020); Decay sugar-rich substrates (Põlme et al. 2020) | Põlme S, et al. 2020. Fungal Diversity 105:1-16 ((https://doi.org/10.1007/s13225-020-00466-2)) |
| Otu20645 | 1 | k__Fungi(100);p__Ascomycota(100);c__Saccharomycetes(100);o__Saccharomycetales(100);f__Saccharomycetales_family_Incertae_sedis(100);g__Candida(100);g__Candida_unclassified(100); | Candida | 13 | Pathotroph-Saprotroph-Symbiotroph | Animal Pathogen-Endophyte-Endosymbiont-Epiphyte-\|Undefined Saprotroph\| | Dimorphic Yeast-Yeast | NULL | Probable | As nectar-tap_saprotroph (Põlme et al. 2020); Decay sugar-rich substrates (Põlme et al. 2020); This asexual genus is polyphyletic and contain species that function very differently depend on what they are. Caution should be taken when interpreting guilds for Candida species. | Manolakaki D, et al. 2010. Virulence. 1: 367-375 ((https://doi.org/10.4161/viru.1.5.12796)); Mekbib SB, et al. 2011. Tropical Plant Pathology 36:233-240 ((https://doi.org/10.1590/S1982-56762011000400004)); Põlme S, et al. 2020. Fungal Diversity 105:1-16 ((https://doi.org/10.1007/s13225-020-00466-2)) |
| Otu20650 | 1 | k__Fungi(100);p__Ascomycota(100);c__Saccharomycetes(100);o__Saccharomycetales(100);f__Saccharomycetales_family_Incertae_sedis(100);g__Candida(100);g__Candida_unclassified(100); | Candida | 13 | Pathotroph-Saprotroph-Symbiotroph | Animal Pathogen-Endophyte-Endosymbiont-Epiphyte-\|Undefined Saprotroph\| | Dimorphic Yeast-Yeast | NULL | Probable | As nectar-tap_saprotroph (Põlme et al. 2020); Decay sugar-rich substrates (Põlme et al. 2020); This asexual genus is polyphyletic and contain species that function very differently depend on what they are. Caution should be taken when interpreting guilds for Candida species. | Manolakaki D, et al. 2010. Virulence. 1: 367-375 ((https://doi.org/10.4161/viru.1.5.12796)); Mekbib SB, et al. 2011. Tropical Plant Pathology 36:233-240 ((https://doi.org/10.1590/S1982-56762011000400004)); Põlme S, et al. 2020. Fungal Diversity 105:1-16 ((https://doi.org/10.1007/s13225-020-00466-2)) |
| Otu20653 | 1 | k__Fungi(100);p__Ascomycota(100);c__Saccharomycetes(100);o__Saccharomycetales(100);f__Metschnikowiaceae(100);g__Metschnikowia(100);g__Metschnikowia_unclassified(100); | Metschnikowia | 13 | Saprotroph | \|Undefined Saprotroph\| | Yeast | NULL | Probable | As nectar-tap_saprotroph (Põlme et al. 2020); Decay sugar-rich substrates (Põlme et al. 2020) | Põlme S, et al. 2020. Fungal Diversity 105:1-16 ((https://doi.org/10.1007/s13225-020-00466-2)) |
| Otu20654 | 1 | k__Fungi(100);p__Ascomycota(100);c__Saccharomycetes(100);o__Saccharomycetales(100);f__Metschnikowiaceae(100);g__Metschnikowia(100);s__Metschnikowia_pulcherrima(100); | Metschnikowia | 13 | Saprotroph | \|Undefined Saprotroph\| | Yeast | NULL | Probable | As nectar-tap_saprotroph (Põlme et al. 2020); Decay sugar-rich substrates (Põlme et al. 2020) | Põlme S, et al. 2020. Fungal Diversity 105:1-16 ((https://doi.org/10.1007/s13225-020-00466-2)) |
| Otu20655 | 1 | k__Fungi(100);p__Ascomycota(100);c__Saccharomycetes(100);o__Saccharomycetales(100);f__Metschnikowiaceae(100);g__Metschnikowia(100);s__Metschnikowia_pulcherrima(100); | Metschnikowia | 13 | Saprotroph | \|Undefined Saprotroph\| | Yeast | NULL | Probable | As nectar-tap_saprotroph (Põlme et al. 2020); Decay sugar-rich substrates (Põlme et al. 2020) | Põlme S, et al. 2020. Fungal Diversity 105:1-16 ((https://doi.org/10.1007/s13225-020-00466-2)) |
| Otu20656 | 1 | k__Fungi(100);p__Ascomycota(100);c__Saccharomycetes(100);o__Saccharomycetales(100);f__Metschnikowiaceae(100);g__Metschnikowia(100);s__Metschnikowia_pulcherrima(100); | Metschnikowia | 13 | Saprotroph | \|Undefined Saprotroph\| | Yeast | NULL | Probable | As nectar-tap_saprotroph (Põlme et al. 2020); Decay sugar-rich substrates (Põlme et al. 2020) | Põlme S, et al. 2020. Fungal Diversity 105:1-16 ((https://doi.org/10.1007/s13225-020-00466-2)) |
| Otu20659 | 1 | k__Fungi(100);p__Ascomycota(100);c__Saccharomycetes(100);o__Saccharomycetales(100);f__Metschnikowiaceae(100);g__Metschnikowia(100);g__Metschnikowia_unclassified(100); | Metschnikowia | 13 | Saprotroph | \|Undefined Saprotroph\| | Yeast | NULL | Probable | As nectar-tap_saprotroph (Põlme et al. 2020); Decay sugar-rich substrates (Põlme et al. 2020) | Põlme S, et al. 2020. Fungal Diversity 105:1-16 ((https://doi.org/10.1007/s13225-020-00466-2)) |
| Otu20660 | 1 | k__Fungi(100);p__Ascomycota(100);c__Saccharomycetes(100);o__Saccharomycetales(100);f__Metschnikowiaceae(100);g__Metschnikowia(100);s__Metschnikowia_pulcherrima(100); | Metschnikowia | 13 | Saprotroph | \|Undefined Saprotroph\| | Yeast | NULL | Probable | As nectar-tap_saprotroph (Põlme et al. 2020); Decay sugar-rich substrates (Põlme et al. 2020) | Põlme S, et al. 2020. Fungal Diversity 105:1-16 ((https://doi.org/10.1007/s13225-020-00466-2)) |
| Otu20661 | 1 | k__Fungi(100);p__Ascomycota(100);c__Saccharomycetes(100);o__Saccharomycetales(100);f__Metschnikowiaceae(100);g__Metschnikowia(100);g__Metschnikowia_unclassified(100); | Metschnikowia | 13 | Saprotroph | \|Undefined Saprotroph\| | Yeast | NULL | Probable | As nectar-tap_saprotroph (Põlme et al. 2020); Decay sugar-rich substrates (Põlme et al. 2020) | Põlme S, et al. 2020. Fungal Diversity 105:1-16 ((https://doi.org/10.1007/s13225-020-00466-2)) |
| Otu20662 | 1 | k__Fungi(100);p__Ascomycota(100);c__Saccharomycetes(100);o__Saccharomycetales(100);f__Metschnikowiaceae(100);g__Metschnikowia(100);s__Metschnikowia_pulcherrima(100); | Metschnikowia | 13 | Saprotroph | \|Undefined Saprotroph\| | Yeast | NULL | Probable | As nectar-tap_saprotroph (Põlme et al. 2020); Decay sugar-rich substrates (Põlme et al. 2020) | Põlme S, et al. 2020. Fungal Diversity 105:1-16 ((https://doi.org/10.1007/s13225-020-00466-2)) |
| Otu20663 | 1 | k__Fungi(100);p__Ascomycota(100);c__Saccharomycetes(100);o__Saccharomycetales(100);f__Metschnikowiaceae(100);g__Metschnikowia(100);s__Metschnikowia_pulcherrima(100); | Metschnikowia | 13 | Saprotroph | \|Undefined Saprotroph\| | Yeast | NULL | Probable | As nectar-tap_saprotroph (Põlme et al. 2020); Decay sugar-rich substrates (Põlme et al. 2020) | Põlme S, et al. 2020. Fungal Diversity 105:1-16 ((https://doi.org/10.1007/s13225-020-00466-2)) |
| Otu20664 | 1 | k__Fungi(100);p__Ascomycota(100);c__Saccharomycetes(100);o__Saccharomycetales(100);f__Metschnikowiaceae(100);g__Metschnikowia(100);s__Metschnikowia_pulcherrima(100); | Metschnikowia | 13 | Saprotroph | \|Undefined Saprotroph\| | Yeast | NULL | Probable | As nectar-tap_saprotroph (Põlme et al. 2020); Decay sugar-rich substrates (Põlme et al. 2020) | Põlme S, et al. 2020. Fungal Diversity 105:1-16 ((https://doi.org/10.1007/s13225-020-00466-2)) |
| Otu20665 | 1 | k__Fungi(100);p__Ascomycota(100);c__Saccharomycetes(100);o__Saccharomycetales(100);f__Metschnikowiaceae(100);g__Metschnikowia(100);s__Metschnikowia_pulcherrima(100); | Metschnikowia | 13 | Saprotroph | \|Undefined Saprotroph\| | Yeast | NULL | Probable | As nectar-tap_saprotroph (Põlme et al. 2020); Decay sugar-rich substrates (Põlme et al. 2020) | Põlme S, et al. 2020. Fungal Diversity 105:1-16 ((https://doi.org/10.1007/s13225-020-00466-2)) |
| Otu20666 | 1 | k__Fungi(100);p__Ascomycota(100);c__Saccharomycetes(100);o__Saccharomycetales(100);f__Metschnikowiaceae(100);g__Metschnikowia(100);s__Metschnikowia_pulcherrima(100); | Metschnikowia | 13 | Saprotroph | \|Undefined Saprotroph\| | Yeast | NULL | Probable | As nectar-tap_saprotroph (Põlme et al. 2020); Decay sugar-rich substrates (Põlme et al. 2020) | Põlme S, et al. 2020. Fungal Diversity 105:1-16 ((https://doi.org/10.1007/s13225-020-00466-2)) |
| Otu20667 | 1 | k__Fungi(100);p__Ascomycota(100);c__Saccharomycetes(100);o__Saccharomycetales(100);f__Metschnikowiaceae(100);g__Metschnikowia(100);s__Metschnikowia_pulcherrima(100); | Metschnikowia | 13 | Saprotroph | \|Undefined Saprotroph\| | Yeast | NULL | Probable | As nectar-tap_saprotroph (Põlme et al. 2020); Decay sugar-rich substrates (Põlme et al. 2020) | Põlme S, et al. 2020. Fungal Diversity 105:1-16 ((https://doi.org/10.1007/s13225-020-00466-2)) |
| Otu20668 | 1 | k__Fungi(100);p__Ascomycota(100);c__Saccharomycetes(100);o__Saccharomycetales(100);f__Metschnikowiaceae(100);g__Metschnikowia(100);g__Metschnikowia_unclassified(100); | Metschnikowia | 13 | Saprotroph | \|Undefined Saprotroph\| | Yeast | NULL | Probable | As nectar-tap_saprotroph (Põlme et al. 2020); Decay sugar-rich substrates (Põlme et al. 2020) | Põlme S, et al. 2020. Fungal Diversity 105:1-16 ((https://doi.org/10.1007/s13225-020-00466-2)) |
| Otu20670 | 1 | k__Fungi(100);p__Ascomycota(100);c__Saccharomycetes(100);o__Saccharomycetales(100);f__Metschnikowiaceae(100);g__Metschnikowia(100);s__Metschnikowia_pulcherrima(100); | Metschnikowia | 13 | Saprotroph | \|Undefined Saprotroph\| | Yeast | NULL | Probable | As nectar-tap_saprotroph (Põlme et al. 2020); Decay sugar-rich substrates (Põlme et al. 2020) | Põlme S, et al. 2020. Fungal Diversity 105:1-16 ((https://doi.org/10.1007/s13225-020-00466-2)) |
| Otu20671 | 1 | k__Fungi(100);p__Ascomycota(100);c__Saccharomycetes(100);o__Saccharomycetales(100);f__Metschnikowiaceae(100);g__Metschnikowia(100);s__Metschnikowia_pulcherrima(100); | Metschnikowia | 13 | Saprotroph | \|Undefined Saprotroph\| | Yeast | NULL | Probable | As nectar-tap_saprotroph (Põlme et al. 2020); Decay sugar-rich substrates (Põlme et al. 2020) | Põlme S, et al. 2020. Fungal Diversity 105:1-16 ((https://doi.org/10.1007/s13225-020-00466-2)) |
| Otu20672 | 1 | k__Fungi(100);p__Ascomycota(100);c__Saccharomycetes(100);o__Saccharomycetales(100);f__Metschnikowiaceae(100);g__Metschnikowia(100);g__Metschnikowia_unclassified(100); | Metschnikowia | 13 | Saprotroph | \|Undefined Saprotroph\| | Yeast | NULL | Probable | As nectar-tap_saprotroph (Põlme et al. 2020); Decay sugar-rich substrates (Põlme et al. 2020) | Põlme S, et al. 2020. Fungal Diversity 105:1-16 ((https://doi.org/10.1007/s13225-020-00466-2)) |
| Otu20673 | 1 | k__Fungi(100);p__Ascomycota(100);c__Saccharomycetes(100);o__Saccharomycetales(100);f__Metschnikowiaceae(100);g__Metschnikowia(100);g__Metschnikowia_unclassified(100); | Metschnikowia | 13 | Saprotroph | \|Undefined Saprotroph\| | Yeast | NULL | Probable | As nectar-tap_saprotroph (Põlme et al. 2020); Decay sugar-rich substrates (Põlme et al. 2020) | Põlme S, et al. 2020. Fungal Diversity 105:1-16 ((https://doi.org/10.1007/s13225-020-00466-2)) |
| Otu20674 | 1 | k__Fungi(100);p__Ascomycota(100);c__Saccharomycetes(100);o__Saccharomycetales(100);f__Metschnikowiaceae(100);g__Metschnikowia(100);g__Metschnikowia_unclassified(100); | Metschnikowia | 13 | Saprotroph | \|Undefined Saprotroph\| | Yeast | NULL | Probable | As nectar-tap_saprotroph (Põlme et al. 2020); Decay sugar-rich substrates (Põlme et al. 2020) | Põlme S, et al. 2020. Fungal Diversity 105:1-16 ((https://doi.org/10.1007/s13225-020-00466-2)) |
| Otu20676 | 1 | k__Fungi(100);p__Ascomycota(100);c__Saccharomycetes(100);o__Saccharomycetales(100);f__Metschnikowiaceae(100);g__Metschnikowia(100);s__Metschnikowia_pulcherrima(100); | Metschnikowia | 13 | Saprotroph | \|Undefined Saprotroph\| | Yeast | NULL | Probable | As nectar-tap_saprotroph (Põlme et al. 2020); Decay sugar-rich substrates (Põlme et al. 2020) | Põlme S, et al. 2020. Fungal Diversity 105:1-16 ((https://doi.org/10.1007/s13225-020-00466-2)) |
| Otu20677 | 1 | k__Fungi(100);p__Ascomycota(100);c__Saccharomycetes(100);o__Saccharomycetales(100);f__Metschnikowiaceae(100);g__Metschnikowia(100);s__Metschnikowia_pulcherrima(100); | Metschnikowia | 13 | Saprotroph | \|Undefined Saprotroph\| | Yeast | NULL | Probable | As nectar-tap_saprotroph (Põlme et al. 2020); Decay sugar-rich substrates (Põlme et al. 2020) | Põlme S, et al. 2020. Fungal Diversity 105:1-16 ((https://doi.org/10.1007/s13225-020-00466-2)) |
| Otu20678 | 1 | k__Fungi(100);p__Ascomycota(100);c__Saccharomycetes(100);o__Saccharomycetales(100);f__Metschnikowiaceae(100);g__Metschnikowia(100);g__Metschnikowia_unclassified(100); | Metschnikowia | 13 | Saprotroph | \|Undefined Saprotroph\| | Yeast | NULL | Probable | As nectar-tap_saprotroph (Põlme et al. 2020); Decay sugar-rich substrates (Põlme et al. 2020) | Põlme S, et al. 2020. Fungal Diversity 105:1-16 ((https://doi.org/10.1007/s13225-020-00466-2)) |
| Otu20679 | 1 | k__Fungi(100);p__Ascomycota(100);c__Saccharomycetes(100);o__Saccharomycetales(100);f__Metschnikowiaceae(100);g__Metschnikowia(100);s__Metschnikowia_pulcherrima(100); | Metschnikowia | 13 | Saprotroph | \|Undefined Saprotroph\| | Yeast | NULL | Probable | As nectar-tap_saprotroph (Põlme et al. 2020); Decay sugar-rich substrates (Põlme et al. 2020) | Põlme S, et al. 2020. Fungal Diversity 105:1-16 ((https://doi.org/10.1007/s13225-020-00466-2)) |
| Otu20682 | 1 | k__Fungi(100);p__Ascomycota(100);c__Saccharomycetes(100);o__Saccharomycetales(100);f__Metschnikowiaceae(100);g__Metschnikowia(100);s__Metschnikowia_pulcherrima(100); | Metschnikowia | 13 | Saprotroph | \|Undefined Saprotroph\| | Yeast | NULL | Probable | As nectar-tap_saprotroph (Põlme et al. 2020); Decay sugar-rich substrates (Põlme et al. 2020) | Põlme S, et al. 2020. Fungal Diversity 105:1-16 ((https://doi.org/10.1007/s13225-020-00466-2)) |
| Otu20683 | 1 | k__Fungi(100);p__Ascomycota(100);c__Saccharomycetes(100);o__Saccharomycetales(100);f__Metschnikowiaceae(100);g__Metschnikowia(100);s__Metschnikowia_pulcherrima(100); | Metschnikowia | 13 | Saprotroph | \|Undefined Saprotroph\| | Yeast | NULL | Probable | As nectar-tap_saprotroph (Põlme et al. 2020); Decay sugar-rich substrates (Põlme et al. 2020) | Põlme S, et al. 2020. Fungal Diversity 105:1-16 ((https://doi.org/10.1007/s13225-020-00466-2)) |
| Otu20684 | 1 | k__Fungi(100);p__Ascomycota(100);c__Saccharomycetes(100);o__Saccharomycetales(100);f__Metschnikowiaceae(100);g__Metschnikowia(100);g__Metschnikowia_unclassified(100); | Metschnikowia | 13 | Saprotroph | \|Undefined Saprotroph\| | Yeast | NULL | Probable | As nectar-tap_saprotroph (Põlme et al. 2020); Decay sugar-rich substrates (Põlme et al. 2020) | Põlme S, et al. 2020. Fungal Diversity 105:1-16 ((https://doi.org/10.1007/s13225-020-00466-2)) |
| Otu20687 | 1 | k__Fungi(100);p__Ascomycota(100);c__Saccharomycetes(100);o__Saccharomycetales(100);f__Metschnikowiaceae(100);g__Metschnikowia(100);g__Metschnikowia_unclassified(100); | Metschnikowia | 13 | Saprotroph | \|Undefined Saprotroph\| | Yeast | NULL | Probable | As nectar-tap_saprotroph (Põlme et al. 2020); Decay sugar-rich substrates (Põlme et al. 2020) | Põlme S, et al. 2020. Fungal Diversity 105:1-16 ((https://doi.org/10.1007/s13225-020-00466-2)) |
| Otu20688 | 1 | k__Fungi(100);p__Ascomycota(100);c__Saccharomycetes(100);o__Saccharomycetales(100);f__Metschnikowiaceae(100);g__Metschnikowia(100);s__Metschnikowia_pulcherrima(100); | Metschnikowia | 13 | Saprotroph | \|Undefined Saprotroph\| | Yeast | NULL | Probable | As nectar-tap_saprotroph (Põlme et al. 2020); Decay sugar-rich substrates (Põlme et al. 2020) | Põlme S, et al. 2020. Fungal Diversity 105:1-16 ((https://doi.org/10.1007/s13225-020-00466-2)) |
| Otu20689 | 1 | k__Fungi(100);p__Ascomycota(100);c__Saccharomycetes(100);o__Saccharomycetales(100);f__Metschnikowiaceae(100);g__Metschnikowia(100);g__Metschnikowia_unclassified(100); | Metschnikowia | 13 | Saprotroph | \|Undefined Saprotroph\| | Yeast | NULL | Probable | As nectar-tap_saprotroph (Põlme et al. 2020); Decay sugar-rich substrates (Põlme et al. 2020) | Põlme S, et al. 2020. Fungal Diversity 105:1-16 ((https://doi.org/10.1007/s13225-020-00466-2)) |

| | | | | | | | | | | | |
|---|---|---|---|---|---|---|---|---|---|---|---|
| Otu20690 | 1 | k__Fungi(100);p__Ascomycota(100);c__Saccharomycetes(100);o__Saccharomycetales(100);f__Metschnikowiaceae(100);g__Metschnikowia(100);g__Metschnikowia_unclassified(100); | Metschnikowia | 13 | Saprotroph | \|Undefined Saprotroph\| | Yeast | NULL | Probable | As nectar-tap_saprotroph (Põlme et al. 2020); Decay sugar-rich substrates (Põlme et al. 2020) | Põlme S, et al. 2020. Fungal Diversity 105:1-16 ((https://doi.org/10.1007/s13225-020-00466-2)) |
| Otu20691 | 1 | k__Fungi(100);p__Ascomycota(100);c__Saccharomycetes(100);o__Saccharomycetales(100);f__Metschnikowiaceae(100);g__Metschnikowia(100);g__Metschnikowia_unclassified(100); | Metschnikowia | 13 | Saprotroph | \|Undefined Saprotroph\| | Yeast | NULL | Probable | As nectar-tap_saprotroph (Põlme et al. 2020); Decay sugar-rich substrates (Põlme et al. 2020) | Põlme S, et al. 2020. Fungal Diversity 105:1-16 ((https://doi.org/10.1007/s13225-020-00466-2)) |
| Otu20692 | 1 | k__Fungi(100);p__Ascomycota(100);c__Saccharomycetes(100);o__Saccharomycetales(100);f__Metschnikowiaceae(100);g__Metschnikowia(100);s__Metschnikowia_pulcherrima(100); | Metschnikowia | 13 | Saprotroph | \|Undefined Saprotroph\| | Yeast | NULL | Probable | As nectar-tap_saprotroph (Põlme et al. 2020); Decay sugar-rich substrates (Põlme et al. 2020) | Põlme S, et al. 2020. Fungal Diversity 105:1-16 ((https://doi.org/10.1007/s13225-020-00466-2)) |
| Otu20693 | 1 | k__Fungi(100);p__Ascomycota(100);c__Saccharomycetes(100);o__Saccharomycetales(100);f__Metschnikowiaceae(100);g__Metschnikowia(100);s__Metschnikowia_pulcherrima(100); | Metschnikowia | 13 | Saprotroph | \|Undefined Saprotroph\| | Yeast | NULL | Probable | As nectar-tap_saprotroph (Põlme et al. 2020); Decay sugar-rich substrates (Põlme et al. 2020) | Põlme S, et al. 2020. Fungal Diversity 105:1-16 ((https://doi.org/10.1007/s13225-020-00466-2)) |
| Otu20694 | 1 | k__Fungi(100);p__Ascomycota(100);c__Saccharomycetes(100);o__Saccharomycetales(100);f__Metschnikowiaceae(100);g__Metschnikowia(100);g__Metschnikowia_unclassified(100); | Metschnikowia | 13 | Saprotroph | \|Undefined Saprotroph\| | Yeast | NULL | Probable | As nectar-tap_saprotroph (Põlme et al. 2020); Decay sugar-rich substrates (Põlme et al. 2020) | Põlme S, et al. 2020. Fungal Diversity 105:1-16 ((https://doi.org/10.1007/s13225-020-00466-2)) |
| Otu20695 | 1 | k__Fungi(100);p__Ascomycota(100);c__Saccharomycetes(100);o__Saccharomycetales(100);f__Metschnikowiaceae(100);g__Metschnikowia(100);s__Metschnikowia_pulcherrima(100); | Metschnikowia | 13 | Saprotroph | \|Undefined Saprotroph\| | Yeast | NULL | Probable | As nectar-tap_saprotroph (Põlme et al. 2020); Decay sugar-rich substrates (Põlme et al. 2020) | Põlme S, et al. 2020. Fungal Diversity 105:1-16 ((https://doi.org/10.1007/s13225-020-00466-2)) |
| Otu20696 | 1 | k__Fungi(100);p__Ascomycota(100);c__Saccharomycetes(100);o__Saccharomycetales(100);f__Metschnikowiaceae(100);g__Metschnikowia(100);g__Metschnikowia_unclassified(100); | Metschnikowia | 13 | Saprotroph | \|Undefined Saprotroph\| | Yeast | NULL | Probable | As nectar-tap_saprotroph (Põlme et al. 2020); Decay sugar-rich substrates (Põlme et al. 2020) | Põlme S, et al. 2020. Fungal Diversity 105:1-16 ((https://doi.org/10.1007/s13225-020-00466-2)) |
| Otu20697 | 1 | k__Fungi(100);p__Basidiomycota(100);c__Microbotryomycetes(100);o__Sporidiobolales(100);f__Sporidiobolales_family_Incertae_sedis(100);g__Sporobolomyces(100);s__Sporobolomyces_lactosus(100); | Sporobolomyces | 13 | Pathotroph-Saprotroph | \|Fungal Parasite\|-Plant Saprotroph | Yeast | NULL | Probable | As litter_saprotroph (Põlme et al. 2020); As mycoparasite (Põlme et al. 2020); Decay fungal material (Põlme et al. 2020) | Tedersoo L, et al. 2014. Science 346:e1256688 ((https://doi.org/10.1126/science.1256688)); Purahong W, et al. 2016. Molecular Ecology 25:4059-4074 ((https://doi.org/10.1111/mec.13739)); Põlme S, et al. 2020. Fungal Diversity 105:1-16 ((https://doi.org/10.1007/s13225-020-00466-2)) |
| Otu20698 | 1 | k__Fungi(100);p__Ascomycota(100);c__Saccharomycetes(100);o__Saccharomycetales(100);f__Metschnikowiaceae(100);g__Metschnikowia(100);s__Metschnikowia_pulcherrima(100); | Metschnikowia | 13 | Saprotroph | \|Undefined Saprotroph\| | Yeast | NULL | Probable | As nectar-tap_saprotroph (Põlme et al. 2020); Decay sugar-rich substrates (Põlme et al. 2020) | Põlme S, et al. 2020. Fungal Diversity 105:1-16 ((https://doi.org/10.1007/s13225-020-00466-2)) |
| Otu20699 | 1 | k__Fungi(100);p__Ascomycota(100);c__Saccharomycetes(100);o__Saccharomycetales(100);f__Metschnikowiaceae(100);g__Metschnikowia(100);s__Metschnikowia_pulcherrima(100); | Metschnikowia | 13 | Saprotroph | \|Undefined Saprotroph\| | Yeast | NULL | Probable | As nectar-tap_saprotroph (Põlme et al. 2020); Decay sugar-rich substrates (Põlme et al. 2020) | Põlme S, et al. 2020. Fungal Diversity 105:1-16 ((https://doi.org/10.1007/s13225-020-00466-2)) |
| Otu20700 | 1 | k__Fungi(100);p__Ascomycota(100);c__Saccharomycetes(100);o__Saccharomycetales(100);f__Metschnikowiaceae(100);g__Metschnikowia(100);s__Metschnikowia_pulcherrima(100); | Metschnikowia | 13 | Saprotroph | \|Undefined Saprotroph\| | Yeast | NULL | Probable | As nectar-tap_saprotroph (Põlme et al. 2020); Decay sugar-rich substrates (Põlme et al. 2020) | Põlme S, et al. 2020. Fungal Diversity 105:1-16 ((https://doi.org/10.1007/s13225-020-00466-2)) |
| Otu20701 | 1 | k__Fungi(100);p__Basidiomycota(100);c__Microbotryomycetes(100);o__Sporidiobolales(100);f__Sporidiobolales_family_Incertae_sedis(100);g__Sporobolomyces(100);s__Sporobolomyces_lactosus(100); | Sporobolomyces | 13 | Pathotroph-Saprotroph | \|Fungal Parasite\|-Plant Saprotroph | Yeast | NULL | Probable | As litter_saprotroph (Põlme et al. 2020); As mycoparasite (Põlme et al. 2020); Decay fungal material (Põlme et al. 2020) | Tedersoo L, et al. 2014. Science 346:e1256688 ((https://doi.org/10.1126/science.1256688)); Purahong W, et al. 2016. Molecular Ecology 25:4059-4074 ((https://doi.org/10.1111/mec.13739)); Põlme S, et al. 2020. Fungal Diversity 105:1-16 ((https://doi.org/10.1007/s13225-020-00466-2)) |
| Otu20702 | 1 | k__Fungi(100);p__Basidiomycota(100);c__Microbotryomycetes(100);o__Sporidiobolales(100);f__Sporidiobolales_family_Incertae_sedis(100);g__Sporobolomyces(100);s__Sporobolomyces_lactosus(100); | Sporobolomyces | 13 | Pathotroph-Saprotroph | \|Fungal Parasite\|-Plant Saprotroph | Yeast | NULL | Probable | As litter_saprotroph (Põlme et al. 2020); As mycoparasite (Põlme et al. 2020); Decay fungal material (Põlme et al. 2020) | Tedersoo L, et al. 2014. Science 346:e1256688 ((https://doi.org/10.1126/science.1256688)); Purahong W, et al. 2016. Molecular Ecology 25:4059-4074 ((https://doi.org/10.1111/mec.13739)); Põlme S, et al. 2020. Fungal Diversity 105:1-16 ((https://doi.org/10.1007/s13225-020-00466-2)) |
| Otu20703 | 1 | k__Fungi(100);p__Ascomycota(100);c__Saccharomycetes(100);o__Saccharomycetales(100);f__Metschnikowiaceae(100);g__Metschnikowia(100);s__Metschnikowia_pulcherrima(100); | Metschnikowia | 13 | Saprotroph | \|Undefined Saprotroph\| | Yeast | NULL | Probable | As nectar-tap_saprotroph (Põlme et al. 2020); Decay sugar-rich substrates (Põlme et al. 2020) | Põlme S, et al. 2020. Fungal Diversity 105:1-16 ((https://doi.org/10.1007/s13225-020-00466-2)) |
| Otu20704 | 1 | k__Fungi(100);p__Ascomycota(100);c__Saccharomycetes(100);o__Saccharomycetales(100);f__Metschnikowiaceae(100);g__Metschnikowia(100);g__Metschnikowia_unclassified(100); | Metschnikowia | 13 | Saprotroph | \|Undefined Saprotroph\| | Yeast | NULL | Probable | As nectar-tap_saprotroph (Põlme et al. 2020); Decay sugar-rich substrates (Põlme et al. 2020) | Põlme S, et al. 2020. Fungal Diversity 105:1-16 ((https://doi.org/10.1007/s13225-020-00466-2)) |
| Otu20705 | 1 | k__Fungi(100);p__Ascomycota(100);c__Saccharomycetes(100);o__Saccharomycetales(100);f__Metschnikowiaceae(100);g__Metschnikowia(100);g__Metschnikowia_unclassified(100); | Metschnikowia | 13 | Saprotroph | \|Undefined Saprotroph\| | Yeast | NULL | Probable | As nectar-tap_saprotroph (Põlme et al. 2020); Decay sugar-rich substrates (Põlme et al. 2020) | Põlme S, et al. 2020. Fungal Diversity 105:1-16 ((https://doi.org/10.1007/s13225-020-00466-2)) |
| Otu20706 | 1 | k__Fungi(100);p__Ascomycota(100);c__Saccharomycetes(100);o__Saccharomycetales(100);f__Metschnikowiaceae(100);g__Metschnikowia(100);s__Metschnikowia_pulcherrima(100); | Metschnikowia | 13 | Saprotroph | \|Undefined Saprotroph\| | Yeast | NULL | Probable | As nectar-tap_saprotroph (Põlme et al. 2020); Decay sugar-rich substrates (Põlme et al. 2020) | Põlme S, et al. 2020. Fungal Diversity 105:1-16 ((https://doi.org/10.1007/s13225-020-00466-2)) |
| Otu20707 | 1 | k__Fungi(100);p__Ascomycota(100);c__Saccharomycetes(100);o__Saccharomycetales(100);f__Metschnikowiaceae(100);g__Metschnikowia(100);g__Metschnikowia_unclassified(100); | Metschnikowia | 13 | Saprotroph | \|Undefined Saprotroph\| | Yeast | NULL | Probable | As nectar-tap_saprotroph (Põlme et al. 2020); Decay sugar-rich substrates (Põlme et al. 2020) | Põlme S, et al. 2020. Fungal Diversity 105:1-16 ((https://doi.org/10.1007/s13225-020-00466-2)) |
| Otu20708 | 1 | k__Fungi(100);p__Ascomycota(100);c__Saccharomycetes(100);o__Saccharomycetales(100);f__Metschnikowiaceae(100);g__Metschnikowia(100);s__Metschnikowia_pulcherrima(100); | Metschnikowia | 13 | Saprotroph | \|Undefined Saprotroph\| | Yeast | NULL | Probable | As nectar-tap_saprotroph (Põlme et al. 2020); Decay sugar-rich substrates (Põlme et al. 2020) | Põlme S, et al. 2020. Fungal Diversity 105:1-16 ((https://doi.org/10.1007/s13225-020-00466-2)) |
| Otu20710 | 1 | k__Fungi(100);p__Ascomycota(100);c__Saccharomycetes(100);o__Saccharomycetales(100);f__Metschnikowiaceae(100);g__Metschnikowia(100);s__Metschnikowia_pulcherrima(100); | Metschnikowia | 13 | Saprotroph | \|Undefined Saprotroph\| | Yeast | NULL | Probable | As nectar-tap_saprotroph (Põlme et al. 2020); Decay sugar-rich substrates (Põlme et al. 2020) | Põlme S, et al. 2020. Fungal Diversity 105:1-16 ((https://doi.org/10.1007/s13225-020-00466-2)) |
| Otu20712 | 1 | k__Fungi(100);p__Ascomycota(100);c__Saccharomycetes(100);o__Saccharomycetales(100);f__Metschnikowiaceae(100);g__Metschnikowia(100);s__Metschnikowia_pulcherrima(100); | Metschnikowia | 13 | Saprotroph | \|Undefined Saprotroph\| | Yeast | NULL | Probable | As nectar-tap_saprotroph (Põlme et al. 2020); Decay sugar-rich substrates (Põlme et al. 2020) | Põlme S, et al. 2020. Fungal Diversity 105:1-16 ((https://doi.org/10.1007/s13225-020-00466-2)) |
| Otu20713 | 1 | k__Fungi(100);p__Ascomycota(100);c__Saccharomycetes(100);o__Saccharomycetales(100);f__Metschnikowiaceae(100);g__Metschnikowia(100);s__Metschnikowia_pulcherrima(100); | Metschnikowia | 13 | Saprotroph | \|Undefined Saprotroph\| | Yeast | NULL | Probable | As nectar-tap_saprotroph (Põlme et al. 2020); Decay sugar-rich substrates (Põlme et al. 2020) | Põlme S, et al. 2020. Fungal Diversity 105:1-16 ((https://doi.org/10.1007/s13225-020-00466-2)) |
| Otu20714 | 1 | k__Fungi(100);p__Ascomycota(100);c__Saccharomycetes(100);o__Saccharomycetales(100);f__Metschnikowiaceae(100);g__Metschnikowia(100);g__Metschnikowia_unclassified(100); | Metschnikowia | 13 | Saprotroph | \|Undefined Saprotroph\| | Yeast | NULL | Probable | As nectar-tap_saprotroph (Põlme et al. 2020); Decay sugar-rich substrates (Põlme et al. 2020) | Põlme S, et al. 2020. Fungal Diversity 105:1-16 ((https://doi.org/10.1007/s13225-020-00466-2)) |
| Otu20715 | 1 | k__Fungi(100);p__Ascomycota(100);c__Saccharomycetes(100);o__Saccharomycetales(100);f__Metschnikowiaceae(100);g__Metschnikowia(100);s__Metschnikowia_pulcherrima(100); | Metschnikowia | 13 | Saprotroph | \|Undefined Saprotroph\| | Yeast | NULL | Probable | As nectar-tap_saprotroph (Põlme et al. 2020); Decay sugar-rich substrates (Põlme et al. 2020) | Põlme S, et al. 2020. Fungal Diversity 105:1-16 ((https://doi.org/10.1007/s13225-020-00466-2)) |
| Otu20716 | 1 | k__Fungi(100);p__Ascomycota(100);c__Saccharomycetes(100);o__Saccharomycetales(100);f__Saccharomycetales_family_Incertae_sedis(100);g__Candida(100);g__Candida_unclassified(100); | Candida | 13 | Pathotroph-Saprotroph-Symbiotroph | Animal Pathogen-Endophyte-Endosymbiont-Epiphyte-\|Undefined Saprotroph\| | Dimorphic Yeast-Yeast | NULL | Probable | As nectar-tap_saprotroph (Põlme et al. 2020); Decay sugar-rich substrates (Põlme et al. 2020); This asexual genus is polyphyletic and contain species that function very differently depend on what they are. Caution should be taken when interpreting guilds for Candida species. | Manolakaki D, et al. 2010. Virulence. 1: 367-375 ((https://doi.org/10.4161/viru.1.5.12796)); Mekbib SB, et al. 2011. Tropical Plant Pathology 36:233-240 ((https://doi.org/10.1590/S1982-56762011000400004)); Põlme S, et al. 2020. Fungal Diversity 105:1-16 ((https://doi.org/10.1007/s13225-020-00466-2)) |
| Otu20719 | 1 | k__Fungi(100);p__Ascomycota(100);c__Saccharomycetes(100);o__Saccharomycetales(100);f__Metschnikowiaceae(100);g__Metschnikowia(100);g__Metschnikowia_unclassified(100); | Metschnikowia | 13 | Saprotroph | \|Undefined Saprotroph\| | Yeast | NULL | Probable | As nectar-tap_saprotroph (Põlme et al. 2020); Decay sugar-rich substrates (Põlme et al. 2020) | Põlme S, et al. 2020. Fungal Diversity 105:1-16 ((https://doi.org/10.1007/s13225-020-00466-2)) |
| Otu20720 | 1 | k__Fungi(100);p__Ascomycota(100);c__Saccharomycetes(100);o__Saccharomycetales(100);f__Metschnikowiaceae(100);g__Metschnikowia(100);s__Metschnikowia_pulcherrima(100); | Metschnikowia | 13 | Saprotroph | \|Undefined Saprotroph\| | Yeast | NULL | Probable | As nectar-tap_saprotroph (Põlme et al. 2020); Decay sugar-rich substrates (Põlme et al. 2020) | Põlme S, et al. 2020. Fungal Diversity 105:1-16 ((https://doi.org/10.1007/s13225-020-00466-2)) |
| Otu20721 | 1 | k__Fungi(100);p__Ascomycota(100);c__Saccharomycetes(100);o__Saccharomycetales(100);f__Metschnikowiaceae(100);g__Metschnikowia(100);s__Metschnikowia_pulcherrima(100); | Metschnikowia | 13 | Saprotroph | \|Undefined Saprotroph\| | Yeast | NULL | Probable | As nectar-tap_saprotroph (Põlme et al. 2020); Decay sugar-rich substrates (Põlme et al. 2020) | Põlme S, et al. 2020. Fungal Diversity 105:1-16 ((https://doi.org/10.1007/s13225-020-00466-2)) |
| Otu20722 | 1 | k__Fungi(100);p__Ascomycota(100);c__Saccharomycetes(100);o__Saccharomycetales(100);f__Metschnikowiaceae(100);g__Metschnikowia(100);g__Metschnikowia_unclassified(100); | Metschnikowia | 13 | Saprotroph | \|Undefined Saprotroph\| | Yeast | NULL | Probable | As nectar-tap_saprotroph (Põlme et al. 2020); Decay sugar-rich substrates (Põlme et al. 2020) | Põlme S, et al. 2020. Fungal Diversity 105:1-16 ((https://doi.org/10.1007/s13225-020-00466-2)) |
| Otu20723 | 1 | k__Fungi(100);p__Ascomycota(100);c__Saccharomycetes(100);o__Saccharomycetales(100);f__Metschnikowiaceae(100);g__Metschnikowia(100);s__Metschnikowia_pulcherrima(100); | Metschnikowia | 13 | Saprotroph | \|Undefined Saprotroph\| | Yeast | NULL | Probable | As nectar-tap_saprotroph (Põlme et al. 2020); Decay sugar-rich substrates (Põlme et al. 2020) | Põlme S, et al. 2020. Fungal Diversity 105:1-16 ((https://doi.org/10.1007/s13225-020-00466-2)) |
| Otu20724 | 1 | k__Fungi(100);p__Basidiomycota(100);c__Microbotryomycetes(100);o__Sporidiobolales(100);f__Sporidiobolales_family_Incertae_sedis(100);g__Sporobolomyces(100);s__Sporobolomyces_lactosus(100); | Sporobolomyces | 13 | Pathotroph-Saprotroph | \|Fungal Parasite\|-Plant Saprotroph | Yeast | NULL | Probable | As litter_saprotroph (Põlme et al. 2020); As mycoparasite (Põlme et al. 2020); Decay fungal material (Põlme et al. 2020) | Tedersoo L, et al. 2014. Science 346:e1256688 ((https://doi.org/10.1126/science.1256688)); Purahong W, et al. 2016. Molecular Ecology 25:4059-4074 ((https://doi.org/10.1111/mec.13739)); Põlme S, et al. 2020. Fungal Diversity 105:1-16 ((https://doi.org/10.1007/s13225-020-00466-2)) |
| Otu20725 | 1 | k__Fungi(100);p__Ascomycota(100);c__Saccharomycetes(100);o__Saccharomycetales(100);f__Metschnikowiaceae(100);g__Metschnikowia(100);s__Metschnikowia_pulcherrima(100); | Metschnikowia | 13 | Saprotroph | \|Undefined Saprotroph\| | Yeast | NULL | Probable | As nectar-tap_saprotroph (Põlme et al. 2020); Decay sugar-rich substrates (Põlme et al. 2020) | Põlme S, et al. 2020. Fungal Diversity 105:1-16 ((https://doi.org/10.1007/s13225-020-00466-2)) |
| Otu20726 | 1 | k__Fungi(100);p__Ascomycota(100);c__Saccharomycetes(100);o__Saccharomycetales(100);f__Metschnikowiaceae(100);g__Metschnikowia(100);s__Metschnikowia_pulcherrima(100); | Metschnikowia | 13 | Saprotroph | \|Undefined Saprotroph\| | Yeast | NULL | Probable | As nectar-tap_saprotroph (Põlme et al. 2020); Decay sugar-rich substrates (Põlme et al. 2020) | Põlme S, et al. 2020. Fungal Diversity 105:1-16 ((https://doi.org/10.1007/s13225-020-00466-2)) |
| Otu20727 | 1 | k__Fungi(100);p__Ascomycota(100);c__Saccharomycetes(100);o__Saccharomycetales(100);f__Metschnikowiaceae(100);g__Metschnikowia(100);s__Metschnikowia_pulcherrima(100); | Metschnikowia | 13 | Saprotroph | \|Undefined Saprotroph\| | Yeast | NULL | Probable | As nectar-tap_saprotroph (Põlme et al. 2020); Decay sugar-rich substrates (Põlme et al. 2020) | Põlme S, et al. 2020. Fungal Diversity 105:1-16 ((https://doi.org/10.1007/s13225-020-00466-2)) |
| Otu20728 | 1 | k__Fungi(100);p__Ascomycota(100);c__Saccharomycetes(100);o__Saccharomycetales(100);f__Metschnikowiaceae(100);g__Metschnikowia(100);g__Metschnikowia_unclassified(100); | Metschnikowia | 13 | Saprotroph | \|Undefined Saprotroph\| | Yeast | NULL | Probable | As nectar-tap_saprotroph (Põlme et al. 2020); Decay sugar-rich substrates (Põlme et al. 2020) | Põlme S, et al. 2020. Fungal Diversity 105:1-16 ((https://doi.org/10.1007/s13225-020-00466-2)) |
| Otu20729 | 1 | k__Fungi(100);p__Ascomycota(100);c__Saccharomycetes(100);o__Saccharomycetales(100);f__Metschnikowiaceae(100);g__Metschnikowia(100);s__Metschnikowia_pulcherrima(100); | Metschnikowia | 13 | Saprotroph | \|Undefined Saprotroph\| | Yeast | NULL | Probable | As nectar-tap_saprotroph (Põlme et al. 2020); Decay sugar-rich substrates (Põlme et al. 2020) | Põlme S, et al. 2020. Fungal Diversity 105:1-16 ((https://doi.org/10.1007/s13225-020-00466-2)) |
| Otu20730 | 1 | k__Fungi(100);p__Ascomycota(100);c__Saccharomycetes(100);o__Saccharomycetales(100);f__Metschnikowiaceae(100);g__Metschnikowia(100);s__Metschnikowia_pulcherrima(100); | Metschnikowia | 13 | Saprotroph | \|Undefined Saprotroph\| | Yeast | NULL | Probable | As nectar-tap_saprotroph (Põlme et al. 2020); Decay sugar-rich substrates (Põlme et al. 2020) | Põlme S, et al. 2020. Fungal Diversity 105:1-16 ((https://doi.org/10.1007/s13225-020-00466-2)) |
| Otu20731 | 1 | k__Fungi(100);p__Ascomycota(100);c__Dothideomycetes(100);o__Dothideomycetes_order_Incertae_sedis(100);f__Pseudeurotiaceae(100);g__Pseudeurotium(100);s__Pseudeurotium_hygrophilum(100); | Pseudeurotium | 13 | Saprotroph | \|Dung Saprotroph\|-Undefined Saprotroph | Microfungus | NULL | Probable | As soil_saprotroph (Põlme et al. 2020); Decay soil, dung (Põlme et al. 2020) | Tedersoo L, et al. 2014. Science 346:e1256688 ((https://doi.org/10.1126/science.1256688)); Põlme S, et al. 2020. Fungal Diversity 105:1-16 ((https://doi.org/10.1007/s13225-020-00466-2)) |
| Otu20732 | 1 | k__Fungi(100);p__Ascomycota(100);c__Saccharomycetes(100);o__Saccharomycetales(100);f__Metschnikowiaceae(100);g__Metschnikowia(100);s__Metschnikowia_pulcherrima(100); | Metschnikowia | 13 | Saprotroph | \|Undefined Saprotroph\| | Yeast | NULL | Probable | As nectar-tap_saprotroph (Põlme et al. 2020); Decay sugar-rich substrates (Põlme et al. 2020) | Põlme S, et al. 2020. Fungal Diversity 105:1-16 ((https://doi.org/10.1007/s13225-020-00466-2)) |
| Otu20733 | 1 | k__Fungi(100);p__Ascomycota(100);c__Saccharomycetes(100);o__Saccharomycetales(100);f__Metschnikowiaceae(100);g__Metschnikowia(100);g__Metschnikowia_unclassified(100); | Metschnikowia | 13 | Saprotroph | \|Undefined Saprotroph\| | Yeast | NULL | Probable | As nectar-tap_saprotroph (Põlme et al. 2020); Decay sugar-rich substrates (Põlme et al. 2020) | Põlme S, et al. 2020. Fungal Diversity 105:1-16 ((https://doi.org/10.1007/s13225-020-00466-2)) |
| Otu20734 | 1 | k__Fungi(100);p__Ascomycota(100);c__Saccharomycetes(100);o__Saccharomycetales(100);f__Saccharomycetales_family_Incertae_sedis(100);g__Candida(100);g__Candida_unclassified(100); | Candida | 13 | Pathotroph-Saprotroph-Symbiotroph | Animal Pathogen-Endophyte-Endosymbiont-Epiphyte-\|Undefined Saprotroph\| | Dimorphic Yeast-Yeast | NULL | Probable | As nectar-tap_saprotroph (Põlme et al. 2020); Decay sugar-rich substrates (Põlme et al. 2020); This asexual genus is polyphyletic and contain species that function very differently depend on what they are. Caution should be taken when interpreting guilds for Candida species. | Manolakaki D, et al. 2010. Virulence. 1: 367-375 ((https://doi.org/10.4161/viru.1.5.12796)); Mekbib SB, et al. 2011. Tropical Plant Pathology 36:233-240 ((https://doi.org/10.1590/S1982-56762011000400004)); Põlme S, et al. 2020. Fungal Diversity 105:1-16 ((https://doi.org/10.1007/s13225-020-00466-2)) |
| Otu20735 | 1 | k__Fungi(100);p__Ascomycota(100);c__Saccharomycetes(100);o__Saccharomycetales(100);f__Metschnikowiaceae(100);g__Metschnikowia(100);s__Metschnikowia_pulcherrima(100); | Metschnikowia | 13 | Saprotroph | \|Undefined Saprotroph\| | Yeast | NULL | Probable | As nectar-tap_saprotroph (Põlme et al. 2020); Decay sugar-rich substrates (Põlme et al. 2020) | Põlme S, et al. 2020. Fungal Diversity 105:1-16 ((https://doi.org/10.1007/s13225-020-00466-2)) |
| Otu20736 | 1 | k__Fungi(100);p__Ascomycota(100);c__Saccharomycetes(100);o__Saccharomycetales(100);f__Metschnikowiaceae(100);g__Metschnikowia(100);g__Metschnikowia_unclassified(100); | Metschnikowia | 13 | Saprotroph | \|Undefined Saprotroph\| | Yeast | NULL | Probable | As nectar-tap_saprotroph (Põlme et al. 2020); Decay sugar-rich substrates (Põlme et al. 2020) | Põlme S, et al. 2020. Fungal Diversity 105:1-16 ((https://doi.org/10.1007/s13225-020-00466-2)) |
| Otu20737 | 1 | k__Fungi(100);p__Ascomycota(100);c__Saccharomycetes(100);o__Saccharomycetales(100);f__Metschnikowiaceae(100);g__Metschnikowia(100);s__Metschnikowia_pulcherrima(100); | Metschnikowia | 13 | Saprotroph | \|Undefined Saprotroph\| | Yeast | NULL | Probable | As nectar-tap_saprotroph (Põlme et al. 2020); Decay sugar-rich substrates (Põlme et al. 2020) | Põlme S, et al. 2020. Fungal Diversity 105:1-16 ((https://doi.org/10.1007/s13225-020-00466-2)) |
| Otu20738 | 1 | k__Fungi(100);p__Ascomycota(100);c__Saccharomycetes(100);o__Saccharomycetales(100);f__Metschnikowiaceae(100);g__Metschnikowia(100);s__Metschnikowia_pulcherrima(100); | Metschnikowia | 13 | Saprotroph | \|Undefined Saprotroph\| | Yeast | NULL | Probable | As nectar-tap_saprotroph (Põlme et al. 2020); Decay sugar-rich substrates (Põlme et al. 2020) | Põlme S, et al. 2020. Fungal Diversity 105:1-16 ((https://doi.org/10.1007/s13225-020-00466-2)) |
| Otu20739 | 1 | k__Fungi(100);p__Ascomycota(100);c__Saccharomycetes(100);o__Saccharomycetales(100);f__Metschnikowiaceae(100);g__Metschnikowia(100);s__Metschnikowia_pulcherrima(100); | Metschnikowia | 13 | Saprotroph | \|Undefined Saprotroph\| | Yeast | NULL | Probable | As nectar-tap_saprotroph (Põlme et al. 2020); Decay sugar-rich substrates (Põlme et al. 2020) | Põlme S, et al. 2020. Fungal Diversity 105:1-16 ((https://doi.org/10.1007/s13225-020-00466-2)) |
| Otu20740 | 1 | k__Fungi(100);p__Ascomycota(100);c__Saccharomycetes(100);o__Saccharomycetales(100);f__Metschnikowiaceae(100);g__Metschnikowia(100);s__Metschnikowia_pulcherrima(100); | Metschnikowia | 13 | Saprotroph | \|Undefined Saprotroph\| | Yeast | NULL | Probable | As nectar-tap_saprotroph (Põlme et al. 2020); Decay sugar-rich substrates (Põlme et al. 2020) | Põlme S, et al. 2020. Fungal Diversity 105:1-16 ((https://doi.org/10.1007/s13225-020-00466-2)) |
| Otu20742 | 1 | k__Fungi(100);p__Ascomycota(100);c__Saccharomycetes(100);o__Saccharomycetales(100);f__Saccharomycetales_family_Incertae_sedis(100);g__Candida(100);g__Candida_unclassified(100); | Candida | 13 | Pathotroph-Saprotroph-Symbiotroph | Animal Pathogen-Endophyte-Endosymbiont-Epiphyte-\|Undefined Saprotroph\| | Dimorphic Yeast-Yeast | NULL | Probable | As nectar-tap_saprotroph (Põlme et al. 2020); Decay sugar-rich substrates (Põlme et al. 2020); This asexual genus is polyphyletic and contain species that function very differently depend on what they are. Caution should be taken when interpreting guilds for Candida species. | Manolakaki D, et al. 2010. Virulence. 1: 367-375 ((https://doi.org/10.4161/viru.1.5.12796)); Mekbib SB, et al. 2011. Tropical Plant Pathology 36:233-240 ((https://doi.org/10.1590/S1982-56762011000400004)); Põlme S, et al. 2020. Fungal Diversity 105:1-16 ((https://doi.org/10.1007/s13225-020-00466-2)) |
| Otu20743 | 1 | k__Fungi(100);p__Ascomycota(100);c__Saccharomycetes(100);o__Saccharomycetales(100);f__Saccharomycetales_family_Incertae_sedis(100);g__Candida(100);g__Candida_unclassified(100); | Candida | 13 | Pathotroph-Saprotroph-Symbiotroph | Animal Pathogen-Endophyte-Endosymbiont-Epiphyte-\|Undefined Saprotroph\| | Dimorphic Yeast-Yeast | NULL | Probable | As nectar-tap_saprotroph (Põlme et al. 2020); Decay sugar-rich substrates (Põlme et al. 2020); This asexual genus is polyphyletic and contain species that function very differently depend on what they are. Caution should be taken when interpreting guilds for Candida species. | Manolakaki D, et al. 2010. Virulence. 1: 367-375 ((https://doi.org/10.4161/viru.1.5.12796)); Mekbib SB, et al. 2011. Tropical Plant Pathology 36:233-240 ((https://doi.org/10.1590/S1982-56762011000400004)); Põlme S, et al. 2020. Fungal Diversity 105:1-16 ((https://doi.org/10.1007/s13225-020-00466-2)) |
| Otu20744 | 1 | k__Fungi(100);p__Ascomycota(100);c__Saccharomycetes(100);o__Saccharomycetales(100);f__Metschnikowiaceae(100);g__Metschnikowia(100);g__Metschnikowia_unclassified(100); | Metschnikowia | 13 | Saprotroph | \|Undefined Saprotroph\| | Yeast | NULL | Probable | As nectar-tap_saprotroph (Põlme et al. 2020); Decay sugar-rich substrates (Põlme et al. 2020) | Põlme S, et al. 2020. Fungal Diversity 105:1-16 ((https://doi.org/10.1007/s13225-020-00466-2)) |
| Otu20745 | 1 | k__Fungi(100);p__Ascomycota(100);c__Saccharomycetes(100);o__Saccharomycetales(100);f__Metschnikowiaceae(100);g__Metschnikowia(100);s__Metschnikowia_pulcherrima(100); | Metschnikowia | 13 | Saprotroph | \|Undefined Saprotroph\| | Yeast | NULL | Probable | As nectar-tap_saprotroph (Põlme et al. 2020); Decay sugar-rich substrates (Põlme et al. 2020) | Põlme S, et al. 2020. Fungal Diversity 105:1-16 ((https://doi.org/10.1007/s13225-020-00466-2)) |
| Otu20747 | 1 | k__Fungi(100);p__Ascomycota(100);c__Saccharomycetes(100);o__Saccharomycetales(100);f__Metschnikowiaceae(100);g__Metschnikowia(100);s__Metschnikowia_pulcherrima(100); | Metschnikowia | 13 | Saprotroph | \|Undefined Saprotroph\| | Yeast | NULL | Probable | As nectar-tap_saprotroph (Põlme et al. 2020); Decay sugar-rich substrates (Põlme et al. 2020) | Põlme S, et al. 2020. Fungal Diversity 105:1-16 ((https://doi.org/10.1007/s13225-020-00466-2)) |
| Otu20748 | 1 | k__Fungi(100);p__Ascomycota(100);c__Saccharomycetes(100);o__Saccharomycetales(100);f__Saccharomycetales_family_Incertae_sedis(100);g__Candida(100);g__Candida_unclassified(100); | Candida | 13 | Pathotroph-Saprotroph-Symbiotroph | Animal Pathogen-Endophyte-Endosymbiont-Epiphyte-\|Undefined Saprotroph\| | Dimorphic Yeast-Yeast | NULL | Probable | As nectar-tap_saprotroph (Põlme et al. 2020); Decay sugar-rich substrates (Põlme et al. 2020); This asexual genus is polyphyletic and contain species that function very differently depend on what they are. Caution should be taken when interpreting guilds for Candida species. | Manolakaki D, et al. 2010. Virulence. 1: 367-375 ((https://doi.org/10.4161/viru.1.5.12796)); Mekbib SB, et al. 2011. Tropical Plant Pathology 36:233-240 ((https://doi.org/10.1590/S1982-56762011000400004)); Põlme S, et al. 2020. Fungal Diversity 105:1-16 ((https://doi.org/10.1007/s13225-020-00466-2)) |
| Otu20749 | 1 | k__Fungi(100);p__Ascomycota(100);c__Saccharomycetes(100);o__Saccharomycetales(100);f__Metschnikowiaceae(100);g__Metschnikowia(100);s__Metschnikowia_pulcherrima(100); | Metschnikowia | 13 | Saprotroph | \|Undefined Saprotroph\| | Yeast | NULL | Probable | As nectar-tap_saprotroph (Põlme et al. 2020); Decay sugar-rich substrates (Põlme et al. 2020) | Põlme S, et al. 2020. Fungal Diversity 105:1-16 ((https://doi.org/10.1007/s13225-020-00466-2)) |
| Otu20750 | 1 | k__Fungi(100);p__Ascomycota(100);c__Saccharomycetes(100);o__Saccharomycetales(100);f__Metschnikowiaceae(100);g__Metschnikowia(100);s__Metschnikowia_pulcherrima(100); | Metschnikowia | 13 | Saprotroph | \|Undefined Saprotroph\| | Yeast | NULL | Probable | As nectar-tap_saprotroph (Põlme et al. 2020); Decay sugar-rich substrates (Põlme et al. 2020) | Põlme S, et al. 2020. Fungal Diversity 105:1-16 ((https://doi.org/10.1007/s13225-020-00466-2)) |
| Otu20752 | 1 | k__Fungi(100);p__Ascomycota(100);c__Saccharomycetes(100);o__Saccharomycetales(100);f__Saccharomycetales_family_Incertae_sedis(100);g__Candida(100);s__Candida_cretensis(100); | Candida | 13 | Pathotroph-Saprotroph-Symbiotroph | Animal Pathogen-Endophyte-Endosymbiont-Epiphyte-\|Undefined Saprotroph\| | Dimorphic Yeast-Yeast | NULL | Probable | As nectar-tap_saprotroph (Põlme et al. 2020); Decay sugar-rich substrates (Põlme et al. 2020); This asexual genus is polyphyletic and contain species that function very differently depend on what they are. Caution should be taken when interpreting guilds for Candida species. | Manolakaki D, et al. 2010. Virulence. 1: 367-375 ((https://doi.org/10.4161/viru.1.5.12796)); Mekbib SB, et al. 2011. Tropical Plant Pathology 36:233-240 ((https://doi.org/10.1590/S1982-56762011000400004)); Põlme S, et al. 2020. Fungal Diversity 105:1-16 ((https://doi.org/10.1007/s13225-020-00466-2)) |
| Otu20753 | 1 | k__Fungi(100);p__Ascomycota(100);c__Saccharomycetes(100);o__Saccharomycetales(100);f__Saccharomycetales_family_Incertae_sedis(100);g__Candida(100);g__Candida_unclassified(100); | Candida | 13 | Pathotroph-Saprotroph-Symbiotroph | Animal Pathogen-Endophyte-Endosymbiont-Epiphyte-\|Undefined Saprotroph\| | Dimorphic Yeast-Yeast | NULL | Probable | As nectar-tap_saprotroph (Põlme et al. 2020); Decay sugar-rich substrates (Põlme et al. 2020); This asexual genus is polyphyletic and contain species that function very differently depend on what they are. Caution should be taken when interpreting guilds for Candida species. | Manolakaki D, et al. 2010. Virulence. 1: 367-375 ((https://doi.org/10.4161/viru.1.5.12796)); Mekbib SB, et al. 2011. Tropical Plant Pathology 36:233-240 ((https://doi.org/10.1590/S1982-56762011000400004)); Põlme S, et al. 2020. Fungal Diversity 105:1-16 ((https://doi.org/10.1007/s13225-020-00466-2)) |
| Otu20754 | 1 | k__Fungi(100);p__Ascomycota(100);c__Saccharomycetes(100);o__Saccharomycetales(100);f__Saccharomycetales_family_Incertae_sedis(100);g__Candida(100);g__Candida_unclassified(100); | Candida | 13 | Pathotroph-Saprotroph-Symbiotroph | Animal Pathogen-Endophyte-Endosymbiont-Epiphyte-\|Undefined Saprotroph\| | Dimorphic Yeast-Yeast | NULL | Probable | As nectar-tap_saprotroph (Põlme et al. 2020); Decay sugar-rich substrates (Põlme et al. 2020); This asexual genus is polyphyletic and contain species that function very differently depend on what they are. Caution should be taken when interpreting guilds for Candida species. | Manolakaki D, et al. 2010. Virulence. 1: 367-375 ((https://doi.org/10.4161/viru.1.5.12796)); Mekbib SB, et al. 2011. Tropical Plant Pathology 36:233-240 ((https://doi.org/10.1590/S1982-56762011000400004)); Põlme S, et al. 2020. Fungal Diversity 105:1-16 ((https://doi.org/10.1007/s13225-020-00466-2)) |
| Otu20755 | 1 | k__Fungi(100);p__Ascomycota(100);c__Saccharomycetes(100);o__Saccharomycetales(100);f__Saccharomycetales_family_Incertae_sedis(100);g__Candida(100);s__Candida_cretensis(100); | Candida | 13 | Pathotroph-Saprotroph-Symbiotroph | Animal Pathogen-Endophyte-Endosymbiont-Epiphyte-\|Undefined Saprotroph\| | Dimorphic Yeast-Yeast | NULL | Probable | As nectar-tap_saprotroph (Põlme et al. 2020); Decay sugar-rich substrates (Põlme et al. 2020); This asexual genus is polyphyletic and contain species that function very differently depend on what they are. Caution should be taken when interpreting guilds for Candida species. | Manolakaki D, et al. 2010. Virulence. 1: 367-375 ((https://doi.org/10.4161/viru.1.5.12796)); Mekbib SB, et al. 2011. Tropical Plant Pathology 36:233-240 ((https://doi.org/10.1590/S1982-56762011000400004)); Põlme S, et al. 2020. Fungal Diversity 105:1-16 ((https://doi.org/10.1007/s13225-020-00466-2)) |
| Otu20756 | 1 | k__Fungi(100);p__Ascomycota(100);c__Saccharomycetes(100);o__Saccharomycetales(100);f__Metschnikowiaceae(100);g__Metschnikowia(100);s__Metschnikowia_pulcherrima(100); | Metschnikowia | 13 | Saprotroph | \|Undefined Saprotroph\| | Yeast | NULL | Probable | As nectar-tap_saprotroph (Põlme et al. 2020); Decay sugar-rich substrates (Põlme et al. 2020) | Põlme S, et al. 2020. Fungal Diversity 105:1-16 ((https://doi.org/10.1007/s13225-020-00466-2)) |
| Otu20757 | 1 | k__Fungi(100);p__Ascomycota(100);c__Saccharomycetes(100);o__Saccharomycetales(100);f__Saccharomycetales_family_Incertae_sedis(100);g__Candida(100);g__Candida_unclassified(100); | Candida | 13 | Pathotroph-Saprotroph-Symbiotroph | Animal Pathogen-Endophyte-Endosymbiont-Epiphyte-\|Undefined Saprotroph\| | Dimorphic Yeast-Yeast | NULL | Probable | As nectar-tap_saprotroph (Põlme et al. 2020); Decay sugar-rich substrates (Põlme et al. 2020); This asexual genus is polyphyletic and contain species that function very differently depend on what they are. Caution should be taken when interpreting guilds for Candida species. | Manolakaki D, et al. 2010. Virulence. 1: 367-375 ((https://doi.org/10.4161/viru.1.5.12796)); Mekbib SB, et al. 2011. Tropical Plant Pathology 36:233-240 ((https://doi.org/10.1590/S1982-56762011000400004)); Põlme S, et al. 2020. Fungal Diversity 105:1-16 ((https://doi.org/10.1007/s13225-020-00466-2)) |
| Otu20758 | 1 | k__Fungi(100);p__Ascomycota(100);c__Saccharomycetes(100);o__Saccharomycetales(100);f__Saccharomycetales_family_Incertae_sedis(100);g__Candida(100);g__Candida_unclassified(100); | Candida | 13 | Pathotroph-Saprotroph-Symbiotroph | Animal Pathogen-Endophyte-Endosymbiont-Epiphyte-\|Undefined Saprotroph\| | Dimorphic Yeast-Yeast | NULL | Probable | As nectar-tap_saprotroph (Põlme et al. 2020); Decay sugar-rich substrates (Põlme et al. 2020); This asexual genus is polyphyletic and contain species that function very differently depend on what they are. Caution should be taken when interpreting guilds for Candida species. | Manolakaki D, et al. 2010. Virulence. 1: 367-375 ((https://doi.org/10.4161/viru.1.5.12796)); Mekbib SB, et al. 2011. Tropical Plant Pathology 36:233-240 ((https://doi.org/10.1590/S1982-56762011000400004)); Põlme S, et al. 2020. Fungal Diversity 105:1-16 ((https://doi.org/10.1007/s13225-020-00466-2)) |
| Otu20759 | 1 | k__Fungi(100);p__Ascomycota(100);c__Saccharomycetes(100);o__Saccharomycetales(100);f__Saccharomycetales_family_Incertae_sedis(100);g__Candida(100);g__Candida_unclassified(100); | Candida | 13 | Pathotroph-Saprotroph-Symbiotroph | Animal Pathogen-Endophyte-Endosymbiont-Epiphyte-\|Undefined Saprotroph\| | Dimorphic Yeast-Yeast | NULL | Probable | As nectar-tap_saprotroph (Põlme et al. 2020); Decay sugar-rich substrates (Põlme et al. 2020); This asexual genus is polyphyletic and contain species that function very differently depend on what they are. Caution should be taken when interpreting guilds for Candida species. | Manolakaki D, et al. 2010. Virulence. 1: 367-375 ((https://doi.org/10.4161/viru.1.5.12796)); Mekbib SB, et al. 2011. Tropical Plant Pathology 36:233-240 ((https://doi.org/10.1590/S1982-56762011000400004)); Põlme S, et al. 2020. Fungal Diversity 105:1-16 ((https://doi.org/10.1007/s13225-020-00466-2)) |
| Otu20761 | 1 | k__Fungi(100);p__Ascomycota(100);c__Saccharomycetes(100);o__Saccharomycetales(100);f__Metschnikowiaceae(100);g__Metschnikowia(100);s__Metschnikowia_pulcherrima(100); | Metschnikowia | 13 | Saprotroph | \|Undefined Saprotroph\| | Yeast | NULL | Probable | As nectar-tap_saprotroph (Põlme et al. 2020); Decay sugar-rich substrates (Põlme et al. 2020) | Põlme S, et al. 2020. Fungal Diversity 105:1-16 ((https://doi.org/10.1007/s13225-020-00466-2)) |
| Otu20762 | 1 | k__Fungi(100);p__Ascomycota(100);c__Saccharomycetes(100);o__Saccharomycetales(100);f__Metschnikowiaceae(100);g__Metschnikowia(100);s__Metschnikowia_pulcherrima(100); | Metschnikowia | 13 | Saprotroph | \|Undefined Saprotroph\| | Yeast | NULL | Probable | As nectar-tap_saprotroph (Põlme et al. 2020); Decay sugar-rich substrates (Põlme et al. 2020) | Põlme S, et al. 2020. Fungal Diversity 105:1-16 ((https://doi.org/10.1007/s13225-020-00466-2)) |
| Otu20763 | 1 | k__Fungi(100);p__Ascomycota(100);c__Saccharomycetes(100);o__Saccharomycetales(100);f__Metschnikowiaceae(100);g__Metschnikowia(100);s__Metschnikowia_pulcherrima(100); | Metschnikowia | 13 | Saprotroph | \|Undefined Saprotroph\| | Yeast | NULL | Probable | As nectar-tap_saprotroph (Põlme et al. 2020); Decay sugar-rich substrates (Põlme et al. 2020) | Põlme S, et al. 2020. Fungal Diversity 105:1-16 ((https://doi.org/10.1007/s13225-020-00466-2)) |
| Otu20764 | 1 | k__Fungi(100);p__Ascomycota(100);c__Saccharomycetes(100);o__Saccharomycetales(100);f__Metschnikowiaceae(100);g__Metschnikowia(100);g__Metschnikowia_unclassified(100); | Metschnikowia | 13 | Saprotroph | \|Undefined Saprotroph\| | Yeast | NULL | Probable | As nectar-tap_saprotroph (Põlme et al. 2020); Decay sugar-rich substrates (Põlme et al. 2020) | Põlme S, et al. 2020. Fungal Diversity 105:1-16 ((https://doi.org/10.1007/s13225-020-00466-2)) |
| Otu20765 | 1 | k__Fungi(100);p__Ascomycota(100);c__Saccharomycetes(100);o__Saccharomycetales(100);f__Metschnikowiaceae(100);g__Metschnikowia(100);s__Metschnikowia_pulcherrima(100); | Metschnikowia | 13 | Saprotroph | \|Undefined Saprotroph\| | Yeast | NULL | Probable | As nectar-tap_saprotroph (Põlme et al. 2020); Decay sugar-rich substrates (Põlme et al. 2020) | Põlme S, et al. 2020. Fungal Diversity 105:1-16 ((https://doi.org/10.1007/s13225-020-00466-2)) |
| Otu20766 | 1 | k__Fungi(100);p__Ascomycota(100);c__Saccharomycetes(100);o__Saccharomycetales(100);f__Metschnikowiaceae(100);g__Metschnikowia(100);s__Metschnikowia_pulcherrima(100); | Metschnikowia | 13 | Saprotroph | \|Undefined Saprotroph\| | Yeast | NULL | Probable | As nectar-tap_saprotroph (Põlme et al. 2020); Decay sugar-rich substrates (Põlme et al. 2020) | Põlme S, et al. 2020. Fungal Diversity 105:1-16 ((https://doi.org/10.1007/s13225-020-00466-2)) |
| Otu20767 | 1 | k__Fungi(100);p__Ascomycota(100);c__Saccharomycetes(100);o__Saccharomycetales(100);f__Metschnikowiaceae(100);g__Metschnikowia(100);s__Metschnikowia_pulcherrima(100); | Metschnikowia | 13 | Saprotroph | \|Undefined Saprotroph\| | Yeast | NULL | Probable | As nectar-tap_saprotroph (Põlme et al. 2020); Decay sugar-rich substrates (Põlme et al. 2020) | Põlme S, et al. 2020. Fungal Diversity 105:1-16 ((https://doi.org/10.1007/s13225-020-00466-2)) |
| Otu20768 | 1 | k__Fungi(100);p__Ascomycota(100);c__Saccharomycetes(100);o__Saccharomycetales(100);f__Metschnikowiaceae(100);g__Metschnikowia(100);s__Metschnikowia_pulcherrima(100); | Metschnikowia | 13 | Saprotroph | \|Undefined Saprotroph\| | Yeast | NULL | Probable | As nectar-tap_saprotroph (Põlme et al. 2020); Decay sugar-rich substrates (Põlme et al. 2020) | Põlme S, et al. 2020. Fungal Diversity 105:1-16 ((https://doi.org/10.1007/s13225-020-00466-2)) |
| Otu20770 | 1 | k__Fungi(100);p__Ascomycota(100);c__Saccharomycetes(100);o__Saccharomycetales(100);f__Metschnikowiaceae(100);g__Metschnikowia(100);s__Metschnikowia_pulcherrima(100); | Metschnikowia | 13 | Saprotroph | \|Undefined Saprotroph\| | Yeast | NULL | Probable | As nectar-tap_saprotroph (Põlme et al. 2020); Decay sugar-rich substrates (Põlme et al. 2020) | Põlme S, et al. 2020. Fungal Diversity 105:1-16 ((https://doi.org/10.1007/s13225-020-00466-2)) |
| Otu20777 | 1 | k__Fungi(100);p__Ascomycota(100);c__Saccharomycetes(100);o__Saccharomycetales(100);f__Metschnikowiaceae(100);g__Metschnikowia(100);s__Metschnikowia_pulcherrima(100); | Metschnikowia | 13 | Saprotroph | \|Undefined Saprotroph\| | Yeast | NULL | Probable | As nectar-tap_saprotroph (Põlme et al. 2020); Decay sugar-rich substrates (Põlme et al. 2020) | Põlme S, et al. 2020. Fungal Diversity 105:1-16 ((https://doi.org/10.1007/s13225-020-00466-2)) |
| Otu20778 | 1 | k__Fungi(100);p__Ascomycota(100);c__Saccharomycetes(100);o__Saccharomycetales(100);f__Metschnikowiaceae(100);g__Metschnikowia(100);s__Metschnikowia_pulcherrima(100); | Metschnikowia | 13 | Saprotroph | \|Undefined Saprotroph\| | Yeast | NULL | Probable | As nectar-tap_saprotroph (Põlme et al. 2020); Decay sugar-rich substrates (Põlme et al. 2020) | Põlme S, et al. 2020. Fungal Diversity 105:1-16 ((https://doi.org/10.1007/s13225-020-00466-2)) |
| Otu20781 | 1 | k__Fungi(100);p__Ascomycota(100);c__Saccharomycetes(100);o__Saccharomycetales(100);f__Metschnikowiaceae(100);g__Metschnikowia(100);s__Metschnikowia_pulcherrima(100); | Metschnikowia | 13 | Saprotroph | \|Undefined Saprotroph\| | Yeast | NULL | Probable | As nectar-tap_saprotroph (Põlme et al. 2020); Decay sugar-rich substrates (Põlme et al. 2020) | Põlme S, et al. 2020. Fungal Diversity 105:1-16 ((https://doi.org/10.1007/s13225-020-00466-2)) |
| Otu20784 | 1 | k__Fungi(100);p__Ascomycota(100);c__Saccharomycetes(100);o__Saccharomycetales(100);f__Metschnikowiaceae(100);g__Metschnikowia(100);s__Metschnikowia_pulcherrima(100); | Metschnikowia | 13 | Saprotroph | \|Undefined Saprotroph\| | Yeast | NULL | Probable | As nectar-tap_saprotroph (Põlme et al. 2020); Decay sugar-rich substrates (Põlme et al. 2020) | Põlme S, et al. 2020. Fungal Diversity 105:1-16 ((https://doi.org/10.1007/s13225-020-00466-2)) |
| Otu20785 | 1 | k__Fungi(100);p__Ascomycota(100);c__Saccharomycetes(100);o__Saccharomycetales(100);f__Metschnikowiaceae(100);g__Metschnikowia(100);s__Metschnikowia_pulcherrima(100); | Metschnikowia | 13 | Saprotroph | \|Undefined Saprotroph\| | Yeast | NULL | Probable | As nectar-tap_saprotroph (Põlme et al. 2020); Decay sugar-rich substrates (Põlme et al. 2020) | Põlme S, et al. 2020. Fungal Diversity 105:1-16 ((https://doi.org/10.1007/s13225-020-00466-2)) |
| Otu20786 | 1 | k__Fungi(100);p__Ascomycota(100);c__Saccharomycetes(100);o__Saccharomycetales(100);f__Metschnikowiaceae(100);g__Metschnikowia(100);s__Metschnikowia_pulcherrima(100); | Metschnikowia | 13 | Saprotroph | \|Undefined Saprotroph\| | Yeast | NULL | Probable | As nectar-tap_saprotroph (Põlme et al. 2020); Decay sugar-rich substrates (Põlme et al. 2020) | Põlme S, et al. 2020. Fungal Diversity 105:1-16 ((https://doi.org/10.1007/s13225-020-00466-2)) |
| Otu20787 | 1 | k__Fungi(100);p__Ascomycota(100);c__Saccharomycetes(100);o__Saccharomycetales(100);f__Metschnikowiaceae(100);g__Metschnikowia(100);s__Metschnikowia_pulcherrima(100); | Metschnikowia | 13 | Saprotroph | \|Undefined Saprotroph\| | Yeast | NULL | Probable | As nectar-tap_saprotroph (Põlme et al. 2020); Decay sugar-rich substrates (Põlme et al. 2020) | Põlme S, et al. 2020. Fungal Diversity 105:1-16 ((https://doi.org/10.1007/s13225-020-00466-2)) |
| Otu20788 | 1 | k__Fungi(100);p__Ascomycota(100);c__Saccharomycetes(100);o__Saccharomycetales(100);f__Metschnikowiaceae(100);g__Metschnikowia(100);s__Metschnikowia_pulcherrima(100); | Metschnikowia | 13 | Saprotroph | \|Undefined Saprotroph\| | Yeast | NULL | Probable | As nectar-tap_saprotroph (Põlme et al. 2020); Decay sugar-rich substrates (Põlme et al. 2020) | Põlme S, et al. 2020. Fungal Diversity 105:1-16 ((https://doi.org/10.1007/s13225-020-00466-2)) |
| Otu20789 | 1 | k__Fungi(100);p__Ascomycota(100);c__Saccharomycetes(100);o__Saccharomycetales(100);f__Metschnikowiaceae(100);g__Metschnikowia(100);s__Metschnikowia_pulcherrima(100); | Metschnikowia | 13 | Saprotroph | \|Undefined Saprotroph\| | Yeast | NULL | Probable | As nectar-tap_saprotroph (Põlme et al. 2020); Decay sugar-rich substrates (Põlme et al. 2020) | Põlme S, et al. 2020. Fungal Diversity 105:1-16 ((https://doi.org/10.1007/s13225-020-00466-2)) |
| Otu20790 | 1 | k__Fungi(100);p__Ascomycota(100);c__Saccharomycetes(100);o__Saccharomycetales(100);f__Metschnikowiaceae(100);g__Metschnikowia(100);s__Metschnikowia_pulcherrima(100); | Metschnikowia | 13 | Saprotroph | \|Undefined Saprotroph\| | Yeast | NULL | Probable | As nectar-tap_saprotroph (Põlme et al. 2020); Decay sugar-rich substrates (Põlme et al. 2020) | Põlme S, et al. 2020. Fungal Diversity 105:1-16 ((https://doi.org/10.1007/s13225-020-00466-2)) |
| Otu20791 | 1 | k__Fungi(100);p__Ascomycota(100);c__Saccharomycetes(100);o__Saccharomycetales(100);f__Metschnikowiaceae(100);g__Metschnikowia(100);s__Metschnikowia_pulcherrima(100); | Metschnikowia | 13 | Saprotroph | \|Undefined Saprotroph\| | Yeast | NULL | Probable | As nectar-tap_saprotroph (Põlme et al. 2020); Decay sugar-rich substrates (Põlme et al. 2020) | Põlme S, et al. 2020. Fungal Diversity 105:1-16 ((https://doi.org/10.1007/s13225-020-00466-2)) |
| Otu20792 | 1 | k__Fungi(100);p__Ascomycota(100);c__Saccharomycetes(100);o__Saccharomycetales(100);f__Metschnikowiaceae(100);g__Metschnikowia(100);g__Metschnikowia_unclassified(100); | Metschnikowia | 13 | Saprotroph | \|Undefined Saprotroph\| | Yeast | NULL | Probable | As nectar-tap_saprotroph (Põlme et al. 2020); Decay sugar-rich substrates (Põlme et al. 2020) | Põlme S, et al. 2020. Fungal Diversity 105:1-16 ((https://doi.org/10.1007/s13225-020-00466-2)) |
| Otu20793 | 1 | k__Fungi(100);p__Ascomycota(100);c__Saccharomycetes(100);o__Saccharomycetales(100);f__Metschnikowiaceae(100);g__Metschnikowia(100);s__Metschnikowia_pulcherrima(100); | Metschnikowia | 13 | Saprotroph | \|Undefined Saprotroph\| | Yeast | NULL | Probable | As nectar-tap_saprotroph (Põlme et al. 2020); Decay sugar-rich substrates (Põlme et al. 2020) | Põlme S, et al. 2020. Fungal Diversity 105:1-16 ((https://doi.org/10.1007/s13225-020-00466-2)) |
| Otu20794 | 1 | k__Fungi(100);p__Ascomycota(100);c__Saccharomycetes(100);o__Saccharomycetales(100);f__Metschnikowiaceae(100);g__Metschnikowia(100);s__Metschnikowia_pulcherrima(100); | Metschnikowia | 13 | Saprotroph | \|Undefined Saprotroph\| | Yeast | NULL | Probable | As nectar-tap_saprotroph (Põlme et al. 2020); Decay sugar-rich substrates (Põlme et al. 2020) | Põlme S, et al. 2020. Fungal Diversity 105:1-16 ((https://doi.org/10.1007/s13225-020-00466-2)) |
| Otu20795 | 1 | k__Fungi(100);p__Ascomycota(100);c__Saccharomycetes(100);o__Saccharomycetales(100);f__Metschnikowiaceae(100);g__Metschnikowia(100);s__Metschnikowia_pulcherrima(100); | Metschnikowia | 13 | Saprotroph | \|Undefined Saprotroph\| | Yeast | NULL | Probable | As nectar-tap_saprotroph (Põlme et al. 2020); Decay sugar-rich substrates (Põlme et al. 2020) | Põlme S, et al. 2020. Fungal Diversity 105:1-16 ((https://doi.org/10.1007/s13225-020-00466-2)) |
| Otu20796 | 1 | k__Fungi(100);p__Ascomycota(100);c__Saccharomycetes(100);o__Saccharomycetales(100);f__Metschnikowiaceae(100);g__Metschnikowia(100);g__Metschnikowia_unclassified(100); | Metschnikowia | 13 | Saprotroph | \|Undefined Saprotroph\| | Yeast | NULL | Probable | As nectar-tap_saprotroph (Põlme et al. 2020); Decay sugar-rich substrates (Põlme et al. 2020) | Põlme S, et al. 2020. Fungal Diversity 105:1-16 ((https://doi.org/10.1007/s13225-020-00466-2)) |
| Otu20797 | 1 | k__Fungi(100);p__Ascomycota(100);c__Saccharomycetes(100);o__Saccharomycetales(100);f__Metschnikowiaceae(100);g__Metschnikowia(100);s__Metschnikowia_sp(100); | Metschnikowia | 13 | Saprotroph | \|Undefined Saprotroph\| | Yeast | NULL | Probable | As nectar-tap_saprotroph (Põlme et al. 2020); Decay sugar-rich substrates (Põlme et al. 2020) | Põlme S, et al. 2020. Fungal Diversity 105:1-16 ((https://doi.org/10.1007/s13225-020-00466-2)) |
| Otu20798 | 1 | k__Fungi(100);p__Ascomycota(100);c__Saccharomycetes(100);o__Saccharomycetales(100);f__Metschnikowiaceae(100);g__Metschnikowia(100);s__Metschnikowia_pulcherrima(100); | Metschnikowia | 13 | Saprotroph | \|Undefined Saprotroph\| | Yeast | NULL | Probable | As nectar-tap_saprotroph (Põlme et al. 2020); Decay sugar-rich substrates (Põlme et al. 2020) | Põlme S, et al. 2020. Fungal Diversity 105:1-16 ((https://doi.org/10.1007/s13225-020-00466-2)) |
| Otu20799 | 1 | k__Fungi(100);p__Ascomycota(100);c__Saccharomycetes(100);o__Saccharomycetales(100);f__Metschnikowiaceae(100);g__Metschnikowia(100);g__Metschnikowia_unclassified(100); | Metschnikowia | 13 | Saprotroph | \|Undefined Saprotroph\| | Yeast | NULL | Probable | As nectar-tap_saprotroph (Põlme et al. 2020); Decay sugar-rich substrates (Põlme et al. 2020) | Põlme S, et al. 2020. Fungal Diversity 105:1-16 ((https://doi.org/10.1007/s13225-020-00466-2)) |
| Otu20800 | 1 | k__Fungi(100);p__Ascomycota(100);c__Saccharomycetes(100);o__Saccharomycetales(100);o__Saccharomycetales_unclassified(100);o__Saccharomycetales_unclassified(100);o__Saccharomycetales_unclassified(100); | Saccharomycetales | 7 | Saprotroph | Undefined Saprotroph | Yeast | NULL | Possible | NULL | Sterkenburg E, et al. 2015. New Phytologist 207:1145-1158 ((https://doi.org/10.1111/nph.13426)) |
| Otu20801 | 1 | k__Fungi(100);p__Ascomycota(100);c__Saccharomycetes(100);o__Saccharomycetales(100);f__Metschnikowiaceae(100);g__Metschnikowia(100);s__Metschnikowia_pulcherrima(100); | Metschnikowia | 13 | Saprotroph | \|Undefined Saprotroph\| | Yeast | NULL | Probable | As nectar-tap_saprotroph (Põlme et al. 2020); Decay sugar-rich substrates (Põlme et al. 2020) | Põlme S, et al. 2020. Fungal Diversity 105:1-16 ((https://doi.org/10.1007/s13225-020-00466-2)) |
| Otu20802 | 1 | k__Fungi(100);p__Ascomycota(100);c__Saccharomycetes(100);o__Saccharomycetales(100);f__Metschnikowiaceae(100);g__Metschnikowia(100);g__Metschnikowia_unclassified(100); | Metschnikowia | 13 | Saprotroph | \|Undefined Saprotroph\| | Yeast | NULL | Probable | As nectar-tap_saprotroph (Põlme et al. 2020); Decay sugar-rich substrates (Põlme et al. 2020) | Põlme S, et al. 2020. Fungal Diversity 105:1-16 ((https://doi.org/10.1007/s13225-020-00466-2)) |
| Otu20804 | 1 | k__Fungi(100);p__Ascomycota(100);c__Saccharomycetes(100);o__Saccharomycetales(100);f__Metschnikowiaceae(100);g__Metschnikowia(100);s__Metschnikowia_pulcherrima(100); | Metschnikowia | 13 | Saprotroph | \|Undefined Saprotroph\| | Yeast | NULL | Probable | As nectar-tap_saprotroph (Põlme et al. 2020); Decay sugar-rich substrates (Põlme et al. 2020) | Põlme S, et al. 2020. Fungal Diversity 105:1-16 ((https://doi.org/10.1007/s13225-020-00466-2)) |
| Otu20806 | 1 | k__Fungi(100);p__Ascomycota(100);c__Saccharomycetes(100);o__Saccharomycetales(100);f__Metschnikowiaceae(100);g__Metschnikowia(100);s__Metschnikowia_pulcherrima(100); | Metschnikowia | 13 | Saprotroph | \|Undefined Saprotroph\| | Yeast | NULL | Probable | As nectar-tap_saprotroph (Põlme et al. 2020); Decay sugar-rich substrates (Põlme et al. 2020) | Põlme S, et al. 2020. Fungal Diversity 105:1-16 ((https://doi.org/10.1007/s13225-020-00466-2)) |
| Otu20808 | 1 | k__Fungi(100);p__Ascomycota(100);c__Saccharomycetes(100);o__Saccharomycetales(100);f__Saccharomycetales_family_Incertae_sedis(100);g__Candida(100);g__Candida_unclassified(100); | Candida | 13 | Pathotroph-Saprotroph-Symbiotroph | Animal Pathogen-Endophyte-Endosymbiont-Epiphyte-\|Undefined Saprotroph\| | Dimorphic Yeast-Yeast | NULL | Probable | As nectar-tap_saprotroph (Põlme et al. 2020); Decay sugar-rich substrates (Põlme et al. 2020); This asexual genus is polyphyletic and contain species that function very differently depend on what they are. Caution should be taken when interpreting guilds for Candida species. | Manolakaki D, et al. 2010. Virulence. 1: 367-375 ((https://doi.org/10.4161/viru.1.5.12796)); Mekbib SB, et al. 2011. Tropical Plant Pathology 36:233-240 ((https://doi.org/10.1590/S1982-56762011000400004)); Põlme S, et al. 2020. Fungal Diversity 105:1-16 ((https://doi.org/10.1007/s13225-020-00466-2)) |
| Otu20813 | 1 | k__Fungi(100);p__Ascomycota(100);c__Saccharomycetes(100);o__Saccharomycetales(100);f__Metschnikowiaceae(100);g__Metschnikowia(100);s__Metschnikowia_pulcherrima(100); | Metschnikowia | 13 | Saprotroph | \|Undefined Saprotroph\| | Yeast | NULL | Probable | As nectar-tap_saprotroph (Põlme et al. 2020); Decay sugar-rich substrates (Põlme et al. 2020) | Põlme S, et al. 2020. Fungal Diversity 105:1-16 ((https://doi.org/10.1007/s13225-020-00466-2)) |
| Otu20814 | 1 | k__Fungi(100);p__Ascomycota(100);c__Saccharomycetes(100);o__Saccharomycetales(100);f__Metschnikowiaceae(100);g__Metschnikowia(100);g__Metschnikowia_unclassified(100); | Metschnikowia | 13 | Saprotroph | \|Undefined Saprotroph\| | Yeast | NULL | Probable | As nectar-tap_saprotroph (Põlme et al. 2020); Decay sugar-rich substrates (Põlme et al. 2020) | Põlme S, et al. 2020. Fungal Diversity 105:1-16 ((https://doi.org/10.1007/s13225-020-00466-2)) |
| Otu20815 | 1 | k__Fungi(100);p__Ascomycota(100);c__Saccharomycetes(100);o__Saccharomycetales(100);f__Metschnikowiaceae(100);g__Metschnikowia(100);s__Metschnikowia_pulcherrima(100); | Metschnikowia | 13 | Saprotroph | \|Undefined Saprotroph\| | Yeast | NULL | Probable | As nectar-tap_saprotroph (Põlme et al. 2020); Decay sugar-rich substrates (Põlme et al. 2020) | Põlme S, et al. 2020. Fungal Diversity 105:1-16 ((https://doi.org/10.1007/s13225-020-00466-2)) |
| Otu20816 | 1 | k__Fungi(100);p__Ascomycota(100);c__Saccharomycetes(100);o__Saccharomycetales(100);f__Metschnikowiaceae(100);g__Metschnikowia(100);s__Metschnikowia_pulcherrima(100); | Metschnikowia | 13 | Saprotroph | \|Undefined Saprotroph\| | Yeast | NULL | Probable | As nectar-tap_saprotroph (Põlme et al. 2020); Decay sugar-rich substrates (Põlme et al. 2020) | Põlme S, et al. 2020. Fungal Diversity 105:1-16 ((https://doi.org/10.1007/s13225-020-00466-2)) |
| Otu20817 | 1 | k__Fungi(100);p__Ascomycota(100);c__Saccharomycetes(100);o__Saccharomycetales(100);f__Metschnikowiaceae(100);g__Metschnikowia(100);s__Metschnikowia_pulcherrima(100); | Metschnikowia | 13 | Saprotroph | \|Undefined Saprotroph\| | Yeast | NULL | Probable | As nectar-tap_saprotroph (Põlme et al. 2020); Decay sugar-rich substrates (Põlme et al. 2020) | Põlme S, et al. 2020. Fungal Diversity 105:1-16 ((https://doi.org/10.1007/s13225-020-00466-2)) |
| Otu20818 | 1 | k__Fungi(100);p__Ascomycota(100);c__Saccharomycetes(100);o__Saccharomycetales(100);f__Metschnikowiaceae(100);g__Metschnikowia(100);s__Metschnikowia_pulcherrima(100); | Metschnikowia | 13 | Saprotroph | \|Undefined Saprotroph\| | Yeast | NULL | Probable | As nectar-tap_saprotroph (Põlme et al. 2020); Decay sugar-rich substrates (Põlme et al. 2020) | Põlme S, et al. 2020. Fungal Diversity 105:1-16 ((https://doi.org/10.1007/s13225-020-00466-2)) |
| Otu20819 | 1 | k__Fungi(100);p__Ascomycota(100);c__Saccharomycetes(100);o__Saccharomycetales(100);f__Metschnikowiaceae(100);g__Metschnikowia(100);g__Metschnikowia_unclassified(100); | Metschnikowia | 13 | Saprotroph | \|Undefined Saprotroph\| | Yeast | NULL | Probable | As nectar-tap_saprotroph (Põlme et al. 2020); Decay sugar-rich substrates (Põlme et al. 2020) | Põlme S, et al. 2020. Fungal Diversity 105:1-16 ((https://doi.org/10.1007/s13225-020-00466-2)) |
| Otu20820 | 1 | k__Fungi(100);p__Ascomycota(100);c__Saccharomycetes(100);o__Saccharomycetales(100);f__Metschnikowiaceae(100);g__Metschnikowia(100);g__Metschnikowia_unclassified(100); | Metschnikowia | 13 | Saprotroph | \|Undefined Saprotroph\| | Yeast | NULL | Probable | As nectar-tap_saprotroph (Põlme et al. 2020); Decay sugar-rich substrates (Põlme et al. 2020) | Põlme S, et al. 2020. Fungal Diversity 105:1-16 ((https://doi.org/10.1007/s13225-020-00466-2)) |
| Otu20822 | 1 | k__Fungi(100);p__Ascomycota(100);c__Saccharomycetes(100);o__Saccharomycetales(100);f__Metschnikowiaceae(100);g__Metschnikowia(100);s__Metschnikowia_pulcherrima(100); | Metschnikowia | 13 | Saprotroph | \|Undefined Saprotroph\| | Yeast | NULL | Probable | As nectar-tap_saprotroph (Põlme et al. 2020); Decay sugar-rich substrates (Põlme et al. 2020) | Põlme S, et al. 2020. Fungal Diversity 105:1-16 ((https://doi.org/10.1007/s13225-020-00466-2)) |
| Otu20823 | 1 | k__Fungi(100);p__Ascomycota(100);c__Saccharomycetes(100);o__Saccharomycetales(100);f__Metschnikowiaceae(100);g__Metschnikowia(100);s__Metschnikowia_pulcherrima(100); | Metschnikowia | 13 | Saprotroph | \|Undefined Saprotroph\| | Yeast | NULL | Probable | As nectar-tap_saprotroph (Põlme et al. 2020); Decay sugar-rich substrates (Põlme et al. 2020) | Põlme S, et al. 2020. Fungal Diversity 105:1-16 ((https://doi.org/10.1007/s13225-020-00466-2)) |
| Otu20824 | 1 | k__Fungi(100);p__Ascomycota(100);c__Saccharomycetes(100);o__Saccharomycetales(100);f__Saccharomycetales_family_Incertae_sedis(100);f__Saccharomycetales_family_Incertae_sedis_unclassified(100);f__Saccharomycetales_family_Incertae_sedis_unclassified(100); | Saccharomycetales | 7 | Saprotroph | Undefined Saprotroph | Yeast | NULL | Possible | NULL | Sterkenburg E, et al. 2015. New Phytologist 207:1145-1158 ((https://doi.org/10.1111/nph.13426)) |
| Otu20825 | 1 | k__Fungi(100);p__Ascomycota(100);c__Saccharomycetes(100);o__Saccharomycetales(100);f__Metschnikowiaceae(100);g__Metschnikowia(100);g__Metschnikowia_unclassified(100); | Metschnikowia | 13 | Saprotroph | \|Undefined Saprotroph\| | Yeast | NULL | Probable | As nectar-tap_saprotroph (Põlme et al. 2020); Decay sugar-rich substrates (Põlme et al. 2020) | Põlme S, et al. 2020. Fungal Diversity 105:1-16 ((https://doi.org/10.1007/s13225-020-00466-2)) |
| Otu20829 | 1 | k__Fungi(100);p__Ascomycota(100);c__Saccharomycetes(100);o__Saccharomycetales(100);f__Metschnikowiaceae(100);g__Metschnikowia(100);s__Metschnikowia_pulcherrima(100); | Metschnikowia | 13 | Saprotroph | \|Undefined Saprotroph\| | Yeast | NULL | Probable | As nectar-tap_saprotroph (Põlme et al. 2020); Decay sugar-rich substrates (Põlme et al. 2020) | Põlme S, et al. 2020. Fungal Diversity 105:1-16 ((https://doi.org/10.1007/s13225-020-00466-2)) |
| Otu20831 | 1 | k__Fungi(100);p__Ascomycota(100);c__Saccharomycetes(100);o__Saccharomycetales(100);f__Metschnikowiaceae(100);g__Metschnikowia(100);s__Metschnikowia_pulcherrima(100); | Metschnikowia | 13 | Saprotroph | \|Undefined Saprotroph\| | Yeast | NULL | Probable | As nectar-tap_saprotroph (Põlme et al. 2020); Decay sugar-rich substrates (Põlme et al. 2020) | Põlme S, et al. 2020. Fungal Diversity 105:1-16 ((https://doi.org/10.1007/s13225-020-00466-2)) |
| Otu20832 | 1 | k__Fungi(100);p__Ascomycota(100);c__Saccharomycetes(100);o__Saccharomycetales(100);f__Metschnikowiaceae(100);g__Metschnikowia(100);s__Metschnikowia_pulcherrima(100); | Metschnikowia | 13 | Saprotroph | \|Undefined Saprotroph\| | Yeast | NULL | Probable | As nectar-tap_saprotroph (Põlme et al. 2020); Decay sugar-rich substrates (Põlme et al. 2020) | Põlme S, et al. 2020. Fungal Diversity 105:1-16 ((https://doi.org/10.1007/s13225-020-00466-2)) |
| Otu20833 | 1 | k__Fungi(100);p__Ascomycota(100);c__Saccharomycetes(100);o__Saccharomycetales(100);f__Metschnikowiaceae(100);g__Metschnikowia(100);s__Metschnikowia_pulcherrima(100); | Metschnikowia | 13 | Saprotroph | \|Undefined Saprotroph\| | Yeast | NULL | Probable | As nectar-tap_saprotroph (Põlme et al. 2020); Decay sugar-rich substrates (Põlme et al. 2020) | Põlme S, et al. 2020. Fungal Diversity 105:1-16 ((https://doi.org/10.1007/s13225-020-00466-2)) |
| Otu20834 | 1 | k__Fungi(100);p__Ascomycota(100);c__Saccharomycetes(100);o__Saccharomycetales(100);f__Metschnikowiaceae(100);g__Metschnikowia(100);g__Metschnikowia_unclassified(100); | Metschnikowia | 13 | Saprotroph | \|Undefined Saprotroph\| | Yeast | NULL | Probable | As nectar-tap_saprotroph (Põlme et al. 2020); Decay sugar-rich substrates (Põlme et al. 2020) | Põlme S, et al. 2020. Fungal Diversity 105:1-16 ((https://doi.org/10.1007/s13225-020-00466-2)) |
| Otu20835 | 1 | k__Fungi(100);p__Ascomycota(100);c__Saccharomycetes(100);o__Saccharomycetales(100);f__Metschnikowiaceae(100);g__Metschnikowia(100);s__Metschnikowia_pulcherrima(100); | Metschnikowia | 13 | Saprotroph | \|Undefined Saprotroph\| | Yeast | NULL | Probable | As nectar-tap_saprotroph (Põlme et al. 2020); Decay sugar-rich substrates (Põlme et al. 2020) | Põlme S, et al. 2020. Fungal Diversity 105:1-16 ((https://doi.org/10.1007/s13225-020-00466-2)) |
| Otu20836 | 1 | k__Fungi(100);p__Ascomycota(100);c__Saccharomycetes(100);o__Saccharomycetales(100);f__Metschnikowiaceae(100);g__Metschnikowia(100);g__Metschnikowia_unclassified(100); | Metschnikowia | 13 | Saprotroph | \|Undefined Saprotroph\| | Yeast | NULL | Probable | As nectar-tap_saprotroph (Põlme et al. 2020); Decay sugar-rich substrates (Põlme et al. 2020) | Põlme S, et al. 2020. Fungal Diversity 105:1-16 ((https://doi.org/10.1007/s13225-020-00466-2)) |
| Otu20837 | 1 | k__Fungi(100);p__Ascomycota(100);c__Saccharomycetes(100);o__Saccharomycetales(100);f__Metschnikowiaceae(100);g__Metschnikowia(100);s__Metschnikowia_pulcherrima(100); | Metschnikowia | 13 | Saprotroph | \|Undefined Saprotroph\| | Yeast | NULL | Probable | As nectar-tap_saprotroph (Põlme et al. 2020); Decay sugar-rich substrates (Põlme et al. 2020) | Põlme S, et al. 2020. Fungal Diversity 105:1-16 ((https://doi.org/10.1007/s13225-020-00466-2)) |
| Otu20838 | 1 | k__Fungi(100);p__Ascomycota(100);c__Saccharomycetes(100);o__Saccharomycetales(100);f__Metschnikowiaceae(100);g__Metschnikowia(100);s__Metschnikowia_pulcherrima(100); | Metschnikowia | 13 | Saprotroph | \|Undefined Saprotroph\| | Yeast | NULL | Probable | As nectar-tap_saprotroph (Põlme et al. 2020); Decay sugar-rich substrates (Põlme et al. 2020) | Põlme S, et al. 2020. Fungal Diversity 105:1-16 ((https://doi.org/10.1007/s13225-020-00466-2)) |
| Otu20839 | 1 | k__Fungi(100);p__Ascomycota(100);c__Saccharomycetes(100);o__Saccharomycetales(100);f__Metschnikowiaceae(100);g__Metschnikowia(100);g__Metschnikowia_unclassified(100); | Metschnikowia | 13 | Saprotroph | \|Undefined Saprotroph\| | Yeast | NULL | Probable | As nectar-tap_saprotroph (Põlme et al. 2020); Decay sugar-rich substrates (Põlme et al. 2020) | Põlme S, et al. 2020. Fungal Diversity 105:1-16 ((https://doi.org/10.1007/s13225-020-00466-2)) |
| Otu20840 | 1 | k__Fungi(100);p__Ascomycota(100);c__Saccharomycetes(100);o__Saccharomycetales(100);f__Metschnikowiaceae(100);g__Metschnikowia(100);s__Metschnikowia_pulcherrima(100); | Metschnikowia | 13 | Saprotroph | \|Undefined Saprotroph\| | Yeast | NULL | Probable | As nectar-tap_saprotroph (Põlme et al. 2020); Decay sugar-rich substrates (Põlme et al. 2020) | Põlme S, et al. 2020. Fungal Diversity 105:1-16 ((https://doi.org/10.1007/s13225-020-00466-2)) |
| Otu20841 | 1 | k__Fungi(100);p__Ascomycota(100);c__Saccharomycetes(100);o__Saccharomycetales(100);f__Metschnikowiaceae(100);g__Metschnikowia(100);s__Metschnikowia_pulcherrima(100); | Metschnikowia | 13 | Saprotroph | \|Undefined Saprotroph\| | Yeast | NULL | Probable | As nectar-tap_saprotroph (Põlme et al. 2020); Decay sugar-rich substrates (Põlme et al. 2020) | Põlme S, et al. 2020. Fungal Diversity 105:1-16 ((https://doi.org/10.1007/s13225-020-00466-2)) |
| Otu20842 | 1 | k__Fungi(100);p__Ascomycota(100);c__Saccharomycetes(100);o__Saccharomycetales(100);f__Metschnikowiaceae(100);g__Metschnikowia(100);s__Metschnikowia_pulcherrima(100); | Metschnikowia | 13 | Saprotroph | \|Undefined Saprotroph\| | Yeast | NULL | Probable | As nectar-tap_saprotroph (Põlme et al. 2020); Decay sugar-rich substrates (Põlme et al. 2020) | Põlme S, et al. 2020. Fungal Diversity 105:1-16 ((https://doi.org/10.1007/s13225-020-00466-2)) |
| Otu20843 | 1 | k__Fungi(100);p__Ascomycota(100);c__Saccharomycetes(100);o__Saccharomycetales(100);f__Metschnikowiaceae(100);g__Metschnikowia(100);s__Metschnikowia_pulcherrima(100); | Metschnikowia | 13 | Saprotroph | \|Undefined Saprotroph\| | Yeast | NULL | Probable | As nectar-tap_saprotroph (Põlme et al. 2020); Decay sugar-rich substrates (Põlme et al. 2020) | Põlme S, et al. 2020. Fungal Diversity 105:1-16 ((https://doi.org/10.1007/s13225-020-00466-2)) |
| Otu20844 | 1 | k__Fungi(100);p__Ascomycota(100);c__Saccharomycetes(100);o__Saccharomycetales(100);f__Metschnikowiaceae(100);g__Metschnikowia(100);g__Metschnikowia_unclassified(100); | Metschnikowia | 13 | Saprotroph | \|Undefined Saprotroph\| | Yeast | NULL | Probable | As nectar-tap_saprotroph (Põlme et al. 2020); Decay sugar-rich substrates (Põlme et al. 2020) | Põlme S, et al. 2020. Fungal Diversity 105:1-16 ((https://doi.org/10.1007/s13225-020-00466-2)) |
| Otu20846 | 1 | k__Fungi(100);p__Ascomycota(100);c__Saccharomycetes(100);o__Saccharomycetales(100);f__Metschnikowiaceae(100);g__Metschnikowia(100);s__Metschnikowia_pulcherrima(100); | Metschnikowia | 13 | Saprotroph | \|Undefined Saprotroph\| | Yeast | NULL | Probable | As nectar-tap_saprotroph (Põlme et al. 2020); Decay sugar-rich substrates (Põlme et al. 2020) | Põlme S, et al. 2020. Fungal Diversity 105:1-16 ((https://doi.org/10.1007/s13225-020-00466-2)) |
| Otu20847 | 1 | k__Fungi(100);p__Ascomycota(100);c__Saccharomycetes(100);o__Saccharomycetales(100);f__Metschnikowiaceae(100);g__Metschnikowia(100);s__Metschnikowia_pulcherrima(100); | Metschnikowia | 13 | Saprotroph | \|Undefined Saprotroph\| | Yeast | NULL | Probable | As nectar-tap_saprotroph (Põlme et al. 2020); Decay sugar-rich substrates (Põlme et al. 2020) | Põlme S, et al. 2020. Fungal Diversity 105:1-16 ((https://doi.org/10.1007/s13225-020-00466-2)) |
| Otu20848 | 1 | k__Fungi(100);p__Ascomycota(100);c__Saccharomycetes(100);o__Saccharomycetales(100);f__Saccharomycetales_family_Incertae_sedis(100);g__Candida(100);g__Candida_unclassified(100); | Candida | 13 | Pathotroph-Saprotroph-Symbiotroph | Animal Pathogen-Endophyte-Endosymbiont-Epiphyte-\|Undefined Saprotroph\| | Dimorphic Yeast-Yeast | NULL | Probable | As nectar-tap_saprotroph (Põlme et al. 2020); Decay sugar-rich substrates (Põlme et al. 2020); This asexual genus is polyphyletic and contain species that function very differently depend on what they are. Caution should be taken when interpreting guilds for Candida species. | Manolakaki D, et al. 2010. Virulence. 1: 367-375 ((https://doi.org/10.4161/viru.1.5.12796)); Mekbib SB, et al. 2011. Tropical Plant Pathology 36:233-240 ((https://doi.org/10.1590/S1982-56762011000400004)); Põlme S, et al. 2020. Fungal Diversity 105:1-16 ((https://doi.org/10.1007/s13225-020-00466-2)) |
| Otu20849 | 1 | k__Fungi(100);p__Ascomycota(100);c__Saccharomycetes(100);o__Saccharomycetales(100);f__Metschnikowiaceae(100);g__Metschnikowia(100);s__Metschnikowia_pulcherrima(100); | Metschnikowia | 13 | Saprotroph | \|Undefined Saprotroph\| | Yeast | NULL | Probable | As nectar-tap_saprotroph (Põlme et al. 2020); Decay sugar-rich substrates (Põlme et al. 2020) | Põlme S, et al. 2020. Fungal Diversity 105:1-16 ((https://doi.org/10.1007/s13225-020-00466-2)) |
| Otu20850 | 1 | k__Fungi(100);p__Ascomycota(100);c__Saccharomycetes(100);o__Saccharomycetales(100);f__Metschnikowiaceae(100);g__Metschnikowia(100);s__Metschnikowia_pulcherrima(100); | Metschnikowia | 13 | Saprotroph | \|Undefined Saprotroph\| | Yeast | NULL | Probable | As nectar-tap_saprotroph (Põlme et al. 2020); Decay sugar-rich substrates (Põlme et al. 2020) | Põlme S, et al. 2020. Fungal Diversity 105:1-16 ((https://doi.org/10.1007/s13225-020-00466-2)) |
| Otu20851 | 1 | k__Fungi(100);p__Ascomycota(100);c__Saccharomycetes(100);o__Saccharomycetales(100);f__Metschnikowiaceae(100);g__Metschnikowia(100);g__Metschnikowia_unclassified(100); | Metschnikowia | 13 | Saprotroph | \|Undefined Saprotroph\| | Yeast | NULL | Probable | As nectar-tap_saprotroph (Põlme et al. 2020); Decay sugar-rich substrates (Põlme et al. 2020) | Põlme S, et al. 2020. Fungal Diversity 105:1-16 ((https://doi.org/10.1007/s13225-020-00466-2)) |
| Otu20852 | 1 | k__Fungi(100);p__Ascomycota(100);c__Saccharomycetes(100);o__Saccharomycetales(100);f__Metschnikowiaceae(100);g__Metschnikowia(100);g__Metschnikowia_unclassified(100); | Metschnikowia | 13 | Saprotroph | \|Undefined Saprotroph\| | Yeast | NULL | Probable | As nectar-tap_saprotroph (Põlme et al. 2020); Decay sugar-rich substrates (Põlme et al. 2020) | Põlme S, et al. 2020. Fungal Diversity 105:1-16 ((https://doi.org/10.1007/s13225-020-00466-2)) |
| Otu20860 | 1 | k__Fungi(100);p__Ascomycota(100);c__Saccharomycetes(100);o__Saccharomycetales(100);f__Metschnikowiaceae(100);g__Metschnikowia(100);s__Metschnikowia_pulcherrima(100); | Metschnikowia | 13 | Saprotroph | \|Undefined Saprotroph\| | Yeast | NULL | Probable | As nectar-tap_saprotroph (Põlme et al. 2020); Decay sugar-rich substrates (Põlme et al. 2020) | Põlme S, et al. 2020. Fungal Diversity 105:1-16 ((https://doi.org/10.1007/s13225-020-00466-2)) |
| Otu20861 | 1 | k__Fungi(100);p__Ascomycota(100);c__Ascomycota_class_Incertae_sedis(100);o__Ascomycota_order_Incertae_sedis(100);f__Ascomycota_family_Incertae_sedis(100);g__Staphylotrichum(100);s__Staphylotrichum_sp_Amy_4(100); | Staphylotrichum | 13 | Saprotroph | \|Undefined Saprotroph\| | Microfungus | NULL | Probable | As soil_saprotroph (Põlme et al. 2020); Decay soil (Põlme et al. 2020) | Tedersoo L, et al. 2014. Science 346:e1256688 ((https://doi.org/10.1126/science.1256688)); Põlme S, et al. 2020. Fungal Diversity 105:1-16 ((https://doi.org/10.1007/s13225-020-00466-2)) |
| Otu20864 | 1 | k__Fungi(100);p__Ascomycota(100);c__Saccharomycetes(100);o__Saccharomycetales(100);f__Metschnikowiaceae(100);g__Metschnikowia(100);g__Metschnikowia_unclassified(100); | Metschnikowia | 13 | Saprotroph | \|Undefined Saprotroph\| | Yeast | NULL | Probable | As nectar-tap_saprotroph (Põlme et al. 2020); Decay sugar-rich substrates (Põlme et al. 2020) | Põlme S, et al. 2020. Fungal Diversity 105:1-16 ((https://doi.org/10.1007/s13225-020-00466-2)) |
| Otu20865 | 1 | k__Fungi(100);p__Ascomycota(100);c__Saccharomycetes(100);o__Saccharomycetales(100);f__Metschnikowiaceae(100);g__Metschnikowia(100);g__Metschnikowia_unclassified(100); | Metschnikowia | 13 | Saprotroph | \|Undefined Saprotroph\| | Yeast | NULL | Probable | As nectar-tap_saprotroph (Põlme et al. 2020); Decay sugar-rich substrates (Põlme et al. 2020) | Põlme S, et al. 2020. Fungal Diversity 105:1-16 ((https://doi.org/10.1007/s13225-020-00466-2)) |
| Otu20866 | 1 | k__Fungi(100);p__Ascomycota(100);c__Saccharomycetes(100);o__Saccharomycetales(100);f__Metschnikowiaceae(100);g__Metschnikowia(100);s__Metschnikowia_pulcherrima(100); | Metschnikowia | 13 | Saprotroph | \|Undefined Saprotroph\| | Yeast | NULL | Probable | As nectar-tap_saprotroph (Põlme et al. 2020); Decay sugar-rich substrates (Põlme et al. 2020) | Põlme S, et al. 2020. Fungal Diversity 105:1-16 ((https://doi.org/10.1007/s13225-020-00466-2)) |
| Otu20867 | 1 | k__Fungi(100);p__Ascomycota(100);c__Saccharomycetes(100);o__Saccharomycetales(100);f__Metschnikowiaceae(100);g__Metschnikowia(100);s__Metschnikowia_pulcherrima(100); | Metschnikowia | 13 | Saprotroph | \|Undefined Saprotroph\| | Yeast | NULL | Probable | As nectar-tap_saprotroph (Põlme et al. 2020); Decay sugar-rich substrates (Põlme et al. 2020) | Põlme S, et al. 2020. Fungal Diversity 105:1-16 ((https://doi.org/10.1007/s13225-020-00466-2)) |
| Otu20868 | 1 | k__Fungi(100);p__Ascomycota(100);c__Saccharomycetes(100);o__Saccharomycetales(100);f__Metschnikowiaceae(100);g__Metschnikowia(100);s__Metschnikowia_pulcherrima(100); | Metschnikowia | 13 | Saprotroph | \|Undefined Saprotroph\| | Yeast | NULL | Probable | As nectar-tap_saprotroph (Põlme et al. 2020); Decay sugar-rich substrates (Põlme et al. 2020) | Põlme S, et al. 2020. Fungal Diversity 105:1-16 ((https://doi.org/10.1007/s13225-020-00466-2)) |
| Otu20869 | 1 | k__Fungi(100);p__Ascomycota(100);c__Saccharomycetes(100);o__Saccharomycetales(100);f__Saccharomycetales_family_Incertae_sedis(100);g__Candida(100);g__Candida_unclassified(100); | Candida | 13 | Pathotroph-Saprotroph-Symbiotroph | Animal Pathogen-Endophyte-Endosymbiont-Epiphyte-\|Undefined Saprotroph\| | Dimorphic Yeast-Yeast | NULL | Probable | As nectar-tap_saprotroph (Põlme et al. 2020); Decay sugar-rich substrates (Põlme et al. 2020); This asexual genus is polyphyletic and contain species that function very differently depend on what they are. Caution should be taken when interpreting guilds for Candida species. | Manolakaki D, et al. 2010. Virulence. 1: 367-375 ((https://doi.org/10.4161/viru.1.5.12796)); Mekbib SB, et al. 2011. Tropical Plant Pathology 36:233-240 ((https://doi.org/10.1590/S1982-56762011000400004)); Põlme S, et al. 2020. Fungal Diversity 105:1-16 ((https://doi.org/10.1007/s13225-020-00466-2)) |
| Otu20871 | 1 | k__Fungi(100);p__Ascomycota(100);c__Saccharomycetes(100);o__Saccharomycetales(100);f__Metschnikowiaceae(100);g__Metschnikowia(100);s__Metschnikowia_pulcherrima(100); | Metschnikowia | 13 | Saprotroph | \|Undefined Saprotroph\| | Yeast | NULL | Probable | As nectar-tap_saprotroph (Põlme et al. 2020); Decay sugar-rich substrates (Põlme et al. 2020) | Põlme S, et al. 2020. Fungal Diversity 105:1-16 ((https://doi.org/10.1007/s13225-020-00466-2)) |
| Otu20872 | 1 | k__Fungi(100);p__Ascomycota(100);c__Saccharomycetes(100);o__Saccharomycetales(100);f__Metschnikowiaceae(100);g__Metschnikowia(100);s__Metschnikowia_pulcherrima(100); | Metschnikowia | 13 | Saprotroph | \|Undefined Saprotroph\| | Yeast | NULL | Probable | As nectar-tap_saprotroph (Põlme et al. 2020); Decay sugar-rich substrates (Põlme et al. 2020) | Põlme S, et al. 2020. Fungal Diversity 105:1-16 ((https://doi.org/10.1007/s13225-020-00466-2)) |
| Otu20873 | 1 | k__Fungi(100);p__Ascomycota(100);c__Saccharomycetes(100);o__Saccharomycetales(100);f__Metschnikowiaceae(100);g__Metschnikowia(100);s__Metschnikowia_pulcherrima(100); | Metschnikowia | 13 | Saprotroph | \|Undefined Saprotroph\| | Yeast | NULL | Probable | As nectar-tap_saprotroph (Põlme et al. 2020); Decay sugar-rich substrates (Põlme et al. 2020) | Põlme S, et al. 2020. Fungal Diversity 105:1-16 ((https://doi.org/10.1007/s13225-020-00466-2)) |
| Otu20874 | 1 | k__Fungi(100);p__Ascomycota(100);c__Saccharomycetes(100);o__Saccharomycetales(100);f__Metschnikowiaceae(100);g__Metschnikowia(100);s__Metschnikowia_pulcherrima(100); | Metschnikowia | 13 | Saprotroph | \|Undefined Saprotroph\| | Yeast | NULL | Probable | As nectar-tap_saprotroph (Põlme et al. 2020); Decay sugar-rich substrates (Põlme et al. 2020) | Põlme S, et al. 2020. Fungal Diversity 105:1-16 ((https://doi.org/10.1007/s13225-020-00466-2)) |
| Otu20875 | 1 | k__Fungi(100);p__Ascomycota(100);c__Saccharomycetes(100);o__Saccharomycetales(100);f__Metschnikowiaceae(100);g__Metschnikowia(100);g__Metschnikowia_unclassified(100); | Metschnikowia | 13 | Saprotroph | \|Undefined Saprotroph\| | Yeast | NULL | Probable | As nectar-tap_saprotroph (Põlme et al. 2020); Decay sugar-rich substrates (Põlme et al. 2020) | Põlme S, et al. 2020. Fungal Diversity 105:1-16 ((https://doi.org/10.1007/s13225-020-00466-2)) |
| Otu20876 | 1 | k__Fungi(100);p__Ascomycota(100);c__Saccharomycetes(100);o__Saccharomycetales(100);f__Metschnikowiaceae(100);g__Metschnikowia(100);s__Metschnikowia_pulcherrima(100); | Metschnikowia | 13 | Saprotroph | \|Undefined Saprotroph\| | Yeast | NULL | Probable | As nectar-tap_saprotroph (Põlme et al. 2020); Decay sugar-rich substrates (Põlme et al. 2020) | Põlme S, et al. 2020. Fungal Diversity 105:1-16 ((https://doi.org/10.1007/s13225-020-00466-2)) |
| Otu20878 | 1 | k__Fungi(100);p__Ascomycota(100);c__Saccharomycetes(100);o__Saccharomycetales(100);f__Metschnikowiaceae(100);g__Metschnikowia(100);s__Metschnikowia_pulcherrima(100); | Metschnikowia | 13 | Saprotroph | \|Undefined Saprotroph\| | Yeast | NULL | Probable | As nectar-tap_saprotroph (Põlme et al. 2020); Decay sugar-rich substrates (Põlme et al. 2020) | Põlme S, et al. 2020. Fungal Diversity 105:1-16 ((https://doi.org/10.1007/s13225-020-00466-2)) |
| Otu20879 | 1 | k__Fungi(100);p__Ascomycota(100);c__Saccharomycetes(100);o__Saccharomycetales(100);f__Metschnikowiaceae(100);g__Metschnikowia(100);g__Metschnikowia_unclassified(100); | Metschnikowia | 13 | Saprotroph | \|Undefined Saprotroph\| | Yeast | NULL | Probable | As nectar-tap_saprotroph (Põlme et al. 2020); Decay sugar-rich substrates (Põlme et al. 2020) | Põlme S, et al. 2020. Fungal Diversity 105:1-16 ((https://doi.org/10.1007/s13225-020-00466-2)) |
| Otu20880 | 1 | k__Fungi(100);p__Ascomycota(100);c__Saccharomycetes(100);o__Saccharomycetales(100);f__Metschnikowiaceae(100);g__Metschnikowia(100);g__Metschnikowia_unclassified(100); | Metschnikowia | 13 | Saprotroph | \|Undefined Saprotroph\| | Yeast | NULL | Probable | As nectar-tap_saprotroph (Põlme et al. 2020); Decay sugar-rich substrates (Põlme et al. 2020) | Põlme S, et al. 2020. Fungal Diversity 105:1-16 ((https://doi.org/10.1007/s13225-020-00466-2)) |
| Otu20881 | 1 | k__Fungi(100);p__Ascomycota(100);c__Saccharomycetes(100);o__Saccharomycetales(100);f__Metschnikowiaceae(100);g__Metschnikowia(100);g__Metschnikowia_unclassified(100); | Metschnikowia | 13 | Saprotroph | \|Undefined Saprotroph\| | Yeast | NULL | Probable | As nectar-tap_saprotroph (Põlme et al. 2020); Decay sugar-rich substrates (Põlme et al. 2020) | Põlme S, et al. 2020. Fungal Diversity 105:1-16 ((https://doi.org/10.1007/s13225-020-00466-2)) |
| Otu20882 | 1 | k__Fungi(100);p__Ascomycota(100);c__Saccharomycetes(100);o__Saccharomycetales(100);f__Metschnikowiaceae(100);g__Metschnikowia(100);s__Metschnikowia_pulcherrima(100); | Metschnikowia | 13 | Saprotroph | \|Undefined Saprotroph\| | Yeast | NULL | Probable | As nectar-tap_saprotroph (Põlme et al. 2020); Decay sugar-rich substrates (Põlme et al. 2020) | Põlme S, et al. 2020. Fungal Diversity 105:1-16 ((https://doi.org/10.1007/s13225-020-00466-2)) |
| Otu20883 | 1 | k__Fungi(100);p__Ascomycota(100);c__Saccharomycetes(100);o__Saccharomycetales(100);f__Metschnikowiaceae(100);g__Metschnikowia(100);s__Metschnikowia_pulcherrima(100); | Metschnikowia | 13 | Saprotroph | \|Undefined Saprotroph\| | Yeast | NULL | Probable | As nectar-tap_saprotroph (Põlme et al. 2020); Decay sugar-rich substrates (Põlme et al. 2020) | Põlme S, et al. 2020. Fungal Diversity 105:1-16 ((https://doi.org/10.1007/s13225-020-00466-2)) |
| Otu20885 | 1 | k__Fungi(100);p__Ascomycota(100);c__Saccharomycetes(100);o__Saccharomycetales(100);f__Metschnikowiaceae(100);g__Metschnikowia(100);g__Metschnikowia_unclassified(100); | Metschnikowia | 13 | Saprotroph | \|Undefined Saprotroph\| | Yeast | NULL | Probable | As nectar-tap_saprotroph (Põlme et al. 2020); Decay sugar-rich substrates (Põlme et al. 2020) | Põlme S, et al. 2020. Fungal Diversity 105:1-16 ((https://doi.org/10.1007/s13225-020-00466-2)) |
| Otu20886 | 1 | k__Fungi(100);p__Ascomycota(100);c__Saccharomycetes(100);o__Saccharomycetales(100);f__Metschnikowiaceae(100);g__Metschnikowia(100);s__Metschnikowia_pulcherrima(100); | Metschnikowia | 13 | Saprotroph | \|Undefined Saprotroph\| | Yeast | NULL | Probable | As nectar-tap_saprotroph (Põlme et al. 2020); Decay sugar-rich substrates (Põlme et al. 2020) | Põlme S, et al. 2020. Fungal Diversity 105:1-16 ((https://doi.org/10.1007/s13225-020-00466-2)) |
| Otu20887 | 1 | k__Fungi(100);p__Ascomycota(100);c__Saccharomycetes(100);o__Saccharomycetales(100);f__Metschnikowiaceae(100);g__Metschnikowia(100);s__Metschnikowia_pulcherrima(100); | Metschnikowia | 13 | Saprotroph | \|Undefined Saprotroph\| | Yeast | NULL | Probable | As nectar-tap_saprotroph (Põlme et al. 2020); Decay sugar-rich substrates (Põlme et al. 2020) | Põlme S, et al. 2020. Fungal Diversity 105:1-16 ((https://doi.org/10.1007/s13225-020-00466-2)) |
| Otu20888 | 1 | k__Fungi(100);p__Ascomycota(100);c__Saccharomycetes(100);o__Saccharomycetales(100);f__Metschnikowiaceae(100);g__Metschnikowia(100);s__Metschnikowia_pulcherrima(100); | Metschnikowia | 13 | Saprotroph | \|Undefined Saprotroph\| | Yeast | NULL | Probable | As nectar-tap_saprotroph (Põlme et al. 2020); Decay sugar-rich substrates (Põlme et al. 2020) | Põlme S, et al. 2020. Fungal Diversity 105:1-16 ((https://doi.org/10.1007/s13225-020-00466-2)) |
| Otu20889 | 1 | k__Fungi(100);p__Ascomycota(100);c__Saccharomycetes(100);o__Saccharomycetales(100);f__Metschnikowiaceae(100);g__Metschnikowia(100);s__Metschnikowia_pulcherrima(100); | Metschnikowia | 13 | Saprotroph | \|Undefined Saprotroph\| | Yeast | NULL | Probable | As nectar-tap_saprotroph (Põlme et al. 2020); Decay sugar-rich substrates (Põlme et al. 2020) | Põlme S, et al. 2020. Fungal Diversity 105:1-16 ((https://doi.org/10.1007/s13225-020-00466-2)) |
| Otu20890 | 1 | k__Fungi(100);p__Ascomycota(100);c__Saccharomycetes(100);o__Saccharomycetales(100);f__Metschnikowiaceae(100);g__Metschnikowia(100);g__Metschnikowia_unclassified(100); | Metschnikowia | 13 | Saprotroph | \|Undefined Saprotroph\| | Yeast | NULL | Probable | As nectar-tap_saprotroph (Põlme et al. 2020); Decay sugar-rich substrates (Põlme et al. 2020) | Põlme S, et al. 2020. Fungal Diversity 105:1-16 ((https://doi.org/10.1007/s13225-020-00466-2)) |
| Otu20891 | 1 | k__Fungi(100);p__Ascomycota(100);c__Saccharomycetes(100);o__Saccharomycetales(100);f__Metschnikowiaceae(100);g__Metschnikowia(100);s__Metschnikowia_pulcherrima(100); | Metschnikowia | 13 | Saprotroph | \|Undefined Saprotroph\| | Yeast | NULL | Probable | As nectar-tap_saprotroph (Põlme et al. 2020); Decay sugar-rich substrates (Põlme et al. 2020) | Põlme S, et al. 2020. Fungal Diversity 105:1-16 ((https://doi.org/10.1007/s13225-020-00466-2)) |
| Otu20892 | 1 | k__Fungi(100);p__Ascomycota(100);c__Saccharomycetes(100);o__Saccharomycetales(100);f__Metschnikowiaceae(100);g__Metschnikowia(100);g__Metschnikowia_unclassified(100); | Metschnikowia | 13 | Saprotroph | \|Undefined Saprotroph\| | Yeast | NULL | Probable | As nectar-tap_saprotroph (Põlme et al. 2020); Decay sugar-rich substrates (Põlme et al. 2020) | Põlme S, et al. 2020. Fungal Diversity 105:1-16 ((https://doi.org/10.1007/s13225-020-00466-2)) |
| Otu20901 | 1 | k__Fungi(100);p__Ascomycota(100);c__Saccharomycetes(100);o__Saccharomycetales(100);f__Metschnikowiaceae(100);g__Metschnikowia(100);s__Metschnikowia_pulcherrima(100); | Metschnikowia | 13 | Saprotroph | \|Undefined Saprotroph\| | Yeast | NULL | Probable | As nectar-tap_saprotroph (Põlme et al. 2020); Decay sugar-rich substrates (Põlme et al. 2020) | Põlme S, et al. 2020. Fungal Diversity 105:1-16 ((https://doi.org/10.1007/s13225-020-00466-2)) |
| Otu20902 | 1 | k__Fungi(100);p__Ascomycota(100);c__Saccharomycetes(100);o__Saccharomycetales(100);f__Metschnikowiaceae(100);g__Metschnikowia(100);s__Metschnikowia_pulcherrima(100); | Metschnikowia | 13 | Saprotroph | \|Undefined Saprotroph\| | Yeast | NULL | Probable | As nectar-tap_saprotroph (Põlme et al. 2020); Decay sugar-rich substrates (Põlme et al. 2020) | Põlme S, et al. 2020. Fungal Diversity 105:1-16 ((https://doi.org/10.1007/s13225-020-00466-2)) |
| Otu20903 | 1 | k__Fungi(100);p__Ascomycota(100);c__Saccharomycetes(100);o__Saccharomycetales(100);f__Metschnikowiaceae(100);g__Metschnikowia(100);g__Metschnikowia_unclassified(100); | Metschnikowia | 13 | Saprotroph | \|Undefined Saprotroph\| | Yeast | NULL | Probable | As nectar-tap_saprotroph (Põlme et al. 2020); Decay sugar-rich substrates (Põlme et al. 2020) | Põlme S, et al. 2020. Fungal Diversity 105:1-16 ((https://doi.org/10.1007/s13225-020-00466-2)) |
| Otu20904 | 1 | k__Fungi(100);p__Ascomycota(100);c__Saccharomycetes(100);o__Saccharomycetales(100);f__Metschnikowiaceae(100);g__Metschnikowia(100);s__Metschnikowia_pulcherrima(100); | Metschnikowia | 13 | Saprotroph | \|Undefined Saprotroph\| | Yeast | NULL | Probable | As nectar-tap_saprotroph (Põlme et al. 2020); Decay sugar-rich substrates (Põlme et al. 2020) | Põlme S, et al. 2020. Fungal Diversity 105:1-16 ((https://doi.org/10.1007/s13225-020-00466-2)) |
| Otu20905 | 1 | k__Fungi(100);p__Ascomycota(100);c__Saccharomycetes(100);o__Saccharomycetales(100);f__Metschnikowiaceae(100);g__Metschnikowia(100);s__Metschnikowia_pulcherrima(100); | Metschnikowia | 13 | Saprotroph | \|Undefined Saprotroph\| | Yeast | NULL | Probable | As nectar-tap_saprotroph (Põlme et al. 2020); Decay sugar-rich substrates (Põlme et al. 2020) | Põlme S, et al. 2020. Fungal Diversity 105:1-16 ((https://doi.org/10.1007/s13225-020-00466-2)) |
| Otu20906 | 1 | k__Fungi(100);p__Ascomycota(100);c__Saccharomycetes(100);o__Saccharomycetales(100);f__Metschnikowiaceae(100);g__Metschnikowia(100);s__Metschnikowia_pulcherrima(100); | Metschnikowia | 13 | Saprotroph | \|Undefined Saprotroph\| | Yeast | NULL | Probable | As nectar-tap_saprotroph (Põlme et al. 2020); Decay sugar-rich substrates (Põlme et al. 2020) | Põlme S, et al. 2020. Fungal Diversity 105:1-16 ((https://doi.org/10.1007/s13225-020-00466-2)) |
| Otu20907 | 1 | k__Fungi(100);p__Ascomycota(100);c__Saccharomycetes(100);o__Saccharomycetales(100);f__Metschnikowiaceae(100);g__Metschnikowia(100);s__Metschnikowia_pulcherrima(100); | Metschnikowia | 13 | Saprotroph | \|Undefined Saprotroph\| | Yeast | NULL | Probable | As nectar-tap_saprotroph (Põlme et al. 2020); Decay sugar-rich substrates (Põlme et al. 2020) | Põlme S, et al. 2020. Fungal Diversity 105:1-16 ((https://doi.org/10.1007/s13225-020-00466-2)) |
| Otu20910 | 1 | k__Fungi(100);p__Ascomycota(100);c__Saccharomycetes(100);o__Saccharomycetales(100);f__Metschnikowiaceae(100);g__Metschnikowia(100);s__Metschnikowia_pulcherrima(100); | Metschnikowia | 13 | Saprotroph | \|Undefined Saprotroph\| | Yeast | NULL | Probable | As nectar-tap_saprotroph (Põlme et al. 2020); Decay sugar-rich substrates (Põlme et al. 2020) | Põlme S, et al. 2020. Fungal Diversity 105:1-16 ((https://doi.org/10.1007/s13225-020-00466-2)) |
| Otu20911 | 1 | k__Fungi(100);p__Ascomycota(100);c__Saccharomycetes(100);o__Saccharomycetales(100);f__Metschnikowiaceae(100);g__Metschnikowia(100);s__Metschnikowia_pulcherrima(100); | Metschnikowia | 13 | Saprotroph | \|Undefined Saprotroph\| | Yeast | NULL | Probable | As nectar-tap_saprotroph (Põlme et al. 2020); Decay sugar-rich substrates (Põlme et al. 2020) | Põlme S, et al. 2020. Fungal Diversity 105:1-16 ((https://doi.org/10.1007/s13225-020-00466-2)) |
| Otu20912 | 1 | k__Fungi(100);p__Ascomycota(100);c__Saccharomycetes(100);o__Saccharomycetales(100);f__Metschnikowiaceae(100);g__Metschnikowia(100);s__Metschnikowia_pulcherrima(100); | Metschnikowia | 13 | Saprotroph | \|Undefined Saprotroph\| | Yeast | NULL | Probable | As nectar-tap_saprotroph (Põlme et al. 2020); Decay sugar-rich substrates (Põlme et al. 2020) | Põlme S, et al. 2020. Fungal Diversity 105:1-16 ((https://doi.org/10.1007/s13225-020-00466-2)) |
| Otu20913 | 1 | k__Fungi(100);p__Ascomycota(100);c__Saccharomycetes(100);o__Saccharomycetales(100);f__Metschnikowiaceae(100);g__Metschnikowia(100);s__Metschnikowia_pulcherrima(100); | Metschnikowia | 13 | Saprotroph | \|Undefined Saprotroph\| | Yeast | NULL | Probable | As nectar-tap_saprotroph (Põlme et al. 2020); Decay sugar-rich substrates (Põlme et al. 2020) | Põlme S, et al. 2020. Fungal Diversity 105:1-16 ((https://doi.org/10.1007/s13225-020-00466-2)) |
| Otu20914 | 1 | k__Fungi(100);p__Ascomycota(100);c__Saccharomycetes(100);o__Saccharomycetales(100);f__Metschnikowiaceae(100);g__Metschnikowia(100);g__Metschnikowia_unclassified(100); | Metschnikowia | 13 | Saprotroph | \|Undefined Saprotroph\| | Yeast | NULL | Probable | As nectar-tap_saprotroph (Põlme et al. 2020); Decay sugar-rich substrates (Põlme et al. 2020) | Põlme S, et al. 2020. Fungal Diversity 105:1-16 ((https://doi.org/10.1007/s13225-020-00466-2)) |
| Otu20916 | 1 | k__Fungi(100);p__Ascomycota(100);c__Saccharomycetes(100);o__Saccharomycetales(100);f__Metschnikowiaceae(100);g__Metschnikowia(100);g__Metschnikowia_unclassified(100); | Metschnikowia | 13 | Saprotroph | \|Undefined Saprotroph\| | Yeast | NULL | Probable | As nectar-tap_saprotroph (Põlme et al. 2020); Decay sugar-rich substrates (Põlme et al. 2020) | Põlme S, et al. 2020. Fungal Diversity 105:1-16 ((https://doi.org/10.1007/s13225-020-00466-2)) |
| Otu20917 | 1 | k__Fungi(100);p__Ascomycota(100);c__Saccharomycetes(100);o__Saccharomycetales(100);o__Saccharomycetales_unclassified(100);o__Saccharomycetales_unclassified(100);o__Saccharomycetales_unclassified(100); | Saccharomycetales | 7 | Saprotroph | Undefined Saprotroph | Yeast | NULL | Possible | NULL | Sterkenburg E, et al. 2015. New Phytologist 207:1145-1158 ((https://doi.org/10.1111/nph.13426)) |
| Otu20918 | 1 | k__Fungi(100);p__Ascomycota(100);c__Saccharomycetes(100);o__Saccharomycetales(100);f__Metschnikowiaceae(100);g__Metschnikowia(100);s__Metschnikowia_pulcherrima(100); | Metschnikowia | 13 | Saprotroph | \|Undefined Saprotroph\| | Yeast | NULL | Probable | As nectar-tap_saprotroph (Põlme et al. 2020); Decay sugar-rich substrates (Põlme et al. 2020) | Põlme S, et al. 2020. Fungal Diversity 105:1-16 ((https://doi.org/10.1007/s13225-020-00466-2)) |
| Otu20919 | 1 | k__Fungi(100);p__Ascomycota(100);c__Ascomycota_class_Incertae_sedis(100);o__Ascomycota_order_Incertae_sedis(100);f__Ascomycota_family_Incertae_sedis(100);g__Zymoseptoria(100);s__Zymoseptoria_verkleyi(100); | Zymoseptoria | 13 | Pathotroph-Saprotroph | \|Plant Pathogen\|-Plant Saprotroph | Microfungus | NULL | Probable | As litter_saprotroph (Põlme et al. 2020); Decay leaf, fruit, seed (Põlme et al. 2020) | Põlme S, et al. 2020. Fungal Diversity 105:1-16 ((https://doi.org/10.1007/s13225-020-00466-2)) |
| Otu20921 | 1 | k__Fungi(100);p__Ascomycota(100);c__Saccharomycetes(100);o__Saccharomycetales(100);f__Metschnikowiaceae(100);g__Metschnikowia(100);s__Metschnikowia_pulcherrima(100); | Metschnikowia | 13 | Saprotroph | \|Undefined Saprotroph\| | Yeast | NULL | Probable | As nectar-tap_saprotroph (Põlme et al. 2020); Decay sugar-rich substrates (Põlme et al. 2020) | Põlme S, et al. 2020. Fungal Diversity 105:1-16 ((https://doi.org/10.1007/s13225-020-00466-2)) |
| Otu20928 | 1 | k__Fungi(100);p__Ascomycota(100);c__Saccharomycetes(100);o__Saccharomycetales(100);f__Metschnikowiaceae(100);g__Metschnikowia(100);g__Metschnikowia_unclassified(100); | Metschnikowia | 13 | Saprotroph | \|Undefined Saprotroph\| | Yeast | NULL | Probable | As nectar-tap_saprotroph (Põlme et al. 2020); Decay sugar-rich substrates (Põlme et al. 2020) | Põlme S, et al. 2020. Fungal Diversity 105:1-16 ((https://doi.org/10.1007/s13225-020-00466-2)) |
| Otu20929 | 1 | k__Fungi(100);p__Ascomycota(100);c__Saccharomycetes(100);o__Saccharomycetales(100);f__Metschnikowiaceae(100);g__Metschnikowia(100);s__Metschnikowia_pulcherrima(100); | Metschnikowia | 13 | Saprotroph | \|Undefined Saprotroph\| | Yeast | NULL | Probable | As nectar-tap_saprotroph (Põlme et al. 2020); Decay sugar-rich substrates (Põlme et al. 2020) | Põlme S, et al. 2020. Fungal Diversity 105:1-16 ((https://doi.org/10.1007/s13225-020-00466-2)) |
| Otu20930 | 1 | k__Fungi(100);p__Ascomycota(100);c__Saccharomycetes(100);o__Saccharomycetales(100);f__Metschnikowiaceae(100);g__Metschnikowia(100);g__Metschnikowia_unclassified(100); | Metschnikowia | 13 | Saprotroph | \|Undefined Saprotroph\| | Yeast | NULL | Probable | As nectar-tap_saprotroph (Põlme et al. 2020); Decay sugar-rich substrates (Põlme et al. 2020) | Põlme S, et al. 2020. Fungal Diversity 105:1-16 ((https://doi.org/10.1007/s13225-020-00466-2)) |
| Otu20931 | 1 | k__Fungi(100);p__Ascomycota(100);c__Saccharomycetes(100);o__Saccharomycetales(100);f__Metschnikowiaceae(100);g__Metschnikowia(100);s__Metschnikowia_pulcherrima(100); | Metschnikowia | 13 | Saprotroph | \|Undefined Saprotroph\| | Yeast | NULL | Probable | As nectar-tap_saprotroph (Põlme et al. 2020); Decay sugar-rich substrates (Põlme et al. 2020) | Põlme S, et al. 2020. Fungal Diversity 105:1-16 ((https://doi.org/10.1007/s13225-020-00466-2)) |
| Otu20932 | 1 | k__Fungi(100);p__Ascomycota(100);c__Dothideomycetes(100);o__Dothideomycetes_order_Incertae_sedis(100);f__Pseudeurotiaceae(100);f__Pseudeurotiaceae_unclassified(100);f__Pseudeurotiaceae_unclassified(100); | Pseudeurotiaceae | 9 | Saprotroph | Plant Saprotroph-Wood Saprotroph | Microfungus | NULL | Probable | Saprobic on woody tissue and rotting vegetation, often isolated from soil (Cannon & Kirk 2007) | Cannon PF, Kirk PM. 2007. Fungal Families of the World. CAB International, Cambridge (ISBN: 978-0851998275) |
| Otu20933 | 1 | k__Fungi(100);p__Ascomycota(100);c__Saccharomycetes(100);o__Saccharomycetales(100);f__Metschnikowiaceae(100);g__Metschnikowia(100);s__Metschnikowia_pulcherrima(100); | Metschnikowia | 13 | Saprotroph | \|Undefined Saprotroph\| | Yeast | NULL | Probable | As nectar-tap_saprotroph (Põlme et al. 2020); Decay sugar-rich substrates (Põlme et al. 2020) | Põlme S, et al. 2020. Fungal Diversity 105:1-16 ((https://doi.org/10.1007/s13225-020-00466-2)) |
| Otu20934 | 1 | k__Fungi(100);p__Ascomycota(100);c__Saccharomycetes(100);o__Saccharomycetales(100);f__Metschnikowiaceae(100);g__Metschnikowia(100);g__Metschnikowia_unclassified(100); | Metschnikowia | 13 | Saprotroph | \|Undefined Saprotroph\| | Yeast | NULL | Probable | As nectar-tap_saprotroph (Põlme et al. 2020); Decay sugar-rich substrates (Põlme et al. 2020) | Põlme S, et al. 2020. Fungal Diversity 105:1-16 ((https://doi.org/10.1007/s13225-020-00466-2)) |
| Otu20937 | 1 | k__Fungi(100);p__Ascomycota(100);c__Saccharomycetes(100);o__Saccharomycetales(100);f__Saccharomycetales_family_Incertae_sedis(100);g__Candida(100);g__Candida_unclassified(100); | Candida | 13 | Pathotroph-Saprotroph-Symbiotroph | Animal Pathogen-Endophyte-Endosymbiont-Epiphyte-\|Undefined Saprotroph\| | Dimorphic Yeast-Yeast | NULL | Probable | As nectar-tap_saprotroph (Põlme et al. 2020); Decay sugar-rich substrates (Põlme et al. 2020); This asexual genus is polyphyletic and contain species that function very differently depend on what they are. Caution should be taken when interpreting guilds for Candida species. | Manolakaki D, et al. 2010. Virulence. 1: 367-375 ((https://doi.org/10.4161/viru.1.5.12796)); Mekbib SB, et al. 2011. Tropical Plant Pathology 36:233-240 ((https://doi.org/10.1590/S1982-56762011000400004)); Põlme S, et al. 2020. Fungal Diversity 105:1-16 ((https://doi.org/10.1007/s13225-020-00466-2)) |
| Otu20938 | 1 | k__Fungi(100);p__Ascomycota(100);c__Saccharomycetes(100);o__Saccharomycetales(100);f__Metschnikowiaceae(100);g__Metschnikowia(100);s__Metschnikowia_pulcherrima(100); | Metschnikowia | 13 | Saprotroph | \|Undefined Saprotroph\| | Yeast | NULL | Probable | As nectar-tap_saprotroph (Põlme et al. 2020); Decay sugar-rich substrates (Põlme et al. 2020) | Põlme S, et al. 2020. Fungal Diversity 105:1-16 ((https://doi.org/10.1007/s13225-020-00466-2)) |
| Otu20939 | 1 | k__Fungi(100);p__Ascomycota(100);c__Saccharomycetes(100);o__Saccharomycetales(100);f__Metschnikowiaceae(100);g__Metschnikowia(100);s__Metschnikowia_pulcherrima(100); | Metschnikowia | 13 | Saprotroph | \|Undefined Saprotroph\| | Yeast | NULL | Probable | As nectar-tap_saprotroph (Põlme et al. 2020); Decay sugar-rich substrates (Põlme et al. 2020) | Põlme S, et al. 2020. Fungal Diversity 105:1-16 ((https://doi.org/10.1007/s13225-020-00466-2)) |
| Otu20940 | 1 | k__Fungi(100);p__Ascomycota(100);c__Saccharomycetes(100);o__Saccharomycetales(100);f__Metschnikowiaceae(100);g__Metschnikowia(100);s__Metschnikowia_pulcherrima(100); | Metschnikowia | 13 | Saprotroph | \|Undefined Saprotroph\| | Yeast | NULL | Probable | As nectar-tap_saprotroph (Põlme et al. 2020); Decay sugar-rich substrates (Põlme et al. 2020) | Põlme S, et al. 2020. Fungal Diversity 105:1-16 ((https://doi.org/10.1007/s13225-020-00466-2)) |

| | | | | | | | | | | | |
|---|---|---|---|---|---|---|---|---|---|---|---|
| Otu20941 | 1 | k__Fungi(100);p__Ascomycota(100);c__Saccharomycetes(100);o__Saccharomycetales(100);f__Metschnikowiaceae(100);g__Metschnikowia(100);s__Metschnikowia_pulcherrima(100); | Metschnikowia | 13 | Saprotroph | \|Undefined Saprotroph\| | Yeast | NULL | Probable | As nectar-tap_saprotroph (Põlme et al. 2020); Decay sugar-rich substrates (Põlme et al. 2020) | Põlme S, et al. 2020. Fungal Diversity 105:1-16 ((https://doi.org/10.1007/s13225-020-00466-2)) |
| Otu20942 | 1 | k__Fungi(100);p__Ascomycota(100);c__Saccharomycetes(100);o__Saccharomycetales(100);f__Metschnikowiaceae(100);g__Metschnikowia(100);s__Metschnikowia_pulcherrima(100); | Metschnikowia | 13 | Saprotroph | \|Undefined Saprotroph\| | Yeast | NULL | Probable | As nectar-tap_saprotroph (Põlme et al. 2020); Decay sugar-rich substrates (Põlme et al. 2020) | Põlme S, et al. 2020. Fungal Diversity 105:1-16 ((https://doi.org/10.1007/s13225-020-00466-2)) |
| Otu20943 | 1 | k__Fungi(100);p__Ascomycota(100);c__Saccharomycetes(100);o__Saccharomycetales(100);f__Metschnikowiaceae(100);g__Metschnikowia(100);s__Metschnikowia_pulcherrima(100); | Metschnikowia | 13 | Saprotroph | \|Undefined Saprotroph\| | Yeast | NULL | Probable | As nectar-tap_saprotroph (Põlme et al. 2020); Decay sugar-rich substrates (Põlme et al. 2020) | Põlme S, et al. 2020. Fungal Diversity 105:1-16 ((https://doi.org/10.1007/s13225-020-00466-2)) |
| Otu20944 | 1 | k__Fungi(100);p__Ascomycota(100);c__Saccharomycetes(100);o__Saccharomycetales(100);f__Metschnikowiaceae(100);g__Metschnikowia(100);s__Metschnikowia_pulcherrima(100); | Metschnikowia | 13 | Saprotroph | \|Undefined Saprotroph\| | Yeast | NULL | Probable | As nectar-tap_saprotroph (Põlme et al. 2020); Decay sugar-rich substrates (Põlme et al. 2020) | Põlme S, et al. 2020. Fungal Diversity 105:1-16 ((https://doi.org/10.1007/s13225-020-00466-2)) |
| Otu20945 | 1 | k__Fungi(100);p__Ascomycota(100);c__Saccharomycetes(100);o__Saccharomycetales(100);f__Metschnikowiaceae(100);g__Metschnikowia(100);s__Metschnikowia_pulcherrima(100); | Metschnikowia | 13 | Saprotroph | \|Undefined Saprotroph\| | Yeast | NULL | Probable | As nectar-tap_saprotroph (Põlme et al. 2020); Decay sugar-rich substrates (Põlme et al. 2020) | Põlme S, et al. 2020. Fungal Diversity 105:1-16 ((https://doi.org/10.1007/s13225-020-00466-2)) |
| Otu20946 | 1 | k__Fungi(100);p__Ascomycota(100);c__Saccharomycetes(100);o__Saccharomycetales(100);f__Metschnikowiaceae(100);g__Metschnikowia(100);s__Metschnikowia_pulcherrima(100); | Metschnikowia | 13 | Saprotroph | \|Undefined Saprotroph\| | Yeast | NULL | Probable | As nectar-tap_saprotroph (Põlme et al. 2020); Decay sugar-rich substrates (Põlme et al. 2020) | Põlme S, et al. 2020. Fungal Diversity 105:1-16 ((https://doi.org/10.1007/s13225-020-00466-2)) |
| Otu20947 | 1 | k__Fungi(100);p__Ascomycota(100);c__Saccharomycetes(100);o__Saccharomycetales(100);f__Metschnikowiaceae(100);g__Metschnikowia(100);s__Metschnikowia_pulcherrima(100); | Metschnikowia | 13 | Saprotroph | \|Undefined Saprotroph\| | Yeast | NULL | Probable | As nectar-tap_saprotroph (Põlme et al. 2020); Decay sugar-rich substrates (Põlme et al. 2020) | Põlme S, et al. 2020. Fungal Diversity 105:1-16 ((https://doi.org/10.1007/s13225-020-00466-2)) |
| Otu20948 | 1 | k__Fungi(100);p__Ascomycota(100);c__Saccharomycetes(100);o__Saccharomycetales(100);f__Metschnikowiaceae(100);g__Metschnikowia(100);s__Metschnikowia_pulcherrima(100); | Metschnikowia | 13 | Saprotroph | \|Undefined Saprotroph\| | Yeast | NULL | Probable | As nectar-tap_saprotroph (Põlme et al. 2020); Decay sugar-rich substrates (Põlme et al. 2020) | Põlme S, et al. 2020. Fungal Diversity 105:1-16 ((https://doi.org/10.1007/s13225-020-00466-2)) |
| Otu20949 | 1 | k__Fungi(100);p__Ascomycota(100);c__Saccharomycetes(100);o__Saccharomycetales(100);f__Metschnikowiaceae(100);g__Metschnikowia(100);s__Metschnikowia_pulcherrima(100); | Metschnikowia | 13 | Saprotroph | \|Undefined Saprotroph\| | Yeast | NULL | Probable | As nectar-tap_saprotroph (Põlme et al. 2020); Decay sugar-rich substrates (Põlme et al. 2020) | Põlme S, et al. 2020. Fungal Diversity 105:1-16 ((https://doi.org/10.1007/s13225-020-00466-2)) |
| Otu20950 | 1 | k__Fungi(100);p__Ascomycota(100);c__Saccharomycetes(100);o__Saccharomycetales(100);f__Metschnikowiaceae(100);g__Metschnikowia(100);s__Metschnikowia_pulcherrima(100); | Metschnikowia | 13 | Saprotroph | \|Undefined Saprotroph\| | Yeast | NULL | Probable | As nectar-tap_saprotroph (Põlme et al. 2020); Decay sugar-rich substrates (Põlme et al. 2020) | Põlme S, et al. 2020. Fungal Diversity 105:1-16 ((https://doi.org/10.1007/s13225-020-00466-2)) |
| Otu20951 | 1 | k__Fungi(100);p__Ascomycota(100);c__Saccharomycetes(100);o__Saccharomycetales(100);f__Metschnikowiaceae(100);g__Metschnikowia(100);s__Metschnikowia_pulcherrima(100); | Metschnikowia | 13 | Saprotroph | \|Undefined Saprotroph\| | Yeast | NULL | Probable | As nectar-tap_saprotroph (Põlme et al. 2020); Decay sugar-rich substrates (Põlme et al. 2020) | Põlme S, et al. 2020. Fungal Diversity 105:1-16 ((https://doi.org/10.1007/s13225-020-00466-2)) |
| Otu20952 | 1 | k__Fungi(100);p__Basidiomycota(100);c__Microbotryomycetes(100);o__Sporidiobolales(100);f__Sporidiobolales_family_Incertae_sedis(100);g__Sporobolomyces(100);s__Sporobolomyces_lactosus(100); | Sporobolomyces | 13 | Pathotroph-Saprotroph | \|Fungal Parasite\|-Plant Saprotroph | Yeast | NULL | Probable | As litter_saprotroph (Põlme et al. 2020); As mycoparasite (Põlme et al. 2020); Decay fungal material (Põlme et al. 2020) | Tedersoo L, et al. 2014. Science 346:e1256688 ((https://doi.org/10.1126/science.1256688)); Purahong W, et al. 2016. Molecular Ecology 25:4059-4074 ((https://doi.org/10.1111/mec.13739)); Põlme S, et al. 2020. Fungal Diversity 105:1-16 ((https://doi.org/10.1007/s13225-020-00466-2)) |
| Otu20953 | 1 | k__Fungi(100);p__Ascomycota(100);c__Saccharomycetes(100);o__Saccharomycetales(100);f__Metschnikowiaceae(100);g__Metschnikowia(100);g__Metschnikowia_unclassified(100); | Metschnikowia | 13 | Saprotroph | \|Undefined Saprotroph\| | Yeast | NULL | Probable | As nectar-tap_saprotroph (Põlme et al. 2020); Decay sugar-rich substrates (Põlme et al. 2020) | Põlme S, et al. 2020. Fungal Diversity 105:1-16 ((https://doi.org/10.1007/s13225-020-00466-2)) |
| Otu20954 | 1 | k__Fungi(100);p__Ascomycota(100);c__Saccharomycetes(100);o__Saccharomycetales(100);f__Metschnikowiaceae(100);g__Metschnikowia(100);g__Metschnikowia_unclassified(100); | Metschnikowia | 13 | Saprotroph | \|Undefined Saprotroph\| | Yeast | NULL | Probable | As nectar-tap_saprotroph (Põlme et al. 2020); Decay sugar-rich substrates (Põlme et al. 2020) | Põlme S, et al. 2020. Fungal Diversity 105:1-16 ((https://doi.org/10.1007/s13225-020-00466-2)) |
| Otu20955 | 1 | k__Fungi(100);p__Ascomycota(100);c__Saccharomycetes(100);o__Saccharomycetales(100);f__Metschnikowiaceae(100);g__Metschnikowia(100);s__Metschnikowia_pulcherrima(100); | Metschnikowia | 13 | Saprotroph | \|Undefined Saprotroph\| | Yeast | NULL | Probable | As nectar-tap_saprotroph (Põlme et al. 2020); Decay sugar-rich substrates (Põlme et al. 2020) | Põlme S, et al. 2020. Fungal Diversity 105:1-16 ((https://doi.org/10.1007/s13225-020-00466-2)) |
| Otu20956 | 1 | k__Fungi(100);p__Ascomycota(100);c__Saccharomycetes(100);o__Saccharomycetales(100);f__Metschnikowiaceae(100);g__Metschnikowia(100);g__Metschnikowia_unclassified(100); | Metschnikowia | 13 | Saprotroph | \|Undefined Saprotroph\| | Yeast | NULL | Probable | As nectar-tap_saprotroph (Põlme et al. 2020); Decay sugar-rich substrates (Põlme et al. 2020) | Põlme S, et al. 2020. Fungal Diversity 105:1-16 ((https://doi.org/10.1007/s13225-020-00466-2)) |
| Otu20957 | 1 | k__Fungi(100);p__Ascomycota(100);c__Saccharomycetes(100);o__Saccharomycetales(100);f__Metschnikowiaceae(100);g__Metschnikowia(100);s__Metschnikowia_pulcherrima(100); | Metschnikowia | 13 | Saprotroph | \|Undefined Saprotroph\| | Yeast | NULL | Probable | As nectar-tap_saprotroph (Põlme et al. 2020); Decay sugar-rich substrates (Põlme et al. 2020) | Põlme S, et al. 2020. Fungal Diversity 105:1-16 ((https://doi.org/10.1007/s13225-020-00466-2)) |
| Otu20959 | 1 | k__Fungi(100);p__Ascomycota(100);c__Saccharomycetes(100);o__Saccharomycetales(100);f__Metschnikowiaceae(100);g__Metschnikowia(100);g__Metschnikowia_unclassified(100); | Metschnikowia | 13 | Saprotroph | \|Undefined Saprotroph\| | Yeast | NULL | Probable | As nectar-tap_saprotroph (Põlme et al. 2020); Decay sugar-rich substrates (Põlme et al. 2020) | Põlme S, et al. 2020. Fungal Diversity 105:1-16 ((https://doi.org/10.1007/s13225-020-00466-2)) |
| Otu20960 | 1 | k__Fungi(100);p__Ascomycota(100);c__Saccharomycetes(100);o__Saccharomycetales(100);f__Metschnikowiaceae(100);g__Metschnikowia(100);g__Metschnikowia_unclassified(100); | Metschnikowia | 13 | Saprotroph | \|Undefined Saprotroph\| | Yeast | NULL | Probable | As nectar-tap_saprotroph (Põlme et al. 2020); Decay sugar-rich substrates (Põlme et al. 2020) | Põlme S, et al. 2020. Fungal Diversity 105:1-16 ((https://doi.org/10.1007/s13225-020-00466-2)) |
| Otu20961 | 1 | k__Fungi(100);p__Ascomycota(100);c__Saccharomycetes(100);o__Saccharomycetales(100);f__Metschnikowiaceae(100);g__Metschnikowia(100);s__Metschnikowia_pulcherrima(100); | Metschnikowia | 13 | Saprotroph | \|Undefined Saprotroph\| | Yeast | NULL | Probable | As nectar-tap_saprotroph (Põlme et al. 2020); Decay sugar-rich substrates (Põlme et al. 2020) | Põlme S, et al. 2020. Fungal Diversity 105:1-16 ((https://doi.org/10.1007/s13225-020-00466-2)) |
| Otu20962 | 1 | k__Fungi(100);p__Ascomycota(100);c__Saccharomycetes(100);o__Saccharomycetales(100);f__Metschnikowiaceae(100);g__Metschnikowia(100);g__Metschnikowia_unclassified(100); | Metschnikowia | 13 | Saprotroph | \|Undefined Saprotroph\| | Yeast | NULL | Probable | As nectar-tap_saprotroph (Põlme et al. 2020); Decay sugar-rich substrates (Põlme et al. 2020) | Põlme S, et al. 2020. Fungal Diversity 105:1-16 ((https://doi.org/10.1007/s13225-020-00466-2)) |
| Otu20963 | 1 | k__Fungi(100);p__Ascomycota(100);c__Saccharomycetes(100);o__Saccharomycetales(100);f__Metschnikowiaceae(100);g__Metschnikowia(100);s__Metschnikowia_pulcherrima(100); | Metschnikowia | 13 | Saprotroph | \|Undefined Saprotroph\| | Yeast | NULL | Probable | As nectar-tap_saprotroph (Põlme et al. 2020); Decay sugar-rich substrates (Põlme et al. 2020) | Põlme S, et al. 2020. Fungal Diversity 105:1-16 ((https://doi.org/10.1007/s13225-020-00466-2)) |
| Otu20964 | 1 | k__Fungi(100);p__Ascomycota(100);c__Saccharomycetes(100);o__Saccharomycetales(100);f__Metschnikowiaceae(100);g__Metschnikowia(100);g__Metschnikowia_unclassified(100); | Metschnikowia | 13 | Saprotroph | \|Undefined Saprotroph\| | Yeast | NULL | Probable | As nectar-tap_saprotroph (Põlme et al. 2020); Decay sugar-rich substrates (Põlme et al. 2020) | Põlme S, et al. 2020. Fungal Diversity 105:1-16 ((https://doi.org/10.1007/s13225-020-00466-2)) |
| Otu20965 | 1 | k__Fungi(100);p__Ascomycota(100);c__Saccharomycetes(100);o__Saccharomycetales(100);f__Metschnikowiaceae(100);g__Metschnikowia(100);g__Metschnikowia_unclassified(100); | Metschnikowia | 13 | Saprotroph | \|Undefined Saprotroph\| | Yeast | NULL | Probable | As nectar-tap_saprotroph (Põlme et al. 2020); Decay sugar-rich substrates (Põlme et al. 2020) | Põlme S, et al. 2020. Fungal Diversity 105:1-16 ((https://doi.org/10.1007/s13225-020-00466-2)) |
| Otu20968 | 1 | k__Fungi(100);p__Ascomycota(100);c__Saccharomycetes(100);o__Saccharomycetales(100);f__Metschnikowiaceae(100);g__Metschnikowia(100);s__Metschnikowia_pulcherrima(100); | Metschnikowia | 13 | Saprotroph | \|Undefined Saprotroph\| | Yeast | NULL | Probable | As nectar-tap_saprotroph (Põlme et al. 2020); Decay sugar-rich substrates (Põlme et al. 2020) | Põlme S, et al. 2020. Fungal Diversity 105:1-16 ((https://doi.org/10.1007/s13225-020-00466-2)) |
| Otu20969 | 1 | k__Fungi(100);p__Ascomycota(100);c__Saccharomycetes(100);o__Saccharomycetales(100);f__Metschnikowiaceae(100);g__Metschnikowia(100);g__Metschnikowia_unclassified(100); | Metschnikowia | 13 | Saprotroph | \|Undefined Saprotroph\| | Yeast | NULL | Probable | As nectar-tap_saprotroph (Põlme et al. 2020); Decay sugar-rich substrates (Põlme et al. 2020) | Põlme S, et al. 2020. Fungal Diversity 105:1-16 ((https://doi.org/10.1007/s13225-020-00466-2)) |
| Otu20970 | 1 | k__Fungi(100);p__Basidiomycota(100);c__Microbotryomycetes(100);o__Sporidiobolales(100);f__Sporidiobolales_family_Incertae_sedis(100);g__Sporobolomyces(100);s__Sporobolomyces_lactosus(100); | Sporobolomyces | 13 | Pathotroph-Saprotroph | \|Fungal Parasite\|-Plant Saprotroph | Yeast | NULL | Probable | As litter_saprotroph (Põlme et al. 2020); As mycoparasite (Põlme et al. 2020); Decay fungal material (Põlme et al. 2020) | Tedersoo L, et al. 2014. Science 346:e1256688 ((https://doi.org/10.1126/science.1256688)); Purahong W, et al. 2016. Molecular Ecology 25:4059-4074 ((https://doi.org/10.1111/mec.13739)); Põlme S, et al. 2020. Fungal Diversity 105:1-16 ((https://doi.org/10.1007/s13225-020-00466-2)) |
| Otu20971 | 1 | k__Fungi(100);p__Ascomycota(100);c__Saccharomycetes(100);o__Saccharomycetales(100);f__Metschnikowiaceae(100);g__Metschnikowia(100);s__Metschnikowia_pulcherrima(100); | Metschnikowia | 13 | Saprotroph | \|Undefined Saprotroph\| | Yeast | NULL | Probable | As nectar-tap_saprotroph (Põlme et al. 2020); Decay sugar-rich substrates (Põlme et al. 2020) | Põlme S, et al. 2020. Fungal Diversity 105:1-16 ((https://doi.org/10.1007/s13225-020-00466-2)) |
| Otu20972 | 1 | k__Fungi(100);p__Ascomycota(100);c__Saccharomycetes(100);o__Saccharomycetales(100);f__Metschnikowiaceae(100);g__Metschnikowia(100);s__Metschnikowia_pulcherrima(100); | Metschnikowia | 13 | Saprotroph | \|Undefined Saprotroph\| | Yeast | NULL | Probable | As nectar-tap_saprotroph (Põlme et al. 2020); Decay sugar-rich substrates (Põlme et al. 2020) | Põlme S, et al. 2020. Fungal Diversity 105:1-16 ((https://doi.org/10.1007/s13225-020-00466-2)) |
| Otu20974 | 1 | k__Fungi(100);p__Ascomycota(100);c__Saccharomycetes(100);o__Saccharomycetales(100);f__Metschnikowiaceae(100);g__Metschnikowia(100);s__Metschnikowia_pulcherrima(100); | Metschnikowia | 13 | Saprotroph | \|Undefined Saprotroph\| | Yeast | NULL | Probable | As nectar-tap_saprotroph (Põlme et al. 2020); Decay sugar-rich substrates (Põlme et al. 2020) | Põlme S, et al. 2020. Fungal Diversity 105:1-16 ((https://doi.org/10.1007/s13225-020-00466-2)) |
| Otu20975 | 1 | k__Fungi(100);p__Ascomycota(100);c__Saccharomycetes(100);o__Saccharomycetales(100);f__Metschnikowiaceae(100);g__Metschnikowia(100);g__Metschnikowia_unclassified(100); | Metschnikowia | 13 | Saprotroph | \|Undefined Saprotroph\| | Yeast | NULL | Probable | As nectar-tap_saprotroph (Põlme et al. 2020); Decay sugar-rich substrates (Põlme et al. 2020) | Põlme S, et al. 2020. Fungal Diversity 105:1-16 ((https://doi.org/10.1007/s13225-020-00466-2)) |
| Otu20976 | 1 | k__Fungi(100);p__Ascomycota(100);c__Saccharomycetes(100);o__Saccharomycetales(100);f__Metschnikowiaceae(100);g__Metschnikowia(100);s__Metschnikowia_pulcherrima(100); | Metschnikowia | 13 | Saprotroph | \|Undefined Saprotroph\| | Yeast | NULL | Probable | As nectar-tap_saprotroph (Põlme et al. 2020); Decay sugar-rich substrates (Põlme et al. 2020) | Põlme S, et al. 2020. Fungal Diversity 105:1-16 ((https://doi.org/10.1007/s13225-020-00466-2)) |
| Otu20977 | 1 | k__Fungi(100);p__Ascomycota(100);c__Saccharomycetes(100);o__Saccharomycetales(100);f__Metschnikowiaceae(100);g__Metschnikowia(100);s__Metschnikowia_pulcherrima(100); | Metschnikowia | 13 | Saprotroph | \|Undefined Saprotroph\| | Yeast | NULL | Probable | As nectar-tap_saprotroph (Põlme et al. 2020); Decay sugar-rich substrates (Põlme et al. 2020) | Põlme S, et al. 2020. Fungal Diversity 105:1-16 ((https://doi.org/10.1007/s13225-020-00466-2)) |
| Otu20979 | 1 | k__Fungi(100);p__Ascomycota(100);c__Saccharomycetes(100);o__Saccharomycetales(100);f__Metschnikowiaceae(100);g__Metschnikowia(100);s__Metschnikowia_pulcherrima(100); | Metschnikowia | 13 | Saprotroph | \|Undefined Saprotroph\| | Yeast | NULL | Probable | As nectar-tap_saprotroph (Põlme et al. 2020); Decay sugar-rich substrates (Põlme et al. 2020) | Põlme S, et al. 2020. Fungal Diversity 105:1-16 ((https://doi.org/10.1007/s13225-020-00466-2)) |
| Otu20982 | 1 | k__Fungi(100);p__Ascomycota(100);c__Saccharomycetes(100);o__Saccharomycetales(100);f__Metschnikowiaceae(100);g__Metschnikowia(100);g__Metschnikowia_unclassified(100); | Metschnikowia | 13 | Saprotroph | \|Undefined Saprotroph\| | Yeast | NULL | Probable | As nectar-tap_saprotroph (Põlme et al. 2020); Decay sugar-rich substrates (Põlme et al. 2020) | Põlme S, et al. 2020. Fungal Diversity 105:1-16 ((https://doi.org/10.1007/s13225-020-00466-2)) |
| Otu20983 | 1 | k__Fungi(100);p__Ascomycota(100);c__Saccharomycetes(100);o__Saccharomycetales(100);f__Metschnikowiaceae(100);g__Metschnikowia(100);g__Metschnikowia_unclassified(100); | Metschnikowia | 13 | Saprotroph | \|Undefined Saprotroph\| | Yeast | NULL | Probable | As nectar-tap_saprotroph (Põlme et al. 2020); Decay sugar-rich substrates (Põlme et al. 2020) | Põlme S, et al. 2020. Fungal Diversity 105:1-16 ((https://doi.org/10.1007/s13225-020-00466-2)) |
| Otu20984 | 1 | k__Fungi(100);p__Ascomycota(100);c__Saccharomycetes(100);o__Saccharomycetales(100);f__Metschnikowiaceae(100);g__Metschnikowia(100);s__Metschnikowia_pulcherrima(100); | Metschnikowia | 13 | Saprotroph | \|Undefined Saprotroph\| | Yeast | NULL | Probable | As nectar-tap_saprotroph (Põlme et al. 2020); Decay sugar-rich substrates (Põlme et al. 2020) | Põlme S, et al. 2020. Fungal Diversity 105:1-16 ((https://doi.org/10.1007/s13225-020-00466-2)) |
| Otu20985 | 1 | k__Fungi(100);p__Basidiomycota(100);c__Microbotryomycetes(100);o__Sporidiobolales(100);f__Sporidiobolales_family_Incertae_sedis(100);g__Sporobolomyces(100);s__Sporobolomyces_lactosus(100); | Sporobolomyces | 13 | Pathotroph-Saprotroph | \|Fungal Parasite\|-Plant Saprotroph | Yeast | NULL | Probable | As litter_saprotroph (Põlme et al. 2020); As mycoparasite (Põlme et al. 2020); Decay fungal material (Põlme et al. 2020) | Tedersoo L, et al. 2014. Science 346:e1256688 ((https://doi.org/10.1126/science.1256688)); Purahong W, et al. 2016. Molecular Ecology 25:4059-4074 ((https://doi.org/10.1111/mec.13739)); Põlme S, et al. 2020. Fungal Diversity 105:1-16 ((https://doi.org/10.1007/s13225-020-00466-2)) |
| Otu20986 | 1 | k__Fungi(100);p__Ascomycota(100);c__Saccharomycetes(100);o__Saccharomycetales(100);f__Metschnikowiaceae(100);g__Metschnikowia(100);s__Metschnikowia_pulcherrima(100); | Metschnikowia | 13 | Saprotroph | \|Undefined Saprotroph\| | Yeast | NULL | Probable | As nectar-tap_saprotroph (Põlme et al. 2020); Decay sugar-rich substrates (Põlme et al. 2020) | Põlme S, et al. 2020. Fungal Diversity 105:1-16 ((https://doi.org/10.1007/s13225-020-00466-2)) |
| Otu20987 | 1 | k__Fungi(100);p__Ascomycota(100);c__Saccharomycetes(100);o__Saccharomycetales(100);f__Metschnikowiaceae(100);g__Metschnikowia(100);s__Metschnikowia_pulcherrima(100); | Metschnikowia | 13 | Saprotroph | \|Undefined Saprotroph\| | Yeast | NULL | Probable | As nectar-tap_saprotroph (Põlme et al. 2020); Decay sugar-rich substrates (Põlme et al. 2020) | Põlme S, et al. 2020. Fungal Diversity 105:1-16 ((https://doi.org/10.1007/s13225-020-00466-2)) |
| Otu20988 | 1 | k__Fungi(100);p__Ascomycota(100);c__Saccharomycetes(100);o__Saccharomycetales(100);f__Metschnikowiaceae(100);g__Metschnikowia(100);g__Metschnikowia_unclassified(100); | Metschnikowia | 13 | Saprotroph | \|Undefined Saprotroph\| | Yeast | NULL | Probable | As nectar-tap_saprotroph (Põlme et al. 2020); Decay sugar-rich substrates (Põlme et al. 2020) | Põlme S, et al. 2020. Fungal Diversity 105:1-16 ((https://doi.org/10.1007/s13225-020-00466-2)) |
| Otu20989 | 1 | k__Fungi(100);p__Ascomycota(100);c__Saccharomycetes(100);o__Saccharomycetales(100);f__Metschnikowiaceae(100);g__Metschnikowia(100);s__Metschnikowia_pulcherrima(100); | Metschnikowia | 13 | Saprotroph | \|Undefined Saprotroph\| | Yeast | NULL | Probable | As nectar-tap_saprotroph (Põlme et al. 2020); Decay sugar-rich substrates (Põlme et al. 2020) | Põlme S, et al. 2020. Fungal Diversity 105:1-16 ((https://doi.org/10.1007/s13225-020-00466-2)) |
| Otu20991 | 1 | k__Fungi(100);p__Ascomycota(100);c__Saccharomycetes(100);o__Saccharomycetales(100);f__Metschnikowiaceae(100);g__Metschnikowia(100);s__Metschnikowia_pulcherrima(100); | Metschnikowia | 13 | Saprotroph | \|Undefined Saprotroph\| | Yeast | NULL | Probable | As nectar-tap_saprotroph (Põlme et al. 2020); Decay sugar-rich substrates (Põlme et al. 2020) | Põlme S, et al. 2020. Fungal Diversity 105:1-16 ((https://doi.org/10.1007/s13225-020-00466-2)) |
| Otu20992 | 1 | k__Fungi(100);p__Ascomycota(100);c__Saccharomycetes(100);o__Saccharomycetales(100);f__Metschnikowiaceae(100);g__Metschnikowia(100);s__Metschnikowia_pulcherrima(100); | Metschnikowia | 13 | Saprotroph | \|Undefined Saprotroph\| | Yeast | NULL | Probable | As nectar-tap_saprotroph (Põlme et al. 2020); Decay sugar-rich substrates (Põlme et al. 2020) | Põlme S, et al. 2020. Fungal Diversity 105:1-16 ((https://doi.org/10.1007/s13225-020-00466-2)) |
| Otu20993 | 1 | k__Fungi(100);p__Ascomycota(100);c__Saccharomycetes(100);o__Saccharomycetales(100);f__Metschnikowiaceae(100);g__Metschnikowia(100);s__Metschnikowia_pulcherrima(100); | Metschnikowia | 13 | Saprotroph | \|Undefined Saprotroph\| | Yeast | NULL | Probable | As nectar-tap_saprotroph (Põlme et al. 2020); Decay sugar-rich substrates (Põlme et al. 2020) | Põlme S, et al. 2020. Fungal Diversity 105:1-16 ((https://doi.org/10.1007/s13225-020-00466-2)) |
| Otu20994 | 1 | k__Fungi(100);p__Ascomycota(100);c__Saccharomycetes(100);o__Saccharomycetales(100);f__Metschnikowiaceae(100);g__Metschnikowia(100);g__Metschnikowia_unclassified(100); | Metschnikowia | 13 | Saprotroph | \|Undefined Saprotroph\| | Yeast | NULL | Probable | As nectar-tap_saprotroph (Põlme et al. 2020); Decay sugar-rich substrates (Põlme et al. 2020) | Põlme S, et al. 2020. Fungal Diversity 105:1-16 ((https://doi.org/10.1007/s13225-020-00466-2)) |
| Otu20996 | 1 | k__Fungi(100);p__Ascomycota(100);c__Saccharomycetes(100);o__Saccharomycetales(100);f__Metschnikowiaceae(100);g__Metschnikowia(100);s__Metschnikowia_pulcherrima(100); | Metschnikowia | 13 | Saprotroph | \|Undefined Saprotroph\| | Yeast | NULL | Probable | As nectar-tap_saprotroph (Põlme et al. 2020); Decay sugar-rich substrates (Põlme et al. 2020) | Põlme S, et al. 2020. Fungal Diversity 105:1-16 ((https://doi.org/10.1007/s13225-020-00466-2)) |
| Otu20997 | 1 | k__Fungi(100);p__Ascomycota(100);c__Saccharomycetes(100);o__Saccharomycetales(100);f__Metschnikowiaceae(100);g__Metschnikowia(100);s__Metschnikowia_pulcherrima(100); | Metschnikowia | 13 | Saprotroph | \|Undefined Saprotroph\| | Yeast | NULL | Probable | As nectar-tap_saprotroph (Põlme et al. 2020); Decay sugar-rich substrates (Põlme et al. 2020) | Põlme S, et al. 2020. Fungal Diversity 105:1-16 ((https://doi.org/10.1007/s13225-020-00466-2)) |
| Otu20998 | 1 | k__Fungi(100);p__Ascomycota(100);c__Saccharomycetes(100);o__Saccharomycetales(100);f__Metschnikowiaceae(100);g__Metschnikowia(100);s__Metschnikowia_pulcherrima(100); | Metschnikowia | 13 | Saprotroph | \|Undefined Saprotroph\| | Yeast | NULL | Probable | As nectar-tap_saprotroph (Põlme et al. 2020); Decay sugar-rich substrates (Põlme et al. 2020) | Põlme S, et al. 2020. Fungal Diversity 105:1-16 ((https://doi.org/10.1007/s13225-020-00466-2)) |
| Otu21000 | 1 | k__Fungi(100);p__Ascomycota(100);c__Saccharomycetes(100);o__Saccharomycetales(100);f__Metschnikowiaceae(100);g__Metschnikowia(100);s__Metschnikowia_pulcherrima(100); | Metschnikowia | 13 | Saprotroph | \|Undefined Saprotroph\| | Yeast | NULL | Probable | As nectar-tap_saprotroph (Põlme et al. 2020); Decay sugar-rich substrates (Põlme et al. 2020) | Põlme S, et al. 2020. Fungal Diversity 105:1-16 ((https://doi.org/10.1007/s13225-020-00466-2)) |
| Otu21001 | 1 | k__Fungi(100);p__Ascomycota(100);c__Saccharomycetes(100);o__Saccharomycetales(100);f__Metschnikowiaceae(100);g__Metschnikowia(100);g__Metschnikowia_unclassified(100); | Metschnikowia | 13 | Saprotroph | \|Undefined Saprotroph\| | Yeast | NULL | Probable | As nectar-tap_saprotroph (Põlme et al. 2020); Decay sugar-rich substrates (Põlme et al. 2020) | Põlme S, et al. 2020. Fungal Diversity 105:1-16 ((https://doi.org/10.1007/s13225-020-00466-2)) |
| Otu21002 | 1 | k__Fungi(100);p__Ascomycota(100);c__Saccharomycetes(100);o__Saccharomycetales(100);f__Metschnikowiaceae(100);g__Metschnikowia(100);s__Metschnikowia_pulcherrima(100); | Metschnikowia | 13 | Saprotroph | \|Undefined Saprotroph\| | Yeast | NULL | Probable | As nectar-tap_saprotroph (Põlme et al. 2020); Decay sugar-rich substrates (Põlme et al. 2020) | Põlme S, et al. 2020. Fungal Diversity 105:1-16 ((https://doi.org/10.1007/s13225-020-00466-2)) |
| Otu21003 | 1 | k__Fungi(100);p__Ascomycota(100);c__Saccharomycetes(100);o__Saccharomycetales(100);f__Metschnikowiaceae(100);g__Metschnikowia(100);s__Metschnikowia_pulcherrima(100); | Metschnikowia | 13 | Saprotroph | \|Undefined Saprotroph\| | Yeast | NULL | Probable | As nectar-tap_saprotroph (Põlme et al. 2020); Decay sugar-rich substrates (Põlme et al. 2020) | Põlme S, et al. 2020. Fungal Diversity 105:1-16 ((https://doi.org/10.1007/s13225-020-00466-2)) |
| Otu21004 | 1 | k__Fungi(100);p__Ascomycota(100);c__Saccharomycetes(100);o__Saccharomycetales(100);f__Metschnikowiaceae(100);g__Metschnikowia(100);g__Metschnikowia_unclassified(100); | Metschnikowia | 13 | Saprotroph | \|Undefined Saprotroph\| | Yeast | NULL | Probable | As nectar-tap_saprotroph (Põlme et al. 2020); Decay sugar-rich substrates (Põlme et al. 2020) | Põlme S, et al. 2020. Fungal Diversity 105:1-16 ((https://doi.org/10.1007/s13225-020-00466-2)) |
| Otu21005 | 1 | k__Fungi(100);p__Basidiomycota(100);c__Microbotryomycetes(100);o__Sporidiobolales(100);f__Sporidiobolales_family_Incertae_sedis(100);g__Sporobolomyces(100);s__Sporobolomyces_lactosus(100); | Sporobolomyces | 13 | Pathotroph-Saprotroph | \|Fungal Parasite\|-Plant Saprotroph | Yeast | NULL | Probable | As litter_saprotroph (Põlme et al. 2020); As mycoparasite (Põlme et al. 2020); Decay fungal material (Põlme et al. 2020) | Tedersoo L, et al. 2014. Science 346:e1256688 ((https://doi.org/10.1126/science.1256688)); Purahong W, et al. 2016. Molecular Ecology 25:4059-4074 ((https://doi.org/10.1111/mec.13739)); Põlme S, et al. 2020. Fungal Diversity 105:1-16 ((https://doi.org/10.1007/s13225-020-00466-2)) |
| Otu21006 | 1 | k__Fungi(100);p__Ascomycota(100);c__Saccharomycetes(100);o__Saccharomycetales(100);f__Metschnikowiaceae(100);g__Metschnikowia(100);s__Metschnikowia_pulcherrima(100); | Metschnikowia | 13 | Saprotroph | \|Undefined Saprotroph\| | Yeast | NULL | Probable | As nectar-tap_saprotroph (Põlme et al. 2020); Decay sugar-rich substrates (Põlme et al. 2020) | Põlme S, et al. 2020. Fungal Diversity 105:1-16 ((https://doi.org/10.1007/s13225-020-00466-2)) |
| Otu21007 | 1 | k__Fungi(100);p__Ascomycota(100);c__Saccharomycetes(100);o__Saccharomycetales(100);f__Metschnikowiaceae(100);g__Metschnikowia(100);g__Metschnikowia_unclassified(100); | Metschnikowia | 13 | Saprotroph | \|Undefined Saprotroph\| | Yeast | NULL | Probable | As nectar-tap_saprotroph (Põlme et al. 2020); Decay sugar-rich substrates (Põlme et al. 2020) | Põlme S, et al. 2020. Fungal Diversity 105:1-16 ((https://doi.org/10.1007/s13225-020-00466-2)) |
| Otu21008 | 1 | k__Fungi(100);p__Ascomycota(100);c__Saccharomycetes(100);o__Saccharomycetales(100);f__Metschnikowiaceae(100);g__Metschnikowia(100);g__Metschnikowia_unclassified(100); | Metschnikowia | 13 | Saprotroph | \|Undefined Saprotroph\| | Yeast | NULL | Probable | As nectar-tap_saprotroph (Põlme et al. 2020); Decay sugar-rich substrates (Põlme et al. 2020) | Põlme S, et al. 2020. Fungal Diversity 105:1-16 ((https://doi.org/10.1007/s13225-020-00466-2)) |
| Otu21009 | 1 | k__Fungi(100);p__Ascomycota(100);c__Saccharomycetes(100);o__Saccharomycetales(100);f__Metschnikowiaceae(100);g__Metschnikowia(100);s__Metschnikowia_pulcherrima(100); | Metschnikowia | 13 | Saprotroph | \|Undefined Saprotroph\| | Yeast | NULL | Probable | As nectar-tap_saprotroph (Põlme et al. 2020); Decay sugar-rich substrates (Põlme et al. 2020) | Põlme S, et al. 2020. Fungal Diversity 105:1-16 ((https://doi.org/10.1007/s13225-020-00466-2)) |
| Otu21010 | 1 | k__Fungi(100);p__Ascomycota(100);c__Saccharomycetes(100);o__Saccharomycetales(100);f__Metschnikowiaceae(100);g__Metschnikowia(100);s__Metschnikowia_pulcherrima(100); | Metschnikowia | 13 | Saprotroph | \|Undefined Saprotroph\| | Yeast | NULL | Probable | As nectar-tap_saprotroph (Põlme et al. 2020); Decay sugar-rich substrates (Põlme et al. 2020) | Põlme S, et al. 2020. Fungal Diversity 105:1-16 ((https://doi.org/10.1007/s13225-020-00466-2)) |
| Otu21011 | 1 | k__Fungi(100);p__Ascomycota(100);c__Saccharomycetes(100);o__Saccharomycetales(100);f__Metschnikowiaceae(100);g__Metschnikowia(100);s__Metschnikowia_pulcherrima(100); | Metschnikowia | 13 | Saprotroph | \|Undefined Saprotroph\| | Yeast | NULL | Probable | As nectar-tap_saprotroph (Põlme et al. 2020); Decay sugar-rich substrates (Põlme et al. 2020) | Põlme S, et al. 2020. Fungal Diversity 105:1-16 ((https://doi.org/10.1007/s13225-020-00466-2)) |
| Otu21012 | 1 | k__Fungi(100);p__Ascomycota(100);c__Saccharomycetes(100);o__Saccharomycetales(100);f__Metschnikowiaceae(100);g__Metschnikowia(100);s__Metschnikowia_pulcherrima(100); | Metschnikowia | 13 | Saprotroph | \|Undefined Saprotroph\| | Yeast | NULL | Probable | As nectar-tap_saprotroph (Põlme et al. 2020); Decay sugar-rich substrates (Põlme et al. 2020) | Põlme S, et al. 2020. Fungal Diversity 105:1-16 ((https://doi.org/10.1007/s13225-020-00466-2)) |
| Otu21013 | 1 | k__Fungi(100);p__Ascomycota(100);c__Saccharomycetes(100);o__Saccharomycetales(100);f__Metschnikowiaceae(100);g__Metschnikowia(100);s__Metschnikowia_pulcherrima(100); | Metschnikowia | 13 | Saprotroph | \|Undefined Saprotroph\| | Yeast | NULL | Probable | As nectar-tap_saprotroph (Põlme et al. 2020); Decay sugar-rich substrates (Põlme et al. 2020) | Põlme S, et al. 2020. Fungal Diversity 105:1-16 ((https://doi.org/10.1007/s13225-020-00466-2)) |
| Otu21014 | 1 | k__Fungi(100);p__Ascomycota(100);c__Saccharomycetes(100);o__Saccharomycetales(100);f__Metschnikowiaceae(100);g__Metschnikowia(100);g__Metschnikowia_unclassified(100); | Metschnikowia | 13 | Saprotroph | \|Undefined Saprotroph\| | Yeast | NULL | Probable | As nectar-tap_saprotroph (Põlme et al. 2020); Decay sugar-rich substrates (Põlme et al. 2020) | Põlme S, et al. 2020. Fungal Diversity 105:1-16 ((https://doi.org/10.1007/s13225-020-00466-2)) |
| Otu21015 | 1 | k__Fungi(100);p__Ascomycota(100);c__Saccharomycetes(100);o__Saccharomycetales(100);f__Metschnikowiaceae(100);g__Metschnikowia(100);s__Metschnikowia_pulcherrima(100); | Metschnikowia | 13 | Saprotroph | \|Undefined Saprotroph\| | Yeast | NULL | Probable | As nectar-tap_saprotroph (Põlme et al. 2020); Decay sugar-rich substrates (Põlme et al. 2020) | Põlme S, et al. 2020. Fungal Diversity 105:1-16 ((https://doi.org/10.1007/s13225-020-00466-2)) |
| Otu21016 | 1 | k__Fungi(100);p__Ascomycota(100);c__Saccharomycetes(100);o__Saccharomycetales(100);f__Metschnikowiaceae(100);g__Metschnikowia(100);g__Metschnikowia_unclassified(100); | Metschnikowia | 13 | Saprotroph | \|Undefined Saprotroph\| | Yeast | NULL | Probable | As nectar-tap_saprotroph (Põlme et al. 2020); Decay sugar-rich substrates (Põlme et al. 2020) | Põlme S, et al. 2020. Fungal Diversity 105:1-16 ((https://doi.org/10.1007/s13225-020-00466-2)) |
| Otu21017 | 1 | k__Fungi(100);p__Ascomycota(100);c__Saccharomycetes(100);o__Saccharomycetales(100);f__Metschnikowiaceae(100);g__Metschnikowia(100);g__Metschnikowia_unclassified(100); | Metschnikowia | 13 | Saprotroph | \|Undefined Saprotroph\| | Yeast | NULL | Probable | As nectar-tap_saprotroph (Põlme et al. 2020); Decay sugar-rich substrates (Põlme et al. 2020) | Põlme S, et al. 2020. Fungal Diversity 105:1-16 ((https://doi.org/10.1007/s13225-020-00466-2)) |
| Otu21018 | 1 | k__Fungi(100);p__Ascomycota(100);c__Saccharomycetes(100);o__Saccharomycetales(100);f__Metschnikowiaceae(100);g__Metschnikowia(100);s__Metschnikowia_pulcherrima(100); | Metschnikowia | 13 | Saprotroph | \|Undefined Saprotroph\| | Yeast | NULL | Probable | As nectar-tap_saprotroph (Põlme et al. 2020); Decay sugar-rich substrates (Põlme et al. 2020) | Põlme S, et al. 2020. Fungal Diversity 105:1-16 ((https://doi.org/10.1007/s13225-020-00466-2)) |
| Otu21019 | 1 | k__Fungi(100);p__Ascomycota(100);c__Saccharomycetes(100);o__Saccharomycetales(100);f__Metschnikowiaceae(100);g__Metschnikowia(100);s__Metschnikowia_pulcherrima(100); | Metschnikowia | 13 | Saprotroph | \|Undefined Saprotroph\| | Yeast | NULL | Probable | As nectar-tap_saprotroph (Põlme et al. 2020); Decay sugar-rich substrates (Põlme et al. 2020) | Põlme S, et al. 2020. Fungal Diversity 105:1-16 ((https://doi.org/10.1007/s13225-020-00466-2)) |
| Otu21022 | 1 | k__Fungi(100);p__Ascomycota(100);c__Saccharomycetes(100);o__Saccharomycetales(100);f__Metschnikowiaceae(100);g__Metschnikowia(100);s__Metschnikowia_pulcherrima(100); | Metschnikowia | 13 | Saprotroph | \|Undefined Saprotroph\| | Yeast | NULL | Probable | As nectar-tap_saprotroph (Põlme et al. 2020); Decay sugar-rich substrates (Põlme et al. 2020) | Põlme S, et al. 2020. Fungal Diversity 105:1-16 ((https://doi.org/10.1007/s13225-020-00466-2)) |
| Otu21023 | 1 | k__Fungi(100);p__Ascomycota(100);c__Saccharomycetes(100);o__Saccharomycetales(100);f__Metschnikowiaceae(100);g__Metschnikowia(100);g__Metschnikowia_unclassified(100); | Metschnikowia | 13 | Saprotroph | \|Undefined Saprotroph\| | Yeast | NULL | Probable | As nectar-tap_saprotroph (Põlme et al. 2020); Decay sugar-rich substrates (Põlme et al. 2020) | Põlme S, et al. 2020. Fungal Diversity 105:1-16 ((https://doi.org/10.1007/s13225-020-00466-2)) |
| Otu21025 | 1 | k__Fungi(100);p__Ascomycota(100);c__Saccharomycetes(100);o__Saccharomycetales(100);f__Metschnikowiaceae(100);g__Metschnikowia(100);s__Metschnikowia_pulcherrima(100); | Metschnikowia | 13 | Saprotroph | \|Undefined Saprotroph\| | Yeast | NULL | Probable | As nectar-tap_saprotroph (Põlme et al. 2020); Decay sugar-rich substrates (Põlme et al. 2020) | Põlme S, et al. 2020. Fungal Diversity 105:1-16 ((https://doi.org/10.1007/s13225-020-00466-2)) |
| Otu21026 | 1 | k__Fungi(100);p__Ascomycota(100);c__Saccharomycetes(100);o__Saccharomycetales(100);f__Metschnikowiaceae(100);g__Metschnikowia(100);s__Metschnikowia_pulcherrima(100); | Metschnikowia | 13 | Saprotroph | \|Undefined Saprotroph\| | Yeast | NULL | Probable | As nectar-tap_saprotroph (Põlme et al. 2020); Decay sugar-rich substrates (Põlme et al. 2020) | Põlme S, et al. 2020. Fungal Diversity 105:1-16 ((https://doi.org/10.1007/s13225-020-00466-2)) |
| Otu21027 | 1 | k__Fungi(100);p__Ascomycota(100);c__Saccharomycetes(100);o__Saccharomycetales(100);f__Metschnikowiaceae(100);g__Metschnikowia(100);s__Metschnikowia_pulcherrima(100); | Metschnikowia | 13 | Saprotroph | \|Undefined Saprotroph\| | Yeast | NULL | Probable | As nectar-tap_saprotroph (Põlme et al. 2020); Decay sugar-rich substrates (Põlme et al. 2020) | Põlme S, et al. 2020. Fungal Diversity 105:1-16 ((https://doi.org/10.1007/s13225-020-00466-2)) |
| Otu21028 | 1 | k__Fungi(100);p__Ascomycota(100);c__Saccharomycetes(100);o__Saccharomycetales(100);f__Metschnikowiaceae(100);g__Metschnikowia(100);s__Metschnikowia_pulcherrima(100); | Metschnikowia | 13 | Saprotroph | \|Undefined Saprotroph\| | Yeast | NULL | Probable | As nectar-tap_saprotroph (Põlme et al. 2020); Decay sugar-rich substrates (Põlme et al. 2020) | Põlme S, et al. 2020. Fungal Diversity 105:1-16 ((https://doi.org/10.1007/s13225-020-00466-2)) |
| Otu21030 | 1 | k__Fungi(100);p__Ascomycota(100);c__Saccharomycetes(100);o__Saccharomycetales(100);f__Metschnikowiaceae(100);g__Metschnikowia(100);g__Metschnikowia_unclassified(100); | Metschnikowia | 13 | Saprotroph | \|Undefined Saprotroph\| | Yeast | NULL | Probable | As nectar-tap_saprotroph (Põlme et al. 2020); Decay sugar-rich substrates (Põlme et al. 2020) | Põlme S, et al. 2020. Fungal Diversity 105:1-16 ((https://doi.org/10.1007/s13225-020-00466-2)) |
| Otu21034 | 1 | k__Fungi(100);p__Ascomycota(100);c__Saccharomycetes(100);o__Saccharomycetales(100);f__Metschnikowiaceae(100);g__Metschnikowia(100);s__Metschnikowia_pulcherrima(100); | Metschnikowia | 13 | Saprotroph | \|Undefined Saprotroph\| | Yeast | NULL | Probable | As nectar-tap_saprotroph (Põlme et al. 2020); Decay sugar-rich substrates (Põlme et al. 2020) | Põlme S, et al. 2020. Fungal Diversity 105:1-16 ((https://doi.org/10.1007/s13225-020-00466-2)) |
| Otu21035 | 1 | k__Fungi(100);p__Basidiomycota(100);c__Microbotryomycetes(100);o__Sporidiobolales(100);f__Sporidiobolales_family_Incertae_sedis(100);g__Sporobolomyces(100);s__Sporobolomyces_lactosus(100); | Sporobolomyces | 13 | Pathotroph-Saprotroph | \|Fungal Parasite\|-Plant Saprotroph | Yeast | NULL | Probable | As litter_saprotroph (Põlme et al. 2020); As mycoparasite (Põlme et al. 2020); Decay fungal material (Põlme et al. 2020) | Tedersoo L, et al. 2014. Science 346:e1256688 ((https://doi.org/10.1126/science.1256688)); Purahong W, et al. 2016. Molecular Ecology 25:4059-4074 ((https://doi.org/10.1111/mec.13739)); Põlme S, et al. 2020. Fungal Diversity 105:1-16 ((https://doi.org/10.1007/s13225-020-00466-2)) |
| Otu21036 | 1 | k__Fungi(100);p__Ascomycota(100);c__Saccharomycetes(100);o__Saccharomycetales(100);f__Metschnikowiaceae(100);g__Metschnikowia(100);g__Metschnikowia_unclassified(100); | Metschnikowia | 13 | Saprotroph | \|Undefined Saprotroph\| | Yeast | NULL | Probable | As nectar-tap_saprotroph (Põlme et al. 2020); Decay sugar-rich substrates (Põlme et al. 2020) | Põlme S, et al. 2020. Fungal Diversity 105:1-16 ((https://doi.org/10.1007/s13225-020-00466-2)) |
| Otu21037 | 1 | k__Fungi(100);p__Ascomycota(100);c__Saccharomycetes(100);o__Saccharomycetales(100);f__Metschnikowiaceae(100);g__Metschnikowia(100);g__Metschnikowia_unclassified(100); | Metschnikowia | 13 | Saprotroph | \|Undefined Saprotroph\| | Yeast | NULL | Probable | As nectar-tap_saprotroph (Põlme et al. 2020); Decay sugar-rich substrates (Põlme et al. 2020) | Põlme S, et al. 2020. Fungal Diversity 105:1-16 ((https://doi.org/10.1007/s13225-020-00466-2)) |
| Otu21039 | 1 | k__Fungi(100);p__Ascomycota(100);c__Saccharomycetes(100);o__Saccharomycetales(100);f__Metschnikowiaceae(100);g__Metschnikowia(100);g__Metschnikowia_unclassified(100); | Metschnikowia | 13 | Saprotroph | \|Undefined Saprotroph\| | Yeast | NULL | Probable | As nectar-tap_saprotroph (Põlme et al. 2020); Decay sugar-rich substrates (Põlme et al. 2020) | Põlme S, et al. 2020. Fungal Diversity 105:1-16 ((https://doi.org/10.1007/s13225-020-00466-2)) |
| Otu21042 | 1 | k__Fungi(100);p__Ascomycota(100);c__Saccharomycetes(100);o__Saccharomycetales(100);f__Metschnikowiaceae(100);g__Metschnikowia(100);s__Metschnikowia_pulcherrima(100); | Metschnikowia | 13 | Saprotroph | \|Undefined Saprotroph\| | Yeast | NULL | Probable | As nectar-tap_saprotroph (Põlme et al. 2020); Decay sugar-rich substrates (Põlme et al. 2020) | Põlme S, et al. 2020. Fungal Diversity 105:1-16 ((https://doi.org/10.1007/s13225-020-00466-2)) |
| Otu21044 | 1 | k__Fungi(100);p__Ascomycota(100);c__Saccharomycetes(100);o__Saccharomycetales(100);f__Metschnikowiaceae(100);g__Metschnikowia(100);s__Metschnikowia_pulcherrima(100); | Metschnikowia | 13 | Saprotroph | \|Undefined Saprotroph\| | Yeast | NULL | Probable | As nectar-tap_saprotroph (Põlme et al. 2020); Decay sugar-rich substrates (Põlme et al. 2020) | Põlme S, et al. 2020. Fungal Diversity 105:1-16 ((https://doi.org/10.1007/s13225-020-00466-2)) |
| Otu21045 | 1 | k__Fungi(100);p__Ascomycota(100);c__Saccharomycetes(100);o__Saccharomycetales(100);f__Metschnikowiaceae(100);g__Metschnikowia(100);s__Metschnikowia_pulcherrima(100); | Metschnikowia | 13 | Saprotroph | \|Undefined Saprotroph\| | Yeast | NULL | Probable | As nectar-tap_saprotroph (Põlme et al. 2020); Decay sugar-rich substrates (Põlme et al. 2020) | Põlme S, et al. 2020. Fungal Diversity 105:1-16 ((https://doi.org/10.1007/s13225-020-00466-2)) |
| Otu21046 | 1 | k__Fungi(100);p__Ascomycota(100);c__Saccharomycetes(100);o__Saccharomycetales(100);f__Saccharomycetales_family_Incertae_sedis(100);g__Candida(100);g__Candida_unclassified(100); | Candida | 13 | Pathotroph-Saprotroph-Symbiotroph | Animal Pathogen-Endophyte-Endosymbiont-Epiphyte-\|Undefined Saprotroph\| | Dimorphic Yeast-Yeast | NULL | Probable | As nectar-tap_saprotroph (Põlme et al. 2020); Decay sugar-rich substrates (Põlme et al. 2020); This asexual genus is polyphyletic and contain species that function very differently depend on what they are. Caution should be taken when interpreting guilds for Candida species. | Manolakaki D, et al. 2010. Virulence. 1: 367-375 ((https://doi.org/10.4161/viru.1.5.12796)); Mekbib SB, et al. 2011. Tropical Plant Pathology 36:233-240 ((https://doi.org/10.1590/S1982-56762011000400004)); Põlme S, et al. 2020. Fungal Diversity 105:1-16 ((https://doi.org/10.1007/s13225-020-00466-2)) |
| Otu21048 | 1 | k__Fungi(100);p__Ascomycota(100);c__Saccharomycetes(100);o__Saccharomycetales(100);f__Saccharomycetales_family_Incertae_sedis(100);f__Saccharomycetales_family_Incertae_sedis_unclassified(100);f__Saccharomycetales_family_Incertae_sedis_unclassified(100); | Saccharomycetales | 7 | Saprotroph | Undefined Saprotroph | Yeast | NULL | Possible | NULL | Sterkenburg E, et al. 2015. New Phytologist 207:1145-1158 ((https://doi.org/10.1111/nph.13426)) |
| Otu21050 | 1 | k__Fungi(100);p__Ascomycota(100);c__Saccharomycetes(100);o__Saccharomycetales(100);f__Saccharomycetales_family_Incertae_sedis(100);g__Candida(100);g__Candida_unclassified(100); | Candida | 13 | Pathotroph-Saprotroph-Symbiotroph | Animal Pathogen-Endophyte-Endosymbiont-Epiphyte-\|Undefined Saprotroph\| | Dimorphic Yeast-Yeast | NULL | Probable | As nectar-tap_saprotroph (Põlme et al. 2020); Decay sugar-rich substrates (Põlme et al. 2020); This asexual genus is polyphyletic and contain species that function very differently depend on what they are. Caution should be taken when interpreting guilds for Candida species. | Manolakaki D, et al. 2010. Virulence. 1: 367-375 ((https://doi.org/10.4161/viru.1.5.12796)); Mekbib SB, et al. 2011. Tropical Plant Pathology 36:233-240 ((https://doi.org/10.1590/S1982-56762011000400004)); Põlme S, et al. 2020. Fungal Diversity 105:1-16 ((https://doi.org/10.1007/s13225-020-00466-2)) |
| Otu21051 | 1 | k__Fungi(100);p__Ascomycota(100);c__Saccharomycetes(100);o__Saccharomycetales(100);o__Saccharomycetales_unclassified(100);o__Saccharomycetales_unclassified(100);o__Saccharomycetales_unclassified(100); | Saccharomycetales | 7 | Saprotroph | Undefined Saprotroph | Yeast | NULL | Possible | NULL | Sterkenburg E, et al. 2015. New Phytologist 207:1145-1158 ((https://doi.org/10.1111/nph.13426)) |
| Otu21052 | 1 | k__Fungi(100);p__Ascomycota(100);c__Saccharomycetes(100);o__Saccharomycetales(100);f__Saccharomycetales_family_Incertae_sedis(100);g__Candida(100);s__Candida_cretensis(100); | Candida | 13 | Pathotroph-Saprotroph-Symbiotroph | Animal Pathogen-Endophyte-Endosymbiont-Epiphyte-\|Undefined Saprotroph\| | Dimorphic Yeast-Yeast | NULL | Probable | As nectar-tap_saprotroph (Põlme et al. 2020); Decay sugar-rich substrates (Põlme et al. 2020); This asexual genus is polyphyletic and contain species that function very differently depend on what they are. Caution should be taken when interpreting guilds for Candida species. | Manolakaki D, et al. 2010. Virulence. 1: 367-375 ((https://doi.org/10.4161/viru.1.5.12796)); Mekbib SB, et al. 2011. Tropical Plant Pathology 36:233-240 ((https://doi.org/10.1590/S1982-56762011000400004)); Põlme S, et al. 2020. Fungal Diversity 105:1-16 ((https://doi.org/10.1007/s13225-020-00466-2)) |
| Otu21059 | 1 | k__Fungi(100);p__Ascomycota(100);c__Saccharomycetes(100);o__Saccharomycetales(100);f__Saccharomycetales_family_Incertae_sedis(100);g__Candida(100);g__Candida_unclassified(100); | Candida | 13 | Pathotroph-Saprotroph-Symbiotroph | Animal Pathogen-Endophyte-Endosymbiont-Epiphyte-\|Undefined Saprotroph\| | Dimorphic Yeast-Yeast | NULL | Probable | As nectar-tap_saprotroph (Põlme et al. 2020); Decay sugar-rich substrates (Põlme et al. 2020); This asexual genus is polyphyletic and contain species that function very differently depend on what they are. Caution should be taken when interpreting guilds for Candida species. | Manolakaki D, et al. 2010. Virulence. 1: 367-375 ((https://doi.org/10.4161/viru.1.5.12796)); Mekbib SB, et al. 2011. Tropical Plant Pathology 36:233-240 ((https://doi.org/10.1590/S1982-56762011000400004)); Põlme S, et al. 2020. Fungal Diversity 105:1-16 ((https://doi.org/10.1007/s13225-020-00466-2)) |
| Otu21060 | 1 | k__Fungi(100);p__Ascomycota(100);c__Saccharomycetes(100);o__Saccharomycetales(100);f__Saccharomycetales_family_Incertae_sedis(100);g__Candida(100);g__Candida_unclassified(100); | Candida | 13 | Pathotroph-Saprotroph-Symbiotroph | Animal Pathogen-Endophyte-Endosymbiont-Epiphyte-\|Undefined Saprotroph\| | Dimorphic Yeast-Yeast | NULL | Probable | As nectar-tap_saprotroph (Põlme et al. 2020); Decay sugar-rich substrates (Põlme et al. 2020); This asexual genus is polyphyletic and contain species that function very differently depend on what they are. Caution should be taken when interpreting guilds for Candida species. | Manolakaki D, et al. 2010. Virulence. 1: 367-375 ((https://doi.org/10.4161/viru.1.5.12796)); Mekbib SB, et al. 2011. Tropical Plant Pathology 36:233-240 ((https://doi.org/10.1590/S1982-56762011000400004)); Põlme S, et al. 2020. Fungal Diversity 105:1-16 ((https://doi.org/10.1007/s13225-020-00466-2)) |
| Otu21062 | 1 | k__Fungi(100);p__Ascomycota(100);c__Saccharomycetes(100);o__Saccharomycetales(100);f__Metschnikowiaceae(100);g__Metschnikowia(100);s__Metschnikowia_pulcherrima(100); | Metschnikowia | 13 | Saprotroph | \|Undefined Saprotroph\| | Yeast | NULL | Probable | As nectar-tap_saprotroph (Põlme et al. 2020); Decay sugar-rich substrates (Põlme et al. 2020) | Põlme S, et al. 2020. Fungal Diversity 105:1-16 ((https://doi.org/10.1007/s13225-020-00466-2)) |
| Otu21064 | 1 | k__Fungi(100);p__Ascomycota(100);c__Saccharomycetes(100);o__Saccharomycetales(100);f__Saccharomycetales_family_Incertae_sedis(100);g__Candida(100);s__Candida_cretensis(100); | Candida | 13 | Pathotroph-Saprotroph-Symbiotroph | Animal Pathogen-Endophyte-Endosymbiont-Epiphyte-\|Undefined Saprotroph\| | Dimorphic Yeast-Yeast | NULL | Probable | As nectar-tap_saprotroph (Põlme et al. 2020); Decay sugar-rich substrates (Põlme et al. 2020); This asexual genus is polyphyletic and contain species that function very differently depend on what they are. Caution should be taken when interpreting guilds for Candida species. | Manolakaki D, et al. 2010. Virulence. 1: 367-375 ((https://doi.org/10.4161/viru.1.5.12796)); Mekbib SB, et al. 2011. Tropical Plant Pathology 36:233-240 ((https://doi.org/10.1590/S1982-56762011000400004)); Põlme S, et al. 2020. Fungal Diversity 105:1-16 ((https://doi.org/10.1007/s13225-020-00466-2)) |
| Otu21065 | 1 | k__Fungi(100);p__Ascomycota(100);c__Saccharomycetes(100);o__Saccharomycetales(100);f__Saccharomycetales_family_Incertae_sedis(100);g__Candida(100);s__Candida_cretensis(100); | Candida | 13 | Pathotroph-Saprotroph-Symbiotroph | Animal Pathogen-Endophyte-Endosymbiont-Epiphyte-\|Undefined Saprotroph\| | Dimorphic Yeast-Yeast | NULL | Probable | As nectar-tap_saprotroph (Põlme et al. 2020); Decay sugar-rich substrates (Põlme et al. 2020); This asexual genus is polyphyletic and contain species that function very differently depend on what they are. Caution should be taken when interpreting guilds for Candida species. | Manolakaki D, et al. 2010. Virulence. 1: 367-375 ((https://doi.org/10.4161/viru.1.5.12796)); Mekbib SB, et al. 2011. Tropical Plant Pathology 36:233-240 ((https://doi.org/10.1590/S1982-56762011000400004)); Põlme S, et al. 2020. Fungal Diversity 105:1-16 ((https://doi.org/10.1007/s13225-020-00466-2)) |
| Otu21066 | 1 | k__Fungi(100);p__Ascomycota(100);c__Saccharomycetes(100);o__Saccharomycetales(100);f__Saccharomycetales_family_Incertae_sedis(100);f__Saccharomycetales_family_Incertae_sedis_unclassified(100);f__Saccharomycetales_family_Incertae_sedis_unclassified(100); | Saccharomycetales | 7 | Saprotroph | Undefined Saprotroph | Yeast | NULL | Possible | NULL | Sterkenburg E, et al. 2015. New Phytologist 207:1145-1158 ((https://doi.org/10.1111/nph.13426)) |
| Otu21068 | 1 | k__Fungi(100);p__Ascomycota(100);c__Saccharomycetes(100);o__Saccharomycetales(100);f__Metschnikowiaceae(100);g__Metschnikowia(100);s__Metschnikowia_pulcherrima(100); | Metschnikowia | 13 | Saprotroph | \|Undefined Saprotroph\| | Yeast | NULL | Probable | As nectar-tap_saprotroph (Põlme et al. 2020); Decay sugar-rich substrates (Põlme et al. 2020) | Põlme S, et al. 2020. Fungal Diversity 105:1-16 ((https://doi.org/10.1007/s13225-020-00466-2)) |
| Otu21069 | 1 | k__Fungi(100);p__Ascomycota(100);c__Saccharomycetes(100);o__Saccharomycetales(100);f__Saccharomycetales_family_Incertae_sedis(100);g__Candida(100);s__Candida_cretensis(100); | Candida | 13 | Pathotroph-Saprotroph-Symbiotroph | Animal Pathogen-Endophyte-Endosymbiont-Epiphyte-\|Undefined Saprotroph\| | Dimorphic Yeast-Yeast | NULL | Probable | As nectar-tap_saprotroph (Põlme et al. 2020); Decay sugar-rich substrates (Põlme et al. 2020); This asexual genus is polyphyletic and contain species that function very differently depend on what they are. Caution should be taken when interpreting guilds for Candida species. | Manolakaki D, et al. 2010. Virulence. 1: 367-375 ((https://doi.org/10.4161/viru.1.5.12796)); Mekbib SB, et al. 2011. Tropical Plant Pathology 36:233-240 ((https://doi.org/10.1590/S1982-56762011000400004)); Põlme S, et al. 2020. Fungal Diversity 105:1-16 ((https://doi.org/10.1007/s13225-020-00466-2)) |
| Otu21070 | 1 | k__Fungi(100);p__Ascomycota(100);c__Saccharomycetes(100);o__Saccharomycetales(100);f__Saccharomycetales_family_Incertae_sedis(100);g__Candida(100);g__Candida_unclassified(100); | Candida | 13 | Pathotroph-Saprotroph-Symbiotroph | Animal Pathogen-Endophyte-Endosymbiont-Epiphyte-\|Undefined Saprotroph\| | Dimorphic Yeast-Yeast | NULL | Probable | As nectar-tap_saprotroph (Põlme et al. 2020); Decay sugar-rich substrates (Põlme et al. 2020); This asexual genus is polyphyletic and contain species that function very differently depend on what they are. Caution should be taken when interpreting guilds for Candida species. | Manolakaki D, et al. 2010. Virulence. 1: 367-375 ((https://doi.org/10.4161/viru.1.5.12796)); Mekbib SB, et al. 2011. Tropical Plant Pathology 36:233-240 ((https://doi.org/10.1590/S1982-56762011000400004)); Põlme S, et al. 2020. Fungal Diversity 105:1-16 ((https://doi.org/10.1007/s13225-020-00466-2)) |
| Otu21073 | 1 | k__Fungi(100);p__Ascomycota(100);c__Saccharomycetes(100);o__Saccharomycetales(100);f__Metschnikowiaceae(100);g__Metschnikowia(100);s__Metschnikowia_pulcherrima(100); | Metschnikowia | 13 | Saprotroph | \|Undefined Saprotroph\| | Yeast | NULL | Probable | As nectar-tap_saprotroph (Põlme et al. 2020); Decay sugar-rich substrates (Põlme et al. 2020) | Põlme S, et al. 2020. Fungal Diversity 105:1-16 ((https://doi.org/10.1007/s13225-020-00466-2)) |
| Otu21074 | 1 | k__Fungi(100);p__Ascomycota(100);c__Saccharomycetes(100);o__Saccharomycetales(100);f__Metschnikowiaceae(100);g__Metschnikowia(100);s__Metschnikowia_pulcherrima(100); | Metschnikowia | 13 | Saprotroph | \|Undefined Saprotroph\| | Yeast | NULL | Probable | As nectar-tap_saprotroph (Põlme et al. 2020); Decay sugar-rich substrates (Põlme et al. 2020) | Põlme S, et al. 2020. Fungal Diversity 105:1-16 ((https://doi.org/10.1007/s13225-020-00466-2)) |
| Otu21075 | 1 | k__Fungi(100);p__Ascomycota(100);c__Saccharomycetes(100);o__Saccharomycetales(100);f__Metschnikowiaceae(100);g__Metschnikowia(100);s__Metschnikowia_pulcherrima(100); | Metschnikowia | 13 | Saprotroph | \|Undefined Saprotroph\| | Yeast | NULL | Probable | As nectar-tap_saprotroph (Põlme et al. 2020); Decay sugar-rich substrates (Põlme et al. 2020) | Põlme S, et al. 2020. Fungal Diversity 105:1-16 ((https://doi.org/10.1007/s13225-020-00466-2)) |
| Otu21076 | 1 | k__Fungi(100);p__Ascomycota(100);c__Saccharomycetes(100);o__Saccharomycetales(100);f__Metschnikowiaceae(100);g__Metschnikowia(100);s__Metschnikowia_pulcherrima(100); | Metschnikowia | 13 | Saprotroph | \|Undefined Saprotroph\| | Yeast | NULL | Probable | As nectar-tap_saprotroph (Põlme et al. 2020); Decay sugar-rich substrates (Põlme et al. 2020) | Põlme S, et al. 2020. Fungal Diversity 105:1-16 ((https://doi.org/10.1007/s13225-020-00466-2)) |
| Otu21077 | 1 | k__Fungi(100);p__Ascomycota(100);c__Saccharomycetes(100);o__Saccharomycetales(100);o__Saccharomycetales_unclassified(100);o__Saccharomycetales_unclassified(100);o__Saccharomycetales_unclassified(100); | Saccharomycetales | 7 | Saprotroph | Undefined Saprotroph | Yeast | NULL | Possible | NULL | Sterkenburg E, et al. 2015. New Phytologist 207:1145-1158 ((https://doi.org/10.1111/nph.13426)) |
| Otu21078 | 1 | k__Fungi(100);p__Ascomycota(100);c__Saccharomycetes(100);o__Saccharomycetales(100);f__Metschnikowiaceae(100);g__Metschnikowia(100);s__Metschnikowia_pulcherrima(100); | Metschnikowia | 13 | Saprotroph | \|Undefined Saprotroph\| | Yeast | NULL | Probable | As nectar-tap_saprotroph (Põlme et al. 2020); Decay sugar-rich substrates (Põlme et al. 2020) | Põlme S, et al. 2020. Fungal Diversity 105:1-16 ((https://doi.org/10.1007/s13225-020-00466-2)) |
| Otu21079 | 1 | k__Fungi(100);p__Ascomycota(100);c__Saccharomycetes(100);o__Saccharomycetales(100);f__Metschnikowiaceae(100);g__Metschnikowia(100);s__Metschnikowia_pulcherrima(100); | Metschnikowia | 13 | Saprotroph | \|Undefined Saprotroph\| | Yeast | NULL | Probable | As nectar-tap_saprotroph (Põlme et al. 2020); Decay sugar-rich substrates (Põlme et al. 2020) | Põlme S, et al. 2020. Fungal Diversity 105:1-16 ((https://doi.org/10.1007/s13225-020-00466-2)) |
| Otu21080 | 1 | k__Fungi(100);p__Ascomycota(100);c__Saccharomycetes(100);o__Saccharomycetales(100);f__Metschnikowiaceae(100);g__Metschnikowia(100);s__Metschnikowia_pulcherrima(100); | Metschnikowia | 13 | Saprotroph | \|Undefined Saprotroph\| | Yeast | NULL | Probable | As nectar-tap_saprotroph (Põlme et al. 2020); Decay sugar-rich substrates (Põlme et al. 2020) | Põlme S, et al. 2020. Fungal Diversity 105:1-16 ((https://doi.org/10.1007/s13225-020-00466-2)) |
| Otu21081 | 1 | k__Fungi(100);p__Ascomycota(100);c__Saccharomycetes(100);o__Saccharomycetales(100);f__Metschnikowiaceae(100);g__Metschnikowia(100);g__Metschnikowia_unclassified(100); | Metschnikowia | 13 | Saprotroph | \|Undefined Saprotroph\| | Yeast | NULL | Probable | As nectar-tap_saprotroph (Põlme et al. 2020); Decay sugar-rich substrates (Põlme et al. 2020) | Põlme S, et al. 2020. Fungal Diversity 105:1-16 ((https://doi.org/10.1007/s13225-020-00466-2)) |
| Otu21082 | 1 | k__Fungi(100);p__Ascomycota(100);c__Saccharomycetes(100);o__Saccharomycetales(100);f__Metschnikowiaceae(100);g__Metschnikowia(100);s__Metschnikowia_pulcherrima(100); | Metschnikowia | 13 | Saprotroph | \|Undefined Saprotroph\| | Yeast | NULL | Probable | As nectar-tap_saprotroph (Põlme et al. 2020); Decay sugar-rich substrates (Põlme et al. 2020) | Põlme S, et al. 2020. Fungal Diversity 105:1-16 ((https://doi.org/10.1007/s13225-020-00466-2)) |
| Otu21083 | 1 | k__Fungi(100);p__Ascomycota(100);c__Saccharomycetes(100);o__Saccharomycetales(100);f__Metschnikowiaceae(100);g__Metschnikowia(100);s__Metschnikowia_pulcherrima(100); | Metschnikowia | 13 | Saprotroph | \|Undefined Saprotroph\| | Yeast | NULL | Probable | As nectar-tap_saprotroph (Põlme et al. 2020); Decay sugar-rich substrates (Põlme et al. 2020) | Põlme S, et al. 2020. Fungal Diversity 105:1-16 ((https://doi.org/10.1007/s13225-020-00466-2)) |
| Otu21084 | 1 | k__Fungi(100);p__Ascomycota(100);c__Saccharomycetes(100);o__Saccharomycetales(100);f__Metschnikowiaceae(100);g__Metschnikowia(100);g__Metschnikowia_unclassified(100); | Metschnikowia | 13 | Saprotroph | \|Undefined Saprotroph\| | Yeast | NULL | Probable | As nectar-tap_saprotroph (Põlme et al. 2020); Decay sugar-rich substrates (Põlme et al. 2020) | Põlme S, et al. 2020. Fungal Diversity 105:1-16 ((https://doi.org/10.1007/s13225-020-00466-2)) |
| Otu21087 | 1 | k__Fungi(100);p__Ascomycota(100);c__Saccharomycetes(100);o__Saccharomycetales(100);f__Saccharomycetales_family_Incertae_sedis(100);g__Candida(100);g__Candida_unclassified(100); | Candida | 13 | Pathotroph-Saprotroph-Symbiotroph | Animal Pathogen-Endophyte-Endosymbiont-Epiphyte-\|Undefined Saprotroph\| | Dimorphic Yeast-Yeast | NULL | Probable | As nectar-tap_saprotroph (Põlme et al. 2020); Decay sugar-rich substrates (Põlme et al. 2020); This asexual genus is polyphyletic and contain species that function very differently depend on what they are. Caution should be taken when interpreting guilds for Candida species. | Manolakaki D, et al. 2010. Virulence. 1: 367-375 ((https://doi.org/10.4161/viru.1.5.12796)); Mekbib SB, et al. 2011. Tropical Plant Pathology 36:233-240 ((https://doi.org/10.1590/S1982-56762011000400004)); Põlme S, et al. 2020. Fungal Diversity 105:1-16 ((https://doi.org/10.1007/s13225-020-00466-2)) |
| Otu21088 | 1 | k__Fungi(100);p__Ascomycota(100);c__Saccharomycetes(100);o__Saccharomycetales(100);f__Saccharomycetales_family_Incertae_sedis(100);g__Candida(100);g__Candida_unclassified(100); | Candida | 13 | Pathotroph-Saprotroph-Symbiotroph | Animal Pathogen-Endophyte-Endosymbiont-Epiphyte-\|Undefined Saprotroph\| | Dimorphic Yeast-Yeast | NULL | Probable | As nectar-tap_saprotroph (Põlme et al. 2020); Decay sugar-rich substrates (Põlme et al. 2020); This asexual genus is polyphyletic and contain species that function very differently depend on what they are. Caution should be taken when interpreting guilds for Candida species. | Manolakaki D, et al. 2010. Virulence. 1: 367-375 ((https://doi.org/10.4161/viru.1.5.12796)); Mekbib SB, et al. 2011. Tropical Plant Pathology 36:233-240 ((https://doi.org/10.1590/S1982-56762011000400004)); Põlme S, et al. 2020. Fungal Diversity 105:1-16 ((https://doi.org/10.1007/s13225-020-00466-2)) |
| Otu21089 | 1 | k__Fungi(100);p__Ascomycota(100);c__Saccharomycetes(100);o__Saccharomycetales(100);o__Saccharomycetales_unclassified(100);o__Saccharomycetales_unclassified(100);o__Saccharomycetales_unclassified(100); | Saccharomycetales | 7 | Saprotroph | Undefined Saprotroph | Yeast | NULL | Possible | NULL | Sterkenburg E, et al. 2015. New Phytologist 207:1145-1158 ((https://doi.org/10.1111/nph.13426)) |
| Otu21090 | 1 | k__Fungi(100);p__Ascomycota(100);c__Saccharomycetes(100);o__Saccharomycetales(100);o__Saccharomycetales_unclassified(100);o__Saccharomycetales_unclassified(100);o__Saccharomycetales_unclassified(100); | Saccharomycetales | 7 | Saprotroph | Undefined Saprotroph | Yeast | NULL | Possible | NULL | Sterkenburg E, et al. 2015. New Phytologist 207:1145-1158 ((https://doi.org/10.1111/nph.13426)) |
| Otu21091 | 1 | k__Fungi(100);p__Ascomycota(100);c__Saccharomycetes(100);o__Saccharomycetales(100);f__Metschnikowiaceae(100);g__Metschnikowia(100);s__Metschnikowia_pulcherrima(100); | Metschnikowia | 13 | Saprotroph | \|Undefined Saprotroph\| | Yeast | NULL | Probable | As nectar-tap_saprotroph (Põlme et al. 2020); Decay sugar-rich substrates (Põlme et al. 2020) | Põlme S, et al. 2020. Fungal Diversity 105:1-16 ((https://doi.org/10.1007/s13225-020-00466-2)) |
| Otu21092 | 1 | k__Fungi(100);p__Ascomycota(100);c__Saccharomycetes(100);o__Saccharomycetales(100);f__Metschnikowiaceae(100);g__Metschnikowia(100);s__Metschnikowia_pulcherrima(100); | Metschnikowia | 13 | Saprotroph | \|Undefined Saprotroph\| | Yeast | NULL | Probable | As nectar-tap_saprotroph (Põlme et al. 2020); Decay sugar-rich substrates (Põlme et al. 2020) | Põlme S, et al. 2020. Fungal Diversity 105:1-16 ((https://doi.org/10.1007/s13225-020-00466-2)) |
| Otu21093 | 1 | k__Fungi(100);p__Ascomycota(100);c__Saccharomycetes(100);o__Saccharomycetales(100);f__Metschnikowiaceae(100);g__Metschnikowia(100);g__Metschnikowia_unclassified(100); | Metschnikowia | 13 | Saprotroph | \|Undefined Saprotroph\| | Yeast | NULL | Probable | As nectar-tap_saprotroph (Põlme et al. 2020); Decay sugar-rich substrates (Põlme et al. 2020) | Põlme S, et al. 2020. Fungal Diversity 105:1-16 ((https://doi.org/10.1007/s13225-020-00466-2)) |
| Otu21096 | 1 | k__Fungi(100);p__Ascomycota(100);c__Saccharomycetes(100);o__Saccharomycetales(100);f__Saccharomycetales_family_Incertae_sedis(100);g__Candida(100);g__Candida_unclassified(100); | Candida | 13 | Pathotroph-Saprotroph-Symbiotroph | Animal Pathogen-Endophyte-Endosymbiont-Epiphyte-\|Undefined Saprotroph\| | Dimorphic Yeast-Yeast | NULL | Probable | As nectar-tap_saprotroph (Põlme et al. 2020); Decay sugar-rich substrates (Põlme et al. 2020); This asexual genus is polyphyletic and contain species that function very differently depend on what they are. Caution should be taken when interpreting guilds for Candida species. | Manolakaki D, et al. 2010. Virulence. 1: 367-375 ((https://doi.org/10.4161/viru.1.5.12796)); Mekbib SB, et al. 2011. Tropical Plant Pathology 36:233-240 ((https://doi.org/10.1590/S1982-56762011000400004)); Põlme S, et al. 2020. Fungal Diversity 105:1-16 ((https://doi.org/10.1007/s13225-020-00466-2)) |
| Otu21097 | 1 | k__Fungi(100);p__Ascomycota(100);c__Saccharomycetes(100);o__Saccharomycetales(100);f__Saccharomycetales_family_Incertae_sedis(100);f__Saccharomycetales_family_Incertae_sedis_unclassified(100);f__Saccharomycetales_family_Incertae_sedis_unclassified(100); | Saccharomycetales | 7 | Saprotroph | Undefined Saprotroph | Yeast | NULL | Possible | NULL | Sterkenburg E, et al. 2015. New Phytologist 207:1145-1158 ((https://doi.org/10.1111/nph.13426)) |
| Otu21098 | 1 | k__Fungi(100);p__Ascomycota(100);c__Saccharomycetes(100);o__Saccharomycetales(100);f__Saccharomycetales_family_Incertae_sedis(100);g__Candida(100);g__Candida_unclassified(100); | Candida | 13 | Pathotroph-Saprotroph-Symbiotroph | Animal Pathogen-Endophyte-Endosymbiont-Epiphyte-\|Undefined Saprotroph\| | Dimorphic Yeast-Yeast | NULL | Probable | As nectar-tap_saprotroph (Põlme et al. 2020); Decay sugar-rich substrates (Põlme et al. 2020); This asexual genus is polyphyletic and contain species that function very differently depend on what they are. Caution should be taken when interpreting guilds for Candida species. | Manolakaki D, et al. 2010. Virulence. 1: 367-375 ((https://doi.org/10.4161/viru.1.5.12796)); Mekbib SB, et al. 2011. Tropical Plant Pathology 36:233-240 ((https://doi.org/10.1590/S1982-56762011000400004)); Põlme S, et al. 2020. Fungal Diversity 105:1-16 ((https://doi.org/10.1007/s13225-020-00466-2)) |
| Otu21104 | 1 | k__Fungi(100);p__Ascomycota(100);c__Saccharomycetes(100);o__Saccharomycetales(100);o__Saccharomycetales_unclassified(100);o__Saccharomycetales_unclassified(100);o__Saccharomycetales_unclassified(100); | Saccharomycetales | 7 | Saprotroph | Undefined Saprotroph | Yeast | NULL | Possible | NULL | Sterkenburg E, et al. 2015. New Phytologist 207:1145-1158 ((https://doi.org/10.1111/nph.13426)) |
| Otu21105 | 1 | k__Fungi(100);p__Ascomycota(100);c__Saccharomycetes(100);o__Saccharomycetales(100);f__Metschnikowiaceae(100);g__Metschnikowia(100);s__Metschnikowia_pulcherrima(100); | Metschnikowia | 13 | Saprotroph | \|Undefined Saprotroph\| | Yeast | NULL | Probable | As nectar-tap_saprotroph (Põlme et al. 2020); Decay sugar-rich substrates (Põlme et al. 2020) | Põlme S, et al. 2020. Fungal Diversity 105:1-16 ((https://doi.org/10.1007/s13225-020-00466-2)) |
| Otu21106 | 1 | k__Fungi(100);p__Ascomycota(100);c__Saccharomycetes(100);o__Saccharomycetales(100);f__Metschnikowiaceae(100);g__Metschnikowia(100);s__Metschnikowia_pulcherrima(100); | Metschnikowia | 13 | Saprotroph | \|Undefined Saprotroph\| | Yeast | NULL | Probable | As nectar-tap_saprotroph (Põlme et al. 2020); Decay sugar-rich substrates (Põlme et al. 2020) | Põlme S, et al. 2020. Fungal Diversity 105:1-16 ((https://doi.org/10.1007/s13225-020-00466-2)) |
| Otu21107 | 1 | k__Fungi(100);p__Ascomycota(100);c__Saccharomycetes(100);o__Saccharomycetales(100);f__Metschnikowiaceae(100);g__Metschnikowia(100);s__Metschnikowia_pulcherrima(100); | Metschnikowia | 13 | Saprotroph | \|Undefined Saprotroph\| | Yeast | NULL | Probable | As nectar-tap_saprotroph (Põlme et al. 2020); Decay sugar-rich substrates (Põlme et al. 2020) | Põlme S, et al. 2020. Fungal Diversity 105:1-16 ((https://doi.org/10.1007/s13225-020-00466-2)) |
| Otu21108 | 1 | k__Fungi(100);p__Ascomycota(100);c__Saccharomycetes(100);o__Saccharomycetales(100);f__Metschnikowiaceae(100);g__Metschnikowia(100);s__Metschnikowia_pulcherrima(100); | Metschnikowia | 13 | Saprotroph | \|Undefined Saprotroph\| | Yeast | NULL | Probable | As nectar-tap_saprotroph (Põlme et al. 2020); Decay sugar-rich substrates (Põlme et al. 2020) | Põlme S, et al. 2020. Fungal Diversity 105:1-16 ((https://doi.org/10.1007/s13225-020-00466-2)) |
| Otu21109 | 1 | k__Fungi(100);p__Ascomycota(100);c__Saccharomycetes(100);o__Saccharomycetales(100);f__Metschnikowiaceae(100);g__Metschnikowia(100);s__Metschnikowia_pulcherrima(100); | Metschnikowia | 13 | Saprotroph | \|Undefined Saprotroph\| | Yeast | NULL | Probable | As nectar-tap_saprotroph (Põlme et al. 2020); Decay sugar-rich substrates (Põlme et al. 2020) | Põlme S, et al. 2020. Fungal Diversity 105:1-16 ((https://doi.org/10.1007/s13225-020-00466-2)) |
| Otu21110 | 1 | k__Fungi(100);p__Ascomycota(100);c__Saccharomycetes(100);o__Saccharomycetales(100);f__Saccharomycetales_family_Incertae_sedis(100);g__Candida(100);g__Candida_unclassified(100); | Candida | 13 | Pathotroph-Saprotroph-Symbiotroph | Animal Pathogen-Endophyte-Endosymbiont-Epiphyte-\|Undefined Saprotroph\| | Dimorphic Yeast-Yeast | NULL | Probable | As nectar-tap_saprotroph (Põlme et al. 2020); Decay sugar-rich substrates (Põlme et al. 2020); This asexual genus is polyphyletic and contain species that function very differently depend on what they are. Caution should be taken when interpreting guilds for Candida species. | Manolakaki D, et al. 2010. Virulence. 1: 367-375 ((https://doi.org/10.4161/viru.1.5.12796)); Mekbib SB, et al. 2011. Tropical Plant Pathology 36:233-240 ((https://doi.org/10.1590/S1982-56762011000400004)); Põlme S, et al. 2020. Fungal Diversity 105:1-16 ((https://doi.org/10.1007/s13225-020-00466-2)) |
| Otu21113 | 1 | k__Fungi(100);p__Ascomycota(100);c__Saccharomycetes(100);o__Saccharomycetales(100);f__Metschnikowiaceae(100);g__Metschnikowia(100);g__Metschnikowia_unclassified(100); | Metschnikowia | 13 | Saprotroph | \|Undefined Saprotroph\| | Yeast | NULL | Probable | As nectar-tap_saprotroph (Põlme et al. 2020); Decay sugar-rich substrates (Põlme et al. 2020) | Põlme S, et al. 2020. Fungal Diversity 105:1-16 ((https://doi.org/10.1007/s13225-020-00466-2)) |
| Otu21114 | 1 | k__Fungi(100);p__Ascomycota(100);c__Saccharomycetes(100);o__Saccharomycetales(100);f__Metschnikowiaceae(100);g__Metschnikowia(100);g__Metschnikowia_unclassified(100); | Metschnikowia | 13 | Saprotroph | \|Undefined Saprotroph\| | Yeast | NULL | Probable | As nectar-tap_saprotroph (Põlme et al. 2020); Decay sugar-rich substrates (Põlme et al. 2020) | Põlme S, et al. 2020. Fungal Diversity 105:1-16 ((https://doi.org/10.1007/s13225-020-00466-2)) |
| Otu21115 | 1 | k__Fungi(100);p__Ascomycota(100);c__Saccharomycetes(100);o__Saccharomycetales(100);f__Metschnikowiaceae(100);g__Metschnikowia(100);s__Metschnikowia_pulcherrima(100); | Metschnikowia | 13 | Saprotroph | \|Undefined Saprotroph\| | Yeast | NULL | Probable | As nectar-tap_saprotroph (Põlme et al. 2020); Decay sugar-rich substrates (Põlme et al. 2020) | Põlme S, et al. 2020. Fungal Diversity 105:1-16 ((https://doi.org/10.1007/s13225-020-00466-2)) |
| Otu21118 | 1 | k__Fungi(100);p__Ascomycota(100);c__Saccharomycetes(100);o__Saccharomycetales(100);f__Saccharomycetales_family_Incertae_sedis(100);g__Candida(100);g__Candida_unclassified(100); | Candida | 13 | Pathotroph-Saprotroph-Symbiotroph | Animal Pathogen-Endophyte-Endosymbiont-Epiphyte-\|Undefined Saprotroph\| | Dimorphic Yeast-Yeast | NULL | Probable | As nectar-tap_saprotroph (Põlme et al. 2020); Decay sugar-rich substrates (Põlme et al. 2020); This asexual genus is polyphyletic and contain species that function very differently depend on what they are. Caution should be taken when interpreting guilds for Candida species. | Manolakaki D, et al. 2010. Virulence. 1: 367-375 ((https://doi.org/10.4161/viru.1.5.12796)); Mekbib SB, et al. 2011. Tropical Plant Pathology 36:233-240 ((https://doi.org/10.1590/S1982-56762011000400004)); Põlme S, et al. 2020. Fungal Diversity 105:1-16 ((https://doi.org/10.1007/s13225-020-00466-2)) |
| Otu21120 | 1 | k__Fungi(100);p__Ascomycota(100);c__Saccharomycetes(100);o__Saccharomycetales(100);o__Saccharomycetales_unclassified(100);o__Saccharomycetales_unclassified(100);o__Saccharomycetales_unclassified(100); | Saccharomycetales | 7 | Saprotroph | Undefined Saprotroph | Yeast | NULL | Possible | NULL | Sterkenburg E, et al. 2015. New Phytologist 207:1145-1158 ((https://doi.org/10.1111/nph.13426)) |
| Otu21122 | 1 | k__Fungi(100);p__Ascomycota(100);c__Saccharomycetes(100);o__Saccharomycetales(100);f__Metschnikowiaceae(100);g__Metschnikowia(100);g__Metschnikowia_unclassified(100); | Metschnikowia | 13 | Saprotroph | \|Undefined Saprotroph\| | Yeast | NULL | Probable | As nectar-tap_saprotroph (Põlme et al. 2020); Decay sugar-rich substrates (Põlme et al. 2020) | Põlme S, et al. 2020. Fungal Diversity 105:1-16 ((https://doi.org/10.1007/s13225-020-00466-2)) |
| Otu21123 | 1 | k__Fungi(100);p__Ascomycota(100);c__Saccharomycetes(100);o__Saccharomycetales(100);f__Metschnikowiaceae(100);g__Metschnikowia(100);s__Metschnikowia_pulcherrima(100); | Metschnikowia | 13 | Saprotroph | \|Undefined Saprotroph\| | Yeast | NULL | Probable | As nectar-tap_saprotroph (Põlme et al. 2020); Decay sugar-rich substrates (Põlme et al. 2020) | Põlme S, et al. 2020. Fungal Diversity 105:1-16 ((https://doi.org/10.1007/s13225-020-00466-2)) |
| Otu21124 | 1 | k__Fungi(100);p__Ascomycota(100);c__Saccharomycetes(100);o__Saccharomycetales(100);f__Saccharomycetales_family_Incertae_sedis(100);g__Candida(100);g__Candida_unclassified(100); | Candida | 13 | Pathotroph-Saprotroph-Symbiotroph | Animal Pathogen-Endophyte-Endosymbiont-Epiphyte-\|Undefined Saprotroph\| | Dimorphic Yeast-Yeast | NULL | Probable | As nectar-tap_saprotroph (Põlme et al. 2020); Decay sugar-rich substrates (Põlme et al. 2020); This asexual genus is polyphyletic and contain species that function very differently depend on what they are. Caution should be taken when interpreting guilds for Candida species. | Manolakaki D, et al. 2010. Virulence. 1: 367-375 ((https://doi.org/10.4161/viru.1.5.12796)); Mekbib SB, et al. 2011. Tropical Plant Pathology 36:233-240 ((https://doi.org/10.1590/S1982-56762011000400004)); Põlme S, et al. 2020. Fungal Diversity 105:1-16 ((https://doi.org/10.1007/s13225-020-00466-2)) |
| Otu21125 | 1 | k__Fungi(100);p__Ascomycota(100);c__Saccharomycetes(100);o__Saccharomycetales(100);f__Saccharomycetales_family_Incertae_sedis(100);f__Saccharomycetales_family_Incertae_sedis_unclassified(100);f__Saccharomycetales_family_Incertae_sedis_unclassified(100); | Saccharomycetales | 7 | Saprotroph | Undefined Saprotroph | Yeast | NULL | Possible | NULL | Sterkenburg E, et al. 2015. New Phytologist 207:1145-1158 ((https://doi.org/10.1111/nph.13426)) |
| Otu21129 | 1 | k__Fungi(100);p__Ascomycota(100);c__Saccharomycetes(100);o__Saccharomycetales(100);o__Saccharomycetales_unclassified(100);o__Saccharomycetales_unclassified(100);o__Saccharomycetales_unclassified(100); | Saccharomycetales | 7 | Saprotroph | Undefined Saprotroph | Yeast | NULL | Possible | NULL | Sterkenburg E, et al. 2015. New Phytologist 207:1145-1158 ((https://doi.org/10.1111/nph.13426)) |
| Otu21130 | 1 | k__Fungi(100);p__Ascomycota(100);c__Saccharomycetes(100);o__Saccharomycetales(100);f__Saccharomycetales_family_Incertae_sedis(100);g__Candida(100);g__Candida_unclassified(100); | Candida | 13 | Pathotroph-Saprotroph-Symbiotroph | Animal Pathogen-Endophyte-Endosymbiont-Epiphyte-\|Undefined Saprotroph\| | Dimorphic Yeast-Yeast | NULL | Probable | As nectar-tap_saprotroph (Põlme et al. 2020); Decay sugar-rich substrates (Põlme et al. 2020); This asexual genus is polyphyletic and contain species that function very differently depend on what they are. Caution should be taken when interpreting guilds for Candida species. | Manolakaki D, et al. 2010. Virulence. 1: 367-375 ((https://doi.org/10.4161/viru.1.5.12796)); Mekbib SB, et al. 2011. Tropical Plant Pathology 36:233-240 ((https://doi.org/10.1590/S1982-56762011000400004)); Põlme S, et al. 2020. Fungal Diversity 105:1-16 ((https://doi.org/10.1007/s13225-020-00466-2)) |
| Otu21134 | 1 | k__Fungi(100);p__Ascomycota(100);c__Saccharomycetes(100);o__Saccharomycetales(100);f__Saccharomycetales_family_Incertae_sedis(100);g__Candida(100);s__Candida_cretensis(100); | Candida | 13 | Pathotroph-Saprotroph-Symbiotroph | Animal Pathogen-Endophyte-Endosymbiont-Epiphyte-\|Undefined Saprotroph\| | Dimorphic Yeast-Yeast | NULL | Probable | As nectar-tap_saprotroph (Põlme et al. 2020); Decay sugar-rich substrates (Põlme et al. 2020); This asexual genus is polyphyletic and contain species that function very differently depend on what they are. Caution should be taken when interpreting guilds for Candida species. | Manolakaki D, et al. 2010. Virulence. 1: 367-375 ((https://doi.org/10.4161/viru.1.5.12796)); Mekbib SB, et al. 2011. Tropical Plant Pathology 36:233-240 ((https://doi.org/10.1590/S1982-56762011000400004)); Põlme S, et al. 2020. Fungal Diversity 105:1-16 ((https://doi.org/10.1007/s13225-020-00466-2)) |
| Otu21135 | 1 | k__Fungi(100);p__Ascomycota(100);c__Saccharomycetes(100);o__Saccharomycetales(100);f__Saccharomycetales_family_Incertae_sedis(100);f__Saccharomycetales_family_Incertae_sedis_unclassified(100);f__Saccharomycetales_family_Incertae_sedis_unclassified(100); | Saccharomycetales | 7 | Saprotroph | Undefined Saprotroph | Yeast | NULL | Possible | NULL | Sterkenburg E, et al. 2015. New Phytologist 207:1145-1158 ((https://doi.org/10.1111/nph.13426)) |
| Otu21136 | 1 | k__Fungi(100);p__Ascomycota(100);c__Saccharomycetes(100);o__Saccharomycetales(100);f__Saccharomycetales_family_Incertae_sedis(100);g__Candida(100);g__Candida_unclassified(100); | Candida | 13 | Pathotroph-Saprotroph-Symbiotroph | Animal Pathogen-Endophyte-Endosymbiont-Epiphyte-\|Undefined Saprotroph\| | Dimorphic Yeast-Yeast | NULL | Probable | As nectar-tap_saprotroph (Põlme et al. 2020); Decay sugar-rich substrates (Põlme et al. 2020); This asexual genus is polyphyletic and contain species that function very differently depend on what they are. Caution should be taken when interpreting guilds for Candida species. | Manolakaki D, et al. 2010. Virulence. 1: 367-375 ((https://doi.org/10.4161/viru.1.5.12796)); Mekbib SB, et al. 2011. Tropical Plant Pathology 36:233-240 ((https://doi.org/10.1590/S1982-56762011000400004)); Põlme S, et al. 2020. Fungal Diversity 105:1-16 ((https://doi.org/10.1007/s13225-020-00466-2)) |
| Otu21138 | 1 | k__Fungi(100);p__Ascomycota(100);c__Saccharomycetes(100);o__Saccharomycetales(100);f__Saccharomycetales_family_Incertae_sedis(100);g__Candida(100);g__Candida_unclassified(100); | Candida | 13 | Pathotroph-Saprotroph-Symbiotroph | Animal Pathogen-Endophyte-Endosymbiont-Epiphyte-\|Undefined Saprotroph\| | Dimorphic Yeast-Yeast | NULL | Probable | As nectar-tap_saprotroph (Põlme et al. 2020); Decay sugar-rich substrates (Põlme et al. 2020); This asexual genus is polyphyletic and contain species that function very differently depend on what they are. Caution should be taken when interpreting guilds for Candida species. | Manolakaki D, et al. 2010. Virulence. 1: 367-375 ((https://doi.org/10.4161/viru.1.5.12796)); Mekbib SB, et al. 2011. Tropical Plant Pathology 36:233-240 ((https://doi.org/10.1590/S1982-56762011000400004)); Põlme S, et al. 2020. Fungal Diversity 105:1-16 ((https://doi.org/10.1007/s13225-020-00466-2)) |
| Otu21141 | 1 | k__Fungi(100);p__Ascomycota(100);c__Saccharomycetes(100);o__Saccharomycetales(100);f__Saccharomycetales_family_Incertae_sedis(100);g__Candida(100);g__Candida_unclassified(100); | Candida | 13 | Pathotroph-Saprotroph-Symbiotroph | Animal Pathogen-Endophyte-Endosymbiont-Epiphyte-\|Undefined Saprotroph\| | Dimorphic Yeast-Yeast | NULL | Probable | As nectar-tap_saprotroph (Põlme et al. 2020); Decay sugar-rich substrates (Põlme et al. 2020); This asexual genus is polyphyletic and contain species that function very differently depend on what they are. Caution should be taken when interpreting guilds for Candida species. | Manolakaki D, et al. 2010. Virulence. 1: 367-375 ((https://doi.org/10.4161/viru.1.5.12796)); Mekbib SB, et al. 2011. Tropical Plant Pathology 36:233-240 ((https://doi.org/10.1590/S1982-56762011000400004)); Põlme S, et al. 2020. Fungal Diversity 105:1-16 ((https://doi.org/10.1007/s13225-020-00466-2)) |
| Otu21146 | 1 | k__Fungi(100);p__Ascomycota(100);c__Saccharomycetes(100);o__Saccharomycetales(100);f__Metschnikowiaceae(100);g__Metschnikowia(100);s__Metschnikowia_pulcherrima(100); | Metschnikowia | 13 | Saprotroph | \|Undefined Saprotroph\| | Yeast | NULL | Probable | As nectar-tap_saprotroph (Põlme et al. 2020); Decay sugar-rich substrates (Põlme et al. 2020) | Põlme S, et al. 2020. Fungal Diversity 105:1-16 ((https://doi.org/10.1007/s13225-020-00466-2)) |
| Otu21147 | 1 | k__Fungi(100);p__Ascomycota(100);c__Saccharomycetes(100);o__Saccharomycetales(100);o__Saccharomycetales_unclassified(100);o__Saccharomycetales_unclassified(100);o__Saccharomycetales_unclassified(100); | Saccharomycetales | 7 | Saprotroph | Undefined Saprotroph | Yeast | NULL | Possible | NULL | Sterkenburg E, et al. 2015. New Phytologist 207:1145-1158 ((https://doi.org/10.1111/nph.13426)) |
| Otu21148 | 1 | k__Fungi(100);p__Ascomycota(100);c__Saccharomycetes(100);o__Saccharomycetales(100);f__Saccharomycetales_family_Incertae_sedis(100);f__Saccharomycetales_family_Incertae_sedis_unclassified(100);f__Saccharomycetales_family_Incertae_sedis_unclassified(100); | Saccharomycetales | 7 | Saprotroph | Undefined Saprotroph | Yeast | NULL | Possible | NULL | Sterkenburg E, et al. 2015. New Phytologist 207:1145-1158 ((https://doi.org/10.1111/nph.13426)) |
| Otu21150 | 1 | k__Fungi(100);p__Ascomycota(100);c__Saccharomycetes(100);o__Saccharomycetales(100);f__Metschnikowiaceae(100);g__Metschnikowia(100);s__Metschnikowia_pulcherrima(100); | Metschnikowia | 13 | Saprotroph | \|Undefined Saprotroph\| | Yeast | NULL | Probable | As nectar-tap_saprotroph (Põlme et al. 2020); Decay sugar-rich substrates (Põlme et al. 2020) | Põlme S, et al. 2020. Fungal Diversity 105:1-16 ((https://doi.org/10.1007/s13225-020-00466-2)) |
| Otu21153 | 1 | k__Fungi(100);p__Ascomycota(100);c__Saccharomycetes(100);o__Saccharomycetales(100);f__Metschnikowiaceae(100);g__Metschnikowia(100);s__Metschnikowia_pulcherrima(100); | Metschnikowia | 13 | Saprotroph | \|Undefined Saprotroph\| | Yeast | NULL | Probable | As nectar-tap_saprotroph (Põlme et al. 2020); Decay sugar-rich substrates (Põlme et al. 2020) | Põlme S, et al. 2020. Fungal Diversity 105:1-16 ((https://doi.org/10.1007/s13225-020-00466-2)) |
| Otu21160 | 1 | k__Fungi(100);p__Ascomycota(100);c__Saccharomycetes(100);o__Saccharomycetales(100);o__Saccharomycetales_unclassified(100);o__Saccharomycetales_unclassified(100);o__Saccharomycetales_unclassified(100); | Saccharomycetales | 7 | Saprotroph | Undefined Saprotroph | Yeast | NULL | Possible | NULL | Sterkenburg E, et al. 2015. New Phytologist 207:1145-1158 ((https://doi.org/10.1111/nph.13426)) |
| Otu21167 | 1 | k__Fungi(100);p__Ascomycota(100);c__Saccharomycetes(100);o__Saccharomycetales(100);f__Metschnikowiaceae(100);g__Metschnikowia(100);g__Metschnikowia_unclassified(100); | Metschnikowia | 13 | Saprotroph | \|Undefined Saprotroph\| | Yeast | NULL | Probable | As nectar-tap_saprotroph (Põlme et al. 2020); Decay sugar-rich substrates (Põlme et al. 2020) | Põlme S, et al. 2020. Fungal Diversity 105:1-16 ((https://doi.org/10.1007/s13225-020-00466-2)) |
| Otu21168 | 1 | k__Fungi(100);p__Ascomycota(100);c__Saccharomycetes(100);o__Saccharomycetales(100);f__Metschnikowiaceae(100);g__Metschnikowia(100);s__Metschnikowia_pulcherrima(100); | Metschnikowia | 13 | Saprotroph | \|Undefined Saprotroph\| | Yeast | NULL | Probable | As nectar-tap_saprotroph (Põlme et al. 2020); Decay sugar-rich substrates (Põlme et al. 2020) | Põlme S, et al. 2020. Fungal Diversity 105:1-16 ((https://doi.org/10.1007/s13225-020-00466-2)) |
| Otu21170 | 1 | k__Fungi(100);p__Ascomycota(100);c__Saccharomycetes(100);o__Saccharomycetales(100);f__Metschnikowiaceae(100);g__Metschnikowia(100);g__Metschnikowia_unclassified(100); | Metschnikowia | 13 | Saprotroph | \|Undefined Saprotroph\| | Yeast | NULL | Probable | As nectar-tap_saprotroph (Põlme et al. 2020); Decay sugar-rich substrates (Põlme et al. 2020) | Põlme S, et al. 2020. Fungal Diversity 105:1-16 ((https://doi.org/10.1007/s13225-020-00466-2)) |
| Otu21171 | 1 | k__Fungi(100);p__Ascomycota(100);c__Saccharomycetes(100);o__Saccharomycetales(100);f__Metschnikowiaceae(100);g__Metschnikowia(100);g__Metschnikowia_unclassified(100); | Metschnikowia | 13 | Saprotroph | \|Undefined Saprotroph\| | Yeast | NULL | Probable | As nectar-tap_saprotroph (Põlme et al. 2020); Decay sugar-rich substrates (Põlme et al. 2020) | Põlme S, et al. 2020. Fungal Diversity 105:1-16 ((https://doi.org/10.1007/s13225-020-00466-2)) |
| Otu21172 | 1 | k__Fungi(100);p__Ascomycota(100);c__Saccharomycetes(100);o__Saccharomycetales(100);f__Metschnikowiaceae(100);g__Metschnikowia(100);s__Metschnikowia_pulcherrima(100); | Metschnikowia | 13 | Saprotroph | \|Undefined Saprotroph\| | Yeast | NULL | Probable | As nectar-tap_saprotroph (Põlme et al. 2020); Decay sugar-rich substrates (Põlme et al. 2020) | Põlme S, et al. 2020. Fungal Diversity 105:1-16 ((https://doi.org/10.1007/s13225-020-00466-2)) |
| Otu21173 | 1 | k__Fungi(100);p__Ascomycota(100);c__Saccharomycetes(100);o__Saccharomycetales(100);f__Metschnikowiaceae(100);g__Metschnikowia(100);s__Metschnikowia_pulcherrima(100); | Metschnikowia | 13 | Saprotroph | \|Undefined Saprotroph\| | Yeast | NULL | Probable | As nectar-tap_saprotroph (Põlme et al. 2020); Decay sugar-rich substrates (Põlme et al. 2020) | Põlme S, et al. 2020. Fungal Diversity 105:1-16 ((https://doi.org/10.1007/s13225-020-00466-2)) |
| Otu21174 | 1 | k__Fungi(100);p__Ascomycota(100);c__Saccharomycetes(100);o__Saccharomycetales(100);f__Metschnikowiaceae(100);g__Metschnikowia(100);s__Metschnikowia_pulcherrima(100); | Metschnikowia | 13 | Saprotroph | \|Undefined Saprotroph\| | Yeast | NULL | Probable | As nectar-tap_saprotroph (Põlme et al. 2020); Decay sugar-rich substrates (Põlme et al. 2020) | Põlme S, et al. 2020. Fungal Diversity 105:1-16 ((https://doi.org/10.1007/s13225-020-00466-2)) |
| Otu21175 | 1 | k__Fungi(100);p__Ascomycota(100);c__Saccharomycetes(100);o__Saccharomycetales(100);f__Metschnikowiaceae(100);g__Metschnikowia(100);s__Metschnikowia_pulcherrima(100); | Metschnikowia | 13 | Saprotroph | \|Undefined Saprotroph\| | Yeast | NULL | Probable | As nectar-tap_saprotroph (Põlme et al. 2020); Decay sugar-rich substrates (Põlme et al. 2020) | Põlme S, et al. 2020. Fungal Diversity 105:1-16 ((https://doi.org/10.1007/s13225-020-00466-2)) |
| Otu21176 | 1 | k__Fungi(100);p__Ascomycota(100);c__Saccharomycetes(100);o__Saccharomycetales(100);f__Saccharomycetales_family_Incertae_sedis(100);f__Saccharomycetales_family_Incertae_sedis_unclassified(100);f__Saccharomycetales_family_Incertae_sedis_unclassified(100); | Saccharomycetales | 7 | Saprotroph | Undefined Saprotroph | Yeast | NULL | Possible | NULL | Sterkenburg E, et al. 2015. New Phytologist 207:1145-1158 ((https://doi.org/10.1111/nph.13426)) |
| Otu21178 | 1 | k__Fungi(100);p__Ascomycota(100);c__Saccharomycetes(100);o__Saccharomycetales(100);f__Metschnikowiaceae(100);g__Metschnikowia(100);s__Metschnikowia_pulcherrima(100); | Metschnikowia | 13 | Saprotroph | \|Undefined Saprotroph\| | Yeast | NULL | Probable | As nectar-tap_saprotroph (Põlme et al. 2020); Decay sugar-rich substrates (Põlme et al. 2020) | Põlme S, et al. 2020. Fungal Diversity 105:1-16 ((https://doi.org/10.1007/s13225-020-00466-2)) |
| Otu21179 | 1 | k__Fungi(100);p__Ascomycota(100);c__Saccharomycetes(100);o__Saccharomycetales(100);o__Saccharomycetales_unclassified(100);o__Saccharomycetales_unclassified(100);o__Saccharomycetales_unclassified(100); | Saccharomycetales | 7 | Saprotroph | Undefined Saprotroph | Yeast | NULL | Possible | NULL | Sterkenburg E, et al. 2015. New Phytologist 207:1145-1158 ((https://doi.org/10.1111/nph.13426)) |
| Otu21180 | 1 | k__Fungi(100);p__Ascomycota(100);c__Saccharomycetes(100);o__Saccharomycetales(100);f__Metschnikowiaceae(100);g__Metschnikowia(100);s__Metschnikowia_pulcherrima(100); | Metschnikowia | 13 | Saprotroph | \|Undefined Saprotroph\| | Yeast | NULL | Probable | As nectar-tap_saprotroph (Põlme et al. 2020); Decay sugar-rich substrates (Põlme et al. 2020) | Põlme S, et al. 2020. Fungal Diversity 105:1-16 ((https://doi.org/10.1007/s13225-020-00466-2)) |
| Otu21181 | 1 | k__Fungi(100);p__Ascomycota(100);c__Saccharomycetes(100);o__Saccharomycetales(100);f__Metschnikowiaceae(100);g__Metschnikowia(100);s__Metschnikowia_pulcherrima(100); | Metschnikowia | 13 | Saprotroph | \|Undefined Saprotroph\| | Yeast | NULL | Probable | As nectar-tap_saprotroph (Põlme et al. 2020); Decay sugar-rich substrates (Põlme et al. 2020) | Põlme S, et al. 2020. Fungal Diversity 105:1-16 ((https://doi.org/10.1007/s13225-020-00466-2)) |
| Otu21184 | 1 | k__Fungi(100);p__Ascomycota(100);c__Saccharomycetes(100);o__Saccharomycetales(100);f__Saccharomycetales_family_Incertae_sedis(100);g__Candida(100);g__Candida_unclassified(100); | Candida | 13 | Pathotroph-Saprotroph-Symbiotroph | Animal Pathogen-Endophyte-Endosymbiont-Epiphyte-\|Undefined Saprotroph\| | Dimorphic Yeast-Yeast | NULL | Probable | As nectar-tap_saprotroph (Põlme et al. 2020); Decay sugar-rich substrates (Põlme et al. 2020); This asexual genus is polyphyletic and contain species that function very differently depend on what they are. Caution should be taken when interpreting guilds for Candida species. | Manolakaki D, et al. 2010. Virulence. 1: 367-375 ((https://doi.org/10.4161/viru.1.5.12796)); Mekbib SB, et al. 2011. Tropical Plant Pathology 36:233-240 ((https://doi.org/10.1590/S1982-56762011000400004)); Põlme S, et al. 2020. Fungal Diversity 105:1-16 ((https://doi.org/10.1007/s13225-020-00466-2)) |
| Otu21185 | 1 | k__Fungi(100);p__Ascomycota(100);c__Saccharomycetes(100);o__Saccharomycetales(100);f__Metschnikowiaceae(100);g__Metschnikowia(100);s__Metschnikowia_pulcherrima(100); | Metschnikowia | 13 | Saprotroph | \|Undefined Saprotroph\| | Yeast | NULL | Probable | As nectar-tap_saprotroph (Põlme et al. 2020); Decay sugar-rich substrates (Põlme et al. 2020) | Põlme S, et al. 2020. Fungal Diversity 105:1-16 ((https://doi.org/10.1007/s13225-020-00466-2)) |
| Otu21186 | 1 | k__Fungi(100);p__Ascomycota(100);c__Saccharomycetes(100);o__Saccharomycetales(100);f__Metschnikowiaceae(100);g__Metschnikowia(100);s__Metschnikowia_pulcherrima(100); | Metschnikowia | 13 | Saprotroph | \|Undefined Saprotroph\| | Yeast | NULL | Probable | As nectar-tap_saprotroph (Põlme et al. 2020); Decay sugar-rich substrates (Põlme et al. 2020) | Põlme S, et al. 2020. Fungal Diversity 105:1-16 ((https://doi.org/10.1007/s13225-020-00466-2)) |
| Otu21187 | 1 | k__Fungi(100);p__Ascomycota(100);c__Saccharomycetes(100);o__Saccharomycetales(100);f__Metschnikowiaceae(100);g__Metschnikowia(100);s__Metschnikowia_pulcherrima(100); | Metschnikowia | 13 | Saprotroph | \|Undefined Saprotroph\| | Yeast | NULL | Probable | As nectar-tap_saprotroph (Põlme et al. 2020); Decay sugar-rich substrates (Põlme et al. 2020) | Põlme S, et al. 2020. Fungal Diversity 105:1-16 ((https://doi.org/10.1007/s13225-020-00466-2)) |
| Otu21188 | 1 | k__Fungi(100);p__Ascomycota(100);c__Saccharomycetes(100);o__Saccharomycetales(100);f__Metschnikowiaceae(100);g__Metschnikowia(100);g__Metschnikowia_unclassified(100); | Metschnikowia | 13 | Saprotroph | \|Undefined Saprotroph\| | Yeast | NULL | Probable | As nectar-tap_saprotroph (Põlme et al. 2020); Decay sugar-rich substrates (Põlme et al. 2020) | Põlme S, et al. 2020. Fungal Diversity 105:1-16 ((https://doi.org/10.1007/s13225-020-00466-2)) |
| Otu21189 | 1 | k__Fungi(100);p__Ascomycota(100);c__Saccharomycetes(100);o__Saccharomycetales(100);f__Metschnikowiaceae(100);g__Metschnikowia(100);g__Metschnikowia_unclassified(100); | Metschnikowia | 13 | Saprotroph | \|Undefined Saprotroph\| | Yeast | NULL | Probable | As nectar-tap_saprotroph (Põlme et al. 2020); Decay sugar-rich substrates (Põlme et al. 2020) | Põlme S, et al. 2020. Fungal Diversity 105:1-16 ((https://doi.org/10.1007/s13225-020-00466-2)) |
| Otu21190 | 1 | k__Fungi(100);p__Ascomycota(100);c__Saccharomycetes(100);o__Saccharomycetales(100);f__Saccharomycetales_family_Incertae_sedis(100);g__Candida(100);s__Candida_cretensis(100); | Candida | 13 | Pathotroph-Saprotroph-Symbiotroph | Animal Pathogen-Endophyte-Endosymbiont-Epiphyte-\|Undefined Saprotroph\| | Dimorphic Yeast-Yeast | NULL | Probable | As nectar-tap_saprotroph (Põlme et al. 2020); Decay sugar-rich substrates (Põlme et al. 2020); This asexual genus is polyphyletic and contain species that function very differently depend on what they are. Caution should be taken when interpreting guilds for Candida species. | Manolakaki D, et al. 2010. Virulence. 1: 367-375 ((https://doi.org/10.4161/viru.1.5.12796)); Mekbib SB, et al. 2011. Tropical Plant Pathology 36:233-240 ((https://doi.org/10.1590/S1982-56762011000400004)); Põlme S, et al. 2020. Fungal Diversity 105:1-16 ((https://doi.org/10.1007/s13225-020-00466-2)) |
| Otu21193 | 1 | k__Fungi(100);p__Ascomycota(100);c__Saccharomycetes(100);o__Saccharomycetales(100);f__Metschnikowiaceae(100);g__Metschnikowia(100);s__Metschnikowia_pulcherrima(100); | Metschnikowia | 13 | Saprotroph | \|Undefined Saprotroph\| | Yeast | NULL | Probable | As nectar-tap_saprotroph (Põlme et al. 2020); Decay sugar-rich substrates (Põlme et al. 2020) | Põlme S, et al. 2020. Fungal Diversity 105:1-16 ((https://doi.org/10.1007/s13225-020-00466-2)) |
| Otu21194 | 1 | k__Fungi(100);p__Ascomycota(100);c__Saccharomycetes(100);o__Saccharomycetales(100);f__Metschnikowiaceae(100);g__Metschnikowia(100);s__Metschnikowia_pulcherrima(100); | Metschnikowia | 13 | Saprotroph | \|Undefined Saprotroph\| | Yeast | NULL | Probable | As nectar-tap_saprotroph (Põlme et al. 2020); Decay sugar-rich substrates (Põlme et al. 2020) | Põlme S, et al. 2020. Fungal Diversity 105:1-16 ((https://doi.org/10.1007/s13225-020-00466-2)) |
| Otu21195 | 1 | k__Fungi(100);p__Ascomycota(100);c__Saccharomycetes(100);o__Saccharomycetales(100);f__Saccharomycetales_family_Incertae_sedis(100);g__Candida(100);g__Candida_unclassified(100); | Candida | 13 | Pathotroph-Saprotroph-Symbiotroph | Animal Pathogen-Endophyte-Endosymbiont-Epiphyte-\|Undefined Saprotroph\| | Dimorphic Yeast-Yeast | NULL | Probable | As nectar-tap_saprotroph (Põlme et al. 2020); Decay sugar-rich substrates (Põlme et al. 2020); This asexual genus is polyphyletic and contain species that function very differently depend on what they are. Caution should be taken when interpreting guilds for Candida species. | Manolakaki D, et al. 2010. Virulence. 1: 367-375 ((https://doi.org/10.4161/viru.1.5.12796)); Mekbib SB, et al. 2011. Tropical Plant Pathology 36:233-240 ((https://doi.org/10.1590/S1982-56762011000400004)); Põlme S, et al. 2020. Fungal Diversity 105:1-16 ((https://doi.org/10.1007/s13225-020-00466-2)) |
| Otu21197 | 1 | k__Fungi(100);p__Ascomycota(100);c__Saccharomycetes(100);o__Saccharomycetales(100);f__Metschnikowiaceae(100);g__Metschnikowia(100);s__Metschnikowia_pulcherrima(100); | Metschnikowia | 13 | Saprotroph | \|Undefined Saprotroph\| | Yeast | NULL | Probable | As nectar-tap_saprotroph (Põlme et al. 2020); Decay sugar-rich substrates (Põlme et al. 2020) | Põlme S, et al. 2020. Fungal Diversity 105:1-16 ((https://doi.org/10.1007/s13225-020-00466-2)) |
| Otu21198 | 1 | k__Fungi(100);p__Ascomycota(100);c__Saccharomycetes(100);o__Saccharomycetales(100);o__Saccharomycetales_unclassified(100);o__Saccharomycetales_unclassified(100);o__Saccharomycetales_unclassified(100); | Saccharomycetales | 7 | Saprotroph | Undefined Saprotroph | Yeast | NULL | Possible | NULL | Sterkenburg E, et al. 2015. New Phytologist 207:1145-1158 ((https://doi.org/10.1111/nph.13426)) |
| Otu21203 | 1 | k__Fungi(100);p__Ascomycota(100);c__Saccharomycetes(100);o__Saccharomycetales(100);f__Metschnikowiaceae(100);g__Metschnikowia(100);g__Metschnikowia_unclassified(100); | Metschnikowia | 13 | Saprotroph | \|Undefined Saprotroph\| | Yeast | NULL | Probable | As nectar-tap_saprotroph (Põlme et al. 2020); Decay sugar-rich substrates (Põlme et al. 2020) | Põlme S, et al. 2020. Fungal Diversity 105:1-16 ((https://doi.org/10.1007/s13225-020-00466-2)) |
| Otu21204 | 1 | k__Fungi(100);p__Ascomycota(100);c__Saccharomycetes(100);o__Saccharomycetales(100);f__Metschnikowiaceae(100);g__Metschnikowia(100);s__Metschnikowia_pulcherrima(100); | Metschnikowia | 13 | Saprotroph | \|Undefined Saprotroph\| | Yeast | NULL | Probable | As nectar-tap_saprotroph (Põlme et al. 2020); Decay sugar-rich substrates (Põlme et al. 2020) | Põlme S, et al. 2020. Fungal Diversity 105:1-16 ((https://doi.org/10.1007/s13225-020-00466-2)) |

| | | | | | | | | | | | |
|---|---|---|---|---|---|---|---|---|---|---|---|
| Otu21205 | 1 | k__Fungi(100);p__Ascomycota(100);c__Saccharomycetes(100);o__Saccharomycetales(100);f__Metschnikowiaceae(100);g__Metschnikowia(100);s__Metschnikowia_pulcherrima(100); | Metschnikowia | 13 | Saprotroph | \|Undefined Saprotroph\| | Yeast | NULL | Probable | As nectar-tap_saprotroph (Põlme et al. 2020); Decay sugar-rich substrates (Põlme et al. 2020) | Põlme S, et al. 2020. Fungal Diversity 105:1-16 ((https://doi.org/10.1007/s13225-020-00466-2)) |
| Otu21206 | 1 | k__Fungi(100);p__Ascomycota(100);c__Saccharomycetes(100);o__Saccharomycetales(100);f__Metschnikowiaceae(100);g__Metschnikowia(100);g__Metschnikowia_unclassified(100); | Metschnikowia | 13 | Saprotroph | \|Undefined Saprotroph\| | Yeast | NULL | Probable | As nectar-tap_saprotroph (Põlme et al. 2020); Decay sugar-rich substrates (Põlme et al. 2020) | Põlme S, et al. 2020. Fungal Diversity 105:1-16 ((https://doi.org/10.1007/s13225-020-00466-2)) |
| Otu21208 | 1 | k__Fungi(100);p__Ascomycota(100);c__Saccharomycetes(100);o__Saccharomycetales(100);f__Saccharomycetales_family_Incertae_sedis(100);g__Candida(100);s__Candida_cretensis(100); | Candida | 13 | Pathotroph-Saprotroph-Symbiotroph | Animal Pathogen-Endophyte-Endosymbiont-Epiphyte-\|Undefined Saprotroph\| | Dimorphic Yeast-Yeast | NULL | Probable | As nectar-tap_saprotroph (Põlme et al. 2020); Decay sugar-rich substrates (Põlme et al. 2020); This asexual genus is polyphyletic and contain species that function very differently depend on what they are. Caution should be taken when interpreting guilds for Candida species. | Manolakaki D, et al. 2010. Virulence. 1: 367-375 ((https://doi.org/10.4161/viru.1.5.12796)); Mekbib SB, et al. 2011. Tropical Plant Pathology 36:233-240 ((https://doi.org/10.1590/S1982-56762011000400004)); Põlme S, et al. 2020. Fungal Diversity 105:1-16 ((https://doi.org/10.1007/s13225-020-00466-2)) |
| Otu21209 | 1 | k__Fungi(100);p__Ascomycota(100);c__Saccharomycetes(100);o__Saccharomycetales(100);f__Saccharomycetales_family_Incertae_sedis(100);g__Candida(100);g__Candida_unclassified(100); | Candida | 13 | Pathotroph-Saprotroph-Symbiotroph | Animal Pathogen-Endophyte-Endosymbiont-Epiphyte-\|Undefined Saprotroph\| | Dimorphic Yeast-Yeast | NULL | Probable | As nectar-tap_saprotroph (Põlme et al. 2020); Decay sugar-rich substrates (Põlme et al. 2020); This asexual genus is polyphyletic and contain species that function very differently depend on what they are. Caution should be taken when interpreting guilds for Candida species. | Manolakaki D, et al. 2010. Virulence. 1: 367-375 ((https://doi.org/10.4161/viru.1.5.12796)); Mekbib SB, et al. 2011. Tropical Plant Pathology 36:233-240 ((https://doi.org/10.1590/S1982-56762011000400004)); Põlme S, et al. 2020. Fungal Diversity 105:1-16 ((https://doi.org/10.1007/s13225-020-00466-2)) |
| Otu21210 | 1 | k__Fungi(100);p__Ascomycota(100);c__Saccharomycetes(100);o__Saccharomycetales(100);f__Saccharomycetales_family_Incertae_sedis(100);f__Saccharomycetales_family_Incertae_sedis_unclassified(100);f__Saccharomycetales_family_Incertae_sedis_unclassified(100); | Saccharomycetales | 7 | Saprotroph | Undefined Saprotroph | Yeast | NULL | Possible | NULL | Sterkenburg E, et al. 2015. New Phytologist 207:1145-1158 ((https://doi.org/10.1111/nph.13426)) |
| Otu21211 | 1 | k__Fungi(100);p__Ascomycota(100);c__Saccharomycetes(100);o__Saccharomycetales(100);f__Saccharomycetales_family_Incertae_sedis(100);g__Candida(100);g__Candida_unclassified(100); | Candida | 13 | Pathotroph-Saprotroph-Symbiotroph | Animal Pathogen-Endophyte-Endosymbiont-Epiphyte-\|Undefined Saprotroph\| | Dimorphic Yeast-Yeast | NULL | Probable | As nectar-tap_saprotroph (Põlme et al. 2020); Decay sugar-rich substrates (Põlme et al. 2020); This asexual genus is polyphyletic and contain species that function very differently depend on what they are. Caution should be taken when interpreting guilds for Candida species. | Manolakaki D, et al. 2010. Virulence. 1: 367-375 ((https://doi.org/10.4161/viru.1.5.12796)); Mekbib SB, et al. 2011. Tropical Plant Pathology 36:233-240 ((https://doi.org/10.1590/S1982-56762011000400004)); Põlme S, et al. 2020. Fungal Diversity 105:1-16 ((https://doi.org/10.1007/s13225-020-00466-2)) |
| Otu21212 | 1 | k__Fungi(100);p__Ascomycota(100);c__Saccharomycetes(100);o__Saccharomycetales(100);f__Saccharomycetales_family_Incertae_sedis(100);g__Candida(100);g__Candida_unclassified(100); | Candida | 13 | Pathotroph-Saprotroph-Symbiotroph | Animal Pathogen-Endophyte-Endosymbiont-Epiphyte-\|Undefined Saprotroph\| | Dimorphic Yeast-Yeast | NULL | Probable | As nectar-tap_saprotroph (Põlme et al. 2020); Decay sugar-rich substrates (Põlme et al. 2020); This asexual genus is polyphyletic and contain species that function very differently depend on what they are. Caution should be taken when interpreting guilds for Candida species. | Manolakaki D, et al. 2010. Virulence. 1: 367-375 ((https://doi.org/10.4161/viru.1.5.12796)); Mekbib SB, et al. 2011. Tropical Plant Pathology 36:233-240 ((https://doi.org/10.1590/S1982-56762011000400004)); Põlme S, et al. 2020. Fungal Diversity 105:1-16 ((https://doi.org/10.1007/s13225-020-00466-2)) |
| Otu21213 | 1 | k__Fungi(100);p__Ascomycota(100);c__Saccharomycetes(100);o__Saccharomycetales(100);o__Saccharomycetales_unclassified(100);o__Saccharomycetales_unclassified(100);o__Saccharomycetales_unclassified(100); | Saccharomycetales | 7 | Saprotroph | Undefined Saprotroph | Yeast | NULL | Possible | NULL | Sterkenburg E, et al. 2015. New Phytologist 207:1145-1158 ((https://doi.org/10.1111/nph.13426)) |
| Otu21215 | 1 | k__Fungi(100);p__Ascomycota(100);c__Saccharomycetes(100);o__Saccharomycetales(100);f__Metschnikowiaceae(100);g__Metschnikowia(100);s__Metschnikowia_pulcherrima(100); | Metschnikowia | 13 | Saprotroph | \|Undefined Saprotroph\| | Yeast | NULL | Probable | As nectar-tap_saprotroph (Põlme et al. 2020); Decay sugar-rich substrates (Põlme et al. 2020) | Põlme S, et al. 2020. Fungal Diversity 105:1-16 ((https://doi.org/10.1007/s13225-020-00466-2)) |
| Otu21216 | 1 | k__Fungi(100);p__Ascomycota(100);c__Saccharomycetes(100);o__Saccharomycetales(100);f__Metschnikowiaceae(100);g__Metschnikowia(100);s__Metschnikowia_pulcherrima(100); | Metschnikowia | 13 | Saprotroph | \|Undefined Saprotroph\| | Yeast | NULL | Probable | As nectar-tap_saprotroph (Põlme et al. 2020); Decay sugar-rich substrates (Põlme et al. 2020) | Põlme S, et al. 2020. Fungal Diversity 105:1-16 ((https://doi.org/10.1007/s13225-020-00466-2)) |
| Otu21217 | 1 | k__Fungi(100);p__Ascomycota(100);c__Saccharomycetes(100);o__Saccharomycetales(100);f__Metschnikowiaceae(100);g__Metschnikowia(100);s__Metschnikowia_pulcherrima(100); | Metschnikowia | 13 | Saprotroph | \|Undefined Saprotroph\| | Yeast | NULL | Probable | As nectar-tap_saprotroph (Põlme et al. 2020); Decay sugar-rich substrates (Põlme et al. 2020) | Põlme S, et al. 2020. Fungal Diversity 105:1-16 ((https://doi.org/10.1007/s13225-020-00466-2)) |
| Otu21218 | 1 | k__Fungi(100);p__Ascomycota(100);c__Saccharomycetes(100);o__Saccharomycetales(100);f__Metschnikowiaceae(100);g__Metschnikowia(100);g__Metschnikowia_unclassified(100); | Metschnikowia | 13 | Saprotroph | \|Undefined Saprotroph\| | Yeast | NULL | Probable | As nectar-tap_saprotroph (Põlme et al. 2020); Decay sugar-rich substrates (Põlme et al. 2020) | Põlme S, et al. 2020. Fungal Diversity 105:1-16 ((https://doi.org/10.1007/s13225-020-00466-2)) |
| Otu21219 | 1 | k__Fungi(100);p__Ascomycota(100);c__Saccharomycetes(100);o__Saccharomycetales(100);f__Metschnikowiaceae(100);g__Metschnikowia(100);s__Metschnikowia_pulcherrima(100); | Metschnikowia | 13 | Saprotroph | \|Undefined Saprotroph\| | Yeast | NULL | Probable | As nectar-tap_saprotroph (Põlme et al. 2020); Decay sugar-rich substrates (Põlme et al. 2020) | Põlme S, et al. 2020. Fungal Diversity 105:1-16 ((https://doi.org/10.1007/s13225-020-00466-2)) |
| Otu21220 | 1 | k__Fungi(100);p__Ascomycota(100);c__Saccharomycetes(100);o__Saccharomycetales(100);f__Metschnikowiaceae(100);g__Metschnikowia(100);s__Metschnikowia_pulcherrima(100); | Metschnikowia | 13 | Saprotroph | \|Undefined Saprotroph\| | Yeast | NULL | Probable | As nectar-tap_saprotroph (Põlme et al. 2020); Decay sugar-rich substrates (Põlme et al. 2020) | Põlme S, et al. 2020. Fungal Diversity 105:1-16 ((https://doi.org/10.1007/s13225-020-00466-2)) |
| Otu21221 | 1 | k__Fungi(100);p__Ascomycota(100);c__Saccharomycetes(100);o__Saccharomycetales(100);f__Metschnikowiaceae(100);g__Metschnikowia(100);s__Metschnikowia_pulcherrima(100); | Metschnikowia | 13 | Saprotroph | \|Undefined Saprotroph\| | Yeast | NULL | Probable | As nectar-tap_saprotroph (Põlme et al. 2020); Decay sugar-rich substrates (Põlme et al. 2020) | Põlme S, et al. 2020. Fungal Diversity 105:1-16 ((https://doi.org/10.1007/s13225-020-00466-2)) |
| Otu21222 | 1 | k__Fungi(100);p__Ascomycota(100);c__Saccharomycetes(100);o__Saccharomycetales(100);f__Metschnikowiaceae(100);g__Metschnikowia(100);s__Metschnikowia_pulcherrima(100); | Metschnikowia | 13 | Saprotroph | \|Undefined Saprotroph\| | Yeast | NULL | Probable | As nectar-tap_saprotroph (Põlme et al. 2020); Decay sugar-rich substrates (Põlme et al. 2020) | Põlme S, et al. 2020. Fungal Diversity 105:1-16 ((https://doi.org/10.1007/s13225-020-00466-2)) |
| Otu21223 | 1 | k__Fungi(100);p__Ascomycota(100);c__Saccharomycetes(100);o__Saccharomycetales(100);f__Metschnikowiaceae(100);g__Metschnikowia(100);g__Metschnikowia_unclassified(100); | Metschnikowia | 13 | Saprotroph | \|Undefined Saprotroph\| | Yeast | NULL | Probable | As nectar-tap_saprotroph (Põlme et al. 2020); Decay sugar-rich substrates (Põlme et al. 2020) | Põlme S, et al. 2020. Fungal Diversity 105:1-16 ((https://doi.org/10.1007/s13225-020-00466-2)) |
| Otu21224 | 1 | k__Fungi(100);p__Ascomycota(100);c__Saccharomycetes(100);o__Saccharomycetales(100);f__Metschnikowiaceae(100);g__Metschnikowia(100);g__Metschnikowia_unclassified(100); | Metschnikowia | 13 | Saprotroph | \|Undefined Saprotroph\| | Yeast | NULL | Probable | As nectar-tap_saprotroph (Põlme et al. 2020); Decay sugar-rich substrates (Põlme et al. 2020) | Põlme S, et al. 2020. Fungal Diversity 105:1-16 ((https://doi.org/10.1007/s13225-020-00466-2)) |
| Otu21225 | 1 | k__Fungi(100);p__Ascomycota(100);c__Saccharomycetes(100);o__Saccharomycetales(100);f__Metschnikowiaceae(100);g__Metschnikowia(100);s__Metschnikowia_pulcherrima(100); | Metschnikowia | 13 | Saprotroph | \|Undefined Saprotroph\| | Yeast | NULL | Probable | As nectar-tap_saprotroph (Põlme et al. 2020); Decay sugar-rich substrates (Põlme et al. 2020) | Põlme S, et al. 2020. Fungal Diversity 105:1-16 ((https://doi.org/10.1007/s13225-020-00466-2)) |
| Otu21226 | 1 | k__Fungi(100);p__Ascomycota(100);c__Saccharomycetes(100);o__Saccharomycetales(100);f__Metschnikowiaceae(100);g__Metschnikowia(100);s__Metschnikowia_pulcherrima(100); | Metschnikowia | 13 | Saprotroph | \|Undefined Saprotroph\| | Yeast | NULL | Probable | As nectar-tap_saprotroph (Põlme et al. 2020); Decay sugar-rich substrates (Põlme et al. 2020) | Põlme S, et al. 2020. Fungal Diversity 105:1-16 ((https://doi.org/10.1007/s13225-020-00466-2)) |
| Otu21228 | 1 | k__Fungi(100);p__Ascomycota(100);c__Saccharomycetes(100);o__Saccharomycetales(100);f__Metschnikowiaceae(100);g__Metschnikowia(100);s__Metschnikowia_pulcherrima(100); | Metschnikowia | 13 | Saprotroph | \|Undefined Saprotroph\| | Yeast | NULL | Probable | As nectar-tap_saprotroph (Põlme et al. 2020); Decay sugar-rich substrates (Põlme et al. 2020) | Põlme S, et al. 2020. Fungal Diversity 105:1-16 ((https://doi.org/10.1007/s13225-020-00466-2)) |
| Otu21229 | 1 | k__Fungi(100);p__Ascomycota(100);c__Saccharomycetes(100);o__Saccharomycetales(100);f__Metschnikowiaceae(100);g__Metschnikowia(100);g__Metschnikowia_unclassified(100); | Metschnikowia | 13 | Saprotroph | \|Undefined Saprotroph\| | Yeast | NULL | Probable | As nectar-tap_saprotroph (Põlme et al. 2020); Decay sugar-rich substrates (Põlme et al. 2020) | Põlme S, et al. 2020. Fungal Diversity 105:1-16 ((https://doi.org/10.1007/s13225-020-00466-2)) |
| Otu21230 | 1 | k__Fungi(100);p__Ascomycota(100);c__Saccharomycetes(100);o__Saccharomycetales(100);f__Metschnikowiaceae(100);g__Metschnikowia(100);s__Metschnikowia_pulcherrima(100); | Metschnikowia | 13 | Saprotroph | \|Undefined Saprotroph\| | Yeast | NULL | Probable | As nectar-tap_saprotroph (Põlme et al. 2020); Decay sugar-rich substrates (Põlme et al. 2020) | Põlme S, et al. 2020. Fungal Diversity 105:1-16 ((https://doi.org/10.1007/s13225-020-00466-2)) |
| Otu21231 | 1 | k__Fungi(100);p__Ascomycota(100);c__Saccharomycetes(100);o__Saccharomycetales(100);f__Metschnikowiaceae(100);g__Metschnikowia(100);s__Metschnikowia_pulcherrima(100); | Metschnikowia | 13 | Saprotroph | \|Undefined Saprotroph\| | Yeast | NULL | Probable | As nectar-tap_saprotroph (Põlme et al. 2020); Decay sugar-rich substrates (Põlme et al. 2020) | Põlme S, et al. 2020. Fungal Diversity 105:1-16 ((https://doi.org/10.1007/s13225-020-00466-2)) |
| Otu21232 | 1 | k__Fungi(100);p__Ascomycota(100);c__Saccharomycetes(100);o__Saccharomycetales(100);f__Metschnikowiaceae(100);g__Metschnikowia(100);s__Metschnikowia_pulcherrima(100); | Metschnikowia | 13 | Saprotroph | \|Undefined Saprotroph\| | Yeast | NULL | Probable | As nectar-tap_saprotroph (Põlme et al. 2020); Decay sugar-rich substrates (Põlme et al. 2020) | Põlme S, et al. 2020. Fungal Diversity 105:1-16 ((https://doi.org/10.1007/s13225-020-00466-2)) |
| Otu21233 | 1 | k__Fungi(100);p__Ascomycota(100);c__Saccharomycetes(100);o__Saccharomycetales(100);f__Metschnikowiaceae(100);g__Metschnikowia(100);s__Metschnikowia_pulcherrima(100); | Metschnikowia | 13 | Saprotroph | \|Undefined Saprotroph\| | Yeast | NULL | Probable | As nectar-tap_saprotroph (Põlme et al. 2020); Decay sugar-rich substrates (Põlme et al. 2020) | Põlme S, et al. 2020. Fungal Diversity 105:1-16 ((https://doi.org/10.1007/s13225-020-00466-2)) |
| Otu21234 | 1 | k__Fungi(100);p__Ascomycota(100);c__Saccharomycetes(100);o__Saccharomycetales(100);f__Metschnikowiaceae(100);g__Metschnikowia(100);s__Metschnikowia_pulcherrima(100); | Metschnikowia | 13 | Saprotroph | \|Undefined Saprotroph\| | Yeast | NULL | Probable | As nectar-tap_saprotroph (Põlme et al. 2020); Decay sugar-rich substrates (Põlme et al. 2020) | Põlme S, et al. 2020. Fungal Diversity 105:1-16 ((https://doi.org/10.1007/s13225-020-00466-2)) |
| Otu21235 | 1 | k__Fungi(100);p__Ascomycota(100);c__Saccharomycetes(100);o__Saccharomycetales(100);f__Metschnikowiaceae(100);g__Metschnikowia(100);s__Metschnikowia_pulcherrima(100); | Metschnikowia | 13 | Saprotroph | \|Undefined Saprotroph\| | Yeast | NULL | Probable | As nectar-tap_saprotroph (Põlme et al. 2020); Decay sugar-rich substrates (Põlme et al. 2020) | Põlme S, et al. 2020. Fungal Diversity 105:1-16 ((https://doi.org/10.1007/s13225-020-00466-2)) |
| Otu21236 | 1 | k__Fungi(100);p__Ascomycota(100);c__Saccharomycetes(100);o__Saccharomycetales(100);f__Metschnikowiaceae(100);g__Metschnikowia(100);s__Metschnikowia_pulcherrima(100); | Metschnikowia | 13 | Saprotroph | \|Undefined Saprotroph\| | Yeast | NULL | Probable | As nectar-tap_saprotroph (Põlme et al. 2020); Decay sugar-rich substrates (Põlme et al. 2020) | Põlme S, et al. 2020. Fungal Diversity 105:1-16 ((https://doi.org/10.1007/s13225-020-00466-2)) |
| Otu21237 | 1 | k__Fungi(100);p__Ascomycota(100);c__Saccharomycetes(100);o__Saccharomycetales(100);f__Metschnikowiaceae(100);g__Metschnikowia(100);g__Metschnikowia_unclassified(100); | Metschnikowia | 13 | Saprotroph | \|Undefined Saprotroph\| | Yeast | NULL | Probable | As nectar-tap_saprotroph (Põlme et al. 2020); Decay sugar-rich substrates (Põlme et al. 2020) | Põlme S, et al. 2020. Fungal Diversity 105:1-16 ((https://doi.org/10.1007/s13225-020-00466-2)) |
| Otu21238 | 1 | k__Fungi(100);p__Ascomycota(100);c__Saccharomycetes(100);o__Saccharomycetales(100);f__Metschnikowiaceae(100);g__Metschnikowia(100);g__Metschnikowia_unclassified(100); | Metschnikowia | 13 | Saprotroph | \|Undefined Saprotroph\| | Yeast | NULL | Probable | As nectar-tap_saprotroph (Põlme et al. 2020); Decay sugar-rich substrates (Põlme et al. 2020) | Põlme S, et al. 2020. Fungal Diversity 105:1-16 ((https://doi.org/10.1007/s13225-020-00466-2)) |
| Otu21239 | 1 | k__Fungi(100);p__Ascomycota(100);c__Saccharomycetes(100);o__Saccharomycetales(100);f__Metschnikowiaceae(100);g__Metschnikowia(100);s__Metschnikowia_pulcherrima(100); | Metschnikowia | 13 | Saprotroph | \|Undefined Saprotroph\| | Yeast | NULL | Probable | As nectar-tap_saprotroph (Põlme et al. 2020); Decay sugar-rich substrates (Põlme et al. 2020) | Põlme S, et al. 2020. Fungal Diversity 105:1-16 ((https://doi.org/10.1007/s13225-020-00466-2)) |
| Otu21240 | 1 | k__Fungi(100);p__Ascomycota(100);c__Saccharomycetes(100);o__Saccharomycetales(100);f__Saccharomycetales_family_Incertae_sedis(100);g__Candida(100);g__Candida_unclassified(100); | Candida | 13 | Pathotroph-Saprotroph-Symbiotroph | Animal Pathogen-Endophyte-Endosymbiont-Epiphyte-\|Undefined Saprotroph\| | Dimorphic Yeast-Yeast | NULL | Probable | As nectar-tap_saprotroph (Põlme et al. 2020); Decay sugar-rich substrates (Põlme et al. 2020); This asexual genus is polyphyletic and contain species that function very differently depend on what they are. Caution should be taken when interpreting guilds for Candida species. | Manolakaki D, et al. 2010. Virulence. 1: 367-375 ((https://doi.org/10.4161/viru.1.5.12796)); Mekbib SB, et al. 2011. Tropical Plant Pathology 36:233-240 ((https://doi.org/10.1590/S1982-56762011000400004)); Põlme S, et al. 2020. Fungal Diversity 105:1-16 ((https://doi.org/10.1007/s13225-020-00466-2)) |
| Otu21241 | 1 | k__Fungi(100);p__Ascomycota(100);c__Saccharomycetes(100);o__Saccharomycetales(100);o__Saccharomycetales_unclassified(100);o__Saccharomycetales_unclassified(100);o__Saccharomycetales_unclassified(100); | Saccharomycetales | 7 | Saprotroph | Undefined Saprotroph | Yeast | NULL | Possible | NULL | Sterkenburg E, et al. 2015. New Phytologist 207:1145-1158 ((https://doi.org/10.1111/nph.13426)) |
| Otu21243 | 1 | k__Fungi(100);p__Ascomycota(100);c__Saccharomycetes(100);o__Saccharomycetales(100);f__Metschnikowiaceae(100);g__Metschnikowia(100);g__Metschnikowia_unclassified(100); | Metschnikowia | 13 | Saprotroph | \|Undefined Saprotroph\| | Yeast | NULL | Probable | As nectar-tap_saprotroph (Põlme et al. 2020); Decay sugar-rich substrates (Põlme et al. 2020) | Põlme S, et al. 2020. Fungal Diversity 105:1-16 ((https://doi.org/10.1007/s13225-020-00466-2)) |
| Otu21245 | 1 | k__Fungi(100);p__Ascomycota(100);c__Saccharomycetes(100);o__Saccharomycetales(100);f__Metschnikowiaceae(100);g__Metschnikowia(100);s__Metschnikowia_pulcherrima(100); | Metschnikowia | 13 | Saprotroph | \|Undefined Saprotroph\| | Yeast | NULL | Probable | As nectar-tap_saprotroph (Põlme et al. 2020); Decay sugar-rich substrates (Põlme et al. 2020) | Põlme S, et al. 2020. Fungal Diversity 105:1-16 ((https://doi.org/10.1007/s13225-020-00466-2)) |
| Otu21246 | 1 | k__Fungi(100);p__Ascomycota(100);c__Saccharomycetes(100);o__Saccharomycetales(100);f__Metschnikowiaceae(100);g__Metschnikowia(100);s__Metschnikowia_pulcherrima(100); | Metschnikowia | 13 | Saprotroph | \|Undefined Saprotroph\| | Yeast | NULL | Probable | As nectar-tap_saprotroph (Põlme et al. 2020); Decay sugar-rich substrates (Põlme et al. 2020) | Põlme S, et al. 2020. Fungal Diversity 105:1-16 ((https://doi.org/10.1007/s13225-020-00466-2)) |
| Otu21247 | 1 | k__Fungi(100);p__Ascomycota(100);c__Saccharomycetes(100);o__Saccharomycetales(100);f__Metschnikowiaceae(100);g__Metschnikowia(100);s__Metschnikowia_pulcherrima(100); | Metschnikowia | 13 | Saprotroph | \|Undefined Saprotroph\| | Yeast | NULL | Probable | As nectar-tap_saprotroph (Põlme et al. 2020); Decay sugar-rich substrates (Põlme et al. 2020) | Põlme S, et al. 2020. Fungal Diversity 105:1-16 ((https://doi.org/10.1007/s13225-020-00466-2)) |
| Otu21248 | 1 | k__Fungi(100);p__Ascomycota(100);c__Saccharomycetes(100);o__Saccharomycetales(100);f__Metschnikowiaceae(100);g__Metschnikowia(100);s__Metschnikowia_pulcherrima(100); | Metschnikowia | 13 | Saprotroph | \|Undefined Saprotroph\| | Yeast | NULL | Probable | As nectar-tap_saprotroph (Põlme et al. 2020); Decay sugar-rich substrates (Põlme et al. 2020) | Põlme S, et al. 2020. Fungal Diversity 105:1-16 ((https://doi.org/10.1007/s13225-020-00466-2)) |
| Otu21249 | 1 | k__Fungi(100);p__Ascomycota(100);c__Saccharomycetes(100);o__Saccharomycetales(100);f__Metschnikowiaceae(100);g__Metschnikowia(100);s__Metschnikowia_pulcherrima(100); | Metschnikowia | 13 | Saprotroph | \|Undefined Saprotroph\| | Yeast | NULL | Probable | As nectar-tap_saprotroph (Põlme et al. 2020); Decay sugar-rich substrates (Põlme et al. 2020) | Põlme S, et al. 2020. Fungal Diversity 105:1-16 ((https://doi.org/10.1007/s13225-020-00466-2)) |
| Otu21251 | 1 | k__Fungi(100);p__Ascomycota(100);c__Saccharomycetes(100);o__Saccharomycetales(100);f__Metschnikowiaceae(100);g__Metschnikowia(100);g__Metschnikowia_unclassified(100); | Metschnikowia | 13 | Saprotroph | \|Undefined Saprotroph\| | Yeast | NULL | Probable | As nectar-tap_saprotroph (Põlme et al. 2020); Decay sugar-rich substrates (Põlme et al. 2020) | Põlme S, et al. 2020. Fungal Diversity 105:1-16 ((https://doi.org/10.1007/s13225-020-00466-2)) |
| Otu21252 | 1 | k__Fungi(100);p__Ascomycota(100);c__Saccharomycetes(100);o__Saccharomycetales(100);f__Saccharomycetales_family_Incertae_sedis(100);g__Candida(100);g__Candida_unclassified(100); | Candida | 13 | Pathotroph-Saprotroph-Symbiotroph | Animal Pathogen-Endophyte-Endosymbiont-Epiphyte-\|Undefined Saprotroph\| | Dimorphic Yeast-Yeast | NULL | Probable | As nectar-tap_saprotroph (Põlme et al. 2020); Decay sugar-rich substrates (Põlme et al. 2020); This asexual genus is polyphyletic and contain species that function very differently depend on what they are. Caution should be taken when interpreting guilds for Candida species. | Manolakaki D, et al. 2010. Virulence. 1: 367-375 ((https://doi.org/10.4161/viru.1.5.12796)); Mekbib SB, et al. 2011. Tropical Plant Pathology 36:233-240 ((https://doi.org/10.1590/S1982-56762011000400004)); Põlme S, et al. 2020. Fungal Diversity 105:1-16 ((https://doi.org/10.1007/s13225-020-00466-2)) |
| Otu21253 | 1 | k__Fungi(100);p__Ascomycota(100);c__Saccharomycetes(100);o__Saccharomycetales(100);f__Saccharomycetales_family_Incertae_sedis(100);g__Candida(100);g__Candida_unclassified(100); | Candida | 13 | Pathotroph-Saprotroph-Symbiotroph | Animal Pathogen-Endophyte-Endosymbiont-Epiphyte-\|Undefined Saprotroph\| | Dimorphic Yeast-Yeast | NULL | Probable | As nectar-tap_saprotroph (Põlme et al. 2020); Decay sugar-rich substrates (Põlme et al. 2020); This asexual genus is polyphyletic and contain species that function very differently depend on what they are. Caution should be taken when interpreting guilds for Candida species. | Manolakaki D, et al. 2010. Virulence. 1: 367-375 ((https://doi.org/10.4161/viru.1.5.12796)); Mekbib SB, et al. 2011. Tropical Plant Pathology 36:233-240 ((https://doi.org/10.1590/S1982-56762011000400004)); Põlme S, et al. 2020. Fungal Diversity 105:1-16 ((https://doi.org/10.1007/s13225-020-00466-2)) |
| Otu21254 | 1 | k__Fungi(100);p__Ascomycota(100);c__Saccharomycetes(100);o__Saccharomycetales(100);f__Metschnikowiaceae(100);g__Metschnikowia(100);g__Metschnikowia_unclassified(100); | Metschnikowia | 13 | Saprotroph | \|Undefined Saprotroph\| | Yeast | NULL | Probable | As nectar-tap_saprotroph (Põlme et al. 2020); Decay sugar-rich substrates (Põlme et al. 2020) | Põlme S, et al. 2020. Fungal Diversity 105:1-16 ((https://doi.org/10.1007/s13225-020-00466-2)) |
| Otu21255 | 1 | k__Fungi(100);p__Ascomycota(100);c__Saccharomycetes(100);o__Saccharomycetales(100);f__Metschnikowiaceae(100);g__Metschnikowia(100);s__Metschnikowia_pulcherrima(100); | Metschnikowia | 13 | Saprotroph | \|Undefined Saprotroph\| | Yeast | NULL | Probable | As nectar-tap_saprotroph (Põlme et al. 2020); Decay sugar-rich substrates (Põlme et al. 2020) | Põlme S, et al. 2020. Fungal Diversity 105:1-16 ((https://doi.org/10.1007/s13225-020-00466-2)) |
| Otu21256 | 1 | k__Fungi(100);p__Ascomycota(100);c__Saccharomycetes(100);o__Saccharomycetales(100);f__Metschnikowiaceae(100);g__Metschnikowia(100);s__Metschnikowia_pulcherrima(100); | Metschnikowia | 13 | Saprotroph | \|Undefined Saprotroph\| | Yeast | NULL | Probable | As nectar-tap_saprotroph (Põlme et al. 2020); Decay sugar-rich substrates (Põlme et al. 2020) | Põlme S, et al. 2020. Fungal Diversity 105:1-16 ((https://doi.org/10.1007/s13225-020-00466-2)) |
| Otu21257 | 1 | k__Fungi(100);p__Ascomycota(100);c__Saccharomycetes(100);o__Saccharomycetales(100);f__Metschnikowiaceae(100);g__Metschnikowia(100);s__Metschnikowia_pulcherrima(100); | Metschnikowia | 13 | Saprotroph | \|Undefined Saprotroph\| | Yeast | NULL | Probable | As nectar-tap_saprotroph (Põlme et al. 2020); Decay sugar-rich substrates (Põlme et al. 2020) | Põlme S, et al. 2020. Fungal Diversity 105:1-16 ((https://doi.org/10.1007/s13225-020-00466-2)) |
| Otu21258 | 1 | k__Fungi(100);p__Ascomycota(100);c__Saccharomycetes(100);o__Saccharomycetales(100);f__Metschnikowiaceae(100);g__Metschnikowia(100);s__Metschnikowia_pulcherrima(100); | Metschnikowia | 13 | Saprotroph | \|Undefined Saprotroph\| | Yeast | NULL | Probable | As nectar-tap_saprotroph (Põlme et al. 2020); Decay sugar-rich substrates (Põlme et al. 2020) | Põlme S, et al. 2020. Fungal Diversity 105:1-16 ((https://doi.org/10.1007/s13225-020-00466-2)) |
| Otu21259 | 1 | k__Fungi(100);p__Ascomycota(100);c__Saccharomycetes(100);o__Saccharomycetales(100);f__Saccharomycetales_family_Incertae_sedis(100);g__Candida(100);g__Candida_unclassified(100); | Candida | 13 | Pathotroph-Saprotroph-Symbiotroph | Animal Pathogen-Endophyte-Endosymbiont-Epiphyte-\|Undefined Saprotroph\| | Dimorphic Yeast-Yeast | NULL | Probable | As nectar-tap_saprotroph (Põlme et al. 2020); Decay sugar-rich substrates (Põlme et al. 2020); This asexual genus is polyphyletic and contain species that function very differently depend on what they are. Caution should be taken when interpreting guilds for Candida species. | Manolakaki D, et al. 2010. Virulence. 1: 367-375 ((https://doi.org/10.4161/viru.1.5.12796)); Mekbib SB, et al. 2011. Tropical Plant Pathology 36:233-240 ((https://doi.org/10.1590/S1982-56762011000400004)); Põlme S, et al. 2020. Fungal Diversity 105:1-16 ((https://doi.org/10.1007/s13225-020-00466-2)) |
| Otu21260 | 1 | k__Fungi(100);p__Ascomycota(100);c__Saccharomycetes(100);o__Saccharomycetales(100);f__Metschnikowiaceae(100);g__Metschnikowia(100);s__Metschnikowia_pulcherrima(100); | Metschnikowia | 13 | Saprotroph | \|Undefined Saprotroph\| | Yeast | NULL | Probable | As nectar-tap_saprotroph (Põlme et al. 2020); Decay sugar-rich substrates (Põlme et al. 2020) | Põlme S, et al. 2020. Fungal Diversity 105:1-16 ((https://doi.org/10.1007/s13225-020-00466-2)) |
| Otu21261 | 1 | k__Fungi(100);p__Ascomycota(100);c__Saccharomycetes(100);o__Saccharomycetales(100);f__Metschnikowiaceae(100);g__Metschnikowia(100);s__Metschnikowia_pulcherrima(100); | Metschnikowia | 13 | Saprotroph | \|Undefined Saprotroph\| | Yeast | NULL | Probable | As nectar-tap_saprotroph (Põlme et al. 2020); Decay sugar-rich substrates (Põlme et al. 2020) | Põlme S, et al. 2020. Fungal Diversity 105:1-16 ((https://doi.org/10.1007/s13225-020-00466-2)) |
| Otu21262 | 1 | k__Fungi(100);p__Ascomycota(100);c__Saccharomycetes(100);o__Saccharomycetales(100);f__Metschnikowiaceae(100);g__Metschnikowia(100);s__Metschnikowia_pulcherrima(100); | Metschnikowia | 13 | Saprotroph | \|Undefined Saprotroph\| | Yeast | NULL | Probable | As nectar-tap_saprotroph (Põlme et al. 2020); Decay sugar-rich substrates (Põlme et al. 2020) | Põlme S, et al. 2020. Fungal Diversity 105:1-16 ((https://doi.org/10.1007/s13225-020-00466-2)) |
| Otu21263 | 1 | k__Fungi(100);p__Ascomycota(100);c__Saccharomycetes(100);o__Saccharomycetales(100);f__Metschnikowiaceae(100);g__Metschnikowia(100);g__Metschnikowia_unclassified(100); | Metschnikowia | 13 | Saprotroph | \|Undefined Saprotroph\| | Yeast | NULL | Probable | As nectar-tap_saprotroph (Põlme et al. 2020); Decay sugar-rich substrates (Põlme et al. 2020) | Põlme S, et al. 2020. Fungal Diversity 105:1-16 ((https://doi.org/10.1007/s13225-020-00466-2)) |
| Otu21264 | 1 | k__Fungi(100);p__Ascomycota(100);c__Saccharomycetes(100);o__Saccharomycetales(100);f__Metschnikowiaceae(100);g__Metschnikowia(100);s__Metschnikowia_pulcherrima(100); | Metschnikowia | 13 | Saprotroph | \|Undefined Saprotroph\| | Yeast | NULL | Probable | As nectar-tap_saprotroph (Põlme et al. 2020); Decay sugar-rich substrates (Põlme et al. 2020) | Põlme S, et al. 2020. Fungal Diversity 105:1-16 ((https://doi.org/10.1007/s13225-020-00466-2)) |
| Otu21265 | 1 | k__Fungi(100);p__Ascomycota(100);c__Saccharomycetes(100);o__Saccharomycetales(100);f__Metschnikowiaceae(100);g__Metschnikowia(100);s__Metschnikowia_pulcherrima(100); | Metschnikowia | 13 | Saprotroph | \|Undefined Saprotroph\| | Yeast | NULL | Probable | As nectar-tap_saprotroph (Põlme et al. 2020); Decay sugar-rich substrates (Põlme et al. 2020) | Põlme S, et al. 2020. Fungal Diversity 105:1-16 ((https://doi.org/10.1007/s13225-020-00466-2)) |
| Otu21266 | 1 | k__Fungi(100);p__Ascomycota(100);c__Saccharomycetes(100);o__Saccharomycetales(100);f__Metschnikowiaceae(100);g__Metschnikowia(100);s__Metschnikowia_pulcherrima(100); | Metschnikowia | 13 | Saprotroph | \|Undefined Saprotroph\| | Yeast | NULL | Probable | As nectar-tap_saprotroph (Põlme et al. 2020); Decay sugar-rich substrates (Põlme et al. 2020) | Põlme S, et al. 2020. Fungal Diversity 105:1-16 ((https://doi.org/10.1007/s13225-020-00466-2)) |
| Otu21267 | 1 | k__Fungi(100);p__Ascomycota(100);c__Saccharomycetes(100);o__Saccharomycetales(100);f__Metschnikowiaceae(100);g__Metschnikowia(100);s__Metschnikowia_pulcherrima(100); | Metschnikowia | 13 | Saprotroph | \|Undefined Saprotroph\| | Yeast | NULL | Probable | As nectar-tap_saprotroph (Põlme et al. 2020); Decay sugar-rich substrates (Põlme et al. 2020) | Põlme S, et al. 2020. Fungal Diversity 105:1-16 ((https://doi.org/10.1007/s13225-020-00466-2)) |
| Otu21268 | 1 | k__Fungi(100);p__Ascomycota(100);c__Saccharomycetes(100);o__Saccharomycetales(100);f__Metschnikowiaceae(100);g__Metschnikowia(100);s__Metschnikowia_pulcherrima(100); | Metschnikowia | 13 | Saprotroph | \|Undefined Saprotroph\| | Yeast | NULL | Probable | As nectar-tap_saprotroph (Põlme et al. 2020); Decay sugar-rich substrates (Põlme et al. 2020) | Põlme S, et al. 2020. Fungal Diversity 105:1-16 ((https://doi.org/10.1007/s13225-020-00466-2)) |
| Otu21269 | 1 | k__Fungi(100);p__Ascomycota(100);c__Saccharomycetes(100);o__Saccharomycetales(100);f__Saccharomycetales_family_Incertae_sedis(100);g__Candida(100);g__Candida_unclassified(100); | Candida | 13 | Pathotroph-Saprotroph-Symbiotroph | Animal Pathogen-Endophyte-Endosymbiont-Epiphyte-\|Undefined Saprotroph\| | Dimorphic Yeast-Yeast | NULL | Probable | As nectar-tap_saprotroph (Põlme et al. 2020); Decay sugar-rich substrates (Põlme et al. 2020); This asexual genus is polyphyletic and contain species that function very differently depend on what they are. Caution should be taken when interpreting guilds for Candida species. | Manolakaki D, et al. 2010. Virulence. 1: 367-375 ((https://doi.org/10.4161/viru.1.5.12796)); Mekbib SB, et al. 2011. Tropical Plant Pathology 36:233-240 ((https://doi.org/10.1590/S1982-56762011000400004)); Põlme S, et al. 2020. Fungal Diversity 105:1-16 ((https://doi.org/10.1007/s13225-020-00466-2)) |
| Otu21271 | 1 | k__Fungi(100);p__Ascomycota(100);c__Saccharomycetes(100);o__Saccharomycetales(100);f__Metschnikowiaceae(100);g__Metschnikowia(100);s__Metschnikowia_pulcherrima(100); | Metschnikowia | 13 | Saprotroph | \|Undefined Saprotroph\| | Yeast | NULL | Probable | As nectar-tap_saprotroph (Põlme et al. 2020); Decay sugar-rich substrates (Põlme et al. 2020) | Põlme S, et al. 2020. Fungal Diversity 105:1-16 ((https://doi.org/10.1007/s13225-020-00466-2)) |
| Otu21272 | 1 | k__Fungi(100);p__Ascomycota(100);c__Saccharomycetes(100);o__Saccharomycetales(100);f__Metschnikowiaceae(100);g__Metschnikowia(100);s__Metschnikowia_pulcherrima(100); | Metschnikowia | 13 | Saprotroph | \|Undefined Saprotroph\| | Yeast | NULL | Probable | As nectar-tap_saprotroph (Põlme et al. 2020); Decay sugar-rich substrates (Põlme et al. 2020) | Põlme S, et al. 2020. Fungal Diversity 105:1-16 ((https://doi.org/10.1007/s13225-020-00466-2)) |
| Otu21273 | 1 | k__Fungi(100);p__Ascomycota(100);c__Saccharomycetes(100);o__Saccharomycetales(100);f__Metschnikowiaceae(100);g__Metschnikowia(100);s__Metschnikowia_pulcherrima(100); | Metschnikowia | 13 | Saprotroph | \|Undefined Saprotroph\| | Yeast | NULL | Probable | As nectar-tap_saprotroph (Põlme et al. 2020); Decay sugar-rich substrates (Põlme et al. 2020) | Põlme S, et al. 2020. Fungal Diversity 105:1-16 ((https://doi.org/10.1007/s13225-020-00466-2)) |
| Otu21277 | 1 | k__Fungi(100);p__Ascomycota(100);c__Saccharomycetes(100);o__Saccharomycetales(100);f__Metschnikowiaceae(100);g__Metschnikowia(100);s__Metschnikowia_pulcherrima(100); | Metschnikowia | 13 | Saprotroph | \|Undefined Saprotroph\| | Yeast | NULL | Probable | As nectar-tap_saprotroph (Põlme et al. 2020); Decay sugar-rich substrates (Põlme et al. 2020) | Põlme S, et al. 2020. Fungal Diversity 105:1-16 ((https://doi.org/10.1007/s13225-020-00466-2)) |
| Otu21280 | 1 | k__Fungi(100);p__Ascomycota(100);c__Saccharomycetes(100);o__Saccharomycetales(100);f__Metschnikowiaceae(100);g__Metschnikowia(100);s__Metschnikowia_pulcherrima(100); | Metschnikowia | 13 | Saprotroph | \|Undefined Saprotroph\| | Yeast | NULL | Probable | As nectar-tap_saprotroph (Põlme et al. 2020); Decay sugar-rich substrates (Põlme et al. 2020) | Põlme S, et al. 2020. Fungal Diversity 105:1-16 ((https://doi.org/10.1007/s13225-020-00466-2)) |
| Otu21282 | 1 | k__Fungi(100);p__Ascomycota(100);c__Saccharomycetes(100);o__Saccharomycetales(100);f__Metschnikowiaceae(100);g__Metschnikowia(100);g__Metschnikowia_unclassified(100); | Metschnikowia | 13 | Saprotroph | \|Undefined Saprotroph\| | Yeast | NULL | Probable | As nectar-tap_saprotroph (Põlme et al. 2020); Decay sugar-rich substrates (Põlme et al. 2020) | Põlme S, et al. 2020. Fungal Diversity 105:1-16 ((https://doi.org/10.1007/s13225-020-00466-2)) |
| Otu21283 | 1 | k__Fungi(100);p__Ascomycota(100);c__Saccharomycetes(100);o__Saccharomycetales(100);f__Metschnikowiaceae(100);g__Metschnikowia(100);s__Metschnikowia_pulcherrima(100); | Metschnikowia | 13 | Saprotroph | \|Undefined Saprotroph\| | Yeast | NULL | Probable | As nectar-tap_saprotroph (Põlme et al. 2020); Decay sugar-rich substrates (Põlme et al. 2020) | Põlme S, et al. 2020. Fungal Diversity 105:1-16 ((https://doi.org/10.1007/s13225-020-00466-2)) |
| Otu21285 | 1 | k__Fungi(100);p__Ascomycota(100);c__Saccharomycetes(100);o__Saccharomycetales(100);f__Metschnikowiaceae(100);g__Metschnikowia(100);s__Metschnikowia_pulcherrima(100); | Metschnikowia | 13 | Saprotroph | \|Undefined Saprotroph\| | Yeast | NULL | Probable | As nectar-tap_saprotroph (Põlme et al. 2020); Decay sugar-rich substrates (Põlme et al. 2020) | Põlme S, et al. 2020. Fungal Diversity 105:1-16 ((https://doi.org/10.1007/s13225-020-00466-2)) |
| Otu21286 | 1 | k__Fungi(100);p__Ascomycota(100);c__Saccharomycetes(100);o__Saccharomycetales(100);f__Metschnikowiaceae(100);g__Metschnikowia(100);g__Metschnikowia_unclassified(100); | Metschnikowia | 13 | Saprotroph | \|Undefined Saprotroph\| | Yeast | NULL | Probable | As nectar-tap_saprotroph (Põlme et al. 2020); Decay sugar-rich substrates (Põlme et al. 2020) | Põlme S, et al. 2020. Fungal Diversity 105:1-16 ((https://doi.org/10.1007/s13225-020-00466-2)) |
| Otu21287 | 1 | k__Fungi(100);p__Ascomycota(100);c__Saccharomycetes(100);o__Saccharomycetales(100);f__Metschnikowiaceae(100);g__Metschnikowia(100);g__Metschnikowia_unclassified(100); | Metschnikowia | 13 | Saprotroph | \|Undefined Saprotroph\| | Yeast | NULL | Probable | As nectar-tap_saprotroph (Põlme et al. 2020); Decay sugar-rich substrates (Põlme et al. 2020) | Põlme S, et al. 2020. Fungal Diversity 105:1-16 ((https://doi.org/10.1007/s13225-020-00466-2)) |
| Otu21288 | 1 | k__Fungi(100);p__Ascomycota(100);c__Saccharomycetes(100);o__Saccharomycetales(100);f__Metschnikowiaceae(100);g__Metschnikowia(100);s__Metschnikowia_pulcherrima(100); | Metschnikowia | 13 | Saprotroph | \|Undefined Saprotroph\| | Yeast | NULL | Probable | As nectar-tap_saprotroph (Põlme et al. 2020); Decay sugar-rich substrates (Põlme et al. 2020) | Põlme S, et al. 2020. Fungal Diversity 105:1-16 ((https://doi.org/10.1007/s13225-020-00466-2)) |
| Otu21289 | 1 | k__Fungi(100);p__Ascomycota(100);c__Saccharomycetes(100);o__Saccharomycetales(100);f__Metschnikowiaceae(100);g__Metschnikowia(100);s__Metschnikowia_pulcherrima(100); | Metschnikowia | 13 | Saprotroph | \|Undefined Saprotroph\| | Yeast | NULL | Probable | As nectar-tap_saprotroph (Põlme et al. 2020); Decay sugar-rich substrates (Põlme et al. 2020) | Põlme S, et al. 2020. Fungal Diversity 105:1-16 ((https://doi.org/10.1007/s13225-020-00466-2)) |
| Otu21290 | 1 | k__Fungi(100);p__Ascomycota(100);c__Saccharomycetes(100);o__Saccharomycetales(100);f__Metschnikowiaceae(100);g__Metschnikowia(100);g__Metschnikowia_unclassified(100); | Metschnikowia | 13 | Saprotroph | \|Undefined Saprotroph\| | Yeast | NULL | Probable | As nectar-tap_saprotroph (Põlme et al. 2020); Decay sugar-rich substrates (Põlme et al. 2020) | Põlme S, et al. 2020. Fungal Diversity 105:1-16 ((https://doi.org/10.1007/s13225-020-00466-2)) |
| Otu21291 | 1 | k__Fungi(100);p__Ascomycota(100);c__Saccharomycetes(100);o__Saccharomycetales(100);f__Metschnikowiaceae(100);g__Metschnikowia(100);s__Metschnikowia_pulcherrima(100); | Metschnikowia | 13 | Saprotroph | \|Undefined Saprotroph\| | Yeast | NULL | Probable | As nectar-tap_saprotroph (Põlme et al. 2020); Decay sugar-rich substrates (Põlme et al. 2020) | Põlme S, et al. 2020. Fungal Diversity 105:1-16 ((https://doi.org/10.1007/s13225-020-00466-2)) |
| Otu21292 | 1 | k__Fungi(100);p__Ascomycota(100);c__Saccharomycetes(100);o__Saccharomycetales(100);f__Metschnikowiaceae(100);g__Metschnikowia(100);s__Metschnikowia_pulcherrima(100); | Metschnikowia | 13 | Saprotroph | \|Undefined Saprotroph\| | Yeast | NULL | Probable | As nectar-tap_saprotroph (Põlme et al. 2020); Decay sugar-rich substrates (Põlme et al. 2020) | Põlme S, et al. 2020. Fungal Diversity 105:1-16 ((https://doi.org/10.1007/s13225-020-00466-2)) |
| Otu21293 | 1 | k__Fungi(100);p__Ascomycota(100);c__Saccharomycetes(100);o__Saccharomycetales(100);f__Metschnikowiaceae(100);g__Metschnikowia(100);g__Metschnikowia_unclassified(100); | Metschnikowia | 13 | Saprotroph | \|Undefined Saprotroph\| | Yeast | NULL | Probable | As nectar-tap_saprotroph (Põlme et al. 2020); Decay sugar-rich substrates (Põlme et al. 2020) | Põlme S, et al. 2020. Fungal Diversity 105:1-16 ((https://doi.org/10.1007/s13225-020-00466-2)) |
| Otu21294 | 1 | k__Fungi(100);p__Ascomycota(100);c__Saccharomycetes(100);o__Saccharomycetales(100);f__Metschnikowiaceae(100);g__Metschnikowia(100);g__Metschnikowia_unclassified(100); | Metschnikowia | 13 | Saprotroph | \|Undefined Saprotroph\| | Yeast | NULL | Probable | As nectar-tap_saprotroph (Põlme et al. 2020); Decay sugar-rich substrates (Põlme et al. 2020) | Põlme S, et al. 2020. Fungal Diversity 105:1-16 ((https://doi.org/10.1007/s13225-020-00466-2)) |
| Otu21295 | 1 | k__Fungi(100);p__Ascomycota(100);c__Saccharomycetes(100);o__Saccharomycetales(100);f__Metschnikowiaceae(100);g__Metschnikowia(100);g__Metschnikowia_unclassified(100); | Metschnikowia | 13 | Saprotroph | \|Undefined Saprotroph\| | Yeast | NULL | Probable | As nectar-tap_saprotroph (Põlme et al. 2020); Decay sugar-rich substrates (Põlme et al. 2020) | Põlme S, et al. 2020. Fungal Diversity 105:1-16 ((https://doi.org/10.1007/s13225-020-00466-2)) |
| Otu21296 | 1 | k__Fungi(100);p__Ascomycota(100);c__Saccharomycetes(100);o__Saccharomycetales(100);f__Metschnikowiaceae(100);g__Metschnikowia(100);s__Metschnikowia_pulcherrima(100); | Metschnikowia | 13 | Saprotroph | \|Undefined Saprotroph\| | Yeast | NULL | Probable | As nectar-tap_saprotroph (Põlme et al. 2020); Decay sugar-rich substrates (Põlme et al. 2020) | Põlme S, et al. 2020. Fungal Diversity 105:1-16 ((https://doi.org/10.1007/s13225-020-00466-2)) |
| Otu21297 | 1 | k__Fungi(100);p__Ascomycota(100);c__Saccharomycetes(100);o__Saccharomycetales(100);f__Metschnikowiaceae(100);g__Metschnikowia(100);s__Metschnikowia_pulcherrima(100); | Metschnikowia | 13 | Saprotroph | \|Undefined Saprotroph\| | Yeast | NULL | Probable | As nectar-tap_saprotroph (Põlme et al. 2020); Decay sugar-rich substrates (Põlme et al. 2020) | Põlme S, et al. 2020. Fungal Diversity 105:1-16 ((https://doi.org/10.1007/s13225-020-00466-2)) |
| Otu21298 | 1 | k__Fungi(100);p__Ascomycota(100);c__Saccharomycetes(100);o__Saccharomycetales(100);f__Metschnikowiaceae(100);g__Metschnikowia(100);s__Metschnikowia_pulcherrima(100); | Metschnikowia | 13 | Saprotroph | \|Undefined Saprotroph\| | Yeast | NULL | Probable | As nectar-tap_saprotroph (Põlme et al. 2020); Decay sugar-rich substrates (Põlme et al. 2020) | Põlme S, et al. 2020. Fungal Diversity 105:1-16 ((https://doi.org/10.1007/s13225-020-00466-2)) |
| Otu21299 | 1 | k__Fungi(100);p__Ascomycota(100);c__Saccharomycetes(100);o__Saccharomycetales(100);f__Metschnikowiaceae(100);g__Metschnikowia(100);s__Metschnikowia_pulcherrima(100); | Metschnikowia | 13 | Saprotroph | \|Undefined Saprotroph\| | Yeast | NULL | Probable | As nectar-tap_saprotroph (Põlme et al. 2020); Decay sugar-rich substrates (Põlme et al. 2020) | Põlme S, et al. 2020. Fungal Diversity 105:1-16 ((https://doi.org/10.1007/s13225-020-00466-2)) |
| Otu21300 | 1 | k__Fungi(100);p__Ascomycota(100);c__Saccharomycetes(100);o__Saccharomycetales(100);f__Metschnikowiaceae(100);g__Metschnikowia(100);s__Metschnikowia_pulcherrima(100); | Metschnikowia | 13 | Saprotroph | \|Undefined Saprotroph\| | Yeast | NULL | Probable | As nectar-tap_saprotroph (Põlme et al. 2020); Decay sugar-rich substrates (Põlme et al. 2020) | Põlme S, et al. 2020. Fungal Diversity 105:1-16 ((https://doi.org/10.1007/s13225-020-00466-2)) |
| Otu21301 | 1 | k__Fungi(100);p__Ascomycota(100);c__Saccharomycetes(100);o__Saccharomycetales(100);f__Metschnikowiaceae(100);g__Metschnikowia(100);s__Metschnikowia_pulcherrima(100); | Metschnikowia | 13 | Saprotroph | \|Undefined Saprotroph\| | Yeast | NULL | Probable | As nectar-tap_saprotroph (Põlme et al. 2020); Decay sugar-rich substrates (Põlme et al. 2020) | Põlme S, et al. 2020. Fungal Diversity 105:1-16 ((https://doi.org/10.1007/s13225-020-00466-2)) |
| Otu21302 | 1 | k__Fungi(100);p__Ascomycota(100);c__Saccharomycetes(100);o__Saccharomycetales(100);f__Metschnikowiaceae(100);g__Metschnikowia(100);s__Metschnikowia_pulcherrima(100); | Metschnikowia | 13 | Saprotroph | \|Undefined Saprotroph\| | Yeast | NULL | Probable | As nectar-tap_saprotroph (Põlme et al. 2020); Decay sugar-rich substrates (Põlme et al. 2020) | Põlme S, et al. 2020. Fungal Diversity 105:1-16 ((https://doi.org/10.1007/s13225-020-00466-2)) |
| Otu21303 | 1 | k__Fungi(100);p__Ascomycota(100);c__Saccharomycetes(100);o__Saccharomycetales(100);f__Metschnikowiaceae(100);g__Metschnikowia(100);s__Metschnikowia_pulcherrima(100); | Metschnikowia | 13 | Saprotroph | \|Undefined Saprotroph\| | Yeast | NULL | Probable | As nectar-tap_saprotroph (Põlme et al. 2020); Decay sugar-rich substrates (Põlme et al. 2020) | Põlme S, et al. 2020. Fungal Diversity 105:1-16 ((https://doi.org/10.1007/s13225-020-00466-2)) |
| Otu21304 | 1 | k__Fungi(100);p__Ascomycota(100);c__Saccharomycetes(100);o__Saccharomycetales(100);f__Metschnikowiaceae(100);g__Metschnikowia(100);g__Metschnikowia_unclassified(100); | Metschnikowia | 13 | Saprotroph | \|Undefined Saprotroph\| | Yeast | NULL | Probable | As nectar-tap_saprotroph (Põlme et al. 2020); Decay sugar-rich substrates (Põlme et al. 2020) | Põlme S, et al. 2020. Fungal Diversity 105:1-16 ((https://doi.org/10.1007/s13225-020-00466-2)) |
| Otu21305 | 1 | k__Fungi(100);p__Ascomycota(100);c__Saccharomycetes(100);o__Saccharomycetales(100);f__Metschnikowiaceae(100);g__Metschnikowia(100);g__Metschnikowia_unclassified(100); | Metschnikowia | 13 | Saprotroph | \|Undefined Saprotroph\| | Yeast | NULL | Probable | As nectar-tap_saprotroph (Põlme et al. 2020); Decay sugar-rich substrates (Põlme et al. 2020) | Põlme S, et al. 2020. Fungal Diversity 105:1-16 ((https://doi.org/10.1007/s13225-020-00466-2)) |
| Otu21306 | 1 | k__Fungi(100);p__Ascomycota(100);c__Saccharomycetes(100);o__Saccharomycetales(100);f__Saccharomycetales_family_Incertae_sedis(100);g__Candida(100);g__Candida_unclassified(100); | Candida | 13 | Pathotroph-Saprotroph-Symbiotroph | Animal Pathogen-Endophyte-Endosymbiont-Epiphyte-\|Undefined Saprotroph\| | Dimorphic Yeast-Yeast | NULL | Probable | As nectar-tap_saprotroph (Põlme et al. 2020); Decay sugar-rich substrates (Põlme et al. 2020); This asexual genus is polyphyletic and contain species that function very differently depend on what they are. Caution should be taken when interpreting guilds for Candida species. | Manolakaki D, et al. 2010. Virulence. 1: 367-375 ((https://doi.org/10.4161/viru.1.5.12796)); Mekbib SB, et al. 2011. Tropical Plant Pathology 36:233-240 ((https://doi.org/10.1590/S1982-56762011000400004)); Põlme S, et al. 2020. Fungal Diversity 105:1-16 ((https://doi.org/10.1007/s13225-020-00466-2)) |
| Otu21307 | 1 | k__Fungi(100);p__Ascomycota(100);c__Saccharomycetes(100);o__Saccharomycetales(100);f__Metschnikowiaceae(100);g__Metschnikowia(100);s__Metschnikowia_pulcherrima(100); | Metschnikowia | 13 | Saprotroph | \|Undefined Saprotroph\| | Yeast | NULL | Probable | As nectar-tap_saprotroph (Põlme et al. 2020); Decay sugar-rich substrates (Põlme et al. 2020) | Põlme S, et al. 2020. Fungal Diversity 105:1-16 ((https://doi.org/10.1007/s13225-020-00466-2)) |
| Otu21308 | 1 | k__Fungi(100);p__Ascomycota(100);c__Saccharomycetes(100);o__Saccharomycetales(100);f__Metschnikowiaceae(100);g__Metschnikowia(100);s__Metschnikowia_pulcherrima(100); | Metschnikowia | 13 | Saprotroph | \|Undefined Saprotroph\| | Yeast | NULL | Probable | As nectar-tap_saprotroph (Põlme et al. 2020); Decay sugar-rich substrates (Põlme et al. 2020) | Põlme S, et al. 2020. Fungal Diversity 105:1-16 ((https://doi.org/10.1007/s13225-020-00466-2)) |
| Otu21309 | 1 | k__Fungi(100);p__Ascomycota(100);c__Saccharomycetes(100);o__Saccharomycetales(100);f__Metschnikowiaceae(100);g__Metschnikowia(100);g__Metschnikowia_unclassified(100); | Metschnikowia | 13 | Saprotroph | \|Undefined Saprotroph\| | Yeast | NULL | Probable | As nectar-tap_saprotroph (Põlme et al. 2020); Decay sugar-rich substrates (Põlme et al. 2020) | Põlme S, et al. 2020. Fungal Diversity 105:1-16 ((https://doi.org/10.1007/s13225-020-00466-2)) |
| Otu21310 | 1 | k__Fungi(100);p__Ascomycota(100);c__Saccharomycetes(100);o__Saccharomycetales(100);f__Metschnikowiaceae(100);g__Metschnikowia(100);s__Metschnikowia_pulcherrima(100); | Metschnikowia | 13 | Saprotroph | \|Undefined Saprotroph\| | Yeast | NULL | Probable | As nectar-tap_saprotroph (Põlme et al. 2020); Decay sugar-rich substrates (Põlme et al. 2020) | Põlme S, et al. 2020. Fungal Diversity 105:1-16 ((https://doi.org/10.1007/s13225-020-00466-2)) |
| Otu21311 | 1 | k__Fungi(100);p__Ascomycota(100);c__Saccharomycetes(100);o__Saccharomycetales(100);f__Metschnikowiaceae(100);g__Metschnikowia(100);g__Metschnikowia_unclassified(100); | Metschnikowia | 13 | Saprotroph | \|Undefined Saprotroph\| | Yeast | NULL | Probable | As nectar-tap_saprotroph (Põlme et al. 2020); Decay sugar-rich substrates (Põlme et al. 2020) | Põlme S, et al. 2020. Fungal Diversity 105:1-16 ((https://doi.org/10.1007/s13225-020-00466-2)) |
| Otu21312 | 1 | k__Fungi(100);p__Ascomycota(100);c__Saccharomycetes(100);o__Saccharomycetales(100);f__Metschnikowiaceae(100);g__Metschnikowia(100);s__Metschnikowia_pulcherrima(100); | Metschnikowia | 13 | Saprotroph | \|Undefined Saprotroph\| | Yeast | NULL | Probable | As nectar-tap_saprotroph (Põlme et al. 2020); Decay sugar-rich substrates (Põlme et al. 2020) | Põlme S, et al. 2020. Fungal Diversity 105:1-16 ((https://doi.org/10.1007/s13225-020-00466-2)) |
| Otu21313 | 1 | k__Fungi(100);p__Ascomycota(100);c__Saccharomycetes(100);o__Saccharomycetales(100);f__Metschnikowiaceae(100);g__Metschnikowia(100);s__Metschnikowia_pulcherrima(100); | Metschnikowia | 13 | Saprotroph | \|Undefined Saprotroph\| | Yeast | NULL | Probable | As nectar-tap_saprotroph (Põlme et al. 2020); Decay sugar-rich substrates (Põlme et al. 2020) | Põlme S, et al. 2020. Fungal Diversity 105:1-16 ((https://doi.org/10.1007/s13225-020-00466-2)) |
| Otu21314 | 1 | k__Fungi(100);p__Ascomycota(100);c__Saccharomycetes(100);o__Saccharomycetales(100);f__Metschnikowiaceae(100);g__Metschnikowia(100);g__Metschnikowia_unclassified(100); | Metschnikowia | 13 | Saprotroph | \|Undefined Saprotroph\| | Yeast | NULL | Probable | As nectar-tap_saprotroph (Põlme et al. 2020); Decay sugar-rich substrates (Põlme et al. 2020) | Põlme S, et al. 2020. Fungal Diversity 105:1-16 ((https://doi.org/10.1007/s13225-020-00466-2)) |
| Otu21315 | 1 | k__Fungi(100);p__Ascomycota(100);c__Saccharomycetes(100);o__Saccharomycetales(100);f__Metschnikowiaceae(100);g__Metschnikowia(100);s__Metschnikowia_pulcherrima(100); | Metschnikowia | 13 | Saprotroph | \|Undefined Saprotroph\| | Yeast | NULL | Probable | As nectar-tap_saprotroph (Põlme et al. 2020); Decay sugar-rich substrates (Põlme et al. 2020) | Põlme S, et al. 2020. Fungal Diversity 105:1-16 ((https://doi.org/10.1007/s13225-020-00466-2)) |
| Otu21316 | 1 | k__Fungi(100);p__Ascomycota(100);c__Saccharomycetes(100);o__Saccharomycetales(100);f__Metschnikowiaceae(100);g__Metschnikowia(100);s__Metschnikowia_pulcherrima(100); | Metschnikowia | 13 | Saprotroph | \|Undefined Saprotroph\| | Yeast | NULL | Probable | As nectar-tap_saprotroph (Põlme et al. 2020); Decay sugar-rich substrates (Põlme et al. 2020) | Põlme S, et al. 2020. Fungal Diversity 105:1-16 ((https://doi.org/10.1007/s13225-020-00466-2)) |
| Otu21317 | 1 | k__Fungi(100);p__Ascomycota(100);c__Saccharomycetes(100);o__Saccharomycetales(100);f__Metschnikowiaceae(100);g__Metschnikowia(100);s__Metschnikowia_pulcherrima(100); | Metschnikowia | 13 | Saprotroph | \|Undefined Saprotroph\| | Yeast | NULL | Probable | As nectar-tap_saprotroph (Põlme et al. 2020); Decay sugar-rich substrates (Põlme et al. 2020) | Põlme S, et al. 2020. Fungal Diversity 105:1-16 ((https://doi.org/10.1007/s13225-020-00466-2)) |
| Otu21318 | 1 | k__Fungi(100);p__Ascomycota(100);c__Saccharomycetes(100);o__Saccharomycetales(100);f__Metschnikowiaceae(100);g__Metschnikowia(100);s__Metschnikowia_pulcherrima(100); | Metschnikowia | 13 | Saprotroph | \|Undefined Saprotroph\| | Yeast | NULL | Probable | As nectar-tap_saprotroph (Põlme et al. 2020); Decay sugar-rich substrates (Põlme et al. 2020) | Põlme S, et al. 2020. Fungal Diversity 105:1-16 ((https://doi.org/10.1007/s13225-020-00466-2)) |
| Otu21319 | 1 | k__Fungi(100);p__Ascomycota(100);c__Saccharomycetes(100);o__Saccharomycetales(100);f__Metschnikowiaceae(100);g__Metschnikowia(100);s__Metschnikowia_pulcherrima(100); | Metschnikowia | 13 | Saprotroph | \|Undefined Saprotroph\| | Yeast | NULL | Probable | As nectar-tap_saprotroph (Põlme et al. 2020); Decay sugar-rich substrates (Põlme et al. 2020) | Põlme S, et al. 2020. Fungal Diversity 105:1-16 ((https://doi.org/10.1007/s13225-020-00466-2)) |
| Otu21321 | 1 | k__Fungi(100);p__Ascomycota(100);c__Saccharomycetes(100);o__Saccharomycetales(100);f__Metschnikowiaceae(100);g__Metschnikowia(100);s__Metschnikowia_pulcherrima(100); | Metschnikowia | 13 | Saprotroph | \|Undefined Saprotroph\| | Yeast | NULL | Probable | As nectar-tap_saprotroph (Põlme et al. 2020); Decay sugar-rich substrates (Põlme et al. 2020) | Põlme S, et al. 2020. Fungal Diversity 105:1-16 ((https://doi.org/10.1007/s13225-020-00466-2)) |
| Otu21323 | 1 | k__Fungi(100);p__Ascomycota(100);c__Saccharomycetes(100);o__Saccharomycetales(100);f__Metschnikowiaceae(100);g__Metschnikowia(100);s__Metschnikowia_pulcherrima(100); | Metschnikowia | 13 | Saprotroph | \|Undefined Saprotroph\| | Yeast | NULL | Probable | As nectar-tap_saprotroph (Põlme et al. 2020); Decay sugar-rich substrates (Põlme et al. 2020) | Põlme S, et al. 2020. Fungal Diversity 105:1-16 ((https://doi.org/10.1007/s13225-020-00466-2)) |
| Otu21324 | 1 | k__Fungi(100);p__Ascomycota(100);c__Saccharomycetes(100);o__Saccharomycetales(100);f__Metschnikowiaceae(100);g__Metschnikowia(100);g__Metschnikowia_unclassified(100); | Metschnikowia | 13 | Saprotroph | \|Undefined Saprotroph\| | Yeast | NULL | Probable | As nectar-tap_saprotroph (Põlme et al. 2020); Decay sugar-rich substrates (Põlme et al. 2020) | Põlme S, et al. 2020. Fungal Diversity 105:1-16 ((https://doi.org/10.1007/s13225-020-00466-2)) |
| Otu21325 | 1 | k__Fungi(100);p__Ascomycota(100);c__Saccharomycetes(100);o__Saccharomycetales(100);f__Metschnikowiaceae(100);g__Metschnikowia(100);s__Metschnikowia_pulcherrima(100); | Metschnikowia | 13 | Saprotroph | \|Undefined Saprotroph\| | Yeast | NULL | Probable | As nectar-tap_saprotroph (Põlme et al. 2020); Decay sugar-rich substrates (Põlme et al. 2020) | Põlme S, et al. 2020. Fungal Diversity 105:1-16 ((https://doi.org/10.1007/s13225-020-00466-2)) |
| Otu21327 | 1 | k__Fungi(100);p__Ascomycota(100);c__Saccharomycetes(100);o__Saccharomycetales(100);f__Saccharomycetales_family_Incertae_sedis(100);g__Candida(100);g__Candida_unclassified(100); | Candida | 13 | Pathotroph-Saprotroph-Symbiotroph | Animal Pathogen-Endophyte-Endosymbiont-Epiphyte-\|Undefined Saprotroph\| | Dimorphic Yeast-Yeast | NULL | Probable | As nectar-tap_saprotroph (Põlme et al. 2020); Decay sugar-rich substrates (Põlme et al. 2020); This asexual genus is polyphyletic and contain species that function very differently depend on what they are. Caution should be taken when interpreting guilds for Candida species. | Manolakaki D, et al. 2010. Virulence. 1: 367-375 ((https://doi.org/10.4161/viru.1.5.12796)); Mekbib SB, et al. 2011. Tropical Plant Pathology 36:233-240 ((https://doi.org/10.1590/S1982-56762011000400004)); Põlme S, et al. 2020. Fungal Diversity 105:1-16 ((https://doi.org/10.1007/s13225-020-00466-2)) |
| Otu21328 | 1 | k__Fungi(100);p__Ascomycota(100);c__Saccharomycetes(100);o__Saccharomycetales(100);f__Metschnikowiaceae(100);g__Metschnikowia(100);s__Metschnikowia_pulcherrima(100); | Metschnikowia | 13 | Saprotroph | \|Undefined Saprotroph\| | Yeast | NULL | Probable | As nectar-tap_saprotroph (Põlme et al. 2020); Decay sugar-rich substrates (Põlme et al. 2020) | Põlme S, et al. 2020. Fungal Diversity 105:1-16 ((https://doi.org/10.1007/s13225-020-00466-2)) |
| Otu21329 | 1 | k__Fungi(100);p__Ascomycota(100);c__Saccharomycetes(100);o__Saccharomycetales(100);f__Metschnikowiaceae(100);g__Metschnikowia(100);s__Metschnikowia_pulcherrima(100); | Metschnikowia | 13 | Saprotroph | \|Undefined Saprotroph\| | Yeast | NULL | Probable | As nectar-tap_saprotroph (Põlme et al. 2020); Decay sugar-rich substrates (Põlme et al. 2020) | Põlme S, et al. 2020. Fungal Diversity 105:1-16 ((https://doi.org/10.1007/s13225-020-00466-2)) |
| Otu21330 | 1 | k__Fungi(100);p__Ascomycota(100);c__Saccharomycetes(100);o__Saccharomycetales(100);f__Metschnikowiaceae(100);g__Metschnikowia(100);g__Metschnikowia_unclassified(100); | Metschnikowia | 13 | Saprotroph | \|Undefined Saprotroph\| | Yeast | NULL | Probable | As nectar-tap_saprotroph (Põlme et al. 2020); Decay sugar-rich substrates (Põlme et al. 2020) | Põlme S, et al. 2020. Fungal Diversity 105:1-16 ((https://doi.org/10.1007/s13225-020-00466-2)) |
| Otu21331 | 1 | k__Fungi(100);p__Ascomycota(100);c__Saccharomycetes(100);o__Saccharomycetales(100);f__Metschnikowiaceae(100);g__Metschnikowia(100);s__Metschnikowia_pulcherrima(100); | Metschnikowia | 13 | Saprotroph | \|Undefined Saprotroph\| | Yeast | NULL | Probable | As nectar-tap_saprotroph (Põlme et al. 2020); Decay sugar-rich substrates (Põlme et al. 2020) | Põlme S, et al. 2020. Fungal Diversity 105:1-16 ((https://doi.org/10.1007/s13225-020-00466-2)) |
| Otu21332 | 1 | k__Fungi(100);p__Ascomycota(100);c__Saccharomycetes(100);o__Saccharomycetales(100);f__Metschnikowiaceae(100);g__Metschnikowia(100);g__Metschnikowia_unclassified(100); | Metschnikowia | 13 | Saprotroph | \|Undefined Saprotroph\| | Yeast | NULL | Probable | As nectar-tap_saprotroph (Põlme et al. 2020); Decay sugar-rich substrates (Põlme et al. 2020) | Põlme S, et al. 2020. Fungal Diversity 105:1-16 ((https://doi.org/10.1007/s13225-020-00466-2)) |
| Otu21333 | 1 | k__Fungi(100);p__Ascomycota(100);c__Saccharomycetes(100);o__Saccharomycetales(100);f__Metschnikowiaceae(100);g__Metschnikowia(100);g__Metschnikowia_unclassified(100); | Metschnikowia | 13 | Saprotroph | \|Undefined Saprotroph\| | Yeast | NULL | Probable | As nectar-tap_saprotroph (Põlme et al. 2020); Decay sugar-rich substrates (Põlme et al. 2020) | Põlme S, et al. 2020. Fungal Diversity 105:1-16 ((https://doi.org/10.1007/s13225-020-00466-2)) |
| Otu21334 | 1 | k__Fungi(100);p__Ascomycota(100);c__Saccharomycetes(100);o__Saccharomycetales(100);f__Metschnikowiaceae(100);g__Metschnikowia(100);s__Metschnikowia_pulcherrima(100); | Metschnikowia | 13 | Saprotroph | \|Undefined Saprotroph\| | Yeast | NULL | Probable | As nectar-tap_saprotroph (Põlme et al. 2020); Decay sugar-rich substrates (Põlme et al. 2020) | Põlme S, et al. 2020. Fungal Diversity 105:1-16 ((https://doi.org/10.1007/s13225-020-00466-2)) |
| Otu21335 | 1 | k__Fungi(100);p__Ascomycota(100);c__Saccharomycetes(100);o__Saccharomycetales(100);f__Metschnikowiaceae(100);g__Metschnikowia(100);s__Metschnikowia_pulcherrima(100); | Metschnikowia | 13 | Saprotroph | \|Undefined Saprotroph\| | Yeast | NULL | Probable | As nectar-tap_saprotroph (Põlme et al. 2020); Decay sugar-rich substrates (Põlme et al. 2020) | Põlme S, et al. 2020. Fungal Diversity 105:1-16 ((https://doi.org/10.1007/s13225-020-00466-2)) |
| Otu21336 | 1 | k__Fungi(100);p__Ascomycota(100);c__Saccharomycetes(100);o__Saccharomycetales(100);f__Metschnikowiaceae(100);g__Metschnikowia(100);s__Metschnikowia_pulcherrima(100); | Metschnikowia | 13 | Saprotroph | \|Undefined Saprotroph\| | Yeast | NULL | Probable | As nectar-tap_saprotroph (Põlme et al. 2020); Decay sugar-rich substrates (Põlme et al. 2020) | Põlme S, et al. 2020. Fungal Diversity 105:1-16 ((https://doi.org/10.1007/s13225-020-00466-2)) |
| Otu21337 | 1 | k__Fungi(100);p__Ascomycota(100);c__Saccharomycetes(100);o__Saccharomycetales(100);f__Metschnikowiaceae(100);g__Metschnikowia(100);g__Metschnikowia_unclassified(100); | Metschnikowia | 13 | Saprotroph | \|Undefined Saprotroph\| | Yeast | NULL | Probable | As nectar-tap_saprotroph (Põlme et al. 2020); Decay sugar-rich substrates (Põlme et al. 2020) | Põlme S, et al. 2020. Fungal Diversity 105:1-16 ((https://doi.org/10.1007/s13225-020-00466-2)) |
| Otu21338 | 1 | k__Fungi(100);p__Ascomycota(100);c__Saccharomycetes(100);o__Saccharomycetales(100);f__Metschnikowiaceae(100);g__Metschnikowia(100);s__Metschnikowia_pulcherrima(100); | Metschnikowia | 13 | Saprotroph | \|Undefined Saprotroph\| | Yeast | NULL | Probable | As nectar-tap_saprotroph (Põlme et al. 2020); Decay sugar-rich substrates (Põlme et al. 2020) | Põlme S, et al. 2020. Fungal Diversity 105:1-16 ((https://doi.org/10.1007/s13225-020-00466-2)) |
| Otu21341 | 1 | k__Fungi(100);p__Ascomycota(100);c__Saccharomycetes(100);o__Saccharomycetales(100);f__Metschnikowiaceae(100);g__Metschnikowia(100);s__Metschnikowia_pulcherrima(100); | Metschnikowia | 13 | Saprotroph | \|Undefined Saprotroph\| | Yeast | NULL | Probable | As nectar-tap_saprotroph (Põlme et al. 2020); Decay sugar-rich substrates (Põlme et al. 2020) | Põlme S, et al. 2020. Fungal Diversity 105:1-16 ((https://doi.org/10.1007/s13225-020-00466-2)) |
| Otu21343 | 1 | k__Fungi(100);p__Ascomycota(100);c__Saccharomycetes(100);o__Saccharomycetales(100);f__Metschnikowiaceae(100);g__Metschnikowia(100);g__Metschnikowia_unclassified(100); | Metschnikowia | 13 | Saprotroph | \|Undefined Saprotroph\| | Yeast | NULL | Probable | As nectar-tap_saprotroph (Põlme et al. 2020); Decay sugar-rich substrates (Põlme et al. 2020) | Põlme S, et al. 2020. Fungal Diversity 105:1-16 ((https://doi.org/10.1007/s13225-020-00466-2)) |
| Otu21344 | 1 | k__Fungi(100);p__Ascomycota(100);c__Saccharomycetes(100);o__Saccharomycetales(100);f__Metschnikowiaceae(100);g__Metschnikowia(100);g__Metschnikowia_unclassified(100); | Metschnikowia | 13 | Saprotroph | \|Undefined Saprotroph\| | Yeast | NULL | Probable | As nectar-tap_saprotroph (Põlme et al. 2020); Decay sugar-rich substrates (Põlme et al. 2020) | Põlme S, et al. 2020. Fungal Diversity 105:1-16 ((https://doi.org/10.1007/s13225-020-00466-2)) |
| Otu21345 | 1 | k__Fungi(100);p__Ascomycota(100);c__Saccharomycetes(100);o__Saccharomycetales(100);f__Saccharomycetales_family_Incertae_sedis(100);g__Candida(100);g__Candida_unclassified(100); | Candida | 13 | Pathotroph-Saprotroph-Symbiotroph | Animal Pathogen-Endophyte-Endosymbiont-Epiphyte-\|Undefined Saprotroph\| | Dimorphic Yeast-Yeast | NULL | Probable | As nectar-tap_saprotroph (Põlme et al. 2020); Decay sugar-rich substrates (Põlme et al. 2020); This asexual genus is polyphyletic and contain species that function very differently depend on what they are. Caution should be taken when interpreting guilds for Candida species. | Manolakaki D, et al. 2010. Virulence. 1: 367-375 ((https://doi.org/10.4161/viru.1.5.12796)); Mekbib SB, et al. 2011. Tropical Plant Pathology 36:233-240 ((https://doi.org/10.1590/S1982-56762011000400004)); Põlme S, et al. 2020. Fungal Diversity 105:1-16 ((https://doi.org/10.1007/s13225-020-00466-2)) |
| Otu21346 | 1 | k__Fungi(100);p__Ascomycota(100);c__Saccharomycetes(100);o__Saccharomycetales(100);f__Metschnikowiaceae(100);g__Metschnikowia(100);g__Metschnikowia_unclassified(100); | Metschnikowia | 13 | Saprotroph | \|Undefined Saprotroph\| | Yeast | NULL | Probable | As nectar-tap_saprotroph (Põlme et al. 2020); Decay sugar-rich substrates (Põlme et al. 2020) | Põlme S, et al. 2020. Fungal Diversity 105:1-16 ((https://doi.org/10.1007/s13225-020-00466-2)) |
| Otu21347 | 1 | k__Fungi(100);p__Ascomycota(100);c__Saccharomycetes(100);o__Saccharomycetales(100);f__Metschnikowiaceae(100);g__Metschnikowia(100);g__Metschnikowia_unclassified(100); | Metschnikowia | 13 | Saprotroph | \|Undefined Saprotroph\| | Yeast | NULL | Probable | As nectar-tap_saprotroph (Põlme et al. 2020); Decay sugar-rich substrates (Põlme et al. 2020) | Põlme S, et al. 2020. Fungal Diversity 105:1-16 ((https://doi.org/10.1007/s13225-020-00466-2)) |
| Otu21348 | 1 | k__Fungi(100);p__Ascomycota(100);c__Saccharomycetes(100);o__Saccharomycetales(100);f__Metschnikowiaceae(100);g__Metschnikowia(100);s__Metschnikowia_pulcherrima(100); | Metschnikowia | 13 | Saprotroph | \|Undefined Saprotroph\| | Yeast | NULL | Probable | As nectar-tap_saprotroph (Põlme et al. 2020); Decay sugar-rich substrates (Põlme et al. 2020) | Põlme S, et al. 2020. Fungal Diversity 105:1-16 ((https://doi.org/10.1007/s13225-020-00466-2)) |
| Otu21349 | 1 | k__Fungi(100);p__Ascomycota(100);c__Saccharomycetes(100);o__Saccharomycetales(100);f__Metschnikowiaceae(100);g__Metschnikowia(100);s__Metschnikowia_pulcherrima(100); | Metschnikowia | 13 | Saprotroph | \|Undefined Saprotroph\| | Yeast | NULL | Probable | As nectar-tap_saprotroph (Põlme et al. 2020); Decay sugar-rich substrates (Põlme et al. 2020) | Põlme S, et al. 2020. Fungal Diversity 105:1-16 ((https://doi.org/10.1007/s13225-020-00466-2)) |
| Otu21350 | 1 | k__Fungi(100);p__Ascomycota(100);c__Saccharomycetes(100);o__Saccharomycetales(100);f__Metschnikowiaceae(100);g__Metschnikowia(100);s__Metschnikowia_pulcherrima(100); | Metschnikowia | 13 | Saprotroph | \|Undefined Saprotroph\| | Yeast | NULL | Probable | As nectar-tap_saprotroph (Põlme et al. 2020); Decay sugar-rich substrates (Põlme et al. 2020) | Põlme S, et al. 2020. Fungal Diversity 105:1-16 ((https://doi.org/10.1007/s13225-020-00466-2)) |
| Otu21351 | 1 | k__Fungi(100);p__Ascomycota(100);c__Saccharomycetes(100);o__Saccharomycetales(100);f__Metschnikowiaceae(100);g__Metschnikowia(100);s__Metschnikowia_pulcherrima(100); | Metschnikowia | 13 | Saprotroph | \|Undefined Saprotroph\| | Yeast | NULL | Probable | As nectar-tap_saprotroph (Põlme et al. 2020); Decay sugar-rich substrates (Põlme et al. 2020) | Põlme S, et al. 2020. Fungal Diversity 105:1-16 ((https://doi.org/10.1007/s13225-020-00466-2)) |
| Otu21352 | 1 | k__Fungi(100);p__Ascomycota(100);c__Saccharomycetes(100);o__Saccharomycetales(100);f__Metschnikowiaceae(100);g__Metschnikowia(100);s__Metschnikowia_pulcherrima(100); | Metschnikowia | 13 | Saprotroph | \|Undefined Saprotroph\| | Yeast | NULL | Probable | As nectar-tap_saprotroph (Põlme et al. 2020); Decay sugar-rich substrates (Põlme et al. 2020) | Põlme S, et al. 2020. Fungal Diversity 105:1-16 ((https://doi.org/10.1007/s13225-020-00466-2)) |
| Otu21353 | 1 | k__Fungi(100);p__Ascomycota(100);c__Saccharomycetes(100);o__Saccharomycetales(100);f__Metschnikowiaceae(100);g__Metschnikowia(100);s__Metschnikowia_pulcherrima(100); | Metschnikowia | 13 | Saprotroph | \|Undefined Saprotroph\| | Yeast | NULL | Probable | As nectar-tap_saprotroph (Põlme et al. 2020); Decay sugar-rich substrates (Põlme et al. 2020) | Põlme S, et al. 2020. Fungal Diversity 105:1-16 ((https://doi.org/10.1007/s13225-020-00466-2)) |
| Otu21354 | 1 | k__Fungi(100);p__Ascomycota(100);c__Saccharomycetes(100);o__Saccharomycetales(100);f__Metschnikowiaceae(100);g__Metschnikowia(100);s__Metschnikowia_pulcherrima(100); | Metschnikowia | 13 | Saprotroph | \|Undefined Saprotroph\| | Yeast | NULL | Probable | As nectar-tap_saprotroph (Põlme et al. 2020); Decay sugar-rich substrates (Põlme et al. 2020) | Põlme S, et al. 2020. Fungal Diversity 105:1-16 ((https://doi.org/10.1007/s13225-020-00466-2)) |
| Otu21355 | 1 | k__Fungi(100);p__Ascomycota(100);c__Saccharomycetes(100);o__Saccharomycetales(100);f__Metschnikowiaceae(100);g__Metschnikowia(100);g__Metschnikowia_unclassified(100); | Metschnikowia | 13 | Saprotroph | \|Undefined Saprotroph\| | Yeast | NULL | Probable | As nectar-tap_saprotroph (Põlme et al. 2020); Decay sugar-rich substrates (Põlme et al. 2020) | Põlme S, et al. 2020. Fungal Diversity 105:1-16 ((https://doi.org/10.1007/s13225-020-00466-2)) |
| Otu21356 | 1 | k__Fungi(100);p__Ascomycota(100);c__Saccharomycetes(100);o__Saccharomycetales(100);f__Metschnikowiaceae(100);g__Metschnikowia(100);s__Metschnikowia_pulcherrima(100); | Metschnikowia | 13 | Saprotroph | \|Undefined Saprotroph\| | Yeast | NULL | Probable | As nectar-tap_saprotroph (Põlme et al. 2020); Decay sugar-rich substrates (Põlme et al. 2020) | Põlme S, et al. 2020. Fungal Diversity 105:1-16 ((https://doi.org/10.1007/s13225-020-00466-2)) |
| Otu21357 | 1 | k__Fungi(100);p__Ascomycota(100);c__Saccharomycetes(100);o__Saccharomycetales(100);f__Metschnikowiaceae(100);g__Metschnikowia(100);s__Metschnikowia_pulcherrima(100); | Metschnikowia | 13 | Saprotroph | \|Undefined Saprotroph\| | Yeast | NULL | Probable | As nectar-tap_saprotroph (Põlme et al. 2020); Decay sugar-rich substrates (Põlme et al. 2020) | Põlme S, et al. 2020. Fungal Diversity 105:1-16 ((https://doi.org/10.1007/s13225-020-00466-2)) |
| Otu21358 | 1 | k__Fungi(100);p__Ascomycota(100);c__Saccharomycetes(100);o__Saccharomycetales(100);f__Metschnikowiaceae(100);g__Metschnikowia(100);s__Metschnikowia_pulcherrima(100); | Metschnikowia | 13 | Saprotroph | \|Undefined Saprotroph\| | Yeast | NULL | Probable | As nectar-tap_saprotroph (Põlme et al. 2020); Decay sugar-rich substrates (Põlme et al. 2020) | Põlme S, et al. 2020. Fungal Diversity 105:1-16 ((https://doi.org/10.1007/s13225-020-00466-2)) |
| Otu21361 | 1 | k__Fungi(100);p__Ascomycota(100);c__Saccharomycetes(100);o__Saccharomycetales(100);f__Metschnikowiaceae(100);g__Metschnikowia(100);s__Metschnikowia_pulcherrima(100); | Metschnikowia | 13 | Saprotroph | \|Undefined Saprotroph\| | Yeast | NULL | Probable | As nectar-tap_saprotroph (Põlme et al. 2020); Decay sugar-rich substrates (Põlme et al. 2020) | Põlme S, et al. 2020. Fungal Diversity 105:1-16 ((https://doi.org/10.1007/s13225-020-00466-2)) |
| Otu21363 | 1 | k__Fungi(100);p__Ascomycota(100);c__Saccharomycetes(100);o__Saccharomycetales(100);f__Metschnikowiaceae(100);g__Metschnikowia(100);g__Metschnikowia_unclassified(100); | Metschnikowia | 13 | Saprotroph | \|Undefined Saprotroph\| | Yeast | NULL | Probable | As nectar-tap_saprotroph (Põlme et al. 2020); Decay sugar-rich substrates (Põlme et al. 2020) | Põlme S, et al. 2020. Fungal Diversity 105:1-16 ((https://doi.org/10.1007/s13225-020-00466-2)) |
| Otu21364 | 1 | k__Fungi(100);p__Ascomycota(100);c__Saccharomycetes(100);o__Saccharomycetales(100);f__Metschnikowiaceae(100);g__Metschnikowia(100);g__Metschnikowia_unclassified(100); | Metschnikowia | 13 | Saprotroph | \|Undefined Saprotroph\| | Yeast | NULL | Probable | As nectar-tap_saprotroph (Põlme et al. 2020); Decay sugar-rich substrates (Põlme et al. 2020) | Põlme S, et al. 2020. Fungal Diversity 105:1-16 ((https://doi.org/10.1007/s13225-020-00466-2)) |
| Otu21365 | 1 | k__Fungi(100);p__Ascomycota(100);c__Saccharomycetes(100);o__Saccharomycetales(100);f__Metschnikowiaceae(100);g__Metschnikowia(100);s__Metschnikowia_pulcherrima(100); | Metschnikowia | 13 | Saprotroph | \|Undefined Saprotroph\| | Yeast | NULL | Probable | As nectar-tap_saprotroph (Põlme et al. 2020); Decay sugar-rich substrates (Põlme et al. 2020) | Põlme S, et al. 2020. Fungal Diversity 105:1-16 ((https://doi.org/10.1007/s13225-020-00466-2)) |
| Otu21366 | 1 | k__Fungi(100);p__Ascomycota(100);c__Saccharomycetes(100);o__Saccharomycetales(100);f__Metschnikowiaceae(100);g__Metschnikowia(100);s__Metschnikowia_pulcherrima(100); | Metschnikowia | 13 | Saprotroph | \|Undefined Saprotroph\| | Yeast | NULL | Probable | As nectar-tap_saprotroph (Põlme et al. 2020); Decay sugar-rich substrates (Põlme et al. 2020) | Põlme S, et al. 2020. Fungal Diversity 105:1-16 ((https://doi.org/10.1007/s13225-020-00466-2)) |
| Otu21367 | 1 | k__Fungi(100);p__Ascomycota(100);c__Saccharomycetes(100);o__Saccharomycetales(100);f__Metschnikowiaceae(100);g__Metschnikowia(100);s__Metschnikowia_pulcherrima(100); | Metschnikowia | 13 | Saprotroph | \|Undefined Saprotroph\| | Yeast | NULL | Probable | As nectar-tap_saprotroph (Põlme et al. 2020); Decay sugar-rich substrates (Põlme et al. 2020) | Põlme S, et al. 2020. Fungal Diversity 105:1-16 ((https://doi.org/10.1007/s13225-020-00466-2)) |
| Otu21368 | 1 | k__Fungi(100);p__Ascomycota(100);c__Saccharomycetes(100);o__Saccharomycetales(100);f__Metschnikowiaceae(100);g__Metschnikowia(100);s__Metschnikowia_pulcherrima(100); | Metschnikowia | 13 | Saprotroph | \|Undefined Saprotroph\| | Yeast | NULL | Probable | As nectar-tap_saprotroph (Põlme et al. 2020); Decay sugar-rich substrates (Põlme et al. 2020) | Põlme S, et al. 2020. Fungal Diversity 105:1-16 ((https://doi.org/10.1007/s13225-020-00466-2)) |
| Otu21369 | 1 | k__Fungi(100);p__Ascomycota(100);c__Saccharomycetes(100);o__Saccharomycetales(100);f__Metschnikowiaceae(100);g__Metschnikowia(100);s__Metschnikowia_pulcherrima(100); | Metschnikowia | 13 | Saprotroph | \|Undefined Saprotroph\| | Yeast | NULL | Probable | As nectar-tap_saprotroph (Põlme et al. 2020); Decay sugar-rich substrates (Põlme et al. 2020) | Põlme S, et al. 2020. Fungal Diversity 105:1-16 ((https://doi.org/10.1007/s13225-020-00466-2)) |
| Otu21370 | 1 | k__Fungi(100);p__Ascomycota(100);c__Saccharomycetes(100);o__Saccharomycetales(100);f__Metschnikowiaceae(100);g__Metschnikowia(100);s__Metschnikowia_pulcherrima(100); | Metschnikowia | 13 | Saprotroph | \|Undefined Saprotroph\| | Yeast | NULL | Probable | As nectar-tap_saprotroph (Põlme et al. 2020); Decay sugar-rich substrates (Põlme et al. 2020) | Põlme S, et al. 2020. Fungal Diversity 105:1-16 ((https://doi.org/10.1007/s13225-020-00466-2)) |
| Otu21372 | 1 | k__Fungi(100);p__Ascomycota(100);c__Saccharomycetes(100);o__Saccharomycetales(100);f__Metschnikowiaceae(100);g__Metschnikowia(100);s__Metschnikowia_pulcherrima(100); | Metschnikowia | 13 | Saprotroph | \|Undefined Saprotroph\| | Yeast | NULL | Probable | As nectar-tap_saprotroph (Põlme et al. 2020); Decay sugar-rich substrates (Põlme et al. 2020) | Põlme S, et al. 2020. Fungal Diversity 105:1-16 ((https://doi.org/10.1007/s13225-020-00466-2)) |
| Otu21373 | 1 | k__Fungi(100);p__Ascomycota(100);c__Saccharomycetes(100);o__Saccharomycetales(100);f__Metschnikowiaceae(100);g__Metschnikowia(100);s__Metschnikowia_pulcherrima(100); | Metschnikowia | 13 | Saprotroph | \|Undefined Saprotroph\| | Yeast | NULL | Probable | As nectar-tap_saprotroph (Põlme et al. 2020); Decay sugar-rich substrates (Põlme et al. 2020) | Põlme S, et al. 2020. Fungal Diversity 105:1-16 ((https://doi.org/10.1007/s13225-020-00466-2)) |
| Otu21374 | 1 | k__Fungi(100);p__Ascomycota(100);c__Saccharomycetes(100);o__Saccharomycetales(100);f__Metschnikowiaceae(100);g__Metschnikowia(100);g__Metschnikowia_unclassified(100); | Metschnikowia | 13 | Saprotroph | \|Undefined Saprotroph\| | Yeast | NULL | Probable | As nectar-tap_saprotroph (Põlme et al. 2020); Decay sugar-rich substrates (Põlme et al. 2020) | Põlme S, et al. 2020. Fungal Diversity 105:1-16 ((https://doi.org/10.1007/s13225-020-00466-2)) |
| Otu21375 | 1 | k__Fungi(100);p__Ascomycota(100);c__Saccharomycetes(100);o__Saccharomycetales(100);f__Metschnikowiaceae(100);g__Metschnikowia(100);s__Metschnikowia_pulcherrima(100); | Metschnikowia | 13 | Saprotroph | \|Undefined Saprotroph\| | Yeast | NULL | Probable | As nectar-tap_saprotroph (Põlme et al. 2020); Decay sugar-rich substrates (Põlme et al. 2020) | Põlme S, et al. 2020. Fungal Diversity 105:1-16 ((https://doi.org/10.1007/s13225-020-00466-2)) |
| Otu21376 | 1 | k__Fungi(100);p__Ascomycota(100);c__Saccharomycetes(100);o__Saccharomycetales(100);f__Metschnikowiaceae(100);g__Metschnikowia(100);s__Metschnikowia_pulcherrima(100); | Metschnikowia | 13 | Saprotroph | \|Undefined Saprotroph\| | Yeast | NULL | Probable | As nectar-tap_saprotroph (Põlme et al. 2020); Decay sugar-rich substrates (Põlme et al. 2020) | Põlme S, et al. 2020. Fungal Diversity 105:1-16 ((https://doi.org/10.1007/s13225-020-00466-2)) |
| Otu21377 | 1 | k__Fungi(100);p__Ascomycota(100);c__Saccharomycetes(100);o__Saccharomycetales(100);f__Metschnikowiaceae(100);g__Metschnikowia(100);s__Metschnikowia_pulcherrima(100); | Metschnikowia | 13 | Saprotroph | \|Undefined Saprotroph\| | Yeast | NULL | Probable | As nectar-tap_saprotroph (Põlme et al. 2020); Decay sugar-rich substrates (Põlme et al. 2020) | Põlme S, et al. 2020. Fungal Diversity 105:1-16 ((https://doi.org/10.1007/s13225-020-00466-2)) |
| Otu21378 | 1 | k__Fungi(100);p__Ascomycota(100);c__Saccharomycetes(100);o__Saccharomycetales(100);f__Metschnikowiaceae(100);g__Metschnikowia(100);s__Metschnikowia_pulcherrima(100); | Metschnikowia | 13 | Saprotroph | \|Undefined Saprotroph\| | Yeast | NULL | Probable | As nectar-tap_saprotroph (Põlme et al. 2020); Decay sugar-rich substrates (Põlme et al. 2020) | Põlme S, et al. 2020. Fungal Diversity 105:1-16 ((https://doi.org/10.1007/s13225-020-00466-2)) |
| Otu21379 | 1 | k__Fungi(100);p__Ascomycota(100);c__Saccharomycetes(100);o__Saccharomycetales(100);f__Metschnikowiaceae(100);g__Metschnikowia(100);s__Metschnikowia_pulcherrima(100); | Metschnikowia | 13 | Saprotroph | \|Undefined Saprotroph\| | Yeast | NULL | Probable | As nectar-tap_saprotroph (Põlme et al. 2020); Decay sugar-rich substrates (Põlme et al. 2020) | Põlme S, et al. 2020. Fungal Diversity 105:1-16 ((https://doi.org/10.1007/s13225-020-00466-2)) |
| Otu21380 | 1 | k__Fungi(100);p__Ascomycota(100);c__Saccharomycetes(100);o__Saccharomycetales(100);f__Metschnikowiaceae(100);g__Metschnikowia(100);s__Metschnikowia_pulcherrima(100); | Metschnikowia | 13 | Saprotroph | \|Undefined Saprotroph\| | Yeast | NULL | Probable | As nectar-tap_saprotroph (Põlme et al. 2020); Decay sugar-rich substrates (Põlme et al. 2020) | Põlme S, et al. 2020. Fungal Diversity 105:1-16 ((https://doi.org/10.1007/s13225-020-00466-2)) |
| Otu21381 | 1 | k__Fungi(100);p__Ascomycota(100);c__Saccharomycetes(100);o__Saccharomycetales(100);f__Metschnikowiaceae(100);g__Metschnikowia(100);g__Metschnikowia_unclassified(100); | Metschnikowia | 13 | Saprotroph | \|Undefined Saprotroph\| | Yeast | NULL | Probable | As nectar-tap_saprotroph (Põlme et al. 2020); Decay sugar-rich substrates (Põlme et al. 2020) | Põlme S, et al. 2020. Fungal Diversity 105:1-16 ((https://doi.org/10.1007/s13225-020-00466-2)) |
| Otu21382 | 1 | k__Fungi(100);p__Ascomycota(100);c__Saccharomycetes(100);o__Saccharomycetales(100);f__Metschnikowiaceae(100);g__Metschnikowia(100);s__Metschnikowia_pulcherrima(100); | Metschnikowia | 13 | Saprotroph | \|Undefined Saprotroph\| | Yeast | NULL | Probable | As nectar-tap_saprotroph (Põlme et al. 2020); Decay sugar-rich substrates (Põlme et al. 2020) | Põlme S, et al. 2020. Fungal Diversity 105:1-16 ((https://doi.org/10.1007/s13225-020-00466-2)) |
| Otu21383 | 1 | k__Fungi(100);p__Ascomycota(100);c__Saccharomycetes(100);o__Saccharomycetales(100);f__Metschnikowiaceae(100);g__Metschnikowia(100);g__Metschnikowia_unclassified(100); | Metschnikowia | 13 | Saprotroph | \|Undefined Saprotroph\| | Yeast | NULL | Probable | As nectar-tap_saprotroph (Põlme et al. 2020); Decay sugar-rich substrates (Põlme et al. 2020) | Põlme S, et al. 2020. Fungal Diversity 105:1-16 ((https://doi.org/10.1007/s13225-020-00466-2)) |
| Otu21384 | 1 | k__Fungi(100);p__Ascomycota(100);c__Saccharomycetes(100);o__Saccharomycetales(100);f__Saccharomycetales_family_Incertae_sedis(100);g__Candida(100);g__Candida_unclassified(100); | Candida | 13 | Pathotroph-Saprotroph-Symbiotroph | Animal Pathogen-Endophyte-Endosymbiont-Epiphyte-\|Undefined Saprotroph\| | Dimorphic Yeast-Yeast | NULL | Probable | As nectar-tap_saprotroph (Põlme et al. 2020); Decay sugar-rich substrates (Põlme et al. 2020); This asexual genus is polyphyletic and contain species that function very differently depend on what they are. Caution should be taken when interpreting guilds for Candida species. | Manolakaki D, et al. 2010. Virulence. 1: 367-375 ((https://doi.org/10.4161/viru.1.5.12796)); Mekbib SB, et al. 2011. Tropical Plant Pathology 36:233-240 ((https://doi.org/10.1590/S1982-56762011000400004)); Põlme S, et al. 2020. Fungal Diversity 105:1-16 ((https://doi.org/10.1007/s13225-020-00466-2)) |
| Otu21387 | 1 | k__Fungi(100);p__Ascomycota(100);c__Saccharomycetes(100);o__Saccharomycetales(100);f__Metschnikowiaceae(100);g__Metschnikowia(100);g__Metschnikowia_unclassified(100); | Metschnikowia | 13 | Saprotroph | \|Undefined Saprotroph\| | Yeast | NULL | Probable | As nectar-tap_saprotroph (Põlme et al. 2020); Decay sugar-rich substrates (Põlme et al. 2020) | Põlme S, et al. 2020. Fungal Diversity 105:1-16 ((https://doi.org/10.1007/s13225-020-00466-2)) |
| Otu21388 | 1 | k__Fungi(100);p__Ascomycota(100);c__Saccharomycetes(100);o__Saccharomycetales(100);f__Saccharomycetales_family_Incertae_sedis(100);g__Candida(100);g__Candida_unclassified(100); | Candida | 13 | Pathotroph-Saprotroph-Symbiotroph | Animal Pathogen-Endophyte-Endosymbiont-Epiphyte-\|Undefined Saprotroph\| | Dimorphic Yeast-Yeast | NULL | Probable | As nectar-tap_saprotroph (Põlme et al. 2020); Decay sugar-rich substrates (Põlme et al. 2020); This asexual genus is polyphyletic and contain species that function very differently depend on what they are. Caution should be taken when interpreting guilds for Candida species. | Manolakaki D, et al. 2010. Virulence. 1: 367-375 ((https://doi.org/10.4161/viru.1.5.12796)); Mekbib SB, et al. 2011. Tropical Plant Pathology 36:233-240 ((https://doi.org/10.1590/S1982-56762011000400004)); Põlme S, et al. 2020. Fungal Diversity 105:1-16 ((https://doi.org/10.1007/s13225-020-00466-2)) |
| Otu21390 | 1 | k__Fungi(100);p__Ascomycota(100);c__Saccharomycetes(100);o__Saccharomycetales(100);f__Metschnikowiaceae(100);g__Metschnikowia(100);s__Metschnikowia_pulcherrima(100); | Metschnikowia | 13 | Saprotroph | \|Undefined Saprotroph\| | Yeast | NULL | Probable | As nectar-tap_saprotroph (Põlme et al. 2020); Decay sugar-rich substrates (Põlme et al. 2020) | Põlme S, et al. 2020. Fungal Diversity 105:1-16 ((https://doi.org/10.1007/s13225-020-00466-2)) |
| Otu21392 | 1 | k__Fungi(100);p__Ascomycota(100);c__Saccharomycetes(100);o__Saccharomycetales(100);f__Metschnikowiaceae(100);g__Metschnikowia(100);g__Metschnikowia_unclassified(100); | Metschnikowia | 13 | Saprotroph | \|Undefined Saprotroph\| | Yeast | NULL | Probable | As nectar-tap_saprotroph (Põlme et al. 2020); Decay sugar-rich substrates (Põlme et al. 2020) | Põlme S, et al. 2020. Fungal Diversity 105:1-16 ((https://doi.org/10.1007/s13225-020-00466-2)) |
| Otu21393 | 1 | k__Fungi(100);p__Ascomycota(100);c__Saccharomycetes(100);o__Saccharomycetales(100);f__Saccharomycetales_family_Incertae_sedis(100);g__Candida(100);s__Candida_cretensis(100); | Candida | 13 | Pathotroph-Saprotroph-Symbiotroph | Animal Pathogen-Endophyte-Endosymbiont-Epiphyte-\|Undefined Saprotroph\| | Dimorphic Yeast-Yeast | NULL | Probable | As nectar-tap_saprotroph (Põlme et al. 2020); Decay sugar-rich substrates (Põlme et al. 2020); This asexual genus is polyphyletic and contain species that function very differently depend on what they are. Caution should be taken when interpreting guilds for Candida species. | Manolakaki D, et al. 2010. Virulence. 1: 367-375 ((https://doi.org/10.4161/viru.1.5.12796)); Mekbib SB, et al. 2011. Tropical Plant Pathology 36:233-240 ((https://doi.org/10.1590/S1982-56762011000400004)); Põlme S, et al. 2020. Fungal Diversity 105:1-16 ((https://doi.org/10.1007/s13225-020-00466-2)) |
| Otu21397 | 1 | k__Fungi(100);p__Ascomycota(100);c__Saccharomycetes(100);o__Saccharomycetales(100);f__Metschnikowiaceae(100);g__Metschnikowia(100);g__Metschnikowia_unclassified(100); | Metschnikowia | 13 | Saprotroph | \|Undefined Saprotroph\| | Yeast | NULL | Probable | As nectar-tap_saprotroph (Põlme et al. 2020); Decay sugar-rich substrates (Põlme et al. 2020) | Põlme S, et al. 2020. Fungal Diversity 105:1-16 ((https://doi.org/10.1007/s13225-020-00466-2)) |
| Otu21398 | 1 | k__Fungi(100);p__Ascomycota(100);c__Saccharomycetes(100);o__Saccharomycetales(100);f__Metschnikowiaceae(100);g__Metschnikowia(100);s__Metschnikowia_pulcherrima(100); | Metschnikowia | 13 | Saprotroph | \|Undefined Saprotroph\| | Yeast | NULL | Probable | As nectar-tap_saprotroph (Põlme et al. 2020); Decay sugar-rich substrates (Põlme et al. 2020) | Põlme S, et al. 2020. Fungal Diversity 105:1-16 ((https://doi.org/10.1007/s13225-020-00466-2)) |
| Otu21399 | 1 | k__Fungi(100);p__Ascomycota(100);c__Saccharomycetes(100);o__Saccharomycetales(100);f__Metschnikowiaceae(100);g__Metschnikowia(100);g__Metschnikowia_unclassified(100); | Metschnikowia | 13 | Saprotroph | \|Undefined Saprotroph\| | Yeast | NULL | Probable | As nectar-tap_saprotroph (Põlme et al. 2020); Decay sugar-rich substrates (Põlme et al. 2020) | Põlme S, et al. 2020. Fungal Diversity 105:1-16 ((https://doi.org/10.1007/s13225-020-00466-2)) |
| Otu21400 | 1 | k__Fungi(100);p__Ascomycota(100);c__Saccharomycetes(100);o__Saccharomycetales(100);o__Saccharomycetales_unclassified(100);o__Saccharomycetales_unclassified(100);o__Saccharomycetales_unclassified(100); | Saccharomycetales | 7 | Saprotroph | Undefined Saprotroph | Yeast | NULL | Possible | NULL | Sterkenburg E, et al. 2015. New Phytologist 207:1145-1158 ((https://doi.org/10.1111/nph.13426)) |
| Otu21401 | 1 | k__Fungi(100);p__Ascomycota(100);c__Saccharomycetes(100);o__Saccharomycetales(100);f__Saccharomycetales_family_Incertae_sedis(100);g__Candida(100);g__Candida_unclassified(100); | Candida | 13 | Pathotroph-Saprotroph-Symbiotroph | Animal Pathogen-Endophyte-Endosymbiont-Epiphyte-\|Undefined Saprotroph\| | Dimorphic Yeast-Yeast | NULL | Probable | As nectar-tap_saprotroph (Põlme et al. 2020); Decay sugar-rich substrates (Põlme et al. 2020); This asexual genus is polyphyletic and contain species that function very differently depend on what they are. Caution should be taken when interpreting guilds for Candida species. | Manolakaki D, et al. 2010. Virulence. 1: 367-375 ((https://doi.org/10.4161/viru.1.5.12796)); Mekbib SB, et al. 2011. Tropical Plant Pathology 36:233-240 ((https://doi.org/10.1590/S1982-56762011000400004)); Põlme S, et al. 2020. Fungal Diversity 105:1-16 ((https://doi.org/10.1007/s13225-020-00466-2)) |
| Otu21403 | 1 | k__Fungi(100);p__Ascomycota(100);c__Saccharomycetes(100);o__Saccharomycetales(100);f__Saccharomycetales_family_Incertae_sedis(100);g__Candida(100);g__Candida_unclassified(100); | Candida | 13 | Pathotroph-Saprotroph-Symbiotroph | Animal Pathogen-Endophyte-Endosymbiont-Epiphyte-\|Undefined Saprotroph\| | Dimorphic Yeast-Yeast | NULL | Probable | As nectar-tap_saprotroph (Põlme et al. 2020); Decay sugar-rich substrates (Põlme et al. 2020); This asexual genus is polyphyletic and contain species that function very differently depend on what they are. Caution should be taken when interpreting guilds for Candida species. | Manolakaki D, et al. 2010. Virulence. 1: 367-375 ((https://doi.org/10.4161/viru.1.5.12796)); Mekbib SB, et al. 2011. Tropical Plant Pathology 36:233-240 ((https://doi.org/10.1590/S1982-56762011000400004)); Põlme S, et al. 2020. Fungal Diversity 105:1-16 ((https://doi.org/10.1007/s13225-020-00466-2)) |
| Otu21407 | 1 | k__Fungi(100);p__Ascomycota(100);c__Saccharomycetes(100);o__Saccharomycetales(100);f__Metschnikowiaceae(100);g__Metschnikowia(100);g__Metschnikowia_unclassified(100); | Metschnikowia | 13 | Saprotroph | \|Undefined Saprotroph\| | Yeast | NULL | Probable | As nectar-tap_saprotroph (Põlme et al. 2020); Decay sugar-rich substrates (Põlme et al. 2020) | Põlme S, et al. 2020. Fungal Diversity 105:1-16 ((https://doi.org/10.1007/s13225-020-00466-2)) |
| Otu21408 | 1 | k__Fungi(100);p__Ascomycota(100);c__Saccharomycetes(100);o__Saccharomycetales(100);f__Metschnikowiaceae(100);g__Metschnikowia(100);s__Metschnikowia_pulcherrima(100); | Metschnikowia | 13 | Saprotroph | \|Undefined Saprotroph\| | Yeast | NULL | Probable | As nectar-tap_saprotroph (Põlme et al. 2020); Decay sugar-rich substrates (Põlme et al. 2020) | Põlme S, et al. 2020. Fungal Diversity 105:1-16 ((https://doi.org/10.1007/s13225-020-00466-2)) |
| Otu21411 | 1 | k__Fungi(100);p__Ascomycota(100);c__Saccharomycetes(100);o__Saccharomycetales(100);f__Metschnikowiaceae(100);g__Metschnikowia(100);s__Metschnikowia_pulcherrima(100); | Metschnikowia | 13 | Saprotroph | \|Undefined Saprotroph\| | Yeast | NULL | Probable | As nectar-tap_saprotroph (Põlme et al. 2020); Decay sugar-rich substrates (Põlme et al. 2020) | Põlme S, et al. 2020. Fungal Diversity 105:1-16 ((https://doi.org/10.1007/s13225-020-00466-2)) |
| Otu21412 | 1 | k__Fungi(100);p__Ascomycota(100);c__Saccharomycetes(100);o__Saccharomycetales(100);f__Metschnikowiaceae(100);g__Metschnikowia(100);s__Metschnikowia_pulcherrima(100); | Metschnikowia | 13 | Saprotroph | \|Undefined Saprotroph\| | Yeast | NULL | Probable | As nectar-tap_saprotroph (Põlme et al. 2020); Decay sugar-rich substrates (Põlme et al. 2020) | Põlme S, et al. 2020. Fungal Diversity 105:1-16 ((https://doi.org/10.1007/s13225-020-00466-2)) |
| Otu21413 | 1 | k__Fungi(100);p__Ascomycota(100);c__Saccharomycetes(100);o__Saccharomycetales(100);f__Metschnikowiaceae(100);g__Metschnikowia(100);g__Metschnikowia_unclassified(100); | Metschnikowia | 13 | Saprotroph | \|Undefined Saprotroph\| | Yeast | NULL | Probable | As nectar-tap_saprotroph (Põlme et al. 2020); Decay sugar-rich substrates (Põlme et al. 2020) | Põlme S, et al. 2020. Fungal Diversity 105:1-16 ((https://doi.org/10.1007/s13225-020-00466-2)) |
| Otu21414 | 1 | k__Fungi(100);p__Ascomycota(100);c__Saccharomycetes(100);o__Saccharomycetales(100);f__Metschnikowiaceae(100);g__Metschnikowia(100);g__Metschnikowia_unclassified(100); | Metschnikowia | 13 | Saprotroph | \|Undefined Saprotroph\| | Yeast | NULL | Probable | As nectar-tap_saprotroph (Põlme et al. 2020); Decay sugar-rich substrates (Põlme et al. 2020) | Põlme S, et al. 2020. Fungal Diversity 105:1-16 ((https://doi.org/10.1007/s13225-020-00466-2)) |
| Otu21415 | 1 | k__Fungi(100);p__Ascomycota(100);c__Saccharomycetes(100);o__Saccharomycetales(100);f__Metschnikowiaceae(100);g__Metschnikowia(100);s__Metschnikowia_pulcherrima(100); | Metschnikowia | 13 | Saprotroph | \|Undefined Saprotroph\| | Yeast | NULL | Probable | As nectar-tap_saprotroph (Põlme et al. 2020); Decay sugar-rich substrates (Põlme et al. 2020) | Põlme S, et al. 2020. Fungal Diversity 105:1-16 ((https://doi.org/10.1007/s13225-020-00466-2)) |
| Otu21416 | 1 | k__Fungi(100);p__Ascomycota(100);c__Saccharomycetes(100);o__Saccharomycetales(100);f__Metschnikowiaceae(100);g__Metschnikowia(100);g__Metschnikowia_unclassified(100); | Metschnikowia | 13 | Saprotroph | \|Undefined Saprotroph\| | Yeast | NULL | Probable | As nectar-tap_saprotroph (Põlme et al. 2020); Decay sugar-rich substrates (Põlme et al. 2020) | Põlme S, et al. 2020. Fungal Diversity 105:1-16 ((https://doi.org/10.1007/s13225-020-00466-2)) |
| Otu21417 | 1 | k__Fungi(100);p__Ascomycota(100);c__Saccharomycetes(100);o__Saccharomycetales(100);f__Metschnikowiaceae(100);g__Metschnikowia(100);s__Metschnikowia_pulcherrima(100); | Metschnikowia | 13 | Saprotroph | \|Undefined Saprotroph\| | Yeast | NULL | Probable | As nectar-tap_saprotroph (Põlme et al. 2020); Decay sugar-rich substrates (Põlme et al. 2020) | Põlme S, et al. 2020. Fungal Diversity 105:1-16 ((https://doi.org/10.1007/s13225-020-00466-2)) |
| Otu21418 | 1 | k__Fungi(100);p__Ascomycota(100);c__Saccharomycetes(100);o__Saccharomycetales(100);f__Metschnikowiaceae(100);g__Metschnikowia(100);s__Metschnikowia_pulcherrima(100); | Metschnikowia | 13 | Saprotroph | \|Undefined Saprotroph\| | Yeast | NULL | Probable | As nectar-tap_saprotroph (Põlme et al. 2020); Decay sugar-rich substrates (Põlme et al. 2020) | Põlme S, et al. 2020. Fungal Diversity 105:1-16 ((https://doi.org/10.1007/s13225-020-00466-2)) |
| Otu21419 | 1 | k__Fungi(100);p__Ascomycota(100);c__Saccharomycetes(100);o__Saccharomycetales(100);f__Metschnikowiaceae(100);g__Metschnikowia(100);g__Metschnikowia_unclassified(100); | Metschnikowia | 13 | Saprotroph | \|Undefined Saprotroph\| | Yeast | NULL | Probable | As nectar-tap_saprotroph (Põlme et al. 2020); Decay sugar-rich substrates (Põlme et al. 2020) | Põlme S, et al. 2020. Fungal Diversity 105:1-16 ((https://doi.org/10.1007/s13225-020-00466-2)) |
| Otu21421 | 1 | k__Fungi(100);p__Ascomycota(100);c__Saccharomycetes(100);o__Saccharomycetales(100);f__Metschnikowiaceae(100);g__Metschnikowia(100);s__Metschnikowia_pulcherrima(100); | Metschnikowia | 13 | Saprotroph | \|Undefined Saprotroph\| | Yeast | NULL | Probable | As nectar-tap_saprotroph (Põlme et al. 2020); Decay sugar-rich substrates (Põlme et al. 2020) | Põlme S, et al. 2020. Fungal Diversity 105:1-16 ((https://doi.org/10.1007/s13225-020-00466-2)) |
| Otu21422 | 1 | k__Fungi(100);p__Ascomycota(100);c__Saccharomycetes(100);o__Saccharomycetales(100);f__Metschnikowiaceae(100);g__Metschnikowia(100);s__Metschnikowia_pulcherrima(100); | Metschnikowia | 13 | Saprotroph | \|Undefined Saprotroph\| | Yeast | NULL | Probable | As nectar-tap_saprotroph (Põlme et al. 2020); Decay sugar-rich substrates (Põlme et al. 2020) | Põlme S, et al. 2020. Fungal Diversity 105:1-16 ((https://doi.org/10.1007/s13225-020-00466-2)) |
| Otu21423 | 1 | k__Fungi(100);p__Ascomycota(100);c__Saccharomycetes(100);o__Saccharomycetales(100);f__Metschnikowiaceae(100);g__Metschnikowia(100);s__Metschnikowia_pulcherrima(100); | Metschnikowia | 13 | Saprotroph | \|Undefined Saprotroph\| | Yeast | NULL | Probable | As nectar-tap_saprotroph (Põlme et al. 2020); Decay sugar-rich substrates (Põlme et al. 2020) | Põlme S, et al. 2020. Fungal Diversity 105:1-16 ((https://doi.org/10.1007/s13225-020-00466-2)) |
| Otu21424 | 1 | k__Fungi(100);p__Ascomycota(100);c__Saccharomycetes(100);o__Saccharomycetales(100);f__Metschnikowiaceae(100);g__Metschnikowia(100);g__Metschnikowia_unclassified(100); | Metschnikowia | 13 | Saprotroph | \|Undefined Saprotroph\| | Yeast | NULL | Probable | As nectar-tap_saprotroph (Põlme et al. 2020); Decay sugar-rich substrates (Põlme et al. 2020) | Põlme S, et al. 2020. Fungal Diversity 105:1-16 ((https://doi.org/10.1007/s13225-020-00466-2)) |
| Otu21425 | 1 | k__Fungi(100);p__Ascomycota(100);c__Saccharomycetes(100);o__Saccharomycetales(100);f__Metschnikowiaceae(100);g__Metschnikowia(100);g__Metschnikowia_unclassified(100); | Metschnikowia | 13 | Saprotroph | \|Undefined Saprotroph\| | Yeast | NULL | Probable | As nectar-tap_saprotroph (Põlme et al. 2020); Decay sugar-rich substrates (Põlme et al. 2020) | Põlme S, et al. 2020. Fungal Diversity 105:1-16 ((https://doi.org/10.1007/s13225-020-00466-2)) |
| Otu21426 | 1 | k__Fungi(100);p__Ascomycota(100);c__Saccharomycetes(100);o__Saccharomycetales(100);f__Metschnikowiaceae(100);g__Metschnikowia(100);s__Metschnikowia_pulcherrima(100); | Metschnikowia | 13 | Saprotroph | \|Undefined Saprotroph\| | Yeast | NULL | Probable | As nectar-tap_saprotroph (Põlme et al. 2020); Decay sugar-rich substrates (Põlme et al. 2020) | Põlme S, et al. 2020. Fungal Diversity 105:1-16 ((https://doi.org/10.1007/s13225-020-00466-2)) |
| Otu21427 | 1 | k__Fungi(100);p__Ascomycota(100);c__Saccharomycetes(100);o__Saccharomycetales(100);f__Metschnikowiaceae(100);g__Metschnikowia(100);s__Metschnikowia_pulcherrima(100); | Metschnikowia | 13 | Saprotroph | \|Undefined Saprotroph\| | Yeast | NULL | Probable | As nectar-tap_saprotroph (Põlme et al. 2020); Decay sugar-rich substrates (Põlme et al. 2020) | Põlme S, et al. 2020. Fungal Diversity 105:1-16 ((https://doi.org/10.1007/s13225-020-00466-2)) |
| Otu21428 | 1 | k__Fungi(100);p__Ascomycota(100);c__Saccharomycetes(100);o__Saccharomycetales(100);f__Metschnikowiaceae(100);g__Metschnikowia(100);s__Metschnikowia_pulcherrima(100); | Metschnikowia | 13 | Saprotroph | \|Undefined Saprotroph\| | Yeast | NULL | Probable | As nectar-tap_saprotroph (Põlme et al. 2020); Decay sugar-rich substrates (Põlme et al. 2020) | Põlme S, et al. 2020. Fungal Diversity 105:1-16 ((https://doi.org/10.1007/s13225-020-00466-2)) |
| Otu21430 | 1 | k__Fungi(100);p__Ascomycota(100);c__Saccharomycetes(100);o__Saccharomycetales(100);f__Metschnikowiaceae(100);g__Metschnikowia(100);s__Metschnikowia_pulcherrima(100); | Metschnikowia | 13 | Saprotroph | \|Undefined Saprotroph\| | Yeast | NULL | Probable | As nectar-tap_saprotroph (Põlme et al. 2020); Decay sugar-rich substrates (Põlme et al. 2020) | Põlme S, et al. 2020. Fungal Diversity 105:1-16 ((https://doi.org/10.1007/s13225-020-00466-2)) |
| Otu21431 | 1 | k__Fungi(100);p__Ascomycota(100);c__Saccharomycetes(100);o__Saccharomycetales(100);f__Metschnikowiaceae(100);g__Metschnikowia(100);s__Metschnikowia_pulcherrima(100); | Metschnikowia | 13 | Saprotroph | \|Undefined Saprotroph\| | Yeast | NULL | Probable | As nectar-tap_saprotroph (Põlme et al. 2020); Decay sugar-rich substrates (Põlme et al. 2020) | Põlme S, et al. 2020. Fungal Diversity 105:1-16 ((https://doi.org/10.1007/s13225-020-00466-2)) |
| Otu21432 | 1 | k__Fungi(100);p__Ascomycota(100);c__Saccharomycetes(100);o__Saccharomycetales(100);f__Metschnikowiaceae(100);g__Metschnikowia(100);s__Metschnikowia_pulcherrima(100); | Metschnikowia | 13 | Saprotroph | \|Undefined Saprotroph\| | Yeast | NULL | Probable | As nectar-tap_saprotroph (Põlme et al. 2020); Decay sugar-rich substrates (Põlme et al. 2020) | Põlme S, et al. 2020. Fungal Diversity 105:1-16 ((https://doi.org/10.1007/s13225-020-00466-2)) |
| Otu21433 | 1 | k__Fungi(100);p__Ascomycota(100);c__Saccharomycetes(100);o__Saccharomycetales(100);f__Metschnikowiaceae(100);g__Metschnikowia(100);s__Metschnikowia_pulcherrima(100); | Metschnikowia | 13 | Saprotroph | \|Undefined Saprotroph\| | Yeast | NULL | Probable | As nectar-tap_saprotroph (Põlme et al. 2020); Decay sugar-rich substrates (Põlme et al. 2020) | Põlme S, et al. 2020. Fungal Diversity 105:1-16 ((https://doi.org/10.1007/s13225-020-00466-2)) |
| Otu21437 | 1 | k__Fungi(100);p__Ascomycota(100);c__Saccharomycetes(100);o__Saccharomycetales(100);f__Metschnikowiaceae(100);g__Metschnikowia(100);s__Metschnikowia_pulcherrima(100); | Metschnikowia | 13 | Saprotroph | \|Undefined Saprotroph\| | Yeast | NULL | Probable | As nectar-tap_saprotroph (Põlme et al. 2020); Decay sugar-rich substrates (Põlme et al. 2020) | Põlme S, et al. 2020. Fungal Diversity 105:1-16 ((https://doi.org/10.1007/s13225-020-00466-2)) |
| Otu21438 | 1 | k__Fungi(100);p__Ascomycota(100);c__Saccharomycetes(100);o__Saccharomycetales(100);f__Metschnikowiaceae(100);g__Metschnikowia(100);s__Metschnikowia_pulcherrima(100); | Metschnikowia | 13 | Saprotroph | \|Undefined Saprotroph\| | Yeast | NULL | Probable | As nectar-tap_saprotroph (Põlme et al. 2020); Decay sugar-rich substrates (Põlme et al. 2020) | Põlme S, et al. 2020. Fungal Diversity 105:1-16 ((https://doi.org/10.1007/s13225-020-00466-2)) |
| Otu21439 | 1 | k__Fungi(100);p__Ascomycota(100);c__Saccharomycetes(100);o__Saccharomycetales(100);f__Metschnikowiaceae(100);g__Metschnikowia(100);g__Metschnikowia_unclassified(100); | Metschnikowia | 13 | Saprotroph | \|Undefined Saprotroph\| | Yeast | NULL | Probable | As nectar-tap_saprotroph (Põlme et al. 2020); Decay sugar-rich substrates (Põlme et al. 2020) | Põlme S, et al. 2020. Fungal Diversity 105:1-16 ((https://doi.org/10.1007/s13225-020-00466-2)) |

| | | | | | | | | | | | |
|---|---|---|---|---|---|---|---|---|---|---|---|
| Otu21440 | 1 | k__Fungi(100);p__Ascomycota(100);c__Saccharomycetes(100);o__Saccharomycetales(100);f__Metschnikowiaceae(100);g__Metschnikowia(100);s__Metschnikowia_pulcherrima(100); | Metschnikowia | 13 | Saprotroph | \|Undefined Saprotroph\| | Yeast | NULL | Probable | As nectar-tap_saprotroph (Põlme et al. 2020); Decay sugar-rich substrates (Põlme et al. 2020) | Põlme S, et al. 2020. Fungal Diversity 105:1-16 ((https://doi.org/10.1007/s13225-020-00466-2)) |
| Otu21441 | 1 | k__Fungi(100);p__Ascomycota(100);c__Saccharomycetes(100);o__Saccharomycetales(100);f__Metschnikowiaceae(100);g__Metschnikowia(100);s__Metschnikowia_pulcherrima(100); | Metschnikowia | 13 | Saprotroph | \|Undefined Saprotroph\| | Yeast | NULL | Probable | As nectar-tap_saprotroph (Põlme et al. 2020); Decay sugar-rich substrates (Põlme et al. 2020) | Põlme S, et al. 2020. Fungal Diversity 105:1-16 ((https://doi.org/10.1007/s13225-020-00466-2)) |
| Otu21442 | 1 | k__Fungi(100);p__Ascomycota(100);c__Saccharomycetes(100);o__Saccharomycetales(100);f__Metschnikowiaceae(100);g__Metschnikowia(100);s__Metschnikowia_pulcherrima(100); | Metschnikowia | 13 | Saprotroph | \|Undefined Saprotroph\| | Yeast | NULL | Probable | As nectar-tap_saprotroph (Põlme et al. 2020); Decay sugar-rich substrates (Põlme et al. 2020) | Põlme S, et al. 2020. Fungal Diversity 105:1-16 ((https://doi.org/10.1007/s13225-020-00466-2)) |
| Otu21443 | 1 | k__Fungi(100);p__Ascomycota(100);c__Saccharomycetes(100);o__Saccharomycetales(100);f__Metschnikowiaceae(100);g__Metschnikowia(100);s__Metschnikowia_pulcherrima(100); | Metschnikowia | 13 | Saprotroph | \|Undefined Saprotroph\| | Yeast | NULL | Probable | As nectar-tap_saprotroph (Põlme et al. 2020); Decay sugar-rich substrates (Põlme et al. 2020) | Põlme S, et al. 2020. Fungal Diversity 105:1-16 ((https://doi.org/10.1007/s13225-020-00466-2)) |
| Otu21444 | 1 | k__Fungi(100);p__Ascomycota(100);c__Saccharomycetes(100);o__Saccharomycetales(100);f__Metschnikowiaceae(100);g__Metschnikowia(100);g__Metschnikowia_unclassified(100); | Metschnikowia | 13 | Saprotroph | \|Undefined Saprotroph\| | Yeast | NULL | Probable | As nectar-tap_saprotroph (Põlme et al. 2020); Decay sugar-rich substrates (Põlme et al. 2020) | Põlme S, et al. 2020. Fungal Diversity 105:1-16 ((https://doi.org/10.1007/s13225-020-00466-2)) |
| Otu21445 | 1 | k__Fungi(100);p__Ascomycota(100);c__Saccharomycetes(100);o__Saccharomycetales(100);f__Metschnikowiaceae(100);g__Metschnikowia(100);s__Metschnikowia_pulcherrima(100); | Metschnikowia | 13 | Saprotroph | \|Undefined Saprotroph\| | Yeast | NULL | Probable | As nectar-tap_saprotroph (Põlme et al. 2020); Decay sugar-rich substrates (Põlme et al. 2020) | Põlme S, et al. 2020. Fungal Diversity 105:1-16 ((https://doi.org/10.1007/s13225-020-00466-2)) |
| Otu21446 | 1 | k__Fungi(100);p__Ascomycota(100);c__Saccharomycetes(100);o__Saccharomycetales(100);f__Metschnikowiaceae(100);g__Metschnikowia(100);s__Metschnikowia_pulcherrima(100); | Metschnikowia | 13 | Saprotroph | \|Undefined Saprotroph\| | Yeast | NULL | Probable | As nectar-tap_saprotroph (Põlme et al. 2020); Decay sugar-rich substrates (Põlme et al. 2020) | Põlme S, et al. 2020. Fungal Diversity 105:1-16 ((https://doi.org/10.1007/s13225-020-00466-2)) |
| Otu21447 | 1 | k__Fungi(100);p__Ascomycota(100);c__Saccharomycetes(100);o__Saccharomycetales(100);f__Metschnikowiaceae(100);g__Metschnikowia(100);s__Metschnikowia_pulcherrima(100); | Metschnikowia | 13 | Saprotroph | \|Undefined Saprotroph\| | Yeast | NULL | Probable | As nectar-tap_saprotroph (Põlme et al. 2020); Decay sugar-rich substrates (Põlme et al. 2020) | Põlme S, et al. 2020. Fungal Diversity 105:1-16 ((https://doi.org/10.1007/s13225-020-00466-2)) |
| Otu21449 | 1 | k__Fungi(100);p__Ascomycota(100);c__Saccharomycetes(100);o__Saccharomycetales(100);o__Saccharomycetales_unclassified(100);o__Saccharomycetales_unclassified(100);o__Saccharomycetales_unclassified(100); | Saccharomycetales | 7 | Saprotroph | Undefined Saprotroph | Yeast | NULL | Possible | NULL | Sterkenburg E, et al. 2015. New Phytologist 207:1145-1158 ((https://doi.org/10.1111/nph.13426)) |
| Otu21450 | 1 | k__Fungi(100);p__Ascomycota(100);c__Saccharomycetes(100);o__Saccharomycetales(100);f__Metschnikowiaceae(100);g__Metschnikowia(100);s__Metschnikowia_pulcherrima(100); | Metschnikowia | 13 | Saprotroph | \|Undefined Saprotroph\| | Yeast | NULL | Probable | As nectar-tap_saprotroph (Põlme et al. 2020); Decay sugar-rich substrates (Põlme et al. 2020) | Põlme S, et al. 2020. Fungal Diversity 105:1-16 ((https://doi.org/10.1007/s13225-020-00466-2)) |
| Otu21452 | 1 | k__Fungi(100);p__Ascomycota(100);c__Saccharomycetes(100);o__Saccharomycetales(100);f__Metschnikowiaceae(100);g__Metschnikowia(100);s__Metschnikowia_pulcherrima(100); | Metschnikowia | 13 | Saprotroph | \|Undefined Saprotroph\| | Yeast | NULL | Probable | As nectar-tap_saprotroph (Põlme et al. 2020); Decay sugar-rich substrates (Põlme et al. 2020) | Põlme S, et al. 2020. Fungal Diversity 105:1-16 ((https://doi.org/10.1007/s13225-020-00466-2)) |
| Otu21453 | 1 | k__Fungi(100);p__Ascomycota(100);c__Saccharomycetes(100);o__Saccharomycetales(100);f__Metschnikowiaceae(100);g__Metschnikowia(100);s__Metschnikowia_pulcherrima(100); | Metschnikowia | 13 | Saprotroph | \|Undefined Saprotroph\| | Yeast | NULL | Probable | As nectar-tap_saprotroph (Põlme et al. 2020); Decay sugar-rich substrates (Põlme et al. 2020) | Põlme S, et al. 2020. Fungal Diversity 105:1-16 ((https://doi.org/10.1007/s13225-020-00466-2)) |
| Otu21454 | 1 | k__Fungi(100);p__Ascomycota(100);c__Saccharomycetes(100);o__Saccharomycetales(100);f__Metschnikowiaceae(100);g__Metschnikowia(100);s__Metschnikowia_pulcherrima(100); | Metschnikowia | 13 | Saprotroph | \|Undefined Saprotroph\| | Yeast | NULL | Probable | As nectar-tap_saprotroph (Põlme et al. 2020); Decay sugar-rich substrates (Põlme et al. 2020) | Põlme S, et al. 2020. Fungal Diversity 105:1-16 ((https://doi.org/10.1007/s13225-020-00466-2)) |
| Otu21455 | 1 | k__Fungi(100);p__Ascomycota(100);c__Saccharomycetes(100);o__Saccharomycetales(100);f__Metschnikowiaceae(100);g__Metschnikowia(100);s__Metschnikowia_pulcherrima(100); | Metschnikowia | 13 | Saprotroph | \|Undefined Saprotroph\| | Yeast | NULL | Probable | As nectar-tap_saprotroph (Põlme et al. 2020); Decay sugar-rich substrates (Põlme et al. 2020) | Põlme S, et al. 2020. Fungal Diversity 105:1-16 ((https://doi.org/10.1007/s13225-020-00466-2)) |
| Otu21456 | 1 | k__Fungi(100);p__Ascomycota(100);c__Saccharomycetes(100);o__Saccharomycetales(100);f__Metschnikowiaceae(100);g__Metschnikowia(100);s__Metschnikowia_pulcherrima(100); | Metschnikowia | 13 | Saprotroph | \|Undefined Saprotroph\| | Yeast | NULL | Probable | As nectar-tap_saprotroph (Põlme et al. 2020); Decay sugar-rich substrates (Põlme et al. 2020) | Põlme S, et al. 2020. Fungal Diversity 105:1-16 ((https://doi.org/10.1007/s13225-020-00466-2)) |
| Otu21457 | 1 | k__Fungi(100);p__Ascomycota(100);c__Saccharomycetes(100);o__Saccharomycetales(100);f__Metschnikowiaceae(100);g__Metschnikowia(100);g__Metschnikowia_unclassified(100); | Metschnikowia | 13 | Saprotroph | \|Undefined Saprotroph\| | Yeast | NULL | Probable | As nectar-tap_saprotroph (Põlme et al. 2020); Decay sugar-rich substrates (Põlme et al. 2020) | Põlme S, et al. 2020. Fungal Diversity 105:1-16 ((https://doi.org/10.1007/s13225-020-00466-2)) |
| Otu21458 | 1 | k__Fungi(100);p__Ascomycota(100);c__Saccharomycetes(100);o__Saccharomycetales(100);f__Metschnikowiaceae(100);g__Metschnikowia(100);s__Metschnikowia_pulcherrima(100); | Metschnikowia | 13 | Saprotroph | \|Undefined Saprotroph\| | Yeast | NULL | Probable | As nectar-tap_saprotroph (Põlme et al. 2020); Decay sugar-rich substrates (Põlme et al. 2020) | Põlme S, et al. 2020. Fungal Diversity 105:1-16 ((https://doi.org/10.1007/s13225-020-00466-2)) |
| Otu21459 | 1 | k__Fungi(100);p__Ascomycota(100);c__Saccharomycetes(100);o__Saccharomycetales(100);f__Metschnikowiaceae(100);g__Metschnikowia(100);g__Metschnikowia_unclassified(100); | Metschnikowia | 13 | Saprotroph | \|Undefined Saprotroph\| | Yeast | NULL | Probable | As nectar-tap_saprotroph (Põlme et al. 2020); Decay sugar-rich substrates (Põlme et al. 2020) | Põlme S, et al. 2020. Fungal Diversity 105:1-16 ((https://doi.org/10.1007/s13225-020-00466-2)) |
| Otu21460 | 1 | k__Fungi(100);p__Ascomycota(100);c__Saccharomycetes(100);o__Saccharomycetales(100);f__Metschnikowiaceae(100);g__Metschnikowia(100);s__Metschnikowia_pulcherrima(100); | Metschnikowia | 13 | Saprotroph | \|Undefined Saprotroph\| | Yeast | NULL | Probable | As nectar-tap_saprotroph (Põlme et al. 2020); Decay sugar-rich substrates (Põlme et al. 2020) | Põlme S, et al. 2020. Fungal Diversity 105:1-16 ((https://doi.org/10.1007/s13225-020-00466-2)) |
| Otu21461 | 1 | k__Fungi(100);p__Ascomycota(100);c__Saccharomycetes(100);o__Saccharomycetales(100);f__Metschnikowiaceae(100);g__Metschnikowia(100);g__Metschnikowia_unclassified(100); | Metschnikowia | 13 | Saprotroph | \|Undefined Saprotroph\| | Yeast | NULL | Probable | As nectar-tap_saprotroph (Põlme et al. 2020); Decay sugar-rich substrates (Põlme et al. 2020) | Põlme S, et al. 2020. Fungal Diversity 105:1-16 ((https://doi.org/10.1007/s13225-020-00466-2)) |
| Otu21465 | 1 | k__Fungi(100);p__Ascomycota(100);c__Dothideomycetes(100);o__Dothideomycetes_order_Incertae_sedis(100);f__Myxotrichaceae(100);g__Oidiodendron(100);g__Oidiodendron_unclassified(100); | Oidiodendron | 13 | Pathotroph-Saprotroph-Symbiotroph | Endophyte-Ericoid Mycorrhizal-\|Undefined Saprotroph\| | Microfungus | NULL | Probable | As root_endophyte (Põlme et al. 2020); As soil_saprotroph (Põlme et al. 2020); Dark Septate Endophyte; Decay soil (Põlme et al. 2020) | Newsham KK. 2011. New Phytologist 190:783-793 ((https://doi.org/10.1111/j.1469-8137.2010.03611.x)) (as dark septate root endophytes); Põlme S, et al. 2020. Fungal Diversity 105:1-16 ((https://doi.org/10.1007/s13225-020-00466-2)) |
| Otu21466 | 1 | k__Fungi(100);p__Ascomycota(100);c__Saccharomycetes(100);o__Saccharomycetales(100);f__Metschnikowiaceae(100);g__Metschnikowia(100);s__Metschnikowia_pulcherrima(100); | Metschnikowia | 13 | Saprotroph | \|Undefined Saprotroph\| | Yeast | NULL | Probable | As nectar-tap_saprotroph (Põlme et al. 2020); Decay sugar-rich substrates (Põlme et al. 2020) | Põlme S, et al. 2020. Fungal Diversity 105:1-16 ((https://doi.org/10.1007/s13225-020-00466-2)) |
| Otu21467 | 1 | k__Fungi(100);p__Ascomycota(100);c__Saccharomycetes(100);o__Saccharomycetales(100);f__Metschnikowiaceae(100);g__Metschnikowia(100);s__Metschnikowia_pulcherrima(100); | Metschnikowia | 13 | Saprotroph | \|Undefined Saprotroph\| | Yeast | NULL | Probable | As nectar-tap_saprotroph (Põlme et al. 2020); Decay sugar-rich substrates (Põlme et al. 2020) | Põlme S, et al. 2020. Fungal Diversity 105:1-16 ((https://doi.org/10.1007/s13225-020-00466-2)) |
| Otu21473 | 1 | k__Fungi(100);p__Ascomycota(100);c__Leotiomycetes(100);o__Helotiales(100);f__Vibrisseaceae(100);g__Phialocephala(100);s__Phialocephala_fusca(100); | Phialocephala | 13 | Saprotroph-Symbiotroph | Endophyte-\|Undefined Saprotroph\| | Microfungus | NULL | Highly Probable | As root_endophyte (Põlme et al. 2020); As root_endophyte_dark_septate (Põlme et al. 2020); As soil_saprotroph (Põlme et al. 2020); Dark Septate Endophyte, known from plant roots.; Decay roots, soil (Põlme et al. 2020) | Newsham KK. 2011. New Phytologist 190:783-793 ((https://doi.org/10.1111/j.1469-8137.2010.03611.x)) (as dark septate root endophytes); Põlme S, et al. 2020. Fungal Diversity 105:1-16 ((https://doi.org/10.1007/s13225-020-00466-2)) |
| Otu21475 | 1 | k__Fungi(100);p__Ascomycota(100);c__Saccharomycetes(100);o__Saccharomycetales(100);f__Metschnikowiaceae(100);g__Metschnikowia(100);s__Metschnikowia_pulcherrima(100); | Metschnikowia | 13 | Saprotroph | \|Undefined Saprotroph\| | Yeast | NULL | Probable | As nectar-tap_saprotroph (Põlme et al. 2020); Decay sugar-rich substrates (Põlme et al. 2020) | Põlme S, et al. 2020. Fungal Diversity 105:1-16 ((https://doi.org/10.1007/s13225-020-00466-2)) |
| Otu21476 | 1 | k__Fungi(100);p__Ascomycota(100);c__Saccharomycetes(100);o__Saccharomycetales(100);f__Metschnikowiaceae(100);g__Metschnikowia(100);s__Metschnikowia_pulcherrima(100); | Metschnikowia | 13 | Saprotroph | \|Undefined Saprotroph\| | Yeast | NULL | Probable | As nectar-tap_saprotroph (Põlme et al. 2020); Decay sugar-rich substrates (Põlme et al. 2020) | Põlme S, et al. 2020. Fungal Diversity 105:1-16 ((https://doi.org/10.1007/s13225-020-00466-2)) |
| Otu21477 | 1 | k__Fungi(100);p__Ascomycota(100);c__Saccharomycetes(100);o__Saccharomycetales(100);f__Metschnikowiaceae(100);g__Metschnikowia(100);g__Metschnikowia_unclassified(100); | Metschnikowia | 13 | Saprotroph | \|Undefined Saprotroph\| | Yeast | NULL | Probable | As nectar-tap_saprotroph (Põlme et al. 2020); Decay sugar-rich substrates (Põlme et al. 2020) | Põlme S, et al. 2020. Fungal Diversity 105:1-16 ((https://doi.org/10.1007/s13225-020-00466-2)) |
| Otu21478 | 1 | k__Fungi(100);p__Ascomycota(100);c__Saccharomycetes(100);o__Saccharomycetales(100);f__Metschnikowiaceae(100);g__Metschnikowia(100);s__Metschnikowia_pulcherrima(100); | Metschnikowia | 13 | Saprotroph | \|Undefined Saprotroph\| | Yeast | NULL | Probable | As nectar-tap_saprotroph (Põlme et al. 2020); Decay sugar-rich substrates (Põlme et al. 2020) | Põlme S, et al. 2020. Fungal Diversity 105:1-16 ((https://doi.org/10.1007/s13225-020-00466-2)) |
| Otu21479 | 1 | k__Fungi(100);p__Ascomycota(100);c__Saccharomycetes(100);o__Saccharomycetales(100);f__Metschnikowiaceae(100);g__Metschnikowia(100);s__Metschnikowia_pulcherrima(100); | Metschnikowia | 13 | Saprotroph | \|Undefined Saprotroph\| | Yeast | NULL | Probable | As nectar-tap_saprotroph (Põlme et al. 2020); Decay sugar-rich substrates (Põlme et al. 2020) | Põlme S, et al. 2020. Fungal Diversity 105:1-16 ((https://doi.org/10.1007/s13225-020-00466-2)) |
| Otu21480 | 1 | k__Fungi(100);p__Ascomycota(100);c__Saccharomycetes(100);o__Saccharomycetales(100);f__Metschnikowiaceae(100);g__Metschnikowia(100);g__Metschnikowia_unclassified(100); | Metschnikowia | 13 | Saprotroph | \|Undefined Saprotroph\| | Yeast | NULL | Probable | As nectar-tap_saprotroph (Põlme et al. 2020); Decay sugar-rich substrates (Põlme et al. 2020) | Põlme S, et al. 2020. Fungal Diversity 105:1-16 ((https://doi.org/10.1007/s13225-020-00466-2)) |
| Otu21481 | 1 | k__Fungi(100);p__Ascomycota(100);c__Saccharomycetes(100);o__Saccharomycetales(100);f__Metschnikowiaceae(100);g__Metschnikowia(100);s__Metschnikowia_pulcherrima(100); | Metschnikowia | 13 | Saprotroph | \|Undefined Saprotroph\| | Yeast | NULL | Probable | As nectar-tap_saprotroph (Põlme et al. 2020); Decay sugar-rich substrates (Põlme et al. 2020) | Põlme S, et al. 2020. Fungal Diversity 105:1-16 ((https://doi.org/10.1007/s13225-020-00466-2)) |
| Otu21482 | 1 | k__Fungi(100);p__Ascomycota(100);c__Saccharomycetes(100);o__Saccharomycetales(100);f__Metschnikowiaceae(100);g__Metschnikowia(100);s__Metschnikowia_pulcherrima(100); | Metschnikowia | 13 | Saprotroph | \|Undefined Saprotroph\| | Yeast | NULL | Probable | As nectar-tap_saprotroph (Põlme et al. 2020); Decay sugar-rich substrates (Põlme et al. 2020) | Põlme S, et al. 2020. Fungal Diversity 105:1-16 ((https://doi.org/10.1007/s13225-020-00466-2)) |
| Otu21483 | 1 | k__Fungi(100);p__Ascomycota(100);c__Saccharomycetes(100);o__Saccharomycetales(100);f__Metschnikowiaceae(100);g__Metschnikowia(100);g__Metschnikowia_unclassified(100); | Metschnikowia | 13 | Saprotroph | \|Undefined Saprotroph\| | Yeast | NULL | Probable | As nectar-tap_saprotroph (Põlme et al. 2020); Decay sugar-rich substrates (Põlme et al. 2020) | Põlme S, et al. 2020. Fungal Diversity 105:1-16 ((https://doi.org/10.1007/s13225-020-00466-2)) |
| Otu21484 | 1 | k__Fungi(100);p__Ascomycota(100);c__Saccharomycetes(100);o__Saccharomycetales(100);f__Metschnikowiaceae(100);g__Metschnikowia(100);s__Metschnikowia_pulcherrima(100); | Metschnikowia | 13 | Saprotroph | \|Undefined Saprotroph\| | Yeast | NULL | Probable | As nectar-tap_saprotroph (Põlme et al. 2020); Decay sugar-rich substrates (Põlme et al. 2020) | Põlme S, et al. 2020. Fungal Diversity 105:1-16 ((https://doi.org/10.1007/s13225-020-00466-2)) |
| Otu21485 | 1 | k__Fungi(100);p__Ascomycota(100);c__Saccharomycetes(100);o__Saccharomycetales(100);f__Metschnikowiaceae(100);g__Metschnikowia(100);g__Metschnikowia_unclassified(100); | Metschnikowia | 13 | Saprotroph | \|Undefined Saprotroph\| | Yeast | NULL | Probable | As nectar-tap_saprotroph (Põlme et al. 2020); Decay sugar-rich substrates (Põlme et al. 2020) | Põlme S, et al. 2020. Fungal Diversity 105:1-16 ((https://doi.org/10.1007/s13225-020-00466-2)) |
| Otu21486 | 1 | k__Fungi(100);p__Ascomycota(100);c__Saccharomycetes(100);o__Saccharomycetales(100);f__Metschnikowiaceae(100);g__Metschnikowia(100);g__Metschnikowia_unclassified(100); | Metschnikowia | 13 | Saprotroph | \|Undefined Saprotroph\| | Yeast | NULL | Probable | As nectar-tap_saprotroph (Põlme et al. 2020); Decay sugar-rich substrates (Põlme et al. 2020) | Põlme S, et al. 2020. Fungal Diversity 105:1-16 ((https://doi.org/10.1007/s13225-020-00466-2)) |
| Otu21487 | 1 | k__Fungi(100);p__Ascomycota(100);c__Saccharomycetes(100);o__Saccharomycetales(100);f__Metschnikowiaceae(100);g__Metschnikowia(100);g__Metschnikowia_unclassified(100); | Metschnikowia | 13 | Saprotroph | \|Undefined Saprotroph\| | Yeast | NULL | Probable | As nectar-tap_saprotroph (Põlme et al. 2020); Decay sugar-rich substrates (Põlme et al. 2020) | Põlme S, et al. 2020. Fungal Diversity 105:1-16 ((https://doi.org/10.1007/s13225-020-00466-2)) |
| Otu21488 | 1 | k__Fungi(100);p__Ascomycota(100);c__Saccharomycetes(100);o__Saccharomycetales(100);f__Metschnikowiaceae(100);g__Metschnikowia(100);s__Metschnikowia_pulcherrima(100); | Metschnikowia | 13 | Saprotroph | \|Undefined Saprotroph\| | Yeast | NULL | Probable | As nectar-tap_saprotroph (Põlme et al. 2020); Decay sugar-rich substrates (Põlme et al. 2020) | Põlme S, et al. 2020. Fungal Diversity 105:1-16 ((https://doi.org/10.1007/s13225-020-00466-2)) |
| Otu21489 | 1 | k__Fungi(100);p__Ascomycota(100);c__Saccharomycetes(100);o__Saccharomycetales(100);f__Metschnikowiaceae(100);g__Metschnikowia(100);s__Metschnikowia_pulcherrima(100); | Metschnikowia | 13 | Saprotroph | \|Undefined Saprotroph\| | Yeast | NULL | Probable | As nectar-tap_saprotroph (Põlme et al. 2020); Decay sugar-rich substrates (Põlme et al. 2020) | Põlme S, et al. 2020. Fungal Diversity 105:1-16 ((https://doi.org/10.1007/s13225-020-00466-2)) |
| Otu21490 | 1 | k__Fungi(100);p__Ascomycota(100);c__Saccharomycetes(100);o__Saccharomycetales(100);f__Metschnikowiaceae(100);g__Metschnikowia(100);s__Metschnikowia_pulcherrima(100); | Metschnikowia | 13 | Saprotroph | \|Undefined Saprotroph\| | Yeast | NULL | Probable | As nectar-tap_saprotroph (Põlme et al. 2020); Decay sugar-rich substrates (Põlme et al. 2020) | Põlme S, et al. 2020. Fungal Diversity 105:1-16 ((https://doi.org/10.1007/s13225-020-00466-2)) |
| Otu21491 | 1 | k__Fungi(100);p__Ascomycota(100);c__Saccharomycetes(100);o__Saccharomycetales(100);f__Metschnikowiaceae(100);g__Metschnikowia(100);s__Metschnikowia_pulcherrima(100); | Metschnikowia | 13 | Saprotroph | \|Undefined Saprotroph\| | Yeast | NULL | Probable | As nectar-tap_saprotroph (Põlme et al. 2020); Decay sugar-rich substrates (Põlme et al. 2020) | Põlme S, et al. 2020. Fungal Diversity 105:1-16 ((https://doi.org/10.1007/s13225-020-00466-2)) |
| Otu21492 | 1 | k__Fungi(100);p__Ascomycota(100);c__Saccharomycetes(100);o__Saccharomycetales(100);f__Metschnikowiaceae(100);g__Metschnikowia(100);s__Metschnikowia_pulcherrima(100); | Metschnikowia | 13 | Saprotroph | \|Undefined Saprotroph\| | Yeast | NULL | Probable | As nectar-tap_saprotroph (Põlme et al. 2020); Decay sugar-rich substrates (Põlme et al. 2020) | Põlme S, et al. 2020. Fungal Diversity 105:1-16 ((https://doi.org/10.1007/s13225-020-00466-2)) |
| Otu21493 | 1 | k__Fungi(100);p__Ascomycota(100);c__Saccharomycetes(100);o__Saccharomycetales(100);f__Metschnikowiaceae(100);g__Metschnikowia(100);s__Metschnikowia_pulcherrima(100); | Metschnikowia | 13 | Saprotroph | \|Undefined Saprotroph\| | Yeast | NULL | Probable | As nectar-tap_saprotroph (Põlme et al. 2020); Decay sugar-rich substrates (Põlme et al. 2020) | Põlme S, et al. 2020. Fungal Diversity 105:1-16 ((https://doi.org/10.1007/s13225-020-00466-2)) |
| Otu21497 | 1 | k__Fungi(100);p__Ascomycota(100);c__Saccharomycetes(100);o__Saccharomycetales(100);f__Saccharomycetales_family_Incertae_sedis(100);g__Candida(100);g__Candida_unclassified(100); | Candida | 13 | Pathotroph-Saprotroph-Symbiotroph | Animal Pathogen-Endophyte-Endosymbiont-Epiphyte-\|Undefined Saprotroph\| | Dimorphic Yeast-Yeast | NULL | Probable | As nectar-tap_saprotroph (Põlme et al. 2020); Decay sugar-rich substrates (Põlme et al. 2020); This asexual genus is polyphyletic and contain species that function very differently depend on what they are. Caution should be taken when interpreting guilds for Candida species. | Manolakaki D, et al. 2010. Virulence. 1: 367-375 ((https://doi.org/10.4161/viru.1.5.12796)); Mekbib SB, et al. 2011. Tropical Plant Pathology 36:233-240 ((https://doi.org/10.1590/S1982-56762011000400004)); Põlme S, et al. 2020. Fungal Diversity 105:1-16 ((https://doi.org/10.1007/s13225-020-00466-2)) |
| Otu21498 | 1 | k__Fungi(100);p__Ascomycota(100);c__Saccharomycetes(100);o__Saccharomycetales(100);f__Metschnikowiaceae(100);g__Metschnikowia(100);g__Metschnikowia_unclassified(100); | Metschnikowia | 13 | Saprotroph | \|Undefined Saprotroph\| | Yeast | NULL | Probable | As nectar-tap_saprotroph (Põlme et al. 2020); Decay sugar-rich substrates (Põlme et al. 2020) | Põlme S, et al. 2020. Fungal Diversity 105:1-16 ((https://doi.org/10.1007/s13225-020-00466-2)) |
| Otu21499 | 1 | k__Fungi(100);p__Ascomycota(100);c__Saccharomycetes(100);o__Saccharomycetales(100);f__Metschnikowiaceae(100);g__Metschnikowia(100);g__Metschnikowia_unclassified(100); | Metschnikowia | 13 | Saprotroph | \|Undefined Saprotroph\| | Yeast | NULL | Probable | As nectar-tap_saprotroph (Põlme et al. 2020); Decay sugar-rich substrates (Põlme et al. 2020) | Põlme S, et al. 2020. Fungal Diversity 105:1-16 ((https://doi.org/10.1007/s13225-020-00466-2)) |
| Otu21500 | 1 | k__Fungi(100);p__Ascomycota(100);c__Saccharomycetes(100);o__Saccharomycetales(100);f__Metschnikowiaceae(100);g__Metschnikowia(100);s__Metschnikowia_pulcherrima(100); | Metschnikowia | 13 | Saprotroph | \|Undefined Saprotroph\| | Yeast | NULL | Probable | As nectar-tap_saprotroph (Põlme et al. 2020); Decay sugar-rich substrates (Põlme et al. 2020) | Põlme S, et al. 2020. Fungal Diversity 105:1-16 ((https://doi.org/10.1007/s13225-020-00466-2)) |
| Otu21501 | 1 | k__Fungi(100);p__Ascomycota(100);c__Saccharomycetes(100);o__Saccharomycetales(100);f__Metschnikowiaceae(100);g__Metschnikowia(100);s__Metschnikowia_pulcherrima(100); | Metschnikowia | 13 | Saprotroph | \|Undefined Saprotroph\| | Yeast | NULL | Probable | As nectar-tap_saprotroph (Põlme et al. 2020); Decay sugar-rich substrates (Põlme et al. 2020) | Põlme S, et al. 2020. Fungal Diversity 105:1-16 ((https://doi.org/10.1007/s13225-020-00466-2)) |
| Otu21503 | 1 | k__Fungi(100);p__Ascomycota(100);c__Saccharomycetes(100);o__Saccharomycetales(100);f__Metschnikowiaceae(100);g__Metschnikowia(100);s__Metschnikowia_pulcherrima(100); | Metschnikowia | 13 | Saprotroph | \|Undefined Saprotroph\| | Yeast | NULL | Probable | As nectar-tap_saprotroph (Põlme et al. 2020); Decay sugar-rich substrates (Põlme et al. 2020) | Põlme S, et al. 2020. Fungal Diversity 105:1-16 ((https://doi.org/10.1007/s13225-020-00466-2)) |
| Otu21504 | 1 | k__Fungi(100);p__Ascomycota(100);c__Saccharomycetes(100);o__Saccharomycetales(100);f__Metschnikowiaceae(100);g__Metschnikowia(100);s__Metschnikowia_pulcherrima(100); | Metschnikowia | 13 | Saprotroph | \|Undefined Saprotroph\| | Yeast | NULL | Probable | As nectar-tap_saprotroph (Põlme et al. 2020); Decay sugar-rich substrates (Põlme et al. 2020) | Põlme S, et al. 2020. Fungal Diversity 105:1-16 ((https://doi.org/10.1007/s13225-020-00466-2)) |
| Otu21505 | 1 | k__Fungi(100);p__Ascomycota(100);c__Saccharomycetes(100);o__Saccharomycetales(100);f__Metschnikowiaceae(100);g__Metschnikowia(100);s__Metschnikowia_pulcherrima(100); | Metschnikowia | 13 | Saprotroph | \|Undefined Saprotroph\| | Yeast | NULL | Probable | As nectar-tap_saprotroph (Põlme et al. 2020); Decay sugar-rich substrates (Põlme et al. 2020) | Põlme S, et al. 2020. Fungal Diversity 105:1-16 ((https://doi.org/10.1007/s13225-020-00466-2)) |
| Otu21506 | 1 | k__Fungi(100);p__Ascomycota(100);c__Saccharomycetes(100);o__Saccharomycetales(100);f__Metschnikowiaceae(100);g__Metschnikowia(100);s__Metschnikowia_pulcherrima(100); | Metschnikowia | 13 | Saprotroph | \|Undefined Saprotroph\| | Yeast | NULL | Probable | As nectar-tap_saprotroph (Põlme et al. 2020); Decay sugar-rich substrates (Põlme et al. 2020) | Põlme S, et al. 2020. Fungal Diversity 105:1-16 ((https://doi.org/10.1007/s13225-020-00466-2)) |
| Otu21510 | 1 | k__Fungi(100);p__Ascomycota(100);c__Saccharomycetes(100);o__Saccharomycetales(100);f__Metschnikowiaceae(100);g__Metschnikowia(100);s__Metschnikowia_pulcherrima(100); | Metschnikowia | 13 | Saprotroph | \|Undefined Saprotroph\| | Yeast | NULL | Probable | As nectar-tap_saprotroph (Põlme et al. 2020); Decay sugar-rich substrates (Põlme et al. 2020) | Põlme S, et al. 2020. Fungal Diversity 105:1-16 ((https://doi.org/10.1007/s13225-020-00466-2)) |
| Otu21511 | 1 | k__Fungi(100);p__Ascomycota(100);c__Saccharomycetes(100);o__Saccharomycetales(100);f__Metschnikowiaceae(100);g__Metschnikowia(100);s__Metschnikowia_pulcherrima(100); | Metschnikowia | 13 | Saprotroph | \|Undefined Saprotroph\| | Yeast | NULL | Probable | As nectar-tap_saprotroph (Põlme et al. 2020); Decay sugar-rich substrates (Põlme et al. 2020) | Põlme S, et al. 2020. Fungal Diversity 105:1-16 ((https://doi.org/10.1007/s13225-020-00466-2)) |
| Otu21512 | 1 | k__Fungi(100);p__Ascomycota(100);c__Saccharomycetes(100);o__Saccharomycetales(100);f__Metschnikowiaceae(100);g__Metschnikowia(100);s__Metschnikowia_pulcherrima(100); | Metschnikowia | 13 | Saprotroph | \|Undefined Saprotroph\| | Yeast | NULL | Probable | As nectar-tap_saprotroph (Põlme et al. 2020); Decay sugar-rich substrates (Põlme et al. 2020) | Põlme S, et al. 2020. Fungal Diversity 105:1-16 ((https://doi.org/10.1007/s13225-020-00466-2)) |
| Otu21513 | 1 | k__Fungi(100);p__Ascomycota(100);c__Saccharomycetes(100);o__Saccharomycetales(100);f__Metschnikowiaceae(100);g__Metschnikowia(100);s__Metschnikowia_pulcherrima(100); | Metschnikowia | 13 | Saprotroph | \|Undefined Saprotroph\| | Yeast | NULL | Probable | As nectar-tap_saprotroph (Põlme et al. 2020); Decay sugar-rich substrates (Põlme et al. 2020) | Põlme S, et al. 2020. Fungal Diversity 105:1-16 ((https://doi.org/10.1007/s13225-020-00466-2)) |
| Otu21514 | 1 | k__Fungi(100);p__Ascomycota(100);c__Saccharomycetes(100);o__Saccharomycetales(100);f__Metschnikowiaceae(100);g__Metschnikowia(100);s__Metschnikowia_pulcherrima(100); | Metschnikowia | 13 | Saprotroph | \|Undefined Saprotroph\| | Yeast | NULL | Probable | As nectar-tap_saprotroph (Põlme et al. 2020); Decay sugar-rich substrates (Põlme et al. 2020) | Põlme S, et al. 2020. Fungal Diversity 105:1-16 ((https://doi.org/10.1007/s13225-020-00466-2)) |
| Otu21516 | 1 | k__Fungi(100);p__Ascomycota(100);c__Saccharomycetes(100);o__Saccharomycetales(100);f__Metschnikowiaceae(100);g__Metschnikowia(100);s__Metschnikowia_pulcherrima(100); | Metschnikowia | 13 | Saprotroph | \|Undefined Saprotroph\| | Yeast | NULL | Probable | As nectar-tap_saprotroph (Põlme et al. 2020); Decay sugar-rich substrates (Põlme et al. 2020) | Põlme S, et al. 2020. Fungal Diversity 105:1-16 ((https://doi.org/10.1007/s13225-020-00466-2)) |
| Otu21517 | 1 | k__Fungi(100);p__Ascomycota(100);c__Saccharomycetes(100);o__Saccharomycetales(100);f__Metschnikowiaceae(100);g__Metschnikowia(100);g__Metschnikowia_unclassified(100); | Metschnikowia | 13 | Saprotroph | \|Undefined Saprotroph\| | Yeast | NULL | Probable | As nectar-tap_saprotroph (Põlme et al. 2020); Decay sugar-rich substrates (Põlme et al. 2020) | Põlme S, et al. 2020. Fungal Diversity 105:1-16 ((https://doi.org/10.1007/s13225-020-00466-2)) |
| Otu21519 | 1 | k__Fungi(100);p__Ascomycota(100);c__Saccharomycetes(100);o__Saccharomycetales(100);f__Metschnikowiaceae(100);g__Metschnikowia(100);s__Metschnikowia_pulcherrima(100); | Metschnikowia | 13 | Saprotroph | \|Undefined Saprotroph\| | Yeast | NULL | Probable | As nectar-tap_saprotroph (Põlme et al. 2020); Decay sugar-rich substrates (Põlme et al. 2020) | Põlme S, et al. 2020. Fungal Diversity 105:1-16 ((https://doi.org/10.1007/s13225-020-00466-2)) |
| Otu21521 | 1 | k__Fungi(100);p__Ascomycota(100);c__Saccharomycetes(100);o__Saccharomycetales(100);f__Metschnikowiaceae(100);g__Metschnikowia(100);s__Metschnikowia_pulcherrima(100); | Metschnikowia | 13 | Saprotroph | \|Undefined Saprotroph\| | Yeast | NULL | Probable | As nectar-tap_saprotroph (Põlme et al. 2020); Decay sugar-rich substrates (Põlme et al. 2020) | Põlme S, et al. 2020. Fungal Diversity 105:1-16 ((https://doi.org/10.1007/s13225-020-00466-2)) |
| Otu21522 | 1 | k__Fungi(100);p__Ascomycota(100);c__Saccharomycetes(100);o__Saccharomycetales(100);f__Metschnikowiaceae(100);g__Metschnikowia(100);s__Metschnikowia_pulcherrima(100); | Metschnikowia | 13 | Saprotroph | \|Undefined Saprotroph\| | Yeast | NULL | Probable | As nectar-tap_saprotroph (Põlme et al. 2020); Decay sugar-rich substrates (Põlme et al. 2020) | Põlme S, et al. 2020. Fungal Diversity 105:1-16 ((https://doi.org/10.1007/s13225-020-00466-2)) |
| Otu21527 | 1 | k__Fungi(100);p__Ascomycota(100);c__Saccharomycetes(100);o__Saccharomycetales(100);f__Metschnikowiaceae(100);g__Metschnikowia(100);s__Metschnikowia_pulcherrima(100); | Metschnikowia | 13 | Saprotroph | \|Undefined Saprotroph\| | Yeast | NULL | Probable | As nectar-tap_saprotroph (Põlme et al. 2020); Decay sugar-rich substrates (Põlme et al. 2020) | Põlme S, et al. 2020. Fungal Diversity 105:1-16 ((https://doi.org/10.1007/s13225-020-00466-2)) |
| Otu21528 | 1 | k__Fungi(100);p__Ascomycota(100);c__Saccharomycetes(100);o__Saccharomycetales(100);f__Metschnikowiaceae(100);g__Metschnikowia(100);s__Metschnikowia_pulcherrima(100); | Metschnikowia | 13 | Saprotroph | \|Undefined Saprotroph\| | Yeast | NULL | Probable | As nectar-tap_saprotroph (Põlme et al. 2020); Decay sugar-rich substrates (Põlme et al. 2020) | Põlme S, et al. 2020. Fungal Diversity 105:1-16 ((https://doi.org/10.1007/s13225-020-00466-2)) |
| Otu21529 | 1 | k__Fungi(100);p__Ascomycota(100);c__Saccharomycetes(100);o__Saccharomycetales(100);f__Metschnikowiaceae(100);g__Metschnikowia(100);s__Metschnikowia_pulcherrima(100); | Metschnikowia | 13 | Saprotroph | \|Undefined Saprotroph\| | Yeast | NULL | Probable | As nectar-tap_saprotroph (Põlme et al. 2020); Decay sugar-rich substrates (Põlme et al. 2020) | Põlme S, et al. 2020. Fungal Diversity 105:1-16 ((https://doi.org/10.1007/s13225-020-00466-2)) |
| Otu21530 | 1 | k__Fungi(100);p__Ascomycota(100);c__Saccharomycetes(100);o__Saccharomycetales(100);f__Metschnikowiaceae(100);g__Metschnikowia(100);s__Metschnikowia_pulcherrima(100); | Metschnikowia | 13 | Saprotroph | \|Undefined Saprotroph\| | Yeast | NULL | Probable | As nectar-tap_saprotroph (Põlme et al. 2020); Decay sugar-rich substrates (Põlme et al. 2020) | Põlme S, et al. 2020. Fungal Diversity 105:1-16 ((https://doi.org/10.1007/s13225-020-00466-2)) |
| Otu21531 | 1 | k__Fungi(100);p__Ascomycota(100);c__Saccharomycetes(100);o__Saccharomycetales(100);f__Metschnikowiaceae(100);g__Metschnikowia(100);s__Metschnikowia_pulcherrima(100); | Metschnikowia | 13 | Saprotroph | \|Undefined Saprotroph\| | Yeast | NULL | Probable | As nectar-tap_saprotroph (Põlme et al. 2020); Decay sugar-rich substrates (Põlme et al. 2020) | Põlme S, et al. 2020. Fungal Diversity 105:1-16 ((https://doi.org/10.1007/s13225-020-00466-2)) |
| Otu21532 | 1 | k__Fungi(100);p__Ascomycota(100);c__Saccharomycetes(100);o__Saccharomycetales(100);f__Metschnikowiaceae(100);g__Metschnikowia(100);g__Metschnikowia_unclassified(100); | Metschnikowia | 13 | Saprotroph | \|Undefined Saprotroph\| | Yeast | NULL | Probable | As nectar-tap_saprotroph (Põlme et al. 2020); Decay sugar-rich substrates (Põlme et al. 2020) | Põlme S, et al. 2020. Fungal Diversity 105:1-16 ((https://doi.org/10.1007/s13225-020-00466-2)) |
| Otu21533 | 1 | k__Fungi(100);p__Ascomycota(100);c__Saccharomycetes(100);o__Saccharomycetales(100);f__Metschnikowiaceae(100);g__Metschnikowia(100);g__Metschnikowia_unclassified(100); | Metschnikowia | 13 | Saprotroph | \|Undefined Saprotroph\| | Yeast | NULL | Probable | As nectar-tap_saprotroph (Põlme et al. 2020); Decay sugar-rich substrates (Põlme et al. 2020) | Põlme S, et al. 2020. Fungal Diversity 105:1-16 ((https://doi.org/10.1007/s13225-020-00466-2)) |
| Otu21534 | 1 | k__Fungi(100);p__Ascomycota(100);c__Saccharomycetes(100);o__Saccharomycetales(100);f__Metschnikowiaceae(100);g__Metschnikowia(100);s__Metschnikowia_pulcherrima(100); | Metschnikowia | 13 | Saprotroph | \|Undefined Saprotroph\| | Yeast | NULL | Probable | As nectar-tap_saprotroph (Põlme et al. 2020); Decay sugar-rich substrates (Põlme et al. 2020) | Põlme S, et al. 2020. Fungal Diversity 105:1-16 ((https://doi.org/10.1007/s13225-020-00466-2)) |
| Otu21535 | 1 | k__Fungi(100);p__Ascomycota(100);c__Saccharomycetes(100);o__Saccharomycetales(100);f__Metschnikowiaceae(100);g__Metschnikowia(100);s__Metschnikowia_pulcherrima(100); | Metschnikowia | 13 | Saprotroph | \|Undefined Saprotroph\| | Yeast | NULL | Probable | As nectar-tap_saprotroph (Põlme et al. 2020); Decay sugar-rich substrates (Põlme et al. 2020) | Põlme S, et al. 2020. Fungal Diversity 105:1-16 ((https://doi.org/10.1007/s13225-020-00466-2)) |
| Otu21537 | 1 | k__Fungi(100);p__Ascomycota(100);c__Saccharomycetes(100);o__Saccharomycetales(100);f__Metschnikowiaceae(100);g__Metschnikowia(100);g__Metschnikowia_unclassified(100); | Metschnikowia | 13 | Saprotroph | \|Undefined Saprotroph\| | Yeast | NULL | Probable | As nectar-tap_saprotroph (Põlme et al. 2020); Decay sugar-rich substrates (Põlme et al. 2020) | Põlme S, et al. 2020. Fungal Diversity 105:1-16 ((https://doi.org/10.1007/s13225-020-00466-2)) |
| Otu21538 | 1 | k__Fungi(100);p__Ascomycota(100);c__Saccharomycetes(100);o__Saccharomycetales(100);o__Saccharomycetales_unclassified(100);o__Saccharomycetales_unclassified(100);o__Saccharomycetales_unclassified(100); | Saccharomycetales | 7 | Saprotroph | Undefined Saprotroph | Yeast | NULL | Possible | NULL | Sterkenburg E, et al. 2015. New Phytologist 207:1145-1158 ((https://doi.org/10.1111/nph.13426)) |
| Otu21540 | 1 | k__Fungi(100);p__Ascomycota(100);c__Saccharomycetes(100);o__Saccharomycetales(100);f__Metschnikowiaceae(100);g__Metschnikowia(100);g__Metschnikowia_unclassified(100); | Metschnikowia | 13 | Saprotroph | \|Undefined Saprotroph\| | Yeast | NULL | Probable | As nectar-tap_saprotroph (Põlme et al. 2020); Decay sugar-rich substrates (Põlme et al. 2020) | Põlme S, et al. 2020. Fungal Diversity 105:1-16 ((https://doi.org/10.1007/s13225-020-00466-2)) |
| Otu21541 | 1 | k__Fungi(100);p__Ascomycota(100);c__Saccharomycetes(100);o__Saccharomycetales(100);f__Metschnikowiaceae(100);g__Metschnikowia(100);s__Metschnikowia_pulcherrima(100); | Metschnikowia | 13 | Saprotroph | \|Undefined Saprotroph\| | Yeast | NULL | Probable | As nectar-tap_saprotroph (Põlme et al. 2020); Decay sugar-rich substrates (Põlme et al. 2020) | Põlme S, et al. 2020. Fungal Diversity 105:1-16 ((https://doi.org/10.1007/s13225-020-00466-2)) |
| Otu21542 | 1 | k__Fungi(100);p__Ascomycota(100);c__Saccharomycetes(100);o__Saccharomycetales(100);f__Metschnikowiaceae(100);g__Metschnikowia(100);s__Metschnikowia_pulcherrima(100); | Metschnikowia | 13 | Saprotroph | \|Undefined Saprotroph\| | Yeast | NULL | Probable | As nectar-tap_saprotroph (Põlme et al. 2020); Decay sugar-rich substrates (Põlme et al. 2020) | Põlme S, et al. 2020. Fungal Diversity 105:1-16 ((https://doi.org/10.1007/s13225-020-00466-2)) |
| Otu21543 | 1 | k__Fungi(100);p__Ascomycota(100);c__Saccharomycetes(100);o__Saccharomycetales(100);f__Metschnikowiaceae(100);g__Metschnikowia(100);s__Metschnikowia_pulcherrima(100); | Metschnikowia | 13 | Saprotroph | \|Undefined Saprotroph\| | Yeast | NULL | Probable | As nectar-tap_saprotroph (Põlme et al. 2020); Decay sugar-rich substrates (Põlme et al. 2020) | Põlme S, et al. 2020. Fungal Diversity 105:1-16 ((https://doi.org/10.1007/s13225-020-00466-2)) |
| Otu21544 | 1 | k__Fungi(100);p__Ascomycota(100);c__Saccharomycetes(100);o__Saccharomycetales(100);f__Metschnikowiaceae(100);g__Metschnikowia(100);s__Metschnikowia_pulcherrima(100); | Metschnikowia | 13 | Saprotroph | \|Undefined Saprotroph\| | Yeast | NULL | Probable | As nectar-tap_saprotroph (Põlme et al. 2020); Decay sugar-rich substrates (Põlme et al. 2020) | Põlme S, et al. 2020. Fungal Diversity 105:1-16 ((https://doi.org/10.1007/s13225-020-00466-2)) |
| Otu21546 | 1 | k__Fungi(100);p__Ascomycota(100);c__Saccharomycetes(100);o__Saccharomycetales(100);f__Metschnikowiaceae(100);g__Metschnikowia(100);s__Metschnikowia_pulcherrima(100); | Metschnikowia | 13 | Saprotroph | \|Undefined Saprotroph\| | Yeast | NULL | Probable | As nectar-tap_saprotroph (Põlme et al. 2020); Decay sugar-rich substrates (Põlme et al. 2020) | Põlme S, et al. 2020. Fungal Diversity 105:1-16 ((https://doi.org/10.1007/s13225-020-00466-2)) |
| Otu21548 | 1 | k__Fungi(100);p__Ascomycota(100);c__Saccharomycetes(100);o__Saccharomycetales(100);f__Metschnikowiaceae(100);g__Metschnikowia(100);s__Metschnikowia_pulcherrima(100); | Metschnikowia | 13 | Saprotroph | \|Undefined Saprotroph\| | Yeast | NULL | Probable | As nectar-tap_saprotroph (Põlme et al. 2020); Decay sugar-rich substrates (Põlme et al. 2020) | Põlme S, et al. 2020. Fungal Diversity 105:1-16 ((https://doi.org/10.1007/s13225-020-00466-2)) |
| Otu21549 | 1 | k__Fungi(100);p__Ascomycota(100);c__Saccharomycetes(100);o__Saccharomycetales(100);f__Metschnikowiaceae(100);g__Metschnikowia(100);s__Metschnikowia_pulcherrima(100); | Metschnikowia | 13 | Saprotroph | \|Undefined Saprotroph\| | Yeast | NULL | Probable | As nectar-tap_saprotroph (Põlme et al. 2020); Decay sugar-rich substrates (Põlme et al. 2020) | Põlme S, et al. 2020. Fungal Diversity 105:1-16 ((https://doi.org/10.1007/s13225-020-00466-2)) |
| Otu21550 | 1 | k__Fungi(100);p__Ascomycota(100);c__Saccharomycetes(100);o__Saccharomycetales(100);f__Metschnikowiaceae(100);g__Metschnikowia(100);s__Metschnikowia_pulcherrima(100); | Metschnikowia | 13 | Saprotroph | \|Undefined Saprotroph\| | Yeast | NULL | Probable | As nectar-tap_saprotroph (Põlme et al. 2020); Decay sugar-rich substrates (Põlme et al. 2020) | Põlme S, et al. 2020. Fungal Diversity 105:1-16 ((https://doi.org/10.1007/s13225-020-00466-2)) |
| Otu21551 | 1 | k__Fungi(100);p__Ascomycota(100);c__Saccharomycetes(100);o__Saccharomycetales(100);o__Saccharomycetales_unclassified(100);o__Saccharomycetales_unclassified(100);o__Saccharomycetales_unclassified(100); | Saccharomycetales | 7 | Saprotroph | Undefined Saprotroph | Yeast | NULL | Possible | NULL | Sterkenburg E, et al. 2015. New Phytologist 207:1145-1158 ((https://doi.org/10.1111/nph.13426)) |
| Otu21552 | 1 | k__Fungi(100);p__Ascomycota(100);c__Saccharomycetes(100);o__Saccharomycetales(100);f__Metschnikowiaceae(100);g__Metschnikowia(100);s__Metschnikowia_pulcherrima(100); | Metschnikowia | 13 | Saprotroph | \|Undefined Saprotroph\| | Yeast | NULL | Probable | As nectar-tap_saprotroph (Põlme et al. 2020); Decay sugar-rich substrates (Põlme et al. 2020) | Põlme S, et al. 2020. Fungal Diversity 105:1-16 ((https://doi.org/10.1007/s13225-020-00466-2)) |
| Otu21553 | 1 | k__Fungi(100);p__Ascomycota(100);c__Saccharomycetes(100);o__Saccharomycetales(100);f__Metschnikowiaceae(100);g__Metschnikowia(100);g__Metschnikowia_unclassified(100); | Metschnikowia | 13 | Saprotroph | \|Undefined Saprotroph\| | Yeast | NULL | Probable | As nectar-tap_saprotroph (Põlme et al. 2020); Decay sugar-rich substrates (Põlme et al. 2020) | Põlme S, et al. 2020. Fungal Diversity 105:1-16 ((https://doi.org/10.1007/s13225-020-00466-2)) |
| Otu21554 | 1 | k__Fungi(100);p__Ascomycota(100);c__Saccharomycetes(100);o__Saccharomycetales(100);f__Metschnikowiaceae(100);g__Metschnikowia(100);g__Metschnikowia_unclassified(100); | Metschnikowia | 13 | Saprotroph | \|Undefined Saprotroph\| | Yeast | NULL | Probable | As nectar-tap_saprotroph (Põlme et al. 2020); Decay sugar-rich substrates (Põlme et al. 2020) | Põlme S, et al. 2020. Fungal Diversity 105:1-16 ((https://doi.org/10.1007/s13225-020-00466-2)) |
| Otu21555 | 1 | k__Fungi(100);p__Ascomycota(100);c__Saccharomycetes(100);o__Saccharomycetales(100);f__Metschnikowiaceae(100);g__Metschnikowia(100);g__Metschnikowia_unclassified(100); | Metschnikowia | 13 | Saprotroph | \|Undefined Saprotroph\| | Yeast | NULL | Probable | As nectar-tap_saprotroph (Põlme et al. 2020); Decay sugar-rich substrates (Põlme et al. 2020) | Põlme S, et al. 2020. Fungal Diversity 105:1-16 ((https://doi.org/10.1007/s13225-020-00466-2)) |
| Otu21557 | 1 | k__Fungi(100);p__Ascomycota(100);c__Saccharomycetes(100);o__Saccharomycetales(100);f__Metschnikowiaceae(100);g__Metschnikowia(100);s__Metschnikowia_pulcherrima(100); | Metschnikowia | 13 | Saprotroph | \|Undefined Saprotroph\| | Yeast | NULL | Probable | As nectar-tap_saprotroph (Põlme et al. 2020); Decay sugar-rich substrates (Põlme et al. 2020) | Põlme S, et al. 2020. Fungal Diversity 105:1-16 ((https://doi.org/10.1007/s13225-020-00466-2)) |
| Otu21558 | 1 | k__Fungi(100);p__Ascomycota(100);c__Saccharomycetes(100);o__Saccharomycetales(100);f__Metschnikowiaceae(100);g__Metschnikowia(100);s__Metschnikowia_pulcherrima(100); | Metschnikowia | 13 | Saprotroph | \|Undefined Saprotroph\| | Yeast | NULL | Probable | As nectar-tap_saprotroph (Põlme et al. 2020); Decay sugar-rich substrates (Põlme et al. 2020) | Põlme S, et al. 2020. Fungal Diversity 105:1-16 ((https://doi.org/10.1007/s13225-020-00466-2)) |
| Otu21559 | 1 | k__Fungi(100);p__Ascomycota(100);c__Saccharomycetes(100);o__Saccharomycetales(100);f__Metschnikowiaceae(100);g__Metschnikowia(100);s__Metschnikowia_pulcherrima(100); | Metschnikowia | 13 | Saprotroph | \|Undefined Saprotroph\| | Yeast | NULL | Probable | As nectar-tap_saprotroph (Põlme et al. 2020); Decay sugar-rich substrates (Põlme et al. 2020) | Põlme S, et al. 2020. Fungal Diversity 105:1-16 ((https://doi.org/10.1007/s13225-020-00466-2)) |
| Otu21560 | 1 | k__Fungi(100);p__Ascomycota(100);c__Saccharomycetes(100);o__Saccharomycetales(100);f__Metschnikowiaceae(100);g__Metschnikowia(100);s__Metschnikowia_pulcherrima(100); | Metschnikowia | 13 | Saprotroph | \|Undefined Saprotroph\| | Yeast | NULL | Probable | As nectar-tap_saprotroph (Põlme et al. 2020); Decay sugar-rich substrates (Põlme et al. 2020) | Põlme S, et al. 2020. Fungal Diversity 105:1-16 ((https://doi.org/10.1007/s13225-020-00466-2)) |
| Otu21561 | 1 | k__Fungi(100);p__Ascomycota(100);c__Saccharomycetes(100);o__Saccharomycetales(100);f__Metschnikowiaceae(100);g__Metschnikowia(100);s__Metschnikowia_pulcherrima(100); | Metschnikowia | 13 | Saprotroph | \|Undefined Saprotroph\| | Yeast | NULL | Probable | As nectar-tap_saprotroph (Põlme et al. 2020); Decay sugar-rich substrates (Põlme et al. 2020) | Põlme S, et al. 2020. Fungal Diversity 105:1-16 ((https://doi.org/10.1007/s13225-020-00466-2)) |
| Otu21562 | 1 | k__Fungi(100);p__Ascomycota(100);c__Saccharomycetes(100);o__Saccharomycetales(100);f__Metschnikowiaceae(100);g__Metschnikowia(100);s__Metschnikowia_pulcherrima(100); | Metschnikowia | 13 | Saprotroph | \|Undefined Saprotroph\| | Yeast | NULL | Probable | As nectar-tap_saprotroph (Põlme et al. 2020); Decay sugar-rich substrates (Põlme et al. 2020) | Põlme S, et al. 2020. Fungal Diversity 105:1-16 ((https://doi.org/10.1007/s13225-020-00466-2)) |
| Otu21563 | 1 | k__Fungi(100);p__Ascomycota(100);c__Saccharomycetes(100);o__Saccharomycetales(100);f__Metschnikowiaceae(100);g__Metschnikowia(100);s__Metschnikowia_pulcherrima(100); | Metschnikowia | 13 | Saprotroph | \|Undefined Saprotroph\| | Yeast | NULL | Probable | As nectar-tap_saprotroph (Põlme et al. 2020); Decay sugar-rich substrates (Põlme et al. 2020) | Põlme S, et al. 2020. Fungal Diversity 105:1-16 ((https://doi.org/10.1007/s13225-020-00466-2)) |
| Otu21564 | 1 | k__Fungi(100);p__Ascomycota(100);c__Saccharomycetes(100);o__Saccharomycetales(100);f__Metschnikowiaceae(100);g__Metschnikowia(100);s__Metschnikowia_pulcherrima(100); | Metschnikowia | 13 | Saprotroph | \|Undefined Saprotroph\| | Yeast | NULL | Probable | As nectar-tap_saprotroph (Põlme et al. 2020); Decay sugar-rich substrates (Põlme et al. 2020) | Põlme S, et al. 2020. Fungal Diversity 105:1-16 ((https://doi.org/10.1007/s13225-020-00466-2)) |
| Otu21565 | 1 | k__Fungi(100);p__Ascomycota(100);c__Saccharomycetes(100);o__Saccharomycetales(100);f__Metschnikowiaceae(100);g__Metschnikowia(100);s__Metschnikowia_pulcherrima(100); | Metschnikowia | 13 | Saprotroph | \|Undefined Saprotroph\| | Yeast | NULL | Probable | As nectar-tap_saprotroph (Põlme et al. 2020); Decay sugar-rich substrates (Põlme et al. 2020) | Põlme S, et al. 2020. Fungal Diversity 105:1-16 ((https://doi.org/10.1007/s13225-020-00466-2)) |
| Otu21566 | 1 | k__Fungi(100);p__Ascomycota(100);c__Saccharomycetes(100);o__Saccharomycetales(100);f__Metschnikowiaceae(100);g__Metschnikowia(100);g__Metschnikowia_unclassified(100); | Metschnikowia | 13 | Saprotroph | \|Undefined Saprotroph\| | Yeast | NULL | Probable | As nectar-tap_saprotroph (Põlme et al. 2020); Decay sugar-rich substrates (Põlme et al. 2020) | Põlme S, et al. 2020. Fungal Diversity 105:1-16 ((https://doi.org/10.1007/s13225-020-00466-2)) |
| Otu21567 | 1 | k__Fungi(100);p__Ascomycota(100);c__Saccharomycetes(100);o__Saccharomycetales(100);f__Metschnikowiaceae(100);g__Metschnikowia(100);g__Metschnikowia_unclassified(100); | Metschnikowia | 13 | Saprotroph | \|Undefined Saprotroph\| | Yeast | NULL | Probable | As nectar-tap_saprotroph (Põlme et al. 2020); Decay sugar-rich substrates (Põlme et al. 2020) | Põlme S, et al. 2020. Fungal Diversity 105:1-16 ((https://doi.org/10.1007/s13225-020-00466-2)) |
| Otu21568 | 1 | k__Fungi(100);p__Ascomycota(100);c__Saccharomycetes(100);o__Saccharomycetales(100);f__Metschnikowiaceae(100);g__Metschnikowia(100);s__Metschnikowia_pulcherrima(100); | Metschnikowia | 13 | Saprotroph | \|Undefined Saprotroph\| | Yeast | NULL | Probable | As nectar-tap_saprotroph (Põlme et al. 2020); Decay sugar-rich substrates (Põlme et al. 2020) | Põlme S, et al. 2020. Fungal Diversity 105:1-16 ((https://doi.org/10.1007/s13225-020-00466-2)) |
| Otu21569 | 1 | k__Fungi(100);p__Ascomycota(100);c__Saccharomycetes(100);o__Saccharomycetales(100);f__Metschnikowiaceae(100);g__Metschnikowia(100);g__Metschnikowia_unclassified(100); | Metschnikowia | 13 | Saprotroph | \|Undefined Saprotroph\| | Yeast | NULL | Probable | As nectar-tap_saprotroph (Põlme et al. 2020); Decay sugar-rich substrates (Põlme et al. 2020) | Põlme S, et al. 2020. Fungal Diversity 105:1-16 ((https://doi.org/10.1007/s13225-020-00466-2)) |
| Otu21570 | 1 | k__Fungi(100);p__Ascomycota(100);c__Saccharomycetes(100);o__Saccharomycetales(100);f__Metschnikowiaceae(100);g__Metschnikowia(100);g__Metschnikowia_unclassified(100); | Metschnikowia | 13 | Saprotroph | \|Undefined Saprotroph\| | Yeast | NULL | Probable | As nectar-tap_saprotroph (Põlme et al. 2020); Decay sugar-rich substrates (Põlme et al. 2020) | Põlme S, et al. 2020. Fungal Diversity 105:1-16 ((https://doi.org/10.1007/s13225-020-00466-2)) |
| Otu21571 | 1 | k__Fungi(100);p__Ascomycota(100);c__Saccharomycetes(100);o__Saccharomycetales(100);f__Metschnikowiaceae(100);g__Metschnikowia(100);g__Metschnikowia_unclassified(100); | Metschnikowia | 13 | Saprotroph | \|Undefined Saprotroph\| | Yeast | NULL | Probable | As nectar-tap_saprotroph (Põlme et al. 2020); Decay sugar-rich substrates (Põlme et al. 2020) | Põlme S, et al. 2020. Fungal Diversity 105:1-16 ((https://doi.org/10.1007/s13225-020-00466-2)) |
| Otu21572 | 1 | k__Fungi(100);p__Ascomycota(100);c__Saccharomycetes(100);o__Saccharomycetales(100);f__Metschnikowiaceae(100);g__Metschnikowia(100);s__Metschnikowia_pulcherrima(100); | Metschnikowia | 13 | Saprotroph | \|Undefined Saprotroph\| | Yeast | NULL | Probable | As nectar-tap_saprotroph (Põlme et al. 2020); Decay sugar-rich substrates (Põlme et al. 2020) | Põlme S, et al. 2020. Fungal Diversity 105:1-16 ((https://doi.org/10.1007/s13225-020-00466-2)) |
| Otu21573 | 1 | k__Fungi(100);p__Ascomycota(100);c__Saccharomycetes(100);o__Saccharomycetales(100);f__Metschnikowiaceae(100);g__Metschnikowia(100);s__Metschnikowia_pulcherrima(100); | Metschnikowia | 13 | Saprotroph | \|Undefined Saprotroph\| | Yeast | NULL | Probable | As nectar-tap_saprotroph (Põlme et al. 2020); Decay sugar-rich substrates (Põlme et al. 2020) | Põlme S, et al. 2020. Fungal Diversity 105:1-16 ((https://doi.org/10.1007/s13225-020-00466-2)) |
| Otu21574 | 1 | k__Fungi(100);p__Ascomycota(100);c__Saccharomycetes(100);o__Saccharomycetales(100);f__Metschnikowiaceae(100);g__Metschnikowia(100);s__Metschnikowia_pulcherrima(100); | Metschnikowia | 13 | Saprotroph | \|Undefined Saprotroph\| | Yeast | NULL | Probable | As nectar-tap_saprotroph (Põlme et al. 2020); Decay sugar-rich substrates (Põlme et al. 2020) | Põlme S, et al. 2020. Fungal Diversity 105:1-16 ((https://doi.org/10.1007/s13225-020-00466-2)) |
| Otu21575 | 1 | k__Fungi(100);p__Ascomycota(100);c__Saccharomycetes(100);o__Saccharomycetales(100);f__Metschnikowiaceae(100);g__Metschnikowia(100);s__Metschnikowia_pulcherrima(100); | Metschnikowia | 13 | Saprotroph | \|Undefined Saprotroph\| | Yeast | NULL | Probable | As nectar-tap_saprotroph (Põlme et al. 2020); Decay sugar-rich substrates (Põlme et al. 2020) | Põlme S, et al. 2020. Fungal Diversity 105:1-16 ((https://doi.org/10.1007/s13225-020-00466-2)) |
| Otu21576 | 1 | k__Fungi(100);p__Ascomycota(100);c__Saccharomycetes(100);o__Saccharomycetales(100);f__Metschnikowiaceae(100);g__Metschnikowia(100);s__Metschnikowia_pulcherrima(100); | Metschnikowia | 13 | Saprotroph | \|Undefined Saprotroph\| | Yeast | NULL | Probable | As nectar-tap_saprotroph (Põlme et al. 2020); Decay sugar-rich substrates (Põlme et al. 2020) | Põlme S, et al. 2020. Fungal Diversity 105:1-16 ((https://doi.org/10.1007/s13225-020-00466-2)) |
| Otu21577 | 1 | k__Fungi(100);p__Ascomycota(100);c__Saccharomycetes(100);o__Saccharomycetales(100);f__Metschnikowiaceae(100);g__Metschnikowia(100);s__Metschnikowia_pulcherrima(100); | Metschnikowia | 13 | Saprotroph | \|Undefined Saprotroph\| | Yeast | NULL | Probable | As nectar-tap_saprotroph (Põlme et al. 2020); Decay sugar-rich substrates (Põlme et al. 2020) | Põlme S, et al. 2020. Fungal Diversity 105:1-16 ((https://doi.org/10.1007/s13225-020-00466-2)) |
| Otu21579 | 1 | k__Fungi(100);p__Ascomycota(100);c__Saccharomycetes(100);o__Saccharomycetales(100);f__Metschnikowiaceae(100);g__Metschnikowia(100);s__Metschnikowia_pulcherrima(100); | Metschnikowia | 13 | Saprotroph | \|Undefined Saprotroph\| | Yeast | NULL | Probable | As nectar-tap_saprotroph (Põlme et al. 2020); Decay sugar-rich substrates (Põlme et al. 2020) | Põlme S, et al. 2020. Fungal Diversity 105:1-16 ((https://doi.org/10.1007/s13225-020-00466-2)) |
| Otu21580 | 1 | k__Fungi(100);p__Ascomycota(100);c__Saccharomycetes(100);o__Saccharomycetales(100);f__Metschnikowiaceae(100);g__Metschnikowia(100);s__Metschnikowia_pulcherrima(100); | Metschnikowia | 13 | Saprotroph | \|Undefined Saprotroph\| | Yeast | NULL | Probable | As nectar-tap_saprotroph (Põlme et al. 2020); Decay sugar-rich substrates (Põlme et al. 2020) | Põlme S, et al. 2020. Fungal Diversity 105:1-16 ((https://doi.org/10.1007/s13225-020-00466-2)) |
| Otu21581 | 1 | k__Fungi(100);p__Ascomycota(100);c__Saccharomycetes(100);o__Saccharomycetales(100);f__Metschnikowiaceae(100);g__Metschnikowia(100);s__Metschnikowia_pulcherrima(100); | Metschnikowia | 13 | Saprotroph | \|Undefined Saprotroph\| | Yeast | NULL | Probable | As nectar-tap_saprotroph (Põlme et al. 2020); Decay sugar-rich substrates (Põlme et al. 2020) | Põlme S, et al. 2020. Fungal Diversity 105:1-16 ((https://doi.org/10.1007/s13225-020-00466-2)) |
| Otu21582 | 1 | k__Fungi(100);p__Ascomycota(100);c__Saccharomycetes(100);o__Saccharomycetales(100);f__Metschnikowiaceae(100);g__Metschnikowia(100);s__Metschnikowia_pulcherrima(100); | Metschnikowia | 13 | Saprotroph | \|Undefined Saprotroph\| | Yeast | NULL | Probable | As nectar-tap_saprotroph (Põlme et al. 2020); Decay sugar-rich substrates (Põlme et al. 2020) | Põlme S, et al. 2020. Fungal Diversity 105:1-16 ((https://doi.org/10.1007/s13225-020-00466-2)) |
| Otu21583 | 1 | k__Fungi(100);p__Ascomycota(100);c__Saccharomycetes(100);o__Saccharomycetales(100);f__Metschnikowiaceae(100);g__Metschnikowia(100);s__Metschnikowia_pulcherrima(100); | Metschnikowia | 13 | Saprotroph | \|Undefined Saprotroph\| | Yeast | NULL | Probable | As nectar-tap_saprotroph (Põlme et al. 2020); Decay sugar-rich substrates (Põlme et al. 2020) | Põlme S, et al. 2020. Fungal Diversity 105:1-16 ((https://doi.org/10.1007/s13225-020-00466-2)) |
| Otu21584 | 1 | k__Fungi(100);p__Ascomycota(100);c__Saccharomycetes(100);o__Saccharomycetales(100);f__Metschnikowiaceae(100);g__Metschnikowia(100);s__Metschnikowia_pulcherrima(100); | Metschnikowia | 13 | Saprotroph | \|Undefined Saprotroph\| | Yeast | NULL | Probable | As nectar-tap_saprotroph (Põlme et al. 2020); Decay sugar-rich substrates (Põlme et al. 2020) | Põlme S, et al. 2020. Fungal Diversity 105:1-16 ((https://doi.org/10.1007/s13225-020-00466-2)) |
| Otu21585 | 1 | k__Fungi(100);p__Ascomycota(100);c__Saccharomycetes(100);o__Saccharomycetales(100);f__Metschnikowiaceae(100);g__Metschnikowia(100);g__Metschnikowia_unclassified(100); | Metschnikowia | 13 | Saprotroph | \|Undefined Saprotroph\| | Yeast | NULL | Probable | As nectar-tap_saprotroph (Põlme et al. 2020); Decay sugar-rich substrates (Põlme et al. 2020) | Põlme S, et al. 2020. Fungal Diversity 105:1-16 ((https://doi.org/10.1007/s13225-020-00466-2)) |
| Otu21586 | 1 | k__Fungi(100);p__Ascomycota(100);c__Saccharomycetes(100);o__Saccharomycetales(100);f__Metschnikowiaceae(100);g__Metschnikowia(100);s__Metschnikowia_pulcherrima(100); | Metschnikowia | 13 | Saprotroph | \|Undefined Saprotroph\| | Yeast | NULL | Probable | As nectar-tap_saprotroph (Põlme et al. 2020); Decay sugar-rich substrates (Põlme et al. 2020) | Põlme S, et al. 2020. Fungal Diversity 105:1-16 ((https://doi.org/10.1007/s13225-020-00466-2)) |
| Otu21587 | 1 | k__Fungi(100);p__Ascomycota(100);c__Saccharomycetes(100);o__Saccharomycetales(100);f__Metschnikowiaceae(100);g__Metschnikowia(100);g__Metschnikowia_unclassified(100); | Metschnikowia | 13 | Saprotroph | \|Undefined Saprotroph\| | Yeast | NULL | Probable | As nectar-tap_saprotroph (Põlme et al. 2020); Decay sugar-rich substrates (Põlme et al. 2020) | Põlme S, et al. 2020. Fungal Diversity 105:1-16 ((https://doi.org/10.1007/s13225-020-00466-2)) |
| Otu21588 | 1 | k__Fungi(100);p__Ascomycota(100);c__Saccharomycetes(100);o__Saccharomycetales(100);f__Metschnikowiaceae(100);g__Metschnikowia(100);s__Metschnikowia_pulcherrima(100); | Metschnikowia | 13 | Saprotroph | \|Undefined Saprotroph\| | Yeast | NULL | Probable | As nectar-tap_saprotroph (Põlme et al. 2020); Decay sugar-rich substrates (Põlme et al. 2020) | Põlme S, et al. 2020. Fungal Diversity 105:1-16 ((https://doi.org/10.1007/s13225-020-00466-2)) |
| Otu21589 | 1 | k__Fungi(100);p__Ascomycota(100);c__Saccharomycetes(100);o__Saccharomycetales(100);f__Metschnikowiaceae(100);g__Metschnikowia(100);s__Metschnikowia_pulcherrima(100); | Metschnikowia | 13 | Saprotroph | \|Undefined Saprotroph\| | Yeast | NULL | Probable | As nectar-tap_saprotroph (Põlme et al. 2020); Decay sugar-rich substrates (Põlme et al. 2020) | Põlme S, et al. 2020. Fungal Diversity 105:1-16 ((https://doi.org/10.1007/s13225-020-00466-2)) |
| Otu21590 | 1 | k__Fungi(100);p__Ascomycota(100);c__Saccharomycetes(100);o__Saccharomycetales(100);f__Metschnikowiaceae(100);g__Metschnikowia(100);s__Metschnikowia_pulcherrima(100); | Metschnikowia | 13 | Saprotroph | \|Undefined Saprotroph\| | Yeast | NULL | Probable | As nectar-tap_saprotroph (Põlme et al. 2020); Decay sugar-rich substrates (Põlme et al. 2020) | Põlme S, et al. 2020. Fungal Diversity 105:1-16 ((https://doi.org/10.1007/s13225-020-00466-2)) |
| Otu21599 | 1 | k__Fungi(100);p__Ascomycota(100);c__Saccharomycetes(100);o__Saccharomycetales(100);f__Metschnikowiaceae(100);g__Metschnikowia(100);s__Metschnikowia_pulcherrima(100); | Metschnikowia | 13 | Saprotroph | \|Undefined Saprotroph\| | Yeast | NULL | Probable | As nectar-tap_saprotroph (Põlme et al. 2020); Decay sugar-rich substrates (Põlme et al. 2020) | Põlme S, et al. 2020. Fungal Diversity 105:1-16 ((https://doi.org/10.1007/s13225-020-00466-2)) |
| Otu21601 | 1 | k__Fungi(100);p__Ascomycota(100);c__Saccharomycetes(100);o__Saccharomycetales(100);f__Metschnikowiaceae(100);g__Metschnikowia(100);s__Metschnikowia_pulcherrima(100); | Metschnikowia | 13 | Saprotroph | \|Undefined Saprotroph\| | Yeast | NULL | Probable | As nectar-tap_saprotroph (Põlme et al. 2020); Decay sugar-rich substrates (Põlme et al. 2020) | Põlme S, et al. 2020. Fungal Diversity 105:1-16 ((https://doi.org/10.1007/s13225-020-00466-2)) |
| Otu21602 | 1 | k__Fungi(100);p__Ascomycota(100);c__Saccharomycetes(100);o__Saccharomycetales(100);f__Metschnikowiaceae(100);g__Metschnikowia(100);s__Metschnikowia_pulcherrima(100); | Metschnikowia | 13 | Saprotroph | \|Undefined Saprotroph\| | Yeast | NULL | Probable | As nectar-tap_saprotroph (Põlme et al. 2020); Decay sugar-rich substrates (Põlme et al. 2020) | Põlme S, et al. 2020. Fungal Diversity 105:1-16 ((https://doi.org/10.1007/s13225-020-00466-2)) |
| Otu21603 | 1 | k__Fungi(100);p__Ascomycota(100);c__Saccharomycetes(100);o__Saccharomycetales(100);f__Metschnikowiaceae(100);g__Metschnikowia(100);s__Metschnikowia_pulcherrima(100); | Metschnikowia | 13 | Saprotroph | \|Undefined Saprotroph\| | Yeast | NULL | Probable | As nectar-tap_saprotroph (Põlme et al. 2020); Decay sugar-rich substrates (Põlme et al. 2020) | Põlme S, et al. 2020. Fungal Diversity 105:1-16 ((https://doi.org/10.1007/s13225-020-00466-2)) |
| Otu21604 | 1 | k__Fungi(100);p__Ascomycota(100);c__Saccharomycetes(100);o__Saccharomycetales(100);f__Metschnikowiaceae(100);g__Metschnikowia(100);s__Metschnikowia_pulcherrima(100); | Metschnikowia | 13 | Saprotroph | \|Undefined Saprotroph\| | Yeast | NULL | Probable | As nectar-tap_saprotroph (Põlme et al. 2020); Decay sugar-rich substrates (Põlme et al. 2020) | Põlme S, et al. 2020. Fungal Diversity 105:1-16 ((https://doi.org/10.1007/s13225-020-00466-2)) |
| Otu21610 | 1 | k__Fungi(100);p__Ascomycota(100);c__Saccharomycetes(100);o__Saccharomycetales(100);f__Metschnikowiaceae(100);g__Metschnikowia(100);g__Metschnikowia_unclassified(100); | Metschnikowia | 13 | Saprotroph | \|Undefined Saprotroph\| | Yeast | NULL | Probable | As nectar-tap_saprotroph (Põlme et al. 2020); Decay sugar-rich substrates (Põlme et al. 2020) | Põlme S, et al. 2020. Fungal Diversity 105:1-16 ((https://doi.org/10.1007/s13225-020-00466-2)) |
| Otu21611 | 1 | k__Fungi(100);p__Ascomycota(100);c__Saccharomycetes(100);o__Saccharomycetales(100);f__Metschnikowiaceae(100);g__Metschnikowia(100);g__Metschnikowia_unclassified(100); | Metschnikowia | 13 | Saprotroph | \|Undefined Saprotroph\| | Yeast | NULL | Probable | As nectar-tap_saprotroph (Põlme et al. 2020); Decay sugar-rich substrates (Põlme et al. 2020) | Põlme S, et al. 2020. Fungal Diversity 105:1-16 ((https://doi.org/10.1007/s13225-020-00466-2)) |
| Otu21612 | 1 | k__Fungi(100);p__Ascomycota(100);c__Saccharomycetes(100);o__Saccharomycetales(100);f__Saccharomycetales_family_Incertae_sedis(100);g__Candida(100);s__Candida_cretensis(100); | Candida | 13 | Pathotroph-Saprotroph-Symbiotroph | Animal Pathogen-Endophyte-Endosymbiont-Epiphyte-\|Undefined Saprotroph\| | Dimorphic Yeast-Yeast | NULL | Probable | As nectar-tap_saprotroph (Põlme et al. 2020); Decay sugar-rich substrates (Põlme et al. 2020); This asexual genus is polyphyletic and contain species that function very differently depend on what they are. Caution should be taken when interpreting guilds for Candida species. | Manolakaki D, et al. 2010. Virulence. 1: 367-375 ((https://doi.org/10.4161/viru.1.5.12796)); Mekbib SB, et al. 2011. Tropical Plant Pathology 36:233-240 ((https://doi.org/10.1590/S1982-56762011000400004)); Põlme S, et al. 2020. Fungal Diversity 105:1-16 ((https://doi.org/10.1007/s13225-020-00466-2)) |
| Otu21613 | 1 | k__Fungi(100);p__Ascomycota(100);c__Saccharomycetes(100);o__Saccharomycetales(100);f__Metschnikowiaceae(100);g__Metschnikowia(100);s__Metschnikowia_pulcherrima(100); | Metschnikowia | 13 | Saprotroph | \|Undefined Saprotroph\| | Yeast | NULL | Probable | As nectar-tap_saprotroph (Põlme et al. 2020); Decay sugar-rich substrates (Põlme et al. 2020) | Põlme S, et al. 2020. Fungal Diversity 105:1-16 ((https://doi.org/10.1007/s13225-020-00466-2)) |
| Otu21614 | 1 | k__Fungi(100);p__Ascomycota(100);c__Saccharomycetes(100);o__Saccharomycetales(100);f__Metschnikowiaceae(100);g__Metschnikowia(100);g__Metschnikowia_unclassified(100); | Metschnikowia | 13 | Saprotroph | \|Undefined Saprotroph\| | Yeast | NULL | Probable | As nectar-tap_saprotroph (Põlme et al. 2020); Decay sugar-rich substrates (Põlme et al. 2020) | Põlme S, et al. 2020. Fungal Diversity 105:1-16 ((https://doi.org/10.1007/s13225-020-00466-2)) |
| Otu21616 | 1 | k__Fungi(100);p__Ascomycota(100);c__Saccharomycetes(100);o__Saccharomycetales(100);f__Metschnikowiaceae(100);g__Metschnikowia(100);s__Metschnikowia_pulcherrima(100); | Metschnikowia | 13 | Saprotroph | \|Undefined Saprotroph\| | Yeast | NULL | Probable | As nectar-tap_saprotroph (Põlme et al. 2020); Decay sugar-rich substrates (Põlme et al. 2020) | Põlme S, et al. 2020. Fungal Diversity 105:1-16 ((https://doi.org/10.1007/s13225-020-00466-2)) |
| Otu21617 | 1 | k__Fungi(100);p__Ascomycota(100);c__Saccharomycetes(100);o__Saccharomycetales(100);f__Metschnikowiaceae(100);g__Metschnikowia(100);s__Metschnikowia_pulcherrima(100); | Metschnikowia | 13 | Saprotroph | \|Undefined Saprotroph\| | Yeast | NULL | Probable | As nectar-tap_saprotroph (Põlme et al. 2020); Decay sugar-rich substrates (Põlme et al. 2020) | Põlme S, et al. 2020. Fungal Diversity 105:1-16 ((https://doi.org/10.1007/s13225-020-00466-2)) |
| Otu21618 | 1 | k__Fungi(100);p__Ascomycota(100);c__Saccharomycetes(100);o__Saccharomycetales(100);f__Metschnikowiaceae(100);g__Metschnikowia(100);s__Metschnikowia_pulcherrima(100); | Metschnikowia | 13 | Saprotroph | \|Undefined Saprotroph\| | Yeast | NULL | Probable | As nectar-tap_saprotroph (Põlme et al. 2020); Decay sugar-rich substrates (Põlme et al. 2020) | Põlme S, et al. 2020. Fungal Diversity 105:1-16 ((https://doi.org/10.1007/s13225-020-00466-2)) |
| Otu21619 | 1 | k__Fungi(100);p__Ascomycota(100);c__Saccharomycetes(100);o__Saccharomycetales(100);f__Metschnikowiaceae(100);g__Metschnikowia(100);g__Metschnikowia_unclassified(100); | Metschnikowia | 13 | Saprotroph | \|Undefined Saprotroph\| | Yeast | NULL | Probable | As nectar-tap_saprotroph (Põlme et al. 2020); Decay sugar-rich substrates (Põlme et al. 2020) | Põlme S, et al. 2020. Fungal Diversity 105:1-16 ((https://doi.org/10.1007/s13225-020-00466-2)) |
| Otu21620 | 1 | k__Fungi(100);p__Ascomycota(100);c__Saccharomycetes(100);o__Saccharomycetales(100);f__Metschnikowiaceae(100);g__Metschnikowia(100);s__Metschnikowia_pulcherrima(100); | Metschnikowia | 13 | Saprotroph | \|Undefined Saprotroph\| | Yeast | NULL | Probable | As nectar-tap_saprotroph (Põlme et al. 2020); Decay sugar-rich substrates (Põlme et al. 2020) | Põlme S, et al. 2020. Fungal Diversity 105:1-16 ((https://doi.org/10.1007/s13225-020-00466-2)) |
| Otu21621 | 1 | k__Fungi(100);p__Ascomycota(100);c__Saccharomycetes(100);o__Saccharomycetales(100);f__Metschnikowiaceae(100);g__Metschnikowia(100);s__Metschnikowia_pulcherrima(100); | Metschnikowia | 13 | Saprotroph | \|Undefined Saprotroph\| | Yeast | NULL | Probable | As nectar-tap_saprotroph (Põlme et al. 2020); Decay sugar-rich substrates (Põlme et al. 2020) | Põlme S, et al. 2020. Fungal Diversity 105:1-16 ((https://doi.org/10.1007/s13225-020-00466-2)) |
| Otu21622 | 1 | k__Fungi(100);p__Ascomycota(100);c__Saccharomycetes(100);o__Saccharomycetales(100);f__Metschnikowiaceae(100);g__Metschnikowia(100);s__Metschnikowia_pulcherrima(100); | Metschnikowia | 13 | Saprotroph | \|Undefined Saprotroph\| | Yeast | NULL | Probable | As nectar-tap_saprotroph (Põlme et al. 2020); Decay sugar-rich substrates (Põlme et al. 2020) | Põlme S, et al. 2020. Fungal Diversity 105:1-16 ((https://doi.org/10.1007/s13225-020-00466-2)) |
| Otu21623 | 1 | k__Fungi(100);p__Ascomycota(100);c__Saccharomycetes(100);o__Saccharomycetales(100);f__Metschnikowiaceae(100);g__Metschnikowia(100);g__Metschnikowia_unclassified(100); | Metschnikowia | 13 | Saprotroph | \|Undefined Saprotroph\| | Yeast | NULL | Probable | As nectar-tap_saprotroph (Põlme et al. 2020); Decay sugar-rich substrates (Põlme et al. 2020) | Põlme S, et al. 2020. Fungal Diversity 105:1-16 ((https://doi.org/10.1007/s13225-020-00466-2)) |
| Otu21624 | 1 | k__Fungi(100);p__Ascomycota(100);c__Saccharomycetes(100);o__Saccharomycetales(100);f__Metschnikowiaceae(100);g__Metschnikowia(100);s__Metschnikowia_pulcherrima(100); | Metschnikowia | 13 | Saprotroph | \|Undefined Saprotroph\| | Yeast | NULL | Probable | As nectar-tap_saprotroph (Põlme et al. 2020); Decay sugar-rich substrates (Põlme et al. 2020) | Põlme S, et al. 2020. Fungal Diversity 105:1-16 ((https://doi.org/10.1007/s13225-020-00466-2)) |
| Otu21625 | 1 | k__Fungi(100);p__Ascomycota(100);c__Saccharomycetes(100);o__Saccharomycetales(100);f__Metschnikowiaceae(100);g__Metschnikowia(100);s__Metschnikowia_pulcherrima(100); | Metschnikowia | 13 | Saprotroph | \|Undefined Saprotroph\| | Yeast | NULL | Probable | As nectar-tap_saprotroph (Põlme et al. 2020); Decay sugar-rich substrates (Põlme et al. 2020) | Põlme S, et al. 2020. Fungal Diversity 105:1-16 ((https://doi.org/10.1007/s13225-020-00466-2)) |
| Otu21626 | 1 | k__Fungi(100);p__Ascomycota(100);c__Saccharomycetes(100);o__Saccharomycetales(100);f__Metschnikowiaceae(100);g__Metschnikowia(100);s__Metschnikowia_pulcherrima(100); | Metschnikowia | 13 | Saprotroph | \|Undefined Saprotroph\| | Yeast | NULL | Probable | As nectar-tap_saprotroph (Põlme et al. 2020); Decay sugar-rich substrates (Põlme et al. 2020) | Põlme S, et al. 2020. Fungal Diversity 105:1-16 ((https://doi.org/10.1007/s13225-020-00466-2)) |
| Otu21627 | 1 | k__Fungi(100);p__Ascomycota(100);c__Saccharomycetes(100);o__Saccharomycetales(100);f__Metschnikowiaceae(100);g__Metschnikowia(100);g__Metschnikowia_unclassified(100); | Metschnikowia | 13 | Saprotroph | \|Undefined Saprotroph\| | Yeast | NULL | Probable | As nectar-tap_saprotroph (Põlme et al. 2020); Decay sugar-rich substrates (Põlme et al. 2020) | Põlme S, et al. 2020. Fungal Diversity 105:1-16 ((https://doi.org/10.1007/s13225-020-00466-2)) |
| Otu21628 | 1 | k__Fungi(100);p__Ascomycota(100);c__Saccharomycetes(100);o__Saccharomycetales(100);f__Metschnikowiaceae(100);g__Metschnikowia(100);s__Metschnikowia_pulcherrima(100); | Metschnikowia | 13 | Saprotroph | \|Undefined Saprotroph\| | Yeast | NULL | Probable | As nectar-tap_saprotroph (Põlme et al. 2020); Decay sugar-rich substrates (Põlme et al. 2020) | Põlme S, et al. 2020. Fungal Diversity 105:1-16 ((https://doi.org/10.1007/s13225-020-00466-2)) |
| Otu21629 | 1 | k__Fungi(100);p__Ascomycota(100);c__Saccharomycetes(100);o__Saccharomycetales(100);f__Metschnikowiaceae(100);g__Metschnikowia(100);g__Metschnikowia_unclassified(100); | Metschnikowia | 13 | Saprotroph | \|Undefined Saprotroph\| | Yeast | NULL | Probable | As nectar-tap_saprotroph (Põlme et al. 2020); Decay sugar-rich substrates (Põlme et al. 2020) | Põlme S, et al. 2020. Fungal Diversity 105:1-16 ((https://doi.org/10.1007/s13225-020-00466-2)) |
| Otu21630 | 1 | k__Fungi(100);p__Ascomycota(100);c__Saccharomycetes(100);o__Saccharomycetales(100);f__Metschnikowiaceae(100);g__Metschnikowia(100);g__Metschnikowia_unclassified(100); | Metschnikowia | 13 | Saprotroph | \|Undefined Saprotroph\| | Yeast | NULL | Probable | As nectar-tap_saprotroph (Põlme et al. 2020); Decay sugar-rich substrates (Põlme et al. 2020) | Põlme S, et al. 2020. Fungal Diversity 105:1-16 ((https://doi.org/10.1007/s13225-020-00466-2)) |
| Otu21631 | 1 | k__Fungi(100);p__Ascomycota(100);c__Saccharomycetes(100);o__Saccharomycetales(100);f__Metschnikowiaceae(100);g__Metschnikowia(100);s__Metschnikowia_pulcherrima(100); | Metschnikowia | 13 | Saprotroph | \|Undefined Saprotroph\| | Yeast | NULL | Probable | As nectar-tap_saprotroph (Põlme et al. 2020); Decay sugar-rich substrates (Põlme et al. 2020) | Põlme S, et al. 2020. Fungal Diversity 105:1-16 ((https://doi.org/10.1007/s13225-020-00466-2)) |
| Otu21632 | 1 | k__Fungi(100);p__Ascomycota(100);c__Saccharomycetes(100);o__Saccharomycetales(100);f__Metschnikowiaceae(100);g__Metschnikowia(100);s__Metschnikowia_pulcherrima(100); | Metschnikowia | 13 | Saprotroph | \|Undefined Saprotroph\| | Yeast | NULL | Probable | As nectar-tap_saprotroph (Põlme et al. 2020); Decay sugar-rich substrates (Põlme et al. 2020) | Põlme S, et al. 2020. Fungal Diversity 105:1-16 ((https://doi.org/10.1007/s13225-020-00466-2)) |
| Otu21633 | 1 | k__Fungi(100);p__Ascomycota(100);c__Saccharomycetes(100);o__Saccharomycetales(100);f__Metschnikowiaceae(100);g__Metschnikowia(100);s__Metschnikowia_pulcherrima(100); | Metschnikowia | 13 | Saprotroph | \|Undefined Saprotroph\| | Yeast | NULL | Probable | As nectar-tap_saprotroph (Põlme et al. 2020); Decay sugar-rich substrates (Põlme et al. 2020) | Põlme S, et al. 2020. Fungal Diversity 105:1-16 ((https://doi.org/10.1007/s13225-020-00466-2)) |
| Otu21634 | 1 | k__Fungi(100);p__Ascomycota(100);c__Saccharomycetes(100);o__Saccharomycetales(100);f__Metschnikowiaceae(100);g__Metschnikowia(100);s__Metschnikowia_pulcherrima(100); | Metschnikowia | 13 | Saprotroph | \|Undefined Saprotroph\| | Yeast | NULL | Probable | As nectar-tap_saprotroph (Põlme et al. 2020); Decay sugar-rich substrates (Põlme et al. 2020) | Põlme S, et al. 2020. Fungal Diversity 105:1-16 ((https://doi.org/10.1007/s13225-020-00466-2)) |
| Otu21635 | 1 | k__Fungi(100);p__Ascomycota(100);c__Saccharomycetes(100);o__Saccharomycetales(100);f__Metschnikowiaceae(100);g__Metschnikowia(100);s__Metschnikowia_pulcherrima(100); | Metschnikowia | 13 | Saprotroph | \|Undefined Saprotroph\| | Yeast | NULL | Probable | As nectar-tap_saprotroph (Põlme et al. 2020); Decay sugar-rich substrates (Põlme et al. 2020) | Põlme S, et al. 2020. Fungal Diversity 105:1-16 ((https://doi.org/10.1007/s13225-020-00466-2)) |
| Otu21636 | 1 | k__Fungi(100);p__Ascomycota(100);c__Saccharomycetes(100);o__Saccharomycetales(100);f__Metschnikowiaceae(100);g__Metschnikowia(100);s__Metschnikowia_pulcherrima(100); | Metschnikowia | 13 | Saprotroph | \|Undefined Saprotroph\| | Yeast | NULL | Probable | As nectar-tap_saprotroph (Põlme et al. 2020); Decay sugar-rich substrates (Põlme et al. 2020) | Põlme S, et al. 2020. Fungal Diversity 105:1-16 ((https://doi.org/10.1007/s13225-020-00466-2)) |
| Otu21637 | 1 | k__Fungi(100);p__Ascomycota(100);c__Saccharomycetes(100);o__Saccharomycetales(100);f__Metschnikowiaceae(100);g__Metschnikowia(100);s__Metschnikowia_pulcherrima(100); | Metschnikowia | 13 | Saprotroph | \|Undefined Saprotroph\| | Yeast | NULL | Probable | As nectar-tap_saprotroph (Põlme et al. 2020); Decay sugar-rich substrates (Põlme et al. 2020) | Põlme S, et al. 2020. Fungal Diversity 105:1-16 ((https://doi.org/10.1007/s13225-020-00466-2)) |
| Otu21638 | 1 | k__Fungi(100);p__Ascomycota(100);c__Saccharomycetes(100);o__Saccharomycetales(100);f__Metschnikowiaceae(100);g__Metschnikowia(100);s__Metschnikowia_pulcherrima(100); | Metschnikowia | 13 | Saprotroph | \|Undefined Saprotroph\| | Yeast | NULL | Probable | As nectar-tap_saprotroph (Põlme et al. 2020); Decay sugar-rich substrates (Põlme et al. 2020) | Põlme S, et al. 2020. Fungal Diversity 105:1-16 ((https://doi.org/10.1007/s13225-020-00466-2)) |
| Otu21639 | 1 | k__Fungi(100);p__Ascomycota(100);c__Saccharomycetes(100);o__Saccharomycetales(100);f__Metschnikowiaceae(100);g__Metschnikowia(100);s__Metschnikowia_pulcherrima(100); | Metschnikowia | 13 | Saprotroph | \|Undefined Saprotroph\| | Yeast | NULL | Probable | As nectar-tap_saprotroph (Põlme et al. 2020); Decay sugar-rich substrates (Põlme et al. 2020) | Põlme S, et al. 2020. Fungal Diversity 105:1-16 ((https://doi.org/10.1007/s13225-020-00466-2)) |
| Otu21640 | 1 | k__Fungi(100);p__Ascomycota(100);c__Saccharomycetes(100);o__Saccharomycetales(100);f__Metschnikowiaceae(100);g__Metschnikowia(100);s__Metschnikowia_pulcherrima(100); | Metschnikowia | 13 | Saprotroph | \|Undefined Saprotroph\| | Yeast | NULL | Probable | As nectar-tap_saprotroph (Põlme et al. 2020); Decay sugar-rich substrates (Põlme et al. 2020) | Põlme S, et al. 2020. Fungal Diversity 105:1-16 ((https://doi.org/10.1007/s13225-020-00466-2)) |
| Otu21641 | 1 | k__Fungi(100);p__Ascomycota(100);c__Saccharomycetes(100);o__Saccharomycetales(100);f__Metschnikowiaceae(100);g__Metschnikowia(100);s__Metschnikowia_pulcherrima(100); | Metschnikowia | 13 | Saprotroph | \|Undefined Saprotroph\| | Yeast | NULL | Probable | As nectar-tap_saprotroph (Põlme et al. 2020); Decay sugar-rich substrates (Põlme et al. 2020) | Põlme S, et al. 2020. Fungal Diversity 105:1-16 ((https://doi.org/10.1007/s13225-020-00466-2)) |
| Otu21642 | 1 | k__Fungi(100);p__Ascomycota(100);c__Saccharomycetes(100);o__Saccharomycetales(100);f__Metschnikowiaceae(100);g__Metschnikowia(100);s__Metschnikowia_pulcherrima(100); | Metschnikowia | 13 | Saprotroph | \|Undefined Saprotroph\| | Yeast | NULL | Probable | As nectar-tap_saprotroph (Põlme et al. 2020); Decay sugar-rich substrates (Põlme et al. 2020) | Põlme S, et al. 2020. Fungal Diversity 105:1-16 ((https://doi.org/10.1007/s13225-020-00466-2)) |
| Otu21643 | 1 | k__Fungi(100);p__Ascomycota(100);c__Saccharomycetes(100);o__Saccharomycetales(100);f__Metschnikowiaceae(100);g__Metschnikowia(100);g__Metschnikowia_unclassified(100); | Metschnikowia | 13 | Saprotroph | \|Undefined Saprotroph\| | Yeast | NULL | Probable | As nectar-tap_saprotroph (Põlme et al. 2020); Decay sugar-rich substrates (Põlme et al. 2020) | Põlme S, et al. 2020. Fungal Diversity 105:1-16 ((https://doi.org/10.1007/s13225-020-00466-2)) |
| Otu21644 | 1 | k__Fungi(100);p__Ascomycota(100);c__Saccharomycetes(100);o__Saccharomycetales(100);f__Metschnikowiaceae(100);g__Metschnikowia(100);g__Metschnikowia_unclassified(100); | Metschnikowia | 13 | Saprotroph | \|Undefined Saprotroph\| | Yeast | NULL | Probable | As nectar-tap_saprotroph (Põlme et al. 2020); Decay sugar-rich substrates (Põlme et al. 2020) | Põlme S, et al. 2020. Fungal Diversity 105:1-16 ((https://doi.org/10.1007/s13225-020-00466-2)) |
| Otu21645 | 1 | k__Fungi(100);p__Ascomycota(100);c__Saccharomycetes(100);o__Saccharomycetales(100);f__Metschnikowiaceae(100);g__Metschnikowia(100);s__Metschnikowia_pulcherrima(100); | Metschnikowia | 13 | Saprotroph | \|Undefined Saprotroph\| | Yeast | NULL | Probable | As nectar-tap_saprotroph (Põlme et al. 2020); Decay sugar-rich substrates (Põlme et al. 2020) | Põlme S, et al. 2020. Fungal Diversity 105:1-16 ((https://doi.org/10.1007/s13225-020-00466-2)) |
| Otu21646 | 1 | k__Fungi(100);p__Ascomycota(100);c__Saccharomycetes(100);o__Saccharomycetales(100);f__Metschnikowiaceae(100);g__Metschnikowia(100);s__Metschnikowia_pulcherrima(100); | Metschnikowia | 13 | Saprotroph | \|Undefined Saprotroph\| | Yeast | NULL | Probable | As nectar-tap_saprotroph (Põlme et al. 2020); Decay sugar-rich substrates (Põlme et al. 2020) | Põlme S, et al. 2020. Fungal Diversity 105:1-16 ((https://doi.org/10.1007/s13225-020-00466-2)) |
| Otu21647 | 1 | k__Fungi(100);p__Ascomycota(100);c__Saccharomycetes(100);o__Saccharomycetales(100);f__Metschnikowiaceae(100);g__Metschnikowia(100);g__Metschnikowia_unclassified(100); | Metschnikowia | 13 | Saprotroph | \|Undefined Saprotroph\| | Yeast | NULL | Probable | As nectar-tap_saprotroph (Põlme et al. 2020); Decay sugar-rich substrates (Põlme et al. 2020) | Põlme S, et al. 2020. Fungal Diversity 105:1-16 ((https://doi.org/10.1007/s13225-020-00466-2)) |
| Otu21649 | 1 | k__Fungi(100);p__Ascomycota(100);c__Saccharomycetes(100);o__Saccharomycetales(100);f__Metschnikowiaceae(100);g__Metschnikowia(100);s__Metschnikowia_pulcherrima(100); | Metschnikowia | 13 | Saprotroph | \|Undefined Saprotroph\| | Yeast | NULL | Probable | As nectar-tap_saprotroph (Põlme et al. 2020); Decay sugar-rich substrates (Põlme et al. 2020) | Põlme S, et al. 2020. Fungal Diversity 105:1-16 ((https://doi.org/10.1007/s13225-020-00466-2)) |
| Otu21650 | 1 | k__Fungi(100);p__Ascomycota(100);c__Saccharomycetes(100);o__Saccharomycetales(100);f__Metschnikowiaceae(100);g__Metschnikowia(100);s__Metschnikowia_pulcherrima(100); | Metschnikowia | 13 | Saprotroph | \|Undefined Saprotroph\| | Yeast | NULL | Probable | As nectar-tap_saprotroph (Põlme et al. 2020); Decay sugar-rich substrates (Põlme et al. 2020) | Põlme S, et al. 2020. Fungal Diversity 105:1-16 ((https://doi.org/10.1007/s13225-020-00466-2)) |
| Otu21651 | 1 | k__Fungi(100);p__Ascomycota(100);c__Saccharomycetes(100);o__Saccharomycetales(100);f__Metschnikowiaceae(100);g__Metschnikowia(100);s__Metschnikowia_pulcherrima(100); | Metschnikowia | 13 | Saprotroph | \|Undefined Saprotroph\| | Yeast | NULL | Probable | As nectar-tap_saprotroph (Põlme et al. 2020); Decay sugar-rich substrates (Põlme et al. 2020) | Põlme S, et al. 2020. Fungal Diversity 105:1-16 ((https://doi.org/10.1007/s13225-020-00466-2)) |
| Otu21652 | 1 | k__Fungi(100);p__Ascomycota(100);c__Saccharomycetes(100);o__Saccharomycetales(100);f__Metschnikowiaceae(100);g__Metschnikowia(100);g__Metschnikowia_unclassified(100); | Metschnikowia | 13 | Saprotroph | \|Undefined Saprotroph\| | Yeast | NULL | Probable | As nectar-tap_saprotroph (Põlme et al. 2020); Decay sugar-rich substrates (Põlme et al. 2020) | Põlme S, et al. 2020. Fungal Diversity 105:1-16 ((https://doi.org/10.1007/s13225-020-00466-2)) |
| Otu21653 | 1 | k__Fungi(100);p__Ascomycota(100);c__Saccharomycetes(100);o__Saccharomycetales(100);f__Metschnikowiaceae(100);g__Metschnikowia(100);g__Metschnikowia_unclassified(100); | Metschnikowia | 13 | Saprotroph | \|Undefined Saprotroph\| | Yeast | NULL | Probable | As nectar-tap_saprotroph (Põlme et al. 2020); Decay sugar-rich substrates (Põlme et al. 2020) | Põlme S, et al. 2020. Fungal Diversity 105:1-16 ((https://doi.org/10.1007/s13225-020-00466-2)) |
| Otu21654 | 1 | k__Fungi(100);p__Ascomycota(100);c__Saccharomycetes(100);o__Saccharomycetales(100);f__Metschnikowiaceae(100);g__Metschnikowia(100);s__Metschnikowia_pulcherrima(100); | Metschnikowia | 13 | Saprotroph | \|Undefined Saprotroph\| | Yeast | NULL | Probable | As nectar-tap_saprotroph (Põlme et al. 2020); Decay sugar-rich substrates (Põlme et al. 2020) | Põlme S, et al. 2020. Fungal Diversity 105:1-16 ((https://doi.org/10.1007/s13225-020-00466-2)) |
| Otu21655 | 1 | k__Fungi(100);p__Ascomycota(100);c__Saccharomycetes(100);o__Saccharomycetales(100);f__Metschnikowiaceae(100);g__Metschnikowia(100);g__Metschnikowia_unclassified(100); | Metschnikowia | 13 | Saprotroph | \|Undefined Saprotroph\| | Yeast | NULL | Probable | As nectar-tap_saprotroph (Põlme et al. 2020); Decay sugar-rich substrates (Põlme et al. 2020) | Põlme S, et al. 2020. Fungal Diversity 105:1-16 ((https://doi.org/10.1007/s13225-020-00466-2)) |
| Otu21656 | 1 | k__Fungi(100);p__Ascomycota(100);c__Saccharomycetes(100);o__Saccharomycetales(100);f__Metschnikowiaceae(100);g__Metschnikowia(100);s__Metschnikowia_pulcherrima(100); | Metschnikowia | 13 | Saprotroph | \|Undefined Saprotroph\| | Yeast | NULL | Probable | As nectar-tap_saprotroph (Põlme et al. 2020); Decay sugar-rich substrates (Põlme et al. 2020) | Põlme S, et al. 2020. Fungal Diversity 105:1-16 ((https://doi.org/10.1007/s13225-020-00466-2)) |
| Otu21657 | 1 | k__Fungi(100);p__Ascomycota(100);c__Saccharomycetes(100);o__Saccharomycetales(100);f__Metschnikowiaceae(100);g__Metschnikowia(100);s__Metschnikowia_pulcherrima(100); | Metschnikowia | 13 | Saprotroph | \|Undefined Saprotroph\| | Yeast | NULL | Probable | As nectar-tap_saprotroph (Põlme et al. 2020); Decay sugar-rich substrates (Põlme et al. 2020) | Põlme S, et al. 2020. Fungal Diversity 105:1-16 ((https://doi.org/10.1007/s13225-020-00466-2)) |
| Otu21658 | 1 | k__Fungi(100);p__Ascomycota(100);c__Saccharomycetes(100);o__Saccharomycetales(100);f__Metschnikowiaceae(100);g__Metschnikowia(100);s__Metschnikowia_pulcherrima(100); | Metschnikowia | 13 | Saprotroph | \|Undefined Saprotroph\| | Yeast | NULL | Probable | As nectar-tap_saprotroph (Põlme et al. 2020); Decay sugar-rich substrates (Põlme et al. 2020) | Põlme S, et al. 2020. Fungal Diversity 105:1-16 ((https://doi.org/10.1007/s13225-020-00466-2)) |
| Otu21659 | 1 | k__Fungi(100);p__Ascomycota(100);c__Saccharomycetes(100);o__Saccharomycetales(100);f__Metschnikowiaceae(100);g__Metschnikowia(100);s__Metschnikowia_pulcherrima(100); | Metschnikowia | 13 | Saprotroph | \|Undefined Saprotroph\| | Yeast | NULL | Probable | As nectar-tap_saprotroph (Põlme et al. 2020); Decay sugar-rich substrates (Põlme et al. 2020) | Põlme S, et al. 2020. Fungal Diversity 105:1-16 ((https://doi.org/10.1007/s13225-020-00466-2)) |
| Otu21660 | 1 | k__Fungi(100);p__Ascomycota(100);c__Saccharomycetes(100);o__Saccharomycetales(100);f__Metschnikowiaceae(100);g__Metschnikowia(100);s__Metschnikowia_pulcherrima(100); | Metschnikowia | 13 | Saprotroph | \|Undefined Saprotroph\| | Yeast | NULL | Probable | As nectar-tap_saprotroph (Põlme et al. 2020); Decay sugar-rich substrates (Põlme et al. 2020) | Põlme S, et al. 2020. Fungal Diversity 105:1-16 ((https://doi.org/10.1007/s13225-020-00466-2)) |
| Otu21661 | 1 | k__Fungi(100);p__Ascomycota(100);c__Saccharomycetes(100);o__Saccharomycetales(100);f__Metschnikowiaceae(100);g__Metschnikowia(100);s__Metschnikowia_pulcherrima(100); | Metschnikowia | 13 | Saprotroph | \|Undefined Saprotroph\| | Yeast | NULL | Probable | As nectar-tap_saprotroph (Põlme et al. 2020); Decay sugar-rich substrates (Põlme et al. 2020) | Põlme S, et al. 2020. Fungal Diversity 105:1-16 ((https://doi.org/10.1007/s13225-020-00466-2)) |
| Otu21662 | 1 | k__Fungi(100);p__Ascomycota(100);c__Saccharomycetes(100);o__Saccharomycetales(100);f__Saccharomycetaceae(100);g__Kazachstania(100);s__Kazachstania_intestinalis(100); | Kazachstania | 13 | Saprotroph | \|Undefined Saprotroph\| | Yeast | NULL | Probable | As nectar-tap_saprotroph (Põlme et al. 2020); Decay sugar-rich substrates (Põlme et al. 2020) | Põlme S, et al. 2020. Fungal Diversity 105:1-16 ((https://doi.org/10.1007/s13225-020-00466-2)) |
| Otu21663 | 1 | k__Fungi(100);p__Ascomycota(100);c__Saccharomycetes(100);o__Saccharomycetales(100);f__Metschnikowiaceae(100);g__Metschnikowia(100);s__Metschnikowia_pulcherrima(100); | Metschnikowia | 13 | Saprotroph | \|Undefined Saprotroph\| | Yeast | NULL | Probable | As nectar-tap_saprotroph (Põlme et al. 2020); Decay sugar-rich substrates (Põlme et al. 2020) | Põlme S, et al. 2020. Fungal Diversity 105:1-16 ((https://doi.org/10.1007/s13225-020-00466-2)) |
| Otu21664 | 1 | k__Fungi(100);p__Ascomycota(100);c__Saccharomycetes(100);o__Saccharomycetales(100);f__Metschnikowiaceae(100);g__Metschnikowia(100);g__Metschnikowia_unclassified(100); | Metschnikowia | 13 | Saprotroph | \|Undefined Saprotroph\| | Yeast | NULL | Probable | As nectar-tap_saprotroph (Põlme et al. 2020); Decay sugar-rich substrates (Põlme et al. 2020) | Põlme S, et al. 2020. Fungal Diversity 105:1-16 ((https://doi.org/10.1007/s13225-020-00466-2)) |
| Otu21665 | 1 | k__Fungi(100);p__Ascomycota(100);c__Saccharomycetes(100);o__Saccharomycetales(100);f__Metschnikowiaceae(100);g__Metschnikowia(100);s__Metschnikowia_pulcherrima(100); | Metschnikowia | 13 | Saprotroph | \|Undefined Saprotroph\| | Yeast | NULL | Probable | As nectar-tap_saprotroph (Põlme et al. 2020); Decay sugar-rich substrates (Põlme et al. 2020) | Põlme S, et al. 2020. Fungal Diversity 105:1-16 ((https://doi.org/10.1007/s13225-020-00466-2)) |
| Otu21666 | 1 | k__Fungi(100);p__Ascomycota(100);c__Saccharomycetes(100);o__Saccharomycetales(100);f__Metschnikowiaceae(100);g__Metschnikowia(100);g__Metschnikowia_unclassified(100); | Metschnikowia | 13 | Saprotroph | \|Undefined Saprotroph\| | Yeast | NULL | Probable | As nectar-tap_saprotroph (Põlme et al. 2020); Decay sugar-rich substrates (Põlme et al. 2020) | Põlme S, et al. 2020. Fungal Diversity 105:1-16 ((https://doi.org/10.1007/s13225-020-00466-2)) |
| Otu21667 | 1 | k__Fungi(100);p__Ascomycota(100);c__Saccharomycetes(100);o__Saccharomycetales(100);f__Metschnikowiaceae(100);g__Metschnikowia(100);s__Metschnikowia_pulcherrima(100); | Metschnikowia | 13 | Saprotroph | \|Undefined Saprotroph\| | Yeast | NULL | Probable | As nectar-tap_saprotroph (Põlme et al. 2020); Decay sugar-rich substrates (Põlme et al. 2020) | Põlme S, et al. 2020. Fungal Diversity 105:1-16 ((https://doi.org/10.1007/s13225-020-00466-2)) |
| Otu21668 | 1 | k__Fungi(100);p__Ascomycota(100);c__Saccharomycetes(100);o__Saccharomycetales(100);f__Metschnikowiaceae(100);g__Metschnikowia(100);s__Metschnikowia_pulcherrima(100); | Metschnikowia | 13 | Saprotroph | \|Undefined Saprotroph\| | Yeast | NULL | Probable | As nectar-tap_saprotroph (Põlme et al. 2020); Decay sugar-rich substrates (Põlme et al. 2020) | Põlme S, et al. 2020. Fungal Diversity 105:1-16 ((https://doi.org/10.1007/s13225-020-00466-2)) |
| Otu21669 | 1 | k__Fungi(100);p__Ascomycota(100);c__Saccharomycetes(100);o__Saccharomycetales(100);f__Metschnikowiaceae(100);g__Metschnikowia(100);s__Metschnikowia_pulcherrima(100); | Metschnikowia | 13 | Saprotroph | \|Undefined Saprotroph\| | Yeast | NULL | Probable | As nectar-tap_saprotroph (Põlme et al. 2020); Decay sugar-rich substrates (Põlme et al. 2020) | Põlme S, et al. 2020. Fungal Diversity 105:1-16 ((https://doi.org/10.1007/s13225-020-00466-2)) |
| Otu21671 | 1 | k__Fungi(100);p__Ascomycota(100);c__Saccharomycetes(100);o__Saccharomycetales(100);f__Metschnikowiaceae(100);g__Metschnikowia(100);s__Metschnikowia_pulcherrima(100); | Metschnikowia | 13 | Saprotroph | \|Undefined Saprotroph\| | Yeast | NULL | Probable | As nectar-tap_saprotroph (Põlme et al. 2020); Decay sugar-rich substrates (Põlme et al. 2020) | Põlme S, et al. 2020. Fungal Diversity 105:1-16 ((https://doi.org/10.1007/s13225-020-00466-2)) |
| Otu21673 | 1 | k__Fungi(100);p__Ascomycota(100);c__Saccharomycetes(100);o__Saccharomycetales(100);f__Metschnikowiaceae(100);g__Metschnikowia(100);g__Metschnikowia_unclassified(100); | Metschnikowia | 13 | Saprotroph | \|Undefined Saprotroph\| | Yeast | NULL | Probable | As nectar-tap_saprotroph (Põlme et al. 2020); Decay sugar-rich substrates (Põlme et al. 2020) | Põlme S, et al. 2020. Fungal Diversity 105:1-16 ((https://doi.org/10.1007/s13225-020-00466-2)) |
| Otu21674 | 1 | k__Fungi(100);p__Ascomycota(100);c__Saccharomycetes(100);o__Saccharomycetales(100);f__Metschnikowiaceae(100);g__Metschnikowia(100);s__Metschnikowia_pulcherrima(100); | Metschnikowia | 13 | Saprotroph | \|Undefined Saprotroph\| | Yeast | NULL | Probable | As nectar-tap_saprotroph (Põlme et al. 2020); Decay sugar-rich substrates (Põlme et al. 2020) | Põlme S, et al. 2020. Fungal Diversity 105:1-16 ((https://doi.org/10.1007/s13225-020-00466-2)) |
| Otu21675 | 1 | k__Fungi(100);p__Ascomycota(100);c__Saccharomycetes(100);o__Saccharomycetales(100);f__Metschnikowiaceae(100);g__Metschnikowia(100);s__Metschnikowia_pulcherrima(100); | Metschnikowia | 13 | Saprotroph | \|Undefined Saprotroph\| | Yeast | NULL | Probable | As nectar-tap_saprotroph (Põlme et al. 2020); Decay sugar-rich substrates (Põlme et al. 2020) | Põlme S, et al. 2020. Fungal Diversity 105:1-16 ((https://doi.org/10.1007/s13225-020-00466-2)) |
| Otu21676 | 1 | k__Fungi(100);p__Ascomycota(100);c__Saccharomycetes(100);o__Saccharomycetales(100);f__Metschnikowiaceae(100);g__Metschnikowia(100);s__Metschnikowia_pulcherrima(100); | Metschnikowia | 13 | Saprotroph | \|Undefined Saprotroph\| | Yeast | NULL | Probable | As nectar-tap_saprotroph (Põlme et al. 2020); Decay sugar-rich substrates (Põlme et al. 2020) | Põlme S, et al. 2020. Fungal Diversity 105:1-16 ((https://doi.org/10.1007/s13225-020-00466-2)) |
| Otu21678 | 1 | k__Fungi(100);p__Ascomycota(100);c__Saccharomycetes(100);o__Saccharomycetales(100);f__Metschnikowiaceae(100);g__Metschnikowia(100);g__Metschnikowia_unclassified(100); | Metschnikowia | 13 | Saprotroph | \|Undefined Saprotroph\| | Yeast | NULL | Probable | As nectar-tap_saprotroph (Põlme et al. 2020); Decay sugar-rich substrates (Põlme et al. 2020) | Põlme S, et al. 2020. Fungal Diversity 105:1-16 ((https://doi.org/10.1007/s13225-020-00466-2)) |
| Otu21680 | 1 | k__Fungi(100);p__Ascomycota(100);c__Saccharomycetes(100);o__Saccharomycetales(100);f__Metschnikowiaceae(100);g__Metschnikowia(100);s__Metschnikowia_pulcherrima(100); | Metschnikowia | 13 | Saprotroph | \|Undefined Saprotroph\| | Yeast | NULL | Probable | As nectar-tap_saprotroph (Põlme et al. 2020); Decay sugar-rich substrates (Põlme et al. 2020) | Põlme S, et al. 2020. Fungal Diversity 105:1-16 ((https://doi.org/10.1007/s13225-020-00466-2)) |
| Otu21681 | 1 | k__Fungi(100);p__Ascomycota(100);c__Saccharomycetes(100);o__Saccharomycetales(100);f__Metschnikowiaceae(100);g__Metschnikowia(100);g__Metschnikowia_unclassified(100); | Metschnikowia | 13 | Saprotroph | \|Undefined Saprotroph\| | Yeast | NULL | Probable | As nectar-tap_saprotroph (Põlme et al. 2020); Decay sugar-rich substrates (Põlme et al. 2020) | Põlme S, et al. 2020. Fungal Diversity 105:1-16 ((https://doi.org/10.1007/s13225-020-00466-2)) |
| Otu21683 | 1 | k__Fungi(100);p__Ascomycota(100);c__Saccharomycetes(100);o__Saccharomycetales(100);f__Metschnikowiaceae(100);g__Metschnikowia(100);s__Metschnikowia_pulcherrima(100); | Metschnikowia | 13 | Saprotroph | \|Undefined Saprotroph\| | Yeast | NULL | Probable | As nectar-tap_saprotroph (Põlme et al. 2020); Decay sugar-rich substrates (Põlme et al. 2020) | Põlme S, et al. 2020. Fungal Diversity 105:1-16 ((https://doi.org/10.1007/s13225-020-00466-2)) |
| Otu21684 | 1 | k__Fungi(100);p__Ascomycota(100);c__Saccharomycetes(100);o__Saccharomycetales(100);f__Metschnikowiaceae(100);g__Metschnikowia(100);s__Metschnikowia_pulcherrima(100); | Metschnikowia | 13 | Saprotroph | \|Undefined Saprotroph\| | Yeast | NULL | Probable | As nectar-tap_saprotroph (Põlme et al. 2020); Decay sugar-rich substrates (Põlme et al. 2020) | Põlme S, et al. 2020. Fungal Diversity 105:1-16 ((https://doi.org/10.1007/s13225-020-00466-2)) |
| Otu21685 | 1 | k__Fungi(100);p__Ascomycota(100);c__Saccharomycetes(100);o__Saccharomycetales(100);f__Metschnikowiaceae(100);g__Metschnikowia(100);s__Metschnikowia_pulcherrima(100); | Metschnikowia | 13 | Saprotroph | \|Undefined Saprotroph\| | Yeast | NULL | Probable | As nectar-tap_saprotroph (Põlme et al. 2020); Decay sugar-rich substrates (Põlme et al. 2020) | Põlme S, et al. 2020. Fungal Diversity 105:1-16 ((https://doi.org/10.1007/s13225-020-00466-2)) |
| Otu21686 | 1 | k__Fungi(100);p__Ascomycota(100);c__Saccharomycetes(100);o__Saccharomycetales(100);f__Metschnikowiaceae(100);g__Metschnikowia(100);s__Metschnikowia_pulcherrima(100); | Metschnikowia | 13 | Saprotroph | \|Undefined Saprotroph\| | Yeast | NULL | Probable | As nectar-tap_saprotroph (Põlme et al. 2020); Decay sugar-rich substrates (Põlme et al. 2020) | Põlme S, et al. 2020. Fungal Diversity 105:1-16 ((https://doi.org/10.1007/s13225-020-00466-2)) |
| Otu21687 | 1 | k__Fungi(100);p__Ascomycota(100);c__Saccharomycetes(100);o__Saccharomycetales(100);f__Metschnikowiaceae(100);g__Metschnikowia(100);s__Metschnikowia_pulcherrima(100); | Metschnikowia | 13 | Saprotroph | \|Undefined Saprotroph\| | Yeast | NULL | Probable | As nectar-tap_saprotroph (Põlme et al. 2020); Decay sugar-rich substrates (Põlme et al. 2020) | Põlme S, et al. 2020. Fungal Diversity 105:1-16 ((https://doi.org/10.1007/s13225-020-00466-2)) |
| Otu21688 | 1 | k__Fungi(100);p__Ascomycota(100);c__Saccharomycetes(100);o__Saccharomycetales(100);f__Metschnikowiaceae(100);g__Metschnikowia(100);s__Metschnikowia_pulcherrima(100); | Metschnikowia | 13 | Saprotroph | \|Undefined Saprotroph\| | Yeast | NULL | Probable | As nectar-tap_saprotroph (Põlme et al. 2020); Decay sugar-rich substrates (Põlme et al. 2020) | Põlme S, et al. 2020. Fungal Diversity 105:1-16 ((https://doi.org/10.1007/s13225-020-00466-2)) |
| Otu21689 | 1 | k__Fungi(100);p__Ascomycota(100);c__Saccharomycetes(100);o__Saccharomycetales(100);f__Metschnikowiaceae(100);g__Metschnikowia(100);s__Metschnikowia_pulcherrima(100); | Metschnikowia | 13 | Saprotroph | \|Undefined Saprotroph\| | Yeast | NULL | Probable | As nectar-tap_saprotroph (Põlme et al. 2020); Decay sugar-rich substrates (Põlme et al. 2020) | Põlme S, et al. 2020. Fungal Diversity 105:1-16 ((https://doi.org/10.1007/s13225-020-00466-2)) |
| Otu21690 | 1 | k__Fungi(100);p__Ascomycota(100);c__Saccharomycetes(100);o__Saccharomycetales(100);f__Metschnikowiaceae(100);g__Metschnikowia(100);s__Metschnikowia_pulcherrima(100); | Metschnikowia | 13 | Saprotroph | \|Undefined Saprotroph\| | Yeast | NULL | Probable | As nectar-tap_saprotroph (Põlme et al. 2020); Decay sugar-rich substrates (Põlme et al. 2020) | Põlme S, et al. 2020. Fungal Diversity 105:1-16 ((https://doi.org/10.1007/s13225-020-00466-2)) |
| Otu21691 | 1 | k__Fungi(100);p__Ascomycota(100);c__Saccharomycetes(100);o__Saccharomycetales(100);f__Metschnikowiaceae(100);g__Metschnikowia(100);g__Metschnikowia_unclassified(100); | Metschnikowia | 13 | Saprotroph | \|Undefined Saprotroph\| | Yeast | NULL | Probable | As nectar-tap_saprotroph (Põlme et al. 2020); Decay sugar-rich substrates (Põlme et al. 2020) | Põlme S, et al. 2020. Fungal Diversity 105:1-16 ((https://doi.org/10.1007/s13225-020-00466-2)) |
| Otu21692 | 1 | k__Fungi(100);p__Ascomycota(100);c__Saccharomycetes(100);o__Saccharomycetales(100);f__Metschnikowiaceae(100);g__Metschnikowia(100);g__Metschnikowia_unclassified(100); | Metschnikowia | 13 | Saprotroph | \|Undefined Saprotroph\| | Yeast | NULL | Probable | As nectar-tap_saprotroph (Põlme et al. 2020); Decay sugar-rich substrates (Põlme et al. 2020) | Põlme S, et al. 2020. Fungal Diversity 105:1-16 ((https://doi.org/10.1007/s13225-020-00466-2)) |
| Otu21693 | 1 | k__Fungi(100);p__Ascomycota(100);c__Saccharomycetes(100);o__Saccharomycetales(100);f__Metschnikowiaceae(100);g__Metschnikowia(100);s__Metschnikowia_pulcherrima(100); | Metschnikowia | 13 | Saprotroph | \|Undefined Saprotroph\| | Yeast | NULL | Probable | As nectar-tap_saprotroph (Põlme et al. 2020); Decay sugar-rich substrates (Põlme et al. 2020) | Põlme S, et al. 2020. Fungal Diversity 105:1-16 ((https://doi.org/10.1007/s13225-020-00466-2)) |
| Otu21694 | 1 | k__Fungi(100);p__Ascomycota(100);c__Saccharomycetes(100);o__Saccharomycetales(100);f__Metschnikowiaceae(100);g__Metschnikowia(100);s__Metschnikowia_pulcherrima(100); | Metschnikowia | 13 | Saprotroph | \|Undefined Saprotroph\| | Yeast | NULL | Probable | As nectar-tap_saprotroph (Põlme et al. 2020); Decay sugar-rich substrates (Põlme et al. 2020) | Põlme S, et al. 2020. Fungal Diversity 105:1-16 ((https://doi.org/10.1007/s13225-020-00466-2)) |
| Otu21696 | 1 | k__Fungi(100);p__Ascomycota(100);c__Saccharomycetes(100);o__Saccharomycetales(100);f__Metschnikowiaceae(100);g__Metschnikowia(100);s__Metschnikowia_pulcherrima(100); | Metschnikowia | 13 | Saprotroph | \|Undefined Saprotroph\| | Yeast | NULL | Probable | As nectar-tap_saprotroph (Põlme et al. 2020); Decay sugar-rich substrates (Põlme et al. 2020) | Põlme S, et al. 2020. Fungal Diversity 105:1-16 ((https://doi.org/10.1007/s13225-020-00466-2)) |
| Otu21697 | 1 | k__Fungi(100);p__Ascomycota(100);c__Saccharomycetes(100);o__Saccharomycetales(100);f__Metschnikowiaceae(100);g__Metschnikowia(100);g__Metschnikowia_unclassified(100); | Metschnikowia | 13 | Saprotroph | \|Undefined Saprotroph\| | Yeast | NULL | Probable | As nectar-tap_saprotroph (Põlme et al. 2020); Decay sugar-rich substrates (Põlme et al. 2020) | Põlme S, et al. 2020. Fungal Diversity 105:1-16 ((https://doi.org/10.1007/s13225-020-00466-2)) |
| Otu21698 | 1 | k__Fungi(100);p__Ascomycota(100);c__Saccharomycetes(100);o__Saccharomycetales(100);f__Metschnikowiaceae(100);g__Metschnikowia(100);g__Metschnikowia_unclassified(100); | Metschnikowia | 13 | Saprotroph | \|Undefined Saprotroph\| | Yeast | NULL | Probable | As nectar-tap_saprotroph (Põlme et al. 2020); Decay sugar-rich substrates (Põlme et al. 2020) | Põlme S, et al. 2020. Fungal Diversity 105:1-16 ((https://doi.org/10.1007/s13225-020-00466-2)) |
| Otu21699 | 1 | k__Fungi(100);p__Ascomycota(100);c__Saccharomycetes(100);o__Saccharomycetales(100);f__Metschnikowiaceae(100);g__Metschnikowia(100);g__Metschnikowia_unclassified(100); | Metschnikowia | 13 | Saprotroph | \|Undefined Saprotroph\| | Yeast | NULL | Probable | As nectar-tap_saprotroph (Põlme et al. 2020); Decay sugar-rich substrates (Põlme et al. 2020) | Põlme S, et al. 2020. Fungal Diversity 105:1-16 ((https://doi.org/10.1007/s13225-020-00466-2)) |
| Otu21700 | 1 | k__Fungi(100);p__Ascomycota(100);c__Saccharomycetes(100);o__Saccharomycetales(100);f__Saccharomycetales_family_Incertae_sedis(100);g__Candida(100);g__Candida_unclassified(100); | Candida | 13 | Pathotroph-Saprotroph-Symbiotroph | Animal Pathogen-Endophyte-Endosymbiont-Epiphyte-\|Undefined Saprotroph\| | Dimorphic Yeast-Yeast | NULL | Probable | As nectar-tap_saprotroph (Põlme et al. 2020); Decay sugar-rich substrates (Põlme et al. 2020); This asexual genus is polyphyletic and contain species that function very differently depend on what they are. Caution should be taken when interpreting guilds for Candida species. | Manolakaki D, et al. 2010. Virulence. 1: 367-375 ((https://doi.org/10.4161/viru.1.5.12796)); Mekbib SB, et al. 2011. Tropical Plant Pathology 36:233-240 ((https://doi.org/10.1590/S1982-56762011000400004)); Põlme S, et al. 2020. Fungal Diversity 105:1-16 ((https://doi.org/10.1007/s13225-020-00466-2)) |
| Otu21701 | 1 | k__Fungi(100);p__Ascomycota(100);c__Saccharomycetes(100);o__Saccharomycetales(100);f__Saccharomycetales_family_Incertae_sedis(100);f__Saccharomycetales_family_Incertae_sedis_unclassified(100);f__Saccharomycetales_family_Incertae_sedis_unclassified(100); | Saccharomycetales | 7 | Saprotroph | Undefined Saprotroph | Yeast | NULL | Possible | NULL | Sterkenburg E, et al. 2015. New Phytologist 207:1145-1158 ((https://doi.org/10.1111/nph.13426)) |
| Otu21702 | 1 | k__Fungi(100);p__Ascomycota(100);c__Saccharomycetes(100);o__Saccharomycetales(100);f__Metschnikowiaceae(100);g__Metschnikowia(100);s__Metschnikowia_pulcherrima(100); | Metschnikowia | 13 | Saprotroph | \|Undefined Saprotroph\| | Yeast | NULL | Probable | As nectar-tap_saprotroph (Põlme et al. 2020); Decay sugar-rich substrates (Põlme et al. 2020) | Põlme S, et al. 2020. Fungal Diversity 105:1-16 ((https://doi.org/10.1007/s13225-020-00466-2)) |
| Otu21703 | 1 | k__Fungi(100);p__Ascomycota(100);c__Saccharomycetes(100);o__Saccharomycetales(100);o__Saccharomycetales_unclassified(100);o__Saccharomycetales_unclassified(100);o__Saccharomycetales_unclassified(100); | Saccharomycetales | 7 | Saprotroph | Undefined Saprotroph | Yeast | NULL | Possible | NULL | Sterkenburg E, et al. 2015. New Phytologist 207:1145-1158 ((https://doi.org/10.1111/nph.13426)) |

| | | | | | | | | | | | |
|---|---|---|---|---|---|---|---|---|---|---|---|
| Otu21704 | 1 | k__Fungi(100);p__Ascomycota(100);c__Saccharomycetes(100);o__Saccharomycetales(100);f__Saccharomycetales_family_Incertae_sedis(100);g__Candida(100);s__Candida_cretensis(100); | Candida | 13 | Pathotroph-Saprotroph-Symbiotroph | Animal Pathogen-Endophyte-Endosymbiont-Epiphyte-\|Undefined Saprotroph\| | Dimorphic Yeast-Yeast | NULL | Probable | As nectar-tap_saprotroph (Põlme et al. 2020); Decay sugar-rich substrates (Põlme et al. 2020); This asexual genus is polyphyletic and contain species that function very differently depend on what they are. Caution should be taken when interpreting guilds for Candida species. | Manolakaki D, et al. 2010. Virulence. 1: 367-375 ((https://doi.org/10.4161/viru.1.5.12796)); Mekbib SB, et al. 2011. Tropical Plant Pathology 36:233-240 ((https://doi.org/10.1590/S1982-56762011000400004)); Põlme S, et al. 2020. Fungal Diversity 105:1-16 ((https://doi.org/10.1007/s13225-020-00466-2)) |
| Otu21707 | 1 | k__Fungi(100);p__Ascomycota(100);c__Saccharomycetes(100);o__Saccharomycetales(100);f__Saccharomycetales_family_Incertae_sedis(100);f__Saccharomycetales_family_Incertae_sedis_unclassified(100);f__Saccharomycetales_family_Incertae_sedis_unclassified(100); | Saccharomycetales | 7 | Saprotroph | Undefined Saprotroph | Yeast | NULL | Possible | NULL | Sterkenburg E, et al. 2015. New Phytologist 207:1145-1158 ((https://doi.org/10.1111/nph.13426)) |
| Otu21708 | 1 | k__Fungi(100);p__Ascomycota(100);c__Saccharomycetes(100);o__Saccharomycetales(100);f__Metschnikowiaceae(100);g__Metschnikowia(100);s__Metschnikowia_pulcherrima(100); | Metschnikowia | 13 | Saprotroph | \|Undefined Saprotroph\| | Yeast | NULL | Probable | As nectar-tap_saprotroph (Põlme et al. 2020); Decay sugar-rich substrates (Põlme et al. 2020) | Põlme S, et al. 2020. Fungal Diversity 105:1-16 ((https://doi.org/10.1007/s13225-020-00466-2)) |
| Otu21709 | 1 | k__Fungi(100);p__Ascomycota(100);c__Saccharomycetes(100);o__Saccharomycetales(100);f__Metschnikowiaceae(100);g__Metschnikowia(100);s__Metschnikowia_pulcherrima(100); | Metschnikowia | 13 | Saprotroph | \|Undefined Saprotroph\| | Yeast | NULL | Probable | As nectar-tap_saprotroph (Põlme et al. 2020); Decay sugar-rich substrates (Põlme et al. 2020) | Põlme S, et al. 2020. Fungal Diversity 105:1-16 ((https://doi.org/10.1007/s13225-020-00466-2)) |
| Otu21714 | 1 | k__Fungi(100);p__Ascomycota(100);c__Saccharomycetes(100);o__Saccharomycetales(100);f__Saccharomycetales_family_Incertae_sedis(100);g__Candida(100);g__Candida_unclassified(100); | Candida | 13 | Pathotroph-Saprotroph-Symbiotroph | Animal Pathogen-Endophyte-Endosymbiont-Epiphyte-\|Undefined Saprotroph\| | Dimorphic Yeast-Yeast | NULL | Probable | As nectar-tap_saprotroph (Põlme et al. 2020); Decay sugar-rich substrates (Põlme et al. 2020); This asexual genus is polyphyletic and contain species that function very differently depend on what they are. Caution should be taken when interpreting guilds for Candida species. | Manolakaki D, et al. 2010. Virulence. 1: 367-375 ((https://doi.org/10.4161/viru.1.5.12796)); Mekbib SB, et al. 2011. Tropical Plant Pathology 36:233-240 ((https://doi.org/10.1590/S1982-56762011000400004)); Põlme S, et al. 2020. Fungal Diversity 105:1-16 ((https://doi.org/10.1007/s13225-020-00466-2)) |
| Otu21715 | 1 | k__Fungi(100);p__Ascomycota(100);c__Saccharomycetes(100);o__Saccharomycetales(100);f__Saccharomycetales_family_Incertae_sedis(100);f__Saccharomycetales_family_Incertae_sedis_unclassified(100);f__Saccharomycetales_family_Incertae_sedis_unclassified(100); | Saccharomycetales | 7 | Saprotroph | Undefined Saprotroph | Yeast | NULL | Possible | NULL | Sterkenburg E, et al. 2015. New Phytologist 207:1145-1158 ((https://doi.org/10.1111/nph.13426)) |
| Otu21716 | 1 | k__Fungi(100);p__Ascomycota(100);c__Saccharomycetes(100);o__Saccharomycetales(100);f__Metschnikowiaceae(100);g__Metschnikowia(100);s__Metschnikowia_pulcherrima(100); | Metschnikowia | 13 | Saprotroph | \|Undefined Saprotroph\| | Yeast | NULL | Probable | As nectar-tap_saprotroph (Põlme et al. 2020); Decay sugar-rich substrates (Põlme et al. 2020) | Põlme S, et al. 2020. Fungal Diversity 105:1-16 ((https://doi.org/10.1007/s13225-020-00466-2)) |
| Otu21717 | 1 | k__Fungi(100);p__Ascomycota(100);c__Saccharomycetes(100);o__Saccharomycetales(100);f__Metschnikowiaceae(100);g__Metschnikowia(100);s__Metschnikowia_pulcherrima(100); | Metschnikowia | 13 | Saprotroph | \|Undefined Saprotroph\| | Yeast | NULL | Probable | As nectar-tap_saprotroph (Põlme et al. 2020); Decay sugar-rich substrates (Põlme et al. 2020) | Põlme S, et al. 2020. Fungal Diversity 105:1-16 ((https://doi.org/10.1007/s13225-020-00466-2)) |
| Otu21718 | 1 | k__Fungi(100);p__Ascomycota(100);c__Saccharomycetes(100);o__Saccharomycetales(100);f__Saccharomycetales_family_Incertae_sedis(100);g__Candida(100);s__Candida_cretensis(100); | Candida | 13 | Pathotroph-Saprotroph-Symbiotroph | Animal Pathogen-Endophyte-Endosymbiont-Epiphyte-\|Undefined Saprotroph\| | Dimorphic Yeast-Yeast | NULL | Probable | As nectar-tap_saprotroph (Põlme et al. 2020); Decay sugar-rich substrates (Põlme et al. 2020); This asexual genus is polyphyletic and contain species that function very differently depend on what they are. Caution should be taken when interpreting guilds for Candida species. | Manolakaki D, et al. 2010. Virulence. 1: 367-375 ((https://doi.org/10.4161/viru.1.5.12796)); Mekbib SB, et al. 2011. Tropical Plant Pathology 36:233-240 ((https://doi.org/10.1590/S1982-56762011000400004)); Põlme S, et al. 2020. Fungal Diversity 105:1-16 ((https://doi.org/10.1007/s13225-020-00466-2)) |
| Otu21719 | 1 | k__Fungi(100);p__Ascomycota(100);c__Saccharomycetes(100);o__Saccharomycetales(100);f__Metschnikowiaceae(100);g__Metschnikowia(100);s__Metschnikowia_pulcherrima(100); | Metschnikowia | 13 | Saprotroph | \|Undefined Saprotroph\| | Yeast | NULL | Probable | As nectar-tap_saprotroph (Põlme et al. 2020); Decay sugar-rich substrates (Põlme et al. 2020) | Põlme S, et al. 2020. Fungal Diversity 105:1-16 ((https://doi.org/10.1007/s13225-020-00466-2)) |
| Otu21721 | 1 | k__Fungi(100);p__Ascomycota(100);c__Saccharomycetes(100);o__Saccharomycetales(100);f__Metschnikowiaceae(100);g__Metschnikowia(100);g__Metschnikowia_unclassified(100); | Metschnikowia | 13 | Saprotroph | \|Undefined Saprotroph\| | Yeast | NULL | Probable | As nectar-tap_saprotroph (Põlme et al. 2020); Decay sugar-rich substrates (Põlme et al. 2020) | Põlme S, et al. 2020. Fungal Diversity 105:1-16 ((https://doi.org/10.1007/s13225-020-00466-2)) |
| Otu21724 | 1 | k__Fungi(100);p__Ascomycota(100);c__Saccharomycetes(100);o__Saccharomycetales(100);f__Metschnikowiaceae(100);g__Metschnikowia(100);s__Metschnikowia_pulcherrima(100); | Metschnikowia | 13 | Saprotroph | \|Undefined Saprotroph\| | Yeast | NULL | Probable | As nectar-tap_saprotroph (Põlme et al. 2020); Decay sugar-rich substrates (Põlme et al. 2020) | Põlme S, et al. 2020. Fungal Diversity 105:1-16 ((https://doi.org/10.1007/s13225-020-00466-2)) |
| Otu21725 | 1 | k__Fungi(100);p__Ascomycota(100);c__Saccharomycetes(100);o__Saccharomycetales(100);f__Saccharomycetales_family_Incertae_sedis(100);g__Candida(100);g__Candida_unclassified(100); | Candida | 13 | Pathotroph-Saprotroph-Symbiotroph | Animal Pathogen-Endophyte-Endosymbiont-Epiphyte-\|Undefined Saprotroph\| | Dimorphic Yeast-Yeast | NULL | Probable | As nectar-tap_saprotroph (Põlme et al. 2020); Decay sugar-rich substrates (Põlme et al. 2020); This asexual genus is polyphyletic and contain species that function very differently depend on what they are. Caution should be taken when interpreting guilds for Candida species. | Manolakaki D, et al. 2010. Virulence. 1: 367-375 ((https://doi.org/10.4161/viru.1.5.12796)); Mekbib SB, et al. 2011. Tropical Plant Pathology 36:233-240 ((https://doi.org/10.1590/S1982-56762011000400004)); Põlme S, et al. 2020. Fungal Diversity 105:1-16 ((https://doi.org/10.1007/s13225-020-00466-2)) |
| Otu21726 | 1 | k__Fungi(100);p__Ascomycota(100);c__Saccharomycetes(100);o__Saccharomycetales(100);f__Metschnikowiaceae(100);g__Metschnikowia(100);s__Metschnikowia_pulcherrima(100); | Metschnikowia | 13 | Saprotroph | \|Undefined Saprotroph\| | Yeast | NULL | Probable | As nectar-tap_saprotroph (Põlme et al. 2020); Decay sugar-rich substrates (Põlme et al. 2020) | Põlme S, et al. 2020. Fungal Diversity 105:1-16 ((https://doi.org/10.1007/s13225-020-00466-2)) |
| Otu21727 | 1 | k__Fungi(100);p__Ascomycota(100);c__Saccharomycetes(100);o__Saccharomycetales(100);f__Metschnikowiaceae(100);g__Metschnikowia(100);g__Metschnikowia_unclassified(100); | Metschnikowia | 13 | Saprotroph | \|Undefined Saprotroph\| | Yeast | NULL | Probable | As nectar-tap_saprotroph (Põlme et al. 2020); Decay sugar-rich substrates (Põlme et al. 2020) | Põlme S, et al. 2020. Fungal Diversity 105:1-16 ((https://doi.org/10.1007/s13225-020-00466-2)) |
| Otu21728 | 1 | k__Fungi(100);p__Ascomycota(100);c__Saccharomycetes(100);o__Saccharomycetales(100);f__Metschnikowiaceae(100);g__Metschnikowia(100);s__Metschnikowia_pulcherrima(100); | Metschnikowia | 13 | Saprotroph | \|Undefined Saprotroph\| | Yeast | NULL | Probable | As nectar-tap_saprotroph (Põlme et al. 2020); Decay sugar-rich substrates (Põlme et al. 2020) | Põlme S, et al. 2020. Fungal Diversity 105:1-16 ((https://doi.org/10.1007/s13225-020-00466-2)) |
| Otu21729 | 1 | k__Fungi(100);p__Ascomycota(100);c__Saccharomycetes(100);o__Saccharomycetales(100);f__Metschnikowiaceae(100);g__Metschnikowia(100);g__Metschnikowia_unclassified(100); | Metschnikowia | 13 | Saprotroph | \|Undefined Saprotroph\| | Yeast | NULL | Probable | As nectar-tap_saprotroph (Põlme et al. 2020); Decay sugar-rich substrates (Põlme et al. 2020) | Põlme S, et al. 2020. Fungal Diversity 105:1-16 ((https://doi.org/10.1007/s13225-020-00466-2)) |
| Otu21730 | 1 | k__Fungi(100);p__Ascomycota(100);c__Saccharomycetes(100);o__Saccharomycetales(100);f__Metschnikowiaceae(100);g__Metschnikowia(100);s__Metschnikowia_pulcherrima(100); | Metschnikowia | 13 | Saprotroph | \|Undefined Saprotroph\| | Yeast | NULL | Probable | As nectar-tap_saprotroph (Põlme et al. 2020); Decay sugar-rich substrates (Põlme et al. 2020) | Põlme S, et al. 2020. Fungal Diversity 105:1-16 ((https://doi.org/10.1007/s13225-020-00466-2)) |
| Otu21732 | 1 | k__Fungi(100);p__Ascomycota(100);c__Saccharomycetes(100);o__Saccharomycetales(100);f__Metschnikowiaceae(100);g__Metschnikowia(100);s__Metschnikowia_pulcherrima(100); | Metschnikowia | 13 | Saprotroph | \|Undefined Saprotroph\| | Yeast | NULL | Probable | As nectar-tap_saprotroph (Põlme et al. 2020); Decay sugar-rich substrates (Põlme et al. 2020) | Põlme S, et al. 2020. Fungal Diversity 105:1-16 ((https://doi.org/10.1007/s13225-020-00466-2)) |
| Otu21734 | 1 | k__Fungi(100);p__Ascomycota(100);c__Saccharomycetes(100);o__Saccharomycetales(100);f__Metschnikowiaceae(100);g__Metschnikowia(100);s__Metschnikowia_pulcherrima(100); | Metschnikowia | 13 | Saprotroph | \|Undefined Saprotroph\| | Yeast | NULL | Probable | As nectar-tap_saprotroph (Põlme et al. 2020); Decay sugar-rich substrates (Põlme et al. 2020) | Põlme S, et al. 2020. Fungal Diversity 105:1-16 ((https://doi.org/10.1007/s13225-020-00466-2)) |
| Otu21735 | 1 | k__Fungi(100);p__Ascomycota(100);c__Saccharomycetes(100);o__Saccharomycetales(100);f__Metschnikowiaceae(100);g__Metschnikowia(100);g__Metschnikowia_unclassified(100); | Metschnikowia | 13 | Saprotroph | \|Undefined Saprotroph\| | Yeast | NULL | Probable | As nectar-tap_saprotroph (Põlme et al. 2020); Decay sugar-rich substrates (Põlme et al. 2020) | Põlme S, et al. 2020. Fungal Diversity 105:1-16 ((https://doi.org/10.1007/s13225-020-00466-2)) |
| Otu21736 | 1 | k__Fungi(100);p__Ascomycota(100);c__Saccharomycetes(100);o__Saccharomycetales(100);f__Metschnikowiaceae(100);g__Metschnikowia(100);g__Metschnikowia_unclassified(100); | Metschnikowia | 13 | Saprotroph | \|Undefined Saprotroph\| | Yeast | NULL | Probable | As nectar-tap_saprotroph (Põlme et al. 2020); Decay sugar-rich substrates (Põlme et al. 2020) | Põlme S, et al. 2020. Fungal Diversity 105:1-16 ((https://doi.org/10.1007/s13225-020-00466-2)) |
| Otu21737 | 1 | k__Fungi(100);p__Ascomycota(100);c__Saccharomycetes(100);o__Saccharomycetales(100);f__Metschnikowiaceae(100);g__Metschnikowia(100);s__Metschnikowia_pulcherrima(100); | Metschnikowia | 13 | Saprotroph | \|Undefined Saprotroph\| | Yeast | NULL | Probable | As nectar-tap_saprotroph (Põlme et al. 2020); Decay sugar-rich substrates (Põlme et al. 2020) | Põlme S, et al. 2020. Fungal Diversity 105:1-16 ((https://doi.org/10.1007/s13225-020-00466-2)) |
| Otu21738 | 1 | k__Fungi(100);p__Ascomycota(100);c__Saccharomycetes(100);o__Saccharomycetales(100);f__Metschnikowiaceae(100);g__Metschnikowia(100);s__Metschnikowia_pulcherrima(100); | Metschnikowia | 13 | Saprotroph | \|Undefined Saprotroph\| | Yeast | NULL | Probable | As nectar-tap_saprotroph (Põlme et al. 2020); Decay sugar-rich substrates (Põlme et al. 2020) | Põlme S, et al. 2020. Fungal Diversity 105:1-16 ((https://doi.org/10.1007/s13225-020-00466-2)) |
| Otu21739 | 1 | k__Fungi(100);p__Ascomycota(100);c__Saccharomycetes(100);o__Saccharomycetales(100);f__Metschnikowiaceae(100);g__Metschnikowia(100);s__Metschnikowia_pulcherrima(100); | Metschnikowia | 13 | Saprotroph | \|Undefined Saprotroph\| | Yeast | NULL | Probable | As nectar-tap_saprotroph (Põlme et al. 2020); Decay sugar-rich substrates (Põlme et al. 2020) | Põlme S, et al. 2020. Fungal Diversity 105:1-16 ((https://doi.org/10.1007/s13225-020-00466-2)) |
| Otu21741 | 1 | k__Fungi(100);p__Ascomycota(100);c__Saccharomycetes(100);o__Saccharomycetales(100);f__Metschnikowiaceae(100);g__Metschnikowia(100);g__Metschnikowia_unclassified(100); | Metschnikowia | 13 | Saprotroph | \|Undefined Saprotroph\| | Yeast | NULL | Probable | As nectar-tap_saprotroph (Põlme et al. 2020); Decay sugar-rich substrates (Põlme et al. 2020) | Põlme S, et al. 2020. Fungal Diversity 105:1-16 ((https://doi.org/10.1007/s13225-020-00466-2)) |
| Otu21742 | 1 | k__Fungi(100);p__Ascomycota(100);c__Saccharomycetes(100);o__Saccharomycetales(100);f__Metschnikowiaceae(100);g__Metschnikowia(100);s__Metschnikowia_pulcherrima(100); | Metschnikowia | 13 | Saprotroph | \|Undefined Saprotroph\| | Yeast | NULL | Probable | As nectar-tap_saprotroph (Põlme et al. 2020); Decay sugar-rich substrates (Põlme et al. 2020) | Põlme S, et al. 2020. Fungal Diversity 105:1-16 ((https://doi.org/10.1007/s13225-020-00466-2)) |
| Otu21743 | 1 | k__Fungi(100);p__Ascomycota(100);c__Saccharomycetes(100);o__Saccharomycetales(100);f__Metschnikowiaceae(100);g__Metschnikowia(100);g__Metschnikowia_unclassified(100); | Metschnikowia | 13 | Saprotroph | \|Undefined Saprotroph\| | Yeast | NULL | Probable | As nectar-tap_saprotroph (Põlme et al. 2020); Decay sugar-rich substrates (Põlme et al. 2020) | Põlme S, et al. 2020. Fungal Diversity 105:1-16 ((https://doi.org/10.1007/s13225-020-00466-2)) |
| Otu21744 | 1 | k__Fungi(100);p__Ascomycota(100);c__Saccharomycetes(100);o__Saccharomycetales(100);f__Metschnikowiaceae(100);g__Metschnikowia(100);s__Metschnikowia_pulcherrima(100); | Metschnikowia | 13 | Saprotroph | \|Undefined Saprotroph\| | Yeast | NULL | Probable | As nectar-tap_saprotroph (Põlme et al. 2020); Decay sugar-rich substrates (Põlme et al. 2020) | Põlme S, et al. 2020. Fungal Diversity 105:1-16 ((https://doi.org/10.1007/s13225-020-00466-2)) |
| Otu21745 | 1 | k__Fungi(100);p__Ascomycota(100);c__Saccharomycetes(100);o__Saccharomycetales(100);f__Metschnikowiaceae(100);g__Metschnikowia(100);s__Metschnikowia_pulcherrima(100); | Metschnikowia | 13 | Saprotroph | \|Undefined Saprotroph\| | Yeast | NULL | Probable | As nectar-tap_saprotroph (Põlme et al. 2020); Decay sugar-rich substrates (Põlme et al. 2020) | Põlme S, et al. 2020. Fungal Diversity 105:1-16 ((https://doi.org/10.1007/s13225-020-00466-2)) |
| Otu21746 | 1 | k__Fungi(100);p__Ascomycota(100);c__Saccharomycetes(100);o__Saccharomycetales(100);f__Metschnikowiaceae(100);g__Metschnikowia(100);s__Metschnikowia_pulcherrima(100); | Metschnikowia | 13 | Saprotroph | \|Undefined Saprotroph\| | Yeast | NULL | Probable | As nectar-tap_saprotroph (Põlme et al. 2020); Decay sugar-rich substrates (Põlme et al. 2020) | Põlme S, et al. 2020. Fungal Diversity 105:1-16 ((https://doi.org/10.1007/s13225-020-00466-2)) |
| Otu21747 | 1 | k__Fungi(100);p__Ascomycota(100);c__Saccharomycetes(100);o__Saccharomycetales(100);f__Metschnikowiaceae(100);g__Metschnikowia(100);s__Metschnikowia_pulcherrima(100); | Metschnikowia | 13 | Saprotroph | \|Undefined Saprotroph\| | Yeast | NULL | Probable | As nectar-tap_saprotroph (Põlme et al. 2020); Decay sugar-rich substrates (Põlme et al. 2020) | Põlme S, et al. 2020. Fungal Diversity 105:1-16 ((https://doi.org/10.1007/s13225-020-00466-2)) |
| Otu21748 | 1 | k__Fungi(100);p__Ascomycota(100);c__Saccharomycetes(100);o__Saccharomycetales(100);f__Metschnikowiaceae(100);g__Metschnikowia(100);g__Metschnikowia_unclassified(100); | Metschnikowia | 13 | Saprotroph | \|Undefined Saprotroph\| | Yeast | NULL | Probable | As nectar-tap_saprotroph (Põlme et al. 2020); Decay sugar-rich substrates (Põlme et al. 2020) | Põlme S, et al. 2020. Fungal Diversity 105:1-16 ((https://doi.org/10.1007/s13225-020-00466-2)) |
| Otu21751 | 1 | k__Fungi(100);p__Ascomycota(100);c__Saccharomycetes(100);o__Saccharomycetales(100);f__Metschnikowiaceae(100);g__Metschnikowia(100);g__Metschnikowia_unclassified(100); | Metschnikowia | 13 | Saprotroph | \|Undefined Saprotroph\| | Yeast | NULL | Probable | As nectar-tap_saprotroph (Põlme et al. 2020); Decay sugar-rich substrates (Põlme et al. 2020) | Põlme S, et al. 2020. Fungal Diversity 105:1-16 ((https://doi.org/10.1007/s13225-020-00466-2)) |
| Otu21752 | 1 | k__Fungi(100);p__Ascomycota(100);c__Saccharomycetes(100);o__Saccharomycetales(100);f__Metschnikowiaceae(100);g__Metschnikowia(100);s__Metschnikowia_pulcherrima(100); | Metschnikowia | 13 | Saprotroph | \|Undefined Saprotroph\| | Yeast | NULL | Probable | As nectar-tap_saprotroph (Põlme et al. 2020); Decay sugar-rich substrates (Põlme et al. 2020) | Põlme S, et al. 2020. Fungal Diversity 105:1-16 ((https://doi.org/10.1007/s13225-020-00466-2)) |
| Otu21753 | 1 | k__Fungi(100);p__Ascomycota(100);c__Saccharomycetes(100);o__Saccharomycetales(100);f__Metschnikowiaceae(100);g__Metschnikowia(100);s__Metschnikowia_pulcherrima(100); | Metschnikowia | 13 | Saprotroph | \|Undefined Saprotroph\| | Yeast | NULL | Probable | As nectar-tap_saprotroph (Põlme et al. 2020); Decay sugar-rich substrates (Põlme et al. 2020) | Põlme S, et al. 2020. Fungal Diversity 105:1-16 ((https://doi.org/10.1007/s13225-020-00466-2)) |
| Otu21754 | 1 | k__Fungi(100);p__Ascomycota(100);c__Saccharomycetes(100);o__Saccharomycetales(100);f__Metschnikowiaceae(100);g__Metschnikowia(100);g__Metschnikowia_unclassified(100); | Metschnikowia | 13 | Saprotroph | \|Undefined Saprotroph\| | Yeast | NULL | Probable | As nectar-tap_saprotroph (Põlme et al. 2020); Decay sugar-rich substrates (Põlme et al. 2020) | Põlme S, et al. 2020. Fungal Diversity 105:1-16 ((https://doi.org/10.1007/s13225-020-00466-2)) |
| Otu21755 | 1 | k__Fungi(100);p__Ascomycota(100);c__Saccharomycetes(100);o__Saccharomycetales(100);f__Metschnikowiaceae(100);g__Metschnikowia(100);s__Metschnikowia_pulcherrima(100); | Metschnikowia | 13 | Saprotroph | \|Undefined Saprotroph\| | Yeast | NULL | Probable | As nectar-tap_saprotroph (Põlme et al. 2020); Decay sugar-rich substrates (Põlme et al. 2020) | Põlme S, et al. 2020. Fungal Diversity 105:1-16 ((https://doi.org/10.1007/s13225-020-00466-2)) |
| Otu21757 | 1 | k__Fungi(100);p__Ascomycota(100);c__Saccharomycetes(100);o__Saccharomycetales(100);f__Metschnikowiaceae(100);g__Metschnikowia(100);s__Metschnikowia_pulcherrima(100); | Metschnikowia | 13 | Saprotroph | \|Undefined Saprotroph\| | Yeast | NULL | Probable | As nectar-tap_saprotroph (Põlme et al. 2020); Decay sugar-rich substrates (Põlme et al. 2020) | Põlme S, et al. 2020. Fungal Diversity 105:1-16 ((https://doi.org/10.1007/s13225-020-00466-2)) |
| Otu21758 | 1 | k__Fungi(100);p__Ascomycota(100);c__Saccharomycetes(100);o__Saccharomycetales(100);f__Metschnikowiaceae(100);g__Metschnikowia(100);g__Metschnikowia_unclassified(100); | Metschnikowia | 13 | Saprotroph | \|Undefined Saprotroph\| | Yeast | NULL | Probable | As nectar-tap_saprotroph (Põlme et al. 2020); Decay sugar-rich substrates (Põlme et al. 2020) | Põlme S, et al. 2020. Fungal Diversity 105:1-16 ((https://doi.org/10.1007/s13225-020-00466-2)) |
| Otu21759 | 1 | k__Fungi(100);p__Ascomycota(100);c__Saccharomycetes(100);o__Saccharomycetales(100);f__Metschnikowiaceae(100);g__Metschnikowia(100);s__Metschnikowia_pulcherrima(100); | Metschnikowia | 13 | Saprotroph | \|Undefined Saprotroph\| | Yeast | NULL | Probable | As nectar-tap_saprotroph (Põlme et al. 2020); Decay sugar-rich substrates (Põlme et al. 2020) | Põlme S, et al. 2020. Fungal Diversity 105:1-16 ((https://doi.org/10.1007/s13225-020-00466-2)) |
| Otu21760 | 1 | k__Fungi(100);p__Ascomycota(100);c__Saccharomycetes(100);o__Saccharomycetales(100);f__Metschnikowiaceae(100);g__Metschnikowia(100);s__Metschnikowia_pulcherrima(100); | Metschnikowia | 13 | Saprotroph | \|Undefined Saprotroph\| | Yeast | NULL | Probable | As nectar-tap_saprotroph (Põlme et al. 2020); Decay sugar-rich substrates (Põlme et al. 2020) | Põlme S, et al. 2020. Fungal Diversity 105:1-16 ((https://doi.org/10.1007/s13225-020-00466-2)) |
| Otu21761 | 1 | k__Fungi(100);p__Ascomycota(100);c__Saccharomycetes(100);o__Saccharomycetales(100);f__Metschnikowiaceae(100);g__Metschnikowia(100);s__Metschnikowia_pulcherrima(100); | Metschnikowia | 13 | Saprotroph | \|Undefined Saprotroph\| | Yeast | NULL | Probable | As nectar-tap_saprotroph (Põlme et al. 2020); Decay sugar-rich substrates (Põlme et al. 2020) | Põlme S, et al. 2020. Fungal Diversity 105:1-16 ((https://doi.org/10.1007/s13225-020-00466-2)) |
| Otu21762 | 1 | k__Fungi(100);p__Ascomycota(100);c__Saccharomycetes(100);o__Saccharomycetales(100);f__Saccharomycetales_family_Incertae_sedis(100);g__Candida(100);g__Candida_unclassified(100); | Candida | 13 | Pathotroph-Saprotroph-Symbiotroph | Animal Pathogen-Endophyte-Endosymbiont-Epiphyte-\|Undefined Saprotroph\| | Dimorphic Yeast-Yeast | NULL | Probable | As nectar-tap_saprotroph (Põlme et al. 2020); Decay sugar-rich substrates (Põlme et al. 2020); This asexual genus is polyphyletic and contain species that function very differently depend on what they are. Caution should be taken when interpreting guilds for Candida species. | Manolakaki D, et al. 2010. Virulence. 1: 367-375 ((https://doi.org/10.4161/viru.1.5.12796)); Mekbib SB, et al. 2011. Tropical Plant Pathology 36:233-240 ((https://doi.org/10.1590/S1982-56762011000400004)); Põlme S, et al. 2020. Fungal Diversity 105:1-16 ((https://doi.org/10.1007/s13225-020-00466-2)) |
| Otu21763 | 1 | k__Fungi(100);p__Ascomycota(100);c__Saccharomycetes(100);o__Saccharomycetales(100);f__Metschnikowiaceae(100);g__Metschnikowia(100);s__Metschnikowia_pulcherrima(100); | Metschnikowia | 13 | Saprotroph | \|Undefined Saprotroph\| | Yeast | NULL | Probable | As nectar-tap_saprotroph (Põlme et al. 2020); Decay sugar-rich substrates (Põlme et al. 2020) | Põlme S, et al. 2020. Fungal Diversity 105:1-16 ((https://doi.org/10.1007/s13225-020-00466-2)) |
| Otu21764 | 1 | k__Fungi(100);p__Ascomycota(100);c__Saccharomycetes(100);o__Saccharomycetales(100);f__Metschnikowiaceae(100);g__Metschnikowia(100);g__Metschnikowia_unclassified(100); | Metschnikowia | 13 | Saprotroph | \|Undefined Saprotroph\| | Yeast | NULL | Probable | As nectar-tap_saprotroph (Põlme et al. 2020); Decay sugar-rich substrates (Põlme et al. 2020) | Põlme S, et al. 2020. Fungal Diversity 105:1-16 ((https://doi.org/10.1007/s13225-020-00466-2)) |
| Otu21766 | 1 | k__Fungi(100);p__Ascomycota(100);c__Saccharomycetes(100);o__Saccharomycetales(100);f__Saccharomycetales_family_Incertae_sedis(100);g__Candida(100);g__Candida_unclassified(100); | Candida | 13 | Pathotroph-Saprotroph-Symbiotroph | Animal Pathogen-Endophyte-Endosymbiont-Epiphyte-\|Undefined Saprotroph\| | Dimorphic Yeast-Yeast | NULL | Probable | As nectar-tap_saprotroph (Põlme et al. 2020); Decay sugar-rich substrates (Põlme et al. 2020); This asexual genus is polyphyletic and contain species that function very differently depend on what they are. Caution should be taken when interpreting guilds for Candida species. | Manolakaki D, et al. 2010. Virulence. 1: 367-375 ((https://doi.org/10.4161/viru.1.5.12796)); Mekbib SB, et al. 2011. Tropical Plant Pathology 36:233-240 ((https://doi.org/10.1590/S1982-56762011000400004)); Põlme S, et al. 2020. Fungal Diversity 105:1-16 ((https://doi.org/10.1007/s13225-020-00466-2)) |
| Otu21767 | 1 | k__Fungi(100);p__Ascomycota(100);c__Saccharomycetes(100);o__Saccharomycetales(100);f__Metschnikowiaceae(100);g__Metschnikowia(100);s__Metschnikowia_pulcherrima(100); | Metschnikowia | 13 | Saprotroph | \|Undefined Saprotroph\| | Yeast | NULL | Probable | As nectar-tap_saprotroph (Põlme et al. 2020); Decay sugar-rich substrates (Põlme et al. 2020) | Põlme S, et al. 2020. Fungal Diversity 105:1-16 ((https://doi.org/10.1007/s13225-020-00466-2)) |
| Otu21768 | 1 | k__Fungi(100);p__Ascomycota(100);c__Saccharomycetes(100);o__Saccharomycetales(100);f__Metschnikowiaceae(100);g__Metschnikowia(100);s__Metschnikowia_pulcherrima(100); | Metschnikowia | 13 | Saprotroph | \|Undefined Saprotroph\| | Yeast | NULL | Probable | As nectar-tap_saprotroph (Põlme et al. 2020); Decay sugar-rich substrates (Põlme et al. 2020) | Põlme S, et al. 2020. Fungal Diversity 105:1-16 ((https://doi.org/10.1007/s13225-020-00466-2)) |
| Otu21770 | 1 | k__Fungi(100);p__Ascomycota(100);c__Saccharomycetes(100);o__Saccharomycetales(100);f__Metschnikowiaceae(100);g__Metschnikowia(100);g__Metschnikowia_unclassified(100); | Metschnikowia | 13 | Saprotroph | \|Undefined Saprotroph\| | Yeast | NULL | Probable | As nectar-tap_saprotroph (Põlme et al. 2020); Decay sugar-rich substrates (Põlme et al. 2020) | Põlme S, et al. 2020. Fungal Diversity 105:1-16 ((https://doi.org/10.1007/s13225-020-00466-2)) |
| Otu21771 | 1 | k__Fungi(100);p__Ascomycota(100);c__Saccharomycetes(100);o__Saccharomycetales(100);f__Metschnikowiaceae(100);g__Metschnikowia(100);g__Metschnikowia_unclassified(100); | Metschnikowia | 13 | Saprotroph | \|Undefined Saprotroph\| | Yeast | NULL | Probable | As nectar-tap_saprotroph (Põlme et al. 2020); Decay sugar-rich substrates (Põlme et al. 2020) | Põlme S, et al. 2020. Fungal Diversity 105:1-16 ((https://doi.org/10.1007/s13225-020-00466-2)) |
| Otu21772 | 1 | k__Fungi(100);p__Ascomycota(100);c__Saccharomycetes(100);o__Saccharomycetales(100);f__Metschnikowiaceae(100);g__Metschnikowia(100);g__Metschnikowia_unclassified(100); | Metschnikowia | 13 | Saprotroph | \|Undefined Saprotroph\| | Yeast | NULL | Probable | As nectar-tap_saprotroph (Põlme et al. 2020); Decay sugar-rich substrates (Põlme et al. 2020) | Põlme S, et al. 2020. Fungal Diversity 105:1-16 ((https://doi.org/10.1007/s13225-020-00466-2)) |
| Otu21773 | 1 | k__Fungi(100);p__Ascomycota(100);c__Saccharomycetes(100);o__Saccharomycetales(100);f__Metschnikowiaceae(100);g__Metschnikowia(100);s__Metschnikowia_pulcherrima(100); | Metschnikowia | 13 | Saprotroph | \|Undefined Saprotroph\| | Yeast | NULL | Probable | As nectar-tap_saprotroph (Põlme et al. 2020); Decay sugar-rich substrates (Põlme et al. 2020) | Põlme S, et al. 2020. Fungal Diversity 105:1-16 ((https://doi.org/10.1007/s13225-020-00466-2)) |
| Otu21774 | 1 | k__Fungi(100);p__Ascomycota(100);c__Saccharomycetes(100);o__Saccharomycetales(100);f__Metschnikowiaceae(100);g__Metschnikowia(100);s__Metschnikowia_pulcherrima(100); | Metschnikowia | 13 | Saprotroph | \|Undefined Saprotroph\| | Yeast | NULL | Probable | As nectar-tap_saprotroph (Põlme et al. 2020); Decay sugar-rich substrates (Põlme et al. 2020) | Põlme S, et al. 2020. Fungal Diversity 105:1-16 ((https://doi.org/10.1007/s13225-020-00466-2)) |
| Otu21776 | 1 | k__Fungi(100);p__Ascomycota(100);c__Saccharomycetes(100);o__Saccharomycetales(100);f__Saccharomycetales_family_Incertae_sedis(100);g__Candida(100);s__Candida_cretensis(100); | Candida | 13 | Pathotroph-Saprotroph-Symbiotroph | Animal Pathogen-Endophyte-Endosymbiont-Epiphyte-\|Undefined Saprotroph\| | Dimorphic Yeast-Yeast | NULL | Probable | As nectar-tap_saprotroph (Põlme et al. 2020); Decay sugar-rich substrates (Põlme et al. 2020); This asexual genus is polyphyletic and contain species that function very differently depend on what they are. Caution should be taken when interpreting guilds for Candida species. | Manolakaki D, et al. 2010. Virulence. 1: 367-375 ((https://doi.org/10.4161/viru.1.5.12796)); Mekbib SB, et al. 2011. Tropical Plant Pathology 36:233-240 ((https://doi.org/10.1590/S1982-56762011000400004)); Põlme S, et al. 2020. Fungal Diversity 105:1-16 ((https://doi.org/10.1007/s13225-020-00466-2)) |
| Otu21786 | 1 | k__Fungi(100);p__Ascomycota(100);c__Saccharomycetes(100);o__Saccharomycetales(100);f__Metschnikowiaceae(100);g__Metschnikowia(100);s__Metschnikowia_pulcherrima(100); | Metschnikowia | 13 | Saprotroph | \|Undefined Saprotroph\| | Yeast | NULL | Probable | As nectar-tap_saprotroph (Põlme et al. 2020); Decay sugar-rich substrates (Põlme et al. 2020) | Põlme S, et al. 2020. Fungal Diversity 105:1-16 ((https://doi.org/10.1007/s13225-020-00466-2)) |
| Otu21787 | 1 | k__Fungi(100);p__Ascomycota(100);c__Saccharomycetes(100);o__Saccharomycetales(100);f__Metschnikowiaceae(100);g__Metschnikowia(100);s__Metschnikowia_pulcherrima(100); | Metschnikowia | 13 | Saprotroph | \|Undefined Saprotroph\| | Yeast | NULL | Probable | As nectar-tap_saprotroph (Põlme et al. 2020); Decay sugar-rich substrates (Põlme et al. 2020) | Põlme S, et al. 2020. Fungal Diversity 105:1-16 ((https://doi.org/10.1007/s13225-020-00466-2)) |
| Otu21788 | 1 | k__Fungi(100);p__Ascomycota(100);c__Saccharomycetes(100);o__Saccharomycetales(100);f__Metschnikowiaceae(100);g__Metschnikowia(100);s__Metschnikowia_pulcherrima(100); | Metschnikowia | 13 | Saprotroph | \|Undefined Saprotroph\| | Yeast | NULL | Probable | As nectar-tap_saprotroph (Põlme et al. 2020); Decay sugar-rich substrates (Põlme et al. 2020) | Põlme S, et al. 2020. Fungal Diversity 105:1-16 ((https://doi.org/10.1007/s13225-020-00466-2)) |
| Otu21789 | 1 | k__Fungi(100);p__Ascomycota(100);c__Saccharomycetes(100);o__Saccharomycetales(100);f__Metschnikowiaceae(100);g__Metschnikowia(100);g__Metschnikowia_unclassified(100); | Metschnikowia | 13 | Saprotroph | \|Undefined Saprotroph\| | Yeast | NULL | Probable | As nectar-tap_saprotroph (Põlme et al. 2020); Decay sugar-rich substrates (Põlme et al. 2020) | Põlme S, et al. 2020. Fungal Diversity 105:1-16 ((https://doi.org/10.1007/s13225-020-00466-2)) |
| Otu21790 | 1 | k__Fungi(100);p__Ascomycota(100);c__Saccharomycetes(100);o__Saccharomycetales(100);f__Metschnikowiaceae(100);g__Metschnikowia(100);s__Metschnikowia_pulcherrima(100); | Metschnikowia | 13 | Saprotroph | \|Undefined Saprotroph\| | Yeast | NULL | Probable | As nectar-tap_saprotroph (Põlme et al. 2020); Decay sugar-rich substrates (Põlme et al. 2020) | Põlme S, et al. 2020. Fungal Diversity 105:1-16 ((https://doi.org/10.1007/s13225-020-00466-2)) |
| Otu21791 | 1 | k__Fungi(100);p__Ascomycota(100);c__Saccharomycetes(100);o__Saccharomycetales(100);f__Metschnikowiaceae(100);g__Metschnikowia(100);s__Metschnikowia_pulcherrima(100); | Metschnikowia | 13 | Saprotroph | \|Undefined Saprotroph\| | Yeast | NULL | Probable | As nectar-tap_saprotroph (Põlme et al. 2020); Decay sugar-rich substrates (Põlme et al. 2020) | Põlme S, et al. 2020. Fungal Diversity 105:1-16 ((https://doi.org/10.1007/s13225-020-00466-2)) |
| Otu21793 | 1 | k__Fungi(100);p__Ascomycota(100);c__Saccharomycetes(100);o__Saccharomycetales(100);f__Metschnikowiaceae(100);g__Metschnikowia(100);g__Metschnikowia_unclassified(100); | Metschnikowia | 13 | Saprotroph | \|Undefined Saprotroph\| | Yeast | NULL | Probable | As nectar-tap_saprotroph (Põlme et al. 2020); Decay sugar-rich substrates (Põlme et al. 2020) | Põlme S, et al. 2020. Fungal Diversity 105:1-16 ((https://doi.org/10.1007/s13225-020-00466-2)) |
| Otu21794 | 1 | k__Fungi(100);p__Ascomycota(100);c__Saccharomycetes(100);o__Saccharomycetales(100);f__Metschnikowiaceae(100);g__Metschnikowia(100);s__Metschnikowia_pulcherrima(100); | Metschnikowia | 13 | Saprotroph | \|Undefined Saprotroph\| | Yeast | NULL | Probable | As nectar-tap_saprotroph (Põlme et al. 2020); Decay sugar-rich substrates (Põlme et al. 2020) | Põlme S, et al. 2020. Fungal Diversity 105:1-16 ((https://doi.org/10.1007/s13225-020-00466-2)) |
| Otu21795 | 1 | k__Fungi(100);p__Ascomycota(100);c__Saccharomycetes(100);o__Saccharomycetales(100);f__Metschnikowiaceae(100);g__Metschnikowia(100);s__Metschnikowia_pulcherrima(100); | Metschnikowia | 13 | Saprotroph | \|Undefined Saprotroph\| | Yeast | NULL | Probable | As nectar-tap_saprotroph (Põlme et al. 2020); Decay sugar-rich substrates (Põlme et al. 2020) | Põlme S, et al. 2020. Fungal Diversity 105:1-16 ((https://doi.org/10.1007/s13225-020-00466-2)) |
| Otu21796 | 1 | k__Fungi(100);p__Ascomycota(100);c__Saccharomycetes(100);o__Saccharomycetales(100);f__Metschnikowiaceae(100);g__Metschnikowia(100);s__Metschnikowia_pulcherrima(100); | Metschnikowia | 13 | Saprotroph | \|Undefined Saprotroph\| | Yeast | NULL | Probable | As nectar-tap_saprotroph (Põlme et al. 2020); Decay sugar-rich substrates (Põlme et al. 2020) | Põlme S, et al. 2020. Fungal Diversity 105:1-16 ((https://doi.org/10.1007/s13225-020-00466-2)) |
| Otu21797 | 1 | k__Fungi(100);p__Ascomycota(100);c__Saccharomycetes(100);o__Saccharomycetales(100);f__Saccharomycetales_family_Incertae_sedis(100);f__Saccharomycetales_family_Incertae_sedis_unclassified(100);f__Saccharomycetales_family_Incertae_sedis_unclassified(100); | Saccharomycetales | 7 | Saprotroph | Undefined Saprotroph | Yeast | NULL | Possible | NULL | Sterkenburg E, et al. 2015. New Phytologist 207:1145-1158 ((https://doi.org/10.1111/nph.13426)) |
| Otu21798 | 1 | k__Fungi(100);p__Ascomycota(100);c__Saccharomycetes(100);o__Saccharomycetales(100);f__Metschnikowiaceae(100);g__Metschnikowia(100);s__Metschnikowia_pulcherrima(100); | Metschnikowia | 13 | Saprotroph | \|Undefined Saprotroph\| | Yeast | NULL | Probable | As nectar-tap_saprotroph (Põlme et al. 2020); Decay sugar-rich substrates (Põlme et al. 2020) | Põlme S, et al. 2020. Fungal Diversity 105:1-16 ((https://doi.org/10.1007/s13225-020-00466-2)) |
| Otu21799 | 1 | k__Fungi(100);p__Ascomycota(100);c__Saccharomycetes(100);o__Saccharomycetales(100);f__Saccharomycetales_family_Incertae_sedis(100);g__Candida(100);g__Candida_unclassified(100); | Candida | 13 | Pathotroph-Saprotroph-Symbiotroph | Animal Pathogen-Endophyte-Endosymbiont-Epiphyte-\|Undefined Saprotroph\| | Dimorphic Yeast-Yeast | NULL | Probable | As nectar-tap_saprotroph (Põlme et al. 2020); Decay sugar-rich substrates (Põlme et al. 2020); This asexual genus is polyphyletic and contain species that function very differently depend on what they are. Caution should be taken when interpreting guilds for Candida species. | Manolakaki D, et al. 2010. Virulence. 1: 367-375 ((https://doi.org/10.4161/viru.1.5.12796)); Mekbib SB, et al. 2011. Tropical Plant Pathology 36:233-240 ((https://doi.org/10.1590/S1982-56762011000400004)); Põlme S, et al. 2020. Fungal Diversity 105:1-16 ((https://doi.org/10.1007/s13225-020-00466-2)) |
| Otu21800 | 1 | k__Fungi(100);p__Ascomycota(100);c__Saccharomycetes(100);o__Saccharomycetales(100);f__Metschnikowiaceae(100);g__Metschnikowia(100);s__Metschnikowia_pulcherrima(100); | Metschnikowia | 13 | Saprotroph | \|Undefined Saprotroph\| | Yeast | NULL | Probable | As nectar-tap_saprotroph (Põlme et al. 2020); Decay sugar-rich substrates (Põlme et al. 2020) | Põlme S, et al. 2020. Fungal Diversity 105:1-16 ((https://doi.org/10.1007/s13225-020-00466-2)) |
| Otu21801 | 1 | k__Fungi(100);p__Ascomycota(100);c__Saccharomycetes(100);o__Saccharomycetales(100);f__Metschnikowiaceae(100);g__Metschnikowia(100);s__Metschnikowia_pulcherrima(100); | Metschnikowia | 13 | Saprotroph | \|Undefined Saprotroph\| | Yeast | NULL | Probable | As nectar-tap_saprotroph (Põlme et al. 2020); Decay sugar-rich substrates (Põlme et al. 2020) | Põlme S, et al. 2020. Fungal Diversity 105:1-16 ((https://doi.org/10.1007/s13225-020-00466-2)) |
| Otu21802 | 1 | k__Fungi(100);p__Ascomycota(100);c__Saccharomycetes(100);o__Saccharomycetales(100);f__Metschnikowiaceae(100);g__Metschnikowia(100);s__Metschnikowia_pulcherrima(100); | Metschnikowia | 13 | Saprotroph | \|Undefined Saprotroph\| | Yeast | NULL | Probable | As nectar-tap_saprotroph (Põlme et al. 2020); Decay sugar-rich substrates (Põlme et al. 2020) | Põlme S, et al. 2020. Fungal Diversity 105:1-16 ((https://doi.org/10.1007/s13225-020-00466-2)) |
| Otu21803 | 1 | k__Fungi(100);p__Ascomycota(100);c__Saccharomycetes(100);o__Saccharomycetales(100);f__Metschnikowiaceae(100);g__Metschnikowia(100);s__Metschnikowia_pulcherrima(100); | Metschnikowia | 13 | Saprotroph | \|Undefined Saprotroph\| | Yeast | NULL | Probable | As nectar-tap_saprotroph (Põlme et al. 2020); Decay sugar-rich substrates (Põlme et al. 2020) | Põlme S, et al. 2020. Fungal Diversity 105:1-16 ((https://doi.org/10.1007/s13225-020-00466-2)) |
| Otu21804 | 1 | k__Fungi(100);p__Ascomycota(100);c__Saccharomycetes(100);o__Saccharomycetales(100);f__Saccharomycetales_family_Incertae_sedis(100);g__Candida(100);g__Candida_unclassified(100); | Candida | 13 | Pathotroph-Saprotroph-Symbiotroph | Animal Pathogen-Endophyte-Endosymbiont-Epiphyte-\|Undefined Saprotroph\| | Dimorphic Yeast-Yeast | NULL | Probable | As nectar-tap_saprotroph (Põlme et al. 2020); Decay sugar-rich substrates (Põlme et al. 2020); This asexual genus is polyphyletic and contain species that function very differently depend on what they are. Caution should be taken when interpreting guilds for Candida species. | Manolakaki D, et al. 2010. Virulence. 1: 367-375 ((https://doi.org/10.4161/viru.1.5.12796)); Mekbib SB, et al. 2011. Tropical Plant Pathology 36:233-240 ((https://doi.org/10.1590/S1982-56762011000400004)); Põlme S, et al. 2020. Fungal Diversity 105:1-16 ((https://doi.org/10.1007/s13225-020-00466-2)) |
| Otu21806 | 1 | k__Fungi(100);p__Ascomycota(100);c__Saccharomycetes(100);o__Saccharomycetales(100);f__Metschnikowiaceae(100);g__Metschnikowia(100);g__Metschnikowia_unclassified(100); | Metschnikowia | 13 | Saprotroph | \|Undefined Saprotroph\| | Yeast | NULL | Probable | As nectar-tap_saprotroph (Põlme et al. 2020); Decay sugar-rich substrates (Põlme et al. 2020) | Põlme S, et al. 2020. Fungal Diversity 105:1-16 ((https://doi.org/10.1007/s13225-020-00466-2)) |
| Otu21807 | 1 | k__Fungi(100);p__Ascomycota(100);c__Saccharomycetes(100);o__Saccharomycetales(100);f__Saccharomycetales_family_Incertae_sedis(100);g__Candida(100);g__Candida_unclassified(100); | Candida | 13 | Pathotroph-Saprotroph-Symbiotroph | Animal Pathogen-Endophyte-Endosymbiont-Epiphyte-\|Undefined Saprotroph\| | Dimorphic Yeast-Yeast | NULL | Probable | As nectar-tap_saprotroph (Põlme et al. 2020); Decay sugar-rich substrates (Põlme et al. 2020); This asexual genus is polyphyletic and contain species that function very differently depend on what they are. Caution should be taken when interpreting guilds for Candida species. | Manolakaki D, et al. 2010. Virulence. 1: 367-375 ((https://doi.org/10.4161/viru.1.5.12796)); Mekbib SB, et al. 2011. Tropical Plant Pathology 36:233-240 ((https://doi.org/10.1590/S1982-56762011000400004)); Põlme S, et al. 2020. Fungal Diversity 105:1-16 ((https://doi.org/10.1007/s13225-020-00466-2)) |
| Otu21808 | 1 | k__Fungi(100);p__Ascomycota(100);c__Saccharomycetes(100);o__Saccharomycetales(100);f__Saccharomycetales_family_Incertae_sedis(100);g__Candida(100);s__Candida_sp_AS23486(100); | Candida | 13 | Pathotroph-Saprotroph-Symbiotroph | Animal Pathogen-Endophyte-Endosymbiont-Epiphyte-\|Undefined Saprotroph\| | Dimorphic Yeast-Yeast | NULL | Probable | As nectar-tap_saprotroph (Põlme et al. 2020); Decay sugar-rich substrates (Põlme et al. 2020); This asexual genus is polyphyletic and contain species that function very differently depend on what they are. Caution should be taken when interpreting guilds for Candida species. | Manolakaki D, et al. 2010. Virulence. 1: 367-375 ((https://doi.org/10.4161/viru.1.5.12796)); Mekbib SB, et al. 2011. Tropical Plant Pathology 36:233-240 ((https://doi.org/10.1590/S1982-56762011000400004)); Põlme S, et al. 2020. Fungal Diversity 105:1-16 ((https://doi.org/10.1007/s13225-020-00466-2)) |
| Otu21811 | 1 | k__Fungi(100);p__Ascomycota(100);c__Saccharomycetes(100);o__Saccharomycetales(100);f__Metschnikowiaceae(100);g__Metschnikowia(100);g__Metschnikowia_unclassified(100); | Metschnikowia | 13 | Saprotroph | \|Undefined Saprotroph\| | Yeast | NULL | Probable | As nectar-tap_saprotroph (Põlme et al. 2020); Decay sugar-rich substrates (Põlme et al. 2020) | Põlme S, et al. 2020. Fungal Diversity 105:1-16 ((https://doi.org/10.1007/s13225-020-00466-2)) |
| Otu21814 | 1 | k__Fungi(100);p__Ascomycota(100);c__Saccharomycetes(100);o__Saccharomycetales(100);f__Saccharomycetales_family_Incertae_sedis(100);f__Saccharomycetales_family_Incertae_sedis_unclassified(100);f__Saccharomycetales_family_Incertae_sedis_unclassified(100); | Saccharomycetales | 7 | Saprotroph | Undefined Saprotroph | Yeast | NULL | Possible | NULL | Sterkenburg E, et al. 2015. New Phytologist 207:1145-1158 ((https://doi.org/10.1111/nph.13426)) |
| Otu21816 | 1 | k__Fungi(100);p__Ascomycota(100);c__Saccharomycetes(100);o__Saccharomycetales(100);f__Saccharomycetales_family_Incertae_sedis(100);g__Candida(100);g__Candida_unclassified(100); | Candida | 13 | Pathotroph-Saprotroph-Symbiotroph | Animal Pathogen-Endophyte-Endosymbiont-Epiphyte-\|Undefined Saprotroph\| | Dimorphic Yeast-Yeast | NULL | Probable | As nectar-tap_saprotroph (Põlme et al. 2020); Decay sugar-rich substrates (Põlme et al. 2020); This asexual genus is polyphyletic and contain species that function very differently depend on what they are. Caution should be taken when interpreting guilds for Candida species. | Manolakaki D, et al. 2010. Virulence. 1: 367-375 ((https://doi.org/10.4161/viru.1.5.12796)); Mekbib SB, et al. 2011. Tropical Plant Pathology 36:233-240 ((https://doi.org/10.1590/S1982-56762011000400004)); Põlme S, et al. 2020. Fungal Diversity 105:1-16 ((https://doi.org/10.1007/s13225-020-00466-2)) |
| Otu21818 | 1 | k__Fungi(100);p__Ascomycota(100);c__Saccharomycetes(100);o__Saccharomycetales(100);f__Metschnikowiaceae(100);g__Metschnikowia(100);g__Metschnikowia_unclassified(100); | Metschnikowia | 13 | Saprotroph | \|Undefined Saprotroph\| | Yeast | NULL | Probable | As nectar-tap_saprotroph (Põlme et al. 2020); Decay sugar-rich substrates (Põlme et al. 2020) | Põlme S, et al. 2020. Fungal Diversity 105:1-16 ((https://doi.org/10.1007/s13225-020-00466-2)) |
| Otu21819 | 1 | k__Fungi(100);p__Ascomycota(100);c__Saccharomycetes(100);o__Saccharomycetales(100);f__Metschnikowiaceae(100);g__Metschnikowia(100);s__Metschnikowia_pulcherrima(100); | Metschnikowia | 13 | Saprotroph | \|Undefined Saprotroph\| | Yeast | NULL | Probable | As nectar-tap_saprotroph (Põlme et al. 2020); Decay sugar-rich substrates (Põlme et al. 2020) | Põlme S, et al. 2020. Fungal Diversity 105:1-16 ((https://doi.org/10.1007/s13225-020-00466-2)) |
| Otu21820 | 1 | k__Fungi(100);p__Ascomycota(100);c__Saccharomycetes(100);o__Saccharomycetales(100);f__Metschnikowiaceae(100);g__Metschnikowia(100);s__Metschnikowia_pulcherrima(100); | Metschnikowia | 13 | Saprotroph | \|Undefined Saprotroph\| | Yeast | NULL | Probable | As nectar-tap_saprotroph (Põlme et al. 2020); Decay sugar-rich substrates (Põlme et al. 2020) | Põlme S, et al. 2020. Fungal Diversity 105:1-16 ((https://doi.org/10.1007/s13225-020-00466-2)) |
| Otu21821 | 1 | k__Fungi(100);p__Ascomycota(100);c__Saccharomycetes(100);o__Saccharomycetales(100);f__Metschnikowiaceae(100);g__Metschnikowia(100);s__Metschnikowia_pulcherrima(100); | Metschnikowia | 13 | Saprotroph | \|Undefined Saprotroph\| | Yeast | NULL | Probable | As nectar-tap_saprotroph (Põlme et al. 2020); Decay sugar-rich substrates (Põlme et al. 2020) | Põlme S, et al. 2020. Fungal Diversity 105:1-16 ((https://doi.org/10.1007/s13225-020-00466-2)) |
| Otu21823 | 1 | k__Fungi(100);p__Ascomycota(100);c__Saccharomycetes(100);o__Saccharomycetales(100);f__Metschnikowiaceae(100);g__Metschnikowia(100);s__Metschnikowia_pulcherrima(100); | Metschnikowia | 13 | Saprotroph | \|Undefined Saprotroph\| | Yeast | NULL | Probable | As nectar-tap_saprotroph (Põlme et al. 2020); Decay sugar-rich substrates (Põlme et al. 2020) | Põlme S, et al. 2020. Fungal Diversity 105:1-16 ((https://doi.org/10.1007/s13225-020-00466-2)) |
| Otu21825 | 1 | k__Fungi(100);p__Ascomycota(100);c__Saccharomycetes(100);o__Saccharomycetales(100);f__Metschnikowiaceae(100);g__Metschnikowia(100);s__Metschnikowia_pulcherrima(100); | Metschnikowia | 13 | Saprotroph | \|Undefined Saprotroph\| | Yeast | NULL | Probable | As nectar-tap_saprotroph (Põlme et al. 2020); Decay sugar-rich substrates (Põlme et al. 2020) | Põlme S, et al. 2020. Fungal Diversity 105:1-16 ((https://doi.org/10.1007/s13225-020-00466-2)) |
| Otu21826 | 1 | k__Fungi(100);p__Ascomycota(100);c__Saccharomycetes(100);o__Saccharomycetales(100);f__Metschnikowiaceae(100);g__Metschnikowia(100);s__Metschnikowia_pulcherrima(100); | Metschnikowia | 13 | Saprotroph | \|Undefined Saprotroph\| | Yeast | NULL | Probable | As nectar-tap_saprotroph (Põlme et al. 2020); Decay sugar-rich substrates (Põlme et al. 2020) | Põlme S, et al. 2020. Fungal Diversity 105:1-16 ((https://doi.org/10.1007/s13225-020-00466-2)) |
| Otu21827 | 1 | k__Fungi(100);p__Ascomycota(100);c__Saccharomycetes(100);o__Saccharomycetales(100);f__Metschnikowiaceae(100);g__Metschnikowia(100);s__Metschnikowia_pulcherrima(100); | Metschnikowia | 13 | Saprotroph | \|Undefined Saprotroph\| | Yeast | NULL | Probable | As nectar-tap_saprotroph (Põlme et al. 2020); Decay sugar-rich substrates (Põlme et al. 2020) | Põlme S, et al. 2020. Fungal Diversity 105:1-16 ((https://doi.org/10.1007/s13225-020-00466-2)) |
| Otu21828 | 1 | k__Fungi(100);p__Ascomycota(100);c__Saccharomycetes(100);o__Saccharomycetales(100);f__Metschnikowiaceae(100);g__Metschnikowia(100);s__Metschnikowia_pulcherrima(100); | Metschnikowia | 13 | Saprotroph | \|Undefined Saprotroph\| | Yeast | NULL | Probable | As nectar-tap_saprotroph (Põlme et al. 2020); Decay sugar-rich substrates (Põlme et al. 2020) | Põlme S, et al. 2020. Fungal Diversity 105:1-16 ((https://doi.org/10.1007/s13225-020-00466-2)) |
| Otu21829 | 1 | k__Fungi(100);p__Ascomycota(100);c__Saccharomycetes(100);o__Saccharomycetales(100);f__Metschnikowiaceae(100);g__Metschnikowia(100);s__Metschnikowia_pulcherrima(100); | Metschnikowia | 13 | Saprotroph | \|Undefined Saprotroph\| | Yeast | NULL | Probable | As nectar-tap_saprotroph (Põlme et al. 2020); Decay sugar-rich substrates (Põlme et al. 2020) | Põlme S, et al. 2020. Fungal Diversity 105:1-16 ((https://doi.org/10.1007/s13225-020-00466-2)) |
| Otu21830 | 1 | k__Fungi(100);p__Ascomycota(100);c__Saccharomycetes(100);o__Saccharomycetales(100);f__Metschnikowiaceae(100);g__Metschnikowia(100);g__Metschnikowia_unclassified(100); | Metschnikowia | 13 | Saprotroph | \|Undefined Saprotroph\| | Yeast | NULL | Probable | As nectar-tap_saprotroph (Põlme et al. 2020); Decay sugar-rich substrates (Põlme et al. 2020) | Põlme S, et al. 2020. Fungal Diversity 105:1-16 ((https://doi.org/10.1007/s13225-020-00466-2)) |
| Otu21831 | 1 | k__Fungi(100);p__Ascomycota(100);c__Saccharomycetes(100);o__Saccharomycetales(100);f__Metschnikowiaceae(100);g__Metschnikowia(100);g__Metschnikowia_unclassified(100); | Metschnikowia | 13 | Saprotroph | \|Undefined Saprotroph\| | Yeast | NULL | Probable | As nectar-tap_saprotroph (Põlme et al. 2020); Decay sugar-rich substrates (Põlme et al. 2020) | Põlme S, et al. 2020. Fungal Diversity 105:1-16 ((https://doi.org/10.1007/s13225-020-00466-2)) |
| Otu21832 | 1 | k__Fungi(100);p__Ascomycota(100);c__Saccharomycetes(100);o__Saccharomycetales(100);f__Metschnikowiaceae(100);g__Metschnikowia(100);g__Metschnikowia_unclassified(100); | Metschnikowia | 13 | Saprotroph | \|Undefined Saprotroph\| | Yeast | NULL | Probable | As nectar-tap_saprotroph (Põlme et al. 2020); Decay sugar-rich substrates (Põlme et al. 2020) | Põlme S, et al. 2020. Fungal Diversity 105:1-16 ((https://doi.org/10.1007/s13225-020-00466-2)) |
| Otu21833 | 1 | k__Fungi(100);p__Ascomycota(100);c__Saccharomycetes(100);o__Saccharomycetales(100);f__Metschnikowiaceae(100);g__Metschnikowia(100);s__Metschnikowia_pulcherrima(100); | Metschnikowia | 13 | Saprotroph | \|Undefined Saprotroph\| | Yeast | NULL | Probable | As nectar-tap_saprotroph (Põlme et al. 2020); Decay sugar-rich substrates (Põlme et al. 2020) | Põlme S, et al. 2020. Fungal Diversity 105:1-16 ((https://doi.org/10.1007/s13225-020-00466-2)) |
| Otu21834 | 1 | k__Fungi(100);p__Ascomycota(100);c__Saccharomycetes(100);o__Saccharomycetales(100);f__Metschnikowiaceae(100);g__Metschnikowia(100);s__Metschnikowia_pulcherrima(100); | Metschnikowia | 13 | Saprotroph | \|Undefined Saprotroph\| | Yeast | NULL | Probable | As nectar-tap_saprotroph (Põlme et al. 2020); Decay sugar-rich substrates (Põlme et al. 2020) | Põlme S, et al. 2020. Fungal Diversity 105:1-16 ((https://doi.org/10.1007/s13225-020-00466-2)) |
| Otu21835 | 1 | k__Fungi(100);p__Ascomycota(100);c__Saccharomycetes(100);o__Saccharomycetales(100);f__Metschnikowiaceae(100);g__Metschnikowia(100);s__Metschnikowia_pulcherrima(100); | Metschnikowia | 13 | Saprotroph | \|Undefined Saprotroph\| | Yeast | NULL | Probable | As nectar-tap_saprotroph (Põlme et al. 2020); Decay sugar-rich substrates (Põlme et al. 2020) | Põlme S, et al. 2020. Fungal Diversity 105:1-16 ((https://doi.org/10.1007/s13225-020-00466-2)) |
| Otu21836 | 1 | k__Fungi(100);p__Ascomycota(100);c__Saccharomycetes(100);o__Saccharomycetales(100);f__Metschnikowiaceae(100);g__Metschnikowia(100);g__Metschnikowia_unclassified(100); | Metschnikowia | 13 | Saprotroph | \|Undefined Saprotroph\| | Yeast | NULL | Probable | As nectar-tap_saprotroph (Põlme et al. 2020); Decay sugar-rich substrates (Põlme et al. 2020) | Põlme S, et al. 2020. Fungal Diversity 105:1-16 ((https://doi.org/10.1007/s13225-020-00466-2)) |
| Otu21837 | 1 | k__Fungi(100);p__Ascomycota(100);c__Saccharomycetes(100);o__Saccharomycetales(100);f__Metschnikowiaceae(100);g__Metschnikowia(100);s__Metschnikowia_pulcherrima(100); | Metschnikowia | 13 | Saprotroph | \|Undefined Saprotroph\| | Yeast | NULL | Probable | As nectar-tap_saprotroph (Põlme et al. 2020); Decay sugar-rich substrates (Põlme et al. 2020) | Põlme S, et al. 2020. Fungal Diversity 105:1-16 ((https://doi.org/10.1007/s13225-020-00466-2)) |
| Otu21838 | 1 | k__Fungi(100);p__Ascomycota(100);c__Saccharomycetes(100);o__Saccharomycetales(100);f__Metschnikowiaceae(100);g__Metschnikowia(100);s__Metschnikowia_pulcherrima(100); | Metschnikowia | 13 | Saprotroph | \|Undefined Saprotroph\| | Yeast | NULL | Probable | As nectar-tap_saprotroph (Põlme et al. 2020); Decay sugar-rich substrates (Põlme et al. 2020) | Põlme S, et al. 2020. Fungal Diversity 105:1-16 ((https://doi.org/10.1007/s13225-020-00466-2)) |
| Otu21839 | 1 | k__Fungi(100);p__Ascomycota(100);c__Saccharomycetes(100);o__Saccharomycetales(100);f__Metschnikowiaceae(100);g__Metschnikowia(100);s__Metschnikowia_pulcherrima(100); | Metschnikowia | 13 | Saprotroph | \|Undefined Saprotroph\| | Yeast | NULL | Probable | As nectar-tap_saprotroph (Põlme et al. 2020); Decay sugar-rich substrates (Põlme et al. 2020) | Põlme S, et al. 2020. Fungal Diversity 105:1-16 ((https://doi.org/10.1007/s13225-020-00466-2)) |
| Otu21845 | 1 | k__Fungi(100);p__Ascomycota(100);c__Saccharomycetes(100);o__Saccharomycetales(100);f__Metschnikowiaceae(100);g__Metschnikowia(100);g__Metschnikowia_unclassified(100); | Metschnikowia | 13 | Saprotroph | \|Undefined Saprotroph\| | Yeast | NULL | Probable | As nectar-tap_saprotroph (Põlme et al. 2020); Decay sugar-rich substrates (Põlme et al. 2020) | Põlme S, et al. 2020. Fungal Diversity 105:1-16 ((https://doi.org/10.1007/s13225-020-00466-2)) |
| Otu21846 | 1 | k__Fungi(100);p__Ascomycota(100);c__Saccharomycetes(100);o__Saccharomycetales(100);f__Metschnikowiaceae(100);g__Metschnikowia(100);s__Metschnikowia_pulcherrima(100); | Metschnikowia | 13 | Saprotroph | \|Undefined Saprotroph\| | Yeast | NULL | Probable | As nectar-tap_saprotroph (Põlme et al. 2020); Decay sugar-rich substrates (Põlme et al. 2020) | Põlme S, et al. 2020. Fungal Diversity 105:1-16 ((https://doi.org/10.1007/s13225-020-00466-2)) |
| Otu21847 | 1 | k__Fungi(100);p__Ascomycota(100);c__Saccharomycetes(100);o__Saccharomycetales(100);f__Metschnikowiaceae(100);g__Metschnikowia(100);s__Metschnikowia_pulcherrima(100); | Metschnikowia | 13 | Saprotroph | \|Undefined Saprotroph\| | Yeast | NULL | Probable | As nectar-tap_saprotroph (Põlme et al. 2020); Decay sugar-rich substrates (Põlme et al. 2020) | Põlme S, et al. 2020. Fungal Diversity 105:1-16 ((https://doi.org/10.1007/s13225-020-00466-2)) |
| Otu21848 | 1 | k__Fungi(100);p__Ascomycota(100);c__Saccharomycetes(100);o__Saccharomycetales(100);f__Metschnikowiaceae(100);g__Metschnikowia(100);s__Metschnikowia_pulcherrima(100); | Metschnikowia | 13 | Saprotroph | \|Undefined Saprotroph\| | Yeast | NULL | Probable | As nectar-tap_saprotroph (Põlme et al. 2020); Decay sugar-rich substrates (Põlme et al. 2020) | Põlme S, et al. 2020. Fungal Diversity 105:1-16 ((https://doi.org/10.1007/s13225-020-00466-2)) |
| Otu21849 | 1 | k__Fungi(100);p__Ascomycota(100);c__Saccharomycetes(100);o__Saccharomycetales(100);f__Metschnikowiaceae(100);g__Metschnikowia(100);s__Metschnikowia_pulcherrima(100); | Metschnikowia | 13 | Saprotroph | \|Undefined Saprotroph\| | Yeast | NULL | Probable | As nectar-tap_saprotroph (Põlme et al. 2020); Decay sugar-rich substrates (Põlme et al. 2020) | Põlme S, et al. 2020. Fungal Diversity 105:1-16 ((https://doi.org/10.1007/s13225-020-00466-2)) |
| Otu21850 | 1 | k__Fungi(100);p__Ascomycota(100);c__Saccharomycetes(100);o__Saccharomycetales(100);f__Metschnikowiaceae(100);g__Metschnikowia(100);s__Metschnikowia_pulcherrima(100); | Metschnikowia | 13 | Saprotroph | \|Undefined Saprotroph\| | Yeast | NULL | Probable | As nectar-tap_saprotroph (Põlme et al. 2020); Decay sugar-rich substrates (Põlme et al. 2020) | Põlme S, et al. 2020. Fungal Diversity 105:1-16 ((https://doi.org/10.1007/s13225-020-00466-2)) |
| Otu21852 | 1 | k__Fungi(100);p__Ascomycota(100);c__Saccharomycetes(100);o__Saccharomycetales(100);f__Saccharomycetales_family_Incertae_sedis(100);g__Candida(100);g__Candida_unclassified(100); | Candida | 13 | Pathotroph-Saprotroph-Symbiotroph | Animal Pathogen-Endophyte-Endosymbiont-Epiphyte-\|Undefined Saprotroph\| | Dimorphic Yeast-Yeast | NULL | Probable | As nectar-tap_saprotroph (Põlme et al. 2020); Decay sugar-rich substrates (Põlme et al. 2020); This asexual genus is polyphyletic and contain species that function very differently depend on what they are. Caution should be taken when interpreting guilds for Candida species. | Manolakaki D, et al. 2010. Virulence. 1: 367-375 ((https://doi.org/10.4161/viru.1.5.12796)); Mekbib SB, et al. 2011. Tropical Plant Pathology 36:233-240 ((https://doi.org/10.1590/S1982-56762011000400004)); Põlme S, et al. 2020. Fungal Diversity 105:1-16 ((https://doi.org/10.1007/s13225-020-00466-2)) |
| Otu21863 | 1 | k__Fungi(100);p__Ascomycota(100);c__Saccharomycetes(100);o__Saccharomycetales(100);f__Metschnikowiaceae(100);g__Metschnikowia(100);s__Metschnikowia_pulcherrima(100); | Metschnikowia | 13 | Saprotroph | \|Undefined Saprotroph\| | Yeast | NULL | Probable | As nectar-tap_saprotroph (Põlme et al. 2020); Decay sugar-rich substrates (Põlme et al. 2020) | Põlme S, et al. 2020. Fungal Diversity 105:1-16 ((https://doi.org/10.1007/s13225-020-00466-2)) |
| Otu21864 | 1 | k__Fungi(100);p__Ascomycota(100);c__Saccharomycetes(100);o__Saccharomycetales(100);f__Metschnikowiaceae(100);g__Metschnikowia(100);s__Metschnikowia_pulcherrima(100); | Metschnikowia | 13 | Saprotroph | \|Undefined Saprotroph\| | Yeast | NULL | Probable | As nectar-tap_saprotroph (Põlme et al. 2020); Decay sugar-rich substrates (Põlme et al. 2020) | Põlme S, et al. 2020. Fungal Diversity 105:1-16 ((https://doi.org/10.1007/s13225-020-00466-2)) |
| Otu21867 | 1 | k__Fungi(100);p__Ascomycota(100);c__Saccharomycetes(100);o__Saccharomycetales(100);f__Metschnikowiaceae(100);g__Metschnikowia(100);s__Metschnikowia_pulcherrima(100); | Metschnikowia | 13 | Saprotroph | \|Undefined Saprotroph\| | Yeast | NULL | Probable | As nectar-tap_saprotroph (Põlme et al. 2020); Decay sugar-rich substrates (Põlme et al. 2020) | Põlme S, et al. 2020. Fungal Diversity 105:1-16 ((https://doi.org/10.1007/s13225-020-00466-2)) |
| Otu21868 | 1 | k__Fungi(100);p__Ascomycota(100);c__Saccharomycetes(100);o__Saccharomycetales(100);f__Metschnikowiaceae(100);g__Metschnikowia(100);s__Metschnikowia_pulcherrima(100); | Metschnikowia | 13 | Saprotroph | \|Undefined Saprotroph\| | Yeast | NULL | Probable | As nectar-tap_saprotroph (Põlme et al. 2020); Decay sugar-rich substrates (Põlme et al. 2020) | Põlme S, et al. 2020. Fungal Diversity 105:1-16 ((https://doi.org/10.1007/s13225-020-00466-2)) |
| Otu21869 | 1 | k__Fungi(100);p__Ascomycota(100);c__Saccharomycetes(100);o__Saccharomycetales(100);f__Metschnikowiaceae(100);g__Metschnikowia(100);s__Metschnikowia_pulcherrima(100); | Metschnikowia | 13 | Saprotroph | \|Undefined Saprotroph\| | Yeast | NULL | Probable | As nectar-tap_saprotroph (Põlme et al. 2020); Decay sugar-rich substrates (Põlme et al. 2020) | Põlme S, et al. 2020. Fungal Diversity 105:1-16 ((https://doi.org/10.1007/s13225-020-00466-2)) |
| Otu21870 | 1 | k__Fungi(100);p__Ascomycota(100);c__Saccharomycetes(100);o__Saccharomycetales(100);f__Metschnikowiaceae(100);g__Metschnikowia(100);s__Metschnikowia_pulcherrima(100); | Metschnikowia | 13 | Saprotroph | \|Undefined Saprotroph\| | Yeast | NULL | Probable | As nectar-tap_saprotroph (Põlme et al. 2020); Decay sugar-rich substrates (Põlme et al. 2020) | Põlme S, et al. 2020. Fungal Diversity 105:1-16 ((https://doi.org/10.1007/s13225-020-00466-2)) |
| Otu21871 | 1 | k__Fungi(100);p__Ascomycota(100);c__Saccharomycetes(100);o__Saccharomycetales(100);f__Metschnikowiaceae(100);g__Metschnikowia(100);g__Metschnikowia_unclassified(100); | Metschnikowia | 13 | Saprotroph | \|Undefined Saprotroph\| | Yeast | NULL | Probable | As nectar-tap_saprotroph (Põlme et al. 2020); Decay sugar-rich substrates (Põlme et al. 2020) | Põlme S, et al. 2020. Fungal Diversity 105:1-16 ((https://doi.org/10.1007/s13225-020-00466-2)) |
| Otu21872 | 1 | k__Fungi(100);p__Ascomycota(100);c__Saccharomycetes(100);o__Saccharomycetales(100);f__Metschnikowiaceae(100);g__Metschnikowia(100);s__Metschnikowia_pulcherrima(100); | Metschnikowia | 13 | Saprotroph | \|Undefined Saprotroph\| | Yeast | NULL | Probable | As nectar-tap_saprotroph (Põlme et al. 2020); Decay sugar-rich substrates (Põlme et al. 2020) | Põlme S, et al. 2020. Fungal Diversity 105:1-16 ((https://doi.org/10.1007/s13225-020-00466-2)) |
| Otu21873 | 1 | k__Fungi(100);p__Ascomycota(100);c__Saccharomycetes(100);o__Saccharomycetales(100);f__Metschnikowiaceae(100);g__Metschnikowia(100);s__Metschnikowia_pulcherrima(100); | Metschnikowia | 13 | Saprotroph | \|Undefined Saprotroph\| | Yeast | NULL | Probable | As nectar-tap_saprotroph (Põlme et al. 2020); Decay sugar-rich substrates (Põlme et al. 2020) | Põlme S, et al. 2020. Fungal Diversity 105:1-16 ((https://doi.org/10.1007/s13225-020-00466-2)) |
| Otu21874 | 1 | k__Fungi(100);p__Ascomycota(100);c__Saccharomycetes(100);o__Saccharomycetales(100);f__Metschnikowiaceae(100);g__Metschnikowia(100);s__Metschnikowia_pulcherrima(100); | Metschnikowia | 13 | Saprotroph | \|Undefined Saprotroph\| | Yeast | NULL | Probable | As nectar-tap_saprotroph (Põlme et al. 2020); Decay sugar-rich substrates (Põlme et al. 2020) | Põlme S, et al. 2020. Fungal Diversity 105:1-16 ((https://doi.org/10.1007/s13225-020-00466-2)) |
| Otu21875 | 1 | k__Fungi(100);p__Ascomycota(100);c__Saccharomycetes(100);o__Saccharomycetales(100);f__Metschnikowiaceae(100);g__Metschnikowia(100);s__Metschnikowia_pulcherrima(100); | Metschnikowia | 13 | Saprotroph | \|Undefined Saprotroph\| | Yeast | NULL | Probable | As nectar-tap_saprotroph (Põlme et al. 2020); Decay sugar-rich substrates (Põlme et al. 2020) | Põlme S, et al. 2020. Fungal Diversity 105:1-16 ((https://doi.org/10.1007/s13225-020-00466-2)) |
| Otu21876 | 1 | k__Fungi(100);p__Ascomycota(100);c__Saccharomycetes(100);o__Saccharomycetales(100);f__Metschnikowiaceae(100);g__Metschnikowia(100);s__Metschnikowia_pulcherrima(100); | Metschnikowia | 13 | Saprotroph | \|Undefined Saprotroph\| | Yeast | NULL | Probable | As nectar-tap_saprotroph (Põlme et al. 2020); Decay sugar-rich substrates (Põlme et al. 2020) | Põlme S, et al. 2020. Fungal Diversity 105:1-16 ((https://doi.org/10.1007/s13225-020-00466-2)) |
| Otu21880 | 1 | k__Fungi(100);p__Ascomycota(100);c__Saccharomycetes(100);o__Saccharomycetales(100);f__Saccharomycetales_family_Incertae_sedis(100);g__Candida(100);g__Candida_unclassified(100); | Candida | 13 | Pathotroph-Saprotroph-Symbiotroph | Animal Pathogen-Endophyte-Endosymbiont-Epiphyte-\|Undefined Saprotroph\| | Dimorphic Yeast-Yeast | NULL | Probable | As nectar-tap_saprotroph (Põlme et al. 2020); Decay sugar-rich substrates (Põlme et al. 2020); This asexual genus is polyphyletic and contain species that function very differently depend on what they are. Caution should be taken when interpreting guilds for Candida species. | Manolakaki D, et al. 2010. Virulence. 1: 367-375 ((https://doi.org/10.4161/viru.1.5.12796)); Mekbib SB, et al. 2011. Tropical Plant Pathology 36:233-240 ((https://doi.org/10.1590/S1982-56762011000400004)); Põlme S, et al. 2020. Fungal Diversity 105:1-16 ((https://doi.org/10.1007/s13225-020-00466-2)) |
| Otu21882 | 1 | k__Fungi(100);p__Ascomycota(100);c__Saccharomycetes(100);o__Saccharomycetales(100);f__Metschnikowiaceae(100);g__Metschnikowia(100);s__Metschnikowia_pulcherrima(100); | Metschnikowia | 13 | Saprotroph | \|Undefined Saprotroph\| | Yeast | NULL | Probable | As nectar-tap_saprotroph (Põlme et al. 2020); Decay sugar-rich substrates (Põlme et al. 2020) | Põlme S, et al. 2020. Fungal Diversity 105:1-16 ((https://doi.org/10.1007/s13225-020-00466-2)) |
| Otu21886 | 1 | k__Fungi(100);p__Ascomycota(100);c__Saccharomycetes(100);o__Saccharomycetales(100);f__Metschnikowiaceae(100);g__Metschnikowia(100);s__Metschnikowia_pulcherrima(100); | Metschnikowia | 13 | Saprotroph | \|Undefined Saprotroph\| | Yeast | NULL | Probable | As nectar-tap_saprotroph (Põlme et al. 2020); Decay sugar-rich substrates (Põlme et al. 2020) | Põlme S, et al. 2020. Fungal Diversity 105:1-16 ((https://doi.org/10.1007/s13225-020-00466-2)) |
| Otu21887 | 1 | k__Fungi(100);p__Ascomycota(100);c__Saccharomycetes(100);o__Saccharomycetales(100);f__Saccharomycetales_family_Incertae_sedis(100);g__Candida(100);s__Candida_cretensis(100); | Candida | 13 | Pathotroph-Saprotroph-Symbiotroph | Animal Pathogen-Endophyte-Endosymbiont-Epiphyte-\|Undefined Saprotroph\| | Dimorphic Yeast-Yeast | NULL | Probable | As nectar-tap_saprotroph (Põlme et al. 2020); Decay sugar-rich substrates (Põlme et al. 2020); This asexual genus is polyphyletic and contain species that function very differently depend on what they are. Caution should be taken when interpreting guilds for Candida species. | Manolakaki D, et al. 2010. Virulence. 1: 367-375 ((https://doi.org/10.4161/viru.1.5.12796)); Mekbib SB, et al. 2011. Tropical Plant Pathology 36:233-240 ((https://doi.org/10.1590/S1982-56762011000400004)); Põlme S, et al. 2020. Fungal Diversity 105:1-16 ((https://doi.org/10.1007/s13225-020-00466-2)) |
| Otu21889 | 1 | k__Fungi(100);p__Ascomycota(100);c__Saccharomycetes(100);o__Saccharomycetales(100);f__Metschnikowiaceae(100);g__Metschnikowia(100);s__Metschnikowia_pulcherrima(100); | Metschnikowia | 13 | Saprotroph | \|Undefined Saprotroph\| | Yeast | NULL | Probable | As nectar-tap_saprotroph (Põlme et al. 2020); Decay sugar-rich substrates (Põlme et al. 2020) | Põlme S, et al. 2020. Fungal Diversity 105:1-16 ((https://doi.org/10.1007/s13225-020-00466-2)) |
| Otu21891 | 1 | k__Fungi(100);p__Ascomycota(100);c__Saccharomycetes(100);o__Saccharomycetales(100);o__Saccharomycetales_unclassified(100);o__Saccharomycetales_unclassified(100);o__Saccharomycetales_unclassified(100); | Saccharomycetales | 7 | Saprotroph | Undefined Saprotroph | Yeast | NULL | Possible | NULL | Sterkenburg E, et al. 2015. New Phytologist 207:1145-1158 ((https://doi.org/10.1111/nph.13426)) |
| Otu21894 | 1 | k__Fungi(100);p__Ascomycota(100);c__Saccharomycetes(100);o__Saccharomycetales(100);f__Saccharomycetales_family_Incertae_sedis(100);g__Candida(100);g__Candida_unclassified(100); | Candida | 13 | Pathotroph-Saprotroph-Symbiotroph | Animal Pathogen-Endophyte-Endosymbiont-Epiphyte-\|Undefined Saprotroph\| | Dimorphic Yeast-Yeast | NULL | Probable | As nectar-tap_saprotroph (Põlme et al. 2020); Decay sugar-rich substrates (Põlme et al. 2020); This asexual genus is polyphyletic and contain species that function very differently depend on what they are. Caution should be taken when interpreting guilds for Candida species. | Manolakaki D, et al. 2010. Virulence. 1: 367-375 ((https://doi.org/10.4161/viru.1.5.12796)); Mekbib SB, et al. 2011. Tropical Plant Pathology 36:233-240 ((https://doi.org/10.1590/S1982-56762011000400004)); Põlme S, et al. 2020. Fungal Diversity 105:1-16 ((https://doi.org/10.1007/s13225-020-00466-2)) |
| Otu21896 | 1 | k__Fungi(100);p__Ascomycota(100);c__Saccharomycetes(100);o__Saccharomycetales(100);f__Metschnikowiaceae(100);g__Metschnikowia(100);s__Metschnikowia_pulcherrima(100); | Metschnikowia | 13 | Saprotroph | \|Undefined Saprotroph\| | Yeast | NULL | Probable | As nectar-tap_saprotroph (Põlme et al. 2020); Decay sugar-rich substrates (Põlme et al. 2020) | Põlme S, et al. 2020. Fungal Diversity 105:1-16 ((https://doi.org/10.1007/s13225-020-00466-2)) |
| Otu21897 | 1 | k__Fungi(100);p__Ascomycota(100);c__Saccharomycetes(100);o__Saccharomycetales(100);f__Saccharomycetales_family_Incertae_sedis(100);g__Candida(100);g__Candida_unclassified(100); | Candida | 13 | Pathotroph-Saprotroph-Symbiotroph | Animal Pathogen-Endophyte-Endosymbiont-Epiphyte-\|Undefined Saprotroph\| | Dimorphic Yeast-Yeast | NULL | Probable | As nectar-tap_saprotroph (Põlme et al. 2020); Decay sugar-rich substrates (Põlme et al. 2020); This asexual genus is polyphyletic and contain species that function very differently depend on what they are. Caution should be taken when interpreting guilds for Candida species. | Manolakaki D, et al. 2010. Virulence. 1: 367-375 ((https://doi.org/10.4161/viru.1.5.12796)); Mekbib SB, et al. 2011. Tropical Plant Pathology 36:233-240 ((https://doi.org/10.1590/S1982-56762011000400004)); Põlme S, et al. 2020. Fungal Diversity 105:1-16 ((https://doi.org/10.1007/s13225-020-00466-2)) |
| Otu21898 | 1 | k__Fungi(100);p__Ascomycota(100);c__Saccharomycetes(100);o__Saccharomycetales(100);f__Metschnikowiaceae(100);g__Metschnikowia(100);s__Metschnikowia_pulcherrima(100); | Metschnikowia | 13 | Saprotroph | \|Undefined Saprotroph\| | Yeast | NULL | Probable | As nectar-tap_saprotroph (Põlme et al. 2020); Decay sugar-rich substrates (Põlme et al. 2020) | Põlme S, et al. 2020. Fungal Diversity 105:1-16 ((https://doi.org/10.1007/s13225-020-00466-2)) |
| Otu21901 | 1 | k__Fungi(100);p__Ascomycota(100);c__Saccharomycetes(100);o__Saccharomycetales(100);f__Metschnikowiaceae(100);g__Metschnikowia(100);g__Metschnikowia_unclassified(100); | Metschnikowia | 13 | Saprotroph | \|Undefined Saprotroph\| | Yeast | NULL | Probable | As nectar-tap_saprotroph (Põlme et al. 2020); Decay sugar-rich substrates (Põlme et al. 2020) | Põlme S, et al. 2020. Fungal Diversity 105:1-16 ((https://doi.org/10.1007/s13225-020-00466-2)) |
| Otu21902 | 1 | k__Fungi(100);p__Ascomycota(100);c__Saccharomycetes(100);o__Saccharomycetales(100);f__Saccharomycetales_family_Incertae_sedis(100);g__Candida(100);g__Candida_unclassified(100); | Candida | 13 | Pathotroph-Saprotroph-Symbiotroph | Animal Pathogen-Endophyte-Endosymbiont-Epiphyte-\|Undefined Saprotroph\| | Dimorphic Yeast-Yeast | NULL | Probable | As nectar-tap_saprotroph (Põlme et al. 2020); Decay sugar-rich substrates (Põlme et al. 2020); This asexual genus is polyphyletic and contain species that function very differently depend on what they are. Caution should be taken when interpreting guilds for Candida species. | Manolakaki D, et al. 2010. Virulence. 1: 367-375 ((https://doi.org/10.4161/viru.1.5.12796)); Mekbib SB, et al. 2011. Tropical Plant Pathology 36:233-240 ((https://doi.org/10.1590/S1982-56762011000400004)); Põlme S, et al. 2020. Fungal Diversity 105:1-16 ((https://doi.org/10.1007/s13225-020-00466-2)) |
| Otu21903 | 1 | k__Fungi(100);p__Ascomycota(100);c__Saccharomycetes(100);o__Saccharomycetales(100);f__Metschnikowiaceae(100);g__Metschnikowia(100);g__Metschnikowia_unclassified(100); | Metschnikowia | 13 | Saprotroph | \|Undefined Saprotroph\| | Yeast | NULL | Probable | As nectar-tap_saprotroph (Põlme et al. 2020); Decay sugar-rich substrates (Põlme et al. 2020) | Põlme S, et al. 2020. Fungal Diversity 105:1-16 ((https://doi.org/10.1007/s13225-020-00466-2)) |
| Otu21904 | 1 | k__Fungi(100);p__Ascomycota(100);c__Saccharomycetes(100);o__Saccharomycetales(100);f__Metschnikowiaceae(100);g__Metschnikowia(100);g__Metschnikowia_unclassified(100); | Metschnikowia | 13 | Saprotroph | \|Undefined Saprotroph\| | Yeast | NULL | Probable | As nectar-tap_saprotroph (Põlme et al. 2020); Decay sugar-rich substrates (Põlme et al. 2020) | Põlme S, et al. 2020. Fungal Diversity 105:1-16 ((https://doi.org/10.1007/s13225-020-00466-2)) |
| Otu21906 | 1 | k__Fungi(100);p__Ascomycota(100);c__Saccharomycetes(100);o__Saccharomycetales(100);f__Metschnikowiaceae(100);g__Metschnikowia(100);s__Metschnikowia_pulcherrima(100); | Metschnikowia | 13 | Saprotroph | \|Undefined Saprotroph\| | Yeast | NULL | Probable | As nectar-tap_saprotroph (Põlme et al. 2020); Decay sugar-rich substrates (Põlme et al. 2020) | Põlme S, et al. 2020. Fungal Diversity 105:1-16 ((https://doi.org/10.1007/s13225-020-00466-2)) |
| Otu21907 | 1 | k__Fungi(100);p__Ascomycota(100);c__Saccharomycetes(100);o__Saccharomycetales(100);f__Metschnikowiaceae(100);g__Metschnikowia(100);g__Metschnikowia_unclassified(100); | Metschnikowia | 13 | Saprotroph | \|Undefined Saprotroph\| | Yeast | NULL | Probable | As nectar-tap_saprotroph (Põlme et al. 2020); Decay sugar-rich substrates (Põlme et al. 2020) | Põlme S, et al. 2020. Fungal Diversity 105:1-16 ((https://doi.org/10.1007/s13225-020-00466-2)) |
| Otu21908 | 1 | k__Fungi(100);p__Ascomycota(100);c__Saccharomycetes(100);o__Saccharomycetales(100);f__Metschnikowiaceae(100);g__Metschnikowia(100);s__Metschnikowia_pulcherrima(100); | Metschnikowia | 13 | Saprotroph | \|Undefined Saprotroph\| | Yeast | NULL | Probable | As nectar-tap_saprotroph (Põlme et al. 2020); Decay sugar-rich substrates (Põlme et al. 2020) | Põlme S, et al. 2020. Fungal Diversity 105:1-16 ((https://doi.org/10.1007/s13225-020-00466-2)) |
| Otu21909 | 1 | k__Fungi(100);p__Ascomycota(100);c__Saccharomycetes(100);o__Saccharomycetales(100);f__Metschnikowiaceae(100);g__Metschnikowia(100);s__Metschnikowia_pulcherrima(100); | Metschnikowia | 13 | Saprotroph | \|Undefined Saprotroph\| | Yeast | NULL | Probable | As nectar-tap_saprotroph (Põlme et al. 2020); Decay sugar-rich substrates (Põlme et al. 2020) | Põlme S, et al. 2020. Fungal Diversity 105:1-16 ((https://doi.org/10.1007/s13225-020-00466-2)) |
| Otu21910 | 1 | k__Fungi(100);p__Ascomycota(100);c__Saccharomycetes(100);o__Saccharomycetales(100);f__Metschnikowiaceae(100);g__Metschnikowia(100);s__Metschnikowia_pulcherrima(100); | Metschnikowia | 13 | Saprotroph | \|Undefined Saprotroph\| | Yeast | NULL | Probable | As nectar-tap_saprotroph (Põlme et al. 2020); Decay sugar-rich substrates (Põlme et al. 2020) | Põlme S, et al. 2020. Fungal Diversity 105:1-16 ((https://doi.org/10.1007/s13225-020-00466-2)) |
| Otu21911 | 1 | k__Fungi(100);p__Ascomycota(100);c__Saccharomycetes(100);o__Saccharomycetales(100);f__Metschnikowiaceae(100);g__Metschnikowia(100);s__Metschnikowia_pulcherrima(100); | Metschnikowia | 13 | Saprotroph | \|Undefined Saprotroph\| | Yeast | NULL | Probable | As nectar-tap_saprotroph (Põlme et al. 2020); Decay sugar-rich substrates (Põlme et al. 2020) | Põlme S, et al. 2020. Fungal Diversity 105:1-16 ((https://doi.org/10.1007/s13225-020-00466-2)) |
| Otu21912 | 1 | k__Fungi(100);p__Ascomycota(100);c__Saccharomycetes(100);o__Saccharomycetales(100);f__Metschnikowiaceae(100);g__Metschnikowia(100);s__Metschnikowia_pulcherrima(100); | Metschnikowia | 13 | Saprotroph | \|Undefined Saprotroph\| | Yeast | NULL | Probable | As nectar-tap_saprotroph (Põlme et al. 2020); Decay sugar-rich substrates (Põlme et al. 2020) | Põlme S, et al. 2020. Fungal Diversity 105:1-16 ((https://doi.org/10.1007/s13225-020-00466-2)) |
| Otu21913 | 1 | k__Fungi(100);p__Ascomycota(100);c__Saccharomycetes(100);o__Saccharomycetales(100);f__Metschnikowiaceae(100);g__Metschnikowia(100);g__Metschnikowia_unclassified(100); | Metschnikowia | 13 | Saprotroph | \|Undefined Saprotroph\| | Yeast | NULL | Probable | As nectar-tap_saprotroph (Põlme et al. 2020); Decay sugar-rich substrates (Põlme et al. 2020) | Põlme S, et al. 2020. Fungal Diversity 105:1-16 ((https://doi.org/10.1007/s13225-020-00466-2)) |
| Otu21914 | 1 | k__Fungi(100);p__Ascomycota(100);c__Saccharomycetes(100);o__Saccharomycetales(100);f__Metschnikowiaceae(100);g__Metschnikowia(100);s__Metschnikowia_pulcherrima(100); | Metschnikowia | 13 | Saprotroph | \|Undefined Saprotroph\| | Yeast | NULL | Probable | As nectar-tap_saprotroph (Põlme et al. 2020); Decay sugar-rich substrates (Põlme et al. 2020) | Põlme S, et al. 2020. Fungal Diversity 105:1-16 ((https://doi.org/10.1007/s13225-020-00466-2)) |
| Otu21915 | 1 | k__Fungi(100);p__Ascomycota(100);c__Saccharomycetes(100);o__Saccharomycetales(100);f__Metschnikowiaceae(100);g__Metschnikowia(100);s__Metschnikowia_pulcherrima(100); | Metschnikowia | 13 | Saprotroph | \|Undefined Saprotroph\| | Yeast | NULL | Probable | As nectar-tap_saprotroph (Põlme et al. 2020); Decay sugar-rich substrates (Põlme et al. 2020) | Põlme S, et al. 2020. Fungal Diversity 105:1-16 ((https://doi.org/10.1007/s13225-020-00466-2)) |
| Otu21916 | 1 | k__Fungi(100);p__Ascomycota(100);c__Saccharomycetes(100);o__Saccharomycetales(100);f__Metschnikowiaceae(100);g__Metschnikowia(100);s__Metschnikowia_pulcherrima(100); | Metschnikowia | 13 | Saprotroph | \|Undefined Saprotroph\| | Yeast | NULL | Probable | As nectar-tap_saprotroph (Põlme et al. 2020); Decay sugar-rich substrates (Põlme et al. 2020) | Põlme S, et al. 2020. Fungal Diversity 105:1-16 ((https://doi.org/10.1007/s13225-020-00466-2)) |
| Otu21917 | 1 | k__Fungi(100);p__Ascomycota(100);c__Saccharomycetes(100);o__Saccharomycetales(100);f__Metschnikowiaceae(100);g__Metschnikowia(100);s__Metschnikowia_pulcherrima(100); | Metschnikowia | 13 | Saprotroph | \|Undefined Saprotroph\| | Yeast | NULL | Probable | As nectar-tap_saprotroph (Põlme et al. 2020); Decay sugar-rich substrates (Põlme et al. 2020) | Põlme S, et al. 2020. Fungal Diversity 105:1-16 ((https://doi.org/10.1007/s13225-020-00466-2)) |
| Otu21918 | 1 | k__Fungi(100);p__Ascomycota(100);c__Saccharomycetes(100);o__Saccharomycetales(100);f__Metschnikowiaceae(100);g__Metschnikowia(100);s__Metschnikowia_pulcherrima(100); | Metschnikowia | 13 | Saprotroph | \|Undefined Saprotroph\| | Yeast | NULL | Probable | As nectar-tap_saprotroph (Põlme et al. 2020); Decay sugar-rich substrates (Põlme et al. 2020) | Põlme S, et al. 2020. Fungal Diversity 105:1-16 ((https://doi.org/10.1007/s13225-020-00466-2)) |
| Otu21919 | 1 | k__Fungi(100);p__Ascomycota(100);c__Saccharomycetes(100);o__Saccharomycetales(100);f__Metschnikowiaceae(100);g__Metschnikowia(100);s__Metschnikowia_pulcherrima(100); | Metschnikowia | 13 | Saprotroph | \|Undefined Saprotroph\| | Yeast | NULL | Probable | As nectar-tap_saprotroph (Põlme et al. 2020); Decay sugar-rich substrates (Põlme et al. 2020) | Põlme S, et al. 2020. Fungal Diversity 105:1-16 ((https://doi.org/10.1007/s13225-020-00466-2)) |
| Otu21920 | 1 | k__Fungi(100);p__Ascomycota(100);c__Saccharomycetes(100);o__Saccharomycetales(100);f__Metschnikowiaceae(100);g__Metschnikowia(100);g__Metschnikowia_unclassified(100); | Metschnikowia | 13 | Saprotroph | \|Undefined Saprotroph\| | Yeast | NULL | Probable | As nectar-tap_saprotroph (Põlme et al. 2020); Decay sugar-rich substrates (Põlme et al. 2020) | Põlme S, et al. 2020. Fungal Diversity 105:1-16 ((https://doi.org/10.1007/s13225-020-00466-2)) |
| Otu21922 | 1 | k__Fungi(100);p__Ascomycota(100);c__Saccharomycetes(100);o__Saccharomycetales(100);f__Metschnikowiaceae(100);g__Metschnikowia(100);g__Metschnikowia_unclassified(100); | Metschnikowia | 13 | Saprotroph | \|Undefined Saprotroph\| | Yeast | NULL | Probable | As nectar-tap_saprotroph (Põlme et al. 2020); Decay sugar-rich substrates (Põlme et al. 2020) | Põlme S, et al. 2020. Fungal Diversity 105:1-16 ((https://doi.org/10.1007/s13225-020-00466-2)) |
| Otu21923 | 1 | k__Fungi(100);p__Ascomycota(100);c__Saccharomycetes(100);o__Saccharomycetales(100);f__Metschnikowiaceae(100);g__Metschnikowia(100);s__Metschnikowia_pulcherrima(100); | Metschnikowia | 13 | Saprotroph | \|Undefined Saprotroph\| | Yeast | NULL | Probable | As nectar-tap_saprotroph (Põlme et al. 2020); Decay sugar-rich substrates (Põlme et al. 2020) | Põlme S, et al. 2020. Fungal Diversity 105:1-16 ((https://doi.org/10.1007/s13225-020-00466-2)) |
| Otu21924 | 1 | k__Fungi(100);p__Ascomycota(100);c__Saccharomycetes(100);o__Saccharomycetales(100);o__Saccharomycetales_unclassified(100);o__Saccharomycetales_unclassified(100);o__Saccharomycetales_unclassified(100); | Saccharomycetales | 7 | Saprotroph | Undefined Saprotroph | Yeast | NULL | Possible | NULL | Sterkenburg E, et al. 2015. New Phytologist 207:1145-1158 ((https://doi.org/10.1111/nph.13426)) |
| Otu21925 | 1 | k__Fungi(100);p__Ascomycota(100);c__Saccharomycetes(100);o__Saccharomycetales(100);f__Saccharomycetales_family_Incertae_sedis(100);g__Candida(100);g__Candida_unclassified(100); | Candida | 13 | Pathotroph-Saprotroph-Symbiotroph | Animal Pathogen-Endophyte-Endosymbiont-Epiphyte-\|Undefined Saprotroph\| | Dimorphic Yeast-Yeast | NULL | Probable | As nectar-tap_saprotroph (Põlme et al. 2020); Decay sugar-rich substrates (Põlme et al. 2020); This asexual genus is polyphyletic and contain species that function very differently depend on what they are. Caution should be taken when interpreting guilds for Candida species. | Manolakaki D, et al. 2010. Virulence. 1: 367-375 ((https://doi.org/10.4161/viru.1.5.12796)); Mekbib SB, et al. 2011. Tropical Plant Pathology 36:233-240 ((https://doi.org/10.1590/S1982-56762011000400004)); Põlme S, et al. 2020. Fungal Diversity 105:1-16 ((https://doi.org/10.1007/s13225-020-00466-2)) |
| Otu21926 | 1 | k__Fungi(100);p__Ascomycota(100);c__Saccharomycetes(100);o__Saccharomycetales(100);f__Metschnikowiaceae(100);g__Metschnikowia(100);s__Metschnikowia_pulcherrima(100); | Metschnikowia | 13 | Saprotroph | \|Undefined Saprotroph\| | Yeast | NULL | Probable | As nectar-tap_saprotroph (Põlme et al. 2020); Decay sugar-rich substrates (Põlme et al. 2020) | Põlme S, et al. 2020. Fungal Diversity 105:1-16 ((https://doi.org/10.1007/s13225-020-00466-2)) |
| Otu21927 | 1 | k__Fungi(100);p__Ascomycota(100);c__Saccharomycetes(100);o__Saccharomycetales(100);f__Metschnikowiaceae(100);g__Metschnikowia(100);s__Metschnikowia_pulcherrima(100); | Metschnikowia | 13 | Saprotroph | \|Undefined Saprotroph\| | Yeast | NULL | Probable | As nectar-tap_saprotroph (Põlme et al. 2020); Decay sugar-rich substrates (Põlme et al. 2020) | Põlme S, et al. 2020. Fungal Diversity 105:1-16 ((https://doi.org/10.1007/s13225-020-00466-2)) |
| Otu21928 | 1 | k__Fungi(100);p__Ascomycota(100);c__Saccharomycetes(100);o__Saccharomycetales(100);f__Saccharomycetales_family_Incertae_sedis(100);g__Candida(100);g__Candida_unclassified(100); | Candida | 13 | Pathotroph-Saprotroph-Symbiotroph | Animal Pathogen-Endophyte-Endosymbiont-Epiphyte-\|Undefined Saprotroph\| | Dimorphic Yeast-Yeast | NULL | Probable | As nectar-tap_saprotroph (Põlme et al. 2020); Decay sugar-rich substrates (Põlme et al. 2020); This asexual genus is polyphyletic and contain species that function very differently depend on what they are. Caution should be taken when interpreting guilds for Candida species. | Manolakaki D, et al. 2010. Virulence. 1: 367-375 ((https://doi.org/10.4161/viru.1.5.12796)); Mekbib SB, et al. 2011. Tropical Plant Pathology 36:233-240 ((https://doi.org/10.1590/S1982-56762011000400004)); Põlme S, et al. 2020. Fungal Diversity 105:1-16 ((https://doi.org/10.1007/s13225-020-00466-2)) |
| Otu21929 | 1 | k__Fungi(100);p__Ascomycota(100);c__Saccharomycetes(100);o__Saccharomycetales(100);f__Metschnikowiaceae(100);g__Metschnikowia(100);g__Metschnikowia_unclassified(100); | Metschnikowia | 13 | Saprotroph | \|Undefined Saprotroph\| | Yeast | NULL | Probable | As nectar-tap_saprotroph (Põlme et al. 2020); Decay sugar-rich substrates (Põlme et al. 2020) | Põlme S, et al. 2020. Fungal Diversity 105:1-16 ((https://doi.org/10.1007/s13225-020-00466-2)) |
| Otu21936 | 1 | k__Fungi(100);p__Ascomycota(100);c__Saccharomycetes(100);o__Saccharomycetales(100);f__Metschnikowiaceae(100);g__Metschnikowia(100);s__Metschnikowia_pulcherrima(100); | Metschnikowia | 13 | Saprotroph | \|Undefined Saprotroph\| | Yeast | NULL | Probable | As nectar-tap_saprotroph (Põlme et al. 2020); Decay sugar-rich substrates (Põlme et al. 2020) | Põlme S, et al. 2020. Fungal Diversity 105:1-16 ((https://doi.org/10.1007/s13225-020-00466-2)) |
| Otu21937 | 1 | k__Fungi(100);p__Ascomycota(100);c__Saccharomycetes(100);o__Saccharomycetales(100);f__Metschnikowiaceae(100);g__Metschnikowia(100);s__Metschnikowia_pulcherrima(100); | Metschnikowia | 13 | Saprotroph | \|Undefined Saprotroph\| | Yeast | NULL | Probable | As nectar-tap_saprotroph (Põlme et al. 2020); Decay sugar-rich substrates (Põlme et al. 2020) | Põlme S, et al. 2020. Fungal Diversity 105:1-16 ((https://doi.org/10.1007/s13225-020-00466-2)) |
| Otu21938 | 1 | k__Fungi(100);p__Ascomycota(100);c__Saccharomycetes(100);o__Saccharomycetales(100);f__Metschnikowiaceae(100);g__Metschnikowia(100);s__Metschnikowia_pulcherrima(100); | Metschnikowia | 13 | Saprotroph | \|Undefined Saprotroph\| | Yeast | NULL | Probable | As nectar-tap_saprotroph (Põlme et al. 2020); Decay sugar-rich substrates (Põlme et al. 2020) | Põlme S, et al. 2020. Fungal Diversity 105:1-16 ((https://doi.org/10.1007/s13225-020-00466-2)) |
| Otu21939 | 1 | k__Fungi(100);p__Ascomycota(100);c__Saccharomycetes(100);o__Saccharomycetales(100);f__Metschnikowiaceae(100);g__Metschnikowia(100);s__Metschnikowia_pulcherrima(100); | Metschnikowia | 13 | Saprotroph | \|Undefined Saprotroph\| | Yeast | NULL | Probable | As nectar-tap_saprotroph (Põlme et al. 2020); Decay sugar-rich substrates (Põlme et al. 2020) | Põlme S, et al. 2020. Fungal Diversity 105:1-16 ((https://doi.org/10.1007/s13225-020-00466-2)) |
| Otu21941 | 1 | k__Fungi(100);p__Ascomycota(100);c__Saccharomycetes(100);o__Saccharomycetales(100);f__Metschnikowiaceae(100);g__Metschnikowia(100);g__Metschnikowia_unclassified(100); | Metschnikowia | 13 | Saprotroph | \|Undefined Saprotroph\| | Yeast | NULL | Probable | As nectar-tap_saprotroph (Põlme et al. 2020); Decay sugar-rich substrates (Põlme et al. 2020) | Põlme S, et al. 2020. Fungal Diversity 105:1-16 ((https://doi.org/10.1007/s13225-020-00466-2)) |
| Otu21942 | 1 | k__Fungi(100);p__Ascomycota(100);c__Saccharomycetes(100);o__Saccharomycetales(100);f__Metschnikowiaceae(100);g__Metschnikowia(100);s__Metschnikowia_pulcherrima(100); | Metschnikowia | 13 | Saprotroph | \|Undefined Saprotroph\| | Yeast | NULL | Probable | As nectar-tap_saprotroph (Põlme et al. 2020); Decay sugar-rich substrates (Põlme et al. 2020) | Põlme S, et al. 2020. Fungal Diversity 105:1-16 ((https://doi.org/10.1007/s13225-020-00466-2)) |
| Otu21943 | 1 | k__Fungi(100);p__Ascomycota(100);c__Saccharomycetes(100);o__Saccharomycetales(100);f__Metschnikowiaceae(100);g__Metschnikowia(100);s__Metschnikowia_pulcherrima(100); | Metschnikowia | 13 | Saprotroph | \|Undefined Saprotroph\| | Yeast | NULL | Probable | As nectar-tap_saprotroph (Põlme et al. 2020); Decay sugar-rich substrates (Põlme et al. 2020) | Põlme S, et al. 2020. Fungal Diversity 105:1-16 ((https://doi.org/10.1007/s13225-020-00466-2)) |
| Otu21944 | 1 | k__Fungi(100);p__Ascomycota(100);c__Saccharomycetes(100);o__Saccharomycetales(100);f__Metschnikowiaceae(100);g__Metschnikowia(100);g__Metschnikowia_unclassified(100); | Metschnikowia | 13 | Saprotroph | \|Undefined Saprotroph\| | Yeast | NULL | Probable | As nectar-tap_saprotroph (Põlme et al. 2020); Decay sugar-rich substrates (Põlme et al. 2020) | Põlme S, et al. 2020. Fungal Diversity 105:1-16 ((https://doi.org/10.1007/s13225-020-00466-2)) |
| Otu21946 | 1 | k__Fungi(100);p__Ascomycota(100);c__Saccharomycetes(100);o__Saccharomycetales(100);f__Metschnikowiaceae(100);g__Metschnikowia(100);g__Metschnikowia_unclassified(100); | Metschnikowia | 13 | Saprotroph | \|Undefined Saprotroph\| | Yeast | NULL | Probable | As nectar-tap_saprotroph (Põlme et al. 2020); Decay sugar-rich substrates (Põlme et al. 2020) | Põlme S, et al. 2020. Fungal Diversity 105:1-16 ((https://doi.org/10.1007/s13225-020-00466-2)) |
| Otu21947 | 1 | k__Fungi(100);p__Ascomycota(100);c__Saccharomycetes(100);o__Saccharomycetales(100);f__Metschnikowiaceae(100);g__Metschnikowia(100);s__Metschnikowia_pulcherrima(100); | Metschnikowia | 13 | Saprotroph | \|Undefined Saprotroph\| | Yeast | NULL | Probable | As nectar-tap_saprotroph (Põlme et al. 2020); Decay sugar-rich substrates (Põlme et al. 2020) | Põlme S, et al. 2020. Fungal Diversity 105:1-16 ((https://doi.org/10.1007/s13225-020-00466-2)) |
| Otu21948 | 1 | k__Fungi(100);p__Ascomycota(100);c__Saccharomycetes(100);o__Saccharomycetales(100);f__Metschnikowiaceae(100);g__Metschnikowia(100);s__Metschnikowia_pulcherrima(100); | Metschnikowia | 13 | Saprotroph | \|Undefined Saprotroph\| | Yeast | NULL | Probable | As nectar-tap_saprotroph (Põlme et al. 2020); Decay sugar-rich substrates (Põlme et al. 2020) | Põlme S, et al. 2020. Fungal Diversity 105:1-16 ((https://doi.org/10.1007/s13225-020-00466-2)) |
| Otu21949 | 1 | k__Fungi(100);p__Ascomycota(100);c__Saccharomycetes(100);o__Saccharomycetales(100);f__Metschnikowiaceae(100);g__Metschnikowia(100);s__Metschnikowia_pulcherrima(100); | Metschnikowia | 13 | Saprotroph | \|Undefined Saprotroph\| | Yeast | NULL | Probable | As nectar-tap_saprotroph (Põlme et al. 2020); Decay sugar-rich substrates (Põlme et al. 2020) | Põlme S, et al. 2020. Fungal Diversity 105:1-16 ((https://doi.org/10.1007/s13225-020-00466-2)) |
| Otu21951 | 1 | k__Fungi(100);p__Ascomycota(100);c__Saccharomycetes(100);o__Saccharomycetales(100);f__Metschnikowiaceae(100);g__Metschnikowia(100);s__Metschnikowia_pulcherrima(100); | Metschnikowia | 13 | Saprotroph | \|Undefined Saprotroph\| | Yeast | NULL | Probable | As nectar-tap_saprotroph (Põlme et al. 2020); Decay sugar-rich substrates (Põlme et al. 2020) | Põlme S, et al. 2020. Fungal Diversity 105:1-16 ((https://doi.org/10.1007/s13225-020-00466-2)) |
| Otu21952 | 1 | k__Fungi(100);p__Ascomycota(100);c__Saccharomycetes(100);o__Saccharomycetales(100);f__Metschnikowiaceae(100);g__Metschnikowia(100);s__Metschnikowia_pulcherrima(100); | Metschnikowia | 13 | Saprotroph | \|Undefined Saprotroph\| | Yeast | NULL | Probable | As nectar-tap_saprotroph (Põlme et al. 2020); Decay sugar-rich substrates (Põlme et al. 2020) | Põlme S, et al. 2020. Fungal Diversity 105:1-16 ((https://doi.org/10.1007/s13225-020-00466-2)) |
| Otu21953 | 1 | k__Fungi(100);p__Ascomycota(100);c__Saccharomycetes(100);o__Saccharomycetales(100);f__Metschnikowiaceae(100);g__Metschnikowia(100);s__Metschnikowia_pulcherrima(100); | Metschnikowia | 13 | Saprotroph | \|Undefined Saprotroph\| | Yeast | NULL | Probable | As nectar-tap_saprotroph (Põlme et al. 2020); Decay sugar-rich substrates (Põlme et al. 2020) | Põlme S, et al. 2020. Fungal Diversity 105:1-16 ((https://doi.org/10.1007/s13225-020-00466-2)) |
| Otu21955 | 1 | k__Fungi(100);p__Ascomycota(100);c__Saccharomycetes(100);o__Saccharomycetales(100);f__Metschnikowiaceae(100);g__Metschnikowia(100);s__Metschnikowia_pulcherrima(100); | Metschnikowia | 13 | Saprotroph | \|Undefined Saprotroph\| | Yeast | NULL | Probable | As nectar-tap_saprotroph (Põlme et al. 2020); Decay sugar-rich substrates (Põlme et al. 2020) | Põlme S, et al. 2020. Fungal Diversity 105:1-16 ((https://doi.org/10.1007/s13225-020-00466-2)) |
| Otu21956 | 1 | k__Fungi(100);p__Ascomycota(100);c__Saccharomycetes(100);o__Saccharomycetales(100);f__Metschnikowiaceae(100);g__Metschnikowia(100);s__Metschnikowia_pulcherrima(100); | Metschnikowia | 13 | Saprotroph | \|Undefined Saprotroph\| | Yeast | NULL | Probable | As nectar-tap_saprotroph (Põlme et al. 2020); Decay sugar-rich substrates (Põlme et al. 2020) | Põlme S, et al. 2020. Fungal Diversity 105:1-16 ((https://doi.org/10.1007/s13225-020-00466-2)) |
| Otu21957 | 1 | k__Fungi(100);p__Ascomycota(100);c__Saccharomycetes(100);o__Saccharomycetales(100);f__Metschnikowiaceae(100);g__Metschnikowia(100);s__Metschnikowia_pulcherrima(100); | Metschnikowia | 13 | Saprotroph | \|Undefined Saprotroph\| | Yeast | NULL | Probable | As nectar-tap_saprotroph (Põlme et al. 2020); Decay sugar-rich substrates (Põlme et al. 2020) | Põlme S, et al. 2020. Fungal Diversity 105:1-16 ((https://doi.org/10.1007/s13225-020-00466-2)) |
| Otu21958 | 1 | k__Fungi(100);p__Ascomycota(100);c__Saccharomycetes(100);o__Saccharomycetales(100);f__Metschnikowiaceae(100);g__Metschnikowia(100);s__Metschnikowia_pulcherrima(100); | Metschnikowia | 13 | Saprotroph | \|Undefined Saprotroph\| | Yeast | NULL | Probable | As nectar-tap_saprotroph (Põlme et al. 2020); Decay sugar-rich substrates (Põlme et al. 2020) | Põlme S, et al. 2020. Fungal Diversity 105:1-16 ((https://doi.org/10.1007/s13225-020-00466-2)) |
| Otu21959 | 1 | k__Fungi(100);p__Ascomycota(100);c__Saccharomycetes(100);o__Saccharomycetales(100);f__Metschnikowiaceae(100);g__Metschnikowia(100);g__Metschnikowia_unclassified(100); | Metschnikowia | 13 | Saprotroph | \|Undefined Saprotroph\| | Yeast | NULL | Probable | As nectar-tap_saprotroph (Põlme et al. 2020); Decay sugar-rich substrates (Põlme et al. 2020) | Põlme S, et al. 2020. Fungal Diversity 105:1-16 ((https://doi.org/10.1007/s13225-020-00466-2)) |
| Otu21960 | 1 | k__Fungi(100);p__Ascomycota(100);c__Saccharomycetes(100);o__Saccharomycetales(100);f__Metschnikowiaceae(100);g__Metschnikowia(100);s__Metschnikowia_pulcherrima(100); | Metschnikowia | 13 | Saprotroph | \|Undefined Saprotroph\| | Yeast | NULL | Probable | As nectar-tap_saprotroph (Põlme et al. 2020); Decay sugar-rich substrates (Põlme et al. 2020) | Põlme S, et al. 2020. Fungal Diversity 105:1-16 ((https://doi.org/10.1007/s13225-020-00466-2)) |
| Otu21961 | 1 | k__Fungi(100);p__Ascomycota(100);c__Saccharomycetes(100);o__Saccharomycetales(100);f__Metschnikowiaceae(100);g__Metschnikowia(100);s__Metschnikowia_pulcherrima(100); | Metschnikowia | 13 | Saprotroph | \|Undefined Saprotroph\| | Yeast | NULL | Probable | As nectar-tap_saprotroph (Põlme et al. 2020); Decay sugar-rich substrates (Põlme et al. 2020) | Põlme S, et al. 2020. Fungal Diversity 105:1-16 ((https://doi.org/10.1007/s13225-020-00466-2)) |
| Otu21962 | 1 | k__Fungi(100);p__Ascomycota(100);c__Saccharomycetes(100);o__Saccharomycetales(100);f__Metschnikowiaceae(100);g__Metschnikowia(100);s__Metschnikowia_pulcherrima(100); | Metschnikowia | 13 | Saprotroph | \|Undefined Saprotroph\| | Yeast | NULL | Probable | As nectar-tap_saprotroph (Põlme et al. 2020); Decay sugar-rich substrates (Põlme et al. 2020) | Põlme S, et al. 2020. Fungal Diversity 105:1-16 ((https://doi.org/10.1007/s13225-020-00466-2)) |
| Otu21963 | 1 | k__Fungi(100);p__Ascomycota(100);c__Saccharomycetes(100);o__Saccharomycetales(100);f__Metschnikowiaceae(100);g__Metschnikowia(100);s__Metschnikowia_pulcherrima(100); | Metschnikowia | 13 | Saprotroph | \|Undefined Saprotroph\| | Yeast | NULL | Probable | As nectar-tap_saprotroph (Põlme et al. 2020); Decay sugar-rich substrates (Põlme et al. 2020) | Põlme S, et al. 2020. Fungal Diversity 105:1-16 ((https://doi.org/10.1007/s13225-020-00466-2)) |
| Otu21964 | 1 | k__Fungi(100);p__Ascomycota(100);c__Saccharomycetes(100);o__Saccharomycetales(100);f__Metschnikowiaceae(100);g__Metschnikowia(100);g__Metschnikowia_unclassified(100); | Metschnikowia | 13 | Saprotroph | \|Undefined Saprotroph\| | Yeast | NULL | Probable | As nectar-tap_saprotroph (Põlme et al. 2020); Decay sugar-rich substrates (Põlme et al. 2020) | Põlme S, et al. 2020. Fungal Diversity 105:1-16 ((https://doi.org/10.1007/s13225-020-00466-2)) |
| Otu21965 | 1 | k__Fungi(100);p__Ascomycota(100);c__Saccharomycetes(100);o__Saccharomycetales(100);f__Metschnikowiaceae(100);g__Metschnikowia(100);s__Metschnikowia_pulcherrima(100); | Metschnikowia | 13 | Saprotroph | \|Undefined Saprotroph\| | Yeast | NULL | Probable | As nectar-tap_saprotroph (Põlme et al. 2020); Decay sugar-rich substrates (Põlme et al. 2020) | Põlme S, et al. 2020. Fungal Diversity 105:1-16 ((https://doi.org/10.1007/s13225-020-00466-2)) |
| Otu21966 | 1 | k__Fungi(100);p__Ascomycota(100);c__Saccharomycetes(100);o__Saccharomycetales(100);f__Metschnikowiaceae(100);g__Metschnikowia(100);g__Metschnikowia_unclassified(100); | Metschnikowia | 13 | Saprotroph | \|Undefined Saprotroph\| | Yeast | NULL | Probable | As nectar-tap_saprotroph (Põlme et al. 2020); Decay sugar-rich substrates (Põlme et al. 2020) | Põlme S, et al. 2020. Fungal Diversity 105:1-16 ((https://doi.org/10.1007/s13225-020-00466-2)) |
| Otu21967 | 1 | k__Fungi(100);p__Ascomycota(100);c__Saccharomycetes(100);o__Saccharomycetales(100);f__Metschnikowiaceae(100);g__Metschnikowia(100);s__Metschnikowia_pulcherrima(100); | Metschnikowia | 13 | Saprotroph | \|Undefined Saprotroph\| | Yeast | NULL | Probable | As nectar-tap_saprotroph (Põlme et al. 2020); Decay sugar-rich substrates (Põlme et al. 2020) | Põlme S, et al. 2020. Fungal Diversity 105:1-16 ((https://doi.org/10.1007/s13225-020-00466-2)) |
| Otu21968 | 1 | k__Fungi(100);p__Ascomycota(100);c__Saccharomycetes(100);o__Saccharomycetales(100);f__Saccharomycetales_family_Incertae_sedis(100);g__Candida(100);g__Candida_unclassified(100); | Candida | 13 | Pathotroph-Saprotroph-Symbiotroph | Animal Pathogen-Endophyte-Endosymbiont-Epiphyte-\|Undefined Saprotroph\| | Dimorphic Yeast-Yeast | NULL | Probable | As nectar-tap_saprotroph (Põlme et al. 2020); Decay sugar-rich substrates (Põlme et al. 2020); This asexual genus is polyphyletic and contain species that function very differently depend on what they are. Caution should be taken when interpreting guilds for Candida species. | Manolakaki D, et al. 2010. Virulence. 1: 367-375 ((https://doi.org/10.4161/viru.1.5.12796)); Mekbib SB, et al. 2011. Tropical Plant Pathology 36:233-240 ((https://doi.org/10.1590/S1982-56762011000400004)); Põlme S, et al. 2020. Fungal Diversity 105:1-16 ((https://doi.org/10.1007/s13225-020-00466-2)) |
| Otu21969 | 1 | k__Fungi(100);p__Ascomycota(100);c__Saccharomycetes(100);o__Saccharomycetales(100);f__Metschnikowiaceae(100);g__Metschnikowia(100);s__Metschnikowia_pulcherrima(100); | Metschnikowia | 13 | Saprotroph | \|Undefined Saprotroph\| | Yeast | NULL | Probable | As nectar-tap_saprotroph (Põlme et al. 2020); Decay sugar-rich substrates (Põlme et al. 2020) | Põlme S, et al. 2020. Fungal Diversity 105:1-16 ((https://doi.org/10.1007/s13225-020-00466-2)) |
| Otu21970 | 1 | k__Fungi(100);p__Ascomycota(100);c__Saccharomycetes(100);o__Saccharomycetales(100);f__Metschnikowiaceae(100);g__Metschnikowia(100);s__Metschnikowia_pulcherrima(100); | Metschnikowia | 13 | Saprotroph | \|Undefined Saprotroph\| | Yeast | NULL | Probable | As nectar-tap_saprotroph (Põlme et al. 2020); Decay sugar-rich substrates (Põlme et al. 2020) | Põlme S, et al. 2020. Fungal Diversity 105:1-16 ((https://doi.org/10.1007/s13225-020-00466-2)) |
| Otu21971 | 1 | k__Fungi(100);p__Ascomycota(100);c__Saccharomycetes(100);o__Saccharomycetales(100);o__Saccharomycetales_unclassified(100);o__Saccharomycetales_unclassified(100);o__Saccharomycetales_unclassified(100); | Saccharomycetales | 7 | Saprotroph | Undefined Saprotroph | Yeast | NULL | Possible | NULL | Sterkenburg E, et al. 2015. New Phytologist 207:1145-1158 ((https://doi.org/10.1111/nph.13426)) |

| | | | | | | | | | | | |
|---|---|---|---|---|---|---|---|---|---|---|---|
| Otu21972 | 1 | k__Fungi(100);p__Ascomycota(100);c__Saccharomycetes(100);o__Saccharomycetales(100);f__Metschnikowiaceae(100);g__Metschnikowia(100);s__Metschnikowia_pulcherrima(100); | Metschnikowia | 13 | Saprotroph | Undefined Saprotroph | Yeast | NULL | Probable | As nectar-tap_saprotroph (Põlme et al. 2020); Decay sugar-rich substrates (Põlme et al. 2020) | Põlme S, et al. 2020. Fungal Diversity 105:1-16 ((https://doi.org/10.1007/s13225-020-00466-2)) |
| Otu21973 | 1 | k__Fungi(100);p__Ascomycota(100);c__Saccharomycetes(100);o__Saccharomycetales(100);f__Metschnikowiaceae(100);g__Metschnikowia(100);s__Metschnikowia_pulcherrima(100); | Metschnikowia | 13 | Saprotroph | Undefined Saprotroph | Yeast | NULL | Probable | As nectar-tap_saprotroph (Põlme et al. 2020); Decay sugar-rich substrates (Põlme et al. 2020) | Põlme S, et al. 2020. Fungal Diversity 105:1-16 ((https://doi.org/10.1007/s13225-020-00466-2)) |
| Otu21974 | 1 | k__Fungi(100);p__Ascomycota(100);c__Saccharomycetes(100);o__Saccharomycetales(100);f__Metschnikowiaceae(100);g__Metschnikowia(100);s__Metschnikowia_pulcherrima(100); | Metschnikowia | 13 | Saprotroph | Undefined Saprotroph | Yeast | NULL | Probable | As nectar-tap_saprotroph (Põlme et al. 2020); Decay sugar-rich substrates (Põlme et al. 2020) | Põlme S, et al. 2020. Fungal Diversity 105:1-16 ((https://doi.org/10.1007/s13225-020-00466-2)) |
| Otu21975 | 1 | k__Fungi(100);p__Ascomycota(100);c__Saccharomycetes(100);o__Saccharomycetales(100);f__Metschnikowiaceae(100);g__Metschnikowia(100);s__Metschnikowia_pulcherrima(100); | Metschnikowia | 13 | Saprotroph | Undefined Saprotroph | Yeast | NULL | Probable | As nectar-tap_saprotroph (Põlme et al. 2020); Decay sugar-rich substrates (Põlme et al. 2020) | Põlme S, et al. 2020. Fungal Diversity 105:1-16 ((https://doi.org/10.1007/s13225-020-00466-2)) |
| Otu21976 | 1 | k__Fungi(100);p__Ascomycota(100);c__Saccharomycetes(100);o__Saccharomycetales(100);f__Metschnikowiaceae(100);g__Metschnikowia(100);g__Metschnikowia_unclassified(100); | Metschnikowia | 13 | Saprotroph | Undefined Saprotroph | Yeast | NULL | Probable | As nectar-tap_saprotroph (Põlme et al. 2020); Decay sugar-rich substrates (Põlme et al. 2020) | Põlme S, et al. 2020. Fungal Diversity 105:1-16 ((https://doi.org/10.1007/s13225-020-00466-2)) |
| Otu21977 | 1 | k__Fungi(100);p__Ascomycota(100);c__Saccharomycetes(100);o__Saccharomycetales(100);f__Saccharomycetales_family_Incertae_sedis(100);g__Candida(100);s__Candida_cretensis(100); | Candida | 13 | Pathotroph-Saprotroph-Symbiotroph | Animal Pathogen-Endophyte-Endosymbiont-Epiphyte-Undefined Saprotroph | Dimorphic Yeast Yeast | NULL | Probable | As nectar-tap_saprotroph (Põlme et al. 2020); Decay sugar-rich substrates (Põlme et al. 2020); This asexual genus is polyphyletic and contain species that function very differently depend on what they are. Caution should be taken when interpreting guilds for Candida species. | Manolakaki D, et al. 2010. Virulence. 1: 367-375 ((https://doi.org/10.4161/viru.1.5.12796)); Mekbib SB, et al. 2011. Tropical Plant Pathology 36:233-240 ((https://doi.org/10.1590/S1982-56762011000400004)); Põlme S, et al. 2020. Fungal Diversity 105:1-16 ((https://doi.org/10.1007/s13225-020-00466-2)) |
| Otu21978 | 1 | k__Fungi(100);p__Ascomycota(100);c__Saccharomycetes(100);o__Saccharomycetales(100);f__Metschnikowiaceae(100);g__Metschnikowia(100);g__Metschnikowia_unclassified(100); | Metschnikowia | 13 | Saprotroph | Undefined Saprotroph | Yeast | NULL | Probable | As nectar-tap_saprotroph (Põlme et al. 2020); Decay sugar-rich substrates (Põlme et al. 2020) | Põlme S, et al. 2020. Fungal Diversity 105:1-16 ((https://doi.org/10.1007/s13225-020-00466-2)) |
| Otu21979 | 1 | k__Fungi(100);p__Ascomycota(100);c__Saccharomycetes(100);o__Saccharomycetales(100);f__Metschnikowiaceae(100);g__Metschnikowia(100);s__Metschnikowia_pulcherrima(100); | Metschnikowia | 13 | Saprotroph | Undefined Saprotroph | Yeast | NULL | Probable | As nectar-tap_saprotroph (Põlme et al. 2020); Decay sugar-rich substrates (Põlme et al. 2020) | Põlme S, et al. 2020. Fungal Diversity 105:1-16 ((https://doi.org/10.1007/s13225-020-00466-2)) |
| Otu21980 | 1 | k__Fungi(100);p__Ascomycota(100);c__Saccharomycetes(100);o__Saccharomycetales(100);f__Metschnikowiaceae(100);g__Metschnikowia(100);s__Metschnikowia_pulcherrima(100); | Metschnikowia | 13 | Saprotroph | Undefined Saprotroph | Yeast | NULL | Probable | As nectar-tap_saprotroph (Põlme et al. 2020); Decay sugar-rich substrates (Põlme et al. 2020) | Põlme S, et al. 2020. Fungal Diversity 105:1-16 ((https://doi.org/10.1007/s13225-020-00466-2)) |
| Otu21982 | 1 | k__Fungi(100);p__Ascomycota(100);c__Saccharomycetes(100);o__Saccharomycetales(100);f__Metschnikowiaceae(100);g__Metschnikowia(100);s__Metschnikowia_pulcherrima(100); | Metschnikowia | 13 | Saprotroph | Undefined Saprotroph | Yeast | NULL | Probable | As nectar-tap_saprotroph (Põlme et al. 2020); Decay sugar-rich substrates (Põlme et al. 2020) | Põlme S, et al. 2020. Fungal Diversity 105:1-16 ((https://doi.org/10.1007/s13225-020-00466-2)) |
| Otu21983 | 1 | k__Fungi(100);p__Ascomycota(100);c__Saccharomycetes(100);o__Saccharomycetales(100);o__Saccharomycetales_unclassified(100);o__Saccharomycetales_unclassified(100);o__Saccharomycetales_unclassified(100); | Saccharomycetales | 7 | Saprotroph | Undefined Saprotroph | Yeast | NULL | Possible | NULL | Sterkenburg E, et al. 2015. New Phytologist 207:1145-1158 ((https://doi.org/10.1111/nph.13426)) |
| Otu21984 | 1 | k__Fungi(100);p__Ascomycota(100);c__Saccharomycetes(100);o__Saccharomycetales(100);f__Metschnikowiaceae(100);g__Metschnikowia(100);g__Metschnikowia_unclassified(100); | Metschnikowia | 13 | Saprotroph | Undefined Saprotroph | Yeast | NULL | Probable | As nectar-tap_saprotroph (Põlme et al. 2020); Decay sugar-rich substrates (Põlme et al. 2020) | Põlme S, et al. 2020. Fungal Diversity 105:1-16 ((https://doi.org/10.1007/s13225-020-00466-2)) |
| Otu21985 | 1 | k__Fungi(100);p__Ascomycota(100);c__Saccharomycetes(100);o__Saccharomycetales(100);f__Metschnikowiaceae(100);g__Metschnikowia(100);s__Metschnikowia_pulcherrima(100); | Metschnikowia | 13 | Saprotroph | Undefined Saprotroph | Yeast | NULL | Probable | As nectar-tap_saprotroph (Põlme et al. 2020); Decay sugar-rich substrates (Põlme et al. 2020) | Põlme S, et al. 2020. Fungal Diversity 105:1-16 ((https://doi.org/10.1007/s13225-020-00466-2)) |
| Otu21986 | 1 | k__Fungi(100);p__Ascomycota(100);c__Saccharomycetes(100);o__Saccharomycetales(100);f__Metschnikowiaceae(100);g__Metschnikowia(100);s__Metschnikowia_pulcherrima(100); | Metschnikowia | 13 | Saprotroph | Undefined Saprotroph | Yeast | NULL | Probable | As nectar-tap_saprotroph (Põlme et al. 2020); Decay sugar-rich substrates (Põlme et al. 2020) | Põlme S, et al. 2020. Fungal Diversity 105:1-16 ((https://doi.org/10.1007/s13225-020-00466-2)) |
| Otu21987 | 1 | k__Fungi(100);p__Ascomycota(100);c__Saccharomycetes(100);o__Saccharomycetales(100);f__Metschnikowiaceae(100);g__Metschnikowia(100);s__Metschnikowia_pulcherrima(100); | Metschnikowia | 13 | Saprotroph | Undefined Saprotroph | Yeast | NULL | Probable | As nectar-tap_saprotroph (Põlme et al. 2020); Decay sugar-rich substrates (Põlme et al. 2020) | Põlme S, et al. 2020. Fungal Diversity 105:1-16 ((https://doi.org/10.1007/s13225-020-00466-2)) |
| Otu21988 | 1 | k__Fungi(100);p__Ascomycota(100);c__Saccharomycetes(100);o__Saccharomycetales(100);f__Metschnikowiaceae(100);g__Metschnikowia(100);s__Metschnikowia_pulcherrima(100); | Metschnikowia | 13 | Saprotroph | Undefined Saprotroph | Yeast | NULL | Probable | As nectar-tap_saprotroph (Põlme et al. 2020); Decay sugar-rich substrates (Põlme et al. 2020) | Põlme S, et al. 2020. Fungal Diversity 105:1-16 ((https://doi.org/10.1007/s13225-020-00466-2)) |
| Otu21990 | 1 | k__Fungi(100);p__Ascomycota(100);c__Saccharomycetes(100);o__Saccharomycetales(100);f__Metschnikowiaceae(100);g__Metschnikowia(100);s__Metschnikowia_pulcherrima(100); | Metschnikowia | 13 | Saprotroph | Undefined Saprotroph | Yeast | NULL | Probable | As nectar-tap_saprotroph (Põlme et al. 2020); Decay sugar-rich substrates (Põlme et al. 2020) | Põlme S, et al. 2020. Fungal Diversity 105:1-16 ((https://doi.org/10.1007/s13225-020-00466-2)) |
| Otu21991 | 1 | k__Fungi(100);p__Ascomycota(100);c__Saccharomycetes(100);o__Saccharomycetales(100);f__Metschnikowiaceae(100);g__Metschnikowia(100);s__Metschnikowia_pulcherrima(100); | Metschnikowia | 13 | Saprotroph | Undefined Saprotroph | Yeast | NULL | Probable | As nectar-tap_saprotroph (Põlme et al. 2020); Decay sugar-rich substrates (Põlme et al. 2020) | Põlme S, et al. 2020. Fungal Diversity 105:1-16 ((https://doi.org/10.1007/s13225-020-00466-2)) |
| Otu21992 | 1 | k__Fungi(100);p__Ascomycota(100);c__Saccharomycetes(100);o__Saccharomycetales(100);f__Saccharomycetales_family_Incertae_sedis(100);g__Candida(100);g__Candida_unclassified(100); | Candida | 13 | Pathotroph-Saprotroph-Symbiotroph | Animal Pathogen-Endophyte-Endosymbiont-Epiphyte-Undefined Saprotroph | Dimorphic Yeast Yeast | NULL | Probable | As nectar-tap_saprotroph (Põlme et al. 2020); Decay sugar-rich substrates (Põlme et al. 2020); This asexual genus is polyphyletic and contain species that function very differently depend on what they are. Caution should be taken when interpreting guilds for Candida species. | Manolakaki D, et al. 2010. Virulence. 1: 367-375 ((https://doi.org/10.4161/viru.1.5.12796)); Mekbib SB, et al. 2011. Tropical Plant Pathology 36:233-240 ((https://doi.org/10.1590/S1982-56762011000400004)); Põlme S, et al. 2020. Fungal Diversity 105:1-16 ((https://doi.org/10.1007/s13225-020-00466-2)) |
| Otu21994 | 1 | k__Fungi(100);p__Ascomycota(100);c__Saccharomycetes(100);o__Saccharomycetales(100);f__Metschnikowiaceae(100);g__Metschnikowia(100);g__Metschnikowia_unclassified(100); | Metschnikowia | 13 | Saprotroph | Undefined Saprotroph | Yeast | NULL | Probable | As nectar-tap_saprotroph (Põlme et al. 2020); Decay sugar-rich substrates (Põlme et al. 2020) | Põlme S, et al. 2020. Fungal Diversity 105:1-16 ((https://doi.org/10.1007/s13225-020-00466-2)) |
| Otu21995 | 1 | k__Fungi(100);p__Ascomycota(100);c__Saccharomycetes(100);o__Saccharomycetales(100);f__Metschnikowiaceae(100);g__Metschnikowia(100);s__Metschnikowia_pulcherrima(100); | Metschnikowia | 13 | Saprotroph | Undefined Saprotroph | Yeast | NULL | Probable | As nectar-tap_saprotroph (Põlme et al. 2020); Decay sugar-rich substrates (Põlme et al. 2020) | Põlme S, et al. 2020. Fungal Diversity 105:1-16 ((https://doi.org/10.1007/s13225-020-00466-2)) |
| Otu21996 | 1 | k__Fungi(100);p__Ascomycota(100);c__Saccharomycetes(100);o__Saccharomycetales(100);f__Saccharomycetales_family_Incertae_sedis(100);g__Candida(100);g__Candida_unclassified(100); | Candida | 13 | Pathotroph-Saprotroph-Symbiotroph | Animal Pathogen-Endophyte-Endosymbiont-Epiphyte-Undefined Saprotroph | Dimorphic Yeast Yeast | NULL | Probable | As nectar-tap_saprotroph (Põlme et al. 2020); Decay sugar-rich substrates (Põlme et al. 2020); This asexual genus is polyphyletic and contain species that function very differently depend on what they are. Caution should be taken when interpreting guilds for Candida species. | Manolakaki D, et al. 2010. Virulence. 1: 367-375 ((https://doi.org/10.4161/viru.1.5.12796)); Mekbib SB, et al. 2011. Tropical Plant Pathology 36:233-240 ((https://doi.org/10.1590/S1982-56762011000400004)); Põlme S, et al. 2020. Fungal Diversity 105:1-16 ((https://doi.org/10.1007/s13225-020-00466-2)) |
| Otu21997 | 1 | k__Fungi(100);p__Ascomycota(100);c__Saccharomycetes(100);o__Saccharomycetales(100);f__Saccharomycetales_family_Incertae_sedis(100);g__Candida(100);g__Candida_unclassified(100); | Candida | 13 | Pathotroph-Saprotroph-Symbiotroph | Animal Pathogen-Endophyte-Endosymbiont-Epiphyte-Undefined Saprotroph | Dimorphic Yeast Yeast | NULL | Probable | As nectar-tap_saprotroph (Põlme et al. 2020); Decay sugar-rich substrates (Põlme et al. 2020); This asexual genus is polyphyletic and contain species that function very differently depend on what they are. Caution should be taken when interpreting guilds for Candida species. | Manolakaki D, et al. 2010. Virulence. 1: 367-375 ((https://doi.org/10.4161/viru.1.5.12796)); Mekbib SB, et al. 2011. Tropical Plant Pathology 36:233-240 ((https://doi.org/10.1590/S1982-56762011000400004)); Põlme S, et al. 2020. Fungal Diversity 105:1-16 ((https://doi.org/10.1007/s13225-020-00466-2)) |
| Otu21998 | 1 | k__Fungi(100);p__Ascomycota(100);c__Saccharomycetes(100);o__Saccharomycetales(100);f__Saccharomycetales_family_Incertae_sedis(100);g__Candida(100);g__Candida_unclassified(100); | Candida | 13 | Pathotroph-Saprotroph-Symbiotroph | Animal Pathogen-Endophyte-Endosymbiont-Epiphyte-Undefined Saprotroph | Dimorphic Yeast Yeast | NULL | Probable | As nectar-tap_saprotroph (Põlme et al. 2020); Decay sugar-rich substrates (Põlme et al. 2020); This asexual genus is polyphyletic and contain species that function very differently depend on what they are. Caution should be taken when interpreting guilds for Candida species. | Manolakaki D, et al. 2010. Virulence. 1: 367-375 ((https://doi.org/10.4161/viru.1.5.12796)); Mekbib SB, et al. 2011. Tropical Plant Pathology 36:233-240 ((https://doi.org/10.1590/S1982-56762011000400004)); Põlme S, et al. 2020. Fungal Diversity 105:1-16 ((https://doi.org/10.1007/s13225-020-00466-2)) |
| Otu22003 | 1 | k__Fungi(100);p__Ascomycota(100);c__Saccharomycetes(100);o__Saccharomycetales(100);f__Metschnikowiaceae(100);g__Metschnikowia(100);s__Metschnikowia_pulcherrima(100); | Metschnikowia | 13 | Saprotroph | Undefined Saprotroph | Yeast | NULL | Probable | As nectar-tap_saprotroph (Põlme et al. 2020); Decay sugar-rich substrates (Põlme et al. 2020) | Põlme S, et al. 2020. Fungal Diversity 105:1-16 ((https://doi.org/10.1007/s13225-020-00466-2)) |
| Otu22005 | 1 | k__Fungi(100);p__Ascomycota(100);c__Saccharomycetes(100);o__Saccharomycetales(100);f__Metschnikowiaceae(100);g__Metschnikowia(100);s__Metschnikowia_pulcherrima(100); | Metschnikowia | 13 | Saprotroph | Undefined Saprotroph | Yeast | NULL | Probable | As nectar-tap_saprotroph (Põlme et al. 2020); Decay sugar-rich substrates (Põlme et al. 2020) | Põlme S, et al. 2020. Fungal Diversity 105:1-16 ((https://doi.org/10.1007/s13225-020-00466-2)) |
| Otu22008 | 1 | k__Fungi(100);p__Ascomycota(100);c__Saccharomycetes(100);o__Saccharomycetales(100);f__Metschnikowiaceae(100);g__Metschnikowia(100);s__Metschnikowia_pulcherrima(100); | Metschnikowia | 13 | Saprotroph | Undefined Saprotroph | Yeast | NULL | Probable | As nectar-tap_saprotroph (Põlme et al. 2020); Decay sugar-rich substrates (Põlme et al. 2020) | Põlme S, et al. 2020. Fungal Diversity 105:1-16 ((https://doi.org/10.1007/s13225-020-00466-2)) |
| Otu22009 | 1 | k__Fungi(100);p__Ascomycota(100);c__Saccharomycetes(100);o__Saccharomycetales(100);f__Metschnikowiaceae(100);g__Metschnikowia(100);s__Metschnikowia_pulcherrima(100); | Metschnikowia | 13 | Saprotroph | Undefined Saprotroph | Yeast | NULL | Probable | As nectar-tap_saprotroph (Põlme et al. 2020); Decay sugar-rich substrates (Põlme et al. 2020) | Põlme S, et al. 2020. Fungal Diversity 105:1-16 ((https://doi.org/10.1007/s13225-020-00466-2)) |
| Otu22010 | 1 | k__Fungi(100);p__Ascomycota(100);c__Saccharomycetes(100);o__Saccharomycetales(100);f__Metschnikowiaceae(100);g__Metschnikowia(100);s__Metschnikowia_pulcherrima(100); | Metschnikowia | 13 | Saprotroph | Undefined Saprotroph | Yeast | NULL | Probable | As nectar-tap_saprotroph (Põlme et al. 2020); Decay sugar-rich substrates (Põlme et al. 2020) | Põlme S, et al. 2020. Fungal Diversity 105:1-16 ((https://doi.org/10.1007/s13225-020-00466-2)) |
| Otu22011 | 1 | k__Fungi(100);p__Ascomycota(100);c__Saccharomycetes(100);o__Saccharomycetales(100);f__Metschnikowiaceae(100);g__Metschnikowia(100);s__Metschnikowia_pulcherrima(100); | Metschnikowia | 13 | Saprotroph | Undefined Saprotroph | Yeast | NULL | Probable | As nectar-tap_saprotroph (Põlme et al. 2020); Decay sugar-rich substrates (Põlme et al. 2020) | Põlme S, et al. 2020. Fungal Diversity 105:1-16 ((https://doi.org/10.1007/s13225-020-00466-2)) |
| Otu22012 | 1 | k__Fungi(100);p__Ascomycota(100);c__Saccharomycetes(100);o__Saccharomycetales(100);f__Metschnikowiaceae(100);g__Metschnikowia(100);g__Metschnikowia_unclassified(100); | Metschnikowia | 13 | Saprotroph | Undefined Saprotroph | Yeast | NULL | Probable | As nectar-tap_saprotroph (Põlme et al. 2020); Decay sugar-rich substrates (Põlme et al. 2020) | Põlme S, et al. 2020. Fungal Diversity 105:1-16 ((https://doi.org/10.1007/s13225-020-00466-2)) |
| Otu22013 | 1 | k__Fungi(100);p__Ascomycota(100);c__Saccharomycetes(100);o__Saccharomycetales(100);f__Metschnikowiaceae(100);g__Metschnikowia(100);s__Metschnikowia_pulcherrima(100); | Metschnikowia | 13 | Saprotroph | Undefined Saprotroph | Yeast | NULL | Probable | As nectar-tap_saprotroph (Põlme et al. 2020); Decay sugar-rich substrates (Põlme et al. 2020) | Põlme S, et al. 2020. Fungal Diversity 105:1-16 ((https://doi.org/10.1007/s13225-020-00466-2)) |
| Otu22014 | 1 | k__Fungi(100);p__Ascomycota(100);c__Saccharomycetes(100);o__Saccharomycetales(100);f__Metschnikowiaceae(100);g__Metschnikowia(100);s__Metschnikowia_pulcherrima(100); | Metschnikowia | 13 | Saprotroph | Undefined Saprotroph | Yeast | NULL | Probable | As nectar-tap_saprotroph (Põlme et al. 2020); Decay sugar-rich substrates (Põlme et al. 2020) | Põlme S, et al. 2020. Fungal Diversity 105:1-16 ((https://doi.org/10.1007/s13225-020-00466-2)) |
| Otu22015 | 1 | k__Fungi(100);p__Ascomycota(100);c__Saccharomycetes(100);o__Saccharomycetales(100);f__Metschnikowiaceae(100);g__Metschnikowia(100);g__Metschnikowia_unclassified(100); | Metschnikowia | 13 | Saprotroph | Undefined Saprotroph | Yeast | NULL | Probable | As nectar-tap_saprotroph (Põlme et al. 2020); Decay sugar-rich substrates (Põlme et al. 2020) | Põlme S, et al. 2020. Fungal Diversity 105:1-16 ((https://doi.org/10.1007/s13225-020-00466-2)) |
| Otu22016 | 1 | k__Fungi(100);p__Ascomycota(100);c__Saccharomycetes(100);o__Saccharomycetales(100);f__Metschnikowiaceae(100);g__Metschnikowia(100);s__Metschnikowia_pulcherrima(100); | Metschnikowia | 13 | Saprotroph | Undefined Saprotroph | Yeast | NULL | Probable | As nectar-tap_saprotroph (Põlme et al. 2020); Decay sugar-rich substrates (Põlme et al. 2020) | Põlme S, et al. 2020. Fungal Diversity 105:1-16 ((https://doi.org/10.1007/s13225-020-00466-2)) |
| Otu22017 | 1 | k__Fungi(100);p__Ascomycota(100);c__Saccharomycetes(100);o__Saccharomycetales(100);f__Metschnikowiaceae(100);g__Metschnikowia(100);s__Metschnikowia_pulcherrima(100); | Metschnikowia | 13 | Saprotroph | Undefined Saprotroph | Yeast | NULL | Probable | As nectar-tap_saprotroph (Põlme et al. 2020); Decay sugar-rich substrates (Põlme et al. 2020) | Põlme S, et al. 2020. Fungal Diversity 105:1-16 ((https://doi.org/10.1007/s13225-020-00466-2)) |
| Otu22018 | 1 | k__Fungi(100);p__Ascomycota(100);c__Saccharomycetes(100);o__Saccharomycetales(100);f__Metschnikowiaceae(100);g__Metschnikowia(100);s__Metschnikowia_pulcherrima(100); | Metschnikowia | 13 | Saprotroph | Undefined Saprotroph | Yeast | NULL | Probable | As nectar-tap_saprotroph (Põlme et al. 2020); Decay sugar-rich substrates (Põlme et al. 2020) | Põlme S, et al. 2020. Fungal Diversity 105:1-16 ((https://doi.org/10.1007/s13225-020-00466-2)) |
| Otu22019 | 1 | k__Fungi(100);p__Ascomycota(100);c__Saccharomycetes(100);o__Saccharomycetales(100);f__Metschnikowiaceae(100);g__Metschnikowia(100);s__Metschnikowia_pulcherrima(100); | Metschnikowia | 13 | Saprotroph | Undefined Saprotroph | Yeast | NULL | Probable | As nectar-tap_saprotroph (Põlme et al. 2020); Decay sugar-rich substrates (Põlme et al. 2020) | Põlme S, et al. 2020. Fungal Diversity 105:1-16 ((https://doi.org/10.1007/s13225-020-00466-2)) |
| Otu22020 | 1 | k__Fungi(100);p__Ascomycota(100);c__Saccharomycetes(100);o__Saccharomycetales(100);f__Metschnikowiaceae(100);g__Metschnikowia(100);s__Metschnikowia_pulcherrima(100); | Metschnikowia | 13 | Saprotroph | Undefined Saprotroph | Yeast | NULL | Probable | As nectar-tap_saprotroph (Põlme et al. 2020); Decay sugar-rich substrates (Põlme et al. 2020) | Põlme S, et al. 2020. Fungal Diversity 105:1-16 ((https://doi.org/10.1007/s13225-020-00466-2)) |
| Otu22021 | 1 | k__Fungi(100);p__Ascomycota(100);c__Saccharomycetes(100);o__Saccharomycetales(100);f__Metschnikowiaceae(100);g__Metschnikowia(100);g__Metschnikowia_unclassified(100); | Metschnikowia | 13 | Saprotroph | Undefined Saprotroph | Yeast | NULL | Probable | As nectar-tap_saprotroph (Põlme et al. 2020); Decay sugar-rich substrates (Põlme et al. 2020) | Põlme S, et al. 2020. Fungal Diversity 105:1-16 ((https://doi.org/10.1007/s13225-020-00466-2)) |
| Otu22022 | 1 | k__Fungi(100);p__Ascomycota(100);c__Saccharomycetes(100);o__Saccharomycetales(100);f__Metschnikowiaceae(100);g__Metschnikowia(100);g__Metschnikowia_unclassified(100); | Metschnikowia | 13 | Saprotroph | Undefined Saprotroph | Yeast | NULL | Probable | As nectar-tap_saprotroph (Põlme et al. 2020); Decay sugar-rich substrates (Põlme et al. 2020) | Põlme S, et al. 2020. Fungal Diversity 105:1-16 ((https://doi.org/10.1007/s13225-020-00466-2)) |
| Otu22030 | 1 | k__Fungi(100);p__Ascomycota(100);c__Saccharomycetes(100);o__Saccharomycetales(100);f__Metschnikowiaceae(100);g__Metschnikowia(100);s__Metschnikowia_pulcherrima(100); | Metschnikowia | 13 | Saprotroph | Undefined Saprotroph | Yeast | NULL | Probable | As nectar-tap_saprotroph (Põlme et al. 2020); Decay sugar-rich substrates (Põlme et al. 2020) | Põlme S, et al. 2020. Fungal Diversity 105:1-16 ((https://doi.org/10.1007/s13225-020-00466-2)) |
| Otu22031 | 1 | k__Fungi(100);p__Ascomycota(100);c__Saccharomycetes(100);o__Saccharomycetales(100);f__Metschnikowiaceae(100);g__Metschnikowia(100);s__Metschnikowia_pulcherrima(100); | Metschnikowia | 13 | Saprotroph | Undefined Saprotroph | Yeast | NULL | Probable | As nectar-tap_saprotroph (Põlme et al. 2020); Decay sugar-rich substrates (Põlme et al. 2020) | Põlme S, et al. 2020. Fungal Diversity 105:1-16 ((https://doi.org/10.1007/s13225-020-00466-2)) |
| Otu22032 | 1 | k__Fungi(100);p__Ascomycota(100);c__Saccharomycetes(100);o__Saccharomycetales(100);f__Metschnikowiaceae(100);g__Metschnikowia(100);s__Metschnikowia_pulcherrima(100); | Metschnikowia | 13 | Saprotroph | Undefined Saprotroph | Yeast | NULL | Probable | As nectar-tap_saprotroph (Põlme et al. 2020); Decay sugar-rich substrates (Põlme et al. 2020) | Põlme S, et al. 2020. Fungal Diversity 105:1-16 ((https://doi.org/10.1007/s13225-020-00466-2)) |
| Otu22034 | 1 | k__Fungi(100);p__Ascomycota(100);c__Saccharomycetes(100);o__Saccharomycetales(100);f__Metschnikowiaceae(100);g__Metschnikowia(100);s__Metschnikowia_pulcherrima(100); | Metschnikowia | 13 | Saprotroph | Undefined Saprotroph | Yeast | NULL | Probable | As nectar-tap_saprotroph (Põlme et al. 2020); Decay sugar-rich substrates (Põlme et al. 2020) | Põlme S, et al. 2020. Fungal Diversity 105:1-16 ((https://doi.org/10.1007/s13225-020-00466-2)) |
| Otu22035 | 1 | k__Fungi(100);p__Ascomycota(100);c__Saccharomycetes(100);o__Saccharomycetales(100);f__Metschnikowiaceae(100);g__Metschnikowia(100);s__Metschnikowia_pulcherrima(100); | Metschnikowia | 13 | Saprotroph | Undefined Saprotroph | Yeast | NULL | Probable | As nectar-tap_saprotroph (Põlme et al. 2020); Decay sugar-rich substrates (Põlme et al. 2020) | Põlme S, et al. 2020. Fungal Diversity 105:1-16 ((https://doi.org/10.1007/s13225-020-00466-2)) |
| Otu22036 | 1 | k__Fungi(100);p__Ascomycota(100);c__Saccharomycetes(100);o__Saccharomycetales(100);f__Saccharomycetales_family_Incertae_sedis(100);f__Saccharomycetales_family_Incertae_sedis_unclassified(100);f__Saccharomycetales_family_Incertae_sedis_unclassified(100); | Saccharomycetales | 7 | Saprotroph | Undefined Saprotroph | Yeast | NULL | Possible | NULL | Sterkenburg E, et al. 2015. New Phytologist 207:1145-1158 ((https://doi.org/10.1111/nph.13426)) |
| Otu22037 | 1 | k__Fungi(100);p__Ascomycota(100);c__Saccharomycetes(100);o__Saccharomycetales(100);f__Metschnikowiaceae(100);g__Metschnikowia(100);g__Metschnikowia_unclassified(100); | Metschnikowia | 13 | Saprotroph | Undefined Saprotroph | Yeast | NULL | Probable | As nectar-tap_saprotroph (Põlme et al. 2020); Decay sugar-rich substrates (Põlme et al. 2020) | Põlme S, et al. 2020. Fungal Diversity 105:1-16 ((https://doi.org/10.1007/s13225-020-00466-2)) |
| Otu22038 | 1 | k__Fungi(100);p__Ascomycota(100);c__Saccharomycetes(100);o__Saccharomycetales(100);f__Saccharomycetales_family_Incertae_sedis(100);g__Candida(100);g__Candida_unclassified(100); | Candida | 13 | Pathotroph-Saprotroph-Symbiotroph | Animal Pathogen-Endophyte-Endosymbiont-Epiphyte-Undefined Saprotroph | Dimorphic Yeast Yeast | NULL | Probable | As nectar-tap_saprotroph (Põlme et al. 2020); Decay sugar-rich substrates (Põlme et al. 2020); This asexual genus is polyphyletic and contain species that function very differently depend on what they are. Caution should be taken when interpreting guilds for Candida species. | Manolakaki D, et al. 2010. Virulence. 1: 367-375 ((https://doi.org/10.4161/viru.1.5.12796)); Mekbib SB, et al. 2011. Tropical Plant Pathology 36:233-240 ((https://doi.org/10.1590/S1982-56762011000400004)); Põlme S, et al. 2020. Fungal Diversity 105:1-16 ((https://doi.org/10.1007/s13225-020-00466-2)) |
| Otu22039 | 1 | k__Fungi(100);p__Ascomycota(100);c__Saccharomycetes(100);o__Saccharomycetales(100);f__Saccharomycetales_family_Incertae_sedis(100);f__Saccharomycetales_family_Incertae_sedis_unclassified(100);f__Saccharomycetales_family_Incertae_sedis_unclassified(100); | Saccharomycetales | 7 | Saprotroph | Undefined Saprotroph | Yeast | NULL | Possible | NULL | Sterkenburg E, et al. 2015. New Phytologist 207:1145-1158 ((https://doi.org/10.1111/nph.13426)) |
| Otu22044 | 1 | k__Fungi(100);p__Ascomycota(100);c__Saccharomycetes(100);o__Saccharomycetales(100);f__Saccharomycetales_family_Incertae_sedis(100);g__Candida(100);g__Candida_unclassified(100); | Candida | 13 | Pathotroph-Saprotroph-Symbiotroph | Animal Pathogen-Endophyte-Endosymbiont-Epiphyte-Undefined Saprotroph | Dimorphic Yeast Yeast | NULL | Probable | As nectar-tap_saprotroph (Põlme et al. 2020); Decay sugar-rich substrates (Põlme et al. 2020); This asexual genus is polyphyletic and contain species that function very differently depend on what they are. Caution should be taken when interpreting guilds for Candida species. | Manolakaki D, et al. 2010. Virulence. 1: 367-375 ((https://doi.org/10.4161/viru.1.5.12796)); Mekbib SB, et al. 2011. Tropical Plant Pathology 36:233-240 ((https://doi.org/10.1590/S1982-56762011000400004)); Põlme S, et al. 2020. Fungal Diversity 105:1-16 ((https://doi.org/10.1007/s13225-020-00466-2)) |
| Otu22046 | 1 | k__Fungi(100);p__Ascomycota(100);c__Saccharomycetes(100);o__Saccharomycetales(100);f__Metschnikowiaceae(100);g__Metschnikowia(100);s__Metschnikowia_pulcherrima(100); | Metschnikowia | 13 | Saprotroph | Undefined Saprotroph | Yeast | NULL | Probable | As nectar-tap_saprotroph (Põlme et al. 2020); Decay sugar-rich substrates (Põlme et al. 2020) | Põlme S, et al. 2020. Fungal Diversity 105:1-16 ((https://doi.org/10.1007/s13225-020-00466-2)) |
| Otu22048 | 1 | k__Fungi(100);p__Ascomycota(100);c__Saccharomycetes(100);o__Saccharomycetales(100);f__Metschnikowiaceae(100);g__Metschnikowia(100);s__Metschnikowia_pulcherrima(100); | Metschnikowia | 13 | Saprotroph | Undefined Saprotroph | Yeast | NULL | Probable | As nectar-tap_saprotroph (Põlme et al. 2020); Decay sugar-rich substrates (Põlme et al. 2020) | Põlme S, et al. 2020. Fungal Diversity 105:1-16 ((https://doi.org/10.1007/s13225-020-00466-2)) |
| Otu22053 | 1 | k__Fungi(100);p__Ascomycota(100);c__Saccharomycetes(100);o__Saccharomycetales(100);f__Metschnikowiaceae(100);g__Metschnikowia(100);s__Metschnikowia_pulcherrima(100); | Metschnikowia | 13 | Saprotroph | Undefined Saprotroph | Yeast | NULL | Probable | As nectar-tap_saprotroph (Põlme et al. 2020); Decay sugar-rich substrates (Põlme et al. 2020) | Põlme S, et al. 2020. Fungal Diversity 105:1-16 ((https://doi.org/10.1007/s13225-020-00466-2)) |
| Otu22054 | 1 | k__Fungi(100);p__Ascomycota(100);c__Saccharomycetes(100);o__Saccharomycetales(100);f__Metschnikowiaceae(100);g__Metschnikowia(100);s__Metschnikowia_pulcherrima(100); | Metschnikowia | 13 | Saprotroph | Undefined Saprotroph | Yeast | NULL | Probable | As nectar-tap_saprotroph (Põlme et al. 2020); Decay sugar-rich substrates (Põlme et al. 2020) | Põlme S, et al. 2020. Fungal Diversity 105:1-16 ((https://doi.org/10.1007/s13225-020-00466-2)) |
| Otu22055 | 1 | k__Fungi(100);p__Ascomycota(100);c__Saccharomycetes(100);o__Saccharomycetales(100);f__Metschnikowiaceae(100);g__Metschnikowia(100);g__Metschnikowia_unclassified(100); | Metschnikowia | 13 | Saprotroph | Undefined Saprotroph | Yeast | NULL | Probable | As nectar-tap_saprotroph (Põlme et al. 2020); Decay sugar-rich substrates (Põlme et al. 2020) | Põlme S, et al. 2020. Fungal Diversity 105:1-16 ((https://doi.org/10.1007/s13225-020-00466-2)) |
| Otu22056 | 1 | k__Fungi(100);p__Ascomycota(100);c__Saccharomycetes(100);o__Saccharomycetales(100);f__Metschnikowiaceae(100);g__Metschnikowia(100);s__Metschnikowia_pulcherrima(100); | Metschnikowia | 13 | Saprotroph | Undefined Saprotroph | Yeast | NULL | Probable | As nectar-tap_saprotroph (Põlme et al. 2020); Decay sugar-rich substrates (Põlme et al. 2020) | Põlme S, et al. 2020. Fungal Diversity 105:1-16 ((https://doi.org/10.1007/s13225-020-00466-2)) |
| Otu22057 | 1 | k__Fungi(100);p__Ascomycota(100);c__Saccharomycetes(100);o__Saccharomycetales(100);f__Metschnikowiaceae(100);g__Metschnikowia(100);s__Metschnikowia_pulcherrima(100); | Metschnikowia | 13 | Saprotroph | Undefined Saprotroph | Yeast | NULL | Probable | As nectar-tap_saprotroph (Põlme et al. 2020); Decay sugar-rich substrates (Põlme et al. 2020) | Põlme S, et al. 2020. Fungal Diversity 105:1-16 ((https://doi.org/10.1007/s13225-020-00466-2)) |
| Otu22058 | 1 | k__Fungi(100);p__Ascomycota(100);c__Saccharomycetes(100);o__Saccharomycetales(100);f__Metschnikowiaceae(100);g__Metschnikowia(100);s__Metschnikowia_pulcherrima(100); | Metschnikowia | 13 | Saprotroph | Undefined Saprotroph | Yeast | NULL | Probable | As nectar-tap_saprotroph (Põlme et al. 2020); Decay sugar-rich substrates (Põlme et al. 2020) | Põlme S, et al. 2020. Fungal Diversity 105:1-16 ((https://doi.org/10.1007/s13225-020-00466-2)) |
| Otu22060 | 1 | k__Fungi(100);p__Ascomycota(100);c__Saccharomycetes(100);o__Saccharomycetales(100);f__Saccharomycetales_family_Incertae_sedis(100);g__Candida(100);s__Candida_cretensis(100); | Candida | 13 | Pathotroph-Saprotroph-Symbiotroph | Animal Pathogen-Endophyte-Endosymbiont-Epiphyte-Undefined Saprotroph | Dimorphic Yeast Yeast | NULL | Probable | As nectar-tap_saprotroph (Põlme et al. 2020); Decay sugar-rich substrates (Põlme et al. 2020); This asexual genus is polyphyletic and contain species that function very differently depend on what they are. Caution should be taken when interpreting guilds for Candida species. | Manolakaki D, et al. 2010. Virulence. 1: 367-375 ((https://doi.org/10.4161/viru.1.5.12796)); Mekbib SB, et al. 2011. Tropical Plant Pathology 36:233-240 ((https://doi.org/10.1590/S1982-56762011000400004)); Põlme S, et al. 2020. Fungal Diversity 105:1-16 ((https://doi.org/10.1007/s13225-020-00466-2)) |
| Otu22061 | 1 | k__Fungi(100);p__Ascomycota(100);c__Saccharomycetes(100);o__Saccharomycetales(100);f__Saccharomycetales_family_Incertae_sedis(100);f__Saccharomycetales_family_Incertae_sedis_unclassified(100);f__Saccharomycetales_family_Incertae_sedis_unclassified(100); | Saccharomycetales | 7 | Saprotroph | Undefined Saprotroph | Yeast | NULL | Possible | NULL | Sterkenburg E, et al. 2015. New Phytologist 207:1145-1158 ((https://doi.org/10.1111/nph.13426)) |
| Otu22063 | 1 | k__Fungi(100);p__Ascomycota(100);c__Saccharomycetes(100);o__Saccharomycetales(100);f__Saccharomycetales_family_Incertae_sedis(100);f__Saccharomycetales_family_Incertae_sedis_unclassified(100);f__Saccharomycetales_family_Incertae_sedis_unclassified(100); | Saccharomycetales | 7 | Saprotroph | Undefined Saprotroph | Yeast | NULL | Possible | NULL | Sterkenburg E, et al. 2015. New Phytologist 207:1145-1158 ((https://doi.org/10.1111/nph.13426)) |
| Otu22065 | 1 | k__Fungi(100);p__Ascomycota(100);c__Saccharomycetes(100);o__Saccharomycetales(100);f__Saccharomycetales_family_Incertae_sedis(100);g__Candida(100);g__Candida_unclassified(100); | Candida | 13 | Pathotroph-Saprotroph-Symbiotroph | Animal Pathogen-Endophyte-Endosymbiont-Epiphyte-Undefined Saprotroph | Dimorphic Yeast Yeast | NULL | Probable | As nectar-tap_saprotroph (Põlme et al. 2020); Decay sugar-rich substrates (Põlme et al. 2020); This asexual genus is polyphyletic and contain species that function very differently depend on what they are. Caution should be taken when interpreting guilds for Candida species. | Manolakaki D, et al. 2010. Virulence. 1: 367-375 ((https://doi.org/10.4161/viru.1.5.12796)); Mekbib SB, et al. 2011. Tropical Plant Pathology 36:233-240 ((https://doi.org/10.1590/S1982-56762011000400004)); Põlme S, et al. 2020. Fungal Diversity 105:1-16 ((https://doi.org/10.1007/s13225-020-00466-2)) |
| Otu22071 | 1 | k__Fungi(100);p__Ascomycota(100);c__Saccharomycetes(100);o__Saccharomycetales(100);f__Saccharomycetales_family_Incertae_sedis(100);f__Saccharomycetales_family_Incertae_sedis_unclassified(100);f__Saccharomycetales_family_Incertae_sedis_unclassified(100); | Saccharomycetales | 7 | Saprotroph | Undefined Saprotroph | Yeast | NULL | Possible | NULL | Sterkenburg E, et al. 2015. New Phytologist 207:1145-1158 ((https://doi.org/10.1111/nph.13426)) |
| Otu22072 | 1 | k__Fungi(100);p__Ascomycota(100);c__Saccharomycetes(100);o__Saccharomycetales(100);f__Saccharomycetales_family_Incertae_sedis(100);f__Saccharomycetales_family_Incertae_sedis_unclassified(100);f__Saccharomycetales_family_Incertae_sedis_unclassified(100); | Saccharomycetales | 7 | Saprotroph | Undefined Saprotroph | Yeast | NULL | Possible | NULL | Sterkenburg E, et al. 2015. New Phytologist 207:1145-1158 ((https://doi.org/10.1111/nph.13426)) |
| Otu22078 | 1 | k__Fungi(100);p__Ascomycota(100);c__Saccharomycetes(100);o__Saccharomycetales(100);f__Metschnikowiaceae(100);g__Metschnikowia(100);s__Metschnikowia_pulcherrima(100); | Metschnikowia | 13 | Saprotroph | Undefined Saprotroph | Yeast | NULL | Probable | As nectar-tap_saprotroph (Põlme et al. 2020); Decay sugar-rich substrates (Põlme et al. 2020) | Põlme S, et al. 2020. Fungal Diversity 105:1-16 ((https://doi.org/10.1007/s13225-020-00466-2)) |
| Otu22079 | 1 | k__Fungi(100);p__Ascomycota(100);c__Saccharomycetes(100);o__Saccharomycetales(100);f__Metschnikowiaceae(100);g__Metschnikowia(100);s__Metschnikowia_pulcherrima(100); | Metschnikowia | 13 | Saprotroph | Undefined Saprotroph | Yeast | NULL | Probable | As nectar-tap_saprotroph (Põlme et al. 2020); Decay sugar-rich substrates (Põlme et al. 2020) | Põlme S, et al. 2020. Fungal Diversity 105:1-16 ((https://doi.org/10.1007/s13225-020-00466-2)) |
| Otu22080 | 1 | k__Fungi(100);p__Ascomycota(100);c__Saccharomycetes(100);o__Saccharomycetales(100);f__Metschnikowiaceae(100);g__Metschnikowia(100);s__Metschnikowia_pulcherrima(100); | Metschnikowia | 13 | Saprotroph | Undefined Saprotroph | Yeast | NULL | Probable | As nectar-tap_saprotroph (Põlme et al. 2020); Decay sugar-rich substrates (Põlme et al. 2020) | Põlme S, et al. 2020. Fungal Diversity 105:1-16 ((https://doi.org/10.1007/s13225-020-00466-2)) |
| Otu22081 | 1 | k__Fungi(100);p__Ascomycota(100);c__Saccharomycetes(100);o__Saccharomycetales(100);f__Metschnikowiaceae(100);g__Metschnikowia(100);s__Metschnikowia_pulcherrima(100); | Metschnikowia | 13 | Saprotroph | Undefined Saprotroph | Yeast | NULL | Probable | As nectar-tap_saprotroph (Põlme et al. 2020); Decay sugar-rich substrates (Põlme et al. 2020) | Põlme S, et al. 2020. Fungal Diversity 105:1-16 ((https://doi.org/10.1007/s13225-020-00466-2)) |
| Otu22082 | 1 | k__Fungi(100);p__Ascomycota(100);c__Saccharomycetes(100);o__Saccharomycetales(100);f__Metschnikowiaceae(100);g__Metschnikowia(100);s__Metschnikowia_pulcherrima(100); | Metschnikowia | 13 | Saprotroph | Undefined Saprotroph | Yeast | NULL | Probable | As nectar-tap_saprotroph (Põlme et al. 2020); Decay sugar-rich substrates (Põlme et al. 2020) | Põlme S, et al. 2020. Fungal Diversity 105:1-16 ((https://doi.org/10.1007/s13225-020-00466-2)) |
| Otu22083 | 1 | k__Fungi(100);p__Ascomycota(100);c__Saccharomycetes(100);o__Saccharomycetales(100);f__Metschnikowiaceae(100);g__Metschnikowia(100);s__Metschnikowia_pulcherrima(100); | Metschnikowia | 13 | Saprotroph | Undefined Saprotroph | Yeast | NULL | Probable | As nectar-tap_saprotroph (Põlme et al. 2020); Decay sugar-rich substrates (Põlme et al. 2020) | Põlme S, et al. 2020. Fungal Diversity 105:1-16 ((https://doi.org/10.1007/s13225-020-00466-2)) |
| Otu22084 | 1 | k__Fungi(100);p__Ascomycota(100);c__Saccharomycetes(100);o__Saccharomycetales(100);f__Saccharomycetales_family_Incertae_sedis(100);g__Candida(100);g__Candida_unclassified(100); | Candida | 13 | Pathotroph-Saprotroph-Symbiotroph | Animal Pathogen-Endophyte-Endosymbiont-Epiphyte-Undefined Saprotroph | Dimorphic Yeast Yeast | NULL | Probable | As nectar-tap_saprotroph (Põlme et al. 2020); Decay sugar-rich substrates (Põlme et al. 2020); This asexual genus is polyphyletic and contain species that function very differently depend on what they are. Caution should be taken when interpreting guilds for Candida species. | Manolakaki D, et al. 2010. Virulence. 1: 367-375 ((https://doi.org/10.4161/viru.1.5.12796)); Mekbib SB, et al. 2011. Tropical Plant Pathology 36:233-240 ((https://doi.org/10.1590/S1982-56762011000400004)); Põlme S, et al. 2020. Fungal Diversity 105:1-16 ((https://doi.org/10.1007/s13225-020-00466-2)) |
| Otu22085 | 1 | k__Fungi(100);p__Ascomycota(100);c__Saccharomycetes(100);o__Saccharomycetales(100);f__Metschnikowiaceae(100);g__Metschnikowia(100);s__Metschnikowia_pulcherrima(100); | Metschnikowia | 13 | Saprotroph | Undefined Saprotroph | Yeast | NULL | Probable | As nectar-tap_saprotroph (Põlme et al. 2020); Decay sugar-rich substrates (Põlme et al. 2020) | Põlme S, et al. 2020. Fungal Diversity 105:1-16 ((https://doi.org/10.1007/s13225-020-00466-2)) |
| Otu22086 | 1 | k__Fungi(100);p__Ascomycota(100);c__Saccharomycetes(100);o__Saccharomycetales(100);f__Metschnikowiaceae(100);g__Metschnikowia(100);s__Metschnikowia_pulcherrima(100); | Metschnikowia | 13 | Saprotroph | Undefined Saprotroph | Yeast | NULL | Probable | As nectar-tap_saprotroph (Põlme et al. 2020); Decay sugar-rich substrates (Põlme et al. 2020) | Põlme S, et al. 2020. Fungal Diversity 105:1-16 ((https://doi.org/10.1007/s13225-020-00466-2)) |
| Otu22087 | 1 | k__Fungi(100);p__Ascomycota(100);c__Saccharomycetes(100);o__Saccharomycetales(100);f__Metschnikowiaceae(100);g__Metschnikowia(100);g__Metschnikowia_unclassified(100); | Metschnikowia | 13 | Saprotroph | Undefined Saprotroph | Yeast | NULL | Probable | As nectar-tap_saprotroph (Põlme et al. 2020); Decay sugar-rich substrates (Põlme et al. 2020) | Põlme S, et al. 2020. Fungal Diversity 105:1-16 ((https://doi.org/10.1007/s13225-020-00466-2)) |
| Otu22088 | 1 | k__Fungi(100);p__Ascomycota(100);c__Saccharomycetes(100);o__Saccharomycetales(100);f__Metschnikowiaceae(100);g__Metschnikowia(100);s__Metschnikowia_pulcherrima(100); | Metschnikowia | 13 | Saprotroph | Undefined Saprotroph | Yeast | NULL | Probable | As nectar-tap_saprotroph (Põlme et al. 2020); Decay sugar-rich substrates (Põlme et al. 2020) | Põlme S, et al. 2020. Fungal Diversity 105:1-16 ((https://doi.org/10.1007/s13225-020-00466-2)) |
| Otu22089 | 1 | k__Fungi(100);p__Ascomycota(100);c__Saccharomycetes(100);o__Saccharomycetales(100);f__Metschnikowiaceae(100);g__Metschnikowia(100);s__Metschnikowia_pulcherrima(100); | Metschnikowia | 13 | Saprotroph | Undefined Saprotroph | Yeast | NULL | Probable | As nectar-tap_saprotroph (Põlme et al. 2020); Decay sugar-rich substrates (Põlme et al. 2020) | Põlme S, et al. 2020. Fungal Diversity 105:1-16 ((https://doi.org/10.1007/s13225-020-00466-2)) |
| Otu22090 | 1 | k__Fungi(100);p__Ascomycota(100);c__Saccharomycetes(100);o__Saccharomycetales(100);f__Metschnikowiaceae(100);g__Metschnikowia(100);s__Metschnikowia_pulcherrima(100); | Metschnikowia | 13 | Saprotroph | Undefined Saprotroph | Yeast | NULL | Probable | As nectar-tap_saprotroph (Põlme et al. 2020); Decay sugar-rich substrates (Põlme et al. 2020) | Põlme S, et al. 2020. Fungal Diversity 105:1-16 ((https://doi.org/10.1007/s13225-020-00466-2)) |
| Otu22091 | 1 | k__Fungi(100);p__Ascomycota(100);c__Saccharomycetes(100);o__Saccharomycetales(100);f__Metschnikowiaceae(100);g__Metschnikowia(100);s__Metschnikowia_pulcherrima(100); | Metschnikowia | 13 | Saprotroph | Undefined Saprotroph | Yeast | NULL | Probable | As nectar-tap_saprotroph (Põlme et al. 2020); Decay sugar-rich substrates (Põlme et al. 2020) | Põlme S, et al. 2020. Fungal Diversity 105:1-16 ((https://doi.org/10.1007/s13225-020-00466-2)) |
| Otu22092 | 1 | k__Fungi(100);p__Ascomycota(100);c__Saccharomycetes(100);o__Saccharomycetales(100);f__Metschnikowiaceae(100);g__Metschnikowia(100);s__Metschnikowia_pulcherrima(100); | Metschnikowia | 13 | Saprotroph | Undefined Saprotroph | Yeast | NULL | Probable | As nectar-tap_saprotroph (Põlme et al. 2020); Decay sugar-rich substrates (Põlme et al. 2020) | Põlme S, et al. 2020. Fungal Diversity 105:1-16 ((https://doi.org/10.1007/s13225-020-00466-2)) |
| Otu22093 | 1 | k__Fungi(100);p__Ascomycota(100);c__Saccharomycetes(100);o__Saccharomycetales(100);f__Metschnikowiaceae(100);g__Metschnikowia(100);s__Metschnikowia_pulcherrima(100); | Metschnikowia | 13 | Saprotroph | Undefined Saprotroph | Yeast | NULL | Probable | As nectar-tap_saprotroph (Põlme et al. 2020); Decay sugar-rich substrates (Põlme et al. 2020) | Põlme S, et al. 2020. Fungal Diversity 105:1-16 ((https://doi.org/10.1007/s13225-020-00466-2)) |
| Otu22095 | 1 | k__Fungi(100);p__Ascomycota(100);c__Saccharomycetes(100);o__Saccharomycetales(100);f__Metschnikowiaceae(100);g__Metschnikowia(100);g__Metschnikowia_unclassified(100); | Metschnikowia | 13 | Saprotroph | Undefined Saprotroph | Yeast | NULL | Probable | As nectar-tap_saprotroph (Põlme et al. 2020); Decay sugar-rich substrates (Põlme et al. 2020) | Põlme S, et al. 2020. Fungal Diversity 105:1-16 ((https://doi.org/10.1007/s13225-020-00466-2)) |
| Otu22096 | 1 | k__Fungi(100);p__Ascomycota(100);c__Saccharomycetes(100);o__Saccharomycetales(100);f__Metschnikowiaceae(100);g__Metschnikowia(100);s__Metschnikowia_pulcherrima(100); | Metschnikowia | 13 | Saprotroph | Undefined Saprotroph | Yeast | NULL | Probable | As nectar-tap_saprotroph (Põlme et al. 2020); Decay sugar-rich substrates (Põlme et al. 2020) | Põlme S, et al. 2020. Fungal Diversity 105:1-16 ((https://doi.org/10.1007/s13225-020-00466-2)) |
| Otu22098 | 1 | k__Fungi(100);p__Ascomycota(100);c__Saccharomycetes(100);o__Saccharomycetales(100);f__Metschnikowiaceae(100);g__Metschnikowia(100);s__Metschnikowia_pulcherrima(100); | Metschnikowia | 13 | Saprotroph | Undefined Saprotroph | Yeast | NULL | Probable | As nectar-tap_saprotroph (Põlme et al. 2020); Decay sugar-rich substrates (Põlme et al. 2020) | Põlme S, et al. 2020. Fungal Diversity 105:1-16 ((https://doi.org/10.1007/s13225-020-00466-2)) |
| Otu22099 | 1 | k__Fungi(100);p__Ascomycota(100);c__Saccharomycetes(100);o__Saccharomycetales(100);f__Metschnikowiaceae(100);g__Metschnikowia(100);s__Metschnikowia_pulcherrima(100); | Metschnikowia | 13 | Saprotroph | Undefined Saprotroph | Yeast | NULL | Probable | As nectar-tap_saprotroph (Põlme et al. 2020); Decay sugar-rich substrates (Põlme et al. 2020) | Põlme S, et al. 2020. Fungal Diversity 105:1-16 ((https://doi.org/10.1007/s13225-020-00466-2)) |
| Otu22100 | 1 | k__Fungi(100);p__Ascomycota(100);c__Saccharomycetes(100);o__Saccharomycetales(100);f__Metschnikowiaceae(100);g__Metschnikowia(100);s__Metschnikowia_pulcherrima(100); | Metschnikowia | 13 | Saprotroph | Undefined Saprotroph | Yeast | NULL | Probable | As nectar-tap_saprotroph (Põlme et al. 2020); Decay sugar-rich substrates (Põlme et al. 2020) | Põlme S, et al. 2020. Fungal Diversity 105:1-16 ((https://doi.org/10.1007/s13225-020-00466-2)) |
| Otu22101 | 1 | k__Fungi(100);p__Ascomycota(100);c__Saccharomycetes(100);o__Saccharomycetales(100);f__Metschnikowiaceae(100);g__Metschnikowia(100);g__Metschnikowia_unclassified(100); | Metschnikowia | 13 | Saprotroph | Undefined Saprotroph | Yeast | NULL | Probable | As nectar-tap_saprotroph (Põlme et al. 2020); Decay sugar-rich substrates (Põlme et al. 2020) | Põlme S, et al. 2020. Fungal Diversity 105:1-16 ((https://doi.org/10.1007/s13225-020-00466-2)) |
| Otu22102 | 1 | k__Fungi(100);p__Ascomycota(100);c__Saccharomycetes(100);o__Saccharomycetales(100);f__Metschnikowiaceae(100);g__Metschnikowia(100);s__Metschnikowia_pulcherrima(100); | Metschnikowia | 13 | Saprotroph | Undefined Saprotroph | Yeast | NULL | Probable | As nectar-tap_saprotroph (Põlme et al. 2020); Decay sugar-rich substrates (Põlme et al. 2020) | Põlme S, et al. 2020. Fungal Diversity 105:1-16 ((https://doi.org/10.1007/s13225-020-00466-2)) |
| Otu22103 | 1 | k__Fungi(100);p__Ascomycota(100);c__Saccharomycetes(100);o__Saccharomycetales(100);f__Metschnikowiaceae(100);g__Metschnikowia(100);g__Metschnikowia_unclassified(100); | Metschnikowia | 13 | Saprotroph | Undefined Saprotroph | Yeast | NULL | Probable | As nectar-tap_saprotroph (Põlme et al. 2020); Decay sugar-rich substrates (Põlme et al. 2020) | Põlme S, et al. 2020. Fungal Diversity 105:1-16 ((https://doi.org/10.1007/s13225-020-00466-2)) |
| Otu22104 | 1 | k__Fungi(100);p__Ascomycota(100);c__Saccharomycetes(100);o__Saccharomycetales(100);f__Metschnikowiaceae(100);g__Metschnikowia(100);s__Metschnikowia_pulcherrima(100); | Metschnikowia | 13 | Saprotroph | Undefined Saprotroph | Yeast | NULL | Probable | As nectar-tap_saprotroph (Põlme et al. 2020); Decay sugar-rich substrates (Põlme et al. 2020) | Põlme S, et al. 2020. Fungal Diversity 105:1-16 ((https://doi.org/10.1007/s13225-020-00466-2)) |
| Otu22105 | 1 | k__Fungi(100);p__Ascomycota(100);c__Saccharomycetes(100);o__Saccharomycetales(100);f__Metschnikowiaceae(100);g__Metschnikowia(100);s__Metschnikowia_pulcherrima(100); | Metschnikowia | 13 | Saprotroph | Undefined Saprotroph | Yeast | NULL | Probable | As nectar-tap_saprotroph (Põlme et al. 2020); Decay sugar-rich substrates (Põlme et al. 2020) | Põlme S, et al. 2020. Fungal Diversity 105:1-16 ((https://doi.org/10.1007/s13225-020-00466-2)) |
| Otu22106 | 1 | k__Fungi(100);p__Ascomycota(100);c__Saccharomycetes(100);o__Saccharomycetales(100);f__Metschnikowiaceae(100);g__Metschnikowia(100);s__Metschnikowia_pulcherrima(100); | Metschnikowia | 13 | Saprotroph | Undefined Saprotroph | Yeast | NULL | Probable | As nectar-tap_saprotroph (Põlme et al. 2020); Decay sugar-rich substrates (Põlme et al. 2020) | Põlme S, et al. 2020. Fungal Diversity 105:1-16 ((https://doi.org/10.1007/s13225-020-00466-2)) |
| Otu22107 | 1 | k__Fungi(100);p__Ascomycota(100);c__Saccharomycetes(100);o__Saccharomycetales(100);f__Metschnikowiaceae(100);g__Metschnikowia(100);g__Metschnikowia_unclassified(100); | Metschnikowia | 13 | Saprotroph | Undefined Saprotroph | Yeast | NULL | Probable | As nectar-tap_saprotroph (Põlme et al. 2020); Decay sugar-rich substrates (Põlme et al. 2020) | Põlme S, et al. 2020. Fungal Diversity 105:1-16 ((https://doi.org/10.1007/s13225-020-00466-2)) |
| Otu22108 | 1 | k__Fungi(100);p__Ascomycota(100);c__Saccharomycetes(100);o__Saccharomycetales(100);f__Metschnikowiaceae(100);g__Metschnikowia(100);s__Metschnikowia_pulcherrima(100); | Metschnikowia | 13 | Saprotroph | Undefined Saprotroph | Yeast | NULL | Probable | As nectar-tap_saprotroph (Põlme et al. 2020); Decay sugar-rich substrates (Põlme et al. 2020) | Põlme S, et al. 2020. Fungal Diversity 105:1-16 ((https://doi.org/10.1007/s13225-020-00466-2)) |
| Otu22109 | 1 | k__Fungi(100);p__Ascomycota(100);c__Saccharomycetes(100);o__Saccharomycetales(100);f__Metschnikowiaceae(100);g__Metschnikowia(100);s__Metschnikowia_pulcherrima(100); | Metschnikowia | 13 | Saprotroph | Undefined Saprotroph | Yeast | NULL | Probable | As nectar-tap_saprotroph (Põlme et al. 2020); Decay sugar-rich substrates (Põlme et al. 2020) | Põlme S, et al. 2020. Fungal Diversity 105:1-16 ((https://doi.org/10.1007/s13225-020-00466-2)) |
| Otu22110 | 1 | k__Fungi(100);p__Ascomycota(100);c__Saccharomycetes(100);o__Saccharomycetales(100);f__Metschnikowiaceae(100);g__Metschnikowia(100);s__Metschnikowia_pulcherrima(100); | Metschnikowia | 13 | Saprotroph | Undefined Saprotroph | Yeast | NULL | Probable | As nectar-tap_saprotroph (Põlme et al. 2020); Decay sugar-rich substrates (Põlme et al. 2020) | Põlme S, et al. 2020. Fungal Diversity 105:1-16 ((https://doi.org/10.1007/s13225-020-00466-2)) |
| Otu22111 | 1 | k__Fungi(100);p__Ascomycota(100);c__Saccharomycetes(100);o__Saccharomycetales(100);f__Metschnikowiaceae(100);g__Metschnikowia(100);s__Metschnikowia_pulcherrima(100); | Metschnikowia | 13 | Saprotroph | Undefined Saprotroph | Yeast | NULL | Probable | As nectar-tap_saprotroph (Põlme et al. 2020); Decay sugar-rich substrates (Põlme et al. 2020) | Põlme S, et al. 2020. Fungal Diversity 105:1-16 ((https://doi.org/10.1007/s13225-020-00466-2)) |
| Otu22112 | 1 | k__Fungi(100);p__Ascomycota(100);c__Saccharomycetes(100);o__Saccharomycetales(100);f__Metschnikowiaceae(100);g__Metschnikowia(100);s__Metschnikowia_pulcherrima(100); | Metschnikowia | 13 | Saprotroph | Undefined Saprotroph | Yeast | NULL | Probable | As nectar-tap_saprotroph (Põlme et al. 2020); Decay sugar-rich substrates (Põlme et al. 2020) | Põlme S, et al. 2020. Fungal Diversity 105:1-16 ((https://doi.org/10.1007/s13225-020-00466-2)) |
| Otu22114 | 1 | k__Fungi(100);p__Ascomycota(100);c__Saccharomycetes(100);o__Saccharomycetales(100);f__Metschnikowiaceae(100);g__Metschnikowia(100);s__Metschnikowia_pulcherrima(100); | Metschnikowia | 13 | Saprotroph | Undefined Saprotroph | Yeast | NULL | Probable | As nectar-tap_saprotroph (Põlme et al. 2020); Decay sugar-rich substrates (Põlme et al. 2020) | Põlme S, et al. 2020. Fungal Diversity 105:1-16 ((https://doi.org/10.1007/s13225-020-00466-2)) |
| Otu22115 | 1 | k__Fungi(100);p__Ascomycota(100);c__Saccharomycetes(100);o__Saccharomycetales(100);f__Metschnikowiaceae(100);g__Metschnikowia(100);g__Metschnikowia_unclassified(100); | Metschnikowia | 13 | Saprotroph | Undefined Saprotroph | Yeast | NULL | Probable | As nectar-tap_saprotroph (Põlme et al. 2020); Decay sugar-rich substrates (Põlme et al. 2020) | Põlme S, et al. 2020. Fungal Diversity 105:1-16 ((https://doi.org/10.1007/s13225-020-00466-2)) |
| Otu22116 | 1 | k__Fungi(100);p__Ascomycota(100);c__Saccharomycetes(100);o__Saccharomycetales(100);f__Metschnikowiaceae(100);g__Metschnikowia(100);g__Metschnikowia_unclassified(100); | Metschnikowia | 13 | Saprotroph | Undefined Saprotroph | Yeast | NULL | Probable | As nectar-tap_saprotroph (Põlme et al. 2020); Decay sugar-rich substrates (Põlme et al. 2020) | Põlme S, et al. 2020. Fungal Diversity 105:1-16 ((https://doi.org/10.1007/s13225-020-00466-2)) |
| Otu22117 | 1 | k__Fungi(100);p__Ascomycota(100);c__Saccharomycetes(100);o__Saccharomycetales(100);f__Metschnikowiaceae(100);g__Metschnikowia(100);s__Metschnikowia_pulcherrima(100); | Metschnikowia | 13 | Saprotroph | Undefined Saprotroph | Yeast | NULL | Probable | As nectar-tap_saprotroph (Põlme et al. 2020); Decay sugar-rich substrates (Põlme et al. 2020) | Põlme S, et al. 2020. Fungal Diversity 105:1-16 ((https://doi.org/10.1007/s13225-020-00466-2)) |
| Otu22118 | 1 | k__Fungi(100);p__Ascomycota(100);c__Saccharomycetes(100);o__Saccharomycetales(100);f__Metschnikowiaceae(100);g__Metschnikowia(100);s__Metschnikowia_pulcherrima(100); | Metschnikowia | 13 | Saprotroph | Undefined Saprotroph | Yeast | NULL | Probable | As nectar-tap_saprotroph (Põlme et al. 2020); Decay sugar-rich substrates (Põlme et al. 2020) | Põlme S, et al. 2020. Fungal Diversity 105:1-16 ((https://doi.org/10.1007/s13225-020-00466-2)) |
| Otu22119 | 1 | k__Fungi(100);p__Ascomycota(100);c__Saccharomycetes(100);o__Saccharomycetales(100);f__Metschnikowiaceae(100);g__Metschnikowia(100);s__Metschnikowia_pulcherrima(100); | Metschnikowia | 13 | Saprotroph | Undefined Saprotroph | Yeast | NULL | Probable | As nectar-tap_saprotroph (Põlme et al. 2020); Decay sugar-rich substrates (Põlme et al. 2020) | Põlme S, et al. 2020. Fungal Diversity 105:1-16 ((https://doi.org/10.1007/s13225-020-00466-2)) |
| Otu22120 | 1 | k__Fungi(100);p__Ascomycota(100);c__Saccharomycetes(100);o__Saccharomycetales(100);f__Metschnikowiaceae(100);g__Metschnikowia(100);g__Metschnikowia_unclassified(100); | Metschnikowia | 13 | Saprotroph | Undefined Saprotroph | Yeast | NULL | Probable | As nectar-tap_saprotroph (Põlme et al. 2020); Decay sugar-rich substrates (Põlme et al. 2020) | Põlme S, et al. 2020. Fungal Diversity 105:1-16 ((https://doi.org/10.1007/s13225-020-00466-2)) |
| Otu22121 | 1 | k__Fungi(100);p__Ascomycota(100);c__Saccharomycetes(100);o__Saccharomycetales(100);f__Metschnikowiaceae(100);g__Metschnikowia(100);s__Metschnikowia_pulcherrima(100); | Metschnikowia | 13 | Saprotroph | Undefined Saprotroph | Yeast | NULL | Probable | As nectar-tap_saprotroph (Põlme et al. 2020); Decay sugar-rich substrates (Põlme et al. 2020) | Põlme S, et al. 2020. Fungal Diversity 105:1-16 ((https://doi.org/10.1007/s13225-020-00466-2)) |
| Otu22122 | 1 | k__Fungi(100);p__Ascomycota(100);c__Saccharomycetes(100);o__Saccharomycetales(100);f__Metschnikowiaceae(100);g__Metschnikowia(100);g__Metschnikowia_unclassified(100); | Metschnikowia | 13 | Saprotroph | Undefined Saprotroph | Yeast | NULL | Probable | As nectar-tap_saprotroph (Põlme et al. 2020); Decay sugar-rich substrates (Põlme et al. 2020) | Põlme S, et al. 2020. Fungal Diversity 105:1-16 ((https://doi.org/10.1007/s13225-020-00466-2)) |
| Otu22123 | 1 | k__Fungi(100);p__Ascomycota(100);c__Saccharomycetes(100);o__Saccharomycetales(100);f__Metschnikowiaceae(100);g__Metschnikowia(100);s__Metschnikowia_pulcherrima(100); | Metschnikowia | 13 | Saprotroph | Undefined Saprotroph | Yeast | NULL | Probable | As nectar-tap_saprotroph (Põlme et al. 2020); Decay sugar-rich substrates (Põlme et al. 2020) | Põlme S, et al. 2020. Fungal Diversity 105:1-16 ((https://doi.org/10.1007/s13225-020-00466-2)) |
| Otu22124 | 1 | k__Fungi(100);p__Ascomycota(100);c__Saccharomycetes(100);o__Saccharomycetales(100);f__Metschnikowiaceae(100);g__Metschnikowia(100);s__Metschnikowia_pulcherrima(100); | Metschnikowia | 13 | Saprotroph | Undefined Saprotroph | Yeast | NULL | Probable | As nectar-tap_saprotroph (Põlme et al. 2020); Decay sugar-rich substrates (Põlme et al. 2020) | Põlme S, et al. 2020. Fungal Diversity 105:1-16 ((https://doi.org/10.1007/s13225-020-00466-2)) |
| Otu22125 | 1 | k__Fungi(100);p__Ascomycota(100);c__Saccharomycetes(100);o__Saccharomycetales(100);f__Metschnikowiaceae(100);g__Metschnikowia(100);s__Metschnikowia_pulcherrima(100); | Metschnikowia | 13 | Saprotroph | Undefined Saprotroph | Yeast | NULL | Probable | As nectar-tap_saprotroph (Põlme et al. 2020); Decay sugar-rich substrates (Põlme et al. 2020) | Põlme S, et al. 2020. Fungal Diversity 105:1-16 ((https://doi.org/10.1007/s13225-020-00466-2)) |
| Otu22126 | 1 | k__Fungi(100);p__Ascomycota(100);c__Saccharomycetes(100);o__Saccharomycetales(100);f__Metschnikowiaceae(100);g__Metschnikowia(100);s__Metschnikowia_pulcherrima(100); | Metschnikowia | 13 | Saprotroph | Undefined Saprotroph | Yeast | NULL | Probable | As nectar-tap_saprotroph (Põlme et al. 2020); Decay sugar-rich substrates (Põlme et al. 2020) | Põlme S, et al. 2020. Fungal Diversity 105:1-16 ((https://doi.org/10.1007/s13225-020-00466-2)) |
| Otu22127 | 1 | k__Fungi(100);p__Ascomycota(100);c__Saccharomycetes(100);o__Saccharomycetales(100);f__Metschnikowiaceae(100);g__Metschnikowia(100);s__Metschnikowia_pulcherrima(100); | Metschnikowia | 13 | Saprotroph | Undefined Saprotroph | Yeast | NULL | Probable | As nectar-tap_saprotroph (Põlme et al. 2020); Decay sugar-rich substrates (Põlme et al. 2020) | Põlme S, et al. 2020. Fungal Diversity 105:1-16 ((https://doi.org/10.1007/s13225-020-00466-2)) |
| Otu22129 | 1 | k__Fungi(100);p__Ascomycota(100);c__Saccharomycetes(100);o__Saccharomycetales(100);f__Metschnikowiaceae(100);g__Metschnikowia(100);s__Metschnikowia_pulcherrima(100); | Metschnikowia | 13 | Saprotroph | Undefined Saprotroph | Yeast | NULL | Probable | As nectar-tap_saprotroph (Põlme et al. 2020); Decay sugar-rich substrates (Põlme et al. 2020) | Põlme S, et al. 2020. Fungal Diversity 105:1-16 ((https://doi.org/10.1007/s13225-020-00466-2)) |
| Otu22131 | 1 | k__Fungi(100);p__Ascomycota(100);c__Saccharomycetes(100);o__Saccharomycetales(100);f__Metschnikowiaceae(100);g__Metschnikowia(100);s__Metschnikowia_pulcherrima(100); | Metschnikowia | 13 | Saprotroph | Undefined Saprotroph | Yeast | NULL | Probable | As nectar-tap_saprotroph (Põlme et al. 2020); Decay sugar-rich substrates (Põlme et al. 2020) | Põlme S, et al. 2020. Fungal Diversity 105:1-16 ((https://doi.org/10.1007/s13225-020-00466-2)) |
| Otu22132 | 1 | k__Fungi(100);p__Ascomycota(100);c__Saccharomycetes(100);o__Saccharomycetales(100);f__Metschnikowiaceae(100);g__Metschnikowia(100);s__Metschnikowia_pulcherrima(100); | Metschnikowia | 13 | Saprotroph | Undefined Saprotroph | Yeast | NULL | Probable | As nectar-tap_saprotroph (Põlme et al. 2020); Decay sugar-rich substrates (Põlme et al. 2020) | Põlme S, et al. 2020. Fungal Diversity 105:1-16 ((https://doi.org/10.1007/s13225-020-00466-2)) |
| Otu22133 | 1 | k__Fungi(100);p__Ascomycota(100);c__Saccharomycetes(100);o__Saccharomycetales(100);f__Metschnikowiaceae(100);g__Metschnikowia(100);s__Metschnikowia_pulcherrima(100); | Metschnikowia | 13 | Saprotroph | Undefined Saprotroph | Yeast | NULL | Probable | As nectar-tap_saprotroph (Põlme et al. 2020); Decay sugar-rich substrates (Põlme et al. 2020) | Põlme S, et al. 2020. Fungal Diversity 105:1-16 ((https://doi.org/10.1007/s13225-020-00466-2)) |
| Otu22134 | 1 | k__Fungi(100);p__Ascomycota(100);c__Saccharomycetes(100);o__Saccharomycetales(100);f__Metschnikowiaceae(100);g__Metschnikowia(100);s__Metschnikowia_pulcherrima(100); | Metschnikowia | 13 | Saprotroph | Undefined Saprotroph | Yeast | NULL | Probable | As nectar-tap_saprotroph (Põlme et al. 2020); Decay sugar-rich substrates (Põlme et al. 2020) | Põlme S, et al. 2020. Fungal Diversity 105:1-16 ((https://doi.org/10.1007/s13225-020-00466-2)) |
| Otu22135 | 1 | k__Fungi(100);p__Ascomycota(100);c__Saccharomycetes(100);o__Saccharomycetales(100);f__Metschnikowiaceae(100);g__Metschnikowia(100);s__Metschnikowia_pulcherrima(100); | Metschnikowia | 13 | Saprotroph | Undefined Saprotroph | Yeast | NULL | Probable | As nectar-tap_saprotroph (Põlme et al. 2020); Decay sugar-rich substrates (Põlme et al. 2020) | Põlme S, et al. 2020. Fungal Diversity 105:1-16 ((https://doi.org/10.1007/s13225-020-00466-2)) |
| Otu22136 | 1 | k__Fungi(100);p__Ascomycota(100);c__Saccharomycetes(100);o__Saccharomycetales(100);f__Metschnikowiaceae(100);g__Metschnikowia(100);s__Metschnikowia_pulcherrima(100); | Metschnikowia | 13 | Saprotroph | Undefined Saprotroph | Yeast | NULL | Probable | As nectar-tap_saprotroph (Põlme et al. 2020); Decay sugar-rich substrates (Põlme et al. 2020) | Põlme S, et al. 2020. Fungal Diversity 105:1-16 ((https://doi.org/10.1007/s13225-020-00466-2)) |
| Otu22138 | 1 | k__Fungi(100);p__Ascomycota(100);c__Saccharomycetes(100);o__Saccharomycetales(100);f__Metschnikowiaceae(100);g__Metschnikowia(100);s__Metschnikowia_pulcherrima(100); | Metschnikowia | 13 | Saprotroph | Undefined Saprotroph | Yeast | NULL | Probable | As nectar-tap_saprotroph (Põlme et al. 2020); Decay sugar-rich substrates (Põlme et al. 2020) | Põlme S, et al. 2020. Fungal Diversity 105:1-16 ((https://doi.org/10.1007/s13225-020-00466-2)) |
| Otu22140 | 1 | k__Fungi(100);p__Ascomycota(100);c__Saccharomycetes(100);o__Saccharomycetales(100);f__Metschnikowiaceae(100);g__Metschnikowia(100);s__Metschnikowia_pulcherrima(100); | Metschnikowia | 13 | Saprotroph | Undefined Saprotroph | Yeast | NULL | Probable | As nectar-tap_saprotroph (Põlme et al. 2020); Decay sugar-rich substrates (Põlme et al. 2020) | Põlme S, et al. 2020. Fungal Diversity 105:1-16 ((https://doi.org/10.1007/s13225-020-00466-2)) |
| Otu22141 | 1 | k__Fungi(100);p__Ascomycota(100);c__Saccharomycetes(100);o__Saccharomycetales(100);f__Metschnikowiaceae(100);g__Metschnikowia(100);s__Metschnikowia_pulcherrima(100); | Metschnikowia | 13 | Saprotroph | Undefined Saprotroph | Yeast | NULL | Probable | As nectar-tap_saprotroph (Põlme et al. 2020); Decay sugar-rich substrates (Põlme et al. 2020) | Põlme S, et al. 2020. Fungal Diversity 105:1-16 ((https://doi.org/10.1007/s13225-020-00466-2)) |
| Otu22145 | 1 | k__Fungi(100);p__Ascomycota(100);c__Saccharomycetes(100);o__Saccharomycetales(100);f__Metschnikowiaceae(100);g__Metschnikowia(100);s__Metschnikowia_pulcherrima(100); | Metschnikowia | 13 | Saprotroph | Undefined Saprotroph | Yeast | NULL | Probable | As nectar-tap_saprotroph (Põlme et al. 2020); Decay sugar-rich substrates (Põlme et al. 2020) | Põlme S, et al. 2020. Fungal Diversity 105:1-16 ((https://doi.org/10.1007/s13225-020-00466-2)) |
| Otu22150 | 1 | k__Fungi(100);p__Ascomycota(100);c__Saccharomycetes(100);o__Saccharomycetales(100);f__Metschnikowiaceae(100);g__Metschnikowia(100);s__Metschnikowia_pulcherrima(100); | Metschnikowia | 13 | Saprotroph | Undefined Saprotroph | Yeast | NULL | Probable | As nectar-tap_saprotroph (Põlme et al. 2020); Decay sugar-rich substrates (Põlme et al. 2020) | Põlme S, et al. 2020. Fungal Diversity 105:1-16 ((https://doi.org/10.1007/s13225-020-00466-2)) |
| Otu22161 | 1 | k__Fungi(100);p__Ascomycota(100);c__Saccharomycetes(100);o__Saccharomycetales(100);f__Metschnikowiaceae(100);g__Metschnikowia(100);g__Metschnikowia_unclassified(100); | Metschnikowia | 13 | Saprotroph | Undefined Saprotroph | Yeast | NULL | Probable | As nectar-tap_saprotroph (Põlme et al. 2020); Decay sugar-rich substrates (Põlme et al. 2020) | Põlme S, et al. 2020. Fungal Diversity 105:1-16 ((https://doi.org/10.1007/s13225-020-00466-2)) |
| Otu22163 | 1 | k__Fungi(100);p__Ascomycota(100);c__Saccharomycetes(100);o__Saccharomycetales(100);f__Metschnikowiaceae(100);g__Metschnikowia(100);s__Metschnikowia_pulcherrima(100); | Metschnikowia | 13 | Saprotroph | Undefined Saprotroph | Yeast | NULL | Probable | As nectar-tap_saprotroph (Põlme et al. 2020); Decay sugar-rich substrates (Põlme et al. 2020) | Põlme S, et al. 2020. Fungal Diversity 105:1-16 ((https://doi.org/10.1007/s13225-020-00466-2)) |
| Otu22164 | 1 | k__Fungi(100);p__Ascomycota(100);c__Saccharomycetes(100);o__Saccharomycetales(100);f__Metschnikowiaceae(100);g__Metschnikowia(100);s__Metschnikowia_pulcherrima(100); | Metschnikowia | 13 | Saprotroph | Undefined Saprotroph | Yeast | NULL | Probable | As nectar-tap_saprotroph (Põlme et al. 2020); Decay sugar-rich substrates (Põlme et al. 2020) | Põlme S, et al. 2020. Fungal Diversity 105:1-16 ((https://doi.org/10.1007/s13225-020-00466-2)) |
| Otu22165 | 1 | k__Fungi(100);p__Ascomycota(100);c__Saccharomycetes(100);o__Saccharomycetales(100);f__Metschnikowiaceae(100);g__Metschnikowia(100);g__Metschnikowia_unclassified(100); | Metschnikowia | 13 | Saprotroph | Undefined Saprotroph | Yeast | NULL | Probable | As nectar-tap_saprotroph (Põlme et al. 2020); Decay sugar-rich substrates (Põlme et al. 2020) | Põlme S, et al. 2020. Fungal Diversity 105:1-16 ((https://doi.org/10.1007/s13225-020-00466-2)) |
| Otu22166 | 1 | k__Fungi(100);p__Ascomycota(100);c__Saccharomycetes(100);o__Saccharomycetales(100);f__Metschnikowiaceae(100);g__Metschnikowia(100);g__Metschnikowia_unclassified(100); | Metschnikowia | 13 | Saprotroph | Undefined Saprotroph | Yeast | NULL | Probable | As nectar-tap_saprotroph (Põlme et al. 2020); Decay sugar-rich substrates (Põlme et al. 2020) | Põlme S, et al. 2020. Fungal Diversity 105:1-16 ((https://doi.org/10.1007/s13225-020-00466-2)) |
| Otu22167 | 1 | k__Fungi(100);p__Ascomycota(100);c__Saccharomycetes(100);o__Saccharomycetales(100);f__Metschnikowiaceae(100);g__Metschnikowia(100);g__Metschnikowia_unclassified(100); | Metschnikowia | 13 | Saprotroph | Undefined Saprotroph | Yeast | NULL | Probable | As nectar-tap_saprotroph (Põlme et al. 2020); Decay sugar-rich substrates (Põlme et al. 2020) | Põlme S, et al. 2020. Fungal Diversity 105:1-16 ((https://doi.org/10.1007/s13225-020-00466-2)) |
| Otu22168 | 1 | k__Fungi(100);p__Ascomycota(100);c__Saccharomycetes(100);o__Saccharomycetales(100);f__Metschnikowiaceae(100);g__Metschnikowia(100);s__Metschnikowia_pulcherrima(100); | Metschnikowia | 13 | Saprotroph | Undefined Saprotroph | Yeast | NULL | Probable | As nectar-tap_saprotroph (Põlme et al. 2020); Decay sugar-rich substrates (Põlme et al. 2020) | Põlme S, et al. 2020. Fungal Diversity 105:1-16 ((https://doi.org/10.1007/s13225-020-00466-2)) |
| Otu22169 | 1 | k__Fungi(100);p__Ascomycota(100);c__Saccharomycetes(100);o__Saccharomycetales(100);f__Metschnikowiaceae(100);g__Metschnikowia(100);s__Metschnikowia_pulcherrima(100); | Metschnikowia | 13 | Saprotroph | Undefined Saprotroph | Yeast | NULL | Probable | As nectar-tap_saprotroph (Põlme et al. 2020); Decay sugar-rich substrates (Põlme et al. 2020) | Põlme S, et al. 2020. Fungal Diversity 105:1-16 ((https://doi.org/10.1007/s13225-020-00466-2)) |
| Otu22170 | 1 | k__Fungi(100);p__Ascomycota(100);c__Saccharomycetes(100);o__Saccharomycetales(100);f__Metschnikowiaceae(100);g__Metschnikowia(100);g__Metschnikowia_unclassified(100); | Metschnikowia | 13 | Saprotroph | Undefined Saprotroph | Yeast | NULL | Probable | As nectar-tap_saprotroph (Põlme et al. 2020); Decay sugar-rich substrates (Põlme et al. 2020) | Põlme S, et al. 2020. Fungal Diversity 105:1-16 ((https://doi.org/10.1007/s13225-020-00466-2)) |
| Otu22171 | 1 | k__Fungi(100);p__Ascomycota(100);c__Saccharomycetes(100);o__Saccharomycetales(100);f__Metschnikowiaceae(100);g__Metschnikowia(100);s__Metschnikowia_pulcherrima(100); | Metschnikowia | 13 | Saprotroph | Undefined Saprotroph | Yeast | NULL | Probable | As nectar-tap_saprotroph (Põlme et al. 2020); Decay sugar-rich substrates (Põlme et al. 2020) | Põlme S, et al. 2020. Fungal Diversity 105:1-16 ((https://doi.org/10.1007/s13225-020-00466-2)) |
| Otu22172 | 1 | k__Fungi(100);p__Ascomycota(100);c__Saccharomycetes(100);o__Saccharomycetales(100);f__Metschnikowiaceae(100);g__Metschnikowia(100);g__Metschnikowia_unclassified(100); | Metschnikowia | 13 | Saprotroph | Undefined Saprotroph | Yeast | NULL | Probable | As nectar-tap_saprotroph (Põlme et al. 2020); Decay sugar-rich substrates (Põlme et al. 2020) | Põlme S, et al. 2020. Fungal Diversity 105:1-16 ((https://doi.org/10.1007/s13225-020-00466-2)) |
| Otu22173 | 1 | k__Fungi(100);p__Ascomycota(100);c__Saccharomycetes(100);o__Saccharomycetales(100);f__Metschnikowiaceae(100);g__Metschnikowia(100);s__Metschnikowia_pulcherrima(100); | Metschnikowia | 13 | Saprotroph | Undefined Saprotroph | Yeast | NULL | Probable | As nectar-tap_saprotroph (Põlme et al. 2020); Decay sugar-rich substrates (Põlme et al. 2020) | Põlme S, et al. 2020. Fungal Diversity 105:1-16 ((https://doi.org/10.1007/s13225-020-00466-2)) |
| Otu22176 | 1 | k__Fungi(100);p__Ascomycota(100);c__Saccharomycetes(100);o__Saccharomycetales(100);f__Metschnikowiaceae(100);g__Metschnikowia(100);g__Metschnikowia_unclassified(100); | Metschnikowia | 13 | Saprotroph | Undefined Saprotroph | Yeast | NULL | Probable | As nectar-tap_saprotroph (Põlme et al. 2020); Decay sugar-rich substrates (Põlme et al. 2020) | Põlme S, et al. 2020. Fungal Diversity 105:1-16 ((https://doi.org/10.1007/s13225-020-00466-2)) |
| Otu22177 | 1 | k__Fungi(100);p__Ascomycota(100);c__Saccharomycetes(100);o__Saccharomycetales(100);f__Metschnikowiaceae(100);g__Metschnikowia(100);s__Metschnikowia_pulcherrima(100); | Metschnikowia | 13 | Saprotroph | Undefined Saprotroph | Yeast | NULL | Probable | As nectar-tap_saprotroph (Põlme et al. 2020); Decay sugar-rich substrates (Põlme et al. 2020) | Põlme S, et al. 2020. Fungal Diversity 105:1-16 ((https://doi.org/10.1007/s13225-020-00466-2)) |
| Otu22178 | 1 | k__Fungi(100);p__Ascomycota(100);c__Saccharomycetes(100);o__Saccharomycetales(100);f__Metschnikowiaceae(100);g__Metschnikowia(100);s__Metschnikowia_pulcherrima(100); | Metschnikowia | 13 | Saprotroph | Undefined Saprotroph | Yeast | NULL | Probable | As nectar-tap_saprotroph (Põlme et al. 2020); Decay sugar-rich substrates (Põlme et al. 2020) | Põlme S, et al. 2020. Fungal Diversity 105:1-16 ((https://doi.org/10.1007/s13225-020-00466-2)) |
| Otu22183 | 1 | k__Fungi(100);p__Ascomycota(100);c__Saccharomycetes(100);o__Saccharomycetales(100);f__Metschnikowiaceae(100);g__Metschnikowia(100);s__Metschnikowia_pulcherrima(100); | Metschnikowia | 13 | Saprotroph | Undefined Saprotroph | Yeast | NULL | Probable | As nectar-tap_saprotroph (Põlme et al. 2020); Decay sugar-rich substrates (Põlme et al. 2020) | Põlme S, et al. 2020. Fungal Diversity 105:1-16 ((https://doi.org/10.1007/s13225-020-00466-2)) |
| Otu22187 | 1 | k__Fungi(100);p__Ascomycota(100);c__Saccharomycetes(100);o__Saccharomycetales(100);f__Metschnikowiaceae(100);g__Metschnikowia(100);s__Metschnikowia_pulcherrima(100); | Metschnikowia | 13 | Saprotroph | Undefined Saprotroph | Yeast | NULL | Probable | As nectar-tap_saprotroph (Põlme et al. 2020); Decay sugar-rich substrates (Põlme et al. 2020) | Põlme S, et al. 2020. Fungal Diversity 105:1-16 ((https://doi.org/10.1007/s13225-020-00466-2)) |
| Otu22189 | 1 | k__Fungi(100);p__Ascomycota(100);c__Saccharomycetes(100);o__Saccharomycetales(100);f__Metschnikowiaceae(100);g__Metschnikowia(100);g__Metschnikowia_unclassified(100); | Metschnikowia | 13 | Saprotroph | Undefined Saprotroph | Yeast | NULL | Probable | As nectar-tap_saprotroph (Põlme et al. 2020); Decay sugar-rich substrates (Põlme et al. 2020) | Põlme S, et al. 2020. Fungal Diversity 105:1-16 ((https://doi.org/10.1007/s13225-020-00466-2)) |
| Otu22190 | 1 | k__Fungi(100);p__Ascomycota(100);c__Saccharomycetes(100);o__Saccharomycetales(100);f__Saccharomycetales_family_Incertae_sedis(100);f__Saccharomycetales_family_Incertae_sedis_unclassified(100);f__Saccharomycetales_family_Incertae_sedis_unclassified(100); | Saccharomycetales | 7 | Saprotroph | Undefined Saprotroph | Yeast | NULL | Possible | NULL | Sterkenburg E, et al. 2015. New Phytologist 207:1145-1158 ((https://doi.org/10.1111/nph.13426)) |
| Otu22192 | 1 | k__Fungi(100);p__Ascomycota(100);c__Saccharomycetes(100);o__Saccharomycetales(100);f__Saccharomycetales_family_Incertae_sedis(100);f__Saccharomycetales_family_Incertae_sedis_unclassified(100);f__Saccharomycetales_family_Incertae_sedis_unclassified(100); | Saccharomycetales | 7 | Saprotroph | Undefined Saprotroph | Yeast | NULL | Possible | NULL | Sterkenburg E, et al. 2015. New Phytologist 207:1145-1158 ((https://doi.org/10.1111/nph.13426)) |
| Otu22195 | 1 | k__Fungi(100);p__Ascomycota(100);c__Saccharomycetes(100);o__Saccharomycetales(100);f__Metschnikowiaceae(100);g__Metschnikowia(100);g__Metschnikowia_unclassified(100); | Metschnikowia | 13 | Saprotroph | Undefined Saprotroph | Yeast | NULL | Probable | As nectar-tap_saprotroph (Põlme et al. 2020); Decay sugar-rich substrates (Põlme et al. 2020) | Põlme S, et al. 2020. Fungal Diversity 105:1-16 ((https://doi.org/10.1007/s13225-020-00466-2)) |
| Otu22196 | 1 | k__Fungi(100);p__Ascomycota(100);c__Saccharomycetes(100);o__Saccharomycetales(100);f__Metschnikowiaceae(100);g__Metschnikowia(100);s__Metschnikowia_pulcherrima(100); | Metschnikowia | 13 | Saprotroph | Undefined Saprotroph | Yeast | NULL | Probable | As nectar-tap_saprotroph (Põlme et al. 2020); Decay sugar-rich substrates (Põlme et al. 2020) | Põlme S, et al. 2020. Fungal Diversity 105:1-16 ((https://doi.org/10.1007/s13225-020-00466-2)) |
| Otu22197 | 1 | k__Fungi(100);p__Ascomycota(100);c__Saccharomycetes(100);o__Saccharomycetales(100);f__Metschnikowiaceae(100);g__Metschnikowia(100);s__Metschnikowia_pulcherrima(100); | Metschnikowia | 13 | Saprotroph | Undefined Saprotroph | Yeast | NULL | Probable | As nectar-tap_saprotroph (Põlme et al. 2020); Decay sugar-rich substrates (Põlme et al. 2020) | Põlme S, et al. 2020. Fungal Diversity 105:1-16 ((https://doi.org/10.1007/s13225-020-00466-2)) |
| Otu22199 | 1 | k__Fungi(100);p__Ascomycota(100);c__Saccharomycetes(100);o__Saccharomycetales(100);f__Metschnikowiaceae(100);g__Metschnikowia(100);g__Metschnikowia_unclassified(100); | Metschnikowia | 13 | Saprotroph | Undefined Saprotroph | Yeast | NULL | Probable | As nectar-tap_saprotroph (Põlme et al. 2020); Decay sugar-rich substrates (Põlme et al. 2020) | Põlme S, et al. 2020. Fungal Diversity 105:1-16 ((https://doi.org/10.1007/s13225-020-00466-2)) |
| Otu22201 | 1 | k__Fungi(100);p__Ascomycota(100);c__Saccharomycetes(100);o__Saccharomycetales(100);f__Metschnikowiaceae(100);g__Metschnikowia(100);s__Metschnikowia_pulcherrima(100); | Metschnikowia | 13 | Saprotroph | Undefined Saprotroph | Yeast | NULL | Probable | As nectar-tap_saprotroph (Põlme et al. 2020); Decay sugar-rich substrates (Põlme et al. 2020) | Põlme S, et al. 2020. Fungal Diversity 105:1-16 ((https://doi.org/10.1007/s13225-020-00466-2)) |
| Otu22203 | 1 | k__Fungi(100);p__Ascomycota(100);c__Saccharomycetes(100);o__Saccharomycetales(100);f__Metschnikowiaceae(100);g__Metschnikowia(100);s__Metschnikowia_pulcherrima(100); | Metschnikowia | 13 | Saprotroph | Undefined Saprotroph | Yeast | NULL | Probable | As nectar-tap_saprotroph (Põlme et al. 2020); Decay sugar-rich substrates (Põlme et al. 2020) | Põlme S, et al. 2020. Fungal Diversity 105:1-16 ((https://doi.org/10.1007/s13225-020-00466-2)) |
| Otu22204 | 1 | k__Fungi(100);p__Ascomycota(100);c__Saccharomycetes(100);o__Saccharomycetales(100);f__Saccharomycetales_family_Incertae_sedis(100);g__Candida(100);s__Candida_cretensis(100); | Candida | 13 | Pathotroph-Saprotroph-Symbiotroph | Animal Pathogen-Endophyte-Endosymbiont-Epiphyte-Undefined Saprotroph | Dimorphic Yeast Yeast | NULL | Probable | As nectar-tap_saprotroph (Põlme et al. 2020); Decay sugar-rich substrates (Põlme et al. 2020); This asexual genus is polyphyletic and contain species that function very differently depend on what they are. Caution should be taken when interpreting guilds for Candida species. | Manolakaki D, et al. 2010. Virulence. 1: 367-375 ((https://doi.org/10.4161/viru.1.5.12796)); Mekbib SB, et al. 2011. Tropical Plant Pathology 36:233-240 ((https://doi.org/10.1590/S1982-56762011000400004)); Põlme S, et al. 2020. Fungal Diversity 105:1-16 ((https://doi.org/10.1007/s13225-020-00466-2)) |
| Otu22205 | 1 | k__Fungi(100);p__Ascomycota(100);c__Saccharomycetes(100);o__Saccharomycetales(100);f__Saccharomycetales_family_Incertae_sedis(100);g__Candida(100);g__Candida_unclassified(100); | Candida | 13 | Pathotroph-Saprotroph-Symbiotroph | Animal Pathogen-Endophyte-Endosymbiont-Epiphyte-Undefined Saprotroph | Dimorphic Yeast Yeast | NULL | Probable | As nectar-tap_saprotroph (Põlme et al. 2020); Decay sugar-rich substrates (Põlme et al. 2020); This asexual genus is polyphyletic and contain species that function very differently depend on what they are. Caution should be taken when interpreting guilds for Candida species. | Manolakaki D, et al. 2010. Virulence. 1: 367-375 ((https://doi.org/10.4161/viru.1.5.12796)); Mekbib SB, et al. 2011. Tropical Plant Pathology 36:233-240 ((https://doi.org/10.1590/S1982-56762011000400004)); Põlme S, et al. 2020. Fungal Diversity 105:1-16 ((https://doi.org/10.1007/s13225-020-00466-2)) |
| Otu22207 | 1 | k__Fungi(100);p__Ascomycota(100);c__Saccharomycetes(100);o__Saccharomycetales(100);f__Metschnikowiaceae(100);g__Metschnikowia(100);s__Metschnikowia_pulcherrima(100); | Metschnikowia | 13 | Saprotroph | Undefined Saprotroph | Yeast | NULL | Probable | As nectar-tap_saprotroph (Põlme et al. 2020); Decay sugar-rich substrates (Põlme et al. 2020) | Põlme S, et al. 2020. Fungal Diversity 105:1-16 ((https://doi.org/10.1007/s13225-020-00466-2)) |
| Otu22208 | 1 | k__Fungi(100);p__Ascomycota(100);c__Saccharomycetes(100);o__Saccharomycetales(100);o__Saccharomycetales_unclassified(100);o__Saccharomycetales_unclassified(100);o__Saccharomycetales_unclassified(100); | Saccharomycetales | 7 | Saprotroph | Undefined Saprotroph | Yeast | NULL | Possible | NULL | Sterkenburg E, et al. 2015. New Phytologist 207:1145-1158 ((https://doi.org/10.1111/nph.13426)) |
| Otu22209 | 1 | k__Fungi(100);p__Ascomycota(100);c__Saccharomycetes(100);o__Saccharomycetales(100);f__Saccharomycetales_family_Incertae_sedis(100);g__Candida(100);s__Candida_sp_AS23486(100); | Candida | 13 | Pathotroph-Saprotroph-Symbiotroph | Animal Pathogen-Endophyte-Endosymbiont-Epiphyte-Undefined Saprotroph | Dimorphic Yeast Yeast | NULL | Probable | As nectar-tap_saprotroph (Põlme et al. 2020); Decay sugar-rich substrates (Põlme et al. 2020); This asexual genus is polyphyletic and contain species that function very differently depend on what they are. Caution should be taken when interpreting guilds for Candida species. | Manolakaki D, et al. 2010. Virulence. 1: 367-375 ((https://doi.org/10.4161/viru.1.5.12796)); Mekbib SB, et al. 2011. Tropical Plant Pathology 36:233-240 ((https://doi.org/10.1590/S1982-56762011000400004)); Põlme S, et al. 2020. Fungal Diversity 105:1-16 ((https://doi.org/10.1007/s13225-020-00466-2)) |
| Otu22210 | 1 | k__Fungi(100);p__Ascomycota(100);c__Saccharomycetes(100);o__Saccharomycetales(100);f__Metschnikowiaceae(100);g__Metschnikowia(100);g__Metschnikowia_unclassified(100); | Metschnikowia | 13 | Saprotroph | Undefined Saprotroph | Yeast | NULL | Probable | As nectar-tap_saprotroph (Põlme et al. 2020); Decay sugar-rich substrates (Põlme et al. 2020) | Põlme S, et al. 2020. Fungal Diversity 105:1-16 ((https://doi.org/10.1007/s13225-020-00466-2)) |
| Otu22211 | 1 | k__Fungi(100);p__Ascomycota(100);c__Saccharomycetes(100);o__Saccharomycetales(100);f__Metschnikowiaceae(100);g__Metschnikowia(100);s__Metschnikowia_pulcherrima(100); | Metschnikowia | 13 | Saprotroph | Undefined Saprotroph | Yeast | NULL | Probable | As nectar-tap_saprotroph (Põlme et al. 2020); Decay sugar-rich substrates (Põlme et al. 2020) | Põlme S, et al. 2020. Fungal Diversity 105:1-16 ((https://doi.org/10.1007/s13225-020-00466-2)) |
| Otu22212 | 1 | k__Fungi(100);p__Ascomycota(100);c__Saccharomycetes(100);o__Saccharomycetales(100);f__Metschnikowiaceae(100);g__Metschnikowia(100);s__Metschnikowia_pulcherrima(100); | Metschnikowia | 13 | Saprotroph | Undefined Saprotroph | Yeast | NULL | Probable | As nectar-tap_saprotroph (Põlme et al. 2020); Decay sugar-rich substrates (Põlme et al. 2020) | Põlme S, et al. 2020. Fungal Diversity 105:1-16 ((https://doi.org/10.1007/s13225-020-00466-2)) |
| Otu22213 | 1 | k__Fungi(100);p__Ascomycota(100);c__Saccharomycetes(100);o__Saccharomycetales(100);f__Metschnikowiaceae(100);g__Metschnikowia(100);s__Metschnikowia_pulcherrima(100); | Metschnikowia | 13 | Saprotroph | Undefined Saprotroph | Yeast | NULL | Probable | As nectar-tap_saprotroph (Põlme et al. 2020); Decay sugar-rich substrates (Põlme et al. 2020) | Põlme S, et al. 2020. Fungal Diversity 105:1-16 ((https://doi.org/10.1007/s13225-020-00466-2)) |
| Otu22214 | 1 | k__Fungi(100);p__Ascomycota(100);c__Saccharomycetes(100);o__Saccharomycetales(100);f__Metschnikowiaceae(100);g__Metschnikowia(100);s__Metschnikowia_pulcherrima(100); | Metschnikowia | 13 | Saprotroph | Undefined Saprotroph | Yeast | NULL | Probable | As nectar-tap_saprotroph (Põlme et al. 2020); Decay sugar-rich substrates (Põlme et al. 2020) | Põlme S, et al. 2020. Fungal Diversity 105:1-16 ((https://doi.org/10.1007/s13225-020-00466-2)) |
| Otu22217 | 1 | k__Fungi(100);p__Ascomycota(100);c__Saccharomycetes(100);o__Saccharomycetales(100);f__Metschnikowiaceae(100);g__Metschnikowia(100);s__Metschnikowia_pulcherrima(100); | Metschnikowia | 13 | Saprotroph | Undefined Saprotroph | Yeast | NULL | Probable | As nectar-tap_saprotroph (Põlme et al. 2020); Decay sugar-rich substrates (Põlme et al. 2020) | Põlme S, et al. 2020. Fungal Diversity 105:1-16 ((https://doi.org/10.1007/s13225-020-00466-2)) |
| Otu22218 | 1 | k__Fungi(100);p__Ascomycota(100);c__Saccharomycetes(100);o__Saccharomycetales(100);f__Metschnikowiaceae(100);g__Metschnikowia(100);s__Metschnikowia_pulcherrima(100); | Metschnikowia | 13 | Saprotroph | Undefined Saprotroph | Yeast | NULL | Probable | As nectar-tap_saprotroph (Põlme et al. 2020); Decay sugar-rich substrates (Põlme et al. 2020) | Põlme S, et al. 2020. Fungal Diversity 105:1-16 ((https://doi.org/10.1007/s13225-020-00466-2)) |
| Otu22219 | 1 | k__Fungi(100);p__Ascomycota(100);c__Saccharomycetes(100);o__Saccharomycetales(100);f__Metschnikowiaceae(100);g__Metschnikowia(100);g__Metschnikowia_unclassified(100); | Metschnikowia | 13 | Saprotroph | Undefined Saprotroph | Yeast | NULL | Probable | As nectar-tap_saprotroph (Põlme et al. 2020); Decay sugar-rich substrates (Põlme et al. 2020) | Põlme S, et al. 2020. Fungal Diversity 105:1-16 ((https://doi.org/10.1007/s13225-020-00466-2)) |
| Otu22220 | 1 | k__Fungi(100);p__Ascomycota(100);c__Saccharomycetes(100);o__Saccharomycetales(100);f__Metschnikowiaceae(100);g__Metschnikowia(100);s__Metschnikowia_pulcherrima(100); | Metschnikowia | 13 | Saprotroph | Undefined Saprotroph | Yeast | NULL | Probable | As nectar-tap_saprotroph (Põlme et al. 2020); Decay sugar-rich substrates (Põlme et al. 2020) | Põlme S, et al. 2020. Fungal Diversity 105:1-16 ((https://doi.org/10.1007/s13225-020-00466-2)) |
| Otu22221 | 1 | k__Fungi(100);p__Ascomycota(100);c__Saccharomycetes(100);o__Saccharomycetales(100);f__Metschnikowiaceae(100);g__Metschnikowia(100);s__Metschnikowia_pulcherrima(100); | Metschnikowia | 13 | Saprotroph | Undefined Saprotroph | Yeast | NULL | Probable | As nectar-tap_saprotroph (Põlme et al. 2020); Decay sugar-rich substrates (Põlme et al. 2020) | Põlme S, et al. 2020. Fungal Diversity 105:1-16 ((https://doi.org/10.1007/s13225-020-00466-2)) |
| Otu22222 | 1 | k__Fungi(100);p__Ascomycota(100);c__Saccharomycetes(100);o__Saccharomycetales(100);f__Saccharomycetales_family_Incertae_sedis(100);g__Candida(100);g__Candida_unclassified(100); | Candida | 13 | Pathotroph-Saprotroph-Symbiotroph | Animal Pathogen-Endophyte-Endosymbiont-Epiphyte-Undefined Saprotroph | Dimorphic Yeast Yeast | NULL | Probable | As nectar-tap_saprotroph (Põlme et al. 2020); Decay sugar-rich substrates (Põlme et al. 2020); This asexual genus is polyphyletic and contain species that function very differently depend on what they are. Caution should be taken when interpreting guilds for Candida species. | Manolakaki D, et al. 2010. Virulence. 1: 367-375 ((https://doi.org/10.4161/viru.1.5.12796)); Mekbib SB, et al. 2011. Tropical Plant Pathology 36:233-240 ((https://doi.org/10.1590/S1982-56762011000400004)); Põlme S, et al. 2020. Fungal Diversity 105:1-16 ((https://doi.org/10.1007/s13225-020-00466-2)) |
| Otu22224 | 1 | k__Fungi(100);p__Ascomycota(100);c__Saccharomycetes(100);o__Saccharomycetales(100);f__Saccharomycetales_family_Incertae_sedis(100);g__Candida(100);s__Candida_sp_AS23486(100); | Candida | 13 | Pathotroph-Saprotroph-Symbiotroph | Animal Pathogen-Endophyte-Endosymbiont-Epiphyte-Undefined Saprotroph | Dimorphic Yeast Yeast | NULL | Probable | As nectar-tap_saprotroph (Põlme et al. 2020); Decay sugar-rich substrates (Põlme et al. 2020); This asexual genus is polyphyletic and contain species that function very differently depend on what they are. Caution should be taken when interpreting guilds for Candida species. | Manolakaki D, et al. 2010. Virulence. 1: 367-375 ((https://doi.org/10.4161/viru.1.5.12796)); Mekbib SB, et al. 2011. Tropical Plant Pathology 36:233-240 ((https://doi.org/10.1590/S1982-56762011000400004)); Põlme S, et al. 2020. Fungal Diversity 105:1-16 ((https://doi.org/10.1007/s13225-020-00466-2)) |
| Otu22227 | 1 | k__Fungi(100);p__Ascomycota(100);c__Saccharomycetes(100);o__Saccharomycetales(100);f__Metschnikowiaceae(100);g__Metschnikowia(100);s__Metschnikowia_pulcherrima(100); | Metschnikowia | 13 | Saprotroph | Undefined Saprotroph | Yeast | NULL | Probable | As nectar-tap_saprotroph (Põlme et al. 2020); Decay sugar-rich substrates (Põlme et al. 2020) | Põlme S, et al. 2020. Fungal Diversity 105:1-16 ((https://doi.org/10.1007/s13225-020-00466-2)) |
| Otu22228 | 1 | k__Fungi(100);p__Ascomycota(100);c__Saccharomycetes(100);o__Saccharomycetales(100);f__Metschnikowiaceae(100);g__Metschnikowia(100);s__Metschnikowia_pulcherrima(100); | Metschnikowia | 13 | Saprotroph | Undefined Saprotroph | Yeast | NULL | Probable | As nectar-tap_saprotroph (Põlme et al. 2020); Decay sugar-rich substrates (Põlme et al. 2020) | Põlme S, et al. 2020. Fungal Diversity 105:1-16 ((https://doi.org/10.1007/s13225-020-00466-2)) |
| Otu22230 | 1 | k__Fungi(100);p__Ascomycota(100);c__Saccharomycetes(100);o__Saccharomycetales(100);f__Metschnikowiaceae(100);g__Metschnikowia(100);g__Metschnikowia_unclassified(100); | Metschnikowia | 13 | Saprotroph | Undefined Saprotroph | Yeast | NULL | Probable | As nectar-tap_saprotroph (Põlme et al. 2020); Decay sugar-rich substrates (Põlme et al. 2020) | Põlme S, et al. 2020. Fungal Diversity 105:1-16 ((https://doi.org/10.1007/s13225-020-00466-2)) |
| Otu22231 | 1 | k__Fungi(100);p__Ascomycota(100);c__Saccharomycetes(100);o__Saccharomycetales(100);f__Metschnikowiaceae(100);g__Metschnikowia(100);s__Metschnikowia_pulcherrima(100); | Metschnikowia | 13 | Saprotroph | Undefined Saprotroph | Yeast | NULL | Probable | As nectar-tap_saprotroph (Põlme et al. 2020); Decay sugar-rich substrates (Põlme et al. 2020) | Põlme S, et al. 2020. Fungal Diversity 105:1-16 ((https://doi.org/10.1007/s13225-020-00466-2)) |
| Otu22232 | 1 | k__Fungi(100);p__Ascomycota(100);c__Saccharomycetes(100);o__Saccharomycetales(100);f__Metschnikowiaceae(100);g__Metschnikowia(100);g__Metschnikowia_unclassified(100); | Metschnikowia | 13 | Saprotroph | Undefined Saprotroph | Yeast | NULL | Probable | As nectar-tap_saprotroph (Põlme et al. 2020); Decay sugar-rich substrates (Põlme et al. 2020) | Põlme S, et al. 2020. Fungal Diversity 105:1-16 ((https://doi.org/10.1007/s13225-020-00466-2)) |
| Otu22233 | 1 | k__Fungi(100);p__Ascomycota(100);c__Saccharomycetes(100);o__Saccharomycetales(100);f__Metschnikowiaceae(100);g__Metschnikowia(100);s__Metschnikowia_pulcherrima(100); | Metschnikowia | 13 | Saprotroph | Undefined Saprotroph | Yeast | NULL | Probable | As nectar-tap_saprotroph (Põlme et al. 2020); Decay sugar-rich substrates (Põlme et al. 2020) | Põlme S, et al. 2020. Fungal Diversity 105:1-16 ((https://doi.org/10.1007/s13225-020-00466-2)) |
| Otu22235 | 1 | k__Fungi(100);p__Ascomycota(100);c__Saccharomycetes(100);o__Saccharomycetales(100);f__Metschnikowiaceae(100);g__Metschnikowia(100);s__Metschnikowia_pulcherrima(100); | Metschnikowia | 13 | Saprotroph | Undefined Saprotroph | Yeast | NULL | Probable | As nectar-tap_saprotroph (Põlme et al. 2020); Decay sugar-rich substrates (Põlme et al. 2020) | Põlme S, et al. 2020. Fungal Diversity 105:1-16 ((https://doi.org/10.1007/s13225-020-00466-2)) |
| Otu22236 | 1 | k__Fungi(100);p__Ascomycota(100);c__Saccharomycetes(100);o__Saccharomycetales(100);f__Metschnikowiaceae(100);g__Metschnikowia(100);s__Metschnikowia_pulcherrima(100); | Metschnikowia | 13 | Saprotroph | Undefined Saprotroph | Yeast | NULL | Probable | As nectar-tap_saprotroph (Põlme et al. 2020); Decay sugar-rich substrates (Põlme et al. 2020) | Põlme S, et al. 2020. Fungal Diversity 105:1-16 ((https://doi.org/10.1007/s13225-020-00466-2)) |
| Otu22237 | 1 | k__Fungi(100);p__Ascomycota(100);c__Saccharomycetes(100);o__Saccharomycetales(100);f__Metschnikowiaceae(100);g__Metschnikowia(100);g__Metschnikowia_unclassified(100); | Metschnikowia | 13 | Saprotroph | Undefined Saprotroph | Yeast | NULL | Probable | As nectar-tap_saprotroph (Põlme et al. 2020); Decay sugar-rich substrates (Põlme et al. 2020) | Põlme S, et al. 2020. Fungal Diversity 105:1-16 ((https://doi.org/10.1007/s13225-020-00466-2)) |
| Otu22242 | 1 | k__Fungi(100);p__Ascomycota(100);c__Saccharomycetes(100);o__Saccharomycetales(100);f__Metschnikowiaceae(100);g__Metschnikowia(100);s__Metschnikowia_pulcherrima(100); | Metschnikowia | 13 | Saprotroph | Undefined Saprotroph | Yeast | NULL | Probable | As nectar-tap_saprotroph (Põlme et al. 2020); Decay sugar-rich substrates (Põlme et al. 2020) | Põlme S, et al. 2020. Fungal Diversity 105:1-16 ((https://doi.org/10.1007/s13225-020-00466-2)) |
| Otu22243 | 1 | k__Fungi(100);p__Ascomycota(100);c__Saccharomycetes(100);o__Saccharomycetales(100);f__Metschnikowiaceae(100);g__Metschnikowia(100);s__Metschnikowia_pulcherrima(100); | Metschnikowia | 13 | Saprotroph | Undefined Saprotroph | Yeast | NULL | Probable | As nectar-tap_saprotroph (Põlme et al. 2020); Decay sugar-rich substrates (Põlme et al. 2020) | Põlme S, et al. 2020. Fungal Diversity 105:1-16 ((https://doi.org/10.1007/s13225-020-00466-2)) |
| Otu22244 | 1 | k__Fungi(100);p__Ascomycota(100);c__Saccharomycetes(100);o__Saccharomycetales(100);f__Metschnikowiaceae(100);g__Metschnikowia(100);s__Metschnikowia_pulcherrima(100); | Metschnikowia | 13 | Saprotroph | Undefined Saprotroph | Yeast | NULL | Probable | As nectar-tap_saprotroph (Põlme et al. 2020); Decay sugar-rich substrates (Põlme et al. 2020) | Põlme S, et al. 2020. Fungal Diversity 105:1-16 ((https://doi.org/10.1007/s13225-020-00466-2)) |
| Otu22245 | 1 | k__Fungi(100);p__Ascomycota(100);c__Saccharomycetes(100);o__Saccharomycetales(100);f__Metschnikowiaceae(100);g__Metschnikowia(100);g__Metschnikowia_unclassified(100); | Metschnikowia | 13 | Saprotroph | Undefined Saprotroph | Yeast | NULL | Probable | As nectar-tap_saprotroph (Põlme et al. 2020); Decay sugar-rich substrates (Põlme et al. 2020) | Põlme S, et al. 2020. Fungal Diversity 105:1-16 ((https://doi.org/10.1007/s13225-020-00466-2)) |
| Otu22246 | 1 | k__Fungi(100);p__Ascomycota(100);c__Saccharomycetes(100);o__Saccharomycetales(100);f__Metschnikowiaceae(100);g__Metschnikowia(100);g__Metschnikowia_unclassified(100); | Metschnikowia | 13 | Saprotroph | Undefined Saprotroph | Yeast | NULL | Probable | As nectar-tap_saprotroph (Põlme et al. 2020); Decay sugar-rich substrates (Põlme et al. 2020) | Põlme S, et al. 2020. Fungal Diversity 105:1-16 ((https://doi.org/10.1007/s13225-020-00466-2)) |
| Otu22247 | 1 | k__Fungi(100);p__Ascomycota(100);c__Saccharomycetes(100);o__Saccharomycetales(100);f__Metschnikowiaceae(100);g__Metschnikowia(100);s__Metschnikowia_pulcherrima(100); | Metschnikowia | 13 | Saprotroph | Undefined Saprotroph | Yeast | NULL | Probable | As nectar-tap_saprotroph (Põlme et al. 2020); Decay sugar-rich substrates (Põlme et al. 2020) | Põlme S, et al. 2020. Fungal Diversity 105:1-16 ((https://doi.org/10.1007/s13225-020-00466-2)) |
| Otu22248 | 1 | k__Fungi(100);p__Ascomycota(100);c__Saccharomycetes(100);o__Saccharomycetales(100);f__Metschnikowiaceae(100);g__Metschnikowia(100);s__Metschnikowia_pulcherrima(100); | Metschnikowia | 13 | Saprotroph | Undefined Saprotroph | Yeast | NULL | Probable | As nectar-tap_saprotroph (Põlme et al. 2020); Decay sugar-rich substrates (Põlme et al. 2020) | Põlme S, et al. 2020. Fungal Diversity 105:1-16 ((https://doi.org/10.1007/s13225-020-00466-2)) |
| Otu22249 | 1 | k__Fungi(100);p__Ascomycota(100);c__Saccharomycetes(100);o__Saccharomycetales(100);f__Metschnikowiaceae(100);g__Metschnikowia(100);s__Metschnikowia_pulcherrima(100); | Metschnikowia | 13 | Saprotroph | Undefined Saprotroph | Yeast | NULL | Probable | As nectar-tap_saprotroph (Põlme et al. 2020); Decay sugar-rich substrates (Põlme et al. 2020) | Põlme S, et al. 2020. Fungal Diversity 105:1-16 ((https://doi.org/10.1007/s13225-020-00466-2)) |
| Otu22252 | 1 | k__Fungi(100);p__Ascomycota(100);c__Saccharomycetes(100);o__Saccharomycetales(100);f__Metschnikowiaceae(100);g__Metschnikowia(100);s__Metschnikowia_pulcherrima(100); | Metschnikowia | 13 | Saprotroph | Undefined Saprotroph | Yeast | NULL | Probable | As nectar-tap_saprotroph (Põlme et al. 2020); Decay sugar-rich substrates (Põlme et al. 2020) | Põlme S, et al. 2020. Fungal Diversity 105:1-16 ((https://doi.org/10.1007/s13225-020-00466-2)) |
| Otu22253 | 1 | k__Fungi(100);p__Ascomycota(100);c__Saccharomycetes(100);o__Saccharomycetales(100);f__Metschnikowiaceae(100);g__Metschnikowia(100);s__Metschnikowia_pulcherrima(100); | Metschnikowia | 13 | Saprotroph | Undefined Saprotroph | Yeast | NULL | Probable | As nectar-tap_saprotroph (Põlme et al. 2020); Decay sugar-rich substrates (Põlme et al. 2020) | Põlme S, et al. 2020. Fungal Diversity 105:1-16 ((https://doi.org/10.1007/s13225-020-00466-2)) |
| Otu22254 | 1 | k__Fungi(100);p__Ascomycota(100);c__Saccharomycetes(100);o__Saccharomycetales(100);f__Metschnikowiaceae(100);g__Metschnikowia(100);g__Metschnikowia_unclassified(100); | Metschnikowia | 13 | Saprotroph | Undefined Saprotroph | Yeast | NULL | Probable | As nectar-tap_saprotroph (Põlme et al. 2020); Decay sugar-rich substrates (Põlme et al. 2020) | Põlme S, et al. 2020. Fungal Diversity 105:1-16 ((https://doi.org/10.1007/s13225-020-00466-2)) |
| Otu22255 | 1 | k__Fungi(100);p__Ascomycota(100);c__Saccharomycetes(100);o__Saccharomycetales(100);f__Metschnikowiaceae(100);g__Metschnikowia(100);g__Metschnikowia_unclassified(100); | Metschnikowia | 13 | Saprotroph | Undefined Saprotroph | Yeast | NULL | Probable | As nectar-tap_saprotroph (Põlme et al. 2020); Decay sugar-rich substrates (Põlme et al. 2020) | Põlme S, et al. 2020. Fungal Diversity 105:1-16 ((https://doi.org/10.1007/s13225-020-00466-2)) |
| Otu22257 | 1 | k__Fungi(100);p__Ascomycota(100);c__Saccharomycetes(100);o__Saccharomycetales(100);f__Metschnikowiaceae(100);g__Metschnikowia(100);s__Metschnikowia_pulcherrima(100); | Metschnikowia | 13 | Saprotroph | Undefined Saprotroph | Yeast | NULL | Probable | As nectar-tap_saprotroph (Põlme et al. 2020); Decay sugar-rich substrates (Põlme et al. 2020) | Põlme S, et al. 2020. Fungal Diversity 105:1-16 ((https://doi.org/10.1007/s13225-020-00466-2)) |
| Otu22258 | 1 | k__Fungi(100);p__Ascomycota(100);c__Saccharomycetes(100);o__Saccharomycetales(100);f__Metschnikowiaceae(100);g__Metschnikowia(100);s__Metschnikowia_pulcherrima(100); | Metschnikowia | 13 | Saprotroph | Undefined Saprotroph | Yeast | NULL | Probable | As nectar-tap_saprotroph (Põlme et al. 2020); Decay sugar-rich substrates (Põlme et al. 2020) | Põlme S, et al. 2020. Fungal Diversity 105:1-16 ((https://doi.org/10.1007/s13225-020-00466-2)) |
| Otu22259 | 1 | k__Fungi(100);p__Ascomycota(100);c__Saccharomycetes(100);o__Saccharomycetales(100);f__Metschnikowiaceae(100);g__Metschnikowia(100);g__Metschnikowia_unclassified(100); | Metschnikowia | 13 | Saprotroph | Undefined Saprotroph | Yeast | NULL | Probable | As nectar-tap_saprotroph (Põlme et al. 2020); Decay sugar-rich substrates (Põlme et al. 2020) | Põlme S, et al. 2020. Fungal Diversity 105:1-16 ((https://doi.org/10.1007/s13225-020-00466-2)) |
| Otu22260 | 1 | k__Fungi(100);p__Ascomycota(100);c__Saccharomycetes(100);o__Saccharomycetales(100);f__Metschnikowiaceae(100);g__Metschnikowia(100);g__Metschnikowia_unclassified(100); | Metschnikowia | 13 | Saprotroph | Undefined Saprotroph | Yeast | NULL | Probable | As nectar-tap_saprotroph (Põlme et al. 2020); Decay sugar-rich substrates (Põlme et al. 2020) | Põlme S, et al. 2020. Fungal Diversity 105:1-16 ((https://doi.org/10.1007/s13225-020-00466-2)) |
| Otu22261 | 1 | k__Fungi(100);p__Ascomycota(100);c__Saccharomycetes(100);o__Saccharomycetales(100);f__Metschnikowiaceae(100);g__Metschnikowia(100);s__Metschnikowia_pulcherrima(100); | Metschnikowia | 13 | Saprotroph | Undefined Saprotroph | Yeast | NULL | Probable | As nectar-tap_saprotroph (Põlme et al. 2020); Decay sugar-rich substrates (Põlme et al. 2020) | Põlme S, et al. 2020. Fungal Diversity 105:1-16 ((https://doi.org/10.1007/s13225-020-00466-2)) |
| Otu22262 | 1 | k__Fungi(100);p__Ascomycota(100);c__Saccharomycetes(100);o__Saccharomycetales(100);f__Metschnikowiaceae(100);g__Metschnikowia(100);s__Metschnikowia_pulcherrima(100); | Metschnikowia | 13 | Saprotroph | Undefined Saprotroph | Yeast | NULL | Probable | As nectar-tap_saprotroph (Põlme et al. 2020); Decay sugar-rich substrates (Põlme et al. 2020) | Põlme S, et al. 2020. Fungal Diversity 105:1-16 ((https://doi.org/10.1007/s13225-020-00466-2)) |
| Otu22263 | 1 | k__Fungi(100);p__Ascomycota(100);c__Saccharomycetes(100);o__Saccharomycetales(100);f__Metschnikowiaceae(100);g__Metschnikowia(100);s__Metschnikowia_pulcherrima(100); | Metschnikowia | 13 | Saprotroph | Undefined Saprotroph | Yeast | NULL | Probable | As nectar-tap_saprotroph (Põlme et al. 2020); Decay sugar-rich substrates (Põlme et al. 2020) | Põlme S, et al. 2020. Fungal Diversity 105:1-16 ((https://doi.org/10.1007/s13225-020-00466-2)) |
| Otu22264 | 1 | k__Fungi(100);p__Ascomycota(100);c__Saccharomycetes(100);o__Saccharomycetales(100);f__Metschnikowiaceae(100);g__Metschnikowia(100);s__Metschnikowia_pulcherrima(100); | Metschnikowia | 13 | Saprotroph | Undefined Saprotroph | Yeast | NULL | Probable | As nectar-tap_saprotroph (Põlme et al. 2020); Decay sugar-rich substrates (Põlme et al. 2020) | Põlme S, et al. 2020. Fungal Diversity 105:1-16 ((https://doi.org/10.1007/s13225-020-00466-2)) |

| | | | | | | | | | | |
|---|---|---|---|---|---|---|---|---|---|---|
| Otu22265 | 1 | k__Fungi(100);p__Ascomycota(100);c__Saccharomycetes(100);o__Saccharomycetales(100);f__Metschnikowiaceae(100);g__Metschnikowia(100);s__Metschnikowia_pulcherrima(100); | Metschnikowia | 13 | Saprotroph | \|Undefined Saprotroph\| | Yeast | NULL | Probable | As nectar-tap_saprotroph (Põlme et al. 2020); Decay sugar-rich substrates (Põlme et al. 2020) | Põlme S, et al. 2020. Fungal Diversity 105:1-16 ((https://doi.org/10.1007/s13225-020-00466-2)) |
| Otu22266 | 1 | k__Fungi(100);p__Ascomycota(100);c__Saccharomycetes(100);o__Saccharomycetales(100);f__Metschnikowiaceae(100);g__Metschnikowia(100);s__Metschnikowia_pulcherrima(100); | Metschnikowia | 13 | Saprotroph | \|Undefined Saprotroph\| | Yeast | NULL | Probable | As nectar-tap_saprotroph (Põlme et al. 2020); Decay sugar-rich substrates (Põlme et al. 2020) | Põlme S, et al. 2020. Fungal Diversity 105:1-16 ((https://doi.org/10.1007/s13225-020-00466-2)) |
| Otu22267 | 1 | k__Fungi(100);p__Ascomycota(100);c__Saccharomycetes(100);o__Saccharomycetales(100);f__Metschnikowiaceae(100);g__Metschnikowia(100);g__Metschnikowia_unclassified(100); | Metschnikowia | 13 | Saprotroph | \|Undefined Saprotroph\| | Yeast | NULL | Probable | As nectar-tap_saprotroph (Põlme et al. 2020); Decay sugar-rich substrates (Põlme et al. 2020) | Põlme S, et al. 2020. Fungal Diversity 105:1-16 ((https://doi.org/10.1007/s13225-020-00466-2)) |
| Otu22268 | 1 | k__Fungi(100);p__Ascomycota(100);c__Saccharomycetes(100);o__Saccharomycetales(100);f__Metschnikowiaceae(100);g__Metschnikowia(100);s__Metschnikowia_pulcherrima(100); | Metschnikowia | 13 | Saprotroph | \|Undefined Saprotroph\| | Yeast | NULL | Probable | As nectar-tap_saprotroph (Põlme et al. 2020); Decay sugar-rich substrates (Põlme et al. 2020) | Põlme S, et al. 2020. Fungal Diversity 105:1-16 ((https://doi.org/10.1007/s13225-020-00466-2)) |
| Otu22269 | 1 | k__Fungi(100);p__Ascomycota(100);c__Saccharomycetes(100);o__Saccharomycetales(100);f__Metschnikowiaceae(100);g__Metschnikowia(100);s__Metschnikowia_pulcherrima(100); | Metschnikowia | 13 | Saprotroph | \|Undefined Saprotroph\| | Yeast | NULL | Probable | As nectar-tap_saprotroph (Põlme et al. 2020); Decay sugar-rich substrates (Põlme et al. 2020) | Põlme S, et al. 2020. Fungal Diversity 105:1-16 ((https://doi.org/10.1007/s13225-020-00466-2)) |
| Otu22270 | 1 | k__Fungi(100);p__Ascomycota(100);c__Saccharomycetes(100);o__Saccharomycetales(100);f__Metschnikowiaceae(100);g__Metschnikowia(100);g__Metschnikowia_unclassified(100); | Metschnikowia | 13 | Saprotroph | \|Undefined Saprotroph\| | Yeast | NULL | Probable | As nectar-tap_saprotroph (Põlme et al. 2020); Decay sugar-rich substrates (Põlme et al. 2020) | Põlme S, et al. 2020. Fungal Diversity 105:1-16 ((https://doi.org/10.1007/s13225-020-00466-2)) |
| Otu22271 | 1 | k__Fungi(100);p__Ascomycota(100);c__Saccharomycetes(100);o__Saccharomycetales(100);f__Metschnikowiaceae(100);g__Metschnikowia(100);s__Metschnikowia_pulcherrima(100); | Metschnikowia | 13 | Saprotroph | \|Undefined Saprotroph\| | Yeast | NULL | Probable | As nectar-tap_saprotroph (Põlme et al. 2020); Decay sugar-rich substrates (Põlme et al. 2020) | Põlme S, et al. 2020. Fungal Diversity 105:1-16 ((https://doi.org/10.1007/s13225-020-00466-2)) |
| Otu22272 | 1 | k__Fungi(100);p__Ascomycota(100);c__Saccharomycetes(100);o__Saccharomycetales(100);f__Metschnikowiaceae(100);g__Metschnikowia(100);g__Metschnikowia_unclassified(100); | Metschnikowia | 13 | Saprotroph | \|Undefined Saprotroph\| | Yeast | NULL | Probable | As nectar-tap_saprotroph (Põlme et al. 2020); Decay sugar-rich substrates (Põlme et al. 2020) | Põlme S, et al. 2020. Fungal Diversity 105:1-16 ((https://doi.org/10.1007/s13225-020-00466-2)) |
| Otu22273 | 1 | k__Fungi(100);p__Ascomycota(100);c__Saccharomycetes(100);o__Saccharomycetales(100);f__Metschnikowiaceae(100);g__Metschnikowia(100);s__Metschnikowia_pulcherrima(100); | Metschnikowia | 13 | Saprotroph | \|Undefined Saprotroph\| | Yeast | NULL | Probable | As nectar-tap_saprotroph (Põlme et al. 2020); Decay sugar-rich substrates (Põlme et al. 2020) | Põlme S, et al. 2020. Fungal Diversity 105:1-16 ((https://doi.org/10.1007/s13225-020-00466-2)) |
| Otu22274 | 1 | k__Fungi(100);p__Ascomycota(100);c__Saccharomycetes(100);o__Saccharomycetales(100);f__Metschnikowiaceae(100);g__Metschnikowia(100);s__Metschnikowia_pulcherrima(100); | Metschnikowia | 13 | Saprotroph | \|Undefined Saprotroph\| | Yeast | NULL | Probable | As nectar-tap_saprotroph (Põlme et al. 2020); Decay sugar-rich substrates (Põlme et al. 2020) | Põlme S, et al. 2020. Fungal Diversity 105:1-16 ((https://doi.org/10.1007/s13225-020-00466-2)) |
| Otu22275 | 1 | k__Fungi(100);p__Ascomycota(100);c__Saccharomycetes(100);o__Saccharomycetales(100);f__Metschnikowiaceae(100);g__Metschnikowia(100);s__Metschnikowia_pulcherrima(100); | Metschnikowia | 13 | Saprotroph | \|Undefined Saprotroph\| | Yeast | NULL | Probable | As nectar-tap_saprotroph (Põlme et al. 2020); Decay sugar-rich substrates (Põlme et al. 2020) | Põlme S, et al. 2020. Fungal Diversity 105:1-16 ((https://doi.org/10.1007/s13225-020-00466-2)) |
| Otu22276 | 1 | k__Fungi(100);p__Ascomycota(100);c__Saccharomycetes(100);o__Saccharomycetales(100);f__Metschnikowiaceae(100);g__Metschnikowia(100);s__Metschnikowia_pulcherrima(100); | Metschnikowia | 13 | Saprotroph | \|Undefined Saprotroph\| | Yeast | NULL | Probable | As nectar-tap_saprotroph (Põlme et al. 2020); Decay sugar-rich substrates (Põlme et al. 2020) | Põlme S, et al. 2020. Fungal Diversity 105:1-16 ((https://doi.org/10.1007/s13225-020-00466-2)) |
| Otu22277 | 1 | k__Fungi(100);p__Ascomycota(100);c__Saccharomycetes(100);o__Saccharomycetales(100);f__Metschnikowiaceae(100);g__Metschnikowia(100);s__Metschnikowia_pulcherrima(100); | Metschnikowia | 13 | Saprotroph | \|Undefined Saprotroph\| | Yeast | NULL | Probable | As nectar-tap_saprotroph (Põlme et al. 2020); Decay sugar-rich substrates (Põlme et al. 2020) | Põlme S, et al. 2020. Fungal Diversity 105:1-16 ((https://doi.org/10.1007/s13225-020-00466-2)) |
| Otu22278 | 1 | k__Fungi(100);p__Ascomycota(100);c__Saccharomycetes(100);o__Saccharomycetales(100);f__Metschnikowiaceae(100);g__Metschnikowia(100);g__Metschnikowia_unclassified(100); | Metschnikowia | 13 | Saprotroph | \|Undefined Saprotroph\| | Yeast | NULL | Probable | As nectar-tap_saprotroph (Põlme et al. 2020); Decay sugar-rich substrates (Põlme et al. 2020) | Põlme S, et al. 2020. Fungal Diversity 105:1-16 ((https://doi.org/10.1007/s13225-020-00466-2)) |
| Otu22279 | 1 | k__Fungi(100);p__Ascomycota(100);c__Saccharomycetes(100);o__Saccharomycetales(100);f__Metschnikowiaceae(100);g__Metschnikowia(100);g__Metschnikowia_unclassified(100); | Metschnikowia | 13 | Saprotroph | \|Undefined Saprotroph\| | Yeast | NULL | Probable | As nectar-tap_saprotroph (Põlme et al. 2020); Decay sugar-rich substrates (Põlme et al. 2020) | Põlme S, et al. 2020. Fungal Diversity 105:1-16 ((https://doi.org/10.1007/s13225-020-00466-2)) |
| Otu22280 | 1 | k__Fungi(100);p__Ascomycota(100);c__Saccharomycetes(100);o__Saccharomycetales(100);f__Metschnikowiaceae(100);g__Metschnikowia(100);s__Metschnikowia_pulcherrima(100); | Metschnikowia | 13 | Saprotroph | \|Undefined Saprotroph\| | Yeast | NULL | Probable | As nectar-tap_saprotroph (Põlme et al. 2020); Decay sugar-rich substrates (Põlme et al. 2020) | Põlme S, et al. 2020. Fungal Diversity 105:1-16 ((https://doi.org/10.1007/s13225-020-00466-2)) |
| Otu22281 | 1 | k__Fungi(100);p__Ascomycota(100);c__Saccharomycetes(100);o__Saccharomycetales(100);f__Metschnikowiaceae(100);g__Metschnikowia(100);s__Metschnikowia_pulcherrima(100); | Metschnikowia | 13 | Saprotroph | \|Undefined Saprotroph\| | Yeast | NULL | Probable | As nectar-tap_saprotroph (Põlme et al. 2020); Decay sugar-rich substrates (Põlme et al. 2020) | Põlme S, et al. 2020. Fungal Diversity 105:1-16 ((https://doi.org/10.1007/s13225-020-00466-2)) |
| Otu22282 | 1 | k__Fungi(100);p__Ascomycota(100);c__Saccharomycetes(100);o__Saccharomycetales(100);f__Metschnikowiaceae(100);g__Metschnikowia(100);s__Metschnikowia_pulcherrima(100); | Metschnikowia | 13 | Saprotroph | \|Undefined Saprotroph\| | Yeast | NULL | Probable | As nectar-tap_saprotroph (Põlme et al. 2020); Decay sugar-rich substrates (Põlme et al. 2020) | Põlme S, et al. 2020. Fungal Diversity 105:1-16 ((https://doi.org/10.1007/s13225-020-00466-2)) |
| Otu22283 | 1 | k__Fungi(100);p__Ascomycota(100);c__Saccharomycetes(100);o__Saccharomycetales(100);f__Metschnikowiaceae(100);g__Metschnikowia(100);s__Metschnikowia_pulcherrima(100); | Metschnikowia | 13 | Saprotroph | \|Undefined Saprotroph\| | Yeast | NULL | Probable | As nectar-tap_saprotroph (Põlme et al. 2020); Decay sugar-rich substrates (Põlme et al. 2020) | Põlme S, et al. 2020. Fungal Diversity 105:1-16 ((https://doi.org/10.1007/s13225-020-00466-2)) |
| Otu22284 | 1 | k__Fungi(100);p__Ascomycota(100);c__Saccharomycetes(100);o__Saccharomycetales(100);f__Metschnikowiaceae(100);g__Metschnikowia(100);g__Metschnikowia_unclassified(100); | Metschnikowia | 13 | Saprotroph | \|Undefined Saprotroph\| | Yeast | NULL | Probable | As nectar-tap_saprotroph (Põlme et al. 2020); Decay sugar-rich substrates (Põlme et al. 2020) | Põlme S, et al. 2020. Fungal Diversity 105:1-16 ((https://doi.org/10.1007/s13225-020-00466-2)) |
| Otu22285 | 1 | k__Fungi(100);p__Ascomycota(100);c__Saccharomycetes(100);o__Saccharomycetales(100);f__Metschnikowiaceae(100);g__Metschnikowia(100);g__Metschnikowia_unclassified(100); | Metschnikowia | 13 | Saprotroph | \|Undefined Saprotroph\| | Yeast | NULL | Probable | As nectar-tap_saprotroph (Põlme et al. 2020); Decay sugar-rich substrates (Põlme et al. 2020) | Põlme S, et al. 2020. Fungal Diversity 105:1-16 ((https://doi.org/10.1007/s13225-020-00466-2)) |
| Otu22286 | 1 | k__Fungi(100);p__Ascomycota(100);c__Saccharomycetes(100);o__Saccharomycetales(100);f__Metschnikowiaceae(100);g__Metschnikowia(100);g__Metschnikowia_unclassified(100); | Metschnikowia | 13 | Saprotroph | \|Undefined Saprotroph\| | Yeast | NULL | Probable | As nectar-tap_saprotroph (Põlme et al. 2020); Decay sugar-rich substrates (Põlme et al. 2020) | Põlme S, et al. 2020. Fungal Diversity 105:1-16 ((https://doi.org/10.1007/s13225-020-00466-2)) |
| Otu22287 | 1 | k__Fungi(100);p__Ascomycota(100);c__Saccharomycetes(100);o__Saccharomycetales(100);f__Metschnikowiaceae(100);g__Metschnikowia(100);s__Metschnikowia_pulcherrima(100); | Metschnikowia | 13 | Saprotroph | \|Undefined Saprotroph\| | Yeast | NULL | Probable | As nectar-tap_saprotroph (Põlme et al. 2020); Decay sugar-rich substrates (Põlme et al. 2020) | Põlme S, et al. 2020. Fungal Diversity 105:1-16 ((https://doi.org/10.1007/s13225-020-00466-2)) |
| Otu22288 | 1 | k__Fungi(100);p__Ascomycota(100);c__Saccharomycetes(100);o__Saccharomycetales(100);f__Metschnikowiaceae(100);g__Metschnikowia(100);g__Metschnikowia_unclassified(100); | Metschnikowia | 13 | Saprotroph | \|Undefined Saprotroph\| | Yeast | NULL | Probable | As nectar-tap_saprotroph (Põlme et al. 2020); Decay sugar-rich substrates (Põlme et al. 2020) | Põlme S, et al. 2020. Fungal Diversity 105:1-16 ((https://doi.org/10.1007/s13225-020-00466-2)) |
| Otu22289 | 1 | k__Fungi(100);p__Ascomycota(100);c__Saccharomycetes(100);o__Saccharomycetales(100);f__Metschnikowiaceae(100);g__Metschnikowia(100);s__Metschnikowia_pulcherrima(100); | Metschnikowia | 13 | Saprotroph | \|Undefined Saprotroph\| | Yeast | NULL | Probable | As nectar-tap_saprotroph (Põlme et al. 2020); Decay sugar-rich substrates (Põlme et al. 2020) | Põlme S, et al. 2020. Fungal Diversity 105:1-16 ((https://doi.org/10.1007/s13225-020-00466-2)) |
| Otu22290 | 1 | k__Fungi(100);p__Ascomycota(100);c__Saccharomycetes(100);o__Saccharomycetales(100);f__Metschnikowiaceae(100);g__Metschnikowia(100);s__Metschnikowia_pulcherrima(100); | Metschnikowia | 13 | Saprotroph | \|Undefined Saprotroph\| | Yeast | NULL | Probable | As nectar-tap_saprotroph (Põlme et al. 2020); Decay sugar-rich substrates (Põlme et al. 2020) | Põlme S, et al. 2020. Fungal Diversity 105:1-16 ((https://doi.org/10.1007/s13225-020-00466-2)) |
| Otu22291 | 1 | k__Fungi(100);p__Ascomycota(100);c__Saccharomycetes(100);o__Saccharomycetales(100);f__Metschnikowiaceae(100);g__Metschnikowia(100);g__Metschnikowia_unclassified(100); | Metschnikowia | 13 | Saprotroph | \|Undefined Saprotroph\| | Yeast | NULL | Probable | As nectar-tap_saprotroph (Põlme et al. 2020); Decay sugar-rich substrates (Põlme et al. 2020) | Põlme S, et al. 2020. Fungal Diversity 105:1-16 ((https://doi.org/10.1007/s13225-020-00466-2)) |
| Otu22292 | 1 | k__Fungi(100);p__Ascomycota(100);c__Saccharomycetes(100);o__Saccharomycetales(100);f__Metschnikowiaceae(100);g__Metschnikowia(100);s__Metschnikowia_pulcherrima(100); | Metschnikowia | 13 | Saprotroph | \|Undefined Saprotroph\| | Yeast | NULL | Probable | As nectar-tap_saprotroph (Põlme et al. 2020); Decay sugar-rich substrates (Põlme et al. 2020) | Põlme S, et al. 2020. Fungal Diversity 105:1-16 ((https://doi.org/10.1007/s13225-020-00466-2)) |
| Otu22294 | 1 | k__Fungi(100);p__Ascomycota(100);c__Saccharomycetes(100);o__Saccharomycetales(100);f__Metschnikowiaceae(100);g__Metschnikowia(100);s__Metschnikowia_pulcherrima(100); | Metschnikowia | 13 | Saprotroph | \|Undefined Saprotroph\| | Yeast | NULL | Probable | As nectar-tap_saprotroph (Põlme et al. 2020); Decay sugar-rich substrates (Põlme et al. 2020) | Põlme S, et al. 2020. Fungal Diversity 105:1-16 ((https://doi.org/10.1007/s13225-020-00466-2)) |
| Otu22296 | 1 | k__Fungi(100);p__Ascomycota(100);c__Saccharomycetes(100);o__Saccharomycetales(100);f__Metschnikowiaceae(100);g__Metschnikowia(100);g__Metschnikowia_unclassified(100); | Metschnikowia | 13 | Saprotroph | \|Undefined Saprotroph\| | Yeast | NULL | Probable | As nectar-tap_saprotroph (Põlme et al. 2020); Decay sugar-rich substrates (Põlme et al. 2020) | Põlme S, et al. 2020. Fungal Diversity 105:1-16 ((https://doi.org/10.1007/s13225-020-00466-2)) |
| Otu22297 | 1 | k__Fungi(100);p__Ascomycota(100);c__Saccharomycetes(100);o__Saccharomycetales(100);f__Metschnikowiaceae(100);g__Metschnikowia(100);s__Metschnikowia_pulcherrima(100); | Metschnikowia | 13 | Saprotroph | \|Undefined Saprotroph\| | Yeast | NULL | Probable | As nectar-tap_saprotroph (Põlme et al. 2020); Decay sugar-rich substrates (Põlme et al. 2020) | Põlme S, et al. 2020. Fungal Diversity 105:1-16 ((https://doi.org/10.1007/s13225-020-00466-2)) |
| Otu22298 | 1 | k__Fungi(100);p__Ascomycota(100);c__Saccharomycetes(100);o__Saccharomycetales(100);f__Metschnikowiaceae(100);g__Metschnikowia(100);g__Metschnikowia_unclassified(100); | Metschnikowia | 13 | Saprotroph | \|Undefined Saprotroph\| | Yeast | NULL | Probable | As nectar-tap_saprotroph (Põlme et al. 2020); Decay sugar-rich substrates (Põlme et al. 2020) | Põlme S, et al. 2020. Fungal Diversity 105:1-16 ((https://doi.org/10.1007/s13225-020-00466-2)) |
| Otu22299 | 1 | k__Fungi(100);p__Ascomycota(100);c__Saccharomycetes(100);o__Saccharomycetales(100);f__Metschnikowiaceae(100);g__Metschnikowia(100);s__Metschnikowia_pulcherrima(100); | Metschnikowia | 13 | Saprotroph | \|Undefined Saprotroph\| | Yeast | NULL | Probable | As nectar-tap_saprotroph (Põlme et al. 2020); Decay sugar-rich substrates (Põlme et al. 2020) | Põlme S, et al. 2020. Fungal Diversity 105:1-16 ((https://doi.org/10.1007/s13225-020-00466-2)) |
| Otu22300 | 1 | k__Fungi(100);p__Ascomycota(100);c__Saccharomycetes(100);o__Saccharomycetales(100);f__Metschnikowiaceae(100);g__Metschnikowia(100);s__Metschnikowia_pulcherrima(100); | Metschnikowia | 13 | Saprotroph | \|Undefined Saprotroph\| | Yeast | NULL | Probable | As nectar-tap_saprotroph (Põlme et al. 2020); Decay sugar-rich substrates (Põlme et al. 2020) | Põlme S, et al. 2020. Fungal Diversity 105:1-16 ((https://doi.org/10.1007/s13225-020-00466-2)) |
| Otu22301 | 1 | k__Fungi(100);p__Ascomycota(100);c__Saccharomycetes(100);o__Saccharomycetales(100);f__Metschnikowiaceae(100);g__Metschnikowia(100);s__Metschnikowia_pulcherrima(100); | Metschnikowia | 13 | Saprotroph | \|Undefined Saprotroph\| | Yeast | NULL | Probable | As nectar-tap_saprotroph (Põlme et al. 2020); Decay sugar-rich substrates (Põlme et al. 2020) | Põlme S, et al. 2020. Fungal Diversity 105:1-16 ((https://doi.org/10.1007/s13225-020-00466-2)) |
| Otu22302 | 1 | k__Fungi(100);p__Ascomycota(100);c__Saccharomycetes(100);o__Saccharomycetales(100);f__Metschnikowiaceae(100);g__Metschnikowia(100);s__Metschnikowia_pulcherrima(100); | Metschnikowia | 13 | Saprotroph | \|Undefined Saprotroph\| | Yeast | NULL | Probable | As nectar-tap_saprotroph (Põlme et al. 2020); Decay sugar-rich substrates (Põlme et al. 2020) | Põlme S, et al. 2020. Fungal Diversity 105:1-16 ((https://doi.org/10.1007/s13225-020-00466-2)) |
| Otu22303 | 1 | k__Fungi(100);p__Ascomycota(100);c__Saccharomycetes(100);o__Saccharomycetales(100);f__Metschnikowiaceae(100);g__Metschnikowia(100);g__Metschnikowia_unclassified(100); | Metschnikowia | 13 | Saprotroph | \|Undefined Saprotroph\| | Yeast | NULL | Probable | As nectar-tap_saprotroph (Põlme et al. 2020); Decay sugar-rich substrates (Põlme et al. 2020) | Põlme S, et al. 2020. Fungal Diversity 105:1-16 ((https://doi.org/10.1007/s13225-020-00466-2)) |
| Otu22305 | 1 | k__Fungi(100);p__Ascomycota(100);c__Saccharomycetes(100);o__Saccharomycetales(100);f__Metschnikowiaceae(100);g__Metschnikowia(100);s__Metschnikowia_pulcherrima(100); | Metschnikowia | 13 | Saprotroph | \|Undefined Saprotroph\| | Yeast | NULL | Probable | As nectar-tap_saprotroph (Põlme et al. 2020); Decay sugar-rich substrates (Põlme et al. 2020) | Põlme S, et al. 2020. Fungal Diversity 105:1-16 ((https://doi.org/10.1007/s13225-020-00466-2)) |
| Otu22306 | 1 | k__Fungi(100);p__Ascomycota(100);c__Saccharomycetes(100);o__Saccharomycetales(100);f__Metschnikowiaceae(100);g__Metschnikowia(100);s__Metschnikowia_pulcherrima(100); | Metschnikowia | 13 | Saprotroph | \|Undefined Saprotroph\| | Yeast | NULL | Probable | As nectar-tap_saprotroph (Põlme et al. 2020); Decay sugar-rich substrates (Põlme et al. 2020) | Põlme S, et al. 2020. Fungal Diversity 105:1-16 ((https://doi.org/10.1007/s13225-020-00466-2)) |
| Otu22307 | 1 | k__Fungi(100);p__Ascomycota(100);c__Saccharomycetes(100);o__Saccharomycetales(100);f__Metschnikowiaceae(100);g__Metschnikowia(100);s__Metschnikowia_pulcherrima(100); | Metschnikowia | 13 | Saprotroph | \|Undefined Saprotroph\| | Yeast | NULL | Probable | As nectar-tap_saprotroph (Põlme et al. 2020); Decay sugar-rich substrates (Põlme et al. 2020) | Põlme S, et al. 2020. Fungal Diversity 105:1-16 ((https://doi.org/10.1007/s13225-020-00466-2)) |
| Otu22308 | 1 | k__Fungi(100);p__Ascomycota(100);c__Saccharomycetes(100);o__Saccharomycetales(100);f__Metschnikowiaceae(100);g__Metschnikowia(100);g__Metschnikowia_unclassified(100); | Metschnikowia | 13 | Saprotroph | \|Undefined Saprotroph\| | Yeast | NULL | Probable | As nectar-tap_saprotroph (Põlme et al. 2020); Decay sugar-rich substrates (Põlme et al. 2020) | Põlme S, et al. 2020. Fungal Diversity 105:1-16 ((https://doi.org/10.1007/s13225-020-00466-2)) |
| Otu22309 | 1 | k__Fungi(100);p__Ascomycota(100);c__Saccharomycetes(100);o__Saccharomycetales(100);f__Metschnikowiaceae(100);g__Metschnikowia(100);g__Metschnikowia_unclassified(100); | Metschnikowia | 13 | Saprotroph | \|Undefined Saprotroph\| | Yeast | NULL | Probable | As nectar-tap_saprotroph (Põlme et al. 2020); Decay sugar-rich substrates (Põlme et al. 2020) | Põlme S, et al. 2020. Fungal Diversity 105:1-16 ((https://doi.org/10.1007/s13225-020-00466-2)) |
| Otu22310 | 1 | k__Fungi(100);p__Ascomycota(100);c__Saccharomycetes(100);o__Saccharomycetales(100);f__Metschnikowiaceae(100);g__Metschnikowia(100);g__Metschnikowia_unclassified(100); | Metschnikowia | 13 | Saprotroph | \|Undefined Saprotroph\| | Yeast | NULL | Probable | As nectar-tap_saprotroph (Põlme et al. 2020); Decay sugar-rich substrates (Põlme et al. 2020) | Põlme S, et al. 2020. Fungal Diversity 105:1-16 ((https://doi.org/10.1007/s13225-020-00466-2)) |
| Otu22311 | 1 | k__Fungi(100);p__Ascomycota(100);c__Saccharomycetes(100);o__Saccharomycetales(100);f__Metschnikowiaceae(100);g__Metschnikowia(100);s__Metschnikowia_pulcherrima(100); | Metschnikowia | 13 | Saprotroph | \|Undefined Saprotroph\| | Yeast | NULL | Probable | As nectar-tap_saprotroph (Põlme et al. 2020); Decay sugar-rich substrates (Põlme et al. 2020) | Põlme S, et al. 2020. Fungal Diversity 105:1-16 ((https://doi.org/10.1007/s13225-020-00466-2)) |
| Otu22312 | 1 | k__Fungi(100);p__Ascomycota(100);c__Saccharomycetes(100);o__Saccharomycetales(100);f__Metschnikowiaceae(100);g__Metschnikowia(100);s__Metschnikowia_pulcherrima(100); | Metschnikowia | 13 | Saprotroph | \|Undefined Saprotroph\| | Yeast | NULL | Probable | As nectar-tap_saprotroph (Põlme et al. 2020); Decay sugar-rich substrates (Põlme et al. 2020) | Põlme S, et al. 2020. Fungal Diversity 105:1-16 ((https://doi.org/10.1007/s13225-020-00466-2)) |
| Otu22313 | 1 | k__Fungi(100);p__Ascomycota(100);c__Saccharomycetes(100);o__Saccharomycetales(100);f__Metschnikowiaceae(100);g__Metschnikowia(100);g__Metschnikowia_unclassified(100); | Metschnikowia | 13 | Saprotroph | \|Undefined Saprotroph\| | Yeast | NULL | Probable | As nectar-tap_saprotroph (Põlme et al. 2020); Decay sugar-rich substrates (Põlme et al. 2020) | Põlme S, et al. 2020. Fungal Diversity 105:1-16 ((https://doi.org/10.1007/s13225-020-00466-2)) |
| Otu22314 | 1 | k__Fungi(100);p__Ascomycota(100);c__Saccharomycetes(100);o__Saccharomycetales(100);f__Metschnikowiaceae(100);g__Metschnikowia(100);s__Metschnikowia_pulcherrima(100); | Metschnikowia | 13 | Saprotroph | \|Undefined Saprotroph\| | Yeast | NULL | Probable | As nectar-tap_saprotroph (Põlme et al. 2020); Decay sugar-rich substrates (Põlme et al. 2020) | Põlme S, et al. 2020. Fungal Diversity 105:1-16 ((https://doi.org/10.1007/s13225-020-00466-2)) |
| Otu22315 | 1 | k__Fungi(100);p__Ascomycota(100);c__Saccharomycetes(100);o__Saccharomycetales(100);f__Metschnikowiaceae(100);g__Metschnikowia(100);s__Metschnikowia_pulcherrima(100); | Metschnikowia | 13 | Saprotroph | \|Undefined Saprotroph\| | Yeast | NULL | Probable | As nectar-tap_saprotroph (Põlme et al. 2020); Decay sugar-rich substrates (Põlme et al. 2020) | Põlme S, et al. 2020. Fungal Diversity 105:1-16 ((https://doi.org/10.1007/s13225-020-00466-2)) |
| Otu22316 | 1 | k__Fungi(100);p__Ascomycota(100);c__Saccharomycetes(100);o__Saccharomycetales(100);f__Metschnikowiaceae(100);g__Metschnikowia(100);g__Metschnikowia_unclassified(100); | Metschnikowia | 13 | Saprotroph | \|Undefined Saprotroph\| | Yeast | NULL | Probable | As nectar-tap_saprotroph (Põlme et al. 2020); Decay sugar-rich substrates (Põlme et al. 2020) | Põlme S, et al. 2020. Fungal Diversity 105:1-16 ((https://doi.org/10.1007/s13225-020-00466-2)) |
| Otu22317 | 1 | k__Fungi(100);p__Ascomycota(100);c__Saccharomycetes(100);o__Saccharomycetales(100);f__Metschnikowiaceae(100);g__Metschnikowia(100);s__Metschnikowia_pulcherrima(100); | Metschnikowia | 13 | Saprotroph | \|Undefined Saprotroph\| | Yeast | NULL | Probable | As nectar-tap_saprotroph (Põlme et al. 2020); Decay sugar-rich substrates (Põlme et al. 2020) | Põlme S, et al. 2020. Fungal Diversity 105:1-16 ((https://doi.org/10.1007/s13225-020-00466-2)) |
| Otu22318 | 1 | k__Fungi(100);p__Ascomycota(100);c__Saccharomycetes(100);o__Saccharomycetales(100);f__Metschnikowiaceae(100);g__Metschnikowia(100);g__Metschnikowia_unclassified(100); | Metschnikowia | 13 | Saprotroph | \|Undefined Saprotroph\| | Yeast | NULL | Probable | As nectar-tap_saprotroph (Põlme et al. 2020); Decay sugar-rich substrates (Põlme et al. 2020) | Põlme S, et al. 2020. Fungal Diversity 105:1-16 ((https://doi.org/10.1007/s13225-020-00466-2)) |
| Otu22319 | 1 | k__Fungi(100);p__Ascomycota(100);c__Saccharomycetes(100);o__Saccharomycetales(100);f__Metschnikowiaceae(100);g__Metschnikowia(100);s__Metschnikowia_pulcherrima(100); | Metschnikowia | 13 | Saprotroph | \|Undefined Saprotroph\| | Yeast | NULL | Probable | As nectar-tap_saprotroph (Põlme et al. 2020); Decay sugar-rich substrates (Põlme et al. 2020) | Põlme S, et al. 2020. Fungal Diversity 105:1-16 ((https://doi.org/10.1007/s13225-020-00466-2)) |
| Otu22320 | 1 | k__Fungi(100);p__Ascomycota(100);c__Saccharomycetes(100);o__Saccharomycetales(100);f__Metschnikowiaceae(100);g__Metschnikowia(100);s__Metschnikowia_pulcherrima(100); | Metschnikowia | 13 | Saprotroph | \|Undefined Saprotroph\| | Yeast | NULL | Probable | As nectar-tap_saprotroph (Põlme et al. 2020); Decay sugar-rich substrates (Põlme et al. 2020) | Põlme S, et al. 2020. Fungal Diversity 105:1-16 ((https://doi.org/10.1007/s13225-020-00466-2)) |
| Otu22321 | 1 | k__Fungi(100);p__Ascomycota(100);c__Saccharomycetes(100);o__Saccharomycetales(100);f__Metschnikowiaceae(100);g__Metschnikowia(100);s__Metschnikowia_pulcherrima(100); | Metschnikowia | 13 | Saprotroph | \|Undefined Saprotroph\| | Yeast | NULL | Probable | As nectar-tap_saprotroph (Põlme et al. 2020); Decay sugar-rich substrates (Põlme et al. 2020) | Põlme S, et al. 2020. Fungal Diversity 105:1-16 ((https://doi.org/10.1007/s13225-020-00466-2)) |
| Otu22322 | 1 | k__Fungi(100);p__Ascomycota(100);c__Saccharomycetes(100);o__Saccharomycetales(100);f__Metschnikowiaceae(100);g__Metschnikowia(100);s__Metschnikowia_pulcherrima(100); | Metschnikowia | 13 | Saprotroph | \|Undefined Saprotroph\| | Yeast | NULL | Probable | As nectar-tap_saprotroph (Põlme et al. 2020); Decay sugar-rich substrates (Põlme et al. 2020) | Põlme S, et al. 2020. Fungal Diversity 105:1-16 ((https://doi.org/10.1007/s13225-020-00466-2)) |
| Otu22323 | 1 | k__Fungi(100);p__Ascomycota(100);c__Saccharomycetes(100);o__Saccharomycetales(100);f__Metschnikowiaceae(100);g__Metschnikowia(100);s__Metschnikowia_pulcherrima(100); | Metschnikowia | 13 | Saprotroph | \|Undefined Saprotroph\| | Yeast | NULL | Probable | As nectar-tap_saprotroph (Põlme et al. 2020); Decay sugar-rich substrates (Põlme et al. 2020) | Põlme S, et al. 2020. Fungal Diversity 105:1-16 ((https://doi.org/10.1007/s13225-020-00466-2)) |
| Otu22324 | 1 | k__Fungi(100);p__Ascomycota(100);c__Saccharomycetes(100);o__Saccharomycetales(100);f__Metschnikowiaceae(100);g__Metschnikowia(100);g__Metschnikowia_unclassified(100); | Metschnikowia | 13 | Saprotroph | \|Undefined Saprotroph\| | Yeast | NULL | Probable | As nectar-tap_saprotroph (Põlme et al. 2020); Decay sugar-rich substrates (Põlme et al. 2020) | Põlme S, et al. 2020. Fungal Diversity 105:1-16 ((https://doi.org/10.1007/s13225-020-00466-2)) |
| Otu22325 | 1 | k__Fungi(100);p__Ascomycota(100);c__Saccharomycetes(100);o__Saccharomycetales(100);f__Metschnikowiaceae(100);g__Metschnikowia(100);s__Metschnikowia_pulcherrima(100); | Metschnikowia | 13 | Saprotroph | \|Undefined Saprotroph\| | Yeast | NULL | Probable | As nectar-tap_saprotroph (Põlme et al. 2020); Decay sugar-rich substrates (Põlme et al. 2020) | Põlme S, et al. 2020. Fungal Diversity 105:1-16 ((https://doi.org/10.1007/s13225-020-00466-2)) |
| Otu22326 | 1 | k__Fungi(100);p__Ascomycota(100);c__Saccharomycetes(100);o__Saccharomycetales(100);f__Metschnikowiaceae(100);g__Metschnikowia(100);s__Metschnikowia_pulcherrima(100); | Metschnikowia | 13 | Saprotroph | \|Undefined Saprotroph\| | Yeast | NULL | Probable | As nectar-tap_saprotroph (Põlme et al. 2020); Decay sugar-rich substrates (Põlme et al. 2020) | Põlme S, et al. 2020. Fungal Diversity 105:1-16 ((https://doi.org/10.1007/s13225-020-00466-2)) |
| Otu22327 | 1 | k__Fungi(100);p__Ascomycota(100);c__Saccharomycetes(100);o__Saccharomycetales(100);f__Metschnikowiaceae(100);g__Metschnikowia(100);g__Metschnikowia_unclassified(100); | Metschnikowia | 13 | Saprotroph | \|Undefined Saprotroph\| | Yeast | NULL | Probable | As nectar-tap_saprotroph (Põlme et al. 2020); Decay sugar-rich substrates (Põlme et al. 2020) | Põlme S, et al. 2020. Fungal Diversity 105:1-16 ((https://doi.org/10.1007/s13225-020-00466-2)) |
| Otu22328 | 1 | k__Fungi(100);p__Ascomycota(100);c__Saccharomycetes(100);o__Saccharomycetales(100);f__Metschnikowiaceae(100);g__Metschnikowia(100);s__Metschnikowia_pulcherrima(100); | Metschnikowia | 13 | Saprotroph | \|Undefined Saprotroph\| | Yeast | NULL | Probable | As nectar-tap_saprotroph (Põlme et al. 2020); Decay sugar-rich substrates (Põlme et al. 2020) | Põlme S, et al. 2020. Fungal Diversity 105:1-16 ((https://doi.org/10.1007/s13225-020-00466-2)) |
| Otu22329 | 1 | k__Fungi(100);p__Ascomycota(100);c__Saccharomycetes(100);o__Saccharomycetales(100);f__Metschnikowiaceae(100);g__Metschnikowia(100);g__Metschnikowia_unclassified(100); | Metschnikowia | 13 | Saprotroph | \|Undefined Saprotroph\| | Yeast | NULL | Probable | As nectar-tap_saprotroph (Põlme et al. 2020); Decay sugar-rich substrates (Põlme et al. 2020) | Põlme S, et al. 2020. Fungal Diversity 105:1-16 ((https://doi.org/10.1007/s13225-020-00466-2)) |
| Otu22330 | 1 | k__Fungi(100);p__Ascomycota(100);c__Saccharomycetes(100);o__Saccharomycetales(100);f__Metschnikowiaceae(100);g__Metschnikowia(100);g__Metschnikowia_unclassified(100); | Metschnikowia | 13 | Saprotroph | \|Undefined Saprotroph\| | Yeast | NULL | Probable | As nectar-tap_saprotroph (Põlme et al. 2020); Decay sugar-rich substrates (Põlme et al. 2020) | Põlme S, et al. 2020. Fungal Diversity 105:1-16 ((https://doi.org/10.1007/s13225-020-00466-2)) |
| Otu22331 | 1 | k__Fungi(100);p__Ascomycota(100);c__Saccharomycetes(100);o__Saccharomycetales(100);f__Metschnikowiaceae(100);g__Metschnikowia(100);s__Metschnikowia_pulcherrima(100); | Metschnikowia | 13 | Saprotroph | \|Undefined Saprotroph\| | Yeast | NULL | Probable | As nectar-tap_saprotroph (Põlme et al. 2020); Decay sugar-rich substrates (Põlme et al. 2020) | Põlme S, et al. 2020. Fungal Diversity 105:1-16 ((https://doi.org/10.1007/s13225-020-00466-2)) |
| Otu22332 | 1 | k__Fungi(100);p__Ascomycota(100);c__Saccharomycetes(100);o__Saccharomycetales(100);f__Metschnikowiaceae(100);g__Metschnikowia(100);s__Metschnikowia_pulcherrima(100); | Metschnikowia | 13 | Saprotroph | \|Undefined Saprotroph\| | Yeast | NULL | Probable | As nectar-tap_saprotroph (Põlme et al. 2020); Decay sugar-rich substrates (Põlme et al. 2020) | Põlme S, et al. 2020. Fungal Diversity 105:1-16 ((https://doi.org/10.1007/s13225-020-00466-2)) |
| Otu22333 | 1 | k__Fungi(100);p__Ascomycota(100);c__Saccharomycetes(100);o__Saccharomycetales(100);f__Metschnikowiaceae(100);g__Metschnikowia(100);s__Metschnikowia_pulcherrima(100); | Metschnikowia | 13 | Saprotroph | \|Undefined Saprotroph\| | Yeast | NULL | Probable | As nectar-tap_saprotroph (Põlme et al. 2020); Decay sugar-rich substrates (Põlme et al. 2020) | Põlme S, et al. 2020. Fungal Diversity 105:1-16 ((https://doi.org/10.1007/s13225-020-00466-2)) |
| Otu22334 | 1 | k__Fungi(100);p__Ascomycota(100);c__Saccharomycetes(100);o__Saccharomycetales(100);f__Metschnikowiaceae(100);g__Metschnikowia(100);g__Metschnikowia_unclassified(100); | Metschnikowia | 13 | Saprotroph | \|Undefined Saprotroph\| | Yeast | NULL | Probable | As nectar-tap_saprotroph (Põlme et al. 2020); Decay sugar-rich substrates (Põlme et al. 2020) | Põlme S, et al. 2020. Fungal Diversity 105:1-16 ((https://doi.org/10.1007/s13225-020-00466-2)) |
| Otu22335 | 1 | k__Fungi(100);p__Ascomycota(100);c__Saccharomycetes(100);o__Saccharomycetales(100);f__Metschnikowiaceae(100);g__Metschnikowia(100);g__Metschnikowia_unclassified(100); | Metschnikowia | 13 | Saprotroph | \|Undefined Saprotroph\| | Yeast | NULL | Probable | As nectar-tap_saprotroph (Põlme et al. 2020); Decay sugar-rich substrates (Põlme et al. 2020) | Põlme S, et al. 2020. Fungal Diversity 105:1-16 ((https://doi.org/10.1007/s13225-020-00466-2)) |
| Otu22336 | 1 | k__Fungi(100);p__Ascomycota(100);c__Saccharomycetes(100);o__Saccharomycetales(100);f__Metschnikowiaceae(100);g__Metschnikowia(100);s__Metschnikowia_pulcherrima(100); | Metschnikowia | 13 | Saprotroph | \|Undefined Saprotroph\| | Yeast | NULL | Probable | As nectar-tap_saprotroph (Põlme et al. 2020); Decay sugar-rich substrates (Põlme et al. 2020) | Põlme S, et al. 2020. Fungal Diversity 105:1-16 ((https://doi.org/10.1007/s13225-020-00466-2)) |
| Otu22337 | 1 | k__Fungi(100);p__Ascomycota(100);c__Saccharomycetes(100);o__Saccharomycetales(100);f__Metschnikowiaceae(100);g__Metschnikowia(100);s__Metschnikowia_pulcherrima(100); | Metschnikowia | 13 | Saprotroph | \|Undefined Saprotroph\| | Yeast | NULL | Probable | As nectar-tap_saprotroph (Põlme et al. 2020); Decay sugar-rich substrates (Põlme et al. 2020) | Põlme S, et al. 2020. Fungal Diversity 105:1-16 ((https://doi.org/10.1007/s13225-020-00466-2)) |
| Otu22338 | 1 | k__Fungi(100);p__Ascomycota(100);c__Saccharomycetes(100);o__Saccharomycetales(100);f__Metschnikowiaceae(100);g__Metschnikowia(100);s__Metschnikowia_pulcherrima(100); | Metschnikowia | 13 | Saprotroph | \|Undefined Saprotroph\| | Yeast | NULL | Probable | As nectar-tap_saprotroph (Põlme et al. 2020); Decay sugar-rich substrates (Põlme et al. 2020) | Põlme S, et al. 2020. Fungal Diversity 105:1-16 ((https://doi.org/10.1007/s13225-020-00466-2)) |
| Otu22339 | 1 | k__Fungi(100);p__Ascomycota(100);c__Saccharomycetes(100);o__Saccharomycetales(100);f__Metschnikowiaceae(100);g__Metschnikowia(100);g__Metschnikowia_unclassified(100); | Metschnikowia | 13 | Saprotroph | \|Undefined Saprotroph\| | Yeast | NULL | Probable | As nectar-tap_saprotroph (Põlme et al. 2020); Decay sugar-rich substrates (Põlme et al. 2020) | Põlme S, et al. 2020. Fungal Diversity 105:1-16 ((https://doi.org/10.1007/s13225-020-00466-2)) |
| Otu22340 | 1 | k__Fungi(100);p__Ascomycota(100);c__Saccharomycetes(100);o__Saccharomycetales(100);f__Metschnikowiaceae(100);g__Metschnikowia(100);s__Metschnikowia_pulcherrima(100); | Metschnikowia | 13 | Saprotroph | \|Undefined Saprotroph\| | Yeast | NULL | Probable | As nectar-tap_saprotroph (Põlme et al. 2020); Decay sugar-rich substrates (Põlme et al. 2020) | Põlme S, et al. 2020. Fungal Diversity 105:1-16 ((https://doi.org/10.1007/s13225-020-00466-2)) |
| Otu22341 | 1 | k__Fungi(100);p__Ascomycota(100);c__Saccharomycetes(100);o__Saccharomycetales(100);f__Metschnikowiaceae(100);g__Metschnikowia(100);s__Metschnikowia_pulcherrima(100); | Metschnikowia | 13 | Saprotroph | \|Undefined Saprotroph\| | Yeast | NULL | Probable | As nectar-tap_saprotroph (Põlme et al. 2020); Decay sugar-rich substrates (Põlme et al. 2020) | Põlme S, et al. 2020. Fungal Diversity 105:1-16 ((https://doi.org/10.1007/s13225-020-00466-2)) |
| Otu22342 | 1 | k__Fungi(100);p__Ascomycota(100);c__Saccharomycetes(100);o__Saccharomycetales(100);f__Metschnikowiaceae(100);g__Metschnikowia(100);g__Metschnikowia_unclassified(100); | Metschnikowia | 13 | Saprotroph | \|Undefined Saprotroph\| | Yeast | NULL | Probable | As nectar-tap_saprotroph (Põlme et al. 2020); Decay sugar-rich substrates (Põlme et al. 2020) | Põlme S, et al. 2020. Fungal Diversity 105:1-16 ((https://doi.org/10.1007/s13225-020-00466-2)) |
| Otu22343 | 1 | k__Fungi(100);p__Ascomycota(100);c__Saccharomycetes(100);o__Saccharomycetales(100);f__Metschnikowiaceae(100);g__Metschnikowia(100);s__Metschnikowia_pulcherrima(100); | Metschnikowia | 13 | Saprotroph | \|Undefined Saprotroph\| | Yeast | NULL | Probable | As nectar-tap_saprotroph (Põlme et al. 2020); Decay sugar-rich substrates (Põlme et al. 2020) | Põlme S, et al. 2020. Fungal Diversity 105:1-16 ((https://doi.org/10.1007/s13225-020-00466-2)) |
| Otu22344 | 1 | k__Fungi(100);p__Ascomycota(100);c__Saccharomycetes(100);o__Saccharomycetales(100);f__Metschnikowiaceae(100);g__Metschnikowia(100);s__Metschnikowia_pulcherrima(100); | Metschnikowia | 13 | Saprotroph | \|Undefined Saprotroph\| | Yeast | NULL | Probable | As nectar-tap_saprotroph (Põlme et al. 2020); Decay sugar-rich substrates (Põlme et al. 2020) | Põlme S, et al. 2020. Fungal Diversity 105:1-16 ((https://doi.org/10.1007/s13225-020-00466-2)) |
| Otu22345 | 1 | k__Fungi(100);p__Ascomycota(100);c__Saccharomycetes(100);o__Saccharomycetales(100);f__Metschnikowiaceae(100);g__Metschnikowia(100);s__Metschnikowia_pulcherrima(100); | Metschnikowia | 13 | Saprotroph | \|Undefined Saprotroph\| | Yeast | NULL | Probable | As nectar-tap_saprotroph (Põlme et al. 2020); Decay sugar-rich substrates (Põlme et al. 2020) | Põlme S, et al. 2020. Fungal Diversity 105:1-16 ((https://doi.org/10.1007/s13225-020-00466-2)) |
| Otu22346 | 1 | k__Fungi(100);p__Ascomycota(100);c__Saccharomycetes(100);o__Saccharomycetales(100);f__Metschnikowiaceae(100);g__Metschnikowia(100);s__Metschnikowia_pulcherrima(100); | Metschnikowia | 13 | Saprotroph | \|Undefined Saprotroph\| | Yeast | NULL | Probable | As nectar-tap_saprotroph (Põlme et al. 2020); Decay sugar-rich substrates (Põlme et al. 2020) | Põlme S, et al. 2020. Fungal Diversity 105:1-16 ((https://doi.org/10.1007/s13225-020-00466-2)) |
| Otu22347 | 1 | k__Fungi(100);p__Ascomycota(100);c__Saccharomycetes(100);o__Saccharomycetales(100);f__Metschnikowiaceae(100);g__Metschnikowia(100);s__Metschnikowia_pulcherrima(100); | Metschnikowia | 13 | Saprotroph | \|Undefined Saprotroph\| | Yeast | NULL | Probable | As nectar-tap_saprotroph (Põlme et al. 2020); Decay sugar-rich substrates (Põlme et al. 2020) | Põlme S, et al. 2020. Fungal Diversity 105:1-16 ((https://doi.org/10.1007/s13225-020-00466-2)) |
| Otu22348 | 1 | k__Fungi(100);p__Ascomycota(100);c__Saccharomycetes(100);o__Saccharomycetales(100);f__Metschnikowiaceae(100);g__Metschnikowia(100);s__Metschnikowia_pulcherrima(100); | Metschnikowia | 13 | Saprotroph | \|Undefined Saprotroph\| | Yeast | NULL | Probable | As nectar-tap_saprotroph (Põlme et al. 2020); Decay sugar-rich substrates (Põlme et al. 2020) | Põlme S, et al. 2020. Fungal Diversity 105:1-16 ((https://doi.org/10.1007/s13225-020-00466-2)) |
| Otu22349 | 1 | k__Fungi(100);p__Ascomycota(100);c__Saccharomycetes(100);o__Saccharomycetales(100);f__Metschnikowiaceae(100);g__Metschnikowia(100);s__Metschnikowia_pulcherrima(100); | Metschnikowia | 13 | Saprotroph | \|Undefined Saprotroph\| | Yeast | NULL | Probable | As nectar-tap_saprotroph (Põlme et al. 2020); Decay sugar-rich substrates (Põlme et al. 2020) | Põlme S, et al. 2020. Fungal Diversity 105:1-16 ((https://doi.org/10.1007/s13225-020-00466-2)) |
| Otu22350 | 1 | k__Fungi(100);p__Ascomycota(100);c__Saccharomycetes(100);o__Saccharomycetales(100);f__Metschnikowiaceae(100);g__Metschnikowia(100);s__Metschnikowia_pulcherrima(100); | Metschnikowia | 13 | Saprotroph | \|Undefined Saprotroph\| | Yeast | NULL | Probable | As nectar-tap_saprotroph (Põlme et al. 2020); Decay sugar-rich substrates (Põlme et al. 2020) | Põlme S, et al. 2020. Fungal Diversity 105:1-16 ((https://doi.org/10.1007/s13225-020-00466-2)) |
| Otu22351 | 1 | k__Fungi(100);p__Ascomycota(100);c__Saccharomycetes(100);o__Saccharomycetales(100);f__Metschnikowiaceae(100);g__Metschnikowia(100);s__Metschnikowia_pulcherrima(100); | Metschnikowia | 13 | Saprotroph | \|Undefined Saprotroph\| | Yeast | NULL | Probable | As nectar-tap_saprotroph (Põlme et al. 2020); Decay sugar-rich substrates (Põlme et al. 2020) | Põlme S, et al. 2020. Fungal Diversity 105:1-16 ((https://doi.org/10.1007/s13225-020-00466-2)) |
| Otu22352 | 1 | k__Fungi(100);p__Ascomycota(100);c__Saccharomycetes(100);o__Saccharomycetales(100);f__Metschnikowiaceae(100);g__Metschnikowia(100);s__Metschnikowia_pulcherrima(100); | Metschnikowia | 13 | Saprotroph | \|Undefined Saprotroph\| | Yeast | NULL | Probable | As nectar-tap_saprotroph (Põlme et al. 2020); Decay sugar-rich substrates (Põlme et al. 2020) | Põlme S, et al. 2020. Fungal Diversity 105:1-16 ((https://doi.org/10.1007/s13225-020-00466-2)) |
| Otu22353 | 1 | k__Fungi(100);p__Ascomycota(100);c__Saccharomycetes(100);o__Saccharomycetales(100);f__Metschnikowiaceae(100);g__Metschnikowia(100);s__Metschnikowia_pulcherrima(100); | Metschnikowia | 13 | Saprotroph | \|Undefined Saprotroph\| | Yeast | NULL | Probable | As nectar-tap_saprotroph (Põlme et al. 2020); Decay sugar-rich substrates (Põlme et al. 2020) | Põlme S, et al. 2020. Fungal Diversity 105:1-16 ((https://doi.org/10.1007/s13225-020-00466-2)) |
| Otu22354 | 1 | k__Fungi(100);p__Ascomycota(100);c__Saccharomycetes(100);o__Saccharomycetales(100);f__Metschnikowiaceae(100);g__Metschnikowia(100);s__Metschnikowia_pulcherrima(100); | Metschnikowia | 13 | Saprotroph | \|Undefined Saprotroph\| | Yeast | NULL | Probable | As nectar-tap_saprotroph (Põlme et al. 2020); Decay sugar-rich substrates (Põlme et al. 2020) | Põlme S, et al. 2020. Fungal Diversity 105:1-16 ((https://doi.org/10.1007/s13225-020-00466-2)) |
| Otu22355 | 1 | k__Fungi(100);p__Ascomycota(100);c__Saccharomycetes(100);o__Saccharomycetales(100);f__Metschnikowiaceae(100);g__Metschnikowia(100);s__Metschnikowia_pulcherrima(100); | Metschnikowia | 13 | Saprotroph | \|Undefined Saprotroph\| | Yeast | NULL | Probable | As nectar-tap_saprotroph (Põlme et al. 2020); Decay sugar-rich substrates (Põlme et al. 2020) | Põlme S, et al. 2020. Fungal Diversity 105:1-16 ((https://doi.org/10.1007/s13225-020-00466-2)) |
| Otu22357 | 1 | k__Fungi(100);p__Ascomycota(100);c__Saccharomycetes(100);o__Saccharomycetales(100);f__Metschnikowiaceae(100);g__Metschnikowia(100);g__Metschnikowia_unclassified(100); | Metschnikowia | 13 | Saprotroph | \|Undefined Saprotroph\| | Yeast | NULL | Probable | As nectar-tap_saprotroph (Põlme et al. 2020); Decay sugar-rich substrates (Põlme et al. 2020) | Põlme S, et al. 2020. Fungal Diversity 105:1-16 ((https://doi.org/10.1007/s13225-020-00466-2)) |
| Otu22358 | 1 | k__Fungi(100);p__Ascomycota(100);c__Saccharomycetes(100);o__Saccharomycetales(100);f__Metschnikowiaceae(100);g__Metschnikowia(100);s__Metschnikowia_pulcherrima(100); | Metschnikowia | 13 | Saprotroph | \|Undefined Saprotroph\| | Yeast | NULL | Probable | As nectar-tap_saprotroph (Põlme et al. 2020); Decay sugar-rich substrates (Põlme et al. 2020) | Põlme S, et al. 2020. Fungal Diversity 105:1-16 ((https://doi.org/10.1007/s13225-020-00466-2)) |
| Otu22359 | 1 | k__Fungi(100);p__Ascomycota(100);c__Saccharomycetes(100);o__Saccharomycetales(100);f__Metschnikowiaceae(100);g__Metschnikowia(100);g__Metschnikowia_unclassified(100); | Metschnikowia | 13 | Saprotroph | \|Undefined Saprotroph\| | Yeast | NULL | Probable | As nectar-tap_saprotroph (Põlme et al. 2020); Decay sugar-rich substrates (Põlme et al. 2020) | Põlme S, et al. 2020. Fungal Diversity 105:1-16 ((https://doi.org/10.1007/s13225-020-00466-2)) |
| Otu22360 | 1 | k__Fungi(100);p__Ascomycota(100);c__Saccharomycetes(100);o__Saccharomycetales(100);f__Metschnikowiaceae(100);g__Metschnikowia(100);s__Metschnikowia_pulcherrima(100); | Metschnikowia | 13 | Saprotroph | \|Undefined Saprotroph\| | Yeast | NULL | Probable | As nectar-tap_saprotroph (Põlme et al. 2020); Decay sugar-rich substrates (Põlme et al. 2020) | Põlme S, et al. 2020. Fungal Diversity 105:1-16 ((https://doi.org/10.1007/s13225-020-00466-2)) |
| Otu22361 | 1 | k__Fungi(100);p__Ascomycota(100);c__Saccharomycetes(100);o__Saccharomycetales(100);f__Metschnikowiaceae(100);g__Metschnikowia(100);g__Metschnikowia_unclassified(100); | Metschnikowia | 13 | Saprotroph | \|Undefined Saprotroph\| | Yeast | NULL | Probable | As nectar-tap_saprotroph (Põlme et al. 2020); Decay sugar-rich substrates (Põlme et al. 2020) | Põlme S, et al. 2020. Fungal Diversity 105:1-16 ((https://doi.org/10.1007/s13225-020-00466-2)) |
| Otu22362 | 1 | k__Fungi(100);p__Ascomycota(100);c__Saccharomycetes(100);o__Saccharomycetales(100);f__Metschnikowiaceae(100);g__Metschnikowia(100);g__Metschnikowia_unclassified(100); | Metschnikowia | 13 | Saprotroph | \|Undefined Saprotroph\| | Yeast | NULL | Probable | As nectar-tap_saprotroph (Põlme et al. 2020); Decay sugar-rich substrates (Põlme et al. 2020) | Põlme S, et al. 2020. Fungal Diversity 105:1-16 ((https://doi.org/10.1007/s13225-020-00466-2)) |
| Otu22363 | 1 | k__Fungi(100);p__Ascomycota(100);c__Saccharomycetes(100);o__Saccharomycetales(100);f__Metschnikowiaceae(100);g__Metschnikowia(100);g__Metschnikowia_unclassified(100); | Metschnikowia | 13 | Saprotroph | \|Undefined Saprotroph\| | Yeast | NULL | Probable | As nectar-tap_saprotroph (Põlme et al. 2020); Decay sugar-rich substrates (Põlme et al. 2020) | Põlme S, et al. 2020. Fungal Diversity 105:1-16 ((https://doi.org/10.1007/s13225-020-00466-2)) |
| Otu22364 | 1 | k__Fungi(100);p__Ascomycota(100);c__Saccharomycetes(100);o__Saccharomycetales(100);f__Metschnikowiaceae(100);g__Metschnikowia(100);g__Metschnikowia_unclassified(100); | Metschnikowia | 13 | Saprotroph | \|Undefined Saprotroph\| | Yeast | NULL | Probable | As nectar-tap_saprotroph (Põlme et al. 2020); Decay sugar-rich substrates (Põlme et al. 2020) | Põlme S, et al. 2020. Fungal Diversity 105:1-16 ((https://doi.org/10.1007/s13225-020-00466-2)) |
| Otu22365 | 1 | k__Fungi(100);p__Ascomycota(100);c__Saccharomycetes(100);o__Saccharomycetales(100);f__Metschnikowiaceae(100);g__Metschnikowia(100);s__Metschnikowia_pulcherrima(100); | Metschnikowia | 13 | Saprotroph | \|Undefined Saprotroph\| | Yeast | NULL | Probable | As nectar-tap_saprotroph (Põlme et al. 2020); Decay sugar-rich substrates (Põlme et al. 2020) | Põlme S, et al. 2020. Fungal Diversity 105:1-16 ((https://doi.org/10.1007/s13225-020-00466-2)) |
| Otu22366 | 1 | k__Fungi(100);p__Ascomycota(100);c__Saccharomycetes(100);o__Saccharomycetales(100);f__Metschnikowiaceae(100);g__Metschnikowia(100);g__Metschnikowia_unclassified(100); | Metschnikowia | 13 | Saprotroph | \|Undefined Saprotroph\| | Yeast | NULL | Probable | As nectar-tap_saprotroph (Põlme et al. 2020); Decay sugar-rich substrates (Põlme et al. 2020) | Põlme S, et al. 2020. Fungal Diversity 105:1-16 ((https://doi.org/10.1007/s13225-020-00466-2)) |
| Otu22367 | 1 | k__Fungi(100);p__Ascomycota(100);c__Saccharomycetes(100);o__Saccharomycetales(100);f__Metschnikowiaceae(100);g__Metschnikowia(100);s__Metschnikowia_pulcherrima(100); | Metschnikowia | 13 | Saprotroph | \|Undefined Saprotroph\| | Yeast | NULL | Probable | As nectar-tap_saprotroph (Põlme et al. 2020); Decay sugar-rich substrates (Põlme et al. 2020) | Põlme S, et al. 2020. Fungal Diversity 105:1-16 ((https://doi.org/10.1007/s13225-020-00466-2)) |
| Otu22368 | 1 | k__Fungi(100);p__Ascomycota(100);c__Saccharomycetes(100);o__Saccharomycetales(100);f__Metschnikowiaceae(100);g__Metschnikowia(100);s__Metschnikowia_pulcherrima(100); | Metschnikowia | 13 | Saprotroph | \|Undefined Saprotroph\| | Yeast | NULL | Probable | As nectar-tap_saprotroph (Põlme et al. 2020); Decay sugar-rich substrates (Põlme et al. 2020) | Põlme S, et al. 2020. Fungal Diversity 105:1-16 ((https://doi.org/10.1007/s13225-020-00466-2)) |
| Otu22369 | 1 | k__Fungi(100);p__Ascomycota(100);c__Saccharomycetes(100);o__Saccharomycetales(100);f__Metschnikowiaceae(100);g__Metschnikowia(100);s__Metschnikowia_pulcherrima(100); | Metschnikowia | 13 | Saprotroph | \|Undefined Saprotroph\| | Yeast | NULL | Probable | As nectar-tap_saprotroph (Põlme et al. 2020); Decay sugar-rich substrates (Põlme et al. 2020) | Põlme S, et al. 2020. Fungal Diversity 105:1-16 ((https://doi.org/10.1007/s13225-020-00466-2)) |
| Otu22370 | 1 | k__Fungi(100);p__Ascomycota(100);c__Saccharomycetes(100);o__Saccharomycetales(100);f__Metschnikowiaceae(100);g__Metschnikowia(100);s__Metschnikowia_pulcherrima(100); | Metschnikowia | 13 | Saprotroph | \|Undefined Saprotroph\| | Yeast | NULL | Probable | As nectar-tap_saprotroph (Põlme et al. 2020); Decay sugar-rich substrates (Põlme et al. 2020) | Põlme S, et al. 2020. Fungal Diversity 105:1-16 ((https://doi.org/10.1007/s13225-020-00466-2)) |
| Otu22371 | 1 | k__Fungi(100);p__Ascomycota(100);c__Saccharomycetes(100);o__Saccharomycetales(100);f__Metschnikowiaceae(100);g__Metschnikowia(100);g__Metschnikowia_unclassified(100); | Metschnikowia | 13 | Saprotroph | \|Undefined Saprotroph\| | Yeast | NULL | Probable | As nectar-tap_saprotroph (Põlme et al. 2020); Decay sugar-rich substrates (Põlme et al. 2020) | Põlme S, et al. 2020. Fungal Diversity 105:1-16 ((https://doi.org/10.1007/s13225-020-00466-2)) |
| Otu22372 | 1 | k__Fungi(100);p__Ascomycota(100);c__Saccharomycetes(100);o__Saccharomycetales(100);f__Metschnikowiaceae(100);g__Metschnikowia(100);g__Metschnikowia_unclassified(100); | Metschnikowia | 13 | Saprotroph | \|Undefined Saprotroph\| | Yeast | NULL | Probable | As nectar-tap_saprotroph (Põlme et al. 2020); Decay sugar-rich substrates (Põlme et al. 2020) | Põlme S, et al. 2020. Fungal Diversity 105:1-16 ((https://doi.org/10.1007/s13225-020-00466-2)) |
| Otu22373 | 1 | k__Fungi(100);p__Ascomycota(100);c__Saccharomycetes(100);o__Saccharomycetales(100);f__Metschnikowiaceae(100);g__Metschnikowia(100);s__Metschnikowia_pulcherrima(100); | Metschnikowia | 13 | Saprotroph | \|Undefined Saprotroph\| | Yeast | NULL | Probable | As nectar-tap_saprotroph (Põlme et al. 2020); Decay sugar-rich substrates (Põlme et al. 2020) | Põlme S, et al. 2020. Fungal Diversity 105:1-16 ((https://doi.org/10.1007/s13225-020-00466-2)) |
| Otu22374 | 1 | k__Fungi(100);p__Ascomycota(100);c__Saccharomycetes(100);o__Saccharomycetales(100);f__Metschnikowiaceae(100);g__Metschnikowia(100);s__Metschnikowia_pulcherrima(100); | Metschnikowia | 13 | Saprotroph | \|Undefined Saprotroph\| | Yeast | NULL | Probable | As nectar-tap_saprotroph (Põlme et al. 2020); Decay sugar-rich substrates (Põlme et al. 2020) | Põlme S, et al. 2020. Fungal Diversity 105:1-16 ((https://doi.org/10.1007/s13225-020-00466-2)) |
| Otu22375 | 1 | k__Fungi(100);p__Ascomycota(100);c__Saccharomycetes(100);o__Saccharomycetales(100);f__Metschnikowiaceae(100);g__Metschnikowia(100);s__Metschnikowia_pulcherrima(100); | Metschnikowia | 13 | Saprotroph | \|Undefined Saprotroph\| | Yeast | NULL | Probable | As nectar-tap_saprotroph (Põlme et al. 2020); Decay sugar-rich substrates (Põlme et al. 2020) | Põlme S, et al. 2020. Fungal Diversity 105:1-16 ((https://doi.org/10.1007/s13225-020-00466-2)) |
| Otu22376 | 1 | k__Fungi(100);p__Ascomycota(100);c__Saccharomycetes(100);o__Saccharomycetales(100);f__Metschnikowiaceae(100);g__Metschnikowia(100);g__Metschnikowia_unclassified(100); | Metschnikowia | 13 | Saprotroph | \|Undefined Saprotroph\| | Yeast | NULL | Probable | As nectar-tap_saprotroph (Põlme et al. 2020); Decay sugar-rich substrates (Põlme et al. 2020) | Põlme S, et al. 2020. Fungal Diversity 105:1-16 ((https://doi.org/10.1007/s13225-020-00466-2)) |
| Otu22377 | 1 | k__Fungi(100);p__Ascomycota(100);c__Saccharomycetes(100);o__Saccharomycetales(100);f__Metschnikowiaceae(100);g__Metschnikowia(100);s__Metschnikowia_pulcherrima(100); | Metschnikowia | 13 | Saprotroph | \|Undefined Saprotroph\| | Yeast | NULL | Probable | As nectar-tap_saprotroph (Põlme et al. 2020); Decay sugar-rich substrates (Põlme et al. 2020) | Põlme S, et al. 2020. Fungal Diversity 105:1-16 ((https://doi.org/10.1007/s13225-020-00466-2)) |
| Otu22378 | 1 | k__Fungi(100);p__Ascomycota(100);c__Saccharomycetes(100);o__Saccharomycetales(100);f__Metschnikowiaceae(100);g__Metschnikowia(100);s__Metschnikowia_pulcherrima(100); | Metschnikowia | 13 | Saprotroph | \|Undefined Saprotroph\| | Yeast | NULL | Probable | As nectar-tap_saprotroph (Põlme et al. 2020); Decay sugar-rich substrates (Põlme et al. 2020) | Põlme S, et al. 2020. Fungal Diversity 105:1-16 ((https://doi.org/10.1007/s13225-020-00466-2)) |
| Otu22379 | 1 | k__Fungi(100);p__Ascomycota(100);c__Saccharomycetes(100);o__Saccharomycetales(100);f__Metschnikowiaceae(100);g__Metschnikowia(100);s__Metschnikowia_pulcherrima(100); | Metschnikowia | 13 | Saprotroph | \|Undefined Saprotroph\| | Yeast | NULL | Probable | As nectar-tap_saprotroph (Põlme et al. 2020); Decay sugar-rich substrates (Põlme et al. 2020) | Põlme S, et al. 2020. Fungal Diversity 105:1-16 ((https://doi.org/10.1007/s13225-020-00466-2)) |
| Otu22380 | 1 | k__Fungi(100);p__Ascomycota(100);c__Saccharomycetes(100);o__Saccharomycetales(100);f__Metschnikowiaceae(100);g__Metschnikowia(100);s__Metschnikowia_pulcherrima(100); | Metschnikowia | 13 | Saprotroph | \|Undefined Saprotroph\| | Yeast | NULL | Probable | As nectar-tap_saprotroph (Põlme et al. 2020); Decay sugar-rich substrates (Põlme et al. 2020) | Põlme S, et al. 2020. Fungal Diversity 105:1-16 ((https://doi.org/10.1007/s13225-020-00466-2)) |
| Otu22381 | 1 | k__Fungi(100);p__Ascomycota(100);c__Saccharomycetes(100);o__Saccharomycetales(100);f__Metschnikowiaceae(100);g__Metschnikowia(100);g__Metschnikowia_unclassified(100); | Metschnikowia | 13 | Saprotroph | \|Undefined Saprotroph\| | Yeast | NULL | Probable | As nectar-tap_saprotroph (Põlme et al. 2020); Decay sugar-rich substrates (Põlme et al. 2020) | Põlme S, et al. 2020. Fungal Diversity 105:1-16 ((https://doi.org/10.1007/s13225-020-00466-2)) |
| Otu22382 | 1 | k__Fungi(100);p__Ascomycota(100);c__Saccharomycetes(100);o__Saccharomycetales(100);f__Metschnikowiaceae(100);g__Metschnikowia(100);s__Metschnikowia_pulcherrima(100); | Metschnikowia | 13 | Saprotroph | \|Undefined Saprotroph\| | Yeast | NULL | Probable | As nectar-tap_saprotroph (Põlme et al. 2020); Decay sugar-rich substrates (Põlme et al. 2020) | Põlme S, et al. 2020. Fungal Diversity 105:1-16 ((https://doi.org/10.1007/s13225-020-00466-2)) |
| Otu22383 | 1 | k__Fungi(100);p__Ascomycota(100);c__Saccharomycetes(100);o__Saccharomycetales(100);f__Metschnikowiaceae(100);g__Metschnikowia(100);g__Metschnikowia_unclassified(100); | Metschnikowia | 13 | Saprotroph | \|Undefined Saprotroph\| | Yeast | NULL | Probable | As nectar-tap_saprotroph (Põlme et al. 2020); Decay sugar-rich substrates (Põlme et al. 2020) | Põlme S, et al. 2020. Fungal Diversity 105:1-16 ((https://doi.org/10.1007/s13225-020-00466-2)) |
| Otu22384 | 1 | k__Fungi(100);p__Ascomycota(100);c__Saccharomycetes(100);o__Saccharomycetales(100);f__Metschnikowiaceae(100);g__Metschnikowia(100);s__Metschnikowia_pulcherrima(100); | Metschnikowia | 13 | Saprotroph | \|Undefined Saprotroph\| | Yeast | NULL | Probable | As nectar-tap_saprotroph (Põlme et al. 2020); Decay sugar-rich substrates (Põlme et al. 2020) | Põlme S, et al. 2020. Fungal Diversity 105:1-16 ((https://doi.org/10.1007/s13225-020-00466-2)) |
| Otu22385 | 1 | k__Fungi(100);p__Ascomycota(100);c__Saccharomycetes(100);o__Saccharomycetales(100);f__Metschnikowiaceae(100);g__Metschnikowia(100);s__Metschnikowia_pulcherrima(100); | Metschnikowia | 13 | Saprotroph | \|Undefined Saprotroph\| | Yeast | NULL | Probable | As nectar-tap_saprotroph (Põlme et al. 2020); Decay sugar-rich substrates (Põlme et al. 2020) | Põlme S, et al. 2020. Fungal Diversity 105:1-16 ((https://doi.org/10.1007/s13225-020-00466-2)) |
| Otu22386 | 1 | k__Fungi(100);p__Ascomycota(100);c__Saccharomycetes(100);o__Saccharomycetales(100);f__Metschnikowiaceae(100);g__Metschnikowia(100);s__Metschnikowia_pulcherrima(100); | Metschnikowia | 13 | Saprotroph | \|Undefined Saprotroph\| | Yeast | NULL | Probable | As nectar-tap_saprotroph (Põlme et al. 2020); Decay sugar-rich substrates (Põlme et al. 2020) | Põlme S, et al. 2020. Fungal Diversity 105:1-16 ((https://doi.org/10.1007/s13225-020-00466-2)) |
| Otu22387 | 1 | k__Fungi(100);p__Ascomycota(100);c__Saccharomycetes(100);o__Saccharomycetales(100);f__Metschnikowiaceae(100);g__Metschnikowia(100);g__Metschnikowia_unclassified(100); | Metschnikowia | 13 | Saprotroph | \|Undefined Saprotroph\| | Yeast | NULL | Probable | As nectar-tap_saprotroph (Põlme et al. 2020); Decay sugar-rich substrates (Põlme et al. 2020) | Põlme S, et al. 2020. Fungal Diversity 105:1-16 ((https://doi.org/10.1007/s13225-020-00466-2)) |
| Otu22388 | 1 | k__Fungi(100);p__Ascomycota(100);c__Saccharomycetes(100);o__Saccharomycetales(100);f__Metschnikowiaceae(100);g__Metschnikowia(100);s__Metschnikowia_pulcherrima(100); | Metschnikowia | 13 | Saprotroph | \|Undefined Saprotroph\| | Yeast | NULL | Probable | As nectar-tap_saprotroph (Põlme et al. 2020); Decay sugar-rich substrates (Põlme et al. 2020) | Põlme S, et al. 2020. Fungal Diversity 105:1-16 ((https://doi.org/10.1007/s13225-020-00466-2)) |
| Otu22389 | 1 | k__Fungi(100);p__Ascomycota(100);c__Saccharomycetes(100);o__Saccharomycetales(100);f__Metschnikowiaceae(100);g__Metschnikowia(100);s__Metschnikowia_pulcherrima(100); | Metschnikowia | 13 | Saprotroph | \|Undefined Saprotroph\| | Yeast | NULL | Probable | As nectar-tap_saprotroph (Põlme et al. 2020); Decay sugar-rich substrates (Põlme et al. 2020) | Põlme S, et al. 2020. Fungal Diversity 105:1-16 ((https://doi.org/10.1007/s13225-020-00466-2)) |
| Otu22391 | 1 | k__Fungi(100);p__Ascomycota(100);c__Saccharomycetes(100);o__Saccharomycetales(100);f__Metschnikowiaceae(100);g__Metschnikowia(100);s__Metschnikowia_pulcherrima(100); | Metschnikowia | 13 | Saprotroph | \|Undefined Saprotroph\| | Yeast | NULL | Probable | As nectar-tap_saprotroph (Põlme et al. 2020); Decay sugar-rich substrates (Põlme et al. 2020) | Põlme S, et al. 2020. Fungal Diversity 105:1-16 ((https://doi.org/10.1007/s13225-020-00466-2)) |
| Otu22392 | 1 | k__Fungi(100);p__Ascomycota(100);c__Saccharomycetes(100);o__Saccharomycetales(100);f__Metschnikowiaceae(100);g__Metschnikowia(100);s__Metschnikowia_pulcherrima(100); | Metschnikowia | 13 | Saprotroph | \|Undefined Saprotroph\| | Yeast | NULL | Probable | As nectar-tap_saprotroph (Põlme et al. 2020); Decay sugar-rich substrates (Põlme et al. 2020) | Põlme S, et al. 2020. Fungal Diversity 105:1-16 ((https://doi.org/10.1007/s13225-020-00466-2)) |
| Otu22393 | 1 | k__Fungi(100);p__Ascomycota(100);c__Saccharomycetes(100);o__Saccharomycetales(100);f__Metschnikowiaceae(100);g__Metschnikowia(100);g__Metschnikowia_unclassified(100); | Metschnikowia | 13 | Saprotroph | \|Undefined Saprotroph\| | Yeast | NULL | Probable | As nectar-tap_saprotroph (Põlme et al. 2020); Decay sugar-rich substrates (Põlme et al. 2020) | Põlme S, et al. 2020. Fungal Diversity 105:1-16 ((https://doi.org/10.1007/s13225-020-00466-2)) |
| Otu22394 | 1 | k__Fungi(100);p__Ascomycota(100);c__Saccharomycetes(100);o__Saccharomycetales(100);f__Metschnikowiaceae(100);g__Metschnikowia(100);s__Metschnikowia_pulcherrima(100); | Metschnikowia | 13 | Saprotroph | \|Undefined Saprotroph\| | Yeast | NULL | Probable | As nectar-tap_saprotroph (Põlme et al. 2020); Decay sugar-rich substrates (Põlme et al. 2020) | Põlme S, et al. 2020. Fungal Diversity 105:1-16 ((https://doi.org/10.1007/s13225-020-00466-2)) |
| Otu22395 | 1 | k__Fungi(100);p__Ascomycota(100);c__Saccharomycetes(100);o__Saccharomycetales(100);f__Metschnikowiaceae(100);g__Metschnikowia(100);s__Metschnikowia_pulcherrima(100); | Metschnikowia | 13 | Saprotroph | \|Undefined Saprotroph\| | Yeast | NULL | Probable | As nectar-tap_saprotroph (Põlme et al. 2020); Decay sugar-rich substrates (Põlme et al. 2020) | Põlme S, et al. 2020. Fungal Diversity 105:1-16 ((https://doi.org/10.1007/s13225-020-00466-2)) |
| Otu22396 | 1 | k__Fungi(100);p__Ascomycota(100);c__Saccharomycetes(100);o__Saccharomycetales(100);f__Metschnikowiaceae(100);g__Metschnikowia(100);g__Metschnikowia_unclassified(100); | Metschnikowia | 13 | Saprotroph | \|Undefined Saprotroph\| | Yeast | NULL | Probable | As nectar-tap_saprotroph (Põlme et al. 2020); Decay sugar-rich substrates (Põlme et al. 2020) | Põlme S, et al. 2020. Fungal Diversity 105:1-16 ((https://doi.org/10.1007/s13225-020-00466-2)) |
| Otu22397 | 1 | k__Fungi(100);p__Ascomycota(100);c__Saccharomycetes(100);o__Saccharomycetales(100);f__Metschnikowiaceae(100);g__Metschnikowia(100);s__Metschnikowia_pulcherrima(100); | Metschnikowia | 13 | Saprotroph | \|Undefined Saprotroph\| | Yeast | NULL | Probable | As nectar-tap_saprotroph (Põlme et al. 2020); Decay sugar-rich substrates (Põlme et al. 2020) | Põlme S, et al. 2020. Fungal Diversity 105:1-16 ((https://doi.org/10.1007/s13225-020-00466-2)) |
| Otu22398 | 1 | k__Fungi(100);p__Ascomycota(100);c__Saccharomycetes(100);o__Saccharomycetales(100);f__Metschnikowiaceae(100);g__Metschnikowia(100);g__Metschnikowia_unclassified(100); | Metschnikowia | 13 | Saprotroph | \|Undefined Saprotroph\| | Yeast | NULL | Probable | As nectar-tap_saprotroph (Põlme et al. 2020); Decay sugar-rich substrates (Põlme et al. 2020) | Põlme S, et al. 2020. Fungal Diversity 105:1-16 ((https://doi.org/10.1007/s13225-020-00466-2)) |
| Otu22399 | 1 | k__Fungi(100);p__Ascomycota(100);c__Saccharomycetes(100);o__Saccharomycetales(100);f__Metschnikowiaceae(100);g__Metschnikowia(100);g__Metschnikowia_unclassified(100); | Metschnikowia | 13 | Saprotroph | \|Undefined Saprotroph\| | Yeast | NULL | Probable | As nectar-tap_saprotroph (Põlme et al. 2020); Decay sugar-rich substrates (Põlme et al. 2020) | Põlme S, et al. 2020. Fungal Diversity 105:1-16 ((https://doi.org/10.1007/s13225-020-00466-2)) |
| Otu22400 | 1 | k__Fungi(100);p__Ascomycota(100);c__Saccharomycetes(100);o__Saccharomycetales(100);f__Metschnikowiaceae(100);g__Metschnikowia(100);s__Metschnikowia_pulcherrima(100); | Metschnikowia | 13 | Saprotroph | \|Undefined Saprotroph\| | Yeast | NULL | Probable | As nectar-tap_saprotroph (Põlme et al. 2020); Decay sugar-rich substrates (Põlme et al. 2020) | Põlme S, et al. 2020. Fungal Diversity 105:1-16 ((https://doi.org/10.1007/s13225-020-00466-2)) |
| Otu22401 | 1 | k__Fungi(100);p__Ascomycota(100);c__Saccharomycetes(100);o__Saccharomycetales(100);f__Metschnikowiaceae(100);g__Metschnikowia(100);s__Metschnikowia_pulcherrima(100); | Metschnikowia | 13 | Saprotroph | \|Undefined Saprotroph\| | Yeast | NULL | Probable | As nectar-tap_saprotroph (Põlme et al. 2020); Decay sugar-rich substrates (Põlme et al. 2020) | Põlme S, et al. 2020. Fungal Diversity 105:1-16 ((https://doi.org/10.1007/s13225-020-00466-2)) |
| Otu22403 | 1 | k__Fungi(100);p__Ascomycota(100);c__Saccharomycetes(100);o__Saccharomycetales(100);f__Metschnikowiaceae(100);g__Metschnikowia(100);g__Metschnikowia_unclassified(100); | Metschnikowia | 13 | Saprotroph | \|Undefined Saprotroph\| | Yeast | NULL | Probable | As nectar-tap_saprotroph (Põlme et al. 2020); Decay sugar-rich substrates (Põlme et al. 2020) | Põlme S, et al. 2020. Fungal Diversity 105:1-16 ((https://doi.org/10.1007/s13225-020-00466-2)) |
| Otu22404 | 1 | k__Fungi(100);p__Ascomycota(100);c__Saccharomycetes(100);o__Saccharomycetales(100);f__Saccharomycetales_family_Incertae_sedis(100);g__Candida(100);g__Candida_unclassified(100); | Candida | 13 | Pathotroph-Saprotroph-Symbiotroph | Animal Pathogen-Endophyte-Endosymbiont-Epiphyte-\|Undefined Saprotroph\| | Dimorphic Yeast-Yeast | NULL | Probable | As nectar-tap_saprotroph (Põlme et al. 2020); Decay sugar-rich substrates (Põlme et al. 2020); This asexual genus is polyphyletic and contain species that function very differently depend on what they are. Caution should be taken when interpreting guilds for Candida species. | Manolakaki D, et al. 2010. Virulence. 1: 367-375 ((https://doi.org/10.4161/viru.1.5.12796)); Mekbib SB, et al. 2011. Tropical Plant Pathology 36:233-240 ((https://doi.org/10.1590/S1982-56762011000400004)); Põlme S, et al. 2020. Fungal Diversity 105:1-16 ((https://doi.org/10.1007/s13225-020-00466-2)) |
| Otu22406 | 1 | k__Fungi(100);p__Ascomycota(100);c__Saccharomycetes(100);o__Saccharomycetales(100);f__Metschnikowiaceae(100);g__Metschnikowia(100);s__Metschnikowia_pulcherrima(100); | Metschnikowia | 13 | Saprotroph | \|Undefined Saprotroph\| | Yeast | NULL | Probable | As nectar-tap_saprotroph (Põlme et al. 2020); Decay sugar-rich substrates (Põlme et al. 2020) | Põlme S, et al. 2020. Fungal Diversity 105:1-16 ((https://doi.org/10.1007/s13225-020-00466-2)) |
| Otu22407 | 1 | k__Fungi(100);p__Ascomycota(100);c__Saccharomycetes(100);o__Saccharomycetales(100);f__Metschnikowiaceae(100);g__Metschnikowia(100);s__Metschnikowia_pulcherrima(100); | Metschnikowia | 13 | Saprotroph | \|Undefined Saprotroph\| | Yeast | NULL | Probable | As nectar-tap_saprotroph (Põlme et al. 2020); Decay sugar-rich substrates (Põlme et al. 2020) | Põlme S, et al. 2020. Fungal Diversity 105:1-16 ((https://doi.org/10.1007/s13225-020-00466-2)) |
| Otu22408 | 1 | k__Fungi(100);p__Ascomycota(100);c__Saccharomycetes(100);o__Saccharomycetales(100);f__Metschnikowiaceae(100);g__Metschnikowia(100);s__Metschnikowia_pulcherrima(100); | Metschnikowia | 13 | Saprotroph | \|Undefined Saprotroph\| | Yeast | NULL | Probable | As nectar-tap_saprotroph (Põlme et al. 2020); Decay sugar-rich substrates (Põlme et al. 2020) | Põlme S, et al. 2020. Fungal Diversity 105:1-16 ((https://doi.org/10.1007/s13225-020-00466-2)) |
| Otu22409 | 1 | k__Fungi(100);p__Ascomycota(100);c__Saccharomycetes(100);o__Saccharomycetales(100);f__Metschnikowiaceae(100);g__Metschnikowia(100);s__Metschnikowia_pulcherrima(100); | Metschnikowia | 13 | Saprotroph | \|Undefined Saprotroph\| | Yeast | NULL | Probable | As nectar-tap_saprotroph (Põlme et al. 2020); Decay sugar-rich substrates (Põlme et al. 2020) | Põlme S, et al. 2020. Fungal Diversity 105:1-16 ((https://doi.org/10.1007/s13225-020-00466-2)) |
| Otu22412 | 1 | k__Fungi(100);p__Ascomycota(100);c__Saccharomycetes(100);o__Saccharomycetales(100);f__Metschnikowiaceae(100);g__Metschnikowia(100);g__Metschnikowia_unclassified(100); | Metschnikowia | 13 | Saprotroph | \|Undefined Saprotroph\| | Yeast | NULL | Probable | As nectar-tap_saprotroph (Põlme et al. 2020); Decay sugar-rich substrates (Põlme et al. 2020) | Põlme S, et al. 2020. Fungal Diversity 105:1-16 ((https://doi.org/10.1007/s13225-020-00466-2)) |
| Otu22413 | 1 | k__Fungi(100);p__Ascomycota(100);c__Saccharomycetes(100);o__Saccharomycetales(100);f__Metschnikowiaceae(100);g__Metschnikowia(100);s__Metschnikowia_pulcherrima(100); | Metschnikowia | 13 | Saprotroph | \|Undefined Saprotroph\| | Yeast | NULL | Probable | As nectar-tap_saprotroph (Põlme et al. 2020); Decay sugar-rich substrates (Põlme et al. 2020) | Põlme S, et al. 2020. Fungal Diversity 105:1-16 ((https://doi.org/10.1007/s13225-020-00466-2)) |
| Otu22414 | 1 | k__Fungi(100);p__Ascomycota(100);c__Saccharomycetes(100);o__Saccharomycetales(100);f__Saccharomycetales_family_Incertae_sedis(100);g__Candida(100);g__Candida_unclassified(100); | Candida | 13 | Pathotroph-Saprotroph-Symbiotroph | Animal Pathogen-Endophyte-Endosymbiont-Epiphyte-\|Undefined Saprotroph\| | Dimorphic Yeast-Yeast | NULL | Probable | As nectar-tap_saprotroph (Põlme et al. 2020); Decay sugar-rich substrates (Põlme et al. 2020); This asexual genus is polyphyletic and contain species that function very differently depend on what they are. Caution should be taken when interpreting guilds for Candida species. | Manolakaki D, et al. 2010. Virulence. 1: 367-375 ((https://doi.org/10.4161/viru.1.5.12796)); Mekbib SB, et al. 2011. Tropical Plant Pathology 36:233-240 ((https://doi.org/10.1590/S1982-56762011000400004)); Põlme S, et al. 2020. Fungal Diversity 105:1-16 ((https://doi.org/10.1007/s13225-020-00466-2)) |
| Otu22415 | 1 | k__Fungi(100);p__Ascomycota(100);c__Saccharomycetes(100);o__Saccharomycetales(100);f__Metschnikowiaceae(100);g__Metschnikowia(100);s__Metschnikowia_pulcherrima(100); | Metschnikowia | 13 | Saprotroph | \|Undefined Saprotroph\| | Yeast | NULL | Probable | As nectar-tap_saprotroph (Põlme et al. 2020); Decay sugar-rich substrates (Põlme et al. 2020) | Põlme S, et al. 2020. Fungal Diversity 105:1-16 ((https://doi.org/10.1007/s13225-020-00466-2)) |
| Otu22416 | 1 | k__Fungi(100);p__Ascomycota(100);c__Saccharomycetes(100);o__Saccharomycetales(100);f__Metschnikowiaceae(100);g__Metschnikowia(100);s__Metschnikowia_pulcherrima(100); | Metschnikowia | 13 | Saprotroph | \|Undefined Saprotroph\| | Yeast | NULL | Probable | As nectar-tap_saprotroph (Põlme et al. 2020); Decay sugar-rich substrates (Põlme et al. 2020) | Põlme S, et al. 2020. Fungal Diversity 105:1-16 ((https://doi.org/10.1007/s13225-020-00466-2)) |
| Otu22417 | 1 | k__Fungi(100);p__Ascomycota(100);c__Saccharomycetes(100);o__Saccharomycetales(100);f__Metschnikowiaceae(100);g__Metschnikowia(100);g__Metschnikowia_unclassified(100); | Metschnikowia | 13 | Saprotroph | \|Undefined Saprotroph\| | Yeast | NULL | Probable | As nectar-tap_saprotroph (Põlme et al. 2020); Decay sugar-rich substrates (Põlme et al. 2020) | Põlme S, et al. 2020. Fungal Diversity 105:1-16 ((https://doi.org/10.1007/s13225-020-00466-2)) |
| Otu22418 | 1 | k__Fungi(100);p__Ascomycota(100);c__Saccharomycetes(100);o__Saccharomycetales(100);f__Metschnikowiaceae(100);g__Metschnikowia(100);s__Metschnikowia_pulcherrima(100); | Metschnikowia | 13 | Saprotroph | \|Undefined Saprotroph\| | Yeast | NULL | Probable | As nectar-tap_saprotroph (Põlme et al. 2020); Decay sugar-rich substrates (Põlme et al. 2020) | Põlme S, et al. 2020. Fungal Diversity 105:1-16 ((https://doi.org/10.1007/s13225-020-00466-2)) |
| Otu22419 | 1 | k__Fungi(100);p__Ascomycota(100);c__Saccharomycetes(100);o__Saccharomycetales(100);f__Metschnikowiaceae(100);g__Metschnikowia(100);s__Metschnikowia_pulcherrima(100); | Metschnikowia | 13 | Saprotroph | \|Undefined Saprotroph\| | Yeast | NULL | Probable | As nectar-tap_saprotroph (Põlme et al. 2020); Decay sugar-rich substrates (Põlme et al. 2020) | Põlme S, et al. 2020. Fungal Diversity 105:1-16 ((https://doi.org/10.1007/s13225-020-00466-2)) |
| Otu22420 | 1 | k__Fungi(100);p__Ascomycota(100);c__Saccharomycetes(100);o__Saccharomycetales(100);f__Metschnikowiaceae(100);g__Metschnikowia(100);s__Metschnikowia_pulcherrima(100); | Metschnikowia | 13 | Saprotroph | \|Undefined Saprotroph\| | Yeast | NULL | Probable | As nectar-tap_saprotroph (Põlme et al. 2020); Decay sugar-rich substrates (Põlme et al. 2020) | Põlme S, et al. 2020. Fungal Diversity 105:1-16 ((https://doi.org/10.1007/s13225-020-00466-2)) |
| Otu22421 | 1 | k__Fungi(100);p__Ascomycota(100);c__Saccharomycetes(100);o__Saccharomycetales(100);f__Metschnikowiaceae(100);g__Metschnikowia(100);s__Metschnikowia_pulcherrima(100); | Metschnikowia | 13 | Saprotroph | \|Undefined Saprotroph\| | Yeast | NULL | Probable | As nectar-tap_saprotroph (Põlme et al. 2020); Decay sugar-rich substrates (Põlme et al. 2020) | Põlme S, et al. 2020. Fungal Diversity 105:1-16 ((https://doi.org/10.1007/s13225-020-00466-2)) |
| Otu22422 | 1 | k__Fungi(100);p__Ascomycota(100);c__Saccharomycetes(100);o__Saccharomycetales(100);f__Metschnikowiaceae(100);g__Metschnikowia(100);s__Metschnikowia_pulcherrima(100); | Metschnikowia | 13 | Saprotroph | \|Undefined Saprotroph\| | Yeast | NULL | Probable | As nectar-tap_saprotroph (Põlme et al. 2020); Decay sugar-rich substrates (Põlme et al. 2020) | Põlme S, et al. 2020. Fungal Diversity 105:1-16 ((https://doi.org/10.1007/s13225-020-00466-2)) |
| Otu22423 | 1 | k__Fungi(100);p__Ascomycota(100);c__Saccharomycetes(100);o__Saccharomycetales(100);f__Metschnikowiaceae(100);g__Metschnikowia(100);g__Metschnikowia_unclassified(100); | Metschnikowia | 13 | Saprotroph | \|Undefined Saprotroph\| | Yeast | NULL | Probable | As nectar-tap_saprotroph (Põlme et al. 2020); Decay sugar-rich substrates (Põlme et al. 2020) | Põlme S, et al. 2020. Fungal Diversity 105:1-16 ((https://doi.org/10.1007/s13225-020-00466-2)) |
| Otu22424 | 1 | k__Fungi(100);p__Ascomycota(100);c__Saccharomycetes(100);o__Saccharomycetales(100);f__Metschnikowiaceae(100);g__Metschnikowia(100);g__Metschnikowia_unclassified(100); | Metschnikowia | 13 | Saprotroph | \|Undefined Saprotroph\| | Yeast | NULL | Probable | As nectar-tap_saprotroph (Põlme et al. 2020); Decay sugar-rich substrates (Põlme et al. 2020) | Põlme S, et al. 2020. Fungal Diversity 105:1-16 ((https://doi.org/10.1007/s13225-020-00466-2)) |
| Otu22428 | 1 | k__Fungi(100);p__Ascomycota(100);c__Saccharomycetes(100);o__Saccharomycetales(100);f__Metschnikowiaceae(100);g__Metschnikowia(100);s__Metschnikowia_pulcherrima(100); | Metschnikowia | 13 | Saprotroph | \|Undefined Saprotroph\| | Yeast | NULL | Probable | As nectar-tap_saprotroph (Põlme et al. 2020); Decay sugar-rich substrates (Põlme et al. 2020) | Põlme S, et al. 2020. Fungal Diversity 105:1-16 ((https://doi.org/10.1007/s13225-020-00466-2)) |
| Otu22429 | 1 | k__Fungi(100);p__Ascomycota(100);c__Saccharomycetes(100);o__Saccharomycetales(100);f__Metschnikowiaceae(100);g__Metschnikowia(100);g__Metschnikowia_unclassified(100); | Metschnikowia | 13 | Saprotroph | \|Undefined Saprotroph\| | Yeast | NULL | Probable | As nectar-tap_saprotroph (Põlme et al. 2020); Decay sugar-rich substrates (Põlme et al. 2020) | Põlme S, et al. 2020. Fungal Diversity 105:1-16 ((https://doi.org/10.1007/s13225-020-00466-2)) |
| Otu22430 | 1 | k__Fungi(100);p__Ascomycota(100);c__Saccharomycetes(100);o__Saccharomycetales(100);f__Metschnikowiaceae(100);g__Metschnikowia(100);g__Metschnikowia_unclassified(100); | Metschnikowia | 13 | Saprotroph | \|Undefined Saprotroph\| | Yeast | NULL | Probable | As nectar-tap_saprotroph (Põlme et al. 2020); Decay sugar-rich substrates (Põlme et al. 2020) | Põlme S, et al. 2020. Fungal Diversity 105:1-16 ((https://doi.org/10.1007/s13225-020-00466-2)) |
| Otu22431 | 1 | k__Fungi(100);p__Ascomycota(100);c__Saccharomycetes(100);o__Saccharomycetales(100);f__Metschnikowiaceae(100);g__Metschnikowia(100);g__Metschnikowia_unclassified(100); | Metschnikowia | 13 | Saprotroph | \|Undefined Saprotroph\| | Yeast | NULL | Probable | As nectar-tap_saprotroph (Põlme et al. 2020); Decay sugar-rich substrates (Põlme et al. 2020) | Põlme S, et al. 2020. Fungal Diversity 105:1-16 ((https://doi.org/10.1007/s13225-020-00466-2)) |
| Otu22432 | 1 | k__Fungi(100);p__Ascomycota(100);c__Saccharomycetes(100);o__Saccharomycetales(100);f__Metschnikowiaceae(100);g__Metschnikowia(100);s__Metschnikowia_pulcherrima(100); | Metschnikowia | 13 | Saprotroph | \|Undefined Saprotroph\| | Yeast | NULL | Probable | As nectar-tap_saprotroph (Põlme et al. 2020); Decay sugar-rich substrates (Põlme et al. 2020) | Põlme S, et al. 2020. Fungal Diversity 105:1-16 ((https://doi.org/10.1007/s13225-020-00466-2)) |
| Otu22433 | 1 | k__Fungi(100);p__Ascomycota(100);c__Saccharomycetes(100);o__Saccharomycetales(100);f__Metschnikowiaceae(100);g__Metschnikowia(100);s__Metschnikowia_pulcherrima(100); | Metschnikowia | 13 | Saprotroph | \|Undefined Saprotroph\| | Yeast | NULL | Probable | As nectar-tap_saprotroph (Põlme et al. 2020); Decay sugar-rich substrates (Põlme et al. 2020) | Põlme S, et al. 2020. Fungal Diversity 105:1-16 ((https://doi.org/10.1007/s13225-020-00466-2)) |
| Otu22434 | 1 | k__Fungi(100);p__Ascomycota(100);c__Saccharomycetes(100);o__Saccharomycetales(100);f__Metschnikowiaceae(100);g__Metschnikowia(100);s__Metschnikowia_pulcherrima(100); | Metschnikowia | 13 | Saprotroph | \|Undefined Saprotroph\| | Yeast | NULL | Probable | As nectar-tap_saprotroph (Põlme et al. 2020); Decay sugar-rich substrates (Põlme et al. 2020) | Põlme S, et al. 2020. Fungal Diversity 105:1-16 ((https://doi.org/10.1007/s13225-020-00466-2)) |
| Otu22435 | 1 | k__Fungi(100);p__Ascomycota(100);c__Saccharomycetes(100);o__Saccharomycetales(100);f__Metschnikowiaceae(100);g__Metschnikowia(100);g__Metschnikowia_unclassified(100); | Metschnikowia | 13 | Saprotroph | \|Undefined Saprotroph\| | Yeast | NULL | Probable | As nectar-tap_saprotroph (Põlme et al. 2020); Decay sugar-rich substrates (Põlme et al. 2020) | Põlme S, et al. 2020. Fungal Diversity 105:1-16 ((https://doi.org/10.1007/s13225-020-00466-2)) |
| Otu22436 | 1 | k__Fungi(100);p__Ascomycota(100);c__Saccharomycetes(100);o__Saccharomycetales(100);f__Metschnikowiaceae(100);g__Metschnikowia(100);s__Metschnikowia_pulcherrima(100); | Metschnikowia | 13 | Saprotroph | \|Undefined Saprotroph\| | Yeast | NULL | Probable | As nectar-tap_saprotroph (Põlme et al. 2020); Decay sugar-rich substrates (Põlme et al. 2020) | Põlme S, et al. 2020. Fungal Diversity 105:1-16 ((https://doi.org/10.1007/s13225-020-00466-2)) |
| Otu22437 | 1 | k__Fungi(100);p__Ascomycota(100);c__Saccharomycetes(100);o__Saccharomycetales(100);f__Metschnikowiaceae(100);g__Metschnikowia(100);s__Metschnikowia_pulcherrima(100); | Metschnikowia | 13 | Saprotroph | \|Undefined Saprotroph\| | Yeast | NULL | Probable | As nectar-tap_saprotroph (Põlme et al. 2020); Decay sugar-rich substrates (Põlme et al. 2020) | Põlme S, et al. 2020. Fungal Diversity 105:1-16 ((https://doi.org/10.1007/s13225-020-00466-2)) |
| Otu22438 | 1 | k__Fungi(100);p__Ascomycota(100);c__Saccharomycetes(100);o__Saccharomycetales(100);f__Metschnikowiaceae(100);g__Metschnikowia(100);s__Metschnikowia_pulcherrima(100); | Metschnikowia | 13 | Saprotroph | \|Undefined Saprotroph\| | Yeast | NULL | Probable | As nectar-tap_saprotroph (Põlme et al. 2020); Decay sugar-rich substrates (Põlme et al. 2020) | Põlme S, et al. 2020. Fungal Diversity 105:1-16 ((https://doi.org/10.1007/s13225-020-00466-2)) |
| Otu22439 | 1 | k__Fungi(100);p__Ascomycota(100);c__Saccharomycetes(100);o__Saccharomycetales(100);f__Metschnikowiaceae(100);g__Metschnikowia(100);s__Metschnikowia_pulcherrima(100); | Metschnikowia | 13 | Saprotroph | \|Undefined Saprotroph\| | Yeast | NULL | Probable | As nectar-tap_saprotroph (Põlme et al. 2020); Decay sugar-rich substrates (Põlme et al. 2020) | Põlme S, et al. 2020. Fungal Diversity 105:1-16 ((https://doi.org/10.1007/s13225-020-00466-2)) |
| Otu22440 | 1 | k__Fungi(100);p__Ascomycota(100);c__Saccharomycetes(100);o__Saccharomycetales(100);f__Metschnikowiaceae(100);g__Metschnikowia(100);s__Metschnikowia_pulcherrima(100); | Metschnikowia | 13 | Saprotroph | \|Undefined Saprotroph\| | Yeast | NULL | Probable | As nectar-tap_saprotroph (Põlme et al. 2020); Decay sugar-rich substrates (Põlme et al. 2020) | Põlme S, et al. 2020. Fungal Diversity 105:1-16 ((https://doi.org/10.1007/s13225-020-00466-2)) |
| Otu22442 | 1 | k__Fungi(100);p__Ascomycota(100);c__Saccharomycetes(100);o__Saccharomycetales(100);f__Metschnikowiaceae(100);g__Metschnikowia(100);s__Metschnikowia_pulcherrima(100); | Metschnikowia | 13 | Saprotroph | \|Undefined Saprotroph\| | Yeast | NULL | Probable | As nectar-tap_saprotroph (Põlme et al. 2020); Decay sugar-rich substrates (Põlme et al. 2020) | Põlme S, et al. 2020. Fungal Diversity 105:1-16 ((https://doi.org/10.1007/s13225-020-00466-2)) |
| Otu22443 | 1 | k__Fungi(100);p__Ascomycota(100);c__Saccharomycetes(100);o__Saccharomycetales(100);f__Metschnikowiaceae(100);g__Metschnikowia(100);s__Metschnikowia_pulcherrima(100); | Metschnikowia | 13 | Saprotroph | \|Undefined Saprotroph\| | Yeast | NULL | Probable | As nectar-tap_saprotroph (Põlme et al. 2020); Decay sugar-rich substrates (Põlme et al. 2020) | Põlme S, et al. 2020. Fungal Diversity 105:1-16 ((https://doi.org/10.1007/s13225-020-00466-2)) |
| Otu22444 | 1 | k__Fungi(100);p__Ascomycota(100);c__Saccharomycetes(100);o__Saccharomycetales(100);f__Metschnikowiaceae(100);g__Metschnikowia(100);s__Metschnikowia_pulcherrima(100); | Metschnikowia | 13 | Saprotroph | \|Undefined Saprotroph\| | Yeast | NULL | Probable | As nectar-tap_saprotroph (Põlme et al. 2020); Decay sugar-rich substrates (Põlme et al. 2020) | Põlme S, et al. 2020. Fungal Diversity 105:1-16 ((https://doi.org/10.1007/s13225-020-00466-2)) |
| Otu22445 | 1 | k__Fungi(100);p__Ascomycota(100);c__Saccharomycetes(100);o__Saccharomycetales(100);f__Metschnikowiaceae(100);g__Metschnikowia(100);g__Metschnikowia_unclassified(100); | Metschnikowia | 13 | Saprotroph | \|Undefined Saprotroph\| | Yeast | NULL | Probable | As nectar-tap_saprotroph (Põlme et al. 2020); Decay sugar-rich substrates (Põlme et al. 2020) | Põlme S, et al. 2020. Fungal Diversity 105:1-16 ((https://doi.org/10.1007/s13225-020-00466-2)) |
| Otu22447 | 1 | k__Fungi(100);p__Ascomycota(100);c__Saccharomycetes(100);o__Saccharomycetales(100);f__Metschnikowiaceae(100);g__Metschnikowia(100);s__Metschnikowia_pulcherrima(100); | Metschnikowia | 13 | Saprotroph | \|Undefined Saprotroph\| | Yeast | NULL | Probable | As nectar-tap_saprotroph (Põlme et al. 2020); Decay sugar-rich substrates (Põlme et al. 2020) | Põlme S, et al. 2020. Fungal Diversity 105:1-16 ((https://doi.org/10.1007/s13225-020-00466-2)) |
| Otu22448 | 1 | k__Fungi(100);p__Ascomycota(100);c__Saccharomycetes(100);o__Saccharomycetales(100);f__Metschnikowiaceae(100);g__Metschnikowia(100);s__Metschnikowia_pulcherrima(100); | Metschnikowia | 13 | Saprotroph | \|Undefined Saprotroph\| | Yeast | NULL | Probable | As nectar-tap_saprotroph (Põlme et al. 2020); Decay sugar-rich substrates (Põlme et al. 2020) | Põlme S, et al. 2020. Fungal Diversity 105:1-16 ((https://doi.org/10.1007/s13225-020-00466-2)) |
| Otu22449 | 1 | k__Fungi(100);p__Ascomycota(100);c__Saccharomycetes(100);o__Saccharomycetales(100);f__Metschnikowiaceae(100);g__Metschnikowia(100);s__Metschnikowia_pulcherrima(100); | Metschnikowia | 13 | Saprotroph | \|Undefined Saprotroph\| | Yeast | NULL | Probable | As nectar-tap_saprotroph (Põlme et al. 2020); Decay sugar-rich substrates (Põlme et al. 2020) | Põlme S, et al. 2020. Fungal Diversity 105:1-16 ((https://doi.org/10.1007/s13225-020-00466-2)) |
| Otu22450 | 1 | k__Fungi(100);p__Ascomycota(100);c__Saccharomycetes(100);o__Saccharomycetales(100);f__Metschnikowiaceae(100);g__Metschnikowia(100);s__Metschnikowia_pulcherrima(100); | Metschnikowia | 13 | Saprotroph | \|Undefined Saprotroph\| | Yeast | NULL | Probable | As nectar-tap_saprotroph (Põlme et al. 2020); Decay sugar-rich substrates (Põlme et al. 2020) | Põlme S, et al. 2020. Fungal Diversity 105:1-16 ((https://doi.org/10.1007/s13225-020-00466-2)) |
| Otu22451 | 1 | k__Fungi(100);p__Ascomycota(100);c__Saccharomycetes(100);o__Saccharomycetales(100);f__Metschnikowiaceae(100);g__Metschnikowia(100);s__Metschnikowia_pulcherrima(100); | Metschnikowia | 13 | Saprotroph | \|Undefined Saprotroph\| | Yeast | NULL | Probable | As nectar-tap_saprotroph (Põlme et al. 2020); Decay sugar-rich substrates (Põlme et al. 2020) | Põlme S, et al. 2020. Fungal Diversity 105:1-16 ((https://doi.org/10.1007/s13225-020-00466-2)) |
| Otu22452 | 1 | k__Fungi(100);p__Ascomycota(100);c__Saccharomycetes(100);o__Saccharomycetales(100);f__Metschnikowiaceae(100);g__Metschnikowia(100);s__Metschnikowia_pulcherrima(100); | Metschnikowia | 13 | Saprotroph | \|Undefined Saprotroph\| | Yeast | NULL | Probable | As nectar-tap_saprotroph (Põlme et al. 2020); Decay sugar-rich substrates (Põlme et al. 2020) | Põlme S, et al. 2020. Fungal Diversity 105:1-16 ((https://doi.org/10.1007/s13225-020-00466-2)) |
| Otu22453 | 1 | k__Fungi(100);p__Ascomycota(100);c__Saccharomycetes(100);o__Saccharomycetales(100);f__Metschnikowiaceae(100);g__Metschnikowia(100);g__Metschnikowia_unclassified(100); | Metschnikowia | 13 | Saprotroph | \|Undefined Saprotroph\| | Yeast | NULL | Probable | As nectar-tap_saprotroph (Põlme et al. 2020); Decay sugar-rich substrates (Põlme et al. 2020) | Põlme S, et al. 2020. Fungal Diversity 105:1-16 ((https://doi.org/10.1007/s13225-020-00466-2)) |
| Otu22454 | 1 | k__Fungi(100);p__Ascomycota(100);c__Saccharomycetes(100);o__Saccharomycetales(100);f__Metschnikowiaceae(100);g__Metschnikowia(100);g__Metschnikowia_unclassified(100); | Metschnikowia | 13 | Saprotroph | \|Undefined Saprotroph\| | Yeast | NULL | Probable | As nectar-tap_saprotroph (Põlme et al. 2020); Decay sugar-rich substrates (Põlme et al. 2020) | Põlme S, et al. 2020. Fungal Diversity 105:1-16 ((https://doi.org/10.1007/s13225-020-00466-2)) |
| Otu22455 | 1 | k__Fungi(100);p__Ascomycota(100);c__Saccharomycetes(100);o__Saccharomycetales(100);f__Metschnikowiaceae(100);g__Metschnikowia(100);s__Metschnikowia_pulcherrima(100); | Metschnikowia | 13 | Saprotroph | \|Undefined Saprotroph\| | Yeast | NULL | Probable | As nectar-tap_saprotroph (Põlme et al. 2020); Decay sugar-rich substrates (Põlme et al. 2020) | Põlme S, et al. 2020. Fungal Diversity 105:1-16 ((https://doi.org/10.1007/s13225-020-00466-2)) |
| Otu22456 | 1 | k__Fungi(100);p__Ascomycota(100);c__Saccharomycetes(100);o__Saccharomycetales(100);f__Metschnikowiaceae(100);g__Metschnikowia(100);s__Metschnikowia_pulcherrima(100); | Metschnikowia | 13 | Saprotroph | \|Undefined Saprotroph\| | Yeast | NULL | Probable | As nectar-tap_saprotroph (Põlme et al. 2020); Decay sugar-rich substrates (Põlme et al. 2020) | Põlme S, et al. 2020. Fungal Diversity 105:1-16 ((https://doi.org/10.1007/s13225-020-00466-2)) |
| Otu22457 | 1 | k__Fungi(100);p__Ascomycota(100);c__Saccharomycetes(100);o__Saccharomycetales(100);f__Metschnikowiaceae(100);g__Metschnikowia(100);g__Metschnikowia_unclassified(100); | Metschnikowia | 13 | Saprotroph | \|Undefined Saprotroph\| | Yeast | NULL | Probable | As nectar-tap_saprotroph (Põlme et al. 2020); Decay sugar-rich substrates (Põlme et al. 2020) | Põlme S, et al. 2020. Fungal Diversity 105:1-16 ((https://doi.org/10.1007/s13225-020-00466-2)) |
| Otu22458 | 1 | k__Fungi(100);p__Ascomycota(100);c__Saccharomycetes(100);o__Saccharomycetales(100);f__Metschnikowiaceae(100);g__Metschnikowia(100);s__Metschnikowia_pulcherrima(100); | Metschnikowia | 13 | Saprotroph | \|Undefined Saprotroph\| | Yeast | NULL | Probable | As nectar-tap_saprotroph (Põlme et al. 2020); Decay sugar-rich substrates (Põlme et al. 2020) | Põlme S, et al. 2020. Fungal Diversity 105:1-16 ((https://doi.org/10.1007/s13225-020-00466-2)) |
| Otu22459 | 1 | k__Fungi(100);p__Ascomycota(100);c__Saccharomycetes(100);o__Saccharomycetales(100);f__Metschnikowiaceae(100);g__Metschnikowia(100);s__Metschnikowia_pulcherrima(100); | Metschnikowia | 13 | Saprotroph | \|Undefined Saprotroph\| | Yeast | NULL | Probable | As nectar-tap_saprotroph (Põlme et al. 2020); Decay sugar-rich substrates (Põlme et al. 2020) | Põlme S, et al. 2020. Fungal Diversity 105:1-16 ((https://doi.org/10.1007/s13225-020-00466-2)) |
| Otu22461 | 1 | k__Fungi(100);p__Ascomycota(100);c__Saccharomycetes(100);o__Saccharomycetales(100);f__Metschnikowiaceae(100);g__Metschnikowia(100);s__Metschnikowia_pulcherrima(100); | Metschnikowia | 13 | Saprotroph | \|Undefined Saprotroph\| | Yeast | NULL | Probable | As nectar-tap_saprotroph (Põlme et al. 2020); Decay sugar-rich substrates (Põlme et al. 2020) | Põlme S, et al. 2020. Fungal Diversity 105:1-16 ((https://doi.org/10.1007/s13225-020-00466-2)) |
| Otu22462 | 1 | k__Fungi(100);p__Ascomycota(100);c__Saccharomycetes(100);o__Saccharomycetales(100);f__Metschnikowiaceae(100);g__Metschnikowia(100);s__Metschnikowia_pulcherrima(100); | Metschnikowia | 13 | Saprotroph | \|Undefined Saprotroph\| | Yeast | NULL | Probable | As nectar-tap_saprotroph (Põlme et al. 2020); Decay sugar-rich substrates (Põlme et al. 2020) | Põlme S, et al. 2020. Fungal Diversity 105:1-16 ((https://doi.org/10.1007/s13225-020-00466-2)) |
| Otu22463 | 1 | k__Fungi(100);p__Ascomycota(100);c__Saccharomycetes(100);o__Saccharomycetales(100);f__Metschnikowiaceae(100);g__Metschnikowia(100);g__Metschnikowia_unclassified(100); | Metschnikowia | 13 | Saprotroph | \|Undefined Saprotroph\| | Yeast | NULL | Probable | As nectar-tap_saprotroph (Põlme et al. 2020); Decay sugar-rich substrates (Põlme et al. 2020) | Põlme S, et al. 2020. Fungal Diversity 105:1-16 ((https://doi.org/10.1007/s13225-020-00466-2)) |
| Otu22464 | 1 | k__Fungi(100);p__Ascomycota(100);c__Saccharomycetes(100);o__Saccharomycetales(100);f__Metschnikowiaceae(100);g__Metschnikowia(100);s__Metschnikowia_pulcherrima(100); | Metschnikowia | 13 | Saprotroph | \|Undefined Saprotroph\| | Yeast | NULL | Probable | As nectar-tap_saprotroph (Põlme et al. 2020); Decay sugar-rich substrates (Põlme et al. 2020) | Põlme S, et al. 2020. Fungal Diversity 105:1-16 ((https://doi.org/10.1007/s13225-020-00466-2)) |
| Otu22465 | 1 | k__Fungi(100);p__Ascomycota(100);c__Saccharomycetes(100);o__Saccharomycetales(100);f__Metschnikowiaceae(100);g__Metschnikowia(100);s__Metschnikowia_pulcherrima(100); | Metschnikowia | 13 | Saprotroph | \|Undefined Saprotroph\| | Yeast | NULL | Probable | As nectar-tap_saprotroph (Põlme et al. 2020); Decay sugar-rich substrates (Põlme et al. 2020) | Põlme S, et al. 2020. Fungal Diversity 105:1-16 ((https://doi.org/10.1007/s13225-020-00466-2)) |
| Otu22466 | 1 | k__Fungi(100);p__Ascomycota(100);c__Saccharomycetes(100);o__Saccharomycetales(100);f__Metschnikowiaceae(100);g__Metschnikowia(100);s__Metschnikowia_pulcherrima(100); | Metschnikowia | 13 | Saprotroph | \|Undefined Saprotroph\| | Yeast | NULL | Probable | As nectar-tap_saprotroph (Põlme et al. 2020); Decay sugar-rich substrates (Põlme et al. 2020) | Põlme S, et al. 2020. Fungal Diversity 105:1-16 ((https://doi.org/10.1007/s13225-020-00466-2)) |
| Otu22469 | 1 | k__Fungi(100);p__Ascomycota(100);c__Saccharomycetes(100);o__Saccharomycetales(100);f__Metschnikowiaceae(100);g__Metschnikowia(100);s__Metschnikowia_pulcherrima(100); | Metschnikowia | 13 | Saprotroph | \|Undefined Saprotroph\| | Yeast | NULL | Probable | As nectar-tap_saprotroph (Põlme et al. 2020); Decay sugar-rich substrates (Põlme et al. 2020) | Põlme S, et al. 2020. Fungal Diversity 105:1-16 ((https://doi.org/10.1007/s13225-020-00466-2)) |
| Otu22470 | 1 | k__Fungi(100);p__Ascomycota(100);c__Saccharomycetes(100);o__Saccharomycetales(100);f__Metschnikowiaceae(100);g__Metschnikowia(100);g__Metschnikowia_unclassified(100); | Metschnikowia | 13 | Saprotroph | \|Undefined Saprotroph\| | Yeast | NULL | Probable | As nectar-tap_saprotroph (Põlme et al. 2020); Decay sugar-rich substrates (Põlme et al. 2020) | Põlme S, et al. 2020. Fungal Diversity 105:1-16 ((https://doi.org/10.1007/s13225-020-00466-2)) |
| Otu22471 | 1 | k__Fungi(100);p__Ascomycota(100);c__Saccharomycetes(100);o__Saccharomycetales(100);f__Metschnikowiaceae(100);g__Metschnikowia(100);s__Metschnikowia_pulcherrima(100); | Metschnikowia | 13 | Saprotroph | \|Undefined Saprotroph\| | Yeast | NULL | Probable | As nectar-tap_saprotroph (Põlme et al. 2020); Decay sugar-rich substrates (Põlme et al. 2020) | Põlme S, et al. 2020. Fungal Diversity 105:1-16 ((https://doi.org/10.1007/s13225-020-00466-2)) |
| Otu22472 | 1 | k__Fungi(100);p__Ascomycota(100);c__Saccharomycetes(100);o__Saccharomycetales(100);f__Metschnikowiaceae(100);g__Metschnikowia(100);s__Metschnikowia_pulcherrima(100); | Metschnikowia | 13 | Saprotroph | \|Undefined Saprotroph\| | Yeast | NULL | Probable | As nectar-tap_saprotroph (Põlme et al. 2020); Decay sugar-rich substrates (Põlme et al. 2020) | Põlme S, et al. 2020. Fungal Diversity 105:1-16 ((https://doi.org/10.1007/s13225-020-00466-2)) |
| Otu22473 | 1 | k__Fungi(100);p__Ascomycota(100);c__Saccharomycetes(100);o__Saccharomycetales(100);f__Metschnikowiaceae(100);g__Metschnikowia(100);s__Metschnikowia_pulcherrima(100); | Metschnikowia | 13 | Saprotroph | \|Undefined Saprotroph\| | Yeast | NULL | Probable | As nectar-tap_saprotroph (Põlme et al. 2020); Decay sugar-rich substrates (Põlme et al. 2020) | Põlme S, et al. 2020. Fungal Diversity 105:1-16 ((https://doi.org/10.1007/s13225-020-00466-2)) |
| Otu22474 | 1 | k__Fungi(100);p__Ascomycota(100);c__Saccharomycetes(100);o__Saccharomycetales(100);f__Metschnikowiaceae(100);g__Metschnikowia(100);g__Metschnikowia_unclassified(100); | Metschnikowia | 13 | Saprotroph | \|Undefined Saprotroph\| | Yeast | NULL | Probable | As nectar-tap_saprotroph (Põlme et al. 2020); Decay sugar-rich substrates (Põlme et al. 2020) | Põlme S, et al. 2020. Fungal Diversity 105:1-16 ((https://doi.org/10.1007/s13225-020-00466-2)) |
| Otu22475 | 1 | k__Fungi(100);p__Ascomycota(100);c__Saccharomycetes(100);o__Saccharomycetales(100);f__Metschnikowiaceae(100);g__Metschnikowia(100);g__Metschnikowia_unclassified(100); | Metschnikowia | 13 | Saprotroph | \|Undefined Saprotroph\| | Yeast | NULL | Probable | As nectar-tap_saprotroph (Põlme et al. 2020); Decay sugar-rich substrates (Põlme et al. 2020) | Põlme S, et al. 2020. Fungal Diversity 105:1-16 ((https://doi.org/10.1007/s13225-020-00466-2)) |
| Otu22476 | 1 | k__Fungi(100);p__Ascomycota(100);c__Saccharomycetes(100);o__Saccharomycetales(100);f__Metschnikowiaceae(100);g__Metschnikowia(100);s__Metschnikowia_pulcherrima(100); | Metschnikowia | 13 | Saprotroph | \|Undefined Saprotroph\| | Yeast | NULL | Probable | As nectar-tap_saprotroph (Põlme et al. 2020); Decay sugar-rich substrates (Põlme et al. 2020) | Põlme S, et al. 2020. Fungal Diversity 105:1-16 ((https://doi.org/10.1007/s13225-020-00466-2)) |
| Otu22477 | 1 | k__Fungi(100);p__Ascomycota(100);c__Saccharomycetes(100);o__Saccharomycetales(100);f__Metschnikowiaceae(100);g__Metschnikowia(100);s__Metschnikowia_pulcherrima(100); | Metschnikowia | 13 | Saprotroph | \|Undefined Saprotroph\| | Yeast | NULL | Probable | As nectar-tap_saprotroph (Põlme et al. 2020); Decay sugar-rich substrates (Põlme et al. 2020) | Põlme S, et al. 2020. Fungal Diversity 105:1-16 ((https://doi.org/10.1007/s13225-020-00466-2)) |
| Otu22478 | 1 | k__Fungi(100);p__Ascomycota(100);c__Saccharomycetes(100);o__Saccharomycetales(100);f__Metschnikowiaceae(100);g__Metschnikowia(100);s__Metschnikowia_pulcherrima(100); | Metschnikowia | 13 | Saprotroph | \|Undefined Saprotroph\| | Yeast | NULL | Probable | As nectar-tap_saprotroph (Põlme et al. 2020); Decay sugar-rich substrates (Põlme et al. 2020) | Põlme S, et al. 2020. Fungal Diversity 105:1-16 ((https://doi.org/10.1007/s13225-020-00466-2)) |
| Otu22479 | 1 | k__Fungi(100);p__Ascomycota(100);c__Saccharomycetes(100);o__Saccharomycetales(100);f__Metschnikowiaceae(100);g__Metschnikowia(100);s__Metschnikowia_pulcherrima(100); | Metschnikowia | 13 | Saprotroph | \|Undefined Saprotroph\| | Yeast | NULL | Probable | As nectar-tap_saprotroph (Põlme et al. 2020); Decay sugar-rich substrates (Põlme et al. 2020) | Põlme S, et al. 2020. Fungal Diversity 105:1-16 ((https://doi.org/10.1007/s13225-020-00466-2)) |
| Otu22480 | 1 | k__Fungi(100);p__Ascomycota(100);c__Saccharomycetes(100);o__Saccharomycetales(100);f__Metschnikowiaceae(100);g__Metschnikowia(100);s__Metschnikowia_pulcherrima(100); | Metschnikowia | 13 | Saprotroph | \|Undefined Saprotroph\| | Yeast | NULL | Probable | As nectar-tap_saprotroph (Põlme et al. 2020); Decay sugar-rich substrates (Põlme et al. 2020) | Põlme S, et al. 2020. Fungal Diversity 105:1-16 ((https://doi.org/10.1007/s13225-020-00466-2)) |
| Otu22481 | 1 | k__Fungi(100);p__Ascomycota(100);c__Saccharomycetes(100);o__Saccharomycetales(100);f__Metschnikowiaceae(100);g__Metschnikowia(100);s__Metschnikowia_pulcherrima(100); | Metschnikowia | 13 | Saprotroph | \|Undefined Saprotroph\| | Yeast | NULL | Probable | As nectar-tap_saprotroph (Põlme et al. 2020); Decay sugar-rich substrates (Põlme et al. 2020) | Põlme S, et al. 2020. Fungal Diversity 105:1-16 ((https://doi.org/10.1007/s13225-020-00466-2)) |
| Otu22482 | 1 | k__Fungi(100);p__Ascomycota(100);c__Saccharomycetes(100);o__Saccharomycetales(100);f__Metschnikowiaceae(100);g__Metschnikowia(100);s__Metschnikowia_pulcherrima(100); | Metschnikowia | 13 | Saprotroph | \|Undefined Saprotroph\| | Yeast | NULL | Probable | As nectar-tap_saprotroph (Põlme et al. 2020); Decay sugar-rich substrates (Põlme et al. 2020) | Põlme S, et al. 2020. Fungal Diversity 105:1-16 ((https://doi.org/10.1007/s13225-020-00466-2)) |
| Otu22483 | 1 | k__Fungi(100);p__Ascomycota(100);c__Saccharomycetes(100);o__Saccharomycetales(100);f__Metschnikowiaceae(100);g__Metschnikowia(100);s__Metschnikowia_pulcherrima(100); | Metschnikowia | 13 | Saprotroph | \|Undefined Saprotroph\| | Yeast | NULL | Probable | As nectar-tap_saprotroph (Põlme et al. 2020); Decay sugar-rich substrates (Põlme et al. 2020) | Põlme S, et al. 2020. Fungal Diversity 105:1-16 ((https://doi.org/10.1007/s13225-020-00466-2)) |
| Otu22484 | 1 | k__Fungi(100);p__Ascomycota(100);c__Saccharomycetes(100);o__Saccharomycetales(100);f__Metschnikowiaceae(100);g__Metschnikowia(100);s__Metschnikowia_pulcherrima(100); | Metschnikowia | 13 | Saprotroph | \|Undefined Saprotroph\| | Yeast | NULL | Probable | As nectar-tap_saprotroph (Põlme et al. 2020); Decay sugar-rich substrates (Põlme et al. 2020) | Põlme S, et al. 2020. Fungal Diversity 105:1-16 ((https://doi.org/10.1007/s13225-020-00466-2)) |
| Otu22485 | 1 | k__Fungi(100);p__Ascomycota(100);c__Saccharomycetes(100);o__Saccharomycetales(100);f__Metschnikowiaceae(100);g__Metschnikowia(100);g__Metschnikowia_unclassified(100); | Metschnikowia | 13 | Saprotroph | \|Undefined Saprotroph\| | Yeast | NULL | Probable | As nectar-tap_saprotroph (Põlme et al. 2020); Decay sugar-rich substrates (Põlme et al. 2020) | Põlme S, et al. 2020. Fungal Diversity 105:1-16 ((https://doi.org/10.1007/s13225-020-00466-2)) |
| Otu22486 | 1 | k__Fungi(100);p__Ascomycota(100);c__Saccharomycetes(100);o__Saccharomycetales(100);f__Metschnikowiaceae(100);g__Metschnikowia(100);s__Metschnikowia_pulcherrima(100); | Metschnikowia | 13 | Saprotroph | \|Undefined Saprotroph\| | Yeast | NULL | Probable | As nectar-tap_saprotroph (Põlme et al. 2020); Decay sugar-rich substrates (Põlme et al. 2020) | Põlme S, et al. 2020. Fungal Diversity 105:1-16 ((https://doi.org/10.1007/s13225-020-00466-2)) |
| Otu22487 | 1 | k__Fungi(100);p__Ascomycota(100);c__Saccharomycetes(100);o__Saccharomycetales(100);f__Metschnikowiaceae(100);g__Metschnikowia(100);s__Metschnikowia_pulcherrima(100); | Metschnikowia | 13 | Saprotroph | \|Undefined Saprotroph\| | Yeast | NULL | Probable | As nectar-tap_saprotroph (Põlme et al. 2020); Decay sugar-rich substrates (Põlme et al. 2020) | Põlme S, et al. 2020. Fungal Diversity 105:1-16 ((https://doi.org/10.1007/s13225-020-00466-2)) |
| Otu22488 | 1 | k__Fungi(100);p__Ascomycota(100);c__Saccharomycetes(100);o__Saccharomycetales(100);f__Metschnikowiaceae(100);g__Metschnikowia(100);s__Metschnikowia_pulcherrima(100); | Metschnikowia | 13 | Saprotroph | \|Undefined Saprotroph\| | Yeast | NULL | Probable | As nectar-tap_saprotroph (Põlme et al. 2020); Decay sugar-rich substrates (Põlme et al. 2020) | Põlme S, et al. 2020. Fungal Diversity 105:1-16 ((https://doi.org/10.1007/s13225-020-00466-2)) |
| Otu22489 | 1 | k__Fungi(100);p__Ascomycota(100);c__Saccharomycetes(100);o__Saccharomycetales(100);f__Metschnikowiaceae(100);g__Metschnikowia(100);s__Metschnikowia_pulcherrima(100); | Metschnikowia | 13 | Saprotroph | \|Undefined Saprotroph\| | Yeast | NULL | Probable | As nectar-tap_saprotroph (Põlme et al. 2020); Decay sugar-rich substrates (Põlme et al. 2020) | Põlme S, et al. 2020. Fungal Diversity 105:1-16 ((https://doi.org/10.1007/s13225-020-00466-2)) |
| Otu22490 | 1 | k__Fungi(100);p__Ascomycota(100);c__Saccharomycetes(100);o__Saccharomycetales(100);f__Metschnikowiaceae(100);g__Metschnikowia(100);s__Metschnikowia_pulcherrima(100); | Metschnikowia | 13 | Saprotroph | \|Undefined Saprotroph\| | Yeast | NULL | Probable | As nectar-tap_saprotroph (Põlme et al. 2020); Decay sugar-rich substrates (Põlme et al. 2020) | Põlme S, et al. 2020. Fungal Diversity 105:1-16 ((https://doi.org/10.1007/s13225-020-00466-2)) |
| Otu22491 | 1 | k__Fungi(100);p__Ascomycota(100);c__Saccharomycetes(100);o__Saccharomycetales(100);f__Metschnikowiaceae(100);g__Metschnikowia(100);g__Metschnikowia_unclassified(100); | Metschnikowia | 13 | Saprotroph | \|Undefined Saprotroph\| | Yeast | NULL | Probable | As nectar-tap_saprotroph (Põlme et al. 2020); Decay sugar-rich substrates (Põlme et al. 2020) | Põlme S, et al. 2020. Fungal Diversity 105:1-16 ((https://doi.org/10.1007/s13225-020-00466-2)) |
| Otu22492 | 1 | k__Fungi(100);p__Ascomycota(100);c__Saccharomycetes(100);o__Saccharomycetales(100);f__Metschnikowiaceae(100);g__Metschnikowia(100);s__Metschnikowia_pulcherrima(100); | Metschnikowia | 13 | Saprotroph | \|Undefined Saprotroph\| | Yeast | NULL | Probable | As nectar-tap_saprotroph (Põlme et al. 2020); Decay sugar-rich substrates (Põlme et al. 2020) | Põlme S, et al. 2020. Fungal Diversity 105:1-16 ((https://doi.org/10.1007/s13225-020-00466-2)) |
| Otu22493 | 1 | k__Fungi(100);p__Ascomycota(100);c__Saccharomycetes(100);o__Saccharomycetales(100);f__Metschnikowiaceae(100);g__Metschnikowia(100);g__Metschnikowia_unclassified(100); | Metschnikowia | 13 | Saprotroph | \|Undefined Saprotroph\| | Yeast | NULL | Probable | As nectar-tap_saprotroph (Põlme et al. 2020); Decay sugar-rich substrates (Põlme et al. 2020) | Põlme S, et al. 2020. Fungal Diversity 105:1-16 ((https://doi.org/10.1007/s13225-020-00466-2)) |
| Otu22496 | 1 | k__Fungi(100);p__Ascomycota(100);c__Saccharomycetes(100);o__Saccharomycetales(100);f__Metschnikowiaceae(100);g__Metschnikowia(100);s__Metschnikowia_pulcherrima(100); | Metschnikowia | 13 | Saprotroph | \|Undefined Saprotroph\| | Yeast | NULL | Probable | As nectar-tap_saprotroph (Põlme et al. 2020); Decay sugar-rich substrates (Põlme et al. 2020) | Põlme S, et al. 2020. Fungal Diversity 105:1-16 ((https://doi.org/10.1007/s13225-020-00466-2)) |
| Otu22497 | 1 | k__Fungi(100);p__Ascomycota(100);c__Saccharomycetes(100);o__Saccharomycetales(100);f__Metschnikowiaceae(100);g__Metschnikowia(100);s__Metschnikowia_pulcherrima(100); | Metschnikowia | 13 | Saprotroph | \|Undefined Saprotroph\| | Yeast | NULL | Probable | As nectar-tap_saprotroph (Põlme et al. 2020); Decay sugar-rich substrates (Põlme et al. 2020) | Põlme S, et al. 2020. Fungal Diversity 105:1-16 ((https://doi.org/10.1007/s13225-020-00466-2)) |

| | | | | | | | | | | | |
|---|---|---|---|---|---|---|---|---|---|---|---|
| Otu22498 | 1 | k__Fungi(100);p__Ascomycota(100);c__Saccharomycetes(100);o__Saccharomycetales(100);f__Metschnikowiaceae(100);g__Metschnikowia(100);s__Metschnikowia_pulcherrima(100); | Metschnikowia | 13 | Saprotroph | \|Undefined Saprotroph\| | Yeast | NULL | Probable | As nectar-tap_saprotroph (Põlme et al. 2020); Decay sugar-rich substrates (Põlme et al. 2020) | Põlme S, et al. 2020. Fungal Diversity 105:1-16 ((https://doi.org/10.1007/s13225-020-00466-2)) |
| Otu22499 | 1 | k__Fungi(100);p__Ascomycota(100);c__Saccharomycetes(100);o__Saccharomycetales(100);f__Metschnikowiaceae(100);g__Metschnikowia(100);s__Metschnikowia_pulcherrima(100); | Metschnikowia | 13 | Saprotroph | \|Undefined Saprotroph\| | Yeast | NULL | Probable | As nectar-tap_saprotroph (Põlme et al. 2020); Decay sugar-rich substrates (Põlme et al. 2020) | Põlme S, et al. 2020. Fungal Diversity 105:1-16 ((https://doi.org/10.1007/s13225-020-00466-2)) |
| Otu22501 | 1 | k__Fungi(100);p__Ascomycota(100);c__Saccharomycetes(100);o__Saccharomycetales(100);f__Metschnikowiaceae(100);g__Metschnikowia(100);g__Metschnikowia_unclassified(100); | Metschnikowia | 13 | Saprotroph | \|Undefined Saprotroph\| | Yeast | NULL | Probable | As nectar-tap_saprotroph (Põlme et al. 2020); Decay sugar-rich substrates (Põlme et al. 2020) | Põlme S, et al. 2020. Fungal Diversity 105:1-16 ((https://doi.org/10.1007/s13225-020-00466-2)) |
| Otu22502 | 1 | k__Fungi(100);p__Ascomycota(100);c__Saccharomycetes(100);o__Saccharomycetales(100);f__Metschnikowiaceae(100);g__Metschnikowia(100);g__Metschnikowia_unclassified(100); | Metschnikowia | 13 | Saprotroph | \|Undefined Saprotroph\| | Yeast | NULL | Probable | As nectar-tap_saprotroph (Põlme et al. 2020); Decay sugar-rich substrates (Põlme et al. 2020) | Põlme S, et al. 2020. Fungal Diversity 105:1-16 ((https://doi.org/10.1007/s13225-020-00466-2)) |
| Otu22503 | 1 | k__Fungi(100);p__Ascomycota(100);c__Saccharomycetes(100);o__Saccharomycetales(100);f__Metschnikowiaceae(100);g__Metschnikowia(100);s__Metschnikowia_pulcherrima(100); | Metschnikowia | 13 | Saprotroph | \|Undefined Saprotroph\| | Yeast | NULL | Probable | As nectar-tap_saprotroph (Põlme et al. 2020); Decay sugar-rich substrates (Põlme et al. 2020) | Põlme S, et al. 2020. Fungal Diversity 105:1-16 ((https://doi.org/10.1007/s13225-020-00466-2)) |
| Otu22504 | 1 | k__Fungi(100);p__Ascomycota(100);c__Saccharomycetes(100);o__Saccharomycetales(100);f__Metschnikowiaceae(100);g__Metschnikowia(100);s__Metschnikowia_pulcherrima(100); | Metschnikowia | 13 | Saprotroph | \|Undefined Saprotroph\| | Yeast | NULL | Probable | As nectar-tap_saprotroph (Põlme et al. 2020); Decay sugar-rich substrates (Põlme et al. 2020) | Põlme S, et al. 2020. Fungal Diversity 105:1-16 ((https://doi.org/10.1007/s13225-020-00466-2)) |
| Otu22505 | 1 | k__Fungi(100);p__Ascomycota(100);c__Saccharomycetes(100);o__Saccharomycetales(100);f__Metschnikowiaceae(100);g__Metschnikowia(100);s__Metschnikowia_pulcherrima(100); | Metschnikowia | 13 | Saprotroph | \|Undefined Saprotroph\| | Yeast | NULL | Probable | As nectar-tap_saprotroph (Põlme et al. 2020); Decay sugar-rich substrates (Põlme et al. 2020) | Põlme S, et al. 2020. Fungal Diversity 105:1-16 ((https://doi.org/10.1007/s13225-020-00466-2)) |
| Otu22506 | 1 | k__Fungi(100);p__Ascomycota(100);c__Saccharomycetes(100);o__Saccharomycetales(100);f__Metschnikowiaceae(100);g__Metschnikowia(100);s__Metschnikowia_pulcherrima(100); | Metschnikowia | 13 | Saprotroph | \|Undefined Saprotroph\| | Yeast | NULL | Probable | As nectar-tap_saprotroph (Põlme et al. 2020); Decay sugar-rich substrates (Põlme et al. 2020) | Põlme S, et al. 2020. Fungal Diversity 105:1-16 ((https://doi.org/10.1007/s13225-020-00466-2)) |
| Otu22507 | 1 | k__Fungi(100);p__Ascomycota(100);c__Saccharomycetes(100);o__Saccharomycetales(100);f__Metschnikowiaceae(100);g__Metschnikowia(100);s__Metschnikowia_pulcherrima(100); | Metschnikowia | 13 | Saprotroph | \|Undefined Saprotroph\| | Yeast | NULL | Probable | As nectar-tap_saprotroph (Põlme et al. 2020); Decay sugar-rich substrates (Põlme et al. 2020) | Põlme S, et al. 2020. Fungal Diversity 105:1-16 ((https://doi.org/10.1007/s13225-020-00466-2)) |
| Otu22508 | 1 | k__Fungi(100);p__Ascomycota(100);c__Saccharomycetes(100);o__Saccharomycetales(100);f__Metschnikowiaceae(100);g__Metschnikowia(100);s__Metschnikowia_pulcherrima(100); | Metschnikowia | 13 | Saprotroph | \|Undefined Saprotroph\| | Yeast | NULL | Probable | As nectar-tap_saprotroph (Põlme et al. 2020); Decay sugar-rich substrates (Põlme et al. 2020) | Põlme S, et al. 2020. Fungal Diversity 105:1-16 ((https://doi.org/10.1007/s13225-020-00466-2)) |
| Otu22509 | 1 | k__Fungi(100);p__Ascomycota(100);c__Saccharomycetes(100);o__Saccharomycetales(100);f__Metschnikowiaceae(100);g__Metschnikowia(100);s__Metschnikowia_pulcherrima(100); | Metschnikowia | 13 | Saprotroph | \|Undefined Saprotroph\| | Yeast | NULL | Probable | As nectar-tap_saprotroph (Põlme et al. 2020); Decay sugar-rich substrates (Põlme et al. 2020) | Põlme S, et al. 2020. Fungal Diversity 105:1-16 ((https://doi.org/10.1007/s13225-020-00466-2)) |
| Otu22511 | 1 | k__Fungi(100);p__Ascomycota(100);c__Saccharomycetes(100);o__Saccharomycetales(100);f__Metschnikowiaceae(100);g__Metschnikowia(100);s__Metschnikowia_pulcherrima(100); | Metschnikowia | 13 | Saprotroph | \|Undefined Saprotroph\| | Yeast | NULL | Probable | As nectar-tap_saprotroph (Põlme et al. 2020); Decay sugar-rich substrates (Põlme et al. 2020) | Põlme S, et al. 2020. Fungal Diversity 105:1-16 ((https://doi.org/10.1007/s13225-020-00466-2)) |
| Otu22512 | 1 | k__Fungi(100);p__Ascomycota(100);c__Saccharomycetes(100);o__Saccharomycetales(100);f__Metschnikowiaceae(100);g__Metschnikowia(100);g__Metschnikowia_unclassified(100); | Metschnikowia | 13 | Saprotroph | \|Undefined Saprotroph\| | Yeast | NULL | Probable | As nectar-tap_saprotroph (Põlme et al. 2020); Decay sugar-rich substrates (Põlme et al. 2020) | Põlme S, et al. 2020. Fungal Diversity 105:1-16 ((https://doi.org/10.1007/s13225-020-00466-2)) |
| Otu22513 | 1 | k__Fungi(100);p__Ascomycota(100);c__Saccharomycetes(100);o__Saccharomycetales(100);f__Metschnikowiaceae(100);g__Metschnikowia(100);s__Metschnikowia_pulcherrima(100); | Metschnikowia | 13 | Saprotroph | \|Undefined Saprotroph\| | Yeast | NULL | Probable | As nectar-tap_saprotroph (Põlme et al. 2020); Decay sugar-rich substrates (Põlme et al. 2020) | Põlme S, et al. 2020. Fungal Diversity 105:1-16 ((https://doi.org/10.1007/s13225-020-00466-2)) |
| Otu22521 | 1 | k__Fungi(100);p__Ascomycota(100);c__Saccharomycetes(100);o__Saccharomycetales(100);f__Metschnikowiaceae(100);g__Metschnikowia(100);s__Metschnikowia_pulcherrima(100); | Metschnikowia | 13 | Saprotroph | \|Undefined Saprotroph\| | Yeast | NULL | Probable | As nectar-tap_saprotroph (Põlme et al. 2020); Decay sugar-rich substrates (Põlme et al. 2020) | Põlme S, et al. 2020. Fungal Diversity 105:1-16 ((https://doi.org/10.1007/s13225-020-00466-2)) |
| Otu22526 | 1 | k__Fungi(100);p__Ascomycota(100);c__Saccharomycetes(100);o__Saccharomycetales(100);f__Metschnikowiaceae(100);g__Metschnikowia(100);s__Metschnikowia_pulcherrima(100); | Metschnikowia | 13 | Saprotroph | \|Undefined Saprotroph\| | Yeast | NULL | Probable | As nectar-tap_saprotroph (Põlme et al. 2020); Decay sugar-rich substrates (Põlme et al. 2020) | Põlme S, et al. 2020. Fungal Diversity 105:1-16 ((https://doi.org/10.1007/s13225-020-00466-2)) |
| Otu22527 | 1 | k__Fungi(100);p__Ascomycota(100);c__Saccharomycetes(100);o__Saccharomycetales(100);f__Metschnikowiaceae(100);g__Metschnikowia(100);g__Metschnikowia_unclassified(100); | Metschnikowia | 13 | Saprotroph | \|Undefined Saprotroph\| | Yeast | NULL | Probable | As nectar-tap_saprotroph (Põlme et al. 2020); Decay sugar-rich substrates (Põlme et al. 2020) | Põlme S, et al. 2020. Fungal Diversity 105:1-16 ((https://doi.org/10.1007/s13225-020-00466-2)) |
| Otu22528 | 1 | k__Fungi(100);p__Ascomycota(100);c__Saccharomycetes(100);o__Saccharomycetales(100);f__Metschnikowiaceae(100);g__Metschnikowia(100);g__Metschnikowia_unclassified(100); | Metschnikowia | 13 | Saprotroph | \|Undefined Saprotroph\| | Yeast | NULL | Probable | As nectar-tap_saprotroph (Põlme et al. 2020); Decay sugar-rich substrates (Põlme et al. 2020) | Põlme S, et al. 2020. Fungal Diversity 105:1-16 ((https://doi.org/10.1007/s13225-020-00466-2)) |
| Otu22529 | 1 | k__Fungi(100);p__Ascomycota(100);c__Saccharomycetes(100);o__Saccharomycetales(100);f__Metschnikowiaceae(100);g__Metschnikowia(100);g__Metschnikowia_unclassified(100); | Metschnikowia | 13 | Saprotroph | \|Undefined Saprotroph\| | Yeast | NULL | Probable | As nectar-tap_saprotroph (Põlme et al. 2020); Decay sugar-rich substrates (Põlme et al. 2020) | Põlme S, et al. 2020. Fungal Diversity 105:1-16 ((https://doi.org/10.1007/s13225-020-00466-2)) |
| Otu22530 | 1 | k__Fungi(100);p__Ascomycota(100);c__Saccharomycetes(100);o__Saccharomycetales(100);f__Metschnikowiaceae(100);g__Metschnikowia(100);s__Metschnikowia_pulcherrima(100); | Metschnikowia | 13 | Saprotroph | \|Undefined Saprotroph\| | Yeast | NULL | Probable | As nectar-tap_saprotroph (Põlme et al. 2020); Decay sugar-rich substrates (Põlme et al. 2020) | Põlme S, et al. 2020. Fungal Diversity 105:1-16 ((https://doi.org/10.1007/s13225-020-00466-2)) |
| Otu22531 | 1 | k__Fungi(100);p__Ascomycota(100);c__Saccharomycetes(100);o__Saccharomycetales(100);f__Metschnikowiaceae(100);g__Metschnikowia(100);s__Metschnikowia_pulcherrima(100); | Metschnikowia | 13 | Saprotroph | \|Undefined Saprotroph\| | Yeast | NULL | Probable | As nectar-tap_saprotroph (Põlme et al. 2020); Decay sugar-rich substrates (Põlme et al. 2020) | Põlme S, et al. 2020. Fungal Diversity 105:1-16 ((https://doi.org/10.1007/s13225-020-00466-2)) |
| Otu22532 | 1 | k__Fungi(100);p__Ascomycota(100);c__Saccharomycetes(100);o__Saccharomycetales(100);f__Metschnikowiaceae(100);g__Metschnikowia(100);s__Metschnikowia_pulcherrima(100); | Metschnikowia | 13 | Saprotroph | \|Undefined Saprotroph\| | Yeast | NULL | Probable | As nectar-tap_saprotroph (Põlme et al. 2020); Decay sugar-rich substrates (Põlme et al. 2020) | Põlme S, et al. 2020. Fungal Diversity 105:1-16 ((https://doi.org/10.1007/s13225-020-00466-2)) |
| Otu22533 | 1 | k__Fungi(100);p__Ascomycota(100);c__Saccharomycetes(100);o__Saccharomycetales(100);f__Metschnikowiaceae(100);g__Metschnikowia(100);s__Metschnikowia_pulcherrima(100); | Metschnikowia | 13 | Saprotroph | \|Undefined Saprotroph\| | Yeast | NULL | Probable | As nectar-tap_saprotroph (Põlme et al. 2020); Decay sugar-rich substrates (Põlme et al. 2020) | Põlme S, et al. 2020. Fungal Diversity 105:1-16 ((https://doi.org/10.1007/s13225-020-00466-2)) |
| Otu22534 | 1 | k__Fungi(100);p__Ascomycota(100);c__Saccharomycetes(100);o__Saccharomycetales(100);f__Metschnikowiaceae(100);g__Metschnikowia(100);s__Metschnikowia_pulcherrima(100); | Metschnikowia | 13 | Saprotroph | \|Undefined Saprotroph\| | Yeast | NULL | Probable | As nectar-tap_saprotroph (Põlme et al. 2020); Decay sugar-rich substrates (Põlme et al. 2020) | Põlme S, et al. 2020. Fungal Diversity 105:1-16 ((https://doi.org/10.1007/s13225-020-00466-2)) |
| Otu22536 | 1 | k__Fungi(100);p__Ascomycota(100);c__Saccharomycetes(100);o__Saccharomycetales(100);f__Metschnikowiaceae(100);g__Metschnikowia(100);s__Metschnikowia_pulcherrima(100); | Metschnikowia | 13 | Saprotroph | \|Undefined Saprotroph\| | Yeast | NULL | Probable | As nectar-tap_saprotroph (Põlme et al. 2020); Decay sugar-rich substrates (Põlme et al. 2020) | Põlme S, et al. 2020. Fungal Diversity 105:1-16 ((https://doi.org/10.1007/s13225-020-00466-2)) |
| Otu22537 | 1 | k__Fungi(100);p__Ascomycota(100);c__Saccharomycetes(100);o__Saccharomycetales(100);f__Metschnikowiaceae(100);g__Metschnikowia(100);s__Metschnikowia_pulcherrima(100); | Metschnikowia | 13 | Saprotroph | \|Undefined Saprotroph\| | Yeast | NULL | Probable | As nectar-tap_saprotroph (Põlme et al. 2020); Decay sugar-rich substrates (Põlme et al. 2020) | Põlme S, et al. 2020. Fungal Diversity 105:1-16 ((https://doi.org/10.1007/s13225-020-00466-2)) |
| Otu22538 | 1 | k__Fungi(100);p__Ascomycota(100);c__Saccharomycetes(100);o__Saccharomycetales(100);f__Metschnikowiaceae(100);g__Metschnikowia(100);s__Metschnikowia_pulcherrima(100); | Metschnikowia | 13 | Saprotroph | \|Undefined Saprotroph\| | Yeast | NULL | Probable | As nectar-tap_saprotroph (Põlme et al. 2020); Decay sugar-rich substrates (Põlme et al. 2020) | Põlme S, et al. 2020. Fungal Diversity 105:1-16 ((https://doi.org/10.1007/s13225-020-00466-2)) |
| Otu22540 | 1 | k__Fungi(100);p__Ascomycota(100);c__Saccharomycetes(100);o__Saccharomycetales(100);f__Metschnikowiaceae(100);g__Metschnikowia(100);g__Metschnikowia_unclassified(100); | Metschnikowia | 13 | Saprotroph | \|Undefined Saprotroph\| | Yeast | NULL | Probable | As nectar-tap_saprotroph (Põlme et al. 2020); Decay sugar-rich substrates (Põlme et al. 2020) | Põlme S, et al. 2020. Fungal Diversity 105:1-16 ((https://doi.org/10.1007/s13225-020-00466-2)) |
| Otu22541 | 1 | k__Fungi(100);p__Ascomycota(100);c__Saccharomycetes(100);o__Saccharomycetales(100);f__Metschnikowiaceae(100);g__Metschnikowia(100);s__Metschnikowia_pulcherrima(100); | Metschnikowia | 13 | Saprotroph | \|Undefined Saprotroph\| | Yeast | NULL | Probable | As nectar-tap_saprotroph (Põlme et al. 2020); Decay sugar-rich substrates (Põlme et al. 2020) | Põlme S, et al. 2020. Fungal Diversity 105:1-16 ((https://doi.org/10.1007/s13225-020-00466-2)) |
| Otu22542 | 1 | k__Fungi(100);p__Ascomycota(100);c__Saccharomycetes(100);o__Saccharomycetales(100);f__Metschnikowiaceae(100);g__Metschnikowia(100);s__Metschnikowia_pulcherrima(100); | Metschnikowia | 13 | Saprotroph | \|Undefined Saprotroph\| | Yeast | NULL | Probable | As nectar-tap_saprotroph (Põlme et al. 2020); Decay sugar-rich substrates (Põlme et al. 2020) | Põlme S, et al. 2020. Fungal Diversity 105:1-16 ((https://doi.org/10.1007/s13225-020-00466-2)) |
| Otu22543 | 1 | k__Fungi(100);p__Ascomycota(100);c__Saccharomycetes(100);o__Saccharomycetales(100);f__Metschnikowiaceae(100);g__Metschnikowia(100);g__Metschnikowia_unclassified(100); | Metschnikowia | 13 | Saprotroph | \|Undefined Saprotroph\| | Yeast | NULL | Probable | As nectar-tap_saprotroph (Põlme et al. 2020); Decay sugar-rich substrates (Põlme et al. 2020) | Põlme S, et al. 2020. Fungal Diversity 105:1-16 ((https://doi.org/10.1007/s13225-020-00466-2)) |
| Otu22544 | 1 | k__Fungi(100);p__Ascomycota(100);c__Saccharomycetes(100);o__Saccharomycetales(100);f__Metschnikowiaceae(100);g__Metschnikowia(100);s__Metschnikowia_pulcherrima(100); | Metschnikowia | 13 | Saprotroph | \|Undefined Saprotroph\| | Yeast | NULL | Probable | As nectar-tap_saprotroph (Põlme et al. 2020); Decay sugar-rich substrates (Põlme et al. 2020) | Põlme S, et al. 2020. Fungal Diversity 105:1-16 ((https://doi.org/10.1007/s13225-020-00466-2)) |
| Otu22545 | 1 | k__Fungi(100);p__Ascomycota(100);c__Saccharomycetes(100);o__Saccharomycetales(100);f__Metschnikowiaceae(100);g__Metschnikowia(100);s__Metschnikowia_pulcherrima(100); | Metschnikowia | 13 | Saprotroph | \|Undefined Saprotroph\| | Yeast | NULL | Probable | As nectar-tap_saprotroph (Põlme et al. 2020); Decay sugar-rich substrates (Põlme et al. 2020) | Põlme S, et al. 2020. Fungal Diversity 105:1-16 ((https://doi.org/10.1007/s13225-020-00466-2)) |
| Otu22546 | 1 | k__Fungi(100);p__Ascomycota(100);c__Saccharomycetes(100);o__Saccharomycetales(100);f__Metschnikowiaceae(100);g__Metschnikowia(100);s__Metschnikowia_pulcherrima(100); | Metschnikowia | 13 | Saprotroph | \|Undefined Saprotroph\| | Yeast | NULL | Probable | As nectar-tap_saprotroph (Põlme et al. 2020); Decay sugar-rich substrates (Põlme et al. 2020) | Põlme S, et al. 2020. Fungal Diversity 105:1-16 ((https://doi.org/10.1007/s13225-020-00466-2)) |
| Otu22547 | 1 | k__Fungi(100);p__Ascomycota(100);c__Saccharomycetes(100);o__Saccharomycetales(100);f__Metschnikowiaceae(100);g__Metschnikowia(100);s__Metschnikowia_pulcherrima(100); | Metschnikowia | 13 | Saprotroph | \|Undefined Saprotroph\| | Yeast | NULL | Probable | As nectar-tap_saprotroph (Põlme et al. 2020); Decay sugar-rich substrates (Põlme et al. 2020) | Põlme S, et al. 2020. Fungal Diversity 105:1-16 ((https://doi.org/10.1007/s13225-020-00466-2)) |
| Otu22548 | 1 | k__Fungi(100);p__Ascomycota(100);c__Saccharomycetes(100);o__Saccharomycetales(100);f__Metschnikowiaceae(100);g__Metschnikowia(100);s__Metschnikowia_pulcherrima(100); | Metschnikowia | 13 | Saprotroph | \|Undefined Saprotroph\| | Yeast | NULL | Probable | As nectar-tap_saprotroph (Põlme et al. 2020); Decay sugar-rich substrates (Põlme et al. 2020) | Põlme S, et al. 2020. Fungal Diversity 105:1-16 ((https://doi.org/10.1007/s13225-020-00466-2)) |
| Otu22549 | 1 | k__Fungi(100);p__Ascomycota(100);c__Saccharomycetes(100);o__Saccharomycetales(100);f__Metschnikowiaceae(100);g__Metschnikowia(100);s__Metschnikowia_pulcherrima(100); | Metschnikowia | 13 | Saprotroph | \|Undefined Saprotroph\| | Yeast | NULL | Probable | As nectar-tap_saprotroph (Põlme et al. 2020); Decay sugar-rich substrates (Põlme et al. 2020) | Põlme S, et al. 2020. Fungal Diversity 105:1-16 ((https://doi.org/10.1007/s13225-020-00466-2)) |
| Otu22550 | 1 | k__Fungi(100);p__Ascomycota(100);c__Saccharomycetes(100);o__Saccharomycetales(100);f__Metschnikowiaceae(100);g__Metschnikowia(100);s__Metschnikowia_pulcherrima(100); | Metschnikowia | 13 | Saprotroph | \|Undefined Saprotroph\| | Yeast | NULL | Probable | As nectar-tap_saprotroph (Põlme et al. 2020); Decay sugar-rich substrates (Põlme et al. 2020) | Põlme S, et al. 2020. Fungal Diversity 105:1-16 ((https://doi.org/10.1007/s13225-020-00466-2)) |
| Otu22551 | 1 | k__Fungi(100);p__Ascomycota(100);c__Saccharomycetes(100);o__Saccharomycetales(100);f__Metschnikowiaceae(100);g__Metschnikowia(100);s__Metschnikowia_pulcherrima(100); | Metschnikowia | 13 | Saprotroph | \|Undefined Saprotroph\| | Yeast | NULL | Probable | As nectar-tap_saprotroph (Põlme et al. 2020); Decay sugar-rich substrates (Põlme et al. 2020) | Põlme S, et al. 2020. Fungal Diversity 105:1-16 ((https://doi.org/10.1007/s13225-020-00466-2)) |
| Otu22552 | 1 | k__Fungi(100);p__Ascomycota(100);c__Saccharomycetes(100);o__Saccharomycetales(100);f__Metschnikowiaceae(100);g__Metschnikowia(100);g__Metschnikowia_unclassified(100); | Metschnikowia | 13 | Saprotroph | \|Undefined Saprotroph\| | Yeast | NULL | Probable | As nectar-tap_saprotroph (Põlme et al. 2020); Decay sugar-rich substrates (Põlme et al. 2020) | Põlme S, et al. 2020. Fungal Diversity 105:1-16 ((https://doi.org/10.1007/s13225-020-00466-2)) |
| Otu22553 | 1 | k__Fungi(100);p__Ascomycota(100);c__Saccharomycetes(100);o__Saccharomycetales(100);f__Metschnikowiaceae(100);g__Metschnikowia(100);s__Metschnikowia_pulcherrima(100); | Metschnikowia | 13 | Saprotroph | \|Undefined Saprotroph\| | Yeast | NULL | Probable | As nectar-tap_saprotroph (Põlme et al. 2020); Decay sugar-rich substrates (Põlme et al. 2020) | Põlme S, et al. 2020. Fungal Diversity 105:1-16 ((https://doi.org/10.1007/s13225-020-00466-2)) |
| Otu22554 | 1 | k__Fungi(100);p__Ascomycota(100);c__Saccharomycetes(100);o__Saccharomycetales(100);o__Saccharomycetales_unclassified(100);o__Saccharomycetales_unclassified(100); | Saccharomycetales | 7 | Saprotroph | Undefined Saprotroph | Yeast | NULL | Possible | NULL | Sterkenburg E, et al. 2015. New Phytologist 207:1145-1158 ((https://doi.org/10.1111/nph.13426)) |
| Otu22556 | 1 | k__Fungi(100);p__Ascomycota(100);c__Saccharomycetes(100);o__Saccharomycetales(100);f__Saccharomycetales_family_Incertae_sedis(100);f__Saccharomycetales_family_Incertae_sedis_unclassified(100);f__Saccharomycetales_family_Incertae_sedis_unclassified(100); | Saccharomycetales | 7 | Saprotroph | Undefined Saprotroph | Yeast | NULL | Possible | NULL | Sterkenburg E, et al. 2015. New Phytologist 207:1145-1158 ((https://doi.org/10.1111/nph.13426)) |
| Otu22561 | 1 | k__Fungi(100);p__Ascomycota(100);c__Saccharomycetes(100);o__Saccharomycetales(100);f__Metschnikowiaceae(100);g__Metschnikowia(100);s__Metschnikowia_pulcherrima(100); | Metschnikowia | 13 | Saprotroph | \|Undefined Saprotroph\| | Yeast | NULL | Probable | As nectar-tap_saprotroph (Põlme et al. 2020); Decay sugar-rich substrates (Põlme et al. 2020) | Põlme S, et al. 2020. Fungal Diversity 105:1-16 ((https://doi.org/10.1007/s13225-020-00466-2)) |
| Otu22564 | 1 | k__Fungi(100);p__Ascomycota(100);c__Saccharomycetes(100);o__Saccharomycetales(100);f__Metschnikowiaceae(100);g__Metschnikowia(100);s__Metschnikowia_pulcherrima(100); | Metschnikowia | 13 | Saprotroph | \|Undefined Saprotroph\| | Yeast | NULL | Probable | As nectar-tap_saprotroph (Põlme et al. 2020); Decay sugar-rich substrates (Põlme et al. 2020) | Põlme S, et al. 2020. Fungal Diversity 105:1-16 ((https://doi.org/10.1007/s13225-020-00466-2)) |
| Otu22565 | 1 | k__Fungi(100);p__Ascomycota(100);c__Saccharomycetes(100);o__Saccharomycetales(100);f__Metschnikowiaceae(100);g__Metschnikowia(100);s__Metschnikowia_pulcherrima(100); | Metschnikowia | 13 | Saprotroph | \|Undefined Saprotroph\| | Yeast | NULL | Probable | As nectar-tap_saprotroph (Põlme et al. 2020); Decay sugar-rich substrates (Põlme et al. 2020) | Põlme S, et al. 2020. Fungal Diversity 105:1-16 ((https://doi.org/10.1007/s13225-020-00466-2)) |
| Otu22566 | 1 | k__Fungi(100);p__Ascomycota(100);c__Saccharomycetes(100);o__Saccharomycetales(100);f__Metschnikowiaceae(100);g__Metschnikowia(100);s__Metschnikowia_pulcherrima(100); | Metschnikowia | 13 | Saprotroph | \|Undefined Saprotroph\| | Yeast | NULL | Probable | As nectar-tap_saprotroph (Põlme et al. 2020); Decay sugar-rich substrates (Põlme et al. 2020) | Põlme S, et al. 2020. Fungal Diversity 105:1-16 ((https://doi.org/10.1007/s13225-020-00466-2)) |
| Otu22567 | 1 | k__Fungi(100);p__Ascomycota(100);c__Saccharomycetes(100);o__Saccharomycetales(100);f__Metschnikowiaceae(100);g__Metschnikowia(100);s__Metschnikowia_pulcherrima(100); | Metschnikowia | 13 | Saprotroph | \|Undefined Saprotroph\| | Yeast | NULL | Probable | As nectar-tap_saprotroph (Põlme et al. 2020); Decay sugar-rich substrates (Põlme et al. 2020) | Põlme S, et al. 2020. Fungal Diversity 105:1-16 ((https://doi.org/10.1007/s13225-020-00466-2)) |
| Otu22568 | 1 | k__Fungi(100);p__Ascomycota(100);c__Saccharomycetes(100);o__Saccharomycetales(100);f__Metschnikowiaceae(100);g__Metschnikowia(100);s__Metschnikowia_pulcherrima(100); | Metschnikowia | 13 | Saprotroph | \|Undefined Saprotroph\| | Yeast | NULL | Probable | As nectar-tap_saprotroph (Põlme et al. 2020); Decay sugar-rich substrates (Põlme et al. 2020) | Põlme S, et al. 2020. Fungal Diversity 105:1-16 ((https://doi.org/10.1007/s13225-020-00466-2)) |
| Otu22574 | 1 | k__Fungi(100);p__Ascomycota(100);c__Saccharomycetes(100);o__Saccharomycetales(100);f__Metschnikowiaceae(100);g__Metschnikowia(100);s__Metschnikowia_pulcherrima(100); | Metschnikowia | 13 | Saprotroph | \|Undefined Saprotroph\| | Yeast | NULL | Probable | As nectar-tap_saprotroph (Põlme et al. 2020); Decay sugar-rich substrates (Põlme et al. 2020) | Põlme S, et al. 2020. Fungal Diversity 105:1-16 ((https://doi.org/10.1007/s13225-020-00466-2)) |
| Otu22575 | 1 | k__Fungi(100);p__Ascomycota(100);c__Saccharomycetes(100);o__Saccharomycetales(100);f__Metschnikowiaceae(100);g__Metschnikowia(100);s__Metschnikowia_pulcherrima(100); | Metschnikowia | 13 | Saprotroph | \|Undefined Saprotroph\| | Yeast | NULL | Probable | As nectar-tap_saprotroph (Põlme et al. 2020); Decay sugar-rich substrates (Põlme et al. 2020) | Põlme S, et al. 2020. Fungal Diversity 105:1-16 ((https://doi.org/10.1007/s13225-020-00466-2)) |
| Otu22576 | 1 | k__Fungi(100);p__Ascomycota(100);c__Saccharomycetes(100);o__Saccharomycetales(100);f__Metschnikowiaceae(100);g__Metschnikowia(100);s__Metschnikowia_pulcherrima(100); | Metschnikowia | 13 | Saprotroph | \|Undefined Saprotroph\| | Yeast | NULL | Probable | As nectar-tap_saprotroph (Põlme et al. 2020); Decay sugar-rich substrates (Põlme et al. 2020) | Põlme S, et al. 2020. Fungal Diversity 105:1-16 ((https://doi.org/10.1007/s13225-020-00466-2)) |
| Otu22577 | 1 | k__Fungi(100);p__Ascomycota(100);c__Saccharomycetes(100);o__Saccharomycetales(100);f__Saccharomycetales_family_Incertae_sedis(100);g__Candida(100);g__Candida_unclassified(100); | Candida | 13 | Pathotroph-Saprotroph-Symbiotroph | Animal Pathogen-Endophyte-Endosymbiont-Epiphyte-\|Undefined Saprotroph\| | Dimorphic Yeast-Yeast | NULL | Probable | As nectar-tap_saprotroph (Põlme et al. 2020); Decay sugar-rich substrates (Põlme et al. 2020); This asexual genus is polyphyletic and contain species that function very differently depend on what they are. Caution should be taken when interpreting guilds for Candida species. | Manolakaki D, et al. 2010. Virulence. 1: 367-375 ((https://doi.org/10.4161/viru.1.5.12796)); Mekbib SB, et al. 2011. Tropical Plant Pathology 36:233-240 ((https://doi.org/10.1590/S1982-56762011000400004)); Põlme S, et al. 2020. Fungal Diversity 105:1-16 ((https://doi.org/10.1007/s13225-020-00466-2)) |
| Otu22578 | 1 | k__Fungi(100);p__Ascomycota(100);c__Saccharomycetes(100);o__Saccharomycetales(100);f__Metschnikowiaceae(100);g__Metschnikowia(100);s__Metschnikowia_pulcherrima(100); | Metschnikowia | 13 | Saprotroph | \|Undefined Saprotroph\| | Yeast | NULL | Probable | As nectar-tap_saprotroph (Põlme et al. 2020); Decay sugar-rich substrates (Põlme et al. 2020) | Põlme S, et al. 2020. Fungal Diversity 105:1-16 ((https://doi.org/10.1007/s13225-020-00466-2)) |
| Otu22582 | 1 | k__Fungi(100);p__Ascomycota(100);c__Saccharomycetes(100);o__Saccharomycetales(100);f__Metschnikowiaceae(100);g__Metschnikowia(100);s__Metschnikowia_pulcherrima(100); | Metschnikowia | 13 | Saprotroph | \|Undefined Saprotroph\| | Yeast | NULL | Probable | As nectar-tap_saprotroph (Põlme et al. 2020); Decay sugar-rich substrates (Põlme et al. 2020) | Põlme S, et al. 2020. Fungal Diversity 105:1-16 ((https://doi.org/10.1007/s13225-020-00466-2)) |
| Otu22583 | 1 | k__Fungi(100);p__Ascomycota(100);c__Saccharomycetes(100);o__Saccharomycetales(100);f__Metschnikowiaceae(100);g__Metschnikowia(100);s__Metschnikowia_pulcherrima(100); | Metschnikowia | 13 | Saprotroph | \|Undefined Saprotroph\| | Yeast | NULL | Probable | As nectar-tap_saprotroph (Põlme et al. 2020); Decay sugar-rich substrates (Põlme et al. 2020) | Põlme S, et al. 2020. Fungal Diversity 105:1-16 ((https://doi.org/10.1007/s13225-020-00466-2)) |
| Otu22584 | 1 | k__Fungi(100);p__Ascomycota(100);c__Saccharomycetes(100);o__Saccharomycetales(100);f__Metschnikowiaceae(100);g__Metschnikowia(100);s__Metschnikowia_pulcherrima(100); | Metschnikowia | 13 | Saprotroph | \|Undefined Saprotroph\| | Yeast | NULL | Probable | As nectar-tap_saprotroph (Põlme et al. 2020); Decay sugar-rich substrates (Põlme et al. 2020) | Põlme S, et al. 2020. Fungal Diversity 105:1-16 ((https://doi.org/10.1007/s13225-020-00466-2)) |
| Otu22585 | 1 | k__Fungi(100);p__Ascomycota(100);c__Saccharomycetes(100);o__Saccharomycetales(100);f__Metschnikowiaceae(100);g__Metschnikowia(100);s__Metschnikowia_pulcherrima(100); | Metschnikowia | 13 | Saprotroph | \|Undefined Saprotroph\| | Yeast | NULL | Probable | As nectar-tap_saprotroph (Põlme et al. 2020); Decay sugar-rich substrates (Põlme et al. 2020) | Põlme S, et al. 2020. Fungal Diversity 105:1-16 ((https://doi.org/10.1007/s13225-020-00466-2)) |
| Otu22586 | 1 | k__Fungi(100);p__Ascomycota(100);c__Saccharomycetes(100);o__Saccharomycetales(100);f__Metschnikowiaceae(100);g__Metschnikowia(100);g__Metschnikowia_unclassified(100); | Metschnikowia | 13 | Saprotroph | \|Undefined Saprotroph\| | Yeast | NULL | Probable | As nectar-tap_saprotroph (Põlme et al. 2020); Decay sugar-rich substrates (Põlme et al. 2020) | Põlme S, et al. 2020. Fungal Diversity 105:1-16 ((https://doi.org/10.1007/s13225-020-00466-2)) |
| Otu22587 | 1 | k__Fungi(100);p__Ascomycota(100);c__Saccharomycetes(100);o__Saccharomycetales(100);f__Metschnikowiaceae(100);g__Metschnikowia(100);s__Metschnikowia_pulcherrima(100); | Metschnikowia | 13 | Saprotroph | \|Undefined Saprotroph\| | Yeast | NULL | Probable | As nectar-tap_saprotroph (Põlme et al. 2020); Decay sugar-rich substrates (Põlme et al. 2020) | Põlme S, et al. 2020. Fungal Diversity 105:1-16 ((https://doi.org/10.1007/s13225-020-00466-2)) |
| Otu22588 | 1 | k__Fungi(100);p__Ascomycota(100);c__Saccharomycetes(100);o__Saccharomycetales(100);f__Metschnikowiaceae(100);g__Metschnikowia(100);s__Metschnikowia_pulcherrima(100); | Metschnikowia | 13 | Saprotroph | \|Undefined Saprotroph\| | Yeast | NULL | Probable | As nectar-tap_saprotroph (Põlme et al. 2020); Decay sugar-rich substrates (Põlme et al. 2020) | Põlme S, et al. 2020. Fungal Diversity 105:1-16 ((https://doi.org/10.1007/s13225-020-00466-2)) |
| Otu22589 | 1 | k__Fungi(100);p__Ascomycota(100);c__Saccharomycetes(100);o__Saccharomycetales(100);f__Metschnikowiaceae(100);g__Metschnikowia(100);s__Metschnikowia_pulcherrima(100); | Metschnikowia | 13 | Saprotroph | \|Undefined Saprotroph\| | Yeast | NULL | Probable | As nectar-tap_saprotroph (Põlme et al. 2020); Decay sugar-rich substrates (Põlme et al. 2020) | Põlme S, et al. 2020. Fungal Diversity 105:1-16 ((https://doi.org/10.1007/s13225-020-00466-2)) |
| Otu22590 | 1 | k__Fungi(100);p__Ascomycota(100);c__Saccharomycetes(100);o__Saccharomycetales(100);f__Metschnikowiaceae(100);g__Metschnikowia(100);s__Metschnikowia_pulcherrima(100); | Metschnikowia | 13 | Saprotroph | \|Undefined Saprotroph\| | Yeast | NULL | Probable | As nectar-tap_saprotroph (Põlme et al. 2020); Decay sugar-rich substrates (Põlme et al. 2020) | Põlme S, et al. 2020. Fungal Diversity 105:1-16 ((https://doi.org/10.1007/s13225-020-00466-2)) |
| Otu22596 | 1 | k__Fungi(100);p__Ascomycota(100);c__Saccharomycetes(100);o__Saccharomycetales(100);f__Saccharomycetales_family_Incertae_sedis(100);g__Candida(100);g__Candida_unclassified(100); | Candida | 13 | Pathotroph-Saprotroph-Symbiotroph | Animal Pathogen-Endophyte-Endosymbiont-Epiphyte-\|Undefined Saprotroph\| | Dimorphic Yeast-Yeast | NULL | Probable | As nectar-tap_saprotroph (Põlme et al. 2020); Decay sugar-rich substrates (Põlme et al. 2020); This asexual genus is polyphyletic and contain species that function very differently depend on what they are. Caution should be taken when interpreting guilds for Candida species. | Manolakaki D, et al. 2010. Virulence. 1: 367-375 ((https://doi.org/10.4161/viru.1.5.12796)); Mekbib SB, et al. 2011. Tropical Plant Pathology 36:233-240 ((https://doi.org/10.1590/S1982-56762011000400004)); Põlme S, et al. 2020. Fungal Diversity 105:1-16 ((https://doi.org/10.1007/s13225-020-00466-2)) |
| Otu22597 | 1 | k__Fungi(100);p__Ascomycota(100);c__Saccharomycetes(100);o__Saccharomycetales(100);f__Metschnikowiaceae(100);g__Metschnikowia(100);s__Metschnikowia_pulcherrima(100); | Metschnikowia | 13 | Saprotroph | \|Undefined Saprotroph\| | Yeast | NULL | Probable | As nectar-tap_saprotroph (Põlme et al. 2020); Decay sugar-rich substrates (Põlme et al. 2020) | Põlme S, et al. 2020. Fungal Diversity 105:1-16 ((https://doi.org/10.1007/s13225-020-00466-2)) |
| Otu22598 | 1 | k__Fungi(100);p__Ascomycota(100);c__Saccharomycetes(100);o__Saccharomycetales(100);f__Metschnikowiaceae(100);g__Metschnikowia(100);s__Metschnikowia_pulcherrima(100); | Metschnikowia | 13 | Saprotroph | \|Undefined Saprotroph\| | Yeast | NULL | Probable | As nectar-tap_saprotroph (Põlme et al. 2020); Decay sugar-rich substrates (Põlme et al. 2020) | Põlme S, et al. 2020. Fungal Diversity 105:1-16 ((https://doi.org/10.1007/s13225-020-00466-2)) |
| Otu22603 | 1 | k__Fungi(100);p__Ascomycota(100);c__Saccharomycetes(100);o__Saccharomycetales(100);f__Metschnikowiaceae(100);g__Metschnikowia(100);g__Metschnikowia_unclassified(100); | Metschnikowia | 13 | Saprotroph | \|Undefined Saprotroph\| | Yeast | NULL | Probable | As nectar-tap_saprotroph (Põlme et al. 2020); Decay sugar-rich substrates (Põlme et al. 2020) | Põlme S, et al. 2020. Fungal Diversity 105:1-16 ((https://doi.org/10.1007/s13225-020-00466-2)) |
| Otu22604 | 1 | k__Fungi(100);p__Ascomycota(100);c__Saccharomycetes(100);o__Saccharomycetales(100);f__Metschnikowiaceae(100);g__Metschnikowia(100);s__Metschnikowia_pulcherrima(100); | Metschnikowia | 13 | Saprotroph | \|Undefined Saprotroph\| | Yeast | NULL | Probable | As nectar-tap_saprotroph (Põlme et al. 2020); Decay sugar-rich substrates (Põlme et al. 2020) | Põlme S, et al. 2020. Fungal Diversity 105:1-16 ((https://doi.org/10.1007/s13225-020-00466-2)) |
| Otu22605 | 1 | k__Fungi(100);p__Ascomycota(100);c__Saccharomycetes(100);o__Saccharomycetales(100);f__Metschnikowiaceae(100);g__Metschnikowia(100);g__Metschnikowia_unclassified(100); | Metschnikowia | 13 | Saprotroph | \|Undefined Saprotroph\| | Yeast | NULL | Probable | As nectar-tap_saprotroph (Põlme et al. 2020); Decay sugar-rich substrates (Põlme et al. 2020) | Põlme S, et al. 2020. Fungal Diversity 105:1-16 ((https://doi.org/10.1007/s13225-020-00466-2)) |
| Otu22606 | 1 | k__Fungi(100);p__Ascomycota(100);c__Saccharomycetes(100);o__Saccharomycetales(100);f__Metschnikowiaceae(100);g__Metschnikowia(100);s__Metschnikowia_pulcherrima(100); | Metschnikowia | 13 | Saprotroph | \|Undefined Saprotroph\| | Yeast | NULL | Probable | As nectar-tap_saprotroph (Põlme et al. 2020); Decay sugar-rich substrates (Põlme et al. 2020) | Põlme S, et al. 2020. Fungal Diversity 105:1-16 ((https://doi.org/10.1007/s13225-020-00466-2)) |
| Otu22607 | 1 | k__Fungi(100);p__Ascomycota(100);c__Saccharomycetes(100);o__Saccharomycetales(100);f__Metschnikowiaceae(100);g__Metschnikowia(100);g__Metschnikowia_unclassified(100); | Metschnikowia | 13 | Saprotroph | \|Undefined Saprotroph\| | Yeast | NULL | Probable | As nectar-tap_saprotroph (Põlme et al. 2020); Decay sugar-rich substrates (Põlme et al. 2020) | Põlme S, et al. 2020. Fungal Diversity 105:1-16 ((https://doi.org/10.1007/s13225-020-00466-2)) |
| Otu22608 | 1 | k__Fungi(100);p__Ascomycota(100);c__Saccharomycetes(100);o__Saccharomycetales(100);f__Metschnikowiaceae(100);g__Metschnikowia(100);s__Metschnikowia_pulcherrima(100); | Metschnikowia | 13 | Saprotroph | \|Undefined Saprotroph\| | Yeast | NULL | Probable | As nectar-tap_saprotroph (Põlme et al. 2020); Decay sugar-rich substrates (Põlme et al. 2020) | Põlme S, et al. 2020. Fungal Diversity 105:1-16 ((https://doi.org/10.1007/s13225-020-00466-2)) |
| Otu22609 | 1 | k__Fungi(100);p__Ascomycota(100);c__Saccharomycetes(100);o__Saccharomycetales(100);f__Metschnikowiaceae(100);g__Metschnikowia(100);s__Metschnikowia_pulcherrima(100); | Metschnikowia | 13 | Saprotroph | \|Undefined Saprotroph\| | Yeast | NULL | Probable | As nectar-tap_saprotroph (Põlme et al. 2020); Decay sugar-rich substrates (Põlme et al. 2020) | Põlme S, et al. 2020. Fungal Diversity 105:1-16 ((https://doi.org/10.1007/s13225-020-00466-2)) |
| Otu22610 | 1 | k__Fungi(100);p__Ascomycota(100);c__Saccharomycetes(100);o__Saccharomycetales(100);f__Metschnikowiaceae(100);g__Metschnikowia(100);g__Metschnikowia_unclassified(100); | Metschnikowia | 13 | Saprotroph | \|Undefined Saprotroph\| | Yeast | NULL | Probable | As nectar-tap_saprotroph (Põlme et al. 2020); Decay sugar-rich substrates (Põlme et al. 2020) | Põlme S, et al. 2020. Fungal Diversity 105:1-16 ((https://doi.org/10.1007/s13225-020-00466-2)) |
| Otu22611 | 1 | k__Fungi(100);p__Ascomycota(100);c__Saccharomycetes(100);o__Saccharomycetales(100);f__Metschnikowiaceae(100);g__Metschnikowia(100);g__Metschnikowia_unclassified(100); | Metschnikowia | 13 | Saprotroph | \|Undefined Saprotroph\| | Yeast | NULL | Probable | As nectar-tap_saprotroph (Põlme et al. 2020); Decay sugar-rich substrates (Põlme et al. 2020) | Põlme S, et al. 2020. Fungal Diversity 105:1-16 ((https://doi.org/10.1007/s13225-020-00466-2)) |
| Otu22612 | 1 | k__Fungi(100);p__Ascomycota(100);c__Saccharomycetes(100);o__Saccharomycetales(100);f__Metschnikowiaceae(100);g__Metschnikowia(100);s__Metschnikowia_pulcherrima(100); | Metschnikowia | 13 | Saprotroph | \|Undefined Saprotroph\| | Yeast | NULL | Probable | As nectar-tap_saprotroph (Põlme et al. 2020); Decay sugar-rich substrates (Põlme et al. 2020) | Põlme S, et al. 2020. Fungal Diversity 105:1-16 ((https://doi.org/10.1007/s13225-020-00466-2)) |
| Otu22613 | 1 | k__Fungi(100);p__Ascomycota(100);c__Saccharomycetes(100);o__Saccharomycetales(100);f__Metschnikowiaceae(100);g__Metschnikowia(100);s__Metschnikowia_pulcherrima(100); | Metschnikowia | 13 | Saprotroph | \|Undefined Saprotroph\| | Yeast | NULL | Probable | As nectar-tap_saprotroph (Põlme et al. 2020); Decay sugar-rich substrates (Põlme et al. 2020) | Põlme S, et al. 2020. Fungal Diversity 105:1-16 ((https://doi.org/10.1007/s13225-020-00466-2)) |
| Otu22614 | 1 | k__Fungi(100);p__Ascomycota(100);c__Saccharomycetes(100);o__Saccharomycetales(100);f__Metschnikowiaceae(100);g__Metschnikowia(100);g__Metschnikowia_unclassified(100); | Metschnikowia | 13 | Saprotroph | \|Undefined Saprotroph\| | Yeast | NULL | Probable | As nectar-tap_saprotroph (Põlme et al. 2020); Decay sugar-rich substrates (Põlme et al. 2020) | Põlme S, et al. 2020. Fungal Diversity 105:1-16 ((https://doi.org/10.1007/s13225-020-00466-2)) |
| Otu22616 | 1 | k__Fungi(100);p__Ascomycota(100);c__Saccharomycetes(100);o__Saccharomycetales(100);f__Metschnikowiaceae(100);g__Metschnikowia(100);s__Metschnikowia_pulcherrima(100); | Metschnikowia | 13 | Saprotroph | \|Undefined Saprotroph\| | Yeast | NULL | Probable | As nectar-tap_saprotroph (Põlme et al. 2020); Decay sugar-rich substrates (Põlme et al. 2020) | Põlme S, et al. 2020. Fungal Diversity 105:1-16 ((https://doi.org/10.1007/s13225-020-00466-2)) |
| Otu22618 | 1 | k__Fungi(100);p__Ascomycota(100);c__Saccharomycetes(100);o__Saccharomycetales(100);f__Metschnikowiaceae(100);g__Metschnikowia(100);s__Metschnikowia_pulcherrima(100); | Metschnikowia | 13 | Saprotroph | \|Undefined Saprotroph\| | Yeast | NULL | Probable | As nectar-tap_saprotroph (Põlme et al. 2020); Decay sugar-rich substrates (Põlme et al. 2020) | Põlme S, et al. 2020. Fungal Diversity 105:1-16 ((https://doi.org/10.1007/s13225-020-00466-2)) |
| Otu22619 | 1 | k__Fungi(100);p__Ascomycota(100);c__Saccharomycetes(100);o__Saccharomycetales(100);f__Metschnikowiaceae(100);g__Metschnikowia(100);s__Metschnikowia_pulcherrima(100); | Metschnikowia | 13 | Saprotroph | \|Undefined Saprotroph\| | Yeast | NULL | Probable | As nectar-tap_saprotroph (Põlme et al. 2020); Decay sugar-rich substrates (Põlme et al. 2020) | Põlme S, et al. 2020. Fungal Diversity 105:1-16 ((https://doi.org/10.1007/s13225-020-00466-2)) |
| Otu22620 | 1 | k__Fungi(100);p__Ascomycota(100);c__Saccharomycetes(100);o__Saccharomycetales(100);f__Metschnikowiaceae(100);g__Metschnikowia(100);s__Metschnikowia_pulcherrima(100); | Metschnikowia | 13 | Saprotroph | \|Undefined Saprotroph\| | Yeast | NULL | Probable | As nectar-tap_saprotroph (Põlme et al. 2020); Decay sugar-rich substrates (Põlme et al. 2020) | Põlme S, et al. 2020. Fungal Diversity 105:1-16 ((https://doi.org/10.1007/s13225-020-00466-2)) |
| Otu22621 | 1 | k__Fungi(100);p__Ascomycota(100);c__Saccharomycetes(100);o__Saccharomycetales(100);f__Metschnikowiaceae(100);g__Metschnikowia(100);s__Metschnikowia_pulcherrima(100); | Metschnikowia | 13 | Saprotroph | \|Undefined Saprotroph\| | Yeast | NULL | Probable | As nectar-tap_saprotroph (Põlme et al. 2020); Decay sugar-rich substrates (Põlme et al. 2020) | Põlme S, et al. 2020. Fungal Diversity 105:1-16 ((https://doi.org/10.1007/s13225-020-00466-2)) |
| Otu22624 | 1 | k__Fungi(100);p__Ascomycota(100);c__Saccharomycetes(100);o__Saccharomycetales(100);f__Metschnikowiaceae(100);g__Metschnikowia(100);g__Metschnikowia_unclassified(100); | Metschnikowia | 13 | Saprotroph | \|Undefined Saprotroph\| | Yeast | NULL | Probable | As nectar-tap_saprotroph (Põlme et al. 2020); Decay sugar-rich substrates (Põlme et al. 2020) | Põlme S, et al. 2020. Fungal Diversity 105:1-16 ((https://doi.org/10.1007/s13225-020-00466-2)) |
| Otu22625 | 1 | k__Fungi(100);p__Ascomycota(100);c__Saccharomycetes(100);o__Saccharomycetales(100);f__Metschnikowiaceae(100);g__Metschnikowia(100);s__Metschnikowia_pulcherrima(100); | Metschnikowia | 13 | Saprotroph | \|Undefined Saprotroph\| | Yeast | NULL | Probable | As nectar-tap_saprotroph (Põlme et al. 2020); Decay sugar-rich substrates (Põlme et al. 2020) | Põlme S, et al. 2020. Fungal Diversity 105:1-16 ((https://doi.org/10.1007/s13225-020-00466-2)) |
| Otu22626 | 1 | k__Fungi(100);p__Ascomycota(100);c__Saccharomycetes(100);o__Saccharomycetales(100);f__Metschnikowiaceae(100);g__Metschnikowia(100);s__Metschnikowia_pulcherrima(100); | Metschnikowia | 13 | Saprotroph | \|Undefined Saprotroph\| | Yeast | NULL | Probable | As nectar-tap_saprotroph (Põlme et al. 2020); Decay sugar-rich substrates (Põlme et al. 2020) | Põlme S, et al. 2020. Fungal Diversity 105:1-16 ((https://doi.org/10.1007/s13225-020-00466-2)) |
| Otu22627 | 1 | k__Fungi(100);p__Ascomycota(100);c__Saccharomycetes(100);o__Saccharomycetales(100);f__Metschnikowiaceae(100);g__Metschnikowia(100);g__Metschnikowia_unclassified(100); | Metschnikowia | 13 | Saprotroph | \|Undefined Saprotroph\| | Yeast | NULL | Probable | As nectar-tap_saprotroph (Põlme et al. 2020); Decay sugar-rich substrates (Põlme et al. 2020) | Põlme S, et al. 2020. Fungal Diversity 105:1-16 ((https://doi.org/10.1007/s13225-020-00466-2)) |
| Otu22628 | 1 | k__Fungi(100);p__Ascomycota(100);c__Saccharomycetes(100);o__Saccharomycetales(100);f__Metschnikowiaceae(100);g__Metschnikowia(100);g__Metschnikowia_unclassified(100); | Metschnikowia | 13 | Saprotroph | \|Undefined Saprotroph\| | Yeast | NULL | Probable | As nectar-tap_saprotroph (Põlme et al. 2020); Decay sugar-rich substrates (Põlme et al. 2020) | Põlme S, et al. 2020. Fungal Diversity 105:1-16 ((https://doi.org/10.1007/s13225-020-00466-2)) |
| Otu22629 | 1 | k__Fungi(100);p__Ascomycota(100);c__Saccharomycetes(100);o__Saccharomycetales(100);f__Metschnikowiaceae(100);g__Metschnikowia(100);s__Metschnikowia_pulcherrima(100); | Metschnikowia | 13 | Saprotroph | \|Undefined Saprotroph\| | Yeast | NULL | Probable | As nectar-tap_saprotroph (Põlme et al. 2020); Decay sugar-rich substrates (Põlme et al. 2020) | Põlme S, et al. 2020. Fungal Diversity 105:1-16 ((https://doi.org/10.1007/s13225-020-00466-2)) |
| Otu22630 | 1 | k__Fungi(100);p__Ascomycota(100);c__Saccharomycetes(100);o__Saccharomycetales(100);f__Metschnikowiaceae(100);g__Metschnikowia(100);s__Metschnikowia_pulcherrima(100); | Metschnikowia | 13 | Saprotroph | \|Undefined Saprotroph\| | Yeast | NULL | Probable | As nectar-tap_saprotroph (Põlme et al. 2020); Decay sugar-rich substrates (Põlme et al. 2020) | Põlme S, et al. 2020. Fungal Diversity 105:1-16 ((https://doi.org/10.1007/s13225-020-00466-2)) |
| Otu22631 | 1 | k__Fungi(100);p__Ascomycota(100);c__Saccharomycetes(100);o__Saccharomycetales(100);f__Metschnikowiaceae(100);g__Metschnikowia(100);g__Metschnikowia_unclassified(100); | Metschnikowia | 13 | Saprotroph | \|Undefined Saprotroph\| | Yeast | NULL | Probable | As nectar-tap_saprotroph (Põlme et al. 2020); Decay sugar-rich substrates (Põlme et al. 2020) | Põlme S, et al. 2020. Fungal Diversity 105:1-16 ((https://doi.org/10.1007/s13225-020-00466-2)) |
| Otu22632 | 1 | k__Fungi(100);p__Ascomycota(100);c__Saccharomycetes(100);o__Saccharomycetales(100);f__Metschnikowiaceae(100);g__Metschnikowia(100);s__Metschnikowia_pulcherrima(100); | Metschnikowia | 13 | Saprotroph | \|Undefined Saprotroph\| | Yeast | NULL | Probable | As nectar-tap_saprotroph (Põlme et al. 2020); Decay sugar-rich substrates (Põlme et al. 2020) | Põlme S, et al. 2020. Fungal Diversity 105:1-16 ((https://doi.org/10.1007/s13225-020-00466-2)) |
| Otu22633 | 1 | k__Fungi(100);p__Ascomycota(100);c__Saccharomycetes(100);o__Saccharomycetales(100);f__Metschnikowiaceae(100);g__Metschnikowia(100);s__Metschnikowia_pulcherrima(100); | Metschnikowia | 13 | Saprotroph | \|Undefined Saprotroph\| | Yeast | NULL | Probable | As nectar-tap_saprotroph (Põlme et al. 2020); Decay sugar-rich substrates (Põlme et al. 2020) | Põlme S, et al. 2020. Fungal Diversity 105:1-16 ((https://doi.org/10.1007/s13225-020-00466-2)) |
| Otu22634 | 1 | k__Fungi(100);p__Ascomycota(100);c__Saccharomycetes(100);o__Saccharomycetales(100);f__Metschnikowiaceae(100);g__Metschnikowia(100);s__Metschnikowia_pulcherrima(100); | Metschnikowia | 13 | Saprotroph | \|Undefined Saprotroph\| | Yeast | NULL | Probable | As nectar-tap_saprotroph (Põlme et al. 2020); Decay sugar-rich substrates (Põlme et al. 2020) | Põlme S, et al. 2020. Fungal Diversity 105:1-16 ((https://doi.org/10.1007/s13225-020-00466-2)) |
| Otu22635 | 1 | k__Fungi(100);p__Ascomycota(100);c__Saccharomycetes(100);o__Saccharomycetales(100);f__Metschnikowiaceae(100);g__Metschnikowia(100);s__Metschnikowia_pulcherrima(100); | Metschnikowia | 13 | Saprotroph | \|Undefined Saprotroph\| | Yeast | NULL | Probable | As nectar-tap_saprotroph (Põlme et al. 2020); Decay sugar-rich substrates (Põlme et al. 2020) | Põlme S, et al. 2020. Fungal Diversity 105:1-16 ((https://doi.org/10.1007/s13225-020-00466-2)) |
| Otu22636 | 1 | k__Fungi(100);p__Ascomycota(100);c__Saccharomycetes(100);o__Saccharomycetales(100);f__Metschnikowiaceae(100);g__Metschnikowia(100);s__Metschnikowia_pulcherrima(100); | Metschnikowia | 13 | Saprotroph | \|Undefined Saprotroph\| | Yeast | NULL | Probable | As nectar-tap_saprotroph (Põlme et al. 2020); Decay sugar-rich substrates (Põlme et al. 2020) | Põlme S, et al. 2020. Fungal Diversity 105:1-16 ((https://doi.org/10.1007/s13225-020-00466-2)) |
| Otu22637 | 1 | k__Fungi(100);p__Ascomycota(100);c__Saccharomycetes(100);o__Saccharomycetales(100);f__Metschnikowiaceae(100);g__Metschnikowia(100);s__Metschnikowia_pulcherrima(100); | Metschnikowia | 13 | Saprotroph | \|Undefined Saprotroph\| | Yeast | NULL | Probable | As nectar-tap_saprotroph (Põlme et al. 2020); Decay sugar-rich substrates (Põlme et al. 2020) | Põlme S, et al. 2020. Fungal Diversity 105:1-16 ((https://doi.org/10.1007/s13225-020-00466-2)) |
| Otu22638 | 1 | k__Fungi(100);p__Ascomycota(100);c__Saccharomycetes(100);o__Saccharomycetales(100);f__Metschnikowiaceae(100);g__Metschnikowia(100);s__Metschnikowia_pulcherrima(100); | Metschnikowia | 13 | Saprotroph | \|Undefined Saprotroph\| | Yeast | NULL | Probable | As nectar-tap_saprotroph (Põlme et al. 2020); Decay sugar-rich substrates (Põlme et al. 2020) | Põlme S, et al. 2020. Fungal Diversity 105:1-16 ((https://doi.org/10.1007/s13225-020-00466-2)) |
| Otu22639 | 1 | k__Fungi(100);p__Ascomycota(100);c__Saccharomycetes(100);o__Saccharomycetales(100);f__Metschnikowiaceae(100);g__Metschnikowia(100);s__Metschnikowia_pulcherrima(100); | Metschnikowia | 13 | Saprotroph | \|Undefined Saprotroph\| | Yeast | NULL | Probable | As nectar-tap_saprotroph (Põlme et al. 2020); Decay sugar-rich substrates (Põlme et al. 2020) | Põlme S, et al. 2020. Fungal Diversity 105:1-16 ((https://doi.org/10.1007/s13225-020-00466-2)) |
| Otu22640 | 1 | k__Fungi(100);p__Ascomycota(100);c__Saccharomycetes(100);o__Saccharomycetales(100);f__Metschnikowiaceae(100);g__Metschnikowia(100);s__Metschnikowia_pulcherrima(100); | Metschnikowia | 13 | Saprotroph | \|Undefined Saprotroph\| | Yeast | NULL | Probable | As nectar-tap_saprotroph (Põlme et al. 2020); Decay sugar-rich substrates (Põlme et al. 2020) | Põlme S, et al. 2020. Fungal Diversity 105:1-16 ((https://doi.org/10.1007/s13225-020-00466-2)) |
| Otu22641 | 1 | k__Fungi(100);p__Ascomycota(100);c__Saccharomycetes(100);o__Saccharomycetales(100);f__Metschnikowiaceae(100);g__Metschnikowia(100);s__Metschnikowia_pulcherrima(100); | Metschnikowia | 13 | Saprotroph | \|Undefined Saprotroph\| | Yeast | NULL | Probable | As nectar-tap_saprotroph (Põlme et al. 2020); Decay sugar-rich substrates (Põlme et al. 2020) | Põlme S, et al. 2020. Fungal Diversity 105:1-16 ((https://doi.org/10.1007/s13225-020-00466-2)) |
| Otu22642 | 1 | k__Fungi(100);p__Ascomycota(100);c__Saccharomycetes(100);o__Saccharomycetales(100);f__Metschnikowiaceae(100);g__Metschnikowia(100);s__Metschnikowia_pulcherrima(100); | Metschnikowia | 13 | Saprotroph | \|Undefined Saprotroph\| | Yeast | NULL | Probable | As nectar-tap_saprotroph (Põlme et al. 2020); Decay sugar-rich substrates (Põlme et al. 2020) | Põlme S, et al. 2020. Fungal Diversity 105:1-16 ((https://doi.org/10.1007/s13225-020-00466-2)) |
| Otu22643 | 1 | k__Fungi(100);p__Ascomycota(100);c__Saccharomycetes(100);o__Saccharomycetales(100);f__Metschnikowiaceae(100);g__Metschnikowia(100);s__Metschnikowia_pulcherrima(100); | Metschnikowia | 13 | Saprotroph | \|Undefined Saprotroph\| | Yeast | NULL | Probable | As nectar-tap_saprotroph (Põlme et al. 2020); Decay sugar-rich substrates (Põlme et al. 2020) | Põlme S, et al. 2020. Fungal Diversity 105:1-16 ((https://doi.org/10.1007/s13225-020-00466-2)) |
| Otu22644 | 1 | k__Fungi(100);p__Ascomycota(100);c__Saccharomycetes(100);o__Saccharomycetales(100);f__Metschnikowiaceae(100);g__Metschnikowia(100);s__Metschnikowia_pulcherrima(100); | Metschnikowia | 13 | Saprotroph | \|Undefined Saprotroph\| | Yeast | NULL | Probable | As nectar-tap_saprotroph (Põlme et al. 2020); Decay sugar-rich substrates (Põlme et al. 2020) | Põlme S, et al. 2020. Fungal Diversity 105:1-16 ((https://doi.org/10.1007/s13225-020-00466-2)) |
| Otu22645 | 1 | k__Fungi(100);p__Ascomycota(100);c__Saccharomycetes(100);o__Saccharomycetales(100);f__Metschnikowiaceae(100);g__Metschnikowia(100);g__Metschnikowia_unclassified(100); | Metschnikowia | 13 | Saprotroph | \|Undefined Saprotroph\| | Yeast | NULL | Probable | As nectar-tap_saprotroph (Põlme et al. 2020); Decay sugar-rich substrates (Põlme et al. 2020) | Põlme S, et al. 2020. Fungal Diversity 105:1-16 ((https://doi.org/10.1007/s13225-020-00466-2)) |
| Otu22646 | 1 | k__Fungi(100);p__Ascomycota(100);c__Saccharomycetes(100);o__Saccharomycetales(100);f__Metschnikowiaceae(100);g__Metschnikowia(100);s__Metschnikowia_pulcherrima(100); | Metschnikowia | 13 | Saprotroph | \|Undefined Saprotroph\| | Yeast | NULL | Probable | As nectar-tap_saprotroph (Põlme et al. 2020); Decay sugar-rich substrates (Põlme et al. 2020) | Põlme S, et al. 2020. Fungal Diversity 105:1-16 ((https://doi.org/10.1007/s13225-020-00466-2)) |
| Otu22647 | 1 | k__Fungi(100);p__Ascomycota(100);c__Saccharomycetes(100);o__Saccharomycetales(100);f__Metschnikowiaceae(100);g__Metschnikowia(100);s__Metschnikowia_pulcherrima(100); | Metschnikowia | 13 | Saprotroph | \|Undefined Saprotroph\| | Yeast | NULL | Probable | As nectar-tap_saprotroph (Põlme et al. 2020); Decay sugar-rich substrates (Põlme et al. 2020) | Põlme S, et al. 2020. Fungal Diversity 105:1-16 ((https://doi.org/10.1007/s13225-020-00466-2)) |
| Otu22648 | 1 | k__Fungi(100);p__Ascomycota(100);c__Saccharomycetes(100);o__Saccharomycetales(100);f__Metschnikowiaceae(100);g__Metschnikowia(100);s__Metschnikowia_pulcherrima(100); | Metschnikowia | 13 | Saprotroph | \|Undefined Saprotroph\| | Yeast | NULL | Probable | As nectar-tap_saprotroph (Põlme et al. 2020); Decay sugar-rich substrates (Põlme et al. 2020) | Põlme S, et al. 2020. Fungal Diversity 105:1-16 ((https://doi.org/10.1007/s13225-020-00466-2)) |
| Otu22650 | 1 | k__Fungi(100);p__Ascomycota(100);c__Saccharomycetes(100);o__Saccharomycetales(100);f__Metschnikowiaceae(100);g__Metschnikowia(100);s__Metschnikowia_pulcherrima(100); | Metschnikowia | 13 | Saprotroph | \|Undefined Saprotroph\| | Yeast | NULL | Probable | As nectar-tap_saprotroph (Põlme et al. 2020); Decay sugar-rich substrates (Põlme et al. 2020) | Põlme S, et al. 2020. Fungal Diversity 105:1-16 ((https://doi.org/10.1007/s13225-020-00466-2)) |
| Otu22651 | 1 | k__Fungi(100);p__Ascomycota(100);c__Saccharomycetes(100);o__Saccharomycetales(100);f__Metschnikowiaceae(100);g__Metschnikowia(100);s__Metschnikowia_pulcherrima(100); | Metschnikowia | 13 | Saprotroph | \|Undefined Saprotroph\| | Yeast | NULL | Probable | As nectar-tap_saprotroph (Põlme et al. 2020); Decay sugar-rich substrates (Põlme et al. 2020) | Põlme S, et al. 2020. Fungal Diversity 105:1-16 ((https://doi.org/10.1007/s13225-020-00466-2)) |
| Otu22653 | 1 | k__Fungi(100);p__Ascomycota(100);c__Saccharomycetes(100);o__Saccharomycetales(100);f__Metschnikowiaceae(100);g__Metschnikowia(100);s__Metschnikowia_pulcherrima(100); | Metschnikowia | 13 | Saprotroph | \|Undefined Saprotroph\| | Yeast | NULL | Probable | As nectar-tap_saprotroph (Põlme et al. 2020); Decay sugar-rich substrates (Põlme et al. 2020) | Põlme S, et al. 2020. Fungal Diversity 105:1-16 ((https://doi.org/10.1007/s13225-020-00466-2)) |
| Otu22654 | 1 | k__Fungi(100);p__Ascomycota(100);c__Saccharomycetes(100);o__Saccharomycetales(100);f__Metschnikowiaceae(100);g__Metschnikowia(100);s__Metschnikowia_pulcherrima(100); | Metschnikowia | 13 | Saprotroph | \|Undefined Saprotroph\| | Yeast | NULL | Probable | As nectar-tap_saprotroph (Põlme et al. 2020); Decay sugar-rich substrates (Põlme et al. 2020) | Põlme S, et al. 2020. Fungal Diversity 105:1-16 ((https://doi.org/10.1007/s13225-020-00466-2)) |
| Otu22655 | 1 | k__Fungi(100);p__Ascomycota(100);c__Saccharomycetes(100);o__Saccharomycetales(100);f__Metschnikowiaceae(100);g__Metschnikowia(100);s__Metschnikowia_pulcherrima(100); | Metschnikowia | 13 | Saprotroph | \|Undefined Saprotroph\| | Yeast | NULL | Probable | As nectar-tap_saprotroph (Põlme et al. 2020); Decay sugar-rich substrates (Põlme et al. 2020) | Põlme S, et al. 2020. Fungal Diversity 105:1-16 ((https://doi.org/10.1007/s13225-020-00466-2)) |
| Otu22656 | 1 | k__Fungi(100);p__Ascomycota(100);c__Saccharomycetes(100);o__Saccharomycetales(100);f__Metschnikowiaceae(100);g__Metschnikowia(100);s__Metschnikowia_pulcherrima(100); | Metschnikowia | 13 | Saprotroph | \|Undefined Saprotroph\| | Yeast | NULL | Probable | As nectar-tap_saprotroph (Põlme et al. 2020); Decay sugar-rich substrates (Põlme et al. 2020) | Põlme S, et al. 2020. Fungal Diversity 105:1-16 ((https://doi.org/10.1007/s13225-020-00466-2)) |
| Otu22682 | 1 | k__Fungi(100);p__Ascomycota(100);c__Saccharomycetes(100);o__Saccharomycetales(100);f__Metschnikowiaceae(100);g__Metschnikowia(100);s__Metschnikowia_pulcherrima(100); | Metschnikowia | 13 | Saprotroph | \|Undefined Saprotroph\| | Yeast | NULL | Probable | As nectar-tap_saprotroph (Põlme et al. 2020); Decay sugar-rich substrates (Põlme et al. 2020) | Põlme S, et al. 2020. Fungal Diversity 105:1-16 ((https://doi.org/10.1007/s13225-020-00466-2)) |
| Otu22683 | 1 | k__Fungi(100);p__Ascomycota(100);c__Saccharomycetes(100);o__Saccharomycetales(100);f__Metschnikowiaceae(100);g__Metschnikowia(100);g__Metschnikowia_unclassified(100); | Metschnikowia | 13 | Saprotroph | \|Undefined Saprotroph\| | Yeast | NULL | Probable | As nectar-tap_saprotroph (Põlme et al. 2020); Decay sugar-rich substrates (Põlme et al. 2020) | Põlme S, et al. 2020. Fungal Diversity 105:1-16 ((https://doi.org/10.1007/s13225-020-00466-2)) |
| Otu22684 | 1 | k__Fungi(100);p__Ascomycota(100);c__Saccharomycetes(100);o__Saccharomycetales(100);f__Metschnikowiaceae(100);g__Metschnikowia(100);s__Metschnikowia_pulcherrima(100); | Metschnikowia | 13 | Saprotroph | \|Undefined Saprotroph\| | Yeast | NULL | Probable | As nectar-tap_saprotroph (Põlme et al. 2020); Decay sugar-rich substrates (Põlme et al. 2020) | Põlme S, et al. 2020. Fungal Diversity 105:1-16 ((https://doi.org/10.1007/s13225-020-00466-2)) |
| Otu22685 | 1 | k__Fungi(100);p__Ascomycota(100);c__Saccharomycetes(100);o__Saccharomycetales(100);f__Metschnikowiaceae(100);g__Metschnikowia(100);s__Metschnikowia_pulcherrima(100); | Metschnikowia | 13 | Saprotroph | \|Undefined Saprotroph\| | Yeast | NULL | Probable | As nectar-tap_saprotroph (Põlme et al. 2020); Decay sugar-rich substrates (Põlme et al. 2020) | Põlme S, et al. 2020. Fungal Diversity 105:1-16 ((https://doi.org/10.1007/s13225-020-00466-2)) |
| Otu22686 | 1 | k__Fungi(100);p__Ascomycota(100);c__Saccharomycetes(100);o__Saccharomycetales(100);f__Metschnikowiaceae(100);g__Metschnikowia(100);g__Metschnikowia_unclassified(100); | Metschnikowia | 13 | Saprotroph | \|Undefined Saprotroph\| | Yeast | NULL | Probable | As nectar-tap_saprotroph (Põlme et al. 2020); Decay sugar-rich substrates (Põlme et al. 2020) | Põlme S, et al. 2020. Fungal Diversity 105:1-16 ((https://doi.org/10.1007/s13225-020-00466-2)) |
| Otu22687 | 1 | k__Fungi(100);p__Ascomycota(100);c__Saccharomycetes(100);o__Saccharomycetales(100);f__Metschnikowiaceae(100);g__Metschnikowia(100);s__Metschnikowia_pulcherrima(100); | Metschnikowia | 13 | Saprotroph | \|Undefined Saprotroph\| | Yeast | NULL | Probable | As nectar-tap_saprotroph (Põlme et al. 2020); Decay sugar-rich substrates (Põlme et al. 2020) | Põlme S, et al. 2020. Fungal Diversity 105:1-16 ((https://doi.org/10.1007/s13225-020-00466-2)) |
| Otu22688 | 1 | k__Fungi(100);p__Ascomycota(100);c__Saccharomycetes(100);o__Saccharomycetales(100);f__Metschnikowiaceae(100);g__Metschnikowia(100);s__Metschnikowia_pulcherrima(100); | Metschnikowia | 13 | Saprotroph | \|Undefined Saprotroph\| | Yeast | NULL | Probable | As nectar-tap_saprotroph (Põlme et al. 2020); Decay sugar-rich substrates (Põlme et al. 2020) | Põlme S, et al. 2020. Fungal Diversity 105:1-16 ((https://doi.org/10.1007/s13225-020-00466-2)) |
| Otu22689 | 1 | k__Fungi(100);p__Ascomycota(100);c__Saccharomycetes(100);o__Saccharomycetales(100);f__Metschnikowiaceae(100);g__Metschnikowia(100);s__Metschnikowia_pulcherrima(100); | Metschnikowia | 13 | Saprotroph | \|Undefined Saprotroph\| | Yeast | NULL | Probable | As nectar-tap_saprotroph (Põlme et al. 2020); Decay sugar-rich substrates (Põlme et al. 2020) | Põlme S, et al. 2020. Fungal Diversity 105:1-16 ((https://doi.org/10.1007/s13225-020-00466-2)) |
| Otu22690 | 1 | k__Fungi(100);p__Ascomycota(100);c__Saccharomycetes(100);o__Saccharomycetales(100);f__Metschnikowiaceae(100);g__Metschnikowia(100);s__Metschnikowia_pulcherrima(100); | Metschnikowia | 13 | Saprotroph | \|Undefined Saprotroph\| | Yeast | NULL | Probable | As nectar-tap_saprotroph (Põlme et al. 2020); Decay sugar-rich substrates (Põlme et al. 2020) | Põlme S, et al. 2020. Fungal Diversity 105:1-16 ((https://doi.org/10.1007/s13225-020-00466-2)) |
| Otu22691 | 1 | k__Fungi(100);p__Ascomycota(100);c__Saccharomycetes(100);o__Saccharomycetales(100);f__Metschnikowiaceae(100);g__Metschnikowia(100);g__Metschnikowia_unclassified(100); | Metschnikowia | 13 | Saprotroph | \|Undefined Saprotroph\| | Yeast | NULL | Probable | As nectar-tap_saprotroph (Põlme et al. 2020); Decay sugar-rich substrates (Põlme et al. 2020) | Põlme S, et al. 2020. Fungal Diversity 105:1-16 ((https://doi.org/10.1007/s13225-020-00466-2)) |
| Otu22692 | 1 | k__Fungi(100);p__Ascomycota(100);c__Saccharomycetes(100);o__Saccharomycetales(100);f__Metschnikowiaceae(100);g__Metschnikowia(100);g__Metschnikowia_unclassified(100); | Metschnikowia | 13 | Saprotroph | \|Undefined Saprotroph\| | Yeast | NULL | Probable | As nectar-tap_saprotroph (Põlme et al. 2020); Decay sugar-rich substrates (Põlme et al. 2020) | Põlme S, et al. 2020. Fungal Diversity 105:1-16 ((https://doi.org/10.1007/s13225-020-00466-2)) |
| Otu22693 | 1 | k__Fungi(100);p__Ascomycota(100);c__Saccharomycetes(100);o__Saccharomycetales(100);f__Metschnikowiaceae(100);g__Metschnikowia(100);s__Metschnikowia_pulcherrima(100); | Metschnikowia | 13 | Saprotroph | \|Undefined Saprotroph\| | Yeast | NULL | Probable | As nectar-tap_saprotroph (Põlme et al. 2020); Decay sugar-rich substrates (Põlme et al. 2020) | Põlme S, et al. 2020. Fungal Diversity 105:1-16 ((https://doi.org/10.1007/s13225-020-00466-2)) |
| Otu22694 | 1 | k__Fungi(100);p__Ascomycota(100);c__Saccharomycetes(100);o__Saccharomycetales(100);f__Metschnikowiaceae(100);g__Metschnikowia(100);s__Metschnikowia_pulcherrima(100); | Metschnikowia | 13 | Saprotroph | \|Undefined Saprotroph\| | Yeast | NULL | Probable | As nectar-tap_saprotroph (Põlme et al. 2020); Decay sugar-rich substrates (Põlme et al. 2020) | Põlme S, et al. 2020. Fungal Diversity 105:1-16 ((https://doi.org/10.1007/s13225-020-00466-2)) |
| Otu22695 | 1 | k__Fungi(100);p__Ascomycota(100);c__Saccharomycetes(100);o__Saccharomycetales(100);f__Metschnikowiaceae(100);g__Metschnikowia(100);s__Metschnikowia_pulcherrima(100); | Metschnikowia | 13 | Saprotroph | \|Undefined Saprotroph\| | Yeast | NULL | Probable | As nectar-tap_saprotroph (Põlme et al. 2020); Decay sugar-rich substrates (Põlme et al. 2020) | Põlme S, et al. 2020. Fungal Diversity 105:1-16 ((https://doi.org/10.1007/s13225-020-00466-2)) |
| Otu22696 | 1 | k__Fungi(100);p__Ascomycota(100);c__Saccharomycetes(100);o__Saccharomycetales(100);f__Metschnikowiaceae(100);g__Metschnikowia(100);g__Metschnikowia_unclassified(100); | Metschnikowia | 13 | Saprotroph | \|Undefined Saprotroph\| | Yeast | NULL | Probable | As nectar-tap_saprotroph (Põlme et al. 2020); Decay sugar-rich substrates (Põlme et al. 2020) | Põlme S, et al. 2020. Fungal Diversity 105:1-16 ((https://doi.org/10.1007/s13225-020-00466-2)) |
| Otu22697 | 1 | k__Fungi(100);p__Ascomycota(100);c__Saccharomycetes(100);o__Saccharomycetales(100);f__Metschnikowiaceae(100);g__Metschnikowia(100);s__Metschnikowia_pulcherrima(100); | Metschnikowia | 13 | Saprotroph | \|Undefined Saprotroph\| | Yeast | NULL | Probable | As nectar-tap_saprotroph (Põlme et al. 2020); Decay sugar-rich substrates (Põlme et al. 2020) | Põlme S, et al. 2020. Fungal Diversity 105:1-16 ((https://doi.org/10.1007/s13225-020-00466-2)) |
| Otu22698 | 1 | k__Fungi(100);p__Ascomycota(100);c__Saccharomycetes(100);o__Saccharomycetales(100);f__Metschnikowiaceae(100);g__Metschnikowia(100);s__Metschnikowia_pulcherrima(100); | Metschnikowia | 13 | Saprotroph | \|Undefined Saprotroph\| | Yeast | NULL | Probable | As nectar-tap_saprotroph (Põlme et al. 2020); Decay sugar-rich substrates (Põlme et al. 2020) | Põlme S, et al. 2020. Fungal Diversity 105:1-16 ((https://doi.org/10.1007/s13225-020-00466-2)) |
| Otu22699 | 1 | k__Fungi(100);p__Ascomycota(100);c__Saccharomycetes(100);o__Saccharomycetales(100);f__Metschnikowiaceae(100);g__Metschnikowia(100);s__Metschnikowia_pulcherrima(100); | Metschnikowia | 13 | Saprotroph | \|Undefined Saprotroph\| | Yeast | NULL | Probable | As nectar-tap_saprotroph (Põlme et al. 2020); Decay sugar-rich substrates (Põlme et al. 2020) | Põlme S, et al. 2020. Fungal Diversity 105:1-16 ((https://doi.org/10.1007/s13225-020-00466-2)) |
| Otu22700 | 1 | k__Fungi(100);p__Ascomycota(100);c__Saccharomycetes(100);o__Saccharomycetales(100);f__Metschnikowiaceae(100);g__Metschnikowia(100);g__Metschnikowia_unclassified(100); | Metschnikowia | 13 | Saprotroph | \|Undefined Saprotroph\| | Yeast | NULL | Probable | As nectar-tap_saprotroph (Põlme et al. 2020); Decay sugar-rich substrates (Põlme et al. 2020) | Põlme S, et al. 2020. Fungal Diversity 105:1-16 ((https://doi.org/10.1007/s13225-020-00466-2)) |
| Otu22701 | 1 | k__Fungi(100);p__Ascomycota(100);c__Saccharomycetes(100);o__Saccharomycetales(100);f__Metschnikowiaceae(100);g__Metschnikowia(100);g__Metschnikowia_unclassified(100); | Metschnikowia | 13 | Saprotroph | \|Undefined Saprotroph\| | Yeast | NULL | Probable | As nectar-tap_saprotroph (Põlme et al. 2020); Decay sugar-rich substrates (Põlme et al. 2020) | Põlme S, et al. 2020. Fungal Diversity 105:1-16 ((https://doi.org/10.1007/s13225-020-00466-2)) |
| Otu22702 | 1 | k__Fungi(100);p__Ascomycota(100);c__Saccharomycetes(100);o__Saccharomycetales(100);f__Metschnikowiaceae(100);g__Metschnikowia(100);s__Metschnikowia_pulcherrima(100); | Metschnikowia | 13 | Saprotroph | \|Undefined Saprotroph\| | Yeast | NULL | Probable | As nectar-tap_saprotroph (Põlme et al. 2020); Decay sugar-rich substrates (Põlme et al. 2020) | Põlme S, et al. 2020. Fungal Diversity 105:1-16 ((https://doi.org/10.1007/s13225-020-00466-2)) |
| Otu22703 | 1 | k__Fungi(100);p__Ascomycota(100);c__Saccharomycetes(100);o__Saccharomycetales(100);f__Metschnikowiaceae(100);g__Metschnikowia(100);s__Metschnikowia_pulcherrima(100); | Metschnikowia | 13 | Saprotroph | \|Undefined Saprotroph\| | Yeast | NULL | Probable | As nectar-tap_saprotroph (Põlme et al. 2020); Decay sugar-rich substrates (Põlme et al. 2020) | Põlme S, et al. 2020. Fungal Diversity 105:1-16 ((https://doi.org/10.1007/s13225-020-00466-2)) |
| Otu22705 | 1 | k__Fungi(100);p__Ascomycota(100);c__Saccharomycetes(100);o__Saccharomycetales(100);f__Metschnikowiaceae(100);g__Metschnikowia(100);s__Metschnikowia_pulcherrima(100); | Metschnikowia | 13 | Saprotroph | \|Undefined Saprotroph\| | Yeast | NULL | Probable | As nectar-tap_saprotroph (Põlme et al. 2020); Decay sugar-rich substrates (Põlme et al. 2020) | Põlme S, et al. 2020. Fungal Diversity 105:1-16 ((https://doi.org/10.1007/s13225-020-00466-2)) |
| Otu22708 | 1 | k__Fungi(100);p__Ascomycota(100);c__Saccharomycetes(100);o__Saccharomycetales(100);f__Metschnikowiaceae(100);g__Metschnikowia(100);s__Metschnikowia_pulcherrima(100); | Metschnikowia | 13 | Saprotroph | \|Undefined Saprotroph\| | Yeast | NULL | Probable | As nectar-tap_saprotroph (Põlme et al. 2020); Decay sugar-rich substrates (Põlme et al. 2020) | Põlme S, et al. 2020. Fungal Diversity 105:1-16 ((https://doi.org/10.1007/s13225-020-00466-2)) |
| Otu22709 | 1 | k__Fungi(100);p__Ascomycota(100);c__Saccharomycetes(100);o__Saccharomycetales(100);f__Metschnikowiaceae(100);g__Metschnikowia(100);s__Metschnikowia_pulcherrima(100); | Metschnikowia | 13 | Saprotroph | \|Undefined Saprotroph\| | Yeast | NULL | Probable | As nectar-tap_saprotroph (Põlme et al. 2020); Decay sugar-rich substrates (Põlme et al. 2020) | Põlme S, et al. 2020. Fungal Diversity 105:1-16 ((https://doi.org/10.1007/s13225-020-00466-2)) |
| Otu22710 | 1 | k__Fungi(100);p__Ascomycota(100);c__Saccharomycetes(100);o__Saccharomycetales(100);f__Metschnikowiaceae(100);g__Metschnikowia(100);s__Metschnikowia_pulcherrima(100); | Metschnikowia | 13 | Saprotroph | \|Undefined Saprotroph\| | Yeast | NULL | Probable | As nectar-tap_saprotroph (Põlme et al. 2020); Decay sugar-rich substrates (Põlme et al. 2020) | Põlme S, et al. 2020. Fungal Diversity 105:1-16 ((https://doi.org/10.1007/s13225-020-00466-2)) |
| Otu22711 | 1 | k__Fungi(100);p__Ascomycota(100);c__Saccharomycetes(100);o__Saccharomycetales(100);f__Metschnikowiaceae(100);g__Metschnikowia(100);s__Metschnikowia_pulcherrima(100); | Metschnikowia | 13 | Saprotroph | \|Undefined Saprotroph\| | Yeast | NULL | Probable | As nectar-tap_saprotroph (Põlme et al. 2020); Decay sugar-rich substrates (Põlme et al. 2020) | Põlme S, et al. 2020. Fungal Diversity 105:1-16 ((https://doi.org/10.1007/s13225-020-00466-2)) |
| Otu22712 | 1 | k__Fungi(100);p__Ascomycota(100);c__Saccharomycetes(100);o__Saccharomycetales(100);f__Metschnikowiaceae(100);g__Metschnikowia(100);s__Metschnikowia_pulcherrima(100); | Metschnikowia | 13 | Saprotroph | \|Undefined Saprotroph\| | Yeast | NULL | Probable | As nectar-tap_saprotroph (Põlme et al. 2020); Decay sugar-rich substrates (Põlme et al. 2020) | Põlme S, et al. 2020. Fungal Diversity 105:1-16 ((https://doi.org/10.1007/s13225-020-00466-2)) |
| Otu22713 | 1 | k__Fungi(100);p__Ascomycota(100);c__Saccharomycetes(100);o__Saccharomycetales(100);f__Metschnikowiaceae(100);g__Metschnikowia(100);s__Metschnikowia_pulcherrima(100); | Metschnikowia | 13 | Saprotroph | \|Undefined Saprotroph\| | Yeast | NULL | Probable | As nectar-tap_saprotroph (Põlme et al. 2020); Decay sugar-rich substrates (Põlme et al. 2020) | Põlme S, et al. 2020. Fungal Diversity 105:1-16 ((https://doi.org/10.1007/s13225-020-00466-2)) |
| Otu22714 | 1 | k__Fungi(100);p__Ascomycota(100);c__Saccharomycetes(100);o__Saccharomycetales(100);f__Metschnikowiaceae(100);g__Metschnikowia(100);s__Metschnikowia_pulcherrima(100); | Metschnikowia | 13 | Saprotroph | \|Undefined Saprotroph\| | Yeast | NULL | Probable | As nectar-tap_saprotroph (Põlme et al. 2020); Decay sugar-rich substrates (Põlme et al. 2020) | Põlme S, et al. 2020. Fungal Diversity 105:1-16 ((https://doi.org/10.1007/s13225-020-00466-2)) |
| Otu22715 | 1 | k__Fungi(100);p__Ascomycota(100);c__Saccharomycetes(100);o__Saccharomycetales(100);f__Metschnikowiaceae(100);g__Metschnikowia(100);g__Metschnikowia_unclassified(100); | Metschnikowia | 13 | Saprotroph | \|Undefined Saprotroph\| | Yeast | NULL | Probable | As nectar-tap_saprotroph (Põlme et al. 2020); Decay sugar-rich substrates (Põlme et al. 2020) | Põlme S, et al. 2020. Fungal Diversity 105:1-16 ((https://doi.org/10.1007/s13225-020-00466-2)) |
| Otu22717 | 1 | k__Fungi(100);p__Ascomycota(100);c__Saccharomycetes(100);o__Saccharomycetales(100);f__Metschnikowiaceae(100);g__Metschnikowia(100);s__Metschnikowia_pulcherrima(100); | Metschnikowia | 13 | Saprotroph | \|Undefined Saprotroph\| | Yeast | NULL | Probable | As nectar-tap_saprotroph (Põlme et al. 2020); Decay sugar-rich substrates (Põlme et al. 2020) | Põlme S, et al. 2020. Fungal Diversity 105:1-16 ((https://doi.org/10.1007/s13225-020-00466-2)) |
| Otu22718 | 1 | k__Fungi(100);p__Ascomycota(100);c__Saccharomycetes(100);o__Saccharomycetales(100);f__Metschnikowiaceae(100);g__Metschnikowia(100);s__Metschnikowia_pulcherrima(100); | Metschnikowia | 13 | Saprotroph | \|Undefined Saprotroph\| | Yeast | NULL | Probable | As nectar-tap_saprotroph (Põlme et al. 2020); Decay sugar-rich substrates (Põlme et al. 2020) | Põlme S, et al. 2020. Fungal Diversity 105:1-16 ((https://doi.org/10.1007/s13225-020-00466-2)) |
| Otu22719 | 1 | k__Fungi(100);p__Ascomycota(100);c__Saccharomycetes(100);o__Saccharomycetales(100);f__Metschnikowiaceae(100);g__Metschnikowia(100);s__Metschnikowia_pulcherrima(100); | Metschnikowia | 13 | Saprotroph | \|Undefined Saprotroph\| | Yeast | NULL | Probable | As nectar-tap_saprotroph (Põlme et al. 2020); Decay sugar-rich substrates (Põlme et al. 2020) | Põlme S, et al. 2020. Fungal Diversity 105:1-16 ((https://doi.org/10.1007/s13225-020-00466-2)) |
| Otu22721 | 1 | k__Fungi(100);p__Ascomycota(100);c__Saccharomycetes(100);o__Saccharomycetales(100);f__Metschnikowiaceae(100);g__Metschnikowia(100);g__Metschnikowia_unclassified(100); | Metschnikowia | 13 | Saprotroph | \|Undefined Saprotroph\| | Yeast | NULL | Probable | As nectar-tap_saprotroph (Põlme et al. 2020); Decay sugar-rich substrates (Põlme et al. 2020) | Põlme S, et al. 2020. Fungal Diversity 105:1-16 ((https://doi.org/10.1007/s13225-020-00466-2)) |
| Otu22722 | 1 | k__Fungi(100);p__Ascomycota(100);c__Saccharomycetes(100);o__Saccharomycetales(100);f__Metschnikowiaceae(100);g__Metschnikowia(100);g__Metschnikowia_unclassified(100); | Metschnikowia | 13 | Saprotroph | \|Undefined Saprotroph\| | Yeast | NULL | Probable | As nectar-tap_saprotroph (Põlme et al. 2020); Decay sugar-rich substrates (Põlme et al. 2020) | Põlme S, et al. 2020. Fungal Diversity 105:1-16 ((https://doi.org/10.1007/s13225-020-00466-2)) |
| Otu22723 | 1 | k__Fungi(100);p__Ascomycota(100);c__Saccharomycetes(100);o__Saccharomycetales(100);f__Metschnikowiaceae(100);g__Metschnikowia(100);s__Metschnikowia_pulcherrima(100); | Metschnikowia | 13 | Saprotroph | \|Undefined Saprotroph\| | Yeast | NULL | Probable | As nectar-tap_saprotroph (Põlme et al. 2020); Decay sugar-rich substrates (Põlme et al. 2020) | Põlme S, et al. 2020. Fungal Diversity 105:1-16 ((https://doi.org/10.1007/s13225-020-00466-2)) |
| Otu22724 | 1 | k__Fungi(100);p__Ascomycota(100);c__Saccharomycetes(100);o__Saccharomycetales(100);f__Metschnikowiaceae(100);g__Metschnikowia(100);g__Metschnikowia_unclassified(100); | Metschnikowia | 13 | Saprotroph | \|Undefined Saprotroph\| | Yeast | NULL | Probable | As nectar-tap_saprotroph (Põlme et al. 2020); Decay sugar-rich substrates (Põlme et al. 2020) | Põlme S, et al. 2020. Fungal Diversity 105:1-16 ((https://doi.org/10.1007/s13225-020-00466-2)) |
| Otu22726 | 1 | k__Fungi(100);p__Ascomycota(100);c__Saccharomycetes(100);o__Saccharomycetales(100);f__Metschnikowiaceae(100);g__Metschnikowia(100);s__Metschnikowia_pulcherrima(100); | Metschnikowia | 13 | Saprotroph | \|Undefined Saprotroph\| | Yeast | NULL | Probable | As nectar-tap_saprotroph (Põlme et al. 2020); Decay sugar-rich substrates (Põlme et al. 2020) | Põlme S, et al. 2020. Fungal Diversity 105:1-16 ((https://doi.org/10.1007/s13225-020-00466-2)) |
| Otu22727 | 1 | k__Fungi(100);p__Ascomycota(100);c__Saccharomycetes(100);o__Saccharomycetales(100);f__Metschnikowiaceae(100);g__Metschnikowia(100);s__Metschnikowia_pulcherrima(100); | Metschnikowia | 13 | Saprotroph | \|Undefined Saprotroph\| | Yeast | NULL | Probable | As nectar-tap_saprotroph (Põlme et al. 2020); Decay sugar-rich substrates (Põlme et al. 2020) | Põlme S, et al. 2020. Fungal Diversity 105:1-16 ((https://doi.org/10.1007/s13225-020-00466-2)) |
| Otu22728 | 1 | k__Fungi(100);p__Ascomycota(100);c__Saccharomycetes(100);o__Saccharomycetales(100);f__Metschnikowiaceae(100);g__Metschnikowia(100);s__Metschnikowia_pulcherrima(100); | Metschnikowia | 13 | Saprotroph | \|Undefined Saprotroph\| | Yeast | NULL | Probable | As nectar-tap_saprotroph (Põlme et al. 2020); Decay sugar-rich substrates (Põlme et al. 2020) | Põlme S, et al. 2020. Fungal Diversity 105:1-16 ((https://doi.org/10.1007/s13225-020-00466-2)) |
| Otu22729 | 1 | k__Fungi(100);p__Ascomycota(100);c__Saccharomycetes(100);o__Saccharomycetales(100);f__Metschnikowiaceae(100);g__Metschnikowia(100);g__Metschnikowia_unclassified(100); | Metschnikowia | 13 | Saprotroph | \|Undefined Saprotroph\| | Yeast | NULL | Probable | As nectar-tap_saprotroph (Põlme et al. 2020); Decay sugar-rich substrates (Põlme et al. 2020) | Põlme S, et al. 2020. Fungal Diversity 105:1-16 ((https://doi.org/10.1007/s13225-020-00466-2)) |
| Otu22730 | 1 | k__Fungi(100);p__Ascomycota(100);c__Saccharomycetes(100);o__Saccharomycetales(100);f__Metschnikowiaceae(100);g__Metschnikowia(100);s__Metschnikowia_pulcherrima(100); | Metschnikowia | 13 | Saprotroph | \|Undefined Saprotroph\| | Yeast | NULL | Probable | As nectar-tap_saprotroph (Põlme et al. 2020); Decay sugar-rich substrates (Põlme et al. 2020) | Põlme S, et al. 2020. Fungal Diversity 105:1-16 ((https://doi.org/10.1007/s13225-020-00466-2)) |
| Otu22731 | 1 | k__Fungi(100);p__Ascomycota(100);c__Saccharomycetes(100);o__Saccharomycetales(100);f__Metschnikowiaceae(100);g__Metschnikowia(100);g__Metschnikowia_unclassified(100); | Metschnikowia | 13 | Saprotroph | \|Undefined Saprotroph\| | Yeast | NULL | Probable | As nectar-tap_saprotroph (Põlme et al. 2020); Decay sugar-rich substrates (Põlme et al. 2020) | Põlme S, et al. 2020. Fungal Diversity 105:1-16 ((https://doi.org/10.1007/s13225-020-00466-2)) |
| Otu22732 | 1 | k__Fungi(100);p__Ascomycota(100);c__Saccharomycetes(100);o__Saccharomycetales(100);f__Metschnikowiaceae(100);g__Metschnikowia(100);s__Metschnikowia_pulcherrima(100); | Metschnikowia | 13 | Saprotroph | \|Undefined Saprotroph\| | Yeast | NULL | Probable | As nectar-tap_saprotroph (Põlme et al. 2020); Decay sugar-rich substrates (Põlme et al. 2020) | Põlme S, et al. 2020. Fungal Diversity 105:1-16 ((https://doi.org/10.1007/s13225-020-00466-2)) |
| Otu22733 | 1 | k__Fungi(100);p__Ascomycota(100);c__Saccharomycetes(100);o__Saccharomycetales(100);f__Metschnikowiaceae(100);g__Metschnikowia(100);s__Metschnikowia_pulcherrima(100); | Metschnikowia | 13 | Saprotroph | \|Undefined Saprotroph\| | Yeast | NULL | Probable | As nectar-tap_saprotroph (Põlme et al. 2020); Decay sugar-rich substrates (Põlme et al. 2020) | Põlme S, et al. 2020. Fungal Diversity 105:1-16 ((https://doi.org/10.1007/s13225-020-00466-2)) |
| Otu22734 | 1 | k__Fungi(100);p__Ascomycota(100);c__Saccharomycetes(100);o__Saccharomycetales(100);f__Metschnikowiaceae(100);g__Metschnikowia(100);g__Metschnikowia_unclassified(100); | Metschnikowia | 13 | Saprotroph | \|Undefined Saprotroph\| | Yeast | NULL | Probable | As nectar-tap_saprotroph (Põlme et al. 2020); Decay sugar-rich substrates (Põlme et al. 2020) | Põlme S, et al. 2020. Fungal Diversity 105:1-16 ((https://doi.org/10.1007/s13225-020-00466-2)) |
| Otu22735 | 1 | k__Fungi(100);p__Ascomycota(100);c__Saccharomycetes(100);o__Saccharomycetales(100);f__Metschnikowiaceae(100);g__Metschnikowia(100);g__Metschnikowia_unclassified(100); | Metschnikowia | 13 | Saprotroph | \|Undefined Saprotroph\| | Yeast | NULL | Probable | As nectar-tap_saprotroph (Põlme et al. 2020); Decay sugar-rich substrates (Põlme et al. 2020) | Põlme S, et al. 2020. Fungal Diversity 105:1-16 ((https://doi.org/10.1007/s13225-020-00466-2)) |
| Otu22736 | 1 | k__Fungi(100);p__Ascomycota(100);c__Saccharomycetes(100);o__Saccharomycetales(100);f__Metschnikowiaceae(100);g__Metschnikowia(100);s__Metschnikowia_pulcherrima(100); | Metschnikowia | 13 | Saprotroph | \|Undefined Saprotroph\| | Yeast | NULL | Probable | As nectar-tap_saprotroph (Põlme et al. 2020); Decay sugar-rich substrates (Põlme et al. 2020) | Põlme S, et al. 2020. Fungal Diversity 105:1-16 ((https://doi.org/10.1007/s13225-020-00466-2)) |
| Otu22737 | 1 | k__Fungi(100);p__Ascomycota(100);c__Saccharomycetes(100);o__Saccharomycetales(100);f__Metschnikowiaceae(100);g__Metschnikowia(100);s__Metschnikowia_pulcherrima(100); | Metschnikowia | 13 | Saprotroph | \|Undefined Saprotroph\| | Yeast | NULL | Probable | As nectar-tap_saprotroph (Põlme et al. 2020); Decay sugar-rich substrates (Põlme et al. 2020) | Põlme S, et al. 2020. Fungal Diversity 105:1-16 ((https://doi.org/10.1007/s13225-020-00466-2)) |
| Otu22738 | 1 | k__Fungi(100);p__Ascomycota(100);c__Saccharomycetes(100);o__Saccharomycetales(100);f__Metschnikowiaceae(100);g__Metschnikowia(100);s__Metschnikowia_pulcherrima(100); | Metschnikowia | 13 | Saprotroph | \|Undefined Saprotroph\| | Yeast | NULL | Probable | As nectar-tap_saprotroph (Põlme et al. 2020); Decay sugar-rich substrates (Põlme et al. 2020) | Põlme S, et al. 2020. Fungal Diversity 105:1-16 ((https://doi.org/10.1007/s13225-020-00466-2)) |
| Otu22739 | 1 | k__Fungi(100);p__Ascomycota(100);c__Saccharomycetes(100);o__Saccharomycetales(100);f__Metschnikowiaceae(100);g__Metschnikowia(100);s__Metschnikowia_pulcherrima(100); | Metschnikowia | 13 | Saprotroph | \|Undefined Saprotroph\| | Yeast | NULL | Probable | As nectar-tap_saprotroph (Põlme et al. 2020); Decay sugar-rich substrates (Põlme et al. 2020) | Põlme S, et al. 2020. Fungal Diversity 105:1-16 ((https://doi.org/10.1007/s13225-020-00466-2)) |
| Otu22740 | 1 | k__Fungi(100);p__Ascomycota(100);c__Saccharomycetes(100);o__Saccharomycetales(100);f__Metschnikowiaceae(100);g__Metschnikowia(100);s__Metschnikowia_pulcherrima(100); | Metschnikowia | 13 | Saprotroph | \|Undefined Saprotroph\| | Yeast | NULL | Probable | As nectar-tap_saprotroph (Põlme et al. 2020); Decay sugar-rich substrates (Põlme et al. 2020) | Põlme S, et al. 2020. Fungal Diversity 105:1-16 ((https://doi.org/10.1007/s13225-020-00466-2)) |
| Otu22741 | 1 | k__Fungi(100);p__Ascomycota(100);c__Saccharomycetes(100);o__Saccharomycetales(100);f__Metschnikowiaceae(100);g__Metschnikowia(100);g__Metschnikowia_unclassified(100); | Metschnikowia | 13 | Saprotroph | \|Undefined Saprotroph\| | Yeast | NULL | Probable | As nectar-tap_saprotroph (Põlme et al. 2020); Decay sugar-rich substrates (Põlme et al. 2020) | Põlme S, et al. 2020. Fungal Diversity 105:1-16 ((https://doi.org/10.1007/s13225-020-00466-2)) |
| Otu22742 | 1 | k__Fungi(100);p__Ascomycota(100);c__Saccharomycetes(100);o__Saccharomycetales(100);f__Metschnikowiaceae(100);g__Metschnikowia(100);g__Metschnikowia_unclassified(100); | Metschnikowia | 13 | Saprotroph | \|Undefined Saprotroph\| | Yeast | NULL | Probable | As nectar-tap_saprotroph (Põlme et al. 2020); Decay sugar-rich substrates (Põlme et al. 2020) | Põlme S, et al. 2020. Fungal Diversity 105:1-16 ((https://doi.org/10.1007/s13225-020-00466-2)) |
| Otu22743 | 1 | k__Fungi(100);p__Ascomycota(100);c__Saccharomycetes(100);o__Saccharomycetales(100);f__Metschnikowiaceae(100);g__Metschnikowia(100);s__Metschnikowia_pulcherrima(100); | Metschnikowia | 13 | Saprotroph | \|Undefined Saprotroph\| | Yeast | NULL | Probable | As nectar-tap_saprotroph (Põlme et al. 2020); Decay sugar-rich substrates (Põlme et al. 2020) | Põlme S, et al. 2020. Fungal Diversity 105:1-16 ((https://doi.org/10.1007/s13225-020-00466-2)) |
| Otu22744 | 1 | k__Fungi(100);p__Ascomycota(100);c__Saccharomycetes(100);o__Saccharomycetales(100);f__Metschnikowiaceae(100);g__Metschnikowia(100);g__Metschnikowia_unclassified(100); | Metschnikowia | 13 | Saprotroph | \|Undefined Saprotroph\| | Yeast | NULL | Probable | As nectar-tap_saprotroph (Põlme et al. 2020); Decay sugar-rich substrates (Põlme et al. 2020) | Põlme S, et al. 2020. Fungal Diversity 105:1-16 ((https://doi.org/10.1007/s13225-020-00466-2)) |
| Otu22745 | 1 | k__Fungi(100);p__Ascomycota(100);c__Saccharomycetes(100);o__Saccharomycetales(100);f__Metschnikowiaceae(100);g__Metschnikowia(100);s__Metschnikowia_pulcherrima(100); | Metschnikowia | 13 | Saprotroph | \|Undefined Saprotroph\| | Yeast | NULL | Probable | As nectar-tap_saprotroph (Põlme et al. 2020); Decay sugar-rich substrates (Põlme et al. 2020) | Põlme S, et al. 2020. Fungal Diversity 105:1-16 ((https://doi.org/10.1007/s13225-020-00466-2)) |
| Otu22746 | 1 | k__Fungi(100);p__Ascomycota(100);c__Saccharomycetes(100);o__Saccharomycetales(100);f__Metschnikowiaceae(100);g__Metschnikowia(100);s__Metschnikowia_pulcherrima(100); | Metschnikowia | 13 | Saprotroph | \|Undefined Saprotroph\| | Yeast | NULL | Probable | As nectar-tap_saprotroph (Põlme et al. 2020); Decay sugar-rich substrates (Põlme et al. 2020) | Põlme S, et al. 2020. Fungal Diversity 105:1-16 ((https://doi.org/10.1007/s13225-020-00466-2)) |
| Otu22747 | 1 | k__Fungi(100);p__Ascomycota(100);c__Saccharomycetes(100);o__Saccharomycetales(100);f__Metschnikowiaceae(100);g__Metschnikowia(100);g__Metschnikowia_unclassified(100); | Metschnikowia | 13 | Saprotroph | \|Undefined Saprotroph\| | Yeast | NULL | Probable | As nectar-tap_saprotroph (Põlme et al. 2020); Decay sugar-rich substrates (Põlme et al. 2020) | Põlme S, et al. 2020. Fungal Diversity 105:1-16 ((https://doi.org/10.1007/s13225-020-00466-2)) |
| Otu22748 | 1 | k__Fungi(100);p__Ascomycota(100);c__Saccharomycetes(100);o__Saccharomycetales(100);f__Metschnikowiaceae(100);g__Metschnikowia(100);s__Metschnikowia_pulcherrima(100); | Metschnikowia | 13 | Saprotroph | \|Undefined Saprotroph\| | Yeast | NULL | Probable | As nectar-tap_saprotroph (Põlme et al. 2020); Decay sugar-rich substrates (Põlme et al. 2020) | Põlme S, et al. 2020. Fungal Diversity 105:1-16 ((https://doi.org/10.1007/s13225-020-00466-2)) |
| Otu22750 | 1 | k__Fungi(100);p__Ascomycota(100);c__Saccharomycetes(100);o__Saccharomycetales(100);f__Metschnikowiaceae(100);g__Metschnikowia(100);s__Metschnikowia_pulcherrima(100); | Metschnikowia | 13 | Saprotroph | \|Undefined Saprotroph\| | Yeast | NULL | Probable | As nectar-tap_saprotroph (Põlme et al. 2020); Decay sugar-rich substrates (Põlme et al. 2020) | Põlme S, et al. 2020. Fungal Diversity 105:1-16 ((https://doi.org/10.1007/s13225-020-00466-2)) |
| Otu22751 | 1 | k__Fungi(100);p__Ascomycota(100);c__Saccharomycetes(100);o__Saccharomycetales(100);f__Metschnikowiaceae(100);g__Metschnikowia(100);s__Metschnikowia_pulcherrima(100); | Metschnikowia | 13 | Saprotroph | \|Undefined Saprotroph\| | Yeast | NULL | Probable | As nectar-tap_saprotroph (Põlme et al. 2020); Decay sugar-rich substrates (Põlme et al. 2020) | Põlme S, et al. 2020. Fungal Diversity 105:1-16 ((https://doi.org/10.1007/s13225-020-00466-2)) |
| Otu22753 | 1 | k__Fungi(100);p__Ascomycota(100);c__Saccharomycetes(100);o__Saccharomycetales(100);f__Metschnikowiaceae(100);g__Metschnikowia(100);s__Metschnikowia_pulcherrima(100); | Metschnikowia | 13 | Saprotroph | \|Undefined Saprotroph\| | Yeast | NULL | Probable | As nectar-tap_saprotroph (Põlme et al. 2020); Decay sugar-rich substrates (Põlme et al. 2020) | Põlme S, et al. 2020. Fungal Diversity 105:1-16 ((https://doi.org/10.1007/s13225-020-00466-2)) |
| Otu22754 | 1 | k__Fungi(100);p__Ascomycota(100);c__Saccharomycetes(100);o__Saccharomycetales(100);f__Metschnikowiaceae(100);g__Metschnikowia(100);s__Metschnikowia_pulcherrima(100); | Metschnikowia | 13 | Saprotroph | \|Undefined Saprotroph\| | Yeast | NULL | Probable | As nectar-tap_saprotroph (Põlme et al. 2020); Decay sugar-rich substrates (Põlme et al. 2020) | Põlme S, et al. 2020. Fungal Diversity 105:1-16 ((https://doi.org/10.1007/s13225-020-00466-2)) |
| Otu22756 | 1 | k__Fungi(100);p__Ascomycota(100);c__Saccharomycetes(100);o__Saccharomycetales(100);f__Metschnikowiaceae(100);g__Metschnikowia(100);s__Metschnikowia_pulcherrima(100); | Metschnikowia | 13 | Saprotroph | \|Undefined Saprotroph\| | Yeast | NULL | Probable | As nectar-tap_saprotroph (Põlme et al. 2020); Decay sugar-rich substrates (Põlme et al. 2020) | Põlme S, et al. 2020. Fungal Diversity 105:1-16 ((https://doi.org/10.1007/s13225-020-00466-2)) |
| Otu22757 | 1 | k__Fungi(100);p__Ascomycota(100);c__Dothideomycetes(100);o__Capnodiales(100);f__Mycosphaerellaceae(100);g__Mycosphaerella(100);s__Mycosphaerella_nyssicola(100); | Mycosphaerella | 13 | Pathotroph-Saprotroph | \|Plant Pathogen\|-Plant Saprotroph | Microfungus | NULL | Probable | As litter_saprotroph (Põlme et al. 2020); Decay leaf, fruit, seed (Põlme et al. 2020) | Tedersoo L, et al. 2014. Science 346:e1256688 ((https://doi.org/10.1126/science.1256688)); Põlme S, et al. 2020. Fungal Diversity 105:1-16 ((https://doi.org/10.1007/s13225-020-00466-2)) |
| Otu22758 | 1 | k__Fungi(100);p__Ascomycota(100);c__Dothideomycetes(100);o__Capnodiales(100);f__Mycosphaerellaceae(100);f__Mycosphaerellaceae_unclassified(100);f__Mycosphaerellaceae_unclassified(100); | Mycosphaerellaceae | 9 | Pathotroph-Saprotroph | Plant Pathogen-Undefined Saprotroph | Microfungus | NULL | Probable | Biotrophic, necrotrophic or saprobic on various plant tissues (Cannon & Kirk 2007) | Cannon PF, Kirk PM. 2007. Fungal Families of the World. CAB International, Cambridge (ISBN: 978-0851998275) |
| Otu22759 | 1 | k__Fungi(100);p__Ascomycota(100);c__Saccharomycetes(100);o__Saccharomycetales(100);f__Metschnikowiaceae(100);g__Metschnikowia(100);s__Metschnikowia_pulcherrima(100); | Metschnikowia | 13 | Saprotroph | \|Undefined Saprotroph\| | Yeast | NULL | Probable | As nectar-tap_saprotroph (Põlme et al. 2020); Decay sugar-rich substrates (Põlme et al. 2020) | Põlme S, et al. 2020. Fungal Diversity 105:1-16 ((https://doi.org/10.1007/s13225-020-00466-2)) |
| Otu22760 | 1 | k__Fungi(100);p__Ascomycota(100);c__Saccharomycetes(100);o__Saccharomycetales(100);f__Metschnikowiaceae(100);g__Metschnikowia(100);g__Metschnikowia_unclassified(100); | Metschnikowia | 13 | Saprotroph | \|Undefined Saprotroph\| | Yeast | NULL | Probable | As nectar-tap_saprotroph (Põlme et al. 2020); Decay sugar-rich substrates (Põlme et al. 2020) | Põlme S, et al. 2020. Fungal Diversity 105:1-16 ((https://doi.org/10.1007/s13225-020-00466-2)) |
| Otu22761 | 1 | k__Fungi(100);p__Ascomycota(100);c__Saccharomycetes(100);o__Saccharomycetales(100);f__Metschnikowiaceae(100);g__Metschnikowia(100);s__Metschnikowia_pulcherrima(100); | Metschnikowia | 13 | Saprotroph | \|Undefined Saprotroph\| | Yeast | NULL | Probable | As nectar-tap_saprotroph (Põlme et al. 2020); Decay sugar-rich substrates (Põlme et al. 2020) | Põlme S, et al. 2020. Fungal Diversity 105:1-16 ((https://doi.org/10.1007/s13225-020-00466-2)) |
| Otu22762 | 1 | k__Fungi(100);p__Ascomycota(100);c__Saccharomycetes(100);o__Saccharomycetales(100);f__Metschnikowiaceae(100);g__Metschnikowia(100);s__Metschnikowia_pulcherrima(100); | Metschnikowia | 13 | Saprotroph | \|Undefined Saprotroph\| | Yeast | NULL | Probable | As nectar-tap_saprotroph (Põlme et al. 2020); Decay sugar-rich substrates (Põlme et al. 2020) | Põlme S, et al. 2020. Fungal Diversity 105:1-16 ((https://doi.org/10.1007/s13225-020-00466-2)) |
| Otu22763 | 1 | k__Fungi(100);p__Ascomycota(100);c__Saccharomycetes(100);o__Saccharomycetales(100);f__Metschnikowiaceae(100);g__Metschnikowia(100);s__Metschnikowia_pulcherrima(100); | Metschnikowia | 13 | Saprotroph | \|Undefined Saprotroph\| | Yeast | NULL | Probable | As nectar-tap_saprotroph (Põlme et al. 2020); Decay sugar-rich substrates (Põlme et al. 2020) | Põlme S, et al. 2020. Fungal Diversity 105:1-16 ((https://doi.org/10.1007/s13225-020-00466-2)) |
| Otu22764 | 1 | k__Fungi(100);p__Ascomycota(100);c__Saccharomycetes(100);o__Saccharomycetales(100);f__Metschnikowiaceae(100);g__Metschnikowia(100);s__Metschnikowia_pulcherrima(100); | Metschnikowia | 13 | Saprotroph | \|Undefined Saprotroph\| | Yeast | NULL | Probable | As nectar-tap_saprotroph (Põlme et al. 2020); Decay sugar-rich substrates (Põlme et al. 2020) | Põlme S, et al. 2020. Fungal Diversity 105:1-16 ((https://doi.org/10.1007/s13225-020-00466-2)) |
| Otu22765 | 1 | k__Fungi(100);p__Ascomycota(100);c__Saccharomycetes(100);o__Saccharomycetales(100);f__Metschnikowiaceae(100);g__Metschnikowia(100);g__Metschnikowia_unclassified(100); | Metschnikowia | 13 | Saprotroph | \|Undefined Saprotroph\| | Yeast | NULL | Probable | As nectar-tap_saprotroph (Põlme et al. 2020); Decay sugar-rich substrates (Põlme et al. 2020) | Põlme S, et al. 2020. Fungal Diversity 105:1-16 ((https://doi.org/10.1007/s13225-020-00466-2)) |
| Otu22766 | 1 | k__Fungi(100);p__Ascomycota(100);c__Saccharomycetes(100);o__Saccharomycetales(100);f__Metschnikowiaceae(100);g__Metschnikowia(100);g__Metschnikowia_unclassified(100); | Metschnikowia | 13 | Saprotroph | \|Undefined Saprotroph\| | Yeast | NULL | Probable | As nectar-tap_saprotroph (Põlme et al. 2020); Decay sugar-rich substrates (Põlme et al. 2020) | Põlme S, et al. 2020. Fungal Diversity 105:1-16 ((https://doi.org/10.1007/s13225-020-00466-2)) |
| Otu22767 | 1 | k__Fungi(100);p__Ascomycota(100);c__Saccharomycetes(100);o__Saccharomycetales(100);f__Metschnikowiaceae(100);g__Metschnikowia(100);s__Metschnikowia_pulcherrima(100); | Metschnikowia | 13 | Saprotroph | \|Undefined Saprotroph\| | Yeast | NULL | Probable | As nectar-tap_saprotroph (Põlme et al. 2020); Decay sugar-rich substrates (Põlme et al. 2020) | Põlme S, et al. 2020. Fungal Diversity 105:1-16 ((https://doi.org/10.1007/s13225-020-00466-2)) |
| Otu22768 | 1 | k__Fungi(100);p__Ascomycota(100);c__Saccharomycetes(100);o__Saccharomycetales(100);f__Metschnikowiaceae(100);g__Metschnikowia(100);s__Metschnikowia_pulcherrima(100); | Metschnikowia | 13 | Saprotroph | \|Undefined Saprotroph\| | Yeast | NULL | Probable | As nectar-tap_saprotroph (Põlme et al. 2020); Decay sugar-rich substrates (Põlme et al. 2020) | Põlme S, et al. 2020. Fungal Diversity 105:1-16 ((https://doi.org/10.1007/s13225-020-00466-2)) |
| Otu22769 | 1 | k__Fungi(100);p__Ascomycota(100);c__Saccharomycetes(100);o__Saccharomycetales(100);f__Metschnikowiaceae(100);g__Metschnikowia(100);s__Metschnikowia_pulcherrima(100); | Metschnikowia | 13 | Saprotroph | \|Undefined Saprotroph\| | Yeast | NULL | Probable | As nectar-tap_saprotroph (Põlme et al. 2020); Decay sugar-rich substrates (Põlme et al. 2020) | Põlme S, et al. 2020. Fungal Diversity 105:1-16 ((https://doi.org/10.1007/s13225-020-00466-2)) |
| Otu22770 | 1 | k__Fungi(100);p__Ascomycota(100);c__Saccharomycetes(100);o__Saccharomycetales(100);f__Metschnikowiaceae(100);g__Metschnikowia(100);s__Metschnikowia_pulcherrima(100); | Metschnikowia | 13 | Saprotroph | \|Undefined Saprotroph\| | Yeast | NULL | Probable | As nectar-tap_saprotroph (Põlme et al. 2020); Decay sugar-rich substrates (Põlme et al. 2020) | Põlme S, et al. 2020. Fungal Diversity 105:1-16 ((https://doi.org/10.1007/s13225-020-00466-2)) |
| Otu22771 | 1 | k__Fungi(100);p__Ascomycota(100);c__Saccharomycetes(100);o__Saccharomycetales(100);f__Metschnikowiaceae(100);g__Metschnikowia(100);s__Metschnikowia_pulcherrima(100); | Metschnikowia | 13 | Saprotroph | \|Undefined Saprotroph\| | Yeast | NULL | Probable | As nectar-tap_saprotroph (Põlme et al. 2020); Decay sugar-rich substrates (Põlme et al. 2020) | Põlme S, et al. 2020. Fungal Diversity 105:1-16 ((https://doi.org/10.1007/s13225-020-00466-2)) |
| Otu22772 | 1 | k__Fungi(100);p__Ascomycota(100);c__Saccharomycetes(100);o__Saccharomycetales(100);f__Metschnikowiaceae(100);g__Metschnikowia(100);s__Metschnikowia_pulcherrima(100); | Metschnikowia | 13 | Saprotroph | \|Undefined Saprotroph\| | Yeast | NULL | Probable | As nectar-tap_saprotroph (Põlme et al. 2020); Decay sugar-rich substrates (Põlme et al. 2020) | Põlme S, et al. 2020. Fungal Diversity 105:1-16 ((https://doi.org/10.1007/s13225-020-00466-2)) |
| Otu22773 | 1 | k__Fungi(100);p__Ascomycota(100);c__Saccharomycetes(100);o__Saccharomycetales(100);f__Metschnikowiaceae(100);g__Metschnikowia(100);s__Metschnikowia_pulcherrima(100); | Metschnikowia | 13 | Saprotroph | \|Undefined Saprotroph\| | Yeast | NULL | Probable | As nectar-tap_saprotroph (Põlme et al. 2020); Decay sugar-rich substrates (Põlme et al. 2020) | Põlme S, et al. 2020. Fungal Diversity 105:1-16 ((https://doi.org/10.1007/s13225-020-00466-2)) |
| Otu22774 | 1 | k__Fungi(100);p__Ascomycota(100);c__Saccharomycetes(100);o__Saccharomycetales(100);f__Metschnikowiaceae(100);g__Metschnikowia(100);s__Metschnikowia_pulcherrima(100); | Metschnikowia | 13 | Saprotroph | \|Undefined Saprotroph\| | Yeast | NULL | Probable | As nectar-tap_saprotroph (Põlme et al. 2020); Decay sugar-rich substrates (Põlme et al. 2020) | Põlme S, et al. 2020. Fungal Diversity 105:1-16 ((https://doi.org/10.1007/s13225-020-00466-2)) |
| Otu22775 | 1 | k__Fungi(100);p__Ascomycota(100);c__Saccharomycetes(100);o__Saccharomycetales(100);f__Metschnikowiaceae(100);g__Metschnikowia(100);g__Metschnikowia_unclassified(100); | Metschnikowia | 13 | Saprotroph | \|Undefined Saprotroph\| | Yeast | NULL | Probable | As nectar-tap_saprotroph (Põlme et al. 2020); Decay sugar-rich substrates (Põlme et al. 2020) | Põlme S, et al. 2020. Fungal Diversity 105:1-16 ((https://doi.org/10.1007/s13225-020-00466-2)) |
| Otu22776 | 1 | k__Fungi(100);p__Ascomycota(100);c__Saccharomycetes(100);o__Saccharomycetales(100);f__Metschnikowiaceae(100);g__Metschnikowia(100);s__Metschnikowia_pulcherrima(100); | Metschnikowia | 13 | Saprotroph | \|Undefined Saprotroph\| | Yeast | NULL | Probable | As nectar-tap_saprotroph (Põlme et al. 2020); Decay sugar-rich substrates (Põlme et al. 2020) | Põlme S, et al. 2020. Fungal Diversity 105:1-16 ((https://doi.org/10.1007/s13225-020-00466-2)) |
| Otu22777 | 1 | k__Fungi(100);p__Ascomycota(100);c__Saccharomycetes(100);o__Saccharomycetales(100);f__Metschnikowiaceae(100);g__Metschnikowia(100);s__Metschnikowia_pulcherrima(100); | Metschnikowia | 13 | Saprotroph | \|Undefined Saprotroph\| | Yeast | NULL | Probable | As nectar-tap_saprotroph (Põlme et al. 2020); Decay sugar-rich substrates (Põlme et al. 2020) | Põlme S, et al. 2020. Fungal Diversity 105:1-16 ((https://doi.org/10.1007/s13225-020-00466-2)) |
| Otu22778 | 1 | k__Fungi(100);p__Ascomycota(100);c__Saccharomycetes(100);o__Saccharomycetales(100);f__Metschnikowiaceae(100);g__Metschnikowia(100);s__Metschnikowia_pulcherrima(100); | Metschnikowia | 13 | Saprotroph | \|Undefined Saprotroph\| | Yeast | NULL | Probable | As nectar-tap_saprotroph (Põlme et al. 2020); Decay sugar-rich substrates (Põlme et al. 2020) | Põlme S, et al. 2020. Fungal Diversity 105:1-16 ((https://doi.org/10.1007/s13225-020-00466-2)) |
| Otu22779 | 1 | k__Fungi(100);p__Ascomycota(100);c__Saccharomycetes(100);o__Saccharomycetales(100);f__Metschnikowiaceae(100);g__Metschnikowia(100);s__Metschnikowia_pulcherrima(100); | Metschnikowia | 13 | Saprotroph | \|Undefined Saprotroph\| | Yeast | NULL | Probable | As nectar-tap_saprotroph (Põlme et al. 2020); Decay sugar-rich substrates (Põlme et al. 2020) | Põlme S, et al. 2020. Fungal Diversity 105:1-16 ((https://doi.org/10.1007/s13225-020-00466-2)) |
| Otu22780 | 1 | k__Fungi(100);p__Ascomycota(100);c__Saccharomycetes(100);o__Saccharomycetales(100);f__Metschnikowiaceae(100);g__Metschnikowia(100);s__Metschnikowia_pulcherrima(100); | Metschnikowia | 13 | Saprotroph | \|Undefined Saprotroph\| | Yeast | NULL | Probable | As nectar-tap_saprotroph (Põlme et al. 2020); Decay sugar-rich substrates (Põlme et al. 2020) | Põlme S, et al. 2020. Fungal Diversity 105:1-16 ((https://doi.org/10.1007/s13225-020-00466-2)) |
| Otu22781 | 1 | k__Fungi(100);p__Ascomycota(100);c__Saccharomycetes(100);o__Saccharomycetales(100);f__Metschnikowiaceae(100);g__Metschnikowia(100);s__Metschnikowia_pulcherrima(100); | Metschnikowia | 13 | Saprotroph | \|Undefined Saprotroph\| | Yeast | NULL | Probable | As nectar-tap_saprotroph (Põlme et al. 2020); Decay sugar-rich substrates (Põlme et al. 2020) | Põlme S, et al. 2020. Fungal Diversity 105:1-16 ((https://doi.org/10.1007/s13225-020-00466-2)) |
| Otu22782 | 1 | k__Fungi(100);p__Ascomycota(100);c__Saccharomycetes(100);o__Saccharomycetales(100);f__Metschnikowiaceae(100);g__Metschnikowia(100);s__Metschnikowia_pulcherrima(100); | Metschnikowia | 13 | Saprotroph | \|Undefined Saprotroph\| | Yeast | NULL | Probable | As nectar-tap_saprotroph (Põlme et al. 2020); Decay sugar-rich substrates (Põlme et al. 2020) | Põlme S, et al. 2020. Fungal Diversity 105:1-16 ((https://doi.org/10.1007/s13225-020-00466-2)) |
| Otu22783 | 1 | k__Fungi(100);p__Ascomycota(100);c__Saccharomycetes(100);o__Saccharomycetales(100);f__Metschnikowiaceae(100);g__Metschnikowia(100);s__Metschnikowia_pulcherrima(100); | Metschnikowia | 13 | Saprotroph | \|Undefined Saprotroph\| | Yeast | NULL | Probable | As nectar-tap_saprotroph (Põlme et al. 2020); Decay sugar-rich substrates (Põlme et al. 2020) | Põlme S, et al. 2020. Fungal Diversity 105:1-16 ((https://doi.org/10.1007/s13225-020-00466-2)) |
| Otu22784 | 1 | k__Fungi(100);p__Ascomycota(100);c__Saccharomycetes(100);o__Saccharomycetales(100);f__Metschnikowiaceae(100);g__Metschnikowia(100);g__Metschnikowia_unclassified(100); | Metschnikowia | 13 | Saprotroph | \|Undefined Saprotroph\| | Yeast | NULL | Probable | As nectar-tap_saprotroph (Põlme et al. 2020); Decay sugar-rich substrates (Põlme et al. 2020) | Põlme S, et al. 2020. Fungal Diversity 105:1-16 ((https://doi.org/10.1007/s13225-020-00466-2)) |
| Otu22785 | 1 | k__Fungi(100);p__Ascomycota(100);c__Saccharomycetes(100);o__Saccharomycetales(100);f__Metschnikowiaceae(100);g__Metschnikowia(100);g__Metschnikowia_unclassified(100); | Metschnikowia | 13 | Saprotroph | \|Undefined Saprotroph\| | Yeast | NULL | Probable | As nectar-tap_saprotroph (Põlme et al. 2020); Decay sugar-rich substrates (Põlme et al. 2020) | Põlme S, et al. 2020. Fungal Diversity 105:1-16 ((https://doi.org/10.1007/s13225-020-00466-2)) |
| Otu22786 | 1 | k__Fungi(100);p__Ascomycota(100);c__Saccharomycetes(100);o__Saccharomycetales(100);f__Metschnikowiaceae(100);g__Metschnikowia(100);s__Metschnikowia_pulcherrima(100); | Metschnikowia | 13 | Saprotroph | \|Undefined Saprotroph\| | Yeast | NULL | Probable | As nectar-tap_saprotroph (Põlme et al. 2020); Decay sugar-rich substrates (Põlme et al. 2020) | Põlme S, et al. 2020. Fungal Diversity 105:1-16 ((https://doi.org/10.1007/s13225-020-00466-2)) |
| Otu22787 | 1 | k__Fungi(100);p__Ascomycota(100);c__Saccharomycetes(100);o__Saccharomycetales(100);f__Metschnikowiaceae(100);g__Metschnikowia(100);g__Metschnikowia_unclassified(100); | Metschnikowia | 13 | Saprotroph | \|Undefined Saprotroph\| | Yeast | NULL | Probable | As nectar-tap_saprotroph (Põlme et al. 2020); Decay sugar-rich substrates (Põlme et al. 2020) | Põlme S, et al. 2020. Fungal Diversity 105:1-16 ((https://doi.org/10.1007/s13225-020-00466-2)) |
| Otu22788 | 1 | k__Fungi(100);p__Ascomycota(100);c__Saccharomycetes(100);o__Saccharomycetales(100);f__Metschnikowiaceae(100);g__Metschnikowia(100);s__Metschnikowia_pulcherrima(100); | Metschnikowia | 13 | Saprotroph | \|Undefined Saprotroph\| | Yeast | NULL | Probable | As nectar-tap_saprotroph (Põlme et al. 2020); Decay sugar-rich substrates (Põlme et al. 2020) | Põlme S, et al. 2020. Fungal Diversity 105:1-16 ((https://doi.org/10.1007/s13225-020-00466-2)) |
| Otu22789 | 1 | k__Fungi(100);p__Ascomycota(100);c__Saccharomycetes(100);o__Saccharomycetales(100);f__Metschnikowiaceae(100);g__Metschnikowia(100);s__Metschnikowia_pulcherrima(100); | Metschnikowia | 13 | Saprotroph | \|Undefined Saprotroph\| | Yeast | NULL | Probable | As nectar-tap_saprotroph (Põlme et al. 2020); Decay sugar-rich substrates (Põlme et al. 2020) | Põlme S, et al. 2020. Fungal Diversity 105:1-16 ((https://doi.org/10.1007/s13225-020-00466-2)) |

| | | | | | | | | | | | |
|---|---|---|---|---|---|---|---|---|---|---|---|
| Otu22790 | 1 | k__Fungi(100);p__Ascomycota(100);c__Saccharomycetes(100);o__Saccharomycetales(100);f__Metschnikowiaceae(100);g__Metschnikowia(100);s__Metschnikowia_pulcherrima(100); | Metschnikowia | 13 | Saprotroph | \|Undefined Saprotroph\| | Yeast | NULL | Probable | As nectar-tap_saprotroph (Põlme et al. 2020); Decay sugar-rich substrates (Põlme et al. 2020) | Põlme S, et al. 2020. Fungal Diversity 105:1-16 ((https://doi.org/10.1007/s13225-020-00466-2)) |
| Otu22794 | 1 | k__Fungi(100);p__Ascomycota(100);c__Saccharomycetes(100);o__Saccharomycetales(100);f__Metschnikowiaceae(100);g__Metschnikowia(100);s__Metschnikowia_pulcherrima(100); | Metschnikowia | 13 | Saprotroph | \|Undefined Saprotroph\| | Yeast | NULL | Probable | As nectar-tap_saprotroph (Põlme et al. 2020); Decay sugar-rich substrates (Põlme et al. 2020) | Põlme S, et al. 2020. Fungal Diversity 105:1-16 ((https://doi.org/10.1007/s13225-020-00466-2)) |
| Otu22795 | 1 | k__Fungi(100);p__Ascomycota(100);c__Saccharomycetes(100);o__Saccharomycetales(100);f__Metschnikowiaceae(100);g__Metschnikowia(100);g__Metschnikowia_unclassified(100); | Metschnikowia | 13 | Saprotroph | \|Undefined Saprotroph\| | Yeast | NULL | Probable | As nectar-tap_saprotroph (Põlme et al. 2020); Decay sugar-rich substrates (Põlme et al. 2020) | Põlme S, et al. 2020. Fungal Diversity 105:1-16 ((https://doi.org/10.1007/s13225-020-00466-2)) |
| Otu22796 | 1 | k__Fungi(100);p__Ascomycota(100);c__Saccharomycetes(100);o__Saccharomycetales(100);f__Metschnikowiaceae(100);g__Metschnikowia(100);g__Metschnikowia_unclassified(100); | Metschnikowia | 13 | Saprotroph | \|Undefined Saprotroph\| | Yeast | NULL | Probable | As nectar-tap_saprotroph (Põlme et al. 2020); Decay sugar-rich substrates (Põlme et al. 2020) | Põlme S, et al. 2020. Fungal Diversity 105:1-16 ((https://doi.org/10.1007/s13225-020-00466-2)) |
| Otu22797 | 1 | k__Fungi(100);p__Ascomycota(100);c__Saccharomycetes(100);o__Saccharomycetales(100);f__Metschnikowiaceae(100);g__Metschnikowia(100);s__Metschnikowia_pulcherrima(100); | Metschnikowia | 13 | Saprotroph | \|Undefined Saprotroph\| | Yeast | NULL | Probable | As nectar-tap_saprotroph (Põlme et al. 2020); Decay sugar-rich substrates (Põlme et al. 2020) | Põlme S, et al. 2020. Fungal Diversity 105:1-16 ((https://doi.org/10.1007/s13225-020-00466-2)) |
| Otu22798 | 1 | k__Fungi(100);p__Ascomycota(100);c__Saccharomycetes(100);o__Saccharomycetales(100);f__Metschnikowiaceae(100);g__Metschnikowia(100);s__Metschnikowia_pulcherrima(100); | Metschnikowia | 13 | Saprotroph | \|Undefined Saprotroph\| | Yeast | NULL | Probable | As nectar-tap_saprotroph (Põlme et al. 2020); Decay sugar-rich substrates (Põlme et al. 2020) | Põlme S, et al. 2020. Fungal Diversity 105:1-16 ((https://doi.org/10.1007/s13225-020-00466-2)) |
| Otu22799 | 1 | k__Fungi(100);p__Ascomycota(100);c__Saccharomycetes(100);o__Saccharomycetales(100);f__Metschnikowiaceae(100);g__Metschnikowia(100);g__Metschnikowia_unclassified(100); | Metschnikowia | 13 | Saprotroph | \|Undefined Saprotroph\| | Yeast | NULL | Probable | As nectar-tap_saprotroph (Põlme et al. 2020); Decay sugar-rich substrates (Põlme et al. 2020) | Põlme S, et al. 2020. Fungal Diversity 105:1-16 ((https://doi.org/10.1007/s13225-020-00466-2)) |
| Otu22801 | 1 | k__Fungi(100);p__Ascomycota(100);c__Saccharomycetes(100);o__Saccharomycetales(100);f__Metschnikowiaceae(100);g__Metschnikowia(100);s__Metschnikowia_pulcherrima(100); | Metschnikowia | 13 | Saprotroph | \|Undefined Saprotroph\| | Yeast | NULL | Probable | As nectar-tap_saprotroph (Põlme et al. 2020); Decay sugar-rich substrates (Põlme et al. 2020) | Põlme S, et al. 2020. Fungal Diversity 105:1-16 ((https://doi.org/10.1007/s13225-020-00466-2)) |
| Otu22802 | 1 | k__Fungi(100);p__Ascomycota(100);c__Saccharomycetes(100);o__Saccharomycetales(100);f__Metschnikowiaceae(100);g__Metschnikowia(100);s__Metschnikowia_pulcherrima(100); | Metschnikowia | 13 | Saprotroph | \|Undefined Saprotroph\| | Yeast | NULL | Probable | As nectar-tap_saprotroph (Põlme et al. 2020); Decay sugar-rich substrates (Põlme et al. 2020) | Põlme S, et al. 2020. Fungal Diversity 105:1-16 ((https://doi.org/10.1007/s13225-020-00466-2)) |
| Otu22804 | 1 | k__Fungi(100);p__Ascomycota(100);c__Saccharomycetes(100);o__Saccharomycetales(100);f__Metschnikowiaceae(100);g__Metschnikowia(100);g__Metschnikowia_unclassified(100); | Metschnikowia | 13 | Saprotroph | \|Undefined Saprotroph\| | Yeast | NULL | Probable | As nectar-tap_saprotroph (Põlme et al. 2020); Decay sugar-rich substrates (Põlme et al. 2020) | Põlme S, et al. 2020. Fungal Diversity 105:1-16 ((https://doi.org/10.1007/s13225-020-00466-2)) |
| Otu22805 | 1 | k__Fungi(100);p__Ascomycota(100);c__Saccharomycetes(100);o__Saccharomycetales(100);f__Metschnikowiaceae(100);g__Metschnikowia(100);g__Metschnikowia_unclassified(100); | Metschnikowia | 13 | Saprotroph | \|Undefined Saprotroph\| | Yeast | NULL | Probable | As nectar-tap_saprotroph (Põlme et al. 2020); Decay sugar-rich substrates (Põlme et al. 2020) | Põlme S, et al. 2020. Fungal Diversity 105:1-16 ((https://doi.org/10.1007/s13225-020-00466-2)) |
| Otu22806 | 1 | k__Fungi(100);p__Ascomycota(100);c__Saccharomycetes(100);o__Saccharomycetales(100);f__Metschnikowiaceae(100);g__Metschnikowia(100);g__Metschnikowia_unclassified(100); | Metschnikowia | 13 | Saprotroph | \|Undefined Saprotroph\| | Yeast | NULL | Probable | As nectar-tap_saprotroph (Põlme et al. 2020); Decay sugar-rich substrates (Põlme et al. 2020) | Põlme S, et al. 2020. Fungal Diversity 105:1-16 ((https://doi.org/10.1007/s13225-020-00466-2)) |
| Otu22807 | 1 | k__Fungi(100);p__Ascomycota(100);c__Saccharomycetes(100);o__Saccharomycetales(100);f__Metschnikowiaceae(100);g__Metschnikowia(100);g__Metschnikowia_unclassified(100); | Metschnikowia | 13 | Saprotroph | \|Undefined Saprotroph\| | Yeast | NULL | Probable | As nectar-tap_saprotroph (Põlme et al. 2020); Decay sugar-rich substrates (Põlme et al. 2020) | Põlme S, et al. 2020. Fungal Diversity 105:1-16 ((https://doi.org/10.1007/s13225-020-00466-2)) |
| Otu22808 | 1 | k__Fungi(100);p__Ascomycota(100);c__Saccharomycetes(100);o__Saccharomycetales(100);f__Metschnikowiaceae(100);g__Metschnikowia(100);s__Metschnikowia_pulcherrima(100); | Metschnikowia | 13 | Saprotroph | \|Undefined Saprotroph\| | Yeast | NULL | Probable | As nectar-tap_saprotroph (Põlme et al. 2020); Decay sugar-rich substrates (Põlme et al. 2020) | Põlme S, et al. 2020. Fungal Diversity 105:1-16 ((https://doi.org/10.1007/s13225-020-00466-2)) |
| Otu22809 | 1 | k__Fungi(100);p__Ascomycota(100);c__Saccharomycetes(100);o__Saccharomycetales(100);f__Metschnikowiaceae(100);g__Metschnikowia(100);g__Metschnikowia_unclassified(100); | Metschnikowia | 13 | Saprotroph | \|Undefined Saprotroph\| | Yeast | NULL | Probable | As nectar-tap_saprotroph (Põlme et al. 2020); Decay sugar-rich substrates (Põlme et al. 2020) | Põlme S, et al. 2020. Fungal Diversity 105:1-16 ((https://doi.org/10.1007/s13225-020-00466-2)) |
| Otu22810 | 1 | k__Fungi(100);p__Ascomycota(100);c__Saccharomycetes(100);o__Saccharomycetales(100);f__Metschnikowiaceae(100);g__Metschnikowia(100);s__Metschnikowia_pulcherrima(100); | Metschnikowia | 13 | Saprotroph | \|Undefined Saprotroph\| | Yeast | NULL | Probable | As nectar-tap_saprotroph (Põlme et al. 2020); Decay sugar-rich substrates (Põlme et al. 2020) | Põlme S, et al. 2020. Fungal Diversity 105:1-16 ((https://doi.org/10.1007/s13225-020-00466-2)) |
| Otu22811 | 1 | k__Fungi(100);p__Ascomycota(100);c__Saccharomycetes(100);o__Saccharomycetales(100);f__Metschnikowiaceae(100);g__Metschnikowia(100);s__Metschnikowia_pulcherrima(100); | Metschnikowia | 13 | Saprotroph | \|Undefined Saprotroph\| | Yeast | NULL | Probable | As nectar-tap_saprotroph (Põlme et al. 2020); Decay sugar-rich substrates (Põlme et al. 2020) | Põlme S, et al. 2020. Fungal Diversity 105:1-16 ((https://doi.org/10.1007/s13225-020-00466-2)) |
| Otu22812 | 1 | k__Fungi(100);p__Ascomycota(100);c__Saccharomycetes(100);o__Saccharomycetales(100);f__Metschnikowiaceae(100);g__Metschnikowia(100);s__Metschnikowia_pulcherrima(100); | Metschnikowia | 13 | Saprotroph | \|Undefined Saprotroph\| | Yeast | NULL | Probable | As nectar-tap_saprotroph (Põlme et al. 2020); Decay sugar-rich substrates (Põlme et al. 2020) | Põlme S, et al. 2020. Fungal Diversity 105:1-16 ((https://doi.org/10.1007/s13225-020-00466-2)) |
| Otu22813 | 1 | k__Fungi(100);p__Ascomycota(100);c__Saccharomycetes(100);o__Saccharomycetales(100);f__Metschnikowiaceae(100);g__Metschnikowia(100);s__Metschnikowia_pulcherrima(100); | Metschnikowia | 13 | Saprotroph | \|Undefined Saprotroph\| | Yeast | NULL | Probable | As nectar-tap_saprotroph (Põlme et al. 2020); Decay sugar-rich substrates (Põlme et al. 2020) | Põlme S, et al. 2020. Fungal Diversity 105:1-16 ((https://doi.org/10.1007/s13225-020-00466-2)) |
| Otu22814 | 1 | k__Fungi(100);p__Ascomycota(100);c__Saccharomycetes(100);o__Saccharomycetales(100);f__Metschnikowiaceae(100);g__Metschnikowia(100);g__Metschnikowia_unclassified(100); | Metschnikowia | 13 | Saprotroph | \|Undefined Saprotroph\| | Yeast | NULL | Probable | As nectar-tap_saprotroph (Põlme et al. 2020); Decay sugar-rich substrates (Põlme et al. 2020) | Põlme S, et al. 2020. Fungal Diversity 105:1-16 ((https://doi.org/10.1007/s13225-020-00466-2)) |
| Otu22816 | 1 | k__Fungi(100);p__Ascomycota(100);c__Saccharomycetes(100);o__Saccharomycetales(100);f__Metschnikowiaceae(100);g__Metschnikowia(100);s__Metschnikowia_pulcherrima(100); | Metschnikowia | 13 | Saprotroph | \|Undefined Saprotroph\| | Yeast | NULL | Probable | As nectar-tap_saprotroph (Põlme et al. 2020); Decay sugar-rich substrates (Põlme et al. 2020) | Põlme S, et al. 2020. Fungal Diversity 105:1-16 ((https://doi.org/10.1007/s13225-020-00466-2)) |
| Otu22817 | 1 | k__Fungi(100);p__Ascomycota(100);c__Saccharomycetes(100);o__Saccharomycetales(100);f__Metschnikowiaceae(100);g__Metschnikowia(100);s__Metschnikowia_pulcherrima(100); | Metschnikowia | 13 | Saprotroph | \|Undefined Saprotroph\| | Yeast | NULL | Probable | As nectar-tap_saprotroph (Põlme et al. 2020); Decay sugar-rich substrates (Põlme et al. 2020) | Põlme S, et al. 2020. Fungal Diversity 105:1-16 ((https://doi.org/10.1007/s13225-020-00466-2)) |
| Otu22818 | 1 | k__Fungi(100);p__Ascomycota(100);c__Saccharomycetes(100);o__Saccharomycetales(100);f__Metschnikowiaceae(100);g__Metschnikowia(100);s__Metschnikowia_pulcherrima(100); | Metschnikowia | 13 | Saprotroph | \|Undefined Saprotroph\| | Yeast | NULL | Probable | As nectar-tap_saprotroph (Põlme et al. 2020); Decay sugar-rich substrates (Põlme et al. 2020) | Põlme S, et al. 2020. Fungal Diversity 105:1-16 ((https://doi.org/10.1007/s13225-020-00466-2)) |
| Otu22819 | 1 | k__Fungi(100);p__Ascomycota(100);c__Saccharomycetes(100);o__Saccharomycetales(100);f__Metschnikowiaceae(100);g__Metschnikowia(100);s__Metschnikowia_pulcherrima(100); | Metschnikowia | 13 | Saprotroph | \|Undefined Saprotroph\| | Yeast | NULL | Probable | As nectar-tap_saprotroph (Põlme et al. 2020); Decay sugar-rich substrates (Põlme et al. 2020) | Põlme S, et al. 2020. Fungal Diversity 105:1-16 ((https://doi.org/10.1007/s13225-020-00466-2)) |
| Otu22820 | 1 | k__Fungi(100);p__Ascomycota(100);c__Saccharomycetes(100);o__Saccharomycetales(100);f__Metschnikowiaceae(100);g__Metschnikowia(100);s__Metschnikowia_pulcherrima(100); | Metschnikowia | 13 | Saprotroph | \|Undefined Saprotroph\| | Yeast | NULL | Probable | As nectar-tap_saprotroph (Põlme et al. 2020); Decay sugar-rich substrates (Põlme et al. 2020) | Põlme S, et al. 2020. Fungal Diversity 105:1-16 ((https://doi.org/10.1007/s13225-020-00466-2)) |
| Otu22821 | 1 | k__Fungi(100);p__Ascomycota(100);c__Saccharomycetes(100);o__Saccharomycetales(100);f__Metschnikowiaceae(100);g__Metschnikowia(100);s__Metschnikowia_pulcherrima(100); | Metschnikowia | 13 | Saprotroph | \|Undefined Saprotroph\| | Yeast | NULL | Probable | As nectar-tap_saprotroph (Põlme et al. 2020); Decay sugar-rich substrates (Põlme et al. 2020) | Põlme S, et al. 2020. Fungal Diversity 105:1-16 ((https://doi.org/10.1007/s13225-020-00466-2)) |
| Otu22822 | 1 | k__Fungi(100);p__Ascomycota(100);c__Saccharomycetes(100);o__Saccharomycetales(100);f__Metschnikowiaceae(100);g__Metschnikowia(100);s__Metschnikowia_pulcherrima(100); | Metschnikowia | 13 | Saprotroph | \|Undefined Saprotroph\| | Yeast | NULL | Probable | As nectar-tap_saprotroph (Põlme et al. 2020); Decay sugar-rich substrates (Põlme et al. 2020) | Põlme S, et al. 2020. Fungal Diversity 105:1-16 ((https://doi.org/10.1007/s13225-020-00466-2)) |
| Otu22823 | 1 | k__Fungi(100);p__Ascomycota(100);c__Saccharomycetes(100);o__Saccharomycetales(100);f__Metschnikowiaceae(100);g__Metschnikowia(100);s__Metschnikowia_pulcherrima(100); | Metschnikowia | 13 | Saprotroph | \|Undefined Saprotroph\| | Yeast | NULL | Probable | As nectar-tap_saprotroph (Põlme et al. 2020); Decay sugar-rich substrates (Põlme et al. 2020) | Põlme S, et al. 2020. Fungal Diversity 105:1-16 ((https://doi.org/10.1007/s13225-020-00466-2)) |
| Otu22824 | 1 | k__Fungi(100);p__Ascomycota(100);c__Saccharomycetes(100);o__Saccharomycetales(100);f__Metschnikowiaceae(100);g__Metschnikowia(100);s__Metschnikowia_pulcherrima(100); | Metschnikowia | 13 | Saprotroph | \|Undefined Saprotroph\| | Yeast | NULL | Probable | As nectar-tap_saprotroph (Põlme et al. 2020); Decay sugar-rich substrates (Põlme et al. 2020) | Põlme S, et al. 2020. Fungal Diversity 105:1-16 ((https://doi.org/10.1007/s13225-020-00466-2)) |
| Otu22825 | 1 | k__Fungi(100);p__Ascomycota(100);c__Saccharomycetes(100);o__Saccharomycetales(100);f__Metschnikowiaceae(100);g__Metschnikowia(100);s__Metschnikowia_pulcherrima(100); | Metschnikowia | 13 | Saprotroph | \|Undefined Saprotroph\| | Yeast | NULL | Probable | As nectar-tap_saprotroph (Põlme et al. 2020); Decay sugar-rich substrates (Põlme et al. 2020) | Põlme S, et al. 2020. Fungal Diversity 105:1-16 ((https://doi.org/10.1007/s13225-020-00466-2)) |
| Otu22826 | 1 | k__Fungi(100);p__Ascomycota(100);c__Saccharomycetes(100);o__Saccharomycetales(100);f__Metschnikowiaceae(100);g__Metschnikowia(100);g__Metschnikowia_unclassified(100); | Metschnikowia | 13 | Saprotroph | \|Undefined Saprotroph\| | Yeast | NULL | Probable | As nectar-tap_saprotroph (Põlme et al. 2020); Decay sugar-rich substrates (Põlme et al. 2020) | Põlme S, et al. 2020. Fungal Diversity 105:1-16 ((https://doi.org/10.1007/s13225-020-00466-2)) |
| Otu22827 | 1 | k__Fungi(100);p__Ascomycota(100);c__Saccharomycetes(100);o__Saccharomycetales(100);f__Metschnikowiaceae(100);g__Metschnikowia(100);g__Metschnikowia_unclassified(100); | Metschnikowia | 13 | Saprotroph | \|Undefined Saprotroph\| | Yeast | NULL | Probable | As nectar-tap_saprotroph (Põlme et al. 2020); Decay sugar-rich substrates (Põlme et al. 2020) | Põlme S, et al. 2020. Fungal Diversity 105:1-16 ((https://doi.org/10.1007/s13225-020-00466-2)) |
| Otu22828 | 1 | k__Fungi(100);p__Ascomycota(100);c__Saccharomycetes(100);o__Saccharomycetales(100);f__Metschnikowiaceae(100);g__Metschnikowia(100);s__Metschnikowia_pulcherrima(100); | Metschnikowia | 13 | Saprotroph | \|Undefined Saprotroph\| | Yeast | NULL | Probable | As nectar-tap_saprotroph (Põlme et al. 2020); Decay sugar-rich substrates (Põlme et al. 2020) | Põlme S, et al. 2020. Fungal Diversity 105:1-16 ((https://doi.org/10.1007/s13225-020-00466-2)) |
| Otu22829 | 1 | k__Fungi(100);p__Ascomycota(100);c__Saccharomycetes(100);o__Saccharomycetales(100);f__Metschnikowiaceae(100);g__Metschnikowia(100);s__Metschnikowia_pulcherrima(100); | Metschnikowia | 13 | Saprotroph | \|Undefined Saprotroph\| | Yeast | NULL | Probable | As nectar-tap_saprotroph (Põlme et al. 2020); Decay sugar-rich substrates (Põlme et al. 2020) | Põlme S, et al. 2020. Fungal Diversity 105:1-16 ((https://doi.org/10.1007/s13225-020-00466-2)) |
| Otu22830 | 1 | k__Fungi(100);p__Ascomycota(100);c__Saccharomycetes(100);o__Saccharomycetales(100);f__Metschnikowiaceae(100);g__Metschnikowia(100);s__Metschnikowia_pulcherrima(100); | Metschnikowia | 13 | Saprotroph | \|Undefined Saprotroph\| | Yeast | NULL | Probable | As nectar-tap_saprotroph (Põlme et al. 2020); Decay sugar-rich substrates (Põlme et al. 2020) | Põlme S, et al. 2020. Fungal Diversity 105:1-16 ((https://doi.org/10.1007/s13225-020-00466-2)) |
| Otu22831 | 1 | k__Fungi(100);p__Ascomycota(100);c__Saccharomycetes(100);o__Saccharomycetales(100);f__Metschnikowiaceae(100);g__Metschnikowia(100);s__Metschnikowia_pulcherrima(100); | Metschnikowia | 13 | Saprotroph | \|Undefined Saprotroph\| | Yeast | NULL | Probable | As nectar-tap_saprotroph (Põlme et al. 2020); Decay sugar-rich substrates (Põlme et al. 2020) | Põlme S, et al. 2020. Fungal Diversity 105:1-16 ((https://doi.org/10.1007/s13225-020-00466-2)) |
| Otu22832 | 1 | k__Fungi(100);p__Ascomycota(100);c__Saccharomycetes(100);o__Saccharomycetales(100);f__Metschnikowiaceae(100);g__Metschnikowia(100);s__Metschnikowia_pulcherrima(100); | Metschnikowia | 13 | Saprotroph | \|Undefined Saprotroph\| | Yeast | NULL | Probable | As nectar-tap_saprotroph (Põlme et al. 2020); Decay sugar-rich substrates (Põlme et al. 2020) | Põlme S, et al. 2020. Fungal Diversity 105:1-16 ((https://doi.org/10.1007/s13225-020-00466-2)) |
| Otu22833 | 1 | k__Fungi(100);p__Ascomycota(100);c__Saccharomycetes(100);o__Saccharomycetales(100);f__Metschnikowiaceae(100);g__Metschnikowia(100);g__Metschnikowia_unclassified(100); | Metschnikowia | 13 | Saprotroph | \|Undefined Saprotroph\| | Yeast | NULL | Probable | As nectar-tap_saprotroph (Põlme et al. 2020); Decay sugar-rich substrates (Põlme et al. 2020) | Põlme S, et al. 2020. Fungal Diversity 105:1-16 ((https://doi.org/10.1007/s13225-020-00466-2)) |
| Otu22834 | 1 | k__Fungi(100);p__Ascomycota(100);c__Saccharomycetes(100);o__Saccharomycetales(100);f__Metschnikowiaceae(100);g__Metschnikowia(100);g__Metschnikowia_unclassified(100); | Metschnikowia | 13 | Saprotroph | \|Undefined Saprotroph\| | Yeast | NULL | Probable | As nectar-tap_saprotroph (Põlme et al. 2020); Decay sugar-rich substrates (Põlme et al. 2020) | Põlme S, et al. 2020. Fungal Diversity 105:1-16 ((https://doi.org/10.1007/s13225-020-00466-2)) |
| Otu22835 | 1 | k__Fungi(100);p__Ascomycota(100);c__Saccharomycetes(100);o__Saccharomycetales(100);f__Metschnikowiaceae(100);g__Metschnikowia(100);g__Metschnikowia_unclassified(100); | Metschnikowia | 13 | Saprotroph | \|Undefined Saprotroph\| | Yeast | NULL | Probable | As nectar-tap_saprotroph (Põlme et al. 2020); Decay sugar-rich substrates (Põlme et al. 2020) | Põlme S, et al. 2020. Fungal Diversity 105:1-16 ((https://doi.org/10.1007/s13225-020-00466-2)) |
| Otu22836 | 1 | k__Fungi(100);p__Ascomycota(100);c__Saccharomycetes(100);o__Saccharomycetales(100);f__Metschnikowiaceae(100);g__Metschnikowia(100);s__Metschnikowia_pulcherrima(100); | Metschnikowia | 13 | Saprotroph | \|Undefined Saprotroph\| | Yeast | NULL | Probable | As nectar-tap_saprotroph (Põlme et al. 2020); Decay sugar-rich substrates (Põlme et al. 2020) | Põlme S, et al. 2020. Fungal Diversity 105:1-16 ((https://doi.org/10.1007/s13225-020-00466-2)) |
| Otu22837 | 1 | k__Fungi(100);p__Ascomycota(100);c__Saccharomycetes(100);o__Saccharomycetales(100);f__Metschnikowiaceae(100);g__Metschnikowia(100);g__Metschnikowia_unclassified(100); | Metschnikowia | 13 | Saprotroph | \|Undefined Saprotroph\| | Yeast | NULL | Probable | As nectar-tap_saprotroph (Põlme et al. 2020); Decay sugar-rich substrates (Põlme et al. 2020) | Põlme S, et al. 2020. Fungal Diversity 105:1-16 ((https://doi.org/10.1007/s13225-020-00466-2)) |
| Otu22838 | 1 | k__Fungi(100);p__Ascomycota(100);c__Saccharomycetes(100);o__Saccharomycetales(100);f__Metschnikowiaceae(100);g__Metschnikowia(100);g__Metschnikowia_unclassified(100); | Metschnikowia | 13 | Saprotroph | \|Undefined Saprotroph\| | Yeast | NULL | Probable | As nectar-tap_saprotroph (Põlme et al. 2020); Decay sugar-rich substrates (Põlme et al. 2020) | Põlme S, et al. 2020. Fungal Diversity 105:1-16 ((https://doi.org/10.1007/s13225-020-00466-2)) |
| Otu22839 | 1 | k__Fungi(100);p__Ascomycota(100);c__Saccharomycetes(100);o__Saccharomycetales(100);f__Metschnikowiaceae(100);g__Metschnikowia(100);g__Metschnikowia_unclassified(100); | Metschnikowia | 13 | Saprotroph | \|Undefined Saprotroph\| | Yeast | NULL | Probable | As nectar-tap_saprotroph (Põlme et al. 2020); Decay sugar-rich substrates (Põlme et al. 2020) | Põlme S, et al. 2020. Fungal Diversity 105:1-16 ((https://doi.org/10.1007/s13225-020-00466-2)) |
| Otu22840 | 1 | k__Fungi(100);p__Ascomycota(100);c__Saccharomycetes(100);o__Saccharomycetales(100);f__Metschnikowiaceae(100);g__Metschnikowia(100);s__Metschnikowia_pulcherrima(100); | Metschnikowia | 13 | Saprotroph | \|Undefined Saprotroph\| | Yeast | NULL | Probable | As nectar-tap_saprotroph (Põlme et al. 2020); Decay sugar-rich substrates (Põlme et al. 2020) | Põlme S, et al. 2020. Fungal Diversity 105:1-16 ((https://doi.org/10.1007/s13225-020-00466-2)) |
| Otu22841 | 1 | k__Fungi(100);p__Ascomycota(100);c__Saccharomycetes(100);o__Saccharomycetales(100);f__Metschnikowiaceae(100);g__Metschnikowia(100);s__Metschnikowia_pulcherrima(100); | Metschnikowia | 13 | Saprotroph | \|Undefined Saprotroph\| | Yeast | NULL | Probable | As nectar-tap_saprotroph (Põlme et al. 2020); Decay sugar-rich substrates (Põlme et al. 2020) | Põlme S, et al. 2020. Fungal Diversity 105:1-16 ((https://doi.org/10.1007/s13225-020-00466-2)) |
| Otu22842 | 1 | k__Fungi(100);p__Ascomycota(100);c__Saccharomycetes(100);o__Saccharomycetales(100);f__Metschnikowiaceae(100);g__Metschnikowia(100);s__Metschnikowia_pulcherrima(100); | Metschnikowia | 13 | Saprotroph | \|Undefined Saprotroph\| | Yeast | NULL | Probable | As nectar-tap_saprotroph (Põlme et al. 2020); Decay sugar-rich substrates (Põlme et al. 2020) | Põlme S, et al. 2020. Fungal Diversity 105:1-16 ((https://doi.org/10.1007/s13225-020-00466-2)) |
| Otu22843 | 1 | k__Fungi(100);p__Ascomycota(100);c__Saccharomycetes(100);o__Saccharomycetales(100);f__Metschnikowiaceae(100);g__Metschnikowia(100);g__Metschnikowia_unclassified(100); | Metschnikowia | 13 | Saprotroph | \|Undefined Saprotroph\| | Yeast | NULL | Probable | As nectar-tap_saprotroph (Põlme et al. 2020); Decay sugar-rich substrates (Põlme et al. 2020) | Põlme S, et al. 2020. Fungal Diversity 105:1-16 ((https://doi.org/10.1007/s13225-020-00466-2)) |
| Otu22844 | 1 | k__Fungi(100);p__Ascomycota(100);c__Saccharomycetes(100);o__Saccharomycetales(100);f__Metschnikowiaceae(100);g__Metschnikowia(100);g__Metschnikowia_unclassified(100); | Metschnikowia | 13 | Saprotroph | \|Undefined Saprotroph\| | Yeast | NULL | Probable | As nectar-tap_saprotroph (Põlme et al. 2020); Decay sugar-rich substrates (Põlme et al. 2020) | Põlme S, et al. 2020. Fungal Diversity 105:1-16 ((https://doi.org/10.1007/s13225-020-00466-2)) |
| Otu22845 | 1 | k__Fungi(100);p__Ascomycota(100);c__Saccharomycetes(100);o__Saccharomycetales(100);f__Metschnikowiaceae(100);g__Metschnikowia(100);g__Metschnikowia_unclassified(100); | Metschnikowia | 13 | Saprotroph | \|Undefined Saprotroph\| | Yeast | NULL | Probable | As nectar-tap_saprotroph (Põlme et al. 2020); Decay sugar-rich substrates (Põlme et al. 2020) | Põlme S, et al. 2020. Fungal Diversity 105:1-16 ((https://doi.org/10.1007/s13225-020-00466-2)) |
| Otu22846 | 1 | k__Fungi(100);p__Ascomycota(100);c__Saccharomycetes(100);o__Saccharomycetales(100);f__Metschnikowiaceae(100);g__Metschnikowia(100);g__Metschnikowia_unclassified(100); | Metschnikowia | 13 | Saprotroph | \|Undefined Saprotroph\| | Yeast | NULL | Probable | As nectar-tap_saprotroph (Põlme et al. 2020); Decay sugar-rich substrates (Põlme et al. 2020) | Põlme S, et al. 2020. Fungal Diversity 105:1-16 ((https://doi.org/10.1007/s13225-020-00466-2)) |
| Otu22847 | 1 | k__Fungi(100);p__Ascomycota(100);c__Saccharomycetes(100);o__Saccharomycetales(100);f__Metschnikowiaceae(100);g__Metschnikowia(100);s__Metschnikowia_pulcherrima(100); | Metschnikowia | 13 | Saprotroph | \|Undefined Saprotroph\| | Yeast | NULL | Probable | As nectar-tap_saprotroph (Põlme et al. 2020); Decay sugar-rich substrates (Põlme et al. 2020) | Põlme S, et al. 2020. Fungal Diversity 105:1-16 ((https://doi.org/10.1007/s13225-020-00466-2)) |
| Otu22848 | 1 | k__Fungi(100);p__Ascomycota(100);c__Saccharomycetes(100);o__Saccharomycetales(100);f__Metschnikowiaceae(100);g__Metschnikowia(100);s__Metschnikowia_pulcherrima(100); | Metschnikowia | 13 | Saprotroph | \|Undefined Saprotroph\| | Yeast | NULL | Probable | As nectar-tap_saprotroph (Põlme et al. 2020); Decay sugar-rich substrates (Põlme et al. 2020) | Põlme S, et al. 2020. Fungal Diversity 105:1-16 ((https://doi.org/10.1007/s13225-020-00466-2)) |
| Otu22849 | 1 | k__Fungi(100);p__Ascomycota(100);c__Saccharomycetes(100);o__Saccharomycetales(100);f__Metschnikowiaceae(100);g__Metschnikowia(100);g__Metschnikowia_unclassified(100); | Metschnikowia | 13 | Saprotroph | \|Undefined Saprotroph\| | Yeast | NULL | Probable | As nectar-tap_saprotroph (Põlme et al. 2020); Decay sugar-rich substrates (Põlme et al. 2020) | Põlme S, et al. 2020. Fungal Diversity 105:1-16 ((https://doi.org/10.1007/s13225-020-00466-2)) |
| Otu22850 | 1 | k__Fungi(100);p__Ascomycota(100);c__Saccharomycetes(100);o__Saccharomycetales(100);f__Metschnikowiaceae(100);g__Metschnikowia(100);g__Metschnikowia_unclassified(100); | Metschnikowia | 13 | Saprotroph | \|Undefined Saprotroph\| | Yeast | NULL | Probable | As nectar-tap_saprotroph (Põlme et al. 2020); Decay sugar-rich substrates (Põlme et al. 2020) | Põlme S, et al. 2020. Fungal Diversity 105:1-16 ((https://doi.org/10.1007/s13225-020-00466-2)) |
| Otu22851 | 1 | k__Fungi(100);p__Ascomycota(100);c__Saccharomycetes(100);o__Saccharomycetales(100);f__Metschnikowiaceae(100);g__Metschnikowia(100);s__Metschnikowia_pulcherrima(100); | Metschnikowia | 13 | Saprotroph | \|Undefined Saprotroph\| | Yeast | NULL | Probable | As nectar-tap_saprotroph (Põlme et al. 2020); Decay sugar-rich substrates (Põlme et al. 2020) | Põlme S, et al. 2020. Fungal Diversity 105:1-16 ((https://doi.org/10.1007/s13225-020-00466-2)) |
| Otu22852 | 1 | k__Fungi(100);p__Ascomycota(100);c__Saccharomycetes(100);o__Saccharomycetales(100);f__Metschnikowiaceae(100);g__Metschnikowia(100);s__Metschnikowia_pulcherrima(100); | Metschnikowia | 13 | Saprotroph | \|Undefined Saprotroph\| | Yeast | NULL | Probable | As nectar-tap_saprotroph (Põlme et al. 2020); Decay sugar-rich substrates (Põlme et al. 2020) | Põlme S, et al. 2020. Fungal Diversity 105:1-16 ((https://doi.org/10.1007/s13225-020-00466-2)) |
| Otu22853 | 1 | k__Fungi(100);p__Ascomycota(100);c__Saccharomycetes(100);o__Saccharomycetales(100);f__Metschnikowiaceae(100);g__Metschnikowia(100);g__Metschnikowia_unclassified(100); | Metschnikowia | 13 | Saprotroph | \|Undefined Saprotroph\| | Yeast | NULL | Probable | As nectar-tap_saprotroph (Põlme et al. 2020); Decay sugar-rich substrates (Põlme et al. 2020) | Põlme S, et al. 2020. Fungal Diversity 105:1-16 ((https://doi.org/10.1007/s13225-020-00466-2)) |
| Otu22854 | 1 | k__Fungi(100);p__Ascomycota(100);c__Saccharomycetes(100);o__Saccharomycetales(100);f__Metschnikowiaceae(100);g__Metschnikowia(100);s__Metschnikowia_pulcherrima(100); | Metschnikowia | 13 | Saprotroph | \|Undefined Saprotroph\| | Yeast | NULL | Probable | As nectar-tap_saprotroph (Põlme et al. 2020); Decay sugar-rich substrates (Põlme et al. 2020) | Põlme S, et al. 2020. Fungal Diversity 105:1-16 ((https://doi.org/10.1007/s13225-020-00466-2)) |
| Otu22855 | 1 | k__Fungi(100);p__Ascomycota(100);c__Saccharomycetes(100);o__Saccharomycetales(100);f__Metschnikowiaceae(100);g__Metschnikowia(100);g__Metschnikowia_unclassified(100); | Metschnikowia | 13 | Saprotroph | \|Undefined Saprotroph\| | Yeast | NULL | Probable | As nectar-tap_saprotroph (Põlme et al. 2020); Decay sugar-rich substrates (Põlme et al. 2020) | Põlme S, et al. 2020. Fungal Diversity 105:1-16 ((https://doi.org/10.1007/s13225-020-00466-2)) |
| Otu22856 | 1 | k__Fungi(100);p__Ascomycota(100);c__Saccharomycetes(100);o__Saccharomycetales(100);f__Metschnikowiaceae(100);g__Metschnikowia(100);s__Metschnikowia_pulcherrima(100); | Metschnikowia | 13 | Saprotroph | \|Undefined Saprotroph\| | Yeast | NULL | Probable | As nectar-tap_saprotroph (Põlme et al. 2020); Decay sugar-rich substrates (Põlme et al. 2020) | Põlme S, et al. 2020. Fungal Diversity 105:1-16 ((https://doi.org/10.1007/s13225-020-00466-2)) |
| Otu22857 | 1 | k__Fungi(100);p__Ascomycota(100);c__Saccharomycetes(100);o__Saccharomycetales(100);f__Metschnikowiaceae(100);g__Metschnikowia(100);s__Metschnikowia_pulcherrima(100); | Metschnikowia | 13 | Saprotroph | \|Undefined Saprotroph\| | Yeast | NULL | Probable | As nectar-tap_saprotroph (Põlme et al. 2020); Decay sugar-rich substrates (Põlme et al. 2020) | Põlme S, et al. 2020. Fungal Diversity 105:1-16 ((https://doi.org/10.1007/s13225-020-00466-2)) |
| Otu22858 | 1 | k__Fungi(100);p__Ascomycota(100);c__Saccharomycetes(100);o__Saccharomycetales(100);f__Metschnikowiaceae(100);g__Metschnikowia(100);g__Metschnikowia_unclassified(100); | Metschnikowia | 13 | Saprotroph | \|Undefined Saprotroph\| | Yeast | NULL | Probable | As nectar-tap_saprotroph (Põlme et al. 2020); Decay sugar-rich substrates (Põlme et al. 2020) | Põlme S, et al. 2020. Fungal Diversity 105:1-16 ((https://doi.org/10.1007/s13225-020-00466-2)) |
| Otu22859 | 1 | k__Fungi(100);p__Ascomycota(100);c__Saccharomycetes(100);o__Saccharomycetales(100);f__Metschnikowiaceae(100);g__Metschnikowia(100);s__Metschnikowia_pulcherrima(100); | Metschnikowia | 13 | Saprotroph | \|Undefined Saprotroph\| | Yeast | NULL | Probable | As nectar-tap_saprotroph (Põlme et al. 2020); Decay sugar-rich substrates (Põlme et al. 2020) | Põlme S, et al. 2020. Fungal Diversity 105:1-16 ((https://doi.org/10.1007/s13225-020-00466-2)) |
| Otu22860 | 1 | k__Fungi(100);p__Ascomycota(100);c__Saccharomycetes(100);o__Saccharomycetales(100);f__Metschnikowiaceae(100);g__Metschnikowia(100);s__Metschnikowia_pulcherrima(100); | Metschnikowia | 13 | Saprotroph | \|Undefined Saprotroph\| | Yeast | NULL | Probable | As nectar-tap_saprotroph (Põlme et al. 2020); Decay sugar-rich substrates (Põlme et al. 2020) | Põlme S, et al. 2020. Fungal Diversity 105:1-16 ((https://doi.org/10.1007/s13225-020-00466-2)) |
| Otu22861 | 1 | k__Fungi(100);p__Ascomycota(100);c__Saccharomycetes(100);o__Saccharomycetales(100);f__Metschnikowiaceae(100);g__Metschnikowia(100);s__Metschnikowia_pulcherrima(100); | Metschnikowia | 13 | Saprotroph | \|Undefined Saprotroph\| | Yeast | NULL | Probable | As nectar-tap_saprotroph (Põlme et al. 2020); Decay sugar-rich substrates (Põlme et al. 2020) | Põlme S, et al. 2020. Fungal Diversity 105:1-16 ((https://doi.org/10.1007/s13225-020-00466-2)) |
| Otu22862 | 1 | k__Fungi(100);p__Ascomycota(100);c__Saccharomycetes(100);o__Saccharomycetales(100);f__Metschnikowiaceae(100);g__Metschnikowia(100);s__Metschnikowia_pulcherrima(100); | Metschnikowia | 13 | Saprotroph | \|Undefined Saprotroph\| | Yeast | NULL | Probable | As nectar-tap_saprotroph (Põlme et al. 2020); Decay sugar-rich substrates (Põlme et al. 2020) | Põlme S, et al. 2020. Fungal Diversity 105:1-16 ((https://doi.org/10.1007/s13225-020-00466-2)) |
| Otu22863 | 1 | k__Fungi(100);p__Ascomycota(100);c__Saccharomycetes(100);o__Saccharomycetales(100);f__Metschnikowiaceae(100);g__Metschnikowia(100);s__Metschnikowia_pulcherrima(100); | Metschnikowia | 13 | Saprotroph | \|Undefined Saprotroph\| | Yeast | NULL | Probable | As nectar-tap_saprotroph (Põlme et al. 2020); Decay sugar-rich substrates (Põlme et al. 2020) | Põlme S, et al. 2020. Fungal Diversity 105:1-16 ((https://doi.org/10.1007/s13225-020-00466-2)) |
| Otu22864 | 1 | k__Fungi(100);p__Ascomycota(100);c__Saccharomycetes(100);o__Saccharomycetales(100);f__Metschnikowiaceae(100);g__Metschnikowia(100);g__Metschnikowia_unclassified(100); | Metschnikowia | 13 | Saprotroph | \|Undefined Saprotroph\| | Yeast | NULL | Probable | As nectar-tap_saprotroph (Põlme et al. 2020); Decay sugar-rich substrates (Põlme et al. 2020) | Põlme S, et al. 2020. Fungal Diversity 105:1-16 ((https://doi.org/10.1007/s13225-020-00466-2)) |
| Otu22865 | 1 | k__Fungi(100);p__Ascomycota(100);c__Saccharomycetes(100);o__Saccharomycetales(100);f__Metschnikowiaceae(100);g__Metschnikowia(100);g__Metschnikowia_unclassified(100); | Metschnikowia | 13 | Saprotroph | \|Undefined Saprotroph\| | Yeast | NULL | Probable | As nectar-tap_saprotroph (Põlme et al. 2020); Decay sugar-rich substrates (Põlme et al. 2020) | Põlme S, et al. 2020. Fungal Diversity 105:1-16 ((https://doi.org/10.1007/s13225-020-00466-2)) |
| Otu22866 | 1 | k__Fungi(100);p__Ascomycota(100);c__Saccharomycetes(100);o__Saccharomycetales(100);f__Metschnikowiaceae(100);g__Metschnikowia(100);s__Metschnikowia_pulcherrima(100); | Metschnikowia | 13 | Saprotroph | \|Undefined Saprotroph\| | Yeast | NULL | Probable | As nectar-tap_saprotroph (Põlme et al. 2020); Decay sugar-rich substrates (Põlme et al. 2020) | Põlme S, et al. 2020. Fungal Diversity 105:1-16 ((https://doi.org/10.1007/s13225-020-00466-2)) |
| Otu22867 | 1 | k__Fungi(100);p__Ascomycota(100);c__Saccharomycetes(100);o__Saccharomycetales(100);f__Metschnikowiaceae(100);g__Metschnikowia(100);s__Metschnikowia_pulcherrima(100); | Metschnikowia | 13 | Saprotroph | \|Undefined Saprotroph\| | Yeast | NULL | Probable | As nectar-tap_saprotroph (Põlme et al. 2020); Decay sugar-rich substrates (Põlme et al. 2020) | Põlme S, et al. 2020. Fungal Diversity 105:1-16 ((https://doi.org/10.1007/s13225-020-00466-2)) |
| Otu22868 | 1 | k__Fungi(100);p__Ascomycota(100);c__Saccharomycetes(100);o__Saccharomycetales(100);f__Metschnikowiaceae(100);g__Metschnikowia(100);s__Metschnikowia_pulcherrima(100); | Metschnikowia | 13 | Saprotroph | \|Undefined Saprotroph\| | Yeast | NULL | Probable | As nectar-tap_saprotroph (Põlme et al. 2020); Decay sugar-rich substrates (Põlme et al. 2020) | Põlme S, et al. 2020. Fungal Diversity 105:1-16 ((https://doi.org/10.1007/s13225-020-00466-2)) |
| Otu22869 | 1 | k__Fungi(100);p__Ascomycota(100);c__Saccharomycetes(100);o__Saccharomycetales(100);f__Metschnikowiaceae(100);g__Metschnikowia(100);s__Metschnikowia_pulcherrima(100); | Metschnikowia | 13 | Saprotroph | \|Undefined Saprotroph\| | Yeast | NULL | Probable | As nectar-tap_saprotroph (Põlme et al. 2020); Decay sugar-rich substrates (Põlme et al. 2020) | Põlme S, et al. 2020. Fungal Diversity 105:1-16 ((https://doi.org/10.1007/s13225-020-00466-2)) |
| Otu22870 | 1 | k__Fungi(100);p__Ascomycota(100);c__Saccharomycetes(100);o__Saccharomycetales(100);f__Metschnikowiaceae(100);g__Metschnikowia(100);s__Metschnikowia_pulcherrima(100); | Metschnikowia | 13 | Saprotroph | \|Undefined Saprotroph\| | Yeast | NULL | Probable | As nectar-tap_saprotroph (Põlme et al. 2020); Decay sugar-rich substrates (Põlme et al. 2020) | Põlme S, et al. 2020. Fungal Diversity 105:1-16 ((https://doi.org/10.1007/s13225-020-00466-2)) |
| Otu22871 | 1 | k__Fungi(100);p__Ascomycota(100);c__Saccharomycetes(100);o__Saccharomycetales(100);f__Metschnikowiaceae(100);g__Metschnikowia(100);g__Metschnikowia_unclassified(100); | Metschnikowia | 13 | Saprotroph | \|Undefined Saprotroph\| | Yeast | NULL | Probable | As nectar-tap_saprotroph (Põlme et al. 2020); Decay sugar-rich substrates (Põlme et al. 2020) | Põlme S, et al. 2020. Fungal Diversity 105:1-16 ((https://doi.org/10.1007/s13225-020-00466-2)) |
| Otu22872 | 1 | k__Fungi(100);p__Ascomycota(100);c__Saccharomycetes(100);o__Saccharomycetales(100);f__Metschnikowiaceae(100);g__Metschnikowia(100);g__Metschnikowia_unclassified(100); | Metschnikowia | 13 | Saprotroph | \|Undefined Saprotroph\| | Yeast | NULL | Probable | As nectar-tap_saprotroph (Põlme et al. 2020); Decay sugar-rich substrates (Põlme et al. 2020) | Põlme S, et al. 2020. Fungal Diversity 105:1-16 ((https://doi.org/10.1007/s13225-020-00466-2)) |
| Otu22873 | 1 | k__Fungi(100);p__Ascomycota(100);c__Saccharomycetes(100);o__Saccharomycetales(100);f__Metschnikowiaceae(100);g__Metschnikowia(100);s__Metschnikowia_pulcherrima(100); | Metschnikowia | 13 | Saprotroph | \|Undefined Saprotroph\| | Yeast | NULL | Probable | As nectar-tap_saprotroph (Põlme et al. 2020); Decay sugar-rich substrates (Põlme et al. 2020) | Põlme S, et al. 2020. Fungal Diversity 105:1-16 ((https://doi.org/10.1007/s13225-020-00466-2)) |
| Otu22874 | 1 | k__Fungi(100);p__Ascomycota(100);c__Saccharomycetes(100);o__Saccharomycetales(100);f__Metschnikowiaceae(100);g__Metschnikowia(100);s__Metschnikowia_pulcherrima(100); | Metschnikowia | 13 | Saprotroph | \|Undefined Saprotroph\| | Yeast | NULL | Probable | As nectar-tap_saprotroph (Põlme et al. 2020); Decay sugar-rich substrates (Põlme et al. 2020) | Põlme S, et al. 2020. Fungal Diversity 105:1-16 ((https://doi.org/10.1007/s13225-020-00466-2)) |
| Otu22875 | 1 | k__Fungi(100);p__Ascomycota(100);c__Saccharomycetes(100);o__Saccharomycetales(100);f__Metschnikowiaceae(100);g__Metschnikowia(100);s__Metschnikowia_pulcherrima(100); | Metschnikowia | 13 | Saprotroph | \|Undefined Saprotroph\| | Yeast | NULL | Probable | As nectar-tap_saprotroph (Põlme et al. 2020); Decay sugar-rich substrates (Põlme et al. 2020) | Põlme S, et al. 2020. Fungal Diversity 105:1-16 ((https://doi.org/10.1007/s13225-020-00466-2)) |
| Otu22876 | 1 | k__Fungi(100);p__Ascomycota(100);c__Saccharomycetes(100);o__Saccharomycetales(100);f__Metschnikowiaceae(100);g__Metschnikowia(100);g__Metschnikowia_unclassified(100); | Metschnikowia | 13 | Saprotroph | \|Undefined Saprotroph\| | Yeast | NULL | Probable | As nectar-tap_saprotroph (Põlme et al. 2020); Decay sugar-rich substrates (Põlme et al. 2020) | Põlme S, et al. 2020. Fungal Diversity 105:1-16 ((https://doi.org/10.1007/s13225-020-00466-2)) |
| Otu22877 | 1 | k__Fungi(100);p__Ascomycota(100);c__Saccharomycetes(100);o__Saccharomycetales(100);f__Metschnikowiaceae(100);g__Metschnikowia(100);g__Metschnikowia_unclassified(100); | Metschnikowia | 13 | Saprotroph | \|Undefined Saprotroph\| | Yeast | NULL | Probable | As nectar-tap_saprotroph (Põlme et al. 2020); Decay sugar-rich substrates (Põlme et al. 2020) | Põlme S, et al. 2020. Fungal Diversity 105:1-16 ((https://doi.org/10.1007/s13225-020-00466-2)) |
| Otu22878 | 1 | k__Fungi(100);p__Ascomycota(100);c__Saccharomycetes(100);o__Saccharomycetales(100);f__Metschnikowiaceae(100);g__Metschnikowia(100);s__Metschnikowia_pulcherrima(100); | Metschnikowia | 13 | Saprotroph | \|Undefined Saprotroph\| | Yeast | NULL | Probable | As nectar-tap_saprotroph (Põlme et al. 2020); Decay sugar-rich substrates (Põlme et al. 2020) | Põlme S, et al. 2020. Fungal Diversity 105:1-16 ((https://doi.org/10.1007/s13225-020-00466-2)) |
| Otu22879 | 1 | k__Fungi(100);p__Ascomycota(100);c__Saccharomycetes(100);o__Saccharomycetales(100);f__Metschnikowiaceae(100);g__Metschnikowia(100);s__Metschnikowia_pulcherrima(100); | Metschnikowia | 13 | Saprotroph | \|Undefined Saprotroph\| | Yeast | NULL | Probable | As nectar-tap_saprotroph (Põlme et al. 2020); Decay sugar-rich substrates (Põlme et al. 2020) | Põlme S, et al. 2020. Fungal Diversity 105:1-16 ((https://doi.org/10.1007/s13225-020-00466-2)) |
| Otu22880 | 1 | k__Fungi(100);p__Ascomycota(100);c__Saccharomycetes(100);o__Saccharomycetales(100);f__Metschnikowiaceae(100);g__Metschnikowia(100);s__Metschnikowia_pulcherrima(100); | Metschnikowia | 13 | Saprotroph | \|Undefined Saprotroph\| | Yeast | NULL | Probable | As nectar-tap_saprotroph (Põlme et al. 2020); Decay sugar-rich substrates (Põlme et al. 2020) | Põlme S, et al. 2020. Fungal Diversity 105:1-16 ((https://doi.org/10.1007/s13225-020-00466-2)) |
| Otu22881 | 1 | k__Fungi(100);p__Ascomycota(100);c__Saccharomycetes(100);o__Saccharomycetales(100);f__Metschnikowiaceae(100);g__Metschnikowia(100);g__Metschnikowia_unclassified(100); | Metschnikowia | 13 | Saprotroph | \|Undefined Saprotroph\| | Yeast | NULL | Probable | As nectar-tap_saprotroph (Põlme et al. 2020); Decay sugar-rich substrates (Põlme et al. 2020) | Põlme S, et al. 2020. Fungal Diversity 105:1-16 ((https://doi.org/10.1007/s13225-020-00466-2)) |
| Otu22882 | 1 | k__Fungi(100);p__Ascomycota(100);c__Saccharomycetes(100);o__Saccharomycetales(100);f__Metschnikowiaceae(100);g__Metschnikowia(100);s__Metschnikowia_pulcherrima(100); | Metschnikowia | 13 | Saprotroph | \|Undefined Saprotroph\| | Yeast | NULL | Probable | As nectar-tap_saprotroph (Põlme et al. 2020); Decay sugar-rich substrates (Põlme et al. 2020) | Põlme S, et al. 2020. Fungal Diversity 105:1-16 ((https://doi.org/10.1007/s13225-020-00466-2)) |
| Otu22883 | 1 | k__Fungi(100);p__Ascomycota(100);c__Saccharomycetes(100);o__Saccharomycetales(100);f__Metschnikowiaceae(100);g__Metschnikowia(100);s__Metschnikowia_pulcherrima(100); | Metschnikowia | 13 | Saprotroph | \|Undefined Saprotroph\| | Yeast | NULL | Probable | As nectar-tap_saprotroph (Põlme et al. 2020); Decay sugar-rich substrates (Põlme et al. 2020) | Põlme S, et al. 2020. Fungal Diversity 105:1-16 ((https://doi.org/10.1007/s13225-020-00466-2)) |
| Otu22884 | 1 | k__Fungi(100);p__Ascomycota(100);c__Saccharomycetes(100);o__Saccharomycetales(100);f__Metschnikowiaceae(100);g__Metschnikowia(100);s__Metschnikowia_pulcherrima(100); | Metschnikowia | 13 | Saprotroph | \|Undefined Saprotroph\| | Yeast | NULL | Probable | As nectar-tap_saprotroph (Põlme et al. 2020); Decay sugar-rich substrates (Põlme et al. 2020) | Põlme S, et al. 2020. Fungal Diversity 105:1-16 ((https://doi.org/10.1007/s13225-020-00466-2)) |
| Otu22885 | 1 | k__Fungi(100);p__Ascomycota(100);c__Saccharomycetes(100);o__Saccharomycetales(100);f__Metschnikowiaceae(100);g__Metschnikowia(100);s__Metschnikowia_pulcherrima(100); | Metschnikowia | 13 | Saprotroph | \|Undefined Saprotroph\| | Yeast | NULL | Probable | As nectar-tap_saprotroph (Põlme et al. 2020); Decay sugar-rich substrates (Põlme et al. 2020) | Põlme S, et al. 2020. Fungal Diversity 105:1-16 ((https://doi.org/10.1007/s13225-020-00466-2)) |
| Otu22886 | 1 | k__Fungi(100);p__Ascomycota(100);c__Saccharomycetes(100);o__Saccharomycetales(100);f__Metschnikowiaceae(100);g__Metschnikowia(100);s__Metschnikowia_pulcherrima(100); | Metschnikowia | 13 | Saprotroph | \|Undefined Saprotroph\| | Yeast | NULL | Probable | As nectar-tap_saprotroph (Põlme et al. 2020); Decay sugar-rich substrates (Põlme et al. 2020) | Põlme S, et al. 2020. Fungal Diversity 105:1-16 ((https://doi.org/10.1007/s13225-020-00466-2)) |
| Otu22887 | 1 | k__Fungi(100);p__Ascomycota(100);c__Saccharomycetes(100);o__Saccharomycetales(100);f__Metschnikowiaceae(100);g__Metschnikowia(100);g__Metschnikowia_unclassified(100); | Metschnikowia | 13 | Saprotroph | \|Undefined Saprotroph\| | Yeast | NULL | Probable | As nectar-tap_saprotroph (Põlme et al. 2020); Decay sugar-rich substrates (Põlme et al. 2020) | Põlme S, et al. 2020. Fungal Diversity 105:1-16 ((https://doi.org/10.1007/s13225-020-00466-2)) |
| Otu22888 | 1 | k__Fungi(100);p__Ascomycota(100);c__Saccharomycetes(100);o__Saccharomycetales(100);f__Metschnikowiaceae(100);g__Metschnikowia(100);s__Metschnikowia_pulcherrima(100); | Metschnikowia | 13 | Saprotroph | \|Undefined Saprotroph\| | Yeast | NULL | Probable | As nectar-tap_saprotroph (Põlme et al. 2020); Decay sugar-rich substrates (Põlme et al. 2020) | Põlme S, et al. 2020. Fungal Diversity 105:1-16 ((https://doi.org/10.1007/s13225-020-00466-2)) |
| Otu22889 | 1 | k__Fungi(100);p__Ascomycota(100);c__Saccharomycetes(100);o__Saccharomycetales(100);f__Metschnikowiaceae(100);g__Metschnikowia(100);g__Metschnikowia_unclassified(100); | Metschnikowia | 13 | Saprotroph | \|Undefined Saprotroph\| | Yeast | NULL | Probable | As nectar-tap_saprotroph (Põlme et al. 2020); Decay sugar-rich substrates (Põlme et al. 2020) | Põlme S, et al. 2020. Fungal Diversity 105:1-16 ((https://doi.org/10.1007/s13225-020-00466-2)) |
| Otu22890 | 1 | k__Fungi(100);p__Ascomycota(100);c__Saccharomycetes(100);o__Saccharomycetales(100);f__Metschnikowiaceae(100);g__Metschnikowia(100);g__Metschnikowia_unclassified(100); | Metschnikowia | 13 | Saprotroph | \|Undefined Saprotroph\| | Yeast | NULL | Probable | As nectar-tap_saprotroph (Põlme et al. 2020); Decay sugar-rich substrates (Põlme et al. 2020) | Põlme S, et al. 2020. Fungal Diversity 105:1-16 ((https://doi.org/10.1007/s13225-020-00466-2)) |
| Otu22891 | 1 | k__Fungi(100);p__Ascomycota(100);c__Saccharomycetes(100);o__Saccharomycetales(100);f__Metschnikowiaceae(100);g__Metschnikowia(100);g__Metschnikowia_unclassified(100); | Metschnikowia | 13 | Saprotroph | \|Undefined Saprotroph\| | Yeast | NULL | Probable | As nectar-tap_saprotroph (Põlme et al. 2020); Decay sugar-rich substrates (Põlme et al. 2020) | Põlme S, et al. 2020. Fungal Diversity 105:1-16 ((https://doi.org/10.1007/s13225-020-00466-2)) |
| Otu22892 | 1 | k__Fungi(100);p__Ascomycota(100);c__Saccharomycetes(100);o__Saccharomycetales(100);f__Metschnikowiaceae(100);g__Metschnikowia(100);s__Metschnikowia_pulcherrima(100); | Metschnikowia | 13 | Saprotroph | \|Undefined Saprotroph\| | Yeast | NULL | Probable | As nectar-tap_saprotroph (Põlme et al. 2020); Decay sugar-rich substrates (Põlme et al. 2020) | Põlme S, et al. 2020. Fungal Diversity 105:1-16 ((https://doi.org/10.1007/s13225-020-00466-2)) |
| Otu22893 | 1 | k__Fungi(100);p__Ascomycota(100);c__Saccharomycetes(100);o__Saccharomycetales(100);f__Metschnikowiaceae(100);g__Metschnikowia(100);s__Metschnikowia_pulcherrima(100); | Metschnikowia | 13 | Saprotroph | \|Undefined Saprotroph\| | Yeast | NULL | Probable | As nectar-tap_saprotroph (Põlme et al. 2020); Decay sugar-rich substrates (Põlme et al. 2020) | Põlme S, et al. 2020. Fungal Diversity 105:1-16 ((https://doi.org/10.1007/s13225-020-00466-2)) |
| Otu22894 | 1 | k__Fungi(100);p__Ascomycota(100);c__Saccharomycetes(100);o__Saccharomycetales(100);f__Metschnikowiaceae(100);g__Metschnikowia(100);g__Metschnikowia_unclassified(100); | Metschnikowia | 13 | Saprotroph | \|Undefined Saprotroph\| | Yeast | NULL | Probable | As nectar-tap_saprotroph (Põlme et al. 2020); Decay sugar-rich substrates (Põlme et al. 2020) | Põlme S, et al. 2020. Fungal Diversity 105:1-16 ((https://doi.org/10.1007/s13225-020-00466-2)) |
| Otu22895 | 1 | k__Fungi(100);p__Ascomycota(100);c__Saccharomycetes(100);o__Saccharomycetales(100);f__Metschnikowiaceae(100);g__Metschnikowia(100);s__Metschnikowia_pulcherrima(100); | Metschnikowia | 13 | Saprotroph | \|Undefined Saprotroph\| | Yeast | NULL | Probable | As nectar-tap_saprotroph (Põlme et al. 2020); Decay sugar-rich substrates (Põlme et al. 2020) | Põlme S, et al. 2020. Fungal Diversity 105:1-16 ((https://doi.org/10.1007/s13225-020-00466-2)) |
| Otu22896 | 1 | k__Fungi(100);p__Ascomycota(100);c__Saccharomycetes(100);o__Saccharomycetales(100);f__Metschnikowiaceae(100);g__Metschnikowia(100);s__Metschnikowia_pulcherrima(100); | Metschnikowia | 13 | Saprotroph | \|Undefined Saprotroph\| | Yeast | NULL | Probable | As nectar-tap_saprotroph (Põlme et al. 2020); Decay sugar-rich substrates (Põlme et al. 2020) | Põlme S, et al. 2020. Fungal Diversity 105:1-16 ((https://doi.org/10.1007/s13225-020-00466-2)) |
| Otu22897 | 1 | k__Fungi(100);p__Ascomycota(100);c__Saccharomycetes(100);o__Saccharomycetales(100);f__Metschnikowiaceae(100);g__Metschnikowia(100);g__Metschnikowia_unclassified(100); | Metschnikowia | 13 | Saprotroph | \|Undefined Saprotroph\| | Yeast | NULL | Probable | As nectar-tap_saprotroph (Põlme et al. 2020); Decay sugar-rich substrates (Põlme et al. 2020) | Põlme S, et al. 2020. Fungal Diversity 105:1-16 ((https://doi.org/10.1007/s13225-020-00466-2)) |
| Otu22898 | 1 | k__Fungi(100);p__Ascomycota(100);c__Saccharomycetes(100);o__Saccharomycetales(100);f__Metschnikowiaceae(100);g__Metschnikowia(100);s__Metschnikowia_pulcherrima(100); | Metschnikowia | 13 | Saprotroph | \|Undefined Saprotroph\| | Yeast | NULL | Probable | As nectar-tap_saprotroph (Põlme et al. 2020); Decay sugar-rich substrates (Põlme et al. 2020) | Põlme S, et al. 2020. Fungal Diversity 105:1-16 ((https://doi.org/10.1007/s13225-020-00466-2)) |
| Otu22899 | 1 | k__Fungi(100);p__Ascomycota(100);c__Saccharomycetes(100);o__Saccharomycetales(100);f__Metschnikowiaceae(100);g__Metschnikowia(100);s__Metschnikowia_pulcherrima(100); | Metschnikowia | 13 | Saprotroph | \|Undefined Saprotroph\| | Yeast | NULL | Probable | As nectar-tap_saprotroph (Põlme et al. 2020); Decay sugar-rich substrates (Põlme et al. 2020) | Põlme S, et al. 2020. Fungal Diversity 105:1-16 ((https://doi.org/10.1007/s13225-020-00466-2)) |
| Otu22900 | 1 | k__Fungi(100);p__Ascomycota(100);c__Saccharomycetes(100);o__Saccharomycetales(100);f__Metschnikowiaceae(100);g__Metschnikowia(100);g__Metschnikowia_unclassified(100); | Metschnikowia | 13 | Saprotroph | \|Undefined Saprotroph\| | Yeast | NULL | Probable | As nectar-tap_saprotroph (Põlme et al. 2020); Decay sugar-rich substrates (Põlme et al. 2020) | Põlme S, et al. 2020. Fungal Diversity 105:1-16 ((https://doi.org/10.1007/s13225-020-00466-2)) |
| Otu22901 | 1 | k__Fungi(100);p__Ascomycota(100);c__Saccharomycetes(100);o__Saccharomycetales(100);f__Metschnikowiaceae(100);g__Metschnikowia(100);g__Metschnikowia_unclassified(100); | Metschnikowia | 13 | Saprotroph | \|Undefined Saprotroph\| | Yeast | NULL | Probable | As nectar-tap_saprotroph (Põlme et al. 2020); Decay sugar-rich substrates (Põlme et al. 2020) | Põlme S, et al. 2020. Fungal Diversity 105:1-16 ((https://doi.org/10.1007/s13225-020-00466-2)) |
| Otu22902 | 1 | k__Fungi(100);p__Ascomycota(100);c__Saccharomycetes(100);o__Saccharomycetales(100);f__Metschnikowiaceae(100);g__Metschnikowia(100);s__Metschnikowia_pulcherrima(100); | Metschnikowia | 13 | Saprotroph | \|Undefined Saprotroph\| | Yeast | NULL | Probable | As nectar-tap_saprotroph (Põlme et al. 2020); Decay sugar-rich substrates (Põlme et al. 2020) | Põlme S, et al. 2020. Fungal Diversity 105:1-16 ((https://doi.org/10.1007/s13225-020-00466-2)) |
| Otu22903 | 1 | k__Fungi(100);p__Ascomycota(100);c__Saccharomycetes(100);o__Saccharomycetales(100);f__Metschnikowiaceae(100);g__Metschnikowia(100);s__Metschnikowia_pulcherrima(100); | Metschnikowia | 13 | Saprotroph | \|Undefined Saprotroph\| | Yeast | NULL | Probable | As nectar-tap_saprotroph (Põlme et al. 2020); Decay sugar-rich substrates (Põlme et al. 2020) | Põlme S, et al. 2020. Fungal Diversity 105:1-16 ((https://doi.org/10.1007/s13225-020-00466-2)) |
| Otu22904 | 1 | k__Fungi(100);p__Ascomycota(100);c__Saccharomycetes(100);o__Saccharomycetales(100);f__Metschnikowiaceae(100);g__Metschnikowia(100);s__Metschnikowia_pulcherrima(100); | Metschnikowia | 13 | Saprotroph | \|Undefined Saprotroph\| | Yeast | NULL | Probable | As nectar-tap_saprotroph (Põlme et al. 2020); Decay sugar-rich substrates (Põlme et al. 2020) | Põlme S, et al. 2020. Fungal Diversity 105:1-16 ((https://doi.org/10.1007/s13225-020-00466-2)) |
| Otu22905 | 1 | k__Fungi(100);p__Ascomycota(100);c__Saccharomycetes(100);o__Saccharomycetales(100);f__Metschnikowiaceae(100);g__Metschnikowia(100);g__Metschnikowia_unclassified(100); | Metschnikowia | 13 | Saprotroph | \|Undefined Saprotroph\| | Yeast | NULL | Probable | As nectar-tap_saprotroph (Põlme et al. 2020); Decay sugar-rich substrates (Põlme et al. 2020) | Põlme S, et al. 2020. Fungal Diversity 105:1-16 ((https://doi.org/10.1007/s13225-020-00466-2)) |
| Otu22906 | 1 | k__Fungi(100);p__Ascomycota(100);c__Saccharomycetes(100);o__Saccharomycetales(100);f__Metschnikowiaceae(100);g__Metschnikowia(100);s__Metschnikowia_pulcherrima(100); | Metschnikowia | 13 | Saprotroph | \|Undefined Saprotroph\| | Yeast | NULL | Probable | As nectar-tap_saprotroph (Põlme et al. 2020); Decay sugar-rich substrates (Põlme et al. 2020) | Põlme S, et al. 2020. Fungal Diversity 105:1-16 ((https://doi.org/10.1007/s13225-020-00466-2)) |
| Otu22907 | 1 | k__Fungi(100);p__Ascomycota(100);c__Saccharomycetes(100);o__Saccharomycetales(100);f__Metschnikowiaceae(100);g__Metschnikowia(100);s__Metschnikowia_pulcherrima(100); | Metschnikowia | 13 | Saprotroph | \|Undefined Saprotroph\| | Yeast | NULL | Probable | As nectar-tap_saprotroph (Põlme et al. 2020); Decay sugar-rich substrates (Põlme et al. 2020) | Põlme S, et al. 2020. Fungal Diversity 105:1-16 ((https://doi.org/10.1007/s13225-020-00466-2)) |
| Otu22908 | 1 | k__Fungi(100);p__Ascomycota(100);c__Saccharomycetes(100);o__Saccharomycetales(100);f__Metschnikowiaceae(100);g__Metschnikowia(100);s__Metschnikowia_pulcherrima(100); | Metschnikowia | 13 | Saprotroph | \|Undefined Saprotroph\| | Yeast | NULL | Probable | As nectar-tap_saprotroph (Põlme et al. 2020); Decay sugar-rich substrates (Põlme et al. 2020) | Põlme S, et al. 2020. Fungal Diversity 105:1-16 ((https://doi.org/10.1007/s13225-020-00466-2)) |
| Otu22909 | 1 | k__Fungi(100);p__Ascomycota(100);c__Saccharomycetes(100);o__Saccharomycetales(100);f__Metschnikowiaceae(100);g__Metschnikowia(100);s__Metschnikowia_pulcherrima(100); | Metschnikowia | 13 | Saprotroph | \|Undefined Saprotroph\| | Yeast | NULL | Probable | As nectar-tap_saprotroph (Põlme et al. 2020); Decay sugar-rich substrates (Põlme et al. 2020) | Põlme S, et al. 2020. Fungal Diversity 105:1-16 ((https://doi.org/10.1007/s13225-020-00466-2)) |
| Otu22910 | 1 | k__Fungi(100);p__Ascomycota(100);c__Saccharomycetes(100);o__Saccharomycetales(100);f__Metschnikowiaceae(100);g__Metschnikowia(100);s__Metschnikowia_pulcherrima(100); | Metschnikowia | 13 | Saprotroph | \|Undefined Saprotroph\| | Yeast | NULL | Probable | As nectar-tap_saprotroph (Põlme et al. 2020); Decay sugar-rich substrates (Põlme et al. 2020) | Põlme S, et al. 2020. Fungal Diversity 105:1-16 ((https://doi.org/10.1007/s13225-020-00466-2)) |
| Otu22911 | 1 | k__Fungi(100);p__Ascomycota(100);c__Saccharomycetes(100);o__Saccharomycetales(100);f__Metschnikowiaceae(100);g__Metschnikowia(100);s__Metschnikowia_pulcherrima(100); | Metschnikowia | 13 | Saprotroph | \|Undefined Saprotroph\| | Yeast | NULL | Probable | As nectar-tap_saprotroph (Põlme et al. 2020); Decay sugar-rich substrates (Põlme et al. 2020) | Põlme S, et al. 2020. Fungal Diversity 105:1-16 ((https://doi.org/10.1007/s13225-020-00466-2)) |
| Otu22912 | 1 | k__Fungi(100);p__Ascomycota(100);c__Saccharomycetes(100);o__Saccharomycetales(100);f__Metschnikowiaceae(100);g__Metschnikowia(100);s__Metschnikowia_pulcherrima(100); | Metschnikowia | 13 | Saprotroph | \|Undefined Saprotroph\| | Yeast | NULL | Probable | As nectar-tap_saprotroph (Põlme et al. 2020); Decay sugar-rich substrates (Põlme et al. 2020) | Põlme S, et al. 2020. Fungal Diversity 105:1-16 ((https://doi.org/10.1007/s13225-020-00466-2)) |
| Otu22913 | 1 | k__Fungi(100);p__Ascomycota(100);c__Saccharomycetes(100);o__Saccharomycetales(100);f__Metschnikowiaceae(100);g__Metschnikowia(100);s__Metschnikowia_pulcherrima(100); | Metschnikowia | 13 | Saprotroph | \|Undefined Saprotroph\| | Yeast | NULL | Probable | As nectar-tap_saprotroph (Põlme et al. 2020); Decay sugar-rich substrates (Põlme et al. 2020) | Põlme S, et al. 2020. Fungal Diversity 105:1-16 ((https://doi.org/10.1007/s13225-020-00466-2)) |
| Otu22914 | 1 | k__Fungi(100);p__Ascomycota(100);c__Saccharomycetes(100);o__Saccharomycetales(100);f__Metschnikowiaceae(100);g__Metschnikowia(100);g__Metschnikowia_unclassified(100); | Metschnikowia | 13 | Saprotroph | \|Undefined Saprotroph\| | Yeast | NULL | Probable | As nectar-tap_saprotroph (Põlme et al. 2020); Decay sugar-rich substrates (Põlme et al. 2020) | Põlme S, et al. 2020. Fungal Diversity 105:1-16 ((https://doi.org/10.1007/s13225-020-00466-2)) |
| Otu22915 | 1 | k__Fungi(100);p__Ascomycota(100);c__Saccharomycetes(100);o__Saccharomycetales(100);f__Metschnikowiaceae(100);g__Metschnikowia(100);s__Metschnikowia_pulcherrima(100); | Metschnikowia | 13 | Saprotroph | \|Undefined Saprotroph\| | Yeast | NULL | Probable | As nectar-tap_saprotroph (Põlme et al. 2020); Decay sugar-rich substrates (Põlme et al. 2020) | Põlme S, et al. 2020. Fungal Diversity 105:1-16 ((https://doi.org/10.1007/s13225-020-00466-2)) |
| Otu22917 | 1 | k__Fungi(100);p__Ascomycota(100);c__Saccharomycetes(100);o__Saccharomycetales(100);f__Metschnikowiaceae(100);g__Metschnikowia(100);s__Metschnikowia_pulcherrima(100); | Metschnikowia | 13 | Saprotroph | \|Undefined Saprotroph\| | Yeast | NULL | Probable | As nectar-tap_saprotroph (Põlme et al. 2020); Decay sugar-rich substrates (Põlme et al. 2020) | Põlme S, et al. 2020. Fungal Diversity 105:1-16 ((https://doi.org/10.1007/s13225-020-00466-2)) |
| Otu22918 | 1 | k__Fungi(100);p__Ascomycota(100);c__Saccharomycetes(100);o__Saccharomycetales(100);f__Metschnikowiaceae(100);g__Metschnikowia(100);s__Metschnikowia_pulcherrima(100); | Metschnikowia | 13 | Saprotroph | \|Undefined Saprotroph\| | Yeast | NULL | Probable | As nectar-tap_saprotroph (Põlme et al. 2020); Decay sugar-rich substrates (Põlme et al. 2020) | Põlme S, et al. 2020. Fungal Diversity 105:1-16 ((https://doi.org/10.1007/s13225-020-00466-2)) |
| Otu22919 | 1 | k__Fungi(100);p__Ascomycota(100);c__Saccharomycetes(100);o__Saccharomycetales(100);f__Metschnikowiaceae(100);g__Metschnikowia(100);s__Metschnikowia_pulcherrima(100); | Metschnikowia | 13 | Saprotroph | \|Undefined Saprotroph\| | Yeast | NULL | Probable | As nectar-tap_saprotroph (Põlme et al. 2020); Decay sugar-rich substrates (Põlme et al. 2020) | Põlme S, et al. 2020. Fungal Diversity 105:1-16 ((https://doi.org/10.1007/s13225-020-00466-2)) |
| Otu22920 | 1 | k__Fungi(100);p__Ascomycota(100);c__Saccharomycetes(100);o__Saccharomycetales(100);f__Metschnikowiaceae(100);g__Metschnikowia(100);g__Metschnikowia_unclassified(100); | Metschnikowia | 13 | Saprotroph | \|Undefined Saprotroph\| | Yeast | NULL | Probable | As nectar-tap_saprotroph (Põlme et al. 2020); Decay sugar-rich substrates (Põlme et al. 2020) | Põlme S, et al. 2020. Fungal Diversity 105:1-16 ((https://doi.org/10.1007/s13225-020-00466-2)) |
| Otu22921 | 1 | k__Fungi(100);p__Ascomycota(100);c__Saccharomycetes(100);o__Saccharomycetales(100);f__Saccharomycetales_family_Incertae_sedis(100);g__Candida(100);g__Candida_unclassified(100); | Candida | 13 | Pathotroph-Saprotroph-Symbiotroph | Animal Pathogen-Endophyte-Endosymbiont-Epiphyte-\|Undefined Saprotroph\| | Dimorphic Yeast-Yeast | NULL | Probable | As nectar-tap_saprotroph (Põlme et al. 2020); Decay sugar-rich substrates (Põlme et al. 2020); This asexual genus is polyphyletic and contain species that function very differently depend on what they are. Caution should be taken when interpreting guilds for Candida species. | Manolakaki D, et al. 2010. Virulence. 1: 367-375 ((https://doi.org/10.4161/viru.1.5.12796)); Mekbib SB, et al. 2011. Tropical Plant Pathology 36:233-240 ((https://doi.org/10.1590/S1982-56762011000400004)); Põlme S, et al. 2020. Fungal Diversity 105:1-16 ((https://doi.org/10.1007/s13225-020-00466-2)) |
| Otu22922 | 1 | k__Fungi(100);p__Ascomycota(100);c__Saccharomycetes(100);o__Saccharomycetales(100);f__Metschnikowiaceae(100);g__Metschnikowia(100);s__Metschnikowia_pulcherrima(100); | Metschnikowia | 13 | Saprotroph | \|Undefined Saprotroph\| | Yeast | NULL | Probable | As nectar-tap_saprotroph (Põlme et al. 2020); Decay sugar-rich substrates (Põlme et al. 2020) | Põlme S, et al. 2020. Fungal Diversity 105:1-16 ((https://doi.org/10.1007/s13225-020-00466-2)) |
| Otu22923 | 1 | k__Fungi(100);p__Ascomycota(100);c__Saccharomycetes(100);o__Saccharomycetales(100);f__Metschnikowiaceae(100);g__Metschnikowia(100);s__Metschnikowia_pulcherrima(100); | Metschnikowia | 13 | Saprotroph | \|Undefined Saprotroph\| | Yeast | NULL | Probable | As nectar-tap_saprotroph (Põlme et al. 2020); Decay sugar-rich substrates (Põlme et al. 2020) | Põlme S, et al. 2020. Fungal Diversity 105:1-16 ((https://doi.org/10.1007/s13225-020-00466-2)) |
| Otu22925 | 1 | k__Fungi(100);p__Ascomycota(100);c__Saccharomycetes(100);o__Saccharomycetales(100);f__Saccharomycetales_family_Incertae_sedis(100);g__Candida(100);s__Candida_cretensis(100); | Candida | 13 | Pathotroph-Saprotroph-Symbiotroph | Animal Pathogen-Endophyte-Endosymbiont-Epiphyte-\|Undefined Saprotroph\| | Dimorphic Yeast-Yeast | NULL | Probable | As nectar-tap_saprotroph (Põlme et al. 2020); Decay sugar-rich substrates (Põlme et al. 2020); This asexual genus is polyphyletic and contain species that function very differently depend on what they are. Caution should be taken when interpreting guilds for Candida species. | Manolakaki D, et al. 2010. Virulence. 1: 367-375 ((https://doi.org/10.4161/viru.1.5.12796)); Mekbib SB, et al. 2011. Tropical Plant Pathology 36:233-240 ((https://doi.org/10.1590/S1982-56762011000400004)); Põlme S, et al. 2020. Fungal Diversity 105:1-16 ((https://doi.org/10.1007/s13225-020-00466-2)) |
| Otu22929 | 1 | k__Fungi(100);p__Ascomycota(100);c__Saccharomycetes(100);o__Saccharomycetales(100);f__Metschnikowiaceae(100);g__Metschnikowia(100);s__Metschnikowia_pulcherrima(100); | Metschnikowia | 13 | Saprotroph | \|Undefined Saprotroph\| | Yeast | NULL | Probable | As nectar-tap_saprotroph (Põlme et al. 2020); Decay sugar-rich substrates (Põlme et al. 2020) | Põlme S, et al. 2020. Fungal Diversity 105:1-16 ((https://doi.org/10.1007/s13225-020-00466-2)) |
| Otu22930 | 1 | k__Fungi(100);p__Ascomycota(100);c__Saccharomycetes(100);o__Saccharomycetales(100);f__Metschnikowiaceae(100);g__Metschnikowia(100);s__Metschnikowia_pulcherrima(100); | Metschnikowia | 13 | Saprotroph | \|Undefined Saprotroph\| | Yeast | NULL | Probable | As nectar-tap_saprotroph (Põlme et al. 2020); Decay sugar-rich substrates (Põlme et al. 2020) | Põlme S, et al. 2020. Fungal Diversity 105:1-16 ((https://doi.org/10.1007/s13225-020-00466-2)) |
| Otu22931 | 1 | k__Fungi(100);p__Ascomycota(100);c__Saccharomycetes(100);o__Saccharomycetales(100);f__Metschnikowiaceae(100);g__Metschnikowia(100);s__Metschnikowia_pulcherrima(100); | Metschnikowia | 13 | Saprotroph | \|Undefined Saprotroph\| | Yeast | NULL | Probable | As nectar-tap_saprotroph (Põlme et al. 2020); Decay sugar-rich substrates (Põlme et al. 2020) | Põlme S, et al. 2020. Fungal Diversity 105:1-16 ((https://doi.org/10.1007/s13225-020-00466-2)) |
| Otu22932 | 1 | k__Fungi(100);p__Ascomycota(100);c__Saccharomycetes(100);o__Saccharomycetales(100);f__Metschnikowiaceae(100);g__Metschnikowia(100);g__Metschnikowia_unclassified(100); | Metschnikowia | 13 | Saprotroph | \|Undefined Saprotroph\| | Yeast | NULL | Probable | As nectar-tap_saprotroph (Põlme et al. 2020); Decay sugar-rich substrates (Põlme et al. 2020) | Põlme S, et al. 2020. Fungal Diversity 105:1-16 ((https://doi.org/10.1007/s13225-020-00466-2)) |
| Otu22933 | 1 | k__Fungi(100);p__Ascomycota(100);c__Saccharomycetes(100);o__Saccharomycetales(100);f__Metschnikowiaceae(100);g__Metschnikowia(100);s__Metschnikowia_pulcherrima(100); | Metschnikowia | 13 | Saprotroph | \|Undefined Saprotroph\| | Yeast | NULL | Probable | As nectar-tap_saprotroph (Põlme et al. 2020); Decay sugar-rich substrates (Põlme et al. 2020) | Põlme S, et al. 2020. Fungal Diversity 105:1-16 ((https://doi.org/10.1007/s13225-020-00466-2)) |
| Otu22934 | 1 | k__Fungi(100);p__Ascomycota(100);c__Saccharomycetes(100);o__Saccharomycetales(100);f__Metschnikowiaceae(100);g__Metschnikowia(100);s__Metschnikowia_pulcherrima(100); | Metschnikowia | 13 | Saprotroph | \|Undefined Saprotroph\| | Yeast | NULL | Probable | As nectar-tap_saprotroph (Põlme et al. 2020); Decay sugar-rich substrates (Põlme et al. 2020) | Põlme S, et al. 2020. Fungal Diversity 105:1-16 ((https://doi.org/10.1007/s13225-020-00466-2)) |
| Otu22935 | 1 | k__Fungi(100);p__Ascomycota(100);c__Saccharomycetes(100);o__Saccharomycetales(100);f__Metschnikowiaceae(100);g__Metschnikowia(100);s__Metschnikowia_pulcherrima(100); | Metschnikowia | 13 | Saprotroph | \|Undefined Saprotroph\| | Yeast | NULL | Probable | As nectar-tap_saprotroph (Põlme et al. 2020); Decay sugar-rich substrates (Põlme et al. 2020) | Põlme S, et al. 2020. Fungal Diversity 105:1-16 ((https://doi.org/10.1007/s13225-020-00466-2)) |
| Otu22936 | 1 | k__Fungi(100);p__Ascomycota(100);c__Saccharomycetes(100);o__Saccharomycetales(100);f__Metschnikowiaceae(100);g__Metschnikowia(100);s__Metschnikowia_pulcherrima(100); | Metschnikowia | 13 | Saprotroph | \|Undefined Saprotroph\| | Yeast | NULL | Probable | As nectar-tap_saprotroph (Põlme et al. 2020); Decay sugar-rich substrates (Põlme et al. 2020) | Põlme S, et al. 2020. Fungal Diversity 105:1-16 ((https://doi.org/10.1007/s13225-020-00466-2)) |
| Otu22937 | 1 | k__Fungi(100);p__Ascomycota(100);c__Saccharomycetes(100);o__Saccharomycetales(100);f__Metschnikowiaceae(100);g__Metschnikowia(100);s__Metschnikowia_pulcherrima(100); | Metschnikowia | 13 | Saprotroph | \|Undefined Saprotroph\| | Yeast | NULL | Probable | As nectar-tap_saprotroph (Põlme et al. 2020); Decay sugar-rich substrates (Põlme et al. 2020) | Põlme S, et al. 2020. Fungal Diversity 105:1-16 ((https://doi.org/10.1007/s13225-020-00466-2)) |
| Otu22938 | 1 | k__Fungi(100);p__Ascomycota(100);c__Saccharomycetes(100);o__Saccharomycetales(100);f__Metschnikowiaceae(100);g__Metschnikowia(100);g__Metschnikowia_unclassified(100); | Metschnikowia | 13 | Saprotroph | \|Undefined Saprotroph\| | Yeast | NULL | Probable | As nectar-tap_saprotroph (Põlme et al. 2020); Decay sugar-rich substrates (Põlme et al. 2020) | Põlme S, et al. 2020. Fungal Diversity 105:1-16 ((https://doi.org/10.1007/s13225-020-00466-2)) |
| Otu22940 | 1 | k__Fungi(100);p__Ascomycota(100);c__Saccharomycetes(100);o__Saccharomycetales(100);f__Metschnikowiaceae(100);g__Metschnikowia(100);s__Metschnikowia_pulcherrima(100); | Metschnikowia | 13 | Saprotroph | \|Undefined Saprotroph\| | Yeast | NULL | Probable | As nectar-tap_saprotroph (Põlme et al. 2020); Decay sugar-rich substrates (Põlme et al. 2020) | Põlme S, et al. 2020. Fungal Diversity 105:1-16 ((https://doi.org/10.1007/s13225-020-00466-2)) |
| Otu22941 | 1 | k__Fungi(100);p__Ascomycota(100);c__Saccharomycetes(100);o__Saccharomycetales(100);f__Metschnikowiaceae(100);g__Metschnikowia(100);s__Metschnikowia_pulcherrima(100); | Metschnikowia | 13 | Saprotroph | \|Undefined Saprotroph\| | Yeast | NULL | Probable | As nectar-tap_saprotroph (Põlme et al. 2020); Decay sugar-rich substrates (Põlme et al. 2020) | Põlme S, et al. 2020. Fungal Diversity 105:1-16 ((https://doi.org/10.1007/s13225-020-00466-2)) |
| Otu22942 | 1 | k__Fungi(100);p__Ascomycota(100);c__Saccharomycetes(100);o__Saccharomycetales(100);f__Metschnikowiaceae(100);g__Metschnikowia(100);s__Metschnikowia_pulcherrima(100); | Metschnikowia | 13 | Saprotroph | \|Undefined Saprotroph\| | Yeast | NULL | Probable | As nectar-tap_saprotroph (Põlme et al. 2020); Decay sugar-rich substrates (Põlme et al. 2020) | Põlme S, et al. 2020. Fungal Diversity 105:1-16 ((https://doi.org/10.1007/s13225-020-00466-2)) |
| Otu22943 | 1 | k__Fungi(100);p__Ascomycota(100);c__Saccharomycetes(100);o__Saccharomycetales(100);f__Metschnikowiaceae(100);g__Metschnikowia(100);s__Metschnikowia_pulcherrima(100); | Metschnikowia | 13 | Saprotroph | \|Undefined Saprotroph\| | Yeast | NULL | Probable | As nectar-tap_saprotroph (Põlme et al. 2020); Decay sugar-rich substrates (Põlme et al. 2020) | Põlme S, et al. 2020. Fungal Diversity 105:1-16 ((https://doi.org/10.1007/s13225-020-00466-2)) |
| Otu22944 | 1 | k__Fungi(100);p__Ascomycota(100);c__Saccharomycetes(100);o__Saccharomycetales(100);f__Metschnikowiaceae(100);g__Metschnikowia(100);s__Metschnikowia_pulcherrima(100); | Metschnikowia | 13 | Saprotroph | \|Undefined Saprotroph\| | Yeast | NULL | Probable | As nectar-tap_saprotroph (Põlme et al. 2020); Decay sugar-rich substrates (Põlme et al. 2020) | Põlme S, et al. 2020. Fungal Diversity 105:1-16 ((https://doi.org/10.1007/s13225-020-00466-2)) |
| Otu22945 | 1 | k__Fungi(100);p__Ascomycota(100);c__Saccharomycetes(100);o__Saccharomycetales(100);f__Metschnikowiaceae(100);g__Metschnikowia(100);s__Metschnikowia_pulcherrima(100); | Metschnikowia | 13 | Saprotroph | \|Undefined Saprotroph\| | Yeast | NULL | Probable | As nectar-tap_saprotroph (Põlme et al. 2020); Decay sugar-rich substrates (Põlme et al. 2020) | Põlme S, et al. 2020. Fungal Diversity 105:1-16 ((https://doi.org/10.1007/s13225-020-00466-2)) |
| Otu22947 | 1 | k__Fungi(100);p__Ascomycota(100);c__Saccharomycetes(100);o__Saccharomycetales(100);f__Saccharomycetales_family_Incertae_sedis(100);g__Candida(100);g__Candida_unclassified(100); | Candida | 13 | Pathotroph-Saprotroph-Symbiotroph | Animal Pathogen-Endophyte-Endosymbiont-Epiphyte-\|Undefined Saprotroph\| | Dimorphic Yeast-Yeast | NULL | Probable | As nectar-tap_saprotroph (Põlme et al. 2020); Decay sugar-rich substrates (Põlme et al. 2020); This asexual genus is polyphyletic and contain species that function very differently depend on what they are. Caution should be taken when interpreting guilds for Candida species. | Manolakaki D, et al. 2010. Virulence. 1: 367-375 ((https://doi.org/10.4161/viru.1.5.12796)); Mekbib SB, et al. 2011. Tropical Plant Pathology 36:233-240 ((https://doi.org/10.1590/S1982-56762011000400004)); Põlme S, et al. 2020. Fungal Diversity 105:1-16 ((https://doi.org/10.1007/s13225-020-00466-2)) |
| Otu22948 | 1 | k__Fungi(100);p__Ascomycota(100);c__Saccharomycetes(100);o__Saccharomycetales(100);f__Metschnikowiaceae(100);g__Metschnikowia(100);g__Metschnikowia_unclassified(100); | Metschnikowia | 13 | Saprotroph | \|Undefined Saprotroph\| | Yeast | NULL | Probable | As nectar-tap_saprotroph (Põlme et al. 2020); Decay sugar-rich substrates (Põlme et al. 2020) | Põlme S, et al. 2020. Fungal Diversity 105:1-16 ((https://doi.org/10.1007/s13225-020-00466-2)) |
| Otu22949 | 1 | k__Fungi(100);p__Ascomycota(100);c__Saccharomycetes(100);o__Saccharomycetales(100);f__Metschnikowiaceae(100);g__Metschnikowia(100);s__Metschnikowia_pulcherrima(100); | Metschnikowia | 13 | Saprotroph | \|Undefined Saprotroph\| | Yeast | NULL | Probable | As nectar-tap_saprotroph (Põlme et al. 2020); Decay sugar-rich substrates (Põlme et al. 2020) | Põlme S, et al. 2020. Fungal Diversity 105:1-16 ((https://doi.org/10.1007/s13225-020-00466-2)) |
| Otu22950 | 1 | k__Fungi(100);p__Ascomycota(100);c__Saccharomycetes(100);o__Saccharomycetales(100);f__Metschnikowiaceae(100);g__Metschnikowia(100);g__Metschnikowia_unclassified(100); | Metschnikowia | 13 | Saprotroph | \|Undefined Saprotroph\| | Yeast | NULL | Probable | As nectar-tap_saprotroph (Põlme et al. 2020); Decay sugar-rich substrates (Põlme et al. 2020) | Põlme S, et al. 2020. Fungal Diversity 105:1-16 ((https://doi.org/10.1007/s13225-020-00466-2)) |
| Otu22951 | 1 | k__Fungi(100);p__Ascomycota(100);c__Saccharomycetes(100);o__Saccharomycetales(100);f__Metschnikowiaceae(100);g__Metschnikowia(100);s__Metschnikowia_pulcherrima(100); | Metschnikowia | 13 | Saprotroph | \|Undefined Saprotroph\| | Yeast | NULL | Probable | As nectar-tap_saprotroph (Põlme et al. 2020); Decay sugar-rich substrates (Põlme et al. 2020) | Põlme S, et al. 2020. Fungal Diversity 105:1-16 ((https://doi.org/10.1007/s13225-020-00466-2)) |
| Otu22952 | 1 | k__Fungi(100);p__Ascomycota(100);c__Saccharomycetes(100);o__Saccharomycetales(100);f__Metschnikowiaceae(100);g__Metschnikowia(100);s__Metschnikowia_pulcherrima(100); | Metschnikowia | 13 | Saprotroph | \|Undefined Saprotroph\| | Yeast | NULL | Probable | As nectar-tap_saprotroph (Põlme et al. 2020); Decay sugar-rich substrates (Põlme et al. 2020) | Põlme S, et al. 2020. Fungal Diversity 105:1-16 ((https://doi.org/10.1007/s13225-020-00466-2)) |
| Otu22953 | 1 | k__Fungi(100);p__Ascomycota(100);c__Saccharomycetes(100);o__Saccharomycetales(100);f__Metschnikowiaceae(100);g__Metschnikowia(100);s__Metschnikowia_pulcherrima(100); | Metschnikowia | 13 | Saprotroph | \|Undefined Saprotroph\| | Yeast | NULL | Probable | As nectar-tap_saprotroph (Põlme et al. 2020); Decay sugar-rich substrates (Põlme et al. 2020) | Põlme S, et al. 2020. Fungal Diversity 105:1-16 ((https://doi.org/10.1007/s13225-020-00466-2)) |
| Otu22954 | 1 | k__Fungi(100);p__Ascomycota(100);c__Saccharomycetes(100);o__Saccharomycetales(100);f__Metschnikowiaceae(100);g__Metschnikowia(100);s__Metschnikowia_pulcherrima(100); | Metschnikowia | 13 | Saprotroph | \|Undefined Saprotroph\| | Yeast | NULL | Probable | As nectar-tap_saprotroph (Põlme et al. 2020); Decay sugar-rich substrates (Põlme et al. 2020) | Põlme S, et al. 2020. Fungal Diversity 105:1-16 ((https://doi.org/10.1007/s13225-020-00466-2)) |
| Otu22955 | 1 | k__Fungi(100);p__Ascomycota(100);c__Saccharomycetes(100);o__Saccharomycetales(100);f__Metschnikowiaceae(100);g__Metschnikowia(100);s__Metschnikowia_pulcherrima(100); | Metschnikowia | 13 | Saprotroph | \|Undefined Saprotroph\| | Yeast | NULL | Probable | As nectar-tap_saprotroph (Põlme et al. 2020); Decay sugar-rich substrates (Põlme et al. 2020) | Põlme S, et al. 2020. Fungal Diversity 105:1-16 ((https://doi.org/10.1007/s13225-020-00466-2)) |
| Otu22956 | 1 | k__Fungi(100);p__Ascomycota(100);c__Saccharomycetes(100);o__Saccharomycetales(100);f__Metschnikowiaceae(100);g__Metschnikowia(100);g__Metschnikowia_unclassified(100); | Metschnikowia | 13 | Saprotroph | \|Undefined Saprotroph\| | Yeast | NULL | Probable | As nectar-tap_saprotroph (Põlme et al. 2020); Decay sugar-rich substrates (Põlme et al. 2020) | Põlme S, et al. 2020. Fungal Diversity 105:1-16 ((https://doi.org/10.1007/s13225-020-00466-2)) |
| Otu22957 | 1 | k__Fungi(100);p__Ascomycota(100);c__Saccharomycetes(100);o__Saccharomycetales(100);f__Metschnikowiaceae(100);g__Metschnikowia(100);s__Metschnikowia_pulcherrima(100); | Metschnikowia | 13 | Saprotroph | \|Undefined Saprotroph\| | Yeast | NULL | Probable | As nectar-tap_saprotroph (Põlme et al. 2020); Decay sugar-rich substrates (Põlme et al. 2020) | Põlme S, et al. 2020. Fungal Diversity 105:1-16 ((https://doi.org/10.1007/s13225-020-00466-2)) |
| Otu22961 | 1 | k__Fungi(100);p__Ascomycota(100);c__Saccharomycetes(100);o__Saccharomycetales(100);f__Metschnikowiaceae(100);g__Metschnikowia(100);s__Metschnikowia_pulcherrima(100); | Metschnikowia | 13 | Saprotroph | \|Undefined Saprotroph\| | Yeast | NULL | Probable | As nectar-tap_saprotroph (Põlme et al. 2020); Decay sugar-rich substrates (Põlme et al. 2020) | Põlme S, et al. 2020. Fungal Diversity 105:1-16 ((https://doi.org/10.1007/s13225-020-00466-2)) |
| Otu22962 | 1 | k__Fungi(100);p__Ascomycota(100);c__Saccharomycetes(100);o__Saccharomycetales(100);f__Metschnikowiaceae(100);g__Metschnikowia(100);s__Metschnikowia_pulcherrima(100); | Metschnikowia | 13 | Saprotroph | \|Undefined Saprotroph\| | Yeast | NULL | Probable | As nectar-tap_saprotroph (Põlme et al. 2020); Decay sugar-rich substrates (Põlme et al. 2020) | Põlme S, et al. 2020. Fungal Diversity 105:1-16 ((https://doi.org/10.1007/s13225-020-00466-2)) |
| Otu22963 | 1 | k__Fungi(100);p__Ascomycota(100);c__Saccharomycetes(100);o__Saccharomycetales(100);f__Metschnikowiaceae(100);g__Metschnikowia(100);s__Metschnikowia_pulcherrima(100); | Metschnikowia | 13 | Saprotroph | \|Undefined Saprotroph\| | Yeast | NULL | Probable | As nectar-tap_saprotroph (Põlme et al. 2020); Decay sugar-rich substrates (Põlme et al. 2020) | Põlme S, et al. 2020. Fungal Diversity 105:1-16 ((https://doi.org/10.1007/s13225-020-00466-2)) |
| Otu22964 | 1 | k__Fungi(100);p__Ascomycota(100);c__Saccharomycetes(100);o__Saccharomycetales(100);f__Metschnikowiaceae(100);g__Metschnikowia(100);s__Metschnikowia_pulcherrima(100); | Metschnikowia | 13 | Saprotroph | \|Undefined Saprotroph\| | Yeast | NULL | Probable | As nectar-tap_saprotroph (Põlme et al. 2020); Decay sugar-rich substrates (Põlme et al. 2020) | Põlme S, et al. 2020. Fungal Diversity 105:1-16 ((https://doi.org/10.1007/s13225-020-00466-2)) |
| Otu22965 | 1 | k__Fungi(100);p__Ascomycota(100);c__Saccharomycetes(100);o__Saccharomycetales(100);f__Metschnikowiaceae(100);g__Metschnikowia(100);g__Metschnikowia_unclassified(100); | Metschnikowia | 13 | Saprotroph | \|Undefined Saprotroph\| | Yeast | NULL | Probable | As nectar-tap_saprotroph (Põlme et al. 2020); Decay sugar-rich substrates (Põlme et al. 2020) | Põlme S, et al. 2020. Fungal Diversity 105:1-16 ((https://doi.org/10.1007/s13225-020-00466-2)) |
| Otu22966 | 1 | k__Fungi(100);p__Ascomycota(100);c__Saccharomycetes(100);o__Saccharomycetales(100);f__Metschnikowiaceae(100);g__Metschnikowia(100);s__Metschnikowia_pulcherrima(100); | Metschnikowia | 13 | Saprotroph | \|Undefined Saprotroph\| | Yeast | NULL | Probable | As nectar-tap_saprotroph (Põlme et al. 2020); Decay sugar-rich substrates (Põlme et al. 2020) | Põlme S, et al. 2020. Fungal Diversity 105:1-16 ((https://doi.org/10.1007/s13225-020-00466-2)) |
| Otu22967 | 1 | k__Fungi(100);p__Ascomycota(100);c__Saccharomycetes(100);o__Saccharomycetales(100);f__Metschnikowiaceae(100);g__Metschnikowia(100);s__Metschnikowia_pulcherrima(100); | Metschnikowia | 13 | Saprotroph | \|Undefined Saprotroph\| | Yeast | NULL | Probable | As nectar-tap_saprotroph (Põlme et al. 2020); Decay sugar-rich substrates (Põlme et al. 2020) | Põlme S, et al. 2020. Fungal Diversity 105:1-16 ((https://doi.org/10.1007/s13225-020-00466-2)) |
| Otu22969 | 1 | k__Fungi(100);p__Ascomycota(100);c__Saccharomycetes(100);o__Saccharomycetales(100);f__Metschnikowiaceae(100);g__Metschnikowia(100);s__Metschnikowia_pulcherrima(100); | Metschnikowia | 13 | Saprotroph | \|Undefined Saprotroph\| | Yeast | NULL | Probable | As nectar-tap_saprotroph (Põlme et al. 2020); Decay sugar-rich substrates (Põlme et al. 2020) | Põlme S, et al. 2020. Fungal Diversity 105:1-16 ((https://doi.org/10.1007/s13225-020-00466-2)) |
| Otu22970 | 1 | k__Fungi(100);p__Ascomycota(100);c__Saccharomycetes(100);o__Saccharomycetales(100);f__Metschnikowiaceae(100);g__Metschnikowia(100);g__Metschnikowia_unclassified(100); | Metschnikowia | 13 | Saprotroph | \|Undefined Saprotroph\| | Yeast | NULL | Probable | As nectar-tap_saprotroph (Põlme et al. 2020); Decay sugar-rich substrates (Põlme et al. 2020) | Põlme S, et al. 2020. Fungal Diversity 105:1-16 ((https://doi.org/10.1007/s13225-020-00466-2)) |
| Otu22971 | 1 | k__Fungi(100);p__Ascomycota(100);c__Saccharomycetes(100);o__Saccharomycetales(100);f__Metschnikowiaceae(100);g__Metschnikowia(100);s__Metschnikowia_pulcherrima(100); | Metschnikowia | 13 | Saprotroph | \|Undefined Saprotroph\| | Yeast | NULL | Probable | As nectar-tap_saprotroph (Põlme et al. 2020); Decay sugar-rich substrates (Põlme et al. 2020) | Põlme S, et al. 2020. Fungal Diversity 105:1-16 ((https://doi.org/10.1007/s13225-020-00466-2)) |
| Otu22972 | 1 | k__Fungi(100);p__Ascomycota(100);c__Saccharomycetes(100);o__Saccharomycetales(100);f__Metschnikowiaceae(100);g__Metschnikowia(100);s__Metschnikowia_pulcherrima(100); | Metschnikowia | 13 | Saprotroph | \|Undefined Saprotroph\| | Yeast | NULL | Probable | As nectar-tap_saprotroph (Põlme et al. 2020); Decay sugar-rich substrates (Põlme et al. 2020) | Põlme S, et al. 2020. Fungal Diversity 105:1-16 ((https://doi.org/10.1007/s13225-020-00466-2)) |
| Otu22973 | 1 | k__Fungi(100);p__Ascomycota(100);c__Saccharomycetes(100);o__Saccharomycetales(100);f__Metschnikowiaceae(100);g__Metschnikowia(100);s__Metschnikowia_pulcherrima(100); | Metschnikowia | 13 | Saprotroph | \|Undefined Saprotroph\| | Yeast | NULL | Probable | As nectar-tap_saprotroph (Põlme et al. 2020); Decay sugar-rich substrates (Põlme et al. 2020) | Põlme S, et al. 2020. Fungal Diversity 105:1-16 ((https://doi.org/10.1007/s13225-020-00466-2)) |
| Otu22974 | 1 | k__Fungi(100);p__Ascomycota(100);c__Saccharomycetes(100);o__Saccharomycetales(100);f__Metschnikowiaceae(100);g__Metschnikowia(100);s__Metschnikowia_pulcherrima(100); | Metschnikowia | 13 | Saprotroph | \|Undefined Saprotroph\| | Yeast | NULL | Probable | As nectar-tap_saprotroph (Põlme et al. 2020); Decay sugar-rich substrates (Põlme et al. 2020) | Põlme S, et al. 2020. Fungal Diversity 105:1-16 ((https://doi.org/10.1007/s13225-020-00466-2)) |
| Otu22975 | 1 | k__Fungi(100);p__Ascomycota(100);c__Saccharomycetes(100);o__Saccharomycetales(100);f__Metschnikowiaceae(100);g__Metschnikowia(100);s__Metschnikowia_pulcherrima(100); | Metschnikowia | 13 | Saprotroph | \|Undefined Saprotroph\| | Yeast | NULL | Probable | As nectar-tap_saprotroph (Põlme et al. 2020); Decay sugar-rich substrates (Põlme et al. 2020) | Põlme S, et al. 2020. Fungal Diversity 105:1-16 ((https://doi.org/10.1007/s13225-020-00466-2)) |
| Otu22976 | 1 | k__Fungi(100);p__Ascomycota(100);c__Saccharomycetes(100);o__Saccharomycetales(100);f__Metschnikowiaceae(100);g__Metschnikowia(100);g__Metschnikowia_unclassified(100); | Metschnikowia | 13 | Saprotroph | \|Undefined Saprotroph\| | Yeast | NULL | Probable | As nectar-tap_saprotroph (Põlme et al. 2020); Decay sugar-rich substrates (Põlme et al. 2020) | Põlme S, et al. 2020. Fungal Diversity 105:1-16 ((https://doi.org/10.1007/s13225-020-00466-2)) |
| Otu22977 | 1 | k__Fungi(100);p__Ascomycota(100);c__Saccharomycetes(100);o__Saccharomycetales(100);f__Metschnikowiaceae(100);g__Metschnikowia(100);s__Metschnikowia_pulcherrima(100); | Metschnikowia | 13 | Saprotroph | \|Undefined Saprotroph\| | Yeast | NULL | Probable | As nectar-tap_saprotroph (Põlme et al. 2020); Decay sugar-rich substrates (Põlme et al. 2020) | Põlme S, et al. 2020. Fungal Diversity 105:1-16 ((https://doi.org/10.1007/s13225-020-00466-2)) |
| Otu22978 | 1 | k__Fungi(100);p__Ascomycota(100);c__Saccharomycetes(100);o__Saccharomycetales(100);f__Metschnikowiaceae(100);g__Metschnikowia(100);g__Metschnikowia_unclassified(100); | Metschnikowia | 13 | Saprotroph | \|Undefined Saprotroph\| | Yeast | NULL | Probable | As nectar-tap_saprotroph (Põlme et al. 2020); Decay sugar-rich substrates (Põlme et al. 2020) | Põlme S, et al. 2020. Fungal Diversity 105:1-16 ((https://doi.org/10.1007/s13225-020-00466-2)) |
| Otu22979 | 1 | k__Fungi(100);p__Ascomycota(100);c__Saccharomycetes(100);o__Saccharomycetales(100);f__Metschnikowiaceae(100);g__Metschnikowia(100);s__Metschnikowia_pulcherrima(100); | Metschnikowia | 13 | Saprotroph | \|Undefined Saprotroph\| | Yeast | NULL | Probable | As nectar-tap_saprotroph (Põlme et al. 2020); Decay sugar-rich substrates (Põlme et al. 2020) | Põlme S, et al. 2020. Fungal Diversity 105:1-16 ((https://doi.org/10.1007/s13225-020-00466-2)) |
| Otu22980 | 1 | k__Fungi(100);p__Ascomycota(100);c__Saccharomycetes(100);o__Saccharomycetales(100);f__Metschnikowiaceae(100);g__Metschnikowia(100);g__Metschnikowia_unclassified(100); | Metschnikowia | 13 | Saprotroph | \|Undefined Saprotroph\| | Yeast | NULL | Probable | As nectar-tap_saprotroph (Põlme et al. 2020); Decay sugar-rich substrates (Põlme et al. 2020) | Põlme S, et al. 2020. Fungal Diversity 105:1-16 ((https://doi.org/10.1007/s13225-020-00466-2)) |
| Otu22981 | 1 | k__Fungi(100);p__Ascomycota(100);c__Saccharomycetes(100);o__Saccharomycetales(100);f__Metschnikowiaceae(100);g__Metschnikowia(100);s__Metschnikowia_pulcherrima(100); | Metschnikowia | 13 | Saprotroph | \|Undefined Saprotroph\| | Yeast | NULL | Probable | As nectar-tap_saprotroph (Põlme et al. 2020); Decay sugar-rich substrates (Põlme et al. 2020) | Põlme S, et al. 2020. Fungal Diversity 105:1-16 ((https://doi.org/10.1007/s13225-020-00466-2)) |
| Otu22982 | 1 | k__Fungi(100);p__Ascomycota(100);c__Saccharomycetes(100);o__Saccharomycetales(100);f__Metschnikowiaceae(100);g__Metschnikowia(100);s__Metschnikowia_pulcherrima(100); | Metschnikowia | 13 | Saprotroph | \|Undefined Saprotroph\| | Yeast | NULL | Probable | As nectar-tap_saprotroph (Põlme et al. 2020); Decay sugar-rich substrates (Põlme et al. 2020) | Põlme S, et al. 2020. Fungal Diversity 105:1-16 ((https://doi.org/10.1007/s13225-020-00466-2)) |
| Otu22983 | 1 | k__Fungi(100);p__Ascomycota(100);c__Saccharomycetes(100);o__Saccharomycetales(100);f__Metschnikowiaceae(100);g__Metschnikowia(100);g__Metschnikowia_unclassified(100); | Metschnikowia | 13 | Saprotroph | \|Undefined Saprotroph\| | Yeast | NULL | Probable | As nectar-tap_saprotroph (Põlme et al. 2020); Decay sugar-rich substrates (Põlme et al. 2020) | Põlme S, et al. 2020. Fungal Diversity 105:1-16 ((https://doi.org/10.1007/s13225-020-00466-2)) |
| Otu22984 | 1 | k__Fungi(100);p__Ascomycota(100);c__Saccharomycetes(100);o__Saccharomycetales(100);f__Metschnikowiaceae(100);g__Metschnikowia(100);s__Metschnikowia_pulcherrima(100); | Metschnikowia | 13 | Saprotroph | \|Undefined Saprotroph\| | Yeast | NULL | Probable | As nectar-tap_saprotroph (Põlme et al. 2020); Decay sugar-rich substrates (Põlme et al. 2020) | Põlme S, et al. 2020. Fungal Diversity 105:1-16 ((https://doi.org/10.1007/s13225-020-00466-2)) |
| Otu22985 | 1 | k__Fungi(100);p__Ascomycota(100);c__Saccharomycetes(100);o__Saccharomycetales(100);f__Metschnikowiaceae(100);g__Metschnikowia(100);s__Metschnikowia_pulcherrima(100); | Metschnikowia | 13 | Saprotroph | \|Undefined Saprotroph\| | Yeast | NULL | Probable | As nectar-tap_saprotroph (Põlme et al. 2020); Decay sugar-rich substrates (Põlme et al. 2020) | Põlme S, et al. 2020. Fungal Diversity 105:1-16 ((https://doi.org/10.1007/s13225-020-00466-2)) |
| Otu22986 | 1 | k__Fungi(100);p__Ascomycota(100);c__Saccharomycetes(100);o__Saccharomycetales(100);f__Metschnikowiaceae(100);g__Metschnikowia(100);g__Metschnikowia_unclassified(100); | Metschnikowia | 13 | Saprotroph | \|Undefined Saprotroph\| | Yeast | NULL | Probable | As nectar-tap_saprotroph (Põlme et al. 2020); Decay sugar-rich substrates (Põlme et al. 2020) | Põlme S, et al. 2020. Fungal Diversity 105:1-16 ((https://doi.org/10.1007/s13225-020-00466-2)) |
| Otu22987 | 1 | k__Fungi(100);p__Ascomycota(100);c__Saccharomycetes(100);o__Saccharomycetales(100);f__Metschnikowiaceae(100);g__Metschnikowia(100);s__Metschnikowia_pulcherrima(100); | Metschnikowia | 13 | Saprotroph | \|Undefined Saprotroph\| | Yeast | NULL | Probable | As nectar-tap_saprotroph (Põlme et al. 2020); Decay sugar-rich substrates (Põlme et al. 2020) | Põlme S, et al. 2020. Fungal Diversity 105:1-16 ((https://doi.org/10.1007/s13225-020-00466-2)) |
| Otu22988 | 1 | k__Fungi(100);p__Ascomycota(100);c__Saccharomycetes(100);o__Saccharomycetales(100);f__Metschnikowiaceae(100);g__Metschnikowia(100);s__Metschnikowia_pulcherrima(100); | Metschnikowia | 13 | Saprotroph | \|Undefined Saprotroph\| | Yeast | NULL | Probable | As nectar-tap_saprotroph (Põlme et al. 2020); Decay sugar-rich substrates (Põlme et al. 2020) | Põlme S, et al. 2020. Fungal Diversity 105:1-16 ((https://doi.org/10.1007/s13225-020-00466-2)) |
| Otu22989 | 1 | k__Fungi(100);p__Ascomycota(100);c__Saccharomycetes(100);o__Saccharomycetales(100);f__Metschnikowiaceae(100);g__Metschnikowia(100);g__Metschnikowia_unclassified(100); | Metschnikowia | 13 | Saprotroph | \|Undefined Saprotroph\| | Yeast | NULL | Probable | As nectar-tap_saprotroph (Põlme et al. 2020); Decay sugar-rich substrates (Põlme et al. 2020) | Põlme S, et al. 2020. Fungal Diversity 105:1-16 ((https://doi.org/10.1007/s13225-020-00466-2)) |
| Otu22990 | 1 | k__Fungi(100);p__Ascomycota(100);c__Saccharomycetes(100);o__Saccharomycetales(100);f__Metschnikowiaceae(100);g__Metschnikowia(100);g__Metschnikowia_unclassified(100); | Metschnikowia | 13 | Saprotroph | \|Undefined Saprotroph\| | Yeast | NULL | Probable | As nectar-tap_saprotroph (Põlme et al. 2020); Decay sugar-rich substrates (Põlme et al. 2020) | Põlme S, et al. 2020. Fungal Diversity 105:1-16 ((https://doi.org/10.1007/s13225-020-00466-2)) |
| Otu22991 | 1 | k__Fungi(100);p__Ascomycota(100);c__Saccharomycetes(100);o__Saccharomycetales(100);f__Metschnikowiaceae(100);g__Metschnikowia(100);s__Metschnikowia_pulcherrima(100); | Metschnikowia | 13 | Saprotroph | \|Undefined Saprotroph\| | Yeast | NULL | Probable | As nectar-tap_saprotroph (Põlme et al. 2020); Decay sugar-rich substrates (Põlme et al. 2020) | Põlme S, et al. 2020. Fungal Diversity 105:1-16 ((https://doi.org/10.1007/s13225-020-00466-2)) |
| Otu22992 | 1 | k__Fungi(100);p__Ascomycota(100);c__Saccharomycetes(100);o__Saccharomycetales(100);f__Metschnikowiaceae(100);g__Metschnikowia(100);g__Metschnikowia_unclassified(100); | Metschnikowia | 13 | Saprotroph | \|Undefined Saprotroph\| | Yeast | NULL | Probable | As nectar-tap_saprotroph (Põlme et al. 2020); Decay sugar-rich substrates (Põlme et al. 2020) | Põlme S, et al. 2020. Fungal Diversity 105:1-16 ((https://doi.org/10.1007/s13225-020-00466-2)) |
| Otu22993 | 1 | k__Fungi(100);p__Ascomycota(100);c__Saccharomycetes(100);o__Saccharomycetales(100);f__Metschnikowiaceae(100);g__Metschnikowia(100);g__Metschnikowia_unclassified(100); | Metschnikowia | 13 | Saprotroph | \|Undefined Saprotroph\| | Yeast | NULL | Probable | As nectar-tap_saprotroph (Põlme et al. 2020); Decay sugar-rich substrates (Põlme et al. 2020) | Põlme S, et al. 2020. Fungal Diversity 105:1-16 ((https://doi.org/10.1007/s13225-020-00466-2)) |
| Otu22994 | 1 | k__Fungi(100);p__Ascomycota(100);c__Saccharomycetes(100);o__Saccharomycetales(100);f__Metschnikowiaceae(100);g__Metschnikowia(100);s__Metschnikowia_pulcherrima(100); | Metschnikowia | 13 | Saprotroph | \|Undefined Saprotroph\| | Yeast | NULL | Probable | As nectar-tap_saprotroph (Põlme et al. 2020); Decay sugar-rich substrates (Põlme et al. 2020) | Põlme S, et al. 2020. Fungal Diversity 105:1-16 ((https://doi.org/10.1007/s13225-020-00466-2)) |
| Otu22995 | 1 | k__Fungi(100);p__Ascomycota(100);c__Saccharomycetes(100);o__Saccharomycetales(100);f__Metschnikowiaceae(100);g__Metschnikowia(100);g__Metschnikowia_unclassified(100); | Metschnikowia | 13 | Saprotroph | \|Undefined Saprotroph\| | Yeast | NULL | Probable | As nectar-tap_saprotroph (Põlme et al. 2020); Decay sugar-rich substrates (Põlme et al. 2020) | Põlme S, et al. 2020. Fungal Diversity 105:1-16 ((https://doi.org/10.1007/s13225-020-00466-2)) |
| Otu22996 | 1 | k__Fungi(100);p__Ascomycota(100);c__Saccharomycetes(100);o__Saccharomycetales(100);f__Metschnikowiaceae(100);g__Metschnikowia(100);s__Metschnikowia_pulcherrima(100); | Metschnikowia | 13 | Saprotroph | \|Undefined Saprotroph\| | Yeast | NULL | Probable | As nectar-tap_saprotroph (Põlme et al. 2020); Decay sugar-rich substrates (Põlme et al. 2020) | Põlme S, et al. 2020. Fungal Diversity 105:1-16 ((https://doi.org/10.1007/s13225-020-00466-2)) |
| Otu22997 | 1 | k__Fungi(100);p__Ascomycota(100);c__Saccharomycetes(100);o__Saccharomycetales(100);f__Metschnikowiaceae(100);g__Metschnikowia(100);s__Metschnikowia_pulcherrima(100); | Metschnikowia | 13 | Saprotroph | \|Undefined Saprotroph\| | Yeast | NULL | Probable | As nectar-tap_saprotroph (Põlme et al. 2020); Decay sugar-rich substrates (Põlme et al. 2020) | Põlme S, et al. 2020. Fungal Diversity 105:1-16 ((https://doi.org/10.1007/s13225-020-00466-2)) |
| Otu22998 | 1 | k__Fungi(100);p__Ascomycota(100);c__Saccharomycetes(100);o__Saccharomycetales(100);f__Metschnikowiaceae(100);g__Metschnikowia(100);s__Metschnikowia_pulcherrima(100); | Metschnikowia | 13 | Saprotroph | \|Undefined Saprotroph\| | Yeast | NULL | Probable | As nectar-tap_saprotroph (Põlme et al. 2020); Decay sugar-rich substrates (Põlme et al. 2020) | Põlme S, et al. 2020. Fungal Diversity 105:1-16 ((https://doi.org/10.1007/s13225-020-00466-2)) |
| Otu22999 | 1 | k__Fungi(100);p__Ascomycota(100);c__Saccharomycetes(100);o__Saccharomycetales(100);f__Metschnikowiaceae(100);g__Metschnikowia(100);s__Metschnikowia_pulcherrima(100); | Metschnikowia | 13 | Saprotroph | \|Undefined Saprotroph\| | Yeast | NULL | Probable | As nectar-tap_saprotroph (Põlme et al. 2020); Decay sugar-rich substrates (Põlme et al. 2020) | Põlme S, et al. 2020. Fungal Diversity 105:1-16 ((https://doi.org/10.1007/s13225-020-00466-2)) |
| Otu23000 | 1 | k__Fungi(100);p__Ascomycota(100);c__Saccharomycetes(100);o__Saccharomycetales(100);f__Metschnikowiaceae(100);g__Metschnikowia(100);s__Metschnikowia_pulcherrima(100); | Metschnikowia | 13 | Saprotroph | \|Undefined Saprotroph\| | Yeast | NULL | Probable | As nectar-tap_saprotroph (Põlme et al. 2020); Decay sugar-rich substrates (Põlme et al. 2020) | Põlme S, et al. 2020. Fungal Diversity 105:1-16 ((https://doi.org/10.1007/s13225-020-00466-2)) |
| Otu23002 | 1 | k__Fungi(100);p__Ascomycota(100);c__Saccharomycetes(100);o__Saccharomycetales(100);f__Metschnikowiaceae(100);g__Metschnikowia(100);s__Metschnikowia_pulcherrima(100); | Metschnikowia | 13 | Saprotroph | \|Undefined Saprotroph\| | Yeast | NULL | Probable | As nectar-tap_saprotroph (Põlme et al. 2020); Decay sugar-rich substrates (Põlme et al. 2020) | Põlme S, et al. 2020. Fungal Diversity 105:1-16 ((https://doi.org/10.1007/s13225-020-00466-2)) |
| Otu23004 | 1 | k__Fungi(100);p__Ascomycota(100);c__Saccharomycetes(100);o__Saccharomycetales(100);f__Metschnikowiaceae(100);g__Metschnikowia(100);g__Metschnikowia_unclassified(100); | Metschnikowia | 13 | Saprotroph | \|Undefined Saprotroph\| | Yeast | NULL | Probable | As nectar-tap_saprotroph (Põlme et al. 2020); Decay sugar-rich substrates (Põlme et al. 2020) | Põlme S, et al. 2020. Fungal Diversity 105:1-16 ((https://doi.org/10.1007/s13225-020-00466-2)) |
| Otu23007 | 1 | k__Fungi(100);p__Ascomycota(100);c__Saccharomycetes(100);o__Saccharomycetales(100);f__Metschnikowiaceae(100);g__Metschnikowia(100);s__Metschnikowia_pulcherrima(100); | Metschnikowia | 13 | Saprotroph | \|Undefined Saprotroph\| | Yeast | NULL | Probable | As nectar-tap_saprotroph (Põlme et al. 2020); Decay sugar-rich substrates (Põlme et al. 2020) | Põlme S, et al. 2020. Fungal Diversity 105:1-16 ((https://doi.org/10.1007/s13225-020-00466-2)) |
| Otu23008 | 1 | k__Fungi(100);p__Ascomycota(100);c__Saccharomycetes(100);o__Saccharomycetales(100);f__Metschnikowiaceae(100);g__Metschnikowia(100);g__Metschnikowia_unclassified(100); | Metschnikowia | 13 | Saprotroph | \|Undefined Saprotroph\| | Yeast | NULL | Probable | As nectar-tap_saprotroph (Põlme et al. 2020); Decay sugar-rich substrates (Põlme et al. 2020) | Põlme S, et al. 2020. Fungal Diversity 105:1-16 ((https://doi.org/10.1007/s13225-020-00466-2)) |
| Otu23009 | 1 | k__Fungi(100);p__Ascomycota(100);c__Saccharomycetes(100);o__Saccharomycetales(100);f__Metschnikowiaceae(100);g__Metschnikowia(100);s__Metschnikowia_pulcherrima(100); | Metschnikowia | 13 | Saprotroph | \|Undefined Saprotroph\| | Yeast | NULL | Probable | As nectar-tap_saprotroph (Põlme et al. 2020); Decay sugar-rich substrates (Põlme et al. 2020) | Põlme S, et al. 2020. Fungal Diversity 105:1-16 ((https://doi.org/10.1007/s13225-020-00466-2)) |
| Otu23010 | 1 | k__Fungi(100);p__Ascomycota(100);c__Saccharomycetes(100);o__Saccharomycetales(100);f__Metschnikowiaceae(100);g__Metschnikowia(100);s__Metschnikowia_pulcherrima(100); | Metschnikowia | 13 | Saprotroph | \|Undefined Saprotroph\| | Yeast | NULL | Probable | As nectar-tap_saprotroph (Põlme et al. 2020); Decay sugar-rich substrates (Põlme et al. 2020) | Põlme S, et al. 2020. Fungal Diversity 105:1-16 ((https://doi.org/10.1007/s13225-020-00466-2)) |
| Otu23011 | 1 | k__Fungi(100);p__Ascomycota(100);c__Saccharomycetes(100);o__Saccharomycetales(100);f__Metschnikowiaceae(100);g__Metschnikowia(100);s__Metschnikowia_pulcherrima(100); | Metschnikowia | 13 | Saprotroph | \|Undefined Saprotroph\| | Yeast | NULL | Probable | As nectar-tap_saprotroph (Põlme et al. 2020); Decay sugar-rich substrates (Põlme et al. 2020) | Põlme S, et al. 2020. Fungal Diversity 105:1-16 ((https://doi.org/10.1007/s13225-020-00466-2)) |
| Otu23012 | 1 | k__Fungi(100);p__Ascomycota(100);c__Saccharomycetes(100);o__Saccharomycetales(100);f__Metschnikowiaceae(100);g__Metschnikowia(100);s__Metschnikowia_pulcherrima(100); | Metschnikowia | 13 | Saprotroph | \|Undefined Saprotroph\| | Yeast | NULL | Probable | As nectar-tap_saprotroph (Põlme et al. 2020); Decay sugar-rich substrates (Põlme et al. 2020) | Põlme S, et al. 2020. Fungal Diversity 105:1-16 ((https://doi.org/10.1007/s13225-020-00466-2)) |
| Otu23013 | 1 | k__Fungi(100);p__Ascomycota(100);c__Saccharomycetes(100);o__Saccharomycetales(100);f__Metschnikowiaceae(100);g__Metschnikowia(100);s__Metschnikowia_pulcherrima(100); | Metschnikowia | 13 | Saprotroph | \|Undefined Saprotroph\| | Yeast | NULL | Probable | As nectar-tap_saprotroph (Põlme et al. 2020); Decay sugar-rich substrates (Põlme et al. 2020) | Põlme S, et al. 2020. Fungal Diversity 105:1-16 ((https://doi.org/10.1007/s13225-020-00466-2)) |
| Otu23014 | 1 | k__Fungi(100);p__Ascomycota(100);c__Saccharomycetes(100);o__Saccharomycetales(100);f__Metschnikowiaceae(100);g__Metschnikowia(100);s__Metschnikowia_pulcherrima(100); | Metschnikowia | 13 | Saprotroph | \|Undefined Saprotroph\| | Yeast | NULL | Probable | As nectar-tap_saprotroph (Põlme et al. 2020); Decay sugar-rich substrates (Põlme et al. 2020) | Põlme S, et al. 2020. Fungal Diversity 105:1-16 ((https://doi.org/10.1007/s13225-020-00466-2)) |
| Otu23015 | 1 | k__Fungi(100);p__Ascomycota(100);c__Saccharomycetes(100);o__Saccharomycetales(100);f__Metschnikowiaceae(100);g__Metschnikowia(100);g__Metschnikowia_unclassified(100); | Metschnikowia | 13 | Saprotroph | \|Undefined Saprotroph\| | Yeast | NULL | Probable | As nectar-tap_saprotroph (Põlme et al. 2020); Decay sugar-rich substrates (Põlme et al. 2020) | Põlme S, et al. 2020. Fungal Diversity 105:1-16 ((https://doi.org/10.1007/s13225-020-00466-2)) |
| Otu23016 | 1 | k__Fungi(100);p__Ascomycota(100);c__Saccharomycetes(100);o__Saccharomycetales(100);f__Metschnikowiaceae(100);g__Metschnikowia(100);s__Metschnikowia_pulcherrima(100); | Metschnikowia | 13 | Saprotroph | \|Undefined Saprotroph\| | Yeast | NULL | Probable | As nectar-tap_saprotroph (Põlme et al. 2020); Decay sugar-rich substrates (Põlme et al. 2020) | Põlme S, et al. 2020. Fungal Diversity 105:1-16 ((https://doi.org/10.1007/s13225-020-00466-2)) |
| Otu23017 | 1 | k__Fungi(100);p__Ascomycota(100);c__Saccharomycetes(100);o__Saccharomycetales(100);f__Metschnikowiaceae(100);g__Metschnikowia(100);s__Metschnikowia_pulcherrima(100); | Metschnikowia | 13 | Saprotroph | \|Undefined Saprotroph\| | Yeast | NULL | Probable | As nectar-tap_saprotroph (Põlme et al. 2020); Decay sugar-rich substrates (Põlme et al. 2020) | Põlme S, et al. 2020. Fungal Diversity 105:1-16 ((https://doi.org/10.1007/s13225-020-00466-2)) |
| Otu23018 | 1 | k__Fungi(100);p__Ascomycota(100);c__Saccharomycetes(100);o__Saccharomycetales(100);f__Metschnikowiaceae(100);g__Metschnikowia(100);s__Metschnikowia_pulcherrima(100); | Metschnikowia | 13 | Saprotroph | \|Undefined Saprotroph\| | Yeast | NULL | Probable | As nectar-tap_saprotroph (Põlme et al. 2020); Decay sugar-rich substrates (Põlme et al. 2020) | Põlme S, et al. 2020. Fungal Diversity 105:1-16 ((https://doi.org/10.1007/s13225-020-00466-2)) |
| Otu23019 | 1 | k__Fungi(100);p__Ascomycota(100);c__Saccharomycetes(100);o__Saccharomycetales(100);f__Metschnikowiaceae(100);g__Metschnikowia(100);s__Metschnikowia_pulcherrima(100); | Metschnikowia | 13 | Saprotroph | \|Undefined Saprotroph\| | Yeast | NULL | Probable | As nectar-tap_saprotroph (Põlme et al. 2020); Decay sugar-rich substrates (Põlme et al. 2020) | Põlme S, et al. 2020. Fungal Diversity 105:1-16 ((https://doi.org/10.1007/s13225-020-00466-2)) |
| Otu23020 | 1 | k__Fungi(100);p__Ascomycota(100);c__Saccharomycetes(100);o__Saccharomycetales(100);f__Metschnikowiaceae(100);g__Metschnikowia(100);s__Metschnikowia_pulcherrima(100); | Metschnikowia | 13 | Saprotroph | \|Undefined Saprotroph\| | Yeast | NULL | Probable | As nectar-tap_saprotroph (Põlme et al. 2020); Decay sugar-rich substrates (Põlme et al. 2020) | Põlme S, et al. 2020. Fungal Diversity 105:1-16 ((https://doi.org/10.1007/s13225-020-00466-2)) |
| Otu23021 | 1 | k__Fungi(100);p__Ascomycota(100);c__Saccharomycetes(100);o__Saccharomycetales(100);f__Metschnikowiaceae(100);g__Metschnikowia(100);s__Metschnikowia_pulcherrima(100); | Metschnikowia | 13 | Saprotroph | \|Undefined Saprotroph\| | Yeast | NULL | Probable | As nectar-tap_saprotroph (Põlme et al. 2020); Decay sugar-rich substrates (Põlme et al. 2020) | Põlme S, et al. 2020. Fungal Diversity 105:1-16 ((https://doi.org/10.1007/s13225-020-00466-2)) |
| Otu23022 | 1 | k__Fungi(100);p__Ascomycota(100);c__Saccharomycetes(100);o__Saccharomycetales(100);f__Metschnikowiaceae(100);g__Metschnikowia(100);s__Metschnikowia_pulcherrima(100); | Metschnikowia | 13 | Saprotroph | \|Undefined Saprotroph\| | Yeast | NULL | Probable | As nectar-tap_saprotroph (Põlme et al. 2020); Decay sugar-rich substrates (Põlme et al. 2020) | Põlme S, et al. 2020. Fungal Diversity 105:1-16 ((https://doi.org/10.1007/s13225-020-00466-2)) |

| | | | | | | | | | | | |
|---|---|---|---|---|---|---|---|---|---|---|---|
| Otu23023 | 1 | k__Fungi(100);p__Ascomycota(100);c__Saccharomycetes(100);o__Saccharomycetales(100);f__Metschnikowiaceae(100);g__Metschnikowia(100);s__Metschnikowia_pulcherrima(100); | Metschnikowia | 13 | Saprotroph | \|Undefined Saprotroph\| | Yeast | NULL | Probable | As nectar-tap_saprotroph (Põlme et al. 2020); Decay sugar-rich substrates (Põlme et al. 2020) | Põlme S, et al. 2020. Fungal Diversity 105:1-16 ((https://doi.org/10.1007/s13225-020-00466-2)) |
| Otu23024 | 1 | k__Fungi(100);p__Ascomycota(100);c__Saccharomycetes(100);o__Saccharomycetales(100);f__Metschnikowiaceae(100);g__Metschnikowia(100);s__Metschnikowia_pulcherrima(100); | Metschnikowia | 13 | Saprotroph | \|Undefined Saprotroph\| | Yeast | NULL | Probable | As nectar-tap_saprotroph (Põlme et al. 2020); Decay sugar-rich substrates (Põlme et al. 2020) | Põlme S, et al. 2020. Fungal Diversity 105:1-16 ((https://doi.org/10.1007/s13225-020-00466-2)) |
| Otu23025 | 1 | k__Fungi(100);p__Ascomycota(100);c__Saccharomycetes(100);o__Saccharomycetales(100);f__Metschnikowiaceae(100);g__Metschnikowia(100);g__Metschnikowia_unclassified(100); | Metschnikowia | 13 | Saprotroph | \|Undefined Saprotroph\| | Yeast | NULL | Probable | As nectar-tap_saprotroph (Põlme et al. 2020); Decay sugar-rich substrates (Põlme et al. 2020) | Põlme S, et al. 2020. Fungal Diversity 105:1-16 ((https://doi.org/10.1007/s13225-020-00466-2)) |
| Otu23026 | 1 | k__Fungi(100);p__Ascomycota(100);c__Saccharomycetes(100);o__Saccharomycetales(100);f__Metschnikowiaceae(100);g__Metschnikowia(100);g__Metschnikowia_unclassified(100); | Metschnikowia | 13 | Saprotroph | \|Undefined Saprotroph\| | Yeast | NULL | Probable | As nectar-tap_saprotroph (Põlme et al. 2020); Decay sugar-rich substrates (Põlme et al. 2020) | Põlme S, et al. 2020. Fungal Diversity 105:1-16 ((https://doi.org/10.1007/s13225-020-00466-2)) |
| Otu23027 | 1 | k__Fungi(100);p__Ascomycota(100);c__Saccharomycetes(100);o__Saccharomycetales(100);f__Metschnikowiaceae(100);g__Metschnikowia(100);g__Metschnikowia_unclassified(100); | Metschnikowia | 13 | Saprotroph | \|Undefined Saprotroph\| | Yeast | NULL | Probable | As nectar-tap_saprotroph (Põlme et al. 2020); Decay sugar-rich substrates (Põlme et al. 2020) | Põlme S, et al. 2020. Fungal Diversity 105:1-16 ((https://doi.org/10.1007/s13225-020-00466-2)) |
| Otu23028 | 1 | k__Fungi(100);p__Ascomycota(100);c__Saccharomycetes(100);o__Saccharomycetales(100);f__Metschnikowiaceae(100);g__Metschnikowia(100);s__Metschnikowia_pulcherrima(100); | Metschnikowia | 13 | Saprotroph | \|Undefined Saprotroph\| | Yeast | NULL | Probable | As nectar-tap_saprotroph (Põlme et al. 2020); Decay sugar-rich substrates (Põlme et al. 2020) | Põlme S, et al. 2020. Fungal Diversity 105:1-16 ((https://doi.org/10.1007/s13225-020-00466-2)) |
| Otu23029 | 1 | k__Fungi(100);p__Ascomycota(100);c__Saccharomycetes(100);o__Saccharomycetales(100);f__Metschnikowiaceae(100);g__Metschnikowia(100);s__Metschnikowia_pulcherrima(100); | Metschnikowia | 13 | Saprotroph | \|Undefined Saprotroph\| | Yeast | NULL | Probable | As nectar-tap_saprotroph (Põlme et al. 2020); Decay sugar-rich substrates (Põlme et al. 2020) | Põlme S, et al. 2020. Fungal Diversity 105:1-16 ((https://doi.org/10.1007/s13225-020-00466-2)) |
| Otu23030 | 1 | k__Fungi(100);p__Ascomycota(100);c__Saccharomycetes(100);o__Saccharomycetales(100);f__Metschnikowiaceae(100);g__Metschnikowia(100);s__Metschnikowia_pulcherrima(100); | Metschnikowia | 13 | Saprotroph | \|Undefined Saprotroph\| | Yeast | NULL | Probable | As nectar-tap_saprotroph (Põlme et al. 2020); Decay sugar-rich substrates (Põlme et al. 2020) | Põlme S, et al. 2020. Fungal Diversity 105:1-16 ((https://doi.org/10.1007/s13225-020-00466-2)) |
| Otu23031 | 1 | k__Fungi(100);p__Ascomycota(100);c__Saccharomycetes(100);o__Saccharomycetales(100);f__Metschnikowiaceae(100);g__Metschnikowia(100);s__Metschnikowia_pulcherrima(100); | Metschnikowia | 13 | Saprotroph | \|Undefined Saprotroph\| | Yeast | NULL | Probable | As nectar-tap_saprotroph (Põlme et al. 2020); Decay sugar-rich substrates (Põlme et al. 2020) | Põlme S, et al. 2020. Fungal Diversity 105:1-16 ((https://doi.org/10.1007/s13225-020-00466-2)) |
| Otu23032 | 1 | k__Fungi(100);p__Ascomycota(100);c__Saccharomycetes(100);o__Saccharomycetales(100);f__Metschnikowiaceae(100);g__Metschnikowia(100);g__Metschnikowia_unclassified(100); | Metschnikowia | 13 | Saprotroph | \|Undefined Saprotroph\| | Yeast | NULL | Probable | As nectar-tap_saprotroph (Põlme et al. 2020); Decay sugar-rich substrates (Põlme et al. 2020) | Põlme S, et al. 2020. Fungal Diversity 105:1-16 ((https://doi.org/10.1007/s13225-020-00466-2)) |
| Otu23033 | 1 | k__Fungi(100);p__Ascomycota(100);c__Saccharomycetes(100);o__Saccharomycetales(100);f__Metschnikowiaceae(100);g__Metschnikowia(100);s__Metschnikowia_pulcherrima(100); | Metschnikowia | 13 | Saprotroph | \|Undefined Saprotroph\| | Yeast | NULL | Probable | As nectar-tap_saprotroph (Põlme et al. 2020); Decay sugar-rich substrates (Põlme et al. 2020) | Põlme S, et al. 2020. Fungal Diversity 105:1-16 ((https://doi.org/10.1007/s13225-020-00466-2)) |
| Otu23034 | 1 | k__Fungi(100);p__Ascomycota(100);c__Saccharomycetes(100);o__Saccharomycetales(100);f__Metschnikowiaceae(100);g__Metschnikowia(100);g__Metschnikowia_unclassified(100); | Metschnikowia | 13 | Saprotroph | \|Undefined Saprotroph\| | Yeast | NULL | Probable | As nectar-tap_saprotroph (Põlme et al. 2020); Decay sugar-rich substrates (Põlme et al. 2020) | Põlme S, et al. 2020. Fungal Diversity 105:1-16 ((https://doi.org/10.1007/s13225-020-00466-2)) |
| Otu23035 | 1 | k__Fungi(100);p__Ascomycota(100);c__Saccharomycetes(100);o__Saccharomycetales(100);f__Metschnikowiaceae(100);g__Metschnikowia(100);g__Metschnikowia_unclassified(100); | Metschnikowia | 13 | Saprotroph | \|Undefined Saprotroph\| | Yeast | NULL | Probable | As nectar-tap_saprotroph (Põlme et al. 2020); Decay sugar-rich substrates (Põlme et al. 2020) | Põlme S, et al. 2020. Fungal Diversity 105:1-16 ((https://doi.org/10.1007/s13225-020-00466-2)) |
| Otu23036 | 1 | k__Fungi(100);p__Ascomycota(100);c__Saccharomycetes(100);o__Saccharomycetales(100);f__Metschnikowiaceae(100);g__Metschnikowia(100);g__Metschnikowia_unclassified(100); | Metschnikowia | 13 | Saprotroph | \|Undefined Saprotroph\| | Yeast | NULL | Probable | As nectar-tap_saprotroph (Põlme et al. 2020); Decay sugar-rich substrates (Põlme et al. 2020) | Põlme S, et al. 2020. Fungal Diversity 105:1-16 ((https://doi.org/10.1007/s13225-020-00466-2)) |
| Otu23037 | 1 | k__Fungi(100);p__Ascomycota(100);c__Saccharomycetes(100);o__Saccharomycetales(100);f__Metschnikowiaceae(100);g__Metschnikowia(100);s__Metschnikowia_pulcherrima(100); | Metschnikowia | 13 | Saprotroph | \|Undefined Saprotroph\| | Yeast | NULL | Probable | As nectar-tap_saprotroph (Põlme et al. 2020); Decay sugar-rich substrates (Põlme et al. 2020) | Põlme S, et al. 2020. Fungal Diversity 105:1-16 ((https://doi.org/10.1007/s13225-020-00466-2)) |
| Otu23039 | 1 | k__Fungi(100);p__Ascomycota(100);c__Saccharomycetes(100);o__Saccharomycetales(100);f__Metschnikowiaceae(100);g__Metschnikowia(100);s__Metschnikowia_pulcherrima(100); | Metschnikowia | 13 | Saprotroph | \|Undefined Saprotroph\| | Yeast | NULL | Probable | As nectar-tap_saprotroph (Põlme et al. 2020); Decay sugar-rich substrates (Põlme et al. 2020) | Põlme S, et al. 2020. Fungal Diversity 105:1-16 ((https://doi.org/10.1007/s13225-020-00466-2)) |
| Otu23040 | 1 | k__Fungi(100);p__Ascomycota(100);c__Saccharomycetes(100);o__Saccharomycetales(100);f__Metschnikowiaceae(100);g__Metschnikowia(100);g__Metschnikowia_unclassified(100); | Metschnikowia | 13 | Saprotroph | \|Undefined Saprotroph\| | Yeast | NULL | Probable | As nectar-tap_saprotroph (Põlme et al. 2020); Decay sugar-rich substrates (Põlme et al. 2020) | Põlme S, et al. 2020. Fungal Diversity 105:1-16 ((https://doi.org/10.1007/s13225-020-00466-2)) |
| Otu23042 | 1 | k__Fungi(100);p__Ascomycota(100);c__Saccharomycetes(100);o__Saccharomycetales(100);f__Metschnikowiaceae(100);g__Metschnikowia(100);s__Metschnikowia_pulcherrima(100); | Metschnikowia | 13 | Saprotroph | \|Undefined Saprotroph\| | Yeast | NULL | Probable | As nectar-tap_saprotroph (Põlme et al. 2020); Decay sugar-rich substrates (Põlme et al. 2020) | Põlme S, et al. 2020. Fungal Diversity 105:1-16 ((https://doi.org/10.1007/s13225-020-00466-2)) |
| Otu23043 | 1 | k__Fungi(100);p__Ascomycota(100);c__Saccharomycetes(100);o__Saccharomycetales(100);f__Metschnikowiaceae(100);g__Metschnikowia(100);s__Metschnikowia_pulcherrima(100); | Metschnikowia | 13 | Saprotroph | \|Undefined Saprotroph\| | Yeast | NULL | Probable | As nectar-tap_saprotroph (Põlme et al. 2020); Decay sugar-rich substrates (Põlme et al. 2020) | Põlme S, et al. 2020. Fungal Diversity 105:1-16 ((https://doi.org/10.1007/s13225-020-00466-2)) |
| Otu23044 | 1 | k__Fungi(100);p__Ascomycota(100);c__Saccharomycetes(100);o__Saccharomycetales(100);f__Metschnikowiaceae(100);g__Metschnikowia(100);g__Metschnikowia_unclassified(100); | Metschnikowia | 13 | Saprotroph | \|Undefined Saprotroph\| | Yeast | NULL | Probable | As nectar-tap_saprotroph (Põlme et al. 2020); Decay sugar-rich substrates (Põlme et al. 2020) | Põlme S, et al. 2020. Fungal Diversity 105:1-16 ((https://doi.org/10.1007/s13225-020-00466-2)) |
| Otu23045 | 1 | k__Fungi(100);p__Ascomycota(100);c__Saccharomycetes(100);o__Saccharomycetales(100);f__Metschnikowiaceae(100);g__Metschnikowia(100);s__Metschnikowia_pulcherrima(100); | Metschnikowia | 13 | Saprotroph | \|Undefined Saprotroph\| | Yeast | NULL | Probable | As nectar-tap_saprotroph (Põlme et al. 2020); Decay sugar-rich substrates (Põlme et al. 2020) | Põlme S, et al. 2020. Fungal Diversity 105:1-16 ((https://doi.org/10.1007/s13225-020-00466-2)) |
| Otu23046 | 1 | k__Fungi(100);p__Ascomycota(100);c__Saccharomycetes(100);o__Saccharomycetales(100);f__Metschnikowiaceae(100);g__Metschnikowia(100);s__Metschnikowia_pulcherrima(100); | Metschnikowia | 13 | Saprotroph | \|Undefined Saprotroph\| | Yeast | NULL | Probable | As nectar-tap_saprotroph (Põlme et al. 2020); Decay sugar-rich substrates (Põlme et al. 2020) | Põlme S, et al. 2020. Fungal Diversity 105:1-16 ((https://doi.org/10.1007/s13225-020-00466-2)) |
| Otu23047 | 1 | k__Fungi(100);p__Ascomycota(100);c__Saccharomycetes(100);o__Saccharomycetales(100);f__Metschnikowiaceae(100);g__Metschnikowia(100);s__Metschnikowia_pulcherrima(100); | Metschnikowia | 13 | Saprotroph | \|Undefined Saprotroph\| | Yeast | NULL | Probable | As nectar-tap_saprotroph (Põlme et al. 2020); Decay sugar-rich substrates (Põlme et al. 2020) | Põlme S, et al. 2020. Fungal Diversity 105:1-16 ((https://doi.org/10.1007/s13225-020-00466-2)) |
| Otu23048 | 1 | k__Fungi(100);p__Ascomycota(100);c__Saccharomycetes(100);o__Saccharomycetales(100);f__Metschnikowiaceae(100);g__Metschnikowia(100);s__Metschnikowia_pulcherrima(100); | Metschnikowia | 13 | Saprotroph | \|Undefined Saprotroph\| | Yeast | NULL | Probable | As nectar-tap_saprotroph (Põlme et al. 2020); Decay sugar-rich substrates (Põlme et al. 2020) | Põlme S, et al. 2020. Fungal Diversity 105:1-16 ((https://doi.org/10.1007/s13225-020-00466-2)) |
| Otu23049 | 1 | k__Fungi(100);p__Ascomycota(100);c__Saccharomycetes(100);o__Saccharomycetales(100);f__Metschnikowiaceae(100);g__Metschnikowia(100);g__Metschnikowia_unclassified(100); | Metschnikowia | 13 | Saprotroph | \|Undefined Saprotroph\| | Yeast | NULL | Probable | As nectar-tap_saprotroph (Põlme et al. 2020); Decay sugar-rich substrates (Põlme et al. 2020) | Põlme S, et al. 2020. Fungal Diversity 105:1-16 ((https://doi.org/10.1007/s13225-020-00466-2)) |
| Otu23050 | 1 | k__Fungi(100);p__Ascomycota(100);c__Saccharomycetes(100);o__Saccharomycetales(100);f__Metschnikowiaceae(100);g__Metschnikowia(100);s__Metschnikowia_pulcherrima(100); | Metschnikowia | 13 | Saprotroph | \|Undefined Saprotroph\| | Yeast | NULL | Probable | As nectar-tap_saprotroph (Põlme et al. 2020); Decay sugar-rich substrates (Põlme et al. 2020) | Põlme S, et al. 2020. Fungal Diversity 105:1-16 ((https://doi.org/10.1007/s13225-020-00466-2)) |
| Otu23051 | 1 | k__Fungi(100);p__Ascomycota(100);c__Saccharomycetes(100);o__Saccharomycetales(100);f__Metschnikowiaceae(100);g__Metschnikowia(100);g__Metschnikowia_unclassified(100); | Metschnikowia | 13 | Saprotroph | \|Undefined Saprotroph\| | Yeast | NULL | Probable | As nectar-tap_saprotroph (Põlme et al. 2020); Decay sugar-rich substrates (Põlme et al. 2020) | Põlme S, et al. 2020. Fungal Diversity 105:1-16 ((https://doi.org/10.1007/s13225-020-00466-2)) |
| Otu23052 | 1 | k__Fungi(100);p__Ascomycota(100);c__Saccharomycetes(100);o__Saccharomycetales(100);f__Metschnikowiaceae(100);g__Metschnikowia(100);g__Metschnikowia_unclassified(100); | Metschnikowia | 13 | Saprotroph | \|Undefined Saprotroph\| | Yeast | NULL | Probable | As nectar-tap_saprotroph (Põlme et al. 2020); Decay sugar-rich substrates (Põlme et al. 2020) | Põlme S, et al. 2020. Fungal Diversity 105:1-16 ((https://doi.org/10.1007/s13225-020-00466-2)) |
| Otu23053 | 1 | k__Fungi(100);p__Ascomycota(100);c__Saccharomycetes(100);o__Saccharomycetales(100);f__Metschnikowiaceae(100);g__Metschnikowia(100);s__Metschnikowia_pulcherrima(100); | Metschnikowia | 13 | Saprotroph | \|Undefined Saprotroph\| | Yeast | NULL | Probable | As nectar-tap_saprotroph (Põlme et al. 2020); Decay sugar-rich substrates (Põlme et al. 2020) | Põlme S, et al. 2020. Fungal Diversity 105:1-16 ((https://doi.org/10.1007/s13225-020-00466-2)) |
| Otu23054 | 1 | k__Fungi(100);p__Ascomycota(100);c__Saccharomycetes(100);o__Saccharomycetales(100);f__Metschnikowiaceae(100);g__Metschnikowia(100);s__Metschnikowia_pulcherrima(100); | Metschnikowia | 13 | Saprotroph | \|Undefined Saprotroph\| | Yeast | NULL | Probable | As nectar-tap_saprotroph (Põlme et al. 2020); Decay sugar-rich substrates (Põlme et al. 2020) | Põlme S, et al. 2020. Fungal Diversity 105:1-16 ((https://doi.org/10.1007/s13225-020-00466-2)) |
| Otu23055 | 1 | k__Fungi(100);p__Ascomycota(100);c__Saccharomycetes(100);o__Saccharomycetales(100);f__Metschnikowiaceae(100);g__Metschnikowia(100);s__Metschnikowia_pulcherrima(100); | Metschnikowia | 13 | Saprotroph | \|Undefined Saprotroph\| | Yeast | NULL | Probable | As nectar-tap_saprotroph (Põlme et al. 2020); Decay sugar-rich substrates (Põlme et al. 2020) | Põlme S, et al. 2020. Fungal Diversity 105:1-16 ((https://doi.org/10.1007/s13225-020-00466-2)) |
| Otu23056 | 1 | k__Fungi(100);p__Ascomycota(100);c__Saccharomycetes(100);o__Saccharomycetales(100);f__Metschnikowiaceae(100);g__Metschnikowia(100);s__Metschnikowia_pulcherrima(100); | Metschnikowia | 13 | Saprotroph | \|Undefined Saprotroph\| | Yeast | NULL | Probable | As nectar-tap_saprotroph (Põlme et al. 2020); Decay sugar-rich substrates (Põlme et al. 2020) | Põlme S, et al. 2020. Fungal Diversity 105:1-16 ((https://doi.org/10.1007/s13225-020-00466-2)) |
| Otu23057 | 1 | k__Fungi(100);p__Ascomycota(100);c__Saccharomycetes(100);o__Saccharomycetales(100);f__Metschnikowiaceae(100);g__Metschnikowia(100);g__Metschnikowia_unclassified(100); | Metschnikowia | 13 | Saprotroph | \|Undefined Saprotroph\| | Yeast | NULL | Probable | As nectar-tap_saprotroph (Põlme et al. 2020); Decay sugar-rich substrates (Põlme et al. 2020) | Põlme S, et al. 2020. Fungal Diversity 105:1-16 ((https://doi.org/10.1007/s13225-020-00466-2)) |
| Otu23058 | 1 | k__Fungi(100);p__Ascomycota(100);c__Saccharomycetes(100);o__Saccharomycetales(100);f__Metschnikowiaceae(100);g__Metschnikowia(100);s__Metschnikowia_pulcherrima(100); | Metschnikowia | 13 | Saprotroph | \|Undefined Saprotroph\| | Yeast | NULL | Probable | As nectar-tap_saprotroph (Põlme et al. 2020); Decay sugar-rich substrates (Põlme et al. 2020) | Põlme S, et al. 2020. Fungal Diversity 105:1-16 ((https://doi.org/10.1007/s13225-020-00466-2)) |
| Otu23059 | 1 | k__Fungi(100);p__Ascomycota(100);c__Saccharomycetes(100);o__Saccharomycetales(100);f__Metschnikowiaceae(100);g__Metschnikowia(100);s__Metschnikowia_pulcherrima(100); | Metschnikowia | 13 | Saprotroph | \|Undefined Saprotroph\| | Yeast | NULL | Probable | As nectar-tap_saprotroph (Põlme et al. 2020); Decay sugar-rich substrates (Põlme et al. 2020) | Põlme S, et al. 2020. Fungal Diversity 105:1-16 ((https://doi.org/10.1007/s13225-020-00466-2)) |
| Otu23060 | 1 | k__Fungi(100);p__Ascomycota(100);c__Saccharomycetes(100);o__Saccharomycetales(100);f__Metschnikowiaceae(100);g__Metschnikowia(100);s__Metschnikowia_pulcherrima(100); | Metschnikowia | 13 | Saprotroph | \|Undefined Saprotroph\| | Yeast | NULL | Probable | As nectar-tap_saprotroph (Põlme et al. 2020); Decay sugar-rich substrates (Põlme et al. 2020) | Põlme S, et al. 2020. Fungal Diversity 105:1-16 ((https://doi.org/10.1007/s13225-020-00466-2)) |
| Otu23061 | 1 | k__Fungi(100);p__Ascomycota(100);c__Saccharomycetes(100);o__Saccharomycetales(100);f__Metschnikowiaceae(100);g__Metschnikowia(100);g__Metschnikowia_unclassified(100); | Metschnikowia | 13 | Saprotroph | \|Undefined Saprotroph\| | Yeast | NULL | Probable | As nectar-tap_saprotroph (Põlme et al. 2020); Decay sugar-rich substrates (Põlme et al. 2020) | Põlme S, et al. 2020. Fungal Diversity 105:1-16 ((https://doi.org/10.1007/s13225-020-00466-2)) |
| Otu23062 | 1 | k__Fungi(100);p__Ascomycota(100);c__Saccharomycetes(100);o__Saccharomycetales(100);f__Metschnikowiaceae(100);g__Metschnikowia(100);s__Metschnikowia_pulcherrima(100); | Metschnikowia | 13 | Saprotroph | \|Undefined Saprotroph\| | Yeast | NULL | Probable | As nectar-tap_saprotroph (Põlme et al. 2020); Decay sugar-rich substrates (Põlme et al. 2020) | Põlme S, et al. 2020. Fungal Diversity 105:1-16 ((https://doi.org/10.1007/s13225-020-00466-2)) |
| Otu23063 | 1 | k__Fungi(100);p__Ascomycota(100);c__Saccharomycetes(100);o__Saccharomycetales(100);f__Metschnikowiaceae(100);g__Metschnikowia(100);s__Metschnikowia_pulcherrima(100); | Metschnikowia | 13 | Saprotroph | \|Undefined Saprotroph\| | Yeast | NULL | Probable | As nectar-tap_saprotroph (Põlme et al. 2020); Decay sugar-rich substrates (Põlme et al. 2020) | Põlme S, et al. 2020. Fungal Diversity 105:1-16 ((https://doi.org/10.1007/s13225-020-00466-2)) |
| Otu23064 | 1 | k__Fungi(100);p__Ascomycota(100);c__Saccharomycetes(100);o__Saccharomycetales(100);f__Metschnikowiaceae(100);g__Metschnikowia(100);s__Metschnikowia_pulcherrima(100); | Metschnikowia | 13 | Saprotroph | \|Undefined Saprotroph\| | Yeast | NULL | Probable | As nectar-tap_saprotroph (Põlme et al. 2020); Decay sugar-rich substrates (Põlme et al. 2020) | Põlme S, et al. 2020. Fungal Diversity 105:1-16 ((https://doi.org/10.1007/s13225-020-00466-2)) |
| Otu23065 | 1 | k__Fungi(100);p__Ascomycota(100);c__Saccharomycetes(100);o__Saccharomycetales(100);f__Metschnikowiaceae(100);g__Metschnikowia(100);s__Metschnikowia_pulcherrima(100); | Metschnikowia | 13 | Saprotroph | \|Undefined Saprotroph\| | Yeast | NULL | Probable | As nectar-tap_saprotroph (Põlme et al. 2020); Decay sugar-rich substrates (Põlme et al. 2020) | Põlme S, et al. 2020. Fungal Diversity 105:1-16 ((https://doi.org/10.1007/s13225-020-00466-2)) |
| Otu23066 | 1 | k__Fungi(100);p__Ascomycota(100);c__Saccharomycetes(100);o__Saccharomycetales(100);f__Metschnikowiaceae(100);g__Metschnikowia(100);s__Metschnikowia_pulcherrima(100); | Metschnikowia | 13 | Saprotroph | \|Undefined Saprotroph\| | Yeast | NULL | Probable | As nectar-tap_saprotroph (Põlme et al. 2020); Decay sugar-rich substrates (Põlme et al. 2020) | Põlme S, et al. 2020. Fungal Diversity 105:1-16 ((https://doi.org/10.1007/s13225-020-00466-2)) |
| Otu23067 | 1 | k__Fungi(100);p__Ascomycota(100);c__Saccharomycetes(100);o__Saccharomycetales(100);f__Metschnikowiaceae(100);g__Metschnikowia(100);s__Metschnikowia_pulcherrima(100); | Metschnikowia | 13 | Saprotroph | \|Undefined Saprotroph\| | Yeast | NULL | Probable | As nectar-tap_saprotroph (Põlme et al. 2020); Decay sugar-rich substrates (Põlme et al. 2020) | Põlme S, et al. 2020. Fungal Diversity 105:1-16 ((https://doi.org/10.1007/s13225-020-00466-2)) |
| Otu23068 | 1 | k__Fungi(100);p__Ascomycota(100);c__Saccharomycetes(100);o__Saccharomycetales(100);f__Metschnikowiaceae(100);g__Metschnikowia(100);g__Metschnikowia_unclassified(100); | Metschnikowia | 13 | Saprotroph | \|Undefined Saprotroph\| | Yeast | NULL | Probable | As nectar-tap_saprotroph (Põlme et al. 2020); Decay sugar-rich substrates (Põlme et al. 2020) | Põlme S, et al. 2020. Fungal Diversity 105:1-16 ((https://doi.org/10.1007/s13225-020-00466-2)) |
| Otu23069 | 1 | k__Fungi(100);p__Ascomycota(100);c__Saccharomycetes(100);o__Saccharomycetales(100);f__Metschnikowiaceae(100);g__Metschnikowia(100);s__Metschnikowia_pulcherrima(100); | Metschnikowia | 13 | Saprotroph | \|Undefined Saprotroph\| | Yeast | NULL | Probable | As nectar-tap_saprotroph (Põlme et al. 2020); Decay sugar-rich substrates (Põlme et al. 2020) | Põlme S, et al. 2020. Fungal Diversity 105:1-16 ((https://doi.org/10.1007/s13225-020-00466-2)) |
| Otu23070 | 1 | k__Fungi(100);p__Ascomycota(100);c__Saccharomycetes(100);o__Saccharomycetales(100);f__Metschnikowiaceae(100);g__Metschnikowia(100);s__Metschnikowia_pulcherrima(100); | Metschnikowia | 13 | Saprotroph | \|Undefined Saprotroph\| | Yeast | NULL | Probable | As nectar-tap_saprotroph (Põlme et al. 2020); Decay sugar-rich substrates (Põlme et al. 2020) | Põlme S, et al. 2020. Fungal Diversity 105:1-16 ((https://doi.org/10.1007/s13225-020-00466-2)) |
| Otu23071 | 1 | k__Fungi(100);p__Ascomycota(100);c__Saccharomycetes(100);o__Saccharomycetales(100);f__Metschnikowiaceae(100);g__Metschnikowia(100);g__Metschnikowia_unclassified(100); | Metschnikowia | 13 | Saprotroph | \|Undefined Saprotroph\| | Yeast | NULL | Probable | As nectar-tap_saprotroph (Põlme et al. 2020); Decay sugar-rich substrates (Põlme et al. 2020) | Põlme S, et al. 2020. Fungal Diversity 105:1-16 ((https://doi.org/10.1007/s13225-020-00466-2)) |
| Otu23072 | 1 | k__Fungi(100);p__Ascomycota(100);c__Saccharomycetes(100);o__Saccharomycetales(100);f__Metschnikowiaceae(100);g__Metschnikowia(100);g__Metschnikowia_unclassified(100); | Metschnikowia | 13 | Saprotroph | \|Undefined Saprotroph\| | Yeast | NULL | Probable | As nectar-tap_saprotroph (Põlme et al. 2020); Decay sugar-rich substrates (Põlme et al. 2020) | Põlme S, et al. 2020. Fungal Diversity 105:1-16 ((https://doi.org/10.1007/s13225-020-00466-2)) |
| Otu23075 | 1 | k__Fungi(100);p__Ascomycota(100);c__Saccharomycetes(100);o__Saccharomycetales(100);f__Metschnikowiaceae(100);g__Metschnikowia(100);g__Metschnikowia_unclassified(100); | Metschnikowia | 13 | Saprotroph | \|Undefined Saprotroph\| | Yeast | NULL | Probable | As nectar-tap_saprotroph (Põlme et al. 2020); Decay sugar-rich substrates (Põlme et al. 2020) | Põlme S, et al. 2020. Fungal Diversity 105:1-16 ((https://doi.org/10.1007/s13225-020-00466-2)) |
| Otu23078 | 1 | k__Fungi(100);p__Ascomycota(100);c__Saccharomycetes(100);o__Saccharomycetales(100);f__Metschnikowiaceae(100);g__Metschnikowia(100);s__Metschnikowia_pulcherrima(100); | Metschnikowia | 13 | Saprotroph | \|Undefined Saprotroph\| | Yeast | NULL | Probable | As nectar-tap_saprotroph (Põlme et al. 2020); Decay sugar-rich substrates (Põlme et al. 2020) | Põlme S, et al. 2020. Fungal Diversity 105:1-16 ((https://doi.org/10.1007/s13225-020-00466-2)) |
| Otu23079 | 1 | k__Fungi(100);p__Ascomycota(100);c__Dothideomycetes(100);o__Capnodiales(100);f__Teratosphaeriaceae(100);g__Devriesia(100);s__Devriesia_sp(100); | Devriesia | 13 | Pathotroph-Saprotroph | \|Plant Pathogen\|-Plant Saprotroph | Microfungus | NULL | Probable | As litter_saprotroph (Põlme et al. 2020); Decay leaf, fruit, seed (Põlme et al. 2020) | Tedersoo L, et al. 2014. Science 346:e1256688 ((https://doi.org/10.1126/science.1256688)); Põlme S, et al. 2020. Fungal Diversity 105:1-16 ((https://doi.org/10.1007/s13225-020-00466-2)) |
| Otu23080 | 1 | k__Fungi(100);p__Ascomycota(100);c__Saccharomycetes(100);o__Saccharomycetales(100);f__Metschnikowiaceae(100);g__Metschnikowia(100);s__Metschnikowia_pulcherrima(100); | Metschnikowia | 13 | Saprotroph | \|Undefined Saprotroph\| | Yeast | NULL | Probable | As nectar-tap_saprotroph (Põlme et al. 2020); Decay sugar-rich substrates (Põlme et al. 2020) | Põlme S, et al. 2020. Fungal Diversity 105:1-16 ((https://doi.org/10.1007/s13225-020-00466-2)) |
| Otu23082 | 1 | k__Fungi(100);p__Ascomycota(100);c__Saccharomycetes(100);o__Saccharomycetales(100);f__Metschnikowiaceae(100);g__Metschnikowia(100);g__Metschnikowia_unclassified(100); | Metschnikowia | 13 | Saprotroph | \|Undefined Saprotroph\| | Yeast | NULL | Probable | As nectar-tap_saprotroph (Põlme et al. 2020); Decay sugar-rich substrates (Põlme et al. 2020) | Põlme S, et al. 2020. Fungal Diversity 105:1-16 ((https://doi.org/10.1007/s13225-020-00466-2)) |
| Otu23083 | 1 | k__Fungi(100);p__Ascomycota(100);c__Saccharomycetes(100);o__Saccharomycetales(100);f__Metschnikowiaceae(100);g__Metschnikowia(100);g__Metschnikowia_unclassified(100); | Metschnikowia | 13 | Saprotroph | \|Undefined Saprotroph\| | Yeast | NULL | Probable | As nectar-tap_saprotroph (Põlme et al. 2020); Decay sugar-rich substrates (Põlme et al. 2020) | Põlme S, et al. 2020. Fungal Diversity 105:1-16 ((https://doi.org/10.1007/s13225-020-00466-2)) |
| Otu23084 | 1 | k__Fungi(100);p__Ascomycota(100);c__Saccharomycetes(100);o__Saccharomycetales(100);f__Metschnikowiaceae(100);g__Metschnikowia(100);s__Metschnikowia_pulcherrima(100); | Metschnikowia | 13 | Saprotroph | \|Undefined Saprotroph\| | Yeast | NULL | Probable | As nectar-tap_saprotroph (Põlme et al. 2020); Decay sugar-rich substrates (Põlme et al. 2020) | Põlme S, et al. 2020. Fungal Diversity 105:1-16 ((https://doi.org/10.1007/s13225-020-00466-2)) |
| Otu23086 | 1 | k__Fungi(100);p__Ascomycota(100);c__Saccharomycetes(100);o__Saccharomycetales(100);f__Metschnikowiaceae(100);g__Metschnikowia(100);s__Metschnikowia_pulcherrima(100); | Metschnikowia | 13 | Saprotroph | \|Undefined Saprotroph\| | Yeast | NULL | Probable | As nectar-tap_saprotroph (Põlme et al. 2020); Decay sugar-rich substrates (Põlme et al. 2020) | Põlme S, et al. 2020. Fungal Diversity 105:1-16 ((https://doi.org/10.1007/s13225-020-00466-2)) |
| Otu23087 | 1 | k__Fungi(100);p__Ascomycota(100);c__Saccharomycetes(100);o__Saccharomycetales(100);f__Metschnikowiaceae(100);g__Metschnikowia(100);s__Metschnikowia_pulcherrima(100); | Metschnikowia | 13 | Saprotroph | \|Undefined Saprotroph\| | Yeast | NULL | Probable | As nectar-tap_saprotroph (Põlme et al. 2020); Decay sugar-rich substrates (Põlme et al. 2020) | Põlme S, et al. 2020. Fungal Diversity 105:1-16 ((https://doi.org/10.1007/s13225-020-00466-2)) |
| Otu23088 | 1 | k__Fungi(100);p__Ascomycota(100);c__Saccharomycetes(100);o__Saccharomycetales(100);f__Metschnikowiaceae(100);g__Metschnikowia(100);s__Metschnikowia_pulcherrima(100); | Metschnikowia | 13 | Saprotroph | \|Undefined Saprotroph\| | Yeast | NULL | Probable | As nectar-tap_saprotroph (Põlme et al. 2020); Decay sugar-rich substrates (Põlme et al. 2020) | Põlme S, et al. 2020. Fungal Diversity 105:1-16 ((https://doi.org/10.1007/s13225-020-00466-2)) |
| Otu23089 | 1 | k__Fungi(100);p__Ascomycota(100);c__Saccharomycetes(100);o__Saccharomycetales(100);f__Metschnikowiaceae(100);g__Metschnikowia(100);s__Metschnikowia_pulcherrima(100); | Metschnikowia | 13 | Saprotroph | \|Undefined Saprotroph\| | Yeast | NULL | Probable | As nectar-tap_saprotroph (Põlme et al. 2020); Decay sugar-rich substrates (Põlme et al. 2020) | Põlme S, et al. 2020. Fungal Diversity 105:1-16 ((https://doi.org/10.1007/s13225-020-00466-2)) |
| Otu23090 | 1 | k__Fungi(100);p__Ascomycota(100);c__Saccharomycetes(100);o__Saccharomycetales(100);f__Metschnikowiaceae(100);g__Metschnikowia(100);s__Metschnikowia_pulcherrima(100); | Metschnikowia | 13 | Saprotroph | \|Undefined Saprotroph\| | Yeast | NULL | Probable | As nectar-tap_saprotroph (Põlme et al. 2020); Decay sugar-rich substrates (Põlme et al. 2020) | Põlme S, et al. 2020. Fungal Diversity 105:1-16 ((https://doi.org/10.1007/s13225-020-00466-2)) |
| Otu23091 | 1 | k__Fungi(100);p__Ascomycota(100);c__Saccharomycetes(100);o__Saccharomycetales(100);f__Metschnikowiaceae(100);g__Metschnikowia(100);g__Metschnikowia_unclassified(100); | Metschnikowia | 13 | Saprotroph | \|Undefined Saprotroph\| | Yeast | NULL | Probable | As nectar-tap_saprotroph (Põlme et al. 2020); Decay sugar-rich substrates (Põlme et al. 2020) | Põlme S, et al. 2020. Fungal Diversity 105:1-16 ((https://doi.org/10.1007/s13225-020-00466-2)) |
| Otu23092 | 1 | k__Fungi(100);p__Ascomycota(100);c__Saccharomycetes(100);o__Saccharomycetales(100);f__Metschnikowiaceae(100);g__Metschnikowia(100);g__Metschnikowia_unclassified(100); | Metschnikowia | 13 | Saprotroph | \|Undefined Saprotroph\| | Yeast | NULL | Probable | As nectar-tap_saprotroph (Põlme et al. 2020); Decay sugar-rich substrates (Põlme et al. 2020) | Põlme S, et al. 2020. Fungal Diversity 105:1-16 ((https://doi.org/10.1007/s13225-020-00466-2)) |
| Otu23093 | 1 | k__Fungi(100);p__Ascomycota(100);c__Saccharomycetes(100);o__Saccharomycetales(100);f__Metschnikowiaceae(100);g__Metschnikowia(100);g__Metschnikowia_unclassified(100); | Metschnikowia | 13 | Saprotroph | \|Undefined Saprotroph\| | Yeast | NULL | Probable | As nectar-tap_saprotroph (Põlme et al. 2020); Decay sugar-rich substrates (Põlme et al. 2020) | Põlme S, et al. 2020. Fungal Diversity 105:1-16 ((https://doi.org/10.1007/s13225-020-00466-2)) |
| Otu23094 | 1 | k__Fungi(100);p__Ascomycota(100);c__Saccharomycetes(100);o__Saccharomycetales(100);f__Metschnikowiaceae(100);g__Metschnikowia(100);g__Metschnikowia_unclassified(100); | Metschnikowia | 13 | Saprotroph | \|Undefined Saprotroph\| | Yeast | NULL | Probable | As nectar-tap_saprotroph (Põlme et al. 2020); Decay sugar-rich substrates (Põlme et al. 2020) | Põlme S, et al. 2020. Fungal Diversity 105:1-16 ((https://doi.org/10.1007/s13225-020-00466-2)) |
| Otu23095 | 1 | k__Fungi(100);p__Ascomycota(100);c__Saccharomycetes(100);o__Saccharomycetales(100);f__Metschnikowiaceae(100);g__Metschnikowia(100);s__Metschnikowia_pulcherrima(100); | Metschnikowia | 13 | Saprotroph | \|Undefined Saprotroph\| | Yeast | NULL | Probable | As nectar-tap_saprotroph (Põlme et al. 2020); Decay sugar-rich substrates (Põlme et al. 2020) | Põlme S, et al. 2020. Fungal Diversity 105:1-16 ((https://doi.org/10.1007/s13225-020-00466-2)) |
| Otu23096 | 1 | k__Fungi(100);p__Ascomycota(100);c__Saccharomycetes(100);o__Saccharomycetales(100);f__Metschnikowiaceae(100);g__Metschnikowia(100);s__Metschnikowia_pulcherrima(100); | Metschnikowia | 13 | Saprotroph | \|Undefined Saprotroph\| | Yeast | NULL | Probable | As nectar-tap_saprotroph (Põlme et al. 2020); Decay sugar-rich substrates (Põlme et al. 2020) | Põlme S, et al. 2020. Fungal Diversity 105:1-16 ((https://doi.org/10.1007/s13225-020-00466-2)) |
| Otu23097 | 1 | k__Fungi(100);p__Ascomycota(100);c__Saccharomycetes(100);o__Saccharomycetales(100);f__Metschnikowiaceae(100);g__Metschnikowia(100);s__Metschnikowia_pulcherrima(100); | Metschnikowia | 13 | Saprotroph | \|Undefined Saprotroph\| | Yeast | NULL | Probable | As nectar-tap_saprotroph (Põlme et al. 2020); Decay sugar-rich substrates (Põlme et al. 2020) | Põlme S, et al. 2020. Fungal Diversity 105:1-16 ((https://doi.org/10.1007/s13225-020-00466-2)) |
| Otu23098 | 1 | k__Fungi(100);p__Ascomycota(100);c__Saccharomycetes(100);o__Saccharomycetales(100);f__Metschnikowiaceae(100);g__Metschnikowia(100);s__Metschnikowia_pulcherrima(100); | Metschnikowia | 13 | Saprotroph | \|Undefined Saprotroph\| | Yeast | NULL | Probable | As nectar-tap_saprotroph (Põlme et al. 2020); Decay sugar-rich substrates (Põlme et al. 2020) | Põlme S, et al. 2020. Fungal Diversity 105:1-16 ((https://doi.org/10.1007/s13225-020-00466-2)) |
| Otu23099 | 1 | k__Fungi(100);p__Ascomycota(100);c__Saccharomycetes(100);o__Saccharomycetales(100);f__Metschnikowiaceae(100);g__Metschnikowia(100);s__Metschnikowia_pulcherrima(100); | Metschnikowia | 13 | Saprotroph | \|Undefined Saprotroph\| | Yeast | NULL | Probable | As nectar-tap_saprotroph (Põlme et al. 2020); Decay sugar-rich substrates (Põlme et al. 2020) | Põlme S, et al. 2020. Fungal Diversity 105:1-16 ((https://doi.org/10.1007/s13225-020-00466-2)) |
| Otu23100 | 1 | k__Fungi(100);p__Ascomycota(100);c__Saccharomycetes(100);o__Saccharomycetales(100);f__Metschnikowiaceae(100);g__Metschnikowia(100);s__Metschnikowia_pulcherrima(100); | Metschnikowia | 13 | Saprotroph | \|Undefined Saprotroph\| | Yeast | NULL | Probable | As nectar-tap_saprotroph (Põlme et al. 2020); Decay sugar-rich substrates (Põlme et al. 2020) | Põlme S, et al. 2020. Fungal Diversity 105:1-16 ((https://doi.org/10.1007/s13225-020-00466-2)) |
| Otu23101 | 1 | k__Fungi(100);p__Ascomycota(100);c__Saccharomycetes(100);o__Saccharomycetales(100);f__Metschnikowiaceae(100);g__Metschnikowia(100);s__Metschnikowia_pulcherrima(100); | Metschnikowia | 13 | Saprotroph | \|Undefined Saprotroph\| | Yeast | NULL | Probable | As nectar-tap_saprotroph (Põlme et al. 2020); Decay sugar-rich substrates (Põlme et al. 2020) | Põlme S, et al. 2020. Fungal Diversity 105:1-16 ((https://doi.org/10.1007/s13225-020-00466-2)) |
| Otu23102 | 1 | k__Fungi(100);p__Ascomycota(100);c__Saccharomycetes(100);o__Saccharomycetales(100);f__Metschnikowiaceae(100);g__Metschnikowia(100);s__Metschnikowia_pulcherrima(100); | Metschnikowia | 13 | Saprotroph | \|Undefined Saprotroph\| | Yeast | NULL | Probable | As nectar-tap_saprotroph (Põlme et al. 2020); Decay sugar-rich substrates (Põlme et al. 2020) | Põlme S, et al. 2020. Fungal Diversity 105:1-16 ((https://doi.org/10.1007/s13225-020-00466-2)) |
| Otu23103 | 1 | k__Fungi(100);p__Ascomycota(100);c__Saccharomycetes(100);o__Saccharomycetales(100);f__Metschnikowiaceae(100);g__Metschnikowia(100);s__Metschnikowia_pulcherrima(100); | Metschnikowia | 13 | Saprotroph | \|Undefined Saprotroph\| | Yeast | NULL | Probable | As nectar-tap_saprotroph (Põlme et al. 2020); Decay sugar-rich substrates (Põlme et al. 2020) | Põlme S, et al. 2020. Fungal Diversity 105:1-16 ((https://doi.org/10.1007/s13225-020-00466-2)) |
| Otu23104 | 1 | k__Fungi(100);p__Ascomycota(100);c__Saccharomycetes(100);o__Saccharomycetales(100);f__Metschnikowiaceae(100);g__Metschnikowia(100);g__Metschnikowia_unclassified(100); | Metschnikowia | 13 | Saprotroph | \|Undefined Saprotroph\| | Yeast | NULL | Probable | As nectar-tap_saprotroph (Põlme et al. 2020); Decay sugar-rich substrates (Põlme et al. 2020) | Põlme S, et al. 2020. Fungal Diversity 105:1-16 ((https://doi.org/10.1007/s13225-020-00466-2)) |
| Otu23105 | 1 | k__Fungi(100);p__Ascomycota(100);c__Saccharomycetes(100);o__Saccharomycetales(100);f__Metschnikowiaceae(100);g__Metschnikowia(100);s__Metschnikowia_pulcherrima(100); | Metschnikowia | 13 | Saprotroph | \|Undefined Saprotroph\| | Yeast | NULL | Probable | As nectar-tap_saprotroph (Põlme et al. 2020); Decay sugar-rich substrates (Põlme et al. 2020) | Põlme S, et al. 2020. Fungal Diversity 105:1-16 ((https://doi.org/10.1007/s13225-020-00466-2)) |
| Otu23106 | 1 | k__Fungi(100);p__Ascomycota(100);c__Saccharomycetes(100);o__Saccharomycetales(100);f__Metschnikowiaceae(100);g__Metschnikowia(100);g__Metschnikowia_unclassified(100); | Metschnikowia | 13 | Saprotroph | \|Undefined Saprotroph\| | Yeast | NULL | Probable | As nectar-tap_saprotroph (Põlme et al. 2020); Decay sugar-rich substrates (Põlme et al. 2020) | Põlme S, et al. 2020. Fungal Diversity 105:1-16 ((https://doi.org/10.1007/s13225-020-00466-2)) |
| Otu23107 | 1 | k__Fungi(100);p__Ascomycota(100);c__Saccharomycetes(100);o__Saccharomycetales(100);f__Metschnikowiaceae(100);g__Metschnikowia(100);s__Metschnikowia_pulcherrima(100); | Metschnikowia | 13 | Saprotroph | \|Undefined Saprotroph\| | Yeast | NULL | Probable | As nectar-tap_saprotroph (Põlme et al. 2020); Decay sugar-rich substrates (Põlme et al. 2020) | Põlme S, et al. 2020. Fungal Diversity 105:1-16 ((https://doi.org/10.1007/s13225-020-00466-2)) |
| Otu23108 | 1 | k__Fungi(100);p__Ascomycota(100);c__Saccharomycetes(100);o__Saccharomycetales(100);f__Metschnikowiaceae(100);g__Metschnikowia(100);s__Metschnikowia_pulcherrima(100); | Metschnikowia | 13 | Saprotroph | \|Undefined Saprotroph\| | Yeast | NULL | Probable | As nectar-tap_saprotroph (Põlme et al. 2020); Decay sugar-rich substrates (Põlme et al. 2020) | Põlme S, et al. 2020. Fungal Diversity 105:1-16 ((https://doi.org/10.1007/s13225-020-00466-2)) |
| Otu23109 | 1 | k__Fungi(100);p__Ascomycota(100);c__Saccharomycetes(100);o__Saccharomycetales(100);f__Metschnikowiaceae(100);g__Metschnikowia(100);s__Metschnikowia_pulcherrima(100); | Metschnikowia | 13 | Saprotroph | \|Undefined Saprotroph\| | Yeast | NULL | Probable | As nectar-tap_saprotroph (Põlme et al. 2020); Decay sugar-rich substrates (Põlme et al. 2020) | Põlme S, et al. 2020. Fungal Diversity 105:1-16 ((https://doi.org/10.1007/s13225-020-00466-2)) |
| Otu23110 | 1 | k__Fungi(100);p__Ascomycota(100);c__Saccharomycetes(100);o__Saccharomycetales(100);f__Metschnikowiaceae(100);g__Metschnikowia(100);s__Metschnikowia_pulcherrima(100); | Metschnikowia | 13 | Saprotroph | \|Undefined Saprotroph\| | Yeast | NULL | Probable | As nectar-tap_saprotroph (Põlme et al. 2020); Decay sugar-rich substrates (Põlme et al. 2020) | Põlme S, et al. 2020. Fungal Diversity 105:1-16 ((https://doi.org/10.1007/s13225-020-00466-2)) |
| Otu23111 | 1 | k__Fungi(100);p__Ascomycota(100);c__Saccharomycetes(100);o__Saccharomycetales(100);f__Saccharomycetales_family_Incertae_sedis(100);g__Candida(100);g__Candida_unclassified(100); | Candida | 13 | Pathotroph-Saprotroph-Symbiotroph | Animal Pathogen-Endophyte-Endosymbiont-Epiphyte-\|Undefined Saprotroph\| | Dimorphic Yeast-Yeast | NULL | Probable | As nectar-tap_saprotroph (Põlme et al. 2020); Decay sugar-rich substrates (Põlme et al. 2020); This asexual genus is polyphyletic and contain species that function very differently depend on what they are. Caution should be taken when interpreting guilds for Candida species. | Manolakaki D, et al. 2010. Virulence. 1: 367-375 ((https://doi.org/10.4161/viru.1.5.12796)); Mekbib SB, et al. 2011. Tropical Plant Pathology 36:233-240 ((https://doi.org/10.1590/S1982-56762011000400004)); Põlme S, et al. 2020. Fungal Diversity 105:1-16 ((https://doi.org/10.1007/s13225-020-00466-2)) |
| Otu23112 | 1 | k__Fungi(100);p__Ascomycota(100);c__Saccharomycetes(100);o__Saccharomycetales(100);f__Metschnikowiaceae(100);g__Metschnikowia(100);s__Metschnikowia_pulcherrima(100); | Metschnikowia | 13 | Saprotroph | \|Undefined Saprotroph\| | Yeast | NULL | Probable | As nectar-tap_saprotroph (Põlme et al. 2020); Decay sugar-rich substrates (Põlme et al. 2020) | Põlme S, et al. 2020. Fungal Diversity 105:1-16 ((https://doi.org/10.1007/s13225-020-00466-2)) |
| Otu23113 | 1 | k__Fungi(100);p__Ascomycota(100);c__Saccharomycetes(100);o__Saccharomycetales(100);f__Metschnikowiaceae(100);g__Metschnikowia(100);s__Metschnikowia_pulcherrima(100); | Metschnikowia | 13 | Saprotroph | \|Undefined Saprotroph\| | Yeast | NULL | Probable | As nectar-tap_saprotroph (Põlme et al. 2020); Decay sugar-rich substrates (Põlme et al. 2020) | Põlme S, et al. 2020. Fungal Diversity 105:1-16 ((https://doi.org/10.1007/s13225-020-00466-2)) |
| Otu23114 | 1 | k__Fungi(100);p__Ascomycota(100);c__Saccharomycetes(100);o__Saccharomycetales(100);f__Metschnikowiaceae(100);g__Metschnikowia(100);s__Metschnikowia_pulcherrima(100); | Metschnikowia | 13 | Saprotroph | \|Undefined Saprotroph\| | Yeast | NULL | Probable | As nectar-tap_saprotroph (Põlme et al. 2020); Decay sugar-rich substrates (Põlme et al. 2020) | Põlme S, et al. 2020. Fungal Diversity 105:1-16 ((https://doi.org/10.1007/s13225-020-00466-2)) |
| Otu23115 | 1 | k__Fungi(100);p__Ascomycota(100);c__Saccharomycetes(100);o__Saccharomycetales(100);f__Metschnikowiaceae(100);g__Metschnikowia(100);g__Metschnikowia_unclassified(100); | Metschnikowia | 13 | Saprotroph | \|Undefined Saprotroph\| | Yeast | NULL | Probable | As nectar-tap_saprotroph (Põlme et al. 2020); Decay sugar-rich substrates (Põlme et al. 2020) | Põlme S, et al. 2020. Fungal Diversity 105:1-16 ((https://doi.org/10.1007/s13225-020-00466-2)) |
| Otu23116 | 1 | k__Fungi(100);p__Ascomycota(100);c__Saccharomycetes(100);o__Saccharomycetales(100);f__Metschnikowiaceae(100);g__Metschnikowia(100);s__Metschnikowia_pulcherrima(100); | Metschnikowia | 13 | Saprotroph | \|Undefined Saprotroph\| | Yeast | NULL | Probable | As nectar-tap_saprotroph (Põlme et al. 2020); Decay sugar-rich substrates (Põlme et al. 2020) | Põlme S, et al. 2020. Fungal Diversity 105:1-16 ((https://doi.org/10.1007/s13225-020-00466-2)) |
| Otu23117 | 1 | k__Fungi(100);p__Ascomycota(100);c__Saccharomycetes(100);o__Saccharomycetales(100);f__Metschnikowiaceae(100);g__Metschnikowia(100);s__Metschnikowia_pulcherrima(100); | Metschnikowia | 13 | Saprotroph | \|Undefined Saprotroph\| | Yeast | NULL | Probable | As nectar-tap_saprotroph (Põlme et al. 2020); Decay sugar-rich substrates (Põlme et al. 2020) | Põlme S, et al. 2020. Fungal Diversity 105:1-16 ((https://doi.org/10.1007/s13225-020-00466-2)) |
| Otu23118 | 1 | k__Fungi(100);p__Ascomycota(100);c__Saccharomycetes(100);o__Saccharomycetales(100);f__Metschnikowiaceae(100);g__Metschnikowia(100);s__Metschnikowia_pulcherrima(100); | Metschnikowia | 13 | Saprotroph | \|Undefined Saprotroph\| | Yeast | NULL | Probable | As nectar-tap_saprotroph (Põlme et al. 2020); Decay sugar-rich substrates (Põlme et al. 2020) | Põlme S, et al. 2020. Fungal Diversity 105:1-16 ((https://doi.org/10.1007/s13225-020-00466-2)) |
| Otu23119 | 1 | k__Fungi(100);p__Ascomycota(100);c__Saccharomycetes(100);o__Saccharomycetales(100);f__Metschnikowiaceae(100);g__Metschnikowia(100);s__Metschnikowia_pulcherrima(100); | Metschnikowia | 13 | Saprotroph | \|Undefined Saprotroph\| | Yeast | NULL | Probable | As nectar-tap_saprotroph (Põlme et al. 2020); Decay sugar-rich substrates (Põlme et al. 2020) | Põlme S, et al. 2020. Fungal Diversity 105:1-16 ((https://doi.org/10.1007/s13225-020-00466-2)) |
| Otu23124 | 1 | k__Fungi(100);p__Ascomycota(100);c__Saccharomycetes(100);o__Saccharomycetales(100);f__Metschnikowiaceae(100);g__Metschnikowia(100);s__Metschnikowia_pulcherrima(100); | Metschnikowia | 13 | Saprotroph | \|Undefined Saprotroph\| | Yeast | NULL | Probable | As nectar-tap_saprotroph (Põlme et al. 2020); Decay sugar-rich substrates (Põlme et al. 2020) | Põlme S, et al. 2020. Fungal Diversity 105:1-16 ((https://doi.org/10.1007/s13225-020-00466-2)) |
| Otu23128 | 1 | k__Fungi(100);p__Ascomycota(100);c__Saccharomycetes(100);o__Saccharomycetales(100);f__Metschnikowiaceae(100);g__Metschnikowia(100);s__Metschnikowia_pulcherrima(100); | Metschnikowia | 13 | Saprotroph | \|Undefined Saprotroph\| | Yeast | NULL | Probable | As nectar-tap_saprotroph (Põlme et al. 2020); Decay sugar-rich substrates (Põlme et al. 2020) | Põlme S, et al. 2020. Fungal Diversity 105:1-16 ((https://doi.org/10.1007/s13225-020-00466-2)) |
| Otu23129 | 1 | k__Fungi(100);p__Ascomycota(100);c__Saccharomycetes(100);o__Saccharomycetales(100);f__Metschnikowiaceae(100);g__Metschnikowia(100);s__Metschnikowia_pulcherrima(100); | Metschnikowia | 13 | Saprotroph | \|Undefined Saprotroph\| | Yeast | NULL | Probable | As nectar-tap_saprotroph (Põlme et al. 2020); Decay sugar-rich substrates (Põlme et al. 2020) | Põlme S, et al. 2020. Fungal Diversity 105:1-16 ((https://doi.org/10.1007/s13225-020-00466-2)) |
| Otu23130 | 1 | k__Fungi(100);p__Ascomycota(100);c__Saccharomycetes(100);o__Saccharomycetales(100);f__Metschnikowiaceae(100);g__Metschnikowia(100);s__Metschnikowia_pulcherrima(100); | Metschnikowia | 13 | Saprotroph | \|Undefined Saprotroph\| | Yeast | NULL | Probable | As nectar-tap_saprotroph (Põlme et al. 2020); Decay sugar-rich substrates (Põlme et al. 2020) | Põlme S, et al. 2020. Fungal Diversity 105:1-16 ((https://doi.org/10.1007/s13225-020-00466-2)) |
| Otu23131 | 1 | k__Fungi(100);p__Ascomycota(100);c__Saccharomycetes(100);o__Saccharomycetales(100);f__Metschnikowiaceae(100);g__Metschnikowia(100);s__Metschnikowia_pulcherrima(100); | Metschnikowia | 13 | Saprotroph | \|Undefined Saprotroph\| | Yeast | NULL | Probable | As nectar-tap_saprotroph (Põlme et al. 2020); Decay sugar-rich substrates (Põlme et al. 2020) | Põlme S, et al. 2020. Fungal Diversity 105:1-16 ((https://doi.org/10.1007/s13225-020-00466-2)) |
| Otu23132 | 1 | k__Fungi(100);p__Ascomycota(100);c__Saccharomycetes(100);o__Saccharomycetales(100);f__Metschnikowiaceae(100);g__Metschnikowia(100);s__Metschnikowia_pulcherrima(100); | Metschnikowia | 13 | Saprotroph | \|Undefined Saprotroph\| | Yeast | NULL | Probable | As nectar-tap_saprotroph (Põlme et al. 2020); Decay sugar-rich substrates (Põlme et al. 2020) | Põlme S, et al. 2020. Fungal Diversity 105:1-16 ((https://doi.org/10.1007/s13225-020-00466-2)) |
| Otu23133 | 1 | k__Fungi(100);p__Ascomycota(100);c__Saccharomycetes(100);o__Saccharomycetales(100);f__Metschnikowiaceae(100);g__Metschnikowia(100);g__Metschnikowia_unclassified(100); | Metschnikowia | 13 | Saprotroph | \|Undefined Saprotroph\| | Yeast | NULL | Probable | As nectar-tap_saprotroph (Põlme et al. 2020); Decay sugar-rich substrates (Põlme et al. 2020) | Põlme S, et al. 2020. Fungal Diversity 105:1-16 ((https://doi.org/10.1007/s13225-020-00466-2)) |
| Otu23134 | 1 | k__Fungi(100);p__Ascomycota(100);c__Saccharomycetes(100);o__Saccharomycetales(100);f__Metschnikowiaceae(100);g__Metschnikowia(100);g__Metschnikowia_unclassified(100); | Metschnikowia | 13 | Saprotroph | \|Undefined Saprotroph\| | Yeast | NULL | Probable | As nectar-tap_saprotroph (Põlme et al. 2020); Decay sugar-rich substrates (Põlme et al. 2020) | Põlme S, et al. 2020. Fungal Diversity 105:1-16 ((https://doi.org/10.1007/s13225-020-00466-2)) |
| Otu23135 | 1 | k__Fungi(100);p__Ascomycota(100);c__Saccharomycetes(100);o__Saccharomycetales(100);f__Metschnikowiaceae(100);g__Metschnikowia(100);s__Metschnikowia_pulcherrima(100); | Metschnikowia | 13 | Saprotroph | \|Undefined Saprotroph\| | Yeast | NULL | Probable | As nectar-tap_saprotroph (Põlme et al. 2020); Decay sugar-rich substrates (Põlme et al. 2020) | Põlme S, et al. 2020. Fungal Diversity 105:1-16 ((https://doi.org/10.1007/s13225-020-00466-2)) |
| Otu23137 | 1 | k__Fungi(100);p__Ascomycota(100);c__Saccharomycetes(100);o__Saccharomycetales(100);f__Metschnikowiaceae(100);g__Metschnikowia(100);s__Metschnikowia_pulcherrima(100); | Metschnikowia | 13 | Saprotroph | \|Undefined Saprotroph\| | Yeast | NULL | Probable | As nectar-tap_saprotroph (Põlme et al. 2020); Decay sugar-rich substrates (Põlme et al. 2020) | Põlme S, et al. 2020. Fungal Diversity 105:1-16 ((https://doi.org/10.1007/s13225-020-00466-2)) |
| Otu23138 | 1 | k__Fungi(100);p__Ascomycota(100);c__Saccharomycetes(100);o__Saccharomycetales(100);f__Metschnikowiaceae(100);g__Metschnikowia(100);s__Metschnikowia_pulcherrima(100); | Metschnikowia | 13 | Saprotroph | \|Undefined Saprotroph\| | Yeast | NULL | Probable | As nectar-tap_saprotroph (Põlme et al. 2020); Decay sugar-rich substrates (Põlme et al. 2020) | Põlme S, et al. 2020. Fungal Diversity 105:1-16 ((https://doi.org/10.1007/s13225-020-00466-2)) |
| Otu23139 | 1 | k__Fungi(100);p__Ascomycota(100);c__Saccharomycetes(100);o__Saccharomycetales(100);f__Saccharomycetales_family_Incertae_sedis(100);g__Candida(100);g__Candida_unclassified(100); | Candida | 13 | Pathotroph-Saprotroph-Symbiotroph | Animal Pathogen-Endophyte-Endosymbiont-Epiphyte-\|Undefined Saprotroph\| | Dimorphic Yeast-Yeast | NULL | Probable | As nectar-tap_saprotroph (Põlme et al. 2020); Decay sugar-rich substrates (Põlme et al. 2020); This asexual genus is polyphyletic and contain species that function very differently depend on what they are. Caution should be taken when interpreting guilds for Candida species. | Manolakaki D, et al. 2010. Virulence. 1: 367-375 ((https://doi.org/10.4161/viru.1.5.12796)); Mekbib SB, et al. 2011. Tropical Plant Pathology 36:233-240 ((https://doi.org/10.1590/S1982-56762011000400004)); Põlme S, et al. 2020. Fungal Diversity 105:1-16 ((https://doi.org/10.1007/s13225-020-00466-2)) |
| Otu23140 | 1 | k__Fungi(100);p__Ascomycota(100);c__Saccharomycetes(100);o__Saccharomycetales(100);f__Metschnikowiaceae(100);g__Metschnikowia(100);g__Metschnikowia_unclassified(100); | Metschnikowia | 13 | Saprotroph | \|Undefined Saprotroph\| | Yeast | NULL | Probable | As nectar-tap_saprotroph (Põlme et al. 2020); Decay sugar-rich substrates (Põlme et al. 2020) | Põlme S, et al. 2020. Fungal Diversity 105:1-16 ((https://doi.org/10.1007/s13225-020-00466-2)) |
| Otu23141 | 1 | k__Fungi(100);p__Ascomycota(100);c__Saccharomycetes(100);o__Saccharomycetales(100);f__Metschnikowiaceae(100);g__Metschnikowia(100);s__Metschnikowia_pulcherrima(100); | Metschnikowia | 13 | Saprotroph | \|Undefined Saprotroph\| | Yeast | NULL | Probable | As nectar-tap_saprotroph (Põlme et al. 2020); Decay sugar-rich substrates (Põlme et al. 2020) | Põlme S, et al. 2020. Fungal Diversity 105:1-16 ((https://doi.org/10.1007/s13225-020-00466-2)) |
| Otu23142 | 1 | k__Fungi(100);p__Ascomycota(100);c__Leotiomycetes(100);o__Helotiales(100);f__Hyaloscyphaceae(100);g__Lachnum(100);s__Lachnum_virgineum(100); | Lachnum | 13 | Saprotroph-Symbiotroph | Endophyte-\|Plant Saprotroph\|-Undefined Saprotroph-Wood Saprotroph | Microfungus | NULL | Probable | As litter_saprotroph (Põlme et al. 2020); As root_endophyte (Põlme et al. 2020); Decay wood, leaf, fruit, seed (Põlme et al. 2020) | Tedersoo L, et al. 2014. Science 346:e1256688 ((https://doi.org/10.1126/science.1256688)); Põlme S, et al. 2020. Fungal Diversity 105:1-16 ((https://doi.org/10.1007/s13225-020-00466-2)) |
| Otu23143 | 1 | k__Fungi(100);p__Ascomycota(100);c__Saccharomycetes(100);o__Saccharomycetales(100);f__Metschnikowiaceae(100);g__Metschnikowia(100);s__Metschnikowia_pulcherrima(100); | Metschnikowia | 13 | Saprotroph | \|Undefined Saprotroph\| | Yeast | NULL | Probable | As nectar-tap_saprotroph (Põlme et al. 2020); Decay sugar-rich substrates (Põlme et al. 2020) | Põlme S, et al. 2020. Fungal Diversity 105:1-16 ((https://doi.org/10.1007/s13225-020-00466-2)) |
| Otu23144 | 1 | k__Fungi(100);p__Ascomycota(100);c__Saccharomycetes(100);o__Saccharomycetales(100);f__Saccharomycetales_family_Incertae_sedis(100);f__Saccharomycetales_family_Incertae_sedis_unclassified(100);f__Saccharomycetales_family_Incertae_sedis_unclassified(100); | Saccharomycetales | 7 | Saprotroph | Undefined Saprotroph | Yeast | NULL | Possible | NULL | Sterkenburg E, et al. 2015. New Phytologist 207:1145-1158 ((https://doi.org/10.1111/nph.13426)) |
| Otu23145 | 1 | k__Fungi(100);p__Ascomycota(100);c__Saccharomycetes(100);o__Saccharomycetales(100);f__Metschnikowiaceae(100);g__Metschnikowia(100);g__Metschnikowia_unclassified(100); | Metschnikowia | 13 | Saprotroph | \|Undefined Saprotroph\| | Yeast | NULL | Probable | As nectar-tap_saprotroph (Põlme et al. 2020); Decay sugar-rich substrates (Põlme et al. 2020) | Põlme S, et al. 2020. Fungal Diversity 105:1-16 ((https://doi.org/10.1007/s13225-020-00466-2)) |
| Otu23146 | 1 | k__Fungi(100);p__Ascomycota(100);c__Saccharomycetes(100);o__Saccharomycetales(100);f__Metschnikowiaceae(100);g__Metschnikowia(100);s__Metschnikowia_pulcherrima(100); | Metschnikowia | 13 | Saprotroph | \|Undefined Saprotroph\| | Yeast | NULL | Probable | As nectar-tap_saprotroph (Põlme et al. 2020); Decay sugar-rich substrates (Põlme et al. 2020) | Põlme S, et al. 2020. Fungal Diversity 105:1-16 ((https://doi.org/10.1007/s13225-020-00466-2)) |
| Otu23147 | 1 | k__Fungi(100);p__Ascomycota(100);c__Saccharomycetes(100);o__Saccharomycetales(100);f__Metschnikowiaceae(100);g__Metschnikowia(100);s__Metschnikowia_pulcherrima(100); | Metschnikowia | 13 | Saprotroph | \|Undefined Saprotroph\| | Yeast | NULL | Probable | As nectar-tap_saprotroph (Põlme et al. 2020); Decay sugar-rich substrates (Põlme et al. 2020) | Põlme S, et al. 2020. Fungal Diversity 105:1-16 ((https://doi.org/10.1007/s13225-020-00466-2)) |
| Otu23148 | 1 | k__Fungi(100);p__Basidiomycota(100);c__Tremellomycetes(100);o__Tremellales(100);f__Tremellales_family_Incertae_sedis(100);g__Cryptococcus(100);s__Cryptococcus_sp_JCM_11356(100); | Cryptococcus | 13 | Pathotroph-Saprotroph-Symbiotroph | Animal Pathogen-Endophyte-Epiphyte-\|Undefined Saprotroph\| | Dimorphic Yeast-Yeast | NULL | Possible | As foliar_endophyte (Põlme et al. 2020); Decay leaf, fruit, seed, soil, dung, animal material (Põlme et al. 2020) | Andrews JH, et al. 1983. Phytopathology 73:228-234 ((https://doi.org/10.1094/Phyto-73-228)); Kurtzman CP, et al. (eds.) 2011. The Yeasts, a Taxonomic Study. Fifth Edition. Vols 1-3. Elsevier, San Diego (ISBN: 9780444521491); Põlme S, et al. 2020. Fungal Diversity 105:1-16 ((https://doi.org/10.1007/s13225-020-00466-2)) |
| Otu23164 | 1 | k__Fungi(100);p__Ascomycota(100);c__Saccharomycetes(100);o__Saccharomycetales(100);f__Metschnikowiaceae(100);g__Metschnikowia(100);g__Metschnikowia_unclassified(100); | Metschnikowia | 13 | Saprotroph | \|Undefined Saprotroph\| | Yeast | NULL | Probable | As nectar-tap_saprotroph (Põlme et al. 2020); Decay sugar-rich substrates (Põlme et al. 2020) | Põlme S, et al. 2020. Fungal Diversity 105:1-16 ((https://doi.org/10.1007/s13225-020-00466-2)) |
| Otu23166 | 1 | k__Fungi(100);p__Ascomycota(100);c__Saccharomycetes(100);o__Saccharomycetales(100);f__Metschnikowiaceae(100);g__Metschnikowia(100);s__Metschnikowia_pulcherrima(100); | Metschnikowia | 13 | Saprotroph | \|Undefined Saprotroph\| | Yeast | NULL | Probable | As nectar-tap_saprotroph (Põlme et al. 2020); Decay sugar-rich substrates (Põlme et al. 2020) | Põlme S, et al. 2020. Fungal Diversity 105:1-16 ((https://doi.org/10.1007/s13225-020-00466-2)) |
| Otu23167 | 1 | k__Fungi(100);p__Ascomycota(100);c__Saccharomycetes(100);o__Saccharomycetales(100);f__Metschnikowiaceae(100);g__Metschnikowia(100);s__Metschnikowia_pulcherrima(100); | Metschnikowia | 13 | Saprotroph | \|Undefined Saprotroph\| | Yeast | NULL | Probable | As nectar-tap_saprotroph (Põlme et al. 2020); Decay sugar-rich substrates (Põlme et al. 2020) | Põlme S, et al. 2020. Fungal Diversity 105:1-16 ((https://doi.org/10.1007/s13225-020-00466-2)) |
| Otu23168 | 1 | k__Fungi(100);p__Ascomycota(100);c__Saccharomycetes(100);o__Saccharomycetales(100);f__Metschnikowiaceae(100);g__Metschnikowia(100);s__Metschnikowia_pulcherrima(100); | Metschnikowia | 13 | Saprotroph | \|Undefined Saprotroph\| | Yeast | NULL | Probable | As nectar-tap_saprotroph (Põlme et al. 2020); Decay sugar-rich substrates (Põlme et al. 2020) | Põlme S, et al. 2020. Fungal Diversity 105:1-16 ((https://doi.org/10.1007/s13225-020-00466-2)) |
| Otu23169 | 1 | k__Fungi(100);p__Ascomycota(100);c__Saccharomycetes(100);o__Saccharomycetales(100);f__Metschnikowiaceae(100);g__Metschnikowia(100);s__Metschnikowia_pulcherrima(100); | Metschnikowia | 13 | Saprotroph | \|Undefined Saprotroph\| | Yeast | NULL | Probable | As nectar-tap_saprotroph (Põlme et al. 2020); Decay sugar-rich substrates (Põlme et al. 2020) | Põlme S, et al. 2020. Fungal Diversity 105:1-16 ((https://doi.org/10.1007/s13225-020-00466-2)) |
| Otu23172 | 1 | k__Fungi(100);p__Ascomycota(100);c__Saccharomycetes(100);o__Saccharomycetales(100);f__Metschnikowiaceae(100);g__Metschnikowia(100);s__Metschnikowia_pulcherrima(100); | Metschnikowia | 13 | Saprotroph | \|Undefined Saprotroph\| | Yeast | NULL | Probable | As nectar-tap_saprotroph (Põlme et al. 2020); Decay sugar-rich substrates (Põlme et al. 2020) | Põlme S, et al. 2020. Fungal Diversity 105:1-16 ((https://doi.org/10.1007/s13225-020-00466-2)) |
| Otu23173 | 1 | k__Fungi(100);p__Ascomycota(100);c__Saccharomycetes(100);o__Saccharomycetales(100);f__Metschnikowiaceae(100);g__Metschnikowia(100);s__Metschnikowia_pulcherrima(100); | Metschnikowia | 13 | Saprotroph | \|Undefined Saprotroph\| | Yeast | NULL | Probable | As nectar-tap_saprotroph (Põlme et al. 2020); Decay sugar-rich substrates (Põlme et al. 2020) | Põlme S, et al. 2020. Fungal Diversity 105:1-16 ((https://doi.org/10.1007/s13225-020-00466-2)) |
| Otu23174 | 1 | k__Fungi(100);p__Ascomycota(100);c__Saccharomycetes(100);o__Saccharomycetales(100);f__Metschnikowiaceae(100);g__Metschnikowia(100);g__Metschnikowia_unclassified(100); | Metschnikowia | 13 | Saprotroph | \|Undefined Saprotroph\| | Yeast | NULL | Probable | As nectar-tap_saprotroph (Põlme et al. 2020); Decay sugar-rich substrates (Põlme et al. 2020) | Põlme S, et al. 2020. Fungal Diversity 105:1-16 ((https://doi.org/10.1007/s13225-020-00466-2)) |
| Otu23176 | 1 | k__Fungi(100);p__Ascomycota(100);c__Dothideomycetes(100);o__Capnodiales(100);f__Mycosphaerellaceae(100);g__Pseudocercosporella(100);s__Pseudocercosporella_chaenomelis(100); | Pseudocercosporella | 13 | Pathotroph | \|Plant Pathogen\| | Microfungus | NULL | Probable | NULL | Tedersoo L, et al. 2014. Science 346:e1256688 ((https://doi.org/10.1126/science.1256688)); Põlme S, et al. 2020. Fungal Diversity 105:1-16 ((https://doi.org/10.1007/s13225-020-00466-2)) |
| Otu23177 | 1 | k__Fungi(100);p__Ascomycota(100);c__Saccharomycetes(100);o__Saccharomycetales(100);f__Metschnikowiaceae(100);g__Metschnikowia(100);s__Metschnikowia_pulcherrima(100); | Metschnikowia | 13 | Saprotroph | \|Undefined Saprotroph\| | Yeast | NULL | Probable | As nectar-tap_saprotroph (Põlme et al. 2020); Decay sugar-rich substrates (Põlme et al. 2020) | Põlme S, et al. 2020. Fungal Diversity 105:1-16 ((https://doi.org/10.1007/s13225-020-00466-2)) |
| Otu23178 | 1 | k__Fungi(100);p__Ascomycota(100);c__Saccharomycetes(100);o__Saccharomycetales(100);f__Saccharomycetales_family_Incertae_sedis(100);g__Candida(100);g__Candida_unclassified(100); | Candida | 13 | Pathotroph-Saprotroph-Symbiotroph | Animal Pathogen-Endophyte-Endosymbiont-Epiphyte-\|Undefined Saprotroph\| | Dimorphic Yeast-Yeast | NULL | Probable | As nectar-tap_saprotroph (Põlme et al. 2020); Decay sugar-rich substrates (Põlme et al. 2020); This asexual genus is polyphyletic and contain species that function very differently depend on what they are. Caution should be taken when interpreting guilds for Candida species. | Manolakaki D, et al. 2010. Virulence. 1: 367-375 ((https://doi.org/10.4161/viru.1.5.12796)); Mekbib SB, et al. 2011. Tropical Plant Pathology 36:233-240 ((https://doi.org/10.1590/S1982-56762011000400004)); Põlme S, et al. 2020. Fungal Diversity 105:1-16 ((https://doi.org/10.1007/s13225-020-00466-2)) |
| Otu23179 | 1 | k__Fungi(100);p__Ascomycota(100);c__Saccharomycetes(100);o__Saccharomycetales(100);f__Metschnikowiaceae(100);g__Metschnikowia(100);s__Metschnikowia_pulcherrima(100); | Metschnikowia | 13 | Saprotroph | \|Undefined Saprotroph\| | Yeast | NULL | Probable | As nectar-tap_saprotroph (Põlme et al. 2020); Decay sugar-rich substrates (Põlme et al. 2020) | Põlme S, et al. 2020. Fungal Diversity 105:1-16 ((https://doi.org/10.1007/s13225-020-00466-2)) |
| Otu23180 | 1 | k__Fungi(100);p__Ascomycota(100);c__Saccharomycetes(100);o__Saccharomycetales(100);f__Metschnikowiaceae(100);g__Metschnikowia(100);g__Metschnikowia_unclassified(100); | Metschnikowia | 13 | Saprotroph | \|Undefined Saprotroph\| | Yeast | NULL | Probable | As nectar-tap_saprotroph (Põlme et al. 2020); Decay sugar-rich substrates (Põlme et al. 2020) | Põlme S, et al. 2020. Fungal Diversity 105:1-16 ((https://doi.org/10.1007/s13225-020-00466-2)) |
| Otu23181 | 1 | k__Fungi(100);p__Ascomycota(100);c__Saccharomycetes(100);o__Saccharomycetales(100);f__Metschnikowiaceae(100);g__Metschnikowia(100);s__Metschnikowia_pulcherrima(100); | Metschnikowia | 13 | Saprotroph | \|Undefined Saprotroph\| | Yeast | NULL | Probable | As nectar-tap_saprotroph (Põlme et al. 2020); Decay sugar-rich substrates (Põlme et al. 2020) | Põlme S, et al. 2020. Fungal Diversity 105:1-16 ((https://doi.org/10.1007/s13225-020-00466-2)) |
| Otu23182 | 1 | k__Fungi(100);p__Ascomycota(100);c__Saccharomycetes(100);o__Saccharomycetales(100);f__Metschnikowiaceae(100);g__Metschnikowia(100);s__Metschnikowia_pulcherrima(100); | Metschnikowia | 13 | Saprotroph | \|Undefined Saprotroph\| | Yeast | NULL | Probable | As nectar-tap_saprotroph (Põlme et al. 2020); Decay sugar-rich substrates (Põlme et al. 2020) | Põlme S, et al. 2020. Fungal Diversity 105:1-16 ((https://doi.org/10.1007/s13225-020-00466-2)) |
| Otu23184 | 1 | k__Fungi(100);p__Ascomycota(100);c__Saccharomycetes(100);o__Saccharomycetales(100);f__Metschnikowiaceae(100);g__Metschnikowia(100);s__Metschnikowia_pulcherrima(100); | Metschnikowia | 13 | Saprotroph | \|Undefined Saprotroph\| | Yeast | NULL | Probable | As nectar-tap_saprotroph (Põlme et al. 2020); Decay sugar-rich substrates (Põlme et al. 2020) | Põlme S, et al. 2020. Fungal Diversity 105:1-16 ((https://doi.org/10.1007/s13225-020-00466-2)) |
| Otu23186 | 1 | k__Fungi(100);p__Ascomycota(100);c__Saccharomycetes(100);o__Saccharomycetales(100);f__Metschnikowiaceae(100);g__Metschnikowia(100);s__Metschnikowia_pulcherrima(100); | Metschnikowia | 13 | Saprotroph | \|Undefined Saprotroph\| | Yeast | NULL | Probable | As nectar-tap_saprotroph (Põlme et al. 2020); Decay sugar-rich substrates (Põlme et al. 2020) | Põlme S, et al. 2020. Fungal Diversity 105:1-16 ((https://doi.org/10.1007/s13225-020-00466-2)) |
| Otu23187 | 1 | k__Fungi(100);p__Ascomycota(100);c__Saccharomycetes(100);o__Saccharomycetales(100);f__Metschnikowiaceae(100);g__Metschnikowia(100);s__Metschnikowia_pulcherrima(100); | Metschnikowia | 13 | Saprotroph | \|Undefined Saprotroph\| | Yeast | NULL | Probable | As nectar-tap_saprotroph (Põlme et al. 2020); Decay sugar-rich substrates (Põlme et al. 2020) | Põlme S, et al. 2020. Fungal Diversity 105:1-16 ((https://doi.org/10.1007/s13225-020-00466-2)) |
| Otu23188 | 1 | k__Fungi(100);p__Ascomycota(100);c__Saccharomycetes(100);o__Saccharomycetales(100);f__Metschnikowiaceae(100);g__Metschnikowia(100);s__Metschnikowia_pulcherrima(100); | Metschnikowia | 13 | Saprotroph | \|Undefined Saprotroph\| | Yeast | NULL | Probable | As nectar-tap_saprotroph (Põlme et al. 2020); Decay sugar-rich substrates (Põlme et al. 2020) | Põlme S, et al. 2020. Fungal Diversity 105:1-16 ((https://doi.org/10.1007/s13225-020-00466-2)) |
| Otu23190 | 1 | k__Fungi(100);p__Ascomycota(100);c__Saccharomycetes(100);o__Saccharomycetales(100);f__Metschnikowiaceae(100);g__Metschnikowia(100);s__Metschnikowia_pulcherrima(100); | Metschnikowia | 13 | Saprotroph | \|Undefined Saprotroph\| | Yeast | NULL | Probable | As nectar-tap_saprotroph (Põlme et al. 2020); Decay sugar-rich substrates (Põlme et al. 2020) | Põlme S, et al. 2020. Fungal Diversity 105:1-16 ((https://doi.org/10.1007/s13225-020-00466-2)) |
| Otu23191 | 1 | k__Fungi(100);p__Ascomycota(100);c__Saccharomycetes(100);o__Saccharomycetales(100);f__Metschnikowiaceae(100);g__Metschnikowia(100);s__Metschnikowia_pulcherrima(100); | Metschnikowia | 13 | Saprotroph | \|Undefined Saprotroph\| | Yeast | NULL | Probable | As nectar-tap_saprotroph (Põlme et al. 2020); Decay sugar-rich substrates (Põlme et al. 2020) | Põlme S, et al. 2020. Fungal Diversity 105:1-16 ((https://doi.org/10.1007/s13225-020-00466-2)) |
| Otu23192 | 1 | k__Fungi(100);p__Ascomycota(100);c__Saccharomycetes(100);o__Saccharomycetales(100);f__Metschnikowiaceae(100);g__Metschnikowia(100);s__Metschnikowia_pulcherrima(100); | Metschnikowia | 13 | Saprotroph | \|Undefined Saprotroph\| | Yeast | NULL | Probable | As nectar-tap_saprotroph (Põlme et al. 2020); Decay sugar-rich substrates (Põlme et al. 2020) | Põlme S, et al. 2020. Fungal Diversity 105:1-16 ((https://doi.org/10.1007/s13225-020-00466-2)) |
| Otu23193 | 1 | k__Fungi(100);p__Ascomycota(100);c__Saccharomycetes(100);o__Saccharomycetales(100);f__Metschnikowiaceae(100);g__Metschnikowia(100);s__Metschnikowia_pulcherrima(100); | Metschnikowia | 13 | Saprotroph | \|Undefined Saprotroph\| | Yeast | NULL | Probable | As nectar-tap_saprotroph (Põlme et al. 2020); Decay sugar-rich substrates (Põlme et al. 2020) | Põlme S, et al. 2020. Fungal Diversity 105:1-16 ((https://doi.org/10.1007/s13225-020-00466-2)) |
| Otu23194 | 1 | k__Fungi(100);p__Ascomycota(100);c__Saccharomycetes(100);o__Saccharomycetales(100);f__Metschnikowiaceae(100);g__Metschnikowia(100);s__Metschnikowia_pulcherrima(100); | Metschnikowia | 13 | Saprotroph | \|Undefined Saprotroph\| | Yeast | NULL | Probable | As nectar-tap_saprotroph (Põlme et al. 2020); Decay sugar-rich substrates (Põlme et al. 2020) | Põlme S, et al. 2020. Fungal Diversity 105:1-16 ((https://doi.org/10.1007/s13225-020-00466-2)) |
| Otu23195 | 1 | k__Fungi(100);p__Ascomycota(100);c__Saccharomycetes(100);o__Saccharomycetales(100);f__Metschnikowiaceae(100);g__Metschnikowia(100);s__Metschnikowia_pulcherrima(100); | Metschnikowia | 13 | Saprotroph | \|Undefined Saprotroph\| | Yeast | NULL | Probable | As nectar-tap_saprotroph (Põlme et al. 2020); Decay sugar-rich substrates (Põlme et al. 2020) | Põlme S, et al. 2020. Fungal Diversity 105:1-16 ((https://doi.org/10.1007/s13225-020-00466-2)) |
| Otu23196 | 1 | k__Fungi(100);p__Ascomycota(100);c__Saccharomycetes(100);o__Saccharomycetales(100);f__Metschnikowiaceae(100);g__Metschnikowia(100);s__Metschnikowia_pulcherrima(100); | Metschnikowia | 13 | Saprotroph | \|Undefined Saprotroph\| | Yeast | NULL | Probable | As nectar-tap_saprotroph (Põlme et al. 2020); Decay sugar-rich substrates (Põlme et al. 2020) | Põlme S, et al. 2020. Fungal Diversity 105:1-16 ((https://doi.org/10.1007/s13225-020-00466-2)) |
| Otu23197 | 1 | k__Fungi(100);p__Ascomycota(100);c__Saccharomycetes(100);o__Saccharomycetales(100);f__Metschnikowiaceae(100);g__Metschnikowia(100);s__Metschnikowia_pulcherrima(100); | Metschnikowia | 13 | Saprotroph | \|Undefined Saprotroph\| | Yeast | NULL | Probable | As nectar-tap_saprotroph (Põlme et al. 2020); Decay sugar-rich substrates (Põlme et al. 2020) | Põlme S, et al. 2020. Fungal Diversity 105:1-16 ((https://doi.org/10.1007/s13225-020-00466-2)) |
| Otu23198 | 1 | k__Fungi(100);p__Ascomycota(100);c__Saccharomycetes(100);o__Saccharomycetales(100);f__Metschnikowiaceae(100);g__Metschnikowia(100);s__Metschnikowia_pulcherrima(100); | Metschnikowia | 13 | Saprotroph | \|Undefined Saprotroph\| | Yeast | NULL | Probable | As nectar-tap_saprotroph (Põlme et al. 2020); Decay sugar-rich substrates (Põlme et al. 2020) | Põlme S, et al. 2020. Fungal Diversity 105:1-16 ((https://doi.org/10.1007/s13225-020-00466-2)) |
| Otu23199 | 1 | k__Fungi(100);p__Ascomycota(100);c__Saccharomycetes(100);o__Saccharomycetales(100);f__Metschnikowiaceae(100);g__Metschnikowia(100);g__Metschnikowia_unclassified(100); | Metschnikowia | 13 | Saprotroph | \|Undefined Saprotroph\| | Yeast | NULL | Probable | As nectar-tap_saprotroph (Põlme et al. 2020); Decay sugar-rich substrates (Põlme et al. 2020) | Põlme S, et al. 2020. Fungal Diversity 105:1-16 ((https://doi.org/10.1007/s13225-020-00466-2)) |
| Otu23200 | 1 | k__Fungi(100);p__Ascomycota(100);c__Saccharomycetes(100);o__Saccharomycetales(100);f__Metschnikowiaceae(100);g__Metschnikowia(100);g__Metschnikowia_unclassified(100); | Metschnikowia | 13 | Saprotroph | \|Undefined Saprotroph\| | Yeast | NULL | Probable | As nectar-tap_saprotroph (Põlme et al. 2020); Decay sugar-rich substrates (Põlme et al. 2020) | Põlme S, et al. 2020. Fungal Diversity 105:1-16 ((https://doi.org/10.1007/s13225-020-00466-2)) |
| Otu23201 | 1 | k__Fungi(100);p__Ascomycota(100);c__Saccharomycetes(100);o__Saccharomycetales(100);f__Metschnikowiaceae(100);g__Metschnikowia(100);s__Metschnikowia_pulcherrima(100); | Metschnikowia | 13 | Saprotroph | \|Undefined Saprotroph\| | Yeast | NULL | Probable | As nectar-tap_saprotroph (Põlme et al. 2020); Decay sugar-rich substrates (Põlme et al. 2020) | Põlme S, et al. 2020. Fungal Diversity 105:1-16 ((https://doi.org/10.1007/s13225-020-00466-2)) |
| Otu23203 | 1 | k__Fungi(100);p__Ascomycota(100);c__Saccharomycetes(100);o__Saccharomycetales(100);f__Metschnikowiaceae(100);g__Metschnikowia(100);s__Metschnikowia_pulcherrima(100); | Metschnikowia | 13 | Saprotroph | \|Undefined Saprotroph\| | Yeast | NULL | Probable | As nectar-tap_saprotroph (Põlme et al. 2020); Decay sugar-rich substrates (Põlme et al. 2020) | Põlme S, et al. 2020. Fungal Diversity 105:1-16 ((https://doi.org/10.1007/s13225-020-00466-2)) |
| Otu23204 | 1 | k__Fungi(100);p__Ascomycota(100);c__Saccharomycetes(100);o__Saccharomycetales(100);f__Metschnikowiaceae(100);g__Metschnikowia(100);g__Metschnikowia_unclassified(100); | Metschnikowia | 13 | Saprotroph | \|Undefined Saprotroph\| | Yeast | NULL | Probable | As nectar-tap_saprotroph (Põlme et al. 2020); Decay sugar-rich substrates (Põlme et al. 2020) | Põlme S, et al. 2020. Fungal Diversity 105:1-16 ((https://doi.org/10.1007/s13225-020-00466-2)) |
| Otu23205 | 1 | k__Fungi(100);p__Ascomycota(100);c__Saccharomycetes(100);o__Saccharomycetales(100);f__Metschnikowiaceae(100);g__Metschnikowia(100);s__Metschnikowia_pulcherrima(100); | Metschnikowia | 13 | Saprotroph | \|Undefined Saprotroph\| | Yeast | NULL | Probable | As nectar-tap_saprotroph (Põlme et al. 2020); Decay sugar-rich substrates (Põlme et al. 2020) | Põlme S, et al. 2020. Fungal Diversity 105:1-16 ((https://doi.org/10.1007/s13225-020-00466-2)) |
| Otu23206 | 1 | k__Fungi(100);p__Ascomycota(100);c__Saccharomycetes(100);o__Saccharomycetales(100);f__Metschnikowiaceae(100);g__Metschnikowia(100);s__Metschnikowia_pulcherrima(100); | Metschnikowia | 13 | Saprotroph | \|Undefined Saprotroph\| | Yeast | NULL | Probable | As nectar-tap_saprotroph (Põlme et al. 2020); Decay sugar-rich substrates (Põlme et al. 2020) | Põlme S, et al. 2020. Fungal Diversity 105:1-16 ((https://doi.org/10.1007/s13225-020-00466-2)) |
| Otu23207 | 1 | k__Fungi(100);p__Ascomycota(100);c__Saccharomycetes(100);o__Saccharomycetales(100);f__Metschnikowiaceae(100);g__Metschnikowia(100);s__Metschnikowia_pulcherrima(100); | Metschnikowia | 13 | Saprotroph | \|Undefined Saprotroph\| | Yeast | NULL | Probable | As nectar-tap_saprotroph (Põlme et al. 2020); Decay sugar-rich substrates (Põlme et al. 2020) | Põlme S, et al. 2020. Fungal Diversity 105:1-16 ((https://doi.org/10.1007/s13225-020-00466-2)) |
| Otu23208 | 1 | k__Fungi(100);p__Ascomycota(100);c__Saccharomycetes(100);o__Saccharomycetales(100);f__Metschnikowiaceae(100);g__Metschnikowia(100);s__Metschnikowia_pulcherrima(100); | Metschnikowia | 13 | Saprotroph | \|Undefined Saprotroph\| | Yeast | NULL | Probable | As nectar-tap_saprotroph (Põlme et al. 2020); Decay sugar-rich substrates (Põlme et al. 2020) | Põlme S, et al. 2020. Fungal Diversity 105:1-16 ((https://doi.org/10.1007/s13225-020-00466-2)) |
| Otu23209 | 1 | k__Fungi(100);p__Ascomycota(100);c__Dothideomycetes(100);o__Dothideomycetes_order_Incertae_sedis(100);f__Pseudeurotiaceae(100);g__Pseudeurotium(100);g__Pseudeurotium_unclassified(100); | Pseudeurotium | 13 | Saprotroph | \|Dung Saprotroph\|-Undefined Saprotroph | Microfungus | NULL | Probable | As soil_saprotroph (Põlme et al. 2020); Decay soil, dung (Põlme et al. 2020) | Tedersoo L, et al. 2014. Science 346:e1256688 ((https://doi.org/10.1126/science.1256688)); Põlme S, et al. 2020. Fungal Diversity 105:1-16 ((https://doi.org/10.1007/s13225-020-00466-2)) |
| Otu23210 | 1 | k__Fungi(100);p__Ascomycota(100);c__Saccharomycetes(100);o__Saccharomycetales(100);f__Metschnikowiaceae(100);g__Metschnikowia(100);s__Metschnikowia_pulcherrima(100); | Metschnikowia | 13 | Saprotroph | \|Undefined Saprotroph\| | Yeast | NULL | Probable | As nectar-tap_saprotroph (Põlme et al. 2020); Decay sugar-rich substrates (Põlme et al. 2020) | Põlme S, et al. 2020. Fungal Diversity 105:1-16 ((https://doi.org/10.1007/s13225-020-00466-2)) |
| Otu23211 | 1 | k__Fungi(100);p__Ascomycota(100);c__Saccharomycetes(100);o__Saccharomycetales(100);f__Metschnikowiaceae(100);g__Metschnikowia(100);g__Metschnikowia_unclassified(100); | Metschnikowia | 13 | Saprotroph | \|Undefined Saprotroph\| | Yeast | NULL | Probable | As nectar-tap_saprotroph (Põlme et al. 2020); Decay sugar-rich substrates (Põlme et al. 2020) | Põlme S, et al. 2020. Fungal Diversity 105:1-16 ((https://doi.org/10.1007/s13225-020-00466-2)) |
| Otu23212 | 1 | k__Fungi(100);p__Ascomycota(100);c__Saccharomycetes(100);o__Saccharomycetales(100);f__Metschnikowiaceae(100);g__Metschnikowia(100);s__Metschnikowia_pulcherrima(100); | Metschnikowia | 13 | Saprotroph | \|Undefined Saprotroph\| | Yeast | NULL | Probable | As nectar-tap_saprotroph (Põlme et al. 2020); Decay sugar-rich substrates (Põlme et al. 2020) | Põlme S, et al. 2020. Fungal Diversity 105:1-16 ((https://doi.org/10.1007/s13225-020-00466-2)) |
| Otu23213 | 1 | k__Fungi(100);p__Ascomycota(100);c__Saccharomycetes(100);o__Saccharomycetales(100);f__Metschnikowiaceae(100);g__Metschnikowia(100);s__Metschnikowia_pulcherrima(100); | Metschnikowia | 13 | Saprotroph | \|Undefined Saprotroph\| | Yeast | NULL | Probable | As nectar-tap_saprotroph (Põlme et al. 2020); Decay sugar-rich substrates (Põlme et al. 2020) | Põlme S, et al. 2020. Fungal Diversity 105:1-16 ((https://doi.org/10.1007/s13225-020-00466-2)) |
| Otu23214 | 1 | k__Fungi(100);p__Ascomycota(100);c__Saccharomycetes(100);o__Saccharomycetales(100);f__Metschnikowiaceae(100);g__Metschnikowia(100);s__Metschnikowia_pulcherrima(100); | Metschnikowia | 13 | Saprotroph | \|Undefined Saprotroph\| | Yeast | NULL | Probable | As nectar-tap_saprotroph (Põlme et al. 2020); Decay sugar-rich substrates (Põlme et al. 2020) | Põlme S, et al. 2020. Fungal Diversity 105:1-16 ((https://doi.org/10.1007/s13225-020-00466-2)) |
| Otu23215 | 1 | k__Fungi(100);p__Ascomycota(100);c__Saccharomycetes(100);o__Saccharomycetales(100);f__Metschnikowiaceae(100);g__Metschnikowia(100);s__Metschnikowia_pulcherrima(100); | Metschnikowia | 13 | Saprotroph | \|Undefined Saprotroph\| | Yeast | NULL | Probable | As nectar-tap_saprotroph (Põlme et al. 2020); Decay sugar-rich substrates (Põlme et al. 2020) | Põlme S, et al. 2020. Fungal Diversity 105:1-16 ((https://doi.org/10.1007/s13225-020-00466-2)) |
| Otu23216 | 1 | k__Fungi(100);p__Ascomycota(100);c__Saccharomycetes(100);o__Saccharomycetales(100);f__Metschnikowiaceae(100);g__Metschnikowia(100);g__Metschnikowia_unclassified(100); | Metschnikowia | 13 | Saprotroph | \|Undefined Saprotroph\| | Yeast | NULL | Probable | As nectar-tap_saprotroph (Põlme et al. 2020); Decay sugar-rich substrates (Põlme et al. 2020) | Põlme S, et al. 2020. Fungal Diversity 105:1-16 ((https://doi.org/10.1007/s13225-020-00466-2)) |
| Otu23217 | 1 | k__Fungi(100);p__Ascomycota(100);c__Saccharomycetes(100);o__Saccharomycetales(100);f__Metschnikowiaceae(100);g__Metschnikowia(100);s__Metschnikowia_pulcherrima(100); | Metschnikowia | 13 | Saprotroph | \|Undefined Saprotroph\| | Yeast | NULL | Probable | As nectar-tap_saprotroph (Põlme et al. 2020); Decay sugar-rich substrates (Põlme et al. 2020) | Põlme S, et al. 2020. Fungal Diversity 105:1-16 ((https://doi.org/10.1007/s13225-020-00466-2)) |
| Otu23218 | 1 | k__Fungi(100);p__Ascomycota(100);c__Saccharomycetes(100);o__Saccharomycetales(100);f__Metschnikowiaceae(100);g__Metschnikowia(100);s__Metschnikowia_pulcherrima(100); | Metschnikowia | 13 | Saprotroph | \|Undefined Saprotroph\| | Yeast | NULL | Probable | As nectar-tap_saprotroph (Põlme et al. 2020); Decay sugar-rich substrates (Põlme et al. 2020) | Põlme S, et al. 2020. Fungal Diversity 105:1-16 ((https://doi.org/10.1007/s13225-020-00466-2)) |
| Otu23219 | 1 | k__Fungi(100);p__Ascomycota(100);c__Saccharomycetes(100);o__Saccharomycetales(100);f__Metschnikowiaceae(100);g__Metschnikowia(100);s__Metschnikowia_pulcherrima(100); | Metschnikowia | 13 | Saprotroph | \|Undefined Saprotroph\| | Yeast | NULL | Probable | As nectar-tap_saprotroph (Põlme et al. 2020); Decay sugar-rich substrates (Põlme et al. 2020) | Põlme S, et al. 2020. Fungal Diversity 105:1-16 ((https://doi.org/10.1007/s13225-020-00466-2)) |
| Otu23220 | 1 | k__Fungi(100);p__Ascomycota(100);c__Saccharomycetes(100);o__Saccharomycetales(100);f__Metschnikowiaceae(100);g__Metschnikowia(100);s__Metschnikowia_pulcherrima(100); | Metschnikowia | 13 | Saprotroph | \|Undefined Saprotroph\| | Yeast | NULL | Probable | As nectar-tap_saprotroph (Põlme et al. 2020); Decay sugar-rich substrates (Põlme et al. 2020) | Põlme S, et al. 2020. Fungal Diversity 105:1-16 ((https://doi.org/10.1007/s13225-020-00466-2)) |
| Otu23221 | 1 | k__Fungi(100);p__Ascomycota(100);c__Saccharomycetes(100);o__Saccharomycetales(100);f__Metschnikowiaceae(100);g__Metschnikowia(100);s__Metschnikowia_pulcherrima(100); | Metschnikowia | 13 | Saprotroph | \|Undefined Saprotroph\| | Yeast | NULL | Probable | As nectar-tap_saprotroph (Põlme et al. 2020); Decay sugar-rich substrates (Põlme et al. 2020) | Põlme S, et al. 2020. Fungal Diversity 105:1-16 ((https://doi.org/10.1007/s13225-020-00466-2)) |
| Otu23222 | 1 | k__Fungi(100);p__Ascomycota(100);c__Saccharomycetes(100);o__Saccharomycetales(100);f__Metschnikowiaceae(100);g__Metschnikowia(100);s__Metschnikowia_pulcherrima(100); | Metschnikowia | 13 | Saprotroph | \|Undefined Saprotroph\| | Yeast | NULL | Probable | As nectar-tap_saprotroph (Põlme et al. 2020); Decay sugar-rich substrates (Põlme et al. 2020) | Põlme S, et al. 2020. Fungal Diversity 105:1-16 ((https://doi.org/10.1007/s13225-020-00466-2)) |
| Otu23223 | 1 | k__Fungi(100);p__Ascomycota(100);c__Saccharomycetes(100);o__Saccharomycetales(100);f__Metschnikowiaceae(100);g__Metschnikowia(100);s__Metschnikowia_pulcherrima(100); | Metschnikowia | 13 | Saprotroph | \|Undefined Saprotroph\| | Yeast | NULL | Probable | As nectar-tap_saprotroph (Põlme et al. 2020); Decay sugar-rich substrates (Põlme et al. 2020) | Põlme S, et al. 2020. Fungal Diversity 105:1-16 ((https://doi.org/10.1007/s13225-020-00466-2)) |
| Otu23224 | 1 | k__Fungi(100);p__Ascomycota(100);c__Saccharomycetes(100);o__Saccharomycetales(100);f__Metschnikowiaceae(100);g__Metschnikowia(100);g__Metschnikowia_unclassified(100); | Metschnikowia | 13 | Saprotroph | \|Undefined Saprotroph\| | Yeast | NULL | Probable | As nectar-tap_saprotroph (Põlme et al. 2020); Decay sugar-rich substrates (Põlme et al. 2020) | Põlme S, et al. 2020. Fungal Diversity 105:1-16 ((https://doi.org/10.1007/s13225-020-00466-2)) |
| Otu23225 | 1 | k__Fungi(100);p__Ascomycota(100);c__Saccharomycetes(100);o__Saccharomycetales(100);f__Metschnikowiaceae(100);g__Metschnikowia(100);s__Metschnikowia_pulcherrima(100); | Metschnikowia | 13 | Saprotroph | \|Undefined Saprotroph\| | Yeast | NULL | Probable | As nectar-tap_saprotroph (Põlme et al. 2020); Decay sugar-rich substrates (Põlme et al. 2020) | Põlme S, et al. 2020. Fungal Diversity 105:1-16 ((https://doi.org/10.1007/s13225-020-00466-2)) |
| Otu23226 | 1 | k__Fungi(100);p__Ascomycota(100);c__Saccharomycetes(100);o__Saccharomycetales(100);f__Metschnikowiaceae(100);g__Metschnikowia(100);s__Metschnikowia_pulcherrima(100); | Metschnikowia | 13 | Saprotroph | \|Undefined Saprotroph\| | Yeast | NULL | Probable | As nectar-tap_saprotroph (Põlme et al. 2020); Decay sugar-rich substrates (Põlme et al. 2020) | Põlme S, et al. 2020. Fungal Diversity 105:1-16 ((https://doi.org/10.1007/s13225-020-00466-2)) |
| Otu23227 | 1 | k__Fungi(100);p__Ascomycota(100);c__Saccharomycetes(100);o__Saccharomycetales(100);f__Metschnikowiaceae(100);g__Metschnikowia(100);s__Metschnikowia_pulcherrima(100); | Metschnikowia | 13 | Saprotroph | \|Undefined Saprotroph\| | Yeast | NULL | Probable | As nectar-tap_saprotroph (Põlme et al. 2020); Decay sugar-rich substrates (Põlme et al. 2020) | Põlme S, et al. 2020. Fungal Diversity 105:1-16 ((https://doi.org/10.1007/s13225-020-00466-2)) |
| Otu23228 | 1 | k__Fungi(100);p__Ascomycota(100);c__Saccharomycetes(100);o__Saccharomycetales(100);f__Metschnikowiaceae(100);g__Metschnikowia(100);s__Metschnikowia_pulcherrima(100); | Metschnikowia | 13 | Saprotroph | \|Undefined Saprotroph\| | Yeast | NULL | Probable | As nectar-tap_saprotroph (Põlme et al. 2020); Decay sugar-rich substrates (Põlme et al. 2020) | Põlme S, et al. 2020. Fungal Diversity 105:1-16 ((https://doi.org/10.1007/s13225-020-00466-2)) |
| Otu23229 | 1 | k__Fungi(100);p__Ascomycota(100);c__Saccharomycetes(100);o__Saccharomycetales(100);f__Metschnikowiaceae(100);g__Metschnikowia(100);s__Metschnikowia_pulcherrima(100); | Metschnikowia | 13 | Saprotroph | \|Undefined Saprotroph\| | Yeast | NULL | Probable | As nectar-tap_saprotroph (Põlme et al. 2020); Decay sugar-rich substrates (Põlme et al. 2020) | Põlme S, et al. 2020. Fungal Diversity 105:1-16 ((https://doi.org/10.1007/s13225-020-00466-2)) |
| Otu23232 | 1 | k__Fungi(100);p__Ascomycota(100);c__Saccharomycetes(100);o__Saccharomycetales(100);f__Metschnikowiaceae(100);g__Metschnikowia(100);s__Metschnikowia_pulcherrima(100); | Metschnikowia | 13 | Saprotroph | \|Undefined Saprotroph\| | Yeast | NULL | Probable | As nectar-tap_saprotroph (Põlme et al. 2020); Decay sugar-rich substrates (Põlme et al. 2020) | Põlme S, et al. 2020. Fungal Diversity 105:1-16 ((https://doi.org/10.1007/s13225-020-00466-2)) |
| Otu23235 | 1 | k__Fungi(100);p__Ascomycota(100);c__Saccharomycetes(100);o__Saccharomycetales(100);f__Metschnikowiaceae(100);g__Metschnikowia(100);s__Metschnikowia_pulcherrima(100); | Metschnikowia | 13 | Saprotroph | \|Undefined Saprotroph\| | Yeast | NULL | Probable | As nectar-tap_saprotroph (Põlme et al. 2020); Decay sugar-rich substrates (Põlme et al. 2020) | Põlme S, et al. 2020. Fungal Diversity 105:1-16 ((https://doi.org/10.1007/s13225-020-00466-2)) |
| Otu23246 | 1 | k__Fungi(100);p__Ascomycota(100);c__Saccharomycetes(100);o__Saccharomycetales(100);f__Saccharomycetales_family_Incertae_sedis(100);g__Candida(100);g__Candida_unclassified(100); | Candida | 13 | Pathotroph-Saprotroph-Symbiotroph | Animal Pathogen-Endophyte-Endosymbiont-Epiphyte-\|Undefined Saprotroph\| | Dimorphic Yeast-Yeast | NULL | Probable | As nectar-tap_saprotroph (Põlme et al. 2020); Decay sugar-rich substrates (Põlme et al. 2020); This asexual genus is polyphyletic and contain species that function very differently depend on what they are. Caution should be taken when interpreting guilds for Candida species. | Manolakaki D, et al. 2010. Virulence. 1: 367-375 ((https://doi.org/10.4161/viru.1.5.12796)); Mekbib SB, et al. 2011. Tropical Plant Pathology 36:233-240 ((https://doi.org/10.1590/S1982-56762011000400004)); Põlme S, et al. 2020. Fungal Diversity 105:1-16 ((https://doi.org/10.1007/s13225-020-00466-2)) |
| Otu23251 | 1 | k__Fungi(100);p__Ascomycota(100);c__Saccharomycetes(100);o__Saccharomycetales(100);f__Metschnikowiaceae(100);g__Metschnikowia(100);s__Metschnikowia_pulcherrima(100); | Metschnikowia | 13 | Saprotroph | \|Undefined Saprotroph\| | Yeast | NULL | Probable | As nectar-tap_saprotroph (Põlme et al. 2020); Decay sugar-rich substrates (Põlme et al. 2020) | Põlme S, et al. 2020. Fungal Diversity 105:1-16 ((https://doi.org/10.1007/s13225-020-00466-2)) |
| Otu23252 | 1 | k__Fungi(100);p__Ascomycota(100);c__Saccharomycetes(100);o__Saccharomycetales(100);f__Metschnikowiaceae(100);g__Metschnikowia(100);g__Metschnikowia_unclassified(100); | Metschnikowia | 13 | Saprotroph | \|Undefined Saprotroph\| | Yeast | NULL | Probable | As nectar-tap_saprotroph (Põlme et al. 2020); Decay sugar-rich substrates (Põlme et al. 2020) | Põlme S, et al. 2020. Fungal Diversity 105:1-16 ((https://doi.org/10.1007/s13225-020-00466-2)) |
| Otu23284 | 1 | k__Fungi(100);p__Ascomycota(100);c__Saccharomycetes(100);o__Saccharomycetales(100);f__Metschnikowiaceae(100);g__Metschnikowia(100);s__Metschnikowia_pulcherrima(100); | Metschnikowia | 13 | Saprotroph | \|Undefined Saprotroph\| | Yeast | NULL | Probable | As nectar-tap_saprotroph (Põlme et al. 2020); Decay sugar-rich substrates (Põlme et al. 2020) | Põlme S, et al. 2020. Fungal Diversity 105:1-16 ((https://doi.org/10.1007/s13225-020-00466-2)) |
| Otu23297 | 1 | k__Fungi(100);p__Ascomycota(100);c__Saccharomycetes(100);o__Saccharomycetales(100);f__Metschnikowiaceae(100);g__Metschnikowia(100);g__Metschnikowia_unclassified(100); | Metschnikowia | 13 | Saprotroph | \|Undefined Saprotroph\| | Yeast | NULL | Probable | As nectar-tap_saprotroph (Põlme et al. 2020); Decay sugar-rich substrates (Põlme et al. 2020) | Põlme S, et al. 2020. Fungal Diversity 105:1-16 ((https://doi.org/10.1007/s13225-020-00466-2)) |
| Otu23306 | 1 | k__Fungi(100);p__Ascomycota(100);c__Saccharomycetes(100);o__Saccharomycetales(100);f__Metschnikowiaceae(100);g__Metschnikowia(100);s__Metschnikowia_pulcherrima(100); | Metschnikowia | 13 | Saprotroph | \|Undefined Saprotroph\| | Yeast | NULL | Probable | As nectar-tap_saprotroph (Põlme et al. 2020); Decay sugar-rich substrates (Põlme et al. 2020) | Põlme S, et al. 2020. Fungal Diversity 105:1-16 ((https://doi.org/10.1007/s13225-020-00466-2)) |
| Otu23307 | 1 | k__Fungi(100);p__Ascomycota(100);c__Saccharomycetes(100);o__Saccharomycetales(100);f__Metschnikowiaceae(100);g__Metschnikowia(100);s__Metschnikowia_pulcherrima(100); | Metschnikowia | 13 | Saprotroph | \|Undefined Saprotroph\| | Yeast | NULL | Probable | As nectar-tap_saprotroph (Põlme et al. 2020); Decay sugar-rich substrates (Põlme et al. 2020) | Põlme S, et al. 2020. Fungal Diversity 105:1-16 ((https://doi.org/10.1007/s13225-020-00466-2)) |
| Otu23310 | 1 | k__Fungi(100);p__Ascomycota(100);c__Saccharomycetes(100);o__Saccharomycetales(100);f__Metschnikowiaceae(100);g__Metschnikowia(100);s__Metschnikowia_pulcherrima(100); | Metschnikowia | 13 | Saprotroph | \|Undefined Saprotroph\| | Yeast | NULL | Probable | As nectar-tap_saprotroph (Põlme et al. 2020); Decay sugar-rich substrates (Põlme et al. 2020) | Põlme S, et al. 2020. Fungal Diversity 105:1-16 ((https://doi.org/10.1007/s13225-020-00466-2)) |
| Otu23320 | 1 | k__Fungi(100);p__Ascomycota(100);c__Saccharomycetes(100);o__Saccharomycetales(100);f__Metschnikowiaceae(100);g__Metschnikowia(100);g__Metschnikowia_unclassified(100); | Metschnikowia | 13 | Saprotroph | \|Undefined Saprotroph\| | Yeast | NULL | Probable | As nectar-tap_saprotroph (Põlme et al. 2020); Decay sugar-rich substrates (Põlme et al. 2020) | Põlme S, et al. 2020. Fungal Diversity 105:1-16 ((https://doi.org/10.1007/s13225-020-00466-2)) |
| Otu23321 | 1 | k__Fungi(100);p__Ascomycota(100);c__Saccharomycetes(100);o__Saccharomycetales(100);f__Metschnikowiaceae(100);g__Metschnikowia(100);s__Metschnikowia_pulcherrima(100); | Metschnikowia | 13 | Saprotroph | \|Undefined Saprotroph\| | Yeast | NULL | Probable | As nectar-tap_saprotroph (Põlme et al. 2020); Decay sugar-rich substrates (Põlme et al. 2020) | Põlme S, et al. 2020. Fungal Diversity 105:1-16 ((https://doi.org/10.1007/s13225-020-00466-2)) |
| Otu23322 | 1 | k__Fungi(100);p__Ascomycota(100);c__Saccharomycetes(100);o__Saccharomycetales(100);f__Metschnikowiaceae(100);g__Metschnikowia(100);s__Metschnikowia_pulcherrima(100); | Metschnikowia | 13 | Saprotroph | \|Undefined Saprotroph\| | Yeast | NULL | Probable | As nectar-tap_saprotroph (Põlme et al. 2020); Decay sugar-rich substrates (Põlme et al. 2020) | Põlme S, et al. 2020. Fungal Diversity 105:1-16 ((https://doi.org/10.1007/s13225-020-00466-2)) |
| Otu23323 | 1 | k__Fungi(100);p__Ascomycota(100);c__Saccharomycetes(100);o__Saccharomycetales(100);f__Metschnikowiaceae(100);g__Metschnikowia(100);s__Metschnikowia_pulcherrima(100); | Metschnikowia | 13 | Saprotroph | \|Undefined Saprotroph\| | Yeast | NULL | Probable | As nectar-tap_saprotroph (Põlme et al. 2020); Decay sugar-rich substrates (Põlme et al. 2020) | Põlme S, et al. 2020. Fungal Diversity 105:1-16 ((https://doi.org/10.1007/s13225-020-00466-2)) |
| Otu23324 | 1 | k__Fungi(100);p__Ascomycota(100);c__Saccharomycetes(100);o__Saccharomycetales(100);f__Metschnikowiaceae(100);g__Metschnikowia(100);s__Metschnikowia_pulcherrima(100); | Metschnikowia | 13 | Saprotroph | \|Undefined Saprotroph\| | Yeast | NULL | Probable | As nectar-tap_saprotroph (Põlme et al. 2020); Decay sugar-rich substrates (Põlme et al. 2020) | Põlme S, et al. 2020. Fungal Diversity 105:1-16 ((https://doi.org/10.1007/s13225-020-00466-2)) |
| Otu23325 | 1 | k__Fungi(100);p__Ascomycota(100);c__Saccharomycetes(100);o__Saccharomycetales(100);f__Metschnikowiaceae(100);g__Metschnikowia(100);s__Metschnikowia_pulcherrima(100); | Metschnikowia | 13 | Saprotroph | \|Undefined Saprotroph\| | Yeast | NULL | Probable | As nectar-tap_saprotroph (Põlme et al. 2020); Decay sugar-rich substrates (Põlme et al. 2020) | Põlme S, et al. 2020. Fungal Diversity 105:1-16 ((https://doi.org/10.1007/s13225-020-00466-2)) |
| Otu23326 | 1 | k__Fungi(100);p__Ascomycota(100);c__Saccharomycetes(100);o__Saccharomycetales(100);f__Metschnikowiaceae(100);g__Metschnikowia(100);s__Metschnikowia_pulcherrima(100); | Metschnikowia | 13 | Saprotroph | \|Undefined Saprotroph\| | Yeast | NULL | Probable | As nectar-tap_saprotroph (Põlme et al. 2020); Decay sugar-rich substrates (Põlme et al. 2020) | Põlme S, et al. 2020. Fungal Diversity 105:1-16 ((https://doi.org/10.1007/s13225-020-00466-2)) |
| Otu23328 | 1 | k__Fungi(100);p__Ascomycota(100);c__Saccharomycetes(100);o__Saccharomycetales(100);f__Metschnikowiaceae(100);g__Metschnikowia(100);s__Metschnikowia_pulcherrima(100); | Metschnikowia | 13 | Saprotroph | \|Undefined Saprotroph\| | Yeast | NULL | Probable | As nectar-tap_saprotroph (Põlme et al. 2020); Decay sugar-rich substrates (Põlme et al. 2020) | Põlme S, et al. 2020. Fungal Diversity 105:1-16 ((https://doi.org/10.1007/s13225-020-00466-2)) |
| Otu23329 | 1 | k__Fungi(100);p__Ascomycota(100);c__Saccharomycetes(100);o__Saccharomycetales(100);f__Metschnikowiaceae(100);g__Metschnikowia(100);s__Metschnikowia_pulcherrima(100); | Metschnikowia | 13 | Saprotroph | \|Undefined Saprotroph\| | Yeast | NULL | Probable | As nectar-tap_saprotroph (Põlme et al. 2020); Decay sugar-rich substrates (Põlme et al. 2020) | Põlme S, et al. 2020. Fungal Diversity 105:1-16 ((https://doi.org/10.1007/s13225-020-00466-2)) |
| Otu23330 | 1 | k__Fungi(100);p__Ascomycota(100);c__Dothideomycetes(100);o__Dothideomycetes_order_Incertae_sedis(100);f__Myxotrichaceae(100);g__Oidiodendron(100);s__Oidiodendron_griseum(100); | Oidiodendron | 13 | Pathotroph-Saprotroph-Symbiotroph | Endophyte-Ericoid Mycorrhizal-\|Undefined Saprotroph\| | Microfungus | NULL | Probable | As root_endophyte (Põlme et al. 2020); As soil_saprotroph (Põlme et al. 2020); Dark Septate Endophyte; Decay soil (Põlme et al. 2020) | Newsham KK. 2011. New Phytologist 190:783-793 ((https://doi.org/10.1111/j.1469-8137.2010.03611.x)) (as dark septate root endophytes); Põlme S, et al. 2020. Fungal Diversity 105:1-16 ((https://doi.org/10.1007/s13225-020-00466-2)) |
| Otu23332 | 1 | k__Fungi(100);p__Ascomycota(100);c__Saccharomycetes(100);o__Saccharomycetales(100);f__Metschnikowiaceae(100);g__Metschnikowia(100);s__Metschnikowia_pulcherrima(100); | Metschnikowia | 13 | Saprotroph | \|Undefined Saprotroph\| | Yeast | NULL | Probable | As nectar-tap_saprotroph (Põlme et al. 2020); Decay sugar-rich substrates (Põlme et al. 2020) | Põlme S, et al. 2020. Fungal Diversity 105:1-16 ((https://doi.org/10.1007/s13225-020-00466-2)) |
| Otu23333 | 1 | k__Fungi(100);p__Ascomycota(100);c__Saccharomycetes(100);o__Saccharomycetales(100);f__Metschnikowiaceae(100);g__Metschnikowia(100);s__Metschnikowia_pulcherrima(100); | Metschnikowia | 13 | Saprotroph | \|Undefined Saprotroph\| | Yeast | NULL | Probable | As nectar-tap_saprotroph (Põlme et al. 2020); Decay sugar-rich substrates (Põlme et al. 2020) | Põlme S, et al. 2020. Fungal Diversity 105:1-16 ((https://doi.org/10.1007/s13225-020-00466-2)) |
| Otu23335 | 1 | k__Fungi(100);p__Ascomycota(100);c__Saccharomycetes(100);o__Saccharomycetales(100);f__Metschnikowiaceae(100);g__Metschnikowia(100);s__Metschnikowia_pulcherrima(100); | Metschnikowia | 13 | Saprotroph | \|Undefined Saprotroph\| | Yeast | NULL | Probable | As nectar-tap_saprotroph (Põlme et al. 2020); Decay sugar-rich substrates (Põlme et al. 2020) | Põlme S, et al. 2020. Fungal Diversity 105:1-16 ((https://doi.org/10.1007/s13225-020-00466-2)) |
| Otu23364 | 1 | k__Fungi(100);p__Ascomycota(100);c__Saccharomycetes(100);o__Saccharomycetales(100);f__Metschnikowiaceae(100);g__Metschnikowia(100);s__Metschnikowia_pulcherrima(100); | Metschnikowia | 13 | Saprotroph | \|Undefined Saprotroph\| | Yeast | NULL | Probable | As nectar-tap_saprotroph (Põlme et al. 2020); Decay sugar-rich substrates (Põlme et al. 2020) | Põlme S, et al. 2020. Fungal Diversity 105:1-16 ((https://doi.org/10.1007/s13225-020-00466-2)) |
| Otu23381 | 1 | k__Fungi(100);p__Ascomycota(100);c__Saccharomycetes(100);o__Saccharomycetales(100);f__Metschnikowiaceae(100);g__Metschnikowia(100);g__Metschnikowia_unclassified(100); | Metschnikowia | 13 | Saprotroph | \|Undefined Saprotroph\| | Yeast | NULL | Probable | As nectar-tap_saprotroph (Põlme et al. 2020); Decay sugar-rich substrates (Põlme et al. 2020) | Põlme S, et al. 2020. Fungal Diversity 105:1-16 ((https://doi.org/10.1007/s13225-020-00466-2)) |
| Otu23383 | 1 | k__Fungi(100);p__Ascomycota(100);c__Saccharomycetes(100);o__Saccharomycetales(100);f__Metschnikowiaceae(100);g__Metschnikowia(100);s__Metschnikowia_pulcherrima(100); | Metschnikowia | 13 | Saprotroph | \|Undefined Saprotroph\| | Yeast | NULL | Probable | As nectar-tap_saprotroph (Põlme et al. 2020); Decay sugar-rich substrates (Põlme et al. 2020) | Põlme S, et al. 2020. Fungal Diversity 105:1-16 ((https://doi.org/10.1007/s13225-020-00466-2)) |
| Otu23407 | 1 | k__Fungi(100);p__Ascomycota(100);c__Saccharomycetes(100);o__Saccharomycetales(100);f__Metschnikowiaceae(100);g__Metschnikowia(100);g__Metschnikowia_unclassified(100); | Metschnikowia | 13 | Saprotroph | \|Undefined Saprotroph\| | Yeast | NULL | Probable | As nectar-tap_saprotroph (Põlme et al. 2020); Decay sugar-rich substrates (Põlme et al. 2020) | Põlme S, et al. 2020. Fungal Diversity 105:1-16 ((https://doi.org/10.1007/s13225-020-00466-2)) |
| Otu23429 | 1 | k__Fungi(100);p__Ascomycota(100);c__Saccharomycetes(100);o__Saccharomycetales(100);f__Metschnikowiaceae(100);g__Metschnikowia(100);s__Metschnikowia_pulcherrima(100); | Metschnikowia | 13 | Saprotroph | \|Undefined Saprotroph\| | Yeast | NULL | Probable | As nectar-tap_saprotroph (Põlme et al. 2020); Decay sugar-rich substrates (Põlme et al. 2020) | Põlme S, et al. 2020. Fungal Diversity 105:1-16 ((https://doi.org/10.1007/s13225-020-00466-2)) |
| Otu23458 | 1 | k__Fungi(100);p__Ascomycota(100);c__Saccharomycetes(100);o__Saccharomycetales(100);f__Metschnikowiaceae(100);g__Metschnikowia(100);s__Metschnikowia_pulcherrima(100); | Metschnikowia | 13 | Saprotroph | \|Undefined Saprotroph\| | Yeast | NULL | Probable | As nectar-tap_saprotroph (Põlme et al. 2020); Decay sugar-rich substrates (Põlme et al. 2020) | Põlme S, et al. 2020. Fungal Diversity 105:1-16 ((https://doi.org/10.1007/s13225-020-00466-2)) |
| Otu23459 | 1 | k__Fungi(100);p__Ascomycota(100);c__Saccharomycetes(100);o__Saccharomycetales(100);f__Metschnikowiaceae(100);g__Metschnikowia(100);s__Metschnikowia_pulcherrima(100); | Metschnikowia | 13 | Saprotroph | \|Undefined Saprotroph\| | Yeast | NULL | Probable | As nectar-tap_saprotroph (Põlme et al. 2020); Decay sugar-rich substrates (Põlme et al. 2020) | Põlme S, et al. 2020. Fungal Diversity 105:1-16 ((https://doi.org/10.1007/s13225-020-00466-2)) |
| Otu23469 | 1 | k__Fungi(100);p__Ascomycota(100);c__Saccharomycetes(100);o__Saccharomycetales(100);f__Metschnikowiaceae(100);g__Metschnikowia(100);s__Metschnikowia_pulcherrima(100); | Metschnikowia | 13 | Saprotroph | \|Undefined Saprotroph\| | Yeast | NULL | Probable | As nectar-tap_saprotroph (Põlme et al. 2020); Decay sugar-rich substrates (Põlme et al. 2020) | Põlme S, et al. 2020. Fungal Diversity 105:1-16 ((https://doi.org/10.1007/s13225-020-00466-2)) |
| Otu23470 | 1 | k__Fungi(100);p__Ascomycota(100);c__Saccharomycetes(100);o__Saccharomycetales(100);f__Metschnikowiaceae(100);g__Metschnikowia(100);s__Metschnikowia_pulcherrima(100); | Metschnikowia | 13 | Saprotroph | \|Undefined Saprotroph\| | Yeast | NULL | Probable | As nectar-tap_saprotroph (Põlme et al. 2020); Decay sugar-rich substrates (Põlme et al. 2020) | Põlme S, et al. 2020. Fungal Diversity 105:1-16 ((https://doi.org/10.1007/s13225-020-00466-2)) |
| Otu23471 | 1 | k__Fungi(100);p__Ascomycota(100);c__Saccharomycetes(100);o__Saccharomycetales(100);f__Metschnikowiaceae(100);g__Metschnikowia(100);s__Metschnikowia_pulcherrima(100); | Metschnikowia | 13 | Saprotroph | \|Undefined Saprotroph\| | Yeast | NULL | Probable | As nectar-tap_saprotroph (Põlme et al. 2020); Decay sugar-rich substrates (Põlme et al. 2020) | Põlme S, et al. 2020. Fungal Diversity 105:1-16 ((https://doi.org/10.1007/s13225-020-00466-2)) |
| Otu23472 | 1 | k__Fungi(100);p__Ascomycota(100);c__Saccharomycetes(100);o__Saccharomycetales(100);f__Saccharomycetales_family_Incertae_sedis(100);f__Saccharomycetales_family_Incertae_sedis_unclassified(100);f__Saccharomycetales_family_Incertae_sedis_unclassified(100); | Saccharomycetales | 7 | Saprotroph | Undefined Saprotroph | Yeast | NULL | Possible | NULL | Sterkenburg E, et al. 2015. New Phytologist 207:1145-1158 ((https://doi.org/10.1111/nph.13426)) |
| Otu23473 | 1 | k__Fungi(100);p__Ascomycota(100);c__Saccharomycetes(100);o__Saccharomycetales(100);f__Metschnikowiaceae(100);g__Metschnikowia(100);g__Metschnikowia_unclassified(100); | Metschnikowia | 13 | Saprotroph | \|Undefined Saprotroph\| | Yeast | NULL | Probable | As nectar-tap_saprotroph (Põlme et al. 2020); Decay sugar-rich substrates (Põlme et al. 2020) | Põlme S, et al. 2020. Fungal Diversity 105:1-16 ((https://doi.org/10.1007/s13225-020-00466-2)) |
| Otu23475 | 1 | k__Fungi(100);p__Ascomycota(100);c__Saccharomycetes(100);o__Saccharomycetales(100);f__Metschnikowiaceae(100);g__Metschnikowia(100);s__Metschnikowia_pulcherrima(100); | Metschnikowia | 13 | Saprotroph | \|Undefined Saprotroph\| | Yeast | NULL | Probable | As nectar-tap_saprotroph (Põlme et al. 2020); Decay sugar-rich substrates (Põlme et al. 2020) | Põlme S, et al. 2020. Fungal Diversity 105:1-16 ((https://doi.org/10.1007/s13225-020-00466-2)) |
| Otu23476 | 1 | k__Fungi(100);p__Ascomycota(100);c__Saccharomycetes(100);o__Saccharomycetales(100);f__Metschnikowiaceae(100);g__Metschnikowia(100);s__Metschnikowia_pulcherrima(100); | Metschnikowia | 13 | Saprotroph | \|Undefined Saprotroph\| | Yeast | NULL | Probable | As nectar-tap_saprotroph (Põlme et al. 2020); Decay sugar-rich substrates (Põlme et al. 2020) | Põlme S, et al. 2020. Fungal Diversity 105:1-16 ((https://doi.org/10.1007/s13225-020-00466-2)) |
| Otu23477 | 1 | k__Fungi(100);p__Ascomycota(100);c__Saccharomycetes(100);o__Saccharomycetales(100);f__Metschnikowiaceae(100);g__Metschnikowia(100);g__Metschnikowia_unclassified(100); | Metschnikowia | 13 | Saprotroph | \|Undefined Saprotroph\| | Yeast | NULL | Probable | As nectar-tap_saprotroph (Põlme et al. 2020); Decay sugar-rich substrates (Põlme et al. 2020) | Põlme S, et al. 2020. Fungal Diversity 105:1-16 ((https://doi.org/10.1007/s13225-020-00466-2)) |

| | | | | | | | | | | | | | |
|---|---|---|---|---|---|---|---|---|---|---|---|---|---|
| Otu23479 | | 1 | k__Fungi(100);p__Ascomycota(100);c__Saccharomycetes(100);o__Saccharomycetales(100);f__Metschnikowiaceae(100);g__Metschnikowia(100);s__Metschnikowia_pulcherrima(100); | Metschnikowia | | 13 | Saprotroph | \|Undefined Saprotroph\| | Yeast | NULL | Probable | As nectar-tap_saprotroph (Põlme et al. 2020); Decay sugar-rich substrates (Põlme et al. 2020) | Põlme S, et al. 2020. Fungal Diversity 105:1-16 ((https://doi.org/10.1007/s13225-020-00466-2)) |
| Otu23480 | | 1 | k__Fungi(100);p__Ascomycota(100);c__Saccharomycetes(100);o__Saccharomycetales(100);f__Metschnikowiaceae(100);g__Metschnikowia(100);s__Metschnikowia_pulcherrima(100); | Metschnikowia | | 13 | Saprotroph | \|Undefined Saprotroph\| | Yeast | NULL | Probable | As nectar-tap_saprotroph (Põlme et al. 2020); Decay sugar-rich substrates (Põlme et al. 2020) | Põlme S, et al. 2020. Fungal Diversity 105:1-16 ((https://doi.org/10.1007/s13225-020-00466-2)) |
| Otu23481 | | 1 | k__Fungi(100);p__Ascomycota(100);c__Saccharomycetes(100);o__Saccharomycetales(100);f__Metschnikowiaceae(100);g__Metschnikowia(100);s__Metschnikowia_pulcherrima(100); | Metschnikowia | | 13 | Saprotroph | \|Undefined Saprotroph\| | Yeast | NULL | Probable | As nectar-tap_saprotroph (Põlme et al. 2020); Decay sugar-rich substrates (Põlme et al. 2020) | Põlme S, et al. 2020. Fungal Diversity 105:1-16 ((https://doi.org/10.1007/s13225-020-00466-2)) |
| Otu23483 | | 1 | k__Fungi(100);p__Ascomycota(100);c__Saccharomycetes(100);o__Saccharomycetales(100);f__Metschnikowiaceae(100);g__Metschnikowia(100);s__Metschnikowia_pulcherrima(100); | Metschnikowia | | 13 | Saprotroph | \|Undefined Saprotroph\| | Yeast | NULL | Probable | As nectar-tap_saprotroph (Põlme et al. 2020); Decay sugar-rich substrates (Põlme et al. 2020) | Põlme S, et al. 2020. Fungal Diversity 105:1-16 ((https://doi.org/10.1007/s13225-020-00466-2)) |
| Otu23485 | | 1 | k__Fungi(100);p__Ascomycota(100);c__Saccharomycetes(100);o__Saccharomycetales(100);f__Metschnikowiaceae(100);g__Metschnikowia(100);g__Metschnikowia_unclassified(100); | Metschnikowia | | 13 | Saprotroph | \|Undefined Saprotroph\| | Yeast | NULL | Probable | As nectar-tap_saprotroph (Põlme et al. 2020); Decay sugar-rich substrates (Põlme et al. 2020) | Põlme S, et al. 2020. Fungal Diversity 105:1-16 ((https://doi.org/10.1007/s13225-020-00466-2)) |
| Otu23486 | | 1 | k__Fungi(100);p__Ascomycota(100);c__Saccharomycetes(100);o__Saccharomycetales(100);f__Metschnikowiaceae(100);g__Metschnikowia(100);s__Metschnikowia_pulcherrima(100); | Metschnikowia | | 13 | Saprotroph | \|Undefined Saprotroph\| | Yeast | NULL | Probable | As nectar-tap_saprotroph (Põlme et al. 2020); Decay sugar-rich substrates (Põlme et al. 2020) | Põlme S, et al. 2020. Fungal Diversity 105:1-16 ((https://doi.org/10.1007/s13225-020-00466-2)) |
| Otu23487 | | 1 | k__Fungi(100);p__Ascomycota(100);c__Saccharomycetes(100);o__Saccharomycetales(100);f__Metschnikowiaceae(100);g__Metschnikowia(100);s__Metschnikowia_pulcherrima(100); | Metschnikowia | | 13 | Saprotroph | \|Undefined Saprotroph\| | Yeast | NULL | Probable | As nectar-tap_saprotroph (Põlme et al. 2020); Decay sugar-rich substrates (Põlme et al. 2020) | Põlme S, et al. 2020. Fungal Diversity 105:1-16 ((https://doi.org/10.1007/s13225-020-00466-2)) |
| Otu23488 | | 1 | k__Fungi(100);p__Ascomycota(100);c__Saccharomycetes(100);o__Saccharomycetales(100);f__Metschnikowiaceae(100);g__Metschnikowia(100);s__Metschnikowia_pulcherrima(100); | Metschnikowia | | 13 | Saprotroph | \|Undefined Saprotroph\| | Yeast | NULL | Probable | As nectar-tap_saprotroph (Põlme et al. 2020); Decay sugar-rich substrates (Põlme et al. 2020) | Põlme S, et al. 2020. Fungal Diversity 105:1-16 ((https://doi.org/10.1007/s13225-020-00466-2)) |
| Otu23489 | | 1 | k__Fungi(100);p__Ascomycota(100);c__Saccharomycetes(100);o__Saccharomycetales(100);f__Metschnikowiaceae(100);g__Metschnikowia(100);s__Metschnikowia_pulcherrima(100); | Metschnikowia | | 13 | Saprotroph | \|Undefined Saprotroph\| | Yeast | NULL | Probable | As nectar-tap_saprotroph (Põlme et al. 2020); Decay sugar-rich substrates (Põlme et al. 2020) | Põlme S, et al. 2020. Fungal Diversity 105:1-16 ((https://doi.org/10.1007/s13225-020-00466-2)) |
| Otu23490 | | 1 | k__Fungi(100);p__Ascomycota(100);c__Saccharomycetes(100);o__Saccharomycetales(100);f__Metschnikowiaceae(100);g__Metschnikowia(100);s__Metschnikowia_pulcherrima(100); | Metschnikowia | | 13 | Saprotroph | \|Undefined Saprotroph\| | Yeast | NULL | Probable | As nectar-tap_saprotroph (Põlme et al. 2020); Decay sugar-rich substrates (Põlme et al. 2020) | Põlme S, et al. 2020. Fungal Diversity 105:1-16 ((https://doi.org/10.1007/s13225-020-00466-2)) |
| Otu23493 | | 1 | k__Fungi(100);p__Ascomycota(100);c__Saccharomycetes(100);o__Saccharomycetales(100);f__Metschnikowiaceae(100);g__Metschnikowia(100);s__Metschnikowia_pulcherrima(100); | Metschnikowia | | 13 | Saprotroph | \|Undefined Saprotroph\| | Yeast | NULL | Probable | As nectar-tap_saprotroph (Põlme et al. 2020); Decay sugar-rich substrates (Põlme et al. 2020) | Põlme S, et al. 2020. Fungal Diversity 105:1-16 ((https://doi.org/10.1007/s13225-020-00466-2)) |
| Otu23494 | | 1 | k__Fungi(100);p__Ascomycota(100);c__Saccharomycetes(100);o__Saccharomycetales(100);f__Metschnikowiaceae(100);g__Metschnikowia(100);s__Metschnikowia_pulcherrima(100); | Metschnikowia | | 13 | Saprotroph | \|Undefined Saprotroph\| | Yeast | NULL | Probable | As nectar-tap_saprotroph (Põlme et al. 2020); Decay sugar-rich substrates (Põlme et al. 2020) | Põlme S, et al. 2020. Fungal Diversity 105:1-16 ((https://doi.org/10.1007/s13225-020-00466-2)) |
| Otu23495 | | 1 | k__Fungi(100);p__Ascomycota(100);c__Saccharomycetes(100);o__Saccharomycetales(100);f__Metschnikowiaceae(100);g__Metschnikowia(100);s__Metschnikowia_pulcherrima(100); | Metschnikowia | | 13 | Saprotroph | \|Undefined Saprotroph\| | Yeast | NULL | Probable | As nectar-tap_saprotroph (Põlme et al. 2020); Decay sugar-rich substrates (Põlme et al. 2020) | Põlme S, et al. 2020. Fungal Diversity 105:1-16 ((https://doi.org/10.1007/s13225-020-00466-2)) |
| Otu23496 | | 1 | k__Fungi(100);p__Ascomycota(100);c__Saccharomycetes(100);o__Saccharomycetales(100);f__Metschnikowiaceae(100);g__Metschnikowia(100);s__Metschnikowia_pulcherrima(100); | Metschnikowia | | 13 | Saprotroph | \|Undefined Saprotroph\| | Yeast | NULL | Probable | As nectar-tap_saprotroph (Põlme et al. 2020); Decay sugar-rich substrates (Põlme et al. 2020) | Põlme S, et al. 2020. Fungal Diversity 105:1-16 ((https://doi.org/10.1007/s13225-020-00466-2)) |
| Otu23497 | | 1 | k__Fungi(100);p__Ascomycota(100);c__Saccharomycetes(100);o__Saccharomycetales(100);f__Metschnikowiaceae(100);g__Metschnikowia(100);g__Metschnikowia_unclassified(100); | Metschnikowia | | 13 | Saprotroph | \|Undefined Saprotroph\| | Yeast | NULL | Probable | As nectar-tap_saprotroph (Põlme et al. 2020); Decay sugar-rich substrates (Põlme et al. 2020) | Põlme S, et al. 2020. Fungal Diversity 105:1-16 ((https://doi.org/10.1007/s13225-020-00466-2)) |
| Otu23498 | | 1 | k__Fungi(100);p__Ascomycota(100);c__Saccharomycetes(100);o__Saccharomycetales(100);f__Metschnikowiaceae(100);g__Metschnikowia(100);s__Metschnikowia_pulcherrima(100); | Metschnikowia | | 13 | Saprotroph | \|Undefined Saprotroph\| | Yeast | NULL | Probable | As nectar-tap_saprotroph (Põlme et al. 2020); Decay sugar-rich substrates (Põlme et al. 2020) | Põlme S, et al. 2020. Fungal Diversity 105:1-16 ((https://doi.org/10.1007/s13225-020-00466-2)) |
| Otu23499 | | 1 | k__Fungi(100);p__Ascomycota(100);c__Saccharomycetes(100);o__Saccharomycetales(100);f__Metschnikowiaceae(100);g__Metschnikowia(100);s__Metschnikowia_pulcherrima(100); | Metschnikowia | | 13 | Saprotroph | \|Undefined Saprotroph\| | Yeast | NULL | Probable | As nectar-tap_saprotroph (Põlme et al. 2020); Decay sugar-rich substrates (Põlme et al. 2020) | Põlme S, et al. 2020. Fungal Diversity 105:1-16 ((https://doi.org/10.1007/s13225-020-00466-2)) |
| Otu23500 | | 1 | k__Fungi(100);p__Ascomycota(100);c__Saccharomycetes(100);o__Saccharomycetales(100);f__Metschnikowiaceae(100);g__Metschnikowia(100);g__Metschnikowia_unclassified(100); | Metschnikowia | | 13 | Saprotroph | \|Undefined Saprotroph\| | Yeast | NULL | Probable | As nectar-tap_saprotroph (Põlme et al. 2020); Decay sugar-rich substrates (Põlme et al. 2020) | Põlme S, et al. 2020. Fungal Diversity 105:1-16 ((https://doi.org/10.1007/s13225-020-00466-2)) |
| Otu23501 | | 1 | k__Fungi(100);p__Ascomycota(100);c__Saccharomycetes(100);o__Saccharomycetales(100);f__Metschnikowiaceae(100);g__Metschnikowia(100);s__Metschnikowia_pulcherrima(100); | Metschnikowia | | 13 | Saprotroph | \|Undefined Saprotroph\| | Yeast | NULL | Probable | As nectar-tap_saprotroph (Põlme et al. 2020); Decay sugar-rich substrates (Põlme et al. 2020) | Põlme S, et al. 2020. Fungal Diversity 105:1-16 ((https://doi.org/10.1007/s13225-020-00466-2)) |
| Otu23502 | | 1 | k__Fungi(100);p__Ascomycota(100);c__Saccharomycetes(100);o__Saccharomycetales(100);f__Metschnikowiaceae(100);g__Metschnikowia(100);g__Metschnikowia_unclassified(100); | Metschnikowia | | 13 | Saprotroph | \|Undefined Saprotroph\| | Yeast | NULL | Probable | As nectar-tap_saprotroph (Põlme et al. 2020); Decay sugar-rich substrates (Põlme et al. 2020) | Põlme S, et al. 2020. Fungal Diversity 105:1-16 ((https://doi.org/10.1007/s13225-020-00466-2)) |
| Otu23508 | | 1 | k__Fungi(100);p__Ascomycota(100);c__Saccharomycetes(100);o__Saccharomycetales(100);f__Metschnikowiaceae(100);g__Metschnikowia(100);g__Metschnikowia_unclassified(100); | Metschnikowia | | 13 | Saprotroph | \|Undefined Saprotroph\| | Yeast | NULL | Probable | As nectar-tap_saprotroph (Põlme et al. 2020); Decay sugar-rich substrates (Põlme et al. 2020) | Põlme S, et al. 2020. Fungal Diversity 105:1-16 ((https://doi.org/10.1007/s13225-020-00466-2)) |
| Otu23509 | | 1 | k__Fungi(100);p__Ascomycota(100);c__Saccharomycetes(100);o__Saccharomycetales(100);f__Metschnikowiaceae(100);g__Metschnikowia(100);s__Metschnikowia_pulcherrima(100); | Metschnikowia | | 13 | Saprotroph | \|Undefined Saprotroph\| | Yeast | NULL | Probable | As nectar-tap_saprotroph (Põlme et al. 2020); Decay sugar-rich substrates (Põlme et al. 2020) | Põlme S, et al. 2020. Fungal Diversity 105:1-16 ((https://doi.org/10.1007/s13225-020-00466-2)) |
| Otu23510 | | 1 | k__Fungi(100);p__Ascomycota(100);c__Saccharomycetes(100);o__Saccharomycetales(100);f__Metschnikowiaceae(100);g__Metschnikowia(100);s__Metschnikowia_pulcherrima(100); | Metschnikowia | | 13 | Saprotroph | \|Undefined Saprotroph\| | Yeast | NULL | Probable | As nectar-tap_saprotroph (Põlme et al. 2020); Decay sugar-rich substrates (Põlme et al. 2020) | Põlme S, et al. 2020. Fungal Diversity 105:1-16 ((https://doi.org/10.1007/s13225-020-00466-2)) |
| Otu23513 | | 1 | k__Fungi(100);p__Ascomycota(100);c__Saccharomycetes(100);o__Saccharomycetales(100);f__Metschnikowiaceae(100);g__Metschnikowia(100);s__Metschnikowia_pulcherrima(100); | Metschnikowia | | 13 | Saprotroph | \|Undefined Saprotroph\| | Yeast | NULL | Probable | As nectar-tap_saprotroph (Põlme et al. 2020); Decay sugar-rich substrates (Põlme et al. 2020) | Põlme S, et al. 2020. Fungal Diversity 105:1-16 ((https://doi.org/10.1007/s13225-020-00466-2)) |
| Otu23514 | | 1 | k__Fungi(100);p__Ascomycota(100);c__Saccharomycetes(100);o__Saccharomycetales(100);f__Metschnikowiaceae(100);g__Metschnikowia(100);s__Metschnikowia_pulcherrima(100); | Metschnikowia | | 13 | Saprotroph | \|Undefined Saprotroph\| | Yeast | NULL | Probable | As nectar-tap_saprotroph (Põlme et al. 2020); Decay sugar-rich substrates (Põlme et al. 2020) | Põlme S, et al. 2020. Fungal Diversity 105:1-16 ((https://doi.org/10.1007/s13225-020-00466-2)) |
| Otu23515 | | 1 | k__Fungi(100);p__Ascomycota(100);c__Saccharomycetes(100);o__Saccharomycetales(100);f__Metschnikowiaceae(100);g__Metschnikowia(100);s__Metschnikowia_pulcherrima(100); | Metschnikowia | | 13 | Saprotroph | \|Undefined Saprotroph\| | Yeast | NULL | Probable | As nectar-tap_saprotroph (Põlme et al. 2020); Decay sugar-rich substrates (Põlme et al. 2020) | Põlme S, et al. 2020. Fungal Diversity 105:1-16 ((https://doi.org/10.1007/s13225-020-00466-2)) |
| Otu23516 | | 1 | k__Fungi(100);p__Ascomycota(100);c__Saccharomycetes(100);o__Saccharomycetales(100);f__Metschnikowiaceae(100);g__Metschnikowia(100);s__Metschnikowia_pulcherrima(100); | Metschnikowia | | 13 | Saprotroph | \|Undefined Saprotroph\| | Yeast | NULL | Probable | As nectar-tap_saprotroph (Põlme et al. 2020); Decay sugar-rich substrates (Põlme et al. 2020) | Põlme S, et al. 2020. Fungal Diversity 105:1-16 ((https://doi.org/10.1007/s13225-020-00466-2)) |
| Otu23520 | | 1 | k__Fungi(100);p__Ascomycota(100);c__Saccharomycetes(100);o__Saccharomycetales(100);f__Metschnikowiaceae(100);g__Metschnikowia(100);g__Metschnikowia_unclassified(100); | Metschnikowia | | 13 | Saprotroph | \|Undefined Saprotroph\| | Yeast | NULL | Probable | As nectar-tap_saprotroph (Põlme et al. 2020); Decay sugar-rich substrates (Põlme et al. 2020) | Põlme S, et al. 2020. Fungal Diversity 105:1-16 ((https://doi.org/10.1007/s13225-020-00466-2)) |
| Otu23521 | | 1 | k__Fungi(100);p__Ascomycota(100);c__Saccharomycetes(100);o__Saccharomycetales(100);f__Metschnikowiaceae(100);g__Metschnikowia(100);s__Metschnikowia_pulcherrima(100); | Metschnikowia | | 13 | Saprotroph | \|Undefined Saprotroph\| | Yeast | NULL | Probable | As nectar-tap_saprotroph (Põlme et al. 2020); Decay sugar-rich substrates (Põlme et al. 2020) | Põlme S, et al. 2020. Fungal Diversity 105:1-16 ((https://doi.org/10.1007/s13225-020-00466-2)) |
| Otu23523 | | 1 | k__Fungi(100);p__Ascomycota(100);c__Saccharomycetes(100);o__Saccharomycetales(100);f__Metschnikowiaceae(100);g__Metschnikowia(100);s__Metschnikowia_pulcherrima(100); | Metschnikowia | | 13 | Saprotroph | \|Undefined Saprotroph\| | Yeast | NULL | Probable | As nectar-tap_saprotroph (Põlme et al. 2020); Decay sugar-rich substrates (Põlme et al. 2020) | Põlme S, et al. 2020. Fungal Diversity 105:1-16 ((https://doi.org/10.1007/s13225-020-00466-2)) |
| Otu23524 | | 1 | k__Fungi(100);p__Ascomycota(100);c__Saccharomycetes(100);o__Saccharomycetales(100);f__Metschnikowiaceae(100);g__Metschnikowia(100);s__Metschnikowia_pulcherrima(100); | Metschnikowia | | 13 | Saprotroph | \|Undefined Saprotroph\| | Yeast | NULL | Probable | As nectar-tap_saprotroph (Põlme et al. 2020); Decay sugar-rich substrates (Põlme et al. 2020) | Põlme S, et al. 2020. Fungal Diversity 105:1-16 ((https://doi.org/10.1007/s13225-020-00466-2)) |
| Otu23525 | | 1 | k__Fungi(100);p__Ascomycota(100);c__Saccharomycetes(100);o__Saccharomycetales(100);f__Metschnikowiaceae(100);g__Metschnikowia(100);g__Metschnikowia_unclassified(100); | Metschnikowia | | 13 | Saprotroph | \|Undefined Saprotroph\| | Yeast | NULL | Probable | As nectar-tap_saprotroph (Põlme et al. 2020); Decay sugar-rich substrates (Põlme et al. 2020) | Põlme S, et al. 2020. Fungal Diversity 105:1-16 ((https://doi.org/10.1007/s13225-020-00466-2)) |
| Otu23526 | | 1 | k__Fungi(100);p__Ascomycota(100);c__Saccharomycetes(100);o__Saccharomycetales(100);f__Metschnikowiaceae(100);g__Metschnikowia(100);s__Metschnikowia_pulcherrima(100); | Metschnikowia | | 13 | Saprotroph | \|Undefined Saprotroph\| | Yeast | NULL | Probable | As nectar-tap_saprotroph (Põlme et al. 2020); Decay sugar-rich substrates (Põlme et al. 2020) | Põlme S, et al. 2020. Fungal Diversity 105:1-16 ((https://doi.org/10.1007/s13225-020-00466-2)) |
| Otu23528 | | 1 | k__Fungi(100);p__Ascomycota(100);c__Saccharomycetes(100);o__Saccharomycetales(100);f__Metschnikowiaceae(100);g__Metschnikowia(100);s__Metschnikowia_pulcherrima(100); | Metschnikowia | | 13 | Saprotroph | \|Undefined Saprotroph\| | Yeast | NULL | Probable | As nectar-tap_saprotroph (Põlme et al. 2020); Decay sugar-rich substrates (Põlme et al. 2020) | Põlme S, et al. 2020. Fungal Diversity 105:1-16 ((https://doi.org/10.1007/s13225-020-00466-2)) |
| Otu23529 | | 1 | k__Fungi(100);p__Ascomycota(100);c__Saccharomycetes(100);o__Saccharomycetales(100);f__Metschnikowiaceae(100);g__Metschnikowia(100);s__Metschnikowia_pulcherrima(100); | Metschnikowia | | 13 | Saprotroph | \|Undefined Saprotroph\| | Yeast | NULL | Probable | As nectar-tap_saprotroph (Põlme et al. 2020); Decay sugar-rich substrates (Põlme et al. 2020) | Põlme S, et al. 2020. Fungal Diversity 105:1-16 ((https://doi.org/10.1007/s13225-020-00466-2)) |
| Otu23530 | | 1 | k__Fungi(100);p__Ascomycota(100);c__Saccharomycetes(100);o__Saccharomycetales(100);f__Metschnikowiaceae(100);g__Metschnikowia(100);s__Metschnikowia_pulcherrima(100); | Metschnikowia | | 13 | Saprotroph | \|Undefined Saprotroph\| | Yeast | NULL | Probable | As nectar-tap_saprotroph (Põlme et al. 2020); Decay sugar-rich substrates (Põlme et al. 2020) | Põlme S, et al. 2020. Fungal Diversity 105:1-16 ((https://doi.org/10.1007/s13225-020-00466-2)) |
| Otu23531 | | 1 | k__Fungi(100);p__Ascomycota(100);c__Saccharomycetes(100);o__Saccharomycetales(100);f__Metschnikowiaceae(100);g__Metschnikowia(100);s__Metschnikowia_pulcherrima(100); | Metschnikowia | | 13 | Saprotroph | \|Undefined Saprotroph\| | Yeast | NULL | Probable | As nectar-tap_saprotroph (Põlme et al. 2020); Decay sugar-rich substrates (Põlme et al. 2020) | Põlme S, et al. 2020. Fungal Diversity 105:1-16 ((https://doi.org/10.1007/s13225-020-00466-2)) |
| Otu23533 | | 1 | k__Fungi(100);p__Ascomycota(100);c__Saccharomycetes(100);o__Saccharomycetales(100);f__Metschnikowiaceae(100);g__Metschnikowia(100);s__Metschnikowia_pulcherrima(100); | Metschnikowia | | 13 | Saprotroph | \|Undefined Saprotroph\| | Yeast | NULL | Probable | As nectar-tap_saprotroph (Põlme et al. 2020); Decay sugar-rich substrates (Põlme et al. 2020) | Põlme S, et al. 2020. Fungal Diversity 105:1-16 ((https://doi.org/10.1007/s13225-020-00466-2)) |
| Otu23538 | | 1 | k__Fungi(100);p__Ascomycota(100);c__Saccharomycetes(100);o__Saccharomycetales(100);f__Metschnikowiaceae(100);g__Metschnikowia(100);s__Metschnikowia_pulcherrima(100); | Metschnikowia | | 13 | Saprotroph | \|Undefined Saprotroph\| | Yeast | NULL | Probable | As nectar-tap_saprotroph (Põlme et al. 2020); Decay sugar-rich substrates (Põlme et al. 2020) | Põlme S, et al. 2020. Fungal Diversity 105:1-16 ((https://doi.org/10.1007/s13225-020-00466-2)) |
| Otu23542 | | 1 | k__Fungi(100);p__Ascomycota(100);c__Saccharomycetes(100);o__Saccharomycetales(100);f__Metschnikowiaceae(100);g__Metschnikowia(100);g__Metschnikowia_unclassified(100); | Metschnikowia | | 13 | Saprotroph | \|Undefined Saprotroph\| | Yeast | NULL | Probable | As nectar-tap_saprotroph (Põlme et al. 2020); Decay sugar-rich substrates (Põlme et al. 2020) | Põlme S, et al. 2020. Fungal Diversity 105:1-16 ((https://doi.org/10.1007/s13225-020-00466-2)) |
| Otu23544 | | 1 | k__Fungi(100);p__Ascomycota(100);c__Saccharomycetes(100);o__Saccharomycetales(100);f__Metschnikowiaceae(100);g__Metschnikowia(100);g__Metschnikowia_unclassified(100); | Metschnikowia | | 13 | Saprotroph | \|Undefined Saprotroph\| | Yeast | NULL | Probable | As nectar-tap_saprotroph (Põlme et al. 2020); Decay sugar-rich substrates (Põlme et al. 2020) | Põlme S, et al. 2020. Fungal Diversity 105:1-16 ((https://doi.org/10.1007/s13225-020-00466-2)) |
| Otu23545 | | 1 | k__Fungi(100);p__Ascomycota(100);c__Saccharomycetes(100);o__Saccharomycetales(100);f__Metschnikowiaceae(100);g__Metschnikowia(100);g__Metschnikowia_unclassified(100); | Metschnikowia | | 13 | Saprotroph | \|Undefined Saprotroph\| | Yeast | NULL | Probable | As nectar-tap_saprotroph (Põlme et al. 2020); Decay sugar-rich substrates (Põlme et al. 2020) | Põlme S, et al. 2020. Fungal Diversity 105:1-16 ((https://doi.org/10.1007/s13225-020-00466-2)) |
| Otu23546 | | 1 | k__Fungi(100);p__Ascomycota(100);c__Saccharomycetes(100);o__Saccharomycetales(100);f__Metschnikowiaceae(100);g__Metschnikowia(100);g__Metschnikowia_unclassified(100); | Metschnikowia | | 13 | Saprotroph | \|Undefined Saprotroph\| | Yeast | NULL | Probable | As nectar-tap_saprotroph (Põlme et al. 2020); Decay sugar-rich substrates (Põlme et al. 2020) | Põlme S, et al. 2020. Fungal Diversity 105:1-16 ((https://doi.org/10.1007/s13225-020-00466-2)) |
| Otu23547 | | 1 | k__Fungi(100);p__Ascomycota(100);c__Saccharomycetes(100);o__Saccharomycetales(100);f__Metschnikowiaceae(100);g__Metschnikowia(100);s__Metschnikowia_pulcherrima(100); | Metschnikowia | | 13 | Saprotroph | \|Undefined Saprotroph\| | Yeast | NULL | Probable | As nectar-tap_saprotroph (Põlme et al. 2020); Decay sugar-rich substrates (Põlme et al. 2020) | Põlme S, et al. 2020. Fungal Diversity 105:1-16 ((https://doi.org/10.1007/s13225-020-00466-2)) |
| Otu23548 | | 1 | k__Fungi(100);p__Ascomycota(100);c__Saccharomycetes(100);o__Saccharomycetales(100);f__Metschnikowiaceae(100);g__Metschnikowia(100);s__Metschnikowia_pulcherrima(100); | Metschnikowia | | 13 | Saprotroph | \|Undefined Saprotroph\| | Yeast | NULL | Probable | As nectar-tap_saprotroph (Põlme et al. 2020); Decay sugar-rich substrates (Põlme et al. 2020) | Põlme S, et al. 2020. Fungal Diversity 105:1-16 ((https://doi.org/10.1007/s13225-020-00466-2)) |
| Otu23549 | | 1 | k__Fungi(100);p__Ascomycota(100);c__Saccharomycetes(100);o__Saccharomycetales(100);f__Metschnikowiaceae(100);g__Metschnikowia(100);s__Metschnikowia_pulcherrima(100); | Metschnikowia | | 13 | Saprotroph | \|Undefined Saprotroph\| | Yeast | NULL | Probable | As nectar-tap_saprotroph (Põlme et al. 2020); Decay sugar-rich substrates (Põlme et al. 2020) | Põlme S, et al. 2020. Fungal Diversity 105:1-16 ((https://doi.org/10.1007/s13225-020-00466-2)) |
| Otu23550 | | 1 | k__Fungi(100);p__Ascomycota(100);c__Saccharomycetes(100);o__Saccharomycetales(100);f__Metschnikowiaceae(100);g__Metschnikowia(100);s__Metschnikowia_pulcherrima(100); | Metschnikowia | | 13 | Saprotroph | \|Undefined Saprotroph\| | Yeast | NULL | Probable | As nectar-tap_saprotroph (Põlme et al. 2020); Decay sugar-rich substrates (Põlme et al. 2020) | Põlme S, et al. 2020. Fungal Diversity 105:1-16 ((https://doi.org/10.1007/s13225-020-00466-2)) |
| Otu23551 | | 1 | k__Fungi(100);p__Ascomycota(100);c__Saccharomycetes(100);o__Saccharomycetales(100);f__Metschnikowiaceae(100);g__Metschnikowia(100);s__Metschnikowia_pulcherrima(100); | Metschnikowia | | 13 | Saprotroph | \|Undefined Saprotroph\| | Yeast | NULL | Probable | As nectar-tap_saprotroph (Põlme et al. 2020); Decay sugar-rich substrates (Põlme et al. 2020) | Põlme S, et al. 2020. Fungal Diversity 105:1-16 ((https://doi.org/10.1007/s13225-020-00466-2)) |
| Otu23552 | | 1 | k__Fungi(100);p__Ascomycota(100);c__Saccharomycetes(100);o__Saccharomycetales(100);f__Metschnikowiaceae(100);g__Metschnikowia(100);s__Metschnikowia_pulcherrima(100); | Metschnikowia | | 13 | Saprotroph | \|Undefined Saprotroph\| | Yeast | NULL | Probable | As nectar-tap_saprotroph (Põlme et al. 2020); Decay sugar-rich substrates (Põlme et al. 2020) | Põlme S, et al. 2020. Fungal Diversity 105:1-16 ((https://doi.org/10.1007/s13225-020-00466-2)) |
| Otu23553 | | 1 | k__Fungi(100);p__Ascomycota(100);c__Saccharomycetes(100);o__Saccharomycetales(100);f__Metschnikowiaceae(100);g__Metschnikowia(100);s__Metschnikowia_pulcherrima(100); | Metschnikowia | | 13 | Saprotroph | \|Undefined Saprotroph\| | Yeast | NULL | Probable | As nectar-tap_saprotroph (Põlme et al. 2020); Decay sugar-rich substrates (Põlme et al. 2020) | Põlme S, et al. 2020. Fungal Diversity 105:1-16 ((https://doi.org/10.1007/s13225-020-00466-2)) |
| Otu23554 | | 1 | k__Fungi(100);p__Ascomycota(100);c__Saccharomycetes(100);o__Saccharomycetales(100);f__Metschnikowiaceae(100);g__Metschnikowia(100);s__Metschnikowia_pulcherrima(100); | Metschnikowia | | 13 | Saprotroph | \|Undefined Saprotroph\| | Yeast | NULL | Probable | As nectar-tap_saprotroph (Põlme et al. 2020); Decay sugar-rich substrates (Põlme et al. 2020) | Põlme S, et al. 2020. Fungal Diversity 105:1-16 ((https://doi.org/10.1007/s13225-020-00466-2)) |
| Otu23555 | | 1 | k__Fungi(100);p__Ascomycota(100);c__Saccharomycetes(100);o__Saccharomycetales(100);f__Metschnikowiaceae(100);g__Metschnikowia(100);s__Metschnikowia_pulcherrima(100); | Metschnikowia | | 13 | Saprotroph | \|Undefined Saprotroph\| | Yeast | NULL | Probable | As nectar-tap_saprotroph (Põlme et al. 2020); Decay sugar-rich substrates (Põlme et al. 2020) | Põlme S, et al. 2020. Fungal Diversity 105:1-16 ((https://doi.org/10.1007/s13225-020-00466-2)) |
| Otu23558 | | 1 | k__Fungi(100);p__Ascomycota(100);c__Saccharomycetes(100);o__Saccharomycetales(100);f__Metschnikowiaceae(100);g__Metschnikowia(100);s__Metschnikowia_pulcherrima(100); | Metschnikowia | | 13 | Saprotroph | \|Undefined Saprotroph\| | Yeast | NULL | Probable | As nectar-tap_saprotroph (Põlme et al. 2020); Decay sugar-rich substrates (Põlme et al. 2020) | Põlme S, et al. 2020. Fungal Diversity 105:1-16 ((https://doi.org/10.1007/s13225-020-00466-2)) |
| Otu23559 | | 1 | k__Fungi(100);p__Ascomycota(100);c__Saccharomycetes(100);o__Saccharomycetales(100);f__Metschnikowiaceae(100);g__Metschnikowia(100);s__Metschnikowia_pulcherrima(100); | Metschnikowia | | 13 | Saprotroph | \|Undefined Saprotroph\| | Yeast | NULL | Probable | As nectar-tap_saprotroph (Põlme et al. 2020); Decay sugar-rich substrates (Põlme et al. 2020) | Põlme S, et al. 2020. Fungal Diversity 105:1-16 ((https://doi.org/10.1007/s13225-020-00466-2)) |
| Otu23560 | | 1 | k__Fungi(100);p__Ascomycota(100);c__Saccharomycetes(100);o__Saccharomycetales(100);f__Metschnikowiaceae(100);g__Metschnikowia(100);s__Metschnikowia_pulcherrima(100); | Metschnikowia | | 13 | Saprotroph | \|Undefined Saprotroph\| | Yeast | NULL | Probable | As nectar-tap_saprotroph (Põlme et al. 2020); Decay sugar-rich substrates (Põlme et al. 2020) | Põlme S, et al. 2020. Fungal Diversity 105:1-16 ((https://doi.org/10.1007/s13225-020-00466-2)) |
| Otu23561 | | 1 | k__Fungi(100);p__Ascomycota(100);c__Saccharomycetes(100);o__Saccharomycetales(100);f__Metschnikowiaceae(100);g__Metschnikowia(100);s__Metschnikowia_pulcherrima(100); | Metschnikowia | | 13 | Saprotroph | \|Undefined Saprotroph\| | Yeast | NULL | Probable | As nectar-tap_saprotroph (Põlme et al. 2020); Decay sugar-rich substrates (Põlme et al. 2020) | Põlme S, et al. 2020. Fungal Diversity 105:1-16 ((https://doi.org/10.1007/s13225-020-00466-2)) |
| Otu23562 | | 1 | k__Fungi(100);p__Ascomycota(100);c__Saccharomycetes(100);o__Saccharomycetales(100);f__Metschnikowiaceae(100);g__Metschnikowia(100);s__Metschnikowia_pulcherrima(100); | Metschnikowia | | 13 | Saprotroph | \|Undefined Saprotroph\| | Yeast | NULL | Probable | As nectar-tap_saprotroph (Põlme et al. 2020); Decay sugar-rich substrates (Põlme et al. 2020) | Põlme S, et al. 2020. Fungal Diversity 105:1-16 ((https://doi.org/10.1007/s13225-020-00466-2)) |
| Otu23563 | | 1 | k__Fungi(100);p__Ascomycota(100);c__Saccharomycetes(100);o__Saccharomycetales(100);f__Metschnikowiaceae(100);g__Metschnikowia(100);s__Metschnikowia_pulcherrima(100); | Metschnikowia | | 13 | Saprotroph | \|Undefined Saprotroph\| | Yeast | NULL | Probable | As nectar-tap_saprotroph (Põlme et al. 2020); Decay sugar-rich substrates (Põlme et al. 2020) | Põlme S, et al. 2020. Fungal Diversity 105:1-16 ((https://doi.org/10.1007/s13225-020-00466-2)) |
| Otu23564 | | 1 | k__Fungi(100);p__Ascomycota(100);c__Saccharomycetes(100);o__Saccharomycetales(100);f__Metschnikowiaceae(100);g__Metschnikowia(100);s__Metschnikowia_pulcherrima(100); | Metschnikowia | | 13 | Saprotroph | \|Undefined Saprotroph\| | Yeast | NULL | Probable | As nectar-tap_saprotroph (Põlme et al. 2020); Decay sugar-rich substrates (Põlme et al. 2020) | Põlme S, et al. 2020. Fungal Diversity 105:1-16 ((https://doi.org/10.1007/s13225-020-00466-2)) |
| Otu23565 | | 1 | k__Fungi(100);p__Ascomycota(100);c__Saccharomycetes(100);o__Saccharomycetales(100);f__Metschnikowiaceae(100);g__Metschnikowia(100);s__Metschnikowia_pulcherrima(100); | Metschnikowia | | 13 | Saprotroph | \|Undefined Saprotroph\| | Yeast | NULL | Probable | As nectar-tap_saprotroph (Põlme et al. 2020); Decay sugar-rich substrates (Põlme et al. 2020) | Põlme S, et al. 2020. Fungal Diversity 105:1-16 ((https://doi.org/10.1007/s13225-020-00466-2)) |
| Otu23566 | | 1 | k__Fungi(100);p__Ascomycota(100);c__Saccharomycetes(100);o__Saccharomycetales(100);f__Metschnikowiaceae(100);g__Metschnikowia(100);s__Metschnikowia_pulcherrima(100); | Metschnikowia | | 13 | Saprotroph | \|Undefined Saprotroph\| | Yeast | NULL | Probable | As nectar-tap_saprotroph (Põlme et al. 2020); Decay sugar-rich substrates (Põlme et al. 2020) | Põlme S, et al. 2020. Fungal Diversity 105:1-16 ((https://doi.org/10.1007/s13225-020-00466-2)) |
| Otu23568 | | 1 | k__Fungi(100);p__Ascomycota(100);c__Saccharomycetes(100);o__Saccharomycetales(100);f__Metschnikowiaceae(100);g__Metschnikowia(100);g__Metschnikowia_unclassified(100); | Metschnikowia | | 13 | Saprotroph | \|Undefined Saprotroph\| | Yeast | NULL | Probable | As nectar-tap_saprotroph (Põlme et al. 2020); Decay sugar-rich substrates (Põlme et al. 2020) | Põlme S, et al. 2020. Fungal Diversity 105:1-16 ((https://doi.org/10.1007/s13225-020-00466-2)) |
| Otu23569 | | 1 | k__Fungi(100);p__Ascomycota(100);c__Saccharomycetes(100);o__Saccharomycetales(100);f__Metschnikowiaceae(100);g__Metschnikowia(100);s__Metschnikowia_pulcherrima(100); | Metschnikowia | | 13 | Saprotroph | \|Undefined Saprotroph\| | Yeast | NULL | Probable | As nectar-tap_saprotroph (Põlme et al. 2020); Decay sugar-rich substrates (Põlme et al. 2020) | Põlme S, et al. 2020. Fungal Diversity 105:1-16 ((https://doi.org/10.1007/s13225-020-00466-2)) |
| Otu23570 | | 1 | k__Fungi(100);p__Ascomycota(100);c__Saccharomycetes(100);o__Saccharomycetales(100);f__Metschnikowiaceae(100);g__Metschnikowia(100);s__Metschnikowia_pulcherrima(100); | Metschnikowia | | 13 | Saprotroph | \|Undefined Saprotroph\| | Yeast | NULL | Probable | As nectar-tap_saprotroph (Põlme et al. 2020); Decay sugar-rich substrates (Põlme et al. 2020) | Põlme S, et al. 2020. Fungal Diversity 105:1-16 ((https://doi.org/10.1007/s13225-020-00466-2)) |
| Otu23571 | | 1 | k__Fungi(100);p__Ascomycota(100);c__Saccharomycetes(100);o__Saccharomycetales(100);f__Metschnikowiaceae(100);g__Metschnikowia(100);s__Metschnikowia_pulcherrima(100); | Metschnikowia | | 13 | Saprotroph | \|Undefined Saprotroph\| | Yeast | NULL | Probable | As nectar-tap_saprotroph (Põlme et al. 2020); Decay sugar-rich substrates (Põlme et al. 2020) | Põlme S, et al. 2020. Fungal Diversity 105:1-16 ((https://doi.org/10.1007/s13225-020-00466-2)) |
| Otu23572 | | 1 | k__Fungi(100);p__Ascomycota(100);c__Saccharomycetes(100);o__Saccharomycetales(100);f__Metschnikowiaceae(100);g__Metschnikowia(100);g__Metschnikowia_unclassified(100); | Metschnikowia | | 13 | Saprotroph | \|Undefined Saprotroph\| | Yeast | NULL | Probable | As nectar-tap_saprotroph (Põlme et al. 2020); Decay sugar-rich substrates (Põlme et al. 2020) | Põlme S, et al. 2020. Fungal Diversity 105:1-16 ((https://doi.org/10.1007/s13225-020-00466-2)) |
| Otu23573 | | 1 | k__Fungi(100);p__Ascomycota(100);c__Saccharomycetes(100);o__Saccharomycetales(100);f__Metschnikowiaceae(100);g__Metschnikowia(100);s__Metschnikowia_pulcherrima(100); | Metschnikowia | | 13 | Saprotroph | \|Undefined Saprotroph\| | Yeast | NULL | Probable | As nectar-tap_saprotroph (Põlme et al. 2020); Decay sugar-rich substrates (Põlme et al. 2020) | Põlme S, et al. 2020. Fungal Diversity 105:1-16 ((https://doi.org/10.1007/s13225-020-00466-2)) |
| Otu23574 | | 1 | k__Fungi(100);p__Ascomycota(100);c__Saccharomycetes(100);o__Saccharomycetales(100);f__Metschnikowiaceae(100);g__Metschnikowia(100);g__Metschnikowia_unclassified(100); | Metschnikowia | | 13 | Saprotroph | \|Undefined Saprotroph\| | Yeast | NULL | Probable | As nectar-tap_saprotroph (Põlme et al. 2020); Decay sugar-rich substrates (Põlme et al. 2020) | Põlme S, et al. 2020. Fungal Diversity 105:1-16 ((https://doi.org/10.1007/s13225-020-00466-2)) |
| Otu23575 | | 1 | k__Fungi(100);p__Ascomycota(100);c__Saccharomycetes(100);o__Saccharomycetales(100);f__Metschnikowiaceae(100);g__Metschnikowia(100);g__Metschnikowia_unclassified(100); | Metschnikowia | | 13 | Saprotroph | \|Undefined Saprotroph\| | Yeast | NULL | Probable | As nectar-tap_saprotroph (Põlme et al. 2020); Decay sugar-rich substrates (Põlme et al. 2020) | Põlme S, et al. 2020. Fungal Diversity 105:1-16 ((https://doi.org/10.1007/s13225-020-00466-2)) |
| Otu23576 | | 1 | k__Fungi(100);p__Ascomycota(100);c__Saccharomycetes(100);o__Saccharomycetales(100);f__Metschnikowiaceae(100);g__Metschnikowia(100);s__Metschnikowia_pulcherrima(100); | Metschnikowia | | 13 | Saprotroph | \|Undefined Saprotroph\| | Yeast | NULL | Probable | As nectar-tap_saprotroph (Põlme et al. 2020); Decay sugar-rich substrates (Põlme et al. 2020) | Põlme S, et al. 2020. Fungal Diversity 105:1-16 ((https://doi.org/10.1007/s13225-020-00466-2)) |
| Otu23577 | | 1 | k__Fungi(100);p__Ascomycota(100);c__Saccharomycetes(100);o__Saccharomycetales(100);f__Metschnikowiaceae(100);g__Metschnikowia(100);g__Metschnikowia_unclassified(100); | Metschnikowia | | 13 | Saprotroph | \|Undefined Saprotroph\| | Yeast | NULL | Probable | As nectar-tap_saprotroph (Põlme et al. 2020); Decay sugar-rich substrates (Põlme et al. 2020) | Põlme S, et al. 2020. Fungal Diversity 105:1-16 ((https://doi.org/10.1007/s13225-020-00466-2)) |
| Otu23578 | | 1 | k__Fungi(100);p__Ascomycota(100);c__Saccharomycetes(100);o__Saccharomycetales(100);f__Metschnikowiaceae(100);g__Metschnikowia(100);s__Metschnikowia_pulcherrima(100); | Metschnikowia | | 13 | Saprotroph | \|Undefined Saprotroph\| | Yeast | NULL | Probable | As nectar-tap_saprotroph (Põlme et al. 2020); Decay sugar-rich substrates (Põlme et al. 2020) | Põlme S, et al. 2020. Fungal Diversity 105:1-16 ((https://doi.org/10.1007/s13225-020-00466-2)) |
| Otu23579 | | 1 | k__Fungi(100);p__Ascomycota(100);c__Saccharomycetes(100);o__Saccharomycetales(100);f__Metschnikowiaceae(100);g__Metschnikowia(100);s__Metschnikowia_pulcherrima(100); | Metschnikowia | | 13 | Saprotroph | \|Undefined Saprotroph\| | Yeast | NULL | Probable | As nectar-tap_saprotroph (Põlme et al. 2020); Decay sugar-rich substrates (Põlme et al. 2020) | Põlme S, et al. 2020. Fungal Diversity 105:1-16 ((https://doi.org/10.1007/s13225-020-00466-2)) |
| Otu23580 | | 1 | k__Fungi(100);p__Ascomycota(100);c__Saccharomycetes(100);o__Saccharomycetales(100);f__Metschnikowiaceae(100);g__Metschnikowia(100);g__Metschnikowia_unclassified(100); | Metschnikowia | | 13 | Saprotroph | \|Undefined Saprotroph\| | Yeast | NULL | Probable | As nectar-tap_saprotroph (Põlme et al. 2020); Decay sugar-rich substrates (Põlme et al. 2020) | Põlme S, et al. 2020. Fungal Diversity 105:1-16 ((https://doi.org/10.1007/s13225-020-00466-2)) |
| Otu23581 | | 1 | k__Fungi(100);p__Ascomycota(100);c__Saccharomycetes(100);o__Saccharomycetales(100);f__Metschnikowiaceae(100);g__Metschnikowia(100);s__Metschnikowia_pulcherrima(100); | Metschnikowia | | 13 | Saprotroph | \|Undefined Saprotroph\| | Yeast | NULL | Probable | As nectar-tap_saprotroph (Põlme et al. 2020); Decay sugar-rich substrates (Põlme et al. 2020) | Põlme S, et al. 2020. Fungal Diversity 105:1-16 ((https://doi.org/10.1007/s13225-020-00466-2)) |
| Otu23585 | | 1 | k__Fungi(100);p__Ascomycota(100);c__Saccharomycetes(100);o__Saccharomycetales(100);f__Metschnikowiaceae(100);g__Metschnikowia(100);g__Metschnikowia_unclassified(100); | Metschnikowia | | 13 | Saprotroph | \|Undefined Saprotroph\| | Yeast | NULL | Probable | As nectar-tap_saprotroph (Põlme et al. 2020); Decay sugar-rich substrates (Põlme et al. 2020) | Põlme S, et al. 2020. Fungal Diversity 105:1-16 ((https://doi.org/10.1007/s13225-020-00466-2)) |
| Otu23586 | | 1 | k__Fungi(100);p__Ascomycota(100);c__Saccharomycetes(100);o__Saccharomycetales(100);f__Metschnikowiaceae(100);g__Metschnikowia(100);s__Metschnikowia_pulcherrima(100); | Metschnikowia | | 13 | Saprotroph | \|Undefined Saprotroph\| | Yeast | NULL | Probable | As nectar-tap_saprotroph (Põlme et al. 2020); Decay sugar-rich substrates (Põlme et al. 2020) | Põlme S, et al. 2020. Fungal Diversity 105:1-16 ((https://doi.org/10.1007/s13225-020-00466-2)) |
| Otu23588 | | 1 | k__Fungi(100);p__Ascomycota(100);c__Saccharomycetes(100);o__Saccharomycetales(100);f__Metschnikowiaceae(100);g__Metschnikowia(100);g__Metschnikowia_unclassified(100); | Metschnikowia | | 13 | Saprotroph | \|Undefined Saprotroph\| | Yeast | NULL | Probable | As nectar-tap_saprotroph (Põlme et al. 2020); Decay sugar-rich substrates (Põlme et al. 2020) | Põlme S, et al. 2020. Fungal Diversity 105:1-16 ((https://doi.org/10.1007/s13225-020-00466-2)) |
| Otu23589 | | 1 | k__Fungi(100);p__Ascomycota(100);c__Saccharomycetes(100);o__Saccharomycetales(100);f__Metschnikowiaceae(100);g__Metschnikowia(100);s__Metschnikowia_pulcherrima(100); | Metschnikowia | | 13 | Saprotroph | \|Undefined Saprotroph\| | Yeast | NULL | Probable | As nectar-tap_saprotroph (Põlme et al. 2020); Decay sugar-rich substrates (Põlme et al. 2020) | Põlme S, et al. 2020. Fungal Diversity 105:1-16 ((https://doi.org/10.1007/s13225-020-00466-2)) |
| Otu23590 | | 1 | k__Fungi(100);p__Ascomycota(100);c__Saccharomycetes(100);o__Saccharomycetales(100);f__Metschnikowiaceae(100);g__Metschnikowia(100);s__Metschnikowia_pulcherrima(100); | Metschnikowia | | 13 | Saprotroph | \|Undefined Saprotroph\| | Yeast | NULL | Probable | As nectar-tap_saprotroph (Põlme et al. 2020); Decay sugar-rich substrates (Põlme et al. 2020) | Põlme S, et al. 2020. Fungal Diversity 105:1-16 ((https://doi.org/10.1007/s13225-020-00466-2)) |
| Otu23591 | | 1 | k__Fungi(100);p__Ascomycota(100);c__Saccharomycetes(100);o__Saccharomycetales(100);f__Metschnikowiaceae(100);g__Metschnikowia(100);g__Metschnikowia_unclassified(100); | Metschnikowia | | 13 | Saprotroph | \|Undefined Saprotroph\| | Yeast | NULL | Probable | As nectar-tap_saprotroph (Põlme et al. 2020); Decay sugar-rich substrates (Põlme et al. 2020) | Põlme S, et al. 2020. Fungal Diversity 105:1-16 ((https://doi.org/10.1007/s13225-020-00466-2)) |
| Otu23592 | | 1 | k__Fungi(100);p__Ascomycota(100);c__Saccharomycetes(100);o__Saccharomycetales(100);f__Metschnikowiaceae(100);g__Metschnikowia(100);s__Metschnikowia_pulcherrima(100); | Metschnikowia | | 13 | Saprotroph | \|Undefined Saprotroph\| | Yeast | NULL | Probable | As nectar-tap_saprotroph (Põlme et al. 2020); Decay sugar-rich substrates (Põlme et al. 2020) | Põlme S, et al. 2020. Fungal Diversity 105:1-16 ((https://doi.org/10.1007/s13225-020-00466-2)) |
| Otu23593 | | 1 | k__Fungi(100);p__Ascomycota(100);c__Saccharomycetes(100);o__Saccharomycetales(100);f__Metschnikowiaceae(100);g__Metschnikowia(100);g__Metschnikowia_unclassified(100); | Metschnikowia | | 13 | Saprotroph | \|Undefined Saprotroph\| | Yeast | NULL | Probable | As nectar-tap_saprotroph (Põlme et al. 2020); Decay sugar-rich substrates (Põlme et al. 2020) | Põlme S, et al. 2020. Fungal Diversity 105:1-16 ((https://doi.org/10.1007/s13225-020-00466-2)) |
| Otu23594 | | 1 | k__Fungi(100);p__Ascomycota(100);c__Saccharomycetes(100);o__Saccharomycetales(100);f__Metschnikowiaceae(100);g__Metschnikowia(100);s__Metschnikowia_pulcherrima(100); | Metschnikowia | | 13 | Saprotroph | \|Undefined Saprotroph\| | Yeast | NULL | Probable | As nectar-tap_saprotroph (Põlme et al. 2020); Decay sugar-rich substrates (Põlme et al. 2020) | Põlme S, et al. 2020. Fungal Diversity 105:1-16 ((https://doi.org/10.1007/s13225-020-00466-2)) |
| Otu23595 | | 1 | k__Fungi(100);p__Ascomycota(100);c__Saccharomycetes(100);o__Saccharomycetales(100);f__Metschnikowiaceae(100);g__Metschnikowia(100);s__Metschnikowia_pulcherrima(100); | Metschnikowia | | 13 | Saprotroph | \|Undefined Saprotroph\| | Yeast | NULL | Probable | As nectar-tap_saprotroph (Põlme et al. 2020); Decay sugar-rich substrates (Põlme et al. 2020) | Põlme S, et al. 2020. Fungal Diversity 105:1-16 ((https://doi.org/10.1007/s13225-020-00466-2)) |
| Otu23596 | | 1 | k__Fungi(100);p__Ascomycota(100);c__Saccharomycetes(100);o__Saccharomycetales(100);f__Metschnikowiaceae(100);g__Metschnikowia(100);g__Metschnikowia_unclassified(100); | Metschnikowia | | 13 | Saprotroph | \|Undefined Saprotroph\| | Yeast | NULL | Probable | As nectar-tap_saprotroph (Põlme et al. 2020); Decay sugar-rich substrates (Põlme et al. 2020) | Põlme S, et al. 2020. Fungal Diversity 105:1-16 ((https://doi.org/10.1007/s13225-020-00466-2)) |
| Otu23597 | | 1 | k__Fungi(100);p__Ascomycota(100);c__Saccharomycetes(100);o__Saccharomycetales(100);f__Metschnikowiaceae(100);g__Metschnikowia(100);s__Metschnikowia_pulcherrima(100); | Metschnikowia | | 13 | Saprotroph | \|Undefined Saprotroph\| | Yeast | NULL | Probable | As nectar-tap_saprotroph (Põlme et al. 2020); Decay sugar-rich substrates (Põlme et al. 2020) | Põlme S, et al. 2020. Fungal Diversity 105:1-16 ((https://doi.org/10.1007/s13225-020-00466-2)) |
| Otu23598 | | 1 | k__Fungi(100);p__Ascomycota(100);c__Saccharomycetes(100);o__Saccharomycetales(100);f__Metschnikowiaceae(100);g__Metschnikowia(100);s__Metschnikowia_pulcherrima(100); | Metschnikowia | | 13 | Saprotroph | \|Undefined Saprotroph\| | Yeast | NULL | Probable | As nectar-tap_saprotroph (Põlme et al. 2020); Decay sugar-rich substrates (Põlme et al. 2020) | Põlme S, et al. 2020. Fungal Diversity 105:1-16 ((https://doi.org/10.1007/s13225-020-00466-2)) |
| Otu23599 | | 1 | k__Fungi(100);p__Ascomycota(100);c__Saccharomycetes(100);o__Saccharomycetales(100);f__Metschnikowiaceae(100);g__Metschnikowia(100);s__Metschnikowia_pulcherrima(100); | Metschnikowia | | 13 | Saprotroph | \|Undefined Saprotroph\| | Yeast | NULL | Probable | As nectar-tap_saprotroph (Põlme et al. 2020); Decay sugar-rich substrates (Põlme et al. 2020) | Põlme S, et al. 2020. Fungal Diversity 105:1-16 ((https://doi.org/10.1007/s13225-020-00466-2)) |
| Otu23601 | | 1 | k__Fungi(100);p__Ascomycota(100);c__Saccharomycetes(100);o__Saccharomycetales(100);f__Metschnikowiaceae(100);g__Metschnikowia(100);s__Metschnikowia_pulcherrima(100); | Metschnikowia | | 13 | Saprotroph | \|Undefined Saprotroph\| | Yeast | NULL | Probable | As nectar-tap_saprotroph (Põlme et al. 2020); Decay sugar-rich substrates (Põlme et al. 2020) | Põlme S, et al. 2020. Fungal Diversity 105:1-16 ((https://doi.org/10.1007/s13225-020-00466-2)) |
| Otu23604 | | 1 | k__Fungi(100);p__Ascomycota(100);c__Saccharomycetes(100);o__Saccharomycetales(100);f__Metschnikowiaceae(100);g__Metschnikowia(100);s__Metschnikowia_pulcherrima(100); | Metschnikowia | | 13 | Saprotroph | \|Undefined Saprotroph\| | Yeast | NULL | Probable | As nectar-tap_saprotroph (Põlme et al. 2020); Decay sugar-rich substrates (Põlme et al. 2020) | Põlme S, et al. 2020. Fungal Diversity 105:1-16 ((https://doi.org/10.1007/s13225-020-00466-2)) |
| Otu23608 | | 1 | k__Fungi(100);p__Ascomycota(100);c__Saccharomycetes(100);o__Saccharomycetales(100);f__Metschnikowiaceae(100);g__Metschnikowia(100);g__Metschnikowia_unclassified(100); | Metschnikowia | | 13 | Saprotroph | \|Undefined Saprotroph\| | Yeast | NULL | Probable | As nectar-tap_saprotroph (Põlme et al. 2020); Decay sugar-rich substrates (Põlme et al. 2020) | Põlme S, et al. 2020. Fungal Diversity 105:1-16 ((https://doi.org/10.1007/s13225-020-00466-2)) |
| Otu23609 | | 1 | k__Fungi(100);p__Ascomycota(100);c__Saccharomycetes(100);o__Saccharomycetales(100);f__Metschnikowiaceae(100);g__Metschnikowia(100);s__Metschnikowia_pulcherrima(100); | Metschnikowia | | 13 | Saprotroph | \|Undefined Saprotroph\| | Yeast | NULL | Probable | As nectar-tap_saprotroph (Põlme et al. 2020); Decay sugar-rich substrates (Põlme et al. 2020) | Põlme S, et al. 2020. Fungal Diversity 105:1-16 ((https://doi.org/10.1007/s13225-020-00466-2)) |
| Otu23612 | | 1 | k__Fungi(100);p__Ascomycota(100);c__Saccharomycetes(100);o__Saccharomycetales(100);f__Metschnikowiaceae(100);g__Metschnikowia(100);s__Metschnikowia_pulcherrima(100); | Metschnikowia | | 13 | Saprotroph | \|Undefined Saprotroph\| | Yeast | NULL | Probable | As nectar-tap_saprotroph (Põlme et al. 2020); Decay sugar-rich substrates (Põlme et al. 2020) | Põlme S, et al. 2020. Fungal Diversity 105:1-16 ((https://doi.org/10.1007/s13225-020-00466-2)) |
| Otu23613 | | 1 | k__Fungi(100);p__Ascomycota(100);c__Saccharomycetes(100);o__Saccharomycetales(100);f__Metschnikowiaceae(100);g__Metschnikowia(100);s__Metschnikowia_pulcherrima(100); | Metschnikowia | | 13 | Saprotroph | \|Undefined Saprotroph\| | Yeast | NULL | Probable | As nectar-tap_saprotroph (Põlme et al. 2020); Decay sugar-rich substrates (Põlme et al. 2020) | Põlme S, et al. 2020. Fungal Diversity 105:1-16 ((https://doi.org/10.1007/s13225-020-00466-2)) |
| Otu23614 | | 1 | k__Fungi(100);p__Ascomycota(100);c__Saccharomycetes(100);o__Saccharomycetales(100);f__Metschnikowiaceae(100);g__Metschnikowia(100);g__Metschnikowia_unclassified(100); | Metschnikowia | | 13 | Saprotroph | \|Undefined Saprotroph\| | Yeast | NULL | Probable | As nectar-tap_saprotroph (Põlme et al. 2020); Decay sugar-rich substrates (Põlme et al. 2020) | Põlme S, et al. 2020. Fungal Diversity 105:1-16 ((https://doi.org/10.1007/s13225-020-00466-2)) |
| Otu23629 | | 1 | k__Fungi(100);p__Ascomycota(100);c__Saccharomycetes(100);o__Saccharomycetales(100);f__Metschnikowiaceae(100);g__Metschnikowia(100);s__Metschnikowia_pulcherrima(100); | Metschnikowia | | 13 | Saprotroph | \|Undefined Saprotroph\| | Yeast | NULL | Probable | As nectar-tap_saprotroph (Põlme et al. 2020); Decay sugar-rich substrates (Põlme et al. 2020) | Põlme S, et al. 2020. Fungal Diversity 105:1-16 ((https://doi.org/10.1007/s13225-020-00466-2)) |
| Otu23630 | | 1 | k__Fungi(100);p__Ascomycota(100);c__Saccharomycetes(100);o__Saccharomycetales(100);f__Metschnikowiaceae(100);g__Metschnikowia(100);s__Metschnikowia_pulcherrima(100); | Metschnikowia | | 13 | Saprotroph | \|Undefined Saprotroph\| | Yeast | NULL | Probable | As nectar-tap_saprotroph (Põlme et al. 2020); Decay sugar-rich substrates (Põlme et al. 2020) | Põlme S, et al. 2020. Fungal Diversity 105:1-16 ((https://doi.org/10.1007/s13225-020-00466-2)) |
| Otu23631 | | 1 | k__Fungi(100);p__Ascomycota(100);c__Saccharomycetes(100);o__Saccharomycetales(100);f__Metschnikowiaceae(100);g__Metschnikowia(100);g__Metschnikowia_unclassified(100); | Metschnikowia | | 13 | Saprotroph | \|Undefined Saprotroph\| | Yeast | NULL | Probable | As nectar-tap_saprotroph (Põlme et al. 2020); Decay sugar-rich substrates (Põlme et al. 2020) | Põlme S, et al. 2020. Fungal Diversity 105:1-16 ((https://doi.org/10.1007/s13225-020-00466-2)) |
| Otu23632 | | 1 | k__Fungi(100);p__Ascomycota(100);c__Saccharomycetes(100);o__Saccharomycetales(100);f__Metschnikowiaceae(100);g__Metschnikowia(100);g__Metschnikowia_unclassified(100); | Metschnikowia | | 13 | Saprotroph | \|Undefined Saprotroph\| | Yeast | NULL | Probable | As nectar-tap_saprotroph (Põlme et al. 2020); Decay sugar-rich substrates (Põlme et al. 2020) | Põlme S, et al. 2020. Fungal Diversity 105:1-16 ((https://doi.org/10.1007/s13225-020-00466-2)) |
| Otu23633 | | 1 | k__Fungi(100);p__Ascomycota(100);c__Saccharomycetes(100);o__Saccharomycetales(100);f__Metschnikowiaceae(100);g__Metschnikowia(100);s__Metschnikowia_pulcherrima(100); | Metschnikowia | | 13 | Saprotroph | \|Undefined Saprotroph\| | Yeast | NULL | Probable | As nectar-tap_saprotroph (Põlme et al. 2020); Decay sugar-rich substrates (Põlme et al. 2020) | Põlme S, et al. 2020. Fungal Diversity 105:1-16 ((https://doi.org/10.1007/s13225-020-00466-2)) |
| Otu23634 | | 1 | k__Fungi(100);p__Ascomycota(100);c__Saccharomycetes(100);o__Saccharomycetales(100);f__Metschnikowiaceae(100);g__Metschnikowia(100);s__Metschnikowia_pulcherrima(100); | Metschnikowia | | 13 | Saprotroph | \|Undefined Saprotroph\| | Yeast | NULL | Probable | As nectar-tap_saprotroph (Põlme et al. 2020); Decay sugar-rich substrates (Põlme et al. 2020) | Põlme S, et al. 2020. Fungal Diversity 105:1-16 ((https://doi.org/10.1007/s13225-020-00466-2)) |
| Otu23635 | | 1 | k__Fungi(100);p__Ascomycota(100);c__Saccharomycetes(100);o__Saccharomycetales(100);f__Metschnikowiaceae(100);g__Metschnikowia(100);s__Metschnikowia_pulcherrima(100); | Metschnikowia | | 13 | Saprotroph | \|Undefined Saprotroph\| | Yeast | NULL | Probable | As nectar-tap_saprotroph (Põlme et al. 2020); Decay sugar-rich substrates (Põlme et al. 2020) | Põlme S, et al. 2020. Fungal Diversity 105:1-16 ((https://doi.org/10.1007/s13225-020-00466-2)) |
| Otu23636 | | 1 | k__Fungi(100);p__Ascomycota(100);c__Saccharomycetes(100);o__Saccharomycetales(100);f__Metschnikowiaceae(100);g__Metschnikowia(100);s__Metschnikowia_pulcherrima(100); | Metschnikowia | | 13 | Saprotroph | \|Undefined Saprotroph\| | Yeast | NULL | Probable | As nectar-tap_saprotroph (Põlme et al. 2020); Decay sugar-rich substrates (Põlme et al. 2020) | Põlme S, et al. 2020. Fungal Diversity 105:1-16 ((https://doi.org/10.1007/s13225-020-00466-2)) |
| Otu23637 | | 1 | k__Fungi(100);p__Ascomycota(100);c__Saccharomycetes(100);o__Saccharomycetales(100);f__Metschnikowiaceae(100);g__Metschnikowia(100);g__Metschnikowia_unclassified(100); | Metschnikowia | | 13 | Saprotroph | \|Undefined Saprotroph\| | Yeast | NULL | Probable | As nectar-tap_saprotroph (Põlme et al. 2020); Decay sugar-rich substrates (Põlme et al. 2020) | Põlme S, et al. 2020. Fungal Diversity 105:1-16 ((https://doi.org/10.1007/s13225-020-00466-2)) |
| Otu23638 | | 1 | k__Fungi(100);p__Ascomycota(100);c__Saccharomycetes(100);o__Saccharomycetales(100);f__Metschnikowiaceae(100);g__Metschnikowia(100);s__Metschnikowia_pulcherrima(100); | Metschnikowia | | 13 | Saprotroph | \|Undefined Saprotroph\| | Yeast | NULL | Probable | As nectar-tap_saprotroph (Põlme et al. 2020); Decay sugar-rich substrates (Põlme et al. 2020) | Põlme S, et al. 2020. Fungal Diversity 105:1-16 ((https://doi.org/10.1007/s13225-020-00466-2)) |
| Otu23639 | | 1 | k__Fungi(100);p__Ascomycota(100);c__Saccharomycetes(100);o__Saccharomycetales(100);f__Metschnikowiaceae(100);g__Metschnikowia(100);s__Metschnikowia_pulcherrima(100); | Metschnikowia | | 13 | Saprotroph | \|Undefined Saprotroph\| | Yeast | NULL | Probable | As nectar-tap_saprotroph (Põlme et al. 2020); Decay sugar-rich substrates (Põlme et al. 2020) | Põlme S, et al. 2020. Fungal Diversity 105:1-16 ((https://doi.org/10.1007/s13225-020-00466-2)) |
| Otu23641 | | 1 | k__Fungi(100);p__Ascomycota(100);c__Saccharomycetes(100);o__Saccharomycetales(100);f__Metschnikowiaceae(100);g__Metschnikowia(100);s__Metschnikowia_pulcherrima(100); | Metschnikowia | | 13 | Saprotroph | \|Undefined Saprotroph\| | Yeast | NULL | Probable | As nectar-tap_saprotroph (Põlme et al. 2020); Decay sugar-rich substrates (Põlme et al. 2020) | Põlme S, et al. 2020. Fungal Diversity 105:1-16 ((https://doi.org/10.1007/s13225-020-00466-2)) |
| Otu23642 | | 1 | k__Fungi(100);p__Ascomycota(100);c__Saccharomycetes(100);o__Saccharomycetales(100);f__Metschnikowiaceae(100);g__Metschnikowia(100);s__Metschnikowia_pulcherrima(100); | Metschnikowia | | 13 | Saprotroph | \|Undefined Saprotroph\| | Yeast | NULL | Probable | As nectar-tap_saprotroph (Põlme et al. 2020); Decay sugar-rich substrates (Põlme et al. 2020) | Põlme S, et al. 2020. Fungal Diversity 105:1-16 ((https://doi.org/10.1007/s13225-020-00466-2)) |
| Otu23643 | | 1 | k__Fungi(100);p__Ascomycota(100);c__Saccharomycetes(100);o__Saccharomycetales(100);f__Metschnikowiaceae(100);g__Metschnikowia(100);s__Metschnikowia_pulcherrima(100); | Metschnikowia | | 13 | Saprotroph | \|Undefined Saprotroph\| | Yeast | NULL | Probable | As nectar-tap_saprotroph (Põlme et al. 2020); Decay sugar-rich substrates (Põlme et al. 2020) | Põlme S, et al. 2020. Fungal Diversity 105:1-16 ((https://doi.org/10.1007/s13225-020-00466-2)) |
| Otu23644 | | 1 | k__Fungi(100);p__Ascomycota(100);c__Saccharomycetes(100);o__Saccharomycetales(100);f__Metschnikowiaceae(100);g__Metschnikowia(100);s__Metschnikowia_pulcherrima(100); | Metschnikowia | | 13 | Saprotroph | \|Undefined Saprotroph\| | Yeast | NULL | Probable | As nectar-tap_saprotroph (Põlme et al. 2020); Decay sugar-rich substrates (Põlme et al. 2020) | Põlme S, et al. 2020. Fungal Diversity 105:1-16 ((https://doi.org/10.1007/s13225-020-00466-2)) |
| Otu23645 | | 1 | k__Fungi(100);p__Ascomycota(100);c__Saccharomycetes(100);o__Saccharomycetales(100);f__Metschnikowiaceae(100);g__Metschnikowia(100);s__Metschnikowia_pulcherrima(100); | Metschnikowia | | 13 | Saprotroph | \|Undefined Saprotroph\| | Yeast | NULL | Probable | As nectar-tap_saprotroph (Põlme et al. 2020); Decay sugar-rich substrates (Põlme et al. 2020) | Põlme S, et al. 2020. Fungal Diversity 105:1-16 ((https://doi.org/10.1007/s13225-020-00466-2)) |
| Otu23646 | | 1 | k__Fungi(100);p__Ascomycota(100);c__Saccharomycetes(100);o__Saccharomycetales(100);f__Metschnikowiaceae(100);g__Metschnikowia(100);s__Metschnikowia_pulcherrima(100); | Metschnikowia | | 13 | Saprotroph | \|Undefined Saprotroph\| | Yeast | NULL | Probable | As nectar-tap_saprotroph (Põlme et al. 2020); Decay sugar-rich substrates (Põlme et al. 2020) | Põlme S, et al. 2020. Fungal Diversity 105:1-16 ((https://doi.org/10.1007/s13225-020-00466-2)) |
| Otu23649 | | 1 | k__Fungi(100);p__Ascomycota(100);c__Saccharomycetes(100);o__Saccharomycetales(100);f__Metschnikowiaceae(100);g__Metschnikowia(100);g__Metschnikowia_unclassified(100); | Metschnikowia | | 13 | Saprotroph | \|Undefined Saprotroph\| | Yeast | NULL | Probable | As nectar-tap_saprotroph (Põlme et al. 2020); Decay sugar-rich substrates (Põlme et al. 2020) | Põlme S, et al. 2020. Fungal Diversity 105:1-16 ((https://doi.org/10.1007/s13225-020-00466-2)) |
| Otu23651 | | 1 | k__Fungi(100);p__Ascomycota(100);c__Saccharomycetes(100);o__Saccharomycetales(100);f__Metschnikowiaceae(100);g__Metschnikowia(100);s__Metschnikowia_pulcherrima(100); | Metschnikowia | | 13 | Saprotroph | \|Undefined Saprotroph\| | Yeast | NULL | Probable | As nectar-tap_saprotroph (Põlme et al. 2020); Decay sugar-rich substrates (Põlme et al. 2020) | Põlme S, et al. 2020. Fungal Diversity 105:1-16 ((https://doi.org/10.1007/s13225-020-00466-2)) |
| Otu23657 | | 1 | k__Fungi(100);p__Ascomycota(100);c__Saccharomycetes(100);o__Saccharomycetales(100);f__Metschnikowiaceae(100);g__Metschnikowia(100);s__Metschnikowia_pulcherrima(100); | Metschnikowia | | 13 | Saprotroph | \|Undefined Saprotroph\| | Yeast | NULL | Probable | As nectar-tap_saprotroph (Põlme et al. 2020); Decay sugar-rich substrates (Põlme et al. 2020) | Põlme S, et al. 2020. Fungal Diversity 105:1-16 ((https://doi.org/10.1007/s13225-020-00466-2)) |
| Otu23658 | | 1 | k__Fungi(100);p__Ascomycota(100);c__Saccharomycetes(100);o__Saccharomycetales(100);f__Metschnikowiaceae(100);g__Metschnikowia(100);s__Metschnikowia_pulcherrima(100); | Metschnikowia | | 13 | Saprotroph | \|Undefined Saprotroph\| | Yeast | NULL | Probable | As nectar-tap_saprotroph (Põlme et al. 2020); Decay sugar-rich substrates (Põlme et al. 2020) | Põlme S, et al. 2020. Fungal Diversity 105:1-16 ((https://doi.org/10.1007/s13225-020-00466-2)) |
| Otu23661 | | 1 | k__Fungi(100);p__Ascomycota(100);c__Saccharomycetes(100);o__Saccharomycetales(100);f__Saccharomycetales_family_Incertae_sedis(100);f__Saccharomycetales_family_Incertae_sedis_unclassified(100);f__Saccharomycetales_family_Incertae_sedis_unclassified(100); | Saccharomycetales | | 7 | Saprotroph | Undefined Saprotroph | Yeast | NULL | Possible | NULL | Sterkenburg E, et al. 2015. New Phytologist 207:1145-1158 ((https://doi.org/10.1111/nph.13426)) |
| Otu23662 | | 1 | k__Fungi(100);p__Ascomycota(100);c__Saccharomycetes(100);o__Saccharomycetales(100);f__Metschnikowiaceae(100);g__Metschnikowia(100);s__Metschnikowia_pulcherrima(100); | Metschnikowia | | 13 | Saprotroph | \|Undefined Saprotroph\| | Yeast | NULL | Probable | As nectar-tap_saprotroph (Põlme et al. 2020); Decay sugar-rich substrates (Põlme et al. 2020) | Põlme S, et al. 2020. Fungal Diversity 105:1-16 ((https://doi.org/10.1007/s13225-020-00466-2)) |
| Otu23665 | | 1 | k__Fungi(100);p__Ascomycota(100);c__Saccharomycetes(100);o__Saccharomycetales(100);f__Metschnikowiaceae(100);g__Metschnikowia(100);s__Metschnikowia_pulcherrima(100); | Metschnikowia | | 13 | Saprotroph | \|Undefined Saprotroph\| | Yeast | NULL | Probable | As nectar-tap_saprotroph (Põlme et al. 2020); Decay sugar-rich substrates (Põlme et al. 2020) | Põlme S, et al. 2020. Fungal Diversity 105:1-16 ((https://doi.org/10.1007/s13225-020-00466-2)) |
| Otu23666 | | 1 | k__Fungi(100);p__Ascomycota(100);c__Saccharomycetes(100);o__Saccharomycetales(100);f__Metschnikowiaceae(100);g__Metschnikowia(100);s__Metschnikowia_pulcherrima(100); | Metschnikowia | | 13 | Saprotroph | \|Undefined Saprotroph\| | Yeast | NULL | Probable | As nectar-tap_saprotroph (Põlme et al. 2020); Decay sugar-rich substrates (Põlme et al. 2020) | Põlme S, et al. 2020. Fungal Diversity 105:1-16 ((https://doi.org/10.1007/s13225-020-00466-2)) |
| Otu23671 | | 1 | k__Fungi(100);p__Ascomycota(100);c__Saccharomycetes(100);o__Saccharomycetales(100);f__Metschnikowiaceae(100);g__Metschnikowia(100);s__Metschnikowia_pulcherrima(100); | Metschnikowia | | 13 | Saprotroph | \|Undefined Saprotroph\| | Yeast | NULL | Probable | As nectar-tap_saprotroph (Põlme et al. 2020); Decay sugar-rich substrates (Põlme et al. 2020) | Põlme S, et al. 2020. Fungal Diversity 105:1-16 ((https://doi.org/10.1007/s13225-020-00466-2)) |
| Otu23672 | | 1 | k__Fungi(100);p__Ascomycota(100);c__Saccharomycetes(100);o__Saccharomycetales(100);f__Metschnikowiaceae(100);g__Metschnikowia(100);s__Metschnikowia_pulcherrima(100); | Metschnikowia | | 13 | Saprotroph | \|Undefined Saprotroph\| | Yeast | NULL | Probable | As nectar-tap_saprotroph (Põlme et al. 2020); Decay sugar-rich substrates (Põlme et al. 2020) | Põlme S, et al. 2020. Fungal Diversity 105:1-16 ((https://doi.org/10.1007/s13225-020-00466-2)) |
| Otu23673 | | 1 | k__Fungi(100);p__Ascomycota(100);c__Saccharomycetes(100);o__Saccharomycetales(100);f__Metschnikowiaceae(100);g__Metschnikowia(100);s__Metschnikowia_pulcherrima(100); | Metschnikowia | | 13 | Saprotroph | \|Undefined Saprotroph\| | Yeast | NULL | Probable | As nectar-tap_saprotroph (Põlme et al. 2020); Decay sugar-rich substrates (Põlme et al. 2020) | Põlme S, et al. 2020. Fungal Diversity 105:1-16 ((https://doi.org/10.1007/s13225-020-00466-2)) |
| Otu23674 | | 1 | k__Fungi(100);p__Ascomycota(100);c__Saccharomycetes(100);o__Saccharomycetales(100);f__Metschnikowiaceae(100);g__Metschnikowia(100);s__Metschnikowia_pulcherrima(100); | Metschnikowia | | 13 | Saprotroph | \|Undefined Saprotroph\| | Yeast | NULL | Probable | As nectar-tap_saprotroph (Põlme et al. 2020); Decay sugar-rich substrates (Põlme et al. 2020) | Põlme S, et al. 2020. Fungal Diversity 105:1-16 ((https://doi.org/10.1007/s13225-020-00466-2)) |
| Otu23675 | | 1 | k__Fungi(100);p__Ascomycota(100);c__Saccharomycetes(100);o__Saccharomycetales(100);f__Metschnikowiaceae(100);g__Metschnikowia(100);g__Metschnikowia_unclassified(100); | Metschnikowia | | 13 | Saprotroph | \|Undefined Saprotroph\| | Yeast | NULL | Probable | As nectar-tap_saprotroph (Põlme et al. 2020); Decay sugar-rich substrates (Põlme et al. 2020) | Põlme S, et al. 2020. Fungal Diversity 105:1-16 ((https://doi.org/10.1007/s13225-020-00466-2)) |
| Otu23676 | | 1 | k__Fungi(100);p__Ascomycota(100);c__Dothideomycetes(100);o__Pleosporales(100);f__Phaeosphaeriaceae(100);g__unclassified_Phaeosphaeriaceae(100);s__Phaeosphaeriaceae_sp(100); | Phaeosphaeriaceae | | 9 | Pathotroph-Saprotroph | Fungal Parasite-Plant Pathogen-Plant Saprotroph | Microfungus | NULL | Probable | Necrotrophic or saprobic on a wide range of plants, especially monocotyledons; a few species parasitic on other fungi (Cannon & Kirk 2007) | Cannon PF, Kirk PM. 2007. Fungal Families of the World. CAB International, Cambridge (ISBN: 978-0851998275) |
| Otu23677 | | 1 | k__Fungi(100);p__Ascomycota(100);c__Saccharomycetes(100);o__Saccharomycetales(100);f__Metschnikowiaceae(100);g__Metschnikowia(100);s__Metschnikowia_pulcherrima(100); | Metschnikowia | | 13 | Saprotroph | \|Undefined Saprotroph\| | Yeast | NULL | Probable | As nectar-tap_saprotroph (Põlme et al. 2020); Decay sugar-rich substrates (Põlme et al. 2020) | Põlme S, et al. 2020. Fungal Diversity 105:1-16 ((https://doi.org/10.1007/s13225-020-00466-2)) |
| Otu23678 | | 1 | k__Fungi(100);p__Ascomycota(100);c__Saccharomycetes(100);o__Saccharomycetales(100);f__Metschnikowiaceae(100);g__Metschnikowia(100);s__Metschnikowia_pulcherrima(100); | Metschnikowia | | 13 | Saprotroph | \|Undefined Saprotroph\| | Yeast | NULL | Probable | As nectar-tap_saprotroph (Põlme et al. 2020); Decay sugar-rich substrates (Põlme et al. 2020) | Põlme S, et al. 2020. Fungal Diversity 105:1-16 ((https://doi.org/10.1007/s13225-020-00466-2)) |
| Otu23679 | | 1 | k__Fungi(100);p__Ascomycota(100);c__Saccharomycetes(100);o__Saccharomycetales(100);f__Metschnikowiaceae(100);g__Metschnikowia(100);s__Metschnikowia_pulcherrima(100); | Metschnikowia | | 13 | Saprotroph | \|Undefined Saprotroph\| | Yeast | NULL | Probable | As nectar-tap_saprotroph (Põlme et al. 2020); Decay sugar-rich substrates (Põlme et al. 2020) | Põlme S, et al. 2020. Fungal Diversity 105:1-16 ((https://doi.org/10.1007/s13225-020-00466-2)) |
| Otu23680 | | 1 | k__Fungi(100);p__Ascomycota(100);c__Saccharomycetes(100);o__Saccharomycetales(100);f__Metschnikowiaceae(100);g__Metschnikowia(100);s__Metschnikowia_pulcherrima(100); | Metschnikowia | | 13 | Saprotroph | \|Undefined Saprotroph\| | Yeast | NULL | Probable | As nectar-tap_saprotroph (Põlme et al. 2020); Decay sugar-rich substrates (Põlme et al. 2020) | Põlme S, et al. 2020. Fungal Diversity 105:1-16 ((https://doi.org/10.1007/s13225-020-00466-2)) |
| Otu23681 | | 1 | k__Fungi(100);p__Ascomycota(100);c__Saccharomycetes(100);o__Saccharomycetales(100);f__Metschnikowiaceae(100);g__Metschnikowia(100);s__Metschnikowia_pulcherrima(100); | Metschnikowia | | 13 | Saprotroph | \|Undefined Saprotroph\| | Yeast | NULL | Probable | As nectar-tap_saprotroph (Põlme et al. 2020); Decay sugar-rich substrates (Põlme et al. 2020) | Põlme S, et al. 2020. Fungal Diversity 105:1-16 ((https://doi.org/10.1007/s13225-020-00466-2)) |
| Otu23686 | | 1 | k__Fungi(100);p__Ascomycota(100);c__Saccharomycetes(100);o__Saccharomycetales(100);f__Metschnikowiaceae(100);g__Metschnikowia(100);s__Metschnikowia_pulcherrima(100); | Metschnikowia | | 13 | Saprotroph | \|Undefined Saprotroph\| | Yeast | NULL | Probable | As nectar-tap_saprotroph (Põlme et al. 2020); Decay sugar-rich substrates (Põlme et al. 2020) | Põlme S, et al. 2020. Fungal Diversity 105:1-16 ((https://doi.org/10.1007/s13225-020-00466-2)) |
| Otu23687 | | 1 | k__Fungi(100);p__Ascomycota(100);c__Saccharomycetes(100);o__Saccharomycetales(100);f__Metschnikowiaceae(100);g__Metschnikowia(100);g__Metschnikowia_unclassified(100); | Metschnikowia | | 13 | Saprotroph | \|Undefined Saprotroph\| | Yeast | NULL | Probable | As nectar-tap_saprotroph (Põlme et al. 2020); Decay sugar-rich substrates (Põlme et al. 2020) | Põlme S, et al. 2020. Fungal Diversity 105:1-16 ((https://doi.org/10.1007/s13225-020-00466-2)) |
| Otu23689 | | 1 | k__Fungi(100);p__Ascomycota(100);c__Saccharomycetes(100);o__Saccharomycetales(100);f__Metschnikowiaceae(100);g__Metschnikowia(100);s__Metschnikowia_pulcherrima(100); | Metschnikowia | | 13 | Saprotroph | \|Undefined Saprotroph\| | Yeast | NULL | Probable | As nectar-tap_saprotroph (Põlme et al. 2020); Decay sugar-rich substrates (Põlme et al. 2020) | Põlme S, et al. 2020. Fungal Diversity 105:1-16 ((https://doi.org/10.1007/s13225-020-00466-2)) |
| Otu23691 | | 1 | k__Fungi(100);p__Ascomycota(100);c__Saccharomycetes(100);o__Saccharomycetales(100);f__Metschnikowiaceae(100);g__Metschnikowia(100);g__Metschnikowia_unclassified(100); | Metschnikowia | | 13 | Saprotroph | \|Undefined Saprotroph\| | Yeast | NULL | Probable | As nectar-tap_saprotroph (Põlme et al. 2020); Decay sugar-rich substrates (Põlme et al. 2020) | Põlme S, et al. 2020. Fungal Diversity 105:1-16 ((https://doi.org/10.1007/s13225-020-00466-2)) |
| Otu23693 | | 1 | k__Fungi(100);p__Ascomycota(100);c__Saccharomycetes(100);o__Saccharomycetales(100);f__Saccharomycetales_family_Incertae_sedis(100);g__Candida(100);g__Candida_unclassified(100); | Candida | | 13 | Pathotroph-Saprotroph-Symbiotroph | Animal Pathogen-Endophyte-Endosymbiont-Epiphyte-\|Undefined Saprotroph\| | Dimorphic Yeast-Yeast | NULL | Probable | As nectar-tap_saprotroph (Põlme et al. 2020); Decay sugar-rich substrates (Põlme et al. 2020); This asexual genus is polyphyletic and contain species that function very differently depend on what they are. Caution should be taken when interpreting guilds for Candida species. | Manolakaki D, et al. 2010. Virulence. 1: 367-375 ((https://doi.org/10.4161/viru.1.5.12796)); Mekbib SB, et al. 2011. Tropical Plant Pathology 36:233-240 ((https://doi.org/10.1590/S1982-56762011000400004)); Põlme S, et al. 2020. Fungal Diversity 105:1-16 ((https://doi.org/10.1007/s13225-020-00466-2)) |
| Otu23694 | | 1 | k__Fungi(100);p__Ascomycota(100);c__Saccharomycetes(100);o__Saccharomycetales(100);f__Saccharomycetales_family_Incertae_sedis(100);g__Candida(100);s__Candida_cretensis(100); | Candida | | 13 | Pathotroph-Saprotroph-Symbiotroph | Animal Pathogen-Endophyte-Endosymbiont-Epiphyte-\|Undefined Saprotroph\| | Dimorphic Yeast-Yeast | NULL | Probable | As nectar-tap_saprotroph (Põlme et al. 2020); Decay sugar-rich substrates (Põlme et al. 2020); This asexual genus is polyphyletic and contain species that function very differently depend on what they are. Caution should be taken when interpreting guilds for Candida species. | Manolakaki D, et al. 2010. Virulence. 1: 367-375 ((https://doi.org/10.4161/viru.1.5.12796)); Mekbib SB, et al. 2011. Tropical Plant Pathology 36:233-240 ((https://doi.org/10.1590/S1982-56762011000400004)); Põlme S, et al. 2020. Fungal Diversity 105:1-16 ((https://doi.org/10.1007/s13225-020-00466-2)) |
| Otu23696 | | 1 | k__Fungi(100);p__Ascomycota(100);c__Saccharomycetes(100);o__Saccharomycetales(100);f__Saccharomycetales_family_Incertae_sedis(100);g__Candida(100);g__Candida_unclassified(100); | Candida | | 13 | Pathotroph-Saprotroph-Symbiotroph | Animal Pathogen-Endophyte-Endosymbiont-Epiphyte-\|Undefined Saprotroph\| | Dimorphic Yeast-Yeast | NULL | Probable | As nectar-tap_saprotroph (Põlme et al. 2020); Decay sugar-rich substrates (Põlme et al. 2020); This asexual genus is polyphyletic and contain species that function very differently depend on what they are. Caution should be taken when interpreting guilds for Candida species. | Manolakaki D, et al. 2010. Virulence. 1: 367-375 ((https://doi.org/10.4161/viru.1.5.12796)); Mekbib SB, et al. 2011. Tropical Plant Pathology 36:233-240 ((https://doi.org/10.1590/S1982-56762011000400004)); Põlme S, et al. 2020. Fungal Diversity 105:1-16 ((https://doi.org/10.1007/s13225-020-00466-2)) |
| Otu23699 | | 1 | k__Fungi(100);p__Ascomycota(100);c__Saccharomycetes(100);o__Saccharomycetales(100);f__Metschnikowiaceae(100);g__Metschnikowia(100);s__Metschnikowia_pulcherrima(100); | Metschnikowia | | 13 | Saprotroph | \|Undefined Saprotroph\| | Yeast | NULL | Probable | As nectar-tap_saprotroph (Põlme et al. 2020); Decay sugar-rich substrates (Põlme et al. 2020) | Põlme S, et al. 2020. Fungal Diversity 105:1-16 ((https://doi.org/10.1007/s13225-020-00466-2)) |
| Otu23700 | | 1 | k__Fungi(100);p__Ascomycota(100);c__Saccharomycetes(100);o__Saccharomycetales(100);f__Metschnikowiaceae(100);g__Metschnikowia(100);s__Metschnikowia_pulcherrima(100); | Metschnikowia | | 13 | Saprotroph | \|Undefined Saprotroph\| | Yeast | NULL | Probable | As nectar-tap_saprotroph (Põlme et al. 2020); Decay sugar-rich substrates (Põlme et al. 2020) | Põlme S, et al. 2020. Fungal Diversity 105:1-16 ((https://doi.org/10.1007/s13225-020-00466-2)) |
| Otu23701 | | 1 | k__Fungi(100);p__Ascomycota(100);c__Saccharomycetes(100);o__Saccharomycetales(100);f__Saccharomycetales_family_Incertae_sedis(100);g__Candida(100);g__Candida_unclassified(100); | Candida | | 13 | Pathotroph-Saprotroph-Symbiotroph | Animal Pathogen-Endophyte-Endosymbiont-Epiphyte-\|Undefined Saprotroph\| | Dimorphic Yeast-Yeast | NULL | Probable | As nectar-tap_saprotroph (Põlme et al. 2020); Decay sugar-rich substrates (Põlme et al. 2020); This asexual genus is polyphyletic and contain species that function very differently depend on what they are. Caution should be taken when interpreting guilds for Candida species. | Manolakaki D, et al. 2010. Virulence. 1: 367-375 ((https://doi.org/10.4161/viru.1.5.12796)); Mekbib SB, et al. 2011. Tropical Plant Pathology 36:233-240 ((https://doi.org/10.1590/S1982-56762011000400004)); Põlme S, et al. 2020. Fungal Diversity 105:1-16 ((https://doi.org/10.1007/s13225-020-00466-2)) |
| Otu23704 | | 1 | k__Fungi(100);p__Ascomycota(100);c__Saccharomycetes(100);o__Saccharomycetales(100);f__Metschnikowiaceae(100);g__Metschnikowia(100);g__Metschnikowia_unclassified(100); | Metschnikowia | | 13 | Saprotroph | \|Undefined Saprotroph\| | Yeast | NULL | Probable | As nectar-tap_saprotroph (Põlme et al. 2020); Decay sugar-rich substrates (Põlme et al. 2020) | Põlme S, et al. 2020. Fungal Diversity 105:1-16 ((https://doi.org/10.1007/s13225-020-00466-2)) |
| Otu23705 | | 1 | k__Fungi(100);p__Ascomycota(100);c__Saccharomycetes(100);o__Saccharomycetales(100);f__Saccharomycetaceae(100);g__Kazachstania(100);s__Kazachstania_intestinalis(100); | Kazachstania | | 13 | Saprotroph | \|Undefined Saprotroph\| | Yeast | NULL | Probable | As nectar-tap_saprotroph (Põlme et al. 2020); Decay sugar-rich substrates (Põlme et al. 2020) | Põlme S, et al. 2020. Fungal Diversity 105:1-16 ((https://doi.org/10.1007/s13225-020-00466-2)) |
| Otu23707 | | 1 | k__Fungi(100);p__Ascomycota(100);c__Saccharomycetes(100);o__Saccharomycetales(100);f__Metschnikowiaceae(100);g__Metschnikowia(100);g__Metschnikowia_unclassified(100); | Metschnikowia | | 13 | Saprotroph | \|Undefined Saprotroph\| | Yeast | NULL | Probable | As nectar-tap_saprotroph (Põlme et al. 2020); Decay sugar-rich substrates (Põlme et al. 2020) | Põlme S, et al. 2020. Fungal Diversity 105:1-16 ((https://doi.org/10.1007/s13225-020-00466-2)) |
| Otu23713 | | 1 | k__Fungi(100);p__Ascomycota(100);c__Saccharomycetes(100);o__Saccharomycetales(100);f__Metschnikowiaceae(100);g__Metschnikowia(100);s__Metschnikowia_pulcherrima(100); | Metschnikowia | | 13 | Saprotroph | \|Undefined Saprotroph\| | Yeast | NULL | Probable | As nectar-tap_saprotroph (Põlme et al. 2020); Decay sugar-rich substrates (Põlme et al. 2020) | Põlme S, et al. 2020. Fungal Diversity 105:1-16 ((https://doi.org/10.1007/s13225-020-00466-2)) |
| Otu23715 | | 1 | k__Fungi(100);p__Ascomycota(100);c__Saccharomycetes(100);o__Saccharomycetales(100);f__Metschnikowiaceae(100);g__Metschnikowia(100);s__Metschnikowia_pulcherrima(100); | Metschnikowia | | 13 | Saprotroph | \|Undefined Saprotroph\| | Yeast | NULL | Probable | As nectar-tap_saprotroph (Põlme et al. 2020); Decay sugar-rich substrates (Põlme et al. 2020) | Põlme S, et al. 2020. Fungal Diversity 105:1-16 ((https://doi.org/10.1007/s13225-020-00466-2)) |
| Otu23717 | | 1 | k__Fungi(100);p__Ascomycota(100);c__Saccharomycetes(100);o__Saccharomycetales(100);f__Metschnikowiaceae(100);g__Metschnikowia(100);s__Metschnikowia_pulcherrima(100); | Metschnikowia | | 13 | Saprotroph | \|Undefined Saprotroph\| | Yeast | NULL | Probable | As nectar-tap_saprotroph (Põlme et al. 2020); Decay sugar-rich substrates (Põlme et al. 2020) | Põlme S, et al. 2020. Fungal Diversity 105:1-16 ((https://doi.org/10.1007/s13225-020-00466-2)) |
| Otu23719 | | 1 | k__Fungi(100);p__Ascomycota(100);c__Saccharomycetes(100);o__Saccharomycetales(100);f__Metschnikowiaceae(100);g__Metschnikowia(100);s__Metschnikowia_pulcherrima(100); | Metschnikowia | | 13 | Saprotroph | \|Undefined Saprotroph\| | Yeast | NULL | Probable | As nectar-tap_saprotroph (Põlme et al. 2020); Decay sugar-rich substrates (Põlme et al. 2020) | Põlme S, et al. 2020. Fungal Diversity 105:1-16 ((https://doi.org/10.1007/s13225-020-00466-2)) |
| Otu23720 | | 1 | k__Fungi(100);p__Ascomycota(100);c__Saccharomycetes(100);o__Saccharomycetales(100);f__Metschnikowiaceae(100);g__Metschnikowia(100);s__Metschnikowia_pulcherrima(100); | Metschnikowia | | 13 | Saprotroph | \|Undefined Saprotroph\| | Yeast | NULL | Probable | As nectar-tap_saprotroph (Põlme et al. 2020); Decay sugar-rich substrates (Põlme et al. 2020) | Põlme S, et al. 2020. Fungal Diversity 105:1-16 ((https://doi.org/10.1007/s13225-020-00466-2)) |
| Otu23721 | | 1 | k__Fungi(100);p__Ascomycota(100);c__Saccharomycetes(100);o__Saccharomycetales(100);f__Metschnikowiaceae(100);g__Metschnikowia(100);s__Metschnikowia_pulcherrima(100); | Metschnikowia | | 13 | Saprotroph | \|Undefined Saprotroph\| | Yeast | NULL | Probable | As nectar-tap_saprotroph (Põlme et al. 2020); Decay sugar-rich substrates (Põlme et al. 2020) | Põlme S, et al. 2020. Fungal Diversity 105:1-16 ((https://doi.org/10.1007/s13225-020-00466-2)) |
| Otu23724 | | 1 | k__Fungi(100);p__Ascomycota(100);c__Saccharomycetes(100);o__Saccharomycetales(100);f__Metschnikowiaceae(100);g__Metschnikowia(100);s__Metschnikowia_pulcherrima(100); | Metschnikowia | | 13 | Saprotroph | \|Undefined Saprotroph\| | Yeast | NULL | Probable | As nectar-tap_saprotroph (Põlme et al. 2020); Decay sugar-rich substrates (Põlme et al. 2020) | Põlme S, et al. 2020. Fungal Diversity 105:1-16 ((https://doi.org/10.1007/s13225-020-00466-2)) |
| Otu23726 | | 1 | k__Fungi(100);p__Ascomycota(100);c__Saccharomycetes(100);o__Saccharomycetales(100);f__Metschnikowiaceae(100);g__Metschnikowia(100);s__Metschnikowia_pulcherrima(100); | Metschnikowia | | 13 | Saprotroph | \|Undefined Saprotroph\| | Yeast | NULL | Probable | As nectar-tap_saprotroph (Põlme et al. 2020); Decay sugar-rich substrates (Põlme et al. 2020) | Põlme S, et al. 2020. Fungal Diversity 105:1-16 ((https://doi.org/10.1007/s13225-020-00466-2)) |
| Otu23727 | | 1 | k__Fungi(100);p__Ascomycota(100);c__Saccharomycetes(100);o__Saccharomycetales(100);f__Metschnikowiaceae(100);g__Metschnikowia(100);s__Metschnikowia_pulcherrima(100); | Metschnikowia | | 13 | Saprotroph | \|Undefined Saprotroph\| | Yeast | NULL | Probable | As nectar-tap_saprotroph (Põlme et al. 2020); Decay sugar-rich substrates (Põlme et al. 2020) | Põlme S, et al. 2020. Fungal Diversity 105:1-16 ((https://doi.org/10.1007/s13225-020-00466-2)) |
| Otu23729 | | 1 | k__Fungi(100);p__Ascomycota(100);c__Saccharomycetes(100);o__Saccharomycetales(100);f__Metschnikowiaceae(100);g__Metschnikowia(100);s__Metschnikowia_pulcherrima(100); | Metschnikowia | | 13 | Saprotroph | \|Undefined Saprotroph\| | Yeast | NULL | Probable | As nectar-tap_saprotroph (Põlme et al. 2020); Decay sugar-rich substrates (Põlme et al. 2020) | Põlme S, et al. 2020. Fungal Diversity 105:1-16 ((https://doi.org/10.1007/s13225-020-00466-2)) |
| Otu23730 | | 1 | k__Fungi(100);p__Ascomycota(100);c__Saccharomycetes(100);o__Saccharomycetales(100);f__Metschnikowiaceae(100);g__Metschnikowia(100);s__Metschnikowia_pulcherrima(100); | Metschnikowia | | 13 | Saprotroph | \|Undefined Saprotroph\| | Yeast | NULL | Probable | As nectar-tap_saprotroph (Põlme et al. 2020); Decay sugar-rich substrates (Põlme et al. 2020) | Põlme S, et al. 2020. Fungal Diversity 105:1-16 ((https://doi.org/10.1007/s13225-020-00466-2)) |
| Otu23731 | | 1 | k__Fungi(100);p__Ascomycota(100);c__Saccharomycetes(100);o__Saccharomycetales(100);f__Metschnikowiaceae(100);g__Metschnikowia(100);g__Metschnikowia_unclassified(100); | Metschnikowia | | 13 | Saprotroph | \|Undefined Saprotroph\| | Yeast | NULL | Probable | As nectar-tap_saprotroph (Põlme et al. 2020); Decay sugar-rich substrates (Põlme et al. 2020) | Põlme S, et al. 2020. Fungal Diversity 105:1-16 ((https://doi.org/10.1007/s13225-020-00466-2)) |
| Otu23732 | | 1 | k__Fungi(100);p__Ascomycota(100);c__Saccharomycetes(100);o__Saccharomycetales(100);f__Metschnikowiaceae(100);g__Metschnikowia(100);s__Metschnikowia_pulcherrima(100); | Metschnikowia | | 13 | Saprotroph | \|Undefined Saprotroph\| | Yeast | NULL | Probable | As nectar-tap_saprotroph (Põlme et al. 2020); Decay sugar-rich substrates (Põlme et al. 2020) | Põlme S, et al. 2020. Fungal Diversity 105:1-16 ((https://doi.org/10.1007/s13225-020-00466-2)) |
| Otu23733 | | 1 | k__Fungi(100);p__Ascomycota(100);c__Saccharomycetes(100);o__Saccharomycetales(100);f__Metschnikowiaceae(100);g__Metschnikowia(100);s__Metschnikowia_pulcherrima(100); | Metschnikowia | | 13 | Saprotroph | \|Undefined Saprotroph\| | Yeast | NULL | Probable | As nectar-tap_saprotroph (Põlme et al. 2020); Decay sugar-rich substrates (Põlme et al. 2020) | Põlme S, et al. 2020. Fungal Diversity 105:1-16 ((https://doi.org/10.1007/s13225-020-00466-2)) |
| Otu23734 | | 1 | k__Fungi(100);p__Ascomycota(100);c__Saccharomycetes(100);o__Saccharomycetales(100);f__Metschnikowiaceae(100);g__Metschnikowia(100);s__Metschnikowia_pulcherrima(100); | Metschnikowia | | 13 | Saprotroph | \|Undefined Saprotroph\| | Yeast | NULL | Probable | As nectar-tap_saprotroph (Põlme et al. 2020); Decay sugar-rich substrates (Põlme et al. 2020) | Põlme S, et al. 2020. Fungal Diversity 105:1-16 ((https://doi.org/10.1007/s13225-020-00466-2)) |
| Otu23735 | | 1 | k__Fungi(100);p__Ascomycota(100);c__Saccharomycetes(100);o__Saccharomycetales(100);f__Metschnikowiaceae(100);g__Metschnikowia(100);s__Metschnikowia_pulcherrima(100); | Metschnikowia | | 13 | Saprotroph | \|Undefined Saprotroph\| | Yeast | NULL | Probable | As nectar-tap_saprotroph (Põlme et al. 2020); Decay sugar-rich substrates (Põlme et al. 2020) | Põlme S, et al. 2020. Fungal Diversity 105:1-16 ((https://doi.org/10.1007/s13225-020-00466-2)) |
| Otu23736 | | 1 | k__Fungi(100);p__Ascomycota(100);c__Saccharomycetes(100);o__Saccharomycetales(100);f__Metschnikowiaceae(100);g__Metschnikowia(100);s__Metschnikowia_pulcherrima(100); | Metschnikowia | | 13 | Saprotroph | \|Undefined Saprotroph\| | Yeast | NULL | Probable | As nectar-tap_saprotroph (Põlme et al. 2020); Decay sugar-rich substrates (Põlme et al. 2020) | Põlme S, et al. 2020. Fungal Diversity 105:1-16 ((https://doi.org/10.1007/s13225-020-00466-2)) |
| Otu23737 | | 1 | k__Fungi(100);p__Ascomycota(100);c__Saccharomycetes(100);o__Saccharomycetales(100);f__Metschnikowiaceae(100);g__Metschnikowia(100);s__Metschnikowia_pulcherrima(100); | Metschnikowia | | 13 | Saprotroph | \|Undefined Saprotroph\| | Yeast | NULL | Probable | As nectar-tap_saprotroph (Põlme et al. 2020); Decay sugar-rich substrates (Põlme et al. 2020) | Põlme S, et al. 2020. Fungal Diversity 105:1-16 ((https://doi.org/10.1007/s13225-020-00466-2)) |
| Otu23738 | | 1 | k__Fungi(100);p__Ascomycota(100);c__Saccharomycetes(100);o__Saccharomycetales(100);f__Metschnikowiaceae(100);g__Metschnikowia(100);s__Metschnikowia_pulcherrima(100); | Metschnikowia | | 13 | Saprotroph | \|Undefined Saprotroph\| | Yeast | NULL | Probable | As nectar-tap_saprotroph (Põlme et al. 2020); Decay sugar-rich substrates (Põlme et al. 2020) | Põlme S, et al. 2020. Fungal Diversity 105:1-16 ((https://doi.org/10.1007/s13225-020-00466-2)) |
| Otu23739 | | 1 | k__Fungi(100);p__Ascomycota(100);c__Saccharomycetes(100);o__Saccharomycetales(100);f__Metschnikowiaceae(100);g__Metschnikowia(100);s__Metschnikowia_pulcherrima(100); | Metschnikowia | | 13 | Saprotroph | \|Undefined Saprotroph\| | Yeast | NULL | Probable | As nectar-tap_saprotroph (Põlme et al. 2020); Decay sugar-rich substrates (Põlme et al. 2020) | Põlme S, et al. 2020. Fungal Diversity 105:1-16 ((https://doi.org/10.1007/s13225-020-00466-2)) |
| Otu23744 | | 1 | k__Fungi(100);p__Ascomycota(100);c__Saccharomycetes(100);o__Saccharomycetales(100);f__Metschnikowiaceae(100);g__Metschnikowia(100);g__Metschnikowia_unclassified(100); | Metschnikowia | | 13 | Saprotroph | \|Undefined Saprotroph\| | Yeast | NULL | Probable | As nectar-tap_saprotroph (Põlme et al. 2020); Decay sugar-rich substrates (Põlme et al. 2020) | Põlme S, et al. 2020. Fungal Diversity 105:1-16 ((https://doi.org/10.1007/s13225-020-00466-2)) |
| Otu23745 | | 1 | k__Fungi(100);p__Ascomycota(100);c__Saccharomycetes(100);o__Saccharomycetales(100);f__Metschnikowiaceae(100);g__Metschnikowia(100);s__Metschnikowia_pulcherrima(100); | Metschnikowia | | 13 | Saprotroph | \|Undefined Saprotroph\| | Yeast | NULL | Probable | As nectar-tap_saprotroph (Põlme et al. 2020); Decay sugar-rich substrates (Põlme et al. 2020) | Põlme S, et al. 2020. Fungal Diversity 105:1-16 ((https://doi.org/10.1007/s13225-020-00466-2)) |
| Otu23746 | | 1 | k__Fungi(100);p__Ascomycota(100);c__Saccharomycetes(100);o__Saccharomycetales(100);f__Metschnikowiaceae(100);g__Metschnikowia(100);s__Metschnikowia_pulcherrima(100); | Metschnikowia | | 13 | Saprotroph | \|Undefined Saprotroph\| | Yeast | NULL | Probable | As nectar-tap_saprotroph (Põlme et al. 2020); Decay sugar-rich substrates (Põlme et al. 2020) | Põlme S, et al. 2020. Fungal Diversity 105:1-16 ((https://doi.org/10.1007/s13225-020-00466-2)) |
| Otu23747 | | 1 | k__Fungi(100);p__Ascomycota(100);c__Saccharomycetes(100);o__Saccharomycetales(100);f__Metschnikowiaceae(100);g__Metschnikowia(100);s__Metschnikowia_pulcherrima(100); | Metschnikowia | | 13 | Saprotroph | \|Undefined Saprotroph\| | Yeast | NULL | Probable | As nectar-tap_saprotroph (Põlme et al. 2020); Decay sugar-rich substrates (Põlme et al. 2020) | Põlme S, et al. 2020. Fungal Diversity 105:1-16 ((https://doi.org/10.1007/s13225-020-00466-2)) |
| Otu23749 | | 1 | k__Fungi(100);p__Ascomycota(100);c__Saccharomycetes(100);o__Saccharomycetales(100);f__Metschnikowiaceae(100);g__Metschnikowia(100);g__Metschnikowia_unclassified(100); | Metschnikowia | | 13 | Saprotroph | \|Undefined Saprotroph\| | Yeast | NULL | Probable | As nectar-tap_saprotroph (Põlme et al. 2020); Decay sugar-rich substrates (Põlme et al. 2020) | Põlme S, et al. 2020. Fungal Diversity 105:1-16 ((https://doi.org/10.1007/s13225-020-00466-2)) |
| Otu23750 | | 1 | k__Fungi(100);p__Ascomycota(100);c__Saccharomycetes(100);o__Saccharomycetales(100);f__Metschnikowiaceae(100);g__Metschnikowia(100);s__Metschnikowia_pulcherrima(100); | Metschnikowia | | 13 | Saprotroph | \|Undefined Saprotroph\| | Yeast | NULL | Probable | As nectar-tap_saprotroph (Põlme et al. 2020); Decay sugar-rich substrates (Põlme et al. 2020) | Põlme S, et al. 2020. Fungal Diversity 105:1-16 ((https://doi.org/10.1007/s13225-020-00466-2)) |
| Otu23751 | | 1 | k__Fungi(100);p__Ascomycota(100);c__Saccharomycetes(100);o__Saccharomycetales(100);f__Metschnikowiaceae(100);g__Metschnikowia(100);s__Metschnikowia_pulcherrima(100); | Metschnikowia | | 13 | Saprotroph | \|Undefined Saprotroph\| | Yeast | NULL | Probable | As nectar-tap_saprotroph (Põlme et al. 2020); Decay sugar-rich substrates (Põlme et al. 2020) | Põlme S, et al. 2020. Fungal Diversity 105:1-16 ((https://doi.org/10.1007/s13225-020-00466-2)) |
| Otu23752 | | 1 | k__Fungi(100);p__Ascomycota(100);c__Saccharomycetes(100);o__Saccharomycetales(100);f__Metschnikowiaceae(100);g__Metschnikowia(100);s__Metschnikowia_pulcherrima(100); | Metschnikowia | | 13 | Saprotroph | \|Undefined Saprotroph\| | Yeast | NULL | Probable | As nectar-tap_saprotroph (Põlme et al. 2020); Decay sugar-rich substrates (Põlme et al. 2020) | Põlme S, et al. 2020. Fungal Diversity 105:1-16 ((https://doi.org/10.1007/s13225-020-00466-2)) |
| Otu23753 | | 1 | k__Fungi(100);p__Ascomycota(100);c__Saccharomycetes(100);o__Saccharomycetales(100);f__Metschnikowiaceae(100);g__Metschnikowia(100);s__Metschnikowia_pulcherrima(100); | Metschnikowia | | 13 | Saprotroph | \|Undefined Saprotroph\| | Yeast | NULL | Probable | As nectar-tap_saprotroph (Põlme et al. 2020); Decay sugar-rich substrates (Põlme et al. 2020) | Põlme S, et al. 2020. Fungal Diversity 105:1-16 ((https://doi.org/10.1007/s13225-020-00466-2)) |
| Otu23754 | | 1 | k__Fungi(100);p__Ascomycota(100);c__Saccharomycetes(100);o__Saccharomycetales(100);f__Metschnikowiaceae(100);g__Metschnikowia(100);s__Metschnikowia_pulcherrima(100); | Metschnikowia | | 13 | Saprotroph | \|Undefined Saprotroph\| | Yeast | NULL | Probable | As nectar-tap_saprotroph (Põlme et al. 2020); Decay sugar-rich substrates (Põlme et al. 2020) | Põlme S, et al. 2020. Fungal Diversity 105:1-16 ((https://doi.org/10.1007/s13225-020-00466-2)) |
| Otu23756 | | 1 | k__Fungi(100);p__Ascomycota(100);c__Saccharomycetes(100);o__Saccharomycetales(100);f__Metschnikowiaceae(100);g__Metschnikowia(100);s__Metschnikowia_pulcherrima(100); | Metschnikowia | | 13 | Saprotroph | \|Undefined Saprotroph\| | Yeast | NULL | Probable | As nectar-tap_saprotroph (Põlme et al. 2020); Decay sugar-rich substrates (Põlme et al. 2020) | Põlme S, et al. 2020. Fungal Diversity 105:1-16 ((https://doi.org/10.1007/s13225-020-00466-2)) |
| Otu23757 | | 1 | k__Fungi(100);p__Ascomycota(100);c__Saccharomycetes(100);o__Saccharomycetales(100);f__Metschnikowiaceae(100);g__Metschnikowia(100);g__Metschnikowia_unclassified(100); | Metschnikowia | | 13 | Saprotroph | \|Undefined Saprotroph\| | Yeast | NULL | Probable | As nectar-tap_saprotroph (Põlme et al. 2020); Decay sugar-rich substrates (Põlme et al. 2020) | Põlme S, et al. 2020. Fungal Diversity 105:1-16 ((https://doi.org/10.1007/s13225-020-00466-2)) |
| Otu23758 | | 1 | k__Fungi(100);p__Ascomycota(100);c__Saccharomycetes(100);o__Saccharomycetales(100);f__Metschnikowiaceae(100);g__Metschnikowia(100);s__Metschnikowia_pulcherrima(100); | Metschnikowia | | 13 | Saprotroph | \|Undefined Saprotroph\| | Yeast | NULL | Probable | As nectar-tap_saprotroph (Põlme et al. 2020); Decay sugar-rich substrates (Põlme et al. 2020) | Põlme S, et al. 2020. Fungal Diversity 105:1-16 ((https://doi.org/10.1007/s13225-020-00466-2)) |
| Otu23761 | | 1 | k__Fungi(100);p__Ascomycota(100);c__Saccharomycetes(100);o__Saccharomycetales(100);f__Metschnikowiaceae(100);g__Metschnikowia(100);s__Metschnikowia_pulcherrima(100); | Metschnikowia | | 13 | Saprotroph | \|Undefined Saprotroph\| | Yeast | NULL | Probable | As nectar-tap_saprotroph (Põlme et al. 2020); Decay sugar-rich substrates (Põlme et al. 2020) | Põlme S, et al. 2020. Fungal Diversity 105:1-16 ((https://doi.org/10.1007/s13225-020-00466-2)) |
| Otu23762 | | 1 | k__Fungi(100);p__Ascomycota(100);c__Saccharomycetes(100);o__Saccharomycetales(100);f__Metschnikowiaceae(100);g__Metschnikowia(100);s__Metschnikowia_pulcherrima(100); | Metschnikowia | | 13 | Saprotroph | \|Undefined Saprotroph\| | Yeast | NULL | Probable | As nectar-tap_saprotroph (Põlme et al. 2020); Decay sugar-rich substrates (Põlme et al. 2020) | Põlme S, et al. 2020. Fungal Diversity 105:1-16 ((https://doi.org/10.1007/s13225-020-00466-2)) |
| Otu23763 | | 1 | k__Fungi(100);p__Ascomycota(100);c__Lecanoromycetes(100);o__Lecanorales(100);f__Lecanoraceae(100);g__Lecanora(100);s__Lecanora_symmicta(100); | Lecanora | | 13 | Symbiotroph | \|Lichenized\| | Thallus | NULL | Highly Probable | NULL | Esslinger TL. 2014. North Dakota State University ((http://www.ndsu.edu/pubweb/~esslinge/chcklst/chcklst7.htm)); Põlme S, et al. 2020. Fungal Diversity 105:1-16 ((https://doi.org/10.1007/s13225-020-00466-2)) |
| Otu23765 | | 1 | k__Fungi(100);p__Ascomycota(100);c__Saccharomycetes(100);o__Saccharomycetales(100);f__Metschnikowiaceae(100);g__Metschnikowia(100);s__Metschnikowia_pulcherrima(100); | Metschnikowia | | 13 | Saprotroph | \|Undefined Saprotroph\| | Yeast | NULL | Probable | As nectar-tap_saprotroph (Põlme et al. 2020); Decay sugar-rich substrates (Põlme et al. 2020) | Põlme S, et al. 2020. Fungal Diversity 105:1-16 ((https://doi.org/10.1007/s13225-020-00466-2)) |
| Otu23766 | | 1 | k__Fungi(100);p__Ascomycota(100);c__Saccharomycetes(100);o__Saccharomycetales(100);f__Metschnikowiaceae(100);g__Metschnikowia(100);g__Metschnikowia_unclassified(100); | Metschnikowia | | 13 | Saprotroph | \|Undefined Saprotroph\| | Yeast | NULL | Probable | As nectar-tap_saprotroph (Põlme et al. 2020); Decay sugar-rich substrates (Põlme et al. 2020) | Põlme S, et al. 2020. Fungal Diversity 105:1-16 ((https://doi.org/10.1007/s13225-020-00466-2)) |
| Otu23767 | | 1 | k__Fungi(100);p__Ascomycota(100);c__Saccharomycetes(100);o__Saccharomycetales(100);f__Metschnikowiaceae(100);g__Metschnikowia(100);s__Metschnikowia_pulcherrima(100); | Metschnikowia | | 13 | Saprotroph | \|Undefined Saprotroph\| | Yeast | NULL | Probable | As nectar-tap_saprotroph (Põlme et al. 2020); Decay sugar-rich substrates (Põlme et al. 2020) | Põlme S, et al. 2020. Fungal Diversity 105:1-16 ((https://doi.org/10.1007/s13225-020-00466-2)) |
| Otu23768 | | 1 | k__Fungi(100);p__Ascomycota(100);c__Saccharomycetes(100);o__Saccharomycetales(100);f__Metschnikowiaceae(100);g__Metschnikowia(100);s__Metschnikowia_pulcherrima(100); | Metschnikowia | | 13 | Saprotroph | \|Undefined Saprotroph\| | Yeast | NULL | Probable | As nectar-tap_saprotroph (Põlme et al. 2020); Decay sugar-rich substrates (Põlme et al. 2020) | Põlme S, et al. 2020. Fungal Diversity 105:1-16 ((https://doi.org/10.1007/s13225-020-00466-2)) |
| Otu23769 | | 1 | k__Fungi(100);p__Ascomycota(100);c__Saccharomycetes(100);o__Saccharomycetales(100);f__Metschnikowiaceae(100);g__Metschnikowia(100);s__Metschnikowia_pulcherrima(100); | Metschnikowia | | 13 | Saprotroph | \|Undefined Saprotroph\| | Yeast | NULL | Probable | As nectar-tap_saprotroph (Põlme et al. 2020); Decay sugar-rich substrates (Põlme et al. 2020) | Põlme S, et al. 2020. Fungal Diversity 105:1-16 ((https://doi.org/10.1007/s13225-020-00466-2)) |
| Otu23770 | | 1 | k__Fungi(100);p__Ascomycota(100);c__Saccharomycetes(100);o__Saccharomycetales(100);f__Metschnikowiaceae(100);g__Metschnikowia(100);s__Metschnikowia_pulcherrima(100); | Metschnikowia | | 13 | Saprotroph | \|Undefined Saprotroph\| | Yeast | NULL | Probable | As nectar-tap_saprotroph (Põlme et al. 2020); Decay sugar-rich substrates (Põlme et al. 2020) | Põlme S, et al. 2020. Fungal Diversity 105:1-16 ((https://doi.org/10.1007/s13225-020-00466-2)) |
| Otu23772 | | 1 | k__Fungi(100);p__Ascomycota(100);c__Saccharomycetes(100);o__Saccharomycetales(100);f__Metschnikowiaceae(100);g__Metschnikowia(100);s__Metschnikowia_pulcherrima(100); | Metschnikowia | | 13 | Saprotroph | \|Undefined Saprotroph\| | Yeast | NULL | Probable | As nectar-tap_saprotroph (Põlme et al. 2020); Decay sugar-rich substrates (Põlme et al. 2020) | Põlme S, et al. 2020. Fungal Diversity 105:1-16 ((https://doi.org/10.1007/s13225-020-00466-2)) |
| Otu23773 | | 1 | k__Fungi(100);p__Ascomycota(100);c__Saccharomycetes(100);o__Saccharomycetales(100);f__Metschnikowiaceae(100);g__Metschnikowia(100);g__Metschnikowia_unclassified(100); | Metschnikowia | | 13 | Saprotroph | \|Undefined Saprotroph\| | Yeast | NULL | Probable | As nectar-tap_saprotroph (Põlme et al. 2020); Decay sugar-rich substrates (Põlme et al. 2020) | Põlme S, et al. 2020. Fungal Diversity 105:1-16 ((https://doi.org/10.1007/s13225-020-00466-2)) |
| Otu23774 | | 1 | k__Fungi(100);p__Ascomycota(100);c__Saccharomycetes(100);o__Saccharomycetales(100);f__Metschnikowiaceae(100);g__Metschnikowia(100);g__Metschnikowia_unclassified(100); | Metschnikowia | | 13 | Saprotroph | \|Undefined Saprotroph\| | Yeast | NULL | Probable | As nectar-tap_saprotroph (Põlme et al. 2020); Decay sugar-rich substrates (Põlme et al. 2020) | Põlme S, et al. 2020. Fungal Diversity 105:1-16 ((https://doi.org/10.1007/s13225-020-00466-2)) |
| Otu23775 | | 1 | k__Fungi(100);p__Ascomycota(100);c__Saccharomycetes(100);o__Saccharomycetales(100);f__Metschnikowiaceae(100);g__Metschnikowia(100);s__Metschnikowia_pulcherrima(100); | Metschnikowia | | 13 | Saprotroph | \|Undefined Saprotroph\| | Yeast | NULL | Probable | As nectar-tap_saprotroph (Põlme et al. 2020); Decay sugar-rich substrates (Põlme et al. 2020) | Põlme S, et al. 2020. Fungal Diversity 105:1-16 ((https://doi.org/10.1007/s13225-020-00466-2)) |
| Otu23776 | | 1 | k__Fungi(100);p__Ascomycota(100);c__Saccharomycetes(100);o__Saccharomycetales(100);f__Metschnikowiaceae(100);g__Metschnikowia(100);s__Metschnikowia_pulcherrima(100); | Metschnikowia | | 13 | Saprotroph | \|Undefined Saprotroph\| | Yeast | NULL | Probable | As nectar-tap_saprotroph (Põlme et al. 2020); Decay sugar-rich substrates (Põlme et al. 2020) | Põlme S, et al. 2020. Fungal Diversity 105:1-16 ((https://doi.org/10.1007/s13225-020-00466-2)) |
| Otu23777 | | 1 | k__Fungi(100);p__Ascomycota(100);c__Saccharomycetes(100);o__Saccharomycetales(100);f__Metschnikowiaceae(100);g__Metschnikowia(100);s__Metschnikowia_pulcherrima(100); | Metschnikowia | | 13 | Saprotroph | \|Undefined Saprotroph\| | Yeast | NULL | Probable | As nectar-tap_saprotroph (Põlme et al. 2020); Decay sugar-rich substrates (Põlme et al. 2020) | Põlme S, et al. 2020. Fungal Diversity 105:1-16 ((https://doi.org/10.1007/s13225-020-00466-2)) |
| Otu23778 | | 1 | k__Fungi(100);p__Ascomycota(100);c__Saccharomycetes(100);o__Saccharomycetales(100);f__Metschnikowiaceae(100);g__Metschnikowia(100);g__Metschnikowia_unclassified(100); | Metschnikowia | | 13 | Saprotroph | \|Undefined Saprotroph\| | Yeast | NULL | Probable | As nectar-tap_saprotroph (Põlme et al. 2020); Decay sugar-rich substrates (Põlme et al. 2020) | Põlme S, et al. 2020. Fungal Diversity 105:1-16 ((https://doi.org/10.1007/s13225-020-00466-2)) |
| Otu23779 | | 1 | k__Fungi(100);p__Ascomycota(100);c__Saccharomycetes(100);o__Saccharomycetales(100);f__Metschnikowiaceae(100);g__Metschnikowia(100);s__Metschnikowia_pulcherrima(100); | Metschnikowia | | 13 | Saprotroph | \|Undefined Saprotroph\| | Yeast | NULL | Probable | As nectar-tap_saprotroph (Põlme et al. 2020); Decay sugar-rich substrates (Põlme et al. 2020) | Põlme S, et al. 2020. Fungal Diversity 105:1-16 ((https://doi.org/10.1007/s13225-020-00466-2)) |
| Otu23780 | | 1 | k__Fungi(100);p__Ascomycota(100);c__Saccharomycetes(100);o__Saccharomycetales(100);f__Metschnikowiaceae(100);g__Metschnikowia(100);g__Metschnikowia_unclassified(100); | Metschnikowia | | 13 | Saprotroph | \|Undefined Saprotroph\| | Yeast | NULL | Probable | As nectar-tap_saprotroph (Põlme et al. 2020); Decay sugar-rich substrates (Põlme et al. 2020) | Põlme S, et al. 2020. Fungal Diversity 105:1-16 ((https://doi.org/10.1007/s13225-020-00466-2)) |
| Otu23781 | | 1 | k__Fungi(100);p__Ascomycota(100);c__Saccharomycetes(100);o__Saccharomycetales(100);f__Metschnikowiaceae(100);g__Metschnikowia(100);g__Metschnikowia_unclassified(100); | Metschnikowia | | 13 | Saprotroph | \|Undefined Saprotroph\| | Yeast | NULL | Probable | As nectar-tap_saprotroph (Põlme et al. 2020); Decay sugar-rich substrates (Põlme et al. 2020) | Põlme S, et al. 2020. Fungal Diversity 105:1-16 ((https://doi.org/10.1007/s13225-020-00466-2)) |
| Otu23783 | | 1 | k__Fungi(100);p__Ascomycota(100);c__Saccharomycetes(100);o__Saccharomycetales(100);f__Metschnikowiaceae(100);g__Metschnikowia(100);s__Metschnikowia_pulcherrima(100); | Metschnikowia | | 13 | Saprotroph | \|Undefined Saprotroph\| | Yeast | NULL | Probable | As nectar-tap_saprotroph (Põlme et al. 2020); Decay sugar-rich substrates (Põlme et al. 2020) | Põlme S, et al. 2020. Fungal Diversity 105:1-16 ((https://doi.org/10.1007/s13225-020-00466-2)) |
| Otu23784 | | 1 | k__Fungi(100);p__Ascomycota(100);c__Saccharomycetes(100);o__Saccharomycetales(100);f__Metschnikowiaceae(100);g__Metschnikowia(100);s__Metschnikowia_pulcherrima(100); | Metschnikowia | | 13 | Saprotroph | \|Undefined Saprotroph\| | Yeast | NULL | Probable | As nectar-tap_saprotroph (Põlme et al. 2020); Decay sugar-rich substrates (Põlme et al. 2020) | Põlme S, et al. 2020. Fungal Diversity 105:1-16 ((https://doi.org/10.1007/s13225-020-00466-2)) |
| Otu23787 | | 1 | k__Fungi(100);p__Ascomycota(100);c__Saccharomycetes(100);o__Saccharomycetales(100);f__Metschnikowiaceae(100);g__Metschnikowia(100);s__Metschnikowia_pulcherrima(100); | Metschnikowia | | 13 | Saprotroph | \|Undefined Saprotroph\| | Yeast | NULL | Probable | As nectar-tap_saprotroph (Põlme et al. 2020); Decay sugar-rich substrates (Põlme et al. 2020) | Põlme S, et al. 2020. Fungal Diversity 105:1-16 ((https://doi.org/10.1007/s13225-020-00466-2)) |
| Otu23789 | | 1 | k__Fungi(100);p__Ascomycota(100);c__Leotiomycetes(100);o__Helotiales(100);f__Helotiales_family_Incertae_sedis(100);g__Scytalidium(100);s__Scytalidium_sp_TMS_2011(100); | Scytalidium | | 13 | Pathotroph-Saprotroph | Animal Parasite-Fungal Parasite-Plant Pathogen-Plant Saprotroph-\|Wood Saprotroph\| | Microfungus | Soft Rot | Probable | As litter_saprotroph (Põlme et al. 2020); As mycoparasite (Põlme et al. 2020); Decay wood, leaf, fruit, seed, soil, animal material (Põlme et al. 2020) | Nilsson T. 1973. Studia Forestalia Suecica 104:1-40 ((https://pub.epsilon.slu.se/5711/)); Costa IPMW, et al. 2012. Mycotaxon 119:e26 ((http://mycotaxon.com/resources/checklists/costa_v119_checklist.pdf)); Tedersoo L, et al. 2014. Science 346:e1256688 ((https://doi.org/10.1126/science.1256688)); Põlme S, et al. 2020. Fungal Diversity 105:1-16 ((https://doi.org/10.1007/s13225-020-00466-2)); The American Phytopathological Society Common Names of Plant Diseases ((http://www.apsnet.org/publications/commonnames/Pages/default.aspx)) |
| Otu23790 | | 1 | k__Fungi(100);p__Ascomycota(100);c__Leotiomycetes(100);o__Helotiales(100);f__Rutstroemiaceae(100);g__Rutstroemia(100);s__Rutstroemia_firma(100); | Rutstroemia | | 13 | Saprotroph | \|Plant Saprotroph\| | Microfungus | NULL | Probable | As litter_saprotroph (Põlme et al. 2020); Decay leaf, fruit, seed (Põlme et al. 2020); Essentially saprobic on various plant parts, especially seeds and fruits, though initial infection may take place in living tissue (Cannon & Kirk 2007) | Cannon PF, Kirk PM. 2007. Fungal Families of the World. CAB International, Cambridge (ISBN: 978-0851998275); Tedersoo L, et al. 2014. Science 346:e1256688 ((https://doi.org/10.1126/science.1256688)); Põlme S, et al. 2020. Fungal Diversity 105:1-16 ((https://doi.org/10.1007/s13225-020-00466-2)) |
| Otu23794 | | 1 | k__Fungi(100);p__Ascomycota(100);c__Saccharomycetes(100);o__Saccharomycetales(100);f__Metschnikowiaceae(100);g__Metschnikowia(100);s__Metschnikowia_pulcherrima(100); | Metschnikowia | | 13 | Saprotroph | \|Undefined Saprotroph\| | Yeast | NULL | Probable | As nectar-tap_saprotroph (Põlme et al. 2020); Decay sugar-rich substrates (Põlme et al. 2020) | Põlme S, et al. 2020. Fungal Diversity 105:1-16 ((https://doi.org/10.1007/s13225-020-00466-2)) |
| Otu23795 | | 1 | k__Fungi(100);p__Ascomycota(100);c__Saccharomycetes(100);o__Saccharomycetales(100);f__Metschnikowiaceae(100);g__Metschnikowia(100);s__Metschnikowia_pulcherrima(100); | Metschnikowia | | 13 | Saprotroph | \|Undefined Saprotroph\| | Yeast | NULL | Probable | As nectar-tap_saprotroph (Põlme et al. 2020); Decay sugar-rich substrates (Põlme et al. 2020) | Põlme S, et al. 2020. Fungal Diversity 105:1-16 ((https://doi.org/10.1007/s13225-020-00466-2)) |
| Otu23796 | | 1 | k__Fungi(100);p__Ascomycota(100);c__Saccharomycetes(100);o__Saccharomycetales(100);f__Metschnikowiaceae(100);g__Metschnikowia(100);g__Metschnikowia_unclassified(100); | Metschnikowia | | 13 | Saprotroph | \|Undefined Saprotroph\| | Yeast | NULL | Probable | As nectar-tap_saprotroph (Põlme et al. 2020); Decay sugar-rich substrates (Põlme et al. 2020) | Põlme S, et al. 2020. Fungal Diversity 105:1-16 ((https://doi.org/10.1007/s13225-020-00466-2)) |
| Otu23797 | | 1 | k__Fungi(100);p__Ascomycota(100);c__Saccharomycetes(100);o__Saccharomycetales(100);f__Metschnikowiaceae(100);g__Metschnikowia(100);s__Metschnikowia_pulcherrima(100); | Metschnikowia | | 13 | Saprotroph | \|Undefined Saprotroph\| | Yeast | NULL | Probable | As nectar-tap_saprotroph (Põlme et al. 2020); Decay sugar-rich substrates (Põlme et al. 2020) | Põlme S, et al. 2020. Fungal Diversity 105:1-16 ((https://doi.org/10.1007/s13225-020-00466-2)) |

| | | | | | | | | | | | |
|---|---|---|---|---|---|---|---|---|---|---|---|
| Otu23799 | 1 | k__Fungi(100);p__Ascomycota(100);c__Saccharomycetes(100);o__Saccharomycetales(100);f__Metschnikowiaceae(100);g__Metschnikowia(100);s__Metschnikowia_pulcherrima(100); | Metschnikowia | 13 | Saprotroph | \|Undefined Saprotroph\| | Yeast | NULL | Probable | As nectar-tap_saprotroph (Põlme et al. 2020); Decay sugar-rich substrates (Põlme et al. 2020) | Põlme S, et al. 2020. Fungal Diversity 105:1-16 ((https://doi.org/10.1007/s13225-020-00466-2)) |
| Otu23802 | 1 | k__Fungi(100);p__Ascomycota(100);c__Saccharomycetes(100);o__Saccharomycetales(100);f__Metschnikowiaceae(100);g__Metschnikowia(100);s__Metschnikowia_pulcherrima(100); | Metschnikowia | 13 | Saprotroph | \|Undefined Saprotroph\| | Yeast | NULL | Probable | As nectar-tap_saprotroph (Põlme et al. 2020); Decay sugar-rich substrates (Põlme et al. 2020) | Põlme S, et al. 2020. Fungal Diversity 105:1-16 ((https://doi.org/10.1007/s13225-020-00466-2)) |
| Otu23803 | 1 | k__Fungi(100);p__Ascomycota(100);c__Saccharomycetes(100);o__Saccharomycetales(100);f__Metschnikowiaceae(100);g__Metschnikowia(100);s__Metschnikowia_pulcherrima(100); | Metschnikowia | 13 | Saprotroph | \|Undefined Saprotroph\| | Yeast | NULL | Probable | As nectar-tap_saprotroph (Põlme et al. 2020); Decay sugar-rich substrates (Põlme et al. 2020) | Põlme S, et al. 2020. Fungal Diversity 105:1-16 ((https://doi.org/10.1007/s13225-020-00466-2)) |
| Otu23804 | 1 | k__Fungi(100);p__Ascomycota(100);c__Saccharomycetes(100);o__Saccharomycetales(100);f__Metschnikowiaceae(100);g__Metschnikowia(100);s__Metschnikowia_pulcherrima(100); | Metschnikowia | 13 | Saprotroph | \|Undefined Saprotroph\| | Yeast | NULL | Probable | As nectar-tap_saprotroph (Põlme et al. 2020); Decay sugar-rich substrates (Põlme et al. 2020) | Põlme S, et al. 2020. Fungal Diversity 105:1-16 ((https://doi.org/10.1007/s13225-020-00466-2)) |
| Otu23805 | 1 | k__Fungi(100);p__Ascomycota(100);c__Saccharomycetes(100);o__Saccharomycetales(100);f__Metschnikowiaceae(100);g__Metschnikowia(100);g__Metschnikowia_unclassified(100); | Metschnikowia | 13 | Saprotroph | \|Undefined Saprotroph\| | Yeast | NULL | Probable | As nectar-tap_saprotroph (Põlme et al. 2020); Decay sugar-rich substrates (Põlme et al. 2020) | Põlme S, et al. 2020. Fungal Diversity 105:1-16 ((https://doi.org/10.1007/s13225-020-00466-2)) |
| Otu23806 | 1 | k__Fungi(100);p__Ascomycota(100);c__Saccharomycetes(100);o__Saccharomycetales(100);f__Metschnikowiaceae(100);g__Metschnikowia(100);s__Metschnikowia_pulcherrima(100); | Metschnikowia | 13 | Saprotroph | \|Undefined Saprotroph\| | Yeast | NULL | Probable | As nectar-tap_saprotroph (Põlme et al. 2020); Decay sugar-rich substrates (Põlme et al. 2020) | Põlme S, et al. 2020. Fungal Diversity 105:1-16 ((https://doi.org/10.1007/s13225-020-00466-2)) |
| Otu23807 | 1 | k__Fungi(100);p__Ascomycota(100);c__Saccharomycetes(100);o__Saccharomycetales(100);f__Metschnikowiaceae(100);g__Metschnikowia(100);g__Metschnikowia_unclassified(100); | Metschnikowia | 13 | Saprotroph | \|Undefined Saprotroph\| | Yeast | NULL | Probable | As nectar-tap_saprotroph (Põlme et al. 2020); Decay sugar-rich substrates (Põlme et al. 2020) | Põlme S, et al. 2020. Fungal Diversity 105:1-16 ((https://doi.org/10.1007/s13225-020-00466-2)) |
| Otu23808 | 1 | k__Fungi(100);p__Ascomycota(100);c__Saccharomycetes(100);o__Saccharomycetales(100);f__Metschnikowiaceae(100);g__Metschnikowia(100);s__Metschnikowia_pulcherrima(100); | Metschnikowia | 13 | Saprotroph | \|Undefined Saprotroph\| | Yeast | NULL | Probable | As nectar-tap_saprotroph (Põlme et al. 2020); Decay sugar-rich substrates (Põlme et al. 2020) | Põlme S, et al. 2020. Fungal Diversity 105:1-16 ((https://doi.org/10.1007/s13225-020-00466-2)) |
| Otu23809 | 1 | k__Fungi(100);p__Ascomycota(100);c__Saccharomycetes(100);o__Saccharomycetales(100);f__Metschnikowiaceae(100);g__Metschnikowia(100);s__Metschnikowia_pulcherrima(100); | Metschnikowia | 13 | Saprotroph | \|Undefined Saprotroph\| | Yeast | NULL | Probable | As nectar-tap_saprotroph (Põlme et al. 2020); Decay sugar-rich substrates (Põlme et al. 2020) | Põlme S, et al. 2020. Fungal Diversity 105:1-16 ((https://doi.org/10.1007/s13225-020-00466-2)) |
| Otu23810 | 1 | k__Fungi(100);p__Ascomycota(100);c__Saccharomycetes(100);o__Saccharomycetales(100);f__Metschnikowiaceae(100);g__Metschnikowia(100);g__Metschnikowia_unclassified(100); | Metschnikowia | 13 | Saprotroph | \|Undefined Saprotroph\| | Yeast | NULL | Probable | As nectar-tap_saprotroph (Põlme et al. 2020); Decay sugar-rich substrates (Põlme et al. 2020) | Põlme S, et al. 2020. Fungal Diversity 105:1-16 ((https://doi.org/10.1007/s13225-020-00466-2)) |
| Otu23811 | 1 | k__Fungi(100);p__Ascomycota(100);c__Saccharomycetes(100);o__Saccharomycetales(100);f__Metschnikowiaceae(100);g__Metschnikowia(100);s__Metschnikowia_pulcherrima(100); | Metschnikowia | 13 | Saprotroph | \|Undefined Saprotroph\| | Yeast | NULL | Probable | As nectar-tap_saprotroph (Põlme et al. 2020); Decay sugar-rich substrates (Põlme et al. 2020) | Põlme S, et al. 2020. Fungal Diversity 105:1-16 ((https://doi.org/10.1007/s13225-020-00466-2)) |
| Otu23812 | 1 | k__Fungi(100);p__Ascomycota(100);c__Saccharomycetes(100);o__Saccharomycetales(100);f__Metschnikowiaceae(100);g__Metschnikowia(100);g__Metschnikowia_unclassified(100); | Metschnikowia | 13 | Saprotroph | \|Undefined Saprotroph\| | Yeast | NULL | Probable | As nectar-tap_saprotroph (Põlme et al. 2020); Decay sugar-rich substrates (Põlme et al. 2020) | Põlme S, et al. 2020. Fungal Diversity 105:1-16 ((https://doi.org/10.1007/s13225-020-00466-2)) |
| Otu23813 | 1 | k__Fungi(100);p__Ascomycota(100);c__Saccharomycetes(100);o__Saccharomycetales(100);f__Metschnikowiaceae(100);g__Metschnikowia(100);s__Metschnikowia_pulcherrima(100); | Metschnikowia | 13 | Saprotroph | \|Undefined Saprotroph\| | Yeast | NULL | Probable | As nectar-tap_saprotroph (Põlme et al. 2020); Decay sugar-rich substrates (Põlme et al. 2020) | Põlme S, et al. 2020. Fungal Diversity 105:1-16 ((https://doi.org/10.1007/s13225-020-00466-2)) |
| Otu23815 | 1 | k__Fungi(100);p__Ascomycota(100);c__Saccharomycetes(100);o__Saccharomycetales(100);f__Metschnikowiaceae(100);g__Metschnikowia(100);g__Metschnikowia_unclassified(100); | Metschnikowia | 13 | Saprotroph | \|Undefined Saprotroph\| | Yeast | NULL | Probable | As nectar-tap_saprotroph (Põlme et al. 2020); Decay sugar-rich substrates (Põlme et al. 2020) | Põlme S, et al. 2020. Fungal Diversity 105:1-16 ((https://doi.org/10.1007/s13225-020-00466-2)) |
| Otu23816 | 1 | k__Fungi(100);p__Ascomycota(100);c__Saccharomycetes(100);o__Saccharomycetales(100);f__Metschnikowiaceae(100);g__Metschnikowia(100);s__Metschnikowia_pulcherrima(100); | Metschnikowia | 13 | Saprotroph | \|Undefined Saprotroph\| | Yeast | NULL | Probable | As nectar-tap_saprotroph (Põlme et al. 2020); Decay sugar-rich substrates (Põlme et al. 2020) | Põlme S, et al. 2020. Fungal Diversity 105:1-16 ((https://doi.org/10.1007/s13225-020-00466-2)) |
| Otu23817 | 1 | k__Fungi(100);p__Ascomycota(100);c__Saccharomycetes(100);o__Saccharomycetales(100);f__Metschnikowiaceae(100);g__Metschnikowia(100);s__Metschnikowia_pulcherrima(100); | Metschnikowia | 13 | Saprotroph | \|Undefined Saprotroph\| | Yeast | NULL | Probable | As nectar-tap_saprotroph (Põlme et al. 2020); Decay sugar-rich substrates (Põlme et al. 2020) | Põlme S, et al. 2020. Fungal Diversity 105:1-16 ((https://doi.org/10.1007/s13225-020-00466-2)) |
| Otu23820 | 1 | k__Fungi(100);p__Ascomycota(100);c__Saccharomycetes(100);o__Saccharomycetales(100);f__Saccharomycetales_family_Incertae_sedis(100);g__Candida(100);s__Candida_sp_AS23486(100); | Candida | 13 | Pathotroph-Saprotroph-Symbiotroph | Animal Pathogen-Endophyte-Endosymbiont-Epiphyte-\|Undefined Saprotroph\| | Dimorphic Yeast-Yeast | NULL | Probable | As nectar-tap_saprotroph (Põlme et al. 2020); Decay sugar-rich substrates (Põlme et al. 2020); This asexual genus is polyphyletic and contain species that function very differently depend on what they are. Caution should be taken when interpreting guilds for Candida species. | Manolakaki D, et al. 2010. Virulence. 1: 367-375 ((https://doi.org/10.4161/viru.1.5.12796)); Mekbib SB, et al. 2011. Tropical Plant Pathology 36:233-240 ((https://doi.org/10.1590/S1982-56762011000400004)); Põlme S, et al. 2020. Fungal Diversity 105:1-16 ((https://doi.org/10.1007/s13225-020-00466-2)) |
| Otu23821 | 1 | k__Fungi(100);p__Ascomycota(100);c__Saccharomycetes(100);o__Saccharomycetales(100);f__Metschnikowiaceae(100);g__Metschnikowia(100);s__Metschnikowia_pulcherrima(100); | Metschnikowia | 13 | Saprotroph | \|Undefined Saprotroph\| | Yeast | NULL | Probable | As nectar-tap_saprotroph (Põlme et al. 2020); Decay sugar-rich substrates (Põlme et al. 2020) | Põlme S, et al. 2020. Fungal Diversity 105:1-16 ((https://doi.org/10.1007/s13225-020-00466-2)) |
| Otu23822 | 1 | k__Fungi(100);p__Ascomycota(100);c__Saccharomycetes(100);o__Saccharomycetales(100);f__Metschnikowiaceae(100);g__Metschnikowia(100);g__Metschnikowia_unclassified(100); | Metschnikowia | 13 | Saprotroph | \|Undefined Saprotroph\| | Yeast | NULL | Probable | As nectar-tap_saprotroph (Põlme et al. 2020); Decay sugar-rich substrates (Põlme et al. 2020) | Põlme S, et al. 2020. Fungal Diversity 105:1-16 ((https://doi.org/10.1007/s13225-020-00466-2)) |
| Otu23823 | 1 | k__Fungi(100);p__Ascomycota(100);c__Saccharomycetes(100);o__Saccharomycetales(100);f__Metschnikowiaceae(100);g__Metschnikowia(100);s__Metschnikowia_pulcherrima(100); | Metschnikowia | 13 | Saprotroph | \|Undefined Saprotroph\| | Yeast | NULL | Probable | As nectar-tap_saprotroph (Põlme et al. 2020); Decay sugar-rich substrates (Põlme et al. 2020) | Põlme S, et al. 2020. Fungal Diversity 105:1-16 ((https://doi.org/10.1007/s13225-020-00466-2)) |
| Otu23824 | 1 | k__Fungi(100);p__Ascomycota(100);c__Saccharomycetes(100);o__Saccharomycetales(100);f__Metschnikowiaceae(100);g__Metschnikowia(100);s__Metschnikowia_pulcherrima(100); | Metschnikowia | 13 | Saprotroph | \|Undefined Saprotroph\| | Yeast | NULL | Probable | As nectar-tap_saprotroph (Põlme et al. 2020); Decay sugar-rich substrates (Põlme et al. 2020) | Põlme S, et al. 2020. Fungal Diversity 105:1-16 ((https://doi.org/10.1007/s13225-020-00466-2)) |
| Otu23830 | 1 | k__Fungi(100);p__Ascomycota(100);c__Saccharomycetes(100);o__Saccharomycetales(100);f__Metschnikowiaceae(100);g__Metschnikowia(100);s__Metschnikowia_pulcherrima(100); | Metschnikowia | 13 | Saprotroph | \|Undefined Saprotroph\| | Yeast | NULL | Probable | As nectar-tap_saprotroph (Põlme et al. 2020); Decay sugar-rich substrates (Põlme et al. 2020) | Põlme S, et al. 2020. Fungal Diversity 105:1-16 ((https://doi.org/10.1007/s13225-020-00466-2)) |
| Otu23832 | 1 | k__Fungi(100);p__Ascomycota(100);c__Saccharomycetes(100);o__Saccharomycetales(100);f__Metschnikowiaceae(100);g__Metschnikowia(100);s__Metschnikowia_pulcherrima(100); | Metschnikowia | 13 | Saprotroph | \|Undefined Saprotroph\| | Yeast | NULL | Probable | As nectar-tap_saprotroph (Põlme et al. 2020); Decay sugar-rich substrates (Põlme et al. 2020) | Põlme S, et al. 2020. Fungal Diversity 105:1-16 ((https://doi.org/10.1007/s13225-020-00466-2)) |
| Otu23835 | 1 | k__Fungi(100);p__Ascomycota(100);c__Saccharomycetes(100);o__Saccharomycetales(100);f__Metschnikowiaceae(100);g__Metschnikowia(100);g__Metschnikowia_unclassified(100); | Metschnikowia | 13 | Saprotroph | \|Undefined Saprotroph\| | Yeast | NULL | Probable | As nectar-tap_saprotroph (Põlme et al. 2020); Decay sugar-rich substrates (Põlme et al. 2020) | Põlme S, et al. 2020. Fungal Diversity 105:1-16 ((https://doi.org/10.1007/s13225-020-00466-2)) |
| Otu23842 | 1 | k__Fungi(100);p__Ascomycota(100);c__Saccharomycetes(100);o__Saccharomycetales(100);f__Metschnikowiaceae(100);g__Metschnikowia(100);s__Metschnikowia_pulcherrima(100); | Metschnikowia | 13 | Saprotroph | \|Undefined Saprotroph\| | Yeast | NULL | Probable | As nectar-tap_saprotroph (Põlme et al. 2020); Decay sugar-rich substrates (Põlme et al. 2020) | Põlme S, et al. 2020. Fungal Diversity 105:1-16 ((https://doi.org/10.1007/s13225-020-00466-2)) |
| Otu23849 | 1 | k__Fungi(100);p__Ascomycota(100);c__Saccharomycetes(100);o__Saccharomycetales(100);f__Metschnikowiaceae(100);g__Metschnikowia(100);s__Metschnikowia_pulcherrima(100); | Metschnikowia | 13 | Saprotroph | \|Undefined Saprotroph\| | Yeast | NULL | Probable | As nectar-tap_saprotroph (Põlme et al. 2020); Decay sugar-rich substrates (Põlme et al. 2020) | Põlme S, et al. 2020. Fungal Diversity 105:1-16 ((https://doi.org/10.1007/s13225-020-00466-2)) |
| Otu23850 | 1 | k__Fungi(100);p__Ascomycota(100);c__Saccharomycetes(100);o__Saccharomycetales(100);f__Metschnikowiaceae(100);g__Metschnikowia(100);g__Metschnikowia_unclassified(100); | Metschnikowia | 13 | Saprotroph | \|Undefined Saprotroph\| | Yeast | NULL | Probable | As nectar-tap_saprotroph (Põlme et al. 2020); Decay sugar-rich substrates (Põlme et al. 2020) | Põlme S, et al. 2020. Fungal Diversity 105:1-16 ((https://doi.org/10.1007/s13225-020-00466-2)) |
| Otu23854 | 1 | k__Fungi(100);p__Ascomycota(100);c__Leotiomycetes(100);o__Helotiales(100);f__Helotiales_family_Incertae_sedis(100);g__Scytalidium(100);s__Scytalidium_sp(100); | Scytalidium | 13 | Pathotroph-Saprotroph | Animal Parasite-Fungal Parasite-Plant Pathogen-Plant Saprotroph-\|Wood Saprotroph\| | Microfungus | Soft Rot | Probable | As litter_saprotroph (Põlme et al. 2020); As mycoparasite (Põlme et al. 2020); Decay wood, leaf, fruit, seed, soil, animal material (Põlme et al. 2020) | Nilsson T. 1973. Studia Forestalia Suecica 104:1-40 ((https://pub.epsilon.slu.se/5711/)); Costa IPMW, et al. 2012. Mycotaxon 119:e26 ((http://mycotaxon.com/resources/checklists/costa_v119_checklist.pdf)); Tedersoo L, et al. 2014. Science 346:e1256688 ((https://doi.org/10.1126/science.1256688)); Põlme S, et al. 2020. Fungal Diversity 105:1-16 ((https://doi.org/10.1007/s13225-020-00466-2)); The American Phytopathological Society Common Names of Plant Diseases ((http://www.apsnet.org/publications/commonnames/Pages/default.aspx)) |
| Otu23856 | 1 | k__Fungi(100);p__Ascomycota(100);c__Saccharomycetes(100);o__Saccharomycetales(100);f__Metschnikowiaceae(100);g__Metschnikowia(100);s__Metschnikowia_pulcherrima(100); | Metschnikowia | 13 | Saprotroph | \|Undefined Saprotroph\| | Yeast | NULL | Probable | As nectar-tap_saprotroph (Põlme et al. 2020); Decay sugar-rich substrates (Põlme et al. 2020) | Põlme S, et al. 2020. Fungal Diversity 105:1-16 ((https://doi.org/10.1007/s13225-020-00466-2)) |
| Otu23859 | 1 | k__Fungi(100);p__Ascomycota(100);c__Saccharomycetes(100);o__Saccharomycetales(100);f__Metschnikowiaceae(100);g__Metschnikowia(100);s__Metschnikowia_pulcherrima(100); | Metschnikowia | 13 | Saprotroph | \|Undefined Saprotroph\| | Yeast | NULL | Probable | As nectar-tap_saprotroph (Põlme et al. 2020); Decay sugar-rich substrates (Põlme et al. 2020) | Põlme S, et al. 2020. Fungal Diversity 105:1-16 ((https://doi.org/10.1007/s13225-020-00466-2)) |
| Otu23867 | 1 | k__Fungi(100);p__Ascomycota(100);c__Saccharomycetes(100);o__Saccharomycetales(100);f__Metschnikowiaceae(100);g__Metschnikowia(100);g__Metschnikowia_unclassified(100); | Metschnikowia | 13 | Saprotroph | \|Undefined Saprotroph\| | Yeast | NULL | Probable | As nectar-tap_saprotroph (Põlme et al. 2020); Decay sugar-rich substrates (Põlme et al. 2020) | Põlme S, et al. 2020. Fungal Diversity 105:1-16 ((https://doi.org/10.1007/s13225-020-00466-2)) |
| Otu23868 | 1 | k__Fungi(100);p__Ascomycota(100);c__Saccharomycetes(100);o__Saccharomycetales(100);f__Metschnikowiaceae(100);g__Metschnikowia(100);s__Metschnikowia_pulcherrima(100); | Metschnikowia | 13 | Saprotroph | \|Undefined Saprotroph\| | Yeast | NULL | Probable | As nectar-tap_saprotroph (Põlme et al. 2020); Decay sugar-rich substrates (Põlme et al. 2020) | Põlme S, et al. 2020. Fungal Diversity 105:1-16 ((https://doi.org/10.1007/s13225-020-00466-2)) |
| Otu23872 | 1 | k__Fungi(100);p__Ascomycota(100);c__Saccharomycetes(100);o__Saccharomycetales(100);f__Metschnikowiaceae(100);g__Metschnikowia(100);s__Metschnikowia_pulcherrima(100); | Metschnikowia | 13 | Saprotroph | \|Undefined Saprotroph\| | Yeast | NULL | Probable | As nectar-tap_saprotroph (Põlme et al. 2020); Decay sugar-rich substrates (Põlme et al. 2020) | Põlme S, et al. 2020. Fungal Diversity 105:1-16 ((https://doi.org/10.1007/s13225-020-00466-2)) |
| Otu23874 | 1 | k__Fungi(100);p__Ascomycota(100);c__Saccharomycetes(100);o__Saccharomycetales(100);f__Metschnikowiaceae(100);g__Metschnikowia(100);s__Metschnikowia_pulcherrima(100); | Metschnikowia | 13 | Saprotroph | \|Undefined Saprotroph\| | Yeast | NULL | Probable | As nectar-tap_saprotroph (Põlme et al. 2020); Decay sugar-rich substrates (Põlme et al. 2020) | Põlme S, et al. 2020. Fungal Diversity 105:1-16 ((https://doi.org/10.1007/s13225-020-00466-2)) |
| Otu23875 | 1 | k__Fungi(100);p__Ascomycota(100);c__Saccharomycetes(100);o__Saccharomycetales(100);f__Metschnikowiaceae(100);g__Metschnikowia(100);g__Metschnikowia_unclassified(100); | Metschnikowia | 13 | Saprotroph | \|Undefined Saprotroph\| | Yeast | NULL | Probable | As nectar-tap_saprotroph (Põlme et al. 2020); Decay sugar-rich substrates (Põlme et al. 2020) | Põlme S, et al. 2020. Fungal Diversity 105:1-16 ((https://doi.org/10.1007/s13225-020-00466-2)) |
| Otu23877 | 1 | k__Fungi(100);p__Ascomycota(100);c__Saccharomycetes(100);o__Saccharomycetales(100);f__Metschnikowiaceae(100);g__Metschnikowia(100);s__Metschnikowia_pulcherrima(100); | Metschnikowia | 13 | Saprotroph | \|Undefined Saprotroph\| | Yeast | NULL | Probable | As nectar-tap_saprotroph (Põlme et al. 2020); Decay sugar-rich substrates (Põlme et al. 2020) | Põlme S, et al. 2020. Fungal Diversity 105:1-16 ((https://doi.org/10.1007/s13225-020-00466-2)) |
| Otu23878 | 1 | k__Fungi(100);p__Ascomycota(100);c__Saccharomycetes(100);o__Saccharomycetales(100);f__Metschnikowiaceae(100);g__Metschnikowia(100);g__Metschnikowia_unclassified(100); | Metschnikowia | 13 | Saprotroph | \|Undefined Saprotroph\| | Yeast | NULL | Probable | As nectar-tap_saprotroph (Põlme et al. 2020); Decay sugar-rich substrates (Põlme et al. 2020) | Põlme S, et al. 2020. Fungal Diversity 105:1-16 ((https://doi.org/10.1007/s13225-020-00466-2)) |
| Otu23880 | 1 | k__Fungi(100);p__Ascomycota(100);c__Saccharomycetes(100);o__Saccharomycetales(100);f__Metschnikowiaceae(100);g__Metschnikowia(100);s__Metschnikowia_pulcherrima(100); | Metschnikowia | 13 | Saprotroph | \|Undefined Saprotroph\| | Yeast | NULL | Probable | As nectar-tap_saprotroph (Põlme et al. 2020); Decay sugar-rich substrates (Põlme et al. 2020) | Põlme S, et al. 2020. Fungal Diversity 105:1-16 ((https://doi.org/10.1007/s13225-020-00466-2)) |
| Otu23882 | 1 | k__Fungi(100);p__Ascomycota(100);c__Saccharomycetes(100);o__Saccharomycetales(100);f__Metschnikowiaceae(100);g__Metschnikowia(100);s__Metschnikowia_pulcherrima(100); | Metschnikowia | 13 | Saprotroph | \|Undefined Saprotroph\| | Yeast | NULL | Probable | As nectar-tap_saprotroph (Põlme et al. 2020); Decay sugar-rich substrates (Põlme et al. 2020) | Põlme S, et al. 2020. Fungal Diversity 105:1-16 ((https://doi.org/10.1007/s13225-020-00466-2)) |
| Otu23883 | 1 | k__Fungi(100);p__Ascomycota(100);c__Saccharomycetes(100);o__Saccharomycetales(100);f__Metschnikowiaceae(100);g__Metschnikowia(100);s__Metschnikowia_pulcherrima(100); | Metschnikowia | 13 | Saprotroph | \|Undefined Saprotroph\| | Yeast | NULL | Probable | As nectar-tap_saprotroph (Põlme et al. 2020); Decay sugar-rich substrates (Põlme et al. 2020) | Põlme S, et al. 2020. Fungal Diversity 105:1-16 ((https://doi.org/10.1007/s13225-020-00466-2)) |
| Otu23884 | 1 | k__Fungi(100);p__Ascomycota(100);c__Saccharomycetes(100);o__Saccharomycetales(100);f__Metschnikowiaceae(100);g__Metschnikowia(100);s__Metschnikowia_pulcherrima(100); | Metschnikowia | 13 | Saprotroph | \|Undefined Saprotroph\| | Yeast | NULL | Probable | As nectar-tap_saprotroph (Põlme et al. 2020); Decay sugar-rich substrates (Põlme et al. 2020) | Põlme S, et al. 2020. Fungal Diversity 105:1-16 ((https://doi.org/10.1007/s13225-020-00466-2)) |
| Otu23885 | 1 | k__Fungi(100);p__Ascomycota(100);c__Saccharomycetes(100);o__Saccharomycetales(100);f__Metschnikowiaceae(100);g__Metschnikowia(100);s__Metschnikowia_pulcherrima(100); | Metschnikowia | 13 | Saprotroph | \|Undefined Saprotroph\| | Yeast | NULL | Probable | As nectar-tap_saprotroph (Põlme et al. 2020); Decay sugar-rich substrates (Põlme et al. 2020) | Põlme S, et al. 2020. Fungal Diversity 105:1-16 ((https://doi.org/10.1007/s13225-020-00466-2)) |
| Otu23886 | 1 | k__Fungi(100);p__Ascomycota(100);c__Saccharomycetes(100);o__Saccharomycetales(100);f__Metschnikowiaceae(100);g__Metschnikowia(100);s__Metschnikowia_pulcherrima(100); | Metschnikowia | 13 | Saprotroph | \|Undefined Saprotroph\| | Yeast | NULL | Probable | As nectar-tap_saprotroph (Põlme et al. 2020); Decay sugar-rich substrates (Põlme et al. 2020) | Põlme S, et al. 2020. Fungal Diversity 105:1-16 ((https://doi.org/10.1007/s13225-020-00466-2)) |
| Otu23897 | 1 | k__Fungi(100);p__Ascomycota(100);c__Saccharomycetes(100);o__Saccharomycetales(100);f__Metschnikowiaceae(100);g__Metschnikowia(100);s__Metschnikowia_pulcherrima(100); | Metschnikowia | 13 | Saprotroph | \|Undefined Saprotroph\| | Yeast | NULL | Probable | As nectar-tap_saprotroph (Põlme et al. 2020); Decay sugar-rich substrates (Põlme et al. 2020) | Põlme S, et al. 2020. Fungal Diversity 105:1-16 ((https://doi.org/10.1007/s13225-020-00466-2)) |
| Otu23899 | 1 | k__Fungi(100);p__Ascomycota(100);c__Saccharomycetes(100);o__Saccharomycetales(100);f__Metschnikowiaceae(100);g__Metschnikowia(100);s__Metschnikowia_pulcherrima(100); | Metschnikowia | 13 | Saprotroph | \|Undefined Saprotroph\| | Yeast | NULL | Probable | As nectar-tap_saprotroph (Põlme et al. 2020); Decay sugar-rich substrates (Põlme et al. 2020) | Põlme S, et al. 2020. Fungal Diversity 105:1-16 ((https://doi.org/10.1007/s13225-020-00466-2)) |
| Otu23900 | 1 | k__Fungi(100);p__Ascomycota(100);c__Saccharomycetes(100);o__Saccharomycetales(100);f__Metschnikowiaceae(100);g__Metschnikowia(100);s__Metschnikowia_pulcherrima(100); | Metschnikowia | 13 | Saprotroph | \|Undefined Saprotroph\| | Yeast | NULL | Probable | As nectar-tap_saprotroph (Põlme et al. 2020); Decay sugar-rich substrates (Põlme et al. 2020) | Põlme S, et al. 2020. Fungal Diversity 105:1-16 ((https://doi.org/10.1007/s13225-020-00466-2)) |
| Otu23906 | 1 | k__Fungi(100);p__Ascomycota(100);c__Saccharomycetes(100);o__Saccharomycetales(100);f__Metschnikowiaceae(100);g__Metschnikowia(100);s__Metschnikowia_pulcherrima(100); | Metschnikowia | 13 | Saprotroph | \|Undefined Saprotroph\| | Yeast | NULL | Probable | As nectar-tap_saprotroph (Põlme et al. 2020); Decay sugar-rich substrates (Põlme et al. 2020) | Põlme S, et al. 2020. Fungal Diversity 105:1-16 ((https://doi.org/10.1007/s13225-020-00466-2)) |
| Otu23909 | 1 | k__Fungi(100);p__Ascomycota(100);c__Saccharomycetes(100);o__Saccharomycetales(100);f__Metschnikowiaceae(100);g__Metschnikowia(100);g__Metschnikowia_unclassified(100); | Metschnikowia | 13 | Saprotroph | \|Undefined Saprotroph\| | Yeast | NULL | Probable | As nectar-tap_saprotroph (Põlme et al. 2020); Decay sugar-rich substrates (Põlme et al. 2020) | Põlme S, et al. 2020. Fungal Diversity 105:1-16 ((https://doi.org/10.1007/s13225-020-00466-2)) |
| Otu23910 | 1 | k__Fungi(100);p__Ascomycota(100);c__Saccharomycetes(100);o__Saccharomycetales(100);f__Metschnikowiaceae(100);g__Metschnikowia(100);s__Metschnikowia_pulcherrima(100); | Metschnikowia | 13 | Saprotroph | \|Undefined Saprotroph\| | Yeast | NULL | Probable | As nectar-tap_saprotroph (Põlme et al. 2020); Decay sugar-rich substrates (Põlme et al. 2020) | Põlme S, et al. 2020. Fungal Diversity 105:1-16 ((https://doi.org/10.1007/s13225-020-00466-2)) |
| Otu23911 | 1 | k__Fungi(100);p__Ascomycota(100);c__Saccharomycetes(100);o__Saccharomycetales(100);f__Saccharomycetaceae(100);g__Kazachstania(100);s__Kazachstania_intestinalis(100); | Kazachstania | 13 | Saprotroph | \|Undefined Saprotroph\| | Yeast | NULL | Probable | As nectar-tap_saprotroph (Põlme et al. 2020); Decay sugar-rich substrates (Põlme et al. 2020) | Põlme S, et al. 2020. Fungal Diversity 105:1-16 ((https://doi.org/10.1007/s13225-020-00466-2)) |
| Otu23912 | 1 | k__Fungi(100);p__Ascomycota(100);c__Saccharomycetes(100);o__Saccharomycetales(100);f__Saccharomycetaceae(100);g__Kazachstania(100);s__Kazachstania_intestinalis(100); | Kazachstania | 13 | Saprotroph | \|Undefined Saprotroph\| | Yeast | NULL | Probable | As nectar-tap_saprotroph (Põlme et al. 2020); Decay sugar-rich substrates (Põlme et al. 2020) | Põlme S, et al. 2020. Fungal Diversity 105:1-16 ((https://doi.org/10.1007/s13225-020-00466-2)) |
| Otu23914 | 1 | k__Fungi(100);p__Ascomycota(100);c__Saccharomycetes(100);o__Saccharomycetales(100);f__Metschnikowiaceae(100);g__Metschnikowia(100);s__Metschnikowia_pulcherrima(100); | Metschnikowia | 13 | Saprotroph | \|Undefined Saprotroph\| | Yeast | NULL | Probable | As nectar-tap_saprotroph (Põlme et al. 2020); Decay sugar-rich substrates (Põlme et al. 2020) | Põlme S, et al. 2020. Fungal Diversity 105:1-16 ((https://doi.org/10.1007/s13225-020-00466-2)) |
| Otu23915 | 1 | k__Fungi(100);p__Ascomycota(100);c__Saccharomycetes(100);o__Saccharomycetales(100);f__Metschnikowiaceae(100);g__Metschnikowia(100);s__Metschnikowia_pulcherrima(100); | Metschnikowia | 13 | Saprotroph | \|Undefined Saprotroph\| | Yeast | NULL | Probable | As nectar-tap_saprotroph (Põlme et al. 2020); Decay sugar-rich substrates (Põlme et al. 2020) | Põlme S, et al. 2020. Fungal Diversity 105:1-16 ((https://doi.org/10.1007/s13225-020-00466-2)) |
| Otu23916 | 1 | k__Fungi(100);p__Ascomycota(100);c__Saccharomycetes(100);o__Saccharomycetales(100);f__Metschnikowiaceae(100);g__Metschnikowia(100);s__Metschnikowia_pulcherrima(100); | Metschnikowia | 13 | Saprotroph | \|Undefined Saprotroph\| | Yeast | NULL | Probable | As nectar-tap_saprotroph (Põlme et al. 2020); Decay sugar-rich substrates (Põlme et al. 2020) | Põlme S, et al. 2020. Fungal Diversity 105:1-16 ((https://doi.org/10.1007/s13225-020-00466-2)) |
| Otu23917 | 1 | k__Fungi(100);p__Ascomycota(100);c__Saccharomycetes(100);o__Saccharomycetales(100);f__Metschnikowiaceae(100);g__Metschnikowia(100);s__Metschnikowia_pulcherrima(100); | Metschnikowia | 13 | Saprotroph | \|Undefined Saprotroph\| | Yeast | NULL | Probable | As nectar-tap_saprotroph (Põlme et al. 2020); Decay sugar-rich substrates (Põlme et al. 2020) | Põlme S, et al. 2020. Fungal Diversity 105:1-16 ((https://doi.org/10.1007/s13225-020-00466-2)) |
| Otu23920 | 1 | k__Fungi(100);p__Ascomycota(100);c__Saccharomycetes(100);o__Saccharomycetales(100);f__Metschnikowiaceae(100);g__Metschnikowia(100);s__Metschnikowia_pulcherrima(100); | Metschnikowia | 13 | Saprotroph | \|Undefined Saprotroph\| | Yeast | NULL | Probable | As nectar-tap_saprotroph (Põlme et al. 2020); Decay sugar-rich substrates (Põlme et al. 2020) | Põlme S, et al. 2020. Fungal Diversity 105:1-16 ((https://doi.org/10.1007/s13225-020-00466-2)) |
| Otu23921 | 1 | k__Fungi(100);p__Ascomycota(100);c__Saccharomycetes(100);o__Saccharomycetales(100);f__Metschnikowiaceae(100);g__Metschnikowia(100);g__Metschnikowia_unclassified(100); | Metschnikowia | 13 | Saprotroph | \|Undefined Saprotroph\| | Yeast | NULL | Probable | As nectar-tap_saprotroph (Põlme et al. 2020); Decay sugar-rich substrates (Põlme et al. 2020) | Põlme S, et al. 2020. Fungal Diversity 105:1-16 ((https://doi.org/10.1007/s13225-020-00466-2)) |
| Otu23923 | 1 | k__Fungi(100);p__Ascomycota(100);c__Saccharomycetes(100);o__Saccharomycetales(100);f__Metschnikowiaceae(100);g__Metschnikowia(100);s__Metschnikowia_pulcherrima(100); | Metschnikowia | 13 | Saprotroph | \|Undefined Saprotroph\| | Yeast | NULL | Probable | As nectar-tap_saprotroph (Põlme et al. 2020); Decay sugar-rich substrates (Põlme et al. 2020) | Põlme S, et al. 2020. Fungal Diversity 105:1-16 ((https://doi.org/10.1007/s13225-020-00466-2)) |
| Otu23924 | 1 | k__Fungi(100);p__Ascomycota(100);c__Saccharomycetes(100);o__Saccharomycetales(100);f__Metschnikowiaceae(100);g__Metschnikowia(100);g__Metschnikowia_unclassified(100); | Metschnikowia | 13 | Saprotroph | \|Undefined Saprotroph\| | Yeast | NULL | Probable | As nectar-tap_saprotroph (Põlme et al. 2020); Decay sugar-rich substrates (Põlme et al. 2020) | Põlme S, et al. 2020. Fungal Diversity 105:1-16 ((https://doi.org/10.1007/s13225-020-00466-2)) |
| Otu23925 | 1 | k__Fungi(100);p__Ascomycota(100);c__Saccharomycetes(100);o__Saccharomycetales(100);f__Metschnikowiaceae(100);g__Metschnikowia(100);g__Metschnikowia_unclassified(100); | Metschnikowia | 13 | Saprotroph | \|Undefined Saprotroph\| | Yeast | NULL | Probable | As nectar-tap_saprotroph (Põlme et al. 2020); Decay sugar-rich substrates (Põlme et al. 2020) | Põlme S, et al. 2020. Fungal Diversity 105:1-16 ((https://doi.org/10.1007/s13225-020-00466-2)) |
| Otu23926 | 1 | k__Fungi(100);p__Ascomycota(100);c__Saccharomycetes(100);o__Saccharomycetales(100);f__Metschnikowiaceae(100);g__Metschnikowia(100);s__Metschnikowia_pulcherrima(100); | Metschnikowia | 13 | Saprotroph | \|Undefined Saprotroph\| | Yeast | NULL | Probable | As nectar-tap_saprotroph (Põlme et al. 2020); Decay sugar-rich substrates (Põlme et al. 2020) | Põlme S, et al. 2020. Fungal Diversity 105:1-16 ((https://doi.org/10.1007/s13225-020-00466-2)) |
| Otu23927 | 1 | k__Fungi(100);p__Ascomycota(100);c__Saccharomycetes(100);o__Saccharomycetales(100);f__Metschnikowiaceae(100);g__Metschnikowia(100);s__Metschnikowia_pulcherrima(100); | Metschnikowia | 13 | Saprotroph | \|Undefined Saprotroph\| | Yeast | NULL | Probable | As nectar-tap_saprotroph (Põlme et al. 2020); Decay sugar-rich substrates (Põlme et al. 2020) | Põlme S, et al. 2020. Fungal Diversity 105:1-16 ((https://doi.org/10.1007/s13225-020-00466-2)) |
| Otu23928 | 1 | k__Fungi(100);p__Ascomycota(100);c__Saccharomycetes(100);o__Saccharomycetales(100);f__Metschnikowiaceae(100);g__Metschnikowia(100);s__Metschnikowia_pulcherrima(100); | Metschnikowia | 13 | Saprotroph | \|Undefined Saprotroph\| | Yeast | NULL | Probable | As nectar-tap_saprotroph (Põlme et al. 2020); Decay sugar-rich substrates (Põlme et al. 2020) | Põlme S, et al. 2020. Fungal Diversity 105:1-16 ((https://doi.org/10.1007/s13225-020-00466-2)) |
| Otu23929 | 1 | k__Fungi(100);p__Ascomycota(100);c__Saccharomycetes(100);o__Saccharomycetales(100);f__Metschnikowiaceae(100);g__Metschnikowia(100);s__Metschnikowia_pulcherrima(100); | Metschnikowia | 13 | Saprotroph | \|Undefined Saprotroph\| | Yeast | NULL | Probable | As nectar-tap_saprotroph (Põlme et al. 2020); Decay sugar-rich substrates (Põlme et al. 2020) | Põlme S, et al. 2020. Fungal Diversity 105:1-16 ((https://doi.org/10.1007/s13225-020-00466-2)) |
| Otu23930 | 1 | k__Fungi(100);p__Ascomycota(100);c__Saccharomycetes(100);o__Saccharomycetales(100);f__Metschnikowiaceae(100);g__Metschnikowia(100);s__Metschnikowia_pulcherrima(100); | Metschnikowia | 13 | Saprotroph | \|Undefined Saprotroph\| | Yeast | NULL | Probable | As nectar-tap_saprotroph (Põlme et al. 2020); Decay sugar-rich substrates (Põlme et al. 2020) | Põlme S, et al. 2020. Fungal Diversity 105:1-16 ((https://doi.org/10.1007/s13225-020-00466-2)) |
| Otu23931 | 1 | k__Fungi(100);p__Ascomycota(100);c__Saccharomycetes(100);o__Saccharomycetales(100);f__Metschnikowiaceae(100);g__Metschnikowia(100);g__Metschnikowia_unclassified(100); | Metschnikowia | 13 | Saprotroph | \|Undefined Saprotroph\| | Yeast | NULL | Probable | As nectar-tap_saprotroph (Põlme et al. 2020); Decay sugar-rich substrates (Põlme et al. 2020) | Põlme S, et al. 2020. Fungal Diversity 105:1-16 ((https://doi.org/10.1007/s13225-020-00466-2)) |
| Otu23932 | 1 | k__Fungi(100);p__Ascomycota(100);c__Saccharomycetes(100);o__Saccharomycetales(100);f__Metschnikowiaceae(100);g__Metschnikowia(100);s__Metschnikowia_pulcherrima(100); | Metschnikowia | 13 | Saprotroph | \|Undefined Saprotroph\| | Yeast | NULL | Probable | As nectar-tap_saprotroph (Põlme et al. 2020); Decay sugar-rich substrates (Põlme et al. 2020) | Põlme S, et al. 2020. Fungal Diversity 105:1-16 ((https://doi.org/10.1007/s13225-020-00466-2)) |
| Otu23933 | 1 | k__Fungi(100);p__Ascomycota(100);c__Saccharomycetes(100);o__Saccharomycetales(100);f__Metschnikowiaceae(100);g__Metschnikowia(100);s__Metschnikowia_pulcherrima(100); | Metschnikowia | 13 | Saprotroph | \|Undefined Saprotroph\| | Yeast | NULL | Probable | As nectar-tap_saprotroph (Põlme et al. 2020); Decay sugar-rich substrates (Põlme et al. 2020) | Põlme S, et al. 2020. Fungal Diversity 105:1-16 ((https://doi.org/10.1007/s13225-020-00466-2)) |
| Otu23934 | 1 | k__Fungi(100);p__Ascomycota(100);c__Saccharomycetes(100);o__Saccharomycetales(100);f__Metschnikowiaceae(100);g__Metschnikowia(100);s__Metschnikowia_pulcherrima(100); | Metschnikowia | 13 | Saprotroph | \|Undefined Saprotroph\| | Yeast | NULL | Probable | As nectar-tap_saprotroph (Põlme et al. 2020); Decay sugar-rich substrates (Põlme et al. 2020) | Põlme S, et al. 2020. Fungal Diversity 105:1-16 ((https://doi.org/10.1007/s13225-020-00466-2)) |
| Otu23935 | 1 | k__Fungi(100);p__Ascomycota(100);c__Saccharomycetes(100);o__Saccharomycetales(100);f__Metschnikowiaceae(100);g__Metschnikowia(100);s__Metschnikowia_pulcherrima(100); | Metschnikowia | 13 | Saprotroph | \|Undefined Saprotroph\| | Yeast | NULL | Probable | As nectar-tap_saprotroph (Põlme et al. 2020); Decay sugar-rich substrates (Põlme et al. 2020) | Põlme S, et al. 2020. Fungal Diversity 105:1-16 ((https://doi.org/10.1007/s13225-020-00466-2)) |
| Otu23937 | 1 | k__Fungi(100);p__Ascomycota(100);c__Saccharomycetes(100);o__Saccharomycetales(100);f__Metschnikowiaceae(100);g__Metschnikowia(100);s__Metschnikowia_pulcherrima(100); | Metschnikowia | 13 | Saprotroph | \|Undefined Saprotroph\| | Yeast | NULL | Probable | As nectar-tap_saprotroph (Põlme et al. 2020); Decay sugar-rich substrates (Põlme et al. 2020) | Põlme S, et al. 2020. Fungal Diversity 105:1-16 ((https://doi.org/10.1007/s13225-020-00466-2)) |
| Otu23938 | 1 | k__Fungi(100);p__Basidiomycota(100);c__Microbotryomycetes(100);o__Sporidiobolales(100);f__Sporidiobolales_family_Incertae_sedis(100);g__Sporobolomyces(100);s__Sporobolomyces_lactosus(100); | Sporobolomyces | 13 | Pathotroph-Saprotroph | \|Fungal Parasite\|-Plant Saprotroph | Yeast | NULL | Probable | As litter_saprotroph (Põlme et al. 2020); As mycoparasite (Põlme et al. 2020); Decay fungal material (Põlme et al. 2020) | Tedersoo L, et al. 2014. Science 346:e1256688 ((https://doi.org/10.1126/science.1256688)); Purahong W, et al. 2016. Molecular Ecology 25:4059-4074 ((https://doi.org/10.1111/mec.13739)); Põlme S, et al. 2020. Fungal Diversity 105:1-16 ((https://doi.org/10.1007/s13225-020-00466-2)) |
| Otu23939 | 1 | k__Fungi(100);p__Basidiomycota(100);c__Microbotryomycetes(100);o__Sporidiobolales(100);f__Sporidiobolales_family_Incertae_sedis(100);g__Sporobolomyces(100);s__Sporobolomyces_lactosus(100); | Sporobolomyces | 13 | Pathotroph-Saprotroph | \|Fungal Parasite\|-Plant Saprotroph | Yeast | NULL | Probable | As litter_saprotroph (Põlme et al. 2020); As mycoparasite (Põlme et al. 2020); Decay fungal material (Põlme et al. 2020) | Tedersoo L, et al. 2014. Science 346:e1256688 ((https://doi.org/10.1126/science.1256688)); Purahong W, et al. 2016. Molecular Ecology 25:4059-4074 ((https://doi.org/10.1111/mec.13739)); Põlme S, et al. 2020. Fungal Diversity 105:1-16 ((https://doi.org/10.1007/s13225-020-00466-2)) |
| Otu23940 | 1 | k__Fungi(100);p__Ascomycota(100);c__Dothideomycetes(100);o__Pleosporales(100);f__Phaeosphaeriaceae(100);g__unclassified_Phaeosphaeriaceae(100);s__Phaeosphaeriaceae_sp(100); | Phaeosphaeriaceae | 9 | Pathotroph-Saprotroph | Fungal Parasite-Plant Pathogen-Plant Saprotroph | Microfungus | NULL | Probable | Necrotrophic or saprobic on a wide range of plants, especially monocotyledons; a few species parasitic on other fungi (Cannon & Kirk 2007) | Cannon PF, Kirk PM. 2007. Fungal Families of the World. CAB International, Cambridge (ISBN: 978-0851998275) |
| Otu23945 | 1 | k__Fungi(100);p__Ascomycota(100);c__Leotiomycetes(100);o__Helotiales(100);f__Helotiales_family_Incertae_sedis(100);g__Scleropezicula(100);s__Scleropezicula_sp(100); | Scleropezicula | 13 | Saprotroph | Undefined Saprotroph-\|Wood Saprotroph\| | Microfungus | NULL | Probable | Decay: wood (Põlme et al. 2020) | Tedersoo L, et al. 2014. Science 346:e1256688 ((https://doi.org/10.1126/science.1256688)); Põlme S, et al. 2020. Fungal Diversity 105:1-16 ((https://doi.org/10.1007/s13225-020-00466-2)) |
| Otu23946 | 1 | k__Fungi(100);p__Ascomycota(100);c__Leotiomycetes(100);o__Helotiales(100);f__Helotiales_family_Incertae_sedis(100);g__Hyphodiscus(100);s__Hyphodiscus_hymeniophilus(100); | Hyphodiscus | 13 | Pathotroph-Saprotroph | \|Fungal Parasite\|-Wood Saprotroph | Microfungus | NULL | Highly Probable | As mycoparasite (Põlme et al. 2020); Decay fungal material (Põlme et al. 2020) | Põlme S, et al. 2020. Fungal Diversity 105:1-16 ((https://doi.org/10.1007/s13225-020-00466-2)); List compiled by John Plischke III ((https://goodmushroom.weebly.com/uploads/5/7/1/7/57176535/fungi_on_fungi_pdf_lq.pdf)) |
| Otu23950 | 1 | k__Fungi(100);p__Ascomycota(100);c__Saccharomycetes(100);o__Saccharomycetales(100);f__Metschnikowiaceae(100);g__Metschnikowia(100);s__Metschnikowia_pulcherrima(100); | Metschnikowia | 13 | Saprotroph | \|Undefined Saprotroph\| | Yeast | NULL | Probable | As nectar-tap_saprotroph (Põlme et al. 2020); Decay sugar-rich substrates (Põlme et al. 2020) | Põlme S, et al. 2020. Fungal Diversity 105:1-16 ((https://doi.org/10.1007/s13225-020-00466-2)) |
| Otu23951 | 1 | k__Fungi(100);p__Ascomycota(100);c__Saccharomycetes(100);o__Saccharomycetales(100);f__Metschnikowiaceae(100);g__Metschnikowia(100);s__Metschnikowia_pulcherrima(100); | Metschnikowia | 13 | Saprotroph | \|Undefined Saprotroph\| | Yeast | NULL | Probable | As nectar-tap_saprotroph (Põlme et al. 2020); Decay sugar-rich substrates (Põlme et al. 2020) | Põlme S, et al. 2020. Fungal Diversity 105:1-16 ((https://doi.org/10.1007/s13225-020-00466-2)) |
| Otu23952 | 1 | k__Fungi(100);p__Ascomycota(100);c__Saccharomycetes(100);o__Saccharomycetales(100);f__Metschnikowiaceae(100);g__Metschnikowia(100);s__Metschnikowia_pulcherrima(100); | Metschnikowia | 13 | Saprotroph | \|Undefined Saprotroph\| | Yeast | NULL | Probable | As nectar-tap_saprotroph (Põlme et al. 2020); Decay sugar-rich substrates (Põlme et al. 2020) | Põlme S, et al. 2020. Fungal Diversity 105:1-16 ((https://doi.org/10.1007/s13225-020-00466-2)) |
| Otu23953 | 1 | k__Fungi(100);p__Ascomycota(100);c__Saccharomycetes(100);o__Saccharomycetales(100);f__Metschnikowiaceae(100);g__Metschnikowia(100);s__Metschnikowia_pulcherrima(100); | Metschnikowia | 13 | Saprotroph | \|Undefined Saprotroph\| | Yeast | NULL | Probable | As nectar-tap_saprotroph (Põlme et al. 2020); Decay sugar-rich substrates (Põlme et al. 2020) | Põlme S, et al. 2020. Fungal Diversity 105:1-16 ((https://doi.org/10.1007/s13225-020-00466-2)) |
| Otu23956 | 1 | k__Fungi(100);p__Ascomycota(100);c__Saccharomycetes(100);o__Saccharomycetales(100);f__Metschnikowiaceae(100);g__Metschnikowia(100);g__Metschnikowia_unclassified(100); | Metschnikowia | 13 | Saprotroph | \|Undefined Saprotroph\| | Yeast | NULL | Probable | As nectar-tap_saprotroph (Põlme et al. 2020); Decay sugar-rich substrates (Põlme et al. 2020) | Põlme S, et al. 2020. Fungal Diversity 105:1-16 ((https://doi.org/10.1007/s13225-020-00466-2)) |
| Otu23957 | 1 | k__Fungi(100);p__Ascomycota(100);c__Saccharomycetes(100);o__Saccharomycetales(100);f__Metschnikowiaceae(100);g__Metschnikowia(100);s__Metschnikowia_pulcherrima(100); | Metschnikowia | 13 | Saprotroph | \|Undefined Saprotroph\| | Yeast | NULL | Probable | As nectar-tap_saprotroph (Põlme et al. 2020); Decay sugar-rich substrates (Põlme et al. 2020) | Põlme S, et al. 2020. Fungal Diversity 105:1-16 ((https://doi.org/10.1007/s13225-020-00466-2)) |
| Otu23958 | 1 | k__Fungi(100);p__Ascomycota(100);c__Saccharomycetes(100);o__Saccharomycetales(100);f__Metschnikowiaceae(100);g__Metschnikowia(100);g__Metschnikowia_unclassified(100); | Metschnikowia | 13 | Saprotroph | \|Undefined Saprotroph\| | Yeast | NULL | Probable | As nectar-tap_saprotroph (Põlme et al. 2020); Decay sugar-rich substrates (Põlme et al. 2020) | Põlme S, et al. 2020. Fungal Diversity 105:1-16 ((https://doi.org/10.1007/s13225-020-00466-2)) |
| Otu23965 | 1 | k__Fungi(100);p__Ascomycota(100);c__Saccharomycetes(100);o__Saccharomycetales(100);f__Metschnikowiaceae(100);g__Metschnikowia(100);s__Metschnikowia_pulcherrima(100); | Metschnikowia | 13 | Saprotroph | \|Undefined Saprotroph\| | Yeast | NULL | Probable | As nectar-tap_saprotroph (Põlme et al. 2020); Decay sugar-rich substrates (Põlme et al. 2020) | Põlme S, et al. 2020. Fungal Diversity 105:1-16 ((https://doi.org/10.1007/s13225-020-00466-2)) |
| Otu23966 | 1 | k__Fungi(100);p__Ascomycota(100);c__Saccharomycetes(100);o__Saccharomycetales(100);f__Metschnikowiaceae(100);g__Metschnikowia(100);g__Metschnikowia_unclassified(100); | Metschnikowia | 13 | Saprotroph | \|Undefined Saprotroph\| | Yeast | NULL | Probable | As nectar-tap_saprotroph (Põlme et al. 2020); Decay sugar-rich substrates (Põlme et al. 2020) | Põlme S, et al. 2020. Fungal Diversity 105:1-16 ((https://doi.org/10.1007/s13225-020-00466-2)) |
| Otu23967 | 1 | k__Fungi(100);p__Ascomycota(100);c__Saccharomycetes(100);o__Saccharomycetales(100);f__Metschnikowiaceae(100);g__Metschnikowia(100);s__Metschnikowia_pulcherrima(100); | Metschnikowia | 13 | Saprotroph | \|Undefined Saprotroph\| | Yeast | NULL | Probable | As nectar-tap_saprotroph (Põlme et al. 2020); Decay sugar-rich substrates (Põlme et al. 2020) | Põlme S, et al. 2020. Fungal Diversity 105:1-16 ((https://doi.org/10.1007/s13225-020-00466-2)) |
| Otu23971 | 1 | k__Fungi(100);p__Ascomycota(100);c__Saccharomycetes(100);o__Saccharomycetales(100);f__Metschnikowiaceae(100);g__Metschnikowia(100);s__Metschnikowia_pulcherrima(100); | Metschnikowia | 13 | Saprotroph | \|Undefined Saprotroph\| | Yeast | NULL | Probable | As nectar-tap_saprotroph (Põlme et al. 2020); Decay sugar-rich substrates (Põlme et al. 2020) | Põlme S, et al. 2020. Fungal Diversity 105:1-16 ((https://doi.org/10.1007/s13225-020-00466-2)) |
| Otu23972 | 1 | k__Fungi(100);p__Ascomycota(100);c__Saccharomycetes(100);o__Saccharomycetales(100);f__Metschnikowiaceae(100);g__Metschnikowia(100);s__Metschnikowia_pulcherrima(100); | Metschnikowia | 13 | Saprotroph | \|Undefined Saprotroph\| | Yeast | NULL | Probable | As nectar-tap_saprotroph (Põlme et al. 2020); Decay sugar-rich substrates (Põlme et al. 2020) | Põlme S, et al. 2020. Fungal Diversity 105:1-16 ((https://doi.org/10.1007/s13225-020-00466-2)) |
| Otu23981 | 1 | k__Fungi(100);p__Ascomycota(100);c__Saccharomycetes(100);o__Saccharomycetales(100);f__Metschnikowiaceae(100);g__Metschnikowia(100);s__Metschnikowia_pulcherrima(100); | Metschnikowia | 13 | Saprotroph | \|Undefined Saprotroph\| | Yeast | NULL | Probable | As nectar-tap_saprotroph (Põlme et al. 2020); Decay sugar-rich substrates (Põlme et al. 2020) | Põlme S, et al. 2020. Fungal Diversity 105:1-16 ((https://doi.org/10.1007/s13225-020-00466-2)) |
| Otu23984 | 1 | k__Fungi(100);p__Ascomycota(100);c__Saccharomycetes(100);o__Saccharomycetales(100);f__Metschnikowiaceae(100);g__Metschnikowia(100);s__Metschnikowia_pulcherrima(100); | Metschnikowia | 13 | Saprotroph | \|Undefined Saprotroph\| | Yeast | NULL | Probable | As nectar-tap_saprotroph (Põlme et al. 2020); Decay sugar-rich substrates (Põlme et al. 2020) | Põlme S, et al. 2020. Fungal Diversity 105:1-16 ((https://doi.org/10.1007/s13225-020-00466-2)) |
| Otu23997 | 1 | k__Fungi(100);p__Ascomycota(100);c__Saccharomycetes(100);o__Saccharomycetales(100);f__Metschnikowiaceae(100);g__Metschnikowia(100);s__Metschnikowia_pulcherrima(100); | Metschnikowia | 13 | Saprotroph | \|Undefined Saprotroph\| | Yeast | NULL | Probable | As nectar-tap_saprotroph (Põlme et al. 2020); Decay sugar-rich substrates (Põlme et al. 2020) | Põlme S, et al. 2020. Fungal Diversity 105:1-16 ((https://doi.org/10.1007/s13225-020-00466-2)) |
| Otu23998 | 1 | k__Fungi(100);p__Ascomycota(100);c__Saccharomycetes(100);o__Saccharomycetales(100);f__Metschnikowiaceae(100);g__Metschnikowia(100);s__Metschnikowia_pulcherrima(100); | Metschnikowia | 13 | Saprotroph | \|Undefined Saprotroph\| | Yeast | NULL | Probable | As nectar-tap_saprotroph (Põlme et al. 2020); Decay sugar-rich substrates (Põlme et al. 2020) | Põlme S, et al. 2020. Fungal Diversity 105:1-16 ((https://doi.org/10.1007/s13225-020-00466-2)) |
| Otu24000 | 1 | k__Fungi(100);p__Ascomycota(100);c__Saccharomycetes(100);o__Saccharomycetales(100);f__Metschnikowiaceae(100);g__Metschnikowia(100);g__Metschnikowia_unclassified(100); | Metschnikowia | 13 | Saprotroph | \|Undefined Saprotroph\| | Yeast | NULL | Probable | As nectar-tap_saprotroph (Põlme et al. 2020); Decay sugar-rich substrates (Põlme et al. 2020) | Põlme S, et al. 2020. Fungal Diversity 105:1-16 ((https://doi.org/10.1007/s13225-020-00466-2)) |
| Otu24001 | 1 | k__Fungi(100);p__Ascomycota(100);c__Saccharomycetes(100);o__Saccharomycetales(100);f__Metschnikowiaceae(100);g__Metschnikowia(100);s__Metschnikowia_pulcherrima(100); | Metschnikowia | 13 | Saprotroph | \|Undefined Saprotroph\| | Yeast | NULL | Probable | As nectar-tap_saprotroph (Põlme et al. 2020); Decay sugar-rich substrates (Põlme et al. 2020) | Põlme S, et al. 2020. Fungal Diversity 105:1-16 ((https://doi.org/10.1007/s13225-020-00466-2)) |
| Otu24002 | 1 | k__Fungi(100);p__Ascomycota(100);c__Saccharomycetes(100);o__Saccharomycetales(100);f__Metschnikowiaceae(100);g__Metschnikowia(100);g__Metschnikowia_unclassified(100); | Metschnikowia | 13 | Saprotroph | \|Undefined Saprotroph\| | Yeast | NULL | Probable | As nectar-tap_saprotroph (Põlme et al. 2020); Decay sugar-rich substrates (Põlme et al. 2020) | Põlme S, et al. 2020. Fungal Diversity 105:1-16 ((https://doi.org/10.1007/s13225-020-00466-2)) |
| Otu24003 | 1 | k__Fungi(100);p__Ascomycota(100);c__Saccharomycetes(100);o__Saccharomycetales(100);f__Metschnikowiaceae(100);g__Metschnikowia(100);g__Metschnikowia_unclassified(100); | Metschnikowia | 13 | Saprotroph | \|Undefined Saprotroph\| | Yeast | NULL | Probable | As nectar-tap_saprotroph (Põlme et al. 2020); Decay sugar-rich substrates (Põlme et al. 2020) | Põlme S, et al. 2020. Fungal Diversity 105:1-16 ((https://doi.org/10.1007/s13225-020-00466-2)) |
| Otu24004 | 1 | k__Fungi(100);p__Ascomycota(100);c__Saccharomycetes(100);o__Saccharomycetales(100);f__Metschnikowiaceae(100);g__Metschnikowia(100);g__Metschnikowia_unclassified(100); | Metschnikowia | 13 | Saprotroph | \|Undefined Saprotroph\| | Yeast | NULL | Probable | As nectar-tap_saprotroph (Põlme et al. 2020); Decay sugar-rich substrates (Põlme et al. 2020) | Põlme S, et al. 2020. Fungal Diversity 105:1-16 ((https://doi.org/10.1007/s13225-020-00466-2)) |
| Otu24005 | 1 | k__Fungi(100);p__Ascomycota(100);c__Saccharomycetes(100);o__Saccharomycetales(100);f__Metschnikowiaceae(100);g__Metschnikowia(100);s__Metschnikowia_pulcherrima(100); | Metschnikowia | 13 | Saprotroph | \|Undefined Saprotroph\| | Yeast | NULL | Probable | As nectar-tap_saprotroph (Põlme et al. 2020); Decay sugar-rich substrates (Põlme et al. 2020) | Põlme S, et al. 2020. Fungal Diversity 105:1-16 ((https://doi.org/10.1007/s13225-020-00466-2)) |
| Otu24006 | 1 | k__Fungi(100);p__Ascomycota(100);c__Saccharomycetes(100);o__Saccharomycetales(100);f__Metschnikowiaceae(100);g__Metschnikowia(100);s__Metschnikowia_pulcherrima(100); | Metschnikowia | 13 | Saprotroph | \|Undefined Saprotroph\| | Yeast | NULL | Probable | As nectar-tap_saprotroph (Põlme et al. 2020); Decay sugar-rich substrates (Põlme et al. 2020) | Põlme S, et al. 2020. Fungal Diversity 105:1-16 ((https://doi.org/10.1007/s13225-020-00466-2)) |
| Otu24007 | 1 | k__Fungi(100);p__Ascomycota(100);c__Saccharomycetes(100);o__Saccharomycetales(100);f__Metschnikowiaceae(100);g__Metschnikowia(100);s__Metschnikowia_pulcherrima(100); | Metschnikowia | 13 | Saprotroph | \|Undefined Saprotroph\| | Yeast | NULL | Probable | As nectar-tap_saprotroph (Põlme et al. 2020); Decay sugar-rich substrates (Põlme et al. 2020) | Põlme S, et al. 2020. Fungal Diversity 105:1-16 ((https://doi.org/10.1007/s13225-020-00466-2)) |
| Otu24008 | 1 | k__Fungi(100);p__Ascomycota(100);c__Saccharomycetes(100);o__Saccharomycetales(100);f__Metschnikowiaceae(100);g__Metschnikowia(100);s__Metschnikowia_pulcherrima(100); | Metschnikowia | 13 | Saprotroph | \|Undefined Saprotroph\| | Yeast | NULL | Probable | As nectar-tap_saprotroph (Põlme et al. 2020); Decay sugar-rich substrates (Põlme et al. 2020) | Põlme S, et al. 2020. Fungal Diversity 105:1-16 ((https://doi.org/10.1007/s13225-020-00466-2)) |
| Otu24009 | 1 | k__Fungi(100);p__Ascomycota(100);c__Saccharomycetes(100);o__Saccharomycetales(100);f__Metschnikowiaceae(100);g__Metschnikowia(100);s__Metschnikowia_pulcherrima(100); | Metschnikowia | 13 | Saprotroph | \|Undefined Saprotroph\| | Yeast | NULL | Probable | As nectar-tap_saprotroph (Põlme et al. 2020); Decay sugar-rich substrates (Põlme et al. 2020) | Põlme S, et al. 2020. Fungal Diversity 105:1-16 ((https://doi.org/10.1007/s13225-020-00466-2)) |
| Otu24010 | 1 | k__Fungi(100);p__Ascomycota(100);c__Saccharomycetes(100);o__Saccharomycetales(100);f__Metschnikowiaceae(100);g__Metschnikowia(100);s__Metschnikowia_pulcherrima(100); | Metschnikowia | 13 | Saprotroph | \|Undefined Saprotroph\| | Yeast | NULL | Probable | As nectar-tap_saprotroph (Põlme et al. 2020); Decay sugar-rich substrates (Põlme et al. 2020) | Põlme S, et al. 2020. Fungal Diversity 105:1-16 ((https://doi.org/10.1007/s13225-020-00466-2)) |
| Otu24012 | 1 | k__Fungi(100);p__Ascomycota(100);c__Saccharomycetes(100);o__Saccharomycetales(100);f__Metschnikowiaceae(100);g__Metschnikowia(100);s__Metschnikowia_pulcherrima(100); | Metschnikowia | 13 | Saprotroph | \|Undefined Saprotroph\| | Yeast | NULL | Probable | As nectar-tap_saprotroph (Põlme et al. 2020); Decay sugar-rich substrates (Põlme et al. 2020) | Põlme S, et al. 2020. Fungal Diversity 105:1-16 ((https://doi.org/10.1007/s13225-020-00466-2)) |
| Otu24013 | 1 | k__Fungi(100);p__Ascomycota(100);c__Saccharomycetes(100);o__Saccharomycetales(100);f__Metschnikowiaceae(100);g__Metschnikowia(100);s__Metschnikowia_pulcherrima(100); | Metschnikowia | 13 | Saprotroph | \|Undefined Saprotroph\| | Yeast | NULL | Probable | As nectar-tap_saprotroph (Põlme et al. 2020); Decay sugar-rich substrates (Põlme et al. 2020) | Põlme S, et al. 2020. Fungal Diversity 105:1-16 ((https://doi.org/10.1007/s13225-020-00466-2)) |
| Otu24015 | 1 | k__Fungi(100);p__Ascomycota(100);c__Saccharomycetes(100);o__Saccharomycetales(100);f__Metschnikowiaceae(100);g__Metschnikowia(100);g__Metschnikowia_unclassified(100); | Metschnikowia | 13 | Saprotroph | \|Undefined Saprotroph\| | Yeast | NULL | Probable | As nectar-tap_saprotroph (Põlme et al. 2020); Decay sugar-rich substrates (Põlme et al. 2020) | Põlme S, et al. 2020. Fungal Diversity 105:1-16 ((https://doi.org/10.1007/s13225-020-00466-2)) |
| Otu24016 | 1 | k__Fungi(100);p__Ascomycota(100);c__Saccharomycetes(100);o__Saccharomycetales(100);f__Metschnikowiaceae(100);g__Metschnikowia(100);s__Metschnikowia_pulcherrima(100); | Metschnikowia | 13 | Saprotroph | \|Undefined Saprotroph\| | Yeast | NULL | Probable | As nectar-tap_saprotroph (Põlme et al. 2020); Decay sugar-rich substrates (Põlme et al. 2020) | Põlme S, et al. 2020. Fungal Diversity 105:1-16 ((https://doi.org/10.1007/s13225-020-00466-2)) |
| Otu24017 | 1 | k__Fungi(100);p__Ascomycota(100);c__Saccharomycetes(100);o__Saccharomycetales(100);f__Metschnikowiaceae(100);g__Metschnikowia(100);s__Metschnikowia_pulcherrima(100); | Metschnikowia | 13 | Saprotroph | \|Undefined Saprotroph\| | Yeast | NULL | Probable | As nectar-tap_saprotroph (Põlme et al. 2020); Decay sugar-rich substrates (Põlme et al. 2020) | Põlme S, et al. 2020. Fungal Diversity 105:1-16 ((https://doi.org/10.1007/s13225-020-00466-2)) |
| Otu24018 | 1 | k__Fungi(100);p__Ascomycota(100);c__Saccharomycetes(100);o__Saccharomycetales(100);f__Metschnikowiaceae(100);g__Metschnikowia(100);s__Metschnikowia_pulcherrima(100); | Metschnikowia | 13 | Saprotroph | \|Undefined Saprotroph\| | Yeast | NULL | Probable | As nectar-tap_saprotroph (Põlme et al. 2020); Decay sugar-rich substrates (Põlme et al. 2020) | Põlme S, et al. 2020. Fungal Diversity 105:1-16 ((https://doi.org/10.1007/s13225-020-00466-2)) |
| Otu24019 | 1 | k__Fungi(100);p__Ascomycota(100);c__Saccharomycetes(100);o__Saccharomycetales(100);f__Metschnikowiaceae(100);g__Metschnikowia(100);g__Metschnikowia_unclassified(100); | Metschnikowia | 13 | Saprotroph | \|Undefined Saprotroph\| | Yeast | NULL | Probable | As nectar-tap_saprotroph (Põlme et al. 2020); Decay sugar-rich substrates (Põlme et al. 2020) | Põlme S, et al. 2020. Fungal Diversity 105:1-16 ((https://doi.org/10.1007/s13225-020-00466-2)) |
| Otu24020 | 1 | k__Fungi(100);p__Ascomycota(100);c__Saccharomycetes(100);o__Saccharomycetales(100);f__Metschnikowiaceae(100);g__Metschnikowia(100);s__Metschnikowia_pulcherrima(100); | Metschnikowia | 13 | Saprotroph | \|Undefined Saprotroph\| | Yeast | NULL | Probable | As nectar-tap_saprotroph (Põlme et al. 2020); Decay sugar-rich substrates (Põlme et al. 2020) | Põlme S, et al. 2020. Fungal Diversity 105:1-16 ((https://doi.org/10.1007/s13225-020-00466-2)) |
| Otu24021 | 1 | k__Fungi(100);p__Ascomycota(100);c__Saccharomycetes(100);o__Saccharomycetales(100);f__Metschnikowiaceae(100);g__Metschnikowia(100);s__Metschnikowia_pulcherrima(100); | Metschnikowia | 13 | Saprotroph | \|Undefined Saprotroph\| | Yeast | NULL | Probable | As nectar-tap_saprotroph (Põlme et al. 2020); Decay sugar-rich substrates (Põlme et al. 2020) | Põlme S, et al. 2020. Fungal Diversity 105:1-16 ((https://doi.org/10.1007/s13225-020-00466-2)) |
| Otu24022 | 1 | k__Fungi(100);p__Ascomycota(100);c__Saccharomycetes(100);o__Saccharomycetales(100);f__Metschnikowiaceae(100);g__Metschnikowia(100);s__Metschnikowia_pulcherrima(100); | Metschnikowia | 13 | Saprotroph | \|Undefined Saprotroph\| | Yeast | NULL | Probable | As nectar-tap_saprotroph (Põlme et al. 2020); Decay sugar-rich substrates (Põlme et al. 2020) | Põlme S, et al. 2020. Fungal Diversity 105:1-16 ((https://doi.org/10.1007/s13225-020-00466-2)) |
| Otu24023 | 1 | k__Fungi(100);p__Ascomycota(100);c__Saccharomycetes(100);o__Saccharomycetales(100);f__Metschnikowiaceae(100);g__Metschnikowia(100);s__Metschnikowia_pulcherrima(100); | Metschnikowia | 13 | Saprotroph | \|Undefined Saprotroph\| | Yeast | NULL | Probable | As nectar-tap_saprotroph (Põlme et al. 2020); Decay sugar-rich substrates (Põlme et al. 2020) | Põlme S, et al. 2020. Fungal Diversity 105:1-16 ((https://doi.org/10.1007/s13225-020-00466-2)) |
| Otu24024 | 1 | k__Fungi(100);p__Ascomycota(100);c__Saccharomycetes(100);o__Saccharomycetales(100);f__Metschnikowiaceae(100);g__Metschnikowia(100);s__Metschnikowia_pulcherrima(100); | Metschnikowia | 13 | Saprotroph | \|Undefined Saprotroph\| | Yeast | NULL | Probable | As nectar-tap_saprotroph (Põlme et al. 2020); Decay sugar-rich substrates (Põlme et al. 2020) | Põlme S, et al. 2020. Fungal Diversity 105:1-16 ((https://doi.org/10.1007/s13225-020-00466-2)) |
| Otu24025 | 1 | k__Fungi(100);p__Ascomycota(100);c__Saccharomycetes(100);o__Saccharomycetales(100);f__Metschnikowiaceae(100);g__Metschnikowia(100);g__Metschnikowia_unclassified(100); | Metschnikowia | 13 | Saprotroph | \|Undefined Saprotroph\| | Yeast | NULL | Probable | As nectar-tap_saprotroph (Põlme et al. 2020); Decay sugar-rich substrates (Põlme et al. 2020) | Põlme S, et al. 2020. Fungal Diversity 105:1-16 ((https://doi.org/10.1007/s13225-020-00466-2)) |
| Otu24026 | 1 | k__Fungi(100);p__Ascomycota(100);c__Saccharomycetes(100);o__Saccharomycetales(100);f__Metschnikowiaceae(100);g__Metschnikowia(100);s__Metschnikowia_pulcherrima(100); | Metschnikowia | 13 | Saprotroph | \|Undefined Saprotroph\| | Yeast | NULL | Probable | As nectar-tap_saprotroph (Põlme et al. 2020); Decay sugar-rich substrates (Põlme et al. 2020) | Põlme S, et al. 2020. Fungal Diversity 105:1-16 ((https://doi.org/10.1007/s13225-020-00466-2)) |
| Otu24029 | 1 | k__Fungi(100);p__Ascomycota(100);c__Saccharomycetes(100);o__Saccharomycetales(100);f__Metschnikowiaceae(100);g__Metschnikowia(100);s__Metschnikowia_pulcherrima(100); | Metschnikowia | 13 | Saprotroph | \|Undefined Saprotroph\| | Yeast | NULL | Probable | As nectar-tap_saprotroph (Põlme et al. 2020); Decay sugar-rich substrates (Põlme et al. 2020) | Põlme S, et al. 2020. Fungal Diversity 105:1-16 ((https://doi.org/10.1007/s13225-020-00466-2)) |
| Otu24041 | 1 | k__Fungi(100);p__Ascomycota(100);c__Saccharomycetes(100);o__Saccharomycetales(100);f__Metschnikowiaceae(100);g__Metschnikowia(100);g__Metschnikowia_unclassified(100); | Metschnikowia | 13 | Saprotroph | \|Undefined Saprotroph\| | Yeast | NULL | Probable | As nectar-tap_saprotroph (Põlme et al. 2020); Decay sugar-rich substrates (Põlme et al. 2020) | Põlme S, et al. 2020. Fungal Diversity 105:1-16 ((https://doi.org/10.1007/s13225-020-00466-2)) |
| Otu24061 | 1 | k__Fungi(100);p__Ascomycota(100);c__Saccharomycetes(100);o__Saccharomycetales(100);f__Metschnikowiaceae(100);g__Metschnikowia(100);s__Metschnikowia_pulcherrima(100); | Metschnikowia | 13 | Saprotroph | \|Undefined Saprotroph\| | Yeast | NULL | Probable | As nectar-tap_saprotroph (Põlme et al. 2020); Decay sugar-rich substrates (Põlme et al. 2020) | Põlme S, et al. 2020. Fungal Diversity 105:1-16 ((https://doi.org/10.1007/s13225-020-00466-2)) |
| Otu24064 | 1 | k__Fungi(100);p__Ascomycota(100);c__Saccharomycetes(100);o__Saccharomycetales(100);f__Metschnikowiaceae(100);g__Metschnikowia(100);s__Metschnikowia_pulcherrima(100); | Metschnikowia | 13 | Saprotroph | \|Undefined Saprotroph\| | Yeast | NULL | Probable | As nectar-tap_saprotroph (Põlme et al. 2020); Decay sugar-rich substrates (Põlme et al. 2020) | Põlme S, et al. 2020. Fungal Diversity 105:1-16 ((https://doi.org/10.1007/s13225-020-00466-2)) |
| Otu24067 | 1 | k__Fungi(100);p__Ascomycota(100);c__Saccharomycetes(100);o__Saccharomycetales(100);f__Metschnikowiaceae(100);g__Metschnikowia(100);g__Metschnikowia_unclassified(100); | Metschnikowia | 13 | Saprotroph | \|Undefined Saprotroph\| | Yeast | NULL | Probable | As nectar-tap_saprotroph (Põlme et al. 2020); Decay sugar-rich substrates (Põlme et al. 2020) | Põlme S, et al. 2020. Fungal Diversity 105:1-16 ((https://doi.org/10.1007/s13225-020-00466-2)) |
| Otu24070 | 1 | k__Fungi(100);p__Ascomycota(100);c__Saccharomycetes(100);o__Saccharomycetales(100);f__Saccharomycetales_family_Incertae_sedis(100);g__Candida(100);g__Candida_unclassified(100); | Candida | 13 | Pathotroph-Saprotroph-Symbiotroph | Animal Pathogen-Endophyte-Endosymbiont-Epiphyte-\|Undefined Saprotroph\| | Dimorphic Yeast-Yeast | NULL | Probable | As nectar-tap_saprotroph (Põlme et al. 2020); Decay sugar-rich substrates (Põlme et al. 2020); This asexual genus is polyphyletic and contain species that function very differently depend on what they are. Caution should be taken when interpreting guilds for Candida species. | Manolakaki D, et al. 2010. Virulence. 1: 367-375 ((https://doi.org/10.4161/viru.1.5.12796)); Mekbib SB, et al. 2011. Tropical Plant Pathology 36:233-240 ((https://doi.org/10.1590/S1982-56762011000400004)); Põlme S, et al. 2020. Fungal Diversity 105:1-16 ((https://doi.org/10.1007/s13225-020-00466-2)) |
| Otu24072 | 1 | k__Fungi(100);p__Ascomycota(100);c__Saccharomycetes(100);o__Saccharomycetales(100);f__Metschnikowiaceae(100);g__Metschnikowia(100);g__Metschnikowia_unclassified(100); | Metschnikowia | 13 | Saprotroph | \|Undefined Saprotroph\| | Yeast | NULL | Probable | As nectar-tap_saprotroph (Põlme et al. 2020); Decay sugar-rich substrates (Põlme et al. 2020) | Põlme S, et al. 2020. Fungal Diversity 105:1-16 ((https://doi.org/10.1007/s13225-020-00466-2)) |
| Otu24075 | 1 | k__Fungi(100);p__Ascomycota(100);c__Saccharomycetes(100);o__Saccharomycetales(100);f__Metschnikowiaceae(100);g__Metschnikowia(100);s__Metschnikowia_pulcherrima(100); | Metschnikowia | 13 | Saprotroph | \|Undefined Saprotroph\| | Yeast | NULL | Probable | As nectar-tap_saprotroph (Põlme et al. 2020); Decay sugar-rich substrates (Põlme et al. 2020) | Põlme S, et al. 2020. Fungal Diversity 105:1-16 ((https://doi.org/10.1007/s13225-020-00466-2)) |
| Otu24077 | 1 | k__Fungi(100);p__Ascomycota(100);c__Saccharomycetes(100);o__Saccharomycetales(100);f__Metschnikowiaceae(100);g__Metschnikowia(100);s__Metschnikowia_pulcherrima(100); | Metschnikowia | 13 | Saprotroph | \|Undefined Saprotroph\| | Yeast | NULL | Probable | As nectar-tap_saprotroph (Põlme et al. 2020); Decay sugar-rich substrates (Põlme et al. 2020) | Põlme S, et al. 2020. Fungal Diversity 105:1-16 ((https://doi.org/10.1007/s13225-020-00466-2)) |
| Otu24078 | 1 | k__Fungi(100);p__Ascomycota(100);c__Saccharomycetes(100);o__Saccharomycetales(100);f__Metschnikowiaceae(100);g__Metschnikowia(100);g__Metschnikowia_unclassified(100); | Metschnikowia | 13 | Saprotroph | \|Undefined Saprotroph\| | Yeast | NULL | Probable | As nectar-tap_saprotroph (Põlme et al. 2020); Decay sugar-rich substrates (Põlme et al. 2020) | Põlme S, et al. 2020. Fungal Diversity 105:1-16 ((https://doi.org/10.1007/s13225-020-00466-2)) |
| Otu24079 | 1 | k__Fungi(100);p__Ascomycota(100);c__Saccharomycetes(100);o__Saccharomycetales(100);f__Metschnikowiaceae(100);g__Metschnikowia(100);g__Metschnikowia_unclassified(100); | Metschnikowia | 13 | Saprotroph | \|Undefined Saprotroph\| | Yeast | NULL | Probable | As nectar-tap_saprotroph (Põlme et al. 2020); Decay sugar-rich substrates (Põlme et al. 2020) | Põlme S, et al. 2020. Fungal Diversity 105:1-16 ((https://doi.org/10.1007/s13225-020-00466-2)) |
| Otu24089 | 1 | k__Fungi(100);p__Ascomycota(100);c__Saccharomycetes(100);o__Saccharomycetales(100);f__Saccharomycetales_family_Incertae_sedis(100);g__Candida(100);g__Candida_unclassified(100); | Candida | 13 | Pathotroph-Saprotroph-Symbiotroph | Animal Pathogen-Endophyte-Endosymbiont-Epiphyte-\|Undefined Saprotroph\| | Dimorphic Yeast-Yeast | NULL | Probable | As nectar-tap_saprotroph (Põlme et al. 2020); Decay sugar-rich substrates (Põlme et al. 2020); This asexual genus is polyphyletic and contain species that function very differently depend on what they are. Caution should be taken when interpreting guilds for Candida species. | Manolakaki D, et al. 2010. Virulence. 1: 367-375 ((https://doi.org/10.4161/viru.1.5.12796)); Mekbib SB, et al. 2011. Tropical Plant Pathology 36:233-240 ((https://doi.org/10.1590/S1982-56762011000400004)); Põlme S, et al. 2020. Fungal Diversity 105:1-16 ((https://doi.org/10.1007/s13225-020-00466-2)) |
| Otu24091 | 1 | k__Fungi(100);p__Ascomycota(100);c__Saccharomycetes(100);o__Saccharomycetales(100);f__Metschnikowiaceae(100);g__Metschnikowia(100);s__Metschnikowia_pulcherrima(100); | Metschnikowia | 13 | Saprotroph | \|Undefined Saprotroph\| | Yeast | NULL | Probable | As nectar-tap_saprotroph (Põlme et al. 2020); Decay sugar-rich substrates (Põlme et al. 2020) | Põlme S, et al. 2020. Fungal Diversity 105:1-16 ((https://doi.org/10.1007/s13225-020-00466-2)) |
| Otu24095 | 1 | k__Fungi(100);p__Ascomycota(100);c__Saccharomycetes(100);o__Saccharomycetales(100);f__Metschnikowiaceae(100);g__Metschnikowia(100);g__Metschnikowia_unclassified(100); | Metschnikowia | 13 | Saprotroph | \|Undefined Saprotroph\| | Yeast | NULL | Probable | As nectar-tap_saprotroph (Põlme et al. 2020); Decay sugar-rich substrates (Põlme et al. 2020) | Põlme S, et al. 2020. Fungal Diversity 105:1-16 ((https://doi.org/10.1007/s13225-020-00466-2)) |
| Otu24100 | 1 | k__Fungi(100);p__Ascomycota(100);c__Saccharomycetes(100);o__Saccharomycetales(100);f__Metschnikowiaceae(100);g__Metschnikowia(100);s__Metschnikowia_pulcherrima(100); | Metschnikowia | 13 | Saprotroph | \|Undefined Saprotroph\| | Yeast | NULL | Probable | As nectar-tap_saprotroph (Põlme et al. 2020); Decay sugar-rich substrates (Põlme et al. 2020) | Põlme S, et al. 2020. Fungal Diversity 105:1-16 ((https://doi.org/10.1007/s13225-020-00466-2)) |
| Otu24101 | 1 | k__Fungi(100);p__Ascomycota(100);c__Saccharomycetes(100);o__Saccharomycetales(100);f__Metschnikowiaceae(100);g__Metschnikowia(100);g__Metschnikowia_unclassified(100); | Metschnikowia | 13 | Saprotroph | \|Undefined Saprotroph\| | Yeast | NULL | Probable | As nectar-tap_saprotroph (Põlme et al. 2020); Decay sugar-rich substrates (Põlme et al. 2020) | Põlme S, et al. 2020. Fungal Diversity 105:1-16 ((https://doi.org/10.1007/s13225-020-00466-2)) |
| Otu24102 | 1 | k__Fungi(100);p__Ascomycota(100);c__Saccharomycetes(100);o__Saccharomycetales(100);f__Metschnikowiaceae(100);g__Metschnikowia(100);g__Metschnikowia_unclassified(100); | Metschnikowia | 13 | Saprotroph | \|Undefined Saprotroph\| | Yeast | NULL | Probable | As nectar-tap_saprotroph (Põlme et al. 2020); Decay sugar-rich substrates (Põlme et al. 2020) | Põlme S, et al. 2020. Fungal Diversity 105:1-16 ((https://doi.org/10.1007/s13225-020-00466-2)) |
| Otu24103 | 1 | k__Fungi(100);p__Ascomycota(100);c__Saccharomycetes(100);o__Saccharomycetales(100);f__Metschnikowiaceae(100);g__Metschnikowia(100);s__Metschnikowia_pulcherrima(100); | Metschnikowia | 13 | Saprotroph | \|Undefined Saprotroph\| | Yeast | NULL | Probable | As nectar-tap_saprotroph (Põlme et al. 2020); Decay sugar-rich substrates (Põlme et al. 2020) | Põlme S, et al. 2020. Fungal Diversity 105:1-16 ((https://doi.org/10.1007/s13225-020-00466-2)) |
| Otu24104 | 1 | k__Fungi(100);p__Ascomycota(100);c__Saccharomycetes(100);o__Saccharomycetales(100);f__Metschnikowiaceae(100);g__Metschnikowia(100);s__Metschnikowia_pulcherrima(100); | Metschnikowia | 13 | Saprotroph | \|Undefined Saprotroph\| | Yeast | NULL | Probable | As nectar-tap_saprotroph (Põlme et al. 2020); Decay sugar-rich substrates (Põlme et al. 2020) | Põlme S, et al. 2020. Fungal Diversity 105:1-16 ((https://doi.org/10.1007/s13225-020-00466-2)) |
| Otu24106 | 1 | k__Fungi(100);p__Ascomycota(100);c__Saccharomycetes(100);o__Saccharomycetales(100);f__Metschnikowiaceae(100);g__Metschnikowia(100);g__Metschnikowia_unclassified(100); | Metschnikowia | 13 | Saprotroph | \|Undefined Saprotroph\| | Yeast | NULL | Probable | As nectar-tap_saprotroph (Põlme et al. 2020); Decay sugar-rich substrates (Põlme et al. 2020) | Põlme S, et al. 2020. Fungal Diversity 105:1-16 ((https://doi.org/10.1007/s13225-020-00466-2)) |
| Otu24107 | 1 | k__Fungi(100);p__Ascomycota(100);c__Saccharomycetes(100);o__Saccharomycetales(100);f__Metschnikowiaceae(100);g__Metschnikowia(100);g__Metschnikowia_unclassified(100); | Metschnikowia | 13 | Saprotroph | \|Undefined Saprotroph\| | Yeast | NULL | Probable | As nectar-tap_saprotroph (Põlme et al. 2020); Decay sugar-rich substrates (Põlme et al. 2020) | Põlme S, et al. 2020. Fungal Diversity 105:1-16 ((https://doi.org/10.1007/s13225-020-00466-2)) |
| Otu24109 | 1 | k__Fungi(100);p__Ascomycota(100);c__Saccharomycetes(100);o__Saccharomycetales(100);f__Metschnikowiaceae(100);g__Metschnikowia(100);s__Metschnikowia_pulcherrima(100); | Metschnikowia | 13 | Saprotroph | \|Undefined Saprotroph\| | Yeast | NULL | Probable | As nectar-tap_saprotroph (Põlme et al. 2020); Decay sugar-rich substrates (Põlme et al. 2020) | Põlme S, et al. 2020. Fungal Diversity 105:1-16 ((https://doi.org/10.1007/s13225-020-00466-2)) |
| Otu24112 | 1 | k__Fungi(100);p__Ascomycota(100);c__Saccharomycetes(100);o__Saccharomycetales(100);f__Metschnikowiaceae(100);g__Metschnikowia(100);s__Metschnikowia_pulcherrima(100); | Metschnikowia | 13 | Saprotroph | \|Undefined Saprotroph\| | Yeast | NULL | Probable | As nectar-tap_saprotroph (Põlme et al. 2020); Decay sugar-rich substrates (Põlme et al. 2020) | Põlme S, et al. 2020. Fungal Diversity 105:1-16 ((https://doi.org/10.1007/s13225-020-00466-2)) |
| Otu24118 | 1 | k__Fungi(100);p__Ascomycota(100);c__Saccharomycetes(100);o__Saccharomycetales(100);f__Metschnikowiaceae(100);g__Metschnikowia(100);g__Metschnikowia_unclassified(100); | Metschnikowia | 13 | Saprotroph | \|Undefined Saprotroph\| | Yeast | NULL | Probable | As nectar-tap_saprotroph (Põlme et al. 2020); Decay sugar-rich substrates (Põlme et al. 2020) | Põlme S, et al. 2020. Fungal Diversity 105:1-16 ((https://doi.org/10.1007/s13225-020-00466-2)) |
| Otu24121 | 1 | k__Fungi(100);p__Ascomycota(100);c__Saccharomycetes(100);o__Saccharomycetales(100);f__Metschnikowiaceae(100);g__Metschnikowia(100);g__Metschnikowia_unclassified(100); | Metschnikowia | 13 | Saprotroph | \|Undefined Saprotroph\| | Yeast | NULL | Probable | As nectar-tap_saprotroph (Põlme et al. 2020); Decay sugar-rich substrates (Põlme et al. 2020) | Põlme S, et al. 2020. Fungal Diversity 105:1-16 ((https://doi.org/10.1007/s13225-020-00466-2)) |
| Otu24122 | 1 | k__Fungi(100);p__Ascomycota(100);c__Saccharomycetes(100);o__Saccharomycetales(100);f__Metschnikowiaceae(100);g__Metschnikowia(100);s__Metschnikowia_pulcherrima(100); | Metschnikowia | 13 | Saprotroph | \|Undefined Saprotroph\| | Yeast | NULL | Probable | As nectar-tap_saprotroph (Põlme et al. 2020); Decay sugar-rich substrates (Põlme et al. 2020) | Põlme S, et al. 2020. Fungal Diversity 105:1-16 ((https://doi.org/10.1007/s13225-020-00466-2)) |
| Otu24123 | 1 | k__Fungi(100);p__Ascomycota(100);c__Saccharomycetes(100);o__Saccharomycetales(100);f__Metschnikowiaceae(100);g__Metschnikowia(100);g__Metschnikowia_unclassified(100); | Metschnikowia | 13 | Saprotroph | \|Undefined Saprotroph\| | Yeast | NULL | Probable | As nectar-tap_saprotroph (Põlme et al. 2020); Decay sugar-rich substrates (Põlme et al. 2020) | Põlme S, et al. 2020. Fungal Diversity 105:1-16 ((https://doi.org/10.1007/s13225-020-00466-2)) |
| Otu24124 | 1 | k__Fungi(100);p__Ascomycota(100);c__Saccharomycetes(100);o__Saccharomycetales(100);f__Metschnikowiaceae(100);g__Metschnikowia(100);g__Metschnikowia_unclassified(100); | Metschnikowia | 13 | Saprotroph | \|Undefined Saprotroph\| | Yeast | NULL | Probable | As nectar-tap_saprotroph (Põlme et al. 2020); Decay sugar-rich substrates (Põlme et al. 2020) | Põlme S, et al. 2020. Fungal Diversity 105:1-16 ((https://doi.org/10.1007/s13225-020-00466-2)) |
| Otu24133 | 1 | k__Fungi(100);p__Ascomycota(100);c__Saccharomycetes(100);o__Saccharomycetales(100);f__Metschnikowiaceae(100);g__Metschnikowia(100);s__Metschnikowia_pulcherrima(100); | Metschnikowia | 13 | Saprotroph | \|Undefined Saprotroph\| | Yeast | NULL | Probable | As nectar-tap_saprotroph (Põlme et al. 2020); Decay sugar-rich substrates (Põlme et al. 2020) | Põlme S, et al. 2020. Fungal Diversity 105:1-16 ((https://doi.org/10.1007/s13225-020-00466-2)) |
| Otu24137 | 1 | k__Fungi(100);p__Ascomycota(100);c__Saccharomycetes(100);o__Saccharomycetales(100);f__Metschnikowiaceae(100);g__Metschnikowia(100);s__Metschnikowia_pulcherrima(100); | Metschnikowia | 13 | Saprotroph | \|Undefined Saprotroph\| | Yeast | NULL | Probable | As nectar-tap_saprotroph (Põlme et al. 2020); Decay sugar-rich substrates (Põlme et al. 2020) | Põlme S, et al. 2020. Fungal Diversity 105:1-16 ((https://doi.org/10.1007/s13225-020-00466-2)) |
| Otu24139 | 1 | k__Fungi(100);p__Ascomycota(100);c__Saccharomycetes(100);o__Saccharomycetales(100);f__Metschnikowiaceae(100);g__Metschnikowia(100);s__Metschnikowia_pulcherrima(100); | Metschnikowia | 13 | Saprotroph | \|Undefined Saprotroph\| | Yeast | NULL | Probable | As nectar-tap_saprotroph (Põlme et al. 2020); Decay sugar-rich substrates (Põlme et al. 2020) | Põlme S, et al. 2020. Fungal Diversity 105:1-16 ((https://doi.org/10.1007/s13225-020-00466-2)) |
| Otu24140 | 1 | k__Fungi(100);p__Ascomycota(100);c__Leotiomycetes(100);o__Helotiales(100);f__Vibrisseaceae(100);g__Phialocephala(100);s__Phialocephala_fusca(100); | Phialocephala | 13 | Saprotroph-Symbiotroph | Endophyte-\|Undefined Saprotroph\| | Microfungus | NULL | Highly Probable | As root_endophyte (Põlme et al. 2020); As root_endophyte_dark_septate (Põlme et al. 2020); As soil_saprotroph (Põlme et al. 2020); Dark Septate Endophyte, known from plant roots.; Decay roots, soil (Põlme et al. 2020) | Newsham KK. 2011. New Phytologist 190:783-793 ((https://doi.org/10.1111/j.1469-8137.2010.03611.x)) (as dark septate root endophytes); Põlme S, et al. 2020. Fungal Diversity 105:1-16 ((https://doi.org/10.1007/s13225-020-00466-2)) |
| Otu24141 | 1 | k__Fungi(100);p__Ascomycota(100);c__Saccharomycetes(100);o__Saccharomycetales(100);f__Metschnikowiaceae(100);g__Metschnikowia(100);g__Metschnikowia_unclassified(100); | Metschnikowia | 13 | Saprotroph | \|Undefined Saprotroph\| | Yeast | NULL | Probable | As nectar-tap_saprotroph (Põlme et al. 2020); Decay sugar-rich substrates (Põlme et al. 2020) | Põlme S, et al. 2020. Fungal Diversity 105:1-16 ((https://doi.org/10.1007/s13225-020-00466-2)) |
| Otu24142 | 1 | k__Fungi(100);p__Ascomycota(100);c__Saccharomycetes(100);o__Saccharomycetales(100);f__Metschnikowiaceae(100);g__Metschnikowia(100);g__Metschnikowia_unclassified(100); | Metschnikowia | 13 | Saprotroph | \|Undefined Saprotroph\| | Yeast | NULL | Probable | As nectar-tap_saprotroph (Põlme et al. 2020); Decay sugar-rich substrates (Põlme et al. 2020) | Põlme S, et al. 2020. Fungal Diversity 105:1-16 ((https://doi.org/10.1007/s13225-020-00466-2)) |
| Otu24143 | 1 | k__Fungi(100);p__Ascomycota(100);c__Saccharomycetes(100);o__Saccharomycetales(100);f__Metschnikowiaceae(100);g__Metschnikowia(100);s__Metschnikowia_pulcherrima(100); | Metschnikowia | 13 | Saprotroph | \|Undefined Saprotroph\| | Yeast | NULL | Probable | As nectar-tap_saprotroph (Põlme et al. 2020); Decay sugar-rich substrates (Põlme et al. 2020) | Põlme S, et al. 2020. Fungal Diversity 105:1-16 ((https://doi.org/10.1007/s13225-020-00466-2)) |
| Otu24144 | 1 | k__Fungi(100);p__Ascomycota(100);c__Saccharomycetes(100);o__Saccharomycetales(100);f__Metschnikowiaceae(100);g__Metschnikowia(100);s__Metschnikowia_pulcherrima(100); | Metschnikowia | 13 | Saprotroph | \|Undefined Saprotroph\| | Yeast | NULL | Probable | As nectar-tap_saprotroph (Põlme et al. 2020); Decay sugar-rich substrates (Põlme et al. 2020) | Põlme S, et al. 2020. Fungal Diversity 105:1-16 ((https://doi.org/10.1007/s13225-020-00466-2)) |
| Otu24145 | 1 | k__Fungi(100);p__Ascomycota(100);c__Saccharomycetes(100);o__Saccharomycetales(100);f__Metschnikowiaceae(100);g__Metschnikowia(100);s__Metschnikowia_pulcherrima(100); | Metschnikowia | 13 | Saprotroph | \|Undefined Saprotroph\| | Yeast | NULL | Probable | As nectar-tap_saprotroph (Põlme et al. 2020); Decay sugar-rich substrates (Põlme et al. 2020) | Põlme S, et al. 2020. Fungal Diversity 105:1-16 ((https://doi.org/10.1007/s13225-020-00466-2)) |
| Otu24146 | 1 | k__Fungi(100);p__Ascomycota(100);c__Saccharomycetes(100);o__Saccharomycetales(100);f__Metschnikowiaceae(100);g__Metschnikowia(100);s__Metschnikowia_pulcherrima(100); | Metschnikowia | 13 | Saprotroph | \|Undefined Saprotroph\| | Yeast | NULL | Probable | As nectar-tap_saprotroph (Põlme et al. 2020); Decay sugar-rich substrates (Põlme et al. 2020) | Põlme S, et al. 2020. Fungal Diversity 105:1-16 ((https://doi.org/10.1007/s13225-020-00466-2)) |
| Otu24147 | 1 | k__Fungi(100);p__Ascomycota(100);c__Saccharomycetes(100);o__Saccharomycetales(100);f__Metschnikowiaceae(100);g__Metschnikowia(100);s__Metschnikowia_pulcherrima(100); | Metschnikowia | 13 | Saprotroph | \|Undefined Saprotroph\| | Yeast | NULL | Probable | As nectar-tap_saprotroph (Põlme et al. 2020); Decay sugar-rich substrates (Põlme et al. 2020) | Põlme S, et al. 2020. Fungal Diversity 105:1-16 ((https://doi.org/10.1007/s13225-020-00466-2)) |
| Otu24149 | 1 | k__Fungi(100);p__Ascomycota(100);c__Saccharomycetes(100);o__Saccharomycetales(100);f__Metschnikowiaceae(100);g__Metschnikowia(100);s__Metschnikowia_pulcherrima(100); | Metschnikowia | 13 | Saprotroph | \|Undefined Saprotroph\| | Yeast | NULL | Probable | As nectar-tap_saprotroph (Põlme et al. 2020); Decay sugar-rich substrates (Põlme et al. 2020) | Põlme S, et al. 2020. Fungal Diversity 105:1-16 ((https://doi.org/10.1007/s13225-020-00466-2)) |
| Otu24150 | 1 | k__Fungi(100);p__Ascomycota(100);c__Saccharomycetes(100);o__Saccharomycetales(100);f__Metschnikowiaceae(100);g__Metschnikowia(100);g__Metschnikowia_unclassified(100); | Metschnikowia | 13 | Saprotroph | \|Undefined Saprotroph\| | Yeast | NULL | Probable | As nectar-tap_saprotroph (Põlme et al. 2020); Decay sugar-rich substrates (Põlme et al. 2020) | Põlme S, et al. 2020. Fungal Diversity 105:1-16 ((https://doi.org/10.1007/s13225-020-00466-2)) |
| Otu24151 | 1 | k__Fungi(100);p__Ascomycota(100);c__Saccharomycetes(100);o__Saccharomycetales(100);f__Metschnikowiaceae(100);g__Metschnikowia(100);s__Metschnikowia_pulcherrima(100); | Metschnikowia | 13 | Saprotroph | \|Undefined Saprotroph\| | Yeast | NULL | Probable | As nectar-tap_saprotroph (Põlme et al. 2020); Decay sugar-rich substrates (Põlme et al. 2020) | Põlme S, et al. 2020. Fungal Diversity 105:1-16 ((https://doi.org/10.1007/s13225-020-00466-2)) |
| Otu24152 | 1 | k__Fungi(100);p__Ascomycota(100);c__Saccharomycetes(100);o__Saccharomycetales(100);f__Metschnikowiaceae(100);g__Metschnikowia(100);s__Metschnikowia_pulcherrima(100); | Metschnikowia | 13 | Saprotroph | \|Undefined Saprotroph\| | Yeast | NULL | Probable | As nectar-tap_saprotroph (Põlme et al. 2020); Decay sugar-rich substrates (Põlme et al. 2020) | Põlme S, et al. 2020. Fungal Diversity 105:1-16 ((https://doi.org/10.1007/s13225-020-00466-2)) |
| Otu24153 | 1 | k__Fungi(100);p__Ascomycota(100);c__Saccharomycetes(100);o__Saccharomycetales(100);f__Metschnikowiaceae(100);g__Metschnikowia(100);s__Metschnikowia_pulcherrima(100); | Metschnikowia | 13 | Saprotroph | \|Undefined Saprotroph\| | Yeast | NULL | Probable | As nectar-tap_saprotroph (Põlme et al. 2020); Decay sugar-rich substrates (Põlme et al. 2020) | Põlme S, et al. 2020. Fungal Diversity 105:1-16 ((https://doi.org/10.1007/s13225-020-00466-2)) |
| Otu24154 | 1 | k__Fungi(100);p__Ascomycota(100);c__Saccharomycetes(100);o__Saccharomycetales(100);f__Metschnikowiaceae(100);g__Metschnikowia(100);g__Metschnikowia_unclassified(100); | Metschnikowia | 13 | Saprotroph | \|Undefined Saprotroph\| | Yeast | NULL | Probable | As nectar-tap_saprotroph (Põlme et al. 2020); Decay sugar-rich substrates (Põlme et al. 2020) | Põlme S, et al. 2020. Fungal Diversity 105:1-16 ((https://doi.org/10.1007/s13225-020-00466-2)) |
| Otu24155 | 1 | k__Fungi(100);p__Ascomycota(100);c__Saccharomycetes(100);o__Saccharomycetales(100);f__Metschnikowiaceae(100);g__Metschnikowia(100);s__Metschnikowia_pulcherrima(100); | Metschnikowia | 13 | Saprotroph | \|Undefined Saprotroph\| | Yeast | NULL | Probable | As nectar-tap_saprotroph (Põlme et al. 2020); Decay sugar-rich substrates (Põlme et al. 2020) | Põlme S, et al. 2020. Fungal Diversity 105:1-16 ((https://doi.org/10.1007/s13225-020-00466-2)) |
| Otu24156 | 1 | k__Fungi(100);p__Ascomycota(100);c__Saccharomycetes(100);o__Saccharomycetales(100);f__Metschnikowiaceae(100);g__Metschnikowia(100);s__Metschnikowia_pulcherrima(100); | Metschnikowia | 13 | Saprotroph | \|Undefined Saprotroph\| | Yeast | NULL | Probable | As nectar-tap_saprotroph (Põlme et al. 2020); Decay sugar-rich substrates (Põlme et al. 2020) | Põlme S, et al. 2020. Fungal Diversity 105:1-16 ((https://doi.org/10.1007/s13225-020-00466-2)) |
| Otu24160 | 1 | k__Fungi(100);p__Ascomycota(100);c__Saccharomycetes(100);o__Saccharomycetales(100);f__Metschnikowiaceae(100);g__Metschnikowia(100);s__Metschnikowia_pulcherrima(100); | Metschnikowia | 13 | Saprotroph | \|Undefined Saprotroph\| | Yeast | NULL | Probable | As nectar-tap_saprotroph (Põlme et al. 2020); Decay sugar-rich substrates (Põlme et al. 2020) | Põlme S, et al. 2020. Fungal Diversity 105:1-16 ((https://doi.org/10.1007/s13225-020-00466-2)) |
| Otu24165 | 1 | k__Fungi(100);p__Ascomycota(100);c__Leotiomycetes(100);o__Helotiales(100);f__Helotiales_family_Incertae_sedis(100);g__Pilidium(100);s__Pilidium_concavum(100); | Pilidium | 13 | Pathotroph | \|Plant Pathogen\| | Microfungus | NULL | Probable | NULL | Tedersoo L, et al. 2014. Science 346:e1256688 ((https://doi.org/10.1126/science.1256688)); Põlme S, et al. 2020. Fungal Diversity 105:1-16 ((https://doi.org/10.1007/s13225-020-00466-2)) |
| Otu24169 | 1 | k__Fungi(100);p__Ascomycota(100);c__Saccharomycetes(100);o__Saccharomycetales(100);o__Saccharomycetales_unclassified(100);o__Saccharomycetales_unclassified(100);o__Saccharomycetales_unclassified(100); | Saccharomycetales | 7 | Saprotroph | Undefined Saprotroph | Yeast | NULL | Possible | NULL | Sterkenburg E, et al. 2015. New Phytologist 207:1145-1158 ((https://doi.org/10.1111/nph.13426)) |
| Otu24171 | 1 | k__Fungi(100);p__Ascomycota(100);c__Saccharomycetes(100);o__Saccharomycetales(100);f__Metschnikowiaceae(100);g__Metschnikowia(100);g__Metschnikowia_unclassified(100); | Metschnikowia | 13 | Saprotroph | \|Undefined Saprotroph\| | Yeast | NULL | Probable | As nectar-tap_saprotroph (Põlme et al. 2020); Decay sugar-rich substrates (Põlme et al. 2020) | Põlme S, et al. 2020. Fungal Diversity 105:1-16 ((https://doi.org/10.1007/s13225-020-00466-2)) |
| Otu24174 | 1 | k__Fungi(100);p__Ascomycota(100);c__Saccharomycetes(100);o__Saccharomycetales(100);f__Saccharomycetales_family_Incertae_sedis(100);g__Candida(100);g__Candida_unclassified(100); | Candida | 13 | Pathotroph-Saprotroph-Symbiotroph | Animal Pathogen-Endophyte-Endosymbiont-Epiphyte-\|Undefined Saprotroph\| | Dimorphic Yeast-Yeast | NULL | Probable | As nectar-tap_saprotroph (Põlme et al. 2020); Decay sugar-rich substrates (Põlme et al. 2020); This asexual genus is polyphyletic and contain species that function very differently depend on what they are. Caution should be taken when interpreting guilds for Candida species. | Manolakaki D, et al. 2010. Virulence. 1: 367-375 ((https://doi.org/10.4161/viru.1.5.12796)); Mekbib SB, et al. 2011. Tropical Plant Pathology 36:233-240 ((https://doi.org/10.1590/S1982-56762011000400004)); Põlme S, et al. 2020. Fungal Diversity 105:1-16 ((https://doi.org/10.1007/s13225-020-00466-2)) |
| Otu24181 | 1 | k__Fungi(100);p__Ascomycota(100);c__Saccharomycetes(100);o__Saccharomycetales(100);f__Metschnikowiaceae(100);g__Metschnikowia(100);s__Metschnikowia_pulcherrima(100); | Metschnikowia | 13 | Saprotroph | \|Undefined Saprotroph\| | Yeast | NULL | Probable | As nectar-tap_saprotroph (Põlme et al. 2020); Decay sugar-rich substrates (Põlme et al. 2020) | Põlme S, et al. 2020. Fungal Diversity 105:1-16 ((https://doi.org/10.1007/s13225-020-00466-2)) |
| Otu24184 | 1 | k__Fungi(100);p__Ascomycota(100);c__Saccharomycetes(100);o__Saccharomycetales(100);f__Metschnikowiaceae(100);g__Metschnikowia(100);s__Metschnikowia_pulcherrima(100); | Metschnikowia | 13 | Saprotroph | \|Undefined Saprotroph\| | Yeast | NULL | Probable | As nectar-tap_saprotroph (Põlme et al. 2020); Decay sugar-rich substrates (Põlme et al. 2020) | Põlme S, et al. 2020. Fungal Diversity 105:1-16 ((https://doi.org/10.1007/s13225-020-00466-2)) |
| Otu24185 | 1 | k__Fungi(100);p__Ascomycota(100);c__Saccharomycetes(100);o__Saccharomycetales(100);f__Metschnikowiaceae(100);g__Metschnikowia(100);s__Metschnikowia_pulcherrima(100); | Metschnikowia | 13 | Saprotroph | \|Undefined Saprotroph\| | Yeast | NULL | Probable | As nectar-tap_saprotroph (Põlme et al. 2020); Decay sugar-rich substrates (Põlme et al. 2020) | Põlme S, et al. 2020. Fungal Diversity 105:1-16 ((https://doi.org/10.1007/s13225-020-00466-2)) |
| Otu24186 | 1 | k__Fungi(100);p__Ascomycota(100);c__Saccharomycetes(100);o__Saccharomycetales(100);f__Metschnikowiaceae(100);g__Metschnikowia(100);s__Metschnikowia_pulcherrima(100); | Metschnikowia | 13 | Saprotroph | \|Undefined Saprotroph\| | Yeast | NULL | Probable | As nectar-tap_saprotroph (Põlme et al. 2020); Decay sugar-rich substrates (Põlme et al. 2020) | Põlme S, et al. 2020. Fungal Diversity 105:1-16 ((https://doi.org/10.1007/s13225-020-00466-2)) |
| Otu24188 | 1 | k__Fungi(100);p__Ascomycota(100);c__Saccharomycetes(100);o__Saccharomycetales(100);f__Metschnikowiaceae(100);g__Metschnikowia(100);s__Metschnikowia_pulcherrima(100); | Metschnikowia | 13 | Saprotroph | \|Undefined Saprotroph\| | Yeast | NULL | Probable | As nectar-tap_saprotroph (Põlme et al. 2020); Decay sugar-rich substrates (Põlme et al. 2020) | Põlme S, et al. 2020. Fungal Diversity 105:1-16 ((https://doi.org/10.1007/s13225-020-00466-2)) |
| Otu24189 | 1 | k__Fungi(100);p__Ascomycota(100);c__Saccharomycetes(100);o__Saccharomycetales(100);f__Metschnikowiaceae(100);g__Metschnikowia(100);s__Metschnikowia_pulcherrima(100); | Metschnikowia | 13 | Saprotroph | \|Undefined Saprotroph\| | Yeast | NULL | Probable | As nectar-tap_saprotroph (Põlme et al. 2020); Decay sugar-rich substrates (Põlme et al. 2020) | Põlme S, et al. 2020. Fungal Diversity 105:1-16 ((https://doi.org/10.1007/s13225-020-00466-2)) |
| Otu24190 | 1 | k__Fungi(100);p__Ascomycota(100);c__Saccharomycetes(100);o__Saccharomycetales(100);f__Metschnikowiaceae(100);g__Metschnikowia(100);g__Metschnikowia_unclassified(100); | Metschnikowia | 13 | Saprotroph | \|Undefined Saprotroph\| | Yeast | NULL | Probable | As nectar-tap_saprotroph (Põlme et al. 2020); Decay sugar-rich substrates (Põlme et al. 2020) | Põlme S, et al. 2020. Fungal Diversity 105:1-16 ((https://doi.org/10.1007/s13225-020-00466-2)) |
| Otu24191 | 1 | k__Fungi(100);p__Ascomycota(100);c__Saccharomycetes(100);o__Saccharomycetales(100);f__Metschnikowiaceae(100);g__Metschnikowia(100);g__Metschnikowia_unclassified(100); | Metschnikowia | 13 | Saprotroph | \|Undefined Saprotroph\| | Yeast | NULL | Probable | As nectar-tap_saprotroph (Põlme et al. 2020); Decay sugar-rich substrates (Põlme et al. 2020) | Põlme S, et al. 2020. Fungal Diversity 105:1-16 ((https://doi.org/10.1007/s13225-020-00466-2)) |
| Otu24192 | 1 | k__Fungi(100);p__Ascomycota(100);c__Saccharomycetes(100);o__Saccharomycetales(100);f__Metschnikowiaceae(100);g__Metschnikowia(100);s__Metschnikowia_pulcherrima(100); | Metschnikowia | 13 | Saprotroph | \|Undefined Saprotroph\| | Yeast | NULL | Probable | As nectar-tap_saprotroph (Põlme et al. 2020); Decay sugar-rich substrates (Põlme et al. 2020) | Põlme S, et al. 2020. Fungal Diversity 105:1-16 ((https://doi.org/10.1007/s13225-020-00466-2)) |
| Otu24193 | 1 | k__Fungi(100);p__Ascomycota(100);c__Saccharomycetes(100);o__Saccharomycetales(100);f__Metschnikowiaceae(100);g__Metschnikowia(100);s__Metschnikowia_pulcherrima(100); | Metschnikowia | 13 | Saprotroph | \|Undefined Saprotroph\| | Yeast | NULL | Probable | As nectar-tap_saprotroph (Põlme et al. 2020); Decay sugar-rich substrates (Põlme et al. 2020) | Põlme S, et al. 2020. Fungal Diversity 105:1-16 ((https://doi.org/10.1007/s13225-020-00466-2)) |
| Otu24194 | 1 | k__Fungi(100);p__Ascomycota(100);c__Saccharomycetes(100);o__Saccharomycetales(100);f__Metschnikowiaceae(100);g__Metschnikowia(100);s__Metschnikowia_pulcherrima(100); | Metschnikowia | 13 | Saprotroph | \|Undefined Saprotroph\| | Yeast | NULL | Probable | As nectar-tap_saprotroph (Põlme et al. 2020); Decay sugar-rich substrates (Põlme et al. 2020) | Põlme S, et al. 2020. Fungal Diversity 105:1-16 ((https://doi.org/10.1007/s13225-020-00466-2)) |
| Otu24195 | 1 | k__Fungi(100);p__Ascomycota(100);c__Saccharomycetes(100);o__Saccharomycetales(100);f__Metschnikowiaceae(100);g__Metschnikowia(100);s__Metschnikowia_pulcherrima(100); | Metschnikowia | 13 | Saprotroph | \|Undefined Saprotroph\| | Yeast | NULL | Probable | As nectar-tap_saprotroph (Põlme et al. 2020); Decay sugar-rich substrates (Põlme et al. 2020) | Põlme S, et al. 2020. Fungal Diversity 105:1-16 ((https://doi.org/10.1007/s13225-020-00466-2)) |
| Otu24196 | 1 | k__Fungi(100);p__Ascomycota(100);c__Saccharomycetes(100);o__Saccharomycetales(100);f__Metschnikowiaceae(100);g__Metschnikowia(100);s__Metschnikowia_pulcherrima(100); | Metschnikowia | 13 | Saprotroph | \|Undefined Saprotroph\| | Yeast | NULL | Probable | As nectar-tap_saprotroph (Põlme et al. 2020); Decay sugar-rich substrates (Põlme et al. 2020) | Põlme S, et al. 2020. Fungal Diversity 105:1-16 ((https://doi.org/10.1007/s13225-020-00466-2)) |
| Otu24197 | 1 | k__Fungi(100);p__Ascomycota(100);c__Saccharomycetes(100);o__Saccharomycetales(100);f__Metschnikowiaceae(100);g__Metschnikowia(100);g__Metschnikowia_unclassified(100); | Metschnikowia | 13 | Saprotroph | \|Undefined Saprotroph\| | Yeast | NULL | Probable | As nectar-tap_saprotroph (Põlme et al. 2020); Decay sugar-rich substrates (Põlme et al. 2020) | Põlme S, et al. 2020. Fungal Diversity 105:1-16 ((https://doi.org/10.1007/s13225-020-00466-2)) |
| Otu24198 | 1 | k__Fungi(100);p__Ascomycota(100);c__Saccharomycetes(100);o__Saccharomycetales(100);f__Metschnikowiaceae(100);g__Metschnikowia(100);s__Metschnikowia_pulcherrima(100); | Metschnikowia | 13 | Saprotroph | \|Undefined Saprotroph\| | Yeast | NULL | Probable | As nectar-tap_saprotroph (Põlme et al. 2020); Decay sugar-rich substrates (Põlme et al. 2020) | Põlme S, et al. 2020. Fungal Diversity 105:1-16 ((https://doi.org/10.1007/s13225-020-00466-2)) |
| Otu24199 | 1 | k__Fungi(100);p__Ascomycota(100);c__Saccharomycetes(100);o__Saccharomycetales(100);f__Metschnikowiaceae(100);g__Metschnikowia(100);g__Metschnikowia_unclassified(100); | Metschnikowia | 13 | Saprotroph | \|Undefined Saprotroph\| | Yeast | NULL | Probable | As nectar-tap_saprotroph (Põlme et al. 2020); Decay sugar-rich substrates (Põlme et al. 2020) | Põlme S, et al. 2020. Fungal Diversity 105:1-16 ((https://doi.org/10.1007/s13225-020-00466-2)) |
| Otu24200 | 1 | k__Fungi(100);p__Ascomycota(100);c__Saccharomycetes(100);o__Saccharomycetales(100);f__Metschnikowiaceae(100);g__Metschnikowia(100);s__Metschnikowia_pulcherrima(100); | Metschnikowia | 13 | Saprotroph | \|Undefined Saprotroph\| | Yeast | NULL | Probable | As nectar-tap_saprotroph (Põlme et al. 2020); Decay sugar-rich substrates (Põlme et al. 2020) | Põlme S, et al. 2020. Fungal Diversity 105:1-16 ((https://doi.org/10.1007/s13225-020-00466-2)) |
| Otu24201 | 1 | k__Fungi(100);p__Ascomycota(100);c__Saccharomycetes(100);o__Saccharomycetales(100);f__Metschnikowiaceae(100);g__Metschnikowia(100);s__Metschnikowia_pulcherrima(100); | Metschnikowia | 13 | Saprotroph | \|Undefined Saprotroph\| | Yeast | NULL | Probable | As nectar-tap_saprotroph (Põlme et al. 2020); Decay sugar-rich substrates (Põlme et al. 2020) | Põlme S, et al. 2020. Fungal Diversity 105:1-16 ((https://doi.org/10.1007/s13225-020-00466-2)) |
| Otu24202 | 1 | k__Fungi(100);p__Ascomycota(100);c__Saccharomycetes(100);o__Saccharomycetales(100);f__Metschnikowiaceae(100);g__Metschnikowia(100);g__Metschnikowia_unclassified(100); | Metschnikowia | 13 | Saprotroph | \|Undefined Saprotroph\| | Yeast | NULL | Probable | As nectar-tap_saprotroph (Põlme et al. 2020); Decay sugar-rich substrates (Põlme et al. 2020) | Põlme S, et al. 2020. Fungal Diversity 105:1-16 ((https://doi.org/10.1007/s13225-020-00466-2)) |
| Otu24203 | 1 | k__Fungi(100);p__Ascomycota(100);c__Saccharomycetes(100);o__Saccharomycetales(100);f__Metschnikowiaceae(100);g__Metschnikowia(100);s__Metschnikowia_pulcherrima(100); | Metschnikowia | 13 | Saprotroph | \|Undefined Saprotroph\| | Yeast | NULL | Probable | As nectar-tap_saprotroph (Põlme et al. 2020); Decay sugar-rich substrates (Põlme et al. 2020) | Põlme S, et al. 2020. Fungal Diversity 105:1-16 ((https://doi.org/10.1007/s13225-020-00466-2)) |
| Otu24204 | 1 | k__Fungi(100);p__Ascomycota(100);c__Saccharomycetes(100);o__Saccharomycetales(100);f__Metschnikowiaceae(100);g__Metschnikowia(100);g__Metschnikowia_unclassified(100); | Metschnikowia | 13 | Saprotroph | \|Undefined Saprotroph\| | Yeast | NULL | Probable | As nectar-tap_saprotroph (Põlme et al. 2020); Decay sugar-rich substrates (Põlme et al. 2020) | Põlme S, et al. 2020. Fungal Diversity 105:1-16 ((https://doi.org/10.1007/s13225-020-00466-2)) |
| Otu24205 | 1 | k__Fungi(100);p__Ascomycota(100);c__Saccharomycetes(100);o__Saccharomycetales(100);f__Metschnikowiaceae(100);g__Metschnikowia(100);s__Metschnikowia_pulcherrima(100); | Metschnikowia | 13 | Saprotroph | \|Undefined Saprotroph\| | Yeast | NULL | Probable | As nectar-tap_saprotroph (Põlme et al. 2020); Decay sugar-rich substrates (Põlme et al. 2020) | Põlme S, et al. 2020. Fungal Diversity 105:1-16 ((https://doi.org/10.1007/s13225-020-00466-2)) |
| Otu24206 | 1 | k__Fungi(100);p__Ascomycota(100);c__Saccharomycetes(100);o__Saccharomycetales(100);f__Metschnikowiaceae(100);g__Metschnikowia(100);s__Metschnikowia_pulcherrima(100); | Metschnikowia | 13 | Saprotroph | \|Undefined Saprotroph\| | Yeast | NULL | Probable | As nectar-tap_saprotroph (Põlme et al. 2020); Decay sugar-rich substrates (Põlme et al. 2020) | Põlme S, et al. 2020. Fungal Diversity 105:1-16 ((https://doi.org/10.1007/s13225-020-00466-2)) |
| Otu24210 | 1 | k__Fungi(100);p__Ascomycota(100);c__Saccharomycetes(100);o__Saccharomycetales(100);f__Metschnikowiaceae(100);g__Metschnikowia(100);s__Metschnikowia_pulcherrima(100); | Metschnikowia | 13 | Saprotroph | \|Undefined Saprotroph\| | Yeast | NULL | Probable | As nectar-tap_saprotroph (Põlme et al. 2020); Decay sugar-rich substrates (Põlme et al. 2020) | Põlme S, et al. 2020. Fungal Diversity 105:1-16 ((https://doi.org/10.1007/s13225-020-00466-2)) |
| Otu24211 | 1 | k__Fungi(100);p__Ascomycota(100);c__Saccharomycetes(100);o__Saccharomycetales(100);f__Metschnikowiaceae(100);g__Metschnikowia(100);s__Metschnikowia_pulcherrima(100); | Metschnikowia | 13 | Saprotroph | \|Undefined Saprotroph\| | Yeast | NULL | Probable | As nectar-tap_saprotroph (Põlme et al. 2020); Decay sugar-rich substrates (Põlme et al. 2020) | Põlme S, et al. 2020. Fungal Diversity 105:1-16 ((https://doi.org/10.1007/s13225-020-00466-2)) |
| Otu24213 | 1 | k__Fungi(100);p__Ascomycota(100);c__Saccharomycetes(100);o__Saccharomycetales(100);f__Metschnikowiaceae(100);g__Metschnikowia(100);g__Metschnikowia_unclassified(100); | Metschnikowia | 13 | Saprotroph | \|Undefined Saprotroph\| | Yeast | NULL | Probable | As nectar-tap_saprotroph (Põlme et al. 2020); Decay sugar-rich substrates (Põlme et al. 2020) | Põlme S, et al. 2020. Fungal Diversity 105:1-16 ((https://doi.org/10.1007/s13225-020-00466-2)) |
| Otu24218 | 1 | k__Fungi(100);p__Ascomycota(100);c__Saccharomycetes(100);o__Saccharomycetales(100);f__Metschnikowiaceae(100);g__Metschnikowia(100);s__Metschnikowia_pulcherrima(100); | Metschnikowia | 13 | Saprotroph | \|Undefined Saprotroph\| | Yeast | NULL | Probable | As nectar-tap_saprotroph (Põlme et al. 2020); Decay sugar-rich substrates (Põlme et al. 2020) | Põlme S, et al. 2020. Fungal Diversity 105:1-16 ((https://doi.org/10.1007/s13225-020-00466-2)) |
| Otu24219 | 1 | k__Fungi(100);p__Ascomycota(100);c__Saccharomycetes(100);o__Saccharomycetales(100);f__Metschnikowiaceae(100);g__Metschnikowia(100);s__Metschnikowia_pulcherrima(100); | Metschnikowia | 13 | Saprotroph | \|Undefined Saprotroph\| | Yeast | NULL | Probable | As nectar-tap_saprotroph (Põlme et al. 2020); Decay sugar-rich substrates (Põlme et al. 2020) | Põlme S, et al. 2020. Fungal Diversity 105:1-16 ((https://doi.org/10.1007/s13225-020-00466-2)) |
| Otu24221 | 1 | k__Fungi(100);p__Ascomycota(100);c__Saccharomycetes(100);o__Saccharomycetales(100);f__Metschnikowiaceae(100);g__Metschnikowia(100);g__Metschnikowia_unclassified(100); | Metschnikowia | 13 | Saprotroph | \|Undefined Saprotroph\| | Yeast | NULL | Probable | As nectar-tap_saprotroph (Põlme et al. 2020); Decay sugar-rich substrates (Põlme et al. 2020) | Põlme S, et al. 2020. Fungal Diversity 105:1-16 ((https://doi.org/10.1007/s13225-020-00466-2)) |
| Otu24223 | 1 | k__Fungi(100);p__Ascomycota(100);c__Saccharomycetes(100);o__Saccharomycetales(100);f__Metschnikowiaceae(100);g__Metschnikowia(100);g__Metschnikowia_unclassified(100); | Metschnikowia | 13 | Saprotroph | \|Undefined Saprotroph\| | Yeast | NULL | Probable | As nectar-tap_saprotroph (Põlme et al. 2020); Decay sugar-rich substrates (Põlme et al. 2020) | Põlme S, et al. 2020. Fungal Diversity 105:1-16 ((https://doi.org/10.1007/s13225-020-00466-2)) |
| Otu24224 | 1 | k__Fungi(100);p__Ascomycota(100);c__Saccharomycetes(100);o__Saccharomycetales(100);f__Metschnikowiaceae(100);g__Metschnikowia(100);s__Metschnikowia_pulcherrima(100); | Metschnikowia | 13 | Saprotroph | \|Undefined Saprotroph\| | Yeast | NULL | Probable | As nectar-tap_saprotroph (Põlme et al. 2020); Decay sugar-rich substrates (Põlme et al. 2020) | Põlme S, et al. 2020. Fungal Diversity 105:1-16 ((https://doi.org/10.1007/s13225-020-00466-2)) |
| Otu24225 | 1 | k__Fungi(100);p__Ascomycota(100);c__Saccharomycetes(100);o__Saccharomycetales(100);f__Metschnikowiaceae(100);g__Metschnikowia(100);g__Metschnikowia_unclassified(100); | Metschnikowia | 13 | Saprotroph | \|Undefined Saprotroph\| | Yeast | NULL | Probable | As nectar-tap_saprotroph (Põlme et al. 2020); Decay sugar-rich substrates (Põlme et al. 2020) | Põlme S, et al. 2020. Fungal Diversity 105:1-16 ((https://doi.org/10.1007/s13225-020-00466-2)) |
| Otu24227 | 1 | k__Fungi(100);p__Ascomycota(100);c__Saccharomycetes(100);o__Saccharomycetales(100);f__Metschnikowiaceae(100);g__Metschnikowia(100);s__Metschnikowia_pulcherrima(100); | Metschnikowia | 13 | Saprotroph | \|Undefined Saprotroph\| | Yeast | NULL | Probable | As nectar-tap_saprotroph (Põlme et al. 2020); Decay sugar-rich substrates (Põlme et al. 2020) | Põlme S, et al. 2020. Fungal Diversity 105:1-16 ((https://doi.org/10.1007/s13225-020-00466-2)) |
| Otu24228 | 1 | k__Fungi(100);p__Ascomycota(100);c__Saccharomycetes(100);o__Saccharomycetales(100);f__Metschnikowiaceae(100);g__Metschnikowia(100);g__Metschnikowia_unclassified(100); | Metschnikowia | 13 | Saprotroph | \|Undefined Saprotroph\| | Yeast | NULL | Probable | As nectar-tap_saprotroph (Põlme et al. 2020); Decay sugar-rich substrates (Põlme et al. 2020) | Põlme S, et al. 2020. Fungal Diversity 105:1-16 ((https://doi.org/10.1007/s13225-020-00466-2)) |

| | | | | | | | | | | | |
|---|---|---|---|---|---|---|---|---|---|---|---|
| Otu24229 | 1 | k__Fungi(100);p__Ascomycota(100);c__Saccharomycetes(100);o__Saccharomycetales(100);f__Metschnikowiaceae(100);g__Metschnikowia(100);s__Metschnikowia_pulcherrima(100); | Metschnikowia | 13 | Saprotroph | \|Undefined Saprotroph\| | Yeast | NULL | Probable | As nectar-tap_saprotroph (Põlme et al. 2020); Decay sugar-rich substrates (Põlme et al. 2020) | Põlme S, et al. 2020. Fungal Diversity 105:1-16 ((https://doi.org/10.1007/s13225-020-00466-2)) |
| Otu24230 | 1 | k__Fungi(100);p__Ascomycota(100);c__Saccharomycetes(100);o__Saccharomycetales(100);f__Metschnikowiaceae(100);g__Metschnikowia(100);g__Metschnikowia_unclassified(100); | Metschnikowia | 13 | Saprotroph | \|Undefined Saprotroph\| | Yeast | NULL | Probable | As nectar-tap_saprotroph (Põlme et al. 2020); Decay sugar-rich substrates (Põlme et al. 2020) | Põlme S, et al. 2020. Fungal Diversity 105:1-16 ((https://doi.org/10.1007/s13225-020-00466-2)) |
| Otu24231 | 1 | k__Fungi(100);p__Ascomycota(100);c__Saccharomycetes(100);o__Saccharomycetales(100);f__Metschnikowiaceae(100);g__Metschnikowia(100);s__Metschnikowia_pulcherrima(100); | Metschnikowia | 13 | Saprotroph | \|Undefined Saprotroph\| | Yeast | NULL | Probable | As nectar-tap_saprotroph (Põlme et al. 2020); Decay sugar-rich substrates (Põlme et al. 2020) | Põlme S, et al. 2020. Fungal Diversity 105:1-16 ((https://doi.org/10.1007/s13225-020-00466-2)) |
| Otu24232 | 1 | k__Fungi(100);p__Ascomycota(100);c__Saccharomycetes(100);o__Saccharomycetales(100);f__Metschnikowiaceae(100);g__Metschnikowia(100);s__Metschnikowia_pulcherrima(100); | Metschnikowia | 13 | Saprotroph | \|Undefined Saprotroph\| | Yeast | NULL | Probable | As nectar-tap_saprotroph (Põlme et al. 2020); Decay sugar-rich substrates (Põlme et al. 2020) | Põlme S, et al. 2020. Fungal Diversity 105:1-16 ((https://doi.org/10.1007/s13225-020-00466-2)) |
| Otu24233 | 1 | k__Fungi(100);p__Ascomycota(100);c__Saccharomycetes(100);o__Saccharomycetales(100);f__Metschnikowiaceae(100);g__Metschnikowia(100);s__Metschnikowia_pulcherrima(100); | Metschnikowia | 13 | Saprotroph | \|Undefined Saprotroph\| | Yeast | NULL | Probable | As nectar-tap_saprotroph (Põlme et al. 2020); Decay sugar-rich substrates (Põlme et al. 2020) | Põlme S, et al. 2020. Fungal Diversity 105:1-16 ((https://doi.org/10.1007/s13225-020-00466-2)) |
| Otu24234 | 1 | k__Fungi(100);p__Ascomycota(100);c__Saccharomycetes(100);o__Saccharomycetales(100);f__Metschnikowiaceae(100);g__Metschnikowia(100);s__Metschnikowia_pulcherrima(100); | Metschnikowia | 13 | Saprotroph | \|Undefined Saprotroph\| | Yeast | NULL | Probable | As nectar-tap_saprotroph (Põlme et al. 2020); Decay sugar-rich substrates (Põlme et al. 2020) | Põlme S, et al. 2020. Fungal Diversity 105:1-16 ((https://doi.org/10.1007/s13225-020-00466-2)) |
| Otu24235 | 1 | k__Fungi(100);p__Ascomycota(100);c__Saccharomycetes(100);o__Saccharomycetales(100);f__Metschnikowiaceae(100);g__Metschnikowia(100);g__Metschnikowia_unclassified(100); | Metschnikowia | 13 | Saprotroph | \|Undefined Saprotroph\| | Yeast | NULL | Probable | As nectar-tap_saprotroph (Põlme et al. 2020); Decay sugar-rich substrates (Põlme et al. 2020) | Põlme S, et al. 2020. Fungal Diversity 105:1-16 ((https://doi.org/10.1007/s13225-020-00466-2)) |
| Otu24236 | 1 | k__Fungi(100);p__Ascomycota(100);c__Saccharomycetes(100);o__Saccharomycetales(100);f__Metschnikowiaceae(100);g__Metschnikowia(100);s__Metschnikowia_pulcherrima(100); | Metschnikowia | 13 | Saprotroph | \|Undefined Saprotroph\| | Yeast | NULL | Probable | As nectar-tap_saprotroph (Põlme et al. 2020); Decay sugar-rich substrates (Põlme et al. 2020) | Põlme S, et al. 2020. Fungal Diversity 105:1-16 ((https://doi.org/10.1007/s13225-020-00466-2)) |
| Otu24237 | 1 | k__Fungi(100);p__Ascomycota(100);c__Saccharomycetes(100);o__Saccharomycetales(100);f__Metschnikowiaceae(100);g__Metschnikowia(100);s__Metschnikowia_pulcherrima(100); | Metschnikowia | 13 | Saprotroph | \|Undefined Saprotroph\| | Yeast | NULL | Probable | As nectar-tap_saprotroph (Põlme et al. 2020); Decay sugar-rich substrates (Põlme et al. 2020) | Põlme S, et al. 2020. Fungal Diversity 105:1-16 ((https://doi.org/10.1007/s13225-020-00466-2)) |
| Otu24238 | 1 | k__Fungi(100);p__Ascomycota(100);c__Saccharomycetes(100);o__Saccharomycetales(100);f__Metschnikowiaceae(100);g__Metschnikowia(100);s__Metschnikowia_pulcherrima(100); | Metschnikowia | 13 | Saprotroph | \|Undefined Saprotroph\| | Yeast | NULL | Probable | As nectar-tap_saprotroph (Põlme et al. 2020); Decay sugar-rich substrates (Põlme et al. 2020) | Põlme S, et al. 2020. Fungal Diversity 105:1-16 ((https://doi.org/10.1007/s13225-020-00466-2)) |
| Otu24239 | 1 | k__Fungi(100);p__Ascomycota(100);c__Saccharomycetes(100);o__Saccharomycetales(100);f__Metschnikowiaceae(100);g__Metschnikowia(100);s__Metschnikowia_pulcherrima(100); | Metschnikowia | 13 | Saprotroph | \|Undefined Saprotroph\| | Yeast | NULL | Probable | As nectar-tap_saprotroph (Põlme et al. 2020); Decay sugar-rich substrates (Põlme et al. 2020) | Põlme S, et al. 2020. Fungal Diversity 105:1-16 ((https://doi.org/10.1007/s13225-020-00466-2)) |
| Otu24240 | 1 | k__Fungi(100);p__Ascomycota(100);c__Saccharomycetes(100);o__Saccharomycetales(100);f__Metschnikowiaceae(100);g__Metschnikowia(100);s__Metschnikowia_pulcherrima(100); | Metschnikowia | 13 | Saprotroph | \|Undefined Saprotroph\| | Yeast | NULL | Probable | As nectar-tap_saprotroph (Põlme et al. 2020); Decay sugar-rich substrates (Põlme et al. 2020) | Põlme S, et al. 2020. Fungal Diversity 105:1-16 ((https://doi.org/10.1007/s13225-020-00466-2)) |
| Otu24251 | 1 | k__Fungi(100);p__Ascomycota(100);c__Saccharomycetes(100);o__Saccharomycetales(100);f__Metschnikowiaceae(100);g__Metschnikowia(100);s__Metschnikowia_pulcherrima(100); | Metschnikowia | 13 | Saprotroph | \|Undefined Saprotroph\| | Yeast | NULL | Probable | As nectar-tap_saprotroph (Põlme et al. 2020); Decay sugar-rich substrates (Põlme et al. 2020) | Põlme S, et al. 2020. Fungal Diversity 105:1-16 ((https://doi.org/10.1007/s13225-020-00466-2)) |
| Otu24252 | 1 | k__Fungi(100);p__Ascomycota(100);c__Saccharomycetes(100);o__Saccharomycetales(100);f__Metschnikowiaceae(100);g__Metschnikowia(100);s__Metschnikowia_pulcherrima(100); | Metschnikowia | 13 | Saprotroph | \|Undefined Saprotroph\| | Yeast | NULL | Probable | As nectar-tap_saprotroph (Põlme et al. 2020); Decay sugar-rich substrates (Põlme et al. 2020) | Põlme S, et al. 2020. Fungal Diversity 105:1-16 ((https://doi.org/10.1007/s13225-020-00466-2)) |
| Otu24255 | 1 | k__Fungi(100);p__Ascomycota(100);c__Saccharomycetes(100);o__Saccharomycetales(100);f__Metschnikowiaceae(100);g__Metschnikowia(100);g__Metschnikowia_unclassified(100); | Metschnikowia | 13 | Saprotroph | \|Undefined Saprotroph\| | Yeast | NULL | Probable | As nectar-tap_saprotroph (Põlme et al. 2020); Decay sugar-rich substrates (Põlme et al. 2020) | Põlme S, et al. 2020. Fungal Diversity 105:1-16 ((https://doi.org/10.1007/s13225-020-00466-2)) |
| Otu24259 | 1 | k__Fungi(100);p__Ascomycota(100);c__Saccharomycetes(100);o__Saccharomycetales(100);f__Metschnikowiaceae(100);g__Metschnikowia(100);s__Metschnikowia_pulcherrima(100); | Metschnikowia | 13 | Saprotroph | \|Undefined Saprotroph\| | Yeast | NULL | Probable | As nectar-tap_saprotroph (Põlme et al. 2020); Decay sugar-rich substrates (Põlme et al. 2020) | Põlme S, et al. 2020. Fungal Diversity 105:1-16 ((https://doi.org/10.1007/s13225-020-00466-2)) |
| Otu24260 | 1 | k__Fungi(100);p__Zygomycota(100);c__Zygomycota_class_Incertae_sedis(100);o__Mucorales(100);f__Mucoraceae(100);g__Mucor(100);s__Mucor_circinelloides_f._griseocyanus(100); | Mucor | 13 | Saprotroph | \|Undefined Saprotroph\| | Microfungus | NULL | Probable | As soil_saprotroph (Põlme et al. 2020); Decay soil (Põlme et al. 2020) | Tedersoo L, et al. 2014. Science 346:e1256688 ((https://doi.org/10.1126/science.1256688)); Põlme S, et al. 2020. Fungal Diversity 105:1-16 ((https://doi.org/10.1007/s13225-020-00466-2)) |
| Otu24263 | 1 | k__Fungi(100);p__Zygomycota(100);c__Zygomycota_class_Incertae_sedis(100);o__Mucorales(100);f__Mucoraceae(100);g__Mucor(100);g__Mucor_unclassified(100); | Mucor | 13 | Saprotroph | \|Undefined Saprotroph\| | Microfungus | NULL | Probable | As soil_saprotroph (Põlme et al. 2020); Decay soil (Põlme et al. 2020) | Tedersoo L, et al. 2014. Science 346:e1256688 ((https://doi.org/10.1126/science.1256688)); Põlme S, et al. 2020. Fungal Diversity 105:1-16 ((https://doi.org/10.1007/s13225-020-00466-2)) |
| Otu24264 | 1 | k__Fungi(100);p__Ascomycota(100);c__Saccharomycetes(100);o__Saccharomycetales(100);f__Saccharomycetaceae(100);g__Kazachstania(100);s__Kazachstania_intestinalis(100); | Kazachstania | 13 | Saprotroph | \|Undefined Saprotroph\| | Yeast | NULL | Probable | As nectar-tap_saprotroph (Põlme et al. 2020); Decay sugar-rich substrates (Põlme et al. 2020) | Põlme S, et al. 2020. Fungal Diversity 105:1-16 ((https://doi.org/10.1007/s13225-020-00466-2)) |
| Otu24270 | 1 | k__Fungi(100);p__Ascomycota(100);c__Saccharomycetes(100);o__Saccharomycetales(100);f__Metschnikowiaceae(100);g__Metschnikowia(100);s__Metschnikowia_pulcherrima(100); | Metschnikowia | 13 | Saprotroph | \|Undefined Saprotroph\| | Yeast | NULL | Probable | As nectar-tap_saprotroph (Põlme et al. 2020); Decay sugar-rich substrates (Põlme et al. 2020) | Põlme S, et al. 2020. Fungal Diversity 105:1-16 ((https://doi.org/10.1007/s13225-020-00466-2)) |
| Otu24272 | 1 | k__Fungi(100);p__Ascomycota(100);c__Saccharomycetes(100);o__Saccharomycetales(100);f__Metschnikowiaceae(100);g__Metschnikowia(100);s__Metschnikowia_pulcherrima(100); | Metschnikowia | 13 | Saprotroph | \|Undefined Saprotroph\| | Yeast | NULL | Probable | As nectar-tap_saprotroph (Põlme et al. 2020); Decay sugar-rich substrates (Põlme et al. 2020) | Põlme S, et al. 2020. Fungal Diversity 105:1-16 ((https://doi.org/10.1007/s13225-020-00466-2)) |
| Otu24280 | 1 | k__Fungi(100);p__Ascomycota(100);c__Saccharomycetes(100);o__Saccharomycetales(100);f__Metschnikowiaceae(100);g__Metschnikowia(100);g__Metschnikowia_unclassified(100); | Metschnikowia | 13 | Saprotroph | \|Undefined Saprotroph\| | Yeast | NULL | Probable | As nectar-tap_saprotroph (Põlme et al. 2020); Decay sugar-rich substrates (Põlme et al. 2020) | Põlme S, et al. 2020. Fungal Diversity 105:1-16 ((https://doi.org/10.1007/s13225-020-00466-2)) |
| Otu24285 | 1 | k__Fungi(100);p__Ascomycota(100);c__Saccharomycetes(100);o__Saccharomycetales(100);f__Metschnikowiaceae(100);g__Metschnikowia(100);g__Metschnikowia_unclassified(100); | Metschnikowia | 13 | Saprotroph | \|Undefined Saprotroph\| | Yeast | NULL | Probable | As nectar-tap_saprotroph (Põlme et al. 2020); Decay sugar-rich substrates (Põlme et al. 2020) | Põlme S, et al. 2020. Fungal Diversity 105:1-16 ((https://doi.org/10.1007/s13225-020-00466-2)) |
| Otu24287 | 1 | k__Fungi(100);p__Ascomycota(100);c__Saccharomycetes(100);o__Saccharomycetales(100);f__Metschnikowiaceae(100);g__Metschnikowia(100);s__Metschnikowia_pulcherrima(100); | Metschnikowia | 13 | Saprotroph | \|Undefined Saprotroph\| | Yeast | NULL | Probable | As nectar-tap_saprotroph (Põlme et al. 2020); Decay sugar-rich substrates (Põlme et al. 2020) | Põlme S, et al. 2020. Fungal Diversity 105:1-16 ((https://doi.org/10.1007/s13225-020-00466-2)) |
| Otu24288 | 1 | k__Fungi(100);p__Ascomycota(100);c__Saccharomycetes(100);o__Saccharomycetales(100);f__Metschnikowiaceae(100);g__Metschnikowia(100);g__Metschnikowia_unclassified(100); | Metschnikowia | 13 | Saprotroph | \|Undefined Saprotroph\| | Yeast | NULL | Probable | As nectar-tap_saprotroph (Põlme et al. 2020); Decay sugar-rich substrates (Põlme et al. 2020) | Põlme S, et al. 2020. Fungal Diversity 105:1-16 ((https://doi.org/10.1007/s13225-020-00466-2)) |
| Otu24289 | 1 | k__Fungi(100);p__Ascomycota(100);c__Saccharomycetes(100);o__Saccharomycetales(100);f__Metschnikowiaceae(100);g__Metschnikowia(100);s__Metschnikowia_pulcherrima(100); | Metschnikowia | 13 | Saprotroph | \|Undefined Saprotroph\| | Yeast | NULL | Probable | As nectar-tap_saprotroph (Põlme et al. 2020); Decay sugar-rich substrates (Põlme et al. 2020) | Põlme S, et al. 2020. Fungal Diversity 105:1-16 ((https://doi.org/10.1007/s13225-020-00466-2)) |
| Otu24291 | 1 | k__Fungi(100);p__Ascomycota(100);c__Saccharomycetes(100);o__Saccharomycetales(100);f__Metschnikowiaceae(100);g__Metschnikowia(100);s__Metschnikowia_pulcherrima(100); | Metschnikowia | 13 | Saprotroph | \|Undefined Saprotroph\| | Yeast | NULL | Probable | As nectar-tap_saprotroph (Põlme et al. 2020); Decay sugar-rich substrates (Põlme et al. 2020) | Põlme S, et al. 2020. Fungal Diversity 105:1-16 ((https://doi.org/10.1007/s13225-020-00466-2)) |
| Otu24305 | 1 | k__Fungi(100);p__Ascomycota(100);c__Saccharomycetes(100);o__Saccharomycetales(100);f__Metschnikowiaceae(100);g__Metschnikowia(100);g__Metschnikowia_unclassified(100); | Metschnikowia | 13 | Saprotroph | \|Undefined Saprotroph\| | Yeast | NULL | Probable | As nectar-tap_saprotroph (Põlme et al. 2020); Decay sugar-rich substrates (Põlme et al. 2020) | Põlme S, et al. 2020. Fungal Diversity 105:1-16 ((https://doi.org/10.1007/s13225-020-00466-2)) |
| Otu24311 | 1 | k__Fungi(100);p__Ascomycota(100);c__Saccharomycetes(100);o__Saccharomycetales(100);f__Metschnikowiaceae(100);g__Metschnikowia(100);s__Metschnikowia_pulcherrima(100); | Metschnikowia | 13 | Saprotroph | \|Undefined Saprotroph\| | Yeast | NULL | Probable | As nectar-tap_saprotroph (Põlme et al. 2020); Decay sugar-rich substrates (Põlme et al. 2020) | Põlme S, et al. 2020. Fungal Diversity 105:1-16 ((https://doi.org/10.1007/s13225-020-00466-2)) |
| Otu24314 | 1 | k__Fungi(100);p__Ascomycota(100);c__Saccharomycetes(100);o__Saccharomycetales(100);f__Metschnikowiaceae(100);g__Metschnikowia(100);g__Metschnikowia_unclassified(100); | Metschnikowia | 13 | Saprotroph | \|Undefined Saprotroph\| | Yeast | NULL | Probable | As nectar-tap_saprotroph (Põlme et al. 2020); Decay sugar-rich substrates (Põlme et al. 2020) | Põlme S, et al. 2020. Fungal Diversity 105:1-16 ((https://doi.org/10.1007/s13225-020-00466-2)) |
| Otu24331 | 1 | k__Fungi(100);p__Ascomycota(100);c__Saccharomycetes(100);o__Saccharomycetales(100);f__Metschnikowiaceae(100);g__Metschnikowia(100);s__Metschnikowia_pulcherrima(100); | Metschnikowia | 13 | Saprotroph | \|Undefined Saprotroph\| | Yeast | NULL | Probable | As nectar-tap_saprotroph (Põlme et al. 2020); Decay sugar-rich substrates (Põlme et al. 2020) | Põlme S, et al. 2020. Fungal Diversity 105:1-16 ((https://doi.org/10.1007/s13225-020-00466-2)) |
| Otu24332 | 1 | k__Fungi(100);p__Ascomycota(100);c__Saccharomycetes(100);o__Saccharomycetales(100);f__Metschnikowiaceae(100);g__Metschnikowia(100);s__Metschnikowia_pulcherrima(100); | Metschnikowia | 13 | Saprotroph | \|Undefined Saprotroph\| | Yeast | NULL | Probable | As nectar-tap_saprotroph (Põlme et al. 2020); Decay sugar-rich substrates (Põlme et al. 2020) | Põlme S, et al. 2020. Fungal Diversity 105:1-16 ((https://doi.org/10.1007/s13225-020-00466-2)) |
| Otu24342 | 1 | k__Fungi(100);p__Ascomycota(100);c__Saccharomycetes(100);o__Saccharomycetales(100);f__Metschnikowiaceae(100);g__Metschnikowia(100);s__Metschnikowia_pulcherrima(100); | Metschnikowia | 13 | Saprotroph | \|Undefined Saprotroph\| | Yeast | NULL | Probable | As nectar-tap_saprotroph (Põlme et al. 2020); Decay sugar-rich substrates (Põlme et al. 2020) | Põlme S, et al. 2020. Fungal Diversity 105:1-16 ((https://doi.org/10.1007/s13225-020-00466-2)) |
| Otu24343 | 1 | k__Fungi(100);p__Ascomycota(100);c__Saccharomycetes(100);o__Saccharomycetales(100);f__Metschnikowiaceae(100);g__Metschnikowia(100);s__Metschnikowia_pulcherrima(100); | Metschnikowia | 13 | Saprotroph | \|Undefined Saprotroph\| | Yeast | NULL | Probable | As nectar-tap_saprotroph (Põlme et al. 2020); Decay sugar-rich substrates (Põlme et al. 2020) | Põlme S, et al. 2020. Fungal Diversity 105:1-16 ((https://doi.org/10.1007/s13225-020-00466-2)) |
| Otu24345 | 1 | k__Fungi(100);p__Basidiomycota(100);c__Microbotryomycetes(100);o__Sporidiobolales(100);f__Sporidiobolales_family_Incertae_sedis(100);g__Sporobolomyces(100);s__Sporobolomyces_lactosus(100); | Sporobolomyces | 13 | Pathotroph-Saprotroph | \|Fungal Parasite\|-Plant Saprotroph | Yeast | NULL | Probable | As litter_saprotroph (Põlme et al. 2020); As mycoparasite (Põlme et al. 2020); Decay fungal material (Põlme et al. 2020) | Tedersoo L, et al. 2014. Science 346:e1256688 ((https://doi.org/10.1126/science.1256688)); Purahong W, et al. 2016. Molecular Ecology 25:4059-4074 ((https://doi.org/10.1111/mec.13739)); Põlme S, et al. 2020. Fungal Diversity 105:1-16 ((https://doi.org/10.1007/s13225-020-00466-2)) |
| Otu24350 | 1 | k__Fungi(100);p__Ascomycota(100);c__Saccharomycetes(100);o__Saccharomycetales(100);f__Metschnikowiaceae(100);g__Metschnikowia(100);s__Metschnikowia_pulcherrima(100); | Metschnikowia | 13 | Saprotroph | \|Undefined Saprotroph\| | Yeast | NULL | Probable | As nectar-tap_saprotroph (Põlme et al. 2020); Decay sugar-rich substrates (Põlme et al. 2020) | Põlme S, et al. 2020. Fungal Diversity 105:1-16 ((https://doi.org/10.1007/s13225-020-00466-2)) |
| Otu24351 | 1 | k__Fungi(100);p__Ascomycota(100);c__Saccharomycetes(100);o__Saccharomycetales(100);f__Metschnikowiaceae(100);g__Metschnikowia(100);g__Metschnikowia_unclassified(100); | Metschnikowia | 13 | Saprotroph | \|Undefined Saprotroph\| | Yeast | NULL | Probable | As nectar-tap_saprotroph (Põlme et al. 2020); Decay sugar-rich substrates (Põlme et al. 2020) | Põlme S, et al. 2020. Fungal Diversity 105:1-16 ((https://doi.org/10.1007/s13225-020-00466-2)) |
| Otu24352 | 1 | k__Fungi(100);p__Ascomycota(100);c__Saccharomycetes(100);o__Saccharomycetales(100);f__Metschnikowiaceae(100);g__Metschnikowia(100);s__Metschnikowia_pulcherrima(100); | Metschnikowia | 13 | Saprotroph | \|Undefined Saprotroph\| | Yeast | NULL | Probable | As nectar-tap_saprotroph (Põlme et al. 2020); Decay sugar-rich substrates (Põlme et al. 2020) | Põlme S, et al. 2020. Fungal Diversity 105:1-16 ((https://doi.org/10.1007/s13225-020-00466-2)) |
| Otu24354 | 1 | k__Fungi(100);p__Ascomycota(100);c__Saccharomycetes(100);o__Saccharomycetales(100);f__Metschnikowiaceae(100);g__Metschnikowia(100);s__Metschnikowia_pulcherrima(100); | Metschnikowia | 13 | Saprotroph | \|Undefined Saprotroph\| | Yeast | NULL | Probable | As nectar-tap_saprotroph (Põlme et al. 2020); Decay sugar-rich substrates (Põlme et al. 2020) | Põlme S, et al. 2020. Fungal Diversity 105:1-16 ((https://doi.org/10.1007/s13225-020-00466-2)) |
| Otu24360 | 1 | k__Fungi(100);p__Ascomycota(100);c__Saccharomycetes(100);o__Saccharomycetales(100);f__Metschnikowiaceae(100);g__Metschnikowia(100);g__Metschnikowia_unclassified(100); | Metschnikowia | 13 | Saprotroph | \|Undefined Saprotroph\| | Yeast | NULL | Probable | As nectar-tap_saprotroph (Põlme et al. 2020); Decay sugar-rich substrates (Põlme et al. 2020) | Põlme S, et al. 2020. Fungal Diversity 105:1-16 ((https://doi.org/10.1007/s13225-020-00466-2)) |
| Otu24362 | 1 | k__Fungi(100);p__Basidiomycota(100);c__Microbotryomycetes(100);o__Sporidiobolales(100);f__Sporidiobolales_family_Incertae_sedis(100);g__Sporobolomyces(100);s__Sporobolomyces_lactosus(100); | Sporobolomyces | 13 | Pathotroph-Saprotroph | \|Fungal Parasite\|-Plant Saprotroph | Yeast | NULL | Probable | As litter_saprotroph (Põlme et al. 2020); As mycoparasite (Põlme et al. 2020); Decay fungal material (Põlme et al. 2020) | Tedersoo L, et al. 2014. Science 346:e1256688 ((https://doi.org/10.1126/science.1256688)); Purahong W, et al. 2016. Molecular Ecology 25:4059-4074 ((https://doi.org/10.1111/mec.13739)); Põlme S, et al. 2020. Fungal Diversity 105:1-16 ((https://doi.org/10.1007/s13225-020-00466-2)) |
| Otu24369 | 1 | k__Fungi(100);p__Ascomycota(100);c__Saccharomycetes(100);o__Saccharomycetales(100);f__Metschnikowiaceae(100);g__Metschnikowia(100);g__Metschnikowia_unclassified(100); | Metschnikowia | 13 | Saprotroph | \|Undefined Saprotroph\| | Yeast | NULL | Probable | As nectar-tap_saprotroph (Põlme et al. 2020); Decay sugar-rich substrates (Põlme et al. 2020) | Põlme S, et al. 2020. Fungal Diversity 105:1-16 ((https://doi.org/10.1007/s13225-020-00466-2)) |
| Otu24370 | 1 | k__Fungi(100);p__Ascomycota(100);c__Saccharomycetes(100);o__Saccharomycetales(100);f__Metschnikowiaceae(100);g__Metschnikowia(100);s__Metschnikowia_pulcherrima(100); | Metschnikowia | 13 | Saprotroph | \|Undefined Saprotroph\| | Yeast | NULL | Probable | As nectar-tap_saprotroph (Põlme et al. 2020); Decay sugar-rich substrates (Põlme et al. 2020) | Põlme S, et al. 2020. Fungal Diversity 105:1-16 ((https://doi.org/10.1007/s13225-020-00466-2)) |
| Otu24371 | 1 | k__Fungi(100);p__Ascomycota(100);c__Saccharomycetes(100);o__Saccharomycetales(100);f__Metschnikowiaceae(100);g__Metschnikowia(100);s__Metschnikowia_pulcherrima(100); | Metschnikowia | 13 | Saprotroph | \|Undefined Saprotroph\| | Yeast | NULL | Probable | As nectar-tap_saprotroph (Põlme et al. 2020); Decay sugar-rich substrates (Põlme et al. 2020) | Põlme S, et al. 2020. Fungal Diversity 105:1-16 ((https://doi.org/10.1007/s13225-020-00466-2)) |
| Otu24373 | 1 | k__Fungi(100);p__Ascomycota(100);c__Saccharomycetes(100);o__Saccharomycetales(100);f__Metschnikowiaceae(100);g__Metschnikowia(100);s__Metschnikowia_pulcherrima(100); | Metschnikowia | 13 | Saprotroph | \|Undefined Saprotroph\| | Yeast | NULL | Probable | As nectar-tap_saprotroph (Põlme et al. 2020); Decay sugar-rich substrates (Põlme et al. 2020) | Põlme S, et al. 2020. Fungal Diversity 105:1-16 ((https://doi.org/10.1007/s13225-020-00466-2)) |
| Otu24374 | 1 | k__Fungi(100);p__Ascomycota(100);c__Saccharomycetes(100);o__Saccharomycetales(100);o__Saccharomycetales_unclassified(100);o__Saccharomycetales_unclassified(100);o__Saccharomycetales_unclassified(100); | Saccharomycetales | 7 | Saprotroph | Undefined Saprotroph | Yeast | NULL | Possible | NULL | Sterkenburg E, et al. 2015. New Phytologist 207:1145-1158 ((https://doi.org/10.1111/nph.13426)) |
| Otu24378 | 1 | k__Fungi(100);p__Ascomycota(100);c__Saccharomycetes(100);o__Saccharomycetales(100);f__Metschnikowiaceae(100);g__Metschnikowia(100);s__Metschnikowia_pulcherrima(100); | Metschnikowia | 13 | Saprotroph | \|Undefined Saprotroph\| | Yeast | NULL | Probable | As nectar-tap_saprotroph (Põlme et al. 2020); Decay sugar-rich substrates (Põlme et al. 2020) | Põlme S, et al. 2020. Fungal Diversity 105:1-16 ((https://doi.org/10.1007/s13225-020-00466-2)) |
| Otu24382 | 1 | k__Fungi(100);p__Ascomycota(100);c__Saccharomycetes(100);o__Saccharomycetales(100);f__Metschnikowiaceae(100);g__Metschnikowia(100);s__Metschnikowia_pulcherrima(100); | Metschnikowia | 13 | Saprotroph | \|Undefined Saprotroph\| | Yeast | NULL | Probable | As nectar-tap_saprotroph (Põlme et al. 2020); Decay sugar-rich substrates (Põlme et al. 2020) | Põlme S, et al. 2020. Fungal Diversity 105:1-16 ((https://doi.org/10.1007/s13225-020-00466-2)) |
| Otu24383 | 1 | k__Fungi(100);p__Ascomycota(100);c__Saccharomycetes(100);o__Saccharomycetales(100);f__Metschnikowiaceae(100);g__Metschnikowia(100);g__Metschnikowia_unclassified(100); | Metschnikowia | 13 | Saprotroph | \|Undefined Saprotroph\| | Yeast | NULL | Probable | As nectar-tap_saprotroph (Põlme et al. 2020); Decay sugar-rich substrates (Põlme et al. 2020) | Põlme S, et al. 2020. Fungal Diversity 105:1-16 ((https://doi.org/10.1007/s13225-020-00466-2)) |
| Otu24386 | 1 | k__Fungi(100);p__Ascomycota(100);c__Saccharomycetes(100);o__Saccharomycetales(100);f__Metschnikowiaceae(100);g__Metschnikowia(100);s__Metschnikowia_pulcherrima(100); | Metschnikowia | 13 | Saprotroph | \|Undefined Saprotroph\| | Yeast | NULL | Probable | As nectar-tap_saprotroph (Põlme et al. 2020); Decay sugar-rich substrates (Põlme et al. 2020) | Põlme S, et al. 2020. Fungal Diversity 105:1-16 ((https://doi.org/10.1007/s13225-020-00466-2)) |
| Otu24387 | 1 | k__Fungi(100);p__Ascomycota(100);c__Saccharomycetes(100);o__Saccharomycetales(100);f__Metschnikowiaceae(100);g__Metschnikowia(100);s__Metschnikowia_pulcherrima(100); | Metschnikowia | 13 | Saprotroph | \|Undefined Saprotroph\| | Yeast | NULL | Probable | As nectar-tap_saprotroph (Põlme et al. 2020); Decay sugar-rich substrates (Põlme et al. 2020) | Põlme S, et al. 2020. Fungal Diversity 105:1-16 ((https://doi.org/10.1007/s13225-020-00466-2)) |
| Otu24388 | 1 | k__Fungi(100);p__Ascomycota(100);c__Saccharomycetes(100);o__Saccharomycetales(100);f__Metschnikowiaceae(100);g__Metschnikowia(100);s__Metschnikowia_pulcherrima(100); | Metschnikowia | 13 | Saprotroph | \|Undefined Saprotroph\| | Yeast | NULL | Probable | As nectar-tap_saprotroph (Põlme et al. 2020); Decay sugar-rich substrates (Põlme et al. 2020) | Põlme S, et al. 2020. Fungal Diversity 105:1-16 ((https://doi.org/10.1007/s13225-020-00466-2)) |
| Otu24389 | 1 | k__Fungi(100);p__Ascomycota(100);c__Saccharomycetes(100);o__Saccharomycetales(100);f__Metschnikowiaceae(100);g__Metschnikowia(100);g__Metschnikowia_unclassified(100); | Metschnikowia | 13 | Saprotroph | \|Undefined Saprotroph\| | Yeast | NULL | Probable | As nectar-tap_saprotroph (Põlme et al. 2020); Decay sugar-rich substrates (Põlme et al. 2020) | Põlme S, et al. 2020. Fungal Diversity 105:1-16 ((https://doi.org/10.1007/s13225-020-00466-2)) |
| Otu24390 | 1 | k__Fungi(100);p__Ascomycota(100);c__Saccharomycetes(100);o__Saccharomycetales(100);f__Metschnikowiaceae(100);g__Metschnikowia(100);g__Metschnikowia_unclassified(100); | Metschnikowia | 13 | Saprotroph | \|Undefined Saprotroph\| | Yeast | NULL | Probable | As nectar-tap_saprotroph (Põlme et al. 2020); Decay sugar-rich substrates (Põlme et al. 2020) | Põlme S, et al. 2020. Fungal Diversity 105:1-16 ((https://doi.org/10.1007/s13225-020-00466-2)) |
| Otu24391 | 1 | k__Fungi(100);p__Ascomycota(100);c__Saccharomycetes(100);o__Saccharomycetales(100);f__Metschnikowiaceae(100);g__Metschnikowia(100);s__Metschnikowia_pulcherrima(100); | Metschnikowia | 13 | Saprotroph | \|Undefined Saprotroph\| | Yeast | NULL | Probable | As nectar-tap_saprotroph (Põlme et al. 2020); Decay sugar-rich substrates (Põlme et al. 2020) | Põlme S, et al. 2020. Fungal Diversity 105:1-16 ((https://doi.org/10.1007/s13225-020-00466-2)) |
| Otu24400 | 1 | k__Fungi(100);p__Ascomycota(100);c__Saccharomycetes(100);o__Saccharomycetales(100);f__Metschnikowiaceae(100);g__Metschnikowia(100);s__Metschnikowia_pulcherrima(100); | Metschnikowia | 13 | Saprotroph | \|Undefined Saprotroph\| | Yeast | NULL | Probable | As nectar-tap_saprotroph (Põlme et al. 2020); Decay sugar-rich substrates (Põlme et al. 2020) | Põlme S, et al. 2020. Fungal Diversity 105:1-16 ((https://doi.org/10.1007/s13225-020-00466-2)) |
| Otu24403 | 1 | k__Fungi(100);p__Ascomycota(100);c__Saccharomycetes(100);o__Saccharomycetales(100);f__Metschnikowiaceae(100);g__Metschnikowia(100);s__Metschnikowia_pulcherrima(100); | Metschnikowia | 13 | Saprotroph | \|Undefined Saprotroph\| | Yeast | NULL | Probable | As nectar-tap_saprotroph (Põlme et al. 2020); Decay sugar-rich substrates (Põlme et al. 2020) | Põlme S, et al. 2020. Fungal Diversity 105:1-16 ((https://doi.org/10.1007/s13225-020-00466-2)) |
| Otu24404 | 1 | k__Fungi(100);p__Ascomycota(100);c__Saccharomycetes(100);o__Saccharomycetales(100);f__Metschnikowiaceae(100);g__Metschnikowia(100);s__Metschnikowia_pulcherrima(100); | Metschnikowia | 13 | Saprotroph | \|Undefined Saprotroph\| | Yeast | NULL | Probable | As nectar-tap_saprotroph (Põlme et al. 2020); Decay sugar-rich substrates (Põlme et al. 2020) | Põlme S, et al. 2020. Fungal Diversity 105:1-16 ((https://doi.org/10.1007/s13225-020-00466-2)) |
| Otu24405 | 1 | k__Fungi(100);p__Ascomycota(100);c__Saccharomycetes(100);o__Saccharomycetales(100);f__Metschnikowiaceae(100);g__Metschnikowia(100);s__Metschnikowia_pulcherrima(100); | Metschnikowia | 13 | Saprotroph | \|Undefined Saprotroph\| | Yeast | NULL | Probable | As nectar-tap_saprotroph (Põlme et al. 2020); Decay sugar-rich substrates (Põlme et al. 2020) | Põlme S, et al. 2020. Fungal Diversity 105:1-16 ((https://doi.org/10.1007/s13225-020-00466-2)) |
| Otu24412 | 1 | k__Fungi(100);p__Ascomycota(100);c__Saccharomycetes(100);o__Saccharomycetales(100);f__Metschnikowiaceae(100);g__Metschnikowia(100);s__Metschnikowia_pulcherrima(100); | Metschnikowia | 13 | Saprotroph | \|Undefined Saprotroph\| | Yeast | NULL | Probable | As nectar-tap_saprotroph (Põlme et al. 2020); Decay sugar-rich substrates (Põlme et al. 2020) | Põlme S, et al. 2020. Fungal Diversity 105:1-16 ((https://doi.org/10.1007/s13225-020-00466-2)) |
| Otu24414 | 1 | k__Fungi(100);p__Ascomycota(100);c__Saccharomycetes(100);o__Saccharomycetales(100);f__Metschnikowiaceae(100);g__Metschnikowia(100);g__Metschnikowia_unclassified(100); | Metschnikowia | 13 | Saprotroph | \|Undefined Saprotroph\| | Yeast | NULL | Probable | As nectar-tap_saprotroph (Põlme et al. 2020); Decay sugar-rich substrates (Põlme et al. 2020) | Põlme S, et al. 2020. Fungal Diversity 105:1-16 ((https://doi.org/10.1007/s13225-020-00466-2)) |
| Otu24415 | 1 | k__Fungi(100);p__Ascomycota(100);c__Saccharomycetes(100);o__Saccharomycetales(100);f__Metschnikowiaceae(100);g__Metschnikowia(100);s__Metschnikowia_pulcherrima(100); | Metschnikowia | 13 | Saprotroph | \|Undefined Saprotroph\| | Yeast | NULL | Probable | As nectar-tap_saprotroph (Põlme et al. 2020); Decay sugar-rich substrates (Põlme et al. 2020) | Põlme S, et al. 2020. Fungal Diversity 105:1-16 ((https://doi.org/10.1007/s13225-020-00466-2)) |
| Otu24416 | 1 | k__Fungi(100);p__Ascomycota(100);c__Saccharomycetes(100);o__Saccharomycetales(100);f__Metschnikowiaceae(100);g__Metschnikowia(100);g__Metschnikowia_unclassified(100); | Metschnikowia | 13 | Saprotroph | \|Undefined Saprotroph\| | Yeast | NULL | Probable | As nectar-tap_saprotroph (Põlme et al. 2020); Decay sugar-rich substrates (Põlme et al. 2020) | Põlme S, et al. 2020. Fungal Diversity 105:1-16 ((https://doi.org/10.1007/s13225-020-00466-2)) |
| Otu24417 | 1 | k__Fungi(100);p__Ascomycota(100);c__Saccharomycetes(100);o__Saccharomycetales(100);f__Metschnikowiaceae(100);g__Metschnikowia(100);s__Metschnikowia_pulcherrima(100); | Metschnikowia | 13 | Saprotroph | \|Undefined Saprotroph\| | Yeast | NULL | Probable | As nectar-tap_saprotroph (Põlme et al. 2020); Decay sugar-rich substrates (Põlme et al. 2020) | Põlme S, et al. 2020. Fungal Diversity 105:1-16 ((https://doi.org/10.1007/s13225-020-00466-2)) |
| Otu24420 | 1 | k__Fungi(100);p__Ascomycota(100);c__Saccharomycetes(100);o__Saccharomycetales(100);f__Metschnikowiaceae(100);g__Metschnikowia(100);g__Metschnikowia_unclassified(100); | Metschnikowia | 13 | Saprotroph | \|Undefined Saprotroph\| | Yeast | NULL | Probable | As nectar-tap_saprotroph (Põlme et al. 2020); Decay sugar-rich substrates (Põlme et al. 2020) | Põlme S, et al. 2020. Fungal Diversity 105:1-16 ((https://doi.org/10.1007/s13225-020-00466-2)) |
| Otu24421 | 1 | k__Fungi(100);p__Ascomycota(100);c__Saccharomycetes(100);o__Saccharomycetales(100);f__Metschnikowiaceae(100);g__Metschnikowia(100);g__Metschnikowia_unclassified(100); | Metschnikowia | 13 | Saprotroph | \|Undefined Saprotroph\| | Yeast | NULL | Probable | As nectar-tap_saprotroph (Põlme et al. 2020); Decay sugar-rich substrates (Põlme et al. 2020) | Põlme S, et al. 2020. Fungal Diversity 105:1-16 ((https://doi.org/10.1007/s13225-020-00466-2)) |
| Otu24422 | 1 | k__Fungi(100);p__Ascomycota(100);c__Saccharomycetes(100);o__Saccharomycetales(100);f__Metschnikowiaceae(100);g__Metschnikowia(100);s__Metschnikowia_pulcherrima(100); | Metschnikowia | 13 | Saprotroph | \|Undefined Saprotroph\| | Yeast | NULL | Probable | As nectar-tap_saprotroph (Põlme et al. 2020); Decay sugar-rich substrates (Põlme et al. 2020) | Põlme S, et al. 2020. Fungal Diversity 105:1-16 ((https://doi.org/10.1007/s13225-020-00466-2)) |
| Otu24423 | 1 | k__Fungi(100);p__Ascomycota(100);c__Saccharomycetes(100);o__Saccharomycetales(100);f__Metschnikowiaceae(100);g__Metschnikowia(100);s__Metschnikowia_pulcherrima(100); | Metschnikowia | 13 | Saprotroph | \|Undefined Saprotroph\| | Yeast | NULL | Probable | As nectar-tap_saprotroph (Põlme et al. 2020); Decay sugar-rich substrates (Põlme et al. 2020) | Põlme S, et al. 2020. Fungal Diversity 105:1-16 ((https://doi.org/10.1007/s13225-020-00466-2)) |
| Otu24424 | 1 | k__Fungi(100);p__Ascomycota(100);c__Saccharomycetes(100);o__Saccharomycetales(100);f__Metschnikowiaceae(100);g__Metschnikowia(100);g__Metschnikowia_unclassified(100); | Metschnikowia | 13 | Saprotroph | \|Undefined Saprotroph\| | Yeast | NULL | Probable | As nectar-tap_saprotroph (Põlme et al. 2020); Decay sugar-rich substrates (Põlme et al. 2020) | Põlme S, et al. 2020. Fungal Diversity 105:1-16 ((https://doi.org/10.1007/s13225-020-00466-2)) |
| Otu24439 | 1 | k__Fungi(100);p__Ascomycota(100);c__Saccharomycetes(100);o__Saccharomycetales(100);f__Metschnikowiaceae(100);g__Metschnikowia(100);s__Metschnikowia_pulcherrima(100); | Metschnikowia | 13 | Saprotroph | \|Undefined Saprotroph\| | Yeast | NULL | Probable | As nectar-tap_saprotroph (Põlme et al. 2020); Decay sugar-rich substrates (Põlme et al. 2020) | Põlme S, et al. 2020. Fungal Diversity 105:1-16 ((https://doi.org/10.1007/s13225-020-00466-2)) |
| Otu24473 | 1 | k__Fungi(100);p__Ascomycota(100);c__Saccharomycetes(100);o__Saccharomycetales(100);f__Metschnikowiaceae(100);g__Metschnikowia(100);s__Metschnikowia_pulcherrima(100); | Metschnikowia | 13 | Saprotroph | \|Undefined Saprotroph\| | Yeast | NULL | Probable | As nectar-tap_saprotroph (Põlme et al. 2020); Decay sugar-rich substrates (Põlme et al. 2020) | Põlme S, et al. 2020. Fungal Diversity 105:1-16 ((https://doi.org/10.1007/s13225-020-00466-2)) |
| Otu24474 | 1 | k__Fungi(100);p__Ascomycota(100);c__Saccharomycetes(100);o__Saccharomycetales(100);f__Metschnikowiaceae(100);g__Metschnikowia(100);s__Metschnikowia_pulcherrima(100); | Metschnikowia | 13 | Saprotroph | \|Undefined Saprotroph\| | Yeast | NULL | Probable | As nectar-tap_saprotroph (Põlme et al. 2020); Decay sugar-rich substrates (Põlme et al. 2020) | Põlme S, et al. 2020. Fungal Diversity 105:1-16 ((https://doi.org/10.1007/s13225-020-00466-2)) |
| Otu24477 | 1 | k__Fungi(100);p__Ascomycota(100);c__Saccharomycetes(100);o__Saccharomycetales(100);f__Metschnikowiaceae(100);g__Metschnikowia(100);g__Metschnikowia_unclassified(100); | Metschnikowia | 13 | Saprotroph | \|Undefined Saprotroph\| | Yeast | NULL | Probable | As nectar-tap_saprotroph (Põlme et al. 2020); Decay sugar-rich substrates (Põlme et al. 2020) | Põlme S, et al. 2020. Fungal Diversity 105:1-16 ((https://doi.org/10.1007/s13225-020-00466-2)) |
| Otu24479 | 1 | k__Fungi(100);p__Ascomycota(100);c__Saccharomycetes(100);o__Saccharomycetales(100);f__Metschnikowiaceae(100);g__Metschnikowia(100);g__Metschnikowia_unclassified(100); | Metschnikowia | 13 | Saprotroph | \|Undefined Saprotroph\| | Yeast | NULL | Probable | As nectar-tap_saprotroph (Põlme et al. 2020); Decay sugar-rich substrates (Põlme et al. 2020) | Põlme S, et al. 2020. Fungal Diversity 105:1-16 ((https://doi.org/10.1007/s13225-020-00466-2)) |
| Otu24480 | 1 | k__Fungi(100);p__Ascomycota(100);c__Saccharomycetes(100);o__Saccharomycetales(100);f__Metschnikowiaceae(100);g__Metschnikowia(100);s__Metschnikowia_pulcherrima(100); | Metschnikowia | 13 | Saprotroph | \|Undefined Saprotroph\| | Yeast | NULL | Probable | As nectar-tap_saprotroph (Põlme et al. 2020); Decay sugar-rich substrates (Põlme et al. 2020) | Põlme S, et al. 2020. Fungal Diversity 105:1-16 ((https://doi.org/10.1007/s13225-020-00466-2)) |
| Otu24483 | 1 | k__Fungi(100);p__Ascomycota(100);c__Saccharomycetes(100);o__Saccharomycetales(100);f__Metschnikowiaceae(100);g__Metschnikowia(100);s__Metschnikowia_pulcherrima(100); | Metschnikowia | 13 | Saprotroph | \|Undefined Saprotroph\| | Yeast | NULL | Probable | As nectar-tap_saprotroph (Põlme et al. 2020); Decay sugar-rich substrates (Põlme et al. 2020) | Põlme S, et al. 2020. Fungal Diversity 105:1-16 ((https://doi.org/10.1007/s13225-020-00466-2)) |
| Otu24484 | 1 | k__Fungi(100);p__Ascomycota(100);c__Saccharomycetes(100);o__Saccharomycetales(100);f__Metschnikowiaceae(100);g__Metschnikowia(100);s__Metschnikowia_pulcherrima(100); | Metschnikowia | 13 | Saprotroph | \|Undefined Saprotroph\| | Yeast | NULL | Probable | As nectar-tap_saprotroph (Põlme et al. 2020); Decay sugar-rich substrates (Põlme et al. 2020) | Põlme S, et al. 2020. Fungal Diversity 105:1-16 ((https://doi.org/10.1007/s13225-020-00466-2)) |
| Otu24485 | 1 | k__Fungi(100);p__Ascomycota(100);c__Saccharomycetes(100);o__Saccharomycetales(100);f__Metschnikowiaceae(100);g__Metschnikowia(100);s__Metschnikowia_pulcherrima(100); | Metschnikowia | 13 | Saprotroph | \|Undefined Saprotroph\| | Yeast | NULL | Probable | As nectar-tap_saprotroph (Põlme et al. 2020); Decay sugar-rich substrates (Põlme et al. 2020) | Põlme S, et al. 2020. Fungal Diversity 105:1-16 ((https://doi.org/10.1007/s13225-020-00466-2)) |
| Otu24486 | 1 | k__Fungi(100);p__Ascomycota(100);c__Saccharomycetes(100);o__Saccharomycetales(100);f__Metschnikowiaceae(100);g__Metschnikowia(100);g__Metschnikowia_unclassified(100); | Metschnikowia | 13 | Saprotroph | \|Undefined Saprotroph\| | Yeast | NULL | Probable | As nectar-tap_saprotroph (Põlme et al. 2020); Decay sugar-rich substrates (Põlme et al. 2020) | Põlme S, et al. 2020. Fungal Diversity 105:1-16 ((https://doi.org/10.1007/s13225-020-00466-2)) |
| Otu24487 | 1 | k__Fungi(100);p__Ascomycota(100);c__Saccharomycetes(100);o__Saccharomycetales(100);f__Metschnikowiaceae(100);g__Metschnikowia(100);s__Metschnikowia_pulcherrima(100); | Metschnikowia | 13 | Saprotroph | \|Undefined Saprotroph\| | Yeast | NULL | Probable | As nectar-tap_saprotroph (Põlme et al. 2020); Decay sugar-rich substrates (Põlme et al. 2020) | Põlme S, et al. 2020. Fungal Diversity 105:1-16 ((https://doi.org/10.1007/s13225-020-00466-2)) |
| Otu24488 | 1 | k__Fungi(100);p__Ascomycota(100);c__Saccharomycetes(100);o__Saccharomycetales(100);f__Metschnikowiaceae(100);g__Metschnikowia(100);g__Metschnikowia_unclassified(100); | Metschnikowia | 13 | Saprotroph | \|Undefined Saprotroph\| | Yeast | NULL | Probable | As nectar-tap_saprotroph (Põlme et al. 2020); Decay sugar-rich substrates (Põlme et al. 2020) | Põlme S, et al. 2020. Fungal Diversity 105:1-16 ((https://doi.org/10.1007/s13225-020-00466-2)) |
| Otu24489 | 1 | k__Fungi(100);p__Ascomycota(100);c__Saccharomycetes(100);o__Saccharomycetales(100);f__Metschnikowiaceae(100);g__Metschnikowia(100);s__Metschnikowia_pulcherrima(100); | Metschnikowia | 13 | Saprotroph | \|Undefined Saprotroph\| | Yeast | NULL | Probable | As nectar-tap_saprotroph (Põlme et al. 2020); Decay sugar-rich substrates (Põlme et al. 2020) | Põlme S, et al. 2020. Fungal Diversity 105:1-16 ((https://doi.org/10.1007/s13225-020-00466-2)) |
| Otu24491 | 1 | k__Fungi(100);p__Ascomycota(100);c__Saccharomycetes(100);o__Saccharomycetales(100);f__Metschnikowiaceae(100);g__Metschnikowia(100);g__Metschnikowia_unclassified(100); | Metschnikowia | 13 | Saprotroph | \|Undefined Saprotroph\| | Yeast | NULL | Probable | As nectar-tap_saprotroph (Põlme et al. 2020); Decay sugar-rich substrates (Põlme et al. 2020) | Põlme S, et al. 2020. Fungal Diversity 105:1-16 ((https://doi.org/10.1007/s13225-020-00466-2)) |
| Otu24492 | 1 | k__Fungi(100);p__Ascomycota(100);c__Saccharomycetes(100);o__Saccharomycetales(100);f__Metschnikowiaceae(100);g__Metschnikowia(100);s__Metschnikowia_pulcherrima(100); | Metschnikowia | 13 | Saprotroph | \|Undefined Saprotroph\| | Yeast | NULL | Probable | As nectar-tap_saprotroph (Põlme et al. 2020); Decay sugar-rich substrates (Põlme et al. 2020) | Põlme S, et al. 2020. Fungal Diversity 105:1-16 ((https://doi.org/10.1007/s13225-020-00466-2)) |
| Otu24495 | 1 | k__Fungi(100);p__Ascomycota(100);c__Saccharomycetes(100);o__Saccharomycetales(100);f__Metschnikowiaceae(100);g__Metschnikowia(100);s__Metschnikowia_pulcherrima(100); | Metschnikowia | 13 | Saprotroph | \|Undefined Saprotroph\| | Yeast | NULL | Probable | As nectar-tap_saprotroph (Põlme et al. 2020); Decay sugar-rich substrates (Põlme et al. 2020) | Põlme S, et al. 2020. Fungal Diversity 105:1-16 ((https://doi.org/10.1007/s13225-020-00466-2)) |
| Otu24496 | 1 | k__Fungi(100);p__Ascomycota(100);c__Saccharomycetes(100);o__Saccharomycetales(100);f__Metschnikowiaceae(100);g__Metschnikowia(100);s__Metschnikowia_pulcherrima(100); | Metschnikowia | 13 | Saprotroph | \|Undefined Saprotroph\| | Yeast | NULL | Probable | As nectar-tap_saprotroph (Põlme et al. 2020); Decay sugar-rich substrates (Põlme et al. 2020) | Põlme S, et al. 2020. Fungal Diversity 105:1-16 ((https://doi.org/10.1007/s13225-020-00466-2)) |
| Otu24498 | 1 | k__Fungi(100);p__Ascomycota(100);c__Saccharomycetes(100);o__Saccharomycetales(100);f__Metschnikowiaceae(100);g__Metschnikowia(100);g__Metschnikowia_unclassified(100); | Metschnikowia | 13 | Saprotroph | \|Undefined Saprotroph\| | Yeast | NULL | Probable | As nectar-tap_saprotroph (Põlme et al. 2020); Decay sugar-rich substrates (Põlme et al. 2020) | Põlme S, et al. 2020. Fungal Diversity 105:1-16 ((https://doi.org/10.1007/s13225-020-00466-2)) |
| Otu24531 | 1 | k__Fungi(100);p__Ascomycota(100);c__Saccharomycetes(100);o__Saccharomycetales(100);f__Metschnikowiaceae(100);g__Metschnikowia(100);g__Metschnikowia_unclassified(100); | Metschnikowia | 13 | Saprotroph | \|Undefined Saprotroph\| | Yeast | NULL | Probable | As nectar-tap_saprotroph (Põlme et al. 2020); Decay sugar-rich substrates (Põlme et al. 2020) | Põlme S, et al. 2020. Fungal Diversity 105:1-16 ((https://doi.org/10.1007/s13225-020-00466-2)) |
| Otu24550 | 1 | k__Fungi(100);p__Ascomycota(100);c__Saccharomycetes(100);o__Saccharomycetales(100);f__Metschnikowiaceae(100);g__Metschnikowia(100);s__Metschnikowia_pulcherrima(100); | Metschnikowia | 13 | Saprotroph | \|Undefined Saprotroph\| | Yeast | NULL | Probable | As nectar-tap_saprotroph (Põlme et al. 2020); Decay sugar-rich substrates (Põlme et al. 2020) | Põlme S, et al. 2020. Fungal Diversity 105:1-16 ((https://doi.org/10.1007/s13225-020-00466-2)) |
| Otu24559 | 1 | k__Fungi(100);p__Ascomycota(100);c__Saccharomycetes(100);o__Saccharomycetales(100);f__Metschnikowiaceae(100);g__Metschnikowia(100);g__Metschnikowia_unclassified(100); | Metschnikowia | 13 | Saprotroph | \|Undefined Saprotroph\| | Yeast | NULL | Probable | As nectar-tap_saprotroph (Põlme et al. 2020); Decay sugar-rich substrates (Põlme et al. 2020) | Põlme S, et al. 2020. Fungal Diversity 105:1-16 ((https://doi.org/10.1007/s13225-020-00466-2)) |
| Otu24561 | 1 | k__Fungi(100);p__Ascomycota(100);c__Saccharomycetes(100);o__Saccharomycetales(100);f__Metschnikowiaceae(100);g__Metschnikowia(100);g__Metschnikowia_unclassified(100); | Metschnikowia | 13 | Saprotroph | \|Undefined Saprotroph\| | Yeast | NULL | Probable | As nectar-tap_saprotroph (Põlme et al. 2020); Decay sugar-rich substrates (Põlme et al. 2020) | Põlme S, et al. 2020. Fungal Diversity 105:1-16 ((https://doi.org/10.1007/s13225-020-00466-2)) |
| Otu24564 | 1 | k__Fungi(100);p__Ascomycota(100);c__Saccharomycetes(100);o__Saccharomycetales(100);f__Metschnikowiaceae(100);g__Metschnikowia(100);s__Metschnikowia_pulcherrima(100); | Metschnikowia | 13 | Saprotroph | \|Undefined Saprotroph\| | Yeast | NULL | Probable | As nectar-tap_saprotroph (Põlme et al. 2020); Decay sugar-rich substrates (Põlme et al. 2020) | Põlme S, et al. 2020. Fungal Diversity 105:1-16 ((https://doi.org/10.1007/s13225-020-00466-2)) |
| Otu24576 | 1 | k__Fungi(100);p__Ascomycota(100);c__Saccharomycetes(100);o__Saccharomycetales(100);f__Metschnikowiaceae(100);g__Metschnikowia(100);s__Metschnikowia_pulcherrima(100); | Metschnikowia | 13 | Saprotroph | \|Undefined Saprotroph\| | Yeast | NULL | Probable | As nectar-tap_saprotroph (Põlme et al. 2020); Decay sugar-rich substrates (Põlme et al. 2020) | Põlme S, et al. 2020. Fungal Diversity 105:1-16 ((https://doi.org/10.1007/s13225-020-00466-2)) |
| Otu24579 | 1 | k__Fungi(100);p__Ascomycota(100);c__Saccharomycetes(100);o__Saccharomycetales(100);f__Metschnikowiaceae(100);g__Metschnikowia(100);s__Metschnikowia_pulcherrima(100); | Metschnikowia | 13 | Saprotroph | \|Undefined Saprotroph\| | Yeast | NULL | Probable | As nectar-tap_saprotroph (Põlme et al. 2020); Decay sugar-rich substrates (Põlme et al. 2020) | Põlme S, et al. 2020. Fungal Diversity 105:1-16 ((https://doi.org/10.1007/s13225-020-00466-2)) |
| Otu24581 | 1 | k__Fungi(100);p__Ascomycota(100);c__Saccharomycetes(100);o__Saccharomycetales(100);f__Metschnikowiaceae(100);g__Metschnikowia(100);s__Metschnikowia_pulcherrima(100); | Metschnikowia | 13 | Saprotroph | \|Undefined Saprotroph\| | Yeast | NULL | Probable | As nectar-tap_saprotroph (Põlme et al. 2020); Decay sugar-rich substrates (Põlme et al. 2020) | Põlme S, et al. 2020. Fungal Diversity 105:1-16 ((https://doi.org/10.1007/s13225-020-00466-2)) |
| Otu24582 | 1 | k__Fungi(100);p__Ascomycota(100);c__Saccharomycetes(100);o__Saccharomycetales(100);f__Metschnikowiaceae(100);g__Metschnikowia(100);s__Metschnikowia_pulcherrima(100); | Metschnikowia | 13 | Saprotroph | \|Undefined Saprotroph\| | Yeast | NULL | Probable | As nectar-tap_saprotroph (Põlme et al. 2020); Decay sugar-rich substrates (Põlme et al. 2020) | Põlme S, et al. 2020. Fungal Diversity 105:1-16 ((https://doi.org/10.1007/s13225-020-00466-2)) |
| Otu24583 | 1 | k__Fungi(100);p__Ascomycota(100);c__Saccharomycetes(100);o__Saccharomycetales(100);f__Metschnikowiaceae(100);g__Metschnikowia(100);s__Metschnikowia_pulcherrima(100); | Metschnikowia | 13 | Saprotroph | \|Undefined Saprotroph\| | Yeast | NULL | Probable | As nectar-tap_saprotroph (Põlme et al. 2020); Decay sugar-rich substrates (Põlme et al. 2020) | Põlme S, et al. 2020. Fungal Diversity 105:1-16 ((https://doi.org/10.1007/s13225-020-00466-2)) |
| Otu24584 | 1 | k__Fungi(100);p__Ascomycota(100);c__Saccharomycetes(100);o__Saccharomycetales(100);f__Metschnikowiaceae(100);g__Metschnikowia(100);g__Metschnikowia_unclassified(100); | Metschnikowia | 13 | Saprotroph | \|Undefined Saprotroph\| | Yeast | NULL | Probable | As nectar-tap_saprotroph (Põlme et al. 2020); Decay sugar-rich substrates (Põlme et al. 2020) | Põlme S, et al. 2020. Fungal Diversity 105:1-16 ((https://doi.org/10.1007/s13225-020-00466-2)) |
| Otu24586 | 1 | k__Fungi(100);p__Ascomycota(100);c__Saccharomycetes(100);o__Saccharomycetales(100);f__Metschnikowiaceae(100);g__Metschnikowia(100);s__Metschnikowia_pulcherrima(100); | Metschnikowia | 13 | Saprotroph | \|Undefined Saprotroph\| | Yeast | NULL | Probable | As nectar-tap_saprotroph (Põlme et al. 2020); Decay sugar-rich substrates (Põlme et al. 2020) | Põlme S, et al. 2020. Fungal Diversity 105:1-16 ((https://doi.org/10.1007/s13225-020-00466-2)) |
| Otu24587 | 1 | k__Fungi(100);p__Ascomycota(100);c__Saccharomycetes(100);o__Saccharomycetales(100);f__Metschnikowiaceae(100);g__Metschnikowia(100);s__Metschnikowia_pulcherrima(100); | Metschnikowia | 13 | Saprotroph | \|Undefined Saprotroph\| | Yeast | NULL | Probable | As nectar-tap_saprotroph (Põlme et al. 2020); Decay sugar-rich substrates (Põlme et al. 2020) | Põlme S, et al. 2020. Fungal Diversity 105:1-16 ((https://doi.org/10.1007/s13225-020-00466-2)) |
| Otu24589 | 1 | k__Fungi(100);p__Ascomycota(100);c__Saccharomycetes(100);o__Saccharomycetales(100);f__Metschnikowiaceae(100);g__Metschnikowia(100);s__Metschnikowia_pulcherrima(100); | Metschnikowia | 13 | Saprotroph | \|Undefined Saprotroph\| | Yeast | NULL | Probable | As nectar-tap_saprotroph (Põlme et al. 2020); Decay sugar-rich substrates (Põlme et al. 2020) | Põlme S, et al. 2020. Fungal Diversity 105:1-16 ((https://doi.org/10.1007/s13225-020-00466-2)) |
| Otu24598 | 1 | k__Fungi(100);p__Ascomycota(100);c__Saccharomycetes(100);o__Saccharomycetales(100);f__Metschnikowiaceae(100);g__Metschnikowia(100);s__Metschnikowia_pulcherrima(100); | Metschnikowia | 13 | Saprotroph | \|Undefined Saprotroph\| | Yeast | NULL | Probable | As nectar-tap_saprotroph (Põlme et al. 2020); Decay sugar-rich substrates (Põlme et al. 2020) | Põlme S, et al. 2020. Fungal Diversity 105:1-16 ((https://doi.org/10.1007/s13225-020-00466-2)) |
| Otu24612 | 1 | k__Fungi(100);p__Ascomycota(100);c__Saccharomycetes(100);o__Saccharomycetales(100);f__Metschnikowiaceae(100);g__Metschnikowia(100);s__Metschnikowia_pulcherrima(100); | Metschnikowia | 13 | Saprotroph | \|Undefined Saprotroph\| | Yeast | NULL | Probable | As nectar-tap_saprotroph (Põlme et al. 2020); Decay sugar-rich substrates (Põlme et al. 2020) | Põlme S, et al. 2020. Fungal Diversity 105:1-16 ((https://doi.org/10.1007/s13225-020-00466-2)) |
| Otu24613 | 1 | k__Fungi(100);p__Ascomycota(100);c__Saccharomycetes(100);o__Saccharomycetales(100);f__Metschnikowiaceae(100);g__Metschnikowia(100);s__Metschnikowia_pulcherrima(100); | Metschnikowia | 13 | Saprotroph | \|Undefined Saprotroph\| | Yeast | NULL | Probable | As nectar-tap_saprotroph (Põlme et al. 2020); Decay sugar-rich substrates (Põlme et al. 2020) | Põlme S, et al. 2020. Fungal Diversity 105:1-16 ((https://doi.org/10.1007/s13225-020-00466-2)) |
| Otu24614 | 1 | k__Fungi(100);p__Ascomycota(100);c__Saccharomycetes(100);o__Saccharomycetales(100);f__Metschnikowiaceae(100);g__Metschnikowia(100);s__Metschnikowia_pulcherrima(100); | Metschnikowia | 13 | Saprotroph | \|Undefined Saprotroph\| | Yeast | NULL | Probable | As nectar-tap_saprotroph (Põlme et al. 2020); Decay sugar-rich substrates (Põlme et al. 2020) | Põlme S, et al. 2020. Fungal Diversity 105:1-16 ((https://doi.org/10.1007/s13225-020-00466-2)) |
| Otu24615 | 1 | k__Fungi(100);p__Ascomycota(100);c__Saccharomycetes(100);o__Saccharomycetales(100);f__Metschnikowiaceae(100);g__Metschnikowia(100);s__Metschnikowia_pulcherrima(100); | Metschnikowia | 13 | Saprotroph | \|Undefined Saprotroph\| | Yeast | NULL | Probable | As nectar-tap_saprotroph (Põlme et al. 2020); Decay sugar-rich substrates (Põlme et al. 2020) | Põlme S, et al. 2020. Fungal Diversity 105:1-16 ((https://doi.org/10.1007/s13225-020-00466-2)) |
| Otu24616 | 1 | k__Fungi(100);p__Ascomycota(100);c__Saccharomycetes(100);o__Saccharomycetales(100);f__Saccharomycetales_family_Incertae_sedis(100);g__Candida(100);g__Candida_unclassified(100); | Candida | 13 | Pathotroph-Saprotroph-Symbiotroph | Animal Pathogen-Endophyte-Endosymbiont-Epiphyte-\|Undefined Saprotroph\| | Dimorphic Yeast-Yeast | NULL | Probable | As nectar-tap_saprotroph (Põlme et al. 2020); Decay sugar-rich substrates (Põlme et al. 2020); This asexual genus is polyphyletic and contain species that function very differently depend on what they are. Caution should be taken when interpreting guilds for Candida species. | Manolakaki D, et al. 2010. Virulence. 1: 367-375 ((https://doi.org/10.4161/viru.1.5.12796)); Mekbib SB, et al. 2011. Tropical Plant Pathology 36:233-240 ((https://doi.org/10.1590/S1982-56762011000400004)); Põlme S, et al. 2020. Fungal Diversity 105:1-16 ((https://doi.org/10.1007/s13225-020-00466-2)) |
| Otu24618 | 1 | k__Fungi(100);p__Ascomycota(100);c__Saccharomycetes(100);o__Saccharomycetales(100);f__Metschnikowiaceae(100);g__Metschnikowia(100);g__Metschnikowia_unclassified(100); | Metschnikowia | 13 | Saprotroph | \|Undefined Saprotroph\| | Yeast | NULL | Probable | As nectar-tap_saprotroph (Põlme et al. 2020); Decay sugar-rich substrates (Põlme et al. 2020) | Põlme S, et al. 2020. Fungal Diversity 105:1-16 ((https://doi.org/10.1007/s13225-020-00466-2)) |
| Otu24619 | 1 | k__Fungi(100);p__Ascomycota(100);c__Saccharomycetes(100);o__Saccharomycetales(100);f__Metschnikowiaceae(100);g__Metschnikowia(100);s__Metschnikowia_pulcherrima(100); | Metschnikowia | 13 | Saprotroph | \|Undefined Saprotroph\| | Yeast | NULL | Probable | As nectar-tap_saprotroph (Põlme et al. 2020); Decay sugar-rich substrates (Põlme et al. 2020) | Põlme S, et al. 2020. Fungal Diversity 105:1-16 ((https://doi.org/10.1007/s13225-020-00466-2)) |
| Otu24620 | 1 | k__Fungi(100);p__Ascomycota(100);c__Saccharomycetes(100);o__Saccharomycetales(100);f__Metschnikowiaceae(100);g__Metschnikowia(100);s__Metschnikowia_pulcherrima(100); | Metschnikowia | 13 | Saprotroph | \|Undefined Saprotroph\| | Yeast | NULL | Probable | As nectar-tap_saprotroph (Põlme et al. 2020); Decay sugar-rich substrates (Põlme et al. 2020) | Põlme S, et al. 2020. Fungal Diversity 105:1-16 ((https://doi.org/10.1007/s13225-020-00466-2)) |
| Otu24621 | 1 | k__Fungi(100);p__Ascomycota(100);c__Saccharomycetes(100);o__Saccharomycetales(100);f__Metschnikowiaceae(100);g__Metschnikowia(100);g__Metschnikowia_unclassified(100); | Metschnikowia | 13 | Saprotroph | \|Undefined Saprotroph\| | Yeast | NULL | Probable | As nectar-tap_saprotroph (Põlme et al. 2020); Decay sugar-rich substrates (Põlme et al. 2020) | Põlme S, et al. 2020. Fungal Diversity 105:1-16 ((https://doi.org/10.1007/s13225-020-00466-2)) |
| Otu24622 | 1 | k__Fungi(100);p__Ascomycota(100);c__Saccharomycetes(100);o__Saccharomycetales(100);f__Metschnikowiaceae(100);g__Metschnikowia(100);g__Metschnikowia_unclassified(100); | Metschnikowia | 13 | Saprotroph | \|Undefined Saprotroph\| | Yeast | NULL | Probable | As nectar-tap_saprotroph (Põlme et al. 2020); Decay sugar-rich substrates (Põlme et al. 2020) | Põlme S, et al. 2020. Fungal Diversity 105:1-16 ((https://doi.org/10.1007/s13225-020-00466-2)) |
| Otu24623 | 1 | k__Fungi(100);p__Ascomycota(100);c__Saccharomycetes(100);o__Saccharomycetales(100);f__Metschnikowiaceae(100);g__Metschnikowia(100);s__Metschnikowia_pulcherrima(100); | Metschnikowia | 13 | Saprotroph | \|Undefined Saprotroph\| | Yeast | NULL | Probable | As nectar-tap_saprotroph (Põlme et al. 2020); Decay sugar-rich substrates (Põlme et al. 2020) | Põlme S, et al. 2020. Fungal Diversity 105:1-16 ((https://doi.org/10.1007/s13225-020-00466-2)) |
| Otu24624 | 1 | k__Fungi(100);p__Ascomycota(100);c__Saccharomycetes(100);o__Saccharomycetales(100);f__Metschnikowiaceae(100);g__Metschnikowia(100);s__Metschnikowia_pulcherrima(100); | Metschnikowia | 13 | Saprotroph | \|Undefined Saprotroph\| | Yeast | NULL | Probable | As nectar-tap_saprotroph (Põlme et al. 2020); Decay sugar-rich substrates (Põlme et al. 2020) | Põlme S, et al. 2020. Fungal Diversity 105:1-16 ((https://doi.org/10.1007/s13225-020-00466-2)) |
| Otu24626 | 1 | k__Fungi(100);p__Ascomycota(100);c__Saccharomycetes(100);o__Saccharomycetales(100);f__Metschnikowiaceae(100);g__Metschnikowia(100);g__Metschnikowia_unclassified(100); | Metschnikowia | 13 | Saprotroph | \|Undefined Saprotroph\| | Yeast | NULL | Probable | As nectar-tap_saprotroph (Põlme et al. 2020); Decay sugar-rich substrates (Põlme et al. 2020) | Põlme S, et al. 2020. Fungal Diversity 105:1-16 ((https://doi.org/10.1007/s13225-020-00466-2)) |
| Otu24627 | 1 | k__Fungi(100);p__Ascomycota(100);c__Saccharomycetes(100);o__Saccharomycetales(100);f__Metschnikowiaceae(100);g__Metschnikowia(100);s__Metschnikowia_pulcherrima(100); | Metschnikowia | 13 | Saprotroph | \|Undefined Saprotroph\| | Yeast | NULL | Probable | As nectar-tap_saprotroph (Põlme et al. 2020); Decay sugar-rich substrates (Põlme et al. 2020) | Põlme S, et al. 2020. Fungal Diversity 105:1-16 ((https://doi.org/10.1007/s13225-020-00466-2)) |
| Otu24628 | 1 | k__Fungi(100);p__Ascomycota(100);c__Saccharomycetes(100);o__Saccharomycetales(100);f__Metschnikowiaceae(100);g__Metschnikowia(100);g__Metschnikowia_unclassified(100); | Metschnikowia | 13 | Saprotroph | \|Undefined Saprotroph\| | Yeast | NULL | Probable | As nectar-tap_saprotroph (Põlme et al. 2020); Decay sugar-rich substrates (Põlme et al. 2020) | Põlme S, et al. 2020. Fungal Diversity 105:1-16 ((https://doi.org/10.1007/s13225-020-00466-2)) |
| Otu24629 | 1 | k__Fungi(100);p__Ascomycota(100);c__Saccharomycetes(100);o__Saccharomycetales(100);f__Metschnikowiaceae(100);g__Metschnikowia(100);s__Metschnikowia_pulcherrima(100); | Metschnikowia | 13 | Saprotroph | \|Undefined Saprotroph\| | Yeast | NULL | Probable | As nectar-tap_saprotroph (Põlme et al. 2020); Decay sugar-rich substrates (Põlme et al. 2020) | Põlme S, et al. 2020. Fungal Diversity 105:1-16 ((https://doi.org/10.1007/s13225-020-00466-2)) |
| Otu24633 | 1 | k__Fungi(100);p__Ascomycota(100);c__Saccharomycetes(100);o__Saccharomycetales(100);f__Metschnikowiaceae(100);g__Metschnikowia(100);s__Metschnikowia_pulcherrima(100); | Metschnikowia | 13 | Saprotroph | \|Undefined Saprotroph\| | Yeast | NULL | Probable | As nectar-tap_saprotroph (Põlme et al. 2020); Decay sugar-rich substrates (Põlme et al. 2020) | Põlme S, et al. 2020. Fungal Diversity 105:1-16 ((https://doi.org/10.1007/s13225-020-00466-2)) |
| Otu24638 | 1 | k__Fungi(100);p__Ascomycota(100);c__Saccharomycetes(100);o__Saccharomycetales(100);f__Metschnikowiaceae(100);g__Metschnikowia(100);s__Metschnikowia_pulcherrima(100); | Metschnikowia | 13 | Saprotroph | \|Undefined Saprotroph\| | Yeast | NULL | Probable | As nectar-tap_saprotroph (Põlme et al. 2020); Decay sugar-rich substrates (Põlme et al. 2020) | Põlme S, et al. 2020. Fungal Diversity 105:1-16 ((https://doi.org/10.1007/s13225-020-00466-2)) |
| Otu24639 | 1 | k__Fungi(100);p__Ascomycota(100);c__Saccharomycetes(100);o__Saccharomycetales(100);f__Metschnikowiaceae(100);g__Metschnikowia(100);g__Metschnikowia_unclassified(100); | Metschnikowia | 13 | Saprotroph | \|Undefined Saprotroph\| | Yeast | NULL | Probable | As nectar-tap_saprotroph (Põlme et al. 2020); Decay sugar-rich substrates (Põlme et al. 2020) | Põlme S, et al. 2020. Fungal Diversity 105:1-16 ((https://doi.org/10.1007/s13225-020-00466-2)) |
| Otu24640 | 1 | k__Fungi(100);p__Ascomycota(100);c__Saccharomycetes(100);o__Saccharomycetales(100);f__Metschnikowiaceae(100);g__Metschnikowia(100);g__Metschnikowia_unclassified(100); | Metschnikowia | 13 | Saprotroph | \|Undefined Saprotroph\| | Yeast | NULL | Probable | As nectar-tap_saprotroph (Põlme et al. 2020); Decay sugar-rich substrates (Põlme et al. 2020) | Põlme S, et al. 2020. Fungal Diversity 105:1-16 ((https://doi.org/10.1007/s13225-020-00466-2)) |
| Otu24641 | 1 | k__Fungi(100);p__Ascomycota(100);c__Saccharomycetes(100);o__Saccharomycetales(100);f__Metschnikowiaceae(100);g__Metschnikowia(100);s__Metschnikowia_pulcherrima(100); | Metschnikowia | 13 | Saprotroph | \|Undefined Saprotroph\| | Yeast | NULL | Probable | As nectar-tap_saprotroph (Põlme et al. 2020); Decay sugar-rich substrates (Põlme et al. 2020) | Põlme S, et al. 2020. Fungal Diversity 105:1-16 ((https://doi.org/10.1007/s13225-020-00466-2)) |
| Otu24642 | 1 | k__Fungi(100);p__Ascomycota(100);c__Saccharomycetes(100);o__Saccharomycetales(100);f__Metschnikowiaceae(100);g__Metschnikowia(100);s__Metschnikowia_pulcherrima(100); | Metschnikowia | 13 | Saprotroph | \|Undefined Saprotroph\| | Yeast | NULL | Probable | As nectar-tap_saprotroph (Põlme et al. 2020); Decay sugar-rich substrates (Põlme et al. 2020) | Põlme S, et al. 2020. Fungal Diversity 105:1-16 ((https://doi.org/10.1007/s13225-020-00466-2)) |
| Otu24643 | 1 | k__Fungi(100);p__Ascomycota(100);c__Saccharomycetes(100);o__Saccharomycetales(100);f__Metschnikowiaceae(100);g__Metschnikowia(100);s__Metschnikowia_pulcherrima(100); | Metschnikowia | 13 | Saprotroph | \|Undefined Saprotroph\| | Yeast | NULL | Probable | As nectar-tap_saprotroph (Põlme et al. 2020); Decay sugar-rich substrates (Põlme et al. 2020) | Põlme S, et al. 2020. Fungal Diversity 105:1-16 ((https://doi.org/10.1007/s13225-020-00466-2)) |
| Otu24648 | 1 | k__Fungi(100);p__Ascomycota(100);c__Saccharomycetes(100);o__Saccharomycetales(100);f__Metschnikowiaceae(100);g__Metschnikowia(100);s__Metschnikowia_pulcherrima(100); | Metschnikowia | 13 | Saprotroph | \|Undefined Saprotroph\| | Yeast | NULL | Probable | As nectar-tap_saprotroph (Põlme et al. 2020); Decay sugar-rich substrates (Põlme et al. 2020) | Põlme S, et al. 2020. Fungal Diversity 105:1-16 ((https://doi.org/10.1007/s13225-020-00466-2)) |
| Otu24649 | 1 | k__Fungi(100);p__Ascomycota(100);c__Saccharomycetes(100);o__Saccharomycetales(100);f__Metschnikowiaceae(100);g__Metschnikowia(100);s__Metschnikowia_pulcherrima(100); | Metschnikowia | 13 | Saprotroph | \|Undefined Saprotroph\| | Yeast | NULL | Probable | As nectar-tap_saprotroph (Põlme et al. 2020); Decay sugar-rich substrates (Põlme et al. 2020) | Põlme S, et al. 2020. Fungal Diversity 105:1-16 ((https://doi.org/10.1007/s13225-020-00466-2)) |
| Otu24650 | 1 | k__Fungi(100);p__Ascomycota(100);c__Saccharomycetes(100);o__Saccharomycetales(100);f__Metschnikowiaceae(100);g__Metschnikowia(100);s__Metschnikowia_pulcherrima(100); | Metschnikowia | 13 | Saprotroph | \|Undefined Saprotroph\| | Yeast | NULL | Probable | As nectar-tap_saprotroph (Põlme et al. 2020); Decay sugar-rich substrates (Põlme et al. 2020) | Põlme S, et al. 2020. Fungal Diversity 105:1-16 ((https://doi.org/10.1007/s13225-020-00466-2)) |
| Otu24652 | 1 | k__Fungi(100);p__Ascomycota(100);c__Saccharomycetes(100);o__Saccharomycetales(100);f__Metschnikowiaceae(100);g__Metschnikowia(100);s__Metschnikowia_pulcherrima(100); | Metschnikowia | 13 | Saprotroph | \|Undefined Saprotroph\| | Yeast | NULL | Probable | As nectar-tap_saprotroph (Põlme et al. 2020); Decay sugar-rich substrates (Põlme et al. 2020) | Põlme S, et al. 2020. Fungal Diversity 105:1-16 ((https://doi.org/10.1007/s13225-020-00466-2)) |
| Otu24653 | 1 | k__Fungi(100);p__Ascomycota(100);c__Saccharomycetes(100);o__Saccharomycetales(100);f__Metschnikowiaceae(100);g__Metschnikowia(100);g__Metschnikowia_unclassified(100); | Metschnikowia | 13 | Saprotroph | \|Undefined Saprotroph\| | Yeast | NULL | Probable | As nectar-tap_saprotroph (Põlme et al. 2020); Decay sugar-rich substrates (Põlme et al. 2020) | Põlme S, et al. 2020. Fungal Diversity 105:1-16 ((https://doi.org/10.1007/s13225-020-00466-2)) |
| Otu24654 | 1 | k__Fungi(100);p__Ascomycota(100);c__Saccharomycetes(100);o__Saccharomycetales(100);f__Metschnikowiaceae(100);g__Metschnikowia(100);s__Metschnikowia_pulcherrima(100); | Metschnikowia | 13 | Saprotroph | \|Undefined Saprotroph\| | Yeast | NULL | Probable | As nectar-tap_saprotroph (Põlme et al. 2020); Decay sugar-rich substrates (Põlme et al. 2020) | Põlme S, et al. 2020. Fungal Diversity 105:1-16 ((https://doi.org/10.1007/s13225-020-00466-2)) |
| Otu24656 | 1 | k__Fungi(100);p__Ascomycota(100);c__Saccharomycetes(100);o__Saccharomycetales(100);f__Metschnikowiaceae(100);g__Metschnikowia(100);s__Metschnikowia_pulcherrima(100); | Metschnikowia | 13 | Saprotroph | \|Undefined Saprotroph\| | Yeast | NULL | Probable | As nectar-tap_saprotroph (Põlme et al. 2020); Decay sugar-rich substrates (Põlme et al. 2020) | Põlme S, et al. 2020. Fungal Diversity 105:1-16 ((https://doi.org/10.1007/s13225-020-00466-2)) |
| Otu24657 | 1 | k__Fungi(100);p__Ascomycota(100);c__Saccharomycetes(100);o__Saccharomycetales(100);f__Metschnikowiaceae(100);g__Metschnikowia(100);s__Metschnikowia_pulcherrima(100); | Metschnikowia | 13 | Saprotroph | \|Undefined Saprotroph\| | Yeast | NULL | Probable | As nectar-tap_saprotroph (Põlme et al. 2020); Decay sugar-rich substrates (Põlme et al. 2020) | Põlme S, et al. 2020. Fungal Diversity 105:1-16 ((https://doi.org/10.1007/s13225-020-00466-2)) |
| Otu24659 | 1 | k__Fungi(100);p__Ascomycota(100);c__Saccharomycetes(100);o__Saccharomycetales(100);f__Metschnikowiaceae(100);g__Metschnikowia(100);g__Metschnikowia_unclassified(100); | Metschnikowia | 13 | Saprotroph | \|Undefined Saprotroph\| | Yeast | NULL | Probable | As nectar-tap_saprotroph (Põlme et al. 2020); Decay sugar-rich substrates (Põlme et al. 2020) | Põlme S, et al. 2020. Fungal Diversity 105:1-16 ((https://doi.org/10.1007/s13225-020-00466-2)) |
| Otu24660 | 1 | k__Fungi(100);p__Ascomycota(100);c__Saccharomycetes(100);o__Saccharomycetales(100);f__Metschnikowiaceae(100);g__Metschnikowia(100);g__Metschnikowia_unclassified(100); | Metschnikowia | 13 | Saprotroph | \|Undefined Saprotroph\| | Yeast | NULL | Probable | As nectar-tap_saprotroph (Põlme et al. 2020); Decay sugar-rich substrates (Põlme et al. 2020) | Põlme S, et al. 2020. Fungal Diversity 105:1-16 ((https://doi.org/10.1007/s13225-020-00466-2)) |
| Otu24661 | 1 | k__Fungi(100);p__Ascomycota(100);c__Saccharomycetes(100);o__Saccharomycetales(100);f__Metschnikowiaceae(100);g__Metschnikowia(100);s__Metschnikowia_pulcherrima(100); | Metschnikowia | 13 | Saprotroph | \|Undefined Saprotroph\| | Yeast | NULL | Probable | As nectar-tap_saprotroph (Põlme et al. 2020); Decay sugar-rich substrates (Põlme et al. 2020) | Põlme S, et al. 2020. Fungal Diversity 105:1-16 ((https://doi.org/10.1007/s13225-020-00466-2)) |
| Otu24662 | 1 | k__Fungi(100);p__Ascomycota(100);c__Saccharomycetes(100);o__Saccharomycetales(100);f__Metschnikowiaceae(100);g__Metschnikowia(100);s__Metschnikowia_pulcherrima(100); | Metschnikowia | 13 | Saprotroph | \|Undefined Saprotroph\| | Yeast | NULL | Probable | As nectar-tap_saprotroph (Põlme et al. 2020); Decay sugar-rich substrates (Põlme et al. 2020) | Põlme S, et al. 2020. Fungal Diversity 105:1-16 ((https://doi.org/10.1007/s13225-020-00466-2)) |
| Otu24663 | 1 | k__Fungi(100);p__Ascomycota(100);c__Saccharomycetes(100);o__Saccharomycetales(100);f__Metschnikowiaceae(100);g__Metschnikowia(100);s__Metschnikowia_pulcherrima(100); | Metschnikowia | 13 | Saprotroph | \|Undefined Saprotroph\| | Yeast | NULL | Probable | As nectar-tap_saprotroph (Põlme et al. 2020); Decay sugar-rich substrates (Põlme et al. 2020) | Põlme S, et al. 2020. Fungal Diversity 105:1-16 ((https://doi.org/10.1007/s13225-020-00466-2)) |
| Otu24665 | 1 | k__Fungi(100);p__Ascomycota(100);c__Saccharomycetes(100);o__Saccharomycetales(100);f__Metschnikowiaceae(100);g__Metschnikowia(100);s__Metschnikowia_pulcherrima(100); | Metschnikowia | 13 | Saprotroph | \|Undefined Saprotroph\| | Yeast | NULL | Probable | As nectar-tap_saprotroph (Põlme et al. 2020); Decay sugar-rich substrates (Põlme et al. 2020) | Põlme S, et al. 2020. Fungal Diversity 105:1-16 ((https://doi.org/10.1007/s13225-020-00466-2)) |
| Otu24666 | 1 | k__Fungi(100);p__Ascomycota(100);c__Saccharomycetes(100);o__Saccharomycetales(100);f__Metschnikowiaceae(100);g__Metschnikowia(100);s__Metschnikowia_pulcherrima(100); | Metschnikowia | 13 | Saprotroph | \|Undefined Saprotroph\| | Yeast | NULL | Probable | As nectar-tap_saprotroph (Põlme et al. 2020); Decay sugar-rich substrates (Põlme et al. 2020) | Põlme S, et al. 2020. Fungal Diversity 105:1-16 ((https://doi.org/10.1007/s13225-020-00466-2)) |
| Otu24667 | 1 | k__Fungi(100);p__Ascomycota(100);c__Saccharomycetes(100);o__Saccharomycetales(100);f__Metschnikowiaceae(100);g__Metschnikowia(100);g__Metschnikowia_unclassified(100); | Metschnikowia | 13 | Saprotroph | \|Undefined Saprotroph\| | Yeast | NULL | Probable | As nectar-tap_saprotroph (Põlme et al. 2020); Decay sugar-rich substrates (Põlme et al. 2020) | Põlme S, et al. 2020. Fungal Diversity 105:1-16 ((https://doi.org/10.1007/s13225-020-00466-2)) |
| Otu24668 | 1 | k__Fungi(100);p__Ascomycota(100);c__Saccharomycetes(100);o__Saccharomycetales(100);f__Metschnikowiaceae(100);g__Metschnikowia(100);s__Metschnikowia_pulcherrima(100); | Metschnikowia | 13 | Saprotroph | \|Undefined Saprotroph\| | Yeast | NULL | Probable | As nectar-tap_saprotroph (Põlme et al. 2020); Decay sugar-rich substrates (Põlme et al. 2020) | Põlme S, et al. 2020. Fungal Diversity 105:1-16 ((https://doi.org/10.1007/s13225-020-00466-2)) |
| Otu24669 | 1 | k__Fungi(100);p__Ascomycota(100);c__Saccharomycetes(100);o__Saccharomycetales(100);f__Metschnikowiaceae(100);g__Metschnikowia(100);s__Metschnikowia_pulcherrima(100); | Metschnikowia | 13 | Saprotroph | \|Undefined Saprotroph\| | Yeast | NULL | Probable | As nectar-tap_saprotroph (Põlme et al. 2020); Decay sugar-rich substrates (Põlme et al. 2020) | Põlme S, et al. 2020. Fungal Diversity 105:1-16 ((https://doi.org/10.1007/s13225-020-00466-2)) |
| Otu24670 | 1 | k__Fungi(100);p__Ascomycota(100);c__Saccharomycetes(100);o__Saccharomycetales(100);f__Metschnikowiaceae(100);g__Metschnikowia(100);s__Metschnikowia_pulcherrima(100); | Metschnikowia | 13 | Saprotroph | \|Undefined Saprotroph\| | Yeast | NULL | Probable | As nectar-tap_saprotroph (Põlme et al. 2020); Decay sugar-rich substrates (Põlme et al. 2020) | Põlme S, et al. 2020. Fungal Diversity 105:1-16 ((https://doi.org/10.1007/s13225-020-00466-2)) |
| Otu24671 | 1 | k__Fungi(100);p__Ascomycota(100);c__Saccharomycetes(100);o__Saccharomycetales(100);f__Metschnikowiaceae(100);g__Metschnikowia(100);s__Metschnikowia_pulcherrima(100); | Metschnikowia | 13 | Saprotroph | \|Undefined Saprotroph\| | Yeast | NULL | Probable | As nectar-tap_saprotroph (Põlme et al. 2020); Decay sugar-rich substrates (Põlme et al. 2020) | Põlme S, et al. 2020. Fungal Diversity 105:1-16 ((https://doi.org/10.1007/s13225-020-00466-2)) |
| Otu24678 | 1 | k__Fungi(100);p__Ascomycota(100);c__Saccharomycetes(100);o__Saccharomycetales(100);f__Metschnikowiaceae(100);g__Metschnikowia(100);g__Metschnikowia_unclassified(100); | Metschnikowia | 13 | Saprotroph | \|Undefined Saprotroph\| | Yeast | NULL | Probable | As nectar-tap_saprotroph (Põlme et al. 2020); Decay sugar-rich substrates (Põlme et al. 2020) | Põlme S, et al. 2020. Fungal Diversity 105:1-16 ((https://doi.org/10.1007/s13225-020-00466-2)) |
| Otu24682 | 1 | k__Fungi(100);p__Ascomycota(100);c__Saccharomycetes(100);o__Saccharomycetales(100);f__Metschnikowiaceae(100);g__Metschnikowia(100);g__Metschnikowia_unclassified(100); | Metschnikowia | 13 | Saprotroph | \|Undefined Saprotroph\| | Yeast | NULL | Probable | As nectar-tap_saprotroph (Põlme et al. 2020); Decay sugar-rich substrates (Põlme et al. 2020) | Põlme S, et al. 2020. Fungal Diversity 105:1-16 ((https://doi.org/10.1007/s13225-020-00466-2)) |
| Otu24697 | 1 | k__Fungi(100);p__Ascomycota(100);c__Saccharomycetes(100);o__Saccharomycetales(100);f__Metschnikowiaceae(100);g__Metschnikowia(100);s__Metschnikowia_pulcherrima(100); | Metschnikowia | 13 | Saprotroph | \|Undefined Saprotroph\| | Yeast | NULL | Probable | As nectar-tap_saprotroph (Põlme et al. 2020); Decay sugar-rich substrates (Põlme et al. 2020) | Põlme S, et al. 2020. Fungal Diversity 105:1-16 ((https://doi.org/10.1007/s13225-020-00466-2)) |
| Otu24698 | 1 | k__Fungi(100);p__Ascomycota(100);c__Saccharomycetes(100);o__Saccharomycetales(100);f__Metschnikowiaceae(100);g__Metschnikowia(100);s__Metschnikowia_pulcherrima(100); | Metschnikowia | 13 | Saprotroph | \|Undefined Saprotroph\| | Yeast | NULL | Probable | As nectar-tap_saprotroph (Põlme et al. 2020); Decay sugar-rich substrates (Põlme et al. 2020) | Põlme S, et al. 2020. Fungal Diversity 105:1-16 ((https://doi.org/10.1007/s13225-020-00466-2)) |
| Otu24704 | 1 | k__Fungi(100);p__Ascomycota(100);c__Saccharomycetes(100);o__Saccharomycetales(100);f__Metschnikowiaceae(100);g__Metschnikowia(100);g__Metschnikowia_unclassified(100); | Metschnikowia | 13 | Saprotroph | \|Undefined Saprotroph\| | Yeast | NULL | Probable | As nectar-tap_saprotroph (Põlme et al. 2020); Decay sugar-rich substrates (Põlme et al. 2020) | Põlme S, et al. 2020. Fungal Diversity 105:1-16 ((https://doi.org/10.1007/s13225-020-00466-2)) |
| Otu24707 | 1 | k__Fungi(100);p__Ascomycota(100);c__Dothideomycetes(100);o__Capnodiales(100);f__Mycosphaerellaceae(100);g__Mycosphaerella(100);s__Mycosphaerella_latebrosa(100); | Mycosphaerella | 13 | Pathotroph-Saprotroph | \|Plant Pathogen\|-Plant Saprotroph | Microfungus | NULL | Probable | As litter_saprotroph (Põlme et al. 2020); Decay leaf, fruit, seed (Põlme et al. 2020) | Tedersoo L, et al. 2014. Science 346:e1256688 ((https://doi.org/10.1126/science.1256688)); Põlme S, et al. 2020. Fungal Diversity 105:1-16 ((https://doi.org/10.1007/s13225-020-00466-2)) |
| Otu24708 | 1 | k__Fungi(100);p__Ascomycota(100);c__Saccharomycetes(100);o__Saccharomycetales(100);f__Metschnikowiaceae(100);g__Metschnikowia(100);s__Metschnikowia_pulcherrima(100); | Metschnikowia | 13 | Saprotroph | \|Undefined Saprotroph\| | Yeast | NULL | Probable | As nectar-tap_saprotroph (Põlme et al. 2020); Decay sugar-rich substrates (Põlme et al. 2020) | Põlme S, et al. 2020. Fungal Diversity 105:1-16 ((https://doi.org/10.1007/s13225-020-00466-2)) |
| Otu24709 | 1 | k__Fungi(100);p__Ascomycota(100);c__Saccharomycetes(100);o__Saccharomycetales(100);f__Metschnikowiaceae(100);g__Metschnikowia(100);g__Metschnikowia_unclassified(100); | Metschnikowia | 13 | Saprotroph | \|Undefined Saprotroph\| | Yeast | NULL | Probable | As nectar-tap_saprotroph (Põlme et al. 2020); Decay sugar-rich substrates (Põlme et al. 2020) | Põlme S, et al. 2020. Fungal Diversity 105:1-16 ((https://doi.org/10.1007/s13225-020-00466-2)) |
| Otu24710 | 1 | k__Fungi(100);p__Ascomycota(100);c__Dothideomycetes(100);o__Capnodiales(100);f__Davidiellaceae(100);g__Cladosporium(100);s__Cladosporium_velox(100); | Cladosporium | 13 | Pathotroph-Saprotroph-Symbiotroph | Animal Pathogen-Endophyte-Lichen Parasite-Plant Pathogen-\|Plant Saprotroph\|-Wood Saprotroph | Microfungus | NULL | Possible | As foliar_endophyte (Põlme et al. 2020); As litter_saprotroph (Põlme et al. 2020); Decay wood, leaf, fruit, seed, soil, dung, animal material (Põlme et al. 2020) | Seehann G, et al. 1975. List of Fungi in Soft-Rot Tissue Tests. Institute fur Holzbiologie und Holzschutz, Bundesforschungsanstalt fur Forst - und Holzwirtshaft, Hamburg - Lohbrugge, Germany; Lawrey JD, Diederich P. 2016. http://www.lichenicolous.net (accessed 24 May 2017) ((http://www.lichenicolous.net)); Põlme S, et al. 2020. Fungal Diversity 105:1-16 ((https://doi.org/10.1007/s13225-020-00466-2)); The American Phytopathological Society Common Names of Plant Diseases ((http://www.apsnet.org/publications/commonnames/Pages/default.aspx)) |
| Otu24711 | 1 | k__Fungi(100);p__Ascomycota(100);c__Saccharomycetes(100);o__Saccharomycetales(100);f__Metschnikowiaceae(100);g__Metschnikowia(100);g__Metschnikowia_unclassified(100); | Metschnikowia | 13 | Saprotroph | \|Undefined Saprotroph\| | Yeast | NULL | Probable | As nectar-tap_saprotroph (Põlme et al. 2020); Decay sugar-rich substrates (Põlme et al. 2020) | Põlme S, et al. 2020. Fungal Diversity 105:1-16 ((https://doi.org/10.1007/s13225-020-00466-2)) |
| Otu24712 | 1 | k__Fungi(100);p__Ascomycota(100);c__Saccharomycetes(100);o__Saccharomycetales(100);f__Metschnikowiaceae(100);g__Metschnikowia(100);g__Metschnikowia_unclassified(100); | Metschnikowia | 13 | Saprotroph | \|Undefined Saprotroph\| | Yeast | NULL | Probable | As nectar-tap_saprotroph (Põlme et al. 2020); Decay sugar-rich substrates (Põlme et al. 2020) | Põlme S, et al. 2020. Fungal Diversity 105:1-16 ((https://doi.org/10.1007/s13225-020-00466-2)) |
| Otu24716 | 1 | k__Fungi(100);p__Ascomycota(100);c__Saccharomycetes(100);o__Saccharomycetales(100);f__Metschnikowiaceae(100);g__Metschnikowia(100);s__Metschnikowia_pulcherrima(100); | Metschnikowia | 13 | Saprotroph | \|Undefined Saprotroph\| | Yeast | NULL | Probable | As nectar-tap_saprotroph (Põlme et al. 2020); Decay sugar-rich substrates (Põlme et al. 2020) | Põlme S, et al. 2020. Fungal Diversity 105:1-16 ((https://doi.org/10.1007/s13225-020-00466-2)) |
| Otu24717 | 1 | k__Fungi(100);p__Ascomycota(100);c__Dothideomycetes(100);o__Capnodiales(100);f__Davidiellaceae(100);g__Cladosporium(100);g__Cladosporium_unclassified(100); | Cladosporium | 13 | Pathotroph-Saprotroph-Symbiotroph | Animal Pathogen-Endophyte-Lichen Parasite-Plant Pathogen-\|Plant Saprotroph\|-Wood Saprotroph | Microfungus | NULL | Possible | As foliar_endophyte (Põlme et al. 2020); As litter_saprotroph (Põlme et al. 2020); Decay wood, leaf, fruit, seed, soil, dung, animal material (Põlme et al. 2020) | Seehann G, et al. 1975. List of Fungi in Soft-Rot Tissue Tests. Institute fur Holzbiologie und Holzschutz, Bundesforschungsanstalt fur Forst - und Holzwirtshaft, Hamburg - Lohbrugge, Germany; Lawrey JD, Diederich P. 2016. http://www.lichenicolous.net (accessed 24 May 2017) ((http://www.lichenicolous.net)); Põlme S, et al. 2020. Fungal Diversity 105:1-16 ((https://doi.org/10.1007/s13225-020-00466-2)); The American Phytopathological Society Common Names of Plant Diseases ((http://www.apsnet.org/publications/commonnames/Pages/default.aspx)) |
| Otu24719 | 1 | k__Fungi(100);p__Ascomycota(100);c__Saccharomycetes(100);o__Saccharomycetales(100);f__Metschnikowiaceae(100);g__Metschnikowia(100);s__Metschnikowia_pulcherrima(100); | Metschnikowia | 13 | Saprotroph | \|Undefined Saprotroph\| | Yeast | NULL | Probable | As nectar-tap_saprotroph (Põlme et al. 2020); Decay sugar-rich substrates (Põlme et al. 2020) | Põlme S, et al. 2020. Fungal Diversity 105:1-16 ((https://doi.org/10.1007/s13225-020-00466-2)) |
| Otu24721 | 1 | k__Fungi(100);p__Ascomycota(100);c__Saccharomycetes(100);o__Saccharomycetales(100);f__Metschnikowiaceae(100);g__Metschnikowia(100);s__Metschnikowia_pulcherrima(100); | Metschnikowia | 13 | Saprotroph | \|Undefined Saprotroph\| | Yeast | NULL | Probable | As nectar-tap_saprotroph (Põlme et al. 2020); Decay sugar-rich substrates (Põlme et al. 2020) | Põlme S, et al. 2020. Fungal Diversity 105:1-16 ((https://doi.org/10.1007/s13225-020-00466-2)) |
| Otu24722 | 1 | k__Fungi(100);p__Ascomycota(100);c__Saccharomycetes(100);o__Saccharomycetales(100);f__Metschnikowiaceae(100);g__Metschnikowia(100);g__Metschnikowia_unclassified(100); | Metschnikowia | 13 | Saprotroph | \|Undefined Saprotroph\| | Yeast | NULL | Probable | As nectar-tap_saprotroph (Põlme et al. 2020); Decay sugar-rich substrates (Põlme et al. 2020) | Põlme S, et al. 2020. Fungal Diversity 105:1-16 ((https://doi.org/10.1007/s13225-020-00466-2)) |
| Otu24723 | 1 | k__Fungi(100);p__Ascomycota(100);c__Saccharomycetes(100);o__Saccharomycetales(100);f__Metschnikowiaceae(100);g__Metschnikowia(100);s__Metschnikowia_pulcherrima(100); | Metschnikowia | 13 | Saprotroph | \|Undefined Saprotroph\| | Yeast | NULL | Probable | As nectar-tap_saprotroph (Põlme et al. 2020); Decay sugar-rich substrates (Põlme et al. 2020) | Põlme S, et al. 2020. Fungal Diversity 105:1-16 ((https://doi.org/10.1007/s13225-020-00466-2)) |
| Otu24724 | 1 | k__Fungi(100);p__Ascomycota(100);c__Saccharomycetes(100);o__Saccharomycetales(100);f__Metschnikowiaceae(100);g__Metschnikowia(100);s__Metschnikowia_pulcherrima(100); | Metschnikowia | 13 | Saprotroph | \|Undefined Saprotroph\| | Yeast | NULL | Probable | As nectar-tap_saprotroph (Põlme et al. 2020); Decay sugar-rich substrates (Põlme et al. 2020) | Põlme S, et al. 2020. Fungal Diversity 105:1-16 ((https://doi.org/10.1007/s13225-020-00466-2)) |
| Otu24725 | 1 | k__Fungi(100);p__Ascomycota(100);c__Saccharomycetes(100);o__Saccharomycetales(100);f__Metschnikowiaceae(100);g__Metschnikowia(100);s__Metschnikowia_pulcherrima(100); | Metschnikowia | 13 | Saprotroph | \|Undefined Saprotroph\| | Yeast | NULL | Probable | As nectar-tap_saprotroph (Põlme et al. 2020); Decay sugar-rich substrates (Põlme et al. 2020) | Põlme S, et al. 2020. Fungal Diversity 105:1-16 ((https://doi.org/10.1007/s13225-020-00466-2)) |
| Otu24726 | 1 | k__Fungi(100);p__Ascomycota(100);c__Saccharomycetes(100);o__Saccharomycetales(100);f__Metschnikowiaceae(100);g__Metschnikowia(100);g__Metschnikowia_unclassified(100); | Metschnikowia | 13 | Saprotroph | \|Undefined Saprotroph\| | Yeast | NULL | Probable | As nectar-tap_saprotroph (Põlme et al. 2020); Decay sugar-rich substrates (Põlme et al. 2020) | Põlme S, et al. 2020. Fungal Diversity 105:1-16 ((https://doi.org/10.1007/s13225-020-00466-2)) |
| Otu24727 | 1 | k__Fungi(100);p__Ascomycota(100);c__Saccharomycetes(100);o__Saccharomycetales(100);f__Metschnikowiaceae(100);g__Metschnikowia(100);g__Metschnikowia_unclassified(100); | Metschnikowia | 13 | Saprotroph | \|Undefined Saprotroph\| | Yeast | NULL | Probable | As nectar-tap_saprotroph (Põlme et al. 2020); Decay sugar-rich substrates (Põlme et al. 2020) | Põlme S, et al. 2020. Fungal Diversity 105:1-16 ((https://doi.org/10.1007/s13225-020-00466-2)) |
| Otu24728 | 1 | k__Fungi(100);p__Ascomycota(100);c__Saccharomycetes(100);o__Saccharomycetales(100);f__Metschnikowiaceae(100);g__Metschnikowia(100);g__Metschnikowia_unclassified(100); | Metschnikowia | 13 | Saprotroph | \|Undefined Saprotroph\| | Yeast | NULL | Probable | As nectar-tap_saprotroph (Põlme et al. 2020); Decay sugar-rich substrates (Põlme et al. 2020) | Põlme S, et al. 2020. Fungal Diversity 105:1-16 ((https://doi.org/10.1007/s13225-020-00466-2)) |
| Otu24729 | 1 | k__Fungi(100);p__Ascomycota(100);c__Saccharomycetes(100);o__Saccharomycetales(100);f__Metschnikowiaceae(100);g__Metschnikowia(100);s__Metschnikowia_pulcherrima(100); | Metschnikowia | 13 | Saprotroph | \|Undefined Saprotroph\| | Yeast | NULL | Probable | As nectar-tap_saprotroph (Põlme et al. 2020); Decay sugar-rich substrates (Põlme et al. 2020) | Põlme S, et al. 2020. Fungal Diversity 105:1-16 ((https://doi.org/10.1007/s13225-020-00466-2)) |
| Otu24730 | 1 | k__Fungi(100);p__Ascomycota(100);c__Saccharomycetes(100);o__Saccharomycetales(100);f__Metschnikowiaceae(100);g__Metschnikowia(100);g__Metschnikowia_unclassified(100); | Metschnikowia | 13 | Saprotroph | \|Undefined Saprotroph\| | Yeast | NULL | Probable | As nectar-tap_saprotroph (Põlme et al. 2020); Decay sugar-rich substrates (Põlme et al. 2020) | Põlme S, et al. 2020. Fungal Diversity 105:1-16 ((https://doi.org/10.1007/s13225-020-00466-2)) |
| Otu24731 | 1 | k__Fungi(100);p__Ascomycota(100);c__Saccharomycetes(100);o__Saccharomycetales(100);f__Metschnikowiaceae(100);g__Metschnikowia(100);s__Metschnikowia_pulcherrima(100); | Metschnikowia | 13 | Saprotroph | \|Undefined Saprotroph\| | Yeast | NULL | Probable | As nectar-tap_saprotroph (Põlme et al. 2020); Decay sugar-rich substrates (Põlme et al. 2020) | Põlme S, et al. 2020. Fungal Diversity 105:1-16 ((https://doi.org/10.1007/s13225-020-00466-2)) |
| Otu24733 | 1 | k__Fungi(100);p__Ascomycota(100);c__Saccharomycetes(100);o__Saccharomycetales(100);f__Metschnikowiaceae(100);g__Metschnikowia(100);s__Metschnikowia_pulcherrima(100); | Metschnikowia | 13 | Saprotroph | \|Undefined Saprotroph\| | Yeast | NULL | Probable | As nectar-tap_saprotroph (Põlme et al. 2020); Decay sugar-rich substrates (Põlme et al. 2020) | Põlme S, et al. 2020. Fungal Diversity 105:1-16 ((https://doi.org/10.1007/s13225-020-00466-2)) |
| Otu24734 | 1 | k__Fungi(100);p__Ascomycota(100);c__Saccharomycetes(100);o__Saccharomycetales(100);f__Metschnikowiaceae(100);g__Metschnikowia(100);s__Metschnikowia_pulcherrima(100); | Metschnikowia | 13 | Saprotroph | \|Undefined Saprotroph\| | Yeast | NULL | Probable | As nectar-tap_saprotroph (Põlme et al. 2020); Decay sugar-rich substrates (Põlme et al. 2020) | Põlme S, et al. 2020. Fungal Diversity 105:1-16 ((https://doi.org/10.1007/s13225-020-00466-2)) |
| Otu24735 | 1 | k__Fungi(100);p__Ascomycota(100);c__Saccharomycetes(100);o__Saccharomycetales(100);f__Metschnikowiaceae(100);g__Metschnikowia(100);g__Metschnikowia_unclassified(100); | Metschnikowia | 13 | Saprotroph | \|Undefined Saprotroph\| | Yeast | NULL | Probable | As nectar-tap_saprotroph (Põlme et al. 2020); Decay sugar-rich substrates (Põlme et al. 2020) | Põlme S, et al. 2020. Fungal Diversity 105:1-16 ((https://doi.org/10.1007/s13225-020-00466-2)) |
| Otu24736 | 1 | k__Fungi(100);p__Ascomycota(100);c__Saccharomycetes(100);o__Saccharomycetales(100);f__Metschnikowiaceae(100);g__Metschnikowia(100);s__Metschnikowia_pulcherrima(100); | Metschnikowia | 13 | Saprotroph | \|Undefined Saprotroph\| | Yeast | NULL | Probable | As nectar-tap_saprotroph (Põlme et al. 2020); Decay sugar-rich substrates (Põlme et al. 2020) | Põlme S, et al. 2020. Fungal Diversity 105:1-16 ((https://doi.org/10.1007/s13225-020-00466-2)) |
| Otu24737 | 1 | k__Fungi(100);p__Ascomycota(100);c__Saccharomycetes(100);o__Saccharomycetales(100);f__Metschnikowiaceae(100);g__Metschnikowia(100);s__Metschnikowia_pulcherrima(100); | Metschnikowia | 13 | Saprotroph | \|Undefined Saprotroph\| | Yeast | NULL | Probable | As nectar-tap_saprotroph (Põlme et al. 2020); Decay sugar-rich substrates (Põlme et al. 2020) | Põlme S, et al. 2020. Fungal Diversity 105:1-16 ((https://doi.org/10.1007/s13225-020-00466-2)) |
| Otu24738 | 1 | k__Fungi(100);p__Ascomycota(100);c__Saccharomycetes(100);o__Saccharomycetales(100);f__Metschnikowiaceae(100);g__Metschnikowia(100);g__Metschnikowia_unclassified(100); | Metschnikowia | 13 | Saprotroph | \|Undefined Saprotroph\| | Yeast | NULL | Probable | As nectar-tap_saprotroph (Põlme et al. 2020); Decay sugar-rich substrates (Põlme et al. 2020) | Põlme S, et al. 2020. Fungal Diversity 105:1-16 ((https://doi.org/10.1007/s13225-020-00466-2)) |
| Otu24739 | 1 | k__Fungi(100);p__Ascomycota(100);c__Saccharomycetes(100);o__Saccharomycetales(100);f__Metschnikowiaceae(100);g__Metschnikowia(100);s__Metschnikowia_pulcherrima(100); | Metschnikowia | 13 | Saprotroph | \|Undefined Saprotroph\| | Yeast | NULL | Probable | As nectar-tap_saprotroph (Põlme et al. 2020); Decay sugar-rich substrates (Põlme et al. 2020) | Põlme S, et al. 2020. Fungal Diversity 105:1-16 ((https://doi.org/10.1007/s13225-020-00466-2)) |
| Otu24740 | 1 | k__Fungi(100);p__Ascomycota(100);c__Saccharomycetes(100);o__Saccharomycetales(100);f__Metschnikowiaceae(100);g__Metschnikowia(100);s__Metschnikowia_pulcherrima(100); | Metschnikowia | 13 | Saprotroph | \|Undefined Saprotroph\| | Yeast | NULL | Probable | As nectar-tap_saprotroph (Põlme et al. 2020); Decay sugar-rich substrates (Põlme et al. 2020) | Põlme S, et al. 2020. Fungal Diversity 105:1-16 ((https://doi.org/10.1007/s13225-020-00466-2)) |
| Otu24741 | 1 | k__Fungi(100);p__Ascomycota(100);c__Saccharomycetes(100);o__Saccharomycetales(100);f__Metschnikowiaceae(100);g__Metschnikowia(100);s__Metschnikowia_pulcherrima(100); | Metschnikowia | 13 | Saprotroph | \|Undefined Saprotroph\| | Yeast | NULL | Probable | As nectar-tap_saprotroph (Põlme et al. 2020); Decay sugar-rich substrates (Põlme et al. 2020) | Põlme S, et al. 2020. Fungal Diversity 105:1-16 ((https://doi.org/10.1007/s13225-020-00466-2)) |
| Otu24742 | 1 | k__Fungi(100);p__Ascomycota(100);c__Saccharomycetes(100);o__Saccharomycetales(100);f__Metschnikowiaceae(100);g__Metschnikowia(100);s__Metschnikowia_pulcherrima(100); | Metschnikowia | 13 | Saprotroph | \|Undefined Saprotroph\| | Yeast | NULL | Probable | As nectar-tap_saprotroph (Põlme et al. 2020); Decay sugar-rich substrates (Põlme et al. 2020) | Põlme S, et al. 2020. Fungal Diversity 105:1-16 ((https://doi.org/10.1007/s13225-020-00466-2)) |
| Otu24743 | 1 | k__Fungi(100);p__Ascomycota(100);c__Saccharomycetes(100);o__Saccharomycetales(100);f__Metschnikowiaceae(100);g__Metschnikowia(100);s__Metschnikowia_pulcherrima(100); | Metschnikowia | 13 | Saprotroph | \|Undefined Saprotroph\| | Yeast | NULL | Probable | As nectar-tap_saprotroph (Põlme et al. 2020); Decay sugar-rich substrates (Põlme et al. 2020) | Põlme S, et al. 2020. Fungal Diversity 105:1-16 ((https://doi.org/10.1007/s13225-020-00466-2)) |
| Otu24749 | 1 | k__Fungi(100);p__Ascomycota(100);c__Saccharomycetes(100);o__Saccharomycetales(100);f__Metschnikowiaceae(100);g__Metschnikowia(100);g__Metschnikowia_unclassified(100); | Metschnikowia | 13 | Saprotroph | \|Undefined Saprotroph\| | Yeast | NULL | Probable | As nectar-tap_saprotroph (Põlme et al. 2020); Decay sugar-rich substrates (Põlme et al. 2020) | Põlme S, et al. 2020. Fungal Diversity 105:1-16 ((https://doi.org/10.1007/s13225-020-00466-2)) |
| Otu24750 | 1 | k__Fungi(100);p__Ascomycota(100);c__Saccharomycetes(100);o__Saccharomycetales(100);f__Metschnikowiaceae(100);g__Metschnikowia(100);s__Metschnikowia_pulcherrima(100); | Metschnikowia | 13 | Saprotroph | \|Undefined Saprotroph\| | Yeast | NULL | Probable | As nectar-tap_saprotroph (Põlme et al. 2020); Decay sugar-rich substrates (Põlme et al. 2020) | Põlme S, et al. 2020. Fungal Diversity 105:1-16 ((https://doi.org/10.1007/s13225-020-00466-2)) |
| Otu24756 | 1 | k__Fungi(100);p__Ascomycota(100);c__Saccharomycetes(100);o__Saccharomycetales(100);f__Metschnikowiaceae(100);g__Metschnikowia(100);g__Metschnikowia_unclassified(100); | Metschnikowia | 13 | Saprotroph | \|Undefined Saprotroph\| | Yeast | NULL | Probable | As nectar-tap_saprotroph (Põlme et al. 2020); Decay sugar-rich substrates (Põlme et al. 2020) | Põlme S, et al. 2020. Fungal Diversity 105:1-16 ((https://doi.org/10.1007/s13225-020-00466-2)) |
| Otu24757 | 1 | k__Fungi(100);p__Ascomycota(100);c__Saccharomycetes(100);o__Saccharomycetales(100);f__Metschnikowiaceae(100);g__Metschnikowia(100);s__Metschnikowia_pulcherrima(100); | Metschnikowia | 13 | Saprotroph | \|Undefined Saprotroph\| | Yeast | NULL | Probable | As nectar-tap_saprotroph (Põlme et al. 2020); Decay sugar-rich substrates (Põlme et al. 2020) | Põlme S, et al. 2020. Fungal Diversity 105:1-16 ((https://doi.org/10.1007/s13225-020-00466-2)) |
| Otu24758 | 1 | k__Fungi(100);p__Ascomycota(100);c__Saccharomycetes(100);o__Saccharomycetales(100);f__Metschnikowiaceae(100);g__Metschnikowia(100);g__Metschnikowia_unclassified(100); | Metschnikowia | 13 | Saprotroph | \|Undefined Saprotroph\| | Yeast | NULL | Probable | As nectar-tap_saprotroph (Põlme et al. 2020); Decay sugar-rich substrates (Põlme et al. 2020) | Põlme S, et al. 2020. Fungal Diversity 105:1-16 ((https://doi.org/10.1007/s13225-020-00466-2)) |
| Otu24760 | 1 | k__Fungi(100);p__Ascomycota(100);c__Saccharomycetes(100);o__Saccharomycetales(100);f__Metschnikowiaceae(100);g__Metschnikowia(100);g__Metschnikowia_unclassified(100); | Metschnikowia | 13 | Saprotroph | \|Undefined Saprotroph\| | Yeast | NULL | Probable | As nectar-tap_saprotroph (Põlme et al. 2020); Decay sugar-rich substrates (Põlme et al. 2020) | Põlme S, et al. 2020. Fungal Diversity 105:1-16 ((https://doi.org/10.1007/s13225-020-00466-2)) |
| Otu24762 | 1 | k__Fungi(100);p__Ascomycota(100);c__Saccharomycetes(100);o__Saccharomycetales(100);f__Metschnikowiaceae(100);g__Metschnikowia(100);g__Metschnikowia_unclassified(100); | Metschnikowia | 13 | Saprotroph | \|Undefined Saprotroph\| | Yeast | NULL | Probable | As nectar-tap_saprotroph (Põlme et al. 2020); Decay sugar-rich substrates (Põlme et al. 2020) | Põlme S, et al. 2020. Fungal Diversity 105:1-16 ((https://doi.org/10.1007/s13225-020-00466-2)) |
| Otu24764 | 1 | k__Fungi(100);p__Ascomycota(100);c__Saccharomycetes(100);o__Saccharomycetales(100);f__Metschnikowiaceae(100);g__Metschnikowia(100);g__Metschnikowia_unclassified(100); | Metschnikowia | 13 | Saprotroph | \|Undefined Saprotroph\| | Yeast | NULL | Probable | As nectar-tap_saprotroph (Põlme et al. 2020); Decay sugar-rich substrates (Põlme et al. 2020) | Põlme S, et al. 2020. Fungal Diversity 105:1-16 ((https://doi.org/10.1007/s13225-020-00466-2)) |
| Otu24765 | 1 | k__Fungi(100);p__Ascomycota(100);c__Saccharomycetes(100);o__Saccharomycetales(100);f__Metschnikowiaceae(100);g__Metschnikowia(100);s__Metschnikowia_pulcherrima(100); | Metschnikowia | 13 | Saprotroph | \|Undefined Saprotroph\| | Yeast | NULL | Probable | As nectar-tap_saprotroph (Põlme et al. 2020); Decay sugar-rich substrates (Põlme et al. 2020) | Põlme S, et al. 2020. Fungal Diversity 105:1-16 ((https://doi.org/10.1007/s13225-020-00466-2)) |
| Otu24766 | 1 | k__Fungi(100);p__Ascomycota(100);c__Saccharomycetes(100);o__Saccharomycetales(100);f__Metschnikowiaceae(100);g__Metschnikowia(100);s__Metschnikowia_pulcherrima(100); | Metschnikowia | 13 | Saprotroph | \|Undefined Saprotroph\| | Yeast | NULL | Probable | As nectar-tap_saprotroph (Põlme et al. 2020); Decay sugar-rich substrates (Põlme et al. 2020) | Põlme S, et al. 2020. Fungal Diversity 105:1-16 ((https://doi.org/10.1007/s13225-020-00466-2)) |
| Otu24767 | 1 | k__Fungi(100);p__Ascomycota(100);c__Saccharomycetes(100);o__Saccharomycetales(100);f__Metschnikowiaceae(100);g__Metschnikowia(100);s__Metschnikowia_pulcherrima(100); | Metschnikowia | 13 | Saprotroph | \|Undefined Saprotroph\| | Yeast | NULL | Probable | As nectar-tap_saprotroph (Põlme et al. 2020); Decay sugar-rich substrates (Põlme et al. 2020) | Põlme S, et al. 2020. Fungal Diversity 105:1-16 ((https://doi.org/10.1007/s13225-020-00466-2)) |
| Otu24768 | 1 | k__Fungi(100);p__Ascomycota(100);c__Saccharomycetes(100);o__Saccharomycetales(100);f__Metschnikowiaceae(100);g__Metschnikowia(100);g__Metschnikowia_unclassified(100); | Metschnikowia | 13 | Saprotroph | \|Undefined Saprotroph\| | Yeast | NULL | Probable | As nectar-tap_saprotroph (Põlme et al. 2020); Decay sugar-rich substrates (Põlme et al. 2020) | Põlme S, et al. 2020. Fungal Diversity 105:1-16 ((https://doi.org/10.1007/s13225-020-00466-2)) |
| Otu24771 | 1 | k__Fungi(100);p__Ascomycota(100);c__Saccharomycetes(100);o__Saccharomycetales(100);f__Metschnikowiaceae(100);g__Metschnikowia(100);s__Metschnikowia_pulcherrima(100); | Metschnikowia | 13 | Saprotroph | \|Undefined Saprotroph\| | Yeast | NULL | Probable | As nectar-tap_saprotroph (Põlme et al. 2020); Decay sugar-rich substrates (Põlme et al. 2020) | Põlme S, et al. 2020. Fungal Diversity 105:1-16 ((https://doi.org/10.1007/s13225-020-00466-2)) |
| Otu24772 | 1 | k__Fungi(100);p__Ascomycota(100);c__Saccharomycetes(100);o__Saccharomycetales(100);f__Metschnikowiaceae(100);g__Metschnikowia(100);s__Metschnikowia_pulcherrima(100); | Metschnikowia | 13 | Saprotroph | \|Undefined Saprotroph\| | Yeast | NULL | Probable | As nectar-tap_saprotroph (Põlme et al. 2020); Decay sugar-rich substrates (Põlme et al. 2020) | Põlme S, et al. 2020. Fungal Diversity 105:1-16 ((https://doi.org/10.1007/s13225-020-00466-2)) |
| Otu24773 | 1 | k__Fungi(100);p__Ascomycota(100);c__Saccharomycetes(100);o__Saccharomycetales(100);f__Metschnikowiaceae(100);g__Metschnikowia(100);s__Metschnikowia_pulcherrima(100); | Metschnikowia | 13 | Saprotroph | \|Undefined Saprotroph\| | Yeast | NULL | Probable | As nectar-tap_saprotroph (Põlme et al. 2020); Decay sugar-rich substrates (Põlme et al. 2020) | Põlme S, et al. 2020. Fungal Diversity 105:1-16 ((https://doi.org/10.1007/s13225-020-00466-2)) |
| Otu24774 | 1 | k__Fungi(100);p__Ascomycota(100);c__Saccharomycetes(100);o__Saccharomycetales(100);f__Metschnikowiaceae(100);g__Metschnikowia(100);s__Metschnikowia_pulcherrima(100); | Metschnikowia | 13 | Saprotroph | \|Undefined Saprotroph\| | Yeast | NULL | Probable | As nectar-tap_saprotroph (Põlme et al. 2020); Decay sugar-rich substrates (Põlme et al. 2020) | Põlme S, et al. 2020. Fungal Diversity 105:1-16 ((https://doi.org/10.1007/s13225-020-00466-2)) |
| Otu24775 | 1 | k__Fungi(100);p__Ascomycota(100);c__Saccharomycetes(100);o__Saccharomycetales(100);f__Metschnikowiaceae(100);g__Metschnikowia(100);s__Metschnikowia_pulcherrima(100); | Metschnikowia | 13 | Saprotroph | \|Undefined Saprotroph\| | Yeast | NULL | Probable | As nectar-tap_saprotroph (Põlme et al. 2020); Decay sugar-rich substrates (Põlme et al. 2020) | Põlme S, et al. 2020. Fungal Diversity 105:1-16 ((https://doi.org/10.1007/s13225-020-00466-2)) |
| Otu24776 | 1 | k__Fungi(100);p__Ascomycota(100);c__Saccharomycetes(100);o__Saccharomycetales(100);f__Metschnikowiaceae(100);g__Metschnikowia(100);g__Metschnikowia_unclassified(100); | Metschnikowia | 13 | Saprotroph | \|Undefined Saprotroph\| | Yeast | NULL | Probable | As nectar-tap_saprotroph (Põlme et al. 2020); Decay sugar-rich substrates (Põlme et al. 2020) | Põlme S, et al. 2020. Fungal Diversity 105:1-16 ((https://doi.org/10.1007/s13225-020-00466-2)) |
| Otu24783 | 1 | k__Fungi(100);p__Ascomycota(100);c__Saccharomycetes(100);o__Saccharomycetales(100);f__Metschnikowiaceae(100);g__Metschnikowia(100);s__Metschnikowia_pulcherrima(100); | Metschnikowia | 13 | Saprotroph | \|Undefined Saprotroph\| | Yeast | NULL | Probable | As nectar-tap_saprotroph (Põlme et al. 2020); Decay sugar-rich substrates (Põlme et al. 2020) | Põlme S, et al. 2020. Fungal Diversity 105:1-16 ((https://doi.org/10.1007/s13225-020-00466-2)) |
| Otu24784 | 1 | k__Fungi(100);p__Ascomycota(100);c__Saccharomycetes(100);o__Saccharomycetales(100);f__Metschnikowiaceae(100);g__Metschnikowia(100);s__Metschnikowia_pulcherrima(100); | Metschnikowia | 13 | Saprotroph | \|Undefined Saprotroph\| | Yeast | NULL | Probable | As nectar-tap_saprotroph (Põlme et al. 2020); Decay sugar-rich substrates (Põlme et al. 2020) | Põlme S, et al. 2020. Fungal Diversity 105:1-16 ((https://doi.org/10.1007/s13225-020-00466-2)) |
| Otu24786 | 1 | k__Fungi(100);p__Ascomycota(100);c__Saccharomycetes(100);o__Saccharomycetales(100);f__Metschnikowiaceae(100);g__Metschnikowia(100);g__Metschnikowia_unclassified(100); | Metschnikowia | 13 | Saprotroph | \|Undefined Saprotroph\| | Yeast | NULL | Probable | As nectar-tap_saprotroph (Põlme et al. 2020); Decay sugar-rich substrates (Põlme et al. 2020) | Põlme S, et al. 2020. Fungal Diversity 105:1-16 ((https://doi.org/10.1007/s13225-020-00466-2)) |
| Otu24790 | 1 | k__Fungi(100);p__Ascomycota(100);c__Saccharomycetes(100);o__Saccharomycetales(100);f__Metschnikowiaceae(100);g__Metschnikowia(100);s__Metschnikowia_pulcherrima(100); | Metschnikowia | 13 | Saprotroph | \|Undefined Saprotroph\| | Yeast | NULL | Probable | As nectar-tap_saprotroph (Põlme et al. 2020); Decay sugar-rich substrates (Põlme et al. 2020) | Põlme S, et al. 2020. Fungal Diversity 105:1-16 ((https://doi.org/10.1007/s13225-020-00466-2)) |
| Otu24792 | 1 | k__Fungi(100);p__Ascomycota(100);c__Saccharomycetes(100);o__Saccharomycetales(100);f__Metschnikowiaceae(100);g__Metschnikowia(100);s__Metschnikowia_pulcherrima(100); | Metschnikowia | 13 | Saprotroph | \|Undefined Saprotroph\| | Yeast | NULL | Probable | As nectar-tap_saprotroph (Põlme et al. 2020); Decay sugar-rich substrates (Põlme et al. 2020) | Põlme S, et al. 2020. Fungal Diversity 105:1-16 ((https://doi.org/10.1007/s13225-020-00466-2)) |
| Otu24793 | 1 | k__Fungi(100);p__Ascomycota(100);c__Saccharomycetes(100);o__Saccharomycetales(100);f__Metschnikowiaceae(100);g__Metschnikowia(100);s__Metschnikowia_pulcherrima(100); | Metschnikowia | 13 | Saprotroph | \|Undefined Saprotroph\| | Yeast | NULL | Probable | As nectar-tap_saprotroph (Põlme et al. 2020); Decay sugar-rich substrates (Põlme et al. 2020) | Põlme S, et al. 2020. Fungal Diversity 105:1-16 ((https://doi.org/10.1007/s13225-020-00466-2)) |

| | | | | | | | | | | | |
|---|---|---|---|---|---|---|---|---|---|---|---|
| Otu24794 | 1 | k__Fungi(100);p__Ascomycota(100);c__Saccharomycetes(100);o__Saccharomycetales(100);f__Metschnikowiaceae(100);g__Metschnikowia(100);s__Metschnikowia_pulcherrima(100); | Metschnikowia | 13 | Saprotroph | \|Undefined Saprotroph\| | Yeast | NULL | Probable | As nectar-tap_saprotroph (Põlme et al. 2020); Decay sugar-rich substrates (Põlme et al. 2020) | Põlme S, et al. 2020. Fungal Diversity 105:1-16 ((https://doi.org/10.1007/s13225-020-00466-2)) |
| Otu24795 | 1 | k__Fungi(100);p__Ascomycota(100);c__Saccharomycetes(100);o__Saccharomycetales(100);f__Metschnikowiaceae(100);g__Metschnikowia(100);s__Metschnikowia_pulcherrima(100); | Metschnikowia | 13 | Saprotroph | \|Undefined Saprotroph\| | Yeast | NULL | Probable | As nectar-tap_saprotroph (Põlme et al. 2020); Decay sugar-rich substrates (Põlme et al. 2020) | Põlme S, et al. 2020. Fungal Diversity 105:1-16 ((https://doi.org/10.1007/s13225-020-00466-2)) |
| Otu24796 | 1 | k__Fungi(100);p__Ascomycota(100);c__Saccharomycetes(100);o__Saccharomycetales(100);f__Metschnikowiaceae(100);g__Metschnikowia(100);s__Metschnikowia_pulcherrima(100); | Metschnikowia | 13 | Saprotroph | \|Undefined Saprotroph\| | Yeast | NULL | Probable | As nectar-tap_saprotroph (Põlme et al. 2020); Decay sugar-rich substrates (Põlme et al. 2020) | Põlme S, et al. 2020. Fungal Diversity 105:1-16 ((https://doi.org/10.1007/s13225-020-00466-2)) |
| Otu24797 | 1 | k__Fungi(100);p__Ascomycota(100);c__Saccharomycetes(100);o__Saccharomycetales(100);f__Metschnikowiaceae(100);g__Metschnikowia(100);g__Metschnikowia_unclassified(100); | Metschnikowia | 13 | Saprotroph | \|Undefined Saprotroph\| | Yeast | NULL | Probable | As nectar-tap_saprotroph (Põlme et al. 2020); Decay sugar-rich substrates (Põlme et al. 2020) | Põlme S, et al. 2020. Fungal Diversity 105:1-16 ((https://doi.org/10.1007/s13225-020-00466-2)) |
| Otu24798 | 1 | k__Fungi(100);p__Ascomycota(100);c__Saccharomycetes(100);o__Saccharomycetales(100);f__Metschnikowiaceae(100);g__Metschnikowia(100);s__Metschnikowia_pulcherrima(100); | Metschnikowia | 13 | Saprotroph | \|Undefined Saprotroph\| | Yeast | NULL | Probable | As nectar-tap_saprotroph (Põlme et al. 2020); Decay sugar-rich substrates (Põlme et al. 2020) | Põlme S, et al. 2020. Fungal Diversity 105:1-16 ((https://doi.org/10.1007/s13225-020-00466-2)) |
| Otu24800 | 1 | k__Fungi(100);p__Ascomycota(100);c__Saccharomycetes(100);o__Saccharomycetales(100);f__Metschnikowiaceae(100);g__Metschnikowia(100);s__Metschnikowia_pulcherrima(100); | Metschnikowia | 13 | Saprotroph | \|Undefined Saprotroph\| | Yeast | NULL | Probable | As nectar-tap_saprotroph (Põlme et al. 2020); Decay sugar-rich substrates (Põlme et al. 2020) | Põlme S, et al. 2020. Fungal Diversity 105:1-16 ((https://doi.org/10.1007/s13225-020-00466-2)) |
| Otu24801 | 1 | k__Fungi(100);p__Ascomycota(100);c__Saccharomycetes(100);o__Saccharomycetales(100);f__Metschnikowiaceae(100);g__Metschnikowia(100);s__Metschnikowia_pulcherrima(100); | Metschnikowia | 13 | Saprotroph | \|Undefined Saprotroph\| | Yeast | NULL | Probable | As nectar-tap_saprotroph (Põlme et al. 2020); Decay sugar-rich substrates (Põlme et al. 2020) | Põlme S, et al. 2020. Fungal Diversity 105:1-16 ((https://doi.org/10.1007/s13225-020-00466-2)) |
| Otu24803 | 1 | k__Fungi(100);p__Ascomycota(100);c__Saccharomycetes(100);o__Saccharomycetales(100);f__Metschnikowiaceae(100);g__Metschnikowia(100);s__Metschnikowia_pulcherrima(100); | Metschnikowia | 13 | Saprotroph | \|Undefined Saprotroph\| | Yeast | NULL | Probable | As nectar-tap_saprotroph (Põlme et al. 2020); Decay sugar-rich substrates (Põlme et al. 2020) | Põlme S, et al. 2020. Fungal Diversity 105:1-16 ((https://doi.org/10.1007/s13225-020-00466-2)) |
| Otu24804 | 1 | k__Fungi(100);p__Ascomycota(100);c__Saccharomycetes(100);o__Saccharomycetales(100);f__Metschnikowiaceae(100);g__Metschnikowia(100);s__Metschnikowia_pulcherrima(100); | Metschnikowia | 13 | Saprotroph | \|Undefined Saprotroph\| | Yeast | NULL | Probable | As nectar-tap_saprotroph (Põlme et al. 2020); Decay sugar-rich substrates (Põlme et al. 2020) | Põlme S, et al. 2020. Fungal Diversity 105:1-16 ((https://doi.org/10.1007/s13225-020-00466-2)) |
| Otu24805 | 1 | k__Fungi(100);p__Ascomycota(100);c__Saccharomycetes(100);o__Saccharomycetales(100);f__Metschnikowiaceae(100);g__Metschnikowia(100);s__Metschnikowia_pulcherrima(100); | Metschnikowia | 13 | Saprotroph | \|Undefined Saprotroph\| | Yeast | NULL | Probable | As nectar-tap_saprotroph (Põlme et al. 2020); Decay sugar-rich substrates (Põlme et al. 2020) | Põlme S, et al. 2020. Fungal Diversity 105:1-16 ((https://doi.org/10.1007/s13225-020-00466-2)) |
| Otu24806 | 1 | k__Fungi(100);p__Ascomycota(100);c__Saccharomycetes(100);o__Saccharomycetales(100);f__Metschnikowiaceae(100);g__Metschnikowia(100);s__Metschnikowia_pulcherrima(100); | Metschnikowia | 13 | Saprotroph | \|Undefined Saprotroph\| | Yeast | NULL | Probable | As nectar-tap_saprotroph (Põlme et al. 2020); Decay sugar-rich substrates (Põlme et al. 2020) | Põlme S, et al. 2020. Fungal Diversity 105:1-16 ((https://doi.org/10.1007/s13225-020-00466-2)) |
| Otu24808 | 1 | k__Fungi(100);p__Ascomycota(100);c__Saccharomycetes(100);o__Saccharomycetales(100);f__Metschnikowiaceae(100);g__Metschnikowia(100);s__Metschnikowia_pulcherrima(100); | Metschnikowia | 13 | Saprotroph | \|Undefined Saprotroph\| | Yeast | NULL | Probable | As nectar-tap_saprotroph (Põlme et al. 2020); Decay sugar-rich substrates (Põlme et al. 2020) | Põlme S, et al. 2020. Fungal Diversity 105:1-16 ((https://doi.org/10.1007/s13225-020-00466-2)) |
| Otu24813 | 1 | k__Fungi(100);p__Ascomycota(100);c__Saccharomycetes(100);o__Saccharomycetales(100);f__Metschnikowiaceae(100);g__Metschnikowia(100);g__Metschnikowia_unclassified(100); | Metschnikowia | 13 | Saprotroph | \|Undefined Saprotroph\| | Yeast | NULL | Probable | As nectar-tap_saprotroph (Põlme et al. 2020); Decay sugar-rich substrates (Põlme et al. 2020) | Põlme S, et al. 2020. Fungal Diversity 105:1-16 ((https://doi.org/10.1007/s13225-020-00466-2)) |
| Otu24814 | 1 | k__Fungi(100);p__Ascomycota(100);c__Saccharomycetes(100);o__Saccharomycetales(100);f__Metschnikowiaceae(100);g__Metschnikowia(100);g__Metschnikowia_unclassified(100); | Metschnikowia | 13 | Saprotroph | \|Undefined Saprotroph\| | Yeast | NULL | Probable | As nectar-tap_saprotroph (Põlme et al. 2020); Decay sugar-rich substrates (Põlme et al. 2020) | Põlme S, et al. 2020. Fungal Diversity 105:1-16 ((https://doi.org/10.1007/s13225-020-00466-2)) |
| Otu24815 | 1 | k__Fungi(100);p__Ascomycota(100);c__Saccharomycetes(100);o__Saccharomycetales(100);f__Metschnikowiaceae(100);g__Metschnikowia(100);s__Metschnikowia_pulcherrima(100); | Metschnikowia | 13 | Saprotroph | \|Undefined Saprotroph\| | Yeast | NULL | Probable | As nectar-tap_saprotroph (Põlme et al. 2020); Decay sugar-rich substrates (Põlme et al. 2020) | Põlme S, et al. 2020. Fungal Diversity 105:1-16 ((https://doi.org/10.1007/s13225-020-00466-2)) |
| Otu24816 | 1 | k__Fungi(100);p__Ascomycota(100);c__Saccharomycetes(100);o__Saccharomycetales(100);f__Metschnikowiaceae(100);g__Metschnikowia(100);s__Metschnikowia_pulcherrima(100); | Metschnikowia | 13 | Saprotroph | \|Undefined Saprotroph\| | Yeast | NULL | Probable | As nectar-tap_saprotroph (Põlme et al. 2020); Decay sugar-rich substrates (Põlme et al. 2020) | Põlme S, et al. 2020. Fungal Diversity 105:1-16 ((https://doi.org/10.1007/s13225-020-00466-2)) |
| Otu24819 | 1 | k__Fungi(100);p__Ascomycota(100);c__Saccharomycetes(100);o__Saccharomycetales(100);f__Metschnikowiaceae(100);g__Metschnikowia(100);s__Metschnikowia_pulcherrima(100); | Metschnikowia | 13 | Saprotroph | \|Undefined Saprotroph\| | Yeast | NULL | Probable | As nectar-tap_saprotroph (Põlme et al. 2020); Decay sugar-rich substrates (Põlme et al. 2020) | Põlme S, et al. 2020. Fungal Diversity 105:1-16 ((https://doi.org/10.1007/s13225-020-00466-2)) |
| Otu24820 | 1 | k__Fungi(100);p__Ascomycota(100);c__Saccharomycetes(100);o__Saccharomycetales(100);f__Metschnikowiaceae(100);g__Metschnikowia(100);s__Metschnikowia_pulcherrima(100); | Metschnikowia | 13 | Saprotroph | \|Undefined Saprotroph\| | Yeast | NULL | Probable | As nectar-tap_saprotroph (Põlme et al. 2020); Decay sugar-rich substrates (Põlme et al. 2020) | Põlme S, et al. 2020. Fungal Diversity 105:1-16 ((https://doi.org/10.1007/s13225-020-00466-2)) |
| Otu24821 | 1 | k__Fungi(100);p__Ascomycota(100);c__Saccharomycetes(100);o__Saccharomycetales(100);f__Metschnikowiaceae(100);g__Metschnikowia(100);s__Metschnikowia_pulcherrima(100); | Metschnikowia | 13 | Saprotroph | \|Undefined Saprotroph\| | Yeast | NULL | Probable | As nectar-tap_saprotroph (Põlme et al. 2020); Decay sugar-rich substrates (Põlme et al. 2020) | Põlme S, et al. 2020. Fungal Diversity 105:1-16 ((https://doi.org/10.1007/s13225-020-00466-2)) |
| Otu24822 | 1 | k__Fungi(100);p__Ascomycota(100);c__Saccharomycetes(100);o__Saccharomycetales(100);f__Metschnikowiaceae(100);g__Metschnikowia(100);s__Metschnikowia_pulcherrima(100); | Metschnikowia | 13 | Saprotroph | \|Undefined Saprotroph\| | Yeast | NULL | Probable | As nectar-tap_saprotroph (Põlme et al. 2020); Decay sugar-rich substrates (Põlme et al. 2020) | Põlme S, et al. 2020. Fungal Diversity 105:1-16 ((https://doi.org/10.1007/s13225-020-00466-2)) |
| Otu24824 | 1 | k__Fungi(100);p__Ascomycota(100);c__Saccharomycetes(100);o__Saccharomycetales(100);f__Metschnikowiaceae(100);g__Metschnikowia(100);s__Metschnikowia_pulcherrima(100); | Metschnikowia | 13 | Saprotroph | \|Undefined Saprotroph\| | Yeast | NULL | Probable | As nectar-tap_saprotroph (Põlme et al. 2020); Decay sugar-rich substrates (Põlme et al. 2020) | Põlme S, et al. 2020. Fungal Diversity 105:1-16 ((https://doi.org/10.1007/s13225-020-00466-2)) |
| Otu24827 | 1 | k__Fungi(100);p__Ascomycota(100);c__Saccharomycetes(100);o__Saccharomycetales(100);f__Metschnikowiaceae(100);g__Metschnikowia(100);s__Metschnikowia_pulcherrima(100); | Metschnikowia | 13 | Saprotroph | \|Undefined Saprotroph\| | Yeast | NULL | Probable | As nectar-tap_saprotroph (Põlme et al. 2020); Decay sugar-rich substrates (Põlme et al. 2020) | Põlme S, et al. 2020. Fungal Diversity 105:1-16 ((https://doi.org/10.1007/s13225-020-00466-2)) |
| Otu24829 | 1 | k__Fungi(100);p__Ascomycota(100);c__Saccharomycetes(100);o__Saccharomycetales(100);f__Metschnikowiaceae(100);g__Metschnikowia(100);s__Metschnikowia_pulcherrima(100); | Metschnikowia | 13 | Saprotroph | \|Undefined Saprotroph\| | Yeast | NULL | Probable | As nectar-tap_saprotroph (Põlme et al. 2020); Decay sugar-rich substrates (Põlme et al. 2020) | Põlme S, et al. 2020. Fungal Diversity 105:1-16 ((https://doi.org/10.1007/s13225-020-00466-2)) |
| Otu24830 | 1 | k__Fungi(100);p__Ascomycota(100);c__Saccharomycetes(100);o__Saccharomycetales(100);f__Metschnikowiaceae(100);g__Metschnikowia(100);s__Metschnikowia_pulcherrima(100); | Metschnikowia | 13 | Saprotroph | \|Undefined Saprotroph\| | Yeast | NULL | Probable | As nectar-tap_saprotroph (Põlme et al. 2020); Decay sugar-rich substrates (Põlme et al. 2020) | Põlme S, et al. 2020. Fungal Diversity 105:1-16 ((https://doi.org/10.1007/s13225-020-00466-2)) |
| Otu24831 | 1 | k__Fungi(100);p__Ascomycota(100);c__Saccharomycetes(100);o__Saccharomycetales(100);f__Metschnikowiaceae(100);g__Metschnikowia(100);g__Metschnikowia_unclassified(100); | Metschnikowia | 13 | Saprotroph | \|Undefined Saprotroph\| | Yeast | NULL | Probable | As nectar-tap_saprotroph (Põlme et al. 2020); Decay sugar-rich substrates (Põlme et al. 2020) | Põlme S, et al. 2020. Fungal Diversity 105:1-16 ((https://doi.org/10.1007/s13225-020-00466-2)) |
| Otu24832 | 1 | k__Fungi(100);p__Ascomycota(100);c__Saccharomycetes(100);o__Saccharomycetales(100);f__Metschnikowiaceae(100);g__Metschnikowia(100);s__Metschnikowia_pulcherrima(100); | Metschnikowia | 13 | Saprotroph | \|Undefined Saprotroph\| | Yeast | NULL | Probable | As nectar-tap_saprotroph (Põlme et al. 2020); Decay sugar-rich substrates (Põlme et al. 2020) | Põlme S, et al. 2020. Fungal Diversity 105:1-16 ((https://doi.org/10.1007/s13225-020-00466-2)) |
| Otu24834 | 1 | k__Fungi(100);p__Ascomycota(100);c__Saccharomycetes(100);o__Saccharomycetales(100);f__Metschnikowiaceae(100);g__Metschnikowia(100);g__Metschnikowia_unclassified(100); | Metschnikowia | 13 | Saprotroph | \|Undefined Saprotroph\| | Yeast | NULL | Probable | As nectar-tap_saprotroph (Põlme et al. 2020); Decay sugar-rich substrates (Põlme et al. 2020) | Põlme S, et al. 2020. Fungal Diversity 105:1-16 ((https://doi.org/10.1007/s13225-020-00466-2)) |
| Otu24835 | 1 | k__Fungi(100);p__Ascomycota(100);c__Saccharomycetes(100);o__Saccharomycetales(100);f__Metschnikowiaceae(100);g__Metschnikowia(100);s__Metschnikowia_pulcherrima(100); | Metschnikowia | 13 | Saprotroph | \|Undefined Saprotroph\| | Yeast | NULL | Probable | As nectar-tap_saprotroph (Põlme et al. 2020); Decay sugar-rich substrates (Põlme et al. 2020) | Põlme S, et al. 2020. Fungal Diversity 105:1-16 ((https://doi.org/10.1007/s13225-020-00466-2)) |
| Otu24838 | 1 | k__Fungi(100);p__Ascomycota(100);c__Saccharomycetes(100);o__Saccharomycetales(100);f__Metschnikowiaceae(100);g__Metschnikowia(100);s__Metschnikowia_pulcherrima(100); | Metschnikowia | 13 | Saprotroph | \|Undefined Saprotroph\| | Yeast | NULL | Probable | As nectar-tap_saprotroph (Põlme et al. 2020); Decay sugar-rich substrates (Põlme et al. 2020) | Põlme S, et al. 2020. Fungal Diversity 105:1-16 ((https://doi.org/10.1007/s13225-020-00466-2)) |
| Otu24842 | 1 | k__Fungi(100);p__Ascomycota(100);c__Saccharomycetes(100);o__Saccharomycetales(100);f__Metschnikowiaceae(100);g__Metschnikowia(100);s__Metschnikowia_pulcherrima(100); | Metschnikowia | 13 | Saprotroph | \|Undefined Saprotroph\| | Yeast | NULL | Probable | As nectar-tap_saprotroph (Põlme et al. 2020); Decay sugar-rich substrates (Põlme et al. 2020) | Põlme S, et al. 2020. Fungal Diversity 105:1-16 ((https://doi.org/10.1007/s13225-020-00466-2)) |
| Otu24844 | 1 | k__Fungi(100);p__Ascomycota(100);c__Saccharomycetes(100);o__Saccharomycetales(100);f__Metschnikowiaceae(100);g__Metschnikowia(100);g__Metschnikowia_unclassified(100); | Metschnikowia | 13 | Saprotroph | \|Undefined Saprotroph\| | Yeast | NULL | Probable | As nectar-tap_saprotroph (Põlme et al. 2020); Decay sugar-rich substrates (Põlme et al. 2020) | Põlme S, et al. 2020. Fungal Diversity 105:1-16 ((https://doi.org/10.1007/s13225-020-00466-2)) |
| Otu24845 | 1 | k__Fungi(100);p__Ascomycota(100);c__Saccharomycetes(100);o__Saccharomycetales(100);f__Metschnikowiaceae(100);g__Metschnikowia(100);s__Metschnikowia_pulcherrima(100); | Metschnikowia | 13 | Saprotroph | \|Undefined Saprotroph\| | Yeast | NULL | Probable | As nectar-tap_saprotroph (Põlme et al. 2020); Decay sugar-rich substrates (Põlme et al. 2020) | Põlme S, et al. 2020. Fungal Diversity 105:1-16 ((https://doi.org/10.1007/s13225-020-00466-2)) |
| Otu24847 | 1 | k__Fungi(100);p__Ascomycota(100);c__Saccharomycetes(100);o__Saccharomycetales(100);f__Metschnikowiaceae(100);g__Metschnikowia(100);g__Metschnikowia_unclassified(100); | Metschnikowia | 13 | Saprotroph | \|Undefined Saprotroph\| | Yeast | NULL | Probable | As nectar-tap_saprotroph (Põlme et al. 2020); Decay sugar-rich substrates (Põlme et al. 2020) | Põlme S, et al. 2020. Fungal Diversity 105:1-16 ((https://doi.org/10.1007/s13225-020-00466-2)) |
| Otu24848 | 1 | k__Fungi(100);p__Ascomycota(100);c__Saccharomycetes(100);o__Saccharomycetales(100);f__Metschnikowiaceae(100);g__Metschnikowia(100);s__Metschnikowia_pulcherrima(100); | Metschnikowia | 13 | Saprotroph | \|Undefined Saprotroph\| | Yeast | NULL | Probable | As nectar-tap_saprotroph (Põlme et al. 2020); Decay sugar-rich substrates (Põlme et al. 2020) | Põlme S, et al. 2020. Fungal Diversity 105:1-16 ((https://doi.org/10.1007/s13225-020-00466-2)) |
| Otu24849 | 1 | k__Fungi(100);p__Ascomycota(100);c__Saccharomycetes(100);o__Saccharomycetales(100);f__Metschnikowiaceae(100);g__Metschnikowia(100);s__Metschnikowia_pulcherrima(100); | Metschnikowia | 13 | Saprotroph | \|Undefined Saprotroph\| | Yeast | NULL | Probable | As nectar-tap_saprotroph (Põlme et al. 2020); Decay sugar-rich substrates (Põlme et al. 2020) | Põlme S, et al. 2020. Fungal Diversity 105:1-16 ((https://doi.org/10.1007/s13225-020-00466-2)) |
| Otu24851 | 1 | k__Fungi(100);p__Ascomycota(100);c__Saccharomycetes(100);o__Saccharomycetales(100);f__Metschnikowiaceae(100);g__Metschnikowia(100);g__Metschnikowia_unclassified(100); | Metschnikowia | 13 | Saprotroph | \|Undefined Saprotroph\| | Yeast | NULL | Probable | As nectar-tap_saprotroph (Põlme et al. 2020); Decay sugar-rich substrates (Põlme et al. 2020) | Põlme S, et al. 2020. Fungal Diversity 105:1-16 ((https://doi.org/10.1007/s13225-020-00466-2)) |
| Otu24852 | 1 | k__Fungi(100);p__Ascomycota(100);c__Saccharomycetes(100);o__Saccharomycetales(100);f__Metschnikowiaceae(100);g__Metschnikowia(100);s__Metschnikowia_pulcherrima(100); | Metschnikowia | 13 | Saprotroph | \|Undefined Saprotroph\| | Yeast | NULL | Probable | As nectar-tap_saprotroph (Põlme et al. 2020); Decay sugar-rich substrates (Põlme et al. 2020) | Põlme S, et al. 2020. Fungal Diversity 105:1-16 ((https://doi.org/10.1007/s13225-020-00466-2)) |
| Otu24853 | 1 | k__Fungi(100);p__Ascomycota(100);c__Saccharomycetes(100);o__Saccharomycetales(100);f__Metschnikowiaceae(100);g__Metschnikowia(100);s__Metschnikowia_pulcherrima(100); | Metschnikowia | 13 | Saprotroph | \|Undefined Saprotroph\| | Yeast | NULL | Probable | As nectar-tap_saprotroph (Põlme et al. 2020); Decay sugar-rich substrates (Põlme et al. 2020) | Põlme S, et al. 2020. Fungal Diversity 105:1-16 ((https://doi.org/10.1007/s13225-020-00466-2)) |
| Otu24854 | 1 | k__Fungi(100);p__Ascomycota(100);c__Saccharomycetes(100);o__Saccharomycetales(100);f__Metschnikowiaceae(100);g__Metschnikowia(100);s__Metschnikowia_pulcherrima(100); | Metschnikowia | 13 | Saprotroph | \|Undefined Saprotroph\| | Yeast | NULL | Probable | As nectar-tap_saprotroph (Põlme et al. 2020); Decay sugar-rich substrates (Põlme et al. 2020) | Põlme S, et al. 2020. Fungal Diversity 105:1-16 ((https://doi.org/10.1007/s13225-020-00466-2)) |
| Otu24863 | 1 | k__Fungi(100);p__Ascomycota(100);c__Saccharomycetes(100);o__Saccharomycetales(100);f__Metschnikowiaceae(100);g__Metschnikowia(100);s__Metschnikowia_pulcherrima(100); | Metschnikowia | 13 | Saprotroph | \|Undefined Saprotroph\| | Yeast | NULL | Probable | As nectar-tap_saprotroph (Põlme et al. 2020); Decay sugar-rich substrates (Põlme et al. 2020) | Põlme S, et al. 2020. Fungal Diversity 105:1-16 ((https://doi.org/10.1007/s13225-020-00466-2)) |
| Otu24865 | 1 | k__Fungi(100);p__Ascomycota(100);c__Saccharomycetes(100);o__Saccharomycetales(100);f__Metschnikowiaceae(100);g__Metschnikowia(100);g__Metschnikowia_unclassified(100); | Metschnikowia | 13 | Saprotroph | \|Undefined Saprotroph\| | Yeast | NULL | Probable | As nectar-tap_saprotroph (Põlme et al. 2020); Decay sugar-rich substrates (Põlme et al. 2020) | Põlme S, et al. 2020. Fungal Diversity 105:1-16 ((https://doi.org/10.1007/s13225-020-00466-2)) |
| Otu24866 | 1 | k__Fungi(100);p__Ascomycota(100);c__Saccharomycetes(100);o__Saccharomycetales(100);f__Metschnikowiaceae(100);g__Metschnikowia(100);s__Metschnikowia_pulcherrima(100); | Metschnikowia | 13 | Saprotroph | \|Undefined Saprotroph\| | Yeast | NULL | Probable | As nectar-tap_saprotroph (Põlme et al. 2020); Decay sugar-rich substrates (Põlme et al. 2020) | Põlme S, et al. 2020. Fungal Diversity 105:1-16 ((https://doi.org/10.1007/s13225-020-00466-2)) |
| Otu24867 | 1 | k__Fungi(100);p__Ascomycota(100);c__Saccharomycetes(100);o__Saccharomycetales(100);f__Metschnikowiaceae(100);g__Metschnikowia(100);s__Metschnikowia_pulcherrima(100); | Metschnikowia | 13 | Saprotroph | \|Undefined Saprotroph\| | Yeast | NULL | Probable | As nectar-tap_saprotroph (Põlme et al. 2020); Decay sugar-rich substrates (Põlme et al. 2020) | Põlme S, et al. 2020. Fungal Diversity 105:1-16 ((https://doi.org/10.1007/s13225-020-00466-2)) |
| Otu24868 | 1 | k__Fungi(100);p__Ascomycota(100);c__Saccharomycetes(100);o__Saccharomycetales(100);f__Metschnikowiaceae(100);g__Metschnikowia(100);s__Metschnikowia_pulcherrima(100); | Metschnikowia | 13 | Saprotroph | \|Undefined Saprotroph\| | Yeast | NULL | Probable | As nectar-tap_saprotroph (Põlme et al. 2020); Decay sugar-rich substrates (Põlme et al. 2020) | Põlme S, et al. 2020. Fungal Diversity 105:1-16 ((https://doi.org/10.1007/s13225-020-00466-2)) |
| Otu24869 | 1 | k__Fungi(100);p__Ascomycota(100);c__Saccharomycetes(100);o__Saccharomycetales(100);f__Metschnikowiaceae(100);g__Metschnikowia(100);s__Metschnikowia_pulcherrima(100); | Metschnikowia | 13 | Saprotroph | \|Undefined Saprotroph\| | Yeast | NULL | Probable | As nectar-tap_saprotroph (Põlme et al. 2020); Decay sugar-rich substrates (Põlme et al. 2020) | Põlme S, et al. 2020. Fungal Diversity 105:1-16 ((https://doi.org/10.1007/s13225-020-00466-2)) |
| Otu24870 | 1 | k__Fungi(100);p__Ascomycota(100);c__Saccharomycetes(100);o__Saccharomycetales(100);f__Metschnikowiaceae(100);g__Metschnikowia(100);s__Metschnikowia_pulcherrima(100); | Metschnikowia | 13 | Saprotroph | \|Undefined Saprotroph\| | Yeast | NULL | Probable | As nectar-tap_saprotroph (Põlme et al. 2020); Decay sugar-rich substrates (Põlme et al. 2020) | Põlme S, et al. 2020. Fungal Diversity 105:1-16 ((https://doi.org/10.1007/s13225-020-00466-2)) |
| Otu24872 | 1 | k__Fungi(100);p__Ascomycota(100);c__Saccharomycetes(100);o__Saccharomycetales(100);f__Metschnikowiaceae(100);g__Metschnikowia(100);s__Metschnikowia_pulcherrima(100); | Metschnikowia | 13 | Saprotroph | \|Undefined Saprotroph\| | Yeast | NULL | Probable | As nectar-tap_saprotroph (Põlme et al. 2020); Decay sugar-rich substrates (Põlme et al. 2020) | Põlme S, et al. 2020. Fungal Diversity 105:1-16 ((https://doi.org/10.1007/s13225-020-00466-2)) |
| Otu24876 | 1 | k__Fungi(100);p__Ascomycota(100);c__Saccharomycetes(100);o__Saccharomycetales(100);f__Metschnikowiaceae(100);g__Metschnikowia(100);g__Metschnikowia_unclassified(100); | Metschnikowia | 13 | Saprotroph | \|Undefined Saprotroph\| | Yeast | NULL | Probable | As nectar-tap_saprotroph (Põlme et al. 2020); Decay sugar-rich substrates (Põlme et al. 2020) | Põlme S, et al. 2020. Fungal Diversity 105:1-16 ((https://doi.org/10.1007/s13225-020-00466-2)) |
| Otu24877 | 1 | k__Fungi(100);p__Ascomycota(100);c__Saccharomycetes(100);o__Saccharomycetales(100);f__Metschnikowiaceae(100);g__Metschnikowia(100);g__Metschnikowia_unclassified(100); | Metschnikowia | 13 | Saprotroph | \|Undefined Saprotroph\| | Yeast | NULL | Probable | As nectar-tap_saprotroph (Põlme et al. 2020); Decay sugar-rich substrates (Põlme et al. 2020) | Põlme S, et al. 2020. Fungal Diversity 105:1-16 ((https://doi.org/10.1007/s13225-020-00466-2)) |
| Otu24878 | 1 | k__Fungi(100);p__Ascomycota(100);c__Saccharomycetes(100);o__Saccharomycetales(100);f__Metschnikowiaceae(100);g__Metschnikowia(100);s__Metschnikowia_pulcherrima(100); | Metschnikowia | 13 | Saprotroph | \|Undefined Saprotroph\| | Yeast | NULL | Probable | As nectar-tap_saprotroph (Põlme et al. 2020); Decay sugar-rich substrates (Põlme et al. 2020) | Põlme S, et al. 2020. Fungal Diversity 105:1-16 ((https://doi.org/10.1007/s13225-020-00466-2)) |
| Otu24879 | 1 | k__Fungi(100);p__Ascomycota(100);c__Saccharomycetes(100);o__Saccharomycetales(100);f__Metschnikowiaceae(100);g__Metschnikowia(100);s__Metschnikowia_pulcherrima(100); | Metschnikowia | 13 | Saprotroph | \|Undefined Saprotroph\| | Yeast | NULL | Probable | As nectar-tap_saprotroph (Põlme et al. 2020); Decay sugar-rich substrates (Põlme et al. 2020) | Põlme S, et al. 2020. Fungal Diversity 105:1-16 ((https://doi.org/10.1007/s13225-020-00466-2)) |
| Otu24880 | 1 | k__Fungi(100);p__Ascomycota(100);c__Saccharomycetes(100);o__Saccharomycetales(100);f__Metschnikowiaceae(100);g__Metschnikowia(100);s__Metschnikowia_pulcherrima(100); | Metschnikowia | 13 | Saprotroph | \|Undefined Saprotroph\| | Yeast | NULL | Probable | As nectar-tap_saprotroph (Põlme et al. 2020); Decay sugar-rich substrates (Põlme et al. 2020) | Põlme S, et al. 2020. Fungal Diversity 105:1-16 ((https://doi.org/10.1007/s13225-020-00466-2)) |
| Otu24881 | 1 | k__Fungi(100);p__Ascomycota(100);c__Saccharomycetes(100);o__Saccharomycetales(100);f__Metschnikowiaceae(100);g__Metschnikowia(100);g__Metschnikowia_unclassified(100); | Metschnikowia | 13 | Saprotroph | \|Undefined Saprotroph\| | Yeast | NULL | Probable | As nectar-tap_saprotroph (Põlme et al. 2020); Decay sugar-rich substrates (Põlme et al. 2020) | Põlme S, et al. 2020. Fungal Diversity 105:1-16 ((https://doi.org/10.1007/s13225-020-00466-2)) |
| Otu24882 | 1 | k__Fungi(100);p__Ascomycota(100);c__Saccharomycetes(100);o__Saccharomycetales(100);f__Metschnikowiaceae(100);g__Metschnikowia(100);g__Metschnikowia_unclassified(100); | Metschnikowia | 13 | Saprotroph | \|Undefined Saprotroph\| | Yeast | NULL | Probable | As nectar-tap_saprotroph (Põlme et al. 2020); Decay sugar-rich substrates (Põlme et al. 2020) | Põlme S, et al. 2020. Fungal Diversity 105:1-16 ((https://doi.org/10.1007/s13225-020-00466-2)) |
| Otu24884 | 1 | k__Fungi(100);p__Ascomycota(100);c__Saccharomycetes(100);o__Saccharomycetales(100);f__Metschnikowiaceae(100);g__Metschnikowia(100);s__Metschnikowia_pulcherrima(100); | Metschnikowia | 13 | Saprotroph | \|Undefined Saprotroph\| | Yeast | NULL | Probable | As nectar-tap_saprotroph (Põlme et al. 2020); Decay sugar-rich substrates (Põlme et al. 2020) | Põlme S, et al. 2020. Fungal Diversity 105:1-16 ((https://doi.org/10.1007/s13225-020-00466-2)) |
| Otu24885 | 1 | k__Fungi(100);p__Ascomycota(100);c__Saccharomycetes(100);o__Saccharomycetales(100);f__Metschnikowiaceae(100);g__Metschnikowia(100);s__Metschnikowia_pulcherrima(100); | Metschnikowia | 13 | Saprotroph | \|Undefined Saprotroph\| | Yeast | NULL | Probable | As nectar-tap_saprotroph (Põlme et al. 2020); Decay sugar-rich substrates (Põlme et al. 2020) | Põlme S, et al. 2020. Fungal Diversity 105:1-16 ((https://doi.org/10.1007/s13225-020-00466-2)) |
| Otu24886 | 1 | k__Fungi(100);p__Ascomycota(100);c__Saccharomycetes(100);o__Saccharomycetales(100);f__Metschnikowiaceae(100);g__Metschnikowia(100);g__Metschnikowia_unclassified(100); | Metschnikowia | 13 | Saprotroph | \|Undefined Saprotroph\| | Yeast | NULL | Probable | As nectar-tap_saprotroph (Põlme et al. 2020); Decay sugar-rich substrates (Põlme et al. 2020) | Põlme S, et al. 2020. Fungal Diversity 105:1-16 ((https://doi.org/10.1007/s13225-020-00466-2)) |
| Otu24887 | 1 | k__Fungi(100);p__Ascomycota(100);c__Saccharomycetes(100);o__Saccharomycetales(100);f__Metschnikowiaceae(100);g__Metschnikowia(100);s__Metschnikowia_pulcherrima(100); | Metschnikowia | 13 | Saprotroph | \|Undefined Saprotroph\| | Yeast | NULL | Probable | As nectar-tap_saprotroph (Põlme et al. 2020); Decay sugar-rich substrates (Põlme et al. 2020) | Põlme S, et al. 2020. Fungal Diversity 105:1-16 ((https://doi.org/10.1007/s13225-020-00466-2)) |
| Otu24889 | 1 | k__Fungi(100);p__Ascomycota(100);c__Saccharomycetes(100);o__Saccharomycetales(100);f__Metschnikowiaceae(100);g__Metschnikowia(100);g__Metschnikowia_unclassified(100); | Metschnikowia | 13 | Saprotroph | \|Undefined Saprotroph\| | Yeast | NULL | Probable | As nectar-tap_saprotroph (Põlme et al. 2020); Decay sugar-rich substrates (Põlme et al. 2020) | Põlme S, et al. 2020. Fungal Diversity 105:1-16 ((https://doi.org/10.1007/s13225-020-00466-2)) |
| Otu24890 | 1 | k__Fungi(100);p__Ascomycota(100);c__Saccharomycetes(100);o__Saccharomycetales(100);f__Metschnikowiaceae(100);g__Metschnikowia(100);s__Metschnikowia_pulcherrima(100); | Metschnikowia | 13 | Saprotroph | \|Undefined Saprotroph\| | Yeast | NULL | Probable | As nectar-tap_saprotroph (Põlme et al. 2020); Decay sugar-rich substrates (Põlme et al. 2020) | Põlme S, et al. 2020. Fungal Diversity 105:1-16 ((https://doi.org/10.1007/s13225-020-00466-2)) |
| Otu24891 | 1 | k__Fungi(100);p__Ascomycota(100);c__Saccharomycetes(100);o__Saccharomycetales(100);f__Metschnikowiaceae(100);g__Metschnikowia(100);s__Metschnikowia_pulcherrima(100); | Metschnikowia | 13 | Saprotroph | \|Undefined Saprotroph\| | Yeast | NULL | Probable | As nectar-tap_saprotroph (Põlme et al. 2020); Decay sugar-rich substrates (Põlme et al. 2020) | Põlme S, et al. 2020. Fungal Diversity 105:1-16 ((https://doi.org/10.1007/s13225-020-00466-2)) |
| Otu24892 | 1 | k__Fungi(100);p__Ascomycota(100);c__Saccharomycetes(100);o__Saccharomycetales(100);f__Metschnikowiaceae(100);g__Metschnikowia(100);s__Metschnikowia_pulcherrima(100); | Metschnikowia | 13 | Saprotroph | \|Undefined Saprotroph\| | Yeast | NULL | Probable | As nectar-tap_saprotroph (Põlme et al. 2020); Decay sugar-rich substrates (Põlme et al. 2020) | Põlme S, et al. 2020. Fungal Diversity 105:1-16 ((https://doi.org/10.1007/s13225-020-00466-2)) |
| Otu24893 | 1 | k__Fungi(100);p__Ascomycota(100);c__Saccharomycetes(100);o__Saccharomycetales(100);f__Metschnikowiaceae(100);g__Metschnikowia(100);s__Metschnikowia_pulcherrima(100); | Metschnikowia | 13 | Saprotroph | \|Undefined Saprotroph\| | Yeast | NULL | Probable | As nectar-tap_saprotroph (Põlme et al. 2020); Decay sugar-rich substrates (Põlme et al. 2020) | Põlme S, et al. 2020. Fungal Diversity 105:1-16 ((https://doi.org/10.1007/s13225-020-00466-2)) |
| Otu24894 | 1 | k__Fungi(100);p__Ascomycota(100);c__Saccharomycetes(100);o__Saccharomycetales(100);f__Metschnikowiaceae(100);g__Metschnikowia(100);s__Metschnikowia_pulcherrima(100); | Metschnikowia | 13 | Saprotroph | \|Undefined Saprotroph\| | Yeast | NULL | Probable | As nectar-tap_saprotroph (Põlme et al. 2020); Decay sugar-rich substrates (Põlme et al. 2020) | Põlme S, et al. 2020. Fungal Diversity 105:1-16 ((https://doi.org/10.1007/s13225-020-00466-2)) |
| Otu24896 | 1 | k__Fungi(100);p__Ascomycota(100);c__Saccharomycetes(100);o__Saccharomycetales(100);f__Metschnikowiaceae(100);g__Metschnikowia(100);s__Metschnikowia_pulcherrima(100); | Metschnikowia | 13 | Saprotroph | \|Undefined Saprotroph\| | Yeast | NULL | Probable | As nectar-tap_saprotroph (Põlme et al. 2020); Decay sugar-rich substrates (Põlme et al. 2020) | Põlme S, et al. 2020. Fungal Diversity 105:1-16 ((https://doi.org/10.1007/s13225-020-00466-2)) |
| Otu24897 | 1 | k__Fungi(100);p__Ascomycota(100);c__Saccharomycetes(100);o__Saccharomycetales(100);f__Metschnikowiaceae(100);g__Metschnikowia(100);s__Metschnikowia_pulcherrima(100); | Metschnikowia | 13 | Saprotroph | \|Undefined Saprotroph\| | Yeast | NULL | Probable | As nectar-tap_saprotroph (Põlme et al. 2020); Decay sugar-rich substrates (Põlme et al. 2020) | Põlme S, et al. 2020. Fungal Diversity 105:1-16 ((https://doi.org/10.1007/s13225-020-00466-2)) |
| Otu24898 | 1 | k__Fungi(100);p__Ascomycota(100);c__Saccharomycetes(100);o__Saccharomycetales(100);f__Metschnikowiaceae(100);g__Metschnikowia(100);s__Metschnikowia_pulcherrima(100); | Metschnikowia | 13 | Saprotroph | \|Undefined Saprotroph\| | Yeast | NULL | Probable | As nectar-tap_saprotroph (Põlme et al. 2020); Decay sugar-rich substrates (Põlme et al. 2020) | Põlme S, et al. 2020. Fungal Diversity 105:1-16 ((https://doi.org/10.1007/s13225-020-00466-2)) |
| Otu24899 | 1 | k__Fungi(100);p__Ascomycota(100);c__Saccharomycetes(100);o__Saccharomycetales(100);f__Metschnikowiaceae(100);g__Metschnikowia(100);s__Metschnikowia_pulcherrima(100); | Metschnikowia | 13 | Saprotroph | \|Undefined Saprotroph\| | Yeast | NULL | Probable | As nectar-tap_saprotroph (Põlme et al. 2020); Decay sugar-rich substrates (Põlme et al. 2020) | Põlme S, et al. 2020. Fungal Diversity 105:1-16 ((https://doi.org/10.1007/s13225-020-00466-2)) |
| Otu24904 | 1 | k__Fungi(100);p__Ascomycota(100);c__Saccharomycetes(100);o__Saccharomycetales(100);f__Metschnikowiaceae(100);g__Metschnikowia(100);s__Metschnikowia_pulcherrima(100); | Metschnikowia | 13 | Saprotroph | \|Undefined Saprotroph\| | Yeast | NULL | Probable | As nectar-tap_saprotroph (Põlme et al. 2020); Decay sugar-rich substrates (Põlme et al. 2020) | Põlme S, et al. 2020. Fungal Diversity 105:1-16 ((https://doi.org/10.1007/s13225-020-00466-2)) |
| Otu24907 | 1 | k__Fungi(100);p__Ascomycota(100);c__Saccharomycetes(100);o__Saccharomycetales(100);f__Metschnikowiaceae(100);g__Metschnikowia(100);g__Metschnikowia_unclassified(100); | Metschnikowia | 13 | Saprotroph | \|Undefined Saprotroph\| | Yeast | NULL | Probable | As nectar-tap_saprotroph (Põlme et al. 2020); Decay sugar-rich substrates (Põlme et al. 2020) | Põlme S, et al. 2020. Fungal Diversity 105:1-16 ((https://doi.org/10.1007/s13225-020-00466-2)) |
| Otu24908 | 1 | k__Fungi(100);p__Ascomycota(100);c__Saccharomycetes(100);o__Saccharomycetales(100);f__Metschnikowiaceae(100);g__Metschnikowia(100);s__Metschnikowia_pulcherrima(100); | Metschnikowia | 13 | Saprotroph | \|Undefined Saprotroph\| | Yeast | NULL | Probable | As nectar-tap_saprotroph (Põlme et al. 2020); Decay sugar-rich substrates (Põlme et al. 2020) | Põlme S, et al. 2020. Fungal Diversity 105:1-16 ((https://doi.org/10.1007/s13225-020-00466-2)) |
| Otu24909 | 1 | k__Fungi(100);p__Ascomycota(100);c__Saccharomycetes(100);o__Saccharomycetales(100);f__Metschnikowiaceae(100);g__Metschnikowia(100);g__Metschnikowia_unclassified(100); | Metschnikowia | 13 | Saprotroph | \|Undefined Saprotroph\| | Yeast | NULL | Probable | As nectar-tap_saprotroph (Põlme et al. 2020); Decay sugar-rich substrates (Põlme et al. 2020) | Põlme S, et al. 2020. Fungal Diversity 105:1-16 ((https://doi.org/10.1007/s13225-020-00466-2)) |
| Otu24910 | 1 | k__Fungi(100);p__Ascomycota(100);c__Saccharomycetes(100);o__Saccharomycetales(100);f__Metschnikowiaceae(100);g__Metschnikowia(100);s__Metschnikowia_pulcherrima(100); | Metschnikowia | 13 | Saprotroph | \|Undefined Saprotroph\| | Yeast | NULL | Probable | As nectar-tap_saprotroph (Põlme et al. 2020); Decay sugar-rich substrates (Põlme et al. 2020) | Põlme S, et al. 2020. Fungal Diversity 105:1-16 ((https://doi.org/10.1007/s13225-020-00466-2)) |
| Otu24911 | 1 | k__Fungi(100);p__Ascomycota(100);c__Saccharomycetes(100);o__Saccharomycetales(100);f__Metschnikowiaceae(100);g__Metschnikowia(100);g__Metschnikowia_unclassified(100); | Metschnikowia | 13 | Saprotroph | \|Undefined Saprotroph\| | Yeast | NULL | Probable | As nectar-tap_saprotroph (Põlme et al. 2020); Decay sugar-rich substrates (Põlme et al. 2020) | Põlme S, et al. 2020. Fungal Diversity 105:1-16 ((https://doi.org/10.1007/s13225-020-00466-2)) |
| Otu24912 | 1 | k__Fungi(100);p__Ascomycota(100);c__Saccharomycetes(100);o__Saccharomycetales(100);f__Metschnikowiaceae(100);g__Metschnikowia(100);g__Metschnikowia_unclassified(100); | Metschnikowia | 13 | Saprotroph | \|Undefined Saprotroph\| | Yeast | NULL | Probable | As nectar-tap_saprotroph (Põlme et al. 2020); Decay sugar-rich substrates (Põlme et al. 2020) | Põlme S, et al. 2020. Fungal Diversity 105:1-16 ((https://doi.org/10.1007/s13225-020-00466-2)) |
| Otu24913 | 1 | k__Fungi(100);p__Ascomycota(100);c__Saccharomycetes(100);o__Saccharomycetales(100);f__Metschnikowiaceae(100);g__Metschnikowia(100);s__Metschnikowia_pulcherrima(100); | Metschnikowia | 13 | Saprotroph | \|Undefined Saprotroph\| | Yeast | NULL | Probable | As nectar-tap_saprotroph (Põlme et al. 2020); Decay sugar-rich substrates (Põlme et al. 2020) | Põlme S, et al. 2020. Fungal Diversity 105:1-16 ((https://doi.org/10.1007/s13225-020-00466-2)) |
| Otu24914 | 1 | k__Fungi(100);p__Ascomycota(100);c__Saccharomycetes(100);o__Saccharomycetales(100);f__Metschnikowiaceae(100);g__Metschnikowia(100);s__Metschnikowia_pulcherrima(100); | Metschnikowia | 13 | Saprotroph | \|Undefined Saprotroph\| | Yeast | NULL | Probable | As nectar-tap_saprotroph (Põlme et al. 2020); Decay sugar-rich substrates (Põlme et al. 2020) | Põlme S, et al. 2020. Fungal Diversity 105:1-16 ((https://doi.org/10.1007/s13225-020-00466-2)) |
| Otu24915 | 1 | k__Fungi(100);p__Ascomycota(100);c__Saccharomycetes(100);o__Saccharomycetales(100);f__Metschnikowiaceae(100);g__Metschnikowia(100);s__Metschnikowia_pulcherrima(100); | Metschnikowia | 13 | Saprotroph | \|Undefined Saprotroph\| | Yeast | NULL | Probable | As nectar-tap_saprotroph (Põlme et al. 2020); Decay sugar-rich substrates (Põlme et al. 2020) | Põlme S, et al. 2020. Fungal Diversity 105:1-16 ((https://doi.org/10.1007/s13225-020-00466-2)) |
| Otu24916 | 1 | k__Fungi(100);p__Ascomycota(100);c__Saccharomycetes(100);o__Saccharomycetales(100);f__Metschnikowiaceae(100);g__Metschnikowia(100);s__Metschnikowia_pulcherrima(100); | Metschnikowia | 13 | Saprotroph | \|Undefined Saprotroph\| | Yeast | NULL | Probable | As nectar-tap_saprotroph (Põlme et al. 2020); Decay sugar-rich substrates (Põlme et al. 2020) | Põlme S, et al. 2020. Fungal Diversity 105:1-16 ((https://doi.org/10.1007/s13225-020-00466-2)) |
| Otu24917 | 1 | k__Fungi(100);p__Ascomycota(100);c__Saccharomycetes(100);o__Saccharomycetales(100);f__Metschnikowiaceae(100);g__Metschnikowia(100);s__Metschnikowia_pulcherrima(100); | Metschnikowia | 13 | Saprotroph | \|Undefined Saprotroph\| | Yeast | NULL | Probable | As nectar-tap_saprotroph (Põlme et al. 2020); Decay sugar-rich substrates (Põlme et al. 2020) | Põlme S, et al. 2020. Fungal Diversity 105:1-16 ((https://doi.org/10.1007/s13225-020-00466-2)) |
| Otu24918 | 1 | k__Fungi(100);p__Ascomycota(100);c__Saccharomycetes(100);o__Saccharomycetales(100);f__Metschnikowiaceae(100);g__Metschnikowia(100);s__Metschnikowia_pulcherrima(100); | Metschnikowia | 13 | Saprotroph | \|Undefined Saprotroph\| | Yeast | NULL | Probable | As nectar-tap_saprotroph (Põlme et al. 2020); Decay sugar-rich substrates (Põlme et al. 2020) | Põlme S, et al. 2020. Fungal Diversity 105:1-16 ((https://doi.org/10.1007/s13225-020-00466-2)) |
| Otu24919 | 1 | k__Fungi(100);p__Ascomycota(100);c__Saccharomycetes(100);o__Saccharomycetales(100);f__Metschnikowiaceae(100);g__Metschnikowia(100);s__Metschnikowia_pulcherrima(100); | Metschnikowia | 13 | Saprotroph | \|Undefined Saprotroph\| | Yeast | NULL | Probable | As nectar-tap_saprotroph (Põlme et al. 2020); Decay sugar-rich substrates (Põlme et al. 2020) | Põlme S, et al. 2020. Fungal Diversity 105:1-16 ((https://doi.org/10.1007/s13225-020-00466-2)) |
| Otu24920 | 1 | k__Fungi(100);p__Ascomycota(100);c__Saccharomycetes(100);o__Saccharomycetales(100);f__Metschnikowiaceae(100);g__Metschnikowia(100);s__Metschnikowia_pulcherrima(100); | Metschnikowia | 13 | Saprotroph | \|Undefined Saprotroph\| | Yeast | NULL | Probable | As nectar-tap_saprotroph (Põlme et al. 2020); Decay sugar-rich substrates (Põlme et al. 2020) | Põlme S, et al. 2020. Fungal Diversity 105:1-16 ((https://doi.org/10.1007/s13225-020-00466-2)) |
| Otu24921 | 1 | k__Fungi(100);p__Ascomycota(100);c__Saccharomycetes(100);o__Saccharomycetales(100);f__Metschnikowiaceae(100);g__Metschnikowia(100);s__Metschnikowia_pulcherrima(100); | Metschnikowia | 13 | Saprotroph | \|Undefined Saprotroph\| | Yeast | NULL | Probable | As nectar-tap_saprotroph (Põlme et al. 2020); Decay sugar-rich substrates (Põlme et al. 2020) | Põlme S, et al. 2020. Fungal Diversity 105:1-16 ((https://doi.org/10.1007/s13225-020-00466-2)) |
| Otu24922 | 1 | k__Fungi(100);p__Ascomycota(100);c__Saccharomycetes(100);o__Saccharomycetales(100);f__Metschnikowiaceae(100);g__Metschnikowia(100);g__Metschnikowia_unclassified(100); | Metschnikowia | 13 | Saprotroph | \|Undefined Saprotroph\| | Yeast | NULL | Probable | As nectar-tap_saprotroph (Põlme et al. 2020); Decay sugar-rich substrates (Põlme et al. 2020) | Põlme S, et al. 2020. Fungal Diversity 105:1-16 ((https://doi.org/10.1007/s13225-020-00466-2)) |
| Otu24923 | 1 | k__Fungi(100);p__Ascomycota(100);c__Saccharomycetes(100);o__Saccharomycetales(100);f__Metschnikowiaceae(100);g__Metschnikowia(100);s__Metschnikowia_pulcherrima(100); | Metschnikowia | 13 | Saprotroph | \|Undefined Saprotroph\| | Yeast | NULL | Probable | As nectar-tap_saprotroph (Põlme et al. 2020); Decay sugar-rich substrates (Põlme et al. 2020) | Põlme S, et al. 2020. Fungal Diversity 105:1-16 ((https://doi.org/10.1007/s13225-020-00466-2)) |
| Otu24924 | 1 | k__Fungi(100);p__Ascomycota(100);c__Saccharomycetes(100);o__Saccharomycetales(100);f__Metschnikowiaceae(100);g__Metschnikowia(100);s__Metschnikowia_pulcherrima(100); | Metschnikowia | 13 | Saprotroph | \|Undefined Saprotroph\| | Yeast | NULL | Probable | As nectar-tap_saprotroph (Põlme et al. 2020); Decay sugar-rich substrates (Põlme et al. 2020) | Põlme S, et al. 2020. Fungal Diversity 105:1-16 ((https://doi.org/10.1007/s13225-020-00466-2)) |
| Otu24925 | 1 | k__Fungi(100);p__Ascomycota(100);c__Saccharomycetes(100);o__Saccharomycetales(100);f__Metschnikowiaceae(100);g__Metschnikowia(100);s__Metschnikowia_pulcherrima(100); | Metschnikowia | 13 | Saprotroph | \|Undefined Saprotroph\| | Yeast | NULL | Probable | As nectar-tap_saprotroph (Põlme et al. 2020); Decay sugar-rich substrates (Põlme et al. 2020) | Põlme S, et al. 2020. Fungal Diversity 105:1-16 ((https://doi.org/10.1007/s13225-020-00466-2)) |
| Otu24926 | 1 | k__Fungi(100);p__Ascomycota(100);c__Saccharomycetes(100);o__Saccharomycetales(100);f__Metschnikowiaceae(100);g__Metschnikowia(100);s__Metschnikowia_pulcherrima(100); | Metschnikowia | 13 | Saprotroph | \|Undefined Saprotroph\| | Yeast | NULL | Probable | As nectar-tap_saprotroph (Põlme et al. 2020); Decay sugar-rich substrates (Põlme et al. 2020) | Põlme S, et al. 2020. Fungal Diversity 105:1-16 ((https://doi.org/10.1007/s13225-020-00466-2)) |
| Otu24927 | 1 | k__Fungi(100);p__Ascomycota(100);c__Saccharomycetes(100);o__Saccharomycetales(100);f__Metschnikowiaceae(100);g__Metschnikowia(100);s__Metschnikowia_pulcherrima(100); | Metschnikowia | 13 | Saprotroph | \|Undefined Saprotroph\| | Yeast | NULL | Probable | As nectar-tap_saprotroph (Põlme et al. 2020); Decay sugar-rich substrates (Põlme et al. 2020) | Põlme S, et al. 2020. Fungal Diversity 105:1-16 ((https://doi.org/10.1007/s13225-020-00466-2)) |
| Otu24928 | 1 | k__Fungi(100);p__Ascomycota(100);c__Saccharomycetes(100);o__Saccharomycetales(100);f__Metschnikowiaceae(100);g__Metschnikowia(100);s__Metschnikowia_pulcherrima(100); | Metschnikowia | 13 | Saprotroph | \|Undefined Saprotroph\| | Yeast | NULL | Probable | As nectar-tap_saprotroph (Põlme et al. 2020); Decay sugar-rich substrates (Põlme et al. 2020) | Põlme S, et al. 2020. Fungal Diversity 105:1-16 ((https://doi.org/10.1007/s13225-020-00466-2)) |
| Otu24929 | 1 | k__Fungi(100);p__Ascomycota(100);c__Saccharomycetes(100);o__Saccharomycetales(100);f__Metschnikowiaceae(100);g__Metschnikowia(100);s__Metschnikowia_pulcherrima(100); | Metschnikowia | 13 | Saprotroph | \|Undefined Saprotroph\| | Yeast | NULL | Probable | As nectar-tap_saprotroph (Põlme et al. 2020); Decay sugar-rich substrates (Põlme et al. 2020) | Põlme S, et al. 2020. Fungal Diversity 105:1-16 ((https://doi.org/10.1007/s13225-020-00466-2)) |
| Otu24950 | 1 | k__Fungi(100);p__Ascomycota(100);c__Saccharomycetes(100);o__Saccharomycetales(100);f__Saccharomycetales_family_Incertae_sedis(100);g__Candida(100);g__Candida_unclassified(100); | Candida | 13 | Pathotroph-Saprotroph-Symbiotroph | Animal Pathogen-Endophyte-Endosymbiont-Epiphyte-\|Undefined Saprotroph\| | Dimorphic Yeast-Yeast | NULL | Probable | As nectar-tap_saprotroph (Põlme et al. 2020); Decay sugar-rich substrates (Põlme et al. 2020); This asexual genus is polyphyletic and contain species that function very differently depend on what they are. Caution should be taken when interpreting guilds for Candida species. | Manolakaki D, et al. 2010. Virulence. 1: 367-375 ((https://doi.org/10.4161/viru.1.5.12796)); Mekbib SB, et al. 2011. Tropical Plant Pathology 36:233-240 ((https://doi.org/10.1590/S1982-56762011000400004)); Põlme S, et al. 2020. Fungal Diversity 105:1-16 ((https://doi.org/10.1007/s13225-020-00466-2)) |
| Otu24954 | 1 | k__Fungi(100);p__Ascomycota(100);c__Saccharomycetes(100);o__Saccharomycetales(100);f__Metschnikowiaceae(100);g__Metschnikowia(100);s__Metschnikowia_pulcherrima(100); | Metschnikowia | 13 | Saprotroph | \|Undefined Saprotroph\| | Yeast | NULL | Probable | As nectar-tap_saprotroph (Põlme et al. 2020); Decay sugar-rich substrates (Põlme et al. 2020) | Põlme S, et al. 2020. Fungal Diversity 105:1-16 ((https://doi.org/10.1007/s13225-020-00466-2)) |
| Otu24958 | 1 | k__Fungi(100);p__Ascomycota(100);c__Saccharomycetes(100);o__Saccharomycetales(100);f__Metschnikowiaceae(100);g__Metschnikowia(100);s__Metschnikowia_pulcherrima(100); | Metschnikowia | 13 | Saprotroph | \|Undefined Saprotroph\| | Yeast | NULL | Probable | As nectar-tap_saprotroph (Põlme et al. 2020); Decay sugar-rich substrates (Põlme et al. 2020) | Põlme S, et al. 2020. Fungal Diversity 105:1-16 ((https://doi.org/10.1007/s13225-020-00466-2)) |
| Otu24959 | 1 | k__Fungi(100);p__Ascomycota(100);c__Saccharomycetes(100);o__Saccharomycetales(100);f__Metschnikowiaceae(100);g__Metschnikowia(100);s__Metschnikowia_pulcherrima(100); | Metschnikowia | 13 | Saprotroph | \|Undefined Saprotroph\| | Yeast | NULL | Probable | As nectar-tap_saprotroph (Põlme et al. 2020); Decay sugar-rich substrates (Põlme et al. 2020) | Põlme S, et al. 2020. Fungal Diversity 105:1-16 ((https://doi.org/10.1007/s13225-020-00466-2)) |
| Otu24960 | 1 | k__Fungi(100);p__Ascomycota(100);c__Saccharomycetes(100);o__Saccharomycetales(100);f__Metschnikowiaceae(100);g__Metschnikowia(100);s__Metschnikowia_pulcherrima(100); | Metschnikowia | 13 | Saprotroph | \|Undefined Saprotroph\| | Yeast | NULL | Probable | As nectar-tap_saprotroph (Põlme et al. 2020); Decay sugar-rich substrates (Põlme et al. 2020) | Põlme S, et al. 2020. Fungal Diversity 105:1-16 ((https://doi.org/10.1007/s13225-020-00466-2)) |
| Otu24961 | 1 | k__Fungi(100);p__Ascomycota(100);c__Saccharomycetes(100);o__Saccharomycetales(100);f__Metschnikowiaceae(100);g__Metschnikowia(100);g__Metschnikowia_unclassified(100); | Metschnikowia | 13 | Saprotroph | \|Undefined Saprotroph\| | Yeast | NULL | Probable | As nectar-tap_saprotroph (Põlme et al. 2020); Decay sugar-rich substrates (Põlme et al. 2020) | Põlme S, et al. 2020. Fungal Diversity 105:1-16 ((https://doi.org/10.1007/s13225-020-00466-2)) |
| Otu24962 | 1 | k__Fungi(100);p__Ascomycota(100);c__Saccharomycetes(100);o__Saccharomycetales(100);f__Metschnikowiaceae(100);g__Metschnikowia(100);g__Metschnikowia_unclassified(100); | Metschnikowia | 13 | Saprotroph | \|Undefined Saprotroph\| | Yeast | NULL | Probable | As nectar-tap_saprotroph (Põlme et al. 2020); Decay sugar-rich substrates (Põlme et al. 2020) | Põlme S, et al. 2020. Fungal Diversity 105:1-16 ((https://doi.org/10.1007/s13225-020-00466-2)) |
| Otu24963 | 1 | k__Fungi(100);p__Ascomycota(100);c__Saccharomycetes(100);o__Saccharomycetales(100);f__Metschnikowiaceae(100);g__Metschnikowia(100);g__Metschnikowia_unclassified(100); | Metschnikowia | 13 | Saprotroph | \|Undefined Saprotroph\| | Yeast | NULL | Probable | As nectar-tap_saprotroph (Põlme et al. 2020); Decay sugar-rich substrates (Põlme et al. 2020) | Põlme S, et al. 2020. Fungal Diversity 105:1-16 ((https://doi.org/10.1007/s13225-020-00466-2)) |
| Otu24964 | 1 | k__Fungi(100);p__Ascomycota(100);c__Saccharomycetes(100);o__Saccharomycetales(100);f__Metschnikowiaceae(100);g__Metschnikowia(100);s__Metschnikowia_pulcherrima(100); | Metschnikowia | 13 | Saprotroph | \|Undefined Saprotroph\| | Yeast | NULL | Probable | As nectar-tap_saprotroph (Põlme et al. 2020); Decay sugar-rich substrates (Põlme et al. 2020) | Põlme S, et al. 2020. Fungal Diversity 105:1-16 ((https://doi.org/10.1007/s13225-020-00466-2)) |
| Otu24965 | 1 | k__Fungi(100);p__Ascomycota(100);c__Saccharomycetes(100);o__Saccharomycetales(100);f__Metschnikowiaceae(100);g__Metschnikowia(100);s__Metschnikowia_pulcherrima(100); | Metschnikowia | 13 | Saprotroph | \|Undefined Saprotroph\| | Yeast | NULL | Probable | As nectar-tap_saprotroph (Põlme et al. 2020); Decay sugar-rich substrates (Põlme et al. 2020) | Põlme S, et al. 2020. Fungal Diversity 105:1-16 ((https://doi.org/10.1007/s13225-020-00466-2)) |
| Otu24966 | 1 | k__Fungi(100);p__Ascomycota(100);c__Saccharomycetes(100);o__Saccharomycetales(100);f__Metschnikowiaceae(100);g__Metschnikowia(100);s__Metschnikowia_pulcherrima(100); | Metschnikowia | 13 | Saprotroph | \|Undefined Saprotroph\| | Yeast | NULL | Probable | As nectar-tap_saprotroph (Põlme et al. 2020); Decay sugar-rich substrates (Põlme et al. 2020) | Põlme S, et al. 2020. Fungal Diversity 105:1-16 ((https://doi.org/10.1007/s13225-020-00466-2)) |
| Otu24967 | 1 | k__Fungi(100);p__Ascomycota(100);c__Saccharomycetes(100);o__Saccharomycetales(100);f__Metschnikowiaceae(100);g__Metschnikowia(100);s__Metschnikowia_pulcherrima(100); | Metschnikowia | 13 | Saprotroph | \|Undefined Saprotroph\| | Yeast | NULL | Probable | As nectar-tap_saprotroph (Põlme et al. 2020); Decay sugar-rich substrates (Põlme et al. 2020) | Põlme S, et al. 2020. Fungal Diversity 105:1-16 ((https://doi.org/10.1007/s13225-020-00466-2)) |
| Otu24970 | 1 | k__Fungi(100);p__Ascomycota(100);c__Saccharomycetes(100);o__Saccharomycetales(100);f__Metschnikowiaceae(100);g__Metschnikowia(100);s__Metschnikowia_pulcherrima(100); | Metschnikowia | 13 | Saprotroph | \|Undefined Saprotroph\| | Yeast | NULL | Probable | As nectar-tap_saprotroph (Põlme et al. 2020); Decay sugar-rich substrates (Põlme et al. 2020) | Põlme S, et al. 2020. Fungal Diversity 105:1-16 ((https://doi.org/10.1007/s13225-020-00466-2)) |
| Otu24971 | 1 | k__Fungi(100);p__Ascomycota(100);c__Saccharomycetes(100);o__Saccharomycetales(100);f__Metschnikowiaceae(100);g__Metschnikowia(100);s__Metschnikowia_pulcherrima(100); | Metschnikowia | 13 | Saprotroph | \|Undefined Saprotroph\| | Yeast | NULL | Probable | As nectar-tap_saprotroph (Põlme et al. 2020); Decay sugar-rich substrates (Põlme et al. 2020) | Põlme S, et al. 2020. Fungal Diversity 105:1-16 ((https://doi.org/10.1007/s13225-020-00466-2)) |
| Otu24986 | 1 | k__Fungi(100);p__Ascomycota(100);c__Saccharomycetes(100);o__Saccharomycetales(100);f__Metschnikowiaceae(100);g__Metschnikowia(100);s__Metschnikowia_pulcherrima(100); | Metschnikowia | 13 | Saprotroph | \|Undefined Saprotroph\| | Yeast | NULL | Probable | As nectar-tap_saprotroph (Põlme et al. 2020); Decay sugar-rich substrates (Põlme et al. 2020) | Põlme S, et al. 2020. Fungal Diversity 105:1-16 ((https://doi.org/10.1007/s13225-020-00466-2)) |
| Otu24988 | 1 | k__Fungi(100);p__Ascomycota(100);c__Saccharomycetes(100);o__Saccharomycetales(100);f__Metschnikowiaceae(100);g__Metschnikowia(100);g__Metschnikowia_unclassified(100); | Metschnikowia | 13 | Saprotroph | \|Undefined Saprotroph\| | Yeast | NULL | Probable | As nectar-tap_saprotroph (Põlme et al. 2020); Decay sugar-rich substrates (Põlme et al. 2020) | Põlme S, et al. 2020. Fungal Diversity 105:1-16 ((https://doi.org/10.1007/s13225-020-00466-2)) |
| Otu24990 | 1 | k__Fungi(100);p__Ascomycota(100);c__Dothideomycetes(100);o__Pleosporales(100);f__Lophiostomataceae(100);g__Lophiostoma(100);g__Lophiostoma_unclassified(100); | Lophiostoma | 13 | Saprotroph | Plant Saprotroph-Undefined Saprotroph-\|Wood Saprotroph\| | Microfungus | NULL | Probable | As litter_saprotroph (Põlme et al. 2020); Decay wood, leaf, fruit, seed (Põlme et al. 2020) | Tedersoo L, et al. 2014. Science 346:e1256688 ((https://doi.org/10.1126/science.1256688)); Põlme S, et al. 2020. Fungal Diversity 105:1-16 ((https://doi.org/10.1007/s13225-020-00466-2)) |
| Otu24991 | 1 | k__Fungi(100);p__Ascomycota(100);c__Saccharomycetes(100);o__Saccharomycetales(100);f__Metschnikowiaceae(100);g__Metschnikowia(100);s__Metschnikowia_pulcherrima(100); | Metschnikowia | 13 | Saprotroph | \|Undefined Saprotroph\| | Yeast | NULL | Probable | As nectar-tap_saprotroph (Põlme et al. 2020); Decay sugar-rich substrates (Põlme et al. 2020) | Põlme S, et al. 2020. Fungal Diversity 105:1-16 ((https://doi.org/10.1007/s13225-020-00466-2)) |
| Otu24992 | 1 | k__Fungi(100);p__Ascomycota(100);c__Saccharomycetes(100);o__Saccharomycetales(100);f__Metschnikowiaceae(100);g__Metschnikowia(100);s__Metschnikowia_pulcherrima(100); | Metschnikowia | 13 | Saprotroph | \|Undefined Saprotroph\| | Yeast | NULL | Probable | As nectar-tap_saprotroph (Põlme et al. 2020); Decay sugar-rich substrates (Põlme et al. 2020) | Põlme S, et al. 2020. Fungal Diversity 105:1-16 ((https://doi.org/10.1007/s13225-020-00466-2)) |
| Otu24993 | 1 | k__Fungi(100);p__Ascomycota(100);c__Saccharomycetes(100);o__Saccharomycetales(100);f__Metschnikowiaceae(100);g__Metschnikowia(100);s__Metschnikowia_pulcherrima(100); | Metschnikowia | 13 | Saprotroph | \|Undefined Saprotroph\| | Yeast | NULL | Probable | As nectar-tap_saprotroph (Põlme et al. 2020); Decay sugar-rich substrates (Põlme et al. 2020) | Põlme S, et al. 2020. Fungal Diversity 105:1-16 ((https://doi.org/10.1007/s13225-020-00466-2)) |
| Otu24999 | 1 | k__Fungi(100);p__Ascomycota(100);c__Saccharomycetes(100);o__Saccharomycetales(100);f__Saccharomycetales_family_Incertae_sedis(100);g__Candida(100);s__Candida_cretensis(100); | Candida | 13 | Pathotroph-Saprotroph-Symbiotroph | Animal Pathogen-Endophyte-Endosymbiont-Epiphyte-\|Undefined Saprotroph\| | Dimorphic Yeast-Yeast | NULL | Probable | As nectar-tap_saprotroph (Põlme et al. 2020); Decay sugar-rich substrates (Põlme et al. 2020); This asexual genus is polyphyletic and contain species that function very differently depend on what they are. Caution should be taken when interpreting guilds for Candida species. | Manolakaki D, et al. 2010. Virulence. 1: 367-375 ((https://doi.org/10.4161/viru.1.5.12796)); Mekbib SB, et al. 2011. Tropical Plant Pathology 36:233-240 ((https://doi.org/10.1590/S1982-56762011000400004)); Põlme S, et al. 2020. Fungal Diversity 105:1-16 ((https://doi.org/10.1007/s13225-020-00466-2)) |
| Otu25001 | 1 | k__Fungi(100);p__Ascomycota(100);c__Saccharomycetes(100);o__Saccharomycetales(100);f__Metschnikowiaceae(100);g__Metschnikowia(100);s__Metschnikowia_pulcherrima(100); | Metschnikowia | 13 | Saprotroph | \|Undefined Saprotroph\| | Yeast | NULL | Probable | As nectar-tap_saprotroph (Põlme et al. 2020); Decay sugar-rich substrates (Põlme et al. 2020) | Põlme S, et al. 2020. Fungal Diversity 105:1-16 ((https://doi.org/10.1007/s13225-020-00466-2)) |
| Otu25002 | 1 | k__Fungi(100);p__Ascomycota(100);c__Saccharomycetes(100);o__Saccharomycetales(100);f__Metschnikowiaceae(100);g__Metschnikowia(100);s__Metschnikowia_pulcherrima(100); | Metschnikowia | 13 | Saprotroph | \|Undefined Saprotroph\| | Yeast | NULL | Probable | As nectar-tap_saprotroph (Põlme et al. 2020); Decay sugar-rich substrates (Põlme et al. 2020) | Põlme S, et al. 2020. Fungal Diversity 105:1-16 ((https://doi.org/10.1007/s13225-020-00466-2)) |
| Otu25004 | 1 | k__Fungi(100);p__Ascomycota(100);c__Saccharomycetes(100);o__Saccharomycetales(100);f__Metschnikowiaceae(100);g__Metschnikowia(100);s__Metschnikowia_pulcherrima(100); | Metschnikowia | 13 | Saprotroph | \|Undefined Saprotroph\| | Yeast | NULL | Probable | As nectar-tap_saprotroph (Põlme et al. 2020); Decay sugar-rich substrates (Põlme et al. 2020) | Põlme S, et al. 2020. Fungal Diversity 105:1-16 ((https://doi.org/10.1007/s13225-020-00466-2)) |
| Otu25006 | 1 | k__Fungi(100);p__Ascomycota(100);c__Saccharomycetes(100);o__Saccharomycetales(100);o__Saccharomycetales_unclassified(100);o__Saccharomycetales_unclassified(100);o__Saccharomycetales_unclassified(100); | Saccharomycetales | 7 | Saprotroph | Undefined Saprotroph | Yeast | NULL | Possible | NULL | Sterkenburg E, et al. 2015. New Phytologist 207:1145-1158 ((https://doi.org/10.1111/nph.13426)) |
| Otu25011 | 1 | k__Fungi(100);p__Ascomycota(100);c__Saccharomycetes(100);o__Saccharomycetales(100);f__Metschnikowiaceae(100);g__Metschnikowia(100);s__Metschnikowia_pulcherrima(100); | Metschnikowia | 13 | Saprotroph | \|Undefined Saprotroph\| | Yeast | NULL | Probable | As nectar-tap_saprotroph (Põlme et al. 2020); Decay sugar-rich substrates (Põlme et al. 2020) | Põlme S, et al. 2020. Fungal Diversity 105:1-16 ((https://doi.org/10.1007/s13225-020-00466-2)) |
| Otu25012 | 1 | k__Fungi(100);p__Ascomycota(100);c__Saccharomycetes(100);o__Saccharomycetales(100);f__Metschnikowiaceae(100);g__Metschnikowia(100);g__Metschnikowia_unclassified(100); | Metschnikowia | 13 | Saprotroph | \|Undefined Saprotroph\| | Yeast | NULL | Probable | As nectar-tap_saprotroph (Põlme et al. 2020); Decay sugar-rich substrates (Põlme et al. 2020) | Põlme S, et al. 2020. Fungal Diversity 105:1-16 ((https://doi.org/10.1007/s13225-020-00466-2)) |
| Otu25013 | 1 | k__Fungi(100);p__Ascomycota(100);c__Saccharomycetes(100);o__Saccharomycetales(100);f__Metschnikowiaceae(100);g__Metschnikowia(100);s__Metschnikowia_pulcherrima(100); | Metschnikowia | 13 | Saprotroph | \|Undefined Saprotroph\| | Yeast | NULL | Probable | As nectar-tap_saprotroph (Põlme et al. 2020); Decay sugar-rich substrates (Põlme et al. 2020) | Põlme S, et al. 2020. Fungal Diversity 105:1-16 ((https://doi.org/10.1007/s13225-020-00466-2)) |
| Otu25014 | 1 | k__Fungi(100);p__Ascomycota(100);c__Saccharomycetes(100);o__Saccharomycetales(100);f__Metschnikowiaceae(100);g__Metschnikowia(100);s__Metschnikowia_pulcherrima(100); | Metschnikowia | 13 | Saprotroph | \|Undefined Saprotroph\| | Yeast | NULL | Probable | As nectar-tap_saprotroph (Põlme et al. 2020); Decay sugar-rich substrates (Põlme et al. 2020) | Põlme S, et al. 2020. Fungal Diversity 105:1-16 ((https://doi.org/10.1007/s13225-020-00466-2)) |
| Otu25015 | 1 | k__Fungi(100);p__Ascomycota(100);c__Saccharomycetes(100);o__Saccharomycetales(100);f__Metschnikowiaceae(100);g__Metschnikowia(100);s__Metschnikowia_pulcherrima(100); | Metschnikowia | 13 | Saprotroph | \|Undefined Saprotroph\| | Yeast | NULL | Probable | As nectar-tap_saprotroph (Põlme et al. 2020); Decay sugar-rich substrates (Põlme et al. 2020) | Põlme S, et al. 2020. Fungal Diversity 105:1-16 ((https://doi.org/10.1007/s13225-020-00466-2)) |
| Otu25016 | 1 | k__Fungi(100);p__Ascomycota(100);c__Saccharomycetes(100);o__Saccharomycetales(100);f__Metschnikowiaceae(100);g__Metschnikowia(100);s__Metschnikowia_pulcherrima(100); | Metschnikowia | 13 | Saprotroph | \|Undefined Saprotroph\| | Yeast | NULL | Probable | As nectar-tap_saprotroph (Põlme et al. 2020); Decay sugar-rich substrates (Põlme et al. 2020) | Põlme S, et al. 2020. Fungal Diversity 105:1-16 ((https://doi.org/10.1007/s13225-020-00466-2)) |
| Otu25017 | 1 | k__Fungi(100);p__Ascomycota(100);c__Saccharomycetes(100);o__Saccharomycetales(100);f__Metschnikowiaceae(100);g__Metschnikowia(100);g__Metschnikowia_unclassified(100); | Metschnikowia | 13 | Saprotroph | \|Undefined Saprotroph\| | Yeast | NULL | Probable | As nectar-tap_saprotroph (Põlme et al. 2020); Decay sugar-rich substrates (Põlme et al. 2020) | Põlme S, et al. 2020. Fungal Diversity 105:1-16 ((https://doi.org/10.1007/s13225-020-00466-2)) |
| Otu25018 | 1 | k__Fungi(100);p__Ascomycota(100);c__Saccharomycetes(100);o__Saccharomycetales(100);f__Metschnikowiaceae(100);g__Metschnikowia(100);s__Metschnikowia_pulcherrima(100); | Metschnikowia | 13 | Saprotroph | \|Undefined Saprotroph\| | Yeast | NULL | Probable | As nectar-tap_saprotroph (Põlme et al. 2020); Decay sugar-rich substrates (Põlme et al. 2020) | Põlme S, et al. 2020. Fungal Diversity 105:1-16 ((https://doi.org/10.1007/s13225-020-00466-2)) |
| Otu25019 | 1 | k__Fungi(100);p__Ascomycota(100);c__Saccharomycetes(100);o__Saccharomycetales(100);f__Metschnikowiaceae(100);g__Metschnikowia(100);s__Metschnikowia_pulcherrima(100); | Metschnikowia | 13 | Saprotroph | \|Undefined Saprotroph\| | Yeast | NULL | Probable | As nectar-tap_saprotroph (Põlme et al. 2020); Decay sugar-rich substrates (Põlme et al. 2020) | Põlme S, et al. 2020. Fungal Diversity 105:1-16 ((https://doi.org/10.1007/s13225-020-00466-2)) |
| Otu25020 | 1 | k__Fungi(100);p__Ascomycota(100);c__Saccharomycetes(100);o__Saccharomycetales(100);f__Metschnikowiaceae(100);g__Metschnikowia(100);g__Metschnikowia_unclassified(100); | Metschnikowia | 13 | Saprotroph | \|Undefined Saprotroph\| | Yeast | NULL | Probable | As nectar-tap_saprotroph (Põlme et al. 2020); Decay sugar-rich substrates (Põlme et al. 2020) | Põlme S, et al. 2020. Fungal Diversity 105:1-16 ((https://doi.org/10.1007/s13225-020-00466-2)) |
| Otu25021 | 1 | k__Fungi(100);p__Ascomycota(100);c__Saccharomycetes(100);o__Saccharomycetales(100);f__Metschnikowiaceae(100);g__Metschnikowia(100);s__Metschnikowia_pulcherrima(100); | Metschnikowia | 13 | Saprotroph | \|Undefined Saprotroph\| | Yeast | NULL | Probable | As nectar-tap_saprotroph (Põlme et al. 2020); Decay sugar-rich substrates (Põlme et al. 2020) | Põlme S, et al. 2020. Fungal Diversity 105:1-16 ((https://doi.org/10.1007/s13225-020-00466-2)) |
| Otu25022 | 1 | k__Fungi(100);p__Ascomycota(100);c__Saccharomycetes(100);o__Saccharomycetales(100);f__Metschnikowiaceae(100);g__Metschnikowia(100);s__Metschnikowia_pulcherrima(100); | Metschnikowia | 13 | Saprotroph | \|Undefined Saprotroph\| | Yeast | NULL | Probable | As nectar-tap_saprotroph (Põlme et al. 2020); Decay sugar-rich substrates (Põlme et al. 2020) | Põlme S, et al. 2020. Fungal Diversity 105:1-16 ((https://doi.org/10.1007/s13225-020-00466-2)) |
| Otu25023 | 1 | k__Fungi(100);p__Ascomycota(100);c__Saccharomycetes(100);o__Saccharomycetales(100);f__Metschnikowiaceae(100);g__Metschnikowia(100);s__Metschnikowia_pulcherrima(100); | Metschnikowia | 13 | Saprotroph | \|Undefined Saprotroph\| | Yeast | NULL | Probable | As nectar-tap_saprotroph (Põlme et al. 2020); Decay sugar-rich substrates (Põlme et al. 2020) | Põlme S, et al. 2020. Fungal Diversity 105:1-16 ((https://doi.org/10.1007/s13225-020-00466-2)) |
| Otu25024 | 1 | k__Fungi(100);p__Ascomycota(100);c__Saccharomycetes(100);o__Saccharomycetales(100);f__Metschnikowiaceae(100);g__Metschnikowia(100);g__Metschnikowia_unclassified(100); | Metschnikowia | 13 | Saprotroph | \|Undefined Saprotroph\| | Yeast | NULL | Probable | As nectar-tap_saprotroph (Põlme et al. 2020); Decay sugar-rich substrates (Põlme et al. 2020) | Põlme S, et al. 2020. Fungal Diversity 105:1-16 ((https://doi.org/10.1007/s13225-020-00466-2)) |
| Otu25025 | 1 | k__Fungi(100);p__Ascomycota(100);c__Saccharomycetes(100);o__Saccharomycetales(100);f__Metschnikowiaceae(100);g__Metschnikowia(100);g__Metschnikowia_unclassified(100); | Metschnikowia | 13 | Saprotroph | \|Undefined Saprotroph\| | Yeast | NULL | Probable | As nectar-tap_saprotroph (Põlme et al. 2020); Decay sugar-rich substrates (Põlme et al. 2020) | Põlme S, et al. 2020. Fungal Diversity 105:1-16 ((https://doi.org/10.1007/s13225-020-00466-2)) |
| Otu25027 | 1 | k__Fungi(100);p__Ascomycota(100);c__Saccharomycetes(100);o__Saccharomycetales(100);f__Metschnikowiaceae(100);g__Metschnikowia(100);s__Metschnikowia_pulcherrima(100); | Metschnikowia | 13 | Saprotroph | \|Undefined Saprotroph\| | Yeast | NULL | Probable | As nectar-tap_saprotroph (Põlme et al. 2020); Decay sugar-rich substrates (Põlme et al. 2020) | Põlme S, et al. 2020. Fungal Diversity 105:1-16 ((https://doi.org/10.1007/s13225-020-00466-2)) |
| Otu25034 | 1 | k__Fungi(100);p__Ascomycota(100);c__Saccharomycetes(100);o__Saccharomycetales(100);f__Saccharomycetales_family_Incertae_sedis(100);g__Candida(100);s__Candida_cretensis(100); | Candida | 13 | Pathotroph-Saprotroph-Symbiotroph | Animal Pathogen-Endophyte-Endosymbiont-Epiphyte-\|Undefined Saprotroph\| | Dimorphic Yeast-Yeast | NULL | Probable | As nectar-tap_saprotroph (Põlme et al. 2020); Decay sugar-rich substrates (Põlme et al. 2020); This asexual genus is polyphyletic and contain species that function very differently depend on what they are. Caution should be taken when interpreting guilds for Candida species. | Manolakaki D, et al. 2010. Virulence. 1: 367-375 ((https://doi.org/10.4161/viru.1.5.12796)); Mekbib SB, et al. 2011. Tropical Plant Pathology 36:233-240 ((https://doi.org/10.1590/S1982-56762011000400004)); Põlme S, et al. 2020. Fungal Diversity 105:1-16 ((https://doi.org/10.1007/s13225-020-00466-2)) |
| Otu25036 | 1 | k__Fungi(100);p__Ascomycota(100);c__Saccharomycetes(100);o__Saccharomycetales(100);f__Metschnikowiaceae(100);g__Metschnikowia(100);s__Metschnikowia_pulcherrima(100); | Metschnikowia | 13 | Saprotroph | \|Undefined Saprotroph\| | Yeast | NULL | Probable | As nectar-tap_saprotroph (Põlme et al. 2020); Decay sugar-rich substrates (Põlme et al. 2020) | Põlme S, et al. 2020. Fungal Diversity 105:1-16 ((https://doi.org/10.1007/s13225-020-00466-2)) |
| Otu25037 | 1 | k__Fungi(100);p__Ascomycota(100);c__Saccharomycetes(100);o__Saccharomycetales(100);f__Metschnikowiaceae(100);g__Metschnikowia(100);g__Metschnikowia_unclassified(100); | Metschnikowia | 13 | Saprotroph | \|Undefined Saprotroph\| | Yeast | NULL | Probable | As nectar-tap_saprotroph (Põlme et al. 2020); Decay sugar-rich substrates (Põlme et al. 2020) | Põlme S, et al. 2020. Fungal Diversity 105:1-16 ((https://doi.org/10.1007/s13225-020-00466-2)) |
| Otu25038 | 1 | k__Fungi(100);p__Ascomycota(100);c__Saccharomycetes(100);o__Saccharomycetales(100);f__Metschnikowiaceae(100);g__Metschnikowia(100);s__Metschnikowia_pulcherrima(100); | Metschnikowia | 13 | Saprotroph | \|Undefined Saprotroph\| | Yeast | NULL | Probable | As nectar-tap_saprotroph (Põlme et al. 2020); Decay sugar-rich substrates (Põlme et al. 2020) | Põlme S, et al. 2020. Fungal Diversity 105:1-16 ((https://doi.org/10.1007/s13225-020-00466-2)) |
| Otu25039 | 1 | k__Fungi(100);p__Ascomycota(100);c__Saccharomycetes(100);o__Saccharomycetales(100);f__Metschnikowiaceae(100);g__Metschnikowia(100);g__Metschnikowia_unclassified(100); | Metschnikowia | 13 | Saprotroph | \|Undefined Saprotroph\| | Yeast | NULL | Probable | As nectar-tap_saprotroph (Põlme et al. 2020); Decay sugar-rich substrates (Põlme et al. 2020) | Põlme S, et al. 2020. Fungal Diversity 105:1-16 ((https://doi.org/10.1007/s13225-020-00466-2)) |
| Otu25040 | 1 | k__Fungi(100);p__Ascomycota(100);c__Saccharomycetes(100);o__Saccharomycetales(100);f__Metschnikowiaceae(100);g__Metschnikowia(100);s__Metschnikowia_pulcherrima(100); | Metschnikowia | 13 | Saprotroph | \|Undefined Saprotroph\| | Yeast | NULL | Probable | As nectar-tap_saprotroph (Põlme et al. 2020); Decay sugar-rich substrates (Põlme et al. 2020) | Põlme S, et al. 2020. Fungal Diversity 105:1-16 ((https://doi.org/10.1007/s13225-020-00466-2)) |
| Otu25041 | 1 | k__Fungi(100);p__Ascomycota(100);c__Saccharomycetes(100);o__Saccharomycetales(100);f__Metschnikowiaceae(100);g__Metschnikowia(100);s__Metschnikowia_pulcherrima(100); | Metschnikowia | 13 | Saprotroph | \|Undefined Saprotroph\| | Yeast | NULL | Probable | As nectar-tap_saprotroph (Põlme et al. 2020); Decay sugar-rich substrates (Põlme et al. 2020) | Põlme S, et al. 2020. Fungal Diversity 105:1-16 ((https://doi.org/10.1007/s13225-020-00466-2)) |
| Otu25042 | 1 | k__Fungi(100);p__Ascomycota(100);c__Saccharomycetes(100);o__Saccharomycetales(100);f__Metschnikowiaceae(100);g__Metschnikowia(100);g__Metschnikowia_unclassified(100); | Metschnikowia | 13 | Saprotroph | \|Undefined Saprotroph\| | Yeast | NULL | Probable | As nectar-tap_saprotroph (Põlme et al. 2020); Decay sugar-rich substrates (Põlme et al. 2020) | Põlme S, et al. 2020. Fungal Diversity 105:1-16 ((https://doi.org/10.1007/s13225-020-00466-2)) |
| Otu25043 | 1 | k__Fungi(100);p__Ascomycota(100);c__Saccharomycetes(100);o__Saccharomycetales(100);f__Metschnikowiaceae(100);g__Metschnikowia(100);s__Metschnikowia_pulcherrima(100); | Metschnikowia | 13 | Saprotroph | \|Undefined Saprotroph\| | Yeast | NULL | Probable | As nectar-tap_saprotroph (Põlme et al. 2020); Decay sugar-rich substrates (Põlme et al. 2020) | Põlme S, et al. 2020. Fungal Diversity 105:1-16 ((https://doi.org/10.1007/s13225-020-00466-2)) |
| Otu25044 | 1 | k__Fungi(100);p__Ascomycota(100);c__Saccharomycetes(100);o__Saccharomycetales(100);f__Metschnikowiaceae(100);g__Metschnikowia(100);s__Metschnikowia_pulcherrima(100); | Metschnikowia | 13 | Saprotroph | \|Undefined Saprotroph\| | Yeast | NULL | Probable | As nectar-tap_saprotroph (Põlme et al. 2020); Decay sugar-rich substrates (Põlme et al. 2020) | Põlme S, et al. 2020. Fungal Diversity 105:1-16 ((https://doi.org/10.1007/s13225-020-00466-2)) |
| Otu25045 | 1 | k__Fungi(100);p__Ascomycota(100);c__Saccharomycetes(100);o__Saccharomycetales(100);f__Metschnikowiaceae(100);g__Metschnikowia(100);g__Metschnikowia_unclassified(100); | Metschnikowia | 13 | Saprotroph | \|Undefined Saprotroph\| | Yeast | NULL | Probable | As nectar-tap_saprotroph (Põlme et al. 2020); Decay sugar-rich substrates (Põlme et al. 2020) | Põlme S, et al. 2020. Fungal Diversity 105:1-16 ((https://doi.org/10.1007/s13225-020-00466-2)) |
| Otu25046 | 1 | k__Fungi(100);p__Ascomycota(100);c__Saccharomycetes(100);o__Saccharomycetales(100);f__Metschnikowiaceae(100);g__Metschnikowia(100);s__Metschnikowia_pulcherrima(100); | Metschnikowia | 13 | Saprotroph | \|Undefined Saprotroph\| | Yeast | NULL | Probable | As nectar-tap_saprotroph (Põlme et al. 2020); Decay sugar-rich substrates (Põlme et al. 2020) | Põlme S, et al. 2020. Fungal Diversity 105:1-16 ((https://doi.org/10.1007/s13225-020-00466-2)) |
| Otu25047 | 1 | k__Fungi(100);p__Ascomycota(100);c__Saccharomycetes(100);o__Saccharomycetales(100);f__Metschnikowiaceae(100);g__Metschnikowia(100);s__Metschnikowia_pulcherrima(100); | Metschnikowia | 13 | Saprotroph | \|Undefined Saprotroph\| | Yeast | NULL | Probable | As nectar-tap_saprotroph (Põlme et al. 2020); Decay sugar-rich substrates (Põlme et al. 2020) | Põlme S, et al. 2020. Fungal Diversity 105:1-16 ((https://doi.org/10.1007/s13225-020-00466-2)) |
| Otu25048 | 1 | k__Fungi(100);p__Ascomycota(100);c__Saccharomycetes(100);o__Saccharomycetales(100);f__Metschnikowiaceae(100);g__Metschnikowia(100);s__Metschnikowia_pulcherrima(100); | Metschnikowia | 13 | Saprotroph | \|Undefined Saprotroph\| | Yeast | NULL | Probable | As nectar-tap_saprotroph (Põlme et al. 2020); Decay sugar-rich substrates (Põlme et al. 2020) | Põlme S, et al. 2020. Fungal Diversity 105:1-16 ((https://doi.org/10.1007/s13225-020-00466-2)) |
| Otu25049 | 1 | k__Fungi(100);p__Ascomycota(100);c__Saccharomycetes(100);o__Saccharomycetales(100);f__Metschnikowiaceae(100);g__Metschnikowia(100);s__Metschnikowia_pulcherrima(100); | Metschnikowia | 13 | Saprotroph | \|Undefined Saprotroph\| | Yeast | NULL | Probable | As nectar-tap_saprotroph (Põlme et al. 2020); Decay sugar-rich substrates (Põlme et al. 2020) | Põlme S, et al. 2020. Fungal Diversity 105:1-16 ((https://doi.org/10.1007/s13225-020-00466-2)) |
| Otu25050 | 1 | k__Fungi(100);p__Ascomycota(100);c__Saccharomycetes(100);o__Saccharomycetales(100);f__Metschnikowiaceae(100);g__Metschnikowia(100);s__Metschnikowia_pulcherrima(100); | Metschnikowia | 13 | Saprotroph | \|Undefined Saprotroph\| | Yeast | NULL | Probable | As nectar-tap_saprotroph (Põlme et al. 2020); Decay sugar-rich substrates (Põlme et al. 2020) | Põlme S, et al. 2020. Fungal Diversity 105:1-16 ((https://doi.org/10.1007/s13225-020-00466-2)) |
| Otu25052 | 1 | k__Fungi(100);p__Ascomycota(100);c__Saccharomycetes(100);o__Saccharomycetales(100);f__Metschnikowiaceae(100);g__Metschnikowia(100);s__Metschnikowia_pulcherrima(100); | Metschnikowia | 13 | Saprotroph | \|Undefined Saprotroph\| | Yeast | NULL | Probable | As nectar-tap_saprotroph (Põlme et al. 2020); Decay sugar-rich substrates (Põlme et al. 2020) | Põlme S, et al. 2020. Fungal Diversity 105:1-16 ((https://doi.org/10.1007/s13225-020-00466-2)) |
| Otu25053 | 1 | k__Fungi(100);p__Ascomycota(100);c__Saccharomycetes(100);o__Saccharomycetales(100);f__Metschnikowiaceae(100);g__Metschnikowia(100);s__Metschnikowia_pulcherrima(100); | Metschnikowia | 13 | Saprotroph | \|Undefined Saprotroph\| | Yeast | NULL | Probable | As nectar-tap_saprotroph (Põlme et al. 2020); Decay sugar-rich substrates (Põlme et al. 2020) | Põlme S, et al. 2020. Fungal Diversity 105:1-16 ((https://doi.org/10.1007/s13225-020-00466-2)) |
| Otu25057 | 1 | k__Fungi(100);p__Ascomycota(100);c__Saccharomycetes(100);o__Saccharomycetales(100);f__Saccharomycetales_family_Incertae_sedis(100);f__Saccharomycetales_family_Incertae_sedis_unclassified(100);f__Saccharomycetales_family_Incertae_sedis_unclassified(100); | Saccharomycetales | 7 | Saprotroph | Undefined Saprotroph | Yeast | NULL | Possible | NULL | Sterkenburg E, et al. 2015. New Phytologist 207:1145-1158 ((https://doi.org/10.1111/nph.13426)) |
| Otu25060 | 1 | k__Fungi(100);p__Ascomycota(100);c__Saccharomycetes(100);o__Saccharomycetales(100);f__Metschnikowiaceae(100);g__Metschnikowia(100);s__Metschnikowia_pulcherrima(100); | Metschnikowia | 13 | Saprotroph | \|Undefined Saprotroph\| | Yeast | NULL | Probable | As nectar-tap_saprotroph (Põlme et al. 2020); Decay sugar-rich substrates (Põlme et al. 2020) | Põlme S, et al. 2020. Fungal Diversity 105:1-16 ((https://doi.org/10.1007/s13225-020-00466-2)) |
| Otu25061 | 1 | k__Fungi(100);p__Ascomycota(100);c__Saccharomycetes(100);o__Saccharomycetales(100);f__Metschnikowiaceae(100);g__Metschnikowia(100);s__Metschnikowia_pulcherrima(100); | Metschnikowia | 13 | Saprotroph | \|Undefined Saprotroph\| | Yeast | NULL | Probable | As nectar-tap_saprotroph (Põlme et al. 2020); Decay sugar-rich substrates (Põlme et al. 2020) | Põlme S, et al. 2020. Fungal Diversity 105:1-16 ((https://doi.org/10.1007/s13225-020-00466-2)) |
| Otu25063 | 1 | k__Fungi(100);p__Ascomycota(100);c__Saccharomycetes(100);o__Saccharomycetales(100);f__Metschnikowiaceae(100);g__Metschnikowia(100);s__Metschnikowia_pulcherrima(100); | Metschnikowia | 13 | Saprotroph | \|Undefined Saprotroph\| | Yeast | NULL | Probable | As nectar-tap_saprotroph (Põlme et al. 2020); Decay sugar-rich substrates (Põlme et al. 2020) | Põlme S, et al. 2020. Fungal Diversity 105:1-16 ((https://doi.org/10.1007/s13225-020-00466-2)) |
| Otu25065 | 1 | k__Fungi(100);p__Ascomycota(100);c__Saccharomycetes(100);o__Saccharomycetales(100);f__Metschnikowiaceae(100);g__Metschnikowia(100);s__Metschnikowia_pulcherrima(100); | Metschnikowia | 13 | Saprotroph | \|Undefined Saprotroph\| | Yeast | NULL | Probable | As nectar-tap_saprotroph (Põlme et al. 2020); Decay sugar-rich substrates (Põlme et al. 2020) | Põlme S, et al. 2020. Fungal Diversity 105:1-16 ((https://doi.org/10.1007/s13225-020-00466-2)) |
| Otu25067 | 1 | k__Fungi(100);p__Ascomycota(100);c__Saccharomycetes(100);o__Saccharomycetales(100);f__Metschnikowiaceae(100);g__Metschnikowia(100);g__Metschnikowia_unclassified(100); | Metschnikowia | 13 | Saprotroph | \|Undefined Saprotroph\| | Yeast | NULL | Probable | As nectar-tap_saprotroph (Põlme et al. 2020); Decay sugar-rich substrates (Põlme et al. 2020) | Põlme S, et al. 2020. Fungal Diversity 105:1-16 ((https://doi.org/10.1007/s13225-020-00466-2)) |
| Otu25070 | 1 | k__Fungi(100);p__Ascomycota(100);c__Saccharomycetes(100);o__Saccharomycetales(100);f__Metschnikowiaceae(100);g__Metschnikowia(100);s__Metschnikowia_pulcherrima(100); | Metschnikowia | 13 | Saprotroph | \|Undefined Saprotroph\| | Yeast | NULL | Probable | As nectar-tap_saprotroph (Põlme et al. 2020); Decay sugar-rich substrates (Põlme et al. 2020) | Põlme S, et al. 2020. Fungal Diversity 105:1-16 ((https://doi.org/10.1007/s13225-020-00466-2)) |
| Otu25073 | 1 | k__Fungi(100);p__Ascomycota(100);c__Saccharomycetes(100);o__Saccharomycetales(100);f__Metschnikowiaceae(100);g__Metschnikowia(100);g__Metschnikowia_unclassified(100); | Metschnikowia | 13 | Saprotroph | \|Undefined Saprotroph\| | Yeast | NULL | Probable | As nectar-tap_saprotroph (Põlme et al. 2020); Decay sugar-rich substrates (Põlme et al. 2020) | Põlme S, et al. 2020. Fungal Diversity 105:1-16 ((https://doi.org/10.1007/s13225-020-00466-2)) |
| Otu25074 | 1 | k__Fungi(100);p__Ascomycota(100);c__Saccharomycetes(100);o__Saccharomycetales(100);f__Metschnikowiaceae(100);g__Metschnikowia(100);s__Metschnikowia_pulcherrima(100); | Metschnikowia | 13 | Saprotroph | \|Undefined Saprotroph\| | Yeast | NULL | Probable | As nectar-tap_saprotroph (Põlme et al. 2020); Decay sugar-rich substrates (Põlme et al. 2020) | Põlme S, et al. 2020. Fungal Diversity 105:1-16 ((https://doi.org/10.1007/s13225-020-00466-2)) |
| Otu25075 | 1 | k__Fungi(100);p__Ascomycota(100);c__Saccharomycetes(100);o__Saccharomycetales(100);f__Metschnikowiaceae(100);g__Metschnikowia(100);g__Metschnikowia_unclassified(100); | Metschnikowia | 13 | Saprotroph | \|Undefined Saprotroph\| | Yeast | NULL | Probable | As nectar-tap_saprotroph (Põlme et al. 2020); Decay sugar-rich substrates (Põlme et al. 2020) | Põlme S, et al. 2020. Fungal Diversity 105:1-16 ((https://doi.org/10.1007/s13225-020-00466-2)) |
| Otu25076 | 1 | k__Fungi(100);p__Ascomycota(100);c__Saccharomycetes(100);o__Saccharomycetales(100);f__Metschnikowiaceae(100);g__Metschnikowia(100);g__Metschnikowia_unclassified(100); | Metschnikowia | 13 | Saprotroph | \|Undefined Saprotroph\| | Yeast | NULL | Probable | As nectar-tap_saprotroph (Põlme et al. 2020); Decay sugar-rich substrates (Põlme et al. 2020) | Põlme S, et al. 2020. Fungal Diversity 105:1-16 ((https://doi.org/10.1007/s13225-020-00466-2)) |
| Otu25077 | 1 | k__Fungi(100);p__Ascomycota(100);c__Saccharomycetes(100);o__Saccharomycetales(100);f__Metschnikowiaceae(100);g__Metschnikowia(100);g__Metschnikowia_unclassified(100); | Metschnikowia | 13 | Saprotroph | \|Undefined Saprotroph\| | Yeast | NULL | Probable | As nectar-tap_saprotroph (Põlme et al. 2020); Decay sugar-rich substrates (Põlme et al. 2020) | Põlme S, et al. 2020. Fungal Diversity 105:1-16 ((https://doi.org/10.1007/s13225-020-00466-2)) |
| Otu25078 | 1 | k__Fungi(100);p__Ascomycota(100);c__Saccharomycetes(100);o__Saccharomycetales(100);f__Metschnikowiaceae(100);g__Metschnikowia(100);g__Metschnikowia_unclassified(100); | Metschnikowia | 13 | Saprotroph | \|Undefined Saprotroph\| | Yeast | NULL | Probable | As nectar-tap_saprotroph (Põlme et al. 2020); Decay sugar-rich substrates (Põlme et al. 2020) | Põlme S, et al. 2020. Fungal Diversity 105:1-16 ((https://doi.org/10.1007/s13225-020-00466-2)) |
| Otu25079 | 1 | k__Fungi(100);p__Ascomycota(100);c__Saccharomycetes(100);o__Saccharomycetales(100);o__Saccharomycetales_unclassified(100);o__Saccharomycetales_unclassified(100);o__Saccharomycetales_unclassified(100); | Saccharomycetales | 7 | Saprotroph | Undefined Saprotroph | Yeast | NULL | Possible | NULL | Sterkenburg E, et al. 2015. New Phytologist 207:1145-1158 ((https://doi.org/10.1111/nph.13426)) |
| Otu25080 | 1 | k__Fungi(100);p__Ascomycota(100);c__Saccharomycetes(100);o__Saccharomycetales(100);f__Metschnikowiaceae(100);g__Metschnikowia(100);s__Metschnikowia_pulcherrima(100); | Metschnikowia | 13 | Saprotroph | \|Undefined Saprotroph\| | Yeast | NULL | Probable | As nectar-tap_saprotroph (Põlme et al. 2020); Decay sugar-rich substrates (Põlme et al. 2020) | Põlme S, et al. 2020. Fungal Diversity 105:1-16 ((https://doi.org/10.1007/s13225-020-00466-2)) |
| Otu25082 | 1 | k__Fungi(100);p__Ascomycota(100);c__Saccharomycetes(100);o__Saccharomycetales(100);f__Saccharomycetales_family_Incertae_sedis(100);f__Saccharomycetales_family_Incertae_sedis_unclassified(100);f__Saccharomycetales_family_Incertae_sedis_unclassified(100); | Saccharomycetales | 7 | Saprotroph | Undefined Saprotroph | Yeast | NULL | Possible | NULL | Sterkenburg E, et al. 2015. New Phytologist 207:1145-1158 ((https://doi.org/10.1111/nph.13426)) |
| Otu25083 | 1 | k__Fungi(100);p__Ascomycota(100);c__Saccharomycetes(100);o__Saccharomycetales(100);f__Metschnikowiaceae(100);g__Metschnikowia(100);g__Metschnikowia_unclassified(100); | Metschnikowia | 13 | Saprotroph | \|Undefined Saprotroph\| | Yeast | NULL | Probable | As nectar-tap_saprotroph (Põlme et al. 2020); Decay sugar-rich substrates (Põlme et al. 2020) | Põlme S, et al. 2020. Fungal Diversity 105:1-16 ((https://doi.org/10.1007/s13225-020-00466-2)) |
| Otu25084 | 1 | k__Fungi(100);p__Ascomycota(100);c__Saccharomycetes(100);o__Saccharomycetales(100);f__Metschnikowiaceae(100);g__Metschnikowia(100);s__Metschnikowia_pulcherrima(100); | Metschnikowia | 13 | Saprotroph | \|Undefined Saprotroph\| | Yeast | NULL | Probable | As nectar-tap_saprotroph (Põlme et al. 2020); Decay sugar-rich substrates (Põlme et al. 2020) | Põlme S, et al. 2020. Fungal Diversity 105:1-16 ((https://doi.org/10.1007/s13225-020-00466-2)) |
| Otu25085 | 1 | k__Fungi(100);p__Ascomycota(100);c__Saccharomycetes(100);o__Saccharomycetales(100);f__Metschnikowiaceae(100);g__Metschnikowia(100);s__Metschnikowia_pulcherrima(100); | Metschnikowia | 13 | Saprotroph | \|Undefined Saprotroph\| | Yeast | NULL | Probable | As nectar-tap_saprotroph (Põlme et al. 2020); Decay sugar-rich substrates (Põlme et al. 2020) | Põlme S, et al. 2020. Fungal Diversity 105:1-16 ((https://doi.org/10.1007/s13225-020-00466-2)) |
| Otu25086 | 1 | k__Fungi(100);p__Ascomycota(100);c__Saccharomycetes(100);o__Saccharomycetales(100);f__Metschnikowiaceae(100);g__Metschnikowia(100);s__Metschnikowia_pulcherrima(100); | Metschnikowia | 13 | Saprotroph | \|Undefined Saprotroph\| | Yeast | NULL | Probable | As nectar-tap_saprotroph (Põlme et al. 2020); Decay sugar-rich substrates (Põlme et al. 2020) | Põlme S, et al. 2020. Fungal Diversity 105:1-16 ((https://doi.org/10.1007/s13225-020-00466-2)) |
| Otu25087 | 1 | k__Fungi(100);p__Ascomycota(100);c__Saccharomycetes(100);o__Saccharomycetales(100);f__Metschnikowiaceae(100);g__Metschnikowia(100);s__Metschnikowia_pulcherrima(100); | Metschnikowia | 13 | Saprotroph | \|Undefined Saprotroph\| | Yeast | NULL | Probable | As nectar-tap_saprotroph (Põlme et al. 2020); Decay sugar-rich substrates (Põlme et al. 2020) | Põlme S, et al. 2020. Fungal Diversity 105:1-16 ((https://doi.org/10.1007/s13225-020-00466-2)) |
| Otu25088 | 1 | k__Fungi(100);p__Ascomycota(100);c__Saccharomycetes(100);o__Saccharomycetales(100);f__Metschnikowiaceae(100);g__Metschnikowia(100);s__Metschnikowia_pulcherrima(100); | Metschnikowia | 13 | Saprotroph | \|Undefined Saprotroph\| | Yeast | NULL | Probable | As nectar-tap_saprotroph (Põlme et al. 2020); Decay sugar-rich substrates (Põlme et al. 2020) | Põlme S, et al. 2020. Fungal Diversity 105:1-16 ((https://doi.org/10.1007/s13225-020-00466-2)) |
| Otu25089 | 1 | k__Fungi(100);p__Ascomycota(100);c__Saccharomycetes(100);o__Saccharomycetales(100);f__Metschnikowiaceae(100);g__Metschnikowia(100);s__Metschnikowia_pulcherrima(100); | Metschnikowia | 13 | Saprotroph | \|Undefined Saprotroph\| | Yeast | NULL | Probable | As nectar-tap_saprotroph (Põlme et al. 2020); Decay sugar-rich substrates (Põlme et al. 2020) | Põlme S, et al. 2020. Fungal Diversity 105:1-16 ((https://doi.org/10.1007/s13225-020-00466-2)) |
| Otu25090 | 1 | k__Fungi(100);p__Ascomycota(100);c__Saccharomycetes(100);o__Saccharomycetales(100);f__Metschnikowiaceae(100);g__Metschnikowia(100);g__Metschnikowia_unclassified(100); | Metschnikowia | 13 | Saprotroph | \|Undefined Saprotroph\| | Yeast | NULL | Probable | As nectar-tap_saprotroph (Põlme et al. 2020); Decay sugar-rich substrates (Põlme et al. 2020) | Põlme S, et al. 2020. Fungal Diversity 105:1-16 ((https://doi.org/10.1007/s13225-020-00466-2)) |
| Otu25091 | 1 | k__Fungi(100);p__Ascomycota(100);c__Saccharomycetes(100);o__Saccharomycetales(100);f__Metschnikowiaceae(100);g__Metschnikowia(100);s__Metschnikowia_pulcherrima(100); | Metschnikowia | 13 | Saprotroph | \|Undefined Saprotroph\| | Yeast | NULL | Probable | As nectar-tap_saprotroph (Põlme et al. 2020); Decay sugar-rich substrates (Põlme et al. 2020) | Põlme S, et al. 2020. Fungal Diversity 105:1-16 ((https://doi.org/10.1007/s13225-020-00466-2)) |
| Otu25092 | 1 | k__Fungi(100);p__Ascomycota(100);c__Saccharomycetes(100);o__Saccharomycetales(100);f__Metschnikowiaceae(100);g__Metschnikowia(100);s__Metschnikowia_pulcherrima(100); | Metschnikowia | 13 | Saprotroph | \|Undefined Saprotroph\| | Yeast | NULL | Probable | As nectar-tap_saprotroph (Põlme et al. 2020); Decay sugar-rich substrates (Põlme et al. 2020) | Põlme S, et al. 2020. Fungal Diversity 105:1-16 ((https://doi.org/10.1007/s13225-020-00466-2)) |
| Otu25093 | 1 | k__Fungi(100);p__Ascomycota(100);c__Saccharomycetes(100);o__Saccharomycetales(100);f__Metschnikowiaceae(100);g__Metschnikowia(100);s__Metschnikowia_pulcherrima(100); | Metschnikowia | 13 | Saprotroph | \|Undefined Saprotroph\| | Yeast | NULL | Probable | As nectar-tap_saprotroph (Põlme et al. 2020); Decay sugar-rich substrates (Põlme et al. 2020) | Põlme S, et al. 2020. Fungal Diversity 105:1-16 ((https://doi.org/10.1007/s13225-020-00466-2)) |
| Otu25094 | 1 | k__Fungi(100);p__Ascomycota(100);c__Saccharomycetes(100);o__Saccharomycetales(100);f__Metschnikowiaceae(100);g__Metschnikowia(100);g__Metschnikowia_unclassified(100); | Metschnikowia | 13 | Saprotroph | \|Undefined Saprotroph\| | Yeast | NULL | Probable | As nectar-tap_saprotroph (Põlme et al. 2020); Decay sugar-rich substrates (Põlme et al. 2020) | Põlme S, et al. 2020. Fungal Diversity 105:1-16 ((https://doi.org/10.1007/s13225-020-00466-2)) |
| Otu25095 | 1 | k__Fungi(100);p__Ascomycota(100);c__Saccharomycetes(100);o__Saccharomycetales(100);f__Metschnikowiaceae(100);g__Metschnikowia(100);s__Metschnikowia_pulcherrima(100); | Metschnikowia | 13 | Saprotroph | \|Undefined Saprotroph\| | Yeast | NULL | Probable | As nectar-tap_saprotroph (Põlme et al. 2020); Decay sugar-rich substrates (Põlme et al. 2020) | Põlme S, et al. 2020. Fungal Diversity 105:1-16 ((https://doi.org/10.1007/s13225-020-00466-2)) |
| Otu25098 | 1 | k__Fungi(100);p__Ascomycota(100);c__Saccharomycetes(100);o__Saccharomycetales(100);f__Metschnikowiaceae(100);g__Metschnikowia(100);g__Metschnikowia_unclassified(100); | Metschnikowia | 13 | Saprotroph | \|Undefined Saprotroph\| | Yeast | NULL | Probable | As nectar-tap_saprotroph (Põlme et al. 2020); Decay sugar-rich substrates (Põlme et al. 2020) | Põlme S, et al. 2020. Fungal Diversity 105:1-16 ((https://doi.org/10.1007/s13225-020-00466-2)) |
| Otu25100 | 1 | k__Fungi(100);p__Ascomycota(100);c__Saccharomycetes(100);o__Saccharomycetales(100);f__Metschnikowiaceae(100);g__Metschnikowia(100);s__Metschnikowia_pulcherrima(100); | Metschnikowia | 13 | Saprotroph | \|Undefined Saprotroph\| | Yeast | NULL | Probable | As nectar-tap_saprotroph (Põlme et al. 2020); Decay sugar-rich substrates (Põlme et al. 2020) | Põlme S, et al. 2020. Fungal Diversity 105:1-16 ((https://doi.org/10.1007/s13225-020-00466-2)) |
| Otu25101 | 1 | k__Fungi(100);p__Ascomycota(100);c__Saccharomycetes(100);o__Saccharomycetales(100);f__Metschnikowiaceae(100);g__Metschnikowia(100);g__Metschnikowia_unclassified(100); | Metschnikowia | 13 | Saprotroph | \|Undefined Saprotroph\| | Yeast | NULL | Probable | As nectar-tap_saprotroph (Põlme et al. 2020); Decay sugar-rich substrates (Põlme et al. 2020) | Põlme S, et al. 2020. Fungal Diversity 105:1-16 ((https://doi.org/10.1007/s13225-020-00466-2)) |
| Otu25102 | 1 | k__Fungi(100);p__Ascomycota(100);c__Saccharomycetes(100);o__Saccharomycetales(100);f__Metschnikowiaceae(100);g__Metschnikowia(100);s__Metschnikowia_pulcherrima(100); | Metschnikowia | 13 | Saprotroph | \|Undefined Saprotroph\| | Yeast | NULL | Probable | As nectar-tap_saprotroph (Põlme et al. 2020); Decay sugar-rich substrates (Põlme et al. 2020) | Põlme S, et al. 2020. Fungal Diversity 105:1-16 ((https://doi.org/10.1007/s13225-020-00466-2)) |
| Otu25103 | 1 | k__Fungi(100);p__Ascomycota(100);c__Saccharomycetes(100);o__Saccharomycetales(100);f__Metschnikowiaceae(100);g__Metschnikowia(100);s__Metschnikowia_pulcherrima(100); | Metschnikowia | 13 | Saprotroph | \|Undefined Saprotroph\| | Yeast | NULL | Probable | As nectar-tap_saprotroph (Põlme et al. 2020); Decay sugar-rich substrates (Põlme et al. 2020) | Põlme S, et al. 2020. Fungal Diversity 105:1-16 ((https://doi.org/10.1007/s13225-020-00466-2)) |
| Otu25104 | 1 | k__Fungi(100);p__Ascomycota(100);c__Saccharomycetes(100);o__Saccharomycetales(100);f__Metschnikowiaceae(100);g__Metschnikowia(100);s__Metschnikowia_pulcherrima(100); | Metschnikowia | 13 | Saprotroph | \|Undefined Saprotroph\| | Yeast | NULL | Probable | As nectar-tap_saprotroph (Põlme et al. 2020); Decay sugar-rich substrates (Põlme et al. 2020) | Põlme S, et al. 2020. Fungal Diversity 105:1-16 ((https://doi.org/10.1007/s13225-020-00466-2)) |
| Otu25105 | 1 | k__Fungi(100);p__Ascomycota(100);c__Saccharomycetes(100);o__Saccharomycetales(100);f__Metschnikowiaceae(100);g__Metschnikowia(100);s__Metschnikowia_pulcherrima(100); | Metschnikowia | 13 | Saprotroph | \|Undefined Saprotroph\| | Yeast | NULL | Probable | As nectar-tap_saprotroph (Põlme et al. 2020); Decay sugar-rich substrates (Põlme et al. 2020) | Põlme S, et al. 2020. Fungal Diversity 105:1-16 ((https://doi.org/10.1007/s13225-020-00466-2)) |
| Otu25106 | 1 | k__Fungi(100);p__Ascomycota(100);c__Saccharomycetes(100);o__Saccharomycetales(100);f__Metschnikowiaceae(100);g__Metschnikowia(100);s__Metschnikowia_pulcherrima(100); | Metschnikowia | 13 | Saprotroph | \|Undefined Saprotroph\| | Yeast | NULL | Probable | As nectar-tap_saprotroph (Põlme et al. 2020); Decay sugar-rich substrates (Põlme et al. 2020) | Põlme S, et al. 2020. Fungal Diversity 105:1-16 ((https://doi.org/10.1007/s13225-020-00466-2)) |
| Otu25107 | 1 | k__Fungi(100);p__Ascomycota(100);c__Saccharomycetes(100);o__Saccharomycetales(100);f__Metschnikowiaceae(100);g__Metschnikowia(100);s__Metschnikowia_pulcherrima(100); | Metschnikowia | 13 | Saprotroph | \|Undefined Saprotroph\| | Yeast | NULL | Probable | As nectar-tap_saprotroph (Põlme et al. 2020); Decay sugar-rich substrates (Põlme et al. 2020) | Põlme S, et al. 2020. Fungal Diversity 105:1-16 ((https://doi.org/10.1007/s13225-020-00466-2)) |
| Otu25108 | 1 | k__Fungi(100);p__Ascomycota(100);c__Saccharomycetes(100);o__Saccharomycetales(100);f__Metschnikowiaceae(100);g__Metschnikowia(100);s__Metschnikowia_pulcherrima(100); | Metschnikowia | 13 | Saprotroph | \|Undefined Saprotroph\| | Yeast | NULL | Probable | As nectar-tap_saprotroph (Põlme et al. 2020); Decay sugar-rich substrates (Põlme et al. 2020) | Põlme S, et al. 2020. Fungal Diversity 105:1-16 ((https://doi.org/10.1007/s13225-020-00466-2)) |
| Otu25109 | 1 | k__Fungi(100);p__Ascomycota(100);c__Saccharomycetes(100);o__Saccharomycetales(100);f__Metschnikowiaceae(100);g__Metschnikowia(100);s__Metschnikowia_pulcherrima(100); | Metschnikowia | 13 | Saprotroph | \|Undefined Saprotroph\| | Yeast | NULL | Probable | As nectar-tap_saprotroph (Põlme et al. 2020); Decay sugar-rich substrates (Põlme et al. 2020) | Põlme S, et al. 2020. Fungal Diversity 105:1-16 ((https://doi.org/10.1007/s13225-020-00466-2)) |
| Otu25110 | 1 | k__Fungi(100);p__Ascomycota(100);c__Saccharomycetes(100);o__Saccharomycetales(100);f__Metschnikowiaceae(100);g__Metschnikowia(100);s__Metschnikowia_pulcherrima(100); | Metschnikowia | 13 | Saprotroph | \|Undefined Saprotroph\| | Yeast | NULL | Probable | As nectar-tap_saprotroph (Põlme et al. 2020); Decay sugar-rich substrates (Põlme et al. 2020) | Põlme S, et al. 2020. Fungal Diversity 105:1-16 ((https://doi.org/10.1007/s13225-020-00466-2)) |
| Otu25111 | 1 | k__Fungi(100);p__Ascomycota(100);c__Saccharomycetes(100);o__Saccharomycetales(100);f__Metschnikowiaceae(100);g__Metschnikowia(100);g__Metschnikowia_unclassified(100); | Metschnikowia | 13 | Saprotroph | \|Undefined Saprotroph\| | Yeast | NULL | Probable | As nectar-tap_saprotroph (Põlme et al. 2020); Decay sugar-rich substrates (Põlme et al. 2020) | Põlme S, et al. 2020. Fungal Diversity 105:1-16 ((https://doi.org/10.1007/s13225-020-00466-2)) |
| Otu25112 | 1 | k__Fungi(100);p__Ascomycota(100);c__Saccharomycetes(100);o__Saccharomycetales(100);f__Metschnikowiaceae(100);g__Metschnikowia(100);s__Metschnikowia_pulcherrima(100); | Metschnikowia | 13 | Saprotroph | \|Undefined Saprotroph\| | Yeast | NULL | Probable | As nectar-tap_saprotroph (Põlme et al. 2020); Decay sugar-rich substrates (Põlme et al. 2020) | Põlme S, et al. 2020. Fungal Diversity 105:1-16 ((https://doi.org/10.1007/s13225-020-00466-2)) |
| Otu25113 | 1 | k__Fungi(100);p__Ascomycota(100);c__Saccharomycetes(100);o__Saccharomycetales(100);f__Metschnikowiaceae(100);g__Metschnikowia(100);s__Metschnikowia_pulcherrima(100); | Metschnikowia | 13 | Saprotroph | \|Undefined Saprotroph\| | Yeast | NULL | Probable | As nectar-tap_saprotroph (Põlme et al. 2020); Decay sugar-rich substrates (Põlme et al. 2020) | Põlme S, et al. 2020. Fungal Diversity 105:1-16 ((https://doi.org/10.1007/s13225-020-00466-2)) |
| Otu25114 | 1 | k__Fungi(100);p__Ascomycota(100);c__Saccharomycetes(100);o__Saccharomycetales(100);f__Metschnikowiaceae(100);g__Metschnikowia(100);g__Metschnikowia_unclassified(100); | Metschnikowia | 13 | Saprotroph | \|Undefined Saprotroph\| | Yeast | NULL | Probable | As nectar-tap_saprotroph (Põlme et al. 2020); Decay sugar-rich substrates (Põlme et al. 2020) | Põlme S, et al. 2020. Fungal Diversity 105:1-16 ((https://doi.org/10.1007/s13225-020-00466-2)) |
| Otu25115 | 1 | k__Fungi(100);p__Ascomycota(100);c__Saccharomycetes(100);o__Saccharomycetales(100);f__Metschnikowiaceae(100);g__Metschnikowia(100);s__Metschnikowia_pulcherrima(100); | Metschnikowia | 13 | Saprotroph | \|Undefined Saprotroph\| | Yeast | NULL | Probable | As nectar-tap_saprotroph (Põlme et al. 2020); Decay sugar-rich substrates (Põlme et al. 2020) | Põlme S, et al. 2020. Fungal Diversity 105:1-16 ((https://doi.org/10.1007/s13225-020-00466-2)) |
| Otu25116 | 1 | k__Fungi(100);p__Ascomycota(100);c__Saccharomycetes(100);o__Saccharomycetales(100);f__Metschnikowiaceae(100);g__Metschnikowia(100);s__Metschnikowia_pulcherrima(100); | Metschnikowia | 13 | Saprotroph | \|Undefined Saprotroph\| | Yeast | NULL | Probable | As nectar-tap_saprotroph (Põlme et al. 2020); Decay sugar-rich substrates (Põlme et al. 2020) | Põlme S, et al. 2020. Fungal Diversity 105:1-16 ((https://doi.org/10.1007/s13225-020-00466-2)) |
| Otu25117 | 1 | k__Fungi(100);p__Ascomycota(100);c__Saccharomycetes(100);o__Saccharomycetales(100);f__Metschnikowiaceae(100);g__Metschnikowia(100);g__Metschnikowia_unclassified(100); | Metschnikowia | 13 | Saprotroph | \|Undefined Saprotroph\| | Yeast | NULL | Probable | As nectar-tap_saprotroph (Põlme et al. 2020); Decay sugar-rich substrates (Põlme et al. 2020) | Põlme S, et al. 2020. Fungal Diversity 105:1-16 ((https://doi.org/10.1007/s13225-020-00466-2)) |
| Otu25118 | 1 | k__Fungi(100);p__Ascomycota(100);c__Saccharomycetes(100);o__Saccharomycetales(100);f__Metschnikowiaceae(100);g__Metschnikowia(100);g__Metschnikowia_unclassified(100); | Metschnikowia | 13 | Saprotroph | \|Undefined Saprotroph\| | Yeast | NULL | Probable | As nectar-tap_saprotroph (Põlme et al. 2020); Decay sugar-rich substrates (Põlme et al. 2020) | Põlme S, et al. 2020. Fungal Diversity 105:1-16 ((https://doi.org/10.1007/s13225-020-00466-2)) |
| Otu25119 | 1 | k__Fungi(100);p__Ascomycota(100);c__Saccharomycetes(100);o__Saccharomycetales(100);f__Metschnikowiaceae(100);g__Metschnikowia(100);s__Metschnikowia_pulcherrima(100); | Metschnikowia | 13 | Saprotroph | \|Undefined Saprotroph\| | Yeast | NULL | Probable | As nectar-tap_saprotroph (Põlme et al. 2020); Decay sugar-rich substrates (Põlme et al. 2020) | Põlme S, et al. 2020. Fungal Diversity 105:1-16 ((https://doi.org/10.1007/s13225-020-00466-2)) |
| Otu25120 | 1 | k__Fungi(100);p__Ascomycota(100);c__Saccharomycetes(100);o__Saccharomycetales(100);f__Metschnikowiaceae(100);g__Metschnikowia(100);s__Metschnikowia_pulcherrima(100); | Metschnikowia | 13 | Saprotroph | \|Undefined Saprotroph\| | Yeast | NULL | Probable | As nectar-tap_saprotroph (Põlme et al. 2020); Decay sugar-rich substrates (Põlme et al. 2020) | Põlme S, et al. 2020. Fungal Diversity 105:1-16 ((https://doi.org/10.1007/s13225-020-00466-2)) |
| Otu25121 | 1 | k__Fungi(100);p__Ascomycota(100);c__Saccharomycetes(100);o__Saccharomycetales(100);f__Metschnikowiaceae(100);g__Metschnikowia(100);s__Metschnikowia_pulcherrima(100); | Metschnikowia | 13 | Saprotroph | \|Undefined Saprotroph\| | Yeast | NULL | Probable | As nectar-tap_saprotroph (Põlme et al. 2020); Decay sugar-rich substrates (Põlme et al. 2020) | Põlme S, et al. 2020. Fungal Diversity 105:1-16 ((https://doi.org/10.1007/s13225-020-00466-2)) |
| Otu25122 | 1 | k__Fungi(100);p__Ascomycota(100);c__Saccharomycetes(100);o__Saccharomycetales(100);f__Metschnikowiaceae(100);g__Metschnikowia(100);s__Metschnikowia_pulcherrima(100); | Metschnikowia | 13 | Saprotroph | \|Undefined Saprotroph\| | Yeast | NULL | Probable | As nectar-tap_saprotroph (Põlme et al. 2020); Decay sugar-rich substrates (Põlme et al. 2020) | Põlme S, et al. 2020. Fungal Diversity 105:1-16 ((https://doi.org/10.1007/s13225-020-00466-2)) |
| Otu25124 | 1 | k__Fungi(100);p__Ascomycota(100);c__Saccharomycetes(100);o__Saccharomycetales(100);f__Metschnikowiaceae(100);g__Metschnikowia(100);s__Metschnikowia_pulcherrima(100); | Metschnikowia | 13 | Saprotroph | \|Undefined Saprotroph\| | Yeast | NULL | Probable | As nectar-tap_saprotroph (Põlme et al. 2020); Decay sugar-rich substrates (Põlme et al. 2020) | Põlme S, et al. 2020. Fungal Diversity 105:1-16 ((https://doi.org/10.1007/s13225-020-00466-2)) |
| Otu25127 | 1 | k__Fungi(100);p__Ascomycota(100);c__Saccharomycetes(100);o__Saccharomycetales(100);f__Metschnikowiaceae(100);g__Metschnikowia(100);g__Metschnikowia_unclassified(100); | Metschnikowia | 13 | Saprotroph | \|Undefined Saprotroph\| | Yeast | NULL | Probable | As nectar-tap_saprotroph (Põlme et al. 2020); Decay sugar-rich substrates (Põlme et al. 2020) | Põlme S, et al. 2020. Fungal Diversity 105:1-16 ((https://doi.org/10.1007/s13225-020-00466-2)) |
| Otu25128 | 1 | k__Fungi(100);p__Ascomycota(100);c__Saccharomycetes(100);o__Saccharomycetales(100);f__Metschnikowiaceae(100);g__Metschnikowia(100);g__Metschnikowia_unclassified(100); | Metschnikowia | 13 | Saprotroph | \|Undefined Saprotroph\| | Yeast | NULL | Probable | As nectar-tap_saprotroph (Põlme et al. 2020); Decay sugar-rich substrates (Põlme et al. 2020) | Põlme S, et al. 2020. Fungal Diversity 105:1-16 ((https://doi.org/10.1007/s13225-020-00466-2)) |
| Otu25129 | 1 | k__Fungi(100);p__Ascomycota(100);c__Saccharomycetes(100);o__Saccharomycetales(100);f__Metschnikowiaceae(100);g__Metschnikowia(100);s__Metschnikowia_pulcherrima(100); | Metschnikowia | 13 | Saprotroph | \|Undefined Saprotroph\| | Yeast | NULL | Probable | As nectar-tap_saprotroph (Põlme et al. 2020); Decay sugar-rich substrates (Põlme et al. 2020) | Põlme S, et al. 2020. Fungal Diversity 105:1-16 ((https://doi.org/10.1007/s13225-020-00466-2)) |
| Otu25130 | 1 | k__Fungi(100);p__Ascomycota(100);c__Saccharomycetes(100);o__Saccharomycetales(100);f__Metschnikowiaceae(100);g__Metschnikowia(100);g__Metschnikowia_unclassified(100); | Metschnikowia | 13 | Saprotroph | \|Undefined Saprotroph\| | Yeast | NULL | Probable | As nectar-tap_saprotroph (Põlme et al. 2020); Decay sugar-rich substrates (Põlme et al. 2020) | Põlme S, et al. 2020. Fungal Diversity 105:1-16 ((https://doi.org/10.1007/s13225-020-00466-2)) |
| Otu25131 | 1 | k__Fungi(100);p__Ascomycota(100);c__Saccharomycetes(100);o__Saccharomycetales(100);f__Metschnikowiaceae(100);g__Metschnikowia(100);s__Metschnikowia_pulcherrima(100); | Metschnikowia | 13 | Saprotroph | \|Undefined Saprotroph\| | Yeast | NULL | Probable | As nectar-tap_saprotroph (Põlme et al. 2020); Decay sugar-rich substrates (Põlme et al. 2020) | Põlme S, et al. 2020. Fungal Diversity 105:1-16 ((https://doi.org/10.1007/s13225-020-00466-2)) |
| Otu25132 | 1 | k__Fungi(100);p__Ascomycota(100);c__Saccharomycetes(100);o__Saccharomycetales(100);f__Metschnikowiaceae(100);g__Metschnikowia(100);g__Metschnikowia_unclassified(100); | Metschnikowia | 13 | Saprotroph | \|Undefined Saprotroph\| | Yeast | NULL | Probable | As nectar-tap_saprotroph (Põlme et al. 2020); Decay sugar-rich substrates (Põlme et al. 2020) | Põlme S, et al. 2020. Fungal Diversity 105:1-16 ((https://doi.org/10.1007/s13225-020-00466-2)) |
| Otu25133 | 1 | k__Fungi(100);p__Ascomycota(100);c__Saccharomycetes(100);o__Saccharomycetales(100);f__Metschnikowiaceae(100);g__Metschnikowia(100);s__Metschnikowia_pulcherrima(100); | Metschnikowia | 13 | Saprotroph | \|Undefined Saprotroph\| | Yeast | NULL | Probable | As nectar-tap_saprotroph (Põlme et al. 2020); Decay sugar-rich substrates (Põlme et al. 2020) | Põlme S, et al. 2020. Fungal Diversity 105:1-16 ((https://doi.org/10.1007/s13225-020-00466-2)) |

| | | | | | | | | | | | |
|---|---|---|---|---|---|---|---|---|---|---|---|
| Otu25134 | 1 | k__Fungi(100);p__Ascomycota(100);c__Saccharomycetes(100);o__Saccharomycetales(100);f__Metschnikowiaceae(100);g__Metschnikowia(100);s__Metschnikowia_pulcherrima(100); | Metschnikowia | 13 | Saprotroph | \|Undefined Saprotroph\| | Yeast | NULL | Probable | As nectar-tap_saprotroph (Põlme et al. 2020); Decay sugar-rich substrates (Põlme et al. 2020) | Põlme S, et al. 2020. Fungal Diversity 105:1-16 ((https://doi.org/10.1007/s13225-020-00466-2)) |
| Otu25136 | 1 | k__Fungi(100);p__Ascomycota(100);c__Saccharomycetes(100);o__Saccharomycetales(100);f__Metschnikowiaceae(100);g__Metschnikowia(100);s__Metschnikowia_pulcherrima(100); | Metschnikowia | 13 | Saprotroph | \|Undefined Saprotroph\| | Yeast | NULL | Probable | As nectar-tap_saprotroph (Põlme et al. 2020); Decay sugar-rich substrates (Põlme et al. 2020) | Põlme S, et al. 2020. Fungal Diversity 105:1-16 ((https://doi.org/10.1007/s13225-020-00466-2)) |
| Otu25137 | 1 | k__Fungi(100);p__Ascomycota(100);c__Saccharomycetes(100);o__Saccharomycetales(100);f__Metschnikowiaceae(100);g__Metschnikowia(100);g__Metschnikowia_unclassified(100); | Metschnikowia | 13 | Saprotroph | \|Undefined Saprotroph\| | Yeast | NULL | Probable | As nectar-tap_saprotroph (Põlme et al. 2020); Decay sugar-rich substrates (Põlme et al. 2020) | Põlme S, et al. 2020. Fungal Diversity 105:1-16 ((https://doi.org/10.1007/s13225-020-00466-2)) |
| Otu25138 | 1 | k__Fungi(100);p__Ascomycota(100);c__Saccharomycetes(100);o__Saccharomycetales(100);f__Metschnikowiaceae(100);g__Metschnikowia(100);s__Metschnikowia_pulcherrima(100); | Metschnikowia | 13 | Saprotroph | \|Undefined Saprotroph\| | Yeast | NULL | Probable | As nectar-tap_saprotroph (Põlme et al. 2020); Decay sugar-rich substrates (Põlme et al. 2020) | Põlme S, et al. 2020. Fungal Diversity 105:1-16 ((https://doi.org/10.1007/s13225-020-00466-2)) |
| Otu25139 | 1 | k__Fungi(100);p__Ascomycota(100);c__Saccharomycetes(100);o__Saccharomycetales(100);f__Metschnikowiaceae(100);g__Metschnikowia(100);s__Metschnikowia_pulcherrima(100); | Metschnikowia | 13 | Saprotroph | \|Undefined Saprotroph\| | Yeast | NULL | Probable | As nectar-tap_saprotroph (Põlme et al. 2020); Decay sugar-rich substrates (Põlme et al. 2020) | Põlme S, et al. 2020. Fungal Diversity 105:1-16 ((https://doi.org/10.1007/s13225-020-00466-2)) |
| Otu25140 | 1 | k__Fungi(100);p__Ascomycota(100);c__Saccharomycetes(100);o__Saccharomycetales(100);f__Metschnikowiaceae(100);g__Metschnikowia(100);s__Metschnikowia_pulcherrima(100); | Metschnikowia | 13 | Saprotroph | \|Undefined Saprotroph\| | Yeast | NULL | Probable | As nectar-tap_saprotroph (Põlme et al. 2020); Decay sugar-rich substrates (Põlme et al. 2020) | Põlme S, et al. 2020. Fungal Diversity 105:1-16 ((https://doi.org/10.1007/s13225-020-00466-2)) |
| Otu25142 | 1 | k__Fungi(100);p__Ascomycota(100);c__Saccharomycetes(100);o__Saccharomycetales(100);f__Metschnikowiaceae(100);g__Metschnikowia(100);s__Metschnikowia_pulcherrima(100); | Metschnikowia | 13 | Saprotroph | \|Undefined Saprotroph\| | Yeast | NULL | Probable | As nectar-tap_saprotroph (Põlme et al. 2020); Decay sugar-rich substrates (Põlme et al. 2020) | Põlme S, et al. 2020. Fungal Diversity 105:1-16 ((https://doi.org/10.1007/s13225-020-00466-2)) |
| Otu25145 | 1 | k__Fungi(100);p__Ascomycota(100);c__Saccharomycetes(100);o__Saccharomycetales(100);f__Metschnikowiaceae(100);g__Metschnikowia(100);s__Metschnikowia_pulcherrima(100); | Metschnikowia | 13 | Saprotroph | \|Undefined Saprotroph\| | Yeast | NULL | Probable | As nectar-tap_saprotroph (Põlme et al. 2020); Decay sugar-rich substrates (Põlme et al. 2020) | Põlme S, et al. 2020. Fungal Diversity 105:1-16 ((https://doi.org/10.1007/s13225-020-00466-2)) |
| Otu25146 | 1 | k__Fungi(100);p__Ascomycota(100);c__Saccharomycetes(100);o__Saccharomycetales(100);f__Metschnikowiaceae(100);g__Metschnikowia(100);g__Metschnikowia_unclassified(100); | Metschnikowia | 13 | Saprotroph | \|Undefined Saprotroph\| | Yeast | NULL | Probable | As nectar-tap_saprotroph (Põlme et al. 2020); Decay sugar-rich substrates (Põlme et al. 2020) | Põlme S, et al. 2020. Fungal Diversity 105:1-16 ((https://doi.org/10.1007/s13225-020-00466-2)) |
| Otu25147 | 1 | k__Fungi(100);p__Ascomycota(100);c__Saccharomycetes(100);o__Saccharomycetales(100);f__Metschnikowiaceae(100);g__Metschnikowia(100);s__Metschnikowia_pulcherrima(100); | Metschnikowia | 13 | Saprotroph | \|Undefined Saprotroph\| | Yeast | NULL | Probable | As nectar-tap_saprotroph (Põlme et al. 2020); Decay sugar-rich substrates (Põlme et al. 2020) | Põlme S, et al. 2020. Fungal Diversity 105:1-16 ((https://doi.org/10.1007/s13225-020-00466-2)) |
| Otu25148 | 1 | k__Fungi(100);p__Ascomycota(100);c__Saccharomycetes(100);o__Saccharomycetales(100);f__Metschnikowiaceae(100);g__Metschnikowia(100);s__Metschnikowia_pulcherrima(100); | Metschnikowia | 13 | Saprotroph | \|Undefined Saprotroph\| | Yeast | NULL | Probable | As nectar-tap_saprotroph (Põlme et al. 2020); Decay sugar-rich substrates (Põlme et al. 2020) | Põlme S, et al. 2020. Fungal Diversity 105:1-16 ((https://doi.org/10.1007/s13225-020-00466-2)) |
| Otu25149 | 1 | k__Fungi(100);p__Ascomycota(100);c__Saccharomycetes(100);o__Saccharomycetales(100);f__Metschnikowiaceae(100);g__Metschnikowia(100);g__Metschnikowia_unclassified(100); | Metschnikowia | 13 | Saprotroph | \|Undefined Saprotroph\| | Yeast | NULL | Probable | As nectar-tap_saprotroph (Põlme et al. 2020); Decay sugar-rich substrates (Põlme et al. 2020) | Põlme S, et al. 2020. Fungal Diversity 105:1-16 ((https://doi.org/10.1007/s13225-020-00466-2)) |
| Otu25150 | 1 | k__Fungi(100);p__Ascomycota(100);c__Saccharomycetes(100);o__Saccharomycetales(100);f__Metschnikowiaceae(100);g__Metschnikowia(100);g__Metschnikowia_unclassified(100); | Metschnikowia | 13 | Saprotroph | \|Undefined Saprotroph\| | Yeast | NULL | Probable | As nectar-tap_saprotroph (Põlme et al. 2020); Decay sugar-rich substrates (Põlme et al. 2020) | Põlme S, et al. 2020. Fungal Diversity 105:1-16 ((https://doi.org/10.1007/s13225-020-00466-2)) |
| Otu25151 | 1 | k__Fungi(100);p__Ascomycota(100);c__Saccharomycetes(100);o__Saccharomycetales(100);f__Metschnikowiaceae(100);g__Metschnikowia(100);s__Metschnikowia_pulcherrima(100); | Metschnikowia | 13 | Saprotroph | \|Undefined Saprotroph\| | Yeast | NULL | Probable | As nectar-tap_saprotroph (Põlme et al. 2020); Decay sugar-rich substrates (Põlme et al. 2020) | Põlme S, et al. 2020. Fungal Diversity 105:1-16 ((https://doi.org/10.1007/s13225-020-00466-2)) |
| Otu25152 | 1 | k__Fungi(100);p__Ascomycota(100);c__Saccharomycetes(100);o__Saccharomycetales(100);f__Metschnikowiaceae(100);g__Metschnikowia(100);s__Metschnikowia_pulcherrima(100); | Metschnikowia | 13 | Saprotroph | \|Undefined Saprotroph\| | Yeast | NULL | Probable | As nectar-tap_saprotroph (Põlme et al. 2020); Decay sugar-rich substrates (Põlme et al. 2020) | Põlme S, et al. 2020. Fungal Diversity 105:1-16 ((https://doi.org/10.1007/s13225-020-00466-2)) |
| Otu25153 | 1 | k__Fungi(100);p__Ascomycota(100);c__Saccharomycetes(100);o__Saccharomycetales(100);f__Metschnikowiaceae(100);g__Metschnikowia(100);s__Metschnikowia_pulcherrima(100); | Metschnikowia | 13 | Saprotroph | \|Undefined Saprotroph\| | Yeast | NULL | Probable | As nectar-tap_saprotroph (Põlme et al. 2020); Decay sugar-rich substrates (Põlme et al. 2020) | Põlme S, et al. 2020. Fungal Diversity 105:1-16 ((https://doi.org/10.1007/s13225-020-00466-2)) |
| Otu25154 | 1 | k__Fungi(100);p__Ascomycota(100);c__Saccharomycetes(100);o__Saccharomycetales(100);f__Metschnikowiaceae(100);g__Metschnikowia(100);s__Metschnikowia_pulcherrima(100); | Metschnikowia | 13 | Saprotroph | \|Undefined Saprotroph\| | Yeast | NULL | Probable | As nectar-tap_saprotroph (Põlme et al. 2020); Decay sugar-rich substrates (Põlme et al. 2020) | Põlme S, et al. 2020. Fungal Diversity 105:1-16 ((https://doi.org/10.1007/s13225-020-00466-2)) |
| Otu25155 | 1 | k__Fungi(100);p__Ascomycota(100);c__Saccharomycetes(100);o__Saccharomycetales(100);f__Metschnikowiaceae(100);g__Metschnikowia(100);s__Metschnikowia_pulcherrima(100); | Metschnikowia | 13 | Saprotroph | \|Undefined Saprotroph\| | Yeast | NULL | Probable | As nectar-tap_saprotroph (Põlme et al. 2020); Decay sugar-rich substrates (Põlme et al. 2020) | Põlme S, et al. 2020. Fungal Diversity 105:1-16 ((https://doi.org/10.1007/s13225-020-00466-2)) |
| Otu25156 | 1 | k__Fungi(100);p__Ascomycota(100);c__Saccharomycetes(100);o__Saccharomycetales(100);f__Metschnikowiaceae(100);g__Metschnikowia(100);s__Metschnikowia_pulcherrima(100); | Metschnikowia | 13 | Saprotroph | \|Undefined Saprotroph\| | Yeast | NULL | Probable | As nectar-tap_saprotroph (Põlme et al. 2020); Decay sugar-rich substrates (Põlme et al. 2020) | Põlme S, et al. 2020. Fungal Diversity 105:1-16 ((https://doi.org/10.1007/s13225-020-00466-2)) |
| Otu25157 | 1 | k__Fungi(100);p__Ascomycota(100);c__Saccharomycetes(100);o__Saccharomycetales(100);f__Metschnikowiaceae(100);g__Metschnikowia(100);s__Metschnikowia_pulcherrima(100); | Metschnikowia | 13 | Saprotroph | \|Undefined Saprotroph\| | Yeast | NULL | Probable | As nectar-tap_saprotroph (Põlme et al. 2020); Decay sugar-rich substrates (Põlme et al. 2020) | Põlme S, et al. 2020. Fungal Diversity 105:1-16 ((https://doi.org/10.1007/s13225-020-00466-2)) |
| Otu25158 | 1 | k__Fungi(100);p__Ascomycota(100);c__Saccharomycetes(100);o__Saccharomycetales(100);f__Metschnikowiaceae(100);g__Metschnikowia(100);s__Metschnikowia_pulcherrima(100); | Metschnikowia | 13 | Saprotroph | \|Undefined Saprotroph\| | Yeast | NULL | Probable | As nectar-tap_saprotroph (Põlme et al. 2020); Decay sugar-rich substrates (Põlme et al. 2020) | Põlme S, et al. 2020. Fungal Diversity 105:1-16 ((https://doi.org/10.1007/s13225-020-00466-2)) |
| Otu25159 | 1 | k__Fungi(100);p__Ascomycota(100);c__Saccharomycetes(100);o__Saccharomycetales(100);f__Metschnikowiaceae(100);g__Metschnikowia(100);s__Metschnikowia_pulcherrima(100); | Metschnikowia | 13 | Saprotroph | \|Undefined Saprotroph\| | Yeast | NULL | Probable | As nectar-tap_saprotroph (Põlme et al. 2020); Decay sugar-rich substrates (Põlme et al. 2020) | Põlme S, et al. 2020. Fungal Diversity 105:1-16 ((https://doi.org/10.1007/s13225-020-00466-2)) |
| Otu25161 | 1 | k__Fungi(100);p__Ascomycota(100);c__Saccharomycetes(100);o__Saccharomycetales(100);f__Metschnikowiaceae(100);g__Metschnikowia(100);g__Metschnikowia_unclassified(100); | Metschnikowia | 13 | Saprotroph | \|Undefined Saprotroph\| | Yeast | NULL | Probable | As nectar-tap_saprotroph (Põlme et al. 2020); Decay sugar-rich substrates (Põlme et al. 2020) | Põlme S, et al. 2020. Fungal Diversity 105:1-16 ((https://doi.org/10.1007/s13225-020-00466-2)) |
| Otu25163 | 1 | k__Fungi(100);p__Ascomycota(100);c__Saccharomycetes(100);o__Saccharomycetales(100);f__Metschnikowiaceae(100);g__Metschnikowia(100);g__Metschnikowia_unclassified(100); | Metschnikowia | 13 | Saprotroph | \|Undefined Saprotroph\| | Yeast | NULL | Probable | As nectar-tap_saprotroph (Põlme et al. 2020); Decay sugar-rich substrates (Põlme et al. 2020) | Põlme S, et al. 2020. Fungal Diversity 105:1-16 ((https://doi.org/10.1007/s13225-020-00466-2)) |
| Otu25164 | 1 | k__Fungi(100);p__Ascomycota(100);c__Saccharomycetes(100);o__Saccharomycetales(100);f__Metschnikowiaceae(100);g__Metschnikowia(100);g__Metschnikowia_unclassified(100); | Metschnikowia | 13 | Saprotroph | \|Undefined Saprotroph\| | Yeast | NULL | Probable | As nectar-tap_saprotroph (Põlme et al. 2020); Decay sugar-rich substrates (Põlme et al. 2020) | Põlme S, et al. 2020. Fungal Diversity 105:1-16 ((https://doi.org/10.1007/s13225-020-00466-2)) |
| Otu25166 | 1 | k__Fungi(100);p__Ascomycota(100);c__Saccharomycetes(100);o__Saccharomycetales(100);f__Metschnikowiaceae(100);g__Metschnikowia(100);s__Metschnikowia_pulcherrima(100); | Metschnikowia | 13 | Saprotroph | \|Undefined Saprotroph\| | Yeast | NULL | Probable | As nectar-tap_saprotroph (Põlme et al. 2020); Decay sugar-rich substrates (Põlme et al. 2020) | Põlme S, et al. 2020. Fungal Diversity 105:1-16 ((https://doi.org/10.1007/s13225-020-00466-2)) |
| Otu25170 | 1 | k__Fungi(100);p__Ascomycota(100);c__Saccharomycetes(100);o__Saccharomycetales(100);f__Metschnikowiaceae(100);g__Metschnikowia(100);s__Metschnikowia_pulcherrima(100); | Metschnikowia | 13 | Saprotroph | \|Undefined Saprotroph\| | Yeast | NULL | Probable | As nectar-tap_saprotroph (Põlme et al. 2020); Decay sugar-rich substrates (Põlme et al. 2020) | Põlme S, et al. 2020. Fungal Diversity 105:1-16 ((https://doi.org/10.1007/s13225-020-00466-2)) |
| Otu25171 | 1 | k__Fungi(100);p__Ascomycota(100);c__Saccharomycetes(100);o__Saccharomycetales(100);f__Metschnikowiaceae(100);g__Metschnikowia(100);s__Metschnikowia_pulcherrima(100); | Metschnikowia | 13 | Saprotroph | \|Undefined Saprotroph\| | Yeast | NULL | Probable | As nectar-tap_saprotroph (Põlme et al. 2020); Decay sugar-rich substrates (Põlme et al. 2020) | Põlme S, et al. 2020. Fungal Diversity 105:1-16 ((https://doi.org/10.1007/s13225-020-00466-2)) |
| Otu25173 | 1 | k__Fungi(100);p__Ascomycota(100);c__Saccharomycetes(100);o__Saccharomycetales(100);f__Metschnikowiaceae(100);g__Metschnikowia(100);s__Metschnikowia_pulcherrima(100); | Metschnikowia | 13 | Saprotroph | \|Undefined Saprotroph\| | Yeast | NULL | Probable | As nectar-tap_saprotroph (Põlme et al. 2020); Decay sugar-rich substrates (Põlme et al. 2020) | Põlme S, et al. 2020. Fungal Diversity 105:1-16 ((https://doi.org/10.1007/s13225-020-00466-2)) |
| Otu25175 | 1 | k__Fungi(100);p__Ascomycota(100);c__Saccharomycetes(100);o__Saccharomycetales(100);f__Metschnikowiaceae(100);g__Metschnikowia(100);s__Metschnikowia_pulcherrima(100); | Metschnikowia | 13 | Saprotroph | \|Undefined Saprotroph\| | Yeast | NULL | Probable | As nectar-tap_saprotroph (Põlme et al. 2020); Decay sugar-rich substrates (Põlme et al. 2020) | Põlme S, et al. 2020. Fungal Diversity 105:1-16 ((https://doi.org/10.1007/s13225-020-00466-2)) |
| Otu25176 | 1 | k__Fungi(100);p__Ascomycota(100);c__Saccharomycetes(100);o__Saccharomycetales(100);f__Metschnikowiaceae(100);g__Metschnikowia(100);s__Metschnikowia_pulcherrima(100); | Metschnikowia | 13 | Saprotroph | \|Undefined Saprotroph\| | Yeast | NULL | Probable | As nectar-tap_saprotroph (Põlme et al. 2020); Decay sugar-rich substrates (Põlme et al. 2020) | Põlme S, et al. 2020. Fungal Diversity 105:1-16 ((https://doi.org/10.1007/s13225-020-00466-2)) |
| Otu25180 | 1 | k__Fungi(100);p__Ascomycota(100);c__Saccharomycetes(100);o__Saccharomycetales(100);f__Metschnikowiaceae(100);g__Metschnikowia(100);s__Metschnikowia_pulcherrima(100); | Metschnikowia | 13 | Saprotroph | \|Undefined Saprotroph\| | Yeast | NULL | Probable | As nectar-tap_saprotroph (Põlme et al. 2020); Decay sugar-rich substrates (Põlme et al. 2020) | Põlme S, et al. 2020. Fungal Diversity 105:1-16 ((https://doi.org/10.1007/s13225-020-00466-2)) |
| Otu25182 | 1 | k__Fungi(100);p__Ascomycota(100);c__Saccharomycetes(100);o__Saccharomycetales(100);f__Metschnikowiaceae(100);g__Metschnikowia(100);g__Metschnikowia_unclassified(100); | Metschnikowia | 13 | Saprotroph | \|Undefined Saprotroph\| | Yeast | NULL | Probable | As nectar-tap_saprotroph (Põlme et al. 2020); Decay sugar-rich substrates (Põlme et al. 2020) | Põlme S, et al. 2020. Fungal Diversity 105:1-16 ((https://doi.org/10.1007/s13225-020-00466-2)) |
| Otu25184 | 1 | k__Fungi(100);p__Ascomycota(100);c__Saccharomycetes(100);o__Saccharomycetales(100);f__Metschnikowiaceae(100);g__Metschnikowia(100);s__Metschnikowia_pulcherrima(100); | Metschnikowia | 13 | Saprotroph | \|Undefined Saprotroph\| | Yeast | NULL | Probable | As nectar-tap_saprotroph (Põlme et al. 2020); Decay sugar-rich substrates (Põlme et al. 2020) | Põlme S, et al. 2020. Fungal Diversity 105:1-16 ((https://doi.org/10.1007/s13225-020-00466-2)) |
| Otu25186 | 1 | k__Fungi(100);p__Ascomycota(100);c__Saccharomycetes(100);o__Saccharomycetales(100);f__Metschnikowiaceae(100);g__Metschnikowia(100);g__Metschnikowia_unclassified(100); | Metschnikowia | 13 | Saprotroph | \|Undefined Saprotroph\| | Yeast | NULL | Probable | As nectar-tap_saprotroph (Põlme et al. 2020); Decay sugar-rich substrates (Põlme et al. 2020) | Põlme S, et al. 2020. Fungal Diversity 105:1-16 ((https://doi.org/10.1007/s13225-020-00466-2)) |
| Otu25187 | 1 | k__Fungi(100);p__Ascomycota(100);c__Saccharomycetes(100);o__Saccharomycetales(100);f__Metschnikowiaceae(100);g__Metschnikowia(100);s__Metschnikowia_pulcherrima(100); | Metschnikowia | 13 | Saprotroph | \|Undefined Saprotroph\| | Yeast | NULL | Probable | As nectar-tap_saprotroph (Põlme et al. 2020); Decay sugar-rich substrates (Põlme et al. 2020) | Põlme S, et al. 2020. Fungal Diversity 105:1-16 ((https://doi.org/10.1007/s13225-020-00466-2)) |
| Otu25188 | 1 | k__Fungi(100);p__Ascomycota(100);c__Saccharomycetes(100);o__Saccharomycetales(100);f__Metschnikowiaceae(100);g__Metschnikowia(100);s__Metschnikowia_pulcherrima(100); | Metschnikowia | 13 | Saprotroph | \|Undefined Saprotroph\| | Yeast | NULL | Probable | As nectar-tap_saprotroph (Põlme et al. 2020); Decay sugar-rich substrates (Põlme et al. 2020) | Põlme S, et al. 2020. Fungal Diversity 105:1-16 ((https://doi.org/10.1007/s13225-020-00466-2)) |
| Otu25189 | 1 | k__Fungi(100);p__Ascomycota(100);c__Saccharomycetes(100);o__Saccharomycetales(100);f__Metschnikowiaceae(100);g__Metschnikowia(100);g__Metschnikowia_unclassified(100); | Metschnikowia | 13 | Saprotroph | \|Undefined Saprotroph\| | Yeast | NULL | Probable | As nectar-tap_saprotroph (Põlme et al. 2020); Decay sugar-rich substrates (Põlme et al. 2020) | Põlme S, et al. 2020. Fungal Diversity 105:1-16 ((https://doi.org/10.1007/s13225-020-00466-2)) |
| Otu25190 | 1 | k__Fungi(100);p__Ascomycota(100);c__Saccharomycetes(100);o__Saccharomycetales(100);f__Metschnikowiaceae(100);g__Metschnikowia(100);s__Metschnikowia_pulcherrima(100); | Metschnikowia | 13 | Saprotroph | \|Undefined Saprotroph\| | Yeast | NULL | Probable | As nectar-tap_saprotroph (Põlme et al. 2020); Decay sugar-rich substrates (Põlme et al. 2020) | Põlme S, et al. 2020. Fungal Diversity 105:1-16 ((https://doi.org/10.1007/s13225-020-00466-2)) |
| Otu25191 | 1 | k__Fungi(100);p__Ascomycota(100);c__Saccharomycetes(100);o__Saccharomycetales(100);f__Metschnikowiaceae(100);g__Metschnikowia(100);s__Metschnikowia_pulcherrima(100); | Metschnikowia | 13 | Saprotroph | \|Undefined Saprotroph\| | Yeast | NULL | Probable | As nectar-tap_saprotroph (Põlme et al. 2020); Decay sugar-rich substrates (Põlme et al. 2020) | Põlme S, et al. 2020. Fungal Diversity 105:1-16 ((https://doi.org/10.1007/s13225-020-00466-2)) |
| Otu25192 | 1 | k__Fungi(100);p__Ascomycota(100);c__Saccharomycetes(100);o__Saccharomycetales(100);f__Metschnikowiaceae(100);g__Metschnikowia(100);g__Metschnikowia_unclassified(100); | Metschnikowia | 13 | Saprotroph | \|Undefined Saprotroph\| | Yeast | NULL | Probable | As nectar-tap_saprotroph (Põlme et al. 2020); Decay sugar-rich substrates (Põlme et al. 2020) | Põlme S, et al. 2020. Fungal Diversity 105:1-16 ((https://doi.org/10.1007/s13225-020-00466-2)) |
| Otu25193 | 1 | k__Fungi(100);p__Ascomycota(100);c__Saccharomycetes(100);o__Saccharomycetales(100);f__Metschnikowiaceae(100);g__Metschnikowia(100);s__Metschnikowia_pulcherrima(100); | Metschnikowia | 13 | Saprotroph | \|Undefined Saprotroph\| | Yeast | NULL | Probable | As nectar-tap_saprotroph (Põlme et al. 2020); Decay sugar-rich substrates (Põlme et al. 2020) | Põlme S, et al. 2020. Fungal Diversity 105:1-16 ((https://doi.org/10.1007/s13225-020-00466-2)) |
| Otu25194 | 1 | k__Fungi(100);p__Ascomycota(100);c__Saccharomycetes(100);o__Saccharomycetales(100);f__Metschnikowiaceae(100);g__Metschnikowia(100);s__Metschnikowia_pulcherrima(100); | Metschnikowia | 13 | Saprotroph | \|Undefined Saprotroph\| | Yeast | NULL | Probable | As nectar-tap_saprotroph (Põlme et al. 2020); Decay sugar-rich substrates (Põlme et al. 2020) | Põlme S, et al. 2020. Fungal Diversity 105:1-16 ((https://doi.org/10.1007/s13225-020-00466-2)) |
| Otu25195 | 1 | k__Fungi(100);p__Ascomycota(100);c__Saccharomycetes(100);o__Saccharomycetales(100);f__Metschnikowiaceae(100);g__Metschnikowia(100);s__Metschnikowia_pulcherrima(100); | Metschnikowia | 13 | Saprotroph | \|Undefined Saprotroph\| | Yeast | NULL | Probable | As nectar-tap_saprotroph (Põlme et al. 2020); Decay sugar-rich substrates (Põlme et al. 2020) | Põlme S, et al. 2020. Fungal Diversity 105:1-16 ((https://doi.org/10.1007/s13225-020-00466-2)) |
| Otu25198 | 1 | k__Fungi(100);p__Ascomycota(100);c__Saccharomycetes(100);o__Saccharomycetales(100);f__Metschnikowiaceae(100);g__Metschnikowia(100);s__Metschnikowia_pulcherrima(100); | Metschnikowia | 13 | Saprotroph | \|Undefined Saprotroph\| | Yeast | NULL | Probable | As nectar-tap_saprotroph (Põlme et al. 2020); Decay sugar-rich substrates (Põlme et al. 2020) | Põlme S, et al. 2020. Fungal Diversity 105:1-16 ((https://doi.org/10.1007/s13225-020-00466-2)) |
| Otu25200 | 1 | k__Fungi(100);p__Ascomycota(100);c__Saccharomycetes(100);o__Saccharomycetales(100);f__Metschnikowiaceae(100);g__Metschnikowia(100);g__Metschnikowia_unclassified(100); | Metschnikowia | 13 | Saprotroph | \|Undefined Saprotroph\| | Yeast | NULL | Probable | As nectar-tap_saprotroph (Põlme et al. 2020); Decay sugar-rich substrates (Põlme et al. 2020) | Põlme S, et al. 2020. Fungal Diversity 105:1-16 ((https://doi.org/10.1007/s13225-020-00466-2)) |
| Otu25203 | 1 | k__Fungi(100);p__Ascomycota(100);c__Saccharomycetes(100);o__Saccharomycetales(100);f__Metschnikowiaceae(100);g__Metschnikowia(100);g__Metschnikowia_unclassified(100); | Metschnikowia | 13 | Saprotroph | \|Undefined Saprotroph\| | Yeast | NULL | Probable | As nectar-tap_saprotroph (Põlme et al. 2020); Decay sugar-rich substrates (Põlme et al. 2020) | Põlme S, et al. 2020. Fungal Diversity 105:1-16 ((https://doi.org/10.1007/s13225-020-00466-2)) |
| Otu25204 | 1 | k__Fungi(100);p__Ascomycota(100);c__Saccharomycetes(100);o__Saccharomycetales(100);f__Metschnikowiaceae(100);g__Metschnikowia(100);s__Metschnikowia_pulcherrima(100); | Metschnikowia | 13 | Saprotroph | \|Undefined Saprotroph\| | Yeast | NULL | Probable | As nectar-tap_saprotroph (Põlme et al. 2020); Decay sugar-rich substrates (Põlme et al. 2020) | Põlme S, et al. 2020. Fungal Diversity 105:1-16 ((https://doi.org/10.1007/s13225-020-00466-2)) |
| Otu25205 | 1 | k__Fungi(100);p__Ascomycota(100);c__Saccharomycetes(100);o__Saccharomycetales(100);f__Metschnikowiaceae(100);g__Metschnikowia(100);s__Metschnikowia_pulcherrima(100); | Metschnikowia | 13 | Saprotroph | \|Undefined Saprotroph\| | Yeast | NULL | Probable | As nectar-tap_saprotroph (Põlme et al. 2020); Decay sugar-rich substrates (Põlme et al. 2020) | Põlme S, et al. 2020. Fungal Diversity 105:1-16 ((https://doi.org/10.1007/s13225-020-00466-2)) |
| Otu25206 | 1 | k__Fungi(100);p__Ascomycota(100);c__Saccharomycetes(100);o__Saccharomycetales(100);f__Metschnikowiaceae(100);g__Metschnikowia(100);g__Metschnikowia_unclassified(100); | Metschnikowia | 13 | Saprotroph | \|Undefined Saprotroph\| | Yeast | NULL | Probable | As nectar-tap_saprotroph (Põlme et al. 2020); Decay sugar-rich substrates (Põlme et al. 2020) | Põlme S, et al. 2020. Fungal Diversity 105:1-16 ((https://doi.org/10.1007/s13225-020-00466-2)) |
| Otu25207 | 1 | k__Fungi(100);p__Ascomycota(100);c__Saccharomycetes(100);o__Saccharomycetales(100);f__Metschnikowiaceae(100);g__Metschnikowia(100);s__Metschnikowia_pulcherrima(100); | Metschnikowia | 13 | Saprotroph | \|Undefined Saprotroph\| | Yeast | NULL | Probable | As nectar-tap_saprotroph (Põlme et al. 2020); Decay sugar-rich substrates (Põlme et al. 2020) | Põlme S, et al. 2020. Fungal Diversity 105:1-16 ((https://doi.org/10.1007/s13225-020-00466-2)) |
| Otu25208 | 1 | k__Fungi(100);p__Ascomycota(100);c__Saccharomycetes(100);o__Saccharomycetales(100);f__Metschnikowiaceae(100);g__Metschnikowia(100);g__Metschnikowia_unclassified(100); | Metschnikowia | 13 | Saprotroph | \|Undefined Saprotroph\| | Yeast | NULL | Probable | As nectar-tap_saprotroph (Põlme et al. 2020); Decay sugar-rich substrates (Põlme et al. 2020) | Põlme S, et al. 2020. Fungal Diversity 105:1-16 ((https://doi.org/10.1007/s13225-020-00466-2)) |
| Otu25209 | 1 | k__Fungi(100);p__Ascomycota(100);c__Saccharomycetes(100);o__Saccharomycetales(100);f__Metschnikowiaceae(100);g__Metschnikowia(100);s__Metschnikowia_pulcherrima(100); | Metschnikowia | 13 | Saprotroph | \|Undefined Saprotroph\| | Yeast | NULL | Probable | As nectar-tap_saprotroph (Põlme et al. 2020); Decay sugar-rich substrates (Põlme et al. 2020) | Põlme S, et al. 2020. Fungal Diversity 105:1-16 ((https://doi.org/10.1007/s13225-020-00466-2)) |
| Otu25210 | 1 | k__Fungi(100);p__Ascomycota(100);c__Saccharomycetes(100);o__Saccharomycetales(100);f__Metschnikowiaceae(100);g__Metschnikowia(100);g__Metschnikowia_unclassified(100); | Metschnikowia | 13 | Saprotroph | \|Undefined Saprotroph\| | Yeast | NULL | Probable | As nectar-tap_saprotroph (Põlme et al. 2020); Decay sugar-rich substrates (Põlme et al. 2020) | Põlme S, et al. 2020. Fungal Diversity 105:1-16 ((https://doi.org/10.1007/s13225-020-00466-2)) |
| Otu25211 | 1 | k__Fungi(100);p__Ascomycota(100);c__Saccharomycetes(100);o__Saccharomycetales(100);f__Metschnikowiaceae(100);g__Metschnikowia(100);s__Metschnikowia_pulcherrima(100); | Metschnikowia | 13 | Saprotroph | \|Undefined Saprotroph\| | Yeast | NULL | Probable | As nectar-tap_saprotroph (Põlme et al. 2020); Decay sugar-rich substrates (Põlme et al. 2020) | Põlme S, et al. 2020. Fungal Diversity 105:1-16 ((https://doi.org/10.1007/s13225-020-00466-2)) |
| Otu25212 | 1 | k__Fungi(100);p__Ascomycota(100);c__Saccharomycetes(100);o__Saccharomycetales(100);f__Metschnikowiaceae(100);g__Metschnikowia(100);s__Metschnikowia_pulcherrima(100); | Metschnikowia | 13 | Saprotroph | \|Undefined Saprotroph\| | Yeast | NULL | Probable | As nectar-tap_saprotroph (Põlme et al. 2020); Decay sugar-rich substrates (Põlme et al. 2020) | Põlme S, et al. 2020. Fungal Diversity 105:1-16 ((https://doi.org/10.1007/s13225-020-00466-2)) |
| Otu25213 | 1 | k__Fungi(100);p__Ascomycota(100);c__Saccharomycetes(100);o__Saccharomycetales(100);f__Metschnikowiaceae(100);g__Metschnikowia(100);s__Metschnikowia_pulcherrima(100); | Metschnikowia | 13 | Saprotroph | \|Undefined Saprotroph\| | Yeast | NULL | Probable | As nectar-tap_saprotroph (Põlme et al. 2020); Decay sugar-rich substrates (Põlme et al. 2020) | Põlme S, et al. 2020. Fungal Diversity 105:1-16 ((https://doi.org/10.1007/s13225-020-00466-2)) |
| Otu25214 | 1 | k__Fungi(100);p__Ascomycota(100);c__Saccharomycetes(100);o__Saccharomycetales(100);f__Metschnikowiaceae(100);g__Metschnikowia(100);g__Metschnikowia_unclassified(100); | Metschnikowia | 13 | Saprotroph | \|Undefined Saprotroph\| | Yeast | NULL | Probable | As nectar-tap_saprotroph (Põlme et al. 2020); Decay sugar-rich substrates (Põlme et al. 2020) | Põlme S, et al. 2020. Fungal Diversity 105:1-16 ((https://doi.org/10.1007/s13225-020-00466-2)) |
| Otu25215 | 1 | k__Fungi(100);p__Ascomycota(100);c__Saccharomycetes(100);o__Saccharomycetales(100);f__Metschnikowiaceae(100);g__Metschnikowia(100);s__Metschnikowia_pulcherrima(100); | Metschnikowia | 13 | Saprotroph | \|Undefined Saprotroph\| | Yeast | NULL | Probable | As nectar-tap_saprotroph (Põlme et al. 2020); Decay sugar-rich substrates (Põlme et al. 2020) | Põlme S, et al. 2020. Fungal Diversity 105:1-16 ((https://doi.org/10.1007/s13225-020-00466-2)) |
| Otu25218 | 1 | k__Fungi(100);p__Ascomycota(100);c__Saccharomycetes(100);o__Saccharomycetales(100);f__Metschnikowiaceae(100);g__Metschnikowia(100);s__Metschnikowia_pulcherrima(100); | Metschnikowia | 13 | Saprotroph | \|Undefined Saprotroph\| | Yeast | NULL | Probable | As nectar-tap_saprotroph (Põlme et al. 2020); Decay sugar-rich substrates (Põlme et al. 2020) | Põlme S, et al. 2020. Fungal Diversity 105:1-16 ((https://doi.org/10.1007/s13225-020-00466-2)) |
| Otu25219 | 1 | k__Fungi(100);p__Ascomycota(100);c__Saccharomycetes(100);o__Saccharomycetales(100);f__Metschnikowiaceae(100);g__Metschnikowia(100);g__Metschnikowia_unclassified(100); | Metschnikowia | 13 | Saprotroph | \|Undefined Saprotroph\| | Yeast | NULL | Probable | As nectar-tap_saprotroph (Põlme et al. 2020); Decay sugar-rich substrates (Põlme et al. 2020) | Põlme S, et al. 2020. Fungal Diversity 105:1-16 ((https://doi.org/10.1007/s13225-020-00466-2)) |
| Otu25220 | 1 | k__Fungi(100);p__Ascomycota(100);c__Saccharomycetes(100);o__Saccharomycetales(100);f__Metschnikowiaceae(100);g__Metschnikowia(100);g__Metschnikowia_unclassified(100); | Metschnikowia | 13 | Saprotroph | \|Undefined Saprotroph\| | Yeast | NULL | Probable | As nectar-tap_saprotroph (Põlme et al. 2020); Decay sugar-rich substrates (Põlme et al. 2020) | Põlme S, et al. 2020. Fungal Diversity 105:1-16 ((https://doi.org/10.1007/s13225-020-00466-2)) |
| Otu25221 | 1 | k__Fungi(100);p__Ascomycota(100);c__Saccharomycetes(100);o__Saccharomycetales(100);f__Metschnikowiaceae(100);g__Metschnikowia(100);g__Metschnikowia_unclassified(100); | Metschnikowia | 13 | Saprotroph | \|Undefined Saprotroph\| | Yeast | NULL | Probable | As nectar-tap_saprotroph (Põlme et al. 2020); Decay sugar-rich substrates (Põlme et al. 2020) | Põlme S, et al. 2020. Fungal Diversity 105:1-16 ((https://doi.org/10.1007/s13225-020-00466-2)) |
| Otu25223 | 1 | k__Fungi(100);p__Ascomycota(100);c__Saccharomycetes(100);o__Saccharomycetales(100);f__Metschnikowiaceae(100);g__Metschnikowia(100);s__Metschnikowia_pulcherrima(100); | Metschnikowia | 13 | Saprotroph | \|Undefined Saprotroph\| | Yeast | NULL | Probable | As nectar-tap_saprotroph (Põlme et al. 2020); Decay sugar-rich substrates (Põlme et al. 2020) | Põlme S, et al. 2020. Fungal Diversity 105:1-16 ((https://doi.org/10.1007/s13225-020-00466-2)) |
| Otu25224 | 1 | k__Fungi(100);p__Ascomycota(100);c__Saccharomycetes(100);o__Saccharomycetales(100);f__Metschnikowiaceae(100);g__Metschnikowia(100);g__Metschnikowia_unclassified(100); | Metschnikowia | 13 | Saprotroph | \|Undefined Saprotroph\| | Yeast | NULL | Probable | As nectar-tap_saprotroph (Põlme et al. 2020); Decay sugar-rich substrates (Põlme et al. 2020) | Põlme S, et al. 2020. Fungal Diversity 105:1-16 ((https://doi.org/10.1007/s13225-020-00466-2)) |
| Otu25227 | 1 | k__Fungi(100);p__Ascomycota(100);c__Saccharomycetes(100);o__Saccharomycetales(100);f__Metschnikowiaceae(100);g__Metschnikowia(100);g__Metschnikowia_unclassified(100); | Metschnikowia | 13 | Saprotroph | \|Undefined Saprotroph\| | Yeast | NULL | Probable | As nectar-tap_saprotroph (Põlme et al. 2020); Decay sugar-rich substrates (Põlme et al. 2020) | Põlme S, et al. 2020. Fungal Diversity 105:1-16 ((https://doi.org/10.1007/s13225-020-00466-2)) |
| Otu25229 | 1 | k__Fungi(100);p__Ascomycota(100);c__Saccharomycetes(100);o__Saccharomycetales(100);f__Metschnikowiaceae(100);g__Metschnikowia(100);s__Metschnikowia_pulcherrima(100); | Metschnikowia | 13 | Saprotroph | \|Undefined Saprotroph\| | Yeast | NULL | Probable | As nectar-tap_saprotroph (Põlme et al. 2020); Decay sugar-rich substrates (Põlme et al. 2020) | Põlme S, et al. 2020. Fungal Diversity 105:1-16 ((https://doi.org/10.1007/s13225-020-00466-2)) |
| Otu25230 | 1 | k__Fungi(100);p__Ascomycota(100);c__Saccharomycetes(100);o__Saccharomycetales(100);f__Metschnikowiaceae(100);g__Metschnikowia(100);g__Metschnikowia_unclassified(100); | Metschnikowia | 13 | Saprotroph | \|Undefined Saprotroph\| | Yeast | NULL | Probable | As nectar-tap_saprotroph (Põlme et al. 2020); Decay sugar-rich substrates (Põlme et al. 2020) | Põlme S, et al. 2020. Fungal Diversity 105:1-16 ((https://doi.org/10.1007/s13225-020-00466-2)) |
| Otu25232 | 1 | k__Fungi(100);p__Ascomycota(100);c__Saccharomycetes(100);o__Saccharomycetales(100);f__Metschnikowiaceae(100);g__Metschnikowia(100);s__Metschnikowia_pulcherrima(100); | Metschnikowia | 13 | Saprotroph | \|Undefined Saprotroph\| | Yeast | NULL | Probable | As nectar-tap_saprotroph (Põlme et al. 2020); Decay sugar-rich substrates (Põlme et al. 2020) | Põlme S, et al. 2020. Fungal Diversity 105:1-16 ((https://doi.org/10.1007/s13225-020-00466-2)) |
| Otu25234 | 1 | k__Fungi(100);p__Ascomycota(100);c__Saccharomycetes(100);o__Saccharomycetales(100);f__Metschnikowiaceae(100);g__Metschnikowia(100);g__Metschnikowia_unclassified(100); | Metschnikowia | 13 | Saprotroph | \|Undefined Saprotroph\| | Yeast | NULL | Probable | As nectar-tap_saprotroph (Põlme et al. 2020); Decay sugar-rich substrates (Põlme et al. 2020) | Põlme S, et al. 2020. Fungal Diversity 105:1-16 ((https://doi.org/10.1007/s13225-020-00466-2)) |
| Otu25235 | 1 | k__Fungi(100);p__Ascomycota(100);c__Saccharomycetes(100);o__Saccharomycetales(100);f__Metschnikowiaceae(100);g__Metschnikowia(100);s__Metschnikowia_pulcherrima(100); | Metschnikowia | 13 | Saprotroph | \|Undefined Saprotroph\| | Yeast | NULL | Probable | As nectar-tap_saprotroph (Põlme et al. 2020); Decay sugar-rich substrates (Põlme et al. 2020) | Põlme S, et al. 2020. Fungal Diversity 105:1-16 ((https://doi.org/10.1007/s13225-020-00466-2)) |
| Otu25239 | 1 | k__Fungi(100);p__Ascomycota(100);c__Saccharomycetes(100);o__Saccharomycetales(100);f__Metschnikowiaceae(100);g__Metschnikowia(100);s__Metschnikowia_pulcherrima(100); | Metschnikowia | 13 | Saprotroph | \|Undefined Saprotroph\| | Yeast | NULL | Probable | As nectar-tap_saprotroph (Põlme et al. 2020); Decay sugar-rich substrates (Põlme et al. 2020) | Põlme S, et al. 2020. Fungal Diversity 105:1-16 ((https://doi.org/10.1007/s13225-020-00466-2)) |
| Otu25240 | 1 | k__Fungi(100);p__Ascomycota(100);c__Saccharomycetes(100);o__Saccharomycetales(100);f__Metschnikowiaceae(100);g__Metschnikowia(100);s__Metschnikowia_pulcherrima(100); | Metschnikowia | 13 | Saprotroph | \|Undefined Saprotroph\| | Yeast | NULL | Probable | As nectar-tap_saprotroph (Põlme et al. 2020); Decay sugar-rich substrates (Põlme et al. 2020) | Põlme S, et al. 2020. Fungal Diversity 105:1-16 ((https://doi.org/10.1007/s13225-020-00466-2)) |
| Otu25243 | 1 | k__Fungi(100);p__Ascomycota(100);c__Saccharomycetes(100);o__Saccharomycetales(100);f__Metschnikowiaceae(100);g__Metschnikowia(100);s__Metschnikowia_pulcherrima(100); | Metschnikowia | 13 | Saprotroph | \|Undefined Saprotroph\| | Yeast | NULL | Probable | As nectar-tap_saprotroph (Põlme et al. 2020); Decay sugar-rich substrates (Põlme et al. 2020) | Põlme S, et al. 2020. Fungal Diversity 105:1-16 ((https://doi.org/10.1007/s13225-020-00466-2)) |
| Otu25245 | 1 | k__Fungi(100);p__Ascomycota(100);c__Saccharomycetes(100);o__Saccharomycetales(100);f__Metschnikowiaceae(100);g__Metschnikowia(100);s__Metschnikowia_pulcherrima(100); | Metschnikowia | 13 | Saprotroph | \|Undefined Saprotroph\| | Yeast | NULL | Probable | As nectar-tap_saprotroph (Põlme et al. 2020); Decay sugar-rich substrates (Põlme et al. 2020) | Põlme S, et al. 2020. Fungal Diversity 105:1-16 ((https://doi.org/10.1007/s13225-020-00466-2)) |
| Otu25246 | 1 | k__Fungi(100);p__Ascomycota(100);c__Saccharomycetes(100);o__Saccharomycetales(100);f__Metschnikowiaceae(100);g__Metschnikowia(100);s__Metschnikowia_pulcherrima(100); | Metschnikowia | 13 | Saprotroph | \|Undefined Saprotroph\| | Yeast | NULL | Probable | As nectar-tap_saprotroph (Põlme et al. 2020); Decay sugar-rich substrates (Põlme et al. 2020) | Põlme S, et al. 2020. Fungal Diversity 105:1-16 ((https://doi.org/10.1007/s13225-020-00466-2)) |
| Otu25247 | 1 | k__Fungi(100);p__Ascomycota(100);c__Saccharomycetes(100);o__Saccharomycetales(100);f__Metschnikowiaceae(100);g__Metschnikowia(100);s__Metschnikowia_pulcherrima(100); | Metschnikowia | 13 | Saprotroph | \|Undefined Saprotroph\| | Yeast | NULL | Probable | As nectar-tap_saprotroph (Põlme et al. 2020); Decay sugar-rich substrates (Põlme et al. 2020) | Põlme S, et al. 2020. Fungal Diversity 105:1-16 ((https://doi.org/10.1007/s13225-020-00466-2)) |
| Otu25248 | 1 | k__Fungi(100);p__Ascomycota(100);c__Saccharomycetes(100);o__Saccharomycetales(100);f__Metschnikowiaceae(100);g__Metschnikowia(100);s__Metschnikowia_pulcherrima(100); | Metschnikowia | 13 | Saprotroph | \|Undefined Saprotroph\| | Yeast | NULL | Probable | As nectar-tap_saprotroph (Põlme et al. 2020); Decay sugar-rich substrates (Põlme et al. 2020) | Põlme S, et al. 2020. Fungal Diversity 105:1-16 ((https://doi.org/10.1007/s13225-020-00466-2)) |
| Otu25249 | 1 | k__Fungi(100);p__Ascomycota(100);c__Saccharomycetes(100);o__Saccharomycetales(100);f__Metschnikowiaceae(100);g__Metschnikowia(100);s__Metschnikowia_pulcherrima(100); | Metschnikowia | 13 | Saprotroph | \|Undefined Saprotroph\| | Yeast | NULL | Probable | As nectar-tap_saprotroph (Põlme et al. 2020); Decay sugar-rich substrates (Põlme et al. 2020) | Põlme S, et al. 2020. Fungal Diversity 105:1-16 ((https://doi.org/10.1007/s13225-020-00466-2)) |
| Otu25250 | 1 | k__Fungi(100);p__Ascomycota(100);c__Saccharomycetes(100);o__Saccharomycetales(100);f__Metschnikowiaceae(100);g__Metschnikowia(100);s__Metschnikowia_pulcherrima(100); | Metschnikowia | 13 | Saprotroph | \|Undefined Saprotroph\| | Yeast | NULL | Probable | As nectar-tap_saprotroph (Põlme et al. 2020); Decay sugar-rich substrates (Põlme et al. 2020) | Põlme S, et al. 2020. Fungal Diversity 105:1-16 ((https://doi.org/10.1007/s13225-020-00466-2)) |
| Otu25251 | 1 | k__Fungi(100);p__Ascomycota(100);c__Saccharomycetes(100);o__Saccharomycetales(100);f__Metschnikowiaceae(100);g__Metschnikowia(100);s__Metschnikowia_pulcherrima(100); | Metschnikowia | 13 | Saprotroph | \|Undefined Saprotroph\| | Yeast | NULL | Probable | As nectar-tap_saprotroph (Põlme et al. 2020); Decay sugar-rich substrates (Põlme et al. 2020) | Põlme S, et al. 2020. Fungal Diversity 105:1-16 ((https://doi.org/10.1007/s13225-020-00466-2)) |
| Otu25252 | 1 | k__Fungi(100);p__Ascomycota(100);c__Saccharomycetes(100);o__Saccharomycetales(100);f__Metschnikowiaceae(100);g__Metschnikowia(100);g__Metschnikowia_unclassified(100); | Metschnikowia | 13 | Saprotroph | \|Undefined Saprotroph\| | Yeast | NULL | Probable | As nectar-tap_saprotroph (Põlme et al. 2020); Decay sugar-rich substrates (Põlme et al. 2020) | Põlme S, et al. 2020. Fungal Diversity 105:1-16 ((https://doi.org/10.1007/s13225-020-00466-2)) |
| Otu25253 | 1 | k__Fungi(100);p__Ascomycota(100);c__Saccharomycetes(100);o__Saccharomycetales(100);f__Metschnikowiaceae(100);g__Metschnikowia(100);g__Metschnikowia_unclassified(100); | Metschnikowia | 13 | Saprotroph | \|Undefined Saprotroph\| | Yeast | NULL | Probable | As nectar-tap_saprotroph (Põlme et al. 2020); Decay sugar-rich substrates (Põlme et al. 2020) | Põlme S, et al. 2020. Fungal Diversity 105:1-16 ((https://doi.org/10.1007/s13225-020-00466-2)) |
| Otu25254 | 1 | k__Fungi(100);p__Ascomycota(100);c__Saccharomycetes(100);o__Saccharomycetales(100);f__Metschnikowiaceae(100);g__Metschnikowia(100);s__Metschnikowia_pulcherrima(100); | Metschnikowia | 13 | Saprotroph | \|Undefined Saprotroph\| | Yeast | NULL | Probable | As nectar-tap_saprotroph (Põlme et al. 2020); Decay sugar-rich substrates (Põlme et al. 2020) | Põlme S, et al. 2020. Fungal Diversity 105:1-16 ((https://doi.org/10.1007/s13225-020-00466-2)) |
| Otu25255 | 1 | k__Fungi(100);p__Ascomycota(100);c__Saccharomycetes(100);o__Saccharomycetales(100);f__Metschnikowiaceae(100);g__Metschnikowia(100);s__Metschnikowia_pulcherrima(100); | Metschnikowia | 13 | Saprotroph | \|Undefined Saprotroph\| | Yeast | NULL | Probable | As nectar-tap_saprotroph (Põlme et al. 2020); Decay sugar-rich substrates (Põlme et al. 2020) | Põlme S, et al. 2020. Fungal Diversity 105:1-16 ((https://doi.org/10.1007/s13225-020-00466-2)) |
| Otu25256 | 1 | k__Fungi(100);p__Ascomycota(100);c__Saccharomycetes(100);o__Saccharomycetales(100);f__Metschnikowiaceae(100);g__Metschnikowia(100);s__Metschnikowia_pulcherrima(100); | Metschnikowia | 13 | Saprotroph | \|Undefined Saprotroph\| | Yeast | NULL | Probable | As nectar-tap_saprotroph (Põlme et al. 2020); Decay sugar-rich substrates (Põlme et al. 2020) | Põlme S, et al. 2020. Fungal Diversity 105:1-16 ((https://doi.org/10.1007/s13225-020-00466-2)) |
| Otu25257 | 1 | k__Fungi(100);p__Ascomycota(100);c__Saccharomycetes(100);o__Saccharomycetales(100);f__Metschnikowiaceae(100);g__Metschnikowia(100);s__Metschnikowia_pulcherrima(100); | Metschnikowia | 13 | Saprotroph | \|Undefined Saprotroph\| | Yeast | NULL | Probable | As nectar-tap_saprotroph (Põlme et al. 2020); Decay sugar-rich substrates (Põlme et al. 2020) | Põlme S, et al. 2020. Fungal Diversity 105:1-16 ((https://doi.org/10.1007/s13225-020-00466-2)) |
| Otu25258 | 1 | k__Fungi(100);p__Ascomycota(100);c__Saccharomycetes(100);o__Saccharomycetales(100);f__Metschnikowiaceae(100);g__Metschnikowia(100);s__Metschnikowia_pulcherrima(100); | Metschnikowia | 13 | Saprotroph | \|Undefined Saprotroph\| | Yeast | NULL | Probable | As nectar-tap_saprotroph (Põlme et al. 2020); Decay sugar-rich substrates (Põlme et al. 2020) | Põlme S, et al. 2020. Fungal Diversity 105:1-16 ((https://doi.org/10.1007/s13225-020-00466-2)) |
| Otu25259 | 1 | k__Fungi(100);p__Ascomycota(100);c__Saccharomycetes(100);o__Saccharomycetales(100);f__Metschnikowiaceae(100);g__Metschnikowia(100);g__Metschnikowia_unclassified(100); | Metschnikowia | 13 | Saprotroph | \|Undefined Saprotroph\| | Yeast | NULL | Probable | As nectar-tap_saprotroph (Põlme et al. 2020); Decay sugar-rich substrates (Põlme et al. 2020) | Põlme S, et al. 2020. Fungal Diversity 105:1-16 ((https://doi.org/10.1007/s13225-020-00466-2)) |
| Otu25260 | 1 | k__Fungi(100);p__Ascomycota(100);c__Saccharomycetes(100);o__Saccharomycetales(100);f__Metschnikowiaceae(100);g__Metschnikowia(100);s__Metschnikowia_pulcherrima(100); | Metschnikowia | 13 | Saprotroph | \|Undefined Saprotroph\| | Yeast | NULL | Probable | As nectar-tap_saprotroph (Põlme et al. 2020); Decay sugar-rich substrates (Põlme et al. 2020) | Põlme S, et al. 2020. Fungal Diversity 105:1-16 ((https://doi.org/10.1007/s13225-020-00466-2)) |
| Otu25261 | 1 | k__Fungi(100);p__Ascomycota(100);c__Saccharomycetes(100);o__Saccharomycetales(100);f__Metschnikowiaceae(100);g__Metschnikowia(100);g__Metschnikowia_unclassified(100); | Metschnikowia | 13 | Saprotroph | \|Undefined Saprotroph\| | Yeast | NULL | Probable | As nectar-tap_saprotroph (Põlme et al. 2020); Decay sugar-rich substrates (Põlme et al. 2020) | Põlme S, et al. 2020. Fungal Diversity 105:1-16 ((https://doi.org/10.1007/s13225-020-00466-2)) |
| Otu25262 | 1 | k__Fungi(100);p__Ascomycota(100);c__Saccharomycetes(100);o__Saccharomycetales(100);f__Metschnikowiaceae(100);g__Metschnikowia(100);s__Metschnikowia_pulcherrima(100); | Metschnikowia | 13 | Saprotroph | \|Undefined Saprotroph\| | Yeast | NULL | Probable | As nectar-tap_saprotroph (Põlme et al. 2020); Decay sugar-rich substrates (Põlme et al. 2020) | Põlme S, et al. 2020. Fungal Diversity 105:1-16 ((https://doi.org/10.1007/s13225-020-00466-2)) |
| Otu25263 | 1 | k__Fungi(100);p__Ascomycota(100);c__Saccharomycetes(100);o__Saccharomycetales(100);f__Metschnikowiaceae(100);g__Metschnikowia(100);s__Metschnikowia_pulcherrima(100); | Metschnikowia | 13 | Saprotroph | \|Undefined Saprotroph\| | Yeast | NULL | Probable | As nectar-tap_saprotroph (Põlme et al. 2020); Decay sugar-rich substrates (Põlme et al. 2020) | Põlme S, et al. 2020. Fungal Diversity 105:1-16 ((https://doi.org/10.1007/s13225-020-00466-2)) |
| Otu25264 | 1 | k__Fungi(100);p__Ascomycota(100);c__Saccharomycetes(100);o__Saccharomycetales(100);f__Metschnikowiaceae(100);g__Metschnikowia(100);s__Metschnikowia_pulcherrima(100); | Metschnikowia | 13 | Saprotroph | \|Undefined Saprotroph\| | Yeast | NULL | Probable | As nectar-tap_saprotroph (Põlme et al. 2020); Decay sugar-rich substrates (Põlme et al. 2020) | Põlme S, et al. 2020. Fungal Diversity 105:1-16 ((https://doi.org/10.1007/s13225-020-00466-2)) |
| Otu25265 | 1 | k__Fungi(100);p__Ascomycota(100);c__Saccharomycetes(100);o__Saccharomycetales(100);f__Metschnikowiaceae(100);g__Metschnikowia(100);s__Metschnikowia_pulcherrima(100); | Metschnikowia | 13 | Saprotroph | \|Undefined Saprotroph\| | Yeast | NULL | Probable | As nectar-tap_saprotroph (Põlme et al. 2020); Decay sugar-rich substrates (Põlme et al. 2020) | Põlme S, et al. 2020. Fungal Diversity 105:1-16 ((https://doi.org/10.1007/s13225-020-00466-2)) |
| Otu25266 | 1 | k__Fungi(100);p__Ascomycota(100);c__Saccharomycetes(100);o__Saccharomycetales(100);f__Metschnikowiaceae(100);g__Metschnikowia(100);s__Metschnikowia_pulcherrima(100); | Metschnikowia | 13 | Saprotroph | \|Undefined Saprotroph\| | Yeast | NULL | Probable | As nectar-tap_saprotroph (Põlme et al. 2020); Decay sugar-rich substrates (Põlme et al. 2020) | Põlme S, et al. 2020. Fungal Diversity 105:1-16 ((https://doi.org/10.1007/s13225-020-00466-2)) |
| Otu25269 | 1 | k__Fungi(100);p__Ascomycota(100);c__Saccharomycetes(100);o__Saccharomycetales(100);f__Metschnikowiaceae(100);g__Metschnikowia(100);s__Metschnikowia_pulcherrima(100); | Metschnikowia | 13 | Saprotroph | \|Undefined Saprotroph\| | Yeast | NULL | Probable | As nectar-tap_saprotroph (Põlme et al. 2020); Decay sugar-rich substrates (Põlme et al. 2020) | Põlme S, et al. 2020. Fungal Diversity 105:1-16 ((https://doi.org/10.1007/s13225-020-00466-2)) |
| Otu25270 | 1 | k__Fungi(100);p__Ascomycota(100);c__Saccharomycetes(100);o__Saccharomycetales(100);f__Metschnikowiaceae(100);g__Metschnikowia(100);g__Metschnikowia_unclassified(100); | Metschnikowia | 13 | Saprotroph | \|Undefined Saprotroph\| | Yeast | NULL | Probable | As nectar-tap_saprotroph (Põlme et al. 2020); Decay sugar-rich substrates (Põlme et al. 2020) | Põlme S, et al. 2020. Fungal Diversity 105:1-16 ((https://doi.org/10.1007/s13225-020-00466-2)) |
| Otu25271 | 1 | k__Fungi(100);p__Ascomycota(100);c__Saccharomycetes(100);o__Saccharomycetales(100);f__Metschnikowiaceae(100);g__Metschnikowia(100);s__Metschnikowia_pulcherrima(100); | Metschnikowia | 13 | Saprotroph | \|Undefined Saprotroph\| | Yeast | NULL | Probable | As nectar-tap_saprotroph (Põlme et al. 2020); Decay sugar-rich substrates (Põlme et al. 2020) | Põlme S, et al. 2020. Fungal Diversity 105:1-16 ((https://doi.org/10.1007/s13225-020-00466-2)) |
| Otu25272 | 1 | k__Fungi(100);p__Ascomycota(100);c__Saccharomycetes(100);o__Saccharomycetales(100);f__Metschnikowiaceae(100);g__Metschnikowia(100);s__Metschnikowia_pulcherrima(100); | Metschnikowia | 13 | Saprotroph | \|Undefined Saprotroph\| | Yeast | NULL | Probable | As nectar-tap_saprotroph (Põlme et al. 2020); Decay sugar-rich substrates (Põlme et al. 2020) | Põlme S, et al. 2020. Fungal Diversity 105:1-16 ((https://doi.org/10.1007/s13225-020-00466-2)) |
| Otu25273 | 1 | k__Fungi(100);p__Ascomycota(100);c__Saccharomycetes(100);o__Saccharomycetales(100);f__Metschnikowiaceae(100);g__Metschnikowia(100);g__Metschnikowia_unclassified(100); | Metschnikowia | 13 | Saprotroph | \|Undefined Saprotroph\| | Yeast | NULL | Probable | As nectar-tap_saprotroph (Põlme et al. 2020); Decay sugar-rich substrates (Põlme et al. 2020) | Põlme S, et al. 2020. Fungal Diversity 105:1-16 ((https://doi.org/10.1007/s13225-020-00466-2)) |
| Otu25274 | 1 | k__Fungi(100);p__Ascomycota(100);c__Saccharomycetes(100);o__Saccharomycetales(100);f__Metschnikowiaceae(100);g__Metschnikowia(100);g__Metschnikowia_unclassified(100); | Metschnikowia | 13 | Saprotroph | \|Undefined Saprotroph\| | Yeast | NULL | Probable | As nectar-tap_saprotroph (Põlme et al. 2020); Decay sugar-rich substrates (Põlme et al. 2020) | Põlme S, et al. 2020. Fungal Diversity 105:1-16 ((https://doi.org/10.1007/s13225-020-00466-2)) |
| Otu25276 | 1 | k__Fungi(100);p__Ascomycota(100);c__Saccharomycetes(100);o__Saccharomycetales(100);f__Metschnikowiaceae(100);g__Metschnikowia(100);s__Metschnikowia_pulcherrima(100); | Metschnikowia | 13 | Saprotroph | \|Undefined Saprotroph\| | Yeast | NULL | Probable | As nectar-tap_saprotroph (Põlme et al. 2020); Decay sugar-rich substrates (Põlme et al. 2020) | Põlme S, et al. 2020. Fungal Diversity 105:1-16 ((https://doi.org/10.1007/s13225-020-00466-2)) |
| Otu25277 | 1 | k__Fungi(100);p__Ascomycota(100);c__Saccharomycetes(100);o__Saccharomycetales(100);f__Metschnikowiaceae(100);g__Metschnikowia(100);s__Metschnikowia_pulcherrima(100); | Metschnikowia | 13 | Saprotroph | \|Undefined Saprotroph\| | Yeast | NULL | Probable | As nectar-tap_saprotroph (Põlme et al. 2020); Decay sugar-rich substrates (Põlme et al. 2020) | Põlme S, et al. 2020. Fungal Diversity 105:1-16 ((https://doi.org/10.1007/s13225-020-00466-2)) |
| Otu25278 | 1 | k__Fungi(100);p__Ascomycota(100);c__Saccharomycetes(100);o__Saccharomycetales(100);f__Metschnikowiaceae(100);g__Metschnikowia(100);s__Metschnikowia_pulcherrima(100); | Metschnikowia | 13 | Saprotroph | \|Undefined Saprotroph\| | Yeast | NULL | Probable | As nectar-tap_saprotroph (Põlme et al. 2020); Decay sugar-rich substrates (Põlme et al. 2020) | Põlme S, et al. 2020. Fungal Diversity 105:1-16 ((https://doi.org/10.1007/s13225-020-00466-2)) |
| Otu25279 | 1 | k__Fungi(100);p__Ascomycota(100);c__Saccharomycetes(100);o__Saccharomycetales(100);f__Metschnikowiaceae(100);g__Metschnikowia(100);g__Metschnikowia_unclassified(100); | Metschnikowia | 13 | Saprotroph | \|Undefined Saprotroph\| | Yeast | NULL | Probable | As nectar-tap_saprotroph (Põlme et al. 2020); Decay sugar-rich substrates (Põlme et al. 2020) | Põlme S, et al. 2020. Fungal Diversity 105:1-16 ((https://doi.org/10.1007/s13225-020-00466-2)) |
| Otu25282 | 1 | k__Fungi(100);p__Ascomycota(100);c__Saccharomycetes(100);o__Saccharomycetales(100);f__Metschnikowiaceae(100);g__Metschnikowia(100);g__Metschnikowia_unclassified(100); | Metschnikowia | 13 | Saprotroph | \|Undefined Saprotroph\| | Yeast | NULL | Probable | As nectar-tap_saprotroph (Põlme et al. 2020); Decay sugar-rich substrates (Põlme et al. 2020) | Põlme S, et al. 2020. Fungal Diversity 105:1-16 ((https://doi.org/10.1007/s13225-020-00466-2)) |
| Otu25283 | 1 | k__Fungi(100);p__Basidiomycota(100);c__Agaricomycetes(100);o__Polyporales(100);f__Meruliaceae(100);g__Scopuloides(100);s__Scopuloides_hydnoides(100); | Scopuloides | 13 | Saprotroph | Undefined Saprotroph-\|Wood Saprotroph\| | Corticioid | White Rot | Probable | As white_rot (Põlme et al. 2020); Decay wood (Põlme et al. 2020) | Tedersoo L, et al. 2014. Science 346:e1256688 ((https://doi.org/10.1126/science.1256688)); Põlme S, et al. 2020. Fungal Diversity 105:1-16 ((https://doi.org/10.1007/s13225-020-00466-2)) |
| Otu25284 | 1 | k__Fungi(100);p__Basidiomycota(100);c__Agaricomycetes(100);o__Agaricales(100);f__Mycenaceae(100);g__Mycena(100);g__Mycena_unclassified(100); | Mycena | 13 | Pathotroph-Saprotroph | Plant Pathogen-\|Plant Saprotroph\|-Undefined Saprotroph-Wood Saprotroph | Agaricoid | NULL | Probable | As litter_saprotroph (Põlme et al. 2020); Decay leaf, fruit, seed (Põlme et al. 2020) | Grgurinovic CA, Simpson JA. 2001. Fungal Diversity 8:97-127 (http://www.fungaldiversity.org/fdp/sfdp/FD_8_97-127.pdf)); Tedersoo L, et al. 2014. Science 346:e1256688 ((https://doi.org/10.1126/science.1256688)); Põlme S, et al. 2020. Fungal Diversity 105:1-16 ((https://doi.org/10.1007/s13225-020-00466-2)); The American Phytopathological Society Common Names of Plant Diseases ((http://www.apsnet.org/publications/commonnames/Pages/default.aspx)) |
| Otu25285 | 1 | k__Fungi(100);p__Ascomycota(100);c__Saccharomycetes(100);o__Saccharomycetales(100);f__Metschnikowiaceae(100);g__Metschnikowia(100);s__Metschnikowia_pulcherrima(100); | Metschnikowia | 13 | Saprotroph | \|Undefined Saprotroph\| | Yeast | NULL | Probable | As nectar-tap_saprotroph (Põlme et al. 2020); Decay sugar-rich substrates (Põlme et al. 2020) | Põlme S, et al. 2020. Fungal Diversity 105:1-16 ((https://doi.org/10.1007/s13225-020-00466-2)) |
| Otu25286 | 1 | k__Fungi(100);p__Ascomycota(100);c__Saccharomycetes(100);o__Saccharomycetales(100);f__Metschnikowiaceae(100);g__Metschnikowia(100);s__Metschnikowia_pulcherrima(100); | Metschnikowia | 13 | Saprotroph | \|Undefined Saprotroph\| | Yeast | NULL | Probable | As nectar-tap_saprotroph (Põlme et al. 2020); Decay sugar-rich substrates (Põlme et al. 2020) | Põlme S, et al. 2020. Fungal Diversity 105:1-16 ((https://doi.org/10.1007/s13225-020-00466-2)) |
| Otu25287 | 1 | k__Fungi(100);p__Ascomycota(100);c__Saccharomycetes(100);o__Saccharomycetales(100);f__Metschnikowiaceae(100);g__Metschnikowia(100);s__Metschnikowia_pulcherrima(100); | Metschnikowia | 13 | Saprotroph | \|Undefined Saprotroph\| | Yeast | NULL | Probable | As nectar-tap_saprotroph (Põlme et al. 2020); Decay sugar-rich substrates (Põlme et al. 2020) | Põlme S, et al. 2020. Fungal Diversity 105:1-16 ((https://doi.org/10.1007/s13225-020-00466-2)) |
| Otu25288 | 1 | k__Fungi(100);p__Ascomycota(100);c__Saccharomycetes(100);o__Saccharomycetales(100);f__Metschnikowiaceae(100);g__Metschnikowia(100);s__Metschnikowia_pulcherrima(100); | Metschnikowia | 13 | Saprotroph | \|Undefined Saprotroph\| | Yeast | NULL | Probable | As nectar-tap_saprotroph (Põlme et al. 2020); Decay sugar-rich substrates (Põlme et al. 2020) | Põlme S, et al. 2020. Fungal Diversity 105:1-16 ((https://doi.org/10.1007/s13225-020-00466-2)) |
| Otu25289 | 1 | k__Fungi(100);p__Ascomycota(100);c__Saccharomycetes(100);o__Saccharomycetales(100);f__Metschnikowiaceae(100);g__Metschnikowia(100);s__Metschnikowia_pulcherrima(100); | Metschnikowia | 13 | Saprotroph | \|Undefined Saprotroph\| | Yeast | NULL | Probable | As nectar-tap_saprotroph (Põlme et al. 2020); Decay sugar-rich substrates (Põlme et al. 2020) | Põlme S, et al. 2020. Fungal Diversity 105:1-16 ((https://doi.org/10.1007/s13225-020-00466-2)) |
| Otu25290 | 1 | k__Fungi(100);p__Ascomycota(100);c__Saccharomycetes(100);o__Saccharomycetales(100);f__Metschnikowiaceae(100);g__Metschnikowia(100);s__Metschnikowia_pulcherrima(100); | Metschnikowia | 13 | Saprotroph | \|Undefined Saprotroph\| | Yeast | NULL | Probable | As nectar-tap_saprotroph (Põlme et al. 2020); Decay sugar-rich substrates (Põlme et al. 2020) | Põlme S, et al. 2020. Fungal Diversity 105:1-16 ((https://doi.org/10.1007/s13225-020-00466-2)) |
| Otu25291 | 1 | k__Fungi(100);p__Ascomycota(100);c__Saccharomycetes(100);o__Saccharomycetales(100);f__Metschnikowiaceae(100);g__Metschnikowia(100);g__Metschnikowia_unclassified(100); | Metschnikowia | 13 | Saprotroph | \|Undefined Saprotroph\| | Yeast | NULL | Probable | As nectar-tap_saprotroph (Põlme et al. 2020); Decay sugar-rich substrates (Põlme et al. 2020) | Põlme S, et al. 2020. Fungal Diversity 105:1-16 ((https://doi.org/10.1007/s13225-020-00466-2)) |
| Otu25292 | 1 | k__Fungi(100);p__Ascomycota(100);c__Saccharomycetes(100);o__Saccharomycetales(100);f__Metschnikowiaceae(100);g__Metschnikowia(100);s__Metschnikowia_pulcherrima(100); | Metschnikowia | 13 | Saprotroph | \|Undefined Saprotroph\| | Yeast | NULL | Probable | As nectar-tap_saprotroph (Põlme et al. 2020); Decay sugar-rich substrates (Põlme et al. 2020) | Põlme S, et al. 2020. Fungal Diversity 105:1-16 ((https://doi.org/10.1007/s13225-020-00466-2)) |
| Otu25293 | 1 | k__Fungi(100);p__Ascomycota(100);c__Saccharomycetes(100);o__Saccharomycetales(100);f__Metschnikowiaceae(100);g__Metschnikowia(100);s__Metschnikowia_pulcherrima(100); | Metschnikowia | 13 | Saprotroph | \|Undefined Saprotroph\| | Yeast | NULL | Probable | As nectar-tap_saprotroph (Põlme et al. 2020); Decay sugar-rich substrates (Põlme et al. 2020) | Põlme S, et al. 2020. Fungal Diversity 105:1-16 ((https://doi.org/10.1007/s13225-020-00466-2)) |
| Otu25294 | 1 | k__Fungi(100);p__Ascomycota(100);c__Saccharomycetes(100);o__Saccharomycetales(100);f__Metschnikowiaceae(100);g__Metschnikowia(100);s__Metschnikowia_pulcherrima(100); | Metschnikowia | 13 | Saprotroph | \|Undefined Saprotroph\| | Yeast | NULL | Probable | As nectar-tap_saprotroph (Põlme et al. 2020); Decay sugar-rich substrates (Põlme et al. 2020) | Põlme S, et al. 2020. Fungal Diversity 105:1-16 ((https://doi.org/10.1007/s13225-020-00466-2)) |
| Otu25295 | 1 | k__Fungi(100);p__Chytridiomycota(100);c__Chytridiomycetes(100);o__Rhizophlyctidales(100);f__Rhizophlyctidaceae(100);g__Rhizophlyctis(100);s__Rhizophlyctis_rosea(100); | Rhizophlyctis | 13 | Pathotroph-Saprotroph | Algal Parasite-\|Plant Saprotroph\|-Undefined Saprotroph | Chytrid-Microfungus | NULL | Probable | As litter_saprotroph (Põlme et al. 2020); Decay leaf, fruit, seed, algal material (Põlme et al. 2020) | James TY, et al. 2006. Nature 443:818-822 ((https://doi.org/10.1038/nature05110)); Põlme S, et al. 2020. Fungal Diversity 105:1-16 ((https://doi.org/10.1007/s13225-020-00466-2)) |
| Otu25296 | 1 | k__Fungi(100);p__Chytridiomycota(100);c__Chytridiomycetes(100);o__Spizellomycetales(100);f__Spizellomycetaceae(100);g__Spizellomyces(100);g__Spizellomyces_unclassified(100); | Spizellomyces | 13 | Pathotroph-Saprotroph | Plant Pathogen-\|Pollen Saprotroph\|-Undefined Parasite-Undefined Saprotroph | Chytrid-Microfungus | NULL | Probable | Decay pollen (Põlme et al. 2020); Pollen degrader; Common in the soil and also as parasites of a range of soil organisms and plants (Cannon & Kirk 2007) | Cannon PF, Kirk PM. 2007. Fungal Families of the World. CAB International, Cambridge (ISBN: 978-0851998275); Tedersoo L, et al. 2014. Science 346:e1256688 ((https://doi.org/10.1126/science.1256688)); Põlme S, et al. 2020. Fungal Diversity 105:1-16 ((https://doi.org/10.1007/s13225-020-00466-2)) |
| Otu25297 | 1 | k__Fungi(100);p__Ascomycota(100);c__Saccharomycetes(100);o__Saccharomycetales(100);f__Metschnikowiaceae(100);g__Metschnikowia(100);g__Metschnikowia_unclassified(100); | Metschnikowia | 13 | Saprotroph | \|Undefined Saprotroph\| | Yeast | NULL | Probable | As nectar-tap_saprotroph (Põlme et al. 2020); Decay sugar-rich substrates (Põlme et al. 2020) | Põlme S, et al. 2020. Fungal Diversity 105:1-16 ((https://doi.org/10.1007/s13225-020-00466-2)) |
| Otu25298 | 1 | k__Fungi(100);p__Ascomycota(100);c__Saccharomycetes(100);o__Saccharomycetales(100);f__Metschnikowiaceae(100);g__Metschnikowia(100);g__Metschnikowia_unclassified(100); | Metschnikowia | 13 | Saprotroph | \|Undefined Saprotroph\| | Yeast | NULL | Probable | As nectar-tap_saprotroph (Põlme et al. 2020); Decay sugar-rich substrates (Põlme et al. 2020) | Põlme S, et al. 2020. Fungal Diversity 105:1-16 ((https://doi.org/10.1007/s13225-020-00466-2)) |
| Otu25300 | 1 | k__Fungi(100);p__Ascomycota(100);c__Saccharomycetes(100);o__Saccharomycetales(100);f__Metschnikowiaceae(100);g__Metschnikowia(100);s__Metschnikowia_pulcherrima(100); | Metschnikowia | 13 | Saprotroph | \|Undefined Saprotroph\| | Yeast | NULL | Probable | As nectar-tap_saprotroph (Põlme et al. 2020); Decay sugar-rich substrates (Põlme et al. 2020) | Põlme S, et al. 2020. Fungal Diversity 105:1-16 ((https://doi.org/10.1007/s13225-020-00466-2)) |
| Otu25301 | 1 | k__Fungi(100);p__Ascomycota(100);c__Saccharomycetes(100);o__Saccharomycetales(100);f__Metschnikowiaceae(100);g__Metschnikowia(100);g__Metschnikowia_unclassified(100); | Metschnikowia | 13 | Saprotroph | \|Undefined Saprotroph\| | Yeast | NULL | Probable | As nectar-tap_saprotroph (Põlme et al. 2020); Decay sugar-rich substrates (Põlme et al. 2020) | Põlme S, et al. 2020. Fungal Diversity 105:1-16 ((https://doi.org/10.1007/s13225-020-00466-2)) |
| Otu25303 | 1 | k__Fungi(100);p__Ascomycota(100);c__Saccharomycetes(100);o__Saccharomycetales(100);f__Metschnikowiaceae(100);g__Metschnikowia(100);s__Metschnikowia_pulcherrima(100); | Metschnikowia | 13 | Saprotroph | \|Undefined Saprotroph\| | Yeast | NULL | Probable | As nectar-tap_saprotroph (Põlme et al. 2020); Decay sugar-rich substrates (Põlme et al. 2020) | Põlme S, et al. 2020. Fungal Diversity 105:1-16 ((https://doi.org/10.1007/s13225-020-00466-2)) |
| Otu25304 | 1 | k__Fungi(100);p__Ascomycota(100);c__Saccharomycetes(100);o__Saccharomycetales(100);f__Metschnikowiaceae(100);g__Metschnikowia(100);g__Metschnikowia_unclassified(100); | Metschnikowia | 13 | Saprotroph | \|Undefined Saprotroph\| | Yeast | NULL | Probable | As nectar-tap_saprotroph (Põlme et al. 2020); Decay sugar-rich substrates (Põlme et al. 2020) | Põlme S, et al. 2020. Fungal Diversity 105:1-16 ((https://doi.org/10.1007/s13225-020-00466-2)) |
| Otu25305 | 1 | k__Fungi(100);p__Ascomycota(100);c__Saccharomycetes(100);o__Saccharomycetales(100);f__Metschnikowiaceae(100);g__Metschnikowia(100);s__Metschnikowia_pulcherrima(100); | Metschnikowia | 13 | Saprotroph | \|Undefined Saprotroph\| | Yeast | NULL | Probable | As nectar-tap_saprotroph (Põlme et al. 2020); Decay sugar-rich substrates (Põlme et al. 2020) | Põlme S, et al. 2020. Fungal Diversity 105:1-16 ((https://doi.org/10.1007/s13225-020-00466-2)) |
| Otu25306 | 1 | k__Fungi(100);p__Ascomycota(100);c__Saccharomycetes(100);o__Saccharomycetales(100);f__Metschnikowiaceae(100);g__Metschnikowia(100);s__Metschnikowia_pulcherrima(100); | Metschnikowia | 13 | Saprotroph | \|Undefined Saprotroph\| | Yeast | NULL | Probable | As nectar-tap_saprotroph (Põlme et al. 2020); Decay sugar-rich substrates (Põlme et al. 2020) | Põlme S, et al. 2020. Fungal Diversity 105:1-16 ((https://doi.org/10.1007/s13225-020-00466-2)) |
| Otu25329 | 1 | k__Fungi(100);p__Ascomycota(100);c__Saccharomycetes(100);o__Saccharomycetales(100);f__Metschnikowiaceae(100);g__Metschnikowia(100);g__Metschnikowia_unclassified(100); | Metschnikowia | 13 | Saprotroph | \|Undefined Saprotroph\| | Yeast | NULL | Probable | As nectar-tap_saprotroph (Põlme et al. 2020); Decay sugar-rich substrates (Põlme et al. 2020) | Põlme S, et al. 2020. Fungal Diversity 105:1-16 ((https://doi.org/10.1007/s13225-020-00466-2)) |
| Otu25330 | 1 | k__Fungi(100);p__Ascomycota(100);c__Saccharomycetes(100);o__Saccharomycetales(100);f__Metschnikowiaceae(100);g__Metschnikowia(100);g__Metschnikowia_unclassified(100); | Metschnikowia | 13 | Saprotroph | \|Undefined Saprotroph\| | Yeast | NULL | Probable | As nectar-tap_saprotroph (Põlme et al. 2020); Decay sugar-rich substrates (Põlme et al. 2020) | Põlme S, et al. 2020. Fungal Diversity 105:1-16 ((https://doi.org/10.1007/s13225-020-00466-2)) |
| Otu25331 | 1 | k__Fungi(100);p__Ascomycota(100);c__Saccharomycetes(100);o__Saccharomycetales(100);f__Metschnikowiaceae(100);g__Metschnikowia(100);s__Metschnikowia_pulcherrima(100); | Metschnikowia | 13 | Saprotroph | \|Undefined Saprotroph\| | Yeast | NULL | Probable | As nectar-tap_saprotroph (Põlme et al. 2020); Decay sugar-rich substrates (Põlme et al. 2020) | Põlme S, et al. 2020. Fungal Diversity 105:1-16 ((https://doi.org/10.1007/s13225-020-00466-2)) |
| Otu25333 | 1 | k__Fungi(100);p__Ascomycota(100);c__Saccharomycetes(100);o__Saccharomycetales(100);f__Metschnikowiaceae(100);g__Metschnikowia(100);s__Metschnikowia_pulcherrima(100); | Metschnikowia | 13 | Saprotroph | \|Undefined Saprotroph\| | Yeast | NULL | Probable | As nectar-tap_saprotroph (Põlme et al. 2020); Decay sugar-rich substrates (Põlme et al. 2020) | Põlme S, et al. 2020. Fungal Diversity 105:1-16 ((https://doi.org/10.1007/s13225-020-00466-2)) |
| Otu25334 | 1 | k__Fungi(100);p__Ascomycota(100);c__Saccharomycetes(100);o__Saccharomycetales(100);f__Metschnikowiaceae(100);g__Metschnikowia(100);g__Metschnikowia_unclassified(100); | Metschnikowia | 13 | Saprotroph | \|Undefined Saprotroph\| | Yeast | NULL | Probable | As nectar-tap_saprotroph (Põlme et al. 2020); Decay sugar-rich substrates (Põlme et al. 2020) | Põlme S, et al. 2020. Fungal Diversity 105:1-16 ((https://doi.org/10.1007/s13225-020-00466-2)) |
| Otu25336 | 1 | k__Fungi(100);p__Ascomycota(100);c__Saccharomycetes(100);o__Saccharomycetales(100);f__Metschnikowiaceae(100);g__Metschnikowia(100);s__Metschnikowia_pulcherrima(100); | Metschnikowia | 13 | Saprotroph | \|Undefined Saprotroph\| | Yeast | NULL | Probable | As nectar-tap_saprotroph (Põlme et al. 2020); Decay sugar-rich substrates (Põlme et al. 2020) | Põlme S, et al. 2020. Fungal Diversity 105:1-16 ((https://doi.org/10.1007/s13225-020-00466-2)) |
| Otu25337 | 1 | k__Fungi(100);p__Ascomycota(100);c__Saccharomycetes(100);o__Saccharomycetales(100);f__Metschnikowiaceae(100);g__Metschnikowia(100);s__Metschnikowia_pulcherrima(100); | Metschnikowia | 13 | Saprotroph | \|Undefined Saprotroph\| | Yeast | NULL | Probable | As nectar-tap_saprotroph (Põlme et al. 2020); Decay sugar-rich substrates (Põlme et al. 2020) | Põlme S, et al. 2020. Fungal Diversity 105:1-16 ((https://doi.org/10.1007/s13225-020-00466-2)) |
| Otu25338 | 1 | k__Fungi(100);p__Ascomycota(100);c__Saccharomycetes(100);o__Saccharomycetales(100);f__Metschnikowiaceae(100);g__Metschnikowia(100);s__Metschnikowia_pulcherrima(100); | Metschnikowia | 13 | Saprotroph | \|Undefined Saprotroph\| | Yeast | NULL | Probable | As nectar-tap_saprotroph (Põlme et al. 2020); Decay sugar-rich substrates (Põlme et al. 2020) | Põlme S, et al. 2020. Fungal Diversity 105:1-16 ((https://doi.org/10.1007/s13225-020-00466-2)) |
| Otu25339 | 1 | k__Fungi(100);p__Ascomycota(100);c__Saccharomycetes(100);o__Saccharomycetales(100);f__Metschnikowiaceae(100);g__Metschnikowia(100);s__Metschnikowia_pulcherrima(100); | Metschnikowia | 13 | Saprotroph | \|Undefined Saprotroph\| | Yeast | NULL | Probable | As nectar-tap_saprotroph (Põlme et al. 2020); Decay sugar-rich substrates (Põlme et al. 2020) | Põlme S, et al. 2020. Fungal Diversity 105:1-16 ((https://doi.org/10.1007/s13225-020-00466-2)) |
| Otu25340 | 1 | k__Fungi(100);p__Ascomycota(100);c__Saccharomycetes(100);o__Saccharomycetales(100);f__Metschnikowiaceae(100);g__Metschnikowia(100);s__Metschnikowia_pulcherrima(100); | Metschnikowia | 13 | Saprotroph | \|Undefined Saprotroph\| | Yeast | NULL | Probable | As nectar-tap_saprotroph (Põlme et al. 2020); Decay sugar-rich substrates (Põlme et al. 2020) | Põlme S, et al. 2020. Fungal Diversity 105:1-16 ((https://doi.org/10.1007/s13225-020-00466-2)) |
| Otu25341 | 1 | k__Fungi(100);p__Ascomycota(100);c__Saccharomycetes(100);o__Saccharomycetales(100);f__Metschnikowiaceae(100);g__Metschnikowia(100);s__Metschnikowia_pulcherrima(100); | Metschnikowia | 13 | Saprotroph | \|Undefined Saprotroph\| | Yeast | NULL | Probable | As nectar-tap_saprotroph (Põlme et al. 2020); Decay sugar-rich substrates (Põlme et al. 2020) | Põlme S, et al. 2020. Fungal Diversity 105:1-16 ((https://doi.org/10.1007/s13225-020-00466-2)) |
| Otu25342 | 1 | k__Fungi(100);p__Ascomycota(100);c__Saccharomycetes(100);o__Saccharomycetales(100);f__Metschnikowiaceae(100);g__Metschnikowia(100);s__Metschnikowia_pulcherrima(100); | Metschnikowia | 13 | Saprotroph | \|Undefined Saprotroph\| | Yeast | NULL | Probable | As nectar-tap_saprotroph (Põlme et al. 2020); Decay sugar-rich substrates (Põlme et al. 2020) | Põlme S, et al. 2020. Fungal Diversity 105:1-16 ((https://doi.org/10.1007/s13225-020-00466-2)) |
| Otu25344 | 1 | k__Fungi(100);p__Ascomycota(100);c__Saccharomycetes(100);o__Saccharomycetales(100);f__Metschnikowiaceae(100);g__Metschnikowia(100);g__Metschnikowia_unclassified(100); | Metschnikowia | 13 | Saprotroph | \|Undefined Saprotroph\| | Yeast | NULL | Probable | As nectar-tap_saprotroph (Põlme et al. 2020); Decay sugar-rich substrates (Põlme et al. 2020) | Põlme S, et al. 2020. Fungal Diversity 105:1-16 ((https://doi.org/10.1007/s13225-020-00466-2)) |
| Otu25345 | 1 | k__Fungi(100);p__Ascomycota(100);c__Saccharomycetes(100);o__Saccharomycetales(100);f__Metschnikowiaceae(100);g__Metschnikowia(100);s__Metschnikowia_pulcherrima(100); | Metschnikowia | 13 | Saprotroph | \|Undefined Saprotroph\| | Yeast | NULL | Probable | As nectar-tap_saprotroph (Põlme et al. 2020); Decay sugar-rich substrates (Põlme et al. 2020) | Põlme S, et al. 2020. Fungal Diversity 105:1-16 ((https://doi.org/10.1007/s13225-020-00466-2)) |
| Otu25346 | 1 | k__Fungi(100);p__Ascomycota(100);c__Saccharomycetes(100);o__Saccharomycetales(100);f__Metschnikowiaceae(100);g__Metschnikowia(100);s__Metschnikowia_pulcherrima(100); | Metschnikowia | 13 | Saprotroph | \|Undefined Saprotroph\| | Yeast | NULL | Probable | As nectar-tap_saprotroph (Põlme et al. 2020); Decay sugar-rich substrates (Põlme et al. 2020) | Põlme S, et al. 2020. Fungal Diversity 105:1-16 ((https://doi.org/10.1007/s13225-020-00466-2)) |
| Otu25349 | 1 | k__Fungi(100);p__Ascomycota(100);c__Saccharomycetes(100);o__Saccharomycetales(100);f__Metschnikowiaceae(100);g__Metschnikowia(100);s__Metschnikowia_pulcherrima(100); | Metschnikowia | 13 | Saprotroph | \|Undefined Saprotroph\| | Yeast | NULL | Probable | As nectar-tap_saprotroph (Põlme et al. 2020); Decay sugar-rich substrates (Põlme et al. 2020) | Põlme S, et al. 2020. Fungal Diversity 105:1-16 ((https://doi.org/10.1007/s13225-020-00466-2)) |
| Otu25350 | 1 | k__Fungi(100);p__Ascomycota(100);c__Saccharomycetes(100);o__Saccharomycetales(100);f__Metschnikowiaceae(100);g__Metschnikowia(100);s__Metschnikowia_pulcherrima(100); | Metschnikowia | 13 | Saprotroph | \|Undefined Saprotroph\| | Yeast | NULL | Probable | As nectar-tap_saprotroph (Põlme et al. 2020); Decay sugar-rich substrates (Põlme et al. 2020) | Põlme S, et al. 2020. Fungal Diversity 105:1-16 ((https://doi.org/10.1007/s13225-020-00466-2)) |
| Otu25351 | 1 | k__Fungi(100);p__Ascomycota(100);c__Saccharomycetes(100);o__Saccharomycetales(100);f__Metschnikowiaceae(100);g__Metschnikowia(100);g__Metschnikowia_unclassified(100); | Metschnikowia | 13 | Saprotroph | \|Undefined Saprotroph\| | Yeast | NULL | Probable | As nectar-tap_saprotroph (Põlme et al. 2020); Decay sugar-rich substrates (Põlme et al. 2020) | Põlme S, et al. 2020. Fungal Diversity 105:1-16 ((https://doi.org/10.1007/s13225-020-00466-2)) |
| Otu25352 | 1 | k__Fungi(100);p__Ascomycota(100);c__Saccharomycetes(100);o__Saccharomycetales(100);f__Metschnikowiaceae(100);g__Metschnikowia(100);s__Metschnikowia_pulcherrima(100); | Metschnikowia | 13 | Saprotroph | \|Undefined Saprotroph\| | Yeast | NULL | Probable | As nectar-tap_saprotroph (Põlme et al. 2020); Decay sugar-rich substrates (Põlme et al. 2020) | Põlme S, et al. 2020. Fungal Diversity 105:1-16 ((https://doi.org/10.1007/s13225-020-00466-2)) |
| Otu25353 | 1 | k__Fungi(100);p__Ascomycota(100);c__Saccharomycetes(100);o__Saccharomycetales(100);f__Metschnikowiaceae(100);g__Metschnikowia(100);s__Metschnikowia_pulcherrima(100); | Metschnikowia | 13 | Saprotroph | \|Undefined Saprotroph\| | Yeast | NULL | Probable | As nectar-tap_saprotroph (Põlme et al. 2020); Decay sugar-rich substrates (Põlme et al. 2020) | Põlme S, et al. 2020. Fungal Diversity 105:1-16 ((https://doi.org/10.1007/s13225-020-00466-2)) |
| Otu25354 | 1 | k__Fungi(100);p__Ascomycota(100);c__Saccharomycetes(100);o__Saccharomycetales(100);f__Metschnikowiaceae(100);g__Metschnikowia(100);g__Metschnikowia_unclassified(100); | Metschnikowia | 13 | Saprotroph | \|Undefined Saprotroph\| | Yeast | NULL | Probable | As nectar-tap_saprotroph (Põlme et al. 2020); Decay sugar-rich substrates (Põlme et al. 2020) | Põlme S, et al. 2020. Fungal Diversity 105:1-16 ((https://doi.org/10.1007/s13225-020-00466-2)) |
| Otu25358 | 1 | k__Fungi(100);p__Ascomycota(100);c__Saccharomycetes(100);o__Saccharomycetales(100);f__Metschnikowiaceae(100);g__Metschnikowia(100);g__Metschnikowia_unclassified(100); | Metschnikowia | 13 | Saprotroph | \|Undefined Saprotroph\| | Yeast | NULL | Probable | As nectar-tap_saprotroph (Põlme et al. 2020); Decay sugar-rich substrates (Põlme et al. 2020) | Põlme S, et al. 2020. Fungal Diversity 105:1-16 ((https://doi.org/10.1007/s13225-020-00466-2)) |
| Otu25359 | 1 | k__Fungi(100);p__Ascomycota(100);c__Saccharomycetes(100);o__Saccharomycetales(100);f__Metschnikowiaceae(100);g__Metschnikowia(100);g__Metschnikowia_unclassified(100); | Metschnikowia | 13 | Saprotroph | \|Undefined Saprotroph\| | Yeast | NULL | Probable | As nectar-tap_saprotroph (Põlme et al. 2020); Decay sugar-rich substrates (Põlme et al. 2020) | Põlme S, et al. 2020. Fungal Diversity 105:1-16 ((https://doi.org/10.1007/s13225-020-00466-2)) |
| Otu25360 | 1 | k__Fungi(100);p__Ascomycota(100);c__Saccharomycetes(100);o__Saccharomycetales(100);f__Metschnikowiaceae(100);g__Metschnikowia(100);s__Metschnikowia_pulcherrima(100); | Metschnikowia | 13 | Saprotroph | \|Undefined Saprotroph\| | Yeast | NULL | Probable | As nectar-tap_saprotroph (Põlme et al. 2020); Decay sugar-rich substrates (Põlme et al. 2020) | Põlme S, et al. 2020. Fungal Diversity 105:1-16 ((https://doi.org/10.1007/s13225-020-00466-2)) |
| Otu25362 | 1 | k__Fungi(100);p__Ascomycota(100);c__Leotiomycetes(100);o__Helotiales(100);f__Helotiales_family_Incertae_sedis(100);g__Rhexocercosporidium(100);g__Rhexocercosporidium_unclassified(100); | Rhexocercosporidium | 13 | Pathotroph-Saprotroph | \|Plant Pathogen\|-Plant Saprotroph | Microfungus | NULL | Probable | As litter_saprotroph (Põlme et al. 2020); As root_pathogen (Põlme et al. 2020); Decay leaf, fruit, seed, roots (Põlme et al. 2020) | Tedersoo L, et al. 2014. Science 346:e1256688 ((https://doi.org/10.1126/science.1256688)); Põlme S, et al. 2020. Fungal Diversity 105:1-16 ((https://doi.org/10.1007/s13225-020-00466-2)) |
| Otu25363 | 1 | k__Fungi(100);p__Ascomycota(100);c__Saccharomycetes(100);o__Saccharomycetales(100);f__Metschnikowiaceae(100);g__Metschnikowia(100);s__Metschnikowia_pulcherrima(100); | Metschnikowia | 13 | Saprotroph | \|Undefined Saprotroph\| | Yeast | NULL | Probable | As nectar-tap_saprotroph (Põlme et al. 2020); Decay sugar-rich substrates (Põlme et al. 2020) | Põlme S, et al. 2020. Fungal Diversity 105:1-16 ((https://doi.org/10.1007/s13225-020-00466-2)) |
| Otu25364 | 1 | k__Fungi(100);p__Ascomycota(100);c__Saccharomycetes(100);o__Saccharomycetales(100);f__Metschnikowiaceae(100);g__Metschnikowia(100);s__Metschnikowia_pulcherrima(100); | Metschnikowia | 13 | Saprotroph | \|Undefined Saprotroph\| | Yeast | NULL | Probable | As nectar-tap_saprotroph (Põlme et al. 2020); Decay sugar-rich substrates (Põlme et al. 2020) | Põlme S, et al. 2020. Fungal Diversity 105:1-16 ((https://doi.org/10.1007/s13225-020-00466-2)) |
| Otu25365 | 1 | k__Fungi(100);p__Ascomycota(100);c__Saccharomycetes(100);o__Saccharomycetales(100);f__Metschnikowiaceae(100);g__Metschnikowia(100);g__Metschnikowia_unclassified(100); | Metschnikowia | 13 | Saprotroph | \|Undefined Saprotroph\| | Yeast | NULL | Probable | As nectar-tap_saprotroph (Põlme et al. 2020); Decay sugar-rich substrates (Põlme et al. 2020) | Põlme S, et al. 2020. Fungal Diversity 105:1-16 ((https://doi.org/10.1007/s13225-020-00466-2)) |
| Otu25366 | 1 | k__Fungi(100);p__Ascomycota(100);c__Saccharomycetes(100);o__Saccharomycetales(100);f__Metschnikowiaceae(100);g__Metschnikowia(100);g__Metschnikowia_unclassified(100); | Metschnikowia | 13 | Saprotroph | \|Undefined Saprotroph\| | Yeast | NULL | Probable | As nectar-tap_saprotroph (Põlme et al. 2020); Decay sugar-rich substrates (Põlme et al. 2020) | Põlme S, et al. 2020. Fungal Diversity 105:1-16 ((https://doi.org/10.1007/s13225-020-00466-2)) |
| Otu25370 | 1 | k__Fungi(100);p__Ascomycota(100);c__Saccharomycetes(100);o__Saccharomycetales(100);f__Metschnikowiaceae(100);g__Metschnikowia(100);s__Metschnikowia_pulcherrima(100); | Metschnikowia | 13 | Saprotroph | \|Undefined Saprotroph\| | Yeast | NULL | Probable | As nectar-tap_saprotroph (Põlme et al. 2020); Decay sugar-rich substrates (Põlme et al. 2020) | Põlme S, et al. 2020. Fungal Diversity 105:1-16 ((https://doi.org/10.1007/s13225-020-00466-2)) |
| Otu25373 | 1 | k__Fungi(100);p__Ascomycota(100);c__Saccharomycetes(100);o__Saccharomycetales(100);f__Metschnikowiaceae(100);g__Metschnikowia(100);s__Metschnikowia_pulcherrima(100); | Metschnikowia | 13 | Saprotroph | \|Undefined Saprotroph\| | Yeast | NULL | Probable | As nectar-tap_saprotroph (Põlme et al. 2020); Decay sugar-rich substrates (Põlme et al. 2020) | Põlme S, et al. 2020. Fungal Diversity 105:1-16 ((https://doi.org/10.1007/s13225-020-00466-2)) |
| Otu25374 | 1 | k__Fungi(100);p__Ascomycota(100);c__Sordariomycetes(100);o__Chaetosphaeriales(100);f__Chaetosphaeriaceae(100);g__Dictyochaeta(100);s__Dictyochaeta_simplex(100); | Dictyochaeta | 13 | Saprotroph | \|Plant Saprotroph\|-Undefined Saprotroph-Wood Saprotroph | Microfungus | NULL | Probable | As litter_saprotroph (Põlme et al. 2020); Decay wood, leaf, fruit, seed (Põlme et al. 2020) | Tedersoo L, et al. 2014. Science 346:e1256688 ((https://doi.org/10.1126/science.1256688)); Põlme S, et al. 2020. Fungal Diversity 105:1-16 ((https://doi.org/10.1007/s13225-020-00466-2)) |
| Otu25375 | 1 | k__Fungi(100);p__Ascomycota(100);c__Saccharomycetes(100);o__Saccharomycetales(100);f__Metschnikowiaceae(100);g__Metschnikowia(100);g__Metschnikowia_unclassified(100); | Metschnikowia | 13 | Saprotroph | \|Undefined Saprotroph\| | Yeast | NULL | Probable | As nectar-tap_saprotroph (Põlme et al. 2020); Decay sugar-rich substrates (Põlme et al. 2020) | Põlme S, et al. 2020. Fungal Diversity 105:1-16 ((https://doi.org/10.1007/s13225-020-00466-2)) |
| Otu25376 | 1 | k__Fungi(100);p__Ascomycota(100);c__Saccharomycetes(100);o__Saccharomycetales(100);f__Metschnikowiaceae(100);g__Metschnikowia(100);s__Metschnikowia_pulcherrima(100); | Metschnikowia | 13 | Saprotroph | \|Undefined Saprotroph\| | Yeast | NULL | Probable | As nectar-tap_saprotroph (Põlme et al. 2020); Decay sugar-rich substrates (Põlme et al. 2020) | Põlme S, et al. 2020. Fungal Diversity 105:1-16 ((https://doi.org/10.1007/s13225-020-00466-2)) |
| Otu25377 | 1 | k__Fungi(100);p__Ascomycota(100);c__Saccharomycetes(100);o__Saccharomycetales(100);f__Metschnikowiaceae(100);g__Metschnikowia(100);s__Metschnikowia_pulcherrima(100); | Metschnikowia | 13 | Saprotroph | \|Undefined Saprotroph\| | Yeast | NULL | Probable | As nectar-tap_saprotroph (Põlme et al. 2020); Decay sugar-rich substrates (Põlme et al. 2020) | Põlme S, et al. 2020. Fungal Diversity 105:1-16 ((https://doi.org/10.1007/s13225-020-00466-2)) |
| Otu25378 | 1 | k__Fungi(100);p__Ascomycota(100);c__Saccharomycetes(100);o__Saccharomycetales(100);f__Metschnikowiaceae(100);g__Metschnikowia(100);s__Metschnikowia_pulcherrima(100); | Metschnikowia | 13 | Saprotroph | \|Undefined Saprotroph\| | Yeast | NULL | Probable | As nectar-tap_saprotroph (Põlme et al. 2020); Decay sugar-rich substrates (Põlme et al. 2020) | Põlme S, et al. 2020. Fungal Diversity 105:1-16 ((https://doi.org/10.1007/s13225-020-00466-2)) |
| Otu25380 | 1 | k__Fungi(100);p__Ascomycota(100);c__Dothideomycetes(100);o__Capnodiales(100);f__Teratosphaeriaceae(100);g__Catenulostroma(100);s__Catenulostroma_hermanusense(100); | Catenulostroma | 13 | Pathotroph | \|Plant Pathogen\| | Microfungus | NULL | Probable | NULL | Tedersoo L, et al. 2014. Science 346:e1256688 ((https://doi.org/10.1126/science.1256688)); Põlme S, et al. 2020. Fungal Diversity 105:1-16 ((https://doi.org/10.1007/s13225-020-00466-2)) |
| Otu25381 | 1 | k__Fungi(100);p__Ascomycota(100);c__Saccharomycetes(100);o__Saccharomycetales(100);f__Metschnikowiaceae(100);g__Metschnikowia(100);s__Metschnikowia_pulcherrima(100); | Metschnikowia | 13 | Saprotroph | \|Undefined Saprotroph\| | Yeast | NULL | Probable | As nectar-tap_saprotroph (Põlme et al. 2020); Decay sugar-rich substrates (Põlme et al. 2020) | Põlme S, et al. 2020. Fungal Diversity 105:1-16 ((https://doi.org/10.1007/s13225-020-00466-2)) |
| Otu25385 | 1 | k__Fungi(100);p__Ascomycota(100);c__Saccharomycetes(100);o__Saccharomycetales(100);f__Metschnikowiaceae(100);g__Metschnikowia(100);s__Metschnikowia_pulcherrima(100); | Metschnikowia | 13 | Saprotroph | \|Undefined Saprotroph\| | Yeast | NULL | Probable | As nectar-tap_saprotroph (Põlme et al. 2020); Decay sugar-rich substrates (Põlme et al. 2020) | Põlme S, et al. 2020. Fungal Diversity 105:1-16 ((https://doi.org/10.1007/s13225-020-00466-2)) |
| Otu25391 | 1 | k__Fungi(100);p__Ascomycota(100);c__Saccharomycetes(100);o__Saccharomycetales(100);f__Metschnikowiaceae(100);g__Metschnikowia(100);s__Metschnikowia_pulcherrima(100); | Metschnikowia | 13 | Saprotroph | \|Undefined Saprotroph\| | Yeast | NULL | Probable | As nectar-tap_saprotroph (Põlme et al. 2020); Decay sugar-rich substrates (Põlme et al. 2020) | Põlme S, et al. 2020. Fungal Diversity 105:1-16 ((https://doi.org/10.1007/s13225-020-00466-2)) |
| Otu25392 | 1 | k__Fungi(100);p__Ascomycota(100);c__Saccharomycetes(100);o__Saccharomycetales(100);f__Metschnikowiaceae(100);g__Metschnikowia(100);s__Metschnikowia_pulcherrima(100); | Metschnikowia | 13 | Saprotroph | \|Undefined Saprotroph\| | Yeast | NULL | Probable | As nectar-tap_saprotroph (Põlme et al. 2020); Decay sugar-rich substrates (Põlme et al. 2020) | Põlme S, et al. 2020. Fungal Diversity 105:1-16 ((https://doi.org/10.1007/s13225-020-00466-2)) |
| Otu25393 | 1 | k__Fungi(100);p__Ascomycota(100);c__Saccharomycetes(100);o__Saccharomycetales(100);f__Metschnikowiaceae(100);g__Metschnikowia(100);s__Metschnikowia_pulcherrima(100); | Metschnikowia | 13 | Saprotroph | \|Undefined Saprotroph\| | Yeast | NULL | Probable | As nectar-tap_saprotroph (Põlme et al. 2020); Decay sugar-rich substrates (Põlme et al. 2020) | Põlme S, et al. 2020. Fungal Diversity 105:1-16 ((https://doi.org/10.1007/s13225-020-00466-2)) |
| Otu25394 | 1 | k__Fungi(100);p__Ascomycota(100);c__Saccharomycetes(100);o__Saccharomycetales(100);f__Metschnikowiaceae(100);g__Metschnikowia(100);s__Metschnikowia_pulcherrima(100); | Metschnikowia | 13 | Saprotroph | \|Undefined Saprotroph\| | Yeast | NULL | Probable | As nectar-tap_saprotroph (Põlme et al. 2020); Decay sugar-rich substrates (Põlme et al. 2020) | Põlme S, et al. 2020. Fungal Diversity 105:1-16 ((https://doi.org/10.1007/s13225-020-00466-2)) |
| Otu25395 | 1 | k__Fungi(100);p__Ascomycota(100);c__Saccharomycetes(100);o__Saccharomycetales(100);f__Metschnikowiaceae(100);g__Metschnikowia(100);g__Metschnikowia_unclassified(100); | Metschnikowia | 13 | Saprotroph | \|Undefined Saprotroph\| | Yeast | NULL | Probable | As nectar-tap_saprotroph (Põlme et al. 2020); Decay sugar-rich substrates (Põlme et al. 2020) | Põlme S, et al. 2020. Fungal Diversity 105:1-16 ((https://doi.org/10.1007/s13225-020-00466-2)) |
| Otu25399 | 1 | k__Fungi(100);p__Ascomycota(100);c__Saccharomycetes(100);o__Saccharomycetales(100);f__Metschnikowiaceae(100);g__Metschnikowia(100);g__Metschnikowia_unclassified(100); | Metschnikowia | 13 | Saprotroph | \|Undefined Saprotroph\| | Yeast | NULL | Probable | As nectar-tap_saprotroph (Põlme et al. 2020); Decay sugar-rich substrates (Põlme et al. 2020) | Põlme S, et al. 2020. Fungal Diversity 105:1-16 ((https://doi.org/10.1007/s13225-020-00466-2)) |
| Otu25400 | 1 | k__Fungi(100);p__Ascomycota(100);c__Saccharomycetes(100);o__Saccharomycetales(100);f__Metschnikowiaceae(100);g__Metschnikowia(100);g__Metschnikowia_unclassified(100); | Metschnikowia | 13 | Saprotroph | \|Undefined Saprotroph\| | Yeast | NULL | Probable | As nectar-tap_saprotroph (Põlme et al. 2020); Decay sugar-rich substrates (Põlme et al. 2020) | Põlme S, et al. 2020. Fungal Diversity 105:1-16 ((https://doi.org/10.1007/s13225-020-00466-2)) |
| Otu25401 | 1 | k__Fungi(100);p__Ascomycota(100);c__Saccharomycetes(100);o__Saccharomycetales(100);f__Metschnikowiaceae(100);g__Metschnikowia(100);g__Metschnikowia_unclassified(100); | Metschnikowia | 13 | Saprotroph | \|Undefined Saprotroph\| | Yeast | NULL | Probable | As nectar-tap_saprotroph (Põlme et al. 2020); Decay sugar-rich substrates (Põlme et al. 2020) | Põlme S, et al. 2020. Fungal Diversity 105:1-16 ((https://doi.org/10.1007/s13225-020-00466-2)) |
| Otu25402 | 1 | k__Fungi(100);p__Ascomycota(100);c__Saccharomycetes(100);o__Saccharomycetales(100);f__Metschnikowiaceae(100);g__Metschnikowia(100);s__Metschnikowia_pulcherrima(100); | Metschnikowia | 13 | Saprotroph | \|Undefined Saprotroph\| | Yeast | NULL | Probable | As nectar-tap_saprotroph (Põlme et al. 2020); Decay sugar-rich substrates (Põlme et al. 2020) | Põlme S, et al. 2020. Fungal Diversity 105:1-16 ((https://doi.org/10.1007/s13225-020-00466-2)) |
| Otu25403 | 1 | k__Fungi(100);p__Ascomycota(100);c__Saccharomycetes(100);o__Saccharomycetales(100);f__Metschnikowiaceae(100);g__Metschnikowia(100);s__Metschnikowia_pulcherrima(100); | Metschnikowia | 13 | Saprotroph | \|Undefined Saprotroph\| | Yeast | NULL | Probable | As nectar-tap_saprotroph (Põlme et al. 2020); Decay sugar-rich substrates (Põlme et al. 2020) | Põlme S, et al. 2020. Fungal Diversity 105:1-16 ((https://doi.org/10.1007/s13225-020-00466-2)) |
| Otu25404 | 1 | k__Fungi(100);p__Ascomycota(100);c__Saccharomycetes(100);o__Saccharomycetales(100);f__Saccharomycetales_family_Incertae_sedis(100);f__Saccharomycetales_family_Incertae_sedis_unclassified(100);f__Saccharomycetales_family_Incertae_sedis_unclassified(100); | Saccharomycetales | 7 | Saprotroph | Undefined Saprotroph | Yeast | NULL | Possible | NULL | Sterkenburg E, et al. 2015. New Phytologist 207:1145-1158 ((https://doi.org/10.1111/nph.13426)) |
| Otu25408 | 1 | k__Fungi(100);p__Ascomycota(100);c__Saccharomycetes(100);o__Saccharomycetales(100);f__Metschnikowiaceae(100);g__Metschnikowia(100);s__Metschnikowia_pulcherrima(100); | Metschnikowia | 13 | Saprotroph | \|Undefined Saprotroph\| | Yeast | NULL | Probable | As nectar-tap_saprotroph (Põlme et al. 2020); Decay sugar-rich substrates (Põlme et al. 2020) | Põlme S, et al. 2020. Fungal Diversity 105:1-16 ((https://doi.org/10.1007/s13225-020-00466-2)) |
| Otu25409 | 1 | k__Fungi(100);p__Ascomycota(100);c__Saccharomycetes(100);o__Saccharomycetales(100);f__Metschnikowiaceae(100);g__Metschnikowia(100);g__Metschnikowia_unclassified(100); | Metschnikowia | 13 | Saprotroph | \|Undefined Saprotroph\| | Yeast | NULL | Probable | As nectar-tap_saprotroph (Põlme et al. 2020); Decay sugar-rich substrates (Põlme et al. 2020) | Põlme S, et al. 2020. Fungal Diversity 105:1-16 ((https://doi.org/10.1007/s13225-020-00466-2)) |
| Otu25410 | 1 | k__Fungi(100);p__Ascomycota(100);c__Saccharomycetes(100);o__Saccharomycetales(100);f__Metschnikowiaceae(100);g__Metschnikowia(100);g__Metschnikowia_unclassified(100); | Metschnikowia | 13 | Saprotroph | \|Undefined Saprotroph\| | Yeast | NULL | Probable | As nectar-tap_saprotroph (Põlme et al. 2020); Decay sugar-rich substrates (Põlme et al. 2020) | Põlme S, et al. 2020. Fungal Diversity 105:1-16 ((https://doi.org/10.1007/s13225-020-00466-2)) |
| Otu25411 | 1 | k__Fungi(100);p__Ascomycota(100);c__Saccharomycetes(100);o__Saccharomycetales(100);f__Metschnikowiaceae(100);g__Metschnikowia(100);s__Metschnikowia_pulcherrima(100); | Metschnikowia | 13 | Saprotroph | \|Undefined Saprotroph\| | Yeast | NULL | Probable | As nectar-tap_saprotroph (Põlme et al. 2020); Decay sugar-rich substrates (Põlme et al. 2020) | Põlme S, et al. 2020. Fungal Diversity 105:1-16 ((https://doi.org/10.1007/s13225-020-00466-2)) |
| Otu25414 | 1 | k__Fungi(100);p__Ascomycota(100);c__Saccharomycetes(100);o__Saccharomycetales(100);f__Metschnikowiaceae(100);g__Metschnikowia(100);g__Metschnikowia_unclassified(100); | Metschnikowia | 13 | Saprotroph | \|Undefined Saprotroph\| | Yeast | NULL | Probable | As nectar-tap_saprotroph (Põlme et al. 2020); Decay sugar-rich substrates (Põlme et al. 2020) | Põlme S, et al. 2020. Fungal Diversity 105:1-16 ((https://doi.org/10.1007/s13225-020-00466-2)) |
| Otu25415 | 1 | k__Fungi(100);p__Ascomycota(100);c__Saccharomycetes(100);o__Saccharomycetales(100);f__Metschnikowiaceae(100);g__Metschnikowia(100);s__Metschnikowia_pulcherrima(100); | Metschnikowia | 13 | Saprotroph | \|Undefined Saprotroph\| | Yeast | NULL | Probable | As nectar-tap_saprotroph (Põlme et al. 2020); Decay sugar-rich substrates (Põlme et al. 2020) | Põlme S, et al. 2020. Fungal Diversity 105:1-16 ((https://doi.org/10.1007/s13225-020-00466-2)) |
| Otu25416 | 1 | k__Fungi(100);p__Ascomycota(100);c__Saccharomycetes(100);o__Saccharomycetales(100);f__Metschnikowiaceae(100);g__Metschnikowia(100);s__Metschnikowia_pulcherrima(100); | Metschnikowia | 13 | Saprotroph | \|Undefined Saprotroph\| | Yeast | NULL | Probable | As nectar-tap_saprotroph (Põlme et al. 2020); Decay sugar-rich substrates (Põlme et al. 2020) | Põlme S, et al. 2020. Fungal Diversity 105:1-16 ((https://doi.org/10.1007/s13225-020-00466-2)) |
| Otu25417 | 1 | k__Fungi(100);p__Ascomycota(100);c__Saccharomycetes(100);o__Saccharomycetales(100);f__Metschnikowiaceae(100);g__Metschnikowia(100);g__Metschnikowia_unclassified(100); | Metschnikowia | 13 | Saprotroph | \|Undefined Saprotroph\| | Yeast | NULL | Probable | As nectar-tap_saprotroph (Põlme et al. 2020); Decay sugar-rich substrates (Põlme et al. 2020) | Põlme S, et al. 2020. Fungal Diversity 105:1-16 ((https://doi.org/10.1007/s13225-020-00466-2)) |
| Otu25418 | 1 | k__Fungi(100);p__Ascomycota(100);c__Saccharomycetes(100);o__Saccharomycetales(100);f__Metschnikowiaceae(100);g__Metschnikowia(100);g__Metschnikowia_unclassified(100); | Metschnikowia | 13 | Saprotroph | \|Undefined Saprotroph\| | Yeast | NULL | Probable | As nectar-tap_saprotroph (Põlme et al. 2020); Decay sugar-rich substrates (Põlme et al. 2020) | Põlme S, et al. 2020. Fungal Diversity 105:1-16 ((https://doi.org/10.1007/s13225-020-00466-2)) |
| Otu25419 | 1 | k__Fungi(100);p__Ascomycota(100);c__Saccharomycetes(100);o__Saccharomycetales(100);f__Metschnikowiaceae(100);g__Metschnikowia(100);s__Metschnikowia_pulcherrima(100); | Metschnikowia | 13 | Saprotroph | \|Undefined Saprotroph\| | Yeast | NULL | Probable | As nectar-tap_saprotroph (Põlme et al. 2020); Decay sugar-rich substrates (Põlme et al. 2020) | Põlme S, et al. 2020. Fungal Diversity 105:1-16 ((https://doi.org/10.1007/s13225-020-00466-2)) |
| Otu25420 | 1 | k__Fungi(100);p__Ascomycota(100);c__Saccharomycetes(100);o__Saccharomycetales(100);f__Metschnikowiaceae(100);g__Metschnikowia(100);s__Metschnikowia_pulcherrima(100); | Metschnikowia | 13 | Saprotroph | \|Undefined Saprotroph\| | Yeast | NULL | Probable | As nectar-tap_saprotroph (Põlme et al. 2020); Decay sugar-rich substrates (Põlme et al. 2020) | Põlme S, et al. 2020. Fungal Diversity 105:1-16 ((https://doi.org/10.1007/s13225-020-00466-2)) |
| Otu25421 | 1 | k__Fungi(100);p__Ascomycota(100);c__Saccharomycetes(100);o__Saccharomycetales(100);f__Metschnikowiaceae(100);g__Metschnikowia(100);g__Metschnikowia_unclassified(100); | Metschnikowia | 13 | Saprotroph | \|Undefined Saprotroph\| | Yeast | NULL | Probable | As nectar-tap_saprotroph (Põlme et al. 2020); Decay sugar-rich substrates (Põlme et al. 2020) | Põlme S, et al. 2020. Fungal Diversity 105:1-16 ((https://doi.org/10.1007/s13225-020-00466-2)) |
| Otu25423 | 1 | k__Fungi(100);p__Ascomycota(100);c__Saccharomycetes(100);o__Saccharomycetales(100);f__Metschnikowiaceae(100);g__Metschnikowia(100);s__Metschnikowia_pulcherrima(100); | Metschnikowia | 13 | Saprotroph | \|Undefined Saprotroph\| | Yeast | NULL | Probable | As nectar-tap_saprotroph (Põlme et al. 2020); Decay sugar-rich substrates (Põlme et al. 2020) | Põlme S, et al. 2020. Fungal Diversity 105:1-16 ((https://doi.org/10.1007/s13225-020-00466-2)) |
| Otu25424 | 1 | k__Fungi(100);p__Ascomycota(100);c__Saccharomycetes(100);o__Saccharomycetales(100);f__Metschnikowiaceae(100);g__Metschnikowia(100);s__Metschnikowia_pulcherrima(100); | Metschnikowia | 13 | Saprotroph | \|Undefined Saprotroph\| | Yeast | NULL | Probable | As nectar-tap_saprotroph (Põlme et al. 2020); Decay sugar-rich substrates (Põlme et al. 2020) | Põlme S, et al. 2020. Fungal Diversity 105:1-16 ((https://doi.org/10.1007/s13225-020-00466-2)) |
| Otu25425 | 1 | k__Fungi(100);p__Ascomycota(100);c__Saccharomycetes(100);o__Saccharomycetales(100);f__Metschnikowiaceae(100);g__Metschnikowia(100);s__Metschnikowia_sp(100); | Metschnikowia | 13 | Saprotroph | \|Undefined Saprotroph\| | Yeast | NULL | Probable | As nectar-tap_saprotroph (Põlme et al. 2020); Decay sugar-rich substrates (Põlme et al. 2020) | Põlme S, et al. 2020. Fungal Diversity 105:1-16 ((https://doi.org/10.1007/s13225-020-00466-2)) |
| Otu25426 | 1 | k__Fungi(100);p__Ascomycota(100);c__Saccharomycetes(100);o__Saccharomycetales(100);f__Metschnikowiaceae(100);g__Metschnikowia(100);g__Metschnikowia_unclassified(100); | Metschnikowia | 13 | Saprotroph | \|Undefined Saprotroph\| | Yeast | NULL | Probable | As nectar-tap_saprotroph (Põlme et al. 2020); Decay sugar-rich substrates (Põlme et al. 2020) | Põlme S, et al. 2020. Fungal Diversity 105:1-16 ((https://doi.org/10.1007/s13225-020-00466-2)) |
| Otu25427 | 1 | k__Fungi(100);p__Ascomycota(100);c__Saccharomycetes(100);o__Saccharomycetales(100);f__Metschnikowiaceae(100);g__Metschnikowia(100);g__Metschnikowia_unclassified(100); | Metschnikowia | 13 | Saprotroph | \|Undefined Saprotroph\| | Yeast | NULL | Probable | As nectar-tap_saprotroph (Põlme et al. 2020); Decay sugar-rich substrates (Põlme et al. 2020) | Põlme S, et al. 2020. Fungal Diversity 105:1-16 ((https://doi.org/10.1007/s13225-020-00466-2)) |
| Otu25428 | 1 | k__Fungi(100);p__Ascomycota(100);c__Saccharomycetes(100);o__Saccharomycetales(100);f__Metschnikowiaceae(100);g__Metschnikowia(100);s__Metschnikowia_pulcherrima(100); | Metschnikowia | 13 | Saprotroph | \|Undefined Saprotroph\| | Yeast | NULL | Probable | As nectar-tap_saprotroph (Põlme et al. 2020); Decay sugar-rich substrates (Põlme et al. 2020) | Põlme S, et al. 2020. Fungal Diversity 105:1-16 ((https://doi.org/10.1007/s13225-020-00466-2)) |
| Otu25429 | 1 | k__Fungi(100);p__Ascomycota(100);c__Saccharomycetes(100);o__Saccharomycetales(100);f__Metschnikowiaceae(100);g__Metschnikowia(100);s__Metschnikowia_pulcherrima(100); | Metschnikowia | 13 | Saprotroph | \|Undefined Saprotroph\| | Yeast | NULL | Probable | As nectar-tap_saprotroph (Põlme et al. 2020); Decay sugar-rich substrates (Põlme et al. 2020) | Põlme S, et al. 2020. Fungal Diversity 105:1-16 ((https://doi.org/10.1007/s13225-020-00466-2)) |
| Otu25430 | 1 | k__Fungi(100);p__Ascomycota(100);c__Saccharomycetes(100);o__Saccharomycetales(100);f__Metschnikowiaceae(100);g__Metschnikowia(100);g__Metschnikowia_unclassified(100); | Metschnikowia | 13 | Saprotroph | \|Undefined Saprotroph\| | Yeast | NULL | Probable | As nectar-tap_saprotroph (Põlme et al. 2020); Decay sugar-rich substrates (Põlme et al. 2020) | Põlme S, et al. 2020. Fungal Diversity 105:1-16 ((https://doi.org/10.1007/s13225-020-00466-2)) |
| Otu25431 | 1 | k__Fungi(100);p__Ascomycota(100);c__Saccharomycetes(100);o__Saccharomycetales(100);f__Metschnikowiaceae(100);g__Metschnikowia(100);s__Metschnikowia_pulcherrima(100); | Metschnikowia | 13 | Saprotroph | \|Undefined Saprotroph\| | Yeast | NULL | Probable | As nectar-tap_saprotroph (Põlme et al. 2020); Decay sugar-rich substrates (Põlme et al. 2020) | Põlme S, et al. 2020. Fungal Diversity 105:1-16 ((https://doi.org/10.1007/s13225-020-00466-2)) |
| Otu25432 | 1 | k__Fungi(100);p__Ascomycota(100);c__Saccharomycetes(100);o__Saccharomycetales(100);f__Metschnikowiaceae(100);g__Metschnikowia(100);s__Metschnikowia_pulcherrima(100); | Metschnikowia | 13 | Saprotroph | \|Undefined Saprotroph\| | Yeast | NULL | Probable | As nectar-tap_saprotroph (Põlme et al. 2020); Decay sugar-rich substrates (Põlme et al. 2020) | Põlme S, et al. 2020. Fungal Diversity 105:1-16 ((https://doi.org/10.1007/s13225-020-00466-2)) |
| Otu25433 | 1 | k__Fungi(100);p__Ascomycota(100);c__Saccharomycetes(100);o__Saccharomycetales(100);f__Metschnikowiaceae(100);g__Metschnikowia(100);s__Metschnikowia_pulcherrima(100); | Metschnikowia | 13 | Saprotroph | \|Undefined Saprotroph\| | Yeast | NULL | Probable | As nectar-tap_saprotroph (Põlme et al. 2020); Decay sugar-rich substrates (Põlme et al. 2020) | Põlme S, et al. 2020. Fungal Diversity 105:1-16 ((https://doi.org/10.1007/s13225-020-00466-2)) |
| Otu25434 | 1 | k__Fungi(100);p__Ascomycota(100);c__Saccharomycetes(100);o__Saccharomycetales(100);f__Metschnikowiaceae(100);g__Metschnikowia(100);s__Metschnikowia_pulcherrima(100); | Metschnikowia | 13 | Saprotroph | \|Undefined Saprotroph\| | Yeast | NULL | Probable | As nectar-tap_saprotroph (Põlme et al. 2020); Decay sugar-rich substrates (Põlme et al. 2020) | Põlme S, et al. 2020. Fungal Diversity 105:1-16 ((https://doi.org/10.1007/s13225-020-00466-2)) |
| Otu25435 | 1 | k__Fungi(100);p__Ascomycota(100);c__Saccharomycetes(100);o__Saccharomycetales(100);f__Metschnikowiaceae(100);g__Metschnikowia(100);s__Metschnikowia_pulcherrima(100); | Metschnikowia | 13 | Saprotroph | \|Undefined Saprotroph\| | Yeast | NULL | Probable | As nectar-tap_saprotroph (Põlme et al. 2020); Decay sugar-rich substrates (Põlme et al. 2020) | Põlme S, et al. 2020. Fungal Diversity 105:1-16 ((https://doi.org/10.1007/s13225-020-00466-2)) |
| Otu25436 | 1 | k__Fungi(100);p__Ascomycota(100);c__Saccharomycetes(100);o__Saccharomycetales(100);f__Metschnikowiaceae(100);g__Metschnikowia(100);s__Metschnikowia_pulcherrima(100); | Metschnikowia | 13 | Saprotroph | \|Undefined Saprotroph\| | Yeast | NULL | Probable | As nectar-tap_saprotroph (Põlme et al. 2020); Decay sugar-rich substrates (Põlme et al. 2020) | Põlme S, et al. 2020. Fungal Diversity 105:1-16 ((https://doi.org/10.1007/s13225-020-00466-2)) |
| Otu25437 | 1 | k__Fungi(100);p__Ascomycota(100);c__Saccharomycetes(100);o__Saccharomycetales(100);f__Metschnikowiaceae(100);g__Metschnikowia(100);s__Metschnikowia_pulcherrima(100); | Metschnikowia | 13 | Saprotroph | \|Undefined Saprotroph\| | Yeast | NULL | Probable | As nectar-tap_saprotroph (Põlme et al. 2020); Decay sugar-rich substrates (Põlme et al. 2020) | Põlme S, et al. 2020. Fungal Diversity 105:1-16 ((https://doi.org/10.1007/s13225-020-00466-2)) |
| Otu25438 | 1 | k__Fungi(100);p__Ascomycota(100);c__Saccharomycetes(100);o__Saccharomycetales(100);f__Metschnikowiaceae(100);g__Metschnikowia(100);s__Metschnikowia_pulcherrima(100); | Metschnikowia | 13 | Saprotroph | \|Undefined Saprotroph\| | Yeast | NULL | Probable | As nectar-tap_saprotroph (Põlme et al. 2020); Decay sugar-rich substrates (Põlme et al. 2020) | Põlme S, et al. 2020. Fungal Diversity 105:1-16 ((https://doi.org/10.1007/s13225-020-00466-2)) |
| Otu25439 | 1 | k__Fungi(100);p__Ascomycota(100);c__Saccharomycetes(100);o__Saccharomycetales(100);f__Metschnikowiaceae(100);g__Metschnikowia(100);s__Metschnikowia_pulcherrima(100); | Metschnikowia | 13 | Saprotroph | \|Undefined Saprotroph\| | Yeast | NULL | Probable | As nectar-tap_saprotroph (Põlme et al. 2020); Decay sugar-rich substrates (Põlme et al. 2020) | Põlme S, et al. 2020. Fungal Diversity 105:1-16 ((https://doi.org/10.1007/s13225-020-00466-2)) |
| Otu25440 | 1 | k__Fungi(100);p__Ascomycota(100);c__Saccharomycetes(100);o__Saccharomycetales(100);f__Metschnikowiaceae(100);g__Metschnikowia(100);s__Metschnikowia_pulcherrima(100); | Metschnikowia | 13 | Saprotroph | \|Undefined Saprotroph\| | Yeast | NULL | Probable | As nectar-tap_saprotroph (Põlme et al. 2020); Decay sugar-rich substrates (Põlme et al. 2020) | Põlme S, et al. 2020. Fungal Diversity 105:1-16 ((https://doi.org/10.1007/s13225-020-00466-2)) |
| Otu25441 | 1 | k__Fungi(100);p__Ascomycota(100);c__Saccharomycetes(100);o__Saccharomycetales(100);f__Metschnikowiaceae(100);g__Metschnikowia(100);s__Metschnikowia_pulcherrima(100); | Metschnikowia | 13 | Saprotroph | \|Undefined Saprotroph\| | Yeast | NULL | Probable | As nectar-tap_saprotroph (Põlme et al. 2020); Decay sugar-rich substrates (Põlme et al. 2020) | Põlme S, et al. 2020. Fungal Diversity 105:1-16 ((https://doi.org/10.1007/s13225-020-00466-2)) |

| | | | | | | | | | | | |
|---|---|---|---|---|---|---|---|---|---|---|---|
| Otu25442 | 1 | k__Fungi(100);p__Ascomycota(100);c__Saccharomycetes(100);o__Saccharomycetales(100);f__Metschnikowiaceae(100);g__Metschnikowia(100);g__Metschnikowia_unclassified(100); | Metschnikowia | 13 | Saprotroph | \|Undefined Saprotroph\| | Yeast | NULL | Probable | As nectar-tap_saprotroph (Põlme et al. 2020); Decay sugar-rich substrates (Põlme et al. 2020) | Põlme S, et al. 2020. Fungal Diversity 105:1-16 ((https://doi.org/10.1007/s13225-020-00466-2)) |
| Otu25443 | 1 | k__Fungi(100);p__Ascomycota(100);c__Saccharomycetes(100);o__Saccharomycetales(100);f__Metschnikowiaceae(100);g__Metschnikowia(100);g__Metschnikowia_unclassified(100); | Metschnikowia | 13 | Saprotroph | \|Undefined Saprotroph\| | Yeast | NULL | Probable | As nectar-tap_saprotroph (Põlme et al. 2020); Decay sugar-rich substrates (Põlme et al. 2020) | Põlme S, et al. 2020. Fungal Diversity 105:1-16 ((https://doi.org/10.1007/s13225-020-00466-2)) |
| Otu25444 | 1 | k__Fungi(100);p__Ascomycota(100);c__Saccharomycetes(100);o__Saccharomycetales(100);f__Metschnikowiaceae(100);g__Metschnikowia(100);s__Metschnikowia_pulcherrima(100); | Metschnikowia | 13 | Saprotroph | \|Undefined Saprotroph\| | Yeast | NULL | Probable | As nectar-tap_saprotroph (Põlme et al. 2020); Decay sugar-rich substrates (Põlme et al. 2020) | Põlme S, et al. 2020. Fungal Diversity 105:1-16 ((https://doi.org/10.1007/s13225-020-00466-2)) |
| Otu25445 | 1 | k__Fungi(100);p__Ascomycota(100);c__Saccharomycetes(100);o__Saccharomycetales(100);f__Metschnikowiaceae(100);g__Metschnikowia(100);s__Metschnikowia_pulcherrima(100); | Metschnikowia | 13 | Saprotroph | \|Undefined Saprotroph\| | Yeast | NULL | Probable | As nectar-tap_saprotroph (Põlme et al. 2020); Decay sugar-rich substrates (Põlme et al. 2020) | Põlme S, et al. 2020. Fungal Diversity 105:1-16 ((https://doi.org/10.1007/s13225-020-00466-2)) |
| Otu25446 | 1 | k__Fungi(100);p__Ascomycota(100);c__Saccharomycetes(100);o__Saccharomycetales(100);f__Metschnikowiaceae(100);g__Metschnikowia(100);s__Metschnikowia_pulcherrima(100); | Metschnikowia | 13 | Saprotroph | \|Undefined Saprotroph\| | Yeast | NULL | Probable | As nectar-tap_saprotroph (Põlme et al. 2020); Decay sugar-rich substrates (Põlme et al. 2020) | Põlme S, et al. 2020. Fungal Diversity 105:1-16 ((https://doi.org/10.1007/s13225-020-00466-2)) |
| Otu25447 | 1 | k__Fungi(100);p__Ascomycota(100);c__Saccharomycetes(100);o__Saccharomycetales(100);f__Metschnikowiaceae(100);g__Metschnikowia(100);g__Metschnikowia_unclassified(100); | Metschnikowia | 13 | Saprotroph | \|Undefined Saprotroph\| | Yeast | NULL | Probable | As nectar-tap_saprotroph (Põlme et al. 2020); Decay sugar-rich substrates (Põlme et al. 2020) | Põlme S, et al. 2020. Fungal Diversity 105:1-16 ((https://doi.org/10.1007/s13225-020-00466-2)) |
| Otu25449 | 1 | k__Fungi(100);p__Ascomycota(100);c__Saccharomycetes(100);o__Saccharomycetales(100);f__Metschnikowiaceae(100);g__Metschnikowia(100);g__Metschnikowia_unclassified(100); | Metschnikowia | 13 | Saprotroph | \|Undefined Saprotroph\| | Yeast | NULL | Probable | As nectar-tap_saprotroph (Põlme et al. 2020); Decay sugar-rich substrates (Põlme et al. 2020) | Põlme S, et al. 2020. Fungal Diversity 105:1-16 ((https://doi.org/10.1007/s13225-020-00466-2)) |
| Otu25450 | 1 | k__Fungi(100);p__Ascomycota(100);c__Saccharomycetes(100);o__Saccharomycetales(100);f__Metschnikowiaceae(100);g__Metschnikowia(100);g__Metschnikowia_unclassified(100); | Metschnikowia | 13 | Saprotroph | \|Undefined Saprotroph\| | Yeast | NULL | Probable | As nectar-tap_saprotroph (Põlme et al. 2020); Decay sugar-rich substrates (Põlme et al. 2020) | Põlme S, et al. 2020. Fungal Diversity 105:1-16 ((https://doi.org/10.1007/s13225-020-00466-2)) |
| Otu25451 | 1 | k__Fungi(100);p__Ascomycota(100);c__Saccharomycetes(100);o__Saccharomycetales(100);f__Metschnikowiaceae(100);g__Metschnikowia(100);g__Metschnikowia_unclassified(100); | Metschnikowia | 13 | Saprotroph | \|Undefined Saprotroph\| | Yeast | NULL | Probable | As nectar-tap_saprotroph (Põlme et al. 2020); Decay sugar-rich substrates (Põlme et al. 2020) | Põlme S, et al. 2020. Fungal Diversity 105:1-16 ((https://doi.org/10.1007/s13225-020-00466-2)) |
| Otu25452 | 1 | k__Fungi(100);p__Ascomycota(100);c__Saccharomycetes(100);o__Saccharomycetales(100);f__Metschnikowiaceae(100);g__Metschnikowia(100);s__Metschnikowia_pulcherrima(100); | Metschnikowia | 13 | Saprotroph | \|Undefined Saprotroph\| | Yeast | NULL | Probable | As nectar-tap_saprotroph (Põlme et al. 2020); Decay sugar-rich substrates (Põlme et al. 2020) | Põlme S, et al. 2020. Fungal Diversity 105:1-16 ((https://doi.org/10.1007/s13225-020-00466-2)) |
| Otu25457 | 1 | k__Fungi(100);p__Ascomycota(100);c__Saccharomycetes(100);o__Saccharomycetales(100);f__Metschnikowiaceae(100);g__Metschnikowia(100);s__Metschnikowia_pulcherrima(100); | Metschnikowia | 13 | Saprotroph | \|Undefined Saprotroph\| | Yeast | NULL | Probable | As nectar-tap_saprotroph (Põlme et al. 2020); Decay sugar-rich substrates (Põlme et al. 2020) | Põlme S, et al. 2020. Fungal Diversity 105:1-16 ((https://doi.org/10.1007/s13225-020-00466-2)) |
| Otu25458 | 1 | k__Fungi(100);p__Ascomycota(100);c__Saccharomycetes(100);o__Saccharomycetales(100);f__Metschnikowiaceae(100);g__Metschnikowia(100);g__Metschnikowia_unclassified(100); | Metschnikowia | 13 | Saprotroph | \|Undefined Saprotroph\| | Yeast | NULL | Probable | As nectar-tap_saprotroph (Põlme et al. 2020); Decay sugar-rich substrates (Põlme et al. 2020) | Põlme S, et al. 2020. Fungal Diversity 105:1-16 ((https://doi.org/10.1007/s13225-020-00466-2)) |
| Otu25462 | 1 | k__Fungi(100);p__Ascomycota(100);c__Saccharomycetes(100);o__Saccharomycetales(100);f__Metschnikowiaceae(100);g__Metschnikowia(100);g__Metschnikowia_unclassified(100); | Metschnikowia | 13 | Saprotroph | \|Undefined Saprotroph\| | Yeast | NULL | Probable | As nectar-tap_saprotroph (Põlme et al. 2020); Decay sugar-rich substrates (Põlme et al. 2020) | Põlme S, et al. 2020. Fungal Diversity 105:1-16 ((https://doi.org/10.1007/s13225-020-00466-2)) |
| Otu25468 | 1 | k__Fungi(100);p__Ascomycota(100);c__Saccharomycetes(100);o__Saccharomycetales(100);f__Metschnikowiaceae(100);g__Metschnikowia(100);s__Metschnikowia_pulcherrima(100); | Metschnikowia | 13 | Saprotroph | \|Undefined Saprotroph\| | Yeast | NULL | Probable | As nectar-tap_saprotroph (Põlme et al. 2020); Decay sugar-rich substrates (Põlme et al. 2020) | Põlme S, et al. 2020. Fungal Diversity 105:1-16 ((https://doi.org/10.1007/s13225-020-00466-2)) |
| Otu25469 | 1 | k__Fungi(100);p__Ascomycota(100);c__Saccharomycetes(100);o__Saccharomycetales(100);f__Metschnikowiaceae(100);g__Metschnikowia(100);g__Metschnikowia_unclassified(100); | Metschnikowia | 13 | Saprotroph | \|Undefined Saprotroph\| | Yeast | NULL | Probable | As nectar-tap_saprotroph (Põlme et al. 2020); Decay sugar-rich substrates (Põlme et al. 2020) | Põlme S, et al. 2020. Fungal Diversity 105:1-16 ((https://doi.org/10.1007/s13225-020-00466-2)) |
| Otu25470 | 1 | k__Fungi(100);p__Ascomycota(100);c__Saccharomycetes(100);o__Saccharomycetales(100);f__Saccharomycetaceae(100);g__Kazachstania(100);s__Kazachstania_intestinalis(100); | Kazachstania | 13 | Saprotroph | \|Undefined Saprotroph\| | Yeast | NULL | Probable | As nectar-tap_saprotroph (Põlme et al. 2020); Decay sugar-rich substrates (Põlme et al. 2020) | Põlme S, et al. 2020. Fungal Diversity 105:1-16 ((https://doi.org/10.1007/s13225-020-00466-2)) |
| Otu25483 | 1 | k__Fungi(100);p__Ascomycota(100);c__Saccharomycetes(100);o__Saccharomycetales(100);f__Saccharomycetales_family_Incertae_sedis(100);g__Candida(100);s__Candida_cretensis(100); | Candida | 13 | Pathotroph-Saprotroph-Symbiotroph | Animal Pathogen-Endophyte-Endosymbiont-Epiphyte-\|Undefined Saprotroph\| | Dimorphic Yeast-Yeast | NULL | Probable | As nectar-tap_saprotroph (Põlme et al. 2020); Decay sugar-rich substrates (Põlme et al. 2020); This asexual genus is polyphyletic and contain species that function very differently depend on what they are. Caution should be taken when interpreting guilds for Candida species. | Manolakaki D, et al. 2010. Virulence. 1: 367-375 ((https://doi.org/10.4161/viru.1.5.12796)); Mekbib SB, et al. 2011. Tropical Plant Pathology 36:233-240 ((https://doi.org/10.1590/S1982-56762011000400004)); Põlme S, et al. 2020. Fungal Diversity 105:1-16 ((https://doi.org/10.1007/s13225-020-00466-2)) |
| Otu25487 | 1 | k__Fungi(100);p__Ascomycota(100);c__Saccharomycetes(100);o__Saccharomycetales(100);f__Metschnikowiaceae(100);g__Metschnikowia(100);s__Metschnikowia_pulcherrima(100); | Metschnikowia | 13 | Saprotroph | \|Undefined Saprotroph\| | Yeast | NULL | Probable | As nectar-tap_saprotroph (Põlme et al. 2020); Decay sugar-rich substrates (Põlme et al. 2020) | Põlme S, et al. 2020. Fungal Diversity 105:1-16 ((https://doi.org/10.1007/s13225-020-00466-2)) |
| Otu25488 | 1 | k__Fungi(100);p__Ascomycota(100);c__Saccharomycetes(100);o__Saccharomycetales(100);f__Metschnikowiaceae(100);g__Metschnikowia(100);s__Metschnikowia_pulcherrima(100); | Metschnikowia | 13 | Saprotroph | \|Undefined Saprotroph\| | Yeast | NULL | Probable | As nectar-tap_saprotroph (Põlme et al. 2020); Decay sugar-rich substrates (Põlme et al. 2020) | Põlme S, et al. 2020. Fungal Diversity 105:1-16 ((https://doi.org/10.1007/s13225-020-00466-2)) |
| Otu25489 | 1 | k__Fungi(100);p__Ascomycota(100);c__Saccharomycetes(100);o__Saccharomycetales(100);f__Metschnikowiaceae(100);g__Metschnikowia(100);g__Metschnikowia_unclassified(100); | Metschnikowia | 13 | Saprotroph | \|Undefined Saprotroph\| | Yeast | NULL | Probable | As nectar-tap_saprotroph (Põlme et al. 2020); Decay sugar-rich substrates (Põlme et al. 2020) | Põlme S, et al. 2020. Fungal Diversity 105:1-16 ((https://doi.org/10.1007/s13225-020-00466-2)) |
| Otu25490 | 1 | k__Fungi(100);p__Ascomycota(100);c__Saccharomycetes(100);o__Saccharomycetales(100);f__Metschnikowiaceae(100);g__Metschnikowia(100);s__Metschnikowia_pulcherrima(100); | Metschnikowia | 13 | Saprotroph | \|Undefined Saprotroph\| | Yeast | NULL | Probable | As nectar-tap_saprotroph (Põlme et al. 2020); Decay sugar-rich substrates (Põlme et al. 2020) | Põlme S, et al. 2020. Fungal Diversity 105:1-16 ((https://doi.org/10.1007/s13225-020-00466-2)) |
| Otu25491 | 1 | k__Fungi(100);p__Ascomycota(100);c__Saccharomycetes(100);o__Saccharomycetales(100);f__Metschnikowiaceae(100);g__Metschnikowia(100);s__Metschnikowia_pulcherrima(100); | Metschnikowia | 13 | Saprotroph | \|Undefined Saprotroph\| | Yeast | NULL | Probable | As nectar-tap_saprotroph (Põlme et al. 2020); Decay sugar-rich substrates (Põlme et al. 2020) | Põlme S, et al. 2020. Fungal Diversity 105:1-16 ((https://doi.org/10.1007/s13225-020-00466-2)) |
| Otu25492 | 1 | k__Fungi(100);p__Ascomycota(100);c__Saccharomycetes(100);o__Saccharomycetales(100);f__Metschnikowiaceae(100);g__Metschnikowia(100);s__Metschnikowia_pulcherrima(100); | Metschnikowia | 13 | Saprotroph | \|Undefined Saprotroph\| | Yeast | NULL | Probable | As nectar-tap_saprotroph (Põlme et al. 2020); Decay sugar-rich substrates (Põlme et al. 2020) | Põlme S, et al. 2020. Fungal Diversity 105:1-16 ((https://doi.org/10.1007/s13225-020-00466-2)) |
| Otu25493 | 1 | k__Fungi(100);p__Ascomycota(100);c__Saccharomycetes(100);o__Saccharomycetales(100);f__Metschnikowiaceae(100);g__Metschnikowia(100);g__Metschnikowia_unclassified(100); | Metschnikowia | 13 | Saprotroph | \|Undefined Saprotroph\| | Yeast | NULL | Probable | As nectar-tap_saprotroph (Põlme et al. 2020); Decay sugar-rich substrates (Põlme et al. 2020) | Põlme S, et al. 2020. Fungal Diversity 105:1-16 ((https://doi.org/10.1007/s13225-020-00466-2)) |
| Otu25501 | 1 | k__Fungi(100);p__Ascomycota(100);c__Saccharomycetes(100);o__Saccharomycetales(100);f__Metschnikowiaceae(100);g__Metschnikowia(100);g__Metschnikowia_unclassified(100); | Metschnikowia | 13 | Saprotroph | \|Undefined Saprotroph\| | Yeast | NULL | Probable | As nectar-tap_saprotroph (Põlme et al. 2020); Decay sugar-rich substrates (Põlme et al. 2020) | Põlme S, et al. 2020. Fungal Diversity 105:1-16 ((https://doi.org/10.1007/s13225-020-00466-2)) |
| Otu25508 | 1 | k__Fungi(100);p__Ascomycota(100);c__Saccharomycetes(100);o__Saccharomycetales(100);f__Saccharomycetales_family_Incertae_sedis(100);g__Candida(100);g__Candida_unclassified(100); | Candida | 13 | Pathotroph-Saprotroph-Symbiotroph | Animal Pathogen-Endophyte-Endosymbiont-Epiphyte-\|Undefined Saprotroph\| | Dimorphic Yeast-Yeast | NULL | Probable | As nectar-tap_saprotroph (Põlme et al. 2020); Decay sugar-rich substrates (Põlme et al. 2020); This asexual genus is polyphyletic and contain species that function very differently depend on what they are. Caution should be taken when interpreting guilds for Candida species. | Manolakaki D, et al. 2010. Virulence. 1: 367-375 ((https://doi.org/10.4161/viru.1.5.12796)); Mekbib SB, et al. 2011. Tropical Plant Pathology 36:233-240 ((https://doi.org/10.1590/S1982-56762011000400004)); Põlme S, et al. 2020. Fungal Diversity 105:1-16 ((https://doi.org/10.1007/s13225-020-00466-2)) |
| Otu25513 | 1 | k__Fungi(100);p__Ascomycota(100);c__Saccharomycetes(100);o__Saccharomycetales(100);f__Metschnikowiaceae(100);g__Metschnikowia(100);g__Metschnikowia_unclassified(100); | Metschnikowia | 13 | Saprotroph | \|Undefined Saprotroph\| | Yeast | NULL | Probable | As nectar-tap_saprotroph (Põlme et al. 2020); Decay sugar-rich substrates (Põlme et al. 2020) | Põlme S, et al. 2020. Fungal Diversity 105:1-16 ((https://doi.org/10.1007/s13225-020-00466-2)) |
| Otu25514 | 1 | k__Fungi(100);p__Ascomycota(100);c__Saccharomycetes(100);o__Saccharomycetales(100);f__Metschnikowiaceae(100);g__Metschnikowia(100);s__Metschnikowia_pulcherrima(100); | Metschnikowia | 13 | Saprotroph | \|Undefined Saprotroph\| | Yeast | NULL | Probable | As nectar-tap_saprotroph (Põlme et al. 2020); Decay sugar-rich substrates (Põlme et al. 2020) | Põlme S, et al. 2020. Fungal Diversity 105:1-16 ((https://doi.org/10.1007/s13225-020-00466-2)) |
| Otu25515 | 1 | k__Fungi(100);p__Ascomycota(100);c__Saccharomycetes(100);o__Saccharomycetales(100);f__Metschnikowiaceae(100);g__Metschnikowia(100);g__Metschnikowia_unclassified(100); | Metschnikowia | 13 | Saprotroph | \|Undefined Saprotroph\| | Yeast | NULL | Probable | As nectar-tap_saprotroph (Põlme et al. 2020); Decay sugar-rich substrates (Põlme et al. 2020) | Põlme S, et al. 2020. Fungal Diversity 105:1-16 ((https://doi.org/10.1007/s13225-020-00466-2)) |
| Otu25516 | 1 | k__Fungi(100);p__Ascomycota(100);c__Saccharomycetes(100);o__Saccharomycetales(100);f__Metschnikowiaceae(100);g__Metschnikowia(100);s__Metschnikowia_pulcherrima(100); | Metschnikowia | 13 | Saprotroph | \|Undefined Saprotroph\| | Yeast | NULL | Probable | As nectar-tap_saprotroph (Põlme et al. 2020); Decay sugar-rich substrates (Põlme et al. 2020) | Põlme S, et al. 2020. Fungal Diversity 105:1-16 ((https://doi.org/10.1007/s13225-020-00466-2)) |
| Otu25517 | 1 | k__Fungi(100);p__Ascomycota(100);c__Saccharomycetes(100);o__Saccharomycetales(100);f__Metschnikowiaceae(100);g__Metschnikowia(100);s__Metschnikowia_pulcherrima(100); | Metschnikowia | 13 | Saprotroph | \|Undefined Saprotroph\| | Yeast | NULL | Probable | As nectar-tap_saprotroph (Põlme et al. 2020); Decay sugar-rich substrates (Põlme et al. 2020) | Põlme S, et al. 2020. Fungal Diversity 105:1-16 ((https://doi.org/10.1007/s13225-020-00466-2)) |
| Otu25518 | 1 | k__Fungi(100);p__Ascomycota(100);c__Saccharomycetes(100);o__Saccharomycetales(100);f__Metschnikowiaceae(100);g__Metschnikowia(100);s__Metschnikowia_pulcherrima(100); | Metschnikowia | 13 | Saprotroph | \|Undefined Saprotroph\| | Yeast | NULL | Probable | As nectar-tap_saprotroph (Põlme et al. 2020); Decay sugar-rich substrates (Põlme et al. 2020) | Põlme S, et al. 2020. Fungal Diversity 105:1-16 ((https://doi.org/10.1007/s13225-020-00466-2)) |
| Otu25520 | 1 | k__Fungi(100);p__Ascomycota(100);c__Saccharomycetes(100);o__Saccharomycetales(100);f__Metschnikowiaceae(100);g__Metschnikowia(100);s__Metschnikowia_pulcherrima(100); | Metschnikowia | 13 | Saprotroph | \|Undefined Saprotroph\| | Yeast | NULL | Probable | As nectar-tap_saprotroph (Põlme et al. 2020); Decay sugar-rich substrates (Põlme et al. 2020) | Põlme S, et al. 2020. Fungal Diversity 105:1-16 ((https://doi.org/10.1007/s13225-020-00466-2)) |
| Otu25521 | 1 | k__Fungi(100);p__Ascomycota(100);c__Saccharomycetes(100);o__Saccharomycetales(100);f__Metschnikowiaceae(100);g__Metschnikowia(100);s__Metschnikowia_pulcherrima(100); | Metschnikowia | 13 | Saprotroph | \|Undefined Saprotroph\| | Yeast | NULL | Probable | As nectar-tap_saprotroph (Põlme et al. 2020); Decay sugar-rich substrates (Põlme et al. 2020) | Põlme S, et al. 2020. Fungal Diversity 105:1-16 ((https://doi.org/10.1007/s13225-020-00466-2)) |
| Otu25522 | 1 | k__Fungi(100);p__Ascomycota(100);c__Saccharomycetes(100);o__Saccharomycetales(100);f__Metschnikowiaceae(100);g__Metschnikowia(100);s__Metschnikowia_pulcherrima(100); | Metschnikowia | 13 | Saprotroph | \|Undefined Saprotroph\| | Yeast | NULL | Probable | As nectar-tap_saprotroph (Põlme et al. 2020); Decay sugar-rich substrates (Põlme et al. 2020) | Põlme S, et al. 2020. Fungal Diversity 105:1-16 ((https://doi.org/10.1007/s13225-020-00466-2)) |
| Otu25524 | 1 | k__Fungi(100);p__Ascomycota(100);c__Saccharomycetes(100);o__Saccharomycetales(100);f__Metschnikowiaceae(100);g__Metschnikowia(100);s__Metschnikowia_pulcherrima(100); | Metschnikowia | 13 | Saprotroph | \|Undefined Saprotroph\| | Yeast | NULL | Probable | As nectar-tap_saprotroph (Põlme et al. 2020); Decay sugar-rich substrates (Põlme et al. 2020) | Põlme S, et al. 2020. Fungal Diversity 105:1-16 ((https://doi.org/10.1007/s13225-020-00466-2)) |
| Otu25525 | 1 | k__Fungi(100);p__Ascomycota(100);c__Saccharomycetes(100);o__Saccharomycetales(100);f__Metschnikowiaceae(100);g__Metschnikowia(100);g__Metschnikowia_unclassified(100); | Metschnikowia | 13 | Saprotroph | \|Undefined Saprotroph\| | Yeast | NULL | Probable | As nectar-tap_saprotroph (Põlme et al. 2020); Decay sugar-rich substrates (Põlme et al. 2020) | Põlme S, et al. 2020. Fungal Diversity 105:1-16 ((https://doi.org/10.1007/s13225-020-00466-2)) |
| Otu25526 | 1 | k__Fungi(100);p__Ascomycota(100);c__Saccharomycetes(100);o__Saccharomycetales(100);f__Metschnikowiaceae(100);g__Metschnikowia(100);g__Metschnikowia_unclassified(100); | Metschnikowia | 13 | Saprotroph | \|Undefined Saprotroph\| | Yeast | NULL | Probable | As nectar-tap_saprotroph (Põlme et al. 2020); Decay sugar-rich substrates (Põlme et al. 2020) | Põlme S, et al. 2020. Fungal Diversity 105:1-16 ((https://doi.org/10.1007/s13225-020-00466-2)) |
| Otu25527 | 1 | k__Fungi(100);p__Ascomycota(100);c__Saccharomycetes(100);o__Saccharomycetales(100);f__Metschnikowiaceae(100);g__Metschnikowia(100);s__Metschnikowia_pulcherrima(100); | Metschnikowia | 13 | Saprotroph | \|Undefined Saprotroph\| | Yeast | NULL | Probable | As nectar-tap_saprotroph (Põlme et al. 2020); Decay sugar-rich substrates (Põlme et al. 2020) | Põlme S, et al. 2020. Fungal Diversity 105:1-16 ((https://doi.org/10.1007/s13225-020-00466-2)) |
| Otu25528 | 1 | k__Fungi(100);p__Ascomycota(100);c__Saccharomycetes(100);o__Saccharomycetales(100);f__Metschnikowiaceae(100);g__Metschnikowia(100);g__Metschnikowia_unclassified(100); | Metschnikowia | 13 | Saprotroph | \|Undefined Saprotroph\| | Yeast | NULL | Probable | As nectar-tap_saprotroph (Põlme et al. 2020); Decay sugar-rich substrates (Põlme et al. 2020) | Põlme S, et al. 2020. Fungal Diversity 105:1-16 ((https://doi.org/10.1007/s13225-020-00466-2)) |
| Otu25530 | 1 | k__Fungi(100);p__Ascomycota(100);c__Saccharomycetes(100);o__Saccharomycetales(100);f__Metschnikowiaceae(100);g__Metschnikowia(100);g__Metschnikowia_unclassified(100); | Metschnikowia | 13 | Saprotroph | \|Undefined Saprotroph\| | Yeast | NULL | Probable | As nectar-tap_saprotroph (Põlme et al. 2020); Decay sugar-rich substrates (Põlme et al. 2020) | Põlme S, et al. 2020. Fungal Diversity 105:1-16 ((https://doi.org/10.1007/s13225-020-00466-2)) |
| Otu25531 | 1 | k__Fungi(100);p__Ascomycota(100);c__Saccharomycetes(100);o__Saccharomycetales(100);f__Metschnikowiaceae(100);g__Metschnikowia(100);g__Metschnikowia_unclassified(100); | Metschnikowia | 13 | Saprotroph | \|Undefined Saprotroph\| | Yeast | NULL | Probable | As nectar-tap_saprotroph (Põlme et al. 2020); Decay sugar-rich substrates (Põlme et al. 2020) | Põlme S, et al. 2020. Fungal Diversity 105:1-16 ((https://doi.org/10.1007/s13225-020-00466-2)) |
| Otu25532 | 1 | k__Fungi(100);p__Ascomycota(100);c__Saccharomycetes(100);o__Saccharomycetales(100);f__Metschnikowiaceae(100);g__Metschnikowia(100);g__Metschnikowia_unclassified(100); | Metschnikowia | 13 | Saprotroph | \|Undefined Saprotroph\| | Yeast | NULL | Probable | As nectar-tap_saprotroph (Põlme et al. 2020); Decay sugar-rich substrates (Põlme et al. 2020) | Põlme S, et al. 2020. Fungal Diversity 105:1-16 ((https://doi.org/10.1007/s13225-020-00466-2)) |
| Otu25534 | 1 | k__Fungi(100);p__Ascomycota(100);c__Saccharomycetes(100);o__Saccharomycetales(100);f__Metschnikowiaceae(100);g__Metschnikowia(100);s__Metschnikowia_pulcherrima(100); | Metschnikowia | 13 | Saprotroph | \|Undefined Saprotroph\| | Yeast | NULL | Probable | As nectar-tap_saprotroph (Põlme et al. 2020); Decay sugar-rich substrates (Põlme et al. 2020) | Põlme S, et al. 2020. Fungal Diversity 105:1-16 ((https://doi.org/10.1007/s13225-020-00466-2)) |
| Otu25535 | 1 | k__Fungi(100);p__Ascomycota(100);c__Saccharomycetes(100);o__Saccharomycetales(100);f__Metschnikowiaceae(100);g__Metschnikowia(100);s__Metschnikowia_pulcherrima(100); | Metschnikowia | 13 | Saprotroph | \|Undefined Saprotroph\| | Yeast | NULL | Probable | As nectar-tap_saprotroph (Põlme et al. 2020); Decay sugar-rich substrates (Põlme et al. 2020) | Põlme S, et al. 2020. Fungal Diversity 105:1-16 ((https://doi.org/10.1007/s13225-020-00466-2)) |
| Otu25536 | 1 | k__Fungi(100);p__Ascomycota(100);c__Saccharomycetes(100);o__Saccharomycetales(100);f__Metschnikowiaceae(100);g__Metschnikowia(100);s__Metschnikowia_pulcherrima(100); | Metschnikowia | 13 | Saprotroph | \|Undefined Saprotroph\| | Yeast | NULL | Probable | As nectar-tap_saprotroph (Põlme et al. 2020); Decay sugar-rich substrates (Põlme et al. 2020) | Põlme S, et al. 2020. Fungal Diversity 105:1-16 ((https://doi.org/10.1007/s13225-020-00466-2)) |
| Otu25537 | 1 | k__Fungi(100);p__Ascomycota(100);c__Saccharomycetes(100);o__Saccharomycetales(100);f__Metschnikowiaceae(100);g__Metschnikowia(100);g__Metschnikowia_unclassified(100); | Metschnikowia | 13 | Saprotroph | \|Undefined Saprotroph\| | Yeast | NULL | Probable | As nectar-tap_saprotroph (Põlme et al. 2020); Decay sugar-rich substrates (Põlme et al. 2020) | Põlme S, et al. 2020. Fungal Diversity 105:1-16 ((https://doi.org/10.1007/s13225-020-00466-2)) |
| Otu25538 | 1 | k__Fungi(100);p__Ascomycota(100);c__Saccharomycetes(100);o__Saccharomycetales(100);f__Metschnikowiaceae(100);g__Metschnikowia(100);s__Metschnikowia_pulcherrima(100); | Metschnikowia | 13 | Saprotroph | \|Undefined Saprotroph\| | Yeast | NULL | Probable | As nectar-tap_saprotroph (Põlme et al. 2020); Decay sugar-rich substrates (Põlme et al. 2020) | Põlme S, et al. 2020. Fungal Diversity 105:1-16 ((https://doi.org/10.1007/s13225-020-00466-2)) |
| Otu25539 | 1 | k__Fungi(100);p__Ascomycota(100);c__Saccharomycetes(100);o__Saccharomycetales(100);f__Metschnikowiaceae(100);g__Metschnikowia(100);g__Metschnikowia_unclassified(100); | Metschnikowia | 13 | Saprotroph | \|Undefined Saprotroph\| | Yeast | NULL | Probable | As nectar-tap_saprotroph (Põlme et al. 2020); Decay sugar-rich substrates (Põlme et al. 2020) | Põlme S, et al. 2020. Fungal Diversity 105:1-16 ((https://doi.org/10.1007/s13225-020-00466-2)) |
| Otu25540 | 1 | k__Fungi(100);p__Ascomycota(100);c__Saccharomycetes(100);o__Saccharomycetales(100);f__Metschnikowiaceae(100);g__Metschnikowia(100);s__Metschnikowia_pulcherrima(100); | Metschnikowia | 13 | Saprotroph | \|Undefined Saprotroph\| | Yeast | NULL | Probable | As nectar-tap_saprotroph (Põlme et al. 2020); Decay sugar-rich substrates (Põlme et al. 2020) | Põlme S, et al. 2020. Fungal Diversity 105:1-16 ((https://doi.org/10.1007/s13225-020-00466-2)) |
| Otu25542 | 1 | k__Fungi(100);p__Ascomycota(100);c__Saccharomycetes(100);o__Saccharomycetales(100);f__Metschnikowiaceae(100);g__Metschnikowia(100);g__Metschnikowia_unclassified(100); | Metschnikowia | 13 | Saprotroph | \|Undefined Saprotroph\| | Yeast | NULL | Probable | As nectar-tap_saprotroph (Põlme et al. 2020); Decay sugar-rich substrates (Põlme et al. 2020) | Põlme S, et al. 2020. Fungal Diversity 105:1-16 ((https://doi.org/10.1007/s13225-020-00466-2)) |
| Otu25543 | 1 | k__Fungi(100);p__Ascomycota(100);c__Saccharomycetes(100);o__Saccharomycetales(100);f__Metschnikowiaceae(100);g__Metschnikowia(100);s__Metschnikowia_pulcherrima(100); | Metschnikowia | 13 | Saprotroph | \|Undefined Saprotroph\| | Yeast | NULL | Probable | As nectar-tap_saprotroph (Põlme et al. 2020); Decay sugar-rich substrates (Põlme et al. 2020) | Põlme S, et al. 2020. Fungal Diversity 105:1-16 ((https://doi.org/10.1007/s13225-020-00466-2)) |
| Otu25545 | 1 | k__Fungi(100);p__Ascomycota(100);c__Saccharomycetes(100);o__Saccharomycetales(100);f__Metschnikowiaceae(100);g__Metschnikowia(100);g__Metschnikowia_unclassified(100); | Metschnikowia | 13 | Saprotroph | \|Undefined Saprotroph\| | Yeast | NULL | Probable | As nectar-tap_saprotroph (Põlme et al. 2020); Decay sugar-rich substrates (Põlme et al. 2020) | Põlme S, et al. 2020. Fungal Diversity 105:1-16 ((https://doi.org/10.1007/s13225-020-00466-2)) |
| Otu25546 | 1 | k__Fungi(100);p__Ascomycota(100);c__Saccharomycetes(100);o__Saccharomycetales(100);f__Metschnikowiaceae(100);g__Metschnikowia(100);s__Metschnikowia_pulcherrima(100); | Metschnikowia | 13 | Saprotroph | \|Undefined Saprotroph\| | Yeast | NULL | Probable | As nectar-tap_saprotroph (Põlme et al. 2020); Decay sugar-rich substrates (Põlme et al. 2020) | Põlme S, et al. 2020. Fungal Diversity 105:1-16 ((https://doi.org/10.1007/s13225-020-00466-2)) |
| Otu25547 | 1 | k__Fungi(100);p__Ascomycota(100);c__Saccharomycetes(100);o__Saccharomycetales(100);f__Metschnikowiaceae(100);g__Metschnikowia(100);s__Metschnikowia_pulcherrima(100); | Metschnikowia | 13 | Saprotroph | \|Undefined Saprotroph\| | Yeast | NULL | Probable | As nectar-tap_saprotroph (Põlme et al. 2020); Decay sugar-rich substrates (Põlme et al. 2020) | Põlme S, et al. 2020. Fungal Diversity 105:1-16 ((https://doi.org/10.1007/s13225-020-00466-2)) |
| Otu25549 | 1 | k__Fungi(100);p__Ascomycota(100);c__Saccharomycetes(100);o__Saccharomycetales(100);f__Metschnikowiaceae(100);g__Metschnikowia(100);s__Metschnikowia_pulcherrima(100); | Metschnikowia | 13 | Saprotroph | \|Undefined Saprotroph\| | Yeast | NULL | Probable | As nectar-tap_saprotroph (Põlme et al. 2020); Decay sugar-rich substrates (Põlme et al. 2020) | Põlme S, et al. 2020. Fungal Diversity 105:1-16 ((https://doi.org/10.1007/s13225-020-00466-2)) |
| Otu25551 | 1 | k__Fungi(100);p__Ascomycota(100);c__Saccharomycetes(100);o__Saccharomycetales(100);f__Metschnikowiaceae(100);g__Metschnikowia(100);s__Metschnikowia_pulcherrima(100); | Metschnikowia | 13 | Saprotroph | \|Undefined Saprotroph\| | Yeast | NULL | Probable | As nectar-tap_saprotroph (Põlme et al. 2020); Decay sugar-rich substrates (Põlme et al. 2020) | Põlme S, et al. 2020. Fungal Diversity 105:1-16 ((https://doi.org/10.1007/s13225-020-00466-2)) |
| Otu25554 | 1 | k__Fungi(100);p__Ascomycota(100);c__Saccharomycetes(100);o__Saccharomycetales(100);f__Metschnikowiaceae(100);g__Metschnikowia(100);s__Metschnikowia_pulcherrima(100); | Metschnikowia | 13 | Saprotroph | \|Undefined Saprotroph\| | Yeast | NULL | Probable | As nectar-tap_saprotroph (Põlme et al. 2020); Decay sugar-rich substrates (Põlme et al. 2020) | Põlme S, et al. 2020. Fungal Diversity 105:1-16 ((https://doi.org/10.1007/s13225-020-00466-2)) |
| Otu25555 | 1 | k__Fungi(100);p__Ascomycota(100);c__Saccharomycetes(100);o__Saccharomycetales(100);f__Metschnikowiaceae(100);g__Metschnikowia(100);g__Metschnikowia_unclassified(100); | Metschnikowia | 13 | Saprotroph | \|Undefined Saprotroph\| | Yeast | NULL | Probable | As nectar-tap_saprotroph (Põlme et al. 2020); Decay sugar-rich substrates (Põlme et al. 2020) | Põlme S, et al. 2020. Fungal Diversity 105:1-16 ((https://doi.org/10.1007/s13225-020-00466-2)) |
| Otu25556 | 1 | k__Fungi(100);p__Ascomycota(100);c__Saccharomycetes(100);o__Saccharomycetales(100);f__Metschnikowiaceae(100);g__Metschnikowia(100);s__Metschnikowia_pulcherrima(100); | Metschnikowia | 13 | Saprotroph | \|Undefined Saprotroph\| | Yeast | NULL | Probable | As nectar-tap_saprotroph (Põlme et al. 2020); Decay sugar-rich substrates (Põlme et al. 2020) | Põlme S, et al. 2020. Fungal Diversity 105:1-16 ((https://doi.org/10.1007/s13225-020-00466-2)) |
| Otu25557 | 1 | k__Fungi(100);p__Ascomycota(100);c__Saccharomycetes(100);o__Saccharomycetales(100);f__Metschnikowiaceae(100);g__Metschnikowia(100);s__Metschnikowia_pulcherrima(100); | Metschnikowia | 13 | Saprotroph | \|Undefined Saprotroph\| | Yeast | NULL | Probable | As nectar-tap_saprotroph (Põlme et al. 2020); Decay sugar-rich substrates (Põlme et al. 2020) | Põlme S, et al. 2020. Fungal Diversity 105:1-16 ((https://doi.org/10.1007/s13225-020-00466-2)) |
| Otu25560 | 1 | k__Fungi(100);p__Ascomycota(100);c__Saccharomycetes(100);o__Saccharomycetales(100);f__Metschnikowiaceae(100);g__Metschnikowia(100);g__Metschnikowia_unclassified(100); | Metschnikowia | 13 | Saprotroph | \|Undefined Saprotroph\| | Yeast | NULL | Probable | As nectar-tap_saprotroph (Põlme et al. 2020); Decay sugar-rich substrates (Põlme et al. 2020) | Põlme S, et al. 2020. Fungal Diversity 105:1-16 ((https://doi.org/10.1007/s13225-020-00466-2)) |
| Otu25561 | 1 | k__Fungi(100);p__Ascomycota(100);c__Saccharomycetes(100);o__Saccharomycetales(100);f__Metschnikowiaceae(100);g__Metschnikowia(100);s__Metschnikowia_pulcherrima(100); | Metschnikowia | 13 | Saprotroph | \|Undefined Saprotroph\| | Yeast | NULL | Probable | As nectar-tap_saprotroph (Põlme et al. 2020); Decay sugar-rich substrates (Põlme et al. 2020) | Põlme S, et al. 2020. Fungal Diversity 105:1-16 ((https://doi.org/10.1007/s13225-020-00466-2)) |
| Otu25562 | 1 | k__Fungi(100);p__Ascomycota(100);c__Saccharomycetes(100);o__Saccharomycetales(100);f__Metschnikowiaceae(100);g__Metschnikowia(100);s__Metschnikowia_pulcherrima(100); | Metschnikowia | 13 | Saprotroph | \|Undefined Saprotroph\| | Yeast | NULL | Probable | As nectar-tap_saprotroph (Põlme et al. 2020); Decay sugar-rich substrates (Põlme et al. 2020) | Põlme S, et al. 2020. Fungal Diversity 105:1-16 ((https://doi.org/10.1007/s13225-020-00466-2)) |
| Otu25563 | 1 | k__Fungi(100);p__Ascomycota(100);c__Saccharomycetes(100);o__Saccharomycetales(100);f__Metschnikowiaceae(100);g__Metschnikowia(100);g__Metschnikowia_unclassified(100); | Metschnikowia | 13 | Saprotroph | \|Undefined Saprotroph\| | Yeast | NULL | Probable | As nectar-tap_saprotroph (Põlme et al. 2020); Decay sugar-rich substrates (Põlme et al. 2020) | Põlme S, et al. 2020. Fungal Diversity 105:1-16 ((https://doi.org/10.1007/s13225-020-00466-2)) |
| Otu25564 | 1 | k__Fungi(100);p__Ascomycota(100);c__Saccharomycetes(100);o__Saccharomycetales(100);f__Metschnikowiaceae(100);g__Metschnikowia(100);s__Metschnikowia_pulcherrima(100); | Metschnikowia | 13 | Saprotroph | \|Undefined Saprotroph\| | Yeast | NULL | Probable | As nectar-tap_saprotroph (Põlme et al. 2020); Decay sugar-rich substrates (Põlme et al. 2020) | Põlme S, et al. 2020. Fungal Diversity 105:1-16 ((https://doi.org/10.1007/s13225-020-00466-2)) |
| Otu25565 | 1 | k__Fungi(100);p__Ascomycota(100);c__Saccharomycetes(100);o__Saccharomycetales(100);f__Metschnikowiaceae(100);g__Metschnikowia(100);s__Metschnikowia_pulcherrima(100); | Metschnikowia | 13 | Saprotroph | \|Undefined Saprotroph\| | Yeast | NULL | Probable | As nectar-tap_saprotroph (Põlme et al. 2020); Decay sugar-rich substrates (Põlme et al. 2020) | Põlme S, et al. 2020. Fungal Diversity 105:1-16 ((https://doi.org/10.1007/s13225-020-00466-2)) |
| Otu25566 | 1 | k__Fungi(100);p__Ascomycota(100);c__Saccharomycetes(100);o__Saccharomycetales(100);f__Metschnikowiaceae(100);g__Metschnikowia(100);g__Metschnikowia_unclassified(100); | Metschnikowia | 13 | Saprotroph | \|Undefined Saprotroph\| | Yeast | NULL | Probable | As nectar-tap_saprotroph (Põlme et al. 2020); Decay sugar-rich substrates (Põlme et al. 2020) | Põlme S, et al. 2020. Fungal Diversity 105:1-16 ((https://doi.org/10.1007/s13225-020-00466-2)) |
| Otu25567 | 1 | k__Fungi(100);p__Ascomycota(100);c__Saccharomycetes(100);o__Saccharomycetales(100);f__Metschnikowiaceae(100);g__Metschnikowia(100);s__Metschnikowia_pulcherrima(100); | Metschnikowia | 13 | Saprotroph | \|Undefined Saprotroph\| | Yeast | NULL | Probable | As nectar-tap_saprotroph (Põlme et al. 2020); Decay sugar-rich substrates (Põlme et al. 2020) | Põlme S, et al. 2020. Fungal Diversity 105:1-16 ((https://doi.org/10.1007/s13225-020-00466-2)) |
| Otu25587 | 1 | k__Fungi(100);p__Ascomycota(100);c__Saccharomycetes(100);o__Saccharomycetales(100);f__Metschnikowiaceae(100);g__Metschnikowia(100);s__Metschnikowia_pulcherrima(100); | Metschnikowia | 13 | Saprotroph | \|Undefined Saprotroph\| | Yeast | NULL | Probable | As nectar-tap_saprotroph (Põlme et al. 2020); Decay sugar-rich substrates (Põlme et al. 2020) | Põlme S, et al. 2020. Fungal Diversity 105:1-16 ((https://doi.org/10.1007/s13225-020-00466-2)) |
| Otu25591 | 1 | k__Fungi(100);p__Ascomycota(100);c__Saccharomycetes(100);o__Saccharomycetales(100);f__Metschnikowiaceae(100);g__Metschnikowia(100);g__Metschnikowia_unclassified(100); | Metschnikowia | 13 | Saprotroph | \|Undefined Saprotroph\| | Yeast | NULL | Probable | As nectar-tap_saprotroph (Põlme et al. 2020); Decay sugar-rich substrates (Põlme et al. 2020) | Põlme S, et al. 2020. Fungal Diversity 105:1-16 ((https://doi.org/10.1007/s13225-020-00466-2)) |
| Otu25593 | 1 | k__Fungi(100);p__Basidiomycota(100);c__Agaricomycetes(100);o__Polyporales(100);f__Ganodermataceae(100);g__Ganoderma(100);g__Ganoderma_unclassified(100); | Ganoderma | 13 | Pathotroph-Saprotroph | \|Plant Pathogen\|-Wood Saprotroph | Polyporoid | White Rot | Highly Probable | As white_rot (Põlme et al. 2020); As wood_pathogen (Põlme et al. 2020); Decay wood (Põlme et al. 2020); Lignicolous, causing sometimes agressive white rots of timber (Cannon & Kirk 2007) | Gilbertson RL, Ryvarden L. 1987-1987. North American Polypores. Fungiflora, Oslo (ISBN: 978-0945345060); Cannon PF, Kirk PM. 2007. Fungal Families of the World. CAB International, Cambridge (ISBN: 978-0851998275); Põlme S, et al. 2020. Fungal Diversity 105:1-16 ((https://doi.org/10.1007/s13225-020-00466-2)) |
| Otu25594 | 1 | k__Fungi(100);p__Ascomycota(100);c__Saccharomycetes(100);o__Saccharomycetales(100);f__Metschnikowiaceae(100);g__Metschnikowia(100);g__Metschnikowia_unclassified(100); | Metschnikowia | 13 | Saprotroph | \|Undefined Saprotroph\| | Yeast | NULL | Probable | As nectar-tap_saprotroph (Põlme et al. 2020); Decay sugar-rich substrates (Põlme et al. 2020) | Põlme S, et al. 2020. Fungal Diversity 105:1-16 ((https://doi.org/10.1007/s13225-020-00466-2)) |
| Otu25608 | 1 | k__Fungi(100);p__Ascomycota(100);c__Saccharomycetes(100);o__Saccharomycetales(100);f__Metschnikowiaceae(100);g__Metschnikowia(100);s__Metschnikowia_pulcherrima(100); | Metschnikowia | 13 | Saprotroph | \|Undefined Saprotroph\| | Yeast | NULL | Probable | As nectar-tap_saprotroph (Põlme et al. 2020); Decay sugar-rich substrates (Põlme et al. 2020) | Põlme S, et al. 2020. Fungal Diversity 105:1-16 ((https://doi.org/10.1007/s13225-020-00466-2)) |
| Otu25609 | 1 | k__Fungi(100);p__Ascomycota(100);c__Saccharomycetes(100);o__Saccharomycetales(100);f__Metschnikowiaceae(100);g__Metschnikowia(100);s__Metschnikowia_pulcherrima(100); | Metschnikowia | 13 | Saprotroph | \|Undefined Saprotroph\| | Yeast | NULL | Probable | As nectar-tap_saprotroph (Põlme et al. 2020); Decay sugar-rich substrates (Põlme et al. 2020) | Põlme S, et al. 2020. Fungal Diversity 105:1-16 ((https://doi.org/10.1007/s13225-020-00466-2)) |
| Otu25610 | 1 | k__Fungi(100);p__Ascomycota(100);c__Saccharomycetes(100);o__Saccharomycetales(100);f__Metschnikowiaceae(100);g__Metschnikowia(100);s__Metschnikowia_pulcherrima(100); | Metschnikowia | 13 | Saprotroph | \|Undefined Saprotroph\| | Yeast | NULL | Probable | As nectar-tap_saprotroph (Põlme et al. 2020); Decay sugar-rich substrates (Põlme et al. 2020) | Põlme S, et al. 2020. Fungal Diversity 105:1-16 ((https://doi.org/10.1007/s13225-020-00466-2)) |
| Otu25611 | 1 | k__Fungi(100);p__Ascomycota(100);c__Saccharomycetes(100);o__Saccharomycetales(100);f__Metschnikowiaceae(100);g__Metschnikowia(100);s__Metschnikowia_pulcherrima(100); | Metschnikowia | 13 | Saprotroph | \|Undefined Saprotroph\| | Yeast | NULL | Probable | As nectar-tap_saprotroph (Põlme et al. 2020); Decay sugar-rich substrates (Põlme et al. 2020) | Põlme S, et al. 2020. Fungal Diversity 105:1-16 ((https://doi.org/10.1007/s13225-020-00466-2)) |
| Otu25614 | 1 | k__Fungi(100);p__Ascomycota(100);c__Saccharomycetes(100);o__Saccharomycetales(100);f__Metschnikowiaceae(100);g__Metschnikowia(100);g__Metschnikowia_unclassified(100); | Metschnikowia | 13 | Saprotroph | \|Undefined Saprotroph\| | Yeast | NULL | Probable | As nectar-tap_saprotroph (Põlme et al. 2020); Decay sugar-rich substrates (Põlme et al. 2020) | Põlme S, et al. 2020. Fungal Diversity 105:1-16 ((https://doi.org/10.1007/s13225-020-00466-2)) |
| Otu25620 | 1 | k__Fungi(100);p__Ascomycota(100);c__Saccharomycetes(100);o__Saccharomycetales(100);f__Metschnikowiaceae(100);g__Metschnikowia(100);g__Metschnikowia_unclassified(100); | Metschnikowia | 13 | Saprotroph | \|Undefined Saprotroph\| | Yeast | NULL | Probable | As nectar-tap_saprotroph (Põlme et al. 2020); Decay sugar-rich substrates (Põlme et al. 2020) | Põlme S, et al. 2020. Fungal Diversity 105:1-16 ((https://doi.org/10.1007/s13225-020-00466-2)) |
| Otu25622 | 1 | k__Fungi(100);p__Ascomycota(100);c__Saccharomycetes(100);o__Saccharomycetales(100);f__Metschnikowiaceae(100);g__Metschnikowia(100);s__Metschnikowia_pulcherrima(100); | Metschnikowia | 13 | Saprotroph | \|Undefined Saprotroph\| | Yeast | NULL | Probable | As nectar-tap_saprotroph (Põlme et al. 2020); Decay sugar-rich substrates (Põlme et al. 2020) | Põlme S, et al. 2020. Fungal Diversity 105:1-16 ((https://doi.org/10.1007/s13225-020-00466-2)) |
| Otu25626 | 1 | k__Fungi(100);p__Ascomycota(100);c__Saccharomycetes(100);o__Saccharomycetales(100);f__Metschnikowiaceae(100);g__Metschnikowia(100);s__Metschnikowia_pulcherrima(100); | Metschnikowia | 13 | Saprotroph | \|Undefined Saprotroph\| | Yeast | NULL | Probable | As nectar-tap_saprotroph (Põlme et al. 2020); Decay sugar-rich substrates (Põlme et al. 2020) | Põlme S, et al. 2020. Fungal Diversity 105:1-16 ((https://doi.org/10.1007/s13225-020-00466-2)) |
| Otu25627 | 1 | k__Fungi(100);p__Ascomycota(100);c__Saccharomycetes(100);o__Saccharomycetales(100);f__Metschnikowiaceae(100);g__Metschnikowia(100);s__Metschnikowia_pulcherrima(100); | Metschnikowia | 13 | Saprotroph | \|Undefined Saprotroph\| | Yeast | NULL | Probable | As nectar-tap_saprotroph (Põlme et al. 2020); Decay sugar-rich substrates (Põlme et al. 2020) | Põlme S, et al. 2020. Fungal Diversity 105:1-16 ((https://doi.org/10.1007/s13225-020-00466-2)) |
| Otu25628 | 1 | k__Fungi(100);p__Ascomycota(100);c__Saccharomycetes(100);o__Saccharomycetales(100);f__Metschnikowiaceae(100);g__Metschnikowia(100);s__Metschnikowia_pulcherrima(100); | Metschnikowia | 13 | Saprotroph | \|Undefined Saprotroph\| | Yeast | NULL | Probable | As nectar-tap_saprotroph (Põlme et al. 2020); Decay sugar-rich substrates (Põlme et al. 2020) | Põlme S, et al. 2020. Fungal Diversity 105:1-16 ((https://doi.org/10.1007/s13225-020-00466-2)) |
| Otu25629 | 1 | k__Fungi(100);p__Ascomycota(100);c__Saccharomycetes(100);o__Saccharomycetales(100);f__Metschnikowiaceae(100);g__Metschnikowia(100);s__Metschnikowia_pulcherrima(100); | Metschnikowia | 13 | Saprotroph | \|Undefined Saprotroph\| | Yeast | NULL | Probable | As nectar-tap_saprotroph (Põlme et al. 2020); Decay sugar-rich substrates (Põlme et al. 2020) | Põlme S, et al. 2020. Fungal Diversity 105:1-16 ((https://doi.org/10.1007/s13225-020-00466-2)) |
| Otu25630 | 1 | k__Fungi(100);p__Ascomycota(100);c__Saccharomycetes(100);o__Saccharomycetales(100);f__Metschnikowiaceae(100);g__Metschnikowia(100);s__Metschnikowia_pulcherrima(100); | Metschnikowia | 13 | Saprotroph | \|Undefined Saprotroph\| | Yeast | NULL | Probable | As nectar-tap_saprotroph (Põlme et al. 2020); Decay sugar-rich substrates (Põlme et al. 2020) | Põlme S, et al. 2020. Fungal Diversity 105:1-16 ((https://doi.org/10.1007/s13225-020-00466-2)) |
| Otu25631 | 1 | k__Fungi(100);p__Ascomycota(100);c__Saccharomycetes(100);o__Saccharomycetales(100);f__Metschnikowiaceae(100);g__Metschnikowia(100);s__Metschnikowia_pulcherrima(100); | Metschnikowia | 13 | Saprotroph | \|Undefined Saprotroph\| | Yeast | NULL | Probable | As nectar-tap_saprotroph (Põlme et al. 2020); Decay sugar-rich substrates (Põlme et al. 2020) | Põlme S, et al. 2020. Fungal Diversity 105:1-16 ((https://doi.org/10.1007/s13225-020-00466-2)) |
| Otu25633 | 1 | k__Fungi(100);p__Ascomycota(100);c__Saccharomycetes(100);o__Saccharomycetales(100);f__Metschnikowiaceae(100);g__Metschnikowia(100);s__Metschnikowia_pulcherrima(100); | Metschnikowia | 13 | Saprotroph | \|Undefined Saprotroph\| | Yeast | NULL | Probable | As nectar-tap_saprotroph (Põlme et al. 2020); Decay sugar-rich substrates (Põlme et al. 2020) | Põlme S, et al. 2020. Fungal Diversity 105:1-16 ((https://doi.org/10.1007/s13225-020-00466-2)) |
| Otu25635 | 1 | k__Fungi(100);p__Ascomycota(100);c__Saccharomycetes(100);o__Saccharomycetales(100);f__Metschnikowiaceae(100);g__Metschnikowia(100);g__Metschnikowia_unclassified(100); | Metschnikowia | 13 | Saprotroph | \|Undefined Saprotroph\| | Yeast | NULL | Probable | As nectar-tap_saprotroph (Põlme et al. 2020); Decay sugar-rich substrates (Põlme et al. 2020) | Põlme S, et al. 2020. Fungal Diversity 105:1-16 ((https://doi.org/10.1007/s13225-020-00466-2)) |
| Otu25636 | 1 | k__Fungi(100);p__Ascomycota(100);c__Saccharomycetes(100);o__Saccharomycetales(100);f__Metschnikowiaceae(100);g__Metschnikowia(100);g__Metschnikowia_unclassified(100); | Metschnikowia | 13 | Saprotroph | \|Undefined Saprotroph\| | Yeast | NULL | Probable | As nectar-tap_saprotroph (Põlme et al. 2020); Decay sugar-rich substrates (Põlme et al. 2020) | Põlme S, et al. 2020. Fungal Diversity 105:1-16 ((https://doi.org/10.1007/s13225-020-00466-2)) |
| Otu25637 | 1 | k__Fungi(100);p__Ascomycota(100);c__Saccharomycetes(100);o__Saccharomycetales(100);f__Metschnikowiaceae(100);g__Metschnikowia(100);g__Metschnikowia_unclassified(100); | Metschnikowia | 13 | Saprotroph | \|Undefined Saprotroph\| | Yeast | NULL | Probable | As nectar-tap_saprotroph (Põlme et al. 2020); Decay sugar-rich substrates (Põlme et al. 2020) | Põlme S, et al. 2020. Fungal Diversity 105:1-16 ((https://doi.org/10.1007/s13225-020-00466-2)) |
| Otu25638 | 1 | k__Fungi(100);p__Ascomycota(100);c__Saccharomycetes(100);o__Saccharomycetales(100);f__Metschnikowiaceae(100);g__Metschnikowia(100);s__Metschnikowia_pulcherrima(100); | Metschnikowia | 13 | Saprotroph | \|Undefined Saprotroph\| | Yeast | NULL | Probable | As nectar-tap_saprotroph (Põlme et al. 2020); Decay sugar-rich substrates (Põlme et al. 2020) | Põlme S, et al. 2020. Fungal Diversity 105:1-16 ((https://doi.org/10.1007/s13225-020-00466-2)) |
| Otu25640 | 1 | k__Fungi(100);p__Ascomycota(100);c__Saccharomycetes(100);o__Saccharomycetales(100);f__Metschnikowiaceae(100);g__Metschnikowia(100);g__Metschnikowia_unclassified(100); | Metschnikowia | 13 | Saprotroph | \|Undefined Saprotroph\| | Yeast | NULL | Probable | As nectar-tap_saprotroph (Põlme et al. 2020); Decay sugar-rich substrates (Põlme et al. 2020) | Põlme S, et al. 2020. Fungal Diversity 105:1-16 ((https://doi.org/10.1007/s13225-020-00466-2)) |
| Otu25641 | 1 | k__Fungi(100);p__Ascomycota(100);c__Saccharomycetes(100);o__Saccharomycetales(100);f__Metschnikowiaceae(100);g__Metschnikowia(100);s__Metschnikowia_pulcherrima(100); | Metschnikowia | 13 | Saprotroph | \|Undefined Saprotroph\| | Yeast | NULL | Probable | As nectar-tap_saprotroph (Põlme et al. 2020); Decay sugar-rich substrates (Põlme et al. 2020) | Põlme S, et al. 2020. Fungal Diversity 105:1-16 ((https://doi.org/10.1007/s13225-020-00466-2)) |
| Otu25642 | 1 | k__Fungi(100);p__Ascomycota(100);c__Saccharomycetes(100);o__Saccharomycetales(100);f__Metschnikowiaceae(100);g__Metschnikowia(100);g__Metschnikowia_unclassified(100); | Metschnikowia | 13 | Saprotroph | \|Undefined Saprotroph\| | Yeast | NULL | Probable | As nectar-tap_saprotroph (Põlme et al. 2020); Decay sugar-rich substrates (Põlme et al. 2020) | Põlme S, et al. 2020. Fungal Diversity 105:1-16 ((https://doi.org/10.1007/s13225-020-00466-2)) |
| Otu25643 | 1 | k__Fungi(100);p__Ascomycota(100);c__Saccharomycetes(100);o__Saccharomycetales(100);f__Metschnikowiaceae(100);g__Metschnikowia(100);g__Metschnikowia_unclassified(100); | Metschnikowia | 13 | Saprotroph | \|Undefined Saprotroph\| | Yeast | NULL | Probable | As nectar-tap_saprotroph (Põlme et al. 2020); Decay sugar-rich substrates (Põlme et al. 2020) | Põlme S, et al. 2020. Fungal Diversity 105:1-16 ((https://doi.org/10.1007/s13225-020-00466-2)) |
| Otu25644 | 1 | k__Fungi(100);p__Ascomycota(100);c__Saccharomycetes(100);o__Saccharomycetales(100);f__Metschnikowiaceae(100);g__Metschnikowia(100);s__Metschnikowia_pulcherrima(100); | Metschnikowia | 13 | Saprotroph | \|Undefined Saprotroph\| | Yeast | NULL | Probable | As nectar-tap_saprotroph (Põlme et al. 2020); Decay sugar-rich substrates (Põlme et al. 2020) | Põlme S, et al. 2020. Fungal Diversity 105:1-16 ((https://doi.org/10.1007/s13225-020-00466-2)) |
| Otu25645 | 1 | k__Fungi(100);p__Ascomycota(100);c__Saccharomycetes(100);o__Saccharomycetales(100);f__Metschnikowiaceae(100);g__Metschnikowia(100);s__Metschnikowia_pulcherrima(100); | Metschnikowia | 13 | Saprotroph | \|Undefined Saprotroph\| | Yeast | NULL | Probable | As nectar-tap_saprotroph (Põlme et al. 2020); Decay sugar-rich substrates (Põlme et al. 2020) | Põlme S, et al. 2020. Fungal Diversity 105:1-16 ((https://doi.org/10.1007/s13225-020-00466-2)) |
| Otu25646 | 1 | k__Fungi(100);p__Ascomycota(100);c__Saccharomycetes(100);o__Saccharomycetales(100);f__Metschnikowiaceae(100);g__Metschnikowia(100);g__Metschnikowia_unclassified(100); | Metschnikowia | 13 | Saprotroph | \|Undefined Saprotroph\| | Yeast | NULL | Probable | As nectar-tap_saprotroph (Põlme et al. 2020); Decay sugar-rich substrates (Põlme et al. 2020) | Põlme S, et al. 2020. Fungal Diversity 105:1-16 ((https://doi.org/10.1007/s13225-020-00466-2)) |
| Otu25651 | 1 | k__Fungi(100);p__Ascomycota(100);c__Saccharomycetes(100);o__Saccharomycetales(100);f__Metschnikowiaceae(100);g__Metschnikowia(100);s__Metschnikowia_pulcherrima(100); | Metschnikowia | 13 | Saprotroph | \|Undefined Saprotroph\| | Yeast | NULL | Probable | As nectar-tap_saprotroph (Põlme et al. 2020); Decay sugar-rich substrates (Põlme et al. 2020) | Põlme S, et al. 2020. Fungal Diversity 105:1-16 ((https://doi.org/10.1007/s13225-020-00466-2)) |
| Otu25653 | 1 | k__Fungi(100);p__Ascomycota(100);c__Saccharomycetes(100);o__Saccharomycetales(100);f__Metschnikowiaceae(100);g__Metschnikowia(100);s__Metschnikowia_pulcherrima(100); | Metschnikowia | 13 | Saprotroph | \|Undefined Saprotroph\| | Yeast | NULL | Probable | As nectar-tap_saprotroph (Põlme et al. 2020); Decay sugar-rich substrates (Põlme et al. 2020) | Põlme S, et al. 2020. Fungal Diversity 105:1-16 ((https://doi.org/10.1007/s13225-020-00466-2)) |
| Otu25654 | 1 | k__Fungi(100);p__Ascomycota(100);c__Saccharomycetes(100);o__Saccharomycetales(100);f__Metschnikowiaceae(100);g__Metschnikowia(100);g__Metschnikowia_unclassified(100); | Metschnikowia | 13 | Saprotroph | \|Undefined Saprotroph\| | Yeast | NULL | Probable | As nectar-tap_saprotroph (Põlme et al. 2020); Decay sugar-rich substrates (Põlme et al. 2020) | Põlme S, et al. 2020. Fungal Diversity 105:1-16 ((https://doi.org/10.1007/s13225-020-00466-2)) |
| Otu25655 | 1 | k__Fungi(100);p__Ascomycota(100);c__Saccharomycetes(100);o__Saccharomycetales(100);f__Metschnikowiaceae(100);g__Metschnikowia(100);g__Metschnikowia_unclassified(100); | Metschnikowia | 13 | Saprotroph | \|Undefined Saprotroph\| | Yeast | NULL | Probable | As nectar-tap_saprotroph (Põlme et al. 2020); Decay sugar-rich substrates (Põlme et al. 2020) | Põlme S, et al. 2020. Fungal Diversity 105:1-16 ((https://doi.org/10.1007/s13225-020-00466-2)) |
| Otu25656 | 1 | k__Fungi(100);p__Ascomycota(100);c__Saccharomycetes(100);o__Saccharomycetales(100);f__Metschnikowiaceae(100);g__Metschnikowia(100);s__Metschnikowia_pulcherrima(100); | Metschnikowia | 13 | Saprotroph | \|Undefined Saprotroph\| | Yeast | NULL | Probable | As nectar-tap_saprotroph (Põlme et al. 2020); Decay sugar-rich substrates (Põlme et al. 2020) | Põlme S, et al. 2020. Fungal Diversity 105:1-16 ((https://doi.org/10.1007/s13225-020-00466-2)) |
| Otu25657 | 1 | k__Fungi(100);p__Ascomycota(100);c__Saccharomycetes(100);o__Saccharomycetales(100);f__Metschnikowiaceae(100);g__Metschnikowia(100);s__Metschnikowia_pulcherrima(100); | Metschnikowia | 13 | Saprotroph | \|Undefined Saprotroph\| | Yeast | NULL | Probable | As nectar-tap_saprotroph (Põlme et al. 2020); Decay sugar-rich substrates (Põlme et al. 2020) | Põlme S, et al. 2020. Fungal Diversity 105:1-16 ((https://doi.org/10.1007/s13225-020-00466-2)) |
| Otu25659 | 1 | k__Fungi(100);p__Ascomycota(100);c__Saccharomycetes(100);o__Saccharomycetales(100);f__Metschnikowiaceae(100);g__Metschnikowia(100);s__Metschnikowia_pulcherrima(100); | Metschnikowia | 13 | Saprotroph | \|Undefined Saprotroph\| | Yeast | NULL | Probable | As nectar-tap_saprotroph (Põlme et al. 2020); Decay sugar-rich substrates (Põlme et al. 2020) | Põlme S, et al. 2020. Fungal Diversity 105:1-16 ((https://doi.org/10.1007/s13225-020-00466-2)) |
| Otu25660 | 1 | k__Fungi(100);p__Ascomycota(100);c__Saccharomycetes(100);o__Saccharomycetales(100);f__Metschnikowiaceae(100);g__Metschnikowia(100);s__Metschnikowia_pulcherrima(100); | Metschnikowia | 13 | Saprotroph | \|Undefined Saprotroph\| | Yeast | NULL | Probable | As nectar-tap_saprotroph (Põlme et al. 2020); Decay sugar-rich substrates (Põlme et al. 2020) | Põlme S, et al. 2020. Fungal Diversity 105:1-16 ((https://doi.org/10.1007/s13225-020-00466-2)) |
| Otu25666 | 1 | k__Fungi(100);p__Ascomycota(100);c__Saccharomycetes(100);o__Saccharomycetales(100);f__Metschnikowiaceae(100);g__Metschnikowia(100);s__Metschnikowia_pulcherrima(100); | Metschnikowia | 13 | Saprotroph | \|Undefined Saprotroph\| | Yeast | NULL | Probable | As nectar-tap_saprotroph (Põlme et al. 2020); Decay sugar-rich substrates (Põlme et al. 2020) | Põlme S, et al. 2020. Fungal Diversity 105:1-16 ((https://doi.org/10.1007/s13225-020-00466-2)) |
| Otu25669 | 1 | k__Fungi(100);p__Ascomycota(100);c__Saccharomycetes(100);o__Saccharomycetales(100);f__Metschnikowiaceae(100);g__Metschnikowia(100);s__Metschnikowia_pulcherrima(100); | Metschnikowia | 13 | Saprotroph | \|Undefined Saprotroph\| | Yeast | NULL | Probable | As nectar-tap_saprotroph (Põlme et al. 2020); Decay sugar-rich substrates (Põlme et al. 2020) | Põlme S, et al. 2020. Fungal Diversity 105:1-16 ((https://doi.org/10.1007/s13225-020-00466-2)) |
| Otu25675 | 1 | k__Fungi(100);p__Ascomycota(100);c__Saccharomycetes(100);o__Saccharomycetales(100);f__Metschnikowiaceae(100);g__Metschnikowia(100);s__Metschnikowia_pulcherrima(100); | Metschnikowia | 13 | Saprotroph | \|Undefined Saprotroph\| | Yeast | NULL | Probable | As nectar-tap_saprotroph (Põlme et al. 2020); Decay sugar-rich substrates (Põlme et al. 2020) | Põlme S, et al. 2020. Fungal Diversity 105:1-16 ((https://doi.org/10.1007/s13225-020-00466-2)) |
| Otu25697 | 1 | k__Fungi(100);p__Ascomycota(100);c__Saccharomycetes(100);o__Saccharomycetales(100);f__Metschnikowiaceae(100);g__Metschnikowia(100);s__Metschnikowia_pulcherrima(100); | Metschnikowia | 13 | Saprotroph | \|Undefined Saprotroph\| | Yeast | NULL | Probable | As nectar-tap_saprotroph (Põlme et al. 2020); Decay sugar-rich substrates (Põlme et al. 2020) | Põlme S, et al. 2020. Fungal Diversity 105:1-16 ((https://doi.org/10.1007/s13225-020-00466-2)) |
| Otu25698 | 1 | k__Fungi(100);p__Ascomycota(100);c__Saccharomycetes(100);o__Saccharomycetales(100);f__Metschnikowiaceae(100);g__Metschnikowia(100);s__Metschnikowia_pulcherrima(100); | Metschnikowia | 13 | Saprotroph | \|Undefined Saprotroph\| | Yeast | NULL | Probable | As nectar-tap_saprotroph (Põlme et al. 2020); Decay sugar-rich substrates (Põlme et al. 2020) | Põlme S, et al. 2020. Fungal Diversity 105:1-16 ((https://doi.org/10.1007/s13225-020-00466-2)) |
| Otu25700 | 1 | k__Fungi(100);p__Ascomycota(100);c__Saccharomycetes(100);o__Saccharomycetales(100);f__Metschnikowiaceae(100);g__Metschnikowia(100);s__Metschnikowia_pulcherrima(100); | Metschnikowia | 13 | Saprotroph | \|Undefined Saprotroph\| | Yeast | NULL | Probable | As nectar-tap_saprotroph (Põlme et al. 2020); Decay sugar-rich substrates (Põlme et al. 2020) | Põlme S, et al. 2020. Fungal Diversity 105:1-16 ((https://doi.org/10.1007/s13225-020-00466-2)) |
| Otu25702 | 1 | k__Fungi(100);p__Ascomycota(100);c__Saccharomycetes(100);o__Saccharomycetales(100);f__Metschnikowiaceae(100);g__Metschnikowia(100);s__Metschnikowia_pulcherrima(100); | Metschnikowia | 13 | Saprotroph | \|Undefined Saprotroph\| | Yeast | NULL | Probable | As nectar-tap_saprotroph (Põlme et al. 2020); Decay sugar-rich substrates (Põlme et al. 2020) | Põlme S, et al. 2020. Fungal Diversity 105:1-16 ((https://doi.org/10.1007/s13225-020-00466-2)) |
| Otu25703 | 1 | k__Fungi(100);p__Ascomycota(100);c__Saccharomycetes(100);o__Saccharomycetales(100);f__Metschnikowiaceae(100);g__Metschnikowia(100);s__Metschnikowia_pulcherrima(100); | Metschnikowia | 13 | Saprotroph | \|Undefined Saprotroph\| | Yeast | NULL | Probable | As nectar-tap_saprotroph (Põlme et al. 2020); Decay sugar-rich substrates (Põlme et al. 2020) | Põlme S, et al. 2020. Fungal Diversity 105:1-16 ((https://doi.org/10.1007/s13225-020-00466-2)) |
| Otu25704 | 1 | k__Fungi(100);p__Ascomycota(100);c__Saccharomycetes(100);o__Saccharomycetales(100);f__Metschnikowiaceae(100);g__Metschnikowia(100);s__Metschnikowia_pulcherrima(100); | Metschnikowia | 13 | Saprotroph | \|Undefined Saprotroph\| | Yeast | NULL | Probable | As nectar-tap_saprotroph (Põlme et al. 2020); Decay sugar-rich substrates (Põlme et al. 2020) | Põlme S, et al. 2020. Fungal Diversity 105:1-16 ((https://doi.org/10.1007/s13225-020-00466-2)) |
| Otu25705 | 1 | k__Fungi(100);p__Ascomycota(100);c__Saccharomycetes(100);o__Saccharomycetales(100);f__Metschnikowiaceae(100);g__Metschnikowia(100);s__Metschnikowia_pulcherrima(100); | Metschnikowia | 13 | Saprotroph | \|Undefined Saprotroph\| | Yeast | NULL | Probable | As nectar-tap_saprotroph (Põlme et al. 2020); Decay sugar-rich substrates (Põlme et al. 2020) | Põlme S, et al. 2020. Fungal Diversity 105:1-16 ((https://doi.org/10.1007/s13225-020-00466-2)) |
| Otu25706 | 1 | k__Fungi(100);p__Ascomycota(100);c__Saccharomycetes(100);o__Saccharomycetales(100);f__Metschnikowiaceae(100);g__Metschnikowia(100);g__Metschnikowia_unclassified(100); | Metschnikowia | 13 | Saprotroph | \|Undefined Saprotroph\| | Yeast | NULL | Probable | As nectar-tap_saprotroph (Põlme et al. 2020); Decay sugar-rich substrates (Põlme et al. 2020) | Põlme S, et al. 2020. Fungal Diversity 105:1-16 ((https://doi.org/10.1007/s13225-020-00466-2)) |
| Otu25707 | 1 | k__Fungi(100);p__Ascomycota(100);c__Saccharomycetes(100);o__Saccharomycetales(100);f__Metschnikowiaceae(100);g__Metschnikowia(100);s__Metschnikowia_pulcherrima(100); | Metschnikowia | 13 | Saprotroph | \|Undefined Saprotroph\| | Yeast | NULL | Probable | As nectar-tap_saprotroph (Põlme et al. 2020); Decay sugar-rich substrates (Põlme et al. 2020) | Põlme S, et al. 2020. Fungal Diversity 105:1-16 ((https://doi.org/10.1007/s13225-020-00466-2)) |
| Otu25708 | 1 | k__Fungi(100);p__Ascomycota(100);c__Saccharomycetes(100);o__Saccharomycetales(100);f__Metschnikowiaceae(100);g__Metschnikowia(100);s__Metschnikowia_pulcherrima(100); | Metschnikowia | 13 | Saprotroph | \|Undefined Saprotroph\| | Yeast | NULL | Probable | As nectar-tap_saprotroph (Põlme et al. 2020); Decay sugar-rich substrates (Põlme et al. 2020) | Põlme S, et al. 2020. Fungal Diversity 105:1-16 ((https://doi.org/10.1007/s13225-020-00466-2)) |
| Otu25709 | 1 | k__Fungi(100);p__Ascomycota(100);c__Saccharomycetes(100);o__Saccharomycetales(100);f__Metschnikowiaceae(100);g__Metschnikowia(100);s__Metschnikowia_pulcherrima(100); | Metschnikowia | 13 | Saprotroph | \|Undefined Saprotroph\| | Yeast | NULL | Probable | As nectar-tap_saprotroph (Põlme et al. 2020); Decay sugar-rich substrates (Põlme et al. 2020) | Põlme S, et al. 2020. Fungal Diversity 105:1-16 ((https://doi.org/10.1007/s13225-020-00466-2)) |
| Otu25710 | 1 | k__Fungi(100);p__Ascomycota(100);c__Saccharomycetes(100);o__Saccharomycetales(100);f__Metschnikowiaceae(100);g__Metschnikowia(100);s__Metschnikowia_pulcherrima(100); | Metschnikowia | 13 | Saprotroph | \|Undefined Saprotroph\| | Yeast | NULL | Probable | As nectar-tap_saprotroph (Põlme et al. 2020); Decay sugar-rich substrates (Põlme et al. 2020) | Põlme S, et al. 2020. Fungal Diversity 105:1-16 ((https://doi.org/10.1007/s13225-020-00466-2)) |
| Otu25711 | 1 | k__Fungi(100);p__Ascomycota(100);c__Saccharomycetes(100);o__Saccharomycetales(100);f__Metschnikowiaceae(100);g__Metschnikowia(100);g__Metschnikowia_unclassified(100); | Metschnikowia | 13 | Saprotroph | \|Undefined Saprotroph\| | Yeast | NULL | Probable | As nectar-tap_saprotroph (Põlme et al. 2020); Decay sugar-rich substrates (Põlme et al. 2020) | Põlme S, et al. 2020. Fungal Diversity 105:1-16 ((https://doi.org/10.1007/s13225-020-00466-2)) |
| Otu25712 | 1 | k__Fungi(100);p__Ascomycota(100);c__Saccharomycetes(100);o__Saccharomycetales(100);f__Metschnikowiaceae(100);g__Metschnikowia(100);g__Metschnikowia_unclassified(100); | Metschnikowia | 13 | Saprotroph | \|Undefined Saprotroph\| | Yeast | NULL | Probable | As nectar-tap_saprotroph (Põlme et al. 2020); Decay sugar-rich substrates (Põlme et al. 2020) | Põlme S, et al. 2020. Fungal Diversity 105:1-16 ((https://doi.org/10.1007/s13225-020-00466-2)) |
| Otu25713 | 1 | k__Fungi(100);p__Ascomycota(100);c__Saccharomycetes(100);o__Saccharomycetales(100);f__Metschnikowiaceae(100);g__Metschnikowia(100);s__Metschnikowia_pulcherrima(100); | Metschnikowia | 13 | Saprotroph | \|Undefined Saprotroph\| | Yeast | NULL | Probable | As nectar-tap_saprotroph (Põlme et al. 2020); Decay sugar-rich substrates (Põlme et al. 2020) | Põlme S, et al. 2020. Fungal Diversity 105:1-16 ((https://doi.org/10.1007/s13225-020-00466-2)) |
| Otu25715 | 1 | k__Fungi(100);p__Ascomycota(100);c__Saccharomycetes(100);o__Saccharomycetales(100);f__Metschnikowiaceae(100);g__Metschnikowia(100);s__Metschnikowia_pulcherrima(100); | Metschnikowia | 13 | Saprotroph | \|Undefined Saprotroph\| | Yeast | NULL | Probable | As nectar-tap_saprotroph (Põlme et al. 2020); Decay sugar-rich substrates (Põlme et al. 2020) | Põlme S, et al. 2020. Fungal Diversity 105:1-16 ((https://doi.org/10.1007/s13225-020-00466-2)) |
| Otu25716 | 1 | k__Fungi(100);p__Ascomycota(100);c__Saccharomycetes(100);o__Saccharomycetales(100);f__Metschnikowiaceae(100);g__Metschnikowia(100);s__Metschnikowia_pulcherrima(100); | Metschnikowia | 13 | Saprotroph | \|Undefined Saprotroph\| | Yeast | NULL | Probable | As nectar-tap_saprotroph (Põlme et al. 2020); Decay sugar-rich substrates (Põlme et al. 2020) | Põlme S, et al. 2020. Fungal Diversity 105:1-16 ((https://doi.org/10.1007/s13225-020-00466-2)) |
| Otu25718 | 1 | k__Fungi(100);p__Ascomycota(100);c__Saccharomycetes(100);o__Saccharomycetales(100);f__Metschnikowiaceae(100);g__Metschnikowia(100);s__Metschnikowia_pulcherrima(100); | Metschnikowia | 13 | Saprotroph | \|Undefined Saprotroph\| | Yeast | NULL | Probable | As nectar-tap_saprotroph (Põlme et al. 2020); Decay sugar-rich substrates (Põlme et al. 2020) | Põlme S, et al. 2020. Fungal Diversity 105:1-16 ((https://doi.org/10.1007/s13225-020-00466-2)) |
| Otu25720 | 1 | k__Fungi(100);p__Ascomycota(100);c__Saccharomycetes(100);o__Saccharomycetales(100);f__Metschnikowiaceae(100);g__Metschnikowia(100);s__Metschnikowia_pulcherrima(100); | Metschnikowia | 13 | Saprotroph | \|Undefined Saprotroph\| | Yeast | NULL | Probable | As nectar-tap_saprotroph (Põlme et al. 2020); Decay sugar-rich substrates (Põlme et al. 2020) | Põlme S, et al. 2020. Fungal Diversity 105:1-16 ((https://doi.org/10.1007/s13225-020-00466-2)) |
| Otu25727 | 1 | k__Fungi(100);p__Ascomycota(100);c__Saccharomycetes(100);o__Saccharomycetales(100);f__Metschnikowiaceae(100);g__Metschnikowia(100);s__Metschnikowia_pulcherrima(100); | Metschnikowia | 13 | Saprotroph | \|Undefined Saprotroph\| | Yeast | NULL | Probable | As nectar-tap_saprotroph (Põlme et al. 2020); Decay sugar-rich substrates (Põlme et al. 2020) | Põlme S, et al. 2020. Fungal Diversity 105:1-16 ((https://doi.org/10.1007/s13225-020-00466-2)) |
| Otu25731 | 1 | k__Fungi(100);p__Ascomycota(100);c__Saccharomycetes(100);o__Saccharomycetales(100);f__Saccharomycetales_family_Incertae_sedis(100);g__Candida(100);g__Candida_unclassified(100); | Candida | 13 | Pathotroph-Saprotroph-Symbiotroph | Animal Pathogen-Endophyte-Endosymbiont-Epiphyte-\|Undefined Saprotroph\| | Dimorphic Yeast-Yeast | NULL | Probable | As nectar-tap_saprotroph (Põlme et al. 2020); Decay sugar-rich substrates (Põlme et al. 2020); This asexual genus is polyphyletic and contain species that function very differently depend on what they are. Caution should be taken when interpreting guilds for Candida species. | Manolakaki D, et al. 2010. Virulence. 1: 367-375 ((https://doi.org/10.4161/viru.1.5.12796)); Mekbib SB, et al. 2011. Tropical Plant Pathology 36:233-240 ((https://doi.org/10.1590/S1982-56762011000400004)); Põlme S, et al. 2020. Fungal Diversity 105:1-16 ((https://doi.org/10.1007/s13225-020-00466-2)) |
| Otu25732 | 1 | k__Fungi(100);p__Ascomycota(100);c__Saccharomycetes(100);o__Saccharomycetales(100);f__Metschnikowiaceae(100);g__Metschnikowia(100);s__Metschnikowia_pulcherrima(100); | Metschnikowia | 13 | Saprotroph | \|Undefined Saprotroph\| | Yeast | NULL | Probable | As nectar-tap_saprotroph (Põlme et al. 2020); Decay sugar-rich substrates (Põlme et al. 2020) | Põlme S, et al. 2020. Fungal Diversity 105:1-16 ((https://doi.org/10.1007/s13225-020-00466-2)) |
| Otu25735 | 1 | k__Fungi(100);p__Ascomycota(100);c__Saccharomycetes(100);o__Saccharomycetales(100);f__Saccharomycetales_family_Incertae_sedis(100);g__Candida(100);g__Candida_unclassified(100); | Candida | 13 | Pathotroph-Saprotroph-Symbiotroph | Animal Pathogen-Endophyte-Endosymbiont-Epiphyte-\|Undefined Saprotroph\| | Dimorphic Yeast-Yeast | NULL | Probable | As nectar-tap_saprotroph (Põlme et al. 2020); Decay sugar-rich substrates (Põlme et al. 2020); This asexual genus is polyphyletic and contain species that function very differently depend on what they are. Caution should be taken when interpreting guilds for Candida species. | Manolakaki D, et al. 2010. Virulence. 1: 367-375 ((https://doi.org/10.4161/viru.1.5.12796)); Mekbib SB, et al. 2011. Tropical Plant Pathology 36:233-240 ((https://doi.org/10.1590/S1982-56762011000400004)); Põlme S, et al. 2020. Fungal Diversity 105:1-16 ((https://doi.org/10.1007/s13225-020-00466-2)) |
| Otu25738 | 1 | k__Fungi(100);p__Ascomycota(100);c__Saccharomycetes(100);o__Saccharomycetales(100);f__Metschnikowiaceae(100);g__Metschnikowia(100);s__Metschnikowia_pulcherrima(100); | Metschnikowia | 13 | Saprotroph | \|Undefined Saprotroph\| | Yeast | NULL | Probable | As nectar-tap_saprotroph (Põlme et al. 2020); Decay sugar-rich substrates (Põlme et al. 2020) | Põlme S, et al. 2020. Fungal Diversity 105:1-16 ((https://doi.org/10.1007/s13225-020-00466-2)) |
| Otu25739 | 1 | k__Fungi(100);p__Ascomycota(100);c__Saccharomycetes(100);o__Saccharomycetales(100);f__Metschnikowiaceae(100);g__Metschnikowia(100);s__Metschnikowia_pulcherrima(100); | Metschnikowia | 13 | Saprotroph | \|Undefined Saprotroph\| | Yeast | NULL | Probable | As nectar-tap_saprotroph (Põlme et al. 2020); Decay sugar-rich substrates (Põlme et al. 2020) | Põlme S, et al. 2020. Fungal Diversity 105:1-16 ((https://doi.org/10.1007/s13225-020-00466-2)) |
| Otu25740 | 1 | k__Fungi(100);p__Ascomycota(100);c__Saccharomycetes(100);o__Saccharomycetales(100);f__Metschnikowiaceae(100);g__Metschnikowia(100);s__Metschnikowia_pulcherrima(100); | Metschnikowia | 13 | Saprotroph | \|Undefined Saprotroph\| | Yeast | NULL | Probable | As nectar-tap_saprotroph (Põlme et al. 2020); Decay sugar-rich substrates (Põlme et al. 2020) | Põlme S, et al. 2020. Fungal Diversity 105:1-16 ((https://doi.org/10.1007/s13225-020-00466-2)) |
| Otu25741 | 1 | k__Fungi(100);p__Ascomycota(100);c__Saccharomycetes(100);o__Saccharomycetales(100);f__Metschnikowiaceae(100);g__Metschnikowia(100);s__Metschnikowia_pulcherrima(100); | Metschnikowia | 13 | Saprotroph | \|Undefined Saprotroph\| | Yeast | NULL | Probable | As nectar-tap_saprotroph (Põlme et al. 2020); Decay sugar-rich substrates (Põlme et al. 2020) | Põlme S, et al. 2020. Fungal Diversity 105:1-16 ((https://doi.org/10.1007/s13225-020-00466-2)) |
| Otu25743 | 1 | k__Fungi(100);p__Ascomycota(100);c__Saccharomycetes(100);o__Saccharomycetales(100);f__Metschnikowiaceae(100);g__Metschnikowia(100);s__Metschnikowia_pulcherrima(100); | Metschnikowia | 13 | Saprotroph | \|Undefined Saprotroph\| | Yeast | NULL | Probable | As nectar-tap_saprotroph (Põlme et al. 2020); Decay sugar-rich substrates (Põlme et al. 2020) | Põlme S, et al. 2020. Fungal Diversity 105:1-16 ((https://doi.org/10.1007/s13225-020-00466-2)) |
| Otu25744 | 1 | k__Fungi(100);p__Ascomycota(100);c__Saccharomycetes(100);o__Saccharomycetales(100);f__Metschnikowiaceae(100);g__Metschnikowia(100);s__Metschnikowia_pulcherrima(100); | Metschnikowia | 13 | Saprotroph | \|Undefined Saprotroph\| | Yeast | NULL | Probable | As nectar-tap_saprotroph (Põlme et al. 2020); Decay sugar-rich substrates (Põlme et al. 2020) | Põlme S, et al. 2020. Fungal Diversity 105:1-16 ((https://doi.org/10.1007/s13225-020-00466-2)) |
| Otu25745 | 1 | k__Fungi(100);p__Ascomycota(100);c__Saccharomycetes(100);o__Saccharomycetales(100);f__Metschnikowiaceae(100);g__Metschnikowia(100);g__Metschnikowia_unclassified(100); | Metschnikowia | 13 | Saprotroph | \|Undefined Saprotroph\| | Yeast | NULL | Probable | As nectar-tap_saprotroph (Põlme et al. 2020); Decay sugar-rich substrates (Põlme et al. 2020) | Põlme S, et al. 2020. Fungal Diversity 105:1-16 ((https://doi.org/10.1007/s13225-020-00466-2)) |
| Otu25746 | 1 | k__Fungi(100);p__Ascomycota(100);c__Saccharomycetes(100);o__Saccharomycetales(100);f__Metschnikowiaceae(100);g__Metschnikowia(100);s__Metschnikowia_pulcherrima(100); | Metschnikowia | 13 | Saprotroph | \|Undefined Saprotroph\| | Yeast | NULL | Probable | As nectar-tap_saprotroph (Põlme et al. 2020); Decay sugar-rich substrates (Põlme et al. 2020) | Põlme S, et al. 2020. Fungal Diversity 105:1-16 ((https://doi.org/10.1007/s13225-020-00466-2)) |
| Otu25747 | 1 | k__Fungi(100);p__Ascomycota(100);c__Saccharomycetes(100);o__Saccharomycetales(100);f__Metschnikowiaceae(100);g__Metschnikowia(100);s__Metschnikowia_pulcherrima(100); | Metschnikowia | 13 | Saprotroph | \|Undefined Saprotroph\| | Yeast | NULL | Probable | As nectar-tap_saprotroph (Põlme et al. 2020); Decay sugar-rich substrates (Põlme et al. 2020) | Põlme S, et al. 2020. Fungal Diversity 105:1-16 ((https://doi.org/10.1007/s13225-020-00466-2)) |
| Otu25753 | 1 | k__Fungi(100);p__Ascomycota(100);c__Saccharomycetes(100);o__Saccharomycetales(100);f__Metschnikowiaceae(100);g__Metschnikowia(100);s__Metschnikowia_pulcherrima(100); | Metschnikowia | 13 | Saprotroph | \|Undefined Saprotroph\| | Yeast | NULL | Probable | As nectar-tap_saprotroph (Põlme et al. 2020); Decay sugar-rich substrates (Põlme et al. 2020) | Põlme S, et al. 2020. Fungal Diversity 105:1-16 ((https://doi.org/10.1007/s13225-020-00466-2)) |
| Otu25755 | 1 | k__Fungi(100);p__Ascomycota(100);c__Saccharomycetes(100);o__Saccharomycetales(100);f__Metschnikowiaceae(100);g__Metschnikowia(100);s__Metschnikowia_pulcherrima(100); | Metschnikowia | 13 | Saprotroph | \|Undefined Saprotroph\| | Yeast | NULL | Probable | As nectar-tap_saprotroph (Põlme et al. 2020); Decay sugar-rich substrates (Põlme et al. 2020) | Põlme S, et al. 2020. Fungal Diversity 105:1-16 ((https://doi.org/10.1007/s13225-020-00466-2)) |
| Otu25756 | 1 | k__Fungi(100);p__Ascomycota(100);c__Saccharomycetes(100);o__Saccharomycetales(100);f__Metschnikowiaceae(100);g__Metschnikowia(100);g__Metschnikowia_unclassified(100); | Metschnikowia | 13 | Saprotroph | \|Undefined Saprotroph\| | Yeast | NULL | Probable | As nectar-tap_saprotroph (Põlme et al. 2020); Decay sugar-rich substrates (Põlme et al. 2020) | Põlme S, et al. 2020. Fungal Diversity 105:1-16 ((https://doi.org/10.1007/s13225-020-00466-2)) |
| Otu25757 | 1 | k__Fungi(100);p__Ascomycota(100);c__Saccharomycetes(100);o__Saccharomycetales(100);f__Metschnikowiaceae(100);g__Metschnikowia(100);g__Metschnikowia_unclassified(100); | Metschnikowia | 13 | Saprotroph | \|Undefined Saprotroph\| | Yeast | NULL | Probable | As nectar-tap_saprotroph (Põlme et al. 2020); Decay sugar-rich substrates (Põlme et al. 2020) | Põlme S, et al. 2020. Fungal Diversity 105:1-16 ((https://doi.org/10.1007/s13225-020-00466-2)) |
| Otu25758 | 1 | k__Fungi(100);p__Ascomycota(100);c__Saccharomycetes(100);o__Saccharomycetales(100);f__Metschnikowiaceae(100);g__Metschnikowia(100);g__Metschnikowia_unclassified(100); | Metschnikowia | 13 | Saprotroph | \|Undefined Saprotroph\| | Yeast | NULL | Probable | As nectar-tap_saprotroph (Põlme et al. 2020); Decay sugar-rich substrates (Põlme et al. 2020) | Põlme S, et al. 2020. Fungal Diversity 105:1-16 ((https://doi.org/10.1007/s13225-020-00466-2)) |
| Otu25759 | 1 | k__Fungi(100);p__Ascomycota(100);c__Saccharomycetes(100);o__Saccharomycetales(100);f__Metschnikowiaceae(100);g__Metschnikowia(100);g__Metschnikowia_unclassified(100); | Metschnikowia | 13 | Saprotroph | \|Undefined Saprotroph\| | Yeast | NULL | Probable | As nectar-tap_saprotroph (Põlme et al. 2020); Decay sugar-rich substrates (Põlme et al. 2020) | Põlme S, et al. 2020. Fungal Diversity 105:1-16 ((https://doi.org/10.1007/s13225-020-00466-2)) |
| Otu25760 | 1 | k__Fungi(100);p__Ascomycota(100);c__Saccharomycetes(100);o__Saccharomycetales(100);f__Metschnikowiaceae(100);g__Metschnikowia(100);s__Metschnikowia_pulcherrima(100); | Metschnikowia | 13 | Saprotroph | \|Undefined Saprotroph\| | Yeast | NULL | Probable | As nectar-tap_saprotroph (Põlme et al. 2020); Decay sugar-rich substrates (Põlme et al. 2020) | Põlme S, et al. 2020. Fungal Diversity 105:1-16 ((https://doi.org/10.1007/s13225-020-00466-2)) |
| Otu25761 | 1 | k__Fungi(100);p__Ascomycota(100);c__Saccharomycetes(100);o__Saccharomycetales(100);f__Metschnikowiaceae(100);g__Metschnikowia(100);s__Metschnikowia_pulcherrima(100); | Metschnikowia | 13 | Saprotroph | \|Undefined Saprotroph\| | Yeast | NULL | Probable | As nectar-tap_saprotroph (Põlme et al. 2020); Decay sugar-rich substrates (Põlme et al. 2020) | Põlme S, et al. 2020. Fungal Diversity 105:1-16 ((https://doi.org/10.1007/s13225-020-00466-2)) |
| Otu25764 | 1 | k__Fungi(100);p__Ascomycota(100);c__Saccharomycetes(100);o__Saccharomycetales(100);f__Metschnikowiaceae(100);g__Metschnikowia(100);s__Metschnikowia_pulcherrima(100); | Metschnikowia | 13 | Saprotroph | \|Undefined Saprotroph\| | Yeast | NULL | Probable | As nectar-tap_saprotroph (Põlme et al. 2020); Decay sugar-rich substrates (Põlme et al. 2020) | Põlme S, et al. 2020. Fungal Diversity 105:1-16 ((https://doi.org/10.1007/s13225-020-00466-2)) |
| Otu25765 | 1 | k__Fungi(100);p__Ascomycota(100);c__Saccharomycetes(100);o__Saccharomycetales(100);f__Metschnikowiaceae(100);g__Metschnikowia(100);s__Metschnikowia_pulcherrima(100); | Metschnikowia | 13 | Saprotroph | \|Undefined Saprotroph\| | Yeast | NULL | Probable | As nectar-tap_saprotroph (Põlme et al. 2020); Decay sugar-rich substrates (Põlme et al. 2020) | Põlme S, et al. 2020. Fungal Diversity 105:1-16 ((https://doi.org/10.1007/s13225-020-00466-2)) |
| Otu25766 | 1 | k__Fungi(100);p__Ascomycota(100);c__Saccharomycetes(100);o__Saccharomycetales(100);f__Metschnikowiaceae(100);g__Metschnikowia(100);s__Metschnikowia_pulcherrima(100); | Metschnikowia | 13 | Saprotroph | \|Undefined Saprotroph\| | Yeast | NULL | Probable | As nectar-tap_saprotroph (Põlme et al. 2020); Decay sugar-rich substrates (Põlme et al. 2020) | Põlme S, et al. 2020. Fungal Diversity 105:1-16 ((https://doi.org/10.1007/s13225-020-00466-2)) |
| Otu25767 | 1 | k__Fungi(100);p__Ascomycota(100);c__Saccharomycetes(100);o__Saccharomycetales(100);f__Metschnikowiaceae(100);g__Metschnikowia(100);g__Metschnikowia_unclassified(100); | Metschnikowia | 13 | Saprotroph | \|Undefined Saprotroph\| | Yeast | NULL | Probable | As nectar-tap_saprotroph (Põlme et al. 2020); Decay sugar-rich substrates (Põlme et al. 2020) | Põlme S, et al. 2020. Fungal Diversity 105:1-16 ((https://doi.org/10.1007/s13225-020-00466-2)) |
| Otu25769 | 1 | k__Fungi(100);p__Ascomycota(100);c__Saccharomycetes(100);o__Saccharomycetales(100);f__Metschnikowiaceae(100);g__Metschnikowia(100);g__Metschnikowia_unclassified(100); | Metschnikowia | 13 | Saprotroph | \|Undefined Saprotroph\| | Yeast | NULL | Probable | As nectar-tap_saprotroph (Põlme et al. 2020); Decay sugar-rich substrates (Põlme et al. 2020) | Põlme S, et al. 2020. Fungal Diversity 105:1-16 ((https://doi.org/10.1007/s13225-020-00466-2)) |
| Otu25774 | 1 | k__Fungi(100);p__Ascomycota(100);c__Saccharomycetes(100);o__Saccharomycetales(100);f__Metschnikowiaceae(100);g__Metschnikowia(100);s__Metschnikowia_pulcherrima(100); | Metschnikowia | 13 | Saprotroph | \|Undefined Saprotroph\| | Yeast | NULL | Probable | As nectar-tap_saprotroph (Põlme et al. 2020); Decay sugar-rich substrates (Põlme et al. 2020) | Põlme S, et al. 2020. Fungal Diversity 105:1-16 ((https://doi.org/10.1007/s13225-020-00466-2)) |
| Otu25775 | 1 | k__Fungi(100);p__Ascomycota(100);c__Saccharomycetes(100);o__Saccharomycetales(100);f__Metschnikowiaceae(100);g__Metschnikowia(100);s__Metschnikowia_pulcherrima(100); | Metschnikowia | 13 | Saprotroph | \|Undefined Saprotroph\| | Yeast | NULL | Probable | As nectar-tap_saprotroph (Põlme et al. 2020); Decay sugar-rich substrates (Põlme et al. 2020) | Põlme S, et al. 2020. Fungal Diversity 105:1-16 ((https://doi.org/10.1007/s13225-020-00466-2)) |
| Otu25777 | 1 | k__Fungi(100);p__Ascomycota(100);c__Saccharomycetes(100);o__Saccharomycetales(100);f__Metschnikowiaceae(100);g__Metschnikowia(100);g__Metschnikowia_unclassified(100); | Metschnikowia | 13 | Saprotroph | \|Undefined Saprotroph\| | Yeast | NULL | Probable | As nectar-tap_saprotroph (Põlme et al. 2020); Decay sugar-rich substrates (Põlme et al. 2020) | Põlme S, et al. 2020. Fungal Diversity 105:1-16 ((https://doi.org/10.1007/s13225-020-00466-2)) |
| Otu25778 | 1 | k__Fungi(100);p__Ascomycota(100);c__Saccharomycetes(100);o__Saccharomycetales(100);f__Metschnikowiaceae(100);g__Metschnikowia(100);s__Metschnikowia_pulcherrima(100); | Metschnikowia | 13 | Saprotroph | \|Undefined Saprotroph\| | Yeast | NULL | Probable | As nectar-tap_saprotroph (Põlme et al. 2020); Decay sugar-rich substrates (Põlme et al. 2020) | Põlme S, et al. 2020. Fungal Diversity 105:1-16 ((https://doi.org/10.1007/s13225-020-00466-2)) |
| Otu25780 | 1 | k__Fungi(100);p__Ascomycota(100);c__Saccharomycetes(100);o__Saccharomycetales(100);f__Metschnikowiaceae(100);g__Metschnikowia(100);s__Metschnikowia_pulcherrima(100); | Metschnikowia | 13 | Saprotroph | \|Undefined Saprotroph\| | Yeast | NULL | Probable | As nectar-tap_saprotroph (Põlme et al. 2020); Decay sugar-rich substrates (Põlme et al. 2020) | Põlme S, et al. 2020. Fungal Diversity 105:1-16 ((https://doi.org/10.1007/s13225-020-00466-2)) |
| Otu25785 | 1 | k__Fungi(100);p__Ascomycota(100);c__Saccharomycetes(100);o__Saccharomycetales(100);f__Metschnikowiaceae(100);g__Metschnikowia(100);s__Metschnikowia_pulcherrima(100); | Metschnikowia | 13 | Saprotroph | \|Undefined Saprotroph\| | Yeast | NULL | Probable | As nectar-tap_saprotroph (Põlme et al. 2020); Decay sugar-rich substrates (Põlme et al. 2020) | Põlme S, et al. 2020. Fungal Diversity 105:1-16 ((https://doi.org/10.1007/s13225-020-00466-2)) |
| Otu25789 | 1 | k__Fungi(100);p__Ascomycota(100);c__Saccharomycetes(100);o__Saccharomycetales(100);f__Metschnikowiaceae(100);g__Metschnikowia(100);g__Metschnikowia_unclassified(100); | Metschnikowia | 13 | Saprotroph | \|Undefined Saprotroph\| | Yeast | NULL | Probable | As nectar-tap_saprotroph (Põlme et al. 2020); Decay sugar-rich substrates (Põlme et al. 2020) | Põlme S, et al. 2020. Fungal Diversity 105:1-16 ((https://doi.org/10.1007/s13225-020-00466-2)) |
| Otu25790 | 1 | k__Fungi(100);p__Ascomycota(100);c__Saccharomycetes(100);o__Saccharomycetales(100);f__Metschnikowiaceae(100);g__Metschnikowia(100);s__Metschnikowia_pulcherrima(100); | Metschnikowia | 13 | Saprotroph | \|Undefined Saprotroph\| | Yeast | NULL | Probable | As nectar-tap_saprotroph (Põlme et al. 2020); Decay sugar-rich substrates (Põlme et al. 2020) | Põlme S, et al. 2020. Fungal Diversity 105:1-16 ((https://doi.org/10.1007/s13225-020-00466-2)) |
| Otu25791 | 1 | k__Fungi(100);p__Ascomycota(100);c__Saccharomycetes(100);o__Saccharomycetales(100);f__Metschnikowiaceae(100);g__Metschnikowia(100);s__Metschnikowia_pulcherrima(100); | Metschnikowia | 13 | Saprotroph | \|Undefined Saprotroph\| | Yeast | NULL | Probable | As nectar-tap_saprotroph (Põlme et al. 2020); Decay sugar-rich substrates (Põlme et al. 2020) | Põlme S, et al. 2020. Fungal Diversity 105:1-16 ((https://doi.org/10.1007/s13225-020-00466-2)) |
| Otu25792 | 1 | k__Fungi(100);p__Ascomycota(100);c__Saccharomycetes(100);o__Saccharomycetales(100);f__Metschnikowiaceae(100);g__Metschnikowia(100);s__Metschnikowia_pulcherrima(100); | Metschnikowia | 13 | Saprotroph | \|Undefined Saprotroph\| | Yeast | NULL | Probable | As nectar-tap_saprotroph (Põlme et al. 2020); Decay sugar-rich substrates (Põlme et al. 2020) | Põlme S, et al. 2020. Fungal Diversity 105:1-16 ((https://doi.org/10.1007/s13225-020-00466-2)) |
| Otu25793 | 1 | k__Fungi(100);p__Ascomycota(100);c__Saccharomycetes(100);o__Saccharomycetales(100);f__Metschnikowiaceae(100);g__Metschnikowia(100);s__Metschnikowia_pulcherrima(100); | Metschnikowia | 13 | Saprotroph | \|Undefined Saprotroph\| | Yeast | NULL | Probable | As nectar-tap_saprotroph (Põlme et al. 2020); Decay sugar-rich substrates (Põlme et al. 2020) | Põlme S, et al. 2020. Fungal Diversity 105:1-16 ((https://doi.org/10.1007/s13225-020-00466-2)) |
| Otu25797 | 1 | k__Fungi(100);p__Ascomycota(100);c__Saccharomycetes(100);o__Saccharomycetales(100);f__Metschnikowiaceae(100);g__Metschnikowia(100);s__Metschnikowia_pulcherrima(100); | Metschnikowia | 13 | Saprotroph | \|Undefined Saprotroph\| | Yeast | NULL | Probable | As nectar-tap_saprotroph (Põlme et al. 2020); Decay sugar-rich substrates (Põlme et al. 2020) | Põlme S, et al. 2020. Fungal Diversity 105:1-16 ((https://doi.org/10.1007/s13225-020-00466-2)) |
| Otu25798 | 1 | k__Fungi(100);p__Ascomycota(100);c__Saccharomycetes(100);o__Saccharomycetales(100);f__Metschnikowiaceae(100);g__Metschnikowia(100);s__Metschnikowia_pulcherrima(100); | Metschnikowia | 13 | Saprotroph | \|Undefined Saprotroph\| | Yeast | NULL | Probable | As nectar-tap_saprotroph (Põlme et al. 2020); Decay sugar-rich substrates (Põlme et al. 2020) | Põlme S, et al. 2020. Fungal Diversity 105:1-16 ((https://doi.org/10.1007/s13225-020-00466-2)) |
| Otu25800 | 1 | k__Fungi(100);p__Ascomycota(100);c__Saccharomycetes(100);o__Saccharomycetales(100);f__Metschnikowiaceae(100);g__Metschnikowia(100);s__Metschnikowia_pulcherrima(100); | Metschnikowia | 13 | Saprotroph | \|Undefined Saprotroph\| | Yeast | NULL | Probable | As nectar-tap_saprotroph (Põlme et al. 2020); Decay sugar-rich substrates (Põlme et al. 2020) | Põlme S, et al. 2020. Fungal Diversity 105:1-16 ((https://doi.org/10.1007/s13225-020-00466-2)) |
| Otu25803 | 1 | k__Fungi(100);p__Ascomycota(100);c__Saccharomycetes(100);o__Saccharomycetales(100);f__Saccharomycetales_family_Incertae_sedis(100);g__Candida(100);g__Candida_unclassified(100); | Candida | 13 | Pathotroph-Saprotroph-Symbiotroph | Animal Pathogen-Endophyte-Endosymbiont-Epiphyte-\|Undefined Saprotroph\| | Dimorphic Yeast-Yeast | NULL | Probable | As nectar-tap_saprotroph (Põlme et al. 2020); Decay sugar-rich substrates (Põlme et al. 2020); This asexual genus is polyphyletic and contain species that function very differently depend on what they are. Caution should be taken when interpreting guilds for Candida species. | Manolakaki D, et al. 2010. Virulence. 1: 367-375 ((https://doi.org/10.4161/viru.1.5.12796)); Mekbib SB, et al. 2011. Tropical Plant Pathology 36:233-240 ((https://doi.org/10.1590/S1982-56762011000400004)); Põlme S, et al. 2020. Fungal Diversity 105:1-16 ((https://doi.org/10.1007/s13225-020-00466-2)) |
| Otu25806 | 1 | k__Fungi(100);p__Ascomycota(100);c__Saccharomycetes(100);o__Saccharomycetales(100);f__Metschnikowiaceae(100);g__Metschnikowia(100);s__Metschnikowia_pulcherrima(100); | Metschnikowia | 13 | Saprotroph | \|Undefined Saprotroph\| | Yeast | NULL | Probable | As nectar-tap_saprotroph (Põlme et al. 2020); Decay sugar-rich substrates (Põlme et al. 2020) | Põlme S, et al. 2020. Fungal Diversity 105:1-16 ((https://doi.org/10.1007/s13225-020-00466-2)) |
| Otu25807 | 1 | k__Fungi(100);p__Ascomycota(100);c__Saccharomycetes(100);o__Saccharomycetales(100);f__Saccharomycetales_family_Incertae_sedis(100);g__Candida(100);g__Candida_unclassified(100); | Candida | 13 | Pathotroph-Saprotroph-Symbiotroph | Animal Pathogen-Endophyte-Endosymbiont-Epiphyte-\|Undefined Saprotroph\| | Dimorphic Yeast-Yeast | NULL | Probable | As nectar-tap_saprotroph (Põlme et al. 2020); Decay sugar-rich substrates (Põlme et al. 2020); This asexual genus is polyphyletic and contain species that function very differently depend on what they are. Caution should be taken when interpreting guilds for Candida species. | Manolakaki D, et al. 2010. Virulence. 1: 367-375 ((https://doi.org/10.4161/viru.1.5.12796)); Mekbib SB, et al. 2011. Tropical Plant Pathology 36:233-240 ((https://doi.org/10.1590/S1982-56762011000400004)); Põlme S, et al. 2020. Fungal Diversity 105:1-16 ((https://doi.org/10.1007/s13225-020-00466-2)) |
| Otu25808 | 1 | k__Fungi(100);p__Ascomycota(100);c__Saccharomycetes(100);o__Saccharomycetales(100);f__Metschnikowiaceae(100);g__Metschnikowia(100);s__Metschnikowia_pulcherrima(100); | Metschnikowia | 13 | Saprotroph | \|Undefined Saprotroph\| | Yeast | NULL | Probable | As nectar-tap_saprotroph (Põlme et al. 2020); Decay sugar-rich substrates (Põlme et al. 2020) | Põlme S, et al. 2020. Fungal Diversity 105:1-16 ((https://doi.org/10.1007/s13225-020-00466-2)) |
| Otu25809 | 1 | k__Fungi(100);p__Ascomycota(100);c__Saccharomycetes(100);o__Saccharomycetales(100);f__Metschnikowiaceae(100);g__Metschnikowia(100);s__Metschnikowia_pulcherrima(100); | Metschnikowia | 13 | Saprotroph | \|Undefined Saprotroph\| | Yeast | NULL | Probable | As nectar-tap_saprotroph (Põlme et al. 2020); Decay sugar-rich substrates (Põlme et al. 2020) | Põlme S, et al. 2020. Fungal Diversity 105:1-16 ((https://doi.org/10.1007/s13225-020-00466-2)) |
| Otu25810 | 1 | k__Fungi(100);p__Ascomycota(100);c__Saccharomycetes(100);o__Saccharomycetales(100);f__Metschnikowiaceae(100);g__Metschnikowia(100);s__Metschnikowia_pulcherrima(100); | Metschnikowia | 13 | Saprotroph | \|Undefined Saprotroph\| | Yeast | NULL | Probable | As nectar-tap_saprotroph (Põlme et al. 2020); Decay sugar-rich substrates (Põlme et al. 2020) | Põlme S, et al. 2020. Fungal Diversity 105:1-16 ((https://doi.org/10.1007/s13225-020-00466-2)) |
| Otu25814 | 1 | k__Fungi(100);p__Ascomycota(100);c__Saccharomycetes(100);o__Saccharomycetales(100);f__Metschnikowiaceae(100);g__Metschnikowia(100);s__Metschnikowia_pulcherrima(100); | Metschnikowia | 13 | Saprotroph | \|Undefined Saprotroph\| | Yeast | NULL | Probable | As nectar-tap_saprotroph (Põlme et al. 2020); Decay sugar-rich substrates (Põlme et al. 2020) | Põlme S, et al. 2020. Fungal Diversity 105:1-16 ((https://doi.org/10.1007/s13225-020-00466-2)) |
| Otu25817 | 1 | k__Fungi(100);p__Ascomycota(100);c__Saccharomycetes(100);o__Saccharomycetales(100);f__Saccharomycetales_family_Incertae_sedis(100);g__Candida(100);g__Candida_unclassified(100); | Candida | 13 | Pathotroph-Saprotroph-Symbiotroph | Animal Pathogen-Endophyte-Endosymbiont-Epiphyte-\|Undefined Saprotroph\| | Dimorphic Yeast-Yeast | NULL | Probable | As nectar-tap_saprotroph (Põlme et al. 2020); Decay sugar-rich substrates (Põlme et al. 2020); This asexual genus is polyphyletic and contain species that function very differently depend on what they are. Caution should be taken when interpreting guilds for Candida species. | Manolakaki D, et al. 2010. Virulence. 1: 367-375 ((https://doi.org/10.4161/viru.1.5.12796)); Mekbib SB, et al. 2011. Tropical Plant Pathology 36:233-240 ((https://doi.org/10.1590/S1982-56762011000400004)); Põlme S, et al. 2020. Fungal Diversity 105:1-16 ((https://doi.org/10.1007/s13225-020-00466-2)) |
| Otu25820 | 1 | k__Fungi(100);p__Ascomycota(100);c__Saccharomycetes(100);o__Saccharomycetales(100);f__Metschnikowiaceae(100);g__Metschnikowia(100);s__Metschnikowia_pulcherrima(100); | Metschnikowia | 13 | Saprotroph | \|Undefined Saprotroph\| | Yeast | NULL | Probable | As nectar-tap_saprotroph (Põlme et al. 2020); Decay sugar-rich substrates (Põlme et al. 2020) | Põlme S, et al. 2020. Fungal Diversity 105:1-16 ((https://doi.org/10.1007/s13225-020-00466-2)) |
| Otu25822 | 1 | k__Fungi(100);p__Ascomycota(100);c__Saccharomycetes(100);o__Saccharomycetales(100);f__Metschnikowiaceae(100);g__Metschnikowia(100);s__Metschnikowia_pulcherrima(100); | Metschnikowia | 13 | Saprotroph | \|Undefined Saprotroph\| | Yeast | NULL | Probable | As nectar-tap_saprotroph (Põlme et al. 2020); Decay sugar-rich substrates (Põlme et al. 2020) | Põlme S, et al. 2020. Fungal Diversity 105:1-16 ((https://doi.org/10.1007/s13225-020-00466-2)) |
| Otu25825 | 1 | k__Fungi(100);p__Ascomycota(100);c__Saccharomycetes(100);o__Saccharomycetales(100);f__Saccharomycetales_family_Incertae_sedis(100);g__Candida(100);g__Candida_unclassified(100); | Candida | 13 | Pathotroph-Saprotroph-Symbiotroph | Animal Pathogen-Endophyte-Endosymbiont-Epiphyte-\|Undefined Saprotroph\| | Dimorphic Yeast-Yeast | NULL | Probable | As nectar-tap_saprotroph (Põlme et al. 2020); Decay sugar-rich substrates (Põlme et al. 2020); This asexual genus is polyphyletic and contain species that function very differently depend on what they are. Caution should be taken when interpreting guilds for Candida species. | Manolakaki D, et al. 2010. Virulence. 1: 367-375 ((https://doi.org/10.4161/viru.1.5.12796)); Mekbib SB, et al. 2011. Tropical Plant Pathology 36:233-240 ((https://doi.org/10.1590/S1982-56762011000400004)); Põlme S, et al. 2020. Fungal Diversity 105:1-16 ((https://doi.org/10.1007/s13225-020-00466-2)) |
| Otu25826 | 1 | k__Fungi(100);p__Ascomycota(100);c__Saccharomycetes(100);o__Saccharomycetales(100);f__Saccharomycetales_family_Incertae_sedis(100);g__Candida(100);s__Candida_cretensis(100); | Candida | 13 | Pathotroph-Saprotroph-Symbiotroph | Animal Pathogen-Endophyte-Endosymbiont-Epiphyte-\|Undefined Saprotroph\| | Dimorphic Yeast-Yeast | NULL | Probable | As nectar-tap_saprotroph (Põlme et al. 2020); Decay sugar-rich substrates (Põlme et al. 2020); This asexual genus is polyphyletic and contain species that function very differently depend on what they are. Caution should be taken when interpreting guilds for Candida species. | Manolakaki D, et al. 2010. Virulence. 1: 367-375 ((https://doi.org/10.4161/viru.1.5.12796)); Mekbib SB, et al. 2011. Tropical Plant Pathology 36:233-240 ((https://doi.org/10.1590/S1982-56762011000400004)); Põlme S, et al. 2020. Fungal Diversity 105:1-16 ((https://doi.org/10.1007/s13225-020-00466-2)) |
| Otu25830 | 1 | k__Fungi(100);p__Ascomycota(100);c__Saccharomycetes(100);o__Saccharomycetales(100);f__Saccharomycetales_family_Incertae_sedis(100);g__Candida(100);g__Candida_unclassified(100); | Candida | 13 | Pathotroph-Saprotroph-Symbiotroph | Animal Pathogen-Endophyte-Endosymbiont-Epiphyte-\|Undefined Saprotroph\| | Dimorphic Yeast-Yeast | NULL | Probable | As nectar-tap_saprotroph (Põlme et al. 2020); Decay sugar-rich substrates (Põlme et al. 2020); This asexual genus is polyphyletic and contain species that function very differently depend on what they are. Caution should be taken when interpreting guilds for Candida species. | Manolakaki D, et al. 2010. Virulence. 1: 367-375 ((https://doi.org/10.4161/viru.1.5.12796)); Mekbib SB, et al. 2011. Tropical Plant Pathology 36:233-240 ((https://doi.org/10.1590/S1982-56762011000400004)); Põlme S, et al. 2020. Fungal Diversity 105:1-16 ((https://doi.org/10.1007/s13225-020-00466-2)) |
| Otu25832 | 1 | k__Fungi(100);p__Ascomycota(100);c__Saccharomycetes(100);o__Saccharomycetales(100);f__Metschnikowiaceae(100);g__Metschnikowia(100);g__Metschnikowia_unclassified(100); | Metschnikowia | 13 | Saprotroph | \|Undefined Saprotroph\| | Yeast | NULL | Probable | As nectar-tap_saprotroph (Põlme et al. 2020); Decay sugar-rich substrates (Põlme et al. 2020) | Põlme S, et al. 2020. Fungal Diversity 105:1-16 ((https://doi.org/10.1007/s13225-020-00466-2)) |
| Otu25833 | 1 | k__Fungi(100);p__Ascomycota(100);c__Saccharomycetes(100);o__Saccharomycetales(100);f__Metschnikowiaceae(100);g__Metschnikowia(100);g__Metschnikowia_unclassified(100); | Metschnikowia | 13 | Saprotroph | \|Undefined Saprotroph\| | Yeast | NULL | Probable | As nectar-tap_saprotroph (Põlme et al. 2020); Decay sugar-rich substrates (Põlme et al. 2020) | Põlme S, et al. 2020. Fungal Diversity 105:1-16 ((https://doi.org/10.1007/s13225-020-00466-2)) |
| Otu25835 | 1 | k__Fungi(100);p__Ascomycota(100);c__Saccharomycetes(100);o__Saccharomycetales(100);f__Metschnikowiaceae(100);g__Metschnikowia(100);s__Metschnikowia_pulcherrima(100); | Metschnikowia | 13 | Saprotroph | \|Undefined Saprotroph\| | Yeast | NULL | Probable | As nectar-tap_saprotroph (Põlme et al. 2020); Decay sugar-rich substrates (Põlme et al. 2020) | Põlme S, et al. 2020. Fungal Diversity 105:1-16 ((https://doi.org/10.1007/s13225-020-00466-2)) |
| Otu25840 | 1 | k__Fungi(100);p__Ascomycota(100);c__Saccharomycetes(100);o__Saccharomycetales(100);f__Metschnikowiaceae(100);g__Metschnikowia(100);s__Metschnikowia_pulcherrima(100); | Metschnikowia | 13 | Saprotroph | \|Undefined Saprotroph\| | Yeast | NULL | Probable | As nectar-tap_saprotroph (Põlme et al. 2020); Decay sugar-rich substrates (Põlme et al. 2020) | Põlme S, et al. 2020. Fungal Diversity 105:1-16 ((https://doi.org/10.1007/s13225-020-00466-2)) |
| Otu25841 | 1 | k__Fungi(100);p__Ascomycota(100);c__Saccharomycetes(100);o__Saccharomycetales(100);f__Metschnikowiaceae(100);g__Metschnikowia(100);s__Metschnikowia_pulcherrima(100); | Metschnikowia | 13 | Saprotroph | \|Undefined Saprotroph\| | Yeast | NULL | Probable | As nectar-tap_saprotroph (Põlme et al. 2020); Decay sugar-rich substrates (Põlme et al. 2020) | Põlme S, et al. 2020. Fungal Diversity 105:1-16 ((https://doi.org/10.1007/s13225-020-00466-2)) |
| Otu25842 | 1 | k__Fungi(100);p__Ascomycota(100);c__Saccharomycetes(100);o__Saccharomycetales(100);f__Metschnikowiaceae(100);g__Metschnikowia(100);s__Metschnikowia_pulcherrima(100); | Metschnikowia | 13 | Saprotroph | \|Undefined Saprotroph\| | Yeast | NULL | Probable | As nectar-tap_saprotroph (Põlme et al. 2020); Decay sugar-rich substrates (Põlme et al. 2020) | Põlme S, et al. 2020. Fungal Diversity 105:1-16 ((https://doi.org/10.1007/s13225-020-00466-2)) |
| Otu25843 | 1 | k__Fungi(100);p__Ascomycota(100);c__Saccharomycetes(100);o__Saccharomycetales(100);f__Metschnikowiaceae(100);g__Metschnikowia(100);s__Metschnikowia_pulcherrima(100); | Metschnikowia | 13 | Saprotroph | \|Undefined Saprotroph\| | Yeast | NULL | Probable | As nectar-tap_saprotroph (Põlme et al. 2020); Decay sugar-rich substrates (Põlme et al. 2020) | Põlme S, et al. 2020. Fungal Diversity 105:1-16 ((https://doi.org/10.1007/s13225-020-00466-2)) |
| Otu25844 | 1 | k__Fungi(100);p__Ascomycota(100);c__Saccharomycetes(100);o__Saccharomycetales(100);f__Metschnikowiaceae(100);g__Metschnikowia(100);g__Metschnikowia_unclassified(100); | Metschnikowia | 13 | Saprotroph | \|Undefined Saprotroph\| | Yeast | NULL | Probable | As nectar-tap_saprotroph (Põlme et al. 2020); Decay sugar-rich substrates (Põlme et al. 2020) | Põlme S, et al. 2020. Fungal Diversity 105:1-16 ((https://doi.org/10.1007/s13225-020-00466-2)) |
| Otu25845 | 1 | k__Fungi(100);p__Ascomycota(100);c__Saccharomycetes(100);o__Saccharomycetales(100);f__Metschnikowiaceae(100);g__Metschnikowia(100);s__Metschnikowia_pulcherrima(100); | Metschnikowia | 13 | Saprotroph | \|Undefined Saprotroph\| | Yeast | NULL | Probable | As nectar-tap_saprotroph (Põlme et al. 2020); Decay sugar-rich substrates (Põlme et al. 2020) | Põlme S, et al. 2020. Fungal Diversity 105:1-16 ((https://doi.org/10.1007/s13225-020-00466-2)) |
| Otu25846 | 1 | k__Fungi(100);p__Ascomycota(100);c__Saccharomycetes(100);o__Saccharomycetales(100);f__Metschnikowiaceae(100);g__Metschnikowia(100);g__Metschnikowia_unclassified(100); | Metschnikowia | 13 | Saprotroph | \|Undefined Saprotroph\| | Yeast | NULL | Probable | As nectar-tap_saprotroph (Põlme et al. 2020); Decay sugar-rich substrates (Põlme et al. 2020) | Põlme S, et al. 2020. Fungal Diversity 105:1-16 ((https://doi.org/10.1007/s13225-020-00466-2)) |
| Otu25847 | 1 | k__Fungi(100);p__Ascomycota(100);c__Saccharomycetes(100);o__Saccharomycetales(100);f__Metschnikowiaceae(100);g__Metschnikowia(100);g__Metschnikowia_unclassified(100); | Metschnikowia | 13 | Saprotroph | \|Undefined Saprotroph\| | Yeast | NULL | Probable | As nectar-tap_saprotroph (Põlme et al. 2020); Decay sugar-rich substrates (Põlme et al. 2020) | Põlme S, et al. 2020. Fungal Diversity 105:1-16 ((https://doi.org/10.1007/s13225-020-00466-2)) |
| Otu25848 | 1 | k__Fungi(100);p__Ascomycota(100);c__Saccharomycetes(100);o__Saccharomycetales(100);f__Metschnikowiaceae(100);g__Metschnikowia(100);s__Metschnikowia_pulcherrima(100); | Metschnikowia | 13 | Saprotroph | \|Undefined Saprotroph\| | Yeast | NULL | Probable | As nectar-tap_saprotroph (Põlme et al. 2020); Decay sugar-rich substrates (Põlme et al. 2020) | Põlme S, et al. 2020. Fungal Diversity 105:1-16 ((https://doi.org/10.1007/s13225-020-00466-2)) |
| Otu25849 | 1 | k__Fungi(100);p__Ascomycota(100);c__Saccharomycetes(100);o__Saccharomycetales(100);f__Metschnikowiaceae(100);g__Metschnikowia(100);s__Metschnikowia_pulcherrima(100); | Metschnikowia | 13 | Saprotroph | \|Undefined Saprotroph\| | Yeast | NULL | Probable | As nectar-tap_saprotroph (Põlme et al. 2020); Decay sugar-rich substrates (Põlme et al. 2020) | Põlme S, et al. 2020. Fungal Diversity 105:1-16 ((https://doi.org/10.1007/s13225-020-00466-2)) |
| Otu25850 | 1 | k__Fungi(100);p__Ascomycota(100);c__Saccharomycetes(100);o__Saccharomycetales(100);f__Metschnikowiaceae(100);g__Metschnikowia(100);s__Metschnikowia_pulcherrima(100); | Metschnikowia | 13 | Saprotroph | \|Undefined Saprotroph\| | Yeast | NULL | Probable | As nectar-tap_saprotroph (Põlme et al. 2020); Decay sugar-rich substrates (Põlme et al. 2020) | Põlme S, et al. 2020. Fungal Diversity 105:1-16 ((https://doi.org/10.1007/s13225-020-00466-2)) |
| Otu25851 | 1 | k__Fungi(100);p__Ascomycota(100);c__Saccharomycetes(100);o__Saccharomycetales(100);f__Metschnikowiaceae(100);g__Metschnikowia(100);s__Metschnikowia_pulcherrima(100); | Metschnikowia | 13 | Saprotroph | \|Undefined Saprotroph\| | Yeast | NULL | Probable | As nectar-tap_saprotroph (Põlme et al. 2020); Decay sugar-rich substrates (Põlme et al. 2020) | Põlme S, et al. 2020. Fungal Diversity 105:1-16 ((https://doi.org/10.1007/s13225-020-00466-2)) |
| Otu25852 | 1 | k__Fungi(100);p__Ascomycota(100);c__Saccharomycetes(100);o__Saccharomycetales(100);f__Metschnikowiaceae(100);g__Metschnikowia(100);g__Metschnikowia_unclassified(100); | Metschnikowia | 13 | Saprotroph | \|Undefined Saprotroph\| | Yeast | NULL | Probable | As nectar-tap_saprotroph (Põlme et al. 2020); Decay sugar-rich substrates (Põlme et al. 2020) | Põlme S, et al. 2020. Fungal Diversity 105:1-16 ((https://doi.org/10.1007/s13225-020-00466-2)) |
| Otu25853 | 1 | k__Fungi(100);p__Ascomycota(100);c__Saccharomycetes(100);o__Saccharomycetales(100);f__Metschnikowiaceae(100);g__Metschnikowia(100);s__Metschnikowia_pulcherrima(100); | Metschnikowia | 13 | Saprotroph | \|Undefined Saprotroph\| | Yeast | NULL | Probable | As nectar-tap_saprotroph (Põlme et al. 2020); Decay sugar-rich substrates (Põlme et al. 2020) | Põlme S, et al. 2020. Fungal Diversity 105:1-16 ((https://doi.org/10.1007/s13225-020-00466-2)) |
| Otu25854 | 1 | k__Fungi(100);p__Ascomycota(100);c__Saccharomycetes(100);o__Saccharomycetales(100);f__Metschnikowiaceae(100);g__Metschnikowia(100);g__Metschnikowia_unclassified(100); | Metschnikowia | 13 | Saprotroph | \|Undefined Saprotroph\| | Yeast | NULL | Probable | As nectar-tap_saprotroph (Põlme et al. 2020); Decay sugar-rich substrates (Põlme et al. 2020) | Põlme S, et al. 2020. Fungal Diversity 105:1-16 ((https://doi.org/10.1007/s13225-020-00466-2)) |
| Otu25855 | 1 | k__Fungi(100);p__Ascomycota(100);c__Saccharomycetes(100);o__Saccharomycetales(100);f__Metschnikowiaceae(100);g__Metschnikowia(100);s__Metschnikowia_pulcherrima(100); | Metschnikowia | 13 | Saprotroph | \|Undefined Saprotroph\| | Yeast | NULL | Probable | As nectar-tap_saprotroph (Põlme et al. 2020); Decay sugar-rich substrates (Põlme et al. 2020) | Põlme S, et al. 2020. Fungal Diversity 105:1-16 ((https://doi.org/10.1007/s13225-020-00466-2)) |
| Otu25856 | 1 | k__Fungi(100);p__Ascomycota(100);c__Saccharomycetes(100);o__Saccharomycetales(100);f__Metschnikowiaceae(100);g__Metschnikowia(100);s__Metschnikowia_pulcherrima(100); | Metschnikowia | 13 | Saprotroph | \|Undefined Saprotroph\| | Yeast | NULL | Probable | As nectar-tap_saprotroph (Põlme et al. 2020); Decay sugar-rich substrates (Põlme et al. 2020) | Põlme S, et al. 2020. Fungal Diversity 105:1-16 ((https://doi.org/10.1007/s13225-020-00466-2)) |

| | | | | | | | | | | | |
|---|---|---|---|---|---|---|---|---|---|---|---|
| Otu25857 | 1 | k__Fungi(100);p__Ascomycota(100);c__Saccharomycetes(100);o__Saccharomycetales(100);f__Metschnikowiaceae(100);g__Metschnikowia(100);g__Metschnikowia_unclassified(100); | Metschnikowia | 13 | Saprotroph | \|Undefined Saprotroph\| | Yeast | NULL | Probable | As nectar-tap_saprotroph (Põlme et al. 2020); Decay sugar-rich substrates (Põlme et al. 2020) | Põlme S, et al. 2020. Fungal Diversity 105:1-16 ((https://doi.org/10.1007/s13225-020-00466-2)) |
| Otu25858 | 1 | k__Fungi(100);p__Ascomycota(100);c__Saccharomycetes(100);o__Saccharomycetales(100);f__Metschnikowiaceae(100);g__Metschnikowia(100);s__Metschnikowia_pulcherrima(100); | Metschnikowia | 13 | Saprotroph | \|Undefined Saprotroph\| | Yeast | NULL | Probable | As nectar-tap_saprotroph (Põlme et al. 2020); Decay sugar-rich substrates (Põlme et al. 2020) | Põlme S, et al. 2020. Fungal Diversity 105:1-16 ((https://doi.org/10.1007/s13225-020-00466-2)) |
| Otu25859 | 1 | k__Fungi(100);p__Ascomycota(100);c__Saccharomycetes(100);o__Saccharomycetales(100);f__Metschnikowiaceae(100);g__Metschnikowia(100);s__Metschnikowia_pulcherrima(100); | Metschnikowia | 13 | Saprotroph | \|Undefined Saprotroph\| | Yeast | NULL | Probable | As nectar-tap_saprotroph (Põlme et al. 2020); Decay sugar-rich substrates (Põlme et al. 2020) | Põlme S, et al. 2020. Fungal Diversity 105:1-16 ((https://doi.org/10.1007/s13225-020-00466-2)) |
| Otu25861 | 1 | k__Fungi(100);p__Ascomycota(100);c__Saccharomycetes(100);o__Saccharomycetales(100);f__Metschnikowiaceae(100);g__Metschnikowia(100);s__Metschnikowia_pulcherrima(100); | Metschnikowia | 13 | Saprotroph | \|Undefined Saprotroph\| | Yeast | NULL | Probable | As nectar-tap_saprotroph (Põlme et al. 2020); Decay sugar-rich substrates (Põlme et al. 2020) | Põlme S, et al. 2020. Fungal Diversity 105:1-16 ((https://doi.org/10.1007/s13225-020-00466-2)) |
| Otu25862 | 1 | k__Fungi(100);p__Ascomycota(100);c__Saccharomycetes(100);o__Saccharomycetales(100);f__Saccharomycetales_family_Incertae_sedis(100);g__Candida(100);g__Candida_unclassified(100); | Candida | 13 | Pathotroph-Saprotroph-Symbiotroph | Animal Pathogen-Endophyte-Endosymbiont-Epiphyte-\|Undefined Saprotroph\| | Dimorphic Yeast-Yeast | NULL | Probable | As nectar-tap_saprotroph (Põlme et al. 2020); Decay sugar-rich substrates (Põlme et al. 2020); This asexual genus is polyphyletic and contain species that function very differently depend on what they are. Caution should be taken when interpreting guilds for Candida species. | Manolakaki D, et al. 2010. Virulence. 1: 367-375 ((https://doi.org/10.4161/viru.1.5.12796)); Mekbib SB, et al. 2011. Tropical Plant Pathology 36:233-240 ((https://doi.org/10.1590/S1982-56762011000400004)); Põlme S, et al. 2020. Fungal Diversity 105:1-16 ((https://doi.org/10.1007/s13225-020-00466-2)) |
| Otu25863 | 1 | k__Fungi(100);p__Ascomycota(100);c__Saccharomycetes(100);o__Saccharomycetales(100);f__Metschnikowiaceae(100);g__Metschnikowia(100);s__Metschnikowia_pulcherrima(100); | Metschnikowia | 13 | Saprotroph | \|Undefined Saprotroph\| | Yeast | NULL | Probable | As nectar-tap_saprotroph (Põlme et al. 2020); Decay sugar-rich substrates (Põlme et al. 2020) | Põlme S, et al. 2020. Fungal Diversity 105:1-16 ((https://doi.org/10.1007/s13225-020-00466-2)) |
| Otu25864 | 1 | k__Fungi(100);p__Ascomycota(100);c__Saccharomycetes(100);o__Saccharomycetales(100);f__Metschnikowiaceae(100);g__Metschnikowia(100);s__Metschnikowia_pulcherrima(100); | Metschnikowia | 13 | Saprotroph | \|Undefined Saprotroph\| | Yeast | NULL | Probable | As nectar-tap_saprotroph (Põlme et al. 2020); Decay sugar-rich substrates (Põlme et al. 2020) | Põlme S, et al. 2020. Fungal Diversity 105:1-16 ((https://doi.org/10.1007/s13225-020-00466-2)) |
| Otu25866 | 1 | k__Fungi(100);p__Ascomycota(100);c__Saccharomycetes(100);o__Saccharomycetales(100);f__Saccharomycetales_family_Incertae_sedis(100);g__Candida(100);g__Candida_unclassified(100); | Candida | 13 | Pathotroph-Saprotroph-Symbiotroph | Animal Pathogen-Endophyte-Endosymbiont-Epiphyte-\|Undefined Saprotroph\| | Dimorphic Yeast-Yeast | NULL | Probable | As nectar-tap_saprotroph (Põlme et al. 2020); Decay sugar-rich substrates (Põlme et al. 2020); This asexual genus is polyphyletic and contain species that function very differently depend on what they are. Caution should be taken when interpreting guilds for Candida species. | Manolakaki D, et al. 2010. Virulence. 1: 367-375 ((https://doi.org/10.4161/viru.1.5.12796)); Mekbib SB, et al. 2011. Tropical Plant Pathology 36:233-240 ((https://doi.org/10.1590/S1982-56762011000400004)); Põlme S, et al. 2020. Fungal Diversity 105:1-16 ((https://doi.org/10.1007/s13225-020-00466-2)) |
| Otu25870 | 1 | k__Fungi(100);p__Ascomycota(100);c__Saccharomycetes(100);o__Saccharomycetales(100);f__Metschnikowiaceae(100);g__Metschnikowia(100);s__Metschnikowia_pulcherrima(100); | Metschnikowia | 13 | Saprotroph | \|Undefined Saprotroph\| | Yeast | NULL | Probable | As nectar-tap_saprotroph (Põlme et al. 2020); Decay sugar-rich substrates (Põlme et al. 2020) | Põlme S, et al. 2020. Fungal Diversity 105:1-16 ((https://doi.org/10.1007/s13225-020-00466-2)) |
| Otu25877 | 1 | k__Fungi(100);p__Ascomycota(100);c__Saccharomycetes(100);o__Saccharomycetales(100);f__Saccharomycetales_family_Incertae_sedis(100);g__Candida(100);s__Candida_cretensis(100); | Candida | 13 | Pathotroph-Saprotroph-Symbiotroph | Animal Pathogen-Endophyte-Endosymbiont-Epiphyte-\|Undefined Saprotroph\| | Dimorphic Yeast-Yeast | NULL | Probable | As nectar-tap_saprotroph (Põlme et al. 2020); Decay sugar-rich substrates (Põlme et al. 2020); This asexual genus is polyphyletic and contain species that function very differently depend on what they are. Caution should be taken when interpreting guilds for Candida species. | Manolakaki D, et al. 2010. Virulence. 1: 367-375 ((https://doi.org/10.4161/viru.1.5.12796)); Mekbib SB, et al. 2011. Tropical Plant Pathology 36:233-240 ((https://doi.org/10.1590/S1982-56762011000400004)); Põlme S, et al. 2020. Fungal Diversity 105:1-16 ((https://doi.org/10.1007/s13225-020-00466-2)) |
| Otu25880 | 1 | k__Fungi(100);p__Ascomycota(100);c__Saccharomycetes(100);o__Saccharomycetales(100);f__Saccharomycetales_family_Incertae_sedis(100);g__Candida(100);s__Candida_cretensis(100); | Candida | 13 | Pathotroph-Saprotroph-Symbiotroph | Animal Pathogen-Endophyte-Endosymbiont-Epiphyte-\|Undefined Saprotroph\| | Dimorphic Yeast-Yeast | NULL | Probable | As nectar-tap_saprotroph (Põlme et al. 2020); Decay sugar-rich substrates (Põlme et al. 2020); This asexual genus is polyphyletic and contain species that function very differently depend on what they are. Caution should be taken when interpreting guilds for Candida species. | Manolakaki D, et al. 2010. Virulence. 1: 367-375 ((https://doi.org/10.4161/viru.1.5.12796)); Mekbib SB, et al. 2011. Tropical Plant Pathology 36:233-240 ((https://doi.org/10.1590/S1982-56762011000400004)); Põlme S, et al. 2020. Fungal Diversity 105:1-16 ((https://doi.org/10.1007/s13225-020-00466-2)) |
| Otu25881 | 1 | k__Fungi(100);p__Ascomycota(100);c__Saccharomycetes(100);o__Saccharomycetales(100);f__Metschnikowiaceae(100);g__Metschnikowia(100);g__Metschnikowia_unclassified(100); | Metschnikowia | 13 | Saprotroph | \|Undefined Saprotroph\| | Yeast | NULL | Probable | As nectar-tap_saprotroph (Põlme et al. 2020); Decay sugar-rich substrates (Põlme et al. 2020) | Põlme S, et al. 2020. Fungal Diversity 105:1-16 ((https://doi.org/10.1007/s13225-020-00466-2)) |
| Otu25884 | 1 | k__Fungi(100);p__Ascomycota(100);c__Saccharomycetes(100);o__Saccharomycetales(100);f__Metschnikowiaceae(100);g__Metschnikowia(100);s__Metschnikowia_pulcherrima(100); | Metschnikowia | 13 | Saprotroph | \|Undefined Saprotroph\| | Yeast | NULL | Probable | As nectar-tap_saprotroph (Põlme et al. 2020); Decay sugar-rich substrates (Põlme et al. 2020) | Põlme S, et al. 2020. Fungal Diversity 105:1-16 ((https://doi.org/10.1007/s13225-020-00466-2)) |
| Otu25900 | 1 | k__Fungi(100);p__Ascomycota(100);c__Saccharomycetes(100);o__Saccharomycetales(100);f__Metschnikowiaceae(100);g__Metschnikowia(100);g__Metschnikowia_unclassified(100); | Metschnikowia | 13 | Saprotroph | \|Undefined Saprotroph\| | Yeast | NULL | Probable | As nectar-tap_saprotroph (Põlme et al. 2020); Decay sugar-rich substrates (Põlme et al. 2020) | Põlme S, et al. 2020. Fungal Diversity 105:1-16 ((https://doi.org/10.1007/s13225-020-00466-2)) |
| Otu25913 | 1 | k__Fungi(100);p__Ascomycota(100);c__Saccharomycetes(100);o__Saccharomycetales(100);f__Metschnikowiaceae(100);g__Metschnikowia(100);s__Metschnikowia_pulcherrima(100); | Metschnikowia | 13 | Saprotroph | \|Undefined Saprotroph\| | Yeast | NULL | Probable | As nectar-tap_saprotroph (Põlme et al. 2020); Decay sugar-rich substrates (Põlme et al. 2020) | Põlme S, et al. 2020. Fungal Diversity 105:1-16 ((https://doi.org/10.1007/s13225-020-00466-2)) |
| Otu25914 | 1 | k__Fungi(100);p__Ascomycota(100);c__Saccharomycetes(100);o__Saccharomycetales(100);f__Metschnikowiaceae(100);g__Metschnikowia(100);s__Metschnikowia_pulcherrima(100); | Metschnikowia | 13 | Saprotroph | \|Undefined Saprotroph\| | Yeast | NULL | Probable | As nectar-tap_saprotroph (Põlme et al. 2020); Decay sugar-rich substrates (Põlme et al. 2020) | Põlme S, et al. 2020. Fungal Diversity 105:1-16 ((https://doi.org/10.1007/s13225-020-00466-2)) |
| Otu25915 | 1 | k__Fungi(100);p__Ascomycota(100);c__Saccharomycetes(100);o__Saccharomycetales(100);f__Metschnikowiaceae(100);g__Metschnikowia(100);s__Metschnikowia_pulcherrima(100); | Metschnikowia | 13 | Saprotroph | \|Undefined Saprotroph\| | Yeast | NULL | Probable | As nectar-tap_saprotroph (Põlme et al. 2020); Decay sugar-rich substrates (Põlme et al. 2020) | Põlme S, et al. 2020. Fungal Diversity 105:1-16 ((https://doi.org/10.1007/s13225-020-00466-2)) |
| Otu25916 | 1 | k__Fungi(100);p__Ascomycota(100);c__Saccharomycetes(100);o__Saccharomycetales(100);f__Metschnikowiaceae(100);g__Metschnikowia(100);s__Metschnikowia_pulcherrima(100); | Metschnikowia | 13 | Saprotroph | \|Undefined Saprotroph\| | Yeast | NULL | Probable | As nectar-tap_saprotroph (Põlme et al. 2020); Decay sugar-rich substrates (Põlme et al. 2020) | Põlme S, et al. 2020. Fungal Diversity 105:1-16 ((https://doi.org/10.1007/s13225-020-00466-2)) |
| Otu25920 | 1 | k__Fungi(100);p__Ascomycota(100);c__Saccharomycetes(100);o__Saccharomycetales(100);f__Metschnikowiaceae(100);g__Metschnikowia(100);s__Metschnikowia_pulcherrima(100); | Metschnikowia | 13 | Saprotroph | \|Undefined Saprotroph\| | Yeast | NULL | Probable | As nectar-tap_saprotroph (Põlme et al. 2020); Decay sugar-rich substrates (Põlme et al. 2020) | Põlme S, et al. 2020. Fungal Diversity 105:1-16 ((https://doi.org/10.1007/s13225-020-00466-2)) |
| Otu25921 | 1 | k__Fungi(100);p__Ascomycota(100);c__Saccharomycetes(100);o__Saccharomycetales(100);f__Metschnikowiaceae(100);g__Metschnikowia(100);g__Metschnikowia_unclassified(100); | Metschnikowia | 13 | Saprotroph | \|Undefined Saprotroph\| | Yeast | NULL | Probable | As nectar-tap_saprotroph (Põlme et al. 2020); Decay sugar-rich substrates (Põlme et al. 2020) | Põlme S, et al. 2020. Fungal Diversity 105:1-16 ((https://doi.org/10.1007/s13225-020-00466-2)) |
| Otu25922 | 1 | k__Fungi(100);p__Ascomycota(100);c__Saccharomycetes(100);o__Saccharomycetales(100);f__Metschnikowiaceae(100);g__Metschnikowia(100);g__Metschnikowia_unclassified(100); | Metschnikowia | 13 | Saprotroph | \|Undefined Saprotroph\| | Yeast | NULL | Probable | As nectar-tap_saprotroph (Põlme et al. 2020); Decay sugar-rich substrates (Põlme et al. 2020) | Põlme S, et al. 2020. Fungal Diversity 105:1-16 ((https://doi.org/10.1007/s13225-020-00466-2)) |
| Otu25923 | 1 | k__Fungi(100);p__Ascomycota(100);c__Saccharomycetes(100);o__Saccharomycetales(100);f__Metschnikowiaceae(100);g__Metschnikowia(100);s__Metschnikowia_pulcherrima(100); | Metschnikowia | 13 | Saprotroph | \|Undefined Saprotroph\| | Yeast | NULL | Probable | As nectar-tap_saprotroph (Põlme et al. 2020); Decay sugar-rich substrates (Põlme et al. 2020) | Põlme S, et al. 2020. Fungal Diversity 105:1-16 ((https://doi.org/10.1007/s13225-020-00466-2)) |
| Otu25924 | 1 | k__Fungi(100);p__Ascomycota(100);c__Saccharomycetes(100);o__Saccharomycetales(100);f__Metschnikowiaceae(100);g__Metschnikowia(100);s__Metschnikowia_pulcherrima(100); | Metschnikowia | 13 | Saprotroph | \|Undefined Saprotroph\| | Yeast | NULL | Probable | As nectar-tap_saprotroph (Põlme et al. 2020); Decay sugar-rich substrates (Põlme et al. 2020) | Põlme S, et al. 2020. Fungal Diversity 105:1-16 ((https://doi.org/10.1007/s13225-020-00466-2)) |
| Otu25925 | 1 | k__Fungi(100);p__Ascomycota(100);c__Saccharomycetes(100);o__Saccharomycetales(100);f__Metschnikowiaceae(100);g__Metschnikowia(100);s__Metschnikowia_pulcherrima(100); | Metschnikowia | 13 | Saprotroph | \|Undefined Saprotroph\| | Yeast | NULL | Probable | As nectar-tap_saprotroph (Põlme et al. 2020); Decay sugar-rich substrates (Põlme et al. 2020) | Põlme S, et al. 2020. Fungal Diversity 105:1-16 ((https://doi.org/10.1007/s13225-020-00466-2)) |
| Otu25927 | 1 | k__Fungi(100);p__Ascomycota(100);c__Saccharomycetes(100);o__Saccharomycetales(100);f__Metschnikowiaceae(100);g__Metschnikowia(100);g__Metschnikowia_unclassified(100); | Metschnikowia | 13 | Saprotroph | \|Undefined Saprotroph\| | Yeast | NULL | Probable | As nectar-tap_saprotroph (Põlme et al. 2020); Decay sugar-rich substrates (Põlme et al. 2020) | Põlme S, et al. 2020. Fungal Diversity 105:1-16 ((https://doi.org/10.1007/s13225-020-00466-2)) |
| Otu25928 | 1 | k__Fungi(100);p__Ascomycota(100);c__Saccharomycetes(100);o__Saccharomycetales(100);f__Metschnikowiaceae(100);g__Metschnikowia(100);g__Metschnikowia_unclassified(100); | Metschnikowia | 13 | Saprotroph | \|Undefined Saprotroph\| | Yeast | NULL | Probable | As nectar-tap_saprotroph (Põlme et al. 2020); Decay sugar-rich substrates (Põlme et al. 2020) | Põlme S, et al. 2020. Fungal Diversity 105:1-16 ((https://doi.org/10.1007/s13225-020-00466-2)) |
| Otu25929 | 1 | k__Fungi(100);p__Ascomycota(100);c__Saccharomycetes(100);o__Saccharomycetales(100);f__Metschnikowiaceae(100);g__Metschnikowia(100);g__Metschnikowia_unclassified(100); | Metschnikowia | 13 | Saprotroph | \|Undefined Saprotroph\| | Yeast | NULL | Probable | As nectar-tap_saprotroph (Põlme et al. 2020); Decay sugar-rich substrates (Põlme et al. 2020) | Põlme S, et al. 2020. Fungal Diversity 105:1-16 ((https://doi.org/10.1007/s13225-020-00466-2)) |
| Otu25930 | 1 | k__Fungi(100);p__Ascomycota(100);c__Saccharomycetes(100);o__Saccharomycetales(100);f__Metschnikowiaceae(100);g__Metschnikowia(100);g__Metschnikowia_unclassified(100); | Metschnikowia | 13 | Saprotroph | \|Undefined Saprotroph\| | Yeast | NULL | Probable | As nectar-tap_saprotroph (Põlme et al. 2020); Decay sugar-rich substrates (Põlme et al. 2020) | Põlme S, et al. 2020. Fungal Diversity 105:1-16 ((https://doi.org/10.1007/s13225-020-00466-2)) |
| Otu25932 | 1 | k__Fungi(100);p__Ascomycota(100);c__Saccharomycetes(100);o__Saccharomycetales(100);f__Metschnikowiaceae(100);g__Metschnikowia(100);s__Metschnikowia_pulcherrima(100); | Metschnikowia | 13 | Saprotroph | \|Undefined Saprotroph\| | Yeast | NULL | Probable | As nectar-tap_saprotroph (Põlme et al. 2020); Decay sugar-rich substrates (Põlme et al. 2020) | Põlme S, et al. 2020. Fungal Diversity 105:1-16 ((https://doi.org/10.1007/s13225-020-00466-2)) |
| Otu25933 | 1 | k__Fungi(100);p__Ascomycota(100);c__Saccharomycetes(100);o__Saccharomycetales(100);f__Metschnikowiaceae(100);g__Metschnikowia(100);s__Metschnikowia_pulcherrima(100); | Metschnikowia | 13 | Saprotroph | \|Undefined Saprotroph\| | Yeast | NULL | Probable | As nectar-tap_saprotroph (Põlme et al. 2020); Decay sugar-rich substrates (Põlme et al. 2020) | Põlme S, et al. 2020. Fungal Diversity 105:1-16 ((https://doi.org/10.1007/s13225-020-00466-2)) |
| Otu25934 | 1 | k__Fungi(100);p__Ascomycota(100);c__Saccharomycetes(100);o__Saccharomycetales(100);f__Metschnikowiaceae(100);g__Metschnikowia(100);g__Metschnikowia_unclassified(100); | Metschnikowia | 13 | Saprotroph | \|Undefined Saprotroph\| | Yeast | NULL | Probable | As nectar-tap_saprotroph (Põlme et al. 2020); Decay sugar-rich substrates (Põlme et al. 2020) | Põlme S, et al. 2020. Fungal Diversity 105:1-16 ((https://doi.org/10.1007/s13225-020-00466-2)) |
| Otu25935 | 1 | k__Fungi(100);p__Ascomycota(100);c__Saccharomycetes(100);o__Saccharomycetales(100);f__Metschnikowiaceae(100);g__Metschnikowia(100);s__Metschnikowia_pulcherrima(100); | Metschnikowia | 13 | Saprotroph | \|Undefined Saprotroph\| | Yeast | NULL | Probable | As nectar-tap_saprotroph (Põlme et al. 2020); Decay sugar-rich substrates (Põlme et al. 2020) | Põlme S, et al. 2020. Fungal Diversity 105:1-16 ((https://doi.org/10.1007/s13225-020-00466-2)) |
| Otu25938 | 1 | k__Fungi(100);p__Ascomycota(100);c__Saccharomycetes(100);o__Saccharomycetales(100);f__Metschnikowiaceae(100);g__Metschnikowia(100);s__Metschnikowia_pulcherrima(100); | Metschnikowia | 13 | Saprotroph | \|Undefined Saprotroph\| | Yeast | NULL | Probable | As nectar-tap_saprotroph (Põlme et al. 2020); Decay sugar-rich substrates (Põlme et al. 2020) | Põlme S, et al. 2020. Fungal Diversity 105:1-16 ((https://doi.org/10.1007/s13225-020-00466-2)) |
| Otu25945 | 1 | k__Fungi(100);p__Ascomycota(100);c__Saccharomycetes(100);o__Saccharomycetales(100);f__Metschnikowiaceae(100);g__Metschnikowia(100);s__Metschnikowia_pulcherrima(100); | Metschnikowia | 13 | Saprotroph | \|Undefined Saprotroph\| | Yeast | NULL | Probable | As nectar-tap_saprotroph (Põlme et al. 2020); Decay sugar-rich substrates (Põlme et al. 2020) | Põlme S, et al. 2020. Fungal Diversity 105:1-16 ((https://doi.org/10.1007/s13225-020-00466-2)) |
| Otu25946 | 1 | k__Fungi(100);p__Ascomycota(100);c__Saccharomycetes(100);o__Saccharomycetales(100);f__Metschnikowiaceae(100);g__Metschnikowia(100);s__Metschnikowia_pulcherrima(100); | Metschnikowia | 13 | Saprotroph | \|Undefined Saprotroph\| | Yeast | NULL | Probable | As nectar-tap_saprotroph (Põlme et al. 2020); Decay sugar-rich substrates (Põlme et al. 2020) | Põlme S, et al. 2020. Fungal Diversity 105:1-16 ((https://doi.org/10.1007/s13225-020-00466-2)) |
| Otu25948 | 1 | k__Fungi(100);p__Ascomycota(100);c__Saccharomycetes(100);o__Saccharomycetales(100);f__Metschnikowiaceae(100);g__Metschnikowia(100);s__Metschnikowia_pulcherrima(100); | Metschnikowia | 13 | Saprotroph | \|Undefined Saprotroph\| | Yeast | NULL | Probable | As nectar-tap_saprotroph (Põlme et al. 2020); Decay sugar-rich substrates (Põlme et al. 2020) | Põlme S, et al. 2020. Fungal Diversity 105:1-16 ((https://doi.org/10.1007/s13225-020-00466-2)) |
| Otu25949 | 1 | k__Fungi(100);p__Ascomycota(100);c__Saccharomycetes(100);o__Saccharomycetales(100);f__Metschnikowiaceae(100);g__Metschnikowia(100);s__Metschnikowia_pulcherrima(100); | Metschnikowia | 13 | Saprotroph | \|Undefined Saprotroph\| | Yeast | NULL | Probable | As nectar-tap_saprotroph (Põlme et al. 2020); Decay sugar-rich substrates (Põlme et al. 2020) | Põlme S, et al. 2020. Fungal Diversity 105:1-16 ((https://doi.org/10.1007/s13225-020-00466-2)) |
| Otu25952 | 1 | k__Fungi(100);p__Ascomycota(100);c__Saccharomycetes(100);o__Saccharomycetales(100);f__Metschnikowiaceae(100);g__Metschnikowia(100);s__Metschnikowia_pulcherrima(100); | Metschnikowia | 13 | Saprotroph | \|Undefined Saprotroph\| | Yeast | NULL | Probable | As nectar-tap_saprotroph (Põlme et al. 2020); Decay sugar-rich substrates (Põlme et al. 2020) | Põlme S, et al. 2020. Fungal Diversity 105:1-16 ((https://doi.org/10.1007/s13225-020-00466-2)) |
| Otu25955 | 1 | k__Fungi(100);p__Ascomycota(100);c__Saccharomycetes(100);o__Saccharomycetales(100);f__Metschnikowiaceae(100);g__Metschnikowia(100);s__Metschnikowia_pulcherrima(100); | Metschnikowia | 13 | Saprotroph | \|Undefined Saprotroph\| | Yeast | NULL | Probable | As nectar-tap_saprotroph (Põlme et al. 2020); Decay sugar-rich substrates (Põlme et al. 2020) | Põlme S, et al. 2020. Fungal Diversity 105:1-16 ((https://doi.org/10.1007/s13225-020-00466-2)) |
| Otu25956 | 1 | k__Fungi(100);p__Ascomycota(100);c__Saccharomycetes(100);o__Saccharomycetales(100);f__Metschnikowiaceae(100);g__Metschnikowia(100);s__Metschnikowia_pulcherrima(100); | Metschnikowia | 13 | Saprotroph | \|Undefined Saprotroph\| | Yeast | NULL | Probable | As nectar-tap_saprotroph (Põlme et al. 2020); Decay sugar-rich substrates (Põlme et al. 2020) | Põlme S, et al. 2020. Fungal Diversity 105:1-16 ((https://doi.org/10.1007/s13225-020-00466-2)) |
| Otu25957 | 1 | k__Fungi(100);p__Ascomycota(100);c__Saccharomycetes(100);o__Saccharomycetales(100);f__Metschnikowiaceae(100);g__Metschnikowia(100);s__Metschnikowia_pulcherrima(100); | Metschnikowia | 13 | Saprotroph | \|Undefined Saprotroph\| | Yeast | NULL | Probable | As nectar-tap_saprotroph (Põlme et al. 2020); Decay sugar-rich substrates (Põlme et al. 2020) | Põlme S, et al. 2020. Fungal Diversity 105:1-16 ((https://doi.org/10.1007/s13225-020-00466-2)) |
| Otu25959 | 1 | k__Fungi(100);p__Ascomycota(100);c__Saccharomycetes(100);o__Saccharomycetales(100);f__Metschnikowiaceae(100);g__Metschnikowia(100);g__Metschnikowia_unclassified(100); | Metschnikowia | 13 | Saprotroph | \|Undefined Saprotroph\| | Yeast | NULL | Probable | As nectar-tap_saprotroph (Põlme et al. 2020); Decay sugar-rich substrates (Põlme et al. 2020) | Põlme S, et al. 2020. Fungal Diversity 105:1-16 ((https://doi.org/10.1007/s13225-020-00466-2)) |
| Otu25960 | 1 | k__Fungi(100);p__Ascomycota(100);c__Saccharomycetes(100);o__Saccharomycetales(100);f__Metschnikowiaceae(100);g__Metschnikowia(100);g__Metschnikowia_unclassified(100); | Metschnikowia | 13 | Saprotroph | \|Undefined Saprotroph\| | Yeast | NULL | Probable | As nectar-tap_saprotroph (Põlme et al. 2020); Decay sugar-rich substrates (Põlme et al. 2020) | Põlme S, et al. 2020. Fungal Diversity 105:1-16 ((https://doi.org/10.1007/s13225-020-00466-2)) |
| Otu25961 | 1 | k__Fungi(100);p__Ascomycota(100);c__Saccharomycetes(100);o__Saccharomycetales(100);f__Metschnikowiaceae(100);g__Metschnikowia(100);g__Metschnikowia_unclassified(100); | Metschnikowia | 13 | Saprotroph | \|Undefined Saprotroph\| | Yeast | NULL | Probable | As nectar-tap_saprotroph (Põlme et al. 2020); Decay sugar-rich substrates (Põlme et al. 2020) | Põlme S, et al. 2020. Fungal Diversity 105:1-16 ((https://doi.org/10.1007/s13225-020-00466-2)) |
| Otu25962 | 1 | k__Fungi(100);p__Ascomycota(100);c__Saccharomycetes(100);o__Saccharomycetales(100);f__Metschnikowiaceae(100);g__Metschnikowia(100);s__Metschnikowia_pulcherrima(100); | Metschnikowia | 13 | Saprotroph | \|Undefined Saprotroph\| | Yeast | NULL | Probable | As nectar-tap_saprotroph (Põlme et al. 2020); Decay sugar-rich substrates (Põlme et al. 2020) | Põlme S, et al. 2020. Fungal Diversity 105:1-16 ((https://doi.org/10.1007/s13225-020-00466-2)) |
| Otu25963 | 1 | k__Fungi(100);p__Ascomycota(100);c__Saccharomycetes(100);o__Saccharomycetales(100);f__Metschnikowiaceae(100);g__Metschnikowia(100);s__Metschnikowia_pulcherrima(100); | Metschnikowia | 13 | Saprotroph | \|Undefined Saprotroph\| | Yeast | NULL | Probable | As nectar-tap_saprotroph (Põlme et al. 2020); Decay sugar-rich substrates (Põlme et al. 2020) | Põlme S, et al. 2020. Fungal Diversity 105:1-16 ((https://doi.org/10.1007/s13225-020-00466-2)) |
| Otu25967 | 1 | k__Fungi(100);p__Ascomycota(100);c__Saccharomycetes(100);o__Saccharomycetales(100);f__Metschnikowiaceae(100);g__Metschnikowia(100);s__Metschnikowia_pulcherrima(100); | Metschnikowia | 13 | Saprotroph | \|Undefined Saprotroph\| | Yeast | NULL | Probable | As nectar-tap_saprotroph (Põlme et al. 2020); Decay sugar-rich substrates (Põlme et al. 2020) | Põlme S, et al. 2020. Fungal Diversity 105:1-16 ((https://doi.org/10.1007/s13225-020-00466-2)) |
| Otu25974 | 1 | k__Fungi(100);p__Ascomycota(100);c__Saccharomycetes(100);o__Saccharomycetales(100);f__Metschnikowiaceae(100);g__Metschnikowia(100);s__Metschnikowia_pulcherrima(100); | Metschnikowia | 13 | Saprotroph | \|Undefined Saprotroph\| | Yeast | NULL | Probable | As nectar-tap_saprotroph (Põlme et al. 2020); Decay sugar-rich substrates (Põlme et al. 2020) | Põlme S, et al. 2020. Fungal Diversity 105:1-16 ((https://doi.org/10.1007/s13225-020-00466-2)) |
| Otu25992 | 1 | k__Fungi(100);p__Ascomycota(100);c__Saccharomycetes(100);o__Saccharomycetales(100);f__Metschnikowiaceae(100);g__Metschnikowia(100);g__Metschnikowia_unclassified(100); | Metschnikowia | 13 | Saprotroph | \|Undefined Saprotroph\| | Yeast | NULL | Probable | As nectar-tap_saprotroph (Põlme et al. 2020); Decay sugar-rich substrates (Põlme et al. 2020) | Põlme S, et al. 2020. Fungal Diversity 105:1-16 ((https://doi.org/10.1007/s13225-020-00466-2)) |
| Otu25994 | 1 | k__Fungi(100);p__Ascomycota(100);c__Saccharomycetes(100);o__Saccharomycetales(100);f__Metschnikowiaceae(100);g__Metschnikowia(100);g__Metschnikowia_unclassified(100); | Metschnikowia | 13 | Saprotroph | \|Undefined Saprotroph\| | Yeast | NULL | Probable | As nectar-tap_saprotroph (Põlme et al. 2020); Decay sugar-rich substrates (Põlme et al. 2020) | Põlme S, et al. 2020. Fungal Diversity 105:1-16 ((https://doi.org/10.1007/s13225-020-00466-2)) |
| Otu25995 | 1 | k__Fungi(100);p__Ascomycota(100);c__Saccharomycetes(100);o__Saccharomycetales(100);f__Saccharomycetales_family_Incertae_sedis(100);g__Candida(100);s__Candida_cretensis(100); | Candida | 13 | Pathotroph-Saprotroph-Symbiotroph | Animal Pathogen-Endophyte-Endosymbiont-Epiphyte-\|Undefined Saprotroph\| | Dimorphic Yeast-Yeast | NULL | Probable | As nectar-tap_saprotroph (Põlme et al. 2020); Decay sugar-rich substrates (Põlme et al. 2020); This asexual genus is polyphyletic and contain species that function very differently depend on what they are. Caution should be taken when interpreting guilds for Candida species. | Manolakaki D, et al. 2010. Virulence. 1: 367-375 ((https://doi.org/10.4161/viru.1.5.12796)); Mekbib SB, et al. 2011. Tropical Plant Pathology 36:233-240 ((https://doi.org/10.1590/S1982-56762011000400004)); Põlme S, et al. 2020. Fungal Diversity 105:1-16 ((https://doi.org/10.1007/s13225-020-00466-2)) |
| Otu25997 | 1 | k__Fungi(100);p__Ascomycota(100);c__Saccharomycetes(100);o__Saccharomycetales(100);f__Saccharomycetales_family_Incertae_sedis(100);g__Candida(100);g__Candida_unclassified(100); | Candida | 13 | Pathotroph-Saprotroph-Symbiotroph | Animal Pathogen-Endophyte-Endosymbiont-Epiphyte-\|Undefined Saprotroph\| | Dimorphic Yeast-Yeast | NULL | Probable | As nectar-tap_saprotroph (Põlme et al. 2020); Decay sugar-rich substrates (Põlme et al. 2020); This asexual genus is polyphyletic and contain species that function very differently depend on what they are. Caution should be taken when interpreting guilds for Candida species. | Manolakaki D, et al. 2010. Virulence. 1: 367-375 ((https://doi.org/10.4161/viru.1.5.12796)); Mekbib SB, et al. 2011. Tropical Plant Pathology 36:233-240 ((https://doi.org/10.1590/S1982-56762011000400004)); Põlme S, et al. 2020. Fungal Diversity 105:1-16 ((https://doi.org/10.1007/s13225-020-00466-2)) |
| Otu25998 | 1 | k__Fungi(100);p__Ascomycota(100);c__Saccharomycetes(100);o__Saccharomycetales(100);f__Metschnikowiaceae(100);g__Metschnikowia(100);s__Metschnikowia_pulcherrima(100); | Metschnikowia | 13 | Saprotroph | \|Undefined Saprotroph\| | Yeast | NULL | Probable | As nectar-tap_saprotroph (Põlme et al. 2020); Decay sugar-rich substrates (Põlme et al. 2020) | Põlme S, et al. 2020. Fungal Diversity 105:1-16 ((https://doi.org/10.1007/s13225-020-00466-2)) |
| Otu25999 | 1 | k__Fungi(100);p__Ascomycota(100);c__Saccharomycetes(100);o__Saccharomycetales(100);o__Saccharomycetales_unclassified(100);o__Saccharomycetales_unclassified(100);o__Saccharomycetales_unclassified(100); | Saccharomycetales | 7 | Saprotroph | Undefined Saprotroph | Yeast | NULL | Possible | NULL | Sterkenburg E, et al. 2015. New Phytologist 207:1145-1158 ((https://doi.org/10.1111/nph.13426)) |
| Otu26001 | 1 | k__Fungi(100);p__Ascomycota(100);c__Saccharomycetes(100);o__Saccharomycetales(100);f__Metschnikowiaceae(100);g__Metschnikowia(100);g__Metschnikowia_unclassified(100); | Metschnikowia | 13 | Saprotroph | \|Undefined Saprotroph\| | Yeast | NULL | Probable | As nectar-tap_saprotroph (Põlme et al. 2020); Decay sugar-rich substrates (Põlme et al. 2020) | Põlme S, et al. 2020. Fungal Diversity 105:1-16 ((https://doi.org/10.1007/s13225-020-00466-2)) |
| Otu26002 | 1 | k__Fungi(100);p__Ascomycota(100);c__Saccharomycetes(100);o__Saccharomycetales(100);f__Metschnikowiaceae(100);g__Metschnikowia(100);s__Metschnikowia_pulcherrima(100); | Metschnikowia | 13 | Saprotroph | \|Undefined Saprotroph\| | Yeast | NULL | Probable | As nectar-tap_saprotroph (Põlme et al. 2020); Decay sugar-rich substrates (Põlme et al. 2020) | Põlme S, et al. 2020. Fungal Diversity 105:1-16 ((https://doi.org/10.1007/s13225-020-00466-2)) |
| Otu26003 | 1 | k__Fungi(100);p__Ascomycota(100);c__Saccharomycetes(100);o__Saccharomycetales(100);f__Metschnikowiaceae(100);g__Metschnikowia(100);s__Metschnikowia_pulcherrima(100); | Metschnikowia | 13 | Saprotroph | \|Undefined Saprotroph\| | Yeast | NULL | Probable | As nectar-tap_saprotroph (Põlme et al. 2020); Decay sugar-rich substrates (Põlme et al. 2020) | Põlme S, et al. 2020. Fungal Diversity 105:1-16 ((https://doi.org/10.1007/s13225-020-00466-2)) |
| Otu26004 | 1 | k__Fungi(100);p__Ascomycota(100);c__Saccharomycetes(100);o__Saccharomycetales(100);f__Metschnikowiaceae(100);g__Metschnikowia(100);g__Metschnikowia_unclassified(100); | Metschnikowia | 13 | Saprotroph | \|Undefined Saprotroph\| | Yeast | NULL | Probable | As nectar-tap_saprotroph (Põlme et al. 2020); Decay sugar-rich substrates (Põlme et al. 2020) | Põlme S, et al. 2020. Fungal Diversity 105:1-16 ((https://doi.org/10.1007/s13225-020-00466-2)) |
| Otu26005 | 1 | k__Fungi(100);p__Ascomycota(100);c__Saccharomycetes(100);o__Saccharomycetales(100);f__Metschnikowiaceae(100);g__Metschnikowia(100);g__Metschnikowia_unclassified(100); | Metschnikowia | 13 | Saprotroph | \|Undefined Saprotroph\| | Yeast | NULL | Probable | As nectar-tap_saprotroph (Põlme et al. 2020); Decay sugar-rich substrates (Põlme et al. 2020) | Põlme S, et al. 2020. Fungal Diversity 105:1-16 ((https://doi.org/10.1007/s13225-020-00466-2)) |
| Otu26006 | 1 | k__Fungi(100);p__Ascomycota(100);c__Saccharomycetes(100);o__Saccharomycetales(100);f__Metschnikowiaceae(100);g__Metschnikowia(100);s__Metschnikowia_pulcherrima(100); | Metschnikowia | 13 | Saprotroph | \|Undefined Saprotroph\| | Yeast | NULL | Probable | As nectar-tap_saprotroph (Põlme et al. 2020); Decay sugar-rich substrates (Põlme et al. 2020) | Põlme S, et al. 2020. Fungal Diversity 105:1-16 ((https://doi.org/10.1007/s13225-020-00466-2)) |
| Otu26007 | 1 | k__Fungi(100);p__Ascomycota(100);c__Saccharomycetes(100);o__Saccharomycetales(100);f__Metschnikowiaceae(100);g__Metschnikowia(100);s__Metschnikowia_pulcherrima(100); | Metschnikowia | 13 | Saprotroph | \|Undefined Saprotroph\| | Yeast | NULL | Probable | As nectar-tap_saprotroph (Põlme et al. 2020); Decay sugar-rich substrates (Põlme et al. 2020) | Põlme S, et al. 2020. Fungal Diversity 105:1-16 ((https://doi.org/10.1007/s13225-020-00466-2)) |
| Otu26009 | 1 | k__Fungi(100);p__Ascomycota(100);c__Saccharomycetes(100);o__Saccharomycetales(100);f__Metschnikowiaceae(100);g__Metschnikowia(100);s__Metschnikowia_pulcherrima(100); | Metschnikowia | 13 | Saprotroph | \|Undefined Saprotroph\| | Yeast | NULL | Probable | As nectar-tap_saprotroph (Põlme et al. 2020); Decay sugar-rich substrates (Põlme et al. 2020) | Põlme S, et al. 2020. Fungal Diversity 105:1-16 ((https://doi.org/10.1007/s13225-020-00466-2)) |
| Otu26011 | 1 | k__Fungi(100);p__Ascomycota(100);c__Saccharomycetes(100);o__Saccharomycetales(100);f__Metschnikowiaceae(100);g__Metschnikowia(100);s__Metschnikowia_pulcherrima(100); | Metschnikowia | 13 | Saprotroph | \|Undefined Saprotroph\| | Yeast | NULL | Probable | As nectar-tap_saprotroph (Põlme et al. 2020); Decay sugar-rich substrates (Põlme et al. 2020) | Põlme S, et al. 2020. Fungal Diversity 105:1-16 ((https://doi.org/10.1007/s13225-020-00466-2)) |
| Otu26013 | 1 | k__Fungi(100);p__Ascomycota(100);c__Saccharomycetes(100);o__Saccharomycetales(100);f__Metschnikowiaceae(100);g__Metschnikowia(100);s__Metschnikowia_pulcherrima(100); | Metschnikowia | 13 | Saprotroph | \|Undefined Saprotroph\| | Yeast | NULL | Probable | As nectar-tap_saprotroph (Põlme et al. 2020); Decay sugar-rich substrates (Põlme et al. 2020) | Põlme S, et al. 2020. Fungal Diversity 105:1-16 ((https://doi.org/10.1007/s13225-020-00466-2)) |
| Otu26016 | 1 | k__Fungi(100);p__Ascomycota(100);c__Saccharomycetes(100);o__Saccharomycetales(100);f__Metschnikowiaceae(100);g__Metschnikowia(100);s__Metschnikowia_pulcherrima(100); | Metschnikowia | 13 | Saprotroph | \|Undefined Saprotroph\| | Yeast | NULL | Probable | As nectar-tap_saprotroph (Põlme et al. 2020); Decay sugar-rich substrates (Põlme et al. 2020) | Põlme S, et al. 2020. Fungal Diversity 105:1-16 ((https://doi.org/10.1007/s13225-020-00466-2)) |
| Otu26017 | 1 | k__Fungi(100);p__Ascomycota(100);c__Saccharomycetes(100);o__Saccharomycetales(100);f__Metschnikowiaceae(100);g__Metschnikowia(100);s__Metschnikowia_pulcherrima(100); | Metschnikowia | 13 | Saprotroph | \|Undefined Saprotroph\| | Yeast | NULL | Probable | As nectar-tap_saprotroph (Põlme et al. 2020); Decay sugar-rich substrates (Põlme et al. 2020) | Põlme S, et al. 2020. Fungal Diversity 105:1-16 ((https://doi.org/10.1007/s13225-020-00466-2)) |
| Otu26018 | 1 | k__Fungi(100);p__Ascomycota(100);c__Saccharomycetes(100);o__Saccharomycetales(100);f__Metschnikowiaceae(100);g__Metschnikowia(100);g__Metschnikowia_unclassified(100); | Metschnikowia | 13 | Saprotroph | \|Undefined Saprotroph\| | Yeast | NULL | Probable | As nectar-tap_saprotroph (Põlme et al. 2020); Decay sugar-rich substrates (Põlme et al. 2020) | Põlme S, et al. 2020. Fungal Diversity 105:1-16 ((https://doi.org/10.1007/s13225-020-00466-2)) |
| Otu26019 | 1 | k__Fungi(100);p__Ascomycota(100);c__Saccharomycetes(100);o__Saccharomycetales(100);f__Metschnikowiaceae(100);g__Metschnikowia(100);g__Metschnikowia_unclassified(100); | Metschnikowia | 13 | Saprotroph | \|Undefined Saprotroph\| | Yeast | NULL | Probable | As nectar-tap_saprotroph (Põlme et al. 2020); Decay sugar-rich substrates (Põlme et al. 2020) | Põlme S, et al. 2020. Fungal Diversity 105:1-16 ((https://doi.org/10.1007/s13225-020-00466-2)) |
| Otu26020 | 1 | k__Fungi(100);p__Ascomycota(100);c__Saccharomycetes(100);o__Saccharomycetales(100);f__Metschnikowiaceae(100);g__Metschnikowia(100);s__Metschnikowia_pulcherrima(100); | Metschnikowia | 13 | Saprotroph | \|Undefined Saprotroph\| | Yeast | NULL | Probable | As nectar-tap_saprotroph (Põlme et al. 2020); Decay sugar-rich substrates (Põlme et al. 2020) | Põlme S, et al. 2020. Fungal Diversity 105:1-16 ((https://doi.org/10.1007/s13225-020-00466-2)) |
| Otu26021 | 1 | k__Fungi(100);p__Ascomycota(100);c__Saccharomycetes(100);o__Saccharomycetales(100);f__Metschnikowiaceae(100);g__Metschnikowia(100);s__Metschnikowia_pulcherrima(100); | Metschnikowia | 13 | Saprotroph | \|Undefined Saprotroph\| | Yeast | NULL | Probable | As nectar-tap_saprotroph (Põlme et al. 2020); Decay sugar-rich substrates (Põlme et al. 2020) | Põlme S, et al. 2020. Fungal Diversity 105:1-16 ((https://doi.org/10.1007/s13225-020-00466-2)) |
| Otu26022 | 1 | k__Fungi(100);p__Ascomycota(100);c__Saccharomycetes(100);o__Saccharomycetales(100);f__Metschnikowiaceae(100);g__Metschnikowia(100);s__Metschnikowia_pulcherrima(100); | Metschnikowia | 13 | Saprotroph | \|Undefined Saprotroph\| | Yeast | NULL | Probable | As nectar-tap_saprotroph (Põlme et al. 2020); Decay sugar-rich substrates (Põlme et al. 2020) | Põlme S, et al. 2020. Fungal Diversity 105:1-16 ((https://doi.org/10.1007/s13225-020-00466-2)) |
| Otu26025 | 1 | k__Fungi(100);p__Ascomycota(100);c__Saccharomycetes(100);o__Saccharomycetales(100);f__Metschnikowiaceae(100);g__Metschnikowia(100);s__Metschnikowia_pulcherrima(100); | Metschnikowia | 13 | Saprotroph | \|Undefined Saprotroph\| | Yeast | NULL | Probable | As nectar-tap_saprotroph (Põlme et al. 2020); Decay sugar-rich substrates (Põlme et al. 2020) | Põlme S, et al. 2020. Fungal Diversity 105:1-16 ((https://doi.org/10.1007/s13225-020-00466-2)) |
| Otu26026 | 1 | k__Fungi(100);p__Ascomycota(100);c__Saccharomycetes(100);o__Saccharomycetales(100);f__Metschnikowiaceae(100);g__Metschnikowia(100);s__Metschnikowia_pulcherrima(100); | Metschnikowia | 13 | Saprotroph | \|Undefined Saprotroph\| | Yeast | NULL | Probable | As nectar-tap_saprotroph (Põlme et al. 2020); Decay sugar-rich substrates (Põlme et al. 2020) | Põlme S, et al. 2020. Fungal Diversity 105:1-16 ((https://doi.org/10.1007/s13225-020-00466-2)) |
| Otu26027 | 1 | k__Fungi(100);p__Ascomycota(100);c__Saccharomycetes(100);o__Saccharomycetales(100);f__Metschnikowiaceae(100);g__Metschnikowia(100);s__Metschnikowia_pulcherrima(100); | Metschnikowia | 13 | Saprotroph | \|Undefined Saprotroph\| | Yeast | NULL | Probable | As nectar-tap_saprotroph (Põlme et al. 2020); Decay sugar-rich substrates (Põlme et al. 2020) | Põlme S, et al. 2020. Fungal Diversity 105:1-16 ((https://doi.org/10.1007/s13225-020-00466-2)) |
| Otu26028 | 1 | k__Fungi(100);p__Ascomycota(100);c__Saccharomycetes(100);o__Saccharomycetales(100);f__Metschnikowiaceae(100);g__Metschnikowia(100);s__Metschnikowia_pulcherrima(100); | Metschnikowia | 13 | Saprotroph | \|Undefined Saprotroph\| | Yeast | NULL | Probable | As nectar-tap_saprotroph (Põlme et al. 2020); Decay sugar-rich substrates (Põlme et al. 2020) | Põlme S, et al. 2020. Fungal Diversity 105:1-16 ((https://doi.org/10.1007/s13225-020-00466-2)) |
| Otu26029 | 1 | k__Fungi(100);p__Ascomycota(100);c__Saccharomycetes(100);o__Saccharomycetales(100);f__Metschnikowiaceae(100);g__Metschnikowia(100);g__Metschnikowia_unclassified(100); | Metschnikowia | 13 | Saprotroph | \|Undefined Saprotroph\| | Yeast | NULL | Probable | As nectar-tap_saprotroph (Põlme et al. 2020); Decay sugar-rich substrates (Põlme et al. 2020) | Põlme S, et al. 2020. Fungal Diversity 105:1-16 ((https://doi.org/10.1007/s13225-020-00466-2)) |
| Otu26030 | 1 | k__Fungi(100);p__Ascomycota(100);c__Saccharomycetes(100);o__Saccharomycetales(100);f__Metschnikowiaceae(100);g__Metschnikowia(100);s__Metschnikowia_pulcherrima(100); | Metschnikowia | 13 | Saprotroph | \|Undefined Saprotroph\| | Yeast | NULL | Probable | As nectar-tap_saprotroph (Põlme et al. 2020); Decay sugar-rich substrates (Põlme et al. 2020) | Põlme S, et al. 2020. Fungal Diversity 105:1-16 ((https://doi.org/10.1007/s13225-020-00466-2)) |
| Otu26031 | 1 | k__Fungi(100);p__Ascomycota(100);c__Saccharomycetes(100);o__Saccharomycetales(100);f__Metschnikowiaceae(100);g__Metschnikowia(100);s__Metschnikowia_pulcherrima(100); | Metschnikowia | 13 | Saprotroph | \|Undefined Saprotroph\| | Yeast | NULL | Probable | As nectar-tap_saprotroph (Põlme et al. 2020); Decay sugar-rich substrates (Põlme et al. 2020) | Põlme S, et al. 2020. Fungal Diversity 105:1-16 ((https://doi.org/10.1007/s13225-020-00466-2)) |
| Otu26032 | 1 | k__Fungi(100);p__Ascomycota(100);c__Saccharomycetes(100);o__Saccharomycetales(100);f__Metschnikowiaceae(100);g__Metschnikowia(100);s__Metschnikowia_pulcherrima(100); | Metschnikowia | 13 | Saprotroph | \|Undefined Saprotroph\| | Yeast | NULL | Probable | As nectar-tap_saprotroph (Põlme et al. 2020); Decay sugar-rich substrates (Põlme et al. 2020) | Põlme S, et al. 2020. Fungal Diversity 105:1-16 ((https://doi.org/10.1007/s13225-020-00466-2)) |
| Otu26033 | 1 | k__Fungi(100);p__Ascomycota(100);c__Saccharomycetes(100);o__Saccharomycetales(100);f__Metschnikowiaceae(100);g__Metschnikowia(100);g__Metschnikowia_unclassified(100); | Metschnikowia | 13 | Saprotroph | \|Undefined Saprotroph\| | Yeast | NULL | Probable | As nectar-tap_saprotroph (Põlme et al. 2020); Decay sugar-rich substrates (Põlme et al. 2020) | Põlme S, et al. 2020. Fungal Diversity 105:1-16 ((https://doi.org/10.1007/s13225-020-00466-2)) |
| Otu26034 | 1 | k__Fungi(100);p__Ascomycota(100);c__Saccharomycetes(100);o__Saccharomycetales(100);f__Metschnikowiaceae(100);g__Metschnikowia(100);s__Metschnikowia_pulcherrima(100); | Metschnikowia | 13 | Saprotroph | \|Undefined Saprotroph\| | Yeast | NULL | Probable | As nectar-tap_saprotroph (Põlme et al. 2020); Decay sugar-rich substrates (Põlme et al. 2020) | Põlme S, et al. 2020. Fungal Diversity 105:1-16 ((https://doi.org/10.1007/s13225-020-00466-2)) |
| Otu26035 | 1 | k__Fungi(100);p__Ascomycota(100);c__Saccharomycetes(100);o__Saccharomycetales(100);f__Saccharomycetales_family_Incertae_sedis(100);g__Candida(100);g__Candida_unclassified(100); | Candida | 13 | Pathotroph-Saprotroph-Symbiotroph | Animal Pathogen-Endophyte-Endosymbiont-Epiphyte-\|Undefined Saprotroph\| | Dimorphic Yeast-Yeast | NULL | Probable | As nectar-tap_saprotroph (Põlme et al. 2020); Decay sugar-rich substrates (Põlme et al. 2020); This asexual genus is polyphyletic and contain species that function very differently depend on what they are. Caution should be taken when interpreting guilds for Candida species. | Manolakaki D, et al. 2010. Virulence. 1: 367-375 ((https://doi.org/10.4161/viru.1.5.12796)); Mekbib SB, et al. 2011. Tropical Plant Pathology 36:233-240 ((https://doi.org/10.1590/S1982-56762011000400004)); Põlme S, et al. 2020. Fungal Diversity 105:1-16 ((https://doi.org/10.1007/s13225-020-00466-2)) |
| Otu26037 | 1 | k__Fungi(100);p__Ascomycota(100);c__Saccharomycetes(100);o__Saccharomycetales(100);o__Saccharomycetales_unclassified(100);o__Saccharomycetales_unclassified(100);o__Saccharomycetales_unclassified(100); | Saccharomycetales | 7 | Saprotroph | Undefined Saprotroph | Yeast | NULL | Possible | NULL | Sterkenburg E, et al. 2015. New Phytologist 207:1145-1158 ((https://doi.org/10.1111/nph.13426)) |
| Otu26038 | 1 | k__Fungi(100);p__Ascomycota(100);c__Saccharomycetes(100);o__Saccharomycetales(100);f__Metschnikowiaceae(100);g__Metschnikowia(100);s__Metschnikowia_pulcherrima(100); | Metschnikowia | 13 | Saprotroph | \|Undefined Saprotroph\| | Yeast | NULL | Probable | As nectar-tap_saprotroph (Põlme et al. 2020); Decay sugar-rich substrates (Põlme et al. 2020) | Põlme S, et al. 2020. Fungal Diversity 105:1-16 ((https://doi.org/10.1007/s13225-020-00466-2)) |
| Otu26039 | 1 | k__Fungi(100);p__Ascomycota(100);c__Saccharomycetes(100);o__Saccharomycetales(100);f__Metschnikowiaceae(100);g__Metschnikowia(100);g__Metschnikowia_unclassified(100); | Metschnikowia | 13 | Saprotroph | \|Undefined Saprotroph\| | Yeast | NULL | Probable | As nectar-tap_saprotroph (Põlme et al. 2020); Decay sugar-rich substrates (Põlme et al. 2020) | Põlme S, et al. 2020. Fungal Diversity 105:1-16 ((https://doi.org/10.1007/s13225-020-00466-2)) |
| Otu26041 | 1 | k__Fungi(100);p__Ascomycota(100);c__Saccharomycetes(100);o__Saccharomycetales(100);f__Metschnikowiaceae(100);g__Metschnikowia(100);s__Metschnikowia_pulcherrima(100); | Metschnikowia | 13 | Saprotroph | \|Undefined Saprotroph\| | Yeast | NULL | Probable | As nectar-tap_saprotroph (Põlme et al. 2020); Decay sugar-rich substrates (Põlme et al. 2020) | Põlme S, et al. 2020. Fungal Diversity 105:1-16 ((https://doi.org/10.1007/s13225-020-00466-2)) |
| Otu26042 | 1 | k__Fungi(100);p__Ascomycota(100);c__Saccharomycetes(100);o__Saccharomycetales(100);f__Metschnikowiaceae(100);g__Metschnikowia(100);s__Metschnikowia_pulcherrima(100); | Metschnikowia | 13 | Saprotroph | \|Undefined Saprotroph\| | Yeast | NULL | Probable | As nectar-tap_saprotroph (Põlme et al. 2020); Decay sugar-rich substrates (Põlme et al. 2020) | Põlme S, et al. 2020. Fungal Diversity 105:1-16 ((https://doi.org/10.1007/s13225-020-00466-2)) |
| Otu26043 | 1 | k__Fungi(100);p__Ascomycota(100);c__Saccharomycetes(100);o__Saccharomycetales(100);f__Metschnikowiaceae(100);g__Metschnikowia(100);g__Metschnikowia_unclassified(100); | Metschnikowia | 13 | Saprotroph | \|Undefined Saprotroph\| | Yeast | NULL | Probable | As nectar-tap_saprotroph (Põlme et al. 2020); Decay sugar-rich substrates (Põlme et al. 2020) | Põlme S, et al. 2020. Fungal Diversity 105:1-16 ((https://doi.org/10.1007/s13225-020-00466-2)) |
| Otu26044 | 1 | k__Fungi(100);p__Ascomycota(100);c__Saccharomycetes(100);o__Saccharomycetales(100);f__Metschnikowiaceae(100);g__Metschnikowia(100);s__Metschnikowia_pulcherrima(100); | Metschnikowia | 13 | Saprotroph | \|Undefined Saprotroph\| | Yeast | NULL | Probable | As nectar-tap_saprotroph (Põlme et al. 2020); Decay sugar-rich substrates (Põlme et al. 2020) | Põlme S, et al. 2020. Fungal Diversity 105:1-16 ((https://doi.org/10.1007/s13225-020-00466-2)) |
| Otu26046 | 1 | k__Fungi(100);p__Ascomycota(100);c__Saccharomycetes(100);o__Saccharomycetales(100);f__Metschnikowiaceae(100);g__Metschnikowia(100);s__Metschnikowia_pulcherrima(100); | Metschnikowia | 13 | Saprotroph | \|Undefined Saprotroph\| | Yeast | NULL | Probable | As nectar-tap_saprotroph (Põlme et al. 2020); Decay sugar-rich substrates (Põlme et al. 2020) | Põlme S, et al. 2020. Fungal Diversity 105:1-16 ((https://doi.org/10.1007/s13225-020-00466-2)) |
| Otu26052 | 1 | k__Fungi(100);p__Ascomycota(100);c__Saccharomycetes(100);o__Saccharomycetales(100);f__Metschnikowiaceae(100);g__Metschnikowia(100);s__Metschnikowia_pulcherrima(100); | Metschnikowia | 13 | Saprotroph | \|Undefined Saprotroph\| | Yeast | NULL | Probable | As nectar-tap_saprotroph (Põlme et al. 2020); Decay sugar-rich substrates (Põlme et al. 2020) | Põlme S, et al. 2020. Fungal Diversity 105:1-16 ((https://doi.org/10.1007/s13225-020-00466-2)) |
| Otu26057 | 1 | k__Fungi(100);p__Ascomycota(100);c__Saccharomycetes(100);o__Saccharomycetales(100);f__Metschnikowiaceae(100);g__Metschnikowia(100);s__Metschnikowia_pulcherrima(100); | Metschnikowia | 13 | Saprotroph | \|Undefined Saprotroph\| | Yeast | NULL | Probable | As nectar-tap_saprotroph (Põlme et al. 2020); Decay sugar-rich substrates (Põlme et al. 2020) | Põlme S, et al. 2020. Fungal Diversity 105:1-16 ((https://doi.org/10.1007/s13225-020-00466-2)) |
| Otu26059 | 1 | k__Fungi(100);p__Ascomycota(100);c__Saccharomycetes(100);o__Saccharomycetales(100);f__Metschnikowiaceae(100);g__Metschnikowia(100);s__Metschnikowia_pulcherrima(100); | Metschnikowia | 13 | Saprotroph | \|Undefined Saprotroph\| | Yeast | NULL | Probable | As nectar-tap_saprotroph (Põlme et al. 2020); Decay sugar-rich substrates (Põlme et al. 2020) | Põlme S, et al. 2020. Fungal Diversity 105:1-16 ((https://doi.org/10.1007/s13225-020-00466-2)) |
| Otu26062 | 1 | k__Fungi(100);p__Ascomycota(100);c__Saccharomycetes(100);o__Saccharomycetales(100);f__Metschnikowiaceae(100);g__Metschnikowia(100);g__Metschnikowia_unclassified(100); | Metschnikowia | 13 | Saprotroph | \|Undefined Saprotroph\| | Yeast | NULL | Probable | As nectar-tap_saprotroph (Põlme et al. 2020); Decay sugar-rich substrates (Põlme et al. 2020) | Põlme S, et al. 2020. Fungal Diversity 105:1-16 ((https://doi.org/10.1007/s13225-020-00466-2)) |
| Otu26065 | 1 | k__Fungi(100);p__Ascomycota(100);c__Saccharomycetes(100);o__Saccharomycetales(100);f__Metschnikowiaceae(100);g__Metschnikowia(100);s__Metschnikowia_pulcherrima(100); | Metschnikowia | 13 | Saprotroph | \|Undefined Saprotroph\| | Yeast | NULL | Probable | As nectar-tap_saprotroph (Põlme et al. 2020); Decay sugar-rich substrates (Põlme et al. 2020) | Põlme S, et al. 2020. Fungal Diversity 105:1-16 ((https://doi.org/10.1007/s13225-020-00466-2)) |
| Otu26071 | 1 | k__Fungi(100);p__Ascomycota(100);c__Saccharomycetes(100);o__Saccharomycetales(100);f__Metschnikowiaceae(100);g__Metschnikowia(100);s__Metschnikowia_pulcherrima(100); | Metschnikowia | 13 | Saprotroph | \|Undefined Saprotroph\| | Yeast | NULL | Probable | As nectar-tap_saprotroph (Põlme et al. 2020); Decay sugar-rich substrates (Põlme et al. 2020) | Põlme S, et al. 2020. Fungal Diversity 105:1-16 ((https://doi.org/10.1007/s13225-020-00466-2)) |
| Otu26072 | 1 | k__Fungi(100);p__Ascomycota(100);c__Saccharomycetes(100);o__Saccharomycetales(100);f__Metschnikowiaceae(100);g__Metschnikowia(100);s__Metschnikowia_pulcherrima(100); | Metschnikowia | 13 | Saprotroph | \|Undefined Saprotroph\| | Yeast | NULL | Probable | As nectar-tap_saprotroph (Põlme et al. 2020); Decay sugar-rich substrates (Põlme et al. 2020) | Põlme S, et al. 2020. Fungal Diversity 105:1-16 ((https://doi.org/10.1007/s13225-020-00466-2)) |
| Otu26073 | 1 | k__Fungi(100);p__Ascomycota(100);c__Saccharomycetes(100);o__Saccharomycetales(100);f__Metschnikowiaceae(100);g__Metschnikowia(100);s__Metschnikowia_pulcherrima(100); | Metschnikowia | 13 | Saprotroph | \|Undefined Saprotroph\| | Yeast | NULL | Probable | As nectar-tap_saprotroph (Põlme et al. 2020); Decay sugar-rich substrates (Põlme et al. 2020) | Põlme S, et al. 2020. Fungal Diversity 105:1-16 ((https://doi.org/10.1007/s13225-020-00466-2)) |
| Otu26074 | 1 | k__Fungi(100);p__Ascomycota(100);c__Saccharomycetes(100);o__Saccharomycetales(100);f__Metschnikowiaceae(100);g__Metschnikowia(100);s__Metschnikowia_pulcherrima(100); | Metschnikowia | 13 | Saprotroph | \|Undefined Saprotroph\| | Yeast | NULL | Probable | As nectar-tap_saprotroph (Põlme et al. 2020); Decay sugar-rich substrates (Põlme et al. 2020) | Põlme S, et al. 2020. Fungal Diversity 105:1-16 ((https://doi.org/10.1007/s13225-020-00466-2)) |
| Otu26075 | 1 | k__Fungi(100);p__Ascomycota(100);c__Saccharomycetes(100);o__Saccharomycetales(100);f__Metschnikowiaceae(100);g__Metschnikowia(100);g__Metschnikowia_unclassified(100); | Metschnikowia | 13 | Saprotroph | \|Undefined Saprotroph\| | Yeast | NULL | Probable | As nectar-tap_saprotroph (Põlme et al. 2020); Decay sugar-rich substrates (Põlme et al. 2020) | Põlme S, et al. 2020. Fungal Diversity 105:1-16 ((https://doi.org/10.1007/s13225-020-00466-2)) |
| Otu26077 | 1 | k__Fungi(100);p__Ascomycota(100);c__Saccharomycetes(100);o__Saccharomycetales(100);f__Metschnikowiaceae(100);g__Metschnikowia(100);s__Metschnikowia_pulcherrima(100); | Metschnikowia | 13 | Saprotroph | \|Undefined Saprotroph\| | Yeast | NULL | Probable | As nectar-tap_saprotroph (Põlme et al. 2020); Decay sugar-rich substrates (Põlme et al. 2020) | Põlme S, et al. 2020. Fungal Diversity 105:1-16 ((https://doi.org/10.1007/s13225-020-00466-2)) |
| Otu26078 | 1 | k__Fungi(100);p__Ascomycota(100);c__Dothideomycetes(100);o__Dothideomycetes_order_Incertae_sedis(100);f__Myxotrichaceae(100);g__Oidiodendron(100);s__Oidiodendron_maius(100); | Oidiodendron | 13 | Pathotroph-Saprotroph-Symbiotroph | Endophyte-Ericoid Mycorrhizal-\|Undefined Saprotroph\| | Microfungus | NULL | Probable | As root_endophyte (Põlme et al. 2020); As soil_saprotroph (Põlme et al. 2020); Dark Septate Endophyte; Decay soil (Põlme et al. 2020) | Newsham KK. 2011. New Phytologist 190:783-793 ((https://doi.org/10.1111/j.1469-8137.2010.03611.x)) (as dark septate root endophytes); Põlme S, et al. 2020. Fungal Diversity 105:1-16 ((https://doi.org/10.1007/s13225-020-00466-2)) |
| Otu26079 | 1 | k__Fungi(100);p__Ascomycota(100);c__Saccharomycetes(100);o__Saccharomycetales(100);f__Metschnikowiaceae(100);g__Metschnikowia(100);s__Metschnikowia_pulcherrima(100); | Metschnikowia | 13 | Saprotroph | \|Undefined Saprotroph\| | Yeast | NULL | Probable | As nectar-tap_saprotroph (Põlme et al. 2020); Decay sugar-rich substrates (Põlme et al. 2020) | Põlme S, et al. 2020. Fungal Diversity 105:1-16 ((https://doi.org/10.1007/s13225-020-00466-2)) |
| Otu26080 | 1 | k__Fungi(100);p__Ascomycota(100);c__Saccharomycetes(100);o__Saccharomycetales(100);f__Metschnikowiaceae(100);g__Metschnikowia(100);g__Metschnikowia_unclassified(100); | Metschnikowia | 13 | Saprotroph | \|Undefined Saprotroph\| | Yeast | NULL | Probable | As nectar-tap_saprotroph (Põlme et al. 2020); Decay sugar-rich substrates (Põlme et al. 2020) | Põlme S, et al. 2020. Fungal Diversity 105:1-16 ((https://doi.org/10.1007/s13225-020-00466-2)) |
| Otu26081 | 1 | k__Fungi(100);p__Ascomycota(100);c__Saccharomycetes(100);o__Saccharomycetales(100);f__Metschnikowiaceae(100);g__Metschnikowia(100);s__Metschnikowia_pulcherrima(100); | Metschnikowia | 13 | Saprotroph | \|Undefined Saprotroph\| | Yeast | NULL | Probable | As nectar-tap_saprotroph (Põlme et al. 2020); Decay sugar-rich substrates (Põlme et al. 2020) | Põlme S, et al. 2020. Fungal Diversity 105:1-16 ((https://doi.org/10.1007/s13225-020-00466-2)) |
| Otu26082 | 1 | k__Fungi(100);p__Ascomycota(100);c__Saccharomycetes(100);o__Saccharomycetales(100);f__Metschnikowiaceae(100);g__Metschnikowia(100);s__Metschnikowia_pulcherrima(100); | Metschnikowia | 13 | Saprotroph | \|Undefined Saprotroph\| | Yeast | NULL | Probable | As nectar-tap_saprotroph (Põlme et al. 2020); Decay sugar-rich substrates (Põlme et al. 2020) | Põlme S, et al. 2020. Fungal Diversity 105:1-16 ((https://doi.org/10.1007/s13225-020-00466-2)) |
| Otu26083 | 1 | k__Fungi(100);p__Ascomycota(100);c__Saccharomycetes(100);o__Saccharomycetales(100);f__Metschnikowiaceae(100);g__Metschnikowia(100);g__Metschnikowia_unclassified(100); | Metschnikowia | 13 | Saprotroph | \|Undefined Saprotroph\| | Yeast | NULL | Probable | As nectar-tap_saprotroph (Põlme et al. 2020); Decay sugar-rich substrates (Põlme et al. 2020) | Põlme S, et al. 2020. Fungal Diversity 105:1-16 ((https://doi.org/10.1007/s13225-020-00466-2)) |
| Otu26084 | 1 | k__Fungi(100);p__Ascomycota(100);c__Saccharomycetes(100);o__Saccharomycetales(100);f__Metschnikowiaceae(100);g__Metschnikowia(100);s__Metschnikowia_pulcherrima(100); | Metschnikowia | 13 | Saprotroph | \|Undefined Saprotroph\| | Yeast | NULL | Probable | As nectar-tap_saprotroph (Põlme et al. 2020); Decay sugar-rich substrates (Põlme et al. 2020) | Põlme S, et al. 2020. Fungal Diversity 105:1-16 ((https://doi.org/10.1007/s13225-020-00466-2)) |
| Otu26086 | 1 | k__Fungi(100);p__Ascomycota(100);c__Saccharomycetes(100);o__Saccharomycetales(100);f__Metschnikowiaceae(100);g__Metschnikowia(100);s__Metschnikowia_pulcherrima(100); | Metschnikowia | 13 | Saprotroph | \|Undefined Saprotroph\| | Yeast | NULL | Probable | As nectar-tap_saprotroph (Põlme et al. 2020); Decay sugar-rich substrates (Põlme et al. 2020) | Põlme S, et al. 2020. Fungal Diversity 105:1-16 ((https://doi.org/10.1007/s13225-020-00466-2)) |
| Otu26087 | 1 | k__Fungi(100);p__Ascomycota(100);c__Saccharomycetes(100);o__Saccharomycetales(100);f__Metschnikowiaceae(100);g__Metschnikowia(100);g__Metschnikowia_unclassified(100); | Metschnikowia | 13 | Saprotroph | \|Undefined Saprotroph\| | Yeast | NULL | Probable | As nectar-tap_saprotroph (Põlme et al. 2020); Decay sugar-rich substrates (Põlme et al. 2020) | Põlme S, et al. 2020. Fungal Diversity 105:1-16 ((https://doi.org/10.1007/s13225-020-00466-2)) |
| Otu26088 | 1 | k__Fungi(100);p__Ascomycota(100);c__Saccharomycetes(100);o__Saccharomycetales(100);f__Metschnikowiaceae(100);g__Metschnikowia(100);s__Metschnikowia_pulcherrima(100); | Metschnikowia | 13 | Saprotroph | \|Undefined Saprotroph\| | Yeast | NULL | Probable | As nectar-tap_saprotroph (Põlme et al. 2020); Decay sugar-rich substrates (Põlme et al. 2020) | Põlme S, et al. 2020. Fungal Diversity 105:1-16 ((https://doi.org/10.1007/s13225-020-00466-2)) |
| Otu26089 | 1 | k__Fungi(100);p__Ascomycota(100);c__Saccharomycetes(100);o__Saccharomycetales(100);f__Metschnikowiaceae(100);g__Metschnikowia(100);s__Metschnikowia_pulcherrima(100); | Metschnikowia | 13 | Saprotroph | \|Undefined Saprotroph\| | Yeast | NULL | Probable | As nectar-tap_saprotroph (Põlme et al. 2020); Decay sugar-rich substrates (Põlme et al. 2020) | Põlme S, et al. 2020. Fungal Diversity 105:1-16 ((https://doi.org/10.1007/s13225-020-00466-2)) |
| Otu26090 | 1 | k__Fungi(100);p__Ascomycota(100);c__Saccharomycetes(100);o__Saccharomycetales(100);f__Metschnikowiaceae(100);g__Metschnikowia(100);s__Metschnikowia_pulcherrima(100); | Metschnikowia | 13 | Saprotroph | \|Undefined Saprotroph\| | Yeast | NULL | Probable | As nectar-tap_saprotroph (Põlme et al. 2020); Decay sugar-rich substrates (Põlme et al. 2020) | Põlme S, et al. 2020. Fungal Diversity 105:1-16 ((https://doi.org/10.1007/s13225-020-00466-2)) |
| Otu26091 | 1 | k__Fungi(100);p__Ascomycota(100);c__Saccharomycetes(100);o__Saccharomycetales(100);f__Metschnikowiaceae(100);g__Metschnikowia(100);s__Metschnikowia_pulcherrima(100); | Metschnikowia | 13 | Saprotroph | \|Undefined Saprotroph\| | Yeast | NULL | Probable | As nectar-tap_saprotroph (Põlme et al. 2020); Decay sugar-rich substrates (Põlme et al. 2020) | Põlme S, et al. 2020. Fungal Diversity 105:1-16 ((https://doi.org/10.1007/s13225-020-00466-2)) |
| Otu26092 | 1 | k__Fungi(100);p__Ascomycota(100);c__Saccharomycetes(100);o__Saccharomycetales(100);f__Metschnikowiaceae(100);g__Metschnikowia(100);s__Metschnikowia_pulcherrima(100); | Metschnikowia | 13 | Saprotroph | \|Undefined Saprotroph\| | Yeast | NULL | Probable | As nectar-tap_saprotroph (Põlme et al. 2020); Decay sugar-rich substrates (Põlme et al. 2020) | Põlme S, et al. 2020. Fungal Diversity 105:1-16 ((https://doi.org/10.1007/s13225-020-00466-2)) |
| Otu26095 | 1 | k__Fungi(100);p__Ascomycota(100);c__Saccharomycetes(100);o__Saccharomycetales(100);f__Metschnikowiaceae(100);g__Metschnikowia(100);s__Metschnikowia_pulcherrima(100); | Metschnikowia | 13 | Saprotroph | \|Undefined Saprotroph\| | Yeast | NULL | Probable | As nectar-tap_saprotroph (Põlme et al. 2020); Decay sugar-rich substrates (Põlme et al. 2020) | Põlme S, et al. 2020. Fungal Diversity 105:1-16 ((https://doi.org/10.1007/s13225-020-00466-2)) |
| Otu26096 | 1 | k__Fungi(100);p__Ascomycota(100);c__Saccharomycetes(100);o__Saccharomycetales(100);f__Metschnikowiaceae(100);g__Metschnikowia(100);g__Metschnikowia_unclassified(100); | Metschnikowia | 13 | Saprotroph | \|Undefined Saprotroph\| | Yeast | NULL | Probable | As nectar-tap_saprotroph (Põlme et al. 2020); Decay sugar-rich substrates (Põlme et al. 2020) | Põlme S, et al. 2020. Fungal Diversity 105:1-16 ((https://doi.org/10.1007/s13225-020-00466-2)) |
| Otu26097 | 1 | k__Fungi(100);p__Ascomycota(100);c__Saccharomycetes(100);o__Saccharomycetales(100);f__Metschnikowiaceae(100);g__Metschnikowia(100);s__Metschnikowia_pulcherrima(100); | Metschnikowia | 13 | Saprotroph | \|Undefined Saprotroph\| | Yeast | NULL | Probable | As nectar-tap_saprotroph (Põlme et al. 2020); Decay sugar-rich substrates (Põlme et al. 2020) | Põlme S, et al. 2020. Fungal Diversity 105:1-16 ((https://doi.org/10.1007/s13225-020-00466-2)) |
| Otu26098 | 1 | k__Fungi(100);p__Ascomycota(100);c__Saccharomycetes(100);o__Saccharomycetales(100);f__Metschnikowiaceae(100);g__Metschnikowia(100);g__Metschnikowia_unclassified(100); | Metschnikowia | 13 | Saprotroph | \|Undefined Saprotroph\| | Yeast | NULL | Probable | As nectar-tap_saprotroph (Põlme et al. 2020); Decay sugar-rich substrates (Põlme et al. 2020) | Põlme S, et al. 2020. Fungal Diversity 105:1-16 ((https://doi.org/10.1007/s13225-020-00466-2)) |
| Otu26099 | 1 | k__Fungi(100);p__Ascomycota(100);c__Saccharomycetes(100);o__Saccharomycetales(100);f__Saccharomycetales_family_Incertae_sedis(100);f__Saccharomycetales_family_Incertae_sedis_unclassified(100);f__Saccharomycetales_family_Incertae_sedis_unclassified(100); | Saccharomycetales | 7 | Saprotroph | Undefined Saprotroph | Yeast | NULL | Possible | NULL | Sterkenburg E, et al. 2015. New Phytologist 207:1145-1158 ((https://doi.org/10.1111/nph.13426)) |
| Otu26100 | 1 | k__Fungi(100);p__Ascomycota(100);c__Saccharomycetes(100);o__Saccharomycetales(100);f__Metschnikowiaceae(100);g__Metschnikowia(100);s__Metschnikowia_pulcherrima(100); | Metschnikowia | 13 | Saprotroph | \|Undefined Saprotroph\| | Yeast | NULL | Probable | As nectar-tap_saprotroph (Põlme et al. 2020); Decay sugar-rich substrates (Põlme et al. 2020) | Põlme S, et al. 2020. Fungal Diversity 105:1-16 ((https://doi.org/10.1007/s13225-020-00466-2)) |
| Otu26103 | 1 | k__Fungi(100);p__Ascomycota(100);c__Saccharomycetes(100);o__Saccharomycetales(100);f__Metschnikowiaceae(100);g__Metschnikowia(100);g__Metschnikowia_unclassified(100); | Metschnikowia | 13 | Saprotroph | \|Undefined Saprotroph\| | Yeast | NULL | Probable | As nectar-tap_saprotroph (Põlme et al. 2020); Decay sugar-rich substrates (Põlme et al. 2020) | Põlme S, et al. 2020. Fungal Diversity 105:1-16 ((https://doi.org/10.1007/s13225-020-00466-2)) |
| Otu26104 | 1 | k__Fungi(100);p__Ascomycota(100);c__Saccharomycetes(100);o__Saccharomycetales(100);f__Metschnikowiaceae(100);g__Metschnikowia(100);s__Metschnikowia_pulcherrima(100); | Metschnikowia | 13 | Saprotroph | \|Undefined Saprotroph\| | Yeast | NULL | Probable | As nectar-tap_saprotroph (Põlme et al. 2020); Decay sugar-rich substrates (Põlme et al. 2020) | Põlme S, et al. 2020. Fungal Diversity 105:1-16 ((https://doi.org/10.1007/s13225-020-00466-2)) |
| Otu26105 | 1 | k__Fungi(100);p__Ascomycota(100);c__Saccharomycetes(100);o__Saccharomycetales(100);f__Metschnikowiaceae(100);g__Metschnikowia(100);s__Metschnikowia_pulcherrima(100); | Metschnikowia | 13 | Saprotroph | \|Undefined Saprotroph\| | Yeast | NULL | Probable | As nectar-tap_saprotroph (Põlme et al. 2020); Decay sugar-rich substrates (Põlme et al. 2020) | Põlme S, et al. 2020. Fungal Diversity 105:1-16 ((https://doi.org/10.1007/s13225-020-00466-2)) |
| Otu26106 | 1 | k__Fungi(100);p__Ascomycota(100);c__Saccharomycetes(100);o__Saccharomycetales(100);f__Metschnikowiaceae(100);g__Metschnikowia(100);s__Metschnikowia_pulcherrima(100); | Metschnikowia | 13 | Saprotroph | \|Undefined Saprotroph\| | Yeast | NULL | Probable | As nectar-tap_saprotroph (Põlme et al. 2020); Decay sugar-rich substrates (Põlme et al. 2020) | Põlme S, et al. 2020. Fungal Diversity 105:1-16 ((https://doi.org/10.1007/s13225-020-00466-2)) |
| Otu26108 | 1 | k__Fungi(100);p__Ascomycota(100);c__Saccharomycetes(100);o__Saccharomycetales(100);f__Metschnikowiaceae(100);g__Metschnikowia(100);s__Metschnikowia_pulcherrima(100); | Metschnikowia | 13 | Saprotroph | \|Undefined Saprotroph\| | Yeast | NULL | Probable | As nectar-tap_saprotroph (Põlme et al. 2020); Decay sugar-rich substrates (Põlme et al. 2020) | Põlme S, et al. 2020. Fungal Diversity 105:1-16 ((https://doi.org/10.1007/s13225-020-00466-2)) |
| Otu26109 | 1 | k__Fungi(100);p__Ascomycota(100);c__Saccharomycetes(100);o__Saccharomycetales(100);f__Metschnikowiaceae(100);g__Metschnikowia(100);s__Metschnikowia_pulcherrima(100); | Metschnikowia | 13 | Saprotroph | \|Undefined Saprotroph\| | Yeast | NULL | Probable | As nectar-tap_saprotroph (Põlme et al. 2020); Decay sugar-rich substrates (Põlme et al. 2020) | Põlme S, et al. 2020. Fungal Diversity 105:1-16 ((https://doi.org/10.1007/s13225-020-00466-2)) |
| Otu26110 | 1 | k__Fungi(100);p__Ascomycota(100);c__Saccharomycetes(100);o__Saccharomycetales(100);f__Metschnikowiaceae(100);g__Metschnikowia(100);s__Metschnikowia_pulcherrima(100); | Metschnikowia | 13 | Saprotroph | \|Undefined Saprotroph\| | Yeast | NULL | Probable | As nectar-tap_saprotroph (Põlme et al. 2020); Decay sugar-rich substrates (Põlme et al. 2020) | Põlme S, et al. 2020. Fungal Diversity 105:1-16 ((https://doi.org/10.1007/s13225-020-00466-2)) |
| Otu26111 | 1 | k__Fungi(100);p__Ascomycota(100);c__Saccharomycetes(100);o__Saccharomycetales(100);f__Metschnikowiaceae(100);g__Metschnikowia(100);s__Metschnikowia_pulcherrima(100); | Metschnikowia | 13 | Saprotroph | \|Undefined Saprotroph\| | Yeast | NULL | Probable | As nectar-tap_saprotroph (Põlme et al. 2020); Decay sugar-rich substrates (Põlme et al. 2020) | Põlme S, et al. 2020. Fungal Diversity 105:1-16 ((https://doi.org/10.1007/s13225-020-00466-2)) |
| Otu26113 | 1 | k__Fungi(100);p__Ascomycota(100);c__Saccharomycetes(100);o__Saccharomycetales(100);f__Metschnikowiaceae(100);g__Metschnikowia(100);s__Metschnikowia_pulcherrima(100); | Metschnikowia | 13 | Saprotroph | \|Undefined Saprotroph\| | Yeast | NULL | Probable | As nectar-tap_saprotroph (Põlme et al. 2020); Decay sugar-rich substrates (Põlme et al. 2020) | Põlme S, et al. 2020. Fungal Diversity 105:1-16 ((https://doi.org/10.1007/s13225-020-00466-2)) |
| Otu26114 | 1 | k__Fungi(100);p__Ascomycota(100);c__Saccharomycetes(100);o__Saccharomycetales(100);f__Metschnikowiaceae(100);g__Metschnikowia(100);g__Metschnikowia_unclassified(100); | Metschnikowia | 13 | Saprotroph | \|Undefined Saprotroph\| | Yeast | NULL | Probable | As nectar-tap_saprotroph (Põlme et al. 2020); Decay sugar-rich substrates (Põlme et al. 2020) | Põlme S, et al. 2020. Fungal Diversity 105:1-16 ((https://doi.org/10.1007/s13225-020-00466-2)) |
| Otu26115 | 1 | k__Fungi(100);p__Ascomycota(100);c__Saccharomycetes(100);o__Saccharomycetales(100);f__Metschnikowiaceae(100);g__Metschnikowia(100);s__Metschnikowia_pulcherrima(100); | Metschnikowia | 13 | Saprotroph | \|Undefined Saprotroph\| | Yeast | NULL | Probable | As nectar-tap_saprotroph (Põlme et al. 2020); Decay sugar-rich substrates (Põlme et al. 2020) | Põlme S, et al. 2020. Fungal Diversity 105:1-16 ((https://doi.org/10.1007/s13225-020-00466-2)) |
| Otu26116 | 1 | k__Fungi(100);p__Ascomycota(100);c__Saccharomycetes(100);o__Saccharomycetales(100);f__Metschnikowiaceae(100);g__Metschnikowia(100);g__Metschnikowia_unclassified(100); | Metschnikowia | 13 | Saprotroph | \|Undefined Saprotroph\| | Yeast | NULL | Probable | As nectar-tap_saprotroph (Põlme et al. 2020); Decay sugar-rich substrates (Põlme et al. 2020) | Põlme S, et al. 2020. Fungal Diversity 105:1-16 ((https://doi.org/10.1007/s13225-020-00466-2)) |
| Otu26117 | 1 | k__Fungi(100);p__Ascomycota(100);c__Saccharomycetes(100);o__Saccharomycetales(100);f__Metschnikowiaceae(100);g__Metschnikowia(100);s__Metschnikowia_pulcherrima(100); | Metschnikowia | 13 | Saprotroph | \|Undefined Saprotroph\| | Yeast | NULL | Probable | As nectar-tap_saprotroph (Põlme et al. 2020); Decay sugar-rich substrates (Põlme et al. 2020) | Põlme S, et al. 2020. Fungal Diversity 105:1-16 ((https://doi.org/10.1007/s13225-020-00466-2)) |
| Otu26118 | 1 | k__Fungi(100);p__Ascomycota(100);c__Saccharomycetes(100);o__Saccharomycetales(100);f__Metschnikowiaceae(100);g__Metschnikowia(100);s__Metschnikowia_pulcherrima(100); | Metschnikowia | 13 | Saprotroph | \|Undefined Saprotroph\| | Yeast | NULL | Probable | As nectar-tap_saprotroph (Põlme et al. 2020); Decay sugar-rich substrates (Põlme et al. 2020) | Põlme S, et al. 2020. Fungal Diversity 105:1-16 ((https://doi.org/10.1007/s13225-020-00466-2)) |
| Otu26121 | 1 | k__Fungi(100);p__Ascomycota(100);c__Saccharomycetes(100);o__Saccharomycetales(100);f__Metschnikowiaceae(100);g__Metschnikowia(100);s__Metschnikowia_pulcherrima(100); | Metschnikowia | 13 | Saprotroph | \|Undefined Saprotroph\| | Yeast | NULL | Probable | As nectar-tap_saprotroph (Põlme et al. 2020); Decay sugar-rich substrates (Põlme et al. 2020) | Põlme S, et al. 2020. Fungal Diversity 105:1-16 ((https://doi.org/10.1007/s13225-020-00466-2)) |
| Otu26123 | 1 | k__Fungi(100);p__Ascomycota(100);c__Saccharomycetes(100);o__Saccharomycetales(100);f__Saccharomycetales_family_Incertae_sedis(100);g__Candida(100);s__Candida_cretensis(100); | Candida | 13 | Pathotroph-Saprotroph-Symbiotroph | Animal Pathogen-Endophyte-Endosymbiont-Epiphyte-\|Undefined Saprotroph\| | Dimorphic Yeast-Yeast | NULL | Probable | As nectar-tap_saprotroph (Põlme et al. 2020); Decay sugar-rich substrates (Põlme et al. 2020); This asexual genus is polyphyletic and contain species that function very differently depend on what they are. Caution should be taken when interpreting guilds for Candida species. | Manolakaki D, et al. 2010. Virulence. 1: 367-375 ((https://doi.org/10.4161/viru.1.5.12796)); Mekbib SB, et al. 2011. Tropical Plant Pathology 36:233-240 ((https://doi.org/10.1590/S1982-56762011000400004)); Põlme S, et al. 2020. Fungal Diversity 105:1-16 ((https://doi.org/10.1007/s13225-020-00466-2)) |
| Otu26125 | 1 | k__Fungi(100);p__Ascomycota(100);c__Saccharomycetes(100);o__Saccharomycetales(100);f__Metschnikowiaceae(100);g__Metschnikowia(100);s__Metschnikowia_pulcherrima(100); | Metschnikowia | 13 | Saprotroph | \|Undefined Saprotroph\| | Yeast | NULL | Probable | As nectar-tap_saprotroph (Põlme et al. 2020); Decay sugar-rich substrates (Põlme et al. 2020) | Põlme S, et al. 2020. Fungal Diversity 105:1-16 ((https://doi.org/10.1007/s13225-020-00466-2)) |
| Otu26130 | 1 | k__Fungi(100);p__Ascomycota(100);c__Saccharomycetes(100);o__Saccharomycetales(100);f__Metschnikowiaceae(100);g__Metschnikowia(100);g__Metschnikowia_unclassified(100); | Metschnikowia | 13 | Saprotroph | \|Undefined Saprotroph\| | Yeast | NULL | Probable | As nectar-tap_saprotroph (Põlme et al. 2020); Decay sugar-rich substrates (Põlme et al. 2020) | Põlme S, et al. 2020. Fungal Diversity 105:1-16 ((https://doi.org/10.1007/s13225-020-00466-2)) |
| Otu26136 | 1 | k__Fungi(100);p__Ascomycota(100);c__Saccharomycetes(100);o__Saccharomycetales(100);f__Metschnikowiaceae(100);g__Metschnikowia(100);g__Metschnikowia_unclassified(100); | Metschnikowia | 13 | Saprotroph | \|Undefined Saprotroph\| | Yeast | NULL | Probable | As nectar-tap_saprotroph (Põlme et al. 2020); Decay sugar-rich substrates (Põlme et al. 2020) | Põlme S, et al. 2020. Fungal Diversity 105:1-16 ((https://doi.org/10.1007/s13225-020-00466-2)) |
| Otu26137 | 1 | k__Fungi(100);p__Ascomycota(100);c__Saccharomycetes(100);o__Saccharomycetales(100);f__Metschnikowiaceae(100);g__Metschnikowia(100);g__Metschnikowia_unclassified(100); | Metschnikowia | 13 | Saprotroph | \|Undefined Saprotroph\| | Yeast | NULL | Probable | As nectar-tap_saprotroph (Põlme et al. 2020); Decay sugar-rich substrates (Põlme et al. 2020) | Põlme S, et al. 2020. Fungal Diversity 105:1-16 ((https://doi.org/10.1007/s13225-020-00466-2)) |
| Otu26138 | 1 | k__Fungi(100);p__Ascomycota(100);c__Saccharomycetes(100);o__Saccharomycetales(100);f__Metschnikowiaceae(100);g__Metschnikowia(100);g__Metschnikowia_unclassified(100); | Metschnikowia | 13 | Saprotroph | \|Undefined Saprotroph\| | Yeast | NULL | Probable | As nectar-tap_saprotroph (Põlme et al. 2020); Decay sugar-rich substrates (Põlme et al. 2020) | Põlme S, et al. 2020. Fungal Diversity 105:1-16 ((https://doi.org/10.1007/s13225-020-00466-2)) |
| Otu26139 | 1 | k__Fungi(100);p__Ascomycota(100);c__Saccharomycetes(100);o__Saccharomycetales(100);f__Metschnikowiaceae(100);g__Metschnikowia(100);g__Metschnikowia_unclassified(100); | Metschnikowia | 13 | Saprotroph | \|Undefined Saprotroph\| | Yeast | NULL | Probable | As nectar-tap_saprotroph (Põlme et al. 2020); Decay sugar-rich substrates (Põlme et al. 2020) | Põlme S, et al. 2020. Fungal Diversity 105:1-16 ((https://doi.org/10.1007/s13225-020-00466-2)) |
| Otu26142 | 1 | k__Fungi(100);p__Ascomycota(100);c__Saccharomycetes(100);o__Saccharomycetales(100);f__Metschnikowiaceae(100);g__Metschnikowia(100);g__Metschnikowia_unclassified(100); | Metschnikowia | 13 | Saprotroph | \|Undefined Saprotroph\| | Yeast | NULL | Probable | As nectar-tap_saprotroph (Põlme et al. 2020); Decay sugar-rich substrates (Põlme et al. 2020) | Põlme S, et al. 2020. Fungal Diversity 105:1-16 ((https://doi.org/10.1007/s13225-020-00466-2)) |
| Otu26143 | 1 | k__Fungi(100);p__Ascomycota(100);c__Saccharomycetes(100);o__Saccharomycetales(100);f__Metschnikowiaceae(100);g__Metschnikowia(100);s__Metschnikowia_pulcherrima(100); | Metschnikowia | 13 | Saprotroph | \|Undefined Saprotroph\| | Yeast | NULL | Probable | As nectar-tap_saprotroph (Põlme et al. 2020); Decay sugar-rich substrates (Põlme et al. 2020) | Põlme S, et al. 2020. Fungal Diversity 105:1-16 ((https://doi.org/10.1007/s13225-020-00466-2)) |
| Otu26144 | 1 | k__Fungi(100);p__Ascomycota(100);c__Saccharomycetes(100);o__Saccharomycetales(100);f__Metschnikowiaceae(100);g__Metschnikowia(100);g__Metschnikowia_unclassified(100); | Metschnikowia | 13 | Saprotroph | \|Undefined Saprotroph\| | Yeast | NULL | Probable | As nectar-tap_saprotroph (Põlme et al. 2020); Decay sugar-rich substrates (Põlme et al. 2020) | Põlme S, et al. 2020. Fungal Diversity 105:1-16 ((https://doi.org/10.1007/s13225-020-00466-2)) |
| Otu26145 | 1 | k__Fungi(100);p__Ascomycota(100);c__Saccharomycetes(100);o__Saccharomycetales(100);f__Metschnikowiaceae(100);g__Metschnikowia(100);s__Metschnikowia_pulcherrima(100); | Metschnikowia | 13 | Saprotroph | \|Undefined Saprotroph\| | Yeast | NULL | Probable | As nectar-tap_saprotroph (Põlme et al. 2020); Decay sugar-rich substrates (Põlme et al. 2020) | Põlme S, et al. 2020. Fungal Diversity 105:1-16 ((https://doi.org/10.1007/s13225-020-00466-2)) |
| Otu26146 | 1 | k__Fungi(100);p__Ascomycota(100);c__Saccharomycetes(100);o__Saccharomycetales(100);f__Metschnikowiaceae(100);g__Metschnikowia(100);s__Metschnikowia_pulcherrima(100); | Metschnikowia | 13 | Saprotroph | \|Undefined Saprotroph\| | Yeast | NULL | Probable | As nectar-tap_saprotroph (Põlme et al. 2020); Decay sugar-rich substrates (Põlme et al. 2020) | Põlme S, et al. 2020. Fungal Diversity 105:1-16 ((https://doi.org/10.1007/s13225-020-00466-2)) |
| Otu26151 | 1 | k__Fungi(100);p__Ascomycota(100);c__Saccharomycetes(100);o__Saccharomycetales(100);f__Metschnikowiaceae(100);g__Metschnikowia(100);s__Metschnikowia_pulcherrima(100); | Metschnikowia | 13 | Saprotroph | \|Undefined Saprotroph\| | Yeast | NULL | Probable | As nectar-tap_saprotroph (Põlme et al. 2020); Decay sugar-rich substrates (Põlme et al. 2020) | Põlme S, et al. 2020. Fungal Diversity 105:1-16 ((https://doi.org/10.1007/s13225-020-00466-2)) |
| Otu26152 | 1 | k__Fungi(100);p__Ascomycota(100);c__Saccharomycetes(100);o__Saccharomycetales(100);f__Metschnikowiaceae(100);g__Metschnikowia(100);g__Metschnikowia_unclassified(100); | Metschnikowia | 13 | Saprotroph | \|Undefined Saprotroph\| | Yeast | NULL | Probable | As nectar-tap_saprotroph (Põlme et al. 2020); Decay sugar-rich substrates (Põlme et al. 2020) | Põlme S, et al. 2020. Fungal Diversity 105:1-16 ((https://doi.org/10.1007/s13225-020-00466-2)) |
| Otu26153 | 1 | k__Fungi(100);p__Ascomycota(100);c__Saccharomycetes(100);o__Saccharomycetales(100);f__Metschnikowiaceae(100);g__Metschnikowia(100);s__Metschnikowia_pulcherrima(100); | Metschnikowia | 13 | Saprotroph | \|Undefined Saprotroph\| | Yeast | NULL | Probable | As nectar-tap_saprotroph (Põlme et al. 2020); Decay sugar-rich substrates (Põlme et al. 2020) | Põlme S, et al. 2020. Fungal Diversity 105:1-16 ((https://doi.org/10.1007/s13225-020-00466-2)) |
| Otu26154 | 1 | k__Fungi(100);p__Ascomycota(100);c__Saccharomycetes(100);o__Saccharomycetales(100);f__Metschnikowiaceae(100);g__Metschnikowia(100);s__Metschnikowia_pulcherrima(100); | Metschnikowia | 13 | Saprotroph | \|Undefined Saprotroph\| | Yeast | NULL | Probable | As nectar-tap_saprotroph (Põlme et al. 2020); Decay sugar-rich substrates (Põlme et al. 2020) | Põlme S, et al. 2020. Fungal Diversity 105:1-16 ((https://doi.org/10.1007/s13225-020-00466-2)) |
| Otu26156 | 1 | k__Fungi(100);p__Ascomycota(100);c__Saccharomycetes(100);o__Saccharomycetales(100);f__Metschnikowiaceae(100);g__Metschnikowia(100);s__Metschnikowia_pulcherrima(100); | Metschnikowia | 13 | Saprotroph | \|Undefined Saprotroph\| | Yeast | NULL | Probable | As nectar-tap_saprotroph (Põlme et al. 2020); Decay sugar-rich substrates (Põlme et al. 2020) | Põlme S, et al. 2020. Fungal Diversity 105:1-16 ((https://doi.org/10.1007/s13225-020-00466-2)) |
| Otu26157 | 1 | k__Fungi(100);p__Ascomycota(100);c__Saccharomycetes(100);o__Saccharomycetales(100);f__Metschnikowiaceae(100);g__Metschnikowia(100);g__Metschnikowia_unclassified(100); | Metschnikowia | 13 | Saprotroph | \|Undefined Saprotroph\| | Yeast | NULL | Probable | As nectar-tap_saprotroph (Põlme et al. 2020); Decay sugar-rich substrates (Põlme et al. 2020) | Põlme S, et al. 2020. Fungal Diversity 105:1-16 ((https://doi.org/10.1007/s13225-020-00466-2)) |
| Otu26158 | 1 | k__Fungi(100);p__Ascomycota(100);c__Saccharomycetes(100);o__Saccharomycetales(100);f__Metschnikowiaceae(100);g__Metschnikowia(100);g__Metschnikowia_unclassified(100); | Metschnikowia | 13 | Saprotroph | \|Undefined Saprotroph\| | Yeast | NULL | Probable | As nectar-tap_saprotroph (Põlme et al. 2020); Decay sugar-rich substrates (Põlme et al. 2020) | Põlme S, et al. 2020. Fungal Diversity 105:1-16 ((https://doi.org/10.1007/s13225-020-00466-2)) |
| Otu26159 | 1 | k__Fungi(100);p__Ascomycota(100);c__Saccharomycetes(100);o__Saccharomycetales(100);f__Metschnikowiaceae(100);g__Metschnikowia(100);s__Metschnikowia_pulcherrima(100); | Metschnikowia | 13 | Saprotroph | \|Undefined Saprotroph\| | Yeast | NULL | Probable | As nectar-tap_saprotroph (Põlme et al. 2020); Decay sugar-rich substrates (Põlme et al. 2020) | Põlme S, et al. 2020. Fungal Diversity 105:1-16 ((https://doi.org/10.1007/s13225-020-00466-2)) |
| Otu26160 | 1 | k__Fungi(100);p__Ascomycota(100);c__Saccharomycetes(100);o__Saccharomycetales(100);f__Metschnikowiaceae(100);g__Metschnikowia(100);s__Metschnikowia_pulcherrima(100); | Metschnikowia | 13 | Saprotroph | \|Undefined Saprotroph\| | Yeast | NULL | Probable | As nectar-tap_saprotroph (Põlme et al. 2020); Decay sugar-rich substrates (Põlme et al. 2020) | Põlme S, et al. 2020. Fungal Diversity 105:1-16 ((https://doi.org/10.1007/s13225-020-00466-2)) |
| Otu26161 | 1 | k__Fungi(100);p__Ascomycota(100);c__Saccharomycetes(100);o__Saccharomycetales(100);f__Metschnikowiaceae(100);g__Metschnikowia(100);g__Metschnikowia_unclassified(100); | Metschnikowia | 13 | Saprotroph | \|Undefined Saprotroph\| | Yeast | NULL | Probable | As nectar-tap_saprotroph (Põlme et al. 2020); Decay sugar-rich substrates (Põlme et al. 2020) | Põlme S, et al. 2020. Fungal Diversity 105:1-16 ((https://doi.org/10.1007/s13225-020-00466-2)) |
| Otu26162 | 1 | k__Fungi(100);p__Ascomycota(100);c__Saccharomycetes(100);o__Saccharomycetales(100);f__Metschnikowiaceae(100);g__Metschnikowia(100);s__Metschnikowia_pulcherrima(100); | Metschnikowia | 13 | Saprotroph | \|Undefined Saprotroph\| | Yeast | NULL | Probable | As nectar-tap_saprotroph (Põlme et al. 2020); Decay sugar-rich substrates (Põlme et al. 2020) | Põlme S, et al. 2020. Fungal Diversity 105:1-16 ((https://doi.org/10.1007/s13225-020-00466-2)) |
| Otu26163 | 1 | k__Fungi(100);p__Ascomycota(100);c__Saccharomycetes(100);o__Saccharomycetales(100);f__Metschnikowiaceae(100);g__Metschnikowia(100);s__Metschnikowia_pulcherrima(100); | Metschnikowia | 13 | Saprotroph | \|Undefined Saprotroph\| | Yeast | NULL | Probable | As nectar-tap_saprotroph (Põlme et al. 2020); Decay sugar-rich substrates (Põlme et al. 2020) | Põlme S, et al. 2020. Fungal Diversity 105:1-16 ((https://doi.org/10.1007/s13225-020-00466-2)) |
| Otu26164 | 1 | k__Fungi(100);p__Ascomycota(100);c__Saccharomycetes(100);o__Saccharomycetales(100);f__Metschnikowiaceae(100);g__Metschnikowia(100);g__Metschnikowia_unclassified(100); | Metschnikowia | 13 | Saprotroph | \|Undefined Saprotroph\| | Yeast | NULL | Probable | As nectar-tap_saprotroph (Põlme et al. 2020); Decay sugar-rich substrates (Põlme et al. 2020) | Põlme S, et al. 2020. Fungal Diversity 105:1-16 ((https://doi.org/10.1007/s13225-020-00466-2)) |
| Otu26165 | 1 | k__Fungi(100);p__Ascomycota(100);c__Saccharomycetes(100);o__Saccharomycetales(100);f__Metschnikowiaceae(100);g__Metschnikowia(100);g__Metschnikowia_unclassified(100); | Metschnikowia | 13 | Saprotroph | \|Undefined Saprotroph\| | Yeast | NULL | Probable | As nectar-tap_saprotroph (Põlme et al. 2020); Decay sugar-rich substrates (Põlme et al. 2020) | Põlme S, et al. 2020. Fungal Diversity 105:1-16 ((https://doi.org/10.1007/s13225-020-00466-2)) |
| Otu26166 | 1 | k__Fungi(100);p__Ascomycota(100);c__Saccharomycetes(100);o__Saccharomycetales(100);f__Metschnikowiaceae(100);g__Metschnikowia(100);s__Metschnikowia_pulcherrima(100); | Metschnikowia | 13 | Saprotroph | \|Undefined Saprotroph\| | Yeast | NULL | Probable | As nectar-tap_saprotroph (Põlme et al. 2020); Decay sugar-rich substrates (Põlme et al. 2020) | Põlme S, et al. 2020. Fungal Diversity 105:1-16 ((https://doi.org/10.1007/s13225-020-00466-2)) |
| Otu26167 | 1 | k__Fungi(100);p__Ascomycota(100);c__Saccharomycetes(100);o__Saccharomycetales(100);f__Metschnikowiaceae(100);g__Metschnikowia(100);s__Metschnikowia_pulcherrima(100); | Metschnikowia | 13 | Saprotroph | \|Undefined Saprotroph\| | Yeast | NULL | Probable | As nectar-tap_saprotroph (Põlme et al. 2020); Decay sugar-rich substrates (Põlme et al. 2020) | Põlme S, et al. 2020. Fungal Diversity 105:1-16 ((https://doi.org/10.1007/s13225-020-00466-2)) |
| Otu26168 | 1 | k__Fungi(100);p__Ascomycota(100);c__Saccharomycetes(100);o__Saccharomycetales(100);f__Metschnikowiaceae(100);g__Metschnikowia(100);s__Metschnikowia_pulcherrima(100); | Metschnikowia | 13 | Saprotroph | \|Undefined Saprotroph\| | Yeast | NULL | Probable | As nectar-tap_saprotroph (Põlme et al. 2020); Decay sugar-rich substrates (Põlme et al. 2020) | Põlme S, et al. 2020. Fungal Diversity 105:1-16 ((https://doi.org/10.1007/s13225-020-00466-2)) |
| Otu26169 | 1 | k__Fungi(100);p__Ascomycota(100);c__Saccharomycetes(100);o__Saccharomycetales(100);f__Metschnikowiaceae(100);g__Metschnikowia(100);s__Metschnikowia_pulcherrima(100); | Metschnikowia | 13 | Saprotroph | \|Undefined Saprotroph\| | Yeast | NULL | Probable | As nectar-tap_saprotroph (Põlme et al. 2020); Decay sugar-rich substrates (Põlme et al. 2020) | Põlme S, et al. 2020. Fungal Diversity 105:1-16 ((https://doi.org/10.1007/s13225-020-00466-2)) |
| Otu26170 | 1 | k__Fungi(100);p__Ascomycota(100);c__Saccharomycetes(100);o__Saccharomycetales(100);f__Metschnikowiaceae(100);g__Metschnikowia(100);g__Metschnikowia_unclassified(100); | Metschnikowia | 13 | Saprotroph | \|Undefined Saprotroph\| | Yeast | NULL | Probable | As nectar-tap_saprotroph (Põlme et al. 2020); Decay sugar-rich substrates (Põlme et al. 2020) | Põlme S, et al. 2020. Fungal Diversity 105:1-16 ((https://doi.org/10.1007/s13225-020-00466-2)) |
| Otu26171 | 1 | k__Fungi(100);p__Ascomycota(100);c__Saccharomycetes(100);o__Saccharomycetales(100);f__Metschnikowiaceae(100);g__Metschnikowia(100);s__Metschnikowia_pulcherrima(100); | Metschnikowia | 13 | Saprotroph | \|Undefined Saprotroph\| | Yeast | NULL | Probable | As nectar-tap_saprotroph (Põlme et al. 2020); Decay sugar-rich substrates (Põlme et al. 2020) | Põlme S, et al. 2020. Fungal Diversity 105:1-16 ((https://doi.org/10.1007/s13225-020-00466-2)) |
| Otu26172 | 1 | k__Fungi(100);p__Ascomycota(100);c__Saccharomycetes(100);o__Saccharomycetales(100);f__Metschnikowiaceae(100);g__Metschnikowia(100);s__Metschnikowia_pulcherrima(100); | Metschnikowia | 13 | Saprotroph | \|Undefined Saprotroph\| | Yeast | NULL | Probable | As nectar-tap_saprotroph (Põlme et al. 2020); Decay sugar-rich substrates (Põlme et al. 2020) | Põlme S, et al. 2020. Fungal Diversity 105:1-16 ((https://doi.org/10.1007/s13225-020-00466-2)) |
| Otu26173 | 1 | k__Fungi(100);p__Ascomycota(100);c__Saccharomycetes(100);o__Saccharomycetales(100);f__Metschnikowiaceae(100);g__Metschnikowia(100);g__Metschnikowia_unclassified(100); | Metschnikowia | 13 | Saprotroph | \|Undefined Saprotroph\| | Yeast | NULL | Probable | As nectar-tap_saprotroph (Põlme et al. 2020); Decay sugar-rich substrates (Põlme et al. 2020) | Põlme S, et al. 2020. Fungal Diversity 105:1-16 ((https://doi.org/10.1007/s13225-020-00466-2)) |
| Otu26174 | 1 | k__Fungi(100);p__Ascomycota(100);c__Saccharomycetes(100);o__Saccharomycetales(100);f__Metschnikowiaceae(100);g__Metschnikowia(100);g__Metschnikowia_unclassified(100); | Metschnikowia | 13 | Saprotroph | \|Undefined Saprotroph\| | Yeast | NULL | Probable | As nectar-tap_saprotroph (Põlme et al. 2020); Decay sugar-rich substrates (Põlme et al. 2020) | Põlme S, et al. 2020. Fungal Diversity 105:1-16 ((https://doi.org/10.1007/s13225-020-00466-2)) |
| Otu26175 | 1 | k__Fungi(100);p__Ascomycota(100);c__Saccharomycetes(100);o__Saccharomycetales(100);f__Metschnikowiaceae(100);g__Metschnikowia(100);s__Metschnikowia_pulcherrima(100); | Metschnikowia | 13 | Saprotroph | \|Undefined Saprotroph\| | Yeast | NULL | Probable | As nectar-tap_saprotroph (Põlme et al. 2020); Decay sugar-rich substrates (Põlme et al. 2020) | Põlme S, et al. 2020. Fungal Diversity 105:1-16 ((https://doi.org/10.1007/s13225-020-00466-2)) |
| Otu26176 | 1 | k__Fungi(100);p__Ascomycota(100);c__Saccharomycetes(100);o__Saccharomycetales(100);f__Metschnikowiaceae(100);g__Metschnikowia(100);s__Metschnikowia_pulcherrima(100); | Metschnikowia | 13 | Saprotroph | \|Undefined Saprotroph\| | Yeast | NULL | Probable | As nectar-tap_saprotroph (Põlme et al. 2020); Decay sugar-rich substrates (Põlme et al. 2020) | Põlme S, et al. 2020. Fungal Diversity 105:1-16 ((https://doi.org/10.1007/s13225-020-00466-2)) |
| Otu26177 | 1 | k__Fungi(100);p__Ascomycota(100);c__Saccharomycetes(100);o__Saccharomycetales(100);f__Metschnikowiaceae(100);g__Metschnikowia(100);s__Metschnikowia_pulcherrima(100); | Metschnikowia | 13 | Saprotroph | \|Undefined Saprotroph\| | Yeast | NULL | Probable | As nectar-tap_saprotroph (Põlme et al. 2020); Decay sugar-rich substrates (Põlme et al. 2020) | Põlme S, et al. 2020. Fungal Diversity 105:1-16 ((https://doi.org/10.1007/s13225-020-00466-2)) |
| Otu26178 | 1 | k__Fungi(100);p__Ascomycota(100);c__Saccharomycetes(100);o__Saccharomycetales(100);f__Metschnikowiaceae(100);g__Metschnikowia(100);s__Metschnikowia_pulcherrima(100); | Metschnikowia | 13 | Saprotroph | \|Undefined Saprotroph\| | Yeast | NULL | Probable | As nectar-tap_saprotroph (Põlme et al. 2020); Decay sugar-rich substrates (Põlme et al. 2020) | Põlme S, et al. 2020. Fungal Diversity 105:1-16 ((https://doi.org/10.1007/s13225-020-00466-2)) |
| Otu26183 | 1 | k__Fungi(100);p__Ascomycota(100);c__Saccharomycetes(100);o__Saccharomycetales(100);f__Metschnikowiaceae(100);g__Metschnikowia(100);s__Metschnikowia_pulcherrima(100); | Metschnikowia | 13 | Saprotroph | \|Undefined Saprotroph\| | Yeast | NULL | Probable | As nectar-tap_saprotroph (Põlme et al. 2020); Decay sugar-rich substrates (Põlme et al. 2020) | Põlme S, et al. 2020. Fungal Diversity 105:1-16 ((https://doi.org/10.1007/s13225-020-00466-2)) |
| Otu26185 | 1 | k__Fungi(100);p__Ascomycota(100);c__Saccharomycetes(100);o__Saccharomycetales(100);f__Metschnikowiaceae(100);g__Metschnikowia(100);g__Metschnikowia_unclassified(100); | Metschnikowia | 13 | Saprotroph | \|Undefined Saprotroph\| | Yeast | NULL | Probable | As nectar-tap_saprotroph (Põlme et al. 2020); Decay sugar-rich substrates (Põlme et al. 2020) | Põlme S, et al. 2020. Fungal Diversity 105:1-16 ((https://doi.org/10.1007/s13225-020-00466-2)) |
| Otu26189 | 1 | k__Fungi(100);p__Ascomycota(100);c__Saccharomycetes(100);o__Saccharomycetales(100);o__Saccharomycetales_unclassified(100);o__Saccharomycetales_unclassified(100);o__Saccharomycetales_unclassified(100); | Saccharomycetales | 7 | Saprotroph | Undefined Saprotroph | Yeast | NULL | Possible | NULL | Sterkenburg E, et al. 2015. New Phytologist 207:1145-1158 ((https://doi.org/10.1111/nph.13426)) |
| Otu26193 | 1 | k__Fungi(100);p__Ascomycota(100);c__Saccharomycetes(100);o__Saccharomycetales(100);f__Metschnikowiaceae(100);g__Metschnikowia(100);s__Metschnikowia_pulcherrima(100); | Metschnikowia | 13 | Saprotroph | \|Undefined Saprotroph\| | Yeast | NULL | Probable | As nectar-tap_saprotroph (Põlme et al. 2020); Decay sugar-rich substrates (Põlme et al. 2020) | Põlme S, et al. 2020. Fungal Diversity 105:1-16 ((https://doi.org/10.1007/s13225-020-00466-2)) |
| Otu26195 | 1 | k__Fungi(100);p__Ascomycota(100);c__Saccharomycetes(100);o__Saccharomycetales(100);f__Metschnikowiaceae(100);g__Metschnikowia(100);s__Metschnikowia_pulcherrima(100); | Metschnikowia | 13 | Saprotroph | \|Undefined Saprotroph\| | Yeast | NULL | Probable | As nectar-tap_saprotroph (Põlme et al. 2020); Decay sugar-rich substrates (Põlme et al. 2020) | Põlme S, et al. 2020. Fungal Diversity 105:1-16 ((https://doi.org/10.1007/s13225-020-00466-2)) |
| Otu26201 | 1 | k__Fungi(100);p__Ascomycota(100);c__Saccharomycetes(100);o__Saccharomycetales(100);f__Metschnikowiaceae(100);g__Metschnikowia(100);s__Metschnikowia_pulcherrima(100); | Metschnikowia | 13 | Saprotroph | \|Undefined Saprotroph\| | Yeast | NULL | Probable | As nectar-tap_saprotroph (Põlme et al. 2020); Decay sugar-rich substrates (Põlme et al. 2020) | Põlme S, et al. 2020. Fungal Diversity 105:1-16 ((https://doi.org/10.1007/s13225-020-00466-2)) |
| Otu26202 | 1 | k__Fungi(100);p__Ascomycota(100);c__Saccharomycetes(100);o__Saccharomycetales(100);f__Metschnikowiaceae(100);g__Metschnikowia(100);s__Metschnikowia_pulcherrima(100); | Metschnikowia | 13 | Saprotroph | \|Undefined Saprotroph\| | Yeast | NULL | Probable | As nectar-tap_saprotroph (Põlme et al. 2020); Decay sugar-rich substrates (Põlme et al. 2020) | Põlme S, et al. 2020. Fungal Diversity 105:1-16 ((https://doi.org/10.1007/s13225-020-00466-2)) |
| Otu26204 | 1 | k__Fungi(100);p__Ascomycota(100);c__Saccharomycetes(100);o__Saccharomycetales(100);f__Metschnikowiaceae(100);g__Metschnikowia(100);g__Metschnikowia_unclassified(100); | Metschnikowia | 13 | Saprotroph | \|Undefined Saprotroph\| | Yeast | NULL | Probable | As nectar-tap_saprotroph (Põlme et al. 2020); Decay sugar-rich substrates (Põlme et al. 2020) | Põlme S, et al. 2020. Fungal Diversity 105:1-16 ((https://doi.org/10.1007/s13225-020-00466-2)) |
| Otu26205 | 1 | k__Fungi(100);p__Ascomycota(100);c__Saccharomycetes(100);o__Saccharomycetales(100);f__Metschnikowiaceae(100);g__Metschnikowia(100);s__Metschnikowia_pulcherrima(100); | Metschnikowia | 13 | Saprotroph | \|Undefined Saprotroph\| | Yeast | NULL | Probable | As nectar-tap_saprotroph (Põlme et al. 2020); Decay sugar-rich substrates (Põlme et al. 2020) | Põlme S, et al. 2020. Fungal Diversity 105:1-16 ((https://doi.org/10.1007/s13225-020-00466-2)) |
| Otu26206 | 1 | k__Fungi(100);p__Ascomycota(100);c__Saccharomycetes(100);o__Saccharomycetales(100);f__Metschnikowiaceae(100);g__Metschnikowia(100);s__Metschnikowia_pulcherrima(100); | Metschnikowia | 13 | Saprotroph | \|Undefined Saprotroph\| | Yeast | NULL | Probable | As nectar-tap_saprotroph (Põlme et al. 2020); Decay sugar-rich substrates (Põlme et al. 2020) | Põlme S, et al. 2020. Fungal Diversity 105:1-16 ((https://doi.org/10.1007/s13225-020-00466-2)) |
| Otu26207 | 1 | k__Fungi(100);p__Ascomycota(100);c__Saccharomycetes(100);o__Saccharomycetales(100);f__Metschnikowiaceae(100);g__Metschnikowia(100);s__Metschnikowia_pulcherrima(100); | Metschnikowia | 13 | Saprotroph | \|Undefined Saprotroph\| | Yeast | NULL | Probable | As nectar-tap_saprotroph (Põlme et al. 2020); Decay sugar-rich substrates (Põlme et al. 2020) | Põlme S, et al. 2020. Fungal Diversity 105:1-16 ((https://doi.org/10.1007/s13225-020-00466-2)) |
| Otu26208 | 1 | k__Fungi(100);p__Ascomycota(100);c__Saccharomycetes(100);o__Saccharomycetales(100);f__Metschnikowiaceae(100);g__Metschnikowia(100);s__Metschnikowia_pulcherrima(100); | Metschnikowia | 13 | Saprotroph | \|Undefined Saprotroph\| | Yeast | NULL | Probable | As nectar-tap_saprotroph (Põlme et al. 2020); Decay sugar-rich substrates (Põlme et al. 2020) | Põlme S, et al. 2020. Fungal Diversity 105:1-16 ((https://doi.org/10.1007/s13225-020-00466-2)) |
| Otu26210 | 1 | k__Fungi(100);p__Ascomycota(100);c__Saccharomycetes(100);o__Saccharomycetales(100);f__Metschnikowiaceae(100);g__Metschnikowia(100);s__Metschnikowia_pulcherrima(100); | Metschnikowia | 13 | Saprotroph | \|Undefined Saprotroph\| | Yeast | NULL | Probable | As nectar-tap_saprotroph (Põlme et al. 2020); Decay sugar-rich substrates (Põlme et al. 2020) | Põlme S, et al. 2020. Fungal Diversity 105:1-16 ((https://doi.org/10.1007/s13225-020-00466-2)) |
| Otu26211 | 1 | k__Fungi(100);p__Ascomycota(100);c__Saccharomycetes(100);o__Saccharomycetales(100);f__Metschnikowiaceae(100);g__Metschnikowia(100);s__Metschnikowia_pulcherrima(100); | Metschnikowia | 13 | Saprotroph | \|Undefined Saprotroph\| | Yeast | NULL | Probable | As nectar-tap_saprotroph (Põlme et al. 2020); Decay sugar-rich substrates (Põlme et al. 2020) | Põlme S, et al. 2020. Fungal Diversity 105:1-16 ((https://doi.org/10.1007/s13225-020-00466-2)) |
| Otu26212 | 1 | k__Fungi(100);p__Ascomycota(100);c__Saccharomycetes(100);o__Saccharomycetales(100);f__Metschnikowiaceae(100);g__Metschnikowia(100);s__Metschnikowia_pulcherrima(100); | Metschnikowia | 13 | Saprotroph | \|Undefined Saprotroph\| | Yeast | NULL | Probable | As nectar-tap_saprotroph (Põlme et al. 2020); Decay sugar-rich substrates (Põlme et al. 2020) | Põlme S, et al. 2020. Fungal Diversity 105:1-16 ((https://doi.org/10.1007/s13225-020-00466-2)) |
| Otu26213 | 1 | k__Fungi(100);p__Ascomycota(100);c__Saccharomycetes(100);o__Saccharomycetales(100);f__Metschnikowiaceae(100);g__Metschnikowia(100);s__Metschnikowia_pulcherrima(100); | Metschnikowia | 13 | Saprotroph | \|Undefined Saprotroph\| | Yeast | NULL | Probable | As nectar-tap_saprotroph (Põlme et al. 2020); Decay sugar-rich substrates (Põlme et al. 2020) | Põlme S, et al. 2020. Fungal Diversity 105:1-16 ((https://doi.org/10.1007/s13225-020-00466-2)) |
| Otu26214 | 1 | k__Fungi(100);p__Ascomycota(100);c__Saccharomycetes(100);o__Saccharomycetales(100);f__Metschnikowiaceae(100);g__Metschnikowia(100);s__Metschnikowia_pulcherrima(100); | Metschnikowia | 13 | Saprotroph | \|Undefined Saprotroph\| | Yeast | NULL | Probable | As nectar-tap_saprotroph (Põlme et al. 2020); Decay sugar-rich substrates (Põlme et al. 2020) | Põlme S, et al. 2020. Fungal Diversity 105:1-16 ((https://doi.org/10.1007/s13225-020-00466-2)) |
| Otu26215 | 1 | k__Fungi(100);p__Ascomycota(100);c__Saccharomycetes(100);o__Saccharomycetales(100);f__Metschnikowiaceae(100);g__Metschnikowia(100);s__Metschnikowia_pulcherrima(100); | Metschnikowia | 13 | Saprotroph | \|Undefined Saprotroph\| | Yeast | NULL | Probable | As nectar-tap_saprotroph (Põlme et al. 2020); Decay sugar-rich substrates (Põlme et al. 2020) | Põlme S, et al. 2020. Fungal Diversity 105:1-16 ((https://doi.org/10.1007/s13225-020-00466-2)) |
| Otu26216 | 1 | k__Fungi(100);p__Ascomycota(100);c__Saccharomycetes(100);o__Saccharomycetales(100);f__Metschnikowiaceae(100);g__Metschnikowia(100);g__Metschnikowia_unclassified(100); | Metschnikowia | 13 | Saprotroph | \|Undefined Saprotroph\| | Yeast | NULL | Probable | As nectar-tap_saprotroph (Põlme et al. 2020); Decay sugar-rich substrates (Põlme et al. 2020) | Põlme S, et al. 2020. Fungal Diversity 105:1-16 ((https://doi.org/10.1007/s13225-020-00466-2)) |
| Otu26217 | 1 | k__Fungi(100);p__Ascomycota(100);c__Saccharomycetes(100);o__Saccharomycetales(100);f__Metschnikowiaceae(100);g__Metschnikowia(100);s__Metschnikowia_pulcherrima(100); | Metschnikowia | 13 | Saprotroph | \|Undefined Saprotroph\| | Yeast | NULL | Probable | As nectar-tap_saprotroph (Põlme et al. 2020); Decay sugar-rich substrates (Põlme et al. 2020) | Põlme S, et al. 2020. Fungal Diversity 105:1-16 ((https://doi.org/10.1007/s13225-020-00466-2)) |
| Otu26218 | 1 | k__Fungi(100);p__Ascomycota(100);c__Saccharomycetes(100);o__Saccharomycetales(100);f__Metschnikowiaceae(100);g__Metschnikowia(100);s__Metschnikowia_pulcherrima(100); | Metschnikowia | 13 | Saprotroph | \|Undefined Saprotroph\| | Yeast | NULL | Probable | As nectar-tap_saprotroph (Põlme et al. 2020); Decay sugar-rich substrates (Põlme et al. 2020) | Põlme S, et al. 2020. Fungal Diversity 105:1-16 ((https://doi.org/10.1007/s13225-020-00466-2)) |
| Otu26219 | 1 | k__Fungi(100);p__Ascomycota(100);c__Saccharomycetes(100);o__Saccharomycetales(100);f__Metschnikowiaceae(100);g__Metschnikowia(100);g__Metschnikowia_unclassified(100); | Metschnikowia | 13 | Saprotroph | \|Undefined Saprotroph\| | Yeast | NULL | Probable | As nectar-tap_saprotroph (Põlme et al. 2020); Decay sugar-rich substrates (Põlme et al. 2020) | Põlme S, et al. 2020. Fungal Diversity 105:1-16 ((https://doi.org/10.1007/s13225-020-00466-2)) |
| Otu26221 | 1 | k__Fungi(100);p__Ascomycota(100);c__Saccharomycetes(100);o__Saccharomycetales(100);f__Metschnikowiaceae(100);g__Metschnikowia(100);g__Metschnikowia_unclassified(100); | Metschnikowia | 13 | Saprotroph | \|Undefined Saprotroph\| | Yeast | NULL | Probable | As nectar-tap_saprotroph (Põlme et al. 2020); Decay sugar-rich substrates (Põlme et al. 2020) | Põlme S, et al. 2020. Fungal Diversity 105:1-16 ((https://doi.org/10.1007/s13225-020-00466-2)) |
| Otu26222 | 1 | k__Fungi(100);p__Ascomycota(100);c__Saccharomycetes(100);o__Saccharomycetales(100);f__Metschnikowiaceae(100);g__Metschnikowia(100);s__Metschnikowia_pulcherrima(100); | Metschnikowia | 13 | Saprotroph | \|Undefined Saprotroph\| | Yeast | NULL | Probable | As nectar-tap_saprotroph (Põlme et al. 2020); Decay sugar-rich substrates (Põlme et al. 2020) | Põlme S, et al. 2020. Fungal Diversity 105:1-16 ((https://doi.org/10.1007/s13225-020-00466-2)) |
| Otu26223 | 1 | k__Fungi(100);p__Ascomycota(100);c__Saccharomycetes(100);o__Saccharomycetales(100);f__Metschnikowiaceae(100);g__Metschnikowia(100);s__Metschnikowia_pulcherrima(100); | Metschnikowia | 13 | Saprotroph | \|Undefined Saprotroph\| | Yeast | NULL | Probable | As nectar-tap_saprotroph (Põlme et al. 2020); Decay sugar-rich substrates (Põlme et al. 2020) | Põlme S, et al. 2020. Fungal Diversity 105:1-16 ((https://doi.org/10.1007/s13225-020-00466-2)) |
| Otu26224 | 1 | k__Fungi(100);p__Ascomycota(100);c__Saccharomycetes(100);o__Saccharomycetales(100);f__Metschnikowiaceae(100);g__Metschnikowia(100);s__Metschnikowia_pulcherrima(100); | Metschnikowia | 13 | Saprotroph | \|Undefined Saprotroph\| | Yeast | NULL | Probable | As nectar-tap_saprotroph (Põlme et al. 2020); Decay sugar-rich substrates (Põlme et al. 2020) | Põlme S, et al. 2020. Fungal Diversity 105:1-16 ((https://doi.org/10.1007/s13225-020-00466-2)) |
| Otu26226 | 1 | k__Fungi(100);p__Ascomycota(100);c__Saccharomycetes(100);o__Saccharomycetales(100);f__Metschnikowiaceae(100);g__Metschnikowia(100);g__Metschnikowia_unclassified(100); | Metschnikowia | 13 | Saprotroph | \|Undefined Saprotroph\| | Yeast | NULL | Probable | As nectar-tap_saprotroph (Põlme et al. 2020); Decay sugar-rich substrates (Põlme et al. 2020) | Põlme S, et al. 2020. Fungal Diversity 105:1-16 ((https://doi.org/10.1007/s13225-020-00466-2)) |
| Otu26227 | 1 | k__Fungi(100);p__Ascomycota(100);c__Saccharomycetes(100);o__Saccharomycetales(100);f__Saccharomycetales_family_Incertae_sedis(100);g__Candida(100);g__Candida_unclassified(100); | Candida | 13 | Pathotroph-Saprotroph-Symbiotroph | Animal Pathogen-Endophyte-Endosymbiont-Epiphyte-\|Undefined Saprotroph\| | Dimorphic Yeast-Yeast | NULL | Probable | As nectar-tap_saprotroph (Põlme et al. 2020); Decay sugar-rich substrates (Põlme et al. 2020); This asexual genus is polyphyletic and contain species that function very differently depend on what they are. Caution should be taken when interpreting guilds for Candida species. | Manolakaki D, et al. 2010. Virulence. 1: 367-375 ((https://doi.org/10.4161/viru.1.5.12796)); Mekbib SB, et al. 2011. Tropical Plant Pathology 36:233-240 ((https://doi.org/10.1590/S1982-56762011000400004)); Põlme S, et al. 2020. Fungal Diversity 105:1-16 ((https://doi.org/10.1007/s13225-020-00466-2)) |

| | | | | | | | | | | | |
|---|---|---|---|---|---|---|---|---|---|---|---|
| Otu26228 | 1 | k__Fungi(100);p__Ascomycota(100);c__Saccharomycetes(100);o__Saccharomycetales(100);f__Metschnikowiaceae(100);g__Metschnikowia(100);s__Metschnikowia_pulcherrima(100); | Metschnikowia | 13 | Saprotroph | \|Undefined Saprotroph\| | Yeast | NULL | Probable | As nectar-tap_saprotroph (Põlme et al. 2020); Decay sugar-rich substrates (Põlme et al. 2020) | Põlme S, et al. 2020. Fungal Diversity 105:1-16 ((https://doi.org/10.1007/s13225-020-00466-2)) |
| Otu26229 | 1 | k__Fungi(100);p__Ascomycota(100);c__Saccharomycetes(100);o__Saccharomycetales(100);f__Metschnikowiaceae(100);g__Metschnikowia(100);s__Metschnikowia_pulcherrima(100); | Metschnikowia | 13 | Saprotroph | \|Undefined Saprotroph\| | Yeast | NULL | Probable | As nectar-tap_saprotroph (Põlme et al. 2020); Decay sugar-rich substrates (Põlme et al. 2020) | Põlme S, et al. 2020. Fungal Diversity 105:1-16 ((https://doi.org/10.1007/s13225-020-00466-2)) |
| Otu26231 | 1 | k__Fungi(100);p__Ascomycota(100);c__Saccharomycetes(100);o__Saccharomycetales(100);f__Metschnikowiaceae(100);g__Metschnikowia(100);g__Metschnikowia_unclassified(100); | Metschnikowia | 13 | Saprotroph | \|Undefined Saprotroph\| | Yeast | NULL | Probable | As nectar-tap_saprotroph (Põlme et al. 2020); Decay sugar-rich substrates (Põlme et al. 2020) | Põlme S, et al. 2020. Fungal Diversity 105:1-16 ((https://doi.org/10.1007/s13225-020-00466-2)) |
| Otu26232 | 1 | k__Fungi(100);p__Ascomycota(100);c__Saccharomycetes(100);o__Saccharomycetales(100);f__Metschnikowiaceae(100);g__Metschnikowia(100);s__Metschnikowia_pulcherrima(100); | Metschnikowia | 13 | Saprotroph | \|Undefined Saprotroph\| | Yeast | NULL | Probable | As nectar-tap_saprotroph (Põlme et al. 2020); Decay sugar-rich substrates (Põlme et al. 2020) | Põlme S, et al. 2020. Fungal Diversity 105:1-16 ((https://doi.org/10.1007/s13225-020-00466-2)) |
| Otu26233 | 1 | k__Fungi(100);p__Ascomycota(100);c__Saccharomycetes(100);o__Saccharomycetales(100);f__Metschnikowiaceae(100);g__Metschnikowia(100);s__Metschnikowia_pulcherrima(100); | Metschnikowia | 13 | Saprotroph | \|Undefined Saprotroph\| | Yeast | NULL | Probable | As nectar-tap_saprotroph (Põlme et al. 2020); Decay sugar-rich substrates (Põlme et al. 2020) | Põlme S, et al. 2020. Fungal Diversity 105:1-16 ((https://doi.org/10.1007/s13225-020-00466-2)) |
| Otu26234 | 1 | k__Fungi(100);p__Ascomycota(100);c__Saccharomycetes(100);o__Saccharomycetales(100);f__Metschnikowiaceae(100);g__Metschnikowia(100);s__Metschnikowia_pulcherrima(100); | Metschnikowia | 13 | Saprotroph | \|Undefined Saprotroph\| | Yeast | NULL | Probable | As nectar-tap_saprotroph (Põlme et al. 2020); Decay sugar-rich substrates (Põlme et al. 2020) | Põlme S, et al. 2020. Fungal Diversity 105:1-16 ((https://doi.org/10.1007/s13225-020-00466-2)) |
| Otu26235 | 1 | k__Fungi(100);p__Ascomycota(100);c__Saccharomycetes(100);o__Saccharomycetales(100);f__Saccharomycetales_family_Incertae_sedis(100);g__Candida(100);g__Candida_unclassified(100); | Candida | 13 | Pathotroph-Saprotroph-Symbiotroph | Animal Pathogen-Endophyte-Endosymbiont-Epiphyte-\|Undefined Saprotroph\| | Dimorphic Yeast-Yeast | NULL | Probable | As nectar-tap_saprotroph (Põlme et al. 2020); Decay sugar-rich substrates (Põlme et al. 2020); This asexual genus is polyphyletic and contain species that function very differently depend on what they are. Caution should be taken when interpreting guilds for Candida species. | Manolakaki D, et al. 2010. Virulence. 1:367-375 ((https://doi.org/10.4161/viru.1.5.12796)); Mekbib SB, et al. 2011. Tropical Plant Pathology 36:233-240 ((https://doi.org/10.1590/S1982-56762011000400004)); Põlme S, et al. 2020. Fungal Diversity 105:1-16 ((https://doi.org/10.1007/s13225-020-00466-2)) |
| Otu26237 | 1 | k__Fungi(100);p__Ascomycota(100);c__Saccharomycetes(100);o__Saccharomycetales(100);f__Metschnikowiaceae(100);g__Metschnikowia(100);s__Metschnikowia_pulcherrima(100); | Metschnikowia | 13 | Saprotroph | \|Undefined Saprotroph\| | Yeast | NULL | Probable | As nectar-tap_saprotroph (Põlme et al. 2020); Decay sugar-rich substrates (Põlme et al. 2020) | Põlme S, et al. 2020. Fungal Diversity 105:1-16 ((https://doi.org/10.1007/s13225-020-00466-2)) |
| Otu26238 | 1 | k__Fungi(100);p__Ascomycota(100);c__Saccharomycetes(100);o__Saccharomycetales(100);f__Metschnikowiaceae(100);g__Metschnikowia(100);s__Metschnikowia_pulcherrima(100); | Metschnikowia | 13 | Saprotroph | \|Undefined Saprotroph\| | Yeast | NULL | Probable | As nectar-tap_saprotroph (Põlme et al. 2020); Decay sugar-rich substrates (Põlme et al. 2020) | Põlme S, et al. 2020. Fungal Diversity 105:1-16 ((https://doi.org/10.1007/s13225-020-00466-2)) |
| Otu26239 | 1 | k__Fungi(100);p__Ascomycota(100);c__Saccharomycetes(100);o__Saccharomycetales(100);f__Metschnikowiaceae(100);g__Metschnikowia(100);s__Metschnikowia_pulcherrima(100); | Metschnikowia | 13 | Saprotroph | \|Undefined Saprotroph\| | Yeast | NULL | Probable | As nectar-tap_saprotroph (Põlme et al. 2020); Decay sugar-rich substrates (Põlme et al. 2020) | Põlme S, et al. 2020. Fungal Diversity 105:1-16 ((https://doi.org/10.1007/s13225-020-00466-2)) |
| Otu26240 | 1 | k__Fungi(100);p__Ascomycota(100);c__Saccharomycetes(100);o__Saccharomycetales(100);f__Metschnikowiaceae(100);g__Metschnikowia(100);s__Metschnikowia_pulcherrima(100); | Metschnikowia | 13 | Saprotroph | \|Undefined Saprotroph\| | Yeast | NULL | Probable | As nectar-tap_saprotroph (Põlme et al. 2020); Decay sugar-rich substrates (Põlme et al. 2020) | Põlme S, et al. 2020. Fungal Diversity 105:1-16 ((https://doi.org/10.1007/s13225-020-00466-2)) |
| Otu26241 | 1 | k__Fungi(100);p__Ascomycota(100);c__Saccharomycetes(100);o__Saccharomycetales(100);f__Metschnikowiaceae(100);g__Metschnikowia(100);g__Metschnikowia_unclassified(100); | Metschnikowia | 13 | Saprotroph | \|Undefined Saprotroph\| | Yeast | NULL | Probable | As nectar-tap_saprotroph (Põlme et al. 2020); Decay sugar-rich substrates (Põlme et al. 2020) | Põlme S, et al. 2020. Fungal Diversity 105:1-16 ((https://doi.org/10.1007/s13225-020-00466-2)) |
| Otu26242 | 1 | k__Fungi(100);p__Ascomycota(100);c__Saccharomycetes(100);o__Saccharomycetales(100);f__Metschnikowiaceae(100);g__Metschnikowia(100);g__Metschnikowia_unclassified(100); | Metschnikowia | 13 | Saprotroph | \|Undefined Saprotroph\| | Yeast | NULL | Probable | As nectar-tap_saprotroph (Põlme et al. 2020); Decay sugar-rich substrates (Põlme et al. 2020) | Põlme S, et al. 2020. Fungal Diversity 105:1-16 ((https://doi.org/10.1007/s13225-020-00466-2)) |
| Otu26243 | 1 | k__Fungi(100);p__Ascomycota(100);c__Saccharomycetes(100);o__Saccharomycetales(100);f__Metschnikowiaceae(100);g__Metschnikowia(100);g__Metschnikowia_unclassified(100); | Metschnikowia | 13 | Saprotroph | \|Undefined Saprotroph\| | Yeast | NULL | Probable | As nectar-tap_saprotroph (Põlme et al. 2020); Decay sugar-rich substrates (Põlme et al. 2020) | Põlme S, et al. 2020. Fungal Diversity 105:1-16 ((https://doi.org/10.1007/s13225-020-00466-2)) |
| Otu26245 | 1 | k__Fungi(100);p__Ascomycota(100);c__Saccharomycetes(100);o__Saccharomycetales(100);f__Metschnikowiaceae(100);g__Metschnikowia(100);s__Metschnikowia_pulcherrima(100); | Metschnikowia | 13 | Saprotroph | \|Undefined Saprotroph\| | Yeast | NULL | Probable | As nectar-tap_saprotroph (Põlme et al. 2020); Decay sugar-rich substrates (Põlme et al. 2020) | Põlme S, et al. 2020. Fungal Diversity 105:1-16 ((https://doi.org/10.1007/s13225-020-00466-2)) |
| Otu26246 | 1 | k__Fungi(100);p__Ascomycota(100);c__Saccharomycetes(100);o__Saccharomycetales(100);f__Metschnikowiaceae(100);g__Metschnikowia(100);s__Metschnikowia_pulcherrima(100); | Metschnikowia | 13 | Saprotroph | \|Undefined Saprotroph\| | Yeast | NULL | Probable | As nectar-tap_saprotroph (Põlme et al. 2020); Decay sugar-rich substrates (Põlme et al. 2020) | Põlme S, et al. 2020. Fungal Diversity 105:1-16 ((https://doi.org/10.1007/s13225-020-00466-2)) |
| Otu26247 | 1 | k__Fungi(100);p__Ascomycota(100);c__Saccharomycetes(100);o__Saccharomycetales(100);f__Metschnikowiaceae(100);g__Metschnikowia(100);s__Metschnikowia_pulcherrima(100); | Metschnikowia | 13 | Saprotroph | \|Undefined Saprotroph\| | Yeast | NULL | Probable | As nectar-tap_saprotroph (Põlme et al. 2020); Decay sugar-rich substrates (Põlme et al. 2020) | Põlme S, et al. 2020. Fungal Diversity 105:1-16 ((https://doi.org/10.1007/s13225-020-00466-2)) |
| Otu26248 | 1 | k__Fungi(100);p__Ascomycota(100);c__Saccharomycetes(100);o__Saccharomycetales(100);f__Metschnikowiaceae(100);g__Metschnikowia(100);g__Metschnikowia_unclassified(100); | Metschnikowia | 13 | Saprotroph | \|Undefined Saprotroph\| | Yeast | NULL | Probable | As nectar-tap_saprotroph (Põlme et al. 2020); Decay sugar-rich substrates (Põlme et al. 2020) | Põlme S, et al. 2020. Fungal Diversity 105:1-16 ((https://doi.org/10.1007/s13225-020-00466-2)) |
| Otu26249 | 1 | k__Fungi(100);p__Ascomycota(100);c__Saccharomycetes(100);o__Saccharomycetales(100);f__Metschnikowiaceae(100);g__Metschnikowia(100);g__Metschnikowia_unclassified(100); | Metschnikowia | 13 | Saprotroph | \|Undefined Saprotroph\| | Yeast | NULL | Probable | As nectar-tap_saprotroph (Põlme et al. 2020); Decay sugar-rich substrates (Põlme et al. 2020) | Põlme S, et al. 2020. Fungal Diversity 105:1-16 ((https://doi.org/10.1007/s13225-020-00466-2)) |
| Otu26250 | 1 | k__Fungi(100);p__Ascomycota(100);c__Saccharomycetes(100);o__Saccharomycetales(100);f__Metschnikowiaceae(100);g__Metschnikowia(100);s__Metschnikowia_pulcherrima(100); | Metschnikowia | 13 | Saprotroph | \|Undefined Saprotroph\| | Yeast | NULL | Probable | As nectar-tap_saprotroph (Põlme et al. 2020); Decay sugar-rich substrates (Põlme et al. 2020) | Põlme S, et al. 2020. Fungal Diversity 105:1-16 ((https://doi.org/10.1007/s13225-020-00466-2)) |
| Otu26251 | 1 | k__Fungi(100);p__Ascomycota(100);c__Saccharomycetes(100);o__Saccharomycetales(100);f__Metschnikowiaceae(100);g__Metschnikowia(100);g__Metschnikowia_unclassified(100); | Metschnikowia | 13 | Saprotroph | \|Undefined Saprotroph\| | Yeast | NULL | Probable | As nectar-tap_saprotroph (Põlme et al. 2020); Decay sugar-rich substrates (Põlme et al. 2020) | Põlme S, et al. 2020. Fungal Diversity 105:1-16 ((https://doi.org/10.1007/s13225-020-00466-2)) |
| Otu26252 | 1 | k__Fungi(100);p__Ascomycota(100);c__Saccharomycetes(100);o__Saccharomycetales(100);f__Metschnikowiaceae(100);g__Metschnikowia(100);g__Metschnikowia_unclassified(100); | Metschnikowia | 13 | Saprotroph | \|Undefined Saprotroph\| | Yeast | NULL | Probable | As nectar-tap_saprotroph (Põlme et al. 2020); Decay sugar-rich substrates (Põlme et al. 2020) | Põlme S, et al. 2020. Fungal Diversity 105:1-16 ((https://doi.org/10.1007/s13225-020-00466-2)) |
| Otu26254 | 1 | k__Fungi(100);p__Ascomycota(100);c__Saccharomycetes(100);o__Saccharomycetales(100);f__Metschnikowiaceae(100);g__Metschnikowia(100);s__Metschnikowia_pulcherrima(100); | Metschnikowia | 13 | Saprotroph | \|Undefined Saprotroph\| | Yeast | NULL | Probable | As nectar-tap_saprotroph (Põlme et al. 2020); Decay sugar-rich substrates (Põlme et al. 2020) | Põlme S, et al. 2020. Fungal Diversity 105:1-16 ((https://doi.org/10.1007/s13225-020-00466-2)) |
| Otu26256 | 1 | k__Fungi(100);p__Ascomycota(100);c__Saccharomycetes(100);o__Saccharomycetales(100);f__Metschnikowiaceae(100);g__Metschnikowia(100);s__Metschnikowia_pulcherrima(100); | Metschnikowia | 13 | Saprotroph | \|Undefined Saprotroph\| | Yeast | NULL | Probable | As nectar-tap_saprotroph (Põlme et al. 2020); Decay sugar-rich substrates (Põlme et al. 2020) | Põlme S, et al. 2020. Fungal Diversity 105:1-16 ((https://doi.org/10.1007/s13225-020-00466-2)) |
| Otu26257 | 1 | k__Fungi(100);p__Ascomycota(100);c__Saccharomycetes(100);o__Saccharomycetales(100);f__Metschnikowiaceae(100);g__Metschnikowia(100);s__Metschnikowia_pulcherrima(100); | Metschnikowia | 13 | Saprotroph | \|Undefined Saprotroph\| | Yeast | NULL | Probable | As nectar-tap_saprotroph (Põlme et al. 2020); Decay sugar-rich substrates (Põlme et al. 2020) | Põlme S, et al. 2020. Fungal Diversity 105:1-16 ((https://doi.org/10.1007/s13225-020-00466-2)) |
| Otu26266 | 1 | k__Fungi(100);p__Ascomycota(100);c__Saccharomycetes(100);o__Saccharomycetales(100);f__Metschnikowiaceae(100);g__Metschnikowia(100);s__Metschnikowia_pulcherrima(100); | Metschnikowia | 13 | Saprotroph | \|Undefined Saprotroph\| | Yeast | NULL | Probable | As nectar-tap_saprotroph (Põlme et al. 2020); Decay sugar-rich substrates (Põlme et al. 2020) | Põlme S, et al. 2020. Fungal Diversity 105:1-16 ((https://doi.org/10.1007/s13225-020-00466-2)) |
| Otu26269 | 1 | k__Fungi(100);p__Ascomycota(100);c__Saccharomycetes(100);o__Saccharomycetales(100);f__Metschnikowiaceae(100);g__Metschnikowia(100);s__Metschnikowia_pulcherrima(100); | Metschnikowia | 13 | Saprotroph | \|Undefined Saprotroph\| | Yeast | NULL | Probable | As nectar-tap_saprotroph (Põlme et al. 2020); Decay sugar-rich substrates (Põlme et al. 2020) | Põlme S, et al. 2020. Fungal Diversity 105:1-16 ((https://doi.org/10.1007/s13225-020-00466-2)) |
| Otu26270 | 1 | k__Fungi(100);p__Ascomycota(100);c__Saccharomycetes(100);o__Saccharomycetales(100);f__Metschnikowiaceae(100);g__Metschnikowia(100);s__Metschnikowia_pulcherrima(100); | Metschnikowia | 13 | Saprotroph | \|Undefined Saprotroph\| | Yeast | NULL | Probable | As nectar-tap_saprotroph (Põlme et al. 2020); Decay sugar-rich substrates (Põlme et al. 2020) | Põlme S, et al. 2020. Fungal Diversity 105:1-16 ((https://doi.org/10.1007/s13225-020-00466-2)) |
| Otu26272 | 1 | k__Fungi(100);p__Ascomycota(100);c__Saccharomycetes(100);o__Saccharomycetales(100);f__Metschnikowiaceae(100);g__Metschnikowia(100);s__Metschnikowia_pulcherrima(100); | Metschnikowia | 13 | Saprotroph | \|Undefined Saprotroph\| | Yeast | NULL | Probable | As nectar-tap_saprotroph (Põlme et al. 2020); Decay sugar-rich substrates (Põlme et al. 2020) | Põlme S, et al. 2020. Fungal Diversity 105:1-16 ((https://doi.org/10.1007/s13225-020-00466-2)) |
| Otu26273 | 1 | k__Fungi(100);p__Ascomycota(100);c__Saccharomycetes(100);o__Saccharomycetales(100);f__Metschnikowiaceae(100);g__Metschnikowia(100);s__Metschnikowia_pulcherrima(100); | Metschnikowia | 13 | Saprotroph | \|Undefined Saprotroph\| | Yeast | NULL | Probable | As nectar-tap_saprotroph (Põlme et al. 2020); Decay sugar-rich substrates (Põlme et al. 2020) | Põlme S, et al. 2020. Fungal Diversity 105:1-16 ((https://doi.org/10.1007/s13225-020-00466-2)) |
| Otu26275 | 1 | k__Fungi(100);p__Ascomycota(100);c__Saccharomycetes(100);o__Saccharomycetales(100);f__Metschnikowiaceae(100);g__Metschnikowia(100);s__Metschnikowia_pulcherrima(100); | Metschnikowia | 13 | Saprotroph | \|Undefined Saprotroph\| | Yeast | NULL | Probable | As nectar-tap_saprotroph (Põlme et al. 2020); Decay sugar-rich substrates (Põlme et al. 2020) | Põlme S, et al. 2020. Fungal Diversity 105:1-16 ((https://doi.org/10.1007/s13225-020-00466-2)) |
| Otu26279 | 1 | k__Fungi(100);p__Ascomycota(100);c__Saccharomycetes(100);o__Saccharomycetales(100);f__Saccharomycetales_family_Incertae_sedis(100);g__Candida(100);s__Candida_stellata(100); | Candida | 13 | Pathotroph-Saprotroph-Symbiotroph | Animal Pathogen-Endophyte-Endosymbiont-Epiphyte-\|Undefined Saprotroph\| | Dimorphic Yeast-Yeast | NULL | Probable | As nectar-tap_saprotroph (Põlme et al. 2020); Decay sugar-rich substrates (Põlme et al. 2020); This asexual genus is polyphyletic and contain species that function very differently depend on what they are. Caution should be taken when interpreting guilds for Candida species. | Manolakaki D, et al. 2010. Virulence. 1:367-375 ((https://doi.org/10.4161/viru.1.5.12796)); Mekbib SB, et al. 2011. Tropical Plant Pathology 36:233-240 ((https://doi.org/10.1590/S1982-56762011000400004)); Põlme S, et al. 2020. Fungal Diversity 105:1-16 ((https://doi.org/10.1007/s13225-020-00466-2)) |
| Otu26281 | 1 | k__Fungi(100);p__Ascomycota(100);c__Saccharomycetes(100);o__Saccharomycetales(100);f__Metschnikowiaceae(100);g__Metschnikowia(100);s__Metschnikowia_pulcherrima(100); | Metschnikowia | 13 | Saprotroph | \|Undefined Saprotroph\| | Yeast | NULL | Probable | As nectar-tap_saprotroph (Põlme et al. 2020); Decay sugar-rich substrates (Põlme et al. 2020) | Põlme S, et al. 2020. Fungal Diversity 105:1-16 ((https://doi.org/10.1007/s13225-020-00466-2)) |
| Otu26282 | 1 | k__Fungi(100);p__Ascomycota(100);c__Saccharomycetes(100);o__Saccharomycetales(100);f__Metschnikowiaceae(100);g__Metschnikowia(100);s__Metschnikowia_pulcherrima(100); | Metschnikowia | 13 | Saprotroph | \|Undefined Saprotroph\| | Yeast | NULL | Probable | As nectar-tap_saprotroph (Põlme et al. 2020); Decay sugar-rich substrates (Põlme et al. 2020) | Põlme S, et al. 2020. Fungal Diversity 105:1-16 ((https://doi.org/10.1007/s13225-020-00466-2)) |
| Otu26283 | 1 | k__Fungi(100);p__Ascomycota(100);c__Saccharomycetes(100);o__Saccharomycetales(100);f__Metschnikowiaceae(100);g__Metschnikowia(100);s__Metschnikowia_pulcherrima(100); | Metschnikowia | 13 | Saprotroph | \|Undefined Saprotroph\| | Yeast | NULL | Probable | As nectar-tap_saprotroph (Põlme et al. 2020); Decay sugar-rich substrates (Põlme et al. 2020) | Põlme S, et al. 2020. Fungal Diversity 105:1-16 ((https://doi.org/10.1007/s13225-020-00466-2)) |
| Otu26287 | 1 | k__Fungi(100);p__Ascomycota(100);c__Dothideomycetes(100);o__Capnodiales(100);f__Mycosphaerellaceae(100);g__Mycosphaerella(100);s__Mycosphaerella_nyssicola(100); | Mycosphaerella | 13 | Pathotroph-Saprotroph | \|Plant Pathogen\|-Plant Saprotroph | Microfungus | NULL | Probable | As litter_saprotroph (Põlme et al. 2020); Decay leaf, fruit, seed (Põlme et al. 2020) | Tedersoo L, et al. 2014. Science 346:e1256688 ((https://doi.org/10.1126/science.1256688)); Põlme S, et al. 2020. Fungal Diversity 105:1-16 ((https://doi.org/10.1007/s13225-020-00466-2)) |
| Otu26297 | 1 | k__Fungi(100);p__Ascomycota(100);c__Saccharomycetes(100);o__Saccharomycetales(100);f__Metschnikowiaceae(100);g__Metschnikowia(100);s__Metschnikowia_pulcherrima(100); | Metschnikowia | 13 | Saprotroph | \|Undefined Saprotroph\| | Yeast | NULL | Probable | As nectar-tap_saprotroph (Põlme et al. 2020); Decay sugar-rich substrates (Põlme et al. 2020) | Põlme S, et al. 2020. Fungal Diversity 105:1-16 ((https://doi.org/10.1007/s13225-020-00466-2)) |
| Otu26308 | 1 | k__Fungi(100);p__Ascomycota(100);c__Saccharomycetes(100);o__Saccharomycetales(100);f__Metschnikowiaceae(100);g__Metschnikowia(100);s__Metschnikowia_pulcherrima(100); | Metschnikowia | 13 | Saprotroph | \|Undefined Saprotroph\| | Yeast | NULL | Probable | As nectar-tap_saprotroph (Põlme et al. 2020); Decay sugar-rich substrates (Põlme et al. 2020) | Põlme S, et al. 2020. Fungal Diversity 105:1-16 ((https://doi.org/10.1007/s13225-020-00466-2)) |
| Otu26312 | 1 | k__Fungi(100);p__Ascomycota(100);c__Saccharomycetes(100);o__Saccharomycetales(100);f__Metschnikowiaceae(100);g__Metschnikowia(100);g__Metschnikowia_unclassified(100); | Metschnikowia | 13 | Saprotroph | \|Undefined Saprotroph\| | Yeast | NULL | Probable | As nectar-tap_saprotroph (Põlme et al. 2020); Decay sugar-rich substrates (Põlme et al. 2020) | Põlme S, et al. 2020. Fungal Diversity 105:1-16 ((https://doi.org/10.1007/s13225-020-00466-2)) |
| Otu26315 | 1 | k__Fungi(100);p__Ascomycota(100);c__Saccharomycetes(100);o__Saccharomycetales(100);f__Metschnikowiaceae(100);g__Metschnikowia(100);s__Metschnikowia_pulcherrima(100); | Metschnikowia | 13 | Saprotroph | \|Undefined Saprotroph\| | Yeast | NULL | Probable | As nectar-tap_saprotroph (Põlme et al. 2020); Decay sugar-rich substrates (Põlme et al. 2020) | Põlme S, et al. 2020. Fungal Diversity 105:1-16 ((https://doi.org/10.1007/s13225-020-00466-2)) |
| Otu26318 | 1 | k__Fungi(100);p__Ascomycota(100);c__Saccharomycetes(100);o__Saccharomycetales(100);f__Metschnikowiaceae(100);g__Metschnikowia(100);s__Metschnikowia_pulcherrima(100); | Metschnikowia | 13 | Saprotroph | \|Undefined Saprotroph\| | Yeast | NULL | Probable | As nectar-tap_saprotroph (Põlme et al. 2020); Decay sugar-rich substrates (Põlme et al. 2020) | Põlme S, et al. 2020. Fungal Diversity 105:1-16 ((https://doi.org/10.1007/s13225-020-00466-2)) |
| Otu26335 | 1 | k__Fungi(100);p__Ascomycota(100);c__Saccharomycetes(100);o__Saccharomycetales(100);f__Metschnikowiaceae(100);g__Metschnikowia(100);s__Metschnikowia_pulcherrima(100); | Metschnikowia | 13 | Saprotroph | \|Undefined Saprotroph\| | Yeast | NULL | Probable | As nectar-tap_saprotroph (Põlme et al. 2020); Decay sugar-rich substrates (Põlme et al. 2020) | Põlme S, et al. 2020. Fungal Diversity 105:1-16 ((https://doi.org/10.1007/s13225-020-00466-2)) |
| Otu26336 | 1 | k__Fungi(100);p__Ascomycota(100);c__Saccharomycetes(100);o__Saccharomycetales(100);f__Metschnikowiaceae(100);g__Metschnikowia(100);s__Metschnikowia_pulcherrima(100); | Metschnikowia | 13 | Saprotroph | \|Undefined Saprotroph\| | Yeast | NULL | Probable | As nectar-tap_saprotroph (Põlme et al. 2020); Decay sugar-rich substrates (Põlme et al. 2020) | Põlme S, et al. 2020. Fungal Diversity 105:1-16 ((https://doi.org/10.1007/s13225-020-00466-2)) |
| Otu26340 | 1 | k__Fungi(100);p__Ascomycota(100);c__Saccharomycetes(100);o__Saccharomycetales(100);f__Metschnikowiaceae(100);g__Metschnikowia(100);g__Metschnikowia_unclassified(100); | Metschnikowia | 13 | Saprotroph | \|Undefined Saprotroph\| | Yeast | NULL | Probable | As nectar-tap_saprotroph (Põlme et al. 2020); Decay sugar-rich substrates (Põlme et al. 2020) | Põlme S, et al. 2020. Fungal Diversity 105:1-16 ((https://doi.org/10.1007/s13225-020-00466-2)) |
| Otu26341 | 1 | k__Fungi(100);p__Ascomycota(100);c__Saccharomycetes(100);o__Saccharomycetales(100);f__Metschnikowiaceae(100);g__Metschnikowia(100);s__Metschnikowia_pulcherrima(100); | Metschnikowia | 13 | Saprotroph | \|Undefined Saprotroph\| | Yeast | NULL | Probable | As nectar-tap_saprotroph (Põlme et al. 2020); Decay sugar-rich substrates (Põlme et al. 2020) | Põlme S, et al. 2020. Fungal Diversity 105:1-16 ((https://doi.org/10.1007/s13225-020-00466-2)) |
| Otu26343 | 1 | k__Fungi(100);p__Ascomycota(100);c__Saccharomycetes(100);o__Saccharomycetales(100);f__Metschnikowiaceae(100);g__Metschnikowia(100);g__Metschnikowia_unclassified(100); | Metschnikowia | 13 | Saprotroph | \|Undefined Saprotroph\| | Yeast | NULL | Probable | As nectar-tap_saprotroph (Põlme et al. 2020); Decay sugar-rich substrates (Põlme et al. 2020) | Põlme S, et al. 2020. Fungal Diversity 105:1-16 ((https://doi.org/10.1007/s13225-020-00466-2)) |
| Otu26344 | 1 | k__Fungi(100);p__Ascomycota(100);c__Saccharomycetes(100);o__Saccharomycetales(100);f__Metschnikowiaceae(100);g__Metschnikowia(100);s__Metschnikowia_pulcherrima(100); | Metschnikowia | 13 | Saprotroph | \|Undefined Saprotroph\| | Yeast | NULL | Probable | As nectar-tap_saprotroph (Põlme et al. 2020); Decay sugar-rich substrates (Põlme et al. 2020) | Põlme S, et al. 2020. Fungal Diversity 105:1-16 ((https://doi.org/10.1007/s13225-020-00466-2)) |
| Otu26348 | 1 | k__Fungi(100);p__Ascomycota(100);c__Saccharomycetes(100);o__Saccharomycetales(100);f__Metschnikowiaceae(100);g__Metschnikowia(100);s__Metschnikowia_pulcherrima(100); | Metschnikowia | 13 | Saprotroph | \|Undefined Saprotroph\| | Yeast | NULL | Probable | As nectar-tap_saprotroph (Põlme et al. 2020); Decay sugar-rich substrates (Põlme et al. 2020) | Põlme S, et al. 2020. Fungal Diversity 105:1-16 ((https://doi.org/10.1007/s13225-020-00466-2)) |
| Otu26355 | 1 | k__Fungi(100);p__Ascomycota(100);c__Saccharomycetes(100);o__Saccharomycetales(100);f__Metschnikowiaceae(100);g__Metschnikowia(100);s__Metschnikowia_pulcherrima(100); | Metschnikowia | 13 | Saprotroph | \|Undefined Saprotroph\| | Yeast | NULL | Probable | As nectar-tap_saprotroph (Põlme et al. 2020); Decay sugar-rich substrates (Põlme et al. 2020) | Põlme S, et al. 2020. Fungal Diversity 105:1-16 ((https://doi.org/10.1007/s13225-020-00466-2)) |
| Otu26357 | 1 | k__Fungi(100);p__Ascomycota(100);c__Saccharomycetes(100);o__Saccharomycetales(100);f__Metschnikowiaceae(100);g__Metschnikowia(100);s__Metschnikowia_pulcherrima(100); | Metschnikowia | 13 | Saprotroph | \|Undefined Saprotroph\| | Yeast | NULL | Probable | As nectar-tap_saprotroph (Põlme et al. 2020); Decay sugar-rich substrates (Põlme et al. 2020) | Põlme S, et al. 2020. Fungal Diversity 105:1-16 ((https://doi.org/10.1007/s13225-020-00466-2)) |
| Otu26358 | 1 | k__Fungi(100);p__Ascomycota(100);c__Saccharomycetes(100);o__Saccharomycetales(100);f__Metschnikowiaceae(100);g__Metschnikowia(100);g__Metschnikowia_unclassified(100); | Metschnikowia | 13 | Saprotroph | \|Undefined Saprotroph\| | Yeast | NULL | Probable | As nectar-tap_saprotroph (Põlme et al. 2020); Decay sugar-rich substrates (Põlme et al. 2020) | Põlme S, et al. 2020. Fungal Diversity 105:1-16 ((https://doi.org/10.1007/s13225-020-00466-2)) |
| Otu26371 | 1 | k__Fungi(100);p__Ascomycota(100);c__Saccharomycetes(100);o__Saccharomycetales(100);f__Metschnikowiaceae(100);g__Metschnikowia(100);s__Metschnikowia_pulcherrima(100); | Metschnikowia | 13 | Saprotroph | \|Undefined Saprotroph\| | Yeast | NULL | Probable | As nectar-tap_saprotroph (Põlme et al. 2020); Decay sugar-rich substrates (Põlme et al. 2020) | Põlme S, et al. 2020. Fungal Diversity 105:1-16 ((https://doi.org/10.1007/s13225-020-00466-2)) |
| Otu26372 | 1 | k__Fungi(100);p__Ascomycota(100);c__Saccharomycetes(100);o__Saccharomycetales(100);f__Metschnikowiaceae(100);g__Metschnikowia(100);s__Metschnikowia_pulcherrima(100); | Metschnikowia | 13 | Saprotroph | \|Undefined Saprotroph\| | Yeast | NULL | Probable | As nectar-tap_saprotroph (Põlme et al. 2020); Decay sugar-rich substrates (Põlme et al. 2020) | Põlme S, et al. 2020. Fungal Diversity 105:1-16 ((https://doi.org/10.1007/s13225-020-00466-2)) |
| Otu26384 | 1 | k__Fungi(100);p__Ascomycota(100);c__Saccharomycetes(100);o__Saccharomycetales(100);f__Metschnikowiaceae(100);g__Metschnikowia(100);g__Metschnikowia_unclassified(100); | Metschnikowia | 13 | Saprotroph | \|Undefined Saprotroph\| | Yeast | NULL | Probable | As nectar-tap_saprotroph (Põlme et al. 2020); Decay sugar-rich substrates (Põlme et al. 2020) | Põlme S, et al. 2020. Fungal Diversity 105:1-16 ((https://doi.org/10.1007/s13225-020-00466-2)) |
| Otu26385 | 1 | k__Fungi(100);p__Ascomycota(100);c__Saccharomycetes(100);o__Saccharomycetales(100);f__Metschnikowiaceae(100);g__Metschnikowia(100);s__Metschnikowia_pulcherrima(100); | Metschnikowia | 13 | Saprotroph | \|Undefined Saprotroph\| | Yeast | NULL | Probable | As nectar-tap_saprotroph (Põlme et al. 2020); Decay sugar-rich substrates (Põlme et al. 2020) | Põlme S, et al. 2020. Fungal Diversity 105:1-16 ((https://doi.org/10.1007/s13225-020-00466-2)) |
| Otu26392 | 1 | k__Fungi(100);p__Ascomycota(100);c__Saccharomycetes(100);o__Saccharomycetales(100);f__Metschnikowiaceae(100);g__Metschnikowia(100);s__Metschnikowia_pulcherrima(100); | Metschnikowia | 13 | Saprotroph | \|Undefined Saprotroph\| | Yeast | NULL | Probable | As nectar-tap_saprotroph (Põlme et al. 2020); Decay sugar-rich substrates (Põlme et al. 2020) | Põlme S, et al. 2020. Fungal Diversity 105:1-16 ((https://doi.org/10.1007/s13225-020-00466-2)) |
| Otu26393 | 1 | k__Fungi(100);p__Ascomycota(100);c__Saccharomycetes(100);o__Saccharomycetales(100);f__Metschnikowiaceae(100);g__Metschnikowia(100);s__Metschnikowia_pulcherrima(100); | Metschnikowia | 13 | Saprotroph | \|Undefined Saprotroph\| | Yeast | NULL | Probable | As nectar-tap_saprotroph (Põlme et al. 2020); Decay sugar-rich substrates (Põlme et al. 2020) | Põlme S, et al. 2020. Fungal Diversity 105:1-16 ((https://doi.org/10.1007/s13225-020-00466-2)) |
| Otu26394 | 1 | k__Fungi(100);p__Ascomycota(100);c__Saccharomycetes(100);o__Saccharomycetales(100);o__Saccharomycetales_unclassified(100);o__Saccharomycetales_unclassified(100);o__Saccharomycetales_unclassified(100); | Saccharomycetales | 7 | Saprotroph | Undefined Saprotroph | Yeast | NULL | Possible | NULL | Sterkenburg E, et al. 2015. New Phytologist 207:1145-1158 ((https://doi.org/10.1111/nph.13426)) |
| Otu26395 | 1 | k__Fungi(100);p__Ascomycota(100);c__Saccharomycetes(100);o__Saccharomycetales(100);f__Metschnikowiaceae(100);g__Metschnikowia(100);s__Metschnikowia_pulcherrima(100); | Metschnikowia | 13 | Saprotroph | \|Undefined Saprotroph\| | Yeast | NULL | Probable | As nectar-tap_saprotroph (Põlme et al. 2020); Decay sugar-rich substrates (Põlme et al. 2020) | Põlme S, et al. 2020. Fungal Diversity 105:1-16 ((https://doi.org/10.1007/s13225-020-00466-2)) |
| Otu26397 | 1 | k__Fungi(100);p__Ascomycota(100);c__Saccharomycetes(100);o__Saccharomycetales(100);f__Metschnikowiaceae(100);g__Metschnikowia(100);s__Metschnikowia_pulcherrima(100); | Metschnikowia | 13 | Saprotroph | \|Undefined Saprotroph\| | Yeast | NULL | Probable | As nectar-tap_saprotroph (Põlme et al. 2020); Decay sugar-rich substrates (Põlme et al. 2020) | Põlme S, et al. 2020. Fungal Diversity 105:1-16 ((https://doi.org/10.1007/s13225-020-00466-2)) |
| Otu26401 | 1 | k__Fungi(100);p__Ascomycota(100);c__Saccharomycetes(100);o__Saccharomycetales(100);f__Metschnikowiaceae(100);g__Metschnikowia(100);g__Metschnikowia_unclassified(100); | Metschnikowia | 13 | Saprotroph | \|Undefined Saprotroph\| | Yeast | NULL | Probable | As nectar-tap_saprotroph (Põlme et al. 2020); Decay sugar-rich substrates (Põlme et al. 2020) | Põlme S, et al. 2020. Fungal Diversity 105:1-16 ((https://doi.org/10.1007/s13225-020-00466-2)) |
| Otu26402 | 1 | k__Fungi(100);p__Ascomycota(100);c__Saccharomycetes(100);o__Saccharomycetales(100);f__Metschnikowiaceae(100);g__Metschnikowia(100);s__Metschnikowia_pulcherrima(100); | Metschnikowia | 13 | Saprotroph | \|Undefined Saprotroph\| | Yeast | NULL | Probable | As nectar-tap_saprotroph (Põlme et al. 2020); Decay sugar-rich substrates (Põlme et al. 2020) | Põlme S, et al. 2020. Fungal Diversity 105:1-16 ((https://doi.org/10.1007/s13225-020-00466-2)) |
| Otu26404 | 1 | k__Fungi(100);p__Ascomycota(100);c__Saccharomycetes(100);o__Saccharomycetales(100);f__Metschnikowiaceae(100);g__Metschnikowia(100);s__Metschnikowia_pulcherrima(100); | Metschnikowia | 13 | Saprotroph | \|Undefined Saprotroph\| | Yeast | NULL | Probable | As nectar-tap_saprotroph (Põlme et al. 2020); Decay sugar-rich substrates (Põlme et al. 2020) | Põlme S, et al. 2020. Fungal Diversity 105:1-16 ((https://doi.org/10.1007/s13225-020-00466-2)) |
| Otu26405 | 1 | k__Fungi(100);p__Ascomycota(100);c__Saccharomycetes(100);o__Saccharomycetales(100);f__Metschnikowiaceae(100);g__Metschnikowia(100);g__Metschnikowia_unclassified(100); | Metschnikowia | 13 | Saprotroph | \|Undefined Saprotroph\| | Yeast | NULL | Probable | As nectar-tap_saprotroph (Põlme et al. 2020); Decay sugar-rich substrates (Põlme et al. 2020) | Põlme S, et al. 2020. Fungal Diversity 105:1-16 ((https://doi.org/10.1007/s13225-020-00466-2)) |
| Otu26407 | 1 | k__Fungi(100);p__Ascomycota(100);c__Saccharomycetes(100);o__Saccharomycetales(100);f__Metschnikowiaceae(100);g__Metschnikowia(100);g__Metschnikowia_unclassified(100); | Metschnikowia | 13 | Saprotroph | \|Undefined Saprotroph\| | Yeast | NULL | Probable | As nectar-tap_saprotroph (Põlme et al. 2020); Decay sugar-rich substrates (Põlme et al. 2020) | Põlme S, et al. 2020. Fungal Diversity 105:1-16 ((https://doi.org/10.1007/s13225-020-00466-2)) |
| Otu26417 | 1 | k__Fungi(100);p__Ascomycota(100);c__Saccharomycetes(100);o__Saccharomycetales(100);f__Metschnikowiaceae(100);g__Metschnikowia(100);s__Metschnikowia_pulcherrima(100); | Metschnikowia | 13 | Saprotroph | \|Undefined Saprotroph\| | Yeast | NULL | Probable | As nectar-tap_saprotroph (Põlme et al. 2020); Decay sugar-rich substrates (Põlme et al. 2020) | Põlme S, et al. 2020. Fungal Diversity 105:1-16 ((https://doi.org/10.1007/s13225-020-00466-2)) |
| Otu26418 | 1 | k__Fungi(100);p__Ascomycota(100);c__Saccharomycetes(100);o__Saccharomycetales(100);f__Metschnikowiaceae(100);g__Metschnikowia(100);g__Metschnikowia_unclassified(100); | Metschnikowia | 13 | Saprotroph | \|Undefined Saprotroph\| | Yeast | NULL | Probable | As nectar-tap_saprotroph (Põlme et al. 2020); Decay sugar-rich substrates (Põlme et al. 2020) | Põlme S, et al. 2020. Fungal Diversity 105:1-16 ((https://doi.org/10.1007/s13225-020-00466-2)) |
| Otu26419 | 1 | k__Fungi(100);p__Ascomycota(100);c__Saccharomycetes(100);o__Saccharomycetales(100);f__Metschnikowiaceae(100);g__Metschnikowia(100);s__Metschnikowia_pulcherrima(100); | Metschnikowia | 13 | Saprotroph | \|Undefined Saprotroph\| | Yeast | NULL | Probable | As nectar-tap_saprotroph (Põlme et al. 2020); Decay sugar-rich substrates (Põlme et al. 2020) | Põlme S, et al. 2020. Fungal Diversity 105:1-16 ((https://doi.org/10.1007/s13225-020-00466-2)) |
| Otu26421 | 1 | k__Fungi(100);p__Ascomycota(100);c__Saccharomycetes(100);o__Saccharomycetales(100);f__Metschnikowiaceae(100);g__Metschnikowia(100);s__Metschnikowia_pulcherrima(100); | Metschnikowia | 13 | Saprotroph | \|Undefined Saprotroph\| | Yeast | NULL | Probable | As nectar-tap_saprotroph (Põlme et al. 2020); Decay sugar-rich substrates (Põlme et al. 2020) | Põlme S, et al. 2020. Fungal Diversity 105:1-16 ((https://doi.org/10.1007/s13225-020-00466-2)) |
| Otu26422 | 1 | k__Fungi(100);p__Ascomycota(100);c__Saccharomycetes(100);o__Saccharomycetales(100);f__Saccharomycetales_family_Incertae_sedis(100);g__Candida(100);g__Candida_unclassified(100); | Candida | 13 | Pathotroph-Saprotroph-Symbiotroph | Animal Pathogen-Endophyte-Endosymbiont-Epiphyte-\|Undefined Saprotroph\| | Dimorphic Yeast-Yeast | NULL | Probable | As nectar-tap_saprotroph (Põlme et al. 2020); Decay sugar-rich substrates (Põlme et al. 2020); This asexual genus is polyphyletic and contain species that function very differently depend on what they are. Caution should be taken when interpreting guilds for Candida species. | Manolakaki D, et al. 2010. Virulence. 1:367-375 ((https://doi.org/10.4161/viru.1.5.12796)); Mekbib SB, et al. 2011. Tropical Plant Pathology 36:233-240 ((https://doi.org/10.1590/S1982-56762011000400004)); Põlme S, et al. 2020. Fungal Diversity 105:1-16 ((https://doi.org/10.1007/s13225-020-00466-2)) |
| Otu26423 | 1 | k__Fungi(100);p__Ascomycota(100);c__Saccharomycetes(100);o__Saccharomycetales(100);f__Metschnikowiaceae(100);g__Metschnikowia(100);s__Metschnikowia_pulcherrima(100); | Metschnikowia | 13 | Saprotroph | \|Undefined Saprotroph\| | Yeast | NULL | Probable | As nectar-tap_saprotroph (Põlme et al. 2020); Decay sugar-rich substrates (Põlme et al. 2020) | Põlme S, et al. 2020. Fungal Diversity 105:1-16 ((https://doi.org/10.1007/s13225-020-00466-2)) |
| Otu26424 | 1 | k__Fungi(100);p__Ascomycota(100);c__Saccharomycetes(100);o__Saccharomycetales(100);f__Metschnikowiaceae(100);g__Metschnikowia(100);g__Metschnikowia_unclassified(100); | Metschnikowia | 13 | Saprotroph | \|Undefined Saprotroph\| | Yeast | NULL | Probable | As nectar-tap_saprotroph (Põlme et al. 2020); Decay sugar-rich substrates (Põlme et al. 2020) | Põlme S, et al. 2020. Fungal Diversity 105:1-16 ((https://doi.org/10.1007/s13225-020-00466-2)) |
| Otu26426 | 1 | k__Fungi(100);p__Ascomycota(100);c__Saccharomycetes(100);o__Saccharomycetales(100);f__Metschnikowiaceae(100);g__Metschnikowia(100);s__Metschnikowia_pulcherrima(100); | Metschnikowia | 13 | Saprotroph | \|Undefined Saprotroph\| | Yeast | NULL | Probable | As nectar-tap_saprotroph (Põlme et al. 2020); Decay sugar-rich substrates (Põlme et al. 2020) | Põlme S, et al. 2020. Fungal Diversity 105:1-16 ((https://doi.org/10.1007/s13225-020-00466-2)) |
| Otu26427 | 1 | k__Fungi(100);p__Ascomycota(100);c__Saccharomycetes(100);o__Saccharomycetales(100);f__Metschnikowiaceae(100);g__Metschnikowia(100);g__Metschnikowia_unclassified(100); | Metschnikowia | 13 | Saprotroph | \|Undefined Saprotroph\| | Yeast | NULL | Probable | As nectar-tap_saprotroph (Põlme et al. 2020); Decay sugar-rich substrates (Põlme et al. 2020) | Põlme S, et al. 2020. Fungal Diversity 105:1-16 ((https://doi.org/10.1007/s13225-020-00466-2)) |
| Otu26428 | 1 | k__Fungi(100);p__Ascomycota(100);c__Saccharomycetes(100);o__Saccharomycetales(100);f__Metschnikowiaceae(100);g__Metschnikowia(100);s__Metschnikowia_pulcherrima(100); | Metschnikowia | 13 | Saprotroph | \|Undefined Saprotroph\| | Yeast | NULL | Probable | As nectar-tap_saprotroph (Põlme et al. 2020); Decay sugar-rich substrates (Põlme et al. 2020) | Põlme S, et al. 2020. Fungal Diversity 105:1-16 ((https://doi.org/10.1007/s13225-020-00466-2)) |
| Otu26429 | 1 | k__Fungi(100);p__Ascomycota(100);c__Saccharomycetes(100);o__Saccharomycetales(100);f__Metschnikowiaceae(100);g__Metschnikowia(100);s__Metschnikowia_pulcherrima(100); | Metschnikowia | 13 | Saprotroph | \|Undefined Saprotroph\| | Yeast | NULL | Probable | As nectar-tap_saprotroph (Põlme et al. 2020); Decay sugar-rich substrates (Põlme et al. 2020) | Põlme S, et al. 2020. Fungal Diversity 105:1-16 ((https://doi.org/10.1007/s13225-020-00466-2)) |
| Otu26430 | 1 | k__Fungi(100);p__Ascomycota(100);c__Saccharomycetes(100);o__Saccharomycetales(100);f__Metschnikowiaceae(100);g__Metschnikowia(100);s__Metschnikowia_pulcherrima(100); | Metschnikowia | 13 | Saprotroph | \|Undefined Saprotroph\| | Yeast | NULL | Probable | As nectar-tap_saprotroph (Põlme et al. 2020); Decay sugar-rich substrates (Põlme et al. 2020) | Põlme S, et al. 2020. Fungal Diversity 105:1-16 ((https://doi.org/10.1007/s13225-020-00466-2)) |
| Otu26431 | 1 | k__Fungi(100);p__Ascomycota(100);c__Saccharomycetes(100);o__Saccharomycetales(100);f__Metschnikowiaceae(100);g__Metschnikowia(100);s__Metschnikowia_pulcherrima(100); | Metschnikowia | 13 | Saprotroph | \|Undefined Saprotroph\| | Yeast | NULL | Probable | As nectar-tap_saprotroph (Põlme et al. 2020); Decay sugar-rich substrates (Põlme et al. 2020) | Põlme S, et al. 2020. Fungal Diversity 105:1-16 ((https://doi.org/10.1007/s13225-020-00466-2)) |
| Otu26432 | 1 | k__Fungi(100);p__Ascomycota(100);c__Saccharomycetes(100);o__Saccharomycetales(100);f__Metschnikowiaceae(100);g__Metschnikowia(100);s__Metschnikowia_pulcherrima(100); | Metschnikowia | 13 | Saprotroph | \|Undefined Saprotroph\| | Yeast | NULL | Probable | As nectar-tap_saprotroph (Põlme et al. 2020); Decay sugar-rich substrates (Põlme et al. 2020) | Põlme S, et al. 2020. Fungal Diversity 105:1-16 ((https://doi.org/10.1007/s13225-020-00466-2)) |
| Otu26433 | 1 | k__Fungi(100);p__Ascomycota(100);c__Saccharomycetes(100);o__Saccharomycetales(100);f__Metschnikowiaceae(100);g__Metschnikowia(100);s__Metschnikowia_pulcherrima(100); | Metschnikowia | 13 | Saprotroph | \|Undefined Saprotroph\| | Yeast | NULL | Probable | As nectar-tap_saprotroph (Põlme et al. 2020); Decay sugar-rich substrates (Põlme et al. 2020) | Põlme S, et al. 2020. Fungal Diversity 105:1-16 ((https://doi.org/10.1007/s13225-020-00466-2)) |
| Otu26434 | 1 | k__Fungi(100);p__Ascomycota(100);c__Saccharomycetes(100);o__Saccharomycetales(100);f__Metschnikowiaceae(100);g__Metschnikowia(100);s__Metschnikowia_pulcherrima(100); | Metschnikowia | 13 | Saprotroph | \|Undefined Saprotroph\| | Yeast | NULL | Probable | As nectar-tap_saprotroph (Põlme et al. 2020); Decay sugar-rich substrates (Põlme et al. 2020) | Põlme S, et al. 2020. Fungal Diversity 105:1-16 ((https://doi.org/10.1007/s13225-020-00466-2)) |
| Otu26435 | 1 | k__Fungi(100);p__Ascomycota(100);c__Saccharomycetes(100);o__Saccharomycetales(100);f__Metschnikowiaceae(100);g__Metschnikowia(100);g__Metschnikowia_unclassified(100); | Metschnikowia | 13 | Saprotroph | \|Undefined Saprotroph\| | Yeast | NULL | Probable | As nectar-tap_saprotroph (Põlme et al. 2020); Decay sugar-rich substrates (Põlme et al. 2020) | Põlme S, et al. 2020. Fungal Diversity 105:1-16 ((https://doi.org/10.1007/s13225-020-00466-2)) |
| Otu26436 | 1 | k__Fungi(100);p__Ascomycota(100);c__Saccharomycetes(100);o__Saccharomycetales(100);f__Metschnikowiaceae(100);g__Metschnikowia(100);s__Metschnikowia_pulcherrima(100); | Metschnikowia | 13 | Saprotroph | \|Undefined Saprotroph\| | Yeast | NULL | Probable | As nectar-tap_saprotroph (Põlme et al. 2020); Decay sugar-rich substrates (Põlme et al. 2020) | Põlme S, et al. 2020. Fungal Diversity 105:1-16 ((https://doi.org/10.1007/s13225-020-00466-2)) |
| Otu26437 | 1 | k__Fungi(100);p__Ascomycota(100);c__Saccharomycetes(100);o__Saccharomycetales(100);f__Metschnikowiaceae(100);g__Metschnikowia(100);g__Metschnikowia_unclassified(100); | Metschnikowia | 13 | Saprotroph | \|Undefined Saprotroph\| | Yeast | NULL | Probable | As nectar-tap_saprotroph (Põlme et al. 2020); Decay sugar-rich substrates (Põlme et al. 2020) | Põlme S, et al. 2020. Fungal Diversity 105:1-16 ((https://doi.org/10.1007/s13225-020-00466-2)) |
| Otu26438 | 1 | k__Fungi(100);p__Ascomycota(100);c__Saccharomycetes(100);o__Saccharomycetales(100);f__Metschnikowiaceae(100);g__Metschnikowia(100);s__Metschnikowia_pulcherrima(100); | Metschnikowia | 13 | Saprotroph | \|Undefined Saprotroph\| | Yeast | NULL | Probable | As nectar-tap_saprotroph (Põlme et al. 2020); Decay sugar-rich substrates (Põlme et al. 2020) | Põlme S, et al. 2020. Fungal Diversity 105:1-16 ((https://doi.org/10.1007/s13225-020-00466-2)) |
| Otu26439 | 1 | k__Fungi(100);p__Ascomycota(100);c__Saccharomycetes(100);o__Saccharomycetales(100);f__Metschnikowiaceae(100);g__Metschnikowia(100);g__Metschnikowia_unclassified(100); | Metschnikowia | 13 | Saprotroph | \|Undefined Saprotroph\| | Yeast | NULL | Probable | As nectar-tap_saprotroph (Põlme et al. 2020); Decay sugar-rich substrates (Põlme et al. 2020) | Põlme S, et al. 2020. Fungal Diversity 105:1-16 ((https://doi.org/10.1007/s13225-020-00466-2)) |
| Otu26440 | 1 | k__Fungi(100);p__Ascomycota(100);c__Saccharomycetes(100);o__Saccharomycetales(100);f__Metschnikowiaceae(100);g__Metschnikowia(100);s__Metschnikowia_pulcherrima(100); | Metschnikowia | 13 | Saprotroph | \|Undefined Saprotroph\| | Yeast | NULL | Probable | As nectar-tap_saprotroph (Põlme et al. 2020); Decay sugar-rich substrates (Põlme et al. 2020) | Põlme S, et al. 2020. Fungal Diversity 105:1-16 ((https://doi.org/10.1007/s13225-020-00466-2)) |
| Otu26441 | 1 | k__Fungi(100);p__Ascomycota(100);c__Saccharomycetes(100);o__Saccharomycetales(100);f__Metschnikowiaceae(100);g__Metschnikowia(100);s__Metschnikowia_pulcherrima(100); | Metschnikowia | 13 | Saprotroph | \|Undefined Saprotroph\| | Yeast | NULL | Probable | As nectar-tap_saprotroph (Põlme et al. 2020); Decay sugar-rich substrates (Põlme et al. 2020) | Põlme S, et al. 2020. Fungal Diversity 105:1-16 ((https://doi.org/10.1007/s13225-020-00466-2)) |
| Otu26442 | 1 | k__Fungi(100);p__Ascomycota(100);c__Saccharomycetes(100);o__Saccharomycetales(100);f__Metschnikowiaceae(100);g__Metschnikowia(100);s__Metschnikowia_pulcherrima(100); | Metschnikowia | 13 | Saprotroph | \|Undefined Saprotroph\| | Yeast | NULL | Probable | As nectar-tap_saprotroph (Põlme et al. 2020); Decay sugar-rich substrates (Põlme et al. 2020) | Põlme S, et al. 2020. Fungal Diversity 105:1-16 ((https://doi.org/10.1007/s13225-020-00466-2)) |
| Otu26443 | 1 | k__Fungi(100);p__Ascomycota(100);c__Saccharomycetes(100);o__Saccharomycetales(100);f__Metschnikowiaceae(100);g__Metschnikowia(100);s__Metschnikowia_pulcherrima(100); | Metschnikowia | 13 | Saprotroph | \|Undefined Saprotroph\| | Yeast | NULL | Probable | As nectar-tap_saprotroph (Põlme et al. 2020); Decay sugar-rich substrates (Põlme et al. 2020) | Põlme S, et al. 2020. Fungal Diversity 105:1-16 ((https://doi.org/10.1007/s13225-020-00466-2)) |
| Otu26444 | 1 | k__Fungi(100);p__Ascomycota(100);c__Saccharomycetes(100);o__Saccharomycetales(100);f__Metschnikowiaceae(100);g__Metschnikowia(100);s__Metschnikowia_pulcherrima(100); | Metschnikowia | 13 | Saprotroph | \|Undefined Saprotroph\| | Yeast | NULL | Probable | As nectar-tap_saprotroph (Põlme et al. 2020); Decay sugar-rich substrates (Põlme et al. 2020) | Põlme S, et al. 2020. Fungal Diversity 105:1-16 ((https://doi.org/10.1007/s13225-020-00466-2)) |
| Otu26445 | 1 | k__Fungi(100);p__Ascomycota(100);c__Saccharomycetes(100);o__Saccharomycetales(100);f__Metschnikowiaceae(100);g__Metschnikowia(100);s__Metschnikowia_pulcherrima(100); | Metschnikowia | 13 | Saprotroph | \|Undefined Saprotroph\| | Yeast | NULL | Probable | As nectar-tap_saprotroph (Põlme et al. 2020); Decay sugar-rich substrates (Põlme et al. 2020) | Põlme S, et al. 2020. Fungal Diversity 105:1-16 ((https://doi.org/10.1007/s13225-020-00466-2)) |
| Otu26446 | 1 | k__Fungi(100);p__Ascomycota(100);c__Saccharomycetes(100);o__Saccharomycetales(100);f__Metschnikowiaceae(100);g__Metschnikowia(100);s__Metschnikowia_pulcherrima(100); | Metschnikowia | 13 | Saprotroph | \|Undefined Saprotroph\| | Yeast | NULL | Probable | As nectar-tap_saprotroph (Põlme et al. 2020); Decay sugar-rich substrates (Põlme et al. 2020) | Põlme S, et al. 2020. Fungal Diversity 105:1-16 ((https://doi.org/10.1007/s13225-020-00466-2)) |
| Otu26447 | 1 | k__Fungi(100);p__Ascomycota(100);c__Saccharomycetes(100);o__Saccharomycetales(100);f__Metschnikowiaceae(100);g__Metschnikowia(100);s__Metschnikowia_pulcherrima(100); | Metschnikowia | 13 | Saprotroph | \|Undefined Saprotroph\| | Yeast | NULL | Probable | As nectar-tap_saprotroph (Põlme et al. 2020); Decay sugar-rich substrates (Põlme et al. 2020) | Põlme S, et al. 2020. Fungal Diversity 105:1-16 ((https://doi.org/10.1007/s13225-020-00466-2)) |
| Otu26448 | 1 | k__Fungi(100);p__Ascomycota(100);c__Saccharomycetes(100);o__Saccharomycetales(100);f__Metschnikowiaceae(100);g__Metschnikowia(100);s__Metschnikowia_pulcherrima(100); | Metschnikowia | 13 | Saprotroph | \|Undefined Saprotroph\| | Yeast | NULL | Probable | As nectar-tap_saprotroph (Põlme et al. 2020); Decay sugar-rich substrates (Põlme et al. 2020) | Põlme S, et al. 2020. Fungal Diversity 105:1-16 ((https://doi.org/10.1007/s13225-020-00466-2)) |
| Otu26450 | 1 | k__Fungi(100);p__Ascomycota(100);c__Saccharomycetes(100);o__Saccharomycetales(100);f__Metschnikowiaceae(100);g__Metschnikowia(100);g__Metschnikowia_unclassified(100); | Metschnikowia | 13 | Saprotroph | \|Undefined Saprotroph\| | Yeast | NULL | Probable | As nectar-tap_saprotroph (Põlme et al. 2020); Decay sugar-rich substrates (Põlme et al. 2020) | Põlme S, et al. 2020. Fungal Diversity 105:1-16 ((https://doi.org/10.1007/s13225-020-00466-2)) |
| Otu26451 | 1 | k__Fungi(100);p__Ascomycota(100);c__Saccharomycetes(100);o__Saccharomycetales(100);f__Metschnikowiaceae(100);g__Metschnikowia(100);s__Metschnikowia_pulcherrima(100); | Metschnikowia | 13 | Saprotroph | \|Undefined Saprotroph\| | Yeast | NULL | Probable | As nectar-tap_saprotroph (Põlme et al. 2020); Decay sugar-rich substrates (Põlme et al. 2020) | Põlme S, et al. 2020. Fungal Diversity 105:1-16 ((https://doi.org/10.1007/s13225-020-00466-2)) |
| Otu26452 | 1 | k__Fungi(100);p__Ascomycota(100);c__Saccharomycetes(100);o__Saccharomycetales(100);f__Metschnikowiaceae(100);g__Metschnikowia(100);s__Metschnikowia_pulcherrima(100); | Metschnikowia | 13 | Saprotroph | \|Undefined Saprotroph\| | Yeast | NULL | Probable | As nectar-tap_saprotroph (Põlme et al. 2020); Decay sugar-rich substrates (Põlme et al. 2020) | Põlme S, et al. 2020. Fungal Diversity 105:1-16 ((https://doi.org/10.1007/s13225-020-00466-2)) |
| Otu26454 | 1 | k__Fungi(100);p__Ascomycota(100);c__Saccharomycetes(100);o__Saccharomycetales(100);f__Metschnikowiaceae(100);g__Metschnikowia(100);s__Metschnikowia_pulcherrima(100); | Metschnikowia | 13 | Saprotroph | \|Undefined Saprotroph\| | Yeast | NULL | Probable | As nectar-tap_saprotroph (Põlme et al. 2020); Decay sugar-rich substrates (Põlme et al. 2020) | Põlme S, et al. 2020. Fungal Diversity 105:1-16 ((https://doi.org/10.1007/s13225-020-00466-2)) |
| Otu26455 | 1 | k__Fungi(100);p__Ascomycota(100);c__Saccharomycetes(100);o__Saccharomycetales(100);f__Metschnikowiaceae(100);g__Metschnikowia(100);s__Metschnikowia_pulcherrima(100); | Metschnikowia | 13 | Saprotroph | \|Undefined Saprotroph\| | Yeast | NULL | Probable | As nectar-tap_saprotroph (Põlme et al. 2020); Decay sugar-rich substrates (Põlme et al. 2020) | Põlme S, et al. 2020. Fungal Diversity 105:1-16 ((https://doi.org/10.1007/s13225-020-00466-2)) |
| Otu26456 | 1 | k__Fungi(100);p__Ascomycota(100);c__Saccharomycetes(100);o__Saccharomycetales(100);f__Metschnikowiaceae(100);g__Metschnikowia(100);s__Metschnikowia_pulcherrima(100); | Metschnikowia | 13 | Saprotroph | \|Undefined Saprotroph\| | Yeast | NULL | Probable | As nectar-tap_saprotroph (Põlme et al. 2020); Decay sugar-rich substrates (Põlme et al. 2020) | Põlme S, et al. 2020. Fungal Diversity 105:1-16 ((https://doi.org/10.1007/s13225-020-00466-2)) |
| Otu26457 | 1 | k__Fungi(100);p__Ascomycota(100);c__Saccharomycetes(100);o__Saccharomycetales(100);f__Metschnikowiaceae(100);g__Metschnikowia(100);g__Metschnikowia_unclassified(100); | Metschnikowia | 13 | Saprotroph | \|Undefined Saprotroph\| | Yeast | NULL | Probable | As nectar-tap_saprotroph (Põlme et al. 2020); Decay sugar-rich substrates (Põlme et al. 2020) | Põlme S, et al. 2020. Fungal Diversity 105:1-16 ((https://doi.org/10.1007/s13225-020-00466-2)) |
| Otu26459 | 1 | k__Fungi(100);p__Basidiomycota(100);c__Tremellomycetes(100);o__Tremellales(100);f__Tremellales_family_Incertae_sedis(100);g__Hannaella(100);g__Hannaella_unclassified(100); | Hannaella | 13 | Pathotroph-Saprotroph | \|Animal Parasite\|-Undefined Saprotroph | Yeast | NULL | Probable | As animal_decomposer (Põlme et al. 2020); Decay animal material (Põlme et al. 2020) | Põlme S, et al. 2020. Fungal Diversity 105:1-16 ((https://doi.org/10.1007/s13225-020-00466-2)) |
| Otu26460 | 1 | k__Fungi(100);p__Ascomycota(100);c__Saccharomycetes(100);o__Saccharomycetales(100);f__Metschnikowiaceae(100);g__Metschnikowia(100);g__Metschnikowia_unclassified(100); | Metschnikowia | 13 | Saprotroph | \|Undefined Saprotroph\| | Yeast | NULL | Probable | As nectar-tap_saprotroph (Põlme et al. 2020); Decay sugar-rich substrates (Põlme et al. 2020) | Põlme S, et al. 2020. Fungal Diversity 105:1-16 ((https://doi.org/10.1007/s13225-020-00466-2)) |
| Otu26461 | 1 | k__Fungi(100);p__Ascomycota(100);c__Saccharomycetes(100);o__Saccharomycetales(100);f__Metschnikowiaceae(100);g__Metschnikowia(100);s__Metschnikowia_pulcherrima(100); | Metschnikowia | 13 | Saprotroph | \|Undefined Saprotroph\| | Yeast | NULL | Probable | As nectar-tap_saprotroph (Põlme et al. 2020); Decay sugar-rich substrates (Põlme et al. 2020) | Põlme S, et al. 2020. Fungal Diversity 105:1-16 ((https://doi.org/10.1007/s13225-020-00466-2)) |
| Otu26463 | 1 | k__Fungi(100);p__Ascomycota(100);c__Saccharomycetes(100);o__Saccharomycetales(100);f__Metschnikowiaceae(100);g__Metschnikowia(100);s__Metschnikowia_pulcherrima(100); | Metschnikowia | 13 | Saprotroph | \|Undefined Saprotroph\| | Yeast | NULL | Probable | As nectar-tap_saprotroph (Põlme et al. 2020); Decay sugar-rich substrates (Põlme et al. 2020) | Põlme S, et al. 2020. Fungal Diversity 105:1-16 ((https://doi.org/10.1007/s13225-020-00466-2)) |
| Otu26464 | 1 | k__Fungi(100);p__Ascomycota(100);c__Saccharomycetes(100);o__Saccharomycetales(100);f__Metschnikowiaceae(100);g__Metschnikowia(100);s__Metschnikowia_pulcherrima(100); | Metschnikowia | 13 | Saprotroph | \|Undefined Saprotroph\| | Yeast | NULL | Probable | As nectar-tap_saprotroph (Põlme et al. 2020); Decay sugar-rich substrates (Põlme et al. 2020) | Põlme S, et al. 2020. Fungal Diversity 105:1-16 ((https://doi.org/10.1007/s13225-020-00466-2)) |
| Otu26465 | 1 | k__Fungi(100);p__Ascomycota(100);c__Saccharomycetes(100);o__Saccharomycetales(100);f__Metschnikowiaceae(100);g__Metschnikowia(100);s__Metschnikowia_pulcherrima(100); | Metschnikowia | 13 | Saprotroph | \|Undefined Saprotroph\| | Yeast | NULL | Probable | As nectar-tap_saprotroph (Põlme et al. 2020); Decay sugar-rich substrates (Põlme et al. 2020) | Põlme S, et al. 2020. Fungal Diversity 105:1-16 ((https://doi.org/10.1007/s13225-020-00466-2)) |
| Otu26466 | 1 | k__Fungi(100);p__Ascomycota(100);c__Saccharomycetes(100);o__Saccharomycetales(100);f__Metschnikowiaceae(100);g__Metschnikowia(100);s__Metschnikowia_pulcherrima(100); | Metschnikowia | 13 | Saprotroph | \|Undefined Saprotroph\| | Yeast | NULL | Probable | As nectar-tap_saprotroph (Põlme et al. 2020); Decay sugar-rich substrates (Põlme et al. 2020) | Põlme S, et al. 2020. Fungal Diversity 105:1-16 ((https://doi.org/10.1007/s13225-020-00466-2)) |
| Otu26467 | 1 | k__Fungi(100);p__Ascomycota(100);c__Saccharomycetes(100);o__Saccharomycetales(100);f__Metschnikowiaceae(100);g__Metschnikowia(100);s__Metschnikowia_pulcherrima(100); | Metschnikowia | 13 | Saprotroph | \|Undefined Saprotroph\| | Yeast | NULL | Probable | As nectar-tap_saprotroph (Põlme et al. 2020); Decay sugar-rich substrates (Põlme et al. 2020) | Põlme S, et al. 2020. Fungal Diversity 105:1-16 ((https://doi.org/10.1007/s13225-020-00466-2)) |
| Otu26496 | 1 | k__Fungi(100);p__Ascomycota(100);c__Saccharomycetes(100);o__Saccharomycetales(100);f__Metschnikowiaceae(100);g__Metschnikowia(100);s__Metschnikowia_pulcherrima(100); | Metschnikowia | 13 | Saprotroph | \|Undefined Saprotroph\| | Yeast | NULL | Probable | As nectar-tap_saprotroph (Põlme et al. 2020); Decay sugar-rich substrates (Põlme et al. 2020) | Põlme S, et al. 2020. Fungal Diversity 105:1-16 ((https://doi.org/10.1007/s13225-020-00466-2)) |
| Otu26498 | 1 | k__Fungi(100);p__Ascomycota(100);c__Saccharomycetes(100);o__Saccharomycetales(100);f__Metschnikowiaceae(100);g__Metschnikowia(100);s__Metschnikowia_pulcherrima(100); | Metschnikowia | 13 | Saprotroph | \|Undefined Saprotroph\| | Yeast | NULL | Probable | As nectar-tap_saprotroph (Põlme et al. 2020); Decay sugar-rich substrates (Põlme et al. 2020) | Põlme S, et al. 2020. Fungal Diversity 105:1-16 ((https://doi.org/10.1007/s13225-020-00466-2)) |
| Otu26499 | 1 | k__Fungi(100);p__Ascomycota(100);c__Saccharomycetes(100);o__Saccharomycetales(100);f__Metschnikowiaceae(100);g__Metschnikowia(100);g__Metschnikowia_unclassified(100); | Metschnikowia | 13 | Saprotroph | \|Undefined Saprotroph\| | Yeast | NULL | Probable | As nectar-tap_saprotroph (Põlme et al. 2020); Decay sugar-rich substrates (Põlme et al. 2020) | Põlme S, et al. 2020. Fungal Diversity 105:1-16 ((https://doi.org/10.1007/s13225-020-00466-2)) |
| Otu26500 | 1 | k__Fungi(100);p__Ascomycota(100);c__Saccharomycetes(100);o__Saccharomycetales(100);f__Metschnikowiaceae(100);g__Metschnikowia(100);g__Metschnikowia_unclassified(100); | Metschnikowia | 13 | Saprotroph | \|Undefined Saprotroph\| | Yeast | NULL | Probable | As nectar-tap_saprotroph (Põlme et al. 2020); Decay sugar-rich substrates (Põlme et al. 2020) | Põlme S, et al. 2020. Fungal Diversity 105:1-16 ((https://doi.org/10.1007/s13225-020-00466-2)) |
| Otu26502 | 1 | k__Fungi(100);p__Ascomycota(100);c__Saccharomycetes(100);o__Saccharomycetales(100);f__Metschnikowiaceae(100);g__Metschnikowia(100);g__Metschnikowia_unclassified(100); | Metschnikowia | 13 | Saprotroph | \|Undefined Saprotroph\| | Yeast | NULL | Probable | As nectar-tap_saprotroph (Põlme et al. 2020); Decay sugar-rich substrates (Põlme et al. 2020) | Põlme S, et al. 2020. Fungal Diversity 105:1-16 ((https://doi.org/10.1007/s13225-020-00466-2)) |
| Otu26503 | 1 | k__Fungi(100);p__Ascomycota(100);c__Saccharomycetes(100);o__Saccharomycetales(100);f__Metschnikowiaceae(100);g__Metschnikowia(100);s__Metschnikowia_pulcherrima(100); | Metschnikowia | 13 | Saprotroph | \|Undefined Saprotroph\| | Yeast | NULL | Probable | As nectar-tap_saprotroph (Põlme et al. 2020); Decay sugar-rich substrates (Põlme et al. 2020) | Põlme S, et al. 2020. Fungal Diversity 105:1-16 ((https://doi.org/10.1007/s13225-020-00466-2)) |
| Otu26504 | 1 | k__Fungi(100);p__Ascomycota(100);c__Saccharomycetes(100);o__Saccharomycetales(100);f__Metschnikowiaceae(100);g__Metschnikowia(100);s__Metschnikowia_pulcherrima(100); | Metschnikowia | 13 | Saprotroph | \|Undefined Saprotroph\| | Yeast | NULL | Probable | As nectar-tap_saprotroph (Põlme et al. 2020); Decay sugar-rich substrates (Põlme et al. 2020) | Põlme S, et al. 2020. Fungal Diversity 105:1-16 ((https://doi.org/10.1007/s13225-020-00466-2)) |
| Otu26505 | 1 | k__Fungi(100);p__Ascomycota(100);c__Saccharomycetes(100);o__Saccharomycetales(100);f__Metschnikowiaceae(100);g__Metschnikowia(100);s__Metschnikowia_pulcherrima(100); | Metschnikowia | 13 | Saprotroph | \|Undefined Saprotroph\| | Yeast | NULL | Probable | As nectar-tap_saprotroph (Põlme et al. 2020); Decay sugar-rich substrates (Põlme et al. 2020) | Põlme S, et al. 2020. Fungal Diversity 105:1-16 ((https://doi.org/10.1007/s13225-020-00466-2)) |
| Otu26506 | 1 | k__Fungi(100);p__Ascomycota(100);c__Saccharomycetes(100);o__Saccharomycetales(100);f__Metschnikowiaceae(100);g__Metschnikowia(100);s__Metschnikowia_pulcherrima(100); | Metschnikowia | 13 | Saprotroph | \|Undefined Saprotroph\| | Yeast | NULL | Probable | As nectar-tap_saprotroph (Põlme et al. 2020); Decay sugar-rich substrates (Põlme et al. 2020) | Põlme S, et al. 2020. Fungal Diversity 105:1-16 ((https://doi.org/10.1007/s13225-020-00466-2)) |
| Otu26507 | 1 | k__Fungi(100);p__Ascomycota(100);c__Saccharomycetes(100);o__Saccharomycetales(100);f__Metschnikowiaceae(100);g__Metschnikowia(100);s__Metschnikowia_pulcherrima(100); | Metschnikowia | 13 | Saprotroph | \|Undefined Saprotroph\| | Yeast | NULL | Probable | As nectar-tap_saprotroph (Põlme et al. 2020); Decay sugar-rich substrates (Põlme et al. 2020) | Põlme S, et al. 2020. Fungal Diversity 105:1-16 ((https://doi.org/10.1007/s13225-020-00466-2)) |
| Otu26508 | 1 | k__Fungi(100);p__Ascomycota(100);c__Sordariomycetes(100);o__Hypocreales(100);f__Nectriaceae(100);g__Gibberella(100);s__Gibberella_intricans(100); | Gibberella | 13 | Pathotroph-Saprotroph-Symbiotroph | Endophyte-\|Plant Pathogen\|-Plant Saprotroph-Undefined Saprotroph | Microfungus | Soft Rot | Probable | As foliar_endophyte (Põlme et al. 2020); As litter_saprotroph (Põlme et al. 2020); Causing damping-off disease (Põlme et al. 2020); Decay leaf, fruit, seed, roots, soil, animal material (Põlme et al. 2020) | Tedersoo L, et al. 2014. Science 346:e1256688 ((https://doi.org/10.1126/science.1256688)); Põlme S, et al. 2020. Fungal Diversity 105:1-16 ((https://doi.org/10.1007/s13225-020-00466-2)) |
| Otu26510 | 1 | k__Fungi(100);p__Ascomycota(100);c__Saccharomycetes(100);o__Saccharomycetales(100);f__Metschnikowiaceae(100);g__Metschnikowia(100);g__Metschnikowia_unclassified(100); | Metschnikowia | 13 | Saprotroph | \|Undefined Saprotroph\| | Yeast | NULL | Probable | As nectar-tap_saprotroph (Põlme et al. 2020); Decay sugar-rich substrates (Põlme et al. 2020) | Põlme S, et al. 2020. Fungal Diversity 105:1-16 ((https://doi.org/10.1007/s13225-020-00466-2)) |
| Otu26511 | 1 | k__Fungi(100);p__Ascomycota(100);c__Saccharomycetes(100);o__Saccharomycetales(100);f__Metschnikowiaceae(100);g__Metschnikowia(100);s__Metschnikowia_pulcherrima(100); | Metschnikowia | 13 | Saprotroph | \|Undefined Saprotroph\| | Yeast | NULL | Probable | As nectar-tap_saprotroph (Põlme et al. 2020); Decay sugar-rich substrates (Põlme et al. 2020) | Põlme S, et al. 2020. Fungal Diversity 105:1-16 ((https://doi.org/10.1007/s13225-020-00466-2)) |
| Otu26512 | 1 | k__Fungi(100);p__Ascomycota(100);c__Saccharomycetes(100);o__Saccharomycetales(100);f__Metschnikowiaceae(100);g__Metschnikowia(100);s__Metschnikowia_pulcherrima(100); | Metschnikowia | 13 | Saprotroph | \|Undefined Saprotroph\| | Yeast | NULL | Probable | As nectar-tap_saprotroph (Põlme et al. 2020); Decay sugar-rich substrates (Põlme et al. 2020) | Põlme S, et al. 2020. Fungal Diversity 105:1-16 ((https://doi.org/10.1007/s13225-020-00466-2)) |
| Otu26514 | 1 | k__Fungi(100);p__Ascomycota(100);c__Saccharomycetes(100);o__Saccharomycetales(100);f__Metschnikowiaceae(100);g__Metschnikowia(100);s__Metschnikowia_pulcherrima(100); | Metschnikowia | 13 | Saprotroph | \|Undefined Saprotroph\| | Yeast | NULL | Probable | As nectar-tap_saprotroph (Põlme et al. 2020); Decay sugar-rich substrates (Põlme et al. 2020) | Põlme S, et al. 2020. Fungal Diversity 105:1-16 ((https://doi.org/10.1007/s13225-020-00466-2)) |
| Otu26515 | 1 | k__Fungi(100);p__Ascomycota(100);c__Saccharomycetes(100);o__Saccharomycetales(100);f__Metschnikowiaceae(100);g__Metschnikowia(100);s__Metschnikowia_pulcherrima(100); | Metschnikowia | 13 | Saprotroph | \|Undefined Saprotroph\| | Yeast | NULL | Probable | As nectar-tap_saprotroph (Põlme et al. 2020); Decay sugar-rich substrates (Põlme et al. 2020) | Põlme S, et al. 2020. Fungal Diversity 105:1-16 ((https://doi.org/10.1007/s13225-020-00466-2)) |
| Otu26516 | 1 | k__Fungi(100);p__Ascomycota(100);c__Saccharomycetes(100);o__Saccharomycetales(100);f__Metschnikowiaceae(100);g__Metschnikowia(100);g__Metschnikowia_unclassified(100); | Metschnikowia | 13 | Saprotroph | \|Undefined Saprotroph\| | Yeast | NULL | Probable | As nectar-tap_saprotroph (Põlme et al. 2020); Decay sugar-rich substrates (Põlme et al. 2020) | Põlme S, et al. 2020. Fungal Diversity 105:1-16 ((https://doi.org/10.1007/s13225-020-00466-2)) |
| Otu26517 | 1 | k__Fungi(100);p__Ascomycota(100);c__Saccharomycetes(100);o__Saccharomycetales(100);f__Metschnikowiaceae(100);g__Metschnikowia(100);g__Metschnikowia_unclassified(100); | Metschnikowia | 13 | Saprotroph | \|Undefined Saprotroph\| | Yeast | NULL | Probable | As nectar-tap_saprotroph (Põlme et al. 2020); Decay sugar-rich substrates (Põlme et al. 2020) | Põlme S, et al. 2020. Fungal Diversity 105:1-16 ((https://doi.org/10.1007/s13225-020-00466-2)) |
| Otu26518 | 1 | k__Fungi(100);p__Ascomycota(100);c__Saccharomycetes(100);o__Saccharomycetales(100);f__Metschnikowiaceae(100);g__Metschnikowia(100);s__Metschnikowia_pulcherrima(100); | Metschnikowia | 13 | Saprotroph | \|Undefined Saprotroph\| | Yeast | NULL | Probable | As nectar-tap_saprotroph (Põlme et al. 2020); Decay sugar-rich substrates (Põlme et al. 2020) | Põlme S, et al. 2020. Fungal Diversity 105:1-16 ((https://doi.org/10.1007/s13225-020-00466-2)) |
| Otu26520 | 1 | k__Fungi(100);p__Ascomycota(100);c__Saccharomycetes(100);o__Saccharomycetales(100);f__Metschnikowiaceae(100);g__Metschnikowia(100);s__Metschnikowia_pulcherrima(100); | Metschnikowia | 13 | Saprotroph | \|Undefined Saprotroph\| | Yeast | NULL | Probable | As nectar-tap_saprotroph (Põlme et al. 2020); Decay sugar-rich substrates (Põlme et al. 2020) | Põlme S, et al. 2020. Fungal Diversity 105:1-16 ((https://doi.org/10.1007/s13225-020-00466-2)) |
| Otu26521 | 1 | k__Fungi(100);p__Ascomycota(100);c__Saccharomycetes(100);o__Saccharomycetales(100);f__Metschnikowiaceae(100);g__Metschnikowia(100);g__Metschnikowia_unclassified(100); | Metschnikowia | 13 | Saprotroph | \|Undefined Saprotroph\| | Yeast | NULL | Probable | As nectar-tap_saprotroph (Põlme et al. 2020); Decay sugar-rich substrates (Põlme et al. 2020) | Põlme S, et al. 2020. Fungal Diversity 105:1-16 ((https://doi.org/10.1007/s13225-020-00466-2)) |
| Otu26523 | 1 | k__Fungi(100);p__Ascomycota(100);c__Saccharomycetes(100);o__Saccharomycetales(100);f__Metschnikowiaceae(100);g__Metschnikowia(100);s__Metschnikowia_pulcherrima(100); | Metschnikowia | 13 | Saprotroph | \|Undefined Saprotroph\| | Yeast | NULL | Probable | As nectar-tap_saprotroph (Põlme et al. 2020); Decay sugar-rich substrates (Põlme et al. 2020) | Põlme S, et al. 2020. Fungal Diversity 105:1-16 ((https://doi.org/10.1007/s13225-020-00466-2)) |
| Otu26524 | 1 | k__Fungi(100);p__Ascomycota(100);c__Saccharomycetes(100);o__Saccharomycetales(100);f__Metschnikowiaceae(100);g__Metschnikowia(100);g__Metschnikowia_unclassified(100); | Metschnikowia | 13 | Saprotroph | \|Undefined Saprotroph\| | Yeast | NULL | Probable | As nectar-tap_saprotroph (Põlme et al. 2020); Decay sugar-rich substrates (Põlme et al. 2020) | Põlme S, et al. 2020. Fungal Diversity 105:1-16 ((https://doi.org/10.1007/s13225-020-00466-2)) |
| Otu26525 | 1 | k__Fungi(100);p__Ascomycota(100);c__Saccharomycetes(100);o__Saccharomycetales(100);f__Metschnikowiaceae(100);g__Metschnikowia(100);s__Metschnikowia_pulcherrima(100); | Metschnikowia | 13 | Saprotroph | \|Undefined Saprotroph\| | Yeast | NULL | Probable | As nectar-tap_saprotroph (Põlme et al. 2020); Decay sugar-rich substrates (Põlme et al. 2020) | Põlme S, et al. 2020. Fungal Diversity 105:1-16 ((https://doi.org/10.1007/s13225-020-00466-2)) |
| Otu26526 | 1 | k__Fungi(100);p__Ascomycota(100);c__Saccharomycetes(100);o__Saccharomycetales(100);f__Metschnikowiaceae(100);g__Metschnikowia(100);s__Metschnikowia_pulcherrima(100); | Metschnikowia | 13 | Saprotroph | \|Undefined Saprotroph\| | Yeast | NULL | Probable | As nectar-tap_saprotroph (Põlme et al. 2020); Decay sugar-rich substrates (Põlme et al. 2020) | Põlme S, et al. 2020. Fungal Diversity 105:1-16 ((https://doi.org/10.1007/s13225-020-00466-2)) |
| Otu26527 | 1 | k__Fungi(100);p__Ascomycota(100);c__Saccharomycetes(100);o__Saccharomycetales(100);o__Saccharomycetales_unclassified(100);o__Saccharomycetales_unclassified(100);o__Saccharomycetales_unclassified(100); | Saccharomycetales | 7 | Saprotroph | Undefined Saprotroph | Yeast | NULL | Possible | NULL | Sterkenburg E, et al. 2015. New Phytologist 207:1145-1158 ((https://doi.org/10.1111/nph.13426)) |
| Otu26528 | 1 | k__Fungi(100);p__Ascomycota(100);c__Saccharomycetes(100);o__Saccharomycetales(100);f__Metschnikowiaceae(100);g__Metschnikowia(100);s__Metschnikowia_pulcherrima(100); | Metschnikowia | 13 | Saprotroph | \|Undefined Saprotroph\| | Yeast | NULL | Probable | As nectar-tap_saprotroph (Põlme et al. 2020); Decay sugar-rich substrates (Põlme et al. 2020) | Põlme S, et al. 2020. Fungal Diversity 105:1-16 ((https://doi.org/10.1007/s13225-020-00466-2)) |
| Otu26529 | 1 | k__Fungi(100);p__Ascomycota(100);c__Saccharomycetes(100);o__Saccharomycetales(100);f__Metschnikowiaceae(100);g__Metschnikowia(100);s__Metschnikowia_pulcherrima(100); | Metschnikowia | 13 | Saprotroph | \|Undefined Saprotroph\| | Yeast | NULL | Probable | As nectar-tap_saprotroph (Põlme et al. 2020); Decay sugar-rich substrates (Põlme et al. 2020) | Põlme S, et al. 2020. Fungal Diversity 105:1-16 ((https://doi.org/10.1007/s13225-020-00466-2)) |
| Otu26530 | 1 | k__Fungi(100);p__Ascomycota(100);c__Saccharomycetes(100);o__Saccharomycetales(100);f__Metschnikowiaceae(100);g__Metschnikowia(100);g__Metschnikowia_unclassified(100); | Metschnikowia | 13 | Saprotroph | \|Undefined Saprotroph\| | Yeast | NULL | Probable | As nectar-tap_saprotroph (Põlme et al. 2020); Decay sugar-rich substrates (Põlme et al. 2020) | Põlme S, et al. 2020. Fungal Diversity 105:1-16 ((https://doi.org/10.1007/s13225-020-00466-2)) |
| Otu26532 | 1 | k__Fungi(100);p__Ascomycota(100);c__Saccharomycetes(100);o__Saccharomycetales(100);f__Metschnikowiaceae(100);g__Metschnikowia(100);g__Metschnikowia_unclassified(100); | Metschnikowia | 13 | Saprotroph | \|Undefined Saprotroph\| | Yeast | NULL | Probable | As nectar-tap_saprotroph (Põlme et al. 2020); Decay sugar-rich substrates (Põlme et al. 2020) | Põlme S, et al. 2020. Fungal Diversity 105:1-16 ((https://doi.org/10.1007/s13225-020-00466-2)) |
| Otu26536 | 1 | k__Fungi(100);p__Ascomycota(100);c__Saccharomycetes(100);o__Saccharomycetales(100);f__Metschnikowiaceae(100);g__Metschnikowia(100);s__Metschnikowia_pulcherrima(100); | Metschnikowia | 13 | Saprotroph | \|Undefined Saprotroph\| | Yeast | NULL | Probable | As nectar-tap_saprotroph (Põlme et al. 2020); Decay sugar-rich substrates (Põlme et al. 2020) | Põlme S, et al. 2020. Fungal Diversity 105:1-16 ((https://doi.org/10.1007/s13225-020-00466-2)) |
| Otu26537 | 1 | k__Fungi(100);p__Ascomycota(100);c__Saccharomycetes(100);o__Saccharomycetales(100);f__Metschnikowiaceae(100);g__Metschnikowia(100);g__Metschnikowia_unclassified(100); | Metschnikowia | 13 | Saprotroph | \|Undefined Saprotroph\| | Yeast | NULL | Probable | As nectar-tap_saprotroph (Põlme et al. 2020); Decay sugar-rich substrates (Põlme et al. 2020) | Põlme S, et al. 2020. Fungal Diversity 105:1-16 ((https://doi.org/10.1007/s13225-020-00466-2)) |
| Otu26538 | 1 | k__Fungi(100);p__Ascomycota(100);c__Saccharomycetes(100);o__Saccharomycetales(100);f__Metschnikowiaceae(100);g__Metschnikowia(100);s__Metschnikowia_pulcherrima(100); | Metschnikowia | 13 | Saprotroph | \|Undefined Saprotroph\| | Yeast | NULL | Probable | As nectar-tap_saprotroph (Põlme et al. 2020); Decay sugar-rich substrates (Põlme et al. 2020) | Põlme S, et al. 2020. Fungal Diversity 105:1-16 ((https://doi.org/10.1007/s13225-020-00466-2)) |
| Otu26557 | 1 | k__Fungi(100);p__Ascomycota(100);c__Dothideomycetes(100);o__Capnodiales(100);f__Mycosphaerellaceae(100);g__Mycosphaerella(100);s__Mycosphaerella_latebrosa(100); | Mycosphaerella | 13 | Pathotroph-Saprotroph | \|Plant Pathogen\|-Plant Saprotroph | Microfungus | NULL | Probable | As litter_saprotroph (Põlme et al. 2020); Decay leaf, fruit, seed (Põlme et al. 2020) | Tedersoo L, et al. 2014. Science 346:e1256688 ((https://doi.org/10.1126/science.1256688)); Põlme S, et al. 2020. Fungal Diversity 105:1-16 ((https://doi.org/10.1007/s13225-020-00466-2)) |
| Otu26558 | 1 | k__Fungi(100);p__Ascomycota(100);c__Saccharomycetes(100);o__Saccharomycetales(100);f__Metschnikowiaceae(100);g__Metschnikowia(100);g__Metschnikowia_unclassified(100); | Metschnikowia | 13 | Saprotroph | \|Undefined Saprotroph\| | Yeast | NULL | Probable | As nectar-tap_saprotroph (Põlme et al. 2020); Decay sugar-rich substrates (Põlme et al. 2020) | Põlme S, et al. 2020. Fungal Diversity 105:1-16 ((https://doi.org/10.1007/s13225-020-00466-2)) |
| Otu26559 | 1 | k__Fungi(100);p__Ascomycota(100);c__Saccharomycetes(100);o__Saccharomycetales(100);f__Metschnikowiaceae(100);g__Metschnikowia(100);g__Metschnikowia_unclassified(100); | Metschnikowia | 13 | Saprotroph | \|Undefined Saprotroph\| | Yeast | NULL | Probable | As nectar-tap_saprotroph (Põlme et al. 2020); Decay sugar-rich substrates (Põlme et al. 2020) | Põlme S, et al. 2020. Fungal Diversity 105:1-16 ((https://doi.org/10.1007/s13225-020-00466-2)) |
| Otu26565 | 1 | k__Fungi(100);p__Ascomycota(100);c__Saccharomycetes(100);o__Saccharomycetales(100);f__Metschnikowiaceae(100);g__Metschnikowia(100);g__Metschnikowia_unclassified(100); | Metschnikowia | 13 | Saprotroph | \|Undefined Saprotroph\| | Yeast | NULL | Probable | As nectar-tap_saprotroph (Põlme et al. 2020); Decay sugar-rich substrates (Põlme et al. 2020) | Põlme S, et al. 2020. Fungal Diversity 105:1-16 ((https://doi.org/10.1007/s13225-020-00466-2)) |
| Otu26566 | 1 | k__Fungi(100);p__Ascomycota(100);c__Saccharomycetes(100);o__Saccharomycetales(100);f__Metschnikowiaceae(100);g__Metschnikowia(100);s__Metschnikowia_pulcherrima(100); | Metschnikowia | 13 | Saprotroph | \|Undefined Saprotroph\| | Yeast | NULL | Probable | As nectar-tap_saprotroph (Põlme et al. 2020); Decay sugar-rich substrates (Põlme et al. 2020) | Põlme S, et al. 2020. Fungal Diversity 105:1-16 ((https://doi.org/10.1007/s13225-020-00466-2)) |
| Otu26567 | 1 | k__Fungi(100);p__Ascomycota(100);c__Saccharomycetes(100);o__Saccharomycetales(100);f__Metschnikowiaceae(100);g__Metschnikowia(100);s__Metschnikowia_pulcherrima(100); | Metschnikowia | 13 | Saprotroph | \|Undefined Saprotroph\| | Yeast | NULL | Probable | As nectar-tap_saprotroph (Põlme et al. 2020); Decay sugar-rich substrates (Põlme et al. 2020) | Põlme S, et al. 2020. Fungal Diversity 105:1-16 ((https://doi.org/10.1007/s13225-020-00466-2)) |
| Otu26568 | 1 | k__Fungi(100);p__Ascomycota(100);c__Saccharomycetes(100);o__Saccharomycetales(100);f__Metschnikowiaceae(100);g__Metschnikowia(100);g__Metschnikowia_unclassified(100); | Metschnikowia | 13 | Saprotroph | \|Undefined Saprotroph\| | Yeast | NULL | Probable | As nectar-tap_saprotroph (Põlme et al. 2020); Decay sugar-rich substrates (Põlme et al. 2020) | Põlme S, et al. 2020. Fungal Diversity 105:1-16 ((https://doi.org/10.1007/s13225-020-00466-2)) |
| Otu26569 | 1 | k__Fungi(100);p__Ascomycota(100);c__Saccharomycetes(100);o__Saccharomycetales(100);f__Metschnikowiaceae(100);g__Metschnikowia(100);s__Metschnikowia_pulcherrima(100); | Metschnikowia | 13 | Saprotroph | \|Undefined Saprotroph\| | Yeast | NULL | Probable | As nectar-tap_saprotroph (Põlme et al. 2020); Decay sugar-rich substrates (Põlme et al. 2020) | Põlme S, et al. 2020. Fungal Diversity 105:1-16 ((https://doi.org/10.1007/s13225-020-00466-2)) |
| Otu26570 | 1 | k__Fungi(100);p__Ascomycota(100);c__Saccharomycetes(100);o__Saccharomycetales(100);f__Metschnikowiaceae(100);g__Metschnikowia(100);s__Metschnikowia_pulcherrima(100); | Metschnikowia | 13 | Saprotroph | \|Undefined Saprotroph\| | Yeast | NULL | Probable | As nectar-tap_saprotroph (Põlme et al. 2020); Decay sugar-rich substrates (Põlme et al. 2020) | Põlme S, et al. 2020. Fungal Diversity 105:1-16 ((https://doi.org/10.1007/s13225-020-00466-2)) |
| Otu26573 | 1 | k__Fungi(100);p__Ascomycota(100);c__Saccharomycetes(100);o__Saccharomycetales(100);f__Metschnikowiaceae(100);g__Metschnikowia(100);s__Metschnikowia_pulcherrima(100); | Metschnikowia | 13 | Saprotroph | \|Undefined Saprotroph\| | Yeast | NULL | Probable | As nectar-tap_saprotroph (Põlme et al. 2020); Decay sugar-rich substrates (Põlme et al. 2020) | Põlme S, et al. 2020. Fungal Diversity 105:1-16 ((https://doi.org/10.1007/s13225-020-00466-2)) |
| Otu26574 | 1 | k__Fungi(100);p__Ascomycota(100);c__Saccharomycetes(100);o__Saccharomycetales(100);f__Metschnikowiaceae(100);g__Metschnikowia(100);s__Metschnikowia_pulcherrima(100); | Metschnikowia | 13 | Saprotroph | \|Undefined Saprotroph\| | Yeast | NULL | Probable | As nectar-tap_saprotroph (Põlme et al. 2020); Decay sugar-rich substrates (Põlme et al. 2020) | Põlme S, et al. 2020. Fungal Diversity 105:1-16 ((https://doi.org/10.1007/s13225-020-00466-2)) |
| Otu26576 | 1 | k__Fungi(100);p__Ascomycota(100);c__Saccharomycetes(100);o__Saccharomycetales(100);f__Metschnikowiaceae(100);g__Metschnikowia(100);g__Metschnikowia_unclassified(100); | Metschnikowia | 13 | Saprotroph | \|Undefined Saprotroph\| | Yeast | NULL | Probable | As nectar-tap_saprotroph (Põlme et al. 2020); Decay sugar-rich substrates (Põlme et al. 2020) | Põlme S, et al. 2020. Fungal Diversity 105:1-16 ((https://doi.org/10.1007/s13225-020-00466-2)) |
| Otu26577 | 1 | k__Fungi(100);p__Ascomycota(100);c__Saccharomycetes(100);o__Saccharomycetales(100);f__Metschnikowiaceae(100);g__Metschnikowia(100);s__Metschnikowia_pulcherrima(100); | Metschnikowia | 13 | Saprotroph | \|Undefined Saprotroph\| | Yeast | NULL | Probable | As nectar-tap_saprotroph (Põlme et al. 2020); Decay sugar-rich substrates (Põlme et al. 2020) | Põlme S, et al. 2020. Fungal Diversity 105:1-16 ((https://doi.org/10.1007/s13225-020-00466-2)) |
| Otu26591 | 1 | k__Fungi(100);p__Ascomycota(100);c__Saccharomycetes(100);o__Saccharomycetales(100);f__Metschnikowiaceae(100);g__Metschnikowia(100);s__Metschnikowia_pulcherrima(100); | Metschnikowia | 13 | Saprotroph | \|Undefined Saprotroph\| | Yeast | NULL | Probable | As nectar-tap_saprotroph (Põlme et al. 2020); Decay sugar-rich substrates (Põlme et al. 2020) | Põlme S, et al. 2020. Fungal Diversity 105:1-16 ((https://doi.org/10.1007/s13225-020-00466-2)) |
| Otu26592 | 1 | k__Fungi(100);p__Ascomycota(100);c__Saccharomycetes(100);o__Saccharomycetales(100);f__Metschnikowiaceae(100);g__Metschnikowia(100);s__Metschnikowia_pulcherrima(100); | Metschnikowia | 13 | Saprotroph | \|Undefined Saprotroph\| | Yeast | NULL | Probable | As nectar-tap_saprotroph (Põlme et al. 2020); Decay sugar-rich substrates (Põlme et al. 2020) | Põlme S, et al. 2020. Fungal Diversity 105:1-16 ((https://doi.org/10.1007/s13225-020-00466-2)) |
| Otu26646 | 1 | k__Fungi(100);p__Ascomycota(100);c__Saccharomycetes(100);o__Saccharomycetales(100);f__Metschnikowiaceae(100);g__Metschnikowia(100);s__Metschnikowia_pulcherrima(100); | Metschnikowia | 13 | Saprotroph | \|Undefined Saprotroph\| | Yeast | NULL | Probable | As nectar-tap_saprotroph (Põlme et al. 2020); Decay sugar-rich substrates (Põlme et al. 2020) | Põlme S, et al. 2020. Fungal Diversity 105:1-16 ((https://doi.org/10.1007/s13225-020-00466-2)) |
| Otu26657 | 1 | k__Fungi(100);p__Ascomycota(100);c__Saccharomycetes(100);o__Saccharomycetales(100);f__Metschnikowiaceae(100);g__Metschnikowia(100);s__Metschnikowia_pulcherrima(100); | Metschnikowia | 13 | Saprotroph | \|Undefined Saprotroph\| | Yeast | NULL | Probable | As nectar-tap_saprotroph (Põlme et al. 2020); Decay sugar-rich substrates (Põlme et al. 2020) | Põlme S, et al. 2020. Fungal Diversity 105:1-16 ((https://doi.org/10.1007/s13225-020-00466-2)) |
| Otu26658 | 1 | k__Fungi(100);p__Ascomycota(100);c__Saccharomycetes(100);o__Saccharomycetales(100);f__Metschnikowiaceae(100);g__Metschnikowia(100);s__Metschnikowia_pulcherrima(100); | Metschnikowia | 13 | Saprotroph | \|Undefined Saprotroph\| | Yeast | NULL | Probable | As nectar-tap_saprotroph (Põlme et al. 2020); Decay sugar-rich substrates (Põlme et al. 2020) | Põlme S, et al. 2020. Fungal Diversity 105:1-16 ((https://doi.org/10.1007/s13225-020-00466-2)) |
| Otu26659 | 1 | k__Fungi(100);p__Ascomycota(100);c__Saccharomycetes(100);o__Saccharomycetales(100);f__Metschnikowiaceae(100);g__Metschnikowia(100);s__Metschnikowia_pulcherrima(100); | Metschnikowia | 13 | Saprotroph | \|Undefined Saprotroph\| | Yeast | NULL | Probable | As nectar-tap_saprotroph (Põlme et al. 2020); Decay sugar-rich substrates (Põlme et al. 2020) | Põlme S, et al. 2020. Fungal Diversity 105:1-16 ((https://doi.org/10.1007/s13225-020-00466-2)) |
| Otu26660 | 1 | k__Fungi(100);p__Ascomycota(100);c__Saccharomycetes(100);o__Saccharomycetales(100);f__Metschnikowiaceae(100);g__Metschnikowia(100);s__Metschnikowia_pulcherrima(100); | Metschnikowia | 13 | Saprotroph | \|Undefined Saprotroph\| | Yeast | NULL | Probable | As nectar-tap_saprotroph (Põlme et al. 2020); Decay sugar-rich substrates (Põlme et al. 2020) | Põlme S, et al. 2020. Fungal Diversity 105:1-16 ((https://doi.org/10.1007/s13225-020-00466-2)) |
| Otu26661 | 1 | k__Fungi(100);p__Ascomycota(100);c__Saccharomycetes(100);o__Saccharomycetales(100);f__Metschnikowiaceae(100);g__Metschnikowia(100);s__Metschnikowia_pulcherrima(100); | Metschnikowia | 13 | Saprotroph | \|Undefined Saprotroph\| | Yeast | NULL | Probable | As nectar-tap_saprotroph (Põlme et al. 2020); Decay sugar-rich substrates (Põlme et al. 2020) | Põlme S, et al. 2020. Fungal Diversity 105:1-16 ((https://doi.org/10.1007/s13225-020-00466-2)) |
| Otu26662 | 1 | k__Fungi(100);p__Ascomycota(100);c__Saccharomycetes(100);o__Saccharomycetales(100);f__Metschnikowiaceae(100);g__Metschnikowia(100);s__Metschnikowia_pulcherrima(100); | Metschnikowia | 13 | Saprotroph | \|Undefined Saprotroph\| | Yeast | NULL | Probable | As nectar-tap_saprotroph (Põlme et al. 2020); Decay sugar-rich substrates (Põlme et al. 2020) | Põlme S, et al. 2020. Fungal Diversity 105:1-16 ((https://doi.org/10.1007/s13225-020-00466-2)) |
| Otu26663 | 1 | k__Fungi(100);p__Ascomycota(100);c__Saccharomycetes(100);o__Saccharomycetales(100);f__Metschnikowiaceae(100);g__Metschnikowia(100);g__Metschnikowia_unclassified(100); | Metschnikowia | 13 | Saprotroph | \|Undefined Saprotroph\| | Yeast | NULL | Probable | As nectar-tap_saprotroph (Põlme et al. 2020); Decay sugar-rich substrates (Põlme et al. 2020) | Põlme S, et al. 2020. Fungal Diversity 105:1-16 ((https://doi.org/10.1007/s13225-020-00466-2)) |
| Otu26664 | 1 | k__Fungi(100);p__Ascomycota(100);c__Saccharomycetes(100);o__Saccharomycetales(100);f__Metschnikowiaceae(100);g__Metschnikowia(100);s__Metschnikowia_pulcherrima(100); | Metschnikowia | 13 | Saprotroph | \|Undefined Saprotroph\| | Yeast | NULL | Probable | As nectar-tap_saprotroph (Põlme et al. 2020); Decay sugar-rich substrates (Põlme et al. 2020) | Põlme S, et al. 2020. Fungal Diversity 105:1-16 ((https://doi.org/10.1007/s13225-020-00466-2)) |
| Otu26665 | 1 | k__Fungi(100);p__Ascomycota(100);c__Saccharomycetes(100);o__Saccharomycetales(100);f__Metschnikowiaceae(100);g__Metschnikowia(100);s__Metschnikowia_pulcherrima(100); | Metschnikowia | 13 | Saprotroph | \|Undefined Saprotroph\| | Yeast | NULL | Probable | As nectar-tap_saprotroph (Põlme et al. 2020); Decay sugar-rich substrates (Põlme et al. 2020) | Põlme S, et al. 2020. Fungal Diversity 105:1-16 ((https://doi.org/10.1007/s13225-020-00466-2)) |
| Otu26666 | 1 | k__Fungi(100);p__Ascomycota(100);c__Saccharomycetes(100);o__Saccharomycetales(100);f__Metschnikowiaceae(100);g__Metschnikowia(100);s__Metschnikowia_pulcherrima(100); | Metschnikowia | 13 | Saprotroph | \|Undefined Saprotroph\| | Yeast | NULL | Probable | As nectar-tap_saprotroph (Põlme et al. 2020); Decay sugar-rich substrates (Põlme et al. 2020) | Põlme S, et al. 2020. Fungal Diversity 105:1-16 ((https://doi.org/10.1007/s13225-020-00466-2)) |
| Otu26668 | 1 | k__Fungi(100);p__Ascomycota(100);c__Saccharomycetes(100);o__Saccharomycetales(100);f__Metschnikowiaceae(100);g__Metschnikowia(100);g__Metschnikowia_unclassified(100); | Metschnikowia | 13 | Saprotroph | \|Undefined Saprotroph\| | Yeast | NULL | Probable | As nectar-tap_saprotroph (Põlme et al. 2020); Decay sugar-rich substrates (Põlme et al. 2020) | Põlme S, et al. 2020. Fungal Diversity 105:1-16 ((https://doi.org/10.1007/s13225-020-00466-2)) |
| Otu26670 | 1 | k__Fungi(100);p__Ascomycota(100);c__Saccharomycetes(100);o__Saccharomycetales(100);f__Metschnikowiaceae(100);g__Metschnikowia(100);s__Metschnikowia_pulcherrima(100); | Metschnikowia | 13 | Saprotroph | \|Undefined Saprotroph\| | Yeast | NULL | Probable | As nectar-tap_saprotroph (Põlme et al. 2020); Decay sugar-rich substrates (Põlme et al. 2020) | Põlme S, et al. 2020. Fungal Diversity 105:1-16 ((https://doi.org/10.1007/s13225-020-00466-2)) |
| Otu26672 | 1 | k__Fungi(100);p__Ascomycota(100);c__Saccharomycetes(100);o__Saccharomycetales(100);f__Metschnikowiaceae(100);g__Metschnikowia(100);s__Metschnikowia_pulcherrima(100); | Metschnikowia | 13 | Saprotroph | \|Undefined Saprotroph\| | Yeast | NULL | Probable | As nectar-tap_saprotroph (Põlme et al. 2020); Decay sugar-rich substrates (Põlme et al. 2020) | Põlme S, et al. 2020. Fungal Diversity 105:1-16 ((https://doi.org/10.1007/s13225-020-00466-2)) |
| Otu26673 | 1 | k__Fungi(100);p__Ascomycota(100);c__Saccharomycetes(100);o__Saccharomycetales(100);f__Metschnikowiaceae(100);g__Metschnikowia(100);s__Metschnikowia_pulcherrima(100); | Metschnikowia | 13 | Saprotroph | \|Undefined Saprotroph\| | Yeast | NULL | Probable | As nectar-tap_saprotroph (Põlme et al. 2020); Decay sugar-rich substrates (Põlme et al. 2020) | Põlme S, et al. 2020. Fungal Diversity 105:1-16 ((https://doi.org/10.1007/s13225-020-00466-2)) |
| Otu26674 | 1 | k__Fungi(100);p__Ascomycota(100);c__Saccharomycetes(100);o__Saccharomycetales(100);f__Metschnikowiaceae(100);g__Metschnikowia(100);g__Metschnikowia_unclassified(100); | Metschnikowia | 13 | Saprotroph | \|Undefined Saprotroph\| | Yeast | NULL | Probable | As nectar-tap_saprotroph (Põlme et al. 2020); Decay sugar-rich substrates (Põlme et al. 2020) | Põlme S, et al. 2020. Fungal Diversity 105:1-16 ((https://doi.org/10.1007/s13225-020-00466-2)) |
| Otu26682 | 1 | k__Fungi(100);p__Ascomycota(100);c__Saccharomycetes(100);o__Saccharomycetales(100);f__Metschnikowiaceae(100);g__Metschnikowia(100);s__Metschnikowia_pulcherrima(100); | Metschnikowia | 13 | Saprotroph | \|Undefined Saprotroph\| | Yeast | NULL | Probable | As nectar-tap_saprotroph (Põlme et al. 2020); Decay sugar-rich substrates (Põlme et al. 2020) | Põlme S, et al. 2020. Fungal Diversity 105:1-16 ((https://doi.org/10.1007/s13225-020-00466-2)) |
| Otu26697 | 1 | k__Fungi(100);p__Ascomycota(100);c__Saccharomycetes(100);o__Saccharomycetales(100);f__Metschnikowiaceae(100);g__Metschnikowia(100);g__Metschnikowia_unclassified(100); | Metschnikowia | 13 | Saprotroph | \|Undefined Saprotroph\| | Yeast | NULL | Probable | As nectar-tap_saprotroph (Põlme et al. 2020); Decay sugar-rich substrates (Põlme et al. 2020) | Põlme S, et al. 2020. Fungal Diversity 105:1-16 ((https://doi.org/10.1007/s13225-020-00466-2)) |
| Otu26700 | 1 | k__Fungi(100);p__Ascomycota(100);c__Saccharomycetes(100);o__Saccharomycetales(100);f__Metschnikowiaceae(100);g__Metschnikowia(100);s__Metschnikowia_pulcherrima(100); | Metschnikowia | 13 | Saprotroph | \|Undefined Saprotroph\| | Yeast | NULL | Probable | As nectar-tap_saprotroph (Põlme et al. 2020); Decay sugar-rich substrates (Põlme et al. 2020) | Põlme S, et al. 2020. Fungal Diversity 105:1-16 ((https://doi.org/10.1007/s13225-020-00466-2)) |
| Otu26707 | 1 | k__Fungi(100);p__Ascomycota(100);c__Saccharomycetes(100);o__Saccharomycetales(100);f__Metschnikowiaceae(100);g__Metschnikowia(100);g__Metschnikowia_unclassified(100); | Metschnikowia | 13 | Saprotroph | \|Undefined Saprotroph\| | Yeast | NULL | Probable | As nectar-tap_saprotroph (Põlme et al. 2020); Decay sugar-rich substrates (Põlme et al. 2020) | Põlme S, et al. 2020. Fungal Diversity 105:1-16 ((https://doi.org/10.1007/s13225-020-00466-2)) |
| Otu26715 | 1 | k__Fungi(100);p__Ascomycota(100);c__Saccharomycetes(100);o__Saccharomycetales(100);f__Metschnikowiaceae(100);g__Metschnikowia(100);g__Metschnikowia_unclassified(100); | Metschnikowia | 13 | Saprotroph | \|Undefined Saprotroph\| | Yeast | NULL | Probable | As nectar-tap_saprotroph (Põlme et al. 2020); Decay sugar-rich substrates (Põlme et al. 2020) | Põlme S, et al. 2020. Fungal Diversity 105:1-16 ((https://doi.org/10.1007/s13225-020-00466-2)) |
| Otu26724 | 1 | k__Fungi(100);p__Ascomycota(100);c__Saccharomycetes(100);o__Saccharomycetales(100);f__Metschnikowiaceae(100);g__Metschnikowia(100);s__Metschnikowia_pulcherrima(100); | Metschnikowia | 13 | Saprotroph | \|Undefined Saprotroph\| | Yeast | NULL | Probable | As nectar-tap_saprotroph (Põlme et al. 2020); Decay sugar-rich substrates (Põlme et al. 2020) | Põlme S, et al. 2020. Fungal Diversity 105:1-16 ((https://doi.org/10.1007/s13225-020-00466-2)) |
| Otu26736 | 1 | k__Fungi(100);p__Ascomycota(100);c__Saccharomycetes(100);o__Saccharomycetales(100);f__Metschnikowiaceae(100);g__Metschnikowia(100);g__Metschnikowia_unclassified(100); | Metschnikowia | 13 | Saprotroph | \|Undefined Saprotroph\| | Yeast | NULL | Probable | As nectar-tap_saprotroph (Põlme et al. 2020); Decay sugar-rich substrates (Põlme et al. 2020) | Põlme S, et al. 2020. Fungal Diversity 105:1-16 ((https://doi.org/10.1007/s13225-020-00466-2)) |
| Otu26738 | 1 | k__Fungi(100);p__Ascomycota(100);c__Saccharomycetes(100);o__Saccharomycetales(100);f__Metschnikowiaceae(100);g__Metschnikowia(100);s__Metschnikowia_pulcherrima(100); | Metschnikowia | 13 | Saprotroph | \|Undefined Saprotroph\| | Yeast | NULL | Probable | As nectar-tap_saprotroph (Põlme et al. 2020); Decay sugar-rich substrates (Põlme et al. 2020) | Põlme S, et al. 2020. Fungal Diversity 105:1-16 ((https://doi.org/10.1007/s13225-020-00466-2)) |
| Otu26742 | 1 | k__Fungi(100);p__Ascomycota(100);c__Saccharomycetes(100);o__Saccharomycetales(100);f__Metschnikowiaceae(100);g__Metschnikowia(100);s__Metschnikowia_pulcherrima(100); | Metschnikowia | 13 | Saprotroph | \|Undefined Saprotroph\| | Yeast | NULL | Probable | As nectar-tap_saprotroph (Põlme et al. 2020); Decay sugar-rich substrates (Põlme et al. 2020) | Põlme S, et al. 2020. Fungal Diversity 105:1-16 ((https://doi.org/10.1007/s13225-020-00466-2)) |
| Otu26745 | 1 | k__Fungi(100);p__Ascomycota(100);c__Dothideomycetes(100);o__Pleosporales(100);f__Leptosphaeriaceae(100);g__Leptosphaeria(100);s__Leptosphaeria_sp(100); | Leptosphaeria | 13 | Pathotroph-Saprotroph | \|Plant Pathogen\|-Wood Saprotroph | Microfungus | NULL | Probable | As wood_pathogen (Põlme et al. 2020); Decay wood (Põlme et al. 2020) | Tedersoo L, et al. 2014. Science 346:e1256688 ((https://doi.org/10.1126/science.1256688)); Põlme S, et al. 2020. Fungal Diversity 105:1-16 ((https://doi.org/10.1007/s13225-020-00466-2)) |
| Otu26747 | 1 | k__Fungi(100);p__Ascomycota(100);c__Saccharomycetes(100);o__Saccharomycetales(100);f__Metschnikowiaceae(100);g__Metschnikowia(100);g__Metschnikowia_unclassified(100); | Metschnikowia | 13 | Saprotroph | \|Undefined Saprotroph\| | Yeast | NULL | Probable | As nectar-tap_saprotroph (Põlme et al. 2020); Decay sugar-rich substrates (Põlme et al. 2020) | Põlme S, et al. 2020. Fungal Diversity 105:1-16 ((https://doi.org/10.1007/s13225-020-00466-2)) |
| Otu26751 | 1 | k__Fungi(100);p__Ascomycota(100);c__Saccharomycetes(100);o__Saccharomycetales(100);f__Metschnikowiaceae(100);g__Metschnikowia(100);g__Metschnikowia_unclassified(100); | Metschnikowia | 13 | Saprotroph | \|Undefined Saprotroph\| | Yeast | NULL | Probable | As nectar-tap_saprotroph (Põlme et al. 2020); Decay sugar-rich substrates (Põlme et al. 2020) | Põlme S, et al. 2020. Fungal Diversity 105:1-16 ((https://doi.org/10.1007/s13225-020-00466-2)) |
| Otu26755 | 1 | k__Fungi(100);p__Ascomycota(100);c__Saccharomycetes(100);o__Saccharomycetales(100);f__Metschnikowiaceae(100);g__Metschnikowia(100);s__Metschnikowia_pulcherrima(100); | Metschnikowia | 13 | Saprotroph | \|Undefined Saprotroph\| | Yeast | NULL | Probable | As nectar-tap_saprotroph (Põlme et al. 2020); Decay sugar-rich substrates (Põlme et al. 2020) | Põlme S, et al. 2020. Fungal Diversity 105:1-16 ((https://doi.org/10.1007/s13225-020-00466-2)) |
| Otu26756 | 1 | k__Fungi(100);p__Ascomycota(100);c__Saccharomycetes(100);o__Saccharomycetales(100);f__Metschnikowiaceae(100);g__Metschnikowia(100);s__Metschnikowia_pulcherrima(100); | Metschnikowia | 13 | Saprotroph | \|Undefined Saprotroph\| | Yeast | NULL | Probable | As nectar-tap_saprotroph (Põlme et al. 2020); Decay sugar-rich substrates (Põlme et al. 2020) | Põlme S, et al. 2020. Fungal Diversity 105:1-16 ((https://doi.org/10.1007/s13225-020-00466-2)) |
| Otu26757 | 1 | k__Fungi(100);p__Ascomycota(100);c__Saccharomycetes(100);o__Saccharomycetales(100);f__Metschnikowiaceae(100);g__Metschnikowia(100);s__Metschnikowia_pulcherrima(100); | Metschnikowia | 13 | Saprotroph | \|Undefined Saprotroph\| | Yeast | NULL | Probable | As nectar-tap_saprotroph (Põlme et al. 2020); Decay sugar-rich substrates (Põlme et al. 2020) | Põlme S, et al. 2020. Fungal Diversity 105:1-16 ((https://doi.org/10.1007/s13225-020-00466-2)) |
| Otu26758 | 1 | k__Fungi(100);p__Ascomycota(100);c__Saccharomycetes(100);o__Saccharomycetales(100);f__Metschnikowiaceae(100);g__Metschnikowia(100);s__Metschnikowia_pulcherrima(100); | Metschnikowia | 13 | Saprotroph | \|Undefined Saprotroph\| | Yeast | NULL | Probable | As nectar-tap_saprotroph (Põlme et al. 2020); Decay sugar-rich substrates (Põlme et al. 2020) | Põlme S, et al. 2020. Fungal Diversity 105:1-16 ((https://doi.org/10.1007/s13225-020-00466-2)) |
| Otu26763 | 1 | k__Fungi(100);p__Ascomycota(100);c__Saccharomycetes(100);o__Saccharomycetales(100);f__Metschnikowiaceae(100);g__Metschnikowia(100);s__Metschnikowia_pulcherrima(100); | Metschnikowia | 13 | Saprotroph | \|Undefined Saprotroph\| | Yeast | NULL | Probable | As nectar-tap_saprotroph (Põlme et al. 2020); Decay sugar-rich substrates (Põlme et al. 2020) | Põlme S, et al. 2020. Fungal Diversity 105:1-16 ((https://doi.org/10.1007/s13225-020-00466-2)) |
| Otu26771 | 1 | k__Fungi(100);p__Ascomycota(100);c__Saccharomycetes(100);o__Saccharomycetales(100);f__Metschnikowiaceae(100);g__Metschnikowia(100);s__Metschnikowia_pulcherrima(100); | Metschnikowia | 13 | Saprotroph | \|Undefined Saprotroph\| | Yeast | NULL | Probable | As nectar-tap_saprotroph (Põlme et al. 2020); Decay sugar-rich substrates (Põlme et al. 2020) | Põlme S, et al. 2020. Fungal Diversity 105:1-16 ((https://doi.org/10.1007/s13225-020-00466-2)) |
| Otu26773 | 1 | k__Fungi(100);p__Ascomycota(100);c__Saccharomycetes(100);o__Saccharomycetales(100);f__Metschnikowiaceae(100);g__Metschnikowia(100);s__Metschnikowia_pulcherrima(100); | Metschnikowia | 13 | Saprotroph | \|Undefined Saprotroph\| | Yeast | NULL | Probable | As nectar-tap_saprotroph (Põlme et al. 2020); Decay sugar-rich substrates (Põlme et al. 2020) | Põlme S, et al. 2020. Fungal Diversity 105:1-16 ((https://doi.org/10.1007/s13225-020-00466-2)) |
| Otu26774 | 1 | k__Fungi(100);p__Ascomycota(100);c__Saccharomycetes(100);o__Saccharomycetales(100);f__Metschnikowiaceae(100);g__Metschnikowia(100);g__Metschnikowia_unclassified(100); | Metschnikowia | 13 | Saprotroph | \|Undefined Saprotroph\| | Yeast | NULL | Probable | As nectar-tap_saprotroph (Põlme et al. 2020); Decay sugar-rich substrates (Põlme et al. 2020) | Põlme S, et al. 2020. Fungal Diversity 105:1-16 ((https://doi.org/10.1007/s13225-020-00466-2)) |
| Otu26777 | 1 | k__Fungi(100);p__Ascomycota(100);c__Saccharomycetes(100);o__Saccharomycetales(100);f__Metschnikowiaceae(100);g__Metschnikowia(100);s__Metschnikowia_pulcherrima(100); | Metschnikowia | 13 | Saprotroph | \|Undefined Saprotroph\| | Yeast | NULL | Probable | As nectar-tap_saprotroph (Põlme et al. 2020); Decay sugar-rich substrates (Põlme et al. 2020) | Põlme S, et al. 2020. Fungal Diversity 105:1-16 ((https://doi.org/10.1007/s13225-020-00466-2)) |
| Otu26778 | 1 | k__Fungi(100);p__Ascomycota(100);c__Saccharomycetes(100);o__Saccharomycetales(100);f__Metschnikowiaceae(100);g__Metschnikowia(100);s__Metschnikowia_pulcherrima(100); | Metschnikowia | 13 | Saprotroph | \|Undefined Saprotroph\| | Yeast | NULL | Probable | As nectar-tap_saprotroph (Põlme et al. 2020); Decay sugar-rich substrates (Põlme et al. 2020) | Põlme S, et al. 2020. Fungal Diversity 105:1-16 ((https://doi.org/10.1007/s13225-020-00466-2)) |
| Otu26785 | 1 | k__Fungi(100);p__Ascomycota(100);c__Saccharomycetes(100);o__Saccharomycetales(100);f__Metschnikowiaceae(100);g__Metschnikowia(100);g__Metschnikowia_unclassified(100); | Metschnikowia | 13 | Saprotroph | \|Undefined Saprotroph\| | Yeast | NULL | Probable | As nectar-tap_saprotroph (Põlme et al. 2020); Decay sugar-rich substrates (Põlme et al. 2020) | Põlme S, et al. 2020. Fungal Diversity 105:1-16 ((https://doi.org/10.1007/s13225-020-00466-2)) |
| Otu26787 | 1 | k__Fungi(100);p__Ascomycota(100);c__Saccharomycetes(100);o__Saccharomycetales(100);f__Metschnikowiaceae(100);g__Metschnikowia(100);s__Metschnikowia_pulcherrima(100); | Metschnikowia | 13 | Saprotroph | \|Undefined Saprotroph\| | Yeast | NULL | Probable | As nectar-tap_saprotroph (Põlme et al. 2020); Decay sugar-rich substrates (Põlme et al. 2020) | Põlme S, et al. 2020. Fungal Diversity 105:1-16 ((https://doi.org/10.1007/s13225-020-00466-2)) |
| Otu26788 | 1 | k__Fungi(100);p__Ascomycota(100);c__Saccharomycetes(100);o__Saccharomycetales(100);f__Metschnikowiaceae(100);g__Metschnikowia(100);s__Metschnikowia_pulcherrima(100); | Metschnikowia | 13 | Saprotroph | \|Undefined Saprotroph\| | Yeast | NULL | Probable | As nectar-tap_saprotroph (Põlme et al. 2020); Decay sugar-rich substrates (Põlme et al. 2020) | Põlme S, et al. 2020. Fungal Diversity 105:1-16 ((https://doi.org/10.1007/s13225-020-00466-2)) |
| Otu26789 | 1 | k__Fungi(100);p__Ascomycota(100);c__Saccharomycetes(100);o__Saccharomycetales(100);f__Metschnikowiaceae(100);g__Metschnikowia(100);s__Metschnikowia_pulcherrima(100); | Metschnikowia | 13 | Saprotroph | \|Undefined Saprotroph\| | Yeast | NULL | Probable | As nectar-tap_saprotroph (Põlme et al. 2020); Decay sugar-rich substrates (Põlme et al. 2020) | Põlme S, et al. 2020. Fungal Diversity 105:1-16 ((https://doi.org/10.1007/s13225-020-00466-2)) |
| Otu26790 | 1 | k__Fungi(100);p__Ascomycota(100);c__Saccharomycetes(100);o__Saccharomycetales(100);f__Metschnikowiaceae(100);g__Metschnikowia(100);g__Metschnikowia_unclassified(100); | Metschnikowia | 13 | Saprotroph | \|Undefined Saprotroph\| | Yeast | NULL | Probable | As nectar-tap_saprotroph (Põlme et al. 2020); Decay sugar-rich substrates (Põlme et al. 2020) | Põlme S, et al. 2020. Fungal Diversity 105:1-16 ((https://doi.org/10.1007/s13225-020-00466-2)) |
| Otu26791 | 1 | k__Fungi(100);p__Ascomycota(100);c__Saccharomycetes(100);o__Saccharomycetales(100);f__Metschnikowiaceae(100);g__Metschnikowia(100);s__Metschnikowia_pulcherrima(100); | Metschnikowia | 13 | Saprotroph | \|Undefined Saprotroph\| | Yeast | NULL | Probable | As nectar-tap_saprotroph (Põlme et al. 2020); Decay sugar-rich substrates (Põlme et al. 2020) | Põlme S, et al. 2020. Fungal Diversity 105:1-16 ((https://doi.org/10.1007/s13225-020-00466-2)) |
| Otu26792 | 1 | k__Fungi(100);p__Ascomycota(100);c__Saccharomycetes(100);o__Saccharomycetales(100);f__Metschnikowiaceae(100);g__Metschnikowia(100);s__Metschnikowia_pulcherrima(100); | Metschnikowia | 13 | Saprotroph | \|Undefined Saprotroph\| | Yeast | NULL | Probable | As nectar-tap_saprotroph (Põlme et al. 2020); Decay sugar-rich substrates (Põlme et al. 2020) | Põlme S, et al. 2020. Fungal Diversity 105:1-16 ((https://doi.org/10.1007/s13225-020-00466-2)) |
| Otu26793 | 1 | k__Fungi(100);p__Ascomycota(100);c__Saccharomycetes(100);o__Saccharomycetales(100);f__Metschnikowiaceae(100);g__Metschnikowia(100);s__Metschnikowia_pulcherrima(100); | Metschnikowia | 13 | Saprotroph | \|Undefined Saprotroph\| | Yeast | NULL | Probable | As nectar-tap_saprotroph (Põlme et al. 2020); Decay sugar-rich substrates (Põlme et al. 2020) | Põlme S, et al. 2020. Fungal Diversity 105:1-16 ((https://doi.org/10.1007/s13225-020-00466-2)) |
| Otu26794 | 1 | k__Fungi(100);p__Ascomycota(100);c__Saccharomycetes(100);o__Saccharomycetales(100);f__Metschnikowiaceae(100);g__Metschnikowia(100);s__Metschnikowia_pulcherrima(100); | Metschnikowia | 13 | Saprotroph | \|Undefined Saprotroph\| | Yeast | NULL | Probable | As nectar-tap_saprotroph (Põlme et al. 2020); Decay sugar-rich substrates (Põlme et al. 2020) | Põlme S, et al. 2020. Fungal Diversity 105:1-16 ((https://doi.org/10.1007/s13225-020-00466-2)) |
| Otu26795 | 1 | k__Fungi(100);p__Ascomycota(100);c__Saccharomycetes(100);o__Saccharomycetales(100);f__Metschnikowiaceae(100);g__Metschnikowia(100);s__Metschnikowia_pulcherrima(100); | Metschnikowia | 13 | Saprotroph | \|Undefined Saprotroph\| | Yeast | NULL | Probable | As nectar-tap_saprotroph (Põlme et al. 2020); Decay sugar-rich substrates (Põlme et al. 2020) | Põlme S, et al. 2020. Fungal Diversity 105:1-16 ((https://doi.org/10.1007/s13225-020-00466-2)) |
| Otu26797 | 1 | k__Fungi(100);p__Ascomycota(100);c__Saccharomycetes(100);o__Saccharomycetales(100);f__Metschnikowiaceae(100);g__Metschnikowia(100);g__Metschnikowia_unclassified(100); | Metschnikowia | 13 | Saprotroph | \|Undefined Saprotroph\| | Yeast | NULL | Probable | As nectar-tap_saprotroph (Põlme et al. 2020); Decay sugar-rich substrates (Põlme et al. 2020) | Põlme S, et al. 2020. Fungal Diversity 105:1-16 ((https://doi.org/10.1007/s13225-020-00466-2)) |
| Otu26798 | 1 | k__Fungi(100);p__Ascomycota(100);c__Saccharomycetes(100);o__Saccharomycetales(100);f__Metschnikowiaceae(100);g__Metschnikowia(100);s__Metschnikowia_pulcherrima(100); | Metschnikowia | 13 | Saprotroph | \|Undefined Saprotroph\| | Yeast | NULL | Probable | As nectar-tap_saprotroph (Põlme et al. 2020); Decay sugar-rich substrates (Põlme et al. 2020) | Põlme S, et al. 2020. Fungal Diversity 105:1-16 ((https://doi.org/10.1007/s13225-020-00466-2)) |
| Otu26801 | 1 | k__Fungi(100);p__Ascomycota(100);c__Saccharomycetes(100);o__Saccharomycetales(100);f__Metschnikowiaceae(100);g__Metschnikowia(100);s__Metschnikowia_pulcherrima(100); | Metschnikowia | 13 | Saprotroph | \|Undefined Saprotroph\| | Yeast | NULL | Probable | As nectar-tap_saprotroph (Põlme et al. 2020); Decay sugar-rich substrates (Põlme et al. 2020) | Põlme S, et al. 2020. Fungal Diversity 105:1-16 ((https://doi.org/10.1007/s13225-020-00466-2)) |
| Otu26802 | 1 | k__Fungi(100);p__Ascomycota(100);c__Saccharomycetes(100);o__Saccharomycetales(100);f__Metschnikowiaceae(100);g__Metschnikowia(100);s__Metschnikowia_pulcherrima(100); | Metschnikowia | 13 | Saprotroph | \|Undefined Saprotroph\| | Yeast | NULL | Probable | As nectar-tap_saprotroph (Põlme et al. 2020); Decay sugar-rich substrates (Põlme et al. 2020) | Põlme S, et al. 2020. Fungal Diversity 105:1-16 ((https://doi.org/10.1007/s13225-020-00466-2)) |

| | | | | | | | | | | | |
|---|---|---|---|---|---|---|---|---|---|---|---|
| Otu26804 | 1 | k__Fungi(100);p__Ascomycota(100);c__Saccharomycetes(100);o__Saccharomycetales(100);f__Metschnikowiaceae(100);g__Metschnikowia(100);s__Metschnikowia_pulcherrima(100); | Metschnikowia | 13 | Saprotroph | \|Undefined Saprotroph\| | Yeast | NULL | Probable | As nectar-tap_saprotroph (Põlme et al. 2020); Decay sugar-rich substrates (Põlme et al. 2020) | Põlme S, et al. 2020. Fungal Diversity 105:1-16 ((https://doi.org/10.1007/s13225-020-00466-2)) |
| Otu26805 | 1 | k__Fungi(100);p__Ascomycota(100);c__Saccharomycetes(100);o__Saccharomycetales(100);f__Metschnikowiaceae(100);g__Metschnikowia(100);s__Metschnikowia_pulcherrima(100); | Metschnikowia | 13 | Saprotroph | \|Undefined Saprotroph\| | Yeast | NULL | Probable | As nectar-tap_saprotroph (Põlme et al. 2020); Decay sugar-rich substrates (Põlme et al. 2020) | Põlme S, et al. 2020. Fungal Diversity 105:1-16 ((https://doi.org/10.1007/s13225-020-00466-2)) |
| Otu26807 | 1 | k__Fungi(100);p__Ascomycota(100);c__Saccharomycetes(100);o__Saccharomycetales(100);f__Metschnikowiaceae(100);g__Metschnikowia(100);s__Metschnikowia_pulcherrima(100); | Metschnikowia | 13 | Saprotroph | \|Undefined Saprotroph\| | Yeast | NULL | Probable | As nectar-tap_saprotroph (Põlme et al. 2020); Decay sugar-rich substrates (Põlme et al. 2020) | Põlme S, et al. 2020. Fungal Diversity 105:1-16 ((https://doi.org/10.1007/s13225-020-00466-2)) |
| Otu26809 | 1 | k__Fungi(100);p__Ascomycota(100);c__Saccharomycetes(100);o__Saccharomycetales(100);f__Metschnikowiaceae(100);g__Metschnikowia(100);g__Metschnikowia_unclassified(100); | Metschnikowia | 13 | Saprotroph | \|Undefined Saprotroph\| | Yeast | NULL | Probable | As nectar-tap_saprotroph (Põlme et al. 2020); Decay sugar-rich substrates (Põlme et al. 2020) | Põlme S, et al. 2020. Fungal Diversity 105:1-16 ((https://doi.org/10.1007/s13225-020-00466-2)) |
| Otu26810 | 1 | k__Fungi(100);p__Ascomycota(100);c__Saccharomycetes(100);o__Saccharomycetales(100);f__Metschnikowiaceae(100);g__Metschnikowia(100);s__Metschnikowia_pulcherrima(100); | Metschnikowia | 13 | Saprotroph | \|Undefined Saprotroph\| | Yeast | NULL | Probable | As nectar-tap_saprotroph (Põlme et al. 2020); Decay sugar-rich substrates (Põlme et al. 2020) | Põlme S, et al. 2020. Fungal Diversity 105:1-16 ((https://doi.org/10.1007/s13225-020-00466-2)) |
| Otu26811 | 1 | k__Fungi(100);p__Ascomycota(100);c__Saccharomycetes(100);o__Saccharomycetales(100);f__Metschnikowiaceae(100);g__Metschnikowia(100);s__Metschnikowia_pulcherrima(100); | Metschnikowia | 13 | Saprotroph | \|Undefined Saprotroph\| | Yeast | NULL | Probable | As nectar-tap_saprotroph (Põlme et al. 2020); Decay sugar-rich substrates (Põlme et al. 2020) | Põlme S, et al. 2020. Fungal Diversity 105:1-16 ((https://doi.org/10.1007/s13225-020-00466-2)) |
| Otu26812 | 1 | k__Fungi(100);p__Ascomycota(100);c__Dothideomycetes(100);o__Venturiales(100);f__Sympoventuriaceae(100);g__unclassified_Sympoventuriaceae(100);s__Sympoventuriaceae_sp(100); | Sympoventuriaceae | 9 | Saprotroph | Undefined Saprotroph | Microfungus | NULL | Probable | Sympoventuriaceae is represented by Sympoventuria, and can be distinguished from species of Venturiales by its saprobic life style (Zhang et al. 2011) | Zhang Y, et al. 2011 Fungal Diversity 51:249-277 ((https://doi.org/10.1007/s13225-011-0141-x)) |
| Otu26814 | 1 | k__Fungi(100);p__Ascomycota(100);c__Dothideomycetes(100);o__Capnodiales(100);f__Mycosphaerellaceae(100);g__Pseudocercospora(100);g__Pseudocercospora_unclassified(100); | Pseudocercospora | 13 | Pathotroph | \|Plant Pathogen\| | Microfungus | NULL | Probable | NULL | Tedersoo L, et al. 2014. Science 346:e1256688 ((https://doi.org/10.1126/science.1256688)); Põlme S, et al. 2020. Fungal Diversity 105:1-16 ((https://doi.org/10.1007/s13225-020-00466-2)) |
| Otu26815 | 1 | k__Fungi(100);p__Ascomycota(100);c__Saccharomycetes(100);o__Saccharomycetales(100);f__Saccharomycetales_family_Incertae_sedis(100);g__Candida(100);g__Candida_unclassified(100); | Candida | 13 | Pathotroph-Saprotroph-Symbiotroph | Animal Pathogen-Endophyte-Endosymbiont-Epiphyte-\|Undefined Saprotroph\| | Dimorphic Yeast-Yeast | NULL | Probable | As nectar-tap_saprotroph (Põlme et al. 2020); Decay sugar-rich substrates (Põlme et al. 2020); This asexual genus is polyphyletic and contain species that function very differently depend on what they are. Caution should be taken when interpreting guilds for Candida species. | Manolakaki D, et al. 2010. Virulence. 1: 367-375 ((https://doi.org/10.4161/viru.1.5.12796)); Mekbib SB, et al. 2011. Tropical Plant Pathology 36:233-240 ((https://doi.org/10.1590/S1982-56762011000400004)); Põlme S, et al. 2020. Fungal Diversity 105:1-16 ((https://doi.org/10.1007/s13225-020-00466-2)) |
| Otu26817 | 1 | k__Fungi(100);p__Ascomycota(100);c__Saccharomycetes(100);o__Saccharomycetales(100);f__Metschnikowiaceae(100);g__Metschnikowia(100);s__Metschnikowia_pulcherrima(100); | Metschnikowia | 13 | Saprotroph | \|Undefined Saprotroph\| | Yeast | NULL | Probable | As nectar-tap_saprotroph (Põlme et al. 2020); Decay sugar-rich substrates (Põlme et al. 2020) | Põlme S, et al. 2020. Fungal Diversity 105:1-16 ((https://doi.org/10.1007/s13225-020-00466-2)) |
| Otu26818 | 1 | k__Fungi(100);p__Ascomycota(100);c__Saccharomycetes(100);o__Saccharomycetales(100);f__Metschnikowiaceae(100);g__Metschnikowia(100);s__Metschnikowia_pulcherrima(100); | Metschnikowia | 13 | Saprotroph | \|Undefined Saprotroph\| | Yeast | NULL | Probable | As nectar-tap_saprotroph (Põlme et al. 2020); Decay sugar-rich substrates (Põlme et al. 2020) | Põlme S, et al. 2020. Fungal Diversity 105:1-16 ((https://doi.org/10.1007/s13225-020-00466-2)) |
| Otu26819 | 1 | k__Fungi(100);p__Ascomycota(100);c__Saccharomycetes(100);o__Saccharomycetales(100);f__Metschnikowiaceae(100);g__Metschnikowia(100);g__Metschnikowia_unclassified(100); | Metschnikowia | 13 | Saprotroph | \|Undefined Saprotroph\| | Yeast | NULL | Probable | As nectar-tap_saprotroph (Põlme et al. 2020); Decay sugar-rich substrates (Põlme et al. 2020) | Põlme S, et al. 2020. Fungal Diversity 105:1-16 ((https://doi.org/10.1007/s13225-020-00466-2)) |
| Otu26822 | 1 | k__Fungi(100);p__Ascomycota(100);c__Saccharomycetes(100);o__Saccharomycetales(100);f__Metschnikowiaceae(100);g__Metschnikowia(100);g__Metschnikowia_unclassified(100); | Metschnikowia | 13 | Saprotroph | \|Undefined Saprotroph\| | Yeast | NULL | Probable | As nectar-tap_saprotroph (Põlme et al. 2020); Decay sugar-rich substrates (Põlme et al. 2020) | Põlme S, et al. 2020. Fungal Diversity 105:1-16 ((https://doi.org/10.1007/s13225-020-00466-2)) |
| Otu26823 | 1 | k__Fungi(100);p__Ascomycota(100);c__Saccharomycetes(100);o__Saccharomycetales(100);f__Saccharomycetaceae(100);g__Kazachstania(100);s__Kazachstania_intestinalis(100); | Kazachstania | 13 | Saprotroph | \|Undefined Saprotroph\| | Yeast | NULL | Probable | As nectar-tap_saprotroph (Põlme et al. 2020); Decay sugar-rich substrates (Põlme et al. 2020) | Põlme S, et al. 2020. Fungal Diversity 105:1-16 ((https://doi.org/10.1007/s13225-020-00466-2)) |
| Otu26824 | 1 | k__Fungi(100);p__Ascomycota(100);c__Saccharomycetes(100);o__Saccharomycetales(100);f__Metschnikowiaceae(100);g__Metschnikowia(100);s__Metschnikowia_pulcherrima(100); | Metschnikowia | 13 | Saprotroph | \|Undefined Saprotroph\| | Yeast | NULL | Probable | As nectar-tap_saprotroph (Põlme et al. 2020); Decay sugar-rich substrates (Põlme et al. 2020) | Põlme S, et al. 2020. Fungal Diversity 105:1-16 ((https://doi.org/10.1007/s13225-020-00466-2)) |
| Otu26825 | 1 | k__Fungi(100);p__Ascomycota(100);c__Saccharomycetes(100);o__Saccharomycetales(100);o__Saccharomycetales_unclassified(100);o__Saccharomycetales_unclassified(100);o__Saccharomycetales_unclassified(100); | Saccharomycetales | 7 | Saprotroph | Undefined Saprotroph | Yeast | NULL | Possible | NULL | Sterkenburg E, et al. 2015. New Phytologist 207:1145-1158 ((https://doi.org/10.1111/nph.13426)) |
| Otu26828 | 1 | k__Fungi(100);p__Ascomycota(100);c__Saccharomycetes(100);o__Saccharomycetales(100);f__Metschnikowiaceae(100);g__Metschnikowia(100);s__Metschnikowia_pulcherrima(100); | Metschnikowia | 13 | Saprotroph | \|Undefined Saprotroph\| | Yeast | NULL | Probable | As nectar-tap_saprotroph (Põlme et al. 2020); Decay sugar-rich substrates (Põlme et al. 2020) | Põlme S, et al. 2020. Fungal Diversity 105:1-16 ((https://doi.org/10.1007/s13225-020-00466-2)) |
| Otu26829 | 1 | k__Fungi(100);p__Ascomycota(100);c__Saccharomycetes(100);o__Saccharomycetales(100);f__Metschnikowiaceae(100);g__Metschnikowia(100);s__Metschnikowia_pulcherrima(100); | Metschnikowia | 13 | Saprotroph | \|Undefined Saprotroph\| | Yeast | NULL | Probable | As nectar-tap_saprotroph (Põlme et al. 2020); Decay sugar-rich substrates (Põlme et al. 2020) | Põlme S, et al. 2020. Fungal Diversity 105:1-16 ((https://doi.org/10.1007/s13225-020-00466-2)) |
| Otu26830 | 1 | k__Fungi(100);p__Ascomycota(100);c__Saccharomycetes(100);o__Saccharomycetales(100);f__Metschnikowiaceae(100);g__Metschnikowia(100);s__Metschnikowia_pulcherrima(100); | Metschnikowia | 13 | Saprotroph | \|Undefined Saprotroph\| | Yeast | NULL | Probable | As nectar-tap_saprotroph (Põlme et al. 2020); Decay sugar-rich substrates (Põlme et al. 2020) | Põlme S, et al. 2020. Fungal Diversity 105:1-16 ((https://doi.org/10.1007/s13225-020-00466-2)) |
| Otu26831 | 1 | k__Fungi(100);p__Ascomycota(100);c__Saccharomycetes(100);o__Saccharomycetales(100);f__Metschnikowiaceae(100);g__Metschnikowia(100);s__Metschnikowia_pulcherrima(100); | Metschnikowia | 13 | Saprotroph | \|Undefined Saprotroph\| | Yeast | NULL | Probable | As nectar-tap_saprotroph (Põlme et al. 2020); Decay sugar-rich substrates (Põlme et al. 2020) | Põlme S, et al. 2020. Fungal Diversity 105:1-16 ((https://doi.org/10.1007/s13225-020-00466-2)) |
| Otu26832 | 1 | k__Fungi(100);p__Ascomycota(100);c__Saccharomycetes(100);o__Saccharomycetales(100);f__Metschnikowiaceae(100);g__Metschnikowia(100);s__Metschnikowia_pulcherrima(100); | Metschnikowia | 13 | Saprotroph | \|Undefined Saprotroph\| | Yeast | NULL | Probable | As nectar-tap_saprotroph (Põlme et al. 2020); Decay sugar-rich substrates (Põlme et al. 2020) | Põlme S, et al. 2020. Fungal Diversity 105:1-16 ((https://doi.org/10.1007/s13225-020-00466-2)) |
| Otu26834 | 1 | k__Fungi(100);p__Ascomycota(100);c__Saccharomycetes(100);o__Saccharomycetales(100);f__Metschnikowiaceae(100);g__Metschnikowia(100);s__Metschnikowia_pulcherrima(100); | Metschnikowia | 13 | Saprotroph | \|Undefined Saprotroph\| | Yeast | NULL | Probable | As nectar-tap_saprotroph (Põlme et al. 2020); Decay sugar-rich substrates (Põlme et al. 2020) | Põlme S, et al. 2020. Fungal Diversity 105:1-16 ((https://doi.org/10.1007/s13225-020-00466-2)) |
| Otu26835 | 1 | k__Fungi(100);p__Ascomycota(100);c__Saccharomycetes(100);o__Saccharomycetales(100);f__Metschnikowiaceae(100);g__Metschnikowia(100);s__Metschnikowia_pulcherrima(100); | Metschnikowia | 13 | Saprotroph | \|Undefined Saprotroph\| | Yeast | NULL | Probable | As nectar-tap_saprotroph (Põlme et al. 2020); Decay sugar-rich substrates (Põlme et al. 2020) | Põlme S, et al. 2020. Fungal Diversity 105:1-16 ((https://doi.org/10.1007/s13225-020-00466-2)) |
| Otu26836 | 1 | k__Fungi(100);p__Ascomycota(100);c__Saccharomycetes(100);o__Saccharomycetales(100);f__Metschnikowiaceae(100);g__Metschnikowia(100);g__Metschnikowia_unclassified(100); | Metschnikowia | 13 | Saprotroph | \|Undefined Saprotroph\| | Yeast | NULL | Probable | As nectar-tap_saprotroph (Põlme et al. 2020); Decay sugar-rich substrates (Põlme et al. 2020) | Põlme S, et al. 2020. Fungal Diversity 105:1-16 ((https://doi.org/10.1007/s13225-020-00466-2)) |
| Otu26838 | 1 | k__Fungi(100);p__Ascomycota(100);c__Sordariomycetes(100);o__Chaetosphaeriales(100);f__Chaetosphaeriaceae(100);g__Chaetosphaeria(100);s__Chaetosphaeria_sp_olrim419(100); | Chaetosphaeria | 13 | Saprotroph-Symbiotroph | Endophyte-\|Plant Saprotroph\|-Wood Saprotroph | Microfungus | NULL | Probable | As foliar_endophyte (Põlme et al. 2020); As litter_saprotroph (Põlme et al. 2020); Decay wood, leaf, fruit, seed (Põlme et al. 2020) | Réblová M, et al. 2006. Mycological Research 110:104-109 ((https://doi.org/10.1016/j.mycres.2005.09.003)); Tedersoo L, et al. 2014. Science 346:e1256688 ((https://doi.org/10.1126/science.1256688)); Põlme S, et al. 2020. Fungal Diversity 105:1-16 ((https://doi.org/10.1007/s13225-020-00466-2)) |
| Otu26839 | 1 | k__Fungi(100);p__Ascomycota(100);c__Saccharomycetes(100);o__Saccharomycetales(100);f__Metschnikowiaceae(100);g__Metschnikowia(100);g__Metschnikowia_unclassified(100); | Metschnikowia | 13 | Saprotroph | \|Undefined Saprotroph\| | Yeast | NULL | Probable | As nectar-tap_saprotroph (Põlme et al. 2020); Decay sugar-rich substrates (Põlme et al. 2020) | Põlme S, et al. 2020. Fungal Diversity 105:1-16 ((https://doi.org/10.1007/s13225-020-00466-2)) |
| Otu26840 | 1 | k__Fungi(100);p__Ascomycota(100);c__Saccharomycetes(100);o__Saccharomycetales(100);f__Metschnikowiaceae(100);g__Metschnikowia(100);s__Metschnikowia_pulcherrima(100); | Metschnikowia | 13 | Saprotroph | \|Undefined Saprotroph\| | Yeast | NULL | Probable | As nectar-tap_saprotroph (Põlme et al. 2020); Decay sugar-rich substrates (Põlme et al. 2020) | Põlme S, et al. 2020. Fungal Diversity 105:1-16 ((https://doi.org/10.1007/s13225-020-00466-2)) |
| Otu26859 | 1 | k__Fungi(100);p__Ascomycota(100);c__Saccharomycetes(100);o__Saccharomycetales(100);f__Saccharomycetales_family_Incertae_sedis(100);g__Candida(100);g__Candida_unclassified(100); | Candida | 13 | Pathotroph-Saprotroph-Symbiotroph | Animal Pathogen-Endophyte-Endosymbiont-Epiphyte-\|Undefined Saprotroph\| | Dimorphic Yeast-Yeast | NULL | Probable | As nectar-tap_saprotroph (Põlme et al. 2020); Decay sugar-rich substrates (Põlme et al. 2020); This asexual genus is polyphyletic and contain species that function very differently depend on what they are. Caution should be taken when interpreting guilds for Candida species. | Manolakaki D, et al. 2010. Virulence. 1: 367-375 ((https://doi.org/10.4161/viru.1.5.12796)); Mekbib SB, et al. 2011. Tropical Plant Pathology 36:233-240 ((https://doi.org/10.1590/S1982-56762011000400004)); Põlme S, et al. 2020. Fungal Diversity 105:1-16 ((https://doi.org/10.1007/s13225-020-00466-2)) |
| Otu26864 | 1 | k__Fungi(100);p__Ascomycota(100);c__Saccharomycetes(100);o__Saccharomycetales(100);f__Metschnikowiaceae(100);g__Metschnikowia(100);s__Metschnikowia_pulcherrima(100); | Metschnikowia | 13 | Saprotroph | \|Undefined Saprotroph\| | Yeast | NULL | Probable | As nectar-tap_saprotroph (Põlme et al. 2020); Decay sugar-rich substrates (Põlme et al. 2020) | Põlme S, et al. 2020. Fungal Diversity 105:1-16 ((https://doi.org/10.1007/s13225-020-00466-2)) |
| Otu26865 | 1 | k__Fungi(100);p__Ascomycota(100);c__Saccharomycetes(100);o__Saccharomycetales(100);f__Metschnikowiaceae(100);g__Metschnikowia(100);s__Metschnikowia_pulcherrima(100); | Metschnikowia | 13 | Saprotroph | \|Undefined Saprotroph\| | Yeast | NULL | Probable | As nectar-tap_saprotroph (Põlme et al. 2020); Decay sugar-rich substrates (Põlme et al. 2020) | Põlme S, et al. 2020. Fungal Diversity 105:1-16 ((https://doi.org/10.1007/s13225-020-00466-2)) |
| Otu26866 | 1 | k__Fungi(100);p__Ascomycota(100);c__Saccharomycetes(100);o__Saccharomycetales(100);f__Metschnikowiaceae(100);g__Metschnikowia(100);s__Metschnikowia_pulcherrima(100); | Metschnikowia | 13 | Saprotroph | \|Undefined Saprotroph\| | Yeast | NULL | Probable | As nectar-tap_saprotroph (Põlme et al. 2020); Decay sugar-rich substrates (Põlme et al. 2020) | Põlme S, et al. 2020. Fungal Diversity 105:1-16 ((https://doi.org/10.1007/s13225-020-00466-2)) |
| Otu26873 | 1 | k__Fungi(100);p__Ascomycota(100);c__Saccharomycetes(100);o__Saccharomycetales(100);f__Metschnikowiaceae(100);g__Metschnikowia(100);g__Metschnikowia_unclassified(100); | Metschnikowia | 13 | Saprotroph | \|Undefined Saprotroph\| | Yeast | NULL | Probable | As nectar-tap_saprotroph (Põlme et al. 2020); Decay sugar-rich substrates (Põlme et al. 2020) | Põlme S, et al. 2020. Fungal Diversity 105:1-16 ((https://doi.org/10.1007/s13225-020-00466-2)) |
| Otu26874 | 1 | k__Fungi(100);p__Ascomycota(100);c__Saccharomycetes(100);o__Saccharomycetales(100);f__Metschnikowiaceae(100);g__Metschnikowia(100);g__Metschnikowia_unclassified(100); | Metschnikowia | 13 | Saprotroph | \|Undefined Saprotroph\| | Yeast | NULL | Probable | As nectar-tap_saprotroph (Põlme et al. 2020); Decay sugar-rich substrates (Põlme et al. 2020) | Põlme S, et al. 2020. Fungal Diversity 105:1-16 ((https://doi.org/10.1007/s13225-020-00466-2)) |
| Otu26875 | 1 | k__Fungi(100);p__Ascomycota(100);c__Saccharomycetes(100);o__Saccharomycetales(100);f__Metschnikowiaceae(100);g__Metschnikowia(100);s__Metschnikowia_pulcherrima(100); | Metschnikowia | 13 | Saprotroph | \|Undefined Saprotroph\| | Yeast | NULL | Probable | As nectar-tap_saprotroph (Põlme et al. 2020); Decay sugar-rich substrates (Põlme et al. 2020) | Põlme S, et al. 2020. Fungal Diversity 105:1-16 ((https://doi.org/10.1007/s13225-020-00466-2)) |
| Otu26876 | 1 | k__Fungi(100);p__Ascomycota(100);c__Saccharomycetes(100);o__Saccharomycetales(100);o__Saccharomycetales_unclassified(100);o__Saccharomycetales_unclassified(100);o__Saccharomycetales_unclassified(100); | Saccharomycetales | 7 | Saprotroph | Undefined Saprotroph | Yeast | NULL | Possible | NULL | Sterkenburg E, et al. 2015. New Phytologist 207:1145-1158 ((https://doi.org/10.1111/nph.13426)) |
| Otu26882 | 1 | k__Fungi(100);p__Ascomycota(100);c__Saccharomycetes(100);o__Saccharomycetales(100);f__Metschnikowiaceae(100);g__Metschnikowia(100);s__Metschnikowia_pulcherrima(100); | Metschnikowia | 13 | Saprotroph | \|Undefined Saprotroph\| | Yeast | NULL | Probable | As nectar-tap_saprotroph (Põlme et al. 2020); Decay sugar-rich substrates (Põlme et al. 2020) | Põlme S, et al. 2020. Fungal Diversity 105:1-16 ((https://doi.org/10.1007/s13225-020-00466-2)) |
| Otu26884 | 1 | k__Fungi(100);p__Ascomycota(100);c__Saccharomycetes(100);o__Saccharomycetales(100);f__Metschnikowiaceae(100);g__Metschnikowia(100);s__Metschnikowia_pulcherrima(100); | Metschnikowia | 13 | Saprotroph | \|Undefined Saprotroph\| | Yeast | NULL | Probable | As nectar-tap_saprotroph (Põlme et al. 2020); Decay sugar-rich substrates (Põlme et al. 2020) | Põlme S, et al. 2020. Fungal Diversity 105:1-16 ((https://doi.org/10.1007/s13225-020-00466-2)) |
| Otu26904 | 1 | k__Fungi(100);p__Ascomycota(100);c__Saccharomycetes(100);o__Saccharomycetales(100);f__Metschnikowiaceae(100);g__Metschnikowia(100);g__Metschnikowia_unclassified(100); | Metschnikowia | 13 | Saprotroph | \|Undefined Saprotroph\| | Yeast | NULL | Probable | As nectar-tap_saprotroph (Põlme et al. 2020); Decay sugar-rich substrates (Põlme et al. 2020) | Põlme S, et al. 2020. Fungal Diversity 105:1-16 ((https://doi.org/10.1007/s13225-020-00466-2)) |
| Otu26905 | 1 | k__Fungi(100);p__Ascomycota(100);c__Saccharomycetes(100);o__Saccharomycetales(100);f__Metschnikowiaceae(100);g__Metschnikowia(100);s__Metschnikowia_pulcherrima(100); | Metschnikowia | 13 | Saprotroph | \|Undefined Saprotroph\| | Yeast | NULL | Probable | As nectar-tap_saprotroph (Põlme et al. 2020); Decay sugar-rich substrates (Põlme et al. 2020) | Põlme S, et al. 2020. Fungal Diversity 105:1-16 ((https://doi.org/10.1007/s13225-020-00466-2)) |
| Otu26906 | 1 | k__Fungi(100);p__Ascomycota(100);c__Sordariomycetes(100);o__Hypocreales(100);f__Nectriaceae(100);g__Fusarium(100);s__Fusarium_tricinctum(100); | Fusarium | 13 | Pathotroph-Saprotroph-Symbiotroph | Animal Pathogen-Endophyte-Lichen Parasite-\|Plant Pathogen\|-Plant Saprotroph-Undefined Saprotroph-Wood Saprotroph | Microfungus | Soft Rot | Possible | As foliar_endophyte (Põlme et al. 2020); As litter_saprotroph (Põlme et al. 2020); Causing damping-off disease (Põlme et al. 2020); Decay leaf, fruit, seed, roots, soil, animal material (Põlme et al. 2020); Likely opportunistic human pathogen (Irinyi et al. 2015) | Duncan CG, Eslyn WE. 1966. Mycologia 58:642-664 ((https://doi.org/10.2307/3757045)); Seehann G, et al. 1975. List of Fungi in Soft-Rot Tissue Tests. Institute fur Holzbiologie und Holzschutz, Bundesforschungsanstalt fur Forst - und Holzwirtschaft, Hamburg - Lohbrugge, Germany; Costa IPMW, et al. 2012. Mycotaxon 119:e26 ((http://mycotaxon.com/resources/checklists/costa_v119_checklist.pdf)); Tedersoo L, et al. 2014. Science 346:e1256688 ((https://doi.org/10.1126/science.1256688)); Irinyi L, et al. 2015. Medical Mycology 53:313-337 ((https://doi.org/10.1093/mmy/myv008)); Schroers, et al. 2016. Mycologia 108:806-819 ((https://doi.org/10.3852/15-255)); Põlme S, et al. 2020. Fungal Diversity 105:1-16 ((https://doi.org/10.1007/s13225-020-00466-2)); The American Phytopathological Society Common Names of Plant Diseases ((http://www.apsnet.org/publications/commonnames/Pages/default.aspx)); Lawrey JD, Diederich P. 2016. http://www.lichenicolous.net (accessed 24 May 2017) ((http://www.lichenicolous.net)) |
| Otu26907 | 1 | k__Fungi(100);p__Ascomycota(100);c__Saccharomycetes(100);o__Saccharomycetales(100);f__Metschnikowiaceae(100);g__Metschnikowia(100);s__Metschnikowia_pulcherrima(100); | Metschnikowia | 13 | Saprotroph | \|Undefined Saprotroph\| | Yeast | NULL | Probable | As nectar-tap_saprotroph (Põlme et al. 2020); Decay sugar-rich substrates (Põlme et al. 2020) | Põlme S, et al. 2020. Fungal Diversity 105:1-16 ((https://doi.org/10.1007/s13225-020-00466-2)) |
| Otu26913 | 1 | k__Fungi(100);p__Ascomycota(100);c__Saccharomycetes(100);o__Saccharomycetales(100);f__Metschnikowiaceae(100);g__Metschnikowia(100);g__Metschnikowia_unclassified(100); | Metschnikowia | 13 | Saprotroph | \|Undefined Saprotroph\| | Yeast | NULL | Probable | As nectar-tap_saprotroph (Põlme et al. 2020); Decay sugar-rich substrates (Põlme et al. 2020) | Põlme S, et al. 2020. Fungal Diversity 105:1-16 ((https://doi.org/10.1007/s13225-020-00466-2)) |
| Otu26914 | 1 | k__Fungi(100);p__Ascomycota(100);c__Saccharomycetes(100);o__Saccharomycetales(100);f__Metschnikowiaceae(100);g__Metschnikowia(100);s__Metschnikowia_pulcherrima(100); | Metschnikowia | 13 | Saprotroph | \|Undefined Saprotroph\| | Yeast | NULL | Probable | As nectar-tap_saprotroph (Põlme et al. 2020); Decay sugar-rich substrates (Põlme et al. 2020) | Põlme S, et al. 2020. Fungal Diversity 105:1-16 ((https://doi.org/10.1007/s13225-020-00466-2)) |
| Otu26916 | 1 | k__Fungi(100);p__Ascomycota(100);c__Saccharomycetes(100);o__Saccharomycetales(100);f__Metschnikowiaceae(100);g__Metschnikowia(100);s__Metschnikowia_pulcherrima(100); | Metschnikowia | 13 | Saprotroph | \|Undefined Saprotroph\| | Yeast | NULL | Probable | As nectar-tap_saprotroph (Põlme et al. 2020); Decay sugar-rich substrates (Põlme et al. 2020) | Põlme S, et al. 2020. Fungal Diversity 105:1-16 ((https://doi.org/10.1007/s13225-020-00466-2)) |
| Otu26917 | 1 | k__Fungi(100);p__Ascomycota(100);c__Saccharomycetes(100);o__Saccharomycetales(100);f__Metschnikowiaceae(100);g__Metschnikowia(100);g__Metschnikowia_unclassified(100); | Metschnikowia | 13 | Saprotroph | \|Undefined Saprotroph\| | Yeast | NULL | Probable | As nectar-tap_saprotroph (Põlme et al. 2020); Decay sugar-rich substrates (Põlme et al. 2020) | Põlme S, et al. 2020. Fungal Diversity 105:1-16 ((https://doi.org/10.1007/s13225-020-00466-2)) |
| Otu26918 | 1 | k__Fungi(100);p__Ascomycota(100);c__Saccharomycetes(100);o__Saccharomycetales(100);f__Metschnikowiaceae(100);g__Metschnikowia(100);s__Metschnikowia_pulcherrima(100); | Metschnikowia | 13 | Saprotroph | \|Undefined Saprotroph\| | Yeast | NULL | Probable | As nectar-tap_saprotroph (Põlme et al. 2020); Decay sugar-rich substrates (Põlme et al. 2020) | Põlme S, et al. 2020. Fungal Diversity 105:1-16 ((https://doi.org/10.1007/s13225-020-00466-2)) |
| Otu26921 | 1 | k__Fungi(100);p__Ascomycota(100);c__Saccharomycetes(100);o__Saccharomycetales(100);f__Metschnikowiaceae(100);g__Metschnikowia(100);g__Metschnikowia_unclassified(100); | Metschnikowia | 13 | Saprotroph | \|Undefined Saprotroph\| | Yeast | NULL | Probable | As nectar-tap_saprotroph (Põlme et al. 2020); Decay sugar-rich substrates (Põlme et al. 2020) | Põlme S, et al. 2020. Fungal Diversity 105:1-16 ((https://doi.org/10.1007/s13225-020-00466-2)) |
| Otu26924 | 1 | k__Fungi(100);p__Ascomycota(100);c__Saccharomycetes(100);o__Saccharomycetales(100);f__Metschnikowiaceae(100);g__Metschnikowia(100);g__Metschnikowia_unclassified(100); | Metschnikowia | 13 | Saprotroph | \|Undefined Saprotroph\| | Yeast | NULL | Probable | As nectar-tap_saprotroph (Põlme et al. 2020); Decay sugar-rich substrates (Põlme et al. 2020) | Põlme S, et al. 2020. Fungal Diversity 105:1-16 ((https://doi.org/10.1007/s13225-020-00466-2)) |
| Otu26929 | 1 | k__Fungi(100);p__Ascomycota(100);c__Saccharomycetes(100);o__Saccharomycetales(100);f__Metschnikowiaceae(100);g__Metschnikowia(100);s__Metschnikowia_pulcherrima(100); | Metschnikowia | 13 | Saprotroph | \|Undefined Saprotroph\| | Yeast | NULL | Probable | As nectar-tap_saprotroph (Põlme et al. 2020); Decay sugar-rich substrates (Põlme et al. 2020) | Põlme S, et al. 2020. Fungal Diversity 105:1-16 ((https://doi.org/10.1007/s13225-020-00466-2)) |
| Otu26930 | 1 | k__Fungi(100);p__Ascomycota(100);c__Saccharomycetes(100);o__Saccharomycetales(100);f__Metschnikowiaceae(100);g__Metschnikowia(100);s__Metschnikowia_pulcherrima(100); | Metschnikowia | 13 | Saprotroph | \|Undefined Saprotroph\| | Yeast | NULL | Probable | As nectar-tap_saprotroph (Põlme et al. 2020); Decay sugar-rich substrates (Põlme et al. 2020) | Põlme S, et al. 2020. Fungal Diversity 105:1-16 ((https://doi.org/10.1007/s13225-020-00466-2)) |
| Otu26931 | 1 | k__Fungi(100);p__Ascomycota(100);c__Saccharomycetes(100);o__Saccharomycetales(100);f__Metschnikowiaceae(100);g__Metschnikowia(100);g__Metschnikowia_unclassified(100); | Metschnikowia | 13 | Saprotroph | \|Undefined Saprotroph\| | Yeast | NULL | Probable | As nectar-tap_saprotroph (Põlme et al. 2020); Decay sugar-rich substrates (Põlme et al. 2020) | Põlme S, et al. 2020. Fungal Diversity 105:1-16 ((https://doi.org/10.1007/s13225-020-00466-2)) |
| Otu26932 | 1 | k__Fungi(100);p__Ascomycota(100);c__Saccharomycetes(100);o__Saccharomycetales(100);f__Metschnikowiaceae(100);g__Metschnikowia(100);s__Metschnikowia_pulcherrima(100); | Metschnikowia | 13 | Saprotroph | \|Undefined Saprotroph\| | Yeast | NULL | Probable | As nectar-tap_saprotroph (Põlme et al. 2020); Decay sugar-rich substrates (Põlme et al. 2020) | Põlme S, et al. 2020. Fungal Diversity 105:1-16 ((https://doi.org/10.1007/s13225-020-00466-2)) |
| Otu26933 | 1 | k__Fungi(100);p__Ascomycota(100);c__Saccharomycetes(100);o__Saccharomycetales(100);f__Metschnikowiaceae(100);g__Metschnikowia(100);s__Metschnikowia_pulcherrima(100); | Metschnikowia | 13 | Saprotroph | \|Undefined Saprotroph\| | Yeast | NULL | Probable | As nectar-tap_saprotroph (Põlme et al. 2020); Decay sugar-rich substrates (Põlme et al. 2020) | Põlme S, et al. 2020. Fungal Diversity 105:1-16 ((https://doi.org/10.1007/s13225-020-00466-2)) |
| Otu26935 | 1 | k__Fungi(100);p__Ascomycota(100);c__Saccharomycetes(100);o__Saccharomycetales(100);f__Metschnikowiaceae(100);g__Metschnikowia(100);g__Metschnikowia_unclassified(100); | Metschnikowia | 13 | Saprotroph | \|Undefined Saprotroph\| | Yeast | NULL | Probable | As nectar-tap_saprotroph (Põlme et al. 2020); Decay sugar-rich substrates (Põlme et al. 2020) | Põlme S, et al. 2020. Fungal Diversity 105:1-16 ((https://doi.org/10.1007/s13225-020-00466-2)) |
| Otu26936 | 1 | k__Fungi(100);p__Ascomycota(100);c__Saccharomycetes(100);o__Saccharomycetales(100);f__Metschnikowiaceae(100);g__Metschnikowia(100);s__Metschnikowia_pulcherrima(100); | Metschnikowia | 13 | Saprotroph | \|Undefined Saprotroph\| | Yeast | NULL | Probable | As nectar-tap_saprotroph (Põlme et al. 2020); Decay sugar-rich substrates (Põlme et al. 2020) | Põlme S, et al. 2020. Fungal Diversity 105:1-16 ((https://doi.org/10.1007/s13225-020-00466-2)) |
| Otu26937 | 1 | k__Fungi(100);p__Ascomycota(100);c__Saccharomycetes(100);o__Saccharomycetales(100);f__Metschnikowiaceae(100);g__Metschnikowia(100);s__Metschnikowia_pulcherrima(100); | Metschnikowia | 13 | Saprotroph | \|Undefined Saprotroph\| | Yeast | NULL | Probable | As nectar-tap_saprotroph (Põlme et al. 2020); Decay sugar-rich substrates (Põlme et al. 2020) | Põlme S, et al. 2020. Fungal Diversity 105:1-16 ((https://doi.org/10.1007/s13225-020-00466-2)) |
| Otu26941 | 1 | k__Fungi(100);p__Ascomycota(100);c__Saccharomycetes(100);o__Saccharomycetales(100);f__Metschnikowiaceae(100);g__Metschnikowia(100);s__Metschnikowia_pulcherrima(100); | Metschnikowia | 13 | Saprotroph | \|Undefined Saprotroph\| | Yeast | NULL | Probable | As nectar-tap_saprotroph (Põlme et al. 2020); Decay sugar-rich substrates (Põlme et al. 2020) | Põlme S, et al. 2020. Fungal Diversity 105:1-16 ((https://doi.org/10.1007/s13225-020-00466-2)) |
| Otu26942 | 1 | k__Fungi(100);p__Ascomycota(100);c__Saccharomycetes(100);o__Saccharomycetales(100);f__Metschnikowiaceae(100);g__Metschnikowia(100);s__Metschnikowia_pulcherrima(100); | Metschnikowia | 13 | Saprotroph | \|Undefined Saprotroph\| | Yeast | NULL | Probable | As nectar-tap_saprotroph (Põlme et al. 2020); Decay sugar-rich substrates (Põlme et al. 2020) | Põlme S, et al. 2020. Fungal Diversity 105:1-16 ((https://doi.org/10.1007/s13225-020-00466-2)) |
| Otu26944 | 1 | k__Fungi(100);p__Ascomycota(100);c__Saccharomycetes(100);o__Saccharomycetales(100);f__Metschnikowiaceae(100);g__Metschnikowia(100);s__Metschnikowia_pulcherrima(100); | Metschnikowia | 13 | Saprotroph | \|Undefined Saprotroph\| | Yeast | NULL | Probable | As nectar-tap_saprotroph (Põlme et al. 2020); Decay sugar-rich substrates (Põlme et al. 2020) | Põlme S, et al. 2020. Fungal Diversity 105:1-16 ((https://doi.org/10.1007/s13225-020-00466-2)) |
| Otu26948 | 1 | k__Fungi(100);p__Ascomycota(100);c__Saccharomycetes(100);o__Saccharomycetales(100);f__Metschnikowiaceae(100);g__Metschnikowia(100);s__Metschnikowia_pulcherrima(100); | Metschnikowia | 13 | Saprotroph | \|Undefined Saprotroph\| | Yeast | NULL | Probable | As nectar-tap_saprotroph (Põlme et al. 2020); Decay sugar-rich substrates (Põlme et al. 2020) | Põlme S, et al. 2020. Fungal Diversity 105:1-16 ((https://doi.org/10.1007/s13225-020-00466-2)) |
| Otu26949 | 1 | k__Fungi(100);p__Ascomycota(100);c__Saccharomycetes(100);o__Saccharomycetales(100);f__Metschnikowiaceae(100);g__Metschnikowia(100);g__Metschnikowia_unclassified(100); | Metschnikowia | 13 | Saprotroph | \|Undefined Saprotroph\| | Yeast | NULL | Probable | As nectar-tap_saprotroph (Põlme et al. 2020); Decay sugar-rich substrates (Põlme et al. 2020) | Põlme S, et al. 2020. Fungal Diversity 105:1-16 ((https://doi.org/10.1007/s13225-020-00466-2)) |
| Otu26965 | 1 | k__Fungi(100);p__Ascomycota(100);c__Saccharomycetes(100);o__Saccharomycetales(100);f__Metschnikowiaceae(100);g__Metschnikowia(100);s__Metschnikowia_pulcherrima(100); | Metschnikowia | 13 | Saprotroph | \|Undefined Saprotroph\| | Yeast | NULL | Probable | As nectar-tap_saprotroph (Põlme et al. 2020); Decay sugar-rich substrates (Põlme et al. 2020) | Põlme S, et al. 2020. Fungal Diversity 105:1-16 ((https://doi.org/10.1007/s13225-020-00466-2)) |
| Otu26966 | 1 | k__Fungi(100);p__Ascomycota(100);c__Saccharomycetes(100);o__Saccharomycetales(100);f__Metschnikowiaceae(100);g__Metschnikowia(100);s__Metschnikowia_pulcherrima(100); | Metschnikowia | 13 | Saprotroph | \|Undefined Saprotroph\| | Yeast | NULL | Probable | As nectar-tap_saprotroph (Põlme et al. 2020); Decay sugar-rich substrates (Põlme et al. 2020) | Põlme S, et al. 2020. Fungal Diversity 105:1-16 ((https://doi.org/10.1007/s13225-020-00466-2)) |
| Otu26974 | 1 | k__Fungi(100);p__Ascomycota(100);c__Saccharomycetes(100);o__Saccharomycetales(100);f__Metschnikowiaceae(100);g__Metschnikowia(100);s__Metschnikowia_pulcherrima(100); | Metschnikowia | 13 | Saprotroph | \|Undefined Saprotroph\| | Yeast | NULL | Probable | As nectar-tap_saprotroph (Põlme et al. 2020); Decay sugar-rich substrates (Põlme et al. 2020) | Põlme S, et al. 2020. Fungal Diversity 105:1-16 ((https://doi.org/10.1007/s13225-020-00466-2)) |
| Otu26975 | 1 | k__Fungi(100);p__Ascomycota(100);c__Saccharomycetes(100);o__Saccharomycetales(100);f__Metschnikowiaceae(100);g__Metschnikowia(100);s__Metschnikowia_pulcherrima(100); | Metschnikowia | 13 | Saprotroph | \|Undefined Saprotroph\| | Yeast | NULL | Probable | As nectar-tap_saprotroph (Põlme et al. 2020); Decay sugar-rich substrates (Põlme et al. 2020) | Põlme S, et al. 2020. Fungal Diversity 105:1-16 ((https://doi.org/10.1007/s13225-020-00466-2)) |
| Otu26976 | 1 | k__Fungi(100);p__Ascomycota(100);c__Saccharomycetes(100);o__Saccharomycetales(100);f__Metschnikowiaceae(100);g__Metschnikowia(100);s__Metschnikowia_pulcherrima(100); | Metschnikowia | 13 | Saprotroph | \|Undefined Saprotroph\| | Yeast | NULL | Probable | As nectar-tap_saprotroph (Põlme et al. 2020); Decay sugar-rich substrates (Põlme et al. 2020) | Põlme S, et al. 2020. Fungal Diversity 105:1-16 ((https://doi.org/10.1007/s13225-020-00466-2)) |
| Otu26977 | 1 | k__Fungi(100);p__Ascomycota(100);c__Saccharomycetes(100);o__Saccharomycetales(100);f__Metschnikowiaceae(100);g__Metschnikowia(100);s__Metschnikowia_pulcherrima(100); | Metschnikowia | 13 | Saprotroph | \|Undefined Saprotroph\| | Yeast | NULL | Probable | As nectar-tap_saprotroph (Põlme et al. 2020); Decay sugar-rich substrates (Põlme et al. 2020) | Põlme S, et al. 2020. Fungal Diversity 105:1-16 ((https://doi.org/10.1007/s13225-020-00466-2)) |
| Otu26978 | 1 | k__Fungi(100);p__Ascomycota(100);c__Saccharomycetes(100);o__Saccharomycetales(100);f__Metschnikowiaceae(100);g__Metschnikowia(100);s__Metschnikowia_pulcherrima(100); | Metschnikowia | 13 | Saprotroph | \|Undefined Saprotroph\| | Yeast | NULL | Probable | As nectar-tap_saprotroph (Põlme et al. 2020); Decay sugar-rich substrates (Põlme et al. 2020) | Põlme S, et al. 2020. Fungal Diversity 105:1-16 ((https://doi.org/10.1007/s13225-020-00466-2)) |
| Otu26979 | 1 | k__Fungi(100);p__Ascomycota(100);c__Saccharomycetes(100);o__Saccharomycetales(100);f__Metschnikowiaceae(100);g__Metschnikowia(100);s__Metschnikowia_pulcherrima(100); | Metschnikowia | 13 | Saprotroph | \|Undefined Saprotroph\| | Yeast | NULL | Probable | As nectar-tap_saprotroph (Põlme et al. 2020); Decay sugar-rich substrates (Põlme et al. 2020) | Põlme S, et al. 2020. Fungal Diversity 105:1-16 ((https://doi.org/10.1007/s13225-020-00466-2)) |
| Otu26980 | 1 | k__Fungi(100);p__Ascomycota(100);c__Saccharomycetes(100);o__Saccharomycetales(100);f__Metschnikowiaceae(100);g__Metschnikowia(100);s__Metschnikowia_pulcherrima(100); | Metschnikowia | 13 | Saprotroph | \|Undefined Saprotroph\| | Yeast | NULL | Probable | As nectar-tap_saprotroph (Põlme et al. 2020); Decay sugar-rich substrates (Põlme et al. 2020) | Põlme S, et al. 2020. Fungal Diversity 105:1-16 ((https://doi.org/10.1007/s13225-020-00466-2)) |
| Otu26981 | 1 | k__Fungi(100);p__Ascomycota(100);c__Saccharomycetes(100);o__Saccharomycetales(100);f__Metschnikowiaceae(100);g__Metschnikowia(100);g__Metschnikowia_unclassified(100); | Metschnikowia | 13 | Saprotroph | \|Undefined Saprotroph\| | Yeast | NULL | Probable | As nectar-tap_saprotroph (Põlme et al. 2020); Decay sugar-rich substrates (Põlme et al. 2020) | Põlme S, et al. 2020. Fungal Diversity 105:1-16 ((https://doi.org/10.1007/s13225-020-00466-2)) |
| Otu26982 | 1 | k__Fungi(100);p__Ascomycota(100);c__Saccharomycetes(100);o__Saccharomycetales(100);f__Metschnikowiaceae(100);g__Metschnikowia(100);g__Metschnikowia_unclassified(100); | Metschnikowia | 13 | Saprotroph | \|Undefined Saprotroph\| | Yeast | NULL | Probable | As nectar-tap_saprotroph (Põlme et al. 2020); Decay sugar-rich substrates (Põlme et al. 2020) | Põlme S, et al. 2020. Fungal Diversity 105:1-16 ((https://doi.org/10.1007/s13225-020-00466-2)) |
| Otu26983 | 1 | k__Fungi(100);p__Ascomycota(100);c__Saccharomycetes(100);o__Saccharomycetales(100);f__Metschnikowiaceae(100);g__Metschnikowia(100);s__Metschnikowia_pulcherrima(100); | Metschnikowia | 13 | Saprotroph | \|Undefined Saprotroph\| | Yeast | NULL | Probable | As nectar-tap_saprotroph (Põlme et al. 2020); Decay sugar-rich substrates (Põlme et al. 2020) | Põlme S, et al. 2020. Fungal Diversity 105:1-16 ((https://doi.org/10.1007/s13225-020-00466-2)) |
| Otu26984 | 1 | k__Fungi(100);p__Ascomycota(100);c__Saccharomycetes(100);o__Saccharomycetales(100);f__Metschnikowiaceae(100);g__Metschnikowia(100);s__Metschnikowia_pulcherrima(100); | Metschnikowia | 13 | Saprotroph | \|Undefined Saprotroph\| | Yeast | NULL | Probable | As nectar-tap_saprotroph (Põlme et al. 2020); Decay sugar-rich substrates (Põlme et al. 2020) | Põlme S, et al. 2020. Fungal Diversity 105:1-16 ((https://doi.org/10.1007/s13225-020-00466-2)) |
| Otu26985 | 1 | k__Fungi(100);p__Ascomycota(100);c__Saccharomycetes(100);o__Saccharomycetales(100);f__Metschnikowiaceae(100);g__Metschnikowia(100);s__Metschnikowia_pulcherrima(100); | Metschnikowia | 13 | Saprotroph | \|Undefined Saprotroph\| | Yeast | NULL | Probable | As nectar-tap_saprotroph (Põlme et al. 2020); Decay sugar-rich substrates (Põlme et al. 2020) | Põlme S, et al. 2020. Fungal Diversity 105:1-16 ((https://doi.org/10.1007/s13225-020-00466-2)) |
| Otu26986 | 1 | k__Fungi(100);p__Ascomycota(100);c__Saccharomycetes(100);o__Saccharomycetales(100);f__Metschnikowiaceae(100);g__Metschnikowia(100);s__Metschnikowia_pulcherrima(100); | Metschnikowia | 13 | Saprotroph | \|Undefined Saprotroph\| | Yeast | NULL | Probable | As nectar-tap_saprotroph (Põlme et al. 2020); Decay sugar-rich substrates (Põlme et al. 2020) | Põlme S, et al. 2020. Fungal Diversity 105:1-16 ((https://doi.org/10.1007/s13225-020-00466-2)) |
| Otu26994 | 1 | k__Fungi(100);p__Ascomycota(100);c__Saccharomycetes(100);o__Saccharomycetales(100);f__Metschnikowiaceae(100);g__Metschnikowia(100);s__Metschnikowia_pulcherrima(100); | Metschnikowia | 13 | Saprotroph | \|Undefined Saprotroph\| | Yeast | NULL | Probable | As nectar-tap_saprotroph (Põlme et al. 2020); Decay sugar-rich substrates (Põlme et al. 2020) | Põlme S, et al. 2020. Fungal Diversity 105:1-16 ((https://doi.org/10.1007/s13225-020-00466-2)) |
| Otu27001 | 1 | k__Fungi(100);p__Ascomycota(100);c__Saccharomycetes(100);o__Saccharomycetales(100);f__Metschnikowiaceae(100);g__Metschnikowia(100);s__Metschnikowia_pulcherrima(100); | Metschnikowia | 13 | Saprotroph | \|Undefined Saprotroph\| | Yeast | NULL | Probable | As nectar-tap_saprotroph (Põlme et al. 2020); Decay sugar-rich substrates (Põlme et al. 2020) | Põlme S, et al. 2020. Fungal Diversity 105:1-16 ((https://doi.org/10.1007/s13225-020-00466-2)) |
| Otu27002 | 1 | k__Fungi(100);p__Ascomycota(100);c__Saccharomycetes(100);o__Saccharomycetales(100);f__Metschnikowiaceae(100);g__Metschnikowia(100);s__Metschnikowia_pulcherrima(100); | Metschnikowia | 13 | Saprotroph | \|Undefined Saprotroph\| | Yeast | NULL | Probable | As nectar-tap_saprotroph (Põlme et al. 2020); Decay sugar-rich substrates (Põlme et al. 2020) | Põlme S, et al. 2020. Fungal Diversity 105:1-16 ((https://doi.org/10.1007/s13225-020-00466-2)) |
| Otu27003 | 1 | k__Fungi(100);p__Ascomycota(100);c__Saccharomycetes(100);o__Saccharomycetales(100);f__Metschnikowiaceae(100);g__Metschnikowia(100);s__Metschnikowia_pulcherrima(100); | Metschnikowia | 13 | Saprotroph | \|Undefined Saprotroph\| | Yeast | NULL | Probable | As nectar-tap_saprotroph (Põlme et al. 2020); Decay sugar-rich substrates (Põlme et al. 2020) | Põlme S, et al. 2020. Fungal Diversity 105:1-16 ((https://doi.org/10.1007/s13225-020-00466-2)) |
| Otu27004 | 1 | k__Fungi(100);p__Ascomycota(100);c__Saccharomycetes(100);o__Saccharomycetales(100);f__Metschnikowiaceae(100);g__Metschnikowia(100);s__Metschnikowia_pulcherrima(100); | Metschnikowia | 13 | Saprotroph | \|Undefined Saprotroph\| | Yeast | NULL | Probable | As nectar-tap_saprotroph (Põlme et al. 2020); Decay sugar-rich substrates (Põlme et al. 2020) | Põlme S, et al. 2020. Fungal Diversity 105:1-16 ((https://doi.org/10.1007/s13225-020-00466-2)) |
| Otu27005 | 1 | k__Fungi(100);p__Ascomycota(100);c__Saccharomycetes(100);o__Saccharomycetales(100);f__Metschnikowiaceae(100);g__Metschnikowia(100);s__Metschnikowia_pulcherrima(100); | Metschnikowia | 13 | Saprotroph | \|Undefined Saprotroph\| | Yeast | NULL | Probable | As nectar-tap_saprotroph (Põlme et al. 2020); Decay sugar-rich substrates (Põlme et al. 2020) | Põlme S, et al. 2020. Fungal Diversity 105:1-16 ((https://doi.org/10.1007/s13225-020-00466-2)) |
| Otu27006 | 1 | k__Fungi(100);p__Ascomycota(100);c__Saccharomycetes(100);o__Saccharomycetales(100);f__Metschnikowiaceae(100);g__Metschnikowia(100);s__Metschnikowia_pulcherrima(100); | Metschnikowia | 13 | Saprotroph | \|Undefined Saprotroph\| | Yeast | NULL | Probable | As nectar-tap_saprotroph (Põlme et al. 2020); Decay sugar-rich substrates (Põlme et al. 2020) | Põlme S, et al. 2020. Fungal Diversity 105:1-16 ((https://doi.org/10.1007/s13225-020-00466-2)) |
| Otu27007 | 1 | k__Fungi(100);p__Ascomycota(100);c__Saccharomycetes(100);o__Saccharomycetales(100);f__Metschnikowiaceae(100);g__Metschnikowia(100);g__Metschnikowia_unclassified(100); | Metschnikowia | 13 | Saprotroph | \|Undefined Saprotroph\| | Yeast | NULL | Probable | As nectar-tap_saprotroph (Põlme et al. 2020); Decay sugar-rich substrates (Põlme et al. 2020) | Põlme S, et al. 2020. Fungal Diversity 105:1-16 ((https://doi.org/10.1007/s13225-020-00466-2)) |
| Otu27008 | 1 | k__Fungi(100);p__Ascomycota(100);c__Saccharomycetes(100);o__Saccharomycetales(100);f__Metschnikowiaceae(100);g__Metschnikowia(100);s__Metschnikowia_pulcherrima(100); | Metschnikowia | 13 | Saprotroph | \|Undefined Saprotroph\| | Yeast | NULL | Probable | As nectar-tap_saprotroph (Põlme et al. 2020); Decay sugar-rich substrates (Põlme et al. 2020) | Põlme S, et al. 2020. Fungal Diversity 105:1-16 ((https://doi.org/10.1007/s13225-020-00466-2)) |
| Otu27010 | 1 | k__Fungi(100);p__Ascomycota(100);c__Saccharomycetes(100);o__Saccharomycetales(100);f__Metschnikowiaceae(100);g__Metschnikowia(100);s__Metschnikowia_pulcherrima(100); | Metschnikowia | 13 | Saprotroph | \|Undefined Saprotroph\| | Yeast | NULL | Probable | As nectar-tap_saprotroph (Põlme et al. 2020); Decay sugar-rich substrates (Põlme et al. 2020) | Põlme S, et al. 2020. Fungal Diversity 105:1-16 ((https://doi.org/10.1007/s13225-020-00466-2)) |
| Otu27015 | 1 | k__Fungi(100);p__Ascomycota(100);c__Saccharomycetes(100);o__Saccharomycetales(100);f__Metschnikowiaceae(100);g__Metschnikowia(100);s__Metschnikowia_pulcherrima(100); | Metschnikowia | 13 | Saprotroph | \|Undefined Saprotroph\| | Yeast | NULL | Probable | As nectar-tap_saprotroph (Põlme et al. 2020); Decay sugar-rich substrates (Põlme et al. 2020) | Põlme S, et al. 2020. Fungal Diversity 105:1-16 ((https://doi.org/10.1007/s13225-020-00466-2)) |
| Otu27016 | 1 | k__Fungi(100);p__Ascomycota(100);c__Saccharomycetes(100);o__Saccharomycetales(100);f__Metschnikowiaceae(100);g__Metschnikowia(100);g__Metschnikowia_unclassified(100); | Metschnikowia | 13 | Saprotroph | \|Undefined Saprotroph\| | Yeast | NULL | Probable | As nectar-tap_saprotroph (Põlme et al. 2020); Decay sugar-rich substrates (Põlme et al. 2020) | Põlme S, et al. 2020. Fungal Diversity 105:1-16 ((https://doi.org/10.1007/s13225-020-00466-2)) |
| Otu27017 | 1 | k__Fungi(100);p__Ascomycota(100);c__Saccharomycetes(100);o__Saccharomycetales(100);f__Metschnikowiaceae(100);g__Metschnikowia(100);g__Metschnikowia_unclassified(100); | Metschnikowia | 13 | Saprotroph | \|Undefined Saprotroph\| | Yeast | NULL | Probable | As nectar-tap_saprotroph (Põlme et al. 2020); Decay sugar-rich substrates (Põlme et al. 2020) | Põlme S, et al. 2020. Fungal Diversity 105:1-16 ((https://doi.org/10.1007/s13225-020-00466-2)) |
| Otu27018 | 1 | k__Fungi(100);p__Ascomycota(100);c__Saccharomycetes(100);o__Saccharomycetales(100);f__Metschnikowiaceae(100);g__Metschnikowia(100);s__Metschnikowia_pulcherrima(100); | Metschnikowia | 13 | Saprotroph | \|Undefined Saprotroph\| | Yeast | NULL | Probable | As nectar-tap_saprotroph (Põlme et al. 2020); Decay sugar-rich substrates (Põlme et al. 2020) | Põlme S, et al. 2020. Fungal Diversity 105:1-16 ((https://doi.org/10.1007/s13225-020-00466-2)) |
| Otu27020 | 1 | k__Fungi(100);p__Ascomycota(100);c__Saccharomycetes(100);o__Saccharomycetales(100);f__Metschnikowiaceae(100);g__Metschnikowia(100);s__Metschnikowia_pulcherrima(100); | Metschnikowia | 13 | Saprotroph | \|Undefined Saprotroph\| | Yeast | NULL | Probable | As nectar-tap_saprotroph (Põlme et al. 2020); Decay sugar-rich substrates (Põlme et al. 2020) | Põlme S, et al. 2020. Fungal Diversity 105:1-16 ((https://doi.org/10.1007/s13225-020-00466-2)) |
| Otu27021 | 1 | k__Fungi(100);p__Ascomycota(100);c__Saccharomycetes(100);o__Saccharomycetales(100);f__Metschnikowiaceae(100);g__Metschnikowia(100);s__Metschnikowia_pulcherrima(100); | Metschnikowia | 13 | Saprotroph | \|Undefined Saprotroph\| | Yeast | NULL | Probable | As nectar-tap_saprotroph (Põlme et al. 2020); Decay sugar-rich substrates (Põlme et al. 2020) | Põlme S, et al. 2020. Fungal Diversity 105:1-16 ((https://doi.org/10.1007/s13225-020-00466-2)) |
| Otu27022 | 1 | k__Fungi(100);p__Ascomycota(100);c__Saccharomycetes(100);o__Saccharomycetales(100);f__Metschnikowiaceae(100);g__Metschnikowia(100);s__Metschnikowia_pulcherrima(100); | Metschnikowia | 13 | Saprotroph | \|Undefined Saprotroph\| | Yeast | NULL | Probable | As nectar-tap_saprotroph (Põlme et al. 2020); Decay sugar-rich substrates (Põlme et al. 2020) | Põlme S, et al. 2020. Fungal Diversity 105:1-16 ((https://doi.org/10.1007/s13225-020-00466-2)) |
| Otu27023 | 1 | k__Fungi(100);p__Ascomycota(100);c__Saccharomycetes(100);o__Saccharomycetales(100);f__Metschnikowiaceae(100);g__Metschnikowia(100);s__Metschnikowia_pulcherrima(100); | Metschnikowia | 13 | Saprotroph | \|Undefined Saprotroph\| | Yeast | NULL | Probable | As nectar-tap_saprotroph (Põlme et al. 2020); Decay sugar-rich substrates (Põlme et al. 2020) | Põlme S, et al. 2020. Fungal Diversity 105:1-16 ((https://doi.org/10.1007/s13225-020-00466-2)) |
| Otu27025 | 1 | k__Fungi(100);p__Ascomycota(100);c__Saccharomycetes(100);o__Saccharomycetales(100);f__Metschnikowiaceae(100);g__Metschnikowia(100);g__Metschnikowia_unclassified(100); | Metschnikowia | 13 | Saprotroph | \|Undefined Saprotroph\| | Yeast | NULL | Probable | As nectar-tap_saprotroph (Põlme et al. 2020); Decay sugar-rich substrates (Põlme et al. 2020) | Põlme S, et al. 2020. Fungal Diversity 105:1-16 ((https://doi.org/10.1007/s13225-020-00466-2)) |
| Otu27026 | 1 | k__Fungi(100);p__Ascomycota(100);c__Saccharomycetes(100);o__Saccharomycetales(100);f__Metschnikowiaceae(100);g__Metschnikowia(100);s__Metschnikowia_pulcherrima(100); | Metschnikowia | 13 | Saprotroph | \|Undefined Saprotroph\| | Yeast | NULL | Probable | As nectar-tap_saprotroph (Põlme et al. 2020); Decay sugar-rich substrates (Põlme et al. 2020) | Põlme S, et al. 2020. Fungal Diversity 105:1-16 ((https://doi.org/10.1007/s13225-020-00466-2)) |
| Otu27027 | 1 | k__Fungi(100);p__Ascomycota(100);c__Saccharomycetes(100);o__Saccharomycetales(100);f__Metschnikowiaceae(100);g__Metschnikowia(100);s__Metschnikowia_pulcherrima(100); | Metschnikowia | 13 | Saprotroph | \|Undefined Saprotroph\| | Yeast | NULL | Probable | As nectar-tap_saprotroph (Põlme et al. 2020); Decay sugar-rich substrates (Põlme et al. 2020) | Põlme S, et al. 2020. Fungal Diversity 105:1-16 ((https://doi.org/10.1007/s13225-020-00466-2)) |
| Otu27032 | 1 | k__Fungi(100);p__Ascomycota(100);c__Saccharomycetes(100);o__Saccharomycetales(100);f__Metschnikowiaceae(100);g__Metschnikowia(100);s__Metschnikowia_pulcherrima(100); | Metschnikowia | 13 | Saprotroph | \|Undefined Saprotroph\| | Yeast | NULL | Probable | As nectar-tap_saprotroph (Põlme et al. 2020); Decay sugar-rich substrates (Põlme et al. 2020) | Põlme S, et al. 2020. Fungal Diversity 105:1-16 ((https://doi.org/10.1007/s13225-020-00466-2)) |
| Otu27035 | 1 | k__Fungi(100);p__Ascomycota(100);c__Saccharomycetes(100);o__Saccharomycetales(100);f__Metschnikowiaceae(100);g__Metschnikowia(100);g__Metschnikowia_unclassified(100); | Metschnikowia | 13 | Saprotroph | \|Undefined Saprotroph\| | Yeast | NULL | Probable | As nectar-tap_saprotroph (Põlme et al. 2020); Decay sugar-rich substrates (Põlme et al. 2020) | Põlme S, et al. 2020. Fungal Diversity 105:1-16 ((https://doi.org/10.1007/s13225-020-00466-2)) |
| Otu27038 | 1 | k__Fungi(100);p__Ascomycota(100);c__Saccharomycetes(100);o__Saccharomycetales(100);f__Metschnikowiaceae(100);g__Metschnikowia(100);g__Metschnikowia_unclassified(100); | Metschnikowia | 13 | Saprotroph | \|Undefined Saprotroph\| | Yeast | NULL | Probable | As nectar-tap_saprotroph (Põlme et al. 2020); Decay sugar-rich substrates (Põlme et al. 2020) | Põlme S, et al. 2020. Fungal Diversity 105:1-16 ((https://doi.org/10.1007/s13225-020-00466-2)) |
| Otu27042 | 1 | k__Fungi(100);p__Ascomycota(100);c__Saccharomycetes(100);o__Saccharomycetales(100);f__Metschnikowiaceae(100);g__Metschnikowia(100);s__Metschnikowia_pulcherrima(100); | Metschnikowia | 13 | Saprotroph | \|Undefined Saprotroph\| | Yeast | NULL | Probable | As nectar-tap_saprotroph (Põlme et al. 2020); Decay sugar-rich substrates (Põlme et al. 2020) | Põlme S, et al. 2020. Fungal Diversity 105:1-16 ((https://doi.org/10.1007/s13225-020-00466-2)) |
| Otu27043 | 1 | k__Fungi(100);p__Ascomycota(100);c__Saccharomycetes(100);o__Saccharomycetales(100);f__Metschnikowiaceae(100);g__Metschnikowia(100);g__Metschnikowia_unclassified(100); | Metschnikowia | 13 | Saprotroph | \|Undefined Saprotroph\| | Yeast | NULL | Probable | As nectar-tap_saprotroph (Põlme et al. 2020); Decay sugar-rich substrates (Põlme et al. 2020) | Põlme S, et al. 2020. Fungal Diversity 105:1-16 ((https://doi.org/10.1007/s13225-020-00466-2)) |
| Otu27044 | 1 | k__Fungi(100);p__Ascomycota(100);c__Saccharomycetes(100);o__Saccharomycetales(100);f__Metschnikowiaceae(100);g__Metschnikowia(100);s__Metschnikowia_pulcherrima(100); | Metschnikowia | 13 | Saprotroph | \|Undefined Saprotroph\| | Yeast | NULL | Probable | As nectar-tap_saprotroph (Põlme et al. 2020); Decay sugar-rich substrates (Põlme et al. 2020) | Põlme S, et al. 2020. Fungal Diversity 105:1-16 ((https://doi.org/10.1007/s13225-020-00466-2)) |
| Otu27045 | 1 | k__Fungi(100);p__Ascomycota(100);c__Saccharomycetes(100);o__Saccharomycetales(100);f__Metschnikowiaceae(100);g__Metschnikowia(100);s__Metschnikowia_pulcherrima(100); | Metschnikowia | 13 | Saprotroph | \|Undefined Saprotroph\| | Yeast | NULL | Probable | As nectar-tap_saprotroph (Põlme et al. 2020); Decay sugar-rich substrates (Põlme et al. 2020) | Põlme S, et al. 2020. Fungal Diversity 105:1-16 ((https://doi.org/10.1007/s13225-020-00466-2)) |
| Otu27047 | 1 | k__Fungi(100);p__Zygomycota(100);c__Zygomycota_class_Incertae_sedis(100);o__Mucorales(100);f__Mucoraceae(100);g__Mucor(100);g__Mucor_unclassified(100); | Mucor | 13 | Saprotroph | \|Undefined Saprotroph\| | Microfungus | NULL | Probable | As soil_saprotroph (Põlme et al. 2020); Decay soil (Põlme et al. 2020) | Tedersoo L, et al. 2014. Science 346:e1256688 ((https://doi.org/10.1126/science.1256688)); Põlme S, et al. 2020. Fungal Diversity 105:1-16 ((https://doi.org/10.1007/s13225-020-00466-2)) |
| Otu27048 | 1 | k__Fungi(100);p__Ascomycota(100);c__Saccharomycetes(100);o__Saccharomycetales(100);f__Metschnikowiaceae(100);g__Metschnikowia(100);s__Metschnikowia_pulcherrima(100); | Metschnikowia | 13 | Saprotroph | \|Undefined Saprotroph\| | Yeast | NULL | Probable | As nectar-tap_saprotroph (Põlme et al. 2020); Decay sugar-rich substrates (Põlme et al. 2020) | Põlme S, et al. 2020. Fungal Diversity 105:1-16 ((https://doi.org/10.1007/s13225-020-00466-2)) |
| Otu27051 | 1 | k__Fungi(100);p__Ascomycota(100);c__Saccharomycetes(100);o__Saccharomycetales(100);f__Saccharomycetaceae(100);g__Kazachstania(100);s__Kazachstania_intestinalis(100); | Kazachstania | 13 | Saprotroph | \|Undefined Saprotroph\| | Yeast | NULL | Probable | As nectar-tap_saprotroph (Põlme et al. 2020); Decay sugar-rich substrates (Põlme et al. 2020) | Põlme S, et al. 2020. Fungal Diversity 105:1-16 ((https://doi.org/10.1007/s13225-020-00466-2)) |
| Otu27052 | 1 | k__Fungi(100);p__Ascomycota(100);c__Saccharomycetes(100);o__Saccharomycetales(100);f__Metschnikowiaceae(100);g__Metschnikowia(100);s__Metschnikowia_pulcherrima(100); | Metschnikowia | 13 | Saprotroph | \|Undefined Saprotroph\| | Yeast | NULL | Probable | As nectar-tap_saprotroph (Põlme et al. 2020); Decay sugar-rich substrates (Põlme et al. 2020) | Põlme S, et al. 2020. Fungal Diversity 105:1-16 ((https://doi.org/10.1007/s13225-020-00466-2)) |
| Otu27054 | 1 | k__Fungi(100);p__Zygomycota(100);c__Zygomycota_class_Incertae_sedis(100);o__Mucorales(100);f__Mucoraceae(100);g__Mucor(100);g__Mucor_unclassified(100); | Mucor | 13 | Saprotroph | \|Undefined Saprotroph\| | Microfungus | NULL | Probable | As soil_saprotroph (Põlme et al. 2020); Decay soil (Põlme et al. 2020) | Tedersoo L, et al. 2014. Science 346:e1256688 ((https://doi.org/10.1126/science.1256688)); Põlme S, et al. 2020. Fungal Diversity 105:1-16 ((https://doi.org/10.1007/s13225-020-00466-2)) |
| Otu27055 | 1 | k__Fungi(100);p__Zygomycota(100);c__Zygomycota_class_Incertae_sedis(100);o__Mucorales(100);f__Mucoraceae(100);g__Mucor(100);s__Mucor_genevensis(100); | Mucor | 13 | Saprotroph | \|Undefined Saprotroph\| | Microfungus | NULL | Probable | As soil_saprotroph (Põlme et al. 2020); Decay soil (Põlme et al. 2020) | Tedersoo L, et al. 2014. Science 346:e1256688 ((https://doi.org/10.1126/science.1256688)); Põlme S, et al. 2020. Fungal Diversity 105:1-16 ((https://doi.org/10.1007/s13225-020-00466-2)) |
| Otu27056 | 1 | k__Fungi(100);p__Zygomycota(100);c__Zygomycota_class_Incertae_sedis(100);o__Mucorales(100);f__Mucoraceae(100);g__Mucor(100);s__Mucor_genevensis(100); | Mucor | 13 | Saprotroph | \|Undefined Saprotroph\| | Microfungus | NULL | Probable | As soil_saprotroph (Põlme et al. 2020); Decay soil (Põlme et al. 2020) | Tedersoo L, et al. 2014. Science 346:e1256688 ((https://doi.org/10.1126/science.1256688)); Põlme S, et al. 2020. Fungal Diversity 105:1-16 ((https://doi.org/10.1007/s13225-020-00466-2)) |
| Otu27057 | 1 | k__Fungi(100);p__Zygomycota(100);c__Zygomycota_class_Incertae_sedis(100);o__Mucorales(100);f__Mucoraceae(100);g__Mucor(100);s__Mucor_genevensis(100); | Mucor | 13 | Saprotroph | \|Undefined Saprotroph\| | Microfungus | NULL | Probable | As soil_saprotroph (Põlme et al. 2020); Decay soil (Põlme et al. 2020) | Tedersoo L, et al. 2014. Science 346:e1256688 ((https://doi.org/10.1126/science.1256688)); Põlme S, et al. 2020. Fungal Diversity 105:1-16 ((https://doi.org/10.1007/s13225-020-00466-2)) |
| Otu27058 | 1 | k__Fungi(100);p__Zygomycota(100);c__Zygomycota_class_Incertae_sedis(100);o__Mucorales(100);f__Mucoraceae(100);g__Mucor(100);s__Mucor_genevensis(100); | Mucor | 13 | Saprotroph | \|Undefined Saprotroph\| | Microfungus | NULL | Probable | As soil_saprotroph (Põlme et al. 2020); Decay soil (Põlme et al. 2020) | Tedersoo L, et al. 2014. Science 346:e1256688 ((https://doi.org/10.1126/science.1256688)); Põlme S, et al. 2020. Fungal Diversity 105:1-16 ((https://doi.org/10.1007/s13225-020-00466-2)) |
| Otu27059 | 1 | k__Fungi(100);p__Zygomycota(100);c__Zygomycota_class_Incertae_sedis(100);o__Mucorales(100);f__Mucoraceae(100);g__Mucor(100);s__Mucor_genevensis(100); | Mucor | 13 | Saprotroph | \|Undefined Saprotroph\| | Microfungus | NULL | Probable | As soil_saprotroph (Põlme et al. 2020); Decay soil (Põlme et al. 2020) | Tedersoo L, et al. 2014. Science 346:e1256688 ((https://doi.org/10.1126/science.1256688)); Põlme S, et al. 2020. Fungal Diversity 105:1-16 ((https://doi.org/10.1007/s13225-020-00466-2)) |
| Otu27060 | 1 | k__Fungi(100);p__Zygomycota(100);c__Zygomycota_class_Incertae_sedis(100);o__Mucorales(100);f__Mucoraceae(100);g__Mucor(100);s__Mucor_genevensis(100); | Mucor | 13 | Saprotroph | \|Undefined Saprotroph\| | Microfungus | NULL | Probable | As soil_saprotroph (Põlme et al. 2020); Decay soil (Põlme et al. 2020) | Tedersoo L, et al. 2014. Science 346:e1256688 ((https://doi.org/10.1126/science.1256688)); Põlme S, et al. 2020. Fungal Diversity 105:1-16 ((https://doi.org/10.1007/s13225-020-00466-2)) |
| Otu27061 | 1 | k__Fungi(100);p__Zygomycota(100);c__Zygomycota_class_Incertae_sedis(100);o__Mucorales(100);f__Mucoraceae(100);g__Mucor(100);s__Mucor_genevensis(100); | Mucor | 13 | Saprotroph | \|Undefined Saprotroph\| | Microfungus | NULL | Probable | As soil_saprotroph (Põlme et al. 2020); Decay soil (Põlme et al. 2020) | Tedersoo L, et al. 2014. Science 346:e1256688 ((https://doi.org/10.1126/science.1256688)); Põlme S, et al. 2020. Fungal Diversity 105:1-16 ((https://doi.org/10.1007/s13225-020-00466-2)) |
| Otu27062 | 1 | k__Fungi(100);p__Zygomycota(100);c__Zygomycota_class_Incertae_sedis(100);o__Mucorales(100);f__Mucoraceae(100);g__Mucor(100);s__Mucor_genevensis(100); | Mucor | 13 | Saprotroph | \|Undefined Saprotroph\| | Microfungus | NULL | Probable | As soil_saprotroph (Põlme et al. 2020); Decay soil (Põlme et al. 2020) | Tedersoo L, et al. 2014. Science 346:e1256688 ((https://doi.org/10.1126/science.1256688)); Põlme S, et al. 2020. Fungal Diversity 105:1-16 ((https://doi.org/10.1007/s13225-020-00466-2)) |
| Otu27063 | 1 | k__Fungi(100);p__Zygomycota(100);c__Zygomycota_class_Incertae_sedis(100);o__Mucorales(100);f__Mucoraceae(100);g__Mucor(100);s__Mucor_genevensis(100); | Mucor | 13 | Saprotroph | \|Undefined Saprotroph\| | Microfungus | NULL | Probable | As soil_saprotroph (Põlme et al. 2020); Decay soil (Põlme et al. 2020) | Tedersoo L, et al. 2014. Science 346:e1256688 ((https://doi.org/10.1126/science.1256688)); Põlme S, et al. 2020. Fungal Diversity 105:1-16 ((https://doi.org/10.1007/s13225-020-00466-2)) |
| Otu27064 | 1 | k__Fungi(100);p__Zygomycota(100);c__Zygomycota_class_Incertae_sedis(100);o__Mucorales(100);f__Mucoraceae(100);g__Mucor(100);s__Mucor_circinelloides_f._lusitanicus(100); | Mucor | 13 | Saprotroph | \|Undefined Saprotroph\| | Microfungus | NULL | Probable | As soil_saprotroph (Põlme et al. 2020); Decay soil (Põlme et al. 2020) | Tedersoo L, et al. 2014. Science 346:e1256688 ((https://doi.org/10.1126/science.1256688)); Põlme S, et al. 2020. Fungal Diversity 105:1-16 ((https://doi.org/10.1007/s13225-020-00466-2)) |
| Otu27065 | 1 | k__Fungi(100);p__Zygomycota(100);c__Zygomycota_class_Incertae_sedis(100);o__Mucorales(100);f__Mucoraceae(100);g__Mucor(100);s__Mucor_circinelloides_f._circinelloides(100); | Mucor | 13 | Saprotroph | \|Undefined Saprotroph\| | Microfungus | NULL | Probable | As soil_saprotroph (Põlme et al. 2020); Decay soil (Põlme et al. 2020) | Tedersoo L, et al. 2014. Science 346:e1256688 ((https://doi.org/10.1126/science.1256688)); Põlme S, et al. 2020. Fungal Diversity 105:1-16 ((https://doi.org/10.1007/s13225-020-00466-2)) |
| Otu27066 | 1 | k__Fungi(100);p__Zygomycota(100);c__Zygomycota_class_Incertae_sedis(100);o__Mucorales(100);f__Mucoraceae(100);g__Mucor(100);g__Mucor_unclassified(100); | Mucor | 13 | Saprotroph | \|Undefined Saprotroph\| | Microfungus | NULL | Probable | As soil_saprotroph (Põlme et al. 2020); Decay soil (Põlme et al. 2020) | Tedersoo L, et al. 2014. Science 346:e1256688 ((https://doi.org/10.1126/science.1256688)); Põlme S, et al. 2020. Fungal Diversity 105:1-16 ((https://doi.org/10.1007/s13225-020-00466-2)) |
| Otu27067 | 1 | k__Fungi(100);p__Zygomycota(100);c__Zygomycota_class_Incertae_sedis(100);o__Mucorales(100);f__Mucoraceae(100);g__Mucor(100);g__Mucor_unclassified(100); | Mucor | 13 | Saprotroph | \|Undefined Saprotroph\| | Microfungus | NULL | Probable | As soil_saprotroph (Põlme et al. 2020); Decay soil (Põlme et al. 2020) | Tedersoo L, et al. 2014. Science 346:e1256688 ((https://doi.org/10.1126/science.1256688)); Põlme S, et al. 2020. Fungal Diversity 105:1-16 ((https://doi.org/10.1007/s13225-020-00466-2)) |
| Otu27068 | 1 | k__Fungi(100);p__Zygomycota(100);c__Zygomycota_class_Incertae_sedis(100);o__Mucorales(100);f__Mucoraceae(100);g__Mucor(100);s__Mucor_genevensis(100); | Mucor | 13 | Saprotroph | \|Undefined Saprotroph\| | Microfungus | NULL | Probable | As soil_saprotroph (Põlme et al. 2020); Decay soil (Põlme et al. 2020) | Tedersoo L, et al. 2014. Science 346:e1256688 ((https://doi.org/10.1126/science.1256688)); Põlme S, et al. 2020. Fungal Diversity 105:1-16 ((https://doi.org/10.1007/s13225-020-00466-2)) |
| Otu27069 | 1 | k__Fungi(100);p__Zygomycota(100);c__Zygomycota_class_Incertae_sedis(100);o__Mucorales(100);f__Mucoraceae(100);g__Mucor(100);g__Mucor_unclassified(100); | Mucor | 13 | Saprotroph | \|Undefined Saprotroph\| | Microfungus | NULL | Probable | As soil_saprotroph (Põlme et al. 2020); Decay soil (Põlme et al. 2020) | Tedersoo L, et al. 2014. Science 346:e1256688 ((https://doi.org/10.1126/science.1256688)); Põlme S, et al. 2020. Fungal Diversity 105:1-16 ((https://doi.org/10.1007/s13225-020-00466-2)) |
| Otu27070 | 1 | k__Fungi(100);p__Zygomycota(100);c__Zygomycota_class_Incertae_sedis(100);o__Mucorales(100);f__Mucoraceae(100);g__Mucor(100);s__Mucor_circinelloides_f._janssenii(100); | Mucor | 13 | Saprotroph | \|Undefined Saprotroph\| | Microfungus | NULL | Probable | As soil_saprotroph (Põlme et al. 2020); Decay soil (Põlme et al. 2020) | Tedersoo L, et al. 2014. Science 346:e1256688 ((https://doi.org/10.1126/science.1256688)); Põlme S, et al. 2020. Fungal Diversity 105:1-16 ((https://doi.org/10.1007/s13225-020-00466-2)) |
| Otu27071 | 1 | k__Fungi(100);p__Basidiomycota(100);c__Tremellomycetes(100);o__Tremellales(100);f__Tremellales_family_Incertae_sedis(100);g__Fellomyces(100);s__Fellomyces_horovitziae(100); | Fellomyces | 13 | Pathotroph-Saprotroph | Fungal Parasite-\|Undefined Saprotroph\| | Yeast | NULL | Probable | As mycoparasite (Põlme et al. 2020) | Põlme S, et al. 2020. Fungal Diversity 105:1-16 ((https://doi.org/10.1007/s13225-020-00466-2)) |
| Otu27072 | 1 | k__Fungi(100);p__Basidiomycota(100);c__Agaricomycetes(100);o__Polyporales(100);f__Polyporaceae(100);g__Skeletocutis(100);s__Skeletocutis_nivea(100); | Skeletocutis | 13 | Saprotroph | \|Wood Saprotroph\| | Polyporoid | White Rot | Highly Probable | As white_rot (Põlme et al. 2020); Decay wood (Põlme et al. 2020) | Gilbertson RL, Ryvarden L. 1987-1987. North American Polypores. Fungiflora, Oslo (ISBN: 978-0945345060); Põlme S, et al. 2020. Fungal Diversity 105:1-16 ((https://doi.org/10.1007/s13225-020-00466-2)) |
| Otu27076 | 1 | k__Fungi(100);p__Zygomycota(100);c__Zygomycota_class_Incertae_sedis(100);o__Mucorales(100);f__Mucoraceae(100);g__Mucor(100);s__Mucor_genevensis(100); | Mucor | 13 | Saprotroph | \|Undefined Saprotroph\| | Microfungus | NULL | Probable | As soil_saprotroph (Põlme et al. 2020); Decay soil (Põlme et al. 2020) | Tedersoo L, et al. 2014. Science 346:e1256688 ((https://doi.org/10.1126/science.1256688)); Põlme S, et al. 2020. Fungal Diversity 105:1-16 ((https://doi.org/10.1007/s13225-020-00466-2)) |
| Otu27077 | 1 | k__Fungi(100);p__Zygomycota(100);c__Zygomycota_class_Incertae_sedis(100);o__Mucorales(100);f__Mucoraceae(100);g__Mucor(100);s__Mucor_microsporus(100); | Mucor | 13 | Saprotroph | \|Undefined Saprotroph\| | Microfungus | NULL | Probable | As soil_saprotroph (Põlme et al. 2020); Decay soil (Põlme et al. 2020) | Tedersoo L, et al. 2014. Science 346:e1256688 ((https://doi.org/10.1126/science.1256688)); Põlme S, et al. 2020. Fungal Diversity 105:1-16 ((https://doi.org/10.1007/s13225-020-00466-2)) |
| Otu27078 | 1 | k__Fungi(100);p__Zygomycota(100);c__Zygomycota_class_Incertae_sedis(100);o__Mucorales(100);f__Mucoraceae(100);g__Mucor(100);g__Mucor_unclassified(100); | Mucor | 13 | Saprotroph | \|Undefined Saprotroph\| | Microfungus | NULL | Probable | As soil_saprotroph (Põlme et al. 2020); Decay soil (Põlme et al. 2020) | Tedersoo L, et al. 2014. Science 346:e1256688 ((https://doi.org/10.1126/science.1256688)); Põlme S, et al. 2020. Fungal Diversity 105:1-16 ((https://doi.org/10.1007/s13225-020-00466-2)) |
| Otu27079 | 1 | k__Fungi(100);p__Zygomycota(100);c__Zygomycota_class_Incertae_sedis(100);o__Mucorales(100);f__Mucoraceae(100);g__Mucor(100);s__Mucor_sp_CS_P/F01(100); | Mucor | 13 | Saprotroph | \|Undefined Saprotroph\| | Microfungus | NULL | Probable | As soil_saprotroph (Põlme et al. 2020); Decay soil (Põlme et al. 2020) | Tedersoo L, et al. 2014. Science 346:e1256688 ((https://doi.org/10.1126/science.1256688)); Põlme S, et al. 2020. Fungal Diversity 105:1-16 ((https://doi.org/10.1007/s13225-020-00466-2)) |
| Otu27081 | 1 | k__Fungi(100);p__Basidiomycota(100);c__Agaricomycetes(100);o__Agaricales(100);o__Agaricales_unclassified(100);o__Agaricales_unclassified(100);o__Agaricales_unclassified(100); | Agaricales | 7 | Pathotroph-Saprotroph-Symbiotroph | Bryophyte Parasite-Dung Saprotroph-Ectomycorrhizal-Fungal Parasite-Leaf Saprotroph-Plant Parasite-Undefined Saprotroph-Wood Saprotroph | Agaricoid-Gasteroid-Microfungus-Secotioid-Yeast | NULL | Possible | Mushrooms and toadstools, Gill fungi, Agarics... lignicolous, sometimes muscicolous or fungicolous, saprobic, mycorrhizal, rarely parasitic on plants or fungi (Kirk et al. 2008) | Kirk PM et al. 2008. Dictionary of the Fungi. Tenth Edition. CAB International, Wallingford (ISBN: 978-0851998268) |
| Otu27087 | 1 | k__Fungi(100);p__Basidiomycota(100);c__Wallemiomycetes(100);o__Wallemiales(100);f__Wallemiaceae(100);g__Wallemia(100);s__Wallemia_sebi(100); | Wallemia | 13 | Saprotroph | \|Undefined Saprotroph\| | Microfungus | NULL | Probable | Decay sugar-rich substrates (Põlme et al. 2020) | Tedersoo L, et al. 2014. Science 346:e1256688 ((https://doi.org/10.1126/science.1256688)); Põlme S, et al. 2020. Fungal Diversity 105:1-16 ((https://doi.org/10.1007/s13225-020-00466-2)) |
| Otu27088 | 1 | k__Fungi(100);p__Ascomycota(100);c__Saccharomycetes(100);o__Saccharomycetales(100);f__Metschnikowiaceae(100);g__Metschnikowia(100);g__Metschnikowia_unclassified(100); | Metschnikowia | 13 | Saprotroph | \|Undefined Saprotroph\| | Yeast | NULL | Probable | As nectar-tap_saprotroph (Põlme et al. 2020); Decay sugar-rich substrates (Põlme et al. 2020) | Põlme S, et al. 2020. Fungal Diversity 105:1-16 ((https://doi.org/10.1007/s13225-020-00466-2)) |
| Otu27091 | 1 | k__Fungi(100);p__Zygomycota(100);c__Zygomycota_class_Incertae_sedis(100);o__Mucorales(100);f__Mucoraceae(100);g__Mucor(100);s__Mucor_genevensis(100); | Mucor | 13 | Saprotroph | \|Undefined Saprotroph\| | Microfungus | NULL | Probable | As soil_saprotroph (Põlme et al. 2020); Decay soil (Põlme et al. 2020) | Tedersoo L, et al. 2014. Science 346:e1256688 ((https://doi.org/10.1126/science.1256688)); Põlme S, et al. 2020. Fungal Diversity 105:1-16 ((https://doi.org/10.1007/s13225-020-00466-2)) |
| Otu27092 | 1 | k__Fungi(100);p__Basidiomycota(100);c__Tremellomycetes(100);o__Trichosporonales(100);f__Trichosporonaceae(100);g__Asterotremella(100);s__Asterotremella_humicola(100); | Asterotremella | 13 | Pathotroph-Saprotroph | Fungal Parasite-Soil Saprotroph | Yeast | NULL | Probable | Isolated from forest soil and murshooms (Liu et al. 2011) | Liu Y-R, et al. 2011. Antonie van Leeuwenhoek 99:643-650 ((https://doi.org/10.1007/s10482-010-9537-y)) |
| Otu27093 | 1 | k__Fungi(100);p__Basidiomycota(100);c__Agaricomycetes(100);o__Agaricales(100);f__Inocybaceae(100);g__Crepidotus(100);s__Crepidotus_applanatus(100); | Crepidotus | 13 | Saprotroph | \|Wood Saprotroph\| | Agaricoid | NULL | Highly Probable | Decay wood (Põlme et al. 2020) | Grgurinovic CA, Simpson JA. 2001. Fungal Diversity 8:97-127 ((http://www.fungaldiversity.org/fdp/sfdp/FD_8_97-127.pdf)); Aime MC, et al. 2005. American Journal of Botany 92:74-82 ((https://doi.org/10.3732/ajb.92.1.74)); Tedersoo L, et al. 2014. Science 346:e1256688 ((https://doi.org/10.1126/science.1256688)); Põlme S, et al. 2020. Fungal Diversity 105:1-16 ((https://doi.org/10.1007/s13225-020-00466-2)) |
| Otu27096 | 1 | k__Fungi(100);p__Zygomycota(100);c__Zygomycota_class_Incertae_sedis(100);o__Mucorales(100);f__Mucoraceae(100);g__Mucor(100);s__Mucor_genevensis(100); | Mucor | 13 | Saprotroph | \|Undefined Saprotroph\| | Microfungus | NULL | Probable | As soil_saprotroph (Põlme et al. 2020); Decay soil (Põlme et al. 2020) | Tedersoo L, et al. 2014. Science 346:e1256688 ((https://doi.org/10.1126/science.1256688)); Põlme S, et al. 2020. Fungal Diversity 105:1-16 ((https://doi.org/10.1007/s13225-020-00466-2)) |
| Otu27097 | 1 | k__Fungi(100);p__Zygomycota(100);c__Zygomycota_class_Incertae_sedis(100);o__Mucorales(100);f__Mucoraceae(100);g__Mucor(100);s__Mucor_genevensis(100); | Mucor | 13 | Saprotroph | \|Undefined Saprotroph\| | Microfungus | NULL | Probable | As soil_saprotroph (Põlme et al. 2020); Decay soil (Põlme et al. 2020) | Tedersoo L, et al. 2014. Science 346:e1256688 ((https://doi.org/10.1126/science.1256688)); Põlme S, et al. 2020. Fungal Diversity 105:1-16 ((https://doi.org/10.1007/s13225-020-00466-2)) |
| Otu27098 | 1 | k__Fungi(100);p__Zygomycota(100);c__Zygomycota_class_Incertae_sedis(100);o__Mucorales(100);f__Mucoraceae(100);g__Mucor(100);s__Mucor_genevensis(100); | Mucor | 13 | Saprotroph | \|Undefined Saprotroph\| | Microfungus | NULL | Probable | As soil_saprotroph (Põlme et al. 2020); Decay soil (Põlme et al. 2020) | Tedersoo L, et al. 2014. Science 346:e1256688 ((https://doi.org/10.1126/science.1256688)); Põlme S, et al. 2020. Fungal Diversity 105:1-16 ((https://doi.org/10.1007/s13225-020-00466-2)) |
| Otu27099 | 1 | k__Fungi(100);p__Zygomycota(100);c__Zygomycota_class_Incertae_sedis(100);o__Mucorales(100);f__Mucoraceae(100);g__Mucor(100);s__Mucor_genevensis(100); | Mucor | 13 | Saprotroph | \|Undefined Saprotroph\| | Microfungus | NULL | Probable | As soil_saprotroph (Põlme et al. 2020); Decay soil (Põlme et al. 2020) | Tedersoo L, et al. 2014. Science 346:e1256688 ((https://doi.org/10.1126/science.1256688)); Põlme S, et al. 2020. Fungal Diversity 105:1-16 ((https://doi.org/10.1007/s13225-020-00466-2)) |
| Otu27100 | 1 | k__Fungi(100);p__Zygomycota(100);c__Zygomycota_class_Incertae_sedis(100);o__Mucorales(100);f__Mucoraceae(100);g__Mucor(100);s__Mucor_genevensis(100); | Mucor | 13 | Saprotroph | \|Undefined Saprotroph\| | Microfungus | NULL | Probable | As soil_saprotroph (Põlme et al. 2020); Decay soil (Põlme et al. 2020) | Tedersoo L, et al. 2014. Science 346:e1256688 ((https://doi.org/10.1126/science.1256688)); Põlme S, et al. 2020. Fungal Diversity 105:1-16 ((https://doi.org/10.1007/s13225-020-00466-2)) |
| Otu27101 | 1 | k__Fungi(100);p__Zygomycota(100);c__Zygomycota_class_Incertae_sedis(100);o__Mucorales(100);f__Mucoraceae(100);g__Mucor(100);s__Mucor_genevensis(100); | Mucor | 13 | Saprotroph | \|Undefined Saprotroph\| | Microfungus | NULL | Probable | As soil_saprotroph (Põlme et al. 2020); Decay soil (Põlme et al. 2020) | Tedersoo L, et al. 2014. Science 346:e1256688 ((https://doi.org/10.1126/science.1256688)); Põlme S, et al. 2020. Fungal Diversity 105:1-16 ((https://doi.org/10.1007/s13225-020-00466-2)) |
| Otu27102 | 1 | k__Fungi(100);p__Zygomycota(100);c__Zygomycota_class_Incertae_sedis(100);o__Mucorales(100);f__Mucoraceae(100);g__Mucor(100);s__Mucor_genevensis(100); | Mucor | 13 | Saprotroph | \|Undefined Saprotroph\| | Microfungus | NULL | Probable | As soil_saprotroph (Põlme et al. 2020); Decay soil (Põlme et al. 2020) | Tedersoo L, et al. 2014. Science 346:e1256688 ((https://doi.org/10.1126/science.1256688)); Põlme S, et al. 2020. Fungal Diversity 105:1-16 ((https://doi.org/10.1007/s13225-020-00466-2)) |
| Otu27103 | 1 | k__Fungi(100);p__Zygomycota(100);c__Zygomycota_class_Incertae_sedis(100);o__Mucorales(100);f__Mucoraceae(100);g__Mucor(100);s__Mucor_genevensis(100); | Mucor | 13 | Saprotroph | \|Undefined Saprotroph\| | Microfungus | NULL | Probable | As soil_saprotroph (Põlme et al. 2020); Decay soil (Põlme et al. 2020) | Tedersoo L, et al. 2014. Science 346:e1256688 ((https://doi.org/10.1126/science.1256688)); Põlme S, et al. 2020. Fungal Diversity 105:1-16 ((https://doi.org/10.1007/s13225-020-00466-2)) |
| Otu27104 | 1 | k__Fungi(100);p__Zygomycota(100);c__Zygomycota_class_Incertae_sedis(100);o__Mucorales(100);f__Mucoraceae(100);g__Mucor(100);s__Mucor_genevensis(100); | Mucor | 13 | Saprotroph | \|Undefined Saprotroph\| | Microfungus | NULL | Probable | As soil_saprotroph (Põlme et al. 2020); Decay soil (Põlme et al. 2020) | Tedersoo L, et al. 2014. Science 346:e1256688 ((https://doi.org/10.1126/science.1256688)); Põlme S, et al. 2020. Fungal Diversity 105:1-16 ((https://doi.org/10.1007/s13225-020-00466-2)) |
| Otu27105 | 1 | k__Fungi(100);p__Zygomycota(100);c__Zygomycota_class_Incertae_sedis(100);o__Mucorales(100);f__Mucoraceae(100);g__Mucor(100);s__Mucor_genevensis(100); | Mucor | 13 | Saprotroph | \|Undefined Saprotroph\| | Microfungus | NULL | Probable | As soil_saprotroph (Põlme et al. 2020); Decay soil (Põlme et al. 2020) | Tedersoo L, et al. 2014. Science 346:e1256688 ((https://doi.org/10.1126/science.1256688)); Põlme S, et al. 2020. Fungal Diversity 105:1-16 ((https://doi.org/10.1007/s13225-020-00466-2)) |
| Otu27106 | 1 | k__Fungi(100);p__Zygomycota(100);c__Zygomycota_class_Incertae_sedis(100);o__Mucorales(100);f__Mucoraceae(100);g__Mucor(100);s__Mucor_genevensis(100); | Mucor | 13 | Saprotroph | \|Undefined Saprotroph\| | Microfungus | NULL | Probable | As soil_saprotroph (Põlme et al. 2020); Decay soil (Põlme et al. 2020) | Tedersoo L, et al. 2014. Science 346:e1256688 ((https://doi.org/10.1126/science.1256688)); Põlme S, et al. 2020. Fungal Diversity 105:1-16 ((https://doi.org/10.1007/s13225-020-00466-2)) |
| Otu27107 | 1 | k__Fungi(100);p__Basidiomycota(100);c__Wallemiomycetes(100);o__Wallemiales(100);f__Wallemiaceae(100);g__Wallemia(100);s__Wallemia_sebi(100); | Wallemia | 13 | Saprotroph | \|Undefined Saprotroph\| | Microfungus | NULL | Probable | Decay sugar-rich substrates (Põlme et al. 2020) | Tedersoo L, et al. 2014. Science 346:e1256688 ((https://doi.org/10.1126/science.1256688)); Põlme S, et al. 2020. Fungal Diversity 105:1-16 ((https://doi.org/10.1007/s13225-020-00466-2)) |
| Otu27114 | 1 | k__Fungi(100);p__Basidiomycota(100);c__Tremellomycetes(100);o__Tremellales(100);f__Tremellales_family_Incertae_sedis(100);g__Cryptococcus(100);s__Cryptococcus_sp_QMW_2009a(100); | Cryptococcus | 13 | Pathotroph-Saprotroph-Symbiotroph | Animal Pathogen-Endophyte-Epiphyte-\|Undefined Saprotroph\| | Dimorphic Yeast-Yeast | NULL | Possible | As foliar_endophyte (Põlme et al. 2020); Decay leaf, fruit, seed, soil, dung, animal material (Põlme et al. 2020) | Andrews JH, et al. 1983. Phytopathology 73:228-234 ((https://doi.org/10.1094/Phyto-73-228)); Kurtzman CP, et al. (eds.) 2011. The Yeasts, a Taxonomic Study. Fifth Edition. Vols 1-3. Elsevier, San Diego (ISBN: 9780444521491); Põlme S, et al. 2020. Fungal Diversity 105:1-16 ((https://doi.org/10.1007/s13225-020-00466-2)) |
| Otu27115 | 1 | k__Fungi(100);p__Basidiomycota(100);c__Tremellomycetes(100);o__Tremellales(100);f__Tremellales_family_Incertae_sedis(100);g__Cryptococcus(100);s__Cryptococcus_sp_QMW_2009a(100); | Cryptococcus | 13 | Pathotroph-Saprotroph-Symbiotroph | Animal Pathogen-Endophyte-Epiphyte-\|Undefined Saprotroph\| | Dimorphic Yeast-Yeast | NULL | Possible | As foliar_endophyte (Põlme et al. 2020); Decay leaf, fruit, seed, soil, dung, animal material (Põlme et al. 2020) | Andrews JH, et al. 1983. Phytopathology 73:228-234 ((https://doi.org/10.1094/Phyto-73-228)); Kurtzman CP, et al. (eds.) 2011. The Yeasts, a Taxonomic Study. Fifth Edition. Vols 1-3. Elsevier, San Diego (ISBN: 9780444521491); Põlme S, et al. 2020. Fungal Diversity 105:1-16 ((https://doi.org/10.1007/s13225-020-00466-2)) |
| Otu27116 | 1 | k__Fungi(100);p__Basidiomycota(100);c__Tremellomycetes(100);o__Tremellales(100);f__Tremellales_family_Incertae_sedis(100);g__Cryptococcus(100);s__Cryptococcus_sp_QMW_2009a(100); | Cryptococcus | 13 | Pathotroph-Saprotroph-Symbiotroph | Animal Pathogen-Endophyte-Epiphyte-\|Undefined Saprotroph\| | Dimorphic Yeast-Yeast | NULL | Possible | As foliar_endophyte (Põlme et al. 2020); Decay leaf, fruit, seed, soil, dung, animal material (Põlme et al. 2020) | Andrews JH, et al. 1983. Phytopathology 73:228-234 ((https://doi.org/10.1094/Phyto-73-228)); Kurtzman CP, et al. (eds.) 2011. The Yeasts, a Taxonomic Study. Fifth Edition. Vols 1-3. Elsevier, San Diego (ISBN: 9780444521491); Põlme S, et al. 2020. Fungal Diversity 105:1-16 ((https://doi.org/10.1007/s13225-020-00466-2)) |
| Otu27119 | 1 | k__Fungi(100);p__Basidiomycota(100);c__Wallemiomycetes(100);o__Wallemiales(100);f__Wallemiaceae(100);g__Wallemia(100);s__Wallemia_sebi(100); | Wallemia | 13 | Saprotroph | \|Undefined Saprotroph\| | Microfungus | NULL | Probable | Decay sugar-rich substrates (Põlme et al. 2020) | Tedersoo L, et al. 2014. Science 346:e1256688 ((https://doi.org/10.1126/science.1256688)); Põlme S, et al. 2020. Fungal Diversity 105:1-16 ((https://doi.org/10.1007/s13225-020-00466-2)) |
| Otu27120 | 1 | k__Fungi(100);p__Basidiomycota(100);c__Wallemiomycetes(100);o__Wallemiales(100);f__Wallemiaceae(100);g__Wallemia(100);s__Wallemia_sebi(100); | Wallemia | 13 | Saprotroph | \|Undefined Saprotroph\| | Microfungus | NULL | Probable | Decay sugar-rich substrates (Põlme et al. 2020) | Tedersoo L, et al. 2014. Science 346:e1256688 ((https://doi.org/10.1126/science.1256688)); Põlme S, et al. 2020. Fungal Diversity 105:1-16 ((https://doi.org/10.1007/s13225-020-00466-2)) |
| Otu27122 | 1 | k__Fungi(100);p__Basidiomycota(100);c__Wallemiomycetes(100);o__Wallemiales(100);f__Wallemiaceae(100);g__Wallemia(100);s__Wallemia_sebi(100); | Wallemia | 13 | Saprotroph | \|Undefined Saprotroph\| | Microfungus | NULL | Probable | Decay sugar-rich substrates (Põlme et al. 2020) | Tedersoo L, et al. 2014. Science 346:e1256688 ((https://doi.org/10.1126/science.1256688)); Põlme S, et al. 2020. Fungal Diversity 105:1-16 ((https://doi.org/10.1007/s13225-020-00466-2)) |
| Otu27123 | 1 | k__Fungi(100);p__Basidiomycota(100);c__Wallemiomycetes(100);o__Wallemiales(100);f__Wallemiaceae(100);g__Wallemia(100);s__Wallemia_sebi(100); | Wallemia | 13 | Saprotroph | \|Undefined Saprotroph\| | Microfungus | NULL | Probable | Decay sugar-rich substrates (Põlme et al. 2020) | Tedersoo L, et al. 2014. Science 346:e1256688 ((https://doi.org/10.1126/science.1256688)); Põlme S, et al. 2020. Fungal Diversity 105:1-16 ((https://doi.org/10.1007/s13225-020-00466-2)) |
| Otu27124 | 1 | k__Fungi(100);p__Basidiomycota(100);c__Wallemiomycetes(100);o__Wallemiales(100);f__Wallemiaceae(100);g__Wallemia(100);s__Wallemia_sebi(100); | Wallemia | 13 | Saprotroph | \|Undefined Saprotroph\| | Microfungus | NULL | Probable | Decay sugar-rich substrates (Põlme et al. 2020) | Tedersoo L, et al. 2014. Science 346:e1256688 ((https://doi.org/10.1126/science.1256688)); Põlme S, et al. 2020. Fungal Diversity 105:1-16 ((https://doi.org/10.1007/s13225-020-00466-2)) |
| Otu27125 | 1 | k__Fungi(100);p__Basidiomycota(100);c__Wallemiomycetes(100);o__Wallemiales(100);f__Wallemiaceae(100);g__Wallemia(100);s__Wallemia_sebi(100); | Wallemia | 13 | Saprotroph | \|Undefined Saprotroph\| | Microfungus | NULL | Probable | Decay sugar-rich substrates (Põlme et al. 2020) | Tedersoo L, et al. 2014. Science 346:e1256688 ((https://doi.org/10.1126/science.1256688)); Põlme S, et al. 2020. Fungal Diversity 105:1-16 ((https://doi.org/10.1007/s13225-020-00466-2)) |
| Otu27126 | 1 | k__Fungi(100);p__Basidiomycota(100);c__Wallemiomycetes(100);o__Wallemiales(100);f__Wallemiaceae(100);g__Wallemia(100);s__Wallemia_sebi(100); | Wallemia | 13 | Saprotroph | \|Undefined Saprotroph\| | Microfungus | NULL | Probable | Decay sugar-rich substrates (Põlme et al. 2020) | Tedersoo L, et al. 2014. Science 346:e1256688 ((https://doi.org/10.1126/science.1256688)); Põlme S, et al. 2020. Fungal Diversity 105:1-16 ((https://doi.org/10.1007/s13225-020-00466-2)) |
| Otu27127 | 1 | k__Fungi(100);p__Basidiomycota(100);c__Wallemiomycetes(100);o__Wallemiales(100);f__Wallemiaceae(100);g__Wallemia(100);s__Wallemia_sebi(100); | Wallemia | 13 | Saprotroph | \|Undefined Saprotroph\| | Microfungus | NULL | Probable | Decay sugar-rich substrates (Põlme et al. 2020) | Tedersoo L, et al. 2014. Science 346:e1256688 ((https://doi.org/10.1126/science.1256688)); Põlme S, et al. 2020. Fungal Diversity 105:1-16 ((https://doi.org/10.1007/s13225-020-00466-2)) |
| Otu27128 | 1 | k__Fungi(100);p__Basidiomycota(100);c__Wallemiomycetes(100);o__Wallemiales(100);f__Wallemiaceae(100);g__Wallemia(100);s__Wallemia_sebi(100); | Wallemia | 13 | Saprotroph | \|Undefined Saprotroph\| | Microfungus | NULL | Probable | Decay sugar-rich substrates (Põlme et al. 2020) | Tedersoo L, et al. 2014. Science 346:e1256688 ((https://doi.org/10.1126/science.1256688)); Põlme S, et al. 2020. Fungal Diversity 105:1-16 ((https://doi.org/10.1007/s13225-020-00466-2)) |
| Otu27129 | 1 | k__Fungi(100);p__Basidiomycota(100);c__Wallemiomycetes(100);o__Wallemiales(100);f__Wallemiaceae(100);g__Wallemia(100);s__Wallemia_sebi(100); | Wallemia | 13 | Saprotroph | \|Undefined Saprotroph\| | Microfungus | NULL | Probable | Decay sugar-rich substrates (Põlme et al. 2020) | Tedersoo L, et al. 2014. Science 346:e1256688 ((https://doi.org/10.1126/science.1256688)); Põlme S, et al. 2020. Fungal Diversity 105:1-16 ((https://doi.org/10.1007/s13225-020-00466-2)) |
| Otu27130 | 1 | k__Fungi(100);p__Basidiomycota(100);c__Wallemiomycetes(100);o__Wallemiales(100);f__Wallemiaceae(100);g__Wallemia(100);s__Wallemia_sebi(100); | Wallemia | 13 | Saprotroph | \|Undefined Saprotroph\| | Microfungus | NULL | Probable | Decay sugar-rich substrates (Põlme et al. 2020) | Tedersoo L, et al. 2014. Science 346:e1256688 ((https://doi.org/10.1126/science.1256688)); Põlme S, et al. 2020. Fungal Diversity 105:1-16 ((https://doi.org/10.1007/s13225-020-00466-2)) |
| Otu27131 | 1 | k__Fungi(100);p__Basidiomycota(100);c__Wallemiomycetes(100);o__Wallemiales(100);f__Wallemiaceae(100);g__Wallemia(100);s__Wallemia_sebi(100); | Wallemia | 13 | Saprotroph | \|Undefined Saprotroph\| | Microfungus | NULL | Probable | Decay sugar-rich substrates (Põlme et al. 2020) | Tedersoo L, et al. 2014. Science 346:e1256688 ((https://doi.org/10.1126/science.1256688)); Põlme S, et al. 2020. Fungal Diversity 105:1-16 ((https://doi.org/10.1007/s13225-020-00466-2)) |
| Otu27132 | 1 | k__Fungi(100);p__Basidiomycota(100);c__Wallemiomycetes(100);o__Wallemiales(100);f__Wallemiaceae(100);g__Wallemia(100);s__Wallemia_sebi(100); | Wallemia | 13 | Saprotroph | \|Undefined Saprotroph\| | Microfungus | NULL | Probable | Decay sugar-rich substrates (Põlme et al. 2020) | Tedersoo L, et al. 2014. Science 346:e1256688 ((https://doi.org/10.1126/science.1256688)); Põlme S, et al. 2020. Fungal Diversity 105:1-16 ((https://doi.org/10.1007/s13225-020-00466-2)) |
| Otu27133 | 1 | k__Fungi(100);p__Basidiomycota(100);c__Wallemiomycetes(100);o__Wallemiales(100);f__Wallemiaceae(100);g__Wallemia(100);s__Wallemia_sebi(100); | Wallemia | 13 | Saprotroph | \|Undefined Saprotroph\| | Microfungus | NULL | Probable | Decay sugar-rich substrates (Põlme et al. 2020) | Tedersoo L, et al. 2014. Science 346:e1256688 ((https://doi.org/10.1126/science.1256688)); Põlme S, et al. 2020. Fungal Diversity 105:1-16 ((https://doi.org/10.1007/s13225-020-00466-2)) |

| | | | | | | | | | | | | | |
|---|---|---|---|---|---|---|---|---|---|---|---|---|---|
| Otu27160 | | 1 | k__Fungi(100);p__Ascomycota(100);c__Sordariomycetes(100);o__Sordariomycetes_order_Incertae_sedis(100);f__Sordariomycetes_family_Incertae_sedis(100);g__Myrmecridium(100);s__Myrmecridium_schulzeri(100); | Myrmecridium | | 13 | Saprotroph | \|Undefined Saprotroph\| | Microfungus | NULL | Probable | As soil_saprotroph (Põlme et al. 2020); Decay soil (Põlme et al. 2020) | Tedersoo L, et al. 2014. Science 346:e1256688 ((https://doi.org/10.1126/science.1256688)); Põlme S, et al. 2020. Fungal Diversity 105:1-16 ((https://doi.org/10.1007/s13225-020-00466-2)) |
| Otu27161 | | 1 | k__Fungi(100);p__Ascomycota(100);c__Saccharomycetes(100);o__Saccharomycetales(100);f__Saccharomycetales_family_Incertae_sedis(100);g__Candida(100);g__Candida_unclassified(100); | Candida | | 13 | Pathotroph-Saprotroph-Symbiotroph | Animal Pathogen-Endophyte-Endosymbiont-Epiphyte-\|Undefined Saprotroph\| | Dimorphic Yeast-Yeast | NULL | Probable | As nectar-tap_saprotroph (Põlme et al. 2020); Decay sugar-rich substrates (Põlme et al. 2020); This asexual genus is polyphyletic and contain species that function very differently depend on what they are. Caution should be taken when interpreting guilds for Candida species. | Manolakaki D, et al. 2010. Virulence. 1: 367-375 ((https://doi.org/10.4161/viru.1.5.12796)); Mekbib SB, et al. 2011. Tropical Plant Pathology 36:233-240 ((https://doi.org/10.1590/S1982-56762011000400004)); Põlme S, et al. 2020. Fungal Diversity 105:1-16 ((https://doi.org/10.1007/s13225-020-00466-2)) |
| Otu27163 | | 1 | k__Fungi(100);p__Ascomycota(100);c__Saccharomycetes(100);o__Saccharomycetales(100);f__Metschnikowiaceae(100);g__Metschnikowia(100);s__Metschnikowia_pulcherrima(100); | Metschnikowia | | 13 | Saprotroph | \|Undefined Saprotroph\| | Yeast | NULL | Probable | As nectar-tap_saprotroph (Põlme et al. 2020); Decay sugar-rich substrates (Põlme et al. 2020) | Põlme S, et al. 2020. Fungal Diversity 105:1-16 ((https://doi.org/10.1007/s13225-020-00466-2)) |
| Otu27164 | | 1 | k__Fungi(100);p__Ascomycota(100);c__Saccharomycetes(100);o__Saccharomycetales(100);f__Metschnikowiaceae(100);g__Metschnikowia(100);s__Metschnikowia_pulcherrima(100); | Metschnikowia | | 13 | Saprotroph | \|Undefined Saprotroph\| | Yeast | NULL | Probable | As nectar-tap_saprotroph (Põlme et al. 2020); Decay sugar-rich substrates (Põlme et al. 2020) | Põlme S, et al. 2020. Fungal Diversity 105:1-16 ((https://doi.org/10.1007/s13225-020-00466-2)) |
| Otu27165 | | 1 | k__Fungi(100);p__Ascomycota(100);c__Saccharomycetes(100);o__Saccharomycetales(100);f__Metschnikowiaceae(100);g__Metschnikowia(100);s__Metschnikowia_pulcherrima(100); | Metschnikowia | | 13 | Saprotroph | \|Undefined Saprotroph\| | Yeast | NULL | Probable | As nectar-tap_saprotroph (Põlme et al. 2020); Decay sugar-rich substrates (Põlme et al. 2020) | Põlme S, et al. 2020. Fungal Diversity 105:1-16 ((https://doi.org/10.1007/s13225-020-00466-2)) |
| Otu27167 | | 1 | k__Fungi(100);p__Ascomycota(100);c__Saccharomycetes(100);o__Saccharomycetales(100);f__Metschnikowiaceae(100);g__Metschnikowia(100);s__Metschnikowia_pulcherrima(100); | Metschnikowia | | 13 | Saprotroph | \|Undefined Saprotroph\| | Yeast | NULL | Probable | As nectar-tap_saprotroph (Põlme et al. 2020); Decay sugar-rich substrates (Põlme et al. 2020) | Põlme S, et al. 2020. Fungal Diversity 105:1-16 ((https://doi.org/10.1007/s13225-020-00466-2)) |
| Otu27169 | | 1 | k__Fungi(100);p__Ascomycota(100);c__Saccharomycetes(100);o__Saccharomycetales(100);f__Metschnikowiaceae(100);g__Metschnikowia(100);g__Metschnikowia_unclassified(100); | Metschnikowia | | 13 | Saprotroph | \|Undefined Saprotroph\| | Yeast | NULL | Probable | As nectar-tap_saprotroph (Põlme et al. 2020); Decay sugar-rich substrates (Põlme et al. 2020) | Põlme S, et al. 2020. Fungal Diversity 105:1-16 ((https://doi.org/10.1007/s13225-020-00466-2)) |
| Otu27171 | | 1 | k__Fungi(100);p__Ascomycota(100);c__Saccharomycetes(100);o__Saccharomycetales(100);f__Metschnikowiaceae(100);g__Metschnikowia(100);s__Metschnikowia_pulcherrima(100); | Metschnikowia | | 13 | Saprotroph | \|Undefined Saprotroph\| | Yeast | NULL | Probable | As nectar-tap_saprotroph (Põlme et al. 2020); Decay sugar-rich substrates (Põlme et al. 2020) | Põlme S, et al. 2020. Fungal Diversity 105:1-16 ((https://doi.org/10.1007/s13225-020-00466-2)) |
| Otu27172 | | 1 | k__Fungi(100);p__Ascomycota(100);c__Saccharomycetes(100);o__Saccharomycetales(100);f__Metschnikowiaceae(100);g__Metschnikowia(100);g__Metschnikowia_unclassified(100); | Metschnikowia | | 13 | Saprotroph | \|Undefined Saprotroph\| | Yeast | NULL | Probable | As nectar-tap_saprotroph (Põlme et al. 2020); Decay sugar-rich substrates (Põlme et al. 2020) | Põlme S, et al. 2020. Fungal Diversity 105:1-16 ((https://doi.org/10.1007/s13225-020-00466-2)) |
| Otu27173 | | 1 | k__Fungi(100);p__Ascomycota(100);c__Saccharomycetes(100);o__Saccharomycetales(100);f__Metschnikowiaceae(100);g__Metschnikowia(100);s__Metschnikowia_pulcherrima(100); | Metschnikowia | | 13 | Saprotroph | \|Undefined Saprotroph\| | Yeast | NULL | Probable | As nectar-tap_saprotroph (Põlme et al. 2020); Decay sugar-rich substrates (Põlme et al. 2020) | Põlme S, et al. 2020. Fungal Diversity 105:1-16 ((https://doi.org/10.1007/s13225-020-00466-2)) |
| Otu27174 | | 1 | k__Fungi(100);p__Ascomycota(100);c__Saccharomycetes(100);o__Saccharomycetales(100);f__Metschnikowiaceae(100);g__Metschnikowia(100);s__Metschnikowia_pulcherrima(100); | Metschnikowia | | 13 | Saprotroph | \|Undefined Saprotroph\| | Yeast | NULL | Probable | As nectar-tap_saprotroph (Põlme et al. 2020); Decay sugar-rich substrates (Põlme et al. 2020) | Põlme S, et al. 2020. Fungal Diversity 105:1-16 ((https://doi.org/10.1007/s13225-020-00466-2)) |
| Otu27175 | | 1 | k__Fungi(100);p__Ascomycota(100);c__Saccharomycetes(100);o__Saccharomycetales(100);f__Metschnikowiaceae(100);g__Metschnikowia(100);s__Metschnikowia_pulcherrima(100); | Metschnikowia | | 13 | Saprotroph | \|Undefined Saprotroph\| | Yeast | NULL | Probable | As nectar-tap_saprotroph (Põlme et al. 2020); Decay sugar-rich substrates (Põlme et al. 2020) | Põlme S, et al. 2020. Fungal Diversity 105:1-16 ((https://doi.org/10.1007/s13225-020-00466-2)) |
| Otu27176 | | 1 | k__Fungi(100);p__Ascomycota(100);c__Saccharomycetes(100);o__Saccharomycetales(100);f__Metschnikowiaceae(100);g__Metschnikowia(100);s__Metschnikowia_pulcherrima(100); | Metschnikowia | | 13 | Saprotroph | \|Undefined Saprotroph\| | Yeast | NULL | Probable | As nectar-tap_saprotroph (Põlme et al. 2020); Decay sugar-rich substrates (Põlme et al. 2020) | Põlme S, et al. 2020. Fungal Diversity 105:1-16 ((https://doi.org/10.1007/s13225-020-00466-2)) |
| Otu27177 | | 1 | k__Fungi(100);p__Ascomycota(100);c__Saccharomycetes(100);o__Saccharomycetales(100);f__Metschnikowiaceae(100);g__Metschnikowia(100);s__Metschnikowia_pulcherrima(100); | Metschnikowia | | 13 | Saprotroph | \|Undefined Saprotroph\| | Yeast | NULL | Probable | As nectar-tap_saprotroph (Põlme et al. 2020); Decay sugar-rich substrates (Põlme et al. 2020) | Põlme S, et al. 2020. Fungal Diversity 105:1-16 ((https://doi.org/10.1007/s13225-020-00466-2)) |
| Otu27178 | | 1 | k__Fungi(100);p__Ascomycota(100);c__Saccharomycetes(100);o__Saccharomycetales(100);f__Metschnikowiaceae(100);g__Metschnikowia(100);g__Metschnikowia_unclassified(100); | Metschnikowia | | 13 | Saprotroph | \|Undefined Saprotroph\| | Yeast | NULL | Probable | As nectar-tap_saprotroph (Põlme et al. 2020); Decay sugar-rich substrates (Põlme et al. 2020) | Põlme S, et al. 2020. Fungal Diversity 105:1-16 ((https://doi.org/10.1007/s13225-020-00466-2)) |
| Otu27179 | | 1 | k__Fungi(100);p__Ascomycota(100);c__Saccharomycetes(100);o__Saccharomycetales(100);f__Metschnikowiaceae(100);g__Metschnikowia(100);s__Metschnikowia_pulcherrima(100); | Metschnikowia | | 13 | Saprotroph | \|Undefined Saprotroph\| | Yeast | NULL | Probable | As nectar-tap_saprotroph (Põlme et al. 2020); Decay sugar-rich substrates (Põlme et al. 2020) | Põlme S, et al. 2020. Fungal Diversity 105:1-16 ((https://doi.org/10.1007/s13225-020-00466-2)) |
| Otu27180 | | 1 | k__Fungi(100);p__Ascomycota(100);c__Saccharomycetes(100);o__Saccharomycetales(100);f__Metschnikowiaceae(100);g__Metschnikowia(100);s__Metschnikowia_pulcherrima(100); | Metschnikowia | | 13 | Saprotroph | \|Undefined Saprotroph\| | Yeast | NULL | Probable | As nectar-tap_saprotroph (Põlme et al. 2020); Decay sugar-rich substrates (Põlme et al. 2020) | Põlme S, et al. 2020. Fungal Diversity 105:1-16 ((https://doi.org/10.1007/s13225-020-00466-2)) |
| Otu27181 | | 1 | k__Fungi(100);p__Ascomycota(100);c__Saccharomycetes(100);o__Saccharomycetales(100);f__Metschnikowiaceae(100);g__Metschnikowia(100);s__Metschnikowia_pulcherrima(100); | Metschnikowia | | 13 | Saprotroph | \|Undefined Saprotroph\| | Yeast | NULL | Probable | As nectar-tap_saprotroph (Põlme et al. 2020); Decay sugar-rich substrates (Põlme et al. 2020) | Põlme S, et al. 2020. Fungal Diversity 105:1-16 ((https://doi.org/10.1007/s13225-020-00466-2)) |
| Otu27182 | | 1 | k__Fungi(100);p__Ascomycota(100);c__Saccharomycetes(100);o__Saccharomycetales(100);f__Metschnikowiaceae(100);g__Metschnikowia(100);g__Metschnikowia_unclassified(100); | Metschnikowia | | 13 | Saprotroph | \|Undefined Saprotroph\| | Yeast | NULL | Probable | As nectar-tap_saprotroph (Põlme et al. 2020); Decay sugar-rich substrates (Põlme et al. 2020) | Põlme S, et al. 2020. Fungal Diversity 105:1-16 ((https://doi.org/10.1007/s13225-020-00466-2)) |
| Otu27183 | | 1 | k__Fungi(100);p__Ascomycota(100);c__Saccharomycetes(100);o__Saccharomycetales(100);f__Metschnikowiaceae(100);g__Metschnikowia(100);s__Metschnikowia_pulcherrima(100); | Metschnikowia | | 13 | Saprotroph | \|Undefined Saprotroph\| | Yeast | NULL | Probable | As nectar-tap_saprotroph (Põlme et al. 2020); Decay sugar-rich substrates (Põlme et al. 2020) | Põlme S, et al. 2020. Fungal Diversity 105:1-16 ((https://doi.org/10.1007/s13225-020-00466-2)) |
| Otu27184 | | 1 | k__Fungi(100);p__Ascomycota(100);c__Saccharomycetes(100);o__Saccharomycetales(100);f__Metschnikowiaceae(100);g__Metschnikowia(100);s__Metschnikowia_pulcherrima(100); | Metschnikowia | | 13 | Saprotroph | \|Undefined Saprotroph\| | Yeast | NULL | Probable | As nectar-tap_saprotroph (Põlme et al. 2020); Decay sugar-rich substrates (Põlme et al. 2020) | Põlme S, et al. 2020. Fungal Diversity 105:1-16 ((https://doi.org/10.1007/s13225-020-00466-2)) |
| Otu27185 | | 1 | k__Fungi(100);p__Ascomycota(100);c__Saccharomycetes(100);o__Saccharomycetales(100);f__Metschnikowiaceae(100);g__Metschnikowia(100);g__Metschnikowia_unclassified(100); | Metschnikowia | | 13 | Saprotroph | \|Undefined Saprotroph\| | Yeast | NULL | Probable | As nectar-tap_saprotroph (Põlme et al. 2020); Decay sugar-rich substrates (Põlme et al. 2020) | Põlme S, et al. 2020. Fungal Diversity 105:1-16 ((https://doi.org/10.1007/s13225-020-00466-2)) |
| Otu27186 | | 1 | k__Fungi(100);p__Ascomycota(100);c__Saccharomycetes(100);o__Saccharomycetales(100);f__Metschnikowiaceae(100);g__Metschnikowia(100);g__Metschnikowia_unclassified(100); | Metschnikowia | | 13 | Saprotroph | \|Undefined Saprotroph\| | Yeast | NULL | Probable | As nectar-tap_saprotroph (Põlme et al. 2020); Decay sugar-rich substrates (Põlme et al. 2020) | Põlme S, et al. 2020. Fungal Diversity 105:1-16 ((https://doi.org/10.1007/s13225-020-00466-2)) |
| Otu27188 | | 1 | k__Fungi(100);p__Ascomycota(100);c__Saccharomycetes(100);o__Saccharomycetales(100);f__Metschnikowiaceae(100);g__Metschnikowia(100);s__Metschnikowia_pulcherrima(100); | Metschnikowia | | 13 | Saprotroph | \|Undefined Saprotroph\| | Yeast | NULL | Probable | As nectar-tap_saprotroph (Põlme et al. 2020); Decay sugar-rich substrates (Põlme et al. 2020) | Põlme S, et al. 2020. Fungal Diversity 105:1-16 ((https://doi.org/10.1007/s13225-020-00466-2)) |
| Otu27189 | | 1 | k__Fungi(100);p__Ascomycota(100);c__Saccharomycetes(100);o__Saccharomycetales(100);f__Metschnikowiaceae(100);g__Metschnikowia(100);s__Metschnikowia_pulcherrima(100); | Metschnikowia | | 13 | Saprotroph | \|Undefined Saprotroph\| | Yeast | NULL | Probable | As nectar-tap_saprotroph (Põlme et al. 2020); Decay sugar-rich substrates (Põlme et al. 2020) | Põlme S, et al. 2020. Fungal Diversity 105:1-16 ((https://doi.org/10.1007/s13225-020-00466-2)) |
| Otu27190 | | 1 | k__Fungi(100);p__Ascomycota(100);c__Saccharomycetes(100);o__Saccharomycetales(100);f__Metschnikowiaceae(100);g__Metschnikowia(100);g__Metschnikowia_unclassified(100); | Metschnikowia | | 13 | Saprotroph | \|Undefined Saprotroph\| | Yeast | NULL | Probable | As nectar-tap_saprotroph (Põlme et al. 2020); Decay sugar-rich substrates (Põlme et al. 2020) | Põlme S, et al. 2020. Fungal Diversity 105:1-16 ((https://doi.org/10.1007/s13225-020-00466-2)) |
| Otu27191 | | 1 | k__Fungi(100);p__Ascomycota(100);c__Saccharomycetes(100);o__Saccharomycetales(100);f__Metschnikowiaceae(100);g__Metschnikowia(100);s__Metschnikowia_pulcherrima(100); | Metschnikowia | | 13 | Saprotroph | \|Undefined Saprotroph\| | Yeast | NULL | Probable | As nectar-tap_saprotroph (Põlme et al. 2020); Decay sugar-rich substrates (Põlme et al. 2020) | Põlme S, et al. 2020. Fungal Diversity 105:1-16 ((https://doi.org/10.1007/s13225-020-00466-2)) |
| Otu27192 | | 1 | k__Fungi(100);p__Basidiomycota(100);c__Tremellomycetes(100);o__Tremellales(100);f__Tremellales_family_Incertae_sedis(100);g__Cryptococcus(100);s__Cryptococcus_sp_JCM_11356(100); | Cryptococcus | | 13 | Pathotroph-Saprotroph-Symbiotroph | Animal Pathogen-Endophyte-Epiphyte-\|Undefined Saprotroph\| | Dimorphic Yeast-Yeast | NULL | Possible | As foliar_endophyte (Põlme et al. 2020); Decay leaf, fruit, seed, soil, dung, animal material (Põlme et al. 2020) | Andrews JH, et al. 1983. Phytopathology 73:228-234 ((https://doi.org/10.1094/Phyto-73-228)); Kurtzman CP, et al. (eds.) 2011. The Yeasts, a Taxonomic Study. Fifth Edition. Vols 1-3. Elsevier, San Diego (ISBN: 9780444521491); Põlme S, et al. 2020. Fungal Diversity 105:1-16 ((https://doi.org/10.1007/s13225-020-00466-2)) |
| Otu27194 | | 1 | k__Fungi(100);p__Ascomycota(100);c__Saccharomycetes(100);o__Saccharomycetales(100);f__Metschnikowiaceae(100);g__Metschnikowia(100);s__Metschnikowia_pulcherrima(100); | Metschnikowia | | 13 | Saprotroph | \|Undefined Saprotroph\| | Yeast | NULL | Probable | As nectar-tap_saprotroph (Põlme et al. 2020); Decay sugar-rich substrates (Põlme et al. 2020) | Põlme S, et al. 2020. Fungal Diversity 105:1-16 ((https://doi.org/10.1007/s13225-020-00466-2)) |
| Otu27195 | | 1 | k__Fungi(100);p__Ascomycota(100);c__Leotiomycetes(100);o__Helotiales(100);f__Hyaloscyphaceae(100);g__Haplographium(100);s__Haplographium_delicatum(100); | Haplographium | | 13 | Saprotroph | \|Undefined Saprotroph\| | Microfungus | NULL | Probable | NULL | Tedersoo L, et al. 2014. Science 346:e1256688 ((https://doi.org/10.1126/science.1256688)); Põlme S, et al. 2020. Fungal Diversity 105:1-16 ((https://doi.org/10.1007/s13225-020-00466-2)) |
| Otu27196 | | 1 | k__Fungi(100);p__Ascomycota(100);c__Saccharomycetes(100);o__Saccharomycetales(100);f__Metschnikowiaceae(100);g__Metschnikowia(100);s__Metschnikowia_pulcherrima(100); | Metschnikowia | | 13 | Saprotroph | \|Undefined Saprotroph\| | Yeast | NULL | Probable | As nectar-tap_saprotroph (Põlme et al. 2020); Decay sugar-rich substrates (Põlme et al. 2020) | Põlme S, et al. 2020. Fungal Diversity 105:1-16 ((https://doi.org/10.1007/s13225-020-00466-2)) |
| Otu27197 | | 1 | k__Fungi(100);p__Ascomycota(100);c__Saccharomycetes(100);o__Saccharomycetales(100);f__Metschnikowiaceae(100);g__Metschnikowia(100);s__Metschnikowia_pulcherrima(100); | Metschnikowia | | 13 | Saprotroph | \|Undefined Saprotroph\| | Yeast | NULL | Probable | As nectar-tap_saprotroph (Põlme et al. 2020); Decay sugar-rich substrates (Põlme et al. 2020) | Põlme S, et al. 2020. Fungal Diversity 105:1-16 ((https://doi.org/10.1007/s13225-020-00466-2)) |
| Otu27198 | | 1 | k__Fungi(100);p__Ascomycota(100);c__Saccharomycetes(100);o__Saccharomycetales(100);f__Metschnikowiaceae(100);g__Metschnikowia(100);s__Metschnikowia_pulcherrima(100); | Metschnikowia | | 13 | Saprotroph | \|Undefined Saprotroph\| | Yeast | NULL | Probable | As nectar-tap_saprotroph (Põlme et al. 2020); Decay sugar-rich substrates (Põlme et al. 2020) | Põlme S, et al. 2020. Fungal Diversity 105:1-16 ((https://doi.org/10.1007/s13225-020-00466-2)) |
| Otu27199 | | 1 | k__Fungi(100);p__Ascomycota(100);c__Saccharomycetes(100);o__Saccharomycetales(100);f__Metschnikowiaceae(100);g__Metschnikowia(100);s__Metschnikowia_pulcherrima(100); | Metschnikowia | | 13 | Saprotroph | \|Undefined Saprotroph\| | Yeast | NULL | Probable | As nectar-tap_saprotroph (Põlme et al. 2020); Decay sugar-rich substrates (Põlme et al. 2020) | Põlme S, et al. 2020. Fungal Diversity 105:1-16 ((https://doi.org/10.1007/s13225-020-00466-2)) |
| Otu27200 | | 1 | k__Fungi(100);p__Ascomycota(100);c__Saccharomycetes(100);o__Saccharomycetales(100);f__Metschnikowiaceae(100);g__Metschnikowia(100);s__Metschnikowia_pulcherrima(100); | Metschnikowia | | 13 | Saprotroph | \|Undefined Saprotroph\| | Yeast | NULL | Probable | As nectar-tap_saprotroph (Põlme et al. 2020); Decay sugar-rich substrates (Põlme et al. 2020) | Põlme S, et al. 2020. Fungal Diversity 105:1-16 ((https://doi.org/10.1007/s13225-020-00466-2)) |
| Otu27204 | | 1 | k__Fungi(100);p__Ascomycota(100);c__Saccharomycetes(100);o__Saccharomycetales(100);f__Metschnikowiaceae(100);g__Metschnikowia(100);s__Metschnikowia_pulcherrima(100); | Metschnikowia | | 13 | Saprotroph | \|Undefined Saprotroph\| | Yeast | NULL | Probable | As nectar-tap_saprotroph (Põlme et al. 2020); Decay sugar-rich substrates (Põlme et al. 2020) | Põlme S, et al. 2020. Fungal Diversity 105:1-16 ((https://doi.org/10.1007/s13225-020-00466-2)) |
| Otu27205 | | 1 | k__Fungi(100);p__Ascomycota(100);c__Saccharomycetes(100);o__Saccharomycetales(100);f__Metschnikowiaceae(100);g__Metschnikowia(100);g__Metschnikowia_unclassified(100); | Metschnikowia | | 13 | Saprotroph | \|Undefined Saprotroph\| | Yeast | NULL | Probable | As nectar-tap_saprotroph (Põlme et al. 2020); Decay sugar-rich substrates (Põlme et al. 2020) | Põlme S, et al. 2020. Fungal Diversity 105:1-16 ((https://doi.org/10.1007/s13225-020-00466-2)) |
| Otu27206 | | 1 | k__Fungi(100);p__Ascomycota(100);c__Saccharomycetes(100);o__Saccharomycetales(100);f__Metschnikowiaceae(100);g__Metschnikowia(100);s__Metschnikowia_pulcherrima(100); | Metschnikowia | | 13 | Saprotroph | \|Undefined Saprotroph\| | Yeast | NULL | Probable | As nectar-tap_saprotroph (Põlme et al. 2020); Decay sugar-rich substrates (Põlme et al. 2020) | Põlme S, et al. 2020. Fungal Diversity 105:1-16 ((https://doi.org/10.1007/s13225-020-00466-2)) |
| Otu27207 | | 1 | k__Fungi(100);p__Ascomycota(100);c__Saccharomycetes(100);o__Saccharomycetales(100);f__Saccharomycetales_family_Incertae_sedis(100);g__Candida(100);s__Candida_cretensis(100); | Candida | | 13 | Pathotroph-Saprotroph-Symbiotroph | Animal Pathogen-Endophyte-Endosymbiont-Epiphyte-\|Undefined Saprotroph\| | Dimorphic Yeast-Yeast | NULL | Probable | As nectar-tap_saprotroph (Põlme et al. 2020); Decay sugar-rich substrates (Põlme et al. 2020); This asexual genus is polyphyletic and contain species that function very differently depend on what they are. Caution should be taken when interpreting guilds for Candida species. | Manolakaki D, et al. 2010. Virulence. 1: 367-375 ((https://doi.org/10.4161/viru.1.5.12796)); Mekbib SB, et al. 2011. Tropical Plant Pathology 36:233-240 ((https://doi.org/10.1590/S1982-56762011000400004)); Põlme S, et al. 2020. Fungal Diversity 105:1-16 ((https://doi.org/10.1007/s13225-020-00466-2)) |
| Otu27211 | | 1 | k__Fungi(100);p__Ascomycota(100);c__Saccharomycetes(100);o__Saccharomycetales(100);f__Saccharomycetales_family_Incertae_sedis(100);g__Candida(100);g__Candida_unclassified(100); | Candida | | 13 | Pathotroph-Saprotroph-Symbiotroph | Animal Pathogen-Endophyte-Endosymbiont-Epiphyte-\|Undefined Saprotroph\| | Dimorphic Yeast-Yeast | NULL | Probable | As nectar-tap_saprotroph (Põlme et al. 2020); Decay sugar-rich substrates (Põlme et al. 2020); This asexual genus is polyphyletic and contain species that function very differently depend on what they are. Caution should be taken when interpreting guilds for Candida species. | Manolakaki D, et al. 2010. Virulence. 1: 367-375 ((https://doi.org/10.4161/viru.1.5.12796)); Mekbib SB, et al. 2011. Tropical Plant Pathology 36:233-240 ((https://doi.org/10.1590/S1982-56762011000400004)); Põlme S, et al. 2020. Fungal Diversity 105:1-16 ((https://doi.org/10.1007/s13225-020-00466-2)) |
| Otu27213 | | 1 | k__Fungi(100);p__Ascomycota(100);c__Saccharomycetes(100);o__Saccharomycetales(100);f__Metschnikowiaceae(100);g__Metschnikowia(100);s__Metschnikowia_pulcherrima(100); | Metschnikowia | | 13 | Saprotroph | \|Undefined Saprotroph\| | Yeast | NULL | Probable | As nectar-tap_saprotroph (Põlme et al. 2020); Decay sugar-rich substrates (Põlme et al. 2020) | Põlme S, et al. 2020. Fungal Diversity 105:1-16 ((https://doi.org/10.1007/s13225-020-00466-2)) |
| Otu27214 | | 1 | k__Fungi(100);p__Ascomycota(100);c__Saccharomycetes(100);o__Saccharomycetales(100);f__Metschnikowiaceae(100);g__Metschnikowia(100);s__Metschnikowia_pulcherrima(100); | Metschnikowia | | 13 | Saprotroph | \|Undefined Saprotroph\| | Yeast | NULL | Probable | As nectar-tap_saprotroph (Põlme et al. 2020); Decay sugar-rich substrates (Põlme et al. 2020) | Põlme S, et al. 2020. Fungal Diversity 105:1-16 ((https://doi.org/10.1007/s13225-020-00466-2)) |
| Otu27215 | | 1 | k__Fungi(100);p__Ascomycota(100);c__Saccharomycetes(100);o__Saccharomycetales(100);f__Metschnikowiaceae(100);g__Metschnikowia(100);s__Metschnikowia_pulcherrima(100); | Metschnikowia | | 13 | Saprotroph | \|Undefined Saprotroph\| | Yeast | NULL | Probable | As nectar-tap_saprotroph (Põlme et al. 2020); Decay sugar-rich substrates (Põlme et al. 2020) | Põlme S, et al. 2020. Fungal Diversity 105:1-16 ((https://doi.org/10.1007/s13225-020-00466-2)) |
| Otu27216 | | 1 | k__Fungi(100);p__Ascomycota(100);c__Saccharomycetes(100);o__Saccharomycetales(100);f__Metschnikowiaceae(100);g__Metschnikowia(100);s__Metschnikowia_pulcherrima(100); | Metschnikowia | | 13 | Saprotroph | \|Undefined Saprotroph\| | Yeast | NULL | Probable | As nectar-tap_saprotroph (Põlme et al. 2020); Decay sugar-rich substrates (Põlme et al. 2020) | Põlme S, et al. 2020. Fungal Diversity 105:1-16 ((https://doi.org/10.1007/s13225-020-00466-2)) |
| Otu27217 | | 1 | k__Fungi(100);p__Ascomycota(100);c__Saccharomycetes(100);o__Saccharomycetales(100);f__Metschnikowiaceae(100);g__Metschnikowia(100);s__Metschnikowia_pulcherrima(100); | Metschnikowia | | 13 | Saprotroph | \|Undefined Saprotroph\| | Yeast | NULL | Probable | As nectar-tap_saprotroph (Põlme et al. 2020); Decay sugar-rich substrates (Põlme et al. 2020) | Põlme S, et al. 2020. Fungal Diversity 105:1-16 ((https://doi.org/10.1007/s13225-020-00466-2)) |
| Otu27218 | | 1 | k__Fungi(100);p__Ascomycota(100);c__Saccharomycetes(100);o__Saccharomycetales(100);f__Metschnikowiaceae(100);g__Metschnikowia(100);g__Metschnikowia_unclassified(100); | Metschnikowia | | 13 | Saprotroph | \|Undefined Saprotroph\| | Yeast | NULL | Probable | As nectar-tap_saprotroph (Põlme et al. 2020); Decay sugar-rich substrates (Põlme et al. 2020) | Põlme S, et al. 2020. Fungal Diversity 105:1-16 ((https://doi.org/10.1007/s13225-020-00466-2)) |
| Otu27220 | | 1 | k__Fungi(100);p__Ascomycota(100);c__Saccharomycetes(100);o__Saccharomycetales(100);f__Metschnikowiaceae(100);g__Metschnikowia(100);s__Metschnikowia_pulcherrima(100); | Metschnikowia | | 13 | Saprotroph | \|Undefined Saprotroph\| | Yeast | NULL | Probable | As nectar-tap_saprotroph (Põlme et al. 2020); Decay sugar-rich substrates (Põlme et al. 2020) | Põlme S, et al. 2020. Fungal Diversity 105:1-16 ((https://doi.org/10.1007/s13225-020-00466-2)) |
| Otu27221 | | 1 | k__Fungi(100);p__Ascomycota(100);c__Saccharomycetes(100);o__Saccharomycetales(100);f__Metschnikowiaceae(100);g__Metschnikowia(100);g__Metschnikowia_unclassified(100); | Metschnikowia | | 13 | Saprotroph | \|Undefined Saprotroph\| | Yeast | NULL | Probable | As nectar-tap_saprotroph (Põlme et al. 2020); Decay sugar-rich substrates (Põlme et al. 2020) | Põlme S, et al. 2020. Fungal Diversity 105:1-16 ((https://doi.org/10.1007/s13225-020-00466-2)) |
| Otu27222 | | 1 | k__Fungi(100);p__Ascomycota(100);c__Saccharomycetes(100);o__Saccharomycetales(100);f__Metschnikowiaceae(100);g__Metschnikowia(100);g__Metschnikowia_unclassified(100); | Metschnikowia | | 13 | Saprotroph | \|Undefined Saprotroph\| | Yeast | NULL | Probable | As nectar-tap_saprotroph (Põlme et al. 2020); Decay sugar-rich substrates (Põlme et al. 2020) | Põlme S, et al. 2020. Fungal Diversity 105:1-16 ((https://doi.org/10.1007/s13225-020-00466-2)) |
| Otu27223 | | 1 | k__Fungi(100);p__Ascomycota(100);c__Saccharomycetes(100);o__Saccharomycetales(100);f__Metschnikowiaceae(100);g__Metschnikowia(100);s__Metschnikowia_pulcherrima(100); | Metschnikowia | | 13 | Saprotroph | \|Undefined Saprotroph\| | Yeast | NULL | Probable | As nectar-tap_saprotroph (Põlme et al. 2020); Decay sugar-rich substrates (Põlme et al. 2020) | Põlme S, et al. 2020. Fungal Diversity 105:1-16 ((https://doi.org/10.1007/s13225-020-00466-2)) |
| Otu27224 | | 1 | k__Fungi(100);p__Ascomycota(100);c__Saccharomycetes(100);o__Saccharomycetales(100);f__Metschnikowiaceae(100);g__Metschnikowia(100);s__Metschnikowia_pulcherrima(100); | Metschnikowia | | 13 | Saprotroph | \|Undefined Saprotroph\| | Yeast | NULL | Probable | As nectar-tap_saprotroph (Põlme et al. 2020); Decay sugar-rich substrates (Põlme et al. 2020) | Põlme S, et al. 2020. Fungal Diversity 105:1-16 ((https://doi.org/10.1007/s13225-020-00466-2)) |
| Otu27225 | | 1 | k__Fungi(100);p__Ascomycota(100);c__Saccharomycetes(100);o__Saccharomycetales(100);f__Metschnikowiaceae(100);g__Metschnikowia(100);s__Metschnikowia_pulcherrima(100); | Metschnikowia | | 13 | Saprotroph | \|Undefined Saprotroph\| | Yeast | NULL | Probable | As nectar-tap_saprotroph (Põlme et al. 2020); Decay sugar-rich substrates (Põlme et al. 2020) | Põlme S, et al. 2020. Fungal Diversity 105:1-16 ((https://doi.org/10.1007/s13225-020-00466-2)) |
| Otu27226 | | 1 | k__Fungi(100);p__Ascomycota(100);c__Saccharomycetes(100);o__Saccharomycetales(100);f__Metschnikowiaceae(100);g__Metschnikowia(100);s__Metschnikowia_pulcherrima(100); | Metschnikowia | | 13 | Saprotroph | \|Undefined Saprotroph\| | Yeast | NULL | Probable | As nectar-tap_saprotroph (Põlme et al. 2020); Decay sugar-rich substrates (Põlme et al. 2020) | Põlme S, et al. 2020. Fungal Diversity 105:1-16 ((https://doi.org/10.1007/s13225-020-00466-2)) |
| Otu27227 | | 1 | k__Fungi(100);p__Ascomycota(100);c__Saccharomycetes(100);o__Saccharomycetales(100);f__Metschnikowiaceae(100);g__Metschnikowia(100);s__Metschnikowia_pulcherrima(100); | Metschnikowia | | 13 | Saprotroph | \|Undefined Saprotroph\| | Yeast | NULL | Probable | As nectar-tap_saprotroph (Põlme et al. 2020); Decay sugar-rich substrates (Põlme et al. 2020) | Põlme S, et al. 2020. Fungal Diversity 105:1-16 ((https://doi.org/10.1007/s13225-020-00466-2)) |
| Otu27228 | | 1 | k__Fungi(100);p__Ascomycota(100);c__Saccharomycetes(100);o__Saccharomycetales(100);f__Metschnikowiaceae(100);g__Metschnikowia(100);s__Metschnikowia_pulcherrima(100); | Metschnikowia | | 13 | Saprotroph | \|Undefined Saprotroph\| | Yeast | NULL | Probable | As nectar-tap_saprotroph (Põlme et al. 2020); Decay sugar-rich substrates (Põlme et al. 2020) | Põlme S, et al. 2020. Fungal Diversity 105:1-16 ((https://doi.org/10.1007/s13225-020-00466-2)) |
| Otu27229 | | 1 | k__Fungi(100);p__Ascomycota(100);c__Saccharomycetes(100);o__Saccharomycetales(100);f__Metschnikowiaceae(100);g__Metschnikowia(100);s__Metschnikowia_pulcherrima(100); | Metschnikowia | | 13 | Saprotroph | \|Undefined Saprotroph\| | Yeast | NULL | Probable | As nectar-tap_saprotroph (Põlme et al. 2020); Decay sugar-rich substrates (Põlme et al. 2020) | Põlme S, et al. 2020. Fungal Diversity 105:1-16 ((https://doi.org/10.1007/s13225-020-00466-2)) |
| Otu27230 | | 1 | k__Fungi(100);p__Ascomycota(100);c__Saccharomycetes(100);o__Saccharomycetales(100);f__Metschnikowiaceae(100);g__Metschnikowia(100);s__Metschnikowia_pulcherrima(100); | Metschnikowia | | 13 | Saprotroph | \|Undefined Saprotroph\| | Yeast | NULL | Probable | As nectar-tap_saprotroph (Põlme et al. 2020); Decay sugar-rich substrates (Põlme et al. 2020) | Põlme S, et al. 2020. Fungal Diversity 105:1-16 ((https://doi.org/10.1007/s13225-020-00466-2)) |
| Otu27232 | | 1 | k__Fungi(100);p__Ascomycota(100);c__Saccharomycetes(100);o__Saccharomycetales(100);f__Metschnikowiaceae(100);g__Metschnikowia(100);g__Metschnikowia_unclassified(100); | Metschnikowia | | 13 | Saprotroph | \|Undefined Saprotroph\| | Yeast | NULL | Probable | As nectar-tap_saprotroph (Põlme et al. 2020); Decay sugar-rich substrates (Põlme et al. 2020) | Põlme S, et al. 2020. Fungal Diversity 105:1-16 ((https://doi.org/10.1007/s13225-020-00466-2)) |
| Otu27233 | | 1 | k__Fungi(100);p__Ascomycota(100);c__Saccharomycetes(100);o__Saccharomycetales(100);f__Metschnikowiaceae(100);g__Metschnikowia(100);s__Metschnikowia_pulcherrima(100); | Metschnikowia | | 13 | Saprotroph | \|Undefined Saprotroph\| | Yeast | NULL | Probable | As nectar-tap_saprotroph (Põlme et al. 2020); Decay sugar-rich substrates (Põlme et al. 2020) | Põlme S, et al. 2020. Fungal Diversity 105:1-16 ((https://doi.org/10.1007/s13225-020-00466-2)) |
| Otu27234 | | 1 | k__Fungi(100);p__Ascomycota(100);c__Saccharomycetes(100);o__Saccharomycetales(100);f__Metschnikowiaceae(100);g__Metschnikowia(100);g__Metschnikowia_unclassified(100); | Metschnikowia | | 13 | Saprotroph | \|Undefined Saprotroph\| | Yeast | NULL | Probable | As nectar-tap_saprotroph (Põlme et al. 2020); Decay sugar-rich substrates (Põlme et al. 2020) | Põlme S, et al. 2020. Fungal Diversity 105:1-16 ((https://doi.org/10.1007/s13225-020-00466-2)) |
| Otu27235 | | 1 | k__Fungi(100);p__Ascomycota(100);c__Saccharomycetes(100);o__Saccharomycetales(100);f__Metschnikowiaceae(100);g__Metschnikowia(100);s__Metschnikowia_pulcherrima(100); | Metschnikowia | | 13 | Saprotroph | \|Undefined Saprotroph\| | Yeast | NULL | Probable | As nectar-tap_saprotroph (Põlme et al. 2020); Decay sugar-rich substrates (Põlme et al. 2020) | Põlme S, et al. 2020. Fungal Diversity 105:1-16 ((https://doi.org/10.1007/s13225-020-00466-2)) |
| Otu27237 | | 1 | k__Fungi(100);p__Ascomycota(100);c__Saccharomycetes(100);o__Saccharomycetales(100);f__Metschnikowiaceae(100);g__Metschnikowia(100);g__Metschnikowia_unclassified(100); | Metschnikowia | | 13 | Saprotroph | \|Undefined Saprotroph\| | Yeast | NULL | Probable | As nectar-tap_saprotroph (Põlme et al. 2020); Decay sugar-rich substrates (Põlme et al. 2020) | Põlme S, et al. 2020. Fungal Diversity 105:1-16 ((https://doi.org/10.1007/s13225-020-00466-2)) |
| Otu27238 | | 1 | k__Fungi(100);p__Ascomycota(100);c__Saccharomycetes(100);o__Saccharomycetales(100);f__Metschnikowiaceae(100);g__Metschnikowia(100);s__Metschnikowia_pulcherrima(100); | Metschnikowia | | 13 | Saprotroph | \|Undefined Saprotroph\| | Yeast | NULL | Probable | As nectar-tap_saprotroph (Põlme et al. 2020); Decay sugar-rich substrates (Põlme et al. 2020) | Põlme S, et al. 2020. Fungal Diversity 105:1-16 ((https://doi.org/10.1007/s13225-020-00466-2)) |
| Otu27239 | | 1 | k__Fungi(100);p__Ascomycota(100);c__Saccharomycetes(100);o__Saccharomycetales(100);f__Metschnikowiaceae(100);g__Metschnikowia(100);s__Metschnikowia_pulcherrima(100); | Metschnikowia | | 13 | Saprotroph | \|Undefined Saprotroph\| | Yeast | NULL | Probable | As nectar-tap_saprotroph (Põlme et al. 2020); Decay sugar-rich substrates (Põlme et al. 2020) | Põlme S, et al. 2020. Fungal Diversity 105:1-16 ((https://doi.org/10.1007/s13225-020-00466-2)) |
| Otu27240 | | 1 | k__Fungi(100);p__Ascomycota(100);c__Saccharomycetes(100);o__Saccharomycetales(100);f__Metschnikowiaceae(100);g__Metschnikowia(100);s__Metschnikowia_pulcherrima(100); | Metschnikowia | | 13 | Saprotroph | \|Undefined Saprotroph\| | Yeast | NULL | Probable | As nectar-tap_saprotroph (Põlme et al. 2020); Decay sugar-rich substrates (Põlme et al. 2020) | Põlme S, et al. 2020. Fungal Diversity 105:1-16 ((https://doi.org/10.1007/s13225-020-00466-2)) |
| Otu27241 | | 1 | k__Fungi(100);p__Ascomycota(100);c__Saccharomycetes(100);o__Saccharomycetales(100);f__Metschnikowiaceae(100);g__Metschnikowia(100);s__Metschnikowia_pulcherrima(100); | Metschnikowia | | 13 | Saprotroph | \|Undefined Saprotroph\| | Yeast | NULL | Probable | As nectar-tap_saprotroph (Põlme et al. 2020); Decay sugar-rich substrates (Põlme et al. 2020) | Põlme S, et al. 2020. Fungal Diversity 105:1-16 ((https://doi.org/10.1007/s13225-020-00466-2)) |
| Otu27242 | | 1 | k__Fungi(100);p__Ascomycota(100);c__Saccharomycetes(100);o__Saccharomycetales(100);f__Metschnikowiaceae(100);g__Metschnikowia(100);g__Metschnikowia_unclassified(100); | Metschnikowia | | 13 | Saprotroph | \|Undefined Saprotroph\| | Yeast | NULL | Probable | As nectar-tap_saprotroph (Põlme et al. 2020); Decay sugar-rich substrates (Põlme et al. 2020) | Põlme S, et al. 2020. Fungal Diversity 105:1-16 ((https://doi.org/10.1007/s13225-020-00466-2)) |
| Otu27243 | | 1 | k__Fungi(100);p__Ascomycota(100);c__Saccharomycetes(100);o__Saccharomycetales(100);f__Metschnikowiaceae(100);g__Metschnikowia(100);s__Metschnikowia_pulcherrima(100); | Metschnikowia | | 13 | Saprotroph | \|Undefined Saprotroph\| | Yeast | NULL | Probable | As nectar-tap_saprotroph (Põlme et al. 2020); Decay sugar-rich substrates (Põlme et al. 2020) | Põlme S, et al. 2020. Fungal Diversity 105:1-16 ((https://doi.org/10.1007/s13225-020-00466-2)) |
| Otu27244 | | 1 | k__Fungi(100);p__Ascomycota(100);c__Saccharomycetes(100);o__Saccharomycetales(100);f__Metschnikowiaceae(100);g__Metschnikowia(100);s__Metschnikowia_pulcherrima(100); | Metschnikowia | | 13 | Saprotroph | \|Undefined Saprotroph\| | Yeast | NULL | Probable | As nectar-tap_saprotroph (Põlme et al. 2020); Decay sugar-rich substrates (Põlme et al. 2020) | Põlme S, et al. 2020. Fungal Diversity 105:1-16 ((https://doi.org/10.1007/s13225-020-00466-2)) |
| Otu27245 | | 1 | k__Fungi(100);p__Ascomycota(100);c__Saccharomycetes(100);o__Saccharomycetales(100);f__Metschnikowiaceae(100);g__Metschnikowia(100);s__Metschnikowia_pulcherrima(100); | Metschnikowia | | 13 | Saprotroph | \|Undefined Saprotroph\| | Yeast | NULL | Probable | As nectar-tap_saprotroph (Põlme et al. 2020); Decay sugar-rich substrates (Põlme et al. 2020) | Põlme S, et al. 2020. Fungal Diversity 105:1-16 ((https://doi.org/10.1007/s13225-020-00466-2)) |
| Otu27248 | | 1 | k__Fungi(100);p__Ascomycota(100);c__Saccharomycetes(100);o__Saccharomycetales(100);f__Metschnikowiaceae(100);g__Metschnikowia(100);s__Metschnikowia_pulcherrima(100); | Metschnikowia | | 13 | Saprotroph | \|Undefined Saprotroph\| | Yeast | NULL | Probable | As nectar-tap_saprotroph (Põlme et al. 2020); Decay sugar-rich substrates (Põlme et al. 2020) | Põlme S, et al. 2020. Fungal Diversity 105:1-16 ((https://doi.org/10.1007/s13225-020-00466-2)) |
| Otu27250 | | 1 | k__Fungi(100);p__Ascomycota(100);c__Saccharomycetes(100);o__Saccharomycetales(100);f__Metschnikowiaceae(100);g__Metschnikowia(100);s__Metschnikowia_pulcherrima(100); | Metschnikowia | | 13 | Saprotroph | \|Undefined Saprotroph\| | Yeast | NULL | Probable | As nectar-tap_saprotroph (Põlme et al. 2020); Decay sugar-rich substrates (Põlme et al. 2020) | Põlme S, et al. 2020. Fungal Diversity 105:1-16 ((https://doi.org/10.1007/s13225-020-00466-2)) |
| Otu27252 | | 1 | k__Fungi(100);p__Ascomycota(100);c__Saccharomycetes(100);o__Saccharomycetales(100);f__Metschnikowiaceae(100);g__Metschnikowia(100);s__Metschnikowia_pulcherrima(100); | Metschnikowia | | 13 | Saprotroph | \|Undefined Saprotroph\| | Yeast | NULL | Probable | As nectar-tap_saprotroph (Põlme et al. 2020); Decay sugar-rich substrates (Põlme et al. 2020) | Põlme S, et al. 2020. Fungal Diversity 105:1-16 ((https://doi.org/10.1007/s13225-020-00466-2)) |
| Otu27253 | | 1 | k__Fungi(100);p__Ascomycota(100);c__Saccharomycetes(100);o__Saccharomycetales(100);f__Metschnikowiaceae(100);g__Metschnikowia(100);s__Metschnikowia_pulcherrima(100); | Metschnikowia | | 13 | Saprotroph | \|Undefined Saprotroph\| | Yeast | NULL | Probable | As nectar-tap_saprotroph (Põlme et al. 2020); Decay sugar-rich substrates (Põlme et al. 2020) | Põlme S, et al. 2020. Fungal Diversity 105:1-16 ((https://doi.org/10.1007/s13225-020-00466-2)) |
| Otu27255 | | 1 | k__Fungi(100);p__Ascomycota(100);c__Saccharomycetes(100);o__Saccharomycetales(100);f__Metschnikowiaceae(100);g__Metschnikowia(100);s__Metschnikowia_pulcherrima(100); | Metschnikowia | | 13 | Saprotroph | \|Undefined Saprotroph\| | Yeast | NULL | Probable | As nectar-tap_saprotroph (Põlme et al. 2020); Decay sugar-rich substrates (Põlme et al. 2020) | Põlme S, et al. 2020. Fungal Diversity 105:1-16 ((https://doi.org/10.1007/s13225-020-00466-2)) |
| Otu27257 | | 1 | k__Fungi(100);p__Ascomycota(100);c__Saccharomycetes(100);o__Saccharomycetales(100);f__Metschnikowiaceae(100);g__Metschnikowia(100);s__Metschnikowia_pulcherrima(100); | Metschnikowia | | 13 | Saprotroph | \|Undefined Saprotroph\| | Yeast | NULL | Probable | As nectar-tap_saprotroph (Põlme et al. 2020); Decay sugar-rich substrates (Põlme et al. 2020) | Põlme S, et al. 2020. Fungal Diversity 105:1-16 ((https://doi.org/10.1007/s13225-020-00466-2)) |
| Otu27258 | | 1 | k__Fungi(100);p__Ascomycota(100);c__Saccharomycetes(100);o__Saccharomycetales(100);f__Metschnikowiaceae(100);g__Metschnikowia(100);g__Metschnikowia_unclassified(100); | Metschnikowia | | 13 | Saprotroph | \|Undefined Saprotroph\| | Yeast | NULL | Probable | As nectar-tap_saprotroph (Põlme et al. 2020); Decay sugar-rich substrates (Põlme et al. 2020) | Põlme S, et al. 2020. Fungal Diversity 105:1-16 ((https://doi.org/10.1007/s13225-020-00466-2)) |
| Otu27259 | | 1 | k__Fungi(100);p__Ascomycota(100);c__Saccharomycetes(100);o__Saccharomycetales(100);f__Metschnikowiaceae(100);g__Metschnikowia(100);g__Metschnikowia_unclassified(100); | Metschnikowia | | 13 | Saprotroph | \|Undefined Saprotroph\| | Yeast | NULL | Probable | As nectar-tap_saprotroph (Põlme et al. 2020); Decay sugar-rich substrates (Põlme et al. 2020) | Põlme S, et al. 2020. Fungal Diversity 105:1-16 ((https://doi.org/10.1007/s13225-020-00466-2)) |
| Otu27260 | | 1 | k__Fungi(100);p__Ascomycota(100);c__Saccharomycetes(100);o__Saccharomycetales(100);f__Metschnikowiaceae(100);g__Metschnikowia(100);s__Metschnikowia_pulcherrima(100); | Metschnikowia | | 13 | Saprotroph | \|Undefined Saprotroph\| | Yeast | NULL | Probable | As nectar-tap_saprotroph (Põlme et al. 2020); Decay sugar-rich substrates (Põlme et al. 2020) | Põlme S, et al. 2020. Fungal Diversity 105:1-16 ((https://doi.org/10.1007/s13225-020-00466-2)) |
| Otu27261 | | 1 | k__Fungi(100);p__Ascomycota(100);c__Saccharomycetes(100);o__Saccharomycetales(100);f__Metschnikowiaceae(100);g__Metschnikowia(100);s__Metschnikowia_pulcherrima(100); | Metschnikowia | | 13 | Saprotroph | \|Undefined Saprotroph\| | Yeast | NULL | Probable | As nectar-tap_saprotroph (Põlme et al. 2020); Decay sugar-rich substrates (Põlme et al. 2020) | Põlme S, et al. 2020. Fungal Diversity 105:1-16 ((https://doi.org/10.1007/s13225-020-00466-2)) |
| Otu27262 | | 1 | k__Fungi(100);p__Ascomycota(100);c__Saccharomycetes(100);o__Saccharomycetales(100);f__Metschnikowiaceae(100);g__Metschnikowia(100);s__Metschnikowia_pulcherrima(100); | Metschnikowia | | 13 | Saprotroph | \|Undefined Saprotroph\| | Yeast | NULL | Probable | As nectar-tap_saprotroph (Põlme et al. 2020); Decay sugar-rich substrates (Põlme et al. 2020) | Põlme S, et al. 2020. Fungal Diversity 105:1-16 ((https://doi.org/10.1007/s13225-020-00466-2)) |
| Otu27263 | | 1 | k__Fungi(100);p__Ascomycota(100);c__Saccharomycetes(100);o__Saccharomycetales(100);f__Metschnikowiaceae(100);g__Metschnikowia(100);s__Metschnikowia_pulcherrima(100); | Metschnikowia | | 13 | Saprotroph | \|Undefined Saprotroph\| | Yeast | NULL | Probable | As nectar-tap_saprotroph (Põlme et al. 2020); Decay sugar-rich substrates (Põlme et al. 2020) | Põlme S, et al. 2020. Fungal Diversity 105:1-16 ((https://doi.org/10.1007/s13225-020-00466-2)) |
| Otu27265 | | 1 | k__Fungi(100);p__Ascomycota(100);c__Saccharomycetes(100);o__Saccharomycetales(100);f__Metschnikowiaceae(100);g__Metschnikowia(100);s__Metschnikowia_pulcherrima(100); | Metschnikowia | | 13 | Saprotroph | \|Undefined Saprotroph\| | Yeast | NULL | Probable | As nectar-tap_saprotroph (Põlme et al. 2020); Decay sugar-rich substrates (Põlme et al. 2020) | Põlme S, et al. 2020. Fungal Diversity 105:1-16 ((https://doi.org/10.1007/s13225-020-00466-2)) |
| Otu27271 | | 1 | k__Fungi(100);p__Ascomycota(100);c__Saccharomycetes(100);o__Saccharomycetales(100);f__Metschnikowiaceae(100);g__Metschnikowia(100);s__Metschnikowia_pulcherrima(100); | Metschnikowia | | 13 | Saprotroph | \|Undefined Saprotroph\| | Yeast | NULL | Probable | As nectar-tap_saprotroph (Põlme et al. 2020); Decay sugar-rich substrates (Põlme et al. 2020) | Põlme S, et al. 2020. Fungal Diversity 105:1-16 ((https://doi.org/10.1007/s13225-020-00466-2)) |
| Otu27277 | | 1 | k__Fungi(100);p__Ascomycota(100);c__Saccharomycetes(100);o__Saccharomycetales(100);f__Metschnikowiaceae(100);g__Metschnikowia(100);s__Metschnikowia_pulcherrima(100); | Metschnikowia | | 13 | Saprotroph | \|Undefined Saprotroph\| | Yeast | NULL | Probable | As nectar-tap_saprotroph (Põlme et al. 2020); Decay sugar-rich substrates (Põlme et al. 2020) | Põlme S, et al. 2020. Fungal Diversity 105:1-16 ((https://doi.org/10.1007/s13225-020-00466-2)) |
| Otu27278 | | 1 | k__Fungi(100);p__Ascomycota(100);c__Saccharomycetes(100);o__Saccharomycetales(100);f__Metschnikowiaceae(100);g__Metschnikowia(100);s__Metschnikowia_pulcherrima(100); | Metschnikowia | | 13 | Saprotroph | \|Undefined Saprotroph\| | Yeast | NULL | Probable | As nectar-tap_saprotroph (Põlme et al. 2020); Decay sugar-rich substrates (Põlme et al. 2020) | Põlme S, et al. 2020. Fungal Diversity 105:1-16 ((https://doi.org/10.1007/s13225-020-00466-2)) |
| Otu27292 | | 1 | k__Fungi(100);p__Ascomycota(100);c__Saccharomycetes(100);o__Saccharomycetales(100);o__Saccharomycetales_unclassified(100);o__Saccharomycetales_unclassified(100);o__Saccharomycetales_unclassified(100); | Saccharomycetales | | 7 | Saprotroph | Undefined Saprotroph | Yeast | NULL | Possible | NULL | Sterkenburg E, et al. 2015. New Phytologist 207:1145-1158 ((https://doi.org/10.1111/nph.13426)) |
| Otu27295 | | 1 | k__Fungi(100);p__Ascomycota(100);c__Saccharomycetes(100);o__Saccharomycetales(100);f__Metschnikowiaceae(100);g__Metschnikowia(100);s__Metschnikowia_pulcherrima(100); | Metschnikowia | | 13 | Saprotroph | \|Undefined Saprotroph\| | Yeast | NULL | Probable | As nectar-tap_saprotroph (Põlme et al. 2020); Decay sugar-rich substrates (Põlme et al. 2020) | Põlme S, et al. 2020. Fungal Diversity 105:1-16 ((https://doi.org/10.1007/s13225-020-00466-2)) |
| Otu27308 | | 1 | k__Fungi(100);p__Ascomycota(100);c__Saccharomycetes(100);o__Saccharomycetales(100);f__Metschnikowiaceae(100);g__Metschnikowia(100);s__Metschnikowia_pulcherrima(100); | Metschnikowia | | 13 | Saprotroph | \|Undefined Saprotroph\| | Yeast | NULL | Probable | As nectar-tap_saprotroph (Põlme et al. 2020); Decay sugar-rich substrates (Põlme et al. 2020) | Põlme S, et al. 2020. Fungal Diversity 105:1-16 ((https://doi.org/10.1007/s13225-020-00466-2)) |
| Otu27319 | | 1 | k__Fungi(100);p__Ascomycota(100);c__Saccharomycetes(100);o__Saccharomycetales(100);f__Metschnikowiaceae(100);g__Metschnikowia(100);g__Metschnikowia_unclassified(100); | Metschnikowia | | 13 | Saprotroph | \|Undefined Saprotroph\| | Yeast | NULL | Probable | As nectar-tap_saprotroph (Põlme et al. 2020); Decay sugar-rich substrates (Põlme et al. 2020) | Põlme S, et al. 2020. Fungal Diversity 105:1-16 ((https://doi.org/10.1007/s13225-020-00466-2)) |
| Otu27320 | | 1 | k__Fungi(100);p__Ascomycota(100);c__Saccharomycetes(100);o__Saccharomycetales(100);f__Metschnikowiaceae(100);g__Metschnikowia(100);s__Metschnikowia_pulcherrima(100); | Metschnikowia | | 13 | Saprotroph | \|Undefined Saprotroph\| | Yeast | NULL | Probable | As nectar-tap_saprotroph (Põlme et al. 2020); Decay sugar-rich substrates (Põlme et al. 2020) | Põlme S, et al. 2020. Fungal Diversity 105:1-16 ((https://doi.org/10.1007/s13225-020-00466-2)) |
| Otu27328 | | 1 | k__Fungi(100);p__Ascomycota(100);c__Saccharomycetes(100);o__Saccharomycetales(100);f__Saccharomycetales_family_Incertae_sedis(100);g__Candida(100);g__Candida_unclassified(100); | Candida | | 13 | Pathotroph-Saprotroph-Symbiotroph | Animal Pathogen-Endophyte-Endosymbiont-Epiphyte-\|Undefined Saprotroph\| | Dimorphic Yeast-Yeast | NULL | Probable | As nectar-tap_saprotroph (Põlme et al. 2020); Decay sugar-rich substrates (Põlme et al. 2020); This asexual genus is polyphyletic and contain species that function very differently depend on what they are. Caution should be taken when interpreting guilds for Candida species. | Manolakaki D, et al. 2010. Virulence. 1: 367-375 ((https://doi.org/10.4161/viru.1.5.12796)); Mekbib SB, et al. 2011. Tropical Plant Pathology 36:233-240 ((https://doi.org/10.1590/S1982-56762011000400004)); Põlme S, et al. 2020. Fungal Diversity 105:1-16 ((https://doi.org/10.1007/s13225-020-00466-2)) |
| Otu27332 | | 1 | k__Fungi(100);p__Ascomycota(100);c__Saccharomycetes(100);o__Saccharomycetales(100);f__Metschnikowiaceae(100);g__Metschnikowia(100);s__Metschnikowia_pulcherrima(100); | Metschnikowia | | 13 | Saprotroph | \|Undefined Saprotroph\| | Yeast | NULL | Probable | As nectar-tap_saprotroph (Põlme et al. 2020); Decay sugar-rich substrates (Põlme et al. 2020) | Põlme S, et al. 2020. Fungal Diversity 105:1-16 ((https://doi.org/10.1007/s13225-020-00466-2)) |
| Otu27336 | | 1 | k__Fungi(100);p__Ascomycota(100);c__Saccharomycetes(100);o__Saccharomycetales(100);f__Metschnikowiaceae(100);g__Metschnikowia(100);s__Metschnikowia_pulcherrima(100); | Metschnikowia | | 13 | Saprotroph | \|Undefined Saprotroph\| | Yeast | NULL | Probable | As nectar-tap_saprotroph (Põlme et al. 2020); Decay sugar-rich substrates (Põlme et al. 2020) | Põlme S, et al. 2020. Fungal Diversity 105:1-16 ((https://doi.org/10.1007/s13225-020-00466-2)) |
| Otu27337 | | 1 | k__Fungi(100);p__Ascomycota(100);c__Saccharomycetes(100);o__Saccharomycetales(100);f__Saccharomycetales_family_Incertae_sedis(100);g__Candida(100);g__Candida_unclassified(100); | Candida | | 13 | Pathotroph-Saprotroph-Symbiotroph | Animal Pathogen-Endophyte-Endosymbiont-Epiphyte-\|Undefined Saprotroph\| | Dimorphic Yeast-Yeast | NULL | Probable | As nectar-tap_saprotroph (Põlme et al. 2020); Decay sugar-rich substrates (Põlme et al. 2020); This asexual genus is polyphyletic and contain species that function very differently depend on what they are. Caution should be taken when interpreting guilds for Candida species. | Manolakaki D, et al. 2010. Virulence. 1: 367-375 ((https://doi.org/10.4161/viru.1.5.12796)); Mekbib SB, et al. 2011. Tropical Plant Pathology 36:233-240 ((https://doi.org/10.1590/S1982-56762011000400004)); Põlme S, et al. 2020. Fungal Diversity 105:1-16 ((https://doi.org/10.1007/s13225-020-00466-2)) |
| Otu27341 | | 1 | k__Fungi(100);p__Ascomycota(100);c__Saccharomycetes(100);o__Saccharomycetales(100);f__Metschnikowiaceae(100);g__Metschnikowia(100);s__Metschnikowia_pulcherrima(100); | Metschnikowia | | 13 | Saprotroph | \|Undefined Saprotroph\| | Yeast | NULL | Probable | As nectar-tap_saprotroph (Põlme et al. 2020); Decay sugar-rich substrates (Põlme et al. 2020) | Põlme S, et al. 2020. Fungal Diversity 105:1-16 ((https://doi.org/10.1007/s13225-020-00466-2)) |
| Otu27343 | | 1 | k__Fungi(100);p__Ascomycota(100);c__Saccharomycetes(100);o__Saccharomycetales(100);f__Saccharomycetales_family_Incertae_sedis(100);g__Candida(100);g__Candida_unclassified(100); | Candida | | 13 | Pathotroph-Saprotroph-Symbiotroph | Animal Pathogen-Endophyte-Endosymbiont-Epiphyte-\|Undefined Saprotroph\| | Dimorphic Yeast-Yeast | NULL | Probable | As nectar-tap_saprotroph (Põlme et al. 2020); Decay sugar-rich substrates (Põlme et al. 2020); This asexual genus is polyphyletic and contain species that function very differently depend on what they are. Caution should be taken when interpreting guilds for Candida species. | Manolakaki D, et al. 2010. Virulence. 1: 367-375 ((https://doi.org/10.4161/viru.1.5.12796)); Mekbib SB, et al. 2011. Tropical Plant Pathology 36:233-240 ((https://doi.org/10.1590/S1982-56762011000400004)); Põlme S, et al. 2020. Fungal Diversity 105:1-16 ((https://doi.org/10.1007/s13225-020-00466-2)) |
| Otu27344 | | 1 | k__Fungi(100);p__Ascomycota(100);c__Saccharomycetes(100);o__Saccharomycetales(100);f__Metschnikowiaceae(100);g__Metschnikowia(100);g__Metschnikowia_unclassified(100); | Metschnikowia | | 13 | Saprotroph | \|Undefined Saprotroph\| | Yeast | NULL | Probable | As nectar-tap_saprotroph (Põlme et al. 2020); Decay sugar-rich substrates (Põlme et al. 2020) | Põlme S, et al. 2020. Fungal Diversity 105:1-16 ((https://doi.org/10.1007/s13225-020-00466-2)) |
| Otu27345 | | 1 | k__Fungi(100);p__Ascomycota(100);c__Saccharomycetes(100);o__Saccharomycetales(100);f__Metschnikowiaceae(100);g__Metschnikowia(100);s__Metschnikowia_pulcherrima(100); | Metschnikowia | | 13 | Saprotroph | \|Undefined Saprotroph\| | Yeast | NULL | Probable | As nectar-tap_saprotroph (Põlme et al. 2020); Decay sugar-rich substrates (Põlme et al. 2020) | Põlme S, et al. 2020. Fungal Diversity 105:1-16 ((https://doi.org/10.1007/s13225-020-00466-2)) |
| Otu27357 | | 1 | k__Fungi(100);p__Ascomycota(100);c__Saccharomycetes(100);o__Saccharomycetales(100);f__Metschnikowiaceae(100);g__Metschnikowia(100);s__Metschnikowia_pulcherrima(100); | Metschnikowia | | 13 | Saprotroph | \|Undefined Saprotroph\| | Yeast | NULL | Probable | As nectar-tap_saprotroph (Põlme et al. 2020); Decay sugar-rich substrates (Põlme et al. 2020) | Põlme S, et al. 2020. Fungal Diversity 105:1-16 ((https://doi.org/10.1007/s13225-020-00466-2)) |
| Otu27358 | | 1 | k__Fungi(100);p__Ascomycota(100);c__Saccharomycetes(100);o__Saccharomycetales(100);f__Metschnikowiaceae(100);g__Metschnikowia(100);g__Metschnikowia_unclassified(100); | Metschnikowia | | 13 | Saprotroph | \|Undefined Saprotroph\| | Yeast | NULL | Probable | As nectar-tap_saprotroph (Põlme et al. 2020); Decay sugar-rich substrates (Põlme et al. 2020) | Põlme S, et al. 2020. Fungal Diversity 105:1-16 ((https://doi.org/10.1007/s13225-020-00466-2)) |
| Otu27360 | | 1 | k__Fungi(100);p__Ascomycota(100);c__Saccharomycetes(100);o__Saccharomycetales(100);f__Saccharomycetales_family_Incertae_sedis(100);g__Candida(100);g__Candida_unclassified(100); | Candida | | 13 | Pathotroph-Saprotroph-Symbiotroph | Animal Pathogen-Endophyte-Endosymbiont-Epiphyte-\|Undefined Saprotroph\| | Dimorphic Yeast-Yeast | NULL | Probable | As nectar-tap_saprotroph (Põlme et al. 2020); Decay sugar-rich substrates (Põlme et al. 2020); This asexual genus is polyphyletic and contain species that function very differently depend on what they are. Caution should be taken when interpreting guilds for Candida species. | Manolakaki D, et al. 2010. Virulence. 1: 367-375 ((https://doi.org/10.4161/viru.1.5.12796)); Mekbib SB, et al. 2011. Tropical Plant Pathology 36:233-240 ((https://doi.org/10.1590/S1982-56762011000400004)); Põlme S, et al. 2020. Fungal Diversity 105:1-16 ((https://doi.org/10.1007/s13225-020-00466-2)) |
| Otu27370 | | 1 | k__Fungi(100);p__Ascomycota(100);c__Saccharomycetes(100);o__Saccharomycetales(100);f__Metschnikowiaceae(100);g__Metschnikowia(100);s__Metschnikowia_pulcherrima(100); | Metschnikowia | | 13 | Saprotroph | \|Undefined Saprotroph\| | Yeast | NULL | Probable | As nectar-tap_saprotroph (Põlme et al. 2020); Decay sugar-rich substrates (Põlme et al. 2020) | Põlme S, et al. 2020. Fungal Diversity 105:1-16 ((https://doi.org/10.1007/s13225-020-00466-2)) |
| Otu27374 | | 1 | k__Fungi(100);p__Ascomycota(100);c__Saccharomycetes(100);o__Saccharomycetales(100);f__Metschnikowiaceae(100);g__Metschnikowia(100);s__Metschnikowia_pulcherrima(100); | Metschnikowia | | 13 | Saprotroph | \|Undefined Saprotroph\| | Yeast | NULL | Probable | As nectar-tap_saprotroph (Põlme et al. 2020); Decay sugar-rich substrates (Põlme et al. 2020) | Põlme S, et al. 2020. Fungal Diversity 105:1-16 ((https://doi.org/10.1007/s13225-020-00466-2)) |
| Otu27376 | | 1 | k__Fungi(100);p__Ascomycota(100);c__Saccharomycetes(100);o__Saccharomycetales(100);f__Metschnikowiaceae(100);g__Metschnikowia(100);g__Metschnikowia_unclassified(100); | Metschnikowia | | 13 | Saprotroph | \|Undefined Saprotroph\| | Yeast | NULL | Probable | As nectar-tap_saprotroph (Põlme et al. 2020); Decay sugar-rich substrates (Põlme et al. 2020) | Põlme S, et al. 2020. Fungal Diversity 105:1-16 ((https://doi.org/10.1007/s13225-020-00466-2)) |
| Otu27378 | | 1 | k__Fungi(100);p__Ascomycota(100);c__Saccharomycetes(100);o__Saccharomycetales(100);f__Metschnikowiaceae(100);g__Metschnikowia(100);g__Metschnikowia_unclassified(100); | Metschnikowia | | 13 | Saprotroph | \|Undefined Saprotroph\| | Yeast | NULL | Probable | As nectar-tap_saprotroph (Põlme et al. 2020); Decay sugar-rich substrates (Põlme et al. 2020) | Põlme S, et al. 2020. Fungal Diversity 105:1-16 ((https://doi.org/10.1007/s13225-020-00466-2)) |
| Otu27379 | | 1 | k__Fungi(100);p__Ascomycota(100);c__Saccharomycetes(100);o__Saccharomycetales(100);f__Metschnikowiaceae(100);g__Metschnikowia(100);s__Metschnikowia_pulcherrima(100); | Metschnikowia | | 13 | Saprotroph | \|Undefined Saprotroph\| | Yeast | NULL | Probable | As nectar-tap_saprotroph (Põlme et al. 2020); Decay sugar-rich substrates (Põlme et al. 2020) | Põlme S, et al. 2020. Fungal Diversity 105:1-16 ((https://doi.org/10.1007/s13225-020-00466-2)) |
| Otu27380 | | 1 | k__Fungi(100);p__Ascomycota(100);c__Saccharomycetes(100);o__Saccharomycetales(100);f__Metschnikowiaceae(100);g__Metschnikowia(100);s__Metschnikowia_pulcherrima(100); | Metschnikowia | | 13 | Saprotroph | \|Undefined Saprotroph\| | Yeast | NULL | Probable | As nectar-tap_saprotroph (Põlme et al. 2020); Decay sugar-rich substrates (Põlme et al. 2020) | Põlme S, et al. 2020. Fungal Diversity 105:1-16 ((https://doi.org/10.1007/s13225-020-00466-2)) |
| Otu27381 | | 1 | k__Fungi(100);p__Ascomycota(100);c__Saccharomycetes(100);o__Saccharomycetales(100);f__Metschnikowiaceae(100);g__Metschnikowia(100);g__Metschnikowia_unclassified(100); | Metschnikowia | | 13 | Saprotroph | \|Undefined Saprotroph\| | Yeast | NULL | Probable | As nectar-tap_saprotroph (Põlme et al. 2020); Decay sugar-rich substrates (Põlme et al. 2020) | Põlme S, et al. 2020. Fungal Diversity 105:1-16 ((https://doi.org/10.1007/s13225-020-00466-2)) |
| Otu27382 | | 1 | k__Fungi(100);p__Ascomycota(100);c__Saccharomycetes(100);o__Saccharomycetales(100);f__Metschnikowiaceae(100);g__Metschnikowia(100);g__Metschnikowia_unclassified(100); | Metschnikowia | | 13 | Saprotroph | \|Undefined Saprotroph\| | Yeast | NULL | Probable | As nectar-tap_saprotroph (Põlme et al. 2020); Decay sugar-rich substrates (Põlme et al. 2020) | Põlme S, et al. 2020. Fungal Diversity 105:1-16 ((https://doi.org/10.1007/s13225-020-00466-2)) |
| Otu27384 | | 1 | k__Fungi(100);p__Ascomycota(100);c__Saccharomycetes(100);o__Saccharomycetales(100);f__Metschnikowiaceae(100);g__Metschnikowia(100);s__Metschnikowia_pulcherrima(100); | Metschnikowia | | 13 | Saprotroph | \|Undefined Saprotroph\| | Yeast | NULL | Probable | As nectar-tap_saprotroph (Põlme et al. 2020); Decay sugar-rich substrates (Põlme et al. 2020) | Põlme S, et al. 2020. Fungal Diversity 105:1-16 ((https://doi.org/10.1007/s13225-020-00466-2)) |
| Otu27385 | | 1 | k__Fungi(100);p__Ascomycota(100);c__Saccharomycetes(100);o__Saccharomycetales(100);f__Metschnikowiaceae(100);g__Metschnikowia(100);g__Metschnikowia_unclassified(100); | Metschnikowia | | 13 | Saprotroph | \|Undefined Saprotroph\| | Yeast | NULL | Probable | As nectar-tap_saprotroph (Põlme et al. 2020); Decay sugar-rich substrates (Põlme et al. 2020) | Põlme S, et al. 2020. Fungal Diversity 105:1-16 ((https://doi.org/10.1007/s13225-020-00466-2)) |
| Otu27387 | | 1 | k__Fungi(100);p__Ascomycota(100);c__Saccharomycetes(100);o__Saccharomycetales(100);f__Metschnikowiaceae(100);g__Metschnikowia(100);g__Metschnikowia_unclassified(100); | Metschnikowia | | 13 | Saprotroph | \|Undefined Saprotroph\| | Yeast | NULL | Probable | As nectar-tap_saprotroph (Põlme et al. 2020); Decay sugar-rich substrates (Põlme et al. 2020) | Põlme S, et al. 2020. Fungal Diversity 105:1-16 ((https://doi.org/10.1007/s13225-020-00466-2)) |
| Otu27388 | | 1 | k__Fungi(100);p__Ascomycota(100);c__Saccharomycetes(100);o__Saccharomycetales(100);f__Metschnikowiaceae(100);g__Metschnikowia(100);s__Metschnikowia_pulcherrima(100); | Metschnikowia | | 13 | Saprotroph | \|Undefined Saprotroph\| | Yeast | NULL | Probable | As nectar-tap_saprotroph (Põlme et al. 2020); Decay sugar-rich substrates (Põlme et al. 2020) | Põlme S, et al. 2020. Fungal Diversity 105:1-16 ((https://doi.org/10.1007/s13225-020-00466-2)) |
| Otu27393 | | 1 | k__Fungi(100);p__Ascomycota(100);c__Saccharomycetes(100);o__Saccharomycetales(100);f__Metschnikowiaceae(100);g__Metschnikowia(100);s__Metschnikowia_pulcherrima(100); | Metschnikowia | | 13 | Saprotroph | \|Undefined Saprotroph\| | Yeast | NULL | Probable | As nectar-tap_saprotroph (Põlme et al. 2020); Decay sugar-rich substrates (Põlme et al. 2020) | Põlme S, et al. 2020. Fungal Diversity 105:1-16 ((https://doi.org/10.1007/s13225-020-00466-2)) |
| Otu27398 | | 1 | k__Fungi(100);p__Ascomycota(100);c__Saccharomycetes(100);o__Saccharomycetales(100);f__Metschnikowiaceae(100);g__Metschnikowia(100);g__Metschnikowia_unclassified(100); | Metschnikowia | | 13 | Saprotroph | \|Undefined Saprotroph\| | Yeast | NULL | Probable | As nectar-tap_saprotroph (Põlme et al. 2020); Decay sugar-rich substrates (Põlme et al. 2020) | Põlme S, et al. 2020. Fungal Diversity 105:1-16 ((https://doi.org/10.1007/s13225-020-00466-2)) |
| Otu27399 | | 1 | k__Fungi(100);p__Ascomycota(100);c__Saccharomycetes(100);o__Saccharomycetales(100);f__Saccharomycetales_family_Incertae_sedis(100);g__Candida(100);g__Candida_unclassified(100); | Candida | | 13 | Pathotroph-Saprotroph-Symbiotroph | Animal Pathogen-Endophyte-Endosymbiont-Epiphyte-\|Undefined Saprotroph\| | Dimorphic Yeast-Yeast | NULL | Probable | As nectar-tap_saprotroph (Põlme et al. 2020); Decay sugar-rich substrates (Põlme et al. 2020); This asexual genus is polyphyletic and contain species that function very differently depend on what they are. Caution should be taken when interpreting guilds for Candida species. | Manolakaki D, et al. 2010. Virulence. 1: 367-375 ((https://doi.org/10.4161/viru.1.5.12796)); Mekbib SB, et al. 2011. Tropical Plant Pathology 36:233-240 ((https://doi.org/10.1590/S1982-56762011000400004)); Põlme S, et al. 2020. Fungal Diversity 105:1-16 ((https://doi.org/10.1007/s13225-020-00466-2)) |
| Otu27409 | | 1 | k__Fungi(100);p__Ascomycota(100);c__Saccharomycetes(100);o__Saccharomycetales(100);f__Metschnikowiaceae(100);g__Metschnikowia(100);s__Metschnikowia_pulcherrima(100); | Metschnikowia | | 13 | Saprotroph | \|Undefined Saprotroph\| | Yeast | NULL | Probable | As nectar-tap_saprotroph (Põlme et al. 2020); Decay sugar-rich substrates (Põlme et al. 2020) | Põlme S, et al. 2020. Fungal Diversity 105:1-16 ((https://doi.org/10.1007/s13225-020-00466-2)) |
| Otu27413 | | 1 | k__Fungi(100);p__Ascomycota(100);c__Saccharomycetes(100);o__Saccharomycetales(100);f__Saccharomycetales_family_Incertae_sedis(100);g__Candida(100);g__Candida_unclassified(100); | Candida | | 13 | Pathotroph-Saprotroph-Symbiotroph | Animal Pathogen-Endophyte-Endosymbiont-Epiphyte-\|Undefined Saprotroph\| | Dimorphic Yeast-Yeast | NULL | Probable | As nectar-tap_saprotroph (Põlme et al. 2020); Decay sugar-rich substrates (Põlme et al. 2020); This asexual genus is polyphyletic and contain species that function very differently depend on what they are. Caution should be taken when interpreting guilds for Candida species. | Manolakaki D, et al. 2010. Virulence. 1: 367-375 ((https://doi.org/10.4161/viru.1.5.12796)); Mekbib SB, et al. 2011. Tropical Plant Pathology 36:233-240 ((https://doi.org/10.1590/S1982-56762011000400004)); Põlme S, et al. 2020. Fungal Diversity 105:1-16 ((https://doi.org/10.1007/s13225-020-00466-2)) |
| Otu27415 | | 1 | k__Fungi(100);p__Ascomycota(100);c__Saccharomycetes(100);o__Saccharomycetales(100);f__Metschnikowiaceae(100);g__Metschnikowia(100);s__Metschnikowia_pulcherrima(100); | Metschnikowia | | 13 | Saprotroph | \|Undefined Saprotroph\| | Yeast | NULL | Probable | As nectar-tap_saprotroph (Põlme et al. 2020); Decay sugar-rich substrates (Põlme et al. 2020) | Põlme S, et al. 2020. Fungal Diversity 105:1-16 ((https://doi.org/10.1007/s13225-020-00466-2)) |
| Otu27416 | | 1 | k__Fungi(100);p__Ascomycota(100);c__Saccharomycetes(100);o__Saccharomycetales(100);f__Metschnikowiaceae(100);g__Metschnikowia(100);s__Metschnikowia_pulcherrima(100); | Metschnikowia | | 13 | Saprotroph | \|Undefined Saprotroph\| | Yeast | NULL | Probable | As nectar-tap_saprotroph (Põlme et al. 2020); Decay sugar-rich substrates (Põlme et al. 2020) | Põlme S, et al. 2020. Fungal Diversity 105:1-16 ((https://doi.org/10.1007/s13225-020-00466-2)) |
| Otu27417 | | 1 | k__Fungi(100);p__Ascomycota(100);c__Saccharomycetes(100);o__Saccharomycetales(100);f__Metschnikowiaceae(100);g__Metschnikowia(100);s__Metschnikowia_pulcherrima(100); | Metschnikowia | | 13 | Saprotroph | \|Undefined Saprotroph\| | Yeast | NULL | Probable | As nectar-tap_saprotroph (Põlme et al. 2020); Decay sugar-rich substrates (Põlme et al. 2020) | Põlme S, et al. 2020. Fungal Diversity 105:1-16 ((https://doi.org/10.1007/s13225-020-00466-2)) |
| Otu27419 | | 1 | k__Fungi(100);p__Ascomycota(100);c__Saccharomycetes(100);o__Saccharomycetales(100);f__Metschnikowiaceae(100);g__Metschnikowia(100);s__Metschnikowia_pulcherrima(100); | Metschnikowia | | 13 | Saprotroph | \|Undefined Saprotroph\| | Yeast | NULL | Probable | As nectar-tap_saprotroph (Põlme et al. 2020); Decay sugar-rich substrates (Põlme et al. 2020) | Põlme S, et al. 2020. Fungal Diversity 105:1-16 ((https://doi.org/10.1007/s13225-020-00466-2)) |
| Otu27423 | | 1 | k__Fungi(100);p__Ascomycota(100);c__Saccharomycetes(100);o__Saccharomycetales(100);f__Saccharomycetales_family_Incertae_sedis(100);g__Candida(100);g__Candida_unclassified(100); | Candida | | 13 | Pathotroph-Saprotroph-Symbiotroph | Animal Pathogen-Endophyte-Endosymbiont-Epiphyte-\|Undefined Saprotroph\| | Dimorphic Yeast-Yeast | NULL | Probable | As nectar-tap_saprotroph (Põlme et al. 2020); Decay sugar-rich substrates (Põlme et al. 2020); This asexual genus is polyphyletic and contain species that function very differently depend on what they are. Caution should be taken when interpreting guilds for Candida species. | Manolakaki D, et al. 2010. Virulence. 1: 367-375 ((https://doi.org/10.4161/viru.1.5.12796)); Mekbib SB, et al. 2011. Tropical Plant Pathology 36:233-240 ((https://doi.org/10.1590/S1982-56762011000400004)); Põlme S, et al. 2020. Fungal Diversity 105:1-16 ((https://doi.org/10.1007/s13225-020-00466-2)) |
| Otu27428 | | 1 | k__Fungi(100);p__Ascomycota(100);c__Saccharomycetes(100);o__Saccharomycetales(100);f__Metschnikowiaceae(100);g__Metschnikowia(100);s__Metschnikowia_pulcherrima(100); | Metschnikowia | | 13 | Saprotroph | \|Undefined Saprotroph\| | Yeast | NULL | Probable | As nectar-tap_saprotroph (Põlme et al. 2020); Decay sugar-rich substrates (Põlme et al. 2020) | Põlme S, et al. 2020. Fungal Diversity 105:1-16 ((https://doi.org/10.1007/s13225-020-00466-2)) |
| Otu27429 | | 1 | k__Fungi(100);p__Basidiomycota(100);c__Microbotryomycetes(100);o__Sporidiobolales(100);f__Sporidiobolales_family_Incertae_sedis(100);g__Sporobolomyces(100);s__Sporobolomyces_lactosus(100); | Sporobolomyces | | 13 | Pathotroph-Saprotroph | \|Fungal Parasite\|-Plant Saprotroph | Yeast | NULL | Probable | As litter_saprotroph (Põlme et al. 2020); As mycoparasite (Põlme et al. 2020); Decay fungal material (Põlme et al. 2020) | Tedersoo L, et al. 2014. Science 346:e1256688 ((https://doi.org/10.1126/science.1256688)); Purahong W, et al. 2016. Molecular Ecology 25:4059-4074 ((https://doi.org/10.1111/mec.13739)); Põlme S, et al. 2020. Fungal Diversity 105:1-16 ((https://doi.org/10.1007/s13225-020-00466-2)) |
| Otu27430 | | 1 | k__Fungi(100);p__Ascomycota(100);c__Saccharomycetes(100);o__Saccharomycetales(100);f__Metschnikowiaceae(100);g__Metschnikowia(100);g__Metschnikowia_unclassified(100); | Metschnikowia | | 13 | Saprotroph | \|Undefined Saprotroph\| | Yeast | NULL | Probable | As nectar-tap_saprotroph (Põlme et al. 2020); Decay sugar-rich substrates (Põlme et al. 2020) | Põlme S, et al. 2020. Fungal Diversity 105:1-16 ((https://doi.org/10.1007/s13225-020-00466-2)) |
| Otu27431 | | 1 | k__Fungi(100);p__Ascomycota(100);c__Saccharomycetes(100);o__Saccharomycetales(100);f__Metschnikowiaceae(100);g__Metschnikowia(100);g__Metschnikowia_unclassified(100); | Metschnikowia | | 13 | Saprotroph | \|Undefined Saprotroph\| | Yeast | NULL | Probable | As nectar-tap_saprotroph (Põlme et al. 2020); Decay sugar-rich substrates (Põlme et al. 2020) | Põlme S, et al. 2020. Fungal Diversity 105:1-16 ((https://doi.org/10.1007/s13225-020-00466-2)) |
| Otu27432 | | 1 | k__Fungi(100);p__Ascomycota(100);c__Saccharomycetes(100);o__Saccharomycetales(100);f__Metschnikowiaceae(100);g__Metschnikowia(100);g__Metschnikowia_unclassified(100); | Metschnikowia | | 13 | Saprotroph | \|Undefined Saprotroph\| | Yeast | NULL | Probable | As nectar-tap_saprotroph (Põlme et al. 2020); Decay sugar-rich substrates (Põlme et al. 2020) | Põlme S, et al. 2020. Fungal Diversity 105:1-16 ((https://doi.org/10.1007/s13225-020-00466-2)) |
| Otu27433 | | 1 | k__Fungi(100);p__Basidiomycota(100);c__Microbotryomycetes(100);o__Sporidiobolales(100);f__Sporidiobolales_family_Incertae_sedis(100);g__Sporobolomyces(100);s__Sporobolomyces_lactosus(100); | Sporobolomyces | | 13 | Pathotroph-Saprotroph | \|Fungal Parasite\|-Plant Saprotroph | Yeast | NULL | Probable | As litter_saprotroph (Põlme et al. 2020); As mycoparasite (Põlme et al. 2020); Decay fungal material (Põlme et al. 2020) | Tedersoo L, et al. 2014. Science 346:e1256688 ((https://doi.org/10.1126/science.1256688)); Purahong W, et al. 2016. Molecular Ecology 25:4059-4074 ((https://doi.org/10.1111/mec.13739)); Põlme S, et al. 2020. Fungal Diversity 105:1-16 ((https://doi.org/10.1007/s13225-020-00466-2)) |
| Otu27436 | | 1 | k__Fungi(100);p__Ascomycota(100);c__Saccharomycetes(100);o__Saccharomycetales(100);f__Metschnikowiaceae(100);g__Metschnikowia(100);g__Metschnikowia_unclassified(100); | Metschnikowia | | 13 | Saprotroph | \|Undefined Saprotroph\| | Yeast | NULL | Probable | As nectar-tap_saprotroph (Põlme et al. 2020); Decay sugar-rich substrates (Põlme et al. 2020) | Põlme S, et al. 2020. Fungal Diversity 105:1-16 ((https://doi.org/10.1007/s13225-020-00466-2)) |
| Otu27438 | | 1 | k__Fungi(100);p__Ascomycota(100);c__Saccharomycetes(100);o__Saccharomycetales(100);f__Metschnikowiaceae(100);g__Metschnikowia(100);s__Metschnikowia_pulcherrima(100); | Metschnikowia | | 13 | Saprotroph | \|Undefined Saprotroph\| | Yeast | NULL | Probable | As nectar-tap_saprotroph (Põlme et al. 2020); Decay sugar-rich substrates (Põlme et al. 2020) | Põlme S, et al. 2020. Fungal Diversity 105:1-16 ((https://doi.org/10.1007/s13225-020-00466-2)) |
| Otu27439 | | 1 | k__Fungi(100);p__Ascomycota(100);c__Saccharomycetes(100);o__Saccharomycetales(100);f__Metschnikowiaceae(100);g__Metschnikowia(100);s__Metschnikowia_pulcherrima(100); | Metschnikowia | | 13 | Saprotroph | \|Undefined Saprotroph\| | Yeast | NULL | Probable | As nectar-tap_saprotroph (Põlme et al. 2020); Decay sugar-rich substrates (Põlme et al. 2020) | Põlme S, et al. 2020. Fungal Diversity 105:1-16 ((https://doi.org/10.1007/s13225-020-00466-2)) |
| Otu27440 | | 1 | k__Fungi(100);p__Ascomycota(100);c__Saccharomycetes(100);o__Saccharomycetales(100);f__Metschnikowiaceae(100);g__Metschnikowia(100);s__Metschnikowia_pulcherrima(100); | Metschnikowia | | 13 | Saprotroph | \|Undefined Saprotroph\| | Yeast | NULL | Probable | As nectar-tap_saprotroph (Põlme et al. 2020); Decay sugar-rich substrates (Põlme et al. 2020) | Põlme S, et al. 2020. Fungal Diversity 105:1-16 ((https://doi.org/10.1007/s13225-020-00466-2)) |
| Otu27454 | | 1 | k__Fungi(100);p__Ascomycota(100);c__Saccharomycetes(100);o__Saccharomycetales(100);f__Saccharomycetales_family_Incertae_sedis(100);f__Saccharomycetales_family_Incertae_sedis_unclassified(100);f__Saccharomycetales_family_Incertae_sedis_unclassified(100); | Saccharomycetales | | 7 | Saprotroph | Undefined Saprotroph | Yeast | NULL | Possible | NULL | Sterkenburg E, et al. 2015. New Phytologist 207:1145-1158 ((https://doi.org/10.1111/nph.13426)) |
| Otu27458 | | 1 | k__Fungi(100);p__Ascomycota(100);c__Saccharomycetes(100);o__Saccharomycetales(100);f__Metschnikowiaceae(100);g__Metschnikowia(100);s__Metschnikowia_pulcherrima(100); | Metschnikowia | | 13 | Saprotroph | \|Undefined Saprotroph\| | Yeast | NULL | Probable | As nectar-tap_saprotroph (Põlme et al. 2020); Decay sugar-rich substrates (Põlme et al. 2020) | Põlme S, et al. 2020. Fungal Diversity 105:1-16 ((https://doi.org/10.1007/s13225-020-00466-2)) |
| Otu27462 | | 1 | k__Fungi(100);p__Ascomycota(100);c__Dothideomycetes(100);o__Pleosporales(100);f__Lophiostomataceae(100);g__Lophiostoma(100);g__Lophiostoma_unclassified(100); | Lophiostoma | | 13 | Saprotroph | Plant Saprotroph-Undefined Saprotroph-\|Wood Saprotroph\| | Microfungus | NULL | Probable | As litter_saprotroph (Põlme et al. 2020); Decay wood, leaf, fruit, seed (Põlme et al. 2020) | Tedersoo L, et al. 2014. Science 346:e1256688 ((https://doi.org/10.1126/science.1256688)); Põlme S, et al. 2020. Fungal Diversity 105:1-16 ((https://doi.org/10.1007/s13225-020-00466-2)) |
| Otu27463 | | 1 | k__Fungi(100);p__Ascomycota(100);c__Saccharomycetes(100);o__Saccharomycetales(100);f__Metschnikowiaceae(100);g__Metschnikowia(100);s__Metschnikowia_pulcherrima(100); | Metschnikowia | | 13 | Saprotroph | \|Undefined Saprotroph\| | Yeast | NULL | Probable | As nectar-tap_saprotroph (Põlme et al. 2020); Decay sugar-rich substrates (Põlme et al. 2020) | Põlme S, et al. 2020. Fungal Diversity 105:1-16 ((https://doi.org/10.1007/s13225-020-00466-2)) |
| Otu27464 | | 1 | k__Fungi(100);p__Ascomycota(100);c__Saccharomycetes(100);o__Saccharomycetales(100);f__Metschnikowiaceae(100);g__Metschnikowia(100);s__Metschnikowia_pulcherrima(100); | Metschnikowia | | 13 | Saprotroph | \|Undefined Saprotroph\| | Yeast | NULL | Probable | As nectar-tap_saprotroph (Põlme et al. 2020); Decay sugar-rich substrates (Põlme et al. 2020) | Põlme S, et al. 2020. Fungal Diversity 105:1-16 ((https://doi.org/10.1007/s13225-020-00466-2)) |
| Otu27469 | | 1 | k__Fungi(100);p__Ascomycota(100);c__Saccharomycetes(100);o__Saccharomycetales(100);f__Saccharomycetales_family_Incertae_sedis(100);g__Candida(100);g__Candida_unclassified(100); | Candida | | 13 | Pathotroph-Saprotroph-Symbiotroph | Animal Pathogen-Endophyte-Endosymbiont-Epiphyte-\|Undefined Saprotroph\| | Dimorphic Yeast-Yeast | NULL | Probable | As nectar-tap_saprotroph (Põlme et al. 2020); Decay sugar-rich substrates (Põlme et al. 2020); This asexual genus is polyphyletic and contain species that function very differently depend on what they are. Caution should be taken when interpreting guilds for Candida species. | Manolakaki D, et al. 2010. Virulence. 1: 367-375 ((https://doi.org/10.4161/viru.1.5.12796)); Mekbib SB, et al. 2011. Tropical Plant Pathology 36:233-240 ((https://doi.org/10.1590/S1982-56762011000400004)); Põlme S, et al. 2020. Fungal Diversity 105:1-16 ((https://doi.org/10.1007/s13225-020-00466-2)) |
| Otu27471 | | 1 | k__Fungi(100);p__Ascomycota(100);c__Saccharomycetes(100);o__Saccharomycetales(100);f__Metschnikowiaceae(100);g__Metschnikowia(100);s__Metschnikowia_pulcherrima(100); | Metschnikowia | | 13 | Saprotroph | \|Undefined Saprotroph\| | Yeast | NULL | Probable | As nectar-tap_saprotroph (Põlme et al. 2020); Decay sugar-rich substrates (Põlme et al. 2020) | Põlme S, et al. 2020. Fungal Diversity 105:1-16 ((https://doi.org/10.1007/s13225-020-00466-2)) |
| Otu27473 | | 1 | k__Fungi(100);p__Ascomycota(100);c__Saccharomycetes(100);o__Saccharomycetales(100);f__Metschnikowiaceae(100);g__Metschnikowia(100);s__Metschnikowia_pulcherrima(100); | Metschnikowia | | 13 | Saprotroph | \|Undefined Saprotroph\| | Yeast | NULL | Probable | As nectar-tap_saprotroph (Põlme et al. 2020); Decay sugar-rich substrates (Põlme et al. 2020) | Põlme S, et al. 2020. Fungal Diversity 105:1-16 ((https://doi.org/10.1007/s13225-020-00466-2)) |
| Otu27474 | | 1 | k__Fungi(100);p__Ascomycota(100);c__Saccharomycetes(100);o__Saccharomycetales(100);f__Metschnikowiaceae(100);g__Metschnikowia(100);s__Metschnikowia_pulcherrima(100); | Metschnikowia | | 13 | Saprotroph | \|Undefined Saprotroph\| | Yeast | NULL | Probable | As nectar-tap_saprotroph (Põlme et al. 2020); Decay sugar-rich substrates (Põlme et al. 2020) | Põlme S, et al. 2020. Fungal Diversity 105:1-16 ((https://doi.org/10.1007/s13225-020-00466-2)) |
| Otu27475 | | 1 | k__Fungi(100);p__Ascomycota(100);c__Saccharomycetes(100);o__Saccharomycetales(100);f__Metschnikowiaceae(100);g__Metschnikowia(100);s__Metschnikowia_pulcherrima(100); | Metschnikowia | | 13 | Saprotroph | \|Undefined Saprotroph\| | Yeast | NULL | Probable | As nectar-tap_saprotroph (Põlme et al. 2020); Decay sugar-rich substrates (Põlme et al. 2020) | Põlme S, et al. 2020. Fungal Diversity 105:1-16 ((https://doi.org/10.1007/s13225-020-00466-2)) |
| Otu27477 | | 1 | k__Fungi(100);p__Ascomycota(100);c__Saccharomycetes(100);o__Saccharomycetales(100);f__Metschnikowiaceae(100);g__Metschnikowia(100);s__Metschnikowia_pulcherrima(100); | Metschnikowia | | 13 | Saprotroph | \|Undefined Saprotroph\| | Yeast | NULL | Probable | As nectar-tap_saprotroph (Põlme et al. 2020); Decay sugar-rich substrates (Põlme et al. 2020) | Põlme S, et al. 2020. Fungal Diversity 105:1-16 ((https://doi.org/10.1007/s13225-020-00466-2)) |
| Otu27478 | | 1 | k__Fungi(100);p__Ascomycota(100);c__Saccharomycetes(100);o__Saccharomycetales(100);f__Metschnikowiaceae(100);g__Metschnikowia(100);s__Metschnikowia_pulcherrima(100); | Metschnikowia | | 13 | Saprotroph | \|Undefined Saprotroph\| | Yeast | NULL | Probable | As nectar-tap_saprotroph (Põlme et al. 2020); Decay sugar-rich substrates (Põlme et al. 2020) | Põlme S, et al. 2020. Fungal Diversity 105:1-16 ((https://doi.org/10.1007/s13225-020-00466-2)) |
| Otu27482 | | 1 | k__Fungi(100);p__Ascomycota(100);c__Saccharomycetes(100);o__Saccharomycetales(100);f__Saccharomycetales_family_Incertae_sedis(100);g__Candida(100);g__Candida_unclassified(100); | Candida | | 13 | Pathotroph-Saprotroph-Symbiotroph | Animal Pathogen-Endophyte-Endosymbiont-Epiphyte-\|Undefined Saprotroph\| | Dimorphic Yeast-Yeast | NULL | Probable | As nectar-tap_saprotroph (Põlme et al. 2020); Decay sugar-rich substrates (Põlme et al. 2020); This asexual genus is polyphyletic and contain species that function very differently depend on what they are. Caution should be taken when interpreting guilds for Candida species. | Manolakaki D, et al. 2010. Virulence. 1: 367-375 ((https://doi.org/10.4161/viru.1.5.12796)); Mekbib SB, et al. 2011. Tropical Plant Pathology 36:233-240 ((https://doi.org/10.1590/S1982-56762011000400004)); Põlme S, et al. 2020. Fungal Diversity 105:1-16 ((https://doi.org/10.1007/s13225-020-00466-2)) |
| Otu27483 | | 1 | k__Fungi(100);p__Ascomycota(100);c__Saccharomycetes(100);o__Saccharomycetales(100);f__Saccharomycetales_family_Incertae_sedis(100);g__Candida(100);g__Candida_unclassified(100); | Candida | | 13 | Pathotroph-Saprotroph-Symbiotroph | Animal Pathogen-Endophyte-Endosymbiont-Epiphyte-\|Undefined Saprotroph\| | Dimorphic Yeast-Yeast | NULL | Probable | As nectar-tap_saprotroph (Põlme et al. 2020); Decay sugar-rich substrates (Põlme et al. 2020); This asexual genus is polyphyletic and contain species that function very differently depend on what they are. Caution should be taken when interpreting guilds for Candida species. | Manolakaki D, et al. 2010. Virulence. 1: 367-375 ((https://doi.org/10.4161/viru.1.5.12796)); Mekbib SB, et al. 2011. Tropical Plant Pathology 36:233-240 ((https://doi.org/10.1590/S1982-56762011000400004)); Põlme S, et al. 2020. Fungal Diversity 105:1-16 ((https://doi.org/10.1007/s13225-020-00466-2)) |
| Otu27488 | | 1 | k__Fungi(100);p__Ascomycota(100);c__Saccharomycetes(100);o__Saccharomycetales(100);f__Metschnikowiaceae(100);g__Metschnikowia(100);s__Metschnikowia_pulcherrima(100); | Metschnikowia | | 13 | Saprotroph | \|Undefined Saprotroph\| | Yeast | NULL | Probable | As nectar-tap_saprotroph (Põlme et al. 2020); Decay sugar-rich substrates (Põlme et al. 2020) | Põlme S, et al. 2020. Fungal Diversity 105:1-16 ((https://doi.org/10.1007/s13225-020-00466-2)) |
| Otu27492 | | 1 | k__Fungi(100);p__Ascomycota(100);c__Saccharomycetes(100);o__Saccharomycetales(100);f__Metschnikowiaceae(100);g__Metschnikowia(100);s__Metschnikowia_pulcherrima(100); | Metschnikowia | | 13 | Saprotroph | \|Undefined Saprotroph\| | Yeast | NULL | Probable | As nectar-tap_saprotroph (Põlme et al. 2020); Decay sugar-rich substrates (Põlme et al. 2020) | Põlme S, et al. 2020. Fungal Diversity 105:1-16 ((https://doi.org/10.1007/s13225-020-00466-2)) |
| Otu27494 | | 1 | k__Fungi(100);p__Ascomycota(100);c__Saccharomycetes(100);o__Saccharomycetales(100);f__Saccharomycetales_family_Incertae_sedis(100);g__Candida(100);g__Candida_unclassified(100); | Candida | | 13 | Pathotroph-Saprotroph-Symbiotroph | Animal Pathogen-Endophyte-Endosymbiont-Epiphyte-\|Undefined Saprotroph\| | Dimorphic Yeast-Yeast | NULL | Probable | As nectar-tap_saprotroph (Põlme et al. 2020); Decay sugar-rich substrates (Põlme et al. 2020); This asexual genus is polyphyletic and contain species that function very differently depend on what they are. Caution should be taken when interpreting guilds for Candida species. | Manolakaki D, et al. 2010. Virulence. 1: 367-375 ((https://doi.org/10.4161/viru.1.5.12796)); Mekbib SB, et al. 2011. Tropical Plant Pathology 36:233-240 ((https://doi.org/10.1590/S1982-56762011000400004)); Põlme S, et al. 2020. Fungal Diversity 105:1-16 ((https://doi.org/10.1007/s13225-020-00466-2)) |
| Otu27496 | | 1 | k__Fungi(100);p__Ascomycota(100);c__Saccharomycetes(100);o__Saccharomycetales(100);f__Saccharomycetales_family_Incertae_sedis(100);g__Candida(100);g__Candida_unclassified(100); | Candida | | 13 | Pathotroph-Saprotroph-Symbiotroph | Animal Pathogen-Endophyte-Endosymbiont-Epiphyte-\|Undefined Saprotroph\| | Dimorphic Yeast-Yeast | NULL | Probable | As nectar-tap_saprotroph (Põlme et al. 2020); Decay sugar-rich substrates (Põlme et al. 2020); This asexual genus is polyphyletic and contain species that function very differently depend on what they are. Caution should be taken when interpreting guilds for Candida species. | Manolakaki D, et al. 2010. Virulence. 1: 367-375 ((https://doi.org/10.4161/viru.1.5.12796)); Mekbib SB, et al. 2011. Tropical Plant Pathology 36:233-240 ((https://doi.org/10.1590/S1982-56762011000400004)); Põlme S, et al. 2020. Fungal Diversity 105:1-16 ((https://doi.org/10.1007/s13225-020-00466-2)) |
| Otu27498 | | 1 | k__Fungi(100);p__Ascomycota(100);c__Saccharomycetes(100);o__Saccharomycetales(100);f__Metschnikowiaceae(100);g__Metschnikowia(100);g__Metschnikowia_unclassified(100); | Metschnikowia | | 13 | Saprotroph | \|Undefined Saprotroph\| | Yeast | NULL | Probable | As nectar-tap_saprotroph (Põlme et al. 2020); Decay sugar-rich substrates (Põlme et al. 2020) | Põlme S, et al. 2020. Fungal Diversity 105:1-16 ((https://doi.org/10.1007/s13225-020-00466-2)) |
| Otu27499 | | 1 | k__Fungi(100);p__Ascomycota(100);c__Saccharomycetes(100);o__Saccharomycetales(100);f__Metschnikowiaceae(100);g__Metschnikowia(100);s__Metschnikowia_pulcherrima(100); | Metschnikowia | | 13 | Saprotroph | \|Undefined Saprotroph\| | Yeast | NULL | Probable | As nectar-tap_saprotroph (Põlme et al. 2020); Decay sugar-rich substrates (Põlme et al. 2020) | Põlme S, et al. 2020. Fungal Diversity 105:1-16 ((https://doi.org/10.1007/s13225-020-00466-2)) |
| Otu27500 | | 1 | k__Fungi(100);p__Ascomycota(100);c__Saccharomycetes(100);o__Saccharomycetales(100);f__Metschnikowiaceae(100);g__Metschnikowia(100);s__Metschnikowia_pulcherrima(100); | Metschnikowia | | 13 | Saprotroph | \|Undefined Saprotroph\| | Yeast | NULL | Probable | As nectar-tap_saprotroph (Põlme et al. 2020); Decay sugar-rich substrates (Põlme et al. 2020) | Põlme S, et al. 2020. Fungal Diversity 105:1-16 ((https://doi.org/10.1007/s13225-020-00466-2)) |
| Otu27501 | | 1 | k__Fungi(100);p__Ascomycota(100);c__Saccharomycetes(100);o__Saccharomycetales(100);f__Metschnikowiaceae(100);g__Metschnikowia(100);s__Metschnikowia_pulcherrima(100); | Metschnikowia | | 13 | Saprotroph | \|Undefined Saprotroph\| | Yeast | NULL | Probable | As nectar-tap_saprotroph (Põlme et al. 2020); Decay sugar-rich substrates (Põlme et al. 2020) | Põlme S, et al. 2020. Fungal Diversity 105:1-16 ((https://doi.org/10.1007/s13225-020-00466-2)) |
| Otu27502 | | 1 | k__Fungi(100);p__Ascomycota(100);c__Saccharomycetes(100);o__Saccharomycetales(100);f__Metschnikowiaceae(100);g__Metschnikowia(100);s__Metschnikowia_pulcherrima(100); | Metschnikowia | | 13 | Saprotroph | \|Undefined Saprotroph\| | Yeast | NULL | Probable | As nectar-tap_saprotroph (Põlme et al. 2020); Decay sugar-rich substrates (Põlme et al. 2020) | Põlme S, et al. 2020. Fungal Diversity 105:1-16 ((https://doi.org/10.1007/s13225-020-00466-2)) |
| Otu27503 | | 1 | k__Fungi(100);p__Ascomycota(100);c__Saccharomycetes(100);o__Saccharomycetales(100);f__Metschnikowiaceae(100);g__Metschnikowia(100);s__Metschnikowia_pulcherrima(100); | Metschnikowia | | 13 | Saprotroph | \|Undefined Saprotroph\| | Yeast | NULL | Probable | As nectar-tap_saprotroph (Põlme et al. 2020); Decay sugar-rich substrates (Põlme et al. 2020) | Põlme S, et al. 2020. Fungal Diversity 105:1-16 ((https://doi.org/10.1007/s13225-020-00466-2)) |
| Otu27504 | | 1 | k__Fungi(100);p__Ascomycota(100);c__Saccharomycetes(100);o__Saccharomycetales(100);f__Metschnikowiaceae(100);g__Metschnikowia(100);s__Metschnikowia_pulcherrima(100); | Metschnikowia | | 13 | Saprotroph | \|Undefined Saprotroph\| | Yeast | NULL | Probable | As nectar-tap_saprotroph (Põlme et al. 2020); Decay sugar-rich substrates (Põlme et al. 2020) | Põlme S, et al. 2020. Fungal Diversity 105:1-16 ((https://doi.org/10.1007/s13225-020-00466-2)) |
| Otu27505 | | 1 | k__Fungi(100);p__Ascomycota(100);c__Saccharomycetes(100);o__Saccharomycetales(100);f__Metschnikowiaceae(100);g__Metschnikowia(100);s__Metschnikowia_pulcherrima(100); | Metschnikowia | | 13 | Saprotroph | \|Undefined Saprotroph\| | Yeast | NULL | Probable | As nectar-tap_saprotroph (Põlme et al. 2020); Decay sugar-rich substrates (Põlme et al. 2020) | Põlme S, et al. 2020. Fungal Diversity 105:1-16 ((https://doi.org/10.1007/s13225-020-00466-2)) |
| Otu27509 | | 1 | k__Fungi(100);p__Ascomycota(100);c__Saccharomycetes(100);o__Saccharomycetales(100);f__Saccharomycetales_family_Incertae_sedis(100);g__Candida(100);s__Candida_cretensis(100); | Candida | | 13 | Pathotroph-Saprotroph-Symbiotroph | Animal Pathogen-Endophyte-Endosymbiont-Epiphyte-\|Undefined Saprotroph\| | Dimorphic Yeast-Yeast | NULL | Probable | As nectar-tap_saprotroph (Põlme et al. 2020); Decay sugar-rich substrates (Põlme et al. 2020); This asexual genus is polyphyletic and contain species that function very differently depend on what they are. Caution should be taken when interpreting guilds for Candida species. | Manolakaki D, et al. 2010. Virulence. 1: 367-375 ((https://doi.org/10.4161/viru.1.5.12796)); Mekbib SB, et al. 2011. Tropical Plant Pathology 36:233-240 ((https://doi.org/10.1590/S1982-56762011000400004)); Põlme S, et al. 2020. Fungal Diversity 105:1-16 ((https://doi.org/10.1007/s13225-020-00466-2)) |
| Otu27513 | | 1 | k__Fungi(100);p__Ascomycota(100);c__Saccharomycetes(100);o__Saccharomycetales(100);f__Saccharomycetales_family_Incertae_sedis(100);g__Candida(100);g__Candida_unclassified(100); | Candida | | 13 | Pathotroph-Saprotroph-Symbiotroph | Animal Pathogen-Endophyte-Endosymbiont-Epiphyte-\|Undefined Saprotroph\| | Dimorphic Yeast-Yeast | NULL | Probable | As nectar-tap_saprotroph (Põlme et al. 2020); Decay sugar-rich substrates (Põlme et al. 2020); This asexual genus is polyphyletic and contain species that function very differently depend on what they are. Caution should be taken when interpreting guilds for Candida species. | Manolakaki D, et al. 2010. Virulence. 1: 367-375 ((https://doi.org/10.4161/viru.1.5.12796)); Mekbib SB, et al. 2011. Tropical Plant Pathology 36:233-240 ((https://doi.org/10.1590/S1982-56762011000400004)); Põlme S, et al. 2020. Fungal Diversity 105:1-16 ((https://doi.org/10.1007/s13225-020-00466-2)) |
| Otu27514 | | 1 | k__Fungi(100);p__Ascomycota(100);c__Saccharomycetes(100);o__Saccharomycetales(100);f__Saccharomycetales_family_Incertae_sedis(100);f__Saccharomycetales_family_Incertae_sedis_unclassified(100);f__Saccharomycetales_family_Incertae_sedis_unclassified(100); | Saccharomycetales | | 7 | Saprotroph | Undefined Saprotroph | Yeast | NULL | Possible | NULL | Sterkenburg E, et al. 2015. New Phytologist 207:1145-1158 ((https://doi.org/10.1111/nph.13426)) |
| Otu27515 | | 1 | k__Fungi(100);p__Ascomycota(100);c__Saccharomycetes(100);o__Saccharomycetales(100);f__Metschnikowiaceae(100);g__Metschnikowia(100);s__Metschnikowia_pulcherrima(100); | Metschnikowia | | 13 | Saprotroph | \|Undefined Saprotroph\| | Yeast | NULL | Probable | As nectar-tap_saprotroph (Põlme et al. 2020); Decay sugar-rich substrates (Põlme et al. 2020) | Põlme S, et al. 2020. Fungal Diversity 105:1-16 ((https://doi.org/10.1007/s13225-020-00466-2)) |
| Otu27517 | | 1 | k__Fungi(100);p__Ascomycota(100);c__Saccharomycetes(100);o__Saccharomycetales(100);f__Metschnikowiaceae(100);g__Metschnikowia(100);s__Metschnikowia_pulcherrima(100); | Metschnikowia | | 13 | Saprotroph | \|Undefined Saprotroph\| | Yeast | NULL | Probable | As nectar-tap_saprotroph (Põlme et al. 2020); Decay sugar-rich substrates (Põlme et al. 2020) | Põlme S, et al. 2020. Fungal Diversity 105:1-16 ((https://doi.org/10.1007/s13225-020-00466-2)) |
| Otu27518 | | 1 | k__Fungi(100);p__Ascomycota(100);c__Saccharomycetes(100);o__Saccharomycetales(100);f__Metschnikowiaceae(100);g__Metschnikowia(100);g__Metschnikowia_unclassified(100); | Metschnikowia | | 13 | Saprotroph | \|Undefined Saprotroph\| | Yeast | NULL | Probable | As nectar-tap_saprotroph (Põlme et al. 2020); Decay sugar-rich substrates (Põlme et al. 2020) | Põlme S, et al. 2020. Fungal Diversity 105:1-16 ((https://doi.org/10.1007/s13225-020-00466-2)) |
| Otu27519 | | 1 | k__Fungi(100);p__Ascomycota(100);c__Saccharomycetes(100);o__Saccharomycetales(100);f__Metschnikowiaceae(100);g__Metschnikowia(100);g__Metschnikowia_unclassified(100); | Metschnikowia | | 13 | Saprotroph | \|Undefined Saprotroph\| | Yeast | NULL | Probable | As nectar-tap_saprotroph (Põlme et al. 2020); Decay sugar-rich substrates (Põlme et al. 2020) | Põlme S, et al. 2020. Fungal Diversity 105:1-16 ((https://doi.org/10.1007/s13225-020-00466-2)) |
| Otu27527 | | 1 | k__Fungi(100);p__Ascomycota(100);c__Saccharomycetes(100);o__Saccharomycetales(100);f__Metschnikowiaceae(100);g__Metschnikowia(100);g__Metschnikowia_unclassified(100); | Metschnikowia | | 13 | Saprotroph | \|Undefined Saprotroph\| | Yeast | NULL | Probable | As nectar-tap_saprotroph (Põlme et al. 2020); Decay sugar-rich substrates (Põlme et al. 2020) | Põlme S, et al. 2020. Fungal Diversity 105:1-16 ((https://doi.org/10.1007/s13225-020-00466-2)) |
| Otu27532 | | 1 | k__Fungi(100);p__Ascomycota(100);c__Saccharomycetes(100);o__Saccharomycetales(100);f__Saccharomycetales_family_Incertae_sedis(100);g__Candida(100);g__Candida_unclassified(100); | Candida | | 13 | Pathotroph-Saprotroph-Symbiotroph | Animal Pathogen-Endophyte-Endosymbiont-Epiphyte-\|Undefined Saprotroph\| | Dimorphic Yeast-Yeast | NULL | Probable | As nectar-tap_saprotroph (Põlme et al. 2020); Decay sugar-rich substrates (Põlme et al. 2020); This asexual genus is polyphyletic and contain species that function very differently depend on what they are. Caution should be taken when interpreting guilds for Candida species. | Manolakaki D, et al. 2010. Virulence. 1: 367-375 ((https://doi.org/10.4161/viru.1.5.12796)); Mekbib SB, et al. 2011. Tropical Plant Pathology 36:233-240 ((https://doi.org/10.1590/S1982-56762011000400004)); Põlme S, et al. 2020. Fungal Diversity 105:1-16 ((https://doi.org/10.1007/s13225-020-00466-2)) |
| Otu27540 | | 1 | k__Fungi(100);p__Ascomycota(100);c__Saccharomycetes(100);o__Saccharomycetales(100);f__Metschnikowiaceae(100);g__Metschnikowia(100);g__Metschnikowia_unclassified(100); | Metschnikowia | | 13 | Saprotroph | \|Undefined Saprotroph\| | Yeast | NULL | Probable | As nectar-tap_saprotroph (Põlme et al. 2020); Decay sugar-rich substrates (Põlme et al. 2020) | Põlme S, et al. 2020. Fungal Diversity 105:1-16 ((https://doi.org/10.1007/s13225-020-00466-2)) |
| Otu27547 | | 1 | k__Fungi(100);p__Ascomycota(100);c__Saccharomycetes(100);o__Saccharomycetales(100);f__Metschnikowiaceae(100);g__Metschnikowia(100);s__Metschnikowia_pulcherrima(100); | Metschnikowia | | 13 | Saprotroph | \|Undefined Saprotroph\| | Yeast | NULL | Probable | As nectar-tap_saprotroph (Põlme et al. 2020); Decay sugar-rich substrates (Põlme et al. 2020) | Põlme S, et al. 2020. Fungal Diversity 105:1-16 ((https://doi.org/10.1007/s13225-020-00466-2)) |
| Otu27552 | | 1 | k__Fungi(100);p__Ascomycota(100);c__Saccharomycetes(100);o__Saccharomycetales(100);f__Metschnikowiaceae(100);g__Metschnikowia(100);s__Metschnikowia_pulcherrima(100); | Metschnikowia | | 13 | Saprotroph | \|Undefined Saprotroph\| | Yeast | NULL | Probable | As nectar-tap_saprotroph (Põlme et al. 2020); Decay sugar-rich substrates (Põlme et al. 2020) | Põlme S, et al. 2020. Fungal Diversity 105:1-16 ((https://doi.org/10.1007/s13225-020-00466-2)) |
| Otu27558 | | 1 | k__Fungi(100);p__Ascomycota(100);c__Saccharomycetes(100);o__Saccharomycetales(100);f__Metschnikowiaceae(100);g__Metschnikowia(100);s__Metschnikowia_pulcherrima(100); | Metschnikowia | | 13 | Saprotroph | \|Undefined Saprotroph\| | Yeast | NULL | Probable | As nectar-tap_saprotroph (Põlme et al. 2020); Decay sugar-rich substrates (Põlme et al. 2020) | Põlme S, et al. 2020. Fungal Diversity 105:1-16 ((https://doi.org/10.1007/s13225-020-00466-2)) |
| Otu27559 | | 1 | k__Fungi(100);p__Ascomycota(100);c__Saccharomycetes(100);o__Saccharomycetales(100);f__Metschnikowiaceae(100);g__Metschnikowia(100);g__Metschnikowia_unclassified(100); | Metschnikowia | | 13 | Saprotroph | \|Undefined Saprotroph\| | Yeast | NULL | Probable | As nectar-tap_saprotroph (Põlme et al. 2020); Decay sugar-rich substrates (Põlme et al. 2020) | Põlme S, et al. 2020. Fungal Diversity 105:1-16 ((https://doi.org/10.1007/s13225-020-00466-2)) |
| Otu27564 | | 1 | k__Fungi(100);p__Ascomycota(100);c__Saccharomycetes(100);o__Saccharomycetales(100);f__Metschnikowiaceae(100);g__Metschnikowia(100);s__Metschnikowia_pulcherrima(100); | Metschnikowia | | 13 | Saprotroph | \|Undefined Saprotroph\| | Yeast | NULL | Probable | As nectar-tap_saprotroph (Põlme et al. 2020); Decay sugar-rich substrates (Põlme et al. 2020) | Põlme S, et al. 2020. Fungal Diversity 105:1-16 ((https://doi.org/10.1007/s13225-020-00466-2)) |
| Otu27565 | | 1 | k__Fungi(100);p__Ascomycota(100);c__Saccharomycetes(100);o__Saccharomycetales(100);f__Metschnikowiaceae(100);g__Metschnikowia(100);s__Metschnikowia_pulcherrima(100); | Metschnikowia | | 13 | Saprotroph | \|Undefined Saprotroph\| | Yeast | NULL | Probable | As nectar-tap_saprotroph (Põlme et al. 2020); Decay sugar-rich substrates (Põlme et al. 2020) | Põlme S, et al. 2020. Fungal Diversity 105:1-16 ((https://doi.org/10.1007/s13225-020-00466-2)) |
| Otu27566 | | 1 | k__Fungi(100);p__Ascomycota(100);c__Saccharomycetes(100);o__Saccharomycetales(100);f__Metschnikowiaceae(100);g__Metschnikowia(100);s__Metschnikowia_pulcherrima(100); | Metschnikowia | | 13 | Saprotroph | \|Undefined Saprotroph\| | Yeast | NULL | Probable | As nectar-tap_saprotroph (Põlme et al. 2020); Decay sugar-rich substrates (Põlme et al. 2020) | Põlme S, et al. 2020. Fungal Diversity 105:1-16 ((https://doi.org/10.1007/s13225-020-00466-2)) |
| Otu27567 | | 1 | k__Fungi(100);p__Ascomycota(100);c__Saccharomycetes(100);o__Saccharomycetales(100);f__Metschnikowiaceae(100);g__Metschnikowia(100);s__Metschnikowia_pulcherrima(100); | Metschnikowia | | 13 | Saprotroph | \|Undefined Saprotroph\| | Yeast | NULL | Probable | As nectar-tap_saprotroph (Põlme et al. 2020); Decay sugar-rich substrates (Põlme et al. 2020) | Põlme S, et al. 2020. Fungal Diversity 105:1-16 ((https://doi.org/10.1007/s13225-020-00466-2)) |
| Otu27568 | | 1 | k__Fungi(100);p__Ascomycota(100);c__Saccharomycetes(100);o__Saccharomycetales(100);f__Metschnikowiaceae(100);g__Metschnikowia(100);s__Metschnikowia_pulcherrima(100); | Metschnikowia | | 13 | Saprotroph | \|Undefined Saprotroph\| | Yeast | NULL | Probable | As nectar-tap_saprotroph (Põlme et al. 2020); Decay sugar-rich substrates (Põlme et al. 2020) | Põlme S, et al. 2020. Fungal Diversity 105:1-16 ((https://doi.org/10.1007/s13225-020-00466-2)) |
| Otu27569 | | 1 | k__Fungi(100);p__Ascomycota(100);c__Saccharomycetes(100);o__Saccharomycetales(100);f__Metschnikowiaceae(100);g__Metschnikowia(100);s__Metschnikowia_pulcherrima(100); | Metschnikowia | | 13 | Saprotroph | \|Undefined Saprotroph\| | Yeast | NULL | Probable | As nectar-tap_saprotroph (Põlme et al. 2020); Decay sugar-rich substrates (Põlme et al. 2020) | Põlme S, et al. 2020. Fungal Diversity 105:1-16 ((https://doi.org/10.1007/s13225-020-00466-2)) |
| Otu27570 | | 1 | k__Fungi(100);p__Ascomycota(100);c__Saccharomycetes(100);o__Saccharomycetales(100);f__Metschnikowiaceae(100);g__Metschnikowia(100);s__Metschnikowia_pulcherrima(100); | Metschnikowia | | 13 | Saprotroph | \|Undefined Saprotroph\| | Yeast | NULL | Probable | As nectar-tap_saprotroph (Põlme et al. 2020); Decay sugar-rich substrates (Põlme et al. 2020) | Põlme S, et al. 2020. Fungal Diversity 105:1-16 ((https://doi.org/10.1007/s13225-020-00466-2)) |
| Otu27573 | | 1 | k__Fungi(100);p__Ascomycota(100);c__Saccharomycetes(100);o__Saccharomycetales(100);f__Metschnikowiaceae(100);g__Metschnikowia(100);s__Metschnikowia_pulcherrima(100); | Metschnikowia | | 13 | Saprotroph | \|Undefined Saprotroph\| | Yeast | NULL | Probable | As nectar-tap_saprotroph (Põlme et al. 2020); Decay sugar-rich substrates (Põlme et al. 2020) | Põlme S, et al. 2020. Fungal Diversity 105:1-16 ((https://doi.org/10.1007/s13225-020-00466-2)) |
| Otu27574 | | 1 | k__Fungi(100);p__Ascomycota(100);c__Saccharomycetes(100);o__Saccharomycetales(100);f__Metschnikowiaceae(100);g__Metschnikowia(100);s__Metschnikowia_pulcherrima(100); | Metschnikowia | | 13 | Saprotroph | \|Undefined Saprotroph\| | Yeast | NULL | Probable | As nectar-tap_saprotroph (Põlme et al. 2020); Decay sugar-rich substrates (Põlme et al. 2020) | Põlme S, et al. 2020. Fungal Diversity 105:1-16 ((https://doi.org/10.1007/s13225-020-00466-2)) |
| Otu27587 | | 1 | k__Fungi(100);p__Ascomycota(100);c__Saccharomycetes(100);o__Saccharomycetales(100);f__Metschnikowiaceae(100);g__Metschnikowia(100);s__Metschnikowia_pulcherrima(100); | Metschnikowia | | 13 | Saprotroph | \|Undefined Saprotroph\| | Yeast | NULL | Probable | As nectar-tap_saprotroph (Põlme et al. 2020); Decay sugar-rich substrates (Põlme et al. 2020) | Põlme S, et al. 2020. Fungal Diversity 105:1-16 ((https://doi.org/10.1007/s13225-020-00466-2)) |
| Otu27593 | | 1 | k__Fungi(100);p__Ascomycota(100);c__Saccharomycetes(100);o__Saccharomycetales(100);f__Metschnikowiaceae(100);g__Metschnikowia(100);g__Metschnikowia_unclassified(100); | Metschnikowia | | 13 | Saprotroph | \|Undefined Saprotroph\| | Yeast | NULL | Probable | As nectar-tap_saprotroph (Põlme et al. 2020); Decay sugar-rich substrates (Põlme et al. 2020) | Põlme S, et al. 2020. Fungal Diversity 105:1-16 ((https://doi.org/10.1007/s13225-020-00466-2)) |

| | | | | | | | | | | | |
|---|---|---|---|---|---|---|---|---|---|---|---|
| Otu27594 | 1 | k__Fungi(100);p__Ascomycota(100);c__Saccharomycetes(100);o__Saccharomycetales(100);f__Metschnikowiaceae(100);g__Metschnikowia(100);s__Metschnikowia_pulcherrima(100); | Metschnikowia | 13 | Saprotroph | \|Undefined Saprotroph\| | Yeast | NULL | Probable | As nectar-tap_saprotroph (Põlme et al. 2020); Decay sugar-rich substrates (Põlme et al. 2020) | Põlme S, et al. 2020. Fungal Diversity 105:1-16 ((https://doi.org/10.1007/s13225-020-00466-2)) |
| Otu27595 | 1 | k__Fungi(100);p__Ascomycota(100);c__Saccharomycetes(100);o__Saccharomycetales(100);f__Metschnikowiaceae(100);g__Metschnikowia(100);s__Metschnikowia_pulcherrima(100); | Metschnikowia | 13 | Saprotroph | \|Undefined Saprotroph\| | Yeast | NULL | Probable | As nectar-tap_saprotroph (Põlme et al. 2020); Decay sugar-rich substrates (Põlme et al. 2020) | Põlme S, et al. 2020. Fungal Diversity 105:1-16 ((https://doi.org/10.1007/s13225-020-00466-2)) |
| Otu27600 | 1 | k__Fungi(100);p__Ascomycota(100);c__Saccharomycetes(100);o__Saccharomycetales(100);f__Metschnikowiaceae(100);g__Metschnikowia(100);g__Metschnikowia_unclassified(100); | Metschnikowia | 13 | Saprotroph | \|Undefined Saprotroph\| | Yeast | NULL | Probable | As nectar-tap_saprotroph (Põlme et al. 2020); Decay sugar-rich substrates (Põlme et al. 2020) | Põlme S, et al. 2020. Fungal Diversity 105:1-16 ((https://doi.org/10.1007/s13225-020-00466-2)) |
| Otu27603 | 1 | k__Fungi(100);p__Ascomycota(100);c__Saccharomycetes(100);o__Saccharomycetales(100);f__Metschnikowiaceae(100);g__Metschnikowia(100);s__Metschnikowia_pulcherrima(100); | Metschnikowia | 13 | Saprotroph | \|Undefined Saprotroph\| | Yeast | NULL | Probable | As nectar-tap_saprotroph (Põlme et al. 2020); Decay sugar-rich substrates (Põlme et al. 2020) | Põlme S, et al. 2020. Fungal Diversity 105:1-16 ((https://doi.org/10.1007/s13225-020-00466-2)) |
| Otu27604 | 1 | k__Fungi(100);p__Ascomycota(100);c__Saccharomycetes(100);o__Saccharomycetales(100);f__Metschnikowiaceae(100);g__Metschnikowia(100);s__Metschnikowia_pulcherrima(100); | Metschnikowia | 13 | Saprotroph | \|Undefined Saprotroph\| | Yeast | NULL | Probable | As nectar-tap_saprotroph (Põlme et al. 2020); Decay sugar-rich substrates (Põlme et al. 2020) | Põlme S, et al. 2020. Fungal Diversity 105:1-16 ((https://doi.org/10.1007/s13225-020-00466-2)) |
| Otu27608 | 1 | k__Fungi(100);p__Ascomycota(100);c__Saccharomycetes(100);o__Saccharomycetales(100);f__Metschnikowiaceae(100);g__Metschnikowia(100);g__Metschnikowia_unclassified(100); | Metschnikowia | 13 | Saprotroph | \|Undefined Saprotroph\| | Yeast | NULL | Probable | As nectar-tap_saprotroph (Põlme et al. 2020); Decay sugar-rich substrates (Põlme et al. 2020) | Põlme S, et al. 2020. Fungal Diversity 105:1-16 ((https://doi.org/10.1007/s13225-020-00466-2)) |
| Otu27609 | 1 | k__Fungi(100);p__Ascomycota(100);c__Saccharomycetes(100);o__Saccharomycetales(100);f__Metschnikowiaceae(100);g__Metschnikowia(100);g__Metschnikowia_unclassified(100); | Metschnikowia | 13 | Saprotroph | \|Undefined Saprotroph\| | Yeast | NULL | Probable | As nectar-tap_saprotroph (Põlme et al. 2020); Decay sugar-rich substrates (Põlme et al. 2020) | Põlme S, et al. 2020. Fungal Diversity 105:1-16 ((https://doi.org/10.1007/s13225-020-00466-2)) |
| Otu27611 | 1 | k__Fungi(100);p__Ascomycota(100);c__Saccharomycetes(100);o__Saccharomycetales(100);f__Metschnikowiaceae(100);g__Metschnikowia(100);g__Metschnikowia_unclassified(100); | Metschnikowia | 13 | Saprotroph | \|Undefined Saprotroph\| | Yeast | NULL | Probable | As nectar-tap_saprotroph (Põlme et al. 2020); Decay sugar-rich substrates (Põlme et al. 2020) | Põlme S, et al. 2020. Fungal Diversity 105:1-16 ((https://doi.org/10.1007/s13225-020-00466-2)) |
| Otu27616 | 1 | k__Fungi(100);p__Ascomycota(100);c__Saccharomycetes(100);o__Saccharomycetales(100);f__Metschnikowiaceae(100);g__Metschnikowia(100);s__Metschnikowia_pulcherrima(100); | Metschnikowia | 13 | Saprotroph | \|Undefined Saprotroph\| | Yeast | NULL | Probable | As nectar-tap_saprotroph (Põlme et al. 2020); Decay sugar-rich substrates (Põlme et al. 2020) | Põlme S, et al. 2020. Fungal Diversity 105:1-16 ((https://doi.org/10.1007/s13225-020-00466-2)) |
| Otu27617 | 1 | k__Fungi(100);p__Ascomycota(100);c__Saccharomycetes(100);o__Saccharomycetales(100);f__Metschnikowiaceae(100);g__Metschnikowia(100);s__Metschnikowia_pulcherrima(100); | Metschnikowia | 13 | Saprotroph | \|Undefined Saprotroph\| | Yeast | NULL | Probable | As nectar-tap_saprotroph (Põlme et al. 2020); Decay sugar-rich substrates (Põlme et al. 2020) | Põlme S, et al. 2020. Fungal Diversity 105:1-16 ((https://doi.org/10.1007/s13225-020-00466-2)) |
| Otu27622 | 1 | k__Fungi(100);p__Ascomycota(100);c__Saccharomycetes(100);o__Saccharomycetales(100);f__Metschnikowiaceae(100);g__Metschnikowia(100);g__Metschnikowia_unclassified(100); | Metschnikowia | 13 | Saprotroph | \|Undefined Saprotroph\| | Yeast | NULL | Probable | As nectar-tap_saprotroph (Põlme et al. 2020); Decay sugar-rich substrates (Põlme et al. 2020) | Põlme S, et al. 2020. Fungal Diversity 105:1-16 ((https://doi.org/10.1007/s13225-020-00466-2)) |
| Otu27624 | 1 | k__Fungi(100);p__Ascomycota(100);c__Saccharomycetes(100);o__Saccharomycetales(100);f__Metschnikowiaceae(100);g__Metschnikowia(100);g__Metschnikowia_unclassified(100); | Metschnikowia | 13 | Saprotroph | \|Undefined Saprotroph\| | Yeast | NULL | Probable | As nectar-tap_saprotroph (Põlme et al. 2020); Decay sugar-rich substrates (Põlme et al. 2020) | Põlme S, et al. 2020. Fungal Diversity 105:1-16 ((https://doi.org/10.1007/s13225-020-00466-2)) |
| Otu27625 | 1 | k__Fungi(100);p__Ascomycota(100);c__Saccharomycetes(100);o__Saccharomycetales(100);f__Metschnikowiaceae(100);g__Metschnikowia(100);s__Metschnikowia_pulcherrima(100); | Metschnikowia | 13 | Saprotroph | \|Undefined Saprotroph\| | Yeast | NULL | Probable | As nectar-tap_saprotroph (Põlme et al. 2020); Decay sugar-rich substrates (Põlme et al. 2020) | Põlme S, et al. 2020. Fungal Diversity 105:1-16 ((https://doi.org/10.1007/s13225-020-00466-2)) |
| Otu27626 | 1 | k__Fungi(100);p__Ascomycota(100);c__Saccharomycetes(100);o__Saccharomycetales(100);f__Metschnikowiaceae(100);g__Metschnikowia(100);s__Metschnikowia_pulcherrima(100); | Metschnikowia | 13 | Saprotroph | \|Undefined Saprotroph\| | Yeast | NULL | Probable | As nectar-tap_saprotroph (Põlme et al. 2020); Decay sugar-rich substrates (Põlme et al. 2020) | Põlme S, et al. 2020. Fungal Diversity 105:1-16 ((https://doi.org/10.1007/s13225-020-00466-2)) |
| Otu27633 | 1 | k__Fungi(100);p__Ascomycota(100);c__Saccharomycetes(100);o__Saccharomycetales(100);f__Metschnikowiaceae(100);g__Metschnikowia(100);s__Metschnikowia_pulcherrima(100); | Metschnikowia | 13 | Saprotroph | \|Undefined Saprotroph\| | Yeast | NULL | Probable | As nectar-tap_saprotroph (Põlme et al. 2020); Decay sugar-rich substrates (Põlme et al. 2020) | Põlme S, et al. 2020. Fungal Diversity 105:1-16 ((https://doi.org/10.1007/s13225-020-00466-2)) |
| Otu27636 | 1 | k__Fungi(100);p__Ascomycota(100);c__Saccharomycetes(100);o__Saccharomycetales(100);f__Metschnikowiaceae(100);g__Metschnikowia(100);s__Metschnikowia_pulcherrima(100); | Metschnikowia | 13 | Saprotroph | \|Undefined Saprotroph\| | Yeast | NULL | Probable | As nectar-tap_saprotroph (Põlme et al. 2020); Decay sugar-rich substrates (Põlme et al. 2020) | Põlme S, et al. 2020. Fungal Diversity 105:1-16 ((https://doi.org/10.1007/s13225-020-00466-2)) |
| Otu27637 | 1 | k__Fungi(100);p__Ascomycota(100);c__Saccharomycetes(100);o__Saccharomycetales(100);f__Metschnikowiaceae(100);g__Metschnikowia(100);g__Metschnikowia_unclassified(100); | Metschnikowia | 13 | Saprotroph | \|Undefined Saprotroph\| | Yeast | NULL | Probable | As nectar-tap_saprotroph (Põlme et al. 2020); Decay sugar-rich substrates (Põlme et al. 2020) | Põlme S, et al. 2020. Fungal Diversity 105:1-16 ((https://doi.org/10.1007/s13225-020-00466-2)) |
| Otu27638 | 1 | k__Fungi(100);p__Ascomycota(100);c__Saccharomycetes(100);o__Saccharomycetales(100);f__Metschnikowiaceae(100);g__Metschnikowia(100);g__Metschnikowia_unclassified(100); | Metschnikowia | 13 | Saprotroph | \|Undefined Saprotroph\| | Yeast | NULL | Probable | As nectar-tap_saprotroph (Põlme et al. 2020); Decay sugar-rich substrates (Põlme et al. 2020) | Põlme S, et al. 2020. Fungal Diversity 105:1-16 ((https://doi.org/10.1007/s13225-020-00466-2)) |
| Otu27640 | 1 | k__Fungi(100);p__Ascomycota(100);c__Saccharomycetes(100);o__Saccharomycetales(100);f__Metschnikowiaceae(100);g__Metschnikowia(100);g__Metschnikowia_unclassified(100); | Metschnikowia | 13 | Saprotroph | \|Undefined Saprotroph\| | Yeast | NULL | Probable | As nectar-tap_saprotroph (Põlme et al. 2020); Decay sugar-rich substrates (Põlme et al. 2020) | Põlme S, et al. 2020. Fungal Diversity 105:1-16 ((https://doi.org/10.1007/s13225-020-00466-2)) |
| Otu27641 | 1 | k__Fungi(100);p__Ascomycota(100);c__Saccharomycetes(100);o__Saccharomycetales(100);f__Metschnikowiaceae(100);g__Metschnikowia(100);g__Metschnikowia_unclassified(100); | Metschnikowia | 13 | Saprotroph | \|Undefined Saprotroph\| | Yeast | NULL | Probable | As nectar-tap_saprotroph (Põlme et al. 2020); Decay sugar-rich substrates (Põlme et al. 2020) | Põlme S, et al. 2020. Fungal Diversity 105:1-16 ((https://doi.org/10.1007/s13225-020-00466-2)) |
| Otu27642 | 1 | k__Fungi(100);p__Ascomycota(100);c__Saccharomycetes(100);o__Saccharomycetales(100);f__Metschnikowiaceae(100);g__Metschnikowia(100);g__Metschnikowia_unclassified(100); | Metschnikowia | 13 | Saprotroph | \|Undefined Saprotroph\| | Yeast | NULL | Probable | As nectar-tap_saprotroph (Põlme et al. 2020); Decay sugar-rich substrates (Põlme et al. 2020) | Põlme S, et al. 2020. Fungal Diversity 105:1-16 ((https://doi.org/10.1007/s13225-020-00466-2)) |
| Otu27643 | 1 | k__Fungi(100);p__Ascomycota(100);c__Saccharomycetes(100);o__Saccharomycetales(100);f__Metschnikowiaceae(100);g__Metschnikowia(100);s__Metschnikowia_pulcherrima(100); | Metschnikowia | 13 | Saprotroph | \|Undefined Saprotroph\| | Yeast | NULL | Probable | As nectar-tap_saprotroph (Põlme et al. 2020); Decay sugar-rich substrates (Põlme et al. 2020) | Põlme S, et al. 2020. Fungal Diversity 105:1-16 ((https://doi.org/10.1007/s13225-020-00466-2)) |
| Otu27646 | 1 | k__Fungi(100);p__Ascomycota(100);c__Saccharomycetes(100);o__Saccharomycetales(100);f__Metschnikowiaceae(100);g__Metschnikowia(100);g__Metschnikowia_unclassified(100); | Metschnikowia | 13 | Saprotroph | \|Undefined Saprotroph\| | Yeast | NULL | Probable | As nectar-tap_saprotroph (Põlme et al. 2020); Decay sugar-rich substrates (Põlme et al. 2020) | Põlme S, et al. 2020. Fungal Diversity 105:1-16 ((https://doi.org/10.1007/s13225-020-00466-2)) |
| Otu27647 | 1 | k__Fungi(100);p__Ascomycota(100);c__Sordariomycetes(100);o__Chaetosphaeriales(100);f__Chaetosphaeriaceae(100);g__Chaetosphaeria(100);s__Chaetosphaeria_sp(100); | Chaetosphaeria | 13 | Saprotroph-Symbiotroph | Endophyte-\|Plant Saprotroph\|-Wood Saprotroph | Microfungus | NULL | Probable | As foliar_endophyte (Põlme et al. 2020); As litter_saprotroph (Põlme et al. 2020); Decay wood, leaf, fruit, seed (Põlme et al. 2020) | Réblová M, et al. 2006. Mycological Research 110:104-109 ((https://doi.org/10.1016/j.mycres.2005.09.003)); Tedersoo L, et al. 2014. Science 346:e1256688 ((https://doi.org/10.1126/science.1256688)); Põlme S, et al. 2020. Fungal Diversity 105:1-16 ((https://doi.org/10.1007/s13225-020-00466-2)) |
| Otu27648 | 1 | k__Fungi(100);p__Ascomycota(100);c__Saccharomycetes(100);o__Saccharomycetales(100);f__Metschnikowiaceae(100);g__Metschnikowia(100);g__Metschnikowia_unclassified(100); | Metschnikowia | 13 | Saprotroph | \|Undefined Saprotroph\| | Yeast | NULL | Probable | As nectar-tap_saprotroph (Põlme et al. 2020); Decay sugar-rich substrates (Põlme et al. 2020) | Põlme S, et al. 2020. Fungal Diversity 105:1-16 ((https://doi.org/10.1007/s13225-020-00466-2)) |
| Otu27649 | 1 | k__Fungi(100);p__Ascomycota(100);c__Saccharomycetes(100);o__Saccharomycetales(100);f__Metschnikowiaceae(100);g__Metschnikowia(100);g__Metschnikowia_unclassified(100); | Metschnikowia | 13 | Saprotroph | \|Undefined Saprotroph\| | Yeast | NULL | Probable | As nectar-tap_saprotroph (Põlme et al. 2020); Decay sugar-rich substrates (Põlme et al. 2020) | Põlme S, et al. 2020. Fungal Diversity 105:1-16 ((https://doi.org/10.1007/s13225-020-00466-2)) |
| Otu27650 | 1 | k__Fungi(100);p__Ascomycota(100);c__Saccharomycetes(100);o__Saccharomycetales(100);f__Metschnikowiaceae(100);g__Metschnikowia(100);s__Metschnikowia_pulcherrima(100); | Metschnikowia | 13 | Saprotroph | \|Undefined Saprotroph\| | Yeast | NULL | Probable | As nectar-tap_saprotroph (Põlme et al. 2020); Decay sugar-rich substrates (Põlme et al. 2020) | Põlme S, et al. 2020. Fungal Diversity 105:1-16 ((https://doi.org/10.1007/s13225-020-00466-2)) |
| Otu27651 | 1 | k__Fungi(100);p__Ascomycota(100);c__Saccharomycetes(100);o__Saccharomycetales(100);f__Metschnikowiaceae(100);g__Metschnikowia(100);g__Metschnikowia_unclassified(100); | Metschnikowia | 13 | Saprotroph | \|Undefined Saprotroph\| | Yeast | NULL | Probable | As nectar-tap_saprotroph (Põlme et al. 2020); Decay sugar-rich substrates (Põlme et al. 2020) | Põlme S, et al. 2020. Fungal Diversity 105:1-16 ((https://doi.org/10.1007/s13225-020-00466-2)) |
| Otu27652 | 1 | k__Fungi(100);p__Ascomycota(100);c__Saccharomycetes(100);o__Saccharomycetales(100);f__Metschnikowiaceae(100);g__Metschnikowia(100);s__Metschnikowia_pulcherrima(100); | Metschnikowia | 13 | Saprotroph | \|Undefined Saprotroph\| | Yeast | NULL | Probable | As nectar-tap_saprotroph (Põlme et al. 2020); Decay sugar-rich substrates (Põlme et al. 2020) | Põlme S, et al. 2020. Fungal Diversity 105:1-16 ((https://doi.org/10.1007/s13225-020-00466-2)) |
| Otu27653 | 1 | k__Fungi(100);p__Ascomycota(100);c__Saccharomycetes(100);o__Saccharomycetales(100);f__Metschnikowiaceae(100);g__Metschnikowia(100);g__Metschnikowia_unclassified(100); | Metschnikowia | 13 | Saprotroph | \|Undefined Saprotroph\| | Yeast | NULL | Probable | As nectar-tap_saprotroph (Põlme et al. 2020); Decay sugar-rich substrates (Põlme et al. 2020) | Põlme S, et al. 2020. Fungal Diversity 105:1-16 ((https://doi.org/10.1007/s13225-020-00466-2)) |
| Otu27655 | 1 | k__Fungi(100);p__Ascomycota(100);c__Saccharomycetes(100);o__Saccharomycetales(100);f__Saccharomycetales_family_Incertae_sedis(100);f__Saccharomycetales_family_Incertae_sedis_unclassified(100);f__Saccharomycetales_family_Incertae_sedis_unclassified(100); | Saccharomycetales | 7 | Saprotroph | Undefined Saprotroph | Yeast | NULL | Possible | NULL | Sterkenburg E, et al. 2015. New Phytologist 207:1145-1158 ((https://doi.org/10.1111/nph.13426)) |
| Otu27656 | 1 | k__Fungi(100);p__Ascomycota(100);c__Saccharomycetes(100);o__Saccharomycetales(100);f__Metschnikowiaceae(100);g__Metschnikowia(100);g__Metschnikowia_unclassified(100); | Metschnikowia | 13 | Saprotroph | \|Undefined Saprotroph\| | Yeast | NULL | Probable | As nectar-tap_saprotroph (Põlme et al. 2020); Decay sugar-rich substrates (Põlme et al. 2020) | Põlme S, et al. 2020. Fungal Diversity 105:1-16 ((https://doi.org/10.1007/s13225-020-00466-2)) |
| Otu27657 | 1 | k__Fungi(100);p__Ascomycota(100);c__Saccharomycetes(100);o__Saccharomycetales(100);f__Metschnikowiaceae(100);g__Metschnikowia(100);g__Metschnikowia_unclassified(100); | Metschnikowia | 13 | Saprotroph | \|Undefined Saprotroph\| | Yeast | NULL | Probable | As nectar-tap_saprotroph (Põlme et al. 2020); Decay sugar-rich substrates (Põlme et al. 2020) | Põlme S, et al. 2020. Fungal Diversity 105:1-16 ((https://doi.org/10.1007/s13225-020-00466-2)) |
| Otu27658 | 1 | k__Fungi(100);p__Ascomycota(100);c__Saccharomycetes(100);o__Saccharomycetales(100);f__Metschnikowiaceae(100);g__Metschnikowia(100);g__Metschnikowia_unclassified(100); | Metschnikowia | 13 | Saprotroph | \|Undefined Saprotroph\| | Yeast | NULL | Probable | As nectar-tap_saprotroph (Põlme et al. 2020); Decay sugar-rich substrates (Põlme et al. 2020) | Põlme S, et al. 2020. Fungal Diversity 105:1-16 ((https://doi.org/10.1007/s13225-020-00466-2)) |
| Otu27660 | 1 | k__Fungi(100);p__Ascomycota(100);c__Saccharomycetes(100);o__Saccharomycetales(100);f__Metschnikowiaceae(100);g__Metschnikowia(100);g__Metschnikowia_unclassified(100); | Metschnikowia | 13 | Saprotroph | \|Undefined Saprotroph\| | Yeast | NULL | Probable | As nectar-tap_saprotroph (Põlme et al. 2020); Decay sugar-rich substrates (Põlme et al. 2020) | Põlme S, et al. 2020. Fungal Diversity 105:1-16 ((https://doi.org/10.1007/s13225-020-00466-2)) |
| Otu27663 | 1 | k__Fungi(100);p__Ascomycota(100);c__Saccharomycetes(100);o__Saccharomycetales(100);f__Metschnikowiaceae(100);g__Metschnikowia(100);g__Metschnikowia_unclassified(100); | Metschnikowia | 13 | Saprotroph | \|Undefined Saprotroph\| | Yeast | NULL | Probable | As nectar-tap_saprotroph (Põlme et al. 2020); Decay sugar-rich substrates (Põlme et al. 2020) | Põlme S, et al. 2020. Fungal Diversity 105:1-16 ((https://doi.org/10.1007/s13225-020-00466-2)) |
| Otu27664 | 1 | k__Fungi(100);p__Ascomycota(100);c__Saccharomycetes(100);o__Saccharomycetales(100);f__Metschnikowiaceae(100);g__Metschnikowia(100);g__Metschnikowia_unclassified(100); | Metschnikowia | 13 | Saprotroph | \|Undefined Saprotroph\| | Yeast | NULL | Probable | As nectar-tap_saprotroph (Põlme et al. 2020); Decay sugar-rich substrates (Põlme et al. 2020) | Põlme S, et al. 2020. Fungal Diversity 105:1-16 ((https://doi.org/10.1007/s13225-020-00466-2)) |
| Otu27665 | 1 | k__Fungi(100);p__Ascomycota(100);c__Saccharomycetes(100);o__Saccharomycetales(100);f__Metschnikowiaceae(100);g__Metschnikowia(100);g__Metschnikowia_unclassified(100); | Metschnikowia | 13 | Saprotroph | \|Undefined Saprotroph\| | Yeast | NULL | Probable | As nectar-tap_saprotroph (Põlme et al. 2020); Decay sugar-rich substrates (Põlme et al. 2020) | Põlme S, et al. 2020. Fungal Diversity 105:1-16 ((https://doi.org/10.1007/s13225-020-00466-2)) |
| Otu27667 | 1 | k__Fungi(100);p__Ascomycota(100);c__Saccharomycetes(100);o__Saccharomycetales(100);f__Metschnikowiaceae(100);g__Metschnikowia(100);g__Metschnikowia_unclassified(100); | Metschnikowia | 13 | Saprotroph | \|Undefined Saprotroph\| | Yeast | NULL | Probable | As nectar-tap_saprotroph (Põlme et al. 2020); Decay sugar-rich substrates (Põlme et al. 2020) | Põlme S, et al. 2020. Fungal Diversity 105:1-16 ((https://doi.org/10.1007/s13225-020-00466-2)) |
| Otu27671 | 1 | k__Fungi(100);p__Ascomycota(100);c__Saccharomycetes(100);o__Saccharomycetales(100);f__Metschnikowiaceae(100);g__Metschnikowia(100);s__Metschnikowia_pulcherrima(100); | Metschnikowia | 13 | Saprotroph | \|Undefined Saprotroph\| | Yeast | NULL | Probable | As nectar-tap_saprotroph (Põlme et al. 2020); Decay sugar-rich substrates (Põlme et al. 2020) | Põlme S, et al. 2020. Fungal Diversity 105:1-16 ((https://doi.org/10.1007/s13225-020-00466-2)) |
| Otu27672 | 1 | k__Fungi(100);p__Ascomycota(100);c__Saccharomycetes(100);o__Saccharomycetales(100);f__Saccharomycetales_family_Incertae_sedis(100);g__Candida(100);g__Candida_unclassified(100); | Candida | 13 | Pathotroph-Saprotroph-Symbiotroph | Animal Pathogen-Endophyte-Endosymbiont-Epiphyte-\|Undefined Saprotroph\| | Dimorphic Yeast-Yeast | NULL | Probable | As nectar-tap_saprotroph (Põlme et al. 2020); Decay sugar-rich substrates (Põlme et al. 2020); This asexual genus is polyphyletic and contain species that function very differently depend on what they are. Caution should be taken when interpreting guilds for Candida species. | Manolakaki D, et al. 2010. Virulence. 1: 367-375 ((https://doi.org/10.4161/viru.1.5.12796)); Mekbib SB, et al. 2011. Tropical Plant Pathology 36:233-240 ((https://doi.org/10.1590/S1982-56762011000400004)); Põlme S, et al. 2020. Fungal Diversity 105:1-16 ((https://doi.org/10.1007/s13225-020-00466-2)) |
| Otu27676 | 1 | k__Fungi(100);p__Ascomycota(100);c__Saccharomycetes(100);o__Saccharomycetales(100);f__Metschnikowiaceae(100);g__Metschnikowia(100);s__Metschnikowia_pulcherrima(100); | Metschnikowia | 13 | Saprotroph | \|Undefined Saprotroph\| | Yeast | NULL | Probable | As nectar-tap_saprotroph (Põlme et al. 2020); Decay sugar-rich substrates (Põlme et al. 2020) | Põlme S, et al. 2020. Fungal Diversity 105:1-16 ((https://doi.org/10.1007/s13225-020-00466-2)) |
| Otu27677 | 1 | k__Fungi(100);p__Ascomycota(100);c__Saccharomycetes(100);o__Saccharomycetales(100);f__Metschnikowiaceae(100);g__Metschnikowia(100);g__Metschnikowia_unclassified(100); | Metschnikowia | 13 | Saprotroph | \|Undefined Saprotroph\| | Yeast | NULL | Probable | As nectar-tap_saprotroph (Põlme et al. 2020); Decay sugar-rich substrates (Põlme et al. 2020) | Põlme S, et al. 2020. Fungal Diversity 105:1-16 ((https://doi.org/10.1007/s13225-020-00466-2)) |
| Otu27679 | 1 | k__Fungi(100);p__Ascomycota(100);c__Saccharomycetes(100);o__Saccharomycetales(100);f__Metschnikowiaceae(100);g__Metschnikowia(100);g__Metschnikowia_unclassified(100); | Metschnikowia | 13 | Saprotroph | \|Undefined Saprotroph\| | Yeast | NULL | Probable | As nectar-tap_saprotroph (Põlme et al. 2020); Decay sugar-rich substrates (Põlme et al. 2020) | Põlme S, et al. 2020. Fungal Diversity 105:1-16 ((https://doi.org/10.1007/s13225-020-00466-2)) |
| Otu27680 | 1 | k__Fungi(100);p__Ascomycota(100);c__Saccharomycetes(100);o__Saccharomycetales(100);f__Saccharomycetales_family_Incertae_sedis(100);g__Candida(100);g__Candida_unclassified(100); | Candida | 13 | Pathotroph-Saprotroph-Symbiotroph | Animal Pathogen-Endophyte-Endosymbiont-Epiphyte-\|Undefined Saprotroph\| | Dimorphic Yeast-Yeast | NULL | Probable | As nectar-tap_saprotroph (Põlme et al. 2020); Decay sugar-rich substrates (Põlme et al. 2020); This asexual genus is polyphyletic and contain species that function very differently depend on what they are. Caution should be taken when interpreting guilds for Candida species. | Manolakaki D, et al. 2010. Virulence. 1: 367-375 ((https://doi.org/10.4161/viru.1.5.12796)); Mekbib SB, et al. 2011. Tropical Plant Pathology 36:233-240 ((https://doi.org/10.1590/S1982-56762011000400004)); Põlme S, et al. 2020. Fungal Diversity 105:1-16 ((https://doi.org/10.1007/s13225-020-00466-2)) |
| Otu27683 | 1 | k__Fungi(100);p__Ascomycota(100);c__Saccharomycetes(100);o__Saccharomycetales(100);f__Metschnikowiaceae(100);g__Metschnikowia(100);s__Metschnikowia_sp(100); | Metschnikowia | 13 | Saprotroph | \|Undefined Saprotroph\| | Yeast | NULL | Probable | As nectar-tap_saprotroph (Põlme et al. 2020); Decay sugar-rich substrates (Põlme et al. 2020) | Põlme S, et al. 2020. Fungal Diversity 105:1-16 ((https://doi.org/10.1007/s13225-020-00466-2)) |
| Otu27686 | 1 | k__Fungi(100);p__Ascomycota(100);c__Saccharomycetes(100);o__Saccharomycetales(100);f__Saccharomycetales_family_Incertae_sedis(100);g__Candida(100);g__Candida_unclassified(100); | Candida | 13 | Pathotroph-Saprotroph-Symbiotroph | Animal Pathogen-Endophyte-Endosymbiont-Epiphyte-\|Undefined Saprotroph\| | Dimorphic Yeast-Yeast | NULL | Probable | As nectar-tap_saprotroph (Põlme et al. 2020); Decay sugar-rich substrates (Põlme et al. 2020); This asexual genus is polyphyletic and contain species that function very differently depend on what they are. Caution should be taken when interpreting guilds for Candida species. | Manolakaki D, et al. 2010. Virulence. 1: 367-375 ((https://doi.org/10.4161/viru.1.5.12796)); Mekbib SB, et al. 2011. Tropical Plant Pathology 36:233-240 ((https://doi.org/10.1590/S1982-56762011000400004)); Põlme S, et al. 2020. Fungal Diversity 105:1-16 ((https://doi.org/10.1007/s13225-020-00466-2)) |
| Otu27689 | 1 | k__Fungi(100);p__Ascomycota(100);c__Saccharomycetes(100);o__Saccharomycetales(100);f__Saccharomycetales_family_Incertae_sedis(100);g__Candida(100);g__Candida_unclassified(100); | Candida | 13 | Pathotroph-Saprotroph-Symbiotroph | Animal Pathogen-Endophyte-Endosymbiont-Epiphyte-\|Undefined Saprotroph\| | Dimorphic Yeast-Yeast | NULL | Probable | As nectar-tap_saprotroph (Põlme et al. 2020); Decay sugar-rich substrates (Põlme et al. 2020); This asexual genus is polyphyletic and contain species that function very differently depend on what they are. Caution should be taken when interpreting guilds for Candida species. | Manolakaki D, et al. 2010. Virulence. 1: 367-375 ((https://doi.org/10.4161/viru.1.5.12796)); Mekbib SB, et al. 2011. Tropical Plant Pathology 36:233-240 ((https://doi.org/10.1590/S1982-56762011000400004)); Põlme S, et al. 2020. Fungal Diversity 105:1-16 ((https://doi.org/10.1007/s13225-020-00466-2)) |
| Otu27690 | 1 | k__Fungi(100);p__Ascomycota(100);c__Saccharomycetes(100);o__Saccharomycetales(100);f__Metschnikowiaceae(100);g__Metschnikowia(100);g__Metschnikowia_unclassified(100); | Metschnikowia | 13 | Saprotroph | \|Undefined Saprotroph\| | Yeast | NULL | Probable | As nectar-tap_saprotroph (Põlme et al. 2020); Decay sugar-rich substrates (Põlme et al. 2020) | Põlme S, et al. 2020. Fungal Diversity 105:1-16 ((https://doi.org/10.1007/s13225-020-00466-2)) |
| Otu27705 | 1 | k__Fungi(100);p__Ascomycota(100);c__Saccharomycetes(100);o__Saccharomycetales(100);f__Metschnikowiaceae(100);g__Metschnikowia(100);g__Metschnikowia_unclassified(100); | Metschnikowia | 13 | Saprotroph | \|Undefined Saprotroph\| | Yeast | NULL | Probable | As nectar-tap_saprotroph (Põlme et al. 2020); Decay sugar-rich substrates (Põlme et al. 2020) | Põlme S, et al. 2020. Fungal Diversity 105:1-16 ((https://doi.org/10.1007/s13225-020-00466-2)) |
| Otu27719 | 1 | k__Fungi(100);p__Basidiomycota(100);c__Microbotryomycetes(100);o__Sporidiobolales(100);f__Sporidiobolales_family_Incertae_sedis(100);g__Sporobolomyces(100);s__Sporobolomyces_lactosus(100); | Sporobolomyces | 13 | Pathotroph-Saprotroph | \|Fungal Parasite\|-Plant Saprotroph | Yeast | NULL | Probable | As litter_saprotroph (Põlme et al. 2020); As mycoparasite (Põlme et al. 2020); Decay fungal material (Põlme et al. 2020) | Tedersoo L, et al. 2014. Science 346:e1256688 ((https://doi.org/10.1126/science.1256688)); Purahong W, et al. 2016. Molecular Ecology 25:4059-4074 ((https://doi.org/10.1111/mec.13739)); Põlme S, et al. 2020. Fungal Diversity 105:1-16 ((https://doi.org/10.1007/s13225-020-00466-2)) |
| Otu27724 | 1 | k__Fungi(100);p__Ascomycota(100);c__Saccharomycetes(100);o__Saccharomycetales(100);f__Metschnikowiaceae(100);g__Metschnikowia(100);s__Metschnikowia_pulcherrima(100); | Metschnikowia | 13 | Saprotroph | \|Undefined Saprotroph\| | Yeast | NULL | Probable | As nectar-tap_saprotroph (Põlme et al. 2020); Decay sugar-rich substrates (Põlme et al. 2020) | Põlme S, et al. 2020. Fungal Diversity 105:1-16 ((https://doi.org/10.1007/s13225-020-00466-2)) |
| Otu27726 | 1 | k__Fungi(100);p__Ascomycota(100);c__Saccharomycetes(100);o__Saccharomycetales(100);f__Metschnikowiaceae(100);g__Metschnikowia(100);g__Metschnikowia_unclassified(100); | Metschnikowia | 13 | Saprotroph | \|Undefined Saprotroph\| | Yeast | NULL | Probable | As nectar-tap_saprotroph (Põlme et al. 2020); Decay sugar-rich substrates (Põlme et al. 2020) | Põlme S, et al. 2020. Fungal Diversity 105:1-16 ((https://doi.org/10.1007/s13225-020-00466-2)) |
| Otu27728 | 1 | k__Fungi(100);p__Ascomycota(100);c__Saccharomycetes(100);o__Saccharomycetales(100);f__Saccharomycetales_family_Incertae_sedis(100);f__Saccharomycetales_family_Incertae_sedis_unclassified(100);f__Saccharomycetales_family_Incertae_sedis_unclassified(100); | Saccharomycetales | 7 | Saprotroph | Undefined Saprotroph | Yeast | NULL | Possible | NULL | Sterkenburg E, et al. 2015. New Phytologist 207:1145-1158 ((https://doi.org/10.1111/nph.13426)) |
| Otu27733 | 1 | k__Fungi(100);p__Ascomycota(100);c__Saccharomycetes(100);o__Saccharomycetales(100);o__Saccharomycetales_unclassified(100);o__Saccharomycetales_unclassified(100);o__Saccharomycetales_unclassified(100); | Saccharomycetales | 7 | Saprotroph | Undefined Saprotroph | Yeast | NULL | Possible | NULL | Sterkenburg E, et al. 2015. New Phytologist 207:1145-1158 ((https://doi.org/10.1111/nph.13426)) |
| Otu27734 | 1 | k__Fungi(100);p__Ascomycota(100);c__Saccharomycetes(100);o__Saccharomycetales(100);f__Metschnikowiaceae(100);g__Metschnikowia(100);g__Metschnikowia_unclassified(100); | Metschnikowia | 13 | Saprotroph | \|Undefined Saprotroph\| | Yeast | NULL | Probable | As nectar-tap_saprotroph (Põlme et al. 2020); Decay sugar-rich substrates (Põlme et al. 2020) | Põlme S, et al. 2020. Fungal Diversity 105:1-16 ((https://doi.org/10.1007/s13225-020-00466-2)) |
| Otu27735 | 1 | k__Fungi(100);p__Basidiomycota(100);c__Microbotryomycetes(100);o__Sporidiobolales(100);f__Sporidiobolales_family_Incertae_sedis(100);g__Sporobolomyces(100);s__Sporobolomyces_lactosus(100); | Sporobolomyces | 13 | Pathotroph-Saprotroph | \|Fungal Parasite\|-Plant Saprotroph | Yeast | NULL | Probable | As litter_saprotroph (Põlme et al. 2020); As mycoparasite (Põlme et al. 2020); Decay fungal material (Põlme et al. 2020) | Tedersoo L, et al. 2014. Science 346:e1256688 ((https://doi.org/10.1126/science.1256688)); Purahong W, et al. 2016. Molecular Ecology 25:4059-4074 ((https://doi.org/10.1111/mec.13739)); Põlme S, et al. 2020. Fungal Diversity 105:1-16 ((https://doi.org/10.1007/s13225-020-00466-2)) |
| Otu27738 | 1 | k__Fungi(100);p__Ascomycota(100);c__Saccharomycetes(100);o__Saccharomycetales(100);f__Metschnikowiaceae(100);g__Metschnikowia(100);g__Metschnikowia_unclassified(100); | Metschnikowia | 13 | Saprotroph | \|Undefined Saprotroph\| | Yeast | NULL | Probable | As nectar-tap_saprotroph (Põlme et al. 2020); Decay sugar-rich substrates (Põlme et al. 2020) | Põlme S, et al. 2020. Fungal Diversity 105:1-16 ((https://doi.org/10.1007/s13225-020-00466-2)) |
| Otu27739 | 1 | k__Fungi(100);p__Ascomycota(100);c__Saccharomycetes(100);o__Saccharomycetales(100);f__Metschnikowiaceae(100);g__Metschnikowia(100);g__Metschnikowia_unclassified(100); | Metschnikowia | 13 | Saprotroph | \|Undefined Saprotroph\| | Yeast | NULL | Probable | As nectar-tap_saprotroph (Põlme et al. 2020); Decay sugar-rich substrates (Põlme et al. 2020) | Põlme S, et al. 2020. Fungal Diversity 105:1-16 ((https://doi.org/10.1007/s13225-020-00466-2)) |
| Otu27742 | 1 | k__Fungi(100);p__Ascomycota(100);c__Saccharomycetes(100);o__Saccharomycetales(100);f__Metschnikowiaceae(100);g__Metschnikowia(100);s__Metschnikowia_pulcherrima(100); | Metschnikowia | 13 | Saprotroph | \|Undefined Saprotroph\| | Yeast | NULL | Probable | As nectar-tap_saprotroph (Põlme et al. 2020); Decay sugar-rich substrates (Põlme et al. 2020) | Põlme S, et al. 2020. Fungal Diversity 105:1-16 ((https://doi.org/10.1007/s13225-020-00466-2)) |
| Otu27745 | 1 | k__Fungi(100);p__Ascomycota(100);c__Saccharomycetes(100);o__Saccharomycetales(100);f__Saccharomycetales_family_Incertae_sedis(100);g__Candida(100);g__Candida_unclassified(100); | Candida | 13 | Pathotroph-Saprotroph-Symbiotroph | Animal Pathogen-Endophyte-Endosymbiont-Epiphyte-\|Undefined Saprotroph\| | Dimorphic Yeast-Yeast | NULL | Probable | As nectar-tap_saprotroph (Põlme et al. 2020); Decay sugar-rich substrates (Põlme et al. 2020); This asexual genus is polyphyletic and contain species that function very differently depend on what they are. Caution should be taken when interpreting guilds for Candida species. | Manolakaki D, et al. 2010. Virulence. 1: 367-375 ((https://doi.org/10.4161/viru.1.5.12796)); Mekbib SB, et al. 2011. Tropical Plant Pathology 36:233-240 ((https://doi.org/10.1590/S1982-56762011000400004)); Põlme S, et al. 2020. Fungal Diversity 105:1-16 ((https://doi.org/10.1007/s13225-020-00466-2)) |
| Otu27746 | 1 | k__Fungi(100);p__Ascomycota(100);c__Saccharomycetes(100);o__Saccharomycetales(100);f__Saccharomycetales_family_Incertae_sedis(100);g__Candida(100);g__Candida_unclassified(100); | Candida | 13 | Pathotroph-Saprotroph-Symbiotroph | Animal Pathogen-Endophyte-Endosymbiont-Epiphyte-\|Undefined Saprotroph\| | Dimorphic Yeast-Yeast | NULL | Probable | As nectar-tap_saprotroph (Põlme et al. 2020); Decay sugar-rich substrates (Põlme et al. 2020); This asexual genus is polyphyletic and contain species that function very differently depend on what they are. Caution should be taken when interpreting guilds for Candida species. | Manolakaki D, et al. 2010. Virulence. 1: 367-375 ((https://doi.org/10.4161/viru.1.5.12796)); Mekbib SB, et al. 2011. Tropical Plant Pathology 36:233-240 ((https://doi.org/10.1590/S1982-56762011000400004)); Põlme S, et al. 2020. Fungal Diversity 105:1-16 ((https://doi.org/10.1007/s13225-020-00466-2)) |
| Otu27747 | 1 | k__Fungi(100);p__Ascomycota(100);c__Saccharomycetes(100);o__Saccharomycetales(100);f__Metschnikowiaceae(100);g__Metschnikowia(100);s__Metschnikowia_pulcherrima(100); | Metschnikowia | 13 | Saprotroph | \|Undefined Saprotroph\| | Yeast | NULL | Probable | As nectar-tap_saprotroph (Põlme et al. 2020); Decay sugar-rich substrates (Põlme et al. 2020) | Põlme S, et al. 2020. Fungal Diversity 105:1-16 ((https://doi.org/10.1007/s13225-020-00466-2)) |
| Otu27751 | 1 | k__Fungi(100);p__Ascomycota(100);c__Saccharomycetes(100);o__Saccharomycetales(100);f__Saccharomycetales_family_Incertae_sedis(100);g__Candida(100);s__Candida_cretensis(100); | Candida | 13 | Pathotroph-Saprotroph-Symbiotroph | Animal Pathogen-Endophyte-Endosymbiont-Epiphyte-\|Undefined Saprotroph\| | Dimorphic Yeast-Yeast | NULL | Probable | As nectar-tap_saprotroph (Põlme et al. 2020); Decay sugar-rich substrates (Põlme et al. 2020); This asexual genus is polyphyletic and contain species that function very differently depend on what they are. Caution should be taken when interpreting guilds for Candida species. | Manolakaki D, et al. 2010. Virulence. 1: 367-375 ((https://doi.org/10.4161/viru.1.5.12796)); Mekbib SB, et al. 2011. Tropical Plant Pathology 36:233-240 ((https://doi.org/10.1590/S1982-56762011000400004)); Põlme S, et al. 2020. Fungal Diversity 105:1-16 ((https://doi.org/10.1007/s13225-020-00466-2)) |
| Otu27753 | 1 | k__Fungi(100);p__Ascomycota(100);c__Saccharomycetes(100);o__Saccharomycetales(100);f__Metschnikowiaceae(100);g__Metschnikowia(100);g__Metschnikowia_unclassified(100); | Metschnikowia | 13 | Saprotroph | \|Undefined Saprotroph\| | Yeast | NULL | Probable | As nectar-tap_saprotroph (Põlme et al. 2020); Decay sugar-rich substrates (Põlme et al. 2020) | Põlme S, et al. 2020. Fungal Diversity 105:1-16 ((https://doi.org/10.1007/s13225-020-00466-2)) |
| Otu27754 | 1 | k__Fungi(100);p__Ascomycota(100);c__Saccharomycetes(100);o__Saccharomycetales(100);f__Metschnikowiaceae(100);g__Metschnikowia(100);g__Metschnikowia_unclassified(100); | Metschnikowia | 13 | Saprotroph | \|Undefined Saprotroph\| | Yeast | NULL | Probable | As nectar-tap_saprotroph (Põlme et al. 2020); Decay sugar-rich substrates (Põlme et al. 2020) | Põlme S, et al. 2020. Fungal Diversity 105:1-16 ((https://doi.org/10.1007/s13225-020-00466-2)) |
| Otu27755 | 1 | k__Fungi(100);p__Ascomycota(100);c__Saccharomycetes(100);o__Saccharomycetales(100);f__Metschnikowiaceae(100);g__Metschnikowia(100);s__Metschnikowia_pulcherrima(100); | Metschnikowia | 13 | Saprotroph | \|Undefined Saprotroph\| | Yeast | NULL | Probable | As nectar-tap_saprotroph (Põlme et al. 2020); Decay sugar-rich substrates (Põlme et al. 2020) | Põlme S, et al. 2020. Fungal Diversity 105:1-16 ((https://doi.org/10.1007/s13225-020-00466-2)) |
| Otu27756 | 1 | k__Fungi(100);p__Ascomycota(100);c__Saccharomycetes(100);o__Saccharomycetales(100);f__Metschnikowiaceae(100);g__Metschnikowia(100);s__Metschnikowia_pulcherrima(100); | Metschnikowia | 13 | Saprotroph | \|Undefined Saprotroph\| | Yeast | NULL | Probable | As nectar-tap_saprotroph (Põlme et al. 2020); Decay sugar-rich substrates (Põlme et al. 2020) | Põlme S, et al. 2020. Fungal Diversity 105:1-16 ((https://doi.org/10.1007/s13225-020-00466-2)) |
| Otu27757 | 1 | k__Fungi(100);p__Ascomycota(100);c__Saccharomycetes(100);o__Saccharomycetales(100);f__Metschnikowiaceae(100);g__Metschnikowia(100);s__Metschnikowia_pulcherrima(100); | Metschnikowia | 13 | Saprotroph | \|Undefined Saprotroph\| | Yeast | NULL | Probable | As nectar-tap_saprotroph (Põlme et al. 2020); Decay sugar-rich substrates (Põlme et al. 2020) | Põlme S, et al. 2020. Fungal Diversity 105:1-16 ((https://doi.org/10.1007/s13225-020-00466-2)) |
| Otu27758 | 1 | k__Fungi(100);p__Ascomycota(100);c__Saccharomycetes(100);o__Saccharomycetales(100);f__Saccharomycetales_family_Incertae_sedis(100);f__Saccharomycetales_family_Incertae_sedis_unclassified(100);f__Saccharomycetales_family_Incertae_sedis_unclassified(100); | Saccharomycetales | 7 | Saprotroph | Undefined Saprotroph | Yeast | NULL | Possible | NULL | Sterkenburg E, et al. 2015. New Phytologist 207:1145-1158 ((https://doi.org/10.1111/nph.13426)) |
| Otu27769 | 1 | k__Fungi(100);p__Ascomycota(100);c__Saccharomycetes(100);o__Saccharomycetales(100);f__Metschnikowiaceae(100);g__Metschnikowia(100);s__Metschnikowia_pulcherrima(100); | Metschnikowia | 13 | Saprotroph | \|Undefined Saprotroph\| | Yeast | NULL | Probable | As nectar-tap_saprotroph (Põlme et al. 2020); Decay sugar-rich substrates (Põlme et al. 2020) | Põlme S, et al. 2020. Fungal Diversity 105:1-16 ((https://doi.org/10.1007/s13225-020-00466-2)) |
| Otu27770 | 1 | k__Fungi(100);p__Ascomycota(100);c__Saccharomycetes(100);o__Saccharomycetales(100);o__Saccharomycetales_unclassified(100);o__Saccharomycetales_unclassified(100);o__Saccharomycetales_unclassified(100); | Saccharomycetales | 7 | Saprotroph | Undefined Saprotroph | Yeast | NULL | Possible | NULL | Sterkenburg E, et al. 2015. New Phytologist 207:1145-1158 ((https://doi.org/10.1111/nph.13426)) |
| Otu27773 | 1 | k__Fungi(100);p__Ascomycota(100);c__Lecanoromycetes(100);o__Teloschistales(100);f__Physciaceae(100);g__Physcia(100);s__Physcia_sp(100); | Physcia | 13 | Symbiotroph | \|Lichenized\| | Thallus | NULL | Highly Probable | NULL | Esslinger TL. 2014. North Dakota State University ((http://www.ndsu.edu/pubweb/~esslinge/chcklst/chcklst7.htm)); Põlme S, et al. 2020. Fungal Diversity 105:1-16 ((https://doi.org/10.1007/s13225-020-00466-2)) |
| Otu27774 | 1 | k__Fungi(100);p__Ascomycota(100);c__Sordariomycetes(100);o__Sordariales(100);f__Chaetomiaceae(100);f__Chaetomiaceae_unclassified(100);f__Chaetomiaceae_unclassified(100); | Chaetomiaceae | 9 | Pathotroph-Saprotroph-Symbiotroph | Animal Pathogen-Dung Saprotroph-Endophyte-Epiphyte-Plant Saprotroph-Wood Saprotroph | Microfungus | Soft Rot | Possible | Saprobic, acting as key decay organisms of plant materials and plant-derived commodities (Cannon & Kirk 2007); Epiphyte detection method-culture (Massimo et al. 2015); Endophyte detection method-culture (Busby et al. 2016, David et al. 2016); Likely opportunistic human pathogen (Irinyi et al. 2015) | Seehann G, et al. 1975. List of Fungi in Soft-Rot Tissue Tests. Institute fur Holzbiologie und Holzschutz, Bundesforschungsanstalt fur Forst - und Holzwirtshaft, Hamburg - Lohbrugge, Germany; Cannon PF, Kirk PM. 2007. Fungal Families of the World. CAB International, Cambridge ((https://isbnsearch.org/isbn/9780851998275)); Bills GF, et al. 2013. Current Opinion in Microbiology 16:549-565 ((https://doi.org/10.1016/j.mib.2013.08.001)); Tedersoo L, et al. 2014. Science 346:e1256688 ((https://doi.org/10.1126/science.1256688)); Irinyi L, et al. 2015. Medical Mycology 53:313-337 ((https://doi.org/10.1093/mmy/myv008)); Massimo NC, et al. 2015. Microbial Ecology 70:61-76 ((https://doi.org/10.1007/s00248-014-0563-6)); Busby PE, et al. 2016. New Phytologist 209:1681-1692 ((https://doi.org/10.1111/nph.13742)); David AS, et al. 2016. Microbial Ecology 71:912-926 ((https://doi.org/10.1007/s00248-015-0712-6)) |
| Otu27795 | 1 | k__Fungi(100);p__Ascomycota(100);c__Saccharomycetes(100);o__Saccharomycetales(100);f__Metschnikowiaceae(100);g__Metschnikowia(100);s__Metschnikowia_pulcherrima(100); | Metschnikowia | 13 | Saprotroph | \|Undefined Saprotroph\| | Yeast | NULL | Probable | As nectar-tap_saprotroph (Põlme et al. 2020); Decay sugar-rich substrates (Põlme et al. 2020) | Põlme S, et al. 2020. Fungal Diversity 105:1-16 ((https://doi.org/10.1007/s13225-020-00466-2)) |
| Otu27797 | 1 | k__Fungi(100);p__Basidiomycota(100);c__Agaricomycetes(100);o__Russulales(100);f__Peniophoraceae(100);f__Peniophoraceae_unclassified(100);f__Peniophoraceae_unclassified(100); | Peniophoraceae | 9 | Pathotroph-Saprotroph | Plant Pathogen-Wood Saprotroph | Corticioid | White Rot | Probable | Most species are saprobic, causing rots of standing and fallen wood. Many species are adapted to xeric conditions, being able to grow on rapidly drying dead attached branches (Cannon & Kirk 2007) | Cannon PF, Kirk PM. 2007. Fungal Families of the World. CAB International, Cambridge (ISBN: 978-0851998275) |
| Otu27806 | 1 | k__Fungi(100);p__Ascomycota(100);c__Saccharomycetes(100);o__Saccharomycetales(100);f__Metschnikowiaceae(100);g__Metschnikowia(100);g__Metschnikowia_unclassified(100); | Metschnikowia | 13 | Saprotroph | \|Undefined Saprotroph\| | Yeast | NULL | Probable | As nectar-tap_saprotroph (Põlme et al. 2020); Decay sugar-rich substrates (Põlme et al. 2020) | Põlme S, et al. 2020. Fungal Diversity 105:1-16 ((https://doi.org/10.1007/s13225-020-00466-2)) |
| Otu27814 | 1 | k__Fungi(100);p__Ascomycota(100);c__Dothideomycetes(100);o__Pleosporales(100);f__Lophiostomataceae(100);g__Lophiostoma(100);g__Lophiostoma_unclassified(100); | Lophiostoma | 13 | Saprotroph | Plant Saprotroph-Undefined Saprotroph-\|Wood Saprotroph\| | Microfungus | NULL | Probable | As litter_saprotroph (Põlme et al. 2020); Decay wood, leaf, fruit, seed (Põlme et al. 2020) | Tedersoo L, et al. 2014. Science 346:e1256688 ((https://doi.org/10.1126/science.1256688)); Põlme S, et al. 2020. Fungal Diversity 105:1-16 ((https://doi.org/10.1007/s13225-020-00466-2)) |
| Otu27827 | 1 | k__Fungi(100);p__Ascomycota(100);c__Saccharomycetes(100);o__Saccharomycetales(100);f__Metschnikowiaceae(100);g__Metschnikowia(100);s__Metschnikowia_pulcherrima(100); | Metschnikowia | 13 | Saprotroph | \|Undefined Saprotroph\| | Yeast | NULL | Probable | As nectar-tap_saprotroph (Põlme et al. 2020); Decay sugar-rich substrates (Põlme et al. 2020) | Põlme S, et al. 2020. Fungal Diversity 105:1-16 ((https://doi.org/10.1007/s13225-020-00466-2)) |
| Otu27828 | 1 | k__Fungi(100);p__Ascomycota(100);c__Saccharomycetes(100);o__Saccharomycetales(100);f__Metschnikowiaceae(100);g__Metschnikowia(100);g__Metschnikowia_unclassified(100); | Metschnikowia | 13 | Saprotroph | \|Undefined Saprotroph\| | Yeast | NULL | Probable | As nectar-tap_saprotroph (Põlme et al. 2020); Decay sugar-rich substrates (Põlme et al. 2020) | Põlme S, et al. 2020. Fungal Diversity 105:1-16 ((https://doi.org/10.1007/s13225-020-00466-2)) |
| Otu27829 | 1 | k__Fungi(100);p__Ascomycota(100);c__Saccharomycetes(100);o__Saccharomycetales(100);f__Metschnikowiaceae(100);g__Metschnikowia(100);s__Metschnikowia_pulcherrima(100); | Metschnikowia | 13 | Saprotroph | \|Undefined Saprotroph\| | Yeast | NULL | Probable | As nectar-tap_saprotroph (Põlme et al. 2020); Decay sugar-rich substrates (Põlme et al. 2020) | Põlme S, et al. 2020. Fungal Diversity 105:1-16 ((https://doi.org/10.1007/s13225-020-00466-2)) |
| Otu27830 | 1 | k__Fungi(100);p__Ascomycota(100);c__Saccharomycetes(100);o__Saccharomycetales(100);f__Metschnikowiaceae(100);g__Metschnikowia(100);s__Metschnikowia_pulcherrima(100); | Metschnikowia | 13 | Saprotroph | \|Undefined Saprotroph\| | Yeast | NULL | Probable | As nectar-tap_saprotroph (Põlme et al. 2020); Decay sugar-rich substrates (Põlme et al. 2020) | Põlme S, et al. 2020. Fungal Diversity 105:1-16 ((https://doi.org/10.1007/s13225-020-00466-2)) |
| Otu27831 | 1 | k__Fungi(100);p__Ascomycota(100);c__Saccharomycetes(100);o__Saccharomycetales(100);f__Metschnikowiaceae(100);g__Metschnikowia(100);s__Metschnikowia_pulcherrima(100); | Metschnikowia | 13 | Saprotroph | \|Undefined Saprotroph\| | Yeast | NULL | Probable | As nectar-tap_saprotroph (Põlme et al. 2020); Decay sugar-rich substrates (Põlme et al. 2020) | Põlme S, et al. 2020. Fungal Diversity 105:1-16 ((https://doi.org/10.1007/s13225-020-00466-2)) |
| Otu27834 | 1 | k__Fungi(100);p__Ascomycota(100);c__Saccharomycetes(100);o__Saccharomycetales(100);f__Metschnikowiaceae(100);g__Metschnikowia(100);s__Metschnikowia_pulcherrima(100); | Metschnikowia | 13 | Saprotroph | \|Undefined Saprotroph\| | Yeast | NULL | Probable | As nectar-tap_saprotroph (Põlme et al. 2020); Decay sugar-rich substrates (Põlme et al. 2020) | Põlme S, et al. 2020. Fungal Diversity 105:1-16 ((https://doi.org/10.1007/s13225-020-00466-2)) |
| Otu27835 | 1 | k__Fungi(100);p__Ascomycota(100);c__Saccharomycetes(100);o__Saccharomycetales(100);f__Metschnikowiaceae(100);g__Metschnikowia(100);s__Metschnikowia_pulcherrima(100); | Metschnikowia | 13 | Saprotroph | \|Undefined Saprotroph\| | Yeast | NULL | Probable | As nectar-tap_saprotroph (Põlme et al. 2020); Decay sugar-rich substrates (Põlme et al. 2020) | Põlme S, et al. 2020. Fungal Diversity 105:1-16 ((https://doi.org/10.1007/s13225-020-00466-2)) |
| Otu27837 | 1 | k__Fungi(100);p__Ascomycota(100);c__Saccharomycetes(100);o__Saccharomycetales(100);f__Metschnikowiaceae(100);g__Metschnikowia(100);s__Metschnikowia_pulcherrima(100); | Metschnikowia | 13 | Saprotroph | \|Undefined Saprotroph\| | Yeast | NULL | Probable | As nectar-tap_saprotroph (Põlme et al. 2020); Decay sugar-rich substrates (Põlme et al. 2020) | Põlme S, et al. 2020. Fungal Diversity 105:1-16 ((https://doi.org/10.1007/s13225-020-00466-2)) |
| Otu27840 | 1 | k__Fungi(100);p__Ascomycota(100);c__Saccharomycetes(100);o__Saccharomycetales(100);f__Metschnikowiaceae(100);g__Metschnikowia(100);s__Metschnikowia_pulcherrima(100); | Metschnikowia | 13 | Saprotroph | \|Undefined Saprotroph\| | Yeast | NULL | Probable | As nectar-tap_saprotroph (Põlme et al. 2020); Decay sugar-rich substrates (Põlme et al. 2020) | Põlme S, et al. 2020. Fungal Diversity 105:1-16 ((https://doi.org/10.1007/s13225-020-00466-2)) |
| Otu27853 | 1 | k__Fungi(100);p__Ascomycota(100);c__Saccharomycetes(100);o__Saccharomycetales(100);f__Metschnikowiaceae(100);g__Metschnikowia(100);s__Metschnikowia_pulcherrima(100); | Metschnikowia | 13 | Saprotroph | \|Undefined Saprotroph\| | Yeast | NULL | Probable | As nectar-tap_saprotroph (Põlme et al. 2020); Decay sugar-rich substrates (Põlme et al. 2020) | Põlme S, et al. 2020. Fungal Diversity 105:1-16 ((https://doi.org/10.1007/s13225-020-00466-2)) |
| Otu27854 | 1 | k__Fungi(100);p__Ascomycota(100);c__Saccharomycetes(100);o__Saccharomycetales(100);f__Metschnikowiaceae(100);g__Metschnikowia(100);g__Metschnikowia_unclassified(100); | Metschnikowia | 13 | Saprotroph | \|Undefined Saprotroph\| | Yeast | NULL | Probable | As nectar-tap_saprotroph (Põlme et al. 2020); Decay sugar-rich substrates (Põlme et al. 2020) | Põlme S, et al. 2020. Fungal Diversity 105:1-16 ((https://doi.org/10.1007/s13225-020-00466-2)) |
| Otu27857 | 1 | k__Fungi(100);p__Ascomycota(100);c__Saccharomycetes(100);o__Saccharomycetales(100);f__Metschnikowiaceae(100);g__Metschnikowia(100);s__Metschnikowia_pulcherrima(100); | Metschnikowia | 13 | Saprotroph | \|Undefined Saprotroph\| | Yeast | NULL | Probable | As nectar-tap_saprotroph (Põlme et al. 2020); Decay sugar-rich substrates (Põlme et al. 2020) | Põlme S, et al. 2020. Fungal Diversity 105:1-16 ((https://doi.org/10.1007/s13225-020-00466-2)) |
| Otu27866 | 1 | k__Fungi(100);p__Ascomycota(100);c__Saccharomycetes(100);o__Saccharomycetales(100);f__Metschnikowiaceae(100);g__Metschnikowia(100);s__Metschnikowia_pulcherrima(100); | Metschnikowia | 13 | Saprotroph | \|Undefined Saprotroph\| | Yeast | NULL | Probable | As nectar-tap_saprotroph (Põlme et al. 2020); Decay sugar-rich substrates (Põlme et al. 2020) | Põlme S, et al. 2020. Fungal Diversity 105:1-16 ((https://doi.org/10.1007/s13225-020-00466-2)) |
| Otu27874 | 1 | k__Fungi(100);p__Ascomycota(100);c__Saccharomycetes(100);o__Saccharomycetales(100);f__Metschnikowiaceae(100);g__Metschnikowia(100);s__Metschnikowia_pulcherrima(100); | Metschnikowia | 13 | Saprotroph | \|Undefined Saprotroph\| | Yeast | NULL | Probable | As nectar-tap_saprotroph (Põlme et al. 2020); Decay sugar-rich substrates (Põlme et al. 2020) | Põlme S, et al. 2020. Fungal Diversity 105:1-16 ((https://doi.org/10.1007/s13225-020-00466-2)) |
| Otu27875 | 1 | k__Fungi(100);p__Ascomycota(100);c__Saccharomycetes(100);o__Saccharomycetales(100);f__Saccharomycetales_family_Incertae_sedis(100);g__Candida(100);g__Candida_unclassified(100); | Candida | 13 | Pathotroph-Saprotroph-Symbiotroph | Animal Pathogen-Endophyte-Endosymbiont-Epiphyte-\|Undefined Saprotroph\| | Dimorphic Yeast-Yeast | NULL | Probable | As nectar-tap_saprotroph (Põlme et al. 2020); Decay sugar-rich substrates (Põlme et al. 2020); This asexual genus is polyphyletic and contain species that function very differently depend on what they are. Caution should be taken when interpreting guilds for Candida species. | Manolakaki D, et al. 2010. Virulence. 1: 367-375 ((https://doi.org/10.4161/viru.1.5.12796)); Mekbib SB, et al. 2011. Tropical Plant Pathology 36:233-240 ((https://doi.org/10.1590/S1982-56762011000400004)); Põlme S, et al. 2020. Fungal Diversity 105:1-16 ((https://doi.org/10.1007/s13225-020-00466-2)) |
| Otu27882 | 1 | k__Fungi(100);p__Basidiomycota(100);c__Agaricomycetes(100);o__Polyporales(100);f__Fomitopsidaceae(100);g__Antrodia(100);s__Antrodia_serialis(100); | Antrodia | 13 | Saprotroph | \|Wood Saprotroph\| | Corticioid | Brown Rot | Highly Probable | As brown-rot (Põlme et al. 2020); Decay wood (Põlme et al. 2020) | Gilbertson RL, Ryvarden L. 1987-1987. North American Polypores. Fungiflora, Oslo (ISBN: 978-0945345060); Põlme S, et al. 2020. Fungal Diversity 105:1-16 ((https://doi.org/10.1007/s13225-020-00466-2)) |
| Otu27884 | 1 | k__Fungi(100);p__Ascomycota(100);c__Saccharomycetes(100);o__Saccharomycetales(100);f__Metschnikowiaceae(100);g__Metschnikowia(100);s__Metschnikowia_pulcherrima(100); | Metschnikowia | 13 | Saprotroph | \|Undefined Saprotroph\| | Yeast | NULL | Probable | As nectar-tap_saprotroph (Põlme et al. 2020); Decay sugar-rich substrates (Põlme et al. 2020) | Põlme S, et al. 2020. Fungal Diversity 105:1-16 ((https://doi.org/10.1007/s13225-020-00466-2)) |
| Otu27887 | 1 | k__Fungi(100);p__Ascomycota(100);c__Saccharomycetes(100);o__Saccharomycetales(100);f__Metschnikowiaceae(100);g__Metschnikowia(100);s__Metschnikowia_pulcherrima(100); | Metschnikowia | 13 | Saprotroph | \|Undefined Saprotroph\| | Yeast | NULL | Probable | As nectar-tap_saprotroph (Põlme et al. 2020); Decay sugar-rich substrates (Põlme et al. 2020) | Põlme S, et al. 2020. Fungal Diversity 105:1-16 ((https://doi.org/10.1007/s13225-020-00466-2)) |
| Otu27891 | 1 | k__Fungi(100);p__Ascomycota(100);c__Saccharomycetes(100);o__Saccharomycetales(100);f__Metschnikowiaceae(100);g__Metschnikowia(100);s__Metschnikowia_pulcherrima(100); | Metschnikowia | 13 | Saprotroph | \|Undefined Saprotroph\| | Yeast | NULL | Probable | As nectar-tap_saprotroph (Põlme et al. 2020); Decay sugar-rich substrates (Põlme et al. 2020) | Põlme S, et al. 2020. Fungal Diversity 105:1-16 ((https://doi.org/10.1007/s13225-020-00466-2)) |
| Otu27896 | 1 | k__Fungi(100);p__Ascomycota(100);c__Saccharomycetes(100);o__Saccharomycetales(100);f__Metschnikowiaceae(100);g__Metschnikowia(100);g__Metschnikowia_unclassified(100); | Metschnikowia | 13 | Saprotroph | \|Undefined Saprotroph\| | Yeast | NULL | Probable | As nectar-tap_saprotroph (Põlme et al. 2020); Decay sugar-rich substrates (Põlme et al. 2020) | Põlme S, et al. 2020. Fungal Diversity 105:1-16 ((https://doi.org/10.1007/s13225-020-00466-2)) |
| Otu27902 | 1 | k__Fungi(100);p__Ascomycota(100);c__Saccharomycetes(100);o__Saccharomycetales(100);f__Metschnikowiaceae(100);g__Metschnikowia(100);s__Metschnikowia_pulcherrima(100); | Metschnikowia | 13 | Saprotroph | \|Undefined Saprotroph\| | Yeast | NULL | Probable | As nectar-tap_saprotroph (Põlme et al. 2020); Decay sugar-rich substrates (Põlme et al. 2020) | Põlme S, et al. 2020. Fungal Diversity 105:1-16 ((https://doi.org/10.1007/s13225-020-00466-2)) |
| Otu27905 | 1 | k__Fungi(100);p__Ascomycota(100);c__Saccharomycetes(100);o__Saccharomycetales(100);f__Metschnikowiaceae(100);g__Metschnikowia(100);s__Metschnikowia_pulcherrima(100); | Metschnikowia | 13 | Saprotroph | \|Undefined Saprotroph\| | Yeast | NULL | Probable | As nectar-tap_saprotroph (Põlme et al. 2020); Decay sugar-rich substrates (Põlme et al. 2020) | Põlme S, et al. 2020. Fungal Diversity 105:1-16 ((https://doi.org/10.1007/s13225-020-00466-2)) |
| Otu27906 | 1 | k__Fungi(100);p__Ascomycota(100);c__Saccharomycetes(100);o__Saccharomycetales(100);f__Metschnikowiaceae(100);g__Metschnikowia(100);s__Metschnikowia_pulcherrima(100); | Metschnikowia | 13 | Saprotroph | \|Undefined Saprotroph\| | Yeast | NULL | Probable | As nectar-tap_saprotroph (Põlme et al. 2020); Decay sugar-rich substrates (Põlme et al. 2020) | Põlme S, et al. 2020. Fungal Diversity 105:1-16 ((https://doi.org/10.1007/s13225-020-00466-2)) |
| Otu27907 | 1 | k__Fungi(100);p__Ascomycota(100);c__Saccharomycetes(100);o__Saccharomycetales(100);f__Metschnikowiaceae(100);g__Metschnikowia(100);s__Metschnikowia_pulcherrima(100); | Metschnikowia | 13 | Saprotroph | \|Undefined Saprotroph\| | Yeast | NULL | Probable | As nectar-tap_saprotroph (Põlme et al. 2020); Decay sugar-rich substrates (Põlme et al. 2020) | Põlme S, et al. 2020. Fungal Diversity 105:1-16 ((https://doi.org/10.1007/s13225-020-00466-2)) |
| Otu27908 | 1 | k__Fungi(100);p__Ascomycota(100);c__Saccharomycetes(100);o__Saccharomycetales(100);f__Metschnikowiaceae(100);g__Metschnikowia(100);s__Metschnikowia_pulcherrima(100); | Metschnikowia | 13 | Saprotroph | \|Undefined Saprotroph\| | Yeast | NULL | Probable | As nectar-tap_saprotroph (Põlme et al. 2020); Decay sugar-rich substrates (Põlme et al. 2020) | Põlme S, et al. 2020. Fungal Diversity 105:1-16 ((https://doi.org/10.1007/s13225-020-00466-2)) |
| Otu27909 | 1 | k__Fungi(100);p__Ascomycota(100);c__Saccharomycetes(100);o__Saccharomycetales(100);f__Metschnikowiaceae(100);g__Metschnikowia(100);s__Metschnikowia_pulcherrima(100); | Metschnikowia | 13 | Saprotroph | \|Undefined Saprotroph\| | Yeast | NULL | Probable | As nectar-tap_saprotroph (Põlme et al. 2020); Decay sugar-rich substrates (Põlme et al. 2020) | Põlme S, et al. 2020. Fungal Diversity 105:1-16 ((https://doi.org/10.1007/s13225-020-00466-2)) |
| Otu27910 | 1 | k__Fungi(100);p__Ascomycota(100);c__Saccharomycetes(100);o__Saccharomycetales(100);f__Metschnikowiaceae(100);g__Metschnikowia(100);g__Metschnikowia_unclassified(100); | Metschnikowia | 13 | Saprotroph | \|Undefined Saprotroph\| | Yeast | NULL | Probable | As nectar-tap_saprotroph (Põlme et al. 2020); Decay sugar-rich substrates (Põlme et al. 2020) | Põlme S, et al. 2020. Fungal Diversity 105:1-16 ((https://doi.org/10.1007/s13225-020-00466-2)) |
| Otu27911 | 1 | k__Fungi(100);p__Ascomycota(100);c__Saccharomycetes(100);o__Saccharomycetales(100);f__Metschnikowiaceae(100);g__Metschnikowia(100);s__Metschnikowia_pulcherrima(100); | Metschnikowia | 13 | Saprotroph | \|Undefined Saprotroph\| | Yeast | NULL | Probable | As nectar-tap_saprotroph (Põlme et al. 2020); Decay sugar-rich substrates (Põlme et al. 2020) | Põlme S, et al. 2020. Fungal Diversity 105:1-16 ((https://doi.org/10.1007/s13225-020-00466-2)) |
| Otu27912 | 1 | k__Fungi(100);p__Ascomycota(100);c__Leotiomycetes(100);o__Helotiales(100);f__Hyaloscyphaceae(100);g__Arachnopeziza(100);g__Arachnopeziza_unclassified(100); | Arachnopeziza | 13 | Saprotroph | Undefined Saprotroph-\|Wood Saprotroph\| | Microfungus-Pezizoid | NULL | Probable | Decay wood (Põlme et al. 2020) | Tedersoo L, et al. 2014. Science 346:e1256688 ((https://doi.org/10.1126/science.1256688)); Põlme S, et al. 2020. Fungal Diversity 105:1-16 ((https://doi.org/10.1007/s13225-020-00466-2)) |
| Otu27913 | 1 | k__Fungi(100);p__Ascomycota(100);c__Saccharomycetes(100);o__Saccharomycetales(100);f__Metschnikowiaceae(100);g__Metschnikowia(100);s__Metschnikowia_pulcherrima(100); | Metschnikowia | 13 | Saprotroph | \|Undefined Saprotroph\| | Yeast | NULL | Probable | As nectar-tap_saprotroph (Põlme et al. 2020); Decay sugar-rich substrates (Põlme et al. 2020) | Põlme S, et al. 2020. Fungal Diversity 105:1-16 ((https://doi.org/10.1007/s13225-020-00466-2)) |
| Otu27921 | 1 | k__Fungi(100);p__Ascomycota(100);c__Saccharomycetes(100);o__Saccharomycetales(100);f__Metschnikowiaceae(100);g__Metschnikowia(100);s__Metschnikowia_pulcherrima(100); | Metschnikowia | 13 | Saprotroph | \|Undefined Saprotroph\| | Yeast | NULL | Probable | As nectar-tap_saprotroph (Põlme et al. 2020); Decay sugar-rich substrates (Põlme et al. 2020) | Põlme S, et al. 2020. Fungal Diversity 105:1-16 ((https://doi.org/10.1007/s13225-020-00466-2)) |
| Otu27923 | 1 | k__Fungi(100);p__Ascomycota(100);c__Saccharomycetes(100);o__Saccharomycetales(100);f__Metschnikowiaceae(100);g__Metschnikowia(100);s__Metschnikowia_pulcherrima(100); | Metschnikowia | 13 | Saprotroph | \|Undefined Saprotroph\| | Yeast | NULL | Probable | As nectar-tap_saprotroph (Põlme et al. 2020); Decay sugar-rich substrates (Põlme et al. 2020) | Põlme S, et al. 2020. Fungal Diversity 105:1-16 ((https://doi.org/10.1007/s13225-020-00466-2)) |
| Otu27926 | 1 | k__Fungi(100);p__Ascomycota(100);c__Saccharomycetes(100);o__Saccharomycetales(100);f__Metschnikowiaceae(100);g__Metschnikowia(100);s__Metschnikowia_pulcherrima(100); | Metschnikowia | 13 | Saprotroph | \|Undefined Saprotroph\| | Yeast | NULL | Probable | As nectar-tap_saprotroph (Põlme et al. 2020); Decay sugar-rich substrates (Põlme et al. 2020) | Põlme S, et al. 2020. Fungal Diversity 105:1-16 ((https://doi.org/10.1007/s13225-020-00466-2)) |
| Otu27927 | 1 | k__Fungi(100);p__Ascomycota(100);c__Saccharomycetes(100);o__Saccharomycetales(100);f__Metschnikowiaceae(100);g__Metschnikowia(100);g__Metschnikowia_unclassified(100); | Metschnikowia | 13 | Saprotroph | \|Undefined Saprotroph\| | Yeast | NULL | Probable | As nectar-tap_saprotroph (Põlme et al. 2020); Decay sugar-rich substrates (Põlme et al. 2020) | Põlme S, et al. 2020. Fungal Diversity 105:1-16 ((https://doi.org/10.1007/s13225-020-00466-2)) |
| Otu27928 | 1 | k__Fungi(100);p__Ascomycota(100);c__Saccharomycetes(100);o__Saccharomycetales(100);f__Metschnikowiaceae(100);g__Metschnikowia(100);s__Metschnikowia_pulcherrima(100); | Metschnikowia | 13 | Saprotroph | \|Undefined Saprotroph\| | Yeast | NULL | Probable | As nectar-tap_saprotroph (Põlme et al. 2020); Decay sugar-rich substrates (Põlme et al. 2020) | Põlme S, et al. 2020. Fungal Diversity 105:1-16 ((https://doi.org/10.1007/s13225-020-00466-2)) |
| Otu27930 | 1 | k__Fungi(100);p__Ascomycota(100);c__Saccharomycetes(100);o__Saccharomycetales(100);f__Metschnikowiaceae(100);g__Metschnikowia(100);s__Metschnikowia_pulcherrima(100); | Metschnikowia | 13 | Saprotroph | \|Undefined Saprotroph\| | Yeast | NULL | Probable | As nectar-tap_saprotroph (Põlme et al. 2020); Decay sugar-rich substrates (Põlme et al. 2020) | Põlme S, et al. 2020. Fungal Diversity 105:1-16 ((https://doi.org/10.1007/s13225-020-00466-2)) |
| Otu27932 | 1 | k__Fungi(100);p__Ascomycota(100);c__Leotiomycetes(100);o__Helotiales(100);f__Helotiales_family_Incertae_sedis(100);g__Leptodontidium(100);s__Leptodontidium_sp_Di124_4(100); | Leptodontidium | 13 | Saprotroph-Symbiotroph | Endophyte-Ericoid Mycorrhizal-\|Plant Saprotroph\| | Microfungus | NULL | Highly Probable | As litter_saprotroph (Põlme et al. 2020); As root_endophyte_dark_septate (Põlme et al. 2020); Dark Septate Endophyte, known from plant roots.; Decay leaf, fruit, seed (Põlme et al. 2020) | Jumpponen A, Trappe J. 1998. New Phytologist 140:295-310 ((https://doi.org/10.1046/j.1469-8137.1998.00265.x)) (as DSE); Taylor DL, et al. 2014. Ecological Monographs 8:43-20 ((https://esajournals.onlinelibrary.wiley.com/doi/10.1890/12-1693.1)); Põlme S, et al. 2020. Fungal Diversity 105:1-16 ((https://doi.org/10.1007/s13225-020-00466-2)) |
| Otu27934 | 1 | k__Fungi(100);p__Ascomycota(100);c__Saccharomycetes(100);o__Saccharomycetales(100);f__Metschnikowiaceae(100);g__Metschnikowia(100);g__Metschnikowia_unclassified(100); | Metschnikowia | 13 | Saprotroph | \|Undefined Saprotroph\| | Yeast | NULL | Probable | As nectar-tap_saprotroph (Põlme et al. 2020); Decay sugar-rich substrates (Põlme et al. 2020) | Põlme S, et al. 2020. Fungal Diversity 105:1-16 ((https://doi.org/10.1007/s13225-020-00466-2)) |
| Otu27935 | 1 | k__Fungi(100);p__Ascomycota(100);c__Saccharomycetes(100);o__Saccharomycetales(100);f__Metschnikowiaceae(100);g__Metschnikowia(100);s__Metschnikowia_pulcherrima(100); | Metschnikowia | 13 | Saprotroph | \|Undefined Saprotroph\| | Yeast | NULL | Probable | As nectar-tap_saprotroph (Põlme et al. 2020); Decay sugar-rich substrates (Põlme et al. 2020) | Põlme S, et al. 2020. Fungal Diversity 105:1-16 ((https://doi.org/10.1007/s13225-020-00466-2)) |
| Otu27936 | 1 | k__Fungi(100);p__Ascomycota(100);c__Saccharomycetes(100);o__Saccharomycetales(100);f__Metschnikowiaceae(100);g__Metschnikowia(100);s__Metschnikowia_pulcherrima(100); | Metschnikowia | 13 | Saprotroph | \|Undefined Saprotroph\| | Yeast | NULL | Probable | As nectar-tap_saprotroph (Põlme et al. 2020); Decay sugar-rich substrates (Põlme et al. 2020) | Põlme S, et al. 2020. Fungal Diversity 105:1-16 ((https://doi.org/10.1007/s13225-020-00466-2)) |
| Otu27937 | 1 | k__Fungi(100);p__Ascomycota(100);c__Saccharomycetes(100);o__Saccharomycetales(100);f__Metschnikowiaceae(100);g__Metschnikowia(100);s__Metschnikowia_pulcherrima(100); | Metschnikowia | 13 | Saprotroph | \|Undefined Saprotroph\| | Yeast | NULL | Probable | As nectar-tap_saprotroph (Põlme et al. 2020); Decay sugar-rich substrates (Põlme et al. 2020) | Põlme S, et al. 2020. Fungal Diversity 105:1-16 ((https://doi.org/10.1007/s13225-020-00466-2)) |
| Otu27939 | 1 | k__Fungi(100);p__Ascomycota(100);c__Saccharomycetes(100);o__Saccharomycetales(100);f__Metschnikowiaceae(100);g__Metschnikowia(100);s__Metschnikowia_pulcherrima(100); | Metschnikowia | 13 | Saprotroph | \|Undefined Saprotroph\| | Yeast | NULL | Probable | As nectar-tap_saprotroph (Põlme et al. 2020); Decay sugar-rich substrates (Põlme et al. 2020) | Põlme S, et al. 2020. Fungal Diversity 105:1-16 ((https://doi.org/10.1007/s13225-020-00466-2)) |
| Otu27947 | 1 | k__Fungi(100);p__Ascomycota(100);c__Saccharomycetes(100);o__Saccharomycetales(100);f__Metschnikowiaceae(100);g__Metschnikowia(100);s__Metschnikowia_pulcherrima(100); | Metschnikowia | 13 | Saprotroph | \|Undefined Saprotroph\| | Yeast | NULL | Probable | As nectar-tap_saprotroph (Põlme et al. 2020); Decay sugar-rich substrates (Põlme et al. 2020) | Põlme S, et al. 2020. Fungal Diversity 105:1-16 ((https://doi.org/10.1007/s13225-020-00466-2)) |
| Otu27953 | 1 | k__Fungi(100);p__Ascomycota(100);c__Saccharomycetes(100);o__Saccharomycetales(100);f__Metschnikowiaceae(100);g__Metschnikowia(100);s__Metschnikowia_pulcherrima(100); | Metschnikowia | 13 | Saprotroph | \|Undefined Saprotroph\| | Yeast | NULL | Probable | As nectar-tap_saprotroph (Põlme et al. 2020); Decay sugar-rich substrates (Põlme et al. 2020) | Põlme S, et al. 2020. Fungal Diversity 105:1-16 ((https://doi.org/10.1007/s13225-020-00466-2)) |
| Otu27959 | 1 | k__Fungi(100);p__Ascomycota(100);c__Saccharomycetes(100);o__Saccharomycetales(100);f__Metschnikowiaceae(100);g__Metschnikowia(100);s__Metschnikowia_pulcherrima(100); | Metschnikowia | 13 | Saprotroph | \|Undefined Saprotroph\| | Yeast | NULL | Probable | As nectar-tap_saprotroph (Põlme et al. 2020); Decay sugar-rich substrates (Põlme et al. 2020) | Põlme S, et al. 2020. Fungal Diversity 105:1-16 ((https://doi.org/10.1007/s13225-020-00466-2)) |
| Otu27962 | 1 | k__Fungi(100);p__Ascomycota(100);c__Saccharomycetes(100);o__Saccharomycetales(100);f__Metschnikowiaceae(100);g__Metschnikowia(100);s__Metschnikowia_pulcherrima(100); | Metschnikowia | 13 | Saprotroph | \|Undefined Saprotroph\| | Yeast | NULL | Probable | As nectar-tap_saprotroph (Põlme et al. 2020); Decay sugar-rich substrates (Põlme et al. 2020) | Põlme S, et al. 2020. Fungal Diversity 105:1-16 ((https://doi.org/10.1007/s13225-020-00466-2)) |
| Otu27963 | 1 | k__Fungi(100);p__Ascomycota(100);c__Saccharomycetes(100);o__Saccharomycetales(100);f__Metschnikowiaceae(100);g__Metschnikowia(100);s__Metschnikowia_pulcherrima(100); | Metschnikowia | 13 | Saprotroph | \|Undefined Saprotroph\| | Yeast | NULL | Probable | As nectar-tap_saprotroph (Põlme et al. 2020); Decay sugar-rich substrates (Põlme et al. 2020) | Põlme S, et al. 2020. Fungal Diversity 105:1-16 ((https://doi.org/10.1007/s13225-020-00466-2)) |
| Otu27964 | 1 | k__Fungi(100);p__Ascomycota(100);c__Saccharomycetes(100);o__Saccharomycetales(100);f__Metschnikowiaceae(100);g__Metschnikowia(100);g__Metschnikowia_unclassified(100); | Metschnikowia | 13 | Saprotroph | \|Undefined Saprotroph\| | Yeast | NULL | Probable | As nectar-tap_saprotroph (Põlme et al. 2020); Decay sugar-rich substrates (Põlme et al. 2020) | Põlme S, et al. 2020. Fungal Diversity 105:1-16 ((https://doi.org/10.1007/s13225-020-00466-2)) |
| Otu27965 | 1 | k__Fungi(100);p__Ascomycota(100);c__Saccharomycetes(100);o__Saccharomycetales(100);f__Metschnikowiaceae(100);g__Metschnikowia(100);s__Metschnikowia_pulcherrima(100); | Metschnikowia | 13 | Saprotroph | \|Undefined Saprotroph\| | Yeast | NULL | Probable | As nectar-tap_saprotroph (Põlme et al. 2020); Decay sugar-rich substrates (Põlme et al. 2020) | Põlme S, et al. 2020. Fungal Diversity 105:1-16 ((https://doi.org/10.1007/s13225-020-00466-2)) |
| Otu27966 | 1 | k__Fungi(100);p__Ascomycota(100);c__Saccharomycetes(100);o__Saccharomycetales(100);f__Metschnikowiaceae(100);g__Metschnikowia(100);s__Metschnikowia_pulcherrima(100); | Metschnikowia | 13 | Saprotroph | \|Undefined Saprotroph\| | Yeast | NULL | Probable | As nectar-tap_saprotroph (Põlme et al. 2020); Decay sugar-rich substrates (Põlme et al. 2020) | Põlme S, et al. 2020. Fungal Diversity 105:1-16 ((https://doi.org/10.1007/s13225-020-00466-2)) |
| Otu27968 | 1 | k__Fungi(100);p__Ascomycota(100);c__Saccharomycetes(100);o__Saccharomycetales(100);f__Metschnikowiaceae(100);g__Metschnikowia(100);s__Metschnikowia_pulcherrima(100); | Metschnikowia | 13 | Saprotroph | \|Undefined Saprotroph\| | Yeast | NULL | Probable | As nectar-tap_saprotroph (Põlme et al. 2020); Decay sugar-rich substrates (Põlme et al. 2020) | Põlme S, et al. 2020. Fungal Diversity 105:1-16 ((https://doi.org/10.1007/s13225-020-00466-2)) |
| Otu27969 | 1 | k__Fungi(100);p__Ascomycota(100);c__Saccharomycetes(100);o__Saccharomycetales(100);f__Metschnikowiaceae(100);g__Metschnikowia(100);s__Metschnikowia_pulcherrima(100); | Metschnikowia | 13 | Saprotroph | \|Undefined Saprotroph\| | Yeast | NULL | Probable | As nectar-tap_saprotroph (Põlme et al. 2020); Decay sugar-rich substrates (Põlme et al. 2020) | Põlme S, et al. 2020. Fungal Diversity 105:1-16 ((https://doi.org/10.1007/s13225-020-00466-2)) |
| Otu27970 | 1 | k__Fungi(100);p__Ascomycota(100);c__Saccharomycetes(100);o__Saccharomycetales(100);f__Metschnikowiaceae(100);g__Metschnikowia(100);g__Metschnikowia_unclassified(100); | Metschnikowia | 13 | Saprotroph | \|Undefined Saprotroph\| | Yeast | NULL | Probable | As nectar-tap_saprotroph (Põlme et al. 2020); Decay sugar-rich substrates (Põlme et al. 2020) | Põlme S, et al. 2020. Fungal Diversity 105:1-16 ((https://doi.org/10.1007/s13225-020-00466-2)) |
| Otu27972 | 1 | k__Fungi(100);p__Ascomycota(100);c__Saccharomycetes(100);o__Saccharomycetales(100);f__Metschnikowiaceae(100);g__Metschnikowia(100);s__Metschnikowia_pulcherrima(100); | Metschnikowia | 13 | Saprotroph | \|Undefined Saprotroph\| | Yeast | NULL | Probable | As nectar-tap_saprotroph (Põlme et al. 2020); Decay sugar-rich substrates (Põlme et al. 2020) | Põlme S, et al. 2020. Fungal Diversity 105:1-16 ((https://doi.org/10.1007/s13225-020-00466-2)) |
| Otu27976 | 1 | k__Fungi(100);p__Ascomycota(100);c__Saccharomycetes(100);o__Saccharomycetales(100);f__Metschnikowiaceae(100);g__Metschnikowia(100);g__Metschnikowia_unclassified(100); | Metschnikowia | 13 | Saprotroph | \|Undefined Saprotroph\| | Yeast | NULL | Probable | As nectar-tap_saprotroph (Põlme et al. 2020); Decay sugar-rich substrates (Põlme et al. 2020) | Põlme S, et al. 2020. Fungal Diversity 105:1-16 ((https://doi.org/10.1007/s13225-020-00466-2)) |
| Otu27978 | 1 | k__Fungi(100);p__Ascomycota(100);c__Saccharomycetes(100);o__Saccharomycetales(100);f__Metschnikowiaceae(100);g__Metschnikowia(100);g__Metschnikowia_unclassified(100); | Metschnikowia | 13 | Saprotroph | \|Undefined Saprotroph\| | Yeast | NULL | Probable | As nectar-tap_saprotroph (Põlme et al. 2020); Decay sugar-rich substrates (Põlme et al. 2020) | Põlme S, et al. 2020. Fungal Diversity 105:1-16 ((https://doi.org/10.1007/s13225-020-00466-2)) |
| Otu27979 | 1 | k__Fungi(100);p__Ascomycota(100);c__Saccharomycetes(100);o__Saccharomycetales(100);f__Metschnikowiaceae(100);g__Metschnikowia(100);s__Metschnikowia_pulcherrima(100); | Metschnikowia | 13 | Saprotroph | \|Undefined Saprotroph\| | Yeast | NULL | Probable | As nectar-tap_saprotroph (Põlme et al. 2020); Decay sugar-rich substrates (Põlme et al. 2020) | Põlme S, et al. 2020. Fungal Diversity 105:1-16 ((https://doi.org/10.1007/s13225-020-00466-2)) |
| Otu27980 | 1 | k__Fungi(100);p__Ascomycota(100);c__Saccharomycetes(100);o__Saccharomycetales(100);f__Metschnikowiaceae(100);g__Metschnikowia(100);g__Metschnikowia_unclassified(100); | Metschnikowia | 13 | Saprotroph | \|Undefined Saprotroph\| | Yeast | NULL | Probable | As nectar-tap_saprotroph (Põlme et al. 2020); Decay sugar-rich substrates (Põlme et al. 2020) | Põlme S, et al. 2020. Fungal Diversity 105:1-16 ((https://doi.org/10.1007/s13225-020-00466-2)) |
| Otu27981 | 1 | k__Fungi(100);p__Ascomycota(100);c__Saccharomycetes(100);o__Saccharomycetales(100);f__Metschnikowiaceae(100);g__Metschnikowia(100);s__Metschnikowia_pulcherrima(100); | Metschnikowia | 13 | Saprotroph | \|Undefined Saprotroph\| | Yeast | NULL | Probable | As nectar-tap_saprotroph (Põlme et al. 2020); Decay sugar-rich substrates (Põlme et al. 2020) | Põlme S, et al. 2020. Fungal Diversity 105:1-16 ((https://doi.org/10.1007/s13225-020-00466-2)) |
| Otu27982 | 1 | k__Fungi(100);p__Ascomycota(100);c__Saccharomycetes(100);o__Saccharomycetales(100);f__Metschnikowiaceae(100);g__Metschnikowia(100);s__Metschnikowia_pulcherrima(100); | Metschnikowia | 13 | Saprotroph | \|Undefined Saprotroph\| | Yeast | NULL | Probable | As nectar-tap_saprotroph (Põlme et al. 2020); Decay sugar-rich substrates (Põlme et al. 2020) | Põlme S, et al. 2020. Fungal Diversity 105:1-16 ((https://doi.org/10.1007/s13225-020-00466-2)) |
| Otu27983 | 1 | k__Fungi(100);p__Ascomycota(100);c__Saccharomycetes(100);o__Saccharomycetales(100);f__Metschnikowiaceae(100);g__Metschnikowia(100);s__Metschnikowia_pulcherrima(100); | Metschnikowia | 13 | Saprotroph | \|Undefined Saprotroph\| | Yeast | NULL | Probable | As nectar-tap_saprotroph (Põlme et al. 2020); Decay sugar-rich substrates (Põlme et al. 2020) | Põlme S, et al. 2020. Fungal Diversity 105:1-16 ((https://doi.org/10.1007/s13225-020-00466-2)) |
| Otu27985 | 1 | k__Fungi(100);p__Ascomycota(100);c__Saccharomycetes(100);o__Saccharomycetales(100);f__Metschnikowiaceae(100);g__Metschnikowia(100);s__Metschnikowia_pulcherrima(100); | Metschnikowia | 13 | Saprotroph | \|Undefined Saprotroph\| | Yeast | NULL | Probable | As nectar-tap_saprotroph (Põlme et al. 2020); Decay sugar-rich substrates (Põlme et al. 2020) | Põlme S, et al. 2020. Fungal Diversity 105:1-16 ((https://doi.org/10.1007/s13225-020-00466-2)) |
| Otu27986 | 1 | k__Fungi(100);p__Ascomycota(100);c__Saccharomycetes(100);o__Saccharomycetales(100);f__Metschnikowiaceae(100);g__Metschnikowia(100);s__Metschnikowia_pulcherrima(100); | Metschnikowia | 13 | Saprotroph | \|Undefined Saprotroph\| | Yeast | NULL | Probable | As nectar-tap_saprotroph (Põlme et al. 2020); Decay sugar-rich substrates (Põlme et al. 2020) | Põlme S, et al. 2020. Fungal Diversity 105:1-16 ((https://doi.org/10.1007/s13225-020-00466-2)) |
| Otu27987 | 1 | k__Fungi(100);p__Ascomycota(100);c__Saccharomycetes(100);o__Saccharomycetales(100);f__Metschnikowiaceae(100);g__Metschnikowia(100);g__Metschnikowia_unclassified(100); | Metschnikowia | 13 | Saprotroph | \|Undefined Saprotroph\| | Yeast | NULL | Probable | As nectar-tap_saprotroph (Põlme et al. 2020); Decay sugar-rich substrates (Põlme et al. 2020) | Põlme S, et al. 2020. Fungal Diversity 105:1-16 ((https://doi.org/10.1007/s13225-020-00466-2)) |
| Otu27988 | 1 | k__Fungi(100);p__Ascomycota(100);c__Saccharomycetes(100);o__Saccharomycetales(100);f__Metschnikowiaceae(100);g__Metschnikowia(100);s__Metschnikowia_pulcherrima(100); | Metschnikowia | 13 | Saprotroph | \|Undefined Saprotroph\| | Yeast | NULL | Probable | As nectar-tap_saprotroph (Põlme et al. 2020); Decay sugar-rich substrates (Põlme et al. 2020) | Põlme S, et al. 2020. Fungal Diversity 105:1-16 ((https://doi.org/10.1007/s13225-020-00466-2)) |
| Otu27994 | 1 | k__Fungi(100);p__Ascomycota(100);c__Saccharomycetes(100);o__Saccharomycetales(100);f__Metschnikowiaceae(100);g__Metschnikowia(100);s__Metschnikowia_pulcherrima(100); | Metschnikowia | 13 | Saprotroph | \|Undefined Saprotroph\| | Yeast | NULL | Probable | As nectar-tap_saprotroph (Põlme et al. 2020); Decay sugar-rich substrates (Põlme et al. 2020) | Põlme S, et al. 2020. Fungal Diversity 105:1-16 ((https://doi.org/10.1007/s13225-020-00466-2)) |
| Otu27996 | 1 | k__Fungi(100);p__Ascomycota(100);c__Saccharomycetes(100);o__Saccharomycetales(100);f__Metschnikowiaceae(100);g__Metschnikowia(100);s__Metschnikowia_pulcherrima(100); | Metschnikowia | 13 | Saprotroph | \|Undefined Saprotroph\| | Yeast | NULL | Probable | As nectar-tap_saprotroph (Põlme et al. 2020); Decay sugar-rich substrates (Põlme et al. 2020) | Põlme S, et al. 2020. Fungal Diversity 105:1-16 ((https://doi.org/10.1007/s13225-020-00466-2)) |
| Otu27999 | 1 | k__Fungi(100);p__Ascomycota(100);c__Saccharomycetes(100);o__Saccharomycetales(100);f__Metschnikowiaceae(100);g__Metschnikowia(100);s__Metschnikowia_pulcherrima(100); | Metschnikowia | 13 | Saprotroph | \|Undefined Saprotroph\| | Yeast | NULL | Probable | As nectar-tap_saprotroph (Põlme et al. 2020); Decay sugar-rich substrates (Põlme et al. 2020) | Põlme S, et al. 2020. Fungal Diversity 105:1-16 ((https://doi.org/10.1007/s13225-020-00466-2)) |
| Otu28000 | 1 | k__Fungi(100);p__Ascomycota(100);c__Saccharomycetes(100);o__Saccharomycetales(100);f__Metschnikowiaceae(100);g__Metschnikowia(100);s__Metschnikowia_pulcherrima(100); | Metschnikowia | 13 | Saprotroph | \|Undefined Saprotroph\| | Yeast | NULL | Probable | As nectar-tap_saprotroph (Põlme et al. 2020); Decay sugar-rich substrates (Põlme et al. 2020) | Põlme S, et al. 2020. Fungal Diversity 105:1-16 ((https://doi.org/10.1007/s13225-020-00466-2)) |
| Otu28001 | 1 | k__Fungi(100);p__Ascomycota(100);c__Saccharomycetes(100);o__Saccharomycetales(100);f__Metschnikowiaceae(100);g__Metschnikowia(100);g__Metschnikowia_unclassified(100); | Metschnikowia | 13 | Saprotroph | \|Undefined Saprotroph\| | Yeast | NULL | Probable | As nectar-tap_saprotroph (Põlme et al. 2020); Decay sugar-rich substrates (Põlme et al. 2020) | Põlme S, et al. 2020. Fungal Diversity 105:1-16 ((https://doi.org/10.1007/s13225-020-00466-2)) |
| Otu28002 | 1 | k__Fungi(100);p__Ascomycota(100);c__Saccharomycetes(100);o__Saccharomycetales(100);f__Metschnikowiaceae(100);g__Metschnikowia(100);s__Metschnikowia_pulcherrima(100); | Metschnikowia | 13 | Saprotroph | \|Undefined Saprotroph\| | Yeast | NULL | Probable | As nectar-tap_saprotroph (Põlme et al. 2020); Decay sugar-rich substrates (Põlme et al. 2020) | Põlme S, et al. 2020. Fungal Diversity 105:1-16 ((https://doi.org/10.1007/s13225-020-00466-2)) |
| Otu28003 | 1 | k__Fungi(100);p__Ascomycota(100);c__Saccharomycetes(100);o__Saccharomycetales(100);f__Metschnikowiaceae(100);g__Metschnikowia(100);s__Metschnikowia_pulcherrima(100); | Metschnikowia | 13 | Saprotroph | \|Undefined Saprotroph\| | Yeast | NULL | Probable | As nectar-tap_saprotroph (Põlme et al. 2020); Decay sugar-rich substrates (Põlme et al. 2020) | Põlme S, et al. 2020. Fungal Diversity 105:1-16 ((https://doi.org/10.1007/s13225-020-00466-2)) |
| Otu28004 | 1 | k__Fungi(100);p__Basidiomycota(100);c__Microbotryomycetes(100);o__Sporidiobolales(100);f__Sporidiobolales_family_Incertae_sedis(100);g__Sporobolomyces(100);s__Sporobolomyces_lactosus(100); | Sporobolomyces | 13 | Pathotroph-Saprotroph | \|Fungal Parasite\|-Plant Saprotroph | Yeast | NULL | Probable | As litter_saprotroph (Põlme et al. 2020); As mycoparasite (Põlme et al. 2020); Decay fungal material (Põlme et al. 2020) | Tedersoo L, et al. 2014. Science 346:e1256688 ((https://doi.org/10.1126/science.1256688)); Purahong W, et al. 2016. Molecular Ecology 25:4059-4074 ((https://doi.org/10.1111/mec.13739)); Põlme S, et al. 2020. Fungal Diversity 105:1-16 ((https://doi.org/10.1007/s13225-020-00466-2)) |
| Otu28005 | 1 | k__Fungi(100);p__Ascomycota(100);c__Saccharomycetes(100);o__Saccharomycetales(100);f__Metschnikowiaceae(100);g__Metschnikowia(100);s__Metschnikowia_pulcherrima(100); | Metschnikowia | 13 | Saprotroph | \|Undefined Saprotroph\| | Yeast | NULL | Probable | As nectar-tap_saprotroph (Põlme et al. 2020); Decay sugar-rich substrates (Põlme et al. 2020) | Põlme S, et al. 2020. Fungal Diversity 105:1-16 ((https://doi.org/10.1007/s13225-020-00466-2)) |
| Otu28007 | 1 | k__Fungi(100);p__Ascomycota(100);c__Saccharomycetes(100);o__Saccharomycetales(100);f__Metschnikowiaceae(100);g__Metschnikowia(100);s__Metschnikowia_pulcherrima(100); | Metschnikowia | 13 | Saprotroph | \|Undefined Saprotroph\| | Yeast | NULL | Probable | As nectar-tap_saprotroph (Põlme et al. 2020); Decay sugar-rich substrates (Põlme et al. 2020) | Põlme S, et al. 2020. Fungal Diversity 105:1-16 ((https://doi.org/10.1007/s13225-020-00466-2)) |
| Otu28016 | 1 | k__Fungi(100);p__Ascomycota(100);c__Dothideomycetes(100);o__Pleosporales(100);f__Cucurbitariaceae(100);g__Curreya(100);g__Curreya_unclassified(100); | Curreya | 13 | Saprotroph | \|Plant Saprotroph\|-Undefined Saprotroph | Microfungus | NULL | Probable | As litter_saprotroph (Põlme et al. 2020); Decay leaf, fruit, seed (Põlme et al. 2020) | Tedersoo L, et al. 2014. Science 346:e1256688 ((https://doi.org/10.1126/science.1256688)); Põlme S, et al. 2020. Fungal Diversity 105:1-16 ((https://doi.org/10.1007/s13225-020-00466-2)) |
| Otu28025 | 1 | k__Fungi(100);p__Ascomycota(100);c__Saccharomycetes(100);o__Saccharomycetales(100);f__Metschnikowiaceae(100);g__Metschnikowia(100);s__Metschnikowia_pulcherrima(100); | Metschnikowia | 13 | Saprotroph | \|Undefined Saprotroph\| | Yeast | NULL | Probable | As nectar-tap_saprotroph (Põlme et al. 2020); Decay sugar-rich substrates (Põlme et al. 2020) | Põlme S, et al. 2020. Fungal Diversity 105:1-16 ((https://doi.org/10.1007/s13225-020-00466-2)) |
| Otu28029 | 1 | k__Fungi(100);p__Ascomycota(100);c__Saccharomycetes(100);o__Saccharomycetales(100);f__Metschnikowiaceae(100);g__Metschnikowia(100);g__Metschnikowia_unclassified(100); | Metschnikowia | 13 | Saprotroph | \|Undefined Saprotroph\| | Yeast | NULL | Probable | As nectar-tap_saprotroph (Põlme et al. 2020); Decay sugar-rich substrates (Põlme et al. 2020) | Põlme S, et al. 2020. Fungal Diversity 105:1-16 ((https://doi.org/10.1007/s13225-020-00466-2)) |
| Otu28046 | 1 | k__Fungi(100);p__Ascomycota(100);c__Dothideomycetes(100);o__Dothideomycetes_order_Incertae_sedis(100);f__Myxotrichaceae(100);g__Oidiodendron(100);s__Oidiodendron_maius(100); | Oidiodendron | 13 | Pathotroph-Saprotroph-Symbiotroph | Endophyte-Ericoid Mycorrhizal-\|Undefined Saprotroph\| | Microfungus | NULL | Probable | As root_endophyte (Põlme et al. 2020); As soil_saprotroph (Põlme et al. 2020); Dark Septate Endophyte; Decay soil (Põlme et al. 2020) | Newsham KK. 2011. New Phytologist 190:783-793 ((https://doi.org/10.1111/j.1469-8137.2010.03611.x)) (as dark septate root endophytes); Põlme S, et al. 2020. Fungal Diversity 105:1-16 ((https://doi.org/10.1007/s13225-020-00466-2)) |
| Otu28047 | 1 | k__Fungi(100);p__Ascomycota(100);c__Saccharomycetes(100);o__Saccharomycetales(100);f__Metschnikowiaceae(100);g__Metschnikowia(100);s__Metschnikowia_pulcherrima(100); | Metschnikowia | 13 | Saprotroph | \|Undefined Saprotroph\| | Yeast | NULL | Probable | As nectar-tap_saprotroph (Põlme et al. 2020); Decay sugar-rich substrates (Põlme et al. 2020) | Põlme S, et al. 2020. Fungal Diversity 105:1-16 ((https://doi.org/10.1007/s13225-020-00466-2)) |
| Otu28049 | 1 | k__Fungi(100);p__Ascomycota(100);c__Saccharomycetes(100);o__Saccharomycetales(100);f__Metschnikowiaceae(100);g__Metschnikowia(100);g__Metschnikowia_unclassified(100); | Metschnikowia | 13 | Saprotroph | \|Undefined Saprotroph\| | Yeast | NULL | Probable | As nectar-tap_saprotroph (Põlme et al. 2020); Decay sugar-rich substrates (Põlme et al. 2020) | Põlme S, et al. 2020. Fungal Diversity 105:1-16 ((https://doi.org/10.1007/s13225-020-00466-2)) |
| Otu28052 | 1 | k__Fungi(100);p__Ascomycota(100);c__Saccharomycetes(100);o__Saccharomycetales(100);f__Metschnikowiaceae(100);g__Metschnikowia(100);g__Metschnikowia_unclassified(100); | Metschnikowia | 13 | Saprotroph | \|Undefined Saprotroph\| | Yeast | NULL | Probable | As nectar-tap_saprotroph (Põlme et al. 2020); Decay sugar-rich substrates (Põlme et al. 2020) | Põlme S, et al. 2020. Fungal Diversity 105:1-16 ((https://doi.org/10.1007/s13225-020-00466-2)) |
| Otu28060 | 1 | k__Fungi(100);p__Ascomycota(100);c__Saccharomycetes(100);o__Saccharomycetales(100);f__Metschnikowiaceae(100);g__Metschnikowia(100);s__Metschnikowia_pulcherrima(100); | Metschnikowia | 13 | Saprotroph | \|Undefined Saprotroph\| | Yeast | NULL | Probable | As nectar-tap_saprotroph (Põlme et al. 2020); Decay sugar-rich substrates (Põlme et al. 2020) | Põlme S, et al. 2020. Fungal Diversity 105:1-16 ((https://doi.org/10.1007/s13225-020-00466-2)) |
| Otu28084 | 1 | k__Fungi(100);p__Ascomycota(100);c__Saccharomycetes(100);o__Saccharomycetales(100);f__Saccharomycetales_family_Incertae_sedis(100);f__Saccharomycetales_family_Incertae_sedis_unclassified(100);f__Saccharomycetales_family_Incertae_sedis_unclassified(100); | Saccharomycetales | 7 | Saprotroph | Undefined Saprotroph | Yeast | NULL | Possible | NULL | Sterkenburg E, et al. 2015. New Phytologist 207:1145-1158 ((https://doi.org/10.1111/nph.13426)) |
| Otu28088 | 1 | k__Fungi(100);p__Ascomycota(100);c__Saccharomycetes(100);o__Saccharomycetales(100);o__Saccharomycetales_unclassified(100);o__Saccharomycetales_unclassified(100);o__Saccharomycetales_unclassified(100); | Saccharomycetales | 7 | Saprotroph | Undefined Saprotroph | Yeast | NULL | Possible | NULL | Sterkenburg E, et al. 2015. New Phytologist 207:1145-1158 ((https://doi.org/10.1111/nph.13426)) |
| Otu28102 | 1 | k__Fungi(100);p__Ascomycota(100);c__Saccharomycetes(100);o__Saccharomycetales(100);f__Saccharomycetales_family_Incertae_sedis(100);f__Saccharomycetales_family_Incertae_sedis_unclassified(100);f__Saccharomycetales_family_Incertae_sedis_unclassified(100); | Saccharomycetales | 7 | Saprotroph | Undefined Saprotroph | Yeast | NULL | Possible | NULL | Sterkenburg E, et al. 2015. New Phytologist 207:1145-1158 ((https://doi.org/10.1111/nph.13426)) |
| Otu28116 | 1 | k__Fungi(100);p__Ascomycota(100);c__Saccharomycetes(100);o__Saccharomycetales(100);f__Saccharomycetales_family_Incertae_sedis(100);g__Candida(100);g__Candida_unclassified(100); | Candida | 13 | Pathotroph-Saprotroph-Symbiotroph | Animal Pathogen-Endophyte-Endosymbiont-Epiphyte-\|Undefined Saprotroph\| | Dimorphic Yeast-Yeast | NULL | Probable | As nectar-tap_saprotroph (Põlme et al. 2020); Decay sugar-rich substrates (Põlme et al. 2020); This asexual genus is polyphyletic and contain species that function very differently depend on what they are. Caution should be taken when interpreting guilds for Candida species. | Manolakaki D, et al. 2010. Virulence. 1: 367-375 ((https://doi.org/10.4161/viru.1.5.12796)); Mekbib SB, et al. 2011. Tropical Plant Pathology 36:233-240 ((https://doi.org/10.1590/S1982-56762011000400004)); Põlme S, et al. 2020. Fungal Diversity 105:1-16 ((https://doi.org/10.1007/s13225-020-00466-2)) |
| Otu28117 | 1 | k__Fungi(100);p__Ascomycota(100);c__Saccharomycetes(100);o__Saccharomycetales(100);f__Metschnikowiaceae(100);g__Metschnikowia(100);s__Metschnikowia_pulcherrima(100); | Metschnikowia | 13 | Saprotroph | \|Undefined Saprotroph\| | Yeast | NULL | Probable | As nectar-tap_saprotroph (Põlme et al. 2020); Decay sugar-rich substrates (Põlme et al. 2020) | Põlme S, et al. 2020. Fungal Diversity 105:1-16 ((https://doi.org/10.1007/s13225-020-00466-2)) |
| Otu28119 | 1 | k__Fungi(100);p__Ascomycota(100);c__Saccharomycetes(100);o__Saccharomycetales(100);f__Metschnikowiaceae(100);g__Metschnikowia(100);s__Metschnikowia_pulcherrima(100); | Metschnikowia | 13 | Saprotroph | \|Undefined Saprotroph\| | Yeast | NULL | Probable | As nectar-tap_saprotroph (Põlme et al. 2020); Decay sugar-rich substrates (Põlme et al. 2020) | Põlme S, et al. 2020. Fungal Diversity 105:1-16 ((https://doi.org/10.1007/s13225-020-00466-2)) |
| Otu28122 | 1 | k__Fungi(100);p__Ascomycota(100);c__Saccharomycetes(100);o__Saccharomycetales(100);f__Metschnikowiaceae(100);g__Metschnikowia(100);g__Metschnikowia_unclassified(100); | Metschnikowia | 13 | Saprotroph | \|Undefined Saprotroph\| | Yeast | NULL | Probable | As nectar-tap_saprotroph (Põlme et al. 2020); Decay sugar-rich substrates (Põlme et al. 2020) | Põlme S, et al. 2020. Fungal Diversity 105:1-16 ((https://doi.org/10.1007/s13225-020-00466-2)) |
| Otu28123 | 1 | k__Fungi(100);p__Ascomycota(100);c__Saccharomycetes(100);o__Saccharomycetales(100);f__Metschnikowiaceae(100);g__Metschnikowia(100);s__Metschnikowia_pulcherrima(100); | Metschnikowia | 13 | Saprotroph | \|Undefined Saprotroph\| | Yeast | NULL | Probable | As nectar-tap_saprotroph (Põlme et al. 2020); Decay sugar-rich substrates (Põlme et al. 2020) | Põlme S, et al. 2020. Fungal Diversity 105:1-16 ((https://doi.org/10.1007/s13225-020-00466-2)) |
| Otu28124 | 1 | k__Fungi(100);p__Ascomycota(100);c__Saccharomycetes(100);o__Saccharomycetales(100);f__Metschnikowiaceae(100);g__Metschnikowia(100);s__Metschnikowia_pulcherrima(100); | Metschnikowia | 13 | Saprotroph | \|Undefined Saprotroph\| | Yeast | NULL | Probable | As nectar-tap_saprotroph (Põlme et al. 2020); Decay sugar-rich substrates (Põlme et al. 2020) | Põlme S, et al. 2020. Fungal Diversity 105:1-16 ((https://doi.org/10.1007/s13225-020-00466-2)) |
| Otu28127 | 1 | k__Fungi(100);p__Ascomycota(100);c__Saccharomycetes(100);o__Saccharomycetales(100);f__Metschnikowiaceae(100);g__Metschnikowia(100);s__Metschnikowia_pulcherrima(100); | Metschnikowia | 13 | Saprotroph | \|Undefined Saprotroph\| | Yeast | NULL | Probable | As nectar-tap_saprotroph (Põlme et al. 2020); Decay sugar-rich substrates (Põlme et al. 2020) | Põlme S, et al. 2020. Fungal Diversity 105:1-16 ((https://doi.org/10.1007/s13225-020-00466-2)) |
| Otu28128 | 1 | k__Fungi(100);p__Basidiomycota(100);c__Microbotryomycetes(100);o__Sporidiobolales(100);f__Sporidiobolales_family_Incertae_sedis(100);g__Rhodotorula(100);s__Rhodotorula_mucilaginosa(100); | Rhodotorula | 13 | Pathotroph-Saprotroph-Symbiotroph | Animal Pathogen-Animal Symbiotroph-Endophyte-Plant Pathogen-\|Undefined Saprotroph\| | Yeast | NULL | Probable | As foliar_endophyte (Põlme et al. 2020); Opportunistic human pathogen | Põlme S, et al. 2020. Fungal Diversity 105:1-16 ((https://doi.org/10.1007/s13225-020-00466-2)); https://www.cdc.gov ((https://www.cdc.gov)) |
| Otu28129 | 1 | k__Fungi(100);p__Basidiomycota(100);c__Microbotryomycetes(100);o__Sporidiobolales(100);f__Sporidiobolales_family_Incertae_sedis(100);g__Rhodotorula(100);s__Rhodotorula_taiwanensis(100); | Rhodotorula | 13 | Pathotroph-Saprotroph-Symbiotroph | Animal Pathogen-Animal Symbiotroph-Endophyte-Plant Pathogen-\|Undefined Saprotroph\| | Yeast | NULL | Probable | As foliar_endophyte (Põlme et al. 2020); Opportunistic human pathogen | Põlme S, et al. 2020. Fungal Diversity 105:1-16 ((https://doi.org/10.1007/s13225-020-00466-2)); https://www.cdc.gov ((https://www.cdc.gov)) |
| Otu28130 | 1 | k__Fungi(100);p__Ascomycota(100);c__Saccharomycetes(100);o__Saccharomycetales(100);f__Metschnikowiaceae(100);g__Metschnikowia(100);s__Metschnikowia_pulcherrima(100); | Metschnikowia | 13 | Saprotroph | \|Undefined Saprotroph\| | Yeast | NULL | Probable | As nectar-tap_saprotroph (Põlme et al. 2020); Decay sugar-rich substrates (Põlme et al. 2020) | Põlme S, et al. 2020. Fungal Diversity 105:1-16 ((https://doi.org/10.1007/s13225-020-00466-2)) |
| Otu28131 | 1 | k__Fungi(100);p__Ascomycota(100);c__Saccharomycetes(100);o__Saccharomycetales(100);f__Metschnikowiaceae(100);g__Metschnikowia(100);s__Metschnikowia_pulcherrima(100); | Metschnikowia | 13 | Saprotroph | \|Undefined Saprotroph\| | Yeast | NULL | Probable | As nectar-tap_saprotroph (Põlme et al. 2020); Decay sugar-rich substrates (Põlme et al. 2020) | Põlme S, et al. 2020. Fungal Diversity 105:1-16 ((https://doi.org/10.1007/s13225-020-00466-2)) |
| Otu28132 | 1 | k__Fungi(100);p__Basidiomycota(100);c__Microbotryomycetes(100);o__Sporidiobolales(100);f__Sporidiobolales_family_Incertae_sedis(100);g__Rhodotorula(100);s__Rhodotorula_mucilaginosa(100); | Rhodotorula | 13 | Pathotroph-Saprotroph-Symbiotroph | Animal Pathogen-Animal Symbiotroph-Endophyte-Plant Pathogen-\|Undefined Saprotroph\| | Yeast | NULL | Probable | As foliar_endophyte (Põlme et al. 2020); Opportunistic human pathogen | Põlme S, et al. 2020. Fungal Diversity 105:1-16 ((https://doi.org/10.1007/s13225-020-00466-2)); https://www.cdc.gov ((https://www.cdc.gov)) |
| Otu28133 | 1 | k__Fungi(100);p__Ascomycota(100);c__Saccharomycetes(100);o__Saccharomycetales(100);f__Saccharomycetaceae(100);g__Saccharomyces(100);g__Saccharomyces_unclassified(100); | Saccharomyces | 13 | Saprotroph | \|Undefined Saprotroph\| | Yeast | NULL | Probable | As nectar-tap_saprotroph (Põlme et al. 2020); Decay sugar-rich substrates (Põlme et al. 2020) | James TY, et al. 2006. Nature 443:818-822 ((https://doi.org/10.1038/nature05110)); Põlme S, et al. 2020. Fungal Diversity 105:1-16 ((https://doi.org/10.1007/s13225-020-00466-2)) |
| Otu28136 | 1 | k__Fungi(100);p__Ascomycota(100);c__Saccharomycetes(100);o__Saccharomycetales(100);o__Saccharomycetales_unclassified(100);o__Saccharomycetales_unclassified(100);o__Saccharomycetales_unclassified(100); | Saccharomycetales | 7 | Saprotroph | Undefined Saprotroph | Yeast | NULL | Possible | NULL | Sterkenburg E, et al. 2015. New Phytologist 207:1145-1158 ((https://doi.org/10.1111/nph.13426)) |
| Otu28137 | 1 | k__Fungi(100);p__Zygomycota(100);c__Zygomycota_class_Incertae_sedis(100);o__Mucorales(100);f__Mucoraceae(100);g__Mucor(100);s__Mucor_genevensis(100); | Mucor | 13 | Saprotroph | \|Undefined Saprotroph\| | Microfungus | NULL | Probable | As soil_saprotroph (Põlme et al. 2020); Decay soil (Põlme et al. 2020) | Tedersoo L, et al. 2014. Science 346:e1256688 ((https://doi.org/10.1126/science.1256688)); Põlme S, et al. 2020. Fungal Diversity 105:1-16 ((https://doi.org/10.1007/s13225-020-00466-2)) |
| Otu28138 | 1 | k__Fungi(100);p__Basidiomycota(100);c__Pucciniomycetes(100);o__Pucciniales(100);f__Pucciniaceae(100);g__Puccinia(100);s__Puccinia_bartholomaei(100); | Puccinia | 13 | Pathotroph | \|Plant Pathogen\| | Rust | NULL | Highly Probable | NULL | Tedersoo L, et al. 2014. Science 346:e1256688 ((https://doi.org/10.1126/science.1256688)); Põlme S, et al. 2020. Fungal Diversity 105:1-16 ((https://doi.org/10.1007/s13225-020-00466-2)) |
| Otu28139 | 1 | k__Fungi(100);p__Basidiomycota(100);c__Wallemiomycetes(100);o__Wallemiales(100);f__Wallemiaceae(100);g__Wallemia(100);s__Wallemia_sebi(100); | Wallemia | 13 | Saprotroph | \|Undefined Saprotroph\| | Microfungus | NULL | Probable | Decay sugar-rich substrates (Põlme et al. 2020) | Tedersoo L, et al. 2014. Science 346:e1256688 ((https://doi.org/10.1126/science.1256688)); Põlme S, et al. 2020. Fungal Diversity 105:1-16 ((https://doi.org/10.1007/s13225-020-00466-2)) |
| Otu28140 | 1 | k__Fungi(100);p__Basidiomycota(100);c__Agaricomycetes(100);o__Hymenochaetales(100);f__Schizoporaceae(100);g__Hyphodontia(100);s__Hyphodontia_aspera(100); | Hyphodontia | 13 | Saprotroph | Undefined Saprotroph-\|Wood Saprotroph\| | Corticioid | White Rot | Probable | As white_rot (Põlme et al. 2020); Decay wood (Põlme et al. 2020) | Tedersoo L, et al. 2014. Science 346:e1256688 ((https://doi.org/10.1126/science.1256688)); Põlme S, et al. 2020. Fungal Diversity 105:1-16 ((https://doi.org/10.1007/s13225-020-00466-2)) |
| Otu28141 | 1 | k__Fungi(100);p__Basidiomycota(100);c__Microbotryomycetes(100);o__Sporidiobolales(100);f__Sporidiobolales_family_Incertae_sedis(100);g__Rhodotorula(100);s__Rhodotorula_nothofagi(100); | Rhodotorula | 13 | Pathotroph-Saprotroph-Symbiotroph | Animal Pathogen-Animal Symbiotroph-Endophyte-Plant Pathogen-\|Undefined Saprotroph\| | Yeast | NULL | Probable | As foliar_endophyte (Põlme et al. 2020); Opportunistic human pathogen | Põlme S, et al. 2020. Fungal Diversity 105:1-16 ((https://doi.org/10.1007/s13225-020-00466-2)); https://www.cdc.gov ((https://www.cdc.gov)) |

| | | | | | | | | | | | |
|---|---|---|---|---|---|---|---|---|---|---|---|
| Otu28142 | 1 | k__Fungi(100);p__Basidiomycota(100);c__Agaricomycetes(100);o__Russulales(100);f__Peniophoraceae(100);f__Peniophoraceae_unclassified(100);f__Peniophoraceae_unclassified(100); | Peniophoraceae | 9 | Pathotroph-Saprotroph | Plant Pathogen-Wood Saprotroph | Corticioid | White Rot | Probable | Most species are saprobic, causing rots of standing and fallen wood. Many species are adapted to xeric conditions, being able to grow on rapidly drying dead attached branches (Cannon & Kirk 2007) | Cannon PF, Kirk PM. 2007. Fungal Families of the World. CAB International, Cambridge (ISBN: 978-0851998275) |
| Otu28143 | 1 | k__Fungi(100);p__Ascomycota(100);c__Saccharomycetes(100);o__Saccharomycetales(100);f__Metschnikowiaceae(100);g__Metschnikowia(100);s__Metschnikowia_pulcherrima(100); | Metschnikowia | 13 | Saprotroph | \|Undefined Saprotroph\| | Yeast | NULL | Probable | As nectar-tap_saprotroph (Põlme et al. 2020); Decay sugar-rich substrates (Põlme et al. 2020) | Põlme S, et al. 2020. Fungal Diversity 105:1-16 ((https://doi.org/10.1007/s13225-020-00466-2)) |
| Otu28148 | 1 | k__Fungi(100);p__Ascomycota(100);c__Saccharomycetes(100);o__Saccharomycetales(100);f__Metschnikowiaceae(100);g__Metschnikowia(100);s__Metschnikowia_pulcherrima(100); | Metschnikowia | 13 | Saprotroph | \|Undefined Saprotroph\| | Yeast | NULL | Probable | As nectar-tap_saprotroph (Põlme et al. 2020); Decay sugar-rich substrates (Põlme et al. 2020) | Põlme S, et al. 2020. Fungal Diversity 105:1-16 ((https://doi.org/10.1007/s13225-020-00466-2)) |
| Otu28150 | 1 | k__Fungi(100);p__Ascomycota(100);c__Saccharomycetes(100);o__Saccharomycetales(100);f__Metschnikowiaceae(100);g__Metschnikowia(100);g__Metschnikowia_unclassified(100); | Metschnikowia | 13 | Saprotroph | \|Undefined Saprotroph\| | Yeast | NULL | Probable | As nectar-tap_saprotroph (Põlme et al. 2020); Decay sugar-rich substrates (Põlme et al. 2020) | Põlme S, et al. 2020. Fungal Diversity 105:1-16 ((https://doi.org/10.1007/s13225-020-00466-2)) |
| Otu28151 | 1 | k__Fungi(100);p__Ascomycota(100);c__Saccharomycetes(100);o__Saccharomycetales(100);f__Metschnikowiaceae(100);g__Metschnikowia(100);s__Metschnikowia_pulcherrima(100); | Metschnikowia | 13 | Saprotroph | \|Undefined Saprotroph\| | Yeast | NULL | Probable | As nectar-tap_saprotroph (Põlme et al. 2020); Decay sugar-rich substrates (Põlme et al. 2020) | Põlme S, et al. 2020. Fungal Diversity 105:1-16 ((https://doi.org/10.1007/s13225-020-00466-2)) |
| Otu28152 | 1 | k__Fungi(100);p__Ascomycota(100);c__Saccharomycetes(100);o__Saccharomycetales(100);f__Metschnikowiaceae(100);g__Metschnikowia(100);g__Metschnikowia_unclassified(100); | Metschnikowia | 13 | Saprotroph | \|Undefined Saprotroph\| | Yeast | NULL | Probable | As nectar-tap_saprotroph (Põlme et al. 2020); Decay sugar-rich substrates (Põlme et al. 2020) | Põlme S, et al. 2020. Fungal Diversity 105:1-16 ((https://doi.org/10.1007/s13225-020-00466-2)) |
| Otu28153 | 1 | k__Fungi(100);p__Ascomycota(100);c__Saccharomycetes(100);o__Saccharomycetales(100);f__Metschnikowiaceae(100);g__Metschnikowia(100);g__Metschnikowia_unclassified(100); | Metschnikowia | 13 | Saprotroph | \|Undefined Saprotroph\| | Yeast | NULL | Probable | As nectar-tap_saprotroph (Põlme et al. 2020); Decay sugar-rich substrates (Põlme et al. 2020) | Põlme S, et al. 2020. Fungal Diversity 105:1-16 ((https://doi.org/10.1007/s13225-020-00466-2)) |
| Otu28154 | 1 | k__Fungi(100);p__Ascomycota(100);c__Saccharomycetes(100);o__Saccharomycetales(100);f__Metschnikowiaceae(100);g__Metschnikowia(100);s__Metschnikowia_pulcherrima(100); | Metschnikowia | 13 | Saprotroph | \|Undefined Saprotroph\| | Yeast | NULL | Probable | As nectar-tap_saprotroph (Põlme et al. 2020); Decay sugar-rich substrates (Põlme et al. 2020) | Põlme S, et al. 2020. Fungal Diversity 105:1-16 ((https://doi.org/10.1007/s13225-020-00466-2)) |
| Otu28155 | 1 | k__Fungi(100);p__Ascomycota(100);c__Saccharomycetes(100);o__Saccharomycetales(100);f__Metschnikowiaceae(100);g__Metschnikowia(100);s__Metschnikowia_pulcherrima(100); | Metschnikowia | 13 | Saprotroph | \|Undefined Saprotroph\| | Yeast | NULL | Probable | As nectar-tap_saprotroph (Põlme et al. 2020); Decay sugar-rich substrates (Põlme et al. 2020) | Põlme S, et al. 2020. Fungal Diversity 105:1-16 ((https://doi.org/10.1007/s13225-020-00466-2)) |
| Otu28157 | 1 | k__Fungi(100);p__Ascomycota(100);c__Saccharomycetes(100);o__Saccharomycetales(100);f__Metschnikowiaceae(100);g__Metschnikowia(100);s__Metschnikowia_pulcherrima(100); | Metschnikowia | 13 | Saprotroph | \|Undefined Saprotroph\| | Yeast | NULL | Probable | As nectar-tap_saprotroph (Põlme et al. 2020); Decay sugar-rich substrates (Põlme et al. 2020) | Põlme S, et al. 2020. Fungal Diversity 105:1-16 ((https://doi.org/10.1007/s13225-020-00466-2)) |
| Otu28158 | 1 | k__Fungi(100);p__Ascomycota(100);c__Saccharomycetes(100);o__Saccharomycetales(100);f__Metschnikowiaceae(100);g__Metschnikowia(100);s__Metschnikowia_pulcherrima(100); | Metschnikowia | 13 | Saprotroph | \|Undefined Saprotroph\| | Yeast | NULL | Probable | As nectar-tap_saprotroph (Põlme et al. 2020); Decay sugar-rich substrates (Põlme et al. 2020) | Põlme S, et al. 2020. Fungal Diversity 105:1-16 ((https://doi.org/10.1007/s13225-020-00466-2)) |
| Otu28170 | 1 | k__Fungi(100);p__Ascomycota(100);c__Saccharomycetes(100);o__Saccharomycetales(100);f__Metschnikowiaceae(100);g__Metschnikowia(100);s__Metschnikowia_pulcherrima(100); | Metschnikowia | 13 | Saprotroph | \|Undefined Saprotroph\| | Yeast | NULL | Probable | As nectar-tap_saprotroph (Põlme et al. 2020); Decay sugar-rich substrates (Põlme et al. 2020) | Põlme S, et al. 2020. Fungal Diversity 105:1-16 ((https://doi.org/10.1007/s13225-020-00466-2)) |
| Otu28171 | 1 | k__Fungi(100);p__Ascomycota(100);c__Saccharomycetes(100);o__Saccharomycetales(100);f__Metschnikowiaceae(100);g__Metschnikowia(100);s__Metschnikowia_pulcherrima(100); | Metschnikowia | 13 | Saprotroph | \|Undefined Saprotroph\| | Yeast | NULL | Probable | As nectar-tap_saprotroph (Põlme et al. 2020); Decay sugar-rich substrates (Põlme et al. 2020) | Põlme S, et al. 2020. Fungal Diversity 105:1-16 ((https://doi.org/10.1007/s13225-020-00466-2)) |
| Otu28172 | 1 | k__Fungi(100);p__Ascomycota(100);c__Saccharomycetes(100);o__Saccharomycetales(100);f__Metschnikowiaceae(100);g__Metschnikowia(100);g__Metschnikowia_unclassified(100); | Metschnikowia | 13 | Saprotroph | \|Undefined Saprotroph\| | Yeast | NULL | Probable | As nectar-tap_saprotroph (Põlme et al. 2020); Decay sugar-rich substrates (Põlme et al. 2020) | Põlme S, et al. 2020. Fungal Diversity 105:1-16 ((https://doi.org/10.1007/s13225-020-00466-2)) |
| Otu28173 | 1 | k__Fungi(100);p__Ascomycota(100);c__Ascomycota_class_Incertae_sedis(100);o__Ascomycota_order_Incertae_sedis(100);f__Ascomycota_family_Incertae_sedis(100);g__Cordana(100);s__Cordana_ellipsoidea(100); | Cordana | 13 | Pathotroph-Saprotroph | \|Plant Pathogen\| | Microfungus | NULL | Probable | NULL | Tedersoo L, et al. 2014. Science 346:e1256688 ((https://doi.org/10.1126/science.1256688)); Põlme S, et al. 2020. Fungal Diversity 105:1-16 ((https://doi.org/10.1007/s13225-020-00466-2)) |
| Otu28174 | 1 | k__Fungi(100);p__Ascomycota(100);c__Saccharomycetes(100);o__Saccharomycetales(100);f__Metschnikowiaceae(100);g__Metschnikowia(100);g__Metschnikowia_unclassified(100); | Metschnikowia | 13 | Saprotroph | \|Undefined Saprotroph\| | Yeast | NULL | Probable | As nectar-tap_saprotroph (Põlme et al. 2020); Decay sugar-rich substrates (Põlme et al. 2020) | Põlme S, et al. 2020. Fungal Diversity 105:1-16 ((https://doi.org/10.1007/s13225-020-00466-2)) |
| Otu28176 | 1 | k__Fungi(100);p__Ascomycota(100);c__Saccharomycetes(100);o__Saccharomycetales(100);f__Metschnikowiaceae(100);g__Metschnikowia(100);s__Metschnikowia_pulcherrima(100); | Metschnikowia | 13 | Saprotroph | \|Undefined Saprotroph\| | Yeast | NULL | Probable | As nectar-tap_saprotroph (Põlme et al. 2020); Decay sugar-rich substrates (Põlme et al. 2020) | Põlme S, et al. 2020. Fungal Diversity 105:1-16 ((https://doi.org/10.1007/s13225-020-00466-2)) |
| Otu28177 | 1 | k__Fungi(100);p__Ascomycota(100);c__Saccharomycetes(100);o__Saccharomycetales(100);f__Metschnikowiaceae(100);g__Metschnikowia(100);g__Metschnikowia_unclassified(100); | Metschnikowia | 13 | Saprotroph | \|Undefined Saprotroph\| | Yeast | NULL | Probable | As nectar-tap_saprotroph (Põlme et al. 2020); Decay sugar-rich substrates (Põlme et al. 2020) | Põlme S, et al. 2020. Fungal Diversity 105:1-16 ((https://doi.org/10.1007/s13225-020-00466-2)) |
| Otu28178 | 1 | k__Fungi(100);p__Ascomycota(100);c__Saccharomycetes(100);o__Saccharomycetales(100);f__Metschnikowiaceae(100);g__Metschnikowia(100);s__Metschnikowia_pulcherrima(100); | Metschnikowia | 13 | Saprotroph | \|Undefined Saprotroph\| | Yeast | NULL | Probable | As nectar-tap_saprotroph (Põlme et al. 2020); Decay sugar-rich substrates (Põlme et al. 2020) | Põlme S, et al. 2020. Fungal Diversity 105:1-16 ((https://doi.org/10.1007/s13225-020-00466-2)) |
| Otu28182 | 1 | k__Fungi(100);p__Ascomycota(100);c__Saccharomycetes(100);o__Saccharomycetales(100);f__Metschnikowiaceae(100);g__Metschnikowia(100);s__Metschnikowia_pulcherrima(100); | Metschnikowia | 13 | Saprotroph | \|Undefined Saprotroph\| | Yeast | NULL | Probable | As nectar-tap_saprotroph (Põlme et al. 2020); Decay sugar-rich substrates (Põlme et al. 2020) | Põlme S, et al. 2020. Fungal Diversity 105:1-16 ((https://doi.org/10.1007/s13225-020-00466-2)) |
| Otu28192 | 1 | k__Fungi(100);p__Ascomycota(100);c__Saccharomycetes(100);o__Saccharomycetales(100);f__Metschnikowiaceae(100);g__Metschnikowia(100);s__Metschnikowia_pulcherrima(100); | Metschnikowia | 13 | Saprotroph | \|Undefined Saprotroph\| | Yeast | NULL | Probable | As nectar-tap_saprotroph (Põlme et al. 2020); Decay sugar-rich substrates (Põlme et al. 2020) | Põlme S, et al. 2020. Fungal Diversity 105:1-16 ((https://doi.org/10.1007/s13225-020-00466-2)) |
| Otu28196 | 1 | k__Fungi(100);p__Ascomycota(100);c__Saccharomycetes(100);o__Saccharomycetales(100);f__Metschnikowiaceae(100);g__Metschnikowia(100);s__Metschnikowia_pulcherrima(100); | Metschnikowia | 13 | Saprotroph | \|Undefined Saprotroph\| | Yeast | NULL | Probable | As nectar-tap_saprotroph (Põlme et al. 2020); Decay sugar-rich substrates (Põlme et al. 2020) | Põlme S, et al. 2020. Fungal Diversity 105:1-16 ((https://doi.org/10.1007/s13225-020-00466-2)) |
| Otu28198 | 1 | k__Fungi(100);p__Ascomycota(100);c__Saccharomycetes(100);o__Saccharomycetales(100);f__Metschnikowiaceae(100);g__Metschnikowia(100);g__Metschnikowia_unclassified(100); | Metschnikowia | 13 | Saprotroph | \|Undefined Saprotroph\| | Yeast | NULL | Probable | As nectar-tap_saprotroph (Põlme et al. 2020); Decay sugar-rich substrates (Põlme et al. 2020) | Põlme S, et al. 2020. Fungal Diversity 105:1-16 ((https://doi.org/10.1007/s13225-020-00466-2)) |
| Otu28202 | 1 | k__Fungi(100);p__Ascomycota(100);c__Saccharomycetes(100);o__Saccharomycetales(100);f__Metschnikowiaceae(100);g__Metschnikowia(100);g__Metschnikowia_unclassified(100); | Metschnikowia | 13 | Saprotroph | \|Undefined Saprotroph\| | Yeast | NULL | Probable | As nectar-tap_saprotroph (Põlme et al. 2020); Decay sugar-rich substrates (Põlme et al. 2020) | Põlme S, et al. 2020. Fungal Diversity 105:1-16 ((https://doi.org/10.1007/s13225-020-00466-2)) |
| Otu28203 | 1 | k__Fungi(100);p__Ascomycota(100);c__Saccharomycetes(100);o__Saccharomycetales(100);f__Metschnikowiaceae(100);g__Metschnikowia(100);s__Metschnikowia_pulcherrima(100); | Metschnikowia | 13 | Saprotroph | \|Undefined Saprotroph\| | Yeast | NULL | Probable | As nectar-tap_saprotroph (Põlme et al. 2020); Decay sugar-rich substrates (Põlme et al. 2020) | Põlme S, et al. 2020. Fungal Diversity 105:1-16 ((https://doi.org/10.1007/s13225-020-00466-2)) |
| Otu28204 | 1 | k__Fungi(100);p__Ascomycota(100);c__Saccharomycetes(100);o__Saccharomycetales(100);f__Metschnikowiaceae(100);g__Metschnikowia(100);s__Metschnikowia_pulcherrima(100); | Metschnikowia | 13 | Saprotroph | \|Undefined Saprotroph\| | Yeast | NULL | Probable | As nectar-tap_saprotroph (Põlme et al. 2020); Decay sugar-rich substrates (Põlme et al. 2020) | Põlme S, et al. 2020. Fungal Diversity 105:1-16 ((https://doi.org/10.1007/s13225-020-00466-2)) |
| Otu28205 | 1 | k__Fungi(100);p__Ascomycota(100);c__Saccharomycetes(100);o__Saccharomycetales(100);f__Metschnikowiaceae(100);g__Metschnikowia(100);s__Metschnikowia_pulcherrima(100); | Metschnikowia | 13 | Saprotroph | \|Undefined Saprotroph\| | Yeast | NULL | Probable | As nectar-tap_saprotroph (Põlme et al. 2020); Decay sugar-rich substrates (Põlme et al. 2020) | Põlme S, et al. 2020. Fungal Diversity 105:1-16 ((https://doi.org/10.1007/s13225-020-00466-2)) |
| Otu28206 | 1 | k__Fungi(100);p__Ascomycota(100);c__Saccharomycetes(100);o__Saccharomycetales(100);f__Metschnikowiaceae(100);g__Metschnikowia(100);s__Metschnikowia_pulcherrima(100); | Metschnikowia | 13 | Saprotroph | \|Undefined Saprotroph\| | Yeast | NULL | Probable | As nectar-tap_saprotroph (Põlme et al. 2020); Decay sugar-rich substrates (Põlme et al. 2020) | Põlme S, et al. 2020. Fungal Diversity 105:1-16 ((https://doi.org/10.1007/s13225-020-00466-2)) |
| Otu28207 | 1 | k__Fungi(100);p__Ascomycota(100);c__Saccharomycetes(100);o__Saccharomycetales(100);f__Metschnikowiaceae(100);g__Metschnikowia(100);g__Metschnikowia_unclassified(100); | Metschnikowia | 13 | Saprotroph | \|Undefined Saprotroph\| | Yeast | NULL | Probable | As nectar-tap_saprotroph (Põlme et al. 2020); Decay sugar-rich substrates (Põlme et al. 2020) | Põlme S, et al. 2020. Fungal Diversity 105:1-16 ((https://doi.org/10.1007/s13225-020-00466-2)) |
| Otu28210 | 1 | k__Fungi(100);p__Ascomycota(100);c__Saccharomycetes(100);o__Saccharomycetales(100);f__Metschnikowiaceae(100);g__Metschnikowia(100);g__Metschnikowia_unclassified(100); | Metschnikowia | 13 | Saprotroph | \|Undefined Saprotroph\| | Yeast | NULL | Probable | As nectar-tap_saprotroph (Põlme et al. 2020); Decay sugar-rich substrates (Põlme et al. 2020) | Põlme S, et al. 2020. Fungal Diversity 105:1-16 ((https://doi.org/10.1007/s13225-020-00466-2)) |
| Otu28211 | 1 | k__Fungi(100);p__Ascomycota(100);c__Saccharomycetes(100);o__Saccharomycetales(100);f__Metschnikowiaceae(100);g__Metschnikowia(100);s__Metschnikowia_pulcherrima(100); | Metschnikowia | 13 | Saprotroph | \|Undefined Saprotroph\| | Yeast | NULL | Probable | As nectar-tap_saprotroph (Põlme et al. 2020); Decay sugar-rich substrates (Põlme et al. 2020) | Põlme S, et al. 2020. Fungal Diversity 105:1-16 ((https://doi.org/10.1007/s13225-020-00466-2)) |
| Otu28214 | 1 | k__Fungi(100);p__Ascomycota(100);c__Saccharomycetes(100);o__Saccharomycetales(100);f__Metschnikowiaceae(100);g__Metschnikowia(100);s__Metschnikowia_pulcherrima(100); | Metschnikowia | 13 | Saprotroph | \|Undefined Saprotroph\| | Yeast | NULL | Probable | As nectar-tap_saprotroph (Põlme et al. 2020); Decay sugar-rich substrates (Põlme et al. 2020) | Põlme S, et al. 2020. Fungal Diversity 105:1-16 ((https://doi.org/10.1007/s13225-020-00466-2)) |
| Otu28216 | 1 | k__Fungi(100);p__Ascomycota(100);c__Saccharomycetes(100);o__Saccharomycetales(100);f__Metschnikowiaceae(100);g__Metschnikowia(100);s__Metschnikowia_pulcherrima(100); | Metschnikowia | 13 | Saprotroph | \|Undefined Saprotroph\| | Yeast | NULL | Probable | As nectar-tap_saprotroph (Põlme et al. 2020); Decay sugar-rich substrates (Põlme et al. 2020) | Põlme S, et al. 2020. Fungal Diversity 105:1-16 ((https://doi.org/10.1007/s13225-020-00466-2)) |
| Otu28217 | 1 | k__Fungi(100);p__Ascomycota(100);c__Saccharomycetes(100);o__Saccharomycetales(100);f__Metschnikowiaceae(100);g__Metschnikowia(100);g__Metschnikowia_unclassified(100); | Metschnikowia | 13 | Saprotroph | \|Undefined Saprotroph\| | Yeast | NULL | Probable | As nectar-tap_saprotroph (Põlme et al. 2020); Decay sugar-rich substrates (Põlme et al. 2020) | Põlme S, et al. 2020. Fungal Diversity 105:1-16 ((https://doi.org/10.1007/s13225-020-00466-2)) |
| Otu28218 | 1 | k__Fungi(100);p__Ascomycota(100);c__Saccharomycetes(100);o__Saccharomycetales(100);f__Metschnikowiaceae(100);g__Metschnikowia(100);g__Metschnikowia_unclassified(100); | Metschnikowia | 13 | Saprotroph | \|Undefined Saprotroph\| | Yeast | NULL | Probable | As nectar-tap_saprotroph (Põlme et al. 2020); Decay sugar-rich substrates (Põlme et al. 2020) | Põlme S, et al. 2020. Fungal Diversity 105:1-16 ((https://doi.org/10.1007/s13225-020-00466-2)) |
| Otu28220 | 1 | k__Fungi(100);p__Ascomycota(100);c__Saccharomycetes(100);o__Saccharomycetales(100);f__Metschnikowiaceae(100);g__Metschnikowia(100);s__Metschnikowia_pulcherrima(100); | Metschnikowia | 13 | Saprotroph | \|Undefined Saprotroph\| | Yeast | NULL | Probable | As nectar-tap_saprotroph (Põlme et al. 2020); Decay sugar-rich substrates (Põlme et al. 2020) | Põlme S, et al. 2020. Fungal Diversity 105:1-16 ((https://doi.org/10.1007/s13225-020-00466-2)) |
| Otu28221 | 1 | k__Fungi(100);p__Ascomycota(100);c__Saccharomycetes(100);o__Saccharomycetales(100);f__Metschnikowiaceae(100);g__Metschnikowia(100);s__Metschnikowia_pulcherrima(100); | Metschnikowia | 13 | Saprotroph | \|Undefined Saprotroph\| | Yeast | NULL | Probable | As nectar-tap_saprotroph (Põlme et al. 2020); Decay sugar-rich substrates (Põlme et al. 2020) | Põlme S, et al. 2020. Fungal Diversity 105:1-16 ((https://doi.org/10.1007/s13225-020-00466-2)) |
| Otu28222 | 1 | k__Fungi(100);p__Basidiomycota(100);c__Agaricomycetes(100);o__Agaricales(100);f__Strophariaceae(100);g__Gymnopilus(100);s__Gymnopilus_picreus(100); | Gymnopilus | 13 | Saprotroph | Plant Saprotroph-\|Wood Saprotroph\| | Agaricoid | White Rot | Probable | [illegible] | [illegible] |
| Otu28223 | 1 | k__Fungi(100);p__Basidiomycota(100);c__Agaricomycetes(100);o__Agaricales(100);f__Cortinariaceae(100);g__Cortinarius(100);g__Cortinarius_unclassified(100); | Cortinarius | 13 | Symbiotroph | \|Ectomycorrhizal\| | Agaricoid | NULL | Highly Probable | NULL | [illegible] |
| Otu28224 | 1 | k__Fungi(100);p__Basidiomycota(100);c__Agaricomycetes(100);o__Agaricales(100);f__Cortinariaceae(100);g__Cortinarius(100);g__Cortinarius_unclassified(100); | Cortinarius | 13 | Symbiotroph | \|Ectomycorrhizal\| | Agaricoid | NULL | Highly Probable | NULL | [illegible] |
| Otu28225 | 1 | k__Fungi(100);p__Basidiomycota(100);c__Agaricomycetes(100);o__Agaricales(100);f__Cortinariaceae(100);g__Cortinarius(100);g__Cortinarius_unclassified(100); | Cortinarius | 13 | Symbiotroph | \|Ectomycorrhizal\| | Agaricoid | NULL | Highly Probable | NULL | [illegible] |
| Otu28226 | 1 | k__Fungi(100);p__Basidiomycota(100);c__Agaricomycetes(100);o__Agaricales(100);f__Agaricaceae(100);g__Echinoderma(100);s__Echinoderma_asperum(100); | Echinoderma | 13 | Saprotroph | \|Undefined Saprotroph\| | Agaricoid | NULL | Probable | [illegible] | [illegible] |
| Otu28229 | 1 | k__Fungi(100);p__Basidiomycota(100);c__Agaricomycetes(100);o__Agaricales(100);f__Bolbitiaceae(100);g__Conocybe(100);s__Conocybe_gigasperma(100); | Conocybe | 13 | Saprotroph | Dung Saprotroph-Plant Saprotroph-\|Undefined Saprotroph\| | Agaricoid | NULL | Probable | [illegible] | [illegible] |
| Otu28230 | 1 | k__Fungi(100);p__Basidiomycota(100);c__Agaricomycetes(100);o__Agaricales(100);f__Psathyrellaceae(100);g__Psathyrella(100);s__Psathyrella_maculata(100); | Psathyrella | 13 | Saprotroph | Plant Saprotroph-\|Wood Saprotroph\| | Agaricoid | NULL | Probable | [illegible] | [illegible] |
| Otu28231 | 1 | k__Fungi(100);p__Basidiomycota(100);c__Agaricomycetes(100);o__Agaricales(100);f__Psathyrellaceae(100);g__Psathyrella(100);s__Psathyrella_romagnesii(100); | Psathyrella | 13 | Saprotroph | Plant Saprotroph-\|Wood Saprotroph\| | Agaricoid | NULL | Probable | [illegible] | [illegible] |
| Otu28232 | 1 | k__Fungi(100);p__Basidiomycota(100);c__Agaricomycetes(100);o__Agaricales(100);f__Psathyrellaceae(100);g__Psathyrella(100);g__Psathyrella_unclassified(100); | Psathyrella | 13 | Saprotroph | Plant Saprotroph-\|Wood Saprotroph\| | Agaricoid | NULL | Probable | [illegible] | [illegible] |
| Otu28233 | 1 | k__Fungi(100);p__Basidiomycota(100);c__Agaricomycetes(100);o__Agaricales(100);o__Agaricales_unclassified(100);o__Agaricales_unclassified(100);o__Agaricales_unclassified(100); | Agaricales | 7 | Pathotroph-Saprotroph-Symbiotroph | Bryophyte Parasite-Dung Saprotroph-Ectomycorrhizal-Fungal Parasite-Leaf Saprotroph-Plant Parasite-Undefined Saprotroph-Wood Saprotroph | Agaricoid-Gasteroid-Microfungus-Secotioid-Yeast | NULL | Possible | [illegible] | [illegible] |
| Otu28234 | 1 | k__Fungi(100);p__Basidiomycota(100);c__Agaricomycetes(100);o__Agaricales(100);o__Agaricales_unclassified(100);o__Agaricales_unclassified(100);o__Agaricales_unclassified(100); | Agaricales | 7 | Pathotroph-Saprotroph-Symbiotroph | Bryophyte Parasite-Dung Saprotroph-Ectomycorrhizal-Fungal Parasite-Leaf Saprotroph-Plant Parasite-Undefined Saprotroph-Wood Saprotroph | Agaricoid-Gasteroid-Microfungus-Secotioid-Yeast | NULL | Possible | [illegible] | [illegible] |
| Otu28235 | 1 | k__Fungi(100);p__Basidiomycota(100);c__Agaricomycetes(100);o__Agaricales(100);f__Tricholomataceae(100);f__Tricholomataceae_unclassified(100);f__Tricholomataceae_unclassified(100); | Tricholomataceae | 9 | Pathotroph-Symbiotroph | Ectomycorrhizal-Fungal Parasite | Agaricoid | NULL | Probable | [illegible] | [illegible] |
| Otu28236 | 1 | k__Fungi(100);p__Basidiomycota(100);c__Agaricomycetes(100);o__Agaricales(100);f__Cortinariaceae(100);g__Cortinarius(100);g__Cortinarius_unclassified(100); | Cortinarius | 13 | Symbiotroph | \|Ectomycorrhizal\| | Agaricoid | NULL | Highly Probable | NULL | [illegible] |
| Otu28237 | 1 | k__Fungi(100);p__Basidiomycota(100);c__Agaricomycetes(100);o__Agaricales(100);f__Strophariaceae(100);g__Naucoria(100);s__Naucoria_celluloderma(100); | Naucoria | 13 | Symbiotroph | \|Ectomycorrhizal\| | Agaricoid | NULL | Highly Probable | NULL | [illegible] |
| Otu28238 | 1 | k__Fungi(100);p__Basidiomycota(100);c__Agaricomycetes(100);o__Agaricales(100);f__Strophariaceae(100);g__Galerina(100);s__Galerina_unicolor_(Fr.)_Singer(100); | Galerina | 13 | Pathotroph-Saprotroph | Bryophyte Parasite-Plant Pathogen-Plant Saprotroph-Undefined Saprotroph-\|Wood Saprotroph\| | Agaricoid | White Rot | Probable | [illegible] | [illegible] |
| Otu28239 | 1 | k__Fungi(100);p__Basidiomycota(100);c__Agaricomycetes(100);o__Agaricales(100);f__Agaricaceae(100);g__Coprinus(100);g__Coprinus_unclassified(100); | Coprinus | 13 | Pathotroph-Saprotroph | \|Dung Saprotroph\|-Plant Pathogen-Plant Saprotroph | Agaricoid | NULL | Highly Probable | [illegible] | [illegible] |
| Otu28240 | 1 | k__Fungi(100);p__Basidiomycota(100);c__Agaricomycetes(100);o__Agaricales(100);f__Strophariaceae(100);g__Psilocybe(100);g__Psilocybe_unclassified(100); | Psilocybe | 13 | Saprotroph | \|Plant Saprotroph\|-Undefined Saprotroph | Agaricoid | Poisonous | Probable | [illegible] | [illegible] |
| Otu28241 | 1 | k__Fungi(100);p__Basidiomycota(100);c__Agaricomycetes(100);o__Agaricales(100);f__Strophariaceae(100);f__Strophariaceae_unclassified(100);f__Strophariaceae_unclassified(100); | Strophariaceae | 9 | Pathotroph-Saprotroph | Dung Saprotroph-Plant Parasite-Soil Saprotroph-Undefined Saprotroph-Wood Saprotroph | Agaricoid | NULL | Probable | [illegible] | [illegible] |
| Otu28242 | 1 | k__Fungi(100);p__Basidiomycota(100);c__Agaricomycetes(100);o__Agaricales(100);f__Strophariaceae(100);f__Strophariaceae_unclassified(100);f__Strophariaceae_unclassified(100); | Strophariaceae | 9 | Pathotroph-Saprotroph | Dung Saprotroph-Plant Parasite-Soil Saprotroph-Undefined Saprotroph-Wood Saprotroph | Agaricoid | NULL | Probable | [illegible] | [illegible] |
| Otu28243 | 1 | k__Fungi(100);p__Basidiomycota(100);c__Agaricomycetes(100);o__Agaricales(100);f__Psathyrellaceae(100);g__Psathyrella(100);g__Psathyrella_unclassified(100); | Psathyrella | 13 | Saprotroph | Plant Saprotroph-\|Wood Saprotroph\| | Agaricoid | NULL | Probable | [illegible] | [illegible] |
| Otu28244 | 1 | k__Fungi(100);p__Basidiomycota(100);c__Agaricomycetes(100);o__Agaricales(100);f__Psathyrellaceae(100);g__Coprinellus(100);g__Coprinellus_unclassified(100); | Coprinellus | 13 | Saprotroph | \|Undefined Saprotroph\| | Agaricoid | NULL | Probable | [illegible] | [illegible] |
| Otu28245 | 1 | k__Fungi(100);p__Basidiomycota(100);c__Agaricomycetes(100);o__Agaricales(100);f__Agaricaceae(100);g__Coprinus(100);s__Coprinus_phaeopunctatus(100); | Coprinus | 13 | Pathotroph-Saprotroph | \|Dung Saprotroph\|-Plant Pathogen-Plant Saprotroph | Agaricoid | NULL | Highly Probable | [illegible] | [illegible] |
| Otu28246 | 1 | k__Fungi(100);p__Basidiomycota(100);c__Agaricomycetes(100);o__Agaricales(100);f__Psathyrellaceae(100);g__Coprinopsis(100);s__Coprinopsis_candidolanata(100); | Coprinopsis | 13 | Saprotroph | \|Undefined Saprotroph\| | Agaricoid | NULL | Probable | [illegible] | [illegible] |
| Otu28247 | 1 | k__Fungi(100);p__Basidiomycota(100);c__Agaricomycetes(100);o__Agaricales(100);f__Lycoperdaceae(100);g__unclassified_Lycoperdaceae(100);s__Lycoperdaceae_sp(100); | Lycoperdaceae | 9 | Saprotroph | Undefined Saprotroph | Gasteroid | NULL | Probable | [illegible] | [illegible] |
| Otu28249 | 1 | k__Fungi(100);p__Basidiomycota(100);c__Agaricomycetes(100);o__Boletales(100);f__Boletaceae(100);g__unclassified_Boletaceae(100);s__Boletaceae_sp(100); | Boletaceae | 9 | Pathotroph-Saprotroph-Symbiotroph | Ectomycorrhizal-Fungal Parasite-Plant Saprotroph-Wood Saprotroph | Boletoid | NULL | Probable | [illegible] | [illegible] |
| Otu28250 | 1 | k__Fungi(100);p__Basidiomycota(100);c__Agaricomycetes(100);o__Trechisporales(100);f__Hydnodontaceae(100);g__Trechispora(100);g__Trechispora_unclassified(100); | Trechispora | 13 | Saprotroph | \|Wood Saprotroph\| | Corticioid | White Rot | Highly Probable | [illegible] | [illegible] |
| Otu28251 | 1 | k__Fungi(100);p__Basidiomycota(100);c__Agaricomycetes(100);o__Trechisporales(100);f__Hydnodontaceae(100);g__Trechispora(100);g__Trechispora_unclassified(100); | Trechispora | 13 | Saprotroph | \|Wood Saprotroph\| | Corticioid | White Rot | Highly Probable | [illegible] | [illegible] |
| Otu28253 | 1 | k__Fungi(100);p__Basidiomycota(100);c__Agaricomycetes(100);o__Agaricales(100);f__Psathyrellaceae(100);g__Coprinellus(100);s__Coprinellus_sp(100); | Coprinellus | 13 | Saprotroph | \|Undefined Saprotroph\| | Agaricoid | NULL | Probable | [illegible] | [illegible] |
| Otu28254 | 1 | k__Fungi(100);p__Basidiomycota(100);c__Agaricomycetes(100);o__Polyporales(100);f__Meruliaceae(100);g__Hypochnicium(100);s__Hypochnicium_sp_DLL2011_2(100); | Hypochnicium | 13 | Saprotroph | Undefined Saprotroph-\|Wood Saprotroph\| | Corticioid | White Rot | Probable | [illegible] | [illegible] |
| Otu28256 | 1 | k__Fungi(100);p__Basidiomycota(100);c__Agaricomycetes(100);o__Hymenochaetales(100);f__Schizoporaceae(100);g__Hyphodontia(100);s__Hyphodontia_radula(100); | Hyphodontia | 13 | Saprotroph | Undefined Saprotroph-\|Wood Saprotroph\| | Corticioid | White Rot | Probable | [illegible] | [illegible] |
| Otu28259 | 1 | k__Fungi(100);p__Basidiomycota(100);c__Agaricomycetes(100);o__Hymenochaetales(100);f__Schizoporaceae(100);g__Schizopora(100);g__Schizopora_unclassified(100); | Schizopora | 13 | Pathotroph-Saprotroph | Plant Pathogen-\|Wood Saprotroph\| | Corticioid | White Rot | Highly Probable | [illegible] | [illegible] |
| Otu28260 | 1 | k__Fungi(100);p__Basidiomycota(100);c__Agaricomycetes(100);o__Agaricales(100);f__Bolbitiaceae(100);g__Conocybe(100);g__Conocybe_unclassified(100); | Conocybe | 13 | Saprotroph | Dung Saprotroph-Plant Saprotroph-\|Undefined Saprotroph\| | Agaricoid | NULL | Probable | [illegible] | [illegible] |
| Otu28261 | 1 | k__Fungi(100);p__Basidiomycota(100);c__Agaricomycetes(100);o__Agaricales(100);f__Cortinariaceae(100);g__Cortinarius(100);g__Cortinarius_unclassified(100); | Cortinarius | 13 | Symbiotroph | \|Ectomycorrhizal\| | Agaricoid | NULL | Highly Probable | NULL | [illegible] |
| Otu28262 | 1 | k__Fungi(100);p__Basidiomycota(100);c__Agaricomycetes(100);o__Boletales(100);f__Boletaceae(100);g__Boletus(100);s__Boletus_sp(100); | Boletus | 13 | Pathotroph-Symbiotroph | \|Ectomycorrhizal\|-Fungal Parasite | Agaricoid-Boletoid | NULL | Highly Probable | NULL | [illegible] |
| Otu28263 | 1 | k__Fungi(100);p__Basidiomycota(100);c__Agaricomycetes(100);o__Agaricales(100);o__Agaricales_unclassified(100);o__Agaricales_unclassified(100);o__Agaricales_unclassified(100); | Agaricales | 7 | Pathotroph-Saprotroph-Symbiotroph | Bryophyte Parasite-Dung Saprotroph-Ectomycorrhizal-Fungal Parasite-Leaf Saprotroph-Plant Parasite-Undefined Saprotroph-Wood Saprotroph | Agaricoid-Gasteroid-Microfungus-Secotioid-Yeast | NULL | Possible | [illegible] | [illegible] |
| Otu28264 | 1 | k__Fungi(100);p__Basidiomycota(100);c__Agaricomycetes(100);o__Agaricales(100);f__Strophariaceae(100);g__Stropharia(100);s__Stropharia_rugosoannulata(100); | Stropharia | 13 | Saprotroph | Dung Saprotroph-Plant Parasite-\|Plant Saprotroph\|-Undefined Saprotroph-Wood Saprotroph | Agaricoid | NULL | Probable | [illegible] | [illegible] |
| Otu28265 | 1 | k__Fungi(100);p__Basidiomycota(100);c__Agaricomycetes(100);o__Agaricales(100);f__Strophariaceae(100);f__Strophariaceae_unclassified(100);f__Strophariaceae_unclassified(100); | Strophariaceae | 9 | Pathotroph-Saprotroph | Dung Saprotroph-Plant Parasite-Soil Saprotroph-Undefined Saprotroph-Wood Saprotroph | Agaricoid | NULL | Probable | [illegible] | [illegible] |
| Otu28266 | 1 | k__Fungi(100);p__Basidiomycota(100);c__Agaricomycetes(100);o__Agaricales(100);f__Strophariaceae(100);g__Stropharia(100);g__Stropharia_unclassified(100); | Stropharia | 13 | Saprotroph | Dung Saprotroph-Plant Parasite-\|Plant Saprotroph\|-Undefined Saprotroph-Wood Saprotroph | Agaricoid | NULL | Probable | [illegible] | [illegible] |
| Otu28267 | 1 | k__Fungi(100);p__Basidiomycota(100);c__Agaricomycetes(100);o__Agaricales(100);f__Cortinariaceae(100);g__Cortinarius(100);g__Cortinarius_unclassified(100); | Cortinarius | 13 | Symbiotroph | \|Ectomycorrhizal\| | Agaricoid | NULL | Highly Probable | NULL | [illegible] |
| Otu28268 | 1 | k__Fungi(100);p__Basidiomycota(100);c__Agaricomycetes(100);o__Agaricales(100);f__Cortinariaceae(100);g__Cortinarius(100);g__Cortinarius_unclassified(100); | Cortinarius | 13 | Symbiotroph | \|Ectomycorrhizal\| | Agaricoid | NULL | Highly Probable | NULL | [illegible] |
| Otu28269 | 1 | k__Fungi(100);p__Basidiomycota(100);c__Agaricomycetes(100);o__Agaricales(100);f__Cortinariaceae(100);g__Cortinarius(100);g__Cortinarius_unclassified(100); | Cortinarius | 13 | Symbiotroph | \|Ectomycorrhizal\| | Agaricoid | NULL | Highly Probable | NULL | [illegible] |
| Otu28270 | 1 | k__Fungi(100);p__Basidiomycota(100);c__Agaricomycetes(100);o__Agaricales(100);f__Cortinariaceae(100);g__Cortinarius(100);g__Cortinarius_unclassified(100); | Cortinarius | 13 | Symbiotroph | \|Ectomycorrhizal\| | Agaricoid | NULL | Highly Probable | NULL | [illegible] |
| Otu28271 | 1 | k__Fungi(100);p__Basidiomycota(100);c__Agaricomycetes(100);o__Agaricales(100);f__Cortinariaceae(100);g__Cortinarius(100);s__Cortinarius_alboviolaceus(100); | Cortinarius | 13 | Symbiotroph | \|Ectomycorrhizal\| | Agaricoid | NULL | Highly Probable | NULL | [illegible] |
| Otu28272 | 1 | k__Fungi(100);p__Basidiomycota(100);c__Agaricomycetes(100);o__Agaricales(100);f__Cortinariaceae(100);g__Cortinarius(100);g__Cortinarius_unclassified(100); | Cortinarius | 13 | Symbiotroph | \|Ectomycorrhizal\| | Agaricoid | NULL | Highly Probable | NULL | [illegible] |
| Otu28273 | 1 | k__Fungi(100);p__Basidiomycota(100);c__Agaricomycetes(100);o__Agaricales(100);f__Cortinariaceae(100);g__Cortinarius(100);g__Cortinarius_unclassified(100); | Cortinarius | 13 | Symbiotroph | \|Ectomycorrhizal\| | Agaricoid | NULL | Highly Probable | NULL | [illegible] |
| Otu28274 | 1 | k__Fungi(100);p__Basidiomycota(100);c__Agaricomycetes(100);o__Agaricales(100);f__Cortinariaceae(100);g__Cortinarius(100);g__Cortinarius_unclassified(100); | Cortinarius | 13 | Symbiotroph | \|Ectomycorrhizal\| | Agaricoid | NULL | Highly Probable | NULL | [illegible] |
| Otu28275 | 1 | k__Fungi(100);p__Basidiomycota(100);c__Agaricomycetes(100);o__Agaricales(100);f__Cortinariaceae(100);g__Cortinarius(100);g__Cortinarius_unclassified(100); | Cortinarius | 13 | Symbiotroph | \|Ectomycorrhizal\| | Agaricoid | NULL | Highly Probable | NULL | [illegible] |
| Otu28276 | 1 | k__Fungi(100);p__Basidiomycota(100);c__Tremellomycetes(100);o__Filobasidiales(100);f__Filobasidiaceae(100);g__Cryptococcus(100);s__Cryptococcus_friedmannii(100); | Cryptococcus | 13 | Pathotroph-Saprotroph-Symbiotroph | Animal Pathogen-Endophyte-Epiphyte-\|Undefined Saprotroph\| | Dimorphic Yeast-Yeast | NULL | Possible | [illegible] | [illegible] |
| Otu28277 | 1 | k__Fungi(100);p__Basidiomycota(100);c__Microbotryomycetes(100);o__Sporidiobolales(100);f__Sporidiobolales_family_Incertae_sedis(100);g__Rhodotorula(100);s__Rhodotorula_graminis(100); | Rhodotorula | 13 | Pathotroph-Saprotroph-Symbiotroph | Animal Pathogen-Animal Symbiotroph-Endophyte-Plant Pathogen-\|Undefined Saprotroph\| | Yeast | NULL | Probable | [illegible] | [illegible] |
| Otu28278 | 1 | k__Fungi(100);p__Basidiomycota(100);c__Agaricomycetes(100);o__Auriculariales(100);f__Auriculariales_family_Incertae_sedis(100);g__Pseudohydnum(100);s__Pseudohydnum_gelatinosum(100); | Pseudohydnum | 13 | Saprotroph | Undefined Saprotroph-\|Wood Saprotroph\| | Corticioid-Tremelloid | NULL | Probable | [illegible] | [illegible] |
| Otu28279 | 1 | k__Fungi(100);p__Basidiomycota(100);c__Agaricomycetes(100);o__Polyporales(100);f__Polyporaceae(100);g__Perenniporia(100);s__Perenniporia_sp(100); | Perenniporia | 13 | Saprotroph | \|Wood Saprotroph\| | Corticioid-Polyporoid | White Rot | Highly Probable | [illegible] | [illegible] |
| Otu28280 | 1 | k__Fungi(100);p__Basidiomycota(100);c__Agaricomycetes(100);o__Agaricales(100);f__Strophariaceae(100);g__Galerina(100);g__Galerina_unclassified(100); | Galerina | 13 | Pathotroph-Saprotroph | Bryophyte Parasite-Plant Pathogen-Plant Saprotroph-Undefined Saprotroph-\|Wood Saprotroph\| | Agaricoid | White Rot | Probable | [illegible] | [illegible] |
| Otu28281 | 1 | k__Fungi(100);p__Basidiomycota(100);c__Agaricomycetes(100);o__Polyporales(100);f__Polyporaceae(100);f__Polyporaceae_unclassified(100);f__Polyporaceae_unclassified(100); | Polyporaceae | 9 | Saprotroph | Wood Saprotroph | Polyporoid | Brown Rot-White | Probable | [illegible] | [illegible] |
| Otu28283 | 1 | k__Fungi(100);p__Basidiomycota(100);c__Microbotryomycetes(100);o__Sporidiobolales(100);f__Sporidiobolales_family_Incertae_sedis(100);g__Rhodotorula(100);s__Rhodotorula_graminis(100); | Rhodotorula | 13 | Pathotroph-Saprotroph-Symbiotroph | Animal Pathogen-Animal Symbiotroph-Endophyte-Plant Pathogen-\|Undefined Saprotroph\| | Yeast | NULL | Probable | [illegible] | [illegible] |
| Otu28284 | 1 | k__Fungi(100);p__Basidiomycota(100);c__Agaricomycetes(100);o__Agaricales(100);o__Agaricales_unclassified(100);o__Agaricales_unclassified(100);o__Agaricales_unclassified(100); | Agaricales | 7 | Pathotroph-Saprotroph-Symbiotroph | Bryophyte Parasite-Dung Saprotroph-Ectomycorrhizal-Fungal Parasite-Leaf Saprotroph-Plant Parasite-Undefined Saprotroph-Wood Saprotroph | Agaricoid-Gasteroid-Microfungus-Secotioid-Yeast | NULL | Possible | [illegible] | [illegible] |
| Otu28285 | 1 | k__Fungi(100);p__Basidiomycota(100);c__Agaricomycetes(100);o__Polyporales(100);f__Meruliaceae(100);g__Steccherinum(100);s__Steccherinum_sp_MLB_2010(100); | Steccherinum | 13 | Saprotroph | Undefined Saprotroph-\|Wood Saprotroph\| | Corticioid | White Rot | Probable | [illegible] | [illegible] |
| Otu28286 | 1 | k__Fungi(100);p__Basidiomycota(100);c__Agaricomycetes(100);o__Agaricales(100);f__Psathyrellaceae(100);g__Coprinopsis(100);s__Coprinopsis_strossmayeri(100); | Coprinopsis | 13 | Saprotroph | \|Undefined Saprotroph\| | Agaricoid | NULL | Probable | [illegible] | [illegible] |
| Otu28289 | 1 | k__Fungi(100);p__Basidiomycota(100);c__Tremellomycetes(100);o__Tremellales(100);f__Tremellales_family_Incertae_sedis(100);g__Cryptococcus(100);s__Cryptococcus_magnus(100); | Cryptococcus | 13 | Pathotroph-Saprotroph-Symbiotroph | Animal Pathogen-Endophyte-Epiphyte-\|Undefined Saprotroph\| | Dimorphic Yeast-Yeast | NULL | Possible | [illegible] | [illegible] |
| Otu28292 | 1 | k__Fungi(100);p__Basidiomycota(100);c__Agaricomycetes(100);o__Boletales(100);f__Tapinellaceae(100);g__Pseudomerulius(100);s__Pseudomerulius_aureus(100); | Pseudomerulius | 13 | Saprotroph | \|Wood Saprotroph\| | Corticioid | Brown Rot | Probable | [illegible] | [illegible] |
| Otu28293 | 1 | k__Fungi(100);p__Basidiomycota(100);c__Microbotryomycetes(100);o__Sporidiobolales(100);f__Sporidiobolales_family_Incertae_sedis(100);g__Rhodotorula(100);s__Rhodotorula_graminis(100); | Rhodotorula | 13 | Pathotroph-Saprotroph-Symbiotroph | Animal Pathogen-Animal Symbiotroph-Endophyte-Plant Pathogen-\|Undefined Saprotroph\| | Yeast | NULL | Probable | [illegible] | [illegible] |
| Otu28294 | 1 | k__Fungi(100);p__Basidiomycota(100);c__Agaricomycetes(100);o__Polyporales(100);f__Polyporaceae(100);g__Perenniporia(100);s__Perenniporia_subacida(100); | Perenniporia | 13 | Saprotroph | \|Wood Saprotroph\| | Corticioid-Polyporoid | White Rot | Highly Probable | [illegible] | [illegible] |
| Otu28295 | 1 | k__Fungi(100);p__Basidiomycota(100);c__Agaricomycetes(100);o__Hymenochaetales(100);f__Schizoporaceae(100);f__Schizoporaceae_unclassified(100);f__Schizoporaceae_unclassified(100); | Schizoporaceae | 9 | Saprotroph | Wood Saprotroph | Corticioid | White Rot | Probable | [illegible] | [illegible] |
| Otu28296 | 1 | k__Fungi(100);p__Basidiomycota(100);c__Agaricomycetes(100);o__Agaricales(100);f__Entolomataceae(100);g__Clitopilus(100);s__Clitopilus_cystidiatus(100); | Clitopilus | 13 | Pathotroph-Saprotroph | Fungal Parasite-\|Plant Saprotroph\|-Undefined Saprotroph | Agaricoid | NULL | Probable | [illegible] | [illegible] |
| Otu28297 | 1 | k__Fungi(100);p__Basidiomycota(100);c__Agaricomycetes(100);o__Polyporales(100);f__Fomitopsidaceae(100);g__unclassified_Fomitopsidaceae(100);s__Fomitopsidaceae_sp(100); | Fomitopsidaceae | 9 | Pathotroph-Saprotroph | Plant Pathogen-Wood Saprotroph | Polyporoid | Brown Rot | Probable | [illegible] | [illegible] |
| Otu28298 | 1 | k__Fungi(100);p__Basidiomycota(100);c__Agaricomycetes(100);o__Polyporales(100);f__Polyporaceae(100);g__Polyporus(100);s__Polyporus_tuberaster(100); | Polyporus | 13 | Pathotroph-Saprotroph | Plant Pathogen-\|Wood Saprotroph\| | Polyporoid | White Rot | Highly Probable | [illegible] | [illegible] |
| Otu28300 | 1 | k__Fungi(100);p__Basidiomycota(100);c__Agaricomycetes(100);o__Agaricales(100);o__Agaricales_unclassified(100);o__Agaricales_unclassified(100);o__Agaricales_unclassified(100); | Agaricales | 7 | Pathotroph-Saprotroph-Symbiotroph | Bryophyte Parasite-Dung Saprotroph-Ectomycorrhizal-Fungal Parasite-Leaf Saprotroph-Plant Parasite-Undefined Saprotroph-Wood Saprotroph | Agaricoid-Gasteroid-Microfungus-Secotioid-Yeast | NULL | Possible | [illegible] | [illegible] |
| Otu28301 | 1 | k__Fungi(100);p__Basidiomycota(100);c__Agaricomycetes(100);o__Polyporales(100);f__Meruliaceae(100);g__Phlebia(100);s__Phlebia_tuberculata(100); | Phlebia | 13 | Saprotroph | \|Wood Saprotroph\| | Corticioid | White Rot | Highly Probable | [illegible] | [illegible] |
| Otu28302 | 1 | k__Fungi(100);p__Basidiomycota(100);c__Agaricomycetes(100);o__Polyporales(100);f__Meruliaceae(100);g__Hyphoderma(100);s__Hyphoderma_praetermissum(100); | Hyphoderma | 13 | Saprotroph | Undefined Saprotroph-\|Wood Saprotroph\| | Corticioid | White Rot | Probable | [illegible] | [illegible] |
| Otu28304 | 1 | k__Fungi(100);p__Basidiomycota(100);c__Agaricomycetes(100);o__Agaricales(100);f__Strophariaceae(100);g__Galerina(100);g__Galerina_unclassified(100); | Galerina | 13 | Pathotroph-Saprotroph | Bryophyte Parasite-Plant Pathogen-Plant Saprotroph-Undefined Saprotroph-\|Wood Saprotroph\| | Agaricoid | White Rot | Probable | [illegible] | [illegible] |
| Otu28305 | 1 | k__Fungi(100);p__Basidiomycota(100);c__Agaricomycetes(100);o__Agaricales(100);f__Cortinariaceae(100);g__Phaeocollybia(100);g__Phaeocollybia_unclassified(100); | Phaeocollybia | 13 | Symbiotroph | \|Ectomycorrhizal\| | Agaricoid | NULL | Possible | NULL | [illegible] |
| Otu28306 | 1 | k__Fungi(100);p__Basidiomycota(100);c__Agaricomycetes(100);o__Agaricales(100);f__Pluteaceae(100);g__Pluteus(100);g__Pluteus_unclassified(100); | Pluteus | 13 | Pathotroph-Saprotroph | Bryophyte Parasite-\|Plant Saprotroph\|-Wood Saprotroph | Agaricoid | White Rot | Probable | [illegible] | [illegible] |
| Otu28307 | 1 | k__Fungi(100);p__Basidiomycota(100);c__Agaricomycetes(100);o__Polyporales(100);f__Meruliaceae(100);g__Hyphoderma(100);s__Hyphoderma_setigerum(100); | Hyphoderma | 13 | Saprotroph | Undefined Saprotroph-\|Wood Saprotroph\| | Corticioid | White Rot | Probable | [illegible] | [illegible] |
| Otu28308 | 1 | k__Fungi(100);p__Basidiomycota(100);c__Agaricomycetes(100);o__Polyporales(100);f__Meruliaceae(100);g__Scopuloides(100);s__Scopuloides_hydnoides(100); | Scopuloides | 13 | Saprotroph | Undefined Saprotroph-\|Wood Saprotroph\| | Corticioid | White Rot | Probable | [illegible] | [illegible] |
| Otu28309 | 1 | k__Fungi(100);p__Basidiomycota(100);c__Agaricomycetes(100);o__Polyporales(100);f__Phanerochaetaceae(100);g__Phanerochaete(100);s__Phanerochaete_rimosa(100); | Phanerochaete | 13 | Saprotroph | \|Wood Saprotroph\| | Corticioid | White Rot | Highly Probable | [illegible] | [illegible] |
| Otu28311 | 1 | k__Fungi(100);p__Basidiomycota(100);c__Agaricomycetes(100);o__Polyporales(100);f__Meruliaceae(100);g__Phlebia(100);s__Phlebia_fuscoatra(100); | Phlebia | 13 | Saprotroph | \|Wood Saprotroph\| | Corticioid | White Rot | Highly Probable | [illegible] | [illegible] |
| Otu28312 | 1 | k__Fungi(100);p__Basidiomycota(100);c__Agaricomycetes(100);o__Polyporales(100);f__Fomitopsidaceae(100);g__Fomitopsis(100);s__Fomitopsis_rosea(100); | Fomitopsis | 13 | Pathotroph-Saprotroph | Plant Pathogen-\|Wood Saprotroph\| | Polyporoid | Brown Rot | Highly Probable | [illegible] | [illegible] |
| Otu28313 | 1 | k__Fungi(100);p__Basidiomycota(100);c__Agaricomycetes(100);o__Polyporales(100);f__Meruliaceae(100);g__Phlebia(100);s__Phlebia_tremellosa(100); | Phlebia | 13 | Saprotroph | \|Wood Saprotroph\| | Corticioid | White Rot | Highly Probable | [illegible] | [illegible] |
| Otu28314 | 1 | k__Fungi(100);p__Basidiomycota(100);c__Agaricomycetes(100);o__Polyporales(100);f__Fomitopsidaceae(100);g__Antrodia(100);s__Antrodia_heteromorpha(100); | Antrodia | 13 | Saprotroph | \|Wood Saprotroph\| | Corticioid | Brown Rot | Highly Probable | [illegible] | [illegible] |
| Otu28316 | 1 | k__Fungi(100);p__Basidiomycota(100);c__Agaricomycetes(100);o__Trechisporales(100);f__Hydnodontaceae(100);g__Trechispora(100);g__Trechispora_unclassified(100); | Trechispora | 13 | Saprotroph | \|Wood Saprotroph\| | Corticioid | White Rot | Highly Probable | [illegible] | [illegible] |
| Otu28318 | 1 | k__Fungi(100);p__Basidiomycota(100);c__Agaricomycetes(100);o__Agaricales(100);f__Entolomataceae(100);g__Entoloma(100);s__Entoloma_infula(100); | Entoloma | 13 | Pathotroph-Saprotroph-Symbiotroph | Ectomycorrhizal-Fungal Parasite-\|Undefined Saprotroph\|-Undefined Symbiotroph | Agaricoid | NULL | Possible | [illegible] | [illegible] |
| Otu28319 | 1 | k__Fungi(100);p__Basidiomycota(100);c__Agaricomycetes(100);o__Polyporales(100);f__Polyporaceae(100);g__Perenniporia(100);s__Perenniporia_sp(100); | Perenniporia | 13 | Saprotroph | \|Wood Saprotroph\| | Corticioid-Polyporoid | White Rot | Highly Probable | [illegible] | [illegible] |
| Otu28322 | 1 | k__Fungi(100);p__Basidiomycota(100);c__Agaricomycetes(100);o__Agaricales(100);f__Tricholomataceae(100);g__Tricholoma(100);g__Tricholoma_unclassified(100); | Tricholoma | 13 | Pathotroph-Symbiotroph | \|Ectomycorrhizal\|-Fungal Parasite | Agaricoid | NULL | Highly Probable | [illegible] | [illegible] |
| Otu28324 | 1 | k__Fungi(100);p__Basidiomycota(100);c__Tremellomycetes(100);o__Filobasidiales(100);f__Filobasidiaceae(100);g__Cryptococcus(100);g__Cryptococcus_unclassified(100); | Cryptococcus | 13 | Pathotroph-Saprotroph-Symbiotroph | Animal Pathogen-Endophyte-Epiphyte-\|Undefined Saprotroph\| | Dimorphic Yeast-Yeast | NULL | Possible | [illegible] | [illegible] |
| Otu28325 | 1 | k__Fungi(100);p__Basidiomycota(100);c__Tremellomycetes(100);o__Filobasidiales(100);f__Filobasidiaceae(100);g__Cryptococcus(100);s__Cryptococcus_friedmannii(100); | Cryptococcus | 13 | Pathotroph-Saprotroph-Symbiotroph | Animal Pathogen-Endophyte-Epiphyte-\|Undefined Saprotroph\| | Dimorphic Yeast-Yeast | NULL | Possible | [illegible] | [illegible] |
| Otu28326 | 1 | k__Fungi(100);p__Basidiomycota(100);c__Tremellomycetes(100);o__Filobasidiales(100);f__Filobasidiaceae(100);g__Cryptococcus(100);s__Cryptococcus_friedmannii(100); | Cryptococcus | 13 | Pathotroph-Saprotroph-Symbiotroph | Animal Pathogen-Endophyte-Epiphyte-\|Undefined Saprotroph\| | Dimorphic Yeast-Yeast | NULL | Possible | [illegible] | [illegible] |
| Otu28327 | 1 | k__Fungi(100);p__Basidiomycota(100);c__Tremellomycetes(100);o__Filobasidiales(100);f__Filobasidiaceae(100);g__Cryptococcus(100);s__Cryptococcus_adeliensis(100); | Cryptococcus | 13 | Pathotroph-Saprotroph-Symbiotroph | Animal Pathogen-Endophyte-Epiphyte-\|Undefined Saprotroph\| | Dimorphic Yeast-Yeast | NULL | Possible | [illegible] | [illegible] |
| Otu28328 | 1 | k__Fungi(100);p__Basidiomycota(100);c__Agaricomycetes(100);o__Agaricales(100);f__Agaricaceae(100);g__Lepiota(100);s__Lepiota_cristata(100); | Lepiota | 13 | Saprotroph | \|Plant Saprotroph\|-Undefined Saprotroph | Agaricoid | NULL | Probable | [illegible] | [illegible] |
| Otu28329 | 1 | k__Fungi(100);p__Basidiomycota(100);c__Agaricomycetes(100);o__Cantharellales(100);f__Ceratobasidiaceae(100);g__Ceratobasidium(100);g__Ceratobasidium_unclassified(100); | Ceratobasidium | 13 | Pathotroph-Saprotroph-Symbiotroph | Ectomycorrhizal-Endomycorrhizal-Orchid Mycorrhizal-\|Plant Pathogen\|-Plant Saprotroph-Undefined Saprotroph | Corticioid-Microfungus | NULL | Possible | [illegible] | [illegible] |
| Otu28330 | 1 | k__Fungi(100);p__Basidiomycota(100);c__Exobasidiomycetes(100);o__Exobasidiales(100);f__Exobasidiaceae(100);g__Exobasidium(100);g__Exobasidium_unclassified(100); | Exobasidium | 13 | Pathotroph | \|Plant Pathogen\| | Leaf Blister-Leaf Gall-Leaf Spot | NULL | Probable | NULL | [illegible] |
| Otu28331 | 1 | k__Fungi(100);p__Basidiomycota(100);c__Exobasidiomycetes(100);o__Exobasidiales(100);f__Exobasidiaceae(100);g__Exobasidium(100);g__Exobasidium_unclassified(100); | Exobasidium | 13 | Pathotroph | \|Plant Pathogen\| | Leaf Blister-Leaf Gall-Leaf Spot | NULL | Probable | NULL | [illegible] |
| Otu28333 | 1 | k__Fungi(100);p__Basidiomycota(100);c__Agaricomycetes(100);o__Cantharellales(100);f__Ceratobasidiaceae(100);g__Ceratobasidium(100);g__Ceratobasidium_unclassified(100); | Ceratobasidium | 13 | Pathotroph-Saprotroph-Symbiotroph | Ectomycorrhizal-Endomycorrhizal-Orchid Mycorrhizal-\|Plant Pathogen\|-Plant Saprotroph-Undefined Saprotroph | Corticioid-Microfungus | NULL | Possible | [illegible] | [illegible] |
| Otu28334 | 1 | k__Fungi(100);p__Basidiomycota(100);c__Agaricomycetes(100);o__Cantharellales(100);f__Ceratobasidiaceae(100);f__Ceratobasidiaceae_unclassified(100);f__Ceratobasidiaceae_unclassified(100); | Ceratobasidiaceae | 9 | Pathotroph-Saprotroph-Symbiotroph | Endomycorrhizal-Plant Pathogen-Undefined Saprotroph | Microfungus | NULL | Possible | [illegible] | [illegible] |
| Otu28335 | 1 | k__Fungi(100);p__Basidiomycota(100);c__Agaricomycetes(100);o__Cantharellales(100);f__Ceratobasidiaceae(100);g__unclassified_Ceratobasidiaceae(100);g__unclassified_Ceratobasidiaceae_unclassified(100); | Ceratobasidiaceae | 9 | Pathotroph-Saprotroph-Symbiotroph | Endomycorrhizal-Plant Pathogen-Undefined Saprotroph | Microfungus | NULL | Possible | [illegible] | [illegible] |
| Otu28336 | 1 | k__Fungi(100);p__Basidiomycota(100);c__Exobasidiomycetes(100);o__Exobasidiales(100);f__Exobasidiaceae(100);g__Exobasidium(100);g__Exobasidium_unclassified(100); | Exobasidium | 13 | Pathotroph | \|Plant Pathogen\| | Leaf Blister-Leaf Gall-Leaf Spot | NULL | Probable | NULL | [illegible] |

| | | | | | | | | | | | |
|---|---|---|---|---|---|---|---|---|---|---|---|
| Otu28337 | 1 | k__Fungi(100);p__Basidiomycota(100);c__Agaricomycetes(100);o__Cantharellales(100);f__Ceratobasidiaceae(100);g__Rhizoctonia(100);s__Rhizoctonia_sp_KW214(100); | Rhizoctonia | 13 | Pathotroph-Saprotroph-Symbiotroph | Ectomycorrhizal-Orchid Mycorrhizal-\|Plant Pathogen\|-Plant Saprotroph-Wood Saprotroph | Corticioid-Microfungus | Soft Rot | Probable | As litter_saprotroph (Põlme et al. 2020); Decay leaf, fruit, seed, roots, soil (Põlme et al. 2020); Only symbiotic (perhaps leaning parasitic) in special cases with orchids, otherwise typically a plant pathogen. | Seehann G, et al. 1975. List of Fungi in Soft-Rot Tissue Tests. Institute fur Holzbiologie und Holzschutz, Bundesforschungsanstalt fur Forst - und Holzwirtshaft, Hamburg - Lohbrugge, Germany; Costa IPMW, et al. 2012. Mycotaxon 119:e26 ((http://mycotaxon.com/resources/checklists/costa_v119_checklist.pdf)); Tedersoo L, et al. 2014. Science 346:e1256688 ((https://doi.org/10.1126/science.1256688)); Põlme S, et al. 2020. Fungal Diversity 105:1-16 ((https://doi.org/10.1007/s13225-020-00466-2)); List compiled by Sara Branco with input from Nhu Nguyen and Else Vellinga; The American Phytopathological Society Common Names of Plant Diseases ((http://www.apsnet.org/publications/commonnames/Pages/default.aspx)) |
| Otu28338 | 1 | k__Fungi(100);p__Basidiomycota(100);c__Agaricomycetes(100);o__Cantharellales(100);f__Ceratobasidiaceae(100);g__Rhizoctonia(100);s__Rhizoctonia_sp_KW214(100); | Rhizoctonia | 13 | Pathotroph-Saprotroph-Symbiotroph | Ectomycorrhizal-Orchid Mycorrhizal-\|Plant Pathogen\|-Plant Saprotroph-Wood Saprotroph | Corticioid-Microfungus | Soft Rot | Probable | As litter_saprotroph (Põlme et al. 2020); Decay leaf, fruit, seed, roots, soil (Põlme et al. 2020); Only symbiotic (perhaps leaning parasitic) in special cases with orchids, otherwise typically a plant pathogen. | Seehann G, et al. 1975. List of Fungi in Soft-Rot Tissue Tests. Institute fur Holzbiologie und Holzschutz, Bundesforschungsanstalt fur Forst - und Holzwirtshaft, Hamburg - Lohbrugge, Germany; Costa IPMW, et al. 2012. Mycotaxon 119:e26 ((http://mycotaxon.com/resources/checklists/costa_v119_checklist.pdf)); Tedersoo L, et al. 2014. Science 346:e1256688 ((https://doi.org/10.1126/science.1256688)); Põlme S, et al. 2020. Fungal Diversity 105:1-16 ((https://doi.org/10.1007/s13225-020-00466-2)); List compiled by Sara Branco with input from Nhu Nguyen and Else Vellinga; The American Phytopathological Society Common Names of Plant Diseases ((http://www.apsnet.org/publications/commonnames/Pages/default.aspx)) |
| Otu28339 | 1 | k__Fungi(100);p__Basidiomycota(100);c__Agaricomycetes(100);o__Cantharellales(100);f__Ceratobasidiaceae(100);g__Ceratobasidium(100);g__Ceratobasidium_unclassified(100); | Ceratobasidium | 13 | Pathotroph-Saprotroph-Symbiotroph | Ectomycorrhizal-Endomycorrhizal-Orchid Mycorrhizal-\|Plant Pathogen\|-Plant Saprotroph-Undefined Saprotroph | Corticioid-Microfungus | NULL | Possible | especially of Orchidaceae but also even bryophytes (Cannon & Kirk 2007); As litter_saprotroph (Põlme et al. 2020); Best known as plant pathogens, but also saprobic. Species are endomycorrhizal with a wide range of plant roots ; Decay leaf, fruit, seed, roots, soil (Põlme et al. 2020) | Cannon PF, Kirk PM. 2007. Fungal Families of the World. CAB International, Cambridge (ISBN: 978-0851998275); Tedersoo L, et al. 2014. Science 346:e1256688 ((https://doi.org/10.1126/science.1256688)); Põlme S, et al. 2020. Fungal Diversity 105:1-16 ((https://doi.org/10.1007/s13225-020-00466-2)) |
| Otu28340 | 1 | k__Fungi(100);p__Basidiomycota(100);c__Agaricomycetes(100);o__Cantharellales(100);f__Ceratobasidiaceae(100);g__Ceratobasidium(100);s__Ceratobasidium_sp(100); | Ceratobasidium | 13 | Pathotroph-Saprotroph-Symbiotroph | Ectomycorrhizal-Endomycorrhizal-Orchid Mycorrhizal-\|Plant Pathogen\|-Plant Saprotroph-Undefined Saprotroph | Corticioid-Microfungus | NULL | Possible | especially of Orchidaceae but also even bryophytes (Cannon & Kirk 2007); As litter_saprotroph (Põlme et al. 2020); Best known as plant pathogens, but also saprobic. Species are endomycorrhizal with a wide range of plant roots ; Decay leaf, fruit, seed, roots, soil (Põlme et al. 2020) | Cannon PF, Kirk PM. 2007. Fungal Families of the World. CAB International, Cambridge (ISBN: 978-0851998275); Tedersoo L, et al. 2014. Science 346:e1256688 ((https://doi.org/10.1126/science.1256688)); Põlme S, et al. 2020. Fungal Diversity 105:1-16 ((https://doi.org/10.1007/s13225-020-00466-2)) |
| Otu28341 | 1 | k__Fungi(100);p__Basidiomycota(100);c__Agaricomycetes(100);o__Polyporales(100);f__Phanerochaetaceae(100);g__Ceriporia(100);g__Ceriporia_unclassified(100); | Ceriporia | 13 | Saprotroph | \|Wood Saprotroph\| | Corticioid | White Rot | Highly Probable | As white_rot (Põlme et al. 2020); Decay wood (Põlme et al. 2020) | Tedersoo L, et al. 2014. Science 346:e1256688 ((https://doi.org/10.1126/science.1256688)); Põlme S, et al. 2020. Fungal Diversity 105:1-16 ((https://doi.org/10.1007/s13225-020-00466-2)) |
| Otu28342 | 1 | k__Fungi(100);p__Basidiomycota(100);c__Agaricomycetes(100);o__Cantharellales(100);f__Ceratobasidiaceae(100);g__Ceratobasidium(100);g__Ceratobasidium_unclassified(100); | Ceratobasidium | 13 | Pathotroph-Saprotroph-Symbiotroph | Ectomycorrhizal-Endomycorrhizal-Orchid Mycorrhizal-\|Plant Pathogen\|-Plant Saprotroph-Undefined Saprotroph | Corticioid-Microfungus | NULL | Possible | especially of Orchidaceae but also even bryophytes (Cannon & Kirk 2007); As litter_saprotroph (Põlme et al. 2020); Best known as plant pathogens, but also saprobic. Species are endomycorrhizal with a wide range of plant roots ; Decay leaf, fruit, seed, roots, soil (Põlme et al. 2020) | Cannon PF, Kirk PM. 2007. Fungal Families of the World. CAB International, Cambridge (ISBN: 978-0851998275); Tedersoo L, et al. 2014. Science 346:e1256688 ((https://doi.org/10.1126/science.1256688)); Põlme S, et al. 2020. Fungal Diversity 105:1-16 ((https://doi.org/10.1007/s13225-020-00466-2)) |
| Otu28343 | 1 | k__Fungi(100);p__Basidiomycota(100);c__Agaricomycetes(100);o__Cantharellales(100);f__Ceratobasidiaceae(100);g__Ceratobasidium(100);s__Ceratobasidium_sp(100); | Ceratobasidium | 13 | Pathotroph-Saprotroph-Symbiotroph | Ectomycorrhizal-Endomycorrhizal-Orchid Mycorrhizal-\|Plant Pathogen\|-Plant Saprotroph-Undefined Saprotroph | Corticioid-Microfungus | NULL | Possible | especially of Orchidaceae but also even bryophytes (Cannon & Kirk 2007); As litter_saprotroph (Põlme et al. 2020); Best known as plant pathogens, but also saprobic. Species are endomycorrhizal with a wide range of plant roots ; Decay leaf, fruit, seed, roots, soil (Põlme et al. 2020) | Cannon PF, Kirk PM. 2007. Fungal Families of the World. CAB International, Cambridge (ISBN: 978-0851998275); Tedersoo L, et al. 2014. Science 346:e1256688 ((https://doi.org/10.1126/science.1256688)); Põlme S, et al. 2020. Fungal Diversity 105:1-16 ((https://doi.org/10.1007/s13225-020-00466-2)) |
| Otu28344 | 1 | k__Fungi(100);p__Basidiomycota(100);c__Agaricomycetes(100);o__Cantharellales(100);f__Ceratobasidiaceae(100);g__Ceratobasidium(100);g__Ceratobasidium_unclassified(100); | Ceratobasidium | 13 | Pathotroph-Saprotroph-Symbiotroph | Ectomycorrhizal-Endomycorrhizal-Orchid Mycorrhizal-\|Plant Pathogen\|-Plant Saprotroph-Undefined Saprotroph | Corticioid-Microfungus | NULL | Possible | especially of Orchidaceae but also even bryophytes (Cannon & Kirk 2007); As litter_saprotroph (Põlme et al. 2020); Best known as plant pathogens, but also saprobic. Species are endomycorrhizal with a wide range of plant roots ; Decay leaf, fruit, seed, roots, soil (Põlme et al. 2020) | Cannon PF, Kirk PM. 2007. Fungal Families of the World. CAB International, Cambridge (ISBN: 978-0851998275); Tedersoo L, et al. 2014. Science 346:e1256688 ((https://doi.org/10.1126/science.1256688)); Põlme S, et al. 2020. Fungal Diversity 105:1-16 ((https://doi.org/10.1007/s13225-020-00466-2)) |
| Otu28345 | 1 | k__Fungi(100);p__Basidiomycota(100);c__Microbotryomycetes(100);o__Sporidiobolales(100);f__Sporidiobolales_family_Incertae_sedis(100);g__Rhodotorula(100);g__Rhodotorula_unclassified(100); | Rhodotorula | 13 | Pathotroph-Saprotroph-Symbiotroph | Animal Pathogen-Animal Symbiotroph-Endophyte-Plant Pathogen-\|Undefined Saprotroph\| | Yeast | NULL | Probable | As foliar_endophyte (Põlme et al. 2020); Opportunistic human pathogen | Põlme S, et al. 2020. Fungal Diversity 105:1-16 ((https://doi.org/10.1007/s13225-020-00466-2)); https://www.cdc.gov ((https://www.cdc.gov)) |
| Otu28346 | 1 | k__Fungi(100);p__Basidiomycota(100);c__Agaricomycetes(100);o__Polyporales(100);f__Ganodermataceae(100);g__Ganoderma(100);g__Ganoderma_unclassified(100); | Ganoderma | 13 | Pathotroph-Saprotroph | \|Plant Pathogen\|-Wood Saprotroph | Polyporoid | White Rot | Highly Probable | As white_rot (Põlme et al. 2020); As wood_pathogen (Põlme et al. 2020); Decay wood (Põlme et al. 2020); Lignicolous, causing sometimes agressive white rots of timber (Cannon & Kirk 2007) | Gilbertson RL, Ryvarden L. 1987-1987. North American Polypores. Fungiflora, Oslo (ISBN: 978-0945345060); Cannon PF, Kirk PM. 2007. Fungal Families of the World. CAB International, Cambridge (ISBN: 978-0851998275); Põlme S, et al. 2020. Fungal Diversity 105:1-16 ((https://doi.org/10.1007/s13225-020-00466-2)) |
| Otu28347 | 1 | k__Fungi(100);p__Basidiomycota(100);c__Tremellomycetes(100);o__Tremellales(100);f__Tremellales_family_Incertae_sedis(100);g__Cryptococcus(100);s__Cryptococcus_magnus(100); | Cryptococcus | 13 | Pathotroph-Saprotroph-Symbiotroph | Animal Pathogen-Endophyte-Epiphyte-\|Undefined Saprotroph\| | Dimorphic Yeast-Yeast | NULL | Possible | As foliar_endophyte (Põlme et al. 2020); Decay leaf, fruit, seed, soil, dung, animal material (Põlme et al. 2020) | Andrews JH, et al. 1983. Phytopathology 73:228-234 ((https://doi.org/10.1094/Phyto-73-228)); Kurtzman CP, et al. (eds.) 2011. The Yeasts, a Taxonomic Study. Fifth Edition. Vols 1-3. Elsevier, San Diego (ISBN: 9780444521491); Põlme S, et al. 2020. Fungal Diversity 105:1-16 ((https://doi.org/10.1007/s13225-020-00466-2)) |
| Otu28348 | 1 | k__Fungi(100);p__Basidiomycota(100);c__Microbotryomycetes(100);o__Sporidiobolales(100);f__Sporidiobolales_family_Incertae_sedis(100);g__Rhodotorula(100);s__Rhodotorula_taiwanensis(100); | Rhodotorula | 13 | Pathotroph-Saprotroph-Symbiotroph | Animal Pathogen-Animal Symbiotroph-Endophyte-Plant Pathogen-\|Undefined Saprotroph\| | Yeast | NULL | Probable | As foliar_endophyte (Põlme et al. 2020); Opportunistic human pathogen | Põlme S, et al. 2020. Fungal Diversity 105:1-16 ((https://doi.org/10.1007/s13225-020-00466-2)); https://www.cdc.gov ((https://www.cdc.gov)) |
| Otu28349 | 1 | k__Fungi(100);p__Basidiomycota(100);c__Agaricomycetes(100);o__Agaricales(100);f__Agaricaceae(100);g__Cystolepiota(100);g__Cystolepiota_unclassified(100); | Cystolepiota | 13 | Saprotroph | \|Undefined Saprotroph\| | Agaricoid | NULL | Probable | As soil_saprotroph (Põlme et al. 2020); Decay soil (Põlme et al. 2020) | Tedersoo L, et al. 2014. Science 346:e1256688 ((https://doi.org/10.1126/science.1256688)); Põlme S, et al. 2020. Fungal Diversity 105:1-16 ((https://doi.org/10.1007/s13225-020-00466-2)) |
| Otu28350 | 1 | k__Fungi(100);p__Basidiomycota(100);c__Agaricomycetes(100);o__Agaricales(100);f__Cortinariaceae(100);g__Cortinarius(100);g__Cortinarius_unclassified(100); | Cortinarius | 13 | Symbiotroph | \|Ectomycorrhizal\| | Agaricoid | NULL | Highly Probable | NULL | Rinaldi AC, et al. 2008. Fungal Diversity 33:1-45 ((http://www.fungaldiversity.org/fdp/sfdp/33-1.pdf)); Tedersoo L, et al. 2010. Mycorrhiza 20:217-263 ((https://doi.org/10.1007/s00572-009-0274-x)); Põlme S, et al. 2020. Fungal Diversity 105:1-16 ((https://doi.org/10.1007/s13225-020-00466-2)); http://mycorrhizas.info/ecmf.html ((http://mycorrhizas.info/ecmf.html)) |
| Otu28351 | 1 | k__Fungi(100);p__Basidiomycota(100);c__Agaricomycetes(100);o__Cantharellales(100);f__Botryobasidiaceae(100);g__Haplotrichum(100);s__Haplotrichum_conspersum(100); | Haplotrichum | 13 | Saprotroph | \|Wood Saprotroph\| | Corticioid | White Rot | Probable | As white_rot (Põlme et al. 2020); Decay wood (Põlme et al. 2020) | Põlme S, et al. 2020. Fungal Diversity 105:1-16 ((https://doi.org/10.1007/s13225-020-00466-2)) |
| Otu28352 | 1 | k__Fungi(100);p__Basidiomycota(100);c__Agaricomycetes(100);o__Polyporales(100);f__Meripilaceae(100);g__Rigidoporus(100);s__Rigidoporus_corticola(100); | Rigidoporus | 13 | Saprotroph | \|Wood Saprotroph\| | Polyporoid | White Rot | Highly Probable | As white_rot (Põlme et al. 2020); Decay wood (Põlme et al. 2020) | Gilbertson RL, Ryvarden L. 1987-1987. North American Polypores. Fungiflora, Oslo (ISBN: 978-0945345060); Põlme S, et al. 2020. Fungal Diversity 105:1-16 ((https://doi.org/10.1007/s13225-020-00466-2)) |
| Otu28355 | 1 | k__Fungi(100);p__Basidiomycota(100);c__Agaricomycetes(100);o__Cantharellales(100);f__Botryobasidiaceae(100);g__Haplotrichum(100);s__Haplotrichum_conspersum(100); | Haplotrichum | 13 | Saprotroph | \|Wood Saprotroph\| | Corticioid | White Rot | Probable | As white_rot (Põlme et al. 2020); Decay wood (Põlme et al. 2020) | Põlme S, et al. 2020. Fungal Diversity 105:1-16 ((https://doi.org/10.1007/s13225-020-00466-2)) |
| Otu28356 | 1 | k__Fungi(100);p__Basidiomycota(100);c__Agaricomycetes(100);o__Polyporales(100);f__Phanerochaetaceae(100);g__Phanerochaete(100);g__Phanerochaete_unclassified(100); | Phanerochaete | 13 | Saprotroph | \|Wood Saprotroph\| | Corticioid | White Rot | Highly Probable | As white_rot (Põlme et al. 2020); Decay wood (Põlme et al. 2020) | Glenn JK, et al. 1983. Biochemical and Biophysical Research Communications 114:1077-1083 ((https://doi.org/10.1016/0006-291x((83))90672-1)); Põlme S, et al. 2020. Fungal Diversity 105:1-16 ((https://doi.org/10.1007/s13225-020-00466-2)) |
| Otu28358 | 1 | k__Fungi(100);p__Basidiomycota(100);c__Microbotryomycetes(100);o__Sporidiobolales(100);f__Sporidiobolales_family_Incertae_sedis(100);g__Rhodotorula(100);s__Rhodotorula_nothofagi(100); | Rhodotorula | 13 | Pathotroph-Saprotroph-Symbiotroph | Animal Pathogen-Animal Symbiotroph-Endophyte-Plant Pathogen-\|Undefined Saprotroph\| | Yeast | NULL | Probable | As foliar_endophyte (Põlme et al. 2020); Opportunistic human pathogen | Põlme S, et al. 2020. Fungal Diversity 105:1-16 ((https://doi.org/10.1007/s13225-020-00466-2)); https://www.cdc.gov ((https://www.cdc.gov)) |
| Otu28359 | 1 | k__Fungi(100);p__Basidiomycota(100);c__Microbotryomycetes(100);o__Sporidiobolales(100);f__Sporidiobolales_family_Incertae_sedis(100);g__Rhodotorula(100);s__Rhodotorula_nothofagi(100); | Rhodotorula | 13 | Pathotroph-Saprotroph-Symbiotroph | Animal Pathogen-Animal Symbiotroph-Endophyte-Plant Pathogen-\|Undefined Saprotroph\| | Yeast | NULL | Probable | As foliar_endophyte (Põlme et al. 2020); Opportunistic human pathogen | Põlme S, et al. 2020. Fungal Diversity 105:1-16 ((https://doi.org/10.1007/s13225-020-00466-2)); https://www.cdc.gov ((https://www.cdc.gov)) |
| Otu28360 | 1 | k__Fungi(100);p__Basidiomycota(100);c__Microbotryomycetes(100);o__Sporidiobolales(100);f__Sporidiobolales_family_Incertae_sedis(100);g__Rhodotorula(100);s__Rhodotorula_nothofagi(100); | Rhodotorula | 13 | Pathotroph-Saprotroph-Symbiotroph | Animal Pathogen-Animal Symbiotroph-Endophyte-Plant Pathogen-\|Undefined Saprotroph\| | Yeast | NULL | Probable | As foliar_endophyte (Põlme et al. 2020); Opportunistic human pathogen | Põlme S, et al. 2020. Fungal Diversity 105:1-16 ((https://doi.org/10.1007/s13225-020-00466-2)); https://www.cdc.gov ((https://www.cdc.gov)) |
| Otu28361 | 1 | k__Fungi(100);p__Basidiomycota(100);c__Microbotryomycetes(100);o__Sporidiobolales(100);f__Sporidiobolales_family_Incertae_sedis(100);g__Rhodotorula(100);s__Rhodotorula_nothofagi(100); | Rhodotorula | 13 | Pathotroph-Saprotroph-Symbiotroph | Animal Pathogen-Animal Symbiotroph-Endophyte-Plant Pathogen-\|Undefined Saprotroph\| | Yeast | NULL | Probable | As foliar_endophyte (Põlme et al. 2020); Opportunistic human pathogen | Põlme S, et al. 2020. Fungal Diversity 105:1-16 ((https://doi.org/10.1007/s13225-020-00466-2)); https://www.cdc.gov ((https://www.cdc.gov)) |
| Otu28362 | 1 | k__Fungi(100);p__Basidiomycota(100);c__Microbotryomycetes(100);o__Microbotryomycetes_order_Incertae_sedis(100);f__Microbotryomycetes_family_Incertae_sedis(100);g__Curvibasidium(100);s__Curvibasidium_cygneicollum(100); | Curvibasidium | 13 | Saprotroph-Symbiotroph | Epiphyte-Plant Saprotroph-Undefined Saprotroph-\|Wood Saprotroph\| | Dimorphic-Microfungus | NULL | Possible | As litter_saprotroph (Põlme et al. 2020); Decay wood, leaf, fruit, seed, soil (Põlme et al. 2020) | Sampaio JP, et al. 2004. International Journal of Systematic and Evolutionary Microbiology 54:1401-1407 ((https://doi.org/10.1099/ijs.0.03037-0)); Põlme S, et al. 2020. Fungal Diversity 105:1-16 ((https://doi.org/10.1007/s13225-020-00466-2)) |
| Otu28363 | 1 | k__Fungi(100);p__Basidiomycota(100);c__Microbotryomycetes(100);o__Microbotryomycetes_order_Incertae_sedis(100);f__Microbotryomycetes_family_Incertae_sedis(100);g__Curvibasidium(100);s__Curvibasidium_cygneicollum(100); | Curvibasidium | 13 | Saprotroph-Symbiotroph | Epiphyte-Plant Saprotroph-Undefined Saprotroph-\|Wood Saprotroph\| | Dimorphic-Microfungus | NULL | Possible | As litter_saprotroph (Põlme et al. 2020); Decay wood, leaf, fruit, seed, soil (Põlme et al. 2020) | Sampaio JP, et al. 2004. International Journal of Systematic and Evolutionary Microbiology 54:1401-1407 ((https://doi.org/10.1099/ijs.0.03037-0)); Põlme S, et al. 2020. Fungal Diversity 105:1-16 ((https://doi.org/10.1007/s13225-020-00466-2)) |
| Otu28364 | 1 | k__Fungi(100);p__Basidiomycota(100);c__Microbotryomycetes(100);o__Sporidiobolales(100);f__Sporidiobolales_family_Incertae_sedis(100);g__Sporobolomyces(100);s__Sporobolomyces_griseoflavus(100); | Sporobolomyces | 13 | Pathotroph-Saprotroph | \|Fungal Parasite\|-Plant Saprotroph | Yeast | NULL | Probable | As litter_saprotroph (Põlme et al. 2020); As mycoparasite (Põlme et al. 2020); Decay fungal material (Põlme et al. 2020) | Tedersoo L, et al. 2014. Science 346:e1256688 ((https://doi.org/10.1126/science.1256688)); Purahong W, et al. 2016. Molecular Ecology 25:4059-4074 ((https://doi.org/10.1111/mec.13739)); Põlme S, et al. 2020. Fungal Diversity 105:1-16 ((https://doi.org/10.1007/s13225-020-00466-2)) |
| Otu28365 | 1 | k__Fungi(100);p__Basidiomycota(100);c__Microbotryomycetes(100);o__Sporidiobolales(100);f__Sporidiobolales_family_Incertae_sedis(100);g__Sporobolomyces(100);s__Sporobolomyces_griseoflavus(100); | Sporobolomyces | 13 | Pathotroph-Saprotroph | \|Fungal Parasite\|-Plant Saprotroph | Yeast | NULL | Probable | As litter_saprotroph (Põlme et al. 2020); As mycoparasite (Põlme et al. 2020); Decay fungal material (Põlme et al. 2020) | Tedersoo L, et al. 2014. Science 346:e1256688 ((https://doi.org/10.1126/science.1256688)); Purahong W, et al. 2016. Molecular Ecology 25:4059-4074 ((https://doi.org/10.1111/mec.13739)); Põlme S, et al. 2020. Fungal Diversity 105:1-16 ((https://doi.org/10.1007/s13225-020-00466-2)) |
| Otu28366 | 1 | k__Fungi(100);p__Basidiomycota(100);c__Microbotryomycetes(100);o__Sporidiobolales(100);f__Sporidiobolales_family_Incertae_sedis(100);g__Sporobolomyces(100);s__Sporobolomyces_griseoflavus(100); | Sporobolomyces | 13 | Pathotroph-Saprotroph | \|Fungal Parasite\|-Plant Saprotroph | Yeast | NULL | Probable | As litter_saprotroph (Põlme et al. 2020); As mycoparasite (Põlme et al. 2020); Decay fungal material (Põlme et al. 2020) | Tedersoo L, et al. 2014. Science 346:e1256688 ((https://doi.org/10.1126/science.1256688)); Purahong W, et al. 2016. Molecular Ecology 25:4059-4074 ((https://doi.org/10.1111/mec.13739)); Põlme S, et al. 2020. Fungal Diversity 105:1-16 ((https://doi.org/10.1007/s13225-020-00466-2)) |
| Otu28367 | 1 | k__Fungi(100);p__Basidiomycota(100);c__Microbotryomycetes(100);o__Microbotryomycetes_order_Incertae_sedis(100);f__Microbotryomycetes_family_Incertae_sedis(100);g__Curvibasidium(100);s__Curvibasidium_cygneicollum(100); | Curvibasidium | 13 | Saprotroph-Symbiotroph | Epiphyte-Plant Saprotroph-Undefined Saprotroph-\|Wood Saprotroph\| | Dimorphic-Microfungus | NULL | Possible | As litter_saprotroph (Põlme et al. 2020); Decay wood, leaf, fruit, seed, soil (Põlme et al. 2020) | Sampaio JP, et al. 2004. International Journal of Systematic and Evolutionary Microbiology 54:1401-1407 ((https://doi.org/10.1099/ijs.0.03037-0)); Põlme S, et al. 2020. Fungal Diversity 105:1-16 ((https://doi.org/10.1007/s13225-020-00466-2)) |
| Otu28368 | 1 | k__Fungi(100);p__Basidiomycota(100);c__Microbotryomycetes(100);o__Microbotryomycetes_order_Incertae_sedis(100);f__Microbotryomycetes_family_Incertae_sedis(100);g__Curvibasidium(100);s__Curvibasidium_cygneicollum(100); | Curvibasidium | 13 | Saprotroph-Symbiotroph | Epiphyte-Plant Saprotroph-Undefined Saprotroph-\|Wood Saprotroph\| | Dimorphic-Microfungus | NULL | Possible | As litter_saprotroph (Põlme et al. 2020); Decay wood, leaf, fruit, seed, soil (Põlme et al. 2020) | Sampaio JP, et al. 2004. International Journal of Systematic and Evolutionary Microbiology 54:1401-1407 ((https://doi.org/10.1099/ijs.0.03037-0)); Põlme S, et al. 2020. Fungal Diversity 105:1-16 ((https://doi.org/10.1007/s13225-020-00466-2)) |
| Otu28369 | 1 | k__Fungi(100);p__Basidiomycota(100);c__Agaricomycetes(100);o__Trechisporales(100);f__Hydnodontaceae(100);g__Trechispora(100);g__Trechispora_unclassified(100); | Trechispora | 13 | Saprotroph | \|Wood Saprotroph\| | Corticioid | White Rot | Highly Probable | As white_rot (Põlme et al. 2020); Decay wood (Põlme et al. 2020) | Gilbertson RL, Ryvarden L. 1987-1987. North American Polypores. Fungiflora, Oslo (ISBN: 978-0945345060); Rinaldi AC, et al. 2008. Fungal Diversity 33:1-45 ((http://www.fungaldiversity.org/fdp/sfdp/33-1.pdf)) (as Uncertain); Tedersoo L, et al. 2010. Mycorrhiza 20:217-263 ((https://doi.org/10.1007/s00572-009-0274-x)) (as not ECM).; Põlme S, et al. 2020. Fungal Diversity 105:1-16 ((https://doi.org/10.1007/s13225-020-00466-2)) |
| Otu28370 | 1 | k__Fungi(100);p__Basidiomycota(100);c__Agaricomycetes(100);o__Corticiales(100);f__Corticiaceae(100);f__Corticiaceae_unclassified(100);f__Corticiaceae_unclassified(100); | Corticiaceae | 9 | Pathotroph-Saprotroph | Lichen Parasite-Plant Pathogen-Wood Saprotroph | Corticioid-Microfungus | NULL | Possible | Saprobes or pathogens on a wide range of mainly woody substrata; a few species are lichenicolous and some are grass pathogens (Cannon & Kirk 2007) | Cannon PF, Kirk PM. 2007. Fungal Families of the World. CAB International, Cambridge (ISBN: 978-0851998275) |
| Otu28371 | 1 | k__Fungi(100);p__Basidiomycota(100);c__Microbotryomycetes(100);o__Sporidiobolales(100);f__Sporidiobolales_family_Incertae_sedis(100);g__Sporobolomyces(100);s__Sporobolomyces_tsugae(100); | Sporobolomyces | 13 | Pathotroph-Saprotroph | \|Fungal Parasite\|-Plant Saprotroph | Yeast | NULL | Probable | As litter_saprotroph (Põlme et al. 2020); As mycoparasite (Põlme et al. 2020); Decay fungal material (Põlme et al. 2020) | Tedersoo L, et al. 2014. Science 346:e1256688 ((https://doi.org/10.1126/science.1256688)); Purahong W, et al. 2016. Molecular Ecology 25:4059-4074 ((https://doi.org/10.1111/mec.13739)); Põlme S, et al. 2020. Fungal Diversity 105:1-16 ((https://doi.org/10.1007/s13225-020-00466-2)) |
| Otu28373 | 1 | k__Fungi(100);p__Basidiomycota(100);c__Microbotryomycetes(100);o__Sporidiobolales(100);f__Sporidiobolales_family_Incertae_sedis(100);g__Sporobolomyces(100);s__Sporobolomyces_inositophilus(100); | Sporobolomyces | 13 | Pathotroph-Saprotroph | \|Fungal Parasite\|-Plant Saprotroph | Yeast | NULL | Probable | As litter_saprotroph (Põlme et al. 2020); As mycoparasite (Põlme et al. 2020); Decay fungal material (Põlme et al. 2020) | Tedersoo L, et al. 2014. Science 346:e1256688 ((https://doi.org/10.1126/science.1256688)); Purahong W, et al. 2016. Molecular Ecology 25:4059-4074 ((https://doi.org/10.1111/mec.13739)); Põlme S, et al. 2020. Fungal Diversity 105:1-16 ((https://doi.org/10.1007/s13225-020-00466-2)) |
| Otu28374 | 1 | k__Fungi(100);p__Basidiomycota(100);c__Microbotryomycetes(100);o__Microbotryomycetes_order_Incertae_sedis(100);f__Microbotryomycetes_family_Incertae_sedis(100);g__Curvibasidium(100);s__Curvibasidium_cygneicollum(100); | Curvibasidium | 13 | Saprotroph-Symbiotroph | Epiphyte-Plant Saprotroph-Undefined Saprotroph-\|Wood Saprotroph\| | Dimorphic-Microfungus | NULL | Possible | As litter_saprotroph (Põlme et al. 2020); Decay wood, leaf, fruit, seed, soil (Põlme et al. 2020) | Sampaio JP, et al. 2004. International Journal of Systematic and Evolutionary Microbiology 54:1401-1407 ((https://doi.org/10.1099/ijs.0.03037-0)); Põlme S, et al. 2020. Fungal Diversity 105:1-16 ((https://doi.org/10.1007/s13225-020-00466-2)) |
| Otu28377 | 1 | k__Fungi(100);p__Basidiomycota(100);c__Agaricomycetes(100);o__Cantharellales(100);f__Ceratobasidiaceae(100);g__Ceratobasidium(100);s__Ceratobasidium_sp_AG_A(100); | Ceratobasidium | 13 | Pathotroph-Saprotroph-Symbiotroph | Ectomycorrhizal-Endomycorrhizal-Orchid Mycorrhizal-\|Plant Pathogen\|-Plant Saprotroph-Undefined Saprotroph | Corticioid-Microfungus | NULL | Possible | especially of Orchidaceae but also even bryophytes (Cannon & Kirk 2007); As litter_saprotroph (Põlme et al. 2020); Best known as plant pathogens, but also saprobic. Species are endomycorrhizal with a wide range of plant roots ; Decay leaf, fruit, seed, roots, soil (Põlme et al. 2020) | Cannon PF, Kirk PM. 2007. Fungal Families of the World. CAB International, Cambridge (ISBN: 978-0851998275); Tedersoo L, et al. 2014. Science 346:e1256688 ((https://doi.org/10.1126/science.1256688)); Põlme S, et al. 2020. Fungal Diversity 105:1-16 ((https://doi.org/10.1007/s13225-020-00466-2)) |
| Otu28379 | 1 | k__Fungi(100);p__Basidiomycota(100);c__Microbotryomycetes(100);o__Sporidiobolales(100);f__Sporidiobolales_family_Incertae_sedis(100);g__Rhodotorula(100);g__Rhodotorula_unclassified(100); | Rhodotorula | 13 | Pathotroph-Saprotroph-Symbiotroph | Animal Pathogen-Animal Symbiotroph-Endophyte-Plant Pathogen-\|Undefined Saprotroph\| | Yeast | NULL | Probable | As foliar_endophyte (Põlme et al. 2020); Opportunistic human pathogen | Põlme S, et al. 2020. Fungal Diversity 105:1-16 ((https://doi.org/10.1007/s13225-020-00466-2)); https://www.cdc.gov ((https://www.cdc.gov)) |
| Otu28380 | 1 | k__Fungi(100);p__Basidiomycota(100);c__Agaricomycetes(100);o__Polyporales(100);f__Meruliaceae(100);g__Steccherinum(100);s__Steccherinum_ochraceum(100); | Steccherinum | 13 | Saprotroph | Undefined Saprotroph-\|Wood Saprotroph\| | Corticioid | White Rot | Probable | As white_rot (Põlme et al. 2020); Decay wood (Põlme et al. 2020) | Tedersoo L, et al. 2014. Science 346:e1256688 ((https://doi.org/10.1126/science.1256688)); Põlme S, et al. 2020. Fungal Diversity 105:1-16 ((https://doi.org/10.1007/s13225-020-00466-2)) |
| Otu28381 | 1 | k__Fungi(100);p__Basidiomycota(100);c__Agaricomycetes(100);o__Polyporales(100);f__Fomitopsidaceae(100);g__Postia(100);s__Postia_stiptica(100); | Postia | 13 | Saprotroph | \|Wood Saprotroph\| | Polyporoid | Brown Rot | Highly Probable | As brown-rot (Põlme et al. 2020); Decay wood (Põlme et al. 2020) | Jordan CR, et al. 1996. Holzforschung - International Journal of the Biology, Chemistry, Physics and Technology of Wood 50:312-318 ((https://doi.org/10.1515/hfsg.1996.50.4.312)); Põlme S, et al. 2020. Fungal Diversity 105:1-16 ((https://doi.org/10.1007/s13225-020-00466-2)) |
| Otu28383 | 1 | k__Fungi(100);p__Basidiomycota(100);c__Agaricomycetes(100);o__Agaricales(100);f__Psathyrellaceae(100);f__Psathyrellaceae_unclassified(100);f__Psathyrellaceae_unclassified(100); | Psathyrellaceae | 9 | Saprotroph | Dung Saprotroph-Plant Saprotroph-Wood Saprotroph | Agaricoid | NULL | Probable | Saprobic, in grass, on dung, rotten wood and similar substrata (Cannon & Kirk 2007) | Cannon PF, Kirk PM. 2007. Fungal Families of the World. CAB International, Cambridge (ISBN: 978-0851998275) |
| Otu28384 | 1 | k__Fungi(100);p__Basidiomycota(100);c__Agaricomycetes(100);o__Trechisporales(100);f__Hydnodontaceae(100);g__Trechispora(100);g__Trechispora_unclassified(100); | Trechispora | 13 | Saprotroph | \|Wood Saprotroph\| | Corticioid | White Rot | Highly Probable | As white_rot (Põlme et al. 2020); Decay wood (Põlme et al. 2020) | Gilbertson RL, Ryvarden L. 1987-1987. North American Polypores. Fungiflora, Oslo (ISBN: 978-0945345060); Rinaldi AC, et al. 2008. Fungal Diversity 33:1-45 ((http://www.fungaldiversity.org/fdp/sfdp/33-1.pdf)) (as Uncertain); Tedersoo L, et al. 2010. Mycorrhiza 20:217-263 ((https://doi.org/10.1007/s00572-009-0274-x)) (as not ECM).; Põlme S, et al. 2020. Fungal Diversity 105:1-16 ((https://doi.org/10.1007/s13225-020-00466-2)) |
| Otu28385 | 1 | k__Fungi(100);p__Basidiomycota(100);c__Agaricomycetes(100);o__Trechisporales(100);f__Hydnodontaceae(100);g__Trechispora(100);g__Trechispora_unclassified(100); | Trechispora | 13 | Saprotroph | \|Wood Saprotroph\| | Corticioid | White Rot | Highly Probable | As white_rot (Põlme et al. 2020); Decay wood (Põlme et al. 2020) | Gilbertson RL, Ryvarden L. 1987-1987. North American Polypores. Fungiflora, Oslo (ISBN: 978-0945345060); Rinaldi AC, et al. 2008. Fungal Diversity 33:1-45 ((http://www.fungaldiversity.org/fdp/sfdp/33-1.pdf)) (as Uncertain); Tedersoo L, et al. 2010. Mycorrhiza 20:217-263 ((https://doi.org/10.1007/s00572-009-0274-x)) (as not ECM).; Põlme S, et al. 2020. Fungal Diversity 105:1-16 ((https://doi.org/10.1007/s13225-020-00466-2)) |
| Otu28386 | 1 | k__Fungi(100);p__Basidiomycota(100);c__Agaricomycetes(100);o__Polyporales(100);f__Polyporaceae(100);g__Amyloporia(100);s__Amyloporia_sinuosa(100); | Amyloporia | 13 | Saprotroph | \|Wood Saprotroph\| | Corticioid | Brown Rot | Probable | As brown-rot (Põlme et al. 2020); Decay wood (Põlme et al. 2020) | Põlme S, et al. 2020. Fungal Diversity 105:1-16 ((https://doi.org/10.1007/s13225-020-00466-2)) |
| Otu28387 | 1 | k__Fungi(100);p__Basidiomycota(100);c__Agaricomycetes(100);o__Polyporales(100);f__Phanerochaetaceae(100);g__Antrodiella(100);s__Antrodiella_faginea(100); | Antrodiella | 13 | Saprotroph | \|Wood Saprotroph\| | Corticioid-Polyporoid | White Rot | Highly Probable | As white_rot (Põlme et al. 2020); Decay wood (Põlme et al. 2020) | Gilbertson RL, Ryvarden L. 1987-1987. North American Polypores. Fungiflora, Oslo (ISBN: 978-0945345060); Põlme S, et al. 2020. Fungal Diversity 105:1-16 ((https://doi.org/10.1007/s13225-020-00466-2)) |
| Otu28388 | 1 | k__Fungi(100);p__Basidiomycota(100);c__Agaricomycetes(100);o__Agaricales(100);f__Strophariaceae(100);g__Kuehneromyces(100);s__Kuehneromyces_rostratus(100); | Kuehneromyces | 13 | Saprotroph | Undefined Saprotroph-\|Wood Saprotroph\| | Agaricoid | White Rot | Probable | As white_rot (Põlme et al. 2020); Decay wood (Põlme et al. 2020) | Tedersoo L, et al. 2014. Science 346:e1256688 ((https://doi.org/10.1126/science.1256688)); Põlme S, et al. 2020. Fungal Diversity 105:1-16 ((https://doi.org/10.1007/s13225-020-00466-2)) |
| Otu28389 | 1 | k__Fungi(100);p__Basidiomycota(100);c__Agaricomycetes(100);o__Agaricales(100);f__Psathyrellaceae(100);g__Coprinellus(100);s__Coprinellus_plagioporus(100); | Coprinellus | 13 | Saprotroph | \|Undefined Saprotroph\| | Agaricoid | NULL | Probable | As soil_saprotroph (Põlme et al. 2020); Decay soil (Põlme et al. 2020) | Tedersoo L, et al. 2014. Science 346:e1256688 ((https://doi.org/10.1126/science.1256688)); Põlme S, et al. 2020. Fungal Diversity 105:1-16 ((https://doi.org/10.1007/s13225-020-00466-2)) |
| Otu28390 | 1 | k__Fungi(100);p__Basidiomycota(100);c__Agaricomycetes(100);o__Trechisporales(100);f__Hydnodontaceae(100);g__Trechispora(100);s__Trechispora_sp(100); | Trechispora | 13 | Saprotroph | \|Wood Saprotroph\| | Corticioid | White Rot | Highly Probable | As white_rot (Põlme et al. 2020); Decay wood (Põlme et al. 2020) | Gilbertson RL, Ryvarden L. 1987-1987. North American Polypores. Fungiflora, Oslo (ISBN: 978-0945345060); Rinaldi AC, et al. 2008. Fungal Diversity 33:1-45 ((http://www.fungaldiversity.org/fdp/sfdp/33-1.pdf)) (as Uncertain); Tedersoo L, et al. 2010. Mycorrhiza 20:217-263 ((https://doi.org/10.1007/s00572-009-0274-x)) (as not ECM).; Põlme S, et al. 2020. Fungal Diversity 105:1-16 ((https://doi.org/10.1007/s13225-020-00466-2)) |
| Otu28391 | 1 | k__Fungi(100);p__Basidiomycota(100);c__Agaricomycetes(100);o__Agaricales(100);f__Psathyrellaceae(100);g__Coprinopsis(100);s__Coprinopsis_pseudofriesii(100); | Coprinopsis | 13 | Saprotroph | \|Undefined Saprotroph\| | Agaricoid | NULL | Probable | As soil_saprotroph (Põlme et al. 2020); Decay soil (Põlme et al. 2020) | Tedersoo L, et al. 2014. Science 346:e1256688 ((https://doi.org/10.1126/science.1256688)); Põlme S, et al. 2020. Fungal Diversity 105:1-16 ((https://doi.org/10.1007/s13225-020-00466-2)) |
| Otu28392 | 1 | k__Fungi(100);p__Basidiomycota(100);c__Agaricomycetes(100);o__Polyporales(100);f__Meruliaceae(100);g__Steccherinum(100);s__Steccherinum_sp(100); | Steccherinum | 13 | Saprotroph | Undefined Saprotroph-\|Wood Saprotroph\| | Corticioid | White Rot | Probable | As white_rot (Põlme et al. 2020); Decay wood (Põlme et al. 2020) | Tedersoo L, et al. 2014. Science 346:e1256688 ((https://doi.org/10.1126/science.1256688)); Põlme S, et al. 2020. Fungal Diversity 105:1-16 ((https://doi.org/10.1007/s13225-020-00466-2)) |
| Otu28393 | 1 | k__Fungi(100);p__Basidiomycota(100);c__Agaricomycetes(100);o__Polyporales(100);f__Phanerochaetaceae(100);g__Phanerochaete(100);s__Phanerochaete_rimosa(100); | Phanerochaete | 13 | Saprotroph | \|Wood Saprotroph\| | Corticioid | White Rot | Highly Probable | As white_rot (Põlme et al. 2020); Decay wood (Põlme et al. 2020) | Glenn JK, et al. 1983. Biochemical and Biophysical Research Communications 114:1077-1083 ((https://doi.org/10.1016/0006-291x((83))90672-1)); Põlme S, et al. 2020. Fungal Diversity 105:1-16 ((https://doi.org/10.1007/s13225-020-00466-2)) |
| Otu28394 | 1 | k__Fungi(100);p__Basidiomycota(100);c__Agaricomycetes(100);o__Polyporales(100);f__Meruliaceae(100);g__Scopuloides(100);s__Scopuloides_hydnoides(100); | Scopuloides | 13 | Saprotroph | Undefined Saprotroph-\|Wood Saprotroph\| | Corticioid | White Rot | Probable | As white_rot (Põlme et al. 2020); Decay wood (Põlme et al. 2020) | Tedersoo L, et al. 2014. Science 346:e1256688 ((https://doi.org/10.1126/science.1256688)); Põlme S, et al. 2020. Fungal Diversity 105:1-16 ((https://doi.org/10.1007/s13225-020-00466-2)) |
| Otu28396 | 1 | k__Fungi(100);p__Basidiomycota(100);c__Agaricomycetes(100);o__Agaricomycetes_order_Incertae_sedis(100);f__Agaricomycetes_family_Incertae_sedis(100);g__Resinicium(100);s__Resinicium_sp(100); | Resinicium | 13 | Saprotroph | Undefined Saprotroph-\|Wood Saprotroph\| | Corticioid | White Rot | Probable | As white_rot (Põlme et al. 2020); Decay wood (Põlme et al. 2020) | Tedersoo L, et al. 2014. Science 346:e1256688 ((https://doi.org/10.1126/science.1256688)); Põlme S, et al. 2020. Fungal Diversity 105:1-16 ((https://doi.org/10.1007/s13225-020-00466-2)) |
| Otu28398 | 1 | k__Fungi(100);p__Basidiomycota(100);c__Agaricomycetes(100);o__Polyporales(100);f__Meruliaceae(100);g__Phlebia(100);s__Phlebia_sp_DABAC9(100); | Phlebia | 13 | Saprotroph | \|Wood Saprotroph\| | Corticioid | White Rot | Highly Probable | As white_rot (Põlme et al. 2020); Decay wood (Põlme et al. 2020) | Kirk PM et al. 2008. Dictionary of the Fungi. Tenth Edition. CAB International, Wallingford (ISBN: 978-0851998268); Põlme S, et al. 2020. Fungal Diversity 105:1-16 ((https://doi.org/10.1007/s13225-020-00466-2)) |
| Otu28399 | 1 | k__Fungi(100);p__Basidiomycota(100);c__Agaricomycetes(100);o__Agaricales(100);f__Strophariaceae(100);g__Kuehneromyces(100);s__Kuehneromyces_rostratus(100); | Kuehneromyces | 13 | Saprotroph | Undefined Saprotroph-\|Wood Saprotroph\| | Agaricoid | White Rot | Probable | As white_rot (Põlme et al. 2020); Decay wood (Põlme et al. 2020) | Tedersoo L, et al. 2014. Science 346:e1256688 ((https://doi.org/10.1126/science.1256688)); Põlme S, et al. 2020. Fungal Diversity 105:1-16 ((https://doi.org/10.1007/s13225-020-00466-2)) |
| Otu28400 | 1 | k__Fungi(100);p__Basidiomycota(100);c__Agaricomycetes(100);o__Polyporales(100);f__Meruliaceae(100);g__Steccherinum(100);s__Steccherinum_ochraceum(100); | Steccherinum | 13 | Saprotroph | Undefined Saprotroph-\|Wood Saprotroph\| | Corticioid | White Rot | Probable | As white_rot (Põlme et al. 2020); Decay wood (Põlme et al. 2020) | Tedersoo L, et al. 2014. Science 346:e1256688 ((https://doi.org/10.1126/science.1256688)); Põlme S, et al. 2020. Fungal Diversity 105:1-16 ((https://doi.org/10.1007/s13225-020-00466-2)) |
| Otu28401 | 1 | k__Fungi(100);p__Basidiomycota(100);c__Agaricomycetes(100);o__Agaricales(100);f__Psathyrellaceae(100);g__Coprinellus(100);g__Coprinellus_unclassified(100); | Coprinellus | 13 | Saprotroph | \|Undefined Saprotroph\| | Agaricoid | NULL | Probable | As soil_saprotroph (Põlme et al. 2020); Decay soil (Põlme et al. 2020) | Tedersoo L, et al. 2014. Science 346:e1256688 ((https://doi.org/10.1126/science.1256688)); Põlme S, et al. 2020. Fungal Diversity 105:1-16 ((https://doi.org/10.1007/s13225-020-00466-2)) |
| Otu28402 | 1 | k__Fungi(100);p__Basidiomycota(100);c__Agaricomycetes(100);o__Agaricales(100);f__Psathyrellaceae(100);g__Coprinopsis(100);s__Coprinopsis_lagopus(100); | Coprinopsis | 13 | Saprotroph | \|Undefined Saprotroph\| | Agaricoid | NULL | Probable | As soil_saprotroph (Põlme et al. 2020); Decay soil (Põlme et al. 2020) | Tedersoo L, et al. 2014. Science 346:e1256688 ((https://doi.org/10.1126/science.1256688)); Põlme S, et al. 2020. Fungal Diversity 105:1-16 ((https://doi.org/10.1007/s13225-020-00466-2)) |
| Otu28404 | 1 | k__Fungi(100);p__Basidiomycota(100);c__Agaricomycetes(100);o__Trechisporales(100);f__Hydnodontaceae(100);g__Trechispora(100);s__Trechispora_sp(100); | Trechispora | 13 | Saprotroph | \|Wood Saprotroph\| | Corticioid | White Rot | Highly Probable | As white_rot (Põlme et al. 2020); Decay wood (Põlme et al. 2020) | Gilbertson RL, Ryvarden L. 1987-1987. North American Polypores. Fungiflora, Oslo (ISBN: 978-0945345060); Rinaldi AC, et al. 2008. Fungal Diversity 33:1-45 ((http://www.fungaldiversity.org/fdp/sfdp/33-1.pdf)) (as Uncertain); Tedersoo L, et al. 2010. Mycorrhiza 20:217-263 ((https://doi.org/10.1007/s00572-009-0274-x)) (as not ECM).; Põlme S, et al. 2020. Fungal Diversity 105:1-16 ((https://doi.org/10.1007/s13225-020-00466-2)) |
| Otu28405 | 1 | k__Fungi(100);p__Basidiomycota(100);c__Agaricomycetes(100);o__Trechisporales(100);f__Hydnodontaceae(100);g__Trechispora(100);s__Trechispora_sp(100); | Trechispora | 13 | Saprotroph | \|Wood Saprotroph\| | Corticioid | White Rot | Highly Probable | As white_rot (Põlme et al. 2020); Decay wood (Põlme et al. 2020) | Gilbertson RL, Ryvarden L. 1987-1987. North American Polypores. Fungiflora, Oslo (ISBN: 978-0945345060); Rinaldi AC, et al. 2008. Fungal Diversity 33:1-45 ((http://www.fungaldiversity.org/fdp/sfdp/33-1.pdf)) (as Uncertain); Tedersoo L, et al. 2010. Mycorrhiza 20:217-263 ((https://doi.org/10.1007/s00572-009-0274-x)) (as not ECM).; Põlme S, et al. 2020. Fungal Diversity 105:1-16 ((https://doi.org/10.1007/s13225-020-00466-2)) |
| Otu28406 | 1 | k__Fungi(100);p__Basidiomycota(100);c__Agaricomycetes(100);o__Trechisporales(100);f__Hydnodontaceae(100);g__Trechispora(100);g__Trechispora_unclassified(100); | Trechispora | 13 | Saprotroph | \|Wood Saprotroph\| | Corticioid | White Rot | Highly Probable | As white_rot (Põlme et al. 2020); Decay wood (Põlme et al. 2020) | Gilbertson RL, Ryvarden L. 1987-1987. North American Polypores. Fungiflora, Oslo (ISBN: 978-0945345060); Rinaldi AC, et al. 2008. Fungal Diversity 33:1-45 ((http://www.fungaldiversity.org/fdp/sfdp/33-1.pdf)) (as Uncertain); Tedersoo L, et al. 2010. Mycorrhiza 20:217-263 ((https://doi.org/10.1007/s00572-009-0274-x)) (as not ECM).; Põlme S, et al. 2020. Fungal Diversity 105:1-16 ((https://doi.org/10.1007/s13225-020-00466-2)) |
| Otu28407 | 1 | k__Fungi(100);p__Basidiomycota(100);c__Agaricomycetes(100);o__Trechisporales(100);f__Hydnodontaceae(100);g__Trechispora(100);s__Trechispora_sp_1_MPM_2013(100); | Trechispora | 13 | Saprotroph | \|Wood Saprotroph\| | Corticioid | White Rot | Highly Probable | As white_rot (Põlme et al. 2020); Decay wood (Põlme et al. 2020) | Gilbertson RL, Ryvarden L. 1987-1987. North American Polypores. Fungiflora, Oslo (ISBN: 978-0945345060); Rinaldi AC, et al. 2008. Fungal Diversity 33:1-45 ((http://www.fungaldiversity.org/fdp/sfdp/33-1.pdf)) (as Uncertain); Tedersoo L, et al. 2010. Mycorrhiza 20:217-263 ((https://doi.org/10.1007/s00572-009-0274-x)) (as not ECM).; Põlme S, et al. 2020. Fungal Diversity 105:1-16 ((https://doi.org/10.1007/s13225-020-00466-2)) |
| Otu28408 | 1 | k__Fungi(100);p__Basidiomycota(100);c__Agaricomycetes(100);o__Trechisporales(100);f__Hydnodontaceae(100);g__Trechispora(100);s__Trechispora_sp_1_MPM_2013(100); | Trechispora | 13 | Saprotroph | \|Wood Saprotroph\| | Corticioid | White Rot | Highly Probable | As white_rot (Põlme et al. 2020); Decay wood (Põlme et al. 2020) | Gilbertson RL, Ryvarden L. 1987-1987. North American Polypores. Fungiflora, Oslo (ISBN: 978-0945345060); Rinaldi AC, et al. 2008. Fungal Diversity 33:1-45 ((http://www.fungaldiversity.org/fdp/sfdp/33-1.pdf)) (as Uncertain); Tedersoo L, et al. 2010. Mycorrhiza 20:217-263 ((https://doi.org/10.1007/s00572-009-0274-x)) (as not ECM).; Põlme S, et al. 2020. Fungal Diversity 105:1-16 ((https://doi.org/10.1007/s13225-020-00466-2)) |
| Otu28409 | 1 | k__Fungi(100);p__Basidiomycota(100);c__Agaricomycetes(100);o__Atheliales(100);f__Atheliaceae(100);g__Piloderma(100);s__Piloderma_olivaceum(100); | Piloderma | 13 | Symbiotroph | \|Ectomycorrhizal\| | Corticioid | NULL | Highly Probable | NULL | Rinaldi AC, et al. 2008. Fungal Diversity 33:1-45 ((http://www.fungaldiversity.org/fdp/sfdp/33-1.pdf)); Tedersoo L, et al. 2010. Mycorrhiza 20:217-263 ((https://doi.org/10.1007/s00572-009-0274-x)); Põlme S, et al. 2020. Fungal Diversity 105:1-16 ((https://doi.org/10.1007/s13225-020-00466-2)); http://mycorrhizas.info/ecmf.html ((http://mycorrhizas.info/ecmf.html)) |
| Otu28410 | 1 | k__Fungi(100);p__Basidiomycota(100);c__Agaricomycetes(100);o__Agaricales(100);f__Psathyrellaceae(100);g__Coprinopsis(100);s__Coprinopsis_calospora(100); | Coprinopsis | 13 | Saprotroph | \|Undefined Saprotroph\| | Agaricoid | NULL | Probable | As soil_saprotroph (Põlme et al. 2020); Decay soil (Põlme et al. 2020) | Tedersoo L, et al. 2014. Science 346:e1256688 ((https://doi.org/10.1126/science.1256688)); Põlme S, et al. 2020. Fungal Diversity 105:1-16 ((https://doi.org/10.1007/s13225-020-00466-2)) |
| Otu28412 | 1 | k__Fungi(100);p__Basidiomycota(100);c__Agaricomycetes(100);o__Agaricales(100);f__Psathyrellaceae(100);g__Coprinellus(100);s__Coprinellus_micaceus(100); | Coprinellus | 13 | Saprotroph | \|Undefined Saprotroph\| | Agaricoid | NULL | Probable | As soil_saprotroph (Põlme et al. 2020); Decay soil (Põlme et al. 2020) | Tedersoo L, et al. 2014. Science 346:e1256688 ((https://doi.org/10.1126/science.1256688)); Põlme S, et al. 2020. Fungal Diversity 105:1-16 ((https://doi.org/10.1007/s13225-020-00466-2)) |
| Otu28413 | 1 | k__Fungi(100);p__Basidiomycota(100);c__Agaricomycetes(100);o__Cantharellales(100);f__Botryobasidiaceae(100);g__unclassified_Botryobasidiaceae(100);s__Botryobasidiaceae_sp(100); | Botryobasidiaceae | 9 | Saprotroph | Undefined Saprotroph-Wood Saprotroph | Corticioid | NULL | Probable | Saprobic, typically on rotten wood (Cannon & Kirk 2007) | Cannon PF, Kirk PM. 2007. Fungal Families of the World. CAB International, Cambridge (ISBN: 978-0851998275) |
| Otu28414 | 1 | k__Fungi(100);p__Basidiomycota(100);c__Agaricomycetes(100);o__Polyporales(100);f__Phanerochaetaceae(100);g__Ceriporia(100);s__Ceriporia_alachuana(100); | Ceriporia | 13 | Saprotroph | \|Wood Saprotroph\| | Corticioid | White Rot | Highly Probable | As white_rot (Põlme et al. 2020); Decay wood (Põlme et al. 2020) | Tedersoo L, et al. 2014. Science 346:e1256688 ((https://doi.org/10.1126/science.1256688)); Põlme S, et al. 2020. Fungal Diversity 105:1-16 ((https://doi.org/10.1007/s13225-020-00466-2)) |
| Otu28416 | 1 | k__Fungi(100);p__Basidiomycota(100);c__Tremellomycetes(100);o__Filobasidiales(100);f__Filobasidiaceae(100);g__Cryptococcus(100);s__Cryptococcus_wieringae(100); | Cryptococcus | 13 | Pathotroph-Saprotroph-Symbiotroph | Animal Pathogen-Endophyte-Epiphyte-\|Undefined Saprotroph\| | Dimorphic Yeast-Yeast | NULL | Possible | As foliar_endophyte (Põlme et al. 2020); Decay leaf, fruit, seed, soil, dung, animal material (Põlme et al. 2020) | Andrews JH, et al. 1983. Phytopathology 73:228-234 ((https://doi.org/10.1094/Phyto-73-228)); Kurtzman CP, et al. (eds.) 2011. The Yeasts, a Taxonomic Study. Fifth Edition. Vols 1-3. Elsevier, San Diego (ISBN: 9780444521491); Põlme S, et al. 2020. Fungal Diversity 105:1-16 ((https://doi.org/10.1007/s13225-020-00466-2)) |
| Otu28417 | 1 | k__Fungi(100);p__Basidiomycota(100);c__Tremellomycetes(100);o__Filobasidiales(100);f__Filobasidiaceae(100);g__Cryptococcus(100);s__Cryptococcus_friedmannii(100); | Cryptococcus | 13 | Pathotroph-Saprotroph-Symbiotroph | Animal Pathogen-Endophyte-Epiphyte-\|Undefined Saprotroph\| | Dimorphic Yeast-Yeast | NULL | Possible | As foliar_endophyte (Põlme et al. 2020); Decay leaf, fruit, seed, soil, dung, animal material (Põlme et al. 2020) | Andrews JH, et al. 1983. Phytopathology 73:228-234 ((https://doi.org/10.1094/Phyto-73-228)); Kurtzman CP, et al. (eds.) 2011. The Yeasts, a Taxonomic Study. Fifth Edition. Vols 1-3. Elsevier, San Diego (ISBN: 9780444521491); Põlme S, et al. 2020. Fungal Diversity 105:1-16 ((https://doi.org/10.1007/s13225-020-00466-2)) |
| Otu28418 | 1 | k__Fungi(100);p__Basidiomycota(100);c__Agaricomycetes(100);o__Polyporales(100);f__Phanerochaetaceae(100);g__Antrodiella(100);g__Antrodiella_unclassified(100); | Antrodiella | 13 | Saprotroph | \|Wood Saprotroph\| | Corticioid-Polyporoid | White Rot | Highly Probable | As white_rot (Põlme et al. 2020); Decay wood (Põlme et al. 2020) | Gilbertson RL, Ryvarden L. 1987-1987. North American Polypores. Fungiflora, Oslo (ISBN: 978-0945345060); Põlme S, et al. 2020. Fungal Diversity 105:1-16 ((https://doi.org/10.1007/s13225-020-00466-2)) |
| Otu28419 | 1 | k__Fungi(100);p__Basidiomycota(100);c__Agaricomycetes(100);o__Polyporales(100);f__Polyporaceae(100);f__Polyporaceae_unclassified(100);f__Polyporaceae_unclassified(100); | Polyporaceae | 9 | Saprotroph | Wood Saprotroph | Polyporoid | Brown Rot-White | Probable | Lignicolous, causing brown or white rot, mostly of broadleaved trees (Cannon & Kirk 2007) | Cannon PF, Kirk PM. 2007. Fungal Families of the World. CAB International, Cambridge (ISBN: 978-0851998275) |
| Otu28420 | 1 | k__Fungi(100);p__Basidiomycota(100);c__Agaricomycetes(100);o__Polyporales(100);f__Polyporaceae(100);g__Perenniporia(100);s__Perenniporia_sp(100); | Perenniporia | 13 | Saprotroph | \|Wood Saprotroph\| | Corticioid-Polyporoid | White Rot | Highly Probable | As white_rot (Põlme et al. 2020); Decay wood (Põlme et al. 2020) | Gilbertson RL, Ryvarden L. 1987-1987. North American Polypores. Fungiflora, Oslo (ISBN: 978-0945345060); Tedersoo L, et al. 2014. Science 346:e1256688 ((https://doi.org/10.1126/science.1256688)); Põlme S, et al. 2020. Fungal Diversity 105:1-16 ((https://doi.org/10.1007/s13225-020-00466-2)) |
| Otu28421 | 1 | k__Fungi(100);p__Basidiomycota(100);c__Agaricomycetes(100);o__Trechisporales(100);f__Hydnodontaceae(100);g__Trechispora(100);s__Trechispora_sp(100); | Trechispora | 13 | Saprotroph | \|Wood Saprotroph\| | Corticioid | White Rot | Highly Probable | As white_rot (Põlme et al. 2020); Decay wood (Põlme et al. 2020) | Gilbertson RL, Ryvarden L. 1987-1987. North American Polypores. Fungiflora, Oslo (ISBN: 978-0945345060); Rinaldi AC, et al. 2008. Fungal Diversity 33:1-45 ((http://www.fungaldiversity.org/fdp/sfdp/33-1.pdf)) (as Uncertain); Tedersoo L, et al. 2010. Mycorrhiza 20:217-263 ((https://doi.org/10.1007/s00572-009-0274-x)) (as not ECM).; Põlme S, et al. 2020. Fungal Diversity 105:1-16 ((https://doi.org/10.1007/s13225-020-00466-2)) |
| Otu28422 | 1 | k__Fungi(100);p__Basidiomycota(100);c__Agaricomycetes(100);o__Hymenochaetales(100);f__Schizoporaceae(100);g__Hyphodontia(100);s__Hyphodontia_sp_DLL2011_1(100); | Hyphodontia | 13 | Saprotroph | Undefined Saprotroph-\|Wood Saprotroph\| | Corticioid | White Rot | Probable | As white_rot (Põlme et al. 2020); Decay wood (Põlme et al. 2020) | Tedersoo L, et al. 2014. Science 346:e1256688 ((https://doi.org/10.1126/science.1256688)); Põlme S, et al. 2020. Fungal Diversity 105:1-16 ((https://doi.org/10.1007/s13225-020-00466-2)) |
| Otu28424 | 1 | k__Fungi(100);p__Basidiomycota(100);c__Microbotryomycetes(100);o__Sporidiobolales(100);f__Sporidiobolales_family_Incertae_sedis(100);g__Rhodotorula(100);s__Rhodotorula_graminis(100); | Rhodotorula | 13 | Pathotroph-Saprotroph-Symbiotroph | Animal Pathogen-Animal Symbiotroph-Endophyte-Plant Pathogen-\|Undefined Saprotroph\| | Yeast | NULL | Probable | As foliar_endophyte (Põlme et al. 2020); Opportunistic human pathogen | Põlme S, et al. 2020. Fungal Diversity 105:1-16 ((https://doi.org/10.1007/s13225-020-00466-2)); https://www.cdc.gov ((https://www.cdc.gov)) |
| Otu28425 | 1 | k__Fungi(100);p__Basidiomycota(100);c__Agaricomycetes(100);o__Hymenochaetales(100);f__Schizoporaceae(100);g__Schizopora(100);s__Schizopora_flavipora(100); | Schizopora | 13 | Pathotroph-Saprotroph | Plant Pathogen-\|Wood Saprotroph\| | Corticioid | White Rot | Highly Probable | As white_rot (Põlme et al. 2020); As wood_pathogen (Põlme et al. 2020); Decay wood (Põlme et al. 2020) | Gilbertson RL, Ryvarden L. 1987-1987. North American Polypores. Fungiflora, Oslo (ISBN: 978-0945345060); Põlme S, et al. 2020. Fungal Diversity 105:1-16 ((https://doi.org/10.1007/s13225-020-00466-2)) |
| Otu28426 | 1 | k__Fungi(100);p__Basidiomycota(100);c__Agaricomycetes(100);o__Polyporales(100);f__Meruliaceae(100);g__Scopuloides(100);s__Scopuloides_hydnoides(100); | Scopuloides | 13 | Saprotroph | Undefined Saprotroph-\|Wood Saprotroph\| | Corticioid | White Rot | Probable | As white_rot (Põlme et al. 2020); Decay wood (Põlme et al. 2020) | Tedersoo L, et al. 2014. Science 346:e1256688 ((https://doi.org/10.1126/science.1256688)); Põlme S, et al. 2020. Fungal Diversity 105:1-16 ((https://doi.org/10.1007/s13225-020-00466-2)) |
| Otu28427 | 1 | k__Fungi(100);p__Basidiomycota(100);c__Agaricomycetes(100);o__Polyporales(100);f__Meruliaceae(100);g__Scopuloides(100);s__Scopuloides_hydnoides(100); | Scopuloides | 13 | Saprotroph | Undefined Saprotroph-\|Wood Saprotroph\| | Corticioid | White Rot | Probable | As white_rot (Põlme et al. 2020); Decay wood (Põlme et al. 2020) | Tedersoo L, et al. 2014. Science 346:e1256688 ((https://doi.org/10.1126/science.1256688)); Põlme S, et al. 2020. Fungal Diversity 105:1-16 ((https://doi.org/10.1007/s13225-020-00466-2)) |
| Otu28428 | 1 | k__Fungi(100);p__Basidiomycota(100);c__Agaricomycetes(100);o__Polyporales(100);f__Meruliaceae(100);g__Junghuhnia(100);s__Junghuhnia_nitida(100); | Junghuhnia | 13 | Saprotroph | \|Wood Saprotroph\| | Corticioid | White Rot | Highly Probable | As white_rot (Põlme et al. 2020); Decay wood (Põlme et al. 2020) | Gilbertson RL, Ryvarden L. 1987-1987. North American Polypores. Fungiflora, Oslo (ISBN: 978-0945345060); Põlme S, et al. 2020. Fungal Diversity 105:1-16 ((https://doi.org/10.1007/s13225-020-00466-2)) |
| Otu28430 | 1 | k__Fungi(100);p__Basidiomycota(100);c__Agaricomycetes(100);o__Polyporales(100);f__Phanerochaetaceae(100);g__Byssomerulius(100);s__Byssomerulius_sp(100); | Byssomerulius | 13 | Saprotroph | Undefined Saprotroph-\|Wood Saprotroph\| | Corticioid | White Rot | Probable | As white_rot (Põlme et al. 2020); Decay wood (Põlme et al. 2020) | Tedersoo L, et al. 2014. Science 346:e1256688 ((https://doi.org/10.1126/science.1256688)); Põlme S, et al. 2020. Fungal Diversity 105:1-16 ((https://doi.org/10.1007/s13225-020-00466-2)) |
| Otu28431 | 1 | k__Fungi(100);p__Basidiomycota(100);c__Agaricomycetes(100);o__Polyporales(100);f__Meruliaceae(100);g__Steccherinum(100);s__Steccherinum_sp_DLL2011_2(100); | Steccherinum | 13 | Saprotroph | Undefined Saprotroph-\|Wood Saprotroph\| | Corticioid | White Rot | Probable | As white_rot (Põlme et al. 2020); Decay wood (Põlme et al. 2020) | Tedersoo L, et al. 2014. Science 346:e1256688 ((https://doi.org/10.1126/science.1256688)); Põlme S, et al. 2020. Fungal Diversity 105:1-16 ((https://doi.org/10.1007/s13225-020-00466-2)) |
| Otu28433 | 1 | k__Fungi(100);p__Basidiomycota(100);c__Microbotryomycetes(100);o__Sporidiobolales(100);f__Sporidiobolales_family_Incertae_sedis(100);g__Rhodotorula(100);s__Rhodotorula_graminis(100); | Rhodotorula | 13 | Pathotroph-Saprotroph-Symbiotroph | Animal Pathogen-Animal Symbiotroph-Endophyte-Plant Pathogen-\|Undefined Saprotroph\| | Yeast | NULL | Probable | As foliar_endophyte (Põlme et al. 2020); Opportunistic human pathogen | Põlme S, et al. 2020. Fungal Diversity 105:1-16 ((https://doi.org/10.1007/s13225-020-00466-2)); https://www.cdc.gov ((https://www.cdc.gov)) |
| Otu28434 | 1 | k__Fungi(100);p__Basidiomycota(100);c__Agaricomycetes(100);o__Cantharellales(100);f__Botryobasidiaceae(100);g__Botryobasidium(100);s__Botryobasidium_subcoronatum(100); | Botryobasidium | 13 | Saprotroph | Undefined Saprotroph-\|Wood Saprotroph\| | Corticioid | White Rot | Probable | As white_rot (Põlme et al. 2020); Decay wood (Põlme et al. 2020) | Tedersoo L, et al. 2014. Science 346:e1256688 ((https://doi.org/10.1126/science.1256688)); Põlme S, et al. 2020. Fungal Diversity 105:1-16 ((https://doi.org/10.1007/s13225-020-00466-2)) |
| Otu28435 | 1 | k__Fungi(100);p__Basidiomycota(100);c__Microbotryomycetes(100);o__Sporidiobolales(100);f__Sporidiobolales_family_Incertae_sedis(100);g__Rhodotorula(100);s__Rhodotorula_taiwanensis(100); | Rhodotorula | 13 | Pathotroph-Saprotroph-Symbiotroph | Animal Pathogen-Animal Symbiotroph-Endophyte-Plant Pathogen-\|Undefined Saprotroph\| | Yeast | NULL | Probable | As foliar_endophyte (Põlme et al. 2020); Opportunistic human pathogen | Põlme S, et al. 2020. Fungal Diversity 105:1-16 ((https://doi.org/10.1007/s13225-020-00466-2)); https://www.cdc.gov ((https://www.cdc.gov)) |
| Otu28436 | 1 | k__Fungi(100);p__Basidiomycota(100);c__Tremellomycetes(100);o__Filobasidiales(100);f__Filobasidiaceae(100);g__Cryptococcus(100);s__Cryptococcus_flavus(100); | Cryptococcus | 13 | Pathotroph-Saprotroph-Symbiotroph | Animal Pathogen-Endophyte-Epiphyte-\|Undefined Saprotroph\| | Dimorphic Yeast-Yeast | NULL | Possible | As foliar_endophyte (Põlme et al. 2020); Decay leaf, fruit, seed, soil, dung, animal material (Põlme et al. 2020) | Andrews JH, et al. 1983. Phytopathology 73:228-234 ((https://doi.org/10.1094/Phyto-73-228)); Kurtzman CP, et al. (eds.) 2011. The Yeasts, a Taxonomic Study. Fifth Edition. Vols 1-3. Elsevier, San Diego (ISBN: 9780444521491); Põlme S, et al. 2020. Fungal Diversity 105:1-16 ((https://doi.org/10.1007/s13225-020-00466-2)) |
| Otu28437 | 1 | k__Fungi(100);p__Basidiomycota(100);c__Microbotryomycetes(100);o__Sporidiobolales(100);f__Sporidiobolales_family_Incertae_sedis(100);g__Rhodotorula(100);s__Rhodotorula_taiwanensis(100); | Rhodotorula | 13 | Pathotroph-Saprotroph-Symbiotroph | Animal Pathogen-Animal Symbiotroph-Endophyte-Plant Pathogen-\|Undefined Saprotroph\| | Yeast | NULL | Probable | As foliar_endophyte (Põlme et al. 2020); Opportunistic human pathogen | Põlme S, et al. 2020. Fungal Diversity 105:1-16 ((https://doi.org/10.1007/s13225-020-00466-2)); https://www.cdc.gov ((https://www.cdc.gov)) |
| Otu28438 | 1 | k__Fungi(100);p__Basidiomycota(100);c__Microbotryomycetes(100);o__Sporidiobolales(100);f__Sporidiobolales_family_Incertae_sedis(100);g__Rhodotorula(100);s__Rhodotorula_taiwanensis(100); | Rhodotorula | 13 | Pathotroph-Saprotroph-Symbiotroph | Animal Pathogen-Animal Symbiotroph-Endophyte-Plant Pathogen-\|Undefined Saprotroph\| | Yeast | NULL | Probable | As foliar_endophyte (Põlme et al. 2020); Opportunistic human pathogen | Põlme S, et al. 2020. Fungal Diversity 105:1-16 ((https://doi.org/10.1007/s13225-020-00466-2)); https://www.cdc.gov ((https://www.cdc.gov)) |
| Otu28439 | 1 | k__Fungi(100);p__Basidiomycota(100);c__Microbotryomycetes(100);o__Sporidiobolales(100);f__Sporidiobolales_family_Incertae_sedis(100);g__Rhodotorula(100);s__Rhodotorula_taiwanensis(100); | Rhodotorula | 13 | Pathotroph-Saprotroph-Symbiotroph | Animal Pathogen-Animal Symbiotroph-Endophyte-Plant Pathogen-\|Undefined Saprotroph\| | Yeast | NULL | Probable | As foliar_endophyte (Põlme et al. 2020); Opportunistic human pathogen | Põlme S, et al. 2020. Fungal Diversity 105:1-16 ((https://doi.org/10.1007/s13225-020-00466-2)); https://www.cdc.gov ((https://www.cdc.gov)) |
| Otu28440 | 1 | k__Fungi(100);p__Basidiomycota(100);c__Microbotryomycetes(100);o__Sporidiobolales(100);f__Sporidiobolales_family_Incertae_sedis(100);g__Rhodotorula(100);s__Rhodotorula_taiwanensis(100); | Rhodotorula | 13 | Pathotroph-Saprotroph-Symbiotroph | Animal Pathogen-Animal Symbiotroph-Endophyte-Plant Pathogen-\|Undefined Saprotroph\| | Yeast | NULL | Probable | As foliar_endophyte (Põlme et al. 2020); Opportunistic human pathogen | Põlme S, et al. 2020. Fungal Diversity 105:1-16 ((https://doi.org/10.1007/s13225-020-00466-2)); https://www.cdc.gov ((https://www.cdc.gov)) |
| Otu28441 | 1 | k__Fungi(100);p__Basidiomycota(100);c__Microbotryomycetes(100);o__Sporidiobolales(100);f__Sporidiobolales_family_Incertae_sedis(100);g__Rhodotorula(100);s__Rhodotorula_taiwanensis(100); | Rhodotorula | 13 | Pathotroph-Saprotroph-Symbiotroph | Animal Pathogen-Animal Symbiotroph-Endophyte-Plant Pathogen-\|Undefined Saprotroph\| | Yeast | NULL | Probable | As foliar_endophyte (Põlme et al. 2020); Opportunistic human pathogen | Põlme S, et al. 2020. Fungal Diversity 105:1-16 ((https://doi.org/10.1007/s13225-020-00466-2)); https://www.cdc.gov ((https://www.cdc.gov)) |
| Otu28442 | 1 | k__Fungi(100);p__Basidiomycota(100);c__Microbotryomycetes(100);o__Sporidiobolales(100);f__Sporidiobolales_family_Incertae_sedis(100);g__Rhodotorula(100);s__Rhodotorula_taiwanensis(100); | Rhodotorula | 13 | Pathotroph-Saprotroph-Symbiotroph | Animal Pathogen-Animal Symbiotroph-Endophyte-Plant Pathogen-\|Undefined Saprotroph\| | Yeast | NULL | Probable | As foliar_endophyte (Põlme et al. 2020); Opportunistic human pathogen | Põlme S, et al. 2020. Fungal Diversity 105:1-16 ((https://doi.org/10.1007/s13225-020-00466-2)); https://www.cdc.gov ((https://www.cdc.gov)) |
| Otu28443 | 1 | k__Fungi(100);p__Basidiomycota(100);c__Microbotryomycetes(100);o__Sporidiobolales(100);f__Sporidiobolales_family_Incertae_sedis(100);g__Rhodotorula(100);s__Rhodotorula_taiwanensis(100); | Rhodotorula | 13 | Pathotroph-Saprotroph-Symbiotroph | Animal Pathogen-Animal Symbiotroph-Endophyte-Plant Pathogen-\|Undefined Saprotroph\| | Yeast | NULL | Probable | As foliar_endophyte (Põlme et al. 2020); Opportunistic human pathogen | Põlme S, et al. 2020. Fungal Diversity 105:1-16 ((https://doi.org/10.1007/s13225-020-00466-2)); https://www.cdc.gov ((https://www.cdc.gov)) |
| Otu28444 | 1 | k__Fungi(100);p__Basidiomycota(100);c__Microbotryomycetes(100);o__Sporidiobolales(100);f__Sporidiobolales_family_Incertae_sedis(100);g__Rhodotorula(100);s__Rhodotorula_taiwanensis(100); | Rhodotorula | 13 | Pathotroph-Saprotroph-Symbiotroph | Animal Pathogen-Animal Symbiotroph-Endophyte-Plant Pathogen-\|Undefined Saprotroph\| | Yeast | NULL | Probable | As foliar_endophyte (Põlme et al. 2020); Opportunistic human pathogen | Põlme S, et al. 2020. Fungal Diversity 105:1-16 ((https://doi.org/10.1007/s13225-020-00466-2)); https://www.cdc.gov ((https://www.cdc.gov)) |
| Otu28445 | 1 | k__Fungi(100);p__Basidiomycota(100);c__Microbotryomycetes(100);o__Sporidiobolales(100);f__Sporidiobolales_family_Incertae_sedis(100);g__Rhodotorula(100);s__Rhodotorula_taiwanensis(100); | Rhodotorula | 13 | Pathotroph-Saprotroph-Symbiotroph | Animal Pathogen-Animal Symbiotroph-Endophyte-Plant Pathogen-\|Undefined Saprotroph\| | Yeast | NULL | Probable | As foliar_endophyte (Põlme et al. 2020); Opportunistic human pathogen | Põlme S, et al. 2020. Fungal Diversity 105:1-16 ((https://doi.org/10.1007/s13225-020-00466-2)); https://www.cdc.gov ((https://www.cdc.gov)) |
| Otu28446 | 1 | k__Fungi(100);p__Basidiomycota(100);c__Microbotryomycetes(100);o__Sporidiobolales(100);f__Sporidiobolales_family_Incertae_sedis(100);g__Rhodotorula(100);s__Rhodotorula_taiwanensis(100); | Rhodotorula | 13 | Pathotroph-Saprotroph-Symbiotroph | Animal Pathogen-Animal Symbiotroph-Endophyte-Plant Pathogen-\|Undefined Saprotroph\| | Yeast | NULL | Probable | As foliar_endophyte (Põlme et al. 2020); Opportunistic human pathogen | Põlme S, et al. 2020. Fungal Diversity 105:1-16 ((https://doi.org/10.1007/s13225-020-00466-2)); https://www.cdc.gov ((https://www.cdc.gov)) |
| Otu28447 | 1 | k__Fungi(100);p__Basidiomycota(100);c__Microbotryomycetes(100);o__Sporidiobolales(100);f__Sporidiobolales_family_Incertae_sedis(100);g__Rhodotorula(100);s__Rhodotorula_taiwanensis(100); | Rhodotorula | 13 | Pathotroph-Saprotroph-Symbiotroph | Animal Pathogen-Animal Symbiotroph-Endophyte-Plant Pathogen-\|Undefined Saprotroph\| | Yeast | NULL | Probable | As foliar_endophyte (Põlme et al. 2020); Opportunistic human pathogen | Põlme S, et al. 2020. Fungal Diversity 105:1-16 ((https://doi.org/10.1007/s13225-020-00466-2)); https://www.cdc.gov ((https://www.cdc.gov)) |
| Otu28448 | 1 | k__Fungi(100);p__Basidiomycota(100);c__Microbotryomycetes(100);o__Sporidiobolales(100);f__Sporidiobolales_family_Incertae_sedis(100);g__Rhodotorula(100);s__Rhodotorula_taiwanensis(100); | Rhodotorula | 13 | Pathotroph-Saprotroph-Symbiotroph | Animal Pathogen-Animal Symbiotroph-Endophyte-Plant Pathogen-\|Undefined Saprotroph\| | Yeast | NULL | Probable | As foliar_endophyte (Põlme et al. 2020); Opportunistic human pathogen | Põlme S, et al. 2020. Fungal Diversity 105:1-16 ((https://doi.org/10.1007/s13225-020-00466-2)); https://www.cdc.gov ((https://www.cdc.gov)) |
| Otu28449 | 1 | k__Fungi(100);p__Basidiomycota(100);c__Microbotryomycetes(100);o__Sporidiobolales(100);f__Sporidiobolales_family_Incertae_sedis(100);g__Rhodotorula(100);s__Rhodotorula_taiwanensis(100); | Rhodotorula | 13 | Pathotroph-Saprotroph-Symbiotroph | Animal Pathogen-Animal Symbiotroph-Endophyte-Plant Pathogen-\|Undefined Saprotroph\| | Yeast | NULL | Probable | As foliar_endophyte (Põlme et al. 2020); Opportunistic human pathogen | Põlme S, et al. 2020. Fungal Diversity 105:1-16 ((https://doi.org/10.1007/s13225-020-00466-2)); https://www.cdc.gov ((https://www.cdc.gov)) |
| Otu28450 | 1 | k__Fungi(100);p__Basidiomycota(100);c__Microbotryomycetes(100);o__Sporidiobolales(100);f__Sporidiobolales_family_Incertae_sedis(100);g__Rhodotorula(100);s__Rhodotorula_taiwanensis(100); | Rhodotorula | 13 | Pathotroph-Saprotroph-Symbiotroph | Animal Pathogen-Animal Symbiotroph-Endophyte-Plant Pathogen-\|Undefined Saprotroph\| | Yeast | NULL | Probable | As foliar_endophyte (Põlme et al. 2020); Opportunistic human pathogen | Põlme S, et al. 2020. Fungal Diversity 105:1-16 ((https://doi.org/10.1007/s13225-020-00466-2)); https://www.cdc.gov ((https://www.cdc.gov)) |
| Otu28451 | 1 | k__Fungi(100);p__Basidiomycota(100);c__Microbotryomycetes(100);o__Sporidiobolales(100);f__Sporidiobolales_family_Incertae_sedis(100);g__Rhodotorula(100);s__Rhodotorula_taiwanensis(100); | Rhodotorula | 13 | Pathotroph-Saprotroph-Symbiotroph | Animal Pathogen-Animal Symbiotroph-Endophyte-Plant Pathogen-\|Undefined Saprotroph\| | Yeast | NULL | Probable | As foliar_endophyte (Põlme et al. 2020); Opportunistic human pathogen | Põlme S, et al. 2020. Fungal Diversity 105:1-16 ((https://doi.org/10.1007/s13225-020-00466-2)); https://www.cdc.gov ((https://www.cdc.gov)) |
| Otu28452 | 1 | k__Fungi(100);p__Basidiomycota(100);c__Microbotryomycetes(100);o__Sporidiobolales(100);f__Sporidiobolales_family_Incertae_sedis(100);g__Rhodotorula(100);s__Rhodotorula_taiwanensis(100); | Rhodotorula | 13 | Pathotroph-Saprotroph-Symbiotroph | Animal Pathogen-Animal Symbiotroph-Endophyte-Plant Pathogen-\|Undefined Saprotroph\| | Yeast | NULL | Probable | As foliar_endophyte (Põlme et al. 2020); Opportunistic human pathogen | Põlme S, et al. 2020. Fungal Diversity 105:1-16 ((https://doi.org/10.1007/s13225-020-00466-2)); https://www.cdc.gov ((https://www.cdc.gov)) |
| Otu28453 | 1 | k__Fungi(100);p__Basidiomycota(100);c__Microbotryomycetes(100);o__Sporidiobolales(100);f__Sporidiobolales_family_Incertae_sedis(100);g__Rhodotorula(100);s__Rhodotorula_mucilaginosa(100); | Rhodotorula | 13 | Pathotroph-Saprotroph-Symbiotroph | Animal Pathogen-Animal Symbiotroph-Endophyte-Plant Pathogen-\|Undefined Saprotroph\| | Yeast | NULL | Probable | As foliar_endophyte (Põlme et al. 2020); Opportunistic human pathogen | Põlme S, et al. 2020. Fungal Diversity 105:1-16 ((https://doi.org/10.1007/s13225-020-00466-2)); https://www.cdc.gov ((https://www.cdc.gov)) |
| Otu28454 | 1 | k__Fungi(100);p__Basidiomycota(100);c__Microbotryomycetes(100);o__Sporidiobolales(100);f__Sporidiobolales_family_Incertae_sedis(100);g__Rhodotorula(100);s__Rhodotorula_mucilaginosa(100); | Rhodotorula | 13 | Pathotroph-Saprotroph-Symbiotroph | Animal Pathogen-Animal Symbiotroph-Endophyte-Plant Pathogen-\|Undefined Saprotroph\| | Yeast | NULL | Probable | As foliar_endophyte (Põlme et al. 2020); Opportunistic human pathogen | Põlme S, et al. 2020. Fungal Diversity 105:1-16 ((https://doi.org/10.1007/s13225-020-00466-2)); https://www.cdc.gov ((https://www.cdc.gov)) |
| Otu28455 | 1 | k__Fungi(100);p__Basidiomycota(100);c__Microbotryomycetes(100);o__Sporidiobolales(100);f__Sporidiobolales_family_Incertae_sedis(100);g__Rhodotorula(100);s__Rhodotorula_mucilaginosa(100); | Rhodotorula | 13 | Pathotroph-Saprotroph-Symbiotroph | Animal Pathogen-Animal Symbiotroph-Endophyte-Plant Pathogen-\|Undefined Saprotroph\| | Yeast | NULL | Probable | As foliar_endophyte (Põlme et al. 2020); Opportunistic human pathogen | Põlme S, et al. 2020. Fungal Diversity 105:1-16 ((https://doi.org/10.1007/s13225-020-00466-2)); https://www.cdc.gov ((https://www.cdc.gov)) |
| Otu28456 | 1 | k__Fungi(100);p__Basidiomycota(100);c__Microbotryomycetes(100);o__Sporidiobolales(100);f__Sporidiobolales_family_Incertae_sedis(100);g__Rhodotorula(100);s__Rhodotorula_mucilaginosa(100); | Rhodotorula | 13 | Pathotroph-Saprotroph-Symbiotroph | Animal Pathogen-Animal Symbiotroph-Endophyte-Plant Pathogen-\|Undefined Saprotroph\| | Yeast | NULL | Probable | As foliar_endophyte (Põlme et al. 2020); Opportunistic human pathogen | Põlme S, et al. 2020. Fungal Diversity 105:1-16 ((https://doi.org/10.1007/s13225-020-00466-2)); https://www.cdc.gov ((https://www.cdc.gov)) |
| Otu28457 | 1 | k__Fungi(100);p__Basidiomycota(100);c__Microbotryomycetes(100);o__Sporidiobolales(100);f__Sporidiobolales_family_Incertae_sedis(100);g__Rhodotorula(100);s__Rhodotorula_mucilaginosa(100); | Rhodotorula | 13 | Pathotroph-Saprotroph-Symbiotroph | Animal Pathogen-Animal Symbiotroph-Endophyte-Plant Pathogen-\|Undefined Saprotroph\| | Yeast | NULL | Probable | As foliar_endophyte (Põlme et al. 2020); Opportunistic human pathogen | Põlme S, et al. 2020. Fungal Diversity 105:1-16 ((https://doi.org/10.1007/s13225-020-00466-2)); https://www.cdc.gov ((https://www.cdc.gov)) |
| Otu28458 | 1 | k__Fungi(100);p__Basidiomycota(100);c__Microbotryomycetes(100);o__Sporidiobolales(100);f__Sporidiobolales_family_Incertae_sedis(100);g__Rhodotorula(100);s__Rhodotorula_mucilaginosa(100); | Rhodotorula | 13 | Pathotroph-Saprotroph-Symbiotroph | Animal Pathogen-Animal Symbiotroph-Endophyte-Plant Pathogen-\|Undefined Saprotroph\| | Yeast | NULL | Probable | As foliar_endophyte (Põlme et al. 2020); Opportunistic human pathogen | Põlme S, et al. 2020. Fungal Diversity 105:1-16 ((https://doi.org/10.1007/s13225-020-00466-2)); https://www.cdc.gov ((https://www.cdc.gov)) |
| Otu28459 | 1 | k__Fungi(100);p__Basidiomycota(100);c__Microbotryomycetes(100);o__Sporidiobolales(100);f__Sporidiobolales_family_Incertae_sedis(100);g__Rhodotorula(100);s__Rhodotorula_mucilaginosa(100); | Rhodotorula | 13 | Pathotroph-Saprotroph-Symbiotroph | Animal Pathogen-Animal Symbiotroph-Endophyte-Plant Pathogen-\|Undefined Saprotroph\| | Yeast | NULL | Probable | As foliar_endophyte (Põlme et al. 2020); Opportunistic human pathogen | Põlme S, et al. 2020. Fungal Diversity 105:1-16 ((https://doi.org/10.1007/s13225-020-00466-2)); https://www.cdc.gov ((https://www.cdc.gov)) |
| Otu28460 | 1 | k__Fungi(100);p__Basidiomycota(100);c__Microbotryomycetes(100);o__Sporidiobolales(100);f__Sporidiobolales_family_Incertae_sedis(100);g__Rhodotorula(100);s__Rhodotorula_mucilaginosa(100); | Rhodotorula | 13 | Pathotroph-Saprotroph-Symbiotroph | Animal Pathogen-Animal Symbiotroph-Endophyte-Plant Pathogen-\|Undefined Saprotroph\| | Yeast | NULL | Probable | As foliar_endophyte (Põlme et al. 2020); Opportunistic human pathogen | Põlme S, et al. 2020. Fungal Diversity 105:1-16 ((https://doi.org/10.1007/s13225-020-00466-2)); https://www.cdc.gov ((https://www.cdc.gov)) |
| Otu28461 | 1 | k__Fungi(100);p__Basidiomycota(100);c__Microbotryomycetes(100);o__Sporidiobolales(100);f__Sporidiobolales_family_Incertae_sedis(100);g__Rhodotorula(100);s__Rhodotorula_mucilaginosa(100); | Rhodotorula | 13 | Pathotroph-Saprotroph-Symbiotroph | Animal Pathogen-Animal Symbiotroph-Endophyte-Plant Pathogen-\|Undefined Saprotroph\| | Yeast | NULL | Probable | As foliar_endophyte (Põlme et al. 2020); Opportunistic human pathogen | Põlme S, et al. 2020. Fungal Diversity 105:1-16 ((https://doi.org/10.1007/s13225-020-00466-2)); https://www.cdc.gov ((https://www.cdc.gov)) |
| Otu28462 | 1 | k__Fungi(100);p__Basidiomycota(100);c__Microbotryomycetes(100);o__Sporidiobolales(100);f__Sporidiobolales_family_Incertae_sedis(100);g__Rhodotorula(100);s__Rhodotorula_taiwanensis(100); | Rhodotorula | 13 | Pathotroph-Saprotroph-Symbiotroph | Animal Pathogen-Animal Symbiotroph-Endophyte-Plant Pathogen-\|Undefined Saprotroph\| | Yeast | NULL | Probable | As foliar_endophyte (Põlme et al. 2020); Opportunistic human pathogen | Põlme S, et al. 2020. Fungal Diversity 105:1-16 ((https://doi.org/10.1007/s13225-020-00466-2)); https://www.cdc.gov ((https://www.cdc.gov)) |
| Otu28463 | 1 | k__Fungi(100);p__Basidiomycota(100);c__Microbotryomycetes(100);o__Sporidiobolales(100);f__Sporidiobolales_family_Incertae_sedis(100);g__Rhodotorula(100);s__Rhodotorula_taiwanensis(100); | Rhodotorula | 13 | Pathotroph-Saprotroph-Symbiotroph | Animal Pathogen-Animal Symbiotroph-Endophyte-Plant Pathogen-\|Undefined Saprotroph\| | Yeast | NULL | Probable | As foliar_endophyte (Põlme et al. 2020); Opportunistic human pathogen | Põlme S, et al. 2020. Fungal Diversity 105:1-16 ((https://doi.org/10.1007/s13225-020-00466-2)); https://www.cdc.gov ((https://www.cdc.gov)) |
| Otu28464 | 1 | k__Fungi(100);p__Basidiomycota(100);c__Microbotryomycetes(100);o__Sporidiobolales(100);f__Sporidiobolales_family_Incertae_sedis(100);g__Rhodotorula(100);s__Rhodotorula_taiwanensis(100); | Rhodotorula | 13 | Pathotroph-Saprotroph-Symbiotroph | Animal Pathogen-Animal Symbiotroph-Endophyte-Plant Pathogen-\|Undefined Saprotroph\| | Yeast | NULL | Probable | As foliar_endophyte (Põlme et al. 2020); Opportunistic human pathogen | Põlme S, et al. 2020. Fungal Diversity 105:1-16 ((https://doi.org/10.1007/s13225-020-00466-2)); https://www.cdc.gov ((https://www.cdc.gov)) |

| | | | | | | | | | | | |
|---|---|---|---|---|---|---|---|---|---|---|---|
| Otu28465 | 1 | k__Fungi(100);p__Basidiomycota(100);c__Microbotryomycetes(100);o__Sporidiobolales(100);f__Sporidiobolales_family_Incertae_sedis(100);g__Rhodotorula(100);s__Rhodotorula_mucilaginosa(100); | Rhodotorula | 13 | Pathotroph-Saprotroph-Symbiotroph | Animal Pathogen-Animal Symbiotroph-Endophyte-Plant Pathogen-\|Undefined Saprotroph\| | Yeast | NULL | Probable | As foliar_endophyte (Põlme et al. 2020); Opportunistic human pathogen | Põlme S, et al. 2020. Fungal Diversity 105:1-16 ((https://doi.org/10.1007/s13225-020-00466-2)); https://www.cdc.gov ((https://www.cdc.gov)) |
| Otu28466 | 1 | k__Fungi(100);p__Basidiomycota(100);c__Microbotryomycetes(100);o__Sporidiobolales(100);f__Sporidiobolales_family_Incertae_sedis(100);g__Rhodotorula(100);s__Rhodotorula_mucilaginosa(100); | Rhodotorula | 13 | Pathotroph-Saprotroph-Symbiotroph | Animal Pathogen-Animal Symbiotroph-Endophyte-Plant Pathogen-\|Undefined Saprotroph\| | Yeast | NULL | Probable | As foliar_endophyte (Põlme et al. 2020); Opportunistic human pathogen | Põlme S, et al. 2020. Fungal Diversity 105:1-16 ((https://doi.org/10.1007/s13225-020-00466-2)); https://www.cdc.gov ((https://www.cdc.gov)) |
| Otu28467 | 1 | k__Fungi(100);p__Basidiomycota(100);c__Microbotryomycetes(100);o__Sporidiobolales(100);f__Sporidiobolales_family_Incertae_sedis(100);g__Rhodotorula(100);s__Rhodotorula_mucilaginosa(100); | Rhodotorula | 13 | Pathotroph-Saprotroph-Symbiotroph | Animal Pathogen-Animal Symbiotroph-Endophyte-Plant Pathogen-\|Undefined Saprotroph\| | Yeast | NULL | Probable | As foliar_endophyte (Põlme et al. 2020); Opportunistic human pathogen | Põlme S, et al. 2020. Fungal Diversity 105:1-16 ((https://doi.org/10.1007/s13225-020-00466-2)); https://www.cdc.gov ((https://www.cdc.gov)) |
| Otu28468 | 1 | k__Fungi(100);p__Basidiomycota(100);c__Microbotryomycetes(100);o__Sporidiobolales(100);f__Sporidiobolales_family_Incertae_sedis(100);g__Rhodotorula(100);g__Rhodotorula_unclassified(100); | Rhodotorula | 13 | Pathotroph-Saprotroph-Symbiotroph | Animal Pathogen-Animal Symbiotroph-Endophyte-Plant Pathogen-\|Undefined Saprotroph\| | Yeast | NULL | Probable | As foliar_endophyte (Põlme et al. 2020); Opportunistic human pathogen | Põlme S, et al. 2020. Fungal Diversity 105:1-16 ((https://doi.org/10.1007/s13225-020-00466-2)); https://www.cdc.gov ((https://www.cdc.gov)) |
| Otu28469 | 1 | k__Fungi(100);p__Basidiomycota(100);c__Microbotryomycetes(100);o__Sporidiobolales(100);f__Sporidiobolales_family_Incertae_sedis(100);g__Rhodotorula(100);s__Rhodotorula_taiwanensis(100); | Rhodotorula | 13 | Pathotroph-Saprotroph-Symbiotroph | Animal Pathogen-Animal Symbiotroph-Endophyte-Plant Pathogen-\|Undefined Saprotroph\| | Yeast | NULL | Probable | As foliar_endophyte (Põlme et al. 2020); Opportunistic human pathogen | Põlme S, et al. 2020. Fungal Diversity 105:1-16 ((https://doi.org/10.1007/s13225-020-00466-2)); https://www.cdc.gov ((https://www.cdc.gov)) |
| Otu28470 | 1 | k__Fungi(100);p__Basidiomycota(100);c__Microbotryomycetes(100);o__Sporidiobolales(100);f__Sporidiobolales_family_Incertae_sedis(100);g__Rhodotorula(100);s__Rhodotorula_mucilaginosa(100); | Rhodotorula | 13 | Pathotroph-Saprotroph-Symbiotroph | Animal Pathogen-Animal Symbiotroph-Endophyte-Plant Pathogen-\|Undefined Saprotroph\| | Yeast | NULL | Probable | As foliar_endophyte (Põlme et al. 2020); Opportunistic human pathogen | Põlme S, et al. 2020. Fungal Diversity 105:1-16 ((https://doi.org/10.1007/s13225-020-00466-2)); https://www.cdc.gov ((https://www.cdc.gov)) |
| Otu28471 | 1 | k__Fungi(100);p__Basidiomycota(100);c__Microbotryomycetes(100);o__Sporidiobolales(100);f__Sporidiobolales_family_Incertae_sedis(100);g__Rhodotorula(100);s__Rhodotorula_taiwanensis(100); | Rhodotorula | 13 | Pathotroph-Saprotroph-Symbiotroph | Animal Pathogen-Animal Symbiotroph-Endophyte-Plant Pathogen-\|Undefined Saprotroph\| | Yeast | NULL | Probable | As foliar_endophyte (Põlme et al. 2020); Opportunistic human pathogen | Põlme S, et al. 2020. Fungal Diversity 105:1-16 ((https://doi.org/10.1007/s13225-020-00466-2)); https://www.cdc.gov ((https://www.cdc.gov)) |
| Otu28472 | 1 | k__Fungi(100);p__Basidiomycota(100);c__Microbotryomycetes(100);o__Sporidiobolales(100);f__Sporidiobolales_family_Incertae_sedis(100);g__Rhodotorula(100);s__Rhodotorula_mucilaginosa(100); | Rhodotorula | 13 | Pathotroph-Saprotroph-Symbiotroph | Animal Pathogen-Animal Symbiotroph-Endophyte-Plant Pathogen-\|Undefined Saprotroph\| | Yeast | NULL | Probable | As foliar_endophyte (Põlme et al. 2020); Opportunistic human pathogen | Põlme S, et al. 2020. Fungal Diversity 105:1-16 ((https://doi.org/10.1007/s13225-020-00466-2)); https://www.cdc.gov ((https://www.cdc.gov)) |
| Otu28473 | 1 | k__Fungi(100);p__Basidiomycota(100);c__Microbotryomycetes(100);o__Sporidiobolales(100);f__Sporidiobolales_family_Incertae_sedis(100);g__Rhodotorula(100);s__Rhodotorula_taiwanensis(100); | Rhodotorula | 13 | Pathotroph-Saprotroph-Symbiotroph | Animal Pathogen-Animal Symbiotroph-Endophyte-Plant Pathogen-\|Undefined Saprotroph\| | Yeast | NULL | Probable | As foliar_endophyte (Põlme et al. 2020); Opportunistic human pathogen | Põlme S, et al. 2020. Fungal Diversity 105:1-16 ((https://doi.org/10.1007/s13225-020-00466-2)); https://www.cdc.gov ((https://www.cdc.gov)) |
| Otu28474 | 1 | k__Fungi(100);p__Basidiomycota(100);c__Microbotryomycetes(100);o__Sporidiobolales(100);f__Sporidiobolales_family_Incertae_sedis(100);g__Rhodotorula(100);s__Rhodotorula_taiwanensis(100); | Rhodotorula | 13 | Pathotroph-Saprotroph-Symbiotroph | Animal Pathogen-Animal Symbiotroph-Endophyte-Plant Pathogen-\|Undefined Saprotroph\| | Yeast | NULL | Probable | As foliar_endophyte (Põlme et al. 2020); Opportunistic human pathogen | Põlme S, et al. 2020. Fungal Diversity 105:1-16 ((https://doi.org/10.1007/s13225-020-00466-2)); https://www.cdc.gov ((https://www.cdc.gov)) |
| Otu28475 | 1 | k__Fungi(100);p__Basidiomycota(100);c__Microbotryomycetes(100);o__Sporidiobolales(100);f__Sporidiobolales_family_Incertae_sedis(100);g__Rhodotorula(100);s__Rhodotorula_taiwanensis(100); | Rhodotorula | 13 | Pathotroph-Saprotroph-Symbiotroph | Animal Pathogen-Animal Symbiotroph-Endophyte-Plant Pathogen-\|Undefined Saprotroph\| | Yeast | NULL | Probable | As foliar_endophyte (Põlme et al. 2020); Opportunistic human pathogen | Põlme S, et al. 2020. Fungal Diversity 105:1-16 ((https://doi.org/10.1007/s13225-020-00466-2)); https://www.cdc.gov ((https://www.cdc.gov)) |
| Otu28476 | 1 | k__Fungi(100);p__Basidiomycota(100);c__Microbotryomycetes(100);o__Sporidiobolales(100);f__Sporidiobolales_family_Incertae_sedis(100);g__Rhodotorula(100);s__Rhodotorula_taiwanensis(100); | Rhodotorula | 13 | Pathotroph-Saprotroph-Symbiotroph | Animal Pathogen-Animal Symbiotroph-Endophyte-Plant Pathogen-\|Undefined Saprotroph\| | Yeast | NULL | Probable | As foliar_endophyte (Põlme et al. 2020); Opportunistic human pathogen | Põlme S, et al. 2020. Fungal Diversity 105:1-16 ((https://doi.org/10.1007/s13225-020-00466-2)); https://www.cdc.gov ((https://www.cdc.gov)) |
| Otu28477 | 1 | k__Fungi(100);p__Basidiomycota(100);c__Microbotryomycetes(100);o__Sporidiobolales(100);f__Sporidiobolales_family_Incertae_sedis(100);g__Rhodotorula(100);s__Rhodotorula_taiwanensis(100); | Rhodotorula | 13 | Pathotroph-Saprotroph-Symbiotroph | Animal Pathogen-Animal Symbiotroph-Endophyte-Plant Pathogen-\|Undefined Saprotroph\| | Yeast | NULL | Probable | As foliar_endophyte (Põlme et al. 2020); Opportunistic human pathogen | Põlme S, et al. 2020. Fungal Diversity 105:1-16 ((https://doi.org/10.1007/s13225-020-00466-2)); https://www.cdc.gov ((https://www.cdc.gov)) |
| Otu28478 | 1 | k__Fungi(100);p__Basidiomycota(100);c__Microbotryomycetes(100);o__Sporidiobolales(100);f__Sporidiobolales_family_Incertae_sedis(100);g__Rhodotorula(100);s__Rhodotorula_mucilaginosa(100); | Rhodotorula | 13 | Pathotroph-Saprotroph-Symbiotroph | Animal Pathogen-Animal Symbiotroph-Endophyte-Plant Pathogen-\|Undefined Saprotroph\| | Yeast | NULL | Probable | As foliar_endophyte (Põlme et al. 2020); Opportunistic human pathogen | Põlme S, et al. 2020. Fungal Diversity 105:1-16 ((https://doi.org/10.1007/s13225-020-00466-2)); https://www.cdc.gov ((https://www.cdc.gov)) |
| Otu28479 | 1 | k__Fungi(100);p__Basidiomycota(100);c__Microbotryomycetes(100);o__Sporidiobolales(100);f__Sporidiobolales_family_Incertae_sedis(100);g__Rhodotorula(100);s__Rhodotorula_mucilaginosa(100); | Rhodotorula | 13 | Pathotroph-Saprotroph-Symbiotroph | Animal Pathogen-Animal Symbiotroph-Endophyte-Plant Pathogen-\|Undefined Saprotroph\| | Yeast | NULL | Probable | As foliar_endophyte (Põlme et al. 2020); Opportunistic human pathogen | Põlme S, et al. 2020. Fungal Diversity 105:1-16 ((https://doi.org/10.1007/s13225-020-00466-2)); https://www.cdc.gov ((https://www.cdc.gov)) |
| Otu28480 | 1 | k__Fungi(100);p__Basidiomycota(100);c__Microbotryomycetes(100);o__Sporidiobolales(100);f__Sporidiobolales_family_Incertae_sedis(100);g__Rhodotorula(100);s__Rhodotorula_taiwanensis(100); | Rhodotorula | 13 | Pathotroph-Saprotroph-Symbiotroph | Animal Pathogen-Animal Symbiotroph-Endophyte-Plant Pathogen-\|Undefined Saprotroph\| | Yeast | NULL | Probable | As foliar_endophyte (Põlme et al. 2020); Opportunistic human pathogen | Põlme S, et al. 2020. Fungal Diversity 105:1-16 ((https://doi.org/10.1007/s13225-020-00466-2)); https://www.cdc.gov ((https://www.cdc.gov)) |
| Otu28481 | 1 | k__Fungi(100);p__Basidiomycota(100);c__Microbotryomycetes(100);o__Sporidiobolales(100);f__Sporidiobolales_family_Incertae_sedis(100);g__Rhodotorula(100);s__Rhodotorula_mucilaginosa(100); | Rhodotorula | 13 | Pathotroph-Saprotroph-Symbiotroph | Animal Pathogen-Animal Symbiotroph-Endophyte-Plant Pathogen-\|Undefined Saprotroph\| | Yeast | NULL | Probable | As foliar_endophyte (Põlme et al. 2020); Opportunistic human pathogen | Põlme S, et al. 2020. Fungal Diversity 105:1-16 ((https://doi.org/10.1007/s13225-020-00466-2)); https://www.cdc.gov ((https://www.cdc.gov)) |
| Otu28482 | 1 | k__Fungi(100);p__Basidiomycota(100);c__Microbotryomycetes(100);o__Sporidiobolales(100);f__Sporidiobolales_family_Incertae_sedis(100);g__Rhodotorula(100);s__Rhodotorula_mucilaginosa(100); | Rhodotorula | 13 | Pathotroph-Saprotroph-Symbiotroph | Animal Pathogen-Animal Symbiotroph-Endophyte-Plant Pathogen-\|Undefined Saprotroph\| | Yeast | NULL | Probable | As foliar_endophyte (Põlme et al. 2020); Opportunistic human pathogen | Põlme S, et al. 2020. Fungal Diversity 105:1-16 ((https://doi.org/10.1007/s13225-020-00466-2)); https://www.cdc.gov ((https://www.cdc.gov)) |
| Otu28483 | 1 | k__Fungi(100);p__Basidiomycota(100);c__Microbotryomycetes(100);o__Sporidiobolales(100);f__Sporidiobolales_family_Incertae_sedis(100);g__Rhodotorula(100);s__Rhodotorula_taiwanensis(100); | Rhodotorula | 13 | Pathotroph-Saprotroph-Symbiotroph | Animal Pathogen-Animal Symbiotroph-Endophyte-Plant Pathogen-\|Undefined Saprotroph\| | Yeast | NULL | Probable | As foliar_endophyte (Põlme et al. 2020); Opportunistic human pathogen | Põlme S, et al. 2020. Fungal Diversity 105:1-16 ((https://doi.org/10.1007/s13225-020-00466-2)); https://www.cdc.gov ((https://www.cdc.gov)) |
| Otu28486 | 1 | k__Fungi(100);p__Basidiomycota(100);c__Microbotryomycetes(100);o__Sporidiobolales(100);f__Sporidiobolales_family_Incertae_sedis(100);g__Rhodotorula(100);s__Rhodotorula_mucilaginosa(100); | Rhodotorula | 13 | Pathotroph-Saprotroph-Symbiotroph | Animal Pathogen-Animal Symbiotroph-Endophyte-Plant Pathogen-\|Undefined Saprotroph\| | Yeast | NULL | Probable | As foliar_endophyte (Põlme et al. 2020); Opportunistic human pathogen | Põlme S, et al. 2020. Fungal Diversity 105:1-16 ((https://doi.org/10.1007/s13225-020-00466-2)); https://www.cdc.gov ((https://www.cdc.gov)) |
| Otu28487 | 1 | k__Fungi(100);p__Basidiomycota(100);c__Microbotryomycetes(100);o__Sporidiobolales(100);f__Sporidiobolales_family_Incertae_sedis(100);g__Rhodotorula(100);s__Rhodotorula_mucilaginosa(100); | Rhodotorula | 13 | Pathotroph-Saprotroph-Symbiotroph | Animal Pathogen-Animal Symbiotroph-Endophyte-Plant Pathogen-\|Undefined Saprotroph\| | Yeast | NULL | Probable | As foliar_endophyte (Põlme et al. 2020); Opportunistic human pathogen | Põlme S, et al. 2020. Fungal Diversity 105:1-16 ((https://doi.org/10.1007/s13225-020-00466-2)); https://www.cdc.gov ((https://www.cdc.gov)) |
| Otu28488 | 1 | k__Fungi(100);p__Basidiomycota(100);c__Agaricomycetes(100);o__Agaricales(100);f__Mycenaceae(100);g__Mycena(100);g__Mycena_unclassified(100); | Mycena | 13 | Pathotroph-Saprotroph | Plant Pathogen-\|Plant Saprotroph\|-Undefined Saprotroph-Wood Saprotroph | Agaricoid | NULL | Probable | As litter_saprotroph (Põlme et al. 2020); Decay leaf, fruit, seed (Põlme et al. 2020) | Grgurinovic CA, Simpson JA. 2001. Fungal Diversity 8:97-127 ((http://www.fungaldiversity.org/fdp/sfdp/FD_8_97-127.pdf)); Tedersoo L, et al. 2014. Science 346:e1256688 ((https://doi.org/10.1126/science.1256688)); Põlme S, et al. 2020. Fungal Diversity 105:1-16 ((https://doi.org/10.1007/s13225-020-00466-2)); The American Phytopathological Society Common Names of Plant Diseases ((http://www.apsnet.org/publications/commonnames/Pages/default.aspx)) |
| Otu28489 | 1 | k__Fungi(100);p__Basidiomycota(100);c__Tremellomycetes(100);o__Tremellales(100);f__Tremellales_family_Incertae_sedis(100);g__Cryptococcus(100);g__Cryptococcus_unclassified(100); | Cryptococcus | 13 | Pathotroph-Saprotroph-Symbiotroph | Animal Pathogen-Endophyte-Epiphyte-\|Undefined Saprotroph\| | Dimorphic Yeast-Yeast | NULL | Possible | As foliar_endophyte (Põlme et al. 2020); Decay leaf, fruit, seed, soil, dung, animal material (Põlme et al. 2020) | Andrews JH, et al. 1983. Phytopathology 73:228-234 ((https://doi.org/10.1094/Phyto-73-228)); Kurtzman CP, et al. (eds.) 2011. The Yeasts, a Taxonomic Study. Fifth Edition. Vols 1-3. Elsevier, San Diego (ISBN: 9780444521491); Põlme S, et al. 2020. Fungal Diversity 105:1-16 ((https://doi.org/10.1007/s13225-020-00466-2)) |
| Otu28490 | 1 | k__Fungi(100);p__Basidiomycota(100);c__Agaricomycetes(100);o__Thelephorales(100);f__Thelephoraceae(100);g__Tomentella(100);s__Tomentella_ellisii(100); | Tomentella | 13 | Symbiotroph | \|Ectomycorrhizal\| | Corticioid | NULL | Highly Probable | NULL | Rinaldi AC, et al. 2008. Fungal Diversity 33:1-45 ((http://www.fungaldiversity.org/fdp/sfdp/33-1.pdf)); Tedersoo L, et al. 2010. Mycorrhiza 20:217-263 ((https://doi.org/10.1007/s00572-009-0274-x)); Põlme S, et al. 2020. Fungal Diversity 105:1-16 ((https://doi.org/10.1007/s13225-020-00466-2)); http://mycorrhizas.info/ecmf.html ((http://mycorrhizas.info/ecmf.html)) |
| Otu28491 | 1 | k__Fungi(100);p__Basidiomycota(100);c__Tremellomycetes(100);o__Tremellales(100);f__Tremellales_family_Incertae_sedis(100);g__Kockovaella(100);g__Kockovaella_unclassified(100); | Kockovaella | 13 | Saprotroph-Symbiotroph | Endophyte-\|Epiphyte\|-Undefined Saprotroph | Yeast | NULL | Probable | As foliar_endophyte (Põlme et al. 2020); Decay leaf, fruit, seed (Põlme et al. 2020); Foliar, isolated from a plant sample collected in a tropical rain forest in Thailand. | Fungsin B, et al. 2002. International Journal of Systematic and Evolutionary Microbiology 52:281-284 ((https://doi.org/10.1099/00207713-52-1-281)); Põlme S, et al. 2020. Fungal Diversity 105:1-16 ((https://doi.org/10.1007/s13225-020-00466-2)) |
| Otu28492 | 1 | k__Fungi(100);p__Basidiomycota(100);c__Tremellomycetes(100);o__Tremellales(100);f__Tremellales_family_Incertae_sedis(100);g__Cryptococcus(100);s__Cryptococcus_aff_amylolyticus_AS_22501(100); | Cryptococcus | 13 | Pathotroph-Saprotroph-Symbiotroph | Animal Pathogen-Endophyte-Epiphyte-\|Undefined Saprotroph\| | Dimorphic Yeast-Yeast | NULL | Possible | As foliar_endophyte (Põlme et al. 2020); Decay leaf, fruit, seed, soil, dung, animal material (Põlme et al. 2020) | Andrews JH, et al. 1983. Phytopathology 73:228-234 ((https://doi.org/10.1094/Phyto-73-228)); Kurtzman CP, et al. (eds.) 2011. The Yeasts, a Taxonomic Study. Fifth Edition. Vols 1-3. Elsevier, San Diego (ISBN: 9780444521491); Põlme S, et al. 2020. Fungal Diversity 105:1-16 ((https://doi.org/10.1007/s13225-020-00466-2)) |
| Otu28495 | 1 | k__Fungi(100);p__Basidiomycota(100);c__Agaricomycetes(100);o__Agaricales(100);f__Mycenaceae(100);g__Mycena(100);s__Mycena_plumipes(100); | Mycena | 13 | Pathotroph-Saprotroph | Plant Pathogen-\|Plant Saprotroph\|-Undefined Saprotroph-Wood Saprotroph | Agaricoid | NULL | Probable | As litter_saprotroph (Põlme et al. 2020); Decay leaf, fruit, seed (Põlme et al. 2020) | Grgurinovic CA, Simpson JA. 2001. Fungal Diversity 8:97-127 ((http://www.fungaldiversity.org/fdp/sfdp/FD_8_97-127.pdf)); Tedersoo L, et al. 2014. Science 346:e1256688 ((https://doi.org/10.1126/science.1256688)); Põlme S, et al. 2020. Fungal Diversity 105:1-16 ((https://doi.org/10.1007/s13225-020-00466-2)); The American Phytopathological Society Common Names of Plant Diseases ((http://www.apsnet.org/publications/commonnames/Pages/default.aspx)) |
| Otu28496 | 1 | k__Fungi(100);p__Basidiomycota(100);c__Agaricomycetes(100);o__Thelephorales(100);f__Thelephoraceae(100);g__Tomentella(100);g__Tomentella_unclassified(100); | Tomentella | 13 | Symbiotroph | \|Ectomycorrhizal\| | Corticioid | NULL | Highly Probable | NULL | Rinaldi AC, et al. 2008. Fungal Diversity 33:1-45 ((http://www.fungaldiversity.org/fdp/sfdp/33-1.pdf)); Tedersoo L, et al. 2010. Mycorrhiza 20:217-263 ((https://doi.org/10.1007/s00572-009-0274-x)); Põlme S, et al. 2020. Fungal Diversity 105:1-16 ((https://doi.org/10.1007/s13225-020-00466-2)); http://mycorrhizas.info/ecmf.html ((http://mycorrhizas.info/ecmf.html)) |
| Otu28499 | 1 | k__Fungi(100);p__Basidiomycota(100);c__Agaricomycetes(100);o__Agaricales(100);f__Schizophyllaceae(100);g__Schizophyllum(100);s__Schizophyllum_commune(100); | Schizophyllum | 13 | Saprotroph | Plant Saprotroph-\|Wood Saprotroph\| | Agaricoid | White Rot | Highly Probable | As litter_saprotroph (Põlme et al. 2020); As white_rot (Põlme et al. 2020); Decay wood, leaf, fruit, seed (Põlme et al. 2020) | Ahmada SS, Miles PG. 1970. Genetical Research 15:19-28 ((https://doi.org/10.1017/S0016672300001336)); Põlme S, et al. 2020. Fungal Diversity 105:1-16 ((https://doi.org/10.1007/s13225-020-00466-2)) |
| Otu28500 | 1 | k__Fungi(100);p__Basidiomycota(100);c__Agaricomycetes(100);o__Agaricales(100);f__Schizophyllaceae(100);g__Schizophyllum(100);s__Schizophyllum_commune(100); | Schizophyllum | 13 | Saprotroph | Plant Saprotroph-\|Wood Saprotroph\| | Agaricoid | White Rot | Highly Probable | As litter_saprotroph (Põlme et al. 2020); As white_rot (Põlme et al. 2020); Decay wood, leaf, fruit, seed (Põlme et al. 2020) | Ahmada SS, Miles PG. 1970. Genetical Research 15:19-28 ((https://doi.org/10.1017/S0016672300001336)); Põlme S, et al. 2020. Fungal Diversity 105:1-16 ((https://doi.org/10.1007/s13225-020-00466-2)) |
| Otu28501 | 1 | k__Fungi(100);p__Basidiomycota(100);c__Agaricomycetes(100);o__Agaricales(100);f__Schizophyllaceae(100);g__Schizophyllum(100);s__Schizophyllum_commune(100); | Schizophyllum | 13 | Saprotroph | Plant Saprotroph-\|Wood Saprotroph\| | Agaricoid | White Rot | Highly Probable | As litter_saprotroph (Põlme et al. 2020); As white_rot (Põlme et al. 2020); Decay wood, leaf, fruit, seed (Põlme et al. 2020) | Ahmada SS, Miles PG. 1970. Genetical Research 15:19-28 ((https://doi.org/10.1017/S0016672300001336)); Põlme S, et al. 2020. Fungal Diversity 105:1-16 ((https://doi.org/10.1007/s13225-020-00466-2)) |
| Otu28502 | 1 | k__Fungi(100);p__Basidiomycota(100);c__Tremellomycetes(100);o__Tremellales(100);f__Tremellales_family_Incertae_sedis(100);g__Cryptococcus(100);s__Cryptococcus_laurentii(100); | Cryptococcus | 13 | Pathotroph-Saprotroph-Symbiotroph | Animal Pathogen-Endophyte-Epiphyte-\|Undefined Saprotroph\| | Dimorphic Yeast-Yeast | NULL | Possible | As foliar_endophyte (Põlme et al. 2020); Decay leaf, fruit, seed, soil, dung, animal material (Põlme et al. 2020) | Andrews JH, et al. 1983. Phytopathology 73:228-234 ((https://doi.org/10.1094/Phyto-73-228)); Kurtzman CP, et al. (eds.) 2011. The Yeasts, a Taxonomic Study. Fifth Edition. Vols 1-3. Elsevier, San Diego (ISBN: 9780444521491); Põlme S, et al. 2020. Fungal Diversity 105:1-16 ((https://doi.org/10.1007/s13225-020-00466-2)) |
| Otu28503 | 1 | k__Fungi(100);p__Basidiomycota(100);c__Agaricomycetes(100);o__Russulales(100);f__Stereaceae(100);g__Stereum(100);s__Stereum_sp_Vega483(100); | Stereum | 13 | Saprotroph | Undefined Saprotroph-\|Wood Saprotroph\| | Polyporoid | White Rot | Probable | As white_rot (Põlme et al. 2020); Decay wood (Põlme et al. 2020) | Tedersoo L, et al. 2014. Science 346:e1256688 ((https://doi.org/10.1126/science.1256688)); Põlme S, et al. 2020. Fungal Diversity 105:1-16 ((https://doi.org/10.1007/s13225-020-00466-2)) |
| Otu28504 | 1 | k__Fungi(100);p__Basidiomycota(100);c__Agaricomycetes(100);o__Agaricales(100);f__Clavariaceae(100);g__unclassified_Clavariaceae(100);s__Clavariaceae_sp(100); | Clavariaceae | 9 | Saprotroph-Symbiotroph | Lichenized-Undefined Saprotroph | Clavarioid | NULL | Probable | Most species are presumed saprobic on the ground or rotten wood, but many Multiclavula species are lichenized with green algae (Cannon & Kirk 2007) | Cannon PF, Kirk PM. 2007. Fungal Families of the World. CAB International, Cambridge (ISBN: 978-0851998275) |
| Otu28505 | 1 | k__Fungi(100);p__Basidiomycota(100);c__Agaricomycetes(100);o__Agaricales(100);f__Mycenaceae(100);g__Mycena(100);g__Mycena_unclassified(100); | Mycena | 13 | Pathotroph-Saprotroph | Plant Pathogen-\|Plant Saprotroph\|-Undefined Saprotroph-Wood Saprotroph | Agaricoid | NULL | Probable | As litter_saprotroph (Põlme et al. 2020); Decay leaf, fruit, seed (Põlme et al. 2020) | Grgurinovic CA, Simpson JA. 2001. Fungal Diversity 8:97-127 ((http://www.fungaldiversity.org/fdp/sfdp/FD_8_97-127.pdf)); Tedersoo L, et al. 2014. Science 346:e1256688 ((https://doi.org/10.1126/science.1256688)); Põlme S, et al. 2020. Fungal Diversity 105:1-16 ((https://doi.org/10.1007/s13225-020-00466-2)); The American Phytopathological Society Common Names of Plant Diseases ((http://www.apsnet.org/publications/commonnames/Pages/default.aspx)) |
| Otu28506 | 1 | k__Fungi(100);p__Basidiomycota(100);c__Tremellomycetes(100);o__Tremellales(100);f__Tremellales_family_Incertae_sedis(100);g__Filobasidiella(100);s__Cryptococcus_sp_Y6_3(100); | Cryptococcus | 13 | Pathotroph-Saprotroph-Symbiotroph | Animal Pathogen-Endophyte-Epiphyte-\|Undefined Saprotroph\| | Dimorphic Yeast-Yeast | NULL | Possible | As foliar_endophyte (Põlme et al. 2020); Decay leaf, fruit, seed, soil, dung, animal material (Põlme et al. 2020) | Andrews JH, et al. 1983. Phytopathology 73:228-234 ((https://doi.org/10.1094/Phyto-73-228)); Kurtzman CP, et al. (eds.) 2011. The Yeasts, a Taxonomic Study. Fifth Edition. Vols 1-3. Elsevier, San Diego (ISBN: 9780444521491); Põlme S, et al. 2020. Fungal Diversity 105:1-16 ((https://doi.org/10.1007/s13225-020-00466-2)) |
| Otu28508 | 1 | k__Fungi(100);p__Basidiomycota(100);c__Agaricomycetes(100);o__Russulales(100);f__Stereaceae(100);g__Gloeocystidiellum(100);g__Gloeocystidiellum_unclassified(100); | Gloeocystidiellum | 13 | Saprotroph | Undefined Saprotroph-\|Wood Saprotroph\| | Corticioid | White Rot | Probable | As white_rot (Põlme et al. 2020); Decay wood (Põlme et al. 2020) | Tedersoo L, et al. 2014. Science 346:e1256688 ((https://doi.org/10.1126/science.1256688)); Põlme S, et al. 2020. Fungal Diversity 105:1-16 ((https://doi.org/10.1007/s13225-020-00466-2)) |
| Otu28509 | 1 | k__Fungi(100);p__Basidiomycota(100);c__Agaricomycetes(100);o__Hymenochaetales(100);f__Hymenochaetaceae(100);g__Hymenochaete(100);g__Hymenochaete_unclassified(100); | Hymenochaete | 13 | Pathotroph-Saprotroph | Plant Pathogen-Undefined Saprotroph-\|Wood Saprotroph\| | Corticioid-Polyporoid | White Rot | Probable | As white_rot (Põlme et al. 2020); As wood_pathogen (Põlme et al. 2020); Decay wood (Põlme et al. 2020) | Tedersoo L, et al. 2014. Science 346:e1256688 ((https://doi.org/10.1126/science.1256688)); Põlme S, et al. 2020. Fungal Diversity 105:1-16 ((https://doi.org/10.1007/s13225-020-00466-2)) |
| Otu28510 | 1 | k__Fungi(100);p__Basidiomycota(100);c__Agaricomycetes(100);o__Russulales(100);f__Stereaceae(100);g__Stereum(100);g__Stereum_unclassified(100); | Stereum | 13 | Saprotroph | Undefined Saprotroph-\|Wood Saprotroph\| | Polyporoid | White Rot | Probable | As white_rot (Põlme et al. 2020); Decay wood (Põlme et al. 2020) | Tedersoo L, et al. 2014. Science 346:e1256688 ((https://doi.org/10.1126/science.1256688)); Põlme S, et al. 2020. Fungal Diversity 105:1-16 ((https://doi.org/10.1007/s13225-020-00466-2)) |
| Otu28511 | 1 | k__Fungi(100);p__Basidiomycota(100);c__Agaricomycetes(100);o__Russulales(100);f__Stereaceae(100);g__Gloeocystidiellum(100);s__Gloeocystidiellum_sp(100); | Gloeocystidiellum | 13 | Saprotroph | Undefined Saprotroph-\|Wood Saprotroph\| | Corticioid | White Rot | Probable | As white_rot (Põlme et al. 2020); Decay wood (Põlme et al. 2020) | Tedersoo L, et al. 2014. Science 346:e1256688 ((https://doi.org/10.1126/science.1256688)); Põlme S, et al. 2020. Fungal Diversity 105:1-16 ((https://doi.org/10.1007/s13225-020-00466-2)) |
| Otu28512 | 1 | k__Fungi(100);p__Basidiomycota(100);c__Tremellomycetes(100);o__Tremellales(100);f__Tremellales_family_Incertae_sedis(100);g__Cryptococcus(100);s__Cryptococcus_sp_TC2_092(100); | Cryptococcus | 13 | Pathotroph-Saprotroph-Symbiotroph | Animal Pathogen-Endophyte-Epiphyte-\|Undefined Saprotroph\| | Dimorphic Yeast-Yeast | NULL | Possible | As foliar_endophyte (Põlme et al. 2020); Decay leaf, fruit, seed, soil, dung, animal material (Põlme et al. 2020) | Andrews JH, et al. 1983. Phytopathology 73:228-234 ((https://doi.org/10.1094/Phyto-73-228)); Kurtzman CP, et al. (eds.) 2011. The Yeasts, a Taxonomic Study. Fifth Edition. Vols 1-3. Elsevier, San Diego (ISBN: 9780444521491); Põlme S, et al. 2020. Fungal Diversity 105:1-16 ((https://doi.org/10.1007/s13225-020-00466-2)) |
| Otu28513 | 1 | k__Fungi(100);p__Basidiomycota(100);c__Agaricomycetes(100);o__Thelephorales(100);f__Thelephoraceae(100);f__Thelephoraceae_unclassified(100);f__Thelephoraceae_unclassified(100); | Thelephoraceae | 9 | Saprotroph-Symbiotroph | Ectomycorrhizal-Undefined Saprotroph | Clavarioid | NULL | Probable | On the ground, on soil, a few on wood. At least some speices ectomycorrhizal (Cannon & Kirk 2007) | Cannon PF, Kirk PM. 2007. Fungal Families of the World. CAB International, Cambridge (ISBN: 978-0851998275) |
| Otu28515 | 1 | k__Fungi(100);p__Basidiomycota(100);c__Microbotryomycetes(100);o__Sporidiobolales(100);f__Sporidiobolales_family_Incertae_sedis(100);g__Rhodotorula(100);s__Rhodotorula_taiwanensis(100); | Rhodotorula | 13 | Pathotroph-Saprotroph-Symbiotroph | Animal Pathogen-Animal Symbiotroph-Endophyte-Plant Pathogen-\|Undefined Saprotroph\| | Yeast | NULL | Probable | As foliar_endophyte (Põlme et al. 2020); Opportunistic human pathogen | Põlme S, et al. 2020. Fungal Diversity 105:1-16 ((https://doi.org/10.1007/s13225-020-00466-2)); https://www.cdc.gov ((https://www.cdc.gov)) |
| Otu28516 | 1 | k__Fungi(100);p__Basidiomycota(100);c__Microbotryomycetes(100);o__Sporidiobolales(100);f__Sporidiobolales_family_Incertae_sedis(100);g__Rhodotorula(100);s__Rhodotorula_taiwanensis(100); | Rhodotorula | 13 | Pathotroph-Saprotroph-Symbiotroph | Animal Pathogen-Animal Symbiotroph-Endophyte-Plant Pathogen-\|Undefined Saprotroph\| | Yeast | NULL | Probable | As foliar_endophyte (Põlme et al. 2020); Opportunistic human pathogen | Põlme S, et al. 2020. Fungal Diversity 105:1-16 ((https://doi.org/10.1007/s13225-020-00466-2)); https://www.cdc.gov ((https://www.cdc.gov)) |
| Otu28518 | 1 | k__Fungi(100);p__Basidiomycota(100);c__Microbotryomycetes(100);o__Sporidiobolales(100);f__Sporidiobolales_family_Incertae_sedis(100);g__Rhodotorula(100);s__Rhodotorula_taiwanensis(100); | Rhodotorula | 13 | Pathotroph-Saprotroph-Symbiotroph | Animal Pathogen-Animal Symbiotroph-Endophyte-Plant Pathogen-\|Undefined Saprotroph\| | Yeast | NULL | Probable | As foliar_endophyte (Põlme et al. 2020); Opportunistic human pathogen | Põlme S, et al. 2020. Fungal Diversity 105:1-16 ((https://doi.org/10.1007/s13225-020-00466-2)); https://www.cdc.gov ((https://www.cdc.gov)) |
| Otu28519 | 1 | k__Fungi(100);p__Basidiomycota(100);c__Microbotryomycetes(100);o__Sporidiobolales(100);f__Sporidiobolales_family_Incertae_sedis(100);g__Rhodotorula(100);g__Rhodotorula_unclassified(100); | Rhodotorula | 13 | Pathotroph-Saprotroph-Symbiotroph | Animal Pathogen-Animal Symbiotroph-Endophyte-Plant Pathogen-\|Undefined Saprotroph\| | Yeast | NULL | Probable | As foliar_endophyte (Põlme et al. 2020); Opportunistic human pathogen | Põlme S, et al. 2020. Fungal Diversity 105:1-16 ((https://doi.org/10.1007/s13225-020-00466-2)); https://www.cdc.gov ((https://www.cdc.gov)) |
| Otu28520 | 1 | k__Fungi(100);p__Basidiomycota(100);c__Microbotryomycetes(100);o__Sporidiobolales(100);f__Sporidiobolales_family_Incertae_sedis(100);g__Rhodotorula(100);s__Rhodotorula_mucilaginosa(100); | Rhodotorula | 13 | Pathotroph-Saprotroph-Symbiotroph | Animal Pathogen-Animal Symbiotroph-Endophyte-Plant Pathogen-\|Undefined Saprotroph\| | Yeast | NULL | Probable | As foliar_endophyte (Põlme et al. 2020); Opportunistic human pathogen | Põlme S, et al. 2020. Fungal Diversity 105:1-16 ((https://doi.org/10.1007/s13225-020-00466-2)); https://www.cdc.gov ((https://www.cdc.gov)) |
| Otu28521 | 1 | k__Fungi(100);p__Basidiomycota(100);c__Microbotryomycetes(100);o__Sporidiobolales(100);f__Sporidiobolales_family_Incertae_sedis(100);g__Rhodotorula(100);s__Rhodotorula_nothofagi(100); | Rhodotorula | 13 | Pathotroph-Saprotroph-Symbiotroph | Animal Pathogen-Animal Symbiotroph-Endophyte-Plant Pathogen-\|Undefined Saprotroph\| | Yeast | NULL | Probable | As foliar_endophyte (Põlme et al. 2020); Opportunistic human pathogen | Põlme S, et al. 2020. Fungal Diversity 105:1-16 ((https://doi.org/10.1007/s13225-020-00466-2)); https://www.cdc.gov ((https://www.cdc.gov)) |
| Otu28522 | 1 | k__Fungi(100);p__Basidiomycota(100);c__Microbotryomycetes(100);o__Sporidiobolales(100);f__Sporidiobolales_family_Incertae_sedis(100);g__Rhodotorula(100);s__Rhodotorula_taiwanensis(100); | Rhodotorula | 13 | Pathotroph-Saprotroph-Symbiotroph | Animal Pathogen-Animal Symbiotroph-Endophyte-Plant Pathogen-\|Undefined Saprotroph\| | Yeast | NULL | Probable | As foliar_endophyte (Põlme et al. 2020); Opportunistic human pathogen | Põlme S, et al. 2020. Fungal Diversity 105:1-16 ((https://doi.org/10.1007/s13225-020-00466-2)); https://www.cdc.gov ((https://www.cdc.gov)) |
| Otu28523 | 1 | k__Fungi(100);p__Basidiomycota(100);c__Agaricomycetes(100);o__Auriculariales(100);f__Auriculariales_family_Incertae_sedis(100);g__Exidia(100);g__Exidia_unclassified(100); | Exidia | 13 | Saprotroph-Symbiotroph | Endophyte-Plant Saprotroph-Undefined Saprotroph-\|Wood Saprotroph\| | Tremelloid | NULL | Probable | As foliar_endophyte (Põlme et al. 2020); As litter_saprotroph (Põlme et al. 2020); Decay wood, leaf, fruit, seed (Põlme et al. 2020) | Tedersoo L, et al. 2014. Science 346:e1256688 ((https://doi.org/10.1126/science.1256688)); David AS, et al. 2016. Microbial Ecology 71:912-926 ((https://doi.org/10.1007/s00248-015-0712-6)); Põlme S, et al. 2020. Fungal Diversity 105:1-16 ((https://doi.org/10.1007/s13225-020-00466-2)) |
| Otu28524 | 1 | k__Fungi(100);p__Basidiomycota(100);c__Microbotryomycetes(100);o__Sporidiobolales(100);f__Sporidiobolales_family_Incertae_sedis(100);g__Rhodotorula(100);s__Rhodotorula_taiwanensis(100); | Rhodotorula | 13 | Pathotroph-Saprotroph-Symbiotroph | Animal Pathogen-Animal Symbiotroph-Endophyte-Plant Pathogen-\|Undefined Saprotroph\| | Yeast | NULL | Probable | As foliar_endophyte (Põlme et al. 2020); Opportunistic human pathogen | Põlme S, et al. 2020. Fungal Diversity 105:1-16 ((https://doi.org/10.1007/s13225-020-00466-2)); https://www.cdc.gov ((https://www.cdc.gov)) |
| Otu28525 | 1 | k__Fungi(100);p__Basidiomycota(100);c__Microbotryomycetes(100);o__Sporidiobolales(100);f__Sporidiobolales_family_Incertae_sedis(100);g__Rhodotorula(100);s__Rhodotorula_mucilaginosa(100); | Rhodotorula | 13 | Pathotroph-Saprotroph-Symbiotroph | Animal Pathogen-Animal Symbiotroph-Endophyte-Plant Pathogen-\|Undefined Saprotroph\| | Yeast | NULL | Probable | As foliar_endophyte (Põlme et al. 2020); Opportunistic human pathogen | Põlme S, et al. 2020. Fungal Diversity 105:1-16 ((https://doi.org/10.1007/s13225-020-00466-2)); https://www.cdc.gov ((https://www.cdc.gov)) |
| Otu28526 | 1 | k__Fungi(100);p__Basidiomycota(100);c__Microbotryomycetes(100);o__Sporidiobolales(100);f__Sporidiobolales_family_Incertae_sedis(100);g__Rhodotorula(100);s__Rhodotorula_taiwanensis(100); | Rhodotorula | 13 | Pathotroph-Saprotroph-Symbiotroph | Animal Pathogen-Animal Symbiotroph-Endophyte-Plant Pathogen-\|Undefined Saprotroph\| | Yeast | NULL | Probable | As foliar_endophyte (Põlme et al. 2020); Opportunistic human pathogen | Põlme S, et al. 2020. Fungal Diversity 105:1-16 ((https://doi.org/10.1007/s13225-020-00466-2)); https://www.cdc.gov ((https://www.cdc.gov)) |
| Otu28527 | 1 | k__Fungi(100);p__Basidiomycota(100);c__Microbotryomycetes(100);o__Sporidiobolales(100);f__Sporidiobolales_family_Incertae_sedis(100);g__Rhodotorula(100);s__Rhodotorula_mucilaginosa(100); | Rhodotorula | 13 | Pathotroph-Saprotroph-Symbiotroph | Animal Pathogen-Animal Symbiotroph-Endophyte-Plant Pathogen-\|Undefined Saprotroph\| | Yeast | NULL | Probable | As foliar_endophyte (Põlme et al. 2020); Opportunistic human pathogen | Põlme S, et al. 2020. Fungal Diversity 105:1-16 ((https://doi.org/10.1007/s13225-020-00466-2)); https://www.cdc.gov ((https://www.cdc.gov)) |
| Otu28528 | 1 | k__Fungi(100);p__Basidiomycota(100);c__Microbotryomycetes(100);o__Sporidiobolales(100);f__Sporidiobolales_family_Incertae_sedis(100);g__Rhodotorula(100);s__Rhodotorula_taiwanensis(100); | Rhodotorula | 13 | Pathotroph-Saprotroph-Symbiotroph | Animal Pathogen-Animal Symbiotroph-Endophyte-Plant Pathogen-\|Undefined Saprotroph\| | Yeast | NULL | Probable | As foliar_endophyte (Põlme et al. 2020); Opportunistic human pathogen | Põlme S, et al. 2020. Fungal Diversity 105:1-16 ((https://doi.org/10.1007/s13225-020-00466-2)); https://www.cdc.gov ((https://www.cdc.gov)) |
| Otu28529 | 1 | k__Fungi(100);p__Basidiomycota(100);c__Microbotryomycetes(100);o__Sporidiobolales(100);f__Sporidiobolales_family_Incertae_sedis(100);g__Rhodotorula(100);s__Rhodotorula_mucilaginosa(100); | Rhodotorula | 13 | Pathotroph-Saprotroph-Symbiotroph | Animal Pathogen-Animal Symbiotroph-Endophyte-Plant Pathogen-\|Undefined Saprotroph\| | Yeast | NULL | Probable | As foliar_endophyte (Põlme et al. 2020); Opportunistic human pathogen | Põlme S, et al. 2020. Fungal Diversity 105:1-16 ((https://doi.org/10.1007/s13225-020-00466-2)); https://www.cdc.gov ((https://www.cdc.gov)) |
| Otu28530 | 1 | k__Fungi(100);p__Basidiomycota(100);c__Microbotryomycetes(100);o__Sporidiobolales(100);f__Sporidiobolales_family_Incertae_sedis(100);g__Rhodotorula(100);s__Rhodotorula_taiwanensis(100); | Rhodotorula | 13 | Pathotroph-Saprotroph-Symbiotroph | Animal Pathogen-Animal Symbiotroph-Endophyte-Plant Pathogen-\|Undefined Saprotroph\| | Yeast | NULL | Probable | As foliar_endophyte (Põlme et al. 2020); Opportunistic human pathogen | Põlme S, et al. 2020. Fungal Diversity 105:1-16 ((https://doi.org/10.1007/s13225-020-00466-2)); https://www.cdc.gov ((https://www.cdc.gov)) |
| Otu28531 | 1 | k__Fungi(100);p__Basidiomycota(100);c__Microbotryomycetes(100);o__Sporidiobolales(100);f__Sporidiobolales_family_Incertae_sedis(100);g__Rhodotorula(100);s__Rhodotorula_mucilaginosa(100); | Rhodotorula | 13 | Pathotroph-Saprotroph-Symbiotroph | Animal Pathogen-Animal Symbiotroph-Endophyte-Plant Pathogen-\|Undefined Saprotroph\| | Yeast | NULL | Probable | As foliar_endophyte (Põlme et al. 2020); Opportunistic human pathogen | Põlme S, et al. 2020. Fungal Diversity 105:1-16 ((https://doi.org/10.1007/s13225-020-00466-2)); https://www.cdc.gov ((https://www.cdc.gov)) |
| Otu28532 | 1 | k__Fungi(100);p__Basidiomycota(100);c__Microbotryomycetes(100);o__Sporidiobolales(100);f__Sporidiobolales_family_Incertae_sedis(100);g__Rhodotorula(100);s__Rhodotorula_taiwanensis(100); | Rhodotorula | 13 | Pathotroph-Saprotroph-Symbiotroph | Animal Pathogen-Animal Symbiotroph-Endophyte-Plant Pathogen-\|Undefined Saprotroph\| | Yeast | NULL | Probable | As foliar_endophyte (Põlme et al. 2020); Opportunistic human pathogen | Põlme S, et al. 2020. Fungal Diversity 105:1-16 ((https://doi.org/10.1007/s13225-020-00466-2)); https://www.cdc.gov ((https://www.cdc.gov)) |
| Otu28533 | 1 | k__Fungi(100);p__Basidiomycota(100);c__Microbotryomycetes(100);o__Sporidiobolales(100);f__Sporidiobolales_family_Incertae_sedis(100);g__Rhodotorula(100);s__Rhodotorula_taiwanensis(100); | Rhodotorula | 13 | Pathotroph-Saprotroph-Symbiotroph | Animal Pathogen-Animal Symbiotroph-Endophyte-Plant Pathogen-\|Undefined Saprotroph\| | Yeast | NULL | Probable | As foliar_endophyte (Põlme et al. 2020); Opportunistic human pathogen | Põlme S, et al. 2020. Fungal Diversity 105:1-16 ((https://doi.org/10.1007/s13225-020-00466-2)); https://www.cdc.gov ((https://www.cdc.gov)) |
| Otu28534 | 1 | k__Fungi(100);p__Basidiomycota(100);c__Microbotryomycetes(100);o__Sporidiobolales(100);f__Sporidiobolales_family_Incertae_sedis(100);g__Rhodotorula(100);s__Rhodotorula_taiwanensis(100); | Rhodotorula | 13 | Pathotroph-Saprotroph-Symbiotroph | Animal Pathogen-Animal Symbiotroph-Endophyte-Plant Pathogen-\|Undefined Saprotroph\| | Yeast | NULL | Probable | As foliar_endophyte (Põlme et al. 2020); Opportunistic human pathogen | Põlme S, et al. 2020. Fungal Diversity 105:1-16 ((https://doi.org/10.1007/s13225-020-00466-2)); https://www.cdc.gov ((https://www.cdc.gov)) |
| Otu28535 | 1 | k__Fungi(100);p__Basidiomycota(100);c__Microbotryomycetes(100);o__Sporidiobolales(100);f__Sporidiobolales_family_Incertae_sedis(100);g__Rhodotorula(100);g__Rhodotorula_unclassified(100); | Rhodotorula | 13 | Pathotroph-Saprotroph-Symbiotroph | Animal Pathogen-Animal Symbiotroph-Endophyte-Plant Pathogen-\|Undefined Saprotroph\| | Yeast | NULL | Probable | As foliar_endophyte (Põlme et al. 2020); Opportunistic human pathogen | Põlme S, et al. 2020. Fungal Diversity 105:1-16 ((https://doi.org/10.1007/s13225-020-00466-2)); https://www.cdc.gov ((https://www.cdc.gov)) |
| Otu28537 | 1 | k__Fungi(100);p__Basidiomycota(100);c__Tremellomycetes(100);o__Tremellales(100);f__Tremellales_family_Incertae_sedis(100);g__Cryptococcus(100);s__Cryptococcus_laurentii(100); | Cryptococcus | 13 | Pathotroph-Saprotroph-Symbiotroph | Animal Pathogen-Endophyte-Epiphyte-\|Undefined Saprotroph\| | Dimorphic Yeast-Yeast | NULL | Possible | As foliar_endophyte (Põlme et al. 2020); Decay leaf, fruit, seed, soil, dung, animal material (Põlme et al. 2020) | Andrews JH, et al. 1983. Phytopathology 73:228-234 ((https://doi.org/10.1094/Phyto-73-228)); Kurtzman CP, et al. (eds.) 2011. The Yeasts, a Taxonomic Study. Fifth Edition. Vols 1-3. Elsevier, San Diego (ISBN: 9780444521491); Põlme S, et al. 2020. Fungal Diversity 105:1-16 ((https://doi.org/10.1007/s13225-020-00466-2)) |
| Otu28539 | 1 | k__Fungi(100);p__Basidiomycota(100);c__Tremellomycetes(100);o__Tremellales(100);f__Tremellales_family_Incertae_sedis(100);g__Bullera(100);s__Bullera_pseudoalba(100); | Bullera | 13 | Saprotroph-Symbiotroph | Endophyte-Epiphyte-Plant Saprotroph-\|Undefined Saprotroph\| | Yeast | NULL | Probable | As foliar_endophyte (Põlme et al. 2020); Isolated from dead leaves of Oryza sativa, Miscanthus sinensis and Sasa sp. in Japan (Nakase & Suzuki 1986; Syn. Udeniomyces puniceus); Foliar, isolated from leaves of Metasolenae candidum (Nakase et al. 2004) | Nakase T, Suzuki M. 1986. Journal of General and Applied Microbiology Tokyo 32:225-240 ((https://doi.org/10.2323/jgam.32.225)); Nakase T, et al. 2004. Systematic and Applied Microbiology 27:558-564 ((https://doi.org/10.1078/0723202041748118)) |
| Otu28540 | 1 | k__Fungi(100);p__Basidiomycota(100);c__Tremellomycetes(100);o__Tremellales(100);f__Tremellales_family_Incertae_sedis(100);g__Cryptococcus(100);s__Cryptococcus_laurentii(100); | Cryptococcus | 13 | Pathotroph-Saprotroph-Symbiotroph | Animal Pathogen-Endophyte-Epiphyte-\|Undefined Saprotroph\| | Dimorphic Yeast-Yeast | NULL | Possible | As foliar_endophyte (Põlme et al. 2020); Decay leaf, fruit, seed, soil, dung, animal material (Põlme et al. 2020) | Andrews JH, et al. 1983. Phytopathology 73:228-234 ((https://doi.org/10.1094/Phyto-73-228)); Kurtzman CP, et al. (eds.) 2011. The Yeasts, a Taxonomic Study. Fifth Edition. Vols 1-3. Elsevier, San Diego (ISBN: 9780444521491); Põlme S, et al. 2020. Fungal Diversity 105:1-16 ((https://doi.org/10.1007/s13225-020-00466-2)) |
| Otu28541 | 1 | k__Fungi(100);p__Basidiomycota(100);c__Microbotryomycetes(100);o__Sporidiobolales(100);f__Sporidiobolales_family_Incertae_sedis(100);g__Rhodotorula(100);s__Rhodotorula_sp(100); | Rhodotorula | 13 | Pathotroph-Saprotroph-Symbiotroph | Animal Pathogen-Animal Symbiotroph-Endophyte-Plant Pathogen-\|Undefined Saprotroph\| | Yeast | NULL | Probable | As foliar_endophyte (Põlme et al. 2020); Opportunistic human pathogen | Põlme S, et al. 2020. Fungal Diversity 105:1-16 ((https://doi.org/10.1007/s13225-020-00466-2)); https://www.cdc.gov ((https://www.cdc.gov)) |
| Otu28545 | 1 | k__Fungi(100);p__Basidiomycota(100);c__Exobasidiomycetes(100);o__Microstromatales(100);f__Microstromataceae(100);g__Microstroma(100);s__Microstroma_juglandis(100); | Microstroma | 13 | Pathotroph | \|Plant Pathogen\| | Dimorphic | NULL | Probable | NULL | Tedersoo L, et al. 2014. Science 346:e1256688 ((https://doi.org/10.1126/science.1256688)); Põlme S, et al. 2020. Fungal Diversity 105:1-16 ((https://doi.org/10.1007/s13225-020-00466-2)) |
| Otu28546 | 1 | k__Fungi(100);p__Basidiomycota(100);c__Microbotryomycetes(100);o__Sporidiobolales(100);f__Sporidiobolales_family_Incertae_sedis(100);g__Rhodotorula(100);s__Rhodotorula_sp(100); | Rhodotorula | 13 | Pathotroph-Saprotroph-Symbiotroph | Animal Pathogen-Animal Symbiotroph-Endophyte-Plant Pathogen-\|Undefined Saprotroph\| | Yeast | NULL | Probable | As foliar_endophyte (Põlme et al. 2020); Opportunistic human pathogen | Põlme S, et al. 2020. Fungal Diversity 105:1-16 ((https://doi.org/10.1007/s13225-020-00466-2)); https://www.cdc.gov ((https://www.cdc.gov)) |
| Otu28547 | 1 | k__Fungi(100);p__Basidiomycota(100);c__Microbotryomycetes(100);o__Sporidiobolales(100);f__Sporidiobolales_family_Incertae_sedis(100);g__Rhodotorula(100);s__Rhodotorula_sp(100); | Rhodotorula | 13 | Pathotroph-Saprotroph-Symbiotroph | Animal Pathogen-Animal Symbiotroph-Endophyte-Plant Pathogen-\|Undefined Saprotroph\| | Yeast | NULL | Probable | As foliar_endophyte (Põlme et al. 2020); Opportunistic human pathogen | Põlme S, et al. 2020. Fungal Diversity 105:1-16 ((https://doi.org/10.1007/s13225-020-00466-2)); https://www.cdc.gov ((https://www.cdc.gov)) |
| Otu28548 | 1 | k__Fungi(100);p__Basidiomycota(100);c__Exobasidiomycetes(100);o__Microstromatales(100);f__Microstromataceae(100);g__Microstroma(100);s__Microstroma_juglandis(100); | Microstroma | 13 | Pathotroph | \|Plant Pathogen\| | Dimorphic | NULL | Probable | NULL | Tedersoo L, et al. 2014. Science 346:e1256688 ((https://doi.org/10.1126/science.1256688)); Põlme S, et al. 2020. Fungal Diversity 105:1-16 ((https://doi.org/10.1007/s13225-020-00466-2)) |
| Otu28551 | 1 | k__Fungi(100);p__Basidiomycota(100);c__Microbotryomycetes(100);o__Sporidiobolales(100);f__Sporidiobolales_family_Incertae_sedis(100);g__Rhodotorula(100);s__Rhodotorula_sp(100); | Rhodotorula | 13 | Pathotroph-Saprotroph-Symbiotroph | Animal Pathogen-Animal Symbiotroph-Endophyte-Plant Pathogen-\|Undefined Saprotroph\| | Yeast | NULL | Probable | As foliar_endophyte (Põlme et al. 2020); Opportunistic human pathogen | Põlme S, et al. 2020. Fungal Diversity 105:1-16 ((https://doi.org/10.1007/s13225-020-00466-2)); https://www.cdc.gov ((https://www.cdc.gov)) |
| Otu28552 | 1 | k__Fungi(100);p__Basidiomycota(100);c__Exobasidiomycetes(100);o__Microstromatales(100);f__Microstromataceae(100);g__Microstroma(100);s__Microstroma_juglandis(100); | Microstroma | 13 | Pathotroph | \|Plant Pathogen\| | Dimorphic | NULL | Probable | NULL | Tedersoo L, et al. 2014. Science 346:e1256688 ((https://doi.org/10.1126/science.1256688)); Põlme S, et al. 2020. Fungal Diversity 105:1-16 ((https://doi.org/10.1007/s13225-020-00466-2)) |
| Otu28553 | 1 | k__Fungi(100);p__Basidiomycota(100);c__Microbotryomycetes(100);o__Sporidiobolales(100);f__Sporidiobolales_family_Incertae_sedis(100);g__Rhodotorula(100);s__Rhodotorula_sp(100); | Rhodotorula | 13 | Pathotroph-Saprotroph-Symbiotroph | Animal Pathogen-Animal Symbiotroph-Endophyte-Plant Pathogen-\|Undefined Saprotroph\| | Yeast | NULL | Probable | As foliar_endophyte (Põlme et al. 2020); Opportunistic human pathogen | Põlme S, et al. 2020. Fungal Diversity 105:1-16 ((https://doi.org/10.1007/s13225-020-00466-2)); https://www.cdc.gov ((https://www.cdc.gov)) |
| Otu28554 | 1 | k__Fungi(100);p__Basidiomycota(100);c__Exobasidiomycetes(100);o__Microstromatales(100);f__Microstromataceae(100);g__Microstroma(100);s__Microstroma_juglandis(100); | Microstroma | 13 | Pathotroph | \|Plant Pathogen\| | Dimorphic | NULL | Probable | NULL | Tedersoo L, et al. 2014. Science 346:e1256688 ((https://doi.org/10.1126/science.1256688)); Põlme S, et al. 2020. Fungal Diversity 105:1-16 ((https://doi.org/10.1007/s13225-020-00466-2)) |
| Otu28555 | 1 | k__Fungi(100);p__Basidiomycota(100);c__Microbotryomycetes(100);o__Sporidiobolales(100);f__Sporidiobolales_family_Incertae_sedis(100);g__Rhodotorula(100);s__Rhodotorula_sp(100); | Rhodotorula | 13 | Pathotroph-Saprotroph-Symbiotroph | Animal Pathogen-Animal Symbiotroph-Endophyte-Plant Pathogen-\|Undefined Saprotroph\| | Yeast | NULL | Probable | As foliar_endophyte (Põlme et al. 2020); Opportunistic human pathogen | Põlme S, et al. 2020. Fungal Diversity 105:1-16 ((https://doi.org/10.1007/s13225-020-00466-2)); https://www.cdc.gov ((https://www.cdc.gov)) |
| Otu28556 | 1 | k__Fungi(100);p__Basidiomycota(100);c__Exobasidiomycetes(100);o__Microstromatales(100);f__Microstromataceae(100);g__Microstroma(100);s__Microstroma_juglandis(100); | Microstroma | 13 | Pathotroph | \|Plant Pathogen\| | Dimorphic | NULL | Probable | NULL | Tedersoo L, et al. 2014. Science 346:e1256688 ((https://doi.org/10.1126/science.1256688)); Põlme S, et al. 2020. Fungal Diversity 105:1-16 ((https://doi.org/10.1007/s13225-020-00466-2)) |
| Otu28557 | 1 | k__Fungi(100);p__Basidiomycota(100);c__Exobasidiomycetes(100);o__Microstromatales(100);f__Microstromataceae(100);g__Microstroma(100);s__Microstroma_juglandis(100); | Microstroma | 13 | Pathotroph | \|Plant Pathogen\| | Dimorphic | NULL | Probable | NULL | Tedersoo L, et al. 2014. Science 346:e1256688 ((https://doi.org/10.1126/science.1256688)); Põlme S, et al. 2020. Fungal Diversity 105:1-16 ((https://doi.org/10.1007/s13225-020-00466-2)) |
| Otu28558 | 1 | k__Fungi(100);p__Basidiomycota(100);c__Microbotryomycetes(100);o__Sporidiobolales(100);f__Sporidiobolales_family_Incertae_sedis(100);g__Rhodotorula(100);s__Rhodotorula_sp(100); | Rhodotorula | 13 | Pathotroph-Saprotroph-Symbiotroph | Animal Pathogen-Animal Symbiotroph-Endophyte-Plant Pathogen-\|Undefined Saprotroph\| | Yeast | NULL | Probable | As foliar_endophyte (Põlme et al. 2020); Opportunistic human pathogen | Põlme S, et al. 2020. Fungal Diversity 105:1-16 ((https://doi.org/10.1007/s13225-020-00466-2)); https://www.cdc.gov ((https://www.cdc.gov)) |
| Otu28559 | 1 | k__Fungi(100);p__Basidiomycota(100);c__Microbotryomycetes(100);o__Sporidiobolales(100);f__Sporidiobolales_family_Incertae_sedis(100);g__Rhodotorula(100);s__Rhodotorula_sp(100); | Rhodotorula | 13 | Pathotroph-Saprotroph-Symbiotroph | Animal Pathogen-Animal Symbiotroph-Endophyte-Plant Pathogen-\|Undefined Saprotroph\| | Yeast | NULL | Probable | As foliar_endophyte (Põlme et al. 2020); Opportunistic human pathogen | Põlme S, et al. 2020. Fungal Diversity 105:1-16 ((https://doi.org/10.1007/s13225-020-00466-2)); https://www.cdc.gov ((https://www.cdc.gov)) |
| Otu28560 | 1 | k__Fungi(100);p__Basidiomycota(100);c__Exobasidiomycetes(100);o__Microstromatales(100);f__Microstromataceae(100);g__Microstroma(100);s__Microstroma_juglandis(100); | Microstroma | 13 | Pathotroph | \|Plant Pathogen\| | Dimorphic | NULL | Probable | NULL | Tedersoo L, et al. 2014. Science 346:e1256688 ((https://doi.org/10.1126/science.1256688)); Põlme S, et al. 2020. Fungal Diversity 105:1-16 ((https://doi.org/10.1007/s13225-020-00466-2)) |
| Otu28561 | 1 | k__Fungi(100);p__Basidiomycota(100);c__Exobasidiomycetes(100);o__Microstromatales(100);f__Microstromataceae(100);g__Microstroma(100);s__Microstroma_juglandis(100); | Microstroma | 13 | Pathotroph | \|Plant Pathogen\| | Dimorphic | NULL | Probable | NULL | Tedersoo L, et al. 2014. Science 346:e1256688 ((https://doi.org/10.1126/science.1256688)); Põlme S, et al. 2020. Fungal Diversity 105:1-16 ((https://doi.org/10.1007/s13225-020-00466-2)) |
| Otu28563 | 1 | k__Fungi(100);p__Basidiomycota(100);c__Exobasidiomycetes(100);o__Microstromatales(100);f__Microstromataceae(100);g__Microstroma(100);s__Microstroma_juglandis(100); | Microstroma | 13 | Pathotroph | \|Plant Pathogen\| | Dimorphic | NULL | Probable | NULL | Tedersoo L, et al. 2014. Science 346:e1256688 ((https://doi.org/10.1126/science.1256688)); Põlme S, et al. 2020. Fungal Diversity 105:1-16 ((https://doi.org/10.1007/s13225-020-00466-2)) |
| Otu28565 | 1 | k__Fungi(100);p__Basidiomycota(100);c__Exobasidiomycetes(100);o__Microstromatales(100);f__Microstromataceae(100);g__Microstroma(100);s__Microstroma_juglandis(100); | Microstroma | 13 | Pathotroph | \|Plant Pathogen\| | Dimorphic | NULL | Probable | NULL | Tedersoo L, et al. 2014. Science 346:e1256688 ((https://doi.org/10.1126/science.1256688)); Põlme S, et al. 2020. Fungal Diversity 105:1-16 ((https://doi.org/10.1007/s13225-020-00466-2)) |
| Otu28568 | 1 | k__Fungi(100);p__Basidiomycota(100);c__Exobasidiomycetes(100);o__Microstromatales(100);f__Microstromataceae(100);g__Microstroma(100);s__Microstroma_juglandis(100); | Microstroma | 13 | Pathotroph | \|Plant Pathogen\| | Dimorphic | NULL | Probable | NULL | Tedersoo L, et al. 2014. Science 346:e1256688 ((https://doi.org/10.1126/science.1256688)); Põlme S, et al. 2020. Fungal Diversity 105:1-16 ((https://doi.org/10.1007/s13225-020-00466-2)) |
| Otu28569 | 1 | k__Fungi(100);p__Basidiomycota(100);c__Agaricomycetes(100);o__Boletales(100);f__Suillaceae(100);g__Suillus(100);s__Suillus_placidus(100); | Suillus | 13 | Symbiotroph | \|Ectomycorrhizal\| | Agaricoid-Boletoid | NULL | Highly Probable | Host - Pinaceae (Põlme et al. 2020) | Rinaldi AC, et al. 2008. Fungal Diversity 33:1-45 ((http://www.fungaldiversity.org/fdp/sfdp/33-1.pdf)); Tedersoo L, et al. 2010. Mycorrhiza 20:217-263 ((https://doi.org/10.1007/s00572-009-0274-x)); Põlme S, et al. 2020. Fungal Diversity 105:1-16 ((https://doi.org/10.1007/s13225-020-00466-2)); http://mycorrhizas.info/ecmf.html ((http://mycorrhizas.info/ecmf.html)) |
| Otu28570 | 1 | k__Fungi(100);p__Basidiomycota(100);c__Agaricomycetes(100);o__Boletales(100);f__Suillaceae(100);g__Suillus(100);s__Suillus_placidus(100); | Suillus | 13 | Symbiotroph | \|Ectomycorrhizal\| | Agaricoid-Boletoid | NULL | Highly Probable | Host - Pinaceae (Põlme et al. 2020) | Rinaldi AC, et al. 2008. Fungal Diversity 33:1-45 ((http://www.fungaldiversity.org/fdp/sfdp/33-1.pdf)); Tedersoo L, et al. 2010. Mycorrhiza 20:217-263 ((https://doi.org/10.1007/s00572-009-0274-x)); Põlme S, et al. 2020. Fungal Diversity 105:1-16 ((https://doi.org/10.1007/s13225-020-00466-2)); http://mycorrhizas.info/ecmf.html ((http://mycorrhizas.info/ecmf.html)) |
| Otu28571 | 1 | k__Fungi(100);p__Basidiomycota(100);c__Agaricomycetes(100);o__Boletales(100);f__Suillaceae(100);g__Suillus(100);s__Suillus_placidus(100); | Suillus | 13 | Symbiotroph | \|Ectomycorrhizal\| | Agaricoid-Boletoid | NULL | Highly Probable | Host - Pinaceae (Põlme et al. 2020) | Rinaldi AC, et al. 2008. Fungal Diversity 33:1-45 ((http://www.fungaldiversity.org/fdp/sfdp/33-1.pdf)); Tedersoo L, et al. 2010. Mycorrhiza 20:217-263 ((https://doi.org/10.1007/s00572-009-0274-x)); Põlme S, et al. 2020. Fungal Diversity 105:1-16 ((https://doi.org/10.1007/s13225-020-00466-2)); http://mycorrhizas.info/ecmf.html ((http://mycorrhizas.info/ecmf.html)) |
| Otu28572 | 1 | k__Fungi(100);p__Basidiomycota(100);c__Agaricomycetes(100);o__Boletales(100);f__Suillaceae(100);g__Suillus(100);s__Suillus_placidus(100); | Suillus | 13 | Symbiotroph | \|Ectomycorrhizal\| | Agaricoid-Boletoid | NULL | Highly Probable | Host - Pinaceae (Põlme et al. 2020) | Rinaldi AC, et al. 2008. Fungal Diversity 33:1-45 ((http://www.fungaldiversity.org/fdp/sfdp/33-1.pdf)); Tedersoo L, et al. 2010. Mycorrhiza 20:217-263 ((https://doi.org/10.1007/s00572-009-0274-x)); Põlme S, et al. 2020. Fungal Diversity 105:1-16 ((https://doi.org/10.1007/s13225-020-00466-2)); http://mycorrhizas.info/ecmf.html ((http://mycorrhizas.info/ecmf.html)) |
| Otu28574 | 1 | k__Fungi(100);p__Basidiomycota(100);c__Tremellomycetes(100);o__Filobasidiales(100);f__Filobasidiaceae(100);g__Cryptococcus(100);s__Cryptococcus_wieringae(100); | Cryptococcus | 13 | Pathotroph-Saprotroph-Symbiotroph | Animal Pathogen-Endophyte-Epiphyte-\|Undefined Saprotroph\| | Dimorphic Yeast-Yeast | NULL | Possible | As foliar_endophyte (Põlme et al. 2020); Decay leaf, fruit, seed, soil, dung, animal material (Põlme et al. 2020) | Andrews JH, et al. 1983. Phytopathology 73:228-234 ((https://doi.org/10.1094/Phyto-73-228)); Kurtzman CP, et al. (eds.) 2011. The Yeasts, a Taxonomic Study. Fifth Edition. Vols 1-3. Elsevier, San Diego (ISBN: 9780444521491); Põlme S, et al. 2020. Fungal Diversity 105:1-16 ((https://doi.org/10.1007/s13225-020-00466-2)) |
| Otu28577 | 1 | k__Fungi(100);p__Basidiomycota(100);c__Exobasidiomycetes(100);o__Microstromatales(100);f__Microstromataceae(100);g__Microstroma(100);s__Microstroma_juglandis(100); | Microstroma | 13 | Pathotroph | \|Plant Pathogen\| | Dimorphic | NULL | Probable | NULL | Tedersoo L, et al. 2014. Science 346:e1256688 ((https://doi.org/10.1126/science.1256688)); Põlme S, et al. 2020. Fungal Diversity 105:1-16 ((https://doi.org/10.1007/s13225-020-00466-2)) |
| Otu28578 | 1 | k__Fungi(100);p__Basidiomycota(100);c__Agaricomycetes(100);o__Polyporales(100);f__Phanerochaetaceae(100);g__Phanerochaete(100);s__Phanerochaete_sp(100); | Phanerochaete | 13 | Saprotroph | \|Wood Saprotroph\| | Corticioid | White Rot | Highly Probable | As white_rot (Põlme et al. 2020); Decay wood (Põlme et al. 2020) | Glenn JK, et al. 1983. Biochemical and Biophysical Research Communications 114:1077-1083 ((https://doi.org/10.1016/0006-291x((83))90672-1)); Põlme S, et al. 2020. Fungal Diversity 105:1-16 ((https://doi.org/10.1007/s13225-020-00466-2)) |
| Otu28579 | 1 | k__Fungi(100);p__Basidiomycota(100);c__Exobasidiomycetes(100);o__Microstromatales(100);f__Microstromataceae(100);g__Microstroma(100);s__Microstroma_juglandis(100); | Microstroma | 13 | Pathotroph | \|Plant Pathogen\| | Dimorphic | NULL | Probable | NULL | Tedersoo L, et al. 2014. Science 346:e1256688 ((https://doi.org/10.1126/science.1256688)); Põlme S, et al. 2020. Fungal Diversity 105:1-16 ((https://doi.org/10.1007/s13225-020-00466-2)) |
| Otu28580 | 1 | k__Fungi(100);p__Basidiomycota(100);c__Microbotryomycetes(100);o__Sporidiobolales(100);f__Sporidiobolales_family_Incertae_sedis(100);g__Rhodotorula(100);s__Rhodotorula_sp(100); | Rhodotorula | 13 | Pathotroph-Saprotroph-Symbiotroph | Animal Pathogen-Animal Symbiotroph-Endophyte-Plant Pathogen-\|Undefined Saprotroph\| | Yeast | NULL | Probable | As foliar_endophyte (Põlme et al. 2020); Opportunistic human pathogen | Põlme S, et al. 2020. Fungal Diversity 105:1-16 ((https://doi.org/10.1007/s13225-020-00466-2)); https://www.cdc.gov ((https://www.cdc.gov)) |
| Otu28581 | 1 | k__Fungi(100);p__Basidiomycota(100);c__Microbotryomycetes(100);o__Sporidiobolales(100);f__Sporidiobolales_family_Incertae_sedis(100);g__Rhodotorula(100);s__Rhodotorula_sp(100); | Rhodotorula | 13 | Pathotroph-Saprotroph-Symbiotroph | Animal Pathogen-Animal Symbiotroph-Endophyte-Plant Pathogen-\|Undefined Saprotroph\| | Yeast | NULL | Probable | As foliar_endophyte (Põlme et al. 2020); Opportunistic human pathogen | Põlme S, et al. 2020. Fungal Diversity 105:1-16 ((https://doi.org/10.1007/s13225-020-00466-2)); https://www.cdc.gov ((https://www.cdc.gov)) |
| Otu28584 | 1 | k__Fungi(100);p__Basidiomycota(100);c__Agaricomycetes(100);o__Cantharellales(100);f__Clavulinaceae(100);f__Clavulinaceae_unclassified(100);f__Clavulinaceae_unclassified(100); | Clavulinaceae | 9 | Saprotroph-Symbiotroph | Ectomycorrhizal-Lichenized-Wood Saprotroph | Clavarioid | NULL | Possible | Some species form ectomycorrhizas, with fruit bodies in leaf litter and on soil. Others are probably saprobes associated with rotten wood (Cannon & Kirk 2007) | Cannon PF, Kirk PM. 2007. Fungal Families of the World. CAB International, Cambridge ((https://isbnsearch.org/isbn/9780851998275)); Nelsen MP, et al. 2007. American Journal of Botany 94:1289-1296 ((https://doi.org/10.3732/ajb.94.8.1289)) |
| Otu28585 | 1 | k__Fungi(100);p__Basidiomycota(100);c__Microbotryomycetes(100);o__Sporidiobolales(100);f__Sporidiobolales_family_Incertae_sedis(100);g__Rhodotorula(100);s__Rhodotorula_sp(100); | Rhodotorula | 13 | Pathotroph-Saprotroph-Symbiotroph | Animal Pathogen-Animal Symbiotroph-Endophyte-Plant Pathogen-\|Undefined Saprotroph\| | Yeast | NULL | Probable | As foliar_endophyte (Põlme et al. 2020); Opportunistic human pathogen | Põlme S, et al. 2020. Fungal Diversity 105:1-16 ((https://doi.org/10.1007/s13225-020-00466-2)); https://www.cdc.gov ((https://www.cdc.gov)) |
| Otu28586 | 1 | k__Fungi(100);p__Basidiomycota(100);c__Exobasidiomycetes(100);o__Microstromatales(100);f__Microstromataceae(100);g__Microstroma(100);s__Microstroma_juglandis(100); | Microstroma | 13 | Pathotroph | \|Plant Pathogen\| | Dimorphic | NULL | Probable | NULL | Tedersoo L, et al. 2014. Science 346:e1256688 ((https://doi.org/10.1126/science.1256688)); Põlme S, et al. 2020. Fungal Diversity 105:1-16 ((https://doi.org/10.1007/s13225-020-00466-2)) |
| Otu28589 | 1 | k__Fungi(100);p__Basidiomycota(100);c__Exobasidiomycetes(100);o__Microstromatales(100);f__Microstromataceae(100);g__Microstroma(100);s__Microstroma_juglandis(100); | Microstroma | 13 | Pathotroph | \|Plant Pathogen\| | Dimorphic | NULL | Probable | NULL | Tedersoo L, et al. 2014. Science 346:e1256688 ((https://doi.org/10.1126/science.1256688)); Põlme S, et al. 2020. Fungal Diversity 105:1-16 ((https://doi.org/10.1007/s13225-020-00466-2)) |
| Otu28590 | 1 | k__Fungi(100);p__Basidiomycota(100);c__Microbotryomycetes(100);o__Sporidiobolales(100);f__Sporidiobolales_family_Incertae_sedis(100);g__Rhodotorula(100);s__Rhodotorula_sp(100); | Rhodotorula | 13 | Pathotroph-Saprotroph-Symbiotroph | Animal Pathogen-Animal Symbiotroph-Endophyte-Plant Pathogen-\|Undefined Saprotroph\| | Yeast | NULL | Probable | As foliar_endophyte (Põlme et al. 2020); Opportunistic human pathogen | Põlme S, et al. 2020. Fungal Diversity 105:1-16 ((https://doi.org/10.1007/s13225-020-00466-2)); https://www.cdc.gov ((https://www.cdc.gov)) |
| Otu28595 | 1 | k__Fungi(100);p__Basidiomycota(100);c__Agaricomycetes(100);o__Polyporales(100);f__Phanerochaetaceae(100);g__Phanerochaete(100);s__Phanerochaete_sp_DLL2011_1(100); | Phanerochaete | 13 | Saprotroph | \|Wood Saprotroph\| | Corticioid | White Rot | Highly Probable | As white_rot (Põlme et al. 2020); Decay wood (Põlme et al. 2020) | Glenn JK, et al. 1983. Biochemical and Biophysical Research Communications 114:1077-1083 ((https://doi.org/10.1016/0006-291x((83))90672-1)); Põlme S, et al. 2020. Fungal Diversity 105:1-16 ((https://doi.org/10.1007/s13225-020-00466-2)) |
| Otu28598 | 1 | k__Fungi(100);p__Basidiomycota(100);c__Microbotryomycetes(100);o__Sporidiobolales(100);f__Sporidiobolales_family_Incertae_sedis(100);g__Rhodotorula(100);s__Rhodotorula_yarrowii(100); | Rhodotorula | 13 | Pathotroph-Saprotroph-Symbiotroph | Animal Pathogen-Animal Symbiotroph-Endophyte-Plant Pathogen-\|Undefined Saprotroph\| | Yeast | NULL | Probable | As foliar_endophyte (Põlme et al. 2020); Opportunistic human pathogen | Põlme S, et al. 2020. Fungal Diversity 105:1-16 ((https://doi.org/10.1007/s13225-020-00466-2)); https://www.cdc.gov ((https://www.cdc.gov)) |
| Otu28601 | 1 | k__Fungi(100);p__Basidiomycota(100);c__Agaricomycetes(100);o__Cantharellales(100);f__Clavulinaceae(100);g__Clavulina(100);g__Clavulina_unclassified(100); | Clavulina | 13 | Symbiotroph | \|Ectomycorrhizal\| | Clavarioid | NULL | Highly Probable | NULL | Tedersoo L, et al. 2014. Science 346:e1256688 ((https://doi.org/10.1126/science.1256688)); Põlme S, et al. 2020. Fungal Diversity 105:1-16 ((https://doi.org/10.1007/s13225-020-00466-2)) |
| Otu28602 | 1 | k__Fungi(100);p__Basidiomycota(100);c__Agaricomycetes(100);o__Cantharellales(100);f__Clavulinaceae(100);g__Clavulina(100);g__Clavulina_unclassified(100); | Clavulina | 13 | Symbiotroph | \|Ectomycorrhizal\| | Clavarioid | NULL | Highly Probable | NULL | Tedersoo L, et al. 2014. Science 346:e1256688 ((https://doi.org/10.1126/science.1256688)); Põlme S, et al. 2020. Fungal Diversity 105:1-16 ((https://doi.org/10.1007/s13225-020-00466-2)) |
| Otu28604 | 1 | k__Fungi(100);p__Basidiomycota(100);c__Agaricomycetes(100);o__Cantharellales(100);f__Clavulinaceae(100);g__Clavulina(100);g__Clavulina_unclassified(100); | Clavulina | 13 | Symbiotroph | \|Ectomycorrhizal\| | Clavarioid | NULL | Highly Probable | NULL | Tedersoo L, et al. 2014. Science 346:e1256688 ((https://doi.org/10.1126/science.1256688)); Põlme S, et al. 2020. Fungal Diversity 105:1-16 ((https://doi.org/10.1007/s13225-020-00466-2)) |
| Otu28605 | 1 | k__Fungi(100);p__Basidiomycota(100);c__Agaricomycetes(100);o__Cantharellales(100);f__Hydnaceae(100);g__Sistotrema(100);s__Sistotrema_brinkmannii(100); | Sistotrema | 13 | Saprotroph-Symbiotroph | Ectomycorrhizal-\|Plant Saprotroph\|-Wood Saprotroph | Corticioid | White Rot | Possible | As litter_saprotroph (Põlme et al. 2020); Decay leaf, fruit, seed (Põlme et al. 2020) | Gilbertson RL, Ryvarden L. 1987-1987. North American Polypores. Fungiflora, Oslo (ISBN: 978-0945345060); Rinaldi AC, et al. 2008. Fungal Diversity 33:1-45 ((http://www.fungaldiversity.org/fdp/sfdp/33-1.pdf)) (as ECM/not ECM); Tedersoo L, et al. 2010. Mycorrhiza 20:217-263 ((https://doi.org/10.1007/s00572-009-0274-x)) (pro parte); Põlme S, et al. 2020. Fungal Diversity 105:1-16 ((https://doi.org/10.1007/s13225-020-00466-2)); http://mycorrhizas.info/ecmf.html ((http://mycorrhizas.info/ecmf.html)) (as ECM/not ECM). |
| Otu28606 | 1 | k__Fungi(100);p__Basidiomycota(100);c__Agaricomycetes(100);o__Polyporales(100);f__Phanerochaetaceae(100);g__Hyphodermella(100);s__Hyphodermella_sp(100); | Hyphodermella | 13 | Saprotroph | Undefined Saprotroph-\|Wood Saprotroph\| | Corticioid | White Rot | Probable | As white_rot (Põlme et al. 2020); Decay wood (Põlme et al. 2020) | Tedersoo L, et al. 2014. Science 346:e1256688 ((https://doi.org/10.1126/science.1256688)); Põlme S, et al. 2020. Fungal Diversity 105:1-16 ((https://doi.org/10.1007/s13225-020-00466-2)) |
| Otu28607 | 1 | k__Fungi(100);p__Basidiomycota(100);c__Agaricomycetes(100);o__Polyporales(100);f__Phanerochaetaceae(100);g__Phanerochaete(100);s__Phanerochaete_stereoides(100); | Phanerochaete | 13 | Saprotroph | \|Wood Saprotroph\| | Corticioid | White Rot | Highly Probable | As white_rot (Põlme et al. 2020); Decay wood (Põlme et al. 2020) | Glenn JK, et al. 1983. Biochemical and Biophysical Research Communications 114:1077-1083 ((https://doi.org/10.1016/0006-291x((83))90672-1)); Põlme S, et al. 2020. Fungal Diversity 105:1-16 ((https://doi.org/10.1007/s13225-020-00466-2)) |
| Otu28608 | 1 | k__Fungi(100);p__Basidiomycota(100);c__Agaricomycetes(100);o__Polyporales(100);f__Phanerochaetaceae(100);g__Phanerochaete(100);s__Phanerochaete_stereoides(100); | Phanerochaete | 13 | Saprotroph | \|Wood Saprotroph\| | Corticioid | White Rot | Highly Probable | As white_rot (Põlme et al. 2020); Decay wood (Põlme et al. 2020) | Glenn JK, et al. 1983. Biochemical and Biophysical Research Communications 114:1077-1083 ((https://doi.org/10.1016/0006-291x((83))90672-1)); Põlme S, et al. 2020. Fungal Diversity 105:1-16 ((https://doi.org/10.1007/s13225-020-00466-2)) |
| Otu28609 | 1 | k__Fungi(100);p__Basidiomycota(100);c__Exobasidiomycetes(100);o__Microstromatales(100);f__Microstromataceae(100);g__Microstroma(100);s__Microstroma_juglandis(100); | Microstroma | 13 | Pathotroph | \|Plant Pathogen\| | Dimorphic | NULL | Probable | NULL | Tedersoo L, et al. 2014. Science 346:e1256688 ((https://doi.org/10.1126/science.1256688)); Põlme S, et al. 2020. Fungal Diversity 105:1-16 ((https://doi.org/10.1007/s13225-020-00466-2)) |
| Otu28610 | 1 | k__Fungi(100);p__Basidiomycota(100);c__Agaricomycetes(100);o__Cantharellales(100);f__Hydnaceae(100);g__Sistotrema(100);s__Sistotrema_brinkmannii(100); | Sistotrema | 13 | Saprotroph-Symbiotroph | Ectomycorrhizal-\|Plant Saprotroph\|-Wood Saprotroph | Corticioid | White Rot | Possible | As litter_saprotroph (Põlme et al. 2020); Decay leaf, fruit, seed (Põlme et al. 2020) | Gilbertson RL, Ryvarden L. 1987-1987. North American Polypores. Fungiflora, Oslo (ISBN: 978-0945345060); Rinaldi AC, et al. 2008. Fungal Diversity 33:1-45 ((http://www.fungaldiversity.org/fdp/sfdp/33-1.pdf)) (as ECM/not ECM); Tedersoo L, et al. 2010. Mycorrhiza 20:217-263 ((https://doi.org/10.1007/s00572-009-0274-x)) (pro parte); Põlme S, et al. 2020. Fungal Diversity 105:1-16 ((https://doi.org/10.1007/s13225-020-00466-2)); http://mycorrhizas.info/ecmf.html ((http://mycorrhizas.info/ecmf.html)) (as ECM/not ECM). |
| Otu28611 | 1 | k__Fungi(100);p__Basidiomycota(100);c__Agaricomycetes(100);o__Cantharellales(100);f__Hydnaceae(100);g__Sistotrema(100);s__Sistotrema_brinkmannii(100); | Sistotrema | 13 | Saprotroph-Symbiotroph | Ectomycorrhizal-\|Plant Saprotroph\|-Wood Saprotroph | Corticioid | White Rot | Possible | As litter_saprotroph (Põlme et al. 2020); Decay leaf, fruit, seed (Põlme et al. 2020) | Gilbertson RL, Ryvarden L. 1987-1987. North American Polypores. Fungiflora, Oslo (ISBN: 978-0945345060); Rinaldi AC, et al. 2008. Fungal Diversity 33:1-45 ((http://www.fungaldiversity.org/fdp/sfdp/33-1.pdf)) (as ECM/not ECM); Tedersoo L, et al. 2010. Mycorrhiza 20:217-263 ((https://doi.org/10.1007/s00572-009-0274-x)) (pro parte); Põlme S, et al. 2020. Fungal Diversity 105:1-16 ((https://doi.org/10.1007/s13225-020-00466-2)); http://mycorrhizas.info/ecmf.html ((http://mycorrhizas.info/ecmf.html)) (as ECM/not ECM). |
| Otu28612 | 1 | k__Fungi(100);p__Basidiomycota(100);c__Ustilaginomycetes(100);o__Ustilaginales(100);f__Ustilaginaceae(100);g__unclassified_Ustilaginaceae(100);s__Ustilaginaceae_sp(100); | Ustilaginaceae | 9 | Pathotroph | Plant Pathogen | Smut | NULL | Probable | Biotrophic in living tissue of Gramineae and Cyperaceae (Cannon & Kirk 2007) | Cannon PF, Kirk PM. 2007. Fungal Families of the World. CAB International, Cambridge (ISBN: 978-0851998275) |
| Otu28613 | 1 | k__Fungi(100);p__Basidiomycota(100);c__Ustilaginomycetes(100);o__Ustilaginales(100);f__Ustilaginaceae(100);g__Pseudozyma(100);s__Pseudozyma_aphidis(100); | Pseudozyma | 13 | Pathotroph-Symbiotroph | Endophyte-\|Epiphyte\|-Fungal Parasite | Yeast | NULL | Probable | As foliar_endophyte (Põlme et al. 2020); As mycoparasite (Põlme et al. 2020); Decay leaf, fruit, seed, fungal material (Põlme et al. 2020) | Põlme S, et al. 2020. Fungal Diversity 105:1-16 ((https://doi.org/10.1007/s13225-020-00466-2)) |
| Otu28614 | 1 | k__Fungi(100);p__Basidiomycota(100);c__Exobasidiomycetes(100);o__Microstromatales(100);f__Microstromataceae(100);g__Microstroma(100);s__Microstroma_juglandis(100); | Microstroma | 13 | Pathotroph | \|Plant Pathogen\| | Dimorphic | NULL | Probable | NULL | Tedersoo L, et al. 2014. Science 346:e1256688 ((https://doi.org/10.1126/science.1256688)); Põlme S, et al. 2020. Fungal Diversity 105:1-16 ((https://doi.org/10.1007/s13225-020-00466-2)) |
| Otu28615 | 1 | k__Fungi(100);p__Basidiomycota(100);c__Microbotryomycetes(100);o__Sporidiobolales(100);f__Sporidiobolales_family_Incertae_sedis(100);g__Rhodotorula(100);s__Rhodotorula_mucilaginosa(100); | Rhodotorula | 13 | Pathotroph-Saprotroph-Symbiotroph | Animal Pathogen-Animal Symbiotroph-Endophyte-Plant Pathogen-\|Undefined Saprotroph\| | Yeast | NULL | Probable | As foliar_endophyte (Põlme et al. 2020); Opportunistic human pathogen | Põlme S, et al. 2020. Fungal Diversity 105:1-16 ((https://doi.org/10.1007/s13225-020-00466-2)); https://www.cdc.gov ((https://www.cdc.gov)) |
| Otu28616 | 1 | k__Fungi(100);p__Basidiomycota(100);c__Microbotryomycetes(100);o__Sporidiobolales(100);f__Sporidiobolales_family_Incertae_sedis(100);g__Rhodotorula(100);s__Rhodotorula_taiwanensis(100); | Rhodotorula | 13 | Pathotroph-Saprotroph-Symbiotroph | Animal Pathogen-Animal Symbiotroph-Endophyte-Plant Pathogen-\|Undefined Saprotroph\| | Yeast | NULL | Probable | As foliar_endophyte (Põlme et al. 2020); Opportunistic human pathogen | Põlme S, et al. 2020. Fungal Diversity 105:1-16 ((https://doi.org/10.1007/s13225-020-00466-2)); https://www.cdc.gov ((https://www.cdc.gov)) |
| Otu28617 | 1 | k__Fungi(100);p__Basidiomycota(100);c__Microbotryomycetes(100);o__Sporidiobolales(100);f__Sporidiobolales_family_Incertae_sedis(100);g__Rhodotorula(100);s__Rhodotorula_mucilaginosa(100); | Rhodotorula | 13 | Pathotroph-Saprotroph-Symbiotroph | Animal Pathogen-Animal Symbiotroph-Endophyte-Plant Pathogen-\|Undefined Saprotroph\| | Yeast | NULL | Probable | As foliar_endophyte (Põlme et al. 2020); Opportunistic human pathogen | Põlme S, et al. 2020. Fungal Diversity 105:1-16 ((https://doi.org/10.1007/s13225-020-00466-2)); https://www.cdc.gov ((https://www.cdc.gov)) |
| Otu28618 | 1 | k__Fungi(100);p__Basidiomycota(100);c__Microbotryomycetes(100);o__Sporidiobolales(100);f__Sporidiobolales_family_Incertae_sedis(100);g__Rhodotorula(100);s__Rhodotorula_mucilaginosa(100); | Rhodotorula | 13 | Pathotroph-Saprotroph-Symbiotroph | Animal Pathogen-Animal Symbiotroph-Endophyte-Plant Pathogen-\|Undefined Saprotroph\| | Yeast | NULL | Probable | As foliar_endophyte (Põlme et al. 2020); Opportunistic human pathogen | Põlme S, et al. 2020. Fungal Diversity 105:1-16 ((https://doi.org/10.1007/s13225-020-00466-2)); https://www.cdc.gov ((https://www.cdc.gov)) |
| Otu28619 | 1 | k__Fungi(100);p__Basidiomycota(100);c__Microbotryomycetes(100);o__Sporidiobolales(100);f__Sporidiobolales_family_Incertae_sedis(100);g__Rhodotorula(100);s__Rhodotorula_mucilaginosa(100); | Rhodotorula | 13 | Pathotroph-Saprotroph-Symbiotroph | Animal Pathogen-Animal Symbiotroph-Endophyte-Plant Pathogen-\|Undefined Saprotroph\| | Yeast | NULL | Probable | As foliar_endophyte (Põlme et al. 2020); Opportunistic human pathogen | Põlme S, et al. 2020. Fungal Diversity 105:1-16 ((https://doi.org/10.1007/s13225-020-00466-2)); https://www.cdc.gov ((https://www.cdc.gov)) |
| Otu28620 | 1 | k__Fungi(100);p__Basidiomycota(100);c__Microbotryomycetes(100);o__Sporidiobolales(100);f__Sporidiobolales_family_Incertae_sedis(100);g__Rhodotorula(100);s__Rhodotorula_taiwanensis(100); | Rhodotorula | 13 | Pathotroph-Saprotroph-Symbiotroph | Animal Pathogen-Animal Symbiotroph-Endophyte-Plant Pathogen-\|Undefined Saprotroph\| | Yeast | NULL | Probable | As foliar_endophyte (Põlme et al. 2020); Opportunistic human pathogen | Põlme S, et al. 2020. Fungal Diversity 105:1-16 ((https://doi.org/10.1007/s13225-020-00466-2)); https://www.cdc.gov ((https://www.cdc.gov)) |

| | | | | | | | | | | | |
|---|---|---|---|---|---|---|---|---|---|---|---|
| Otu28624 | 1 | k__Fungi(100);p__Basidiomycota(100);c__Exobasidiomycetes(100);o__Microstromatales(100);f__Microstromataceae(100);g__Microstroma(100);s__Microstroma_juglandis(100); | Microstroma | 13 | Pathotroph | \|Plant Pathogen\| | Dimorphic | NULL | Probable | NULL | Tedersoo L, et al. 2014. Science 346:e1256688 ((https://doi.org/10.1126/science.1256688)); Põlme S, et al. 2020. Fungal Diversity 105:1-16 ((https://doi.org/10.1007/s13225-020-00466-2)) |
| Otu28625 | 1 | k__Fungi(100);p__Basidiomycota(100);c__Exobasidiomycetes(100);o__Microstromatales(100);f__Microstromataceae(100);g__Microstroma(100);s__Microstroma_juglandis(100); | Microstroma | 13 | Pathotroph | \|Plant Pathogen\| | Dimorphic | NULL | Probable | NULL | Tedersoo L, et al. 2014. Science 346:e1256688 ((https://doi.org/10.1126/science.1256688)); Põlme S, et al. 2020. Fungal Diversity 105:1-16 ((https://doi.org/10.1007/s13225-020-00466-2)) |
| Otu28627 | 1 | k__Fungi(100);p__Basidiomycota(100);c__Exobasidiomycetes(100);o__Microstromatales(100);f__Microstromataceae(100);g__Microstroma(100);s__Microstroma_juglandis(100); | Microstroma | 13 | Pathotroph | \|Plant Pathogen\| | Dimorphic | NULL | Probable | NULL | Tedersoo L, et al. 2014. Science 346:e1256688 ((https://doi.org/10.1126/science.1256688)); Põlme S, et al. 2020. Fungal Diversity 105:1-16 ((https://doi.org/10.1007/s13225-020-00466-2)) |
| Otu28628 | 1 | k__Fungi(100);p__Basidiomycota(100);c__Exobasidiomycetes(100);o__Microstromatales(100);f__Microstromataceae(100);g__Microstroma(100);s__Microstroma_juglandis(100); | Microstroma | 13 | Pathotroph | \|Plant Pathogen\| | Dimorphic | NULL | Probable | NULL | Tedersoo L, et al. 2014. Science 346:e1256688 ((https://doi.org/10.1126/science.1256688)); Põlme S, et al. 2020. Fungal Diversity 105:1-16 ((https://doi.org/10.1007/s13225-020-00466-2)) |
| Otu28630 | 1 | k__Fungi(100);p__Basidiomycota(100);c__Microbotryomycetes(100);o__Sporidiobolales(100);f__Sporidiobolales_family_Incertae_sedis(100);g__Rhodotorula(100);s__Rhodotorula_sp(100); | Rhodotorula | 13 | Pathotroph-Saprotroph-Symbiotroph | Animal Pathogen-Animal Symbiotroph-Endophyte-Plant Pathogen-\|Undefined Saprotroph\| | Yeast | NULL | Probable | As foliar_endophyte (Põlme et al. 2020); Opportunistic human pathogen | Põlme S, et al. 2020. Fungal Diversity 105:1-16 ((https://doi.org/10.1007/s13225-020-00466-2)); https://www.cdc.gov ((https://www.cdc.gov)) |
| Otu28631 | 1 | k__Fungi(100);p__Basidiomycota(100);c__Microbotryomycetes(100);o__Sporidiobolales(100);f__Sporidiobolales_family_Incertae_sedis(100);g__Rhodotorula(100);s__Rhodotorula_sp(100); | Rhodotorula | 13 | Pathotroph-Saprotroph-Symbiotroph | Animal Pathogen-Animal Symbiotroph-Endophyte-Plant Pathogen-\|Undefined Saprotroph\| | Yeast | NULL | Probable | As foliar_endophyte (Põlme et al. 2020); Opportunistic human pathogen | Põlme S, et al. 2020. Fungal Diversity 105:1-16 ((https://doi.org/10.1007/s13225-020-00466-2)); https://www.cdc.gov ((https://www.cdc.gov)) |
| Otu28632 | 1 | k__Fungi(100);p__Basidiomycota(100);c__Microbotryomycetes(100);o__Sporidiobolales(100);f__Sporidiobolales_family_Incertae_sedis(100);g__Rhodotorula(100);s__Rhodotorula_sp(100); | Rhodotorula | 13 | Pathotroph-Saprotroph-Symbiotroph | Animal Pathogen-Animal Symbiotroph-Endophyte-Plant Pathogen-\|Undefined Saprotroph\| | Yeast | NULL | Probable | As foliar_endophyte (Põlme et al. 2020); Opportunistic human pathogen | Põlme S, et al. 2020. Fungal Diversity 105:1-16 ((https://doi.org/10.1007/s13225-020-00466-2)); https://www.cdc.gov ((https://www.cdc.gov)) |
| Otu28633 | 1 | k__Fungi(100);p__Basidiomycota(100);c__Exobasidiomycetes(100);o__Microstromatales(100);f__Microstromataceae(100);g__Microstroma(100);s__Microstroma_juglandis(100); | Microstroma | 13 | Pathotroph | \|Plant Pathogen\| | Dimorphic | NULL | Probable | NULL | Tedersoo L, et al. 2014. Science 346:e1256688 ((https://doi.org/10.1126/science.1256688)); Põlme S, et al. 2020. Fungal Diversity 105:1-16 ((https://doi.org/10.1007/s13225-020-00466-2)) |
| Otu28634 | 1 | k__Fungi(100);p__Basidiomycota(100);c__Microbotryomycetes(100);o__Sporidiobolales(100);f__Sporidiobolales_family_Incertae_sedis(100);g__Rhodotorula(100);s__Rhodotorula_nothofagi(100); | Rhodotorula | 13 | Pathotroph-Saprotroph-Symbiotroph | Animal Pathogen-Animal Symbiotroph-Endophyte-Plant Pathogen-\|Undefined Saprotroph\| | Yeast | NULL | Probable | As foliar_endophyte (Põlme et al. 2020); Opportunistic human pathogen | Põlme S, et al. 2020. Fungal Diversity 105:1-16 ((https://doi.org/10.1007/s13225-020-00466-2)); https://www.cdc.gov ((https://www.cdc.gov)) |
| Otu28636 | 1 | k__Fungi(100);p__Basidiomycota(100);c__Exobasidiomycetes(100);o__Microstromatales(100);f__Microstromataceae(100);g__Microstroma(100);s__Microstroma_juglandis(100); | Microstroma | 13 | Pathotroph | \|Plant Pathogen\| | Dimorphic | NULL | Probable | NULL | Tedersoo L, et al. 2014. Science 346:e1256688 ((https://doi.org/10.1126/science.1256688)); Põlme S, et al. 2020. Fungal Diversity 105:1-16 ((https://doi.org/10.1007/s13225-020-00466-2)) |
| Otu28637 | 1 | k__Fungi(100);p__Basidiomycota(100);c__Exobasidiomycetes(100);o__Microstromatales(100);f__Microstromataceae(100);g__Microstroma(100);s__Microstroma_juglandis(100); | Microstroma | 13 | Pathotroph | \|Plant Pathogen\| | Dimorphic | NULL | Probable | NULL | Tedersoo L, et al. 2014. Science 346:e1256688 ((https://doi.org/10.1126/science.1256688)); Põlme S, et al. 2020. Fungal Diversity 105:1-16 ((https://doi.org/10.1007/s13225-020-00466-2)) |
| Otu28638 | 1 | k__Fungi(100);p__Basidiomycota(100);c__Microbotryomycetes(100);o__Sporidiobolales(100);f__Sporidiobolales_family_Incertae_sedis(100);g__Rhodotorula(100);s__Rhodotorula_sp(100); | Rhodotorula | 13 | Pathotroph-Saprotroph-Symbiotroph | Animal Pathogen-Animal Symbiotroph-Endophyte-Plant Pathogen-\|Undefined Saprotroph\| | Yeast | NULL | Probable | As foliar_endophyte (Põlme et al. 2020); Opportunistic human pathogen | Põlme S, et al. 2020. Fungal Diversity 105:1-16 ((https://doi.org/10.1007/s13225-020-00466-2)); https://www.cdc.gov ((https://www.cdc.gov)) |
| Otu28639 | 1 | k__Fungi(100);p__Basidiomycota(100);c__Microbotryomycetes(100);o__Sporidiobolales(100);f__Sporidiobolales_family_Incertae_sedis(100);g__Rhodotorula(100);s__Rhodotorula_sp(100); | Rhodotorula | 13 | Pathotroph-Saprotroph-Symbiotroph | Animal Pathogen-Animal Symbiotroph-Endophyte-Plant Pathogen-\|Undefined Saprotroph\| | Yeast | NULL | Probable | As foliar_endophyte (Põlme et al. 2020); Opportunistic human pathogen | Põlme S, et al. 2020. Fungal Diversity 105:1-16 ((https://doi.org/10.1007/s13225-020-00466-2)); https://www.cdc.gov ((https://www.cdc.gov)) |
| Otu28640 | 1 | k__Fungi(100);p__Basidiomycota(100);c__Microbotryomycetes(100);o__Sporidiobolales(100);f__Sporidiobolales_family_Incertae_sedis(100);g__Rhodotorula(100);s__Rhodotorula_sp(100); | Rhodotorula | 13 | Pathotroph-Saprotroph-Symbiotroph | Animal Pathogen-Animal Symbiotroph-Endophyte-Plant Pathogen-\|Undefined Saprotroph\| | Yeast | NULL | Probable | As foliar_endophyte (Põlme et al. 2020); Opportunistic human pathogen | Põlme S, et al. 2020. Fungal Diversity 105:1-16 ((https://doi.org/10.1007/s13225-020-00466-2)); https://www.cdc.gov ((https://www.cdc.gov)) |
| Otu28641 | 1 | k__Fungi(100);p__Basidiomycota(100);c__Exobasidiomycetes(100);o__Microstromatales(100);f__Microstromataceae(100);g__Microstroma(100);s__Microstroma_juglandis(100); | Microstroma | 13 | Pathotroph | \|Plant Pathogen\| | Dimorphic | NULL | Probable | NULL | Tedersoo L, et al. 2014. Science 346:e1256688 ((https://doi.org/10.1126/science.1256688)); Põlme S, et al. 2020. Fungal Diversity 105:1-16 ((https://doi.org/10.1007/s13225-020-00466-2)) |
| Otu28644 | 1 | k__Fungi(100);p__Basidiomycota(100);c__Exobasidiomycetes(100);o__Microstromatales(100);f__Microstromataceae(100);g__Microstroma(100);s__Microstroma_juglandis(100); | Microstroma | 13 | Pathotroph | \|Plant Pathogen\| | Dimorphic | NULL | Probable | NULL | Tedersoo L, et al. 2014. Science 346:e1256688 ((https://doi.org/10.1126/science.1256688)); Põlme S, et al. 2020. Fungal Diversity 105:1-16 ((https://doi.org/10.1007/s13225-020-00466-2)) |
| Otu28645 | 1 | k__Fungi(100);p__Basidiomycota(100);c__Agaricomycetes(100);o__Cantharellales(100);f__Clavulinaceae(100);f__Clavulinaceae_unclassified(100);f__Clavulinaceae_unclassified(100); | Clavulinaceae | 9 | Saprotroph-Symbiotroph | Ectomycorrhizal-Lichenized-Wood Saprotroph | Clavarioid | NULL | Possible | Some species form ectomycorrhizas, with fruit bodies in leaf litter and on soil. Others are probably saprobes associated with rotten wood (Cannon & Kirk 2007) | Cannon PF, Kirk PM. 2007. Fungal Families of the World. CAB International, Cambridge ((https://isbnsearch.org/isbn/9780851998275)); Nelsen MP, et al. 2007. American Journal of Botany 94:1289-1296 ((https://doi.org/10.3732/ajb.94.8.1289)) |
| Otu28646 | 1 | k__Fungi(100);p__Basidiomycota(100);c__Agaricomycetes(100);o__Polyporales(100);f__Phanerochaetaceae(100);g__Phanerochaete(100);s__Phanerochaete_stereoides(100); | Phanerochaete | 13 | Saprotroph | \|Wood Saprotroph\| | Corticioid | White Rot | Highly Probable | As white_rot (Põlme et al. 2020); Decay wood (Põlme et al. 2020) | Glenn JK, et al. 1983. Biochemical and Biophysical Research Communications 114:1077-1083 ((https://doi.org/10.1016/0006-291x((83)) 90672-1)); Põlme S, et al. 2020. Fungal Diversity 105:1-16 ((https://doi.org/10.1007/s13225-020-00466-2)) |
| Otu28648 | 1 | k__Fungi(100);p__Ascomycota(100);c__Saccharomycetes(100);o__Saccharomycetales(100);f__Metschnikowiaceae(100);g__Metschnikowia(100);g__Metschnikowia_unclassified(100); | Metschnikowia | 13 | Saprotroph | \|Undefined Saprotroph\| | Yeast | NULL | Probable | As nectar-tap_saprotroph (Põlme et al. 2020); Decay sugar-rich substrates (Põlme et al. 2020) | Põlme S, et al. 2020. Fungal Diversity 105:1-16 ((https://doi.org/10.1007/s13225-020-00466-2)) |
| Otu28649 | 1 | k__Fungi(100);p__Ascomycota(100);c__Saccharomycetes(100);o__Saccharomycetales(100);f__Metschnikowiaceae(100);g__Metschnikowia(100);s__Metschnikowia_pulcherrima(100); | Metschnikowia | 13 | Saprotroph | \|Undefined Saprotroph\| | Yeast | NULL | Probable | As nectar-tap_saprotroph (Põlme et al. 2020); Decay sugar-rich substrates (Põlme et al. 2020) | Põlme S, et al. 2020. Fungal Diversity 105:1-16 ((https://doi.org/10.1007/s13225-020-00466-2)) |
| Otu28650 | 1 | k__Fungi(100);p__Ascomycota(100);c__Saccharomycetes(100);o__Saccharomycetales(100);f__Metschnikowiaceae(100);g__Metschnikowia(100);s__Metschnikowia_pulcherrima(100); | Metschnikowia | 13 | Saprotroph | \|Undefined Saprotroph\| | Yeast | NULL | Probable | As nectar-tap_saprotroph (Põlme et al. 2020); Decay sugar-rich substrates (Põlme et al. 2020) | Põlme S, et al. 2020. Fungal Diversity 105:1-16 ((https://doi.org/10.1007/s13225-020-00466-2)) |
| Otu28661 | 1 | k__Fungi(100);p__Ascomycota(100);c__Saccharomycetes(100);o__Saccharomycetales(100);f__Metschnikowiaceae(100);g__Metschnikowia(100);g__Metschnikowia_unclassified(100); | Metschnikowia | 13 | Saprotroph | \|Undefined Saprotroph\| | Yeast | NULL | Probable | As nectar-tap_saprotroph (Põlme et al. 2020); Decay sugar-rich substrates (Põlme et al. 2020) | Põlme S, et al. 2020. Fungal Diversity 105:1-16 ((https://doi.org/10.1007/s13225-020-00466-2)) |
| Otu28675 | 1 | k__Fungi(100);p__Ascomycota(100);c__Saccharomycetes(100);o__Saccharomycetales(100);f__Metschnikowiaceae(100);g__Metschnikowia(100);s__Metschnikowia_pulcherrima(100); | Metschnikowia | 13 | Saprotroph | \|Undefined Saprotroph\| | Yeast | NULL | Probable | As nectar-tap_saprotroph (Põlme et al. 2020); Decay sugar-rich substrates (Põlme et al. 2020) | Põlme S, et al. 2020. Fungal Diversity 105:1-16 ((https://doi.org/10.1007/s13225-020-00466-2)) |
| Otu28677 | 1 | k__Fungi(100);p__Ascomycota(100);c__Saccharomycetes(100);o__Saccharomycetales(100);f__Metschnikowiaceae(100);g__Metschnikowia(100);s__Metschnikowia_pulcherrima(100); | Metschnikowia | 13 | Saprotroph | \|Undefined Saprotroph\| | Yeast | NULL | Probable | As nectar-tap_saprotroph (Põlme et al. 2020); Decay sugar-rich substrates (Põlme et al. 2020) | Põlme S, et al. 2020. Fungal Diversity 105:1-16 ((https://doi.org/10.1007/s13225-020-00466-2)) |
| Otu28680 | 1 | k__Fungi(100);p__Ascomycota(100);c__Saccharomycetes(100);o__Saccharomycetales(100);f__Metschnikowiaceae(100);g__Metschnikowia(100);s__Metschnikowia_pulcherrima(100); | Metschnikowia | 13 | Saprotroph | \|Undefined Saprotroph\| | Yeast | NULL | Probable | As nectar-tap_saprotroph (Põlme et al. 2020); Decay sugar-rich substrates (Põlme et al. 2020) | Põlme S, et al. 2020. Fungal Diversity 105:1-16 ((https://doi.org/10.1007/s13225-020-00466-2)) |
| Otu28681 | 1 | k__Fungi(100);p__Ascomycota(100);c__Saccharomycetes(100);o__Saccharomycetales(100);f__Metschnikowiaceae(100);g__Metschnikowia(100);s__Metschnikowia_pulcherrima(100); | Metschnikowia | 13 | Saprotroph | \|Undefined Saprotroph\| | Yeast | NULL | Probable | As nectar-tap_saprotroph (Põlme et al. 2020); Decay sugar-rich substrates (Põlme et al. 2020) | Põlme S, et al. 2020. Fungal Diversity 105:1-16 ((https://doi.org/10.1007/s13225-020-00466-2)) |
| Otu28683 | 1 | k__Fungi(100);p__Ascomycota(100);c__Saccharomycetes(100);o__Saccharomycetales(100);f__Metschnikowiaceae(100);g__Metschnikowia(100);s__Metschnikowia_pulcherrima(100); | Metschnikowia | 13 | Saprotroph | \|Undefined Saprotroph\| | Yeast | NULL | Probable | As nectar-tap_saprotroph (Põlme et al. 2020); Decay sugar-rich substrates (Põlme et al. 2020) | Põlme S, et al. 2020. Fungal Diversity 105:1-16 ((https://doi.org/10.1007/s13225-020-00466-2)) |
| Otu28684 | 1 | k__Fungi(100);p__Ascomycota(100);c__Saccharomycetes(100);o__Saccharomycetales(100);f__Metschnikowiaceae(100);g__Metschnikowia(100);g__Metschnikowia_unclassified(100); | Metschnikowia | 13 | Saprotroph | \|Undefined Saprotroph\| | Yeast | NULL | Probable | As nectar-tap_saprotroph (Põlme et al. 2020); Decay sugar-rich substrates (Põlme et al. 2020) | Põlme S, et al. 2020. Fungal Diversity 105:1-16 ((https://doi.org/10.1007/s13225-020-00466-2)) |
| Otu28687 | 1 | k__Fungi(100);p__Ascomycota(100);c__Saccharomycetes(100);o__Saccharomycetales(100);f__Metschnikowiaceae(100);g__Metschnikowia(100);s__Metschnikowia_pulcherrima(100); | Metschnikowia | 13 | Saprotroph | \|Undefined Saprotroph\| | Yeast | NULL | Probable | As nectar-tap_saprotroph (Põlme et al. 2020); Decay sugar-rich substrates (Põlme et al. 2020) | Põlme S, et al. 2020. Fungal Diversity 105:1-16 ((https://doi.org/10.1007/s13225-020-00466-2)) |
| Otu28691 | 1 | k__Fungi(100);p__Ascomycota(100);c__Saccharomycetes(100);o__Saccharomycetales(100);f__Metschnikowiaceae(100);g__Metschnikowia(100);s__Metschnikowia_pulcherrima(100); | Metschnikowia | 13 | Saprotroph | \|Undefined Saprotroph\| | Yeast | NULL | Probable | As nectar-tap_saprotroph (Põlme et al. 2020); Decay sugar-rich substrates (Põlme et al. 2020) | Põlme S, et al. 2020. Fungal Diversity 105:1-16 ((https://doi.org/10.1007/s13225-020-00466-2)) |
| Otu28692 | 1 | k__Fungi(100);p__Ascomycota(100);c__Saccharomycetes(100);o__Saccharomycetales(100);f__Metschnikowiaceae(100);g__Metschnikowia(100);s__Metschnikowia_pulcherrima(100); | Metschnikowia | 13 | Saprotroph | \|Undefined Saprotroph\| | Yeast | NULL | Probable | As nectar-tap_saprotroph (Põlme et al. 2020); Decay sugar-rich substrates (Põlme et al. 2020) | Põlme S, et al. 2020. Fungal Diversity 105:1-16 ((https://doi.org/10.1007/s13225-020-00466-2)) |
| Otu28693 | 1 | k__Fungi(100);p__Ascomycota(100);c__Saccharomycetes(100);o__Saccharomycetales(100);f__Metschnikowiaceae(100);g__Metschnikowia(100);g__Metschnikowia_unclassified(100); | Metschnikowia | 13 | Saprotroph | \|Undefined Saprotroph\| | Yeast | NULL | Probable | As nectar-tap_saprotroph (Põlme et al. 2020); Decay sugar-rich substrates (Põlme et al. 2020) | Põlme S, et al. 2020. Fungal Diversity 105:1-16 ((https://doi.org/10.1007/s13225-020-00466-2)) |
| Otu28694 | 1 | k__Fungi(100);p__Ascomycota(100);c__Saccharomycetes(100);o__Saccharomycetales(100);f__Metschnikowiaceae(100);g__Metschnikowia(100);s__Metschnikowia_pulcherrima(100); | Metschnikowia | 13 | Saprotroph | \|Undefined Saprotroph\| | Yeast | NULL | Probable | As nectar-tap_saprotroph (Põlme et al. 2020); Decay sugar-rich substrates (Põlme et al. 2020) | Põlme S, et al. 2020. Fungal Diversity 105:1-16 ((https://doi.org/10.1007/s13225-020-00466-2)) |
| Otu28695 | 1 | k__Fungi(100);p__Ascomycota(100);c__Saccharomycetes(100);o__Saccharomycetales(100);f__Metschnikowiaceae(100);g__Metschnikowia(100);g__Metschnikowia_unclassified(100); | Metschnikowia | 13 | Saprotroph | \|Undefined Saprotroph\| | Yeast | NULL | Probable | As nectar-tap_saprotroph (Põlme et al. 2020); Decay sugar-rich substrates (Põlme et al. 2020) | Põlme S, et al. 2020. Fungal Diversity 105:1-16 ((https://doi.org/10.1007/s13225-020-00466-2)) |
| Otu28696 | 1 | k__Fungi(100);p__Ascomycota(100);c__Saccharomycetes(100);o__Saccharomycetales(100);f__Metschnikowiaceae(100);g__Metschnikowia(100);s__Metschnikowia_pulcherrima(100); | Metschnikowia | 13 | Saprotroph | \|Undefined Saprotroph\| | Yeast | NULL | Probable | As nectar-tap_saprotroph (Põlme et al. 2020); Decay sugar-rich substrates (Põlme et al. 2020) | Põlme S, et al. 2020. Fungal Diversity 105:1-16 ((https://doi.org/10.1007/s13225-020-00466-2)) |
| Otu28697 | 1 | k__Fungi(100);p__Ascomycota(100);c__Saccharomycetes(100);o__Saccharomycetales(100);f__Metschnikowiaceae(100);g__Metschnikowia(100);g__Metschnikowia_unclassified(100); | Metschnikowia | 13 | Saprotroph | \|Undefined Saprotroph\| | Yeast | NULL | Probable | As nectar-tap_saprotroph (Põlme et al. 2020); Decay sugar-rich substrates (Põlme et al. 2020) | Põlme S, et al. 2020. Fungal Diversity 105:1-16 ((https://doi.org/10.1007/s13225-020-00466-2)) |
| Otu28698 | 1 | k__Fungi(100);p__Basidiomycota(100);c__Microbotryomycetes(100);o__Sporidiobolales(100);f__Sporidiobolales_family_Incertae_sedis(100);g__Rhodotorula(100);s__Rhodotorula_sp(100); | Rhodotorula | 13 | Pathotroph-Saprotroph-Symbiotroph | Animal Pathogen-Animal Symbiotroph-Endophyte-Plant Pathogen-\|Undefined Saprotroph\| | Yeast | NULL | Probable | As foliar_endophyte (Põlme et al. 2020); Opportunistic human pathogen | Põlme S, et al. 2020. Fungal Diversity 105:1-16 ((https://doi.org/10.1007/s13225-020-00466-2)); https://www.cdc.gov ((https://www.cdc.gov)) |
| Otu28699 | 1 | k__Fungi(100);p__Basidiomycota(100);c__Agaricomycetes(100);o__Russulales(100);f__Peniophoraceae(100);g__Peniophora(100);g__Peniophora_unclassified(100); | Peniophora | 13 | Pathotroph-Saprotroph | Plant Pathogen-\|Wood Saprotroph\| | Corticioid | White Rot | Probable | As white_rot (Põlme et al. 2020); As wood_pathogen (Põlme et al. 2020); Decay wood (Põlme et al. 2020) | Tedersoo L, et al. 2014. Science 346:e1256688 ((https://doi.org/10.1126/science.1256688)); Põlme S, et al. 2020. Fungal Diversity 105:1-16 ((https://doi.org/10.1007/s13225-020-00466-2)) |
| Otu28700 | 1 | k__Fungi(100);p__Basidiomycota(100);c__Agaricomycetes(100);o__Russulales(100);f__Peniophoraceae(100);g__Peniophora(100);g__Peniophora_unclassified(100); | Peniophora | 13 | Pathotroph-Saprotroph | Plant Pathogen-\|Wood Saprotroph\| | Corticioid | White Rot | Probable | As white_rot (Põlme et al. 2020); As wood_pathogen (Põlme et al. 2020); Decay wood (Põlme et al. 2020) | Tedersoo L, et al. 2014. Science 346:e1256688 ((https://doi.org/10.1126/science.1256688)); Põlme S, et al. 2020. Fungal Diversity 105:1-16 ((https://doi.org/10.1007/s13225-020-00466-2)) |
| Otu28729 | 1 | k__Fungi(100);p__Ascomycota(100);c__Saccharomycetes(100);o__Saccharomycetales(100);f__Metschnikowiaceae(100);g__Metschnikowia(100);g__Metschnikowia_unclassified(100); | Metschnikowia | 13 | Saprotroph | \|Undefined Saprotroph\| | Yeast | NULL | Probable | As nectar-tap_saprotroph (Põlme et al. 2020); Decay sugar-rich substrates (Põlme et al. 2020) | Põlme S, et al. 2020. Fungal Diversity 105:1-16 ((https://doi.org/10.1007/s13225-020-00466-2)) |
| Otu28741 | 1 | k__Fungi(100);p__Ascomycota(100);c__Saccharomycetes(100);o__Saccharomycetales(100);f__Metschnikowiaceae(100);g__Metschnikowia(100);s__Metschnikowia_pulcherrima(100); | Metschnikowia | 13 | Saprotroph | \|Undefined Saprotroph\| | Yeast | NULL | Probable | As nectar-tap_saprotroph (Põlme et al. 2020); Decay sugar-rich substrates (Põlme et al. 2020) | Põlme S, et al. 2020. Fungal Diversity 105:1-16 ((https://doi.org/10.1007/s13225-020-00466-2)) |
| Otu28754 | 1 | k__Fungi(100);p__Ascomycota(100);c__Saccharomycetes(100);o__Saccharomycetales(100);f__Metschnikowiaceae(100);g__Metschnikowia(100);s__Metschnikowia_pulcherrima(100); | Metschnikowia | 13 | Saprotroph | \|Undefined Saprotroph\| | Yeast | NULL | Probable | As nectar-tap_saprotroph (Põlme et al. 2020); Decay sugar-rich substrates (Põlme et al. 2020) | Põlme S, et al. 2020. Fungal Diversity 105:1-16 ((https://doi.org/10.1007/s13225-020-00466-2)) |
| Otu28755 | 1 | k__Fungi(100);p__Ascomycota(100);c__Saccharomycetes(100);o__Saccharomycetales(100);f__Metschnikowiaceae(100);g__Metschnikowia(100);s__Metschnikowia_pulcherrima(100); | Metschnikowia | 13 | Saprotroph | \|Undefined Saprotroph\| | Yeast | NULL | Probable | As nectar-tap_saprotroph (Põlme et al. 2020); Decay sugar-rich substrates (Põlme et al. 2020) | Põlme S, et al. 2020. Fungal Diversity 105:1-16 ((https://doi.org/10.1007/s13225-020-00466-2)) |
| Otu28756 | 1 | k__Fungi(100);p__Ascomycota(100);c__Saccharomycetes(100);o__Saccharomycetales(100);f__Metschnikowiaceae(100);g__Metschnikowia(100);s__Metschnikowia_pulcherrima(100); | Metschnikowia | 13 | Saprotroph | \|Undefined Saprotroph\| | Yeast | NULL | Probable | As nectar-tap_saprotroph (Põlme et al. 2020); Decay sugar-rich substrates (Põlme et al. 2020) | Põlme S, et al. 2020. Fungal Diversity 105:1-16 ((https://doi.org/10.1007/s13225-020-00466-2)) |
| Otu28760 | 1 | k__Fungi(100);p__Ascomycota(100);c__Saccharomycetes(100);o__Saccharomycetales(100);f__Metschnikowiaceae(100);g__Metschnikowia(100);g__Metschnikowia_unclassified(100); | Metschnikowia | 13 | Saprotroph | \|Undefined Saprotroph\| | Yeast | NULL | Probable | As nectar-tap_saprotroph (Põlme et al. 2020); Decay sugar-rich substrates (Põlme et al. 2020) | Põlme S, et al. 2020. Fungal Diversity 105:1-16 ((https://doi.org/10.1007/s13225-020-00466-2)) |
| Otu28761 | 1 | k__Fungi(100);p__Ascomycota(100);c__Saccharomycetes(100);o__Saccharomycetales(100);f__Metschnikowiaceae(100);g__Metschnikowia(100);g__Metschnikowia_unclassified(100); | Metschnikowia | 13 | Saprotroph | \|Undefined Saprotroph\| | Yeast | NULL | Probable | As nectar-tap_saprotroph (Põlme et al. 2020); Decay sugar-rich substrates (Põlme et al. 2020) | Põlme S, et al. 2020. Fungal Diversity 105:1-16 ((https://doi.org/10.1007/s13225-020-00466-2)) |
| Otu28764 | 1 | k__Fungi(100);p__Ascomycota(100);c__Saccharomycetes(100);o__Saccharomycetales(100);f__Metschnikowiaceae(100);g__Metschnikowia(100);s__Metschnikowia_pulcherrima(100); | Metschnikowia | 13 | Saprotroph | \|Undefined Saprotroph\| | Yeast | NULL | Probable | As nectar-tap_saprotroph (Põlme et al. 2020); Decay sugar-rich substrates (Põlme et al. 2020) | Põlme S, et al. 2020. Fungal Diversity 105:1-16 ((https://doi.org/10.1007/s13225-020-00466-2)) |
| Otu28765 | 1 | k__Fungi(100);p__Ascomycota(100);c__Saccharomycetes(100);o__Saccharomycetales(100);f__Metschnikowiaceae(100);g__Metschnikowia(100);s__Metschnikowia_pulcherrima(100); | Metschnikowia | 13 | Saprotroph | \|Undefined Saprotroph\| | Yeast | NULL | Probable | As nectar-tap_saprotroph (Põlme et al. 2020); Decay sugar-rich substrates (Põlme et al. 2020) | Põlme S, et al. 2020. Fungal Diversity 105:1-16 ((https://doi.org/10.1007/s13225-020-00466-2)) |
| Otu28767 | 1 | k__Fungi(100);p__Ascomycota(100);c__Saccharomycetes(100);o__Saccharomycetales(100);f__Metschnikowiaceae(100);g__Metschnikowia(100);s__Metschnikowia_pulcherrima(100); | Metschnikowia | 13 | Saprotroph | \|Undefined Saprotroph\| | Yeast | NULL | Probable | As nectar-tap_saprotroph (Põlme et al. 2020); Decay sugar-rich substrates (Põlme et al. 2020) | Põlme S, et al. 2020. Fungal Diversity 105:1-16 ((https://doi.org/10.1007/s13225-020-00466-2)) |
| Otu28768 | 1 | k__Fungi(100);p__Ascomycota(100);c__Sordariomycetes(100);o__Hypocreales(100);f__Ophiocordycipitaceae(100);f__Ophiocordycipitaceae_unclassified(100);f__Ophiocordycipitaceae_unclassified(100); | Ophiocordycipitaceae | 9 | Pathotroph-Symbiotroph | Endophyte-Insect Pathogen | Microfungus | NULL | Probable | A family of parasitic fungi (Sung et al. 2007) | Sung GH, et al. 2007. Studies in Mycology 57:5-59 ((https://doi.org/10.3114/sim.2007.57.01)); Costa IPMW, et al. 2012. Mycotaxon 119: 425 ((http://mycotaxon.com/resources/checklists/costa_v119_checklist.pdf)) |
| Otu28769 | 1 | k__Fungi(100);p__Ascomycota(100);c__Saccharomycetes(100);o__Saccharomycetales(100);f__Metschnikowiaceae(100);g__Metschnikowia(100);s__Metschnikowia_pulcherrima(100); | Metschnikowia | 13 | Saprotroph | \|Undefined Saprotroph\| | Yeast | NULL | Probable | As nectar-tap_saprotroph (Põlme et al. 2020); Decay sugar-rich substrates (Põlme et al. 2020) | Põlme S, et al. 2020. Fungal Diversity 105:1-16 ((https://doi.org/10.1007/s13225-020-00466-2)) |
| Otu28770 | 1 | k__Fungi(100);p__Ascomycota(100);c__Saccharomycetes(100);o__Saccharomycetales(100);f__Metschnikowiaceae(100);g__Metschnikowia(100);s__Metschnikowia_pulcherrima(100); | Metschnikowia | 13 | Saprotroph | \|Undefined Saprotroph\| | Yeast | NULL | Probable | As nectar-tap_saprotroph (Põlme et al. 2020); Decay sugar-rich substrates (Põlme et al. 2020) | Põlme S, et al. 2020. Fungal Diversity 105:1-16 ((https://doi.org/10.1007/s13225-020-00466-2)) |
| Otu28771 | 1 | k__Fungi(100);p__Ascomycota(100);c__Saccharomycetes(100);o__Saccharomycetales(100);f__Metschnikowiaceae(100);g__Metschnikowia(100);g__Metschnikowia_unclassified(100); | Metschnikowia | 13 | Saprotroph | \|Undefined Saprotroph\| | Yeast | NULL | Probable | As nectar-tap_saprotroph (Põlme et al. 2020); Decay sugar-rich substrates (Põlme et al. 2020) | Põlme S, et al. 2020. Fungal Diversity 105:1-16 ((https://doi.org/10.1007/s13225-020-00466-2)) |
| Otu28772 | 1 | k__Fungi(100);p__Ascomycota(100);c__Saccharomycetes(100);o__Saccharomycetales(100);f__Metschnikowiaceae(100);g__Metschnikowia(100);g__Metschnikowia_unclassified(100); | Metschnikowia | 13 | Saprotroph | \|Undefined Saprotroph\| | Yeast | NULL | Probable | As nectar-tap_saprotroph (Põlme et al. 2020); Decay sugar-rich substrates (Põlme et al. 2020) | Põlme S, et al. 2020. Fungal Diversity 105:1-16 ((https://doi.org/10.1007/s13225-020-00466-2)) |
| Otu28774 | 1 | k__Fungi(100);p__Ascomycota(100);c__Saccharomycetes(100);o__Saccharomycetales(100);f__Metschnikowiaceae(100);g__Metschnikowia(100);s__Metschnikowia_pulcherrima(100); | Metschnikowia | 13 | Saprotroph | \|Undefined Saprotroph\| | Yeast | NULL | Probable | As nectar-tap_saprotroph (Põlme et al. 2020); Decay sugar-rich substrates (Põlme et al. 2020) | Põlme S, et al. 2020. Fungal Diversity 105:1-16 ((https://doi.org/10.1007/s13225-020-00466-2)) |
| Otu28776 | 1 | k__Fungi(100);p__Ascomycota(100);c__Saccharomycetes(100);o__Saccharomycetales(100);f__Metschnikowiaceae(100);g__Metschnikowia(100);g__Metschnikowia_unclassified(100); | Metschnikowia | 13 | Saprotroph | \|Undefined Saprotroph\| | Yeast | NULL | Probable | As nectar-tap_saprotroph (Põlme et al. 2020); Decay sugar-rich substrates (Põlme et al. 2020) | Põlme S, et al. 2020. Fungal Diversity 105:1-16 ((https://doi.org/10.1007/s13225-020-00466-2)) |
| Otu28777 | 1 | k__Fungi(100);p__Ascomycota(100);c__Saccharomycetes(100);o__Saccharomycetales(100);f__Metschnikowiaceae(100);g__Metschnikowia(100);g__Metschnikowia_unclassified(100); | Metschnikowia | 13 | Saprotroph | \|Undefined Saprotroph\| | Yeast | NULL | Probable | As nectar-tap_saprotroph (Põlme et al. 2020); Decay sugar-rich substrates (Põlme et al. 2020) | Põlme S, et al. 2020. Fungal Diversity 105:1-16 ((https://doi.org/10.1007/s13225-020-00466-2)) |
| Otu28778 | 1 | k__Fungi(100);p__Ascomycota(100);c__Saccharomycetes(100);o__Saccharomycetales(100);f__Metschnikowiaceae(100);g__Metschnikowia(100);g__Metschnikowia_unclassified(100); | Metschnikowia | 13 | Saprotroph | \|Undefined Saprotroph\| | Yeast | NULL | Probable | As nectar-tap_saprotroph (Põlme et al. 2020); Decay sugar-rich substrates (Põlme et al. 2020) | Põlme S, et al. 2020. Fungal Diversity 105:1-16 ((https://doi.org/10.1007/s13225-020-00466-2)) |
| Otu28779 | 1 | k__Fungi(100);p__Ascomycota(100);c__Saccharomycetes(100);o__Saccharomycetales(100);f__Metschnikowiaceae(100);g__Metschnikowia(100);g__Metschnikowia_unclassified(100); | Metschnikowia | 13 | Saprotroph | \|Undefined Saprotroph\| | Yeast | NULL | Probable | As nectar-tap_saprotroph (Põlme et al. 2020); Decay sugar-rich substrates (Põlme et al. 2020) | Põlme S, et al. 2020. Fungal Diversity 105:1-16 ((https://doi.org/10.1007/s13225-020-00466-2)) |
| Otu28780 | 1 | k__Fungi(100);p__Ascomycota(100);c__Saccharomycetes(100);o__Saccharomycetales(100);f__Metschnikowiaceae(100);g__Metschnikowia(100);s__Metschnikowia_pulcherrima(100); | Metschnikowia | 13 | Saprotroph | \|Undefined Saprotroph\| | Yeast | NULL | Probable | As nectar-tap_saprotroph (Põlme et al. 2020); Decay sugar-rich substrates (Põlme et al. 2020) | Põlme S, et al. 2020. Fungal Diversity 105:1-16 ((https://doi.org/10.1007/s13225-020-00466-2)) |
| Otu28782 | 1 | k__Fungi(100);p__Ascomycota(100);c__Eurotiomycetes(100);o__Eurotiales(100);f__Trichocomaceae(100);g__Penicillium(100);g__Penicillium_unclassified(100); | Penicillium | 13 | Pathotroph-Saprotroph-Symbiotroph | Animal Parasite-Dung Saprotroph-Endophyte-Fungal Parasite-Plant Pathogen-\|Undefined Saprotroph\|-Wood Saprotroph | Microfungus | Soft Rot | Highly Probable | As foliar_endophyte (Põlme et al. 2020); As mycoparasite (Põlme et al. 2020); Decay wood (Põlme et al. 2020) | Duncan CG, Eslyn WE. 1966. Mycologia 58:642-654 [illegible] |
| Otu28786 | 1 | k__Fungi(100);p__Ascomycota(100);c__Saccharomycetes(100);o__Saccharomycetales(100);f__Metschnikowiaceae(100);g__Metschnikowia(100);g__Metschnikowia_unclassified(100); | Metschnikowia | 13 | Saprotroph | \|Undefined Saprotroph\| | Yeast | NULL | Probable | As nectar-tap_saprotroph (Põlme et al. 2020); Decay sugar-rich substrates (Põlme et al. 2020) | Põlme S, et al. 2020. Fungal Diversity 105:1-16 ((https://doi.org/10.1007/s13225-020-00466-2)) |
| Otu28787 | 1 | k__Fungi(100);p__Ascomycota(100);c__Saccharomycetes(100);o__Saccharomycetales(100);f__Metschnikowiaceae(100);g__Metschnikowia(100);s__Metschnikowia_pulcherrima(100); | Metschnikowia | 13 | Saprotroph | \|Undefined Saprotroph\| | Yeast | NULL | Probable | As nectar-tap_saprotroph (Põlme et al. 2020); Decay sugar-rich substrates (Põlme et al. 2020) | Põlme S, et al. 2020. Fungal Diversity 105:1-16 ((https://doi.org/10.1007/s13225-020-00466-2)) |
| Otu28790 | 1 | k__Fungi(100);p__Ascomycota(100);c__Saccharomycetes(100);o__Saccharomycetales(100);f__Metschnikowiaceae(100);g__Metschnikowia(100);s__Metschnikowia_pulcherrima(100); | Metschnikowia | 13 | Saprotroph | \|Undefined Saprotroph\| | Yeast | NULL | Probable | As nectar-tap_saprotroph (Põlme et al. 2020); Decay sugar-rich substrates (Põlme et al. 2020) | Põlme S, et al. 2020. Fungal Diversity 105:1-16 ((https://doi.org/10.1007/s13225-020-00466-2)) |
| Otu28791 | 1 | k__Fungi(100);p__Ascomycota(100);c__Saccharomycetes(100);o__Saccharomycetales(100);f__Metschnikowiaceae(100);g__Metschnikowia(100);s__Metschnikowia_pulcherrima(100); | Metschnikowia | 13 | Saprotroph | \|Undefined Saprotroph\| | Yeast | NULL | Probable | As nectar-tap_saprotroph (Põlme et al. 2020); Decay sugar-rich substrates (Põlme et al. 2020) | Põlme S, et al. 2020. Fungal Diversity 105:1-16 ((https://doi.org/10.1007/s13225-020-00466-2)) |
| Otu28793 | 1 | k__Fungi(100);p__Ascomycota(100);c__Saccharomycetes(100);o__Saccharomycetales(100);f__Metschnikowiaceae(100);g__Metschnikowia(100);g__Metschnikowia_unclassified(100); | Metschnikowia | 13 | Saprotroph | \|Undefined Saprotroph\| | Yeast | NULL | Probable | As nectar-tap_saprotroph (Põlme et al. 2020); Decay sugar-rich substrates (Põlme et al. 2020) | Põlme S, et al. 2020. Fungal Diversity 105:1-16 ((https://doi.org/10.1007/s13225-020-00466-2)) |
| Otu28794 | 1 | k__Fungi(100);p__Ascomycota(100);c__Saccharomycetes(100);o__Saccharomycetales(100);f__Metschnikowiaceae(100);g__Metschnikowia(100);s__Metschnikowia_pulcherrima(100); | Metschnikowia | 13 | Saprotroph | \|Undefined Saprotroph\| | Yeast | NULL | Probable | As nectar-tap_saprotroph (Põlme et al. 2020); Decay sugar-rich substrates (Põlme et al. 2020) | Põlme S, et al. 2020. Fungal Diversity 105:1-16 ((https://doi.org/10.1007/s13225-020-00466-2)) |
| Otu28798 | 1 | k__Fungi(100);p__Ascomycota(100);c__Saccharomycetes(100);o__Saccharomycetales(100);f__Metschnikowiaceae(100);g__Metschnikowia(100);g__Metschnikowia_unclassified(100); | Metschnikowia | 13 | Saprotroph | \|Undefined Saprotroph\| | Yeast | NULL | Probable | As nectar-tap_saprotroph (Põlme et al. 2020); Decay sugar-rich substrates (Põlme et al. 2020) | Põlme S, et al. 2020. Fungal Diversity 105:1-16 ((https://doi.org/10.1007/s13225-020-00466-2)) |
| Otu28799 | 1 | k__Fungi(100);p__Ascomycota(100);c__Saccharomycetes(100);o__Saccharomycetales(100);f__Metschnikowiaceae(100);g__Metschnikowia(100);s__Metschnikowia_pulcherrima(100); | Metschnikowia | 13 | Saprotroph | \|Undefined Saprotroph\| | Yeast | NULL | Probable | As nectar-tap_saprotroph (Põlme et al. 2020); Decay sugar-rich substrates (Põlme et al. 2020) | Põlme S, et al. 2020. Fungal Diversity 105:1-16 ((https://doi.org/10.1007/s13225-020-00466-2)) |
| Otu28802 | 1 | k__Fungi(100);p__Basidiomycota(100);c__Agaricomycetes(100);o__Phallales(100);f__Phallaceae(100);g__Phallus(100);g__Phallus_unclassified(100); | Phallus | 13 | Saprotroph | Plant Saprotroph-\|Undefined Saprotroph\|-Wood Saprotroph | Phalloid | White Rot | Probable | As soil_saprotroph (Põlme et al. 2020). Common name - stinkhorn; Decay soil (Põlme et al. 2020); Saprobic, on soil or decaying wood, with spores dispersed by insects that are attracted to the fruit bodies by their distinctive foetid smell (Cannon & Kirk 2007) | Dix NJ, Carney JWG. 1995. Transactions of the British Mycological Society 85:514-520 [illegible] |
| Otu28804 | 1 | k__Fungi(100);p__Ascomycota(100);c__Saccharomycetes(100);o__Saccharomycetales(100);f__Metschnikowiaceae(100);g__Metschnikowia(100);s__Metschnikowia_pulcherrima(100); | Metschnikowia | 13 | Saprotroph | \|Undefined Saprotroph\| | Yeast | NULL | Probable | As nectar-tap_saprotroph (Põlme et al. 2020); Decay sugar-rich substrates (Põlme et al. 2020) | Põlme S, et al. 2020. Fungal Diversity 105:1-16 ((https://doi.org/10.1007/s13225-020-00466-2)) |
| Otu28808 | 1 | k__Fungi(100);p__Ascomycota(100);c__Saccharomycetes(100);o__Saccharomycetales(100);f__Metschnikowiaceae(100);g__Metschnikowia(100);s__Metschnikowia_pulcherrima(100); | Metschnikowia | 13 | Saprotroph | \|Undefined Saprotroph\| | Yeast | NULL | Probable | As nectar-tap_saprotroph (Põlme et al. 2020); Decay sugar-rich substrates (Põlme et al. 2020) | Põlme S, et al. 2020. Fungal Diversity 105:1-16 ((https://doi.org/10.1007/s13225-020-00466-2)) |
| Otu28809 | 1 | k__Fungi(100);p__Ascomycota(100);c__Saccharomycetes(100);o__Saccharomycetales(100);f__Metschnikowiaceae(100);g__Metschnikowia(100);g__Metschnikowia_unclassified(100); | Metschnikowia | 13 | Saprotroph | \|Undefined Saprotroph\| | Yeast | NULL | Probable | As nectar-tap_saprotroph (Põlme et al. 2020); Decay sugar-rich substrates (Põlme et al. 2020) | Põlme S, et al. 2020. Fungal Diversity 105:1-16 ((https://doi.org/10.1007/s13225-020-00466-2)) |
| Otu28811 | 1 | k__Fungi(100);p__Ascomycota(100);c__Saccharomycetes(100);o__Saccharomycetales(100);f__Metschnikowiaceae(100);g__Metschnikowia(100);s__Metschnikowia_pulcherrima(100); | Metschnikowia | 13 | Saprotroph | \|Undefined Saprotroph\| | Yeast | NULL | Probable | As nectar-tap_saprotroph (Põlme et al. 2020); Decay sugar-rich substrates (Põlme et al. 2020) | Põlme S, et al. 2020. Fungal Diversity 105:1-16 ((https://doi.org/10.1007/s13225-020-00466-2)) |
| Otu28812 | 1 | k__Fungi(100);p__Ascomycota(100);c__Saccharomycetes(100);o__Saccharomycetales(100);f__Metschnikowiaceae(100);g__Metschnikowia(100);s__Metschnikowia_pulcherrima(100); | Metschnikowia | 13 | Saprotroph | \|Undefined Saprotroph\| | Yeast | NULL | Probable | As nectar-tap_saprotroph (Põlme et al. 2020); Decay sugar-rich substrates (Põlme et al. 2020) | Põlme S, et al. 2020. Fungal Diversity 105:1-16 ((https://doi.org/10.1007/s13225-020-00466-2)) |
| Otu28813 | 1 | k__Fungi(100);p__Ascomycota(100);c__Saccharomycetes(100);o__Saccharomycetales(100);f__Metschnikowiaceae(100);g__Metschnikowia(100);s__Metschnikowia_pulcherrima(100); | Metschnikowia | 13 | Saprotroph | \|Undefined Saprotroph\| | Yeast | NULL | Probable | As nectar-tap_saprotroph (Põlme et al. 2020); Decay sugar-rich substrates (Põlme et al. 2020) | Põlme S, et al. 2020. Fungal Diversity 105:1-16 ((https://doi.org/10.1007/s13225-020-00466-2)) |
| Otu28814 | 1 | k__Fungi(100);p__Ascomycota(100);c__Saccharomycetes(100);o__Saccharomycetales(100);f__Metschnikowiaceae(100);g__Metschnikowia(100);g__Metschnikowia_unclassified(100); | Metschnikowia | 13 | Saprotroph | \|Undefined Saprotroph\| | Yeast | NULL | Probable | As nectar-tap_saprotroph (Põlme et al. 2020); Decay sugar-rich substrates (Põlme et al. 2020) | Põlme S, et al. 2020. Fungal Diversity 105:1-16 ((https://doi.org/10.1007/s13225-020-00466-2)) |
| Otu28815 | 1 | k__Fungi(100);p__Ascomycota(100);c__Saccharomycetes(100);o__Saccharomycetales(100);f__Metschnikowiaceae(100);g__Metschnikowia(100);s__Metschnikowia_pulcherrima(100); | Metschnikowia | 13 | Saprotroph | \|Undefined Saprotroph\| | Yeast | NULL | Probable | As nectar-tap_saprotroph (Põlme et al. 2020); Decay sugar-rich substrates (Põlme et al. 2020) | Põlme S, et al. 2020. Fungal Diversity 105:1-16 ((https://doi.org/10.1007/s13225-020-00466-2)) |
| Otu28816 | 1 | k__Fungi(100);p__Ascomycota(100);c__Saccharomycetes(100);o__Saccharomycetales(100);f__Metschnikowiaceae(100);g__Metschnikowia(100);s__Metschnikowia_pulcherrima(100); | Metschnikowia | 13 | Saprotroph | \|Undefined Saprotroph\| | Yeast | NULL | Probable | As nectar-tap_saprotroph (Põlme et al. 2020); Decay sugar-rich substrates (Põlme et al. 2020) | Põlme S, et al. 2020. Fungal Diversity 105:1-16 ((https://doi.org/10.1007/s13225-020-00466-2)) |
| Otu28817 | 1 | k__Fungi(100);p__Ascomycota(100);c__Saccharomycetes(100);o__Saccharomycetales(100);f__Metschnikowiaceae(100);g__Metschnikowia(100);g__Metschnikowia_unclassified(100); | Metschnikowia | 13 | Saprotroph | \|Undefined Saprotroph\| | Yeast | NULL | Probable | As nectar-tap_saprotroph (Põlme et al. 2020); Decay sugar-rich substrates (Põlme et al. 2020) | Põlme S, et al. 2020. Fungal Diversity 105:1-16 ((https://doi.org/10.1007/s13225-020-00466-2)) |
| Otu28818 | 1 | k__Fungi(100);p__Ascomycota(100);c__Saccharomycetes(100);o__Saccharomycetales(100);f__Metschnikowiaceae(100);g__Metschnikowia(100);s__Metschnikowia_pulcherrima(100); | Metschnikowia | 13 | Saprotroph | \|Undefined Saprotroph\| | Yeast | NULL | Probable | As nectar-tap_saprotroph (Põlme et al. 2020); Decay sugar-rich substrates (Põlme et al. 2020) | Põlme S, et al. 2020. Fungal Diversity 105:1-16 ((https://doi.org/10.1007/s13225-020-00466-2)) |
| Otu28819 | 1 | k__Fungi(100);p__Ascomycota(100);c__Saccharomycetes(100);o__Saccharomycetales(100);f__Metschnikowiaceae(100);g__Metschnikowia(100);g__Metschnikowia_unclassified(100); | Metschnikowia | 13 | Saprotroph | \|Undefined Saprotroph\| | Yeast | NULL | Probable | As nectar-tap_saprotroph (Põlme et al. 2020); Decay sugar-rich substrates (Põlme et al. 2020) | Põlme S, et al. 2020. Fungal Diversity 105:1-16 ((https://doi.org/10.1007/s13225-020-00466-2)) |
| Otu28820 | 1 | k__Fungi(100);p__Ascomycota(100);c__Saccharomycetes(100);o__Saccharomycetales(100);f__Metschnikowiaceae(100);g__Metschnikowia(100);s__Metschnikowia_pulcherrima(100); | Metschnikowia | 13 | Saprotroph | \|Undefined Saprotroph\| | Yeast | NULL | Probable | As nectar-tap_saprotroph (Põlme et al. 2020); Decay sugar-rich substrates (Põlme et al. 2020) | Põlme S, et al. 2020. Fungal Diversity 105:1-16 ((https://doi.org/10.1007/s13225-020-00466-2)) |
| Otu28821 | 1 | k__Fungi(100);p__Ascomycota(100);c__Saccharomycetes(100);o__Saccharomycetales(100);f__Metschnikowiaceae(100);g__Metschnikowia(100);s__Metschnikowia_pulcherrima(100); | Metschnikowia | 13 | Saprotroph | \|Undefined Saprotroph\| | Yeast | NULL | Probable | As nectar-tap_saprotroph (Põlme et al. 2020); Decay sugar-rich substrates (Põlme et al. 2020) | Põlme S, et al. 2020. Fungal Diversity 105:1-16 ((https://doi.org/10.1007/s13225-020-00466-2)) |
| Otu28822 | 1 | k__Fungi(100);p__Ascomycota(100);c__Saccharomycetes(100);o__Saccharomycetales(100);f__Metschnikowiaceae(100);g__Metschnikowia(100);g__Metschnikowia_unclassified(100); | Metschnikowia | 13 | Saprotroph | \|Undefined Saprotroph\| | Yeast | NULL | Probable | As nectar-tap_saprotroph (Põlme et al. 2020); Decay sugar-rich substrates (Põlme et al. 2020) | Põlme S, et al. 2020. Fungal Diversity 105:1-16 ((https://doi.org/10.1007/s13225-020-00466-2)) |
| Otu28823 | 1 | k__Fungi(100);p__Ascomycota(100);c__Saccharomycetes(100);o__Saccharomycetales(100);f__Metschnikowiaceae(100);g__Metschnikowia(100);s__Metschnikowia_pulcherrima(100); | Metschnikowia | 13 | Saprotroph | \|Undefined Saprotroph\| | Yeast | NULL | Probable | As nectar-tap_saprotroph (Põlme et al. 2020); Decay sugar-rich substrates (Põlme et al. 2020) | Põlme S, et al. 2020. Fungal Diversity 105:1-16 ((https://doi.org/10.1007/s13225-020-00466-2)) |
| Otu28824 | 1 | k__Fungi(100);p__Ascomycota(100);c__Saccharomycetes(100);o__Saccharomycetales(100);f__Metschnikowiaceae(100);g__Metschnikowia(100);g__Metschnikowia_unclassified(100); | Metschnikowia | 13 | Saprotroph | \|Undefined Saprotroph\| | Yeast | NULL | Probable | As nectar-tap_saprotroph (Põlme et al. 2020); Decay sugar-rich substrates (Põlme et al. 2020) | Põlme S, et al. 2020. Fungal Diversity 105:1-16 ((https://doi.org/10.1007/s13225-020-00466-2)) |
| Otu28825 | 1 | k__Fungi(100);p__Ascomycota(100);c__Saccharomycetes(100);o__Saccharomycetales(100);f__Metschnikowiaceae(100);g__Metschnikowia(100);s__Metschnikowia_pulcherrima(100); | Metschnikowia | 13 | Saprotroph | \|Undefined Saprotroph\| | Yeast | NULL | Probable | As nectar-tap_saprotroph (Põlme et al. 2020); Decay sugar-rich substrates (Põlme et al. 2020) | Põlme S, et al. 2020. Fungal Diversity 105:1-16 ((https://doi.org/10.1007/s13225-020-00466-2)) |
| Otu28826 | 1 | k__Fungi(100);p__Ascomycota(100);c__Saccharomycetes(100);o__Saccharomycetales(100);f__Metschnikowiaceae(100);g__Metschnikowia(100);s__Metschnikowia_pulcherrima(100); | Metschnikowia | 13 | Saprotroph | \|Undefined Saprotroph\| | Yeast | NULL | Probable | As nectar-tap_saprotroph (Põlme et al. 2020); Decay sugar-rich substrates (Põlme et al. 2020) | Põlme S, et al. 2020. Fungal Diversity 105:1-16 ((https://doi.org/10.1007/s13225-020-00466-2)) |
| Otu28827 | 1 | k__Fungi(100);p__Ascomycota(100);c__Saccharomycetes(100);o__Saccharomycetales(100);f__Metschnikowiaceae(100);g__Metschnikowia(100);s__Metschnikowia_pulcherrima(100); | Metschnikowia | 13 | Saprotroph | \|Undefined Saprotroph\| | Yeast | NULL | Probable | As nectar-tap_saprotroph (Põlme et al. 2020); Decay sugar-rich substrates (Põlme et al. 2020) | Põlme S, et al. 2020. Fungal Diversity 105:1-16 ((https://doi.org/10.1007/s13225-020-00466-2)) |
| Otu28828 | 1 | k__Fungi(100);p__Ascomycota(100);c__Saccharomycetes(100);o__Saccharomycetales(100);f__Metschnikowiaceae(100);g__Metschnikowia(100);g__Metschnikowia_unclassified(100); | Metschnikowia | 13 | Saprotroph | \|Undefined Saprotroph\| | Yeast | NULL | Probable | As nectar-tap_saprotroph (Põlme et al. 2020); Decay sugar-rich substrates (Põlme et al. 2020) | Põlme S, et al. 2020. Fungal Diversity 105:1-16 ((https://doi.org/10.1007/s13225-020-00466-2)) |
| Otu28829 | 1 | k__Fungi(100);p__Ascomycota(100);c__Saccharomycetes(100);o__Saccharomycetales(100);f__Metschnikowiaceae(100);g__Metschnikowia(100);s__Metschnikowia_pulcherrima(100); | Metschnikowia | 13 | Saprotroph | \|Undefined Saprotroph\| | Yeast | NULL | Probable | As nectar-tap_saprotroph (Põlme et al. 2020); Decay sugar-rich substrates (Põlme et al. 2020) | Põlme S, et al. 2020. Fungal Diversity 105:1-16 ((https://doi.org/10.1007/s13225-020-00466-2)) |
| Otu28830 | 1 | k__Fungi(100);p__Ascomycota(100);c__Saccharomycetes(100);o__Saccharomycetales(100);f__Metschnikowiaceae(100);g__Metschnikowia(100);s__Metschnikowia_pulcherrima(100); | Metschnikowia | 13 | Saprotroph | \|Undefined Saprotroph\| | Yeast | NULL | Probable | As nectar-tap_saprotroph (Põlme et al. 2020); Decay sugar-rich substrates (Põlme et al. 2020) | Põlme S, et al. 2020. Fungal Diversity 105:1-16 ((https://doi.org/10.1007/s13225-020-00466-2)) |
| Otu28831 | 1 | k__Fungi(100);p__Ascomycota(100);c__Saccharomycetes(100);o__Saccharomycetales(100);f__Metschnikowiaceae(100);g__Metschnikowia(100);g__Metschnikowia_unclassified(100); | Metschnikowia | 13 | Saprotroph | \|Undefined Saprotroph\| | Yeast | NULL | Probable | As nectar-tap_saprotroph (Põlme et al. 2020); Decay sugar-rich substrates (Põlme et al. 2020) | Põlme S, et al. 2020. Fungal Diversity 105:1-16 ((https://doi.org/10.1007/s13225-020-00466-2)) |
| Otu28832 | 1 | k__Fungi(100);p__Ascomycota(100);c__Saccharomycetes(100);o__Saccharomycetales(100);f__Metschnikowiaceae(100);g__Metschnikowia(100);s__Metschnikowia_pulcherrima(100); | Metschnikowia | 13 | Saprotroph | \|Undefined Saprotroph\| | Yeast | NULL | Probable | As nectar-tap_saprotroph (Põlme et al. 2020); Decay sugar-rich substrates (Põlme et al. 2020) | Põlme S, et al. 2020. Fungal Diversity 105:1-16 ((https://doi.org/10.1007/s13225-020-00466-2)) |
| Otu28833 | 1 | k__Fungi(100);p__Ascomycota(100);c__Saccharomycetes(100);o__Saccharomycetales(100);f__Metschnikowiaceae(100);g__Metschnikowia(100);s__Metschnikowia_pulcherrima(100); | Metschnikowia | 13 | Saprotroph | \|Undefined Saprotroph\| | Yeast | NULL | Probable | As nectar-tap_saprotroph (Põlme et al. 2020); Decay sugar-rich substrates (Põlme et al. 2020) | Põlme S, et al. 2020. Fungal Diversity 105:1-16 ((https://doi.org/10.1007/s13225-020-00466-2)) |
| Otu28834 | 1 | k__Fungi(100);p__Ascomycota(100);c__Saccharomycetes(100);o__Saccharomycetales(100);f__Metschnikowiaceae(100);g__Metschnikowia(100);s__Metschnikowia_pulcherrima(100); | Metschnikowia | 13 | Saprotroph | \|Undefined Saprotroph\| | Yeast | NULL | Probable | As nectar-tap_saprotroph (Põlme et al. 2020); Decay sugar-rich substrates (Põlme et al. 2020) | Põlme S, et al. 2020. Fungal Diversity 105:1-16 ((https://doi.org/10.1007/s13225-020-00466-2)) |
| Otu28835 | 1 | k__Fungi(100);p__Ascomycota(100);c__Saccharomycetes(100);o__Saccharomycetales(100);f__Metschnikowiaceae(100);g__Metschnikowia(100);s__Metschnikowia_pulcherrima(100); | Metschnikowia | 13 | Saprotroph | \|Undefined Saprotroph\| | Yeast | NULL | Probable | As nectar-tap_saprotroph (Põlme et al. 2020); Decay sugar-rich substrates (Põlme et al. 2020) | Põlme S, et al. 2020. Fungal Diversity 105:1-16 ((https://doi.org/10.1007/s13225-020-00466-2)) |
| Otu28836 | 1 | k__Fungi(100);p__Ascomycota(100);c__Saccharomycetes(100);o__Saccharomycetales(100);f__Metschnikowiaceae(100);g__Metschnikowia(100);s__Metschnikowia_pulcherrima(100); | Metschnikowia | 13 | Saprotroph | \|Undefined Saprotroph\| | Yeast | NULL | Probable | As nectar-tap_saprotroph (Põlme et al. 2020); Decay sugar-rich substrates (Põlme et al. 2020) | Põlme S, et al. 2020. Fungal Diversity 105:1-16 ((https://doi.org/10.1007/s13225-020-00466-2)) |
| Otu28837 | 1 | k__Fungi(100);p__Ascomycota(100);c__Saccharomycetes(100);o__Saccharomycetales(100);f__Metschnikowiaceae(100);g__Metschnikowia(100);s__Metschnikowia_pulcherrima(100); | Metschnikowia | 13 | Saprotroph | \|Undefined Saprotroph\| | Yeast | NULL | Probable | As nectar-tap_saprotroph (Põlme et al. 2020); Decay sugar-rich substrates (Põlme et al. 2020) | Põlme S, et al. 2020. Fungal Diversity 105:1-16 ((https://doi.org/10.1007/s13225-020-00466-2)) |
| Otu28839 | 1 | k__Fungi(100);p__Ascomycota(100);c__Saccharomycetes(100);o__Saccharomycetales(100);f__Metschnikowiaceae(100);g__Metschnikowia(100);s__Metschnikowia_pulcherrima(100); | Metschnikowia | 13 | Saprotroph | \|Undefined Saprotroph\| | Yeast | NULL | Probable | As nectar-tap_saprotroph (Põlme et al. 2020); Decay sugar-rich substrates (Põlme et al. 2020) | Põlme S, et al. 2020. Fungal Diversity 105:1-16 ((https://doi.org/10.1007/s13225-020-00466-2)) |
| Otu28843 | 1 | k__Fungi(100);p__Ascomycota(100);c__Saccharomycetes(100);o__Saccharomycetales(100);f__Metschnikowiaceae(100);g__Metschnikowia(100);s__Metschnikowia_pulcherrima(100); | Metschnikowia | 13 | Saprotroph | \|Undefined Saprotroph\| | Yeast | NULL | Probable | As nectar-tap_saprotroph (Põlme et al. 2020); Decay sugar-rich substrates (Põlme et al. 2020) | Põlme S, et al. 2020. Fungal Diversity 105:1-16 ((https://doi.org/10.1007/s13225-020-00466-2)) |
| Otu28844 | 1 | k__Fungi(100);p__Ascomycota(100);c__Saccharomycetes(100);o__Saccharomycetales(100);f__Metschnikowiaceae(100);g__Metschnikowia(100);s__Metschnikowia_pulcherrima(100); | Metschnikowia | 13 | Saprotroph | \|Undefined Saprotroph\| | Yeast | NULL | Probable | As nectar-tap_saprotroph (Põlme et al. 2020); Decay sugar-rich substrates (Põlme et al. 2020) | Põlme S, et al. 2020. Fungal Diversity 105:1-16 ((https://doi.org/10.1007/s13225-020-00466-2)) |
| Otu28845 | 1 | k__Fungi(100);p__Ascomycota(100);c__Saccharomycetes(100);o__Saccharomycetales(100);f__Metschnikowiaceae(100);g__Metschnikowia(100);s__Metschnikowia_pulcherrima(100); | Metschnikowia | 13 | Saprotroph | \|Undefined Saprotroph\| | Yeast | NULL | Probable | As nectar-tap_saprotroph (Põlme et al. 2020); Decay sugar-rich substrates (Põlme et al. 2020) | Põlme S, et al. 2020. Fungal Diversity 105:1-16 ((https://doi.org/10.1007/s13225-020-00466-2)) |
| Otu28846 | 1 | k__Fungi(100);p__Ascomycota(100);c__Saccharomycetes(100);o__Saccharomycetales(100);f__Metschnikowiaceae(100);g__Metschnikowia(100);s__Metschnikowia_pulcherrima(100); | Metschnikowia | 13 | Saprotroph | \|Undefined Saprotroph\| | Yeast | NULL | Probable | As nectar-tap_saprotroph (Põlme et al. 2020); Decay sugar-rich substrates (Põlme et al. 2020) | Põlme S, et al. 2020. Fungal Diversity 105:1-16 ((https://doi.org/10.1007/s13225-020-00466-2)) |
| Otu28848 | 1 | k__Fungi(100);p__Ascomycota(100);c__Saccharomycetes(100);o__Saccharomycetales(100);f__Metschnikowiaceae(100);g__Metschnikowia(100);s__Metschnikowia_pulcherrima(100); | Metschnikowia | 13 | Saprotroph | \|Undefined Saprotroph\| | Yeast | NULL | Probable | As nectar-tap_saprotroph (Põlme et al. 2020); Decay sugar-rich substrates (Põlme et al. 2020) | Põlme S, et al. 2020. Fungal Diversity 105:1-16 ((https://doi.org/10.1007/s13225-020-00466-2)) |
| Otu28850 | 1 | k__Fungi(100);p__Ascomycota(100);c__Saccharomycetes(100);o__Saccharomycetales(100);f__Metschnikowiaceae(100);g__Metschnikowia(100);s__Metschnikowia_pulcherrima(100); | Metschnikowia | 13 | Saprotroph | \|Undefined Saprotroph\| | Yeast | NULL | Probable | As nectar-tap_saprotroph (Põlme et al. 2020); Decay sugar-rich substrates (Põlme et al. 2020) | Põlme S, et al. 2020. Fungal Diversity 105:1-16 ((https://doi.org/10.1007/s13225-020-00466-2)) |
| Otu28851 | 1 | k__Fungi(100);p__Ascomycota(100);c__Saccharomycetes(100);o__Saccharomycetales(100);f__Metschnikowiaceae(100);g__Metschnikowia(100);s__Metschnikowia_pulcherrima(100); | Metschnikowia | 13 | Saprotroph | \|Undefined Saprotroph\| | Yeast | NULL | Probable | As nectar-tap_saprotroph (Põlme et al. 2020); Decay sugar-rich substrates (Põlme et al. 2020) | Põlme S, et al. 2020. Fungal Diversity 105:1-16 ((https://doi.org/10.1007/s13225-020-00466-2)) |
| Otu28853 | 1 | k__Fungi(100);p__Ascomycota(100);c__Saccharomycetes(100);o__Saccharomycetales(100);f__Metschnikowiaceae(100);g__Metschnikowia(100);g__Metschnikowia_unclassified(100); | Metschnikowia | 13 | Saprotroph | \|Undefined Saprotroph\| | Yeast | NULL | Probable | As nectar-tap_saprotroph (Põlme et al. 2020); Decay sugar-rich substrates (Põlme et al. 2020) | Põlme S, et al. 2020. Fungal Diversity 105:1-16 ((https://doi.org/10.1007/s13225-020-00466-2)) |
| Otu28856 | 1 | k__Fungi(100);p__Ascomycota(100);c__Saccharomycetes(100);o__Saccharomycetales(100);f__Metschnikowiaceae(100);g__Metschnikowia(100);s__Metschnikowia_pulcherrima(100); | Metschnikowia | 13 | Saprotroph | \|Undefined Saprotroph\| | Yeast | NULL | Probable | As nectar-tap_saprotroph (Põlme et al. 2020); Decay sugar-rich substrates (Põlme et al. 2020) | Põlme S, et al. 2020. Fungal Diversity 105:1-16 ((https://doi.org/10.1007/s13225-020-00466-2)) |
| Otu28858 | 1 | k__Fungi(100);p__Ascomycota(100);c__Saccharomycetes(100);o__Saccharomycetales(100);f__Metschnikowiaceae(100);g__Metschnikowia(100);s__Metschnikowia_pulcherrima(100); | Metschnikowia | 13 | Saprotroph | \|Undefined Saprotroph\| | Yeast | NULL | Probable | As nectar-tap_saprotroph (Põlme et al. 2020); Decay sugar-rich substrates (Põlme et al. 2020) | Põlme S, et al. 2020. Fungal Diversity 105:1-16 ((https://doi.org/10.1007/s13225-020-00466-2)) |
| Otu28859 | 1 | k__Fungi(100);p__Ascomycota(100);c__Saccharomycetes(100);o__Saccharomycetales(100);f__Metschnikowiaceae(100);g__Metschnikowia(100);s__Metschnikowia_pulcherrima(100); | Metschnikowia | 13 | Saprotroph | \|Undefined Saprotroph\| | Yeast | NULL | Probable | As nectar-tap_saprotroph (Põlme et al. 2020); Decay sugar-rich substrates (Põlme et al. 2020) | Põlme S, et al. 2020. Fungal Diversity 105:1-16 ((https://doi.org/10.1007/s13225-020-00466-2)) |
| Otu28861 | 1 | k__Fungi(100);p__Zygomycota(100);c__Zygomycota_class_Incertae_sedis(100);o__Mortierellales(100);f__Mortierellaceae(100);g__Mortierella(100);s__Mortierella_umbellata(100); | Mortierella | 13 | Saprotroph-Symbiotroph | Endophyte-Plant Saprotroph-\|Undefined Saprotroph\| | Microfungus | NULL | Probable | As chitinolytic (Põlme et al. 2020); As soil_saprotroph (Põlme et al. 2020); Decay roots, soil (Põlme et al. 2020); Saprobic in soil and thus frequently isolated from soil and material contaminated with soil such as hay or rotten silage and spoil tips (Cannon & Kirk 2007) | Cannon PF, Kirk PM. 2007. Fungal Families of the World. CAB International, Cambridge (ISBN: 978-0851998275); Tedersoo L, et al. 2014. Science 346:e1256688 ((https://doi.org/10.1126/science.1256688)); Purahong W, et al. 2016. Molecular Ecology 25:4059-4074 ((https://doi.org/10.1111/mec.13739)); Põlme S, et al. 2020. Fungal Diversity 105:1-16 ((https://doi.org/10.1007/s13225-020-00466-2)) |
| Otu28863 | 1 | k__Fungi(100);p__Zygomycota(100);c__Zygomycota_class_Incertae_sedis(100);o__Mortierellales(100);f__Mortierellaceae(100);g__Mortierella(100);s__Mortierella_umbellata(100); | Mortierella | 13 | Saprotroph-Symbiotroph | Endophyte-Plant Saprotroph-\|Undefined Saprotroph\| | Microfungus | NULL | Probable | As chitinolytic (Põlme et al. 2020); As soil_saprotroph (Põlme et al. 2020); Decay roots, soil (Põlme et al. 2020); Saprobic in soil and thus frequently isolated from soil and material contaminated with soil such as hay or rotten silage and spoil tips (Cannon & Kirk 2007) | Cannon PF, Kirk PM. 2007. Fungal Families of the World. CAB International, Cambridge (ISBN: 978-0851998275); Tedersoo L, et al. 2014. Science 346:e1256688 ((https://doi.org/10.1126/science.1256688)); Purahong W, et al. 2016. Molecular Ecology 25:4059-4074 ((https://doi.org/10.1111/mec.13739)); Põlme S, et al. 2020. Fungal Diversity 105:1-16 ((https://doi.org/10.1007/s13225-020-00466-2)) |
| Otu28864 | 1 | k__Fungi(100);p__Zygomycota(100);c__Zygomycota_class_Incertae_sedis(100);o__Mortierellales(100);f__Mortierellaceae(100);g__Mortierella(100);s__Mortierella_sp_18_M_4(100); | Mortierella | 13 | Saprotroph-Symbiotroph | Endophyte-Plant Saprotroph-\|Undefined Saprotroph\| | Microfungus | NULL | Probable | As chitinolytic (Põlme et al. 2020); As soil_saprotroph (Põlme et al. 2020); Decay roots, soil (Põlme et al. 2020); Saprobic in soil and thus frequently isolated from soil and material contaminated with soil such as hay or rotten silage and spoil tips (Cannon & Kirk 2007) | Cannon PF, Kirk PM. 2007. Fungal Families of the World. CAB International, Cambridge (ISBN: 978-0851998275); Tedersoo L, et al. 2014. Science 346:e1256688 ((https://doi.org/10.1126/science.1256688)); Purahong W, et al. 2016. Molecular Ecology 25:4059-4074 ((https://doi.org/10.1111/mec.13739)); Põlme S, et al. 2020. Fungal Diversity 105:1-16 ((https://doi.org/10.1007/s13225-020-00466-2)) |
| Otu28865 | 1 | k__Fungi(100);p__Zygomycota(100);c__Zygomycota_class_Incertae_sedis(100);o__Mortierellales(100);f__Mortierellaceae(100);g__Mortierella(100);s__Mortierella_gemmifera(100); | Mortierella | 13 | Saprotroph-Symbiotroph | Endophyte-Plant Saprotroph-\|Undefined Saprotroph\| | Microfungus | NULL | Probable | As chitinolytic (Põlme et al. 2020); As soil_saprotroph (Põlme et al. 2020); Decay roots, soil (Põlme et al. 2020); Saprobic in soil and thus frequently isolated from soil and material contaminated with soil such as hay or rotten silage and spoil tips (Cannon & Kirk 2007) | Cannon PF, Kirk PM. 2007. Fungal Families of the World. CAB International, Cambridge (ISBN: 978-0851998275); Tedersoo L, et al. 2014. Science 346:e1256688 ((https://doi.org/10.1126/science.1256688)); Purahong W, et al. 2016. Molecular Ecology 25:4059-4074 ((https://doi.org/10.1111/mec.13739)); Põlme S, et al. 2020. Fungal Diversity 105:1-16 ((https://doi.org/10.1007/s13225-020-00466-2)) |
| Otu28866 | 1 | k__Fungi(100);p__Zygomycota(100);c__Zygomycota_class_Incertae_sedis(100);o__Mortierellales(100);f__Mortierellaceae(100);g__Mortierella(100);g__Mortierella_unclassified(100); | Mortierella | 13 | Saprotroph-Symbiotroph | Endophyte-Plant Saprotroph-\|Undefined Saprotroph\| | Microfungus | NULL | Probable | As chitinolytic (Põlme et al. 2020); As soil_saprotroph (Põlme et al. 2020); Decay roots, soil (Põlme et al. 2020); Saprobic in soil and thus frequently isolated from soil and material contaminated with soil such as hay or rotten silage and spoil tips (Cannon & Kirk 2007) | Cannon PF, Kirk PM. 2007. Fungal Families of the World. CAB International, Cambridge (ISBN: 978-0851998275); Tedersoo L, et al. 2014. Science 346:e1256688 ((https://doi.org/10.1126/science.1256688)); Purahong W, et al. 2016. Molecular Ecology 25:4059-4074 ((https://doi.org/10.1111/mec.13739)); Põlme S, et al. 2020. Fungal Diversity 105:1-16 ((https://doi.org/10.1007/s13225-020-00466-2)) |
| Otu28867 | 1 | k__Fungi(100);p__Zygomycota(100);c__Zygomycota_class_Incertae_sedis(100);o__Mortierellales(100);f__Mortierellaceae(100);g__Mortierella(100);g__Mortierella_unclassified(100); | Mortierella | 13 | Saprotroph-Symbiotroph | Endophyte-Plant Saprotroph-\|Undefined Saprotroph\| | Microfungus | NULL | Probable | As chitinolytic (Põlme et al. 2020); As soil_saprotroph (Põlme et al. 2020); Decay roots, soil (Põlme et al. 2020); Saprobic in soil and thus frequently isolated from soil and material contaminated with soil such as hay or rotten silage and spoil tips (Cannon & Kirk 2007) | Cannon PF, Kirk PM. 2007. Fungal Families of the World. CAB International, Cambridge (ISBN: 978-0851998275); Tedersoo L, et al. 2014. Science 346:e1256688 ((https://doi.org/10.1126/science.1256688)); Purahong W, et al. 2016. Molecular Ecology 25:4059-4074 ((https://doi.org/10.1111/mec.13739)); Põlme S, et al. 2020. Fungal Diversity 105:1-16 ((https://doi.org/10.1007/s13225-020-00466-2)) |
| Otu28868 | 1 | k__Fungi(100);p__Zygomycota(100);c__Zygomycota_class_Incertae_sedis(100);o__Mortierellales(100);f__Mortierellaceae(100);g__Mortierella(100);g__Mortierella_unclassified(100); | Mortierella | 13 | Saprotroph-Symbiotroph | Endophyte-Plant Saprotroph-\|Undefined Saprotroph\| | Microfungus | NULL | Probable | As chitinolytic (Põlme et al. 2020); As soil_saprotroph (Põlme et al. 2020); Decay roots, soil (Põlme et al. 2020); Saprobic in soil and thus frequently isolated from soil and material contaminated with soil such as hay or rotten silage and spoil tips (Cannon & Kirk 2007) | Cannon PF, Kirk PM. 2007. Fungal Families of the World. CAB International, Cambridge (ISBN: 978-0851998275); Tedersoo L, et al. 2014. Science 346:e1256688 ((https://doi.org/10.1126/science.1256688)); Purahong W, et al. 2016. Molecular Ecology 25:4059-4074 ((https://doi.org/10.1111/mec.13739)); Põlme S, et al. 2020. Fungal Diversity 105:1-16 ((https://doi.org/10.1007/s13225-020-00466-2)) |
| Otu28869 | 1 | k__Fungi(100);p__Zygomycota(100);c__Zygomycota_class_Incertae_sedis(100);o__Mortierellales(100);f__Mortierellaceae(100);g__Mortierella(100);g__Mortierella_unclassified(100); | Mortierella | 13 | Saprotroph-Symbiotroph | Endophyte-Plant Saprotroph-\|Undefined Saprotroph\| | Microfungus | NULL | Probable | As chitinolytic (Põlme et al. 2020); As soil_saprotroph (Põlme et al. 2020); Decay roots, soil (Põlme et al. 2020); Saprobic in soil and thus frequently isolated from soil and material contaminated with soil such as hay or rotten silage and spoil tips (Cannon & Kirk 2007) | Cannon PF, Kirk PM. 2007. Fungal Families of the World. CAB International, Cambridge (ISBN: 978-0851998275); Tedersoo L, et al. 2014. Science 346:e1256688 ((https://doi.org/10.1126/science.1256688)); Purahong W, et al. 2016. Molecular Ecology 25:4059-4074 ((https://doi.org/10.1111/mec.13739)); Põlme S, et al. 2020. Fungal Diversity 105:1-16 ((https://doi.org/10.1007/s13225-020-00466-2)) |
| Otu28871 | 1 | k__Fungi(100);p__Zygomycota(100);c__Zygomycota_class_Incertae_sedis(100);o__Mortierellales(100);f__Mortierellaceae(100);g__Mortierella(100);g__Mortierella_unclassified(100); | Mortierella | 13 | Saprotroph-Symbiotroph | Endophyte-Plant Saprotroph-\|Undefined Saprotroph\| | Microfungus | NULL | Probable | As chitinolytic (Põlme et al. 2020); As soil_saprotroph (Põlme et al. 2020); Decay roots, soil (Põlme et al. 2020); Saprobic in soil and thus frequently isolated from soil and material contaminated with soil such as hay or rotten silage and spoil tips (Cannon & Kirk 2007) | Cannon PF, Kirk PM. 2007. Fungal Families of the World. CAB International, Cambridge (ISBN: 978-0851998275); Tedersoo L, et al. 2014. Science 346:e1256688 ((https://doi.org/10.1126/science.1256688)); Purahong W, et al. 2016. Molecular Ecology 25:4059-4074 ((https://doi.org/10.1111/mec.13739)); Põlme S, et al. 2020. Fungal Diversity 105:1-16 ((https://doi.org/10.1007/s13225-020-00466-2)) |
| Otu28874 | 1 | k__Fungi(100);p__Zygomycota(100);c__Zygomycota_class_Incertae_sedis(100);o__Mortierellales(100);f__Mortierellaceae(100);g__Mortierella(100);s__Mortierella_umbellata(100); | Mortierella | 13 | Saprotroph-Symbiotroph | Endophyte-Plant Saprotroph-\|Undefined Saprotroph\| | Microfungus | NULL | Probable | As chitinolytic (Põlme et al. 2020); As soil_saprotroph (Põlme et al. 2020); Decay roots, soil (Põlme et al. 2020); Saprobic in soil and thus frequently isolated from soil and material contaminated with soil such as hay or rotten silage and spoil tips (Cannon & Kirk 2007) | Cannon PF, Kirk PM. 2007. Fungal Families of the World. CAB International, Cambridge (ISBN: 978-0851998275); Tedersoo L, et al. 2014. Science 346:e1256688 ((https://doi.org/10.1126/science.1256688)); Purahong W, et al. 2016. Molecular Ecology 25:4059-4074 ((https://doi.org/10.1111/mec.13739)); Põlme S, et al. 2020. Fungal Diversity 105:1-16 ((https://doi.org/10.1007/s13225-020-00466-2)) |
| Otu28881 | 1 | k__Fungi(100);p__Zygomycota(100);c__Zygomycota_class_Incertae_sedis(100);o__Mortierellales(100);f__Mortierellaceae(100);g__Mortierella(100);g__Mortierella_unclassified(100); | Mortierella | 13 | Saprotroph-Symbiotroph | Endophyte-Plant Saprotroph-\|Undefined Saprotroph\| | Microfungus | NULL | Probable | As chitinolytic (Põlme et al. 2020); As soil_saprotroph (Põlme et al. 2020); Decay roots, soil (Põlme et al. 2020); Saprobic in soil and thus frequently isolated from soil and material contaminated with soil such as hay or rotten silage and spoil tips (Cannon & Kirk 2007) | Cannon PF, Kirk PM. 2007. Fungal Families of the World. CAB International, Cambridge (ISBN: 978-0851998275); Tedersoo L, et al. 2014. Science 346:e1256688 ((https://doi.org/10.1126/science.1256688)); Purahong W, et al. 2016. Molecular Ecology 25:4059-4074 ((https://doi.org/10.1111/mec.13739)); Põlme S, et al. 2020. Fungal Diversity 105:1-16 ((https://doi.org/10.1007/s13225-020-00466-2)) |
| Otu28884 | 1 | k__Fungi(100);p__Ascomycota(100);c__Eurotiomycetes(100);o__Chaetothyriales(100);f__Chaetothyriaceae(100);g__Cyphellophora(100);s__Cyphellophora_sp(100); | Cyphellophora | 13 | Pathotroph-Saprotroph | Animal Parasite-Animal Pathogen-\|Plant Saprotroph\|-Undefined Saprotroph | Dimorphic-Facultative Yeast-Microfungus | NULL | Probable | As litter_saprotroph (Põlme et al. 2020); Consult the current literature for updated names on this group that included the Black Yeasts; Some species are likely opportunistic human pathogen (Irinyi et al. 2015); Decay leaf, fruit, seed, animal material (Põlme et al. 2020) | Tedersoo L, et al. 2014. Science 346:e1256688 ((https://doi.org/10.1126/science.1256688)); Irinyi L, et al. 2015. Medical Mycology 53:313-337 ((https://doi.org/10.1093/mmy/myv008)); Põlme S, et al. 2020. Fungal Diversity 105:1-16 ((https://doi.org/10.1007/s13225-020-00466-2)) |
| Otu28887 | 1 | k__Fungi(100);p__Zygomycota(100);c__Zygomycota_class_Incertae_sedis(100);o__Kickxellales(100);f__Kickxellaceae(100);g__Ramicandelaber(100);s__Ramicandelaber_sp(100); | Ramicandelaber | 13 | Saprotroph | \|Undefined Saprotroph\| | Microfungus | NULL | Probable | As soil_saprotroph (Põlme et al. 2020); Decay soil (Põlme et al. 2020) | Tedersoo L, et al. 2014. Science 346:e1256688 ((https://doi.org/10.1126/science.1256688)); Põlme S, et al. 2020. Fungal Diversity 105:1-16 ((https://doi.org/10.1007/s13225-020-00466-2)) |
| Otu28890 | 1 | k__Fungi(100);p__Zygomycota(100);c__Zygomycota_class_Incertae_sedis(100);o__Mucorales(100);f__Cunninghamellaceae(100);g__Gongronella(100);s__Gongronella_sp_TL_2013(100); | Gongronella | 13 | Saprotroph | \|Undefined Saprotroph\| | Microfungus | NULL | Probable | As soil_saprotroph (Põlme et al. 2020); Decay soil (Põlme et al. 2020) | Tedersoo L, et al. 2014. Science 346:e1256688 ((https://doi.org/10.1126/science.1256688)); Põlme S, et al. 2020. Fungal Diversity 105:1-16 ((https://doi.org/10.1007/s13225-020-00466-2)) |
| Otu28891 | 1 | k__Fungi(100);p__Zygomycota(100);c__Zygomycota_class_Incertae_sedis(100);o__Mucorales(100);f__Backusellaceae(100);g__Backusella(100);s__Backusella_recurva(100); | Backusella | 13 | Saprotroph | Dung Saprotroph-\|Undefined Saprotroph\| | Microfungus-Sporophores | NULL | Probable | As soil_saprotroph (Põlme et al. 2020); Decay soil, dung (Põlme et al. 2020) | Walther G, et al. 2013. Persoonia 30:11-47 ((https://doi.org/10.3767/003158513X665070)); Põlme S, et al. 2020. Fungal Diversity 105:1-16 ((https://doi.org/10.1007/s13225-020-00466-2)) |
| Otu28892 | 1 | k__Fungi(100);p__Zygomycota(100);c__Zygomycota_class_Incertae_sedis(100);o__Mucorales(100);f__Backusellaceae(100);g__Backusella(100);s__Backusella_recurva(100); | Backusella | 13 | Saprotroph | Dung Saprotroph-\|Undefined Saprotroph\| | Microfungus-Sporophores | NULL | Probable | As soil_saprotroph (Põlme et al. 2020); Decay soil, dung (Põlme et al. 2020) | Walther G, et al. 2013. Persoonia 30:11-47 ((https://doi.org/10.3767/003158513X665070)); Põlme S, et al. 2020. Fungal Diversity 105:1-16 ((https://doi.org/10.1007/s13225-020-00466-2)) |
| Otu28893 | 1 | k__Fungi(100);p__Basidiomycota(100);c__Microbotryomycetes(100);o__Sporidiobolales(100);f__Sporidiobolales_family_Incertae_sedis(100);g__Sporobolomyces(100);s__Sporobolomyces_symmetricus(100); | Sporobolomyces | 13 | Pathotroph-Saprotroph | \|Fungal Parasite\|-Plant Saprotroph | Yeast | NULL | Probable | As litter_saprotroph (Põlme et al. 2020); As mycoparasite (Põlme et al. 2020); Decay fungal material (Põlme et al. 2020) | Tedersoo L, et al. 2014. Science 346:e1256688 ((https://doi.org/10.1126/science.1256688)); Purahong W, et al. 2016. Molecular Ecology 25:4059-4074 ((https://doi.org/10.1111/mec.13739)); Põlme S, et al. 2020. Fungal Diversity 105:1-16 ((https://doi.org/10.1007/s13225-020-00466-2)) |
| Otu28894 | 1 | k__Fungi(100);p__Basidiomycota(100);c__Microbotryomycetes(100);o__Sporidiobolales(100);f__Sporidiobolales_family_Incertae_sedis(100);g__Sporobolomyces(100);g__Sporobolomyces_unclassified(100); | Sporobolomyces | 13 | Pathotroph-Saprotroph | \|Fungal Parasite\|-Plant Saprotroph | Yeast | NULL | Probable | As litter_saprotroph (Põlme et al. 2020); As mycoparasite (Põlme et al. 2020); Decay fungal material (Põlme et al. 2020) | Tedersoo L, et al. 2014. Science 346:e1256688 ((https://doi.org/10.1126/science.1256688)); Purahong W, et al. 2016. Molecular Ecology 25:4059-4074 ((https://doi.org/10.1111/mec.13739)); Põlme S, et al. 2020. Fungal Diversity 105:1-16 ((https://doi.org/10.1007/s13225-020-00466-2)) |
| Otu28896 | 1 | k__Fungi(100);p__Basidiomycota(100);c__Agaricostilbomycetes(100);o__Agaricostilbales(100);f__Agaricostilbaceae(100);g__Sterigmatomyces(100);s__Sterigmatomyces_halophilus(100); | Sterigmatomyces | 13 | Saprotroph | \|Undefined Saprotroph\| | Dimorphic | NULL | Probable | NULL | Põlme S, et al. 2020. Fungal Diversity 105:1-16 ((https://doi.org/10.1007/s13225-020-00466-2)) |
| Otu28897 | 1 | k__Fungi(100);p__Zygomycota(100);c__Zygomycota_class_Incertae_sedis(100);o__Kickxellales(100);f__Kickxellaceae(100);g__Ramicandelaber(100);s__Ramicandelaber_sp(100); | Ramicandelaber | 13 | Saprotroph | \|Undefined Saprotroph\| | Microfungus | NULL | Probable | As soil_saprotroph (Põlme et al. 2020); Decay soil (Põlme et al. 2020) | Tedersoo L, et al. 2014. Science 346:e1256688 ((https://doi.org/10.1126/science.1256688)); Põlme S, et al. 2020. Fungal Diversity 105:1-16 ((https://doi.org/10.1007/s13225-020-00466-2)) |
| Otu28898 | 1 | k__Fungi(100);p__Zygomycota(100);c__Zygomycota_class_Incertae_sedis(100);o__Kickxellales(100);f__Kickxellaceae(100);g__Ramicandelaber(100);s__Ramicandelaber_sp(100); | Ramicandelaber | 13 | Saprotroph | \|Undefined Saprotroph\| | Microfungus | NULL | Probable | As soil_saprotroph (Põlme et al. 2020); Decay soil (Põlme et al. 2020) | Tedersoo L, et al. 2014. Science 346:e1256688 ((https://doi.org/10.1126/science.1256688)); Põlme S, et al. 2020. Fungal Diversity 105:1-16 ((https://doi.org/10.1007/s13225-020-00466-2)) |
| Otu28899 | 1 | k__Fungi(100);p__Zygomycota(100);c__Zygomycota_class_Incertae_sedis(100);o__Kickxellales(100);f__Kickxellaceae(100);g__Ramicandelaber(100);s__Ramicandelaber_sp(100); | Ramicandelaber | 13 | Saprotroph | \|Undefined Saprotroph\| | Microfungus | NULL | Probable | As soil_saprotroph (Põlme et al. 2020); Decay soil (Põlme et al. 2020) | Tedersoo L, et al. 2014. Science 346:e1256688 ((https://doi.org/10.1126/science.1256688)); Põlme S, et al. 2020. Fungal Diversity 105:1-16 ((https://doi.org/10.1007/s13225-020-00466-2)) |
| Otu28901 | 1 | k__Fungi(100);p__Zygomycota(100);c__Zygomycota_class_Incertae_sedis(100);o__Kickxellales(100);f__Kickxellaceae(100);g__Ramicandelaber(100);s__Ramicandelaber_sp(100); | Ramicandelaber | 13 | Saprotroph | \|Undefined Saprotroph\| | Microfungus | NULL | Probable | As soil_saprotroph (Põlme et al. 2020); Decay soil (Põlme et al. 2020) | Tedersoo L, et al. 2014. Science 346:e1256688 ((https://doi.org/10.1126/science.1256688)); Põlme S, et al. 2020. Fungal Diversity 105:1-16 ((https://doi.org/10.1007/s13225-020-00466-2)) |
| Otu28903 | 1 | k__Fungi(100);p__Zygomycota(100);c__Zygomycota_class_Incertae_sedis(100);o__Mucorales(100);f__Lentamycetaceae(100);g__Lentamyces(100);s__Lentamyces_parricida(100); | Lentamyces | 13 | Pathotroph-Symbiotroph | Animal Symbiotroph-\|Fungal Parasite\| | Microfungus | NULL | Probable | Animal gut fungus; As mycoparasite (Põlme et al. 2020); Decay fungal material (Põlme et al. 2020) | Tedersoo L, et al. 2014. Science 346:e1256688 ((https://doi.org/10.1126/science.1256688)); Põlme S, et al. 2020. Fungal Diversity 105:1-16 ((https://doi.org/10.1007/s13225-020-00466-2)) |
| Otu28904 | 1 | k__Fungi(100);p__Zygomycota(100);c__Zygomycota_class_Incertae_sedis(100);o__Mucorales(100);f__Cunninghamellaceae(100);g__Absidia(100);g__Absidia_unclassified(100); | Absidia | 13 | Saprotroph | \|Undefined Saprotroph\| | Microfungus | NULL | Probable | As soil_saprotroph (Põlme et al. 2020); Decay soil (Põlme et al. 2020) | Tedersoo L, et al. 2014. Science 346:e1256688 ((https://doi.org/10.1126/science.1256688)); Põlme S, et al. 2020. Fungal Diversity 105:1-16 ((https://doi.org/10.1007/s13225-020-00466-2)) |
| Otu28906 | 1 | k__Fungi(100);p__Ascomycota(100);c__Saccharomycetes(100);o__Saccharomycetales(100);f__Metschnikowiaceae(100);g__Metschnikowia(100);s__Metschnikowia_pulcherrima(100); | Metschnikowia | 13 | Saprotroph | \|Undefined Saprotroph\| | Yeast | NULL | Probable | As nectar-tap_saprotroph (Põlme et al. 2020); Decay sugar-rich substrates (Põlme et al. 2020) | Põlme S, et al. 2020. Fungal Diversity 105:1-16 ((https://doi.org/10.1007/s13225-020-00466-2)) |
| Otu28907 | 1 | k__Fungi(100);p__Ascomycota(100);c__Saccharomycetes(100);o__Saccharomycetales(100);f__Metschnikowiaceae(100);g__Metschnikowia(100);s__Metschnikowia_pulcherrima(100); | Metschnikowia | 13 | Saprotroph | \|Undefined Saprotroph\| | Yeast | NULL | Probable | As nectar-tap_saprotroph (Põlme et al. 2020); Decay sugar-rich substrates (Põlme et al. 2020) | Põlme S, et al. 2020. Fungal Diversity 105:1-16 ((https://doi.org/10.1007/s13225-020-00466-2)) |
| Otu28909 | 1 | k__Fungi(100);p__Zygomycota(100);c__Zygomycota_class_Incertae_sedis(100);o__Mortierellales(100);f__Mortierellaceae(100);g__Mortierella(100);s__Mortierella_sp_18_M_4(100); | Mortierella | 13 | Saprotroph-Symbiotroph | Endophyte-Plant Saprotroph-\|Undefined Saprotroph\| | Microfungus | NULL | Probable | As chitinolytic (Põlme et al. 2020); As soil_saprotroph (Põlme et al. 2020); Decay roots, soil (Põlme et al. 2020); Saprobic in soil and thus frequently isolated from soil and material contaminated with soil such as hay or rotten silage and spoil tips (Cannon & Kirk 2007) | Cannon PF, Kirk PM. 2007. Fungal Families of the World. CAB International, Cambridge (ISBN: 978-0851998275); Tedersoo L, et al. 2014. Science 346:e1256688 ((https://doi.org/10.1126/science.1256688)); Purahong W, et al. 2016. Molecular Ecology 25:4059-4074 ((https://doi.org/10.1111/mec.13739)); Põlme S, et al. 2020. Fungal Diversity 105:1-16 ((https://doi.org/10.1007/s13225-020-00466-2)) |
| Otu28910 | 1 | k__Fungi(100);p__Zygomycota(100);c__Zygomycota_class_Incertae_sedis(100);o__Mortierellales(100);f__Mortierellaceae(100);g__Mortierella(100);g__Mortierella_unclassified(100); | Mortierella | 13 | Saprotroph-Symbiotroph | Endophyte-Plant Saprotroph-\|Undefined Saprotroph\| | Microfungus | NULL | Probable | As chitinolytic (Põlme et al. 2020); As soil_saprotroph (Põlme et al. 2020); Decay roots, soil (Põlme et al. 2020); Saprobic in soil and thus frequently isolated from soil and material contaminated with soil such as hay or rotten silage and spoil tips (Cannon & Kirk 2007) | Cannon PF, Kirk PM. 2007. Fungal Families of the World. CAB International, Cambridge (ISBN: 978-0851998275); Tedersoo L, et al. 2014. Science 346:e1256688 ((https://doi.org/10.1126/science.1256688)); Purahong W, et al. 2016. Molecular Ecology 25:4059-4074 ((https://doi.org/10.1111/mec.13739)); Põlme S, et al. 2020. Fungal Diversity 105:1-16 ((https://doi.org/10.1007/s13225-020-00466-2)) |
| Otu28911 | 1 | k__Fungi(100);p__Zygomycota(100);c__Zygomycota_class_Incertae_sedis(100);o__Mortierellales(100);f__Mortierellaceae(100);g__Mortierella(100);s__Mortierella_rishikesha(100); | Mortierella | 13 | Saprotroph-Symbiotroph | Endophyte-Plant Saprotroph-\|Undefined Saprotroph\| | Microfungus | NULL | Probable | As chitinolytic (Põlme et al. 2020); As soil_saprotroph (Põlme et al. 2020); Decay roots, soil (Põlme et al. 2020); Saprobic in soil and thus frequently isolated from soil and material contaminated with soil such as hay or rotten silage and spoil tips (Cannon & Kirk 2007) | Cannon PF, Kirk PM. 2007. Fungal Families of the World. CAB International, Cambridge (ISBN: 978-0851998275); Tedersoo L, et al. 2014. Science 346:e1256688 ((https://doi.org/10.1126/science.1256688)); Purahong W, et al. 2016. Molecular Ecology 25:4059-4074 ((https://doi.org/10.1111/mec.13739)); Põlme S, et al. 2020. Fungal Diversity 105:1-16 ((https://doi.org/10.1007/s13225-020-00466-2)) |
| Otu28912 | 1 | k__Fungi(100);p__Zygomycota(100);c__Zygomycota_class_Incertae_sedis(100);o__Mortierellales(100);f__Mortierellaceae(100);g__Mortierella(100);g__Mortierella_unclassified(100); | Mortierella | 13 | Saprotroph-Symbiotroph | Endophyte-Plant Saprotroph-\|Undefined Saprotroph\| | Microfungus | NULL | Probable | As chitinolytic (Põlme et al. 2020); As soil_saprotroph (Põlme et al. 2020); Decay roots, soil (Põlme et al. 2020); Saprobic in soil and thus frequently isolated from soil and material contaminated with soil such as hay or rotten silage and spoil tips (Cannon & Kirk 2007) | Cannon PF, Kirk PM. 2007. Fungal Families of the World. CAB International, Cambridge (ISBN: 978-0851998275); Tedersoo L, et al. 2014. Science 346:e1256688 ((https://doi.org/10.1126/science.1256688)); Purahong W, et al. 2016. Molecular Ecology 25:4059-4074 ((https://doi.org/10.1111/mec.13739)); Põlme S, et al. 2020. Fungal Diversity 105:1-16 ((https://doi.org/10.1007/s13225-020-00466-2)) |
| Otu28913 | 1 | k__Fungi(100);p__Ascomycota(100);c__Sordariomycetes(100);o__Melanosporales(100);f__Ceratostomataceae(100);g__Harzia(100);s__Harzia_acremonioides(100); | Harzia | 13 | Saprotroph | Plant Saprotroph-Undefined Saprotroph-\|Wood Saprotroph\| | Microfungus | NULL | Probable | As litter_saprotroph (Põlme et al. 2020); Decay wood, leaf, fruit, seed (Põlme et al. 2020) | Tedersoo L, et al. 2014. Science 346:e1256688 ((https://doi.org/10.1126/science.1256688)); Põlme S, et al. 2020. Fungal Diversity 105:1-16 ((https://doi.org/10.1007/s13225-020-00466-2)) |
| Otu28915 | 1 | k__Fungi(100);p__Basidiomycota(100);c__Pucciniomycetes(100);o__Pucciniales(100);f__Melampsoraceae(100);g__Melampsora(100);s__Melampsora_abietis_canadensis(100); | Melampsora | 13 | Pathotroph | \|Plant Pathogen\| | Rust | NULL | Probable | A prominent group of species produces aecia on conifer needles and telia or leaves of Salicaceae (Cannon & Kirk 2007); Biotrophic, heteroecious or autoecious | Cannon PF, Kirk PM. 2007. Fungal Families of the World. CAB International, Cambridge (ISBN: 978-0851998275); Tedersoo L, et al. 2014. Science 346:e1256688 ((https://doi.org/10.1126/science.1256688)); Põlme S, et al. 2020. Fungal Diversity 105:1-16 ((https://doi.org/10.1007/s13225-020-00466-2)) |
| Otu28916 | 1 | k__Fungi(100);p__Basidiomycota(100);c__Pucciniomycetes(100);o__Pucciniales(100);f__Melampsoraceae(100);g__Melampsora(100);s__Melampsora_sp_hyb8_3_1(100); | Melampsora | 13 | Pathotroph | \|Plant Pathogen\| | Rust | NULL | Probable | A prominent group of species produces aecia on conifer needles and telia or leaves of Salicaceae (Cannon & Kirk 2007); Biotrophic, heteroecious or autoecious | Cannon PF, Kirk PM. 2007. Fungal Families of the World. CAB International, Cambridge (ISBN: 978-0851998275); Tedersoo L, et al. 2014. Science 346:e1256688 ((https://doi.org/10.1126/science.1256688)); Põlme S, et al. 2020. Fungal Diversity 105:1-16 ((https://doi.org/10.1007/s13225-020-00466-2)) |
| Otu28917 | 1 | k__Fungi(100);p__Basidiomycota(100);c__Pucciniomycetes(100);o__Pucciniales(100);f__Melampsoraceae(100);g__Melampsora(100);s__Melampsora_abietis_canadensis(100); | Melampsora | 13 | Pathotroph | \|Plant Pathogen\| | Rust | NULL | Probable | A prominent group of species produces aecia on conifer needles and telia or leaves of Salicaceae (Cannon & Kirk 2007); Biotrophic, heteroecious or autoecious | Cannon PF, Kirk PM. 2007. Fungal Families of the World. CAB International, Cambridge (ISBN: 978-0851998275); Tedersoo L, et al. 2014. Science 346:e1256688 ((https://doi.org/10.1126/science.1256688)); Põlme S, et al. 2020. Fungal Diversity 105:1-16 ((https://doi.org/10.1007/s13225-020-00466-2)) |
| Otu28918 | 1 | k__Fungi(100);p__Basidiomycota(100);c__Pucciniomycetes(100);o__Pucciniales(100);f__Melampsoraceae(100);g__Melampsora(100);s__Melampsora_abietis_canadensis(100); | Melampsora | 13 | Pathotroph | \|Plant Pathogen\| | Rust | NULL | Probable | A prominent group of species produces aecia on conifer needles and telia or leaves of Salicaceae (Cannon & Kirk 2007); Biotrophic, heteroecious or autoecious | Cannon PF, Kirk PM. 2007. Fungal Families of the World. CAB International, Cambridge (ISBN: 978-0851998275); Tedersoo L, et al. 2014. Science 346:e1256688 ((https://doi.org/10.1126/science.1256688)); Põlme S, et al. 2020. Fungal Diversity 105:1-16 ((https://doi.org/10.1007/s13225-020-00466-2)) |
| Otu28920 | 1 | k__Fungi(100);p__Basidiomycota(100);c__Agaricomycetes(100);o__Cantharellales(100);f__Tulasnellaceae(100);g__Tulasnella(100);s__Tulasnella_sp_184(100); | Tulasnella | 13 | Saprotroph-Symbiotroph | Ectomycorrhizal-\|Plant Saprotroph\|-Undefined Saprotroph | Corticioid | NULL | Possible | As litter_saprotroph (Põlme et al. 2020); Decay leaf, fruit, seed (Põlme et al. 2020) | Rinaldi AC, et al. 2008. Fungal Diversity 33:1-45 ((http://www.fungaldiversity.org/fdp/sfdp/33-1.pdf)); Tedersoo L, et al. 2010. Mycorrhiza 20:217-263 ((https://doi.org/10.1007/s00572-009-0274-x)) (pro parte); Põlme S, et al. 2020. Fungal Diversity 105:1-16 ((https://doi.org/10.1007/s13225-020-00466-2)) |
| Otu28922 | 1 | k__Fungi(100);p__Ascomycota(100);c__Saccharomycetes(100);o__Saccharomycetales(100);f__Saccharomycetaceae(100);g__Kazachstania(100);s__Kazachstania_intestinalis(100); | Kazachstania | 13 | Saprotroph | \|Undefined Saprotroph\| | Yeast | NULL | Probable | As nectar-tap_saprotroph (Põlme et al. 2020); Decay sugar-rich substrates (Põlme et al. 2020) | Põlme S, et al. 2020. Fungal Diversity 105:1-16 ((https://doi.org/10.1007/s13225-020-00466-2)) |
| Otu28923 | 1 | k__Fungi(100);p__Basidiomycota(100);c__Exobasidiomycetes(100);o__Microstromatales(100);f__Microstromataceae(100);g__Microstroma(100);s__Microstroma_juglandis(100); | Microstroma | 13 | Pathotroph | \|Plant Pathogen\| | Dimorphic | NULL | Probable | NULL | Tedersoo L, et al. 2014. Science 346:e1256688 ((https://doi.org/10.1126/science.1256688)); Põlme S, et al. 2020. Fungal Diversity 105:1-16 ((https://doi.org/10.1007/s13225-020-00466-2)) |
| Otu28925 | 1 | k__Fungi(100);p__Ascomycota(100);c__Sordariomycetes(100);o__Melanosporales(100);f__Ceratostomataceae(100);g__Harzia(100);s__Harzia_acremonioides(100); | Harzia | 13 | Saprotroph | Plant Saprotroph-Undefined Saprotroph-\|Wood Saprotroph\| | Microfungus | NULL | Probable | As litter_saprotroph (Põlme et al. 2020); Decay wood, leaf, fruit, seed (Põlme et al. 2020) | Tedersoo L, et al. 2014. Science 346:e1256688 ((https://doi.org/10.1126/science.1256688)); Põlme S, et al. 2020. Fungal Diversity 105:1-16 ((https://doi.org/10.1007/s13225-020-00466-2)) |
| Otu28926 | 1 | k__Fungi(100);p__Ascomycota(100);c__Sordariomycetes(100);o__Melanosporales(100);f__Ceratostomataceae(100);g__Harzia(100);s__Harzia_acremonioides(100); | Harzia | 13 | Saprotroph | Plant Saprotroph-Undefined Saprotroph-\|Wood Saprotroph\| | Microfungus | NULL | Probable | As litter_saprotroph (Põlme et al. 2020); Decay wood, leaf, fruit, seed (Põlme et al. 2020) | Tedersoo L, et al. 2014. Science 346:e1256688 ((https://doi.org/10.1126/science.1256688)); Põlme S, et al. 2020. Fungal Diversity 105:1-16 ((https://doi.org/10.1007/s13225-020-00466-2)) |
| Otu28928 | 1 | k__Fungi(100);p__Ascomycota(100);c__Sordariomycetes(100);o__Melanosporales(100);f__Ceratostomataceae(100);g__Harzia(100);s__Harzia_acremonioides(100); | Harzia | 13 | Saprotroph | Plant Saprotroph-Undefined Saprotroph-\|Wood Saprotroph\| | Microfungus | NULL | Probable | As litter_saprotroph (Põlme et al. 2020); Decay wood, leaf, fruit, seed (Põlme et al. 2020) | Tedersoo L, et al. 2014. Science 346:e1256688 ((https://doi.org/10.1126/science.1256688)); Põlme S, et al. 2020. Fungal Diversity 105:1-16 ((https://doi.org/10.1007/s13225-020-00466-2)) |

| | | | | | | | | | | | |
|---|---|---|---|---|---|---|---|---|---|---|---|
| Otu28930 | 1 | k__Fungi(100);p__Basidiomycota(100);c__Agaricostilbomycetes(100);o__Agaricostilbales(100);f__Agaricostilbaceae(100);g__Bensingtonia(100);g__Bensingtonia_unclassified(100); | Bensingtonia | 13 | Saprotroph | \|Undefined Saprotroph\| | Dimorphic-Facultative Yeast | NULL | Highly Probable | NULL | Kurtzman CP, et al. (eds.) 2011. The Yeasts, a Taxonomic Study. Fifth Edition. Vols 1-3. Elsevier, San Diego (ISBN: 9780444521491); Põlme S, et al. 2020. Fungal Diversity 105:1-16 ((https://doi.org/10.1007/s13225-020-00466-2)) |
| Otu28931 | 1 | k__Fungi(100);p__Basidiomycota(100);c__Agaricomycetes(100);o__Hymenochaetales(100);f__Schizoporaceae(100);g__Hyphodontia(100);s__Hyphodontia_subalutacea(100); | Hyphodontia | 13 | Saprotroph | Undefined Saprotroph-\|Wood Saprotroph\| | Corticioid | White Rot | Probable | As white_rot (Põlme et al. 2020); Decay wood (Põlme et al. 2020) | Tedersoo L, et al. 2014. Science 346:e1256688 ((https://doi.org/10.1126/science.1256688)); Põlme S, et al. 2020. Fungal Diversity 105:1-16 ((https://doi.org/10.1007/s13225-020-00466-2)) |
| Otu28932 | 1 | k__Fungi(100);p__Basidiomycota(100);c__Tremellomycetes(100);o__Filobasidiales(100);f__Filobasidiaceae(100);g__Cryptococcus(100);s__Cryptococcus_terricola(100); | Cryptococcus | 13 | Pathotroph-Saprotroph-Symbiotroph | Animal Pathogen-Endophyte-Epiphyte-\|Undefined Saprotroph\| | Dimorphic Yeast-Yeast | NULL | Possible | As foliar_endophyte (Põlme et al. 2020); Decay leaf, fruit, seed, soil, dung, animal material (Põlme et al. 2020) | Andrews JH, et al. 1983. Phytopathology 73:228-234 ((https://doi.org/10.1094/Phyto-73-228)); Kurtzman CP, et al. (eds.) 2011. The Yeasts, a Taxonomic Study. Fifth Edition. Vols 1-3. Elsevier, San Diego (ISBN: 9780444521491); Põlme S, et al. 2020. Fungal Diversity 105:1-16 ((https://doi.org/10.1007/s13225-020-00466-2)) |
| Otu28937 | 1 | k__Fungi(100);p__Basidiomycota(100);c__Exobasidiomycetes(100);o__Exobasidiomycetes_order_Incertae_sedis(100);f__Exobasidiomycetes_family_Incertae_sedis(100);g__Tilletiopsis(100);s__Tilletiopsis_fulvescens(100); | Tilletiopsis | 13 | Pathotroph-Saprotroph | Endophyte-Fungal Parasite-\|Plant Pathogen\|-Plant Saprotroph-Undefined Saprotroph | Microfungus-Yeast | NULL | Probable | As litter_saprotroph (Põlme et al. 2020); As mycoparasite (Põlme et al. 2020); Decay leaf, fruit, seed (Põlme et al. 2020); Isolated from leaves...dead leaves (Nyland 1950) | Nyland G. 1950. Mycologia 42:487-496 ((ttps://doi.org/10.1080/00275514.1950.12017853)); Põlme S, et al. 2020. Fungal Diversity 105:1-16 ((https://doi.org/10.1007/s13225-020-00466-2)) |
| Otu28939 | 1 | k__Fungi(100);p__Zygomycota(100);c__Zygomycota_class_Incertae_sedis(100);o__Mortierellales(100);f__Mortierellaceae(100);f__Mortierellaceae_unclassified(100);f__Mortierellaceae_unclassified(100); | Mortierellaceae | 9 | Saprotroph-Symbiotroph | Endophyte-Litter Saprotroph-Soil Saprotroph-Undefined Saprotroph | Microfungus | NULL | Possible | Saprobic in soil and thus frequenctly isolated from soil and material contaminated with soil such as hay or rotten silage and spoil tips (Cannon & Kirk 2007) | Cannon PF, Kirk PM. 2007. Fungal Families of the World. CAB International, Cambridge (ISBN: 978-0851998275); Purahong W, et al. 2016. Molecular Ecology 25:4059-4074 ((https://doi.org/10.1111/mec.13739)) |
| Otu28940 | 1 | k__Fungi(100);p__Ascomycota(100);c__Saccharomycetes(100);o__Saccharomycetales(100);f__Metschnikowiaceae(100);g__Metschnikowia(100);g__Metschnikowia_unclassified(100); | Metschnikowia | 13 | Saprotroph | \|Undefined Saprotroph\| | Yeast | NULL | Probable | As nectar-tap_saprotroph (Põlme et al. 2020); Decay sugar-rich substrates (Põlme et al. 2020) | Põlme S, et al. 2020. Fungal Diversity 105:1-16 ((https://doi.org/10.1007/s13225-020-00466-2)) |
| Otu28945 | 1 | k__Fungi(100);p__Ascomycota(100);c__Orbiliomycetes(100);o__Orbiliales(100);f__Orbiliaceae(100);g__Drechslerella(100);s__Drechslerella_sp_ZY_2011a(100); | Drechslerella | 13 | Pathotroph-Saprotroph | \|Animal Parasite\|-Nematophagous-Wood Saprotroph | Microfungus | Nematode-trappi | Probable | Decay wood, leaf, fruit, seed, soil, animal material (Põlme et al. 2020) | Tedersoo L, et al. 2014. Science 346:e1256688 ((https://doi.org/10.1126/science.1256688)); Yang C-T, et al. 2020. Proceedings of the National Academy of Sciences 117:6762-6770 ((https://doi.org/10.1073/pnas.1919726117)) |
| Otu28953 | 1 | k__Fungi(100);p__Basidiomycota(100);c__Agaricomycetes(100);o__Agaricales(100);o__Agaricales_unclassified(100);o__Agaricales_unclassified(100);o__Agaricales_unclassified(100); | Agaricales | 7 | Pathotroph-Saprotroph-Symbiotroph | Bryophyte Parasite-Dung Saprotroph-Ectomycorrhizal-Fungal Parasite-Leaf Saprotroph-Plant Parasite-Undefined Saprotroph-Wood Saprotroph | Agaricoid-Gasteroid-Microfungus-Secotioid-Yeast | NULL | Possible | Mushrooms and toadstools, Gill fungi, Agarics... lignicolous, sometimes muscicolous or fungicolous, saprobic, mycorrhizal, rarely parasitic on plants or fungi (Kirk et al. 2008) | Kirk PM et al. 2008. Dictionary of the Fungi. Tenth Edition. CAB International, Wallingford (ISBN: 978-0851998268) |
| Otu28955 | 1 | k__Fungi(100);p__Ascomycota(100);c__Saccharomycetes(100);o__Saccharomycetales(100);f__Metschnikowiaceae(100);g__Metschnikowia(100);s__Metschnikowia_pulcherrima(100); | Metschnikowia | 13 | Saprotroph | \|Undefined Saprotroph\| | Yeast | NULL | Probable | As nectar-tap_saprotroph (Põlme et al. 2020); Decay sugar-rich substrates (Põlme et al. 2020) | Põlme S, et al. 2020. Fungal Diversity 105:1-16 ((https://doi.org/10.1007/s13225-020-00466-2)) |
| Otu28958 | 1 | k__Fungi(100);p__Ascomycota(100);c__Saccharomycetes(100);o__Saccharomycetales(100);f__Saccharomycetales_family_Incertae_sedis(100);g__Candida(100);g__Candida_unclassified(100); | Candida | 13 | Pathotroph-Saprotroph-Symbiotroph | Animal Pathogen-Endophyte-Endosymbiont-Epiphyte-\|Undefined Saprotroph\| | Dimorphic Yeast-Yeast | NULL | Probable | As nectar-tap_saprotroph (Põlme et al. 2020); Decay sugar-rich substrates (Põlme et al. 2020); This asexual genus is polyphyletic and contain species that function very differently depend on what they are. Caution should be taken when interpreting guilds for Candida species. | Manolakaki D, et al. 2010. Virulence. 1: 367-375 ((https://doi.org/10.4161/viru.1.5.12796)); Mekbib SB, et al. 2011. Tropical Plant Pathology 36:233-240 ((https://doi.org/10.1590/S1982-56762011000400004)); Põlme S, et al. 2020. Fungal Diversity 105:1-16 ((https://doi.org/10.1007/s13225-020-00466-2)) |
| Otu28966 | 1 | k__Fungi(100);p__Basidiomycota(100);c__Exobasidiomycetes(100);o__Exobasidiomycetes_order_Incertae_sedis(100);f__Exobasidiomycetes_family_Incertae_sedis(100);g__Tilletiopsis(100);s__Tilletiopsis_pallescens(100); | Tilletiopsis | 13 | Pathotroph-Saprotroph | Endophyte-Fungal Parasite-\|Plant Pathogen\|-Plant Saprotroph-Undefined Saprotroph | Microfungus-Yeast | NULL | Probable | As litter_saprotroph (Põlme et al. 2020); As mycoparasite (Põlme et al. 2020); Decay leaf, fruit, seed (Põlme et al. 2020); Isolated from leaves...dead leaves (Nyland 1950) | Nyland G. 1950. Mycologia 42:487-496 ((ttps://doi.org/10.1080/00275514.1950.12017853)); Põlme S, et al. 2020. Fungal Diversity 105:1-16 ((https://doi.org/10.1007/s13225-020-00466-2)) |
| Otu28969 | 1 | k__Fungi(100);p__Zygomycota(100);c__Zygomycota_class_Incertae_sedis(100);o__Mortierellales(100);f__Mortierellaceae(100);f__Mortierellaceae_unclassified(100);f__Mortierellaceae_unclassified(100); | Mortierellaceae | 9 | Saprotroph-Symbiotroph | Endophyte-Litter Saprotroph-Soil Saprotroph-Undefined Saprotroph | Microfungus | NULL | Possible | Saprobic in soil and thus frequenctly isolated from soil and material contaminated with soil such as hay or rotten silage and spoil tips (Cannon & Kirk 2007) | Cannon PF, Kirk PM. 2007. Fungal Families of the World. CAB International, Cambridge (ISBN: 978-0851998275); Purahong W, et al. 2016. Molecular Ecology 25:4059-4074 ((https://doi.org/10.1111/mec.13739)) |
| Otu28970 | 1 | k__Fungi(100);p__Ascomycota(100);c__Taphrinomycetes(100);o__Taphrinales(100);f__Taphrinaceae(100);f__Taphrinaceae_unclassified(100);f__Taphrinaceae_unclassified(100); | Taphrinaceae | 9 | Pathotroph | Plant Pathogen | Microfungus | NULL | Probable | Biotrophic on plants, usually causing hyperplasia (galls, witches broom) or lesions (Cannon & Kirk 2007) | Cannon PF, Kirk PM. 2007. Fungal Families of the World. CAB International, Cambridge (ISBN: 978-0851998275) |
| Otu28993 | 1 | k__Fungi(100);p__Ascomycota(100);c__Saccharomycetes(100);o__Saccharomycetales(100);f__Metschnikowiaceae(100);g__Metschnikowia(100);g__Metschnikowia_unclassified(100); | Metschnikowia | 13 | Saprotroph | \|Undefined Saprotroph\| | Yeast | NULL | Probable | As nectar-tap_saprotroph (Põlme et al. 2020); Decay sugar-rich substrates (Põlme et al. 2020) | Põlme S, et al. 2020. Fungal Diversity 105:1-16 ((https://doi.org/10.1007/s13225-020-00466-2)) |
| Otu29003 | 1 | k__Fungi(100);p__Ascomycota(100);c__Saccharomycetes(100);o__Saccharomycetales(100);f__Metschnikowiaceae(100);g__Metschnikowia(100);g__Metschnikowia_unclassified(100); | Metschnikowia | 13 | Saprotroph | \|Undefined Saprotroph\| | Yeast | NULL | Probable | As nectar-tap_saprotroph (Põlme et al. 2020); Decay sugar-rich substrates (Põlme et al. 2020) | Põlme S, et al. 2020. Fungal Diversity 105:1-16 ((https://doi.org/10.1007/s13225-020-00466-2)) |
| Otu29004 | 1 | k__Fungi(100);p__Ascomycota(100);c__Saccharomycetes(100);o__Saccharomycetales(100);f__Metschnikowiaceae(100);g__Metschnikowia(100);s__Metschnikowia_pulcherrima(100); | Metschnikowia | 13 | Saprotroph | \|Undefined Saprotroph\| | Yeast | NULL | Probable | As nectar-tap_saprotroph (Põlme et al. 2020); Decay sugar-rich substrates (Põlme et al. 2020) | Põlme S, et al. 2020. Fungal Diversity 105:1-16 ((https://doi.org/10.1007/s13225-020-00466-2)) |
| Otu29008 | 1 | k__Fungi(100);p__Zygomycota(100);c__Zygomycota_class_Incertae_sedis(100);o__Mucorales(100);f__Backusellaceae(100);g__Backusella(100);s__Backusella_circina(100); | Backusella | 13 | Saprotroph | Dung Saprotroph-\|Undefined Saprotroph\| | Microfungus-Sporophores | NULL | Probable | As soil_saprotroph (Põlme et al. 2020); Decay soil, dung (Põlme et al. 2020) | Walther G, et al. 2013. Persoonia 30:11-47 ((https://doi.org/10.3767/003158513X665070)); Põlme S, et al. 2020. Fungal Diversity 105:1-16 ((https://doi.org/10.1007/s13225-020-00466-2)) |
| Otu29017 | 1 | k__Fungi(100);p__Basidiomycota(100);c__Agaricomycetes(100);o__Hymenochaetales(100);f__Hymenochaetaceae(100);g__Phellinus(100);s__Phellinus_alni(100); | Phellinus | 13 | Pathotroph-Saprotroph | \|Plant Pathogen\|-Wood Saprotroph | Polyporoid | White Rot | Highly Probable | As white_rot (Põlme et al. 2020); As wood_pathogen (Põlme et al. 2020); Decay wood (Põlme et al. 2020) | Gilbertson RL, Ryvarden L. 1987-1987. North American Polypores. Fungiflora, Oslo (ISBN: 978-0945345060); Põlme S, et al. 2020. Fungal Diversity 105:1-16 ((https://doi.org/10.1007/s13225-020-00466-2)) |
| Otu29019 | 1 | k__Fungi(100);p__Basidiomycota(100);c__Agaricomycetes(100);o__Agaricales(100);f__Inocybaceae(100);g__Crepidotus(100);s__Crepidotus_applanatus(100); | Crepidotus | 13 | Saprotroph | \|Wood Saprotroph\| | Agaricoid | NULL | Highly Probable | Decay wood (Põlme et al. 2020) | Grgurinovic CA, Simpson JA. 2001. Fungal Diversity 8:97-127 ((http://www.fungaldiversity.org/fdp/sfdp/FD_8_97-127.pdf)); Aime MC, et al. 2005. American Journal of Botany 92:74-82 ((https://doi.org/10.3732/ajb.92.1.74)); Tedersoo L, et al. 2014. Science 346:e1256688 ((https://doi.org/10.1126/science.1256688)); Põlme S, et al. 2020. Fungal Diversity 105:1-16 ((https://doi.org/10.1007/s13225-020-00466-2)) |
| Otu29020 | 1 | k__Fungi(100);p__Basidiomycota(100);c__Microbotryomycetes(100);o__Microbotryales(100);f__Microbotryaceae(100);g__Microbotryum(100);s__Microbotryum_cordae(100); | Microbotryum | 13 | Pathotroph | \|Plant Pathogen\| | Smut | NULL | Probable | NULL | Tedersoo L, et al. 2014. Science 346:e1256688 ((https://doi.org/10.1126/science.1256688)); Põlme S, et al. 2020. Fungal Diversity 105:1-16 ((https://doi.org/10.1007/s13225-020-00466-2)) |
| Otu29021 | 1 | k__Fungi(100);p__Basidiomycota(100);c__Agaricomycetes(100);o__Agaricales(100);f__Pterulaceae(100);g__Aphanobasidium(100);s__Aphanobasidium_pseudotsugae(100); | Aphanobasidium | 13 | Saprotroph | Undefined Saprotroph-\|Wood Saprotroph\| | Corticioid | White Rot | Probable | As white_rot (Põlme et al. 2020); Decay wood (Põlme et al. 2020) | Tedersoo L, et al. 2014. Science 346:e1256688 ((https://doi.org/10.1126/science.1256688)); Põlme S, et al. 2020. Fungal Diversity 105:1-16 ((https://doi.org/10.1007/s13225-020-00466-2)) |
| Otu29022 | 1 | k__Fungi(100);p__Basidiomycota(100);c__Exobasidiomycetes(100);o__Exobasidiomycetes_order_Incertae_sedis(100);f__Exobasidiomycetes_family_Incertae_sedis(100);g__Meira(100);s__Meira_geulakonigii(100); | Meira | 13 | Pathotroph | \|Animal Parasite\|-Animal Pathogen-Plant Pathogen | Dimorphic | NULL | Probable | An anamorphic fungus isolated from cadavers of the citrus rust mite (P. oleivora) infesting grapefruit (C. paradisi) leaves (Boekhout et al. 2003); Plant pathogen (Põlme et al. 2020); Similar to the yeast-like fungi classified in the Ustilaginales, such as Pseudozyma species (Boekhout et al. 2003) | Boekhout T, et al. 2003. International Journal of Systematic and Evolutionary Microbiology 53:1655-1664 ((https://doi.org/10.1099/ijs.0.02434-0)); Põlme S, et al. 2020. Fungal Diversity 105:1-16 ((https://doi.org/10.1007/s13225-020-00466-2)) |
| Otu29023 | 1 | k__Fungi(100);p__Basidiomycota(100);c__Agaricomycetes(100);o__Gomphales(100);f__Gomphaceae(100);f__Gomphaceae_unclassified(100);f__Gomphaceae_unclassified(100); | Gomphaceae | 9 | Saprotroph-Symbiotroph | Ectomycorrhizal-Undefined Saprotroph-Wood Saprotroph | Clavarioid-Gomphoid | NULL | Probable | Terrestrial or lignicolous, either saprobic or forming ectomycorrhizal relationships with woody plants (Cannon & Kirk 2007) | Cannon PF, Kirk PM. 2007. Fungal Families of the World. CAB International, Cambridge (ISBN: 978-0851998275) |
| Otu29024 | 1 | k__Fungi(100);p__Basidiomycota(100);c__Agaricomycetes(100);o__Agaricales(100);f__Inocybaceae(100);g__Crepidotus(100);s__Crepidotus_mollis(100); | Crepidotus | 13 | Saprotroph | \|Wood Saprotroph\| | Agaricoid | NULL | Highly Probable | Decay wood (Põlme et al. 2020) | Grgurinovic CA, Simpson JA. 2001. Fungal Diversity 8:97-127 ((http://www.fungaldiversity.org/fdp/sfdp/FD_8_97-127.pdf)); Aime MC, et al. 2005. American Journal of Botany 92:74-82 ((https://doi.org/10.3732/ajb.92.1.74)); Tedersoo L, et al. 2014. Science 346:e1256688 ((https://doi.org/10.1126/science.1256688)); Põlme S, et al. 2020. Fungal Diversity 105:1-16 ((https://doi.org/10.1007/s13225-020-00466-2)) |
| Otu29025 | 1 | k__Fungi(100);p__Basidiomycota(100);c__Agaricomycetes(100);o__Gomphales(100);f__Lentariaceae(100);g__Kavinia(100);s__Kavinia_alboviridis(100); | Kavinia | 13 | Saprotroph | Undefined Saprotroph-\|Wood Saprotroph\| | Clavarioid | White Rot | Probable | As white_rot (Põlme et al. 2020); Decay wood (Põlme et al. 2020) | Tedersoo L, et al. 2014. Science 346:e1256688 ((https://doi.org/10.1126/science.1256688)); Põlme S, et al. 2020. Fungal Diversity 105:1-16 ((https://doi.org/10.1007/s13225-020-00466-2)) |
| Otu29026 | 1 | k__Fungi(100);p__Basidiomycota(100);c__Agaricomycetes(100);o__Gomphales(100);f__Gomphaceae(100);g__Ramaria(100);s__Ramaria_sandaracina(100); | Ramaria | 13 | Saprotroph-Symbiotroph | \|Ectomycorrhizal\|-Plant Saprotroph | Clavarioid | NULL | Possible | As litter_saprotroph (Põlme et al. 2020) | Rinaldi AC, et al. 2008. Fungal Diversity 33:1-45 ((http://www.fungaldiversity.org/fdp/sfdp/33-1.pdf)) (as ECM/not ECM); Tedersoo L, et al. 2010. Mycorrhiza 20:217-263 ((10.1007/s00572-009-0274-x)) (pro parte); Põlme S, et al. 2020. Fungal Diversity 105:1-16 ((https://doi.org/10.1007/s13225-020-00466-2)); http://mycorrhizas.info/ecmf.html ((http://mycorrhizas.info/ecmf.html)) (as ECM) |
| Otu29027 | 1 | k__Fungi(100);p__Basidiomycota(100);c__Agaricomycetes(100);o__Gomphales(100);f__Gomphaceae(100);g__Ramaria(100);s__Ramaria_maculatipes(100); | Ramaria | 13 | Saprotroph-Symbiotroph | \|Ectomycorrhizal\|-Plant Saprotroph | Clavarioid | NULL | Possible | As litter_saprotroph (Põlme et al. 2020) | Rinaldi AC, et al. 2008. Fungal Diversity 33:1-45 ((http://www.fungaldiversity.org/fdp/sfdp/33-1.pdf)) (as ECM/not ECM); Tedersoo L, et al. 2010. Mycorrhiza 20:217-263 ((10.1007/s00572-009-0274-x)) (pro parte); Põlme S, et al. 2020. Fungal Diversity 105:1-16 ((https://doi.org/10.1007/s13225-020-00466-2)); http://mycorrhizas.info/ecmf.html ((http://mycorrhizas.info/ecmf.html)) (as ECM) |
| Otu29028 | 1 | k__Fungi(100);p__Basidiomycota(100);c__Agaricomycetes(100);o__Agaricales(100);f__Agaricaceae(100);g__Agaricus(100);s__Agaricus_bitorquis(100); | Agaricus | 13 | Saprotroph | \|Undefined Saprotroph\| | Agaricoid-Gasteroid-Secotioid | NULL | Probable | As soil_saprotroph (Põlme et al. 2020); Decay soil (Põlme et al. 2020) | Tedersoo L, et al. 2014. Science 346:e1256688 ((https://doi.org/10.1126/science.1256688)); Põlme S, et al. 2020. Fungal Diversity 105:1-16 ((https://doi.org/10.1007/s13225-020-00466-2)) |
| Otu29029 | 1 | k__Fungi(100);p__Basidiomycota(100);c__Agaricomycetes(100);o__Agaricales(100);o__Agaricales_unclassified(100);o__Agaricales_unclassified(100);o__Agaricales_unclassified(100); | Agaricales | 7 | Pathotroph-Saprotroph-Symbiotroph | Bryophyte Parasite-Dung Saprotroph-Ectomycorrhizal-Fungal Parasite-Leaf Saprotroph-Plant Parasite-Undefined Saprotroph-Wood Saprotroph | Agaricoid-Gasteroid-Microfungus-Secotioid-Yeast | NULL | Possible | Mushrooms and toadstools, Gill fungi, Agarics... lignicolous, sometimes muscicolous or fungicolous, saprobic, mycorrhizal, rarely parasitic on plants or fungi (Kirk et al. 2008) | Kirk PM et al. 2008. Dictionary of the Fungi. Tenth Edition. CAB International, Wallingford (ISBN: 978-0851998268) |
| Otu29030 | 1 | k__Fungi(100);p__Basidiomycota(100);c__Agaricomycetes(100);o__Agaricales(100);f__Agaricaceae(100);g__Agaricus(100);g__Agaricus_unclassified(100); | Agaricus | 13 | Saprotroph | \|Undefined Saprotroph\| | Agaricoid-Gasteroid-Secotioid | NULL | Probable | As soil_saprotroph (Põlme et al. 2020); Decay soil (Põlme et al. 2020) | Tedersoo L, et al. 2014. Science 346:e1256688 ((https://doi.org/10.1126/science.1256688)); Põlme S, et al. 2020. Fungal Diversity 105:1-16 ((https://doi.org/10.1007/s13225-020-00466-2)) |
| Otu29031 | 1 | k__Fungi(100);p__Basidiomycota(100);c__Agaricomycetes(100);o__Agaricales(100);f__Inocybaceae(100);g__Crepidotus(100);s__Crepidotus_sphaerosporus(100); | Crepidotus | 13 | Saprotroph | \|Wood Saprotroph\| | Agaricoid | NULL | Highly Probable | Decay wood (Põlme et al. 2020) | Grgurinovic CA, Simpson JA. 2001. Fungal Diversity 8:97-127 ((http://www.fungaldiversity.org/fdp/sfdp/FD_8_97-127.pdf)); Aime MC, et al. 2005. American Journal of Botany 92:74-82 ((https://doi.org/10.3732/ajb.92.1.74)); Tedersoo L, et al. 2014. Science 346:e1256688 ((https://doi.org/10.1126/science.1256688)); Põlme S, et al. 2020. Fungal Diversity 105:1-16 ((https://doi.org/10.1007/s13225-020-00466-2)) |
| Otu29032 | 1 | k__Fungi(100);p__Basidiomycota(100);c__Agaricomycetes(100);o__Agaricales(100);f__Inocybaceae(100);g__Crepidotus(100);s__Crepidotus_mollis(100); | Crepidotus | 13 | Saprotroph | \|Wood Saprotroph\| | Agaricoid | NULL | Highly Probable | Decay wood (Põlme et al. 2020) | Grgurinovic CA, Simpson JA. 2001. Fungal Diversity 8:97-127 ((http://www.fungaldiversity.org/fdp/sfdp/FD_8_97-127.pdf)); Aime MC, et al. 2005. American Journal of Botany 92:74-82 ((https://doi.org/10.3732/ajb.92.1.74)); Tedersoo L, et al. 2014. Science 346:e1256688 ((https://doi.org/10.1126/science.1256688)); Põlme S, et al. 2020. Fungal Diversity 105:1-16 ((https://doi.org/10.1007/s13225-020-00466-2)) |
| Otu29035 | 1 | k__Fungi(100);p__Basidiomycota(100);c__Agaricomycetes(100);o__Gomphales(100);f__Gomphaceae(100);g__Ramaria(100);s__Ramaria_stricta(100); | Ramaria | 13 | Saprotroph-Symbiotroph | \|Ectomycorrhizal\|-Plant Saprotroph | Clavarioid | NULL | Possible | As litter_saprotroph (Põlme et al. 2020) | Rinaldi AC, et al. 2008. Fungal Diversity 33:1-45 ((http://www.fungaldiversity.org/fdp/sfdp/33-1.pdf)) (as ECM/not ECM); Tedersoo L, et al. 2010. Mycorrhiza 20:217-263 ((10.1007/s00572-009-0274-x)) (pro parte); Põlme S, et al. 2020. Fungal Diversity 105:1-16 ((https://doi.org/10.1007/s13225-020-00466-2)); http://mycorrhizas.info/ecmf.html ((http://mycorrhizas.info/ecmf.html)) (as ECM) |
| Otu29036 | 1 | k__Fungi(100);p__Basidiomycota(100);c__Agaricomycetes(100);o__Polyporales(100);f__Meruliaceae(100);g__Phlebia(100);s__Phlebia_sp_olrim964(100); | Phlebia | 13 | Saprotroph | \|Wood Saprotroph\| | Corticioid | White Rot | Highly Probable | As white_rot (Põlme et al. 2020); Decay wood (Põlme et al. 2020) | Kirk PM et al. 2008. Dictionary of the Fungi. Tenth Edition. CAB International, Wallingford (ISBN: 978-0851998268); Põlme S, et al. 2020. Fungal Diversity 105:1-16 ((https://doi.org/10.1007/s13225-020-00466-2)) |
| Otu29037 | 1 | k__Fungi(100);p__Basidiomycota(100);c__Agaricomycetes(100);o__Phallales(100);f__Phallaceae(100);g__Mutinus(100);s__Mutinus_sp_MA_2013(100); | Mutinus | 13 | Saprotroph | Plant Saprotroph-\|Undefined Saprotroph\| | Phalloid | NULL | Probable | stinkhorn; As soil_saprotroph (Põlme et al. 2020); Common name ; Decay soil (Põlme et al. 2020) | Grgurinovic CA, Simpson JA. 2001. Fungal Diversity 8:97-127 ((http://www.fungaldiversity.org/fdp/sfdp/FD_8_97-127.pdf)); Tedersoo L, et al. 2014. Science 346:e1256688 ((https://doi.org/10.1126/science.1256688)); Põlme S, et al. 2020. Fungal Diversity 105:1-16 ((https://doi.org/10.1007/s13225-020-00466-2)) |
| Otu29038 | 1 | k__Fungi(100);p__Basidiomycota(100);c__Agaricomycetes(100);o__Auriculariales(100);f__Auriculariales_family_Incertae_sedis(100);g__Heterochaete(100);s__Heterochaete_shearii(100); | Heterochaete | 13 | Saprotroph | Undefined Saprotroph-\|Wood Saprotroph\| | Corticioid | NULL | Probable | Decay wood (Põlme et al. 2020) | Tedersoo L, et al. 2014. Science 346:e1256688 ((https://doi.org/10.1126/science.1256688)); Põlme S, et al. 2020. Fungal Diversity 105:1-16 ((https://doi.org/10.1007/s13225-020-00466-2)) |
| Otu29040 | 1 | k__Fungi(100);p__Zygomycota(100);c__Zygomycota_class_Incertae_sedis(100);o__Mucorales(100);f__Mucoraceae(100);g__Mucor(100);s__Mucor_sp_JJP_2009a(100); | Mucor | 13 | Saprotroph | \|Undefined Saprotroph\| | Microfungus | NULL | Probable | As soil_saprotroph (Põlme et al. 2020); Decay soil (Põlme et al. 2020) | Tedersoo L, et al. 2014. Science 346:e1256688 ((https://doi.org/10.1126/science.1256688)); Põlme S, et al. 2020. Fungal Diversity 105:1-16 ((https://doi.org/10.1007/s13225-020-00466-2)) |
| Otu29041 | 1 | k__Fungi(100);p__Zygomycota(100);c__Zygomycota_class_Incertae_sedis(100);o__Mucorales(100);o__Mucorales_unclassified(100);o__Mucorales_unclassified(100);o__Mucorales_unclassified(100); | Mucorales | 7 | Saprotroph | Undefined Saprotroph | Microfungus | NULL | Possible | NULL | Sterkenburg E, et al. 2015. New Phytologist 207:1145-1158 ((https://doi.org/10.1111/nph.13426)) |
| Otu29042 | 1 | k__Fungi(100);p__Zygomycota(100);c__Zygomycota_class_Incertae_sedis(100);o__Mucorales(100);f__Mucoraceae(100);g__Mucor(100);g__Mucor_unclassified(100); | Mucor | 13 | Saprotroph | \|Undefined Saprotroph\| | Microfungus | NULL | Probable | As soil_saprotroph (Põlme et al. 2020); Decay soil (Põlme et al. 2020) | Tedersoo L, et al. 2014. Science 346:e1256688 ((https://doi.org/10.1126/science.1256688)); Põlme S, et al. 2020. Fungal Diversity 105:1-16 ((https://doi.org/10.1007/s13225-020-00466-2)) |
| Otu29046 | 1 | k__Fungi(100);p__Zygomycota(100);c__Zygomycota_class_Incertae_sedis(100);o__Mucorales(100);f__Mucoraceae(100);g__Mucor(100);s__Mucor_mucedo(100); | Mucor | 13 | Saprotroph | \|Undefined Saprotroph\| | Microfungus | NULL | Probable | As soil_saprotroph (Põlme et al. 2020); Decay soil (Põlme et al. 2020) | Tedersoo L, et al. 2014. Science 346:e1256688 ((https://doi.org/10.1126/science.1256688)); Põlme S, et al. 2020. Fungal Diversity 105:1-16 ((https://doi.org/10.1007/s13225-020-00466-2)) |
| Otu29047 | 1 | k__Fungi(100);p__Ascomycota(100);c__Saccharomycetes(100);o__Saccharomycetales(100);f__Metschnikowiaceae(100);g__Metschnikowia(100);g__Metschnikowia_unclassified(100); | Metschnikowia | 13 | Saprotroph | \|Undefined Saprotroph\| | Yeast | NULL | Probable | As nectar-tap_saprotroph (Põlme et al. 2020); Decay sugar-rich substrates (Põlme et al. 2020) | Põlme S, et al. 2020. Fungal Diversity 105:1-16 ((https://doi.org/10.1007/s13225-020-00466-2)) |
| Otu29055 | 1 | k__Fungi(100);p__Ascomycota(100);c__Saccharomycetes(100);o__Saccharomycetales(100);f__Metschnikowiaceae(100);g__Metschnikowia(100);s__Metschnikowia_pulcherrima(100); | Metschnikowia | 13 | Saprotroph | \|Undefined Saprotroph\| | Yeast | NULL | Probable | As nectar-tap_saprotroph (Põlme et al. 2020); Decay sugar-rich substrates (Põlme et al. 2020) | Põlme S, et al. 2020. Fungal Diversity 105:1-16 ((https://doi.org/10.1007/s13225-020-00466-2)) |
| Otu29058 | 1 | k__Fungi(100);p__Ascomycota(100);c__Saccharomycetes(100);o__Saccharomycetales(100);f__Metschnikowiaceae(100);g__Metschnikowia(100);s__Metschnikowia_pulcherrima(100); | Metschnikowia | 13 | Saprotroph | \|Undefined Saprotroph\| | Yeast | NULL | Probable | As nectar-tap_saprotroph (Põlme et al. 2020); Decay sugar-rich substrates (Põlme et al. 2020) | Põlme S, et al. 2020. Fungal Diversity 105:1-16 ((https://doi.org/10.1007/s13225-020-00466-2)) |
| Otu29059 | 1 | k__Fungi(100);p__Basidiomycota(100);c__Agaricomycetes(100);o__Thelephorales(100);f__Thelephoraceae(100);g__Tomentella(100);s__Tomentella_stuposa(100); | Tomentella | 13 | Symbiotroph | \|Ectomycorrhizal\| | Corticioid | NULL | Highly Probable | NULL | Rinaldi AC, et al. 2008. Fungal Diversity 33:1-45 ((http://www.fungaldiversity.org/fdp/sfdp/33-1.pdf)); Tedersoo L, et al. 2010. Mycorrhiza 20:217-263 ((https://doi.org/10.1007/s00572-009-0274-x)); Põlme S, et al. 2020. Fungal Diversity 105:1-16 ((https://doi.org/10.1007/s13225-020-00466-2)); http://mycorrhizas.info/ecmf.html ((http://mycorrhizas.info/ecmf.html)) |
| Otu29060 | 1 | k__Fungi(100);p__Zygomycota(100);c__Zygomycota_class_Incertae_sedis(100);o__Mucorales(100);f__Mucoraceae(100);g__Mucor(100);s__Mucor_mucedo(100); | Mucor | 13 | Saprotroph | \|Undefined Saprotroph\| | Microfungus | NULL | Probable | As soil_saprotroph (Põlme et al. 2020); Decay soil (Põlme et al. 2020) | Tedersoo L, et al. 2014. Science 346:e1256688 ((https://doi.org/10.1126/science.1256688)); Põlme S, et al. 2020. Fungal Diversity 105:1-16 ((https://doi.org/10.1007/s13225-020-00466-2)) |
| Otu29061 | 1 | k__Fungi(100);p__Zygomycota(100);c__Zygomycota_class_Incertae_sedis(100);o__Mucorales(100);f__Mucoraceae(100);g__Mucor(100);s__Mucor_mucedo(100); | Mucor | 13 | Saprotroph | \|Undefined Saprotroph\| | Microfungus | NULL | Probable | As soil_saprotroph (Põlme et al. 2020); Decay soil (Põlme et al. 2020) | Tedersoo L, et al. 2014. Science 346:e1256688 ((https://doi.org/10.1126/science.1256688)); Põlme S, et al. 2020. Fungal Diversity 105:1-16 ((https://doi.org/10.1007/s13225-020-00466-2)) |
| Otu29062 | 1 | k__Fungi(100);p__Basidiomycota(100);c__Agaricomycetes(100);o__Agaricales(100);f__Inocybaceae(100);g__Inocybe(100);s__Inocybe_acuta(100); | Inocybe | 13 | Symbiotroph | \|Ectomycorrhizal\| | Agaricoid | NULL | Highly Probable | Sensu stricto (Matheny 2009). | Rinaldi AC, et al. 2008. Fungal Diversity 33:1-45 ((http://www.fungaldiversity.org/fdp/sfdp/33-1.pdf)); Matheny PB. 2009. McIlvainea 18:11-21 ((https://namyco.org/docs/3_Matheny.pdf)); Tedersoo L, et al. 2010. Mycorrhiza 20:217-263 ((https://doi.org/10.1007/s00572-009-0274-x)); Põlme S, et al. 2020. Fungal Diversity 105:1-16 ((https://doi.org/10.1007/s13225-020-00466-2)); http://mycorrhizas.info/ecmf.html ((http://mycorrhizas.info/ecmf.html)) |
| Otu29063 | 1 | k__Fungi(100);p__Basidiomycota(100);c__Microbotryomycetes(100);o__Sporidiobolales(100);f__Sporidiobolales_family_Incertae_sedis(100);g__Sporobolomyces(100);s__Sporobolomyces_coprosmicola(100); | Sporobolomyces | 13 | Pathotroph-Saprotroph | \|Fungal Parasite\|-Plant Saprotroph | Yeast | NULL | Probable | As litter_saprotroph (Põlme et al. 2020); As mycoparasite (Põlme et al. 2020); Decay fungal material (Põlme et al. 2020) | Tedersoo L, et al. 2014. Science 346:e1256688 ((https://doi.org/10.1126/science.1256688)); Purahong W, et al. 2016. Molecular Ecology 25:4059-4074 ((https://doi.org/10.1111/mec.13739)); Põlme S, et al. 2020. Fungal Diversity 105:1-16 ((https://doi.org/10.1007/s13225-020-00466-2)) |
| Otu29065 | 1 | k__Fungi(100);p__Basidiomycota(100);c__Microbotryomycetes(100);o__Sporidiobolales(100);f__Sporidiobolales_family_Incertae_sedis(100);g__Sporobolomyces(100);g__Sporobolomyces_unclassified(100); | Sporobolomyces | 13 | Pathotroph-Saprotroph | \|Fungal Parasite\|-Plant Saprotroph | Yeast | NULL | Probable | As litter_saprotroph (Põlme et al. 2020); As mycoparasite (Põlme et al. 2020); Decay fungal material (Põlme et al. 2020) | Tedersoo L, et al. 2014. Science 346:e1256688 ((https://doi.org/10.1126/science.1256688)); Purahong W, et al. 2016. Molecular Ecology 25:4059-4074 ((https://doi.org/10.1111/mec.13739)); Põlme S, et al. 2020. Fungal Diversity 105:1-16 ((https://doi.org/10.1007/s13225-020-00466-2)) |
| Otu29069 | 1 | k__Fungi(100);p__Basidiomycota(100);c__Microbotryomycetes(100);o__Sporidiobolales(100);f__Sporidiobolales_family_Incertae_sedis(100);g__Sporobolomyces(100);g__Sporobolomyces_unclassified(100); | Sporobolomyces | 13 | Pathotroph-Saprotroph | \|Fungal Parasite\|-Plant Saprotroph | Yeast | NULL | Probable | As litter_saprotroph (Põlme et al. 2020); As mycoparasite (Põlme et al. 2020); Decay fungal material (Põlme et al. 2020) | Tedersoo L, et al. 2014. Science 346:e1256688 ((https://doi.org/10.1126/science.1256688)); Purahong W, et al. 2016. Molecular Ecology 25:4059-4074 ((https://doi.org/10.1111/mec.13739)); Põlme S, et al. 2020. Fungal Diversity 105:1-16 ((https://doi.org/10.1007/s13225-020-00466-2)) |
| Otu29070 | 1 | k__Fungi(100);p__Basidiomycota(100);c__Microbotryomycetes(100);o__Sporidiobolales(100);f__Sporidiobolales_family_Incertae_sedis(100);g__Sporobolomyces(100);s__Sporobolomyces_ruberrimus(100); | Sporobolomyces | 13 | Pathotroph-Saprotroph | \|Fungal Parasite\|-Plant Saprotroph | Yeast | NULL | Probable | As litter_saprotroph (Põlme et al. 2020); As mycoparasite (Põlme et al. 2020); Decay fungal material (Põlme et al. 2020) | Tedersoo L, et al. 2014. Science 346:e1256688 ((https://doi.org/10.1126/science.1256688)); Purahong W, et al. 2016. Molecular Ecology 25:4059-4074 ((https://doi.org/10.1111/mec.13739)); Põlme S, et al. 2020. Fungal Diversity 105:1-16 ((https://doi.org/10.1007/s13225-020-00466-2)) |
| Otu29073 | 1 | k__Fungi(100);p__Ascomycota(100);c__Sordariomycetes(100);o__Hypocreales(100);o__Hypocreales_unclassified(100);o__Hypocreales_unclassified(100);o__Hypocreales_unclassified(100); | Hypocreales | 7 | Saprotroph | Undefined Saprotroph | Microfungus | NULL | Possible | NULL | Sterkenburg E, et al. 2015. New Phytologist 207:1145-1158 ((https://doi.org/10.1111/nph.13426)) |
| Otu29074 | 1 | k__Fungi(100);p__Ascomycota(100);c__Sordariomycetes(100);o__Hypocreales(100);o__Hypocreales_unclassified(100);o__Hypocreales_unclassified(100);o__Hypocreales_unclassified(100); | Hypocreales | 7 | Saprotroph | Undefined Saprotroph | Microfungus | NULL | Possible | NULL | Sterkenburg E, et al. 2015. New Phytologist 207:1145-1158 ((https://doi.org/10.1111/nph.13426)) |
| Otu29082 | 1 | k__Fungi(100);p__Ascomycota(100);c__Sordariomycetes(100);o__Hypocreales(100);o__Hypocreales_unclassified(100);o__Hypocreales_unclassified(100);o__Hypocreales_unclassified(100); | Hypocreales | 7 | Saprotroph | Undefined Saprotroph | Microfungus | NULL | Possible | NULL | Sterkenburg E, et al. 2015. New Phytologist 207:1145-1158 ((https://doi.org/10.1111/nph.13426)) |
| Otu29084 | 1 | k__Fungi(100);p__Ascomycota(100);c__Sordariomycetes(100);o__Hypocreales(100);o__Hypocreales_unclassified(100);o__Hypocreales_unclassified(100);o__Hypocreales_unclassified(100); | Hypocreales | 7 | Saprotroph | Undefined Saprotroph | Microfungus | NULL | Possible | NULL | Sterkenburg E, et al. 2015. New Phytologist 207:1145-1158 ((https://doi.org/10.1111/nph.13426)) |
| Otu29096 | 1 | k__Fungi(100);p__Ascomycota(100);c__Dothideomycetes(100);o__Dothideales(100);f__Dothideaceae(100);g__Endoconidioma(100);s__Endoconidioma_populi(100); | Endoconidioma | 13 | Saprotroph | Undefined Saprotroph-\|Wood Saprotroph\| | Microfungus | NULL | Probable | Decay: wood (Põlme et al. 2020) | Tedersoo L, et al. 2014. Science 346:e1256688 ((https://doi.org/10.1126/science.1256688)); Põlme S, et al. 2020. Fungal Diversity 105:1-16 ((https://doi.org/10.1007/s13225-020-00466-2)) |
| Otu29098 | 1 | k__Fungi(100);p__Ascomycota(100);c__Dothideomycetes(100);o__Dothideales(100);f__Dothioraceae(100);g__Sydowia(100);s__Sydowia_sp(100); | Sydowia | 13 | Pathotroph-Saprotroph | Plant Pathogen-Undefined Saprotroph-\|Wood Saprotroph\| | Microfungus | NULL | Probable | Decay: wood, leaf, fruit, seed (Põlme et al. 2020) | Tedersoo L, et al. 2014. Science 346:e1256688 ((https://doi.org/10.1126/science.1256688)); Põlme S, et al. 2020. Fungal Diversity 105:1-16 ((https://doi.org/10.1007/s13225-020-00466-2)) |
| Otu29100 | 1 | k__Fungi(100);p__Ascomycota(100);c__Sordariomycetes(100);o__Hypocreales(100);o__Hypocreales_unclassified(100);o__Hypocreales_unclassified(100);o__Hypocreales_unclassified(100); | Hypocreales | 7 | Saprotroph | Undefined Saprotroph | Microfungus | NULL | Possible | NULL | Sterkenburg E, et al. 2015. New Phytologist 207:1145-1158 ((https://doi.org/10.1111/nph.13426)) |
| Otu29102 | 1 | k__Fungi(100);p__Ascomycota(100);c__Sordariomycetes(100);o__Hypocreales(100);o__Hypocreales_unclassified(100);o__Hypocreales_unclassified(100);o__Hypocreales_unclassified(100); | Hypocreales | 7 | Saprotroph | Undefined Saprotroph | Microfungus | NULL | Possible | NULL | Sterkenburg E, et al. 2015. New Phytologist 207:1145-1158 ((https://doi.org/10.1111/nph.13426)) |
| Otu29105 | 1 | k__Fungi(100);p__Ascomycota(100);c__Sordariomycetes(100);o__Hypocreales(100);o__Hypocreales_unclassified(100);o__Hypocreales_unclassified(100);o__Hypocreales_unclassified(100); | Hypocreales | 7 | Saprotroph | Undefined Saprotroph | Microfungus | NULL | Possible | NULL | Sterkenburg E, et al. 2015. New Phytologist 207:1145-1158 ((https://doi.org/10.1111/nph.13426)) |
| Otu29106 | 1 | k__Fungi(100);p__Ascomycota(100);c__Sordariomycetes(100);o__Hypocreales(100);o__Hypocreales_unclassified(100);o__Hypocreales_unclassified(100);o__Hypocreales_unclassified(100); | Hypocreales | 7 | Saprotroph | Undefined Saprotroph | Microfungus | NULL | Possible | NULL | Sterkenburg E, et al. 2015. New Phytologist 207:1145-1158 ((https://doi.org/10.1111/nph.13426)) |
| Otu29108 | 1 | k__Fungi(100);p__Ascomycota(100);c__Sordariomycetes(100);o__Hypocreales(100);o__Hypocreales_unclassified(100);o__Hypocreales_unclassified(100);o__Hypocreales_unclassified(100); | Hypocreales | 7 | Saprotroph | Undefined Saprotroph | Microfungus | NULL | Possible | NULL | Sterkenburg E, et al. 2015. New Phytologist 207:1145-1158 ((https://doi.org/10.1111/nph.13426)) |
| Otu29109 | 1 | k__Fungi(100);p__Ascomycota(100);c__Sordariomycetes(100);o__Hypocreales(100);o__Hypocreales_unclassified(100);o__Hypocreales_unclassified(100);o__Hypocreales_unclassified(100); | Hypocreales | 7 | Saprotroph | Undefined Saprotroph | Microfungus | NULL | Possible | NULL | Sterkenburg E, et al. 2015. New Phytologist 207:1145-1158 ((https://doi.org/10.1111/nph.13426)) |
| Otu29110 | 1 | k__Fungi(100);p__Ascomycota(100);c__Sordariomycetes(100);o__Hypocreales(100);o__Hypocreales_unclassified(100);o__Hypocreales_unclassified(100);o__Hypocreales_unclassified(100); | Hypocreales | 7 | Saprotroph | Undefined Saprotroph | Microfungus | NULL | Possible | NULL | Sterkenburg E, et al. 2015. New Phytologist 207:1145-1158 ((https://doi.org/10.1111/nph.13426)) |
| Otu29112 | 1 | k__Fungi(100);p__Ascomycota(100);c__Sordariomycetes(100);o__Hypocreales(100);o__Hypocreales_unclassified(100);o__Hypocreales_unclassified(100);o__Hypocreales_unclassified(100); | Hypocreales | 7 | Saprotroph | Undefined Saprotroph | Microfungus | NULL | Possible | NULL | Sterkenburg E, et al. 2015. New Phytologist 207:1145-1158 ((https://doi.org/10.1111/nph.13426)) |
| Otu29118 | 1 | k__Fungi(100);p__Ascomycota(100);c__Sordariomycetes(100);o__Hypocreales(100);o__Hypocreales_unclassified(100);o__Hypocreales_unclassified(100);o__Hypocreales_unclassified(100); | Hypocreales | 7 | Saprotroph | Undefined Saprotroph | Microfungus | NULL | Possible | NULL | Sterkenburg E, et al. 2015. New Phytologist 207:1145-1158 ((https://doi.org/10.1111/nph.13426)) |
| Otu29119 | 1 | k__Fungi(100);p__Ascomycota(100);c__Sordariomycetes(100);o__Hypocreales(100);o__Hypocreales_unclassified(100);o__Hypocreales_unclassified(100);o__Hypocreales_unclassified(100); | Hypocreales | 7 | Saprotroph | Undefined Saprotroph | Microfungus | NULL | Possible | NULL | Sterkenburg E, et al. 2015. New Phytologist 207:1145-1158 ((https://doi.org/10.1111/nph.13426)) |
| Otu29122 | 1 | k__Fungi(100);p__Ascomycota(100);c__Sordariomycetes(100);o__Hypocreales(100);o__Hypocreales_unclassified(100);o__Hypocreales_unclassified(100);o__Hypocreales_unclassified(100); | Hypocreales | 7 | Saprotroph | Undefined Saprotroph | Microfungus | NULL | Possible | NULL | Sterkenburg E, et al. 2015. New Phytologist 207:1145-1158 ((https://doi.org/10.1111/nph.13426)) |
| Otu29124 | 1 | k__Fungi(100);p__Ascomycota(100);c__Sordariomycetes(100);o__Hypocreales(100);o__Hypocreales_unclassified(100);o__Hypocreales_unclassified(100);o__Hypocreales_unclassified(100); | Hypocreales | 7 | Saprotroph | Undefined Saprotroph | Microfungus | NULL | Possible | NULL | Sterkenburg E, et al. 2015. New Phytologist 207:1145-1158 ((https://doi.org/10.1111/nph.13426)) |
| Otu29126 | 1 | k__Fungi(100);p__Ascomycota(100);c__Saccharomycetes(100);o__Saccharomycetales(100);f__Saccharomycetaceae(100);g__Saccharomyces(100);s__Saccharomyces_kudriavzevii(100); | Saccharomyces | 13 | Saprotroph | \|Undefined Saprotroph\| | Yeast | NULL | Probable | As nectar-tap_saprotroph (Põlme et al. 2020); Decay sugar-rich substrates (Põlme et al. 2020) | James TY, et al. 2006. Nature 443:818-822 ((https://doi.org/10.1038/nature05110)); Põlme S, et al. 2020. Fungal Diversity 105:1-16 ((https://doi.org/10.1007/s13225-020-00466-2)) |
| Otu29127 | 1 | k__Fungi(100);p__Ascomycota(100);c__Saccharomycetes(100);o__Saccharomycetales(100);f__Saccharomycetaceae(100);g__Saccharomyces(100);s__Saccharomyces_kudriavzevii(100); | Saccharomyces | 13 | Saprotroph | \|Undefined Saprotroph\| | Yeast | NULL | Probable | As nectar-tap_saprotroph (Põlme et al. 2020); Decay sugar-rich substrates (Põlme et al. 2020) | James TY, et al. 2006. Nature 443:818-822 ((https://doi.org/10.1038/nature05110)); Põlme S, et al. 2020. Fungal Diversity 105:1-16 ((https://doi.org/10.1007/s13225-020-00466-2)) |
| Otu29128 | 1 | k__Fungi(100);p__Ascomycota(100);c__Saccharomycetes(100);o__Saccharomycetales(100);f__Saccharomycetaceae(100);g__Saccharomyces(100);g__Saccharomyces_unclassified(100); | Saccharomyces | 13 | Saprotroph | \|Undefined Saprotroph\| | Yeast | NULL | Probable | As nectar-tap_saprotroph (Põlme et al. 2020); Decay sugar-rich substrates (Põlme et al. 2020) | James TY, et al. 2006. Nature 443:818-822 ((https://doi.org/10.1038/nature05110)); Põlme S, et al. 2020. Fungal Diversity 105:1-16 ((https://doi.org/10.1007/s13225-020-00466-2)) |
| Otu29129 | 1 | k__Fungi(100);p__Ascomycota(100);c__Saccharomycetes(100);o__Saccharomycetales(100);f__Saccharomycetaceae(100);g__Saccharomyces(100);g__Saccharomyces_unclassified(100); | Saccharomyces | 13 | Saprotroph | \|Undefined Saprotroph\| | Yeast | NULL | Probable | As nectar-tap_saprotroph (Põlme et al. 2020); Decay sugar-rich substrates (Põlme et al. 2020) | James TY, et al. 2006. Nature 443:818-822 ((https://doi.org/10.1038/nature05110)); Põlme S, et al. 2020. Fungal Diversity 105:1-16 ((https://doi.org/10.1007/s13225-020-00466-2)) |
| Otu29130 | 1 | k__Fungi(100);p__Ascomycota(100);c__Saccharomycetes(100);o__Saccharomycetales(100);f__Saccharomycetaceae(100);g__Saccharomyces(100);s__Saccharomyces_kudriavzevii(100); | Saccharomyces | 13 | Saprotroph | \|Undefined Saprotroph\| | Yeast | NULL | Probable | As nectar-tap_saprotroph (Põlme et al. 2020); Decay sugar-rich substrates (Põlme et al. 2020) | James TY, et al. 2006. Nature 443:818-822 ((https://doi.org/10.1038/nature05110)); Põlme S, et al. 2020. Fungal Diversity 105:1-16 ((https://doi.org/10.1007/s13225-020-00466-2)) |
| Otu29131 | 1 | k__Fungi(100);p__Ascomycota(100);c__Saccharomycetes(100);o__Saccharomycetales(100);f__Saccharomycetaceae(100);g__Saccharomyces(100);g__Saccharomyces_unclassified(100); | Saccharomyces | 13 | Saprotroph | \|Undefined Saprotroph\| | Yeast | NULL | Probable | As nectar-tap_saprotroph (Põlme et al. 2020); Decay sugar-rich substrates (Põlme et al. 2020) | James TY, et al. 2006. Nature 443:818-822 ((https://doi.org/10.1038/nature05110)); Põlme S, et al. 2020. Fungal Diversity 105:1-16 ((https://doi.org/10.1007/s13225-020-00466-2)) |
| Otu29132 | 1 | k__Fungi(100);p__Ascomycota(100);c__Saccharomycetes(100);o__Saccharomycetales(100);f__Saccharomycetaceae(100);g__Saccharomyces(100);g__Saccharomyces_unclassified(100); | Saccharomyces | 13 | Saprotroph | \|Undefined Saprotroph\| | Yeast | NULL | Probable | As nectar-tap_saprotroph (Põlme et al. 2020); Decay sugar-rich substrates (Põlme et al. 2020) | James TY, et al. 2006. Nature 443:818-822 ((https://doi.org/10.1038/nature05110)); Põlme S, et al. 2020. Fungal Diversity 105:1-16 ((https://doi.org/10.1007/s13225-020-00466-2)) |
| Otu29133 | 1 | k__Fungi(100);p__Ascomycota(100);c__Saccharomycetes(100);o__Saccharomycetales(100);f__Saccharomycetaceae(100);g__Saccharomyces(100);s__Saccharomyces_kudriavzevii(100); | Saccharomyces | 13 | Saprotroph | \|Undefined Saprotroph\| | Yeast | NULL | Probable | As nectar-tap_saprotroph (Põlme et al. 2020); Decay sugar-rich substrates (Põlme et al. 2020) | James TY, et al. 2006. Nature 443:818-822 ((https://doi.org/10.1038/nature05110)); Põlme S, et al. 2020. Fungal Diversity 105:1-16 ((https://doi.org/10.1007/s13225-020-00466-2)) |
| Otu29134 | 1 | k__Fungi(100);p__Ascomycota(100);c__Saccharomycetes(100);o__Saccharomycetales(100);f__Saccharomycetaceae(100);g__Saccharomyces(100);s__Saccharomyces_cerevisiae(100); | Saccharomyces | 13 | Saprotroph | \|Undefined Saprotroph\| | Yeast | NULL | Probable | As nectar-tap_saprotroph (Põlme et al. 2020); Decay sugar-rich substrates (Põlme et al. 2020) | James TY, et al. 2006. Nature 443:818-822 ((https://doi.org/10.1038/nature05110)); Põlme S, et al. 2020. Fungal Diversity 105:1-16 ((https://doi.org/10.1007/s13225-020-00466-2)) |
| Otu29135 | 1 | k__Fungi(100);p__Ascomycota(100);c__Saccharomycetes(100);o__Saccharomycetales(100);f__Saccharomycetaceae(100);g__Saccharomyces(100);g__Saccharomyces_unclassified(100); | Saccharomyces | 13 | Saprotroph | \|Undefined Saprotroph\| | Yeast | NULL | Probable | As nectar-tap_saprotroph (Põlme et al. 2020); Decay sugar-rich substrates (Põlme et al. 2020) | James TY, et al. 2006. Nature 443:818-822 ((https://doi.org/10.1038/nature05110)); Põlme S, et al. 2020. Fungal Diversity 105:1-16 ((https://doi.org/10.1007/s13225-020-00466-2)) |
| Otu29136 | 1 | k__Fungi(100);p__Ascomycota(100);c__Saccharomycetes(100);o__Saccharomycetales(100);f__Saccharomycetaceae(100);g__Saccharomyces(100);s__Saccharomyces_kudriavzevii(100); | Saccharomyces | 13 | Saprotroph | \|Undefined Saprotroph\| | Yeast | NULL | Probable | As nectar-tap_saprotroph (Põlme et al. 2020); Decay sugar-rich substrates (Põlme et al. 2020) | James TY, et al. 2006. Nature 443:818-822 ((https://doi.org/10.1038/nature05110)); Põlme S, et al. 2020. Fungal Diversity 105:1-16 ((https://doi.org/10.1007/s13225-020-00466-2)) |
| Otu29137 | 1 | k__Fungi(100);p__Ascomycota(100);c__Saccharomycetes(100);o__Saccharomycetales(100);f__Saccharomycetaceae(100);g__Saccharomyces(100);s__Saccharomyces_kudriavzevii(100); | Saccharomyces | 13 | Saprotroph | \|Undefined Saprotroph\| | Yeast | NULL | Probable | As nectar-tap_saprotroph (Põlme et al. 2020); Decay sugar-rich substrates (Põlme et al. 2020) | James TY, et al. 2006. Nature 443:818-822 ((https://doi.org/10.1038/nature05110)); Põlme S, et al. 2020. Fungal Diversity 105:1-16 ((https://doi.org/10.1007/s13225-020-00466-2)) |
| Otu29138 | 1 | k__Fungi(100);p__Ascomycota(100);c__Saccharomycetes(100);o__Saccharomycetales(100);f__Saccharomycetaceae(100);g__Saccharomyces(100);g__Saccharomyces_unclassified(100); | Saccharomyces | 13 | Saprotroph | \|Undefined Saprotroph\| | Yeast | NULL | Probable | As nectar-tap_saprotroph (Põlme et al. 2020); Decay sugar-rich substrates (Põlme et al. 2020) | James TY, et al. 2006. Nature 443:818-822 ((https://doi.org/10.1038/nature05110)); Põlme S, et al. 2020. Fungal Diversity 105:1-16 ((https://doi.org/10.1007/s13225-020-00466-2)) |
| Otu29139 | 1 | k__Fungi(100);p__Ascomycota(100);c__Saccharomycetes(100);o__Saccharomycetales(100);f__Saccharomycetaceae(100);g__Saccharomyces(100);g__Saccharomyces_unclassified(100); | Saccharomyces | 13 | Saprotroph | \|Undefined Saprotroph\| | Yeast | NULL | Probable | As nectar-tap_saprotroph (Põlme et al. 2020); Decay sugar-rich substrates (Põlme et al. 2020) | James TY, et al. 2006. Nature 443:818-822 ((https://doi.org/10.1038/nature05110)); Põlme S, et al. 2020. Fungal Diversity 105:1-16 ((https://doi.org/10.1007/s13225-020-00466-2)) |
| Otu29140 | 1 | k__Fungi(100);p__Ascomycota(100);c__Saccharomycetes(100);o__Saccharomycetales(100);f__Saccharomycetaceae(100);g__Saccharomyces(100);g__Saccharomyces_unclassified(100); | Saccharomyces | 13 | Saprotroph | \|Undefined Saprotroph\| | Yeast | NULL | Probable | As nectar-tap_saprotroph (Põlme et al. 2020); Decay sugar-rich substrates (Põlme et al. 2020) | James TY, et al. 2006. Nature 443:818-822 ((https://doi.org/10.1038/nature05110)); Põlme S, et al. 2020. Fungal Diversity 105:1-16 ((https://doi.org/10.1007/s13225-020-00466-2)) |
| Otu29141 | 1 | k__Fungi(100);p__Ascomycota(100);c__Saccharomycetes(100);o__Saccharomycetales(100);f__Saccharomycetaceae(100);g__Saccharomyces(100);s__Saccharomyces_kudriavzevii(100); | Saccharomyces | 13 | Saprotroph | \|Undefined Saprotroph\| | Yeast | NULL | Probable | As nectar-tap_saprotroph (Põlme et al. 2020); Decay sugar-rich substrates (Põlme et al. 2020) | James TY, et al. 2006. Nature 443:818-822 ((https://doi.org/10.1038/nature05110)); Põlme S, et al. 2020. Fungal Diversity 105:1-16 ((https://doi.org/10.1007/s13225-020-00466-2)) |
| Otu29142 | 1 | k__Fungi(100);p__Ascomycota(100);c__Saccharomycetes(100);o__Saccharomycetales(100);f__Saccharomycetaceae(100);g__Saccharomyces(100);s__Saccharomyces_kudriavzevii(100); | Saccharomyces | 13 | Saprotroph | \|Undefined Saprotroph\| | Yeast | NULL | Probable | As nectar-tap_saprotroph (Põlme et al. 2020); Decay sugar-rich substrates (Põlme et al. 2020) | James TY, et al. 2006. Nature 443:818-822 ((https://doi.org/10.1038/nature05110)); Põlme S, et al. 2020. Fungal Diversity 105:1-16 ((https://doi.org/10.1007/s13225-020-00466-2)) |
| Otu29143 | 1 | k__Fungi(100);p__Ascomycota(100);c__Saccharomycetes(100);o__Saccharomycetales(100);f__Saccharomycetaceae(100);g__Torulaspora(100);s__Torulaspora_delbrueckii(100); | Torulaspora | 13 | Pathotroph-Saprotroph | Animal Parasite-\|Undefined Saprotroph\| | Yeast | NULL | Probable | As nectar-tap_saprotroph (Põlme et al. 2020); Decay animal material, sugar-rich substrates (Põlme et al. 2020) | Põlme S, et al. 2020. Fungal Diversity 105:1-16 ((https://doi.org/10.1007/s13225-020-00466-2)) |
| Otu29144 | 1 | k__Fungi(100);p__Ascomycota(100);c__Saccharomycetes(100);o__Saccharomycetales(100);f__Saccharomycetaceae(100);g__Saccharomyces(100);s__Saccharomyces_kudriavzevii(100); | Saccharomyces | 13 | Saprotroph | \|Undefined Saprotroph\| | Yeast | NULL | Probable | As nectar-tap_saprotroph (Põlme et al. 2020); Decay sugar-rich substrates (Põlme et al. 2020) | James TY, et al. 2006. Nature 443:818-822 ((https://doi.org/10.1038/nature05110)); Põlme S, et al. 2020. Fungal Diversity 105:1-16 ((https://doi.org/10.1007/s13225-020-00466-2)) |
| Otu29145 | 1 | k__Fungi(100);p__Ascomycota(100);c__Saccharomycetes(100);o__Saccharomycetales(100);f__Saccharomycetaceae(100);g__Torulaspora(100);s__Torulaspora_delbrueckii(100); | Torulaspora | 13 | Pathotroph-Saprotroph | Animal Parasite-\|Undefined Saprotroph\| | Yeast | NULL | Probable | As nectar-tap_saprotroph (Põlme et al. 2020); Decay animal material, sugar-rich substrates (Põlme et al. 2020) | Põlme S, et al. 2020. Fungal Diversity 105:1-16 ((https://doi.org/10.1007/s13225-020-00466-2)) |
| Otu29146 | 1 | k__Fungi(100);p__Ascomycota(100);c__Saccharomycetes(100);o__Saccharomycetales(100);f__Saccharomycetaceae(100);g__Saccharomyces(100);s__Saccharomyces_kudriavzevii(100); | Saccharomyces | 13 | Saprotroph | \|Undefined Saprotroph\| | Yeast | NULL | Probable | As nectar-tap_saprotroph (Põlme et al. 2020); Decay sugar-rich substrates (Põlme et al. 2020) | James TY, et al. 2006. Nature 443:818-822 ((https://doi.org/10.1038/nature05110)); Põlme S, et al. 2020. Fungal Diversity 105:1-16 ((https://doi.org/10.1007/s13225-020-00466-2)) |
| Otu29147 | 1 | k__Fungi(100);p__Ascomycota(100);c__Lecanoromycetes(100);o__Lecanorales(100);f__Ramalinaceae(100);f__Ramalinaceae_unclassified(100);f__Ramalinaceae_unclassified(100); | Ramalinaceae | 9 | Symbiotroph | Lichenized | Thallus | NULL | Probable | Lichenized with green algae, on rocks , branches etc., often forming extensive carpets of thalli; often prominent in maritime communities (Cannon & Kirk 2007) | Cannon PF, Kirk PM. 2007. Fungal Families of the World. CAB International, Cambridge (ISBN: 978-0851998275) |
| Otu29148 | 1 | k__Fungi(100);p__Ascomycota(100);c__Saccharomycetes(100);o__Saccharomycetales(100);f__Saccharomycetaceae(100);g__Saccharomyces(100);s__Saccharomyces_kudriavzevii(100); | Saccharomyces | 13 | Saprotroph | \|Undefined Saprotroph\| | Yeast | NULL | Probable | As nectar-tap_saprotroph (Põlme et al. 2020); Decay sugar-rich substrates (Põlme et al. 2020) | James TY, et al. 2006. Nature 443:818-822 ((https://doi.org/10.1038/nature05110)); Põlme S, et al. 2020. Fungal Diversity 105:1-16 ((https://doi.org/10.1007/s13225-020-00466-2)) |
| Otu29149 | 1 | k__Fungi(100);p__Ascomycota(100);c__Saccharomycetes(100);o__Saccharomycetales(100);f__Saccharomycetaceae(100);g__Saccharomyces(100);s__Saccharomyces_kudriavzevii(100); | Saccharomyces | 13 | Saprotroph | \|Undefined Saprotroph\| | Yeast | NULL | Probable | As nectar-tap_saprotroph (Põlme et al. 2020); Decay sugar-rich substrates (Põlme et al. 2020) | James TY, et al. 2006. Nature 443:818-822 ((https://doi.org/10.1038/nature05110)); Põlme S, et al. 2020. Fungal Diversity 105:1-16 ((https://doi.org/10.1007/s13225-020-00466-2)) |
| Otu29150 | 1 | k__Fungi(100);p__Ascomycota(100);c__Saccharomycetes(100);o__Saccharomycetales(100);f__Saccharomycetaceae(100);g__Torulaspora(100);s__Torulaspora_pretoriensis(100); | Torulaspora | 13 | Pathotroph-Saprotroph | Animal Parasite-\|Undefined Saprotroph\| | Yeast | NULL | Probable | As nectar-tap_saprotroph (Põlme et al. 2020); Decay animal material, sugar-rich substrates (Põlme et al. 2020) | Põlme S, et al. 2020. Fungal Diversity 105:1-16 ((https://doi.org/10.1007/s13225-020-00466-2)) |
| Otu29151 | 1 | k__Fungi(100);p__Ascomycota(100);c__Saccharomycetes(100);o__Saccharomycetales(100);f__Saccharomycetaceae(100);g__Saccharomyces(100);g__Saccharomyces_unclassified(100); | Saccharomyces | 13 | Saprotroph | \|Undefined Saprotroph\| | Yeast | NULL | Probable | As nectar-tap_saprotroph (Põlme et al. 2020); Decay sugar-rich substrates (Põlme et al. 2020) | James TY, et al. 2006. Nature 443:818-822 ((https://doi.org/10.1038/nature05110)); Põlme S, et al. 2020. Fungal Diversity 105:1-16 ((https://doi.org/10.1007/s13225-020-00466-2)) |
| Otu29152 | 1 | k__Fungi(100);p__Ascomycota(100);c__Saccharomycetes(100);o__Saccharomycetales(100);f__Saccharomycetaceae(100);g__Saccharomyces(100);s__Saccharomyces_kudriavzevii(100); | Saccharomyces | 13 | Saprotroph | \|Undefined Saprotroph\| | Yeast | NULL | Probable | As nectar-tap_saprotroph (Põlme et al. 2020); Decay sugar-rich substrates (Põlme et al. 2020) | James TY, et al. 2006. Nature 443:818-822 ((https://doi.org/10.1038/nature05110)); Põlme S, et al. 2020. Fungal Diversity 105:1-16 ((https://doi.org/10.1007/s13225-020-00466-2)) |
| Otu29153 | 1 | k__Fungi(100);p__Ascomycota(100);c__Saccharomycetes(100);o__Saccharomycetales(100);f__Saccharomycetaceae(100);g__Torulaspora(100);s__Torulaspora_delbrueckii(100); | Torulaspora | 13 | Pathotroph-Saprotroph | Animal Parasite-\|Undefined Saprotroph\| | Yeast | NULL | Probable | As nectar-tap_saprotroph (Põlme et al. 2020); Decay animal material, sugar-rich substrates (Põlme et al. 2020) | Põlme S, et al. 2020. Fungal Diversity 105:1-16 ((https://doi.org/10.1007/s13225-020-00466-2)) |
| Otu29155 | 1 | k__Fungi(100);p__Ascomycota(100);c__Dothideomycetes(100);o__Dothideales(100);f__Dothideaceae(100);g__Endoconidioma(100);s__Endoconidioma_populi(100); | Endoconidioma | 13 | Saprotroph | Undefined Saprotroph-\|Wood Saprotroph\| | Microfungus | NULL | Probable | Decay: wood (Põlme et al. 2020) | Tedersoo L, et al. 2014. Science 346:e1256688 ((https://doi.org/10.1126/science.1256688)); Põlme S, et al. 2020. Fungal Diversity 105:1-16 ((https://doi.org/10.1007/s13225-020-00466-2)) |
| Otu29156 | 1 | k__Fungi(100);p__Ascomycota(100);c__Dothideomycetes(100);o__Myriangiales(100);f__Elsinoaceae(100);g__Sphaceloma(100);s__Sphaceloma_erythrinae(100); | Sphaceloma | 13 | Pathotroph | \|Plant Pathogen\| | Microfungus | NULL | Probable | NULL | Tedersoo L, et al. 2014. Science 346:e1256688 ((https://doi.org/10.1126/science.1256688)); Põlme S, et al. 2020. Fungal Diversity 105:1-16 ((https://doi.org/10.1007/s13225-020-00466-2)) |
| Otu29157 | 1 | k__Fungi(100);p__Ascomycota(100);c__Dothideomycetes(100);o__Myriangiales(100);f__Elsinoaceae(100);g__Elsinoe(100);g__Elsinoe_unclassified(100); | Elsinoe | 13 | Pathotroph-Saprotroph | \|Plant Pathogen\|-Plant Saprotroph | Microfungus | NULL | Probable | As litter_saprotroph (Põlme et al. 2020); Decay leaf, fruit, seed (Põlme et al. 2020) | Tedersoo L, et al. 2014. Science 346:e1256688 ((https://doi.org/10.1126/science.1256688)); Põlme S, et al. 2020. Fungal Diversity 105:1-16 ((https://doi.org/10.1007/s13225-020-00466-2)) |
| Otu29158 | 1 | k__Fungi(100);p__Ascomycota(100);c__Dothideomycetes(100);o__Myriangiales(100);f__Elsinoaceae(100);g__Elsinoe(100);g__Elsinoe_unclassified(100); | Elsinoe | 13 | Pathotroph-Saprotroph | \|Plant Pathogen\|-Plant Saprotroph | Microfungus | NULL | Probable | As litter_saprotroph (Põlme et al. 2020); Decay leaf, fruit, seed (Põlme et al. 2020) | Tedersoo L, et al. 2014. Science 346:e1256688 ((https://doi.org/10.1126/science.1256688)); Põlme S, et al. 2020. Fungal Diversity 105:1-16 ((https://doi.org/10.1007/s13225-020-00466-2)) |
| Otu29159 | 1 | k__Fungi(100);p__Ascomycota(100);c__Dothideomycetes(100);o__Myriangiales(100);f__Elsinoaceae(100);g__unclassified_Elsinoaceae(100);s__Elsinoaceae_sp(100); | Elsinoaceae | 9 | Pathotroph | Plant Pathogen | Microfungus | NULL | Probable | Biotrophic or necrotrophic on leaves, stems and fruits of a wide range of plants (Cannon & Kirk 2007) | Cannon PF, Kirk PM. 2007. Fungal Families of the World. CAB International, Cambridge (ISBN: 978-0851998275) |
| Otu29161 | 1 | k__Fungi(100);p__Ascomycota(100);c__Dothideomycetes(100);o__Dothideales(100);f__Dothideaceae(100);g__Endoconidioma(100);s__Endoconidioma_populi(100); | Endoconidioma | 13 | Saprotroph | Undefined Saprotroph-\|Wood Saprotroph\| | Microfungus | NULL | Probable | Decay: wood (Põlme et al. 2020) | Tedersoo L, et al. 2014. Science 346:e1256688 ((https://doi.org/10.1126/science.1256688)); Põlme S, et al. 2020. Fungal Diversity 105:1-16 ((https://doi.org/10.1007/s13225-020-00466-2)) |
| Otu29164 | 1 | k__Fungi(100);p__Ascomycota(100);c__Dothideomycetes(100);o__Dothideomycetes_order_Incertae_sedis(100);f__Dothideomycetes_family_Incertae_sedis(100);g__Leptospora(100);s__Leptospora_rubella(100); | Leptospora | 13 | Saprotroph | \|Plant Saprotroph\| | Microfungus | NULL | Probable | As litter_saprotroph (Põlme et al. 2020); Decay leaf, fruit, seed (Põlme et al. 2020) | Põlme S, et al. 2020. Fungal Diversity 105:1-16 ((https://doi.org/10.1007/s13225-020-00466-2)) |
| Otu29165 | 1 | k__Fungi(100);p__Ascomycota(100);c__Sordariomycetes(100);o__Hypocreales(100);o__Hypocreales_unclassified(100);o__Hypocreales_unclassified(100);o__Hypocreales_unclassified(100); | Hypocreales | 7 | Saprotroph | Undefined Saprotroph | Microfungus | NULL | Possible | NULL | Sterkenburg E, et al. 2015. New Phytologist 207:1145-1158 ((https://doi.org/10.1111/nph.13426)) |
| Otu29168 | 1 | k__Fungi(100);p__Ascomycota(100);c__Saccharomycetes(100);o__Saccharomycetales(100);f__Saccharomycetaceae(100);g__Saccharomyces(100);g__Saccharomyces_unclassified(100); | Saccharomyces | 13 | Saprotroph | \|Undefined Saprotroph\| | Yeast | NULL | Probable | As nectar-tap_saprotroph (Põlme et al. 2020); Decay sugar-rich substrates (Põlme et al. 2020) | James TY, et al. 2006. Nature 443:818-822 ((https://doi.org/10.1038/nature05110)); Põlme S, et al. 2020. Fungal Diversity 105:1-16 ((https://doi.org/10.1007/s13225-020-00466-2)) |
| Otu29169 | 1 | k__Fungi(100);p__Ascomycota(100);c__Saccharomycetes(100);o__Saccharomycetales(100);f__Saccharomycetaceae(100);g__Saccharomyces(100);s__Saccharomyces_kudriavzevii(100); | Saccharomyces | 13 | Saprotroph | \|Undefined Saprotroph\| | Yeast | NULL | Probable | As nectar-tap_saprotroph (Põlme et al. 2020); Decay sugar-rich substrates (Põlme et al. 2020) | James TY, et al. 2006. Nature 443:818-822 ((https://doi.org/10.1038/nature05110)); Põlme S, et al. 2020. Fungal Diversity 105:1-16 ((https://doi.org/10.1007/s13225-020-00466-2)) |
| Otu29170 | 1 | k__Fungi(100);p__Ascomycota(100);c__Saccharomycetes(100);o__Saccharomycetales(100);f__Saccharomycetaceae(100);g__Zygosaccharomyces(100);s__Zygosaccharomyces_bailii(100); | Zygosaccharomyces | 13 | Pathotroph-Saprotroph | Plant Pathogen-\|Undefined Saprotroph\| | Facultative Yeast-Yeast | NULL | Probable | As nectar-tap_saprotroph (Põlme et al. 2020); Decay sugar-rich substrates (Põlme et al. 2020) | Kurtzman CP, et al. (eds.) 2011. The Yeasts, a Taxonomic Study. Fifth Edition. Vols 1-3. Elsevier, San Diego (ISBN: 9780444521491); Põlme S, et al. 2020. Fungal Diversity 105:1-16 ((https://doi.org/10.1007/s13225-020-00466-2)); The American Phytopathological Society Common Names of Plant Diseases ((http://www.apsnet.org/publications/commonnames/Pages/default.aspx)) |
| Otu29171 | 1 | k__Fungi(100);p__Ascomycota(100);c__Saccharomycetes(100);o__Saccharomycetales(100);f__Saccharomycetaceae(100);g__Torulaspora(100);s__Torulaspora_delbrueckii(100); | Torulaspora | 13 | Pathotroph-Saprotroph | Animal Parasite-\|Undefined Saprotroph\| | Yeast | NULL | Probable | As nectar-tap_saprotroph (Põlme et al. 2020); Decay animal material, sugar-rich substrates (Põlme et al. 2020) | Põlme S, et al. 2020. Fungal Diversity 105:1-16 ((https://doi.org/10.1007/s13225-020-00466-2)) |
| Otu29172 | 1 | k__Fungi(100);p__Ascomycota(100);c__Saccharomycetes(100);o__Saccharomycetales(100);f__Saccharomycetaceae(100);g__Saccharomyces(100);g__Saccharomyces_unclassified(100); | Saccharomyces | 13 | Saprotroph | \|Undefined Saprotroph\| | Yeast | NULL | Probable | As nectar-tap_saprotroph (Põlme et al. 2020); Decay sugar-rich substrates (Põlme et al. 2020) | James TY, et al. 2006. Nature 443:818-822 ((https://doi.org/10.1038/nature05110)); Põlme S, et al. 2020. Fungal Diversity 105:1-16 ((https://doi.org/10.1007/s13225-020-00466-2)) |
| Otu29173 | 1 | k__Fungi(100);p__Ascomycota(100);c__Saccharomycetes(100);o__Saccharomycetales(100);f__Saccharomycetaceae(100);g__Saccharomyces(100);g__Saccharomyces_unclassified(100); | Saccharomyces | 13 | Saprotroph | \|Undefined Saprotroph\| | Yeast | NULL | Probable | As nectar-tap_saprotroph (Põlme et al. 2020); Decay sugar-rich substrates (Põlme et al. 2020) | James TY, et al. 2006. Nature 443:818-822 ((https://doi.org/10.1038/nature05110)); Põlme S, et al. 2020. Fungal Diversity 105:1-16 ((https://doi.org/10.1007/s13225-020-00466-2)) |
| Otu29174 | 1 | k__Fungi(100);p__Ascomycota(100);c__Saccharomycetes(100);o__Saccharomycetales(100);f__Saccharomycetaceae(100);g__Saccharomyces(100);g__Saccharomyces_unclassified(100); | Saccharomyces | 13 | Saprotroph | \|Undefined Saprotroph\| | Yeast | NULL | Probable | As nectar-tap_saprotroph (Põlme et al. 2020); Decay sugar-rich substrates (Põlme et al. 2020) | James TY, et al. 2006. Nature 443:818-822 ((https://doi.org/10.1038/nature05110)); Põlme S, et al. 2020. Fungal Diversity 105:1-16 ((https://doi.org/10.1007/s13225-020-00466-2)) |
| Otu29175 | 1 | k__Fungi(100);p__Ascomycota(100);c__Saccharomycetes(100);o__Saccharomycetales(100);f__Saccharomycetaceae(100);g__Saccharomyces(100);g__Saccharomyces_unclassified(100); | Saccharomyces | 13 | Saprotroph | \|Undefined Saprotroph\| | Yeast | NULL | Probable | As nectar-tap_saprotroph (Põlme et al. 2020); Decay sugar-rich substrates (Põlme et al. 2020) | James TY, et al. 2006. Nature 443:818-822 ((https://doi.org/10.1038/nature05110)); Põlme S, et al. 2020. Fungal Diversity 105:1-16 ((https://doi.org/10.1007/s13225-020-00466-2)) |
| Otu29176 | 1 | k__Fungi(100);p__Ascomycota(100);c__Saccharomycetes(100);o__Saccharomycetales(100);f__Saccharomycetaceae(100);g__Saccharomyces(100);s__Saccharomyces_kudriavzevii(100); | Saccharomyces | 13 | Saprotroph | \|Undefined Saprotroph\| | Yeast | NULL | Probable | As nectar-tap_saprotroph (Põlme et al. 2020); Decay sugar-rich substrates (Põlme et al. 2020) | James TY, et al. 2006. Nature 443:818-822 ((https://doi.org/10.1038/nature05110)); Põlme S, et al. 2020. Fungal Diversity 105:1-16 ((https://doi.org/10.1007/s13225-020-00466-2)) |
| Otu29177 | 1 | k__Fungi(100);p__Ascomycota(100);c__Saccharomycetes(100);o__Saccharomycetales(100);f__Saccharomycetaceae(100);g__Saccharomyces(100);g__Saccharomyces_unclassified(100); | Saccharomyces | 13 | Saprotroph | \|Undefined Saprotroph\| | Yeast | NULL | Probable | As nectar-tap_saprotroph (Põlme et al. 2020); Decay sugar-rich substrates (Põlme et al. 2020) | James TY, et al. 2006. Nature 443:818-822 ((https://doi.org/10.1038/nature05110)); Põlme S, et al. 2020. Fungal Diversity 105:1-16 ((https://doi.org/10.1007/s13225-020-00466-2)) |
| Otu29178 | 1 | k__Fungi(100);p__Ascomycota(100);c__Saccharomycetes(100);o__Saccharomycetales(100);f__Saccharomycetaceae(100);g__Saccharomyces(100);s__Saccharomyces_eubayanus(100); | Saccharomyces | 13 | Saprotroph | \|Undefined Saprotroph\| | Yeast | NULL | Probable | As nectar-tap_saprotroph (Põlme et al. 2020); Decay sugar-rich substrates (Põlme et al. 2020) | James TY, et al. 2006. Nature 443:818-822 ((https://doi.org/10.1038/nature05110)); Põlme S, et al. 2020. Fungal Diversity 105:1-16 ((https://doi.org/10.1007/s13225-020-00466-2)) |
| Otu29180 | 1 | k__Fungi(100);p__Ascomycota(100);c__Sordariomycetes(100);o__Hypocreales(100);o__Hypocreales_unclassified(100);o__Hypocreales_unclassified(100);o__Hypocreales_unclassified(100); | Hypocreales | 7 | Saprotroph | Undefined Saprotroph | Microfungus | NULL | Possible | NULL | Sterkenburg E, et al. 2015. New Phytologist 207:1145-1158 ((https://doi.org/10.1111/nph.13426)) |
| Otu29181 | 1 | k__Fungi(100);p__Ascomycota(100);c__Sordariomycetes(100);o__Hypocreales(100);o__Hypocreales_unclassified(100);o__Hypocreales_unclassified(100);o__Hypocreales_unclassified(100); | Hypocreales | 7 | Saprotroph | Undefined Saprotroph | Microfungus | NULL | Possible | NULL | Sterkenburg E, et al. 2015. New Phytologist 207:1145-1158 ((https://doi.org/10.1111/nph.13426)) |
| Otu29193 | 1 | k__Fungi(100);p__Ascomycota(100);c__Sordariomycetes(100);o__Hypocreales(100);f__Ophiocordycipitaceae(100);f__Ophiocordycipitaceae_unclassified(100);f__Ophiocordycipitaceae_unclassified(100); | Ophiocordycipitaceae | 9 | Pathotroph-Symbiotroph | Endophyte-Insect Pathogen | Microfungus | NULL | Probable | A family of parasitic fungi (Sung et al. 2007) | Sung GH, et al. 2007. Studies in Mycology 57:5-59 ((https://doi.org/10.3114/sim.2007.57.01)); Costa IPMW, et al. 2012. Mycotaxon 119: e26 ((http://mycotaxon.com/resources/checklists/costa_v119_checklist.pdf)) |
| Otu29206 | 1 | k__Fungi(100);p__Ascomycota(100);c__Sordariomycetes(100);o__Hypocreales(100);f__Cordycipitaceae(100);g__Beauveria(100);g__Beauveria_unclassified(100); | Beauveria | 13 | Pathotroph-Saprotroph-Symbiotroph | \|Animal Parasite\|-Animal Pathogen-Endophyte-Undefined Saprotroph | Microfungus | NULL | Probable | As animal_decomposer (Põlme et al. 2020); As foliar_endophyte (Põlme et al. 2020); Decay animal material (Põlme et al. 2020) | Tedersoo L, et al. 2014. Science 346:e1256688 ((https://doi.org/10.1126/science.1256688)); Põlme S, et al. 2020. Fungal Diversity 105:1-16 ((https://doi.org/10.1007/s13225-020-00466-2)) |
| Otu29218 | 1 | k__Fungi(100);p__Ascomycota(100);c__Sordariomycetes(100);o__Hypocreales(100);f__Cordycipitaceae(100);g__Lecanicillium(100);s__Lecanicillium_sp(100); | Lecanicillium | 13 | Pathotroph-Saprotroph-Symbiotroph | \|Animal Parasite\|-Animal Pathogen-Endophyte-Undefined Saprotroph | Microfungus | NULL | Probable | As animal_decomposer (Põlme et al. 2020); As foliar_endophyte (Põlme et al. 2020); Decay animal material (Põlme et al. 2020) | Tedersoo L, et al. 2014. Science 346:e1256688 ((https://doi.org/10.1126/science.1256688)); Põlme S, et al. 2020. Fungal Diversity 105:1-16 ((https://doi.org/10.1007/s13225-020-00466-2)) |
| Otu29220 | 1 | k__Fungi(100);p__Ascomycota(100);c__Sordariomycetes(100);o__Hypocreales(100);f__Clavicipitaceae(100);g__Metarhiziopsis(100);s__Metarhiziopsis_microspora(100); | Metarhiziopsis | 13 | Pathotroph-Saprotroph | \|Animal Parasite\|-Undefined Saprotroph | Microfungus | NULL | Probable | As animal_decomposer (Põlme et al. 2020); Decay animal material (Põlme et al. 2020) | Põlme S, et al. 2020. Fungal Diversity 105:1-16 ((https://doi.org/10.1007/s13225-020-00466-2)) |
| Otu29223 | 1 | k__Fungi(100);p__Ascomycota(100);c__Sordariomycetes(100);o__Hypocreales(100);f__Cordycipitaceae(100);g__Simplicillium(100);g__Simplicillium_unclassified(100); | Simplicillium | 13 | Pathotroph-Saprotroph | \|Animal Parasite\|-Animal Pathogen-Fungal Parasite-Undefined Saprotroph | Microfungus | NULL | Probable | As animal_decomposer (Põlme et al. 2020); As mycoparasite (Põlme et al. 2020); Decay animal material (Põlme et al. 2020) | Tedersoo L, et al. 2014. Science 346:e1256688 ((https://doi.org/10.1126/science.1256688)); Põlme S, et al. 2020. Fungal Diversity 105:1-16 ((https://doi.org/10.1007/s13225-020-00466-2)) |
| Otu29224 | 1 | k__Fungi(100);p__Ascomycota(100);c__Sordariomycetes(100);o__Hypocreales(100);f__Cordycipitaceae(100);g__Lecanicillium(100);g__Lecanicillium_unclassified(100); | Lecanicillium | 13 | Pathotroph-Saprotroph-Symbiotroph | \|Animal Parasite\|-Animal Pathogen-Endophyte-Undefined Saprotroph | Microfungus | NULL | Probable | As animal_decomposer (Põlme et al. 2020); As foliar_endophyte (Põlme et al. 2020); Decay animal material (Põlme et al. 2020) | Tedersoo L, et al. 2014. Science 346:e1256688 ((https://doi.org/10.1126/science.1256688)); Põlme S, et al. 2020. Fungal Diversity 105:1-16 ((https://doi.org/10.1007/s13225-020-00466-2)) |
| Otu29225 | 1 | k__Fungi(100);p__Ascomycota(100);c__Sordariomycetes(100);o__Hypocreales(100);f__Hypocreales_family_Incertae_sedis(100);g__Acremonium(100);s__Acremonium_cavaraeanum(100); | Acremonium | 13 | Pathotroph-Saprotroph-Symbiotroph | Animal Pathogen-Endophyte-Fungal Parasite-Plant Pathogen-\|Undefined Saprotroph\|-Wood Saprotroph | Microfungus | Soft Rot | Possible | As foliar_endophyte (Põlme et al. 2020); culture (Hanada et al. 2010, Del Olmo; Decay leaf, fruit, seed, soil, dung, animal material (Põlme et al. 2020); Endophyte detection method; Ruiz & Arnold 2014); Likely opportunistic human pathogen (Irinyi et al. 2015) | Duncan CG, Eslyn WE. 1966. Mycologia 58:642-645 ((https://doi.org/10.2307/3757045)); Nilsson T. 1973. Studia Forestalia Suecica 104: 1-40 ((https://pub.epsilon.slu.se/5711/)); Worrall JJ, et al. 1997. Mycologia 89:199-219 ((https://doi.org/10.2307/3761073)); Hanada RE, et al. 2010. Fungal Biology 114:901-910 ((https://doi.org/10.1016/j.funbio.2010.08.006)); Del Olmo-Ruiz M, Arnold AE. 2014. Mycologia 106: 8-21 ((https://doi.org/10.3852/13-098)); Irinyi L, et al. 2015. Medical Mycology 53313-337 ((https://doi.org/10.1093/mmy/myv008)); Põlme S, et al. 2020. Fungal Diversity 105:1-16 ((https://doi.org/10.1007/s13225-020-00466-2)) |
| Otu29229 | 1 | k__Fungi(100);p__Ascomycota(100);c__Sordariomycetes(100);o__Ophiostomatales(100);f__Ophiostomataceae(100);g__Pesotum(100);s__Pesotum_australi(100); | Pesotum | 13 | Pathotroph-Saprotroph | \|Plant Pathogen\|-Wood Saprotroph | Microfungus | NULL | Probable | As wood_pathogen (Põlme et al. 2020); Decay wood (Põlme et al. 2020) | Tedersoo L, et al. 2014. Science 346:e1256688 ((https://doi.org/10.1126/science.1256688)); Põlme S, et al. 2020. Fungal Diversity 105:1-16 ((https://doi.org/10.1007/s13225-020-00466-2)) |
| Otu29231 | 1 | k__Fungi(100);p__Ascomycota(100);c__Sordariomycetes(100);o__Hypocreales(100);o__Hypocreales_unclassified(100);o__Hypocreales_unclassified(100);o__Hypocreales_unclassified(100); | Hypocreales | 7 | Saprotroph | Undefined Saprotroph | Microfungus | NULL | Possible | NULL | Sterkenburg E, et al. 2015. New Phytologist 207:1145-1158 ((https://doi.org/10.1111/nph.13426)) |
| Otu29232 | 1 | k__Fungi(100);p__Ascomycota(100);c__Sordariomycetes(100);o__Hypocreales(100);f__Ophiocordycipitaceae(100);g__Hirsutella(100);s__Hirsutella_gregis(100); | Hirsutella | 13 | Pathotroph-Saprotroph | \|Animal Parasite\|-Animal Pathogen-Undefined Saprotroph | Microfungus | NULL | Probable | As animal_decomposer (Põlme et al. 2020); Decay animal material (Põlme et al. 2020) | Tedersoo L, et al. 2014. Science 346:e1256688 ((https://doi.org/10.1126/science.1256688)); Põlme S, et al. 2020. Fungal Diversity 105:1-16 ((https://doi.org/10.1007/s13225-020-00466-2)) |
| Otu29244 | 1 | k__Fungi(100);p__Ascomycota(100);c__Sordariomycetes(100);o__Hypocreales(100);o__Hypocreales_unclassified(100);o__Hypocreales_unclassified(100);o__Hypocreales_unclassified(100); | Hypocreales | 7 | Saprotroph | Undefined Saprotroph | Microfungus | NULL | Possible | NULL | Sterkenburg E, et al. 2015. New Phytologist 207:1145-1158 ((https://doi.org/10.1111/nph.13426)) |
| Otu29245 | 1 | k__Fungi(100);p__Ascomycota(100);c__Sordariomycetes(100);o__Hypocreales(100);o__Hypocreales_unclassified(100);o__Hypocreales_unclassified(100);o__Hypocreales_unclassified(100); | Hypocreales | 7 | Saprotroph | Undefined Saprotroph | Microfungus | NULL | Possible | NULL | Sterkenburg E, et al. 2015. New Phytologist 207:1145-1158 ((https://doi.org/10.1111/nph.13426)) |
| Otu29247 | 1 | k__Fungi(100);p__Ascomycota(100);c__Sordariomycetes(100);o__Hypocreales(100);f__Hypocreaceae(100);g__Hypocrea(100);s__Hypocrea_thelephoricola(100); | Hypocrea | 13 | Pathotroph | Endophyte-Fungal Parasite | Microfungus | NULL | Probable | NULL | Tedersoo L, et al. 2014. Science 346:e1256688 ((https://doi.org/10.1126/science.1256688)); Christian N, et al. 2016. Microbial Ecology 72: 621-632 ((https://doi.org/10.1007/s00248-016-0804-y)) |
| Otu29248 | 1 | k__Fungi(100);p__Ascomycota(100);c__Sordariomycetes(100);o__Hypocreales(100);o__Hypocreales_unclassified(100);o__Hypocreales_unclassified(100);o__Hypocreales_unclassified(100); | Hypocreales | 7 | Saprotroph | Undefined Saprotroph | Microfungus | NULL | Possible | NULL | Sterkenburg E, et al. 2015. New Phytologist 207:1145-1158 ((https://doi.org/10.1111/nph.13426)) |
| Otu29270 | 1 | k__Fungi(100);p__Ascomycota(100);c__Sordariomycetes(100);o__Hypocreales(100);f__Ophiocordycipitaceae(100);g__Purpureocillium(100);g__Purpureocillium_unclassified(100); | Purpureocillium | 13 | Pathotroph-Saprotroph | \|Animal Parasite\|-Fungal Parasite-Undefined Saprotroph | Microfungus | NULL | Probable | As animal_decomposer (Põlme et al. 2020); As keratinolytic (Põlme et al. 2020); Decay animal material (Põlme et al. 2020) | Tedersoo L, et al. 2014. Science 346:e1256688 ((https://doi.org/10.1126/science.1256688)); Põlme S, et al. 2020. Fungal Diversity 105:1-16 ((https://doi.org/10.1007/s13225-020-00466-2)) |
| Otu29393 | 1 | k__Fungi(100);p__Basidiomycota(100);c__Tremellomycetes(100);o__Tremellales(100);f__Tremellales_family_Incertae_sedis(100);g__Cryptococcus(100);s__Cryptococcus_laurentii(100); | Cryptococcus | 13 | Pathotroph-Saprotroph-Symbiotroph | Animal Pathogen-Endophyte-Epiphyte-\|Undefined Saprotroph\| | Dimorphic Yeast-Yeast | NULL | Possible | As foliar_endophyte (Põlme et al. 2020); Decay leaf, fruit, seed, soil, dung, animal material (Põlme et al. 2020) | Andrews JH, et al. 1983. Phytopathology 73:228-234 ((https://doi.org/10.1094/Phyto-73-228)); Kurtzman CP, et al. (eds.) 2011. The Yeasts, a Taxonomic Study. Fifth Edition. Vols 1-3. Elsevier, San Diego (ISBN: 9780444521491); Põlme S, et al. 2020. Fungal Diversity 105:1-16 ((https://doi.org/10.1007/s13225-020-00466-2)) |
| Otu29394 | 1 | k__Fungi(100);p__Zygomycota(100);c__Zygomycota_class_Incertae_sedis(100);o__Mucorales(100);f__Mucoraceae(100);g__Mucor(100);s__Mucor_genevensis(100); | Mucor | 13 | Saprotroph | \|Undefined Saprotroph\| | Microfungus | NULL | Probable | As soil_saprotroph (Põlme et al. 2020); Decay soil (Põlme et al. 2020) | Tedersoo L, et al. 2014. Science 346:e1256688 ((https://doi.org/10.1126/science.1256688)); Põlme S, et al. 2020. Fungal Diversity 105:1-16 ((https://doi.org/10.1007/s13225-020-00466-2)) |
| Otu29395 | 1 | k__Fungi(100);p__Ascomycota(100);c__Saccharomycetes(100);o__Saccharomycetales(100);f__Saccharomycetaceae(100);g__Kazachstania(100);s__Kazachstania_intestinalis(100); | Kazachstania | 13 | Saprotroph | \|Undefined Saprotroph\| | Yeast | NULL | Probable | As nectar-tap_saprotroph (Põlme et al. 2020); Decay sugar-rich substrates (Põlme et al. 2020) | Põlme S, et al. 2020. Fungal Diversity 105:1-16 ((https://doi.org/10.1007/s13225-020-00466-2)) |
| Otu29404 | 1 | k__Fungi(100);p__Ascomycota(100);c__Eurotiomycetes(100);o__Eurotiales(100);f__Trichocomaceae(100);g__Paecilomyces(100);g__Paecilomyces_unclassified(100); | Paecilomyces | 13 | Pathotroph-Saprotroph-Symbiotroph | Clavicipitaceous \|Endophyte\|-Plant Pathogen-Plant Saprotroph-Undefined Saprotroph-Wood Saprotroph | Microfungus | Soft Rot | Probable | As foliar_endophyte (Põlme et al. 2020); As litter_saprotroph (Põlme et al. 2020); culture (Lopez et al. 2014); Decay leaf, fruit, seed, animal material (Põlme et al. 2020); Endophyte detection method | Seehann G, et al. 1975. List of Fungi in Soft-Rot Tissue Tests. Institute fur Holzbiologie und Holzschutz, Bundesforschungsanstalt fur Forst - und Holzwirtshaft, Hamburg - Lohbrugge, Germany; Costa IPMW, et al. 2012. Mycotaxon 119:e26 ((http://mycotaxon.com/resources/checklists/costa_v119_checklist.pdf)); Tedersoo L, et al. 2014. Science 346:e1256688 ((https://doi.org/10.1126/science.1256688)); Põlme S, et al. 2020. Fungal Diversity 105:1-16 ((https://doi.org/10.1007/s13225-020-00466-2)); The American Phytopathological Society Common Names of Plant Diseases ((http://www.apsnet.org/publications/commonnames/Pages/default.aspx)); Lopez DC, et al. 2014. Plos One 9:e103891 ((https://doi.org/10.1371/journal.pone.0103891)) |

| | | | | | | | | | | | |
|---|---|---|---|---|---|---|---|---|---|---|---|
| Otu29405 | 1 | k__Fungi(100);p__Ascomycota(100);c__Sordariomycetes(100);o__Hypocreales(100);o__Hypocreales_unclassified(100);o__Hypocreales_unclassified(100);o__Hypocreales_unclassified(100); | Hypocreales | 7 | Saprotroph | Undefined Saprotroph | Microfungus | NULL | Possible | NULL | [illegible] |
| Otu29407 | 1 | k__Fungi(100);p__Ascomycota(100);c__Orbiliomycetes(100);o__Orbiliales(100);f__Orbiliaceae(100);f__Orbiliaceae_unclassified(100);f__Orbiliaceae_unclassified(100); | Orbiliaceae | 9 | Saprotroph | Wood Saprotroph | Microfungus | NULL | Probable | Saprobic, found especially on wood in both wet and dry habitats. Many species are nematophagous, forming adhesive knobs, nets or ring-like traps in the presence of a food source; perhaps a strategy for life in low-nitrogen environments. There have been reports of associations with algae for a few species, but an established lichen association appears not to exist (Cannon & Kirk 2007) | [illegible] |
| Otu29411 | 1 | k__Fungi(100);p__Ascomycota(100);c__Sordariomycetes(100);o__Xylariales(100);f__Amphisphaeriaceae(100);g__Monochaetia(100);g__Monochaetia_unclassified(100); | Monochaetia | 13 | Pathotroph | \|Plant Pathogen\| | Microfungus | NULL | Probable | NULL | [illegible] |
| Otu29412 | 1 | k__Fungi(100);p__Ascomycota(100);c__Sordariomycetes(100);o__Sordariomycetes_order_Incertae_sedis(100);f__Annulatascaceae(100);g__Rhodoveronaea(100);s__Rhodoveronaea_varioseptata(100); | Rhodoveronaea | 13 | Saprotroph | \|Undefined Saprotroph\| | Microfungus | NULL | Probable | NULL | [illegible] |
| Otu29414 | 1 | k__Fungi(100);p__Ascomycota(100);c__Sordariomycetes(100);o__Xylariales(100);f__Xylariales_family_Incertae_sedis(100);g__Microdochium(100);s__Microdochium_sp(100); | Microdochium | 13 | Pathotroph-Symbiotroph | Endophyte-\|Plant Pathogen\| | Microfungus | NULL | Possible | As foliar_endophyte (Põlme et al. 2020); Dark Septate Endophyte; Decay leaf, fruit, seed (Põlme et al. 2020) | [illegible] |
| Otu29415 | 1 | k__Fungi(100);p__Ascomycota(100);c__Sordariomycetes(100);o__Xylariales(100);f__Amphisphaeriaceae(100);f__Amphisphaeriaceae_unclassified(100);f__Amphisphaeriaceae_unclassified(100); | Amphisphaeriaceae | 9 | Saprotroph | Undefined Saprotroph | Microfungus | NULL | Possible | Saprobes or weak parasites of leaves, stems, etc., often appearing as secondary colonizers following damage to plants by other agents (Cannon & Kirk 2007) | [illegible] |
| Otu29416 | 1 | k__Fungi(100);p__Ascomycota(100);c__Sordariomycetes(100);o__Xylariales(100);f__Xylariaceae(100);g__Rosellinia(100);s__Rosellinia_abscondita(100); | Rosellinia | 13 | Pathotroph-Saprotroph | \|Plant Pathogen\|-Undefined Saprotroph-Wood Saprotroph | Microfungus | NULL | Probable | As wood_pathogen (Põlme et al. 2020); Decay wood (Põlme et al. 2020); Host - angiosperms (Põlme et al. 2020) | [illegible] |
| Otu29417 | 1 | k__Fungi(100);p__Ascomycota(100);c__Sordariomycetes(100);o__Xylariales(100);f__Xylariaceae(100);f__Xylariaceae_unclassified(100);f__Xylariaceae_unclassified(100); | Xylariaceae | 9 | Pathotroph-Saprotroph-Symbiotroph | Dung Saprotroph-Endophyte-Plant Pathogen-Undefined Saprotroph | Xylarioid | NULL | Probable | Saprobes or weak pathogens, many endophytes, mainly in wood and bark, occasionally in soil and dung (Cannon & Kirk 2007) | [illegible] |
| Otu29418 | 1 | k__Fungi(100);p__Ascomycota(100);c__Eurotiomycetes(100);o__Mycocaliciales(100);f__Mycocaliciaceae(100);g__Mycocalicium(100);s__Mycocalicium_subtile(100); | Mycocalicium | 13 | Pathotroph-Saprotroph-Symbiotroph | Lichen Parasite-Lichenized-\|Undefined Saprotroph\| | Calicioid-Thallus | NULL | Possible | As resin_saprotroph (Põlme et al. 2020); licenized taxa; Lichenized and non | [illegible] |
| Otu29420 | 1 | k__Fungi(100);p__Ascomycota(100);c__Dothideomycetes(100);o__Pleosporales(100);f__Pleosporales_family_Incertae_sedis(100);g__Periconia(100);g__Periconia_unclassified(100); | Periconia | 13 | Pathotroph-Saprotroph-Symbiotroph | Endophyte-\|Plant Pathogen\|-Wood Saprotroph | Microfungus | Soft Rot | Possible | Asteraceae, Musaceae (Costa et al. 2012); As foliar_endophyte (Põlme et al. 2020); Dark Septate Endophyte, known from plant roots (Mandyam et al. 2010); Endophyte Host ; Decay leaf, fruit, seed (Põlme et al. 2020) | [illegible] |
| Otu29423 | 1 | k__Fungi(100);p__Ascomycota(100);c__Sordariomycetes(100);o__Lulworthiales(100);f__Lulworthiaceae(100);g__Zalerion(100);s__Zalerion_arboricola(100); | Zalerion | 13 | Pathotroph-Saprotroph | Plant Pathogen-\|Plant Saprotroph\|-Undefined Saprotroph | Microfungus | NULL | Probable | As litter_saprotroph (Põlme et al. 2020); Decay leaf, fruit, seed (Põlme et al. 2020) | [illegible] |
| Otu29425 | 1 | k__Fungi(100);p__Ascomycota(100);c__Dothideomycetes(100);o__Pleosporales(100);f__Didymosphaeriaceae(100);g__Roussoella(100);g__Roussoella_unclassified(100); | Roussoella | 13 | Saprotroph | Undefined Saprotroph-\|Wood Saprotroph\| | Microfungus | NULL | Probable | Decay wood (Põlme et al. 2020) | [illegible] |
| Otu29426 | 1 | k__Fungi(100);p__Ascomycota(100);c__Sordariomycetes(100);o__Sordariales(100);f__Lasiosphaeriaceae(100);g__Cladorrhinum(100);s__Cladorrhinum_sp(100); | Cladorrhinum | 13 | Pathotroph-Saprotroph-Symbiotroph | Animal Pathogen-Dung Saprotroph-Endophyte-Plant Pathogen-Plant Saprotroph-\|Undefined Saprotroph\|-Wood Saprotroph | Microfungus | Soft Rot | Probable | As foliar_endophyte (Põlme et al. 2020); Decay leaf, fruit, seed (Põlme et al. 2020); Reported in soils (including agricultural) as saprotrophs on dung or plant material, or in roots as endophytes, while some species are associated with human and animal opportunistic diseases (Carmaran et al.2015) | [illegible] |
| Otu29427 | 1 | k__Fungi(100);p__Ascomycota(100);c__Sordariomycetes(100);o__Hypocreales(100);f__Nectriaceae(100);g__Fusarium(100);s__Fusarium_tricinctum(100); | Fusarium | 13 | Pathotroph-Saprotroph-Symbiotroph | Animal Pathogen-Endophyte-Lichen Parasite-\|Plant Pathogen\|-Plant Saprotroph-Undefined Saprotroph-Wood Saprotroph | Microfungus | Soft Rot | Possible | As foliar_endophyte (Põlme et al. 2020); As litter_saprotroph (Põlme et al. 2020); Causing damping-off disease (Põlme et al. 2020); Decay leaf, fruit, seed, roots, soil, animal material (Põlme et al. 2020); Likely opportunistic human pathogen (Irinyi et al. 2015) | [illegible] |
| Otu29428 | 1 | k__Fungi(100);p__Ascomycota(100);c__Dothideomycetes(100);o__Pleosporales(100);f__Pleosporaceae(100);f__Pleosporaceae_unclassified(100);f__Pleosporaceae_unclassified(100); | Pleosporaceae | 9 | Pathotroph-Saprotroph | Endophyte-Lichen Parasite-Plant Pathogen-Undefined Saprotroph | Microfungus | NULL | Probable | Necrotrophic pathogens and saprobes, especially associated with grasses. Many species are seed-borne (Cannon & Kirk 2007) | [illegible] |
| Otu29429 | 1 | k__Fungi(100);p__Ascomycota(100);c__Eurotiomycetes(100);o__Onygenales(100);f__Arthrodermataceae(100);f__Arthrodermataceae_unclassified(100);f__Arthrodermataceae_unclassified(100); | Arthrodermataceae | 9 | Pathotroph-Saprotroph | Animal Pathogen-Undefined Saprotroph | NULL | NULL | Probable | Principally affecting mammals (including humans) - Colonies present in substrata containing keratin, such as soil and birds nests - Dermatophyte fungi, causing ringworm and tinea of keratinous tissue e.g. hair, nails, and skin - Anamorphorphs-Chrysosporium, Microsporum, Trichophyton (Cannon & Kirk 2007) | [illegible] |
| Otu29430 | 1 | k__Fungi(100);p__Ascomycota(100);c__Eurotiomycetes(100);o__Mycocaliciales(100);f__Mycocaliciaceae(100);g__Mycocalicium(100);s__Mycocalicium_subtile(100); | Mycocalicium | 13 | Pathotroph-Saprotroph-Symbiotroph | Lichen Parasite-Lichenized-\|Undefined Saprotroph\| | Calicioid-Thallus | NULL | Possible | As resin_saprotroph (Põlme et al. 2020); licenized taxa; Lichenized and non | [illegible] |
| Otu29431 | 1 | k__Fungi(100);p__Ascomycota(100);c__Sordariomycetes(100);o__Xylariales(100);f__Xylariaceae(100);g__Anthostomella(100);s__Anthostomella_sp(100); | Anthostomella | 13 | Saprotroph-Symbiotroph | \|Endophyte\|-Plant Saprotroph-Undefined Saprotroph | Microfungus | NULL | Probable | As foliar_endophyte (Põlme et al. 2020); As litter_saprotroph (Põlme et al. 2020); Decay leaf, fruit, seed (Põlme et al. 2020) | [illegible] |
| Otu29432 | 1 | k__Fungi(100);p__Ascomycota(100);c__Eurotiomycetes(100);o__Onygenales(100);f__Ajellomycetaceae(100);g__Emmonsia(100);s__Emmonsia_crescens(100); | Emmonsia | 13 | Pathotroph-Saprotroph | Animal Parasite-\|Undefined Saprotroph\| | Dimorphic-Microfungus | NULL | Probable | As soil_saprotroph (Põlme et al. 2020); Decay soil, animal material (Põlme et al. 2020) | [illegible] |
| Otu29433 | 1 | k__Fungi(100);p__Ascomycota(100);c__Eurotiomycetes(100);o__Eurotiales(100);f__Trichocomaceae(100);g__Eupenicillium(100);s__Eupenicillium_cinnamopurpureum(100); | Eupenicillium | 13 | Saprotroph | Undefined Saprotroph | NULL | NULL | Probable | NULL | [illegible] |
| Otu29434 | 1 | k__Fungi(100);p__Ascomycota(100);c__Eurotiomycetes(100);o__Eurotiales(100);f__Trichocomaceae(100);g__Penicillium(100);g__Penicillium_unclassified(100); | Penicillium | 13 | Pathotroph-Saprotroph-Symbiotroph | Animal Parasite-Dung Saprotroph-Endophyte-Fungal Parasite-Plant Pathogen-\|Undefined Saprotroph\|-Wood Saprotroph | Microfungus | Soft Rot | Highly Probable | As foliar_endophyte (Põlme et al. 2020); As mycoparasite (Põlme et al. 2020); Decay soil (Põlme et al. 2020) | [illegible] |
| Otu29435 | 1 | k__Fungi(100);p__Ascomycota(100);c__Eurotiomycetes(100);o__Eurotiales(100);f__Trichocomaceae(100);g__Aspergillus(100);s__Aspergillus_sp_F55(100); | Aspergillus | 13 | Pathotroph-Saprotroph-Symbiotroph | Animal Pathogen-Endophyte-Plant Saprotroph-\|Undefined Saprotroph\|-Wood Saprotroph | Microfungus | NULL | Probable | As foliar_endophyte (Põlme et al. 2020); Decay leaf, fruit, seed, soil, dung, animal material (Põlme et al. 2020) | [illegible] |
| Otu29436 | 1 | k__Fungi(100);p__Ascomycota(100);c__Eurotiomycetes(100);o__Eurotiales(100);f__Trichocomaceae(100);g__Penicillium(100);g__Penicillium_unclassified(100); | Penicillium | 13 | Pathotroph-Saprotroph-Symbiotroph | Animal Parasite-Dung Saprotroph-Endophyte-Fungal Parasite-Plant Pathogen-\|Undefined Saprotroph\|-Wood Saprotroph | Microfungus | Soft Rot | Highly Probable | As foliar_endophyte (Põlme et al. 2020); As mycoparasite (Põlme et al. 2020); Decay soil (Põlme et al. 2020) | [illegible] |
| Otu29437 | 1 | k__Fungi(100);p__Ascomycota(100);c__Eurotiomycetes(100);o__Eurotiales(100);f__Trichocomaceae(100);g__Penicillium(100);g__Penicillium_unclassified(100); | Penicillium | 13 | Pathotroph-Saprotroph-Symbiotroph | Animal Parasite-Dung Saprotroph-Endophyte-Fungal Parasite-Plant Pathogen-\|Undefined Saprotroph\|-Wood Saprotroph | Microfungus | Soft Rot | Highly Probable | As foliar_endophyte (Põlme et al. 2020); As mycoparasite (Põlme et al. 2020); Decay soil (Põlme et al. 2020) | [illegible] |
| Otu29440 | 1 | k__Fungi(100);p__Ascomycota(100);c__Ascomycota_class_Incertae_sedis(100);o__Ascomycota_order_Incertae_sedis(100);f__Ascomycota_family_Incertae_sedis(100);g__Myriodontium(100);s__Myriodontium_keratinophilum(100); | Myriodontium | 13 | Saprotroph | \|Undefined Saprotroph\| | Microfungus | NULL | Probable | As animal_decomposer (Põlme et al. 2020); As keratinolytic (Põlme et al. 2020); As soil_saprotroph (Põlme et al. 2020); Decay soil, animal material (Põlme et al. 2020) | [illegible] |
| Otu29441 | 1 | k__Fungi(100);p__Ascomycota(100);c__Eurotiomycetes(100);o__Eurotiales(100);f__Trichocomaceae(100);g__Aspergillus(100);g__Aspergillus_unclassified(100); | Aspergillus | 13 | Pathotroph-Saprotroph-Symbiotroph | Animal Pathogen-Endophyte-Plant Saprotroph-\|Undefined Saprotroph\|-Wood Saprotroph | Microfungus | NULL | Probable | As foliar_endophyte (Põlme et al. 2020); Decay leaf, fruit, seed, soil, dung, animal material (Põlme et al. 2020) | [illegible] |
| Otu29445 | 1 | k__Fungi(100);p__Zygomycota(100);c__Zygomycota_class_Incertae_sedis(100);o__Mortierellales(100);f__Mortierellaceae(100);g__Mortierella(100);s__Mortierella_sp_WD32A(100); | Mortierella | 13 | Saprotroph-Symbiotroph | Endophyte-Plant Saprotroph-\|Undefined Saprotroph\| | Microfungus | NULL | Probable | As chitinolytic (Põlme et al. 2020); As soil_saprotroph (Põlme et al. 2020); Decay roots, soil (Põlme et al. 2020); Saprobic in soil and thus frequenctly isolated from soil and material contaminated with soil such as hay or rotten silage and spoil tips (Cannon & Kirk 2007) | [illegible] |
| Otu29446 | 1 | k__Fungi(100);p__Zygomycota(100);c__Zygomycota_class_Incertae_sedis(100);o__Mucorales(100);f__Umbelopsidaceae(100);g__Umbelopsis(100);s__Umbelopsis_ramanniana(100); | Umbelopsis | 13 | Saprotroph | \|Undefined Saprotroph\| | Microfungus | NULL | Probable | As soil_saprotroph (Põlme et al. 2020); Decay roots, soil (Põlme et al. 2020) | [illegible] |
| Otu29447 | 1 | k__Fungi(100);p__Zygomycota(100);c__Zygomycota_class_Incertae_sedis(100);o__Mucorales(100);f__Umbelopsidaceae(100);g__Umbelopsis(100);s__Umbelopsis_ramanniana(100); | Umbelopsis | 13 | Saprotroph | \|Undefined Saprotroph\| | Microfungus | NULL | Probable | As soil_saprotroph (Põlme et al. 2020); Decay roots, soil (Põlme et al. 2020) | [illegible] |
| Otu29448 | 1 | k__Fungi(100);p__Zygomycota(100);c__Zygomycota_class_Incertae_sedis(100);o__Mucorales(100);f__Umbelopsidaceae(100);g__Umbelopsis(100);s__Umbelopsis_isabellina(100); | Umbelopsis | 13 | Saprotroph | \|Undefined Saprotroph\| | Microfungus | NULL | Probable | As soil_saprotroph (Põlme et al. 2020); Decay roots, soil (Põlme et al. 2020) | [illegible] |
| Otu29449 | 1 | k__Fungi(100);p__Zygomycota(100);c__Zygomycota_class_Incertae_sedis(100);o__Mucorales(100);f__Umbelopsidaceae(100);g__Umbelopsis(100);s__Umbelopsis_isabellina(100); | Umbelopsis | 13 | Saprotroph | \|Undefined Saprotroph\| | Microfungus | NULL | Probable | As soil_saprotroph (Põlme et al. 2020); Decay roots, soil (Põlme et al. 2020) | [illegible] |
| Otu29452 | 1 | k__Fungi(100);p__Ascomycota(100);c__Saccharomycetes(100);o__Saccharomycetales(100);o__Saccharomycetales_unclassified(100);o__Saccharomycetales_unclassified(100);o__Saccharomycetales_unclassified(100); | Saccharomycetales | 7 | Saprotroph | Undefined Saprotroph | Yeast | NULL | Possible | NULL | [illegible] |
| Otu29453 | 1 | k__Fungi(100);p__Zygomycota(100);c__Zygomycota_class_Incertae_sedis(100);o__Mucorales(100);f__Umbelopsidaceae(100);g__Umbelopsis(100);s__Umbelopsis_ramanniana(100); | Umbelopsis | 13 | Saprotroph | \|Undefined Saprotroph\| | Microfungus | NULL | Probable | As soil_saprotroph (Põlme et al. 2020); Decay roots, soil (Põlme et al. 2020) | [illegible] |
| Otu29454 | 1 | k__Fungi(100);p__Ascomycota(100);c__Leotiomycetes(100);o__Helotiales(100);f__Vibrisseaceae(100);g__Phialocephala(100);s__Phialocephala_sp_CM16s1(100); | Phialocephala | 13 | Saprotroph-Symbiotroph | Endophyte-\|Undefined Saprotroph\| | Microfungus | NULL | Highly Probable | As root_endophyte (Põlme et al. 2020); As root_endophyte_dark_septate (Põlme et al. 2020); As soil_saprotroph (Põlme et al. 2020); Dark Septate Endophyte, known from plant roots.; Decay roots, soil (Põlme et al. 2020) | [illegible] |
| Otu29455 | 1 | k__Fungi(100);p__Ascomycota(100);c__Sordariomycetes(100);o__Hypocreales(100);f__Cordycipitaceae(100);g__Lecanicillium(100);s__Lecanicillium_fusisporum(100); | Lecanicillium | 13 | Pathotroph-Saprotroph-Symbiotroph | \|Animal Parasite\|-Animal Pathogen-Endophyte-Undefined Saprotroph | Microfungus | NULL | Probable | As animal_decomposer (Põlme et al. 2020); As foliar_endophyte (Põlme et al. 2020); Decay animal material (Põlme et al. 2020) | [illegible] |
| Otu29456 | 1 | k__Fungi(100);p__Ascomycota(100);c__Eurotiomycetes(100);o__Eurotiales(100);f__Trichocomaceae(100);g__Penicillium(100);s__Penicillium_olsonii(100); | Penicillium | 13 | Pathotroph-Saprotroph-Symbiotroph | Animal Parasite-Dung Saprotroph-Endophyte-Fungal Parasite-Plant Pathogen-\|Undefined Saprotroph\|-Wood Saprotroph | Microfungus | Soft Rot | Highly Probable | As foliar_endophyte (Põlme et al. 2020); As mycoparasite (Põlme et al. 2020); Decay soil (Põlme et al. 2020) | [illegible] |
| Otu29457 | 1 | k__Fungi(100);p__Ascomycota(100);c__Sordariomycetes(100);o__Hypocreales(100);f__Hypocreales_family_Incertae_sedis(100);g__Acremonium(100);s__Acremonium_sp_24_OA_2013(100); | Acremonium | 13 | Pathotroph-Saprotroph-Symbiotroph | Animal Pathogen-Endophyte-Fungal Parasite-Plant Pathogen-\|Undefined Saprotroph\|-Wood Saprotroph | Microfungus | Soft Rot | Possible | As foliar_endophyte (Põlme et al. 2020); culture (Hanada et al. 2010, Del Olmo; Decay leaf, fruit, seed, soil, dung, animal material (Põlme et al. 2020); Endophyte detection method; Ruiz & Arnold 2014); Likely opportunistic human pathogen (Irinyi et al. 2015) | [illegible] |
| Otu29458 | 1 | k__Fungi(100);p__Zygomycota(100);c__Zygomycota_class_Incertae_sedis(100);o__Mortierellales(100);f__Mortierellaceae(100);g__Mortierella(100);g__Mortierella_unclassified(100); | Mortierella | 13 | Saprotroph-Symbiotroph | Endophyte-Plant Saprotroph-\|Undefined Saprotroph\| | Microfungus | NULL | Probable | As chitinolytic (Põlme et al. 2020); As soil_saprotroph (Põlme et al. 2020); Decay roots, soil (Põlme et al. 2020); Saprobic in soil and thus frequenctly isolated from soil and material contaminated with soil such as hay or rotten silage and spoil tips (Cannon & Kirk 2007) | [illegible] |
| Otu29459 | 1 | k__Fungi(100);p__Ascomycota(100);c__Saccharomycetes(100);o__Saccharomycetales(100);f__Saccharomycetaceae(100);g__Kazachstania(100);s__Kazachstania_intestinalis(100); | Kazachstania | 13 | Saprotroph | \|Undefined Saprotroph\| | Yeast | NULL | Probable | As nectar-tap_saprotroph (Põlme et al. 2020); Decay sugar-rich substrates (Põlme et al. 2020) | [illegible] |
| Otu29460 | 1 | k__Fungi(100);p__Ascomycota(100);c__Sordariomycetes(100);o__Hypocreales(100);f__Cordycipitaceae(100);g__Gibellula(100);s__Gibellula_sp_WK_2013(100); | Gibellula | 13 | Pathotroph | \|Animal Parasite\|-Animal Pathogen-Arthropod Parasite | Microfungus | NULL | Probable | Decay animal material (Põlme et al. 2020); Host - spiders (Põlme et al. 2020) | [illegible] |
| Otu29462 | 1 | k__Fungi(100);p__Ascomycota(100);c__Saccharomycetes(100);o__Saccharomycetales(100);f__Saccharomycetaceae(100);g__Kazachstania(100);s__Kazachstania_intestinalis(100); | Kazachstania | 13 | Saprotroph | \|Undefined Saprotroph\| | Yeast | NULL | Probable | As nectar-tap_saprotroph (Põlme et al. 2020); Decay sugar-rich substrates (Põlme et al. 2020) | [illegible] |
| Otu29472 | 1 | k__Fungi(100);p__Ascomycota(100);c__Sordariomycetes(100);o__Xylariales(100);f__Xylariaceae(100);g__Anthostomella(100);s__Anthostomella_sp(100); | Anthostomella | 13 | Saprotroph-Symbiotroph | \|Endophyte\|-Plant Saprotroph-Undefined Saprotroph | Microfungus | NULL | Probable | As foliar_endophyte (Põlme et al. 2020); As litter_saprotroph (Põlme et al. 2020); Decay leaf, fruit, seed (Põlme et al. 2020) | [illegible] |
| Otu29474 | 1 | k__Fungi(100);p__Ascomycota(100);c__Sordariomycetes(100);o__Xylariales(100);f__Xylariaceae(100);g__Hypoxylon(100);s__Hypoxylon_intermedium(100); | Hypoxylon | 13 | Saprotroph-Saprotroph-Symbiotroph | Endophyte-Undefined Saprotroph-\|Wood Saprotroph\| | Corticioid-Microfungus-Xylarioid | NULL | Probable | As foliar_endophyte (Põlme et al. 2020); Decay wood, leaf, fruit, seed (Põlme et al. 2020); Host - angiosperms (Põlme et al. 2020) | [illegible] |
| Otu29475 | 1 | k__Fungi(100);p__Ascomycota(100);c__Sordariomycetes(100);o__Xylariales(100);f__Xylariaceae(100);g__Hypoxylon(100);s__Hypoxylon_papillatum(100); | Hypoxylon | 13 | Saprotroph-Saprotroph-Symbiotroph | Endophyte-Undefined Saprotroph-\|Wood Saprotroph\| | Corticioid-Microfungus-Xylarioid | NULL | Probable | As foliar_endophyte (Põlme et al. 2020); Decay wood, leaf, fruit, seed (Põlme et al. 2020); Host - angiosperms (Põlme et al. 2020) | [illegible] |
| Otu29476 | 1 | k__Fungi(100);p__Ascomycota(100);c__Sordariomycetes(100);o__Xylariales(100);f__Amphisphaeriaceae(100);g__Pestalotiopsis(100);g__Pestalotiopsis_unclassified(100); | Pestalotiopsis | 13 | Pathotroph | \|Plant Pathogen\| | Microfungus | NULL | Probable | NULL | [illegible] |
| Otu29477 | 1 | k__Fungi(100);p__Ascomycota(100);c__Sordariomycetes(100);o__Xylariales(100);f__Xylariaceae(100);f__Xylariaceae_unclassified(100);f__Xylariaceae_unclassified(100); | Xylariaceae | 9 | Pathotroph-Saprotroph-Symbiotroph | Dung Saprotroph-Endophyte-Plant Pathogen-Undefined Saprotroph | Xylarioid | NULL | Probable | Saprobes or weak pathogens, many endophytes, mainly in wood and bark, occasionally in soil and dung (Cannon & Kirk 2007) | [illegible] |
| Otu29480 | 1 | k__Fungi(100);p__Ascomycota(100);c__Sordariomycetes(100);o__Microascales(100);f__Microascaceae(100);g__Microascus(100);g__Microascus_unclassified(100); | Microascus | 13 | Pathotroph-Saprotroph-Symbiotroph | Animal Pathogen-Dung Saprotroph-Endophyte-Plant Pathogen-Plant Saprotroph-\|Undefined Saprotroph\| | Microfungus | NULL | Possible | As foliar_endophyte (Põlme et al. 2020); As litter_saprotroph (Põlme et al. 2020); As soil_saprotroph (Põlme et al. 2020); Decay leaf, fruit, seed, soil (Põlme et al. 2020) | [illegible] |
| Otu29488 | 1 | k__Fungi(100);p__Ascomycota(100);c__Sordariomycetes(100);o__Sordariomycetes_order_Incertae_sedis(100);f__Sordariomycetes_family_Incertae_sedis(100);g__Xylomelasma(100);s__Xylomelasma_sp_ZK29/08(100); | Xylomelasma | 13 | Saprotroph | Undefined Saprotroph-\|Wood Saprotroph\| | Microfungus | NULL | Probable | Decay wood (Põlme et al. 2020) | [illegible] |
| Otu29489 | 1 | k__Fungi(100);p__Ascomycota(100);c__Sordariomycetes(100);o__Sordariomycetes_order_Incertae_sedis(100);f__Sordariomycetes_family_Incertae_sedis(100);g__Xylomelasma(100);s__Xylomelasma_sp_ZK29/08(100); | Xylomelasma | 13 | Saprotroph | Undefined Saprotroph-\|Wood Saprotroph\| | Microfungus | NULL | Probable | Decay wood (Põlme et al. 2020) | [illegible] |
| Otu29491 | 1 | k__Fungi(100);p__Ascomycota(100);c__Sordariomycetes(100);o__Hypocreales(100);f__Nectriaceae(100);g__Fusarium(100);s__Fusarium_tricinctum(100); | Fusarium | 13 | Pathotroph-Saprotroph-Symbiotroph | Animal Pathogen-Endophyte-Lichen Parasite-\|Plant Pathogen\|-Plant Saprotroph-Undefined Saprotroph-Wood Saprotroph | Microfungus | Soft Rot | Possible | As foliar_endophyte (Põlme et al. 2020); As litter_saprotroph (Põlme et al. 2020); Causing damping-off disease (Põlme et al. 2020); Decay leaf, fruit, seed, roots, soil, animal material (Põlme et al. 2020); Likely opportunistic human pathogen (Irinyi et al. 2015) | [illegible] |
| Otu29492 | 1 | k__Fungi(100);p__Basidiomycota(100);c__Agaricomycetes(100);o__Agaricales(100);f__Bolbitiaceae(100);g__Conocybe(100);s__Conocybe_watlingii(100); | Conocybe | 13 | Saprotroph | Dung Saprotroph-Plant Saprotroph-\|Undefined Saprotroph\| | Agaricoid | NULL | Probable | As litter_saprotroph (Põlme et al. 2020); As soil_saprotroph (Põlme et al. 2020); Decay soil (Põlme et al. 2020) | [illegible] |
| Otu29493 | 1 | k__Fungi(100);p__Ascomycota(100);c__Sordariomycetes(100);o__Xylariales(100);f__Amphisphaeriaceae(100);g__Monochaetia(100);g__Monochaetia_unclassified(100); | Monochaetia | 13 | Pathotroph | \|Plant Pathogen\| | Microfungus | NULL | Probable | NULL | [illegible] |
| Otu29494 | 1 | k__Fungi(100);p__Chytridiomycota(100);c__Chytridiomycetes(100);o__Spizellomycetales(100);f__Spizellomycetaceae(100);g__unclassified_Spizellomycetaceae(100);s__Spizellomycetaceae_sp(100); | Spizellomycetaceae | 9 | Pathotroph | Plant Pathogen-Undefined Parasite-Undefined Saprotroph | Microfungus | NULL | Probable | Common in the soil and also as parasites of a range of soil organisms and plants (Cannon & Kirk 2007) | [illegible] |
| Otu29495 | 1 | k__Fungi(100);p__Chytridiomycota(100);c__Chytridiomycetes(100);o__Spizellomycetales(100);f__Spizellomycetaceae(100);g__unclassified_Spizellomycetaceae(100);s__Spizellomycetaceae_sp(100); | Spizellomycetaceae | 9 | Pathotroph | Plant Pathogen-Undefined Parasite-Undefined Saprotroph | Microfungus | NULL | Probable | Common in the soil and also as parasites of a range of soil organisms and plants (Cannon & Kirk 2007) | [illegible] |
| Otu29496 | 1 | k__Fungi(100);p__Chytridiomycota(100);c__Chytridiomycetes(100);o__Rhizophydiales(100);f__Rhizophydiaceae(100);g__Rhizophydium(100);s__Rhizophydium_sp(100); | Rhizophydium | 13 | Pathotroph-Pathotroph-Saprotroph | \|Algal Parasite\|-Plant Pathogen-Plant Saprotroph-Protist Parasite-Undefined Saprotroph | Chytrid-Microfungus | NULL | Probable | As animal_decomposer (Põlme et al. 2020); As keratinolytic (Põlme et al. 2020); As litter_saprotroph (Põlme et al. 2020); Decay leaf, fruit, seed, algal material, fungal material, protist material (Põlme et al. 2020) | [illegible] |
| Otu29497 | 1 | k__Fungi(100);p__Zygomycota(100);c__Zygomycota_class_Incertae_sedis(100);o__Mortierellales(100);f__Mortierellaceae(100);g__unclassified_Mortierellaceae(100);s__Mortierellaceae_sp(100); | Mortierellaceae | 9 | Saprotroph-Symbiotroph | Endophyte-Litter Saprotroph-Soil Saprotroph-Undefined Saprotroph | Microfungus | NULL | Possible | Saprobic in soil and thus frequenctly isolated from soil and material contaminated with soil such as hay or rotten silage and spoil tips (Cannon & Kirk 2007) | [illegible] |
| Otu29498 | 1 | k__Fungi(100);p__Zygomycota(100);c__Zygomycota_class_Incertae_sedis(100);o__Mortierellales(100);f__Mortierellaceae(100);f__Mortierellaceae_unclassified(100);f__Mortierellaceae_unclassified(100); | Mortierellaceae | 9 | Saprotroph-Symbiotroph | Endophyte-Litter Saprotroph-Soil Saprotroph-Undefined Saprotroph | Microfungus | NULL | Possible | Saprobic in soil and thus frequenctly isolated from soil and material contaminated with soil such as hay or rotten silage and spoil tips (Cannon & Kirk 2007) | [illegible] |
| Otu29499 | 1 | k__Fungi(100);p__Zygomycota(100);c__Zygomycota_class_Incertae_sedis(100);o__Mortierellales(100);f__Mortierellaceae(100);g__unclassified_Mortierellaceae(100);s__Mortierellaceae_sp(100); | Mortierellaceae | 9 | Saprotroph-Symbiotroph | Endophyte-Litter Saprotroph-Soil Saprotroph-Undefined Saprotroph | Microfungus | NULL | Possible | Saprobic in soil and thus frequenctly isolated from soil and material contaminated with soil such as hay or rotten silage and spoil tips (Cannon & Kirk 2007) | [illegible] |
| Otu29500 | 1 | k__Fungi(100);p__Zygomycota(100);c__Zygomycota_class_Incertae_sedis(100);o__Mortierellales(100);f__Mortierellaceae(100);g__unclassified_Mortierellaceae(100);s__Mortierellaceae_sp(100); | Mortierellaceae | 9 | Saprotroph-Symbiotroph | Endophyte-Litter Saprotroph-Soil Saprotroph-Undefined Saprotroph | Microfungus | NULL | Possible | Saprobic in soil and thus frequenctly isolated from soil and material contaminated with soil such as hay or rotten silage and spoil tips (Cannon & Kirk 2007) | [illegible] |
| Otu29501 | 1 | k__Fungi(100);p__Zygomycota(100);c__Zygomycota_class_Incertae_sedis(100);o__Mortierellales(100);f__Mortierellaceae(100);g__unclassified_Mortierellaceae(100);s__Mortierellaceae_sp(100); | Mortierellaceae | 9 | Saprotroph-Symbiotroph | Endophyte-Litter Saprotroph-Soil Saprotroph-Undefined Saprotroph | Microfungus | NULL | Possible | Saprobic in soil and thus frequenctly isolated from soil and material contaminated with soil such as hay or rotten silage and spoil tips (Cannon & Kirk 2007) | [illegible] |
| Otu29502 | 1 | k__Fungi(100);p__Zygomycota(100);c__Zygomycota_class_Incertae_sedis(100);o__Mortierellales(100);f__Mortierellaceae(100);g__unclassified_Mortierellaceae(100);s__Mortierellaceae_sp(100); | Mortierellaceae | 9 | Saprotroph-Symbiotroph | Endophyte-Litter Saprotroph-Soil Saprotroph-Undefined Saprotroph | Microfungus | NULL | Possible | Saprobic in soil and thus frequenctly isolated from soil and material contaminated with soil such as hay or rotten silage and spoil tips (Cannon & Kirk 2007) | [illegible] |
| Otu29503 | 1 | k__Fungi(100);p__Zygomycota(100);c__Zygomycota_class_Incertae_sedis(100);o__Mortierellales(100);f__Mortierellaceae(100);g__unclassified_Mortierellaceae(100);s__Mortierellaceae_sp(100); | Mortierellaceae | 9 | Saprotroph-Symbiotroph | Endophyte-Litter Saprotroph-Soil Saprotroph-Undefined Saprotroph | Microfungus | NULL | Possible | Saprobic in soil and thus frequenctly isolated from soil and material contaminated with soil such as hay or rotten silage and spoil tips (Cannon & Kirk 2007) | [illegible] |
| Otu29504 | 1 | k__Fungi(100);p__Zygomycota(100);c__Zygomycota_class_Incertae_sedis(100);o__Mortierellales(100);f__Mortierellaceae(100);g__unclassified_Mortierellaceae(100);s__Mortierellaceae_sp(100); | Mortierellaceae | 9 | Saprotroph-Symbiotroph | Endophyte-Litter Saprotroph-Soil Saprotroph-Undefined Saprotroph | Microfungus | NULL | Possible | Saprobic in soil and thus frequenctly isolated from soil and material contaminated with soil such as hay or rotten silage and spoil tips (Cannon & Kirk 2007) | [illegible] |
| Otu29505 | 1 | k__Fungi(100);p__Zygomycota(100);c__Zygomycota_class_Incertae_sedis(100);o__Mortierellales(100);f__Mortierellaceae(100);g__unclassified_Mortierellaceae(100);s__Mortierellaceae_sp(100); | Mortierellaceae | 9 | Saprotroph-Symbiotroph | Endophyte-Litter Saprotroph-Soil Saprotroph-Undefined Saprotroph | Microfungus | NULL | Possible | Saprobic in soil and thus frequenctly isolated from soil and material contaminated with soil such as hay or rotten silage and spoil tips (Cannon & Kirk 2007) | [illegible] |
| Otu29506 | 1 | k__Fungi(100);p__Zygomycota(100);c__Zygomycota_class_Incertae_sedis(100);o__Mortierellales(100);f__Mortierellaceae(100);g__unclassified_Mortierellaceae(100);s__Mortierellaceae_sp(100); | Mortierellaceae | 9 | Saprotroph-Symbiotroph | Endophyte-Litter Saprotroph-Soil Saprotroph-Undefined Saprotroph | Microfungus | NULL | Possible | Saprobic in soil and thus frequenctly isolated from soil and material contaminated with soil such as hay or rotten silage and spoil tips (Cannon & Kirk 2007) | [illegible] |
| Otu29507 | 1 | k__Fungi(100);p__Zygomycota(100);c__Zygomycota_class_Incertae_sedis(100);o__Basidiobolales(100);f__Basidiobolaceae(100);g__Basidiobolus(100);s__Basidiobolus_ranarum(100); | Basidiobolus | 13 | Pathotroph-Saprotroph-Symbiotroph | Animal Pathogen-Animal Symbiotroph-\|Undefined Saprotroph\| | Microfungus | NULL | Probable | Cause of subcutaneous zygomycosis in humans (Cannon & Kirk 2007); Basidiobolus ranarum is commonly present in decaying fruit and vegetable matter, and as a commensal in the intestinal tract of frogs, toads, and lizards ; Decay leaf, fruit, seed, soil, dung, animal material (Põlme et al. 2020) | [illegible] |
| Otu29509 | 1 | k__Fungi(100);p__Zygomycota(100);c__Zygomycota_class_Incertae_sedis(100);o__Basidiobolales(100);f__Basidiobolaceae(100);g__Basidiobolus(100);g__Basidiobolus_unclassified(100); | Basidiobolus | 13 | Pathotroph-Saprotroph-Symbiotroph | Animal Pathogen-Animal Symbiotroph-\|Undefined Saprotroph\| | Microfungus | NULL | Probable | Cause of subcutaneous zygomycosis in humans (Cannon & Kirk 2007); Basidiobolus ranarum is commonly present in decaying fruit and vegetable matter, and as a commensal in the intestinal tract of frogs, toads, and lizards ; Decay leaf, fruit, seed, soil, dung, animal material (Põlme et al. 2020) | [illegible] |
| Otu29511 | 1 | k__Fungi(100);p__Zygomycota(100);c__Zygomycota_class_Incertae_sedis(100);o__Mortierellales(100);f__Mortierellaceae(100);g__unclassified_Mortierellaceae(100);s__Mortierellaceae_sp(100); | Mortierellaceae | 9 | Saprotroph-Symbiotroph | Endophyte-Litter Saprotroph-Soil Saprotroph-Undefined Saprotroph | Microfungus | NULL | Possible | Saprobic in soil and thus frequenctly isolated from soil and material contaminated with soil such as hay or rotten silage and spoil tips (Cannon & Kirk 2007) | [illegible] |
| Otu29512 | 1 | k__Fungi(100);p__Zygomycota(100);c__Zygomycota_class_Incertae_sedis(100);o__Mortierellales(100);f__Mortierellaceae(100);g__Mortierella(100);g__Mortierella_unclassified(100); | Mortierella | 13 | Saprotroph-Symbiotroph | Endophyte-Plant Saprotroph-\|Undefined Saprotroph\| | Microfungus | NULL | Probable | As chitinolytic (Põlme et al. 2020); As soil_saprotroph (Põlme et al. 2020); Decay roots, soil (Põlme et al. 2020); Saprobic in soil and thus frequenctly isolated from soil and material contaminated with soil such as hay or rotten silage and spoil tips (Cannon & Kirk 2007) | [illegible] |
| Otu29516 | 1 | k__Fungi(100);p__Chytridiomycota(100);c__Chytridiomycetes(100);o__Rhizophlyctidales(100);f__Rhizophlyctidaceae(100);g__Rhizophlyctis(100);s__Rhizophlyctis_rosea(100); | Rhizophlyctis | 13 | Pathotroph-Saprotroph | Algal Parasite-\|Plant Saprotroph\|-Undefined Saprotroph | Chytrid-Microfungus | NULL | Probable | As litter_saprotroph (Põlme et al. 2020); Decay leaf, fruit, seed, algal material (Põlme et al. 2020) | [illegible] |
| Otu29517 | 1 | k__Fungi(100);p__Basidiomycota(100);c__Tremellomycetes(100);o__Filobasidiales(100);f__Filobasidiaceae(100);g__Cryptococcus(100);g__Cryptococcus_unclassified(100); | Cryptococcus | 13 | Pathotroph-Saprotroph-Symbiotroph | Animal Pathogen-Endophyte-Epiphyte-\|Undefined Saprotroph\| | Dimorphic Yeast-Yeast | NULL | Possible | As foliar_endophyte (Põlme et al. 2020); Decay leaf, fruit, seed, soil, dung, animal material (Põlme et al. 2020) | [illegible] |
| Otu29524 | 1 | k__Fungi(100);p__Chytridiomycota(100);c__Chytridiomycetes(100);o__Rhizophlyctidales(100);f__Rhizophlyctidaceae(100);g__Rhizophlyctis(100);s__Rhizophlyctis_rosea(100); | Rhizophlyctis | 13 | Pathotroph-Saprotroph | Algal Parasite-\|Plant Saprotroph\|-Undefined Saprotroph | Chytrid-Microfungus | NULL | Probable | As litter_saprotroph (Põlme et al. 2020); Decay leaf, fruit, seed, algal material (Põlme et al. 2020) | [illegible] |
| Otu29526 | 1 | k__Fungi(100);p__Basidiomycota(100);c__Microbotryomycetes(100);o__Sporidiobolales(100);f__Sporidiobolales_family_Incertae_sedis(100);g__Rhodotorula(100);s__Rhodotorula_sp(100); | Rhodotorula | 13 | Pathotroph-Saprotroph-Symbiotroph | Animal Pathogen-Animal Symbiotroph-Endophyte-Plant Pathogen-\|Undefined Saprotroph\| | Yeast | NULL | Probable | As foliar_endophyte (Põlme et al. 2020); Opportunistic human pathogen | [illegible] |
| Otu29528 | 1 | k__Fungi(100);p__Basidiomycota(100);c__Agaricomycetes(100);o__Hymenochaetales(100);f__Schizoporaceae(100);f__Schizoporaceae_unclassified(100);f__Schizoporaceae_unclassified(100); | Schizoporaceae | 9 | Saprotroph | Wood Saprotroph | Corticioid | White Rot | Probable | Saprobic, causing white rot of standing and fallen wood of coniferous and broadleaved trees, rarely on herbaceous stems (Cannon & Kirk 2007) | [illegible] |
| Otu29529 | 1 | k__Fungi(100);p__Ascomycota(100);c__Sordariomycetes(100);o__Diaporthales(100);f__Diaporthaceae(100);f__Diaporthaceae_unclassified(100);f__Diaporthaceae_unclassified(100); | Diaporthaceae | 9 | Pathotroph-Saprotroph-Symbiotroph | Endophyte-Plant Pathogen-Plant Saprotroph | Microfungus | NULL | Probable | Most species are weak pathogens or endophytes in leaf or stem tissue, growing saprobically after the plant dies (Cannon & Kirk 2007) | [illegible] |
| Otu29530 | 1 | k__Fungi(100);p__Ascomycota(100);c__Sordariomycetes(100);o__Xylariales(100);f__Amphisphaeriaceae(100);g__Truncatella(100);s__Truncatella_spadicea(100); | Truncatella | 13 | Pathotroph | \|Plant Pathogen\| | Microfungus | NULL | Probable | NULL | [illegible] |
| Otu29531 | 1 | k__Fungi(100);p__Basidiomycota(100);c__Microbotryomycetes(100);o__Sporidiobolales(100);f__Sporidiobolales_family_Incertae_sedis(100);g__Rhodotorula(100);s__Rhodotorula_mucilaginosa(100); | Rhodotorula | 13 | Pathotroph-Saprotroph-Symbiotroph | Animal Pathogen-Animal Symbiotroph-Endophyte-Plant Pathogen-\|Undefined Saprotroph\| | Yeast | NULL | Probable | As foliar_endophyte (Põlme et al. 2020); Opportunistic human pathogen | [illegible] |
| Otu29532 | 1 | k__Fungi(100);p__Chytridiomycota(100);c__Chytridiomycetes(100);o__Rhizophlyctidales(100);f__Rhizophlyctidaceae(100);g__Rhizophlyctis(100);s__Rhizophlyctis_rosea(100); | Rhizophlyctis | 13 | Pathotroph-Saprotroph | Algal Parasite-\|Plant Saprotroph\|-Undefined Saprotroph | Chytrid-Microfungus | NULL | Probable | As litter_saprotroph (Põlme et al. 2020); Decay leaf, fruit, seed, algal material (Põlme et al. 2020) | [illegible] |
| Otu29533 | 1 | k__Fungi(100);p__Ascomycota(100);c__Dothideomycetes(100);o__Pleosporales(100);f__Sporormiaceae(100);g__Preussia(100);g__Preussia_unclassified(100); | Preussia | 13 | Saprotroph | \|Dung Saprotroph\|-Undefined Saprotroph | Microfungus | NULL | Probable | Decay dung (Põlme et al. 2020) | [illegible] |
| Otu29534 | 1 | k__Fungi(100);p__Ascomycota(100);c__Sordariomycetes(100);o__Hypocreales(100);f__Hypocreaceae(100);g__Hypocrea(100);g__Hypocrea_unclassified(100); | Hypocrea | 13 | Pathotroph | Endophyte-Fungal Parasite | Microfungus | NULL | Probable | NULL | [illegible] |
| Otu29535 | 1 | k__Fungi(100);p__Ascomycota(100);c__Ascomycota_class_Incertae_sedis(100);o__Ascomycota_order_Incertae_sedis(100);f__Ascomycota_family_Incertae_sedis(100);g__Parasympodiella(100);s__Parasympodiella_elongata(100); | Parasympodiella | 13 | Saprotroph | Undefined Saprotroph-\|Wood Saprotroph\| | Microfungus | NULL | Probable | Decay wood (Põlme et al. 2020) | [illegible] |
| Otu29538 | 1 | k__Fungi(100);p__Chytridiomycota(100);c__Chytridiomycetes(100);o__Rhizophlyctidales(100);f__Rhizophlyctidaceae(100);g__Rhizophlyctis(100);s__Rhizophlyctis_rosea(100); | Rhizophlyctis | 13 | Pathotroph-Saprotroph | Algal Parasite-\|Plant Saprotroph\|-Undefined Saprotroph | Chytrid-Microfungus | NULL | Probable | As litter_saprotroph (Põlme et al. 2020); Decay leaf, fruit, seed, algal material (Põlme et al. 2020) | [illegible] |
| Otu29539 | 1 | k__Fungi(100);p__Zygomycota(100);c__Zygomycota_class_Incertae_sedis(100);o__Mortierellales(100);f__Mortierellaceae(100);g__Mortierella(100);s__Mortierella_reticulata(100); | Mortierella | 13 | Saprotroph-Symbiotroph | Endophyte-Plant Saprotroph-\|Undefined Saprotroph\| | Microfungus | NULL | Probable | As chitinolytic (Põlme et al. 2020); As soil_saprotroph (Põlme et al. 2020); Decay roots, soil (Põlme et al. 2020); Saprobic in soil and thus frequenctly isolated from soil and material contaminated with soil such as hay or rotten silage and spoil tips (Cannon & Kirk 2007) | [illegible] |
| Otu29540 | 1 | k__Fungi(100);p__Zygomycota(100);c__Zygomycota_class_Incertae_sedis(100);o__Mortierellales(100);f__Mortierellaceae(100);g__unclassified_Mortierellaceae(100);s__Mortierellaceae_sp(100); | Mortierellaceae | 9 | Saprotroph-Symbiotroph | Endophyte-Litter Saprotroph-Soil Saprotroph-Undefined Saprotroph | Microfungus | NULL | Possible | Saprobic in soil and thus frequenctly isolated from soil and material contaminated with soil such as hay or rotten silage and spoil tips (Cannon & Kirk 2007) | [illegible] |
| Otu29541 | 1 | k__Fungi(100);p__Zygomycota(100);c__Zygomycota_class_Incertae_sedis(100);o__Mortierellales(100);f__Mortierellaceae(100);g__unclassified_Mortierellaceae(100);s__Mortierellaceae_sp(100); | Mortierellaceae | 9 | Saprotroph-Symbiotroph | Endophyte-Litter Saprotroph-Soil Saprotroph-Undefined Saprotroph | Microfungus | NULL | Possible | Saprobic in soil and thus frequenctly isolated from soil and material contaminated with soil such as hay or rotten silage and spoil tips (Cannon & Kirk 2007) | [illegible] |
| Otu29542 | 1 | k__Fungi(100);p__Zygomycota(100);c__Zygomycota_class_Incertae_sedis(100);o__Basidiobolales(100);f__Basidiobolaceae(100);g__Basidiobolus(100);s__Basidiobolus_ranarum(100); | Basidiobolus | 13 | Pathotroph-Saprotroph-Symbiotroph | Animal Pathogen-Animal Symbiotroph-\|Undefined Saprotroph\| | Microfungus | NULL | Probable | Cause of subcutaneous zygomycosis in humans (Cannon & Kirk 2007); Basidiobolus ranarum is commonly present in decaying fruit and vegetable matter, and as a commensal in the intestinal tract of frogs, toads, and lizards ; Decay leaf, fruit, seed, soil, dung, animal material (Põlme et al. 2020) | [illegible] |
| Otu29543 | 1 | k__Fungi(100);p__Zygomycota(100);c__Zygomycota_class_Incertae_sedis(100);o__Basidiobolales(100);f__Basidiobolaceae(100);g__Basidiobolus(100);s__Basidiobolus_ranarum(100); | Basidiobolus | 13 | Pathotroph-Saprotroph-Symbiotroph | Animal Pathogen-Animal Symbiotroph-\|Undefined Saprotroph\| | Microfungus | NULL | Probable | Cause of subcutaneous zygomycosis in humans (Cannon & Kirk 2007); Basidiobolus ranarum is commonly present in decaying fruit and vegetable matter, and as a commensal in the intestinal tract of frogs, toads, and lizards ; Decay leaf, fruit, seed, soil, dung, animal material (Põlme et al. 2020) | [illegible] |
| Otu29544 | 1 | k__Fungi(100);p__Zygomycota(100);c__Zygomycota_class_Incertae_sedis(100);o__Basidiobolales(100);f__Basidiobolaceae(100);g__Basidiobolus(100);g__Basidiobolus_unclassified(100); | Basidiobolus | 13 | Pathotroph-Saprotroph-Symbiotroph | Animal Pathogen-Animal Symbiotroph-\|Undefined Saprotroph\| | Microfungus | NULL | Probable | Cause of subcutaneous zygomycosis in humans (Cannon & Kirk 2007); Basidiobolus ranarum is commonly present in decaying fruit and vegetable matter, and as a commensal in the intestinal tract of frogs, toads, and lizards ; Decay leaf, fruit, seed, soil, dung, animal material (Põlme et al. 2020) | [illegible] |
| Otu29545 | 1 | k__Fungi(100);p__Zygomycota(100);c__Zygomycota_class_Incertae_sedis(100);o__Mortierellales(100);f__Mortierellaceae(100);g__Mortierella(100);s__Mortierella_rostafinskii(100); | Mortierella | 13 | Saprotroph-Symbiotroph | Endophyte-Plant Saprotroph-\|Undefined Saprotroph\| | Microfungus | NULL | Probable | As chitinolytic (Põlme et al. 2020); As soil_saprotroph (Põlme et al. 2020); Decay roots, soil (Põlme et al. 2020); Saprobic in soil and thus frequenctly isolated from soil and material contaminated with soil such as hay or rotten silage and spoil tips (Cannon & Kirk 2007) | [illegible] |
| Otu29546 | 1 | k__Fungi(100);p__Zygomycota(100);c__Zygomycota_class_Incertae_sedis(100);o__Mortierellales(100);f__Mortierellaceae(100);g__unclassified_Mortierellaceae(100);s__Mortierellaceae_sp(100); | Mortierellaceae | 9 | Saprotroph-Symbiotroph | Endophyte-Litter Saprotroph-Soil Saprotroph-Undefined Saprotroph | Microfungus | NULL | Possible | Saprobic in soil and thus frequenctly isolated from soil and material contaminated with soil such as hay or rotten silage and spoil tips (Cannon & Kirk 2007) | [illegible] |
| Otu29547 | 1 | k__Fungi(100);p__Zygomycota(100);c__Zygomycota_class_Incertae_sedis(100);o__Mortierellales(100);f__Mortierellaceae(100);g__unclassified_Mortierellaceae(100);s__Mortierellaceae_sp(100); | Mortierellaceae | 9 | Saprotroph-Symbiotroph | Endophyte-Litter Saprotroph-Soil Saprotroph-Undefined Saprotroph | Microfungus | NULL | Possible | Saprobic in soil and thus frequenctly isolated from soil and material contaminated with soil such as hay or rotten silage and spoil tips (Cannon & Kirk 2007) | [illegible] |
| Otu29548 | 1 | k__Fungi(100);p__Zygomycota(100);c__Zygomycota_class_Incertae_sedis(100);o__Mortierellales(100);f__Mortierellaceae(100);g__unclassified_Mortierellaceae(100);s__Mortierellaceae_sp(100); | Mortierellaceae | 9 | Saprotroph-Symbiotroph | Endophyte-Litter Saprotroph-Soil Saprotroph-Undefined Saprotroph | Microfungus | NULL | Possible | Saprobic in soil and thus frequenctly isolated from soil and material contaminated with soil such as hay or rotten silage and spoil tips (Cannon & Kirk 2007) | [illegible] |
| Otu29549 | 1 | k__Fungi(100);p__Zygomycota(100);c__Zygomycota_class_Incertae_sedis(100);o__Mortierellales(100);f__Mortierellaceae(100);g__unclassified_Mortierellaceae(100);s__Mortierellaceae_sp(100); | Mortierellaceae | 9 | Saprotroph-Symbiotroph | Endophyte-Litter Saprotroph-Soil Saprotroph-Undefined Saprotroph | Microfungus | NULL | Possible | Saprobic in soil and thus frequenctly isolated from soil and material contaminated with soil such as hay or rotten silage and spoil tips (Cannon & Kirk 2007) | [illegible] |
| Otu29550 | 1 | k__Fungi(100);p__Ascomycota(100);c__Dothideomycetes(100);o__Pleosporales(100);f__Tubeufiaceae(100);f__Tubeufiaceae_unclassified(100);f__Tubeufiaceae_unclassified(100); | Tubeufiaceae | 9 | Pathotroph-Saprotroph | Animal Pathogen-Undefined Saprotroph | NULL | NULL | Probable | Biotrophic or saprobic, especially on scale insects or other fungi. The anamorphic forms are fequently aero-aquatic (Cannon & Kirk 2007) | [illegible] |
| Otu29552 | 1 | k__Fungi(100);p__Ascomycota(100);c__Sordariomycetes(100);o__Hypocreales(100);o__Hypocreales_unclassified(100);o__Hypocreales_unclassified(100);o__Hypocreales_unclassified(100); | Hypocreales | 7 | Saprotroph | Undefined Saprotroph | Microfungus | NULL | Possible | NULL | [illegible] |
| Otu29554 | 1 | k__Fungi(100);p__Ascomycota(100);c__Dothideomycetes(100);o__Pleosporales(100);f__Venturiaceae(100);g__Fusicladium(100);s__Fusicladium_cordae(100); | Fusicladium | 13 | Pathotroph-Symbiotroph | Endophyte-Lichen Parasite-\|Plant Pathogen\| | Microfungus | NULL | Probable | NULL | [illegible] |
| Otu29558 | 1 | k__Fungi(100);p__Ascomycota(100);c__Ascomycota_class_Incertae_sedis(100);o__Ascomycota_order_Incertae_sedis(100);f__Ascomycota_family_Incertae_sedis(100);g__Triscelophorus(100);s__Triscelophorus_cf_acuminatus_UMB_55810(100); | Triscelophorus | 13 | Saprotroph-Symbiotroph | \|Endophyte\|-Plant Saprotroph | Microfungus | NULL | Highly Probable | Rubiaceae, Marattiaceae, Rhizophoraceae, Thelypteridaceae; As foliar_endophyte (Põlme et al. 2020); As litter_saprotroph (Põlme et al. 2020); Decay leaf, fruit, seed (Põlme et al. 2020); Host | [illegible] |
| Otu29559 | 1 | k__Fungi(100);p__Ascomycota(100);c__Pezizomycetes(100);o__Pezizales(100);f__Pezizaceae(100);g__Hydnobolites(100);s__Hydnobolites_sp_11_RH_2012(100); | Hydnobolites | 13 | Symbiotroph | \|Ectomycorrhizal\| | Gasteroid | Hypogeous | Highly Probable | NULL | [illegible] |
| Otu29561 | 1 | k__Fungi(100);p__Ascomycota(100);c__Dothideomycetes(100);o__Pleosporales(100);f__Lophiostomataceae(100);g__Lophiostoma(100);s__Lophiostoma_arundinis(100); | Lophiostoma | 13 | Saprotroph | Plant Saprotroph-Undefined Saprotroph-\|Wood Saprotroph\| | Microfungus | NULL | Probable | As litter_saprotroph (Põlme et al. 2020); Decay wood, leaf, fruit, seed (Põlme et al. 2020) | [illegible] |
| Otu29563 | 1 | k__Fungi(100);p__Ascomycota(100);c__Eurotiomycetes(100);o__Chaetothyriales(100);f__Herpotrichiellaceae(100);g__Exophiala(100);g__Exophiala_unclassified(100); | Exophiala | 13 | Pathotroph-Saprotroph-Symbiotroph | \|Animal Parasite\|-Animal Pathogen-Endophyte-Plant Saprotroph-Undefined Saprotroph | Dimorphic-Facultative Yeast | NULL | Probable | As litter_saprotroph (Põlme et al. 2020); As root_endophyte_dark_septate (Põlme et al. 2020); Decay leaf, fruit, seed, animal material (Põlme et al. 2020) | [illegible] |
| Otu29568 | 1 | k__Fungi(100);p__Ascomycota(100);c__Eurotiomycetes(100);o__Chaetothyriales(100);f__Herpotrichiellaceae(100);g__Cladophialophora(100);g__Cladophialophora_unclassified(100); | Cladophialophora | 13 | Pathotroph-Saprotroph-Symbiotroph | Animal Pathogen-Endophyte-Fungal Parasite-\|Undefined Saprotroph\| | Dimorphic-Microfungus | NULL | Probable | As mycoparasite (Põlme et al. 2020); As root_endophyte_dark_septate (Põlme et al. 2020); As soil_saprotroph (Põlme et al. 2020); Decay soil (Põlme et al. 2020) | [illegible] |
| Otu29569 | 1 | k__Fungi(100);p__Ascomycota(100);c__Eurotiomycetes(100);o__Chaetothyriales(100);f__Herpotrichiellaceae(100);g__Rhinocladiella(100);g__Rhinocladiella_unclassified(100); | Rhinocladiella | 13 | Pathotroph-Saprotroph-Symbiotroph | Animal Parasite-Endophyte-\|Undefined Saprotroph\|-Wood Saprotroph | Dimorphic-Facultative Yeast | Soft Rot | Probable | As root_endophyte_dark_septate (Põlme et al. 2020); As soil_saprotroph (Põlme et al. 2020); Decay soil, animal material (Põlme et al. 2020) | [illegible] |
| Otu29570 | 1 | k__Fungi(100);p__Ascomycota(100);c__Eurotiomycetes(100);o__Chaetothyriales(100);f__Herpotrichiellaceae(100);f__Herpotrichiellaceae_unclassified(100);f__Herpotrichiellaceae_unclassified(100); | Herpotrichiellaceae | 9 | Pathotroph-Saprotroph | Animal Pathogen-Fungal Parasite-Undefined Saprotroph | Facultative Yeast-Microfungus | NULL | Probable | Mostly saprobic on plants or other fungi, isolated from soil etc., and from medical sources (Cannon & Kirk 2007) | [illegible] |
| Otu29571 | 1 | k__Fungi(100);p__Ascomycota(100);c__Eurotiomycetes(100);o__Chaetothyriales(100);f__Herpotrichiellaceae(100);f__Herpotrichiellaceae_unclassified(100);f__Herpotrichiellaceae_unclassified(100); | Herpotrichiellaceae | 9 | Pathotroph-Saprotroph | Animal Pathogen-Fungal Parasite-Undefined Saprotroph | Facultative Yeast-Microfungus | NULL | Probable | Mostly saprobic on plants or other fungi, isolated from soil etc., and from medical sources (Cannon & Kirk 2007) | [illegible] |
| Otu29572 | 1 | k__Fungi(100);p__Ascomycota(100);c__Eurotiomycetes(100);o__Chaetothyriales(100);f__Herpotrichiellaceae(100);g__Cladophialophora(100);s__Cladophialophora_sp_olrim407(100); | Cladophialophora | 13 | Pathotroph-Saprotroph-Symbiotroph | Animal Pathogen-Endophyte-Fungal Parasite-\|Undefined Saprotroph\| | Dimorphic-Microfungus | NULL | Probable | As mycoparasite (Põlme et al. 2020); As root_endophyte_dark_septate (Põlme et al. 2020); As soil_saprotroph (Põlme et al. 2020); Decay soil (Põlme et al. 2020) | [illegible] |
| Otu29574 | 1 | k__Fungi(100);p__Ascomycota(100);c__Eurotiomycetes(100);o__Chaetothyriales(100);f__Herpotrichiellaceae(100);g__Cladophialophora(100);s__Cladophialophora_sp_LS_2013(100); | Cladophialophora | 13 | Pathotroph-Saprotroph-Symbiotroph | Animal Pathogen-Endophyte-Fungal Parasite-\|Undefined Saprotroph\| | Dimorphic-Microfungus | NULL | Probable | As mycoparasite (Põlme et al. 2020); As root_endophyte_dark_septate (Põlme et al. 2020); As soil_saprotroph (Põlme et al. 2020); Decay soil (Põlme et al. 2020) | [illegible] |
| Otu29575 | 1 | k__Fungi(100);p__Ascomycota(100);c__Eurotiomycetes(100);o__Chaetothyriales(100);f__Herpotrichiellaceae(100);g__Cladophialophora(100);s__Cladophialophora_chaetospira(100); | Cladophialophora | 13 | Pathotroph-Saprotroph-Symbiotroph | Animal Pathogen-Endophyte-Fungal Parasite-\|Undefined Saprotroph\| | Dimorphic-Microfungus | NULL | Probable | As mycoparasite (Põlme et al. 2020); As root_endophyte_dark_septate (Põlme et al. 2020); As soil_saprotroph (Põlme et al. 2020); Decay soil (Põlme et al. 2020) | [illegible] |
| Otu29576 | 1 | k__Fungi(100);p__Ascomycota(100);c__Eurotiomycetes(100);o__Chaetothyriales(100);f__Herpotrichiellaceae(100);g__Exophiala(100);g__Exophiala_unclassified(100); | Exophiala | 13 | Pathotroph-Saprotroph-Symbiotroph | \|Animal Parasite\|-Animal Pathogen-Endophyte-Plant Saprotroph-Undefined Saprotroph | Dimorphic-Facultative Yeast | NULL | Probable | As litter_saprotroph (Põlme et al. 2020); As root_endophyte_dark_septate (Põlme et al. 2020); Decay leaf, fruit, seed, animal material (Põlme et al. 2020) | [illegible] |

| | | | | | | | | | | | |
|---|---|---|---|---|---|---|---|---|---|---|---|
| Otu29579 | 1 | k__Fungi(100);p__Ascomycota(100);c__Sordariomycetes(100);o__Hypocreales(100);f__Nectriaceae(100);g__Fusarium(100);s__Fusarium_tricinctum(100); | Fusarium | 13 | Pathotroph-Saprotroph-Symbiotroph | Animal Pathogen-Endophyte-Lichen Parasite-\|Plant Pathogen\|-Plant Saprotroph-Undefined Saprotroph-Wood Saprotroph | Microfungus | Soft Rot | Possible | As foliar_endophyte (Põlme et al. 2020); As litter_saprotroph (Põlme et al. 2020); Causing damping-off disease (Põlme et al. 2020); Decay leaf, fruit, seed, roots, soil, animal material (Põlme et al. 2020); Likely opportunistic human pathogen (Irinyi et al. 2015) | [illegible] |
| Otu29580 | 1 | k__Fungi(100);p__Ascomycota(100);c__Sordariomycetes(100);o__Hypocreales(100);f__Nectriaceae(100);g__Fusarium(100);g__Fusarium_unclassified(100); | Fusarium | 13 | Pathotroph-Saprotroph-Symbiotroph | Animal Pathogen-Endophyte-Lichen Parasite-\|Plant Pathogen\|-Plant Saprotroph-Undefined Saprotroph-Wood Saprotroph | Microfungus | Soft Rot | Possible | As foliar_endophyte (Põlme et al. 2020); As litter_saprotroph (Põlme et al. 2020); Causing damping-off disease (Põlme et al. 2020); Decay leaf, fruit, seed, roots, soil, animal material (Põlme et al. 2020); Likely opportunistic human pathogen (Irinyi et al. 2015) | [illegible] |
| Otu29582 | 1 | k__Fungi(100);p__Chytridiomycota(100);c__Chytridiomycetes(100);o__Spizellomycetales(100);f__Spizellomycetaceae(100);f__Spizellomycetaceae_unclassified(100);f__Spizellomycetaceae_unclassified(100); | Spizellomycetaceae | 9 | Pathotroph | Plant Pathogen-Undefined Parasite-Undefined Saprotroph | Microfungus | NULL | Probable | Common in the soil and also as parasites of a range of soil organisms and plants (Cannon & Kirk 2007) | [illegible] |
| Otu29583 | 1 | k__Fungi(100);p__Chytridiomycota(100);c__Chytridiomycetes(100);o__Spizellomycetales(100);f__Spizellomycetaceae(100);g__Spizellomyces(100);g__Spizellomyces_unclassified(100); | Spizellomyces | 13 | Pathotroph-Saprotroph | Plant Pathogen-\|Pollen Saprotroph\|-Undefined Parasite-Undefined Saprotroph | Chytrid-Microfungus | NULL | Probable | Decay pollen (Põlme et al. 2020); Pollen degrader; Common in the soil and also as parasites of a range of soil organisms and plants (Cannon & Kirk 2007) | [illegible] |
| Otu29584 | 1 | k__Fungi(100);p__Chytridiomycota(100);c__Chytridiomycetes(100);o__Spizellomycetales(100);f__Spizellomycetaceae(100);f__Spizellomycetaceae_unclassified(100);f__Spizellomycetaceae_unclassified(100); | Spizellomycetaceae | 9 | Pathotroph | Plant Pathogen-Undefined Parasite-Undefined Saprotroph | Microfungus | NULL | Probable | Common in the soil and also as parasites of a range of soil organisms and plants (Cannon & Kirk 2007) | [illegible] |
| Otu29586 | 1 | k__Fungi(100);p__Basidiomycota(100);c__Microbotryomycetes(100);o__Sporidiobolales(100);f__Sporidiobolales_family_Incertae_sedis(100);g__Rhodotorula(100);s__Rhodotorula_mucilaginosa(100); | Rhodotorula | 13 | Pathotroph-Saprotroph-Symbiotroph | Animal Pathogen-Animal Symbiotroph-Endophyte-Plant Pathogen-\|Undefined Saprotroph\| | Yeast | NULL | Probable | As foliar_endophyte (Põlme et al. 2020); Opportunistic human pathogen | [illegible] |
| Otu29587 | 1 | k__Fungi(100);p__Basidiomycota(100);c__Microbotryomycetes(100);o__Sporidiobolales(100);f__Sporidiobolales_family_Incertae_sedis(100);g__Rhodotorula(100);s__Rhodotorula_taiwanensis(100); | Rhodotorula | 13 | Pathotroph-Saprotroph-Symbiotroph | Animal Pathogen-Animal Symbiotroph-Endophyte-Plant Pathogen-\|Undefined Saprotroph\| | Yeast | NULL | Probable | As foliar_endophyte (Põlme et al. 2020); Opportunistic human pathogen | [illegible] |
| Otu29588 | 1 | k__Fungi(100);p__Ascomycota(100);c__Sordariomycetes(100);o__Sordariomycetes_order_Incertae_sedis(100);f__Glomerellaceae(100);g__Colletotrichum(100);g__Colletotrichum_unclassified(100); | Colletotrichum | 13 | Pathotroph-Saprotroph-Symbiotroph | Endophyte-\|Plant Pathogen\|-Plant Saprotroph | Microfungus | NULL | Probable | As foliar_endophyte (Põlme et al. 2020); As litter_saprotroph (Põlme et al. 2020); Decay leaf, fruit, seed (Põlme et al. 2020) | [illegible] |
| Otu29590 | 1 | k__Fungi(100);p__Basidiomycota(100);c__Microbotryomycetes(100);o__Sporidiobolales(100);f__Sporidiobolales_family_Incertae_sedis(100);g__Rhodotorula(100);s__Rhodotorula_taiwanensis(100); | Rhodotorula | 13 | Pathotroph-Saprotroph-Symbiotroph | Animal Pathogen-Animal Symbiotroph-Endophyte-Plant Pathogen-\|Undefined Saprotroph\| | Yeast | NULL | Probable | As foliar_endophyte (Põlme et al. 2020); Opportunistic human pathogen | [illegible] |
| Otu29597 | 1 | k__Fungi(100);p__Zygomycota(100);c__Zygomycota_class_Incertae_sedis(100);o__Mortierellales(100);f__Mortierellaceae(100);g__Mortierella(100);g__Mortierella_unclassified(100); | Mortierella | 13 | Saprotroph-Symbiotroph | Endophyte-Plant Saprotroph-\|Undefined Saprotroph\| | Microfungus | NULL | Probable | As chitinolytic (Põlme et al. 2020); As soil_saprotroph (Põlme et al. 2020); Decay roots, soil (Põlme et al. 2020); Saprobic in soil and thus frequently isolated from soil and material contaminated with soil such as hay or rotten silage and spoil tips (Cannon & Kirk 2007) | [illegible] |
| Otu29600 | 1 | k__Fungi(100);p__Ascomycota(100);c__Pezizomycetes(100);o__Pezizales(100);f__Pyronemataceae(100);g__Trichophaea(100);s__Trichophaea_woolhopeia(100); | Trichophaea | 13 | Saprotroph-Symbiotroph | Dung Saprotroph-\|Ectomycorrhizal\|-Undefined Saprotroph | Microfungus-Pezizoid | NULL | Probable | As soil_saprotroph (Põlme et al. 2020) | [illegible] |
| Otu29601 | 1 | k__Fungi(100);p__Ascomycota(100);c__Pezizomycetes(100);o__Pezizales(100);f__Sarcosomataceae(100);f__Sarcosomataceae_unclassified(100);f__Sarcosomataceae_unclassified(100); | Sarcosomataceae | 9 | Saprotroph | Wood Saprotroph | Pezizoid | NULL | Probable | Saprobic on rotten, sometimes buried, wood (Cannon & Kirk 2007) | [illegible] |
| Otu29604 | 1 | k__Fungi(100);p__Basidiomycota(100);c__Agaricomycetes(100);o__Thelephorales(100);f__Thelephoraceae(100);g__Tomentella(100);s__Tomentella_terrestris(100); | Tomentella | 13 | Symbiotroph | \|Ectomycorrhizal\| | Corticioid | NULL | Highly Probable | NULL | [illegible] |
| Otu29605 | 1 | k__Fungi(100);p__Basidiomycota(100);c__Agaricomycetes(100);o__Thelephorales(100);f__Thelephoraceae(100);g__Tomentella(100);g__Tomentella_unclassified(100); | Tomentella | 13 | Symbiotroph | \|Ectomycorrhizal\| | Corticioid | NULL | Highly Probable | NULL | [illegible] |
| Otu29606 | 1 | k__Fungi(100);p__Basidiomycota(100);c__Agaricomycetes(100);o__Russulales(100);f__Peniophoraceae(100);f__Peniophoraceae_unclassified(100);f__Peniophoraceae_unclassified(100); | Peniophoraceae | 9 | Pathotroph-Saprotroph | Plant Pathogen-Wood Saprotroph | Corticioid | White Rot | Probable | Most species are saprobic, causing rots of standing and fallen wood. Many species are adapted to xeric conditions, being able to grow on rapidly drying dead attached branches (Cannon & Kirk 2007) | [illegible] |
| Otu29607 | 1 | k__Fungi(100);p__Basidiomycota(100);c__Agaricomycetes(100);o__Russulales(100);f__Russulaceae(100);g__Russula(100);s__Russula_brunneoviolacea(100); | Russula | 13 | Symbiotroph | \|Ectomycorrhizal\| | Agaricoid | NULL | Highly Probable | NULL | [illegible] |
| Otu29608 | 1 | k__Fungi(100);p__Basidiomycota(100);c__Agaricomycetes(100);o__Russulales(100);f__Peniophoraceae(100);g__Peniophora(100);g__Peniophora_unclassified(100); | Peniophora | 13 | Pathotroph-Saprotroph | Plant Pathogen-\|Wood Saprotroph\| | Corticioid | White Rot | Probable | As white_rot (Põlme et al. 2020); As wood_pathogen (Põlme et al. 2020); Decay wood (Põlme et al. 2020) | [illegible] |
| Otu29609 | 1 | k__Fungi(100);p__Basidiomycota(100);c__Agaricomycetes(100);o__Russulales(100);f__Peniophoraceae(100);f__Peniophoraceae_unclassified(100);f__Peniophoraceae_unclassified(100); | Peniophoraceae | 9 | Pathotroph-Saprotroph | Plant Pathogen-Wood Saprotroph | Corticioid | White Rot | Probable | Most species are saprobic, causing rots of standing and fallen wood. Many species are adapted to xeric conditions, being able to grow on rapidly drying dead attached branches (Cannon & Kirk 2007) | [illegible] |
| Otu29610 | 1 | k__Fungi(100);p__Basidiomycota(100);c__Agaricomycetes(100);o__Russulales(100);f__Peniophoraceae(100);g__Peniophora(100);g__Peniophora_unclassified(100); | Peniophora | 13 | Pathotroph-Saprotroph | Plant Pathogen-\|Wood Saprotroph\| | Corticioid | White Rot | Probable | As white_rot (Põlme et al. 2020); As wood_pathogen (Põlme et al. 2020); Decay wood (Põlme et al. 2020) | [illegible] |
| Otu29611 | 1 | k__Fungi(100);p__Basidiomycota(100);c__Agaricomycetes(100);o__Russulales(100);f__Peniophoraceae(100);g__Dendrophora(100);s__Dendrophora_sp(100); | Dendrophora | 13 | Saprotroph | Undefined Saprotroph-\|Wood Saprotroph\| | Corticioid | White Rot | Probable | As white_rot (Põlme et al. 2020); Decay wood (Põlme et al. 2020) | [illegible] |
| Otu29612 | 1 | k__Fungi(100);p__Basidiomycota(100);c__Agaricomycetes(100);o__Russulales(100);f__Peniophoraceae(100);f__Peniophoraceae_unclassified(100);f__Peniophoraceae_unclassified(100); | Peniophoraceae | 9 | Pathotroph-Saprotroph | Plant Pathogen-Wood Saprotroph | Corticioid | White Rot | Probable | Most species are saprobic, causing rots of standing and fallen wood. Many species are adapted to xeric conditions, being able to grow on rapidly drying dead attached branches (Cannon & Kirk 2007) | [illegible] |
| Otu29613 | 1 | k__Fungi(100);p__Basidiomycota(100);c__Agaricomycetes(100);o__Russulales(100);f__Peniophoraceae(100);f__Peniophoraceae_unclassified(100);f__Peniophoraceae_unclassified(100); | Peniophoraceae | 9 | Pathotroph-Saprotroph | Plant Pathogen-Wood Saprotroph | Corticioid | White Rot | Probable | Most species are saprobic, causing rots of standing and fallen wood. Many species are adapted to xeric conditions, being able to grow on rapidly drying dead attached branches (Cannon & Kirk 2007) | [illegible] |
| Otu29614 | 1 | k__Fungi(100);p__Basidiomycota(100);c__Agaricomycetes(100);o__Russulales(100);f__Peniophoraceae(100);g__Peniophora(100);g__Peniophora_unclassified(100); | Peniophora | 13 | Pathotroph-Saprotroph | Plant Pathogen-\|Wood Saprotroph\| | Corticioid | White Rot | Probable | As white_rot (Põlme et al. 2020); As wood_pathogen (Põlme et al. 2020); Decay wood (Põlme et al. 2020) | [illegible] |
| Otu29615 | 1 | k__Fungi(100);p__Basidiomycota(100);c__Agaricomycetes(100);o__Russulales(100);f__Peniophoraceae(100);g__Peniophora(100);s__Peniophora_sp_2_RG_2014(100); | Peniophora | 13 | Pathotroph-Saprotroph | Plant Pathogen-\|Wood Saprotroph\| | Corticioid | White Rot | Probable | As white_rot (Põlme et al. 2020); As wood_pathogen (Põlme et al. 2020); Decay wood (Põlme et al. 2020) | [illegible] |
| Otu29616 | 1 | k__Fungi(100);p__Basidiomycota(100);c__Agaricomycetes(100);o__Russulales(100);f__Peniophoraceae(100);g__Peniophora(100);g__Peniophora_unclassified(100); | Peniophora | 13 | Pathotroph-Saprotroph | Plant Pathogen-\|Wood Saprotroph\| | Corticioid | White Rot | Probable | As white_rot (Põlme et al. 2020); As wood_pathogen (Põlme et al. 2020); Decay wood (Põlme et al. 2020) | [illegible] |
| Otu29618 | 1 | k__Fungi(100);p__Basidiomycota(100);c__Agaricomycetes(100);o__Russulales(100);f__Peniophoraceae(100);f__Peniophoraceae_unclassified(100);f__Peniophoraceae_unclassified(100); | Peniophoraceae | 9 | Pathotroph-Saprotroph | Plant Pathogen-Wood Saprotroph | Corticioid | White Rot | Probable | Most species are saprobic, causing rots of standing and fallen wood. Many species are adapted to xeric conditions, being able to grow on rapidly drying dead attached branches (Cannon & Kirk 2007) | [illegible] |
| Otu29619 | 1 | k__Fungi(100);p__Basidiomycota(100);c__Agaricomycetes(100);o__Russulales(100);f__Peniophoraceae(100);g__Peniophora(100);g__Peniophora_unclassified(100); | Peniophora | 13 | Pathotroph-Saprotroph | Plant Pathogen-\|Wood Saprotroph\| | Corticioid | White Rot | Probable | As white_rot (Põlme et al. 2020); As wood_pathogen (Põlme et al. 2020); Decay wood (Põlme et al. 2020) | [illegible] |
| Otu29620 | 1 | k__Fungi(100);p__Basidiomycota(100);c__Agaricomycetes(100);o__Russulales(100);f__Russulaceae(100);g__Russula(100);s__Russula_puellaris(100); | Russula | 13 | Symbiotroph | \|Ectomycorrhizal\| | Agaricoid | NULL | Highly Probable | NULL | [illegible] |
| Otu29621 | 1 | k__Fungi(100);p__Basidiomycota(100);c__Agaricomycetes(100);o__Russulales(100);f__Russulaceae(100);g__Lactifluus(100);g__Lactifluus_unclassified(100); | Lactifluus | 13 | Symbiotroph | \|Ectomycorrhizal\| | Agaricoid | NULL | Highly Probable | NULL | [illegible] |
| Otu29622 | 1 | k__Fungi(100);p__Basidiomycota(100);c__Agaricomycetes(100);o__Thelephorales(100);f__Thelephoraceae(100);g__unclassified_Thelephoraceae(100);s__Thelephoraceae_sp(100); | Thelephoraceae | 9 | Saprotroph-Symbiotroph | Ectomycorrhizal-Undefined Saprotroph | Clavarioid | NULL | Probable | On the ground, on soil, a few on wood. At least some speices ectomycorrhizal (Cannon & Kirk 2007) | [illegible] |
| Otu29623 | 1 | k__Fungi(100);p__Basidiomycota(100);c__Agaricomycetes(100);o__Russulales(100);f__Russulaceae(100);g__Russula(100);s__Russula_sp_FH_HES04F1SY(100); | Russula | 13 | Symbiotroph | \|Ectomycorrhizal\| | Agaricoid | NULL | Highly Probable | NULL | [illegible] |
| Otu29624 | 1 | k__Fungi(100);p__Basidiomycota(100);c__Exobasidiomycetes(100);o__Microstromatales(100);f__Microstromataceae(100);g__Microstroma(100);s__Microstroma_juglandis(100); | Microstroma | 13 | Pathotroph | \|Plant Pathogen\| | Dimorphic | NULL | Probable | NULL | [illegible] |
| Otu29625 | 1 | k__Fungi(100);p__Basidiomycota(100);c__Microbotryomycetes(100);o__Sporidiobolales(100);f__Sporidiobolales_family_Incertae_sedis(100);g__Rhodotorula(100);s__Rhodotorula_taiwanensis(100); | Rhodotorula | 13 | Pathotroph-Saprotroph-Symbiotroph | Animal Pathogen-Animal Symbiotroph-Endophyte-Plant Pathogen-\|Undefined Saprotroph\| | Yeast | NULL | Probable | As foliar_endophyte (Põlme et al. 2020); Opportunistic human pathogen | [illegible] |
| Otu29628 | 1 | k__Fungi(100);p__Basidiomycota(100);c__Agaricostilbomycetes(100);o__Agaricostilbales(100);f__Agaricostilbaceae(100);g__Bensingtonia(100);g__Bensingtonia_unclassified(100); | Bensingtonia | 13 | Saprotroph | \|Undefined Saprotroph\| | Dimorphic-Facultative Yeast | NULL | Highly Probable | NULL | [illegible] |
| Otu29630 | 1 | k__Fungi(100);p__Basidiomycota(100);c__Agaricomycetes(100);o__Agaricales(100);f__Pleurotaceae(100);g__Hohenbuehelia(100);s__Hohenbuehelia_reniformis(100); | Hohenbuehelia | 13 | Pathotroph-Saprotroph | Nematophagous-Undefined Saprotroph-\|Wood Saprotroph\| | Agaricoid | White Rot | Probable | As white_rot (Põlme et al. 2020); Decay wood, animal material (Põlme et al. 2020) | [illegible] |
| Otu29631 | 1 | k__Fungi(100);p__Basidiomycota(100);c__Agaricomycetes(100);o__Thelephorales(100);f__Thelephoraceae(100);g__Tomentella(100);s__Tomentella_sp(100); | Tomentella | 13 | Symbiotroph | \|Ectomycorrhizal\| | Corticioid | NULL | Highly Probable | NULL | [illegible] |
| Otu29632 | 1 | k__Fungi(100);p__Basidiomycota(100);c__Agaricomycetes(100);o__Thelephorales(100);f__Thelephoraceae(100);g__Tomentella(100);s__Tomentella_badia(100); | Tomentella | 13 | Symbiotroph | \|Ectomycorrhizal\| | Corticioid | NULL | Highly Probable | NULL | [illegible] |
| Otu29635 | 1 | k__Fungi(100);p__Basidiomycota(100);c__Tremellomycetes(100);o__Tremellales(100);f__Tremellales_family_Incertae_sedis(100);f__Tremellales_family_Incertae_sedis_unclassified(100);f__Tremellales_family_Incertae_sedis_unclassified(100); | Tremellales | 7 | Pathotroph-Saprotroph-Symbiotroph | Fungal Parasite-Undefined Saprotroph | Tremelloid-Yeast | NULL | Possible | The order contains both teleomorphic and anamorphic species, most of the latter being yeasts | [illegible] |
| Otu29636 | 1 | k__Fungi(100);p__Basidiomycota(100);c__Dacrymycetes(100);o__Dacrymycetales(100);f__Dacrymycetaceae(100);g__Cerinosterus(100);s__Cerinosterus_sp(100); | Cerinosterus | 13 | Saprotroph | \|Undefined Saprotroph\| | Microfungus | Brown Rot | Probable | NULL | [illegible] |
| Otu29639 | 1 | k__Fungi(100);p__Basidiomycota(100);c__Agaricomycetes(100);o__Hymenochaetales(100);f__Hymenochaetaceae(100);g__Inonotus(100);s__Inonotus_obliquus(100); | Inonotus | 13 | Pathotroph-Saprotroph | Plant Pathogen-\|Wood Saprotroph\| | Polyporoid | White Rot | Highly Probable | As white_rot (Põlme et al. 2020); As wood_pathogen (Põlme et al. 2020); Decay wood (Põlme et al. 2020) | [illegible] |
| Otu29640 | 1 | k__Fungi(100);p__Basidiomycota(100);c__Agaricomycetes(100);o__Hymenochaetales(100);f__Hymenochaetaceae(100);g__Fomitiporia(100);s__Fomitiporia_punctata(100); | Fomitiporia | 13 | Pathotroph-Saprotroph | \|Plant Pathogen\|-Wood Saprotroph | Polyporoid | White Rot | Probable | As white_rot (Põlme et al. 2020); As wood_pathogen (Põlme et al. 2020); Decay wood (Põlme et al. 2020) | [illegible] |
| Otu29641 | 1 | k__Fungi(100);p__Basidiomycota(100);c__Agaricomycetes(100);o__Russulales(100);f__Auriscalpiaceae(100);g__Artomyces(100);s__Artomyces_pyxidatus(100); | Artomyces | 13 | Saprotroph | Undefined Saprotroph-\|Wood Saprotroph\| | Clavarioid | White Rot | Probable | As white_rot (Põlme et al. 2020); Decay wood (Põlme et al. 2020) | [illegible] |
| Otu29642 | 1 | k__Fungi(100);p__Basidiomycota(100);c__Agaricomycetes(100);o__Boletales(100);f__Boletaceae(100);g__Leccinum(100);s__Leccinum_schistophilum(100); | Leccinum | 13 | Symbiotroph | \|Ectomycorrhizal\| | Agaricoid-Boletoid | NULL | Highly Probable | NULL | [illegible] |
| Otu29643 | 1 | k__Fungi(100);p__Basidiomycota(100);c__Cystobasidiomycetes(100);o__Erythrobasidiales(100);f__Erythrobasidiales_family_Incertae_sedis(100);g__Bannoa(100);s__Bannoa_hahajimensis(100); | Bannoa | 13 | Saprotroph | \|Plant Saprotroph\|-Undefined Saprotroph | Dimorphic-Yeast | NULL | Possible | As litter_saprotroph (Põlme et al. 2020); Decay leaf, fruit, seed (Põlme et al. 2020); Symbiotroph isolated from dead leaves (Hamamoto et al. 2002) | [illegible] |
| Otu29644 | 1 | k__Fungi(100);p__Basidiomycota(100);c__Agaricomycetes(100);o__Hymenochaetales(100);f__Hymenochaetaceae(100);g__Inocutis(100);s__Inocutis_dryophilus(100); | Inocutis | 13 | Pathotroph-Saprotroph | Plant Pathogen-Undefined Saprotroph-\|Wood Saprotroph\| | Polyporoid | White Rot | Probable | As white_rot (Põlme et al. 2020); As wood_pathogen (Põlme et al. 2020); Decay wood (Põlme et al. 2020) | [illegible] |
| Otu29645 | 1 | k__Fungi(100);p__Basidiomycota(100);c__Agaricomycetes(100);o__Agaricales(100);f__Pleurotaceae(100);g__Pleurotus(100);s__Pleurotus_ostreatus(100); | Pleurotus | 13 | Pathotroph-Saprotroph | Endophyte-Nematophagous-Plant Pathogen-\|Wood Saprotroph\| | Agaricoid | White Rot | Probable | As white_rot (Põlme et al. 2020); As wood_pathogen (Põlme et al. 2020); Decay wood, animal material (Põlme et al. 2020); This genus is known to be nematophagous, capturing and consuming nematodes. | [illegible] |
| Otu29649 | 1 | k__Fungi(100);p__Basidiomycota(100);c__Agaricomycetes(100);o__Russulales(100);f__Hericiaceae(100);g__Hericium(100);s__Hericium_erinaceum(100); | Hericium | 13 | Saprotroph-Pathotroph-Saprotroph | Plant Pathogen-Undefined Saprotroph-\|Wood Saprotroph\| | Hydnoid-Polyporoid | White Rot | Probable | As root_pathogen (Põlme et al. 2020); As white_rot (Põlme et al. 2020); Decay wood, roots (Põlme et al. 2020) | [illegible] |
| Otu29650 | 1 | k__Fungi(100);p__Basidiomycota(100);c__Agaricomycetes(100);o__Russulales(100);f__Russulaceae(100);g__Lactarius(100);s__Lactarius_quietus(100); | Lactarius | 13 | Symbiotroph | \|Ectomycorrhizal\| | Agaricoid | NULL | Highly Probable | milk; cap; Common name | [illegible] |
| Otu29652 | 1 | k__Fungi(100);p__Basidiomycota(100);c__Agaricomycetes(100);o__Russulales(100);f__Russulaceae(100);g__Lactarius(100);s__Lactarius_tabidus(100); | Lactarius | 13 | Symbiotroph | \|Ectomycorrhizal\| | Agaricoid | NULL | Highly Probable | milk; cap; Common name | [illegible] |
| Otu29653 | 1 | k__Fungi(100);p__Basidiomycota(100);c__Agaricomycetes(100);o__Russulales(100);f__Russulaceae(100);g__Lactarius(100);g__Lactarius_unclassified(100); | Lactarius | 13 | Symbiotroph | \|Ectomycorrhizal\| | Agaricoid | NULL | Highly Probable | milk; cap; Common name | [illegible] |
| Otu29656 | 1 | k__Fungi(100);p__Basidiomycota(100);c__Agaricomycetes(100);o__Russulales(100);f__Russulaceae(100);g__Russula(100);s__Russula_ionochlora(100); | Russula | 13 | Symbiotroph | \|Ectomycorrhizal\| | Agaricoid | NULL | Highly Probable | NULL | [illegible] |
| Otu29657 | 1 | k__Fungi(100);p__Basidiomycota(100);c__Pucciniomycetes(100);o__Pucciniales(100);f__Pucciniastraceae(100);g__Thekopsora(100);g__Thekopsora_unclassified(100); | Thekopsora | 13 | Pathotroph | \|Plant Pathogen\| | Rust | NULL | Probable | NULL | [illegible] |
| Otu29658 | 1 | k__Fungi(100);p__Basidiomycota(100);c__Agaricomycetes(100);o__Agaricales(100);f__Niaceae(100);g__Merismodes(100);s__Merismodes_fasciculata(100); | Merismodes | 13 | Saprotroph | Undefined Saprotroph-\|Wood Saprotroph\| | Cyphelloid | NULL | Probable | Decay wood (Põlme et al. 2020) | [illegible] |
| Otu29665 | 1 | k__Fungi(100);p__Basidiomycota(100);c__Agaricomycetes(100);o__Agaricales(100);o__Agaricales_unclassified(100);o__Agaricales_unclassified(100);o__Agaricales_unclassified(100); | Agaricales | 7 | Pathotroph-Saprotroph-Symbiotroph | Bryophyte Parasite-Dung Saprotroph-Ectomycorrhizal-Fungal Parasite-Leaf Saprotroph-Plant Parasite-Undefined Saprotroph-Wood Saprotroph | Agaricoid-Gasteroid-Microfungus-Secotioid-Yeast | NULL | Possible | Mushrooms and toadstools, Gill fungi, Agarics... lignicolous, sometimes muscicolous or fungicolous, saprobic, mycorrhizal, rarely parasitic on plants or fungi (Kirk et al. 2008) | [illegible] |
| Otu29670 | 1 | k__Fungi(100);p__Basidiomycota(100);c__Pucciniomycetes(100);o__Pucciniales(100);f__Pucciniaceae(100);g__Gymnosporangium(100);s__Gymnosporangium_globosum(100); | Gymnosporangium | 13 | Pathotroph | \|Plant Pathogen\| | Rust-Telial Horns | NULL | Probable | NULL | [illegible] |
| Otu29672 | 1 | k__Fungi(100);p__Basidiomycota(100);c__Agaricomycetes(100);o__Russulales(100);f__Auriscalpiaceae(100);g__Lentinellus(100);s__Lentinellus_ursinus(100); | Lentinellus | 13 | Pathotroph-Saprotroph | Plant Pathogen-Undefined Saprotroph-\|Wood Saprotroph\| | Agaricoid | White Rot | Probable | As white_rot (Põlme et al. 2020); As wood_pathogen (Põlme et al. 2020); Decay wood (Põlme et al. 2020) | [illegible] |
| Otu29673 | 1 | k__Fungi(100);p__Basidiomycota(100);c__Agaricomycetes(100);o__Russulales(100);f__Auriscalpiaceae(100);g__Lentinellus(100);s__Lentinellus_ursinus(100); | Lentinellus | 13 | Pathotroph-Saprotroph | Plant Pathogen-Undefined Saprotroph-\|Wood Saprotroph\| | Agaricoid | White Rot | Probable | As white_rot (Põlme et al. 2020); As wood_pathogen (Põlme et al. 2020); Decay wood (Põlme et al. 2020) | [illegible] |
| Otu29674 | 1 | k__Fungi(100);p__Basidiomycota(100);c__Agaricomycetes(100);o__Russulales(100);f__Russulaceae(100);g__Russula(100);s__Russula_badia(100); | Russula | 13 | Symbiotroph | \|Ectomycorrhizal\| | Agaricoid | NULL | Highly Probable | NULL | [illegible] |
| Otu29675 | 1 | k__Fungi(100);p__Basidiomycota(100);c__Agaricomycetes(100);o__Russulales(100);f__Russulaceae(100);g__Russula(100);g__Russula_unclassified(100); | Russula | 13 | Symbiotroph | \|Ectomycorrhizal\| | Agaricoid | NULL | Highly Probable | NULL | [illegible] |
| Otu29677 | 1 | k__Fungi(100);p__Basidiomycota(100);c__Microbotryomycetes(100);o__Sporidiobolales(100);f__Sporidiobolales_family_Incertae_sedis(100);g__Rhodotorula(100);s__Rhodotorula_taiwanensis(100); | Rhodotorula | 13 | Pathotroph-Saprotroph-Symbiotroph | Animal Pathogen-Animal Symbiotroph-Endophyte-Plant Pathogen-\|Undefined Saprotroph\| | Yeast | NULL | Probable | As foliar_endophyte (Põlme et al. 2020); Opportunistic human pathogen | [illegible] |
| Otu29678 | 1 | k__Fungi(100);p__Basidiomycota(100);c__Agaricostilbomycetes(100);o__Agaricostilbales(100);f__Chionosphaeraceae(100);g__Kurtzmanomyces(100);g__Kurtzmanomyces_unclassified(100); | Kurtzmanomyces | 13 | Saprotroph | \|Undefined Saprotroph\| | Dimorphic | NULL | Probable | NULL | [illegible] |
| Otu29679 | 1 | k__Fungi(100);p__Basidiomycota(100);c__Microbotryomycetes(100);o__Sporidiobolales(100);f__Sporidiobolales_family_Incertae_sedis(100);g__Rhodotorula(100);s__Rhodotorula_taiwanensis(100); | Rhodotorula | 13 | Pathotroph-Saprotroph-Symbiotroph | Animal Pathogen-Animal Symbiotroph-Endophyte-Plant Pathogen-\|Undefined Saprotroph\| | Yeast | NULL | Probable | As foliar_endophyte (Põlme et al. 2020); Opportunistic human pathogen | [illegible] |
| Otu29680 | 1 | k__Fungi(100);p__Basidiomycota(100);c__Agaricomycetes(100);o__Trechisporales(100);f__Hydnodontaceae(100);g__Trechispora(100);g__Trechispora_unclassified(100); | Trechispora | 13 | Saprotroph | \|Wood Saprotroph\| | Corticioid | White Rot | Highly Probable | As white_rot (Põlme et al. 2020); Decay wood (Põlme et al. 2020) | [illegible] |
| Otu29681 | 1 | k__Fungi(100);p__Basidiomycota(100);c__Agaricomycetes(100);o__Agaricales(100);f__Cortinariaceae(100);g__Cortinarius(100);s__Cortinarius_tirolianus(100); | Cortinarius | 13 | Symbiotroph | \|Ectomycorrhizal\| | Agaricoid | NULL | Highly Probable | NULL | [illegible] |
| Otu29682 | 1 | k__Fungi(100);p__Basidiomycota(100);c__Microbotryomycetes(100);o__Sporidiobolales(100);f__Sporidiobolales_family_Incertae_sedis(100);g__Rhodotorula(100);s__Rhodotorula_taiwanensis(100); | Rhodotorula | 13 | Pathotroph-Saprotroph-Symbiotroph | Animal Pathogen-Animal Symbiotroph-Endophyte-Plant Pathogen-\|Undefined Saprotroph\| | Yeast | NULL | Probable | As foliar_endophyte (Põlme et al. 2020); Opportunistic human pathogen | [illegible] |
| Otu29683 | 1 | k__Fungi(100);p__Basidiomycota(100);c__Agaricomycetes(100);o__Agaricales(100);f__Bolbitiaceae(100);g__Conocybe(100);g__Conocybe_unclassified(100); | Conocybe | 13 | Saprotroph | Dung Saprotroph-Plant Saprotroph-\|Undefined Saprotroph\| | Agaricoid | NULL | Probable | As litter_saprotroph (Põlme et al. 2020); As soil_saprotroph (Põlme et al. 2020); Decay soil (Põlme et al. 2020) | [illegible] |
| Otu29685 | 1 | k__Fungi(100);p__Basidiomycota(100);c__Microbotryomycetes(100);o__Sporidiobolales(100);f__Sporidiobolales_family_Incertae_sedis(100);g__Rhodotorula(100);g__Rhodotorula_unclassified(100); | Rhodotorula | 13 | Pathotroph-Saprotroph-Symbiotroph | Animal Pathogen-Animal Symbiotroph-Endophyte-Plant Pathogen-\|Undefined Saprotroph\| | Yeast | NULL | Probable | As foliar_endophyte (Põlme et al. 2020); Opportunistic human pathogen | [illegible] |
| Otu29686 | 1 | k__Fungi(100);p__Basidiomycota(100);c__Microbotryomycetes(100);o__Sporidiobolales(100);f__Sporidiobolales_family_Incertae_sedis(100);g__Rhodotorula(100);s__Rhodotorula_lamellibrachiae(100); | Rhodotorula | 13 | Pathotroph-Saprotroph-Symbiotroph | Animal Pathogen-Animal Symbiotroph-Endophyte-Plant Pathogen-\|Undefined Saprotroph\| | Yeast | NULL | Probable | As foliar_endophyte (Põlme et al. 2020); Opportunistic human pathogen | [illegible] |
| Otu29687 | 1 | k__Fungi(100);p__Basidiomycota(100);c__Cystobasidiomycetes(100);o__Erythrobasidiales(100);f__Erythrobasidiales_family_Incertae_sedis(100);g__Erythrobasidium(100);s__Erythrobasidium_hasegawianum(100); | Erythrobasidium | 13 | Pathotroph-Saprotroph | \|Fungal Parasite\|-Undefined Saprotroph | Yeast | NULL | Possible | As mycoparasite (Põlme et al. 2020); Decay fungal material (Põlme et al. 2020) | [illegible] |
| Otu29688 | 1 | k__Fungi(100);p__Basidiomycota(100);c__Microbotryomycetes(100);o__Sporidiobolales(100);f__Sporidiobolales_family_Incertae_sedis(100);g__Rhodotorula(100);g__Rhodotorula_unclassified(100); | Rhodotorula | 13 | Pathotroph-Saprotroph-Symbiotroph | Animal Pathogen-Animal Symbiotroph-Endophyte-Plant Pathogen-\|Undefined Saprotroph\| | Yeast | NULL | Probable | As foliar_endophyte (Põlme et al. 2020); Opportunistic human pathogen | [illegible] |
| Otu29689 | 1 | k__Fungi(100);p__Basidiomycota(100);c__Microbotryomycetes(100);o__Sporidiobolales(100);f__Sporidiobolales_family_Incertae_sedis(100);g__Sporobolomyces(100);s__Sporobolomyces_ogasawarensis(100); | Sporobolomyces | 13 | Pathotroph-Saprotroph | \|Fungal Parasite\|-Plant Saprotroph | Yeast | NULL | Probable | As litter_saprotroph (Põlme et al. 2020); As mycoparasite (Põlme et al. 2020); Decay fungal material (Põlme et al. 2020) | [illegible] |
| Otu29690 | 1 | k__Fungi(100);p__Basidiomycota(100);c__Microbotryomycetes(100);o__Sporidiobolales(100);f__Sporidiobolales_family_Incertae_sedis(100);g__Rhodotorula(100);s__Rhodotorula_lactosa(100); | Rhodotorula | 13 | Pathotroph-Saprotroph-Symbiotroph | Animal Pathogen-Animal Symbiotroph-Endophyte-Plant Pathogen-\|Undefined Saprotroph\| | Yeast | NULL | Probable | As foliar_endophyte (Põlme et al. 2020); Opportunistic human pathogen | [illegible] |
| Otu29694 | 1 | k__Fungi(100);p__Basidiomycota(100);c__Tremellomycetes(100);o__Cystofilobasidiales(100);f__Cystofilobasidiaceae(100);g__Udeniomyces(100);s__Udeniomyces_pyricola(100); | Udeniomyces | 13 | Saprotroph-Symbiotroph | Epiphyte-Plant Saprotroph-\|Undefined Saprotroph\| | Yeast | NULL | Probable | Isolated from the pear phylloplane (Stadelmann 1975); Isolated from dead leaves of Oryza sativa, Miscanthus sinensis and Sasa sp. in Japan (Nakase & Suzuki 1986; Bullera spp.) | [illegible] |
| Otu29695 | 1 | k__Fungi(100);p__Basidiomycota(100);c__Agaricomycetes(100);o__Polyporales(100);f__Xenasmataceae(100);g__Xenasmatella(100);s__Xenasmatella_sp(100); | Xenasmatella | 13 | Saprotroph | Undefined Saprotroph-\|Wood Saprotroph\| | Corticioid | White Rot | Probable | As white_rot (Põlme et al. 2020); Decay wood (Põlme et al. 2020) | [illegible] |
| Otu29696 | 1 | k__Fungi(100);p__Ascomycota(100);c__Eurotiomycetes(100);o__Chaetothyriales(100);f__Herpotrichiellaceae(100);f__Herpotrichiellaceae_unclassified(100);f__Herpotrichiellaceae_unclassified(100); | Herpotrichiellaceae | 9 | Pathotroph-Saprotroph | Animal Pathogen-Fungal Parasite-Undefined Saprotroph | Facultative Yeast-Microfungus | NULL | Probable | Mostly saprobic on plants or other fungi, isolated from soil etc., and from medical sources (Cannon & Kirk 2007) | [illegible] |
| Otu29697 | 1 | k__Fungi(100);p__Basidiomycota(100);c__Agaricomycetes(100);o__Polyporales(100);f__Meruliaceae(100);g__Phlebia(100);s__Phlebia_subserialis(100); | Phlebia | 13 | Saprotroph | \|Wood Saprotroph\| | Corticioid | White Rot | Highly Probable | As white_rot (Põlme et al. 2020); Decay wood (Põlme et al. 2020) | [illegible] |
| Otu29698 | 1 | k__Fungi(100);p__Basidiomycota(100);c__Agaricomycetes(100);o__Polyporales(100);f__Meruliaceae(100);g__Irpex(100);s__Irpex_sp_JSL_2009a(100); | Irpex | 13 | Saprotroph | \|Wood Saprotroph\| | Corticioid | White Rot | Highly Probable | As white_rot (Põlme et al. 2020); Decay wood (Põlme et al. 2020); White rot of dead hardwoods, rarely on conifers | [illegible] |
| Otu29699 | 1 | k__Fungi(100);p__Basidiomycota(100);c__Agaricomycetes(100);o__Polyporales(100);f__Meruliaceae(100);g__Irpex(100);s__Irpex_sp_JSL_2009a(100); | Irpex | 13 | Saprotroph | \|Wood Saprotroph\| | Corticioid | White Rot | Highly Probable | As white_rot (Põlme et al. 2020); Decay wood (Põlme et al. 2020); White rot of dead hardwoods, rarely on conifers | [illegible] |
| Otu29700 | 1 | k__Fungi(100);p__Basidiomycota(100);c__Agaricomycetes(100);o__Agaricales(100);f__Pluteaceae(100);g__Pluteus(100);s__Pluteus_oreibatus(100); | Pluteus | 13 | Pathotroph-Saprotroph | Bryophyte Parasite-\|Plant Saprotroph\|-Wood Saprotroph | Agaricoid | White Rot | Probable | As litter_saprotroph (Põlme et al. 2020); As white_rot (Põlme et al. 2020); Decay leaf, fruit, seed (Põlme et al. 2020) | [illegible] |
| Otu29701 | 1 | k__Fungi(100);p__Basidiomycota(100);c__Pucciniomycetes(100);o__Pucciniales(100);f__Melampsoraceae(100);g__Melampsora(100);s__Melampsora_sp_Mel_G(100); | Melampsora | 13 | Pathotroph | \|Plant Pathogen\| | Rust | NULL | Probable | A prominent group of species produces aecia on conifer needles and telia or leaves of Salicaceae (Cannon & Kirk 2007); Biotrophic, heteroecious or autoecious | [illegible] |
| Otu29703 | 1 | k__Fungi(100);p__Basidiomycota(100);c__Pucciniomycetes(100);o__Pucciniales(100);f__Coleosporiaceae(100);g__Chrysomyxa(100);s__Chrysomyxa_cassandrae(100); | Chrysomyxa | 13 | Pathotroph | \|Plant Pathogen\| | Rust | NULL | Probable | NULL | [illegible] |
| Otu29704 | 1 | k__Fungi(100);p__Basidiomycota(100);c__Tremellomycetes(100);o__Tremellales(100);f__Tremellales_family_Incertae_sedis(100);f__Tremellales_family_Incertae_sedis_unclassified(100);f__Tremellales_family_Incertae_sedis_unclassified(100); | Tremellales | 7 | Pathotroph-Saprotroph-Symbiotroph | Fungal Parasite-Undefined Saprotroph | Tremelloid-Yeast | NULL | Possible | The order contains both teleomorphic and anamorphic species, most of the latter being yeasts | [illegible] |
| Otu29705 | 1 | k__Fungi(100);p__Basidiomycota(100);c__Pucciniomycetes(100);o__Pucciniales(100);f__Pucciniaceae(100);g__Puccinia(100);s__Puccinia_bartholomaei(100); | Puccinia | 13 | Pathotroph | \|Plant Pathogen\| | Rust | NULL | Highly Probable | NULL | [illegible] |
| Otu29706 | 1 | k__Fungi(100);p__Basidiomycota(100);c__Pucciniomycetes(100);o__Pucciniales(100);f__Pucciniaceae(100);g__Puccinia(100);s__Puccinia_bartholomaei(100); | Puccinia | 13 | Pathotroph | \|Plant Pathogen\| | Rust | NULL | Highly Probable | NULL | [illegible] |
| Otu29707 | 1 | k__Fungi(100);p__Basidiomycota(100);c__Pucciniomycetes(100);o__Pucciniales(100);f__Pucciniaceae(100);g__Gymnosporangium(100);g__Gymnosporangium_unclassified(100); | Gymnosporangium | 13 | Pathotroph | \|Plant Pathogen\| | Rust-Telial Horns | NULL | Probable | NULL | [illegible] |
| Otu29708 | 1 | k__Fungi(100);p__Basidiomycota(100);c__Pucciniomycetes(100);o__Pucciniales(100);f__Pucciniaceae(100);g__Uromyces(100);g__Uromyces_unclassified(100); | Uromyces | 13 | Pathotroph | \|Plant Pathogen\| | Rust | NULL | Probable | NULL | [illegible] |
| Otu29711 | 1 | k__Fungi(100);p__Ascomycota(100);c__Saccharomycetes(100);o__Saccharomycetales(100);f__Saccharomycetaceae(100);g__Kazachstania(100);s__Kazachstania_intestinalis(100); | Kazachstania | 13 | Saprotroph | \|Undefined Saprotroph\| | Yeast | NULL | Probable | As nectar-tap_saprotroph (Põlme et al. 2020); Decay sugar-rich substrates (Põlme et al. 2020) | [illegible] |
| Otu29716 | 1 | k__Fungi(100);p__Ascomycota(100);c__Pezizomycetes(100);o__Pezizales(100);f__Pezizaceae(100);g__Terfezia(100);g__Terfezia_unclassified(100); | Terfezia | 13 | Symbiotroph | \|Ectomycorrhizal\| | Gasteroid | Hypogeous | Highly Probable | desert false truffle; Common name | [illegible] |
| Otu29717 | 1 | k__Fungi(100);p__Ascomycota(100);c__Sordariomycetes(100);o__Hypocreales(100);f__Hypocreales_family_Incertae_sedis(100);g__Stachybotrys(100);s__Stachybotrys_kampalensis(100); | Stachybotrys | 13 | Pathotroph-Saprotroph | Plant Pathogen-Plant Saprotroph-\|Wood Saprotroph\| | Microfungus | Soft Rot | Highly Probable | As litter_saprotroph (Põlme et al. 2020); Decay wood, leaf, fruit, seed (Põlme et al. 2020) | [illegible] |
| Otu29718 | 1 | k__Fungi(100);p__Ascomycota(100);c__Sordariomycetes(100);o__Diaporthales(100);f__Gnomoniaceae(100);g__Cryptosporella(100);s__Cryptosporella_betulae(100); | Cryptosporella | 13 | Pathotroph-Saprotroph | \|Plant Pathogen\|-Undefined Saprotroph | Microfungus | NULL | Probable | NULL | [illegible] |
| Otu29720 | 1 | k__Fungi(100);p__Ascomycota(100);c__Sordariomycetes(100);o__Microascales(100);f__Microascaceae(100);g__Scedosporium(100);s__Scedosporium_prolificans(100); | Scedosporium | 13 | Pathotroph-Saprotroph | Animal Parasite-\|Undefined Saprotroph\| | Microfungus | NULL | Probable | As soil_saprotroph (Põlme et al. 2020); Decay soil, animal material (Põlme et al. 2020) | [illegible] |
| Otu29721 | 1 | k__Fungi(100);p__Ascomycota(100);c__Sordariomycetes(100);o__Microascales(100);f__Microascaceae(100);g__unclassified_Microascaceae(100);s__Microascaceae_sp(100); | Microascaceae | 9 | Pathotroph-Saprotroph-Symbiotroph | Animal Pathogen-Endophyte-Plant Pathogen-Undefined Saprotroph | Microfungus | NULL | Possible | Saprobic in soil or rotting vegetation, some species heat-tolerant and associatated with silage and compost, a few associated with humans and animals (Cannon & Kirk 2007) | [illegible] |
| Otu29722 | 1 | k__Fungi(100);p__Ascomycota(100);c__Sordariomycetes(100);o__Microascales(100);f__Microascaceae(100);g__Petriella(100);s__Petriella_guttulata(100); | Petriella | 13 | Saprotroph | \|Dung Saprotroph\|-Undefined Saprotroph | Microfungus | NULL | Probable | As soil_saprotroph (Põlme et al. 2020); Decay soil, dung (Põlme et al. 2020) | [illegible] |
| Otu29723 | 1 | k__Fungi(100);p__Ascomycota(100);c__Saccharomycetes(100);o__Saccharomycetales(100);f__Saccharomycetaceae(100);g__Kazachstania(100);s__Kazachstania_intestinalis(100); | Kazachstania | 13 | Saprotroph | \|Undefined Saprotroph\| | Yeast | NULL | Probable | As nectar-tap_saprotroph (Põlme et al. 2020); Decay sugar-rich substrates (Põlme et al. 2020) | [illegible] |
| Otu29724 | 1 | k__Fungi(100);p__Ascomycota(100);c__Sordariomycetes(100);o__Hypocreales(100);f__Cordycipitaceae(100);g__Gibellula(100);s__Gibellula_sp_WK_2013(100); | Gibellula | 13 | Pathotroph | \|Animal Parasite\|-Animal Pathogen-Arthropod Parasite | Microfungus | NULL | Probable | Decay animal material (Põlme et al. 2020); Host - spiders (Põlme et al. 2020) | [illegible] |
| Otu29725 | 1 | k__Fungi(100);p__Ascomycota(100);c__Sordariomycetes(100);o__Hypocreales(100);o__Hypocreales_unclassified(100);o__Hypocreales_unclassified(100);o__Hypocreales_unclassified(100); | Hypocreales | 7 | Saprotroph | Undefined Saprotroph | Microfungus | NULL | Possible | NULL | [illegible] |
| Otu29727 | 1 | k__Fungi(100);p__Ascomycota(100);c__Sordariomycetes(100);o__Microascales(100);f__Microascaceae(100);g__Microascus(100);s__Microascus_manginii(100); | Microascus | 13 | Pathotroph-Saprotroph-Symbiotroph | Animal Pathogen-Dung Saprotroph-Endophyte-Plant Pathogen-Plant Saprotroph-\|Undefined Saprotroph\| | Microfungus | NULL | Possible | As foliar_endophyte (Põlme et al. 2020); As litter_saprotroph (Põlme et al. 2020); As soil_saprotroph (Põlme et al. 2020); Decay leaf, fruit, seed, soil (Põlme et al. 2020) | [illegible] |
| Otu29728 | 1 | k__Fungi(100);p__Ascomycota(100);c__Sordariomycetes(100);o__Microascales(100);f__Microascaceae(100);g__Scopulariopsis(100);s__Scopulariopsis_sp_A_4_1(100); | Scopulariopsis | 13 | Pathotroph-Saprotroph-Symbiotroph | Dung Saprotroph-Endophyte-Plant Pathogen-Plant Saprotroph-\|Undefined Saprotroph\|-Wood Saprotroph | Microfungus | Soft Rot | Highly Probable | As foliar_endophyte (Põlme et al. 2020); As litter_saprotroph (Põlme et al. 2020); As soil_saprotroph (Põlme et al. 2020); Decay leaf, fruit, seed, soil (Põlme et al. 2020) | [illegible] |
| Otu29732 | 1 | k__Fungi(100);p__Ascomycota(100);c__Sordariomycetes(100);o__Diaporthales(100);f__Melanconidaceae(100);g__Melanconiella(100);s__Melanconiella_flavovirens(100); | Melanconiella | 13 | Saprotroph | Undefined Saprotroph-\|Wood Saprotroph\| | Microfungus | NULL | Probable | Decay wood (Põlme et al. 2020) | [illegible] |
| Otu29733 | 1 | k__Fungi(100);p__Ascomycota(100);c__Sordariomycetes(100);o__Hypocreales(100);f__unclassified_Hypocreales(100);g__unclassified_Hypocreales(100);s__Hypocreales_sp(100); | Hypocreales | 7 | Saprotroph | Undefined Saprotroph | Microfungus | NULL | Possible | NULL | [illegible] |
| Otu29734 | 1 | k__Fungi(100);p__Ascomycota(100);c__Sordariomycetes(100);o__Hypocreales(100);o__Hypocreales_unclassified(100);o__Hypocreales_unclassified(100);o__Hypocreales_unclassified(100); | Hypocreales | 7 | Saprotroph | Undefined Saprotroph | Microfungus | NULL | Possible | NULL | [illegible] |
| Otu29735 | 1 | k__Fungi(100);p__Ascomycota(100);c__Sordariomycetes(100);o__Hypocreales(100);o__Hypocreales_unclassified(100);o__Hypocreales_unclassified(100);o__Hypocreales_unclassified(100); | Hypocreales | 7 | Saprotroph | Undefined Saprotroph | Microfungus | NULL | Possible | NULL | [illegible] |
| Otu29736 | 1 | k__Fungi(100);p__Ascomycota(100);c__Saccharomycetes(100);o__Saccharomycetales(100);f__Saccharomycetaceae(100);g__Kazachstania(100);s__Kazachstania_intestinalis(100); | Kazachstania | 13 | Saprotroph | \|Undefined Saprotroph\| | Yeast | NULL | Probable | As nectar-tap_saprotroph (Põlme et al. 2020); Decay sugar-rich substrates (Põlme et al. 2020) | [illegible] |
| Otu29737 | 1 | k__Fungi(100);p__Ascomycota(100);c__Saccharomycetes(100);o__Saccharomycetales(100);f__Saccharomycetaceae(100);g__Kazachstania(100);s__Kazachstania_intestinalis(100); | Kazachstania | 13 | Saprotroph | \|Undefined Saprotroph\| | Yeast | NULL | Probable | As nectar-tap_saprotroph (Põlme et al. 2020); Decay sugar-rich substrates (Põlme et al. 2020) | [illegible] |
| Otu29738 | 1 | k__Fungi(100);p__Ascomycota(100);c__Saccharomycetes(100);o__Saccharomycetales(100);f__Saccharomycetaceae(100);g__Kazachstania(100);s__Kazachstania_intestinalis(100); | Kazachstania | 13 | Saprotroph | \|Undefined Saprotroph\| | Yeast | NULL | Probable | As nectar-tap_saprotroph (Põlme et al. 2020); Decay sugar-rich substrates (Põlme et al. 2020) | [illegible] |
| Otu29739 | 1 | k__Fungi(100);p__Ascomycota(100);c__Saccharomycetes(100);o__Saccharomycetales(100);f__Saccharomycetaceae(100);g__Kazachstania(100);s__Kazachstania_intestinalis(100); | Kazachstania | 13 | Saprotroph | \|Undefined Saprotroph\| | Yeast | NULL | Probable | As nectar-tap_saprotroph (Põlme et al. 2020); Decay sugar-rich substrates (Põlme et al. 2020) | [illegible] |
| Otu29740 | 1 | k__Fungi(100);p__Ascomycota(100);c__Saccharomycetes(100);o__Saccharomycetales(100);f__Saccharomycetaceae(100);g__Kazachstania(100);s__Kazachstania_intestinalis(100); | Kazachstania | 13 | Saprotroph | \|Undefined Saprotroph\| | Yeast | NULL | Probable | As nectar-tap_saprotroph (Põlme et al. 2020); Decay sugar-rich substrates (Põlme et al. 2020) | [illegible] |
| Otu29741 | 1 | k__Fungi(100);p__Ascomycota(100);c__Saccharomycetes(100);o__Saccharomycetales(100);f__Saccharomycetaceae(100);g__Kazachstania(100);s__Kazachstania_intestinalis(100); | Kazachstania | 13 | Saprotroph | \|Undefined Saprotroph\| | Yeast | NULL | Probable | As nectar-tap_saprotroph (Põlme et al. 2020); Decay sugar-rich substrates (Põlme et al. 2020) | [illegible] |
| Otu29742 | 1 | k__Fungi(100);p__Ascomycota(100);c__Saccharomycetes(100);o__Saccharomycetales(100);f__Saccharomycetaceae(100);g__Kazachstania(100);s__Kazachstania_intestinalis(100); | Kazachstania | 13 | Saprotroph | \|Undefined Saprotroph\| | Yeast | NULL | Probable | As nectar-tap_saprotroph (Põlme et al. 2020); Decay sugar-rich substrates (Põlme et al. 2020) | [illegible] |
| Otu29743 | 1 | k__Fungi(100);p__Ascomycota(100);c__Saccharomycetes(100);o__Saccharomycetales(100);f__Saccharomycetaceae(100);g__Kazachstania(100);s__Kazachstania_intestinalis(100); | Kazachstania | 13 | Saprotroph | \|Undefined Saprotroph\| | Yeast | NULL | Probable | As nectar-tap_saprotroph (Põlme et al. 2020); Decay sugar-rich substrates (Põlme et al. 2020) | [illegible] |
| Otu29744 | 1 | k__Fungi(100);p__Ascomycota(100);c__Saccharomycetes(100);o__Saccharomycetales(100);f__Saccharomycetaceae(100);g__Kazachstania(100);s__Kazachstania_intestinalis(100); | Kazachstania | 13 | Saprotroph | \|Undefined Saprotroph\| | Yeast | NULL | Probable | As nectar-tap_saprotroph (Põlme et al. 2020); Decay sugar-rich substrates (Põlme et al. 2020) | [illegible] |
| Otu29745 | 1 | k__Fungi(100);p__Ascomycota(100);c__Saccharomycetes(100);o__Saccharomycetales(100);f__Saccharomycetaceae(100);g__Kazachstania(100);s__Kazachstania_intestinalis(100); | Kazachstania | 13 | Saprotroph | \|Undefined Saprotroph\| | Yeast | NULL | Probable | As nectar-tap_saprotroph (Põlme et al. 2020); Decay sugar-rich substrates (Põlme et al. 2020) | [illegible] |
| Otu29746 | 1 | k__Fungi(100);p__Ascomycota(100);c__Saccharomycetes(100);o__Saccharomycetales(100);f__Saccharomycetaceae(100);g__Kazachstania(100);s__Kazachstania_intestinalis(100); | Kazachstania | 13 | Saprotroph | \|Undefined Saprotroph\| | Yeast | NULL | Probable | As nectar-tap_saprotroph (Põlme et al. 2020); Decay sugar-rich substrates (Põlme et al. 2020) | [illegible] |
| Otu29747 | 1 | k__Fungi(100);p__Ascomycota(100);c__Saccharomycetes(100);o__Saccharomycetales(100);f__Saccharomycetaceae(100);g__Lachancea(100);s__Lachancea_waltii(100); | Lachancea | 13 | Pathotroph-Saprotroph | Animal Parasite-Animal Pathogen-\|Undefined Saprotroph\| | Yeast | NULL | Possible | As nectar-tap_saprotroph (Põlme et al. 2020); Decay animal material, sugar-rich substrates (Põlme et al. 2020); Likely opportunistic human pathogen (Irinyi et al. 2015; see L. fermentati) | [illegible] |
| Otu29748 | 1 | k__Fungi(100);p__Ascomycota(100);c__Saccharomycetes(100);o__Saccharomycetales(100);f__Saccharomycetaceae(100);g__Kazachstania(100);s__Kazachstania_intestinalis(100); | Kazachstania | 13 | Saprotroph | \|Undefined Saprotroph\| | Yeast | NULL | Probable | As nectar-tap_saprotroph (Põlme et al. 2020); Decay sugar-rich substrates (Põlme et al. 2020) | [illegible] |
| Otu29750 | 1 | k__Fungi(100);p__Ascomycota(100);c__Saccharomycetes(100);o__Saccharomycetales(100);f__Saccharomycetaceae(100);g__Kazachstania(100);s__Kazachstania_intestinalis(100); | Kazachstania | 13 | Saprotroph | \|Undefined Saprotroph\| | Yeast | NULL | Probable | As nectar-tap_saprotroph (Põlme et al. 2020); Decay sugar-rich substrates (Põlme et al. 2020) | [illegible] |
| Otu29751 | 1 | k__Fungi(100);p__Ascomycota(100);c__Saccharomycetes(100);o__Saccharomycetales(100);f__Saccharomycetaceae(100);g__Kazachstania(100);s__Kazachstania_intestinalis(100); | Kazachstania | 13 | Saprotroph | \|Undefined Saprotroph\| | Yeast | NULL | Probable | As nectar-tap_saprotroph (Põlme et al. 2020); Decay sugar-rich substrates (Põlme et al. 2020) | [illegible] |
| Otu29752 | 1 | k__Fungi(100);p__Ascomycota(100);c__Saccharomycetes(100);o__Saccharomycetales(100);f__Saccharomycetaceae(100);g__Kazachstania(100);s__Kazachstania_intestinalis(100); | Kazachstania | 13 | Saprotroph | \|Undefined Saprotroph\| | Yeast | NULL | Probable | As nectar-tap_saprotroph (Põlme et al. 2020); Decay sugar-rich substrates (Põlme et al. 2020) | [illegible] |
| Otu29753 | 1 | k__Fungi(100);p__Ascomycota(100);c__Saccharomycetes(100);o__Saccharomycetales(100);f__Saccharomycetaceae(100);g__Kazachstania(100);s__Kazachstania_intestinalis(100); | Kazachstania | 13 | Saprotroph | \|Undefined Saprotroph\| | Yeast | NULL | Probable | As nectar-tap_saprotroph (Põlme et al. 2020); Decay sugar-rich substrates (Põlme et al. 2020) | [illegible] |
| Otu29754 | 1 | k__Fungi(100);p__Basidiomycota(100);c__Agaricomycetes(100);o__Polyporales(100);f__Meruliaceae(100);g__Phlebia(100);s__Phlebia_sp_olrim964(100); | Phlebia | 13 | Saprotroph | \|Wood Saprotroph\| | Corticioid | White Rot | Highly Probable | As white_rot (Põlme et al. 2020); Decay wood (Põlme et al. 2020) | [illegible] |
| Otu29755 | 1 | k__Fungi(100);p__Ascomycota(100);c__Saccharomycetes(100);o__Saccharomycetales(100);f__Saccharomycetaceae(100);g__Kazachstania(100);s__Kazachstania_intestinalis(100); | Kazachstania | 13 | Saprotroph | \|Undefined Saprotroph\| | Yeast | NULL | Probable | As nectar-tap_saprotroph (Põlme et al. 2020); Decay sugar-rich substrates (Põlme et al. 2020) | [illegible] |
| Otu29756 | 1 | k__Fungi(100);p__Ascomycota(100);c__Saccharomycetes(100);o__Saccharomycetales(100);f__Saccharomycetaceae(100);g__Kazachstania(100);s__Kazachstania_intestinalis(100); | Kazachstania | 13 | Saprotroph | \|Undefined Saprotroph\| | Yeast | NULL | Probable | As nectar-tap_saprotroph (Põlme et al. 2020); Decay sugar-rich substrates (Põlme et al. 2020) | [illegible] |

| | | | | | | | | | | | |
|---|---|---|---|---|---|---|---|---|---|---|---|
| Otu29757 | 1 | k__Fungi(100);p__Ascomycota(100);c__Saccharomycetes(100);o__Saccharomycetales(100);f__Saccharomycetaceae(100);g__Lachancea(100);s__Lachancea_thermotolerans(100); | Lachancea | 13 | Pathotroph-Saprotroph | Animal Parasite-Animal Pathogen-\|Undefined Saprotroph\| | Yeast | NULL | Possible | As nectar-tap_saprotroph (Põlme et al. 2020); Decay animal material, sugar-rich substrates (Põlme et al. 2020); Likely opportunistic human pathogen (Irinyi et al. 2015; see L. fermentati) | Irinyi L, et al. 2015. Medical Mycology 53:313-337 ((https://doi.org/10.1093/mmy/myv008)); Põlme S, et al. 2020. Fungal Diversity 105:1-16 ((https://doi.org/10.1007/s13225-020-00466-2)) |
| Otu29758 | 1 | k__Fungi(100);p__Ascomycota(100);c__Saccharomycetes(100);o__Saccharomycetales(100);f__Saccharomycetaceae(100);g__Kazachstania(100);s__Kazachstania_intestinalis(100); | Kazachstania | 13 | Saprotroph | \|Undefined Saprotroph\| | Yeast | NULL | Probable | As nectar-tap_saprotroph (Põlme et al. 2020); Decay sugar-rich substrates (Põlme et al. 2020) | Põlme S, et al. 2020. Fungal Diversity 105:1-16 ((https://doi.org/10.1007/s13225-020-00466-2)) |
| Otu29759 | 1 | k__Fungi(100);p__Ascomycota(100);c__Saccharomycetes(100);o__Saccharomycetales(100);f__Saccharomycetaceae(100);g__Kazachstania(100);s__Kazachstania_intestinalis(100); | Kazachstania | 13 | Saprotroph | \|Undefined Saprotroph\| | Yeast | NULL | Probable | As nectar-tap_saprotroph (Põlme et al. 2020); Decay sugar-rich substrates (Põlme et al. 2020) | Põlme S, et al. 2020. Fungal Diversity 105:1-16 ((https://doi.org/10.1007/s13225-020-00466-2)) |
| Otu29760 | 1 | k__Fungi(100);p__Ascomycota(100);c__Saccharomycetes(100);o__Saccharomycetales(100);f__Saccharomycetaceae(100);g__Kazachstania(100);s__Kazachstania_intestinalis(100); | Kazachstania | 13 | Saprotroph | \|Undefined Saprotroph\| | Yeast | NULL | Probable | As nectar-tap_saprotroph (Põlme et al. 2020); Decay sugar-rich substrates (Põlme et al. 2020) | Põlme S, et al. 2020. Fungal Diversity 105:1-16 ((https://doi.org/10.1007/s13225-020-00466-2)) |
| Otu29761 | 1 | k__Fungi(100);p__Ascomycota(100);c__Saccharomycetes(100);o__Saccharomycetales(100);f__Saccharomycetaceae(100);g__Kazachstania(100);s__Kazachstania_intestinalis(100); | Kazachstania | 13 | Saprotroph | \|Undefined Saprotroph\| | Yeast | NULL | Probable | As nectar-tap_saprotroph (Põlme et al. 2020); Decay sugar-rich substrates (Põlme et al. 2020) | Põlme S, et al. 2020. Fungal Diversity 105:1-16 ((https://doi.org/10.1007/s13225-020-00466-2)) |
| Otu29762 | 1 | k__Fungi(100);p__Ascomycota(100);c__Sordariomycetes(100);o__Boliniales(100);f__Boliniaceae(100);g__unclassified_Boliniaceae(100);s__Boliniaceae_sp(100); | Boliniaceae | 9 | Saprotroph | Wood Saprotroph | Microfungus | NULL | Probable | Saprobic on and in wood and bark (Cannon & Kirk 2007) | Cannon PF, Kirk PM. 2007. Fungal Families of the World. CAB International, Cambridge (ISBN: 978-0851998275) |
| Otu29763 | 1 | k__Fungi(100);p__Ascomycota(100);c__Sordariomycetes(100);o__Hypocreales(100);o__Hypocreales_unclassified(100);o__Hypocreales_unclassified(100);o__Hypocreales_unclassified(100); | Hypocreales | 7 | Saprotroph | Undefined Saprotroph | Microfungus | NULL | Possible | NULL | Sterkenburg E, et al. 2015. New Phytologist 207:1145-1158 ((https://doi.org/10.1111/nph.13426)) |
| Otu29764 | 1 | k__Fungi(100);p__Ascomycota(100);c__Eurotiomycetes(100);o__Eurotiales(100);f__Trichocomaceae(100);g__Paecilomyces(100);s__Paecilomyces_niphetodes(100); | Paecilomyces | 13 | Pathotroph-Saprotroph-Symbiotroph | Clavicipitaceous \|Endophyte\|-Plant Pathogen-Plant Saprotroph-Undefined Saprotroph-Wood Saprotroph | Microfungus | Soft Rot | Probable | As foliar_endophyte (Põlme et al. 2020); As litter_saprotroph (Põlme et al. 2020); culture (Lopez et al. 2014); Decay leaf, fruit, seed, animal material (Põlme et al. 2020); Endophyte detection method | Seehann G, et al. 1975. List of Fungi in Soft-Rot Tissue Tests. Institute fur Holzbiologie und Holzschutz, Bundesforschungsanstalt fur Forst - und Holzwirtshaft, Hamburg - Lohbrugge, Germany; Costa IPMW, et al. 2012. Mycotaxon 119:e26 ((http://mycotaxon.com/resources/checklists/costa_v119_checklist.pdf)); Tedersoo L, et al. 2014. Science 346:e1256688 ((https://doi.org/10.1126/science.1256688)); Põlme S, et al. 2020. Fungal Diversity 105:1-16 ((https://doi.org/10.1007/s13225-020-00466-2)); The American Phytopathological Society Common Names of Plant Diseases ((http://www.apsnet.org/publications/commonnames/Pages/default.aspx)); Lopez DC, et al. 2014. Plos One 9:e103891 ((https://doi.org/10.1371/journal.pone.0103891)) |
| Otu29765 | 1 | k__Fungi(100);p__Ascomycota(100);c__Sordariomycetes(100);o__Hypocreales(100);f__Hypocreales_family_Incertae_sedis(100);g__Acremonium(100);g__Acremonium_unclassified(100); | Acremonium | 13 | Pathotroph-Saprotroph-Symbiotroph | Animal Pathogen-Endophyte-Fungal Parasite-Plant Pathogen-\|Undefined Saprotroph\|-Wood Saprotroph | Microfungus | Soft Rot | Possible | As foliar_endophyte (Põlme et al. 2020); culture (Hanada et al. 2010, Del Olmo; Decay leaf, fruit, seed, soil, dung, animal material (Põlme et al. 2020); Endophyte detection method; Ruiz & Arnold 2014); Likely opportunistic human pathogen (Irinyi et al. 2015) | Duncan CG, Eslyn WE. 1966. Mycologia 58:642-645 ((https://doi.org/10.2307/3757045)); Nilsson T. 1973. Studia Forestalia Suecica 104: 1-40 ((https://pub.epsilon.slu.se/5711/)); Worrall JJ, et al. 1997. Mycologia 89:199-219 ((https://doi.org/10.2307/3761073)); Hanada RE, et al. 2010. Fungal Biology 114:901-910 ((https://doi.org/10.1016/j.funbio.2010.08.006)); Del Olmo-Ruiz M, Arnold AE. 2014. Mycologia 106: 8-21 ((https://doi.org/10.3852/13-098)); Irinyi L, et al. 2015. Medical Mycology 53313-337 ((https://doi.org/10.1093/mmy/myv008)); Põlme S, et al. 2020. Fungal Diversity 105:1-16 ((https://doi.org/10.1007/s13225-020-00466-2)) |
| Otu29766 | 1 | k__Fungi(100);p__Ascomycota(100);c__Sordariomycetes(100);o__Hypocreales(100);o__Hypocreales_unclassified(100);o__Hypocreales_unclassified(100);o__Hypocreales_unclassified(100); | Hypocreales | 7 | Saprotroph | Undefined Saprotroph | Microfungus | NULL | Possible | NULL | Sterkenburg E, et al. 2015. New Phytologist 207:1145-1158 ((https://doi.org/10.1111/nph.13426)) |
| Otu29767 | 1 | k__Fungi(100);p__Ascomycota(100);c__Sordariomycetes(100);o__Hypocreales(100);f__Bionectriaceae(100);g__Bionectria(100);s__Bionectria_sp(100); | Bionectria | 13 | Pathotroph-Saprotroph | Fungal Parasite-Plant Pathogen-Plant Saprotroph-Undefined Saprotroph-\|Wood Saprotroph\| | Microfungus | NULL | Probable | As litter_saprotroph (Põlme et al. 2020); As mycoparasite (Põlme et al. 2020); Decay wood, leaf, fruit, seed, fungal material, animal material (Põlme et al. 2020) | Tedersoo L, et al. 2014. Science 346:e1256688 ((https://doi.org/10.1126/science.1256688)); Põlme S, et al. 2020. Fungal Diversity 105:1-16 ((https://doi.org/10.1007/s13225-020-00466-2)) |
| Otu29768 | 1 | k__Fungi(100);p__Ascomycota(100);c__Sordariomycetes(100);o__Sordariomycetes_order_Incertae_sedis(100);f__Sordariomycetes_family_Incertae_sedis(100);g__Eucasphaeria(100);s__Eucasphaeria_capensis(100); | Eucasphaeria | 13 | Pathotroph-Saprotroph | \|Plant Pathogen\|-Plant Saprotroph-Undefined Saprotroph | Microfungus | NULL | Probable | As litter_saprotroph (Põlme et al. 2020); Decay leaf, fruit, seed (Põlme et al. 2020) | Tedersoo L, et al. 2014. Science 346:e1256688 ((https://doi.org/10.1126/science.1256688)); Põlme S, et al. 2020. Fungal Diversity 105:1-16 ((https://doi.org/10.1007/s13225-020-00466-2)) |
| Otu29770 | 1 | k__Fungi(100);p__Ascomycota(100);c__Sordariomycetes(100);o__Hypocreales(100);o__Hypocreales_unclassified(100);o__Hypocreales_unclassified(100);o__Hypocreales_unclassified(100); | Hypocreales | 7 | Saprotroph | Undefined Saprotroph | Microfungus | NULL | Possible | NULL | Sterkenburg E, et al. 2015. New Phytologist 207:1145-1158 ((https://doi.org/10.1111/nph.13426)) |
| Otu29772 | 1 | k__Fungi(100);p__Ascomycota(100);c__Sordariomycetes(100);o__Hypocreales(100);f__Hypocreaceae(100);g__Hypocrea(100);s__Hypocrea_voglmayrii(100); | Hypocrea | 13 | Pathotroph | Endophyte-Fungal Parasite | Microfungus | NULL | Probable | NULL | Tedersoo L, et al. 2014. Science 346:e1256688 ((https://doi.org/10.1126/science.1256688)); Christian N, et al. 2016. Microbial Ecology 72: 621-632 ((https://doi.org/10.1007/s00248-016-0804-y)) |
| Otu29774 | 1 | k__Fungi(100);p__Ascomycota(100);c__Sordariomycetes(100);o__Boliniales(100);f__Boliniaceae(100);f__Boliniaceae_unclassified(100);f__Boliniaceae_unclassified(100); | Boliniaceae | 9 | Saprotroph | Wood Saprotroph | Microfungus | NULL | Probable | Saprobic on and in wood and bark (Cannon & Kirk 2007) | Cannon PF, Kirk PM. 2007. Fungal Families of the World. CAB International, Cambridge (ISBN: 978-0851998275) |
| Otu29777 | 1 | k__Fungi(100);p__Ascomycota(100);c__Sordariomycetes(100);o__Hypocreales(100);f__Hypocreales_family_Incertae_sedis(100);f__Hypocreales_family_Incertae_sedis_unclassified(100);f__Hypocreales_family_Incertae_sedis_unclassified(100); | Hypocreales | 7 | Saprotroph | Undefined Saprotroph | Microfungus | NULL | Possible | NULL | Sterkenburg E, et al. 2015. New Phytologist 207:1145-1158 ((https://doi.org/10.1111/nph.13426)) |
| Otu29779 | 1 | k__Fungi(100);p__Ascomycota(100);c__Sordariomycetes(100);o__Hypocreales(100);f__Hypocreaceae(100);g__Hypocrea(100);s__Hypocrea_moravica(100); | Hypocrea | 13 | Pathotroph | Endophyte-Fungal Parasite | Microfungus | NULL | Probable | NULL | Tedersoo L, et al. 2014. Science 346:e1256688 ((https://doi.org/10.1126/science.1256688)); Christian N, et al. 2016. Microbial Ecology 72: 621-632 ((https://doi.org/10.1007/s00248-016-0804-y)) |
| Otu29780 | 1 | k__Fungi(100);p__Ascomycota(100);c__Sordariomycetes(100);o__Hypocreales(100);o__Hypocreales_unclassified(100);o__Hypocreales_unclassified(100);o__Hypocreales_unclassified(100); | Hypocreales | 7 | Saprotroph | Undefined Saprotroph | Microfungus | NULL | Possible | NULL | Sterkenburg E, et al. 2015. New Phytologist 207:1145-1158 ((https://doi.org/10.1111/nph.13426)) |
| Otu29781 | 1 | k__Fungi(100);p__Ascomycota(100);c__Sordariomycetes(100);o__Hypocreales(100);f__Hypocreales_family_Incertae_sedis(100);g__Geosmithia(100);s__Geosmithia_sp_2_FR_2014(100); | Geosmithia | 13 | Saprotroph | \|Undefined Saprotroph\| | Microfungus | NULL | Probable | Decay leaf, fruit, seed, soil, dung, animal material (Põlme et al. 2020) | Põlme S, et al. 2020. Fungal Diversity 105:1-16 ((https://doi.org/10.1007/s13225-020-00466-2)) |
| Otu29782 | 1 | k__Fungi(100);p__Ascomycota(100);c__Sordariomycetes(100);o__Hypocreales(100);o__Hypocreales_unclassified(100);o__Hypocreales_unclassified(100);o__Hypocreales_unclassified(100); | Hypocreales | 7 | Saprotroph | Undefined Saprotroph | Microfungus | NULL | Possible | NULL | Sterkenburg E, et al. 2015. New Phytologist 207:1145-1158 ((https://doi.org/10.1111/nph.13426)) |
| Otu29785 | 1 | k__Fungi(100);p__Ascomycota(100);c__Sordariomycetes(100);o__Hypocreales(100);o__Hypocreales_unclassified(100);o__Hypocreales_unclassified(100);o__Hypocreales_unclassified(100); | Hypocreales | 7 | Saprotroph | Undefined Saprotroph | Microfungus | NULL | Possible | NULL | Sterkenburg E, et al. 2015. New Phytologist 207:1145-1158 ((https://doi.org/10.1111/nph.13426)) |
| Otu29791 | 1 | k__Fungi(100);p__Ascomycota(100);c__Sordariomycetes(100);o__Hypocreales(100);f__Hypocreales_family_Incertae_sedis(100);g__Acremonium(100);g__Acremonium_unclassified(100); | Acremonium | 13 | Pathotroph-Saprotroph-Symbiotroph | Animal Pathogen-Endophyte-Fungal Parasite-Plant Pathogen-\|Undefined Saprotroph\|-Wood Saprotroph | Microfungus | Soft Rot | Possible | As foliar_endophyte (Põlme et al. 2020); culture (Hanada et al. 2010, Del Olmo; Decay leaf, fruit, seed, soil, dung, animal material (Põlme et al. 2020); Endophyte detection method; Ruiz & Arnold 2014); Likely opportunistic human pathogen (Irinyi et al. 2015) | Duncan CG, Eslyn WE. 1966. Mycologia 58:642-645 ((https://doi.org/10.2307/3757045)); Nilsson T. 1973. Studia Forestalia Suecica 104: 1-40 ((https://pub.epsilon.slu.se/5711/)); Worrall JJ, et al. 1997. Mycologia 89:199-219 ((https://doi.org/10.2307/3761073)); Hanada RE, et al. 2010. Fungal Biology 114:901-910 ((https://doi.org/10.1016/j.funbio.2010.08.006)); Del Olmo-Ruiz M, Arnold AE. 2014. Mycologia 106: 8-21 ((https://doi.org/10.3852/13-098)); Irinyi L, et al. 2015. Medical Mycology 53313-337 ((https://doi.org/10.1093/mmy/myv008)); Põlme S, et al. 2020. Fungal Diversity 105:1-16 ((https://doi.org/10.1007/s13225-020-00466-2)) |
| Otu29792 | 1 | k__Fungi(100);p__Ascomycota(100);c__Sordariomycetes(100);o__Hypocreales(100);f__Nectriaceae(100);g__Volutella(100);s__Volutella_sp(100); | Volutella | 13 | Pathotroph | \|Plant Pathogen\| | Microfungus | NULL | Probable | NULL | Tedersoo L, et al. 2014. Science 346:e1256688 ((https://doi.org/10.1126/science.1256688)); Põlme S, et al. 2020. Fungal Diversity 105:1-16 ((https://doi.org/10.1007/s13225-020-00466-2)) |
| Otu29793 | 1 | k__Fungi(100);p__Ascomycota(100);c__Sordariomycetes(100);o__Hypocreales(100);o__Hypocreales_unclassified(100);o__Hypocreales_unclassified(100);o__Hypocreales_unclassified(100); | Hypocreales | 7 | Saprotroph | Undefined Saprotroph | Microfungus | NULL | Possible | NULL | Sterkenburg E, et al. 2015. New Phytologist 207:1145-1158 ((https://doi.org/10.1111/nph.13426)) |
| Otu29796 | 1 | k__Fungi(100);p__Ascomycota(100);c__Sordariomycetes(100);o__Hypocreales(100);f__Nectriaceae(100);f__Nectriaceae_unclassified(100);f__Nectriaceae_unclassified(100); | Nectriaceae | 9 | Pathotroph-Saprotroph-Symbiotroph | Animal Pathogen-Endophyte-Fungal Parasite-Lichen Parasite-Plant Pathogen-Wood Saprotroph | Microfungus | NULL | Possible | Fusarium species are amongst the economically significant plant pathogens (Cannon & Kirk 2007) | Cannon PF, Kirk PM. 2007. Fungal Families of the World. CAB International, Cambridge (ISBN: 978-0851998275); Hirooka Y, et al. 2012. Studies in Mycology 711-210 ((https://doi.org/10.3114/sim0001)); Tedersoo L, et al. 2014. Science 346:e1256688 ((https://doi.org/10.1126/science.1256688)); The American Phytopathological Society Common Names of Plant Diseases ((http://www.apsnet.org/publications/commonnames/Pages/default.aspx)); https://www.cdc.gov ((https://www.cdc.gov)) |
| Otu29798 | 1 | k__Fungi(100);p__Ascomycota(100);c__Sordariomycetes(100);o__Hypocreales(100);f__Nectriaceae(100);g__Pleonectria(100);g__Pleonectria_unclassified(100); | Pleonectria | 13 | Saprotroph | \|Wood Saprotroph\| | Microfungus | NULL | Probable | Decay wood (Põlme et al. 2020) | Põlme S, et al. 2020. Fungal Diversity 105:1-16 ((https://doi.org/10.1007/s13225-020-00466-2)) |
| Otu29800 | 1 | k__Fungi(100);p__Ascomycota(100);c__Sordariomycetes(100);o__Hypocreales(100);f__Hypocreales_family_Incertae_sedis(100);g__Acremonium(100);s__Acremonium_implicatum(100); | Acremonium | 13 | Pathotroph-Saprotroph-Symbiotroph | Animal Pathogen-Endophyte-Fungal Parasite-Plant Pathogen-\|Undefined Saprotroph\|-Wood Saprotroph | Microfungus | Soft Rot | Possible | As foliar_endophyte (Põlme et al. 2020); culture (Hanada et al. 2010, Del Olmo; Decay leaf, fruit, seed, soil, dung, animal material (Põlme et al. 2020); Endophyte detection method; Ruiz & Arnold 2014); Likely opportunistic human pathogen (Irinyi et al. 2015) | Duncan CG, Eslyn WE. 1966. Mycologia 58:642-645 ((https://doi.org/10.2307/3757045)); Nilsson T. 1973. Studia Forestalia Suecica 104: 1-40 ((https://pub.epsilon.slu.se/5711/)); Worrall JJ, et al. 1997. Mycologia 89:199-219 ((https://doi.org/10.2307/3761073)); Hanada RE, et al. 2010. Fungal Biology 114:901-910 ((https://doi.org/10.1016/j.funbio.2010.08.006)); Del Olmo-Ruiz M, Arnold AE. 2014. Mycologia 106: 8-21 ((https://doi.org/10.3852/13-098)); Irinyi L, et al. 2015. Medical Mycology 53313-337 ((https://doi.org/10.1093/mmy/myv008)); Põlme S, et al. 2020. Fungal Diversity 105:1-16 ((https://doi.org/10.1007/s13225-020-00466-2)) |
| Otu29802 | 1 | k__Fungi(100);p__Ascomycota(100);c__Saccharomycetes(100);o__Saccharomycetales(100);f__Saccharomycetaceae(100);g__Kazachstania(100);s__Kazachstania_intestinalis(100); | Kazachstania | 13 | Saprotroph | \|Undefined Saprotroph\| | Yeast | NULL | Probable | As nectar-tap_saprotroph (Põlme et al. 2020); Decay sugar-rich substrates (Põlme et al. 2020) | Põlme S, et al. 2020. Fungal Diversity 105:1-16 ((https://doi.org/10.1007/s13225-020-00466-2)) |
| Otu29804 | 1 | k__Fungi(100);p__Ascomycota(100);c__Sordariomycetes(100);o__Diaporthales(100);f__Diaporthales_family_Incertae_sedis(100);g__Ophiognomonia(100);s__Ophiognomonia_sp_CBP23C(100); | Ophiognomonia | 13 | Pathotroph | \|Plant Pathogen\| | Microfungus | NULL | Highly Probable | See Marssoniella juglandis, Ophiognomonia leptostyla, and Syn. Gnomonia leptostyla; Causes leaf blotch and spots, also known as walnut anthracnose or walnut black spot (Sogonov et al. 2008) | Sogonov MV, et al. 2008. Studies in Mycology 62:1-79 ((https://www.studiesinmycology.org/sim/Sim62/03_Leaf-inhabiting_genera_of_the_Gnomoniaceae,_Diaporthales.pdf)); Põlme S, et al. 2020. Fungal Diversity 105:1-16 ((https://doi.org/10.1007/s13225-020-00466-2)) |
| Otu29805 | 1 | k__Fungi(100);p__Ascomycota(100);c__Sordariomycetes(100);o__Calosphaeriales(100);f__Calosphaeriaceae(100);g__Jattaea(100);s__Jattaea_prunicola(100); | Jattaea | 13 | Saprotroph | \|Undefined Saprotroph\| | Microfungus | NULL | Probable | NULL | Tedersoo L, et al. 2014. Science 346:e1256688 ((https://doi.org/10.1126/science.1256688)); Põlme S, et al. 2020. Fungal Diversity 105:1-16 ((https://doi.org/10.1007/s13225-020-00466-2)) |
| Otu29806 | 1 | k__Fungi(100);p__Ascomycota(100);c__Sordariomycetes(100);o__Hypocreales(100);o__Hypocreales_unclassified(100);o__Hypocreales_unclassified(100);o__Hypocreales_unclassified(100); | Hypocreales | 7 | Saprotroph | Undefined Saprotroph | Microfungus | NULL | Possible | NULL | Sterkenburg E, et al. 2015. New Phytologist 207:1145-1158 ((https://doi.org/10.1111/nph.13426)) |
| Otu29807 | 1 | k__Fungi(100);p__Ascomycota(100);c__Sordariomycetes(100);o__Hypocreales(100);f__Clavicipitaceae(100);g__Metarhiziopsis(100);s__Metarhiziopsis_microspora(100); | Metarhiziopsis | 13 | Pathotroph-Saprotroph | \|Animal Parasite\|-Undefined Saprotroph | Microfungus | NULL | Probable | As animal_decomposer (Põlme et al. 2020); Decay animal material (Põlme et al. 2020) | Põlme S, et al. 2020. Fungal Diversity 105:1-16 ((https://doi.org/10.1007/s13225-020-00466-2)) |
| Otu29808 | 1 | k__Fungi(100);p__Ascomycota(100);c__Eurotiomycetes(100);o__Eurotiales(100);f__Trichocomaceae(100);g__Paecilomyces(100);s__Paecilomyces_lilacinus(100); | Paecilomyces | 13 | Pathotroph-Saprotroph-Symbiotroph | Clavicipitaceous \|Endophyte\|-Plant Pathogen-Plant Saprotroph-Undefined Saprotroph-Wood Saprotroph | Microfungus | Soft Rot | Probable | As foliar_endophyte (Põlme et al. 2020); As litter_saprotroph (Põlme et al. 2020); culture (Lopez et al. 2014); Decay leaf, fruit, seed, animal material (Põlme et al. 2020); Endophyte detection method | Seehann G, et al. 1975. List of Fungi in Soft-Rot Tissue Tests. Institute fur Holzbiologie und Holzschutz, Bundesforschungsanstalt fur Forst - und Holzwirtshaft, Hamburg - Lohbrugge, Germany; Costa IPMW, et al. 2012. Mycotaxon 119:e26 ((http://mycotaxon.com/resources/checklists/costa_v119_checklist.pdf)); Tedersoo L, et al. 2014. Science 346:e1256688 ((https://doi.org/10.1126/science.1256688)); Põlme S, et al. 2020. Fungal Diversity 105:1-16 ((https://doi.org/10.1007/s13225-020-00466-2)); The American Phytopathological Society Common Names of Plant Diseases ((http://www.apsnet.org/publications/commonnames/Pages/default.aspx)); Lopez DC, et al. 2014. Plos One 9:e103891 ((https://doi.org/10.1371/journal.pone.0103891)) |
| Otu29809 | 1 | k__Fungi(100);p__Ascomycota(100);c__Sordariomycetes(100);o__Hypocreales(100);f__Nectriaceae(100);g__Gibberella(100);s__Gibberella_intricans(100); | Gibberella | 13 | Pathotroph-Saprotroph-Symbiotroph | Endophyte-\|Plant Pathogen\|-Plant Saprotroph-Undefined Saprotroph | Microfungus | Soft Rot | Probable | As foliar_endophyte (Põlme et al. 2020); As litter_saprotroph (Põlme et al. 2020); Causing damping-off disease (Põlme et al. 2020); Decay leaf, fruit, seed, roots, soil, animal material (Põlme et al. 2020) | Tedersoo L, et al. 2014. Science 346:e1256688 ((https://doi.org/10.1126/science.1256688)); Põlme S, et al. 2020. Fungal Diversity 105:1-16 ((https://doi.org/10.1007/s13225-020-00466-2)) |
| Otu29810 | 1 | k__Fungi(100);p__Ascomycota(100);c__Sordariomycetes(100);o__Hypocreales(100);f__Ophiocordycipitaceae(100);g__Elaphocordyceps(100);g__Elaphocordyceps_unclassified(100); | Elaphocordyceps | 13 | Pathotroph | Fungal Parasite | Secotioid-Spathuloid | NULL | Highly Probable | NULL | Samuels GJ. 2007. Self-published; Sung G-H, et al. 2007. Studies in Mycology 57:5-59 ((https://www.studiesinmycology.org/index.php/issue/59-studies-in-mycology-no-57)); List compiled by John Plischke III ((https://goodmushroom.weebly.com/uploads/5/7/1/7/57176535/fungi_on_fungi_pdf_lq.pdf)) |
| Otu29811 | 1 | k__Fungi(100);p__Ascomycota(100);c__Sordariomycetes(100);o__Hypocreales(100);f__Hypocreaceae(100);g__Hypomyces(100);g__Hypomyces_unclassified(100); | Hypomyces | 13 | Pathotroph | \|Fungal Parasite\| | Microfungus | NULL | Highly Probable | As mycoparasite (Põlme et al. 2020); Decay fungal material (Põlme et al. 2020); Sexual form with asexual forms spread across multiple genera | Põlme S, et al. 2020. Fungal Diversity 105:1-16 ((https://doi.org/10.1007/s13225-020-00466-2)); List compiled by John Plischke III ((https://goodmushroom.weebly.com/uploads/5/7/1/7/57176535/fungi_on_fungi_pdf_lq.pdf)) |
| Otu29812 | 1 | k__Fungi(100);p__Ascomycota(100);c__Sordariomycetes(100);o__Hypocreales(100);f__Nectriaceae(100);f__Nectriaceae_unclassified(100);f__Nectriaceae_unclassified(100); | Nectriaceae | 9 | Pathotroph-Saprotroph-Symbiotroph | Animal Pathogen-Endophyte-Fungal Parasite-Lichen Parasite-Plant Pathogen-Wood Saprotroph | Microfungus | NULL | Possible | Fusarium species are amongst the economically significant plant pathogens (Cannon & Kirk 2007) | Cannon PF, Kirk PM. 2007. Fungal Families of the World. CAB International, Cambridge (ISBN: 978-0851998275); Hirooka Y, et al. 2012. Studies in Mycology 711-210 ((https://doi.org/10.3114/sim0001)); Tedersoo L, et al. 2014. Science 346:e1256688 ((https://doi.org/10.1126/science.1256688)); The American Phytopathological Society Common Names of Plant Diseases ((http://www.apsnet.org/publications/commonnames/Pages/default.aspx)); https://www.cdc.gov ((https://www.cdc.gov)) |
| Otu29813 | 1 | k__Fungi(100);p__Ascomycota(100);c__Sordariomycetes(100);o__Hypocreales(100);f__Clavicipitaceae(100);g__Pochonia(100);s__Pochonia_bulbillosa(100); | Pochonia | 13 | Pathotroph-Saprotroph | \|Animal Parasite\|-Animal Pathogen-Undefined Saprotroph | Microfungus | NULL | Probable | As animal_decomposer (Põlme et al. 2020); Decay animal material (Põlme et al. 2020) | Tedersoo L, et al. 2014. Science 346:e1256688 ((https://doi.org/10.1126/science.1256688)); Põlme S, et al. 2020. Fungal Diversity 105:1-16 ((https://doi.org/10.1007/s13225-020-00466-2)) |
| Otu29814 | 1 | k__Fungi(100);p__Ascomycota(100);c__Sordariomycetes(100);o__Hypocreales(100);f__Hypocreales_family_Incertae_sedis(100);g__Acremonium(100);s__Acremonium_sp(100); | Acremonium | 13 | Pathotroph-Saprotroph-Symbiotroph | Animal Pathogen-Endophyte-Fungal Parasite-Plant Pathogen-\|Undefined Saprotroph\|-Wood Saprotroph | Microfungus | Soft Rot | Possible | As foliar_endophyte (Põlme et al. 2020); culture (Hanada et al. 2010, Del Olmo; Decay leaf, fruit, seed, soil, dung, animal material (Põlme et al. 2020); Endophyte detection method; Ruiz & Arnold 2014); Likely opportunistic human pathogen (Irinyi et al. 2015) | Duncan CG, Eslyn WE. 1966. Mycologia 58:642-645 ((https://doi.org/10.2307/3757045)); Nilsson T. 1973. Studia Forestalia Suecica 104: 1-40 ((https://pub.epsilon.slu.se/5711/)); Worrall JJ, et al. 1997. Mycologia 89:199-219 ((https://doi.org/10.2307/3761073)); Hanada RE, et al. 2010. Fungal Biology 114:901-910 ((https://doi.org/10.1016/j.funbio.2010.08.006)); Del Olmo-Ruiz M, Arnold AE. 2014. Mycologia 106: 8-21 ((https://doi.org/10.3852/13-098)); Irinyi L, et al. 2015. Medical Mycology 53313-337 ((https://doi.org/10.1093/mmy/myv008)); Põlme S, et al. 2020. Fungal Diversity 105:1-16 ((https://doi.org/10.1007/s13225-020-00466-2)) |
| Otu29818 | 1 | k__Fungi(100);p__Ascomycota(100);c__Lecanoromycetes(100);o__Ostropales(100);f__Stictidaceae(100);f__Stictidaceae_unclassified(100);f__Stictidaceae_unclassified(100); | Stictidaceae | 9 | Symbiotroph | Lichenized-Wood Saprotroph | Microfungus-Thallus | NULL | Probable | Saprobic on wood and stems, a few species lichenized or lignicolous (Cannon & Kirk 2007) | Cannon PF, Kirk PM. 2007. Fungal Families of the World. CAB International, Cambridge (ISBN: 978-0851998275) |
| Otu29822 | 1 | k__Fungi(100);p__Ascomycota(100);c__Pezizomycetes(100);o__Pezizales(100);f__Pyronemataceae(100);g__Tarzetta(100);s__Tarzetta_sp(100); | Tarzetta | 13 | Symbiotroph | \|Ectomycorrhizal\| | Microfungus-Pezizoid | NULL | Highly Probable | NULL | Rinaldi AC, et al. 2008. Fungal Diversity 33:1-45 ((http://www.fungaldiversity.org/fdp/sfdp/33-1.pdf)); Tedersoo L, et al. 2010. Mycorrhiza 20:217-263 ((https://doi.org/10.1007/s00572-009-0274-x)); Põlme S, et al. 2020. Fungal Diversity 105:1-16 ((https://doi.org/10.1007/s13225-020-00466-2)) |
| Otu29823 | 1 | k__Fungi(100);p__Ascomycota(100);c__Eurotiomycetes(100);o__Eurotiales(100);f__Trichocomaceae(100);g__Aspergillus(100);g__Aspergillus_unclassified(100); | Aspergillus | 13 | Pathotroph-Saprotroph-Symbiotroph | Animal Pathogen-Endophyte-Plant Saprotroph-\|Undefined Saprotroph\|-Wood Saprotroph | Microfungus | NULL | Probable | As foliar_endophyte (Põlme et al. 2020); Decay leaf, fruit, seed, soil, dung, animal material (Põlme et al. 2020) | Seehann G, et al. 1975. List of Fungi in Soft-Rot Tissue Tests. Institute fur Holzbiologie und Holzschutz, Bundesforschungsanstalt fur Forst - und Holzwirtshaft, Hamburg - Lohbrugge, Germany; Tedersoo L, et al. 2014. Science 346:e1256688 ((https://doi.org/10.1126/science.1256688)); Irinyi L, et al. 2015. Medical Mycology 53:313-337 ((https://doi.org/10.1093/mmy/myv008)); Põlme S, et al. 2020. Fungal Diversity 105:1-16 ((https://doi.org/10.1007/s13225-020-00466-2)); Massimo NC, et al. 2015. Microbial Ecology 70:61-76 ((https://doi.org/10.1007/s00248-014-0563-6)); The American Phytopathological Society Common Names of Plant Diseases ((http://www.apsnet.org/publications/commonnames/Pages/default.aspx)); https://www.cdc.gov ((https://www.cdc.gov)) |
| Otu29825 | 1 | k__Fungi(100);p__Ascomycota(100);c__Taphrinomycetes(100);o__Taphrinales(100);f__Taphrinaceae(100);g__Taphrina(100);g__Taphrina_unclassified(100); | Taphrina | 13 | Pathotroph | \|Plant Pathogen\| | Dimorphic-Microfungus | NULL | Probable | NULL | Põlme S, et al. 2020. Fungal Diversity 105:1-16 ((https://doi.org/10.1007/s13225-020-00466-2)); The American Phytopathological Society Common Names of Plant Diseases ((http://www.apsnet.org/publications/commonnames/Pages/default.aspx)) |
| Otu29826 | 1 | k__Fungi(100);p__Ascomycota(100);c__Taphrinomycetes(100);o__Taphrinales(100);f__Taphrinaceae(100);g__Taphrina(100);g__Taphrina_unclassified(100); | Taphrina | 13 | Pathotroph | \|Plant Pathogen\| | Dimorphic-Microfungus | NULL | Probable | NULL | Põlme S, et al. 2020. Fungal Diversity 105:1-16 ((https://doi.org/10.1007/s13225-020-00466-2)); The American Phytopathological Society Common Names of Plant Diseases ((http://www.apsnet.org/publications/commonnames/Pages/default.aspx)) |
| Otu29827 | 1 | k__Fungi(100);p__Ascomycota(100);c__Taphrinomycetes(100);o__Taphrinales(100);f__Taphrinaceae(100);f__Taphrinaceae_unclassified(100);f__Taphrinaceae_unclassified(100); | Taphrinaceae | 9 | Pathotroph | Plant Pathogen | Microfungus | NULL | Probable | Biotrophic on plants, usually causing hyperplasia (galls, witches broom) or lesions (Cannon & Kirk 2007) | Cannon PF, Kirk PM. 2007. Fungal Families of the World. CAB International, Cambridge (ISBN: 978-0851998275) |
| Otu29828 | 1 | k__Fungi(100);p__Ascomycota(100);c__Taphrinomycetes(100);o__Taphrinales(100);f__Taphrinaceae(100);f__Taphrinaceae_unclassified(100);f__Taphrinaceae_unclassified(100); | Taphrinaceae | 9 | Pathotroph | Plant Pathogen | Microfungus | NULL | Probable | Biotrophic on plants, usually causing hyperplasia (galls, witches broom) or lesions (Cannon & Kirk 2007) | Cannon PF, Kirk PM. 2007. Fungal Families of the World. CAB International, Cambridge (ISBN: 978-0851998275) |
| Otu29829 | 1 | k__Fungi(100);p__Ascomycota(100);c__Saccharomycetes(100);o__Saccharomycetales(100);f__Saccharomycetaceae(100);g__Kazachstania(100);s__Kazachstania_intestinalis(100); | Kazachstania | 13 | Saprotroph | \|Undefined Saprotroph\| | Yeast | NULL | Probable | As nectar-tap_saprotroph (Põlme et al. 2020); Decay sugar-rich substrates (Põlme et al. 2020) | Põlme S, et al. 2020. Fungal Diversity 105:1-16 ((https://doi.org/10.1007/s13225-020-00466-2)) |
| Otu29831 | 1 | k__Fungi(100);p__Ascomycota(100);c__Taphrinomycetes(100);o__Taphrinales(100);f__Taphrinaceae(100);g__Taphrina(100);s__Taphrina_confusa(100); | Taphrina | 13 | Pathotroph | \|Plant Pathogen\| | Dimorphic-Microfungus | NULL | Probable | NULL | Põlme S, et al. 2020. Fungal Diversity 105:1-16 ((https://doi.org/10.1007/s13225-020-00466-2)); The American Phytopathological Society Common Names of Plant Diseases ((http://www.apsnet.org/publications/commonnames/Pages/default.aspx)) |
| Otu29833 | 1 | k__Fungi(100);p__Ascomycota(100);c__Taphrinomycetes(100);o__Taphrinales(100);f__Taphrinaceae(100);g__Taphrina(100);g__Taphrina_unclassified(100); | Taphrina | 13 | Pathotroph | \|Plant Pathogen\| | Dimorphic-Microfungus | NULL | Probable | NULL | Põlme S, et al. 2020. Fungal Diversity 105:1-16 ((https://doi.org/10.1007/s13225-020-00466-2)); The American Phytopathological Society Common Names of Plant Diseases ((http://www.apsnet.org/publications/commonnames/Pages/default.aspx)) |
| Otu29836 | 1 | k__Fungi(100);p__Ascomycota(100);c__Taphrinomycetes(100);o__Taphrinales(100);f__Taphrinaceae(100);f__Taphrinaceae_unclassified(100);f__Taphrinaceae_unclassified(100); | Taphrinaceae | 9 | Pathotroph | Plant Pathogen | Microfungus | NULL | Probable | Biotrophic on plants, usually causing hyperplasia (galls, witches broom) or lesions (Cannon & Kirk 2007) | Cannon PF, Kirk PM. 2007. Fungal Families of the World. CAB International, Cambridge (ISBN: 978-0851998275) |
| Otu29838 | 1 | k__Fungi(100);p__Ascomycota(100);c__Taphrinomycetes(100);o__Taphrinales(100);f__Taphrinaceae(100);g__Taphrina(100);g__Taphrina_unclassified(100); | Taphrina | 13 | Pathotroph | \|Plant Pathogen\| | Dimorphic-Microfungus | NULL | Probable | NULL | Põlme S, et al. 2020. Fungal Diversity 105:1-16 ((https://doi.org/10.1007/s13225-020-00466-2)); The American Phytopathological Society Common Names of Plant Diseases ((http://www.apsnet.org/publications/commonnames/Pages/default.aspx)) |
| Otu29839 | 1 | k__Fungi(100);p__Ascomycota(100);c__Taphrinomycetes(100);o__Taphrinales(100);f__Taphrinaceae(100);g__Taphrina(100);g__Taphrina_unclassified(100); | Taphrina | 13 | Pathotroph | \|Plant Pathogen\| | Dimorphic-Microfungus | NULL | Probable | NULL | Põlme S, et al. 2020. Fungal Diversity 105:1-16 ((https://doi.org/10.1007/s13225-020-00466-2)); The American Phytopathological Society Common Names of Plant Diseases ((http://www.apsnet.org/publications/commonnames/Pages/default.aspx)) |
| Otu29841 | 1 | k__Fungi(100);p__Ascomycota(100);c__Taphrinomycetes(100);o__Taphrinales(100);f__Taphrinaceae(100);g__Taphrina(100);g__Taphrina_unclassified(100); | Taphrina | 13 | Pathotroph | \|Plant Pathogen\| | Dimorphic-Microfungus | NULL | Probable | NULL | Põlme S, et al. 2020. Fungal Diversity 105:1-16 ((https://doi.org/10.1007/s13225-020-00466-2)); The American Phytopathological Society Common Names of Plant Diseases ((http://www.apsnet.org/publications/commonnames/Pages/default.aspx)) |
| Otu29843 | 1 | k__Fungi(100);p__Ascomycota(100);c__Eurotiomycetes(100);o__Chaetothyriales(100);f__Herpotrichiellaceae(100);g__Cladophialophora(100);g__Cladophialophora_unclassified(100); | Cladophialophora | 13 | Pathotroph-Saprotroph-Symbiotroph | Animal Pathogen-Endophyte-Fungal Parasite-\|Undefined Saprotroph\| | Dimorphic-Microfungus | NULL | Probable | As mycoparasite (Põlme et al. 2020); As root_endophyte_dark_septate (Põlme et al. 2020); As soil_saprotroph (Põlme et al. 2020); Decay soil (Põlme et al. 2020) | James TY, et al. 2006. Nature 443:818-822 ((https://doi.org/10.1038/nature05110)); Põlme S, et al. 2020. Fungal Diversity 105:1-16 ((https://doi.org/10.1007/s13225-020-00466-2)) |
| Otu29845 | 1 | k__Fungi(100);p__Ascomycota(100);c__Eurotiomycetes(100);o__Chaetothyriales(100);f__Herpotrichiellaceae(100);g__Phaeomoniella(100);g__Phaeomoniella_unclassified(100); | Phaeomoniella | 13 | Pathotroph | \|Plant Pathogen\| | Dimorphic-Microfungus; Facultative Yeast (Tedersoo et | NULL | Highly Probable | Pathotroph isolated from necrosis in Prunus wood (Damm et al. 2010) | Damm U, et al. 2010. Persoonia 24:60-80 ((http://doi.org/10.3767/003158510X500705)); Tedersoo L, et al. 2014. Science 346:e1256688 ((https://doi.org/10.1126/science.1256688)); Põlme S, et al. 2020. Fungal Diversity 105:1-16 ((https://doi.org/10.1007/s13225-020-00466-2)) |
| Otu29846 | 1 | k__Fungi(100);p__Ascomycota(100);c__Eurotiomycetes(100);o__Chaetothyriales(100);f__Herpotrichiellaceae(100);g__Cladophialophora(100);s__Cladophialophora_chaetospira(100); | Cladophialophora | 13 | Pathotroph-Saprotroph-Symbiotroph | Animal Pathogen-Endophyte-Fungal Parasite-\|Undefined Saprotroph\| | Dimorphic-Microfungus | NULL | Probable | As mycoparasite (Põlme et al. 2020); As root_endophyte_dark_septate (Põlme et al. 2020); As soil_saprotroph (Põlme et al. 2020); Decay soil (Põlme et al. 2020) | James TY, et al. 2006. Nature 443:818-822 ((https://doi.org/10.1038/nature05110)); Põlme S, et al. 2020. Fungal Diversity 105:1-16 ((https://doi.org/10.1007/s13225-020-00466-2)) |
| Otu29847 | 1 | k__Fungi(100);p__Ascomycota(100);c__Pezizomycetes(100);o__Pezizales(100);f__Pezizaceae(100);g__Peziza(100);s__Peziza_varia(100); | Peziza | 13 | Saprotroph-Symbiotroph | Dung Saprotroph-Ectomycorrhizal-Endophyte-Plant Saprotroph-\|Undefined Saprotroph\| | Pezizoid | NULL | Possible | As foliar_endophyte (Põlme et al. 2020); As soil_saprotroph (Põlme et al. 2020); Decay soil (Põlme et al. 2020); Saprobic or ectomycorrhizal, on or in soil, leafmould etc., some species associated with dung or fire sites (Cannon & Kirk 2007) | Cannon PF, Kirk PM. 2007. Fungal Families of the World. CAB International, Cambridge (ISBN: 978-0851998275); Rinaldi AC, et al. 2008. Fungal Diversity 33:1-45 ((http://www.fungaldiversity.org/fdp/sfdp/33-1.pdf)); Tedersoo L, et al. 2010. Mycorrhiza 20:217-263 ((https://doi.org/10.1007/s00572-009-0274-x)) (pro parte); Põlme S, et al. 2020. Fungal Diversity 105:1-16 ((https://doi.org/10.1007/s13225-020-00466-2)) |
| Otu29848 | 1 | k__Fungi(100);p__Ascomycota(100);c__Dothideomycetes(100);o__Pleosporales(100);f__Pleosporaceae(100);g__Alternaria(100);g__Alternaria_unclassified(100); | Alternaria | 13 | Pathotroph-Saprotroph-Symbiotroph | Animal Pathogen-Endophyte-\|Plant Pathogen\|-Plant Saprotroph-Wood Saprotroph | Microfungus | Soft Rot | Possible | As foliar_endophyte (Põlme et al. 2020); As litter_saprotroph (Põlme et al. 2020); Decay leaf, fruit, seed (Põlme et al. 2020) | Duncan CG, Eslyn WE. 1966. Mycologia 58:642-664 ((https://doi.org/10.2307/3757045)); Seehann G, et al. 1975. List of Fungi in Soft-Rot Tissue Tests. Institute fur Holzbiologie und Holzschutz, Bundesforschungsanstalt fur Forst - und Holzwirtshaft, Hamburg - Lohbrugge, Germany ((https://www.irg-wp.com/irgdocs/details.php?d6493033-c9ea-40f3-8e3e-5697f32d640d)); Tedersoo L, et al. 2014. Science 346: e1256688 ((https://doi.org/10.1126/science.1256688)); Põlme S, et al. 2020. Fungal Diversity 105:1-16 ((https://doi.org/10.1007/s13225-020-00466-2)) |
| Otu29849 | 1 | k__Fungi(100);p__Basidiomycota(100);c__Microbotryomycetes(100);o__Sporidiobolales(100);f__Sporidiobolales_family_Incertae_sedis(100);g__Rhodotorula(100);s__Rhodotorula_taiwanensis(100); | Rhodotorula | 13 | Pathotroph-Saprotroph-Symbiotroph | Animal Pathogen-Animal Symbiotroph-Endophyte-Plant Pathogen-\|Undefined Saprotroph\| | Yeast | NULL | Probable | As foliar_endophyte (Põlme et al. 2020); Opportunistic human pathogen | Põlme S, et al. 2020. Fungal Diversity 105:1-16 ((https://doi.org/10.1007/s13225-020-00466-2)); https://www.cdc.gov ((https://www.cdc.gov)) |
| Otu29850 | 1 | k__Fungi(100);p__Ascomycota(100);c__Dothideomycetes(100);o__Pleosporales(100);f__Pleosporaceae(100);f__Pleosporaceae_unclassified(100);f__Pleosporaceae_unclassified(100); | Pleosporaceae | 9 | Pathotroph-Saprotroph | Endophyte-Lichen Parasite-Plant Pathogen-Undefined Saprotroph | Microfungus | NULL | Probable | Necrotrophic pathogens and saprobes, especially associated with grasses. Many species are seed-borne (Cannon & Kirk 2007) | Cannon PF, Kirk PM. 2007. Fungal Families of the World. CAB International, Cambridge (ISBN: 978-0851998275); David AS, et al. 2016. Microbial Ecology 71:912-926 ((https://doi.org/10.1007/s00248-015-0712-6)); Lawrey JD, Diederich P. 2016. http://www.lichenicolous.net (accessed 24 May 2017) ((http://www.lichenicolous.net)) |
| Otu29851 | 1 | k__Fungi(100);p__Ascomycota(100);c__Dothideomycetes(100);o__Pleosporales(100);f__Pleosporaceae(100);f__Pleosporaceae_unclassified(100);f__Pleosporaceae_unclassified(100); | Pleosporaceae | 9 | Pathotroph-Saprotroph | Endophyte-Lichen Parasite-Plant Pathogen-Undefined Saprotroph | Microfungus | NULL | Probable | Necrotrophic pathogens and saprobes, especially associated with grasses. Many species are seed-borne (Cannon & Kirk 2007) | Cannon PF, Kirk PM. 2007. Fungal Families of the World. CAB International, Cambridge (ISBN: 978-0851998275); David AS, et al. 2016. Microbial Ecology 71:912-926 ((https://doi.org/10.1007/s00248-015-0712-6)); Lawrey JD, Diederich P. 2016. http://www.lichenicolous.net (accessed 24 May 2017) ((http://www.lichenicolous.net)) |
| Otu29852 | 1 | k__Fungi(100);p__Ascomycota(100);c__Dothideomycetes(100);o__Pleosporales(100);f__Pleosporaceae(100);g__Alternaria(100);g__Alternaria_unclassified(100); | Alternaria | 13 | Pathotroph-Saprotroph-Symbiotroph | Animal Pathogen-Endophyte-\|Plant Pathogen\|-Plant Saprotroph-Wood Saprotroph | Microfungus | Soft Rot | Possible | As foliar_endophyte (Põlme et al. 2020); As litter_saprotroph (Põlme et al. 2020); Decay leaf, fruit, seed (Põlme et al. 2020) | Duncan CG, Eslyn WE. 1966. Mycologia 58:642-664 ((https://doi.org/10.2307/3757045)); Seehann G, et al. 1975. List of Fungi in Soft-Rot Tissue Tests. Institute fur Holzbiologie und Holzschutz, Bundesforschungsanstalt fur Forst - und Holzwirtshaft, Hamburg - Lohbrugge, Germany ((https://www.irg-wp.com/irgdocs/details.php?d6493033-c9ea-40f3-8e3e-5697f32d640d)); Tedersoo L, et al. 2014. Science 346: e1256688 ((https://doi.org/10.1126/science.1256688)); Põlme S, et al. 2020. Fungal Diversity 105:1-16 ((https://doi.org/10.1007/s13225-020-00466-2)) |
| Otu29854 | 1 | k__Fungi(100);p__Ascomycota(100);c__Eurotiomycetes(100);o__Chaetothyriales(100);f__Herpotrichiellaceae(100);g__Phaeomoniella(100);s__Phaeomoniella_niveniae(100); | Phaeomoniella | 13 | Pathotroph | \|Plant Pathogen\| | Dimorphic-Microfungus; Facultative Yeast (Tedersoo et | NULL | Highly Probable | Pathotroph isolated from necrosis in Prunus wood (Damm et al. 2010) | Damm U, et al. 2010. Persoonia 24:60-80 ((http://doi.org/10.3767/003158510X500705)); Tedersoo L, et al. 2014. Science 346:e1256688 ((https://doi.org/10.1126/science.1256688)); Põlme S, et al. 2020. Fungal Diversity 105:1-16 ((https://doi.org/10.1007/s13225-020-00466-2)) |
| Otu29855 | 1 | k__Fungi(100);p__Ascomycota(100);c__Saccharomycetes(100);o__Saccharomycetales(100);o__Saccharomycetales_unclassified(100);o__Saccharomycetales_unclassified(100);o__Saccharomycetales_unclassified(100); | Saccharomycetales | 7 | Saprotroph | Undefined Saprotroph | Yeast | NULL | Possible | NULL | Sterkenburg E, et al. 2015. New Phytologist 207:1145-1158 ((https://doi.org/10.1111/nph.13426)) |
| Otu29857 | 1 | k__Fungi(100);p__Ascomycota(100);c__Eurotiomycetes(100);o__Chaetothyriales(100);f__Herpotrichiellaceae(100);g__Exophiala(100);s__Exophiala_attenuata(100); | Exophiala | 13 | Pathotroph-Saprotroph-Symbiotroph | \|Animal Parasite\|-Animal Pathogen-Endophyte-Plant Saprotroph-Undefined Saprotroph | Dimorphic-Facultative Yeast | NULL | Probable | As litter_saprotroph (Põlme et al. 2020); As root_endophyte_dark_septate (Põlme et al. 2020); Decay leaf, fruit, seed, animal material (Põlme et al. 2020) | Tedersoo L, et al. 2014. Science 346:e1256688 ((https://doi.org/10.1126/science.1256688)), Nagy LG, et al. 2014. Nature Communications 5:e4471 ((https://doi.org/10.1038/ncomms5471)); Põlme S, et al. 2020. Fungal Diversity 105:1-16 ((https://doi.org/10.1007/s13225-020-00466-2)) |
| Otu29859 | 1 | k__Fungi(100);p__Ascomycota(100);c__Dothideomycetes(100);o__Capnodiales(100);f__Davidiellaceae(100);g__Cladosporium(100);s__Cladosporium_sp_CBS_55279(100); | Cladosporium | 13 | Pathotroph-Saprotroph-Symbiotroph | Animal Pathogen-Endophyte-Lichen Parasite-Plant Pathogen-\|Plant Saprotroph\|-Wood Saprotroph | Microfungus | NULL | Possible | As foliar_endophyte (Põlme et al. 2020); As litter_saprotroph (Põlme et al. 2020); Decay wood, leaf, fruit, seed, soil, dung, animal material (Põlme et al. 2020) | Seehann G, et al. 1975. List of Fungi in Soft-Rot Tissue Tests. Institute fur Holzbiologie und Holzschutz, Bundesforschungsanstalt fur Forst - und Holzwirtshaft, Hamburg - Lohbrugge, Germany; Lawrey JD, Diederich P. 2016. http://www.lichenicolous.net (accessed 24 May 2017) ((http://www.lichenicolous.net)); Põlme S, et al. 2020. Fungal Diversity 105:1-16 ((https://doi.org/10.1007/s13225-020-00466-2)); The American Phytopathological Society Common Names of Plant Diseases ((http://www.apsnet.org/publications/commonnames/Pages/default.aspx)) |
| Otu29862 | 1 | k__Fungi(100);p__Ascomycota(100);c__Leotiomycetes(100);o__Helotiales(100);f__Vibrisseaceae(100);f__Vibrisseaceae_unclassified(100);f__Vibrisseaceae_unclassified(100); | Vibrisseaceae | 9 | Saprotroph-Symbiotroph | Endophyte-Undefined Saprotroph-Wood Saprotroph | Microfungus | NULL | Possible | Saprobic, usually on wood and bark immersed in fresh water. Some species may be root endophytes (Cannon & Kirk 2007) | Cannon PF, Kirk PM. 2007. Fungal Families of the World. CAB International, Cambridge (ISBN: 978-0851998275) |
| Otu29863 | 1 | k__Fungi(100);p__Ascomycota(100);c__Saccharomycetes(100);o__Saccharomycetales(100);f__Saccharomycetaceae(100);g__Kazachstania(100);s__Kazachstania_intestinalis(100); | Kazachstania | 13 | Saprotroph | \|Undefined Saprotroph\| | Yeast | NULL | Probable | As nectar-tap_saprotroph (Põlme et al. 2020); Decay sugar-rich substrates (Põlme et al. 2020) | Põlme S, et al. 2020. Fungal Diversity 105:1-16 ((https://doi.org/10.1007/s13225-020-00466-2)) |
| Otu29864 | 1 | k__Fungi(100);p__Ascomycota(100);c__Saccharomycetes(100);o__Saccharomycetales(100);f__Saccharomycetaceae(100);g__Kazachstania(100);s__Kazachstania_intestinalis(100); | Kazachstania | 13 | Saprotroph | \|Undefined Saprotroph\| | Yeast | NULL | Probable | As nectar-tap_saprotroph (Põlme et al. 2020); Decay sugar-rich substrates (Põlme et al. 2020) | Põlme S, et al. 2020. Fungal Diversity 105:1-16 ((https://doi.org/10.1007/s13225-020-00466-2)) |
| Otu29865 | 1 | k__Fungi(100);p__Ascomycota(100);c__Saccharomycetes(100);o__Saccharomycetales(100);f__Saccharomycetaceae(100);g__Kazachstania(100);s__Kazachstania_intestinalis(100); | Kazachstania | 13 | Saprotroph | \|Undefined Saprotroph\| | Yeast | NULL | Probable | As nectar-tap_saprotroph (Põlme et al. 2020); Decay sugar-rich substrates (Põlme et al. 2020) | Põlme S, et al. 2020. Fungal Diversity 105:1-16 ((https://doi.org/10.1007/s13225-020-00466-2)) |
| Otu29866 | 1 | k__Fungi(100);p__Ascomycota(100);c__Saccharomycetes(100);o__Saccharomycetales(100);f__Saccharomycetaceae(100);g__Kazachstania(100);s__Kazachstania_intestinalis(100); | Kazachstania | 13 | Saprotroph | \|Undefined Saprotroph\| | Yeast | NULL | Probable | As nectar-tap_saprotroph (Põlme et al. 2020); Decay sugar-rich substrates (Põlme et al. 2020) | Põlme S, et al. 2020. Fungal Diversity 105:1-16 ((https://doi.org/10.1007/s13225-020-00466-2)) |
| Otu29867 | 1 | k__Fungi(100);p__Ascomycota(100);c__Saccharomycetes(100);o__Saccharomycetales(100);f__Saccharomycetaceae(100);g__Kazachstania(100);s__Kazachstania_intestinalis(100); | Kazachstania | 13 | Saprotroph | \|Undefined Saprotroph\| | Yeast | NULL | Probable | As nectar-tap_saprotroph (Põlme et al. 2020); Decay sugar-rich substrates (Põlme et al. 2020) | Põlme S, et al. 2020. Fungal Diversity 105:1-16 ((https://doi.org/10.1007/s13225-020-00466-2)) |
| Otu29868 | 1 | k__Fungi(100);p__Ascomycota(100);c__Saccharomycetes(100);o__Saccharomycetales(100);f__Saccharomycetaceae(100);g__Kazachstania(100);s__Kazachstania_intestinalis(100); | Kazachstania | 13 | Saprotroph | \|Undefined Saprotroph\| | Yeast | NULL | Probable | As nectar-tap_saprotroph (Põlme et al. 2020); Decay sugar-rich substrates (Põlme et al. 2020) | Põlme S, et al. 2020. Fungal Diversity 105:1-16 ((https://doi.org/10.1007/s13225-020-00466-2)) |
| Otu29869 | 1 | k__Fungi(100);p__Ascomycota(100);c__Saccharomycetes(100);o__Saccharomycetales(100);f__Saccharomycetaceae(100);g__Kazachstania(100);s__Kazachstania_intestinalis(100); | Kazachstania | 13 | Saprotroph | \|Undefined Saprotroph\| | Yeast | NULL | Probable | As nectar-tap_saprotroph (Põlme et al. 2020); Decay sugar-rich substrates (Põlme et al. 2020) | Põlme S, et al. 2020. Fungal Diversity 105:1-16 ((https://doi.org/10.1007/s13225-020-00466-2)) |
| Otu29870 | 1 | k__Fungi(100);p__Ascomycota(100);c__Saccharomycetes(100);o__Saccharomycetales(100);f__Saccharomycetaceae(100);g__Kazachstania(100);s__Kazachstania_intestinalis(100); | Kazachstania | 13 | Saprotroph | \|Undefined Saprotroph\| | Yeast | NULL | Probable | As nectar-tap_saprotroph (Põlme et al. 2020); Decay sugar-rich substrates (Põlme et al. 2020) | Põlme S, et al. 2020. Fungal Diversity 105:1-16 ((https://doi.org/10.1007/s13225-020-00466-2)) |
| Otu29871 | 1 | k__Fungi(100);p__Ascomycota(100);c__Saccharomycetes(100);o__Saccharomycetales(100);f__Saccharomycetaceae(100);g__Kazachstania(100);s__Kazachstania_intestinalis(100); | Kazachstania | 13 | Saprotroph | \|Undefined Saprotroph\| | Yeast | NULL | Probable | As nectar-tap_saprotroph (Põlme et al. 2020); Decay sugar-rich substrates (Põlme et al. 2020) | Põlme S, et al. 2020. Fungal Diversity 105:1-16 ((https://doi.org/10.1007/s13225-020-00466-2)) |
| Otu29872 | 1 | k__Fungi(100);p__Ascomycota(100);c__Saccharomycetes(100);o__Saccharomycetales(100);f__Saccharomycetaceae(100);g__Lachancea(100);s__Lachancea_thermotolerans(100); | Lachancea | 13 | Pathotroph-Saprotroph | Animal Parasite-Animal Pathogen-\|Undefined Saprotroph\| | Yeast | NULL | Possible | As nectar-tap_saprotroph (Põlme et al. 2020); Decay animal material, sugar-rich substrates (Põlme et al. 2020); Likely opportunistic human pathogen (Irinyi et al. 2015; see L. fermentati) | Irinyi L, et al. 2015. Medical Mycology 53:313-337 ((https://doi.org/10.1093/mmy/myv008)); Põlme S, et al. 2020. Fungal Diversity 105:1-16 ((https://doi.org/10.1007/s13225-020-00466-2)) |
| Otu29873 | 1 | k__Fungi(100);p__Ascomycota(100);c__Saccharomycetes(100);o__Saccharomycetales(100);f__Saccharomycetaceae(100);g__Lachancea(100);s__Lachancea_thermotolerans(100); | Lachancea | 13 | Pathotroph-Saprotroph | Animal Parasite-Animal Pathogen-\|Undefined Saprotroph\| | Yeast | NULL | Possible | As nectar-tap_saprotroph (Põlme et al. 2020); Decay animal material, sugar-rich substrates (Põlme et al. 2020); Likely opportunistic human pathogen (Irinyi et al. 2015; see L. fermentati) | Irinyi L, et al. 2015. Medical Mycology 53:313-337 ((https://doi.org/10.1093/mmy/myv008)); Põlme S, et al. 2020. Fungal Diversity 105:1-16 ((https://doi.org/10.1007/s13225-020-00466-2)) |
| Otu29874 | 1 | k__Fungi(100);p__Ascomycota(100);c__Saccharomycetes(100);o__Saccharomycetales(100);f__Saccharomycetaceae(100);g__Kazachstania(100);s__Kazachstania_intestinalis(100); | Kazachstania | 13 | Saprotroph | \|Undefined Saprotroph\| | Yeast | NULL | Probable | As nectar-tap_saprotroph (Põlme et al. 2020); Decay sugar-rich substrates (Põlme et al. 2020) | Põlme S, et al. 2020. Fungal Diversity 105:1-16 ((https://doi.org/10.1007/s13225-020-00466-2)) |
| Otu29875 | 1 | k__Fungi(100);p__Ascomycota(100);c__Saccharomycetes(100);o__Saccharomycetales(100);f__Saccharomycetaceae(100);g__Kazachstania(100);s__Kazachstania_intestinalis(100); | Kazachstania | 13 | Saprotroph | \|Undefined Saprotroph\| | Yeast | NULL | Probable | As nectar-tap_saprotroph (Põlme et al. 2020); Decay sugar-rich substrates (Põlme et al. 2020) | Põlme S, et al. 2020. Fungal Diversity 105:1-16 ((https://doi.org/10.1007/s13225-020-00466-2)) |
| Otu29876 | 1 | k__Fungi(100);p__Ascomycota(100);c__Saccharomycetes(100);o__Saccharomycetales(100);f__Saccharomycetaceae(100);g__Kazachstania(100);s__Kazachstania_intestinalis(100); | Kazachstania | 13 | Saprotroph | \|Undefined Saprotroph\| | Yeast | NULL | Probable | As nectar-tap_saprotroph (Põlme et al. 2020); Decay sugar-rich substrates (Põlme et al. 2020) | Põlme S, et al. 2020. Fungal Diversity 105:1-16 ((https://doi.org/10.1007/s13225-020-00466-2)) |
| Otu29877 | 1 | k__Fungi(100);p__Ascomycota(100);c__Saccharomycetes(100);o__Saccharomycetales(100);f__Saccharomycetaceae(100);g__Kazachstania(100);s__Kazachstania_intestinalis(100); | Kazachstania | 13 | Saprotroph | \|Undefined Saprotroph\| | Yeast | NULL | Probable | As nectar-tap_saprotroph (Põlme et al. 2020); Decay sugar-rich substrates (Põlme et al. 2020) | Põlme S, et al. 2020. Fungal Diversity 105:1-16 ((https://doi.org/10.1007/s13225-020-00466-2)) |
| Otu29878 | 1 | k__Fungi(100);p__Ascomycota(100);c__Saccharomycetes(100);o__Saccharomycetales(100);f__Saccharomycetaceae(100);g__Kazachstania(100);s__Kazachstania_intestinalis(100); | Kazachstania | 13 | Saprotroph | \|Undefined Saprotroph\| | Yeast | NULL | Probable | As nectar-tap_saprotroph (Põlme et al. 2020); Decay sugar-rich substrates (Põlme et al. 2020) | Põlme S, et al. 2020. Fungal Diversity 105:1-16 ((https://doi.org/10.1007/s13225-020-00466-2)) |
| Otu29879 | 1 | k__Fungi(100);p__Ascomycota(100);c__Saccharomycetes(100);o__Saccharomycetales(100);f__Saccharomycetaceae(100);g__Kazachstania(100);s__Kazachstania_intestinalis(100); | Kazachstania | 13 | Saprotroph | \|Undefined Saprotroph\| | Yeast | NULL | Probable | As nectar-tap_saprotroph (Põlme et al. 2020); Decay sugar-rich substrates (Põlme et al. 2020) | Põlme S, et al. 2020. Fungal Diversity 105:1-16 ((https://doi.org/10.1007/s13225-020-00466-2)) |
| Otu29880 | 1 | k__Fungi(100);p__Ascomycota(100);c__Saccharomycetes(100);o__Saccharomycetales(100);f__Saccharomycetaceae(100);g__Lachancea(100);s__Lachancea_thermotolerans(100); | Lachancea | 13 | Pathotroph-Saprotroph | Animal Parasite-Animal Pathogen-\|Undefined Saprotroph\| | Yeast | NULL | Possible | As nectar-tap_saprotroph (Põlme et al. 2020); Decay animal material, sugar-rich substrates (Põlme et al. 2020); Likely opportunistic human pathogen (Irinyi et al. 2015; see L. fermentati) | Irinyi L, et al. 2015. Medical Mycology 53:313-337 ((https://doi.org/10.1093/mmy/myv008)); Põlme S, et al. 2020. Fungal Diversity 105:1-16 ((https://doi.org/10.1007/s13225-020-00466-2)) |
| Otu29881 | 1 | k__Fungi(100);p__Ascomycota(100);c__Saccharomycetes(100);o__Saccharomycetales(100);f__Saccharomycetaceae(100);g__Kazachstania(100);s__Kazachstania_intestinalis(100); | Kazachstania | 13 | Saprotroph | \|Undefined Saprotroph\| | Yeast | NULL | Probable | As nectar-tap_saprotroph (Põlme et al. 2020); Decay sugar-rich substrates (Põlme et al. 2020) | Põlme S, et al. 2020. Fungal Diversity 105:1-16 ((https://doi.org/10.1007/s13225-020-00466-2)) |
| Otu29882 | 1 | k__Fungi(100);p__Ascomycota(100);c__Saccharomycetes(100);o__Saccharomycetales(100);f__Saccharomycetaceae(100);g__Kazachstania(100);s__Kazachstania_intestinalis(100); | Kazachstania | 13 | Saprotroph | \|Undefined Saprotroph\| | Yeast | NULL | Probable | As nectar-tap_saprotroph (Põlme et al. 2020); Decay sugar-rich substrates (Põlme et al. 2020) | Põlme S, et al. 2020. Fungal Diversity 105:1-16 ((https://doi.org/10.1007/s13225-020-00466-2)) |
| Otu29883 | 1 | k__Fungi(100);p__Ascomycota(100);c__Saccharomycetes(100);o__Saccharomycetales(100);f__Saccharomycetaceae(100);g__Kazachstania(100);s__Kazachstania_intestinalis(100); | Kazachstania | 13 | Saprotroph | \|Undefined Saprotroph\| | Yeast | NULL | Probable | As nectar-tap_saprotroph (Põlme et al. 2020); Decay sugar-rich substrates (Põlme et al. 2020) | Põlme S, et al. 2020. Fungal Diversity 105:1-16 ((https://doi.org/10.1007/s13225-020-00466-2)) |
| Otu29884 | 1 | k__Fungi(100);p__Ascomycota(100);c__Saccharomycetes(100);o__Saccharomycetales(100);f__Saccharomycetaceae(100);g__Kazachstania(100);s__Kazachstania_intestinalis(100); | Kazachstania | 13 | Saprotroph | \|Undefined Saprotroph\| | Yeast | NULL | Probable | As nectar-tap_saprotroph (Põlme et al. 2020); Decay sugar-rich substrates (Põlme et al. 2020) | Põlme S, et al. 2020. Fungal Diversity 105:1-16 ((https://doi.org/10.1007/s13225-020-00466-2)) |
| Otu29885 | 1 | k__Fungi(100);p__Ascomycota(100);c__Saccharomycetes(100);o__Saccharomycetales(100);f__Saccharomycetaceae(100);g__Kazachstania(100);s__Kazachstania_intestinalis(100); | Kazachstania | 13 | Saprotroph | \|Undefined Saprotroph\| | Yeast | NULL | Probable | As nectar-tap_saprotroph (Põlme et al. 2020); Decay sugar-rich substrates (Põlme et al. 2020) | Põlme S, et al. 2020. Fungal Diversity 105:1-16 ((https://doi.org/10.1007/s13225-020-00466-2)) |
| Otu29886 | 1 | k__Fungi(100);p__Ascomycota(100);c__Saccharomycetes(100);o__Saccharomycetales(100);f__Saccharomycetaceae(100);g__Kazachstania(100);s__Kazachstania_intestinalis(100); | Kazachstania | 13 | Saprotroph | \|Undefined Saprotroph\| | Yeast | NULL | Probable | As nectar-tap_saprotroph (Põlme et al. 2020); Decay sugar-rich substrates (Põlme et al. 2020) | Põlme S, et al. 2020. Fungal Diversity 105:1-16 ((https://doi.org/10.1007/s13225-020-00466-2)) |
| Otu29887 | 1 | k__Fungi(100);p__Ascomycota(100);c__Saccharomycetes(100);o__Saccharomycetales(100);f__Saccharomycetaceae(100);g__Kazachstania(100);s__Kazachstania_intestinalis(100); | Kazachstania | 13 | Saprotroph | \|Undefined Saprotroph\| | Yeast | NULL | Probable | As nectar-tap_saprotroph (Põlme et al. 2020); Decay sugar-rich substrates (Põlme et al. 2020) | Põlme S, et al. 2020. Fungal Diversity 105:1-16 ((https://doi.org/10.1007/s13225-020-00466-2)) |
| Otu29888 | 1 | k__Fungi(100);p__Ascomycota(100);c__Saccharomycetes(100);o__Saccharomycetales(100);f__Saccharomycetaceae(100);g__Lachancea(100);s__Lachancea_thermotolerans(100); | Lachancea | 13 | Pathotroph-Saprotroph | Animal Parasite-Animal Pathogen-\|Undefined Saprotroph\| | Yeast | NULL | Possible | As nectar-tap_saprotroph (Põlme et al. 2020); Decay animal material, sugar-rich substrates (Põlme et al. 2020); Likely opportunistic human pathogen (Irinyi et al. 2015; see L. fermentati) | Irinyi L, et al. 2015. Medical Mycology 53:313-337 ((https://doi.org/10.1093/mmy/myv008)); Põlme S, et al. 2020. Fungal Diversity 105:1-16 ((https://doi.org/10.1007/s13225-020-00466-2)) |
| Otu29889 | 1 | k__Fungi(100);p__Ascomycota(100);c__Saccharomycetes(100);o__Saccharomycetales(100);f__Saccharomycetaceae(100);g__Lachancea(100);s__Lachancea_thermotolerans(100); | Lachancea | 13 | Pathotroph-Saprotroph | Animal Parasite-Animal Pathogen-\|Undefined Saprotroph\| | Yeast | NULL | Possible | As nectar-tap_saprotroph (Põlme et al. 2020); Decay animal material, sugar-rich substrates (Põlme et al. 2020); Likely opportunistic human pathogen (Irinyi et al. 2015; see L. fermentati) | Irinyi L, et al. 2015. Medical Mycology 53:313-337 ((https://doi.org/10.1093/mmy/myv008)); Põlme S, et al. 2020. Fungal Diversity 105:1-16 ((https://doi.org/10.1007/s13225-020-00466-2)) |
| Otu29890 | 1 | k__Fungi(100);p__Ascomycota(100);c__Saccharomycetes(100);o__Saccharomycetales(100);f__Saccharomycetaceae(100);g__Lachancea(100);s__Lachancea_thermotolerans(100); | Lachancea | 13 | Pathotroph-Saprotroph | Animal Parasite-Animal Pathogen-\|Undefined Saprotroph\| | Yeast | NULL | Possible | As nectar-tap_saprotroph (Põlme et al. 2020); Decay animal material, sugar-rich substrates (Põlme et al. 2020); Likely opportunistic human pathogen (Irinyi et al. 2015; see L. fermentati) | Irinyi L, et al. 2015. Medical Mycology 53:313-337 ((https://doi.org/10.1093/mmy/myv008)); Põlme S, et al. 2020. Fungal Diversity 105:1-16 ((https://doi.org/10.1007/s13225-020-00466-2)) |
| Otu29891 | 1 | k__Fungi(100);p__Ascomycota(100);c__Saccharomycetes(100);o__Saccharomycetales(100);f__Saccharomycetaceae(100);g__Lachancea(100);s__Lachancea_thermotolerans(100); | Lachancea | 13 | Pathotroph-Saprotroph | Animal Parasite-Animal Pathogen-\|Undefined Saprotroph\| | Yeast | NULL | Possible | As nectar-tap_saprotroph (Põlme et al. 2020); Decay animal material, sugar-rich substrates (Põlme et al. 2020); Likely opportunistic human pathogen (Irinyi et al. 2015; see L. fermentati) | Irinyi L, et al. 2015. Medical Mycology 53:313-337 ((https://doi.org/10.1093/mmy/myv008)); Põlme S, et al. 2020. Fungal Diversity 105:1-16 ((https://doi.org/10.1007/s13225-020-00466-2)) |
| Otu29892 | 1 | k__Fungi(100);p__Ascomycota(100);c__Saccharomycetes(100);o__Saccharomycetales(100);f__Saccharomycetaceae(100);g__Kazachstania(100);s__Kazachstania_intestinalis(100); | Kazachstania | 13 | Saprotroph | \|Undefined Saprotroph\| | Yeast | NULL | Probable | As nectar-tap_saprotroph (Põlme et al. 2020); Decay sugar-rich substrates (Põlme et al. 2020) | Põlme S, et al. 2020. Fungal Diversity 105:1-16 ((https://doi.org/10.1007/s13225-020-00466-2)) |
| Otu29893 | 1 | k__Fungi(100);p__Ascomycota(100);c__Saccharomycetes(100);o__Saccharomycetales(100);f__Saccharomycetaceae(100);g__Kazachstania(100);s__Kazachstania_intestinalis(100); | Kazachstania | 13 | Saprotroph | \|Undefined Saprotroph\| | Yeast | NULL | Probable | As nectar-tap_saprotroph (Põlme et al. 2020); Decay sugar-rich substrates (Põlme et al. 2020) | Põlme S, et al. 2020. Fungal Diversity 105:1-16 ((https://doi.org/10.1007/s13225-020-00466-2)) |
| Otu29894 | 1 | k__Fungi(100);p__Ascomycota(100);c__Saccharomycetes(100);o__Saccharomycetales(100);f__Saccharomycetaceae(100);g__Kazachstania(100);s__Kazachstania_intestinalis(100); | Kazachstania | 13 | Saprotroph | \|Undefined Saprotroph\| | Yeast | NULL | Probable | As nectar-tap_saprotroph (Põlme et al. 2020); Decay sugar-rich substrates (Põlme et al. 2020) | Põlme S, et al. 2020. Fungal Diversity 105:1-16 ((https://doi.org/10.1007/s13225-020-00466-2)) |
| Otu29895 | 1 | k__Fungi(100);p__Ascomycota(100);c__Saccharomycetes(100);o__Saccharomycetales(100);f__Saccharomycetaceae(100);g__Kazachstania(100);s__Kazachstania_intestinalis(100); | Kazachstania | 13 | Saprotroph | \|Undefined Saprotroph\| | Yeast | NULL | Probable | As nectar-tap_saprotroph (Põlme et al. 2020); Decay sugar-rich substrates (Põlme et al. 2020) | Põlme S, et al. 2020. Fungal Diversity 105:1-16 ((https://doi.org/10.1007/s13225-020-00466-2)) |
| Otu29896 | 1 | k__Fungi(100);p__Ascomycota(100);c__Saccharomycetes(100);o__Saccharomycetales(100);f__Pichiaceae(100);g__Pichia(100);s__Pichia_sp_LCF_18(100); | Pichia | 13 | Pathotroph-Saprotroph-Symbiotroph | Animal Parasite-Animal Pathogen-Animal Symbiotroph-Plant Pathogen-\|Undefined Saprotroph\| | Yeast | NULL | Probable | As nectar-tap_saprotroph (Põlme et al. 2020); Decay animal material, sugar-rich substrates (Põlme et al. 2020); Found in wood and digestive systems of the beetle Odontotaenius disjunctus; Likely opportunistic human pathogen (Irinyi et al. 2015) | Kurtzman CP, et al. (eds.) 2011. The Yeasts, a Taxonomic Study. Fifth Edition. Vols 1-3. Elsevier, San Diego (ISBN: 9780444521491); Irinyi L, et al. 2015. Medical Mycology 53:313-337 ((https://doi.org/10.1093/mmy/myv008)); Põlme S, et al. 2020. Fungal Diversity 105:1-16 ((https://doi.org/10.1007/s13225-020-00466-2)); The American Phytopathological Society Common Names of Plant Diseases ((https://www.apsnet.org/edcenter/resources/commonnames/Pages/default.aspx)); https://www.cdc.gov ((https://www.cdc.gov)) |
| Otu29897 | 1 | k__Fungi(100);p__Ascomycota(100);c__Taphrinomycetes(100);o__Taphrinales(100);f__Taphrinaceae(100);g__unclassified_Taphrinaceae(100);s__Taphrinaceae_sp(100); | Taphrinaceae | 9 | Pathotroph | Plant Pathogen | Microfungus | NULL | Probable | Biotrophic on plants, usually causing hyperplasia (galls, witches broom) or lesions (Cannon & Kirk 2007) | Cannon PF, Kirk PM. 2007. Fungal Families of the World. CAB International, Cambridge (ISBN: 978-0851998275) |
| Otu29899 | 1 | k__Fungi(100);p__Ascomycota(100);c__Eurotiomycetes(100);o__Chaetothyriales(100);f__Chaetothyriaceae(100);g__Cyphellophora(100);s__Cyphellophora_europaea(100); | Cyphellophora | 13 | Pathotroph-Saprotroph | Animal Parasite-Animal Pathogen-\|Plant Saprotroph\|-Undefined Saprotroph | Dimorphic-Facultative Yeast-Microfungus | NULL | Probable | As litter_saprotroph (Põlme et al. 2020); Consult the current literature for updated names on this group that included the Black Yeasts; Some species are likely opportunistic human pathogen (Irinyi et al. 2015); Decay leaf, fruit, seed, animal material (Põlme et al. 2020) | Tedersoo L, et al. 2014. Science 346:e1256688 ((https://doi.org/10.1126/science.1256688)); Irinyi L, et al. 2015. Medical Mycology 53:313-337 ((https://doi.org/10.1093/mmy/myv008)); Põlme S, et al. 2020. Fungal Diversity 105:1-16 ((https://doi.org/10.1007/s13225-020-00466-2)) |
| Otu29902 | 1 | k__Fungi(100);p__Ascomycota(100);c__Eurotiomycetes(100);o__Eurotiales(100);f__Trichocomaceae(100);g__Penicillium(100);g__Penicillium_unclassified(100); | Penicillium | 13 | Pathotroph-Saprotroph-Symbiotroph | Animal Parasite-Dung Saprotroph-Endophyte-Fungal Parasite-Plant Pathogen-\|Undefined Saprotroph\|-Wood Saprotroph | Microfungus | Soft Rot | Highly Probable | As foliar_endophyte (Põlme et al. 2020); As mycoparasite (Põlme et al. 2020); Decay soil (Põlme et al. 2020) | Duncan CG, Eslyn WE. 1966. Mycologia 58:642-664 ((https://doi.org/10.2307/3757045)); Seehann G, et al. 1975. List of Fungi in Soft-Rot Tissue Tests. Institute fur Holzbiologie und Holzschutz, Bundesforschungsanstalt fur Forst - und Holzwirtshaft, Hamburg - Lohbrugge, Germany ((https://www.irg-wp.com/irgdocs/details.php?d6493033-c9ea-40f3-8e3e-5697f32d640d)); Costa IPMW, et al. 2012. Mycotaxon 119:e26 ((http://mycotaxon.com/resources/checklists/costa_v119_checklist.pdf)); Bills GF, et al. 2013. Current Opinion in Microbiology 16: 549-565 ((https://doi.org/10.1016/j.mib.2013.08.001)); Tedersoo L, et al. 2014. Science 346:e1256688 ((https://doi.org/10.1126/science.1256688)); Põlme S, et al. 2020. Fungal Diversity 105:1-16 ((https://doi.org/10.1007/s13225-020-00466-2)) |
| Otu29904 | 1 | k__Fungi(100);p__Ascomycota(100);c__Eurotiomycetes(100);o__Chaetothyriales(100);f__Herpotrichiellaceae(100);g__unclassified_Herpotrichiellaceae(100);s__Herpotrichiellaceae_sp(100); | Herpotrichiellaceae | 9 | Pathotroph-Saprotroph | Animal Pathogen-Fungal Parasite-Undefined Saprotroph | Facultative Yeast-Microfungus | NULL | Probable | Mostly saprobic on plants or other fungi, isolated from soil etc., and from medical sources (Cannon & Kirk 2007) | Cannon PF, Kirk PM. 2007. Fungal Families of the World. CAB International, Cambridge (ISBN: 978-0851998275) |
| Otu29906 | 1 | k__Fungi(100);p__Ascomycota(100);c__Saccharomycetes(100);o__Saccharomycetales(100);f__Saccharomycetaceae(100);g__Kazachstania(100);s__Kazachstania_intestinalis(100); | Kazachstania | 13 | Saprotroph | \|Undefined Saprotroph\| | Yeast | NULL | Probable | As nectar-tap_saprotroph (Põlme et al. 2020); Decay sugar-rich substrates (Põlme et al. 2020) | Põlme S, et al. 2020. Fungal Diversity 105:1-16 ((https://doi.org/10.1007/s13225-020-00466-2)) |
| Otu29907 | 1 | k__Fungi(100);p__Ascomycota(100);c__Saccharomycetes(100);o__Saccharomycetales(100);f__Saccharomycetaceae(100);g__Kazachstania(100);s__Kazachstania_intestinalis(100); | Kazachstania | 13 | Saprotroph | \|Undefined Saprotroph\| | Yeast | NULL | Probable | As nectar-tap_saprotroph (Põlme et al. 2020); Decay sugar-rich substrates (Põlme et al. 2020) | Põlme S, et al. 2020. Fungal Diversity 105:1-16 ((https://doi.org/10.1007/s13225-020-00466-2)) |
| Otu29909 | 1 | k__Fungi(100);p__Ascomycota(100);c__Saccharomycetes(100);o__Saccharomycetales(100);f__Saccharomycetaceae(100);g__Kazachstania(100);s__Kazachstania_intestinalis(100); | Kazachstania | 13 | Saprotroph | \|Undefined Saprotroph\| | Yeast | NULL | Probable | As nectar-tap_saprotroph (Põlme et al. 2020); Decay sugar-rich substrates (Põlme et al. 2020) | Põlme S, et al. 2020. Fungal Diversity 105:1-16 ((https://doi.org/10.1007/s13225-020-00466-2)) |
| Otu29910 | 1 | k__Fungi(100);p__Ascomycota(100);c__Saccharomycetes(100);o__Saccharomycetales(100);f__Saccharomycetaceae(100);g__Kazachstania(100);s__Kazachstania_intestinalis(100); | Kazachstania | 13 | Saprotroph | \|Undefined Saprotroph\| | Yeast | NULL | Probable | As nectar-tap_saprotroph (Põlme et al. 2020); Decay sugar-rich substrates (Põlme et al. 2020) | Põlme S, et al. 2020. Fungal Diversity 105:1-16 ((https://doi.org/10.1007/s13225-020-00466-2)) |
| Otu29911 | 1 | k__Fungi(100);p__Ascomycota(100);c__Saccharomycetes(100);o__Saccharomycetales(100);f__Saccharomycetaceae(100);g__Kazachstania(100);s__Kazachstania_intestinalis(100); | Kazachstania | 13 | Saprotroph | \|Undefined Saprotroph\| | Yeast | NULL | Probable | As nectar-tap_saprotroph (Põlme et al. 2020); Decay sugar-rich substrates (Põlme et al. 2020) | Põlme S, et al. 2020. Fungal Diversity 105:1-16 ((https://doi.org/10.1007/s13225-020-00466-2)) |
| Otu29912 | 1 | k__Fungi(100);p__Ascomycota(100);c__Saccharomycetes(100);o__Saccharomycetales(100);f__Saccharomycetaceae(100);g__Kazachstania(100);s__Kazachstania_intestinalis(100); | Kazachstania | 13 | Saprotroph | \|Undefined Saprotroph\| | Yeast | NULL | Probable | As nectar-tap_saprotroph (Põlme et al. 2020); Decay sugar-rich substrates (Põlme et al. 2020) | Põlme S, et al. 2020. Fungal Diversity 105:1-16 ((https://doi.org/10.1007/s13225-020-00466-2)) |
| Otu29913 | 1 | k__Fungi(100);p__Ascomycota(100);c__Saccharomycetes(100);o__Saccharomycetales(100);f__Saccharomycetaceae(100);g__Lachancea(100);s__Lachancea_thermotolerans(100); | Lachancea | 13 | Pathotroph-Saprotroph | Animal Parasite-Animal Pathogen-\|Undefined Saprotroph\| | Yeast | NULL | Possible | As nectar-tap_saprotroph (Põlme et al. 2020); Decay animal material, sugar-rich substrates (Põlme et al. 2020); Likely opportunistic human pathogen (Irinyi et al. 2015; see L. fermentati) | Irinyi L, et al. 2015. Medical Mycology 53:313-337 ((https://doi.org/10.1093/mmy/myv008)); Põlme S, et al. 2020. Fungal Diversity 105:1-16 ((https://doi.org/10.1007/s13225-020-00466-2)) |
| Otu29914 | 1 | k__Fungi(100);p__Ascomycota(100);c__Saccharomycetes(100);o__Saccharomycetales(100);f__Saccharomycetaceae(100);g__Kazachstania(100);s__Kazachstania_intestinalis(100); | Kazachstania | 13 | Saprotroph | \|Undefined Saprotroph\| | Yeast | NULL | Probable | As nectar-tap_saprotroph (Põlme et al. 2020); Decay sugar-rich substrates (Põlme et al. 2020) | Põlme S, et al. 2020. Fungal Diversity 105:1-16 ((https://doi.org/10.1007/s13225-020-00466-2)) |
| Otu29915 | 1 | k__Fungi(100);p__Ascomycota(100);c__Saccharomycetes(100);o__Saccharomycetales(100);f__Saccharomycetaceae(100);g__Kazachstania(100);s__Kazachstania_intestinalis(100); | Kazachstania | 13 | Saprotroph | \|Undefined Saprotroph\| | Yeast | NULL | Probable | As nectar-tap_saprotroph (Põlme et al. 2020); Decay sugar-rich substrates (Põlme et al. 2020) | Põlme S, et al. 2020. Fungal Diversity 105:1-16 ((https://doi.org/10.1007/s13225-020-00466-2)) |
| Otu29916 | 1 | k__Fungi(100);p__Ascomycota(100);c__Saccharomycetes(100);o__Saccharomycetales(100);f__Saccharomycetaceae(100);g__Lachancea(100);s__Lachancea_thermotolerans(100); | Lachancea | 13 | Pathotroph-Saprotroph | Animal Parasite-Animal Pathogen-\|Undefined Saprotroph\| | Yeast | NULL | Possible | As nectar-tap_saprotroph (Põlme et al. 2020); Decay animal material, sugar-rich substrates (Põlme et al. 2020); Likely opportunistic human pathogen (Irinyi et al. 2015; see L. fermentati) | Irinyi L, et al. 2015. Medical Mycology 53:313-337 ((https://doi.org/10.1093/mmy/myv008)); Põlme S, et al. 2020. Fungal Diversity 105:1-16 ((https://doi.org/10.1007/s13225-020-00466-2)) |
| Otu29917 | 1 | k__Fungi(100);p__Ascomycota(100);c__Saccharomycetes(100);o__Saccharomycetales(100);f__Saccharomycetaceae(100);g__Kazachstania(100);s__Kazachstania_intestinalis(100); | Kazachstania | 13 | Saprotroph | \|Undefined Saprotroph\| | Yeast | NULL | Probable | As nectar-tap_saprotroph (Põlme et al. 2020); Decay sugar-rich substrates (Põlme et al. 2020) | Põlme S, et al. 2020. Fungal Diversity 105:1-16 ((https://doi.org/10.1007/s13225-020-00466-2)) |
| Otu29918 | 1 | k__Fungi(100);p__Ascomycota(100);c__Saccharomycetes(100);o__Saccharomycetales(100);f__Saccharomycetaceae(100);g__Kazachstania(100);s__Kazachstania_intestinalis(100); | Kazachstania | 13 | Saprotroph | \|Undefined Saprotroph\| | Yeast | NULL | Probable | As nectar-tap_saprotroph (Põlme et al. 2020); Decay sugar-rich substrates (Põlme et al. 2020) | Põlme S, et al. 2020. Fungal Diversity 105:1-16 ((https://doi.org/10.1007/s13225-020-00466-2)) |
| Otu29919 | 1 | k__Fungi(100);p__Ascomycota(100);c__Saccharomycetes(100);o__Saccharomycetales(100);f__Saccharomycetaceae(100);g__Kazachstania(100);s__Kazachstania_intestinalis(100); | Kazachstania | 13 | Saprotroph | \|Undefined Saprotroph\| | Yeast | NULL | Probable | As nectar-tap_saprotroph (Põlme et al. 2020); Decay sugar-rich substrates (Põlme et al. 2020) | Põlme S, et al. 2020. Fungal Diversity 105:1-16 ((https://doi.org/10.1007/s13225-020-00466-2)) |
| Otu29920 | 1 | k__Fungi(100);p__Ascomycota(100);c__Saccharomycetes(100);o__Saccharomycetales(100);f__Saccharomycetaceae(100);g__Kazachstania(100);s__Kazachstania_intestinalis(100); | Kazachstania | 13 | Saprotroph | \|Undefined Saprotroph\| | Yeast | NULL | Probable | As nectar-tap_saprotroph (Põlme et al. 2020); Decay sugar-rich substrates (Põlme et al. 2020) | Põlme S, et al. 2020. Fungal Diversity 105:1-16 ((https://doi.org/10.1007/s13225-020-00466-2)) |
| Otu29921 | 1 | k__Fungi(100);p__Ascomycota(100);c__Saccharomycetes(100);o__Saccharomycetales(100);f__Metschnikowiaceae(100);g__Metschnikowia(100);s__Metschnikowia_pulcherrima(100); | Metschnikowia | 13 | Saprotroph | \|Undefined Saprotroph\| | Yeast | NULL | Probable | As nectar-tap_saprotroph (Põlme et al. 2020); Decay sugar-rich substrates (Põlme et al. 2020) | Põlme S, et al. 2020. Fungal Diversity 105:1-16 ((https://doi.org/10.1007/s13225-020-00466-2)) |
| Otu29923 | 1 | k__Fungi(100);p__Ascomycota(100);c__Saccharomycetes(100);o__Saccharomycetales(100);f__Metschnikowiaceae(100);g__Metschnikowia(100);s__Metschnikowia_pulcherrima(100); | Metschnikowia | 13 | Saprotroph | \|Undefined Saprotroph\| | Yeast | NULL | Probable | As nectar-tap_saprotroph (Põlme et al. 2020); Decay sugar-rich substrates (Põlme et al. 2020) | Põlme S, et al. 2020. Fungal Diversity 105:1-16 ((https://doi.org/10.1007/s13225-020-00466-2)) |
| Otu29924 | 1 | k__Fungi(100);p__Ascomycota(100);c__Saccharomycetes(100);o__Saccharomycetales(100);f__Metschnikowiaceae(100);g__Metschnikowia(100);s__Metschnikowia_pulcherrima(100); | Metschnikowia | 13 | Saprotroph | \|Undefined Saprotroph\| | Yeast | NULL | Probable | As nectar-tap_saprotroph (Põlme et al. 2020); Decay sugar-rich substrates (Põlme et al. 2020) | Põlme S, et al. 2020. Fungal Diversity 105:1-16 ((https://doi.org/10.1007/s13225-020-00466-2)) |
| Otu29925 | 1 | k__Fungi(100);p__Ascomycota(100);c__Saccharomycetes(100);o__Saccharomycetales(100);f__Metschnikowiaceae(100);g__Metschnikowia(100);s__Metschnikowia_pulcherrima(100); | Metschnikowia | 13 | Saprotroph | \|Undefined Saprotroph\| | Yeast | NULL | Probable | As nectar-tap_saprotroph (Põlme et al. 2020); Decay sugar-rich substrates (Põlme et al. 2020) | Põlme S, et al. 2020. Fungal Diversity 105:1-16 ((https://doi.org/10.1007/s13225-020-00466-2)) |
| Otu29927 | 1 | k__Fungi(100);p__Ascomycota(100);c__Saccharomycetes(100);o__Saccharomycetales(100);f__Metschnikowiaceae(100);g__Metschnikowia(100);g__Metschnikowia_unclassified(100); | Metschnikowia | 13 | Saprotroph | \|Undefined Saprotroph\| | Yeast | NULL | Probable | As nectar-tap_saprotroph (Põlme et al. 2020); Decay sugar-rich substrates (Põlme et al. 2020) | Põlme S, et al. 2020. Fungal Diversity 105:1-16 ((https://doi.org/10.1007/s13225-020-00466-2)) |
| Otu29930 | 1 | k__Fungi(100);p__Ascomycota(100);c__Dothideomycetes(100);o__Dothideomycetes_order_Incertae_sedis(100);f__Pseudeurotiaceae(100);g__Pseudeurotium(100);g__Pseudeurotium_unclassified(100); | Pseudeurotium | 13 | Saprotroph | \|Dung Saprotroph\|-Undefined Saprotroph | Microfungus | NULL | Probable | As soil_saprotroph (Põlme et al. 2020); Decay soil, dung (Põlme et al. 2020) | Tedersoo L, et al. 2014. Science 346:e1256688 ((https://doi.org/10.1126/science.1256688)); Põlme S, et al. 2020. Fungal Diversity 105:1-16 ((https://doi.org/10.1007/s13225-020-00466-2)) |
| Otu29931 | 1 | k__Fungi(100);p__Ascomycota(100);c__Saccharomycetes(100);o__Saccharomycetales(100);f__Metschnikowiaceae(100);g__Metschnikowia(100);s__Metschnikowia_pulcherrima(100); | Metschnikowia | 13 | Saprotroph | \|Undefined Saprotroph\| | Yeast | NULL | Probable | As nectar-tap_saprotroph (Põlme et al. 2020); Decay sugar-rich substrates (Põlme et al. 2020) | Põlme S, et al. 2020. Fungal Diversity 105:1-16 ((https://doi.org/10.1007/s13225-020-00466-2)) |
| Otu29932 | 1 | k__Fungi(100);p__Ascomycota(100);c__Saccharomycetes(100);o__Saccharomycetales(100);f__Metschnikowiaceae(100);g__Metschnikowia(100);s__Metschnikowia_pulcherrima(100); | Metschnikowia | 13 | Saprotroph | \|Undefined Saprotroph\| | Yeast | NULL | Probable | As nectar-tap_saprotroph (Põlme et al. 2020); Decay sugar-rich substrates (Põlme et al. 2020) | Põlme S, et al. 2020. Fungal Diversity 105:1-16 ((https://doi.org/10.1007/s13225-020-00466-2)) |
| Otu29934 | 1 | k__Fungi(100);p__Ascomycota(100);c__Saccharomycetes(100);o__Saccharomycetales(100);f__Metschnikowiaceae(100);g__Metschnikowia(100);g__Metschnikowia_unclassified(100); | Metschnikowia | 13 | Saprotroph | \|Undefined Saprotroph\| | Yeast | NULL | Probable | As nectar-tap_saprotroph (Põlme et al. 2020); Decay sugar-rich substrates (Põlme et al. 2020) | Põlme S, et al. 2020. Fungal Diversity 105:1-16 ((https://doi.org/10.1007/s13225-020-00466-2)) |
| Otu29936 | 1 | k__Fungi(100);p__Ascomycota(100);c__Saccharomycetes(100);o__Saccharomycetales(100);f__Metschnikowiaceae(100);g__Metschnikowia(100);g__Metschnikowia_unclassified(100); | Metschnikowia | 13 | Saprotroph | \|Undefined Saprotroph\| | Yeast | NULL | Probable | As nectar-tap_saprotroph (Põlme et al. 2020); Decay sugar-rich substrates (Põlme et al. 2020) | Põlme S, et al. 2020. Fungal Diversity 105:1-16 ((https://doi.org/10.1007/s13225-020-00466-2)) |
| Otu29937 | 1 | k__Fungi(100);p__Ascomycota(100);c__Saccharomycetes(100);o__Saccharomycetales(100);f__Metschnikowiaceae(100);g__Metschnikowia(100);s__Metschnikowia_pulcherrima(100); | Metschnikowia | 13 | Saprotroph | \|Undefined Saprotroph\| | Yeast | NULL | Probable | As nectar-tap_saprotroph (Põlme et al. 2020); Decay sugar-rich substrates (Põlme et al. 2020) | Põlme S, et al. 2020. Fungal Diversity 105:1-16 ((https://doi.org/10.1007/s13225-020-00466-2)) |
| Otu29938 | 1 | k__Fungi(100);p__Ascomycota(100);c__Saccharomycetes(100);o__Saccharomycetales(100);f__Metschnikowiaceae(100);g__Metschnikowia(100);s__Metschnikowia_pulcherrima(100); | Metschnikowia | 13 | Saprotroph | \|Undefined Saprotroph\| | Yeast | NULL | Probable | As nectar-tap_saprotroph (Põlme et al. 2020); Decay sugar-rich substrates (Põlme et al. 2020) | Põlme S, et al. 2020. Fungal Diversity 105:1-16 ((https://doi.org/10.1007/s13225-020-00466-2)) |
| Otu29940 | 1 | k__Fungi(100);p__Ascomycota(100);c__Saccharomycetes(100);o__Saccharomycetales(100);f__Metschnikowiaceae(100);g__Metschnikowia(100);s__Metschnikowia_pulcherrima(100); | Metschnikowia | 13 | Saprotroph | \|Undefined Saprotroph\| | Yeast | NULL | Probable | As nectar-tap_saprotroph (Põlme et al. 2020); Decay sugar-rich substrates (Põlme et al. 2020) | Põlme S, et al. 2020. Fungal Diversity 105:1-16 ((https://doi.org/10.1007/s13225-020-00466-2)) |
| Otu29941 | 1 | k__Fungi(100);p__Ascomycota(100);c__Saccharomycetes(100);o__Saccharomycetales(100);f__Metschnikowiaceae(100);g__Metschnikowia(100);s__Metschnikowia_pulcherrima(100); | Metschnikowia | 13 | Saprotroph | \|Undefined Saprotroph\| | Yeast | NULL | Probable | As nectar-tap_saprotroph (Põlme et al. 2020); Decay sugar-rich substrates (Põlme et al. 2020) | Põlme S, et al. 2020. Fungal Diversity 105:1-16 ((https://doi.org/10.1007/s13225-020-00466-2)) |
| Otu29942 | 1 | k__Fungi(100);p__Ascomycota(100);c__Saccharomycetes(100);o__Saccharomycetales(100);f__Metschnikowiaceae(100);g__Metschnikowia(100);s__Metschnikowia_sp(100); | Metschnikowia | 13 | Saprotroph | \|Undefined Saprotroph\| | Yeast | NULL | Probable | As nectar-tap_saprotroph (Põlme et al. 2020); Decay sugar-rich substrates (Põlme et al. 2020) | Põlme S, et al. 2020. Fungal Diversity 105:1-16 ((https://doi.org/10.1007/s13225-020-00466-2)) |
| Otu29943 | 1 | k__Fungi(100);p__Ascomycota(100);c__Saccharomycetes(100);o__Saccharomycetales(100);f__Metschnikowiaceae(100);g__Metschnikowia(100);s__Metschnikowia_pulcherrima(100); | Metschnikowia | 13 | Saprotroph | \|Undefined Saprotroph\| | Yeast | NULL | Probable | As nectar-tap_saprotroph (Põlme et al. 2020); Decay sugar-rich substrates (Põlme et al. 2020) | Põlme S, et al. 2020. Fungal Diversity 105:1-16 ((https://doi.org/10.1007/s13225-020-00466-2)) |
| Otu29948 | 1 | k__Fungi(100);p__Ascomycota(100);c__Saccharomycetes(100);o__Saccharomycetales(100);f__Metschnikowiaceae(100);g__Metschnikowia(100);s__Metschnikowia_pulcherrima(100); | Metschnikowia | 13 | Saprotroph | \|Undefined Saprotroph\| | Yeast | NULL | Probable | As nectar-tap_saprotroph (Põlme et al. 2020); Decay sugar-rich substrates (Põlme et al. 2020) | Põlme S, et al. 2020. Fungal Diversity 105:1-16 ((https://doi.org/10.1007/s13225-020-00466-2)) |
| Otu29950 | 1 | k__Fungi(100);p__Ascomycota(100);c__Saccharomycetes(100);o__Saccharomycetales(100);f__Metschnikowiaceae(100);g__Metschnikowia(100);s__Metschnikowia_pulcherrima(100); | Metschnikowia | 13 | Saprotroph | \|Undefined Saprotroph\| | Yeast | NULL | Probable | As nectar-tap_saprotroph (Põlme et al. 2020); Decay sugar-rich substrates (Põlme et al. 2020) | Põlme S, et al. 2020. Fungal Diversity 105:1-16 ((https://doi.org/10.1007/s13225-020-00466-2)) |
| Otu29957 | 1 | k__Fungi(100);p__Ascomycota(100);c__Dothideomycetes(100);o__Capnodiales(100);f__Mycosphaerellaceae(100);f__Mycosphaerellaceae_unclassified(100);f__Mycosphaerellaceae_unclassified(100); | Mycosphaerellaceae | 9 | Pathotroph-Saprotroph | Plant Pathogen-Undefined Saprotroph | Microfungus | NULL | Probable | Biotrophic, necrotrophic or saprobic on various plant tissues (Cannon & Kirk 2007) | Cannon PF, Kirk PM. 2007. Fungal Families of the World. CAB International, Cambridge (ISBN: 978-0851998275) |
| Otu29960 | 1 | k__Fungi(100);p__Basidiomycota(100);c__Microbotryomycetes(100);o__Sporidiobolales(100);f__Sporidiobolales_family_Incertae_sedis(100);g__Sporobolomyces(100);s__Sporobolomyces_lactosus(100); | Sporobolomyces | 13 | Pathotroph-Saprotroph | \|Fungal Parasite\|-Plant Saprotroph | Yeast | NULL | Probable | As litter_saprotroph (Põlme et al. 2020); As mycoparasite (Põlme et al. 2020); Decay fungal material (Põlme et al. 2020) | Tedersoo L, et al. 2014. Science 346:e1256688 ((https://doi.org/10.1126/science.1256688)); Purahong W, et al. 2016. Molecular Ecology 25:4059-4074 ((https://doi.org/10.1111/mec.13739)); Põlme S, et al. 2020. Fungal Diversity 105:1-16 ((https://doi.org/10.1007/s13225-020-00466-2)) |
| Otu29969 | 1 | k__Fungi(100);p__Ascomycota(100);c__Saccharomycetes(100);o__Saccharomycetales(100);f__Metschnikowiaceae(100);g__Metschnikowia(100);s__Metschnikowia_pulcherrima(100); | Metschnikowia | 13 | Saprotroph | \|Undefined Saprotroph\| | Yeast | NULL | Probable | As nectar-tap_saprotroph (Põlme et al. 2020); Decay sugar-rich substrates (Põlme et al. 2020) | Põlme S, et al. 2020. Fungal Diversity 105:1-16 ((https://doi.org/10.1007/s13225-020-00466-2)) |

| | | | | | | | | | | | |
|---|---|---|---|---|---|---|---|---|---|---|---|
| Otu29973 | 1 | k__Fungi(100);p__Ascomycota(100);c__Saccharomycetes(100);o__Saccharomycetales(100);f__Metschnikowiaceae(100);g__Metschnikowia(100);s__Metschnikowia_pulcherrima(100); | Metschnikowia | 13 | Saprotroph | \|Undefined Saprotroph\| | Yeast | NULL | Probable | As nectar-tap_saprotroph (Põlme et al. 2020); Decay sugar-rich substrates (Põlme et al. 2020) | Põlme S, et al. 2020. Fungal Diversity 105:1-16 ((https://doi.org/10.1007/s13225-020-00466-2)) |
| Otu29974 | 1 | k__Fungi(100);p__Ascomycota(100);c__Saccharomycetes(100);o__Saccharomycetales(100);f__Metschnikowiaceae(100);g__Metschnikowia(100);g__Metschnikowia_unclassified(100); | Metschnikowia | 13 | Saprotroph | \|Undefined Saprotroph\| | Yeast | NULL | Probable | As nectar-tap_saprotroph (Põlme et al. 2020); Decay sugar-rich substrates (Põlme et al. 2020) | Põlme S, et al. 2020. Fungal Diversity 105:1-16 ((https://doi.org/10.1007/s13225-020-00466-2)) |
| Otu29975 | 1 | k__Fungi(100);p__Ascomycota(100);c__Saccharomycetes(100);o__Saccharomycetales(100);f__Metschnikowiaceae(100);g__Metschnikowia(100);s__Metschnikowia_pulcherrima(100); | Metschnikowia | 13 | Saprotroph | \|Undefined Saprotroph\| | Yeast | NULL | Probable | As nectar-tap_saprotroph (Põlme et al. 2020); Decay sugar-rich substrates (Põlme et al. 2020) | Põlme S, et al. 2020. Fungal Diversity 105:1-16 ((https://doi.org/10.1007/s13225-020-00466-2)) |
| Otu29978 | 1 | k__Fungi(100);p__Ascomycota(100);c__Saccharomycetes(100);o__Saccharomycetales(100);f__Saccharomycetales_family_Incertae_sedis(100);g__Candida(100);g__Candida_unclassified(100); | Candida | 13 | Pathotroph-Saprotroph-Symbiotroph | Animal Pathogen-Endophyte-Endosymbiont-Epiphyte-\|Undefined Saprotroph\| | Dimorphic Yeast-Yeast | NULL | Probable | As nectar-tap_saprotroph (Põlme et al. 2020); Decay sugar-rich substrates (Põlme et al. 2020); This asexual genus is polyphyletic and contain species that function very differently depend on what they are. Caution should be taken when interpreting guilds for Candida species. | Manolakaki D, et al. 2010. Virulence. 1: 367-375 ((https://doi.org/10.4161/viru.1.5.12796)); Mekbib SB, et al. 2011. Tropical Plant Pathology 36:233-240 ((https://doi.org/10.1590/S1982-56762011000400004)); Põlme S, et al. 2020. Fungal Diversity 105:1-16 ((https://doi.org/10.1007/s13225-020-00466-2)) |
| Otu29979 | 1 | k__Fungi(100);p__Ascomycota(100);c__Saccharomycetes(100);o__Saccharomycetales(100);f__Saccharomycetales_family_Incertae_sedis(100);g__Candida(100);g__Candida_unclassified(100); | Candida | 13 | Pathotroph-Saprotroph-Symbiotroph | Animal Pathogen-Endophyte-Endosymbiont-Epiphyte-\|Undefined Saprotroph\| | Dimorphic Yeast-Yeast | NULL | Probable | As nectar-tap_saprotroph (Põlme et al. 2020); Decay sugar-rich substrates (Põlme et al. 2020); This asexual genus is polyphyletic and contain species that function very differently depend on what they are. Caution should be taken when interpreting guilds for Candida species. | Manolakaki D, et al. 2010. Virulence. 1: 367-375 ((https://doi.org/10.4161/viru.1.5.12796)); Mekbib SB, et al. 2011. Tropical Plant Pathology 36:233-240 ((https://doi.org/10.1590/S1982-56762011000400004)); Põlme S, et al. 2020. Fungal Diversity 105:1-16 ((https://doi.org/10.1007/s13225-020-00466-2)) |
| Otu29981 | 1 | k__Fungi(100);p__Ascomycota(100);c__Saccharomycetes(100);o__Saccharomycetales(100);f__Saccharomycetales_family_Incertae_sedis(100);g__Candida(100);s__Candida_cretensis(100); | Candida | 13 | Pathotroph-Saprotroph-Symbiotroph | Animal Pathogen-Endophyte-Endosymbiont-Epiphyte-\|Undefined Saprotroph\| | Dimorphic Yeast-Yeast | NULL | Probable | As nectar-tap_saprotroph (Põlme et al. 2020); Decay sugar-rich substrates (Põlme et al. 2020); This asexual genus is polyphyletic and contain species that function very differently depend on what they are. Caution should be taken when interpreting guilds for Candida species. | Manolakaki D, et al. 2010. Virulence. 1: 367-375 ((https://doi.org/10.4161/viru.1.5.12796)); Mekbib SB, et al. 2011. Tropical Plant Pathology 36:233-240 ((https://doi.org/10.1590/S1982-56762011000400004)); Põlme S, et al. 2020. Fungal Diversity 105:1-16 ((https://doi.org/10.1007/s13225-020-00466-2)) |
| Otu29984 | 1 | k__Fungi(100);p__Ascomycota(100);c__Saccharomycetes(100);o__Saccharomycetales(100);f__Saccharomycetales_family_Incertae_sedis(100);g__Candida(100);g__Candida_unclassified(100); | Candida | 13 | Pathotroph-Saprotroph-Symbiotroph | Animal Pathogen-Endophyte-Endosymbiont-Epiphyte-\|Undefined Saprotroph\| | Dimorphic Yeast-Yeast | NULL | Probable | As nectar-tap_saprotroph (Põlme et al. 2020); Decay sugar-rich substrates (Põlme et al. 2020); This asexual genus is polyphyletic and contain species that function very differently depend on what they are. Caution should be taken when interpreting guilds for Candida species. | Manolakaki D, et al. 2010. Virulence. 1: 367-375 ((https://doi.org/10.4161/viru.1.5.12796)); Mekbib SB, et al. 2011. Tropical Plant Pathology 36:233-240 ((https://doi.org/10.1590/S1982-56762011000400004)); Põlme S, et al. 2020. Fungal Diversity 105:1-16 ((https://doi.org/10.1007/s13225-020-00466-2)) |
| Otu29988 | 1 | k__Fungi(100);p__Ascomycota(100);c__Saccharomycetes(100);o__Saccharomycetales(100);f__Metschnikowiaceae(100);g__Metschnikowia(100);s__Metschnikowia_pulcherrima(100); | Metschnikowia | 13 | Saprotroph | \|Undefined Saprotroph\| | Yeast | NULL | Probable | As nectar-tap_saprotroph (Põlme et al. 2020); Decay sugar-rich substrates (Põlme et al. 2020) | Põlme S, et al. 2020. Fungal Diversity 105:1-16 ((https://doi.org/10.1007/s13225-020-00466-2)) |
| Otu29990 | 1 | k__Fungi(100);p__Ascomycota(100);c__Saccharomycetes(100);o__Saccharomycetales(100);f__Metschnikowiaceae(100);g__Metschnikowia(100);g__Metschnikowia_unclassified(100); | Metschnikowia | 13 | Saprotroph | \|Undefined Saprotroph\| | Yeast | NULL | Probable | As nectar-tap_saprotroph (Põlme et al. 2020); Decay sugar-rich substrates (Põlme et al. 2020) | Põlme S, et al. 2020. Fungal Diversity 105:1-16 ((https://doi.org/10.1007/s13225-020-00466-2)) |
| Otu29992 | 1 | k__Fungi(100);p__Ascomycota(100);c__Saccharomycetes(100);o__Saccharomycetales(100);f__Metschnikowiaceae(100);g__Metschnikowia(100);s__Metschnikowia_pulcherrima(100); | Metschnikowia | 13 | Saprotroph | \|Undefined Saprotroph\| | Yeast | NULL | Probable | As nectar-tap_saprotroph (Põlme et al. 2020); Decay sugar-rich substrates (Põlme et al. 2020) | Põlme S, et al. 2020. Fungal Diversity 105:1-16 ((https://doi.org/10.1007/s13225-020-00466-2)) |
| Otu29994 | 1 | k__Fungi(100);p__Ascomycota(100);c__Saccharomycetes(100);o__Saccharomycetales(100);f__Metschnikowiaceae(100);g__Metschnikowia(100);g__Metschnikowia_unclassified(100); | Metschnikowia | 13 | Saprotroph | \|Undefined Saprotroph\| | Yeast | NULL | Probable | As nectar-tap_saprotroph (Põlme et al. 2020); Decay sugar-rich substrates (Põlme et al. 2020) | Põlme S, et al. 2020. Fungal Diversity 105:1-16 ((https://doi.org/10.1007/s13225-020-00466-2)) |
| Otu29995 | 1 | k__Fungi(100);p__Ascomycota(100);c__Saccharomycetes(100);o__Saccharomycetales(100);f__Metschnikowiaceae(100);g__Metschnikowia(100);s__Metschnikowia_pulcherrima(100); | Metschnikowia | 13 | Saprotroph | \|Undefined Saprotroph\| | Yeast | NULL | Probable | As nectar-tap_saprotroph (Põlme et al. 2020); Decay sugar-rich substrates (Põlme et al. 2020) | Põlme S, et al. 2020. Fungal Diversity 105:1-16 ((https://doi.org/10.1007/s13225-020-00466-2)) |
| Otu29996 | 1 | k__Fungi(100);p__Ascomycota(100);c__Saccharomycetes(100);o__Saccharomycetales(100);f__Metschnikowiaceae(100);g__Metschnikowia(100);s__Metschnikowia_pulcherrima(100); | Metschnikowia | 13 | Saprotroph | \|Undefined Saprotroph\| | Yeast | NULL | Probable | As nectar-tap_saprotroph (Põlme et al. 2020); Decay sugar-rich substrates (Põlme et al. 2020) | Põlme S, et al. 2020. Fungal Diversity 105:1-16 ((https://doi.org/10.1007/s13225-020-00466-2)) |
| Otu29997 | 1 | k__Fungi(100);p__Ascomycota(100);c__Saccharomycetes(100);o__Saccharomycetales(100);f__Saccharomycetales_family_Incertae_sedis(100);g__Candida(100);g__Candida_unclassified(100); | Candida | 13 | Pathotroph-Saprotroph-Symbiotroph | Animal Pathogen-Endophyte-Endosymbiont-Epiphyte-\|Undefined Saprotroph\| | Dimorphic Yeast-Yeast | NULL | Probable | As nectar-tap_saprotroph (Põlme et al. 2020); Decay sugar-rich substrates (Põlme et al. 2020); This asexual genus is polyphyletic and contain species that function very differently depend on what they are. Caution should be taken when interpreting guilds for Candida species. | Manolakaki D, et al. 2010. Virulence. 1: 367-375 ((https://doi.org/10.4161/viru.1.5.12796)); Mekbib SB, et al. 2011. Tropical Plant Pathology 36:233-240 ((https://doi.org/10.1590/S1982-56762011000400004)); Põlme S, et al. 2020. Fungal Diversity 105:1-16 ((https://doi.org/10.1007/s13225-020-00466-2)) |
| Otu29999 | 1 | k__Fungi(100);p__Ascomycota(100);c__Saccharomycetes(100);o__Saccharomycetales(100);f__Metschnikowiaceae(100);g__Metschnikowia(100);s__Metschnikowia_pulcherrima(100); | Metschnikowia | 13 | Saprotroph | \|Undefined Saprotroph\| | Yeast | NULL | Probable | As nectar-tap_saprotroph (Põlme et al. 2020); Decay sugar-rich substrates (Põlme et al. 2020) | Põlme S, et al. 2020. Fungal Diversity 105:1-16 ((https://doi.org/10.1007/s13225-020-00466-2)) |
| Otu30000 | 1 | k__Fungi(100);p__Ascomycota(100);c__Saccharomycetes(100);o__Saccharomycetales(100);f__Metschnikowiaceae(100);g__Metschnikowia(100);s__Metschnikowia_pulcherrima(100); | Metschnikowia | 13 | Saprotroph | \|Undefined Saprotroph\| | Yeast | NULL | Probable | As nectar-tap_saprotroph (Põlme et al. 2020); Decay sugar-rich substrates (Põlme et al. 2020) | Põlme S, et al. 2020. Fungal Diversity 105:1-16 ((https://doi.org/10.1007/s13225-020-00466-2)) |
| Otu30001 | 1 | k__Fungi(100);p__Ascomycota(100);c__Saccharomycetes(100);o__Saccharomycetales(100);f__Metschnikowiaceae(100);g__Metschnikowia(100);s__Metschnikowia_pulcherrima(100); | Metschnikowia | 13 | Saprotroph | \|Undefined Saprotroph\| | Yeast | NULL | Probable | As nectar-tap_saprotroph (Põlme et al. 2020); Decay sugar-rich substrates (Põlme et al. 2020) | Põlme S, et al. 2020. Fungal Diversity 105:1-16 ((https://doi.org/10.1007/s13225-020-00466-2)) |
| Otu30011 | 1 | k__Fungi(100);p__Ascomycota(100);c__Saccharomycetes(100);o__Saccharomycetales(100);f__Saccharomycetales_family_Incertae_sedis(100);g__Candida(100);g__Candida_unclassified(100); | Candida | 13 | Pathotroph-Saprotroph-Symbiotroph | Animal Pathogen-Endophyte-Endosymbiont-Epiphyte-\|Undefined Saprotroph\| | Dimorphic Yeast-Yeast | NULL | Probable | As nectar-tap_saprotroph (Põlme et al. 2020); Decay sugar-rich substrates (Põlme et al. 2020); This asexual genus is polyphyletic and contain species that function very differently depend on what they are. Caution should be taken when interpreting guilds for Candida species. | Manolakaki D, et al. 2010. Virulence. 1: 367-375 ((https://doi.org/10.4161/viru.1.5.12796)); Mekbib SB, et al. 2011. Tropical Plant Pathology 36:233-240 ((https://doi.org/10.1590/S1982-56762011000400004)); Põlme S, et al. 2020. Fungal Diversity 105:1-16 ((https://doi.org/10.1007/s13225-020-00466-2)) |
| Otu30015 | 1 | k__Fungi(100);p__Ascomycota(100);c__Saccharomycetes(100);o__Saccharomycetales(100);f__Metschnikowiaceae(100);g__Metschnikowia(100);s__Metschnikowia_pulcherrima(100); | Metschnikowia | 13 | Saprotroph | \|Undefined Saprotroph\| | Yeast | NULL | Probable | As nectar-tap_saprotroph (Põlme et al. 2020); Decay sugar-rich substrates (Põlme et al. 2020) | Põlme S, et al. 2020. Fungal Diversity 105:1-16 ((https://doi.org/10.1007/s13225-020-00466-2)) |
| Otu30016 | 1 | k__Fungi(100);p__Ascomycota(100);c__Saccharomycetes(100);o__Saccharomycetales(100);f__Saccharomycetales_family_Incertae_sedis(100);g__Candida(100);g__Candida_unclassified(100); | Candida | 13 | Pathotroph-Saprotroph-Symbiotroph | Animal Pathogen-Endophyte-Endosymbiont-Epiphyte-\|Undefined Saprotroph\| | Dimorphic Yeast-Yeast | NULL | Probable | As nectar-tap_saprotroph (Põlme et al. 2020); Decay sugar-rich substrates (Põlme et al. 2020); This asexual genus is polyphyletic and contain species that function very differently depend on what they are. Caution should be taken when interpreting guilds for Candida species. | Manolakaki D, et al. 2010. Virulence. 1: 367-375 ((https://doi.org/10.4161/viru.1.5.12796)); Mekbib SB, et al. 2011. Tropical Plant Pathology 36:233-240 ((https://doi.org/10.1590/S1982-56762011000400004)); Põlme S, et al. 2020. Fungal Diversity 105:1-16 ((https://doi.org/10.1007/s13225-020-00466-2)) |
| Otu30017 | 1 | k__Fungi(100);p__Ascomycota(100);c__Saccharomycetes(100);o__Saccharomycetales(100);f__Metschnikowiaceae(100);g__Metschnikowia(100);s__Metschnikowia_pulcherrima(100); | Metschnikowia | 13 | Saprotroph | \|Undefined Saprotroph\| | Yeast | NULL | Probable | As nectar-tap_saprotroph (Põlme et al. 2020); Decay sugar-rich substrates (Põlme et al. 2020) | Põlme S, et al. 2020. Fungal Diversity 105:1-16 ((https://doi.org/10.1007/s13225-020-00466-2)) |
| Otu30019 | 1 | k__Fungi(100);p__Ascomycota(100);c__Saccharomycetes(100);o__Saccharomycetales(100);f__Metschnikowiaceae(100);g__Metschnikowia(100);s__Metschnikowia_pulcherrima(100); | Metschnikowia | 13 | Saprotroph | \|Undefined Saprotroph\| | Yeast | NULL | Probable | As nectar-tap_saprotroph (Põlme et al. 2020); Decay sugar-rich substrates (Põlme et al. 2020) | Põlme S, et al. 2020. Fungal Diversity 105:1-16 ((https://doi.org/10.1007/s13225-020-00466-2)) |
| Otu30028 | 1 | k__Fungi(100);p__Ascomycota(100);c__Saccharomycetes(100);o__Saccharomycetales(100);f__Metschnikowiaceae(100);g__Metschnikowia(100);g__Metschnikowia_unclassified(100); | Metschnikowia | 13 | Saprotroph | \|Undefined Saprotroph\| | Yeast | NULL | Probable | As nectar-tap_saprotroph (Põlme et al. 2020); Decay sugar-rich substrates (Põlme et al. 2020) | Põlme S, et al. 2020. Fungal Diversity 105:1-16 ((https://doi.org/10.1007/s13225-020-00466-2)) |
| Otu30029 | 1 | k__Fungi(100);p__Ascomycota(100);c__Saccharomycetes(100);o__Saccharomycetales(100);f__Saccharomycetales_family_Incertae_sedis(100);g__Candida(100);g__Candida_unclassified(100); | Candida | 13 | Pathotroph-Saprotroph-Symbiotroph | Animal Pathogen-Endophyte-Endosymbiont-Epiphyte-\|Undefined Saprotroph\| | Dimorphic Yeast-Yeast | NULL | Probable | As nectar-tap_saprotroph (Põlme et al. 2020); Decay sugar-rich substrates (Põlme et al. 2020); This asexual genus is polyphyletic and contain species that function very differently depend on what they are. Caution should be taken when interpreting guilds for Candida species. | Manolakaki D, et al. 2010. Virulence. 1: 367-375 ((https://doi.org/10.4161/viru.1.5.12796)); Mekbib SB, et al. 2011. Tropical Plant Pathology 36:233-240 ((https://doi.org/10.1590/S1982-56762011000400004)); Põlme S, et al. 2020. Fungal Diversity 105:1-16 ((https://doi.org/10.1007/s13225-020-00466-2)) |
| Otu30030 | 1 | k__Fungi(100);p__Ascomycota(100);c__Saccharomycetes(100);o__Saccharomycetales(100);f__Saccharomycetales_family_Incertae_sedis(100);g__Candida(100);g__Candida_unclassified(100); | Candida | 13 | Pathotroph-Saprotroph-Symbiotroph | Animal Pathogen-Endophyte-Endosymbiont-Epiphyte-\|Undefined Saprotroph\| | Dimorphic Yeast-Yeast | NULL | Probable | As nectar-tap_saprotroph (Põlme et al. 2020); Decay sugar-rich substrates (Põlme et al. 2020); This asexual genus is polyphyletic and contain species that function very differently depend on what they are. Caution should be taken when interpreting guilds for Candida species. | Manolakaki D, et al. 2010. Virulence. 1: 367-375 ((https://doi.org/10.4161/viru.1.5.12796)); Mekbib SB, et al. 2011. Tropical Plant Pathology 36:233-240 ((https://doi.org/10.1590/S1982-56762011000400004)); Põlme S, et al. 2020. Fungal Diversity 105:1-16 ((https://doi.org/10.1007/s13225-020-00466-2)) |
| Otu30033 | 1 | k__Fungi(100);p__Ascomycota(100);c__Saccharomycetes(100);o__Saccharomycetales(100);f__Saccharomycetales_family_Incertae_sedis(100);f__Saccharomycetales_family_Incertae_sedis_unclassified(100);f__Saccharomycetales_family_Incertae_sedis_unclassified(100); | Saccharomycetales | 7 | Saprotroph | Undefined Saprotroph | Yeast | NULL | Possible | NULL | Starkenburg S, et al. 2015. New Phytologist 207:1145-1158 ((https://doi.org/10.1111/nph.13426)) |
| Otu30035 | 1 | k__Fungi(100);p__Ascomycota(100);c__Saccharomycetes(100);o__Saccharomycetales(100);f__Saccharomycetales_family_Incertae_sedis(100);g__Candida(100);g__Candida_unclassified(100); | Candida | 13 | Pathotroph-Saprotroph-Symbiotroph | Animal Pathogen-Endophyte-Endosymbiont-Epiphyte-\|Undefined Saprotroph\| | Dimorphic Yeast-Yeast | NULL | Probable | As nectar-tap_saprotroph (Põlme et al. 2020); Decay sugar-rich substrates (Põlme et al. 2020); This asexual genus is polyphyletic and contain species that function very differently depend on what they are. Caution should be taken when interpreting guilds for Candida species. | Manolakaki D, et al. 2010. Virulence. 1: 367-375 ((https://doi.org/10.4161/viru.1.5.12796)); Mekbib SB, et al. 2011. Tropical Plant Pathology 36:233-240 ((https://doi.org/10.1590/S1982-56762011000400004)); Põlme S, et al. 2020. Fungal Diversity 105:1-16 ((https://doi.org/10.1007/s13225-020-00466-2)) |
| Otu30041 | 1 | k__Fungi(100);p__Ascomycota(100);c__Saccharomycetes(100);o__Saccharomycetales(100);f__Metschnikowiaceae(100);g__Metschnikowia(100);g__Metschnikowia_unclassified(100); | Metschnikowia | 13 | Saprotroph | \|Undefined Saprotroph\| | Yeast | NULL | Probable | As nectar-tap_saprotroph (Põlme et al. 2020); Decay sugar-rich substrates (Põlme et al. 2020) | Põlme S, et al. 2020. Fungal Diversity 105:1-16 ((https://doi.org/10.1007/s13225-020-00466-2)) |
| Otu30043 | 1 | k__Fungi(100);p__Ascomycota(100);c__Saccharomycetes(100);o__Saccharomycetales(100);f__Metschnikowiaceae(100);g__Metschnikowia(100);s__Metschnikowia_pulcherrima(100); | Metschnikowia | 13 | Saprotroph | \|Undefined Saprotroph\| | Yeast | NULL | Probable | As nectar-tap_saprotroph (Põlme et al. 2020); Decay sugar-rich substrates (Põlme et al. 2020) | Põlme S, et al. 2020. Fungal Diversity 105:1-16 ((https://doi.org/10.1007/s13225-020-00466-2)) |
| Otu30045 | 1 | k__Fungi(100);p__Ascomycota(100);c__Saccharomycetes(100);o__Saccharomycetales(100);f__Metschnikowiaceae(100);g__Metschnikowia(100);g__Metschnikowia_unclassified(100); | Metschnikowia | 13 | Saprotroph | \|Undefined Saprotroph\| | Yeast | NULL | Probable | As nectar-tap_saprotroph (Põlme et al. 2020); Decay sugar-rich substrates (Põlme et al. 2020) | Põlme S, et al. 2020. Fungal Diversity 105:1-16 ((https://doi.org/10.1007/s13225-020-00466-2)) |
| Otu30052 | 1 | k__Fungi(100);p__Ascomycota(100);c__Saccharomycetes(100);o__Saccharomycetales(100);f__Metschnikowiaceae(100);g__Metschnikowia(100);s__Metschnikowia_pulcherrima(100); | Metschnikowia | 13 | Saprotroph | \|Undefined Saprotroph\| | Yeast | NULL | Probable | As nectar-tap_saprotroph (Põlme et al. 2020); Decay sugar-rich substrates (Põlme et al. 2020) | Põlme S, et al. 2020. Fungal Diversity 105:1-16 ((https://doi.org/10.1007/s13225-020-00466-2)) |
| Otu30053 | 1 | k__Fungi(100);p__Ascomycota(100);c__Saccharomycetes(100);o__Saccharomycetales(100);f__Metschnikowiaceae(100);g__Metschnikowia(100);s__Metschnikowia_pulcherrima(100); | Metschnikowia | 13 | Saprotroph | \|Undefined Saprotroph\| | Yeast | NULL | Probable | As nectar-tap_saprotroph (Põlme et al. 2020); Decay sugar-rich substrates (Põlme et al. 2020) | Põlme S, et al. 2020. Fungal Diversity 105:1-16 ((https://doi.org/10.1007/s13225-020-00466-2)) |
| Otu30054 | 1 | k__Fungi(100);p__Ascomycota(100);c__Saccharomycetes(100);o__Saccharomycetales(100);f__Metschnikowiaceae(100);g__Metschnikowia(100);s__Metschnikowia_pulcherrima(100); | Metschnikowia | 13 | Saprotroph | \|Undefined Saprotroph\| | Yeast | NULL | Probable | As nectar-tap_saprotroph (Põlme et al. 2020); Decay sugar-rich substrates (Põlme et al. 2020) | Põlme S, et al. 2020. Fungal Diversity 105:1-16 ((https://doi.org/10.1007/s13225-020-00466-2)) |
| Otu30057 | 1 | k__Fungi(100);p__Ascomycota(100);c__Saccharomycetes(100);o__Saccharomycetales(100);f__Metschnikowiaceae(100);g__Metschnikowia(100);g__Metschnikowia_unclassified(100); | Metschnikowia | 13 | Saprotroph | \|Undefined Saprotroph\| | Yeast | NULL | Probable | As nectar-tap_saprotroph (Põlme et al. 2020); Decay sugar-rich substrates (Põlme et al. 2020) | Põlme S, et al. 2020. Fungal Diversity 105:1-16 ((https://doi.org/10.1007/s13225-020-00466-2)) |
| Otu30059 | 1 | k__Fungi(100);p__Ascomycota(100);c__Saccharomycetes(100);o__Saccharomycetales(100);f__Metschnikowiaceae(100);g__Metschnikowia(100);s__Metschnikowia_pulcherrima(100); | Metschnikowia | 13 | Saprotroph | \|Undefined Saprotroph\| | Yeast | NULL | Probable | As nectar-tap_saprotroph (Põlme et al. 2020); Decay sugar-rich substrates (Põlme et al. 2020) | Põlme S, et al. 2020. Fungal Diversity 105:1-16 ((https://doi.org/10.1007/s13225-020-00466-2)) |
| Otu30061 | 1 | k__Fungi(100);p__Ascomycota(100);c__Saccharomycetes(100);o__Saccharomycetales(100);f__Metschnikowiaceae(100);g__Metschnikowia(100);s__Metschnikowia_pulcherrima(100); | Metschnikowia | 13 | Saprotroph | \|Undefined Saprotroph\| | Yeast | NULL | Probable | As nectar-tap_saprotroph (Põlme et al. 2020); Decay sugar-rich substrates (Põlme et al. 2020) | Põlme S, et al. 2020. Fungal Diversity 105:1-16 ((https://doi.org/10.1007/s13225-020-00466-2)) |
| Otu30062 | 1 | k__Fungi(100);p__Ascomycota(100);c__Saccharomycetes(100);o__Saccharomycetales(100);f__Metschnikowiaceae(100);g__Metschnikowia(100);g__Metschnikowia_unclassified(100); | Metschnikowia | 13 | Saprotroph | \|Undefined Saprotroph\| | Yeast | NULL | Probable | As nectar-tap_saprotroph (Põlme et al. 2020); Decay sugar-rich substrates (Põlme et al. 2020) | Põlme S, et al. 2020. Fungal Diversity 105:1-16 ((https://doi.org/10.1007/s13225-020-00466-2)) |
| Otu30063 | 1 | k__Fungi(100);p__Ascomycota(100);c__Saccharomycetes(100);o__Saccharomycetales(100);f__Metschnikowiaceae(100);g__Metschnikowia(100);g__Metschnikowia_unclassified(100); | Metschnikowia | 13 | Saprotroph | \|Undefined Saprotroph\| | Yeast | NULL | Probable | As nectar-tap_saprotroph (Põlme et al. 2020); Decay sugar-rich substrates (Põlme et al. 2020) | Põlme S, et al. 2020. Fungal Diversity 105:1-16 ((https://doi.org/10.1007/s13225-020-00466-2)) |
| Otu30064 | 1 | k__Fungi(100);p__Ascomycota(100);c__Saccharomycetes(100);o__Saccharomycetales(100);f__Saccharomycetales_family_Incertae_sedis(100);g__Candida(100);g__Candida_unclassified(100); | Candida | 13 | Pathotroph-Saprotroph-Symbiotroph | Animal Pathogen-Endophyte-Endosymbiont-Epiphyte-\|Undefined Saprotroph\| | Dimorphic Yeast-Yeast | NULL | Probable | As nectar-tap_saprotroph (Põlme et al. 2020); Decay sugar-rich substrates (Põlme et al. 2020); This asexual genus is polyphyletic and contain species that function very differently depend on what they are. Caution should be taken when interpreting guilds for Candida species. | Manolakaki D, et al. 2010. Virulence. 1: 367-375 ((https://doi.org/10.4161/viru.1.5.12796)); Mekbib SB, et al. 2011. Tropical Plant Pathology 36:233-240 ((https://doi.org/10.1590/S1982-56762011000400004)); Põlme S, et al. 2020. Fungal Diversity 105:1-16 ((https://doi.org/10.1007/s13225-020-00466-2)) |
| Otu30065 | 1 | k__Fungi(100);p__Ascomycota(100);c__Saccharomycetes(100);o__Saccharomycetales(100);f__Metschnikowiaceae(100);g__Metschnikowia(100);g__Metschnikowia_unclassified(100); | Metschnikowia | 13 | Saprotroph | \|Undefined Saprotroph\| | Yeast | NULL | Probable | As nectar-tap_saprotroph (Põlme et al. 2020); Decay sugar-rich substrates (Põlme et al. 2020) | Põlme S, et al. 2020. Fungal Diversity 105:1-16 ((https://doi.org/10.1007/s13225-020-00466-2)) |
| Otu30066 | 1 | k__Fungi(100);p__Ascomycota(100);c__Saccharomycetes(100);o__Saccharomycetales(100);f__Saccharomycetales_family_Incertae_sedis(100);f__Saccharomycetales_family_Incertae_sedis_unclassified(100);f__Saccharomycetales_family_Incertae_sedis_unclassified(100); | Saccharomycetales | 7 | Saprotroph | Undefined Saprotroph | Yeast | NULL | Possible | NULL | Starkenburg S, et al. 2015. New Phytologist 207:1145-1158 ((https://doi.org/10.1111/nph.13426)) |
| Otu30067 | 1 | k__Fungi(100);p__Ascomycota(100);c__Saccharomycetes(100);o__Saccharomycetales(100);f__Saccharomycetales_family_Incertae_sedis(100);g__Candida(100);g__Candida_unclassified(100); | Candida | 13 | Pathotroph-Saprotroph-Symbiotroph | Animal Pathogen-Endophyte-Endosymbiont-Epiphyte-\|Undefined Saprotroph\| | Dimorphic Yeast-Yeast | NULL | Probable | As nectar-tap_saprotroph (Põlme et al. 2020); Decay sugar-rich substrates (Põlme et al. 2020); This asexual genus is polyphyletic and contain species that function very differently depend on what they are. Caution should be taken when interpreting guilds for Candida species. | Manolakaki D, et al. 2010. Virulence. 1: 367-375 ((https://doi.org/10.4161/viru.1.5.12796)); Mekbib SB, et al. 2011. Tropical Plant Pathology 36:233-240 ((https://doi.org/10.1590/S1982-56762011000400004)); Põlme S, et al. 2020. Fungal Diversity 105:1-16 ((https://doi.org/10.1007/s13225-020-00466-2)) |
| Otu30068 | 1 | k__Fungi(100);p__Ascomycota(100);c__Saccharomycetes(100);o__Saccharomycetales(100);f__Metschnikowiaceae(100);g__Metschnikowia(100);s__Metschnikowia_pulcherrima(100); | Metschnikowia | 13 | Saprotroph | \|Undefined Saprotroph\| | Yeast | NULL | Probable | As nectar-tap_saprotroph (Põlme et al. 2020); Decay sugar-rich substrates (Põlme et al. 2020) | Põlme S, et al. 2020. Fungal Diversity 105:1-16 ((https://doi.org/10.1007/s13225-020-00466-2)) |
| Otu30073 | 1 | k__Fungi(100);p__Ascomycota(100);c__Saccharomycetes(100);o__Saccharomycetales(100);f__Metschnikowiaceae(100);g__Metschnikowia(100);s__Metschnikowia_pulcherrima(100); | Metschnikowia | 13 | Saprotroph | \|Undefined Saprotroph\| | Yeast | NULL | Probable | As nectar-tap_saprotroph (Põlme et al. 2020); Decay sugar-rich substrates (Põlme et al. 2020) | Põlme S, et al. 2020. Fungal Diversity 105:1-16 ((https://doi.org/10.1007/s13225-020-00466-2)) |
| Otu30075 | 1 | k__Fungi(100);p__Ascomycota(100);c__Saccharomycetes(100);o__Saccharomycetales(100);f__Metschnikowiaceae(100);g__Metschnikowia(100);g__Metschnikowia_unclassified(100); | Metschnikowia | 13 | Saprotroph | \|Undefined Saprotroph\| | Yeast | NULL | Probable | As nectar-tap_saprotroph (Põlme et al. 2020); Decay sugar-rich substrates (Põlme et al. 2020) | Põlme S, et al. 2020. Fungal Diversity 105:1-16 ((https://doi.org/10.1007/s13225-020-00466-2)) |
| Otu30076 | 1 | k__Fungi(100);p__Ascomycota(100);c__Saccharomycetes(100);o__Saccharomycetales(100);f__Metschnikowiaceae(100);g__Metschnikowia(100);s__Metschnikowia_pulcherrima(100); | Metschnikowia | 13 | Saprotroph | \|Undefined Saprotroph\| | Yeast | NULL | Probable | As nectar-tap_saprotroph (Põlme et al. 2020); Decay sugar-rich substrates (Põlme et al. 2020) | Põlme S, et al. 2020. Fungal Diversity 105:1-16 ((https://doi.org/10.1007/s13225-020-00466-2)) |
| Otu30078 | 1 | k__Fungi(100);p__Ascomycota(100);c__Saccharomycetes(100);o__Saccharomycetales(100);f__Metschnikowiaceae(100);g__Metschnikowia(100);g__Metschnikowia_unclassified(100); | Metschnikowia | 13 | Saprotroph | \|Undefined Saprotroph\| | Yeast | NULL | Probable | As nectar-tap_saprotroph (Põlme et al. 2020); Decay sugar-rich substrates (Põlme et al. 2020) | Põlme S, et al. 2020. Fungal Diversity 105:1-16 ((https://doi.org/10.1007/s13225-020-00466-2)) |
| Otu30079 | 1 | k__Fungi(100);p__Ascomycota(100);c__Saccharomycetes(100);o__Saccharomycetales(100);f__Metschnikowiaceae(100);g__Metschnikowia(100);s__Metschnikowia_pulcherrima(100); | Metschnikowia | 13 | Saprotroph | \|Undefined Saprotroph\| | Yeast | NULL | Probable | As nectar-tap_saprotroph (Põlme et al. 2020); Decay sugar-rich substrates (Põlme et al. 2020) | Põlme S, et al. 2020. Fungal Diversity 105:1-16 ((https://doi.org/10.1007/s13225-020-00466-2)) |
| Otu30080 | 1 | k__Fungi(100);p__Ascomycota(100);c__Saccharomycetes(100);o__Saccharomycetales(100);f__Metschnikowiaceae(100);g__Metschnikowia(100);s__Metschnikowia_pulcherrima(100); | Metschnikowia | 13 | Saprotroph | \|Undefined Saprotroph\| | Yeast | NULL | Probable | As nectar-tap_saprotroph (Põlme et al. 2020); Decay sugar-rich substrates (Põlme et al. 2020) | Põlme S, et al. 2020. Fungal Diversity 105:1-16 ((https://doi.org/10.1007/s13225-020-00466-2)) |
| Otu30082 | 1 | k__Fungi(100);p__Ascomycota(100);c__Saccharomycetes(100);o__Saccharomycetales(100);f__Saccharomycetales_family_Incertae_sedis(100);g__Candida(100);g__Candida_unclassified(100); | Candida | 13 | Pathotroph-Saprotroph-Symbiotroph | Animal Pathogen-Endophyte-Endosymbiont-Epiphyte-\|Undefined Saprotroph\| | Dimorphic Yeast-Yeast | NULL | Probable | As nectar-tap_saprotroph (Põlme et al. 2020); Decay sugar-rich substrates (Põlme et al. 2020); This asexual genus is polyphyletic and contain species that function very differently depend on what they are. Caution should be taken when interpreting guilds for Candida species. | Manolakaki D, et al. 2010. Virulence. 1: 367-375 ((https://doi.org/10.4161/viru.1.5.12796)); Mekbib SB, et al. 2011. Tropical Plant Pathology 36:233-240 ((https://doi.org/10.1590/S1982-56762011000400004)); Põlme S, et al. 2020. Fungal Diversity 105:1-16 ((https://doi.org/10.1007/s13225-020-00466-2)) |
| Otu30084 | 1 | k__Fungi(100);p__Ascomycota(100);c__Saccharomycetes(100);o__Saccharomycetales(100);f__Saccharomycetales_family_Incertae_sedis(100);g__Candida(100);g__Candida_unclassified(100); | Candida | 13 | Pathotroph-Saprotroph-Symbiotroph | Animal Pathogen-Endophyte-Endosymbiont-Epiphyte-\|Undefined Saprotroph\| | Dimorphic Yeast-Yeast | NULL | Probable | As nectar-tap_saprotroph (Põlme et al. 2020); Decay sugar-rich substrates (Põlme et al. 2020); This asexual genus is polyphyletic and contain species that function very differently depend on what they are. Caution should be taken when interpreting guilds for Candida species. | Manolakaki D, et al. 2010. Virulence. 1: 367-375 ((https://doi.org/10.4161/viru.1.5.12796)); Mekbib SB, et al. 2011. Tropical Plant Pathology 36:233-240 ((https://doi.org/10.1590/S1982-56762011000400004)); Põlme S, et al. 2020. Fungal Diversity 105:1-16 ((https://doi.org/10.1007/s13225-020-00466-2)) |
| Otu30087 | 1 | k__Fungi(100);p__Ascomycota(100);c__Saccharomycetes(100);o__Saccharomycetales(100);f__Metschnikowiaceae(100);g__Metschnikowia(100);s__Metschnikowia_pulcherrima(100); | Metschnikowia | 13 | Saprotroph | \|Undefined Saprotroph\| | Yeast | NULL | Probable | As nectar-tap_saprotroph (Põlme et al. 2020); Decay sugar-rich substrates (Põlme et al. 2020) | Põlme S, et al. 2020. Fungal Diversity 105:1-16 ((https://doi.org/10.1007/s13225-020-00466-2)) |
| Otu30088 | 1 | k__Fungi(100);p__Ascomycota(100);c__Saccharomycetes(100);o__Saccharomycetales(100);f__Metschnikowiaceae(100);g__Metschnikowia(100);s__Metschnikowia_pulcherrima(100); | Metschnikowia | 13 | Saprotroph | \|Undefined Saprotroph\| | Yeast | NULL | Probable | As nectar-tap_saprotroph (Põlme et al. 2020); Decay sugar-rich substrates (Põlme et al. 2020) | Põlme S, et al. 2020. Fungal Diversity 105:1-16 ((https://doi.org/10.1007/s13225-020-00466-2)) |
| Otu30090 | 1 | k__Fungi(100);p__Basidiomycota(100);c__Microbotryomycetes(100);o__Sporidiobolales(100);f__Sporidiobolales_family_Incertae_sedis(100);g__Sporobolomyces(100);s__Sporobolomyces_lactosus(100); | Sporobolomyces | 13 | Pathotroph-Saprotroph | \|Fungal Parasite\|-Plant Saprotroph | Yeast | NULL | Probable | As litter_saprotroph (Põlme et al. 2020); As mycoparasite (Põlme et al. 2020); Decay fungal material (Põlme et al. 2020) | Tedersoo L, et al. 2014. Science 346:e1256688 ((https://doi.org/10.1126/science.1256688)); Purahong W, et al. 2016. Molecular Ecology 25:4059-4074 ((https://doi.org/10.1111/mec.13739)); Põlme S, et al. 2020. Fungal Diversity 105:1-16 ((https://doi.org/10.1007/s13225-020-00466-2)) |
| Otu30091 | 1 | k__Fungi(100);p__Ascomycota(100);c__Saccharomycetes(100);o__Saccharomycetales(100);f__Metschnikowiaceae(100);g__Metschnikowia(100);s__Metschnikowia_pulcherrima(100); | Metschnikowia | 13 | Saprotroph | \|Undefined Saprotroph\| | Yeast | NULL | Probable | As nectar-tap_saprotroph (Põlme et al. 2020); Decay sugar-rich substrates (Põlme et al. 2020) | Põlme S, et al. 2020. Fungal Diversity 105:1-16 ((https://doi.org/10.1007/s13225-020-00466-2)) |
| Otu30092 | 1 | k__Fungi(100);p__Ascomycota(100);c__Saccharomycetes(100);o__Saccharomycetales(100);f__Metschnikowiaceae(100);g__Metschnikowia(100);s__Metschnikowia_pulcherrima(100); | Metschnikowia | 13 | Saprotroph | \|Undefined Saprotroph\| | Yeast | NULL | Probable | As nectar-tap_saprotroph (Põlme et al. 2020); Decay sugar-rich substrates (Põlme et al. 2020) | Põlme S, et al. 2020. Fungal Diversity 105:1-16 ((https://doi.org/10.1007/s13225-020-00466-2)) |
| Otu30093 | 1 | k__Fungi(100);p__Ascomycota(100);c__Saccharomycetes(100);o__Saccharomycetales(100);f__Metschnikowiaceae(100);g__Metschnikowia(100);g__Metschnikowia_unclassified(100); | Metschnikowia | 13 | Saprotroph | \|Undefined Saprotroph\| | Yeast | NULL | Probable | As nectar-tap_saprotroph (Põlme et al. 2020); Decay sugar-rich substrates (Põlme et al. 2020) | Põlme S, et al. 2020. Fungal Diversity 105:1-16 ((https://doi.org/10.1007/s13225-020-00466-2)) |
| Otu30095 | 1 | k__Fungi(100);p__Ascomycota(100);c__Saccharomycetes(100);o__Saccharomycetales(100);f__Metschnikowiaceae(100);g__Metschnikowia(100);s__Metschnikowia_pulcherrima(100); | Metschnikowia | 13 | Saprotroph | \|Undefined Saprotroph\| | Yeast | NULL | Probable | As nectar-tap_saprotroph (Põlme et al. 2020); Decay sugar-rich substrates (Põlme et al. 2020) | Põlme S, et al. 2020. Fungal Diversity 105:1-16 ((https://doi.org/10.1007/s13225-020-00466-2)) |
| Otu30096 | 1 | k__Fungi(100);p__Basidiomycota(100);c__Microbotryomycetes(100);o__Sporidiobolales(100);f__Sporidiobolales_family_Incertae_sedis(100);g__Sporobolomyces(100);s__Sporobolomyces_lactosus(100); | Sporobolomyces | 13 | Pathotroph-Saprotroph | \|Fungal Parasite\|-Plant Saprotroph | Yeast | NULL | Probable | As litter_saprotroph (Põlme et al. 2020); As mycoparasite (Põlme et al. 2020); Decay fungal material (Põlme et al. 2020) | Tedersoo L, et al. 2014. Science 346:e1256688 ((https://doi.org/10.1126/science.1256688)); Purahong W, et al. 2016. Molecular Ecology 25:4059-4074 ((https://doi.org/10.1111/mec.13739)); Põlme S, et al. 2020. Fungal Diversity 105:1-16 ((https://doi.org/10.1007/s13225-020-00466-2)) |
| Otu30097 | 1 | k__Fungi(100);p__Ascomycota(100);c__Saccharomycetes(100);o__Saccharomycetales(100);f__Metschnikowiaceae(100);g__Metschnikowia(100);s__Metschnikowia_pulcherrima(100); | Metschnikowia | 13 | Saprotroph | \|Undefined Saprotroph\| | Yeast | NULL | Probable | As nectar-tap_saprotroph (Põlme et al. 2020); Decay sugar-rich substrates (Põlme et al. 2020) | Põlme S, et al. 2020. Fungal Diversity 105:1-16 ((https://doi.org/10.1007/s13225-020-00466-2)) |
| Otu30099 | 1 | k__Fungi(100);p__Basidiomycota(100);c__Microbotryomycetes(100);o__Sporidiobolales(100);f__Sporidiobolales_family_Incertae_sedis(100);g__Sporobolomyces(100);s__Sporobolomyces_lactosus(100); | Sporobolomyces | 13 | Pathotroph-Saprotroph | \|Fungal Parasite\|-Plant Saprotroph | Yeast | NULL | Probable | As litter_saprotroph (Põlme et al. 2020); As mycoparasite (Põlme et al. 2020); Decay fungal material (Põlme et al. 2020) | Tedersoo L, et al. 2014. Science 346:e1256688 ((https://doi.org/10.1126/science.1256688)); Purahong W, et al. 2016. Molecular Ecology 25:4059-4074 ((https://doi.org/10.1111/mec.13739)); Põlme S, et al. 2020. Fungal Diversity 105:1-16 ((https://doi.org/10.1007/s13225-020-00466-2)) |
| Otu30101 | 1 | k__Fungi(100);p__Ascomycota(100);c__Saccharomycetes(100);o__Saccharomycetales(100);f__Metschnikowiaceae(100);g__Metschnikowia(100);s__Metschnikowia_pulcherrima(100); | Metschnikowia | 13 | Saprotroph | \|Undefined Saprotroph\| | Yeast | NULL | Probable | As nectar-tap_saprotroph (Põlme et al. 2020); Decay sugar-rich substrates (Põlme et al. 2020) | Põlme S, et al. 2020. Fungal Diversity 105:1-16 ((https://doi.org/10.1007/s13225-020-00466-2)) |
| Otu30102 | 1 | k__Fungi(100);p__Ascomycota(100);c__Saccharomycetes(100);o__Saccharomycetales(100);f__Metschnikowiaceae(100);g__Metschnikowia(100);s__Metschnikowia_pulcherrima(100); | Metschnikowia | 13 | Saprotroph | \|Undefined Saprotroph\| | Yeast | NULL | Probable | As nectar-tap_saprotroph (Põlme et al. 2020); Decay sugar-rich substrates (Põlme et al. 2020) | Põlme S, et al. 2020. Fungal Diversity 105:1-16 ((https://doi.org/10.1007/s13225-020-00466-2)) |
| Otu30103 | 1 | k__Fungi(100);p__Ascomycota(100);c__Saccharomycetes(100);o__Saccharomycetales(100);f__Metschnikowiaceae(100);g__Metschnikowia(100);g__Metschnikowia_unclassified(100); | Metschnikowia | 13 | Saprotroph | \|Undefined Saprotroph\| | Yeast | NULL | Probable | As nectar-tap_saprotroph (Põlme et al. 2020); Decay sugar-rich substrates (Põlme et al. 2020) | Põlme S, et al. 2020. Fungal Diversity 105:1-16 ((https://doi.org/10.1007/s13225-020-00466-2)) |
| Otu30104 | 1 | k__Fungi(100);p__Ascomycota(100);c__Saccharomycetes(100);o__Saccharomycetales(100);f__Metschnikowiaceae(100);g__Metschnikowia(100);g__Metschnikowia_unclassified(100); | Metschnikowia | 13 | Saprotroph | \|Undefined Saprotroph\| | Yeast | NULL | Probable | As nectar-tap_saprotroph (Põlme et al. 2020); Decay sugar-rich substrates (Põlme et al. 2020) | Põlme S, et al. 2020. Fungal Diversity 105:1-16 ((https://doi.org/10.1007/s13225-020-00466-2)) |
| Otu30105 | 1 | k__Fungi(100);p__Ascomycota(100);c__Saccharomycetes(100);o__Saccharomycetales(100);f__Metschnikowiaceae(100);g__Metschnikowia(100);g__Metschnikowia_unclassified(100); | Metschnikowia | 13 | Saprotroph | \|Undefined Saprotroph\| | Yeast | NULL | Probable | As nectar-tap_saprotroph (Põlme et al. 2020); Decay sugar-rich substrates (Põlme et al. 2020) | Põlme S, et al. 2020. Fungal Diversity 105:1-16 ((https://doi.org/10.1007/s13225-020-00466-2)) |
| Otu30109 | 1 | k__Fungi(100);p__Ascomycota(100);c__Saccharomycetes(100);o__Saccharomycetales(100);f__Metschnikowiaceae(100);g__Metschnikowia(100);s__Metschnikowia_pulcherrima(100); | Metschnikowia | 13 | Saprotroph | \|Undefined Saprotroph\| | Yeast | NULL | Probable | As nectar-tap_saprotroph (Põlme et al. 2020); Decay sugar-rich substrates (Põlme et al. 2020) | Põlme S, et al. 2020. Fungal Diversity 105:1-16 ((https://doi.org/10.1007/s13225-020-00466-2)) |
| Otu30110 | 1 | k__Fungi(100);p__Ascomycota(100);c__Saccharomycetes(100);o__Saccharomycetales(100);f__Metschnikowiaceae(100);g__Metschnikowia(100);s__Metschnikowia_pulcherrima(100); | Metschnikowia | 13 | Saprotroph | \|Undefined Saprotroph\| | Yeast | NULL | Probable | As nectar-tap_saprotroph (Põlme et al. 2020); Decay sugar-rich substrates (Põlme et al. 2020) | Põlme S, et al. 2020. Fungal Diversity 105:1-16 ((https://doi.org/10.1007/s13225-020-00466-2)) |
| Otu30111 | 1 | k__Fungi(100);p__Ascomycota(100);c__Saccharomycetes(100);o__Saccharomycetales(100);f__Metschnikowiaceae(100);g__Metschnikowia(100);s__Metschnikowia_pulcherrima(100); | Metschnikowia | 13 | Saprotroph | \|Undefined Saprotroph\| | Yeast | NULL | Probable | As nectar-tap_saprotroph (Põlme et al. 2020); Decay sugar-rich substrates (Põlme et al. 2020) | Põlme S, et al. 2020. Fungal Diversity 105:1-16 ((https://doi.org/10.1007/s13225-020-00466-2)) |
| Otu30113 | 1 | k__Fungi(100);p__Ascomycota(100);c__Saccharomycetes(100);o__Saccharomycetales(100);f__Saccharomycetales_family_Incertae_sedis(100);g__Candida(100);g__Candida_unclassified(100); | Candida | 13 | Pathotroph-Saprotroph-Symbiotroph | Animal Pathogen-Endophyte-Endosymbiont-Epiphyte-\|Undefined Saprotroph\| | Dimorphic Yeast-Yeast | NULL | Probable | As nectar-tap_saprotroph (Põlme et al. 2020); Decay sugar-rich substrates (Põlme et al. 2020); This asexual genus is polyphyletic and contain species that function very differently depend on what they are. Caution should be taken when interpreting guilds for Candida species. | Manolakaki D, et al. 2010. Virulence. 1: 367-375 ((https://doi.org/10.4161/viru.1.5.12796)); Mekbib SB, et al. 2011. Tropical Plant Pathology 36:233-240 ((https://doi.org/10.1590/S1982-56762011000400004)); Põlme S, et al. 2020. Fungal Diversity 105:1-16 ((https://doi.org/10.1007/s13225-020-00466-2)) |
| Otu30114 | 1 | k__Fungi(100);p__Ascomycota(100);c__Saccharomycetes(100);o__Saccharomycetales(100);f__Metschnikowiaceae(100);g__Metschnikowia(100);g__Metschnikowia_unclassified(100); | Metschnikowia | 13 | Saprotroph | \|Undefined Saprotroph\| | Yeast | NULL | Probable | As nectar-tap_saprotroph (Põlme et al. 2020); Decay sugar-rich substrates (Põlme et al. 2020) | Põlme S, et al. 2020. Fungal Diversity 105:1-16 ((https://doi.org/10.1007/s13225-020-00466-2)) |
| Otu30116 | 1 | k__Fungi(100);p__Ascomycota(100);c__Saccharomycetes(100);o__Saccharomycetales(100);f__Metschnikowiaceae(100);g__Metschnikowia(100);s__Metschnikowia_pulcherrima(100); | Metschnikowia | 13 | Saprotroph | \|Undefined Saprotroph\| | Yeast | NULL | Probable | As nectar-tap_saprotroph (Põlme et al. 2020); Decay sugar-rich substrates (Põlme et al. 2020) | Põlme S, et al. 2020. Fungal Diversity 105:1-16 ((https://doi.org/10.1007/s13225-020-00466-2)) |
| Otu30118 | 1 | k__Fungi(100);p__Ascomycota(100);c__Saccharomycetes(100);o__Saccharomycetales(100);f__Saccharomycetales_family_Incertae_sedis(100);g__Candida(100);g__Candida_unclassified(100); | Candida | 13 | Pathotroph-Saprotroph-Symbiotroph | Animal Pathogen-Endophyte-Endosymbiont-Epiphyte-\|Undefined Saprotroph\| | Dimorphic Yeast-Yeast | NULL | Probable | As nectar-tap_saprotroph (Põlme et al. 2020); Decay sugar-rich substrates (Põlme et al. 2020); This asexual genus is polyphyletic and contain species that function very differently depend on what they are. Caution should be taken when interpreting guilds for Candida species. | Manolakaki D, et al. 2010. Virulence. 1: 367-375 ((https://doi.org/10.4161/viru.1.5.12796)); Mekbib SB, et al. 2011. Tropical Plant Pathology 36:233-240 ((https://doi.org/10.1590/S1982-56762011000400004)); Põlme S, et al. 2020. Fungal Diversity 105:1-16 ((https://doi.org/10.1007/s13225-020-00466-2)) |
| Otu30119 | 1 | k__Fungi(100);p__Ascomycota(100);c__Saccharomycetes(100);o__Saccharomycetales(100);f__Saccharomycetales_family_Incertae_sedis(100);g__Candida(100);g__Candida_unclassified(100); | Candida | 13 | Pathotroph-Saprotroph-Symbiotroph | Animal Pathogen-Endophyte-Endosymbiont-Epiphyte-\|Undefined Saprotroph\| | Dimorphic Yeast-Yeast | NULL | Probable | As nectar-tap_saprotroph (Põlme et al. 2020); Decay sugar-rich substrates (Põlme et al. 2020); This asexual genus is polyphyletic and contain species that function very differently depend on what they are. Caution should be taken when interpreting guilds for Candida species. | Manolakaki D, et al. 2010. Virulence. 1: 367-375 ((https://doi.org/10.4161/viru.1.5.12796)); Mekbib SB, et al. 2011. Tropical Plant Pathology 36:233-240 ((https://doi.org/10.1590/S1982-56762011000400004)); Põlme S, et al. 2020. Fungal Diversity 105:1-16 ((https://doi.org/10.1007/s13225-020-00466-2)) |
| Otu30120 | 1 | k__Fungi(100);p__Ascomycota(100);c__Saccharomycetes(100);o__Saccharomycetales(100);f__Saccharomycetales_family_Incertae_sedis(100);g__Candida(100);s__Candida_cretensis(100); | Candida | 13 | Pathotroph-Saprotroph-Symbiotroph | Animal Pathogen-Endophyte-Endosymbiont-Epiphyte-\|Undefined Saprotroph\| | Dimorphic Yeast-Yeast | NULL | Probable | As nectar-tap_saprotroph (Põlme et al. 2020); Decay sugar-rich substrates (Põlme et al. 2020); This asexual genus is polyphyletic and contain species that function very differently depend on what they are. Caution should be taken when interpreting guilds for Candida species. | Manolakaki D, et al. 2010. Virulence. 1: 367-375 ((https://doi.org/10.4161/viru.1.5.12796)); Mekbib SB, et al. 2011. Tropical Plant Pathology 36:233-240 ((https://doi.org/10.1590/S1982-56762011000400004)); Põlme S, et al. 2020. Fungal Diversity 105:1-16 ((https://doi.org/10.1007/s13225-020-00466-2)) |
| Otu30122 | 1 | k__Fungi(100);p__Ascomycota(100);c__Saccharomycetes(100);o__Saccharomycetales(100);o__Saccharomycetales_unclassified(100);o__Saccharomycetales_unclassified(100);o__Saccharomycetales_unclassified(100); | Saccharomycetales | 7 | Saprotroph | Undefined Saprotroph | Yeast | NULL | Possible | NULL | Starkenburg S, et al. 2015. New Phytologist 207:1145-1158 ((https://doi.org/10.1111/nph.13426)) |
| Otu30123 | 1 | k__Fungi(100);p__Ascomycota(100);c__Saccharomycetes(100);o__Saccharomycetales(100);f__Saccharomycetales_family_Incertae_sedis(100);g__Candida(100);g__Candida_unclassified(100); | Candida | 13 | Pathotroph-Saprotroph-Symbiotroph | Animal Pathogen-Endophyte-Endosymbiont-Epiphyte-\|Undefined Saprotroph\| | Dimorphic Yeast-Yeast | NULL | Probable | As nectar-tap_saprotroph (Põlme et al. 2020); Decay sugar-rich substrates (Põlme et al. 2020); This asexual genus is polyphyletic and contain species that function very differently depend on what they are. Caution should be taken when interpreting guilds for Candida species. | Manolakaki D, et al. 2010. Virulence. 1: 367-375 ((https://doi.org/10.4161/viru.1.5.12796)); Mekbib SB, et al. 2011. Tropical Plant Pathology 36:233-240 ((https://doi.org/10.1590/S1982-56762011000400004)); Põlme S, et al. 2020. Fungal Diversity 105:1-16 ((https://doi.org/10.1007/s13225-020-00466-2)) |
| Otu30125 | 1 | k__Fungi(100);p__Ascomycota(100);c__Saccharomycetes(100);o__Saccharomycetales(100);f__Saccharomycetales_family_Incertae_sedis(100);g__Candida(100);g__Candida_unclassified(100); | Candida | 13 | Pathotroph-Saprotroph-Symbiotroph | Animal Pathogen-Endophyte-Endosymbiont-Epiphyte-\|Undefined Saprotroph\| | Dimorphic Yeast-Yeast | NULL | Probable | As nectar-tap_saprotroph (Põlme et al. 2020); Decay sugar-rich substrates (Põlme et al. 2020); This asexual genus is polyphyletic and contain species that function very differently depend on what they are. Caution should be taken when interpreting guilds for Candida species. | Manolakaki D, et al. 2010. Virulence. 1: 367-375 ((https://doi.org/10.4161/viru.1.5.12796)); Mekbib SB, et al. 2011. Tropical Plant Pathology 36:233-240 ((https://doi.org/10.1590/S1982-56762011000400004)); Põlme S, et al. 2020. Fungal Diversity 105:1-16 ((https://doi.org/10.1007/s13225-020-00466-2)) |
| Otu30127 | 1 | k__Fungi(100);p__Ascomycota(100);c__Saccharomycetes(100);o__Saccharomycetales(100);f__Saccharomycetales_family_Incertae_sedis(100);f__Saccharomycetales_family_Incertae_sedis_unclassified(100);f__Saccharomycetales_family_Incertae_sedis_unclassified(100); | Saccharomycetales | 7 | Saprotroph | Undefined Saprotroph | Yeast | NULL | Possible | NULL | Starkenburg S, et al. 2015. New Phytologist 207:1145-1158 ((https://doi.org/10.1111/nph.13426)) |
| Otu30128 | 1 | k__Fungi(100);p__Ascomycota(100);c__Saccharomycetes(100);o__Saccharomycetales(100);o__Saccharomycetales_unclassified(100);o__Saccharomycetales_unclassified(100);o__Saccharomycetales_unclassified(100); | Saccharomycetales | 7 | Saprotroph | Undefined Saprotroph | Yeast | NULL | Possible | NULL | Starkenburg S, et al. 2015. New Phytologist 207:1145-1158 ((https://doi.org/10.1111/nph.13426)) |
| Otu30132 | 1 | k__Fungi(100);p__Ascomycota(100);c__Saccharomycetes(100);o__Saccharomycetales(100);f__Metschnikowiaceae(100);g__Metschnikowia(100);g__Metschnikowia_unclassified(100); | Metschnikowia | 13 | Saprotroph | \|Undefined Saprotroph\| | Yeast | NULL | Probable | As nectar-tap_saprotroph (Põlme et al. 2020); Decay sugar-rich substrates (Põlme et al. 2020) | Põlme S, et al. 2020. Fungal Diversity 105:1-16 ((https://doi.org/10.1007/s13225-020-00466-2)) |
| Otu30133 | 1 | k__Fungi(100);p__Ascomycota(100);c__Saccharomycetes(100);o__Saccharomycetales(100);f__Saccharomycetales_family_Incertae_sedis(100);f__Saccharomycetales_family_Incertae_sedis_unclassified(100);f__Saccharomycetales_family_Incertae_sedis_unclassified(100); | Saccharomycetales | 7 | Saprotroph | Undefined Saprotroph | Yeast | NULL | Possible | NULL | Starkenburg S, et al. 2015. New Phytologist 207:1145-1158 ((https://doi.org/10.1111/nph.13426)) |
| Otu30134 | 1 | k__Fungi(100);p__Ascomycota(100);c__Saccharomycetes(100);o__Saccharomycetales(100);f__Metschnikowiaceae(100);g__Metschnikowia(100);s__Metschnikowia_pulcherrima(100); | Metschnikowia | 13 | Saprotroph | \|Undefined Saprotroph\| | Yeast | NULL | Probable | As nectar-tap_saprotroph (Põlme et al. 2020); Decay sugar-rich substrates (Põlme et al. 2020) | Põlme S, et al. 2020. Fungal Diversity 105:1-16 ((https://doi.org/10.1007/s13225-020-00466-2)) |
| Otu30136 | 1 | k__Fungi(100);p__Ascomycota(100);c__Saccharomycetes(100);o__Saccharomycetales(100);f__Metschnikowiaceae(100);g__Metschnikowia(100);g__Metschnikowia_unclassified(100); | Metschnikowia | 13 | Saprotroph | \|Undefined Saprotroph\| | Yeast | NULL | Probable | As nectar-tap_saprotroph (Põlme et al. 2020); Decay sugar-rich substrates (Põlme et al. 2020) | Põlme S, et al. 2020. Fungal Diversity 105:1-16 ((https://doi.org/10.1007/s13225-020-00466-2)) |
| Otu30137 | 1 | k__Fungi(100);p__Ascomycota(100);c__Saccharomycetes(100);o__Saccharomycetales(100);f__Metschnikowiaceae(100);g__Metschnikowia(100);g__Metschnikowia_unclassified(100); | Metschnikowia | 13 | Saprotroph | \|Undefined Saprotroph\| | Yeast | NULL | Probable | As nectar-tap_saprotroph (Põlme et al. 2020); Decay sugar-rich substrates (Põlme et al. 2020) | Põlme S, et al. 2020. Fungal Diversity 105:1-16 ((https://doi.org/10.1007/s13225-020-00466-2)) |
| Otu30138 | 1 | k__Fungi(100);p__Ascomycota(100);c__Saccharomycetes(100);o__Saccharomycetales(100);f__Metschnikowiaceae(100);g__Metschnikowia(100);s__Metschnikowia_pulcherrima(100); | Metschnikowia | 13 | Saprotroph | \|Undefined Saprotroph\| | Yeast | NULL | Probable | As nectar-tap_saprotroph (Põlme et al. 2020); Decay sugar-rich substrates (Põlme et al. 2020) | Põlme S, et al. 2020. Fungal Diversity 105:1-16 ((https://doi.org/10.1007/s13225-020-00466-2)) |
| Otu30139 | 1 | k__Fungi(100);p__Ascomycota(100);c__Saccharomycetes(100);o__Saccharomycetales(100);f__Metschnikowiaceae(100);g__Metschnikowia(100);s__Metschnikowia_pulcherrima(100); | Metschnikowia | 13 | Saprotroph | \|Undefined Saprotroph\| | Yeast | NULL | Probable | As nectar-tap_saprotroph (Põlme et al. 2020); Decay sugar-rich substrates (Põlme et al. 2020) | Põlme S, et al. 2020. Fungal Diversity 105:1-16 ((https://doi.org/10.1007/s13225-020-00466-2)) |
| Otu30140 | 1 | k__Fungi(100);p__Ascomycota(100);c__Saccharomycetes(100);o__Saccharomycetales(100);f__Metschnikowiaceae(100);g__Metschnikowia(100);g__Metschnikowia_unclassified(100); | Metschnikowia | 13 | Saprotroph | \|Undefined Saprotroph\| | Yeast | NULL | Probable | As nectar-tap_saprotroph (Põlme et al. 2020); Decay sugar-rich substrates (Põlme et al. 2020) | Põlme S, et al. 2020. Fungal Diversity 105:1-16 ((https://doi.org/10.1007/s13225-020-00466-2)) |
| Otu30141 | 1 | k__Fungi(100);p__Ascomycota(100);c__Saccharomycetes(100);o__Saccharomycetales(100);f__Metschnikowiaceae(100);g__Metschnikowia(100);s__Metschnikowia_pulcherrima(100); | Metschnikowia | 13 | Saprotroph | \|Undefined Saprotroph\| | Yeast | NULL | Probable | As nectar-tap_saprotroph (Põlme et al. 2020); Decay sugar-rich substrates (Põlme et al. 2020) | Põlme S, et al. 2020. Fungal Diversity 105:1-16 ((https://doi.org/10.1007/s13225-020-00466-2)) |
| Otu30142 | 1 | k__Fungi(100);p__Ascomycota(100);c__Saccharomycetes(100);o__Saccharomycetales(100);f__Metschnikowiaceae(100);g__Metschnikowia(100);s__Metschnikowia_pulcherrima(100); | Metschnikowia | 13 | Saprotroph | \|Undefined Saprotroph\| | Yeast | NULL | Probable | As nectar-tap_saprotroph (Põlme et al. 2020); Decay sugar-rich substrates (Põlme et al. 2020) | Põlme S, et al. 2020. Fungal Diversity 105:1-16 ((https://doi.org/10.1007/s13225-020-00466-2)) |
| Otu30143 | 1 | k__Fungi(100);p__Ascomycota(100);c__Saccharomycetes(100);o__Saccharomycetales(100);f__Metschnikowiaceae(100);g__Metschnikowia(100);s__Metschnikowia_pulcherrima(100); | Metschnikowia | 13 | Saprotroph | \|Undefined Saprotroph\| | Yeast | NULL | Probable | As nectar-tap_saprotroph (Põlme et al. 2020); Decay sugar-rich substrates (Põlme et al. 2020) | Põlme S, et al. 2020. Fungal Diversity 105:1-16 ((https://doi.org/10.1007/s13225-020-00466-2)) |
| Otu30144 | 1 | k__Fungi(100);p__Ascomycota(100);c__Saccharomycetes(100);o__Saccharomycetales(100);f__Metschnikowiaceae(100);g__Metschnikowia(100);s__Metschnikowia_pulcherrima(100); | Metschnikowia | 13 | Saprotroph | \|Undefined Saprotroph\| | Yeast | NULL | Probable | As nectar-tap_saprotroph (Põlme et al. 2020); Decay sugar-rich substrates (Põlme et al. 2020) | Põlme S, et al. 2020. Fungal Diversity 105:1-16 ((https://doi.org/10.1007/s13225-020-00466-2)) |
| Otu30145 | 1 | k__Fungi(100);p__Ascomycota(100);c__Saccharomycetes(100);o__Saccharomycetales(100);f__Metschnikowiaceae(100);g__Metschnikowia(100);s__Metschnikowia_pulcherrima(100); | Metschnikowia | 13 | Saprotroph | \|Undefined Saprotroph\| | Yeast | NULL | Probable | As nectar-tap_saprotroph (Põlme et al. 2020); Decay sugar-rich substrates (Põlme et al. 2020) | Põlme S, et al. 2020. Fungal Diversity 105:1-16 ((https://doi.org/10.1007/s13225-020-00466-2)) |
| Otu30146 | 1 | k__Fungi(100);p__Ascomycota(100);c__Saccharomycetes(100);o__Saccharomycetales(100);f__Metschnikowiaceae(100);g__Metschnikowia(100);s__Metschnikowia_pulcherrima(100); | Metschnikowia | 13 | Saprotroph | \|Undefined Saprotroph\| | Yeast | NULL | Probable | As nectar-tap_saprotroph (Põlme et al. 2020); Decay sugar-rich substrates (Põlme et al. 2020) | Põlme S, et al. 2020. Fungal Diversity 105:1-16 ((https://doi.org/10.1007/s13225-020-00466-2)) |
| Otu30148 | 1 | k__Fungi(100);p__Ascomycota(100);c__Saccharomycetes(100);o__Saccharomycetales(100);f__Metschnikowiaceae(100);g__Metschnikowia(100);g__Metschnikowia_unclassified(100); | Metschnikowia | 13 | Saprotroph | \|Undefined Saprotroph\| | Yeast | NULL | Probable | As nectar-tap_saprotroph (Põlme et al. 2020); Decay sugar-rich substrates (Põlme et al. 2020) | Põlme S, et al. 2020. Fungal Diversity 105:1-16 ((https://doi.org/10.1007/s13225-020-00466-2)) |
| Otu30149 | 1 | k__Fungi(100);p__Ascomycota(100);c__Saccharomycetes(100);o__Saccharomycetales(100);f__Metschnikowiaceae(100);g__Metschnikowia(100);s__Metschnikowia_pulcherrima(100); | Metschnikowia | 13 | Saprotroph | \|Undefined Saprotroph\| | Yeast | NULL | Probable | As nectar-tap_saprotroph (Põlme et al. 2020); Decay sugar-rich substrates (Põlme et al. 2020) | Põlme S, et al. 2020. Fungal Diversity 105:1-16 ((https://doi.org/10.1007/s13225-020-00466-2)) |
| Otu30150 | 1 | k__Fungi(100);p__Ascomycota(100);c__Saccharomycetes(100);o__Saccharomycetales(100);f__Metschnikowiaceae(100);g__Metschnikowia(100);g__Metschnikowia_unclassified(100); | Metschnikowia | 13 | Saprotroph | \|Undefined Saprotroph\| | Yeast | NULL | Probable | As nectar-tap_saprotroph (Põlme et al. 2020); Decay sugar-rich substrates (Põlme et al. 2020) | Põlme S, et al. 2020. Fungal Diversity 105:1-16 ((https://doi.org/10.1007/s13225-020-00466-2)) |
| Otu30151 | 1 | k__Fungi(100);p__Ascomycota(100);c__Saccharomycetes(100);o__Saccharomycetales(100);f__Metschnikowiaceae(100);g__Metschnikowia(100);g__Metschnikowia_unclassified(100); | Metschnikowia | 13 | Saprotroph | \|Undefined Saprotroph\| | Yeast | NULL | Probable | As nectar-tap_saprotroph (Põlme et al. 2020); Decay sugar-rich substrates (Põlme et al. 2020) | Põlme S, et al. 2020. Fungal Diversity 105:1-16 ((https://doi.org/10.1007/s13225-020-00466-2)) |
| Otu30152 | 1 | k__Fungi(100);p__Ascomycota(100);c__Saccharomycetes(100);o__Saccharomycetales(100);f__Metschnikowiaceae(100);g__Metschnikowia(100);s__Metschnikowia_pulcherrima(100); | Metschnikowia | 13 | Saprotroph | \|Undefined Saprotroph\| | Yeast | NULL | Probable | As nectar-tap_saprotroph (Põlme et al. 2020); Decay sugar-rich substrates (Põlme et al. 2020) | Põlme S, et al. 2020. Fungal Diversity 105:1-16 ((https://doi.org/10.1007/s13225-020-00466-2)) |
| Otu30154 | 1 | k__Fungi(100);p__Ascomycota(100);c__Saccharomycetes(100);o__Saccharomycetales(100);f__Metschnikowiaceae(100);g__Metschnikowia(100);s__Metschnikowia_pulcherrima(100); | Metschnikowia | 13 | Saprotroph | \|Undefined Saprotroph\| | Yeast | NULL | Probable | As nectar-tap_saprotroph (Põlme et al. 2020); Decay sugar-rich substrates (Põlme et al. 2020) | Põlme S, et al. 2020. Fungal Diversity 105:1-16 ((https://doi.org/10.1007/s13225-020-00466-2)) |
| Otu30155 | 1 | k__Fungi(100);p__Ascomycota(100);c__Saccharomycetes(100);o__Saccharomycetales(100);f__Metschnikowiaceae(100);g__Metschnikowia(100);g__Metschnikowia_unclassified(100); | Metschnikowia | 13 | Saprotroph | \|Undefined Saprotroph\| | Yeast | NULL | Probable | As nectar-tap_saprotroph (Põlme et al. 2020); Decay sugar-rich substrates (Põlme et al. 2020) | Põlme S, et al. 2020. Fungal Diversity 105:1-16 ((https://doi.org/10.1007/s13225-020-00466-2)) |
| Otu30156 | 1 | k__Fungi(100);p__Ascomycota(100);c__Saccharomycetes(100);o__Saccharomycetales(100);f__Metschnikowiaceae(100);g__Metschnikowia(100);g__Metschnikowia_unclassified(100); | Metschnikowia | 13 | Saprotroph | \|Undefined Saprotroph\| | Yeast | NULL | Probable | As nectar-tap_saprotroph (Põlme et al. 2020); Decay sugar-rich substrates (Põlme et al. 2020) | Põlme S, et al. 2020. Fungal Diversity 105:1-16 ((https://doi.org/10.1007/s13225-020-00466-2)) |
| Otu30158 | 1 | k__Fungi(100);p__Ascomycota(100);c__Saccharomycetes(100);o__Saccharomycetales(100);f__Metschnikowiaceae(100);g__Metschnikowia(100);s__Metschnikowia_pulcherrima(100); | Metschnikowia | 13 | Saprotroph | \|Undefined Saprotroph\| | Yeast | NULL | Probable | As nectar-tap_saprotroph (Põlme et al. 2020); Decay sugar-rich substrates (Põlme et al. 2020) | Põlme S, et al. 2020. Fungal Diversity 105:1-16 ((https://doi.org/10.1007/s13225-020-00466-2)) |
| Otu30159 | 1 | k__Fungi(100);p__Ascomycota(100);c__Saccharomycetes(100);o__Saccharomycetales(100);f__Metschnikowiaceae(100);g__Metschnikowia(100);g__Metschnikowia_unclassified(100); | Metschnikowia | 13 | Saprotroph | \|Undefined Saprotroph\| | Yeast | NULL | Probable | As nectar-tap_saprotroph (Põlme et al. 2020); Decay sugar-rich substrates (Põlme et al. 2020) | Põlme S, et al. 2020. Fungal Diversity 105:1-16 ((https://doi.org/10.1007/s13225-020-00466-2)) |
| Otu30161 | 1 | k__Fungi(100);p__Ascomycota(100);c__Saccharomycetes(100);o__Saccharomycetales(100);f__Metschnikowiaceae(100);g__Metschnikowia(100);g__Metschnikowia_unclassified(100); | Metschnikowia | 13 | Saprotroph | \|Undefined Saprotroph\| | Yeast | NULL | Probable | As nectar-tap_saprotroph (Põlme et al. 2020); Decay sugar-rich substrates (Põlme et al. 2020) | Põlme S, et al. 2020. Fungal Diversity 105:1-16 ((https://doi.org/10.1007/s13225-020-00466-2)) |
| Otu30162 | 1 | k__Fungi(100);p__Ascomycota(100);c__Saccharomycetes(100);o__Saccharomycetales(100);f__Metschnikowiaceae(100);g__Metschnikowia(100);g__Metschnikowia_unclassified(100); | Metschnikowia | 13 | Saprotroph | \|Undefined Saprotroph\| | Yeast | NULL | Probable | As nectar-tap_saprotroph (Põlme et al. 2020); Decay sugar-rich substrates (Põlme et al. 2020) | Põlme S, et al. 2020. Fungal Diversity 105:1-16 ((https://doi.org/10.1007/s13225-020-00466-2)) |
| Otu30163 | 1 | k__Fungi(100);p__Ascomycota(100);c__Saccharomycetes(100);o__Saccharomycetales(100);f__Metschnikowiaceae(100);g__Metschnikowia(100);s__Metschnikowia_pulcherrima(100); | Metschnikowia | 13 | Saprotroph | \|Undefined Saprotroph\| | Yeast | NULL | Probable | As nectar-tap_saprotroph (Põlme et al. 2020); Decay sugar-rich substrates (Põlme et al. 2020) | Põlme S, et al. 2020. Fungal Diversity 105:1-16 ((https://doi.org/10.1007/s13225-020-00466-2)) |
| Otu30164 | 1 | k__Fungi(100);p__Ascomycota(100);c__Saccharomycetes(100);o__Saccharomycetales(100);f__Metschnikowiaceae(100);g__Metschnikowia(100);g__Metschnikowia_unclassified(100); | Metschnikowia | 13 | Saprotroph | \|Undefined Saprotroph\| | Yeast | NULL | Probable | As nectar-tap_saprotroph (Põlme et al. 2020); Decay sugar-rich substrates (Põlme et al. 2020) | Põlme S, et al. 2020. Fungal Diversity 105:1-16 ((https://doi.org/10.1007/s13225-020-00466-2)) |
| Otu30165 | 1 | k__Fungi(100);p__Ascomycota(100);c__Saccharomycetes(100);o__Saccharomycetales(100);f__Metschnikowiaceae(100);g__Metschnikowia(100);s__Metschnikowia_pulcherrima(100); | Metschnikowia | 13 | Saprotroph | \|Undefined Saprotroph\| | Yeast | NULL | Probable | As nectar-tap_saprotroph (Põlme et al. 2020); Decay sugar-rich substrates (Põlme et al. 2020) | Põlme S, et al. 2020. Fungal Diversity 105:1-16 ((https://doi.org/10.1007/s13225-020-00466-2)) |
| Otu30166 | 1 | k__Fungi(100);p__Ascomycota(100);c__Saccharomycetes(100);o__Saccharomycetales(100);f__Metschnikowiaceae(100);g__Metschnikowia(100);g__Metschnikowia_unclassified(100); | Metschnikowia | 13 | Saprotroph | \|Undefined Saprotroph\| | Yeast | NULL | Probable | As nectar-tap_saprotroph (Põlme et al. 2020); Decay sugar-rich substrates (Põlme et al. 2020) | Põlme S, et al. 2020. Fungal Diversity 105:1-16 ((https://doi.org/10.1007/s13225-020-00466-2)) |
| Otu30167 | 1 | k__Fungi(100);p__Ascomycota(100);c__Saccharomycetes(100);o__Saccharomycetales(100);f__Metschnikowiaceae(100);g__Metschnikowia(100);s__Metschnikowia_pulcherrima(100); | Metschnikowia | 13 | Saprotroph | \|Undefined Saprotroph\| | Yeast | NULL | Probable | As nectar-tap_saprotroph (Põlme et al. 2020); Decay sugar-rich substrates (Põlme et al. 2020) | Põlme S, et al. 2020. Fungal Diversity 105:1-16 ((https://doi.org/10.1007/s13225-020-00466-2)) |
| Otu30168 | 1 | k__Fungi(100);p__Ascomycota(100);c__Saccharomycetes(100);o__Saccharomycetales(100);f__Metschnikowiaceae(100);g__Metschnikowia(100);s__Metschnikowia_pulcherrima(100); | Metschnikowia | 13 | Saprotroph | \|Undefined Saprotroph\| | Yeast | NULL | Probable | As nectar-tap_saprotroph (Põlme et al. 2020); Decay sugar-rich substrates (Põlme et al. 2020) | Põlme S, et al. 2020. Fungal Diversity 105:1-16 ((https://doi.org/10.1007/s13225-020-00466-2)) |
| Otu30169 | 1 | k__Fungi(100);p__Ascomycota(100);c__Saccharomycetes(100);o__Saccharomycetales(100);f__Metschnikowiaceae(100);g__Metschnikowia(100);s__Metschnikowia_pulcherrima(100); | Metschnikowia | 13 | Saprotroph | \|Undefined Saprotroph\| | Yeast | NULL | Probable | As nectar-tap_saprotroph (Põlme et al. 2020); Decay sugar-rich substrates (Põlme et al. 2020) | Põlme S, et al. 2020. Fungal Diversity 105:1-16 ((https://doi.org/10.1007/s13225-020-00466-2)) |
| Otu30170 | 1 | k__Fungi(100);p__Ascomycota(100);c__Saccharomycetes(100);o__Saccharomycetales(100);f__Metschnikowiaceae(100);g__Metschnikowia(100);s__Metschnikowia_pulcherrima(100); | Metschnikowia | 13 | Saprotroph | \|Undefined Saprotroph\| | Yeast | NULL | Probable | As nectar-tap_saprotroph (Põlme et al. 2020); Decay sugar-rich substrates (Põlme et al. 2020) | Põlme S, et al. 2020. Fungal Diversity 105:1-16 ((https://doi.org/10.1007/s13225-020-00466-2)) |
| Otu30171 | 1 | k__Fungi(100);p__Ascomycota(100);c__Saccharomycetes(100);o__Saccharomycetales(100);f__Metschnikowiaceae(100);g__Metschnikowia(100);s__Metschnikowia_pulcherrima(100); | Metschnikowia | 13 | Saprotroph | \|Undefined Saprotroph\| | Yeast | NULL | Probable | As nectar-tap_saprotroph (Põlme et al. 2020); Decay sugar-rich substrates (Põlme et al. 2020) | Põlme S, et al. 2020. Fungal Diversity 105:1-16 ((https://doi.org/10.1007/s13225-020-00466-2)) |
| Otu30172 | 1 | k__Fungi(100);p__Ascomycota(100);c__Saccharomycetes(100);o__Saccharomycetales(100);f__Metschnikowiaceae(100);g__Metschnikowia(100);g__Metschnikowia_unclassified(100); | Metschnikowia | 13 | Saprotroph | \|Undefined Saprotroph\| | Yeast | NULL | Probable | As nectar-tap_saprotroph (Põlme et al. 2020); Decay sugar-rich substrates (Põlme et al. 2020) | Põlme S, et al. 2020. Fungal Diversity 105:1-16 ((https://doi.org/10.1007/s13225-020-00466-2)) |
| Otu30173 | 1 | k__Fungi(100);p__Ascomycota(100);c__Saccharomycetes(100);o__Saccharomycetales(100);f__Metschnikowiaceae(100);g__Metschnikowia(100);g__Metschnikowia_unclassified(100); | Metschnikowia | 13 | Saprotroph | \|Undefined Saprotroph\| | Yeast | NULL | Probable | As nectar-tap_saprotroph (Põlme et al. 2020); Decay sugar-rich substrates (Põlme et al. 2020) | Põlme S, et al. 2020. Fungal Diversity 105:1-16 ((https://doi.org/10.1007/s13225-020-00466-2)) |
| Otu30174 | 1 | k__Fungi(100);p__Ascomycota(100);c__Saccharomycetes(100);o__Saccharomycetales(100);f__Metschnikowiaceae(100);g__Metschnikowia(100);s__Metschnikowia_pulcherrima(100); | Metschnikowia | 13 | Saprotroph | \|Undefined Saprotroph\| | Yeast | NULL | Probable | As nectar-tap_saprotroph (Põlme et al. 2020); Decay sugar-rich substrates (Põlme et al. 2020) | Põlme S, et al. 2020. Fungal Diversity 105:1-16 ((https://doi.org/10.1007/s13225-020-00466-2)) |
| Otu30175 | 1 | k__Fungi(100);p__Ascomycota(100);c__Saccharomycetes(100);o__Saccharomycetales(100);f__Metschnikowiaceae(100);g__Metschnikowia(100);g__Metschnikowia_unclassified(100); | Metschnikowia | 13 | Saprotroph | \|Undefined Saprotroph\| | Yeast | NULL | Probable | As nectar-tap_saprotroph (Põlme et al. 2020); Decay sugar-rich substrates (Põlme et al. 2020) | Põlme S, et al. 2020. Fungal Diversity 105:1-16 ((https://doi.org/10.1007/s13225-020-00466-2)) |
| Otu30176 | 1 | k__Fungi(100);p__Ascomycota(100);c__Saccharomycetes(100);o__Saccharomycetales(100);f__Metschnikowiaceae(100);g__Metschnikowia(100);s__Metschnikowia_pulcherrima(100); | Metschnikowia | 13 | Saprotroph | \|Undefined Saprotroph\| | Yeast | NULL | Probable | As nectar-tap_saprotroph (Põlme et al. 2020); Decay sugar-rich substrates (Põlme et al. 2020) | Põlme S, et al. 2020. Fungal Diversity 105:1-16 ((https://doi.org/10.1007/s13225-020-00466-2)) |
| Otu30177 | 1 | k__Fungi(100);p__Ascomycota(100);c__Saccharomycetes(100);o__Saccharomycetales(100);f__Metschnikowiaceae(100);g__Metschnikowia(100);s__Metschnikowia_pulcherrima(100); | Metschnikowia | 13 | Saprotroph | \|Undefined Saprotroph\| | Yeast | NULL | Probable | As nectar-tap_saprotroph (Põlme et al. 2020); Decay sugar-rich substrates (Põlme et al. 2020) | Põlme S, et al. 2020. Fungal Diversity 105:1-16 ((https://doi.org/10.1007/s13225-020-00466-2)) |
| Otu30178 | 1 | k__Fungi(100);p__Ascomycota(100);c__Saccharomycetes(100);o__Saccharomycetales(100);f__Metschnikowiaceae(100);g__Metschnikowia(100);s__Metschnikowia_pulcherrima(100); | Metschnikowia | 13 | Saprotroph | \|Undefined Saprotroph\| | Yeast | NULL | Probable | As nectar-tap_saprotroph (Põlme et al. 2020); Decay sugar-rich substrates (Põlme et al. 2020) | Põlme S, et al. 2020. Fungal Diversity 105:1-16 ((https://doi.org/10.1007/s13225-020-00466-2)) |
| Otu30179 | 1 | k__Fungi(100);p__Ascomycota(100);c__Saccharomycetes(100);o__Saccharomycetales(100);f__Metschnikowiaceae(100);g__Metschnikowia(100);g__Metschnikowia_unclassified(100); | Metschnikowia | 13 | Saprotroph | \|Undefined Saprotroph\| | Yeast | NULL | Probable | As nectar-tap_saprotroph (Põlme et al. 2020); Decay sugar-rich substrates (Põlme et al. 2020) | Põlme S, et al. 2020. Fungal Diversity 105:1-16 ((https://doi.org/10.1007/s13225-020-00466-2)) |
| Otu30180 | 1 | k__Fungi(100);p__Ascomycota(100);c__Saccharomycetes(100);o__Saccharomycetales(100);f__Metschnikowiaceae(100);g__Metschnikowia(100);s__Metschnikowia_pulcherrima(100); | Metschnikowia | 13 | Saprotroph | \|Undefined Saprotroph\| | Yeast | NULL | Probable | As nectar-tap_saprotroph (Põlme et al. 2020); Decay sugar-rich substrates (Põlme et al. 2020) | Põlme S, et al. 2020. Fungal Diversity 105:1-16 ((https://doi.org/10.1007/s13225-020-00466-2)) |
| Otu30181 | 1 | k__Fungi(100);p__Ascomycota(100);c__Saccharomycetes(100);o__Saccharomycetales(100);f__Metschnikowiaceae(100);g__Metschnikowia(100);s__Metschnikowia_pulcherrima(100); | Metschnikowia | 13 | Saprotroph | \|Undefined Saprotroph\| | Yeast | NULL | Probable | As nectar-tap_saprotroph (Põlme et al. 2020); Decay sugar-rich substrates (Põlme et al. 2020) | Põlme S, et al. 2020. Fungal Diversity 105:1-16 ((https://doi.org/10.1007/s13225-020-00466-2)) |
| Otu30182 | 1 | k__Fungi(100);p__Ascomycota(100);c__Saccharomycetes(100);o__Saccharomycetales(100);f__Metschnikowiaceae(100);g__Metschnikowia(100);g__Metschnikowia_unclassified(100); | Metschnikowia | 13 | Saprotroph | \|Undefined Saprotroph\| | Yeast | NULL | Probable | As nectar-tap_saprotroph (Põlme et al. 2020); Decay sugar-rich substrates (Põlme et al. 2020) | Põlme S, et al. 2020. Fungal Diversity 105:1-16 ((https://doi.org/10.1007/s13225-020-00466-2)) |
| Otu30183 | 1 | k__Fungi(100);p__Ascomycota(100);c__Saccharomycetes(100);o__Saccharomycetales(100);f__Metschnikowiaceae(100);g__Metschnikowia(100);s__Metschnikowia_pulcherrima(100); | Metschnikowia | 13 | Saprotroph | \|Undefined Saprotroph\| | Yeast | NULL | Probable | As nectar-tap_saprotroph (Põlme et al. 2020); Decay sugar-rich substrates (Põlme et al. 2020) | Põlme S, et al. 2020. Fungal Diversity 105:1-16 ((https://doi.org/10.1007/s13225-020-00466-2)) |
| Otu30184 | 1 | k__Fungi(100);p__Ascomycota(100);c__Saccharomycetes(100);o__Saccharomycetales(100);f__Metschnikowiaceae(100);g__Metschnikowia(100);g__Metschnikowia_unclassified(100); | Metschnikowia | 13 | Saprotroph | \|Undefined Saprotroph\| | Yeast | NULL | Probable | As nectar-tap_saprotroph (Põlme et al. 2020); Decay sugar-rich substrates (Põlme et al. 2020) | Põlme S, et al. 2020. Fungal Diversity 105:1-16 ((https://doi.org/10.1007/s13225-020-00466-2)) |
| Otu30185 | 1 | k__Fungi(100);p__Ascomycota(100);c__Saccharomycetes(100);o__Saccharomycetales(100);f__Metschnikowiaceae(100);g__Metschnikowia(100);s__Metschnikowia_pulcherrima(100); | Metschnikowia | 13 | Saprotroph | \|Undefined Saprotroph\| | Yeast | NULL | Probable | As nectar-tap_saprotroph (Põlme et al. 2020); Decay sugar-rich substrates (Põlme et al. 2020) | Põlme S, et al. 2020. Fungal Diversity 105:1-16 ((https://doi.org/10.1007/s13225-020-00466-2)) |
| Otu30186 | 1 | k__Fungi(100);p__Ascomycota(100);c__Saccharomycetes(100);o__Saccharomycetales(100);f__Metschnikowiaceae(100);g__Metschnikowia(100);s__Metschnikowia_pulcherrima(100); | Metschnikowia | 13 | Saprotroph | \|Undefined Saprotroph\| | Yeast | NULL | Probable | As nectar-tap_saprotroph (Põlme et al. 2020); Decay sugar-rich substrates (Põlme et al. 2020) | Põlme S, et al. 2020. Fungal Diversity 105:1-16 ((https://doi.org/10.1007/s13225-020-00466-2)) |
| Otu30187 | 1 | k__Fungi(100);p__Ascomycota(100);c__Saccharomycetes(100);o__Saccharomycetales(100);f__Metschnikowiaceae(100);g__Metschnikowia(100);s__Metschnikowia_pulcherrima(100); | Metschnikowia | 13 | Saprotroph | \|Undefined Saprotroph\| | Yeast | NULL | Probable | As nectar-tap_saprotroph (Põlme et al. 2020); Decay sugar-rich substrates (Põlme et al. 2020) | Põlme S, et al. 2020. Fungal Diversity 105:1-16 ((https://doi.org/10.1007/s13225-020-00466-2)) |
| Otu30188 | 1 | k__Fungi(100);p__Ascomycota(100);c__Saccharomycetes(100);o__Saccharomycetales(100);f__Metschnikowiaceae(100);g__Metschnikowia(100);s__Metschnikowia_pulcherrima(100); | Metschnikowia | 13 | Saprotroph | \|Undefined Saprotroph\| | Yeast | NULL | Probable | As nectar-tap_saprotroph (Põlme et al. 2020); Decay sugar-rich substrates (Põlme et al. 2020) | Põlme S, et al. 2020. Fungal Diversity 105:1-16 ((https://doi.org/10.1007/s13225-020-00466-2)) |
| Otu30189 | 1 | k__Fungi(100);p__Ascomycota(100);c__Saccharomycetes(100);o__Saccharomycetales(100);f__Metschnikowiaceae(100);g__Metschnikowia(100);s__Metschnikowia_pulcherrima(100); | Metschnikowia | 13 | Saprotroph | \|Undefined Saprotroph\| | Yeast | NULL | Probable | As nectar-tap_saprotroph (Põlme et al. 2020); Decay sugar-rich substrates (Põlme et al. 2020) | Põlme S, et al. 2020. Fungal Diversity 105:1-16 ((https://doi.org/10.1007/s13225-020-00466-2)) |
| Otu30190 | 1 | k__Fungi(100);p__Ascomycota(100);c__Saccharomycetes(100);o__Saccharomycetales(100);f__Metschnikowiaceae(100);g__Metschnikowia(100);g__Metschnikowia_unclassified(100); | Metschnikowia | 13 | Saprotroph | \|Undefined Saprotroph\| | Yeast | NULL | Probable | As nectar-tap_saprotroph (Põlme et al. 2020); Decay sugar-rich substrates (Põlme et al. 2020) | Põlme S, et al. 2020. Fungal Diversity 105:1-16 ((https://doi.org/10.1007/s13225-020-00466-2)) |
| Otu30192 | 1 | k__Fungi(100);p__Ascomycota(100);c__Saccharomycetes(100);o__Saccharomycetales(100);f__Metschnikowiaceae(100);g__Metschnikowia(100);s__Metschnikowia_pulcherrima(100); | Metschnikowia | 13 | Saprotroph | \|Undefined Saprotroph\| | Yeast | NULL | Probable | As nectar-tap_saprotroph (Põlme et al. 2020); Decay sugar-rich substrates (Põlme et al. 2020) | Põlme S, et al. 2020. Fungal Diversity 105:1-16 ((https://doi.org/10.1007/s13225-020-00466-2)) |
| Otu30193 | 1 | k__Fungi(100);p__Ascomycota(100);c__Saccharomycetes(100);o__Saccharomycetales(100);f__Metschnikowiaceae(100);g__Metschnikowia(100);g__Metschnikowia_unclassified(100); | Metschnikowia | 13 | Saprotroph | \|Undefined Saprotroph\| | Yeast | NULL | Probable | As nectar-tap_saprotroph (Põlme et al. 2020); Decay sugar-rich substrates (Põlme et al. 2020) | Põlme S, et al. 2020. Fungal Diversity 105:1-16 ((https://doi.org/10.1007/s13225-020-00466-2)) |
| Otu30194 | 1 | k__Fungi(100);p__Ascomycota(100);c__Saccharomycetes(100);o__Saccharomycetales(100);f__Metschnikowiaceae(100);g__Metschnikowia(100);s__Metschnikowia_pulcherrima(100); | Metschnikowia | 13 | Saprotroph | \|Undefined Saprotroph\| | Yeast | NULL | Probable | As nectar-tap_saprotroph (Põlme et al. 2020); Decay sugar-rich substrates (Põlme et al. 2020) | Põlme S, et al. 2020. Fungal Diversity 105:1-16 ((https://doi.org/10.1007/s13225-020-00466-2)) |
| Otu30195 | 1 | k__Fungi(100);p__Basidiomycota(100);c__Microbotryomycetes(100);o__Sporidiobolales(100);f__Sporidiobolales_family_Incertae_sedis(100);g__Rhodotorula(100);s__Rhodotorula_taiwanensis(100); | Rhodotorula | 13 | Pathotroph-Saprotroph-Symbiotroph | Animal Pathogen-Animal Symbiotroph-Endophyte-Plant Pathogen-\|Undefined Saprotroph\| | Yeast | NULL | Probable | As foliar_endophyte (Põlme et al. 2020); Opportunistic human pathogen | Põlme S, et al. 2020. Fungal Diversity 105:1-16 ((https://doi.org/10.1007/s13225-020-00466-2)); https://www.cdc.gov ((https://www.cdc.gov)) |
| Otu30196 | 1 | k__Fungi(100);p__Ascomycota(100);c__Saccharomycetes(100);o__Saccharomycetales(100);f__Metschnikowiaceae(100);g__Metschnikowia(100);s__Metschnikowia_pulcherrima(100); | Metschnikowia | 13 | Saprotroph | \|Undefined Saprotroph\| | Yeast | NULL | Probable | As nectar-tap_saprotroph (Põlme et al. 2020); Decay sugar-rich substrates (Põlme et al. 2020) | Põlme S, et al. 2020. Fungal Diversity 105:1-16 ((https://doi.org/10.1007/s13225-020-00466-2)) |
| Otu30197 | 1 | k__Fungi(100);p__Ascomycota(100);c__Saccharomycetes(100);o__Saccharomycetales(100);f__Metschnikowiaceae(100);g__Metschnikowia(100);s__Metschnikowia_pulcherrima(100); | Metschnikowia | 13 | Saprotroph | \|Undefined Saprotroph\| | Yeast | NULL | Probable | As nectar-tap_saprotroph (Põlme et al. 2020); Decay sugar-rich substrates (Põlme et al. 2020) | Põlme S, et al. 2020. Fungal Diversity 105:1-16 ((https://doi.org/10.1007/s13225-020-00466-2)) |
| Otu30199 | 1 | k__Fungi(100);p__Ascomycota(100);c__Saccharomycetes(100);o__Saccharomycetales(100);f__Metschnikowiaceae(100);g__Metschnikowia(100);s__Metschnikowia_pulcherrima(100); | Metschnikowia | 13 | Saprotroph | \|Undefined Saprotroph\| | Yeast | NULL | Probable | As nectar-tap_saprotroph (Põlme et al. 2020); Decay sugar-rich substrates (Põlme et al. 2020) | Põlme S, et al. 2020. Fungal Diversity 105:1-16 ((https://doi.org/10.1007/s13225-020-00466-2)) |
| Otu30200 | 1 | k__Fungi(100);p__Ascomycota(100);c__Saccharomycetes(100);o__Saccharomycetales(100);f__Metschnikowiaceae(100);g__Metschnikowia(100);g__Metschnikowia_unclassified(100); | Metschnikowia | 13 | Saprotroph | \|Undefined Saprotroph\| | Yeast | NULL | Probable | As nectar-tap_saprotroph (Põlme et al. 2020); Decay sugar-rich substrates (Põlme et al. 2020) | Põlme S, et al. 2020. Fungal Diversity 105:1-16 ((https://doi.org/10.1007/s13225-020-00466-2)) |
| Otu30201 | 1 | k__Fungi(100);p__Ascomycota(100);c__Saccharomycetes(100);o__Saccharomycetales(100);f__Metschnikowiaceae(100);g__Metschnikowia(100);s__Metschnikowia_pulcherrima(100); | Metschnikowia | 13 | Saprotroph | \|Undefined Saprotroph\| | Yeast | NULL | Probable | As nectar-tap_saprotroph (Põlme et al. 2020); Decay sugar-rich substrates (Põlme et al. 2020) | Põlme S, et al. 2020. Fungal Diversity 105:1-16 ((https://doi.org/10.1007/s13225-020-00466-2)) |
| Otu30202 | 1 | k__Fungi(100);p__Ascomycota(100);c__Saccharomycetes(100);o__Saccharomycetales(100);f__Metschnikowiaceae(100);g__Metschnikowia(100);s__Metschnikowia_pulcherrima(100); | Metschnikowia | 13 | Saprotroph | \|Undefined Saprotroph\| | Yeast | NULL | Probable | As nectar-tap_saprotroph (Põlme et al. 2020); Decay sugar-rich substrates (Põlme et al. 2020) | Põlme S, et al. 2020. Fungal Diversity 105:1-16 ((https://doi.org/10.1007/s13225-020-00466-2)) |
| Otu30203 | 1 | k__Fungi(100);p__Ascomycota(100);c__Saccharomycetes(100);o__Saccharomycetales(100);f__Metschnikowiaceae(100);g__Metschnikowia(100);s__Metschnikowia_pulcherrima(100); | Metschnikowia | 13 | Saprotroph | \|Undefined Saprotroph\| | Yeast | NULL | Probable | As nectar-tap_saprotroph (Põlme et al. 2020); Decay sugar-rich substrates (Põlme et al. 2020) | Põlme S, et al. 2020. Fungal Diversity 105:1-16 ((https://doi.org/10.1007/s13225-020-00466-2)) |
| Otu30204 | 1 | k__Fungi(100);p__Ascomycota(100);c__Saccharomycetes(100);o__Saccharomycetales(100);f__Metschnikowiaceae(100);g__Metschnikowia(100);s__Metschnikowia_pulcherrima(100); | Metschnikowia | 13 | Saprotroph | \|Undefined Saprotroph\| | Yeast | NULL | Probable | As nectar-tap_saprotroph (Põlme et al. 2020); Decay sugar-rich substrates (Põlme et al. 2020) | Põlme S, et al. 2020. Fungal Diversity 105:1-16 ((https://doi.org/10.1007/s13225-020-00466-2)) |
| Otu30205 | 1 | k__Fungi(100);p__Ascomycota(100);c__Saccharomycetes(100);o__Saccharomycetales(100);f__Metschnikowiaceae(100);g__Metschnikowia(100);g__Metschnikowia_unclassified(100); | Metschnikowia | 13 | Saprotroph | \|Undefined Saprotroph\| | Yeast | NULL | Probable | As nectar-tap_saprotroph (Põlme et al. 2020); Decay sugar-rich substrates (Põlme et al. 2020) | Põlme S, et al. 2020. Fungal Diversity 105:1-16 ((https://doi.org/10.1007/s13225-020-00466-2)) |
| Otu30207 | 1 | k__Fungi(100);p__Ascomycota(100);c__Saccharomycetes(100);o__Saccharomycetales(100);f__Metschnikowiaceae(100);g__Metschnikowia(100);g__Metschnikowia_unclassified(100); | Metschnikowia | 13 | Saprotroph | \|Undefined Saprotroph\| | Yeast | NULL | Probable | As nectar-tap_saprotroph (Põlme et al. 2020); Decay sugar-rich substrates (Põlme et al. 2020) | Põlme S, et al. 2020. Fungal Diversity 105:1-16 ((https://doi.org/10.1007/s13225-020-00466-2)) |
| Otu30208 | 1 | k__Fungi(100);p__Ascomycota(100);c__Saccharomycetes(100);o__Saccharomycetales(100);f__Metschnikowiaceae(100);g__Metschnikowia(100);g__Metschnikowia_unclassified(100); | Metschnikowia | 13 | Saprotroph | \|Undefined Saprotroph\| | Yeast | NULL | Probable | As nectar-tap_saprotroph (Põlme et al. 2020); Decay sugar-rich substrates (Põlme et al. 2020) | Põlme S, et al. 2020. Fungal Diversity 105:1-16 ((https://doi.org/10.1007/s13225-020-00466-2)) |
| Otu30209 | 1 | k__Fungi(100);p__Ascomycota(100);c__Saccharomycetes(100);o__Saccharomycetales(100);f__Metschnikowiaceae(100);g__Metschnikowia(100);s__Metschnikowia_pulcherrima(100); | Metschnikowia | 13 | Saprotroph | \|Undefined Saprotroph\| | Yeast | NULL | Probable | As nectar-tap_saprotroph (Põlme et al. 2020); Decay sugar-rich substrates (Põlme et al. 2020) | Põlme S, et al. 2020. Fungal Diversity 105:1-16 ((https://doi.org/10.1007/s13225-020-00466-2)) |
| Otu30210 | 1 | k__Fungi(100);p__Ascomycota(100);c__Saccharomycetes(100);o__Saccharomycetales(100);f__Metschnikowiaceae(100);g__Metschnikowia(100);g__Metschnikowia_unclassified(100); | Metschnikowia | 13 | Saprotroph | \|Undefined Saprotroph\| | Yeast | NULL | Probable | As nectar-tap_saprotroph (Põlme et al. 2020); Decay sugar-rich substrates (Põlme et al. 2020) | Põlme S, et al. 2020. Fungal Diversity 105:1-16 ((https://doi.org/10.1007/s13225-020-00466-2)) |
| Otu30211 | 1 | k__Fungi(100);p__Ascomycota(100);c__Saccharomycetes(100);o__Saccharomycetales(100);f__Metschnikowiaceae(100);g__Metschnikowia(100);s__Metschnikowia_pulcherrima(100); | Metschnikowia | 13 | Saprotroph | \|Undefined Saprotroph\| | Yeast | NULL | Probable | As nectar-tap_saprotroph (Põlme et al. 2020); Decay sugar-rich substrates (Põlme et al. 2020) | Põlme S, et al. 2020. Fungal Diversity 105:1-16 ((https://doi.org/10.1007/s13225-020-00466-2)) |
| Otu30212 | 1 | k__Fungi(100);p__Ascomycota(100);c__Saccharomycetes(100);o__Saccharomycetales(100);f__Metschnikowiaceae(100);g__Metschnikowia(100);s__Metschnikowia_pulcherrima(100); | Metschnikowia | 13 | Saprotroph | \|Undefined Saprotroph\| | Yeast | NULL | Probable | As nectar-tap_saprotroph (Põlme et al. 2020); Decay sugar-rich substrates (Põlme et al. 2020) | Põlme S, et al. 2020. Fungal Diversity 105:1-16 ((https://doi.org/10.1007/s13225-020-00466-2)) |
| Otu30213 | 1 | k__Fungi(100);p__Ascomycota(100);c__Saccharomycetes(100);o__Saccharomycetales(100);f__Metschnikowiaceae(100);g__Metschnikowia(100);s__Metschnikowia_pulcherrima(100); | Metschnikowia | 13 | Saprotroph | \|Undefined Saprotroph\| | Yeast | NULL | Probable | As nectar-tap_saprotroph (Põlme et al. 2020); Decay sugar-rich substrates (Põlme et al. 2020) | Põlme S, et al. 2020. Fungal Diversity 105:1-16 ((https://doi.org/10.1007/s13225-020-00466-2)) |
| Otu30214 | 1 | k__Fungi(100);p__Ascomycota(100);c__Saccharomycetes(100);o__Saccharomycetales(100);f__Metschnikowiaceae(100);g__Metschnikowia(100);s__Metschnikowia_pulcherrima(100); | Metschnikowia | 13 | Saprotroph | \|Undefined Saprotroph\| | Yeast | NULL | Probable | As nectar-tap_saprotroph (Põlme et al. 2020); Decay sugar-rich substrates (Põlme et al. 2020) | Põlme S, et al. 2020. Fungal Diversity 105:1-16 ((https://doi.org/10.1007/s13225-020-00466-2)) |
| Otu30215 | 1 | k__Fungi(100);p__Ascomycota(100);c__Saccharomycetes(100);o__Saccharomycetales(100);f__Metschnikowiaceae(100);g__Metschnikowia(100);s__Metschnikowia_pulcherrima(100); | Metschnikowia | 13 | Saprotroph | \|Undefined Saprotroph\| | Yeast | NULL | Probable | As nectar-tap_saprotroph (Põlme et al. 2020); Decay sugar-rich substrates (Põlme et al. 2020) | Põlme S, et al. 2020. Fungal Diversity 105:1-16 ((https://doi.org/10.1007/s13225-020-00466-2)) |
| Otu30216 | 1 | k__Fungi(100);p__Ascomycota(100);c__Saccharomycetes(100);o__Saccharomycetales(100);f__Metschnikowiaceae(100);g__Metschnikowia(100);s__Metschnikowia_pulcherrima(100); | Metschnikowia | 13 | Saprotroph | \|Undefined Saprotroph\| | Yeast | NULL | Probable | As nectar-tap_saprotroph (Põlme et al. 2020); Decay sugar-rich substrates (Põlme et al. 2020) | Põlme S, et al. 2020. Fungal Diversity 105:1-16 ((https://doi.org/10.1007/s13225-020-00466-2)) |
| Otu30217 | 1 | k__Fungi(100);p__Ascomycota(100);c__Saccharomycetes(100);o__Saccharomycetales(100);f__Metschnikowiaceae(100);g__Metschnikowia(100);s__Metschnikowia_pulcherrima(100); | Metschnikowia | 13 | Saprotroph | \|Undefined Saprotroph\| | Yeast | NULL | Probable | As nectar-tap_saprotroph (Põlme et al. 2020); Decay sugar-rich substrates (Põlme et al. 2020) | Põlme S, et al. 2020. Fungal Diversity 105:1-16 ((https://doi.org/10.1007/s13225-020-00466-2)) |
| Otu30218 | 1 | k__Fungi(100);p__Ascomycota(100);c__Saccharomycetes(100);o__Saccharomycetales(100);f__Metschnikowiaceae(100);g__Metschnikowia(100);s__Metschnikowia_pulcherrima(100); | Metschnikowia | 13 | Saprotroph | \|Undefined Saprotroph\| | Yeast | NULL | Probable | As nectar-tap_saprotroph (Põlme et al. 2020); Decay sugar-rich substrates (Põlme et al. 2020) | Põlme S, et al. 2020. Fungal Diversity 105:1-16 ((https://doi.org/10.1007/s13225-020-00466-2)) |
| Otu30219 | 1 | k__Fungi(100);p__Ascomycota(100);c__Saccharomycetes(100);o__Saccharomycetales(100);f__Metschnikowiaceae(100);g__Metschnikowia(100);s__Metschnikowia_pulcherrima(100); | Metschnikowia | 13 | Saprotroph | \|Undefined Saprotroph\| | Yeast | NULL | Probable | As nectar-tap_saprotroph (Põlme et al. 2020); Decay sugar-rich substrates (Põlme et al. 2020) | Põlme S, et al. 2020. Fungal Diversity 105:1-16 ((https://doi.org/10.1007/s13225-020-00466-2)) |
| Otu30220 | 1 | k__Fungi(100);p__Ascomycota(100);c__Saccharomycetes(100);o__Saccharomycetales(100);f__Metschnikowiaceae(100);g__Metschnikowia(100);s__Metschnikowia_pulcherrima(100); | Metschnikowia | 13 | Saprotroph | \|Undefined Saprotroph\| | Yeast | NULL | Probable | As nectar-tap_saprotroph (Põlme et al. 2020); Decay sugar-rich substrates (Põlme et al. 2020) | Põlme S, et al. 2020. Fungal Diversity 105:1-16 ((https://doi.org/10.1007/s13225-020-00466-2)) |
| Otu30221 | 1 | k__Fungi(100);p__Ascomycota(100);c__Saccharomycetes(100);o__Saccharomycetales(100);f__Metschnikowiaceae(100);g__Metschnikowia(100);s__Metschnikowia_pulcherrima(100); | Metschnikowia | 13 | Saprotroph | \|Undefined Saprotroph\| | Yeast | NULL | Probable | As nectar-tap_saprotroph (Põlme et al. 2020); Decay sugar-rich substrates (Põlme et al. 2020) | Põlme S, et al. 2020. Fungal Diversity 105:1-16 ((https://doi.org/10.1007/s13225-020-00466-2)) |
| Otu30222 | 1 | k__Fungi(100);p__Ascomycota(100);c__Saccharomycetes(100);o__Saccharomycetales(100);f__Metschnikowiaceae(100);g__Metschnikowia(100);s__Metschnikowia_pulcherrima(100); | Metschnikowia | 13 | Saprotroph | \|Undefined Saprotroph\| | Yeast | NULL | Probable | As nectar-tap_saprotroph (Põlme et al. 2020); Decay sugar-rich substrates (Põlme et al. 2020) | Põlme S, et al. 2020. Fungal Diversity 105:1-16 ((https://doi.org/10.1007/s13225-020-00466-2)) |
| Otu30223 | 1 | k__Fungi(100);p__Ascomycota(100);c__Saccharomycetes(100);o__Saccharomycetales(100);f__Metschnikowiaceae(100);g__Metschnikowia(100);s__Metschnikowia_pulcherrima(100); | Metschnikowia | 13 | Saprotroph | \|Undefined Saprotroph\| | Yeast | NULL | Probable | As nectar-tap_saprotroph (Põlme et al. 2020); Decay sugar-rich substrates (Põlme et al. 2020) | Põlme S, et al. 2020. Fungal Diversity 105:1-16 ((https://doi.org/10.1007/s13225-020-00466-2)) |
| Otu30224 | 1 | k__Fungi(100);p__Ascomycota(100);c__Saccharomycetes(100);o__Saccharomycetales(100);f__Metschnikowiaceae(100);g__Metschnikowia(100);s__Metschnikowia_pulcherrima(100); | Metschnikowia | 13 | Saprotroph | \|Undefined Saprotroph\| | Yeast | NULL | Probable | As nectar-tap_saprotroph (Põlme et al. 2020); Decay sugar-rich substrates (Põlme et al. 2020) | Põlme S, et al. 2020. Fungal Diversity 105:1-16 ((https://doi.org/10.1007/s13225-020-00466-2)) |
| Otu30225 | 1 | k__Fungi(100);p__Ascomycota(100);c__Saccharomycetes(100);o__Saccharomycetales(100);f__Metschnikowiaceae(100);g__Metschnikowia(100);s__Metschnikowia_pulcherrima(100); | Metschnikowia | 13 | Saprotroph | \|Undefined Saprotroph\| | Yeast | NULL | Probable | As nectar-tap_saprotroph (Põlme et al. 2020); Decay sugar-rich substrates (Põlme et al. 2020) | Põlme S, et al. 2020. Fungal Diversity 105:1-16 ((https://doi.org/10.1007/s13225-020-00466-2)) |
| Otu30226 | 1 | k__Fungi(100);p__Ascomycota(100);c__Saccharomycetes(100);o__Saccharomycetales(100);f__Metschnikowiaceae(100);g__Metschnikowia(100);s__Metschnikowia_pulcherrima(100); | Metschnikowia | 13 | Saprotroph | \|Undefined Saprotroph\| | Yeast | NULL | Probable | As nectar-tap_saprotroph (Põlme et al. 2020); Decay sugar-rich substrates (Põlme et al. 2020) | Põlme S, et al. 2020. Fungal Diversity 105:1-16 ((https://doi.org/10.1007/s13225-020-00466-2)) |
| Otu30227 | 1 | k__Fungi(100);p__Ascomycota(100);c__Saccharomycetes(100);o__Saccharomycetales(100);f__Metschnikowiaceae(100);g__Metschnikowia(100);s__Metschnikowia_pulcherrima(100); | Metschnikowia | 13 | Saprotroph | \|Undefined Saprotroph\| | Yeast | NULL | Probable | As nectar-tap_saprotroph (Põlme et al. 2020); Decay sugar-rich substrates (Põlme et al. 2020) | Põlme S, et al. 2020. Fungal Diversity 105:1-16 ((https://doi.org/10.1007/s13225-020-00466-2)) |
| Otu30229 | 1 | k__Fungi(100);p__Ascomycota(100);c__Saccharomycetes(100);o__Saccharomycetales(100);f__Metschnikowiaceae(100);g__Metschnikowia(100);s__Metschnikowia_pulcherrima(100); | Metschnikowia | 13 | Saprotroph | \|Undefined Saprotroph\| | Yeast | NULL | Probable | As nectar-tap_saprotroph (Põlme et al. 2020); Decay sugar-rich substrates (Põlme et al. 2020) | Põlme S, et al. 2020. Fungal Diversity 105:1-16 ((https://doi.org/10.1007/s13225-020-00466-2)) |
| Otu30230 | 1 | k__Fungi(100);p__Ascomycota(100);c__Saccharomycetes(100);o__Saccharomycetales(100);f__Metschnikowiaceae(100);g__Metschnikowia(100);g__Metschnikowia_unclassified(100); | Metschnikowia | 13 | Saprotroph | \|Undefined Saprotroph\| | Yeast | NULL | Probable | As nectar-tap_saprotroph (Põlme et al. 2020); Decay sugar-rich substrates (Põlme et al. 2020) | Põlme S, et al. 2020. Fungal Diversity 105:1-16 ((https://doi.org/10.1007/s13225-020-00466-2)) |
| Otu30231 | 1 | k__Fungi(100);p__Ascomycota(100);c__Saccharomycetes(100);o__Saccharomycetales(100);f__Metschnikowiaceae(100);g__Metschnikowia(100);s__Metschnikowia_pulcherrima(100); | Metschnikowia | 13 | Saprotroph | \|Undefined Saprotroph\| | Yeast | NULL | Probable | As nectar-tap_saprotroph (Põlme et al. 2020); Decay sugar-rich substrates (Põlme et al. 2020) | Põlme S, et al. 2020. Fungal Diversity 105:1-16 ((https://doi.org/10.1007/s13225-020-00466-2)) |
| Otu30232 | 1 | k__Fungi(100);p__Ascomycota(100);c__Saccharomycetes(100);o__Saccharomycetales(100);f__Metschnikowiaceae(100);g__Metschnikowia(100);s__Metschnikowia_pulcherrima(100); | Metschnikowia | 13 | Saprotroph | \|Undefined Saprotroph\| | Yeast | NULL | Probable | As nectar-tap_saprotroph (Põlme et al. 2020); Decay sugar-rich substrates (Põlme et al. 2020) | Põlme S, et al. 2020. Fungal Diversity 105:1-16 ((https://doi.org/10.1007/s13225-020-00466-2)) |
| Otu30233 | 1 | k__Fungi(100);p__Ascomycota(100);c__Saccharomycetes(100);o__Saccharomycetales(100);f__Metschnikowiaceae(100);g__Metschnikowia(100);s__Metschnikowia_pulcherrima(100); | Metschnikowia | 13 | Saprotroph | \|Undefined Saprotroph\| | Yeast | NULL | Probable | As nectar-tap_saprotroph (Põlme et al. 2020); Decay sugar-rich substrates (Põlme et al. 2020) | Põlme S, et al. 2020. Fungal Diversity 105:1-16 ((https://doi.org/10.1007/s13225-020-00466-2)) |
| Otu30234 | 1 | k__Fungi(100);p__Ascomycota(100);c__Saccharomycetes(100);o__Saccharomycetales(100);f__Metschnikowiaceae(100);g__Metschnikowia(100);s__Metschnikowia_pulcherrima(100); | Metschnikowia | 13 | Saprotroph | \|Undefined Saprotroph\| | Yeast | NULL | Probable | As nectar-tap_saprotroph (Põlme et al. 2020); Decay sugar-rich substrates (Põlme et al. 2020) | Põlme S, et al. 2020. Fungal Diversity 105:1-16 ((https://doi.org/10.1007/s13225-020-00466-2)) |
| Otu30235 | 1 | k__Fungi(100);p__Ascomycota(100);c__Saccharomycetes(100);o__Saccharomycetales(100);f__Metschnikowiaceae(100);g__Metschnikowia(100);s__Metschnikowia_pulcherrima(100); | Metschnikowia | 13 | Saprotroph | \|Undefined Saprotroph\| | Yeast | NULL | Probable | As nectar-tap_saprotroph (Põlme et al. 2020); Decay sugar-rich substrates (Põlme et al. 2020) | Põlme S, et al. 2020. Fungal Diversity 105:1-16 ((https://doi.org/10.1007/s13225-020-00466-2)) |
| Otu30236 | 1 | k__Fungi(100);p__Ascomycota(100);c__Saccharomycetes(100);o__Saccharomycetales(100);f__Metschnikowiaceae(100);g__Metschnikowia(100);s__Metschnikowia_pulcherrima(100); | Metschnikowia | 13 | Saprotroph | \|Undefined Saprotroph\| | Yeast | NULL | Probable | As nectar-tap_saprotroph (Põlme et al. 2020); Decay sugar-rich substrates (Põlme et al. 2020) | Põlme S, et al. 2020. Fungal Diversity 105:1-16 ((https://doi.org/10.1007/s13225-020-00466-2)) |
| Otu30237 | 1 | k__Fungi(100);p__Ascomycota(100);c__Saccharomycetes(100);o__Saccharomycetales(100);f__Metschnikowiaceae(100);g__Metschnikowia(100);s__Metschnikowia_pulcherrima(100); | Metschnikowia | 13 | Saprotroph | \|Undefined Saprotroph\| | Yeast | NULL | Probable | As nectar-tap_saprotroph (Põlme et al. 2020); Decay sugar-rich substrates (Põlme et al. 2020) | Põlme S, et al. 2020. Fungal Diversity 105:1-16 ((https://doi.org/10.1007/s13225-020-00466-2)) |
| Otu30238 | 1 | k__Fungi(100);p__Ascomycota(100);c__Saccharomycetes(100);o__Saccharomycetales(100);f__Metschnikowiaceae(100);g__Metschnikowia(100);s__Metschnikowia_pulcherrima(100); | Metschnikowia | 13 | Saprotroph | \|Undefined Saprotroph\| | Yeast | NULL | Probable | As nectar-tap_saprotroph (Põlme et al. 2020); Decay sugar-rich substrates (Põlme et al. 2020) | Põlme S, et al. 2020. Fungal Diversity 105:1-16 ((https://doi.org/10.1007/s13225-020-00466-2)) |
| Otu30239 | 1 | k__Fungi(100);p__Ascomycota(100);c__Saccharomycetes(100);o__Saccharomycetales(100);f__Metschnikowiaceae(100);g__Metschnikowia(100);s__Metschnikowia_pulcherrima(100); | Metschnikowia | 13 | Saprotroph | \|Undefined Saprotroph\| | Yeast | NULL | Probable | As nectar-tap_saprotroph (Põlme et al. 2020); Decay sugar-rich substrates (Põlme et al. 2020) | Põlme S, et al. 2020. Fungal Diversity 105:1-16 ((https://doi.org/10.1007/s13225-020-00466-2)) |
| Otu30242 | 1 | k__Fungi(100);p__Ascomycota(100);c__Saccharomycetes(100);o__Saccharomycetales(100);o__Saccharomycetales_unclassified(100);o__Saccharomycetales_unclassified(100);o__Saccharomycetales_unclassified(100); | Saccharomycetales | 7 | Saprotroph | Undefined Saprotroph | Yeast | NULL | Possible | NULL | Starkenburg S, et al. 2015. New Phytologist 207:1145-1158 ((https://doi.org/10.1111/nph.13426)) |
| Otu30245 | 1 | k__Fungi(100);p__Ascomycota(100);c__Saccharomycetes(100);o__Saccharomycetales(100);f__Metschnikowiaceae(100);g__Metschnikowia(100);s__Metschnikowia_pulcherrima(100); | Metschnikowia | 13 | Saprotroph | \|Undefined Saprotroph\| | Yeast | NULL | Probable | As nectar-tap_saprotroph (Põlme et al. 2020); Decay sugar-rich substrates (Põlme et al. 2020) | Põlme S, et al. 2020. Fungal Diversity 105:1-16 ((https://doi.org/10.1007/s13225-020-00466-2)) |
| Otu30246 | 1 | k__Fungi(100);p__Ascomycota(100);c__Saccharomycetes(100);o__Saccharomycetales(100);f__Metschnikowiaceae(100);g__Metschnikowia(100);s__Metschnikowia_pulcherrima(100); | Metschnikowia | 13 | Saprotroph | \|Undefined Saprotroph\| | Yeast | NULL | Probable | As nectar-tap_saprotroph (Põlme et al. 2020); Decay sugar-rich substrates (Põlme et al. 2020) | Põlme S, et al. 2020. Fungal Diversity 105:1-16 ((https://doi.org/10.1007/s13225-020-00466-2)) |
| Otu30247 | 1 | k__Fungi(100);p__Ascomycota(100);c__Saccharomycetes(100);o__Saccharomycetales(100);f__Metschnikowiaceae(100);g__Metschnikowia(100);g__Metschnikowia_unclassified(100); | Metschnikowia | 13 | Saprotroph | \|Undefined Saprotroph\| | Yeast | NULL | Probable | As nectar-tap_saprotroph (Põlme et al. 2020); Decay sugar-rich substrates (Põlme et al. 2020) | Põlme S, et al. 2020. Fungal Diversity 105:1-16 ((https://doi.org/10.1007/s13225-020-00466-2)) |
| Otu30248 | 1 | k__Fungi(100);p__Ascomycota(100);c__Saccharomycetes(100);o__Saccharomycetales(100);f__Metschnikowiaceae(100);g__Metschnikowia(100);s__Metschnikowia_pulcherrima(100); | Metschnikowia | 13 | Saprotroph | \|Undefined Saprotroph\| | Yeast | NULL | Probable | As nectar-tap_saprotroph (Põlme et al. 2020); Decay sugar-rich substrates (Põlme et al. 2020) | Põlme S, et al. 2020. Fungal Diversity 105:1-16 ((https://doi.org/10.1007/s13225-020-00466-2)) |

| | | | | | | | | | | | |
|---|---|---|---|---|---|---|---|---|---|---|---|
| Otu30249 | 1 | k__Fungi(100);p__Ascomycota(100);c__Saccharomycetes(100);o__Saccharomycetales(100);f__Metschnikowiaceae(100);g__Metschnikowia(100);s__Metschnikowia_pulcherrima(100); | Metschnikowia | 13 | Saprotroph | \|Undefined Saprotroph\| | Yeast | NULL | Probable | As nectar-tap_saprotroph (Põlme et al. 2020); Decay sugar-rich substrates (Põlme et al. 2020) | Põlme S, et al. 2020. Fungal Diversity 105:1-16 ((https://doi.org/10.1007/s13225-020-00466-2)) |
| Otu30250 | 1 | k__Fungi(100);p__Ascomycota(100);c__Saccharomycetes(100);o__Saccharomycetales(100);f__Metschnikowiaceae(100);g__Metschnikowia(100);s__Metschnikowia_pulcherrima(100); | Metschnikowia | 13 | Saprotroph | \|Undefined Saprotroph\| | Yeast | NULL | Probable | As nectar-tap_saprotroph (Põlme et al. 2020); Decay sugar-rich substrates (Põlme et al. 2020) | Põlme S, et al. 2020. Fungal Diversity 105:1-16 ((https://doi.org/10.1007/s13225-020-00466-2)) |
| Otu30251 | 1 | k__Fungi(100);p__Ascomycota(100);c__Saccharomycetes(100);o__Saccharomycetales(100);f__Metschnikowiaceae(100);g__Metschnikowia(100);s__Metschnikowia_pulcherrima(100); | Metschnikowia | 13 | Saprotroph | \|Undefined Saprotroph\| | Yeast | NULL | Probable | As nectar-tap_saprotroph (Põlme et al. 2020); Decay sugar-rich substrates (Põlme et al. 2020) | Põlme S, et al. 2020. Fungal Diversity 105:1-16 ((https://doi.org/10.1007/s13225-020-00466-2)) |
| Otu30252 | 1 | k__Fungi(100);p__Ascomycota(100);c__Saccharomycetes(100);o__Saccharomycetales(100);f__Metschnikowiaceae(100);g__Metschnikowia(100);s__Metschnikowia_pulcherrima(100); | Metschnikowia | 13 | Saprotroph | \|Undefined Saprotroph\| | Yeast | NULL | Probable | As nectar-tap_saprotroph (Põlme et al. 2020); Decay sugar-rich substrates (Põlme et al. 2020) | Põlme S, et al. 2020. Fungal Diversity 105:1-16 ((https://doi.org/10.1007/s13225-020-00466-2)) |
| Otu30254 | 1 | k__Fungi(100);p__Ascomycota(100);c__Saccharomycetes(100);o__Saccharomycetales(100);f__Metschnikowiaceae(100);g__Metschnikowia(100);s__Metschnikowia_pulcherrima(100); | Metschnikowia | 13 | Saprotroph | \|Undefined Saprotroph\| | Yeast | NULL | Probable | As nectar-tap_saprotroph (Põlme et al. 2020); Decay sugar-rich substrates (Põlme et al. 2020) | Põlme S, et al. 2020. Fungal Diversity 105:1-16 ((https://doi.org/10.1007/s13225-020-00466-2)) |
| Otu30257 | 1 | k__Fungi(100);p__Basidiomycota(100);c__Microbotryomycetes(100);o__Sporidiobolales(100);f__Sporidiobolales_family_Incertae_sedis(100);g__Sporobolomyces(100);s__Sporobolomyces_lactosus(100); | Sporobolomyces | 13 | Pathotroph-Saprotroph | \|Fungal Parasite\|-Plant Saprotroph | Yeast | NULL | Probable | As litter_saprotroph (Põlme et al. 2020); As mycoparasite (Põlme et al. 2020); Decay fungal material (Põlme et al. 2020) | Tedersoo L, et al. 2014. Science 346:e1256688 ((https://doi.org/10.1126/science.1256688)); Purahong W, et al. 2016. Molecular Ecology 25:4059-4074 ((https://doi.org/10.1111/mec.13739)); Põlme S, et al. 2020. Fungal Diversity 105:1-16 ((https://doi.org/10.1007/s13225-020-00466-2)) |
| Otu30258 | 1 | k__Fungi(100);p__Ascomycota(100);c__Saccharomycetes(100);o__Saccharomycetales(100);f__Metschnikowiaceae(100);g__Metschnikowia(100);s__Metschnikowia_pulcherrima(100); | Metschnikowia | 13 | Saprotroph | \|Undefined Saprotroph\| | Yeast | NULL | Probable | As nectar-tap_saprotroph (Põlme et al. 2020); Decay sugar-rich substrates (Põlme et al. 2020) | Põlme S, et al. 2020. Fungal Diversity 105:1-16 ((https://doi.org/10.1007/s13225-020-00466-2)) |
| Otu30261 | 1 | k__Fungi(100);p__Ascomycota(100);c__Saccharomycetes(100);o__Saccharomycetales(100);f__Saccharomycetales_family_Incertae_sedis(100);g__Candida(100);g__Candida_unclassified(100); | Candida | 13 | Pathotroph-Saprotroph-Symbiotroph | Animal Pathogen-Endophyte-Endosymbiont-Epiphyte-\|Undefined Saprotroph\| | Dimorphic Yeast-Yeast | NULL | Probable | As nectar-tap_saprotroph (Põlme et al. 2020); Decay sugar-rich substrates (Põlme et al. 2020); This asexual genus is polyphyletic and contain species that function very differently depend on what they are. Caution should be taken when interpreting guilds for Candida species. | Manolakaki D, et al. 2010. Virulence. 1: 367-375 ((https://doi.org/10.4161/viru.1.5.12796)); Mekbib SB, et al. 2011. Tropical Plant Pathology 36:233-240 ((https://doi.org/10.1590/S1982-56762011000400004)); Põlme S, et al. 2020. Fungal Diversity 105:1-16 ((https://doi.org/10.1007/s13225-020-00466-2)) |
| Otu30263 | 1 | k__Fungi(100);p__Ascomycota(100);c__Saccharomycetes(100);o__Saccharomycetales(100);f__Saccharomycetales_family_Incertae_sedis(100);g__Candida(100);g__Candida_unclassified(100); | Candida | 13 | Pathotroph-Saprotroph-Symbiotroph | Animal Pathogen-Endophyte-Endosymbiont-Epiphyte-\|Undefined Saprotroph\| | Dimorphic Yeast-Yeast | NULL | Probable | As nectar-tap_saprotroph (Põlme et al. 2020); Decay sugar-rich substrates (Põlme et al. 2020); This asexual genus is polyphyletic and contain species that function very differently depend on what they are. Caution should be taken when interpreting guilds for Candida species. | Manolakaki D, et al. 2010. Virulence. 1: 367-375 ((https://doi.org/10.4161/viru.1.5.12796)); Mekbib SB, et al. 2011. Tropical Plant Pathology 36:233-240 ((https://doi.org/10.1590/S1982-56762011000400004)); Põlme S, et al. 2020. Fungal Diversity 105:1-16 ((https://doi.org/10.1007/s13225-020-00466-2)) |
| Otu30265 | 1 | k__Fungi(100);p__Ascomycota(100);c__Saccharomycetes(100);o__Saccharomycetales(100);f__Metschnikowiaceae(100);g__Metschnikowia(100);s__Metschnikowia_pulcherrima(100); | Metschnikowia | 13 | Saprotroph | \|Undefined Saprotroph\| | Yeast | NULL | Probable | As nectar-tap_saprotroph (Põlme et al. 2020); Decay sugar-rich substrates (Põlme et al. 2020) | Põlme S, et al. 2020. Fungal Diversity 105:1-16 ((https://doi.org/10.1007/s13225-020-00466-2)) |
| Otu30269 | 1 | k__Fungi(100);p__Ascomycota(100);c__Sordariomycetes(100);o__Sordariales(100);f__Chaetomiaceae(100);g__Humicola(100);s__Humicola_grisea_var._grisea(100); | Humicola | 13 | Pathotroph-Saprotroph | Plant Pathogen-Undefined Saprotroph-\|Wood Saprotroph\| | Microfungus | Soft Rot | Probable | Decay wood (Põlme et al. 2020) | Seehann G, et al. 1975. List of Fungi in Soft-Rot Tissue Tests. Institute fur Holzbiologie und Holzschutz, Bundesforschungsanstalt fur Forst - und Holzwirtshaft, Hamburg - Lohbrugge, Germany; Costa IPMW, et al. 2012. Mycotaxon 119:e26 ((http://mycotaxon.com/resources/checklists/costa_v119_checklist.pdf)); Tedersoo L, et al. 2014. Science 346:e1256688 ((https://doi.org/10.1126/science.1256688)); Põlme S, et al. 2020. Fungal Diversity 105:1-16 ((https://doi.org/10.1007/s13225-020-00466-2)) |
| Otu30270 | 1 | k__Fungi(100);p__Ascomycota(100);c__Saccharomycetes(100);o__Saccharomycetales(100);f__Metschnikowiaceae(100);g__Metschnikowia(100);g__Metschnikowia_unclassified(100); | Metschnikowia | 13 | Saprotroph | \|Undefined Saprotroph\| | Yeast | NULL | Probable | As nectar-tap_saprotroph (Põlme et al. 2020); Decay sugar-rich substrates (Põlme et al. 2020) | Põlme S, et al. 2020. Fungal Diversity 105:1-16 ((https://doi.org/10.1007/s13225-020-00466-2)) |
| Otu30271 | 1 | k__Fungi(100);p__Ascomycota(100);c__Saccharomycetes(100);o__Saccharomycetales(100);f__Metschnikowiaceae(100);g__Metschnikowia(100);s__Metschnikowia_pulcherrima(100); | Metschnikowia | 13 | Saprotroph | \|Undefined Saprotroph\| | Yeast | NULL | Probable | As nectar-tap_saprotroph (Põlme et al. 2020); Decay sugar-rich substrates (Põlme et al. 2020) | Põlme S, et al. 2020. Fungal Diversity 105:1-16 ((https://doi.org/10.1007/s13225-020-00466-2)) |
| Otu30272 | 1 | k__Fungi(100);p__Ascomycota(100);c__Saccharomycetes(100);o__Saccharomycetales(100);f__Metschnikowiaceae(100);g__Metschnikowia(100);s__Metschnikowia_pulcherrima(100); | Metschnikowia | 13 | Saprotroph | \|Undefined Saprotroph\| | Yeast | NULL | Probable | As nectar-tap_saprotroph (Põlme et al. 2020); Decay sugar-rich substrates (Põlme et al. 2020) | Põlme S, et al. 2020. Fungal Diversity 105:1-16 ((https://doi.org/10.1007/s13225-020-00466-2)) |
| Otu30273 | 1 | k__Fungi(100);p__Ascomycota(100);c__Saccharomycetes(100);o__Saccharomycetales(100);f__Metschnikowiaceae(100);g__Metschnikowia(100);s__Metschnikowia_pulcherrima(100); | Metschnikowia | 13 | Saprotroph | \|Undefined Saprotroph\| | Yeast | NULL | Probable | As nectar-tap_saprotroph (Põlme et al. 2020); Decay sugar-rich substrates (Põlme et al. 2020) | Põlme S, et al. 2020. Fungal Diversity 105:1-16 ((https://doi.org/10.1007/s13225-020-00466-2)) |
| Otu30276 | 1 | k__Fungi(100);p__Ascomycota(100);c__Saccharomycetes(100);o__Saccharomycetales(100);o__Saccharomycetales_unclassified(100);o__Saccharomycetales_unclassified(100);o__Saccharomycetales_unclassified(100); | Saccharomycetales | 7 | Saprotroph | Undefined Saprotroph | Yeast | NULL | Possible | NULL | Sterkenburg E, et al. 2015. New Phytologist 207:1145-1158 ((https://doi.org/10.1111/nph.13426)) |
| Otu30279 | 1 | k__Fungi(100);p__Ascomycota(100);c__Saccharomycetes(100);o__Saccharomycetales(100);f__Saccharomycetales_family_Incertae_sedis(100);g__Candida(100);g__Candida_unclassified(100); | Candida | 13 | Pathotroph-Saprotroph-Symbiotroph | Animal Pathogen-Endophyte-Endosymbiont-Epiphyte-\|Undefined Saprotroph\| | Dimorphic Yeast-Yeast | NULL | Probable | As nectar-tap_saprotroph (Põlme et al. 2020); Decay sugar-rich substrates (Põlme et al. 2020); This asexual genus is polyphyletic and contain species that function very differently depend on what they are. Caution should be taken when interpreting guilds for Candida species. | Manolakaki D, et al. 2010. Virulence. 1: 367-375 ((https://doi.org/10.4161/viru.1.5.12796)); Mekbib SB, et al. 2011. Tropical Plant Pathology 36:233-240 ((https://doi.org/10.1590/S1982-56762011000400004)); Põlme S, et al. 2020. Fungal Diversity 105:1-16 ((https://doi.org/10.1007/s13225-020-00466-2)) |
| Otu30281 | 1 | k__Fungi(100);p__Ascomycota(100);c__Saccharomycetes(100);o__Saccharomycetales(100);f__Saccharomycetales_family_Incertae_sedis(100);g__Candida(100);g__Candida_unclassified(100); | Candida | 13 | Pathotroph-Saprotroph-Symbiotroph | Animal Pathogen-Endophyte-Endosymbiont-Epiphyte-\|Undefined Saprotroph\| | Dimorphic Yeast-Yeast | NULL | Probable | As nectar-tap_saprotroph (Põlme et al. 2020); Decay sugar-rich substrates (Põlme et al. 2020); This asexual genus is polyphyletic and contain species that function very differently depend on what they are. Caution should be taken when interpreting guilds for Candida species. | Manolakaki D, et al. 2010. Virulence. 1: 367-375 ((https://doi.org/10.4161/viru.1.5.12796)); Mekbib SB, et al. 2011. Tropical Plant Pathology 36:233-240 ((https://doi.org/10.1590/S1982-56762011000400004)); Põlme S, et al. 2020. Fungal Diversity 105:1-16 ((https://doi.org/10.1007/s13225-020-00466-2)) |
| Otu30282 | 1 | k__Fungi(100);p__Ascomycota(100);c__Saccharomycetes(100);o__Saccharomycetales(100);o__Saccharomycetales_unclassified(100);o__Saccharomycetales_unclassified(100);o__Saccharomycetales_unclassified(100); | Saccharomycetales | 7 | Saprotroph | Undefined Saprotroph | Yeast | NULL | Possible | NULL | Sterkenburg E, et al. 2015. New Phytologist 207:1145-1158 ((https://doi.org/10.1111/nph.13426)) |
| Otu30284 | 1 | k__Fungi(100);p__Ascomycota(100);c__Saccharomycetes(100);o__Saccharomycetales(100);f__Metschnikowiaceae(100);g__Metschnikowia(100);s__Metschnikowia_pulcherrima(100); | Metschnikowia | 13 | Saprotroph | \|Undefined Saprotroph\| | Yeast | NULL | Probable | As nectar-tap_saprotroph (Põlme et al. 2020); Decay sugar-rich substrates (Põlme et al. 2020) | Põlme S, et al. 2020. Fungal Diversity 105:1-16 ((https://doi.org/10.1007/s13225-020-00466-2)) |
| Otu30287 | 1 | k__Fungi(100);p__Ascomycota(100);c__Saccharomycetes(100);o__Saccharomycetales(100);f__Saccharomycetales_family_Incertae_sedis(100);g__Candida(100);g__Candida_unclassified(100); | Candida | 13 | Pathotroph-Saprotroph-Symbiotroph | Animal Pathogen-Endophyte-Endosymbiont-Epiphyte-\|Undefined Saprotroph\| | Dimorphic Yeast-Yeast | NULL | Probable | As nectar-tap_saprotroph (Põlme et al. 2020); Decay sugar-rich substrates (Põlme et al. 2020); This asexual genus is polyphyletic and contain species that function very differently depend on what they are. Caution should be taken when interpreting guilds for Candida species. | Manolakaki D, et al. 2010. Virulence. 1: 367-375 ((https://doi.org/10.4161/viru.1.5.12796)); Mekbib SB, et al. 2011. Tropical Plant Pathology 36:233-240 ((https://doi.org/10.1590/S1982-56762011000400004)); Põlme S, et al. 2020. Fungal Diversity 105:1-16 ((https://doi.org/10.1007/s13225-020-00466-2)) |
| Otu30288 | 1 | k__Fungi(100);p__Ascomycota(100);c__Saccharomycetes(100);o__Saccharomycetales(100);f__Metschnikowiaceae(100);g__Metschnikowia(100);s__Metschnikowia_pulcherrima(100); | Metschnikowia | 13 | Saprotroph | \|Undefined Saprotroph\| | Yeast | NULL | Probable | As nectar-tap_saprotroph (Põlme et al. 2020); Decay sugar-rich substrates (Põlme et al. 2020) | Põlme S, et al. 2020. Fungal Diversity 105:1-16 ((https://doi.org/10.1007/s13225-020-00466-2)) |
| Otu30290 | 1 | k__Fungi(100);p__Ascomycota(100);c__Saccharomycetes(100);o__Saccharomycetales(100);f__Metschnikowiaceae(100);g__Metschnikowia(100);s__Metschnikowia_pulcherrima(100); | Metschnikowia | 13 | Saprotroph | \|Undefined Saprotroph\| | Yeast | NULL | Probable | As nectar-tap_saprotroph (Põlme et al. 2020); Decay sugar-rich substrates (Põlme et al. 2020) | Põlme S, et al. 2020. Fungal Diversity 105:1-16 ((https://doi.org/10.1007/s13225-020-00466-2)) |
| Otu30291 | 1 | k__Fungi(100);p__Ascomycota(100);c__Saccharomycetes(100);o__Saccharomycetales(100);f__Saccharomycetales_family_Incertae_sedis(100);g__Candida(100);g__Candida_unclassified(100); | Candida | 13 | Pathotroph-Saprotroph-Symbiotroph | Animal Pathogen-Endophyte-Endosymbiont-Epiphyte-\|Undefined Saprotroph\| | Dimorphic Yeast-Yeast | NULL | Probable | As nectar-tap_saprotroph (Põlme et al. 2020); Decay sugar-rich substrates (Põlme et al. 2020); This asexual genus is polyphyletic and contain species that function very differently depend on what they are. Caution should be taken when interpreting guilds for Candida species. | Manolakaki D, et al. 2010. Virulence. 1: 367-375 ((https://doi.org/10.4161/viru.1.5.12796)); Mekbib SB, et al. 2011. Tropical Plant Pathology 36:233-240 ((https://doi.org/10.1590/S1982-56762011000400004)); Põlme S, et al. 2020. Fungal Diversity 105:1-16 ((https://doi.org/10.1007/s13225-020-00466-2)) |
| Otu30293 | 1 | k__Fungi(100);p__Ascomycota(100);c__Saccharomycetes(100);o__Saccharomycetales(100);f__Metschnikowiaceae(100);g__Metschnikowia(100);g__Metschnikowia_unclassified(100); | Metschnikowia | 13 | Saprotroph | \|Undefined Saprotroph\| | Yeast | NULL | Probable | As nectar-tap_saprotroph (Põlme et al. 2020); Decay sugar-rich substrates (Põlme et al. 2020) | Põlme S, et al. 2020. Fungal Diversity 105:1-16 ((https://doi.org/10.1007/s13225-020-00466-2)) |
| Otu30294 | 1 | k__Fungi(100);p__Ascomycota(100);c__Saccharomycetes(100);o__Saccharomycetales(100);f__Metschnikowiaceae(100);g__Metschnikowia(100);s__Metschnikowia_pulcherrima(100); | Metschnikowia | 13 | Saprotroph | \|Undefined Saprotroph\| | Yeast | NULL | Probable | As nectar-tap_saprotroph (Põlme et al. 2020); Decay sugar-rich substrates (Põlme et al. 2020) | Põlme S, et al. 2020. Fungal Diversity 105:1-16 ((https://doi.org/10.1007/s13225-020-00466-2)) |
| Otu30295 | 1 | k__Fungi(100);p__Ascomycota(100);c__Saccharomycetes(100);o__Saccharomycetales(100);f__Metschnikowiaceae(100);g__Metschnikowia(100);s__Metschnikowia_pulcherrima(100); | Metschnikowia | 13 | Saprotroph | \|Undefined Saprotroph\| | Yeast | NULL | Probable | As nectar-tap_saprotroph (Põlme et al. 2020); Decay sugar-rich substrates (Põlme et al. 2020) | Põlme S, et al. 2020. Fungal Diversity 105:1-16 ((https://doi.org/10.1007/s13225-020-00466-2)) |
| Otu30296 | 1 | k__Fungi(100);p__Ascomycota(100);c__Saccharomycetes(100);o__Saccharomycetales(100);f__Metschnikowiaceae(100);g__Metschnikowia(100);s__Metschnikowia_pulcherrima(100); | Metschnikowia | 13 | Saprotroph | \|Undefined Saprotroph\| | Yeast | NULL | Probable | As nectar-tap_saprotroph (Põlme et al. 2020); Decay sugar-rich substrates (Põlme et al. 2020) | Põlme S, et al. 2020. Fungal Diversity 105:1-16 ((https://doi.org/10.1007/s13225-020-00466-2)) |
| Otu30297 | 1 | k__Fungi(100);p__Ascomycota(100);c__Saccharomycetes(100);o__Saccharomycetales(100);f__Metschnikowiaceae(100);g__Metschnikowia(100);s__Metschnikowia_pulcherrima(100); | Metschnikowia | 13 | Saprotroph | \|Undefined Saprotroph\| | Yeast | NULL | Probable | As nectar-tap_saprotroph (Põlme et al. 2020); Decay sugar-rich substrates (Põlme et al. 2020) | Põlme S, et al. 2020. Fungal Diversity 105:1-16 ((https://doi.org/10.1007/s13225-020-00466-2)) |
| Otu30298 | 1 | k__Fungi(100);p__Ascomycota(100);c__Saccharomycetes(100);o__Saccharomycetales(100);f__Metschnikowiaceae(100);g__Metschnikowia(100);s__Metschnikowia_pulcherrima(100); | Metschnikowia | 13 | Saprotroph | \|Undefined Saprotroph\| | Yeast | NULL | Probable | As nectar-tap_saprotroph (Põlme et al. 2020); Decay sugar-rich substrates (Põlme et al. 2020) | Põlme S, et al. 2020. Fungal Diversity 105:1-16 ((https://doi.org/10.1007/s13225-020-00466-2)) |
| Otu30299 | 1 | k__Fungi(100);p__Ascomycota(100);c__Saccharomycetes(100);o__Saccharomycetales(100);f__Metschnikowiaceae(100);g__Metschnikowia(100);s__Metschnikowia_pulcherrima(100); | Metschnikowia | 13 | Saprotroph | \|Undefined Saprotroph\| | Yeast | NULL | Probable | As nectar-tap_saprotroph (Põlme et al. 2020); Decay sugar-rich substrates (Põlme et al. 2020) | Põlme S, et al. 2020. Fungal Diversity 105:1-16 ((https://doi.org/10.1007/s13225-020-00466-2)) |
| Otu30300 | 1 | k__Fungi(100);p__Ascomycota(100);c__Saccharomycetes(100);o__Saccharomycetales(100);f__Metschnikowiaceae(100);g__Metschnikowia(100);s__Metschnikowia_pulcherrima(100); | Metschnikowia | 13 | Saprotroph | \|Undefined Saprotroph\| | Yeast | NULL | Probable | As nectar-tap_saprotroph (Põlme et al. 2020); Decay sugar-rich substrates (Põlme et al. 2020) | Põlme S, et al. 2020. Fungal Diversity 105:1-16 ((https://doi.org/10.1007/s13225-020-00466-2)) |
| Otu30301 | 1 | k__Fungi(100);p__Ascomycota(100);c__Saccharomycetes(100);o__Saccharomycetales(100);f__Metschnikowiaceae(100);g__Metschnikowia(100);s__Metschnikowia_pulcherrima(100); | Metschnikowia | 13 | Saprotroph | \|Undefined Saprotroph\| | Yeast | NULL | Probable | As nectar-tap_saprotroph (Põlme et al. 2020); Decay sugar-rich substrates (Põlme et al. 2020) | Põlme S, et al. 2020. Fungal Diversity 105:1-16 ((https://doi.org/10.1007/s13225-020-00466-2)) |
| Otu30302 | 1 | k__Fungi(100);p__Ascomycota(100);c__Saccharomycetes(100);o__Saccharomycetales(100);f__Metschnikowiaceae(100);g__Metschnikowia(100);s__Metschnikowia_pulcherrima(100); | Metschnikowia | 13 | Saprotroph | \|Undefined Saprotroph\| | Yeast | NULL | Probable | As nectar-tap_saprotroph (Põlme et al. 2020); Decay sugar-rich substrates (Põlme et al. 2020) | Põlme S, et al. 2020. Fungal Diversity 105:1-16 ((https://doi.org/10.1007/s13225-020-00466-2)) |
| Otu30303 | 1 | k__Fungi(100);p__Ascomycota(100);c__Saccharomycetes(100);o__Saccharomycetales(100);f__Metschnikowiaceae(100);g__Metschnikowia(100);s__Metschnikowia_pulcherrima(100); | Metschnikowia | 13 | Saprotroph | \|Undefined Saprotroph\| | Yeast | NULL | Probable | As nectar-tap_saprotroph (Põlme et al. 2020); Decay sugar-rich substrates (Põlme et al. 2020) | Põlme S, et al. 2020. Fungal Diversity 105:1-16 ((https://doi.org/10.1007/s13225-020-00466-2)) |
| Otu30304 | 1 | k__Fungi(100);p__Ascomycota(100);c__Saccharomycetes(100);o__Saccharomycetales(100);f__Metschnikowiaceae(100);g__Metschnikowia(100);s__Metschnikowia_pulcherrima(100); | Metschnikowia | 13 | Saprotroph | \|Undefined Saprotroph\| | Yeast | NULL | Probable | As nectar-tap_saprotroph (Põlme et al. 2020); Decay sugar-rich substrates (Põlme et al. 2020) | Põlme S, et al. 2020. Fungal Diversity 105:1-16 ((https://doi.org/10.1007/s13225-020-00466-2)) |
| Otu30305 | 1 | k__Fungi(100);p__Ascomycota(100);c__Saccharomycetes(100);o__Saccharomycetales(100);f__Metschnikowiaceae(100);g__Metschnikowia(100);s__Metschnikowia_pulcherrima(100); | Metschnikowia | 13 | Saprotroph | \|Undefined Saprotroph\| | Yeast | NULL | Probable | As nectar-tap_saprotroph (Põlme et al. 2020); Decay sugar-rich substrates (Põlme et al. 2020) | Põlme S, et al. 2020. Fungal Diversity 105:1-16 ((https://doi.org/10.1007/s13225-020-00466-2)) |
| Otu30306 | 1 | k__Fungi(100);p__Ascomycota(100);c__Saccharomycetes(100);o__Saccharomycetales(100);f__Metschnikowiaceae(100);g__Metschnikowia(100);s__Metschnikowia_pulcherrima(100); | Metschnikowia | 13 | Saprotroph | \|Undefined Saprotroph\| | Yeast | NULL | Probable | As nectar-tap_saprotroph (Põlme et al. 2020); Decay sugar-rich substrates (Põlme et al. 2020) | Põlme S, et al. 2020. Fungal Diversity 105:1-16 ((https://doi.org/10.1007/s13225-020-00466-2)) |
| Otu30307 | 1 | k__Fungi(100);p__Ascomycota(100);c__Saccharomycetes(100);o__Saccharomycetales(100);f__Metschnikowiaceae(100);g__Metschnikowia(100);s__Metschnikowia_pulcherrima(100); | Metschnikowia | 13 | Saprotroph | \|Undefined Saprotroph\| | Yeast | NULL | Probable | As nectar-tap_saprotroph (Põlme et al. 2020); Decay sugar-rich substrates (Põlme et al. 2020) | Põlme S, et al. 2020. Fungal Diversity 105:1-16 ((https://doi.org/10.1007/s13225-020-00466-2)) |
| Otu30308 | 1 | k__Fungi(100);p__Ascomycota(100);c__Saccharomycetes(100);o__Saccharomycetales(100);f__Metschnikowiaceae(100);g__Metschnikowia(100);s__Metschnikowia_pulcherrima(100); | Metschnikowia | 13 | Saprotroph | \|Undefined Saprotroph\| | Yeast | NULL | Probable | As nectar-tap_saprotroph (Põlme et al. 2020); Decay sugar-rich substrates (Põlme et al. 2020) | Põlme S, et al. 2020. Fungal Diversity 105:1-16 ((https://doi.org/10.1007/s13225-020-00466-2)) |
| Otu30309 | 1 | k__Fungi(100);p__Ascomycota(100);c__Saccharomycetes(100);o__Saccharomycetales(100);f__Metschnikowiaceae(100);g__Metschnikowia(100);s__Metschnikowia_pulcherrima(100); | Metschnikowia | 13 | Saprotroph | \|Undefined Saprotroph\| | Yeast | NULL | Probable | As nectar-tap_saprotroph (Põlme et al. 2020); Decay sugar-rich substrates (Põlme et al. 2020) | Põlme S, et al. 2020. Fungal Diversity 105:1-16 ((https://doi.org/10.1007/s13225-020-00466-2)) |
| Otu30310 | 1 | k__Fungi(100);p__Ascomycota(100);c__Saccharomycetes(100);o__Saccharomycetales(100);f__Metschnikowiaceae(100);g__Metschnikowia(100);s__Metschnikowia_pulcherrima(100); | Metschnikowia | 13 | Saprotroph | \|Undefined Saprotroph\| | Yeast | NULL | Probable | As nectar-tap_saprotroph (Põlme et al. 2020); Decay sugar-rich substrates (Põlme et al. 2020) | Põlme S, et al. 2020. Fungal Diversity 105:1-16 ((https://doi.org/10.1007/s13225-020-00466-2)) |
| Otu30311 | 1 | k__Fungi(100);p__Ascomycota(100);c__Saccharomycetes(100);o__Saccharomycetales(100);f__Metschnikowiaceae(100);g__Metschnikowia(100);s__Metschnikowia_pulcherrima(100); | Metschnikowia | 13 | Saprotroph | \|Undefined Saprotroph\| | Yeast | NULL | Probable | As nectar-tap_saprotroph (Põlme et al. 2020); Decay sugar-rich substrates (Põlme et al. 2020) | Põlme S, et al. 2020. Fungal Diversity 105:1-16 ((https://doi.org/10.1007/s13225-020-00466-2)) |
| Otu30312 | 1 | k__Fungi(100);p__Ascomycota(100);c__Saccharomycetes(100);o__Saccharomycetales(100);f__Metschnikowiaceae(100);g__Metschnikowia(100);g__Metschnikowia_unclassified(100); | Metschnikowia | 13 | Saprotroph | \|Undefined Saprotroph\| | Yeast | NULL | Probable | As nectar-tap_saprotroph (Põlme et al. 2020); Decay sugar-rich substrates (Põlme et al. 2020) | Põlme S, et al. 2020. Fungal Diversity 105:1-16 ((https://doi.org/10.1007/s13225-020-00466-2)) |
| Otu30313 | 1 | k__Fungi(100);p__Ascomycota(100);c__Saccharomycetes(100);o__Saccharomycetales(100);f__Metschnikowiaceae(100);g__Metschnikowia(100);s__Metschnikowia_pulcherrima(100); | Metschnikowia | 13 | Saprotroph | \|Undefined Saprotroph\| | Yeast | NULL | Probable | As nectar-tap_saprotroph (Põlme et al. 2020); Decay sugar-rich substrates (Põlme et al. 2020) | Põlme S, et al. 2020. Fungal Diversity 105:1-16 ((https://doi.org/10.1007/s13225-020-00466-2)) |
| Otu30314 | 1 | k__Fungi(100);p__Ascomycota(100);c__Saccharomycetes(100);o__Saccharomycetales(100);f__Metschnikowiaceae(100);g__Metschnikowia(100);g__Metschnikowia_unclassified(100); | Metschnikowia | 13 | Saprotroph | \|Undefined Saprotroph\| | Yeast | NULL | Probable | As nectar-tap_saprotroph (Põlme et al. 2020); Decay sugar-rich substrates (Põlme et al. 2020) | Põlme S, et al. 2020. Fungal Diversity 105:1-16 ((https://doi.org/10.1007/s13225-020-00466-2)) |
| Otu30315 | 1 | k__Fungi(100);p__Ascomycota(100);c__Saccharomycetes(100);o__Saccharomycetales(100);f__Metschnikowiaceae(100);g__Metschnikowia(100);g__Metschnikowia_unclassified(100); | Metschnikowia | 13 | Saprotroph | \|Undefined Saprotroph\| | Yeast | NULL | Probable | As nectar-tap_saprotroph (Põlme et al. 2020); Decay sugar-rich substrates (Põlme et al. 2020) | Põlme S, et al. 2020. Fungal Diversity 105:1-16 ((https://doi.org/10.1007/s13225-020-00466-2)) |
| Otu30316 | 1 | k__Fungi(100);p__Ascomycota(100);c__Saccharomycetes(100);o__Saccharomycetales(100);f__Metschnikowiaceae(100);g__Metschnikowia(100);g__Metschnikowia_unclassified(100); | Metschnikowia | 13 | Saprotroph | \|Undefined Saprotroph\| | Yeast | NULL | Probable | As nectar-tap_saprotroph (Põlme et al. 2020); Decay sugar-rich substrates (Põlme et al. 2020) | Põlme S, et al. 2020. Fungal Diversity 105:1-16 ((https://doi.org/10.1007/s13225-020-00466-2)) |
| Otu30319 | 1 | k__Fungi(100);p__Ascomycota(100);c__Saccharomycetes(100);o__Saccharomycetales(100);f__Metschnikowiaceae(100);g__Metschnikowia(100);s__Metschnikowia_pulcherrima(100); | Metschnikowia | 13 | Saprotroph | \|Undefined Saprotroph\| | Yeast | NULL | Probable | As nectar-tap_saprotroph (Põlme et al. 2020); Decay sugar-rich substrates (Põlme et al. 2020) | Põlme S, et al. 2020. Fungal Diversity 105:1-16 ((https://doi.org/10.1007/s13225-020-00466-2)) |
| Otu30320 | 1 | k__Fungi(100);p__Ascomycota(100);c__Saccharomycetes(100);o__Saccharomycetales(100);f__Metschnikowiaceae(100);g__Metschnikowia(100);g__Metschnikowia_unclassified(100); | Metschnikowia | 13 | Saprotroph | \|Undefined Saprotroph\| | Yeast | NULL | Probable | As nectar-tap_saprotroph (Põlme et al. 2020); Decay sugar-rich substrates (Põlme et al. 2020) | Põlme S, et al. 2020. Fungal Diversity 105:1-16 ((https://doi.org/10.1007/s13225-020-00466-2)) |
| Otu30323 | 1 | k__Fungi(100);p__Ascomycota(100);c__Saccharomycetes(100);o__Saccharomycetales(100);f__Saccharomycetales_family_Incertae_sedis(100);g__Candida(100);s__Candida_cretensis(100); | Candida | 13 | Pathotroph-Saprotroph-Symbiotroph | Animal Pathogen-Endophyte-Endosymbiont-Epiphyte-\|Undefined Saprotroph\| | Dimorphic Yeast-Yeast | NULL | Probable | As nectar-tap_saprotroph (Põlme et al. 2020); Decay sugar-rich substrates (Põlme et al. 2020); This asexual genus is polyphyletic and contain species that function very differently depend on what they are. Caution should be taken when interpreting guilds for Candida species. | Manolakaki D, et al. 2010. Virulence. 1: 367-375 ((https://doi.org/10.4161/viru.1.5.12796)); Mekbib SB, et al. 2011. Tropical Plant Pathology 36:233-240 ((https://doi.org/10.1590/S1982-56762011000400004)); Põlme S, et al. 2020. Fungal Diversity 105:1-16 ((https://doi.org/10.1007/s13225-020-00466-2)) |
| Otu30324 | 1 | k__Fungi(100);p__Ascomycota(100);c__Saccharomycetes(100);o__Saccharomycetales(100);f__Metschnikowiaceae(100);g__Metschnikowia(100);g__Metschnikowia_unclassified(100); | Metschnikowia | 13 | Saprotroph | \|Undefined Saprotroph\| | Yeast | NULL | Probable | As nectar-tap_saprotroph (Põlme et al. 2020); Decay sugar-rich substrates (Põlme et al. 2020) | Põlme S, et al. 2020. Fungal Diversity 105:1-16 ((https://doi.org/10.1007/s13225-020-00466-2)) |
| Otu30325 | 1 | k__Fungi(100);p__Ascomycota(100);c__Saccharomycetes(100);o__Saccharomycetales(100);f__Metschnikowiaceae(100);g__Metschnikowia(100);s__Metschnikowia_pulcherrima(100); | Metschnikowia | 13 | Saprotroph | \|Undefined Saprotroph\| | Yeast | NULL | Probable | As nectar-tap_saprotroph (Põlme et al. 2020); Decay sugar-rich substrates (Põlme et al. 2020) | Põlme S, et al. 2020. Fungal Diversity 105:1-16 ((https://doi.org/10.1007/s13225-020-00466-2)) |
| Otu30326 | 1 | k__Fungi(100);p__Ascomycota(100);c__Saccharomycetes(100);o__Saccharomycetales(100);f__Metschnikowiaceae(100);g__Metschnikowia(100);s__Metschnikowia_pulcherrima(100); | Metschnikowia | 13 | Saprotroph | \|Undefined Saprotroph\| | Yeast | NULL | Probable | As nectar-tap_saprotroph (Põlme et al. 2020); Decay sugar-rich substrates (Põlme et al. 2020) | Põlme S, et al. 2020. Fungal Diversity 105:1-16 ((https://doi.org/10.1007/s13225-020-00466-2)) |
| Otu30327 | 1 | k__Fungi(100);p__Ascomycota(100);c__Ascomycota_class_Incertae_sedis(100);o__Ascomycota_order_Incertae_sedis(100);f__Ascomycota_family_Incertae_sedis(100);g__Seifertia(100);s__Seifertia_azaleae(100); | Seifertia | 13 | Pathotroph | \|Plant Pathogen\| | Microfungus | NULL | Probable | NULL | Põlme S, et al. 2020. Fungal Diversity 105:1-16 ((https://doi.org/10.1007/s13225-020-00466-2)) |
| Otu30328 | 1 | k__Fungi(100);p__Ascomycota(100);c__Saccharomycetes(100);o__Saccharomycetales(100);f__Metschnikowiaceae(100);g__Metschnikowia(100);s__Metschnikowia_pulcherrima(100); | Metschnikowia | 13 | Saprotroph | \|Undefined Saprotroph\| | Yeast | NULL | Probable | As nectar-tap_saprotroph (Põlme et al. 2020); Decay sugar-rich substrates (Põlme et al. 2020) | Põlme S, et al. 2020. Fungal Diversity 105:1-16 ((https://doi.org/10.1007/s13225-020-00466-2)) |
| Otu30329 | 1 | k__Fungi(100);p__Ascomycota(100);c__Saccharomycetes(100);o__Saccharomycetales(100);f__Metschnikowiaceae(100);g__Metschnikowia(100);g__Metschnikowia_unclassified(100); | Metschnikowia | 13 | Saprotroph | \|Undefined Saprotroph\| | Yeast | NULL | Probable | As nectar-tap_saprotroph (Põlme et al. 2020); Decay sugar-rich substrates (Põlme et al. 2020) | Põlme S, et al. 2020. Fungal Diversity 105:1-16 ((https://doi.org/10.1007/s13225-020-00466-2)) |
| Otu30330 | 1 | k__Fungi(100);p__Ascomycota(100);c__Saccharomycetes(100);o__Saccharomycetales(100);f__Metschnikowiaceae(100);g__Metschnikowia(100);s__Metschnikowia_pulcherrima(100); | Metschnikowia | 13 | Saprotroph | \|Undefined Saprotroph\| | Yeast | NULL | Probable | As nectar-tap_saprotroph (Põlme et al. 2020); Decay sugar-rich substrates (Põlme et al. 2020) | Põlme S, et al. 2020. Fungal Diversity 105:1-16 ((https://doi.org/10.1007/s13225-020-00466-2)) |
| Otu30331 | 1 | k__Fungi(100);p__Ascomycota(100);c__Saccharomycetes(100);o__Saccharomycetales(100);f__Metschnikowiaceae(100);g__Metschnikowia(100);s__Metschnikowia_pulcherrima(100); | Metschnikowia | 13 | Saprotroph | \|Undefined Saprotroph\| | Yeast | NULL | Probable | As nectar-tap_saprotroph (Põlme et al. 2020); Decay sugar-rich substrates (Põlme et al. 2020) | Põlme S, et al. 2020. Fungal Diversity 105:1-16 ((https://doi.org/10.1007/s13225-020-00466-2)) |
| Otu30332 | 1 | k__Fungi(100);p__Ascomycota(100);c__Saccharomycetes(100);o__Saccharomycetales(100);f__Metschnikowiaceae(100);g__Metschnikowia(100);g__Metschnikowia_unclassified(100); | Metschnikowia | 13 | Saprotroph | \|Undefined Saprotroph\| | Yeast | NULL | Probable | As nectar-tap_saprotroph (Põlme et al. 2020); Decay sugar-rich substrates (Põlme et al. 2020) | Põlme S, et al. 2020. Fungal Diversity 105:1-16 ((https://doi.org/10.1007/s13225-020-00466-2)) |
| Otu30333 | 1 | k__Fungi(100);p__Ascomycota(100);c__Saccharomycetes(100);o__Saccharomycetales(100);f__Metschnikowiaceae(100);g__Metschnikowia(100);s__Metschnikowia_pulcherrima(100); | Metschnikowia | 13 | Saprotroph | \|Undefined Saprotroph\| | Yeast | NULL | Probable | As nectar-tap_saprotroph (Põlme et al. 2020); Decay sugar-rich substrates (Põlme et al. 2020) | Põlme S, et al. 2020. Fungal Diversity 105:1-16 ((https://doi.org/10.1007/s13225-020-00466-2)) |
| Otu30334 | 1 | k__Fungi(100);p__Ascomycota(100);c__Saccharomycetes(100);o__Saccharomycetales(100);f__Metschnikowiaceae(100);g__Metschnikowia(100);s__Metschnikowia_pulcherrima(100); | Metschnikowia | 13 | Saprotroph | \|Undefined Saprotroph\| | Yeast | NULL | Probable | As nectar-tap_saprotroph (Põlme et al. 2020); Decay sugar-rich substrates (Põlme et al. 2020) | Põlme S, et al. 2020. Fungal Diversity 105:1-16 ((https://doi.org/10.1007/s13225-020-00466-2)) |
| Otu30335 | 1 | k__Fungi(100);p__Ascomycota(100);c__Saccharomycetes(100);o__Saccharomycetales(100);f__Metschnikowiaceae(100);g__Metschnikowia(100);s__Metschnikowia_pulcherrima(100); | Metschnikowia | 13 | Saprotroph | \|Undefined Saprotroph\| | Yeast | NULL | Probable | As nectar-tap_saprotroph (Põlme et al. 2020); Decay sugar-rich substrates (Põlme et al. 2020) | Põlme S, et al. 2020. Fungal Diversity 105:1-16 ((https://doi.org/10.1007/s13225-020-00466-2)) |
| Otu30336 | 1 | k__Fungi(100);p__Ascomycota(100);c__Saccharomycetes(100);o__Saccharomycetales(100);f__Metschnikowiaceae(100);g__Metschnikowia(100);s__Metschnikowia_pulcherrima(100); | Metschnikowia | 13 | Saprotroph | \|Undefined Saprotroph\| | Yeast | NULL | Probable | As nectar-tap_saprotroph (Põlme et al. 2020); Decay sugar-rich substrates (Põlme et al. 2020) | Põlme S, et al. 2020. Fungal Diversity 105:1-16 ((https://doi.org/10.1007/s13225-020-00466-2)) |
| Otu30337 | 1 | k__Fungi(100);p__Ascomycota(100);c__Saccharomycetes(100);o__Saccharomycetales(100);f__Metschnikowiaceae(100);g__Metschnikowia(100);s__Metschnikowia_pulcherrima(100); | Metschnikowia | 13 | Saprotroph | \|Undefined Saprotroph\| | Yeast | NULL | Probable | As nectar-tap_saprotroph (Põlme et al. 2020); Decay sugar-rich substrates (Põlme et al. 2020) | Põlme S, et al. 2020. Fungal Diversity 105:1-16 ((https://doi.org/10.1007/s13225-020-00466-2)) |
| Otu30338 | 1 | k__Fungi(100);p__Ascomycota(100);c__Saccharomycetes(100);o__Saccharomycetales(100);f__Metschnikowiaceae(100);g__Metschnikowia(100);s__Metschnikowia_pulcherrima(100); | Metschnikowia | 13 | Saprotroph | \|Undefined Saprotroph\| | Yeast | NULL | Probable | As nectar-tap_saprotroph (Põlme et al. 2020); Decay sugar-rich substrates (Põlme et al. 2020) | Põlme S, et al. 2020. Fungal Diversity 105:1-16 ((https://doi.org/10.1007/s13225-020-00466-2)) |
| Otu30339 | 1 | k__Fungi(100);p__Ascomycota(100);c__Saccharomycetes(100);o__Saccharomycetales(100);f__Metschnikowiaceae(100);g__Metschnikowia(100);s__Metschnikowia_pulcherrima(100); | Metschnikowia | 13 | Saprotroph | \|Undefined Saprotroph\| | Yeast | NULL | Probable | As nectar-tap_saprotroph (Põlme et al. 2020); Decay sugar-rich substrates (Põlme et al. 2020) | Põlme S, et al. 2020. Fungal Diversity 105:1-16 ((https://doi.org/10.1007/s13225-020-00466-2)) |
| Otu30342 | 1 | k__Fungi(100);p__Ascomycota(100);c__Saccharomycetes(100);o__Saccharomycetales(100);f__Metschnikowiaceae(100);g__Metschnikowia(100);s__Metschnikowia_pulcherrima(100); | Metschnikowia | 13 | Saprotroph | \|Undefined Saprotroph\| | Yeast | NULL | Probable | As nectar-tap_saprotroph (Põlme et al. 2020); Decay sugar-rich substrates (Põlme et al. 2020) | Põlme S, et al. 2020. Fungal Diversity 105:1-16 ((https://doi.org/10.1007/s13225-020-00466-2)) |
| Otu30343 | 1 | k__Fungi(100);p__Ascomycota(100);c__Saccharomycetes(100);o__Saccharomycetales(100);f__Metschnikowiaceae(100);g__Metschnikowia(100);s__Metschnikowia_pulcherrima(100); | Metschnikowia | 13 | Saprotroph | \|Undefined Saprotroph\| | Yeast | NULL | Probable | As nectar-tap_saprotroph (Põlme et al. 2020); Decay sugar-rich substrates (Põlme et al. 2020) | Põlme S, et al. 2020. Fungal Diversity 105:1-16 ((https://doi.org/10.1007/s13225-020-00466-2)) |
| Otu30345 | 1 | k__Fungi(100);p__Ascomycota(100);c__Saccharomycetes(100);o__Saccharomycetales(100);f__Metschnikowiaceae(100);g__Metschnikowia(100);s__Metschnikowia_pulcherrima(100); | Metschnikowia | 13 | Saprotroph | \|Undefined Saprotroph\| | Yeast | NULL | Probable | As nectar-tap_saprotroph (Põlme et al. 2020); Decay sugar-rich substrates (Põlme et al. 2020) | Põlme S, et al. 2020. Fungal Diversity 105:1-16 ((https://doi.org/10.1007/s13225-020-00466-2)) |
| Otu30354 | 1 | k__Fungi(100);p__Ascomycota(100);c__Saccharomycetes(100);o__Saccharomycetales(100);f__Metschnikowiaceae(100);g__Metschnikowia(100);s__Metschnikowia_pulcherrima(100); | Metschnikowia | 13 | Saprotroph | \|Undefined Saprotroph\| | Yeast | NULL | Probable | As nectar-tap_saprotroph (Põlme et al. 2020); Decay sugar-rich substrates (Põlme et al. 2020) | Põlme S, et al. 2020. Fungal Diversity 105:1-16 ((https://doi.org/10.1007/s13225-020-00466-2)) |
| Otu30355 | 1 | k__Fungi(100);p__Ascomycota(100);c__Saccharomycetes(100);o__Saccharomycetales(100);f__Metschnikowiaceae(100);g__Metschnikowia(100);s__Metschnikowia_pulcherrima(100); | Metschnikowia | 13 | Saprotroph | \|Undefined Saprotroph\| | Yeast | NULL | Probable | As nectar-tap_saprotroph (Põlme et al. 2020); Decay sugar-rich substrates (Põlme et al. 2020) | Põlme S, et al. 2020. Fungal Diversity 105:1-16 ((https://doi.org/10.1007/s13225-020-00466-2)) |
| Otu30356 | 1 | k__Fungi(100);p__Ascomycota(100);c__Saccharomycetes(100);o__Saccharomycetales(100);f__Metschnikowiaceae(100);g__Metschnikowia(100);s__Metschnikowia_pulcherrima(100); | Metschnikowia | 13 | Saprotroph | \|Undefined Saprotroph\| | Yeast | NULL | Probable | As nectar-tap_saprotroph (Põlme et al. 2020); Decay sugar-rich substrates (Põlme et al. 2020) | Põlme S, et al. 2020. Fungal Diversity 105:1-16 ((https://doi.org/10.1007/s13225-020-00466-2)) |
| Otu30358 | 1 | k__Fungi(100);p__Ascomycota(100);c__Saccharomycetes(100);o__Saccharomycetales(100);f__Metschnikowiaceae(100);g__Metschnikowia(100);s__Metschnikowia_pulcherrima(100); | Metschnikowia | 13 | Saprotroph | \|Undefined Saprotroph\| | Yeast | NULL | Probable | As nectar-tap_saprotroph (Põlme et al. 2020); Decay sugar-rich substrates (Põlme et al. 2020) | Põlme S, et al. 2020. Fungal Diversity 105:1-16 ((https://doi.org/10.1007/s13225-020-00466-2)) |
| Otu30359 | 1 | k__Fungi(100);p__Ascomycota(100);c__Saccharomycetes(100);o__Saccharomycetales(100);f__Metschnikowiaceae(100);g__Metschnikowia(100);g__Metschnikowia_unclassified(100); | Metschnikowia | 13 | Saprotroph | \|Undefined Saprotroph\| | Yeast | NULL | Probable | As nectar-tap_saprotroph (Põlme et al. 2020); Decay sugar-rich substrates (Põlme et al. 2020) | Põlme S, et al. 2020. Fungal Diversity 105:1-16 ((https://doi.org/10.1007/s13225-020-00466-2)) |
| Otu30361 | 1 | k__Fungi(100);p__Ascomycota(100);c__Saccharomycetes(100);o__Saccharomycetales(100);f__Metschnikowiaceae(100);g__Metschnikowia(100);s__Metschnikowia_pulcherrima(100); | Metschnikowia | 13 | Saprotroph | \|Undefined Saprotroph\| | Yeast | NULL | Probable | As nectar-tap_saprotroph (Põlme et al. 2020); Decay sugar-rich substrates (Põlme et al. 2020) | Põlme S, et al. 2020. Fungal Diversity 105:1-16 ((https://doi.org/10.1007/s13225-020-00466-2)) |
| Otu30365 | 1 | k__Fungi(100);p__Ascomycota(100);c__Saccharomycetes(100);o__Saccharomycetales(100);f__Metschnikowiaceae(100);g__Metschnikowia(100);g__Metschnikowia_unclassified(100); | Metschnikowia | 13 | Saprotroph | \|Undefined Saprotroph\| | Yeast | NULL | Probable | As nectar-tap_saprotroph (Põlme et al. 2020); Decay sugar-rich substrates (Põlme et al. 2020) | Põlme S, et al. 2020. Fungal Diversity 105:1-16 ((https://doi.org/10.1007/s13225-020-00466-2)) |
| Otu30366 | 1 | k__Fungi(100);p__Ascomycota(100);c__Saccharomycetes(100);o__Saccharomycetales(100);f__Metschnikowiaceae(100);g__Metschnikowia(100);g__Metschnikowia_unclassified(100); | Metschnikowia | 13 | Saprotroph | \|Undefined Saprotroph\| | Yeast | NULL | Probable | As nectar-tap_saprotroph (Põlme et al. 2020); Decay sugar-rich substrates (Põlme et al. 2020) | Põlme S, et al. 2020. Fungal Diversity 105:1-16 ((https://doi.org/10.1007/s13225-020-00466-2)) |
| Otu30367 | 1 | k__Fungi(100);p__Ascomycota(100);c__Saccharomycetes(100);o__Saccharomycetales(100);f__Metschnikowiaceae(100);g__Metschnikowia(100);s__Metschnikowia_pulcherrima(100); | Metschnikowia | 13 | Saprotroph | \|Undefined Saprotroph\| | Yeast | NULL | Probable | As nectar-tap_saprotroph (Põlme et al. 2020); Decay sugar-rich substrates (Põlme et al. 2020) | Põlme S, et al. 2020. Fungal Diversity 105:1-16 ((https://doi.org/10.1007/s13225-020-00466-2)) |
| Otu30368 | 1 | k__Fungi(100);p__Ascomycota(100);c__Saccharomycetes(100);o__Saccharomycetales(100);f__Metschnikowiaceae(100);g__Metschnikowia(100);s__Metschnikowia_pulcherrima(100); | Metschnikowia | 13 | Saprotroph | \|Undefined Saprotroph\| | Yeast | NULL | Probable | As nectar-tap_saprotroph (Põlme et al. 2020); Decay sugar-rich substrates (Põlme et al. 2020) | Põlme S, et al. 2020. Fungal Diversity 105:1-16 ((https://doi.org/10.1007/s13225-020-00466-2)) |
| Otu30369 | 1 | k__Fungi(100);p__Ascomycota(100);c__Saccharomycetes(100);o__Saccharomycetales(100);f__Saccharomycetales_family_Incertae_sedis(100);g__Candida(100);g__Candida_unclassified(100); | Candida | 13 | Pathotroph-Saprotroph-Symbiotroph | Animal Pathogen-Endophyte-Endosymbiont-Epiphyte-\|Undefined Saprotroph\| | Dimorphic Yeast-Yeast | NULL | Probable | As nectar-tap_saprotroph (Põlme et al. 2020); Decay sugar-rich substrates (Põlme et al. 2020); This asexual genus is polyphyletic and contain species that function very differently depend on what they are. Caution should be taken when interpreting guilds for Candida species. | Manolakaki D, et al. 2010. Virulence. 1: 367-375 ((https://doi.org/10.4161/viru.1.5.12796)); Mekbib SB, et al. 2011. Tropical Plant Pathology 36:233-240 ((https://doi.org/10.1590/S1982-56762011000400004)); Põlme S, et al. 2020. Fungal Diversity 105:1-16 ((https://doi.org/10.1007/s13225-020-00466-2)) |
| Otu30370 | 1 | k__Fungi(100);p__Ascomycota(100);c__Saccharomycetes(100);o__Saccharomycetales(100);f__Saccharomycetales_family_Incertae_sedis(100);g__Candida(100);g__Candida_unclassified(100); | Candida | 13 | Pathotroph-Saprotroph-Symbiotroph | Animal Pathogen-Endophyte-Endosymbiont-Epiphyte-\|Undefined Saprotroph\| | Dimorphic Yeast-Yeast | NULL | Probable | As nectar-tap_saprotroph (Põlme et al. 2020); Decay sugar-rich substrates (Põlme et al. 2020); This asexual genus is polyphyletic and contain species that function very differently depend on what they are. Caution should be taken when interpreting guilds for Candida species. | Manolakaki D, et al. 2010. Virulence. 1: 367-375 ((https://doi.org/10.4161/viru.1.5.12796)); Mekbib SB, et al. 2011. Tropical Plant Pathology 36:233-240 ((https://doi.org/10.1590/S1982-56762011000400004)); Põlme S, et al. 2020. Fungal Diversity 105:1-16 ((https://doi.org/10.1007/s13225-020-00466-2)) |
| Otu30371 | 1 | k__Fungi(100);p__Ascomycota(100);c__Saccharomycetes(100);o__Saccharomycetales(100);f__Metschnikowiaceae(100);g__Metschnikowia(100);s__Metschnikowia_pulcherrima(100); | Metschnikowia | 13 | Saprotroph | \|Undefined Saprotroph\| | Yeast | NULL | Probable | As nectar-tap_saprotroph (Põlme et al. 2020); Decay sugar-rich substrates (Põlme et al. 2020) | Põlme S, et al. 2020. Fungal Diversity 105:1-16 ((https://doi.org/10.1007/s13225-020-00466-2)) |
| Otu30372 | 1 | k__Fungi(100);p__Ascomycota(100);c__Saccharomycetes(100);o__Saccharomycetales(100);f__Metschnikowiaceae(100);g__Metschnikowia(100);s__Metschnikowia_pulcherrima(100); | Metschnikowia | 13 | Saprotroph | \|Undefined Saprotroph\| | Yeast | NULL | Probable | As nectar-tap_saprotroph (Põlme et al. 2020); Decay sugar-rich substrates (Põlme et al. 2020) | Põlme S, et al. 2020. Fungal Diversity 105:1-16 ((https://doi.org/10.1007/s13225-020-00466-2)) |
| Otu30373 | 1 | k__Fungi(100);p__Ascomycota(100);c__Saccharomycetes(100);o__Saccharomycetales(100);f__Metschnikowiaceae(100);g__Metschnikowia(100);s__Metschnikowia_pulcherrima(100); | Metschnikowia | 13 | Saprotroph | \|Undefined Saprotroph\| | Yeast | NULL | Probable | As nectar-tap_saprotroph (Põlme et al. 2020); Decay sugar-rich substrates (Põlme et al. 2020) | Põlme S, et al. 2020. Fungal Diversity 105:1-16 ((https://doi.org/10.1007/s13225-020-00466-2)) |
| Otu30376 | 1 | k__Fungi(100);p__Ascomycota(100);c__Dothideomycetes(100);o__Pleosporales(100);f__Cucurbitariaceae(100);g__Curreya(100);g__Curreya_unclassified(100); | Curreya | 13 | Saprotroph | \|Plant Saprotroph\|-Undefined Saprotroph | Microfungus | NULL | Probable | As litter_saprotroph (Põlme et al. 2020); Decay leaf, fruit, seed (Põlme et al. 2020) | Tedersoo L, et al. 2014. Science 346:e1256688 ((https://doi.org/10.1126/science.1256688)); Põlme S, et al. 2020. Fungal Diversity 105:1-16 ((https://doi.org/10.1007/s13225-020-00466-2)) |
| Otu30387 | 1 | k__Fungi(100);p__Ascomycota(100);c__Saccharomycetes(100);o__Saccharomycetales(100);f__Metschnikowiaceae(100);g__Metschnikowia(100);g__Metschnikowia_unclassified(100); | Metschnikowia | 13 | Saprotroph | \|Undefined Saprotroph\| | Yeast | NULL | Probable | As nectar-tap_saprotroph (Põlme et al. 2020); Decay sugar-rich substrates (Põlme et al. 2020) | Põlme S, et al. 2020. Fungal Diversity 105:1-16 ((https://doi.org/10.1007/s13225-020-00466-2)) |
| Otu30389 | 1 | k__Fungi(100);p__Ascomycota(100);c__Eurotiomycetes(100);o__Eurotiales(100);f__Trichocomaceae(100);g__Talaromyces(100);s__Talaromyces_aurantiacus(100); | Talaromyces | 13 | Pathotroph-Saprotroph | Animal Parasite-\|Undefined Saprotroph\| | Microfungus | NULL | Probable | Decay leaf, fruit, seed, soil, animal material (Põlme et al. 2020) | Tedersoo L, et al. 2014. Science 346:e1256688 ((https://doi.org/10.1126/science.1256688)); Põlme S, et al. 2020. Fungal Diversity 105:1-16 ((https://doi.org/10.1007/s13225-020-00466-2)) |
| Otu30392 | 1 | k__Fungi(100);p__Ascomycota(100);c__Saccharomycetes(100);o__Saccharomycetales(100);f__Metschnikowiaceae(100);g__Metschnikowia(100);s__Metschnikowia_pulcherrima(100); | Metschnikowia | 13 | Saprotroph | \|Undefined Saprotroph\| | Yeast | NULL | Probable | As nectar-tap_saprotroph (Põlme et al. 2020); Decay sugar-rich substrates (Põlme et al. 2020) | Põlme S, et al. 2020. Fungal Diversity 105:1-16 ((https://doi.org/10.1007/s13225-020-00466-2)) |
| Otu30396 | 1 | k__Fungi(100);p__Ascomycota(100);c__Saccharomycetes(100);o__Saccharomycetales(100);f__Metschnikowiaceae(100);g__Metschnikowia(100);s__Metschnikowia_pulcherrima(100); | Metschnikowia | 13 | Saprotroph | \|Undefined Saprotroph\| | Yeast | NULL | Probable | As nectar-tap_saprotroph (Põlme et al. 2020); Decay sugar-rich substrates (Põlme et al. 2020) | Põlme S, et al. 2020. Fungal Diversity 105:1-16 ((https://doi.org/10.1007/s13225-020-00466-2)) |
| Otu30397 | 1 | k__Fungi(100);p__Ascomycota(100);c__Saccharomycetes(100);o__Saccharomycetales(100);f__Metschnikowiaceae(100);g__Metschnikowia(100);g__Metschnikowia_unclassified(100); | Metschnikowia | 13 | Saprotroph | \|Undefined Saprotroph\| | Yeast | NULL | Probable | As nectar-tap_saprotroph (Põlme et al. 2020); Decay sugar-rich substrates (Põlme et al. 2020) | Põlme S, et al. 2020. Fungal Diversity 105:1-16 ((https://doi.org/10.1007/s13225-020-00466-2)) |
| Otu30412 | 1 | k__Fungi(100);p__Ascomycota(100);c__Saccharomycetes(100);o__Saccharomycetales(100);f__Saccharomycetales_family_Incertae_sedis(100);g__Candida(100);g__Candida_unclassified(100); | Candida | 13 | Pathotroph-Saprotroph-Symbiotroph | Animal Pathogen-Endophyte-Endosymbiont-Epiphyte-\|Undefined Saprotroph\| | Dimorphic Yeast-Yeast | NULL | Probable | As nectar-tap_saprotroph (Põlme et al. 2020); Decay sugar-rich substrates (Põlme et al. 2020); This asexual genus is polyphyletic and contain species that function very differently depend on what they are. Caution should be taken when interpreting guilds for Candida species. | Manolakaki D, et al. 2010. Virulence. 1: 367-375 ((https://doi.org/10.4161/viru.1.5.12796)); Mekbib SB, et al. 2011. Tropical Plant Pathology 36:233-240 ((https://doi.org/10.1590/S1982-56762011000400004)); Põlme S, et al. 2020. Fungal Diversity 105:1-16 ((https://doi.org/10.1007/s13225-020-00466-2)) |
| Otu30413 | 1 | k__Fungi(100);p__Ascomycota(100);c__Saccharomycetes(100);o__Saccharomycetales(100);f__Saccharomycetales_family_Incertae_sedis(100);f__Saccharomycetales_family_Incertae_sedis_unclassified(100);f__Saccharomycetales_family_Incertae_sedis_unclassified(100); | Saccharomycetales | 7 | Saprotroph | Undefined Saprotroph | Yeast | NULL | Possible | NULL | Sterkenburg E, et al. 2015. New Phytologist 207:1145-1158 ((https://doi.org/10.1111/nph.13426)) |
| Otu30414 | 1 | k__Fungi(100);p__Ascomycota(100);c__Saccharomycetes(100);o__Saccharomycetales(100);f__Metschnikowiaceae(100);g__Metschnikowia(100);g__Metschnikowia_unclassified(100); | Metschnikowia | 13 | Saprotroph | \|Undefined Saprotroph\| | Yeast | NULL | Probable | As nectar-tap_saprotroph (Põlme et al. 2020); Decay sugar-rich substrates (Põlme et al. 2020) | Põlme S, et al. 2020. Fungal Diversity 105:1-16 ((https://doi.org/10.1007/s13225-020-00466-2)) |
| Otu30421 | 1 | k__Fungi(100);p__Ascomycota(100);c__Saccharomycetes(100);o__Saccharomycetales(100);f__Metschnikowiaceae(100);g__Metschnikowia(100);g__Metschnikowia_unclassified(100); | Metschnikowia | 13 | Saprotroph | \|Undefined Saprotroph\| | Yeast | NULL | Probable | As nectar-tap_saprotroph (Põlme et al. 2020); Decay sugar-rich substrates (Põlme et al. 2020) | Põlme S, et al. 2020. Fungal Diversity 105:1-16 ((https://doi.org/10.1007/s13225-020-00466-2)) |
| Otu30422 | 1 | k__Fungi(100);p__Ascomycota(100);c__Saccharomycetes(100);o__Saccharomycetales(100);f__Metschnikowiaceae(100);g__Metschnikowia(100);s__Metschnikowia_pulcherrima(100); | Metschnikowia | 13 | Saprotroph | \|Undefined Saprotroph\| | Yeast | NULL | Probable | As nectar-tap_saprotroph (Põlme et al. 2020); Decay sugar-rich substrates (Põlme et al. 2020) | Põlme S, et al. 2020. Fungal Diversity 105:1-16 ((https://doi.org/10.1007/s13225-020-00466-2)) |
| Otu30424 | 1 | k__Fungi(100);p__Ascomycota(100);c__Saccharomycetes(100);o__Saccharomycetales(100);f__Saccharomycetales_family_Incertae_sedis(100);f__Saccharomycetales_family_Incertae_sedis_unclassified(100);f__Saccharomycetales_family_Incertae_sedis_unclassified(100); | Saccharomycetales | 7 | Saprotroph | Undefined Saprotroph | Yeast | NULL | Possible | NULL | Sterkenburg E, et al. 2015. New Phytologist 207:1145-1158 ((https://doi.org/10.1111/nph.13426)) |
| Otu30427 | 1 | k__Fungi(100);p__Ascomycota(100);c__Saccharomycetes(100);o__Saccharomycetales(100);f__Metschnikowiaceae(100);g__Metschnikowia(100);s__Metschnikowia_pulcherrima(100); | Metschnikowia | 13 | Saprotroph | \|Undefined Saprotroph\| | Yeast | NULL | Probable | As nectar-tap_saprotroph (Põlme et al. 2020); Decay sugar-rich substrates (Põlme et al. 2020) | Põlme S, et al. 2020. Fungal Diversity 105:1-16 ((https://doi.org/10.1007/s13225-020-00466-2)) |
| Otu30428 | 1 | k__Fungi(100);p__Ascomycota(100);c__Saccharomycetes(100);o__Saccharomycetales(100);f__Metschnikowiaceae(100);g__Metschnikowia(100);s__Metschnikowia_pulcherrima(100); | Metschnikowia | 13 | Saprotroph | \|Undefined Saprotroph\| | Yeast | NULL | Probable | As nectar-tap_saprotroph (Põlme et al. 2020); Decay sugar-rich substrates (Põlme et al. 2020) | Põlme S, et al. 2020. Fungal Diversity 105:1-16 ((https://doi.org/10.1007/s13225-020-00466-2)) |
| Otu30429 | 1 | k__Fungi(100);p__Ascomycota(100);c__Saccharomycetes(100);o__Saccharomycetales(100);f__Saccharomycetales_family_Incertae_sedis(100);f__Saccharomycetales_family_Incertae_sedis_unclassified(100);f__Saccharomycetales_family_Incertae_sedis_unclassified(100); | Saccharomycetales | 7 | Saprotroph | Undefined Saprotroph | Yeast | NULL | Possible | NULL | Sterkenburg E, et al. 2015. New Phytologist 207:1145-1158 ((https://doi.org/10.1111/nph.13426)) |
| Otu30430 | 1 | k__Fungi(100);p__Ascomycota(100);c__Saccharomycetes(100);o__Saccharomycetales(100);f__Saccharomycetales_family_Incertae_sedis(100);g__Candida(100);s__Candida_cretensis(100); | Candida | 13 | Pathotroph-Saprotroph-Symbiotroph | Animal Pathogen-Endophyte-Endosymbiont-Epiphyte-\|Undefined Saprotroph\| | Dimorphic Yeast-Yeast | NULL | Probable | As nectar-tap_saprotroph (Põlme et al. 2020); Decay sugar-rich substrates (Põlme et al. 2020); This asexual genus is polyphyletic and contain species that function very differently depend on what they are. Caution should be taken when interpreting guilds for Candida species. | Manolakaki D, et al. 2010. Virulence. 1: 367-375 ((https://doi.org/10.4161/viru.1.5.12796)); Mekbib SB, et al. 2011. Tropical Plant Pathology 36:233-240 ((https://doi.org/10.1590/S1982-56762011000400004)); Põlme S, et al. 2020. Fungal Diversity 105:1-16 ((https://doi.org/10.1007/s13225-020-00466-2)) |
| Otu30432 | 1 | k__Fungi(100);p__Ascomycota(100);c__Saccharomycetes(100);o__Saccharomycetales(100);f__Metschnikowiaceae(100);g__Metschnikowia(100);s__Metschnikowia_pulcherrima(100); | Metschnikowia | 13 | Saprotroph | \|Undefined Saprotroph\| | Yeast | NULL | Probable | As nectar-tap_saprotroph (Põlme et al. 2020); Decay sugar-rich substrates (Põlme et al. 2020) | Põlme S, et al. 2020. Fungal Diversity 105:1-16 ((https://doi.org/10.1007/s13225-020-00466-2)) |
| Otu30433 | 1 | k__Fungi(100);p__Ascomycota(100);c__Saccharomycetes(100);o__Saccharomycetales(100);f__Metschnikowiaceae(100);g__Metschnikowia(100);g__Metschnikowia_unclassified(100); | Metschnikowia | 13 | Saprotroph | \|Undefined Saprotroph\| | Yeast | NULL | Probable | As nectar-tap_saprotroph (Põlme et al. 2020); Decay sugar-rich substrates (Põlme et al. 2020) | Põlme S, et al. 2020. Fungal Diversity 105:1-16 ((https://doi.org/10.1007/s13225-020-00466-2)) |
| Otu30434 | 1 | k__Fungi(100);p__Ascomycota(100);c__Saccharomycetes(100);o__Saccharomycetales(100);f__Metschnikowiaceae(100);g__Metschnikowia(100);s__Metschnikowia_pulcherrima(100); | Metschnikowia | 13 | Saprotroph | \|Undefined Saprotroph\| | Yeast | NULL | Probable | As nectar-tap_saprotroph (Põlme et al. 2020); Decay sugar-rich substrates (Põlme et al. 2020) | Põlme S, et al. 2020. Fungal Diversity 105:1-16 ((https://doi.org/10.1007/s13225-020-00466-2)) |
| Otu30435 | 1 | k__Fungi(100);p__Ascomycota(100);c__Saccharomycetes(100);o__Saccharomycetales(100);f__Metschnikowiaceae(100);g__Metschnikowia(100);s__Metschnikowia_pulcherrima(100); | Metschnikowia | 13 | Saprotroph | \|Undefined Saprotroph\| | Yeast | NULL | Probable | As nectar-tap_saprotroph (Põlme et al. 2020); Decay sugar-rich substrates (Põlme et al. 2020) | Põlme S, et al. 2020. Fungal Diversity 105:1-16 ((https://doi.org/10.1007/s13225-020-00466-2)) |
| Otu30436 | 1 | k__Fungi(100);p__Ascomycota(100);c__Saccharomycetes(100);o__Saccharomycetales(100);f__Metschnikowiaceae(100);g__Metschnikowia(100);g__Metschnikowia_unclassified(100); | Metschnikowia | 13 | Saprotroph | \|Undefined Saprotroph\| | Yeast | NULL | Probable | As nectar-tap_saprotroph (Põlme et al. 2020); Decay sugar-rich substrates (Põlme et al. 2020) | Põlme S, et al. 2020. Fungal Diversity 105:1-16 ((https://doi.org/10.1007/s13225-020-00466-2)) |
| Otu30437 | 1 | k__Fungi(100);p__Ascomycota(100);c__Saccharomycetes(100);o__Saccharomycetales(100);f__Metschnikowiaceae(100);g__Metschnikowia(100);s__Metschnikowia_pulcherrima(100); | Metschnikowia | 13 | Saprotroph | \|Undefined Saprotroph\| | Yeast | NULL | Probable | As nectar-tap_saprotroph (Põlme et al. 2020); Decay sugar-rich substrates (Põlme et al. 2020) | Põlme S, et al. 2020. Fungal Diversity 105:1-16 ((https://doi.org/10.1007/s13225-020-00466-2)) |
| Otu30438 | 1 | k__Fungi(100);p__Ascomycota(100);c__Saccharomycetes(100);o__Saccharomycetales(100);f__Metschnikowiaceae(100);g__Metschnikowia(100);g__Metschnikowia_unclassified(100); | Metschnikowia | 13 | Saprotroph | \|Undefined Saprotroph\| | Yeast | NULL | Probable | As nectar-tap_saprotroph (Põlme et al. 2020); Decay sugar-rich substrates (Põlme et al. 2020) | Põlme S, et al. 2020. Fungal Diversity 105:1-16 ((https://doi.org/10.1007/s13225-020-00466-2)) |
| Otu30441 | 1 | k__Fungi(100);p__Ascomycota(100);c__Saccharomycetes(100);o__Saccharomycetales(100);f__Metschnikowiaceae(100);g__Metschnikowia(100);s__Metschnikowia_pulcherrima(100); | Metschnikowia | 13 | Saprotroph | \|Undefined Saprotroph\| | Yeast | NULL | Probable | As nectar-tap_saprotroph (Põlme et al. 2020); Decay sugar-rich substrates (Põlme et al. 2020) | Põlme S, et al. 2020. Fungal Diversity 105:1-16 ((https://doi.org/10.1007/s13225-020-00466-2)) |
| Otu30443 | 1 | k__Fungi(100);p__Ascomycota(100);c__Saccharomycetes(100);o__Saccharomycetales(100);f__Metschnikowiaceae(100);g__Metschnikowia(100);s__Metschnikowia_pulcherrima(100); | Metschnikowia | 13 | Saprotroph | \|Undefined Saprotroph\| | Yeast | NULL | Probable | As nectar-tap_saprotroph (Põlme et al. 2020); Decay sugar-rich substrates (Põlme et al. 2020) | Põlme S, et al. 2020. Fungal Diversity 105:1-16 ((https://doi.org/10.1007/s13225-020-00466-2)) |
| Otu30444 | 1 | k__Fungi(100);p__Ascomycota(100);c__Saccharomycetes(100);o__Saccharomycetales(100);f__Metschnikowiaceae(100);g__Metschnikowia(100);g__Metschnikowia_unclassified(100); | Metschnikowia | 13 | Saprotroph | \|Undefined Saprotroph\| | Yeast | NULL | Probable | As nectar-tap_saprotroph (Põlme et al. 2020); Decay sugar-rich substrates (Põlme et al. 2020) | Põlme S, et al. 2020. Fungal Diversity 105:1-16 ((https://doi.org/10.1007/s13225-020-00466-2)) |
| Otu30445 | 1 | k__Fungi(100);p__Ascomycota(100);c__Saccharomycetes(100);o__Saccharomycetales(100);f__Metschnikowiaceae(100);g__Metschnikowia(100);g__Metschnikowia_unclassified(100); | Metschnikowia | 13 | Saprotroph | \|Undefined Saprotroph\| | Yeast | NULL | Probable | As nectar-tap_saprotroph (Põlme et al. 2020); Decay sugar-rich substrates (Põlme et al. 2020) | Põlme S, et al. 2020. Fungal Diversity 105:1-16 ((https://doi.org/10.1007/s13225-020-00466-2)) |
| Otu30446 | 1 | k__Fungi(100);p__Ascomycota(100);c__Saccharomycetes(100);o__Saccharomycetales(100);o__Saccharomycetales_unclassified(100);o__Saccharomycetales_unclassified(100);o__Saccharomycetales_unclassified(100); | Saccharomycetales | 7 | Saprotroph | Undefined Saprotroph | Yeast | NULL | Possible | NULL | Sterkenburg E, et al. 2015. New Phytologist 207:1145-1158 ((https://doi.org/10.1111/nph.13426)) |
| Otu30447 | 1 | k__Fungi(100);p__Ascomycota(100);c__Saccharomycetes(100);o__Saccharomycetales(100);f__Metschnikowiaceae(100);g__Metschnikowia(100);s__Metschnikowia_pulcherrima(100); | Metschnikowia | 13 | Saprotroph | \|Undefined Saprotroph\| | Yeast | NULL | Probable | As nectar-tap_saprotroph (Põlme et al. 2020); Decay sugar-rich substrates (Põlme et al. 2020) | Põlme S, et al. 2020. Fungal Diversity 105:1-16 ((https://doi.org/10.1007/s13225-020-00466-2)) |
| Otu30448 | 1 | k__Fungi(100);p__Ascomycota(100);c__Saccharomycetes(100);o__Saccharomycetales(100);f__Metschnikowiaceae(100);g__Metschnikowia(100);s__Metschnikowia_pulcherrima(100); | Metschnikowia | 13 | Saprotroph | \|Undefined Saprotroph\| | Yeast | NULL | Probable | As nectar-tap_saprotroph (Põlme et al. 2020); Decay sugar-rich substrates (Põlme et al. 2020) | Põlme S, et al. 2020. Fungal Diversity 105:1-16 ((https://doi.org/10.1007/s13225-020-00466-2)) |
| Otu30449 | 1 | k__Fungi(100);p__Ascomycota(100);c__Saccharomycetes(100);o__Saccharomycetales(100);f__Metschnikowiaceae(100);g__Metschnikowia(100);s__Metschnikowia_pulcherrima(100); | Metschnikowia | 13 | Saprotroph | \|Undefined Saprotroph\| | Yeast | NULL | Probable | As nectar-tap_saprotroph (Põlme et al. 2020); Decay sugar-rich substrates (Põlme et al. 2020) | Põlme S, et al. 2020. Fungal Diversity 105:1-16 ((https://doi.org/10.1007/s13225-020-00466-2)) |
| Otu30450 | 1 | k__Fungi(100);p__Ascomycota(100);c__Saccharomycetes(100);o__Saccharomycetales(100);f__Metschnikowiaceae(100);g__Metschnikowia(100);s__Metschnikowia_pulcherrima(100); | Metschnikowia | 13 | Saprotroph | \|Undefined Saprotroph\| | Yeast | NULL | Probable | As nectar-tap_saprotroph (Põlme et al. 2020); Decay sugar-rich substrates (Põlme et al. 2020) | Põlme S, et al. 2020. Fungal Diversity 105:1-16 ((https://doi.org/10.1007/s13225-020-00466-2)) |
| Otu30452 | 1 | k__Fungi(100);p__Ascomycota(100);c__Saccharomycetes(100);o__Saccharomycetales(100);f__Metschnikowiaceae(100);g__Metschnikowia(100);s__Metschnikowia_pulcherrima(100); | Metschnikowia | 13 | Saprotroph | \|Undefined Saprotroph\| | Yeast | NULL | Probable | As nectar-tap_saprotroph (Põlme et al. 2020); Decay sugar-rich substrates (Põlme et al. 2020) | Põlme S, et al. 2020. Fungal Diversity 105:1-16 ((https://doi.org/10.1007/s13225-020-00466-2)) |
| Otu30454 | 1 | k__Fungi(100);p__Ascomycota(100);c__Saccharomycetes(100);o__Saccharomycetales(100);f__Metschnikowiaceae(100);g__Metschnikowia(100);s__Metschnikowia_pulcherrima(100); | Metschnikowia | 13 | Saprotroph | \|Undefined Saprotroph\| | Yeast | NULL | Probable | As nectar-tap_saprotroph (Põlme et al. 2020); Decay sugar-rich substrates (Põlme et al. 2020) | Põlme S, et al. 2020. Fungal Diversity 105:1-16 ((https://doi.org/10.1007/s13225-020-00466-2)) |
| Otu30455 | 1 | k__Fungi(100);p__Ascomycota(100);c__Saccharomycetes(100);o__Saccharomycetales(100);f__Metschnikowiaceae(100);g__Metschnikowia(100);s__Metschnikowia_pulcherrima(100); | Metschnikowia | 13 | Saprotroph | \|Undefined Saprotroph\| | Yeast | NULL | Probable | As nectar-tap_saprotroph (Põlme et al. 2020); Decay sugar-rich substrates (Põlme et al. 2020) | Põlme S, et al. 2020. Fungal Diversity 105:1-16 ((https://doi.org/10.1007/s13225-020-00466-2)) |
| Otu30456 | 1 | k__Fungi(100);p__Ascomycota(100);c__Saccharomycetes(100);o__Saccharomycetales(100);f__Metschnikowiaceae(100);g__Metschnikowia(100);g__Metschnikowia_unclassified(100); | Metschnikowia | 13 | Saprotroph | \|Undefined Saprotroph\| | Yeast | NULL | Probable | As nectar-tap_saprotroph (Põlme et al. 2020); Decay sugar-rich substrates (Põlme et al. 2020) | Põlme S, et al. 2020. Fungal Diversity 105:1-16 ((https://doi.org/10.1007/s13225-020-00466-2)) |
| Otu30457 | 1 | k__Fungi(100);p__Ascomycota(100);c__Saccharomycetes(100);o__Saccharomycetales(100);f__Metschnikowiaceae(100);g__Metschnikowia(100);g__Metschnikowia_unclassified(100); | Metschnikowia | 13 | Saprotroph | \|Undefined Saprotroph\| | Yeast | NULL | Probable | As nectar-tap_saprotroph (Põlme et al. 2020); Decay sugar-rich substrates (Põlme et al. 2020) | Põlme S, et al. 2020. Fungal Diversity 105:1-16 ((https://doi.org/10.1007/s13225-020-00466-2)) |
| Otu30458 | 1 | k__Fungi(100);p__Ascomycota(100);c__Saccharomycetes(100);o__Saccharomycetales(100);f__Metschnikowiaceae(100);g__Metschnikowia(100);s__Metschnikowia_pulcherrima(100); | Metschnikowia | 13 | Saprotroph | \|Undefined Saprotroph\| | Yeast | NULL | Probable | As nectar-tap_saprotroph (Põlme et al. 2020); Decay sugar-rich substrates (Põlme et al. 2020) | Põlme S, et al. 2020. Fungal Diversity 105:1-16 ((https://doi.org/10.1007/s13225-020-00466-2)) |
| Otu30459 | 1 | k__Fungi(100);p__Ascomycota(100);c__Saccharomycetes(100);o__Saccharomycetales(100);f__Metschnikowiaceae(100);g__Metschnikowia(100);s__Metschnikowia_pulcherrima(100); | Metschnikowia | 13 | Saprotroph | \|Undefined Saprotroph\| | Yeast | NULL | Probable | As nectar-tap_saprotroph (Põlme et al. 2020); Decay sugar-rich substrates (Põlme et al. 2020) | Põlme S, et al. 2020. Fungal Diversity 105:1-16 ((https://doi.org/10.1007/s13225-020-00466-2)) |
| Otu30460 | 1 | k__Fungi(100);p__Ascomycota(100);c__Saccharomycetes(100);o__Saccharomycetales(100);f__Metschnikowiaceae(100);g__Metschnikowia(100);g__Metschnikowia_unclassified(100); | Metschnikowia | 13 | Saprotroph | \|Undefined Saprotroph\| | Yeast | NULL | Probable | As nectar-tap_saprotroph (Põlme et al. 2020); Decay sugar-rich substrates (Põlme et al. 2020) | Põlme S, et al. 2020. Fungal Diversity 105:1-16 ((https://doi.org/10.1007/s13225-020-00466-2)) |
| Otu30461 | 1 | k__Fungi(100);p__Ascomycota(100);c__Saccharomycetes(100);o__Saccharomycetales(100);f__Metschnikowiaceae(100);g__Metschnikowia(100);s__Metschnikowia_pulcherrima(100); | Metschnikowia | 13 | Saprotroph | \|Undefined Saprotroph\| | Yeast | NULL | Probable | As nectar-tap_saprotroph (Põlme et al. 2020); Decay sugar-rich substrates (Põlme et al. 2020) | Põlme S, et al. 2020. Fungal Diversity 105:1-16 ((https://doi.org/10.1007/s13225-020-00466-2)) |
| Otu30462 | 1 | k__Fungi(100);p__Ascomycota(100);c__Saccharomycetes(100);o__Saccharomycetales(100);f__Metschnikowiaceae(100);g__Metschnikowia(100);g__Metschnikowia_unclassified(100); | Metschnikowia | 13 | Saprotroph | \|Undefined Saprotroph\| | Yeast | NULL | Probable | As nectar-tap_saprotroph (Põlme et al. 2020); Decay sugar-rich substrates (Põlme et al. 2020) | Põlme S, et al. 2020. Fungal Diversity 105:1-16 ((https://doi.org/10.1007/s13225-020-00466-2)) |
| Otu30463 | 1 | k__Fungi(100);p__Ascomycota(100);c__Saccharomycetes(100);o__Saccharomycetales(100);f__Metschnikowiaceae(100);g__Metschnikowia(100);g__Metschnikowia_unclassified(100); | Metschnikowia | 13 | Saprotroph | \|Undefined Saprotroph\| | Yeast | NULL | Probable | As nectar-tap_saprotroph (Põlme et al. 2020); Decay sugar-rich substrates (Põlme et al. 2020) | Põlme S, et al. 2020. Fungal Diversity 105:1-16 ((https://doi.org/10.1007/s13225-020-00466-2)) |
| Otu30465 | 1 | k__Fungi(100);p__Ascomycota(100);c__Saccharomycetes(100);o__Saccharomycetales(100);o__Saccharomycetales_unclassified(100);o__Saccharomycetales_unclassified(100);o__Saccharomycetales_unclassified(100); | Saccharomycetales | 7 | Saprotroph | Undefined Saprotroph | Yeast | NULL | Possible | NULL | Sterkenburg E, et al. 2015. New Phytologist 207:1145-1158 ((https://doi.org/10.1111/nph.13426)) |
| Otu30467 | 1 | k__Fungi(100);p__Ascomycota(100);c__Saccharomycetes(100);o__Saccharomycetales(100);f__Metschnikowiaceae(100);g__Metschnikowia(100);g__Metschnikowia_unclassified(100); | Metschnikowia | 13 | Saprotroph | \|Undefined Saprotroph\| | Yeast | NULL | Probable | As nectar-tap_saprotroph (Põlme et al. 2020); Decay sugar-rich substrates (Põlme et al. 2020) | Põlme S, et al. 2020. Fungal Diversity 105:1-16 ((https://doi.org/10.1007/s13225-020-00466-2)) |
| Otu30468 | 1 | k__Fungi(100);p__Ascomycota(100);c__Saccharomycetes(100);o__Saccharomycetales(100);f__Metschnikowiaceae(100);g__Metschnikowia(100);s__Metschnikowia_pulcherrima(100); | Metschnikowia | 13 | Saprotroph | \|Undefined Saprotroph\| | Yeast | NULL | Probable | As nectar-tap_saprotroph (Põlme et al. 2020); Decay sugar-rich substrates (Põlme et al. 2020) | Põlme S, et al. 2020. Fungal Diversity 105:1-16 ((https://doi.org/10.1007/s13225-020-00466-2)) |
| Otu30469 | 1 | k__Fungi(100);p__Ascomycota(100);c__Saccharomycetes(100);o__Saccharomycetales(100);f__Metschnikowiaceae(100);g__Metschnikowia(100);g__Metschnikowia_unclassified(100); | Metschnikowia | 13 | Saprotroph | \|Undefined Saprotroph\| | Yeast | NULL | Probable | As nectar-tap_saprotroph (Põlme et al. 2020); Decay sugar-rich substrates (Põlme et al. 2020) | Põlme S, et al. 2020. Fungal Diversity 105:1-16 ((https://doi.org/10.1007/s13225-020-00466-2)) |
| Otu30470 | 1 | k__Fungi(100);p__Ascomycota(100);c__Saccharomycetes(100);o__Saccharomycetales(100);f__Metschnikowiaceae(100);g__Metschnikowia(100);s__Metschnikowia_pulcherrima(100); | Metschnikowia | 13 | Saprotroph | \|Undefined Saprotroph\| | Yeast | NULL | Probable | As nectar-tap_saprotroph (Põlme et al. 2020); Decay sugar-rich substrates (Põlme et al. 2020) | Põlme S, et al. 2020. Fungal Diversity 105:1-16 ((https://doi.org/10.1007/s13225-020-00466-2)) |
| Otu30471 | 1 | k__Fungi(100);p__Ascomycota(100);c__Saccharomycetes(100);o__Saccharomycetales(100);f__Metschnikowiaceae(100);g__Metschnikowia(100);s__Metschnikowia_pulcherrima(100); | Metschnikowia | 13 | Saprotroph | \|Undefined Saprotroph\| | Yeast | NULL | Probable | As nectar-tap_saprotroph (Põlme et al. 2020); Decay sugar-rich substrates (Põlme et al. 2020) | Põlme S, et al. 2020. Fungal Diversity 105:1-16 ((https://doi.org/10.1007/s13225-020-00466-2)) |
| Otu30472 | 1 | k__Fungi(100);p__Ascomycota(100);c__Saccharomycetes(100);o__Saccharomycetales(100);f__Metschnikowiaceae(100);g__Metschnikowia(100);s__Metschnikowia_pulcherrima(100); | Metschnikowia | 13 | Saprotroph | \|Undefined Saprotroph\| | Yeast | NULL | Probable | As nectar-tap_saprotroph (Põlme et al. 2020); Decay sugar-rich substrates (Põlme et al. 2020) | Põlme S, et al. 2020. Fungal Diversity 105:1-16 ((https://doi.org/10.1007/s13225-020-00466-2)) |
| Otu30473 | 1 | k__Fungi(100);p__Ascomycota(100);c__Saccharomycetes(100);o__Saccharomycetales(100);f__Metschnikowiaceae(100);g__Metschnikowia(100);g__Metschnikowia_unclassified(100); | Metschnikowia | 13 | Saprotroph | \|Undefined Saprotroph\| | Yeast | NULL | Probable | As nectar-tap_saprotroph (Põlme et al. 2020); Decay sugar-rich substrates (Põlme et al. 2020) | Põlme S, et al. 2020. Fungal Diversity 105:1-16 ((https://doi.org/10.1007/s13225-020-00466-2)) |
| Otu30474 | 1 | k__Fungi(100);p__Ascomycota(100);c__Saccharomycetes(100);o__Saccharomycetales(100);f__Metschnikowiaceae(100);g__Metschnikowia(100);s__Metschnikowia_pulcherrima(100); | Metschnikowia | 13 | Saprotroph | \|Undefined Saprotroph\| | Yeast | NULL | Probable | As nectar-tap_saprotroph (Põlme et al. 2020); Decay sugar-rich substrates (Põlme et al. 2020) | Põlme S, et al. 2020. Fungal Diversity 105:1-16 ((https://doi.org/10.1007/s13225-020-00466-2)) |
| Otu30475 | 1 | k__Fungi(100);p__Ascomycota(100);c__Saccharomycetes(100);o__Saccharomycetales(100);f__Metschnikowiaceae(100);g__Metschnikowia(100);s__Metschnikowia_pulcherrima(100); | Metschnikowia | 13 | Saprotroph | \|Undefined Saprotroph\| | Yeast | NULL | Probable | As nectar-tap_saprotroph (Põlme et al. 2020); Decay sugar-rich substrates (Põlme et al. 2020) | Põlme S, et al. 2020. Fungal Diversity 105:1-16 ((https://doi.org/10.1007/s13225-020-00466-2)) |
| Otu30477 | 1 | k__Fungi(100);p__Ascomycota(100);c__Saccharomycetes(100);o__Saccharomycetales(100);f__Metschnikowiaceae(100);g__Metschnikowia(100);s__Metschnikowia_pulcherrima(100); | Metschnikowia | 13 | Saprotroph | \|Undefined Saprotroph\| | Yeast | NULL | Probable | As nectar-tap_saprotroph (Põlme et al. 2020); Decay sugar-rich substrates (Põlme et al. 2020) | Põlme S, et al. 2020. Fungal Diversity 105:1-16 ((https://doi.org/10.1007/s13225-020-00466-2)) |
| Otu30478 | 1 | k__Fungi(100);p__Ascomycota(100);c__Saccharomycetes(100);o__Saccharomycetales(100);f__Metschnikowiaceae(100);g__Metschnikowia(100);g__Metschnikowia_unclassified(100); | Metschnikowia | 13 | Saprotroph | \|Undefined Saprotroph\| | Yeast | NULL | Probable | As nectar-tap_saprotroph (Põlme et al. 2020); Decay sugar-rich substrates (Põlme et al. 2020) | Põlme S, et al. 2020. Fungal Diversity 105:1-16 ((https://doi.org/10.1007/s13225-020-00466-2)) |
| Otu30479 | 1 | k__Fungi(100);p__Ascomycota(100);c__Saccharomycetes(100);o__Saccharomycetales(100);f__Metschnikowiaceae(100);g__Metschnikowia(100);s__Metschnikowia_pulcherrima(100); | Metschnikowia | 13 | Saprotroph | \|Undefined Saprotroph\| | Yeast | NULL | Probable | As nectar-tap_saprotroph (Põlme et al. 2020); Decay sugar-rich substrates (Põlme et al. 2020) | Põlme S, et al. 2020. Fungal Diversity 105:1-16 ((https://doi.org/10.1007/s13225-020-00466-2)) |
| Otu30480 | 1 | k__Fungi(100);p__Ascomycota(100);c__Saccharomycetes(100);o__Saccharomycetales(100);f__Metschnikowiaceae(100);g__Metschnikowia(100);s__Metschnikowia_pulcherrima(100); | Metschnikowia | 13 | Saprotroph | \|Undefined Saprotroph\| | Yeast | NULL | Probable | As nectar-tap_saprotroph (Põlme et al. 2020); Decay sugar-rich substrates (Põlme et al. 2020) | Põlme S, et al. 2020. Fungal Diversity 105:1-16 ((https://doi.org/10.1007/s13225-020-00466-2)) |
| Otu30481 | 1 | k__Fungi(100);p__Ascomycota(100);c__Saccharomycetes(100);o__Saccharomycetales(100);f__Metschnikowiaceae(100);g__Metschnikowia(100);s__Metschnikowia_pulcherrima(100); | Metschnikowia | 13 | Saprotroph | \|Undefined Saprotroph\| | Yeast | NULL | Probable | As nectar-tap_saprotroph (Põlme et al. 2020); Decay sugar-rich substrates (Põlme et al. 2020) | Põlme S, et al. 2020. Fungal Diversity 105:1-16 ((https://doi.org/10.1007/s13225-020-00466-2)) |
| Otu30482 | 1 | k__Fungi(100);p__Ascomycota(100);c__Saccharomycetes(100);o__Saccharomycetales(100);f__Metschnikowiaceae(100);g__Metschnikowia(100);g__Metschnikowia_unclassified(100); | Metschnikowia | 13 | Saprotroph | \|Undefined Saprotroph\| | Yeast | NULL | Probable | As nectar-tap_saprotroph (Põlme et al. 2020); Decay sugar-rich substrates (Põlme et al. 2020) | Põlme S, et al. 2020. Fungal Diversity 105:1-16 ((https://doi.org/10.1007/s13225-020-00466-2)) |
| Otu30483 | 1 | k__Fungi(100);p__Ascomycota(100);c__Saccharomycetes(100);o__Saccharomycetales(100);f__Metschnikowiaceae(100);g__Metschnikowia(100);g__Metschnikowia_unclassified(100); | Metschnikowia | 13 | Saprotroph | \|Undefined Saprotroph\| | Yeast | NULL | Probable | As nectar-tap_saprotroph (Põlme et al. 2020); Decay sugar-rich substrates (Põlme et al. 2020) | Põlme S, et al. 2020. Fungal Diversity 105:1-16 ((https://doi.org/10.1007/s13225-020-00466-2)) |
| Otu30484 | 1 | k__Fungi(100);p__Ascomycota(100);c__Saccharomycetes(100);o__Saccharomycetales(100);f__Metschnikowiaceae(100);g__Metschnikowia(100);s__Metschnikowia_pulcherrima(100); | Metschnikowia | 13 | Saprotroph | \|Undefined Saprotroph\| | Yeast | NULL | Probable | As nectar-tap_saprotroph (Põlme et al. 2020); Decay sugar-rich substrates (Põlme et al. 2020) | Põlme S, et al. 2020. Fungal Diversity 105:1-16 ((https://doi.org/10.1007/s13225-020-00466-2)) |
| Otu30485 | 1 | k__Fungi(100);p__Ascomycota(100);c__Saccharomycetes(100);o__Saccharomycetales(100);f__Metschnikowiaceae(100);g__Metschnikowia(100);g__Metschnikowia_unclassified(100); | Metschnikowia | 13 | Saprotroph | \|Undefined Saprotroph\| | Yeast | NULL | Probable | As nectar-tap_saprotroph (Põlme et al. 2020); Decay sugar-rich substrates (Põlme et al. 2020) | Põlme S, et al. 2020. Fungal Diversity 105:1-16 ((https://doi.org/10.1007/s13225-020-00466-2)) |
| Otu30486 | 1 | k__Fungi(100);p__Ascomycota(100);c__Saccharomycetes(100);o__Saccharomycetales(100);f__Metschnikowiaceae(100);g__Metschnikowia(100);s__Metschnikowia_pulcherrima(100); | Metschnikowia | 13 | Saprotroph | \|Undefined Saprotroph\| | Yeast | NULL | Probable | As nectar-tap_saprotroph (Põlme et al. 2020); Decay sugar-rich substrates (Põlme et al. 2020) | Põlme S, et al. 2020. Fungal Diversity 105:1-16 ((https://doi.org/10.1007/s13225-020-00466-2)) |
| Otu30487 | 1 | k__Fungi(100);p__Ascomycota(100);c__Saccharomycetes(100);o__Saccharomycetales(100);f__Metschnikowiaceae(100);g__Metschnikowia(100);s__Metschnikowia_pulcherrima(100); | Metschnikowia | 13 | Saprotroph | \|Undefined Saprotroph\| | Yeast | NULL | Probable | As nectar-tap_saprotroph (Põlme et al. 2020); Decay sugar-rich substrates (Põlme et al. 2020) | Põlme S, et al. 2020. Fungal Diversity 105:1-16 ((https://doi.org/10.1007/s13225-020-00466-2)) |
| Otu30488 | 1 | k__Fungi(100);p__Ascomycota(100);c__Saccharomycetes(100);o__Saccharomycetales(100);f__Metschnikowiaceae(100);g__Metschnikowia(100);g__Metschnikowia_unclassified(100); | Metschnikowia | 13 | Saprotroph | \|Undefined Saprotroph\| | Yeast | NULL | Probable | As nectar-tap_saprotroph (Põlme et al. 2020); Decay sugar-rich substrates (Põlme et al. 2020) | Põlme S, et al. 2020. Fungal Diversity 105:1-16 ((https://doi.org/10.1007/s13225-020-00466-2)) |
| Otu30489 | 1 | k__Fungi(100);p__Ascomycota(100);c__Saccharomycetes(100);o__Saccharomycetales(100);f__Metschnikowiaceae(100);g__Metschnikowia(100);s__Metschnikowia_pulcherrima(100); | Metschnikowia | 13 | Saprotroph | \|Undefined Saprotroph\| | Yeast | NULL | Probable | As nectar-tap_saprotroph (Põlme et al. 2020); Decay sugar-rich substrates (Põlme et al. 2020) | Põlme S, et al. 2020. Fungal Diversity 105:1-16 ((https://doi.org/10.1007/s13225-020-00466-2)) |
| Otu30490 | 1 | k__Fungi(100);p__Ascomycota(100);c__Saccharomycetes(100);o__Saccharomycetales(100);f__Metschnikowiaceae(100);g__Metschnikowia(100);s__Metschnikowia_pulcherrima(100); | Metschnikowia | 13 | Saprotroph | \|Undefined Saprotroph\| | Yeast | NULL | Probable | As nectar-tap_saprotroph (Põlme et al. 2020); Decay sugar-rich substrates (Põlme et al. 2020) | Põlme S, et al. 2020. Fungal Diversity 105:1-16 ((https://doi.org/10.1007/s13225-020-00466-2)) |
| Otu30491 | 1 | k__Fungi(100);p__Ascomycota(100);c__Saccharomycetes(100);o__Saccharomycetales(100);f__Metschnikowiaceae(100);g__Metschnikowia(100);s__Metschnikowia_pulcherrima(100); | Metschnikowia | 13 | Saprotroph | \|Undefined Saprotroph\| | Yeast | NULL | Probable | As nectar-tap_saprotroph (Põlme et al. 2020); Decay sugar-rich substrates (Põlme et al. 2020) | Põlme S, et al. 2020. Fungal Diversity 105:1-16 ((https://doi.org/10.1007/s13225-020-00466-2)) |
| Otu30492 | 1 | k__Fungi(100);p__Ascomycota(100);c__Saccharomycetes(100);o__Saccharomycetales(100);f__Metschnikowiaceae(100);g__Metschnikowia(100);s__Metschnikowia_pulcherrima(100); | Metschnikowia | 13 | Saprotroph | \|Undefined Saprotroph\| | Yeast | NULL | Probable | As nectar-tap_saprotroph (Põlme et al. 2020); Decay sugar-rich substrates (Põlme et al. 2020) | Põlme S, et al. 2020. Fungal Diversity 105:1-16 ((https://doi.org/10.1007/s13225-020-00466-2)) |
| Otu30493 | 1 | k__Fungi(100);p__Ascomycota(100);c__Saccharomycetes(100);o__Saccharomycetales(100);f__Metschnikowiaceae(100);g__Metschnikowia(100);s__Metschnikowia_pulcherrima(100); | Metschnikowia | 13 | Saprotroph | \|Undefined Saprotroph\| | Yeast | NULL | Probable | As nectar-tap_saprotroph (Põlme et al. 2020); Decay sugar-rich substrates (Põlme et al. 2020) | Põlme S, et al. 2020. Fungal Diversity 105:1-16 ((https://doi.org/10.1007/s13225-020-00466-2)) |
| Otu30494 | 1 | k__Fungi(100);p__Ascomycota(100);c__Saccharomycetes(100);o__Saccharomycetales(100);f__Metschnikowiaceae(100);g__Metschnikowia(100);s__Metschnikowia_pulcherrima(100); | Metschnikowia | 13 | Saprotroph | \|Undefined Saprotroph\| | Yeast | NULL | Probable | As nectar-tap_saprotroph (Põlme et al. 2020); Decay sugar-rich substrates (Põlme et al. 2020) | Põlme S, et al. 2020. Fungal Diversity 105:1-16 ((https://doi.org/10.1007/s13225-020-00466-2)) |
| Otu30495 | 1 | k__Fungi(100);p__Ascomycota(100);c__Saccharomycetes(100);o__Saccharomycetales(100);f__Metschnikowiaceae(100);g__Metschnikowia(100);s__Metschnikowia_pulcherrima(100); | Metschnikowia | 13 | Saprotroph | \|Undefined Saprotroph\| | Yeast | NULL | Probable | As nectar-tap_saprotroph (Põlme et al. 2020); Decay sugar-rich substrates (Põlme et al. 2020) | Põlme S, et al. 2020. Fungal Diversity 105:1-16 ((https://doi.org/10.1007/s13225-020-00466-2)) |
| Otu30496 | 1 | k__Fungi(100);p__Ascomycota(100);c__Saccharomycetes(100);o__Saccharomycetales(100);f__Metschnikowiaceae(100);g__Metschnikowia(100);s__Metschnikowia_pulcherrima(100); | Metschnikowia | 13 | Saprotroph | \|Undefined Saprotroph\| | Yeast | NULL | Probable | As nectar-tap_saprotroph (Põlme et al. 2020); Decay sugar-rich substrates (Põlme et al. 2020) | Põlme S, et al. 2020. Fungal Diversity 105:1-16 ((https://doi.org/10.1007/s13225-020-00466-2)) |
| Otu30497 | 1 | k__Fungi(100);p__Ascomycota(100);c__Saccharomycetes(100);o__Saccharomycetales(100);f__Metschnikowiaceae(100);g__Metschnikowia(100);s__Metschnikowia_pulcherrima(100); | Metschnikowia | 13 | Saprotroph | \|Undefined Saprotroph\| | Yeast | NULL | Probable | As nectar-tap_saprotroph (Põlme et al. 2020); Decay sugar-rich substrates (Põlme et al. 2020) | Põlme S, et al. 2020. Fungal Diversity 105:1-16 ((https://doi.org/10.1007/s13225-020-00466-2)) |
| Otu30498 | 1 | k__Fungi(100);p__Ascomycota(100);c__Saccharomycetes(100);o__Saccharomycetales(100);f__Metschnikowiaceae(100);g__Metschnikowia(100);s__Metschnikowia_pulcherrima(100); | Metschnikowia | 13 | Saprotroph | \|Undefined Saprotroph\| | Yeast | NULL | Probable | As nectar-tap_saprotroph (Põlme et al. 2020); Decay sugar-rich substrates (Põlme et al. 2020) | Põlme S, et al. 2020. Fungal Diversity 105:1-16 ((https://doi.org/10.1007/s13225-020-00466-2)) |
| Otu30499 | 1 | k__Fungi(100);p__Ascomycota(100);c__Saccharomycetes(100);o__Saccharomycetales(100);f__Metschnikowiaceae(100);g__Metschnikowia(100);s__Metschnikowia_pulcherrima(100); | Metschnikowia | 13 | Saprotroph | \|Undefined Saprotroph\| | Yeast | NULL | Probable | As nectar-tap_saprotroph (Põlme et al. 2020); Decay sugar-rich substrates (Põlme et al. 2020) | Põlme S, et al. 2020. Fungal Diversity 105:1-16 ((https://doi.org/10.1007/s13225-020-00466-2)) |
| Otu30500 | 1 | k__Fungi(100);p__Ascomycota(100);c__Saccharomycetes(100);o__Saccharomycetales(100);f__Metschnikowiaceae(100);g__Metschnikowia(100);g__Metschnikowia_unclassified(100); | Metschnikowia | 13 | Saprotroph | \|Undefined Saprotroph\| | Yeast | NULL | Probable | As nectar-tap_saprotroph (Põlme et al. 2020); Decay sugar-rich substrates (Põlme et al. 2020) | Põlme S, et al. 2020. Fungal Diversity 105:1-16 ((https://doi.org/10.1007/s13225-020-00466-2)) |
| Otu30501 | 1 | k__Fungi(100);p__Ascomycota(100);c__Saccharomycetes(100);o__Saccharomycetales(100);f__Metschnikowiaceae(100);g__Metschnikowia(100);s__Metschnikowia_pulcherrima(100); | Metschnikowia | 13 | Saprotroph | \|Undefined Saprotroph\| | Yeast | NULL | Probable | As nectar-tap_saprotroph (Põlme et al. 2020); Decay sugar-rich substrates (Põlme et al. 2020) | Põlme S, et al. 2020. Fungal Diversity 105:1-16 ((https://doi.org/10.1007/s13225-020-00466-2)) |
| Otu30502 | 1 | k__Fungi(100);p__Ascomycota(100);c__Saccharomycetes(100);o__Saccharomycetales(100);f__Metschnikowiaceae(100);g__Metschnikowia(100);s__Metschnikowia_pulcherrima(100); | Metschnikowia | 13 | Saprotroph | \|Undefined Saprotroph\| | Yeast | NULL | Probable | As nectar-tap_saprotroph (Põlme et al. 2020); Decay sugar-rich substrates (Põlme et al. 2020) | Põlme S, et al. 2020. Fungal Diversity 105:1-16 ((https://doi.org/10.1007/s13225-020-00466-2)) |
| Otu30508 | 1 | k__Fungi(100);p__Ascomycota(100);c__Saccharomycetes(100);o__Saccharomycetales(100);f__Metschnikowiaceae(100);g__Metschnikowia(100);s__Metschnikowia_pulcherrima(100); | Metschnikowia | 13 | Saprotroph | \|Undefined Saprotroph\| | Yeast | NULL | Probable | As nectar-tap_saprotroph (Põlme et al. 2020); Decay sugar-rich substrates (Põlme et al. 2020) | Põlme S, et al. 2020. Fungal Diversity 105:1-16 ((https://doi.org/10.1007/s13225-020-00466-2)) |
| Otu30510 | 1 | k__Fungi(100);p__Ascomycota(100);c__Saccharomycetes(100);o__Saccharomycetales(100);f__Metschnikowiaceae(100);g__Metschnikowia(100);s__Metschnikowia_pulcherrima(100); | Metschnikowia | 13 | Saprotroph | \|Undefined Saprotroph\| | Yeast | NULL | Probable | As nectar-tap_saprotroph (Põlme et al. 2020); Decay sugar-rich substrates (Põlme et al. 2020) | Põlme S, et al. 2020. Fungal Diversity 105:1-16 ((https://doi.org/10.1007/s13225-020-00466-2)) |
| Otu30512 | 1 | k__Fungi(100);p__Ascomycota(100);c__Saccharomycetes(100);o__Saccharomycetales(100);f__Metschnikowiaceae(100);g__Metschnikowia(100);s__Metschnikowia_pulcherrima(100); | Metschnikowia | 13 | Saprotroph | \|Undefined Saprotroph\| | Yeast | NULL | Probable | As nectar-tap_saprotroph (Põlme et al. 2020); Decay sugar-rich substrates (Põlme et al. 2020) | Põlme S, et al. 2020. Fungal Diversity 105:1-16 ((https://doi.org/10.1007/s13225-020-00466-2)) |
| Otu30514 | 1 | k__Fungi(100);p__Ascomycota(100);c__Saccharomycetes(100);o__Saccharomycetales(100);f__Metschnikowiaceae(100);g__Metschnikowia(100);g__Metschnikowia_unclassified(100); | Metschnikowia | 13 | Saprotroph | \|Undefined Saprotroph\| | Yeast | NULL | Probable | As nectar-tap_saprotroph (Põlme et al. 2020); Decay sugar-rich substrates (Põlme et al. 2020) | Põlme S, et al. 2020. Fungal Diversity 105:1-16 ((https://doi.org/10.1007/s13225-020-00466-2)) |
| Otu30515 | 1 | k__Fungi(100);p__Ascomycota(100);c__Saccharomycetes(100);o__Saccharomycetales(100);f__Metschnikowiaceae(100);g__Metschnikowia(100);s__Metschnikowia_pulcherrima(100); | Metschnikowia | 13 | Saprotroph | \|Undefined Saprotroph\| | Yeast | NULL | Probable | As nectar-tap_saprotroph (Põlme et al. 2020); Decay sugar-rich substrates (Põlme et al. 2020) | Põlme S, et al. 2020. Fungal Diversity 105:1-16 ((https://doi.org/10.1007/s13225-020-00466-2)) |
| Otu30516 | 1 | k__Fungi(100);p__Ascomycota(100);c__Saccharomycetes(100);o__Saccharomycetales(100);f__Metschnikowiaceae(100);g__Metschnikowia(100);s__Metschnikowia_pulcherrima(100); | Metschnikowia | 13 | Saprotroph | \|Undefined Saprotroph\| | Yeast | NULL | Probable | As nectar-tap_saprotroph (Põlme et al. 2020); Decay sugar-rich substrates (Põlme et al. 2020) | Põlme S, et al. 2020. Fungal Diversity 105:1-16 ((https://doi.org/10.1007/s13225-020-00466-2)) |
| Otu30517 | 1 | k__Fungi(100);p__Ascomycota(100);c__Saccharomycetes(100);o__Saccharomycetales(100);f__Metschnikowiaceae(100);g__Metschnikowia(100);g__Metschnikowia_unclassified(100); | Metschnikowia | 13 | Saprotroph | \|Undefined Saprotroph\| | Yeast | NULL | Probable | As nectar-tap_saprotroph (Põlme et al. 2020); Decay sugar-rich substrates (Põlme et al. 2020) | Põlme S, et al. 2020. Fungal Diversity 105:1-16 ((https://doi.org/10.1007/s13225-020-00466-2)) |
| Otu30518 | 1 | k__Fungi(100);p__Ascomycota(100);c__Saccharomycetes(100);o__Saccharomycetales(100);f__Metschnikowiaceae(100);g__Metschnikowia(100);s__Metschnikowia_pulcherrima(100); | Metschnikowia | 13 | Saprotroph | \|Undefined Saprotroph\| | Yeast | NULL | Probable | As nectar-tap_saprotroph (Põlme et al. 2020); Decay sugar-rich substrates (Põlme et al. 2020) | Põlme S, et al. 2020. Fungal Diversity 105:1-16 ((https://doi.org/10.1007/s13225-020-00466-2)) |
| Otu30520 | 1 | k__Fungi(100);p__Ascomycota(100);c__Saccharomycetes(100);o__Saccharomycetales(100);f__Metschnikowiaceae(100);g__Metschnikowia(100);s__Metschnikowia_pulcherrima(100); | Metschnikowia | 13 | Saprotroph | \|Undefined Saprotroph\| | Yeast | NULL | Probable | As nectar-tap_saprotroph (Põlme et al. 2020); Decay sugar-rich substrates (Põlme et al. 2020) | Põlme S, et al. 2020. Fungal Diversity 105:1-16 ((https://doi.org/10.1007/s13225-020-00466-2)) |
| Otu30532 | 1 | k__Fungi(100);p__Ascomycota(100);c__Dothideomycetes(100);o__Capnodiales(100);f__Mycosphaerellaceae(100);g__Mycosphaerella(100);s__Mycosphaerella_nyssicola(100); | Mycosphaerella | 13 | Pathotroph-Saprotroph | \|Plant Pathogen\|-Plant Saprotroph | Microfungus | NULL | Probable | As litter_saprotroph (Põlme et al. 2020); Decay leaf, fruit, seed (Põlme et al. 2020) | Tedersoo L, et al. 2014. Science 346:e1256688 ((https://doi.org/10.1126/science.1256688)); Põlme S, et al. 2020. Fungal Diversity 105:1-16 ((https://doi.org/10.1007/s13225-020-00466-2)) |
| Otu30533 | 1 | k__Fungi(100);p__Ascomycota(100);c__Saccharomycetes(100);o__Saccharomycetales(100);f__Metschnikowiaceae(100);g__Metschnikowia(100);s__Metschnikowia_pulcherrima(100); | Metschnikowia | 13 | Saprotroph | \|Undefined Saprotroph\| | Yeast | NULL | Probable | As nectar-tap_saprotroph (Põlme et al. 2020); Decay sugar-rich substrates (Põlme et al. 2020) | Põlme S, et al. 2020. Fungal Diversity 105:1-16 ((https://doi.org/10.1007/s13225-020-00466-2)) |
| Otu30534 | 1 | k__Fungi(100);p__Ascomycota(100);c__Saccharomycetes(100);o__Saccharomycetales(100);f__Metschnikowiaceae(100);g__Metschnikowia(100);s__Metschnikowia_pulcherrima(100); | Metschnikowia | 13 | Saprotroph | \|Undefined Saprotroph\| | Yeast | NULL | Probable | As nectar-tap_saprotroph (Põlme et al. 2020); Decay sugar-rich substrates (Põlme et al. 2020) | Põlme S, et al. 2020. Fungal Diversity 105:1-16 ((https://doi.org/10.1007/s13225-020-00466-2)) |
| Otu30535 | 1 | k__Fungi(100);p__Ascomycota(100);c__Saccharomycetes(100);o__Saccharomycetales(100);f__Metschnikowiaceae(100);g__Metschnikowia(100);g__Metschnikowia_unclassified(100); | Metschnikowia | 13 | Saprotroph | \|Undefined Saprotroph\| | Yeast | NULL | Probable | As nectar-tap_saprotroph (Põlme et al. 2020); Decay sugar-rich substrates (Põlme et al. 2020) | Põlme S, et al. 2020. Fungal Diversity 105:1-16 ((https://doi.org/10.1007/s13225-020-00466-2)) |
| Otu30536 | 1 | k__Fungi(100);p__Ascomycota(100);c__Saccharomycetes(100);o__Saccharomycetales(100);f__Metschnikowiaceae(100);g__Metschnikowia(100);s__Metschnikowia_pulcherrima(100); | Metschnikowia | 13 | Saprotroph | \|Undefined Saprotroph\| | Yeast | NULL | Probable | As nectar-tap_saprotroph (Põlme et al. 2020); Decay sugar-rich substrates (Põlme et al. 2020) | Põlme S, et al. 2020. Fungal Diversity 105:1-16 ((https://doi.org/10.1007/s13225-020-00466-2)) |
| Otu30537 | 1 | k__Fungi(100);p__Ascomycota(100);c__Saccharomycetes(100);o__Saccharomycetales(100);f__Metschnikowiaceae(100);g__Metschnikowia(100);s__Metschnikowia_pulcherrima(100); | Metschnikowia | 13 | Saprotroph | \|Undefined Saprotroph\| | Yeast | NULL | Probable | As nectar-tap_saprotroph (Põlme et al. 2020); Decay sugar-rich substrates (Põlme et al. 2020) | Põlme S, et al. 2020. Fungal Diversity 105:1-16 ((https://doi.org/10.1007/s13225-020-00466-2)) |
| Otu30538 | 1 | k__Fungi(100);p__Ascomycota(100);c__Saccharomycetes(100);o__Saccharomycetales(100);f__Metschnikowiaceae(100);g__Metschnikowia(100);g__Metschnikowia_unclassified(100); | Metschnikowia | 13 | Saprotroph | \|Undefined Saprotroph\| | Yeast | NULL | Probable | As nectar-tap_saprotroph (Põlme et al. 2020); Decay sugar-rich substrates (Põlme et al. 2020) | Põlme S, et al. 2020. Fungal Diversity 105:1-16 ((https://doi.org/10.1007/s13225-020-00466-2)) |
| Otu30539 | 1 | k__Fungi(100);p__Ascomycota(100);c__Saccharomycetes(100);o__Saccharomycetales(100);f__Metschnikowiaceae(100);g__Metschnikowia(100);g__Metschnikowia_unclassified(100); | Metschnikowia | 13 | Saprotroph | \|Undefined Saprotroph\| | Yeast | NULL | Probable | As nectar-tap_saprotroph (Põlme et al. 2020); Decay sugar-rich substrates (Põlme et al. 2020) | Põlme S, et al. 2020. Fungal Diversity 105:1-16 ((https://doi.org/10.1007/s13225-020-00466-2)) |
| Otu30540 | 1 | k__Fungi(100);p__Ascomycota(100);c__Saccharomycetes(100);o__Saccharomycetales(100);f__Metschnikowiaceae(100);g__Metschnikowia(100);s__Metschnikowia_pulcherrima(100); | Metschnikowia | 13 | Saprotroph | \|Undefined Saprotroph\| | Yeast | NULL | Probable | As nectar-tap_saprotroph (Põlme et al. 2020); Decay sugar-rich substrates (Põlme et al. 2020) | Põlme S, et al. 2020. Fungal Diversity 105:1-16 ((https://doi.org/10.1007/s13225-020-00466-2)) |
| Otu30541 | 1 | k__Fungi(100);p__Ascomycota(100);c__Saccharomycetes(100);o__Saccharomycetales(100);f__Metschnikowiaceae(100);g__Metschnikowia(100);s__Metschnikowia_pulcherrima(100); | Metschnikowia | 13 | Saprotroph | \|Undefined Saprotroph\| | Yeast | NULL | Probable | As nectar-tap_saprotroph (Põlme et al. 2020); Decay sugar-rich substrates (Põlme et al. 2020) | Põlme S, et al. 2020. Fungal Diversity 105:1-16 ((https://doi.org/10.1007/s13225-020-00466-2)) |
| Otu30542 | 1 | k__Fungi(100);p__Ascomycota(100);c__Saccharomycetes(100);o__Saccharomycetales(100);f__Metschnikowiaceae(100);g__Metschnikowia(100);s__Metschnikowia_pulcherrima(100); | Metschnikowia | 13 | Saprotroph | \|Undefined Saprotroph\| | Yeast | NULL | Probable | As nectar-tap_saprotroph (Põlme et al. 2020); Decay sugar-rich substrates (Põlme et al. 2020) | Põlme S, et al. 2020. Fungal Diversity 105:1-16 ((https://doi.org/10.1007/s13225-020-00466-2)) |
| Otu30543 | 1 | k__Fungi(100);p__Ascomycota(100);c__Saccharomycetes(100);o__Saccharomycetales(100);f__Metschnikowiaceae(100);g__Metschnikowia(100);s__Metschnikowia_pulcherrima(100); | Metschnikowia | 13 | Saprotroph | \|Undefined Saprotroph\| | Yeast | NULL | Probable | As nectar-tap_saprotroph (Põlme et al. 2020); Decay sugar-rich substrates (Põlme et al. 2020) | Põlme S, et al. 2020. Fungal Diversity 105:1-16 ((https://doi.org/10.1007/s13225-020-00466-2)) |
| Otu30544 | 1 | k__Fungi(100);p__Ascomycota(100);c__Saccharomycetes(100);o__Saccharomycetales(100);f__Metschnikowiaceae(100);g__Metschnikowia(100);s__Metschnikowia_pulcherrima(100); | Metschnikowia | 13 | Saprotroph | \|Undefined Saprotroph\| | Yeast | NULL | Probable | As nectar-tap_saprotroph (Põlme et al. 2020); Decay sugar-rich substrates (Põlme et al. 2020) | Põlme S, et al. 2020. Fungal Diversity 105:1-16 ((https://doi.org/10.1007/s13225-020-00466-2)) |
| Otu30545 | 1 | k__Fungi(100);p__Ascomycota(100);c__Saccharomycetes(100);o__Saccharomycetales(100);f__Metschnikowiaceae(100);g__Metschnikowia(100);g__Metschnikowia_unclassified(100); | Metschnikowia | 13 | Saprotroph | \|Undefined Saprotroph\| | Yeast | NULL | Probable | As nectar-tap_saprotroph (Põlme et al. 2020); Decay sugar-rich substrates (Põlme et al. 2020) | Põlme S, et al. 2020. Fungal Diversity 105:1-16 ((https://doi.org/10.1007/s13225-020-00466-2)) |
| Otu30547 | 1 | k__Fungi(100);p__Ascomycota(100);c__Saccharomycetes(100);o__Saccharomycetales(100);f__Metschnikowiaceae(100);g__Metschnikowia(100);g__Metschnikowia_unclassified(100); | Metschnikowia | 13 | Saprotroph | \|Undefined Saprotroph\| | Yeast | NULL | Probable | As nectar-tap_saprotroph (Põlme et al. 2020); Decay sugar-rich substrates (Põlme et al. 2020) | Põlme S, et al. 2020. Fungal Diversity 105:1-16 ((https://doi.org/10.1007/s13225-020-00466-2)) |
| Otu30548 | 1 | k__Fungi(100);p__Ascomycota(100);c__Saccharomycetes(100);o__Saccharomycetales(100);f__Metschnikowiaceae(100);g__Metschnikowia(100);s__Metschnikowia_pulcherrima(100); | Metschnikowia | 13 | Saprotroph | \|Undefined Saprotroph\| | Yeast | NULL | Probable | As nectar-tap_saprotroph (Põlme et al. 2020); Decay sugar-rich substrates (Põlme et al. 2020) | Põlme S, et al. 2020. Fungal Diversity 105:1-16 ((https://doi.org/10.1007/s13225-020-00466-2)) |
| Otu30549 | 1 | k__Fungi(100);p__Ascomycota(100);c__Saccharomycetes(100);o__Saccharomycetales(100);f__Metschnikowiaceae(100);g__Metschnikowia(100);s__Metschnikowia_pulcherrima(100); | Metschnikowia | 13 | Saprotroph | \|Undefined Saprotroph\| | Yeast | NULL | Probable | As nectar-tap_saprotroph (Põlme et al. 2020); Decay sugar-rich substrates (Põlme et al. 2020) | Põlme S, et al. 2020. Fungal Diversity 105:1-16 ((https://doi.org/10.1007/s13225-020-00466-2)) |
| Otu30550 | 1 | k__Fungi(100);p__Ascomycota(100);c__Saccharomycetes(100);o__Saccharomycetales(100);f__Metschnikowiaceae(100);g__Metschnikowia(100);s__Metschnikowia_pulcherrima(100); | Metschnikowia | 13 | Saprotroph | \|Undefined Saprotroph\| | Yeast | NULL | Probable | As nectar-tap_saprotroph (Põlme et al. 2020); Decay sugar-rich substrates (Põlme et al. 2020) | Põlme S, et al. 2020. Fungal Diversity 105:1-16 ((https://doi.org/10.1007/s13225-020-00466-2)) |
| Otu30551 | 1 | k__Fungi(100);p__Ascomycota(100);c__Saccharomycetes(100);o__Saccharomycetales(100);f__Metschnikowiaceae(100);g__Metschnikowia(100);s__Metschnikowia_pulcherrima(100); | Metschnikowia | 13 | Saprotroph | \|Undefined Saprotroph\| | Yeast | NULL | Probable | As nectar-tap_saprotroph (Põlme et al. 2020); Decay sugar-rich substrates (Põlme et al. 2020) | Põlme S, et al. 2020. Fungal Diversity 105:1-16 ((https://doi.org/10.1007/s13225-020-00466-2)) |
| Otu30552 | 1 | k__Fungi(100);p__Ascomycota(100);c__Saccharomycetes(100);o__Saccharomycetales(100);f__Metschnikowiaceae(100);g__Metschnikowia(100);s__Metschnikowia_pulcherrima(100); | Metschnikowia | 13 | Saprotroph | \|Undefined Saprotroph\| | Yeast | NULL | Probable | As nectar-tap_saprotroph (Põlme et al. 2020); Decay sugar-rich substrates (Põlme et al. 2020) | Põlme S, et al. 2020. Fungal Diversity 105:1-16 ((https://doi.org/10.1007/s13225-020-00466-2)) |
| Otu30553 | 1 | k__Fungi(100);p__Ascomycota(100);c__Saccharomycetes(100);o__Saccharomycetales(100);f__Metschnikowiaceae(100);g__Metschnikowia(100);s__Metschnikowia_pulcherrima(100); | Metschnikowia | 13 | Saprotroph | \|Undefined Saprotroph\| | Yeast | NULL | Probable | As nectar-tap_saprotroph (Põlme et al. 2020); Decay sugar-rich substrates (Põlme et al. 2020) | Põlme S, et al. 2020. Fungal Diversity 105:1-16 ((https://doi.org/10.1007/s13225-020-00466-2)) |
| Otu30555 | 1 | k__Fungi(100);p__Ascomycota(100);c__Saccharomycetes(100);o__Saccharomycetales(100);f__Metschnikowiaceae(100);g__Metschnikowia(100);s__Metschnikowia_pulcherrima(100); | Metschnikowia | 13 | Saprotroph | \|Undefined Saprotroph\| | Yeast | NULL | Probable | As nectar-tap_saprotroph (Põlme et al. 2020); Decay sugar-rich substrates (Põlme et al. 2020) | Põlme S, et al. 2020. Fungal Diversity 105:1-16 ((https://doi.org/10.1007/s13225-020-00466-2)) |
| Otu30556 | 1 | k__Fungi(100);p__Ascomycota(100);c__Saccharomycetes(100);o__Saccharomycetales(100);f__Metschnikowiaceae(100);g__Metschnikowia(100);s__Metschnikowia_pulcherrima(100); | Metschnikowia | 13 | Saprotroph | \|Undefined Saprotroph\| | Yeast | NULL | Probable | As nectar-tap_saprotroph (Põlme et al. 2020); Decay sugar-rich substrates (Põlme et al. 2020) | Põlme S, et al. 2020. Fungal Diversity 105:1-16 ((https://doi.org/10.1007/s13225-020-00466-2)) |

| | | | | | | | | | | | |
|---|---|---|---|---|---|---|---|---|---|---|---|
| Otu30557 | 1 | k__Fungi(100);p__Ascomycota(100);c__Saccharomycetes(100);o__Saccharomycetales(100);f__Metschnikowiaceae(100);g__Metschnikowia(100);s__Metschnikowia_pulcherrima(100); | Metschnikowia | 13 | Saprotroph | \|Undefined Saprotroph\| | Yeast | NULL | Probable | As nectar-tap_saprotroph (Põlme et al. 2020); Decay sugar-rich substrates (Põlme et al. 2020) | Põlme S, et al. 2020. Fungal Diversity 105:1-16 ((https://doi.org/10.1007/s13225-020-00466-2)) |
| Otu30558 | 1 | k__Fungi(100);p__Ascomycota(100);c__Saccharomycetes(100);o__Saccharomycetales(100);f__Metschnikowiaceae(100);g__Metschnikowia(100);s__Metschnikowia_pulcherrima(100); | Metschnikowia | 13 | Saprotroph | \|Undefined Saprotroph\| | Yeast | NULL | Probable | As nectar-tap_saprotroph (Põlme et al. 2020); Decay sugar-rich substrates (Põlme et al. 2020) | Põlme S, et al. 2020. Fungal Diversity 105:1-16 ((https://doi.org/10.1007/s13225-020-00466-2)) |
| Otu30559 | 1 | k__Fungi(100);p__Ascomycota(100);c__Saccharomycetes(100);o__Saccharomycetales(100);f__Metschnikowiaceae(100);g__Metschnikowia(100);g__Metschnikowia_unclassified(100); | Metschnikowia | 13 | Saprotroph | \|Undefined Saprotroph\| | Yeast | NULL | Probable | As nectar-tap_saprotroph (Põlme et al. 2020); Decay sugar-rich substrates (Põlme et al. 2020) | Põlme S, et al. 2020. Fungal Diversity 105:1-16 ((https://doi.org/10.1007/s13225-020-00466-2)) |
| Otu30563 | 1 | k__Fungi(100);p__Ascomycota(100);c__Saccharomycetes(100);o__Saccharomycetales(100);f__Metschnikowiaceae(100);g__Metschnikowia(100);s__Metschnikowia_pulcherrima(100); | Metschnikowia | 13 | Saprotroph | \|Undefined Saprotroph\| | Yeast | NULL | Probable | As nectar-tap_saprotroph (Põlme et al. 2020); Decay sugar-rich substrates (Põlme et al. 2020) | Põlme S, et al. 2020. Fungal Diversity 105:1-16 ((https://doi.org/10.1007/s13225-020-00466-2)) |
| Otu30564 | 1 | k__Fungi(100);p__Ascomycota(100);c__Saccharomycetes(100);o__Saccharomycetales(100);f__Metschnikowiaceae(100);g__Metschnikowia(100);s__Metschnikowia_pulcherrima(100); | Metschnikowia | 13 | Saprotroph | \|Undefined Saprotroph\| | Yeast | NULL | Probable | As nectar-tap_saprotroph (Põlme et al. 2020); Decay sugar-rich substrates (Põlme et al. 2020) | Põlme S, et al. 2020. Fungal Diversity 105:1-16 ((https://doi.org/10.1007/s13225-020-00466-2)) |
| Otu30565 | 1 | k__Fungi(100);p__Ascomycota(100);c__Saccharomycetes(100);o__Saccharomycetales(100);f__Saccharomycetales_family_Incertae_sedis(100);f__Saccharomycetales_family_Incertae_sedis_unclassified(100);f__Saccharomycetales_family_Incertae_sedis_unclassified(100); | Saccharomycetales | 7 | Saprotroph | Undefined Saprotroph | Yeast | NULL | Possible | NULL | Sterkenburg E, et al. 2015. New Phytologist 207:1145-1158 ((https://doi.org/10.1111/nph.13426)) |
| Otu30570 | 1 | k__Fungi(100);p__Ascomycota(100);c__Saccharomycetes(100);o__Saccharomycetales(100);f__Saccharomycetales_family_Incertae_sedis(100);g__Candida(100);s__Candida_cretensis(100); | Candida | 13 | Pathotroph-Saprotroph-Symbiotroph | Animal Pathogen-Endophyte-Endosymbiont-Epiphyte-\|Undefined Saprotroph\| | Dimorphic Yeast-Yeast | NULL | Probable | As nectar-tap_saprotroph (Põlme et al. 2020); Decay sugar-rich substrates (Põlme et al. 2020); This asexual genus is polyphyletic and contain species that function very differently depend on what they are. Caution should be taken when interpreting guilds for Candida species. | Manolakaki D, et al. 2010. Virulence. 1: 367-375 ((https://doi.org/10.4161/viru.1.5.12796)); Mekbib SB, et al. 2011. Tropical Plant Pathology 36:233-240 ((https://doi.org/10.1590/S1982-56762011000400004)); Põlme S, et al. 2020. Fungal Diversity 105:1-16 ((https://doi.org/10.1007/s13225-020-00466-2)) |
| Otu30572 | 1 | k__Fungi(100);p__Ascomycota(100);c__Saccharomycetes(100);o__Saccharomycetales(100);f__Saccharomycetales_family_Incertae_sedis(100);g__Candida(100);g__Candida_unclassified(100); | Candida | 13 | Pathotroph-Saprotroph-Symbiotroph | Animal Pathogen-Endophyte-Endosymbiont-Epiphyte-\|Undefined Saprotroph\| | Dimorphic Yeast-Yeast | NULL | Probable | As nectar-tap_saprotroph (Põlme et al. 2020); Decay sugar-rich substrates (Põlme et al. 2020); This asexual genus is polyphyletic and contain species that function very differently depend on what they are. Caution should be taken when interpreting guilds for Candida species. | Manolakaki D, et al. 2010. Virulence. 1: 367-375 ((https://doi.org/10.4161/viru.1.5.12796)); Mekbib SB, et al. 2011. Tropical Plant Pathology 36:233-240 ((https://doi.org/10.1590/S1982-56762011000400004)); Põlme S, et al. 2020. Fungal Diversity 105:1-16 ((https://doi.org/10.1007/s13225-020-00466-2)) |
| Otu30573 | 1 | k__Fungi(100);p__Ascomycota(100);c__Saccharomycetes(100);o__Saccharomycetales(100);f__Metschnikowiaceae(100);g__Metschnikowia(100);s__Metschnikowia_pulcherrima(100); | Metschnikowia | 13 | Saprotroph | \|Undefined Saprotroph\| | Yeast | NULL | Probable | As nectar-tap_saprotroph (Põlme et al. 2020); Decay sugar-rich substrates (Põlme et al. 2020) | Põlme S, et al. 2020. Fungal Diversity 105:1-16 ((https://doi.org/10.1007/s13225-020-00466-2)) |
| Otu30574 | 1 | k__Fungi(100);p__Ascomycota(100);c__Saccharomycetes(100);o__Saccharomycetales(100);f__Metschnikowiaceae(100);g__Metschnikowia(100);g__Metschnikowia_unclassified(100); | Metschnikowia | 13 | Saprotroph | \|Undefined Saprotroph\| | Yeast | NULL | Probable | As nectar-tap_saprotroph (Põlme et al. 2020); Decay sugar-rich substrates (Põlme et al. 2020) | Põlme S, et al. 2020. Fungal Diversity 105:1-16 ((https://doi.org/10.1007/s13225-020-00466-2)) |
| Otu30575 | 1 | k__Fungi(100);p__Ascomycota(100);c__Saccharomycetes(100);o__Saccharomycetales(100);f__Metschnikowiaceae(100);g__Metschnikowia(100);s__Metschnikowia_pulcherrima(100); | Metschnikowia | 13 | Saprotroph | \|Undefined Saprotroph\| | Yeast | NULL | Probable | As nectar-tap_saprotroph (Põlme et al. 2020); Decay sugar-rich substrates (Põlme et al. 2020) | Põlme S, et al. 2020. Fungal Diversity 105:1-16 ((https://doi.org/10.1007/s13225-020-00466-2)) |
| Otu30576 | 1 | k__Fungi(100);p__Ascomycota(100);c__Saccharomycetes(100);o__Saccharomycetales(100);f__Metschnikowiaceae(100);g__Metschnikowia(100);s__Metschnikowia_pulcherrima(100); | Metschnikowia | 13 | Saprotroph | \|Undefined Saprotroph\| | Yeast | NULL | Probable | As nectar-tap_saprotroph (Põlme et al. 2020); Decay sugar-rich substrates (Põlme et al. 2020) | Põlme S, et al. 2020. Fungal Diversity 105:1-16 ((https://doi.org/10.1007/s13225-020-00466-2)) |
| Otu30577 | 1 | k__Fungi(100);p__Ascomycota(100);c__Saccharomycetes(100);o__Saccharomycetales(100);f__Metschnikowiaceae(100);g__Metschnikowia(100);s__Metschnikowia_pulcherrima(100); | Metschnikowia | 13 | Saprotroph | \|Undefined Saprotroph\| | Yeast | NULL | Probable | As nectar-tap_saprotroph (Põlme et al. 2020); Decay sugar-rich substrates (Põlme et al. 2020) | Põlme S, et al. 2020. Fungal Diversity 105:1-16 ((https://doi.org/10.1007/s13225-020-00466-2)) |
| Otu30578 | 1 | k__Fungi(100);p__Ascomycota(100);c__Saccharomycetes(100);o__Saccharomycetales(100);f__Metschnikowiaceae(100);g__Metschnikowia(100);s__Metschnikowia_pulcherrima(100); | Metschnikowia | 13 | Saprotroph | \|Undefined Saprotroph\| | Yeast | NULL | Probable | As nectar-tap_saprotroph (Põlme et al. 2020); Decay sugar-rich substrates (Põlme et al. 2020) | Põlme S, et al. 2020. Fungal Diversity 105:1-16 ((https://doi.org/10.1007/s13225-020-00466-2)) |
| Otu30579 | 1 | k__Fungi(100);p__Ascomycota(100);c__Saccharomycetes(100);o__Saccharomycetales(100);f__Metschnikowiaceae(100);g__Metschnikowia(100);g__Metschnikowia_unclassified(100); | Metschnikowia | 13 | Saprotroph | \|Undefined Saprotroph\| | Yeast | NULL | Probable | As nectar-tap_saprotroph (Põlme et al. 2020); Decay sugar-rich substrates (Põlme et al. 2020) | Põlme S, et al. 2020. Fungal Diversity 105:1-16 ((https://doi.org/10.1007/s13225-020-00466-2)) |
| Otu30580 | 1 | k__Fungi(100);p__Ascomycota(100);c__Saccharomycetes(100);o__Saccharomycetales(100);f__Metschnikowiaceae(100);g__Metschnikowia(100);s__Metschnikowia_pulcherrima(100); | Metschnikowia | 13 | Saprotroph | \|Undefined Saprotroph\| | Yeast | NULL | Probable | As nectar-tap_saprotroph (Põlme et al. 2020); Decay sugar-rich substrates (Põlme et al. 2020) | Põlme S, et al. 2020. Fungal Diversity 105:1-16 ((https://doi.org/10.1007/s13225-020-00466-2)) |
| Otu30581 | 1 | k__Fungi(100);p__Ascomycota(100);c__Saccharomycetes(100);o__Saccharomycetales(100);f__Metschnikowiaceae(100);g__Metschnikowia(100);s__Metschnikowia_pulcherrima(100); | Metschnikowia | 13 | Saprotroph | \|Undefined Saprotroph\| | Yeast | NULL | Probable | As nectar-tap_saprotroph (Põlme et al. 2020); Decay sugar-rich substrates (Põlme et al. 2020) | Põlme S, et al. 2020. Fungal Diversity 105:1-16 ((https://doi.org/10.1007/s13225-020-00466-2)) |
| Otu30582 | 1 | k__Fungi(100);p__Ascomycota(100);c__Saccharomycetes(100);o__Saccharomycetales(100);f__Metschnikowiaceae(100);g__Metschnikowia(100);g__Metschnikowia_unclassified(100); | Metschnikowia | 13 | Saprotroph | \|Undefined Saprotroph\| | Yeast | NULL | Probable | As nectar-tap_saprotroph (Põlme et al. 2020); Decay sugar-rich substrates (Põlme et al. 2020) | Põlme S, et al. 2020. Fungal Diversity 105:1-16 ((https://doi.org/10.1007/s13225-020-00466-2)) |
| Otu30583 | 1 | k__Fungi(100);p__Ascomycota(100);c__Saccharomycetes(100);o__Saccharomycetales(100);f__Metschnikowiaceae(100);g__Metschnikowia(100);s__Metschnikowia_pulcherrima(100); | Metschnikowia | 13 | Saprotroph | \|Undefined Saprotroph\| | Yeast | NULL | Probable | As nectar-tap_saprotroph (Põlme et al. 2020); Decay sugar-rich substrates (Põlme et al. 2020) | Põlme S, et al. 2020. Fungal Diversity 105:1-16 ((https://doi.org/10.1007/s13225-020-00466-2)) |
| Otu30584 | 1 | k__Fungi(100);p__Ascomycota(100);c__Saccharomycetes(100);o__Saccharomycetales(100);f__Metschnikowiaceae(100);g__Metschnikowia(100);g__Metschnikowia_unclassified(100); | Metschnikowia | 13 | Saprotroph | \|Undefined Saprotroph\| | Yeast | NULL | Probable | As nectar-tap_saprotroph (Põlme et al. 2020); Decay sugar-rich substrates (Põlme et al. 2020) | Põlme S, et al. 2020. Fungal Diversity 105:1-16 ((https://doi.org/10.1007/s13225-020-00466-2)) |
| Otu30585 | 1 | k__Fungi(100);p__Ascomycota(100);c__Saccharomycetes(100);o__Saccharomycetales(100);f__Metschnikowiaceae(100);g__Metschnikowia(100);g__Metschnikowia_unclassified(100); | Metschnikowia | 13 | Saprotroph | \|Undefined Saprotroph\| | Yeast | NULL | Probable | As nectar-tap_saprotroph (Põlme et al. 2020); Decay sugar-rich substrates (Põlme et al. 2020) | Põlme S, et al. 2020. Fungal Diversity 105:1-16 ((https://doi.org/10.1007/s13225-020-00466-2)) |
| Otu30586 | 1 | k__Fungi(100);p__Ascomycota(100);c__Saccharomycetes(100);o__Saccharomycetales(100);f__Saccharomycetales_family_Incertae_sedis(100);f__Saccharomycetales_family_Incertae_sedis_unclassified(100);f__Saccharomycetales_family_Incertae_sedis_unclassified(100); | Saccharomycetales | 7 | Saprotroph | Undefined Saprotroph | Yeast | NULL | Possible | NULL | Sterkenburg E, et al. 2015. New Phytologist 207:1145-1158 ((https://doi.org/10.1111/nph.13426)) |
| Otu30589 | 1 | k__Fungi(100);p__Ascomycota(100);c__Saccharomycetes(100);o__Saccharomycetales(100);o__Saccharomycetales_unclassified(100);o__Saccharomycetales_unclassified(100);o__Saccharomycetales_unclassified(100); | Saccharomycetales | 7 | Saprotroph | Undefined Saprotroph | Yeast | NULL | Possible | NULL | Sterkenburg E, et al. 2015. New Phytologist 207:1145-1158 ((https://doi.org/10.1111/nph.13426)) |
| Otu30590 | 1 | k__Fungi(100);p__Ascomycota(100);c__Saccharomycetes(100);o__Saccharomycetales(100);f__Saccharomycetales_family_Incertae_sedis(100);g__Candida(100);s__Candida_cretensis(100); | Candida | 13 | Pathotroph-Saprotroph-Symbiotroph | Animal Pathogen-Endophyte-Endosymbiont-Epiphyte-\|Undefined Saprotroph\| | Dimorphic Yeast-Yeast | NULL | Probable | As nectar-tap_saprotroph (Põlme et al. 2020); Decay sugar-rich substrates (Põlme et al. 2020); This asexual genus is polyphyletic and contain species that function very differently depend on what they are. Caution should be taken when interpreting guilds for Candida species. | Manolakaki D, et al. 2010. Virulence. 1: 367-375 ((https://doi.org/10.4161/viru.1.5.12796)); Mekbib SB, et al. 2011. Tropical Plant Pathology 36:233-240 ((https://doi.org/10.1590/S1982-56762011000400004)); Põlme S, et al. 2020. Fungal Diversity 105:1-16 ((https://doi.org/10.1007/s13225-020-00466-2)) |
| Otu30591 | 1 | k__Fungi(100);p__Ascomycota(100);c__Saccharomycetes(100);o__Saccharomycetales(100);f__Saccharomycetales_family_Incertae_sedis(100);f__Saccharomycetales_family_Incertae_sedis_unclassified(100);f__Saccharomycetales_family_Incertae_sedis_unclassified(100); | Saccharomycetales | 7 | Saprotroph | Undefined Saprotroph | Yeast | NULL | Possible | NULL | Sterkenburg E, et al. 2015. New Phytologist 207:1145-1158 ((https://doi.org/10.1111/nph.13426)) |
| Otu30594 | 1 | k__Fungi(100);p__Ascomycota(100);c__Saccharomycetes(100);o__Saccharomycetales(100);f__Saccharomycetales_family_Incertae_sedis(100);g__Candida(100);g__Candida_unclassified(100); | Candida | 13 | Pathotroph-Saprotroph-Symbiotroph | Animal Pathogen-Endophyte-Endosymbiont-Epiphyte-\|Undefined Saprotroph\| | Dimorphic Yeast-Yeast | NULL | Probable | As nectar-tap_saprotroph (Põlme et al. 2020); Decay sugar-rich substrates (Põlme et al. 2020); This asexual genus is polyphyletic and contain species that function very differently depend on what they are. Caution should be taken when interpreting guilds for Candida species. | Manolakaki D, et al. 2010. Virulence. 1: 367-375 ((https://doi.org/10.4161/viru.1.5.12796)); Mekbib SB, et al. 2011. Tropical Plant Pathology 36:233-240 ((https://doi.org/10.1590/S1982-56762011000400004)); Põlme S, et al. 2020. Fungal Diversity 105:1-16 ((https://doi.org/10.1007/s13225-020-00466-2)) |
| Otu30595 | 1 | k__Fungi(100);p__Ascomycota(100);c__Saccharomycetes(100);o__Saccharomycetales(100);f__Saccharomycetales_family_Incertae_sedis(100);g__Candida(100);g__Candida_unclassified(100); | Candida | 13 | Pathotroph-Saprotroph-Symbiotroph | Animal Pathogen-Endophyte-Endosymbiont-Epiphyte-\|Undefined Saprotroph\| | Dimorphic Yeast-Yeast | NULL | Probable | As nectar-tap_saprotroph (Põlme et al. 2020); Decay sugar-rich substrates (Põlme et al. 2020); This asexual genus is polyphyletic and contain species that function very differently depend on what they are. Caution should be taken when interpreting guilds for Candida species. | Manolakaki D, et al. 2010. Virulence. 1: 367-375 ((https://doi.org/10.4161/viru.1.5.12796)); Mekbib SB, et al. 2011. Tropical Plant Pathology 36:233-240 ((https://doi.org/10.1590/S1982-56762011000400004)); Põlme S, et al. 2020. Fungal Diversity 105:1-16 ((https://doi.org/10.1007/s13225-020-00466-2)) |
| Otu30599 | 1 | k__Fungi(100);p__Ascomycota(100);c__Saccharomycetes(100);o__Saccharomycetales(100);f__Metschnikowiaceae(100);g__Metschnikowia(100);g__Metschnikowia_unclassified(100); | Metschnikowia | 13 | Saprotroph | \|Undefined Saprotroph\| | Yeast | NULL | Probable | As nectar-tap_saprotroph (Põlme et al. 2020); Decay sugar-rich substrates (Põlme et al. 2020) | Põlme S, et al. 2020. Fungal Diversity 105:1-16 ((https://doi.org/10.1007/s13225-020-00466-2)) |
| Otu30600 | 1 | k__Fungi(100);p__Ascomycota(100);c__Saccharomycetes(100);o__Saccharomycetales(100);f__Saccharomycetales_family_Incertae_sedis(100);g__Candida(100);s__Candida_cretensis(100); | Candida | 13 | Pathotroph-Saprotroph-Symbiotroph | Animal Pathogen-Endophyte-Endosymbiont-Epiphyte-\|Undefined Saprotroph\| | Dimorphic Yeast-Yeast | NULL | Probable | As nectar-tap_saprotroph (Põlme et al. 2020); Decay sugar-rich substrates (Põlme et al. 2020); This asexual genus is polyphyletic and contain species that function very differently depend on what they are. Caution should be taken when interpreting guilds for Candida species. | Manolakaki D, et al. 2010. Virulence. 1: 367-375 ((https://doi.org/10.4161/viru.1.5.12796)); Mekbib SB, et al. 2011. Tropical Plant Pathology 36:233-240 ((https://doi.org/10.1590/S1982-56762011000400004)); Põlme S, et al. 2020. Fungal Diversity 105:1-16 ((https://doi.org/10.1007/s13225-020-00466-2)) |
| Otu30603 | 1 | k__Fungi(100);p__Ascomycota(100);c__Saccharomycetes(100);o__Saccharomycetales(100);o__Saccharomycetales_unclassified(100);o__Saccharomycetales_unclassified(100);o__Saccharomycetales_unclassified(100); | Saccharomycetales | 7 | Saprotroph | Undefined Saprotroph | Yeast | NULL | Possible | NULL | Sterkenburg E, et al. 2015. New Phytologist 207:1145-1158 ((https://doi.org/10.1111/nph.13426)) |
| Otu30604 | 1 | k__Fungi(100);p__Ascomycota(100);c__Saccharomycetes(100);o__Saccharomycetales(100);f__Saccharomycetales_family_Incertae_sedis(100);g__Candida(100);s__Candida_cretensis(100); | Candida | 13 | Pathotroph-Saprotroph-Symbiotroph | Animal Pathogen-Endophyte-Endosymbiont-Epiphyte-\|Undefined Saprotroph\| | Dimorphic Yeast-Yeast | NULL | Probable | As nectar-tap_saprotroph (Põlme et al. 2020); Decay sugar-rich substrates (Põlme et al. 2020); This asexual genus is polyphyletic and contain species that function very differently depend on what they are. Caution should be taken when interpreting guilds for Candida species. | Manolakaki D, et al. 2010. Virulence. 1: 367-375 ((https://doi.org/10.4161/viru.1.5.12796)); Mekbib SB, et al. 2011. Tropical Plant Pathology 36:233-240 ((https://doi.org/10.1590/S1982-56762011000400004)); Põlme S, et al. 2020. Fungal Diversity 105:1-16 ((https://doi.org/10.1007/s13225-020-00466-2)) |
| Otu30605 | 1 | k__Fungi(100);p__Ascomycota(100);c__Saccharomycetes(100);o__Saccharomycetales(100);f__Saccharomycetales_family_Incertae_sedis(100);g__Candida(100);g__Candida_unclassified(100); | Candida | 13 | Pathotroph-Saprotroph-Symbiotroph | Animal Pathogen-Endophyte-Endosymbiont-Epiphyte-\|Undefined Saprotroph\| | Dimorphic Yeast-Yeast | NULL | Probable | As nectar-tap_saprotroph (Põlme et al. 2020); Decay sugar-rich substrates (Põlme et al. 2020); This asexual genus is polyphyletic and contain species that function very differently depend on what they are. Caution should be taken when interpreting guilds for Candida species. | Manolakaki D, et al. 2010. Virulence. 1: 367-375 ((https://doi.org/10.4161/viru.1.5.12796)); Mekbib SB, et al. 2011. Tropical Plant Pathology 36:233-240 ((https://doi.org/10.1590/S1982-56762011000400004)); Põlme S, et al. 2020. Fungal Diversity 105:1-16 ((https://doi.org/10.1007/s13225-020-00466-2)) |
| Otu30607 | 1 | k__Fungi(100);p__Ascomycota(100);c__Saccharomycetes(100);o__Saccharomycetales(100);f__Saccharomycetales_family_Incertae_sedis(100);g__Candida(100);s__Candida_cretensis(100); | Candida | 13 | Pathotroph-Saprotroph-Symbiotroph | Animal Pathogen-Endophyte-Endosymbiont-Epiphyte-\|Undefined Saprotroph\| | Dimorphic Yeast-Yeast | NULL | Probable | As nectar-tap_saprotroph (Põlme et al. 2020); Decay sugar-rich substrates (Põlme et al. 2020); This asexual genus is polyphyletic and contain species that function very differently depend on what they are. Caution should be taken when interpreting guilds for Candida species. | Manolakaki D, et al. 2010. Virulence. 1: 367-375 ((https://doi.org/10.4161/viru.1.5.12796)); Mekbib SB, et al. 2011. Tropical Plant Pathology 36:233-240 ((https://doi.org/10.1590/S1982-56762011000400004)); Põlme S, et al. 2020. Fungal Diversity 105:1-16 ((https://doi.org/10.1007/s13225-020-00466-2)) |
| Otu30608 | 1 | k__Fungi(100);p__Ascomycota(100);c__Saccharomycetes(100);o__Saccharomycetales(100);f__Saccharomycetales_family_Incertae_sedis(100);g__Candida(100);g__Candida_unclassified(100); | Candida | 13 | Pathotroph-Saprotroph-Symbiotroph | Animal Pathogen-Endophyte-Endosymbiont-Epiphyte-\|Undefined Saprotroph\| | Dimorphic Yeast-Yeast | NULL | Probable | As nectar-tap_saprotroph (Põlme et al. 2020); Decay sugar-rich substrates (Põlme et al. 2020); This asexual genus is polyphyletic and contain species that function very differently depend on what they are. Caution should be taken when interpreting guilds for Candida species. | Manolakaki D, et al. 2010. Virulence. 1: 367-375 ((https://doi.org/10.4161/viru.1.5.12796)); Mekbib SB, et al. 2011. Tropical Plant Pathology 36:233-240 ((https://doi.org/10.1590/S1982-56762011000400004)); Põlme S, et al. 2020. Fungal Diversity 105:1-16 ((https://doi.org/10.1007/s13225-020-00466-2)) |
| Otu30609 | 1 | k__Fungi(100);p__Ascomycota(100);c__Saccharomycetes(100);o__Saccharomycetales(100);f__Saccharomycetales_family_Incertae_sedis(100);f__Saccharomycetales_family_Incertae_sedis_unclassified(100);f__Saccharomycetales_family_Incertae_sedis_unclassified(100); | Saccharomycetales | 7 | Saprotroph | Undefined Saprotroph | Yeast | NULL | Possible | NULL | Sterkenburg E, et al. 2015. New Phytologist 207:1145-1158 ((https://doi.org/10.1111/nph.13426)) |
| Otu30611 | 1 | k__Fungi(100);p__Ascomycota(100);c__Saccharomycetes(100);o__Saccharomycetales(100);f__Metschnikowiaceae(100);g__Metschnikowia(100);s__Metschnikowia_pulcherrima(100); | Metschnikowia | 13 | Saprotroph | \|Undefined Saprotroph\| | Yeast | NULL | Probable | As nectar-tap_saprotroph (Põlme et al. 2020); Decay sugar-rich substrates (Põlme et al. 2020) | Põlme S, et al. 2020. Fungal Diversity 105:1-16 ((https://doi.org/10.1007/s13225-020-00466-2)) |
| Otu30612 | 1 | k__Fungi(100);p__Ascomycota(100);c__Saccharomycetes(100);o__Saccharomycetales(100);f__Metschnikowiaceae(100);g__Metschnikowia(100);g__Metschnikowia_unclassified(100); | Metschnikowia | 13 | Saprotroph | \|Undefined Saprotroph\| | Yeast | NULL | Probable | As nectar-tap_saprotroph (Põlme et al. 2020); Decay sugar-rich substrates (Põlme et al. 2020) | Põlme S, et al. 2020. Fungal Diversity 105:1-16 ((https://doi.org/10.1007/s13225-020-00466-2)) |
| Otu30613 | 1 | k__Fungi(100);p__Ascomycota(100);c__Saccharomycetes(100);o__Saccharomycetales(100);f__Saccharomycetales_family_Incertae_sedis(100);g__Candida(100);g__Candida_unclassified(100); | Candida | 13 | Pathotroph-Saprotroph-Symbiotroph | Animal Pathogen-Endophyte-Endosymbiont-Epiphyte-\|Undefined Saprotroph\| | Dimorphic Yeast-Yeast | NULL | Probable | As nectar-tap_saprotroph (Põlme et al. 2020); Decay sugar-rich substrates (Põlme et al. 2020); This asexual genus is polyphyletic and contain species that function very differently depend on what they are. Caution should be taken when interpreting guilds for Candida species. | Manolakaki D, et al. 2010. Virulence. 1: 367-375 ((https://doi.org/10.4161/viru.1.5.12796)); Mekbib SB, et al. 2011. Tropical Plant Pathology 36:233-240 ((https://doi.org/10.1590/S1982-56762011000400004)); Põlme S, et al. 2020. Fungal Diversity 105:1-16 ((https://doi.org/10.1007/s13225-020-00466-2)) |
| Otu30617 | 1 | k__Fungi(100);p__Ascomycota(100);c__Saccharomycetes(100);o__Saccharomycetales(100);f__Metschnikowiaceae(100);g__Metschnikowia(100);s__Metschnikowia_pulcherrima(100); | Metschnikowia | 13 | Saprotroph | \|Undefined Saprotroph\| | Yeast | NULL | Probable | As nectar-tap_saprotroph (Põlme et al. 2020); Decay sugar-rich substrates (Põlme et al. 2020) | Põlme S, et al. 2020. Fungal Diversity 105:1-16 ((https://doi.org/10.1007/s13225-020-00466-2)) |
| Otu30620 | 1 | k__Fungi(100);p__Ascomycota(100);c__Saccharomycetes(100);o__Saccharomycetales(100);f__Saccharomycetales_family_Incertae_sedis(100);g__Candida(100);s__Candida_cretensis(100); | Candida | 13 | Pathotroph-Saprotroph-Symbiotroph | Animal Pathogen-Endophyte-Endosymbiont-Epiphyte-\|Undefined Saprotroph\| | Dimorphic Yeast-Yeast | NULL | Probable | As nectar-tap_saprotroph (Põlme et al. 2020); Decay sugar-rich substrates (Põlme et al. 2020); This asexual genus is polyphyletic and contain species that function very differently depend on what they are. Caution should be taken when interpreting guilds for Candida species. | Manolakaki D, et al. 2010. Virulence. 1: 367-375 ((https://doi.org/10.4161/viru.1.5.12796)); Mekbib SB, et al. 2011. Tropical Plant Pathology 36:233-240 ((https://doi.org/10.1590/S1982-56762011000400004)); Põlme S, et al. 2020. Fungal Diversity 105:1-16 ((https://doi.org/10.1007/s13225-020-00466-2)) |
| Otu30625 | 1 | k__Fungi(100);p__Ascomycota(100);c__Saccharomycetes(100);o__Saccharomycetales(100);f__Metschnikowiaceae(100);g__Metschnikowia(100);s__Metschnikowia_pulcherrima(100); | Metschnikowia | 13 | Saprotroph | \|Undefined Saprotroph\| | Yeast | NULL | Probable | As nectar-tap_saprotroph (Põlme et al. 2020); Decay sugar-rich substrates (Põlme et al. 2020) | Põlme S, et al. 2020. Fungal Diversity 105:1-16 ((https://doi.org/10.1007/s13225-020-00466-2)) |
| Otu30627 | 1 | k__Fungi(100);p__Ascomycota(100);c__Saccharomycetes(100);o__Saccharomycetales(100);f__Metschnikowiaceae(100);g__Metschnikowia(100);s__Metschnikowia_pulcherrima(100); | Metschnikowia | 13 | Saprotroph | \|Undefined Saprotroph\| | Yeast | NULL | Probable | As nectar-tap_saprotroph (Põlme et al. 2020); Decay sugar-rich substrates (Põlme et al. 2020) | Põlme S, et al. 2020. Fungal Diversity 105:1-16 ((https://doi.org/10.1007/s13225-020-00466-2)) |
| Otu30629 | 1 | k__Fungi(100);p__Ascomycota(100);c__Saccharomycetes(100);o__Saccharomycetales(100);f__Saccharomycetales_family_Incertae_sedis(100);g__Candida(100);g__Candida_unclassified(100); | Candida | 13 | Pathotroph-Saprotroph-Symbiotroph | Animal Pathogen-Endophyte-Endosymbiont-Epiphyte-\|Undefined Saprotroph\| | Dimorphic Yeast-Yeast | NULL | Probable | As nectar-tap_saprotroph (Põlme et al. 2020); Decay sugar-rich substrates (Põlme et al. 2020); This asexual genus is polyphyletic and contain species that function very differently depend on what they are. Caution should be taken when interpreting guilds for Candida species. | Manolakaki D, et al. 2010. Virulence. 1: 367-375 ((https://doi.org/10.4161/viru.1.5.12796)); Mekbib SB, et al. 2011. Tropical Plant Pathology 36:233-240 ((https://doi.org/10.1590/S1982-56762011000400004)); Põlme S, et al. 2020. Fungal Diversity 105:1-16 ((https://doi.org/10.1007/s13225-020-00466-2)) |
| Otu30630 | 1 | k__Fungi(100);p__Ascomycota(100);c__Saccharomycetes(100);o__Saccharomycetales(100);f__Metschnikowiaceae(100);g__Metschnikowia(100);g__Metschnikowia_unclassified(100); | Metschnikowia | 13 | Saprotroph | \|Undefined Saprotroph\| | Yeast | NULL | Probable | As nectar-tap_saprotroph (Põlme et al. 2020); Decay sugar-rich substrates (Põlme et al. 2020) | Põlme S, et al. 2020. Fungal Diversity 105:1-16 ((https://doi.org/10.1007/s13225-020-00466-2)) |
| Otu30631 | 1 | k__Fungi(100);p__Ascomycota(100);c__Saccharomycetes(100);o__Saccharomycetales(100);f__Metschnikowiaceae(100);g__Metschnikowia(100);g__Metschnikowia_unclassified(100); | Metschnikowia | 13 | Saprotroph | \|Undefined Saprotroph\| | Yeast | NULL | Probable | As nectar-tap_saprotroph (Põlme et al. 2020); Decay sugar-rich substrates (Põlme et al. 2020) | Põlme S, et al. 2020. Fungal Diversity 105:1-16 ((https://doi.org/10.1007/s13225-020-00466-2)) |
| Otu30632 | 1 | k__Fungi(100);p__Ascomycota(100);c__Saccharomycetes(100);o__Saccharomycetales(100);f__Metschnikowiaceae(100);g__Metschnikowia(100);g__Metschnikowia_unclassified(100); | Metschnikowia | 13 | Saprotroph | \|Undefined Saprotroph\| | Yeast | NULL | Probable | As nectar-tap_saprotroph (Põlme et al. 2020); Decay sugar-rich substrates (Põlme et al. 2020) | Põlme S, et al. 2020. Fungal Diversity 105:1-16 ((https://doi.org/10.1007/s13225-020-00466-2)) |
| Otu30633 | 1 | k__Fungi(100);p__Ascomycota(100);c__Saccharomycetes(100);o__Saccharomycetales(100);f__Metschnikowiaceae(100);g__Metschnikowia(100);s__Metschnikowia_pulcherrima(100); | Metschnikowia | 13 | Saprotroph | \|Undefined Saprotroph\| | Yeast | NULL | Probable | As nectar-tap_saprotroph (Põlme et al. 2020); Decay sugar-rich substrates (Põlme et al. 2020) | Põlme S, et al. 2020. Fungal Diversity 105:1-16 ((https://doi.org/10.1007/s13225-020-00466-2)) |
| Otu30635 | 1 | k__Fungi(100);p__Ascomycota(100);c__Saccharomycetes(100);o__Saccharomycetales(100);f__Metschnikowiaceae(100);g__Metschnikowia(100);g__Metschnikowia_unclassified(100); | Metschnikowia | 13 | Saprotroph | \|Undefined Saprotroph\| | Yeast | NULL | Probable | As nectar-tap_saprotroph (Põlme et al. 2020); Decay sugar-rich substrates (Põlme et al. 2020) | Põlme S, et al. 2020. Fungal Diversity 105:1-16 ((https://doi.org/10.1007/s13225-020-00466-2)) |
| Otu30636 | 1 | k__Fungi(100);p__Ascomycota(100);c__Saccharomycetes(100);o__Saccharomycetales(100);f__Metschnikowiaceae(100);g__Metschnikowia(100);g__Metschnikowia_unclassified(100); | Metschnikowia | 13 | Saprotroph | \|Undefined Saprotroph\| | Yeast | NULL | Probable | As nectar-tap_saprotroph (Põlme et al. 2020); Decay sugar-rich substrates (Põlme et al. 2020) | Põlme S, et al. 2020. Fungal Diversity 105:1-16 ((https://doi.org/10.1007/s13225-020-00466-2)) |
| Otu30637 | 1 | k__Fungi(100);p__Ascomycota(100);c__Saccharomycetes(100);o__Saccharomycetales(100);f__Metschnikowiaceae(100);g__Metschnikowia(100);s__Metschnikowia_pulcherrima(100); | Metschnikowia | 13 | Saprotroph | \|Undefined Saprotroph\| | Yeast | NULL | Probable | As nectar-tap_saprotroph (Põlme et al. 2020); Decay sugar-rich substrates (Põlme et al. 2020) | Põlme S, et al. 2020. Fungal Diversity 105:1-16 ((https://doi.org/10.1007/s13225-020-00466-2)) |
| Otu30639 | 1 | k__Fungi(100);p__Ascomycota(100);c__Saccharomycetes(100);o__Saccharomycetales(100);f__Metschnikowiaceae(100);g__Metschnikowia(100);g__Metschnikowia_unclassified(100); | Metschnikowia | 13 | Saprotroph | \|Undefined Saprotroph\| | Yeast | NULL | Probable | As nectar-tap_saprotroph (Põlme et al. 2020); Decay sugar-rich substrates (Põlme et al. 2020) | Põlme S, et al. 2020. Fungal Diversity 105:1-16 ((https://doi.org/10.1007/s13225-020-00466-2)) |
| Otu30640 | 1 | k__Fungi(100);p__Ascomycota(100);c__Saccharomycetes(100);o__Saccharomycetales(100);f__Metschnikowiaceae(100);g__Metschnikowia(100);s__Metschnikowia_pulcherrima(100); | Metschnikowia | 13 | Saprotroph | \|Undefined Saprotroph\| | Yeast | NULL | Probable | As nectar-tap_saprotroph (Põlme et al. 2020); Decay sugar-rich substrates (Põlme et al. 2020) | Põlme S, et al. 2020. Fungal Diversity 105:1-16 ((https://doi.org/10.1007/s13225-020-00466-2)) |
| Otu30641 | 1 | k__Fungi(100);p__Ascomycota(100);c__Saccharomycetes(100);o__Saccharomycetales(100);f__Metschnikowiaceae(100);g__Metschnikowia(100);s__Metschnikowia_pulcherrima(100); | Metschnikowia | 13 | Saprotroph | \|Undefined Saprotroph\| | Yeast | NULL | Probable | As nectar-tap_saprotroph (Põlme et al. 2020); Decay sugar-rich substrates (Põlme et al. 2020) | Põlme S, et al. 2020. Fungal Diversity 105:1-16 ((https://doi.org/10.1007/s13225-020-00466-2)) |
| Otu30642 | 1 | k__Fungi(100);p__Ascomycota(100);c__Saccharomycetes(100);o__Saccharomycetales(100);f__Metschnikowiaceae(100);g__Metschnikowia(100);g__Metschnikowia_unclassified(100); | Metschnikowia | 13 | Saprotroph | \|Undefined Saprotroph\| | Yeast | NULL | Probable | As nectar-tap_saprotroph (Põlme et al. 2020); Decay sugar-rich substrates (Põlme et al. 2020) | Põlme S, et al. 2020. Fungal Diversity 105:1-16 ((https://doi.org/10.1007/s13225-020-00466-2)) |
| Otu30643 | 1 | k__Fungi(100);p__Ascomycota(100);c__Saccharomycetes(100);o__Saccharomycetales(100);f__Metschnikowiaceae(100);g__Metschnikowia(100);g__Metschnikowia_unclassified(100); | Metschnikowia | 13 | Saprotroph | \|Undefined Saprotroph\| | Yeast | NULL | Probable | As nectar-tap_saprotroph (Põlme et al. 2020); Decay sugar-rich substrates (Põlme et al. 2020) | Põlme S, et al. 2020. Fungal Diversity 105:1-16 ((https://doi.org/10.1007/s13225-020-00466-2)) |
| Otu30644 | 1 | k__Fungi(100);p__Ascomycota(100);c__Saccharomycetes(100);o__Saccharomycetales(100);f__Metschnikowiaceae(100);g__Metschnikowia(100);s__Metschnikowia_pulcherrima(100); | Metschnikowia | 13 | Saprotroph | \|Undefined Saprotroph\| | Yeast | NULL | Probable | As nectar-tap_saprotroph (Põlme et al. 2020); Decay sugar-rich substrates (Põlme et al. 2020) | Põlme S, et al. 2020. Fungal Diversity 105:1-16 ((https://doi.org/10.1007/s13225-020-00466-2)) |
| Otu30645 | 1 | k__Fungi(100);p__Ascomycota(100);c__Saccharomycetes(100);o__Saccharomycetales(100);f__Metschnikowiaceae(100);g__Metschnikowia(100);s__Metschnikowia_pulcherrima(100); | Metschnikowia | 13 | Saprotroph | \|Undefined Saprotroph\| | Yeast | NULL | Probable | As nectar-tap_saprotroph (Põlme et al. 2020); Decay sugar-rich substrates (Põlme et al. 2020) | Põlme S, et al. 2020. Fungal Diversity 105:1-16 ((https://doi.org/10.1007/s13225-020-00466-2)) |
| Otu30646 | 1 | k__Fungi(100);p__Ascomycota(100);c__Saccharomycetes(100);o__Saccharomycetales(100);f__Metschnikowiaceae(100);g__Metschnikowia(100);g__Metschnikowia_unclassified(100); | Metschnikowia | 13 | Saprotroph | \|Undefined Saprotroph\| | Yeast | NULL | Probable | As nectar-tap_saprotroph (Põlme et al. 2020); Decay sugar-rich substrates (Põlme et al. 2020) | Põlme S, et al. 2020. Fungal Diversity 105:1-16 ((https://doi.org/10.1007/s13225-020-00466-2)) |
| Otu30647 | 1 | k__Fungi(100);p__Ascomycota(100);c__Saccharomycetes(100);o__Saccharomycetales(100);f__Metschnikowiaceae(100);g__Metschnikowia(100);s__Metschnikowia_pulcherrima(100); | Metschnikowia | 13 | Saprotroph | \|Undefined Saprotroph\| | Yeast | NULL | Probable | As nectar-tap_saprotroph (Põlme et al. 2020); Decay sugar-rich substrates (Põlme et al. 2020) | Põlme S, et al. 2020. Fungal Diversity 105:1-16 ((https://doi.org/10.1007/s13225-020-00466-2)) |
| Otu30648 | 1 | k__Fungi(100);p__Ascomycota(100);c__Saccharomycetes(100);o__Saccharomycetales(100);f__Metschnikowiaceae(100);g__Metschnikowia(100);s__Metschnikowia_pulcherrima(100); | Metschnikowia | 13 | Saprotroph | \|Undefined Saprotroph\| | Yeast | NULL | Probable | As nectar-tap_saprotroph (Põlme et al. 2020); Decay sugar-rich substrates (Põlme et al. 2020) | Põlme S, et al. 2020. Fungal Diversity 105:1-16 ((https://doi.org/10.1007/s13225-020-00466-2)) |
| Otu30649 | 1 | k__Fungi(100);p__Ascomycota(100);c__Saccharomycetes(100);o__Saccharomycetales(100);f__Metschnikowiaceae(100);g__Metschnikowia(100);g__Metschnikowia_unclassified(100); | Metschnikowia | 13 | Saprotroph | \|Undefined Saprotroph\| | Yeast | NULL | Probable | As nectar-tap_saprotroph (Põlme et al. 2020); Decay sugar-rich substrates (Põlme et al. 2020) | Põlme S, et al. 2020. Fungal Diversity 105:1-16 ((https://doi.org/10.1007/s13225-020-00466-2)) |
| Otu30651 | 1 | k__Fungi(100);p__Ascomycota(100);c__Saccharomycetes(100);o__Saccharomycetales(100);f__Metschnikowiaceae(100);g__Metschnikowia(100);s__Metschnikowia_pulcherrima(100); | Metschnikowia | 13 | Saprotroph | \|Undefined Saprotroph\| | Yeast | NULL | Probable | As nectar-tap_saprotroph (Põlme et al. 2020); Decay sugar-rich substrates (Põlme et al. 2020) | Põlme S, et al. 2020. Fungal Diversity 105:1-16 ((https://doi.org/10.1007/s13225-020-00466-2)) |
| Otu30653 | 1 | k__Fungi(100);p__Ascomycota(100);c__Saccharomycetes(100);o__Saccharomycetales(100);f__Metschnikowiaceae(100);g__Metschnikowia(100);s__Metschnikowia_pulcherrima(100); | Metschnikowia | 13 | Saprotroph | \|Undefined Saprotroph\| | Yeast | NULL | Probable | As nectar-tap_saprotroph (Põlme et al. 2020); Decay sugar-rich substrates (Põlme et al. 2020) | Põlme S, et al. 2020. Fungal Diversity 105:1-16 ((https://doi.org/10.1007/s13225-020-00466-2)) |
| Otu30654 | 1 | k__Fungi(100);p__Ascomycota(100);c__Saccharomycetes(100);o__Saccharomycetales(100);f__Metschnikowiaceae(100);g__Metschnikowia(100);s__Metschnikowia_pulcherrima(100); | Metschnikowia | 13 | Saprotroph | \|Undefined Saprotroph\| | Yeast | NULL | Probable | As nectar-tap_saprotroph (Põlme et al. 2020); Decay sugar-rich substrates (Põlme et al. 2020) | Põlme S, et al. 2020. Fungal Diversity 105:1-16 ((https://doi.org/10.1007/s13225-020-00466-2)) |
| Otu30655 | 1 | k__Fungi(100);p__Ascomycota(100);c__Saccharomycetes(100);o__Saccharomycetales(100);f__Metschnikowiaceae(100);g__Metschnikowia(100);s__Metschnikowia_pulcherrima(100); | Metschnikowia | 13 | Saprotroph | \|Undefined Saprotroph\| | Yeast | NULL | Probable | As nectar-tap_saprotroph (Põlme et al. 2020); Decay sugar-rich substrates (Põlme et al. 2020) | Põlme S, et al. 2020. Fungal Diversity 105:1-16 ((https://doi.org/10.1007/s13225-020-00466-2)) |
| Otu30656 | 1 | k__Fungi(100);p__Ascomycota(100);c__Saccharomycetes(100);o__Saccharomycetales(100);o__Saccharomycetales_unclassified(100);o__Saccharomycetales_unclassified(100);o__Saccharomycetales_unclassified(100); | Saccharomycetales | 7 | Saprotroph | Undefined Saprotroph | Yeast | NULL | Possible | NULL | Sterkenburg E, et al. 2015. New Phytologist 207:1145-1158 ((https://doi.org/10.1111/nph.13426)) |
| Otu30657 | 1 | k__Fungi(100);p__Ascomycota(100);c__Saccharomycetes(100);o__Saccharomycetales(100);f__Metschnikowiaceae(100);g__Metschnikowia(100);s__Metschnikowia_pulcherrima(100); | Metschnikowia | 13 | Saprotroph | \|Undefined Saprotroph\| | Yeast | NULL | Probable | As nectar-tap_saprotroph (Põlme et al. 2020); Decay sugar-rich substrates (Põlme et al. 2020) | Põlme S, et al. 2020. Fungal Diversity 105:1-16 ((https://doi.org/10.1007/s13225-020-00466-2)) |
| Otu30658 | 1 | k__Fungi(100);p__Ascomycota(100);c__Saccharomycetes(100);o__Saccharomycetales(100);f__Metschnikowiaceae(100);g__Metschnikowia(100);g__Metschnikowia_unclassified(100); | Metschnikowia | 13 | Saprotroph | \|Undefined Saprotroph\| | Yeast | NULL | Probable | As nectar-tap_saprotroph (Põlme et al. 2020); Decay sugar-rich substrates (Põlme et al. 2020) | Põlme S, et al. 2020. Fungal Diversity 105:1-16 ((https://doi.org/10.1007/s13225-020-00466-2)) |
| Otu30660 | 1 | k__Fungi(100);p__Ascomycota(100);c__Saccharomycetes(100);o__Saccharomycetales(100);f__Metschnikowiaceae(100);g__Metschnikowia(100);g__Metschnikowia_unclassified(100); | Metschnikowia | 13 | Saprotroph | \|Undefined Saprotroph\| | Yeast | NULL | Probable | As nectar-tap_saprotroph (Põlme et al. 2020); Decay sugar-rich substrates (Põlme et al. 2020) | Põlme S, et al. 2020. Fungal Diversity 105:1-16 ((https://doi.org/10.1007/s13225-020-00466-2)) |
| Otu30661 | 1 | k__Fungi(100);p__Ascomycota(100);c__Saccharomycetes(100);o__Saccharomycetales(100);f__Metschnikowiaceae(100);g__Metschnikowia(100);s__Metschnikowia_pulcherrima(100); | Metschnikowia | 13 | Saprotroph | \|Undefined Saprotroph\| | Yeast | NULL | Probable | As nectar-tap_saprotroph (Põlme et al. 2020); Decay sugar-rich substrates (Põlme et al. 2020) | Põlme S, et al. 2020. Fungal Diversity 105:1-16 ((https://doi.org/10.1007/s13225-020-00466-2)) |
| Otu30662 | 1 | k__Fungi(100);p__Ascomycota(100);c__Saccharomycetes(100);o__Saccharomycetales(100);f__Metschnikowiaceae(100);g__Metschnikowia(100);s__Metschnikowia_pulcherrima(100); | Metschnikowia | 13 | Saprotroph | \|Undefined Saprotroph\| | Yeast | NULL | Probable | As nectar-tap_saprotroph (Põlme et al. 2020); Decay sugar-rich substrates (Põlme et al. 2020) | Põlme S, et al. 2020. Fungal Diversity 105:1-16 ((https://doi.org/10.1007/s13225-020-00466-2)) |
| Otu30665 | 1 | k__Fungi(100);p__Ascomycota(100);c__Saccharomycetes(100);o__Saccharomycetales(100);f__Saccharomycetales_family_Incertae_sedis(100);g__Candida(100);g__Candida_unclassified(100); | Candida | 13 | Pathotroph-Saprotroph-Symbiotroph | Animal Pathogen-Endophyte-Endosymbiont-Epiphyte-\|Undefined Saprotroph\| | Dimorphic Yeast-Yeast | NULL | Probable | As nectar-tap_saprotroph (Põlme et al. 2020); Decay sugar-rich substrates (Põlme et al. 2020); This asexual genus is polyphyletic and contain species that function very differently depend on what they are. Caution should be taken when interpreting guilds for Candida species. | Manolakaki D, et al. 2010. Virulence. 1: 367-375 ((https://doi.org/10.4161/viru.1.5.12796)); Mekbib SB, et al. 2011. Tropical Plant Pathology 36:233-240 ((https://doi.org/10.1590/S1982-56762011000400004)); Põlme S, et al. 2020. Fungal Diversity 105:1-16 ((https://doi.org/10.1007/s13225-020-00466-2)) |
| Otu30666 | 1 | k__Fungi(100);p__Ascomycota(100);c__Saccharomycetes(100);o__Saccharomycetales(100);o__Saccharomycetales_unclassified(100);o__Saccharomycetales_unclassified(100);o__Saccharomycetales_unclassified(100); | Saccharomycetales | 7 | Saprotroph | Undefined Saprotroph | Yeast | NULL | Possible | NULL | Sterkenburg E, et al. 2015. New Phytologist 207:1145-1158 ((https://doi.org/10.1111/nph.13426)) |
| Otu30667 | 1 | k__Fungi(100);p__Ascomycota(100);c__Saccharomycetes(100);o__Saccharomycetales(100);f__Saccharomycetales_family_Incertae_sedis(100);g__Candida(100);g__Candida_unclassified(100); | Candida | 13 | Pathotroph-Saprotroph-Symbiotroph | Animal Pathogen-Endophyte-Endosymbiont-Epiphyte-\|Undefined Saprotroph\| | Dimorphic Yeast-Yeast | NULL | Probable | As nectar-tap_saprotroph (Põlme et al. 2020); Decay sugar-rich substrates (Põlme et al. 2020); This asexual genus is polyphyletic and contain species that function very differently depend on what they are. Caution should be taken when interpreting guilds for Candida species. | Manolakaki D, et al. 2010. Virulence. 1: 367-375 ((https://doi.org/10.4161/viru.1.5.12796)); Mekbib SB, et al. 2011. Tropical Plant Pathology 36:233-240 ((https://doi.org/10.1590/S1982-56762011000400004)); Põlme S, et al. 2020. Fungal Diversity 105:1-16 ((https://doi.org/10.1007/s13225-020-00466-2)) |
| Otu30670 | 1 | k__Fungi(100);p__Ascomycota(100);c__Saccharomycetes(100);o__Saccharomycetales(100);f__Saccharomycetales_family_Incertae_sedis(100);g__Candida(100);g__Candida_unclassified(100); | Candida | 13 | Pathotroph-Saprotroph-Symbiotroph | Animal Pathogen-Endophyte-Endosymbiont-Epiphyte-\|Undefined Saprotroph\| | Dimorphic Yeast-Yeast | NULL | Probable | As nectar-tap_saprotroph (Põlme et al. 2020); Decay sugar-rich substrates (Põlme et al. 2020); This asexual genus is polyphyletic and contain species that function very differently depend on what they are. Caution should be taken when interpreting guilds for Candida species. | Manolakaki D, et al. 2010. Virulence. 1: 367-375 ((https://doi.org/10.4161/viru.1.5.12796)); Mekbib SB, et al. 2011. Tropical Plant Pathology 36:233-240 ((https://doi.org/10.1590/S1982-56762011000400004)); Põlme S, et al. 2020. Fungal Diversity 105:1-16 ((https://doi.org/10.1007/s13225-020-00466-2)) |
| Otu30677 | 1 | k__Fungi(100);p__Ascomycota(100);c__Saccharomycetes(100);o__Saccharomycetales(100);f__Saccharomycetales_family_Incertae_sedis(100);g__Candida(100);g__Candida_unclassified(100); | Candida | 13 | Pathotroph-Saprotroph-Symbiotroph | Animal Pathogen-Endophyte-Endosymbiont-Epiphyte-\|Undefined Saprotroph\| | Dimorphic Yeast-Yeast | NULL | Probable | As nectar-tap_saprotroph (Põlme et al. 2020); Decay sugar-rich substrates (Põlme et al. 2020); This asexual genus is polyphyletic and contain species that function very differently depend on what they are. Caution should be taken when interpreting guilds for Candida species. | Manolakaki D, et al. 2010. Virulence. 1: 367-375 ((https://doi.org/10.4161/viru.1.5.12796)); Mekbib SB, et al. 2011. Tropical Plant Pathology 36:233-240 ((https://doi.org/10.1590/S1982-56762011000400004)); Põlme S, et al. 2020. Fungal Diversity 105:1-16 ((https://doi.org/10.1007/s13225-020-00466-2)) |
| Otu30679 | 1 | k__Fungi(100);p__Ascomycota(100);c__Saccharomycetes(100);o__Saccharomycetales(100);f__Saccharomycetales_family_Incertae_sedis(100);g__Candida(100);s__Candida_cretensis(100); | Candida | 13 | Pathotroph-Saprotroph-Symbiotroph | Animal Pathogen-Endophyte-Endosymbiont-Epiphyte-\|Undefined Saprotroph\| | Dimorphic Yeast-Yeast | NULL | Probable | As nectar-tap_saprotroph (Põlme et al. 2020); Decay sugar-rich substrates (Põlme et al. 2020); This asexual genus is polyphyletic and contain species that function very differently depend on what they are. Caution should be taken when interpreting guilds for Candida species. | Manolakaki D, et al. 2010. Virulence. 1: 367-375 ((https://doi.org/10.4161/viru.1.5.12796)); Mekbib SB, et al. 2011. Tropical Plant Pathology 36:233-240 ((https://doi.org/10.1590/S1982-56762011000400004)); Põlme S, et al. 2020. Fungal Diversity 105:1-16 ((https://doi.org/10.1007/s13225-020-00466-2)) |
| Otu30680 | 1 | k__Fungi(100);p__Ascomycota(100);c__Saccharomycetes(100);o__Saccharomycetales(100);f__Saccharomycetales_family_Incertae_sedis(100);g__Candida(100);g__Candida_unclassified(100); | Candida | 13 | Pathotroph-Saprotroph-Symbiotroph | Animal Pathogen-Endophyte-Endosymbiont-Epiphyte-\|Undefined Saprotroph\| | Dimorphic Yeast-Yeast | NULL | Probable | As nectar-tap_saprotroph (Põlme et al. 2020); Decay sugar-rich substrates (Põlme et al. 2020); This asexual genus is polyphyletic and contain species that function very differently depend on what they are. Caution should be taken when interpreting guilds for Candida species. | Manolakaki D, et al. 2010. Virulence. 1: 367-375 ((https://doi.org/10.4161/viru.1.5.12796)); Mekbib SB, et al. 2011. Tropical Plant Pathology 36:233-240 ((https://doi.org/10.1590/S1982-56762011000400004)); Põlme S, et al. 2020. Fungal Diversity 105:1-16 ((https://doi.org/10.1007/s13225-020-00466-2)) |
| Otu30681 | 1 | k__Fungi(100);p__Ascomycota(100);c__Saccharomycetes(100);o__Saccharomycetales(100);o__Saccharomycetales_unclassified(100);o__Saccharomycetales_unclassified(100);o__Saccharomycetales_unclassified(100); | Saccharomycetales | 7 | Saprotroph | Undefined Saprotroph | Yeast | NULL | Possible | NULL | Sterkenburg E, et al. 2015. New Phytologist 207:1145-1158 ((https://doi.org/10.1111/nph.13426)) |
| Otu30684 | 1 | k__Fungi(100);p__Ascomycota(100);c__Saccharomycetes(100);o__Saccharomycetales(100);f__Saccharomycetales_family_Incertae_sedis(100);g__Candida(100);s__Candida_cretensis(100); | Candida | 13 | Pathotroph-Saprotroph-Symbiotroph | Animal Pathogen-Endophyte-Endosymbiont-Epiphyte-\|Undefined Saprotroph\| | Dimorphic Yeast-Yeast | NULL | Probable | As nectar-tap_saprotroph (Põlme et al. 2020); Decay sugar-rich substrates (Põlme et al. 2020); This asexual genus is polyphyletic and contain species that function very differently depend on what they are. Caution should be taken when interpreting guilds for Candida species. | Manolakaki D, et al. 2010. Virulence. 1: 367-375 ((https://doi.org/10.4161/viru.1.5.12796)); Mekbib SB, et al. 2011. Tropical Plant Pathology 36:233-240 ((https://doi.org/10.1590/S1982-56762011000400004)); Põlme S, et al. 2020. Fungal Diversity 105:1-16 ((https://doi.org/10.1007/s13225-020-00466-2)) |
| Otu30685 | 1 | k__Fungi(100);p__Ascomycota(100);c__Saccharomycetes(100);o__Saccharomycetales(100);o__Saccharomycetales_unclassified(100);o__Saccharomycetales_unclassified(100);o__Saccharomycetales_unclassified(100); | Saccharomycetales | 7 | Saprotroph | Undefined Saprotroph | Yeast | NULL | Possible | NULL | Sterkenburg E, et al. 2015. New Phytologist 207:1145-1158 ((https://doi.org/10.1111/nph.13426)) |
| Otu30686 | 1 | k__Fungi(100);p__Ascomycota(100);c__Saccharomycetes(100);o__Saccharomycetales(100);f__Metschnikowiaceae(100);g__Metschnikowia(100);g__Metschnikowia_unclassified(100); | Metschnikowia | 13 | Saprotroph | \|Undefined Saprotroph\| | Yeast | NULL | Probable | As nectar-tap_saprotroph (Põlme et al. 2020); Decay sugar-rich substrates (Põlme et al. 2020) | Põlme S, et al. 2020. Fungal Diversity 105:1-16 ((https://doi.org/10.1007/s13225-020-00466-2)) |
| Otu30688 | 1 | k__Fungi(100);p__Ascomycota(100);c__Saccharomycetes(100);o__Saccharomycetales(100);f__Saccharomycetales_family_Incertae_sedis(100);g__Candida(100);g__Candida_unclassified(100); | Candida | 13 | Pathotroph-Saprotroph-Symbiotroph | Animal Pathogen-Endophyte-Endosymbiont-Epiphyte-\|Undefined Saprotroph\| | Dimorphic Yeast-Yeast | NULL | Probable | As nectar-tap_saprotroph (Põlme et al. 2020); Decay sugar-rich substrates (Põlme et al. 2020); This asexual genus is polyphyletic and contain species that function very differently depend on what they are. Caution should be taken when interpreting guilds for Candida species. | Manolakaki D, et al. 2010. Virulence. 1: 367-375 ((https://doi.org/10.4161/viru.1.5.12796)); Mekbib SB, et al. 2011. Tropical Plant Pathology 36:233-240 ((https://doi.org/10.1590/S1982-56762011000400004)); Põlme S, et al. 2020. Fungal Diversity 105:1-16 ((https://doi.org/10.1007/s13225-020-00466-2)) |
| Otu30689 | 1 | k__Fungi(100);p__Ascomycota(100);c__Saccharomycetes(100);o__Saccharomycetales(100);f__Saccharomycetales_family_Incertae_sedis(100);f__Saccharomycetales_family_Incertae_sedis_unclassified(100);f__Saccharomycetales_family_Incertae_sedis_unclassified(100); | Saccharomycetales | 7 | Saprotroph | Undefined Saprotroph | Yeast | NULL | Possible | NULL | Sterkenburg E, et al. 2015. New Phytologist 207:1145-1158 ((https://doi.org/10.1111/nph.13426)) |
| Otu30694 | 1 | k__Fungi(100);p__Ascomycota(100);c__Saccharomycetes(100);o__Saccharomycetales(100);f__Metschnikowiaceae(100);g__Metschnikowia(100);g__Metschnikowia_unclassified(100); | Metschnikowia | 13 | Saprotroph | \|Undefined Saprotroph\| | Yeast | NULL | Probable | As nectar-tap_saprotroph (Põlme et al. 2020); Decay sugar-rich substrates (Põlme et al. 2020) | Põlme S, et al. 2020. Fungal Diversity 105:1-16 ((https://doi.org/10.1007/s13225-020-00466-2)) |
| Otu30696 | 1 | k__Fungi(100);p__Ascomycota(100);c__Saccharomycetes(100);o__Saccharomycetales(100);f__Metschnikowiaceae(100);g__Metschnikowia(100);s__Metschnikowia_pulcherrima(100); | Metschnikowia | 13 | Saprotroph | \|Undefined Saprotroph\| | Yeast | NULL | Probable | As nectar-tap_saprotroph (Põlme et al. 2020); Decay sugar-rich substrates (Põlme et al. 2020) | Põlme S, et al. 2020. Fungal Diversity 105:1-16 ((https://doi.org/10.1007/s13225-020-00466-2)) |
| Otu30706 | 1 | k__Fungi(100);p__Ascomycota(100);c__Saccharomycetes(100);o__Saccharomycetales(100);f__Saccharomycetales_family_Incertae_sedis(100);g__Candida(100);g__Candida_unclassified(100); | Candida | 13 | Pathotroph-Saprotroph-Symbiotroph | Animal Pathogen-Endophyte-Endosymbiont-Epiphyte-\|Undefined Saprotroph\| | Dimorphic Yeast-Yeast | NULL | Probable | As nectar-tap_saprotroph (Põlme et al. 2020); Decay sugar-rich substrates (Põlme et al. 2020); This asexual genus is polyphyletic and contain species that function very differently depend on what they are. Caution should be taken when interpreting guilds for Candida species. | Manolakaki D, et al. 2010. Virulence. 1: 367-375 ((https://doi.org/10.4161/viru.1.5.12796)); Mekbib SB, et al. 2011. Tropical Plant Pathology 36:233-240 ((https://doi.org/10.1590/S1982-56762011000400004)); Põlme S, et al. 2020. Fungal Diversity 105:1-16 ((https://doi.org/10.1007/s13225-020-00466-2)) |
| Otu30708 | 1 | k__Fungi(100);p__Ascomycota(100);c__Saccharomycetes(100);o__Saccharomycetales(100);f__Metschnikowiaceae(100);g__Metschnikowia(100);s__Metschnikowia_pulcherrima(100); | Metschnikowia | 13 | Saprotroph | \|Undefined Saprotroph\| | Yeast | NULL | Probable | As nectar-tap_saprotroph (Põlme et al. 2020); Decay sugar-rich substrates (Põlme et al. 2020) | Põlme S, et al. 2020. Fungal Diversity 105:1-16 ((https://doi.org/10.1007/s13225-020-00466-2)) |
| Otu30709 | 1 | k__Fungi(100);p__Ascomycota(100);c__Saccharomycetes(100);o__Saccharomycetales(100);f__Metschnikowiaceae(100);g__Metschnikowia(100);g__Metschnikowia_unclassified(100); | Metschnikowia | 13 | Saprotroph | \|Undefined Saprotroph\| | Yeast | NULL | Probable | As nectar-tap_saprotroph (Põlme et al. 2020); Decay sugar-rich substrates (Põlme et al. 2020) | Põlme S, et al. 2020. Fungal Diversity 105:1-16 ((https://doi.org/10.1007/s13225-020-00466-2)) |
| Otu30711 | 1 | k__Fungi(100);p__Ascomycota(100);c__Saccharomycetes(100);o__Saccharomycetales(100);f__Saccharomycetales_family_Incertae_sedis(100);g__Candida(100);g__Candida_unclassified(100); | Candida | 13 | Pathotroph-Saprotroph-Symbiotroph | Animal Pathogen-Endophyte-Endosymbiont-Epiphyte-\|Undefined Saprotroph\| | Dimorphic Yeast-Yeast | NULL | Probable | As nectar-tap_saprotroph (Põlme et al. 2020); Decay sugar-rich substrates (Põlme et al. 2020); This asexual genus is polyphyletic and contain species that function very differently depend on what they are. Caution should be taken when interpreting guilds for Candida species. | Manolakaki D, et al. 2010. Virulence. 1: 367-375 ((https://doi.org/10.4161/viru.1.5.12796)); Mekbib SB, et al. 2011. Tropical Plant Pathology 36:233-240 ((https://doi.org/10.1590/S1982-56762011000400004)); Põlme S, et al. 2020. Fungal Diversity 105:1-16 ((https://doi.org/10.1007/s13225-020-00466-2)) |
| Otu30712 | 1 | k__Fungi(100);p__Ascomycota(100);c__Saccharomycetes(100);o__Saccharomycetales(100);f__Saccharomycetales_family_Incertae_sedis(100);f__Saccharomycetales_family_Incertae_sedis_unclassified(100);f__Saccharomycetales_family_Incertae_sedis_unclassified(100); | Saccharomycetales | 7 | Saprotroph | Undefined Saprotroph | Yeast | NULL | Possible | NULL | Sterkenburg E, et al. 2015. New Phytologist 207:1145-1158 ((https://doi.org/10.1111/nph.13426)) |
| Otu30714 | 1 | k__Fungi(100);p__Ascomycota(100);c__Saccharomycetes(100);o__Saccharomycetales(100);f__Saccharomycetales_family_Incertae_sedis(100);g__Candida(100);g__Candida_unclassified(100); | Candida | 13 | Pathotroph-Saprotroph-Symbiotroph | Animal Pathogen-Endophyte-Endosymbiont-Epiphyte-\|Undefined Saprotroph\| | Dimorphic Yeast-Yeast | NULL | Probable | As nectar-tap_saprotroph (Põlme et al. 2020); Decay sugar-rich substrates (Põlme et al. 2020); This asexual genus is polyphyletic and contain species that function very differently depend on what they are. Caution should be taken when interpreting guilds for Candida species. | Manolakaki D, et al. 2010. Virulence. 1: 367-375 ((https://doi.org/10.4161/viru.1.5.12796)); Mekbib SB, et al. 2011. Tropical Plant Pathology 36:233-240 ((https://doi.org/10.1590/S1982-56762011000400004)); Põlme S, et al. 2020. Fungal Diversity 105:1-16 ((https://doi.org/10.1007/s13225-020-00466-2)) |
| Otu30716 | 1 | k__Fungi(100);p__Ascomycota(100);c__Saccharomycetes(100);o__Saccharomycetales(100);o__Saccharomycetales_unclassified(100);o__Saccharomycetales_unclassified(100);o__Saccharomycetales_unclassified(100); | Saccharomycetales | 7 | Saprotroph | Undefined Saprotroph | Yeast | NULL | Possible | NULL | Sterkenburg E, et al. 2015. New Phytologist 207:1145-1158 ((https://doi.org/10.1111/nph.13426)) |
| Otu30717 | 1 | k__Fungi(100);p__Ascomycota(100);c__Saccharomycetes(100);o__Saccharomycetales(100);f__Saccharomycetales_family_Incertae_sedis(100);g__Candida(100);s__Candida_cretensis(100); | Candida | 13 | Pathotroph-Saprotroph-Symbiotroph | Animal Pathogen-Endophyte-Endosymbiont-Epiphyte-\|Undefined Saprotroph\| | Dimorphic Yeast-Yeast | NULL | Probable | As nectar-tap_saprotroph (Põlme et al. 2020); Decay sugar-rich substrates (Põlme et al. 2020); This asexual genus is polyphyletic and contain species that function very differently depend on what they are. Caution should be taken when interpreting guilds for Candida species. | Manolakaki D, et al. 2010. Virulence. 1: 367-375 ((https://doi.org/10.4161/viru.1.5.12796)); Mekbib SB, et al. 2011. Tropical Plant Pathology 36:233-240 ((https://doi.org/10.1590/S1982-56762011000400004)); Põlme S, et al. 2020. Fungal Diversity 105:1-16 ((https://doi.org/10.1007/s13225-020-00466-2)) |
| Otu30718 | 1 | k__Fungi(100);p__Ascomycota(100);c__Saccharomycetes(100);o__Saccharomycetales(100);f__Metschnikowiaceae(100);g__Metschnikowia(100);s__Metschnikowia_pulcherrima(100); | Metschnikowia | 13 | Saprotroph | \|Undefined Saprotroph\| | Yeast | NULL | Probable | As nectar-tap_saprotroph (Põlme et al. 2020); Decay sugar-rich substrates (Põlme et al. 2020) | Põlme S, et al. 2020. Fungal Diversity 105:1-16 ((https://doi.org/10.1007/s13225-020-00466-2)) |
| Otu30719 | 1 | k__Fungi(100);p__Ascomycota(100);c__Saccharomycetes(100);o__Saccharomycetales(100);o__Saccharomycetales_unclassified(100);o__Saccharomycetales_unclassified(100);o__Saccharomycetales_unclassified(100); | Saccharomycetales | 7 | Saprotroph | Undefined Saprotroph | Yeast | NULL | Possible | NULL | Sterkenburg E, et al. 2015. New Phytologist 207:1145-1158 ((https://doi.org/10.1111/nph.13426)) |
| Otu30720 | 1 | k__Fungi(100);p__Ascomycota(100);c__Saccharomycetes(100);o__Saccharomycetales(100);f__Saccharomycetales_family_Incertae_sedis(100);f__Saccharomycetales_family_Incertae_sedis_unclassified(100);f__Saccharomycetales_family_Incertae_sedis_unclassified(100); | Saccharomycetales | 7 | Saprotroph | Undefined Saprotroph | Yeast | NULL | Possible | NULL | Sterkenburg E, et al. 2015. New Phytologist 207:1145-1158 ((https://doi.org/10.1111/nph.13426)) |
| Otu30721 | 1 | k__Fungi(100);p__Ascomycota(100);c__Saccharomycetes(100);o__Saccharomycetales(100);f__Metschnikowiaceae(100);g__Metschnikowia(100);s__Metschnikowia_pulcherrima(100); | Metschnikowia | 13 | Saprotroph | \|Undefined Saprotroph\| | Yeast | NULL | Probable | As nectar-tap_saprotroph (Põlme et al. 2020); Decay sugar-rich substrates (Põlme et al. 2020) | Põlme S, et al. 2020. Fungal Diversity 105:1-16 ((https://doi.org/10.1007/s13225-020-00466-2)) |
| Otu30723 | 1 | k__Fungi(100);p__Ascomycota(100);c__Saccharomycetes(100);o__Saccharomycetales(100);f__Saccharomycetales_family_Incertae_sedis(100);g__Candida(100);g__Candida_unclassified(100); | Candida | 13 | Pathotroph-Saprotroph-Symbiotroph | Animal Pathogen-Endophyte-Endosymbiont-Epiphyte-\|Undefined Saprotroph\| | Dimorphic Yeast-Yeast | NULL | Probable | As nectar-tap_saprotroph (Põlme et al. 2020); Decay sugar-rich substrates (Põlme et al. 2020); This asexual genus is polyphyletic and contain species that function very differently depend on what they are. Caution should be taken when interpreting guilds for Candida species. | Manolakaki D, et al. 2010. Virulence. 1: 367-375 ((https://doi.org/10.4161/viru.1.5.12796)); Mekbib SB, et al. 2011. Tropical Plant Pathology 36:233-240 ((https://doi.org/10.1590/S1982-56762011000400004)); Põlme S, et al. 2020. Fungal Diversity 105:1-16 ((https://doi.org/10.1007/s13225-020-00466-2)) |
| Otu30724 | 1 | k__Fungi(100);p__Ascomycota(100);c__Saccharomycetes(100);o__Saccharomycetales(100);f__Saccharomycetales_family_Incertae_sedis(100);g__Candida(100);g__Candida_unclassified(100); | Candida | 13 | Pathotroph-Saprotroph-Symbiotroph | Animal Pathogen-Endophyte-Endosymbiont-Epiphyte-\|Undefined Saprotroph\| | Dimorphic Yeast-Yeast | NULL | Probable | As nectar-tap_saprotroph (Põlme et al. 2020); Decay sugar-rich substrates (Põlme et al. 2020); This asexual genus is polyphyletic and contain species that function very differently depend on what they are. Caution should be taken when interpreting guilds for Candida species. | Manolakaki D, et al. 2010. Virulence. 1: 367-375 ((https://doi.org/10.4161/viru.1.5.12796)); Mekbib SB, et al. 2011. Tropical Plant Pathology 36:233-240 ((https://doi.org/10.1590/S1982-56762011000400004)); Põlme S, et al. 2020. Fungal Diversity 105:1-16 ((https://doi.org/10.1007/s13225-020-00466-2)) |
| Otu30726 | 1 | k__Fungi(100);p__Ascomycota(100);c__Saccharomycetes(100);o__Saccharomycetales(100);f__Saccharomycetales_family_Incertae_sedis(100);g__Candida(100);g__Candida_unclassified(100); | Candida | 13 | Pathotroph-Saprotroph-Symbiotroph | Animal Pathogen-Endophyte-Endosymbiont-Epiphyte-\|Undefined Saprotroph\| | Dimorphic Yeast-Yeast | NULL | Probable | As nectar-tap_saprotroph (Põlme et al. 2020); Decay sugar-rich substrates (Põlme et al. 2020); This asexual genus is polyphyletic and contain species that function very differently depend on what they are. Caution should be taken when interpreting guilds for Candida species. | Manolakaki D, et al. 2010. Virulence. 1: 367-375 ((https://doi.org/10.4161/viru.1.5.12796)); Mekbib SB, et al. 2011. Tropical Plant Pathology 36:233-240 ((https://doi.org/10.1590/S1982-56762011000400004)); Põlme S, et al. 2020. Fungal Diversity 105:1-16 ((https://doi.org/10.1007/s13225-020-00466-2)) |
| Otu30731 | 1 | k__Fungi(100);p__Ascomycota(100);c__Saccharomycetes(100);o__Saccharomycetales(100);f__Metschnikowiaceae(100);g__Metschnikowia(100);g__Metschnikowia_unclassified(100); | Metschnikowia | 13 | Saprotroph | \|Undefined Saprotroph\| | Yeast | NULL | Probable | As nectar-tap_saprotroph (Põlme et al. 2020); Decay sugar-rich substrates (Põlme et al. 2020) | Põlme S, et al. 2020. Fungal Diversity 105:1-16 ((https://doi.org/10.1007/s13225-020-00466-2)) |
| Otu30732 | 1 | k__Fungi(100);p__Ascomycota(100);c__Saccharomycetes(100);o__Saccharomycetales(100);f__Metschnikowiaceae(100);g__Metschnikowia(100);s__Metschnikowia_pulcherrima(100); | Metschnikowia | 13 | Saprotroph | \|Undefined Saprotroph\| | Yeast | NULL | Probable | As nectar-tap_saprotroph (Põlme et al. 2020); Decay sugar-rich substrates (Põlme et al. 2020) | Põlme S, et al. 2020. Fungal Diversity 105:1-16 ((https://doi.org/10.1007/s13225-020-00466-2)) |
| Otu30733 | 1 | k__Fungi(100);p__Ascomycota(100);c__Saccharomycetes(100);o__Saccharomycetales(100);o__Saccharomycetales_unclassified(100);o__Saccharomycetales_unclassified(100);o__Saccharomycetales_unclassified(100); | Saccharomycetales | 7 | Saprotroph | Undefined Saprotroph | Yeast | NULL | Possible | NULL | Sterkenburg E, et al. 2015. New Phytologist 207:1145-1158 ((https://doi.org/10.1111/nph.13426)) |
| Otu30734 | 1 | k__Fungi(100);p__Ascomycota(100);c__Saccharomycetes(100);o__Saccharomycetales(100);f__Saccharomycetales_family_Incertae_sedis(100);g__Candida(100);g__Candida_unclassified(100); | Candida | 13 | Pathotroph-Saprotroph-Symbiotroph | Animal Pathogen-Endophyte-Endosymbiont-Epiphyte-\|Undefined Saprotroph\| | Dimorphic Yeast-Yeast | NULL | Probable | As nectar-tap_saprotroph (Põlme et al. 2020); Decay sugar-rich substrates (Põlme et al. 2020); This asexual genus is polyphyletic and contain species that function very differently depend on what they are. Caution should be taken when interpreting guilds for Candida species. | Manolakaki D, et al. 2010. Virulence. 1: 367-375 ((https://doi.org/10.4161/viru.1.5.12796)); Mekbib SB, et al. 2011. Tropical Plant Pathology 36:233-240 ((https://doi.org/10.1590/S1982-56762011000400004)); Põlme S, et al. 2020. Fungal Diversity 105:1-16 ((https://doi.org/10.1007/s13225-020-00466-2)) |
| Otu30735 | 1 | k__Fungi(100);p__Ascomycota(100);c__Saccharomycetes(100);o__Saccharomycetales(100);f__Metschnikowiaceae(100);g__Metschnikowia(100);g__Metschnikowia_unclassified(100); | Metschnikowia | 13 | Saprotroph | \|Undefined Saprotroph\| | Yeast | NULL | Probable | As nectar-tap_saprotroph (Põlme et al. 2020); Decay sugar-rich substrates (Põlme et al. 2020) | Põlme S, et al. 2020. Fungal Diversity 105:1-16 ((https://doi.org/10.1007/s13225-020-00466-2)) |
| Otu30736 | 1 | k__Fungi(100);p__Ascomycota(100);c__Saccharomycetes(100);o__Saccharomycetales(100);f__Metschnikowiaceae(100);g__Metschnikowia(100);s__Metschnikowia_pulcherrima(100); | Metschnikowia | 13 | Saprotroph | \|Undefined Saprotroph\| | Yeast | NULL | Probable | As nectar-tap_saprotroph (Põlme et al. 2020); Decay sugar-rich substrates (Põlme et al. 2020) | Põlme S, et al. 2020. Fungal Diversity 105:1-16 ((https://doi.org/10.1007/s13225-020-00466-2)) |
| Otu30737 | 1 | k__Fungi(100);p__Ascomycota(100);c__Saccharomycetes(100);o__Saccharomycetales(100);f__Metschnikowiaceae(100);g__Metschnikowia(100);s__Metschnikowia_pulcherrima(100); | Metschnikowia | 13 | Saprotroph | \|Undefined Saprotroph\| | Yeast | NULL | Probable | As nectar-tap_saprotroph (Põlme et al. 2020); Decay sugar-rich substrates (Põlme et al. 2020) | Põlme S, et al. 2020. Fungal Diversity 105:1-16 ((https://doi.org/10.1007/s13225-020-00466-2)) |
| Otu30738 | 1 | k__Fungi(100);p__Ascomycota(100);c__Saccharomycetes(100);o__Saccharomycetales(100);f__Saccharomycetales_family_Incertae_sedis(100);g__Candida(100);s__Candida_cretensis(100); | Candida | 13 | Pathotroph-Saprotroph-Symbiotroph | Animal Pathogen-Endophyte-Endosymbiont-Epiphyte-\|Undefined Saprotroph\| | Dimorphic Yeast-Yeast | NULL | Probable | As nectar-tap_saprotroph (Põlme et al. 2020); Decay sugar-rich substrates (Põlme et al. 2020); This asexual genus is polyphyletic and contain species that function very differently depend on what they are. Caution should be taken when interpreting guilds for Candida species. | Manolakaki D, et al. 2010. Virulence. 1: 367-375 ((https://doi.org/10.4161/viru.1.5.12796)); Mekbib SB, et al. 2011. Tropical Plant Pathology 36:233-240 ((https://doi.org/10.1590/S1982-56762011000400004)); Põlme S, et al. 2020. Fungal Diversity 105:1-16 ((https://doi.org/10.1007/s13225-020-00466-2)) |
| Otu30739 | 1 | k__Fungi(100);p__Ascomycota(100);c__Saccharomycetes(100);o__Saccharomycetales(100);f__Saccharomycetales_family_Incertae_sedis(100);g__Candida(100);g__Candida_unclassified(100); | Candida | 13 | Pathotroph-Saprotroph-Symbiotroph | Animal Pathogen-Endophyte-Endosymbiont-Epiphyte-\|Undefined Saprotroph\| | Dimorphic Yeast-Yeast | NULL | Probable | As nectar-tap_saprotroph (Põlme et al. 2020); Decay sugar-rich substrates (Põlme et al. 2020); This asexual genus is polyphyletic and contain species that function very differently depend on what they are. Caution should be taken when interpreting guilds for Candida species. | Manolakaki D, et al. 2010. Virulence. 1: 367-375 ((https://doi.org/10.4161/viru.1.5.12796)); Mekbib SB, et al. 2011. Tropical Plant Pathology 36:233-240 ((https://doi.org/10.1590/S1982-56762011000400004)); Põlme S, et al. 2020. Fungal Diversity 105:1-16 ((https://doi.org/10.1007/s13225-020-00466-2)) |
| Otu30742 | 1 | k__Fungi(100);p__Ascomycota(100);c__Saccharomycetes(100);o__Saccharomycetales(100);f__Metschnikowiaceae(100);g__Metschnikowia(100);g__Metschnikowia_unclassified(100); | Metschnikowia | 13 | Saprotroph | \|Undefined Saprotroph\| | Yeast | NULL | Probable | As nectar-tap_saprotroph (Põlme et al. 2020); Decay sugar-rich substrates (Põlme et al. 2020) | Põlme S, et al. 2020. Fungal Diversity 105:1-16 ((https://doi.org/10.1007/s13225-020-00466-2)) |
| Otu30743 | 1 | k__Fungi(100);p__Ascomycota(100);c__Saccharomycetes(100);o__Saccharomycetales(100);f__Saccharomycetales_family_Incertae_sedis(100);g__Candida(100);g__Candida_unclassified(100); | Candida | 13 | Pathotroph-Saprotroph-Symbiotroph | Animal Pathogen-Endophyte-Endosymbiont-Epiphyte-\|Undefined Saprotroph\| | Dimorphic Yeast-Yeast | NULL | Probable | As nectar-tap_saprotroph (Põlme et al. 2020); Decay sugar-rich substrates (Põlme et al. 2020); This asexual genus is polyphyletic and contain species that function very differently depend on what they are. Caution should be taken when interpreting guilds for Candida species. | Manolakaki D, et al. 2010. Virulence. 1: 367-375 ((https://doi.org/10.4161/viru.1.5.12796)); Mekbib SB, et al. 2011. Tropical Plant Pathology 36:233-240 ((https://doi.org/10.1590/S1982-56762011000400004)); Põlme S, et al. 2020. Fungal Diversity 105:1-16 ((https://doi.org/10.1007/s13225-020-00466-2)) |
| Otu30748 | 1 | k__Fungi(100);p__Ascomycota(100);c__Saccharomycetes(100);o__Saccharomycetales(100);f__Metschnikowiaceae(100);g__Metschnikowia(100);g__Metschnikowia_unclassified(100); | Metschnikowia | 13 | Saprotroph | \|Undefined Saprotroph\| | Yeast | NULL | Probable | As nectar-tap_saprotroph (Põlme et al. 2020); Decay sugar-rich substrates (Põlme et al. 2020) | Põlme S, et al. 2020. Fungal Diversity 105:1-16 ((https://doi.org/10.1007/s13225-020-00466-2)) |
| Otu30750 | 1 | k__Fungi(100);p__Ascomycota(100);c__Saccharomycetes(100);o__Saccharomycetales(100);o__Saccharomycetales_unclassified(100);o__Saccharomycetales_unclassified(100);o__Saccharomycetales_unclassified(100); | Saccharomycetales | 7 | Saprotroph | Undefined Saprotroph | Yeast | NULL | Possible | NULL | Sterkenburg E, et al. 2015. New Phytologist 207:1145-1158 ((https://doi.org/10.1111/nph.13426)) |
| Otu30751 | 1 | k__Fungi(100);p__Ascomycota(100);c__Saccharomycetes(100);o__Saccharomycetales(100);f__Saccharomycetales_family_Incertae_sedis(100);g__Candida(100);g__Candida_unclassified(100); | Candida | 13 | Pathotroph-Saprotroph-Symbiotroph | Animal Pathogen-Endophyte-Endosymbiont-Epiphyte-\|Undefined Saprotroph\| | Dimorphic Yeast-Yeast | NULL | Probable | As nectar-tap_saprotroph (Põlme et al. 2020); Decay sugar-rich substrates (Põlme et al. 2020); This asexual genus is polyphyletic and contain species that function very differently depend on what they are. Caution should be taken when interpreting guilds for Candida species. | Manolakaki D, et al. 2010. Virulence. 1: 367-375 ((https://doi.org/10.4161/viru.1.5.12796)); Mekbib SB, et al. 2011. Tropical Plant Pathology 36:233-240 ((https://doi.org/10.1590/S1982-56762011000400004)); Põlme S, et al. 2020. Fungal Diversity 105:1-16 ((https://doi.org/10.1007/s13225-020-00466-2)) |
| Otu30752 | 1 | k__Fungi(100);p__Ascomycota(100);c__Saccharomycetes(100);o__Saccharomycetales(100);f__Metschnikowiaceae(100);g__Metschnikowia(100);g__Metschnikowia_unclassified(100); | Metschnikowia | 13 | Saprotroph | \|Undefined Saprotroph\| | Yeast | NULL | Probable | As nectar-tap_saprotroph (Põlme et al. 2020); Decay sugar-rich substrates (Põlme et al. 2020) | Põlme S, et al. 2020. Fungal Diversity 105:1-16 ((https://doi.org/10.1007/s13225-020-00466-2)) |
| Otu30753 | 1 | k__Fungi(100);p__Ascomycota(100);c__Saccharomycetes(100);o__Saccharomycetales(100);f__Metschnikowiaceae(100);g__Metschnikowia(100);s__Metschnikowia_pulcherrima(100); | Metschnikowia | 13 | Saprotroph | \|Undefined Saprotroph\| | Yeast | NULL | Probable | As nectar-tap_saprotroph (Põlme et al. 2020); Decay sugar-rich substrates (Põlme et al. 2020) | Põlme S, et al. 2020. Fungal Diversity 105:1-16 ((https://doi.org/10.1007/s13225-020-00466-2)) |
| Otu30754 | 1 | k__Fungi(100);p__Ascomycota(100);c__Saccharomycetes(100);o__Saccharomycetales(100);f__Metschnikowiaceae(100);g__Metschnikowia(100);s__Metschnikowia_pulcherrima(100); | Metschnikowia | 13 | Saprotroph | \|Undefined Saprotroph\| | Yeast | NULL | Probable | As nectar-tap_saprotroph (Põlme et al. 2020); Decay sugar-rich substrates (Põlme et al. 2020) | Põlme S, et al. 2020. Fungal Diversity 105:1-16 ((https://doi.org/10.1007/s13225-020-00466-2)) |
| Otu30755 | 1 | k__Fungi(100);p__Ascomycota(100);c__Saccharomycetes(100);o__Saccharomycetales(100);f__Metschnikowiaceae(100);g__Metschnikowia(100);s__Metschnikowia_pulcherrima(100); | Metschnikowia | 13 | Saprotroph | \|Undefined Saprotroph\| | Yeast | NULL | Probable | As nectar-tap_saprotroph (Põlme et al. 2020); Decay sugar-rich substrates (Põlme et al. 2020) | Põlme S, et al. 2020. Fungal Diversity 105:1-16 ((https://doi.org/10.1007/s13225-020-00466-2)) |
| Otu30756 | 1 | k__Fungi(100);p__Ascomycota(100);c__Saccharomycetes(100);o__Saccharomycetales(100);f__Metschnikowiaceae(100);g__Metschnikowia(100);s__Metschnikowia_pulcherrima(100); | Metschnikowia | 13 | Saprotroph | \|Undefined Saprotroph\| | Yeast | NULL | Probable | As nectar-tap_saprotroph (Põlme et al. 2020); Decay sugar-rich substrates (Põlme et al. 2020) | Põlme S, et al. 2020. Fungal Diversity 105:1-16 ((https://doi.org/10.1007/s13225-020-00466-2)) |
| Otu30758 | 1 | k__Fungi(100);p__Ascomycota(100);c__Saccharomycetes(100);o__Saccharomycetales(100);f__Metschnikowiaceae(100);g__Metschnikowia(100);s__Metschnikowia_pulcherrima(100); | Metschnikowia | 13 | Saprotroph | \|Undefined Saprotroph\| | Yeast | NULL | Probable | As nectar-tap_saprotroph (Põlme et al. 2020); Decay sugar-rich substrates (Põlme et al. 2020) | Põlme S, et al. 2020. Fungal Diversity 105:1-16 ((https://doi.org/10.1007/s13225-020-00466-2)) |
| Otu30759 | 1 | k__Fungi(100);p__Ascomycota(100);c__Saccharomycetes(100);o__Saccharomycetales(100);f__Metschnikowiaceae(100);g__Metschnikowia(100);s__Metschnikowia_pulcherrima(100); | Metschnikowia | 13 | Saprotroph | \|Undefined Saprotroph\| | Yeast | NULL | Probable | As nectar-tap_saprotroph (Põlme et al. 2020); Decay sugar-rich substrates (Põlme et al. 2020) | Põlme S, et al. 2020. Fungal Diversity 105:1-16 ((https://doi.org/10.1007/s13225-020-00466-2)) |
| Otu30761 | 1 | k__Fungi(100);p__Ascomycota(100);c__Saccharomycetes(100);o__Saccharomycetales(100);f__Metschnikowiaceae(100);g__Metschnikowia(100);g__Metschnikowia_unclassified(100); | Metschnikowia | 13 | Saprotroph | \|Undefined Saprotroph\| | Yeast | NULL | Probable | As nectar-tap_saprotroph (Põlme et al. 2020); Decay sugar-rich substrates (Põlme et al. 2020) | Põlme S, et al. 2020. Fungal Diversity 105:1-16 ((https://doi.org/10.1007/s13225-020-00466-2)) |
| Otu30763 | 1 | k__Fungi(100);p__Ascomycota(100);c__Saccharomycetes(100);o__Saccharomycetales(100);f__Metschnikowiaceae(100);g__Metschnikowia(100);g__Metschnikowia_unclassified(100); | Metschnikowia | 13 | Saprotroph | \|Undefined Saprotroph\| | Yeast | NULL | Probable | As nectar-tap_saprotroph (Põlme et al. 2020); Decay sugar-rich substrates (Põlme et al. 2020) | Põlme S, et al. 2020. Fungal Diversity 105:1-16 ((https://doi.org/10.1007/s13225-020-00466-2)) |
| Otu30764 | 1 | k__Fungi(100);p__Ascomycota(100);c__Saccharomycetes(100);o__Saccharomycetales(100);f__Metschnikowiaceae(100);g__Metschnikowia(100);s__Metschnikowia_pulcherrima(100); | Metschnikowia | 13 | Saprotroph | \|Undefined Saprotroph\| | Yeast | NULL | Probable | As nectar-tap_saprotroph (Põlme et al. 2020); Decay sugar-rich substrates (Põlme et al. 2020) | Põlme S, et al. 2020. Fungal Diversity 105:1-16 ((https://doi.org/10.1007/s13225-020-00466-2)) |
| Otu30765 | 1 | k__Fungi(100);p__Ascomycota(100);c__Saccharomycetes(100);o__Saccharomycetales(100);f__Metschnikowiaceae(100);g__Metschnikowia(100);s__Metschnikowia_pulcherrima(100); | Metschnikowia | 13 | Saprotroph | \|Undefined Saprotroph\| | Yeast | NULL | Probable | As nectar-tap_saprotroph (Põlme et al. 2020); Decay sugar-rich substrates (Põlme et al. 2020) | Põlme S, et al. 2020. Fungal Diversity 105:1-16 ((https://doi.org/10.1007/s13225-020-00466-2)) |
| Otu30766 | 1 | k__Fungi(100);p__Ascomycota(100);c__Saccharomycetes(100);o__Saccharomycetales(100);f__Metschnikowiaceae(100);g__Metschnikowia(100);s__Metschnikowia_pulcherrima(100); | Metschnikowia | 13 | Saprotroph | \|Undefined Saprotroph\| | Yeast | NULL | Probable | As nectar-tap_saprotroph (Põlme et al. 2020); Decay sugar-rich substrates (Põlme et al. 2020) | Põlme S, et al. 2020. Fungal Diversity 105:1-16 ((https://doi.org/10.1007/s13225-020-00466-2)) |
| Otu30767 | 1 | k__Fungi(100);p__Ascomycota(100);c__Saccharomycetes(100);o__Saccharomycetales(100);f__Metschnikowiaceae(100);g__Metschnikowia(100);g__Metschnikowia_unclassified(100); | Metschnikowia | 13 | Saprotroph | \|Undefined Saprotroph\| | Yeast | NULL | Probable | As nectar-tap_saprotroph (Põlme et al. 2020); Decay sugar-rich substrates (Põlme et al. 2020) | Põlme S, et al. 2020. Fungal Diversity 105:1-16 ((https://doi.org/10.1007/s13225-020-00466-2)) |
| Otu30768 | 1 | k__Fungi(100);p__Ascomycota(100);c__Saccharomycetes(100);o__Saccharomycetales(100);f__Metschnikowiaceae(100);g__Metschnikowia(100);s__Metschnikowia_pulcherrima(100); | Metschnikowia | 13 | Saprotroph | \|Undefined Saprotroph\| | Yeast | NULL | Probable | As nectar-tap_saprotroph (Põlme et al. 2020); Decay sugar-rich substrates (Põlme et al. 2020) | Põlme S, et al. 2020. Fungal Diversity 105:1-16 ((https://doi.org/10.1007/s13225-020-00466-2)) |
| Otu30769 | 1 | k__Fungi(100);p__Ascomycota(100);c__Saccharomycetes(100);o__Saccharomycetales(100);f__Metschnikowiaceae(100);g__Metschnikowia(100);s__Metschnikowia_pulcherrima(100); | Metschnikowia | 13 | Saprotroph | \|Undefined Saprotroph\| | Yeast | NULL | Probable | As nectar-tap_saprotroph (Põlme et al. 2020); Decay sugar-rich substrates (Põlme et al. 2020) | Põlme S, et al. 2020. Fungal Diversity 105:1-16 ((https://doi.org/10.1007/s13225-020-00466-2)) |
| Otu30771 | 1 | k__Fungi(100);p__Ascomycota(100);c__Saccharomycetes(100);o__Saccharomycetales(100);f__Metschnikowiaceae(100);g__Metschnikowia(100);s__Metschnikowia_pulcherrima(100); | Metschnikowia | 13 | Saprotroph | \|Undefined Saprotroph\| | Yeast | NULL | Probable | As nectar-tap_saprotroph (Põlme et al. 2020); Decay sugar-rich substrates (Põlme et al. 2020) | Põlme S, et al. 2020. Fungal Diversity 105:1-16 ((https://doi.org/10.1007/s13225-020-00466-2)) |
| Otu30772 | 1 | k__Fungi(100);p__Ascomycota(100);c__Saccharomycetes(100);o__Saccharomycetales(100);f__Metschnikowiaceae(100);g__Metschnikowia(100);s__Metschnikowia_pulcherrima(100); | Metschnikowia | 13 | Saprotroph | \|Undefined Saprotroph\| | Yeast | NULL | Probable | As nectar-tap_saprotroph (Põlme et al. 2020); Decay sugar-rich substrates (Põlme et al. 2020) | Põlme S, et al. 2020. Fungal Diversity 105:1-16 ((https://doi.org/10.1007/s13225-020-00466-2)) |
| Otu30773 | 1 | k__Fungi(100);p__Ascomycota(100);c__Saccharomycetes(100);o__Saccharomycetales(100);f__Metschnikowiaceae(100);g__Metschnikowia(100);s__Metschnikowia_pulcherrima(100); | Metschnikowia | 13 | Saprotroph | \|Undefined Saprotroph\| | Yeast | NULL | Probable | As nectar-tap_saprotroph (Põlme et al. 2020); Decay sugar-rich substrates (Põlme et al. 2020) | Põlme S, et al. 2020. Fungal Diversity 105:1-16 ((https://doi.org/10.1007/s13225-020-00466-2)) |
| Otu30774 | 1 | k__Fungi(100);p__Ascomycota(100);c__Saccharomycetes(100);o__Saccharomycetales(100);f__Metschnikowiaceae(100);g__Metschnikowia(100);g__Metschnikowia_unclassified(100); | Metschnikowia | 13 | Saprotroph | \|Undefined Saprotroph\| | Yeast | NULL | Probable | As nectar-tap_saprotroph (Põlme et al. 2020); Decay sugar-rich substrates (Põlme et al. 2020) | Põlme S, et al. 2020. Fungal Diversity 105:1-16 ((https://doi.org/10.1007/s13225-020-00466-2)) |
| Otu30775 | 1 | k__Fungi(100);p__Ascomycota(100);c__Saccharomycetes(100);o__Saccharomycetales(100);f__Metschnikowiaceae(100);g__Metschnikowia(100);s__Metschnikowia_pulcherrima(100); | Metschnikowia | 13 | Saprotroph | \|Undefined Saprotroph\| | Yeast | NULL | Probable | As nectar-tap_saprotroph (Põlme et al. 2020); Decay sugar-rich substrates (Põlme et al. 2020) | Põlme S, et al. 2020. Fungal Diversity 105:1-16 ((https://doi.org/10.1007/s13225-020-00466-2)) |
| Otu30776 | 1 | k__Fungi(100);p__Ascomycota(100);c__Saccharomycetes(100);o__Saccharomycetales(100);f__Metschnikowiaceae(100);g__Metschnikowia(100);s__Metschnikowia_pulcherrima(100); | Metschnikowia | 13 | Saprotroph | \|Undefined Saprotroph\| | Yeast | NULL | Probable | As nectar-tap_saprotroph (Põlme et al. 2020); Decay sugar-rich substrates (Põlme et al. 2020) | Põlme S, et al. 2020. Fungal Diversity 105:1-16 ((https://doi.org/10.1007/s13225-020-00466-2)) |
| Otu30777 | 1 | k__Fungi(100);p__Ascomycota(100);c__Saccharomycetes(100);o__Saccharomycetales(100);f__Metschnikowiaceae(100);g__Metschnikowia(100);s__Metschnikowia_pulcherrima(100); | Metschnikowia | 13 | Saprotroph | \|Undefined Saprotroph\| | Yeast | NULL | Probable | As nectar-tap_saprotroph (Põlme et al. 2020); Decay sugar-rich substrates (Põlme et al. 2020) | Põlme S, et al. 2020. Fungal Diversity 105:1-16 ((https://doi.org/10.1007/s13225-020-00466-2)) |
| Otu30778 | 1 | k__Fungi(100);p__Ascomycota(100);c__Saccharomycetes(100);o__Saccharomycetales(100);f__Metschnikowiaceae(100);g__Metschnikowia(100);s__Metschnikowia_pulcherrima(100); | Metschnikowia | 13 | Saprotroph | \|Undefined Saprotroph\| | Yeast | NULL | Probable | As nectar-tap_saprotroph (Põlme et al. 2020); Decay sugar-rich substrates (Põlme et al. 2020) | Põlme S, et al. 2020. Fungal Diversity 105:1-16 ((https://doi.org/10.1007/s13225-020-00466-2)) |
| Otu30779 | 1 | k__Fungi(100);p__Ascomycota(100);c__Saccharomycetes(100);o__Saccharomycetales(100);f__Metschnikowiaceae(100);g__Metschnikowia(100);s__Metschnikowia_pulcherrima(100); | Metschnikowia | 13 | Saprotroph | \|Undefined Saprotroph\| | Yeast | NULL | Probable | As nectar-tap_saprotroph (Põlme et al. 2020); Decay sugar-rich substrates (Põlme et al. 2020) | Põlme S, et al. 2020. Fungal Diversity 105:1-16 ((https://doi.org/10.1007/s13225-020-00466-2)) |
| Otu30780 | 1 | k__Fungi(100);p__Ascomycota(100);c__Saccharomycetes(100);o__Saccharomycetales(100);f__Metschnikowiaceae(100);g__Metschnikowia(100);s__Metschnikowia_pulcherrima(100); | Metschnikowia | 13 | Saprotroph | \|Undefined Saprotroph\| | Yeast | NULL | Probable | As nectar-tap_saprotroph (Põlme et al. 2020); Decay sugar-rich substrates (Põlme et al. 2020) | Põlme S, et al. 2020. Fungal Diversity 105:1-16 ((https://doi.org/10.1007/s13225-020-00466-2)) |
| Otu30781 | 1 | k__Fungi(100);p__Ascomycota(100);c__Saccharomycetes(100);o__Saccharomycetales(100);f__Metschnikowiaceae(100);g__Metschnikowia(100);g__Metschnikowia_unclassified(100); | Metschnikowia | 13 | Saprotroph | \|Undefined Saprotroph\| | Yeast | NULL | Probable | As nectar-tap_saprotroph (Põlme et al. 2020); Decay sugar-rich substrates (Põlme et al. 2020) | Põlme S, et al. 2020. Fungal Diversity 105:1-16 ((https://doi.org/10.1007/s13225-020-00466-2)) |
| Otu30782 | 1 | k__Fungi(100);p__Ascomycota(100);c__Saccharomycetes(100);o__Saccharomycetales(100);f__Metschnikowiaceae(100);g__Metschnikowia(100);s__Metschnikowia_pulcherrima(100); | Metschnikowia | 13 | Saprotroph | \|Undefined Saprotroph\| | Yeast | NULL | Probable | As nectar-tap_saprotroph (Põlme et al. 2020); Decay sugar-rich substrates (Põlme et al. 2020) | Põlme S, et al. 2020. Fungal Diversity 105:1-16 ((https://doi.org/10.1007/s13225-020-00466-2)) |
| Otu30783 | 1 | k__Fungi(100);p__Ascomycota(100);c__Saccharomycetes(100);o__Saccharomycetales(100);f__Metschnikowiaceae(100);g__Metschnikowia(100);s__Metschnikowia_pulcherrima(100); | Metschnikowia | 13 | Saprotroph | \|Undefined Saprotroph\| | Yeast | NULL | Probable | As nectar-tap_saprotroph (Põlme et al. 2020); Decay sugar-rich substrates (Põlme et al. 2020) | Põlme S, et al. 2020. Fungal Diversity 105:1-16 ((https://doi.org/10.1007/s13225-020-00466-2)) |
| Otu30784 | 1 | k__Fungi(100);p__Ascomycota(100);c__Saccharomycetes(100);o__Saccharomycetales(100);f__Metschnikowiaceae(100);g__Metschnikowia(100);s__Metschnikowia_pulcherrima(100); | Metschnikowia | 13 | Saprotroph | \|Undefined Saprotroph\| | Yeast | NULL | Probable | As nectar-tap_saprotroph (Põlme et al. 2020); Decay sugar-rich substrates (Põlme et al. 2020) | Põlme S, et al. 2020. Fungal Diversity 105:1-16 ((https://doi.org/10.1007/s13225-020-00466-2)) |
| Otu30785 | 1 | k__Fungi(100);p__Ascomycota(100);c__Saccharomycetes(100);o__Saccharomycetales(100);f__Metschnikowiaceae(100);g__Metschnikowia(100);s__Metschnikowia_pulcherrima(100); | Metschnikowia | 13 | Saprotroph | \|Undefined Saprotroph\| | Yeast | NULL | Probable | As nectar-tap_saprotroph (Põlme et al. 2020); Decay sugar-rich substrates (Põlme et al. 2020) | Põlme S, et al. 2020. Fungal Diversity 105:1-16 ((https://doi.org/10.1007/s13225-020-00466-2)) |
| Otu30786 | 1 | k__Fungi(100);p__Ascomycota(100);c__Saccharomycetes(100);o__Saccharomycetales(100);f__Metschnikowiaceae(100);g__Metschnikowia(100);g__Metschnikowia_unclassified(100); | Metschnikowia | 13 | Saprotroph | \|Undefined Saprotroph\| | Yeast | NULL | Probable | As nectar-tap_saprotroph (Põlme et al. 2020); Decay sugar-rich substrates (Põlme et al. 2020) | Põlme S, et al. 2020. Fungal Diversity 105:1-16 ((https://doi.org/10.1007/s13225-020-00466-2)) |
| Otu30787 | 1 | k__Fungi(100);p__Ascomycota(100);c__Saccharomycetes(100);o__Saccharomycetales(100);f__Metschnikowiaceae(100);g__Metschnikowia(100);s__Metschnikowia_pulcherrima(100); | Metschnikowia | 13 | Saprotroph | \|Undefined Saprotroph\| | Yeast | NULL | Probable | As nectar-tap_saprotroph (Põlme et al. 2020); Decay sugar-rich substrates (Põlme et al. 2020) | Põlme S, et al. 2020. Fungal Diversity 105:1-16 ((https://doi.org/10.1007/s13225-020-00466-2)) |
| Otu30788 | 1 | k__Fungi(100);p__Ascomycota(100);c__Saccharomycetes(100);o__Saccharomycetales(100);f__Metschnikowiaceae(100);g__Metschnikowia(100);g__Metschnikowia_unclassified(100); | Metschnikowia | 13 | Saprotroph | \|Undefined Saprotroph\| | Yeast | NULL | Probable | As nectar-tap_saprotroph (Põlme et al. 2020); Decay sugar-rich substrates (Põlme et al. 2020) | Põlme S, et al. 2020. Fungal Diversity 105:1-16 ((https://doi.org/10.1007/s13225-020-00466-2)) |
| Otu30790 | 1 | k__Fungi(100);p__Ascomycota(100);c__Saccharomycetes(100);o__Saccharomycetales(100);f__Metschnikowiaceae(100);g__Metschnikowia(100);s__Metschnikowia_pulcherrima(100); | Metschnikowia | 13 | Saprotroph | \|Undefined Saprotroph\| | Yeast | NULL | Probable | As nectar-tap_saprotroph (Põlme et al. 2020); Decay sugar-rich substrates (Põlme et al. 2020) | Põlme S, et al. 2020. Fungal Diversity 105:1-16 ((https://doi.org/10.1007/s13225-020-00466-2)) |
| Otu30791 | 1 | k__Fungi(100);p__Ascomycota(100);c__Saccharomycetes(100);o__Saccharomycetales(100);f__Metschnikowiaceae(100);g__Metschnikowia(100);g__Metschnikowia_unclassified(100); | Metschnikowia | 13 | Saprotroph | \|Undefined Saprotroph\| | Yeast | NULL | Probable | As nectar-tap_saprotroph (Põlme et al. 2020); Decay sugar-rich substrates (Põlme et al. 2020) | Põlme S, et al. 2020. Fungal Diversity 105:1-16 ((https://doi.org/10.1007/s13225-020-00466-2)) |
| Otu30792 | 1 | k__Fungi(100);p__Ascomycota(100);c__Saccharomycetes(100);o__Saccharomycetales(100);f__Metschnikowiaceae(100);g__Metschnikowia(100);g__Metschnikowia_unclassified(100); | Metschnikowia | 13 | Saprotroph | \|Undefined Saprotroph\| | Yeast | NULL | Probable | As nectar-tap_saprotroph (Põlme et al. 2020); Decay sugar-rich substrates (Põlme et al. 2020) | Põlme S, et al. 2020. Fungal Diversity 105:1-16 ((https://doi.org/10.1007/s13225-020-00466-2)) |
| Otu30793 | 1 | k__Fungi(100);p__Ascomycota(100);c__Saccharomycetes(100);o__Saccharomycetales(100);f__Metschnikowiaceae(100);g__Metschnikowia(100);g__Metschnikowia_unclassified(100); | Metschnikowia | 13 | Saprotroph | \|Undefined Saprotroph\| | Yeast | NULL | Probable | As nectar-tap_saprotroph (Põlme et al. 2020); Decay sugar-rich substrates (Põlme et al. 2020) | Põlme S, et al. 2020. Fungal Diversity 105:1-16 ((https://doi.org/10.1007/s13225-020-00466-2)) |
| Otu30794 | 1 | k__Fungi(100);p__Ascomycota(100);c__Saccharomycetes(100);o__Saccharomycetales(100);f__Metschnikowiaceae(100);g__Metschnikowia(100);s__Metschnikowia_pulcherrima(100); | Metschnikowia | 13 | Saprotroph | \|Undefined Saprotroph\| | Yeast | NULL | Probable | As nectar-tap_saprotroph (Põlme et al. 2020); Decay sugar-rich substrates (Põlme et al. 2020) | Põlme S, et al. 2020. Fungal Diversity 105:1-16 ((https://doi.org/10.1007/s13225-020-00466-2)) |
| Otu30795 | 1 | k__Fungi(100);p__Ascomycota(100);c__Saccharomycetes(100);o__Saccharomycetales(100);f__Metschnikowiaceae(100);g__Metschnikowia(100);s__Metschnikowia_pulcherrima(100); | Metschnikowia | 13 | Saprotroph | \|Undefined Saprotroph\| | Yeast | NULL | Probable | As nectar-tap_saprotroph (Põlme et al. 2020); Decay sugar-rich substrates (Põlme et al. 2020) | Põlme S, et al. 2020. Fungal Diversity 105:1-16 ((https://doi.org/10.1007/s13225-020-00466-2)) |
| Otu30796 | 1 | k__Fungi(100);p__Ascomycota(100);c__Saccharomycetes(100);o__Saccharomycetales(100);f__Metschnikowiaceae(100);g__Metschnikowia(100);s__Metschnikowia_pulcherrima(100); | Metschnikowia | 13 | Saprotroph | \|Undefined Saprotroph\| | Yeast | NULL | Probable | As nectar-tap_saprotroph (Põlme et al. 2020); Decay sugar-rich substrates (Põlme et al. 2020) | Põlme S, et al. 2020. Fungal Diversity 105:1-16 ((https://doi.org/10.1007/s13225-020-00466-2)) |
| Otu30797 | 1 | k__Fungi(100);p__Ascomycota(100);c__Saccharomycetes(100);o__Saccharomycetales(100);f__Metschnikowiaceae(100);g__Metschnikowia(100);g__Metschnikowia_unclassified(100); | Metschnikowia | 13 | Saprotroph | \|Undefined Saprotroph\| | Yeast | NULL | Probable | As nectar-tap_saprotroph (Põlme et al. 2020); Decay sugar-rich substrates (Põlme et al. 2020) | Põlme S, et al. 2020. Fungal Diversity 105:1-16 ((https://doi.org/10.1007/s13225-020-00466-2)) |
| Otu30798 | 1 | k__Fungi(100);p__Ascomycota(100);c__Saccharomycetes(100);o__Saccharomycetales(100);f__Metschnikowiaceae(100);g__Metschnikowia(100);s__Metschnikowia_pulcherrima(100); | Metschnikowia | 13 | Saprotroph | \|Undefined Saprotroph\| | Yeast | NULL | Probable | As nectar-tap_saprotroph (Põlme et al. 2020); Decay sugar-rich substrates (Põlme et al. 2020) | Põlme S, et al. 2020. Fungal Diversity 105:1-16 ((https://doi.org/10.1007/s13225-020-00466-2)) |
| Otu30799 | 1 | k__Fungi(100);p__Ascomycota(100);c__Saccharomycetes(100);o__Saccharomycetales(100);f__Metschnikowiaceae(100);g__Metschnikowia(100);g__Metschnikowia_unclassified(100); | Metschnikowia | 13 | Saprotroph | \|Undefined Saprotroph\| | Yeast | NULL | Probable | As nectar-tap_saprotroph (Põlme et al. 2020); Decay sugar-rich substrates (Põlme et al. 2020) | Põlme S, et al. 2020. Fungal Diversity 105:1-16 ((https://doi.org/10.1007/s13225-020-00466-2)) |
| Otu30800 | 1 | k__Fungi(100);p__Ascomycota(100);c__Saccharomycetes(100);o__Saccharomycetales(100);f__Metschnikowiaceae(100);g__Metschnikowia(100);s__Metschnikowia_pulcherrima(100); | Metschnikowia | 13 | Saprotroph | \|Undefined Saprotroph\| | Yeast | NULL | Probable | As nectar-tap_saprotroph (Põlme et al. 2020); Decay sugar-rich substrates (Põlme et al. 2020) | Põlme S, et al. 2020. Fungal Diversity 105:1-16 ((https://doi.org/10.1007/s13225-020-00466-2)) |
| Otu30801 | 1 | k__Fungi(100);p__Ascomycota(100);c__Saccharomycetes(100);o__Saccharomycetales(100);f__Metschnikowiaceae(100);g__Metschnikowia(100);s__Metschnikowia_pulcherrima(100); | Metschnikowia | 13 | Saprotroph | \|Undefined Saprotroph\| | Yeast | NULL | Probable | As nectar-tap_saprotroph (Põlme et al. 2020); Decay sugar-rich substrates (Põlme et al. 2020) | Põlme S, et al. 2020. Fungal Diversity 105:1-16 ((https://doi.org/10.1007/s13225-020-00466-2)) |
| Otu30802 | 1 | k__Fungi(100);p__Ascomycota(100);c__Saccharomycetes(100);o__Saccharomycetales(100);f__Metschnikowiaceae(100);g__Metschnikowia(100);s__Metschnikowia_pulcherrima(100); | Metschnikowia | 13 | Saprotroph | \|Undefined Saprotroph\| | Yeast | NULL | Probable | As nectar-tap_saprotroph (Põlme et al. 2020); Decay sugar-rich substrates (Põlme et al. 2020) | Põlme S, et al. 2020. Fungal Diversity 105:1-16 ((https://doi.org/10.1007/s13225-020-00466-2)) |
| Otu30803 | 1 | k__Fungi(100);p__Ascomycota(100);c__Saccharomycetes(100);o__Saccharomycetales(100);f__Metschnikowiaceae(100);g__Metschnikowia(100);s__Metschnikowia_pulcherrima(100); | Metschnikowia | 13 | Saprotroph | \|Undefined Saprotroph\| | Yeast | NULL | Probable | As nectar-tap_saprotroph (Põlme et al. 2020); Decay sugar-rich substrates (Põlme et al. 2020) | Põlme S, et al. 2020. Fungal Diversity 105:1-16 ((https://doi.org/10.1007/s13225-020-00466-2)) |
| Otu30804 | 1 | k__Fungi(100);p__Ascomycota(100);c__Saccharomycetes(100);o__Saccharomycetales(100);f__Metschnikowiaceae(100);g__Metschnikowia(100);s__Metschnikowia_pulcherrima(100); | Metschnikowia | 13 | Saprotroph | \|Undefined Saprotroph\| | Yeast | NULL | Probable | As nectar-tap_saprotroph (Põlme et al. 2020); Decay sugar-rich substrates (Põlme et al. 2020) | Põlme S, et al. 2020. Fungal Diversity 105:1-16 ((https://doi.org/10.1007/s13225-020-00466-2)) |
| Otu30805 | 1 | k__Fungi(100);p__Ascomycota(100);c__Saccharomycetes(100);o__Saccharomycetales(100);f__Metschnikowiaceae(100);g__Metschnikowia(100);s__Metschnikowia_pulcherrima(100); | Metschnikowia | 13 | Saprotroph | \|Undefined Saprotroph\| | Yeast | NULL | Probable | As nectar-tap_saprotroph (Põlme et al. 2020); Decay sugar-rich substrates (Põlme et al. 2020) | Põlme S, et al. 2020. Fungal Diversity 105:1-16 ((https://doi.org/10.1007/s13225-020-00466-2)) |
| Otu30806 | 1 | k__Fungi(100);p__Ascomycota(100);c__Saccharomycetes(100);o__Saccharomycetales(100);f__Metschnikowiaceae(100);g__Metschnikowia(100);s__Metschnikowia_pulcherrima(100); | Metschnikowia | 13 | Saprotroph | \|Undefined Saprotroph\| | Yeast | NULL | Probable | As nectar-tap_saprotroph (Põlme et al. 2020); Decay sugar-rich substrates (Põlme et al. 2020) | Põlme S, et al. 2020. Fungal Diversity 105:1-16 ((https://doi.org/10.1007/s13225-020-00466-2)) |
| Otu30808 | 1 | k__Fungi(100);p__Ascomycota(100);c__Saccharomycetes(100);o__Saccharomycetales(100);f__Metschnikowiaceae(100);g__Metschnikowia(100);s__Metschnikowia_pulcherrima(100); | Metschnikowia | 13 | Saprotroph | \|Undefined Saprotroph\| | Yeast | NULL | Probable | As nectar-tap_saprotroph (Põlme et al. 2020); Decay sugar-rich substrates (Põlme et al. 2020) | Põlme S, et al. 2020. Fungal Diversity 105:1-16 ((https://doi.org/10.1007/s13225-020-00466-2)) |
| Otu30811 | 1 | k__Fungi(100);p__Ascomycota(100);c__Saccharomycetes(100);o__Saccharomycetales(100);f__Metschnikowiaceae(100);g__Metschnikowia(100);s__Metschnikowia_pulcherrima(100); | Metschnikowia | 13 | Saprotroph | \|Undefined Saprotroph\| | Yeast | NULL | Probable | As nectar-tap_saprotroph (Põlme et al. 2020); Decay sugar-rich substrates (Põlme et al. 2020) | Põlme S, et al. 2020. Fungal Diversity 105:1-16 ((https://doi.org/10.1007/s13225-020-00466-2)) |
| Otu30812 | 1 | k__Fungi(100);p__Ascomycota(100);c__Saccharomycetes(100);o__Saccharomycetales(100);f__Metschnikowiaceae(100);g__Metschnikowia(100);s__Metschnikowia_pulcherrima(100); | Metschnikowia | 13 | Saprotroph | \|Undefined Saprotroph\| | Yeast | NULL | Probable | As nectar-tap_saprotroph (Põlme et al. 2020); Decay sugar-rich substrates (Põlme et al. 2020) | Põlme S, et al. 2020. Fungal Diversity 105:1-16 ((https://doi.org/10.1007/s13225-020-00466-2)) |
| Otu30814 | 1 | k__Fungi(100);p__Ascomycota(100);c__Saccharomycetes(100);o__Saccharomycetales(100);f__Metschnikowiaceae(100);g__Metschnikowia(100);s__Metschnikowia_pulcherrima(100); | Metschnikowia | 13 | Saprotroph | \|Undefined Saprotroph\| | Yeast | NULL | Probable | As nectar-tap_saprotroph (Põlme et al. 2020); Decay sugar-rich substrates (Põlme et al. 2020) | Põlme S, et al. 2020. Fungal Diversity 105:1-16 ((https://doi.org/10.1007/s13225-020-00466-2)) |
| Otu30815 | 1 | k__Fungi(100);p__Ascomycota(100);c__Saccharomycetes(100);o__Saccharomycetales(100);f__Metschnikowiaceae(100);g__Metschnikowia(100);s__Metschnikowia_pulcherrima(100); | Metschnikowia | 13 | Saprotroph | \|Undefined Saprotroph\| | Yeast | NULL | Probable | As nectar-tap_saprotroph (Põlme et al. 2020); Decay sugar-rich substrates (Põlme et al. 2020) | Põlme S, et al. 2020. Fungal Diversity 105:1-16 ((https://doi.org/10.1007/s13225-020-00466-2)) |
| Otu30817 | 1 | k__Fungi(100);p__Ascomycota(100);c__Saccharomycetes(100);o__Saccharomycetales(100);f__Metschnikowiaceae(100);g__Metschnikowia(100);s__Metschnikowia_pulcherrima(100); | Metschnikowia | 13 | Saprotroph | \|Undefined Saprotroph\| | Yeast | NULL | Probable | As nectar-tap_saprotroph (Põlme et al. 2020); Decay sugar-rich substrates (Põlme et al. 2020) | Põlme S, et al. 2020. Fungal Diversity 105:1-16 ((https://doi.org/10.1007/s13225-020-00466-2)) |

| | | | | | | | | | | | |
|---|---|---|---|---|---|---|---|---|---|---|---|
| Otu30818 | 1 | k__Fungi(100);p__Ascomycota(100);c__Saccharomycetes(100);o__Saccharomycetales(100);f__Metschnikowiaceae(100);g__Metschnikowia(100);s__Metschnikowia_pulcherrima(100); | Metschnikowia | 13 | Saprotroph | \|Undefined Saprotroph\| | Yeast | NULL | Probable | As nectar-tap_saprotroph (Põlme et al. 2020); Decay sugar-rich substrates (Põlme et al. 2020) | Põlme S, et al. 2020. Fungal Diversity 105:1-16 ((https://doi.org/10.1007/s13225-020-00466-2)) |
| Otu30819 | 1 | k__Fungi(100);p__Ascomycota(100);c__Saccharomycetes(100);o__Saccharomycetales(100);f__Metschnikowiaceae(100);g__Metschnikowia(100);s__Metschnikowia_pulcherrima(100); | Metschnikowia | 13 | Saprotroph | \|Undefined Saprotroph\| | Yeast | NULL | Probable | As nectar-tap_saprotroph (Põlme et al. 2020); Decay sugar-rich substrates (Põlme et al. 2020) | Põlme S, et al. 2020. Fungal Diversity 105:1-16 ((https://doi.org/10.1007/s13225-020-00466-2)) |
| Otu30820 | 1 | k__Fungi(100);p__Ascomycota(100);c__Saccharomycetes(100);o__Saccharomycetales(100);f__Metschnikowiaceae(100);g__Metschnikowia(100);s__Metschnikowia_pulcherrima(100); | Metschnikowia | 13 | Saprotroph | \|Undefined Saprotroph\| | Yeast | NULL | Probable | As nectar-tap_saprotroph (Põlme et al. 2020); Decay sugar-rich substrates (Põlme et al. 2020) | Põlme S, et al. 2020. Fungal Diversity 105:1-16 ((https://doi.org/10.1007/s13225-020-00466-2)) |
| Otu30821 | 1 | k__Fungi(100);p__Ascomycota(100);c__Saccharomycetes(100);o__Saccharomycetales(100);f__Metschnikowiaceae(100);g__Metschnikowia(100);s__Metschnikowia_pulcherrima(100); | Metschnikowia | 13 | Saprotroph | \|Undefined Saprotroph\| | Yeast | NULL | Probable | As nectar-tap_saprotroph (Põlme et al. 2020); Decay sugar-rich substrates (Põlme et al. 2020) | Põlme S, et al. 2020. Fungal Diversity 105:1-16 ((https://doi.org/10.1007/s13225-020-00466-2)) |
| Otu30822 | 1 | k__Fungi(100);p__Ascomycota(100);c__Saccharomycetes(100);o__Saccharomycetales(100);f__Metschnikowiaceae(100);g__Metschnikowia(100);g__Metschnikowia_unclassified(100); | Metschnikowia | 13 | Saprotroph | \|Undefined Saprotroph\| | Yeast | NULL | Probable | As nectar-tap_saprotroph (Põlme et al. 2020); Decay sugar-rich substrates (Põlme et al. 2020) | Põlme S, et al. 2020. Fungal Diversity 105:1-16 ((https://doi.org/10.1007/s13225-020-00466-2)) |
| Otu30823 | 1 | k__Fungi(100);p__Ascomycota(100);c__Saccharomycetes(100);o__Saccharomycetales(100);f__Metschnikowiaceae(100);g__Metschnikowia(100);g__Metschnikowia_unclassified(100); | Metschnikowia | 13 | Saprotroph | \|Undefined Saprotroph\| | Yeast | NULL | Probable | As nectar-tap_saprotroph (Põlme et al. 2020); Decay sugar-rich substrates (Põlme et al. 2020) | Põlme S, et al. 2020. Fungal Diversity 105:1-16 ((https://doi.org/10.1007/s13225-020-00466-2)) |
| Otu30824 | 1 | k__Fungi(100);p__Ascomycota(100);c__Saccharomycetes(100);o__Saccharomycetales(100);f__Metschnikowiaceae(100);g__Metschnikowia(100);s__Metschnikowia_pulcherrima(100); | Metschnikowia | 13 | Saprotroph | \|Undefined Saprotroph\| | Yeast | NULL | Probable | As nectar-tap_saprotroph (Põlme et al. 2020); Decay sugar-rich substrates (Põlme et al. 2020) | Põlme S, et al. 2020. Fungal Diversity 105:1-16 ((https://doi.org/10.1007/s13225-020-00466-2)) |
| Otu30825 | 1 | k__Fungi(100);p__Ascomycota(100);c__Saccharomycetes(100);o__Saccharomycetales(100);f__Metschnikowiaceae(100);g__Metschnikowia(100);g__Metschnikowia_unclassified(100); | Metschnikowia | 13 | Saprotroph | \|Undefined Saprotroph\| | Yeast | NULL | Probable | As nectar-tap_saprotroph (Põlme et al. 2020); Decay sugar-rich substrates (Põlme et al. 2020) | Põlme S, et al. 2020. Fungal Diversity 105:1-16 ((https://doi.org/10.1007/s13225-020-00466-2)) |
| Otu30826 | 1 | k__Fungi(100);p__Ascomycota(100);c__Saccharomycetes(100);o__Saccharomycetales(100);f__Metschnikowiaceae(100);g__Metschnikowia(100);s__Metschnikowia_pulcherrima(100); | Metschnikowia | 13 | Saprotroph | \|Undefined Saprotroph\| | Yeast | NULL | Probable | As nectar-tap_saprotroph (Põlme et al. 2020); Decay sugar-rich substrates (Põlme et al. 2020) | Põlme S, et al. 2020. Fungal Diversity 105:1-16 ((https://doi.org/10.1007/s13225-020-00466-2)) |
| Otu30827 | 1 | k__Fungi(100);p__Ascomycota(100);c__Saccharomycetes(100);o__Saccharomycetales(100);f__Metschnikowiaceae(100);g__Metschnikowia(100);s__Metschnikowia_pulcherrima(100); | Metschnikowia | 13 | Saprotroph | \|Undefined Saprotroph\| | Yeast | NULL | Probable | As nectar-tap_saprotroph (Põlme et al. 2020); Decay sugar-rich substrates (Põlme et al. 2020) | Põlme S, et al. 2020. Fungal Diversity 105:1-16 ((https://doi.org/10.1007/s13225-020-00466-2)) |
| Otu30828 | 1 | k__Fungi(100);p__Ascomycota(100);c__Saccharomycetes(100);o__Saccharomycetales(100);f__Metschnikowiaceae(100);g__Metschnikowia(100);s__Metschnikowia_pulcherrima(100); | Metschnikowia | 13 | Saprotroph | \|Undefined Saprotroph\| | Yeast | NULL | Probable | As nectar-tap_saprotroph (Põlme et al. 2020); Decay sugar-rich substrates (Põlme et al. 2020) | Põlme S, et al. 2020. Fungal Diversity 105:1-16 ((https://doi.org/10.1007/s13225-020-00466-2)) |
| Otu30829 | 1 | k__Fungi(100);p__Ascomycota(100);c__Saccharomycetes(100);o__Saccharomycetales(100);f__Metschnikowiaceae(100);g__Metschnikowia(100);s__Metschnikowia_pulcherrima(100); | Metschnikowia | 13 | Saprotroph | \|Undefined Saprotroph\| | Yeast | NULL | Probable | As nectar-tap_saprotroph (Põlme et al. 2020); Decay sugar-rich substrates (Põlme et al. 2020) | Põlme S, et al. 2020. Fungal Diversity 105:1-16 ((https://doi.org/10.1007/s13225-020-00466-2)) |
| Otu30830 | 1 | k__Fungi(100);p__Ascomycota(100);c__Saccharomycetes(100);o__Saccharomycetales(100);f__Metschnikowiaceae(100);g__Metschnikowia(100);s__Metschnikowia_pulcherrima(100); | Metschnikowia | 13 | Saprotroph | \|Undefined Saprotroph\| | Yeast | NULL | Probable | As nectar-tap_saprotroph (Põlme et al. 2020); Decay sugar-rich substrates (Põlme et al. 2020) | Põlme S, et al. 2020. Fungal Diversity 105:1-16 ((https://doi.org/10.1007/s13225-020-00466-2)) |
| Otu30831 | 1 | k__Fungi(100);p__Ascomycota(100);c__Saccharomycetes(100);o__Saccharomycetales(100);f__Metschnikowiaceae(100);g__Metschnikowia(100);s__Metschnikowia_pulcherrima(100); | Metschnikowia | 13 | Saprotroph | \|Undefined Saprotroph\| | Yeast | NULL | Probable | As nectar-tap_saprotroph (Põlme et al. 2020); Decay sugar-rich substrates (Põlme et al. 2020) | Põlme S, et al. 2020. Fungal Diversity 105:1-16 ((https://doi.org/10.1007/s13225-020-00466-2)) |
| Otu30832 | 1 | k__Fungi(100);p__Ascomycota(100);c__Saccharomycetes(100);o__Saccharomycetales(100);f__Metschnikowiaceae(100);g__Metschnikowia(100);g__Metschnikowia_unclassified(100); | Metschnikowia | 13 | Saprotroph | \|Undefined Saprotroph\| | Yeast | NULL | Probable | As nectar-tap_saprotroph (Põlme et al. 2020); Decay sugar-rich substrates (Põlme et al. 2020) | Põlme S, et al. 2020. Fungal Diversity 105:1-16 ((https://doi.org/10.1007/s13225-020-00466-2)) |
| Otu30833 | 1 | k__Fungi(100);p__Ascomycota(100);c__Saccharomycetes(100);o__Saccharomycetales(100);f__Metschnikowiaceae(100);g__Metschnikowia(100);g__Metschnikowia_unclassified(100); | Metschnikowia | 13 | Saprotroph | \|Undefined Saprotroph\| | Yeast | NULL | Probable | As nectar-tap_saprotroph (Põlme et al. 2020); Decay sugar-rich substrates (Põlme et al. 2020) | Põlme S, et al. 2020. Fungal Diversity 105:1-16 ((https://doi.org/10.1007/s13225-020-00466-2)) |
| Otu30834 | 1 | k__Fungi(100);p__Ascomycota(100);c__Saccharomycetes(100);o__Saccharomycetales(100);f__Metschnikowiaceae(100);g__Metschnikowia(100);s__Metschnikowia_pulcherrima(100); | Metschnikowia | 13 | Saprotroph | \|Undefined Saprotroph\| | Yeast | NULL | Probable | As nectar-tap_saprotroph (Põlme et al. 2020); Decay sugar-rich substrates (Põlme et al. 2020) | Põlme S, et al. 2020. Fungal Diversity 105:1-16 ((https://doi.org/10.1007/s13225-020-00466-2)) |
| Otu30835 | 1 | k__Fungi(100);p__Ascomycota(100);c__Saccharomycetes(100);o__Saccharomycetales(100);f__Metschnikowiaceae(100);g__Metschnikowia(100);g__Metschnikowia_unclassified(100); | Metschnikowia | 13 | Saprotroph | \|Undefined Saprotroph\| | Yeast | NULL | Probable | As nectar-tap_saprotroph (Põlme et al. 2020); Decay sugar-rich substrates (Põlme et al. 2020) | Põlme S, et al. 2020. Fungal Diversity 105:1-16 ((https://doi.org/10.1007/s13225-020-00466-2)) |
| Otu30836 | 1 | k__Fungi(100);p__Ascomycota(100);c__Saccharomycetes(100);o__Saccharomycetales(100);f__Metschnikowiaceae(100);g__Metschnikowia(100);s__Metschnikowia_pulcherrima(100); | Metschnikowia | 13 | Saprotroph | \|Undefined Saprotroph\| | Yeast | NULL | Probable | As nectar-tap_saprotroph (Põlme et al. 2020); Decay sugar-rich substrates (Põlme et al. 2020) | Põlme S, et al. 2020. Fungal Diversity 105:1-16 ((https://doi.org/10.1007/s13225-020-00466-2)) |
| Otu30838 | 1 | k__Fungi(100);p__Ascomycota(100);c__Saccharomycetes(100);o__Saccharomycetales(100);f__Metschnikowiaceae(100);g__Metschnikowia(100);s__Metschnikowia_pulcherrima(100); | Metschnikowia | 13 | Saprotroph | \|Undefined Saprotroph\| | Yeast | NULL | Probable | As nectar-tap_saprotroph (Põlme et al. 2020); Decay sugar-rich substrates (Põlme et al. 2020) | Põlme S, et al. 2020. Fungal Diversity 105:1-16 ((https://doi.org/10.1007/s13225-020-00466-2)) |
| Otu30839 | 1 | k__Fungi(100);p__Ascomycota(100);c__Saccharomycetes(100);o__Saccharomycetales(100);f__Metschnikowiaceae(100);g__Metschnikowia(100);s__Metschnikowia_pulcherrima(100); | Metschnikowia | 13 | Saprotroph | \|Undefined Saprotroph\| | Yeast | NULL | Probable | As nectar-tap_saprotroph (Põlme et al. 2020); Decay sugar-rich substrates (Põlme et al. 2020) | Põlme S, et al. 2020. Fungal Diversity 105:1-16 ((https://doi.org/10.1007/s13225-020-00466-2)) |
| Otu30840 | 1 | k__Fungi(100);p__Ascomycota(100);c__Saccharomycetes(100);o__Saccharomycetales(100);f__Metschnikowiaceae(100);g__Metschnikowia(100);s__Metschnikowia_pulcherrima(100); | Metschnikowia | 13 | Saprotroph | \|Undefined Saprotroph\| | Yeast | NULL | Probable | As nectar-tap_saprotroph (Põlme et al. 2020); Decay sugar-rich substrates (Põlme et al. 2020) | Põlme S, et al. 2020. Fungal Diversity 105:1-16 ((https://doi.org/10.1007/s13225-020-00466-2)) |
| Otu30842 | 1 | k__Fungi(100);p__Ascomycota(100);c__Saccharomycetes(100);o__Saccharomycetales(100);f__Metschnikowiaceae(100);g__Metschnikowia(100);s__Metschnikowia_pulcherrima(100); | Metschnikowia | 13 | Saprotroph | \|Undefined Saprotroph\| | Yeast | NULL | Probable | As nectar-tap_saprotroph (Põlme et al. 2020); Decay sugar-rich substrates (Põlme et al. 2020) | Põlme S, et al. 2020. Fungal Diversity 105:1-16 ((https://doi.org/10.1007/s13225-020-00466-2)) |
| Otu30843 | 1 | k__Fungi(100);p__Ascomycota(100);c__Saccharomycetes(100);o__Saccharomycetales(100);f__Metschnikowiaceae(100);g__Metschnikowia(100);g__Metschnikowia_unclassified(100); | Metschnikowia | 13 | Saprotroph | \|Undefined Saprotroph\| | Yeast | NULL | Probable | As nectar-tap_saprotroph (Põlme et al. 2020); Decay sugar-rich substrates (Põlme et al. 2020) | Põlme S, et al. 2020. Fungal Diversity 105:1-16 ((https://doi.org/10.1007/s13225-020-00466-2)) |
| Otu30844 | 1 | k__Fungi(100);p__Ascomycota(100);c__Saccharomycetes(100);o__Saccharomycetales(100);f__Metschnikowiaceae(100);g__Metschnikowia(100);s__Metschnikowia_pulcherrima(100); | Metschnikowia | 13 | Saprotroph | \|Undefined Saprotroph\| | Yeast | NULL | Probable | As nectar-tap_saprotroph (Põlme et al. 2020); Decay sugar-rich substrates (Põlme et al. 2020) | Põlme S, et al. 2020. Fungal Diversity 105:1-16 ((https://doi.org/10.1007/s13225-020-00466-2)) |
| Otu30845 | 1 | k__Fungi(100);p__Ascomycota(100);c__Leotiomycetes(100);o__Helotiales(100);f__Phacidiaceae(100);g__Phacidium(100);s__Phacidium_lacerum(100); | Phacidium | 13 | Pathotroph-Symbiotroph | Endophyte-\|Plant Pathogen\| | Microfungus | NULL | Probable | As foliar_endophyte (Põlme et al. 2020); Decay leaf, fruit, seed (Põlme et al. 2020) | Tedersoo L, et al. 2014. Science 346:e1256688 ((https://doi.org/10.1126/science.1256688)); Põlme S, et al. 2020. Fungal Diversity 105:1-16 ((https://doi.org/10.1007/s13225-020-00466-2)) |
| Otu30846 | 1 | k__Fungi(100);p__Ascomycota(100);c__Saccharomycetes(100);o__Saccharomycetales(100);f__Metschnikowiaceae(100);g__Metschnikowia(100);s__Metschnikowia_pulcherrima(100); | Metschnikowia | 13 | Saprotroph | \|Undefined Saprotroph\| | Yeast | NULL | Probable | As nectar-tap_saprotroph (Põlme et al. 2020); Decay sugar-rich substrates (Põlme et al. 2020) | Põlme S, et al. 2020. Fungal Diversity 105:1-16 ((https://doi.org/10.1007/s13225-020-00466-2)) |
| Otu30847 | 1 | k__Fungi(100);p__Ascomycota(100);c__Saccharomycetes(100);o__Saccharomycetales(100);f__Metschnikowiaceae(100);g__Metschnikowia(100);s__Metschnikowia_pulcherrima(100); | Metschnikowia | 13 | Saprotroph | \|Undefined Saprotroph\| | Yeast | NULL | Probable | As nectar-tap_saprotroph (Põlme et al. 2020); Decay sugar-rich substrates (Põlme et al. 2020) | Põlme S, et al. 2020. Fungal Diversity 105:1-16 ((https://doi.org/10.1007/s13225-020-00466-2)) |
| Otu30848 | 1 | k__Fungi(100);p__Ascomycota(100);c__Saccharomycetes(100);o__Saccharomycetales(100);f__Metschnikowiaceae(100);g__Metschnikowia(100);s__Metschnikowia_pulcherrima(100); | Metschnikowia | 13 | Saprotroph | \|Undefined Saprotroph\| | Yeast | NULL | Probable | As nectar-tap_saprotroph (Põlme et al. 2020); Decay sugar-rich substrates (Põlme et al. 2020) | Põlme S, et al. 2020. Fungal Diversity 105:1-16 ((https://doi.org/10.1007/s13225-020-00466-2)) |
| Otu30849 | 1 | k__Fungi(100);p__Ascomycota(100);c__Saccharomycetes(100);o__Saccharomycetales(100);f__Metschnikowiaceae(100);g__Metschnikowia(100);s__Metschnikowia_pulcherrima(100); | Metschnikowia | 13 | Saprotroph | \|Undefined Saprotroph\| | Yeast | NULL | Probable | As nectar-tap_saprotroph (Põlme et al. 2020); Decay sugar-rich substrates (Põlme et al. 2020) | Põlme S, et al. 2020. Fungal Diversity 105:1-16 ((https://doi.org/10.1007/s13225-020-00466-2)) |
| Otu30850 | 1 | k__Fungi(100);p__Ascomycota(100);c__Saccharomycetes(100);o__Saccharomycetales(100);f__Metschnikowiaceae(100);g__Metschnikowia(100);s__Metschnikowia_pulcherrima(100); | Metschnikowia | 13 | Saprotroph | \|Undefined Saprotroph\| | Yeast | NULL | Probable | As nectar-tap_saprotroph (Põlme et al. 2020); Decay sugar-rich substrates (Põlme et al. 2020) | Põlme S, et al. 2020. Fungal Diversity 105:1-16 ((https://doi.org/10.1007/s13225-020-00466-2)) |
| Otu30851 | 1 | k__Fungi(100);p__Ascomycota(100);c__Saccharomycetes(100);o__Saccharomycetales(100);f__Metschnikowiaceae(100);g__Metschnikowia(100);s__Metschnikowia_pulcherrima(100); | Metschnikowia | 13 | Saprotroph | \|Undefined Saprotroph\| | Yeast | NULL | Probable | As nectar-tap_saprotroph (Põlme et al. 2020); Decay sugar-rich substrates (Põlme et al. 2020) | Põlme S, et al. 2020. Fungal Diversity 105:1-16 ((https://doi.org/10.1007/s13225-020-00466-2)) |
| Otu30852 | 1 | k__Fungi(100);p__Ascomycota(100);c__Saccharomycetes(100);o__Saccharomycetales(100);f__Metschnikowiaceae(100);g__Metschnikowia(100);g__Metschnikowia_unclassified(100); | Metschnikowia | 13 | Saprotroph | \|Undefined Saprotroph\| | Yeast | NULL | Probable | As nectar-tap_saprotroph (Põlme et al. 2020); Decay sugar-rich substrates (Põlme et al. 2020) | Põlme S, et al. 2020. Fungal Diversity 105:1-16 ((https://doi.org/10.1007/s13225-020-00466-2)) |
| Otu30853 | 1 | k__Fungi(100);p__Ascomycota(100);c__Saccharomycetes(100);o__Saccharomycetales(100);f__Metschnikowiaceae(100);g__Metschnikowia(100);s__Metschnikowia_pulcherrima(100); | Metschnikowia | 13 | Saprotroph | \|Undefined Saprotroph\| | Yeast | NULL | Probable | As nectar-tap_saprotroph (Põlme et al. 2020); Decay sugar-rich substrates (Põlme et al. 2020) | Põlme S, et al. 2020. Fungal Diversity 105:1-16 ((https://doi.org/10.1007/s13225-020-00466-2)) |
| Otu30854 | 1 | k__Fungi(100);p__Ascomycota(100);c__Saccharomycetes(100);o__Saccharomycetales(100);f__Metschnikowiaceae(100);g__Metschnikowia(100);s__Metschnikowia_pulcherrima(100); | Metschnikowia | 13 | Saprotroph | \|Undefined Saprotroph\| | Yeast | NULL | Probable | As nectar-tap_saprotroph (Põlme et al. 2020); Decay sugar-rich substrates (Põlme et al. 2020) | Põlme S, et al. 2020. Fungal Diversity 105:1-16 ((https://doi.org/10.1007/s13225-020-00466-2)) |
| Otu30855 | 1 | k__Fungi(100);p__Ascomycota(100);c__Saccharomycetes(100);o__Saccharomycetales(100);f__Metschnikowiaceae(100);g__Metschnikowia(100);s__Metschnikowia_pulcherrima(100); | Metschnikowia | 13 | Saprotroph | \|Undefined Saprotroph\| | Yeast | NULL | Probable | As nectar-tap_saprotroph (Põlme et al. 2020); Decay sugar-rich substrates (Põlme et al. 2020) | Põlme S, et al. 2020. Fungal Diversity 105:1-16 ((https://doi.org/10.1007/s13225-020-00466-2)) |
| Otu30856 | 1 | k__Fungi(100);p__Ascomycota(100);c__Saccharomycetes(100);o__Saccharomycetales(100);f__Metschnikowiaceae(100);g__Metschnikowia(100);g__Metschnikowia_unclassified(100); | Metschnikowia | 13 | Saprotroph | \|Undefined Saprotroph\| | Yeast | NULL | Probable | As nectar-tap_saprotroph (Põlme et al. 2020); Decay sugar-rich substrates (Põlme et al. 2020) | Põlme S, et al. 2020. Fungal Diversity 105:1-16 ((https://doi.org/10.1007/s13225-020-00466-2)) |
| Otu30857 | 1 | k__Fungi(100);p__Ascomycota(100);c__Saccharomycetes(100);o__Saccharomycetales(100);f__Metschnikowiaceae(100);g__Metschnikowia(100);s__Metschnikowia_pulcherrima(100); | Metschnikowia | 13 | Saprotroph | \|Undefined Saprotroph\| | Yeast | NULL | Probable | As nectar-tap_saprotroph (Põlme et al. 2020); Decay sugar-rich substrates (Põlme et al. 2020) | Põlme S, et al. 2020. Fungal Diversity 105:1-16 ((https://doi.org/10.1007/s13225-020-00466-2)) |
| Otu30858 | 1 | k__Fungi(100);p__Ascomycota(100);c__Saccharomycetes(100);o__Saccharomycetales(100);f__Metschnikowiaceae(100);g__Metschnikowia(100);s__Metschnikowia_pulcherrima(100); | Metschnikowia | 13 | Saprotroph | \|Undefined Saprotroph\| | Yeast | NULL | Probable | As nectar-tap_saprotroph (Põlme et al. 2020); Decay sugar-rich substrates (Põlme et al. 2020) | Põlme S, et al. 2020. Fungal Diversity 105:1-16 ((https://doi.org/10.1007/s13225-020-00466-2)) |
| Otu30859 | 1 | k__Fungi(100);p__Ascomycota(100);c__Saccharomycetes(100);o__Saccharomycetales(100);f__Metschnikowiaceae(100);g__Metschnikowia(100);s__Metschnikowia_pulcherrima(100); | Metschnikowia | 13 | Saprotroph | \|Undefined Saprotroph\| | Yeast | NULL | Probable | As nectar-tap_saprotroph (Põlme et al. 2020); Decay sugar-rich substrates (Põlme et al. 2020) | Põlme S, et al. 2020. Fungal Diversity 105:1-16 ((https://doi.org/10.1007/s13225-020-00466-2)) |
| Otu30860 | 1 | k__Fungi(100);p__Ascomycota(100);c__Saccharomycetes(100);o__Saccharomycetales(100);f__Metschnikowiaceae(100);g__Metschnikowia(100);g__Metschnikowia_unclassified(100); | Metschnikowia | 13 | Saprotroph | \|Undefined Saprotroph\| | Yeast | NULL | Probable | As nectar-tap_saprotroph (Põlme et al. 2020); Decay sugar-rich substrates (Põlme et al. 2020) | Põlme S, et al. 2020. Fungal Diversity 105:1-16 ((https://doi.org/10.1007/s13225-020-00466-2)) |
| Otu30861 | 1 | k__Fungi(100);p__Ascomycota(100);c__Saccharomycetes(100);o__Saccharomycetales(100);f__Metschnikowiaceae(100);g__Metschnikowia(100);s__Metschnikowia_pulcherrima(100); | Metschnikowia | 13 | Saprotroph | \|Undefined Saprotroph\| | Yeast | NULL | Probable | As nectar-tap_saprotroph (Põlme et al. 2020); Decay sugar-rich substrates (Põlme et al. 2020) | Põlme S, et al. 2020. Fungal Diversity 105:1-16 ((https://doi.org/10.1007/s13225-020-00466-2)) |
| Otu30862 | 1 | k__Fungi(100);p__Ascomycota(100);c__Saccharomycetes(100);o__Saccharomycetales(100);f__Metschnikowiaceae(100);g__Metschnikowia(100);s__Metschnikowia_pulcherrima(100); | Metschnikowia | 13 | Saprotroph | \|Undefined Saprotroph\| | Yeast | NULL | Probable | As nectar-tap_saprotroph (Põlme et al. 2020); Decay sugar-rich substrates (Põlme et al. 2020) | Põlme S, et al. 2020. Fungal Diversity 105:1-16 ((https://doi.org/10.1007/s13225-020-00466-2)) |
| Otu30863 | 1 | k__Fungi(100);p__Ascomycota(100);c__Saccharomycetes(100);o__Saccharomycetales(100);f__Metschnikowiaceae(100);g__Metschnikowia(100);g__Metschnikowia_unclassified(100); | Metschnikowia | 13 | Saprotroph | \|Undefined Saprotroph\| | Yeast | NULL | Probable | As nectar-tap_saprotroph (Põlme et al. 2020); Decay sugar-rich substrates (Põlme et al. 2020) | Põlme S, et al. 2020. Fungal Diversity 105:1-16 ((https://doi.org/10.1007/s13225-020-00466-2)) |
| Otu30864 | 1 | k__Fungi(100);p__Ascomycota(100);c__Saccharomycetes(100);o__Saccharomycetales(100);f__Metschnikowiaceae(100);g__Metschnikowia(100);g__Metschnikowia_unclassified(100); | Metschnikowia | 13 | Saprotroph | \|Undefined Saprotroph\| | Yeast | NULL | Probable | As nectar-tap_saprotroph (Põlme et al. 2020); Decay sugar-rich substrates (Põlme et al. 2020) | Põlme S, et al. 2020. Fungal Diversity 105:1-16 ((https://doi.org/10.1007/s13225-020-00466-2)) |
| Otu30865 | 1 | k__Fungi(100);p__Ascomycota(100);c__Saccharomycetes(100);o__Saccharomycetales(100);f__Metschnikowiaceae(100);g__Metschnikowia(100);s__Metschnikowia_pulcherrima(100); | Metschnikowia | 13 | Saprotroph | \|Undefined Saprotroph\| | Yeast | NULL | Probable | As nectar-tap_saprotroph (Põlme et al. 2020); Decay sugar-rich substrates (Põlme et al. 2020) | Põlme S, et al. 2020. Fungal Diversity 105:1-16 ((https://doi.org/10.1007/s13225-020-00466-2)) |
| Otu30866 | 1 | k__Fungi(100);p__Ascomycota(100);c__Saccharomycetes(100);o__Saccharomycetales(100);f__Metschnikowiaceae(100);g__Metschnikowia(100);s__Metschnikowia_pulcherrima(100); | Metschnikowia | 13 | Saprotroph | \|Undefined Saprotroph\| | Yeast | NULL | Probable | As nectar-tap_saprotroph (Põlme et al. 2020); Decay sugar-rich substrates (Põlme et al. 2020) | Põlme S, et al. 2020. Fungal Diversity 105:1-16 ((https://doi.org/10.1007/s13225-020-00466-2)) |
| Otu30867 | 1 | k__Fungi(100);p__Ascomycota(100);c__Saccharomycetes(100);o__Saccharomycetales(100);f__Metschnikowiaceae(100);g__Metschnikowia(100);s__Metschnikowia_pulcherrima(100); | Metschnikowia | 13 | Saprotroph | \|Undefined Saprotroph\| | Yeast | NULL | Probable | As nectar-tap_saprotroph (Põlme et al. 2020); Decay sugar-rich substrates (Põlme et al. 2020) | Põlme S, et al. 2020. Fungal Diversity 105:1-16 ((https://doi.org/10.1007/s13225-020-00466-2)) |
| Otu30868 | 1 | k__Fungi(100);p__Ascomycota(100);c__Saccharomycetes(100);o__Saccharomycetales(100);f__Metschnikowiaceae(100);g__Metschnikowia(100);g__Metschnikowia_unclassified(100); | Metschnikowia | 13 | Saprotroph | \|Undefined Saprotroph\| | Yeast | NULL | Probable | As nectar-tap_saprotroph (Põlme et al. 2020); Decay sugar-rich substrates (Põlme et al. 2020) | Põlme S, et al. 2020. Fungal Diversity 105:1-16 ((https://doi.org/10.1007/s13225-020-00466-2)) |
| Otu30869 | 1 | k__Fungi(100);p__Ascomycota(100);c__Saccharomycetes(100);o__Saccharomycetales(100);f__Metschnikowiaceae(100);g__Metschnikowia(100);s__Metschnikowia_pulcherrima(100); | Metschnikowia | 13 | Saprotroph | \|Undefined Saprotroph\| | Yeast | NULL | Probable | As nectar-tap_saprotroph (Põlme et al. 2020); Decay sugar-rich substrates (Põlme et al. 2020) | Põlme S, et al. 2020. Fungal Diversity 105:1-16 ((https://doi.org/10.1007/s13225-020-00466-2)) |
| Otu30870 | 1 | k__Fungi(100);p__Ascomycota(100);c__Saccharomycetes(100);o__Saccharomycetales(100);f__Metschnikowiaceae(100);g__Metschnikowia(100);s__Metschnikowia_pulcherrima(100); | Metschnikowia | 13 | Saprotroph | \|Undefined Saprotroph\| | Yeast | NULL | Probable | As nectar-tap_saprotroph (Põlme et al. 2020); Decay sugar-rich substrates (Põlme et al. 2020) | Põlme S, et al. 2020. Fungal Diversity 105:1-16 ((https://doi.org/10.1007/s13225-020-00466-2)) |
| Otu30871 | 1 | k__Fungi(100);p__Ascomycota(100);c__Saccharomycetes(100);o__Saccharomycetales(100);f__Metschnikowiaceae(100);g__Metschnikowia(100);s__Metschnikowia_pulcherrima(100); | Metschnikowia | 13 | Saprotroph | \|Undefined Saprotroph\| | Yeast | NULL | Probable | As nectar-tap_saprotroph (Põlme et al. 2020); Decay sugar-rich substrates (Põlme et al. 2020) | Põlme S, et al. 2020. Fungal Diversity 105:1-16 ((https://doi.org/10.1007/s13225-020-00466-2)) |
| Otu30872 | 1 | k__Fungi(100);p__Ascomycota(100);c__Saccharomycetes(100);o__Saccharomycetales(100);f__Metschnikowiaceae(100);g__Metschnikowia(100);s__Metschnikowia_pulcherrima(100); | Metschnikowia | 13 | Saprotroph | \|Undefined Saprotroph\| | Yeast | NULL | Probable | As nectar-tap_saprotroph (Põlme et al. 2020); Decay sugar-rich substrates (Põlme et al. 2020) | Põlme S, et al. 2020. Fungal Diversity 105:1-16 ((https://doi.org/10.1007/s13225-020-00466-2)) |
| Otu30873 | 1 | k__Fungi(100);p__Ascomycota(100);c__Saccharomycetes(100);o__Saccharomycetales(100);f__Metschnikowiaceae(100);g__Metschnikowia(100);g__Metschnikowia_unclassified(100); | Metschnikowia | 13 | Saprotroph | \|Undefined Saprotroph\| | Yeast | NULL | Probable | As nectar-tap_saprotroph (Põlme et al. 2020); Decay sugar-rich substrates (Põlme et al. 2020) | Põlme S, et al. 2020. Fungal Diversity 105:1-16 ((https://doi.org/10.1007/s13225-020-00466-2)) |
| Otu30874 | 1 | k__Fungi(100);p__Ascomycota(100);c__Saccharomycetes(100);o__Saccharomycetales(100);f__Metschnikowiaceae(100);g__Metschnikowia(100);g__Metschnikowia_unclassified(100); | Metschnikowia | 13 | Saprotroph | \|Undefined Saprotroph\| | Yeast | NULL | Probable | As nectar-tap_saprotroph (Põlme et al. 2020); Decay sugar-rich substrates (Põlme et al. 2020) | Põlme S, et al. 2020. Fungal Diversity 105:1-16 ((https://doi.org/10.1007/s13225-020-00466-2)) |
| Otu30875 | 1 | k__Fungi(100);p__Ascomycota(100);c__Saccharomycetes(100);o__Saccharomycetales(100);f__Metschnikowiaceae(100);g__Metschnikowia(100);s__Metschnikowia_pulcherrima(100); | Metschnikowia | 13 | Saprotroph | \|Undefined Saprotroph\| | Yeast | NULL | Probable | As nectar-tap_saprotroph (Põlme et al. 2020); Decay sugar-rich substrates (Põlme et al. 2020) | Põlme S, et al. 2020. Fungal Diversity 105:1-16 ((https://doi.org/10.1007/s13225-020-00466-2)) |
| Otu30876 | 1 | k__Fungi(100);p__Ascomycota(100);c__Saccharomycetes(100);o__Saccharomycetales(100);f__Metschnikowiaceae(100);g__Metschnikowia(100);s__Metschnikowia_pulcherrima(100); | Metschnikowia | 13 | Saprotroph | \|Undefined Saprotroph\| | Yeast | NULL | Probable | As nectar-tap_saprotroph (Põlme et al. 2020); Decay sugar-rich substrates (Põlme et al. 2020) | Põlme S, et al. 2020. Fungal Diversity 105:1-16 ((https://doi.org/10.1007/s13225-020-00466-2)) |
| Otu30877 | 1 | k__Fungi(100);p__Ascomycota(100);c__Saccharomycetes(100);o__Saccharomycetales(100);f__Metschnikowiaceae(100);g__Metschnikowia(100);g__Metschnikowia_unclassified(100); | Metschnikowia | 13 | Saprotroph | \|Undefined Saprotroph\| | Yeast | NULL | Probable | As nectar-tap_saprotroph (Põlme et al. 2020); Decay sugar-rich substrates (Põlme et al. 2020) | Põlme S, et al. 2020. Fungal Diversity 105:1-16 ((https://doi.org/10.1007/s13225-020-00466-2)) |
| Otu30878 | 1 | k__Fungi(100);p__Ascomycota(100);c__Saccharomycetes(100);o__Saccharomycetales(100);f__Metschnikowiaceae(100);g__Metschnikowia(100);s__Metschnikowia_pulcherrima(100); | Metschnikowia | 13 | Saprotroph | \|Undefined Saprotroph\| | Yeast | NULL | Probable | As nectar-tap_saprotroph (Põlme et al. 2020); Decay sugar-rich substrates (Põlme et al. 2020) | Põlme S, et al. 2020. Fungal Diversity 105:1-16 ((https://doi.org/10.1007/s13225-020-00466-2)) |
| Otu30879 | 1 | k__Fungi(100);p__Ascomycota(100);c__Saccharomycetes(100);o__Saccharomycetales(100);f__Metschnikowiaceae(100);g__Metschnikowia(100);s__Metschnikowia_pulcherrima(100); | Metschnikowia | 13 | Saprotroph | \|Undefined Saprotroph\| | Yeast | NULL | Probable | As nectar-tap_saprotroph (Põlme et al. 2020); Decay sugar-rich substrates (Põlme et al. 2020) | Põlme S, et al. 2020. Fungal Diversity 105:1-16 ((https://doi.org/10.1007/s13225-020-00466-2)) |
| Otu30880 | 1 | k__Fungi(100);p__Ascomycota(100);c__Saccharomycetes(100);o__Saccharomycetales(100);f__Metschnikowiaceae(100);g__Metschnikowia(100);g__Metschnikowia_unclassified(100); | Metschnikowia | 13 | Saprotroph | \|Undefined Saprotroph\| | Yeast | NULL | Probable | As nectar-tap_saprotroph (Põlme et al. 2020); Decay sugar-rich substrates (Põlme et al. 2020) | Põlme S, et al. 2020. Fungal Diversity 105:1-16 ((https://doi.org/10.1007/s13225-020-00466-2)) |
| Otu30881 | 1 | k__Fungi(100);p__Ascomycota(100);c__Saccharomycetes(100);o__Saccharomycetales(100);f__Metschnikowiaceae(100);g__Metschnikowia(100);s__Metschnikowia_pulcherrima(100); | Metschnikowia | 13 | Saprotroph | \|Undefined Saprotroph\| | Yeast | NULL | Probable | As nectar-tap_saprotroph (Põlme et al. 2020); Decay sugar-rich substrates (Põlme et al. 2020) | Põlme S, et al. 2020. Fungal Diversity 105:1-16 ((https://doi.org/10.1007/s13225-020-00466-2)) |
| Otu30882 | 1 | k__Fungi(100);p__Ascomycota(100);c__Saccharomycetes(100);o__Saccharomycetales(100);f__Metschnikowiaceae(100);g__Metschnikowia(100);s__Metschnikowia_pulcherrima(100); | Metschnikowia | 13 | Saprotroph | \|Undefined Saprotroph\| | Yeast | NULL | Probable | As nectar-tap_saprotroph (Põlme et al. 2020); Decay sugar-rich substrates (Põlme et al. 2020) | Põlme S, et al. 2020. Fungal Diversity 105:1-16 ((https://doi.org/10.1007/s13225-020-00466-2)) |
| Otu30883 | 1 | k__Fungi(100);p__Ascomycota(100);c__Saccharomycetes(100);o__Saccharomycetales(100);f__Metschnikowiaceae(100);g__Metschnikowia(100);s__Metschnikowia_pulcherrima(100); | Metschnikowia | 13 | Saprotroph | \|Undefined Saprotroph\| | Yeast | NULL | Probable | As nectar-tap_saprotroph (Põlme et al. 2020); Decay sugar-rich substrates (Põlme et al. 2020) | Põlme S, et al. 2020. Fungal Diversity 105:1-16 ((https://doi.org/10.1007/s13225-020-00466-2)) |
| Otu30884 | 1 | k__Fungi(100);p__Ascomycota(100);c__Saccharomycetes(100);o__Saccharomycetales(100);f__Metschnikowiaceae(100);g__Metschnikowia(100);g__Metschnikowia_unclassified(100); | Metschnikowia | 13 | Saprotroph | \|Undefined Saprotroph\| | Yeast | NULL | Probable | As nectar-tap_saprotroph (Põlme et al. 2020); Decay sugar-rich substrates (Põlme et al. 2020) | Põlme S, et al. 2020. Fungal Diversity 105:1-16 ((https://doi.org/10.1007/s13225-020-00466-2)) |
| Otu30886 | 1 | k__Fungi(100);p__Ascomycota(100);c__Saccharomycetes(100);o__Saccharomycetales(100);f__Metschnikowiaceae(100);g__Metschnikowia(100);s__Metschnikowia_pulcherrima(100); | Metschnikowia | 13 | Saprotroph | \|Undefined Saprotroph\| | Yeast | NULL | Probable | As nectar-tap_saprotroph (Põlme et al. 2020); Decay sugar-rich substrates (Põlme et al. 2020) | Põlme S, et al. 2020. Fungal Diversity 105:1-16 ((https://doi.org/10.1007/s13225-020-00466-2)) |
| Otu30892 | 1 | k__Fungi(100);p__Ascomycota(100);c__Saccharomycetes(100);o__Saccharomycetales(100);f__Metschnikowiaceae(100);g__Metschnikowia(100);s__Metschnikowia_pulcherrima(100); | Metschnikowia | 13 | Saprotroph | \|Undefined Saprotroph\| | Yeast | NULL | Probable | As nectar-tap_saprotroph (Põlme et al. 2020); Decay sugar-rich substrates (Põlme et al. 2020) | Põlme S, et al. 2020. Fungal Diversity 105:1-16 ((https://doi.org/10.1007/s13225-020-00466-2)) |
| Otu30895 | 1 | k__Fungi(100);p__Ascomycota(100);c__Saccharomycetes(100);o__Saccharomycetales(100);f__Metschnikowiaceae(100);g__Metschnikowia(100);g__Metschnikowia_unclassified(100); | Metschnikowia | 13 | Saprotroph | \|Undefined Saprotroph\| | Yeast | NULL | Probable | As nectar-tap_saprotroph (Põlme et al. 2020); Decay sugar-rich substrates (Põlme et al. 2020) | Põlme S, et al. 2020. Fungal Diversity 105:1-16 ((https://doi.org/10.1007/s13225-020-00466-2)) |
| Otu30896 | 1 | k__Fungi(100);p__Ascomycota(100);c__Saccharomycetes(100);o__Saccharomycetales(100);f__Metschnikowiaceae(100);g__Metschnikowia(100);s__Metschnikowia_pulcherrima(100); | Metschnikowia | 13 | Saprotroph | \|Undefined Saprotroph\| | Yeast | NULL | Probable | As nectar-tap_saprotroph (Põlme et al. 2020); Decay sugar-rich substrates (Põlme et al. 2020) | Põlme S, et al. 2020. Fungal Diversity 105:1-16 ((https://doi.org/10.1007/s13225-020-00466-2)) |
| Otu30898 | 1 | k__Fungi(100);p__Ascomycota(100);c__Saccharomycetes(100);o__Saccharomycetales(100);f__Metschnikowiaceae(100);g__Metschnikowia(100);g__Metschnikowia_unclassified(100); | Metschnikowia | 13 | Saprotroph | \|Undefined Saprotroph\| | Yeast | NULL | Probable | As nectar-tap_saprotroph (Põlme et al. 2020); Decay sugar-rich substrates (Põlme et al. 2020) | Põlme S, et al. 2020. Fungal Diversity 105:1-16 ((https://doi.org/10.1007/s13225-020-00466-2)) |
| Otu30899 | 1 | k__Fungi(100);p__Ascomycota(100);c__Saccharomycetes(100);o__Saccharomycetales(100);f__Metschnikowiaceae(100);g__Metschnikowia(100);s__Metschnikowia_pulcherrima(100); | Metschnikowia | 13 | Saprotroph | \|Undefined Saprotroph\| | Yeast | NULL | Probable | As nectar-tap_saprotroph (Põlme et al. 2020); Decay sugar-rich substrates (Põlme et al. 2020) | Põlme S, et al. 2020. Fungal Diversity 105:1-16 ((https://doi.org/10.1007/s13225-020-00466-2)) |
| Otu30900 | 1 | k__Fungi(100);p__Ascomycota(100);c__Saccharomycetes(100);o__Saccharomycetales(100);f__Metschnikowiaceae(100);g__Metschnikowia(100);g__Metschnikowia_unclassified(100); | Metschnikowia | 13 | Saprotroph | \|Undefined Saprotroph\| | Yeast | NULL | Probable | As nectar-tap_saprotroph (Põlme et al. 2020); Decay sugar-rich substrates (Põlme et al. 2020) | Põlme S, et al. 2020. Fungal Diversity 105:1-16 ((https://doi.org/10.1007/s13225-020-00466-2)) |
| Otu30901 | 1 | k__Fungi(100);p__Ascomycota(100);c__Saccharomycetes(100);o__Saccharomycetales(100);f__Metschnikowiaceae(100);g__Metschnikowia(100);s__Metschnikowia_pulcherrima(100); | Metschnikowia | 13 | Saprotroph | \|Undefined Saprotroph\| | Yeast | NULL | Probable | As nectar-tap_saprotroph (Põlme et al. 2020); Decay sugar-rich substrates (Põlme et al. 2020) | Põlme S, et al. 2020. Fungal Diversity 105:1-16 ((https://doi.org/10.1007/s13225-020-00466-2)) |
| Otu30902 | 1 | k__Fungi(100);p__Ascomycota(100);c__Saccharomycetes(100);o__Saccharomycetales(100);f__Metschnikowiaceae(100);g__Metschnikowia(100);s__Metschnikowia_pulcherrima(100); | Metschnikowia | 13 | Saprotroph | \|Undefined Saprotroph\| | Yeast | NULL | Probable | As nectar-tap_saprotroph (Põlme et al. 2020); Decay sugar-rich substrates (Põlme et al. 2020) | Põlme S, et al. 2020. Fungal Diversity 105:1-16 ((https://doi.org/10.1007/s13225-020-00466-2)) |
| Otu30904 | 1 | k__Fungi(100);p__Ascomycota(100);c__Saccharomycetes(100);o__Saccharomycetales(100);f__Metschnikowiaceae(100);g__Metschnikowia(100);s__Metschnikowia_pulcherrima(100); | Metschnikowia | 13 | Saprotroph | \|Undefined Saprotroph\| | Yeast | NULL | Probable | As nectar-tap_saprotroph (Põlme et al. 2020); Decay sugar-rich substrates (Põlme et al. 2020) | Põlme S, et al. 2020. Fungal Diversity 105:1-16 ((https://doi.org/10.1007/s13225-020-00466-2)) |
| Otu30913 | 1 | k__Fungi(100);p__Ascomycota(100);c__Saccharomycetes(100);o__Saccharomycetales(100);f__Metschnikowiaceae(100);g__Metschnikowia(100);s__Metschnikowia_pulcherrima(100); | Metschnikowia | 13 | Saprotroph | \|Undefined Saprotroph\| | Yeast | NULL | Probable | As nectar-tap_saprotroph (Põlme et al. 2020); Decay sugar-rich substrates (Põlme et al. 2020) | Põlme S, et al. 2020. Fungal Diversity 105:1-16 ((https://doi.org/10.1007/s13225-020-00466-2)) |
| Otu30914 | 1 | k__Fungi(100);p__Ascomycota(100);c__Saccharomycetes(100);o__Saccharomycetales(100);f__Metschnikowiaceae(100);g__Metschnikowia(100);s__Metschnikowia_pulcherrima(100); | Metschnikowia | 13 | Saprotroph | \|Undefined Saprotroph\| | Yeast | NULL | Probable | As nectar-tap_saprotroph (Põlme et al. 2020); Decay sugar-rich substrates (Põlme et al. 2020) | Põlme S, et al. 2020. Fungal Diversity 105:1-16 ((https://doi.org/10.1007/s13225-020-00466-2)) |
| Otu30919 | 1 | k__Fungi(100);p__Ascomycota(100);c__Saccharomycetes(100);o__Saccharomycetales(100);f__Metschnikowiaceae(100);g__Metschnikowia(100);s__Metschnikowia_pulcherrima(100); | Metschnikowia | 13 | Saprotroph | \|Undefined Saprotroph\| | Yeast | NULL | Probable | As nectar-tap_saprotroph (Põlme et al. 2020); Decay sugar-rich substrates (Põlme et al. 2020) | Põlme S, et al. 2020. Fungal Diversity 105:1-16 ((https://doi.org/10.1007/s13225-020-00466-2)) |
| Otu30925 | 1 | k__Fungi(100);p__Ascomycota(100);c__Saccharomycetes(100);o__Saccharomycetales(100);f__Metschnikowiaceae(100);g__Metschnikowia(100);s__Metschnikowia_pulcherrima(100); | Metschnikowia | 13 | Saprotroph | \|Undefined Saprotroph\| | Yeast | NULL | Probable | As nectar-tap_saprotroph (Põlme et al. 2020); Decay sugar-rich substrates (Põlme et al. 2020) | Põlme S, et al. 2020. Fungal Diversity 105:1-16 ((https://doi.org/10.1007/s13225-020-00466-2)) |
| Otu30928 | 1 | k__Fungi(100);p__Ascomycota(100);c__Saccharomycetes(100);o__Saccharomycetales(100);f__Metschnikowiaceae(100);g__Metschnikowia(100);s__Metschnikowia_pulcherrima(100); | Metschnikowia | 13 | Saprotroph | \|Undefined Saprotroph\| | Yeast | NULL | Probable | As nectar-tap_saprotroph (Põlme et al. 2020); Decay sugar-rich substrates (Põlme et al. 2020) | Põlme S, et al. 2020. Fungal Diversity 105:1-16 ((https://doi.org/10.1007/s13225-020-00466-2)) |
| Otu30929 | 1 | k__Fungi(100);p__Ascomycota(100);c__Saccharomycetes(100);o__Saccharomycetales(100);f__Metschnikowiaceae(100);g__Metschnikowia(100);s__Metschnikowia_pulcherrima(100); | Metschnikowia | 13 | Saprotroph | \|Undefined Saprotroph\| | Yeast | NULL | Probable | As nectar-tap_saprotroph (Põlme et al. 2020); Decay sugar-rich substrates (Põlme et al. 2020) | Põlme S, et al. 2020. Fungal Diversity 105:1-16 ((https://doi.org/10.1007/s13225-020-00466-2)) |
| Otu30930 | 1 | k__Fungi(100);p__Ascomycota(100);c__Saccharomycetes(100);o__Saccharomycetales(100);f__Metschnikowiaceae(100);g__Metschnikowia(100);g__Metschnikowia_unclassified(100); | Metschnikowia | 13 | Saprotroph | \|Undefined Saprotroph\| | Yeast | NULL | Probable | As nectar-tap_saprotroph (Põlme et al. 2020); Decay sugar-rich substrates (Põlme et al. 2020) | Põlme S, et al. 2020. Fungal Diversity 105:1-16 ((https://doi.org/10.1007/s13225-020-00466-2)) |
| Otu30931 | 1 | k__Fungi(100);p__Ascomycota(100);c__Saccharomycetes(100);o__Saccharomycetales(100);f__Metschnikowiaceae(100);g__Metschnikowia(100);s__Metschnikowia_pulcherrima(100); | Metschnikowia | 13 | Saprotroph | \|Undefined Saprotroph\| | Yeast | NULL | Probable | As nectar-tap_saprotroph (Põlme et al. 2020); Decay sugar-rich substrates (Põlme et al. 2020) | Põlme S, et al. 2020. Fungal Diversity 105:1-16 ((https://doi.org/10.1007/s13225-020-00466-2)) |
| Otu30932 | 1 | k__Fungi(100);p__Basidiomycota(100);c__Microbotryomycetes(100);o__Sporidiobolales(100);f__Sporidiobolales_family_Incertae_sedis(100);g__Sporobolomyces(100);s__Sporobolomyces_lactosus(100); | Sporobolomyces | 13 | Pathotroph-Saprotroph | \|Fungal Parasite\|-Plant Saprotroph | Yeast | NULL | Probable | As litter_saprotroph (Põlme et al. 2020); As mycoparasite (Põlme et al. 2020); Decay fungal material (Põlme et al. 2020) | Tedersoo L, et al. 2014. Science 346:e1256688 ((https://doi.org/10.1126/science.1256688)); Purahong W, et al. 2016. Molecular Ecology 25:4059-4074 ((https://doi.org/10.1111/mec.13739)); Põlme S, et al. 2020. Fungal Diversity 105:1-16 ((https://doi.org/10.1007/s13225-020-00466-2)) |
| Otu30933 | 1 | k__Fungi(100);p__Ascomycota(100);c__Saccharomycetes(100);o__Saccharomycetales(100);f__Metschnikowiaceae(100);g__Metschnikowia(100);s__Metschnikowia_pulcherrima(100); | Metschnikowia | 13 | Saprotroph | \|Undefined Saprotroph\| | Yeast | NULL | Probable | As nectar-tap_saprotroph (Põlme et al. 2020); Decay sugar-rich substrates (Põlme et al. 2020) | Põlme S, et al. 2020. Fungal Diversity 105:1-16 ((https://doi.org/10.1007/s13225-020-00466-2)) |
| Otu30936 | 1 | k__Fungi(100);p__Ascomycota(100);c__Sordariomycetes(100);o__Hypocreales(100);f__Nectriaceae(100);f__Nectriaceae_unclassified(100);f__Nectriaceae_unclassified(100); | Nectriaceae | 9 | Pathotroph-Saprotroph-Symbiotroph | Animal Pathogen-Endophyte-Fungal Parasite-Lichen Parasite-Plant Pathogen-Wood Saprotroph | Microfungus | NULL | Possible | Fusarium species are amongst the economically significant plant pathogens (Cannon & Kirk 2007) | Cannon PF, Kirk PM. 2007. Fungal Families of the World. CAB International, Cambridge (ISBN: 978-0851998275); Hirooka Y, et al. 2012. Studies in Mycology 71:1-210 ((https://doi.org/10.3114/sim0001)); Tedersoo L, et al. 2014. Science 346:e1256688 ((https://doi.org/10.1126/science.1256688)); The American Phytopathological Society Common Names of Plant Diseases ((http://www.apsnet.org/publications/commonnames/Pages/default.aspx)); https://www.cdc.gov ((https://www.cdc.gov)) |
| Otu30937 | 1 | k__Fungi(100);p__Ascomycota(100);c__Dothideomycetes(100);o__Pleosporales(100);f__Pleosporales_family_Incertae_sedis(100);g__Paraphoma(100);g__Paraphoma_unclassified(100); | Paraphoma | 13 | Pathotroph-Saprotroph | \|Plant Pathogen\|-Plant Saprotroph-Undefined Saprotroph-Wood Saprotroph | Microfungus | NULL | Probable | As wood_pathogen (Põlme et al. 2020); Associated with leaf and crown infection of pyrethrum (Moslemi et al. 2017); Decay wood (Põlme et al. 2020) | Quaedvlieg W, et al. 2013. Studies in Mycology 75:307-390 ((https://doi.org/10.3114/sim0017)); Moslemi A, et al. 2017. Plant Pathology 67:124-135 ((https://doi.org/10.1111/ppa.12719)); Põlme S, et al. 2020. Fungal Diversity 105:1-16 ((https://doi.org/10.1007/s13225-020-00466-2)) |
| Otu30939 | 1 | k__Fungi(100);p__Ascomycota(100);c__Leotiomycetes(100);o__Helotiales(100);f__Rutstroemiaceae(100);f__Rutstroemiaceae_unclassified(100);f__Rutstroemiaceae_unclassified(100); | Rutstroemiaceae | 9 | Saprotroph | Plant Saprotroph | Microfungus | NULL | Probable | Essentially saprobic on various plant parts, especially seeds and fruits, though initial infection may take place in living tissue (Cannon & Kirk 2007) | Cannon PF, Kirk PM. 2007. Fungal Families of the World. CAB International, Cambridge (ISBN: 978-0851998275) |
| Otu30940 | 1 | k__Fungi(100);p__Ascomycota(100);c__Dothideomycetes(100);o__Capnodiales(100);f__Davidiellaceae(100);g__Cladosporium(100);g__Cladosporium_unclassified(100); | Cladosporium | 13 | Pathotroph-Saprotroph-Symbiotroph | Animal Pathogen-Endophyte-Lichen Parasite-Plant Pathogen-\|Plant Saprotroph\|-Wood Saprotroph | Microfungus | NULL | Possible | As foliar_endophyte (Põlme et al. 2020); As litter_saprotroph (Põlme et al. 2020); Decay wood, leaf, fruit, seed, soil, dung, animal material (Põlme et al. 2020) | Seehann G, et al. 1975. List of Fungi in Soft-Rot Tissue Tests. Institute fur Holzbiologie und Holzschutz, Bundesforschungsanstalt fur Forst - und Holzwirtshaft, Hamburg - Lohbrugge, Germany; Lawrey JD, Diederich P. 2016. http://www.lichenicolous.net (accessed 24 May 2017) ((http://www.lichenicolous.net)); Põlme S, et al. 2020. Fungal Diversity 105:1-16 ((https://doi.org/10.1007/s13225-020-00466-2)); The American Phytopathological Society Common Names of Plant Diseases ((http://www.apsnet.org/publications/commonnames/Pages/default.aspx)) |
| Otu30941 | 1 | k__Fungi(100);p__Basidiomycota(100);c__Tremellomycetes(100);o__Tremellales(100);f__Tremellales_family_Incertae_sedis(100);g__Derxomyces(100);g__Derxomyces_unclassified(100); | Derxomyces | 13 | Saprotroph-Symbiotroph | Epiphyte-\|Undefined Saprotroph\| | Dimorphic | NULL | Probable | NULL | Põlme S, et al. 2020. Fungal Diversity 105:1-16 ((https://doi.org/10.1007/s13225-020-00466-2)) |
| Otu30942 | 1 | k__Fungi(100);p__Ascomycota(100);c__Saccharomycetes(100);o__Saccharomycetales(100);f__Metschnikowiaceae(100);g__Metschnikowia(100);s__Metschnikowia_pulcherrima(100); | Metschnikowia | 13 | Saprotroph | \|Undefined Saprotroph\| | Yeast | NULL | Probable | As nectar-tap_saprotroph (Põlme et al. 2020); Decay sugar-rich substrates (Põlme et al. 2020) | Põlme S, et al. 2020. Fungal Diversity 105:1-16 ((https://doi.org/10.1007/s13225-020-00466-2)) |
| Otu30943 | 1 | k__Fungi(100);p__Ascomycota(100);c__Saccharomycetes(100);o__Saccharomycetales(100);f__Metschnikowiaceae(100);g__Metschnikowia(100);s__Metschnikowia_pulcherrima(100); | Metschnikowia | 13 | Saprotroph | \|Undefined Saprotroph\| | Yeast | NULL | Probable | As nectar-tap_saprotroph (Põlme et al. 2020); Decay sugar-rich substrates (Põlme et al. 2020) | Põlme S, et al. 2020. Fungal Diversity 105:1-16 ((https://doi.org/10.1007/s13225-020-00466-2)) |
| Otu30944 | 1 | k__Fungi(100);p__Ascomycota(100);c__Saccharomycetes(100);o__Saccharomycetales(100);f__Metschnikowiaceae(100);g__Metschnikowia(100);g__Metschnikowia_unclassified(100); | Metschnikowia | 13 | Saprotroph | \|Undefined Saprotroph\| | Yeast | NULL | Probable | As nectar-tap_saprotroph (Põlme et al. 2020); Decay sugar-rich substrates (Põlme et al. 2020) | Põlme S, et al. 2020. Fungal Diversity 105:1-16 ((https://doi.org/10.1007/s13225-020-00466-2)) |
| Otu30945 | 1 | k__Fungi(100);p__Ascomycota(100);c__Saccharomycetes(100);o__Saccharomycetales(100);f__Metschnikowiaceae(100);g__Metschnikowia(100);s__Metschnikowia_pulcherrima(100); | Metschnikowia | 13 | Saprotroph | \|Undefined Saprotroph\| | Yeast | NULL | Probable | As nectar-tap_saprotroph (Põlme et al. 2020); Decay sugar-rich substrates (Põlme et al. 2020) | Põlme S, et al. 2020. Fungal Diversity 105:1-16 ((https://doi.org/10.1007/s13225-020-00466-2)) |
| Otu30946 | 1 | k__Fungi(100);p__Ascomycota(100);c__Saccharomycetes(100);o__Saccharomycetales(100);f__Metschnikowiaceae(100);g__Metschnikowia(100);s__Metschnikowia_pulcherrima(100); | Metschnikowia | 13 | Saprotroph | \|Undefined Saprotroph\| | Yeast | NULL | Probable | As nectar-tap_saprotroph (Põlme et al. 2020); Decay sugar-rich substrates (Põlme et al. 2020) | Põlme S, et al. 2020. Fungal Diversity 105:1-16 ((https://doi.org/10.1007/s13225-020-00466-2)) |
| Otu30947 | 1 | k__Fungi(100);p__Ascomycota(100);c__Saccharomycetes(100);o__Saccharomycetales(100);f__Metschnikowiaceae(100);g__Metschnikowia(100);s__Metschnikowia_pulcherrima(100); | Metschnikowia | 13 | Saprotroph | \|Undefined Saprotroph\| | Yeast | NULL | Probable | As nectar-tap_saprotroph (Põlme et al. 2020); Decay sugar-rich substrates (Põlme et al. 2020) | Põlme S, et al. 2020. Fungal Diversity 105:1-16 ((https://doi.org/10.1007/s13225-020-00466-2)) |
| Otu30949 | 1 | k__Fungi(100);p__Ascomycota(100);c__Saccharomycetes(100);o__Saccharomycetales(100);f__Metschnikowiaceae(100);g__Metschnikowia(100);s__Metschnikowia_pulcherrima(100); | Metschnikowia | 13 | Saprotroph | \|Undefined Saprotroph\| | Yeast | NULL | Probable | As nectar-tap_saprotroph (Põlme et al. 2020); Decay sugar-rich substrates (Põlme et al. 2020) | Põlme S, et al. 2020. Fungal Diversity 105:1-16 ((https://doi.org/10.1007/s13225-020-00466-2)) |
| Otu30950 | 1 | k__Fungi(100);p__Ascomycota(100);c__Saccharomycetes(100);o__Saccharomycetales(100);f__Metschnikowiaceae(100);g__Metschnikowia(100);s__Metschnikowia_pulcherrima(100); | Metschnikowia | 13 | Saprotroph | \|Undefined Saprotroph\| | Yeast | NULL | Probable | As nectar-tap_saprotroph (Põlme et al. 2020); Decay sugar-rich substrates (Põlme et al. 2020) | Põlme S, et al. 2020. Fungal Diversity 105:1-16 ((https://doi.org/10.1007/s13225-020-00466-2)) |
| Otu30951 | 1 | k__Fungi(100);p__Ascomycota(100);c__Saccharomycetes(100);o__Saccharomycetales(100);f__Metschnikowiaceae(100);g__Metschnikowia(100);s__Metschnikowia_pulcherrima(100); | Metschnikowia | 13 | Saprotroph | \|Undefined Saprotroph\| | Yeast | NULL | Probable | As nectar-tap_saprotroph (Põlme et al. 2020); Decay sugar-rich substrates (Põlme et al. 2020) | Põlme S, et al. 2020. Fungal Diversity 105:1-16 ((https://doi.org/10.1007/s13225-020-00466-2)) |
| Otu30952 | 1 | k__Fungi(100);p__Ascomycota(100);c__Saccharomycetes(100);o__Saccharomycetales(100);f__Metschnikowiaceae(100);g__Metschnikowia(100);s__Metschnikowia_pulcherrima(100); | Metschnikowia | 13 | Saprotroph | \|Undefined Saprotroph\| | Yeast | NULL | Probable | As nectar-tap_saprotroph (Põlme et al. 2020); Decay sugar-rich substrates (Põlme et al. 2020) | Põlme S, et al. 2020. Fungal Diversity 105:1-16 ((https://doi.org/10.1007/s13225-020-00466-2)) |
| Otu30953 | 1 | k__Fungi(100);p__Ascomycota(100);c__Saccharomycetes(100);o__Saccharomycetales(100);f__Metschnikowiaceae(100);g__Metschnikowia(100);s__Metschnikowia_pulcherrima(100); | Metschnikowia | 13 | Saprotroph | \|Undefined Saprotroph\| | Yeast | NULL | Probable | As nectar-tap_saprotroph (Põlme et al. 2020); Decay sugar-rich substrates (Põlme et al. 2020) | Põlme S, et al. 2020. Fungal Diversity 105:1-16 ((https://doi.org/10.1007/s13225-020-00466-2)) |
| Otu30954 | 1 | k__Fungi(100);p__Ascomycota(100);c__Saccharomycetes(100);o__Saccharomycetales(100);f__Metschnikowiaceae(100);g__Metschnikowia(100);s__Metschnikowia_pulcherrima(100); | Metschnikowia | 13 | Saprotroph | \|Undefined Saprotroph\| | Yeast | NULL | Probable | As nectar-tap_saprotroph (Põlme et al. 2020); Decay sugar-rich substrates (Põlme et al. 2020) | Põlme S, et al. 2020. Fungal Diversity 105:1-16 ((https://doi.org/10.1007/s13225-020-00466-2)) |
| Otu30955 | 1 | k__Fungi(100);p__Ascomycota(100);c__Saccharomycetes(100);o__Saccharomycetales(100);f__Metschnikowiaceae(100);g__Metschnikowia(100);s__Metschnikowia_pulcherrima(100); | Metschnikowia | 13 | Saprotroph | \|Undefined Saprotroph\| | Yeast | NULL | Probable | As nectar-tap_saprotroph (Põlme et al. 2020); Decay sugar-rich substrates (Põlme et al. 2020) | Põlme S, et al. 2020. Fungal Diversity 105:1-16 ((https://doi.org/10.1007/s13225-020-00466-2)) |
| Otu30956 | 1 | k__Fungi(100);p__Ascomycota(100);c__Saccharomycetes(100);o__Saccharomycetales(100);f__Metschnikowiaceae(100);g__Metschnikowia(100);g__Metschnikowia_unclassified(100); | Metschnikowia | 13 | Saprotroph | \|Undefined Saprotroph\| | Yeast | NULL | Probable | As nectar-tap_saprotroph (Põlme et al. 2020); Decay sugar-rich substrates (Põlme et al. 2020) | Põlme S, et al. 2020. Fungal Diversity 105:1-16 ((https://doi.org/10.1007/s13225-020-00466-2)) |
| Otu30957 | 1 | k__Fungi(100);p__Ascomycota(100);c__Saccharomycetes(100);o__Saccharomycetales(100);f__Metschnikowiaceae(100);g__Metschnikowia(100);g__Metschnikowia_unclassified(100); | Metschnikowia | 13 | Saprotroph | \|Undefined Saprotroph\| | Yeast | NULL | Probable | As nectar-tap_saprotroph (Põlme et al. 2020); Decay sugar-rich substrates (Põlme et al. 2020) | Põlme S, et al. 2020. Fungal Diversity 105:1-16 ((https://doi.org/10.1007/s13225-020-00466-2)) |
| Otu30959 | 1 | k__Fungi(100);p__Ascomycota(100);c__Saccharomycetes(100);o__Saccharomycetales(100);f__Metschnikowiaceae(100);g__Metschnikowia(100);g__Metschnikowia_unclassified(100); | Metschnikowia | 13 | Saprotroph | \|Undefined Saprotroph\| | Yeast | NULL | Probable | As nectar-tap_saprotroph (Põlme et al. 2020); Decay sugar-rich substrates (Põlme et al. 2020) | Põlme S, et al. 2020. Fungal Diversity 105:1-16 ((https://doi.org/10.1007/s13225-020-00466-2)) |
| Otu30961 | 1 | k__Fungi(100);p__Ascomycota(100);c__Saccharomycetes(100);o__Saccharomycetales(100);f__Metschnikowiaceae(100);g__Metschnikowia(100);s__Metschnikowia_pulcherrima(100); | Metschnikowia | 13 | Saprotroph | \|Undefined Saprotroph\| | Yeast | NULL | Probable | As nectar-tap_saprotroph (Põlme et al. 2020); Decay sugar-rich substrates (Põlme et al. 2020) | Põlme S, et al. 2020. Fungal Diversity 105:1-16 ((https://doi.org/10.1007/s13225-020-00466-2)) |
| Otu30962 | 1 | k__Fungi(100);p__Ascomycota(100);c__Saccharomycetes(100);o__Saccharomycetales(100);f__Metschnikowiaceae(100);g__Metschnikowia(100);s__Metschnikowia_pulcherrima(100); | Metschnikowia | 13 | Saprotroph | \|Undefined Saprotroph\| | Yeast | NULL | Probable | As nectar-tap_saprotroph (Põlme et al. 2020); Decay sugar-rich substrates (Põlme et al. 2020) | Põlme S, et al. 2020. Fungal Diversity 105:1-16 ((https://doi.org/10.1007/s13225-020-00466-2)) |
| Otu30963 | 1 | k__Fungi(100);p__Ascomycota(100);c__Saccharomycetes(100);o__Saccharomycetales(100);f__Metschnikowiaceae(100);g__Metschnikowia(100);g__Metschnikowia_unclassified(100); | Metschnikowia | 13 | Saprotroph | \|Undefined Saprotroph\| | Yeast | NULL | Probable | As nectar-tap_saprotroph (Põlme et al. 2020); Decay sugar-rich substrates (Põlme et al. 2020) | Põlme S, et al. 2020. Fungal Diversity 105:1-16 ((https://doi.org/10.1007/s13225-020-00466-2)) |
| Otu30964 | 1 | k__Fungi(100);p__Ascomycota(100);c__Saccharomycetes(100);o__Saccharomycetales(100);f__Metschnikowiaceae(100);g__Metschnikowia(100);s__Metschnikowia_pulcherrima(100); | Metschnikowia | 13 | Saprotroph | \|Undefined Saprotroph\| | Yeast | NULL | Probable | As nectar-tap_saprotroph (Põlme et al. 2020); Decay sugar-rich substrates (Põlme et al. 2020) | Põlme S, et al. 2020. Fungal Diversity 105:1-16 ((https://doi.org/10.1007/s13225-020-00466-2)) |
| Otu30965 | 1 | k__Fungi(100);p__Ascomycota(100);c__Saccharomycetes(100);o__Saccharomycetales(100);f__Metschnikowiaceae(100);g__Metschnikowia(100);s__Metschnikowia_pulcherrima(100); | Metschnikowia | 13 | Saprotroph | \|Undefined Saprotroph\| | Yeast | NULL | Probable | As nectar-tap_saprotroph (Põlme et al. 2020); Decay sugar-rich substrates (Põlme et al. 2020) | Põlme S, et al. 2020. Fungal Diversity 105:1-16 ((https://doi.org/10.1007/s13225-020-00466-2)) |
| Otu30966 | 1 | k__Fungi(100);p__Ascomycota(100);c__Saccharomycetes(100);o__Saccharomycetales(100);f__Metschnikowiaceae(100);g__Metschnikowia(100);s__Metschnikowia_pulcherrima(100); | Metschnikowia | 13 | Saprotroph | \|Undefined Saprotroph\| | Yeast | NULL | Probable | As nectar-tap_saprotroph (Põlme et al. 2020); Decay sugar-rich substrates (Põlme et al. 2020) | Põlme S, et al. 2020. Fungal Diversity 105:1-16 ((https://doi.org/10.1007/s13225-020-00466-2)) |
| Otu30967 | 1 | k__Fungi(100);p__Ascomycota(100);c__Saccharomycetes(100);o__Saccharomycetales(100);f__Metschnikowiaceae(100);g__Metschnikowia(100);s__Metschnikowia_pulcherrima(100); | Metschnikowia | 13 | Saprotroph | \|Undefined Saprotroph\| | Yeast | NULL | Probable | As nectar-tap_saprotroph (Põlme et al. 2020); Decay sugar-rich substrates (Põlme et al. 2020) | Põlme S, et al. 2020. Fungal Diversity 105:1-16 ((https://doi.org/10.1007/s13225-020-00466-2)) |
| Otu30968 | 1 | k__Fungi(100);p__Ascomycota(100);c__Saccharomycetes(100);o__Saccharomycetales(100);f__Metschnikowiaceae(100);g__Metschnikowia(100);s__Metschnikowia_pulcherrima(100); | Metschnikowia | 13 | Saprotroph | \|Undefined Saprotroph\| | Yeast | NULL | Probable | As nectar-tap_saprotroph (Põlme et al. 2020); Decay sugar-rich substrates (Põlme et al. 2020) | Põlme S, et al. 2020. Fungal Diversity 105:1-16 ((https://doi.org/10.1007/s13225-020-00466-2)) |
| Otu30969 | 1 | k__Fungi(100);p__Ascomycota(100);c__Saccharomycetes(100);o__Saccharomycetales(100);f__Metschnikowiaceae(100);g__Metschnikowia(100);s__Metschnikowia_pulcherrima(100); | Metschnikowia | 13 | Saprotroph | \|Undefined Saprotroph\| | Yeast | NULL | Probable | As nectar-tap_saprotroph (Põlme et al. 2020); Decay sugar-rich substrates (Põlme et al. 2020) | Põlme S, et al. 2020. Fungal Diversity 105:1-16 ((https://doi.org/10.1007/s13225-020-00466-2)) |
| Otu30971 | 1 | k__Fungi(100);p__Ascomycota(100);c__Saccharomycetes(100);o__Saccharomycetales(100);f__Metschnikowiaceae(100);g__Metschnikowia(100);s__Metschnikowia_pulcherrima(100); | Metschnikowia | 13 | Saprotroph | \|Undefined Saprotroph\| | Yeast | NULL | Probable | As nectar-tap_saprotroph (Põlme et al. 2020); Decay sugar-rich substrates (Põlme et al. 2020) | Põlme S, et al. 2020. Fungal Diversity 105:1-16 ((https://doi.org/10.1007/s13225-020-00466-2)) |
| Otu30972 | 1 | k__Fungi(100);p__Ascomycota(100);c__Saccharomycetes(100);o__Saccharomycetales(100);f__Metschnikowiaceae(100);g__Metschnikowia(100);g__Metschnikowia_unclassified(100); | Metschnikowia | 13 | Saprotroph | \|Undefined Saprotroph\| | Yeast | NULL | Probable | As nectar-tap_saprotroph (Põlme et al. 2020); Decay sugar-rich substrates (Põlme et al. 2020) | Põlme S, et al. 2020. Fungal Diversity 105:1-16 ((https://doi.org/10.1007/s13225-020-00466-2)) |
| Otu30973 | 1 | k__Fungi(100);p__Ascomycota(100);c__Saccharomycetes(100);o__Saccharomycetales(100);f__Metschnikowiaceae(100);g__Metschnikowia(100);s__Metschnikowia_pulcherrima(100); | Metschnikowia | 13 | Saprotroph | \|Undefined Saprotroph\| | Yeast | NULL | Probable | As nectar-tap_saprotroph (Põlme et al. 2020); Decay sugar-rich substrates (Põlme et al. 2020) | Põlme S, et al. 2020. Fungal Diversity 105:1-16 ((https://doi.org/10.1007/s13225-020-00466-2)) |
| Otu30974 | 1 | k__Fungi(100);p__Ascomycota(100);c__Saccharomycetes(100);o__Saccharomycetales(100);f__Metschnikowiaceae(100);g__Metschnikowia(100);s__Metschnikowia_pulcherrima(100); | Metschnikowia | 13 | Saprotroph | \|Undefined Saprotroph\| | Yeast | NULL | Probable | As nectar-tap_saprotroph (Põlme et al. 2020); Decay sugar-rich substrates (Põlme et al. 2020) | Põlme S, et al. 2020. Fungal Diversity 105:1-16 ((https://doi.org/10.1007/s13225-020-00466-2)) |
| Otu30975 | 1 | k__Fungi(100);p__Ascomycota(100);c__Saccharomycetes(100);o__Saccharomycetales(100);f__Metschnikowiaceae(100);g__Metschnikowia(100);g__Metschnikowia_unclassified(100); | Metschnikowia | 13 | Saprotroph | \|Undefined Saprotroph\| | Yeast | NULL | Probable | As nectar-tap_saprotroph (Põlme et al. 2020); Decay sugar-rich substrates (Põlme et al. 2020) | Põlme S, et al. 2020. Fungal Diversity 105:1-16 ((https://doi.org/10.1007/s13225-020-00466-2)) |
| Otu30976 | 1 | k__Fungi(100);p__Ascomycota(100);c__Saccharomycetes(100);o__Saccharomycetales(100);f__Metschnikowiaceae(100);g__Metschnikowia(100);g__Metschnikowia_unclassified(100); | Metschnikowia | 13 | Saprotroph | \|Undefined Saprotroph\| | Yeast | NULL | Probable | As nectar-tap_saprotroph (Põlme et al. 2020); Decay sugar-rich substrates (Põlme et al. 2020) | Põlme S, et al. 2020. Fungal Diversity 105:1-16 ((https://doi.org/10.1007/s13225-020-00466-2)) |
| Otu30977 | 1 | k__Fungi(100);p__Ascomycota(100);c__Saccharomycetes(100);o__Saccharomycetales(100);f__Metschnikowiaceae(100);g__Metschnikowia(100);g__Metschnikowia_unclassified(100); | Metschnikowia | 13 | Saprotroph | \|Undefined Saprotroph\| | Yeast | NULL | Probable | As nectar-tap_saprotroph (Põlme et al. 2020); Decay sugar-rich substrates (Põlme et al. 2020) | Põlme S, et al. 2020. Fungal Diversity 105:1-16 ((https://doi.org/10.1007/s13225-020-00466-2)) |
| Otu30978 | 1 | k__Fungi(100);p__Ascomycota(100);c__Saccharomycetes(100);o__Saccharomycetales(100);f__Metschnikowiaceae(100);g__Metschnikowia(100);s__Metschnikowia_pulcherrima(100); | Metschnikowia | 13 | Saprotroph | \|Undefined Saprotroph\| | Yeast | NULL | Probable | As nectar-tap_saprotroph (Põlme et al. 2020); Decay sugar-rich substrates (Põlme et al. 2020) | Põlme S, et al. 2020. Fungal Diversity 105:1-16 ((https://doi.org/10.1007/s13225-020-00466-2)) |
| Otu30979 | 1 | k__Fungi(100);p__Ascomycota(100);c__Saccharomycetes(100);o__Saccharomycetales(100);f__Metschnikowiaceae(100);g__Metschnikowia(100);s__Metschnikowia_pulcherrima(100); | Metschnikowia | 13 | Saprotroph | \|Undefined Saprotroph\| | Yeast | NULL | Probable | As nectar-tap_saprotroph (Põlme et al. 2020); Decay sugar-rich substrates (Põlme et al. 2020) | Põlme S, et al. 2020. Fungal Diversity 105:1-16 ((https://doi.org/10.1007/s13225-020-00466-2)) |
| Otu30980 | 1 | k__Fungi(100);p__Ascomycota(100);c__Saccharomycetes(100);o__Saccharomycetales(100);f__Metschnikowiaceae(100);g__Metschnikowia(100);s__Metschnikowia_pulcherrima(100); | Metschnikowia | 13 | Saprotroph | \|Undefined Saprotroph\| | Yeast | NULL | Probable | As nectar-tap_saprotroph (Põlme et al. 2020); Decay sugar-rich substrates (Põlme et al. 2020) | Põlme S, et al. 2020. Fungal Diversity 105:1-16 ((https://doi.org/10.1007/s13225-020-00466-2)) |
| Otu30981 | 1 | k__Fungi(100);p__Ascomycota(100);c__Saccharomycetes(100);o__Saccharomycetales(100);f__Metschnikowiaceae(100);g__Metschnikowia(100);s__Metschnikowia_pulcherrima(100); | Metschnikowia | 13 | Saprotroph | \|Undefined Saprotroph\| | Yeast | NULL | Probable | As nectar-tap_saprotroph (Põlme et al. 2020); Decay sugar-rich substrates (Põlme et al. 2020) | Põlme S, et al. 2020. Fungal Diversity 105:1-16 ((https://doi.org/10.1007/s13225-020-00466-2)) |
| Otu30982 | 1 | k__Fungi(100);p__Ascomycota(100);c__Saccharomycetes(100);o__Saccharomycetales(100);f__Metschnikowiaceae(100);g__Metschnikowia(100);s__Metschnikowia_pulcherrima(100); | Metschnikowia | 13 | Saprotroph | \|Undefined Saprotroph\| | Yeast | NULL | Probable | As nectar-tap_saprotroph (Põlme et al. 2020); Decay sugar-rich substrates (Põlme et al. 2020) | Põlme S, et al. 2020. Fungal Diversity 105:1-16 ((https://doi.org/10.1007/s13225-020-00466-2)) |
| Otu30983 | 1 | k__Fungi(100);p__Ascomycota(100);c__Saccharomycetes(100);o__Saccharomycetales(100);f__Metschnikowiaceae(100);g__Metschnikowia(100);s__Metschnikowia_pulcherrima(100); | Metschnikowia | 13 | Saprotroph | \|Undefined Saprotroph\| | Yeast | NULL | Probable | As nectar-tap_saprotroph (Põlme et al. 2020); Decay sugar-rich substrates (Põlme et al. 2020) | Põlme S, et al. 2020. Fungal Diversity 105:1-16 ((https://doi.org/10.1007/s13225-020-00466-2)) |
| Otu30984 | 1 | k__Fungi(100);p__Ascomycota(100);c__Saccharomycetes(100);o__Saccharomycetales(100);f__Metschnikowiaceae(100);g__Metschnikowia(100);g__Metschnikowia_unclassified(100); | Metschnikowia | 13 | Saprotroph | \|Undefined Saprotroph\| | Yeast | NULL | Probable | As nectar-tap_saprotroph (Põlme et al. 2020); Decay sugar-rich substrates (Põlme et al. 2020) | Põlme S, et al. 2020. Fungal Diversity 105:1-16 ((https://doi.org/10.1007/s13225-020-00466-2)) |
| Otu30986 | 1 | k__Fungi(100);p__Ascomycota(100);c__Saccharomycetes(100);o__Saccharomycetales(100);f__Metschnikowiaceae(100);g__Metschnikowia(100);s__Metschnikowia_pulcherrima(100); | Metschnikowia | 13 | Saprotroph | \|Undefined Saprotroph\| | Yeast | NULL | Probable | As nectar-tap_saprotroph (Põlme et al. 2020); Decay sugar-rich substrates (Põlme et al. 2020) | Põlme S, et al. 2020. Fungal Diversity 105:1-16 ((https://doi.org/10.1007/s13225-020-00466-2)) |
| Otu30987 | 1 | k__Fungi(100);p__Ascomycota(100);c__Saccharomycetes(100);o__Saccharomycetales(100);f__Metschnikowiaceae(100);g__Metschnikowia(100);s__Metschnikowia_pulcherrima(100); | Metschnikowia | 13 | Saprotroph | \|Undefined Saprotroph\| | Yeast | NULL | Probable | As nectar-tap_saprotroph (Põlme et al. 2020); Decay sugar-rich substrates (Põlme et al. 2020) | Põlme S, et al. 2020. Fungal Diversity 105:1-16 ((https://doi.org/10.1007/s13225-020-00466-2)) |
| Otu30988 | 1 | k__Fungi(100);p__Ascomycota(100);c__Saccharomycetes(100);o__Saccharomycetales(100);f__Metschnikowiaceae(100);g__Metschnikowia(100);g__Metschnikowia_unclassified(100); | Metschnikowia | 13 | Saprotroph | \|Undefined Saprotroph\| | Yeast | NULL | Probable | As nectar-tap_saprotroph (Põlme et al. 2020); Decay sugar-rich substrates (Põlme et al. 2020) | Põlme S, et al. 2020. Fungal Diversity 105:1-16 ((https://doi.org/10.1007/s13225-020-00466-2)) |
| Otu30998 | 1 | k__Fungi(100);p__Ascomycota(100);c__Saccharomycetes(100);o__Saccharomycetales(100);f__Metschnikowiaceae(100);g__Metschnikowia(100);s__Metschnikowia_pulcherrima(100); | Metschnikowia | 13 | Saprotroph | \|Undefined Saprotroph\| | Yeast | NULL | Probable | As nectar-tap_saprotroph (Põlme et al. 2020); Decay sugar-rich substrates (Põlme et al. 2020) | Põlme S, et al. 2020. Fungal Diversity 105:1-16 ((https://doi.org/10.1007/s13225-020-00466-2)) |
| Otu31008 | 1 | k__Fungi(100);p__Ascomycota(100);c__Saccharomycetes(100);o__Saccharomycetales(100);f__Metschnikowiaceae(100);g__Metschnikowia(100);s__Metschnikowia_pulcherrima(100); | Metschnikowia | 13 | Saprotroph | \|Undefined Saprotroph\| | Yeast | NULL | Probable | As nectar-tap_saprotroph (Põlme et al. 2020); Decay sugar-rich substrates (Põlme et al. 2020) | Põlme S, et al. 2020. Fungal Diversity 105:1-16 ((https://doi.org/10.1007/s13225-020-00466-2)) |
| Otu31009 | 1 | k__Fungi(100);p__Ascomycota(100);c__Saccharomycetes(100);o__Saccharomycetales(100);f__Metschnikowiaceae(100);g__Metschnikowia(100);s__Metschnikowia_pulcherrima(100); | Metschnikowia | 13 | Saprotroph | \|Undefined Saprotroph\| | Yeast | NULL | Probable | As nectar-tap_saprotroph (Põlme et al. 2020); Decay sugar-rich substrates (Põlme et al. 2020) | Põlme S, et al. 2020. Fungal Diversity 105:1-16 ((https://doi.org/10.1007/s13225-020-00466-2)) |
| Otu31011 | 1 | k__Fungi(100);p__Ascomycota(100);c__Saccharomycetes(100);o__Saccharomycetales(100);f__Metschnikowiaceae(100);g__Metschnikowia(100);g__Metschnikowia_unclassified(100); | Metschnikowia | 13 | Saprotroph | \|Undefined Saprotroph\| | Yeast | NULL | Probable | As nectar-tap_saprotroph (Põlme et al. 2020); Decay sugar-rich substrates (Põlme et al. 2020) | Põlme S, et al. 2020. Fungal Diversity 105:1-16 ((https://doi.org/10.1007/s13225-020-00466-2)) |
| Otu31012 | 1 | k__Fungi(100);p__Ascomycota(100);c__Saccharomycetes(100);o__Saccharomycetales(100);f__Metschnikowiaceae(100);g__Metschnikowia(100);g__Metschnikowia_unclassified(100); | Metschnikowia | 13 | Saprotroph | \|Undefined Saprotroph\| | Yeast | NULL | Probable | As nectar-tap_saprotroph (Põlme et al. 2020); Decay sugar-rich substrates (Põlme et al. 2020) | Põlme S, et al. 2020. Fungal Diversity 105:1-16 ((https://doi.org/10.1007/s13225-020-00466-2)) |
| Otu31014 | 1 | k__Fungi(100);p__Ascomycota(100);c__Saccharomycetes(100);o__Saccharomycetales(100);f__Metschnikowiaceae(100);g__Metschnikowia(100);s__Metschnikowia_pulcherrima(100); | Metschnikowia | 13 | Saprotroph | \|Undefined Saprotroph\| | Yeast | NULL | Probable | As nectar-tap_saprotroph (Põlme et al. 2020); Decay sugar-rich substrates (Põlme et al. 2020) | Põlme S, et al. 2020. Fungal Diversity 105:1-16 ((https://doi.org/10.1007/s13225-020-00466-2)) |
| Otu31015 | 1 | k__Fungi(100);p__Ascomycota(100);c__Saccharomycetes(100);o__Saccharomycetales(100);f__Metschnikowiaceae(100);g__Metschnikowia(100);g__Metschnikowia_unclassified(100); | Metschnikowia | 13 | Saprotroph | \|Undefined Saprotroph\| | Yeast | NULL | Probable | As nectar-tap_saprotroph (Põlme et al. 2020); Decay sugar-rich substrates (Põlme et al. 2020) | Põlme S, et al. 2020. Fungal Diversity 105:1-16 ((https://doi.org/10.1007/s13225-020-00466-2)) |
| Otu31016 | 1 | k__Fungi(100);p__Ascomycota(100);c__Saccharomycetes(100);o__Saccharomycetales(100);f__Metschnikowiaceae(100);g__Metschnikowia(100);s__Metschnikowia_pulcherrima(100); | Metschnikowia | 13 | Saprotroph | \|Undefined Saprotroph\| | Yeast | NULL | Probable | As nectar-tap_saprotroph (Põlme et al. 2020); Decay sugar-rich substrates (Põlme et al. 2020) | Põlme S, et al. 2020. Fungal Diversity 105:1-16 ((https://doi.org/10.1007/s13225-020-00466-2)) |
| Otu31017 | 1 | k__Fungi(100);p__Ascomycota(100);c__Saccharomycetes(100);o__Saccharomycetales(100);f__Metschnikowiaceae(100);g__Metschnikowia(100);s__Metschnikowia_pulcherrima(100); | Metschnikowia | 13 | Saprotroph | \|Undefined Saprotroph\| | Yeast | NULL | Probable | As nectar-tap_saprotroph (Põlme et al. 2020); Decay sugar-rich substrates (Põlme et al. 2020) | Põlme S, et al. 2020. Fungal Diversity 105:1-16 ((https://doi.org/10.1007/s13225-020-00466-2)) |
| Otu31018 | 1 | k__Fungi(100);p__Ascomycota(100);c__Saccharomycetes(100);o__Saccharomycetales(100);f__Metschnikowiaceae(100);g__Metschnikowia(100);s__Metschnikowia_pulcherrima(100); | Metschnikowia | 13 | Saprotroph | \|Undefined Saprotroph\| | Yeast | NULL | Probable | As nectar-tap_saprotroph (Põlme et al. 2020); Decay sugar-rich substrates (Põlme et al. 2020) | Põlme S, et al. 2020. Fungal Diversity 105:1-16 ((https://doi.org/10.1007/s13225-020-00466-2)) |
| Otu31019 | 1 | k__Fungi(100);p__Ascomycota(100);c__Saccharomycetes(100);o__Saccharomycetales(100);f__Metschnikowiaceae(100);g__Metschnikowia(100);g__Metschnikowia_unclassified(100); | Metschnikowia | 13 | Saprotroph | \|Undefined Saprotroph\| | Yeast | NULL | Probable | As nectar-tap_saprotroph (Põlme et al. 2020); Decay sugar-rich substrates (Põlme et al. 2020) | Põlme S, et al. 2020. Fungal Diversity 105:1-16 ((https://doi.org/10.1007/s13225-020-00466-2)) |
| Otu31020 | 1 | k__Fungi(100);p__Ascomycota(100);c__Saccharomycetes(100);o__Saccharomycetales(100);f__Metschnikowiaceae(100);g__Metschnikowia(100);s__Metschnikowia_pulcherrima(100); | Metschnikowia | 13 | Saprotroph | \|Undefined Saprotroph\| | Yeast | NULL | Probable | As nectar-tap_saprotroph (Põlme et al. 2020); Decay sugar-rich substrates (Põlme et al. 2020) | Põlme S, et al. 2020. Fungal Diversity 105:1-16 ((https://doi.org/10.1007/s13225-020-00466-2)) |
| Otu31021 | 1 | k__Fungi(100);p__Ascomycota(100);c__Saccharomycetes(100);o__Saccharomycetales(100);f__Metschnikowiaceae(100);g__Metschnikowia(100);s__Metschnikowia_pulcherrima(100); | Metschnikowia | 13 | Saprotroph | \|Undefined Saprotroph\| | Yeast | NULL | Probable | As nectar-tap_saprotroph (Põlme et al. 2020); Decay sugar-rich substrates (Põlme et al. 2020) | Põlme S, et al. 2020. Fungal Diversity 105:1-16 ((https://doi.org/10.1007/s13225-020-00466-2)) |
| Otu31022 | 1 | k__Fungi(100);p__Ascomycota(100);c__Saccharomycetes(100);o__Saccharomycetales(100);f__Metschnikowiaceae(100);g__Metschnikowia(100);s__Metschnikowia_pulcherrima(100); | Metschnikowia | 13 | Saprotroph | \|Undefined Saprotroph\| | Yeast | NULL | Probable | As nectar-tap_saprotroph (Põlme et al. 2020); Decay sugar-rich substrates (Põlme et al. 2020) | Põlme S, et al. 2020. Fungal Diversity 105:1-16 ((https://doi.org/10.1007/s13225-020-00466-2)) |
| Otu31023 | 1 | k__Fungi(100);p__Ascomycota(100);c__Saccharomycetes(100);o__Saccharomycetales(100);f__Metschnikowiaceae(100);g__Metschnikowia(100);s__Metschnikowia_pulcherrima(100); | Metschnikowia | 13 | Saprotroph | \|Undefined Saprotroph\| | Yeast | NULL | Probable | As nectar-tap_saprotroph (Põlme et al. 2020); Decay sugar-rich substrates (Põlme et al. 2020) | Põlme S, et al. 2020. Fungal Diversity 105:1-16 ((https://doi.org/10.1007/s13225-020-00466-2)) |
| Otu31028 | 1 | k__Fungi(100);p__Ascomycota(100);c__Saccharomycetes(100);o__Saccharomycetales(100);f__Metschnikowiaceae(100);g__Metschnikowia(100);s__Metschnikowia_pulcherrima(100); | Metschnikowia | 13 | Saprotroph | \|Undefined Saprotroph\| | Yeast | NULL | Probable | As nectar-tap_saprotroph (Põlme et al. 2020); Decay sugar-rich substrates (Põlme et al. 2020) | Põlme S, et al. 2020. Fungal Diversity 105:1-16 ((https://doi.org/10.1007/s13225-020-00466-2)) |
| Otu31029 | 1 | k__Fungi(100);p__Ascomycota(100);c__Saccharomycetes(100);o__Saccharomycetales(100);f__Metschnikowiaceae(100);g__Metschnikowia(100);g__Metschnikowia_unclassified(100); | Metschnikowia | 13 | Saprotroph | \|Undefined Saprotroph\| | Yeast | NULL | Probable | As nectar-tap_saprotroph (Põlme et al. 2020); Decay sugar-rich substrates (Põlme et al. 2020) | Põlme S, et al. 2020. Fungal Diversity 105:1-16 ((https://doi.org/10.1007/s13225-020-00466-2)) |
| Otu31030 | 1 | k__Fungi(100);p__Ascomycota(100);c__Saccharomycetes(100);o__Saccharomycetales(100);f__Metschnikowiaceae(100);g__Metschnikowia(100);s__Metschnikowia_pulcherrima(100); | Metschnikowia | 13 | Saprotroph | \|Undefined Saprotroph\| | Yeast | NULL | Probable | As nectar-tap_saprotroph (Põlme et al. 2020); Decay sugar-rich substrates (Põlme et al. 2020) | Põlme S, et al. 2020. Fungal Diversity 105:1-16 ((https://doi.org/10.1007/s13225-020-00466-2)) |
| Otu31031 | 1 | k__Fungi(100);p__Ascomycota(100);c__Saccharomycetes(100);o__Saccharomycetales(100);f__Metschnikowiaceae(100);g__Metschnikowia(100);s__Metschnikowia_pulcherrima(100); | Metschnikowia | 13 | Saprotroph | \|Undefined Saprotroph\| | Yeast | NULL | Probable | As nectar-tap_saprotroph (Põlme et al. 2020); Decay sugar-rich substrates (Põlme et al. 2020) | Põlme S, et al. 2020. Fungal Diversity 105:1-16 ((https://doi.org/10.1007/s13225-020-00466-2)) |
| Otu31033 | 1 | k__Fungi(100);p__Ascomycota(100);c__Saccharomycetes(100);o__Saccharomycetales(100);f__Metschnikowiaceae(100);g__Metschnikowia(100);g__Metschnikowia_unclassified(100); | Metschnikowia | 13 | Saprotroph | \|Undefined Saprotroph\| | Yeast | NULL | Probable | As nectar-tap_saprotroph (Põlme et al. 2020); Decay sugar-rich substrates (Põlme et al. 2020) | Põlme S, et al. 2020. Fungal Diversity 105:1-16 ((https://doi.org/10.1007/s13225-020-00466-2)) |
| Otu31034 | 1 | k__Fungi(100);p__Ascomycota(100);c__Saccharomycetes(100);o__Saccharomycetales(100);f__Metschnikowiaceae(100);g__Metschnikowia(100);s__Metschnikowia_pulcherrima(100); | Metschnikowia | 13 | Saprotroph | \|Undefined Saprotroph\| | Yeast | NULL | Probable | As nectar-tap_saprotroph (Põlme et al. 2020); Decay sugar-rich substrates (Põlme et al. 2020) | Põlme S, et al. 2020. Fungal Diversity 105:1-16 ((https://doi.org/10.1007/s13225-020-00466-2)) |
| Otu31035 | 1 | k__Fungi(100);p__Ascomycota(100);c__Saccharomycetes(100);o__Saccharomycetales(100);f__Metschnikowiaceae(100);g__Metschnikowia(100);s__Metschnikowia_pulcherrima(100); | Metschnikowia | 13 | Saprotroph | \|Undefined Saprotroph\| | Yeast | NULL | Probable | As nectar-tap_saprotroph (Põlme et al. 2020); Decay sugar-rich substrates (Põlme et al. 2020) | Põlme S, et al. 2020. Fungal Diversity 105:1-16 ((https://doi.org/10.1007/s13225-020-00466-2)) |
| Otu31036 | 1 | k__Fungi(100);p__Ascomycota(100);c__Saccharomycetes(100);o__Saccharomycetales(100);f__Metschnikowiaceae(100);g__Metschnikowia(100);g__Metschnikowia_unclassified(100); | Metschnikowia | 13 | Saprotroph | \|Undefined Saprotroph\| | Yeast | NULL | Probable | As nectar-tap_saprotroph (Põlme et al. 2020); Decay sugar-rich substrates (Põlme et al. 2020) | Põlme S, et al. 2020. Fungal Diversity 105:1-16 ((https://doi.org/10.1007/s13225-020-00466-2)) |
| Otu31037 | 1 | k__Fungi(100);p__Ascomycota(100);c__Saccharomycetes(100);o__Saccharomycetales(100);f__Metschnikowiaceae(100);g__Metschnikowia(100);g__Metschnikowia_unclassified(100); | Metschnikowia | 13 | Saprotroph | \|Undefined Saprotroph\| | Yeast | NULL | Probable | As nectar-tap_saprotroph (Põlme et al. 2020); Decay sugar-rich substrates (Põlme et al. 2020) | Põlme S, et al. 2020. Fungal Diversity 105:1-16 ((https://doi.org/10.1007/s13225-020-00466-2)) |
| Otu31038 | 1 | k__Fungi(100);p__Ascomycota(100);c__Saccharomycetes(100);o__Saccharomycetales(100);f__Metschnikowiaceae(100);g__Metschnikowia(100);s__Metschnikowia_pulcherrima(100); | Metschnikowia | 13 | Saprotroph | \|Undefined Saprotroph\| | Yeast | NULL | Probable | As nectar-tap_saprotroph (Põlme et al. 2020); Decay sugar-rich substrates (Põlme et al. 2020) | Põlme S, et al. 2020. Fungal Diversity 105:1-16 ((https://doi.org/10.1007/s13225-020-00466-2)) |
| Otu31039 | 1 | k__Fungi(100);p__Ascomycota(100);c__Saccharomycetes(100);o__Saccharomycetales(100);f__Metschnikowiaceae(100);g__Metschnikowia(100);s__Metschnikowia_pulcherrima(100); | Metschnikowia | 13 | Saprotroph | \|Undefined Saprotroph\| | Yeast | NULL | Probable | As nectar-tap_saprotroph (Põlme et al. 2020); Decay sugar-rich substrates (Põlme et al. 2020) | Põlme S, et al. 2020. Fungal Diversity 105:1-16 ((https://doi.org/10.1007/s13225-020-00466-2)) |
| Otu31040 | 1 | k__Fungi(100);p__Ascomycota(100);c__Saccharomycetes(100);o__Saccharomycetales(100);f__Metschnikowiaceae(100);g__Metschnikowia(100);s__Metschnikowia_pulcherrima(100); | Metschnikowia | 13 | Saprotroph | \|Undefined Saprotroph\| | Yeast | NULL | Probable | As nectar-tap_saprotroph (Põlme et al. 2020); Decay sugar-rich substrates (Põlme et al. 2020) | Põlme S, et al. 2020. Fungal Diversity 105:1-16 ((https://doi.org/10.1007/s13225-020-00466-2)) |
| Otu31041 | 1 | k__Fungi(100);p__Ascomycota(100);c__Saccharomycetes(100);o__Saccharomycetales(100);f__Metschnikowiaceae(100);g__Metschnikowia(100);s__Metschnikowia_pulcherrima(100); | Metschnikowia | 13 | Saprotroph | \|Undefined Saprotroph\| | Yeast | NULL | Probable | As nectar-tap_saprotroph (Põlme et al. 2020); Decay sugar-rich substrates (Põlme et al. 2020) | Põlme S, et al. 2020. Fungal Diversity 105:1-16 ((https://doi.org/10.1007/s13225-020-00466-2)) |
| Otu31042 | 1 | k__Fungi(100);p__Ascomycota(100);c__Saccharomycetes(100);o__Saccharomycetales(100);f__Metschnikowiaceae(100);g__Metschnikowia(100);g__Metschnikowia_unclassified(100); | Metschnikowia | 13 | Saprotroph | \|Undefined Saprotroph\| | Yeast | NULL | Probable | As nectar-tap_saprotroph (Põlme et al. 2020); Decay sugar-rich substrates (Põlme et al. 2020) | Põlme S, et al. 2020. Fungal Diversity 105:1-16 ((https://doi.org/10.1007/s13225-020-00466-2)) |
| Otu31043 | 1 | k__Fungi(100);p__Ascomycota(100);c__Saccharomycetes(100);o__Saccharomycetales(100);f__Metschnikowiaceae(100);g__Metschnikowia(100);g__Metschnikowia_unclassified(100); | Metschnikowia | 13 | Saprotroph | \|Undefined Saprotroph\| | Yeast | NULL | Probable | As nectar-tap_saprotroph (Põlme et al. 2020); Decay sugar-rich substrates (Põlme et al. 2020) | Põlme S, et al. 2020. Fungal Diversity 105:1-16 ((https://doi.org/10.1007/s13225-020-00466-2)) |
| Otu31044 | 1 | k__Fungi(100);p__Ascomycota(100);c__Saccharomycetes(100);o__Saccharomycetales(100);f__Metschnikowiaceae(100);g__Metschnikowia(100);s__Metschnikowia_pulcherrima(100); | Metschnikowia | 13 | Saprotroph | \|Undefined Saprotroph\| | Yeast | NULL | Probable | As nectar-tap_saprotroph (Põlme et al. 2020); Decay sugar-rich substrates (Põlme et al. 2020) | Põlme S, et al. 2020. Fungal Diversity 105:1-16 ((https://doi.org/10.1007/s13225-020-00466-2)) |
| Otu31046 | 1 | k__Fungi(100);p__Ascomycota(100);c__Saccharomycetes(100);o__Saccharomycetales(100);f__Metschnikowiaceae(100);g__Metschnikowia(100);s__Metschnikowia_pulcherrima(100); | Metschnikowia | 13 | Saprotroph | \|Undefined Saprotroph\| | Yeast | NULL | Probable | As nectar-tap_saprotroph (Põlme et al. 2020); Decay sugar-rich substrates (Põlme et al. 2020) | Põlme S, et al. 2020. Fungal Diversity 105:1-16 ((https://doi.org/10.1007/s13225-020-00466-2)) |
| Otu31047 | 1 | k__Fungi(100);p__Ascomycota(100);c__Saccharomycetes(100);o__Saccharomycetales(100);f__Metschnikowiaceae(100);g__Metschnikowia(100);s__Metschnikowia_pulcherrima(100); | Metschnikowia | 13 | Saprotroph | \|Undefined Saprotroph\| | Yeast | NULL | Probable | As nectar-tap_saprotroph (Põlme et al. 2020); Decay sugar-rich substrates (Põlme et al. 2020) | Põlme S, et al. 2020. Fungal Diversity 105:1-16 ((https://doi.org/10.1007/s13225-020-00466-2)) |
| Otu31048 | 1 | k__Fungi(100);p__Ascomycota(100);c__Saccharomycetes(100);o__Saccharomycetales(100);f__Metschnikowiaceae(100);g__Metschnikowia(100);s__Metschnikowia_pulcherrima(100); | Metschnikowia | 13 | Saprotroph | \|Undefined Saprotroph\| | Yeast | NULL | Probable | As nectar-tap_saprotroph (Põlme et al. 2020); Decay sugar-rich substrates (Põlme et al. 2020) | Põlme S, et al. 2020. Fungal Diversity 105:1-16 ((https://doi.org/10.1007/s13225-020-00466-2)) |
| Otu31049 | 1 | k__Fungi(100);p__Ascomycota(100);c__Saccharomycetes(100);o__Saccharomycetales(100);f__Metschnikowiaceae(100);g__Metschnikowia(100);s__Metschnikowia_pulcherrima(100); | Metschnikowia | 13 | Saprotroph | \|Undefined Saprotroph\| | Yeast | NULL | Probable | As nectar-tap_saprotroph (Põlme et al. 2020); Decay sugar-rich substrates (Põlme et al. 2020) | Põlme S, et al. 2020. Fungal Diversity 105:1-16 ((https://doi.org/10.1007/s13225-020-00466-2)) |
| Otu31050 | 1 | k__Fungi(100);p__Ascomycota(100);c__Saccharomycetes(100);o__Saccharomycetales(100);f__Metschnikowiaceae(100);g__Metschnikowia(100);s__Metschnikowia_pulcherrima(100); | Metschnikowia | 13 | Saprotroph | \|Undefined Saprotroph\| | Yeast | NULL | Probable | As nectar-tap_saprotroph (Põlme et al. 2020); Decay sugar-rich substrates (Põlme et al. 2020) | Põlme S, et al. 2020. Fungal Diversity 105:1-16 ((https://doi.org/10.1007/s13225-020-00466-2)) |
| Otu31051 | 1 | k__Fungi(100);p__Ascomycota(100);c__Saccharomycetes(100);o__Saccharomycetales(100);f__Metschnikowiaceae(100);g__Metschnikowia(100);s__Metschnikowia_pulcherrima(100); | Metschnikowia | 13 | Saprotroph | \|Undefined Saprotroph\| | Yeast | NULL | Probable | As nectar-tap_saprotroph (Põlme et al. 2020); Decay sugar-rich substrates (Põlme et al. 2020) | Põlme S, et al. 2020. Fungal Diversity 105:1-16 ((https://doi.org/10.1007/s13225-020-00466-2)) |
| Otu31052 | 1 | k__Fungi(100);p__Ascomycota(100);c__Saccharomycetes(100);o__Saccharomycetales(100);f__Metschnikowiaceae(100);g__Metschnikowia(100);s__Metschnikowia_pulcherrima(100); | Metschnikowia | 13 | Saprotroph | \|Undefined Saprotroph\| | Yeast | NULL | Probable | As nectar-tap_saprotroph (Põlme et al. 2020); Decay sugar-rich substrates (Põlme et al. 2020) | Põlme S, et al. 2020. Fungal Diversity 105:1-16 ((https://doi.org/10.1007/s13225-020-00466-2)) |
| Otu31053 | 1 | k__Fungi(100);p__Ascomycota(100);c__Saccharomycetes(100);o__Saccharomycetales(100);f__Metschnikowiaceae(100);g__Metschnikowia(100);s__Metschnikowia_pulcherrima(100); | Metschnikowia | 13 | Saprotroph | \|Undefined Saprotroph\| | Yeast | NULL | Probable | As nectar-tap_saprotroph (Põlme et al. 2020); Decay sugar-rich substrates (Põlme et al. 2020) | Põlme S, et al. 2020. Fungal Diversity 105:1-16 ((https://doi.org/10.1007/s13225-020-00466-2)) |
| Otu31054 | 1 | k__Fungi(100);p__Ascomycota(100);c__Saccharomycetes(100);o__Saccharomycetales(100);f__Metschnikowiaceae(100);g__Metschnikowia(100);s__Metschnikowia_pulcherrima(100); | Metschnikowia | 13 | Saprotroph | \|Undefined Saprotroph\| | Yeast | NULL | Probable | As nectar-tap_saprotroph (Põlme et al. 2020); Decay sugar-rich substrates (Põlme et al. 2020) | Põlme S, et al. 2020. Fungal Diversity 105:1-16 ((https://doi.org/10.1007/s13225-020-00466-2)) |
| Otu31055 | 1 | k__Fungi(100);p__Ascomycota(100);c__Saccharomycetes(100);o__Saccharomycetales(100);f__Metschnikowiaceae(100);g__Metschnikowia(100);s__Metschnikowia_pulcherrima(100); | Metschnikowia | 13 | Saprotroph | \|Undefined Saprotroph\| | Yeast | NULL | Probable | As nectar-tap_saprotroph (Põlme et al. 2020); Decay sugar-rich substrates (Põlme et al. 2020) | Põlme S, et al. 2020. Fungal Diversity 105:1-16 ((https://doi.org/10.1007/s13225-020-00466-2)) |
| Otu31056 | 1 | k__Fungi(100);p__Ascomycota(100);c__Saccharomycetes(100);o__Saccharomycetales(100);f__Metschnikowiaceae(100);g__Metschnikowia(100);s__Metschnikowia_pulcherrima(100); | Metschnikowia | 13 | Saprotroph | \|Undefined Saprotroph\| | Yeast | NULL | Probable | As nectar-tap_saprotroph (Põlme et al. 2020); Decay sugar-rich substrates (Põlme et al. 2020) | Põlme S, et al. 2020. Fungal Diversity 105:1-16 ((https://doi.org/10.1007/s13225-020-00466-2)) |
| Otu31057 | 1 | k__Fungi(100);p__Ascomycota(100);c__Saccharomycetes(100);o__Saccharomycetales(100);f__Metschnikowiaceae(100);g__Metschnikowia(100);s__Metschnikowia_pulcherrima(100); | Metschnikowia | 13 | Saprotroph | \|Undefined Saprotroph\| | Yeast | NULL | Probable | As nectar-tap_saprotroph (Põlme et al. 2020); Decay sugar-rich substrates (Põlme et al. 2020) | Põlme S, et al. 2020. Fungal Diversity 105:1-16 ((https://doi.org/10.1007/s13225-020-00466-2)) |
| Otu31058 | 1 | k__Fungi(100);p__Ascomycota(100);c__Saccharomycetes(100);o__Saccharomycetales(100);f__Metschnikowiaceae(100);g__Metschnikowia(100);g__Metschnikowia_unclassified(100); | Metschnikowia | 13 | Saprotroph | \|Undefined Saprotroph\| | Yeast | NULL | Probable | As nectar-tap_saprotroph (Põlme et al. 2020); Decay sugar-rich substrates (Põlme et al. 2020) | Põlme S, et al. 2020. Fungal Diversity 105:1-16 ((https://doi.org/10.1007/s13225-020-00466-2)) |
| Otu31059 | 1 | k__Fungi(100);p__Ascomycota(100);c__Saccharomycetes(100);o__Saccharomycetales(100);f__Metschnikowiaceae(100);g__Metschnikowia(100);s__Metschnikowia_pulcherrima(100); | Metschnikowia | 13 | Saprotroph | \|Undefined Saprotroph\| | Yeast | NULL | Probable | As nectar-tap_saprotroph (Põlme et al. 2020); Decay sugar-rich substrates (Põlme et al. 2020) | Põlme S, et al. 2020. Fungal Diversity 105:1-16 ((https://doi.org/10.1007/s13225-020-00466-2)) |
| Otu31060 | 1 | k__Fungi(100);p__Ascomycota(100);c__Saccharomycetes(100);o__Saccharomycetales(100);f__Metschnikowiaceae(100);g__Metschnikowia(100);g__Metschnikowia_unclassified(100); | Metschnikowia | 13 | Saprotroph | \|Undefined Saprotroph\| | Yeast | NULL | Probable | As nectar-tap_saprotroph (Põlme et al. 2020); Decay sugar-rich substrates (Põlme et al. 2020) | Põlme S, et al. 2020. Fungal Diversity 105:1-16 ((https://doi.org/10.1007/s13225-020-00466-2)) |
| Otu31061 | 1 | k__Fungi(100);p__Ascomycota(100);c__Saccharomycetes(100);o__Saccharomycetales(100);f__Metschnikowiaceae(100);g__Metschnikowia(100);s__Metschnikowia_pulcherrima(100); | Metschnikowia | 13 | Saprotroph | \|Undefined Saprotroph\| | Yeast | NULL | Probable | As nectar-tap_saprotroph (Põlme et al. 2020); Decay sugar-rich substrates (Põlme et al. 2020) | Põlme S, et al. 2020. Fungal Diversity 105:1-16 ((https://doi.org/10.1007/s13225-020-00466-2)) |
| Otu31062 | 1 | k__Fungi(100);p__Ascomycota(100);c__Saccharomycetes(100);o__Saccharomycetales(100);f__Metschnikowiaceae(100);g__Metschnikowia(100);s__Metschnikowia_pulcherrima(100); | Metschnikowia | 13 | Saprotroph | \|Undefined Saprotroph\| | Yeast | NULL | Probable | As nectar-tap_saprotroph (Põlme et al. 2020); Decay sugar-rich substrates (Põlme et al. 2020) | Põlme S, et al. 2020. Fungal Diversity 105:1-16 ((https://doi.org/10.1007/s13225-020-00466-2)) |
| Otu31063 | 1 | k__Fungi(100);p__Ascomycota(100);c__Saccharomycetes(100);o__Saccharomycetales(100);f__Metschnikowiaceae(100);g__Metschnikowia(100);s__Metschnikowia_pulcherrima(100); | Metschnikowia | 13 | Saprotroph | \|Undefined Saprotroph\| | Yeast | NULL | Probable | As nectar-tap_saprotroph (Põlme et al. 2020); Decay sugar-rich substrates (Põlme et al. 2020) | Põlme S, et al. 2020. Fungal Diversity 105:1-16 ((https://doi.org/10.1007/s13225-020-00466-2)) |
| Otu31066 | 1 | k__Fungi(100);p__Ascomycota(100);c__Saccharomycetes(100);o__Saccharomycetales(100);f__Metschnikowiaceae(100);g__Metschnikowia(100);s__Metschnikowia_pulcherrima(100); | Metschnikowia | 13 | Saprotroph | \|Undefined Saprotroph\| | Yeast | NULL | Probable | As nectar-tap_saprotroph (Põlme et al. 2020); Decay sugar-rich substrates (Põlme et al. 2020) | Põlme S, et al. 2020. Fungal Diversity 105:1-16 ((https://doi.org/10.1007/s13225-020-00466-2)) |
| Otu31068 | 1 | k__Fungi(100);p__Ascomycota(100);c__Saccharomycetes(100);o__Saccharomycetales(100);f__Metschnikowiaceae(100);g__Metschnikowia(100);s__Metschnikowia_pulcherrima(100); | Metschnikowia | 13 | Saprotroph | \|Undefined Saprotroph\| | Yeast | NULL | Probable | As nectar-tap_saprotroph (Põlme et al. 2020); Decay sugar-rich substrates (Põlme et al. 2020) | Põlme S, et al. 2020. Fungal Diversity 105:1-16 ((https://doi.org/10.1007/s13225-020-00466-2)) |
| Otu31069 | 1 | k__Fungi(100);p__Ascomycota(100);c__Saccharomycetes(100);o__Saccharomycetales(100);f__Metschnikowiaceae(100);g__Metschnikowia(100);g__Metschnikowia_unclassified(100); | Metschnikowia | 13 | Saprotroph | \|Undefined Saprotroph\| | Yeast | NULL | Probable | As nectar-tap_saprotroph (Põlme et al. 2020); Decay sugar-rich substrates (Põlme et al. 2020) | Põlme S, et al. 2020. Fungal Diversity 105:1-16 ((https://doi.org/10.1007/s13225-020-00466-2)) |
| Otu31077 | 1 | k__Fungi(100);p__Ascomycota(100);c__Saccharomycetes(100);o__Saccharomycetales(100);f__Metschnikowiaceae(100);g__Metschnikowia(100);s__Metschnikowia_pulcherrima(100); | Metschnikowia | 13 | Saprotroph | \|Undefined Saprotroph\| | Yeast | NULL | Probable | As nectar-tap_saprotroph (Põlme et al. 2020); Decay sugar-rich substrates (Põlme et al. 2020) | Põlme S, et al. 2020. Fungal Diversity 105:1-16 ((https://doi.org/10.1007/s13225-020-00466-2)) |
| Otu31079 | 1 | k__Fungi(100);p__Ascomycota(100);c__Saccharomycetes(100);o__Saccharomycetales(100);f__Metschnikowiaceae(100);g__Metschnikowia(100);g__Metschnikowia_unclassified(100); | Metschnikowia | 13 | Saprotroph | \|Undefined Saprotroph\| | Yeast | NULL | Probable | As nectar-tap_saprotroph (Põlme et al. 2020); Decay sugar-rich substrates (Põlme et al. 2020) | Põlme S, et al. 2020. Fungal Diversity 105:1-16 ((https://doi.org/10.1007/s13225-020-00466-2)) |
| Otu31080 | 1 | k__Fungi(100);p__Ascomycota(100);c__Saccharomycetes(100);o__Saccharomycetales(100);f__Metschnikowiaceae(100);g__Metschnikowia(100);g__Metschnikowia_unclassified(100); | Metschnikowia | 13 | Saprotroph | \|Undefined Saprotroph\| | Yeast | NULL | Probable | As nectar-tap_saprotroph (Põlme et al. 2020); Decay sugar-rich substrates (Põlme et al. 2020) | Põlme S, et al. 2020. Fungal Diversity 105:1-16 ((https://doi.org/10.1007/s13225-020-00466-2)) |
| Otu31082 | 1 | k__Fungi(100);p__Ascomycota(100);c__Saccharomycetes(100);o__Saccharomycetales(100);f__Metschnikowiaceae(100);g__Metschnikowia(100);g__Metschnikowia_unclassified(100); | Metschnikowia | 13 | Saprotroph | \|Undefined Saprotroph\| | Yeast | NULL | Probable | As nectar-tap_saprotroph (Põlme et al. 2020); Decay sugar-rich substrates (Põlme et al. 2020) | Põlme S, et al. 2020. Fungal Diversity 105:1-16 ((https://doi.org/10.1007/s13225-020-00466-2)) |
| Otu31083 | 1 | k__Fungi(100);p__Ascomycota(100);c__Saccharomycetes(100);o__Saccharomycetales(100);f__Metschnikowiaceae(100);g__Metschnikowia(100);s__Metschnikowia_pulcherrima(100); | Metschnikowia | 13 | Saprotroph | \|Undefined Saprotroph\| | Yeast | NULL | Probable | As nectar-tap_saprotroph (Põlme et al. 2020); Decay sugar-rich substrates (Põlme et al. 2020) | Põlme S, et al. 2020. Fungal Diversity 105:1-16 ((https://doi.org/10.1007/s13225-020-00466-2)) |
| Otu31084 | 1 | k__Fungi(100);p__Ascomycota(100);c__Saccharomycetes(100);o__Saccharomycetales(100);f__Metschnikowiaceae(100);g__Metschnikowia(100);s__Metschnikowia_pulcherrima(100); | Metschnikowia | 13 | Saprotroph | \|Undefined Saprotroph\| | Yeast | NULL | Probable | As nectar-tap_saprotroph (Põlme et al. 2020); Decay sugar-rich substrates (Põlme et al. 2020) | Põlme S, et al. 2020. Fungal Diversity 105:1-16 ((https://doi.org/10.1007/s13225-020-00466-2)) |
| Otu31085 | 1 | k__Fungi(100);p__Ascomycota(100);c__Saccharomycetes(100);o__Saccharomycetales(100);f__Metschnikowiaceae(100);g__Metschnikowia(100);s__Metschnikowia_pulcherrima(100); | Metschnikowia | 13 | Saprotroph | \|Undefined Saprotroph\| | Yeast | NULL | Probable | As nectar-tap_saprotroph (Põlme et al. 2020); Decay sugar-rich substrates (Põlme et al. 2020) | Põlme S, et al. 2020. Fungal Diversity 105:1-16 ((https://doi.org/10.1007/s13225-020-00466-2)) |
| Otu31086 | 1 | k__Fungi(100);p__Ascomycota(100);c__Saccharomycetes(100);o__Saccharomycetales(100);f__Metschnikowiaceae(100);g__Metschnikowia(100);g__Metschnikowia_unclassified(100); | Metschnikowia | 13 | Saprotroph | \|Undefined Saprotroph\| | Yeast | NULL | Probable | As nectar-tap_saprotroph (Põlme et al. 2020); Decay sugar-rich substrates (Põlme et al. 2020) | Põlme S, et al. 2020. Fungal Diversity 105:1-16 ((https://doi.org/10.1007/s13225-020-00466-2)) |
| Otu31087 | 1 | k__Fungi(100);p__Ascomycota(100);c__Saccharomycetes(100);o__Saccharomycetales(100);f__Metschnikowiaceae(100);g__Metschnikowia(100);s__Metschnikowia_pulcherrima(100); | Metschnikowia | 13 | Saprotroph | \|Undefined Saprotroph\| | Yeast | NULL | Probable | As nectar-tap_saprotroph (Põlme et al. 2020); Decay sugar-rich substrates (Põlme et al. 2020) | Põlme S, et al. 2020. Fungal Diversity 105:1-16 ((https://doi.org/10.1007/s13225-020-00466-2)) |
| Otu31091 | 1 | k__Fungi(100);p__Ascomycota(100);c__Saccharomycetes(100);o__Saccharomycetales(100);f__Metschnikowiaceae(100);g__Metschnikowia(100);g__Metschnikowia_unclassified(100); | Metschnikowia | 13 | Saprotroph | \|Undefined Saprotroph\| | Yeast | NULL | Probable | As nectar-tap_saprotroph (Põlme et al. 2020); Decay sugar-rich substrates (Põlme et al. 2020) | Põlme S, et al. 2020. Fungal Diversity 105:1-16 ((https://doi.org/10.1007/s13225-020-00466-2)) |
| Otu31092 | 1 | k__Fungi(100);p__Ascomycota(100);c__Saccharomycetes(100);o__Saccharomycetales(100);f__Metschnikowiaceae(100);g__Metschnikowia(100);s__Metschnikowia_pulcherrima(100); | Metschnikowia | 13 | Saprotroph | \|Undefined Saprotroph\| | Yeast | NULL | Probable | As nectar-tap_saprotroph (Põlme et al. 2020); Decay sugar-rich substrates (Põlme et al. 2020) | Põlme S, et al. 2020. Fungal Diversity 105:1-16 ((https://doi.org/10.1007/s13225-020-00466-2)) |
| Otu31093 | 1 | k__Fungi(100);p__Ascomycota(100);c__Saccharomycetes(100);o__Saccharomycetales(100);f__Metschnikowiaceae(100);g__Metschnikowia(100);g__Metschnikowia_unclassified(100); | Metschnikowia | 13 | Saprotroph | \|Undefined Saprotroph\| | Yeast | NULL | Probable | As nectar-tap_saprotroph (Põlme et al. 2020); Decay sugar-rich substrates (Põlme et al. 2020) | Põlme S, et al. 2020. Fungal Diversity 105:1-16 ((https://doi.org/10.1007/s13225-020-00466-2)) |
| Otu31094 | 1 | k__Fungi(100);p__Ascomycota(100);c__Saccharomycetes(100);o__Saccharomycetales(100);f__Metschnikowiaceae(100);g__Metschnikowia(100);s__Metschnikowia_pulcherrima(100); | Metschnikowia | 13 | Saprotroph | \|Undefined Saprotroph\| | Yeast | NULL | Probable | As nectar-tap_saprotroph (Põlme et al. 2020); Decay sugar-rich substrates (Põlme et al. 2020) | Põlme S, et al. 2020. Fungal Diversity 105:1-16 ((https://doi.org/10.1007/s13225-020-00466-2)) |
| Otu31095 | 1 | k__Fungi(100);p__Ascomycota(100);c__Saccharomycetes(100);o__Saccharomycetales(100);f__Metschnikowiaceae(100);g__Metschnikowia(100);s__Metschnikowia_pulcherrima(100); | Metschnikowia | 13 | Saprotroph | \|Undefined Saprotroph\| | Yeast | NULL | Probable | As nectar-tap_saprotroph (Põlme et al. 2020); Decay sugar-rich substrates (Põlme et al. 2020) | Põlme S, et al. 2020. Fungal Diversity 105:1-16 ((https://doi.org/10.1007/s13225-020-00466-2)) |
| Otu31099 | 1 | k__Fungi(100);p__Ascomycota(100);c__Saccharomycetes(100);o__Saccharomycetales(100);f__Metschnikowiaceae(100);g__Metschnikowia(100);s__Metschnikowia_pulcherrima(100); | Metschnikowia | 13 | Saprotroph | \|Undefined Saprotroph\| | Yeast | NULL | Probable | As nectar-tap_saprotroph (Põlme et al. 2020); Decay sugar-rich substrates (Põlme et al. 2020) | Põlme S, et al. 2020. Fungal Diversity 105:1-16 ((https://doi.org/10.1007/s13225-020-00466-2)) |
| Otu31100 | 1 | k__Fungi(100);p__Ascomycota(100);c__Saccharomycetes(100);o__Saccharomycetales(100);f__Metschnikowiaceae(100);g__Metschnikowia(100);s__Metschnikowia_pulcherrima(100); | Metschnikowia | 13 | Saprotroph | \|Undefined Saprotroph\| | Yeast | NULL | Probable | As nectar-tap_saprotroph (Põlme et al. 2020); Decay sugar-rich substrates (Põlme et al. 2020) | Põlme S, et al. 2020. Fungal Diversity 105:1-16 ((https://doi.org/10.1007/s13225-020-00466-2)) |
| Otu31104 | 1 | k__Fungi(100);p__Ascomycota(100);c__Saccharomycetes(100);o__Saccharomycetales(100);f__Metschnikowiaceae(100);g__Metschnikowia(100);s__Metschnikowia_pulcherrima(100); | Metschnikowia | 13 | Saprotroph | \|Undefined Saprotroph\| | Yeast | NULL | Probable | As nectar-tap_saprotroph (Põlme et al. 2020); Decay sugar-rich substrates (Põlme et al. 2020) | Põlme S, et al. 2020. Fungal Diversity 105:1-16 ((https://doi.org/10.1007/s13225-020-00466-2)) |
| Otu31110 | 1 | k__Fungi(100);p__Ascomycota(100);c__Saccharomycetes(100);o__Saccharomycetales(100);f__Metschnikowiaceae(100);g__Metschnikowia(100);s__Metschnikowia_pulcherrima(100); | Metschnikowia | 13 | Saprotroph | \|Undefined Saprotroph\| | Yeast | NULL | Probable | As nectar-tap_saprotroph (Põlme et al. 2020); Decay sugar-rich substrates (Põlme et al. 2020) | Põlme S, et al. 2020. Fungal Diversity 105:1-16 ((https://doi.org/10.1007/s13225-020-00466-2)) |
| Otu31116 | 1 | k__Fungi(100);p__Ascomycota(100);c__Saccharomycetes(100);o__Saccharomycetales(100);f__Metschnikowiaceae(100);g__Metschnikowia(100);s__Metschnikowia_pulcherrima(100); | Metschnikowia | 13 | Saprotroph | \|Undefined Saprotroph\| | Yeast | NULL | Probable | As nectar-tap_saprotroph (Põlme et al. 2020); Decay sugar-rich substrates (Põlme et al. 2020) | Põlme S, et al. 2020. Fungal Diversity 105:1-16 ((https://doi.org/10.1007/s13225-020-00466-2)) |
| Otu31117 | 1 | k__Fungi(100);p__Ascomycota(100);c__Saccharomycetes(100);o__Saccharomycetales(100);f__Metschnikowiaceae(100);g__Metschnikowia(100);s__Metschnikowia_pulcherrima(100); | Metschnikowia | 13 | Saprotroph | \|Undefined Saprotroph\| | Yeast | NULL | Probable | As nectar-tap_saprotroph (Põlme et al. 2020); Decay sugar-rich substrates (Põlme et al. 2020) | Põlme S, et al. 2020. Fungal Diversity 105:1-16 ((https://doi.org/10.1007/s13225-020-00466-2)) |
| Otu31118 | 1 | k__Fungi(100);p__Ascomycota(100);c__Saccharomycetes(100);o__Saccharomycetales(100);f__Metschnikowiaceae(100);g__Metschnikowia(100);g__Metschnikowia_unclassified(100); | Metschnikowia | 13 | Saprotroph | \|Undefined Saprotroph\| | Yeast | NULL | Probable | As nectar-tap_saprotroph (Põlme et al. 2020); Decay sugar-rich substrates (Põlme et al. 2020) | Põlme S, et al. 2020. Fungal Diversity 105:1-16 ((https://doi.org/10.1007/s13225-020-00466-2)) |
| Otu31119 | 1 | k__Fungi(100);p__Ascomycota(100);c__Saccharomycetes(100);o__Saccharomycetales(100);f__Metschnikowiaceae(100);g__Metschnikowia(100);s__Metschnikowia_pulcherrima(100); | Metschnikowia | 13 | Saprotroph | \|Undefined Saprotroph\| | Yeast | NULL | Probable | As nectar-tap_saprotroph (Põlme et al. 2020); Decay sugar-rich substrates (Põlme et al. 2020) | Põlme S, et al. 2020. Fungal Diversity 105:1-16 ((https://doi.org/10.1007/s13225-020-00466-2)) |
| Otu31120 | 1 | k__Fungi(100);p__Ascomycota(100);c__Saccharomycetes(100);o__Saccharomycetales(100);f__Metschnikowiaceae(100);g__Metschnikowia(100);s__Metschnikowia_pulcherrima(100); | Metschnikowia | 13 | Saprotroph | \|Undefined Saprotroph\| | Yeast | NULL | Probable | As nectar-tap_saprotroph (Põlme et al. 2020); Decay sugar-rich substrates (Põlme et al. 2020) | Põlme S, et al. 2020. Fungal Diversity 105:1-16 ((https://doi.org/10.1007/s13225-020-00466-2)) |
| Otu31121 | 1 | k__Fungi(100);p__Ascomycota(100);c__Saccharomycetes(100);o__Saccharomycetales(100);f__Metschnikowiaceae(100);g__Metschnikowia(100);s__Metschnikowia_pulcherrima(100); | Metschnikowia | 13 | Saprotroph | \|Undefined Saprotroph\| | Yeast | NULL | Probable | As nectar-tap_saprotroph (Põlme et al. 2020); Decay sugar-rich substrates (Põlme et al. 2020) | Põlme S, et al. 2020. Fungal Diversity 105:1-16 ((https://doi.org/10.1007/s13225-020-00466-2)) |

| | | | | | | | | | | | | |
|---|---|---|---|---|---|---|---|---|---|---|---|---|
| Otu31403 | | 1 | k__Fungi(100);p__Ascomycota(100);c__Saccharomycetes(100);o__Saccharomycetales(100);f__Metschnikowiaceae(100);g__Metschnikowia(100);g__Metschnikowia_unclassified(100); | Metschnikowia | 13 | Saprotroph | \|Undefined Saprotroph\| | Yeast | NULL | Probable | As nectar-tap_saprotroph (Põlme et al. 2020); Decay sugar-rich substrates (Põlme et al. 2020) | Põlme S, et al. 2020. Fungal Diversity 105:1-16 ((https://doi.org/10.1007/s13225-020-00466-2)) |
| Otu31404 | | 1 | k__Fungi(100);p__Ascomycota(100);c__Saccharomycetes(100);o__Saccharomycetales(100);f__Metschnikowiaceae(100);g__Metschnikowia(100);g__Metschnikowia_unclassified(100); | Metschnikowia | 13 | Saprotroph | \|Undefined Saprotroph\| | Yeast | NULL | Probable | As nectar-tap_saprotroph (Põlme et al. 2020); Decay sugar-rich substrates (Põlme et al. 2020) | Põlme S, et al. 2020. Fungal Diversity 105:1-16 ((https://doi.org/10.1007/s13225-020-00466-2)) |
| Otu31405 | | 1 | k__Fungi(100);p__Ascomycota(100);c__Saccharomycetes(100);o__Saccharomycetales(100);f__Metschnikowiaceae(100);g__Metschnikowia(100);g__Metschnikowia_unclassified(100); | Metschnikowia | 13 | Saprotroph | \|Undefined Saprotroph\| | Yeast | NULL | Probable | As nectar-tap_saprotroph (Põlme et al. 2020); Decay sugar-rich substrates (Põlme et al. 2020) | Põlme S, et al. 2020. Fungal Diversity 105:1-16 ((https://doi.org/10.1007/s13225-020-00466-2)) |
| Otu31406 | | 1 | k__Fungi(100);p__Ascomycota(100);c__Saccharomycetes(100);o__Saccharomycetales(100);f__Metschnikowiaceae(100);g__Metschnikowia(100);s__Metschnikowia_pulcherrima(100); | Metschnikowia | 13 | Saprotroph | \|Undefined Saprotroph\| | Yeast | NULL | Probable | As nectar-tap_saprotroph (Põlme et al. 2020); Decay sugar-rich substrates (Põlme et al. 2020) | Põlme S, et al. 2020. Fungal Diversity 105:1-16 ((https://doi.org/10.1007/s13225-020-00466-2)) |
| Otu31407 | | 1 | k__Fungi(100);p__Ascomycota(100);c__Saccharomycetes(100);o__Saccharomycetales(100);f__Metschnikowiaceae(100);g__Metschnikowia(100);s__Metschnikowia_pulcherrima(100); | Metschnikowia | 13 | Saprotroph | \|Undefined Saprotroph\| | Yeast | NULL | Probable | As nectar-tap_saprotroph (Põlme et al. 2020); Decay sugar-rich substrates (Põlme et al. 2020) | Põlme S, et al. 2020. Fungal Diversity 105:1-16 ((https://doi.org/10.1007/s13225-020-00466-2)) |
| Otu31408 | | 1 | k__Fungi(100);p__Ascomycota(100);c__Saccharomycetes(100);o__Saccharomycetales(100);f__Metschnikowiaceae(100);g__Metschnikowia(100);s__Metschnikowia_pulcherrima(100); | Metschnikowia | 13 | Saprotroph | \|Undefined Saprotroph\| | Yeast | NULL | Probable | As nectar-tap_saprotroph (Põlme et al. 2020); Decay sugar-rich substrates (Põlme et al. 2020) | Põlme S, et al. 2020. Fungal Diversity 105:1-16 ((https://doi.org/10.1007/s13225-020-00466-2)) |
| Otu31409 | | 1 | k__Fungi(100);p__Ascomycota(100);c__Saccharomycetes(100);o__Saccharomycetales(100);f__Metschnikowiaceae(100);g__Metschnikowia(100);g__Metschnikowia_unclassified(100); | Metschnikowia | 13 | Saprotroph | \|Undefined Saprotroph\| | Yeast | NULL | Probable | As nectar-tap_saprotroph (Põlme et al. 2020); Decay sugar-rich substrates (Põlme et al. 2020) | Põlme S, et al. 2020. Fungal Diversity 105:1-16 ((https://doi.org/10.1007/s13225-020-00466-2)) |
| Otu31411 | | 1 | k__Fungi(100);p__Ascomycota(100);c__Saccharomycetes(100);o__Saccharomycetales(100);f__Metschnikowiaceae(100);g__Metschnikowia(100);g__Metschnikowia_unclassified(100); | Metschnikowia | 13 | Saprotroph | \|Undefined Saprotroph\| | Yeast | NULL | Probable | As nectar-tap_saprotroph (Põlme et al. 2020); Decay sugar-rich substrates (Põlme et al. 2020) | Põlme S, et al. 2020. Fungal Diversity 105:1-16 ((https://doi.org/10.1007/s13225-020-00466-2)) |
| Otu31412 | | 1 | k__Fungi(100);p__Ascomycota(100);c__Saccharomycetes(100);o__Saccharomycetales(100);f__Metschnikowiaceae(100);g__Metschnikowia(100);s__Metschnikowia_pulcherrima(100); | Metschnikowia | 13 | Saprotroph | \|Undefined Saprotroph\| | Yeast | NULL | Probable | As nectar-tap_saprotroph (Põlme et al. 2020); Decay sugar-rich substrates (Põlme et al. 2020) | Põlme S, et al. 2020. Fungal Diversity 105:1-16 ((https://doi.org/10.1007/s13225-020-00466-2)) |
| Otu31413 | | 1 | k__Fungi(100);p__Ascomycota(100);c__Saccharomycetes(100);o__Saccharomycetales(100);f__Metschnikowiaceae(100);g__Metschnikowia(100);s__Metschnikowia_pulcherrima(100); | Metschnikowia | 13 | Saprotroph | \|Undefined Saprotroph\| | Yeast | NULL | Probable | As nectar-tap_saprotroph (Põlme et al. 2020); Decay sugar-rich substrates (Põlme et al. 2020) | Põlme S, et al. 2020. Fungal Diversity 105:1-16 ((https://doi.org/10.1007/s13225-020-00466-2)) |
| Otu31414 | | 1 | k__Fungi(100);p__Ascomycota(100);c__Saccharomycetes(100);o__Saccharomycetales(100);f__Metschnikowiaceae(100);g__Metschnikowia(100);s__Metschnikowia_pulcherrima(100); | Metschnikowia | 13 | Saprotroph | \|Undefined Saprotroph\| | Yeast | NULL | Probable | As nectar-tap_saprotroph (Põlme et al. 2020); Decay sugar-rich substrates (Põlme et al. 2020) | Põlme S, et al. 2020. Fungal Diversity 105:1-16 ((https://doi.org/10.1007/s13225-020-00466-2)) |
| Otu31415 | | 1 | k__Fungi(100);p__Ascomycota(100);c__Saccharomycetes(100);o__Saccharomycetales(100);f__Metschnikowiaceae(100);g__Metschnikowia(100);s__Metschnikowia_pulcherrima(100); | Metschnikowia | 13 | Saprotroph | \|Undefined Saprotroph\| | Yeast | NULL | Probable | As nectar-tap_saprotroph (Põlme et al. 2020); Decay sugar-rich substrates (Põlme et al. 2020) | Põlme S, et al. 2020. Fungal Diversity 105:1-16 ((https://doi.org/10.1007/s13225-020-00466-2)) |
| Otu31416 | | 1 | k__Fungi(100);p__Ascomycota(100);c__Saccharomycetes(100);o__Saccharomycetales(100);f__Metschnikowiaceae(100);g__Metschnikowia(100);s__Metschnikowia_pulcherrima(100); | Metschnikowia | 13 | Saprotroph | \|Undefined Saprotroph\| | Yeast | NULL | Probable | As nectar-tap_saprotroph (Põlme et al. 2020); Decay sugar-rich substrates (Põlme et al. 2020) | Põlme S, et al. 2020. Fungal Diversity 105:1-16 ((https://doi.org/10.1007/s13225-020-00466-2)) |
| Otu31419 | | 1 | k__Fungi(100);p__Ascomycota(100);c__Saccharomycetes(100);o__Saccharomycetales(100);f__Metschnikowiaceae(100);g__Metschnikowia(100);g__Metschnikowia_unclassified(100); | Metschnikowia | 13 | Saprotroph | \|Undefined Saprotroph\| | Yeast | NULL | Probable | As nectar-tap_saprotroph (Põlme et al. 2020); Decay sugar-rich substrates (Põlme et al. 2020) | Põlme S, et al. 2020. Fungal Diversity 105:1-16 ((https://doi.org/10.1007/s13225-020-00466-2)) |
| Otu31420 | | 1 | k__Fungi(100);p__Ascomycota(100);c__Dothideomycetes(100);o__Pleosporales(100);f__Venturiaceae(100);g__Venturia(100);g__Venturia_unclassified(100); | Venturia | 13 | Pathotroph-Pathotroph-Saprotroph | \|Plant Pathogen\|-Plant Saprotroph | Microfungus | NULL | Probable | As litter_saprotroph (Põlme et al. 2020); Decay leaf, fruit, seed (Põlme et al. 2020) | Tedersoo L, et al. 2014. Science 346:e1256688 ((https://doi.org/10.1126/science.1256688)); Põlme S, et al. 2020. Fungal Diversity 105:1-16 ((https://doi.org/10.1007/s13225-020-00466-2)) |
| Otu31421 | | 1 | k__Fungi(100);p__Ascomycota(100);c__Saccharomycetes(100);o__Saccharomycetales(100);f__Metschnikowiaceae(100);g__Metschnikowia(100);g__Metschnikowia_unclassified(100); | Metschnikowia | 13 | Saprotroph | \|Undefined Saprotroph\| | Yeast | NULL | Probable | As nectar-tap_saprotroph (Põlme et al. 2020); Decay sugar-rich substrates (Põlme et al. 2020) | Põlme S, et al. 2020. Fungal Diversity 105:1-16 ((https://doi.org/10.1007/s13225-020-00466-2)) |
| Otu31423 | | 1 | k__Fungi(100);p__Ascomycota(100);c__Saccharomycetes(100);o__Saccharomycetales(100);f__Metschnikowiaceae(100);g__Metschnikowia(100);s__Metschnikowia_pulcherrima(100); | Metschnikowia | 13 | Saprotroph | \|Undefined Saprotroph\| | Yeast | NULL | Probable | As nectar-tap_saprotroph (Põlme et al. 2020); Decay sugar-rich substrates (Põlme et al. 2020) | Põlme S, et al. 2020. Fungal Diversity 105:1-16 ((https://doi.org/10.1007/s13225-020-00466-2)) |
| Otu31424 | | 1 | k__Fungi(100);p__Ascomycota(100);c__Saccharomycetes(100);o__Saccharomycetales(100);f__Metschnikowiaceae(100);g__Metschnikowia(100);s__Metschnikowia_pulcherrima(100); | Metschnikowia | 13 | Saprotroph | \|Undefined Saprotroph\| | Yeast | NULL | Probable | As nectar-tap_saprotroph (Põlme et al. 2020); Decay sugar-rich substrates (Põlme et al. 2020) | Põlme S, et al. 2020. Fungal Diversity 105:1-16 ((https://doi.org/10.1007/s13225-020-00466-2)) |
| Otu31425 | | 1 | k__Fungi(100);p__Ascomycota(100);c__Saccharomycetes(100);o__Saccharomycetales(100);f__Metschnikowiaceae(100);g__Metschnikowia(100);s__Metschnikowia_pulcherrima(100); | Metschnikowia | 13 | Saprotroph | \|Undefined Saprotroph\| | Yeast | NULL | Probable | As nectar-tap_saprotroph (Põlme et al. 2020); Decay sugar-rich substrates (Põlme et al. 2020) | Põlme S, et al. 2020. Fungal Diversity 105:1-16 ((https://doi.org/10.1007/s13225-020-00466-2)) |
| Otu31426 | | 1 | k__Fungi(100);p__Ascomycota(100);c__Saccharomycetes(100);o__Saccharomycetales(100);f__Metschnikowiaceae(100);g__Metschnikowia(100);s__Metschnikowia_pulcherrima(100); | Metschnikowia | 13 | Saprotroph | \|Undefined Saprotroph\| | Yeast | NULL | Probable | As nectar-tap_saprotroph (Põlme et al. 2020); Decay sugar-rich substrates (Põlme et al. 2020) | Põlme S, et al. 2020. Fungal Diversity 105:1-16 ((https://doi.org/10.1007/s13225-020-00466-2)) |
| Otu31427 | | 1 | k__Fungi(100);p__Ascomycota(100);c__Saccharomycetes(100);o__Saccharomycetales(100);f__Metschnikowiaceae(100);g__Metschnikowia(100);g__Metschnikowia_unclassified(100); | Metschnikowia | 13 | Saprotroph | \|Undefined Saprotroph\| | Yeast | NULL | Probable | As nectar-tap_saprotroph (Põlme et al. 2020); Decay sugar-rich substrates (Põlme et al. 2020) | Põlme S, et al. 2020. Fungal Diversity 105:1-16 ((https://doi.org/10.1007/s13225-020-00466-2)) |
| Otu31428 | | 1 | k__Fungi(100);p__Ascomycota(100);c__Saccharomycetes(100);o__Saccharomycetales(100);f__Metschnikowiaceae(100);g__Metschnikowia(100);s__Metschnikowia_pulcherrima(100); | Metschnikowia | 13 | Saprotroph | \|Undefined Saprotroph\| | Yeast | NULL | Probable | As nectar-tap_saprotroph (Põlme et al. 2020); Decay sugar-rich substrates (Põlme et al. 2020) | Põlme S, et al. 2020. Fungal Diversity 105:1-16 ((https://doi.org/10.1007/s13225-020-00466-2)) |
| Otu31432 | | 1 | k__Fungi(100);p__Ascomycota(100);c__Saccharomycetes(100);o__Saccharomycetales(100);f__Metschnikowiaceae(100);g__Metschnikowia(100);s__Metschnikowia_pulcherrima(100); | Metschnikowia | 13 | Saprotroph | \|Undefined Saprotroph\| | Yeast | NULL | Probable | As nectar-tap_saprotroph (Põlme et al. 2020); Decay sugar-rich substrates (Põlme et al. 2020) | Põlme S, et al. 2020. Fungal Diversity 105:1-16 ((https://doi.org/10.1007/s13225-020-00466-2)) |
| Otu31437 | | 1 | k__Fungi(100);p__Ascomycota(100);c__Saccharomycetes(100);o__Saccharomycetales(100);f__Metschnikowiaceae(100);g__Metschnikowia(100);s__Metschnikowia_pulcherrima(100); | Metschnikowia | 13 | Saprotroph | \|Undefined Saprotroph\| | Yeast | NULL | Probable | As nectar-tap_saprotroph (Põlme et al. 2020); Decay sugar-rich substrates (Põlme et al. 2020) | Põlme S, et al. 2020. Fungal Diversity 105:1-16 ((https://doi.org/10.1007/s13225-020-00466-2)) |
| Otu31439 | | 1 | k__Fungi(100);p__Ascomycota(100);c__Saccharomycetes(100);o__Saccharomycetales(100);f__Metschnikowiaceae(100);g__Metschnikowia(100);s__Metschnikowia_pulcherrima(100); | Metschnikowia | 13 | Saprotroph | \|Undefined Saprotroph\| | Yeast | NULL | Probable | As nectar-tap_saprotroph (Põlme et al. 2020); Decay sugar-rich substrates (Põlme et al. 2020) | Põlme S, et al. 2020. Fungal Diversity 105:1-16 ((https://doi.org/10.1007/s13225-020-00466-2)) |
| Otu31440 | | 1 | k__Fungi(100);p__Ascomycota(100);c__Saccharomycetes(100);o__Saccharomycetales(100);f__Metschnikowiaceae(100);g__Metschnikowia(100);s__Metschnikowia_pulcherrima(100); | Metschnikowia | 13 | Saprotroph | \|Undefined Saprotroph\| | Yeast | NULL | Probable | As nectar-tap_saprotroph (Põlme et al. 2020); Decay sugar-rich substrates (Põlme et al. 2020) | Põlme S, et al. 2020. Fungal Diversity 105:1-16 ((https://doi.org/10.1007/s13225-020-00466-2)) |
| Otu31441 | | 1 | k__Fungi(100);p__Ascomycota(100);c__Saccharomycetes(100);o__Saccharomycetales(100);f__Metschnikowiaceae(100);g__Metschnikowia(100);g__Metschnikowia_unclassified(100); | Metschnikowia | 13 | Saprotroph | \|Undefined Saprotroph\| | Yeast | NULL | Probable | As nectar-tap_saprotroph (Põlme et al. 2020); Decay sugar-rich substrates (Põlme et al. 2020) | Põlme S, et al. 2020. Fungal Diversity 105:1-16 ((https://doi.org/10.1007/s13225-020-00466-2)) |
| Otu31442 | | 1 | k__Fungi(100);p__Ascomycota(100);c__Saccharomycetes(100);o__Saccharomycetales(100);f__Metschnikowiaceae(100);g__Metschnikowia(100);s__Metschnikowia_pulcherrima(100); | Metschnikowia | 13 | Saprotroph | \|Undefined Saprotroph\| | Yeast | NULL | Probable | As nectar-tap_saprotroph (Põlme et al. 2020); Decay sugar-rich substrates (Põlme et al. 2020) | Põlme S, et al. 2020. Fungal Diversity 105:1-16 ((https://doi.org/10.1007/s13225-020-00466-2)) |
| Otu31443 | | 1 | k__Fungi(100);p__Ascomycota(100);c__Saccharomycetes(100);o__Saccharomycetales(100);f__Metschnikowiaceae(100);g__Metschnikowia(100);s__Metschnikowia_pulcherrima(100); | Metschnikowia | 13 | Saprotroph | \|Undefined Saprotroph\| | Yeast | NULL | Probable | As nectar-tap_saprotroph (Põlme et al. 2020); Decay sugar-rich substrates (Põlme et al. 2020) | Põlme S, et al. 2020. Fungal Diversity 105:1-16 ((https://doi.org/10.1007/s13225-020-00466-2)) |
| Otu31444 | | 1 | k__Fungi(100);p__Ascomycota(100);c__Saccharomycetes(100);o__Saccharomycetales(100);f__Metschnikowiaceae(100);g__Metschnikowia(100);s__Metschnikowia_pulcherrima(100); | Metschnikowia | 13 | Saprotroph | \|Undefined Saprotroph\| | Yeast | NULL | Probable | As nectar-tap_saprotroph (Põlme et al. 2020); Decay sugar-rich substrates (Põlme et al. 2020) | Põlme S, et al. 2020. Fungal Diversity 105:1-16 ((https://doi.org/10.1007/s13225-020-00466-2)) |
| Otu31445 | | 1 | k__Fungi(100);p__Ascomycota(100);c__Saccharomycetes(100);o__Saccharomycetales(100);f__Metschnikowiaceae(100);g__Metschnikowia(100);g__Metschnikowia_unclassified(100); | Metschnikowia | 13 | Saprotroph | \|Undefined Saprotroph\| | Yeast | NULL | Probable | As nectar-tap_saprotroph (Põlme et al. 2020); Decay sugar-rich substrates (Põlme et al. 2020) | Põlme S, et al. 2020. Fungal Diversity 105:1-16 ((https://doi.org/10.1007/s13225-020-00466-2)) |
| Otu31446 | | 1 | k__Fungi(100);p__Ascomycota(100);c__Saccharomycetes(100);o__Saccharomycetales(100);f__Metschnikowiaceae(100);g__Metschnikowia(100);s__Metschnikowia_pulcherrima(100); | Metschnikowia | 13 | Saprotroph | \|Undefined Saprotroph\| | Yeast | NULL | Probable | As nectar-tap_saprotroph (Põlme et al. 2020); Decay sugar-rich substrates (Põlme et al. 2020) | Põlme S, et al. 2020. Fungal Diversity 105:1-16 ((https://doi.org/10.1007/s13225-020-00466-2)) |
| Otu31447 | | 1 | k__Fungi(100);p__Ascomycota(100);c__Saccharomycetes(100);o__Saccharomycetales(100);f__Metschnikowiaceae(100);g__Metschnikowia(100);g__Metschnikowia_unclassified(100); | Metschnikowia | 13 | Saprotroph | \|Undefined Saprotroph\| | Yeast | NULL | Probable | As nectar-tap_saprotroph (Põlme et al. 2020); Decay sugar-rich substrates (Põlme et al. 2020) | Põlme S, et al. 2020. Fungal Diversity 105:1-16 ((https://doi.org/10.1007/s13225-020-00466-2)) |
| Otu31450 | | 1 | k__Fungi(100);p__Ascomycota(100);c__Saccharomycetes(100);o__Saccharomycetales(100);f__Metschnikowiaceae(100);g__Metschnikowia(100);s__Metschnikowia_pulcherrima(100); | Metschnikowia | 13 | Saprotroph | \|Undefined Saprotroph\| | Yeast | NULL | Probable | As nectar-tap_saprotroph (Põlme et al. 2020); Decay sugar-rich substrates (Põlme et al. 2020) | Põlme S, et al. 2020. Fungal Diversity 105:1-16 ((https://doi.org/10.1007/s13225-020-00466-2)) |
| Otu31451 | | 1 | k__Fungi(100);p__Ascomycota(100);c__Saccharomycetes(100);o__Saccharomycetales(100);f__Metschnikowiaceae(100);g__Metschnikowia(100);s__Metschnikowia_pulcherrima(100); | Metschnikowia | 13 | Saprotroph | \|Undefined Saprotroph\| | Yeast | NULL | Probable | As nectar-tap_saprotroph (Põlme et al. 2020); Decay sugar-rich substrates (Põlme et al. 2020) | Põlme S, et al. 2020. Fungal Diversity 105:1-16 ((https://doi.org/10.1007/s13225-020-00466-2)) |
| Otu31452 | | 1 | k__Fungi(100);p__Ascomycota(100);c__Saccharomycetes(100);o__Saccharomycetales(100);f__Metschnikowiaceae(100);g__Metschnikowia(100);s__Metschnikowia_pulcherrima(100); | Metschnikowia | 13 | Saprotroph | \|Undefined Saprotroph\| | Yeast | NULL | Probable | As nectar-tap_saprotroph (Põlme et al. 2020); Decay sugar-rich substrates (Põlme et al. 2020) | Põlme S, et al. 2020. Fungal Diversity 105:1-16 ((https://doi.org/10.1007/s13225-020-00466-2)) |
| Otu31462 | | 1 | k__Fungi(100);p__Ascomycota(100);c__Saccharomycetes(100);o__Saccharomycetales(100);f__Metschnikowiaceae(100);g__Metschnikowia(100);s__Metschnikowia_pulcherrima(100); | Metschnikowia | 13 | Saprotroph | \|Undefined Saprotroph\| | Yeast | NULL | Probable | As nectar-tap_saprotroph (Põlme et al. 2020); Decay sugar-rich substrates (Põlme et al. 2020) | Põlme S, et al. 2020. Fungal Diversity 105:1-16 ((https://doi.org/10.1007/s13225-020-00466-2)) |
| Otu31463 | | 1 | k__Fungi(100);p__Ascomycota(100);c__Saccharomycetes(100);o__Saccharomycetales(100);f__Metschnikowiaceae(100);g__Metschnikowia(100);s__Metschnikowia_pulcherrima(100); | Metschnikowia | 13 | Saprotroph | \|Undefined Saprotroph\| | Yeast | NULL | Probable | As nectar-tap_saprotroph (Põlme et al. 2020); Decay sugar-rich substrates (Põlme et al. 2020) | Põlme S, et al. 2020. Fungal Diversity 105:1-16 ((https://doi.org/10.1007/s13225-020-00466-2)) |
| Otu31464 | | 1 | k__Fungi(100);p__Ascomycota(100);c__Saccharomycetes(100);o__Saccharomycetales(100);f__Metschnikowiaceae(100);g__Metschnikowia(100);s__Metschnikowia_pulcherrima(100); | Metschnikowia | 13 | Saprotroph | \|Undefined Saprotroph\| | Yeast | NULL | Probable | As nectar-tap_saprotroph (Põlme et al. 2020); Decay sugar-rich substrates (Põlme et al. 2020) | Põlme S, et al. 2020. Fungal Diversity 105:1-16 ((https://doi.org/10.1007/s13225-020-00466-2)) |
| Otu31467 | | 1 | k__Fungi(100);p__Ascomycota(100);c__Saccharomycetes(100);o__Saccharomycetales(100);f__Metschnikowiaceae(100);g__Metschnikowia(100);g__Metschnikowia_unclassified(100); | Metschnikowia | 13 | Saprotroph | \|Undefined Saprotroph\| | Yeast | NULL | Probable | As nectar-tap_saprotroph (Põlme et al. 2020); Decay sugar-rich substrates (Põlme et al. 2020) | Põlme S, et al. 2020. Fungal Diversity 105:1-16 ((https://doi.org/10.1007/s13225-020-00466-2)) |
| Otu31468 | | 1 | k__Fungi(100);p__Ascomycota(100);c__Saccharomycetes(100);o__Saccharomycetales(100);f__Metschnikowiaceae(100);g__Metschnikowia(100);s__Metschnikowia_pulcherrima(100); | Metschnikowia | 13 | Saprotroph | \|Undefined Saprotroph\| | Yeast | NULL | Probable | As nectar-tap_saprotroph (Põlme et al. 2020); Decay sugar-rich substrates (Põlme et al. 2020) | Põlme S, et al. 2020. Fungal Diversity 105:1-16 ((https://doi.org/10.1007/s13225-020-00466-2)) |
| Otu31469 | | 1 | k__Fungi(100);p__Ascomycota(100);c__Saccharomycetes(100);o__Saccharomycetales(100);f__Metschnikowiaceae(100);g__Metschnikowia(100);g__Metschnikowia_unclassified(100); | Metschnikowia | 13 | Saprotroph | \|Undefined Saprotroph\| | Yeast | NULL | Probable | As nectar-tap_saprotroph (Põlme et al. 2020); Decay sugar-rich substrates (Põlme et al. 2020) | Põlme S, et al. 2020. Fungal Diversity 105:1-16 ((https://doi.org/10.1007/s13225-020-00466-2)) |
| Otu31470 | | 1 | k__Fungi(100);p__Ascomycota(100);c__Saccharomycetes(100);o__Saccharomycetales(100);f__Metschnikowiaceae(100);g__Metschnikowia(100);s__Metschnikowia_pulcherrima(100); | Metschnikowia | 13 | Saprotroph | \|Undefined Saprotroph\| | Yeast | NULL | Probable | As nectar-tap_saprotroph (Põlme et al. 2020); Decay sugar-rich substrates (Põlme et al. 2020) | Põlme S, et al. 2020. Fungal Diversity 105:1-16 ((https://doi.org/10.1007/s13225-020-00466-2)) |
| Otu31471 | | 1 | k__Fungi(100);p__Ascomycota(100);c__Saccharomycetes(100);o__Saccharomycetales(100);f__Metschnikowiaceae(100);g__Metschnikowia(100);s__Metschnikowia_pulcherrima(100); | Metschnikowia | 13 | Saprotroph | \|Undefined Saprotroph\| | Yeast | NULL | Probable | As nectar-tap_saprotroph (Põlme et al. 2020); Decay sugar-rich substrates (Põlme et al. 2020) | Põlme S, et al. 2020. Fungal Diversity 105:1-16 ((https://doi.org/10.1007/s13225-020-00466-2)) |
| Otu31472 | | 1 | k__Fungi(100);p__Ascomycota(100);c__Saccharomycetes(100);o__Saccharomycetales(100);f__Metschnikowiaceae(100);g__Metschnikowia(100);s__Metschnikowia_pulcherrima(100); | Metschnikowia | 13 | Saprotroph | \|Undefined Saprotroph\| | Yeast | NULL | Probable | As nectar-tap_saprotroph (Põlme et al. 2020); Decay sugar-rich substrates (Põlme et al. 2020) | Põlme S, et al. 2020. Fungal Diversity 105:1-16 ((https://doi.org/10.1007/s13225-020-00466-2)) |
| Otu31473 | | 1 | k__Fungi(100);p__Ascomycota(100);c__Saccharomycetes(100);o__Saccharomycetales(100);f__Metschnikowiaceae(100);g__Metschnikowia(100);s__Metschnikowia_pulcherrima(100); | Metschnikowia | 13 | Saprotroph | \|Undefined Saprotroph\| | Yeast | NULL | Probable | As nectar-tap_saprotroph (Põlme et al. 2020); Decay sugar-rich substrates (Põlme et al. 2020) | Põlme S, et al. 2020. Fungal Diversity 105:1-16 ((https://doi.org/10.1007/s13225-020-00466-2)) |
| Otu31474 | | 1 | k__Fungi(100);p__Ascomycota(100);c__Saccharomycetes(100);o__Saccharomycetales(100);f__Metschnikowiaceae(100);g__Metschnikowia(100);g__Metschnikowia_unclassified(100); | Metschnikowia | 13 | Saprotroph | \|Undefined Saprotroph\| | Yeast | NULL | Probable | As nectar-tap_saprotroph (Põlme et al. 2020); Decay sugar-rich substrates (Põlme et al. 2020) | Põlme S, et al. 2020. Fungal Diversity 105:1-16 ((https://doi.org/10.1007/s13225-020-00466-2)) |
| Otu31475 | | 1 | k__Fungi(100);p__Ascomycota(100);c__Saccharomycetes(100);o__Saccharomycetales(100);f__Metschnikowiaceae(100);g__Metschnikowia(100);g__Metschnikowia_unclassified(100); | Metschnikowia | 13 | Saprotroph | \|Undefined Saprotroph\| | Yeast | NULL | Probable | As nectar-tap_saprotroph (Põlme et al. 2020); Decay sugar-rich substrates (Põlme et al. 2020) | Põlme S, et al. 2020. Fungal Diversity 105:1-16 ((https://doi.org/10.1007/s13225-020-00466-2)) |
| Otu31476 | | 1 | k__Fungi(100);p__Ascomycota(100);c__Saccharomycetes(100);o__Saccharomycetales(100);f__Metschnikowiaceae(100);g__Metschnikowia(100);s__Metschnikowia_pulcherrima(100); | Metschnikowia | 13 | Saprotroph | \|Undefined Saprotroph\| | Yeast | NULL | Probable | As nectar-tap_saprotroph (Põlme et al. 2020); Decay sugar-rich substrates (Põlme et al. 2020) | Põlme S, et al. 2020. Fungal Diversity 105:1-16 ((https://doi.org/10.1007/s13225-020-00466-2)) |
| Otu31477 | | 1 | k__Fungi(100);p__Ascomycota(100);c__Saccharomycetes(100);o__Saccharomycetales(100);f__Metschnikowiaceae(100);g__Metschnikowia(100);s__Metschnikowia_pulcherrima(100); | Metschnikowia | 13 | Saprotroph | \|Undefined Saprotroph\| | Yeast | NULL | Probable | As nectar-tap_saprotroph (Põlme et al. 2020); Decay sugar-rich substrates (Põlme et al. 2020) | Põlme S, et al. 2020. Fungal Diversity 105:1-16 ((https://doi.org/10.1007/s13225-020-00466-2)) |
| Otu31478 | | 1 | k__Fungi(100);p__Ascomycota(100);c__Saccharomycetes(100);o__Saccharomycetales(100);f__Metschnikowiaceae(100);g__Metschnikowia(100);s__Metschnikowia_pulcherrima(100); | Metschnikowia | 13 | Saprotroph | \|Undefined Saprotroph\| | Yeast | NULL | Probable | As nectar-tap_saprotroph (Põlme et al. 2020); Decay sugar-rich substrates (Põlme et al. 2020) | Põlme S, et al. 2020. Fungal Diversity 105:1-16 ((https://doi.org/10.1007/s13225-020-00466-2)) |
| Otu31479 | | 1 | k__Fungi(100);p__Ascomycota(100);c__Saccharomycetes(100);o__Saccharomycetales(100);f__Metschnikowiaceae(100);g__Metschnikowia(100);s__Metschnikowia_pulcherrima(100); | Metschnikowia | 13 | Saprotroph | \|Undefined Saprotroph\| | Yeast | NULL | Probable | As nectar-tap_saprotroph (Põlme et al. 2020); Decay sugar-rich substrates (Põlme et al. 2020) | Põlme S, et al. 2020. Fungal Diversity 105:1-16 ((https://doi.org/10.1007/s13225-020-00466-2)) |
| Otu31480 | | 1 | k__Fungi(100);p__Ascomycota(100);c__Saccharomycetes(100);o__Saccharomycetales(100);f__Metschnikowiaceae(100);g__Metschnikowia(100);s__Metschnikowia_pulcherrima(100); | Metschnikowia | 13 | Saprotroph | \|Undefined Saprotroph\| | Yeast | NULL | Probable | As nectar-tap_saprotroph (Põlme et al. 2020); Decay sugar-rich substrates (Põlme et al. 2020) | Põlme S, et al. 2020. Fungal Diversity 105:1-16 ((https://doi.org/10.1007/s13225-020-00466-2)) |
| Otu31481 | | 1 | k__Fungi(100);p__Ascomycota(100);c__Saccharomycetes(100);o__Saccharomycetales(100);f__Metschnikowiaceae(100);g__Metschnikowia(100);s__Metschnikowia_pulcherrima(100); | Metschnikowia | 13 | Saprotroph | \|Undefined Saprotroph\| | Yeast | NULL | Probable | As nectar-tap_saprotroph (Põlme et al. 2020); Decay sugar-rich substrates (Põlme et al. 2020) | Põlme S, et al. 2020. Fungal Diversity 105:1-16 ((https://doi.org/10.1007/s13225-020-00466-2)) |
| Otu31482 | | 1 | k__Fungi(100);p__Ascomycota(100);c__Saccharomycetes(100);o__Saccharomycetales(100);f__Metschnikowiaceae(100);g__Metschnikowia(100);s__Metschnikowia_pulcherrima(100); | Metschnikowia | 13 | Saprotroph | \|Undefined Saprotroph\| | Yeast | NULL | Probable | As nectar-tap_saprotroph (Põlme et al. 2020); Decay sugar-rich substrates (Põlme et al. 2020) | Põlme S, et al. 2020. Fungal Diversity 105:1-16 ((https://doi.org/10.1007/s13225-020-00466-2)) |
| Otu31483 | | 1 | k__Fungi(100);p__Ascomycota(100);c__Saccharomycetes(100);o__Saccharomycetales(100);f__Metschnikowiaceae(100);g__Metschnikowia(100);g__Metschnikowia_unclassified(100); | Metschnikowia | 13 | Saprotroph | \|Undefined Saprotroph\| | Yeast | NULL | Probable | As nectar-tap_saprotroph (Põlme et al. 2020); Decay sugar-rich substrates (Põlme et al. 2020) | Põlme S, et al. 2020. Fungal Diversity 105:1-16 ((https://doi.org/10.1007/s13225-020-00466-2)) |
| Otu31484 | | 1 | k__Fungi(100);p__Ascomycota(100);c__Saccharomycetes(100);o__Saccharomycetales(100);f__Metschnikowiaceae(100);g__Metschnikowia(100);s__Metschnikowia_pulcherrima(100); | Metschnikowia | 13 | Saprotroph | \|Undefined Saprotroph\| | Yeast | NULL | Probable | As nectar-tap_saprotroph (Põlme et al. 2020); Decay sugar-rich substrates (Põlme et al. 2020) | Põlme S, et al. 2020. Fungal Diversity 105:1-16 ((https://doi.org/10.1007/s13225-020-00466-2)) |
| Otu31485 | | 1 | k__Fungi(100);p__Ascomycota(100);c__Saccharomycetes(100);o__Saccharomycetales(100);f__Metschnikowiaceae(100);g__Metschnikowia(100);s__Metschnikowia_pulcherrima(100); | Metschnikowia | 13 | Saprotroph | \|Undefined Saprotroph\| | Yeast | NULL | Probable | As nectar-tap_saprotroph (Põlme et al. 2020); Decay sugar-rich substrates (Põlme et al. 2020) | Põlme S, et al. 2020. Fungal Diversity 105:1-16 ((https://doi.org/10.1007/s13225-020-00466-2)) |
| Otu31486 | | 1 | k__Fungi(100);p__Ascomycota(100);c__Saccharomycetes(100);o__Saccharomycetales(100);f__Metschnikowiaceae(100);g__Metschnikowia(100);s__Metschnikowia_pulcherrima(100); | Metschnikowia | 13 | Saprotroph | \|Undefined Saprotroph\| | Yeast | NULL | Probable | As nectar-tap_saprotroph (Põlme et al. 2020); Decay sugar-rich substrates (Põlme et al. 2020) | Põlme S, et al. 2020. Fungal Diversity 105:1-16 ((https://doi.org/10.1007/s13225-020-00466-2)) |
| Otu31487 | | 1 | k__Fungi(100);p__Ascomycota(100);c__Saccharomycetes(100);o__Saccharomycetales(100);f__Metschnikowiaceae(100);g__Metschnikowia(100);s__Metschnikowia_pulcherrima(100); | Metschnikowia | 13 | Saprotroph | \|Undefined Saprotroph\| | Yeast | NULL | Probable | As nectar-tap_saprotroph (Põlme et al. 2020); Decay sugar-rich substrates (Põlme et al. 2020) | Põlme S, et al. 2020. Fungal Diversity 105:1-16 ((https://doi.org/10.1007/s13225-020-00466-2)) |
| Otu31488 | | 1 | k__Fungi(100);p__Ascomycota(100);c__Saccharomycetes(100);o__Saccharomycetales(100);f__Metschnikowiaceae(100);g__Metschnikowia(100);s__Metschnikowia_pulcherrima(100); | Metschnikowia | 13 | Saprotroph | \|Undefined Saprotroph\| | Yeast | NULL | Probable | As nectar-tap_saprotroph (Põlme et al. 2020); Decay sugar-rich substrates (Põlme et al. 2020) | Põlme S, et al. 2020. Fungal Diversity 105:1-16 ((https://doi.org/10.1007/s13225-020-00466-2)) |
| Otu31489 | | 1 | k__Fungi(100);p__Ascomycota(100);c__Saccharomycetes(100);o__Saccharomycetales(100);f__Metschnikowiaceae(100);g__Metschnikowia(100);g__Metschnikowia_unclassified(100); | Metschnikowia | 13 | Saprotroph | \|Undefined Saprotroph\| | Yeast | NULL | Probable | As nectar-tap_saprotroph (Põlme et al. 2020); Decay sugar-rich substrates (Põlme et al. 2020) | Põlme S, et al. 2020. Fungal Diversity 105:1-16 ((https://doi.org/10.1007/s13225-020-00466-2)) |
| Otu31491 | | 1 | k__Fungi(100);p__Ascomycota(100);c__Saccharomycetes(100);o__Saccharomycetales(100);f__Metschnikowiaceae(100);g__Metschnikowia(100);s__Metschnikowia_pulcherrima(100); | Metschnikowia | 13 | Saprotroph | \|Undefined Saprotroph\| | Yeast | NULL | Probable | As nectar-tap_saprotroph (Põlme et al. 2020); Decay sugar-rich substrates (Põlme et al. 2020) | Põlme S, et al. 2020. Fungal Diversity 105:1-16 ((https://doi.org/10.1007/s13225-020-00466-2)) |
| Otu31493 | | 1 | k__Fungi(100);p__Ascomycota(100);c__Saccharomycetes(100);o__Saccharomycetales(100);f__Metschnikowiaceae(100);g__Metschnikowia(100);g__Metschnikowia_unclassified(100); | Metschnikowia | 13 | Saprotroph | \|Undefined Saprotroph\| | Yeast | NULL | Probable | As nectar-tap_saprotroph (Põlme et al. 2020); Decay sugar-rich substrates (Põlme et al. 2020) | Põlme S, et al. 2020. Fungal Diversity 105:1-16 ((https://doi.org/10.1007/s13225-020-00466-2)) |
| Otu31494 | | 1 | k__Fungi(100);p__Ascomycota(100);c__Saccharomycetes(100);o__Saccharomycetales(100);f__Metschnikowiaceae(100);g__Metschnikowia(100);s__Metschnikowia_pulcherrima(100); | Metschnikowia | 13 | Saprotroph | \|Undefined Saprotroph\| | Yeast | NULL | Probable | As nectar-tap_saprotroph (Põlme et al. 2020); Decay sugar-rich substrates (Põlme et al. 2020) | Põlme S, et al. 2020. Fungal Diversity 105:1-16 ((https://doi.org/10.1007/s13225-020-00466-2)) |
| Otu31496 | | 1 | k__Fungi(100);p__Ascomycota(100);c__Saccharomycetes(100);o__Saccharomycetales(100);f__Metschnikowiaceae(100);g__Metschnikowia(100);s__Metschnikowia_pulcherrima(100); | Metschnikowia | 13 | Saprotroph | \|Undefined Saprotroph\| | Yeast | NULL | Probable | As nectar-tap_saprotroph (Põlme et al. 2020); Decay sugar-rich substrates (Põlme et al. 2020) | Põlme S, et al. 2020. Fungal Diversity 105:1-16 ((https://doi.org/10.1007/s13225-020-00466-2)) |
| Otu31497 | | 1 | k__Fungi(100);p__Ascomycota(100);c__Saccharomycetes(100);o__Saccharomycetales(100);f__Metschnikowiaceae(100);g__Metschnikowia(100);s__Metschnikowia_pulcherrima(100); | Metschnikowia | 13 | Saprotroph | \|Undefined Saprotroph\| | Yeast | NULL | Probable | As nectar-tap_saprotroph (Põlme et al. 2020); Decay sugar-rich substrates (Põlme et al. 2020) | Põlme S, et al. 2020. Fungal Diversity 105:1-16 ((https://doi.org/10.1007/s13225-020-00466-2)) |
| Otu31498 | | 1 | k__Fungi(100);p__Ascomycota(100);c__Saccharomycetes(100);o__Saccharomycetales(100);f__Metschnikowiaceae(100);g__Metschnikowia(100);g__Metschnikowia_unclassified(100); | Metschnikowia | 13 | Saprotroph | \|Undefined Saprotroph\| | Yeast | NULL | Probable | As nectar-tap_saprotroph (Põlme et al. 2020); Decay sugar-rich substrates (Põlme et al. 2020) | Põlme S, et al. 2020. Fungal Diversity 105:1-16 ((https://doi.org/10.1007/s13225-020-00466-2)) |
| Otu31499 | | 1 | k__Fungi(100);p__Ascomycota(100);c__Saccharomycetes(100);o__Saccharomycetales(100);f__Metschnikowiaceae(100);g__Metschnikowia(100);s__Metschnikowia_pulcherrima(100); | Metschnikowia | 13 | Saprotroph | \|Undefined Saprotroph\| | Yeast | NULL | Probable | As nectar-tap_saprotroph (Põlme et al. 2020); Decay sugar-rich substrates (Põlme et al. 2020) | Põlme S, et al. 2020. Fungal Diversity 105:1-16 ((https://doi.org/10.1007/s13225-020-00466-2)) |
| Otu31500 | | 1 | k__Fungi(100);p__Ascomycota(100);c__Saccharomycetes(100);o__Saccharomycetales(100);f__Metschnikowiaceae(100);g__Metschnikowia(100);s__Metschnikowia_pulcherrima(100); | Metschnikowia | 13 | Saprotroph | \|Undefined Saprotroph\| | Yeast | NULL | Probable | As nectar-tap_saprotroph (Põlme et al. 2020); Decay sugar-rich substrates (Põlme et al. 2020) | Põlme S, et al. 2020. Fungal Diversity 105:1-16 ((https://doi.org/10.1007/s13225-020-00466-2)) |
| Otu31501 | | 1 | k__Fungi(100);p__Ascomycota(100);c__Saccharomycetes(100);o__Saccharomycetales(100);f__Metschnikowiaceae(100);g__Metschnikowia(100);s__Metschnikowia_pulcherrima(100); | Metschnikowia | 13 | Saprotroph | \|Undefined Saprotroph\| | Yeast | NULL | Probable | As nectar-tap_saprotroph (Põlme et al. 2020); Decay sugar-rich substrates (Põlme et al. 2020) | Põlme S, et al. 2020. Fungal Diversity 105:1-16 ((https://doi.org/10.1007/s13225-020-00466-2)) |
| Otu31502 | | 1 | k__Fungi(100);p__Ascomycota(100);c__Saccharomycetes(100);o__Saccharomycetales(100);f__Metschnikowiaceae(100);g__Metschnikowia(100);s__Metschnikowia_pulcherrima(100); | Metschnikowia | 13 | Saprotroph | \|Undefined Saprotroph\| | Yeast | NULL | Probable | As nectar-tap_saprotroph (Põlme et al. 2020); Decay sugar-rich substrates (Põlme et al. 2020) | Põlme S, et al. 2020. Fungal Diversity 105:1-16 ((https://doi.org/10.1007/s13225-020-00466-2)) |
| Otu31503 | | 1 | k__Fungi(100);p__Ascomycota(100);c__Saccharomycetes(100);o__Saccharomycetales(100);f__Metschnikowiaceae(100);g__Metschnikowia(100);g__Metschnikowia_unclassified(100); | Metschnikowia | 13 | Saprotroph | \|Undefined Saprotroph\| | Yeast | NULL | Probable | As nectar-tap_saprotroph (Põlme et al. 2020); Decay sugar-rich substrates (Põlme et al. 2020) | Põlme S, et al. 2020. Fungal Diversity 105:1-16 ((https://doi.org/10.1007/s13225-020-00466-2)) |
| Otu31504 | | 1 | k__Fungi(100);p__Ascomycota(100);c__Saccharomycetes(100);o__Saccharomycetales(100);f__Metschnikowiaceae(100);g__Metschnikowia(100);s__Metschnikowia_pulcherrima(100); | Metschnikowia | 13 | Saprotroph | \|Undefined Saprotroph\| | Yeast | NULL | Probable | As nectar-tap_saprotroph (Põlme et al. 2020); Decay sugar-rich substrates (Põlme et al. 2020) | Põlme S, et al. 2020. Fungal Diversity 105:1-16 ((https://doi.org/10.1007/s13225-020-00466-2)) |
| Otu31505 | | 1 | k__Fungi(100);p__Ascomycota(100);c__Saccharomycetes(100);o__Saccharomycetales(100);f__Metschnikowiaceae(100);g__Metschnikowia(100);s__Metschnikowia_pulcherrima(100); | Metschnikowia | 13 | Saprotroph | \|Undefined Saprotroph\| | Yeast | NULL | Probable | As nectar-tap_saprotroph (Põlme et al. 2020); Decay sugar-rich substrates (Põlme et al. 2020) | Põlme S, et al. 2020. Fungal Diversity 105:1-16 ((https://doi.org/10.1007/s13225-020-00466-2)) |
| Otu31506 | | 1 | k__Fungi(100);p__Ascomycota(100);c__Saccharomycetes(100);o__Saccharomycetales(100);f__Metschnikowiaceae(100);g__Metschnikowia(100);s__Metschnikowia_pulcherrima(100); | Metschnikowia | 13 | Saprotroph | \|Undefined Saprotroph\| | Yeast | NULL | Probable | As nectar-tap_saprotroph (Põlme et al. 2020); Decay sugar-rich substrates (Põlme et al. 2020) | Põlme S, et al. 2020. Fungal Diversity 105:1-16 ((https://doi.org/10.1007/s13225-020-00466-2)) |
| Otu31507 | | 1 | k__Fungi(100);p__Ascomycota(100);c__Saccharomycetes(100);o__Saccharomycetales(100);f__Metschnikowiaceae(100);g__Metschnikowia(100);s__Metschnikowia_pulcherrima(100); | Metschnikowia | 13 | Saprotroph | \|Undefined Saprotroph\| | Yeast | NULL | Probable | As nectar-tap_saprotroph (Põlme et al. 2020); Decay sugar-rich substrates (Põlme et al. 2020) | Põlme S, et al. 2020. Fungal Diversity 105:1-16 ((https://doi.org/10.1007/s13225-020-00466-2)) |
| Otu31508 | | 1 | k__Fungi(100);p__Ascomycota(100);c__Saccharomycetes(100);o__Saccharomycetales(100);f__Metschnikowiaceae(100);g__Metschnikowia(100);s__Metschnikowia_pulcherrima(100); | Metschnikowia | 13 | Saprotroph | \|Undefined Saprotroph\| | Yeast | NULL | Probable | As nectar-tap_saprotroph (Põlme et al. 2020); Decay sugar-rich substrates (Põlme et al. 2020) | Põlme S, et al. 2020. Fungal Diversity 105:1-16 ((https://doi.org/10.1007/s13225-020-00466-2)) |
| Otu31509 | | 1 | k__Fungi(100);p__Ascomycota(100);c__Saccharomycetes(100);o__Saccharomycetales(100);f__Metschnikowiaceae(100);g__Metschnikowia(100);s__Metschnikowia_pulcherrima(100); | Metschnikowia | 13 | Saprotroph | \|Undefined Saprotroph\| | Yeast | NULL | Probable | As nectar-tap_saprotroph (Põlme et al. 2020); Decay sugar-rich substrates (Põlme et al. 2020) | Põlme S, et al. 2020. Fungal Diversity 105:1-16 ((https://doi.org/10.1007/s13225-020-00466-2)) |
| Otu31510 | | 1 | k__Fungi(100);p__Ascomycota(100);c__Saccharomycetes(100);o__Saccharomycetales(100);f__Metschnikowiaceae(100);g__Metschnikowia(100);s__Metschnikowia_pulcherrima(100); | Metschnikowia | 13 | Saprotroph | \|Undefined Saprotroph\| | Yeast | NULL | Probable | As nectar-tap_saprotroph (Põlme et al. 2020); Decay sugar-rich substrates (Põlme et al. 2020) | Põlme S, et al. 2020. Fungal Diversity 105:1-16 ((https://doi.org/10.1007/s13225-020-00466-2)) |
| Otu31511 | | 1 | k__Fungi(100);p__Ascomycota(100);c__Saccharomycetes(100);o__Saccharomycetales(100);f__Metschnikowiaceae(100);g__Metschnikowia(100);s__Metschnikowia_pulcherrima(100); | Metschnikowia | 13 | Saprotroph | \|Undefined Saprotroph\| | Yeast | NULL | Probable | As nectar-tap_saprotroph (Põlme et al. 2020); Decay sugar-rich substrates (Põlme et al. 2020) | Põlme S, et al. 2020. Fungal Diversity 105:1-16 ((https://doi.org/10.1007/s13225-020-00466-2)) |
| Otu31512 | | 1 | k__Fungi(100);p__Ascomycota(100);c__Saccharomycetes(100);o__Saccharomycetales(100);f__Metschnikowiaceae(100);g__Metschnikowia(100);s__Metschnikowia_pulcherrima(100); | Metschnikowia | 13 | Saprotroph | \|Undefined Saprotroph\| | Yeast | NULL | Probable | As nectar-tap_saprotroph (Põlme et al. 2020); Decay sugar-rich substrates (Põlme et al. 2020) | Põlme S, et al. 2020. Fungal Diversity 105:1-16 ((https://doi.org/10.1007/s13225-020-00466-2)) |
| Otu31513 | | 1 | k__Fungi(100);p__Ascomycota(100);c__Saccharomycetes(100);o__Saccharomycetales(100);f__Metschnikowiaceae(100);g__Metschnikowia(100);s__Metschnikowia_pulcherrima(100); | Metschnikowia | 13 | Saprotroph | \|Undefined Saprotroph\| | Yeast | NULL | Probable | As nectar-tap_saprotroph (Põlme et al. 2020); Decay sugar-rich substrates (Põlme et al. 2020) | Põlme S, et al. 2020. Fungal Diversity 105:1-16 ((https://doi.org/10.1007/s13225-020-00466-2)) |
| Otu31514 | | 1 | k__Fungi(100);p__Ascomycota(100);c__Saccharomycetes(100);o__Saccharomycetales(100);f__Metschnikowiaceae(100);g__Metschnikowia(100);s__Metschnikowia_pulcherrima(100); | Metschnikowia | 13 | Saprotroph | \|Undefined Saprotroph\| | Yeast | NULL | Probable | As nectar-tap_saprotroph (Põlme et al. 2020); Decay sugar-rich substrates (Põlme et al. 2020) | Põlme S, et al. 2020. Fungal Diversity 105:1-16 ((https://doi.org/10.1007/s13225-020-00466-2)) |
| Otu31515 | | 1 | k__Fungi(100);p__Ascomycota(100);c__Saccharomycetes(100);o__Saccharomycetales(100);f__Metschnikowiaceae(100);g__Metschnikowia(100);g__Metschnikowia_unclassified(100); | Metschnikowia | 13 | Saprotroph | \|Undefined Saprotroph\| | Yeast | NULL | Probable | As nectar-tap_saprotroph (Põlme et al. 2020); Decay sugar-rich substrates (Põlme et al. 2020) | Põlme S, et al. 2020. Fungal Diversity 105:1-16 ((https://doi.org/10.1007/s13225-020-00466-2)) |
| Otu31516 | | 1 | k__Fungi(100);p__Ascomycota(100);c__Saccharomycetes(100);o__Saccharomycetales(100);f__Metschnikowiaceae(100);g__Metschnikowia(100);s__Metschnikowia_pulcherrima(100); | Metschnikowia | 13 | Saprotroph | \|Undefined Saprotroph\| | Yeast | NULL | Probable | As nectar-tap_saprotroph (Põlme et al. 2020); Decay sugar-rich substrates (Põlme et al. 2020) | Põlme S, et al. 2020. Fungal Diversity 105:1-16 ((https://doi.org/10.1007/s13225-020-00466-2)) |
| Otu31517 | | 1 | k__Fungi(100);p__Ascomycota(100);c__Saccharomycetes(100);o__Saccharomycetales(100);f__Metschnikowiaceae(100);g__Metschnikowia(100);g__Metschnikowia_unclassified(100); | Metschnikowia | 13 | Saprotroph | \|Undefined Saprotroph\| | Yeast | NULL | Probable | As nectar-tap_saprotroph (Põlme et al. 2020); Decay sugar-rich substrates (Põlme et al. 2020) | Põlme S, et al. 2020. Fungal Diversity 105:1-16 ((https://doi.org/10.1007/s13225-020-00466-2)) |
| Otu31518 | | 1 | k__Fungi(100);p__Ascomycota(100);c__Saccharomycetes(100);o__Saccharomycetales(100);f__Metschnikowiaceae(100);g__Metschnikowia(100);s__Metschnikowia_pulcherrima(100); | Metschnikowia | 13 | Saprotroph | \|Undefined Saprotroph\| | Yeast | NULL | Probable | As nectar-tap_saprotroph (Põlme et al. 2020); Decay sugar-rich substrates (Põlme et al. 2020) | Põlme S, et al. 2020. Fungal Diversity 105:1-16 ((https://doi.org/10.1007/s13225-020-00466-2)) |
| Otu31519 | | 1 | k__Fungi(100);p__Ascomycota(100);c__Saccharomycetes(100);o__Saccharomycetales(100);f__Metschnikowiaceae(100);g__Metschnikowia(100);s__Metschnikowia_pulcherrima(100); | Metschnikowia | 13 | Saprotroph | \|Undefined Saprotroph\| | Yeast | NULL | Probable | As nectar-tap_saprotroph (Põlme et al. 2020); Decay sugar-rich substrates (Põlme et al. 2020) | Põlme S, et al. 2020. Fungal Diversity 105:1-16 ((https://doi.org/10.1007/s13225-020-00466-2)) |
| Otu31520 | | 1 | k__Fungi(100);p__Ascomycota(100);c__Saccharomycetes(100);o__Saccharomycetales(100);f__Metschnikowiaceae(100);g__Metschnikowia(100);g__Metschnikowia_unclassified(100); | Metschnikowia | 13 | Saprotroph | \|Undefined Saprotroph\| | Yeast | NULL | Probable | As nectar-tap_saprotroph (Põlme et al. 2020); Decay sugar-rich substrates (Põlme et al. 2020) | Põlme S, et al. 2020. Fungal Diversity 105:1-16 ((https://doi.org/10.1007/s13225-020-00466-2)) |
| Otu31521 | | 1 | k__Fungi(100);p__Ascomycota(100);c__Saccharomycetes(100);o__Saccharomycetales(100);f__Metschnikowiaceae(100);g__Metschnikowia(100);s__Metschnikowia_pulcherrima(100); | Metschnikowia | 13 | Saprotroph | \|Undefined Saprotroph\| | Yeast | NULL | Probable | As nectar-tap_saprotroph (Põlme et al. 2020); Decay sugar-rich substrates (Põlme et al. 2020) | Põlme S, et al. 2020. Fungal Diversity 105:1-16 ((https://doi.org/10.1007/s13225-020-00466-2)) |
| Otu31522 | | 1 | k__Fungi(100);p__Ascomycota(100);c__Saccharomycetes(100);o__Saccharomycetales(100);f__Metschnikowiaceae(100);g__Metschnikowia(100);s__Metschnikowia_pulcherrima(100); | Metschnikowia | 13 | Saprotroph | \|Undefined Saprotroph\| | Yeast | NULL | Probable | As nectar-tap_saprotroph (Põlme et al. 2020); Decay sugar-rich substrates (Põlme et al. 2020) | Põlme S, et al. 2020. Fungal Diversity 105:1-16 ((https://doi.org/10.1007/s13225-020-00466-2)) |
| Otu31524 | | 1 | k__Fungi(100);p__Ascomycota(100);c__Saccharomycetes(100);o__Saccharomycetales(100);f__Metschnikowiaceae(100);g__Metschnikowia(100);g__Metschnikowia_unclassified(100); | Metschnikowia | 13 | Saprotroph | \|Undefined Saprotroph\| | Yeast | NULL | Probable | As nectar-tap_saprotroph (Põlme et al. 2020); Decay sugar-rich substrates (Põlme et al. 2020) | Põlme S, et al. 2020. Fungal Diversity 105:1-16 ((https://doi.org/10.1007/s13225-020-00466-2)) |
| Otu31525 | | 1 | k__Fungi(100);p__Ascomycota(100);c__Saccharomycetes(100);o__Saccharomycetales(100);f__Metschnikowiaceae(100);g__Metschnikowia(100);s__Metschnikowia_pulcherrima(100); | Metschnikowia | 13 | Saprotroph | \|Undefined Saprotroph\| | Yeast | NULL | Probable | As nectar-tap_saprotroph (Põlme et al. 2020); Decay sugar-rich substrates (Põlme et al. 2020) | Põlme S, et al. 2020. Fungal Diversity 105:1-16 ((https://doi.org/10.1007/s13225-020-00466-2)) |
| Otu31527 | | 1 | k__Fungi(100);p__Ascomycota(100);c__Saccharomycetes(100);o__Saccharomycetales(100);f__Metschnikowiaceae(100);g__Metschnikowia(100);s__Metschnikowia_pulcherrima(100); | Metschnikowia | 13 | Saprotroph | \|Undefined Saprotroph\| | Yeast | NULL | Probable | As nectar-tap_saprotroph (Põlme et al. 2020); Decay sugar-rich substrates (Põlme et al. 2020) | Põlme S, et al. 2020. Fungal Diversity 105:1-16 ((https://doi.org/10.1007/s13225-020-00466-2)) |
| Otu31528 | | 1 | k__Fungi(100);p__Ascomycota(100);c__Saccharomycetes(100);o__Saccharomycetales(100);f__Metschnikowiaceae(100);g__Metschnikowia(100);g__Metschnikowia_unclassified(100); | Metschnikowia | 13 | Saprotroph | \|Undefined Saprotroph\| | Yeast | NULL | Probable | As nectar-tap_saprotroph (Põlme et al. 2020); Decay sugar-rich substrates (Põlme et al. 2020) | Põlme S, et al. 2020. Fungal Diversity 105:1-16 ((https://doi.org/10.1007/s13225-020-00466-2)) |
| Otu31529 | | 1 | k__Fungi(100);p__Ascomycota(100);c__Saccharomycetes(100);o__Saccharomycetales(100);f__Metschnikowiaceae(100);g__Metschnikowia(100);s__Metschnikowia_pulcherrima(100); | Metschnikowia | 13 | Saprotroph | \|Undefined Saprotroph\| | Yeast | NULL | Probable | As nectar-tap_saprotroph (Põlme et al. 2020); Decay sugar-rich substrates (Põlme et al. 2020) | Põlme S, et al. 2020. Fungal Diversity 105:1-16 ((https://doi.org/10.1007/s13225-020-00466-2)) |
| Otu31530 | | 1 | k__Fungi(100);p__Ascomycota(100);c__Saccharomycetes(100);o__Saccharomycetales(100);f__Metschnikowiaceae(100);g__Metschnikowia(100);s__Metschnikowia_pulcherrima(100); | Metschnikowia | 13 | Saprotroph | \|Undefined Saprotroph\| | Yeast | NULL | Probable | As nectar-tap_saprotroph (Põlme et al. 2020); Decay sugar-rich substrates (Põlme et al. 2020) | Põlme S, et al. 2020. Fungal Diversity 105:1-16 ((https://doi.org/10.1007/s13225-020-00466-2)) |
| Otu31531 | | 1 | k__Fungi(100);p__Ascomycota(100);c__Saccharomycetes(100);o__Saccharomycetales(100);f__Metschnikowiaceae(100);g__Metschnikowia(100);s__Metschnikowia_pulcherrima(100); | Metschnikowia | 13 | Saprotroph | \|Undefined Saprotroph\| | Yeast | NULL | Probable | As nectar-tap_saprotroph (Põlme et al. 2020); Decay sugar-rich substrates (Põlme et al. 2020) | Põlme S, et al. 2020. Fungal Diversity 105:1-16 ((https://doi.org/10.1007/s13225-020-00466-2)) |
| Otu31532 | | 1 | k__Fungi(100);p__Ascomycota(100);c__Saccharomycetes(100);o__Saccharomycetales(100);f__Metschnikowiaceae(100);g__Metschnikowia(100);s__Metschnikowia_pulcherrima(100); | Metschnikowia | 13 | Saprotroph | \|Undefined Saprotroph\| | Yeast | NULL | Probable | As nectar-tap_saprotroph (Põlme et al. 2020); Decay sugar-rich substrates (Põlme et al. 2020) | Põlme S, et al. 2020. Fungal Diversity 105:1-16 ((https://doi.org/10.1007/s13225-020-00466-2)) |
| Otu31533 | | 1 | k__Fungi(100);p__Ascomycota(100);c__Saccharomycetes(100);o__Saccharomycetales(100);f__Metschnikowiaceae(100);g__Metschnikowia(100);s__Metschnikowia_pulcherrima(100); | Metschnikowia | 13 | Saprotroph | \|Undefined Saprotroph\| | Yeast | NULL | Probable | As nectar-tap_saprotroph (Põlme et al. 2020); Decay sugar-rich substrates (Põlme et al. 2020) | Põlme S, et al. 2020. Fungal Diversity 105:1-16 ((https://doi.org/10.1007/s13225-020-00466-2)) |
| Otu31534 | | 1 | k__Fungi(100);p__Ascomycota(100);c__Saccharomycetes(100);o__Saccharomycetales(100);f__Metschnikowiaceae(100);g__Metschnikowia(100);g__Metschnikowia_unclassified(100); | Metschnikowia | 13 | Saprotroph | \|Undefined Saprotroph\| | Yeast | NULL | Probable | As nectar-tap_saprotroph (Põlme et al. 2020); Decay sugar-rich substrates (Põlme et al. 2020) | Põlme S, et al. 2020. Fungal Diversity 105:1-16 ((https://doi.org/10.1007/s13225-020-00466-2)) |
| Otu31535 | | 1 | k__Fungi(100);p__Ascomycota(100);c__Saccharomycetes(100);o__Saccharomycetales(100);f__Metschnikowiaceae(100);g__Metschnikowia(100);g__Metschnikowia_unclassified(100); | Metschnikowia | 13 | Saprotroph | \|Undefined Saprotroph\| | Yeast | NULL | Probable | As nectar-tap_saprotroph (Põlme et al. 2020); Decay sugar-rich substrates (Põlme et al. 2020) | Põlme S, et al. 2020. Fungal Diversity 105:1-16 ((https://doi.org/10.1007/s13225-020-00466-2)) |
| Otu31536 | | 1 | k__Fungi(100);p__Ascomycota(100);c__Saccharomycetes(100);o__Saccharomycetales(100);f__Metschnikowiaceae(100);g__Metschnikowia(100);s__Metschnikowia_pulcherrima(100); | Metschnikowia | 13 | Saprotroph | \|Undefined Saprotroph\| | Yeast | NULL | Probable | As nectar-tap_saprotroph (Põlme et al. 2020); Decay sugar-rich substrates (Põlme et al. 2020) | Põlme S, et al. 2020. Fungal Diversity 105:1-16 ((https://doi.org/10.1007/s13225-020-00466-2)) |
| Otu31537 | | 1 | k__Fungi(100);p__Ascomycota(100);c__Saccharomycetes(100);o__Saccharomycetales(100);f__Metschnikowiaceae(100);g__Metschnikowia(100);s__Metschnikowia_pulcherrima(100); | Metschnikowia | 13 | Saprotroph | \|Undefined Saprotroph\| | Yeast | NULL | Probable | As nectar-tap_saprotroph (Põlme et al. 2020); Decay sugar-rich substrates (Põlme et al. 2020) | Põlme S, et al. 2020. Fungal Diversity 105:1-16 ((https://doi.org/10.1007/s13225-020-00466-2)) |
| Otu31538 | | 1 | k__Fungi(100);p__Ascomycota(100);c__Saccharomycetes(100);o__Saccharomycetales(100);f__Metschnikowiaceae(100);g__Metschnikowia(100);g__Metschnikowia_unclassified(100); | Metschnikowia | 13 | Saprotroph | \|Undefined Saprotroph\| | Yeast | NULL | Probable | As nectar-tap_saprotroph (Põlme et al. 2020); Decay sugar-rich substrates (Põlme et al. 2020) | Põlme S, et al. 2020. Fungal Diversity 105:1-16 ((https://doi.org/10.1007/s13225-020-00466-2)) |
| Otu31540 | | 1 | k__Fungi(100);p__Ascomycota(100);c__Saccharomycetes(100);o__Saccharomycetales(100);o__Saccharomycetales_unclassified(100);o__Saccharomycetales_unclassified(100);o__Saccharomycetales_unclassified(100); | Saccharomycetales | 7 | Saprotroph | Undefined Saprotroph | Yeast | NULL | Possible | NULL | Sterkenburg E, et al. 2015. New Phytologist 207:1145-1158 ((https://doi.org/10.1111/nph.13426)) |
| Otu31541 | | 1 | k__Fungi(100);p__Ascomycota(100);c__Saccharomycetes(100);o__Saccharomycetales(100);f__Metschnikowiaceae(100);g__Metschnikowia(100);s__Metschnikowia_pulcherrima(100); | Metschnikowia | 13 | Saprotroph | \|Undefined Saprotroph\| | Yeast | NULL | Probable | As nectar-tap_saprotroph (Põlme et al. 2020); Decay sugar-rich substrates (Põlme et al. 2020) | Põlme S, et al. 2020. Fungal Diversity 105:1-16 ((https://doi.org/10.1007/s13225-020-00466-2)) |
| Otu31542 | | 1 | k__Fungi(100);p__Ascomycota(100);c__Saccharomycetes(100);o__Saccharomycetales(100);f__Metschnikowiaceae(100);g__Metschnikowia(100);g__Metschnikowia_unclassified(100); | Metschnikowia | 13 | Saprotroph | \|Undefined Saprotroph\| | Yeast | NULL | Probable | As nectar-tap_saprotroph (Põlme et al. 2020); Decay sugar-rich substrates (Põlme et al. 2020) | Põlme S, et al. 2020. Fungal Diversity 105:1-16 ((https://doi.org/10.1007/s13225-020-00466-2)) |
| Otu31543 | | 1 | k__Fungi(100);p__Ascomycota(100);c__Saccharomycetes(100);o__Saccharomycetales(100);f__Metschnikowiaceae(100);g__Metschnikowia(100);g__Metschnikowia_unclassified(100); | Metschnikowia | 13 | Saprotroph | \|Undefined Saprotroph\| | Yeast | NULL | Probable | As nectar-tap_saprotroph (Põlme et al. 2020); Decay sugar-rich substrates (Põlme et al. 2020) | Põlme S, et al. 2020. Fungal Diversity 105:1-16 ((https://doi.org/10.1007/s13225-020-00466-2)) |
| Otu31544 | | 1 | k__Fungi(100);p__Basidiomycota(100);c__Microbotryomycetes(100);o__Sporidiobolales(100);f__Sporidiobolales_family_Incertae_sedis(100);g__Sporobolomyces(100);s__Sporobolomyces_lactosus(100); | Sporobolomyces | 13 | Pathotroph-Saprotroph | \|Fungal Parasite\|-Plant Saprotroph | Yeast | NULL | Probable | As litter_saprotroph (Põlme et al. 2020); As mycoparasite (Põlme et al. 2020); Decay fungal material (Põlme et al. 2020) | Tedersoo L, et al. 2014. Science 346:e1256688 ((https://doi.org/10.1126/science.1256688)); Purahong W, et al. 2016. Molecular Ecology 25:4059-4074 ((https://doi.org/10.1111/mec.13739)); Põlme S, et al. 2020. Fungal Diversity 105:1-16 ((https://doi.org/10.1007/s13225-020-00466-2)) |
| Otu31545 | | 1 | k__Fungi(100);p__Ascomycota(100);c__Saccharomycetes(100);o__Saccharomycetales(100);f__Metschnikowiaceae(100);g__Metschnikowia(100);s__Metschnikowia_pulcherrima(100); | Metschnikowia | 13 | Saprotroph | \|Undefined Saprotroph\| | Yeast | NULL | Probable | As nectar-tap_saprotroph (Põlme et al. 2020); Decay sugar-rich substrates (Põlme et al. 2020) | Põlme S, et al. 2020. Fungal Diversity 105:1-16 ((https://doi.org/10.1007/s13225-020-00466-2)) |
| Otu31546 | | 1 | k__Fungi(100);p__Ascomycota(100);c__Saccharomycetes(100);o__Saccharomycetales(100);f__Metschnikowiaceae(100);g__Metschnikowia(100);g__Metschnikowia_unclassified(100); | Metschnikowia | 13 | Saprotroph | \|Undefined Saprotroph\| | Yeast | NULL | Probable | As nectar-tap_saprotroph (Põlme et al. 2020); Decay sugar-rich substrates (Põlme et al. 2020) | Põlme S, et al. 2020. Fungal Diversity 105:1-16 ((https://doi.org/10.1007/s13225-020-00466-2)) |
| Otu31547 | | 1 | k__Fungi(100);p__Ascomycota(100);c__Saccharomycetes(100);o__Saccharomycetales(100);f__Metschnikowiaceae(100);g__Metschnikowia(100);s__Metschnikowia_pulcherrima(100); | Metschnikowia | 13 | Saprotroph | \|Undefined Saprotroph\| | Yeast | NULL | Probable | As nectar-tap_saprotroph (Põlme et al. 2020); Decay sugar-rich substrates (Põlme et al. 2020) | Põlme S, et al. 2020. Fungal Diversity 105:1-16 ((https://doi.org/10.1007/s13225-020-00466-2)) |
| Otu31548 | | 1 | k__Fungi(100);p__Ascomycota(100);c__Saccharomycetes(100);o__Saccharomycetales(100);f__Metschnikowiaceae(100);g__Metschnikowia(100);s__Metschnikowia_pulcherrima(100); | Metschnikowia | 13 | Saprotroph | \|Undefined Saprotroph\| | Yeast | NULL | Probable | As nectar-tap_saprotroph (Põlme et al. 2020); Decay sugar-rich substrates (Põlme et al. 2020) | Põlme S, et al. 2020. Fungal Diversity 105:1-16 ((https://doi.org/10.1007/s13225-020-00466-2)) |
| Otu31549 | | 1 | k__Fungi(100);p__Ascomycota(100);c__Saccharomycetes(100);o__Saccharomycetales(100);f__Metschnikowiaceae(100);g__Metschnikowia(100);s__Metschnikowia_pulcherrima(100); | Metschnikowia | 13 | Saprotroph | \|Undefined Saprotroph\| | Yeast | NULL | Probable | As nectar-tap_saprotroph (Põlme et al. 2020); Decay sugar-rich substrates (Põlme et al. 2020) | Põlme S, et al. 2020. Fungal Diversity 105:1-16 ((https://doi.org/10.1007/s13225-020-00466-2)) |
| Otu31550 | | 1 | k__Fungi(100);p__Ascomycota(100);c__Saccharomycetes(100);o__Saccharomycetales(100);f__Metschnikowiaceae(100);g__Metschnikowia(100);s__Metschnikowia_pulcherrima(100); | Metschnikowia | 13 | Saprotroph | \|Undefined Saprotroph\| | Yeast | NULL | Probable | As nectar-tap_saprotroph (Põlme et al. 2020); Decay sugar-rich substrates (Põlme et al. 2020) | Põlme S, et al. 2020. Fungal Diversity 105:1-16 ((https://doi.org/10.1007/s13225-020-00466-2)) |
| Otu31551 | | 1 | k__Fungi(100);p__Ascomycota(100);c__Saccharomycetes(100);o__Saccharomycetales(100);f__Metschnikowiaceae(100);g__Metschnikowia(100);s__Metschnikowia_pulcherrima(100); | Metschnikowia | 13 | Saprotroph | \|Undefined Saprotroph\| | Yeast | NULL | Probable | As nectar-tap_saprotroph (Põlme et al. 2020); Decay sugar-rich substrates (Põlme et al. 2020) | Põlme S, et al. 2020. Fungal Diversity 105:1-16 ((https://doi.org/10.1007/s13225-020-00466-2)) |
| Otu31552 | | 1 | k__Fungi(100);p__Ascomycota(100);c__Saccharomycetes(100);o__Saccharomycetales(100);f__Metschnikowiaceae(100);g__Metschnikowia(100);s__Metschnikowia_pulcherrima(100); | Metschnikowia | 13 | Saprotroph | \|Undefined Saprotroph\| | Yeast | NULL | Probable | As nectar-tap_saprotroph (Põlme et al. 2020); Decay sugar-rich substrates (Põlme et al. 2020) | Põlme S, et al. 2020. Fungal Diversity 105:1-16 ((https://doi.org/10.1007/s13225-020-00466-2)) |
| Otu31553 | | 1 | k__Fungi(100);p__Ascomycota(100);c__Saccharomycetes(100);o__Saccharomycetales(100);f__Metschnikowiaceae(100);g__Metschnikowia(100);s__Metschnikowia_pulcherrima(100); | Metschnikowia | 13 | Saprotroph | \|Undefined Saprotroph\| | Yeast | NULL | Probable | As nectar-tap_saprotroph (Põlme et al. 2020); Decay sugar-rich substrates (Põlme et al. 2020) | Põlme S, et al. 2020. Fungal Diversity 105:1-16 ((https://doi.org/10.1007/s13225-020-00466-2)) |
| Otu31554 | | 1 | k__Fungi(100);p__Ascomycota(100);c__Saccharomycetes(100);o__Saccharomycetales(100);f__Metschnikowiaceae(100);g__Metschnikowia(100);s__Metschnikowia_pulcherrima(100); | Metschnikowia | 13 | Saprotroph | \|Undefined Saprotroph\| | Yeast | NULL | Probable | As nectar-tap_saprotroph (Põlme et al. 2020); Decay sugar-rich substrates (Põlme et al. 2020) | Põlme S, et al. 2020. Fungal Diversity 105:1-16 ((https://doi.org/10.1007/s13225-020-00466-2)) |
| Otu31555 | | 1 | k__Fungi(100);p__Ascomycota(100);c__Saccharomycetes(100);o__Saccharomycetales(100);f__Metschnikowiaceae(100);g__Metschnikowia(100);s__Metschnikowia_pulcherrima(100); | Metschnikowia | 13 | Saprotroph | \|Undefined Saprotroph\| | Yeast | NULL | Probable | As nectar-tap_saprotroph (Põlme et al. 2020); Decay sugar-rich substrates (Põlme et al. 2020) | Põlme S, et al. 2020. Fungal Diversity 105:1-16 ((https://doi.org/10.1007/s13225-020-00466-2)) |
| Otu31556 | | 1 | k__Fungi(100);p__Ascomycota(100);c__Saccharomycetes(100);o__Saccharomycetales(100);f__Metschnikowiaceae(100);g__Metschnikowia(100);s__Metschnikowia_pulcherrima(100); | Metschnikowia | 13 | Saprotroph | \|Undefined Saprotroph\| | Yeast | NULL | Probable | As nectar-tap_saprotroph (Põlme et al. 2020); Decay sugar-rich substrates (Põlme et al. 2020) | Põlme S, et al. 2020. Fungal Diversity 105:1-16 ((https://doi.org/10.1007/s13225-020-00466-2)) |
| Otu31557 | | 1 | k__Fungi(100);p__Ascomycota(100);c__Saccharomycetes(100);o__Saccharomycetales(100);f__Metschnikowiaceae(100);g__Metschnikowia(100);g__Metschnikowia_unclassified(100); | Metschnikowia | 13 | Saprotroph | \|Undefined Saprotroph\| | Yeast | NULL | Probable | As nectar-tap_saprotroph (Põlme et al. 2020); Decay sugar-rich substrates (Põlme et al. 2020) | Põlme S, et al. 2020. Fungal Diversity 105:1-16 ((https://doi.org/10.1007/s13225-020-00466-2)) |
| Otu31558 | | 1 | k__Fungi(100);p__Ascomycota(100);c__Saccharomycetes(100);o__Saccharomycetales(100);f__Metschnikowiaceae(100);g__Metschnikowia(100);s__Metschnikowia_pulcherrima(100); | Metschnikowia | 13 | Saprotroph | \|Undefined Saprotroph\| | Yeast | NULL | Probable | As nectar-tap_saprotroph (Põlme et al. 2020); Decay sugar-rich substrates (Põlme et al. 2020) | Põlme S, et al. 2020. Fungal Diversity 105:1-16 ((https://doi.org/10.1007/s13225-020-00466-2)) |
| Otu31559 | | 1 | k__Fungi(100);p__Ascomycota(100);c__Saccharomycetes(100);o__Saccharomycetales(100);f__Metschnikowiaceae(100);g__Metschnikowia(100);s__Metschnikowia_pulcherrima(100); | Metschnikowia | 13 | Saprotroph | \|Undefined Saprotroph\| | Yeast | NULL | Probable | As nectar-tap_saprotroph (Põlme et al. 2020); Decay sugar-rich substrates (Põlme et al. 2020) | Põlme S, et al. 2020. Fungal Diversity 105:1-16 ((https://doi.org/10.1007/s13225-020-00466-2)) |
| Otu31560 | | 1 | k__Fungi(100);p__Ascomycota(100);c__Saccharomycetes(100);o__Saccharomycetales(100);f__Metschnikowiaceae(100);g__Metschnikowia(100);s__Metschnikowia_pulcherrima(100); | Metschnikowia | 13 | Saprotroph | \|Undefined Saprotroph\| | Yeast | NULL | Probable | As nectar-tap_saprotroph (Põlme et al. 2020); Decay sugar-rich substrates (Põlme et al. 2020) | Põlme S, et al. 2020. Fungal Diversity 105:1-16 ((https://doi.org/10.1007/s13225-020-00466-2)) |
| Otu31561 | | 1 | k__Fungi(100);p__Ascomycota(100);c__Saccharomycetes(100);o__Saccharomycetales(100);f__Metschnikowiaceae(100);g__Metschnikowia(100);s__Metschnikowia_pulcherrima(100); | Metschnikowia | 13 | Saprotroph | \|Undefined Saprotroph\| | Yeast | NULL | Probable | As nectar-tap_saprotroph (Põlme et al. 2020); Decay sugar-rich substrates (Põlme et al. 2020) | Põlme S, et al. 2020. Fungal Diversity 105:1-16 ((https://doi.org/10.1007/s13225-020-00466-2)) |
| Otu31562 | | 1 | k__Fungi(100);p__Ascomycota(100);c__Saccharomycetes(100);o__Saccharomycetales(100);f__Metschnikowiaceae(100);g__Metschnikowia(100);g__Metschnikowia_unclassified(100); | Metschnikowia | 13 | Saprotroph | \|Undefined Saprotroph\| | Yeast | NULL | Probable | As nectar-tap_saprotroph (Põlme et al. 2020); Decay sugar-rich substrates (Põlme et al. 2020) | Põlme S, et al. 2020. Fungal Diversity 105:1-16 ((https://doi.org/10.1007/s13225-020-00466-2)) |
| Otu31563 | | 1 | k__Fungi(100);p__Ascomycota(100);c__Saccharomycetes(100);o__Saccharomycetales(100);f__Metschnikowiaceae(100);g__Metschnikowia(100);g__Metschnikowia_unclassified(100); | Metschnikowia | 13 | Saprotroph | \|Undefined Saprotroph\| | Yeast | NULL | Probable | As nectar-tap_saprotroph (Põlme et al. 2020); Decay sugar-rich substrates (Põlme et al. 2020) | Põlme S, et al. 2020. Fungal Diversity 105:1-16 ((https://doi.org/10.1007/s13225-020-00466-2)) |
| Otu31564 | | 1 | k__Fungi(100);p__Ascomycota(100);c__Saccharomycetes(100);o__Saccharomycetales(100);f__Metschnikowiaceae(100);g__Metschnikowia(100);s__Metschnikowia_pulcherrima(100); | Metschnikowia | 13 | Saprotroph | \|Undefined Saprotroph\| | Yeast | NULL | Probable | As nectar-tap_saprotroph (Põlme et al. 2020); Decay sugar-rich substrates (Põlme et al. 2020) | Põlme S, et al. 2020. Fungal Diversity 105:1-16 ((https://doi.org/10.1007/s13225-020-00466-2)) |
| Otu31565 | | 1 | k__Fungi(100);p__Ascomycota(100);c__Saccharomycetes(100);o__Saccharomycetales(100);f__Metschnikowiaceae(100);g__Metschnikowia(100);s__Metschnikowia_pulcherrima(100); | Metschnikowia | 13 | Saprotroph | \|Undefined Saprotroph\| | Yeast | NULL | Probable | As nectar-tap_saprotroph (Põlme et al. 2020); Decay sugar-rich substrates (Põlme et al. 2020) | Põlme S, et al. 2020. Fungal Diversity 105:1-16 ((https://doi.org/10.1007/s13225-020-00466-2)) |
| Otu31566 | | 1 | k__Fungi(100);p__Ascomycota(100);c__Saccharomycetes(100);o__Saccharomycetales(100);f__Metschnikowiaceae(100);g__Metschnikowia(100);g__Metschnikowia_unclassified(100); | Metschnikowia | 13 | Saprotroph | \|Undefined Saprotroph\| | Yeast | NULL | Probable | As nectar-tap_saprotroph (Põlme et al. 2020); Decay sugar-rich substrates (Põlme et al. 2020) | Põlme S, et al. 2020. Fungal Diversity 105:1-16 ((https://doi.org/10.1007/s13225-020-00466-2)) |
| Otu31567 | | 1 | k__Fungi(100);p__Ascomycota(100);c__Saccharomycetes(100);o__Saccharomycetales(100);f__Metschnikowiaceae(100);g__Metschnikowia(100);s__Metschnikowia_pulcherrima(100); | Metschnikowia | 13 | Saprotroph | \|Undefined Saprotroph\| | Yeast | NULL | Probable | As nectar-tap_saprotroph (Põlme et al. 2020); Decay sugar-rich substrates (Põlme et al. 2020) | Põlme S, et al. 2020. Fungal Diversity 105:1-16 ((https://doi.org/10.1007/s13225-020-00466-2)) |
| Otu31569 | | 1 | k__Fungi(100);p__Ascomycota(100);c__Saccharomycetes(100);o__Saccharomycetales(100);f__Metschnikowiaceae(100);g__Metschnikowia(100);s__Metschnikowia_pulcherrima(100); | Metschnikowia | 13 | Saprotroph | \|Undefined Saprotroph\| | Yeast | NULL | Probable | As nectar-tap_saprotroph (Põlme et al. 2020); Decay sugar-rich substrates (Põlme et al. 2020) | Põlme S, et al. 2020. Fungal Diversity 105:1-16 ((https://doi.org/10.1007/s13225-020-00466-2)) |
| Otu31570 | | 1 | k__Fungi(100);p__Ascomycota(100);c__Saccharomycetes(100);o__Saccharomycetales(100);f__Metschnikowiaceae(100);g__Metschnikowia(100);s__Metschnikowia_pulcherrima(100); | Metschnikowia | 13 | Saprotroph | \|Undefined Saprotroph\| | Yeast | NULL | Probable | As nectar-tap_saprotroph (Põlme et al. 2020); Decay sugar-rich substrates (Põlme et al. 2020) | Põlme S, et al. 2020. Fungal Diversity 105:1-16 ((https://doi.org/10.1007/s13225-020-00466-2)) |
| Otu31571 | | 1 | k__Fungi(100);p__Ascomycota(100);c__Saccharomycetes(100);o__Saccharomycetales(100);f__Metschnikowiaceae(100);g__Metschnikowia(100);s__Metschnikowia_pulcherrima(100); | Metschnikowia | 13 | Saprotroph | \|Undefined Saprotroph\| | Yeast | NULL | Probable | As nectar-tap_saprotroph (Põlme et al. 2020); Decay sugar-rich substrates (Põlme et al. 2020) | Põlme S, et al. 2020. Fungal Diversity 105:1-16 ((https://doi.org/10.1007/s13225-020-00466-2)) |
| Otu31572 | | 1 | k__Fungi(100);p__Ascomycota(100);c__Saccharomycetes(100);o__Saccharomycetales(100);f__Metschnikowiaceae(100);g__Metschnikowia(100);s__Metschnikowia_pulcherrima(100); | Metschnikowia | 13 | Saprotroph | \|Undefined Saprotroph\| | Yeast | NULL | Probable | As nectar-tap_saprotroph (Põlme et al. 2020); Decay sugar-rich substrates (Põlme et al. 2020) | Põlme S, et al. 2020. Fungal Diversity 105:1-16 ((https://doi.org/10.1007/s13225-020-00466-2)) |
| Otu31573 | | 1 | k__Fungi(100);p__Ascomycota(100);c__Saccharomycetes(100);o__Saccharomycetales(100);f__Metschnikowiaceae(100);g__Metschnikowia(100);g__Metschnikowia_unclassified(100); | Metschnikowia | 13 | Saprotroph | \|Undefined Saprotroph\| | Yeast | NULL | Probable | As nectar-tap_saprotroph (Põlme et al. 2020); Decay sugar-rich substrates (Põlme et al. 2020) | Põlme S, et al. 2020. Fungal Diversity 105:1-16 ((https://doi.org/10.1007/s13225-020-00466-2)) |
| Otu31574 | | 1 | k__Fungi(100);p__Ascomycota(100);c__Saccharomycetes(100);o__Saccharomycetales(100);f__Metschnikowiaceae(100);g__Metschnikowia(100);s__Metschnikowia_pulcherrima(100); | Metschnikowia | 13 | Saprotroph | \|Undefined Saprotroph\| | Yeast | NULL | Probable | As nectar-tap_saprotroph (Põlme et al. 2020); Decay sugar-rich substrates (Põlme et al. 2020) | Põlme S, et al. 2020. Fungal Diversity 105:1-16 ((https://doi.org/10.1007/s13225-020-00466-2)) |
| Otu31575 | | 1 | k__Fungi(100);p__Ascomycota(100);c__Saccharomycetes(100);o__Saccharomycetales(100);f__Metschnikowiaceae(100);g__Metschnikowia(100);s__Metschnikowia_pulcherrima(100); | Metschnikowia | 13 | Saprotroph | \|Undefined Saprotroph\| | Yeast | NULL | Probable | As nectar-tap_saprotroph (Põlme et al. 2020); Decay sugar-rich substrates (Põlme et al. 2020) | Põlme S, et al. 2020. Fungal Diversity 105:1-16 ((https://doi.org/10.1007/s13225-020-00466-2)) |
| Otu31576 | | 1 | k__Fungi(100);p__Ascomycota(100);c__Saccharomycetes(100);o__Saccharomycetales(100);f__Metschnikowiaceae(100);g__Metschnikowia(100);s__Metschnikowia_pulcherrima(100); | Metschnikowia | 13 | Saprotroph | \|Undefined Saprotroph\| | Yeast | NULL | Probable | As nectar-tap_saprotroph (Põlme et al. 2020); Decay sugar-rich substrates (Põlme et al. 2020) | Põlme S, et al. 2020. Fungal Diversity 105:1-16 ((https://doi.org/10.1007/s13225-020-00466-2)) |
| Otu31577 | | 1 | k__Fungi(100);p__Ascomycota(100);c__Saccharomycetes(100);o__Saccharomycetales(100);f__Metschnikowiaceae(100);g__Metschnikowia(100);g__Metschnikowia_unclassified(100); | Metschnikowia | 13 | Saprotroph | \|Undefined Saprotroph\| | Yeast | NULL | Probable | As nectar-tap_saprotroph (Põlme et al. 2020); Decay sugar-rich substrates (Põlme et al. 2020) | Põlme S, et al. 2020. Fungal Diversity 105:1-16 ((https://doi.org/10.1007/s13225-020-00466-2)) |
| Otu31578 | | 1 | k__Fungi(100);p__Ascomycota(100);c__Saccharomycetes(100);o__Saccharomycetales(100);f__Metschnikowiaceae(100);g__Metschnikowia(100);g__Metschnikowia_unclassified(100); | Metschnikowia | 13 | Saprotroph | \|Undefined Saprotroph\| | Yeast | NULL | Probable | As nectar-tap_saprotroph (Põlme et al. 2020); Decay sugar-rich substrates (Põlme et al. 2020) | Põlme S, et al. 2020. Fungal Diversity 105:1-16 ((https://doi.org/10.1007/s13225-020-00466-2)) |
| Otu31579 | | 1 | k__Fungi(100);p__Ascomycota(100);c__Saccharomycetes(100);o__Saccharomycetales(100);f__Metschnikowiaceae(100);g__Metschnikowia(100);g__Metschnikowia_unclassified(100); | Metschnikowia | 13 | Saprotroph | \|Undefined Saprotroph\| | Yeast | NULL | Probable | As nectar-tap_saprotroph (Põlme et al. 2020); Decay sugar-rich substrates (Põlme et al. 2020) | Põlme S, et al. 2020. Fungal Diversity 105:1-16 ((https://doi.org/10.1007/s13225-020-00466-2)) |
| Otu31580 | | 1 | k__Fungi(100);p__Ascomycota(100);c__Saccharomycetes(100);o__Saccharomycetales(100);f__Metschnikowiaceae(100);g__Metschnikowia(100);g__Metschnikowia_unclassified(100); | Metschnikowia | 13 | Saprotroph | \|Undefined Saprotroph\| | Yeast | NULL | Probable | As nectar-tap_saprotroph (Põlme et al. 2020); Decay sugar-rich substrates (Põlme et al. 2020) | Põlme S, et al. 2020. Fungal Diversity 105:1-16 ((https://doi.org/10.1007/s13225-020-00466-2)) |
| Otu31581 | | 1 | k__Fungi(100);p__Ascomycota(100);c__Saccharomycetes(100);o__Saccharomycetales(100);f__Metschnikowiaceae(100);g__Metschnikowia(100);g__Metschnikowia_unclassified(100); | Metschnikowia | 13 | Saprotroph | \|Undefined Saprotroph\| | Yeast | NULL | Probable | As nectar-tap_saprotroph (Põlme et al. 2020); Decay sugar-rich substrates (Põlme et al. 2020) | Põlme S, et al. 2020. Fungal Diversity 105:1-16 ((https://doi.org/10.1007/s13225-020-00466-2)) |
| Otu31582 | | 1 | k__Fungi(100);p__Ascomycota(100);c__Saccharomycetes(100);o__Saccharomycetales(100);f__Metschnikowiaceae(100);g__Metschnikowia(100);s__Metschnikowia_pulcherrima(100); | Metschnikowia | 13 | Saprotroph | \|Undefined Saprotroph\| | Yeast | NULL | Probable | As nectar-tap_saprotroph (Põlme et al. 2020); Decay sugar-rich substrates (Põlme et al. 2020) | Põlme S, et al. 2020. Fungal Diversity 105:1-16 ((https://doi.org/10.1007/s13225-020-00466-2)) |
| Otu31583 | | 1 | k__Fungi(100);p__Ascomycota(100);c__Saccharomycetes(100);o__Saccharomycetales(100);f__Metschnikowiaceae(100);g__Metschnikowia(100);g__Metschnikowia_unclassified(100); | Metschnikowia | 13 | Saprotroph | \|Undefined Saprotroph\| | Yeast | NULL | Probable | As nectar-tap_saprotroph (Põlme et al. 2020); Decay sugar-rich substrates (Põlme et al. 2020) | Põlme S, et al. 2020. Fungal Diversity 105:1-16 ((https://doi.org/10.1007/s13225-020-00466-2)) |
| Otu31584 | | 1 | k__Fungi(100);p__Ascomycota(100);c__Saccharomycetes(100);o__Saccharomycetales(100);f__Metschnikowiaceae(100);g__Metschnikowia(100);s__Metschnikowia_pulcherrima(100); | Metschnikowia | 13 | Saprotroph | \|Undefined Saprotroph\| | Yeast | NULL | Probable | As nectar-tap_saprotroph (Põlme et al. 2020); Decay sugar-rich substrates (Põlme et al. 2020) | Põlme S, et al. 2020. Fungal Diversity 105:1-16 ((https://doi.org/10.1007/s13225-020-00466-2)) |
| Otu31586 | | 1 | k__Fungi(100);p__Ascomycota(100);c__Saccharomycetes(100);o__Saccharomycetales(100);f__Metschnikowiaceae(100);g__Metschnikowia(100);s__Metschnikowia_pulcherrima(100); | Metschnikowia | 13 | Saprotroph | \|Undefined Saprotroph\| | Yeast | NULL | Probable | As nectar-tap_saprotroph (Põlme et al. 2020); Decay sugar-rich substrates (Põlme et al. 2020) | Põlme S, et al. 2020. Fungal Diversity 105:1-16 ((https://doi.org/10.1007/s13225-020-00466-2)) |
| Otu31587 | | 1 | k__Fungi(100);p__Ascomycota(100);c__Saccharomycetes(100);o__Saccharomycetales(100);f__Metschnikowiaceae(100);g__Metschnikowia(100);g__Metschnikowia_unclassified(100); | Metschnikowia | 13 | Saprotroph | \|Undefined Saprotroph\| | Yeast | NULL | Probable | As nectar-tap_saprotroph (Põlme et al. 2020); Decay sugar-rich substrates (Põlme et al. 2020) | Põlme S, et al. 2020. Fungal Diversity 105:1-16 ((https://doi.org/10.1007/s13225-020-00466-2)) |
| Otu31588 | | 1 | k__Fungi(100);p__Ascomycota(100);c__Saccharomycetes(100);o__Saccharomycetales(100);f__Metschnikowiaceae(100);g__Metschnikowia(100);s__Metschnikowia_pulcherrima(100); | Metschnikowia | 13 | Saprotroph | \|Undefined Saprotroph\| | Yeast | NULL | Probable | As nectar-tap_saprotroph (Põlme et al. 2020); Decay sugar-rich substrates (Põlme et al. 2020) | Põlme S, et al. 2020. Fungal Diversity 105:1-16 ((https://doi.org/10.1007/s13225-020-00466-2)) |
| Otu31589 | | 1 | k__Fungi(100);p__Ascomycota(100);c__Saccharomycetes(100);o__Saccharomycetales(100);f__Metschnikowiaceae(100);g__Metschnikowia(100);g__Metschnikowia_unclassified(100); | Metschnikowia | 13 | Saprotroph | \|Undefined Saprotroph\| | Yeast | NULL | Probable | As nectar-tap_saprotroph (Põlme et al. 2020); Decay sugar-rich substrates (Põlme et al. 2020) | Põlme S, et al. 2020. Fungal Diversity 105:1-16 ((https://doi.org/10.1007/s13225-020-00466-2)) |
| Otu31590 | | 1 | k__Fungi(100);p__Ascomycota(100);c__Saccharomycetes(100);o__Saccharomycetales(100);f__Metschnikowiaceae(100);g__Metschnikowia(100);g__Metschnikowia_unclassified(100); | Metschnikowia | 13 | Saprotroph | \|Undefined Saprotroph\| | Yeast | NULL | Probable | As nectar-tap_saprotroph (Põlme et al. 2020); Decay sugar-rich substrates (Põlme et al. 2020) | Põlme S, et al. 2020. Fungal Diversity 105:1-16 ((https://doi.org/10.1007/s13225-020-00466-2)) |
| Otu31591 | | 1 | k__Fungi(100);p__Ascomycota(100);c__Saccharomycetes(100);o__Saccharomycetales(100);f__Metschnikowiaceae(100);g__Metschnikowia(100);g__Metschnikowia_unclassified(100); | Metschnikowia | 13 | Saprotroph | \|Undefined Saprotroph\| | Yeast | NULL | Probable | As nectar-tap_saprotroph (Põlme et al. 2020); Decay sugar-rich substrates (Põlme et al. 2020) | Põlme S, et al. 2020. Fungal Diversity 105:1-16 ((https://doi.org/10.1007/s13225-020-00466-2)) |
| Otu31592 | | 1 | k__Fungi(100);p__Ascomycota(100);c__Saccharomycetes(100);o__Saccharomycetales(100);f__Metschnikowiaceae(100);g__Metschnikowia(100);s__Metschnikowia_pulcherrima(100); | Metschnikowia | 13 | Saprotroph | \|Undefined Saprotroph\| | Yeast | NULL | Probable | As nectar-tap_saprotroph (Põlme et al. 2020); Decay sugar-rich substrates (Põlme et al. 2020) | Põlme S, et al. 2020. Fungal Diversity 105:1-16 ((https://doi.org/10.1007/s13225-020-00466-2)) |
| Otu31593 | | 1 | k__Fungi(100);p__Ascomycota(100);c__Saccharomycetes(100);o__Saccharomycetales(100);f__Metschnikowiaceae(100);g__Metschnikowia(100);s__Metschnikowia_pulcherrima(100); | Metschnikowia | 13 | Saprotroph | \|Undefined Saprotroph\| | Yeast | NULL | Probable | As nectar-tap_saprotroph (Põlme et al. 2020); Decay sugar-rich substrates (Põlme et al. 2020) | Põlme S, et al. 2020. Fungal Diversity 105:1-16 ((https://doi.org/10.1007/s13225-020-00466-2)) |
| Otu31594 | | 1 | k__Fungi(100);p__Ascomycota(100);c__Saccharomycetes(100);o__Saccharomycetales(100);f__Metschnikowiaceae(100);g__Metschnikowia(100);g__Metschnikowia_unclassified(100); | Metschnikowia | 13 | Saprotroph | \|Undefined Saprotroph\| | Yeast | NULL | Probable | As nectar-tap_saprotroph (Põlme et al. 2020); Decay sugar-rich substrates (Põlme et al. 2020) | Põlme S, et al. 2020. Fungal Diversity 105:1-16 ((https://doi.org/10.1007/s13225-020-00466-2)) |
| Otu31597 | | 1 | k__Fungi(100);p__Ascomycota(100);c__Saccharomycetes(100);o__Saccharomycetales(100);f__Metschnikowiaceae(100);g__Metschnikowia(100);s__Metschnikowia_pulcherrima(100); | Metschnikowia | 13 | Saprotroph | \|Undefined Saprotroph\| | Yeast | NULL | Probable | As nectar-tap_saprotroph (Põlme et al. 2020); Decay sugar-rich substrates (Põlme et al. 2020) | Põlme S, et al. 2020. Fungal Diversity 105:1-16 ((https://doi.org/10.1007/s13225-020-00466-2)) |
| Otu31598 | | 1 | k__Fungi(100);p__Ascomycota(100);c__Saccharomycetes(100);o__Saccharomycetales(100);f__Metschnikowiaceae(100);g__Metschnikowia(100);s__Metschnikowia_pulcherrima(100); | Metschnikowia | 13 | Saprotroph | \|Undefined Saprotroph\| | Yeast | NULL | Probable | As nectar-tap_saprotroph (Põlme et al. 2020); Decay sugar-rich substrates (Põlme et al. 2020) | Põlme S, et al. 2020. Fungal Diversity 105:1-16 ((https://doi.org/10.1007/s13225-020-00466-2)) |
| Otu31599 | | 1 | k__Fungi(100);p__Ascomycota(100);c__Saccharomycetes(100);o__Saccharomycetales(100);f__Metschnikowiaceae(100);g__Metschnikowia(100);g__Metschnikowia_unclassified(100); | Metschnikowia | 13 | Saprotroph | \|Undefined Saprotroph\| | Yeast | NULL | Probable | As nectar-tap_saprotroph (Põlme et al. 2020); Decay sugar-rich substrates (Põlme et al. 2020) | Põlme S, et al. 2020. Fungal Diversity 105:1-16 ((https://doi.org/10.1007/s13225-020-00466-2)) |
| Otu31600 | | 1 | k__Fungi(100);p__Ascomycota(100);c__Saccharomycetes(100);o__Saccharomycetales(100);f__Metschnikowiaceae(100);g__Metschnikowia(100);s__Metschnikowia_pulcherrima(100); | Metschnikowia | 13 | Saprotroph | \|Undefined Saprotroph\| | Yeast | NULL | Probable | As nectar-tap_saprotroph (Põlme et al. 2020); Decay sugar-rich substrates (Põlme et al. 2020) | Põlme S, et al. 2020. Fungal Diversity 105:1-16 ((https://doi.org/10.1007/s13225-020-00466-2)) |
| Otu31601 | | 1 | k__Fungi(100);p__Ascomycota(100);c__Saccharomycetes(100);o__Saccharomycetales(100);f__Metschnikowiaceae(100);g__Metschnikowia(100);s__Metschnikowia_pulcherrima(100); | Metschnikowia | 13 | Saprotroph | \|Undefined Saprotroph\| | Yeast | NULL | Probable | As nectar-tap_saprotroph (Põlme et al. 2020); Decay sugar-rich substrates (Põlme et al. 2020) | Põlme S, et al. 2020. Fungal Diversity 105:1-16 ((https://doi.org/10.1007/s13225-020-00466-2)) |
| Otu31603 | | 1 | k__Fungi(100);p__Ascomycota(100);c__Saccharomycetes(100);o__Saccharomycetales(100);f__Metschnikowiaceae(100);g__Metschnikowia(100);s__Metschnikowia_pulcherrima(100); | Metschnikowia | 13 | Saprotroph | \|Undefined Saprotroph\| | Yeast | NULL | Probable | As nectar-tap_saprotroph (Põlme et al. 2020); Decay sugar-rich substrates (Põlme et al. 2020) | Põlme S, et al. 2020. Fungal Diversity 105:1-16 ((https://doi.org/10.1007/s13225-020-00466-2)) |
| Otu31604 | | 1 | k__Fungi(100);p__Ascomycota(100);c__Saccharomycetes(100);o__Saccharomycetales(100);f__Metschnikowiaceae(100);g__Metschnikowia(100);s__Metschnikowia_pulcherrima(100); | Metschnikowia | 13 | Saprotroph | \|Undefined Saprotroph\| | Yeast | NULL | Probable | As nectar-tap_saprotroph (Põlme et al. 2020); Decay sugar-rich substrates (Põlme et al. 2020) | Põlme S, et al. 2020. Fungal Diversity 105:1-16 ((https://doi.org/10.1007/s13225-020-00466-2)) |
| Otu31605 | | 1 | k__Fungi(100);p__Ascomycota(100);c__Saccharomycetes(100);o__Saccharomycetales(100);f__Metschnikowiaceae(100);g__Metschnikowia(100);g__Metschnikowia_unclassified(100); | Metschnikowia | 13 | Saprotroph | \|Undefined Saprotroph\| | Yeast | NULL | Probable | As nectar-tap_saprotroph (Põlme et al. 2020); Decay sugar-rich substrates (Põlme et al. 2020) | Põlme S, et al. 2020. Fungal Diversity 105:1-16 ((https://doi.org/10.1007/s13225-020-00466-2)) |
| Otu31606 | | 1 | k__Fungi(100);p__Ascomycota(100);c__Saccharomycetes(100);o__Saccharomycetales(100);f__Metschnikowiaceae(100);g__Metschnikowia(100);s__Metschnikowia_pulcherrima(100); | Metschnikowia | 13 | Saprotroph | \|Undefined Saprotroph\| | Yeast | NULL | Probable | As nectar-tap_saprotroph (Põlme et al. 2020); Decay sugar-rich substrates (Põlme et al. 2020) | Põlme S, et al. 2020. Fungal Diversity 105:1-16 ((https://doi.org/10.1007/s13225-020-00466-2)) |
| Otu31607 | | 1 | k__Fungi(100);p__Ascomycota(100);c__Saccharomycetes(100);o__Saccharomycetales(100);f__Metschnikowiaceae(100);g__Metschnikowia(100);s__Metschnikowia_pulcherrima(100); | Metschnikowia | 13 | Saprotroph | \|Undefined Saprotroph\| | Yeast | NULL | Probable | As nectar-tap_saprotroph (Põlme et al. 2020); Decay sugar-rich substrates (Põlme et al. 2020) | Põlme S, et al. 2020. Fungal Diversity 105:1-16 ((https://doi.org/10.1007/s13225-020-00466-2)) |
| Otu31608 | | 1 | k__Fungi(100);p__Ascomycota(100);c__Saccharomycetes(100);o__Saccharomycetales(100);f__Metschnikowiaceae(100);g__Metschnikowia(100);s__Metschnikowia_pulcherrima(100); | Metschnikowia | 13 | Saprotroph | \|Undefined Saprotroph\| | Yeast | NULL | Probable | As nectar-tap_saprotroph (Põlme et al. 2020); Decay sugar-rich substrates (Põlme et al. 2020) | Põlme S, et al. 2020. Fungal Diversity 105:1-16 ((https://doi.org/10.1007/s13225-020-00466-2)) |
| Otu31609 | | 1 | k__Fungi(100);p__Ascomycota(100);c__Saccharomycetes(100);o__Saccharomycetales(100);f__Metschnikowiaceae(100);g__Metschnikowia(100);g__Metschnikowia_unclassified(100); | Metschnikowia | 13 | Saprotroph | \|Undefined Saprotroph\| | Yeast | NULL | Probable | As nectar-tap_saprotroph (Põlme et al. 2020); Decay sugar-rich substrates (Põlme et al. 2020) | Põlme S, et al. 2020. Fungal Diversity 105:1-16 ((https://doi.org/10.1007/s13225-020-00466-2)) |
| Otu31610 | | 1 | k__Fungi(100);p__Ascomycota(100);c__Saccharomycetes(100);o__Saccharomycetales(100);f__Metschnikowiaceae(100);g__Metschnikowia(100);s__Metschnikowia_pulcherrima(100); | Metschnikowia | 13 | Saprotroph | \|Undefined Saprotroph\| | Yeast | NULL | Probable | As nectar-tap_saprotroph (Põlme et al. 2020); Decay sugar-rich substrates (Põlme et al. 2020) | Põlme S, et al. 2020. Fungal Diversity 105:1-16 ((https://doi.org/10.1007/s13225-020-00466-2)) |
| Otu31611 | | 1 | k__Fungi(100);p__Ascomycota(100);c__Saccharomycetes(100);o__Saccharomycetales(100);f__Metschnikowiaceae(100);g__Metschnikowia(100);g__Metschnikowia_unclassified(100); | Metschnikowia | 13 | Saprotroph | \|Undefined Saprotroph\| | Yeast | NULL | Probable | As nectar-tap_saprotroph (Põlme et al. 2020); Decay sugar-rich substrates (Põlme et al. 2020) | Põlme S, et al. 2020. Fungal Diversity 105:1-16 ((https://doi.org/10.1007/s13225-020-00466-2)) |
| Otu31612 | | 1 | k__Fungi(100);p__Ascomycota(100);c__Saccharomycetes(100);o__Saccharomycetales(100);f__Metschnikowiaceae(100);g__Metschnikowia(100);s__Metschnikowia_pulcherrima(100); | Metschnikowia | 13 | Saprotroph | \|Undefined Saprotroph\| | Yeast | NULL | Probable | As nectar-tap_saprotroph (Põlme et al. 2020); Decay sugar-rich substrates (Põlme et al. 2020) | Põlme S, et al. 2020. Fungal Diversity 105:1-16 ((https://doi.org/10.1007/s13225-020-00466-2)) |
| Otu31613 | | 1 | k__Fungi(100);p__Ascomycota(100);c__Saccharomycetes(100);o__Saccharomycetales(100);f__Metschnikowiaceae(100);g__Metschnikowia(100);s__Metschnikowia_pulcherrima(100); | Metschnikowia | 13 | Saprotroph | \|Undefined Saprotroph\| | Yeast | NULL | Probable | As nectar-tap_saprotroph (Põlme et al. 2020); Decay sugar-rich substrates (Põlme et al. 2020) | Põlme S, et al. 2020. Fungal Diversity 105:1-16 ((https://doi.org/10.1007/s13225-020-00466-2)) |
| Otu31614 | | 1 | k__Fungi(100);p__Ascomycota(100);c__Saccharomycetes(100);o__Saccharomycetales(100);f__Metschnikowiaceae(100);g__Metschnikowia(100);s__Metschnikowia_pulcherrima(100); | Metschnikowia | 13 | Saprotroph | \|Undefined Saprotroph\| | Yeast | NULL | Probable | As nectar-tap_saprotroph (Põlme et al. 2020); Decay sugar-rich substrates (Põlme et al. 2020) | Põlme S, et al. 2020. Fungal Diversity 105:1-16 ((https://doi.org/10.1007/s13225-020-00466-2)) |
| Otu31615 | | 1 | k__Fungi(100);p__Ascomycota(100);c__Saccharomycetes(100);o__Saccharomycetales(100);f__Metschnikowiaceae(100);g__Metschnikowia(100);g__Metschnikowia_unclassified(100); | Metschnikowia | 13 | Saprotroph | \|Undefined Saprotroph\| | Yeast | NULL | Probable | As nectar-tap_saprotroph (Põlme et al. 2020); Decay sugar-rich substrates (Põlme et al. 2020) | Põlme S, et al. 2020. Fungal Diversity 105:1-16 ((https://doi.org/10.1007/s13225-020-00466-2)) |
| Otu31616 | | 1 | k__Fungi(100);p__Ascomycota(100);c__Saccharomycetes(100);o__Saccharomycetales(100);f__Metschnikowiaceae(100);g__Metschnikowia(100);g__Metschnikowia_unclassified(100); | Metschnikowia | 13 | Saprotroph | \|Undefined Saprotroph\| | Yeast | NULL | Probable | As nectar-tap_saprotroph (Põlme et al. 2020); Decay sugar-rich substrates (Põlme et al. 2020) | Põlme S, et al. 2020. Fungal Diversity 105:1-16 ((https://doi.org/10.1007/s13225-020-00466-2)) |
| Otu31617 | | 1 | k__Fungi(100);p__Ascomycota(100);c__Saccharomycetes(100);o__Saccharomycetales(100);f__Metschnikowiaceae(100);g__Metschnikowia(100);s__Metschnikowia_pulcherrima(100); | Metschnikowia | 13 | Saprotroph | \|Undefined Saprotroph\| | Yeast | NULL | Probable | As nectar-tap_saprotroph (Põlme et al. 2020); Decay sugar-rich substrates (Põlme et al. 2020) | Põlme S, et al. 2020. Fungal Diversity 105:1-16 ((https://doi.org/10.1007/s13225-020-00466-2)) |
| Otu31618 | | 1 | k__Fungi(100);p__Ascomycota(100);c__Saccharomycetes(100);o__Saccharomycetales(100);f__Metschnikowiaceae(100);g__Metschnikowia(100);g__Metschnikowia_unclassified(100); | Metschnikowia | 13 | Saprotroph | \|Undefined Saprotroph\| | Yeast | NULL | Probable | As nectar-tap_saprotroph (Põlme et al. 2020); Decay sugar-rich substrates (Põlme et al. 2020) | Põlme S, et al. 2020. Fungal Diversity 105:1-16 ((https://doi.org/10.1007/s13225-020-00466-2)) |
| Otu31619 | | 1 | k__Fungi(100);p__Ascomycota(100);c__Saccharomycetes(100);o__Saccharomycetales(100);f__Metschnikowiaceae(100);g__Metschnikowia(100);g__Metschnikowia_unclassified(100); | Metschnikowia | 13 | Saprotroph | \|Undefined Saprotroph\| | Yeast | NULL | Probable | As nectar-tap_saprotroph (Põlme et al. 2020); Decay sugar-rich substrates (Põlme et al. 2020) | Põlme S, et al. 2020. Fungal Diversity 105:1-16 ((https://doi.org/10.1007/s13225-020-00466-2)) |
| Otu31620 | | 1 | k__Fungi(100);p__Ascomycota(100);c__Saccharomycetes(100);o__Saccharomycetales(100);f__Metschnikowiaceae(100);g__Metschnikowia(100);s__Metschnikowia_pulcherrima(100); | Metschnikowia | 13 | Saprotroph | \|Undefined Saprotroph\| | Yeast | NULL | Probable | As nectar-tap_saprotroph (Põlme et al. 2020); Decay sugar-rich substrates (Põlme et al. 2020) | Põlme S, et al. 2020. Fungal Diversity 105:1-16 ((https://doi.org/10.1007/s13225-020-00466-2)) |
| Otu31621 | | 1 | k__Fungi(100);p__Ascomycota(100);c__Saccharomycetes(100);o__Saccharomycetales(100);f__Metschnikowiaceae(100);g__Metschnikowia(100);g__Metschnikowia_unclassified(100); | Metschnikowia | 13 | Saprotroph | \|Undefined Saprotroph\| | Yeast | NULL | Probable | As nectar-tap_saprotroph (Põlme et al. 2020); Decay sugar-rich substrates (Põlme et al. 2020) | Põlme S, et al. 2020. Fungal Diversity 105:1-16 ((https://doi.org/10.1007/s13225-020-00466-2)) |
| Otu31622 | | 1 | k__Fungi(100);p__Ascomycota(100);c__Saccharomycetes(100);o__Saccharomycetales(100);f__Metschnikowiaceae(100);g__Metschnikowia(100);s__Metschnikowia_pulcherrima(100); | Metschnikowia | 13 | Saprotroph | \|Undefined Saprotroph\| | Yeast | NULL | Probable | As nectar-tap_saprotroph (Põlme et al. 2020); Decay sugar-rich substrates (Põlme et al. 2020) | Põlme S, et al. 2020. Fungal Diversity 105:1-16 ((https://doi.org/10.1007/s13225-020-00466-2)) |
| Otu31623 | | 1 | k__Fungi(100);p__Ascomycota(100);c__Saccharomycetes(100);o__Saccharomycetales(100);f__Metschnikowiaceae(100);g__Metschnikowia(100);s__Metschnikowia_pulcherrima(100); | Metschnikowia | 13 | Saprotroph | \|Undefined Saprotroph\| | Yeast | NULL | Probable | As nectar-tap_saprotroph (Põlme et al. 2020); Decay sugar-rich substrates (Põlme et al. 2020) | Põlme S, et al. 2020. Fungal Diversity 105:1-16 ((https://doi.org/10.1007/s13225-020-00466-2)) |
| Otu31624 | | 1 | k__Fungi(100);p__Ascomycota(100);c__Saccharomycetes(100);o__Saccharomycetales(100);f__Metschnikowiaceae(100);g__Metschnikowia(100);g__Metschnikowia_unclassified(100); | Metschnikowia | 13 | Saprotroph | \|Undefined Saprotroph\| | Yeast | NULL | Probable | As nectar-tap_saprotroph (Põlme et al. 2020); Decay sugar-rich substrates (Põlme et al. 2020) | Põlme S, et al. 2020. Fungal Diversity 105:1-16 ((https://doi.org/10.1007/s13225-020-00466-2)) |
| Otu31625 | | 1 | k__Fungi(100);p__Ascomycota(100);c__Saccharomycetes(100);o__Saccharomycetales(100);f__Metschnikowiaceae(100);g__Metschnikowia(100);g__Metschnikowia_unclassified(100); | Metschnikowia | 13 | Saprotroph | \|Undefined Saprotroph\| | Yeast | NULL | Probable | As nectar-tap_saprotroph (Põlme et al. 2020); Decay sugar-rich substrates (Põlme et al. 2020) | Põlme S, et al. 2020. Fungal Diversity 105:1-16 ((https://doi.org/10.1007/s13225-020-00466-2)) |
| Otu31626 | | 1 | k__Fungi(100);p__Ascomycota(100);c__Saccharomycetes(100);o__Saccharomycetales(100);f__Metschnikowiaceae(100);g__Metschnikowia(100);s__Metschnikowia_pulcherrima(100); | Metschnikowia | 13 | Saprotroph | \|Undefined Saprotroph\| | Yeast | NULL | Probable | As nectar-tap_saprotroph (Põlme et al. 2020); Decay sugar-rich substrates (Põlme et al. 2020) | Põlme S, et al. 2020. Fungal Diversity 105:1-16 ((https://doi.org/10.1007/s13225-020-00466-2)) |
| Otu31627 | | 1 | k__Fungi(100);p__Ascomycota(100);c__Saccharomycetes(100);o__Saccharomycetales(100);f__Metschnikowiaceae(100);g__Metschnikowia(100);s__Metschnikowia_pulcherrima(100); | Metschnikowia | 13 | Saprotroph | \|Undefined Saprotroph\| | Yeast | NULL | Probable | As nectar-tap_saprotroph (Põlme et al. 2020); Decay sugar-rich substrates (Põlme et al. 2020) | Põlme S, et al. 2020. Fungal Diversity 105:1-16 ((https://doi.org/10.1007/s13225-020-00466-2)) |
| Otu31628 | | 1 | k__Fungi(100);p__Ascomycota(100);c__Saccharomycetes(100);o__Saccharomycetales(100);f__Metschnikowiaceae(100);g__Metschnikowia(100);s__Metschnikowia_pulcherrima(100); | Metschnikowia | 13 | Saprotroph | \|Undefined Saprotroph\| | Yeast | NULL | Probable | As nectar-tap_saprotroph (Põlme et al. 2020); Decay sugar-rich substrates (Põlme et al. 2020) | Põlme S, et al. 2020. Fungal Diversity 105:1-16 ((https://doi.org/10.1007/s13225-020-00466-2)) |
| Otu31629 | | 1 | k__Fungi(100);p__Ascomycota(100);c__Saccharomycetes(100);o__Saccharomycetales(100);f__Metschnikowiaceae(100);g__Metschnikowia(100);g__Metschnikowia_unclassified(100); | Metschnikowia | 13 | Saprotroph | \|Undefined Saprotroph\| | Yeast | NULL | Probable | As nectar-tap_saprotroph (Põlme et al. 2020); Decay sugar-rich substrates (Põlme et al. 2020) | Põlme S, et al. 2020. Fungal Diversity 105:1-16 ((https://doi.org/10.1007/s13225-020-00466-2)) |
| Otu31630 | | 1 | k__Fungi(100);p__Ascomycota(100);c__Saccharomycetes(100);o__Saccharomycetales(100);f__Metschnikowiaceae(100);g__Metschnikowia(100);g__Metschnikowia_unclassified(100); | Metschnikowia | 13 | Saprotroph | \|Undefined Saprotroph\| | Yeast | NULL | Probable | As nectar-tap_saprotroph (Põlme et al. 2020); Decay sugar-rich substrates (Põlme et al. 2020) | Põlme S, et al. 2020. Fungal Diversity 105:1-16 ((https://doi.org/10.1007/s13225-020-00466-2)) |
| Otu31631 | | 1 | k__Fungi(100);p__Ascomycota(100);c__Saccharomycetes(100);o__Saccharomycetales(100);f__Metschnikowiaceae(100);g__Metschnikowia(100);g__Metschnikowia_unclassified(100); | Metschnikowia | 13 | Saprotroph | \|Undefined Saprotroph\| | Yeast | NULL | Probable | As nectar-tap_saprotroph (Põlme et al. 2020); Decay sugar-rich substrates (Põlme et al. 2020) | Põlme S, et al. 2020. Fungal Diversity 105:1-16 ((https://doi.org/10.1007/s13225-020-00466-2)) |
| Otu31632 | | 1 | k__Fungi(100);p__Ascomycota(100);c__Saccharomycetes(100);o__Saccharomycetales(100);f__Metschnikowiaceae(100);g__Metschnikowia(100);g__Metschnikowia_unclassified(100); | Metschnikowia | 13 | Saprotroph | \|Undefined Saprotroph\| | Yeast | NULL | Probable | As nectar-tap_saprotroph (Põlme et al. 2020); Decay sugar-rich substrates (Põlme et al. 2020) | Põlme S, et al. 2020. Fungal Diversity 105:1-16 ((https://doi.org/10.1007/s13225-020-00466-2)) |
| Otu31633 | | 1 | k__Fungi(100);p__Ascomycota(100);c__Saccharomycetes(100);o__Saccharomycetales(100);f__Metschnikowiaceae(100);g__Metschnikowia(100);g__Metschnikowia_unclassified(100); | Metschnikowia | 13 | Saprotroph | \|Undefined Saprotroph\| | Yeast | NULL | Probable | As nectar-tap_saprotroph (Põlme et al. 2020); Decay sugar-rich substrates (Põlme et al. 2020) | Põlme S, et al. 2020. Fungal Diversity 105:1-16 ((https://doi.org/10.1007/s13225-020-00466-2)) |
| Otu31634 | | 1 | k__Fungi(100);p__Ascomycota(100);c__Saccharomycetes(100);o__Saccharomycetales(100);f__Metschnikowiaceae(100);g__Metschnikowia(100);s__Metschnikowia_pulcherrima(100); | Metschnikowia | 13 | Saprotroph | \|Undefined Saprotroph\| | Yeast | NULL | Probable | As nectar-tap_saprotroph (Põlme et al. 2020); Decay sugar-rich substrates (Põlme et al. 2020) | Põlme S, et al. 2020. Fungal Diversity 105:1-16 ((https://doi.org/10.1007/s13225-020-00466-2)) |
| Otu31635 | | 1 | k__Fungi(100);p__Ascomycota(100);c__Saccharomycetes(100);o__Saccharomycetales(100);f__Metschnikowiaceae(100);g__Metschnikowia(100);s__Metschnikowia_pulcherrima(100); | Metschnikowia | 13 | Saprotroph | \|Undefined Saprotroph\| | Yeast | NULL | Probable | As nectar-tap_saprotroph (Põlme et al. 2020); Decay sugar-rich substrates (Põlme et al. 2020) | Põlme S, et al. 2020. Fungal Diversity 105:1-16 ((https://doi.org/10.1007/s13225-020-00466-2)) |
| Otu31636 | | 1 | k__Fungi(100);p__Ascomycota(100);c__Saccharomycetes(100);o__Saccharomycetales(100);f__Metschnikowiaceae(100);g__Metschnikowia(100);g__Metschnikowia_unclassified(100); | Metschnikowia | 13 | Saprotroph | \|Undefined Saprotroph\| | Yeast | NULL | Probable | As nectar-tap_saprotroph (Põlme et al. 2020); Decay sugar-rich substrates (Põlme et al. 2020) | Põlme S, et al. 2020. Fungal Diversity 105:1-16 ((https://doi.org/10.1007/s13225-020-00466-2)) |
| Otu31637 | | 1 | k__Fungi(100);p__Ascomycota(100);c__Saccharomycetes(100);o__Saccharomycetales(100);f__Metschnikowiaceae(100);g__Metschnikowia(100);s__Metschnikowia_pulcherrima(100); | Metschnikowia | 13 | Saprotroph | \|Undefined Saprotroph\| | Yeast | NULL | Probable | As nectar-tap_saprotroph (Põlme et al. 2020); Decay sugar-rich substrates (Põlme et al. 2020) | Põlme S, et al. 2020. Fungal Diversity 105:1-16 ((https://doi.org/10.1007/s13225-020-00466-2)) |
| Otu31638 | | 1 | k__Fungi(100);p__Ascomycota(100);c__Saccharomycetes(100);o__Saccharomycetales(100);f__Metschnikowiaceae(100);g__Metschnikowia(100);g__Metschnikowia_unclassified(100); | Metschnikowia | 13 | Saprotroph | \|Undefined Saprotroph\| | Yeast | NULL | Probable | As nectar-tap_saprotroph (Põlme et al. 2020); Decay sugar-rich substrates (Põlme et al. 2020) | Põlme S, et al. 2020. Fungal Diversity 105:1-16 ((https://doi.org/10.1007/s13225-020-00466-2)) |
| Otu31639 | | 1 | k__Fungi(100);p__Ascomycota(100);c__Saccharomycetes(100);o__Saccharomycetales(100);f__Metschnikowiaceae(100);g__Metschnikowia(100);g__Metschnikowia_unclassified(100); | Metschnikowia | 13 | Saprotroph | \|Undefined Saprotroph\| | Yeast | NULL | Probable | As nectar-tap_saprotroph (Põlme et al. 2020); Decay sugar-rich substrates (Põlme et al. 2020) | Põlme S, et al. 2020. Fungal Diversity 105:1-16 ((https://doi.org/10.1007/s13225-020-00466-2)) |
| Otu31640 | | 1 | k__Fungi(100);p__Ascomycota(100);c__Saccharomycetes(100);o__Saccharomycetales(100);f__Metschnikowiaceae(100);g__Metschnikowia(100);s__Metschnikowia_pulcherrima(100); | Metschnikowia | 13 | Saprotroph | \|Undefined Saprotroph\| | Yeast | NULL | Probable | As nectar-tap_saprotroph (Põlme et al. 2020); Decay sugar-rich substrates (Põlme et al. 2020) | Põlme S, et al. 2020. Fungal Diversity 105:1-16 ((https://doi.org/10.1007/s13225-020-00466-2)) |
| Otu31641 | | 1 | k__Fungi(100);p__Ascomycota(100);c__Saccharomycetes(100);o__Saccharomycetales(100);f__Metschnikowiaceae(100);g__Metschnikowia(100);g__Metschnikowia_unclassified(100); | Metschnikowia | 13 | Saprotroph | \|Undefined Saprotroph\| | Yeast | NULL | Probable | As nectar-tap_saprotroph (Põlme et al. 2020); Decay sugar-rich substrates (Põlme et al. 2020) | Põlme S, et al. 2020. Fungal Diversity 105:1-16 ((https://doi.org/10.1007/s13225-020-00466-2)) |
| Otu31642 | | 1 | k__Fungi(100);p__Ascomycota(100);c__Saccharomycetes(100);o__Saccharomycetales(100);f__Metschnikowiaceae(100);g__Metschnikowia(100);s__Metschnikowia_pulcherrima(100); | Metschnikowia | 13 | Saprotroph | \|Undefined Saprotroph\| | Yeast | NULL | Probable | As nectar-tap_saprotroph (Põlme et al. 2020); Decay sugar-rich substrates (Põlme et al. 2020) | Põlme S, et al. 2020. Fungal Diversity 105:1-16 ((https://doi.org/10.1007/s13225-020-00466-2)) |
| Otu31643 | | 1 | k__Fungi(100);p__Ascomycota(100);c__Saccharomycetes(100);o__Saccharomycetales(100);f__Metschnikowiaceae(100);g__Metschnikowia(100);s__Metschnikowia_pulcherrima(100); | Metschnikowia | 13 | Saprotroph | \|Undefined Saprotroph\| | Yeast | NULL | Probable | As nectar-tap_saprotroph (Põlme et al. 2020); Decay sugar-rich substrates (Põlme et al. 2020) | Põlme S, et al. 2020. Fungal Diversity 105:1-16 ((https://doi.org/10.1007/s13225-020-00466-2)) |
| Otu31644 | | 1 | k__Fungi(100);p__Ascomycota(100);c__Saccharomycetes(100);o__Saccharomycetales(100);f__Metschnikowiaceae(100);g__Metschnikowia(100);s__Metschnikowia_pulcherrima(100); | Metschnikowia | 13 | Saprotroph | \|Undefined Saprotroph\| | Yeast | NULL | Probable | As nectar-tap_saprotroph (Põlme et al. 2020); Decay sugar-rich substrates (Põlme et al. 2020) | Põlme S, et al. 2020. Fungal Diversity 105:1-16 ((https://doi.org/10.1007/s13225-020-00466-2)) |
| Otu31645 | | 1 | k__Fungi(100);p__Ascomycota(100);c__Saccharomycetes(100);o__Saccharomycetales(100);f__Metschnikowiaceae(100);g__Metschnikowia(100);g__Metschnikowia_unclassified(100); | Metschnikowia | 13 | Saprotroph | \|Undefined Saprotroph\| | Yeast | NULL | Probable | As nectar-tap_saprotroph (Põlme et al. 2020); Decay sugar-rich substrates (Põlme et al. 2020) | Põlme S, et al. 2020. Fungal Diversity 105:1-16 ((https://doi.org/10.1007/s13225-020-00466-2)) |
| Otu31646 | | 1 | k__Fungi(100);p__Ascomycota(100);c__Saccharomycetes(100);o__Saccharomycetales(100);f__Metschnikowiaceae(100);g__Metschnikowia(100);s__Metschnikowia_pulcherrima(100); | Metschnikowia | 13 | Saprotroph | \|Undefined Saprotroph\| | Yeast | NULL | Probable | As nectar-tap_saprotroph (Põlme et al. 2020); Decay sugar-rich substrates (Põlme et al. 2020) | Põlme S, et al. 2020. Fungal Diversity 105:1-16 ((https://doi.org/10.1007/s13225-020-00466-2)) |
| Otu31647 | | 1 | k__Fungi(100);p__Ascomycota(100);c__Saccharomycetes(100);o__Saccharomycetales(100);f__Metschnikowiaceae(100);g__Metschnikowia(100);s__Metschnikowia_pulcherrima(100); | Metschnikowia | 13 | Saprotroph | \|Undefined Saprotroph\| | Yeast | NULL | Probable | As nectar-tap_saprotroph (Põlme et al. 2020); Decay sugar-rich substrates (Põlme et al. 2020) | Põlme S, et al. 2020. Fungal Diversity 105:1-16 ((https://doi.org/10.1007/s13225-020-00466-2)) |
| Otu31648 | | 1 | k__Fungi(100);p__Ascomycota(100);c__Saccharomycetes(100);o__Saccharomycetales(100);f__Metschnikowiaceae(100);g__Metschnikowia(100);g__Metschnikowia_unclassified(100); | Metschnikowia | 13 | Saprotroph | \|Undefined Saprotroph\| | Yeast | NULL | Probable | As nectar-tap_saprotroph (Põlme et al. 2020); Decay sugar-rich substrates (Põlme et al. 2020) | Põlme S, et al. 2020. Fungal Diversity 105:1-16 ((https://doi.org/10.1007/s13225-020-00466-2)) |
| Otu31649 | | 1 | k__Fungi(100);p__Ascomycota(100);c__Saccharomycetes(100);o__Saccharomycetales(100);f__Metschnikowiaceae(100);g__Metschnikowia(100);s__Metschnikowia_pulcherrima(100); | Metschnikowia | 13 | Saprotroph | \|Undefined Saprotroph\| | Yeast | NULL | Probable | As nectar-tap_saprotroph (Põlme et al. 2020); Decay sugar-rich substrates (Põlme et al. 2020) | Põlme S, et al. 2020. Fungal Diversity 105:1-16 ((https://doi.org/10.1007/s13225-020-00466-2)) |
| Otu31652 | | 1 | k__Fungi(100);p__Ascomycota(100);c__Saccharomycetes(100);o__Saccharomycetales(100);f__Metschnikowiaceae(100);g__Metschnikowia(100);g__Metschnikowia_unclassified(100); | Metschnikowia | 13 | Saprotroph | \|Undefined Saprotroph\| | Yeast | NULL | Probable | As nectar-tap_saprotroph (Põlme et al. 2020); Decay sugar-rich substrates (Põlme et al. 2020) | Põlme S, et al. 2020. Fungal Diversity 105:1-16 ((https://doi.org/10.1007/s13225-020-00466-2)) |
| Otu31654 | | 1 | k__Fungi(100);p__Ascomycota(100);c__Saccharomycetes(100);o__Saccharomycetales(100);f__Metschnikowiaceae(100);g__Metschnikowia(100);s__Metschnikowia_pulcherrima(100); | Metschnikowia | 13 | Saprotroph | \|Undefined Saprotroph\| | Yeast | NULL | Probable | As nectar-tap_saprotroph (Põlme et al. 2020); Decay sugar-rich substrates (Põlme et al. 2020) | Põlme S, et al. 2020. Fungal Diversity 105:1-16 ((https://doi.org/10.1007/s13225-020-00466-2)) |
| Otu31655 | | 1 | k__Fungi(100);p__Ascomycota(100);c__Saccharomycetes(100);o__Saccharomycetales(100);f__Metschnikowiaceae(100);g__Metschnikowia(100);s__Metschnikowia_pulcherrima(100); | Metschnikowia | 13 | Saprotroph | \|Undefined Saprotroph\| | Yeast | NULL | Probable | As nectar-tap_saprotroph (Põlme et al. 2020); Decay sugar-rich substrates (Põlme et al. 2020) | Põlme S, et al. 2020. Fungal Diversity 105:1-16 ((https://doi.org/10.1007/s13225-020-00466-2)) |

| | | | | | | | | | | | |
|---|---|---|---|---|---|---|---|---|---|---|---|
| Otu31656 | 1 | k__Fungi(100);p__Ascomycota(100);c__Saccharomycetes(100);o__Saccharomycetales(100);f__Metschnikowiaceae(100);g__Metschnikowia(100);g__Metschnikowia_unclassified(100); | Metschnikowia | 13 | Saprotroph | \|Undefined Saprotroph\| | Yeast | NULL | Probable | As nectar-tap_saprotroph (Põlme et al. 2020); Decay sugar-rich substrates (Põlme et al. 2020) | Põlme S, et al. 2020. Fungal Diversity 105:1-16 ((https://doi.org/10.1007/s13225-020-00466-2)) |
| Otu31657 | 1 | k__Fungi(100);p__Ascomycota(100);c__Saccharomycetes(100);o__Saccharomycetales(100);f__Metschnikowiaceae(100);g__Metschnikowia(100);s__Metschnikowia_pulcherrima(100); | Metschnikowia | 13 | Saprotroph | \|Undefined Saprotroph\| | Yeast | NULL | Probable | As nectar-tap_saprotroph (Põlme et al. 2020); Decay sugar-rich substrates (Põlme et al. 2020) | Põlme S, et al. 2020. Fungal Diversity 105:1-16 ((https://doi.org/10.1007/s13225-020-00466-2)) |
| Otu31658 | 1 | k__Fungi(100);p__Ascomycota(100);c__Saccharomycetes(100);o__Saccharomycetales(100);f__Metschnikowiaceae(100);g__Metschnikowia(100);s__Metschnikowia_pulcherrima(100); | Metschnikowia | 13 | Saprotroph | \|Undefined Saprotroph\| | Yeast | NULL | Probable | As nectar-tap_saprotroph (Põlme et al. 2020); Decay sugar-rich substrates (Põlme et al. 2020) | Põlme S, et al. 2020. Fungal Diversity 105:1-16 ((https://doi.org/10.1007/s13225-020-00466-2)) |
| Otu31659 | 1 | k__Fungi(100);p__Ascomycota(100);c__Saccharomycetes(100);o__Saccharomycetales(100);f__Metschnikowiaceae(100);g__Metschnikowia(100);g__Metschnikowia_unclassified(100); | Metschnikowia | 13 | Saprotroph | \|Undefined Saprotroph\| | Yeast | NULL | Probable | As nectar-tap_saprotroph (Põlme et al. 2020); Decay sugar-rich substrates (Põlme et al. 2020) | Põlme S, et al. 2020. Fungal Diversity 105:1-16 ((https://doi.org/10.1007/s13225-020-00466-2)) |
| Otu31660 | 1 | k__Fungi(100);p__Ascomycota(100);c__Saccharomycetes(100);o__Saccharomycetales(100);f__Metschnikowiaceae(100);g__Metschnikowia(100);s__Metschnikowia_pulcherrima(100); | Metschnikowia | 13 | Saprotroph | \|Undefined Saprotroph\| | Yeast | NULL | Probable | As nectar-tap_saprotroph (Põlme et al. 2020); Decay sugar-rich substrates (Põlme et al. 2020) | Põlme S, et al. 2020. Fungal Diversity 105:1-16 ((https://doi.org/10.1007/s13225-020-00466-2)) |
| Otu31661 | 1 | k__Fungi(100);p__Ascomycota(100);c__Saccharomycetes(100);o__Saccharomycetales(100);f__Metschnikowiaceae(100);g__Metschnikowia(100);s__Metschnikowia_pulcherrima(100); | Metschnikowia | 13 | Saprotroph | \|Undefined Saprotroph\| | Yeast | NULL | Probable | As nectar-tap_saprotroph (Põlme et al. 2020); Decay sugar-rich substrates (Põlme et al. 2020) | Põlme S, et al. 2020. Fungal Diversity 105:1-16 ((https://doi.org/10.1007/s13225-020-00466-2)) |
| Otu31662 | 1 | k__Fungi(100);p__Ascomycota(100);c__Saccharomycetes(100);o__Saccharomycetales(100);f__Metschnikowiaceae(100);g__Metschnikowia(100);s__Metschnikowia_pulcherrima(100); | Metschnikowia | 13 | Saprotroph | \|Undefined Saprotroph\| | Yeast | NULL | Probable | As nectar-tap_saprotroph (Põlme et al. 2020); Decay sugar-rich substrates (Põlme et al. 2020) | Põlme S, et al. 2020. Fungal Diversity 105:1-16 ((https://doi.org/10.1007/s13225-020-00466-2)) |
| Otu31663 | 1 | k__Fungi(100);p__Ascomycota(100);c__Saccharomycetes(100);o__Saccharomycetales(100);f__Metschnikowiaceae(100);g__Metschnikowia(100);s__Metschnikowia_pulcherrima(100); | Metschnikowia | 13 | Saprotroph | \|Undefined Saprotroph\| | Yeast | NULL | Probable | As nectar-tap_saprotroph (Põlme et al. 2020); Decay sugar-rich substrates (Põlme et al. 2020) | Põlme S, et al. 2020. Fungal Diversity 105:1-16 ((https://doi.org/10.1007/s13225-020-00466-2)) |
| Otu31664 | 1 | k__Fungi(100);p__Ascomycota(100);c__Saccharomycetes(100);o__Saccharomycetales(100);f__Metschnikowiaceae(100);g__Metschnikowia(100);s__Metschnikowia_pulcherrima(100); | Metschnikowia | 13 | Saprotroph | \|Undefined Saprotroph\| | Yeast | NULL | Probable | As nectar-tap_saprotroph (Põlme et al. 2020); Decay sugar-rich substrates (Põlme et al. 2020) | Põlme S, et al. 2020. Fungal Diversity 105:1-16 ((https://doi.org/10.1007/s13225-020-00466-2)) |
| Otu31665 | 1 | k__Fungi(100);p__Ascomycota(100);c__Saccharomycetes(100);o__Saccharomycetales(100);f__Metschnikowiaceae(100);g__Metschnikowia(100);g__Metschnikowia_unclassified(100); | Metschnikowia | 13 | Saprotroph | \|Undefined Saprotroph\| | Yeast | NULL | Probable | As nectar-tap_saprotroph (Põlme et al. 2020); Decay sugar-rich substrates (Põlme et al. 2020) | Põlme S, et al. 2020. Fungal Diversity 105:1-16 ((https://doi.org/10.1007/s13225-020-00466-2)) |
| Otu31666 | 1 | k__Fungi(100);p__Ascomycota(100);c__Saccharomycetes(100);o__Saccharomycetales(100);f__Metschnikowiaceae(100);g__Metschnikowia(100);g__Metschnikowia_unclassified(100); | Metschnikowia | 13 | Saprotroph | \|Undefined Saprotroph\| | Yeast | NULL | Probable | As nectar-tap_saprotroph (Põlme et al. 2020); Decay sugar-rich substrates (Põlme et al. 2020) | Põlme S, et al. 2020. Fungal Diversity 105:1-16 ((https://doi.org/10.1007/s13225-020-00466-2)) |
| Otu31667 | 1 | k__Fungi(100);p__Ascomycota(100);c__Saccharomycetes(100);o__Saccharomycetales(100);f__Metschnikowiaceae(100);g__Metschnikowia(100);g__Metschnikowia_unclassified(100); | Metschnikowia | 13 | Saprotroph | \|Undefined Saprotroph\| | Yeast | NULL | Probable | As nectar-tap_saprotroph (Põlme et al. 2020); Decay sugar-rich substrates (Põlme et al. 2020) | Põlme S, et al. 2020. Fungal Diversity 105:1-16 ((https://doi.org/10.1007/s13225-020-00466-2)) |
| Otu31668 | 1 | k__Fungi(100);p__Ascomycota(100);c__Saccharomycetes(100);o__Saccharomycetales(100);f__Metschnikowiaceae(100);g__Metschnikowia(100);s__Metschnikowia_pulcherrima(100); | Metschnikowia | 13 | Saprotroph | \|Undefined Saprotroph\| | Yeast | NULL | Probable | As nectar-tap_saprotroph (Põlme et al. 2020); Decay sugar-rich substrates (Põlme et al. 2020) | Põlme S, et al. 2020. Fungal Diversity 105:1-16 ((https://doi.org/10.1007/s13225-020-00466-2)) |
| Otu31669 | 1 | k__Fungi(100);p__Ascomycota(100);c__Saccharomycetes(100);o__Saccharomycetales(100);f__Metschnikowiaceae(100);g__Metschnikowia(100);s__Metschnikowia_pulcherrima(100); | Metschnikowia | 13 | Saprotroph | \|Undefined Saprotroph\| | Yeast | NULL | Probable | As nectar-tap_saprotroph (Põlme et al. 2020); Decay sugar-rich substrates (Põlme et al. 2020) | Põlme S, et al. 2020. Fungal Diversity 105:1-16 ((https://doi.org/10.1007/s13225-020-00466-2)) |
| Otu31670 | 1 | k__Fungi(100);p__Ascomycota(100);c__Saccharomycetes(100);o__Saccharomycetales(100);f__Metschnikowiaceae(100);g__Metschnikowia(100);s__Metschnikowia_pulcherrima(100); | Metschnikowia | 13 | Saprotroph | \|Undefined Saprotroph\| | Yeast | NULL | Probable | As nectar-tap_saprotroph (Põlme et al. 2020); Decay sugar-rich substrates (Põlme et al. 2020) | Põlme S, et al. 2020. Fungal Diversity 105:1-16 ((https://doi.org/10.1007/s13225-020-00466-2)) |
| Otu31671 | 1 | k__Fungi(100);p__Ascomycota(100);c__Saccharomycetes(100);o__Saccharomycetales(100);f__Metschnikowiaceae(100);g__Metschnikowia(100);g__Metschnikowia_unclassified(100); | Metschnikowia | 13 | Saprotroph | \|Undefined Saprotroph\| | Yeast | NULL | Probable | As nectar-tap_saprotroph (Põlme et al. 2020); Decay sugar-rich substrates (Põlme et al. 2020) | Põlme S, et al. 2020. Fungal Diversity 105:1-16 ((https://doi.org/10.1007/s13225-020-00466-2)) |
| Otu31672 | 1 | k__Fungi(100);p__Ascomycota(100);c__Saccharomycetes(100);o__Saccharomycetales(100);f__Metschnikowiaceae(100);g__Metschnikowia(100);s__Metschnikowia_pulcherrima(100); | Metschnikowia | 13 | Saprotroph | \|Undefined Saprotroph\| | Yeast | NULL | Probable | As nectar-tap_saprotroph (Põlme et al. 2020); Decay sugar-rich substrates (Põlme et al. 2020) | Põlme S, et al. 2020. Fungal Diversity 105:1-16 ((https://doi.org/10.1007/s13225-020-00466-2)) |
| Otu31673 | 1 | k__Fungi(100);p__Ascomycota(100);c__Saccharomycetes(100);o__Saccharomycetales(100);f__Metschnikowiaceae(100);g__Metschnikowia(100);g__Metschnikowia_unclassified(100); | Metschnikowia | 13 | Saprotroph | \|Undefined Saprotroph\| | Yeast | NULL | Probable | As nectar-tap_saprotroph (Põlme et al. 2020); Decay sugar-rich substrates (Põlme et al. 2020) | Põlme S, et al. 2020. Fungal Diversity 105:1-16 ((https://doi.org/10.1007/s13225-020-00466-2)) |
| Otu31674 | 1 | k__Fungi(100);p__Ascomycota(100);c__Saccharomycetes(100);o__Saccharomycetales(100);f__Metschnikowiaceae(100);g__Metschnikowia(100);g__Metschnikowia_unclassified(100); | Metschnikowia | 13 | Saprotroph | \|Undefined Saprotroph\| | Yeast | NULL | Probable | As nectar-tap_saprotroph (Põlme et al. 2020); Decay sugar-rich substrates (Põlme et al. 2020) | Põlme S, et al. 2020. Fungal Diversity 105:1-16 ((https://doi.org/10.1007/s13225-020-00466-2)) |
| Otu31675 | 1 | k__Fungi(100);p__Ascomycota(100);c__Saccharomycetes(100);o__Saccharomycetales(100);f__Metschnikowiaceae(100);g__Metschnikowia(100);g__Metschnikowia_unclassified(100); | Metschnikowia | 13 | Saprotroph | \|Undefined Saprotroph\| | Yeast | NULL | Probable | As nectar-tap_saprotroph (Põlme et al. 2020); Decay sugar-rich substrates (Põlme et al. 2020) | Põlme S, et al. 2020. Fungal Diversity 105:1-16 ((https://doi.org/10.1007/s13225-020-00466-2)) |
| Otu31676 | 1 | k__Fungi(100);p__Ascomycota(100);c__Saccharomycetes(100);o__Saccharomycetales(100);f__Metschnikowiaceae(100);g__Metschnikowia(100);s__Metschnikowia_pulcherrima(100); | Metschnikowia | 13 | Saprotroph | \|Undefined Saprotroph\| | Yeast | NULL | Probable | As nectar-tap_saprotroph (Põlme et al. 2020); Decay sugar-rich substrates (Põlme et al. 2020) | Põlme S, et al. 2020. Fungal Diversity 105:1-16 ((https://doi.org/10.1007/s13225-020-00466-2)) |
| Otu31679 | 1 | k__Fungi(100);p__Ascomycota(100);c__Saccharomycetes(100);o__Saccharomycetales(100);f__Metschnikowiaceae(100);g__Metschnikowia(100);s__Metschnikowia_pulcherrima(100); | Metschnikowia | 13 | Saprotroph | \|Undefined Saprotroph\| | Yeast | NULL | Probable | As nectar-tap_saprotroph (Põlme et al. 2020); Decay sugar-rich substrates (Põlme et al. 2020) | Põlme S, et al. 2020. Fungal Diversity 105:1-16 ((https://doi.org/10.1007/s13225-020-00466-2)) |
| Otu31680 | 1 | k__Fungi(100);p__Ascomycota(100);c__Saccharomycetes(100);o__Saccharomycetales(100);f__Metschnikowiaceae(100);g__Metschnikowia(100);s__Metschnikowia_pulcherrima(100); | Metschnikowia | 13 | Saprotroph | \|Undefined Saprotroph\| | Yeast | NULL | Probable | As nectar-tap_saprotroph (Põlme et al. 2020); Decay sugar-rich substrates (Põlme et al. 2020) | Põlme S, et al. 2020. Fungal Diversity 105:1-16 ((https://doi.org/10.1007/s13225-020-00466-2)) |
| Otu31681 | 1 | k__Fungi(100);p__Ascomycota(100);c__Saccharomycetes(100);o__Saccharomycetales(100);f__Saccharomycetales_family_Incertae_sedis(100);g__Candida(100);g__Candida_unclassified(100); | Candida | 13 | Pathotroph-Saprotroph-Symbiotroph | Animal Pathogen-Endophyte-Endosymbiont-Epiphyte-\|Undefined Saprotroph\| | Dimorphic Yeast-Yeast | NULL | Probable | As nectar-tap_saprotroph (Põlme et al. 2020); Decay sugar-rich substrates (Põlme et al. 2020); This asexual genus is polyphyletic and contain species that function very differently depend on what they are. Caution should be taken when interpreting guilds for Candida species. | Manolakaki D, et al. 2010. Virulence. 1: 367-375 ((https://doi.org/10.4161/viru.1.5.12796)); Mekbib SB, et al. 2011. Tropical Plant Pathology 36:233-240 ((https://doi.org/10.1590/S1982-56762011000400004)); Põlme S, et al. 2020. Fungal Diversity 105:1-16 ((https://doi.org/10.1007/s13225-020-00466-2)) |
| Otu31683 | 1 | k__Fungi(100);p__Ascomycota(100);c__Saccharomycetes(100);o__Saccharomycetales(100);f__Metschnikowiaceae(100);g__Metschnikowia(100);s__Metschnikowia_pulcherrima(100); | Metschnikowia | 13 | Saprotroph | \|Undefined Saprotroph\| | Yeast | NULL | Probable | As nectar-tap_saprotroph (Põlme et al. 2020); Decay sugar-rich substrates (Põlme et al. 2020) | Põlme S, et al. 2020. Fungal Diversity 105:1-16 ((https://doi.org/10.1007/s13225-020-00466-2)) |
| Otu31684 | 1 | k__Fungi(100);p__Ascomycota(100);c__Saccharomycetes(100);o__Saccharomycetales(100);f__Metschnikowiaceae(100);g__Metschnikowia(100);s__Metschnikowia_pulcherrima(100); | Metschnikowia | 13 | Saprotroph | \|Undefined Saprotroph\| | Yeast | NULL | Probable | As nectar-tap_saprotroph (Põlme et al. 2020); Decay sugar-rich substrates (Põlme et al. 2020) | Põlme S, et al. 2020. Fungal Diversity 105:1-16 ((https://doi.org/10.1007/s13225-020-00466-2)) |
| Otu31685 | 1 | k__Fungi(100);p__Ascomycota(100);c__Saccharomycetes(100);o__Saccharomycetales(100);f__Metschnikowiaceae(100);g__Metschnikowia(100);s__Metschnikowia_pulcherrima(100); | Metschnikowia | 13 | Saprotroph | \|Undefined Saprotroph\| | Yeast | NULL | Probable | As nectar-tap_saprotroph (Põlme et al. 2020); Decay sugar-rich substrates (Põlme et al. 2020) | Põlme S, et al. 2020. Fungal Diversity 105:1-16 ((https://doi.org/10.1007/s13225-020-00466-2)) |
| Otu31686 | 1 | k__Fungi(100);p__Ascomycota(100);c__Saccharomycetes(100);o__Saccharomycetales(100);f__Metschnikowiaceae(100);g__Metschnikowia(100);s__Metschnikowia_pulcherrima(100); | Metschnikowia | 13 | Saprotroph | \|Undefined Saprotroph\| | Yeast | NULL | Probable | As nectar-tap_saprotroph (Põlme et al. 2020); Decay sugar-rich substrates (Põlme et al. 2020) | Põlme S, et al. 2020. Fungal Diversity 105:1-16 ((https://doi.org/10.1007/s13225-020-00466-2)) |
| Otu31687 | 1 | k__Fungi(100);p__Ascomycota(100);c__Saccharomycetes(100);o__Saccharomycetales(100);f__Metschnikowiaceae(100);g__Metschnikowia(100);s__Metschnikowia_pulcherrima(100); | Metschnikowia | 13 | Saprotroph | \|Undefined Saprotroph\| | Yeast | NULL | Probable | As nectar-tap_saprotroph (Põlme et al. 2020); Decay sugar-rich substrates (Põlme et al. 2020) | Põlme S, et al. 2020. Fungal Diversity 105:1-16 ((https://doi.org/10.1007/s13225-020-00466-2)) |
| Otu31689 | 1 | k__Fungi(100);p__Ascomycota(100);c__Saccharomycetes(100);o__Saccharomycetales(100);f__Metschnikowiaceae(100);g__Metschnikowia(100);g__Metschnikowia_unclassified(100); | Metschnikowia | 13 | Saprotroph | \|Undefined Saprotroph\| | Yeast | NULL | Probable | As nectar-tap_saprotroph (Põlme et al. 2020); Decay sugar-rich substrates (Põlme et al. 2020) | Põlme S, et al. 2020. Fungal Diversity 105:1-16 ((https://doi.org/10.1007/s13225-020-00466-2)) |
| Otu31690 | 1 | k__Fungi(100);p__Ascomycota(100);c__Saccharomycetes(100);o__Saccharomycetales(100);f__Metschnikowiaceae(100);g__Metschnikowia(100);g__Metschnikowia_unclassified(100); | Metschnikowia | 13 | Saprotroph | \|Undefined Saprotroph\| | Yeast | NULL | Probable | As nectar-tap_saprotroph (Põlme et al. 2020); Decay sugar-rich substrates (Põlme et al. 2020) | Põlme S, et al. 2020. Fungal Diversity 105:1-16 ((https://doi.org/10.1007/s13225-020-00466-2)) |
| Otu31691 | 1 | k__Fungi(100);p__Ascomycota(100);c__Saccharomycetes(100);o__Saccharomycetales(100);f__Metschnikowiaceae(100);g__Metschnikowia(100);g__Metschnikowia_unclassified(100); | Metschnikowia | 13 | Saprotroph | \|Undefined Saprotroph\| | Yeast | NULL | Probable | As nectar-tap_saprotroph (Põlme et al. 2020); Decay sugar-rich substrates (Põlme et al. 2020) | Põlme S, et al. 2020. Fungal Diversity 105:1-16 ((https://doi.org/10.1007/s13225-020-00466-2)) |
| Otu31692 | 1 | k__Fungi(100);p__Ascomycota(100);c__Saccharomycetes(100);o__Saccharomycetales(100);f__Metschnikowiaceae(100);g__Metschnikowia(100);g__Metschnikowia_unclassified(100); | Metschnikowia | 13 | Saprotroph | \|Undefined Saprotroph\| | Yeast | NULL | Probable | As nectar-tap_saprotroph (Põlme et al. 2020); Decay sugar-rich substrates (Põlme et al. 2020) | Põlme S, et al. 2020. Fungal Diversity 105:1-16 ((https://doi.org/10.1007/s13225-020-00466-2)) |
| Otu31693 | 1 | k__Fungi(100);p__Ascomycota(100);c__Saccharomycetes(100);o__Saccharomycetales(100);f__Metschnikowiaceae(100);g__Metschnikowia(100);s__Metschnikowia_pulcherrima(100); | Metschnikowia | 13 | Saprotroph | \|Undefined Saprotroph\| | Yeast | NULL | Probable | As nectar-tap_saprotroph (Põlme et al. 2020); Decay sugar-rich substrates (Põlme et al. 2020) | Põlme S, et al. 2020. Fungal Diversity 105:1-16 ((https://doi.org/10.1007/s13225-020-00466-2)) |
| Otu31694 | 1 | k__Fungi(100);p__Ascomycota(100);c__Saccharomycetes(100);o__Saccharomycetales(100);f__Metschnikowiaceae(100);g__Metschnikowia(100);s__Metschnikowia_pulcherrima(100); | Metschnikowia | 13 | Saprotroph | \|Undefined Saprotroph\| | Yeast | NULL | Probable | As nectar-tap_saprotroph (Põlme et al. 2020); Decay sugar-rich substrates (Põlme et al. 2020) | Põlme S, et al. 2020. Fungal Diversity 105:1-16 ((https://doi.org/10.1007/s13225-020-00466-2)) |
| Otu31695 | 1 | k__Fungi(100);p__Ascomycota(100);c__Saccharomycetes(100);o__Saccharomycetales(100);f__Metschnikowiaceae(100);g__Metschnikowia(100);s__Metschnikowia_pulcherrima(100); | Metschnikowia | 13 | Saprotroph | \|Undefined Saprotroph\| | Yeast | NULL | Probable | As nectar-tap_saprotroph (Põlme et al. 2020); Decay sugar-rich substrates (Põlme et al. 2020) | Põlme S, et al. 2020. Fungal Diversity 105:1-16 ((https://doi.org/10.1007/s13225-020-00466-2)) |
| Otu31696 | 1 | k__Fungi(100);p__Ascomycota(100);c__Saccharomycetes(100);o__Saccharomycetales(100);f__Metschnikowiaceae(100);g__Metschnikowia(100);s__Metschnikowia_pulcherrima(100); | Metschnikowia | 13 | Saprotroph | \|Undefined Saprotroph\| | Yeast | NULL | Probable | As nectar-tap_saprotroph (Põlme et al. 2020); Decay sugar-rich substrates (Põlme et al. 2020) | Põlme S, et al. 2020. Fungal Diversity 105:1-16 ((https://doi.org/10.1007/s13225-020-00466-2)) |
| Otu31697 | 1 | k__Fungi(100);p__Ascomycota(100);c__Saccharomycetes(100);o__Saccharomycetales(100);f__Metschnikowiaceae(100);g__Metschnikowia(100);s__Metschnikowia_pulcherrima(100); | Metschnikowia | 13 | Saprotroph | \|Undefined Saprotroph\| | Yeast | NULL | Probable | As nectar-tap_saprotroph (Põlme et al. 2020); Decay sugar-rich substrates (Põlme et al. 2020) | Põlme S, et al. 2020. Fungal Diversity 105:1-16 ((https://doi.org/10.1007/s13225-020-00466-2)) |
| Otu31698 | 1 | k__Fungi(100);p__Ascomycota(100);c__Saccharomycetes(100);o__Saccharomycetales(100);f__Metschnikowiaceae(100);g__Metschnikowia(100);s__Metschnikowia_pulcherrima(100); | Metschnikowia | 13 | Saprotroph | \|Undefined Saprotroph\| | Yeast | NULL | Probable | As nectar-tap_saprotroph (Põlme et al. 2020); Decay sugar-rich substrates (Põlme et al. 2020) | Põlme S, et al. 2020. Fungal Diversity 105:1-16 ((https://doi.org/10.1007/s13225-020-00466-2)) |
| Otu31699 | 1 | k__Fungi(100);p__Ascomycota(100);c__Saccharomycetes(100);o__Saccharomycetales(100);f__Metschnikowiaceae(100);g__Metschnikowia(100);g__Metschnikowia_unclassified(100); | Metschnikowia | 13 | Saprotroph | \|Undefined Saprotroph\| | Yeast | NULL | Probable | As nectar-tap_saprotroph (Põlme et al. 2020); Decay sugar-rich substrates (Põlme et al. 2020) | Põlme S, et al. 2020. Fungal Diversity 105:1-16 ((https://doi.org/10.1007/s13225-020-00466-2)) |
| Otu31700 | 1 | k__Fungi(100);p__Ascomycota(100);c__Saccharomycetes(100);o__Saccharomycetales(100);f__Metschnikowiaceae(100);g__Metschnikowia(100);s__Metschnikowia_pulcherrima(100); | Metschnikowia | 13 | Saprotroph | \|Undefined Saprotroph\| | Yeast | NULL | Probable | As nectar-tap_saprotroph (Põlme et al. 2020); Decay sugar-rich substrates (Põlme et al. 2020) | Põlme S, et al. 2020. Fungal Diversity 105:1-16 ((https://doi.org/10.1007/s13225-020-00466-2)) |
| Otu31701 | 1 | k__Fungi(100);p__Ascomycota(100);c__Saccharomycetes(100);o__Saccharomycetales(100);f__Metschnikowiaceae(100);g__Metschnikowia(100);s__Metschnikowia_pulcherrima(100); | Metschnikowia | 13 | Saprotroph | \|Undefined Saprotroph\| | Yeast | NULL | Probable | As nectar-tap_saprotroph (Põlme et al. 2020); Decay sugar-rich substrates (Põlme et al. 2020) | Põlme S, et al. 2020. Fungal Diversity 105:1-16 ((https://doi.org/10.1007/s13225-020-00466-2)) |
| Otu31702 | 1 | k__Fungi(100);p__Ascomycota(100);c__Saccharomycetes(100);o__Saccharomycetales(100);f__Metschnikowiaceae(100);g__Metschnikowia(100);g__Metschnikowia_unclassified(100); | Metschnikowia | 13 | Saprotroph | \|Undefined Saprotroph\| | Yeast | NULL | Probable | As nectar-tap_saprotroph (Põlme et al. 2020); Decay sugar-rich substrates (Põlme et al. 2020) | Põlme S, et al. 2020. Fungal Diversity 105:1-16 ((https://doi.org/10.1007/s13225-020-00466-2)) |
| Otu31703 | 1 | k__Fungi(100);p__Ascomycota(100);c__Saccharomycetes(100);o__Saccharomycetales(100);f__Metschnikowiaceae(100);g__Metschnikowia(100);s__Metschnikowia_pulcherrima(100); | Metschnikowia | 13 | Saprotroph | \|Undefined Saprotroph\| | Yeast | NULL | Probable | As nectar-tap_saprotroph (Põlme et al. 2020); Decay sugar-rich substrates (Põlme et al. 2020) | Põlme S, et al. 2020. Fungal Diversity 105:1-16 ((https://doi.org/10.1007/s13225-020-00466-2)) |
| Otu31704 | 1 | k__Fungi(100);p__Ascomycota(100);c__Saccharomycetes(100);o__Saccharomycetales(100);f__Metschnikowiaceae(100);g__Metschnikowia(100);s__Metschnikowia_pulcherrima(100); | Metschnikowia | 13 | Saprotroph | \|Undefined Saprotroph\| | Yeast | NULL | Probable | As nectar-tap_saprotroph (Põlme et al. 2020); Decay sugar-rich substrates (Põlme et al. 2020) | Põlme S, et al. 2020. Fungal Diversity 105:1-16 ((https://doi.org/10.1007/s13225-020-00466-2)) |
| Otu31705 | 1 | k__Fungi(100);p__Ascomycota(100);c__Saccharomycetes(100);o__Saccharomycetales(100);f__Metschnikowiaceae(100);g__Metschnikowia(100);s__Metschnikowia_pulcherrima(100); | Metschnikowia | 13 | Saprotroph | \|Undefined Saprotroph\| | Yeast | NULL | Probable | As nectar-tap_saprotroph (Põlme et al. 2020); Decay sugar-rich substrates (Põlme et al. 2020) | Põlme S, et al. 2020. Fungal Diversity 105:1-16 ((https://doi.org/10.1007/s13225-020-00466-2)) |
| Otu31706 | 1 | k__Fungi(100);p__Ascomycota(100);c__Saccharomycetes(100);o__Saccharomycetales(100);f__Metschnikowiaceae(100);g__Metschnikowia(100);s__Metschnikowia_pulcherrima(100); | Metschnikowia | 13 | Saprotroph | \|Undefined Saprotroph\| | Yeast | NULL | Probable | As nectar-tap_saprotroph (Põlme et al. 2020); Decay sugar-rich substrates (Põlme et al. 2020) | Põlme S, et al. 2020. Fungal Diversity 105:1-16 ((https://doi.org/10.1007/s13225-020-00466-2)) |
| Otu31707 | 1 | k__Fungi(100);p__Ascomycota(100);c__Saccharomycetes(100);o__Saccharomycetales(100);f__Metschnikowiaceae(100);g__Metschnikowia(100);s__Metschnikowia_pulcherrima(100); | Metschnikowia | 13 | Saprotroph | \|Undefined Saprotroph\| | Yeast | NULL | Probable | As nectar-tap_saprotroph (Põlme et al. 2020); Decay sugar-rich substrates (Põlme et al. 2020) | Põlme S, et al. 2020. Fungal Diversity 105:1-16 ((https://doi.org/10.1007/s13225-020-00466-2)) |
| Otu31708 | 1 | k__Fungi(100);p__Ascomycota(100);c__Saccharomycetes(100);o__Saccharomycetales(100);f__Metschnikowiaceae(100);g__Metschnikowia(100);s__Metschnikowia_pulcherrima(100); | Metschnikowia | 13 | Saprotroph | \|Undefined Saprotroph\| | Yeast | NULL | Probable | As nectar-tap_saprotroph (Põlme et al. 2020); Decay sugar-rich substrates (Põlme et al. 2020) | Põlme S, et al. 2020. Fungal Diversity 105:1-16 ((https://doi.org/10.1007/s13225-020-00466-2)) |
| Otu31709 | 1 | k__Fungi(100);p__Ascomycota(100);c__Saccharomycetes(100);o__Saccharomycetales(100);f__Metschnikowiaceae(100);g__Metschnikowia(100);s__Metschnikowia_pulcherrima(100); | Metschnikowia | 13 | Saprotroph | \|Undefined Saprotroph\| | Yeast | NULL | Probable | As nectar-tap_saprotroph (Põlme et al. 2020); Decay sugar-rich substrates (Põlme et al. 2020) | Põlme S, et al. 2020. Fungal Diversity 105:1-16 ((https://doi.org/10.1007/s13225-020-00466-2)) |
| Otu31710 | 1 | k__Fungi(100);p__Ascomycota(100);c__Saccharomycetes(100);o__Saccharomycetales(100);f__Metschnikowiaceae(100);g__Metschnikowia(100);g__Metschnikowia_unclassified(100); | Metschnikowia | 13 | Saprotroph | \|Undefined Saprotroph\| | Yeast | NULL | Probable | As nectar-tap_saprotroph (Põlme et al. 2020); Decay sugar-rich substrates (Põlme et al. 2020) | Põlme S, et al. 2020. Fungal Diversity 105:1-16 ((https://doi.org/10.1007/s13225-020-00466-2)) |
| Otu31711 | 1 | k__Fungi(100);p__Ascomycota(100);c__Saccharomycetes(100);o__Saccharomycetales(100);f__Metschnikowiaceae(100);g__Metschnikowia(100);s__Metschnikowia_pulcherrima(100); | Metschnikowia | 13 | Saprotroph | \|Undefined Saprotroph\| | Yeast | NULL | Probable | As nectar-tap_saprotroph (Põlme et al. 2020); Decay sugar-rich substrates (Põlme et al. 2020) | Põlme S, et al. 2020. Fungal Diversity 105:1-16 ((https://doi.org/10.1007/s13225-020-00466-2)) |
| Otu31712 | 1 | k__Fungi(100);p__Ascomycota(100);c__Saccharomycetes(100);o__Saccharomycetales(100);f__Metschnikowiaceae(100);g__Metschnikowia(100);s__Metschnikowia_pulcherrima(100); | Metschnikowia | 13 | Saprotroph | \|Undefined Saprotroph\| | Yeast | NULL | Probable | As nectar-tap_saprotroph (Põlme et al. 2020); Decay sugar-rich substrates (Põlme et al. 2020) | Põlme S, et al. 2020. Fungal Diversity 105:1-16 ((https://doi.org/10.1007/s13225-020-00466-2)) |
| Otu31713 | 1 | k__Fungi(100);p__Ascomycota(100);c__Saccharomycetes(100);o__Saccharomycetales(100);f__Metschnikowiaceae(100);g__Metschnikowia(100);g__Metschnikowia_unclassified(100); | Metschnikowia | 13 | Saprotroph | \|Undefined Saprotroph\| | Yeast | NULL | Probable | As nectar-tap_saprotroph (Põlme et al. 2020); Decay sugar-rich substrates (Põlme et al. 2020) | Põlme S, et al. 2020. Fungal Diversity 105:1-16 ((https://doi.org/10.1007/s13225-020-00466-2)) |
| Otu31714 | 1 | k__Fungi(100);p__Ascomycota(100);c__Saccharomycetes(100);o__Saccharomycetales(100);f__Metschnikowiaceae(100);g__Metschnikowia(100);s__Metschnikowia_pulcherrima(100); | Metschnikowia | 13 | Saprotroph | \|Undefined Saprotroph\| | Yeast | NULL | Probable | As nectar-tap_saprotroph (Põlme et al. 2020); Decay sugar-rich substrates (Põlme et al. 2020) | Põlme S, et al. 2020. Fungal Diversity 105:1-16 ((https://doi.org/10.1007/s13225-020-00466-2)) |
| Otu31715 | 1 | k__Fungi(100);p__Ascomycota(100);c__Saccharomycetes(100);o__Saccharomycetales(100);f__Metschnikowiaceae(100);g__Metschnikowia(100);s__Metschnikowia_pulcherrima(100); | Metschnikowia | 13 | Saprotroph | \|Undefined Saprotroph\| | Yeast | NULL | Probable | As nectar-tap_saprotroph (Põlme et al. 2020); Decay sugar-rich substrates (Põlme et al. 2020) | Põlme S, et al. 2020. Fungal Diversity 105:1-16 ((https://doi.org/10.1007/s13225-020-00466-2)) |
| Otu31716 | 1 | k__Fungi(100);p__Ascomycota(100);c__Saccharomycetes(100);o__Saccharomycetales(100);f__Metschnikowiaceae(100);g__Metschnikowia(100);g__Metschnikowia_unclassified(100); | Metschnikowia | 13 | Saprotroph | \|Undefined Saprotroph\| | Yeast | NULL | Probable | As nectar-tap_saprotroph (Põlme et al. 2020); Decay sugar-rich substrates (Põlme et al. 2020) | Põlme S, et al. 2020. Fungal Diversity 105:1-16 ((https://doi.org/10.1007/s13225-020-00466-2)) |
| Otu31717 | 1 | k__Fungi(100);p__Ascomycota(100);c__Saccharomycetes(100);o__Saccharomycetales(100);f__Metschnikowiaceae(100);g__Metschnikowia(100);s__Metschnikowia_pulcherrima(100); | Metschnikowia | 13 | Saprotroph | \|Undefined Saprotroph\| | Yeast | NULL | Probable | As nectar-tap_saprotroph (Põlme et al. 2020); Decay sugar-rich substrates (Põlme et al. 2020) | Põlme S, et al. 2020. Fungal Diversity 105:1-16 ((https://doi.org/10.1007/s13225-020-00466-2)) |
| Otu31718 | 1 | k__Fungi(100);p__Ascomycota(100);c__Saccharomycetes(100);o__Saccharomycetales(100);f__Metschnikowiaceae(100);g__Metschnikowia(100);s__Metschnikowia_pulcherrima(100); | Metschnikowia | 13 | Saprotroph | \|Undefined Saprotroph\| | Yeast | NULL | Probable | As nectar-tap_saprotroph (Põlme et al. 2020); Decay sugar-rich substrates (Põlme et al. 2020) | Põlme S, et al. 2020. Fungal Diversity 105:1-16 ((https://doi.org/10.1007/s13225-020-00466-2)) |
| Otu31719 | 1 | k__Fungi(100);p__Ascomycota(100);c__Saccharomycetes(100);o__Saccharomycetales(100);f__Metschnikowiaceae(100);g__Metschnikowia(100);s__Metschnikowia_pulcherrima(100); | Metschnikowia | 13 | Saprotroph | \|Undefined Saprotroph\| | Yeast | NULL | Probable | As nectar-tap_saprotroph (Põlme et al. 2020); Decay sugar-rich substrates (Põlme et al. 2020) | Põlme S, et al. 2020. Fungal Diversity 105:1-16 ((https://doi.org/10.1007/s13225-020-00466-2)) |
| Otu31720 | 1 | k__Fungi(100);p__Ascomycota(100);c__Saccharomycetes(100);o__Saccharomycetales(100);f__Metschnikowiaceae(100);g__Metschnikowia(100);g__Metschnikowia_unclassified(100); | Metschnikowia | 13 | Saprotroph | \|Undefined Saprotroph\| | Yeast | NULL | Probable | As nectar-tap_saprotroph (Põlme et al. 2020); Decay sugar-rich substrates (Põlme et al. 2020) | Põlme S, et al. 2020. Fungal Diversity 105:1-16 ((https://doi.org/10.1007/s13225-020-00466-2)) |
| Otu31721 | 1 | k__Fungi(100);p__Ascomycota(100);c__Saccharomycetes(100);o__Saccharomycetales(100);f__Metschnikowiaceae(100);g__Metschnikowia(100);s__Metschnikowia_pulcherrima(100); | Metschnikowia | 13 | Saprotroph | \|Undefined Saprotroph\| | Yeast | NULL | Probable | As nectar-tap_saprotroph (Põlme et al. 2020); Decay sugar-rich substrates (Põlme et al. 2020) | Põlme S, et al. 2020. Fungal Diversity 105:1-16 ((https://doi.org/10.1007/s13225-020-00466-2)) |
| Otu31722 | 1 | k__Fungi(100);p__Ascomycota(100);c__Saccharomycetes(100);o__Saccharomycetales(100);f__Metschnikowiaceae(100);g__Metschnikowia(100);s__Metschnikowia_pulcherrima(100); | Metschnikowia | 13 | Saprotroph | \|Undefined Saprotroph\| | Yeast | NULL | Probable | As nectar-tap_saprotroph (Põlme et al. 2020); Decay sugar-rich substrates (Põlme et al. 2020) | Põlme S, et al. 2020. Fungal Diversity 105:1-16 ((https://doi.org/10.1007/s13225-020-00466-2)) |
| Otu31723 | 1 | k__Fungi(100);p__Ascomycota(100);c__Saccharomycetes(100);o__Saccharomycetales(100);f__Metschnikowiaceae(100);g__Metschnikowia(100);s__Metschnikowia_pulcherrima(100); | Metschnikowia | 13 | Saprotroph | \|Undefined Saprotroph\| | Yeast | NULL | Probable | As nectar-tap_saprotroph (Põlme et al. 2020); Decay sugar-rich substrates (Põlme et al. 2020) | Põlme S, et al. 2020. Fungal Diversity 105:1-16 ((https://doi.org/10.1007/s13225-020-00466-2)) |
| Otu31726 | 1 | k__Fungi(100);p__Ascomycota(100);c__Saccharomycetes(100);o__Saccharomycetales(100);f__Metschnikowiaceae(100);g__Metschnikowia(100);s__Metschnikowia_pulcherrima(100); | Metschnikowia | 13 | Saprotroph | \|Undefined Saprotroph\| | Yeast | NULL | Probable | As nectar-tap_saprotroph (Põlme et al. 2020); Decay sugar-rich substrates (Põlme et al. 2020) | Põlme S, et al. 2020. Fungal Diversity 105:1-16 ((https://doi.org/10.1007/s13225-020-00466-2)) |
| Otu31727 | 1 | k__Fungi(100);p__Ascomycota(100);c__Saccharomycetes(100);o__Saccharomycetales(100);f__Metschnikowiaceae(100);g__Metschnikowia(100);s__Metschnikowia_pulcherrima(100); | Metschnikowia | 13 | Saprotroph | \|Undefined Saprotroph\| | Yeast | NULL | Probable | As nectar-tap_saprotroph (Põlme et al. 2020); Decay sugar-rich substrates (Põlme et al. 2020) | Põlme S, et al. 2020. Fungal Diversity 105:1-16 ((https://doi.org/10.1007/s13225-020-00466-2)) |
| Otu31728 | 1 | k__Fungi(100);p__Ascomycota(100);c__Saccharomycetes(100);o__Saccharomycetales(100);f__Metschnikowiaceae(100);g__Metschnikowia(100);s__Metschnikowia_pulcherrima(100); | Metschnikowia | 13 | Saprotroph | \|Undefined Saprotroph\| | Yeast | NULL | Probable | As nectar-tap_saprotroph (Põlme et al. 2020); Decay sugar-rich substrates (Põlme et al. 2020) | Põlme S, et al. 2020. Fungal Diversity 105:1-16 ((https://doi.org/10.1007/s13225-020-00466-2)) |
| Otu31730 | 1 | k__Fungi(100);p__Ascomycota(100);c__Saccharomycetes(100);o__Saccharomycetales(100);f__Metschnikowiaceae(100);g__Metschnikowia(100);s__Metschnikowia_pulcherrima(100); | Metschnikowia | 13 | Saprotroph | \|Undefined Saprotroph\| | Yeast | NULL | Probable | As nectar-tap_saprotroph (Põlme et al. 2020); Decay sugar-rich substrates (Põlme et al. 2020) | Põlme S, et al. 2020. Fungal Diversity 105:1-16 ((https://doi.org/10.1007/s13225-020-00466-2)) |
| Otu31733 | 1 | k__Fungi(100);p__Ascomycota(100);c__Saccharomycetes(100);o__Saccharomycetales(100);f__Metschnikowiaceae(100);g__Metschnikowia(100);g__Metschnikowia_unclassified(100); | Metschnikowia | 13 | Saprotroph | \|Undefined Saprotroph\| | Yeast | NULL | Probable | As nectar-tap_saprotroph (Põlme et al. 2020); Decay sugar-rich substrates (Põlme et al. 2020) | Põlme S, et al. 2020. Fungal Diversity 105:1-16 ((https://doi.org/10.1007/s13225-020-00466-2)) |
| Otu31739 | 1 | k__Fungi(100);p__Ascomycota(100);c__Saccharomycetes(100);o__Saccharomycetales(100);f__Metschnikowiaceae(100);g__Metschnikowia(100);s__Metschnikowia_pulcherrima(100); | Metschnikowia | 13 | Saprotroph | \|Undefined Saprotroph\| | Yeast | NULL | Probable | As nectar-tap_saprotroph (Põlme et al. 2020); Decay sugar-rich substrates (Põlme et al. 2020) | Põlme S, et al. 2020. Fungal Diversity 105:1-16 ((https://doi.org/10.1007/s13225-020-00466-2)) |
| Otu31741 | 1 | k__Fungi(100);p__Ascomycota(100);c__Saccharomycetes(100);o__Saccharomycetales(100);f__Metschnikowiaceae(100);g__Metschnikowia(100);g__Metschnikowia_unclassified(100); | Metschnikowia | 13 | Saprotroph | \|Undefined Saprotroph\| | Yeast | NULL | Probable | As nectar-tap_saprotroph (Põlme et al. 2020); Decay sugar-rich substrates (Põlme et al. 2020) | Põlme S, et al. 2020. Fungal Diversity 105:1-16 ((https://doi.org/10.1007/s13225-020-00466-2)) |
| Otu31742 | 1 | k__Fungi(100);p__Ascomycota(100);c__Saccharomycetes(100);o__Saccharomycetales(100);f__Metschnikowiaceae(100);g__Metschnikowia(100);g__Metschnikowia_unclassified(100); | Metschnikowia | 13 | Saprotroph | \|Undefined Saprotroph\| | Yeast | NULL | Probable | As nectar-tap_saprotroph (Põlme et al. 2020); Decay sugar-rich substrates (Põlme et al. 2020) | Põlme S, et al. 2020. Fungal Diversity 105:1-16 ((https://doi.org/10.1007/s13225-020-00466-2)) |
| Otu31743 | 1 | k__Fungi(100);p__Ascomycota(100);c__Saccharomycetes(100);o__Saccharomycetales(100);f__Metschnikowiaceae(100);g__Metschnikowia(100);s__Metschnikowia_pulcherrima(100); | Metschnikowia | 13 | Saprotroph | \|Undefined Saprotroph\| | Yeast | NULL | Probable | As nectar-tap_saprotroph (Põlme et al. 2020); Decay sugar-rich substrates (Põlme et al. 2020) | Põlme S, et al. 2020. Fungal Diversity 105:1-16 ((https://doi.org/10.1007/s13225-020-00466-2)) |
| Otu31744 | 1 | k__Fungi(100);p__Ascomycota(100);c__Saccharomycetes(100);o__Saccharomycetales(100);f__Metschnikowiaceae(100);g__Metschnikowia(100);g__Metschnikowia_unclassified(100); | Metschnikowia | 13 | Saprotroph | \|Undefined Saprotroph\| | Yeast | NULL | Probable | As nectar-tap_saprotroph (Põlme et al. 2020); Decay sugar-rich substrates (Põlme et al. 2020) | Põlme S, et al. 2020. Fungal Diversity 105:1-16 ((https://doi.org/10.1007/s13225-020-00466-2)) |
| Otu31745 | 1 | k__Fungi(100);p__Ascomycota(100);c__Saccharomycetes(100);o__Saccharomycetales(100);f__Metschnikowiaceae(100);g__Metschnikowia(100);s__Metschnikowia_pulcherrima(100); | Metschnikowia | 13 | Saprotroph | \|Undefined Saprotroph\| | Yeast | NULL | Probable | As nectar-tap_saprotroph (Põlme et al. 2020); Decay sugar-rich substrates (Põlme et al. 2020) | Põlme S, et al. 2020. Fungal Diversity 105:1-16 ((https://doi.org/10.1007/s13225-020-00466-2)) |
| Otu31747 | 1 | k__Fungi(100);p__Ascomycota(100);c__Saccharomycetes(100);o__Saccharomycetales(100);f__Metschnikowiaceae(100);g__Metschnikowia(100);s__Metschnikowia_pulcherrima(100); | Metschnikowia | 13 | Saprotroph | \|Undefined Saprotroph\| | Yeast | NULL | Probable | As nectar-tap_saprotroph (Põlme et al. 2020); Decay sugar-rich substrates (Põlme et al. 2020) | Põlme S, et al. 2020. Fungal Diversity 105:1-16 ((https://doi.org/10.1007/s13225-020-00466-2)) |
| Otu31749 | 1 | k__Fungi(100);p__Ascomycota(100);c__Saccharomycetes(100);o__Saccharomycetales(100);f__Metschnikowiaceae(100);g__Metschnikowia(100);s__Metschnikowia_pulcherrima(100); | Metschnikowia | 13 | Saprotroph | \|Undefined Saprotroph\| | Yeast | NULL | Probable | As nectar-tap_saprotroph (Põlme et al. 2020); Decay sugar-rich substrates (Põlme et al. 2020) | Põlme S, et al. 2020. Fungal Diversity 105:1-16 ((https://doi.org/10.1007/s13225-020-00466-2)) |
| Otu31750 | 1 | k__Fungi(100);p__Ascomycota(100);c__Saccharomycetes(100);o__Saccharomycetales(100);f__Metschnikowiaceae(100);g__Metschnikowia(100);s__Metschnikowia_pulcherrima(100); | Metschnikowia | 13 | Saprotroph | \|Undefined Saprotroph\| | Yeast | NULL | Probable | As nectar-tap_saprotroph (Põlme et al. 2020); Decay sugar-rich substrates (Põlme et al. 2020) | Põlme S, et al. 2020. Fungal Diversity 105:1-16 ((https://doi.org/10.1007/s13225-020-00466-2)) |
| Otu31751 | 1 | k__Fungi(100);p__Ascomycota(100);c__Saccharomycetes(100);o__Saccharomycetales(100);f__Saccharomycetales_family_Incertae_sedis(100);f__Saccharomycetales_family_Incertae_sedis_unclassified(100);f__Saccharomycetales_family_Incertae_sedis_unclassified(100); | Saccharomycetales | 7 | Saprotroph | Undefined Saprotroph | Yeast | NULL | Possible | NULL | Sterkenburg E, et al. 2015. New Phytologist 207:1145-1158 ((https://doi.org/10.1111/nph.13426)) |
| Otu31752 | 1 | k__Fungi(100);p__Ascomycota(100);c__Saccharomycetes(100);o__Saccharomycetales(100);f__Saccharomycetales_family_Incertae_sedis(100);g__Candida(100);g__Candida_unclassified(100); | Candida | 13 | Pathotroph-Saprotroph-Symbiotroph | Animal Pathogen-Endophyte-Endosymbiont-Epiphyte-\|Undefined Saprotroph\| | Dimorphic Yeast-Yeast | NULL | Probable | As nectar-tap_saprotroph (Põlme et al. 2020); Decay sugar-rich substrates (Põlme et al. 2020); This asexual genus is polyphyletic and contain species that function very differently depend on what they are. Caution should be taken when interpreting guilds for Candida species. | Manolakaki D, et al. 2010. Virulence. 1: 367-375 ((https://doi.org/10.4161/viru.1.5.12796)); Mekbib SB, et al. 2011. Tropical Plant Pathology 36:233-240 ((https://doi.org/10.1590/S1982-56762011000400004)); Põlme S, et al. 2020. Fungal Diversity 105:1-16 ((https://doi.org/10.1007/s13225-020-00466-2)) |
| Otu31753 | 1 | k__Fungi(100);p__Ascomycota(100);c__Saccharomycetes(100);o__Saccharomycetales(100);f__Saccharomycetales_family_Incertae_sedis(100);g__Candida(100);s__Candida_cretensis(100); | Candida | 13 | Pathotroph-Saprotroph-Symbiotroph | Animal Pathogen-Endophyte-Endosymbiont-Epiphyte-\|Undefined Saprotroph\| | Dimorphic Yeast-Yeast | NULL | Probable | As nectar-tap_saprotroph (Põlme et al. 2020); Decay sugar-rich substrates (Põlme et al. 2020); This asexual genus is polyphyletic and contain species that function very differently depend on what they are. Caution should be taken when interpreting guilds for Candida species. | Manolakaki D, et al. 2010. Virulence. 1: 367-375 ((https://doi.org/10.4161/viru.1.5.12796)); Mekbib SB, et al. 2011. Tropical Plant Pathology 36:233-240 ((https://doi.org/10.1590/S1982-56762011000400004)); Põlme S, et al. 2020. Fungal Diversity 105:1-16 ((https://doi.org/10.1007/s13225-020-00466-2)) |
| Otu31754 | 1 | k__Fungi(100);p__Ascomycota(100);c__Saccharomycetes(100);o__Saccharomycetales(100);f__Saccharomycetales_family_Incertae_sedis(100);f__Saccharomycetales_family_Incertae_sedis_unclassified(100);f__Saccharomycetales_family_Incertae_sedis_unclassified(100); | Saccharomycetales | 7 | Saprotroph | Undefined Saprotroph | Yeast | NULL | Possible | NULL | Sterkenburg E, et al. 2015. New Phytologist 207:1145-1158 ((https://doi.org/10.1111/nph.13426)) |
| Otu31755 | 1 | k__Fungi(100);p__Ascomycota(100);c__Saccharomycetes(100);o__Saccharomycetales(100);f__Metschnikowiaceae(100);g__Metschnikowia(100);g__Metschnikowia_unclassified(100); | Metschnikowia | 13 | Saprotroph | \|Undefined Saprotroph\| | Yeast | NULL | Probable | As nectar-tap_saprotroph (Põlme et al. 2020); Decay sugar-rich substrates (Põlme et al. 2020) | Põlme S, et al. 2020. Fungal Diversity 105:1-16 ((https://doi.org/10.1007/s13225-020-00466-2)) |
| Otu31758 | 1 | k__Fungi(100);p__Ascomycota(100);c__Saccharomycetes(100);o__Saccharomycetales(100);f__Metschnikowiaceae(100);g__Metschnikowia(100);g__Metschnikowia_unclassified(100); | Metschnikowia | 13 | Saprotroph | \|Undefined Saprotroph\| | Yeast | NULL | Probable | As nectar-tap_saprotroph (Põlme et al. 2020); Decay sugar-rich substrates (Põlme et al. 2020) | Põlme S, et al. 2020. Fungal Diversity 105:1-16 ((https://doi.org/10.1007/s13225-020-00466-2)) |
| Otu31760 | 1 | k__Fungi(100);p__Ascomycota(100);c__Dothideomycetes(100);o__Capnodiales(100);f__Mycosphaerellaceae(100);g__Mycosphaerella(100);s__Mycosphaerella_latebrosa(100); | Mycosphaerella | 13 | Pathotroph-Saprotroph | \|Plant Pathogen\|-Plant Saprotroph | Microfungus | NULL | Probable | As litter_saprotroph (Põlme et al. 2020); Decay leaf, fruit, seed (Põlme et al. 2020) | Tedersoo L, et al. 2014. Science 346:e1256688 ((https://doi.org/10.1126/science.1256688)); Põlme S, et al. 2020. Fungal Diversity 105:1-16 ((https://doi.org/10.1007/s13225-020-00466-2)) |
| Otu31761 | 1 | k__Fungi(100);p__Ascomycota(100);c__Saccharomycetes(100);o__Saccharomycetales(100);f__Metschnikowiaceae(100);g__Metschnikowia(100);s__Metschnikowia_pulcherrima(100); | Metschnikowia | 13 | Saprotroph | \|Undefined Saprotroph\| | Yeast | NULL | Probable | As nectar-tap_saprotroph (Põlme et al. 2020); Decay sugar-rich substrates (Põlme et al. 2020) | Põlme S, et al. 2020. Fungal Diversity 105:1-16 ((https://doi.org/10.1007/s13225-020-00466-2)) |
| Otu31762 | 1 | k__Fungi(100);p__Ascomycota(100);c__Saccharomycetes(100);o__Saccharomycetales(100);f__Saccharomycetales_family_Incertae_sedis(100);g__Candida(100);s__Candida_cretensis(100); | Candida | 13 | Pathotroph-Saprotroph-Symbiotroph | Animal Pathogen-Endophyte-Endosymbiont-Epiphyte-\|Undefined Saprotroph\| | Dimorphic Yeast-Yeast | NULL | Probable | As nectar-tap_saprotroph (Põlme et al. 2020); Decay sugar-rich substrates (Põlme et al. 2020); This asexual genus is polyphyletic and contain species that function very differently depend on what they are. Caution should be taken when interpreting guilds for Candida species. | Manolakaki D, et al. 2010. Virulence. 1: 367-375 ((https://doi.org/10.4161/viru.1.5.12796)); Mekbib SB, et al. 2011. Tropical Plant Pathology 36:233-240 ((https://doi.org/10.1590/S1982-56762011000400004)); Põlme S, et al. 2020. Fungal Diversity 105:1-16 ((https://doi.org/10.1007/s13225-020-00466-2)) |
| Otu31765 | 1 | k__Fungi(100);p__Ascomycota(100);c__Saccharomycetes(100);o__Saccharomycetales(100);f__Metschnikowiaceae(100);g__Metschnikowia(100);s__Metschnikowia_pulcherrima(100); | Metschnikowia | 13 | Saprotroph | \|Undefined Saprotroph\| | Yeast | NULL | Probable | As nectar-tap_saprotroph (Põlme et al. 2020); Decay sugar-rich substrates (Põlme et al. 2020) | Põlme S, et al. 2020. Fungal Diversity 105:1-16 ((https://doi.org/10.1007/s13225-020-00466-2)) |
| Otu31766 | 1 | k__Fungi(100);p__Ascomycota(100);c__Saccharomycetes(100);o__Saccharomycetales(100);f__Metschnikowiaceae(100);g__Metschnikowia(100);g__Metschnikowia_unclassified(100); | Metschnikowia | 13 | Saprotroph | \|Undefined Saprotroph\| | Yeast | NULL | Probable | As nectar-tap_saprotroph (Põlme et al. 2020); Decay sugar-rich substrates (Põlme et al. 2020) | Põlme S, et al. 2020. Fungal Diversity 105:1-16 ((https://doi.org/10.1007/s13225-020-00466-2)) |
| Otu31767 | 1 | k__Fungi(100);p__Ascomycota(100);c__Saccharomycetes(100);o__Saccharomycetales(100);f__Metschnikowiaceae(100);g__Metschnikowia(100);s__Metschnikowia_pulcherrima(100); | Metschnikowia | 13 | Saprotroph | \|Undefined Saprotroph\| | Yeast | NULL | Probable | As nectar-tap_saprotroph (Põlme et al. 2020); Decay sugar-rich substrates (Põlme et al. 2020) | Põlme S, et al. 2020. Fungal Diversity 105:1-16 ((https://doi.org/10.1007/s13225-020-00466-2)) |
| Otu31769 | 1 | k__Fungi(100);p__Ascomycota(100);c__Saccharomycetes(100);o__Saccharomycetales(100);f__Metschnikowiaceae(100);g__Metschnikowia(100);g__Metschnikowia_unclassified(100); | Metschnikowia | 13 | Saprotroph | \|Undefined Saprotroph\| | Yeast | NULL | Probable | As nectar-tap_saprotroph (Põlme et al. 2020); Decay sugar-rich substrates (Põlme et al. 2020) | Põlme S, et al. 2020. Fungal Diversity 105:1-16 ((https://doi.org/10.1007/s13225-020-00466-2)) |
| Otu31770 | 1 | k__Fungi(100);p__Ascomycota(100);c__Saccharomycetes(100);o__Saccharomycetales(100);f__Metschnikowiaceae(100);g__Metschnikowia(100);g__Metschnikowia_unclassified(100); | Metschnikowia | 13 | Saprotroph | \|Undefined Saprotroph\| | Yeast | NULL | Probable | As nectar-tap_saprotroph (Põlme et al. 2020); Decay sugar-rich substrates (Põlme et al. 2020) | Põlme S, et al. 2020. Fungal Diversity 105:1-16 ((https://doi.org/10.1007/s13225-020-00466-2)) |
| Otu31771 | 1 | k__Fungi(100);p__Ascomycota(100);c__Saccharomycetes(100);o__Saccharomycetales(100);f__Metschnikowiaceae(100);g__Metschnikowia(100);s__Metschnikowia_pulcherrima(100); | Metschnikowia | 13 | Saprotroph | \|Undefined Saprotroph\| | Yeast | NULL | Probable | As nectar-tap_saprotroph (Põlme et al. 2020); Decay sugar-rich substrates (Põlme et al. 2020) | Põlme S, et al. 2020. Fungal Diversity 105:1-16 ((https://doi.org/10.1007/s13225-020-00466-2)) |
| Otu31772 | 1 | k__Fungi(100);p__Ascomycota(100);c__Saccharomycetes(100);o__Saccharomycetales(100);f__Metschnikowiaceae(100);g__Metschnikowia(100);s__Metschnikowia_pulcherrima(100); | Metschnikowia | 13 | Saprotroph | \|Undefined Saprotroph\| | Yeast | NULL | Probable | As nectar-tap_saprotroph (Põlme et al. 2020); Decay sugar-rich substrates (Põlme et al. 2020) | Põlme S, et al. 2020. Fungal Diversity 105:1-16 ((https://doi.org/10.1007/s13225-020-00466-2)) |
| Otu31773 | 1 | k__Fungi(100);p__Ascomycota(100);c__Saccharomycetes(100);o__Saccharomycetales(100);f__Metschnikowiaceae(100);g__Metschnikowia(100);s__Metschnikowia_pulcherrima(100); | Metschnikowia | 13 | Saprotroph | \|Undefined Saprotroph\| | Yeast | NULL | Probable | As nectar-tap_saprotroph (Põlme et al. 2020); Decay sugar-rich substrates (Põlme et al. 2020) | Põlme S, et al. 2020. Fungal Diversity 105:1-16 ((https://doi.org/10.1007/s13225-020-00466-2)) |
| Otu31774 | 1 | k__Fungi(100);p__Ascomycota(100);c__Saccharomycetes(100);o__Saccharomycetales(100);f__Metschnikowiaceae(100);g__Metschnikowia(100);s__Metschnikowia_pulcherrima(100); | Metschnikowia | 13 | Saprotroph | \|Undefined Saprotroph\| | Yeast | NULL | Probable | As nectar-tap_saprotroph (Põlme et al. 2020); Decay sugar-rich substrates (Põlme et al. 2020) | Põlme S, et al. 2020. Fungal Diversity 105:1-16 ((https://doi.org/10.1007/s13225-020-00466-2)) |
| Otu31775 | 1 | k__Fungi(100);p__Ascomycota(100);c__Saccharomycetes(100);o__Saccharomycetales(100);f__Metschnikowiaceae(100);g__Metschnikowia(100);s__Metschnikowia_pulcherrima(100); | Metschnikowia | 13 | Saprotroph | \|Undefined Saprotroph\| | Yeast | NULL | Probable | As nectar-tap_saprotroph (Põlme et al. 2020); Decay sugar-rich substrates (Põlme et al. 2020) | Põlme S, et al. 2020. Fungal Diversity 105:1-16 ((https://doi.org/10.1007/s13225-020-00466-2)) |
| Otu31776 | 1 | k__Fungi(100);p__Ascomycota(100);c__Saccharomycetes(100);o__Saccharomycetales(100);f__Metschnikowiaceae(100);g__Metschnikowia(100);g__Metschnikowia_unclassified(100); | Metschnikowia | 13 | Saprotroph | \|Undefined Saprotroph\| | Yeast | NULL | Probable | As nectar-tap_saprotroph (Põlme et al. 2020); Decay sugar-rich substrates (Põlme et al. 2020) | Põlme S, et al. 2020. Fungal Diversity 105:1-16 ((https://doi.org/10.1007/s13225-020-00466-2)) |
| Otu31777 | 1 | k__Fungi(100);p__Ascomycota(100);c__Saccharomycetes(100);o__Saccharomycetales(100);f__Metschnikowiaceae(100);g__Metschnikowia(100);g__Metschnikowia_unclassified(100); | Metschnikowia | 13 | Saprotroph | \|Undefined Saprotroph\| | Yeast | NULL | Probable | As nectar-tap_saprotroph (Põlme et al. 2020); Decay sugar-rich substrates (Põlme et al. 2020) | Põlme S, et al. 2020. Fungal Diversity 105:1-16 ((https://doi.org/10.1007/s13225-020-00466-2)) |
| Otu31778 | 1 | k__Fungi(100);p__Ascomycota(100);c__Saccharomycetes(100);o__Saccharomycetales(100);f__Metschnikowiaceae(100);g__Metschnikowia(100);g__Metschnikowia_unclassified(100); | Metschnikowia | 13 | Saprotroph | \|Undefined Saprotroph\| | Yeast | NULL | Probable | As nectar-tap_saprotroph (Põlme et al. 2020); Decay sugar-rich substrates (Põlme et al. 2020) | Põlme S, et al. 2020. Fungal Diversity 105:1-16 ((https://doi.org/10.1007/s13225-020-00466-2)) |
| Otu31780 | 1 | k__Fungi(100);p__Ascomycota(100);c__Saccharomycetes(100);o__Saccharomycetales(100);f__Metschnikowiaceae(100);g__Metschnikowia(100);g__Metschnikowia_unclassified(100); | Metschnikowia | 13 | Saprotroph | \|Undefined Saprotroph\| | Yeast | NULL | Probable | As nectar-tap_saprotroph (Põlme et al. 2020); Decay sugar-rich substrates (Põlme et al. 2020) | Põlme S, et al. 2020. Fungal Diversity 105:1-16 ((https://doi.org/10.1007/s13225-020-00466-2)) |
| Otu31781 | 1 | k__Fungi(100);p__Ascomycota(100);c__Saccharomycetes(100);o__Saccharomycetales(100);f__Metschnikowiaceae(100);g__Metschnikowia(100);g__Metschnikowia_unclassified(100); | Metschnikowia | 13 | Saprotroph | \|Undefined Saprotroph\| | Yeast | NULL | Probable | As nectar-tap_saprotroph (Põlme et al. 2020); Decay sugar-rich substrates (Põlme et al. 2020) | Põlme S, et al. 2020. Fungal Diversity 105:1-16 ((https://doi.org/10.1007/s13225-020-00466-2)) |
| Otu31782 | 1 | k__Fungi(100);p__Ascomycota(100);c__Saccharomycetes(100);o__Saccharomycetales(100);f__Metschnikowiaceae(100);g__Metschnikowia(100);s__Metschnikowia_pulcherrima(100); | Metschnikowia | 13 | Saprotroph | \|Undefined Saprotroph\| | Yeast | NULL | Probable | As nectar-tap_saprotroph (Põlme et al. 2020); Decay sugar-rich substrates (Põlme et al. 2020) | Põlme S, et al. 2020. Fungal Diversity 105:1-16 ((https://doi.org/10.1007/s13225-020-00466-2)) |
| Otu31783 | 1 | k__Fungi(100);p__Ascomycota(100);c__Saccharomycetes(100);o__Saccharomycetales(100);f__Metschnikowiaceae(100);g__Metschnikowia(100);s__Metschnikowia_pulcherrima(100); | Metschnikowia | 13 | Saprotroph | \|Undefined Saprotroph\| | Yeast | NULL | Probable | As nectar-tap_saprotroph (Põlme et al. 2020); Decay sugar-rich substrates (Põlme et al. 2020) | Põlme S, et al. 2020. Fungal Diversity 105:1-16 ((https://doi.org/10.1007/s13225-020-00466-2)) |
| Otu31784 | 1 | k__Fungi(100);p__Ascomycota(100);c__Saccharomycetes(100);o__Saccharomycetales(100);f__Metschnikowiaceae(100);g__Metschnikowia(100);s__Metschnikowia_pulcherrima(100); | Metschnikowia | 13 | Saprotroph | \|Undefined Saprotroph\| | Yeast | NULL | Probable | As nectar-tap_saprotroph (Põlme et al. 2020); Decay sugar-rich substrates (Põlme et al. 2020) | Põlme S, et al. 2020. Fungal Diversity 105:1-16 ((https://doi.org/10.1007/s13225-020-00466-2)) |
| Otu31785 | 1 | k__Fungi(100);p__Ascomycota(100);c__Saccharomycetes(100);o__Saccharomycetales(100);f__Metschnikowiaceae(100);g__Metschnikowia(100);g__Metschnikowia_unclassified(100); | Metschnikowia | 13 | Saprotroph | \|Undefined Saprotroph\| | Yeast | NULL | Probable | As nectar-tap_saprotroph (Põlme et al. 2020); Decay sugar-rich substrates (Põlme et al. 2020) | Põlme S, et al. 2020. Fungal Diversity 105:1-16 ((https://doi.org/10.1007/s13225-020-00466-2)) |
| Otu31786 | 1 | k__Fungi(100);p__Ascomycota(100);c__Saccharomycetes(100);o__Saccharomycetales(100);f__Saccharomycetales_family_Incertae_sedis(100);g__Candida(100);g__Candida_unclassified(100); | Candida | 13 | Pathotroph-Saprotroph-Symbiotroph | Animal Pathogen-Endophyte-Endosymbiont-Epiphyte-\|Undefined Saprotroph\| | Dimorphic Yeast-Yeast | NULL | Probable | As nectar-tap_saprotroph (Põlme et al. 2020); Decay sugar-rich substrates (Põlme et al. 2020); This asexual genus is polyphyletic and contain species that function very differently depend on what they are. Caution should be taken when interpreting guilds for Candida species. | Manolakaki D, et al. 2010. Virulence. 1: 367-375 ((https://doi.org/10.4161/viru.1.5.12796)); Mekbib SB, et al. 2011. Tropical Plant Pathology 36:233-240 ((https://doi.org/10.1590/S1982-56762011000400004)); Põlme S, et al. 2020. Fungal Diversity 105:1-16 ((https://doi.org/10.1007/s13225-020-00466-2)) |
| Otu31788 | 1 | k__Fungi(100);p__Ascomycota(100);c__Saccharomycetes(100);o__Saccharomycetales(100);f__Saccharomycetales_family_Incertae_sedis(100);g__Candida(100);g__Candida_unclassified(100); | Candida | 13 | Pathotroph-Saprotroph-Symbiotroph | Animal Pathogen-Endophyte-Endosymbiont-Epiphyte-\|Undefined Saprotroph\| | Dimorphic Yeast-Yeast | NULL | Probable | As nectar-tap_saprotroph (Põlme et al. 2020); Decay sugar-rich substrates (Põlme et al. 2020); This asexual genus is polyphyletic and contain species that function very differently depend on what they are. Caution should be taken when interpreting guilds for Candida species. | Manolakaki D, et al. 2010. Virulence. 1: 367-375 ((https://doi.org/10.4161/viru.1.5.12796)); Mekbib SB, et al. 2011. Tropical Plant Pathology 36:233-240 ((https://doi.org/10.1590/S1982-56762011000400004)); Põlme S, et al. 2020. Fungal Diversity 105:1-16 ((https://doi.org/10.1007/s13225-020-00466-2)) |
| Otu31791 | 1 | k__Fungi(100);p__Ascomycota(100);c__Saccharomycetes(100);o__Saccharomycetales(100);f__Metschnikowiaceae(100);g__Metschnikowia(100);s__Metschnikowia_pulcherrima(100); | Metschnikowia | 13 | Saprotroph | \|Undefined Saprotroph\| | Yeast | NULL | Probable | As nectar-tap_saprotroph (Põlme et al. 2020); Decay sugar-rich substrates (Põlme et al. 2020) | Põlme S, et al. 2020. Fungal Diversity 105:1-16 ((https://doi.org/10.1007/s13225-020-00466-2)) |
| Otu31792 | 1 | k__Fungi(100);p__Ascomycota(100);c__Saccharomycetes(100);o__Saccharomycetales(100);f__Metschnikowiaceae(100);g__Metschnikowia(100);g__Metschnikowia_unclassified(100); | Metschnikowia | 13 | Saprotroph | \|Undefined Saprotroph\| | Yeast | NULL | Probable | As nectar-tap_saprotroph (Põlme et al. 2020); Decay sugar-rich substrates (Põlme et al. 2020) | Põlme S, et al. 2020. Fungal Diversity 105:1-16 ((https://doi.org/10.1007/s13225-020-00466-2)) |
| Otu31793 | 1 | k__Fungi(100);p__Ascomycota(100);c__Saccharomycetes(100);o__Saccharomycetales(100);f__Metschnikowiaceae(100);g__Metschnikowia(100);s__Metschnikowia_pulcherrima(100); | Metschnikowia | 13 | Saprotroph | \|Undefined Saprotroph\| | Yeast | NULL | Probable | As nectar-tap_saprotroph (Põlme et al. 2020); Decay sugar-rich substrates (Põlme et al. 2020) | Põlme S, et al. 2020. Fungal Diversity 105:1-16 ((https://doi.org/10.1007/s13225-020-00466-2)) |
| Otu31794 | 1 | k__Fungi(100);p__Ascomycota(100);c__Saccharomycetes(100);o__Saccharomycetales(100);f__Metschnikowiaceae(100);g__Metschnikowia(100);s__Metschnikowia_pulcherrima(100); | Metschnikowia | 13 | Saprotroph | \|Undefined Saprotroph\| | Yeast | NULL | Probable | As nectar-tap_saprotroph (Põlme et al. 2020); Decay sugar-rich substrates (Põlme et al. 2020) | Põlme S, et al. 2020. Fungal Diversity 105:1-16 ((https://doi.org/10.1007/s13225-020-00466-2)) |
| Otu31795 | 1 | k__Fungi(100);p__Ascomycota(100);c__Saccharomycetes(100);o__Saccharomycetales(100);f__Saccharomycetales_family_Incertae_sedis(100);g__Candida(100);g__Candida_unclassified(100); | Candida | 13 | Pathotroph-Saprotroph-Symbiotroph | Animal Pathogen-Endophyte-Endosymbiont-Epiphyte-\|Undefined Saprotroph\| | Dimorphic Yeast-Yeast | NULL | Probable | As nectar-tap_saprotroph (Põlme et al. 2020); Decay sugar-rich substrates (Põlme et al. 2020); This asexual genus is polyphyletic and contain species that function very differently depend on what they are. Caution should be taken when interpreting guilds for Candida species. | Manolakaki D, et al. 2010. Virulence. 1: 367-375 ((https://doi.org/10.4161/viru.1.5.12796)); Mekbib SB, et al. 2011. Tropical Plant Pathology 36:233-240 ((https://doi.org/10.1590/S1982-56762011000400004)); Põlme S, et al. 2020. Fungal Diversity 105:1-16 ((https://doi.org/10.1007/s13225-020-00466-2)) |
| Otu31796 | 1 | k__Fungi(100);p__Ascomycota(100);c__Saccharomycetes(100);o__Saccharomycetales(100);f__Metschnikowiaceae(100);g__Metschnikowia(100);s__Metschnikowia_pulcherrima(100); | Metschnikowia | 13 | Saprotroph | \|Undefined Saprotroph\| | Yeast | NULL | Probable | As nectar-tap_saprotroph (Põlme et al. 2020); Decay sugar-rich substrates (Põlme et al. 2020) | Põlme S, et al. 2020. Fungal Diversity 105:1-16 ((https://doi.org/10.1007/s13225-020-00466-2)) |
| Otu31798 | 1 | k__Fungi(100);p__Ascomycota(100);c__Saccharomycetes(100);o__Saccharomycetales(100);f__Metschnikowiaceae(100);g__Metschnikowia(100);s__Metschnikowia_pulcherrima(100); | Metschnikowia | 13 | Saprotroph | \|Undefined Saprotroph\| | Yeast | NULL | Probable | As nectar-tap_saprotroph (Põlme et al. 2020); Decay sugar-rich substrates (Põlme et al. 2020) | Põlme S, et al. 2020. Fungal Diversity 105:1-16 ((https://doi.org/10.1007/s13225-020-00466-2)) |
| Otu31800 | 1 | k__Fungi(100);p__Ascomycota(100);c__Saccharomycetes(100);o__Saccharomycetales(100);f__Metschnikowiaceae(100);g__Metschnikowia(100);s__Metschnikowia_pulcherrima(100); | Metschnikowia | 13 | Saprotroph | \|Undefined Saprotroph\| | Yeast | NULL | Probable | As nectar-tap_saprotroph (Põlme et al. 2020); Decay sugar-rich substrates (Põlme et al. 2020) | Põlme S, et al. 2020. Fungal Diversity 105:1-16 ((https://doi.org/10.1007/s13225-020-00466-2)) |
| Otu31801 | 1 | k__Fungi(100);p__Ascomycota(100);c__Saccharomycetes(100);o__Saccharomycetales(100);f__Metschnikowiaceae(100);g__Metschnikowia(100);s__Metschnikowia_pulcherrima(100); | Metschnikowia | 13 | Saprotroph | \|Undefined Saprotroph\| | Yeast | NULL | Probable | As nectar-tap_saprotroph (Põlme et al. 2020); Decay sugar-rich substrates (Põlme et al. 2020) | Põlme S, et al. 2020. Fungal Diversity 105:1-16 ((https://doi.org/10.1007/s13225-020-00466-2)) |
| Otu31802 | 1 | k__Fungi(100);p__Ascomycota(100);c__Saccharomycetes(100);o__Saccharomycetales(100);f__Metschnikowiaceae(100);g__Metschnikowia(100);g__Metschnikowia_unclassified(100); | Metschnikowia | 13 | Saprotroph | \|Undefined Saprotroph\| | Yeast | NULL | Probable | As nectar-tap_saprotroph (Põlme et al. 2020); Decay sugar-rich substrates (Põlme et al. 2020) | Põlme S, et al. 2020. Fungal Diversity 105:1-16 ((https://doi.org/10.1007/s13225-020-00466-2)) |
| Otu31803 | 1 | k__Fungi(100);p__Ascomycota(100);c__Saccharomycetes(100);o__Saccharomycetales(100);f__Metschnikowiaceae(100);g__Metschnikowia(100);g__Metschnikowia_unclassified(100); | Metschnikowia | 13 | Saprotroph | \|Undefined Saprotroph\| | Yeast | NULL | Probable | As nectar-tap_saprotroph (Põlme et al. 2020); Decay sugar-rich substrates (Põlme et al. 2020) | Põlme S, et al. 2020. Fungal Diversity 105:1-16 ((https://doi.org/10.1007/s13225-020-00466-2)) |
| Otu31804 | 1 | k__Fungi(100);p__Ascomycota(100);c__Saccharomycetes(100);o__Saccharomycetales(100);o__Saccharomycetales_unclassified(100);o__Saccharomycetales_unclassified(100);o__Saccharomycetales_unclassified(100); | Saccharomycetales | 7 | Saprotroph | Undefined Saprotroph | Yeast | NULL | Possible | NULL | Sterkenburg E, et al. 2015. New Phytologist 207:1145-1158 ((https://doi.org/10.1111/nph.13426)) |
| Otu31805 | 1 | k__Fungi(100);p__Ascomycota(100);c__Saccharomycetes(100);o__Saccharomycetales(100);f__Metschnikowiaceae(100);g__Metschnikowia(100);s__Metschnikowia_pulcherrima(100); | Metschnikowia | 13 | Saprotroph | \|Undefined Saprotroph\| | Yeast | NULL | Probable | As nectar-tap_saprotroph (Põlme et al. 2020); Decay sugar-rich substrates (Põlme et al. 2020) | Põlme S, et al. 2020. Fungal Diversity 105:1-16 ((https://doi.org/10.1007/s13225-020-00466-2)) |
| Otu31806 | 1 | k__Fungi(100);p__Ascomycota(100);c__Saccharomycetes(100);o__Saccharomycetales(100);f__Saccharomycetales_family_Incertae_sedis(100);g__Candida(100);g__Candida_unclassified(100); | Candida | 13 | Pathotroph-Saprotroph-Symbiotroph | Animal Pathogen-Endophyte-Endosymbiont-Epiphyte-\|Undefined Saprotroph\| | Dimorphic Yeast-Yeast | NULL | Probable | As nectar-tap_saprotroph (Põlme et al. 2020); Decay sugar-rich substrates (Põlme et al. 2020); This asexual genus is polyphyletic and contain species that function very differently depend on what they are. Caution should be taken when interpreting guilds for Candida species. | Manolakaki D, et al. 2010. Virulence. 1: 367-375 ((https://doi.org/10.4161/viru.1.5.12796)); Mekbib SB, et al. 2011. Tropical Plant Pathology 36:233-240 ((https://doi.org/10.1590/S1982-56762011000400004)); Põlme S, et al. 2020. Fungal Diversity 105:1-16 ((https://doi.org/10.1007/s13225-020-00466-2)) |
| Otu31807 | 1 | k__Fungi(100);p__Ascomycota(100);c__Saccharomycetes(100);o__Saccharomycetales(100);f__Metschnikowiaceae(100);g__Metschnikowia(100);s__Metschnikowia_pulcherrima(100); | Metschnikowia | 13 | Saprotroph | \|Undefined Saprotroph\| | Yeast | NULL | Probable | As nectar-tap_saprotroph (Põlme et al. 2020); Decay sugar-rich substrates (Põlme et al. 2020) | Põlme S, et al. 2020. Fungal Diversity 105:1-16 ((https://doi.org/10.1007/s13225-020-00466-2)) |
| Otu31810 | 1 | k__Fungi(100);p__Ascomycota(100);c__Saccharomycetes(100);o__Saccharomycetales(100);f__Saccharomycetales_family_Incertae_sedis(100);f__Saccharomycetales_family_Incertae_sedis_unclassified(100);f__Saccharomycetales_family_Incertae_sedis_unclassified(100); | Saccharomycetales | 7 | Saprotroph | Undefined Saprotroph | Yeast | NULL | Possible | NULL | Sterkenburg E, et al. 2015. New Phytologist 207:1145-1158 ((https://doi.org/10.1111/nph.13426)) |
| Otu31811 | 1 | k__Fungi(100);p__Ascomycota(100);c__Saccharomycetes(100);o__Saccharomycetales(100);f__Saccharomycetales_family_Incertae_sedis(100);f__Saccharomycetales_family_Incertae_sedis_unclassified(100);f__Saccharomycetales_family_Incertae_sedis_unclassified(100); | Saccharomycetales | 7 | Saprotroph | Undefined Saprotroph | Yeast | NULL | Possible | NULL | Sterkenburg E, et al. 2015. New Phytologist 207:1145-1158 ((https://doi.org/10.1111/nph.13426)) |
| Otu31812 | 1 | k__Fungi(100);p__Ascomycota(100);c__Saccharomycetes(100);o__Saccharomycetales(100);f__Metschnikowiaceae(100);g__Metschnikowia(100);s__Metschnikowia_pulcherrima(100); | Metschnikowia | 13 | Saprotroph | \|Undefined Saprotroph\| | Yeast | NULL | Probable | As nectar-tap_saprotroph (Põlme et al. 2020); Decay sugar-rich substrates (Põlme et al. 2020) | Põlme S, et al. 2020. Fungal Diversity 105:1-16 ((https://doi.org/10.1007/s13225-020-00466-2)) |
| Otu31815 | 1 | k__Fungi(100);p__Ascomycota(100);c__Saccharomycetes(100);o__Saccharomycetales(100);f__Metschnikowiaceae(100);g__Metschnikowia(100);s__Metschnikowia_pulcherrima(100); | Metschnikowia | 13 | Saprotroph | \|Undefined Saprotroph\| | Yeast | NULL | Probable | As nectar-tap_saprotroph (Põlme et al. 2020); Decay sugar-rich substrates (Põlme et al. 2020) | Põlme S, et al. 2020. Fungal Diversity 105:1-16 ((https://doi.org/10.1007/s13225-020-00466-2)) |
| Otu31821 | 1 | k__Fungi(100);p__Ascomycota(100);c__Saccharomycetes(100);o__Saccharomycetales(100);f__Metschnikowiaceae(100);g__Metschnikowia(100);s__Metschnikowia_pulcherrima(100); | Metschnikowia | 13 | Saprotroph | \|Undefined Saprotroph\| | Yeast | NULL | Probable | As nectar-tap_saprotroph (Põlme et al. 2020); Decay sugar-rich substrates (Põlme et al. 2020) | Põlme S, et al. 2020. Fungal Diversity 105:1-16 ((https://doi.org/10.1007/s13225-020-00466-2)) |
| Otu31822 | 1 | k__Fungi(100);p__Ascomycota(100);c__Saccharomycetes(100);o__Saccharomycetales(100);f__Metschnikowiaceae(100);g__Metschnikowia(100);s__Metschnikowia_pulcherrima(100); | Metschnikowia | 13 | Saprotroph | \|Undefined Saprotroph\| | Yeast | NULL | Probable | As nectar-tap_saprotroph (Põlme et al. 2020); Decay sugar-rich substrates (Põlme et al. 2020) | Põlme S, et al. 2020. Fungal Diversity 105:1-16 ((https://doi.org/10.1007/s13225-020-00466-2)) |
| Otu31823 | 1 | k__Fungi(100);p__Ascomycota(100);c__Saccharomycetes(100);o__Saccharomycetales(100);f__Metschnikowiaceae(100);g__Metschnikowia(100);s__Metschnikowia_pulcherrima(100); | Metschnikowia | 13 | Saprotroph | \|Undefined Saprotroph\| | Yeast | NULL | Probable | As nectar-tap_saprotroph (Põlme et al. 2020); Decay sugar-rich substrates (Põlme et al. 2020) | Põlme S, et al. 2020. Fungal Diversity 105:1-16 ((https://doi.org/10.1007/s13225-020-00466-2)) |
| Otu31824 | 1 | k__Fungi(100);p__Ascomycota(100);c__Saccharomycetes(100);o__Saccharomycetales(100);f__Metschnikowiaceae(100);g__Metschnikowia(100);g__Metschnikowia_unclassified(100); | Metschnikowia | 13 | Saprotroph | \|Undefined Saprotroph\| | Yeast | NULL | Probable | As nectar-tap_saprotroph (Põlme et al. 2020); Decay sugar-rich substrates (Põlme et al. 2020) | Põlme S, et al. 2020. Fungal Diversity 105:1-16 ((https://doi.org/10.1007/s13225-020-00466-2)) |
| Otu31828 | 1 | k__Fungi(100);p__Ascomycota(100);c__Saccharomycetes(100);o__Saccharomycetales(100);f__Metschnikowiaceae(100);g__Metschnikowia(100);g__Metschnikowia_unclassified(100); | Metschnikowia | 13 | Saprotroph | \|Undefined Saprotroph\| | Yeast | NULL | Probable | As nectar-tap_saprotroph (Põlme et al. 2020); Decay sugar-rich substrates (Põlme et al. 2020) | Põlme S, et al. 2020. Fungal Diversity 105:1-16 ((https://doi.org/10.1007/s13225-020-00466-2)) |
| Otu31829 | 1 | k__Fungi(100);p__Ascomycota(100);c__Saccharomycetes(100);o__Saccharomycetales(100);f__Metschnikowiaceae(100);g__Metschnikowia(100);s__Metschnikowia_pulcherrima(100); | Metschnikowia | 13 | Saprotroph | \|Undefined Saprotroph\| | Yeast | NULL | Probable | As nectar-tap_saprotroph (Põlme et al. 2020); Decay sugar-rich substrates (Põlme et al. 2020) | Põlme S, et al. 2020. Fungal Diversity 105:1-16 ((https://doi.org/10.1007/s13225-020-00466-2)) |
| Otu31830 | 1 | k__Fungi(100);p__Ascomycota(100);c__Saccharomycetes(100);o__Saccharomycetales(100);f__Metschnikowiaceae(100);g__Metschnikowia(100);g__Metschnikowia_unclassified(100); | Metschnikowia | 13 | Saprotroph | \|Undefined Saprotroph\| | Yeast | NULL | Probable | As nectar-tap_saprotroph (Põlme et al. 2020); Decay sugar-rich substrates (Põlme et al. 2020) | Põlme S, et al. 2020. Fungal Diversity 105:1-16 ((https://doi.org/10.1007/s13225-020-00466-2)) |
| Otu31831 | 1 | k__Fungi(100);p__Ascomycota(100);c__Saccharomycetes(100);o__Saccharomycetales(100);f__Metschnikowiaceae(100);g__Metschnikowia(100);s__Metschnikowia_pulcherrima(100); | Metschnikowia | 13 | Saprotroph | \|Undefined Saprotroph\| | Yeast | NULL | Probable | As nectar-tap_saprotroph (Põlme et al. 2020); Decay sugar-rich substrates (Põlme et al. 2020) | Põlme S, et al. 2020. Fungal Diversity 105:1-16 ((https://doi.org/10.1007/s13225-020-00466-2)) |
| Otu31832 | 1 | k__Fungi(100);p__Ascomycota(100);c__Saccharomycetes(100);o__Saccharomycetales(100);f__Metschnikowiaceae(100);g__Metschnikowia(100);s__Metschnikowia_pulcherrima(100); | Metschnikowia | 13 | Saprotroph | \|Undefined Saprotroph\| | Yeast | NULL | Probable | As nectar-tap_saprotroph (Põlme et al. 2020); Decay sugar-rich substrates (Põlme et al. 2020) | Põlme S, et al. 2020. Fungal Diversity 105:1-16 ((https://doi.org/10.1007/s13225-020-00466-2)) |
| Otu31833 | 1 | k__Fungi(100);p__Ascomycota(100);c__Saccharomycetes(100);o__Saccharomycetales(100);o__Saccharomycetales_unclassified(100);o__Saccharomycetales_unclassified(100);o__Saccharomycetales_unclassified(100); | Saccharomycetales | 7 | Saprotroph | Undefined Saprotroph | Yeast | NULL | Possible | NULL | Sterkenburg E, et al. 2015. New Phytologist 207:1145-1158 ((https://doi.org/10.1111/nph.13426)) |
| Otu31834 | 1 | k__Fungi(100);p__Ascomycota(100);c__Saccharomycetes(100);o__Saccharomycetales(100);f__Metschnikowiaceae(100);g__Metschnikowia(100);s__Metschnikowia_pulcherrima(100); | Metschnikowia | 13 | Saprotroph | \|Undefined Saprotroph\| | Yeast | NULL | Probable | As nectar-tap_saprotroph (Põlme et al. 2020); Decay sugar-rich substrates (Põlme et al. 2020) | Põlme S, et al. 2020. Fungal Diversity 105:1-16 ((https://doi.org/10.1007/s13225-020-00466-2)) |
| Otu31836 | 1 | k__Fungi(100);p__Ascomycota(100);c__Saccharomycetes(100);o__Saccharomycetales(100);f__Metschnikowiaceae(100);g__Metschnikowia(100);s__Metschnikowia_pulcherrima(100); | Metschnikowia | 13 | Saprotroph | \|Undefined Saprotroph\| | Yeast | NULL | Probable | As nectar-tap_saprotroph (Põlme et al. 2020); Decay sugar-rich substrates (Põlme et al. 2020) | Põlme S, et al. 2020. Fungal Diversity 105:1-16 ((https://doi.org/10.1007/s13225-020-00466-2)) |
| Otu31837 | 1 | k__Fungi(100);p__Ascomycota(100);c__Saccharomycetes(100);o__Saccharomycetales(100);f__Metschnikowiaceae(100);g__Metschnikowia(100);s__Metschnikowia_pulcherrima(100); | Metschnikowia | 13 | Saprotroph | \|Undefined Saprotroph\| | Yeast | NULL | Probable | As nectar-tap_saprotroph (Põlme et al. 2020); Decay sugar-rich substrates (Põlme et al. 2020) | Põlme S, et al. 2020. Fungal Diversity 105:1-16 ((https://doi.org/10.1007/s13225-020-00466-2)) |
| Otu31842 | 1 | k__Fungi(100);p__Ascomycota(100);c__Saccharomycetes(100);o__Saccharomycetales(100);f__Metschnikowiaceae(100);g__Metschnikowia(100);s__Metschnikowia_pulcherrima(100); | Metschnikowia | 13 | Saprotroph | \|Undefined Saprotroph\| | Yeast | NULL | Probable | As nectar-tap_saprotroph (Põlme et al. 2020); Decay sugar-rich substrates (Põlme et al. 2020) | Põlme S, et al. 2020. Fungal Diversity 105:1-16 ((https://doi.org/10.1007/s13225-020-00466-2)) |
| Otu31846 | 1 | k__Fungi(100);p__Ascomycota(100);c__Saccharomycetes(100);o__Saccharomycetales(100);f__Metschnikowiaceae(100);g__Metschnikowia(100);g__Metschnikowia_unclassified(100); | Metschnikowia | 13 | Saprotroph | \|Undefined Saprotroph\| | Yeast | NULL | Probable | As nectar-tap_saprotroph (Põlme et al. 2020); Decay sugar-rich substrates (Põlme et al. 2020) | Põlme S, et al. 2020. Fungal Diversity 105:1-16 ((https://doi.org/10.1007/s13225-020-00466-2)) |
| Otu31847 | 1 | k__Fungi(100);p__Ascomycota(100);c__Saccharomycetes(100);o__Saccharomycetales(100);f__Metschnikowiaceae(100);g__Metschnikowia(100);s__Metschnikowia_pulcherrima(100); | Metschnikowia | 13 | Saprotroph | \|Undefined Saprotroph\| | Yeast | NULL | Probable | As nectar-tap_saprotroph (Põlme et al. 2020); Decay sugar-rich substrates (Põlme et al. 2020) | Põlme S, et al. 2020. Fungal Diversity 105:1-16 ((https://doi.org/10.1007/s13225-020-00466-2)) |
| Otu31849 | 1 | k__Fungi(100);p__Ascomycota(100);c__Saccharomycetes(100);o__Saccharomycetales(100);f__Metschnikowiaceae(100);g__Metschnikowia(100);s__Metschnikowia_pulcherrima(100); | Metschnikowia | 13 | Saprotroph | \|Undefined Saprotroph\| | Yeast | NULL | Probable | As nectar-tap_saprotroph (Põlme et al. 2020); Decay sugar-rich substrates (Põlme et al. 2020) | Põlme S, et al. 2020. Fungal Diversity 105:1-16 ((https://doi.org/10.1007/s13225-020-00466-2)) |
| Otu31850 | 1 | k__Fungi(100);p__Ascomycota(100);c__Saccharomycetes(100);o__Saccharomycetales(100);f__Metschnikowiaceae(100);g__Metschnikowia(100);s__Metschnikowia_pulcherrima(100); | Metschnikowia | 13 | Saprotroph | \|Undefined Saprotroph\| | Yeast | NULL | Probable | As nectar-tap_saprotroph (Põlme et al. 2020); Decay sugar-rich substrates (Põlme et al. 2020) | Põlme S, et al. 2020. Fungal Diversity 105:1-16 ((https://doi.org/10.1007/s13225-020-00466-2)) |
| Otu31853 | 1 | k__Fungi(100);p__Ascomycota(100);c__Saccharomycetes(100);o__Saccharomycetales(100);f__Metschnikowiaceae(100);g__Metschnikowia(100);g__Metschnikowia_unclassified(100); | Metschnikowia | 13 | Saprotroph | \|Undefined Saprotroph\| | Yeast | NULL | Probable | As nectar-tap_saprotroph (Põlme et al. 2020); Decay sugar-rich substrates (Põlme et al. 2020) | Põlme S, et al. 2020. Fungal Diversity 105:1-16 ((https://doi.org/10.1007/s13225-020-00466-2)) |
| Otu31858 | 1 | k__Fungi(100);p__Ascomycota(100);c__Saccharomycetes(100);o__Saccharomycetales(100);f__Metschnikowiaceae(100);g__Metschnikowia(100);s__Metschnikowia_pulcherrima(100); | Metschnikowia | 13 | Saprotroph | \|Undefined Saprotroph\| | Yeast | NULL | Probable | As nectar-tap_saprotroph (Põlme et al. 2020); Decay sugar-rich substrates (Põlme et al. 2020) | Põlme S, et al. 2020. Fungal Diversity 105:1-16 ((https://doi.org/10.1007/s13225-020-00466-2)) |
| Otu31864 | 1 | k__Fungi(100);p__Ascomycota(100);c__Saccharomycetes(100);o__Saccharomycetales(100);f__Metschnikowiaceae(100);g__Metschnikowia(100);s__Metschnikowia_pulcherrima(100); | Metschnikowia | 13 | Saprotroph | \|Undefined Saprotroph\| | Yeast | NULL | Probable | As nectar-tap_saprotroph (Põlme et al. 2020); Decay sugar-rich substrates (Põlme et al. 2020) | Põlme S, et al. 2020. Fungal Diversity 105:1-16 ((https://doi.org/10.1007/s13225-020-00466-2)) |
| Otu31867 | 1 | k__Fungi(100);p__Ascomycota(100);c__Saccharomycetes(100);o__Saccharomycetales(100);f__Metschnikowiaceae(100);g__Metschnikowia(100);s__Metschnikowia_pulcherrima(100); | Metschnikowia | 13 | Saprotroph | \|Undefined Saprotroph\| | Yeast | NULL | Probable | As nectar-tap_saprotroph (Põlme et al. 2020); Decay sugar-rich substrates (Põlme et al. 2020) | Põlme S, et al. 2020. Fungal Diversity 105:1-16 ((https://doi.org/10.1007/s13225-020-00466-2)) |
| Otu31882 | 1 | k__Fungi(100);p__Ascomycota(100);c__Saccharomycetes(100);o__Saccharomycetales(100);f__Metschnikowiaceae(100);g__Metschnikowia(100);s__Metschnikowia_pulcherrima(100); | Metschnikowia | 13 | Saprotroph | \|Undefined Saprotroph\| | Yeast | NULL | Probable | As nectar-tap_saprotroph (Põlme et al. 2020); Decay sugar-rich substrates (Põlme et al. 2020) | Põlme S, et al. 2020. Fungal Diversity 105:1-16 ((https://doi.org/10.1007/s13225-020-00466-2)) |
| Otu31885 | 1 | k__Fungi(100);p__Ascomycota(100);c__Saccharomycetes(100);o__Saccharomycetales(100);f__Metschnikowiaceae(100);g__Metschnikowia(100);s__Metschnikowia_pulcherrima(100); | Metschnikowia | 13 | Saprotroph | \|Undefined Saprotroph\| | Yeast | NULL | Probable | As nectar-tap_saprotroph (Põlme et al. 2020); Decay sugar-rich substrates (Põlme et al. 2020) | Põlme S, et al. 2020. Fungal Diversity 105:1-16 ((https://doi.org/10.1007/s13225-020-00466-2)) |
| Otu31887 | 1 | k__Fungi(100);p__Ascomycota(100);c__Saccharomycetes(100);o__Saccharomycetales(100);f__Metschnikowiaceae(100);g__Metschnikowia(100);g__Metschnikowia_unclassified(100); | Metschnikowia | 13 | Saprotroph | \|Undefined Saprotroph\| | Yeast | NULL | Probable | As nectar-tap_saprotroph (Põlme et al. 2020); Decay sugar-rich substrates (Põlme et al. 2020) | Põlme S, et al. 2020. Fungal Diversity 105:1-16 ((https://doi.org/10.1007/s13225-020-00466-2)) |
| Otu31890 | 1 | k__Fungi(100);p__Ascomycota(100);c__Saccharomycetes(100);o__Saccharomycetales(100);f__Metschnikowiaceae(100);g__Metschnikowia(100);s__Metschnikowia_pulcherrima(100); | Metschnikowia | 13 | Saprotroph | \|Undefined Saprotroph\| | Yeast | NULL | Probable | As nectar-tap_saprotroph (Põlme et al. 2020); Decay sugar-rich substrates (Põlme et al. 2020) | Põlme S, et al. 2020. Fungal Diversity 105:1-16 ((https://doi.org/10.1007/s13225-020-00466-2)) |
| Otu31902 | 1 | k__Fungi(100);p__Ascomycota(100);c__Saccharomycetes(100);o__Saccharomycetales(100);f__Metschnikowiaceae(100);g__Metschnikowia(100);g__Metschnikowia_unclassified(100); | Metschnikowia | 13 | Saprotroph | \|Undefined Saprotroph\| | Yeast | NULL | Probable | As nectar-tap_saprotroph (Põlme et al. 2020); Decay sugar-rich substrates (Põlme et al. 2020) | Põlme S, et al. 2020. Fungal Diversity 105:1-16 ((https://doi.org/10.1007/s13225-020-00466-2)) |
| Otu31903 | 1 | k__Fungi(100);p__Ascomycota(100);c__Saccharomycetes(100);o__Saccharomycetales(100);f__Metschnikowiaceae(100);g__Metschnikowia(100);s__Metschnikowia_pulcherrima(100); | Metschnikowia | 13 | Saprotroph | \|Undefined Saprotroph\| | Yeast | NULL | Probable | As nectar-tap_saprotroph (Põlme et al. 2020); Decay sugar-rich substrates (Põlme et al. 2020) | Põlme S, et al. 2020. Fungal Diversity 105:1-16 ((https://doi.org/10.1007/s13225-020-00466-2)) |
| Otu31904 | 1 | k__Fungi(100);p__Ascomycota(100);c__Saccharomycetes(100);o__Saccharomycetales(100);f__Metschnikowiaceae(100);g__Metschnikowia(100);s__Metschnikowia_pulcherrima(100); | Metschnikowia | 13 | Saprotroph | \|Undefined Saprotroph\| | Yeast | NULL | Probable | As nectar-tap_saprotroph (Põlme et al. 2020); Decay sugar-rich substrates (Põlme et al. 2020) | Põlme S, et al. 2020. Fungal Diversity 105:1-16 ((https://doi.org/10.1007/s13225-020-00466-2)) |
| Otu31907 | 1 | k__Fungi(100);p__Ascomycota(100);c__Saccharomycetes(100);o__Saccharomycetales(100);f__Metschnikowiaceae(100);g__Metschnikowia(100);s__Metschnikowia_pulcherrima(100); | Metschnikowia | 13 | Saprotroph | \|Undefined Saprotroph\| | Yeast | NULL | Probable | As nectar-tap_saprotroph (Põlme et al. 2020); Decay sugar-rich substrates (Põlme et al. 2020) | Põlme S, et al. 2020. Fungal Diversity 105:1-16 ((https://doi.org/10.1007/s13225-020-00466-2)) |
| Otu31909 | 1 | k__Fungi(100);p__Ascomycota(100);c__Saccharomycetes(100);o__Saccharomycetales(100);f__Metschnikowiaceae(100);g__Metschnikowia(100);s__Metschnikowia_pulcherrima(100); | Metschnikowia | 13 | Saprotroph | \|Undefined Saprotroph\| | Yeast | NULL | Probable | As nectar-tap_saprotroph (Põlme et al. 2020); Decay sugar-rich substrates (Põlme et al. 2020) | Põlme S, et al. 2020. Fungal Diversity 105:1-16 ((https://doi.org/10.1007/s13225-020-00466-2)) |
| Otu31910 | 1 | k__Fungi(100);p__Ascomycota(100);c__Saccharomycetes(100);o__Saccharomycetales(100);f__Metschnikowiaceae(100);g__Metschnikowia(100);s__Metschnikowia_pulcherrima(100); | Metschnikowia | 13 | Saprotroph | \|Undefined Saprotroph\| | Yeast | NULL | Probable | As nectar-tap_saprotroph (Põlme et al. 2020); Decay sugar-rich substrates (Põlme et al. 2020) | Põlme S, et al. 2020. Fungal Diversity 105:1-16 ((https://doi.org/10.1007/s13225-020-00466-2)) |
| Otu31932 | 1 | k__Fungi(100);p__Ascomycota(100);c__Saccharomycetes(100);o__Saccharomycetales(100);f__Metschnikowiaceae(100);g__Metschnikowia(100);s__Metschnikowia_pulcherrima(100); | Metschnikowia | 13 | Saprotroph | \|Undefined Saprotroph\| | Yeast | NULL | Probable | As nectar-tap_saprotroph (Põlme et al. 2020); Decay sugar-rich substrates (Põlme et al. 2020) | Põlme S, et al. 2020. Fungal Diversity 105:1-16 ((https://doi.org/10.1007/s13225-020-00466-2)) |
| Otu31933 | 1 | k__Fungi(100);p__Ascomycota(100);c__Saccharomycetes(100);o__Saccharomycetales(100);f__Metschnikowiaceae(100);g__Metschnikowia(100);s__Metschnikowia_pulcherrima(100); | Metschnikowia | 13 | Saprotroph | \|Undefined Saprotroph\| | Yeast | NULL | Probable | As nectar-tap_saprotroph (Põlme et al. 2020); Decay sugar-rich substrates (Põlme et al. 2020) | Põlme S, et al. 2020. Fungal Diversity 105:1-16 ((https://doi.org/10.1007/s13225-020-00466-2)) |
| Otu31934 | 1 | k__Fungi(100);p__Ascomycota(100);c__Saccharomycetes(100);o__Saccharomycetales(100);f__Metschnikowiaceae(100);g__Metschnikowia(100);s__Metschnikowia_pulcherrima(100); | Metschnikowia | 13 | Saprotroph | \|Undefined Saprotroph\| | Yeast | NULL | Probable | As nectar-tap_saprotroph (Põlme et al. 2020); Decay sugar-rich substrates (Põlme et al. 2020) | Põlme S, et al. 2020. Fungal Diversity 105:1-16 ((https://doi.org/10.1007/s13225-020-00466-2)) |
| Otu31935 | 1 | k__Fungi(100);p__Ascomycota(100);c__Saccharomycetes(100);o__Saccharomycetales(100);f__Metschnikowiaceae(100);g__Metschnikowia(100);g__Metschnikowia_unclassified(100); | Metschnikowia | 13 | Saprotroph | \|Undefined Saprotroph\| | Yeast | NULL | Probable | As nectar-tap_saprotroph (Põlme et al. 2020); Decay sugar-rich substrates (Põlme et al. 2020) | Põlme S, et al. 2020. Fungal Diversity 105:1-16 ((https://doi.org/10.1007/s13225-020-00466-2)) |
| Otu31940 | 1 | k__Fungi(100);p__Ascomycota(100);c__Saccharomycetes(100);o__Saccharomycetales(100);f__Metschnikowiaceae(100);g__Metschnikowia(100);s__Metschnikowia_pulcherrima(100); | Metschnikowia | 13 | Saprotroph | \|Undefined Saprotroph\| | Yeast | NULL | Probable | As nectar-tap_saprotroph (Põlme et al. 2020); Decay sugar-rich substrates (Põlme et al. 2020) | Põlme S, et al. 2020. Fungal Diversity 105:1-16 ((https://doi.org/10.1007/s13225-020-00466-2)) |
| Otu31941 | 1 | k__Fungi(100);p__Ascomycota(100);c__Saccharomycetes(100);o__Saccharomycetales(100);f__Metschnikowiaceae(100);g__Metschnikowia(100);g__Metschnikowia_unclassified(100); | Metschnikowia | 13 | Saprotroph | \|Undefined Saprotroph\| | Yeast | NULL | Probable | As nectar-tap_saprotroph (Põlme et al. 2020); Decay sugar-rich substrates (Põlme et al. 2020) | Põlme S, et al. 2020. Fungal Diversity 105:1-16 ((https://doi.org/10.1007/s13225-020-00466-2)) |
| Otu31942 | 1 | k__Fungi(100);p__Ascomycota(100);c__Saccharomycetes(100);o__Saccharomycetales(100);f__Metschnikowiaceae(100);g__Metschnikowia(100);s__Metschnikowia_pulcherrima(100); | Metschnikowia | 13 | Saprotroph | \|Undefined Saprotroph\| | Yeast | NULL | Probable | As nectar-tap_saprotroph (Põlme et al. 2020); Decay sugar-rich substrates (Põlme et al. 2020) | Põlme S, et al. 2020. Fungal Diversity 105:1-16 ((https://doi.org/10.1007/s13225-020-00466-2)) |
| Otu31943 | 1 | k__Fungi(100);p__Ascomycota(100);c__Saccharomycetes(100);o__Saccharomycetales(100);f__Metschnikowiaceae(100);g__Metschnikowia(100);s__Metschnikowia_pulcherrima(100); | Metschnikowia | 13 | Saprotroph | \|Undefined Saprotroph\| | Yeast | NULL | Probable | As nectar-tap_saprotroph (Põlme et al. 2020); Decay sugar-rich substrates (Põlme et al. 2020) | Põlme S, et al. 2020. Fungal Diversity 105:1-16 ((https://doi.org/10.1007/s13225-020-00466-2)) |
| Otu31944 | 1 | k__Fungi(100);p__Ascomycota(100);c__Saccharomycetes(100);o__Saccharomycetales(100);f__Metschnikowiaceae(100);g__Metschnikowia(100);s__Metschnikowia_pulcherrima(100); | Metschnikowia | 13 | Saprotroph | \|Undefined Saprotroph\| | Yeast | NULL | Probable | As nectar-tap_saprotroph (Põlme et al. 2020); Decay sugar-rich substrates (Põlme et al. 2020) | Põlme S, et al. 2020. Fungal Diversity 105:1-16 ((https://doi.org/10.1007/s13225-020-00466-2)) |
| Otu31945 | 1 | k__Fungi(100);p__Ascomycota(100);c__Saccharomycetes(100);o__Saccharomycetales(100);f__Metschnikowiaceae(100);g__Metschnikowia(100);s__Metschnikowia_pulcherrima(100); | Metschnikowia | 13 | Saprotroph | \|Undefined Saprotroph\| | Yeast | NULL | Probable | As nectar-tap_saprotroph (Põlme et al. 2020); Decay sugar-rich substrates (Põlme et al. 2020) | Põlme S, et al. 2020. Fungal Diversity 105:1-16 ((https://doi.org/10.1007/s13225-020-00466-2)) |
| Otu31946 | 1 | k__Fungi(100);p__Ascomycota(100);c__Saccharomycetes(100);o__Saccharomycetales(100);f__Metschnikowiaceae(100);g__Metschnikowia(100);s__Metschnikowia_pulcherrima(100); | Metschnikowia | 13 | Saprotroph | \|Undefined Saprotroph\| | Yeast | NULL | Probable | As nectar-tap_saprotroph (Põlme et al. 2020); Decay sugar-rich substrates (Põlme et al. 2020) | Põlme S, et al. 2020. Fungal Diversity 105:1-16 ((https://doi.org/10.1007/s13225-020-00466-2)) |
| Otu31947 | 1 | k__Fungi(100);p__Ascomycota(100);c__Saccharomycetes(100);o__Saccharomycetales(100);f__Metschnikowiaceae(100);g__Metschnikowia(100);s__Metschnikowia_pulcherrima(100); | Metschnikowia | 13 | Saprotroph | \|Undefined Saprotroph\| | Yeast | NULL | Probable | As nectar-tap_saprotroph (Põlme et al. 2020); Decay sugar-rich substrates (Põlme et al. 2020) | Põlme S, et al. 2020. Fungal Diversity 105:1-16 ((https://doi.org/10.1007/s13225-020-00466-2)) |
| Otu31948 | 1 | k__Fungi(100);p__Ascomycota(100);c__Saccharomycetes(100);o__Saccharomycetales(100);f__Metschnikowiaceae(100);g__Metschnikowia(100);s__Metschnikowia_pulcherrima(100); | Metschnikowia | 13 | Saprotroph | \|Undefined Saprotroph\| | Yeast | NULL | Probable | As nectar-tap_saprotroph (Põlme et al. 2020); Decay sugar-rich substrates (Põlme et al. 2020) | Põlme S, et al. 2020. Fungal Diversity 105:1-16 ((https://doi.org/10.1007/s13225-020-00466-2)) |
| Otu31949 | 1 | k__Fungi(100);p__Ascomycota(100);c__Saccharomycetes(100);o__Saccharomycetales(100);f__Metschnikowiaceae(100);g__Metschnikowia(100);g__Metschnikowia_unclassified(100); | Metschnikowia | 13 | Saprotroph | \|Undefined Saprotroph\| | Yeast | NULL | Probable | As nectar-tap_saprotroph (Põlme et al. 2020); Decay sugar-rich substrates (Põlme et al. 2020) | Põlme S, et al. 2020. Fungal Diversity 105:1-16 ((https://doi.org/10.1007/s13225-020-00466-2)) |
| Otu31950 | 1 | k__Fungi(100);p__Ascomycota(100);c__Saccharomycetes(100);o__Saccharomycetales(100);f__Metschnikowiaceae(100);g__Metschnikowia(100);s__Metschnikowia_pulcherrima(100); | Metschnikowia | 13 | Saprotroph | \|Undefined Saprotroph\| | Yeast | NULL | Probable | As nectar-tap_saprotroph (Põlme et al. 2020); Decay sugar-rich substrates (Põlme et al. 2020) | Põlme S, et al. 2020. Fungal Diversity 105:1-16 ((https://doi.org/10.1007/s13225-020-00466-2)) |
| Otu31951 | 1 | k__Fungi(100);p__Ascomycota(100);c__Saccharomycetes(100);o__Saccharomycetales(100);f__Metschnikowiaceae(100);g__Metschnikowia(100);s__Metschnikowia_pulcherrima(100); | Metschnikowia | 13 | Saprotroph | \|Undefined Saprotroph\| | Yeast | NULL | Probable | As nectar-tap_saprotroph (Põlme et al. 2020); Decay sugar-rich substrates (Põlme et al. 2020) | Põlme S, et al. 2020. Fungal Diversity 105:1-16 ((https://doi.org/10.1007/s13225-020-00466-2)) |
| Otu31952 | 1 | k__Fungi(100);p__Ascomycota(100);c__Saccharomycetes(100);o__Saccharomycetales(100);f__Metschnikowiaceae(100);g__Metschnikowia(100);s__Metschnikowia_pulcherrima(100); | Metschnikowia | 13 | Saprotroph | \|Undefined Saprotroph\| | Yeast | NULL | Probable | As nectar-tap_saprotroph (Põlme et al. 2020); Decay sugar-rich substrates (Põlme et al. 2020) | Põlme S, et al. 2020. Fungal Diversity 105:1-16 ((https://doi.org/10.1007/s13225-020-00466-2)) |
| Otu31953 | 1 | k__Fungi(100);p__Ascomycota(100);c__Saccharomycetes(100);o__Saccharomycetales(100);f__Metschnikowiaceae(100);g__Metschnikowia(100);s__Metschnikowia_pulcherrima(100); | Metschnikowia | 13 | Saprotroph | \|Undefined Saprotroph\| | Yeast | NULL | Probable | As nectar-tap_saprotroph (Põlme et al. 2020); Decay sugar-rich substrates (Põlme et al. 2020) | Põlme S, et al. 2020. Fungal Diversity 105:1-16 ((https://doi.org/10.1007/s13225-020-00466-2)) |
| Otu31955 | 1 | k__Fungi(100);p__Ascomycota(100);c__Saccharomycetes(100);o__Saccharomycetales(100);f__Metschnikowiaceae(100);g__Metschnikowia(100);g__Metschnikowia_unclassified(100); | Metschnikowia | 13 | Saprotroph | \|Undefined Saprotroph\| | Yeast | NULL | Probable | As nectar-tap_saprotroph (Põlme et al. 2020); Decay sugar-rich substrates (Põlme et al. 2020) | Põlme S, et al. 2020. Fungal Diversity 105:1-16 ((https://doi.org/10.1007/s13225-020-00466-2)) |
| Otu31957 | 1 | k__Fungi(100);p__Ascomycota(100);c__Saccharomycetes(100);o__Saccharomycetales(100);f__Metschnikowiaceae(100);g__Metschnikowia(100);g__Metschnikowia_unclassified(100); | Metschnikowia | 13 | Saprotroph | \|Undefined Saprotroph\| | Yeast | NULL | Probable | As nectar-tap_saprotroph (Põlme et al. 2020); Decay sugar-rich substrates (Põlme et al. 2020) | Põlme S, et al. 2020. Fungal Diversity 105:1-16 ((https://doi.org/10.1007/s13225-020-00466-2)) |
| Otu31958 | 1 | k__Fungi(100);p__Ascomycota(100);c__Saccharomycetes(100);o__Saccharomycetales(100);f__Metschnikowiaceae(100);g__Metschnikowia(100);s__Metschnikowia_pulcherrima(100); | Metschnikowia | 13 | Saprotroph | \|Undefined Saprotroph\| | Yeast | NULL | Probable | As nectar-tap_saprotroph (Põlme et al. 2020); Decay sugar-rich substrates (Põlme et al. 2020) | Põlme S, et al. 2020. Fungal Diversity 105:1-16 ((https://doi.org/10.1007/s13225-020-00466-2)) |
| Otu31959 | 1 | k__Fungi(100);p__Ascomycota(100);c__Saccharomycetes(100);o__Saccharomycetales(100);f__Metschnikowiaceae(100);g__Metschnikowia(100);s__Metschnikowia_pulcherrima(100); | Metschnikowia | 13 | Saprotroph | \|Undefined Saprotroph\| | Yeast | NULL | Probable | As nectar-tap_saprotroph (Põlme et al. 2020); Decay sugar-rich substrates (Põlme et al. 2020) | Põlme S, et al. 2020. Fungal Diversity 105:1-16 ((https://doi.org/10.1007/s13225-020-00466-2)) |
| Otu31961 | 1 | k__Fungi(100);p__Ascomycota(100);c__Saccharomycetes(100);o__Saccharomycetales(100);f__Metschnikowiaceae(100);g__Metschnikowia(100);s__Metschnikowia_pulcherrima(100); | Metschnikowia | 13 | Saprotroph | \|Undefined Saprotroph\| | Yeast | NULL | Probable | As nectar-tap_saprotroph (Põlme et al. 2020); Decay sugar-rich substrates (Põlme et al. 2020) | Põlme S, et al. 2020. Fungal Diversity 105:1-16 ((https://doi.org/10.1007/s13225-020-00466-2)) |
| Otu31963 | 1 | k__Fungi(100);p__Ascomycota(100);c__Saccharomycetes(100);o__Saccharomycetales(100);f__Metschnikowiaceae(100);g__Metschnikowia(100);g__Metschnikowia_unclassified(100); | Metschnikowia | 13 | Saprotroph | \|Undefined Saprotroph\| | Yeast | NULL | Probable | As nectar-tap_saprotroph (Põlme et al. 2020); Decay sugar-rich substrates (Põlme et al. 2020) | Põlme S, et al. 2020. Fungal Diversity 105:1-16 ((https://doi.org/10.1007/s13225-020-00466-2)) |
| Otu31964 | 1 | k__Fungi(100);p__Ascomycota(100);c__Saccharomycetes(100);o__Saccharomycetales(100);f__Metschnikowiaceae(100);g__Metschnikowia(100);s__Metschnikowia_pulcherrima(100); | Metschnikowia | 13 | Saprotroph | \|Undefined Saprotroph\| | Yeast | NULL | Probable | As nectar-tap_saprotroph (Põlme et al. 2020); Decay sugar-rich substrates (Põlme et al. 2020) | Põlme S, et al. 2020. Fungal Diversity 105:1-16 ((https://doi.org/10.1007/s13225-020-00466-2)) |
| Otu31965 | 1 | k__Fungi(100);p__Ascomycota(100);c__Saccharomycetes(100);o__Saccharomycetales(100);f__Metschnikowiaceae(100);g__Metschnikowia(100);s__Metschnikowia_pulcherrima(100); | Metschnikowia | 13 | Saprotroph | \|Undefined Saprotroph\| | Yeast | NULL | Probable | As nectar-tap_saprotroph (Põlme et al. 2020); Decay sugar-rich substrates (Põlme et al. 2020) | Põlme S, et al. 2020. Fungal Diversity 105:1-16 ((https://doi.org/10.1007/s13225-020-00466-2)) |
| Otu31966 | 1 | k__Fungi(100);p__Ascomycota(100);c__Saccharomycetes(100);o__Saccharomycetales(100);f__Metschnikowiaceae(100);g__Metschnikowia(100);s__Metschnikowia_pulcherrima(100); | Metschnikowia | 13 | Saprotroph | \|Undefined Saprotroph\| | Yeast | NULL | Probable | As nectar-tap_saprotroph (Põlme et al. 2020); Decay sugar-rich substrates (Põlme et al. 2020) | Põlme S, et al. 2020. Fungal Diversity 105:1-16 ((https://doi.org/10.1007/s13225-020-00466-2)) |
| Otu31967 | 1 | k__Fungi(100);p__Ascomycota(100);c__Saccharomycetes(100);o__Saccharomycetales(100);f__Metschnikowiaceae(100);g__Metschnikowia(100);g__Metschnikowia_unclassified(100); | Metschnikowia | 13 | Saprotroph | \|Undefined Saprotroph\| | Yeast | NULL | Probable | As nectar-tap_saprotroph (Põlme et al. 2020); Decay sugar-rich substrates (Põlme et al. 2020) | Põlme S, et al. 2020. Fungal Diversity 105:1-16 ((https://doi.org/10.1007/s13225-020-00466-2)) |
| Otu31968 | 1 | k__Fungi(100);p__Ascomycota(100);c__Saccharomycetes(100);o__Saccharomycetales(100);f__Metschnikowiaceae(100);g__Metschnikowia(100);s__Metschnikowia_pulcherrima(100); | Metschnikowia | 13 | Saprotroph | \|Undefined Saprotroph\| | Yeast | NULL | Probable | As nectar-tap_saprotroph (Põlme et al. 2020); Decay sugar-rich substrates (Põlme et al. 2020) | Põlme S, et al. 2020. Fungal Diversity 105:1-16 ((https://doi.org/10.1007/s13225-020-00466-2)) |
| Otu31969 | 1 | k__Fungi(100);p__Ascomycota(100);c__Saccharomycetes(100);o__Saccharomycetales(100);f__Metschnikowiaceae(100);g__Metschnikowia(100);s__Metschnikowia_pulcherrima(100); | Metschnikowia | 13 | Saprotroph | \|Undefined Saprotroph\| | Yeast | NULL | Probable | As nectar-tap_saprotroph (Põlme et al. 2020); Decay sugar-rich substrates (Põlme et al. 2020) | Põlme S, et al. 2020. Fungal Diversity 105:1-16 ((https://doi.org/10.1007/s13225-020-00466-2)) |
| Otu31970 | 1 | k__Fungi(100);p__Ascomycota(100);c__Saccharomycetes(100);o__Saccharomycetales(100);f__Metschnikowiaceae(100);g__Metschnikowia(100);g__Metschnikowia_unclassified(100); | Metschnikowia | 13 | Saprotroph | \|Undefined Saprotroph\| | Yeast | NULL | Probable | As nectar-tap_saprotroph (Põlme et al. 2020); Decay sugar-rich substrates (Põlme et al. 2020) | Põlme S, et al. 2020. Fungal Diversity 105:1-16 ((https://doi.org/10.1007/s13225-020-00466-2)) |
| Otu31971 | 1 | k__Fungi(100);p__Ascomycota(100);c__Saccharomycetes(100);o__Saccharomycetales(100);f__Metschnikowiaceae(100);g__Metschnikowia(100);s__Metschnikowia_pulcherrima(100); | Metschnikowia | 13 | Saprotroph | \|Undefined Saprotroph\| | Yeast | NULL | Probable | As nectar-tap_saprotroph (Põlme et al. 2020); Decay sugar-rich substrates (Põlme et al. 2020) | Põlme S, et al. 2020. Fungal Diversity 105:1-16 ((https://doi.org/10.1007/s13225-020-00466-2)) |
| Otu31972 | 1 | k__Fungi(100);p__Ascomycota(100);c__Saccharomycetes(100);o__Saccharomycetales(100);f__Metschnikowiaceae(100);g__Metschnikowia(100);s__Metschnikowia_pulcherrima(100); | Metschnikowia | 13 | Saprotroph | \|Undefined Saprotroph\| | Yeast | NULL | Probable | As nectar-tap_saprotroph (Põlme et al. 2020); Decay sugar-rich substrates (Põlme et al. 2020) | Põlme S, et al. 2020. Fungal Diversity 105:1-16 ((https://doi.org/10.1007/s13225-020-00466-2)) |
| Otu31973 | 1 | k__Fungi(100);p__Ascomycota(100);c__Saccharomycetes(100);o__Saccharomycetales(100);f__Metschnikowiaceae(100);g__Metschnikowia(100);s__Metschnikowia_pulcherrima(100); | Metschnikowia | 13 | Saprotroph | \|Undefined Saprotroph\| | Yeast | NULL | Probable | As nectar-tap_saprotroph (Põlme et al. 2020); Decay sugar-rich substrates (Põlme et al. 2020) | Põlme S, et al. 2020. Fungal Diversity 105:1-16 ((https://doi.org/10.1007/s13225-020-00466-2)) |
| Otu31974 | 1 | k__Fungi(100);p__Ascomycota(100);c__Saccharomycetes(100);o__Saccharomycetales(100);f__Metschnikowiaceae(100);g__Metschnikowia(100);s__Metschnikowia_pulcherrima(100); | Metschnikowia | 13 | Saprotroph | \|Undefined Saprotroph\| | Yeast | NULL | Probable | As nectar-tap_saprotroph (Põlme et al. 2020); Decay sugar-rich substrates (Põlme et al. 2020) | Põlme S, et al. 2020. Fungal Diversity 105:1-16 ((https://doi.org/10.1007/s13225-020-00466-2)) |
| Otu31975 | 1 | k__Fungi(100);p__Ascomycota(100);c__Saccharomycetes(100);o__Saccharomycetales(100);f__Metschnikowiaceae(100);g__Metschnikowia(100);s__Metschnikowia_pulcherrima(100); | Metschnikowia | 13 | Saprotroph | \|Undefined Saprotroph\| | Yeast | NULL | Probable | As nectar-tap_saprotroph (Põlme et al. 2020); Decay sugar-rich substrates (Põlme et al. 2020) | Põlme S, et al. 2020. Fungal Diversity 105:1-16 ((https://doi.org/10.1007/s13225-020-00466-2)) |
| Otu31976 | 1 | k__Fungi(100);p__Ascomycota(100);c__Saccharomycetes(100);o__Saccharomycetales(100);f__Metschnikowiaceae(100);g__Metschnikowia(100);g__Metschnikowia_unclassified(100); | Metschnikowia | 13 | Saprotroph | \|Undefined Saprotroph\| | Yeast | NULL | Probable | As nectar-tap_saprotroph (Põlme et al. 2020); Decay sugar-rich substrates (Põlme et al. 2020) | Põlme S, et al. 2020. Fungal Diversity 105:1-16 ((https://doi.org/10.1007/s13225-020-00466-2)) |
| Otu31977 | 1 | k__Fungi(100);p__Ascomycota(100);c__Saccharomycetes(100);o__Saccharomycetales(100);f__Metschnikowiaceae(100);g__Metschnikowia(100);g__Metschnikowia_unclassified(100); | Metschnikowia | 13 | Saprotroph | \|Undefined Saprotroph\| | Yeast | NULL | Probable | As nectar-tap_saprotroph (Põlme et al. 2020); Decay sugar-rich substrates (Põlme et al. 2020) | Põlme S, et al. 2020. Fungal Diversity 105:1-16 ((https://doi.org/10.1007/s13225-020-00466-2)) |
| Otu31978 | 1 | k__Fungi(100);p__Ascomycota(100);c__Saccharomycetes(100);o__Saccharomycetales(100);f__Metschnikowiaceae(100);g__Metschnikowia(100);s__Metschnikowia_pulcherrima(100); | Metschnikowia | 13 | Saprotroph | \|Undefined Saprotroph\| | Yeast | NULL | Probable | As nectar-tap_saprotroph (Põlme et al. 2020); Decay sugar-rich substrates (Põlme et al. 2020) | Põlme S, et al. 2020. Fungal Diversity 105:1-16 ((https://doi.org/10.1007/s13225-020-00466-2)) |
| Otu31979 | 1 | k__Fungi(100);p__Ascomycota(100);c__Saccharomycetes(100);o__Saccharomycetales(100);f__Metschnikowiaceae(100);g__Metschnikowia(100);s__Metschnikowia_pulcherrima(100); | Metschnikowia | 13 | Saprotroph | \|Undefined Saprotroph\| | Yeast | NULL | Probable | As nectar-tap_saprotroph (Põlme et al. 2020); Decay sugar-rich substrates (Põlme et al. 2020) | Põlme S, et al. 2020. Fungal Diversity 105:1-16 ((https://doi.org/10.1007/s13225-020-00466-2)) |
| Otu31980 | 1 | k__Fungi(100);p__Ascomycota(100);c__Saccharomycetes(100);o__Saccharomycetales(100);f__Metschnikowiaceae(100);g__Metschnikowia(100);s__Metschnikowia_pulcherrima(100); | Metschnikowia | 13 | Saprotroph | \|Undefined Saprotroph\| | Yeast | NULL | Probable | As nectar-tap_saprotroph (Põlme et al. 2020); Decay sugar-rich substrates (Põlme et al. 2020) | Põlme S, et al. 2020. Fungal Diversity 105:1-16 ((https://doi.org/10.1007/s13225-020-00466-2)) |
| Otu31981 | 1 | k__Fungi(100);p__Ascomycota(100);c__Saccharomycetes(100);o__Saccharomycetales(100);f__Metschnikowiaceae(100);g__Metschnikowia(100);g__Metschnikowia_unclassified(100); | Metschnikowia | 13 | Saprotroph | \|Undefined Saprotroph\| | Yeast | NULL | Probable | As nectar-tap_saprotroph (Põlme et al. 2020); Decay sugar-rich substrates (Põlme et al. 2020) | Põlme S, et al. 2020. Fungal Diversity 105:1-16 ((https://doi.org/10.1007/s13225-020-00466-2)) |
| Otu31982 | 1 | k__Fungi(100);p__Ascomycota(100);c__Saccharomycetes(100);o__Saccharomycetales(100);f__Metschnikowiaceae(100);g__Metschnikowia(100);g__Metschnikowia_unclassified(100); | Metschnikowia | 13 | Saprotroph | \|Undefined Saprotroph\| | Yeast | NULL | Probable | As nectar-tap_saprotroph (Põlme et al. 2020); Decay sugar-rich substrates (Põlme et al. 2020) | Põlme S, et al. 2020. Fungal Diversity 105:1-16 ((https://doi.org/10.1007/s13225-020-00466-2)) |
| Otu31983 | 1 | k__Fungi(100);p__Ascomycota(100);c__Saccharomycetes(100);o__Saccharomycetales(100);f__Metschnikowiaceae(100);g__Metschnikowia(100);s__Metschnikowia_pulcherrima(100); | Metschnikowia | 13 | Saprotroph | \|Undefined Saprotroph\| | Yeast | NULL | Probable | As nectar-tap_saprotroph (Põlme et al. 2020); Decay sugar-rich substrates (Põlme et al. 2020) | Põlme S, et al. 2020. Fungal Diversity 105:1-16 ((https://doi.org/10.1007/s13225-020-00466-2)) |
| Otu31984 | 1 | k__Fungi(100);p__Ascomycota(100);c__Saccharomycetes(100);o__Saccharomycetales(100);f__Metschnikowiaceae(100);g__Metschnikowia(100);s__Metschnikowia_pulcherrima(100); | Metschnikowia | 13 | Saprotroph | \|Undefined Saprotroph\| | Yeast | NULL | Probable | As nectar-tap_saprotroph (Põlme et al. 2020); Decay sugar-rich substrates (Põlme et al. 2020) | Põlme S, et al. 2020. Fungal Diversity 105:1-16 ((https://doi.org/10.1007/s13225-020-00466-2)) |
| Otu31985 | 1 | k__Fungi(100);p__Ascomycota(100);c__Saccharomycetes(100);o__Saccharomycetales(100);f__Metschnikowiaceae(100);g__Metschnikowia(100);g__Metschnikowia_unclassified(100); | Metschnikowia | 13 | Saprotroph | \|Undefined Saprotroph\| | Yeast | NULL | Probable | As nectar-tap_saprotroph (Põlme et al. 2020); Decay sugar-rich substrates (Põlme et al. 2020) | Põlme S, et al. 2020. Fungal Diversity 105:1-16 ((https://doi.org/10.1007/s13225-020-00466-2)) |

| | | | | | | | | | | | | |
|---|---|---|---|---|---|---|---|---|---|---|---|---|
| Otu31986 | 1 | k__Fungi(100);p__Ascomycota(100);c__Saccharomycetes(100);o__Saccharomycetales(100);f__Metschnikowiaceae(100);g__Metschnikowia(100);s__Metschnikowia_pulcherrima(100); | Metschnikowia | 13 | Saprotroph | \|Undefined Saprotroph\| | Yeast | NULL | Probable | As nectar-tap_saprotroph (Põlme et al. 2020); Decay sugar-rich substrates (Põlme et al. 2020) | | Põlme S, et al. 2020. Fungal Diversity 105:1-16 ((https://doi.org/10.1007/s13225-020-00466-2)) |
| Otu31987 | 1 | k__Fungi(100);p__Ascomycota(100);c__Saccharomycetes(100);o__Saccharomycetales(100);f__Metschnikowiaceae(100);g__Metschnikowia(100);s__Metschnikowia_pulcherrima(100); | Metschnikowia | 13 | Saprotroph | \|Undefined Saprotroph\| | Yeast | NULL | Probable | As nectar-tap_saprotroph (Põlme et al. 2020); Decay sugar-rich substrates (Põlme et al. 2020) | | Põlme S, et al. 2020. Fungal Diversity 105:1-16 ((https://doi.org/10.1007/s13225-020-00466-2)) |
| Otu31988 | 1 | k__Fungi(100);p__Ascomycota(100);c__Saccharomycetes(100);o__Saccharomycetales(100);f__Metschnikowiaceae(100);g__Metschnikowia(100);g__Metschnikowia_unclassified(100); | Metschnikowia | 13 | Saprotroph | \|Undefined Saprotroph\| | Yeast | NULL | Probable | As nectar-tap_saprotroph (Põlme et al. 2020); Decay sugar-rich substrates (Põlme et al. 2020) | | Põlme S, et al. 2020. Fungal Diversity 105:1-16 ((https://doi.org/10.1007/s13225-020-00466-2)) |
| Otu31989 | 1 | k__Fungi(100);p__Ascomycota(100);c__Saccharomycetes(100);o__Saccharomycetales(100);f__Metschnikowiaceae(100);g__Metschnikowia(100);s__Metschnikowia_pulcherrima(100); | Metschnikowia | 13 | Saprotroph | \|Undefined Saprotroph\| | Yeast | NULL | Probable | As nectar-tap_saprotroph (Põlme et al. 2020); Decay sugar-rich substrates (Põlme et al. 2020) | | Põlme S, et al. 2020. Fungal Diversity 105:1-16 ((https://doi.org/10.1007/s13225-020-00466-2)) |
| Otu31990 | 1 | k__Fungi(100);p__Ascomycota(100);c__Saccharomycetes(100);o__Saccharomycetales(100);f__Metschnikowiaceae(100);g__Metschnikowia(100);s__Metschnikowia_pulcherrima(100); | Metschnikowia | 13 | Saprotroph | \|Undefined Saprotroph\| | Yeast | NULL | Probable | As nectar-tap_saprotroph (Põlme et al. 2020); Decay sugar-rich substrates (Põlme et al. 2020) | | Põlme S, et al. 2020. Fungal Diversity 105:1-16 ((https://doi.org/10.1007/s13225-020-00466-2)) |
| Otu31991 | 1 | k__Fungi(100);p__Ascomycota(100);c__Saccharomycetes(100);o__Saccharomycetales(100);f__Metschnikowiaceae(100);g__Metschnikowia(100);s__Metschnikowia_pulcherrima(100); | Metschnikowia | 13 | Saprotroph | \|Undefined Saprotroph\| | Yeast | NULL | Probable | As nectar-tap_saprotroph (Põlme et al. 2020); Decay sugar-rich substrates (Põlme et al. 2020) | | Põlme S, et al. 2020. Fungal Diversity 105:1-16 ((https://doi.org/10.1007/s13225-020-00466-2)) |
| Otu31992 | 1 | k__Fungi(100);p__Ascomycota(100);c__Saccharomycetes(100);o__Saccharomycetales(100);f__Metschnikowiaceae(100);g__Metschnikowia(100);s__Metschnikowia_pulcherrima(100); | Metschnikowia | 13 | Saprotroph | \|Undefined Saprotroph\| | Yeast | NULL | Probable | As nectar-tap_saprotroph (Põlme et al. 2020); Decay sugar-rich substrates (Põlme et al. 2020) | | Põlme S, et al. 2020. Fungal Diversity 105:1-16 ((https://doi.org/10.1007/s13225-020-00466-2)) |
| Otu31995 | 1 | k__Fungi(100);p__Ascomycota(100);c__Saccharomycetes(100);o__Saccharomycetales(100);f__Metschnikowiaceae(100);g__Metschnikowia(100);s__Metschnikowia_pulcherrima(100); | Metschnikowia | 13 | Saprotroph | \|Undefined Saprotroph\| | Yeast | NULL | Probable | As nectar-tap_saprotroph (Põlme et al. 2020); Decay sugar-rich substrates (Põlme et al. 2020) | | Põlme S, et al. 2020. Fungal Diversity 105:1-16 ((https://doi.org/10.1007/s13225-020-00466-2)) |
| Otu31996 | 1 | k__Fungi(100);p__Ascomycota(100);c__Saccharomycetes(100);o__Saccharomycetales(100);f__Metschnikowiaceae(100);g__Metschnikowia(100);s__Metschnikowia_pulcherrima(100); | Metschnikowia | 13 | Saprotroph | \|Undefined Saprotroph\| | Yeast | NULL | Probable | As nectar-tap_saprotroph (Põlme et al. 2020); Decay sugar-rich substrates (Põlme et al. 2020) | | Põlme S, et al. 2020. Fungal Diversity 105:1-16 ((https://doi.org/10.1007/s13225-020-00466-2)) |
| Otu31997 | 1 | k__Fungi(100);p__Ascomycota(100);c__Saccharomycetes(100);o__Saccharomycetales(100);f__Metschnikowiaceae(100);g__Metschnikowia(100);s__Metschnikowia_pulcherrima(100); | Metschnikowia | 13 | Saprotroph | \|Undefined Saprotroph\| | Yeast | NULL | Probable | As nectar-tap_saprotroph (Põlme et al. 2020); Decay sugar-rich substrates (Põlme et al. 2020) | | Põlme S, et al. 2020. Fungal Diversity 105:1-16 ((https://doi.org/10.1007/s13225-020-00466-2)) |
| Otu31998 | 1 | k__Fungi(100);p__Ascomycota(100);c__Saccharomycetes(100);o__Saccharomycetales(100);f__Metschnikowiaceae(100);g__Metschnikowia(100);s__Metschnikowia_pulcherrima(100); | Metschnikowia | 13 | Saprotroph | \|Undefined Saprotroph\| | Yeast | NULL | Probable | As nectar-tap_saprotroph (Põlme et al. 2020); Decay sugar-rich substrates (Põlme et al. 2020) | | Põlme S, et al. 2020. Fungal Diversity 105:1-16 ((https://doi.org/10.1007/s13225-020-00466-2)) |
| Otu31999 | 1 | k__Fungi(100);p__Ascomycota(100);c__Saccharomycetes(100);o__Saccharomycetales(100);f__Metschnikowiaceae(100);g__Metschnikowia(100);g__Metschnikowia_unclassified(100); | Metschnikowia | 13 | Saprotroph | \|Undefined Saprotroph\| | Yeast | NULL | Probable | As nectar-tap_saprotroph (Põlme et al. 2020); Decay sugar-rich substrates (Põlme et al. 2020) | | Põlme S, et al. 2020. Fungal Diversity 105:1-16 ((https://doi.org/10.1007/s13225-020-00466-2)) |
| Otu32000 | 1 | k__Fungi(100);p__Ascomycota(100);c__Saccharomycetes(100);o__Saccharomycetales(100);f__Metschnikowiaceae(100);g__Metschnikowia(100);g__Metschnikowia_unclassified(100); | Metschnikowia | 13 | Saprotroph | \|Undefined Saprotroph\| | Yeast | NULL | Probable | As nectar-tap_saprotroph (Põlme et al. 2020); Decay sugar-rich substrates (Põlme et al. 2020) | | Põlme S, et al. 2020. Fungal Diversity 105:1-16 ((https://doi.org/10.1007/s13225-020-00466-2)) |
| Otu32001 | 1 | k__Fungi(100);p__Ascomycota(100);c__Saccharomycetes(100);o__Saccharomycetales(100);f__Metschnikowiaceae(100);g__Metschnikowia(100);s__Metschnikowia_pulcherrima(100); | Metschnikowia | 13 | Saprotroph | \|Undefined Saprotroph\| | Yeast | NULL | Probable | As nectar-tap_saprotroph (Põlme et al. 2020); Decay sugar-rich substrates (Põlme et al. 2020) | | Põlme S, et al. 2020. Fungal Diversity 105:1-16 ((https://doi.org/10.1007/s13225-020-00466-2)) |
| Otu32002 | 1 | k__Fungi(100);p__Ascomycota(100);c__Saccharomycetes(100);o__Saccharomycetales(100);f__Metschnikowiaceae(100);g__Metschnikowia(100);g__Metschnikowia_unclassified(100); | Metschnikowia | 13 | Saprotroph | \|Undefined Saprotroph\| | Yeast | NULL | Probable | As nectar-tap_saprotroph (Põlme et al. 2020); Decay sugar-rich substrates (Põlme et al. 2020) | | Põlme S, et al. 2020. Fungal Diversity 105:1-16 ((https://doi.org/10.1007/s13225-020-00466-2)) |
| Otu32004 | 1 | k__Fungi(100);p__Ascomycota(100);c__Saccharomycetes(100);o__Saccharomycetales(100);f__Metschnikowiaceae(100);g__Metschnikowia(100);s__Metschnikowia_pulcherrima(100); | Metschnikowia | 13 | Saprotroph | \|Undefined Saprotroph\| | Yeast | NULL | Probable | As nectar-tap_saprotroph (Põlme et al. 2020); Decay sugar-rich substrates (Põlme et al. 2020) | | Põlme S, et al. 2020. Fungal Diversity 105:1-16 ((https://doi.org/10.1007/s13225-020-00466-2)) |
| Otu32005 | 1 | k__Fungi(100);p__Ascomycota(100);c__Saccharomycetes(100);o__Saccharomycetales(100);f__Metschnikowiaceae(100);g__Metschnikowia(100);s__Metschnikowia_pulcherrima(100); | Metschnikowia | 13 | Saprotroph | \|Undefined Saprotroph\| | Yeast | NULL | Probable | As nectar-tap_saprotroph (Põlme et al. 2020); Decay sugar-rich substrates (Põlme et al. 2020) | | Põlme S, et al. 2020. Fungal Diversity 105:1-16 ((https://doi.org/10.1007/s13225-020-00466-2)) |
| Otu32006 | 1 | k__Fungi(100);p__Ascomycota(100);c__Saccharomycetes(100);o__Saccharomycetales(100);f__Metschnikowiaceae(100);g__Metschnikowia(100);s__Metschnikowia_pulcherrima(100); | Metschnikowia | 13 | Saprotroph | \|Undefined Saprotroph\| | Yeast | NULL | Probable | As nectar-tap_saprotroph (Põlme et al. 2020); Decay sugar-rich substrates (Põlme et al. 2020) | | Põlme S, et al. 2020. Fungal Diversity 105:1-16 ((https://doi.org/10.1007/s13225-020-00466-2)) |
| Otu32011 | 1 | k__Fungi(100);p__Ascomycota(100);c__Saccharomycetes(100);o__Saccharomycetales(100);f__Metschnikowiaceae(100);g__Metschnikowia(100);s__Metschnikowia_pulcherrima(100); | Metschnikowia | 13 | Saprotroph | \|Undefined Saprotroph\| | Yeast | NULL | Probable | As nectar-tap_saprotroph (Põlme et al. 2020); Decay sugar-rich substrates (Põlme et al. 2020) | | Põlme S, et al. 2020. Fungal Diversity 105:1-16 ((https://doi.org/10.1007/s13225-020-00466-2)) |
| Otu32012 | 1 | k__Fungi(100);p__Ascomycota(100);c__Saccharomycetes(100);o__Saccharomycetales(100);f__Metschnikowiaceae(100);g__Metschnikowia(100);s__Metschnikowia_pulcherrima(100); | Metschnikowia | 13 | Saprotroph | \|Undefined Saprotroph\| | Yeast | NULL | Probable | As nectar-tap_saprotroph (Põlme et al. 2020); Decay sugar-rich substrates (Põlme et al. 2020) | | Põlme S, et al. 2020. Fungal Diversity 105:1-16 ((https://doi.org/10.1007/s13225-020-00466-2)) |
| Otu32014 | 1 | k__Fungi(100);p__Ascomycota(100);c__Saccharomycetes(100);o__Saccharomycetales(100);f__Metschnikowiaceae(100);g__Metschnikowia(100);s__Metschnikowia_pulcherrima(100); | Metschnikowia | 13 | Saprotroph | \|Undefined Saprotroph\| | Yeast | NULL | Probable | As nectar-tap_saprotroph (Põlme et al. 2020); Decay sugar-rich substrates (Põlme et al. 2020) | | Põlme S, et al. 2020. Fungal Diversity 105:1-16 ((https://doi.org/10.1007/s13225-020-00466-2)) |
| Otu32015 | 1 | k__Fungi(100);p__Ascomycota(100);c__Saccharomycetes(100);o__Saccharomycetales(100);f__Metschnikowiaceae(100);g__Metschnikowia(100);s__Metschnikowia_pulcherrima(100); | Metschnikowia | 13 | Saprotroph | \|Undefined Saprotroph\| | Yeast | NULL | Probable | As nectar-tap_saprotroph (Põlme et al. 2020); Decay sugar-rich substrates (Põlme et al. 2020) | | Põlme S, et al. 2020. Fungal Diversity 105:1-16 ((https://doi.org/10.1007/s13225-020-00466-2)) |
| Otu32016 | 1 | k__Fungi(100);p__Ascomycota(100);c__Saccharomycetes(100);o__Saccharomycetales(100);f__Metschnikowiaceae(100);g__Metschnikowia(100);g__Metschnikowia_unclassified(100); | Metschnikowia | 13 | Saprotroph | \|Undefined Saprotroph\| | Yeast | NULL | Probable | As nectar-tap_saprotroph (Põlme et al. 2020); Decay sugar-rich substrates (Põlme et al. 2020) | | Põlme S, et al. 2020. Fungal Diversity 105:1-16 ((https://doi.org/10.1007/s13225-020-00466-2)) |
| Otu32017 | 1 | k__Fungi(100);p__Ascomycota(100);c__Saccharomycetes(100);o__Saccharomycetales(100);f__Metschnikowiaceae(100);g__Metschnikowia(100);s__Metschnikowia_pulcherrima(100); | Metschnikowia | 13 | Saprotroph | \|Undefined Saprotroph\| | Yeast | NULL | Probable | As nectar-tap_saprotroph (Põlme et al. 2020); Decay sugar-rich substrates (Põlme et al. 2020) | | Põlme S, et al. 2020. Fungal Diversity 105:1-16 ((https://doi.org/10.1007/s13225-020-00466-2)) |
| Otu32018 | 1 | k__Fungi(100);p__Ascomycota(100);c__Saccharomycetes(100);o__Saccharomycetales(100);f__Metschnikowiaceae(100);g__Metschnikowia(100);s__Metschnikowia_pulcherrima(100); | Metschnikowia | 13 | Saprotroph | \|Undefined Saprotroph\| | Yeast | NULL | Probable | As nectar-tap_saprotroph (Põlme et al. 2020); Decay sugar-rich substrates (Põlme et al. 2020) | | Põlme S, et al. 2020. Fungal Diversity 105:1-16 ((https://doi.org/10.1007/s13225-020-00466-2)) |
| Otu32020 | 1 | k__Fungi(100);p__Ascomycota(100);c__Saccharomycetes(100);o__Saccharomycetales(100);f__Metschnikowiaceae(100);g__Metschnikowia(100);s__Metschnikowia_pulcherrima(100); | Metschnikowia | 13 | Saprotroph | \|Undefined Saprotroph\| | Yeast | NULL | Probable | As nectar-tap_saprotroph (Põlme et al. 2020); Decay sugar-rich substrates (Põlme et al. 2020) | | Põlme S, et al. 2020. Fungal Diversity 105:1-16 ((https://doi.org/10.1007/s13225-020-00466-2)) |
| Otu32021 | 1 | k__Fungi(100);p__Ascomycota(100);c__Saccharomycetes(100);o__Saccharomycetales(100);f__Metschnikowiaceae(100);g__Metschnikowia(100);g__Metschnikowia_unclassified(100); | Metschnikowia | 13 | Saprotroph | \|Undefined Saprotroph\| | Yeast | NULL | Probable | As nectar-tap_saprotroph (Põlme et al. 2020); Decay sugar-rich substrates (Põlme et al. 2020) | | Põlme S, et al. 2020. Fungal Diversity 105:1-16 ((https://doi.org/10.1007/s13225-020-00466-2)) |
| Otu32022 | 1 | k__Fungi(100);p__Ascomycota(100);c__Saccharomycetes(100);o__Saccharomycetales(100);f__Metschnikowiaceae(100);g__Metschnikowia(100);s__Metschnikowia_pulcherrima(100); | Metschnikowia | 13 | Saprotroph | \|Undefined Saprotroph\| | Yeast | NULL | Probable | As nectar-tap_saprotroph (Põlme et al. 2020); Decay sugar-rich substrates (Põlme et al. 2020) | | Põlme S, et al. 2020. Fungal Diversity 105:1-16 ((https://doi.org/10.1007/s13225-020-00466-2)) |
| Otu32023 | 1 | k__Fungi(100);p__Ascomycota(100);c__Saccharomycetes(100);o__Saccharomycetales(100);f__Metschnikowiaceae(100);g__Metschnikowia(100);g__Metschnikowia_unclassified(100); | Metschnikowia | 13 | Saprotroph | \|Undefined Saprotroph\| | Yeast | NULL | Probable | As nectar-tap_saprotroph (Põlme et al. 2020); Decay sugar-rich substrates (Põlme et al. 2020) | | Põlme S, et al. 2020. Fungal Diversity 105:1-16 ((https://doi.org/10.1007/s13225-020-00466-2)) |
| Otu32024 | 1 | k__Fungi(100);p__Ascomycota(100);c__Saccharomycetes(100);o__Saccharomycetales(100);f__Metschnikowiaceae(100);g__Metschnikowia(100);s__Metschnikowia_pulcherrima(100); | Metschnikowia | 13 | Saprotroph | \|Undefined Saprotroph\| | Yeast | NULL | Probable | As nectar-tap_saprotroph (Põlme et al. 2020); Decay sugar-rich substrates (Põlme et al. 2020) | | Põlme S, et al. 2020. Fungal Diversity 105:1-16 ((https://doi.org/10.1007/s13225-020-00466-2)) |
| Otu32025 | 1 | k__Fungi(100);p__Ascomycota(100);c__Saccharomycetes(100);o__Saccharomycetales(100);f__Metschnikowiaceae(100);g__Metschnikowia(100);s__Metschnikowia_pulcherrima(100); | Metschnikowia | 13 | Saprotroph | \|Undefined Saprotroph\| | Yeast | NULL | Probable | As nectar-tap_saprotroph (Põlme et al. 2020); Decay sugar-rich substrates (Põlme et al. 2020) | | Põlme S, et al. 2020. Fungal Diversity 105:1-16 ((https://doi.org/10.1007/s13225-020-00466-2)) |
| Otu32026 | 1 | k__Fungi(100);p__Ascomycota(100);c__Saccharomycetes(100);o__Saccharomycetales(100);f__Metschnikowiaceae(100);g__Metschnikowia(100);g__Metschnikowia_unclassified(100); | Metschnikowia | 13 | Saprotroph | \|Undefined Saprotroph\| | Yeast | NULL | Probable | As nectar-tap_saprotroph (Põlme et al. 2020); Decay sugar-rich substrates (Põlme et al. 2020) | | Põlme S, et al. 2020. Fungal Diversity 105:1-16 ((https://doi.org/10.1007/s13225-020-00466-2)) |
| Otu32027 | 1 | k__Fungi(100);p__Ascomycota(100);c__Saccharomycetes(100);o__Saccharomycetales(100);f__Metschnikowiaceae(100);g__Metschnikowia(100);s__Metschnikowia_pulcherrima(100); | Metschnikowia | 13 | Saprotroph | \|Undefined Saprotroph\| | Yeast | NULL | Probable | As nectar-tap_saprotroph (Põlme et al. 2020); Decay sugar-rich substrates (Põlme et al. 2020) | | Põlme S, et al. 2020. Fungal Diversity 105:1-16 ((https://doi.org/10.1007/s13225-020-00466-2)) |
| Otu32028 | 1 | k__Fungi(100);p__Ascomycota(100);c__Saccharomycetes(100);o__Saccharomycetales(100);f__Saccharomycetales_family_Incertae_sedis(100);g__Candida(100);g__Candida_unclassified(100); | Candida | 13 | Pathotroph-Saprotroph-Symbiotroph | Animal Pathogen-Endophyte-Endosymbiont-Epiphyte-\|Undefined Saprotroph\| | Dimorphic Yeast-Yeast | NULL | Probable | As nectar-tap_saprotroph (Põlme et al. 2020); Decay sugar-rich substrates (Põlme et al. 2020); This asexual genus is polyphyletic and contain species that function very differently depend on what they are. Caution should be taken when interpreting guilds for Candida species. | | Manolakaki D, et al. 2010. Virulence. 1: 367-375 ((https://doi.org/10.4161/viru.1.5.12796)); Mekbib SB, et al. 2011. Tropical Plant Pathology 36:233-240 ((https://doi.org/10.1590/S1982-56762011000400004)); Põlme S, et al. 2020. Fungal Diversity 105:1-16 ((https://doi.org/10.1007/s13225-020-00466-2)) |
| Otu32029 | 1 | k__Fungi(100);p__Ascomycota(100);c__Saccharomycetes(100);o__Saccharomycetales(100);f__Metschnikowiaceae(100);g__Metschnikowia(100);g__Metschnikowia_unclassified(100); | Metschnikowia | 13 | Saprotroph | \|Undefined Saprotroph\| | Yeast | NULL | Probable | As nectar-tap_saprotroph (Põlme et al. 2020); Decay sugar-rich substrates (Põlme et al. 2020) | | Põlme S, et al. 2020. Fungal Diversity 105:1-16 ((https://doi.org/10.1007/s13225-020-00466-2)) |
| Otu32030 | 1 | k__Fungi(100);p__Ascomycota(100);c__Saccharomycetes(100);o__Saccharomycetales(100);f__Metschnikowiaceae(100);g__Metschnikowia(100);s__Metschnikowia_pulcherrima(100); | Metschnikowia | 13 | Saprotroph | \|Undefined Saprotroph\| | Yeast | NULL | Probable | As nectar-tap_saprotroph (Põlme et al. 2020); Decay sugar-rich substrates (Põlme et al. 2020) | | Põlme S, et al. 2020. Fungal Diversity 105:1-16 ((https://doi.org/10.1007/s13225-020-00466-2)) |
| Otu32031 | 1 | k__Fungi(100);p__Ascomycota(100);c__Saccharomycetes(100);o__Saccharomycetales(100);f__Metschnikowiaceae(100);g__Metschnikowia(100);s__Metschnikowia_pulcherrima(100); | Metschnikowia | 13 | Saprotroph | \|Undefined Saprotroph\| | Yeast | NULL | Probable | As nectar-tap_saprotroph (Põlme et al. 2020); Decay sugar-rich substrates (Põlme et al. 2020) | | Põlme S, et al. 2020. Fungal Diversity 105:1-16 ((https://doi.org/10.1007/s13225-020-00466-2)) |
| Otu32032 | 1 | k__Fungi(100);p__Ascomycota(100);c__Saccharomycetes(100);o__Saccharomycetales(100);f__Metschnikowiaceae(100);g__Metschnikowia(100);s__Metschnikowia_pulcherrima(100); | Metschnikowia | 13 | Saprotroph | \|Undefined Saprotroph\| | Yeast | NULL | Probable | As nectar-tap_saprotroph (Põlme et al. 2020); Decay sugar-rich substrates (Põlme et al. 2020) | | Põlme S, et al. 2020. Fungal Diversity 105:1-16 ((https://doi.org/10.1007/s13225-020-00466-2)) |
| Otu32034 | 1 | k__Fungi(100);p__Ascomycota(100);c__Saccharomycetes(100);o__Saccharomycetales(100);f__Metschnikowiaceae(100);g__Metschnikowia(100);g__Metschnikowia_unclassified(100); | Metschnikowia | 13 | Saprotroph | \|Undefined Saprotroph\| | Yeast | NULL | Probable | As nectar-tap_saprotroph (Põlme et al. 2020); Decay sugar-rich substrates (Põlme et al. 2020) | | Põlme S, et al. 2020. Fungal Diversity 105:1-16 ((https://doi.org/10.1007/s13225-020-00466-2)) |
| Otu32035 | 1 | k__Fungi(100);p__Ascomycota(100);c__Saccharomycetes(100);o__Saccharomycetales(100);f__Metschnikowiaceae(100);g__Metschnikowia(100);s__Metschnikowia_pulcherrima(100); | Metschnikowia | 13 | Saprotroph | \|Undefined Saprotroph\| | Yeast | NULL | Probable | As nectar-tap_saprotroph (Põlme et al. 2020); Decay sugar-rich substrates (Põlme et al. 2020) | | Põlme S, et al. 2020. Fungal Diversity 105:1-16 ((https://doi.org/10.1007/s13225-020-00466-2)) |
| Otu32036 | 1 | k__Fungi(100);p__Ascomycota(100);c__Saccharomycetes(100);o__Saccharomycetales(100);f__Metschnikowiaceae(100);g__Metschnikowia(100);s__Metschnikowia_pulcherrima(100); | Metschnikowia | 13 | Saprotroph | \|Undefined Saprotroph\| | Yeast | NULL | Probable | As nectar-tap_saprotroph (Põlme et al. 2020); Decay sugar-rich substrates (Põlme et al. 2020) | | Põlme S, et al. 2020. Fungal Diversity 105:1-16 ((https://doi.org/10.1007/s13225-020-00466-2)) |
| Otu32037 | 1 | k__Fungi(100);p__Ascomycota(100);c__Saccharomycetes(100);o__Saccharomycetales(100);f__Metschnikowiaceae(100);g__Metschnikowia(100);g__Metschnikowia_unclassified(100); | Metschnikowia | 13 | Saprotroph | \|Undefined Saprotroph\| | Yeast | NULL | Probable | As nectar-tap_saprotroph (Põlme et al. 2020); Decay sugar-rich substrates (Põlme et al. 2020) | | Põlme S, et al. 2020. Fungal Diversity 105:1-16 ((https://doi.org/10.1007/s13225-020-00466-2)) |
| Otu32038 | 1 | k__Fungi(100);p__Ascomycota(100);c__Saccharomycetes(100);o__Saccharomycetales(100);f__Metschnikowiaceae(100);g__Metschnikowia(100);s__Metschnikowia_pulcherrima(100); | Metschnikowia | 13 | Saprotroph | \|Undefined Saprotroph\| | Yeast | NULL | Probable | As nectar-tap_saprotroph (Põlme et al. 2020); Decay sugar-rich substrates (Põlme et al. 2020) | | Põlme S, et al. 2020. Fungal Diversity 105:1-16 ((https://doi.org/10.1007/s13225-020-00466-2)) |
| Otu32039 | 1 | k__Fungi(100);p__Ascomycota(100);c__Saccharomycetes(100);o__Saccharomycetales(100);f__Metschnikowiaceae(100);g__Metschnikowia(100);s__Metschnikowia_pulcherrima(100); | Metschnikowia | 13 | Saprotroph | \|Undefined Saprotroph\| | Yeast | NULL | Probable | As nectar-tap_saprotroph (Põlme et al. 2020); Decay sugar-rich substrates (Põlme et al. 2020) | | Põlme S, et al. 2020. Fungal Diversity 105:1-16 ((https://doi.org/10.1007/s13225-020-00466-2)) |
| Otu32040 | 1 | k__Fungi(100);p__Ascomycota(100);c__Saccharomycetes(100);o__Saccharomycetales(100);f__Metschnikowiaceae(100);g__Metschnikowia(100);g__Metschnikowia_unclassified(100); | Metschnikowia | 13 | Saprotroph | \|Undefined Saprotroph\| | Yeast | NULL | Probable | As nectar-tap_saprotroph (Põlme et al. 2020); Decay sugar-rich substrates (Põlme et al. 2020) | | Põlme S, et al. 2020. Fungal Diversity 105:1-16 ((https://doi.org/10.1007/s13225-020-00466-2)) |
| Otu32042 | 1 | k__Fungi(100);p__Ascomycota(100);c__Saccharomycetes(100);o__Saccharomycetales(100);f__Metschnikowiaceae(100);g__Metschnikowia(100);g__Metschnikowia_unclassified(100); | Metschnikowia | 13 | Saprotroph | \|Undefined Saprotroph\| | Yeast | NULL | Probable | As nectar-tap_saprotroph (Põlme et al. 2020); Decay sugar-rich substrates (Põlme et al. 2020) | | Põlme S, et al. 2020. Fungal Diversity 105:1-16 ((https://doi.org/10.1007/s13225-020-00466-2)) |
| Otu32043 | 1 | k__Fungi(100);p__Ascomycota(100);c__Saccharomycetes(100);o__Saccharomycetales(100);f__Metschnikowiaceae(100);g__Metschnikowia(100);s__Metschnikowia_pulcherrima(100); | Metschnikowia | 13 | Saprotroph | \|Undefined Saprotroph\| | Yeast | NULL | Probable | As nectar-tap_saprotroph (Põlme et al. 2020); Decay sugar-rich substrates (Põlme et al. 2020) | | Põlme S, et al. 2020. Fungal Diversity 105:1-16 ((https://doi.org/10.1007/s13225-020-00466-2)) |
| Otu32044 | 1 | k__Fungi(100);p__Ascomycota(100);c__Saccharomycetes(100);o__Saccharomycetales(100);f__Metschnikowiaceae(100);g__Metschnikowia(100);s__Metschnikowia_pulcherrima(100); | Metschnikowia | 13 | Saprotroph | \|Undefined Saprotroph\| | Yeast | NULL | Probable | As nectar-tap_saprotroph (Põlme et al. 2020); Decay sugar-rich substrates (Põlme et al. 2020) | | Põlme S, et al. 2020. Fungal Diversity 105:1-16 ((https://doi.org/10.1007/s13225-020-00466-2)) |
| Otu32045 | 1 | k__Fungi(100);p__Ascomycota(100);c__Saccharomycetes(100);o__Saccharomycetales(100);f__Metschnikowiaceae(100);g__Metschnikowia(100);g__Metschnikowia_unclassified(100); | Metschnikowia | 13 | Saprotroph | \|Undefined Saprotroph\| | Yeast | NULL | Probable | As nectar-tap_saprotroph (Põlme et al. 2020); Decay sugar-rich substrates (Põlme et al. 2020) | | Põlme S, et al. 2020. Fungal Diversity 105:1-16 ((https://doi.org/10.1007/s13225-020-00466-2)) |
| Otu32046 | 1 | k__Fungi(100);p__Ascomycota(100);c__Saccharomycetes(100);o__Saccharomycetales(100);f__Metschnikowiaceae(100);g__Metschnikowia(100);g__Metschnikowia_unclassified(100); | Metschnikowia | 13 | Saprotroph | \|Undefined Saprotroph\| | Yeast | NULL | Probable | As nectar-tap_saprotroph (Põlme et al. 2020); Decay sugar-rich substrates (Põlme et al. 2020) | | Põlme S, et al. 2020. Fungal Diversity 105:1-16 ((https://doi.org/10.1007/s13225-020-00466-2)) |
| Otu32058 | 1 | k__Fungi(100);p__Ascomycota(100);c__Saccharomycetes(100);o__Saccharomycetales(100);f__Metschnikowiaceae(100);g__Metschnikowia(100);s__Metschnikowia_pulcherrima(100); | Metschnikowia | 13 | Saprotroph | \|Undefined Saprotroph\| | Yeast | NULL | Probable | As nectar-tap_saprotroph (Põlme et al. 2020); Decay sugar-rich substrates (Põlme et al. 2020) | | Põlme S, et al. 2020. Fungal Diversity 105:1-16 ((https://doi.org/10.1007/s13225-020-00466-2)) |
| Otu32061 | 1 | k__Fungi(100);p__Ascomycota(100);c__Saccharomycetes(100);o__Saccharomycetales(100);f__Metschnikowiaceae(100);g__Metschnikowia(100);s__Metschnikowia_pulcherrima(100); | Metschnikowia | 13 | Saprotroph | \|Undefined Saprotroph\| | Yeast | NULL | Probable | As nectar-tap_saprotroph (Põlme et al. 2020); Decay sugar-rich substrates (Põlme et al. 2020) | | Põlme S, et al. 2020. Fungal Diversity 105:1-16 ((https://doi.org/10.1007/s13225-020-00466-2)) |
| Otu32063 | 1 | k__Fungi(100);p__Ascomycota(100);c__Leotiomycetes(100);o__Helotiales(100);f__Vibrisseaceae(100);g__Phialocephala(100);s__Phialocephala_fusca(100); | Phialocephala | 13 | Saprotroph-Symbiotroph | Endophyte-\|Undefined Saprotroph\| | Microfungus | NULL | Highly Probable | As root_endophyte (Põlme et al. 2020); As root_endophyte_dark_septate (Põlme et al. 2020); As soil_saprotroph (Põlme et al. 2020); Dark Septate Endophyte, known from plant roots.; Decay roots, soil (Põlme et al. 2020) | | Newsham KK. 2011. New Phytologist 190:783-793 ((https://doi.org/10.1111/j.1469-8137.2010.03611.x)) (as dark septate root endophytes); Põlme S, et al. 2020. Fungal Diversity 105:1-16 ((https://doi.org/10.1007/s13225-020-00466-2)) |
| Otu32065 | 1 | k__Fungi(100);p__Ascomycota(100);c__Saccharomycetes(100);o__Saccharomycetales(100);f__Saccharomycetales_family_Incertae_sedis(100);f__Saccharomycetales_family_Incertae_sedis_unclassified(100);f__Saccharomycetales_family_Incertae_sedis_unclassified(100); | Saccharomycetales | 7 | Saprotroph | Undefined Saprotroph | Yeast | NULL | Possible | NULL | | Sterkenburg E, et al. 2015. New Phytologist 207:1145-1158 ((https://doi.org/10.1111/nph.13426)) |
| Otu32070 | 1 | k__Fungi(100);p__Ascomycota(100);c__Saccharomycetes(100);o__Saccharomycetales(100);f__Metschnikowiaceae(100);g__Metschnikowia(100);g__Metschnikowia_unclassified(100); | Metschnikowia | 13 | Saprotroph | \|Undefined Saprotroph\| | Yeast | NULL | Probable | As nectar-tap_saprotroph (Põlme et al. 2020); Decay sugar-rich substrates (Põlme et al. 2020) | | Põlme S, et al. 2020. Fungal Diversity 105:1-16 ((https://doi.org/10.1007/s13225-020-00466-2)) |
| Otu32072 | 1 | k__Fungi(100);p__Ascomycota(100);c__Saccharomycetes(100);o__Saccharomycetales(100);f__Saccharomycetales_family_Incertae_sedis(100);g__Candida(100);g__Candida_unclassified(100); | Candida | 13 | Pathotroph-Saprotroph-Symbiotroph | Animal Pathogen-Endophyte-Endosymbiont-Epiphyte-\|Undefined Saprotroph\| | Dimorphic Yeast-Yeast | NULL | Probable | As nectar-tap_saprotroph (Põlme et al. 2020); Decay sugar-rich substrates (Põlme et al. 2020); This asexual genus is polyphyletic and contain species that function very differently depend on what they are. Caution should be taken when interpreting guilds for Candida species. | | Manolakaki D, et al. 2010. Virulence. 1: 367-375 ((https://doi.org/10.4161/viru.1.5.12796)); Mekbib SB, et al. 2011. Tropical Plant Pathology 36:233-240 ((https://doi.org/10.1590/S1982-56762011000400004)); Põlme S, et al. 2020. Fungal Diversity 105:1-16 ((https://doi.org/10.1007/s13225-020-00466-2)) |
| Otu32078 | 1 | k__Fungi(100);p__Ascomycota(100);c__Saccharomycetes(100);o__Saccharomycetales(100);f__Metschnikowiaceae(100);g__Metschnikowia(100);s__Metschnikowia_pulcherrima(100); | Metschnikowia | 13 | Saprotroph | \|Undefined Saprotroph\| | Yeast | NULL | Probable | As nectar-tap_saprotroph (Põlme et al. 2020); Decay sugar-rich substrates (Põlme et al. 2020) | | Põlme S, et al. 2020. Fungal Diversity 105:1-16 ((https://doi.org/10.1007/s13225-020-00466-2)) |
| Otu32079 | 1 | k__Fungi(100);p__Ascomycota(100);c__Saccharomycetes(100);o__Saccharomycetales(100);f__Metschnikowiaceae(100);g__Metschnikowia(100);s__Metschnikowia_pulcherrima(100); | Metschnikowia | 13 | Saprotroph | \|Undefined Saprotroph\| | Yeast | NULL | Probable | As nectar-tap_saprotroph (Põlme et al. 2020); Decay sugar-rich substrates (Põlme et al. 2020) | | Põlme S, et al. 2020. Fungal Diversity 105:1-16 ((https://doi.org/10.1007/s13225-020-00466-2)) |
| Otu32080 | 1 | k__Fungi(100);p__Ascomycota(100);c__Saccharomycetes(100);o__Saccharomycetales(100);f__Metschnikowiaceae(100);g__Metschnikowia(100);s__Metschnikowia_pulcherrima(100); | Metschnikowia | 13 | Saprotroph | \|Undefined Saprotroph\| | Yeast | NULL | Probable | As nectar-tap_saprotroph (Põlme et al. 2020); Decay sugar-rich substrates (Põlme et al. 2020) | | Põlme S, et al. 2020. Fungal Diversity 105:1-16 ((https://doi.org/10.1007/s13225-020-00466-2)) |
| Otu32096 | 1 | k__Fungi(100);p__Ascomycota(100);c__Saccharomycetes(100);o__Saccharomycetales(100);f__Metschnikowiaceae(100);g__Metschnikowia(100);g__Metschnikowia_unclassified(100); | Metschnikowia | 13 | Saprotroph | \|Undefined Saprotroph\| | Yeast | NULL | Probable | As nectar-tap_saprotroph (Põlme et al. 2020); Decay sugar-rich substrates (Põlme et al. 2020) | | Põlme S, et al. 2020. Fungal Diversity 105:1-16 ((https://doi.org/10.1007/s13225-020-00466-2)) |
| Otu32100 | 1 | k__Fungi(100);p__Ascomycota(100);c__Saccharomycetes(100);o__Saccharomycetales(100);f__Metschnikowiaceae(100);g__Metschnikowia(100);s__Metschnikowia_pulcherrima(100); | Metschnikowia | 13 | Saprotroph | \|Undefined Saprotroph\| | Yeast | NULL | Probable | As nectar-tap_saprotroph (Põlme et al. 2020); Decay sugar-rich substrates (Põlme et al. 2020) | | Põlme S, et al. 2020. Fungal Diversity 105:1-16 ((https://doi.org/10.1007/s13225-020-00466-2)) |
| Otu32101 | 1 | k__Fungi(100);p__Ascomycota(100);c__Saccharomycetes(100);o__Saccharomycetales(100);f__Metschnikowiaceae(100);g__Metschnikowia(100);s__Metschnikowia_pulcherrima(100); | Metschnikowia | 13 | Saprotroph | \|Undefined Saprotroph\| | Yeast | NULL | Probable | As nectar-tap_saprotroph (Põlme et al. 2020); Decay sugar-rich substrates (Põlme et al. 2020) | | Põlme S, et al. 2020. Fungal Diversity 105:1-16 ((https://doi.org/10.1007/s13225-020-00466-2)) |
| Otu32102 | 1 | k__Fungi(100);p__Ascomycota(100);c__Saccharomycetes(100);o__Saccharomycetales(100);f__Metschnikowiaceae(100);g__Metschnikowia(100);s__Metschnikowia_pulcherrima(100); | Metschnikowia | 13 | Saprotroph | \|Undefined Saprotroph\| | Yeast | NULL | Probable | As nectar-tap_saprotroph (Põlme et al. 2020); Decay sugar-rich substrates (Põlme et al. 2020) | | Põlme S, et al. 2020. Fungal Diversity 105:1-16 ((https://doi.org/10.1007/s13225-020-00466-2)) |
| Otu32103 | 1 | k__Fungi(100);p__Ascomycota(100);c__Saccharomycetes(100);o__Saccharomycetales(100);f__Metschnikowiaceae(100);g__Metschnikowia(100);s__Metschnikowia_pulcherrima(100); | Metschnikowia | 13 | Saprotroph | \|Undefined Saprotroph\| | Yeast | NULL | Probable | As nectar-tap_saprotroph (Põlme et al. 2020); Decay sugar-rich substrates (Põlme et al. 2020) | | Põlme S, et al. 2020. Fungal Diversity 105:1-16 ((https://doi.org/10.1007/s13225-020-00466-2)) |
| Otu32104 | 1 | k__Fungi(100);p__Ascomycota(100);c__Saccharomycetes(100);o__Saccharomycetales(100);f__Metschnikowiaceae(100);g__Metschnikowia(100);s__Metschnikowia_pulcherrima(100); | Metschnikowia | 13 | Saprotroph | \|Undefined Saprotroph\| | Yeast | NULL | Probable | As nectar-tap_saprotroph (Põlme et al. 2020); Decay sugar-rich substrates (Põlme et al. 2020) | | Põlme S, et al. 2020. Fungal Diversity 105:1-16 ((https://doi.org/10.1007/s13225-020-00466-2)) |
| Otu32105 | 1 | k__Fungi(100);p__Ascomycota(100);c__Saccharomycetes(100);o__Saccharomycetales(100);f__Metschnikowiaceae(100);g__Metschnikowia(100);s__Metschnikowia_pulcherrima(100); | Metschnikowia | 13 | Saprotroph | \|Undefined Saprotroph\| | Yeast | NULL | Probable | As nectar-tap_saprotroph (Põlme et al. 2020); Decay sugar-rich substrates (Põlme et al. 2020) | | Põlme S, et al. 2020. Fungal Diversity 105:1-16 ((https://doi.org/10.1007/s13225-020-00466-2)) |
| Otu32106 | 1 | k__Fungi(100);p__Ascomycota(100);c__Saccharomycetes(100);o__Saccharomycetales(100);f__Metschnikowiaceae(100);g__Metschnikowia(100);s__Metschnikowia_pulcherrima(100); | Metschnikowia | 13 | Saprotroph | \|Undefined Saprotroph\| | Yeast | NULL | Probable | As nectar-tap_saprotroph (Põlme et al. 2020); Decay sugar-rich substrates (Põlme et al. 2020) | | Põlme S, et al. 2020. Fungal Diversity 105:1-16 ((https://doi.org/10.1007/s13225-020-00466-2)) |
| Otu32107 | 1 | k__Fungi(100);p__Ascomycota(100);c__Saccharomycetes(100);o__Saccharomycetales(100);f__Metschnikowiaceae(100);g__Metschnikowia(100);s__Metschnikowia_pulcherrima(100); | Metschnikowia | 13 | Saprotroph | \|Undefined Saprotroph\| | Yeast | NULL | Probable | As nectar-tap_saprotroph (Põlme et al. 2020); Decay sugar-rich substrates (Põlme et al. 2020) | | Põlme S, et al. 2020. Fungal Diversity 105:1-16 ((https://doi.org/10.1007/s13225-020-00466-2)) |
| Otu32108 | 1 | k__Fungi(100);p__Ascomycota(100);c__Saccharomycetes(100);o__Saccharomycetales(100);f__Metschnikowiaceae(100);g__Metschnikowia(100);g__Metschnikowia_unclassified(100); | Metschnikowia | 13 | Saprotroph | \|Undefined Saprotroph\| | Yeast | NULL | Probable | As nectar-tap_saprotroph (Põlme et al. 2020); Decay sugar-rich substrates (Põlme et al. 2020) | | Põlme S, et al. 2020. Fungal Diversity 105:1-16 ((https://doi.org/10.1007/s13225-020-00466-2)) |
| Otu32109 | 1 | k__Fungi(100);p__Ascomycota(100);c__Saccharomycetes(100);o__Saccharomycetales(100);f__Metschnikowiaceae(100);g__Metschnikowia(100);s__Metschnikowia_pulcherrima(100); | Metschnikowia | 13 | Saprotroph | \|Undefined Saprotroph\| | Yeast | NULL | Probable | As nectar-tap_saprotroph (Põlme et al. 2020); Decay sugar-rich substrates (Põlme et al. 2020) | | Põlme S, et al. 2020. Fungal Diversity 105:1-16 ((https://doi.org/10.1007/s13225-020-00466-2)) |
| Otu32111 | 1 | k__Fungi(100);p__Ascomycota(100);c__Saccharomycetes(100);o__Saccharomycetales(100);f__Metschnikowiaceae(100);g__Metschnikowia(100);s__Metschnikowia_pulcherrima(100); | Metschnikowia | 13 | Saprotroph | \|Undefined Saprotroph\| | Yeast | NULL | Probable | As nectar-tap_saprotroph (Põlme et al. 2020); Decay sugar-rich substrates (Põlme et al. 2020) | | Põlme S, et al. 2020. Fungal Diversity 105:1-16 ((https://doi.org/10.1007/s13225-020-00466-2)) |
| Otu32112 | 1 | k__Fungi(100);p__Ascomycota(100);c__Saccharomycetes(100);o__Saccharomycetales(100);f__Metschnikowiaceae(100);g__Metschnikowia(100);s__Metschnikowia_pulcherrima(100); | Metschnikowia | 13 | Saprotroph | \|Undefined Saprotroph\| | Yeast | NULL | Probable | As nectar-tap_saprotroph (Põlme et al. 2020); Decay sugar-rich substrates (Põlme et al. 2020) | | Põlme S, et al. 2020. Fungal Diversity 105:1-16 ((https://doi.org/10.1007/s13225-020-00466-2)) |
| Otu32113 | 1 | k__Fungi(100);p__Ascomycota(100);c__Saccharomycetes(100);o__Saccharomycetales(100);f__Metschnikowiaceae(100);g__Metschnikowia(100);s__Metschnikowia_pulcherrima(100); | Metschnikowia | 13 | Saprotroph | \|Undefined Saprotroph\| | Yeast | NULL | Probable | As nectar-tap_saprotroph (Põlme et al. 2020); Decay sugar-rich substrates (Põlme et al. 2020) | | Põlme S, et al. 2020. Fungal Diversity 105:1-16 ((https://doi.org/10.1007/s13225-020-00466-2)) |
| Otu32114 | 1 | k__Fungi(100);p__Ascomycota(100);c__Saccharomycetes(100);o__Saccharomycetales(100);f__Metschnikowiaceae(100);g__Metschnikowia(100);s__Metschnikowia_pulcherrima(100); | Metschnikowia | 13 | Saprotroph | \|Undefined Saprotroph\| | Yeast | NULL | Probable | As nectar-tap_saprotroph (Põlme et al. 2020); Decay sugar-rich substrates (Põlme et al. 2020) | | Põlme S, et al. 2020. Fungal Diversity 105:1-16 ((https://doi.org/10.1007/s13225-020-00466-2)) |
| Otu32115 | 1 | k__Fungi(100);p__Ascomycota(100);c__Saccharomycetes(100);o__Saccharomycetales(100);f__Metschnikowiaceae(100);g__Metschnikowia(100);s__Metschnikowia_pulcherrima(100); | Metschnikowia | 13 | Saprotroph | \|Undefined Saprotroph\| | Yeast | NULL | Probable | As nectar-tap_saprotroph (Põlme et al. 2020); Decay sugar-rich substrates (Põlme et al. 2020) | | Põlme S, et al. 2020. Fungal Diversity 105:1-16 ((https://doi.org/10.1007/s13225-020-00466-2)) |
| Otu32116 | 1 | k__Fungi(100);p__Ascomycota(100);c__Saccharomycetes(100);o__Saccharomycetales(100);f__Metschnikowiaceae(100);g__Metschnikowia(100);s__Metschnikowia_pulcherrima(100); | Metschnikowia | 13 | Saprotroph | \|Undefined Saprotroph\| | Yeast | NULL | Probable | As nectar-tap_saprotroph (Põlme et al. 2020); Decay sugar-rich substrates (Põlme et al. 2020) | | Põlme S, et al. 2020. Fungal Diversity 105:1-16 ((https://doi.org/10.1007/s13225-020-00466-2)) |
| Otu32119 | 1 | k__Fungi(100);p__Ascomycota(100);c__Saccharomycetes(100);o__Saccharomycetales(100);f__Metschnikowiaceae(100);g__Metschnikowia(100);g__Metschnikowia_unclassified(100); | Metschnikowia | 13 | Saprotroph | \|Undefined Saprotroph\| | Yeast | NULL | Probable | As nectar-tap_saprotroph (Põlme et al. 2020); Decay sugar-rich substrates (Põlme et al. 2020) | | Põlme S, et al. 2020. Fungal Diversity 105:1-16 ((https://doi.org/10.1007/s13225-020-00466-2)) |
| Otu32120 | 1 | k__Fungi(100);p__Ascomycota(100);c__Saccharomycetes(100);o__Saccharomycetales(100);f__Metschnikowiaceae(100);g__Metschnikowia(100);s__Metschnikowia_pulcherrima(100); | Metschnikowia | 13 | Saprotroph | \|Undefined Saprotroph\| | Yeast | NULL | Probable | As nectar-tap_saprotroph (Põlme et al. 2020); Decay sugar-rich substrates (Põlme et al. 2020) | | Põlme S, et al. 2020. Fungal Diversity 105:1-16 ((https://doi.org/10.1007/s13225-020-00466-2)) |
| Otu32123 | 1 | k__Fungi(100);p__Ascomycota(100);c__Saccharomycetes(100);o__Saccharomycetales(100);f__Metschnikowiaceae(100);g__Metschnikowia(100);s__Metschnikowia_pulcherrima(100); | Metschnikowia | 13 | Saprotroph | \|Undefined Saprotroph\| | Yeast | NULL | Probable | As nectar-tap_saprotroph (Põlme et al. 2020); Decay sugar-rich substrates (Põlme et al. 2020) | | Põlme S, et al. 2020. Fungal Diversity 105:1-16 ((https://doi.org/10.1007/s13225-020-00466-2)) |
| Otu32127 | 1 | k__Fungi(100);p__Ascomycota(100);c__Saccharomycetes(100);o__Saccharomycetales(100);f__Metschnikowiaceae(100);g__Metschnikowia(100);g__Metschnikowia_unclassified(100); | Metschnikowia | 13 | Saprotroph | \|Undefined Saprotroph\| | Yeast | NULL | Probable | As nectar-tap_saprotroph (Põlme et al. 2020); Decay sugar-rich substrates (Põlme et al. 2020) | | Põlme S, et al. 2020. Fungal Diversity 105:1-16 ((https://doi.org/10.1007/s13225-020-00466-2)) |
| Otu32128 | 1 | k__Fungi(100);p__Ascomycota(100);c__Saccharomycetes(100);o__Saccharomycetales(100);f__Metschnikowiaceae(100);g__Metschnikowia(100);s__Metschnikowia_pulcherrima(100); | Metschnikowia | 13 | Saprotroph | \|Undefined Saprotroph\| | Yeast | NULL | Probable | As nectar-tap_saprotroph (Põlme et al. 2020); Decay sugar-rich substrates (Põlme et al. 2020) | | Põlme S, et al. 2020. Fungal Diversity 105:1-16 ((https://doi.org/10.1007/s13225-020-00466-2)) |
| Otu32129 | 1 | k__Fungi(100);p__Ascomycota(100);c__Saccharomycetes(100);o__Saccharomycetales(100);f__Metschnikowiaceae(100);g__Metschnikowia(100);s__Metschnikowia_pulcherrima(100); | Metschnikowia | 13 | Saprotroph | \|Undefined Saprotroph\| | Yeast | NULL | Probable | As nectar-tap_saprotroph (Põlme et al. 2020); Decay sugar-rich substrates (Põlme et al. 2020) | | Põlme S, et al. 2020. Fungal Diversity 105:1-16 ((https://doi.org/10.1007/s13225-020-00466-2)) |
| Otu32136 | 1 | k__Fungi(100);p__Ascomycota(100);c__Saccharomycetes(100);o__Saccharomycetales(100);f__Metschnikowiaceae(100);g__Metschnikowia(100);s__Metschnikowia_pulcherrima(100); | Metschnikowia | 13 | Saprotroph | \|Undefined Saprotroph\| | Yeast | NULL | Probable | As nectar-tap_saprotroph (Põlme et al. 2020); Decay sugar-rich substrates (Põlme et al. 2020) | | Põlme S, et al. 2020. Fungal Diversity 105:1-16 ((https://doi.org/10.1007/s13225-020-00466-2)) |
| Otu32137 | 1 | k__Fungi(100);p__Ascomycota(100);c__Saccharomycetes(100);o__Saccharomycetales(100);f__Metschnikowiaceae(100);g__Metschnikowia(100);s__Metschnikowia_pulcherrima(100); | Metschnikowia | 13 | Saprotroph | \|Undefined Saprotroph\| | Yeast | NULL | Probable | As nectar-tap_saprotroph (Põlme et al. 2020); Decay sugar-rich substrates (Põlme et al. 2020) | | Põlme S, et al. 2020. Fungal Diversity 105:1-16 ((https://doi.org/10.1007/s13225-020-00466-2)) |
| Otu32138 | 1 | k__Fungi(100);p__Ascomycota(100);c__Saccharomycetes(100);o__Saccharomycetales(100);f__Metschnikowiaceae(100);g__Metschnikowia(100);s__Metschnikowia_pulcherrima(100); | Metschnikowia | 13 | Saprotroph | \|Undefined Saprotroph\| | Yeast | NULL | Probable | As nectar-tap_saprotroph (Põlme et al. 2020); Decay sugar-rich substrates (Põlme et al. 2020) | | Põlme S, et al. 2020. Fungal Diversity 105:1-16 ((https://doi.org/10.1007/s13225-020-00466-2)) |
| Otu32139 | 1 | k__Fungi(100);p__Ascomycota(100);c__Saccharomycetes(100);o__Saccharomycetales(100);f__Metschnikowiaceae(100);g__Metschnikowia(100);g__Metschnikowia_unclassified(100); | Metschnikowia | 13 | Saprotroph | \|Undefined Saprotroph\| | Yeast | NULL | Probable | As nectar-tap_saprotroph (Põlme et al. 2020); Decay sugar-rich substrates (Põlme et al. 2020) | | Põlme S, et al. 2020. Fungal Diversity 105:1-16 ((https://doi.org/10.1007/s13225-020-00466-2)) |
| Otu32141 | 1 | k__Fungi(100);p__Ascomycota(100);c__Saccharomycetes(100);o__Saccharomycetales(100);f__Metschnikowiaceae(100);g__Metschnikowia(100);s__Metschnikowia_pulcherrima(100); | Metschnikowia | 13 | Saprotroph | \|Undefined Saprotroph\| | Yeast | NULL | Probable | As nectar-tap_saprotroph (Põlme et al. 2020); Decay sugar-rich substrates (Põlme et al. 2020) | | Põlme S, et al. 2020. Fungal Diversity 105:1-16 ((https://doi.org/10.1007/s13225-020-00466-2)) |
| Otu32142 | 1 | k__Fungi(100);p__Ascomycota(100);c__Saccharomycetes(100);o__Saccharomycetales(100);f__Metschnikowiaceae(100);g__Metschnikowia(100);s__Metschnikowia_pulcherrima(100); | Metschnikowia | 13 | Saprotroph | \|Undefined Saprotroph\| | Yeast | NULL | Probable | As nectar-tap_saprotroph (Põlme et al. 2020); Decay sugar-rich substrates (Põlme et al. 2020) | | Põlme S, et al. 2020. Fungal Diversity 105:1-16 ((https://doi.org/10.1007/s13225-020-00466-2)) |
| Otu32143 | 1 | k__Fungi(100);p__Ascomycota(100);c__Saccharomycetes(100);o__Saccharomycetales(100);f__Metschnikowiaceae(100);g__Metschnikowia(100);s__Metschnikowia_pulcherrima(100); | Metschnikowia | 13 | Saprotroph | \|Undefined Saprotroph\| | Yeast | NULL | Probable | As nectar-tap_saprotroph (Põlme et al. 2020); Decay sugar-rich substrates (Põlme et al. 2020) | | Põlme S, et al. 2020. Fungal Diversity 105:1-16 ((https://doi.org/10.1007/s13225-020-00466-2)) |
| Otu32144 | 1 | k__Fungi(100);p__Ascomycota(100);c__Saccharomycetes(100);o__Saccharomycetales(100);f__Metschnikowiaceae(100);g__Metschnikowia(100);s__Metschnikowia_pulcherrima(100); | Metschnikowia | 13 | Saprotroph | \|Undefined Saprotroph\| | Yeast | NULL | Probable | As nectar-tap_saprotroph (Põlme et al. 2020); Decay sugar-rich substrates (Põlme et al. 2020) | | Põlme S, et al. 2020. Fungal Diversity 105:1-16 ((https://doi.org/10.1007/s13225-020-00466-2)) |
| Otu32145 | 1 | k__Fungi(100);p__Ascomycota(100);c__Saccharomycetes(100);o__Saccharomycetales(100);f__Metschnikowiaceae(100);g__Metschnikowia(100);s__Metschnikowia_pulcherrima(100); | Metschnikowia | 13 | Saprotroph | \|Undefined Saprotroph\| | Yeast | NULL | Probable | As nectar-tap_saprotroph (Põlme et al. 2020); Decay sugar-rich substrates (Põlme et al. 2020) | | Põlme S, et al. 2020. Fungal Diversity 105:1-16 ((https://doi.org/10.1007/s13225-020-00466-2)) |
| Otu32146 | 1 | k__Fungi(100);p__Ascomycota(100);c__Saccharomycetes(100);o__Saccharomycetales(100);f__Saccharomycetales_family_Incertae_sedis(100);g__Candida(100);g__Candida_unclassified(100); | Candida | 13 | Pathotroph-Saprotroph-Symbiotroph | Animal Pathogen-Endophyte-Endosymbiont-Epiphyte-\|Undefined Saprotroph\| | Dimorphic Yeast-Yeast | NULL | Probable | As nectar-tap_saprotroph (Põlme et al. 2020); Decay sugar-rich substrates (Põlme et al. 2020); This asexual genus is polyphyletic and contain species that function very differently depend on what they are. Caution should be taken when interpreting guilds for Candida species. | | Manolakaki D, et al. 2010. Virulence. 1: 367-375 ((https://doi.org/10.4161/viru.1.5.12796)); Mekbib SB, et al. 2011. Tropical Plant Pathology 36:233-240 ((https://doi.org/10.1590/S1982-56762011000400004)); Põlme S, et al. 2020. Fungal Diversity 105:1-16 ((https://doi.org/10.1007/s13225-020-00466-2)) |
| Otu32148 | 1 | k__Fungi(100);p__Ascomycota(100);c__Saccharomycetes(100);o__Saccharomycetales(100);f__Metschnikowiaceae(100);g__Metschnikowia(100);s__Metschnikowia_pulcherrima(100); | Metschnikowia | 13 | Saprotroph | \|Undefined Saprotroph\| | Yeast | NULL | Probable | As nectar-tap_saprotroph (Põlme et al. 2020); Decay sugar-rich substrates (Põlme et al. 2020) | | Põlme S, et al. 2020. Fungal Diversity 105:1-16 ((https://doi.org/10.1007/s13225-020-00466-2)) |
| Otu32150 | 1 | k__Fungi(100);p__Ascomycota(100);c__Saccharomycetes(100);o__Saccharomycetales(100);f__Metschnikowiaceae(100);g__Metschnikowia(100);s__Metschnikowia_pulcherrima(100); | Metschnikowia | 13 | Saprotroph | \|Undefined Saprotroph\| | Yeast | NULL | Probable | As nectar-tap_saprotroph (Põlme et al. 2020); Decay sugar-rich substrates (Põlme et al. 2020) | | Põlme S, et al. 2020. Fungal Diversity 105:1-16 ((https://doi.org/10.1007/s13225-020-00466-2)) |
| Otu32151 | 1 | k__Fungi(100);p__Ascomycota(100);c__Saccharomycetes(100);o__Saccharomycetales(100);f__Metschnikowiaceae(100);g__Metschnikowia(100);s__Metschnikowia_pulcherrima(100); | Metschnikowia | 13 | Saprotroph | \|Undefined Saprotroph\| | Yeast | NULL | Probable | As nectar-tap_saprotroph (Põlme et al. 2020); Decay sugar-rich substrates (Põlme et al. 2020) | | Põlme S, et al. 2020. Fungal Diversity 105:1-16 ((https://doi.org/10.1007/s13225-020-00466-2)) |
| Otu32152 | 1 | k__Fungi(100);p__Ascomycota(100);c__Saccharomycetes(100);o__Saccharomycetales(100);f__Metschnikowiaceae(100);g__Metschnikowia(100);s__Metschnikowia_pulcherrima(100); | Metschnikowia | 13 | Saprotroph | \|Undefined Saprotroph\| | Yeast | NULL | Probable | As nectar-tap_saprotroph (Põlme et al. 2020); Decay sugar-rich substrates (Põlme et al. 2020) | | Põlme S, et al. 2020. Fungal Diversity 105:1-16 ((https://doi.org/10.1007/s13225-020-00466-2)) |
| Otu32153 | 1 | k__Fungi(100);p__Ascomycota(100);c__Saccharomycetes(100);o__Saccharomycetales(100);f__Metschnikowiaceae(100);g__Metschnikowia(100);s__Metschnikowia_pulcherrima(100); | Metschnikowia | 13 | Saprotroph | \|Undefined Saprotroph\| | Yeast | NULL | Probable | As nectar-tap_saprotroph (Põlme et al. 2020); Decay sugar-rich substrates (Põlme et al. 2020) | | Põlme S, et al. 2020. Fungal Diversity 105:1-16 ((https://doi.org/10.1007/s13225-020-00466-2)) |
| Otu32154 | 1 | k__Fungi(100);p__Ascomycota(100);c__Saccharomycetes(100);o__Saccharomycetales(100);f__Metschnikowiaceae(100);g__Metschnikowia(100);s__Metschnikowia_pulcherrima(100); | Metschnikowia | 13 | Saprotroph | \|Undefined Saprotroph\| | Yeast | NULL | Probable | As nectar-tap_saprotroph (Põlme et al. 2020); Decay sugar-rich substrates (Põlme et al. 2020) | | Põlme S, et al. 2020. Fungal Diversity 105:1-16 ((https://doi.org/10.1007/s13225-020-00466-2)) |
| Otu32155 | 1 | k__Fungi(100);p__Ascomycota(100);c__Saccharomycetes(100);o__Saccharomycetales(100);f__Metschnikowiaceae(100);g__Metschnikowia(100);s__Metschnikowia_pulcherrima(100); | Metschnikowia | 13 | Saprotroph | \|Undefined Saprotroph\| | Yeast | NULL | Probable | As nectar-tap_saprotroph (Põlme et al. 2020); Decay sugar-rich substrates (Põlme et al. 2020) | | Põlme S, et al. 2020. Fungal Diversity 105:1-16 ((https://doi.org/10.1007/s13225-020-00466-2)) |
| Otu32156 | 1 | k__Fungi(100);p__Ascomycota(100);c__Saccharomycetes(100);o__Saccharomycetales(100);f__Metschnikowiaceae(100);g__Metschnikowia(100);g__Metschnikowia_unclassified(100); | Metschnikowia | 13 | Saprotroph | \|Undefined Saprotroph\| | Yeast | NULL | Probable | As nectar-tap_saprotroph (Põlme et al. 2020); Decay sugar-rich substrates (Põlme et al. 2020) | | Põlme S, et al. 2020. Fungal Diversity 105:1-16 ((https://doi.org/10.1007/s13225-020-00466-2)) |
| Otu32157 | 1 | k__Fungi(100);p__Ascomycota(100);c__Saccharomycetes(100);o__Saccharomycetales(100);f__Metschnikowiaceae(100);g__Metschnikowia(100);s__Metschnikowia_pulcherrima(100); | Metschnikowia | 13 | Saprotroph | \|Undefined Saprotroph\| | Yeast | NULL | Probable | As nectar-tap_saprotroph (Põlme et al. 2020); Decay sugar-rich substrates (Põlme et al. 2020) | | Põlme S, et al. 2020. Fungal Diversity 105:1-16 ((https://doi.org/10.1007/s13225-020-00466-2)) |
| Otu32158 | 1 | k__Fungi(100);p__Ascomycota(100);c__Saccharomycetes(100);o__Saccharomycetales(100);f__Metschnikowiaceae(100);g__Metschnikowia(100);s__Metschnikowia_pulcherrima(100); | Metschnikowia | 13 | Saprotroph | \|Undefined Saprotroph\| | Yeast | NULL | Probable | As nectar-tap_saprotroph (Põlme et al. 2020); Decay sugar-rich substrates (Põlme et al. 2020) | | Põlme S, et al. 2020. Fungal Diversity 105:1-16 ((https://doi.org/10.1007/s13225-020-00466-2)) |
| Otu32159 | 1 | k__Fungi(100);p__Ascomycota(100);c__Saccharomycetes(100);o__Saccharomycetales(100);f__Metschnikowiaceae(100);g__Metschnikowia(100);s__Metschnikowia_pulcherrima(100); | Metschnikowia | 13 | Saprotroph | \|Undefined Saprotroph\| | Yeast | NULL | Probable | As nectar-tap_saprotroph (Põlme et al. 2020); Decay sugar-rich substrates (Põlme et al. 2020) | | Põlme S, et al. 2020. Fungal Diversity 105:1-16 ((https://doi.org/10.1007/s13225-020-00466-2)) |
| Otu32160 | 1 | k__Fungi(100);p__Ascomycota(100);c__Saccharomycetes(100);o__Saccharomycetales(100);f__Metschnikowiaceae(100);g__Metschnikowia(100);s__Metschnikowia_pulcherrima(100); | Metschnikowia | 13 | Saprotroph | \|Undefined Saprotroph\| | Yeast | NULL | Probable | As nectar-tap_saprotroph (Põlme et al. 2020); Decay sugar-rich substrates (Põlme et al. 2020) | | Põlme S, et al. 2020. Fungal Diversity 105:1-16 ((https://doi.org/10.1007/s13225-020-00466-2)) |
| Otu32161 | 1 | k__Fungi(100);p__Ascomycota(100);c__Saccharomycetes(100);o__Saccharomycetales(100);f__Metschnikowiaceae(100);g__Metschnikowia(100);g__Metschnikowia_unclassified(100); | Metschnikowia | 13 | Saprotroph | \|Undefined Saprotroph\| | Yeast | NULL | Probable | As nectar-tap_saprotroph (Põlme et al. 2020); Decay sugar-rich substrates (Põlme et al. 2020) | | Põlme S, et al. 2020. Fungal Diversity 105:1-16 ((https://doi.org/10.1007/s13225-020-00466-2)) |
| Otu32163 | 1 | k__Fungi(100);p__Ascomycota(100);c__Saccharomycetes(100);o__Saccharomycetales(100);f__Metschnikowiaceae(100);g__Metschnikowia(100);g__Metschnikowia_unclassified(100); | Metschnikowia | 13 | Saprotroph | \|Undefined Saprotroph\| | Yeast | NULL | Probable | As nectar-tap_saprotroph (Põlme et al. 2020); Decay sugar-rich substrates (Põlme et al. 2020) | | Põlme S, et al. 2020. Fungal Diversity 105:1-16 ((https://doi.org/10.1007/s13225-020-00466-2)) |
| Otu32164 | 1 | k__Fungi(100);p__Ascomycota(100);c__Saccharomycetes(100);o__Saccharomycetales(100);f__Metschnikowiaceae(100);g__Metschnikowia(100);s__Metschnikowia_pulcherrima(100); | Metschnikowia | 13 | Saprotroph | \|Undefined Saprotroph\| | Yeast | NULL | Probable | As nectar-tap_saprotroph (Põlme et al. 2020); Decay sugar-rich substrates (Põlme et al. 2020) | | Põlme S, et al. 2020. Fungal Diversity 105:1-16 ((https://doi.org/10.1007/s13225-020-00466-2)) |
| Otu32165 | 1 | k__Fungi(100);p__Ascomycota(100);c__Saccharomycetes(100);o__Saccharomycetales(100);f__Metschnikowiaceae(100);g__Metschnikowia(100);s__Metschnikowia_pulcherrima(100); | Metschnikowia | 13 | Saprotroph | \|Undefined Saprotroph\| | Yeast | NULL | Probable | As nectar-tap_saprotroph (Põlme et al. 2020); Decay sugar-rich substrates (Põlme et al. 2020) | | Põlme S, et al. 2020. Fungal Diversity 105:1-16 ((https://doi.org/10.1007/s13225-020-00466-2)) |
| Otu32166 | 1 | k__Fungi(100);p__Ascomycota(100);c__Saccharomycetes(100);o__Saccharomycetales(100);f__Metschnikowiaceae(100);g__Metschnikowia(100);s__Metschnikowia_pulcherrima(100); | Metschnikowia | 13 | Saprotroph | \|Undefined Saprotroph\| | Yeast | NULL | Probable | As nectar-tap_saprotroph (Põlme et al. 2020); Decay sugar-rich substrates (Põlme et al. 2020) | | Põlme S, et al. 2020. Fungal Diversity 105:1-16 ((https://doi.org/10.1007/s13225-020-00466-2)) |
| Otu32167 | 1 | k__Fungi(100);p__Ascomycota(100);c__Saccharomycetes(100);o__Saccharomycetales(100);f__Metschnikowiaceae(100);g__Metschnikowia(100);s__Metschnikowia_pulcherrima(100); | Metschnikowia | 13 | Saprotroph | \|Undefined Saprotroph\| | Yeast | NULL | Probable | As nectar-tap_saprotroph (Põlme et al. 2020); Decay sugar-rich substrates (Põlme et al. 2020) | | Põlme S, et al. 2020. Fungal Diversity 105:1-16 ((https://doi.org/10.1007/s13225-020-00466-2)) |
| Otu32168 | 1 | k__Fungi(100);p__Ascomycota(100);c__Saccharomycetes(100);o__Saccharomycetales(100);f__Metschnikowiaceae(100);g__Metschnikowia(100);s__Metschnikowia_pulcherrima(100); | Metschnikowia | 13 | Saprotroph | \|Undefined Saprotroph\| | Yeast | NULL | Probable | As nectar-tap_saprotroph (Põlme et al. 2020); Decay sugar-rich substrates (Põlme et al. 2020) | | Põlme S, et al. 2020. Fungal Diversity 105:1-16 ((https://doi.org/10.1007/s13225-020-00466-2)) |
| Otu32169 | 1 | k__Fungi(100);p__Ascomycota(100);c__Saccharomycetes(100);o__Saccharomycetales(100);f__Metschnikowiaceae(100);g__Metschnikowia(100);s__Metschnikowia_pulcherrima(100); | Metschnikowia | 13 | Saprotroph | \|Undefined Saprotroph\| | Yeast | NULL | Probable | As nectar-tap_saprotroph (Põlme et al. 2020); Decay sugar-rich substrates (Põlme et al. 2020) | | Põlme S, et al. 2020. Fungal Diversity 105:1-16 ((https://doi.org/10.1007/s13225-020-00466-2)) |
| Otu32170 | 1 | k__Fungi(100);p__Ascomycota(100);c__Saccharomycetes(100);o__Saccharomycetales(100);f__Metschnikowiaceae(100);g__Metschnikowia(100);s__Metschnikowia_pulcherrima(100); | Metschnikowia | 13 | Saprotroph | \|Undefined Saprotroph\| | Yeast | NULL | Probable | As nectar-tap_saprotroph (Põlme et al. 2020); Decay sugar-rich substrates (Põlme et al. 2020) | | Põlme S, et al. 2020. Fungal Diversity 105:1-16 ((https://doi.org/10.1007/s13225-020-00466-2)) |
| Otu32171 | 1 | k__Fungi(100);p__Ascomycota(100);c__Saccharomycetes(100);o__Saccharomycetales(100);f__Saccharomycetales_family_Incertae_sedis(100);f__Saccharomycetales_family_Incertae_sedis_unclassified(100);f__Saccharomycetales_family_Incertae_sedis_unclassified(100); | Saccharomycetales | 7 | Saprotroph | Undefined Saprotroph | Yeast | NULL | Possible | NULL | | Sterkenburg E, et al. 2015. New Phytologist 207:1145-1158 ((https://doi.org/10.1111/nph.13426)) |
| Otu32172 | 1 | k__Fungi(100);p__Ascomycota(100);c__Saccharomycetes(100);o__Saccharomycetales(100);f__Metschnikowiaceae(100);g__Metschnikowia(100);g__Metschnikowia_unclassified(100); | Metschnikowia | 13 | Saprotroph | \|Undefined Saprotroph\| | Yeast | NULL | Probable | As nectar-tap_saprotroph (Põlme et al. 2020); Decay sugar-rich substrates (Põlme et al. 2020) | | Põlme S, et al. 2020. Fungal Diversity 105:1-16 ((https://doi.org/10.1007/s13225-020-00466-2)) |
| Otu32173 | 1 | k__Fungi(100);p__Ascomycota(100);c__Saccharomycetes(100);o__Saccharomycetales(100);f__Metschnikowiaceae(100);g__Metschnikowia(100);s__Metschnikowia_pulcherrima(100); | Metschnikowia | 13 | Saprotroph | \|Undefined Saprotroph\| | Yeast | NULL | Probable | As nectar-tap_saprotroph (Põlme et al. 2020); Decay sugar-rich substrates (Põlme et al. 2020) | | Põlme S, et al. 2020. Fungal Diversity 105:1-16 ((https://doi.org/10.1007/s13225-020-00466-2)) |
| Otu32174 | 1 | k__Fungi(100);p__Ascomycota(100);c__Saccharomycetes(100);o__Saccharomycetales(100);f__Metschnikowiaceae(100);g__Metschnikowia(100);s__Metschnikowia_pulcherrima(100); | Metschnikowia | 13 | Saprotroph | \|Undefined Saprotroph\| | Yeast | NULL | Probable | As nectar-tap_saprotroph (Põlme et al. 2020); Decay sugar-rich substrates (Põlme et al. 2020) | | Põlme S, et al. 2020. Fungal Diversity 105:1-16 ((https://doi.org/10.1007/s13225-020-00466-2)) |
| Otu32175 | 1 | k__Fungi(100);p__Ascomycota(100);c__Saccharomycetes(100);o__Saccharomycetales(100);f__Metschnikowiaceae(100);g__Metschnikowia(100);s__Metschnikowia_pulcherrima(100); | Metschnikowia | 13 | Saprotroph | \|Undefined Saprotroph\| | Yeast | NULL | Probable | As nectar-tap_saprotroph (Põlme et al. 2020); Decay sugar-rich substrates (Põlme et al. 2020) | | Põlme S, et al. 2020. Fungal Diversity 105:1-16 ((https://doi.org/10.1007/s13225-020-00466-2)) |
| Otu32177 | 1 | k__Fungi(100);p__Ascomycota(100);c__Saccharomycetes(100);o__Saccharomycetales(100);f__Metschnikowiaceae(100);g__Metschnikowia(100);s__Metschnikowia_pulcherrima(100); | Metschnikowia | 13 | Saprotroph | \|Undefined Saprotroph\| | Yeast | NULL | Probable | As nectar-tap_saprotroph (Põlme et al. 2020); Decay sugar-rich substrates (Põlme et al. 2020) | | Põlme S, et al. 2020. Fungal Diversity 105:1-16 ((https://doi.org/10.1007/s13225-020-00466-2)) |
| Otu32178 | 1 | k__Fungi(100);p__Ascomycota(100);c__Saccharomycetes(100);o__Saccharomycetales(100);f__Metschnikowiaceae(100);g__Metschnikowia(100);g__Metschnikowia_unclassified(100); | Metschnikowia | 13 | Saprotroph | \|Undefined Saprotroph\| | Yeast | NULL | Probable | As nectar-tap_saprotroph (Põlme et al. 2020); Decay sugar-rich substrates (Põlme et al. 2020) | | Põlme S, et al. 2020. Fungal Diversity 105:1-16 ((https://doi.org/10.1007/s13225-020-00466-2)) |
| Otu32179 | 1 | k__Fungi(100);p__Ascomycota(100);c__Saccharomycetes(100);o__Saccharomycetales(100);f__Metschnikowiaceae(100);g__Metschnikowia(100);s__Metschnikowia_pulcherrima(100); | Metschnikowia | 13 | Saprotroph | \|Undefined Saprotroph\| | Yeast | NULL | Probable | As nectar-tap_saprotroph (Põlme et al. 2020); Decay sugar-rich substrates (Põlme et al. 2020) | | Põlme S, et al. 2020. Fungal Diversity 105:1-16 ((https://doi.org/10.1007/s13225-020-00466-2)) |
| Otu32180 | 1 | k__Fungi(100);p__Ascomycota(100);c__Saccharomycetes(100);o__Saccharomycetales(100);f__Metschnikowiaceae(100);g__Metschnikowia(100);s__Metschnikowia_pulcherrima(100); | Metschnikowia | 13 | Saprotroph | \|Undefined Saprotroph\| | Yeast | NULL | Probable | As nectar-tap_saprotroph (Põlme et al. 2020); Decay sugar-rich substrates (Põlme et al. 2020) | | Põlme S, et al. 2020. Fungal Diversity 105:1-16 ((https://doi.org/10.1007/s13225-020-00466-2)) |
| Otu32181 | 1 | k__Fungi(100);p__Ascomycota(100);c__Saccharomycetes(100);o__Saccharomycetales(100);f__Metschnikowiaceae(100);g__Metschnikowia(100);s__Metschnikowia_pulcherrima(100); | Metschnikowia | 13 | Saprotroph | \|Undefined Saprotroph\| | Yeast | NULL | Probable | As nectar-tap_saprotroph (Põlme et al. 2020); Decay sugar-rich substrates (Põlme et al. 2020) | | Põlme S, et al. 2020. Fungal Diversity 105:1-16 ((https://doi.org/10.1007/s13225-020-00466-2)) |
| Otu32184 | 1 | k__Fungi(100);p__Ascomycota(100);c__Saccharomycetes(100);o__Saccharomycetales(100);f__Metschnikowiaceae(100);g__Metschnikowia(100);g__Metschnikowia_unclassified(100); | Metschnikowia | 13 | Saprotroph | \|Undefined Saprotroph\| | Yeast | NULL | Probable | As nectar-tap_saprotroph (Põlme et al. 2020); Decay sugar-rich substrates (Põlme et al. 2020) | | Põlme S, et al. 2020. Fungal Diversity 105:1-16 ((https://doi.org/10.1007/s13225-020-00466-2)) |
| Otu32185 | 1 | k__Fungi(100);p__Ascomycota(100);c__Saccharomycetes(100);o__Saccharomycetales(100);f__Metschnikowiaceae(100);g__Metschnikowia(100);s__Metschnikowia_pulcherrima(100); | Metschnikowia | 13 | Saprotroph | \|Undefined Saprotroph\| | Yeast | NULL | Probable | As nectar-tap_saprotroph (Põlme et al. 2020); Decay sugar-rich substrates (Põlme et al. 2020) | | Põlme S, et al. 2020. Fungal Diversity 105:1-16 ((https://doi.org/10.1007/s13225-020-00466-2)) |
| Otu32186 | 1 | k__Fungi(100);p__Ascomycota(100);c__Saccharomycetes(100);o__Saccharomycetales(100);f__Metschnikowiaceae(100);g__Metschnikowia(100);s__Metschnikowia_pulcherrima(100); | Metschnikowia | 13 | Saprotroph | \|Undefined Saprotroph\| | Yeast | NULL | Probable | As nectar-tap_saprotroph (Põlme et al. 2020); Decay sugar-rich substrates (Põlme et al. 2020) | | Põlme S, et al. 2020. Fungal Diversity 105:1-16 ((https://doi.org/10.1007/s13225-020-00466-2)) |
| Otu32187 | 1 | k__Fungi(100);p__Ascomycota(100);c__Saccharomycetes(100);o__Saccharomycetales(100);f__Metschnikowiaceae(100);g__Metschnikowia(100);s__Metschnikowia_pulcherrima(100); | Metschnikowia | 13 | Saprotroph | \|Undefined Saprotroph\| | Yeast | NULL | Probable | As nectar-tap_saprotroph (Põlme et al. 2020); Decay sugar-rich substrates (Põlme et al. 2020) | | Põlme S, et al. 2020. Fungal Diversity 105:1-16 ((https://doi.org/10.1007/s13225-020-00466-2)) |
| Otu32188 | 1 | k__Fungi(100);p__Ascomycota(100);c__Saccharomycetes(100);o__Saccharomycetales(100);f__Metschnikowiaceae(100);g__Metschnikowia(100);s__Metschnikowia_pulcherrima(100); | Metschnikowia | 13 | Saprotroph | \|Undefined Saprotroph\| | Yeast | NULL | Probable | As nectar-tap_saprotroph (Põlme et al. 2020); Decay sugar-rich substrates (Põlme et al. 2020) | | Põlme S, et al. 2020. Fungal Diversity 105:1-16 ((https://doi.org/10.1007/s13225-020-00466-2)) |
| Otu32190 | 1 | k__Fungi(100);p__Ascomycota(100);c__Saccharomycetes(100);o__Saccharomycetales(100);f__Metschnikowiaceae(100);g__Metschnikowia(100);s__Metschnikowia_pulcherrima(100); | Metschnikowia | 13 | Saprotroph | \|Undefined Saprotroph\| | Yeast | NULL | Probable | As nectar-tap_saprotroph (Põlme et al. 2020); Decay sugar-rich substrates (Põlme et al. 2020) | | Põlme S, et al. 2020. Fungal Diversity 105:1-16 ((https://doi.org/10.1007/s13225-020-00466-2)) |
| Otu32191 | 1 | k__Fungi(100);p__Ascomycota(100);c__Saccharomycetes(100);o__Saccharomycetales(100);f__Metschnikowiaceae(100);g__Metschnikowia(100);s__Metschnikowia_pulcherrima(100); | Metschnikowia | 13 | Saprotroph | \|Undefined Saprotroph\| | Yeast | NULL | Probable | As nectar-tap_saprotroph (Põlme et al. 2020); Decay sugar-rich substrates (Põlme et al. 2020) | | Põlme S, et al. 2020. Fungal Diversity 105:1-16 ((https://doi.org/10.1007/s13225-020-00466-2)) |
| Otu32192 | 1 | k__Fungi(100);p__Ascomycota(100);c__Saccharomycetes(100);o__Saccharomycetales(100);f__Metschnikowiaceae(100);g__Metschnikowia(100);s__Metschnikowia_pulcherrima(100); | Metschnikowia | 13 | Saprotroph | \|Undefined Saprotroph\| | Yeast | NULL | Probable | As nectar-tap_saprotroph (Põlme et al. 2020); Decay sugar-rich substrates (Põlme et al. 2020) | | Põlme S, et al. 2020. Fungal Diversity 105:1-16 ((https://doi.org/10.1007/s13225-020-00466-2)) |
| Otu32193 | 1 | k__Fungi(100);p__Ascomycota(100);c__Saccharomycetes(100);o__Saccharomycetales(100);f__Metschnikowiaceae(100);g__Metschnikowia(100);s__Metschnikowia_pulcherrima(100); | Metschnikowia | 13 | Saprotroph | \|Undefined Saprotroph\| | Yeast | NULL | Probable | As nectar-tap_saprotroph (Põlme et al. 2020); Decay sugar-rich substrates (Põlme et al. 2020) | | Põlme S, et al. 2020. Fungal Diversity 105:1-16 ((https://doi.org/10.1007/s13225-020-00466-2)) |
| Otu32194 | 1 | k__Fungi(100);p__Ascomycota(100);c__Saccharomycetes(100);o__Saccharomycetales(100);f__Metschnikowiaceae(100);g__Metschnikowia(100);s__Metschnikowia_pulcherrima(100); | Metschnikowia | 13 | Saprotroph | \|Undefined Saprotroph\| | Yeast | NULL | Probable | As nectar-tap_saprotroph (Põlme et al. 2020); Decay sugar-rich substrates (Põlme et al. 2020) | | Põlme S, et al. 2020. Fungal Diversity 105:1-16 ((https://doi.org/10.1007/s13225-020-00466-2)) |
| Otu32195 | 1 | k__Fungi(100);p__Ascomycota(100);c__Saccharomycetes(100);o__Saccharomycetales(100);f__Metschnikowiaceae(100);g__Metschnikowia(100);s__Metschnikowia_pulcherrima(100); | Metschnikowia | 13 | Saprotroph | \|Undefined Saprotroph\| | Yeast | NULL | Probable | As nectar-tap_saprotroph (Põlme et al. 2020); Decay sugar-rich substrates (Põlme et al. 2020) | | Põlme S, et al. 2020. Fungal Diversity 105:1-16 ((https://doi.org/10.1007/s13225-020-00466-2)) |
| Otu32196 | 1 | k__Fungi(100);p__Ascomycota(100);c__Saccharomycetes(100);o__Saccharomycetales(100);f__Metschnikowiaceae(100);g__Metschnikowia(100);g__Metschnikowia_unclassified(100); | Metschnikowia | 13 | Saprotroph | \|Undefined Saprotroph\| | Yeast | NULL | Probable | As nectar-tap_saprotroph (Põlme et al. 2020); Decay sugar-rich substrates (Põlme et al. 2020) | | Põlme S, et al. 2020. Fungal Diversity 105:1-16 ((https://doi.org/10.1007/s13225-020-00466-2)) |
| Otu32197 | 1 | k__Fungi(100);p__Ascomycota(100);c__Saccharomycetes(100);o__Saccharomycetales(100);f__Metschnikowiaceae(100);g__Metschnikowia(100);g__Metschnikowia_unclassified(100); | Metschnikowia | 13 | Saprotroph | \|Undefined Saprotroph\| | Yeast | NULL | Probable | As nectar-tap_saprotroph (Põlme et al. 2020); Decay sugar-rich substrates (Põlme et al. 2020) | | Põlme S, et al. 2020. Fungal Diversity 105:1-16 ((https://doi.org/10.1007/s13225-020-00466-2)) |
| Otu32198 | 1 | k__Fungi(100);p__Ascomycota(100);c__Dothideomycetes(100);o__Capnodiales(100);f__Davidiellaceae(100);g__Cladosporium(100);g__Cladosporium_unclassified(100); | Cladosporium | 13 | Pathotroph-Saprotroph-Symbiotroph | Animal Pathogen-Endophyte-Lichen Parasite-Plant Pathogen-\|Plant Saprotroph\|-Wood Saprotroph | Microfungus | NULL | Possible | As foliar_endophyte (Põlme et al. 2020); As litter_saprotroph (Põlme et al. 2020); Decay wood, leaf, fruit, seed, soil, dung, animal material (Põlme et al. 2020) | | Seehann G, et al. 1975. List of Fungi in Soft-Rot Tissue Tests. Institute fur Holzbiologie und Holzschutz, Bundesforschungsanstalt fur Forst - und Holzwirtshaft, Hamburg - Lohbrugge, Germany; Lawrey JD, Diederich P. 2016. http://www.lichenicolous.net (accessed 24 May 2017) ((http://www.lichenicolous.net)); Põlme S, et al. 2020. Fungal Diversity 105:1-16 ((https://doi.org/10.1007/s13225-020-00466-2)); The American Phytopathological Society Common Names of Plant Diseases ((http://www.apsnet.org/publications/commonnames/Pages/default.aspx)) |
| Otu32199 | 1 | k__Fungi(100);p__Ascomycota(100);c__Saccharomycetes(100);o__Saccharomycetales(100);f__Metschnikowiaceae(100);g__Metschnikowia(100);s__Metschnikowia_pulcherrima(100); | Metschnikowia | 13 | Saprotroph | \|Undefined Saprotroph\| | Yeast | NULL | Probable | As nectar-tap_saprotroph (Põlme et al. 2020); Decay sugar-rich substrates (Põlme et al. 2020) | | Põlme S, et al. 2020. Fungal Diversity 105:1-16 ((https://doi.org/10.1007/s13225-020-00466-2)) |
| Otu32200 | 1 | k__Fungi(100);p__Ascomycota(100);c__Saccharomycetes(100);o__Saccharomycetales(100);f__Metschnikowiaceae(100);g__Metschnikowia(100);s__Metschnikowia_pulcherrima(100); | Metschnikowia | 13 | Saprotroph | \|Undefined Saprotroph\| | Yeast | NULL | Probable | As nectar-tap_saprotroph (Põlme et al. 2020); Decay sugar-rich substrates (Põlme et al. 2020) | | Põlme S, et al. 2020. Fungal Diversity 105:1-16 ((https://doi.org/10.1007/s13225-020-00466-2)) |
| Otu32201 | 1 | k__Fungi(100);p__Ascomycota(100);c__Saccharomycetes(100);o__Saccharomycetales(100);f__Metschnikowiaceae(100);g__Metschnikowia(100);s__Metschnikowia_pulcherrima(100); | Metschnikowia | 13 | Saprotroph | \|Undefined Saprotroph\| | Yeast | NULL | Probable | As nectar-tap_saprotroph (Põlme et al. 2020); Decay sugar-rich substrates (Põlme et al. 2020) | | Põlme S, et al. 2020. Fungal Diversity 105:1-16 ((https://doi.org/10.1007/s13225-020-00466-2)) |
| Otu32202 | 1 | k__Fungi(100);p__Ascomycota(100);c__Saccharomycetes(100);o__Saccharomycetales(100);f__Metschnikowiaceae(100);g__Metschnikowia(100);s__Metschnikowia_pulcherrima(100); | Metschnikowia | 13 | Saprotroph | \|Undefined Saprotroph\| | Yeast | NULL | Probable | As nectar-tap_saprotroph (Põlme et al. 2020); Decay sugar-rich substrates (Põlme et al. 2020) | | Põlme S, et al. 2020. Fungal Diversity 105:1-16 ((https://doi.org/10.1007/s13225-020-00466-2)) |
| Otu32203 | 1 | k__Fungi(100);p__Ascomycota(100);c__Saccharomycetes(100);o__Saccharomycetales(100);f__Metschnikowiaceae(100);g__Metschnikowia(100);s__Metschnikowia_pulcherrima(100); | Metschnikowia | 13 | Saprotroph | \|Undefined Saprotroph\| | Yeast | NULL | Probable | As nectar-tap_saprotroph (Põlme et al. 2020); Decay sugar-rich substrates (Põlme et al. 2020) | | Põlme S, et al. 2020. Fungal Diversity 105:1-16 ((https://doi.org/10.1007/s13225-020-00466-2)) |
| Otu32204 | 1 | k__Fungi(100);p__Ascomycota(100);c__Saccharomycetes(100);o__Saccharomycetales(100);f__Metschnikowiaceae(100);g__Metschnikowia(100);s__Metschnikowia_pulcherrima(100); | Metschnikowia | 13 | Saprotroph | \|Undefined Saprotroph\| | Yeast | NULL | Probable | As nectar-tap_saprotroph (Põlme et al. 2020); Decay sugar-rich substrates (Põlme et al. 2020) | | Põlme S, et al. 2020. Fungal Diversity 105:1-16 ((https://doi.org/10.1007/s13225-020-00466-2)) |
| Otu32205 | 1 | k__Fungi(100);p__Ascomycota(100);c__Saccharomycetes(100);o__Saccharomycetales(100);f__Metschnikowiaceae(100);g__Metschnikowia(100);s__Metschnikowia_pulcherrima(100); | Metschnikowia | 13 | Saprotroph | \|Undefined Saprotroph\| | Yeast | NULL | Probable | As nectar-tap_saprotroph (Põlme et al. 2020); Decay sugar-rich substrates (Põlme et al. 2020) | | Põlme S, et al. 2020. Fungal Diversity 105:1-16 ((https://doi.org/10.1007/s13225-020-00466-2)) |
| Otu32206 | 1 | k__Fungi(100);p__Ascomycota(100);c__Saccharomycetes(100);o__Saccharomycetales(100);f__Metschnikowiaceae(100);g__Metschnikowia(100);s__Metschnikowia_pulcherrima(100); | Metschnikowia | 13 | Saprotroph | \|Undefined Saprotroph\| | Yeast | NULL | Probable | As nectar-tap_saprotroph (Põlme et al. 2020); Decay sugar-rich substrates (Põlme et al. 2020) | | Põlme S, et al. 2020. Fungal Diversity 105:1-16 ((https://doi.org/10.1007/s13225-020-00466-2)) |
| Otu32210 | 1 | k__Fungi(100);p__Ascomycota(100);c__Saccharomycetes(100);o__Saccharomycetales(100);f__Metschnikowiaceae(100);g__Metschnikowia(100);g__Metschnikowia_unclassified(100); | Metschnikowia | 13 | Saprotroph | \|Undefined Saprotroph\| | Yeast | NULL | Probable | As nectar-tap_saprotroph (Põlme et al. 2020); Decay sugar-rich substrates (Põlme et al. 2020) | | Põlme S, et al. 2020. Fungal Diversity 105:1-16 ((https://doi.org/10.1007/s13225-020-00466-2)) |
| Otu32211 | 1 | k__Fungi(100);p__Ascomycota(100);c__Saccharomycetes(100);o__Saccharomycetales(100);f__Metschnikowiaceae(100);g__Metschnikowia(100);s__Metschnikowia_pulcherrima(100); | Metschnikowia | 13 | Saprotroph | \|Undefined Saprotroph\| | Yeast | NULL | Probable | As nectar-tap_saprotroph (Põlme et al. 2020); Decay sugar-rich substrates (Põlme et al. 2020) | | Põlme S, et al. 2020. Fungal Diversity 105:1-16 ((https://doi.org/10.1007/s13225-020-00466-2)) |
| Otu32221 | 1 | k__Fungi(100);p__Ascomycota(100);c__Saccharomycetes(100);o__Saccharomycetales(100);f__Saccharomycetales_family_Incertae_sedis(100);f__Saccharomycetales_family_Incertae_sedis_unclassified(100);f__Saccharomycetales_family_Incertae_sedis_unclassified(100); | Saccharomycetales | 7 | Saprotroph | Undefined Saprotroph | Yeast | NULL | Possible | NULL | | Sterkenburg E, et al. 2015. New Phytologist 207:1145-1158 ((https://doi.org/10.1111/nph.13426)) |
| Otu32234 | 1 | k__Fungi(100);p__Ascomycota(100);c__Saccharomycetes(100);o__Saccharomycetales(100);f__Metschnikowiaceae(100);g__Metschnikowia(100);s__Metschnikowia_pulcherrima(100); | Metschnikowia | 13 | Saprotroph | \|Undefined Saprotroph\| | Yeast | NULL | Probable | As nectar-tap_saprotroph (Põlme et al. 2020); Decay sugar-rich substrates (Põlme et al. 2020) | | Põlme S, et al. 2020. Fungal Diversity 105:1-16 ((https://doi.org/10.1007/s13225-020-00466-2)) |
| Otu32235 | 1 | k__Fungi(100);p__Ascomycota(100);c__Saccharomycetes(100);o__Saccharomycetales(100);f__Metschnikowiaceae(100);g__Metschnikowia(100);s__Metschnikowia_pulcherrima(100); | Metschnikowia | 13 | Saprotroph | \|Undefined Saprotroph\| | Yeast | NULL | Probable | As nectar-tap_saprotroph (Põlme et al. 2020); Decay sugar-rich substrates (Põlme et al. 2020) | | Põlme S, et al. 2020. Fungal Diversity 105:1-16 ((https://doi.org/10.1007/s13225-020-00466-2)) |
| Otu32236 | 1 | k__Fungi(100);p__Ascomycota(100);c__Saccharomycetes(100);o__Saccharomycetales(100);f__Metschnikowiaceae(100);g__Metschnikowia(100);s__Metschnikowia_pulcherrima(100); | Metschnikowia | 13 | Saprotroph | \|Undefined Saprotroph\| | Yeast | NULL | Probable | As nectar-tap_saprotroph (Põlme et al. 2020); Decay sugar-rich substrates (Põlme et al. 2020) | | Põlme S, et al. 2020. Fungal Diversity 105:1-16 ((https://doi.org/10.1007/s13225-020-00466-2)) |
| Otu32238 | 1 | k__Fungi(100);p__Ascomycota(100);c__Saccharomycetes(100);o__Saccharomycetales(100);f__Metschnikowiaceae(100);g__Metschnikowia(100);s__Metschnikowia_pulcherrima(100); | Metschnikowia | 13 | Saprotroph | \|Undefined Saprotroph\| | Yeast | NULL | Probable | As nectar-tap_saprotroph (Põlme et al. 2020); Decay sugar-rich substrates (Põlme et al. 2020) | | Põlme S, et al. 2020. Fungal Diversity 105:1-16 ((https://doi.org/10.1007/s13225-020-00466-2)) |
| Otu32239 | 1 | k__Fungi(100);p__Ascomycota(100);c__Saccharomycetes(100);o__Saccharomycetales(100);f__Metschnikowiaceae(100);g__Metschnikowia(100);s__Metschnikowia_pulcherrima(100); | Metschnikowia | 13 | Saprotroph | \|Undefined Saprotroph\| | Yeast | NULL | Probable | As nectar-tap_saprotroph (Põlme et al. 2020); Decay sugar-rich substrates (Põlme et al. 2020) | | Põlme S, et al. 2020. Fungal Diversity 105:1-16 ((https://doi.org/10.1007/s13225-020-00466-2)) |
| Otu32248 | 1 | k__Fungi(100);p__Ascomycota(100);c__Saccharomycetes(100);o__Saccharomycetales(100);f__Metschnikowiaceae(100);g__Metschnikowia(100);s__Metschnikowia_pulcherrima(100); | Metschnikowia | 13 | Saprotroph | \|Undefined Saprotroph\| | Yeast | NULL | Probable | As nectar-tap_saprotroph (Põlme et al. 2020); Decay sugar-rich substrates (Põlme et al. 2020) | | Põlme S, et al. 2020. Fungal Diversity 105:1-16 ((https://doi.org/10.1007/s13225-020-00466-2)) |
| Otu32259 | 1 | k__Fungi(100);p__Ascomycota(100);c__Saccharomycetes(100);o__Saccharomycetales(100);f__Metschnikowiaceae(100);g__Metschnikowia(100);s__Metschnikowia_pulcherrima(100); | Metschnikowia | 13 | Saprotroph | \|Undefined Saprotroph\| | Yeast | NULL | Probable | As nectar-tap_saprotroph (Põlme et al. 2020); Decay sugar-rich substrates (Põlme et al. 2020) | | Põlme S, et al. 2020. Fungal Diversity 105:1-16 ((https://doi.org/10.1007/s13225-020-00466-2)) |
| Otu32261 | 1 | k__Fungi(100);p__Ascomycota(100);c__Saccharomycetes(100);o__Saccharomycetales(100);f__Metschnikowiaceae(100);g__Metschnikowia(100);g__Metschnikowia_unclassified(100); | Metschnikowia | 13 | Saprotroph | \|Undefined Saprotroph\| | Yeast | NULL | Probable | As nectar-tap_saprotroph (Põlme et al. 2020); Decay sugar-rich substrates (Põlme et al. 2020) | | Põlme S, et al. 2020. Fungal Diversity 105:1-16 ((https://doi.org/10.1007/s13225-020-00466-2)) |
| Otu32264 | 1 | k__Fungi(100);p__Ascomycota(100);c__Saccharomycetes(100);o__Saccharomycetales(100);f__Saccharomycetales_family_Incertae_sedis(100);g__Candida(100);g__Candida_unclassified(100); | Candida | 13 | Pathotroph-Saprotroph-Symbiotroph | Animal Pathogen-Endophyte-Endosymbiont-Epiphyte-\|Undefined Saprotroph\| | Dimorphic Yeast-Yeast | NULL | Probable | As nectar-tap_saprotroph (Põlme et al. 2020); Decay sugar-rich substrates (Põlme et al. 2020); This asexual genus is polyphyletic and contain species that function very differently depend on what they are. Caution should be taken when interpreting guilds for Candida species. | | Manolakaki D, et al. 2010. Virulence. 1: 367-375 ((https://doi.org/10.4161/viru.1.5.12796)); Mekbib SB, et al. 2011. Tropical Plant Pathology 36:233-240 ((https://doi.org/10.1590/S1982-56762011000400004)); Põlme S, et al. 2020. Fungal Diversity 105:1-16 ((https://doi.org/10.1007/s13225-020-00466-2)) |
| Otu32267 | 1 | k__Fungi(100);p__Ascomycota(100);c__Saccharomycetes(100);o__Saccharomycetales(100);f__Saccharomycetales_family_Incertae_sedis(100);g__Candida(100);g__Candida_unclassified(100); | Candida | 13 | Pathotroph-Saprotroph-Symbiotroph | Animal Pathogen-Endophyte-Endosymbiont-Epiphyte-\|Undefined Saprotroph\| | Dimorphic Yeast-Yeast | NULL | Probable | As nectar-tap_saprotroph (Põlme et al. 2020); Decay sugar-rich substrates (Põlme et al. 2020); This asexual genus is polyphyletic and contain species that function very differently depend on what they are. Caution should be taken when interpreting guilds for Candida species. | | Manolakaki D, et al. 2010. Virulence. 1: 367-375 ((https://doi.org/10.4161/viru.1.5.12796)); Mekbib SB, et al. 2011. Tropical Plant Pathology 36:233-240 ((https://doi.org/10.1590/S1982-56762011000400004)); Põlme S, et al. 2020. Fungal Diversity 105:1-16 ((https://doi.org/10.1007/s13225-020-00466-2)) |
| Otu32270 | 1 | k__Fungi(100);p__Ascomycota(100);c__Saccharomycetes(100);o__Saccharomycetales(100);f__Saccharomycetales_family_Incertae_sedis(100);g__Candida(100);g__Candida_unclassified(100); | Candida | 13 | Pathotroph-Saprotroph-Symbiotroph | Animal Pathogen-Endophyte-Endosymbiont-Epiphyte-\|Undefined Saprotroph\| | Dimorphic Yeast-Yeast | NULL | Probable | As nectar-tap_saprotroph (Põlme et al. 2020); Decay sugar-rich substrates (Põlme et al. 2020); This asexual genus is polyphyletic and contain species that function very differently depend on what they are. Caution should be taken when interpreting guilds for Candida species. | | Manolakaki D, et al. 2010. Virulence. 1: 367-375 ((https://doi.org/10.4161/viru.1.5.12796)); Mekbib SB, et al. 2011. Tropical Plant Pathology 36:233-240 ((https://doi.org/10.1590/S1982-56762011000400004)); Põlme S, et al. 2020. Fungal Diversity 105:1-16 ((https://doi.org/10.1007/s13225-020-00466-2)) |
| Otu32272 | 1 | k__Fungi(100);p__Ascomycota(100);c__Saccharomycetes(100);o__Saccharomycetales(100);f__Metschnikowiaceae(100);g__Metschnikowia(100);s__Metschnikowia_pulcherrima(100); | Metschnikowia | 13 | Saprotroph | \|Undefined Saprotroph\| | Yeast | NULL | Probable | As nectar-tap_saprotroph (Põlme et al. 2020); Decay sugar-rich substrates (Põlme et al. 2020) | | Põlme S, et al. 2020. Fungal Diversity 105:1-16 ((https://doi.org/10.1007/s13225-020-00466-2)) |
| Otu32273 | 1 | k__Fungi(100);p__Ascomycota(100);c__Saccharomycetes(100);o__Saccharomycetales(100);f__Saccharomycetales_family_Incertae_sedis(100);g__Candida(100);g__Candida_unclassified(100); | Candida | 13 | Pathotroph-Saprotroph-Symbiotroph | Animal Pathogen-Endophyte-Endosymbiont-Epiphyte-\|Undefined Saprotroph\| | Dimorphic Yeast-Yeast | NULL | Probable | As nectar-tap_saprotroph (Põlme et al. 2020); Decay sugar-rich substrates (Põlme et al. 2020); This asexual genus is polyphyletic and contain species that function very differently depend on what they are. Caution should be taken when interpreting guilds for Candida species. | | Manolakaki D, et al. 2010. Virulence. 1: 367-375 ((https://doi.org/10.4161/viru.1.5.12796)); Mekbib SB, et al. 2011. Tropical Plant Pathology 36:233-240 ((https://doi.org/10.1590/S1982-56762011000400004)); Põlme S, et al. 2020. Fungal Diversity 105:1-16 ((https://doi.org/10.1007/s13225-020-00466-2)) |
| Otu32275 | 1 | k__Fungi(100);p__Ascomycota(100);c__Saccharomycetes(100);o__Saccharomycetales(100);f__Saccharomycetales_family_Incertae_sedis(100);g__Candida(100);g__Candida_unclassified(100); | Candida | 13 | Pathotroph-Saprotroph-Symbiotroph | Animal Pathogen-Endophyte-Endosymbiont-Epiphyte-\|Undefined Saprotroph\| | Dimorphic Yeast-Yeast | NULL | Probable | As nectar-tap_saprotroph (Põlme et al. 2020); Decay sugar-rich substrates (Põlme et al. 2020); This asexual genus is polyphyletic and contain species that function very differently depend on what they are. Caution should be taken when interpreting guilds for Candida species. | | Manolakaki D, et al. 2010. Virulence. 1: 367-375 ((https://doi.org/10.4161/viru.1.5.12796)); Mekbib SB, et al. 2011. Tropical Plant Pathology 36:233-240 ((https://doi.org/10.1590/S1982-56762011000400004)); Põlme S, et al. 2020. Fungal Diversity 105:1-16 ((https://doi.org/10.1007/s13225-020-00466-2)) |
| Otu32276 | 1 | k__Fungi(100);p__Ascomycota(100);c__Saccharomycetes(100);o__Saccharomycetales(100);o__Saccharomycetales_unclassified(100);o__Saccharomycetales_unclassified(100);o__Saccharomycetales_unclassified(100); | Saccharomycetales | 7 | Saprotroph | Undefined Saprotroph | Yeast | NULL | Possible | NULL | | Sterkenburg E, et al. 2015. New Phytologist 207:1145-1158 ((https://doi.org/10.1111/nph.13426)) |
| Otu32277 | 1 | k__Fungi(100);p__Ascomycota(100);c__Saccharomycetes(100);o__Saccharomycetales(100);f__Saccharomycetales_family_Incertae_sedis(100);g__Candida(100);g__Candida_unclassified(100); | Candida | 13 | Pathotroph-Saprotroph-Symbiotroph | Animal Pathogen-Endophyte-Endosymbiont-Epiphyte-\|Undefined Saprotroph\| | Dimorphic Yeast-Yeast | NULL | Probable | As nectar-tap_saprotroph (Põlme et al. 2020); Decay sugar-rich substrates (Põlme et al. 2020); This asexual genus is polyphyletic and contain species that function very differently depend on what they are. Caution should be taken when interpreting guilds for Candida species. | | Manolakaki D, et al. 2010. Virulence. 1: 367-375 ((https://doi.org/10.4161/viru.1.5.12796)); Mekbib SB, et al. 2011. Tropical Plant Pathology 36:233-240 ((https://doi.org/10.1590/S1982-56762011000400004)); Põlme S, et al. 2020. Fungal Diversity 105:1-16 ((https://doi.org/10.1007/s13225-020-00466-2)) |
| Otu32281 | 1 | k__Fungi(100);p__Ascomycota(100);c__Saccharomycetes(100);o__Saccharomycetales(100);f__Metschnikowiaceae(100);g__Metschnikowia(100);s__Metschnikowia_pulcherrima(100); | Metschnikowia | 13 | Saprotroph | \|Undefined Saprotroph\| | Yeast | NULL | Probable | As nectar-tap_saprotroph (Põlme et al. 2020); Decay sugar-rich substrates (Põlme et al. 2020) | | Põlme S, et al. 2020. Fungal Diversity 105:1-16 ((https://doi.org/10.1007/s13225-020-00466-2)) |
| Otu32282 | 1 | k__Fungi(100);p__Ascomycota(100);c__Saccharomycetes(100);o__Saccharomycetales(100);f__Metschnikowiaceae(100);g__Metschnikowia(100);g__Metschnikowia_unclassified(100); | Metschnikowia | 13 | Saprotroph | \|Undefined Saprotroph\| | Yeast | NULL | Probable | As nectar-tap_saprotroph (Põlme et al. 2020); Decay sugar-rich substrates (Põlme et al. 2020) | | Põlme S, et al. 2020. Fungal Diversity 105:1-16 ((https://doi.org/10.1007/s13225-020-00466-2)) |
| Otu32283 | 1 | k__Fungi(100);p__Ascomycota(100);c__Saccharomycetes(100);o__Saccharomycetales(100);f__Metschnikowiaceae(100);g__Metschnikowia(100);s__Metschnikowia_pulcherrima(100); | Metschnikowia | 13 | Saprotroph | \|Undefined Saprotroph\| | Yeast | NULL | Probable | As nectar-tap_saprotroph (Põlme et al. 2020); Decay sugar-rich substrates (Põlme et al. 2020) | | Põlme S, et al. 2020. Fungal Diversity 105:1-16 ((https://doi.org/10.1007/s13225-020-00466-2)) |
| Otu32284 | 1 | k__Fungi(100);p__Ascomycota(100);c__Saccharomycetes(100);o__Saccharomycetales(100);f__Metschnikowiaceae(100);g__Metschnikowia(100);s__Metschnikowia_pulcherrima(100); | Metschnikowia | 13 | Saprotroph | \|Undefined Saprotroph\| | Yeast | NULL | Probable | As nectar-tap_saprotroph (Põlme et al. 2020); Decay sugar-rich substrates (Põlme et al. 2020) | | Põlme S, et al. 2020. Fungal Diversity 105:1-16 ((https://doi.org/10.1007/s13225-020-00466-2)) |
| Otu32285 | 1 | k__Fungi(100);p__Ascomycota(100);c__Saccharomycetes(100);o__Saccharomycetales(100);f__Metschnikowiaceae(100);g__Metschnikowia(100);s__Metschnikowia_pulcherrima(100); | Metschnikowia | 13 | Saprotroph | \|Undefined Saprotroph\| | Yeast | NULL | Probable | As nectar-tap_saprotroph (Põlme et al. 2020); Decay sugar-rich substrates (Põlme et al. 2020) | | Põlme S, et al. 2020. Fungal Diversity 105:1-16 ((https://doi.org/10.1007/s13225-020-00466-2)) |
| Otu32286 | 1 | k__Fungi(100);p__Ascomycota(100);c__Saccharomycetes(100);o__Saccharomycetales(100);f__Saccharomycetales_family_Incertae_sedis(100);g__Candida(100);g__Candida_unclassified(100); | Candida | 13 | Pathotroph-Saprotroph-Symbiotroph | Animal Pathogen-Endophyte-Endosymbiont-Epiphyte-\|Undefined Saprotroph\| | Dimorphic Yeast-Yeast | NULL | Probable | As nectar-tap_saprotroph (Põlme et al. 2020); Decay sugar-rich substrates (Põlme et al. 2020); This asexual genus is polyphyletic and contain species that function very differently depend on what they are. Caution should be taken when interpreting guilds for Candida species. | | Manolakaki D, et al. 2010. Virulence. 1: 367-375 ((https://doi.org/10.4161/viru.1.5.12796)); Mekbib SB, et al. 2011. Tropical Plant Pathology 36:233-240 ((https://doi.org/10.1590/S1982-56762011000400004)); Põlme S, et al. 2020. Fungal Diversity 105:1-16 ((https://doi.org/10.1007/s13225-020-00466-2)) |
| Otu32289 | 1 | k__Fungi(100);p__Ascomycota(100);c__Saccharomycetes(100);o__Saccharomycetales(100);f__Metschnikowiaceae(100);g__Metschnikowia(100);g__Metschnikowia_unclassified(100); | Metschnikowia | 13 | Saprotroph | \|Undefined Saprotroph\| | Yeast | NULL | Probable | As nectar-tap_saprotroph (Põlme et al. 2020); Decay sugar-rich substrates (Põlme et al. 2020) | | Põlme S, et al. 2020. Fungal Diversity 105:1-16 ((https://doi.org/10.1007/s13225-020-00466-2)) |
| Otu32290 | 1 | k__Fungi(100);p__Ascomycota(100);c__Saccharomycetes(100);o__Saccharomycetales(100);f__Metschnikowiaceae(100);g__Metschnikowia(100);s__Metschnikowia_pulcherrima(100); | Metschnikowia | 13 | Saprotroph | \|Undefined Saprotroph\| | Yeast | NULL | Probable | As nectar-tap_saprotroph (Põlme et al. 2020); Decay sugar-rich substrates (Põlme et al. 2020) | | Põlme S, et al. 2020. Fungal Diversity 105:1-16 ((https://doi.org/10.1007/s13225-020-00466-2)) |
| Otu32291 | 1 | k__Fungi(100);p__Ascomycota(100);c__Saccharomycetes(100);o__Saccharomycetales(100);f__Metschnikowiaceae(100);g__Metschnikowia(100);s__Metschnikowia_pulcherrima(100); | Metschnikowia | 13 | Saprotroph | \|Undefined Saprotroph\| | Yeast | NULL | Probable | As nectar-tap_saprotroph (Põlme et al. 2020); Decay sugar-rich substrates (Põlme et al. 2020) | | Põlme S, et al. 2020. Fungal Diversity 105:1-16 ((https://doi.org/10.1007/s13225-020-00466-2)) |
| Otu32292 | 1 | k__Fungi(100);p__Ascomycota(100);c__Saccharomycetes(100);o__Saccharomycetales(100);f__Metschnikowiaceae(100);g__Metschnikowia(100);g__Metschnikowia_unclassified(100); | Metschnikowia | 13 | Saprotroph | \|Undefined Saprotroph\| | Yeast | NULL | Probable | As nectar-tap_saprotroph (Põlme et al. 2020); Decay sugar-rich substrates (Põlme et al. 2020) | | Põlme S, et al. 2020. Fungal Diversity 105:1-16 ((https://doi.org/10.1007/s13225-020-00466-2)) |
| Otu32293 | 1 | k__Fungi(100);p__Ascomycota(100);c__Saccharomycetes(100);o__Saccharomycetales(100);f__Metschnikowiaceae(100);g__Metschnikowia(100);g__Metschnikowia_unclassified(100); | Metschnikowia | 13 | Saprotroph | \|Undefined Saprotroph\| | Yeast | NULL | Probable | As nectar-tap_saprotroph (Põlme et al. 2020); Decay sugar-rich substrates (Põlme et al. 2020) | | Põlme S, et al. 2020. Fungal Diversity 105:1-16 ((https://doi.org/10.1007/s13225-020-00466-2)) |
| Otu32294 | 1 | k__Fungi(100);p__Ascomycota(100);c__Saccharomycetes(100);o__Saccharomycetales(100);f__Metschnikowiaceae(100);g__Metschnikowia(100);s__Metschnikowia_pulcherrima(100); | Metschnikowia | 13 | Saprotroph | \|Undefined Saprotroph\| | Yeast | NULL | Probable | As nectar-tap_saprotroph (Põlme et al. 2020); Decay sugar-rich substrates (Põlme et al. 2020) | | Põlme S, et al. 2020. Fungal Diversity 105:1-16 ((https://doi.org/10.1007/s13225-020-00466-2)) |
| Otu32295 | 1 | k__Fungi(100);p__Ascomycota(100);c__Saccharomycetes(100);o__Saccharomycetales(100);f__Saccharomycetales_family_Incertae_sedis(100);g__Candida(100);g__Candida_unclassified(100); | Candida | 13 | Pathotroph-Saprotroph-Symbiotroph | Animal Pathogen-Endophyte-Endosymbiont-Epiphyte-\|Undefined Saprotroph\| | Dimorphic Yeast-Yeast | NULL | Probable | As nectar-tap_saprotroph (Põlme et al. 2020); Decay sugar-rich substrates (Põlme et al. 2020); This asexual genus is polyphyletic and contain species that function very differently depend on what they are. Caution should be taken when interpreting guilds for Candida species. | | Manolakaki D, et al. 2010. Virulence. 1: 367-375 ((https://doi.org/10.4161/viru.1.5.12796)); Mekbib SB, et al. 2011. Tropical Plant Pathology 36:233-240 ((https://doi.org/10.1590/S1982-56762011000400004)); Põlme S, et al. 2020. Fungal Diversity 105:1-16 ((https://doi.org/10.1007/s13225-020-00466-2)) |
| Otu32297 | 1 | k__Fungi(100);p__Ascomycota(100);c__Saccharomycetes(100);o__Saccharomycetales(100);f__Metschnikowiaceae(100);g__Metschnikowia(100);s__Metschnikowia_pulcherrima(100); | Metschnikowia | 13 | Saprotroph | \|Undefined Saprotroph\| | Yeast | NULL | Probable | As nectar-tap_saprotroph (Põlme et al. 2020); Decay sugar-rich substrates (Põlme et al. 2020) | | Põlme S, et al. 2020. Fungal Diversity 105:1-16 ((https://doi.org/10.1007/s13225-020-00466-2)) |
| Otu32298 | 1 | k__Fungi(100);p__Ascomycota(100);c__Saccharomycetes(100);o__Saccharomycetales(100);f__Saccharomycetales_family_Incertae_sedis(100);g__Candida(100);g__Candida_unclassified(100); | Candida | 13 | Pathotroph-Saprotroph-Symbiotroph | Animal Pathogen-Endophyte-Endosymbiont-Epiphyte-\|Undefined Saprotroph\| | Dimorphic Yeast-Yeast | NULL | Probable | As nectar-tap_saprotroph (Põlme et al. 2020); Decay sugar-rich substrates (Põlme et al. 2020); This asexual genus is polyphyletic and contain species that function very differently depend on what they are. Caution should be taken when interpreting guilds for Candida species. | | Manolakaki D, et al. 2010. Virulence. 1: 367-375 ((https://doi.org/10.4161/viru.1.5.12796)); Mekbib SB, et al. 2011. Tropical Plant Pathology 36:233-240 ((https://doi.org/10.1590/S1982-56762011000400004)); Põlme S, et al. 2020. Fungal Diversity 105:1-16 ((https://doi.org/10.1007/s13225-020-00466-2)) |
| Otu32299 | 1 | k__Fungi(100);p__Ascomycota(100);c__Saccharomycetes(100);o__Saccharomycetales(100);f__Metschnikowiaceae(100);g__Metschnikowia(100);s__Metschnikowia_pulcherrima(100); | Metschnikowia | 13 | Saprotroph | \|Undefined Saprotroph\| | Yeast | NULL | Probable | As nectar-tap_saprotroph (Põlme et al. 2020); Decay sugar-rich substrates (Põlme et al. 2020) | | Põlme S, et al. 2020. Fungal Diversity 105:1-16 ((https://doi.org/10.1007/s13225-020-00466-2)) |
| Otu32302 | 1 | k__Fungi(100);p__Ascomycota(100);c__Saccharomycetes(100);o__Saccharomycetales(100);o__Saccharomycetales_unclassified(100);o__Saccharomycetales_unclassified(100);o__Saccharomycetales_unclassified(100); | Saccharomycetales | 7 | Saprotroph | Undefined Saprotroph | Yeast | NULL | Possible | NULL | | Sterkenburg E, et al. 2015. New Phytologist 207:1145-1158 ((https://doi.org/10.1111/nph.13426)) |
| Otu32305 | 1 | k__Fungi(100);p__Ascomycota(100);c__Saccharomycetes(100);o__Saccharomycetales(100);f__Saccharomycetales_family_Incertae_sedis(100);g__Candida(100);s__Candida_cretensis(100); | Candida | 13 | Pathotroph-Saprotroph-Symbiotroph | Animal Pathogen-Endophyte-Endosymbiont-Epiphyte-\|Undefined Saprotroph\| | Dimorphic Yeast-Yeast | NULL | Probable | As nectar-tap_saprotroph (Põlme et al. 2020); Decay sugar-rich substrates (Põlme et al. 2020); This asexual genus is polyphyletic and contain species that function very differently depend on what they are. Caution should be taken when interpreting guilds for Candida species. | | Manolakaki D, et al. 2010. Virulence. 1: 367-375 ((https://doi.org/10.4161/viru.1.5.12796)); Mekbib SB, et al. 2011. Tropical Plant Pathology 36:233-240 ((https://doi.org/10.1590/S1982-56762011000400004)); Põlme S, et al. 2020. Fungal Diversity 105:1-16 ((https://doi.org/10.1007/s13225-020-00466-2)) |
| Otu32306 | 1 | k__Fungi(100);p__Ascomycota(100);c__Saccharomycetes(100);o__Saccharomycetales(100);o__Saccharomycetales_unclassified(100);o__Saccharomycetales_unclassified(100);o__Saccharomycetales_unclassified(100); | Saccharomycetales | 7 | Saprotroph | Undefined Saprotroph | Yeast | NULL | Possible | NULL | | Sterkenburg E, et al. 2015. New Phytologist 207:1145-1158 ((https://doi.org/10.1111/nph.13426)) |
| Otu32307 | 1 | k__Fungi(100);p__Ascomycota(100);c__Saccharomycetes(100);o__Saccharomycetales(100);o__Saccharomycetales_unclassified(100);o__Saccharomycetales_unclassified(100);o__Saccharomycetales_unclassified(100); | Saccharomycetales | 7 | Saprotroph | Undefined Saprotroph | Yeast | NULL | Possible | NULL | | Sterkenburg E, et al. 2015. New Phytologist 207:1145-1158 ((https://doi.org/10.1111/nph.13426)) |
| Otu32308 | 1 | k__Fungi(100);p__Ascomycota(100);c__Saccharomycetes(100);o__Saccharomycetales(100);f__Saccharomycetales_family_Incertae_sedis(100);g__Candida(100);g__Candida_unclassified(100); | Candida | 13 | Pathotroph-Saprotroph-Symbiotroph | Animal Pathogen-Endophyte-Endosymbiont-Epiphyte-\|Undefined Saprotroph\| | Dimorphic Yeast-Yeast | NULL | Probable | As nectar-tap_saprotroph (Põlme et al. 2020); Decay sugar-rich substrates (Põlme et al. 2020); This asexual genus is polyphyletic and contain species that function very differently depend on what they are. Caution should be taken when interpreting guilds for Candida species. | | Manolakaki D, et al. 2010. Virulence. 1: 367-375 ((https://doi.org/10.4161/viru.1.5.12796)); Mekbib SB, et al. 2011. Tropical Plant Pathology 36:233-240 ((https://doi.org/10.1590/S1982-56762011000400004)); Põlme S, et al. 2020. Fungal Diversity 105:1-16 ((https://doi.org/10.1007/s13225-020-00466-2)) |
| Otu32310 | 1 | k__Fungi(100);p__Ascomycota(100);c__Saccharomycetes(100);o__Saccharomycetales(100);f__Metschnikowiaceae(100);g__Metschnikowia(100);s__Metschnikowia_pulcherrima(100); | Metschnikowia | 13 | Saprotroph | \|Undefined Saprotroph\| | Yeast | NULL | Probable | As nectar-tap_saprotroph (Põlme et al. 2020); Decay sugar-rich substrates (Põlme et al. 2020) | | Põlme S, et al. 2020. Fungal Diversity 105:1-16 ((https://doi.org/10.1007/s13225-020-00466-2)) |
| Otu32311 | 1 | k__Fungi(100);p__Ascomycota(100);c__Saccharomycetes(100);o__Saccharomycetales(100);f__Saccharomycetales_family_Incertae_sedis(100);g__Candida(100);g__Candida_unclassified(100); | Candida | 13 | Pathotroph-Saprotroph-Symbiotroph | Animal Pathogen-Endophyte-Endosymbiont-Epiphyte-\|Undefined Saprotroph\| | Dimorphic Yeast-Yeast | NULL | Probable | As nectar-tap_saprotroph (Põlme et al. 2020); Decay sugar-rich substrates (Põlme et al. 2020); This asexual genus is polyphyletic and contain species that function very differently depend on what they are. Caution should be taken when interpreting guilds for Candida species. | | Manolakaki D, et al. 2010. Virulence. 1: 367-375 ((https://doi.org/10.4161/viru.1.5.12796)); Mekbib SB, et al. 2011. Tropical Plant Pathology 36:233-240 ((https://doi.org/10.1590/S1982-56762011000400004)); Põlme S, et al. 2020. Fungal Diversity 105:1-16 ((https://doi.org/10.1007/s13225-020-00466-2)) |
| Otu32312 | 1 | k__Fungi(100);p__Ascomycota(100);c__Saccharomycetes(100);o__Saccharomycetales(100);f__Metschnikowiaceae(100);g__Metschnikowia(100);s__Metschnikowia_pulcherrima(100); | Metschnikowia | 13 | Saprotroph | \|Undefined Saprotroph\| | Yeast | NULL | Probable | As nectar-tap_saprotroph (Põlme et al. 2020); Decay sugar-rich substrates (Põlme et al. 2020) | | Põlme S, et al. 2020. Fungal Diversity 105:1-16 ((https://doi.org/10.1007/s13225-020-00466-2)) |
| Otu32313 | 1 | k__Fungi(100);p__Ascomycota(100);c__Saccharomycetes(100);o__Saccharomycetales(100);f__Metschnikowiaceae(100);g__Metschnikowia(100);s__Metschnikowia_pulcherrima(100); | Metschnikowia | 13 | Saprotroph | \|Undefined Saprotroph\| | Yeast | NULL | Probable | As nectar-tap_saprotroph (Põlme et al. 2020); Decay sugar-rich substrates (Põlme et al. 2020) | | Põlme S, et al. 2020. Fungal Diversity 105:1-16 ((https://doi.org/10.1007/s13225-020-00466-2)) |
| Otu32314 | 1 | k__Fungi(100);p__Ascomycota(100);c__Saccharomycetes(100);o__Saccharomycetales(100);f__Metschnikowiaceae(100);g__Metschnikowia(100);s__Metschnikowia_pulcherrima(100); | Metschnikowia | 13 | Saprotroph | \|Undefined Saprotroph\| | Yeast | NULL | Probable | As nectar-tap_saprotroph (Põlme et al. 2020); Decay sugar-rich substrates (Põlme et al. 2020) | | Põlme S, et al. 2020. Fungal Diversity 105:1-16 ((https://doi.org/10.1007/s13225-020-00466-2)) |
| Otu32315 | 1 | k__Fungi(100);p__Ascomycota(100);c__Saccharomycetes(100);o__Saccharomycetales(100);f__Metschnikowiaceae(100);g__Metschnikowia(100);s__Metschnikowia_pulcherrima(100); | Metschnikowia | 13 | Saprotroph | \|Undefined Saprotroph\| | Yeast | NULL | Probable | As nectar-tap_saprotroph (Põlme et al. 2020); Decay sugar-rich substrates (Põlme et al. 2020) | | Põlme S, et al. 2020. Fungal Diversity 105:1-16 ((https://doi.org/10.1007/s13225-020-00466-2)) |
| Otu32316 | 1 | k__Fungi(100);p__Ascomycota(100);c__Saccharomycetes(100);o__Saccharomycetales(100);f__Metschnikowiaceae(100);g__Metschnikowia(100);s__Metschnikowia_pulcherrima(100); | Metschnikowia | 13 | Saprotroph | \|Undefined Saprotroph\| | Yeast | NULL | Probable | As nectar-tap_saprotroph (Põlme et al. 2020); Decay sugar-rich substrates (Põlme et al. 2020) | | Põlme S, et al. 2020. Fungal Diversity 105:1-16 ((https://doi.org/10.1007/s13225-020-00466-2)) |

| | | | | | | | | | | | |
|---|---|---|---|---|---|---|---|---|---|---|---|
| Otu32317 | 1 | k__Fungi(100);p__Ascomycota(100);c__Saccharomycetes(100);o__Saccharomycetales(100);f__Metschnikowiaceae(100);g__Metschnikowia(100);s__Metschnikowia_pulcherrima(100); | Metschnikowia | 13 | Saprotroph | \|Undefined Saprotroph\| | Yeast | NULL | Probable | As nectar-tap_saprotroph (Põlme et al. 2020); Decay sugar-rich substrates (Põlme et al. 2020) | Põlme S, et al. 2020. Fungal Diversity 105:1-16 ((https://doi.org/10.1007/s13225-020-00466-2)) |
| Otu32318 | 1 | k__Fungi(100);p__Ascomycota(100);c__Saccharomycetes(100);o__Saccharomycetales(100);f__Metschnikowiaceae(100);g__Metschnikowia(100);s__Metschnikowia_pulcherrima(100); | Metschnikowia | 13 | Saprotroph | \|Undefined Saprotroph\| | Yeast | NULL | Probable | As nectar-tap_saprotroph (Põlme et al. 2020); Decay sugar-rich substrates (Põlme et al. 2020) | Põlme S, et al. 2020. Fungal Diversity 105:1-16 ((https://doi.org/10.1007/s13225-020-00466-2)) |
| Otu32319 | 1 | k__Fungi(100);p__Ascomycota(100);c__Saccharomycetes(100);o__Saccharomycetales(100);f__Metschnikowiaceae(100);g__Metschnikowia(100);s__Metschnikowia_pulcherrima(100); | Metschnikowia | 13 | Saprotroph | \|Undefined Saprotroph\| | Yeast | NULL | Probable | As nectar-tap_saprotroph (Põlme et al. 2020); Decay sugar-rich substrates (Põlme et al. 2020) | Põlme S, et al. 2020. Fungal Diversity 105:1-16 ((https://doi.org/10.1007/s13225-020-00466-2)) |
| Otu32320 | 1 | k__Fungi(100);p__Ascomycota(100);c__Saccharomycetes(100);o__Saccharomycetales(100);f__Metschnikowiaceae(100);g__Metschnikowia(100);s__Metschnikowia_pulcherrima(100); | Metschnikowia | 13 | Saprotroph | \|Undefined Saprotroph\| | Yeast | NULL | Probable | As nectar-tap_saprotroph (Põlme et al. 2020); Decay sugar-rich substrates (Põlme et al. 2020) | Põlme S, et al. 2020. Fungal Diversity 105:1-16 ((https://doi.org/10.1007/s13225-020-00466-2)) |
| Otu32322 | 1 | k__Fungi(100);p__Ascomycota(100);c__Saccharomycetes(100);o__Saccharomycetales(100);f__Metschnikowiaceae(100);g__Metschnikowia(100);g__Metschnikowia_unclassified(100); | Metschnikowia | 13 | Saprotroph | \|Undefined Saprotroph\| | Yeast | NULL | Probable | As nectar-tap_saprotroph (Põlme et al. 2020); Decay sugar-rich substrates (Põlme et al. 2020) | Põlme S, et al. 2020. Fungal Diversity 105:1-16 ((https://doi.org/10.1007/s13225-020-00466-2)) |
| Otu32323 | 1 | k__Fungi(100);p__Ascomycota(100);c__Saccharomycetes(100);o__Saccharomycetales(100);f__Metschnikowiaceae(100);g__Metschnikowia(100);s__Metschnikowia_pulcherrima(100); | Metschnikowia | 13 | Saprotroph | \|Undefined Saprotroph\| | Yeast | NULL | Probable | As nectar-tap_saprotroph (Põlme et al. 2020); Decay sugar-rich substrates (Põlme et al. 2020) | Põlme S, et al. 2020. Fungal Diversity 105:1-16 ((https://doi.org/10.1007/s13225-020-00466-2)) |
| Otu32324 | 1 | k__Fungi(100);p__Ascomycota(100);c__Saccharomycetes(100);o__Saccharomycetales(100);f__Metschnikowiaceae(100);g__Metschnikowia(100);s__Metschnikowia_pulcherrima(100); | Metschnikowia | 13 | Saprotroph | \|Undefined Saprotroph\| | Yeast | NULL | Probable | As nectar-tap_saprotroph (Põlme et al. 2020); Decay sugar-rich substrates (Põlme et al. 2020) | Põlme S, et al. 2020. Fungal Diversity 105:1-16 ((https://doi.org/10.1007/s13225-020-00466-2)) |
| Otu32325 | 1 | k__Fungi(100);p__Ascomycota(100);c__Saccharomycetes(100);o__Saccharomycetales(100);f__Metschnikowiaceae(100);g__Metschnikowia(100);s__Metschnikowia_pulcherrima(100); | Metschnikowia | 13 | Saprotroph | \|Undefined Saprotroph\| | Yeast | NULL | Probable | As nectar-tap_saprotroph (Põlme et al. 2020); Decay sugar-rich substrates (Põlme et al. 2020) | Põlme S, et al. 2020. Fungal Diversity 105:1-16 ((https://doi.org/10.1007/s13225-020-00466-2)) |
| Otu32326 | 1 | k__Fungi(100);p__Ascomycota(100);c__Saccharomycetes(100);o__Saccharomycetales(100);f__Metschnikowiaceae(100);g__Metschnikowia(100);s__Metschnikowia_pulcherrima(100); | Metschnikowia | 13 | Saprotroph | \|Undefined Saprotroph\| | Yeast | NULL | Probable | As nectar-tap_saprotroph (Põlme et al. 2020); Decay sugar-rich substrates (Põlme et al. 2020) | Põlme S, et al. 2020. Fungal Diversity 105:1-16 ((https://doi.org/10.1007/s13225-020-00466-2)) |
| Otu32336 | 1 | k__Fungi(100);p__Ascomycota(100);c__Saccharomycetes(100);o__Saccharomycetales(100);f__Metschnikowiaceae(100);g__Metschnikowia(100);s__Metschnikowia_pulcherrima(100); | Metschnikowia | 13 | Saprotroph | \|Undefined Saprotroph\| | Yeast | NULL | Probable | As nectar-tap_saprotroph (Põlme et al. 2020); Decay sugar-rich substrates (Põlme et al. 2020) | Põlme S, et al. 2020. Fungal Diversity 105:1-16 ((https://doi.org/10.1007/s13225-020-00466-2)) |
| Otu32337 | 1 | k__Fungi(100);p__Ascomycota(100);c__Saccharomycetes(100);o__Saccharomycetales(100);f__Saccharomycetales_family_Incertae_sedis(100);g__Candida(100);s__Candida_cretensis(100); | Candida | 13 | Pathotroph-Saprotroph-Symbiotroph | Animal Pathogen-Endophyte-Endosymbiont-Epiphyte-\|Undefined Saprotroph\| | Dimorphic Yeast-Yeast | NULL | Probable | As nectar-tap_saprotroph (Põlme et al. 2020); Decay sugar-rich substrates (Põlme et al. 2020); This asexual genus is polyphyletic and contain species that function very differently depend on what they are. Caution should be taken when interpreting guilds for Candida species. | Manolakaki D, et al. 2010. Virulence. 1: 367-375 ((https://doi.org/10.4161/viru.1.5.12796)); Mekbib SB, et al. 2011. Tropical Plant Pathology 36:233-240 ((https://doi.org/10.1590/S1982-56762011000400004)); Põlme S, et al. 2020. Fungal Diversity 105:1-16 ((https://doi.org/10.1007/s13225-020-00466-2)) |
| Otu32339 | 1 | k__Fungi(100);p__Ascomycota(100);c__Saccharomycetes(100);o__Saccharomycetales(100);f__Metschnikowiaceae(100);g__Metschnikowia(100);g__Metschnikowia_unclassified(100); | Metschnikowia | 13 | Saprotroph | \|Undefined Saprotroph\| | Yeast | NULL | Probable | As nectar-tap_saprotroph (Põlme et al. 2020); Decay sugar-rich substrates (Põlme et al. 2020) | Põlme S, et al. 2020. Fungal Diversity 105:1-16 ((https://doi.org/10.1007/s13225-020-00466-2)) |
| Otu32340 | 1 | k__Fungi(100);p__Ascomycota(100);c__Saccharomycetes(100);o__Saccharomycetales(100);f__Metschnikowiaceae(100);g__Metschnikowia(100);g__Metschnikowia_unclassified(100); | Metschnikowia | 13 | Saprotroph | \|Undefined Saprotroph\| | Yeast | NULL | Probable | As nectar-tap_saprotroph (Põlme et al. 2020); Decay sugar-rich substrates (Põlme et al. 2020) | Põlme S, et al. 2020. Fungal Diversity 105:1-16 ((https://doi.org/10.1007/s13225-020-00466-2)) |
| Otu32341 | 1 | k__Fungi(100);p__Ascomycota(100);c__Saccharomycetes(100);o__Saccharomycetales(100);f__Metschnikowiaceae(100);g__Metschnikowia(100);s__Metschnikowia_pulcherrima(100); | Metschnikowia | 13 | Saprotroph | \|Undefined Saprotroph\| | Yeast | NULL | Probable | As nectar-tap_saprotroph (Põlme et al. 2020); Decay sugar-rich substrates (Põlme et al. 2020) | Põlme S, et al. 2020. Fungal Diversity 105:1-16 ((https://doi.org/10.1007/s13225-020-00466-2)) |
| Otu32342 | 1 | k__Fungi(100);p__Ascomycota(100);c__Saccharomycetes(100);o__Saccharomycetales(100);f__Metschnikowiaceae(100);g__Metschnikowia(100);s__Metschnikowia_pulcherrima(100); | Metschnikowia | 13 | Saprotroph | \|Undefined Saprotroph\| | Yeast | NULL | Probable | As nectar-tap_saprotroph (Põlme et al. 2020); Decay sugar-rich substrates (Põlme et al. 2020) | Põlme S, et al. 2020. Fungal Diversity 105:1-16 ((https://doi.org/10.1007/s13225-020-00466-2)) |
| Otu32343 | 1 | k__Fungi(100);p__Ascomycota(100);c__Saccharomycetes(100);o__Saccharomycetales(100);f__Metschnikowiaceae(100);g__Metschnikowia(100);g__Metschnikowia_unclassified(100); | Metschnikowia | 13 | Saprotroph | \|Undefined Saprotroph\| | Yeast | NULL | Probable | As nectar-tap_saprotroph (Põlme et al. 2020); Decay sugar-rich substrates (Põlme et al. 2020) | Põlme S, et al. 2020. Fungal Diversity 105:1-16 ((https://doi.org/10.1007/s13225-020-00466-2)) |
| Otu32344 | 1 | k__Fungi(100);p__Ascomycota(100);c__Saccharomycetes(100);o__Saccharomycetales(100);f__Metschnikowiaceae(100);g__Metschnikowia(100);s__Metschnikowia_pulcherrima(100); | Metschnikowia | 13 | Saprotroph | \|Undefined Saprotroph\| | Yeast | NULL | Probable | As nectar-tap_saprotroph (Põlme et al. 2020); Decay sugar-rich substrates (Põlme et al. 2020) | Põlme S, et al. 2020. Fungal Diversity 105:1-16 ((https://doi.org/10.1007/s13225-020-00466-2)) |
| Otu32345 | 1 | k__Fungi(100);p__Ascomycota(100);c__Saccharomycetes(100);o__Saccharomycetales(100);f__Metschnikowiaceae(100);g__Metschnikowia(100);g__Metschnikowia_unclassified(100); | Metschnikowia | 13 | Saprotroph | \|Undefined Saprotroph\| | Yeast | NULL | Probable | As nectar-tap_saprotroph (Põlme et al. 2020); Decay sugar-rich substrates (Põlme et al. 2020) | Põlme S, et al. 2020. Fungal Diversity 105:1-16 ((https://doi.org/10.1007/s13225-020-00466-2)) |
| Otu32346 | 1 | k__Fungi(100);p__Ascomycota(100);c__Saccharomycetes(100);o__Saccharomycetales(100);f__Metschnikowiaceae(100);g__Metschnikowia(100);s__Metschnikowia_pulcherrima(100); | Metschnikowia | 13 | Saprotroph | \|Undefined Saprotroph\| | Yeast | NULL | Probable | As nectar-tap_saprotroph (Põlme et al. 2020); Decay sugar-rich substrates (Põlme et al. 2020) | Põlme S, et al. 2020. Fungal Diversity 105:1-16 ((https://doi.org/10.1007/s13225-020-00466-2)) |
| Otu32347 | 1 | k__Fungi(100);p__Ascomycota(100);c__Saccharomycetes(100);o__Saccharomycetales(100);f__Metschnikowiaceae(100);g__Metschnikowia(100);s__Metschnikowia_pulcherrima(100); | Metschnikowia | 13 | Saprotroph | \|Undefined Saprotroph\| | Yeast | NULL | Probable | As nectar-tap_saprotroph (Põlme et al. 2020); Decay sugar-rich substrates (Põlme et al. 2020) | Põlme S, et al. 2020. Fungal Diversity 105:1-16 ((https://doi.org/10.1007/s13225-020-00466-2)) |
| Otu32349 | 1 | k__Fungi(100);p__Ascomycota(100);c__Saccharomycetes(100);o__Saccharomycetales(100);f__Metschnikowiaceae(100);g__Metschnikowia(100);g__Metschnikowia_unclassified(100); | Metschnikowia | 13 | Saprotroph | \|Undefined Saprotroph\| | Yeast | NULL | Probable | As nectar-tap_saprotroph (Põlme et al. 2020); Decay sugar-rich substrates (Põlme et al. 2020) | Põlme S, et al. 2020. Fungal Diversity 105:1-16 ((https://doi.org/10.1007/s13225-020-00466-2)) |
| Otu32350 | 1 | k__Fungi(100);p__Ascomycota(100);c__Saccharomycetes(100);o__Saccharomycetales(100);f__Metschnikowiaceae(100);g__Metschnikowia(100);g__Metschnikowia_unclassified(100); | Metschnikowia | 13 | Saprotroph | \|Undefined Saprotroph\| | Yeast | NULL | Probable | As nectar-tap_saprotroph (Põlme et al. 2020); Decay sugar-rich substrates (Põlme et al. 2020) | Põlme S, et al. 2020. Fungal Diversity 105:1-16 ((https://doi.org/10.1007/s13225-020-00466-2)) |
| Otu32351 | 1 | k__Fungi(100);p__Ascomycota(100);c__Saccharomycetes(100);o__Saccharomycetales(100);f__Metschnikowiaceae(100);g__Metschnikowia(100);s__Metschnikowia_pulcherrima(100); | Metschnikowia | 13 | Saprotroph | \|Undefined Saprotroph\| | Yeast | NULL | Probable | As nectar-tap_saprotroph (Põlme et al. 2020); Decay sugar-rich substrates (Põlme et al. 2020) | Põlme S, et al. 2020. Fungal Diversity 105:1-16 ((https://doi.org/10.1007/s13225-020-00466-2)) |
| Otu32353 | 1 | k__Fungi(100);p__Ascomycota(100);c__Saccharomycetes(100);o__Saccharomycetales(100);f__Metschnikowiaceae(100);g__Metschnikowia(100);s__Metschnikowia_pulcherrima(100); | Metschnikowia | 13 | Saprotroph | \|Undefined Saprotroph\| | Yeast | NULL | Probable | As nectar-tap_saprotroph (Põlme et al. 2020); Decay sugar-rich substrates (Põlme et al. 2020) | Põlme S, et al. 2020. Fungal Diversity 105:1-16 ((https://doi.org/10.1007/s13225-020-00466-2)) |
| Otu32354 | 1 | k__Fungi(100);p__Ascomycota(100);c__Saccharomycetes(100);o__Saccharomycetales(100);f__Metschnikowiaceae(100);g__Metschnikowia(100);s__Metschnikowia_pulcherrima(100); | Metschnikowia | 13 | Saprotroph | \|Undefined Saprotroph\| | Yeast | NULL | Probable | As nectar-tap_saprotroph (Põlme et al. 2020); Decay sugar-rich substrates (Põlme et al. 2020) | Põlme S, et al. 2020. Fungal Diversity 105:1-16 ((https://doi.org/10.1007/s13225-020-00466-2)) |
| Otu32355 | 1 | k__Fungi(100);p__Ascomycota(100);c__Saccharomycetes(100);o__Saccharomycetales(100);f__Metschnikowiaceae(100);g__Metschnikowia(100);s__Metschnikowia_pulcherrima(100); | Metschnikowia | 13 | Saprotroph | \|Undefined Saprotroph\| | Yeast | NULL | Probable | As nectar-tap_saprotroph (Põlme et al. 2020); Decay sugar-rich substrates (Põlme et al. 2020) | Põlme S, et al. 2020. Fungal Diversity 105:1-16 ((https://doi.org/10.1007/s13225-020-00466-2)) |
| Otu32356 | 1 | k__Fungi(100);p__Ascomycota(100);c__Saccharomycetes(100);o__Saccharomycetales(100);f__Metschnikowiaceae(100);g__Metschnikowia(100);s__Metschnikowia_pulcherrima(100); | Metschnikowia | 13 | Saprotroph | \|Undefined Saprotroph\| | Yeast | NULL | Probable | As nectar-tap_saprotroph (Põlme et al. 2020); Decay sugar-rich substrates (Põlme et al. 2020) | Põlme S, et al. 2020. Fungal Diversity 105:1-16 ((https://doi.org/10.1007/s13225-020-00466-2)) |
| Otu32357 | 1 | k__Fungi(100);p__Ascomycota(100);c__Saccharomycetes(100);o__Saccharomycetales(100);f__Saccharomycetales_family_Incertae_sedis(100);g__Candida(100);g__Candida_unclassified(100); | Candida | 13 | Pathotroph-Saprotroph-Symbiotroph | Animal Pathogen-Endophyte-Endosymbiont-Epiphyte-\|Undefined Saprotroph\| | Dimorphic Yeast-Yeast | NULL | Probable | As nectar-tap_saprotroph (Põlme et al. 2020); Decay sugar-rich substrates (Põlme et al. 2020); This asexual genus is polyphyletic and contain species that function very differently depend on what they are. Caution should be taken when interpreting guilds for Candida species. | Manolakaki D, et al. 2010. Virulence. 1: 367-375 ((https://doi.org/10.4161/viru.1.5.12796)); Mekbib SB, et al. 2011. Tropical Plant Pathology 36:233-240 ((https://doi.org/10.1590/S1982-56762011000400004)); Põlme S, et al. 2020. Fungal Diversity 105:1-16 ((https://doi.org/10.1007/s13225-020-00466-2)) |
| Otu32359 | 1 | k__Fungi(100);p__Ascomycota(100);c__Saccharomycetes(100);o__Saccharomycetales(100);f__Metschnikowiaceae(100);g__Metschnikowia(100);s__Metschnikowia_pulcherrima(100); | Metschnikowia | 13 | Saprotroph | \|Undefined Saprotroph\| | Yeast | NULL | Probable | As nectar-tap_saprotroph (Põlme et al. 2020); Decay sugar-rich substrates (Põlme et al. 2020) | Põlme S, et al. 2020. Fungal Diversity 105:1-16 ((https://doi.org/10.1007/s13225-020-00466-2)) |
| Otu32360 | 1 | k__Fungi(100);p__Ascomycota(100);c__Saccharomycetes(100);o__Saccharomycetales(100);f__Saccharomycetales_family_Incertae_sedis(100);g__Candida(100);g__Candida_unclassified(100); | Candida | 13 | Pathotroph-Saprotroph-Symbiotroph | Animal Pathogen-Endophyte-Endosymbiont-Epiphyte-\|Undefined Saprotroph\| | Dimorphic Yeast-Yeast | NULL | Probable | As nectar-tap_saprotroph (Põlme et al. 2020); Decay sugar-rich substrates (Põlme et al. 2020); This asexual genus is polyphyletic and contain species that function very differently depend on what they are. Caution should be taken when interpreting guilds for Candida species. | Manolakaki D, et al. 2010. Virulence. 1: 367-375 ((https://doi.org/10.4161/viru.1.5.12796)); Mekbib SB, et al. 2011. Tropical Plant Pathology 36:233-240 ((https://doi.org/10.1590/S1982-56762011000400004)); Põlme S, et al. 2020. Fungal Diversity 105:1-16 ((https://doi.org/10.1007/s13225-020-00466-2)) |
| Otu32361 | 1 | k__Fungi(100);p__Ascomycota(100);c__Saccharomycetes(100);o__Saccharomycetales(100);f__Metschnikowiaceae(100);g__Metschnikowia(100);s__Metschnikowia_pulcherrima(100); | Metschnikowia | 13 | Saprotroph | \|Undefined Saprotroph\| | Yeast | NULL | Probable | As nectar-tap_saprotroph (Põlme et al. 2020); Decay sugar-rich substrates (Põlme et al. 2020) | Põlme S, et al. 2020. Fungal Diversity 105:1-16 ((https://doi.org/10.1007/s13225-020-00466-2)) |
| Otu32362 | 1 | k__Fungi(100);p__Ascomycota(100);c__Saccharomycetes(100);o__Saccharomycetales(100);f__Saccharomycetales_family_Incertae_sedis(100);g__Candida(100);s__Candida_cretensis(100); | Candida | 13 | Pathotroph-Saprotroph-Symbiotroph | Animal Pathogen-Endophyte-Endosymbiont-Epiphyte-\|Undefined Saprotroph\| | Dimorphic Yeast-Yeast | NULL | Probable | As nectar-tap_saprotroph (Põlme et al. 2020); Decay sugar-rich substrates (Põlme et al. 2020); This asexual genus is polyphyletic and contain species that function very differently depend on what they are. Caution should be taken when interpreting guilds for Candida species. | Manolakaki D, et al. 2010. Virulence. 1: 367-375 ((https://doi.org/10.4161/viru.1.5.12796)); Mekbib SB, et al. 2011. Tropical Plant Pathology 36:233-240 ((https://doi.org/10.1590/S1982-56762011000400004)); Põlme S, et al. 2020. Fungal Diversity 105:1-16 ((https://doi.org/10.1007/s13225-020-00466-2)) |
| Otu32363 | 1 | k__Fungi(100);p__Ascomycota(100);c__Saccharomycetes(100);o__Saccharomycetales(100);f__Metschnikowiaceae(100);g__Metschnikowia(100);s__Metschnikowia_pulcherrima(100); | Metschnikowia | 13 | Saprotroph | \|Undefined Saprotroph\| | Yeast | NULL | Probable | As nectar-tap_saprotroph (Põlme et al. 2020); Decay sugar-rich substrates (Põlme et al. 2020) | Põlme S, et al. 2020. Fungal Diversity 105:1-16 ((https://doi.org/10.1007/s13225-020-00466-2)) |
| Otu32364 | 1 | k__Fungi(100);p__Ascomycota(100);c__Saccharomycetes(100);o__Saccharomycetales(100);f__Metschnikowiaceae(100);g__Metschnikowia(100);g__Metschnikowia_unclassified(100); | Metschnikowia | 13 | Saprotroph | \|Undefined Saprotroph\| | Yeast | NULL | Probable | As nectar-tap_saprotroph (Põlme et al. 2020); Decay sugar-rich substrates (Põlme et al. 2020) | Põlme S, et al. 2020. Fungal Diversity 105:1-16 ((https://doi.org/10.1007/s13225-020-00466-2)) |
| Otu32365 | 1 | k__Fungi(100);p__Ascomycota(100);c__Saccharomycetes(100);o__Saccharomycetales(100);f__Metschnikowiaceae(100);g__Metschnikowia(100);g__Metschnikowia_unclassified(100); | Metschnikowia | 13 | Saprotroph | \|Undefined Saprotroph\| | Yeast | NULL | Probable | As nectar-tap_saprotroph (Põlme et al. 2020); Decay sugar-rich substrates (Põlme et al. 2020) | Põlme S, et al. 2020. Fungal Diversity 105:1-16 ((https://doi.org/10.1007/s13225-020-00466-2)) |
| Otu32366 | 1 | k__Fungi(100);p__Ascomycota(100);c__Saccharomycetes(100);o__Saccharomycetales(100);f__Saccharomycetales_family_Incertae_sedis(100);g__Candida(100);g__Candida_unclassified(100); | Candida | 13 | Pathotroph-Saprotroph-Symbiotroph | Animal Pathogen-Endophyte-Endosymbiont-Epiphyte-\|Undefined Saprotroph\| | Dimorphic Yeast-Yeast | NULL | Probable | As nectar-tap_saprotroph (Põlme et al. 2020); Decay sugar-rich substrates (Põlme et al. 2020); This asexual genus is polyphyletic and contain species that function very differently depend on what they are. Caution should be taken when interpreting guilds for Candida species. | Manolakaki D, et al. 2010. Virulence. 1: 367-375 ((https://doi.org/10.4161/viru.1.5.12796)); Mekbib SB, et al. 2011. Tropical Plant Pathology 36:233-240 ((https://doi.org/10.1590/S1982-56762011000400004)); Põlme S, et al. 2020. Fungal Diversity 105:1-16 ((https://doi.org/10.1007/s13225-020-00466-2)) |
| Otu32367 | 1 | k__Fungi(100);p__Ascomycota(100);c__Saccharomycetes(100);o__Saccharomycetales(100);f__Metschnikowiaceae(100);g__Metschnikowia(100);s__Metschnikowia_pulcherrima(100); | Metschnikowia | 13 | Saprotroph | \|Undefined Saprotroph\| | Yeast | NULL | Probable | As nectar-tap_saprotroph (Põlme et al. 2020); Decay sugar-rich substrates (Põlme et al. 2020) | Põlme S, et al. 2020. Fungal Diversity 105:1-16 ((https://doi.org/10.1007/s13225-020-00466-2)) |
| Otu32368 | 1 | k__Fungi(100);p__Ascomycota(100);c__Saccharomycetes(100);o__Saccharomycetales(100);f__Saccharomycetales_family_Incertae_sedis(100);g__Candida(100);g__Candida_unclassified(100); | Candida | 13 | Pathotroph-Saprotroph-Symbiotroph | Animal Pathogen-Endophyte-Endosymbiont-Epiphyte-\|Undefined Saprotroph\| | Dimorphic Yeast-Yeast | NULL | Probable | As nectar-tap_saprotroph (Põlme et al. 2020); Decay sugar-rich substrates (Põlme et al. 2020); This asexual genus is polyphyletic and contain species that function very differently depend on what they are. Caution should be taken when interpreting guilds for Candida species. | Manolakaki D, et al. 2010. Virulence. 1: 367-375 ((https://doi.org/10.4161/viru.1.5.12796)); Mekbib SB, et al. 2011. Tropical Plant Pathology 36:233-240 ((https://doi.org/10.1590/S1982-56762011000400004)); Põlme S, et al. 2020. Fungal Diversity 105:1-16 ((https://doi.org/10.1007/s13225-020-00466-2)) |
| Otu32369 | 1 | k__Fungi(100);p__Ascomycota(100);c__Saccharomycetes(100);o__Saccharomycetales(100);o__Saccharomycetales_unclassified(100);o__Saccharomycetales_unclassified(100);o__Saccharomycetales_unclassified(100); | Saccharomycetales | 7 | Saprotroph | Undefined Saprotroph | Yeast | NULL | Possible | NULL | Sterkenburg E, et al. 2015. New Phytologist 207:1145-1158 ((https://doi.org/10.1111/nph.13426)) |
| Otu32370 | 1 | k__Fungi(100);p__Ascomycota(100);c__Saccharomycetes(100);o__Saccharomycetales(100);f__Metschnikowiaceae(100);g__Metschnikowia(100);g__Metschnikowia_unclassified(100); | Metschnikowia | 13 | Saprotroph | \|Undefined Saprotroph\| | Yeast | NULL | Probable | As nectar-tap_saprotroph (Põlme et al. 2020); Decay sugar-rich substrates (Põlme et al. 2020) | Põlme S, et al. 2020. Fungal Diversity 105:1-16 ((https://doi.org/10.1007/s13225-020-00466-2)) |
| Otu32373 | 1 | k__Fungi(100);p__Ascomycota(100);c__Saccharomycetes(100);o__Saccharomycetales(100);f__Saccharomycetales_family_Incertae_sedis(100);g__Candida(100);g__Candida_unclassified(100); | Candida | 13 | Pathotroph-Saprotroph-Symbiotroph | Animal Pathogen-Endophyte-Endosymbiont-Epiphyte-\|Undefined Saprotroph\| | Dimorphic Yeast-Yeast | NULL | Probable | As nectar-tap_saprotroph (Põlme et al. 2020); Decay sugar-rich substrates (Põlme et al. 2020); This asexual genus is polyphyletic and contain species that function very differently depend on what they are. Caution should be taken when interpreting guilds for Candida species. | Manolakaki D, et al. 2010. Virulence. 1: 367-375 ((https://doi.org/10.4161/viru.1.5.12796)); Mekbib SB, et al. 2011. Tropical Plant Pathology 36:233-240 ((https://doi.org/10.1590/S1982-56762011000400004)); Põlme S, et al. 2020. Fungal Diversity 105:1-16 ((https://doi.org/10.1007/s13225-020-00466-2)) |
| Otu32375 | 1 | k__Fungi(100);p__Ascomycota(100);c__Saccharomycetes(100);o__Saccharomycetales(100);f__Saccharomycetales_family_Incertae_sedis(100);f__Saccharomycetales_family_Incertae_sedis_unclassified(100);f__Saccharomycetales_family_Incertae_sedis_unclassified(100); | Saccharomycetales | 7 | Saprotroph | Undefined Saprotroph | Yeast | NULL | Possible | NULL | Sterkenburg E, et al. 2015. New Phytologist 207:1145-1158 ((https://doi.org/10.1111/nph.13426)) |
| Otu32378 | 1 | k__Fungi(100);p__Ascomycota(100);c__Saccharomycetes(100);o__Saccharomycetales(100);f__Metschnikowiaceae(100);g__Metschnikowia(100);g__Metschnikowia_unclassified(100); | Metschnikowia | 13 | Saprotroph | \|Undefined Saprotroph\| | Yeast | NULL | Probable | As nectar-tap_saprotroph (Põlme et al. 2020); Decay sugar-rich substrates (Põlme et al. 2020) | Põlme S, et al. 2020. Fungal Diversity 105:1-16 ((https://doi.org/10.1007/s13225-020-00466-2)) |
| Otu32379 | 1 | k__Fungi(100);p__Ascomycota(100);c__Saccharomycetes(100);o__Saccharomycetales(100);f__Saccharomycetales_family_Incertae_sedis(100);g__Candida(100);g__Candida_unclassified(100); | Candida | 13 | Pathotroph-Saprotroph-Symbiotroph | Animal Pathogen-Endophyte-Endosymbiont-Epiphyte-\|Undefined Saprotroph\| | Dimorphic Yeast-Yeast | NULL | Probable | As nectar-tap_saprotroph (Põlme et al. 2020); Decay sugar-rich substrates (Põlme et al. 2020); This asexual genus is polyphyletic and contain species that function very differently depend on what they are. Caution should be taken when interpreting guilds for Candida species. | Manolakaki D, et al. 2010. Virulence. 1: 367-375 ((https://doi.org/10.4161/viru.1.5.12796)); Mekbib SB, et al. 2011. Tropical Plant Pathology 36:233-240 ((https://doi.org/10.1590/S1982-56762011000400004)); Põlme S, et al. 2020. Fungal Diversity 105:1-16 ((https://doi.org/10.1007/s13225-020-00466-2)) |
| Otu32381 | 1 | k__Fungi(100);p__Ascomycota(100);c__Saccharomycetes(100);o__Saccharomycetales(100);f__Saccharomycetales_family_Incertae_sedis(100);g__Candida(100);g__Candida_unclassified(100); | Candida | 13 | Pathotroph-Saprotroph-Symbiotroph | Animal Pathogen-Endophyte-Endosymbiont-Epiphyte-\|Undefined Saprotroph\| | Dimorphic Yeast-Yeast | NULL | Probable | As nectar-tap_saprotroph (Põlme et al. 2020); Decay sugar-rich substrates (Põlme et al. 2020); This asexual genus is polyphyletic and contain species that function very differently depend on what they are. Caution should be taken when interpreting guilds for Candida species. | Manolakaki D, et al. 2010. Virulence. 1: 367-375 ((https://doi.org/10.4161/viru.1.5.12796)); Mekbib SB, et al. 2011. Tropical Plant Pathology 36:233-240 ((https://doi.org/10.1590/S1982-56762011000400004)); Põlme S, et al. 2020. Fungal Diversity 105:1-16 ((https://doi.org/10.1007/s13225-020-00466-2)) |
| Otu32382 | 1 | k__Fungi(100);p__Ascomycota(100);c__Saccharomycetes(100);o__Saccharomycetales(100);f__Metschnikowiaceae(100);g__Metschnikowia(100);s__Metschnikowia_pulcherrima(100); | Metschnikowia | 13 | Saprotroph | \|Undefined Saprotroph\| | Yeast | NULL | Probable | As nectar-tap_saprotroph (Põlme et al. 2020); Decay sugar-rich substrates (Põlme et al. 2020) | Põlme S, et al. 2020. Fungal Diversity 105:1-16 ((https://doi.org/10.1007/s13225-020-00466-2)) |
| Otu32384 | 1 | k__Fungi(100);p__Ascomycota(100);c__Saccharomycetes(100);o__Saccharomycetales(100);f__Saccharomycetales_family_Incertae_sedis(100);g__Candida(100);s__Candida_cretensis(100); | Candida | 13 | Pathotroph-Saprotroph-Symbiotroph | Animal Pathogen-Endophyte-Endosymbiont-Epiphyte-\|Undefined Saprotroph\| | Dimorphic Yeast-Yeast | NULL | Probable | As nectar-tap_saprotroph (Põlme et al. 2020); Decay sugar-rich substrates (Põlme et al. 2020); This asexual genus is polyphyletic and contain species that function very differently depend on what they are. Caution should be taken when interpreting guilds for Candida species. | Manolakaki D, et al. 2010. Virulence. 1: 367-375 ((https://doi.org/10.4161/viru.1.5.12796)); Mekbib SB, et al. 2011. Tropical Plant Pathology 36:233-240 ((https://doi.org/10.1590/S1982-56762011000400004)); Põlme S, et al. 2020. Fungal Diversity 105:1-16 ((https://doi.org/10.1007/s13225-020-00466-2)) |
| Otu32385 | 1 | k__Fungi(100);p__Ascomycota(100);c__Saccharomycetes(100);o__Saccharomycetales(100);f__Metschnikowiaceae(100);g__Metschnikowia(100);g__Metschnikowia_unclassified(100); | Metschnikowia | 13 | Saprotroph | \|Undefined Saprotroph\| | Yeast | NULL | Probable | As nectar-tap_saprotroph (Põlme et al. 2020); Decay sugar-rich substrates (Põlme et al. 2020) | Põlme S, et al. 2020. Fungal Diversity 105:1-16 ((https://doi.org/10.1007/s13225-020-00466-2)) |
| Otu32386 | 1 | k__Fungi(100);p__Ascomycota(100);c__Saccharomycetes(100);o__Saccharomycetales(100);f__Metschnikowiaceae(100);g__Metschnikowia(100);s__Metschnikowia_pulcherrima(100); | Metschnikowia | 13 | Saprotroph | \|Undefined Saprotroph\| | Yeast | NULL | Probable | As nectar-tap_saprotroph (Põlme et al. 2020); Decay sugar-rich substrates (Põlme et al. 2020) | Põlme S, et al. 2020. Fungal Diversity 105:1-16 ((https://doi.org/10.1007/s13225-020-00466-2)) |
| Otu32387 | 1 | k__Fungi(100);p__Ascomycota(100);c__Saccharomycetes(100);o__Saccharomycetales(100);f__Metschnikowiaceae(100);g__Metschnikowia(100);g__Metschnikowia_unclassified(100); | Metschnikowia | 13 | Saprotroph | \|Undefined Saprotroph\| | Yeast | NULL | Probable | As nectar-tap_saprotroph (Põlme et al. 2020); Decay sugar-rich substrates (Põlme et al. 2020) | Põlme S, et al. 2020. Fungal Diversity 105:1-16 ((https://doi.org/10.1007/s13225-020-00466-2)) |
| Otu32388 | 1 | k__Fungi(100);p__Ascomycota(100);c__Saccharomycetes(100);o__Saccharomycetales(100);f__Saccharomycetales_family_Incertae_sedis(100);g__Candida(100);g__Candida_unclassified(100); | Candida | 13 | Pathotroph-Saprotroph-Symbiotroph | Animal Pathogen-Endophyte-Endosymbiont-Epiphyte-\|Undefined Saprotroph\| | Dimorphic Yeast-Yeast | NULL | Probable | As nectar-tap_saprotroph (Põlme et al. 2020); Decay sugar-rich substrates (Põlme et al. 2020); This asexual genus is polyphyletic and contain species that function very differently depend on what they are. Caution should be taken when interpreting guilds for Candida species. | Manolakaki D, et al. 2010. Virulence. 1: 367-375 ((https://doi.org/10.4161/viru.1.5.12796)); Mekbib SB, et al. 2011. Tropical Plant Pathology 36:233-240 ((https://doi.org/10.1590/S1982-56762011000400004)); Põlme S, et al. 2020. Fungal Diversity 105:1-16 ((https://doi.org/10.1007/s13225-020-00466-2)) |
| Otu32389 | 1 | k__Fungi(100);p__Ascomycota(100);c__Saccharomycetes(100);o__Saccharomycetales(100);f__Saccharomycetales_family_Incertae_sedis(100);g__Candida(100);g__Candida_unclassified(100); | Candida | 13 | Pathotroph-Saprotroph-Symbiotroph | Animal Pathogen-Endophyte-Endosymbiont-Epiphyte-\|Undefined Saprotroph\| | Dimorphic Yeast-Yeast | NULL | Probable | As nectar-tap_saprotroph (Põlme et al. 2020); Decay sugar-rich substrates (Põlme et al. 2020); This asexual genus is polyphyletic and contain species that function very differently depend on what they are. Caution should be taken when interpreting guilds for Candida species. | Manolakaki D, et al. 2010. Virulence. 1: 367-375 ((https://doi.org/10.4161/viru.1.5.12796)); Mekbib SB, et al. 2011. Tropical Plant Pathology 36:233-240 ((https://doi.org/10.1590/S1982-56762011000400004)); Põlme S, et al. 2020. Fungal Diversity 105:1-16 ((https://doi.org/10.1007/s13225-020-00466-2)) |
| Otu32390 | 1 | k__Fungi(100);p__Ascomycota(100);c__Saccharomycetes(100);o__Saccharomycetales(100);f__Saccharomycetales_family_Incertae_sedis(100);g__Candida(100);g__Candida_unclassified(100); | Candida | 13 | Pathotroph-Saprotroph-Symbiotroph | Animal Pathogen-Endophyte-Endosymbiont-Epiphyte-\|Undefined Saprotroph\| | Dimorphic Yeast-Yeast | NULL | Probable | As nectar-tap_saprotroph (Põlme et al. 2020); Decay sugar-rich substrates (Põlme et al. 2020); This asexual genus is polyphyletic and contain species that function very differently depend on what they are. Caution should be taken when interpreting guilds for Candida species. | Manolakaki D, et al. 2010. Virulence. 1: 367-375 ((https://doi.org/10.4161/viru.1.5.12796)); Mekbib SB, et al. 2011. Tropical Plant Pathology 36:233-240 ((https://doi.org/10.1590/S1982-56762011000400004)); Põlme S, et al. 2020. Fungal Diversity 105:1-16 ((https://doi.org/10.1007/s13225-020-00466-2)) |
| Otu32391 | 1 | k__Fungi(100);p__Ascomycota(100);c__Saccharomycetes(100);o__Saccharomycetales(100);f__Saccharomycetales_family_Incertae_sedis(100);g__Candida(100);g__Candida_unclassified(100); | Candida | 13 | Pathotroph-Saprotroph-Symbiotroph | Animal Pathogen-Endophyte-Endosymbiont-Epiphyte-\|Undefined Saprotroph\| | Dimorphic Yeast-Yeast | NULL | Probable | As nectar-tap_saprotroph (Põlme et al. 2020); Decay sugar-rich substrates (Põlme et al. 2020); This asexual genus is polyphyletic and contain species that function very differently depend on what they are. Caution should be taken when interpreting guilds for Candida species. | Manolakaki D, et al. 2010. Virulence. 1: 367-375 ((https://doi.org/10.4161/viru.1.5.12796)); Mekbib SB, et al. 2011. Tropical Plant Pathology 36:233-240 ((https://doi.org/10.1590/S1982-56762011000400004)); Põlme S, et al. 2020. Fungal Diversity 105:1-16 ((https://doi.org/10.1007/s13225-020-00466-2)) |
| Otu32395 | 1 | k__Fungi(100);p__Ascomycota(100);c__Saccharomycetes(100);o__Saccharomycetales(100);o__Saccharomycetales_unclassified(100);o__Saccharomycetales_unclassified(100);o__Saccharomycetales_unclassified(100); | Saccharomycetales | 7 | Saprotroph | Undefined Saprotroph | Yeast | NULL | Possible | NULL | Sterkenburg E, et al. 2015. New Phytologist 207:1145-1158 ((https://doi.org/10.1111/nph.13426)) |
| Otu32397 | 1 | k__Fungi(100);p__Basidiomycota(100);c__Tremellomycetes(100);o__Tremellales(100);f__Tremellales_family_Incertae_sedis(100);g__Hannaella(100);s__Hannaella_oryzae(100); | Hannaella | 13 | Pathotroph-Saprotroph | \|Animal Parasite\|-Undefined Saprotroph | Yeast | NULL | Probable | As animal_decomposer (Põlme et al. 2020); Decay animal material (Põlme et al. 2020) | Põlme S, et al. 2020. Fungal Diversity 105:1-16 ((https://doi.org/10.1007/s13225-020-00466-2)) |
| Otu32405 | 1 | k__Fungi(100);p__Ascomycota(100);c__Saccharomycetes(100);o__Saccharomycetales(100);f__Saccharomycetales_family_Incertae_sedis(100);g__Candida(100);g__Candida_unclassified(100); | Candida | 13 | Pathotroph-Saprotroph-Symbiotroph | Animal Pathogen-Endophyte-Endosymbiont-Epiphyte-\|Undefined Saprotroph\| | Dimorphic Yeast-Yeast | NULL | Probable | As nectar-tap_saprotroph (Põlme et al. 2020); Decay sugar-rich substrates (Põlme et al. 2020); This asexual genus is polyphyletic and contain species that function very differently depend on what they are. Caution should be taken when interpreting guilds for Candida species. | Manolakaki D, et al. 2010. Virulence. 1: 367-375 ((https://doi.org/10.4161/viru.1.5.12796)); Mekbib SB, et al. 2011. Tropical Plant Pathology 36:233-240 ((https://doi.org/10.1590/S1982-56762011000400004)); Põlme S, et al. 2020. Fungal Diversity 105:1-16 ((https://doi.org/10.1007/s13225-020-00466-2)) |
| Otu32409 | 1 | k__Fungi(100);p__Ascomycota(100);c__Saccharomycetes(100);o__Saccharomycetales(100);f__Metschnikowiaceae(100);g__Metschnikowia(100);s__Metschnikowia_pulcherrima(100); | Metschnikowia | 13 | Saprotroph | \|Undefined Saprotroph\| | Yeast | NULL | Probable | As nectar-tap_saprotroph (Põlme et al. 2020); Decay sugar-rich substrates (Põlme et al. 2020) | Põlme S, et al. 2020. Fungal Diversity 105:1-16 ((https://doi.org/10.1007/s13225-020-00466-2)) |
| Otu32411 | 1 | k__Fungi(100);p__Ascomycota(100);c__Saccharomycetes(100);o__Saccharomycetales(100);f__Metschnikowiaceae(100);g__Metschnikowia(100);s__Metschnikowia_pulcherrima(100); | Metschnikowia | 13 | Saprotroph | \|Undefined Saprotroph\| | Yeast | NULL | Probable | As nectar-tap_saprotroph (Põlme et al. 2020); Decay sugar-rich substrates (Põlme et al. 2020) | Põlme S, et al. 2020. Fungal Diversity 105:1-16 ((https://doi.org/10.1007/s13225-020-00466-2)) |
| Otu32412 | 1 | k__Fungi(100);p__Ascomycota(100);c__Saccharomycetes(100);o__Saccharomycetales(100);f__Metschnikowiaceae(100);g__Metschnikowia(100);s__Metschnikowia_pulcherrima(100); | Metschnikowia | 13 | Saprotroph | \|Undefined Saprotroph\| | Yeast | NULL | Probable | As nectar-tap_saprotroph (Põlme et al. 2020); Decay sugar-rich substrates (Põlme et al. 2020) | Põlme S, et al. 2020. Fungal Diversity 105:1-16 ((https://doi.org/10.1007/s13225-020-00466-2)) |
| Otu32413 | 1 | k__Fungi(100);p__Ascomycota(100);c__Saccharomycetes(100);o__Saccharomycetales(100);f__Metschnikowiaceae(100);g__Metschnikowia(100);s__Metschnikowia_pulcherrima(100); | Metschnikowia | 13 | Saprotroph | \|Undefined Saprotroph\| | Yeast | NULL | Probable | As nectar-tap_saprotroph (Põlme et al. 2020); Decay sugar-rich substrates (Põlme et al. 2020) | Põlme S, et al. 2020. Fungal Diversity 105:1-16 ((https://doi.org/10.1007/s13225-020-00466-2)) |
| Otu32415 | 1 | k__Fungi(100);p__Ascomycota(100);c__Saccharomycetes(100);o__Saccharomycetales(100);f__Saccharomycetales_family_Incertae_sedis(100);g__Candida(100);s__Candida_cretensis(100); | Candida | 13 | Pathotroph-Saprotroph-Symbiotroph | Animal Pathogen-Endophyte-Endosymbiont-Epiphyte-\|Undefined Saprotroph\| | Dimorphic Yeast-Yeast | NULL | Probable | As nectar-tap_saprotroph (Põlme et al. 2020); Decay sugar-rich substrates (Põlme et al. 2020); This asexual genus is polyphyletic and contain species that function very differently depend on what they are. Caution should be taken when interpreting guilds for Candida species. | Manolakaki D, et al. 2010. Virulence. 1: 367-375 ((https://doi.org/10.4161/viru.1.5.12796)); Mekbib SB, et al. 2011. Tropical Plant Pathology 36:233-240 ((https://doi.org/10.1590/S1982-56762011000400004)); Põlme S, et al. 2020. Fungal Diversity 105:1-16 ((https://doi.org/10.1007/s13225-020-00466-2)) |
| Otu32418 | 1 | k__Fungi(100);p__Ascomycota(100);c__Saccharomycetes(100);o__Saccharomycetales(100);o__Saccharomycetales_unclassified(100);o__Saccharomycetales_unclassified(100);o__Saccharomycetales_unclassified(100); | Saccharomycetales | 7 | Saprotroph | Undefined Saprotroph | Yeast | NULL | Possible | NULL | Sterkenburg E, et al. 2015. New Phytologist 207:1145-1158 ((https://doi.org/10.1111/nph.13426)) |
| Otu32420 | 1 | k__Fungi(100);p__Ascomycota(100);c__Saccharomycetes(100);o__Saccharomycetales(100);f__Metschnikowiaceae(100);g__Metschnikowia(100);s__Metschnikowia_pulcherrima(100); | Metschnikowia | 13 | Saprotroph | \|Undefined Saprotroph\| | Yeast | NULL | Probable | As nectar-tap_saprotroph (Põlme et al. 2020); Decay sugar-rich substrates (Põlme et al. 2020) | Põlme S, et al. 2020. Fungal Diversity 105:1-16 ((https://doi.org/10.1007/s13225-020-00466-2)) |
| Otu32423 | 1 | k__Fungi(100);p__Ascomycota(100);c__Saccharomycetes(100);o__Saccharomycetales(100);f__Saccharomycetales_family_Incertae_sedis(100);g__Candida(100);g__Candida_unclassified(100); | Candida | 13 | Pathotroph-Saprotroph-Symbiotroph | Animal Pathogen-Endophyte-Endosymbiont-Epiphyte-\|Undefined Saprotroph\| | Dimorphic Yeast-Yeast | NULL | Probable | As nectar-tap_saprotroph (Põlme et al. 2020); Decay sugar-rich substrates (Põlme et al. 2020); This asexual genus is polyphyletic and contain species that function very differently depend on what they are. Caution should be taken when interpreting guilds for Candida species. | Manolakaki D, et al. 2010. Virulence. 1: 367-375 ((https://doi.org/10.4161/viru.1.5.12796)); Mekbib SB, et al. 2011. Tropical Plant Pathology 36:233-240 ((https://doi.org/10.1590/S1982-56762011000400004)); Põlme S, et al. 2020. Fungal Diversity 105:1-16 ((https://doi.org/10.1007/s13225-020-00466-2)) |
| Otu32424 | 1 | k__Fungi(100);p__Ascomycota(100);c__Saccharomycetes(100);o__Saccharomycetales(100);f__Saccharomycetales_family_Incertae_sedis(100);g__Candida(100);g__Candida_unclassified(100); | Candida | 13 | Pathotroph-Saprotroph-Symbiotroph | Animal Pathogen-Endophyte-Endosymbiont-Epiphyte-\|Undefined Saprotroph\| | Dimorphic Yeast-Yeast | NULL | Probable | As nectar-tap_saprotroph (Põlme et al. 2020); Decay sugar-rich substrates (Põlme et al. 2020); This asexual genus is polyphyletic and contain species that function very differently depend on what they are. Caution should be taken when interpreting guilds for Candida species. | Manolakaki D, et al. 2010. Virulence. 1: 367-375 ((https://doi.org/10.4161/viru.1.5.12796)); Mekbib SB, et al. 2011. Tropical Plant Pathology 36:233-240 ((https://doi.org/10.1590/S1982-56762011000400004)); Põlme S, et al. 2020. Fungal Diversity 105:1-16 ((https://doi.org/10.1007/s13225-020-00466-2)) |
| Otu32428 | 1 | k__Fungi(100);p__Ascomycota(100);c__Saccharomycetes(100);o__Saccharomycetales(100);f__Metschnikowiaceae(100);g__Metschnikowia(100);g__Metschnikowia_unclassified(100); | Metschnikowia | 13 | Saprotroph | \|Undefined Saprotroph\| | Yeast | NULL | Probable | As nectar-tap_saprotroph (Põlme et al. 2020); Decay sugar-rich substrates (Põlme et al. 2020) | Põlme S, et al. 2020. Fungal Diversity 105:1-16 ((https://doi.org/10.1007/s13225-020-00466-2)) |
| Otu32429 | 1 | k__Fungi(100);p__Ascomycota(100);c__Saccharomycetes(100);o__Saccharomycetales(100);f__Metschnikowiaceae(100);g__Metschnikowia(100);s__Metschnikowia_pulcherrima(100); | Metschnikowia | 13 | Saprotroph | \|Undefined Saprotroph\| | Yeast | NULL | Probable | As nectar-tap_saprotroph (Põlme et al. 2020); Decay sugar-rich substrates (Põlme et al. 2020) | Põlme S, et al. 2020. Fungal Diversity 105:1-16 ((https://doi.org/10.1007/s13225-020-00466-2)) |
| Otu32430 | 1 | k__Fungi(100);p__Ascomycota(100);c__Saccharomycetes(100);o__Saccharomycetales(100);f__Metschnikowiaceae(100);g__Metschnikowia(100);g__Metschnikowia_unclassified(100); | Metschnikowia | 13 | Saprotroph | \|Undefined Saprotroph\| | Yeast | NULL | Probable | As nectar-tap_saprotroph (Põlme et al. 2020); Decay sugar-rich substrates (Põlme et al. 2020) | Põlme S, et al. 2020. Fungal Diversity 105:1-16 ((https://doi.org/10.1007/s13225-020-00466-2)) |
| Otu32431 | 1 | k__Fungi(100);p__Ascomycota(100);c__Saccharomycetes(100);o__Saccharomycetales(100);f__Metschnikowiaceae(100);g__Metschnikowia(100);s__Metschnikowia_pulcherrima(100); | Metschnikowia | 13 | Saprotroph | \|Undefined Saprotroph\| | Yeast | NULL | Probable | As nectar-tap_saprotroph (Põlme et al. 2020); Decay sugar-rich substrates (Põlme et al. 2020) | Põlme S, et al. 2020. Fungal Diversity 105:1-16 ((https://doi.org/10.1007/s13225-020-00466-2)) |
| Otu32433 | 1 | k__Fungi(100);p__Ascomycota(100);c__Saccharomycetes(100);o__Saccharomycetales(100);f__Metschnikowiaceae(100);g__Metschnikowia(100);s__Metschnikowia_pulcherrima(100); | Metschnikowia | 13 | Saprotroph | \|Undefined Saprotroph\| | Yeast | NULL | Probable | As nectar-tap_saprotroph (Põlme et al. 2020); Decay sugar-rich substrates (Põlme et al. 2020) | Põlme S, et al. 2020. Fungal Diversity 105:1-16 ((https://doi.org/10.1007/s13225-020-00466-2)) |
| Otu32434 | 1 | k__Fungi(100);p__Ascomycota(100);c__Saccharomycetes(100);o__Saccharomycetales(100);f__Metschnikowiaceae(100);g__Metschnikowia(100);g__Metschnikowia_unclassified(100); | Metschnikowia | 13 | Saprotroph | \|Undefined Saprotroph\| | Yeast | NULL | Probable | As nectar-tap_saprotroph (Põlme et al. 2020); Decay sugar-rich substrates (Põlme et al. 2020) | Põlme S, et al. 2020. Fungal Diversity 105:1-16 ((https://doi.org/10.1007/s13225-020-00466-2)) |
| Otu32435 | 1 | k__Fungi(100);p__Ascomycota(100);c__Saccharomycetes(100);o__Saccharomycetales(100);f__Saccharomycetales_family_Incertae_sedis(100);g__Candida(100);g__Candida_unclassified(100); | Candida | 13 | Pathotroph-Saprotroph-Symbiotroph | Animal Pathogen-Endophyte-Endosymbiont-Epiphyte-\|Undefined Saprotroph\| | Dimorphic Yeast-Yeast | NULL | Probable | As nectar-tap_saprotroph (Põlme et al. 2020); Decay sugar-rich substrates (Põlme et al. 2020); This asexual genus is polyphyletic and contain species that function very differently depend on what they are. Caution should be taken when interpreting guilds for Candida species. | Manolakaki D, et al. 2010. Virulence. 1: 367-375 ((https://doi.org/10.4161/viru.1.5.12796)); Mekbib SB, et al. 2011. Tropical Plant Pathology 36:233-240 ((https://doi.org/10.1590/S1982-56762011000400004)); Põlme S, et al. 2020. Fungal Diversity 105:1-16 ((https://doi.org/10.1007/s13225-020-00466-2)) |
| Otu32437 | 1 | k__Fungi(100);p__Basidiomycota(100);c__Tremellomycetes(100);o__Tremellales(100);f__Tremellales_family_Incertae_sedis(100);f__Tremellales_family_Incertae_sedis_unclassified(100);f__Tremellales_family_Incertae_sedis_unclassified(100); | Tremellales | 7 | Pathotroph-Saprotroph-Symbiotroph | Fungal Parasite-Undefined Saprotroph | Tremelloid-Yeast | NULL | Possible | The order contains both teleomorphic and anamorphic species, most of the latter being yeasts | Sterkenburg E, et al. 2015. New Phytologist 207:1145-1158 ((https://doi.org/10.1111/nph.13426)) |
| Otu32438 | 1 | k__Fungi(100);p__Ascomycota(100);c__Saccharomycetes(100);o__Saccharomycetales(100);f__Metschnikowiaceae(100);g__Metschnikowia(100);s__Metschnikowia_pulcherrima(100); | Metschnikowia | 13 | Saprotroph | \|Undefined Saprotroph\| | Yeast | NULL | Probable | As nectar-tap_saprotroph (Põlme et al. 2020); Decay sugar-rich substrates (Põlme et al. 2020) | Põlme S, et al. 2020. Fungal Diversity 105:1-16 ((https://doi.org/10.1007/s13225-020-00466-2)) |
| Otu32439 | 1 | k__Fungi(100);p__Ascomycota(100);c__Saccharomycetes(100);o__Saccharomycetales(100);f__Saccharomycetales_family_Incertae_sedis(100);g__Candida(100);s__Candida_cretensis(100); | Candida | 13 | Pathotroph-Saprotroph-Symbiotroph | Animal Pathogen-Endophyte-Endosymbiont-Epiphyte-\|Undefined Saprotroph\| | Dimorphic Yeast-Yeast | NULL | Probable | As nectar-tap_saprotroph (Põlme et al. 2020); Decay sugar-rich substrates (Põlme et al. 2020); This asexual genus is polyphyletic and contain species that function very differently depend on what they are. Caution should be taken when interpreting guilds for Candida species. | Manolakaki D, et al. 2010. Virulence. 1: 367-375 ((https://doi.org/10.4161/viru.1.5.12796)); Mekbib SB, et al. 2011. Tropical Plant Pathology 36:233-240 ((https://doi.org/10.1590/S1982-56762011000400004)); Põlme S, et al. 2020. Fungal Diversity 105:1-16 ((https://doi.org/10.1007/s13225-020-00466-2)) |
| Otu32441 | 1 | k__Fungi(100);p__Ascomycota(100);c__Saccharomycetes(100);o__Saccharomycetales(100);f__Metschnikowiaceae(100);g__Metschnikowia(100);s__Metschnikowia_pulcherrima(100); | Metschnikowia | 13 | Saprotroph | \|Undefined Saprotroph\| | Yeast | NULL | Probable | As nectar-tap_saprotroph (Põlme et al. 2020); Decay sugar-rich substrates (Põlme et al. 2020) | Põlme S, et al. 2020. Fungal Diversity 105:1-16 ((https://doi.org/10.1007/s13225-020-00466-2)) |
| Otu32442 | 1 | k__Fungi(100);p__Ascomycota(100);c__Saccharomycetes(100);o__Saccharomycetales(100);f__Saccharomycetales_family_Incertae_sedis(100);g__Candida(100);g__Candida_unclassified(100); | Candida | 13 | Pathotroph-Saprotroph-Symbiotroph | Animal Pathogen-Endophyte-Endosymbiont-Epiphyte-\|Undefined Saprotroph\| | Dimorphic Yeast-Yeast | NULL | Probable | As nectar-tap_saprotroph (Põlme et al. 2020); Decay sugar-rich substrates (Põlme et al. 2020); This asexual genus is polyphyletic and contain species that function very differently depend on what they are. Caution should be taken when interpreting guilds for Candida species. | Manolakaki D, et al. 2010. Virulence. 1: 367-375 ((https://doi.org/10.4161/viru.1.5.12796)); Mekbib SB, et al. 2011. Tropical Plant Pathology 36:233-240 ((https://doi.org/10.1590/S1982-56762011000400004)); Põlme S, et al. 2020. Fungal Diversity 105:1-16 ((https://doi.org/10.1007/s13225-020-00466-2)) |
| Otu32443 | 1 | k__Fungi(100);p__Ascomycota(100);c__Saccharomycetes(100);o__Saccharomycetales(100);f__Saccharomycetales_family_Incertae_sedis(100);g__Candida(100);s__Candida_cretensis(100); | Candida | 13 | Pathotroph-Saprotroph-Symbiotroph | Animal Pathogen-Endophyte-Endosymbiont-Epiphyte-\|Undefined Saprotroph\| | Dimorphic Yeast-Yeast | NULL | Probable | As nectar-tap_saprotroph (Põlme et al. 2020); Decay sugar-rich substrates (Põlme et al. 2020); This asexual genus is polyphyletic and contain species that function very differently depend on what they are. Caution should be taken when interpreting guilds for Candida species. | Manolakaki D, et al. 2010. Virulence. 1: 367-375 ((https://doi.org/10.4161/viru.1.5.12796)); Mekbib SB, et al. 2011. Tropical Plant Pathology 36:233-240 ((https://doi.org/10.1590/S1982-56762011000400004)); Põlme S, et al. 2020. Fungal Diversity 105:1-16 ((https://doi.org/10.1007/s13225-020-00466-2)) |
| Otu32446 | 1 | k__Fungi(100);p__Ascomycota(100);c__Saccharomycetes(100);o__Saccharomycetales(100);f__Saccharomycetales_family_Incertae_sedis(100);g__Candida(100);g__Candida_unclassified(100); | Candida | 13 | Pathotroph-Saprotroph-Symbiotroph | Animal Pathogen-Endophyte-Endosymbiont-Epiphyte-\|Undefined Saprotroph\| | Dimorphic Yeast-Yeast | NULL | Probable | As nectar-tap_saprotroph (Põlme et al. 2020); Decay sugar-rich substrates (Põlme et al. 2020); This asexual genus is polyphyletic and contain species that function very differently depend on what they are. Caution should be taken when interpreting guilds for Candida species. | Manolakaki D, et al. 2010. Virulence. 1: 367-375 ((https://doi.org/10.4161/viru.1.5.12796)); Mekbib SB, et al. 2011. Tropical Plant Pathology 36:233-240 ((https://doi.org/10.1590/S1982-56762011000400004)); Põlme S, et al. 2020. Fungal Diversity 105:1-16 ((https://doi.org/10.1007/s13225-020-00466-2)) |
| Otu32449 | 1 | k__Fungi(100);p__Ascomycota(100);c__Saccharomycetes(100);o__Saccharomycetales(100);f__Saccharomycetales_family_Incertae_sedis(100);f__Saccharomycetales_family_Incertae_sedis_unclassified(100);f__Saccharomycetales_family_Incertae_sedis_unclassified(100); | Saccharomycetales | 7 | Saprotroph | Undefined Saprotroph | Yeast | NULL | Possible | NULL | Sterkenburg E, et al. 2015. New Phytologist 207:1145-1158 ((https://doi.org/10.1111/nph.13426)) |
| Otu32452 | 1 | k__Fungi(100);p__Ascomycota(100);c__Saccharomycetes(100);o__Saccharomycetales(100);o__Saccharomycetales_unclassified(100);o__Saccharomycetales_unclassified(100);o__Saccharomycetales_unclassified(100); | Saccharomycetales | 7 | Saprotroph | Undefined Saprotroph | Yeast | NULL | Possible | NULL | Sterkenburg E, et al. 2015. New Phytologist 207:1145-1158 ((https://doi.org/10.1111/nph.13426)) |
| Otu32453 | 1 | k__Fungi(100);p__Ascomycota(100);c__Saccharomycetes(100);o__Saccharomycetales(100);f__Saccharomycetales_family_Incertae_sedis(100);g__Candida(100);g__Candida_unclassified(100); | Candida | 13 | Pathotroph-Saprotroph-Symbiotroph | Animal Pathogen-Endophyte-Endosymbiont-Epiphyte-\|Undefined Saprotroph\| | Dimorphic Yeast-Yeast | NULL | Probable | As nectar-tap_saprotroph (Põlme et al. 2020); Decay sugar-rich substrates (Põlme et al. 2020); This asexual genus is polyphyletic and contain species that function very differently depend on what they are. Caution should be taken when interpreting guilds for Candida species. | Manolakaki D, et al. 2010. Virulence. 1: 367-375 ((https://doi.org/10.4161/viru.1.5.12796)); Mekbib SB, et al. 2011. Tropical Plant Pathology 36:233-240 ((https://doi.org/10.1590/S1982-56762011000400004)); Põlme S, et al. 2020. Fungal Diversity 105:1-16 ((https://doi.org/10.1007/s13225-020-00466-2)) |
| Otu32454 | 1 | k__Fungi(100);p__Ascomycota(100);c__Saccharomycetes(100);o__Saccharomycetales(100);f__Saccharomycetales_family_Incertae_sedis(100);g__Candida(100);s__Candida_cretensis(100); | Candida | 13 | Pathotroph-Saprotroph-Symbiotroph | Animal Pathogen-Endophyte-Endosymbiont-Epiphyte-\|Undefined Saprotroph\| | Dimorphic Yeast-Yeast | NULL | Probable | As nectar-tap_saprotroph (Põlme et al. 2020); Decay sugar-rich substrates (Põlme et al. 2020); This asexual genus is polyphyletic and contain species that function very differently depend on what they are. Caution should be taken when interpreting guilds for Candida species. | Manolakaki D, et al. 2010. Virulence. 1: 367-375 ((https://doi.org/10.4161/viru.1.5.12796)); Mekbib SB, et al. 2011. Tropical Plant Pathology 36:233-240 ((https://doi.org/10.1590/S1982-56762011000400004)); Põlme S, et al. 2020. Fungal Diversity 105:1-16 ((https://doi.org/10.1007/s13225-020-00466-2)) |
| Otu32455 | 1 | k__Fungi(100);p__Ascomycota(100);c__Saccharomycetes(100);o__Saccharomycetales(100);f__Saccharomycetales_family_Incertae_sedis(100);g__Candida(100);g__Candida_unclassified(100); | Candida | 13 | Pathotroph-Saprotroph-Symbiotroph | Animal Pathogen-Endophyte-Endosymbiont-Epiphyte-\|Undefined Saprotroph\| | Dimorphic Yeast-Yeast | NULL | Probable | As nectar-tap_saprotroph (Põlme et al. 2020); Decay sugar-rich substrates (Põlme et al. 2020); This asexual genus is polyphyletic and contain species that function very differently depend on what they are. Caution should be taken when interpreting guilds for Candida species. | Manolakaki D, et al. 2010. Virulence. 1: 367-375 ((https://doi.org/10.4161/viru.1.5.12796)); Mekbib SB, et al. 2011. Tropical Plant Pathology 36:233-240 ((https://doi.org/10.1590/S1982-56762011000400004)); Põlme S, et al. 2020. Fungal Diversity 105:1-16 ((https://doi.org/10.1007/s13225-020-00466-2)) |
| Otu32458 | 1 | k__Fungi(100);p__Ascomycota(100);c__Saccharomycetes(100);o__Saccharomycetales(100);f__Metschnikowiaceae(100);g__Metschnikowia(100);s__Metschnikowia_pulcherrima(100); | Metschnikowia | 13 | Saprotroph | \|Undefined Saprotroph\| | Yeast | NULL | Probable | As nectar-tap_saprotroph (Põlme et al. 2020); Decay sugar-rich substrates (Põlme et al. 2020) | Põlme S, et al. 2020. Fungal Diversity 105:1-16 ((https://doi.org/10.1007/s13225-020-00466-2)) |
| Otu32461 | 1 | k__Fungi(100);p__Ascomycota(100);c__Saccharomycetes(100);o__Saccharomycetales(100);f__Saccharomycetales_family_Incertae_sedis(100);g__Candida(100);g__Candida_unclassified(100); | Candida | 13 | Pathotroph-Saprotroph-Symbiotroph | Animal Pathogen-Endophyte-Endosymbiont-Epiphyte-\|Undefined Saprotroph\| | Dimorphic Yeast-Yeast | NULL | Probable | As nectar-tap_saprotroph (Põlme et al. 2020); Decay sugar-rich substrates (Põlme et al. 2020); This asexual genus is polyphyletic and contain species that function very differently depend on what they are. Caution should be taken when interpreting guilds for Candida species. | Manolakaki D, et al. 2010. Virulence. 1: 367-375 ((https://doi.org/10.4161/viru.1.5.12796)); Mekbib SB, et al. 2011. Tropical Plant Pathology 36:233-240 ((https://doi.org/10.1590/S1982-56762011000400004)); Põlme S, et al. 2020. Fungal Diversity 105:1-16 ((https://doi.org/10.1007/s13225-020-00466-2)) |
| Otu32462 | 1 | k__Fungi(100);p__Ascomycota(100);c__Saccharomycetes(100);o__Saccharomycetales(100);f__Saccharomycetales_family_Incertae_sedis(100);g__Candida(100);g__Candida_unclassified(100); | Candida | 13 | Pathotroph-Saprotroph-Symbiotroph | Animal Pathogen-Endophyte-Endosymbiont-Epiphyte-\|Undefined Saprotroph\| | Dimorphic Yeast-Yeast | NULL | Probable | As nectar-tap_saprotroph (Põlme et al. 2020); Decay sugar-rich substrates (Põlme et al. 2020); This asexual genus is polyphyletic and contain species that function very differently depend on what they are. Caution should be taken when interpreting guilds for Candida species. | Manolakaki D, et al. 2010. Virulence. 1: 367-375 ((https://doi.org/10.4161/viru.1.5.12796)); Mekbib SB, et al. 2011. Tropical Plant Pathology 36:233-240 ((https://doi.org/10.1590/S1982-56762011000400004)); Põlme S, et al. 2020. Fungal Diversity 105:1-16 ((https://doi.org/10.1007/s13225-020-00466-2)) |
| Otu32463 | 1 | k__Fungi(100);p__Ascomycota(100);c__Saccharomycetes(100);o__Saccharomycetales(100);f__Saccharomycetales_family_Incertae_sedis(100);g__Candida(100);g__Candida_unclassified(100); | Candida | 13 | Pathotroph-Saprotroph-Symbiotroph | Animal Pathogen-Endophyte-Endosymbiont-Epiphyte-\|Undefined Saprotroph\| | Dimorphic Yeast-Yeast | NULL | Probable | As nectar-tap_saprotroph (Põlme et al. 2020); Decay sugar-rich substrates (Põlme et al. 2020); This asexual genus is polyphyletic and contain species that function very differently depend on what they are. Caution should be taken when interpreting guilds for Candida species. | Manolakaki D, et al. 2010. Virulence. 1: 367-375 ((https://doi.org/10.4161/viru.1.5.12796)); Mekbib SB, et al. 2011. Tropical Plant Pathology 36:233-240 ((https://doi.org/10.1590/S1982-56762011000400004)); Põlme S, et al. 2020. Fungal Diversity 105:1-16 ((https://doi.org/10.1007/s13225-020-00466-2)) |
| Otu32464 | 1 | k__Fungi(100);p__Ascomycota(100);c__Saccharomycetes(100);o__Saccharomycetales(100);f__Metschnikowiaceae(100);g__Metschnikowia(100);s__Metschnikowia_pulcherrima(100); | Metschnikowia | 13 | Saprotroph | \|Undefined Saprotroph\| | Yeast | NULL | Probable | As nectar-tap_saprotroph (Põlme et al. 2020); Decay sugar-rich substrates (Põlme et al. 2020) | Põlme S, et al. 2020. Fungal Diversity 105:1-16 ((https://doi.org/10.1007/s13225-020-00466-2)) |
| Otu32465 | 1 | k__Fungi(100);p__Ascomycota(100);c__Saccharomycetes(100);o__Saccharomycetales(100);f__Metschnikowiaceae(100);g__Metschnikowia(100);g__Metschnikowia_unclassified(100); | Metschnikowia | 13 | Saprotroph | \|Undefined Saprotroph\| | Yeast | NULL | Probable | As nectar-tap_saprotroph (Põlme et al. 2020); Decay sugar-rich substrates (Põlme et al. 2020) | Põlme S, et al. 2020. Fungal Diversity 105:1-16 ((https://doi.org/10.1007/s13225-020-00466-2)) |
| Otu32466 | 1 | k__Fungi(100);p__Ascomycota(100);c__Saccharomycetes(100);o__Saccharomycetales(100);f__Metschnikowiaceae(100);g__Metschnikowia(100);s__Metschnikowia_pulcherrima(100); | Metschnikowia | 13 | Saprotroph | \|Undefined Saprotroph\| | Yeast | NULL | Probable | As nectar-tap_saprotroph (Põlme et al. 2020); Decay sugar-rich substrates (Põlme et al. 2020) | Põlme S, et al. 2020. Fungal Diversity 105:1-16 ((https://doi.org/10.1007/s13225-020-00466-2)) |
| Otu32467 | 1 | k__Fungi(100);p__Ascomycota(100);c__Saccharomycetes(100);o__Saccharomycetales(100);f__Metschnikowiaceae(100);g__Metschnikowia(100);s__Metschnikowia_pulcherrima(100); | Metschnikowia | 13 | Saprotroph | \|Undefined Saprotroph\| | Yeast | NULL | Probable | As nectar-tap_saprotroph (Põlme et al. 2020); Decay sugar-rich substrates (Põlme et al. 2020) | Põlme S, et al. 2020. Fungal Diversity 105:1-16 ((https://doi.org/10.1007/s13225-020-00466-2)) |
| Otu32468 | 1 | k__Fungi(100);p__Ascomycota(100);c__Saccharomycetes(100);o__Saccharomycetales(100);f__Saccharomycetales_family_Incertae_sedis(100);g__Candida(100);g__Candida_unclassified(100); | Candida | 13 | Pathotroph-Saprotroph-Symbiotroph | Animal Pathogen-Endophyte-Endosymbiont-Epiphyte-\|Undefined Saprotroph\| | Dimorphic Yeast-Yeast | NULL | Probable | As nectar-tap_saprotroph (Põlme et al. 2020); Decay sugar-rich substrates (Põlme et al. 2020); This asexual genus is polyphyletic and contain species that function very differently depend on what they are. Caution should be taken when interpreting guilds for Candida species. | Manolakaki D, et al. 2010. Virulence. 1: 367-375 ((https://doi.org/10.4161/viru.1.5.12796)); Mekbib SB, et al. 2011. Tropical Plant Pathology 36:233-240 ((https://doi.org/10.1590/S1982-56762011000400004)); Põlme S, et al. 2020. Fungal Diversity 105:1-16 ((https://doi.org/10.1007/s13225-020-00466-2)) |
| Otu32469 | 1 | k__Fungi(100);p__Ascomycota(100);c__Saccharomycetes(100);o__Saccharomycetales(100);f__Saccharomycetales_family_Incertae_sedis(100);g__Candida(100);g__Candida_unclassified(100); | Candida | 13 | Pathotroph-Saprotroph-Symbiotroph | Animal Pathogen-Endophyte-Endosymbiont-Epiphyte-\|Undefined Saprotroph\| | Dimorphic Yeast-Yeast | NULL | Probable | As nectar-tap_saprotroph (Põlme et al. 2020); Decay sugar-rich substrates (Põlme et al. 2020); This asexual genus is polyphyletic and contain species that function very differently depend on what they are. Caution should be taken when interpreting guilds for Candida species. | Manolakaki D, et al. 2010. Virulence. 1: 367-375 ((https://doi.org/10.4161/viru.1.5.12796)); Mekbib SB, et al. 2011. Tropical Plant Pathology 36:233-240 ((https://doi.org/10.1590/S1982-56762011000400004)); Põlme S, et al. 2020. Fungal Diversity 105:1-16 ((https://doi.org/10.1007/s13225-020-00466-2)) |
| Otu32470 | 1 | k__Fungi(100);p__Ascomycota(100);c__Saccharomycetes(100);o__Saccharomycetales(100);f__Saccharomycetales_family_Incertae_sedis(100);g__Candida(100);g__Candida_unclassified(100); | Candida | 13 | Pathotroph-Saprotroph-Symbiotroph | Animal Pathogen-Endophyte-Endosymbiont-Epiphyte-\|Undefined Saprotroph\| | Dimorphic Yeast-Yeast | NULL | Probable | As nectar-tap_saprotroph (Põlme et al. 2020); Decay sugar-rich substrates (Põlme et al. 2020); This asexual genus is polyphyletic and contain species that function very differently depend on what they are. Caution should be taken when interpreting guilds for Candida species. | Manolakaki D, et al. 2010. Virulence. 1: 367-375 ((https://doi.org/10.4161/viru.1.5.12796)); Mekbib SB, et al. 2011. Tropical Plant Pathology 36:233-240 ((https://doi.org/10.1590/S1982-56762011000400004)); Põlme S, et al. 2020. Fungal Diversity 105:1-16 ((https://doi.org/10.1007/s13225-020-00466-2)) |
| Otu32475 | 1 | k__Fungi(100);p__Ascomycota(100);c__Saccharomycetes(100);o__Saccharomycetales(100);f__Metschnikowiaceae(100);g__Metschnikowia(100);s__Metschnikowia_pulcherrima(100); | Metschnikowia | 13 | Saprotroph | \|Undefined Saprotroph\| | Yeast | NULL | Probable | As nectar-tap_saprotroph (Põlme et al. 2020); Decay sugar-rich substrates (Põlme et al. 2020) | Põlme S, et al. 2020. Fungal Diversity 105:1-16 ((https://doi.org/10.1007/s13225-020-00466-2)) |
| Otu32479 | 1 | k__Fungi(100);p__Ascomycota(100);c__Saccharomycetes(100);o__Saccharomycetales(100);f__Metschnikowiaceae(100);g__Metschnikowia(100);s__Metschnikowia_pulcherrima(100); | Metschnikowia | 13 | Saprotroph | \|Undefined Saprotroph\| | Yeast | NULL | Probable | As nectar-tap_saprotroph (Põlme et al. 2020); Decay sugar-rich substrates (Põlme et al. 2020) | Põlme S, et al. 2020. Fungal Diversity 105:1-16 ((https://doi.org/10.1007/s13225-020-00466-2)) |
| Otu32481 | 1 | k__Fungi(100);p__Ascomycota(100);c__Saccharomycetes(100);o__Saccharomycetales(100);f__Metschnikowiaceae(100);g__Metschnikowia(100);s__Metschnikowia_pulcherrima(100); | Metschnikowia | 13 | Saprotroph | \|Undefined Saprotroph\| | Yeast | NULL | Probable | As nectar-tap_saprotroph (Põlme et al. 2020); Decay sugar-rich substrates (Põlme et al. 2020) | Põlme S, et al. 2020. Fungal Diversity 105:1-16 ((https://doi.org/10.1007/s13225-020-00466-2)) |
| Otu32483 | 1 | k__Fungi(100);p__Ascomycota(100);c__Saccharomycetes(100);o__Saccharomycetales(100);f__Metschnikowiaceae(100);g__Metschnikowia(100);g__Metschnikowia_unclassified(100); | Metschnikowia | 13 | Saprotroph | \|Undefined Saprotroph\| | Yeast | NULL | Probable | As nectar-tap_saprotroph (Põlme et al. 2020); Decay sugar-rich substrates (Põlme et al. 2020) | Põlme S, et al. 2020. Fungal Diversity 105:1-16 ((https://doi.org/10.1007/s13225-020-00466-2)) |
| Otu32484 | 1 | k__Fungi(100);p__Ascomycota(100);c__Saccharomycetes(100);o__Saccharomycetales(100);f__Metschnikowiaceae(100);g__Metschnikowia(100);s__Metschnikowia_pulcherrima(100); | Metschnikowia | 13 | Saprotroph | \|Undefined Saprotroph\| | Yeast | NULL | Probable | As nectar-tap_saprotroph (Põlme et al. 2020); Decay sugar-rich substrates (Põlme et al. 2020) | Põlme S, et al. 2020. Fungal Diversity 105:1-16 ((https://doi.org/10.1007/s13225-020-00466-2)) |
| Otu32488 | 1 | k__Fungi(100);p__Ascomycota(100);c__Saccharomycetes(100);o__Saccharomycetales(100);f__Metschnikowiaceae(100);g__Metschnikowia(100);s__Metschnikowia_pulcherrima(100); | Metschnikowia | 13 | Saprotroph | \|Undefined Saprotroph\| | Yeast | NULL | Probable | As nectar-tap_saprotroph (Põlme et al. 2020); Decay sugar-rich substrates (Põlme et al. 2020) | Põlme S, et al. 2020. Fungal Diversity 105:1-16 ((https://doi.org/10.1007/s13225-020-00466-2)) |
| Otu32489 | 1 | k__Fungi(100);p__Ascomycota(100);c__Saccharomycetes(100);o__Saccharomycetales(100);o__Saccharomycetales_unclassified(100);o__Saccharomycetales_unclassified(100);o__Saccharomycetales_unclassified(100); | Saccharomycetales | 7 | Saprotroph | Undefined Saprotroph | Yeast | NULL | Possible | NULL | Sterkenburg E, et al. 2015. New Phytologist 207:1145-1158 ((https://doi.org/10.1111/nph.13426)) |
| Otu32491 | 1 | k__Fungi(100);p__Ascomycota(100);c__Sordariomycetes(100);o__Hypocreales(100);f__Nectriaceae(100);g__Gibberella(100);s__Gibberella_intricans(100); | Gibberella | 13 | Pathotroph-Saprotroph-Symbiotroph | Endophyte-\|Plant Pathogen\|-Plant Saprotroph-Undefined Saprotroph | Microfungus | Soft Rot | Probable | As foliar_endophyte (Põlme et al. 2020); As litter_saprotroph (Põlme et al. 2020); Causing damping-off disease (Põlme et al. 2020); Decay leaf, fruit, seed, roots, soil, animal material (Põlme et al. 2020) | Tedersoo L, et al. 2014. Science 346:e1256688 ((https://doi.org/10.1126/science.1256688)); Põlme S, et al. 2020. Fungal Diversity 105:1-16 ((https://doi.org/10.1007/s13225-020-00466-2)) |
| Otu32492 | 1 | k__Fungi(100);p__Ascomycota(100);c__Saccharomycetes(100);o__Saccharomycetales(100);f__Metschnikowiaceae(100);g__Metschnikowia(100);s__Metschnikowia_pulcherrima(100); | Metschnikowia | 13 | Saprotroph | \|Undefined Saprotroph\| | Yeast | NULL | Probable | As nectar-tap_saprotroph (Põlme et al. 2020); Decay sugar-rich substrates (Põlme et al. 2020) | Põlme S, et al. 2020. Fungal Diversity 105:1-16 ((https://doi.org/10.1007/s13225-020-00466-2)) |
| Otu32493 | 1 | k__Fungi(100);p__Ascomycota(100);c__Saccharomycetes(100);o__Saccharomycetales(100);f__Metschnikowiaceae(100);g__Metschnikowia(100);s__Metschnikowia_pulcherrima(100); | Metschnikowia | 13 | Saprotroph | \|Undefined Saprotroph\| | Yeast | NULL | Probable | As nectar-tap_saprotroph (Põlme et al. 2020); Decay sugar-rich substrates (Põlme et al. 2020) | Põlme S, et al. 2020. Fungal Diversity 105:1-16 ((https://doi.org/10.1007/s13225-020-00466-2)) |
| Otu32494 | 1 | k__Fungi(100);p__Ascomycota(100);c__Saccharomycetes(100);o__Saccharomycetales(100);f__Metschnikowiaceae(100);g__Metschnikowia(100);s__Metschnikowia_pulcherrima(100); | Metschnikowia | 13 | Saprotroph | \|Undefined Saprotroph\| | Yeast | NULL | Probable | As nectar-tap_saprotroph (Põlme et al. 2020); Decay sugar-rich substrates (Põlme et al. 2020) | Põlme S, et al. 2020. Fungal Diversity 105:1-16 ((https://doi.org/10.1007/s13225-020-00466-2)) |
| Otu32495 | 1 | k__Fungi(100);p__Ascomycota(100);c__Saccharomycetes(100);o__Saccharomycetales(100);f__Metschnikowiaceae(100);g__Metschnikowia(100);s__Metschnikowia_pulcherrima(100); | Metschnikowia | 13 | Saprotroph | \|Undefined Saprotroph\| | Yeast | NULL | Probable | As nectar-tap_saprotroph (Põlme et al. 2020); Decay sugar-rich substrates (Põlme et al. 2020) | Põlme S, et al. 2020. Fungal Diversity 105:1-16 ((https://doi.org/10.1007/s13225-020-00466-2)) |
| Otu32497 | 1 | k__Fungi(100);p__Ascomycota(100);c__Saccharomycetes(100);o__Saccharomycetales(100);f__Metschnikowiaceae(100);g__Metschnikowia(100);s__Metschnikowia_pulcherrima(100); | Metschnikowia | 13 | Saprotroph | \|Undefined Saprotroph\| | Yeast | NULL | Probable | As nectar-tap_saprotroph (Põlme et al. 2020); Decay sugar-rich substrates (Põlme et al. 2020) | Põlme S, et al. 2020. Fungal Diversity 105:1-16 ((https://doi.org/10.1007/s13225-020-00466-2)) |
| Otu32498 | 1 | k__Fungi(100);p__Ascomycota(100);c__Saccharomycetes(100);o__Saccharomycetales(100);f__Metschnikowiaceae(100);g__Metschnikowia(100);g__Metschnikowia_unclassified(100); | Metschnikowia | 13 | Saprotroph | \|Undefined Saprotroph\| | Yeast | NULL | Probable | As nectar-tap_saprotroph (Põlme et al. 2020); Decay sugar-rich substrates (Põlme et al. 2020) | Põlme S, et al. 2020. Fungal Diversity 105:1-16 ((https://doi.org/10.1007/s13225-020-00466-2)) |
| Otu32499 | 1 | k__Fungi(100);p__Ascomycota(100);c__Saccharomycetes(100);o__Saccharomycetales(100);f__Metschnikowiaceae(100);g__Metschnikowia(100);s__Metschnikowia_pulcherrima(100); | Metschnikowia | 13 | Saprotroph | \|Undefined Saprotroph\| | Yeast | NULL | Probable | As nectar-tap_saprotroph (Põlme et al. 2020); Decay sugar-rich substrates (Põlme et al. 2020) | Põlme S, et al. 2020. Fungal Diversity 105:1-16 ((https://doi.org/10.1007/s13225-020-00466-2)) |
| Otu32500 | 1 | k__Fungi(100);p__Ascomycota(100);c__Saccharomycetes(100);o__Saccharomycetales(100);f__Saccharomycetales_family_Incertae_sedis(100);g__Candida(100);g__Candida_unclassified(100); | Candida | 13 | Pathotroph-Saprotroph-Symbiotroph | Animal Pathogen-Endophyte-Endosymbiont-Epiphyte-\|Undefined Saprotroph\| | Dimorphic Yeast-Yeast | NULL | Probable | As nectar-tap_saprotroph (Põlme et al. 2020); Decay sugar-rich substrates (Põlme et al. 2020); This asexual genus is polyphyletic and contain species that function very differently depend on what they are. Caution should be taken when interpreting guilds for Candida species. | Manolakaki D, et al. 2010. Virulence. 1: 367-375 ((https://doi.org/10.4161/viru.1.5.12796)); Mekbib SB, et al. 2011. Tropical Plant Pathology 36:233-240 ((https://doi.org/10.1590/S1982-56762011000400004)); Põlme S, et al. 2020. Fungal Diversity 105:1-16 ((https://doi.org/10.1007/s13225-020-00466-2)) |
| Otu32501 | 1 | k__Fungi(100);p__Ascomycota(100);c__Saccharomycetes(100);o__Saccharomycetales(100);f__Metschnikowiaceae(100);g__Metschnikowia(100);s__Metschnikowia_pulcherrima(100); | Metschnikowia | 13 | Saprotroph | \|Undefined Saprotroph\| | Yeast | NULL | Probable | As nectar-tap_saprotroph (Põlme et al. 2020); Decay sugar-rich substrates (Põlme et al. 2020) | Põlme S, et al. 2020. Fungal Diversity 105:1-16 ((https://doi.org/10.1007/s13225-020-00466-2)) |
| Otu32502 | 1 | k__Fungi(100);p__Ascomycota(100);c__Saccharomycetes(100);o__Saccharomycetales(100);f__Metschnikowiaceae(100);g__Metschnikowia(100);s__Metschnikowia_pulcherrima(100); | Metschnikowia | 13 | Saprotroph | \|Undefined Saprotroph\| | Yeast | NULL | Probable | As nectar-tap_saprotroph (Põlme et al. 2020); Decay sugar-rich substrates (Põlme et al. 2020) | Põlme S, et al. 2020. Fungal Diversity 105:1-16 ((https://doi.org/10.1007/s13225-020-00466-2)) |
| Otu32503 | 1 | k__Fungi(100);p__Ascomycota(100);c__Saccharomycetes(100);o__Saccharomycetales(100);f__Metschnikowiaceae(100);g__Metschnikowia(100);s__Metschnikowia_pulcherrima(100); | Metschnikowia | 13 | Saprotroph | \|Undefined Saprotroph\| | Yeast | NULL | Probable | As nectar-tap_saprotroph (Põlme et al. 2020); Decay sugar-rich substrates (Põlme et al. 2020) | Põlme S, et al. 2020. Fungal Diversity 105:1-16 ((https://doi.org/10.1007/s13225-020-00466-2)) |
| Otu32504 | 1 | k__Fungi(100);p__Ascomycota(100);c__Saccharomycetes(100);o__Saccharomycetales(100);f__Metschnikowiaceae(100);g__Metschnikowia(100);s__Metschnikowia_pulcherrima(100); | Metschnikowia | 13 | Saprotroph | \|Undefined Saprotroph\| | Yeast | NULL | Probable | As nectar-tap_saprotroph (Põlme et al. 2020); Decay sugar-rich substrates (Põlme et al. 2020) | Põlme S, et al. 2020. Fungal Diversity 105:1-16 ((https://doi.org/10.1007/s13225-020-00466-2)) |
| Otu32506 | 1 | k__Fungi(100);p__Ascomycota(100);c__Saccharomycetes(100);o__Saccharomycetales(100);f__Metschnikowiaceae(100);g__Metschnikowia(100);s__Metschnikowia_pulcherrima(100); | Metschnikowia | 13 | Saprotroph | \|Undefined Saprotroph\| | Yeast | NULL | Probable | As nectar-tap_saprotroph (Põlme et al. 2020); Decay sugar-rich substrates (Põlme et al. 2020) | Põlme S, et al. 2020. Fungal Diversity 105:1-16 ((https://doi.org/10.1007/s13225-020-00466-2)) |
| Otu32507 | 1 | k__Fungi(100);p__Ascomycota(100);c__Saccharomycetes(100);o__Saccharomycetales(100);f__Metschnikowiaceae(100);g__Metschnikowia(100);g__Metschnikowia_unclassified(100); | Metschnikowia | 13 | Saprotroph | \|Undefined Saprotroph\| | Yeast | NULL | Probable | As nectar-tap_saprotroph (Põlme et al. 2020); Decay sugar-rich substrates (Põlme et al. 2020) | Põlme S, et al. 2020. Fungal Diversity 105:1-16 ((https://doi.org/10.1007/s13225-020-00466-2)) |
| Otu32508 | 1 | k__Fungi(100);p__Ascomycota(100);c__Saccharomycetes(100);o__Saccharomycetales(100);f__Metschnikowiaceae(100);g__Metschnikowia(100);s__Metschnikowia_pulcherrima(100); | Metschnikowia | 13 | Saprotroph | \|Undefined Saprotroph\| | Yeast | NULL | Probable | As nectar-tap_saprotroph (Põlme et al. 2020); Decay sugar-rich substrates (Põlme et al. 2020) | Põlme S, et al. 2020. Fungal Diversity 105:1-16 ((https://doi.org/10.1007/s13225-020-00466-2)) |
| Otu32509 | 1 | k__Fungi(100);p__Ascomycota(100);c__Saccharomycetes(100);o__Saccharomycetales(100);f__Metschnikowiaceae(100);g__Metschnikowia(100);s__Metschnikowia_pulcherrima(100); | Metschnikowia | 13 | Saprotroph | \|Undefined Saprotroph\| | Yeast | NULL | Probable | As nectar-tap_saprotroph (Põlme et al. 2020); Decay sugar-rich substrates (Põlme et al. 2020) | Põlme S, et al. 2020. Fungal Diversity 105:1-16 ((https://doi.org/10.1007/s13225-020-00466-2)) |
| Otu32510 | 1 | k__Fungi(100);p__Ascomycota(100);c__Saccharomycetes(100);o__Saccharomycetales(100);f__Metschnikowiaceae(100);g__Metschnikowia(100);s__Metschnikowia_pulcherrima(100); | Metschnikowia | 13 | Saprotroph | \|Undefined Saprotroph\| | Yeast | NULL | Probable | As nectar-tap_saprotroph (Põlme et al. 2020); Decay sugar-rich substrates (Põlme et al. 2020) | Põlme S, et al. 2020. Fungal Diversity 105:1-16 ((https://doi.org/10.1007/s13225-020-00466-2)) |
| Otu32511 | 1 | k__Fungi(100);p__Ascomycota(100);c__Saccharomycetes(100);o__Saccharomycetales(100);f__Metschnikowiaceae(100);g__Metschnikowia(100);s__Metschnikowia_pulcherrima(100); | Metschnikowia | 13 | Saprotroph | \|Undefined Saprotroph\| | Yeast | NULL | Probable | As nectar-tap_saprotroph (Põlme et al. 2020); Decay sugar-rich substrates (Põlme et al. 2020) | Põlme S, et al. 2020. Fungal Diversity 105:1-16 ((https://doi.org/10.1007/s13225-020-00466-2)) |
| Otu32512 | 1 | k__Fungi(100);p__Ascomycota(100);c__Saccharomycetes(100);o__Saccharomycetales(100);f__Metschnikowiaceae(100);g__Metschnikowia(100);s__Metschnikowia_pulcherrima(100); | Metschnikowia | 13 | Saprotroph | \|Undefined Saprotroph\| | Yeast | NULL | Probable | As nectar-tap_saprotroph (Põlme et al. 2020); Decay sugar-rich substrates (Põlme et al. 2020) | Põlme S, et al. 2020. Fungal Diversity 105:1-16 ((https://doi.org/10.1007/s13225-020-00466-2)) |
| Otu32513 | 1 | k__Fungi(100);p__Ascomycota(100);c__Saccharomycetes(100);o__Saccharomycetales(100);f__Metschnikowiaceae(100);g__Metschnikowia(100);s__Metschnikowia_pulcherrima(100); | Metschnikowia | 13 | Saprotroph | \|Undefined Saprotroph\| | Yeast | NULL | Probable | As nectar-tap_saprotroph (Põlme et al. 2020); Decay sugar-rich substrates (Põlme et al. 2020) | Põlme S, et al. 2020. Fungal Diversity 105:1-16 ((https://doi.org/10.1007/s13225-020-00466-2)) |
| Otu32514 | 1 | k__Fungi(100);p__Ascomycota(100);c__Saccharomycetes(100);o__Saccharomycetales(100);f__Metschnikowiaceae(100);g__Metschnikowia(100);s__Metschnikowia_pulcherrima(100); | Metschnikowia | 13 | Saprotroph | \|Undefined Saprotroph\| | Yeast | NULL | Probable | As nectar-tap_saprotroph (Põlme et al. 2020); Decay sugar-rich substrates (Põlme et al. 2020) | Põlme S, et al. 2020. Fungal Diversity 105:1-16 ((https://doi.org/10.1007/s13225-020-00466-2)) |
| Otu32515 | 1 | k__Fungi(100);p__Ascomycota(100);c__Saccharomycetes(100);o__Saccharomycetales(100);f__Metschnikowiaceae(100);g__Metschnikowia(100);s__Metschnikowia_pulcherrima(100); | Metschnikowia | 13 | Saprotroph | \|Undefined Saprotroph\| | Yeast | NULL | Probable | As nectar-tap_saprotroph (Põlme et al. 2020); Decay sugar-rich substrates (Põlme et al. 2020) | Põlme S, et al. 2020. Fungal Diversity 105:1-16 ((https://doi.org/10.1007/s13225-020-00466-2)) |
| Otu32516 | 1 | k__Fungi(100);p__Ascomycota(100);c__Saccharomycetes(100);o__Saccharomycetales(100);f__Metschnikowiaceae(100);g__Metschnikowia(100);s__Metschnikowia_pulcherrima(100); | Metschnikowia | 13 | Saprotroph | \|Undefined Saprotroph\| | Yeast | NULL | Probable | As nectar-tap_saprotroph (Põlme et al. 2020); Decay sugar-rich substrates (Põlme et al. 2020) | Põlme S, et al. 2020. Fungal Diversity 105:1-16 ((https://doi.org/10.1007/s13225-020-00466-2)) |
| Otu32517 | 1 | k__Fungi(100);p__Ascomycota(100);c__Saccharomycetes(100);o__Saccharomycetales(100);f__Metschnikowiaceae(100);g__Metschnikowia(100);g__Metschnikowia_unclassified(100); | Metschnikowia | 13 | Saprotroph | \|Undefined Saprotroph\| | Yeast | NULL | Probable | As nectar-tap_saprotroph (Põlme et al. 2020); Decay sugar-rich substrates (Põlme et al. 2020) | Põlme S, et al. 2020. Fungal Diversity 105:1-16 ((https://doi.org/10.1007/s13225-020-00466-2)) |
| Otu32518 | 1 | k__Fungi(100);p__Ascomycota(100);c__Saccharomycetes(100);o__Saccharomycetales(100);f__Metschnikowiaceae(100);g__Metschnikowia(100);s__Metschnikowia_pulcherrima(100); | Metschnikowia | 13 | Saprotroph | \|Undefined Saprotroph\| | Yeast | NULL | Probable | As nectar-tap_saprotroph (Põlme et al. 2020); Decay sugar-rich substrates (Põlme et al. 2020) | Põlme S, et al. 2020. Fungal Diversity 105:1-16 ((https://doi.org/10.1007/s13225-020-00466-2)) |
| Otu32519 | 1 | k__Fungi(100);p__Ascomycota(100);c__Saccharomycetes(100);o__Saccharomycetales(100);f__Metschnikowiaceae(100);g__Metschnikowia(100);g__Metschnikowia_unclassified(100); | Metschnikowia | 13 | Saprotroph | \|Undefined Saprotroph\| | Yeast | NULL | Probable | As nectar-tap_saprotroph (Põlme et al. 2020); Decay sugar-rich substrates (Põlme et al. 2020) | Põlme S, et al. 2020. Fungal Diversity 105:1-16 ((https://doi.org/10.1007/s13225-020-00466-2)) |
| Otu32520 | 1 | k__Fungi(100);p__Ascomycota(100);c__Saccharomycetes(100);o__Saccharomycetales(100);f__Metschnikowiaceae(100);g__Metschnikowia(100);s__Metschnikowia_pulcherrima(100); | Metschnikowia | 13 | Saprotroph | \|Undefined Saprotroph\| | Yeast | NULL | Probable | As nectar-tap_saprotroph (Põlme et al. 2020); Decay sugar-rich substrates (Põlme et al. 2020) | Põlme S, et al. 2020. Fungal Diversity 105:1-16 ((https://doi.org/10.1007/s13225-020-00466-2)) |
| Otu32525 | 1 | k__Fungi(100);p__Ascomycota(100);c__Saccharomycetes(100);o__Saccharomycetales(100);o__Saccharomycetales_unclassified(100);o__Saccharomycetales_unclassified(100);o__Saccharomycetales_unclassified(100); | Saccharomycetales | 7 | Saprotroph | Undefined Saprotroph | Yeast | NULL | Possible | NULL | Sterkenburg E, et al. 2015. New Phytologist 207:1145-1158 ((https://doi.org/10.1111/nph.13426)) |
| Otu32526 | 1 | k__Fungi(100);p__Ascomycota(100);c__Saccharomycetes(100);o__Saccharomycetales(100);f__Metschnikowiaceae(100);g__Metschnikowia(100);s__Metschnikowia_pulcherrima(100); | Metschnikowia | 13 | Saprotroph | \|Undefined Saprotroph\| | Yeast | NULL | Probable | As nectar-tap_saprotroph (Põlme et al. 2020); Decay sugar-rich substrates (Põlme et al. 2020) | Põlme S, et al. 2020. Fungal Diversity 105:1-16 ((https://doi.org/10.1007/s13225-020-00466-2)) |
| Otu32528 | 1 | k__Fungi(100);p__Ascomycota(100);c__Saccharomycetes(100);o__Saccharomycetales(100);f__Metschnikowiaceae(100);g__Metschnikowia(100);s__Metschnikowia_pulcherrima(100); | Metschnikowia | 13 | Saprotroph | \|Undefined Saprotroph\| | Yeast | NULL | Probable | As nectar-tap_saprotroph (Põlme et al. 2020); Decay sugar-rich substrates (Põlme et al. 2020) | Põlme S, et al. 2020. Fungal Diversity 105:1-16 ((https://doi.org/10.1007/s13225-020-00466-2)) |
| Otu32529 | 1 | k__Fungi(100);p__Ascomycota(100);c__Saccharomycetes(100);o__Saccharomycetales(100);f__Metschnikowiaceae(100);g__Metschnikowia(100);g__Metschnikowia_unclassified(100); | Metschnikowia | 13 | Saprotroph | \|Undefined Saprotroph\| | Yeast | NULL | Probable | As nectar-tap_saprotroph (Põlme et al. 2020); Decay sugar-rich substrates (Põlme et al. 2020) | Põlme S, et al. 2020. Fungal Diversity 105:1-16 ((https://doi.org/10.1007/s13225-020-00466-2)) |
| Otu32530 | 1 | k__Fungi(100);p__Ascomycota(100);c__Saccharomycetes(100);o__Saccharomycetales(100);f__Metschnikowiaceae(100);g__Metschnikowia(100);s__Metschnikowia_pulcherrima(100); | Metschnikowia | 13 | Saprotroph | \|Undefined Saprotroph\| | Yeast | NULL | Probable | As nectar-tap_saprotroph (Põlme et al. 2020); Decay sugar-rich substrates (Põlme et al. 2020) | Põlme S, et al. 2020. Fungal Diversity 105:1-16 ((https://doi.org/10.1007/s13225-020-00466-2)) |
| Otu32531 | 1 | k__Fungi(100);p__Ascomycota(100);c__Saccharomycetes(100);o__Saccharomycetales(100);f__Metschnikowiaceae(100);g__Metschnikowia(100);s__Metschnikowia_pulcherrima(100); | Metschnikowia | 13 | Saprotroph | \|Undefined Saprotroph\| | Yeast | NULL | Probable | As nectar-tap_saprotroph (Põlme et al. 2020); Decay sugar-rich substrates (Põlme et al. 2020) | Põlme S, et al. 2020. Fungal Diversity 105:1-16 ((https://doi.org/10.1007/s13225-020-00466-2)) |
| Otu32532 | 1 | k__Fungi(100);p__Ascomycota(100);c__Saccharomycetes(100);o__Saccharomycetales(100);f__Metschnikowiaceae(100);g__Metschnikowia(100);g__Metschnikowia_unclassified(100); | Metschnikowia | 13 | Saprotroph | \|Undefined Saprotroph\| | Yeast | NULL | Probable | As nectar-tap_saprotroph (Põlme et al. 2020); Decay sugar-rich substrates (Põlme et al. 2020) | Põlme S, et al. 2020. Fungal Diversity 105:1-16 ((https://doi.org/10.1007/s13225-020-00466-2)) |
| Otu32533 | 1 | k__Fungi(100);p__Ascomycota(100);c__Saccharomycetes(100);o__Saccharomycetales(100);f__Metschnikowiaceae(100);g__Metschnikowia(100);s__Metschnikowia_pulcherrima(100); | Metschnikowia | 13 | Saprotroph | \|Undefined Saprotroph\| | Yeast | NULL | Probable | As nectar-tap_saprotroph (Põlme et al. 2020); Decay sugar-rich substrates (Põlme et al. 2020) | Põlme S, et al. 2020. Fungal Diversity 105:1-16 ((https://doi.org/10.1007/s13225-020-00466-2)) |
| Otu32534 | 1 | k__Fungi(100);p__Ascomycota(100);c__Saccharomycetes(100);o__Saccharomycetales(100);f__Metschnikowiaceae(100);g__Metschnikowia(100);s__Metschnikowia_pulcherrima(100); | Metschnikowia | 13 | Saprotroph | \|Undefined Saprotroph\| | Yeast | NULL | Probable | As nectar-tap_saprotroph (Põlme et al. 2020); Decay sugar-rich substrates (Põlme et al. 2020) | Põlme S, et al. 2020. Fungal Diversity 105:1-16 ((https://doi.org/10.1007/s13225-020-00466-2)) |
| Otu32535 | 1 | k__Fungi(100);p__Ascomycota(100);c__Saccharomycetes(100);o__Saccharomycetales(100);f__Metschnikowiaceae(100);g__Metschnikowia(100);g__Metschnikowia_unclassified(100); | Metschnikowia | 13 | Saprotroph | \|Undefined Saprotroph\| | Yeast | NULL | Probable | As nectar-tap_saprotroph (Põlme et al. 2020); Decay sugar-rich substrates (Põlme et al. 2020) | Põlme S, et al. 2020. Fungal Diversity 105:1-16 ((https://doi.org/10.1007/s13225-020-00466-2)) |
| Otu32536 | 1 | k__Fungi(100);p__Ascomycota(100);c__Saccharomycetes(100);o__Saccharomycetales(100);f__Metschnikowiaceae(100);g__Metschnikowia(100);s__Metschnikowia_pulcherrima(100); | Metschnikowia | 13 | Saprotroph | \|Undefined Saprotroph\| | Yeast | NULL | Probable | As nectar-tap_saprotroph (Põlme et al. 2020); Decay sugar-rich substrates (Põlme et al. 2020) | Põlme S, et al. 2020. Fungal Diversity 105:1-16 ((https://doi.org/10.1007/s13225-020-00466-2)) |
| Otu32537 | 1 | k__Fungi(100);p__Ascomycota(100);c__Saccharomycetes(100);o__Saccharomycetales(100);f__Metschnikowiaceae(100);g__Metschnikowia(100);s__Metschnikowia_pulcherrima(100); | Metschnikowia | 13 | Saprotroph | \|Undefined Saprotroph\| | Yeast | NULL | Probable | As nectar-tap_saprotroph (Põlme et al. 2020); Decay sugar-rich substrates (Põlme et al. 2020) | Põlme S, et al. 2020. Fungal Diversity 105:1-16 ((https://doi.org/10.1007/s13225-020-00466-2)) |
| Otu32538 | 1 | k__Fungi(100);p__Ascomycota(100);c__Saccharomycetes(100);o__Saccharomycetales(100);f__Metschnikowiaceae(100);g__Metschnikowia(100);g__Metschnikowia_unclassified(100); | Metschnikowia | 13 | Saprotroph | \|Undefined Saprotroph\| | Yeast | NULL | Probable | As nectar-tap_saprotroph (Põlme et al. 2020); Decay sugar-rich substrates (Põlme et al. 2020) | Põlme S, et al. 2020. Fungal Diversity 105:1-16 ((https://doi.org/10.1007/s13225-020-00466-2)) |
| Otu32540 | 1 | k__Fungi(100);p__Ascomycota(100);c__Saccharomycetes(100);o__Saccharomycetales(100);f__Metschnikowiaceae(100);g__Metschnikowia(100);s__Metschnikowia_pulcherrima(100); | Metschnikowia | 13 | Saprotroph | \|Undefined Saprotroph\| | Yeast | NULL | Probable | As nectar-tap_saprotroph (Põlme et al. 2020); Decay sugar-rich substrates (Põlme et al. 2020) | Põlme S, et al. 2020. Fungal Diversity 105:1-16 ((https://doi.org/10.1007/s13225-020-00466-2)) |
| Otu32541 | 1 | k__Fungi(100);p__Ascomycota(100);c__Saccharomycetes(100);o__Saccharomycetales(100);o__Saccharomycetales_unclassified(100);o__Saccharomycetales_unclassified(100);o__Saccharomycetales_unclassified(100); | Saccharomycetales | 7 | Saprotroph | Undefined Saprotroph | Yeast | NULL | Possible | NULL | Sterkenburg E, et al. 2015. New Phytologist 207:1145-1158 ((https://doi.org/10.1111/nph.13426)) |
| Otu32542 | 1 | k__Fungi(100);p__Ascomycota(100);c__Saccharomycetes(100);o__Saccharomycetales(100);f__Saccharomycetales_family_Incertae_sedis(100);f__Saccharomycetales_family_Incertae_sedis_unclassified(100);f__Saccharomycetales_family_Incertae_sedis_unclassified(100); | Saccharomycetales | 7 | Saprotroph | Undefined Saprotroph | Yeast | NULL | Possible | NULL | Sterkenburg E, et al. 2015. New Phytologist 207:1145-1158 ((https://doi.org/10.1111/nph.13426)) |
| Otu32547 | 1 | k__Fungi(100);p__Ascomycota(100);c__Saccharomycetes(100);o__Saccharomycetales(100);f__Metschnikowiaceae(100);g__Metschnikowia(100);s__Metschnikowia_pulcherrima(100); | Metschnikowia | 13 | Saprotroph | \|Undefined Saprotroph\| | Yeast | NULL | Probable | As nectar-tap_saprotroph (Põlme et al. 2020); Decay sugar-rich substrates (Põlme et al. 2020) | Põlme S, et al. 2020. Fungal Diversity 105:1-16 ((https://doi.org/10.1007/s13225-020-00466-2)) |
| Otu32548 | 1 | k__Fungi(100);p__Ascomycota(100);c__Saccharomycetes(100);o__Saccharomycetales(100);f__Metschnikowiaceae(100);g__Metschnikowia(100);g__Metschnikowia_unclassified(100); | Metschnikowia | 13 | Saprotroph | \|Undefined Saprotroph\| | Yeast | NULL | Probable | As nectar-tap_saprotroph (Põlme et al. 2020); Decay sugar-rich substrates (Põlme et al. 2020) | Põlme S, et al. 2020. Fungal Diversity 105:1-16 ((https://doi.org/10.1007/s13225-020-00466-2)) |
| Otu32549 | 1 | k__Fungi(100);p__Ascomycota(100);c__Saccharomycetes(100);o__Saccharomycetales(100);f__Metschnikowiaceae(100);g__Metschnikowia(100);g__Metschnikowia_unclassified(100); | Metschnikowia | 13 | Saprotroph | \|Undefined Saprotroph\| | Yeast | NULL | Probable | As nectar-tap_saprotroph (Põlme et al. 2020); Decay sugar-rich substrates (Põlme et al. 2020) | Põlme S, et al. 2020. Fungal Diversity 105:1-16 ((https://doi.org/10.1007/s13225-020-00466-2)) |
| Otu32552 | 1 | k__Fungi(100);p__Ascomycota(100);c__Saccharomycetes(100);o__Saccharomycetales(100);f__Metschnikowiaceae(100);g__Metschnikowia(100);s__Metschnikowia_pulcherrima(100); | Metschnikowia | 13 | Saprotroph | \|Undefined Saprotroph\| | Yeast | NULL | Probable | As nectar-tap_saprotroph (Põlme et al. 2020); Decay sugar-rich substrates (Põlme et al. 2020) | Põlme S, et al. 2020. Fungal Diversity 105:1-16 ((https://doi.org/10.1007/s13225-020-00466-2)) |
| Otu32553 | 1 | k__Fungi(100);p__Ascomycota(100);c__Saccharomycetes(100);o__Saccharomycetales(100);f__Metschnikowiaceae(100);g__Metschnikowia(100);g__Metschnikowia_unclassified(100); | Metschnikowia | 13 | Saprotroph | \|Undefined Saprotroph\| | Yeast | NULL | Probable | As nectar-tap_saprotroph (Põlme et al. 2020); Decay sugar-rich substrates (Põlme et al. 2020) | Põlme S, et al. 2020. Fungal Diversity 105:1-16 ((https://doi.org/10.1007/s13225-020-00466-2)) |
| Otu32554 | 1 | k__Fungi(100);p__Ascomycota(100);c__Saccharomycetes(100);o__Saccharomycetales(100);f__Metschnikowiaceae(100);g__Metschnikowia(100);g__Metschnikowia_unclassified(100); | Metschnikowia | 13 | Saprotroph | \|Undefined Saprotroph\| | Yeast | NULL | Probable | As nectar-tap_saprotroph (Põlme et al. 2020); Decay sugar-rich substrates (Põlme et al. 2020) | Põlme S, et al. 2020. Fungal Diversity 105:1-16 ((https://doi.org/10.1007/s13225-020-00466-2)) |
| [illegible] | [illegible] | [illegible] | [illegible] | [illegible] | [illegible] | [illegible] | [illegible] | [illegible] | [illegible] | [illegible] | [illegible] |
| [illegible] | [illegible] | [illegible] | [illegible] | [illegible] | [illegible] | [illegible] | [illegible] | [illegible] | [illegible] | [illegible] | [illegible] |
| [illegible] | [illegible] | [illegible] | [illegible] | [illegible] | [illegible] | [illegible] | [illegible] | [illegible] | [illegible] | [illegible] | [illegible] |
| [illegible] | [illegible] | [illegible] | [illegible] | [illegible] | [illegible] | [illegible] | [illegible] | [illegible] | [illegible] | [illegible] | [illegible] |
| [illegible] | [illegible] | [illegible] | [illegible] | [illegible] | [illegible] | [illegible] | [illegible] | [illegible] | [illegible] | [illegible] | [illegible] |
| [illegible] | [illegible] | [illegible] | [illegible] | [illegible] | [illegible] | [illegible] | [illegible] | [illegible] | [illegible] | [illegible] | [illegible] |
| [illegible] | [illegible] | [illegible] | [illegible] | [illegible] | [illegible] | [illegible] | [illegible] | [illegible] | [illegible] | [illegible] | [illegible] |
| [illegible] | [illegible] | [illegible] | [illegible] | [illegible] | [illegible] | [illegible] | [illegible] | [illegible] | [illegible] | [illegible] | [illegible] |
| [illegible] | [illegible] | [illegible] | [illegible] | [illegible] | [illegible] | [illegible] | [illegible] | [illegible] | [illegible] | [illegible] | [illegible] |
| [illegible] | [illegible] | [illegible] | [illegible] | [illegible] | [illegible] | [illegible] | [illegible] | [illegible] | [illegible] | [illegible] | [illegible] |
| [illegible] | [illegible] | [illegible] | [illegible] | [illegible] | [illegible] | [illegible] | [illegible] | [illegible] | [illegible] | [illegible] | [illegible] |
| [illegible] | [illegible] | [illegible] | [illegible] | [illegible] | [illegible] | [illegible] | [illegible] | [illegible] | [illegible] | [illegible] | [illegible] |
| [illegible] | [illegible] | [illegible] | [illegible] | [illegible] | [illegible] | [illegible] | [illegible] | [illegible] | [illegible] | [illegible] | [illegible] |
| [illegible] | [illegible] | [illegible] | [illegible] | [illegible] | [illegible] | [illegible] | [illegible] | [illegible] | [illegible] | [illegible] | [illegible] |

| | | | | | | | | | | | |
|---|---|---|---|---|---|---|---|---|---|---|---|
| Otu32579 | 1 | k__Fungi(100);p__Ascomycota(100);c__Saccharomycetes(100);o__Saccharomycetales(100);f__Metschnikowiaceae(100);g__Metschnikowia(100);s__Metschnikowia_pulcherrima(100); | Metschnikowia | 13 | Saprotroph | \|Undefined Saprotroph\| | Yeast | NULL | Probable | As nectar-tap_saprotroph (Põlme et al. 2020); Decay sugar-rich substrates (Põlme et al. 2020) | Põlme S, et al. 2020. Fungal Diversity 105:1-16 ((https://doi.org/10.1007/s13225-020-00466-2)) |
| Otu32580 | 1 | k__Fungi(100);p__Ascomycota(100);c__Saccharomycetes(100);o__Saccharomycetales(100);f__Metschnikowiaceae(100);g__Metschnikowia(100);s__Metschnikowia_pulcherrima(100); | Metschnikowia | 13 | Saprotroph | \|Undefined Saprotroph\| | Yeast | NULL | Probable | As nectar-tap_saprotroph (Põlme et al. 2020); Decay sugar-rich substrates (Põlme et al. 2020) | Põlme S, et al. 2020. Fungal Diversity 105:1-16 ((https://doi.org/10.1007/s13225-020-00466-2)) |
| Otu32581 | 1 | k__Fungi(100);p__Ascomycota(100);c__Saccharomycetes(100);o__Saccharomycetales(100);f__Metschnikowiaceae(100);g__Metschnikowia(100);s__Metschnikowia_pulcherrima(100); | Metschnikowia | 13 | Saprotroph | \|Undefined Saprotroph\| | Yeast | NULL | Probable | As nectar-tap_saprotroph (Põlme et al. 2020); Decay sugar-rich substrates (Põlme et al. 2020) | Põlme S, et al. 2020. Fungal Diversity 105:1-16 ((https://doi.org/10.1007/s13225-020-00466-2)) |
| Otu32582 | 1 | k__Fungi(100);p__Ascomycota(100);c__Saccharomycetes(100);o__Saccharomycetales(100);f__Metschnikowiaceae(100);g__Metschnikowia(100);g__Metschnikowia_unclassified(100); | Metschnikowia | 13 | Saprotroph | \|Undefined Saprotroph\| | Yeast | NULL | Probable | As nectar-tap_saprotroph (Põlme et al. 2020); Decay sugar-rich substrates (Põlme et al. 2020) | Põlme S, et al. 2020. Fungal Diversity 105:1-16 ((https://doi.org/10.1007/s13225-020-00466-2)) |
| Otu32583 | 1 | k__Fungi(100);p__Ascomycota(100);c__Saccharomycetes(100);o__Saccharomycetales(100);f__Metschnikowiaceae(100);g__Metschnikowia(100);g__Metschnikowia_unclassified(100); | Metschnikowia | 13 | Saprotroph | \|Undefined Saprotroph\| | Yeast | NULL | Probable | As nectar-tap_saprotroph (Põlme et al. 2020); Decay sugar-rich substrates (Põlme et al. 2020) | Põlme S, et al. 2020. Fungal Diversity 105:1-16 ((https://doi.org/10.1007/s13225-020-00466-2)) |
| Otu32586 | 1 | k__Fungi(100);p__Ascomycota(100);c__Saccharomycetes(100);o__Saccharomycetales(100);f__Metschnikowiaceae(100);g__Metschnikowia(100);s__Metschnikowia_pulcherrima(100); | Metschnikowia | 13 | Saprotroph | \|Undefined Saprotroph\| | Yeast | NULL | Probable | As nectar-tap_saprotroph (Põlme et al. 2020); Decay sugar-rich substrates (Põlme et al. 2020) | Põlme S, et al. 2020. Fungal Diversity 105:1-16 ((https://doi.org/10.1007/s13225-020-00466-2)) |
| Otu32587 | 1 | k__Fungi(100);p__Ascomycota(100);c__Saccharomycetes(100);o__Saccharomycetales(100);f__Metschnikowiaceae(100);g__Metschnikowia(100);s__Metschnikowia_pulcherrima(100); | Metschnikowia | 13 | Saprotroph | \|Undefined Saprotroph\| | Yeast | NULL | Probable | As nectar-tap_saprotroph (Põlme et al. 2020); Decay sugar-rich substrates (Põlme et al. 2020) | Põlme S, et al. 2020. Fungal Diversity 105:1-16 ((https://doi.org/10.1007/s13225-020-00466-2)) |
| Otu32588 | 1 | k__Fungi(100);p__Ascomycota(100);c__Saccharomycetes(100);o__Saccharomycetales(100);f__Metschnikowiaceae(100);g__Metschnikowia(100);s__Metschnikowia_pulcherrima(100); | Metschnikowia | 13 | Saprotroph | \|Undefined Saprotroph\| | Yeast | NULL | Probable | As nectar-tap_saprotroph (Põlme et al. 2020); Decay sugar-rich substrates (Põlme et al. 2020) | Põlme S, et al. 2020. Fungal Diversity 105:1-16 ((https://doi.org/10.1007/s13225-020-00466-2)) |
| Otu32589 | 1 | k__Fungi(100);p__Ascomycota(100);c__Saccharomycetes(100);o__Saccharomycetales(100);f__Metschnikowiaceae(100);g__Metschnikowia(100);s__Metschnikowia_pulcherrima(100); | Metschnikowia | 13 | Saprotroph | \|Undefined Saprotroph\| | Yeast | NULL | Probable | As nectar-tap_saprotroph (Põlme et al. 2020); Decay sugar-rich substrates (Põlme et al. 2020) | Põlme S, et al. 2020. Fungal Diversity 105:1-16 ((https://doi.org/10.1007/s13225-020-00466-2)) |
| Otu32590 | 1 | k__Fungi(100);p__Ascomycota(100);c__Saccharomycetes(100);o__Saccharomycetales(100);f__Metschnikowiaceae(100);g__Metschnikowia(100);s__Metschnikowia_pulcherrima(100); | Metschnikowia | 13 | Saprotroph | \|Undefined Saprotroph\| | Yeast | NULL | Probable | As nectar-tap_saprotroph (Põlme et al. 2020); Decay sugar-rich substrates (Põlme et al. 2020) | Põlme S, et al. 2020. Fungal Diversity 105:1-16 ((https://doi.org/10.1007/s13225-020-00466-2)) |
| Otu32592 | 1 | k__Fungi(100);p__Ascomycota(100);c__Saccharomycetes(100);o__Saccharomycetales(100);f__Metschnikowiaceae(100);g__Metschnikowia(100);s__Metschnikowia_pulcherrima(100); | Metschnikowia | 13 | Saprotroph | \|Undefined Saprotroph\| | Yeast | NULL | Probable | As nectar-tap_saprotroph (Põlme et al. 2020); Decay sugar-rich substrates (Põlme et al. 2020) | Põlme S, et al. 2020. Fungal Diversity 105:1-16 ((https://doi.org/10.1007/s13225-020-00466-2)) |
| Otu32594 | 1 | k__Fungi(100);p__Ascomycota(100);c__Saccharomycetes(100);o__Saccharomycetales(100);f__Metschnikowiaceae(100);g__Metschnikowia(100);s__Metschnikowia_pulcherrima(100); | Metschnikowia | 13 | Saprotroph | \|Undefined Saprotroph\| | Yeast | NULL | Probable | As nectar-tap_saprotroph (Põlme et al. 2020); Decay sugar-rich substrates (Põlme et al. 2020) | Põlme S, et al. 2020. Fungal Diversity 105:1-16 ((https://doi.org/10.1007/s13225-020-00466-2)) |
| Otu32596 | 1 | k__Fungi(100);p__Ascomycota(100);c__Saccharomycetes(100);o__Saccharomycetales(100);f__Metschnikowiaceae(100);g__Metschnikowia(100);s__Metschnikowia_pulcherrima(100); | Metschnikowia | 13 | Saprotroph | \|Undefined Saprotroph\| | Yeast | NULL | Probable | As nectar-tap_saprotroph (Põlme et al. 2020); Decay sugar-rich substrates (Põlme et al. 2020) | Põlme S, et al. 2020. Fungal Diversity 105:1-16 ((https://doi.org/10.1007/s13225-020-00466-2)) |
| Otu32599 | 1 | k__Fungi(100);p__Ascomycota(100);c__Saccharomycetes(100);o__Saccharomycetales(100);f__Metschnikowiaceae(100);g__Metschnikowia(100);s__Metschnikowia_pulcherrima(100); | Metschnikowia | 13 | Saprotroph | \|Undefined Saprotroph\| | Yeast | NULL | Probable | As nectar-tap_saprotroph (Põlme et al. 2020); Decay sugar-rich substrates (Põlme et al. 2020) | Põlme S, et al. 2020. Fungal Diversity 105:1-16 ((https://doi.org/10.1007/s13225-020-00466-2)) |
| Otu32600 | 1 | k__Fungi(100);p__Ascomycota(100);c__Saccharomycetes(100);o__Saccharomycetales(100);f__Metschnikowiaceae(100);g__Metschnikowia(100);s__Metschnikowia_pulcherrima(100); | Metschnikowia | 13 | Saprotroph | \|Undefined Saprotroph\| | Yeast | NULL | Probable | As nectar-tap_saprotroph (Põlme et al. 2020); Decay sugar-rich substrates (Põlme et al. 2020) | Põlme S, et al. 2020. Fungal Diversity 105:1-16 ((https://doi.org/10.1007/s13225-020-00466-2)) |
| Otu32601 | 1 | k__Fungi(100);p__Ascomycota(100);c__Saccharomycetes(100);o__Saccharomycetales(100);f__Metschnikowiaceae(100);g__Metschnikowia(100);s__Metschnikowia_pulcherrima(100); | Metschnikowia | 13 | Saprotroph | \|Undefined Saprotroph\| | Yeast | NULL | Probable | As nectar-tap_saprotroph (Põlme et al. 2020); Decay sugar-rich substrates (Põlme et al. 2020) | Põlme S, et al. 2020. Fungal Diversity 105:1-16 ((https://doi.org/10.1007/s13225-020-00466-2)) |
| Otu32602 | 1 | k__Fungi(100);p__Ascomycota(100);c__Saccharomycetes(100);o__Saccharomycetales(100);f__Metschnikowiaceae(100);g__Metschnikowia(100);s__Metschnikowia_pulcherrima(100); | Metschnikowia | 13 | Saprotroph | \|Undefined Saprotroph\| | Yeast | NULL | Probable | As nectar-tap_saprotroph (Põlme et al. 2020); Decay sugar-rich substrates (Põlme et al. 2020) | Põlme S, et al. 2020. Fungal Diversity 105:1-16 ((https://doi.org/10.1007/s13225-020-00466-2)) |
| Otu32603 | 1 | k__Fungi(100);p__Ascomycota(100);c__Saccharomycetes(100);o__Saccharomycetales(100);f__Metschnikowiaceae(100);g__Metschnikowia(100);s__Metschnikowia_pulcherrima(100); | Metschnikowia | 13 | Saprotroph | \|Undefined Saprotroph\| | Yeast | NULL | Probable | As nectar-tap_saprotroph (Põlme et al. 2020); Decay sugar-rich substrates (Põlme et al. 2020) | Põlme S, et al. 2020. Fungal Diversity 105:1-16 ((https://doi.org/10.1007/s13225-020-00466-2)) |
| Otu32604 | 1 | k__Fungi(100);p__Ascomycota(100);c__Saccharomycetes(100);o__Saccharomycetales(100);f__Metschnikowiaceae(100);g__Metschnikowia(100);s__Metschnikowia_pulcherrima(100); | Metschnikowia | 13 | Saprotroph | \|Undefined Saprotroph\| | Yeast | NULL | Probable | As nectar-tap_saprotroph (Põlme et al. 2020); Decay sugar-rich substrates (Põlme et al. 2020) | Põlme S, et al. 2020. Fungal Diversity 105:1-16 ((https://doi.org/10.1007/s13225-020-00466-2)) |
| Otu32605 | 1 | k__Fungi(100);p__Ascomycota(100);c__Saccharomycetes(100);o__Saccharomycetales(100);f__Metschnikowiaceae(100);g__Metschnikowia(100);g__Metschnikowia_unclassified(100); | Metschnikowia | 13 | Saprotroph | \|Undefined Saprotroph\| | Yeast | NULL | Probable | As nectar-tap_saprotroph (Põlme et al. 2020); Decay sugar-rich substrates (Põlme et al. 2020) | Põlme S, et al. 2020. Fungal Diversity 105:1-16 ((https://doi.org/10.1007/s13225-020-00466-2)) |
| Otu32608 | 1 | k__Fungi(100);p__Ascomycota(100);c__Saccharomycetes(100);o__Saccharomycetales(100);f__Metschnikowiaceae(100);g__Metschnikowia(100);s__Metschnikowia_pulcherrima(100); | Metschnikowia | 13 | Saprotroph | \|Undefined Saprotroph\| | Yeast | NULL | Probable | As nectar-tap_saprotroph (Põlme et al. 2020); Decay sugar-rich substrates (Põlme et al. 2020) | Põlme S, et al. 2020. Fungal Diversity 105:1-16 ((https://doi.org/10.1007/s13225-020-00466-2)) |
| Otu32609 | 1 | k__Fungi(100);p__Ascomycota(100);c__Saccharomycetes(100);o__Saccharomycetales(100);f__Metschnikowiaceae(100);g__Metschnikowia(100);g__Metschnikowia_unclassified(100); | Metschnikowia | 13 | Saprotroph | \|Undefined Saprotroph\| | Yeast | NULL | Probable | As nectar-tap_saprotroph (Põlme et al. 2020); Decay sugar-rich substrates (Põlme et al. 2020) | Põlme S, et al. 2020. Fungal Diversity 105:1-16 ((https://doi.org/10.1007/s13225-020-00466-2)) |
| Otu32610 | 1 | k__Fungi(100);p__Ascomycota(100);c__Saccharomycetes(100);o__Saccharomycetales(100);f__Metschnikowiaceae(100);g__Metschnikowia(100);s__Metschnikowia_pulcherrima(100); | Metschnikowia | 13 | Saprotroph | \|Undefined Saprotroph\| | Yeast | NULL | Probable | As nectar-tap_saprotroph (Põlme et al. 2020); Decay sugar-rich substrates (Põlme et al. 2020) | Põlme S, et al. 2020. Fungal Diversity 105:1-16 ((https://doi.org/10.1007/s13225-020-00466-2)) |
| Otu32611 | 1 | k__Fungi(100);p__Ascomycota(100);c__Saccharomycetes(100);o__Saccharomycetales(100);f__Metschnikowiaceae(100);g__Metschnikowia(100);g__Metschnikowia_unclassified(100); | Metschnikowia | 13 | Saprotroph | \|Undefined Saprotroph\| | Yeast | NULL | Probable | As nectar-tap_saprotroph (Põlme et al. 2020); Decay sugar-rich substrates (Põlme et al. 2020) | Põlme S, et al. 2020. Fungal Diversity 105:1-16 ((https://doi.org/10.1007/s13225-020-00466-2)) |
| Otu32612 | 1 | k__Fungi(100);p__Ascomycota(100);c__Saccharomycetes(100);o__Saccharomycetales(100);f__Metschnikowiaceae(100);g__Metschnikowia(100);s__Metschnikowia_pulcherrima(100); | Metschnikowia | 13 | Saprotroph | \|Undefined Saprotroph\| | Yeast | NULL | Probable | As nectar-tap_saprotroph (Põlme et al. 2020); Decay sugar-rich substrates (Põlme et al. 2020) | Põlme S, et al. 2020. Fungal Diversity 105:1-16 ((https://doi.org/10.1007/s13225-020-00466-2)) |
| Otu32613 | 1 | k__Fungi(100);p__Ascomycota(100);c__Saccharomycetes(100);o__Saccharomycetales(100);f__Metschnikowiaceae(100);g__Metschnikowia(100);s__Metschnikowia_pulcherrima(100); | Metschnikowia | 13 | Saprotroph | \|Undefined Saprotroph\| | Yeast | NULL | Probable | As nectar-tap_saprotroph (Põlme et al. 2020); Decay sugar-rich substrates (Põlme et al. 2020) | Põlme S, et al. 2020. Fungal Diversity 105:1-16 ((https://doi.org/10.1007/s13225-020-00466-2)) |
| Otu32614 | 1 | k__Fungi(100);p__Ascomycota(100);c__Saccharomycetes(100);o__Saccharomycetales(100);f__Metschnikowiaceae(100);g__Metschnikowia(100);s__Metschnikowia_pulcherrima(100); | Metschnikowia | 13 | Saprotroph | \|Undefined Saprotroph\| | Yeast | NULL | Probable | As nectar-tap_saprotroph (Põlme et al. 2020); Decay sugar-rich substrates (Põlme et al. 2020) | Põlme S, et al. 2020. Fungal Diversity 105:1-16 ((https://doi.org/10.1007/s13225-020-00466-2)) |
| Otu32615 | 1 | k__Fungi(100);p__Ascomycota(100);c__Saccharomycetes(100);o__Saccharomycetales(100);f__Metschnikowiaceae(100);g__Metschnikowia(100);s__Metschnikowia_pulcherrima(100); | Metschnikowia | 13 | Saprotroph | \|Undefined Saprotroph\| | Yeast | NULL | Probable | As nectar-tap_saprotroph (Põlme et al. 2020); Decay sugar-rich substrates (Põlme et al. 2020) | Põlme S, et al. 2020. Fungal Diversity 105:1-16 ((https://doi.org/10.1007/s13225-020-00466-2)) |
| Otu32616 | 1 | k__Fungi(100);p__Ascomycota(100);c__Saccharomycetes(100);o__Saccharomycetales(100);f__Metschnikowiaceae(100);g__Metschnikowia(100);s__Metschnikowia_pulcherrima(100); | Metschnikowia | 13 | Saprotroph | \|Undefined Saprotroph\| | Yeast | NULL | Probable | As nectar-tap_saprotroph (Põlme et al. 2020); Decay sugar-rich substrates (Põlme et al. 2020) | Põlme S, et al. 2020. Fungal Diversity 105:1-16 ((https://doi.org/10.1007/s13225-020-00466-2)) |
| Otu32617 | 1 | k__Fungi(100);p__Ascomycota(100);c__Saccharomycetes(100);o__Saccharomycetales(100);f__Metschnikowiaceae(100);g__Metschnikowia(100);s__Metschnikowia_pulcherrima(100); | Metschnikowia | 13 | Saprotroph | \|Undefined Saprotroph\| | Yeast | NULL | Probable | As nectar-tap_saprotroph (Põlme et al. 2020); Decay sugar-rich substrates (Põlme et al. 2020) | Põlme S, et al. 2020. Fungal Diversity 105:1-16 ((https://doi.org/10.1007/s13225-020-00466-2)) |
| Otu32618 | 1 | k__Fungi(100);p__Ascomycota(100);c__Saccharomycetes(100);o__Saccharomycetales(100);f__Metschnikowiaceae(100);g__Metschnikowia(100);s__Metschnikowia_pulcherrima(100); | Metschnikowia | 13 | Saprotroph | \|Undefined Saprotroph\| | Yeast | NULL | Probable | As nectar-tap_saprotroph (Põlme et al. 2020); Decay sugar-rich substrates (Põlme et al. 2020) | Põlme S, et al. 2020. Fungal Diversity 105:1-16 ((https://doi.org/10.1007/s13225-020-00466-2)) |
| Otu32619 | 1 | k__Fungi(100);p__Ascomycota(100);c__Saccharomycetes(100);o__Saccharomycetales(100);f__Metschnikowiaceae(100);g__Metschnikowia(100);s__Metschnikowia_pulcherrima(100); | Metschnikowia | 13 | Saprotroph | \|Undefined Saprotroph\| | Yeast | NULL | Probable | As nectar-tap_saprotroph (Põlme et al. 2020); Decay sugar-rich substrates (Põlme et al. 2020) | Põlme S, et al. 2020. Fungal Diversity 105:1-16 ((https://doi.org/10.1007/s13225-020-00466-2)) |
| Otu32620 | 1 | k__Fungi(100);p__Ascomycota(100);c__Saccharomycetes(100);o__Saccharomycetales(100);f__Metschnikowiaceae(100);g__Metschnikowia(100);s__Metschnikowia_pulcherrima(100); | Metschnikowia | 13 | Saprotroph | \|Undefined Saprotroph\| | Yeast | NULL | Probable | As nectar-tap_saprotroph (Põlme et al. 2020); Decay sugar-rich substrates (Põlme et al. 2020) | Põlme S, et al. 2020. Fungal Diversity 105:1-16 ((https://doi.org/10.1007/s13225-020-00466-2)) |
| Otu32621 | 1 | k__Fungi(100);p__Ascomycota(100);c__Saccharomycetes(100);o__Saccharomycetales(100);f__Metschnikowiaceae(100);g__Metschnikowia(100);s__Metschnikowia_pulcherrima(100); | Metschnikowia | 13 | Saprotroph | \|Undefined Saprotroph\| | Yeast | NULL | Probable | As nectar-tap_saprotroph (Põlme et al. 2020); Decay sugar-rich substrates (Põlme et al. 2020) | Põlme S, et al. 2020. Fungal Diversity 105:1-16 ((https://doi.org/10.1007/s13225-020-00466-2)) |
| Otu32622 | 1 | k__Fungi(100);p__Ascomycota(100);c__Saccharomycetes(100);o__Saccharomycetales(100);f__Metschnikowiaceae(100);g__Metschnikowia(100);s__Metschnikowia_pulcherrima(100); | Metschnikowia | 13 | Saprotroph | \|Undefined Saprotroph\| | Yeast | NULL | Probable | As nectar-tap_saprotroph (Põlme et al. 2020); Decay sugar-rich substrates (Põlme et al. 2020) | Põlme S, et al. 2020. Fungal Diversity 105:1-16 ((https://doi.org/10.1007/s13225-020-00466-2)) |
| Otu32623 | 1 | k__Fungi(100);p__Ascomycota(100);c__Saccharomycetes(100);o__Saccharomycetales(100);f__Metschnikowiaceae(100);g__Metschnikowia(100);g__Metschnikowia_unclassified(100); | Metschnikowia | 13 | Saprotroph | \|Undefined Saprotroph\| | Yeast | NULL | Probable | As nectar-tap_saprotroph (Põlme et al. 2020); Decay sugar-rich substrates (Põlme et al. 2020) | Põlme S, et al. 2020. Fungal Diversity 105:1-16 ((https://doi.org/10.1007/s13225-020-00466-2)) |
| Otu32624 | 1 | k__Fungi(100);p__Ascomycota(100);c__Saccharomycetes(100);o__Saccharomycetales(100);f__Metschnikowiaceae(100);g__Metschnikowia(100);g__Metschnikowia_unclassified(100); | Metschnikowia | 13 | Saprotroph | \|Undefined Saprotroph\| | Yeast | NULL | Probable | As nectar-tap_saprotroph (Põlme et al. 2020); Decay sugar-rich substrates (Põlme et al. 2020) | Põlme S, et al. 2020. Fungal Diversity 105:1-16 ((https://doi.org/10.1007/s13225-020-00466-2)) |
| Otu32625 | 1 | k__Fungi(100);p__Ascomycota(100);c__Saccharomycetes(100);o__Saccharomycetales(100);f__Metschnikowiaceae(100);g__Metschnikowia(100);s__Metschnikowia_pulcherrima(100); | Metschnikowia | 13 | Saprotroph | \|Undefined Saprotroph\| | Yeast | NULL | Probable | As nectar-tap_saprotroph (Põlme et al. 2020); Decay sugar-rich substrates (Põlme et al. 2020) | Põlme S, et al. 2020. Fungal Diversity 105:1-16 ((https://doi.org/10.1007/s13225-020-00466-2)) |
| Otu32626 | 1 | k__Fungi(100);p__Ascomycota(100);c__Saccharomycetes(100);o__Saccharomycetales(100);f__Metschnikowiaceae(100);g__Metschnikowia(100);s__Metschnikowia_pulcherrima(100); | Metschnikowia | 13 | Saprotroph | \|Undefined Saprotroph\| | Yeast | NULL | Probable | As nectar-tap_saprotroph (Põlme et al. 2020); Decay sugar-rich substrates (Põlme et al. 2020) | Põlme S, et al. 2020. Fungal Diversity 105:1-16 ((https://doi.org/10.1007/s13225-020-00466-2)) |
| Otu32627 | 1 | k__Fungi(100);p__Ascomycota(100);c__Saccharomycetes(100);o__Saccharomycetales(100);f__Metschnikowiaceae(100);g__Metschnikowia(100);s__Metschnikowia_pulcherrima(100); | Metschnikowia | 13 | Saprotroph | \|Undefined Saprotroph\| | Yeast | NULL | Probable | As nectar-tap_saprotroph (Põlme et al. 2020); Decay sugar-rich substrates (Põlme et al. 2020) | Põlme S, et al. 2020. Fungal Diversity 105:1-16 ((https://doi.org/10.1007/s13225-020-00466-2)) |
| Otu32629 | 1 | k__Fungi(100);p__Ascomycota(100);c__Saccharomycetes(100);o__Saccharomycetales(100);f__Metschnikowiaceae(100);g__Metschnikowia(100);g__Metschnikowia_unclassified(100); | Metschnikowia | 13 | Saprotroph | \|Undefined Saprotroph\| | Yeast | NULL | Probable | As nectar-tap_saprotroph (Põlme et al. 2020); Decay sugar-rich substrates (Põlme et al. 2020) | Põlme S, et al. 2020. Fungal Diversity 105:1-16 ((https://doi.org/10.1007/s13225-020-00466-2)) |
| Otu32630 | 1 | k__Fungi(100);p__Ascomycota(100);c__Saccharomycetes(100);o__Saccharomycetales(100);f__Metschnikowiaceae(100);g__Metschnikowia(100);g__Metschnikowia_unclassified(100); | Metschnikowia | 13 | Saprotroph | \|Undefined Saprotroph\| | Yeast | NULL | Probable | As nectar-tap_saprotroph (Põlme et al. 2020); Decay sugar-rich substrates (Põlme et al. 2020) | Põlme S, et al. 2020. Fungal Diversity 105:1-16 ((https://doi.org/10.1007/s13225-020-00466-2)) |
| Otu32631 | 1 | k__Fungi(100);p__Ascomycota(100);c__Saccharomycetes(100);o__Saccharomycetales(100);f__Saccharomycetales_family_Incertae_sedis(100);f__Saccharomycetales_family_Incertae_sedis_unclassified(100);f__Saccharomycetales_family_Incertae_sedis_unclassified(100); | Saccharomycetales | 7 | Saprotroph | Undefined Saprotroph | Yeast | NULL | Possible | NULL | Sterkenburg E, et al. 2015. New Phytologist 207:1145-1158 ((https://doi.org/10.1111/nph.13426)) |
| Otu32635 | 1 | k__Fungi(100);p__Ascomycota(100);c__Saccharomycetes(100);o__Saccharomycetales(100);f__Metschnikowiaceae(100);g__Metschnikowia(100);g__Metschnikowia_unclassified(100); | Metschnikowia | 13 | Saprotroph | \|Undefined Saprotroph\| | Yeast | NULL | Probable | As nectar-tap_saprotroph (Põlme et al. 2020); Decay sugar-rich substrates (Põlme et al. 2020) | Põlme S, et al. 2020. Fungal Diversity 105:1-16 ((https://doi.org/10.1007/s13225-020-00466-2)) |
| Otu32636 | 1 | k__Fungi(100);p__Ascomycota(100);c__Saccharomycetes(100);o__Saccharomycetales(100);f__Metschnikowiaceae(100);g__Metschnikowia(100);g__Metschnikowia_unclassified(100); | Metschnikowia | 13 | Saprotroph | \|Undefined Saprotroph\| | Yeast | NULL | Probable | As nectar-tap_saprotroph (Põlme et al. 2020); Decay sugar-rich substrates (Põlme et al. 2020) | Põlme S, et al. 2020. Fungal Diversity 105:1-16 ((https://doi.org/10.1007/s13225-020-00466-2)) |
| Otu32637 | 1 | k__Fungi(100);p__Ascomycota(100);c__Saccharomycetes(100);o__Saccharomycetales(100);f__Metschnikowiaceae(100);g__Metschnikowia(100);s__Metschnikowia_pulcherrima(100); | Metschnikowia | 13 | Saprotroph | \|Undefined Saprotroph\| | Yeast | NULL | Probable | As nectar-tap_saprotroph (Põlme et al. 2020); Decay sugar-rich substrates (Põlme et al. 2020) | Põlme S, et al. 2020. Fungal Diversity 105:1-16 ((https://doi.org/10.1007/s13225-020-00466-2)) |
| Otu32638 | 1 | k__Fungi(100);p__Ascomycota(100);c__Saccharomycetes(100);o__Saccharomycetales(100);f__Metschnikowiaceae(100);g__Metschnikowia(100);g__Metschnikowia_unclassified(100); | Metschnikowia | 13 | Saprotroph | \|Undefined Saprotroph\| | Yeast | NULL | Probable | As nectar-tap_saprotroph (Põlme et al. 2020); Decay sugar-rich substrates (Põlme et al. 2020) | Põlme S, et al. 2020. Fungal Diversity 105:1-16 ((https://doi.org/10.1007/s13225-020-00466-2)) |
| Otu32639 | 1 | k__Fungi(100);p__Ascomycota(100);c__Saccharomycetes(100);o__Saccharomycetales(100);f__Metschnikowiaceae(100);g__Metschnikowia(100);s__Metschnikowia_pulcherrima(100); | Metschnikowia | 13 | Saprotroph | \|Undefined Saprotroph\| | Yeast | NULL | Probable | As nectar-tap_saprotroph (Põlme et al. 2020); Decay sugar-rich substrates (Põlme et al. 2020) | Põlme S, et al. 2020. Fungal Diversity 105:1-16 ((https://doi.org/10.1007/s13225-020-00466-2)) |
| Otu32640 | 1 | k__Fungi(100);p__Ascomycota(100);c__Saccharomycetes(100);o__Saccharomycetales(100);f__Metschnikowiaceae(100);g__Metschnikowia(100);s__Metschnikowia_pulcherrima(100); | Metschnikowia | 13 | Saprotroph | \|Undefined Saprotroph\| | Yeast | NULL | Probable | As nectar-tap_saprotroph (Põlme et al. 2020); Decay sugar-rich substrates (Põlme et al. 2020) | Põlme S, et al. 2020. Fungal Diversity 105:1-16 ((https://doi.org/10.1007/s13225-020-00466-2)) |
| Otu32642 | 1 | k__Fungi(100);p__Ascomycota(100);c__Saccharomycetes(100);o__Saccharomycetales(100);f__Metschnikowiaceae(100);g__Metschnikowia(100);g__Metschnikowia_unclassified(100); | Metschnikowia | 13 | Saprotroph | \|Undefined Saprotroph\| | Yeast | NULL | Probable | As nectar-tap_saprotroph (Põlme et al. 2020); Decay sugar-rich substrates (Põlme et al. 2020) | Põlme S, et al. 2020. Fungal Diversity 105:1-16 ((https://doi.org/10.1007/s13225-020-00466-2)) |
| Otu32643 | 1 | k__Fungi(100);p__Ascomycota(100);c__Saccharomycetes(100);o__Saccharomycetales(100);f__Metschnikowiaceae(100);g__Metschnikowia(100);g__Metschnikowia_unclassified(100); | Metschnikowia | 13 | Saprotroph | \|Undefined Saprotroph\| | Yeast | NULL | Probable | As nectar-tap_saprotroph (Põlme et al. 2020); Decay sugar-rich substrates (Põlme et al. 2020) | Põlme S, et al. 2020. Fungal Diversity 105:1-16 ((https://doi.org/10.1007/s13225-020-00466-2)) |
| Otu32645 | 1 | k__Fungi(100);p__Ascomycota(100);c__Saccharomycetes(100);o__Saccharomycetales(100);f__Metschnikowiaceae(100);g__Metschnikowia(100);g__Metschnikowia_unclassified(100); | Metschnikowia | 13 | Saprotroph | \|Undefined Saprotroph\| | Yeast | NULL | Probable | As nectar-tap_saprotroph (Põlme et al. 2020); Decay sugar-rich substrates (Põlme et al. 2020) | Põlme S, et al. 2020. Fungal Diversity 105:1-16 ((https://doi.org/10.1007/s13225-020-00466-2)) |
| Otu32646 | 1 | k__Fungi(100);p__Ascomycota(100);c__Saccharomycetes(100);o__Saccharomycetales(100);f__Metschnikowiaceae(100);g__Metschnikowia(100);s__Metschnikowia_pulcherrima(100); | Metschnikowia | 13 | Saprotroph | \|Undefined Saprotroph\| | Yeast | NULL | Probable | As nectar-tap_saprotroph (Põlme et al. 2020); Decay sugar-rich substrates (Põlme et al. 2020) | Põlme S, et al. 2020. Fungal Diversity 105:1-16 ((https://doi.org/10.1007/s13225-020-00466-2)) |
| Otu32647 | 1 | k__Fungi(100);p__Ascomycota(100);c__Saccharomycetes(100);o__Saccharomycetales(100);f__Metschnikowiaceae(100);g__Metschnikowia(100);s__Metschnikowia_pulcherrima(100); | Metschnikowia | 13 | Saprotroph | \|Undefined Saprotroph\| | Yeast | NULL | Probable | As nectar-tap_saprotroph (Põlme et al. 2020); Decay sugar-rich substrates (Põlme et al. 2020) | Põlme S, et al. 2020. Fungal Diversity 105:1-16 ((https://doi.org/10.1007/s13225-020-00466-2)) |
| Otu32648 | 1 | k__Fungi(100);p__Ascomycota(100);c__Saccharomycetes(100);o__Saccharomycetales(100);f__Metschnikowiaceae(100);g__Metschnikowia(100);s__Metschnikowia_pulcherrima(100); | Metschnikowia | 13 | Saprotroph | \|Undefined Saprotroph\| | Yeast | NULL | Probable | As nectar-tap_saprotroph (Põlme et al. 2020); Decay sugar-rich substrates (Põlme et al. 2020) | Põlme S, et al. 2020. Fungal Diversity 105:1-16 ((https://doi.org/10.1007/s13225-020-00466-2)) |
| Otu32649 | 1 | k__Fungi(100);p__Ascomycota(100);c__Saccharomycetes(100);o__Saccharomycetales(100);f__Metschnikowiaceae(100);g__Metschnikowia(100);g__Metschnikowia_unclassified(100); | Metschnikowia | 13 | Saprotroph | \|Undefined Saprotroph\| | Yeast | NULL | Probable | As nectar-tap_saprotroph (Põlme et al. 2020); Decay sugar-rich substrates (Põlme et al. 2020) | Põlme S, et al. 2020. Fungal Diversity 105:1-16 ((https://doi.org/10.1007/s13225-020-00466-2)) |
| Otu32651 | 1 | k__Fungi(100);p__Ascomycota(100);c__Saccharomycetes(100);o__Saccharomycetales(100);f__Metschnikowiaceae(100);g__Metschnikowia(100);s__Metschnikowia_pulcherrima(100); | Metschnikowia | 13 | Saprotroph | \|Undefined Saprotroph\| | Yeast | NULL | Probable | As nectar-tap_saprotroph (Põlme et al. 2020); Decay sugar-rich substrates (Põlme et al. 2020) | Põlme S, et al. 2020. Fungal Diversity 105:1-16 ((https://doi.org/10.1007/s13225-020-00466-2)) |
| Otu32653 | 1 | k__Fungi(100);p__Ascomycota(100);c__Saccharomycetes(100);o__Saccharomycetales(100);f__Metschnikowiaceae(100);g__Metschnikowia(100);s__Metschnikowia_pulcherrima(100); | Metschnikowia | 13 | Saprotroph | \|Undefined Saprotroph\| | Yeast | NULL | Probable | As nectar-tap_saprotroph (Põlme et al. 2020); Decay sugar-rich substrates (Põlme et al. 2020) | Põlme S, et al. 2020. Fungal Diversity 105:1-16 ((https://doi.org/10.1007/s13225-020-00466-2)) |
| Otu32655 | 1 | k__Fungi(100);p__Ascomycota(100);c__Saccharomycetes(100);o__Saccharomycetales(100);f__Metschnikowiaceae(100);g__Metschnikowia(100);s__Metschnikowia_pulcherrima(100); | Metschnikowia | 13 | Saprotroph | \|Undefined Saprotroph\| | Yeast | NULL | Probable | As nectar-tap_saprotroph (Põlme et al. 2020); Decay sugar-rich substrates (Põlme et al. 2020) | Põlme S, et al. 2020. Fungal Diversity 105:1-16 ((https://doi.org/10.1007/s13225-020-00466-2)) |
| Otu32656 | 1 | k__Fungi(100);p__Ascomycota(100);c__Saccharomycetes(100);o__Saccharomycetales(100);f__Metschnikowiaceae(100);g__Metschnikowia(100);s__Metschnikowia_pulcherrima(100); | Metschnikowia | 13 | Saprotroph | \|Undefined Saprotroph\| | Yeast | NULL | Probable | As nectar-tap_saprotroph (Põlme et al. 2020); Decay sugar-rich substrates (Põlme et al. 2020) | Põlme S, et al. 2020. Fungal Diversity 105:1-16 ((https://doi.org/10.1007/s13225-020-00466-2)) |
| Otu32657 | 1 | k__Fungi(100);p__Ascomycota(100);c__Saccharomycetes(100);o__Saccharomycetales(100);f__Metschnikowiaceae(100);g__Metschnikowia(100);g__Metschnikowia_unclassified(100); | Metschnikowia | 13 | Saprotroph | \|Undefined Saprotroph\| | Yeast | NULL | Probable | As nectar-tap_saprotroph (Põlme et al. 2020); Decay sugar-rich substrates (Põlme et al. 2020) | Põlme S, et al. 2020. Fungal Diversity 105:1-16 ((https://doi.org/10.1007/s13225-020-00466-2)) |
| Otu32658 | 1 | k__Fungi(100);p__Ascomycota(100);c__Saccharomycetes(100);o__Saccharomycetales(100);f__Metschnikowiaceae(100);g__Metschnikowia(100);s__Metschnikowia_pulcherrima(100); | Metschnikowia | 13 | Saprotroph | \|Undefined Saprotroph\| | Yeast | NULL | Probable | As nectar-tap_saprotroph (Põlme et al. 2020); Decay sugar-rich substrates (Põlme et al. 2020) | Põlme S, et al. 2020. Fungal Diversity 105:1-16 ((https://doi.org/10.1007/s13225-020-00466-2)) |
| Otu32659 | 1 | k__Fungi(100);p__Ascomycota(100);c__Saccharomycetes(100);o__Saccharomycetales(100);f__Metschnikowiaceae(100);g__Metschnikowia(100);s__Metschnikowia_pulcherrima(100); | Metschnikowia | 13 | Saprotroph | \|Undefined Saprotroph\| | Yeast | NULL | Probable | As nectar-tap_saprotroph (Põlme et al. 2020); Decay sugar-rich substrates (Põlme et al. 2020) | Põlme S, et al. 2020. Fungal Diversity 105:1-16 ((https://doi.org/10.1007/s13225-020-00466-2)) |
| Otu32660 | 1 | k__Fungi(100);p__Ascomycota(100);c__Saccharomycetes(100);o__Saccharomycetales(100);f__Metschnikowiaceae(100);g__Metschnikowia(100);s__Metschnikowia_pulcherrima(100); | Metschnikowia | 13 | Saprotroph | \|Undefined Saprotroph\| | Yeast | NULL | Probable | As nectar-tap_saprotroph (Põlme et al. 2020); Decay sugar-rich substrates (Põlme et al. 2020) | Põlme S, et al. 2020. Fungal Diversity 105:1-16 ((https://doi.org/10.1007/s13225-020-00466-2)) |
| Otu32661 | 1 | k__Fungi(100);p__Ascomycota(100);c__Saccharomycetes(100);o__Saccharomycetales(100);f__Metschnikowiaceae(100);g__Metschnikowia(100);s__Metschnikowia_pulcherrima(100); | Metschnikowia | 13 | Saprotroph | \|Undefined Saprotroph\| | Yeast | NULL | Probable | As nectar-tap_saprotroph (Põlme et al. 2020); Decay sugar-rich substrates (Põlme et al. 2020) | Põlme S, et al. 2020. Fungal Diversity 105:1-16 ((https://doi.org/10.1007/s13225-020-00466-2)) |
| Otu32662 | 1 | k__Fungi(100);p__Ascomycota(100);c__Saccharomycetes(100);o__Saccharomycetales(100);f__Metschnikowiaceae(100);g__Metschnikowia(100);g__Metschnikowia_unclassified(100); | Metschnikowia | 13 | Saprotroph | \|Undefined Saprotroph\| | Yeast | NULL | Probable | As nectar-tap_saprotroph (Põlme et al. 2020); Decay sugar-rich substrates (Põlme et al. 2020) | Põlme S, et al. 2020. Fungal Diversity 105:1-16 ((https://doi.org/10.1007/s13225-020-00466-2)) |
| Otu32663 | 1 | k__Fungi(100);p__Ascomycota(100);c__Saccharomycetes(100);o__Saccharomycetales(100);f__Metschnikowiaceae(100);g__Metschnikowia(100);g__Metschnikowia_unclassified(100); | Metschnikowia | 13 | Saprotroph | \|Undefined Saprotroph\| | Yeast | NULL | Probable | As nectar-tap_saprotroph (Põlme et al. 2020); Decay sugar-rich substrates (Põlme et al. 2020) | Põlme S, et al. 2020. Fungal Diversity 105:1-16 ((https://doi.org/10.1007/s13225-020-00466-2)) |
| Otu32664 | 1 | k__Fungi(100);p__Ascomycota(100);c__Saccharomycetes(100);o__Saccharomycetales(100);f__Metschnikowiaceae(100);g__Metschnikowia(100);g__Metschnikowia_unclassified(100); | Metschnikowia | 13 | Saprotroph | \|Undefined Saprotroph\| | Yeast | NULL | Probable | As nectar-tap_saprotroph (Põlme et al. 2020); Decay sugar-rich substrates (Põlme et al. 2020) | Põlme S, et al. 2020. Fungal Diversity 105:1-16 ((https://doi.org/10.1007/s13225-020-00466-2)) |
| Otu32666 | 1 | k__Fungi(100);p__Ascomycota(100);c__Saccharomycetes(100);o__Saccharomycetales(100);f__Metschnikowiaceae(100);g__Metschnikowia(100);s__Metschnikowia_pulcherrima(100); | Metschnikowia | 13 | Saprotroph | \|Undefined Saprotroph\| | Yeast | NULL | Probable | As nectar-tap_saprotroph (Põlme et al. 2020); Decay sugar-rich substrates (Põlme et al. 2020) | Põlme S, et al. 2020. Fungal Diversity 105:1-16 ((https://doi.org/10.1007/s13225-020-00466-2)) |
| Otu32667 | 1 | k__Fungi(100);p__Ascomycota(100);c__Saccharomycetes(100);o__Saccharomycetales(100);f__Metschnikowiaceae(100);g__Metschnikowia(100);g__Metschnikowia_unclassified(100); | Metschnikowia | 13 | Saprotroph | \|Undefined Saprotroph\| | Yeast | NULL | Probable | As nectar-tap_saprotroph (Põlme et al. 2020); Decay sugar-rich substrates (Põlme et al. 2020) | Põlme S, et al. 2020. Fungal Diversity 105:1-16 ((https://doi.org/10.1007/s13225-020-00466-2)) |
| Otu32668 | 1 | k__Fungi(100);p__Ascomycota(100);c__Saccharomycetes(100);o__Saccharomycetales(100);f__Metschnikowiaceae(100);g__Metschnikowia(100);g__Metschnikowia_unclassified(100); | Metschnikowia | 13 | Saprotroph | \|Undefined Saprotroph\| | Yeast | NULL | Probable | As nectar-tap_saprotroph (Põlme et al. 2020); Decay sugar-rich substrates (Põlme et al. 2020) | Põlme S, et al. 2020. Fungal Diversity 105:1-16 ((https://doi.org/10.1007/s13225-020-00466-2)) |
| Otu32669 | 1 | k__Fungi(100);p__Ascomycota(100);c__Saccharomycetes(100);o__Saccharomycetales(100);f__Metschnikowiaceae(100);g__Metschnikowia(100);s__Metschnikowia_pulcherrima(100); | Metschnikowia | 13 | Saprotroph | \|Undefined Saprotroph\| | Yeast | NULL | Probable | As nectar-tap_saprotroph (Põlme et al. 2020); Decay sugar-rich substrates (Põlme et al. 2020) | Põlme S, et al. 2020. Fungal Diversity 105:1-16 ((https://doi.org/10.1007/s13225-020-00466-2)) |
| Otu32670 | 1 | k__Fungi(100);p__Ascomycota(100);c__Saccharomycetes(100);o__Saccharomycetales(100);f__Metschnikowiaceae(100);g__Metschnikowia(100);s__Metschnikowia_pulcherrima(100); | Metschnikowia | 13 | Saprotroph | \|Undefined Saprotroph\| | Yeast | NULL | Probable | As nectar-tap_saprotroph (Põlme et al. 2020); Decay sugar-rich substrates (Põlme et al. 2020) | Põlme S, et al. 2020. Fungal Diversity 105:1-16 ((https://doi.org/10.1007/s13225-020-00466-2)) |
| Otu32674 | 1 | k__Fungi(100);p__Ascomycota(100);c__Saccharomycetes(100);o__Saccharomycetales(100);f__Metschnikowiaceae(100);g__Metschnikowia(100);s__Metschnikowia_pulcherrima(100); | Metschnikowia | 13 | Saprotroph | \|Undefined Saprotroph\| | Yeast | NULL | Probable | As nectar-tap_saprotroph (Põlme et al. 2020); Decay sugar-rich substrates (Põlme et al. 2020) | Põlme S, et al. 2020. Fungal Diversity 105:1-16 ((https://doi.org/10.1007/s13225-020-00466-2)) |
| Otu32678 | 1 | k__Fungi(100);p__Ascomycota(100);c__Saccharomycetes(100);o__Saccharomycetales(100);f__Metschnikowiaceae(100);g__Metschnikowia(100);g__Metschnikowia_unclassified(100); | Metschnikowia | 13 | Saprotroph | \|Undefined Saprotroph\| | Yeast | NULL | Probable | As nectar-tap_saprotroph (Põlme et al. 2020); Decay sugar-rich substrates (Põlme et al. 2020) | Põlme S, et al. 2020. Fungal Diversity 105:1-16 ((https://doi.org/10.1007/s13225-020-00466-2)) |
| Otu32679 | 1 | k__Fungi(100);p__Ascomycota(100);c__Saccharomycetes(100);o__Saccharomycetales(100);f__Metschnikowiaceae(100);g__Metschnikowia(100);s__Metschnikowia_pulcherrima(100); | Metschnikowia | 13 | Saprotroph | \|Undefined Saprotroph\| | Yeast | NULL | Probable | As nectar-tap_saprotroph (Põlme et al. 2020); Decay sugar-rich substrates (Põlme et al. 2020) | Põlme S, et al. 2020. Fungal Diversity 105:1-16 ((https://doi.org/10.1007/s13225-020-00466-2)) |
| Otu32680 | 1 | k__Fungi(100);p__Ascomycota(100);c__Saccharomycetes(100);o__Saccharomycetales(100);f__Metschnikowiaceae(100);g__Metschnikowia(100);s__Metschnikowia_pulcherrima(100); | Metschnikowia | 13 | Saprotroph | \|Undefined Saprotroph\| | Yeast | NULL | Probable | As nectar-tap_saprotroph (Põlme et al. 2020); Decay sugar-rich substrates (Põlme et al. 2020) | Põlme S, et al. 2020. Fungal Diversity 105:1-16 ((https://doi.org/10.1007/s13225-020-00466-2)) |
| Otu32681 | 1 | k__Fungi(100);p__Ascomycota(100);c__Saccharomycetes(100);o__Saccharomycetales(100);f__Metschnikowiaceae(100);g__Metschnikowia(100);s__Metschnikowia_pulcherrima(100); | Metschnikowia | 13 | Saprotroph | \|Undefined Saprotroph\| | Yeast | NULL | Probable | As nectar-tap_saprotroph (Põlme et al. 2020); Decay sugar-rich substrates (Põlme et al. 2020) | Põlme S, et al. 2020. Fungal Diversity 105:1-16 ((https://doi.org/10.1007/s13225-020-00466-2)) |
| Otu32682 | 1 | k__Fungi(100);p__Ascomycota(100);c__Saccharomycetes(100);o__Saccharomycetales(100);f__Metschnikowiaceae(100);g__Metschnikowia(100);s__Metschnikowia_pulcherrima(100); | Metschnikowia | 13 | Saprotroph | \|Undefined Saprotroph\| | Yeast | NULL | Probable | As nectar-tap_saprotroph (Põlme et al. 2020); Decay sugar-rich substrates (Põlme et al. 2020) | Põlme S, et al. 2020. Fungal Diversity 105:1-16 ((https://doi.org/10.1007/s13225-020-00466-2)) |
| Otu32683 | 1 | k__Fungi(100);p__Ascomycota(100);c__Saccharomycetes(100);o__Saccharomycetales(100);f__Metschnikowiaceae(100);g__Metschnikowia(100);g__Metschnikowia_unclassified(100); | Metschnikowia | 13 | Saprotroph | \|Undefined Saprotroph\| | Yeast | NULL | Probable | As nectar-tap_saprotroph (Põlme et al. 2020); Decay sugar-rich substrates (Põlme et al. 2020) | Põlme S, et al. 2020. Fungal Diversity 105:1-16 ((https://doi.org/10.1007/s13225-020-00466-2)) |
| Otu32684 | 1 | k__Fungi(100);p__Ascomycota(100);c__Saccharomycetes(100);o__Saccharomycetales(100);f__Saccharomycetales_family_Incertae_sedis(100);g__Candida(100);g__Candida_unclassified(100); | Candida | 13 | Pathotroph-Saprotroph-Symbiotroph | Animal Pathogen-Endophyte-Endosymbiont-Epiphyte-\|Undefined Saprotroph\| | Dimorphic Yeast-Yeast | NULL | Probable | As nectar-tap_saprotroph (Põlme et al. 2020); Decay sugar-rich substrates (Põlme et al. 2020); This asexual genus is polyphyletic and contain species that function very differently depend on what they are. Caution should be taken when interpreting guilds for Candida species. | Manolakaki D, et al. 2010. Virulence. 1: 367-375 ((https://doi.org/10.4161/viru.1.5.12796)); Mekbib SB, et al. 2011. Tropical Plant Pathology 36:233-240 ((https://doi.org/10.1590/S1982-56762011000400004)); Põlme S, et al. 2020. Fungal Diversity 105:1-16 ((https://doi.org/10.1007/s13225-020-00466-2)) |
| Otu32688 | 1 | k__Fungi(100);p__Ascomycota(100);c__Saccharomycetes(100);o__Saccharomycetales(100);f__Metschnikowiaceae(100);g__Metschnikowia(100);g__Metschnikowia_unclassified(100); | Metschnikowia | 13 | Saprotroph | \|Undefined Saprotroph\| | Yeast | NULL | Probable | As nectar-tap_saprotroph (Põlme et al. 2020); Decay sugar-rich substrates (Põlme et al. 2020) | Põlme S, et al. 2020. Fungal Diversity 105:1-16 ((https://doi.org/10.1007/s13225-020-00466-2)) |
| Otu32689 | 1 | k__Fungi(100);p__Ascomycota(100);c__Saccharomycetes(100);o__Saccharomycetales(100);f__Metschnikowiaceae(100);g__Metschnikowia(100);s__Metschnikowia_pulcherrima(100); | Metschnikowia | 13 | Saprotroph | \|Undefined Saprotroph\| | Yeast | NULL | Probable | As nectar-tap_saprotroph (Põlme et al. 2020); Decay sugar-rich substrates (Põlme et al. 2020) | Põlme S, et al. 2020. Fungal Diversity 105:1-16 ((https://doi.org/10.1007/s13225-020-00466-2)) |
| Otu32690 | 1 | k__Fungi(100);p__Ascomycota(100);c__Saccharomycetes(100);o__Saccharomycetales(100);f__Metschnikowiaceae(100);g__Metschnikowia(100);s__Metschnikowia_pulcherrima(100); | Metschnikowia | 13 | Saprotroph | \|Undefined Saprotroph\| | Yeast | NULL | Probable | As nectar-tap_saprotroph (Põlme et al. 2020); Decay sugar-rich substrates (Põlme et al. 2020) | Põlme S, et al. 2020. Fungal Diversity 105:1-16 ((https://doi.org/10.1007/s13225-020-00466-2)) |
| Otu32691 | 1 | k__Fungi(100);p__Ascomycota(100);c__Saccharomycetes(100);o__Saccharomycetales(100);f__Metschnikowiaceae(100);g__Metschnikowia(100);s__Metschnikowia_pulcherrima(100); | Metschnikowia | 13 | Saprotroph | \|Undefined Saprotroph\| | Yeast | NULL | Probable | As nectar-tap_saprotroph (Põlme et al. 2020); Decay sugar-rich substrates (Põlme et al. 2020) | Põlme S, et al. 2020. Fungal Diversity 105:1-16 ((https://doi.org/10.1007/s13225-020-00466-2)) |
| Otu32693 | 1 | k__Fungi(100);p__Ascomycota(100);c__Saccharomycetes(100);o__Saccharomycetales(100);f__Metschnikowiaceae(100);g__Metschnikowia(100);g__Metschnikowia_unclassified(100); | Metschnikowia | 13 | Saprotroph | \|Undefined Saprotroph\| | Yeast | NULL | Probable | As nectar-tap_saprotroph (Põlme et al. 2020); Decay sugar-rich substrates (Põlme et al. 2020) | Põlme S, et al. 2020. Fungal Diversity 105:1-16 ((https://doi.org/10.1007/s13225-020-00466-2)) |
| Otu32694 | 1 | k__Fungi(100);p__Ascomycota(100);c__Saccharomycetes(100);o__Saccharomycetales(100);f__Metschnikowiaceae(100);g__Metschnikowia(100);g__Metschnikowia_unclassified(100); | Metschnikowia | 13 | Saprotroph | \|Undefined Saprotroph\| | Yeast | NULL | Probable | As nectar-tap_saprotroph (Põlme et al. 2020); Decay sugar-rich substrates (Põlme et al. 2020) | Põlme S, et al. 2020. Fungal Diversity 105:1-16 ((https://doi.org/10.1007/s13225-020-00466-2)) |
| Otu32697 | 1 | k__Fungi(100);p__Ascomycota(100);c__Saccharomycetes(100);o__Saccharomycetales(100);f__Metschnikowiaceae(100);g__Metschnikowia(100);g__Metschnikowia_unclassified(100); | Metschnikowia | 13 | Saprotroph | \|Undefined Saprotroph\| | Yeast | NULL | Probable | As nectar-tap_saprotroph (Põlme et al. 2020); Decay sugar-rich substrates (Põlme et al. 2020) | Põlme S, et al. 2020. Fungal Diversity 105:1-16 ((https://doi.org/10.1007/s13225-020-00466-2)) |
| Otu32701 | 1 | k__Fungi(100);p__Ascomycota(100);c__Saccharomycetes(100);o__Saccharomycetales(100);f__Metschnikowiaceae(100);g__Metschnikowia(100);s__Metschnikowia_pulcherrima(100); | Metschnikowia | 13 | Saprotroph | \|Undefined Saprotroph\| | Yeast | NULL | Probable | As nectar-tap_saprotroph (Põlme et al. 2020); Decay sugar-rich substrates (Põlme et al. 2020) | Põlme S, et al. 2020. Fungal Diversity 105:1-16 ((https://doi.org/10.1007/s13225-020-00466-2)) |
| Otu32704 | 1 | k__Fungi(100);p__Ascomycota(100);c__Saccharomycetes(100);o__Saccharomycetales(100);f__Metschnikowiaceae(100);g__Metschnikowia(100);s__Metschnikowia_pulcherrima(100); | Metschnikowia | 13 | Saprotroph | \|Undefined Saprotroph\| | Yeast | NULL | Probable | As nectar-tap_saprotroph (Põlme et al. 2020); Decay sugar-rich substrates (Põlme et al. 2020) | Põlme S, et al. 2020. Fungal Diversity 105:1-16 ((https://doi.org/10.1007/s13225-020-00466-2)) |
| Otu32706 | 1 | k__Fungi(100);p__Ascomycota(100);c__Saccharomycetes(100);o__Saccharomycetales(100);f__Metschnikowiaceae(100);g__Metschnikowia(100);g__Metschnikowia_unclassified(100); | Metschnikowia | 13 | Saprotroph | \|Undefined Saprotroph\| | Yeast | NULL | Probable | As nectar-tap_saprotroph (Põlme et al. 2020); Decay sugar-rich substrates (Põlme et al. 2020) | Põlme S, et al. 2020. Fungal Diversity 105:1-16 ((https://doi.org/10.1007/s13225-020-00466-2)) |
| Otu32710 | 1 | k__Fungi(100);p__Ascomycota(100);c__Saccharomycetes(100);o__Saccharomycetales(100);f__Saccharomycetales_family_Incertae_sedis(100);g__Candida(100);s__Candida_cretensis(100); | Candida | 13 | Pathotroph-Saprotroph-Symbiotroph | Animal Pathogen-Endophyte-Endosymbiont-Epiphyte-\|Undefined Saprotroph\| | Dimorphic Yeast-Yeast | NULL | Probable | As nectar-tap_saprotroph (Põlme et al. 2020); Decay sugar-rich substrates (Põlme et al. 2020); This asexual genus is polyphyletic and contain species that function very differently depend on what they are. Caution should be taken when interpreting guilds for Candida species. | Manolakaki D, et al. 2010. Virulence. 1: 367-375 ((https://doi.org/10.4161/viru.1.5.12796)); Mekbib SB, et al. 2011. Tropical Plant Pathology 36:233-240 ((https://doi.org/10.1590/S1982-56762011000400004)); Põlme S, et al. 2020. Fungal Diversity 105:1-16 ((https://doi.org/10.1007/s13225-020-00466-2)) |
| Otu32711 | 1 | k__Fungi(100);p__Ascomycota(100);c__Saccharomycetes(100);o__Saccharomycetales(100);f__Metschnikowiaceae(100);g__Metschnikowia(100);s__Metschnikowia_pulcherrima(100); | Metschnikowia | 13 | Saprotroph | \|Undefined Saprotroph\| | Yeast | NULL | Probable | As nectar-tap_saprotroph (Põlme et al. 2020); Decay sugar-rich substrates (Põlme et al. 2020) | Põlme S, et al. 2020. Fungal Diversity 105:1-16 ((https://doi.org/10.1007/s13225-020-00466-2)) |
| Otu32712 | 1 | k__Fungi(100);p__Ascomycota(100);c__Saccharomycetes(100);o__Saccharomycetales(100);f__Metschnikowiaceae(100);g__Metschnikowia(100);s__Metschnikowia_pulcherrima(100); | Metschnikowia | 13 | Saprotroph | \|Undefined Saprotroph\| | Yeast | NULL | Probable | As nectar-tap_saprotroph (Põlme et al. 2020); Decay sugar-rich substrates (Põlme et al. 2020) | Põlme S, et al. 2020. Fungal Diversity 105:1-16 ((https://doi.org/10.1007/s13225-020-00466-2)) |
| Otu32713 | 1 | k__Fungi(100);p__Ascomycota(100);c__Saccharomycetes(100);o__Saccharomycetales(100);f__Metschnikowiaceae(100);g__Metschnikowia(100);s__Metschnikowia_pulcherrima(100); | Metschnikowia | 13 | Saprotroph | \|Undefined Saprotroph\| | Yeast | NULL | Probable | As nectar-tap_saprotroph (Põlme et al. 2020); Decay sugar-rich substrates (Põlme et al. 2020) | Põlme S, et al. 2020. Fungal Diversity 105:1-16 ((https://doi.org/10.1007/s13225-020-00466-2)) |
| Otu32714 | 1 | k__Fungi(100);p__Ascomycota(100);c__Saccharomycetes(100);o__Saccharomycetales(100);f__Metschnikowiaceae(100);g__Metschnikowia(100);g__Metschnikowia_unclassified(100); | Metschnikowia | 13 | Saprotroph | \|Undefined Saprotroph\| | Yeast | NULL | Probable | As nectar-tap_saprotroph (Põlme et al. 2020); Decay sugar-rich substrates (Põlme et al. 2020) | Põlme S, et al. 2020. Fungal Diversity 105:1-16 ((https://doi.org/10.1007/s13225-020-00466-2)) |
| Otu32715 | 1 | k__Fungi(100);p__Ascomycota(100);c__Saccharomycetes(100);o__Saccharomycetales(100);f__Metschnikowiaceae(100);g__Metschnikowia(100);g__Metschnikowia_unclassified(100); | Metschnikowia | 13 | Saprotroph | \|Undefined Saprotroph\| | Yeast | NULL | Probable | As nectar-tap_saprotroph (Põlme et al. 2020); Decay sugar-rich substrates (Põlme et al. 2020) | Põlme S, et al. 2020. Fungal Diversity 105:1-16 ((https://doi.org/10.1007/s13225-020-00466-2)) |
| Otu32716 | 1 | k__Fungi(100);p__Ascomycota(100);c__Saccharomycetes(100);o__Saccharomycetales(100);f__Metschnikowiaceae(100);g__Metschnikowia(100);s__Metschnikowia_pulcherrima(100); | Metschnikowia | 13 | Saprotroph | \|Undefined Saprotroph\| | Yeast | NULL | Probable | As nectar-tap_saprotroph (Põlme et al. 2020); Decay sugar-rich substrates (Põlme et al. 2020) | Põlme S, et al. 2020. Fungal Diversity 105:1-16 ((https://doi.org/10.1007/s13225-020-00466-2)) |
| Otu32717 | 1 | k__Fungi(100);p__Ascomycota(100);c__Saccharomycetes(100);o__Saccharomycetales(100);f__Saccharomycetales_family_Incertae_sedis(100);g__Candida(100);g__Candida_unclassified(100); | Candida | 13 | Pathotroph-Saprotroph-Symbiotroph | Animal Pathogen-Endophyte-Endosymbiont-Epiphyte-\|Undefined Saprotroph\| | Dimorphic Yeast-Yeast | NULL | Probable | As nectar-tap_saprotroph (Põlme et al. 2020); Decay sugar-rich substrates (Põlme et al. 2020); This asexual genus is polyphyletic and contain species that function very differently depend on what they are. Caution should be taken when interpreting guilds for Candida species. | Manolakaki D, et al. 2010. Virulence. 1: 367-375 ((https://doi.org/10.4161/viru.1.5.12796)); Mekbib SB, et al. 2011. Tropical Plant Pathology 36:233-240 ((https://doi.org/10.1590/S1982-56762011000400004)); Põlme S, et al. 2020. Fungal Diversity 105:1-16 ((https://doi.org/10.1007/s13225-020-00466-2)) |
| Otu32718 | 1 | k__Fungi(100);p__Ascomycota(100);c__Saccharomycetes(100);o__Saccharomycetales(100);f__Metschnikowiaceae(100);g__Metschnikowia(100);g__Metschnikowia_unclassified(100); | Metschnikowia | 13 | Saprotroph | \|Undefined Saprotroph\| | Yeast | NULL | Probable | As nectar-tap_saprotroph (Põlme et al. 2020); Decay sugar-rich substrates (Põlme et al. 2020) | Põlme S, et al. 2020. Fungal Diversity 105:1-16 ((https://doi.org/10.1007/s13225-020-00466-2)) |
| Otu32719 | 1 | k__Fungi(100);p__Ascomycota(100);c__Saccharomycetes(100);o__Saccharomycetales(100);f__Metschnikowiaceae(100);g__Metschnikowia(100);s__Metschnikowia_pulcherrima(100); | Metschnikowia | 13 | Saprotroph | \|Undefined Saprotroph\| | Yeast | NULL | Probable | As nectar-tap_saprotroph (Põlme et al. 2020); Decay sugar-rich substrates (Põlme et al. 2020) | Põlme S, et al. 2020. Fungal Diversity 105:1-16 ((https://doi.org/10.1007/s13225-020-00466-2)) |
| Otu32721 | 1 | k__Fungi(100);p__Ascomycota(100);c__Saccharomycetes(100);o__Saccharomycetales(100);f__Metschnikowiaceae(100);g__Metschnikowia(100);g__Metschnikowia_unclassified(100); | Metschnikowia | 13 | Saprotroph | \|Undefined Saprotroph\| | Yeast | NULL | Probable | As nectar-tap_saprotroph (Põlme et al. 2020); Decay sugar-rich substrates (Põlme et al. 2020) | Põlme S, et al. 2020. Fungal Diversity 105:1-16 ((https://doi.org/10.1007/s13225-020-00466-2)) |
| Otu32722 | 1 | k__Fungi(100);p__Ascomycota(100);c__Saccharomycetes(100);o__Saccharomycetales(100);f__Metschnikowiaceae(100);g__Metschnikowia(100);s__Metschnikowia_pulcherrima(100); | Metschnikowia | 13 | Saprotroph | \|Undefined Saprotroph\| | Yeast | NULL | Probable | As nectar-tap_saprotroph (Põlme et al. 2020); Decay sugar-rich substrates (Põlme et al. 2020) | Põlme S, et al. 2020. Fungal Diversity 105:1-16 ((https://doi.org/10.1007/s13225-020-00466-2)) |
| Otu32733 | 1 | k__Fungi(100);p__Ascomycota(100);c__Saccharomycetes(100);o__Saccharomycetales(100);f__Metschnikowiaceae(100);g__Metschnikowia(100);s__Metschnikowia_pulcherrima(100); | Metschnikowia | 13 | Saprotroph | \|Undefined Saprotroph\| | Yeast | NULL | Probable | As nectar-tap_saprotroph (Põlme et al. 2020); Decay sugar-rich substrates (Põlme et al. 2020) | Põlme S, et al. 2020. Fungal Diversity 105:1-16 ((https://doi.org/10.1007/s13225-020-00466-2)) |
| Otu32737 | 1 | k__Fungi(100);p__Ascomycota(100);c__Saccharomycetes(100);o__Saccharomycetales(100);f__Metschnikowiaceae(100);g__Metschnikowia(100);s__Metschnikowia_pulcherrima(100); | Metschnikowia | 13 | Saprotroph | \|Undefined Saprotroph\| | Yeast | NULL | Probable | As nectar-tap_saprotroph (Põlme et al. 2020); Decay sugar-rich substrates (Põlme et al. 2020) | Põlme S, et al. 2020. Fungal Diversity 105:1-16 ((https://doi.org/10.1007/s13225-020-00466-2)) |
| Otu32738 | 1 | k__Fungi(100);p__Ascomycota(100);c__Saccharomycetes(100);o__Saccharomycetales(100);f__Metschnikowiaceae(100);g__Metschnikowia(100);s__Metschnikowia_pulcherrima(100); | Metschnikowia | 13 | Saprotroph | \|Undefined Saprotroph\| | Yeast | NULL | Probable | As nectar-tap_saprotroph (Põlme et al. 2020); Decay sugar-rich substrates (Põlme et al. 2020) | Põlme S, et al. 2020. Fungal Diversity 105:1-16 ((https://doi.org/10.1007/s13225-020-00466-2)) |
| Otu32739 | 1 | k__Fungi(100);p__Ascomycota(100);c__Saccharomycetes(100);o__Saccharomycetales(100);f__Metschnikowiaceae(100);g__Metschnikowia(100);s__Metschnikowia_pulcherrima(100); | Metschnikowia | 13 | Saprotroph | \|Undefined Saprotroph\| | Yeast | NULL | Probable | As nectar-tap_saprotroph (Põlme et al. 2020); Decay sugar-rich substrates (Põlme et al. 2020) | Põlme S, et al. 2020. Fungal Diversity 105:1-16 ((https://doi.org/10.1007/s13225-020-00466-2)) |
| Otu32740 | 1 | k__Fungi(100);p__Ascomycota(100);c__Saccharomycetes(100);o__Saccharomycetales(100);f__Metschnikowiaceae(100);g__Metschnikowia(100);s__Metschnikowia_pulcherrima(100); | Metschnikowia | 13 | Saprotroph | \|Undefined Saprotroph\| | Yeast | NULL | Probable | As nectar-tap_saprotroph (Põlme et al. 2020); Decay sugar-rich substrates (Põlme et al. 2020) | Põlme S, et al. 2020. Fungal Diversity 105:1-16 ((https://doi.org/10.1007/s13225-020-00466-2)) |
| Otu32741 | 1 | k__Fungi(100);p__Ascomycota(100);c__Saccharomycetes(100);o__Saccharomycetales(100);f__Metschnikowiaceae(100);g__Metschnikowia(100);g__Metschnikowia_unclassified(100); | Metschnikowia | 13 | Saprotroph | \|Undefined Saprotroph\| | Yeast | NULL | Probable | As nectar-tap_saprotroph (Põlme et al. 2020); Decay sugar-rich substrates (Põlme et al. 2020) | Põlme S, et al. 2020. Fungal Diversity 105:1-16 ((https://doi.org/10.1007/s13225-020-00466-2)) |
| Otu32742 | 1 | k__Fungi(100);p__Ascomycota(100);c__Saccharomycetes(100);o__Saccharomycetales(100);f__Metschnikowiaceae(100);g__Metschnikowia(100);g__Metschnikowia_unclassified(100); | Metschnikowia | 13 | Saprotroph | \|Undefined Saprotroph\| | Yeast | NULL | Probable | As nectar-tap_saprotroph (Põlme et al. 2020); Decay sugar-rich substrates (Põlme et al. 2020) | Põlme S, et al. 2020. Fungal Diversity 105:1-16 ((https://doi.org/10.1007/s13225-020-00466-2)) |
| Otu32743 | 1 | k__Fungi(100);p__Ascomycota(100);c__Saccharomycetes(100);o__Saccharomycetales(100);f__Metschnikowiaceae(100);g__Metschnikowia(100);g__Metschnikowia_unclassified(100); | Metschnikowia | 13 | Saprotroph | \|Undefined Saprotroph\| | Yeast | NULL | Probable | As nectar-tap_saprotroph (Põlme et al. 2020); Decay sugar-rich substrates (Põlme et al. 2020) | Põlme S, et al. 2020. Fungal Diversity 105:1-16 ((https://doi.org/10.1007/s13225-020-00466-2)) |
| Otu32744 | 1 | k__Fungi(100);p__Ascomycota(100);c__Saccharomycetes(100);o__Saccharomycetales(100);f__Metschnikowiaceae(100);g__Metschnikowia(100);g__Metschnikowia_unclassified(100); | Metschnikowia | 13 | Saprotroph | \|Undefined Saprotroph\| | Yeast | NULL | Probable | As nectar-tap_saprotroph (Põlme et al. 2020); Decay sugar-rich substrates (Põlme et al. 2020) | Põlme S, et al. 2020. Fungal Diversity 105:1-16 ((https://doi.org/10.1007/s13225-020-00466-2)) |
| Otu32747 | 1 | k__Fungi(100);p__Ascomycota(100);c__Saccharomycetes(100);o__Saccharomycetales(100);f__Metschnikowiaceae(100);g__Metschnikowia(100);g__Metschnikowia_unclassified(100); | Metschnikowia | 13 | Saprotroph | \|Undefined Saprotroph\| | Yeast | NULL | Probable | As nectar-tap_saprotroph (Põlme et al. 2020); Decay sugar-rich substrates (Põlme et al. 2020) | Põlme S, et al. 2020. Fungal Diversity 105:1-16 ((https://doi.org/10.1007/s13225-020-00466-2)) |
| Otu32749 | 1 | k__Fungi(100);p__Ascomycota(100);c__Saccharomycetes(100);o__Saccharomycetales(100);f__Metschnikowiaceae(100);g__Metschnikowia(100);s__Metschnikowia_pulcherrima(100); | Metschnikowia | 13 | Saprotroph | \|Undefined Saprotroph\| | Yeast | NULL | Probable | As nectar-tap_saprotroph (Põlme et al. 2020); Decay sugar-rich substrates (Põlme et al. 2020) | Põlme S, et al. 2020. Fungal Diversity 105:1-16 ((https://doi.org/10.1007/s13225-020-00466-2)) |
| Otu32751 | 1 | k__Fungi(100);p__Ascomycota(100);c__Saccharomycetes(100);o__Saccharomycetales(100);f__Metschnikowiaceae(100);g__Metschnikowia(100);g__Metschnikowia_unclassified(100); | Metschnikowia | 13 | Saprotroph | \|Undefined Saprotroph\| | Yeast | NULL | Probable | As nectar-tap_saprotroph (Põlme et al. 2020); Decay sugar-rich substrates (Põlme et al. 2020) | Põlme S, et al. 2020. Fungal Diversity 105:1-16 ((https://doi.org/10.1007/s13225-020-00466-2)) |
| Otu32752 | 1 | k__Fungi(100);p__Ascomycota(100);c__Saccharomycetes(100);o__Saccharomycetales(100);f__Metschnikowiaceae(100);g__Metschnikowia(100);s__Metschnikowia_pulcherrima(100); | Metschnikowia | 13 | Saprotroph | \|Undefined Saprotroph\| | Yeast | NULL | Probable | As nectar-tap_saprotroph (Põlme et al. 2020); Decay sugar-rich substrates (Põlme et al. 2020) | Põlme S, et al. 2020. Fungal Diversity 105:1-16 ((https://doi.org/10.1007/s13225-020-00466-2)) |
| Otu32756 | 1 | k__Fungi(100);p__Ascomycota(100);c__Saccharomycetes(100);o__Saccharomycetales(100);f__Metschnikowiaceae(100);g__Metschnikowia(100);s__Metschnikowia_pulcherrima(100); | Metschnikowia | 13 | Saprotroph | \|Undefined Saprotroph\| | Yeast | NULL | Probable | As nectar-tap_saprotroph (Põlme et al. 2020); Decay sugar-rich substrates (Põlme et al. 2020) | Põlme S, et al. 2020. Fungal Diversity 105:1-16 ((https://doi.org/10.1007/s13225-020-00466-2)) |
| Otu32757 | 1 | k__Fungi(100);p__Ascomycota(100);c__Saccharomycetes(100);o__Saccharomycetales(100);f__Metschnikowiaceae(100);g__Metschnikowia(100);s__Metschnikowia_pulcherrima(100); | Metschnikowia | 13 | Saprotroph | \|Undefined Saprotroph\| | Yeast | NULL | Probable | As nectar-tap_saprotroph (Põlme et al. 2020); Decay sugar-rich substrates (Põlme et al. 2020) | Põlme S, et al. 2020. Fungal Diversity 105:1-16 ((https://doi.org/10.1007/s13225-020-00466-2)) |
| Otu32760 | 1 | k__Fungi(100);p__Ascomycota(100);c__Saccharomycetes(100);o__Saccharomycetales(100);f__Metschnikowiaceae(100);g__Metschnikowia(100);s__Metschnikowia_pulcherrima(100); | Metschnikowia | 13 | Saprotroph | \|Undefined Saprotroph\| | Yeast | NULL | Probable | As nectar-tap_saprotroph (Põlme et al. 2020); Decay sugar-rich substrates (Põlme et al. 2020) | Põlme S, et al. 2020. Fungal Diversity 105:1-16 ((https://doi.org/10.1007/s13225-020-00466-2)) |
| Otu32761 | 1 | k__Fungi(100);p__Ascomycota(100);c__Saccharomycetes(100);o__Saccharomycetales(100);f__Metschnikowiaceae(100);g__Metschnikowia(100);s__Metschnikowia_pulcherrima(100); | Metschnikowia | 13 | Saprotroph | \|Undefined Saprotroph\| | Yeast | NULL | Probable | As nectar-tap_saprotroph (Põlme et al. 2020); Decay sugar-rich substrates (Põlme et al. 2020) | Põlme S, et al. 2020. Fungal Diversity 105:1-16 ((https://doi.org/10.1007/s13225-020-00466-2)) |
| Otu32763 | 1 | k__Fungi(100);p__Ascomycota(100);c__Saccharomycetes(100);o__Saccharomycetales(100);f__Metschnikowiaceae(100);g__Metschnikowia(100);s__Metschnikowia_pulcherrima(100); | Metschnikowia | 13 | Saprotroph | \|Undefined Saprotroph\| | Yeast | NULL | Probable | As nectar-tap_saprotroph (Põlme et al. 2020); Decay sugar-rich substrates (Põlme et al. 2020) | Põlme S, et al. 2020. Fungal Diversity 105:1-16 ((https://doi.org/10.1007/s13225-020-00466-2)) |
| Otu32767 | 1 | k__Fungi(100);p__Ascomycota(100);c__Saccharomycetes(100);o__Saccharomycetales(100);o__Saccharomycetales_unclassified(100);o__Saccharomycetales_unclassified(100);o__Saccharomycetales_unclassified(100); | Saccharomycetales | 7 | Saprotroph | Undefined Saprotroph | Yeast | NULL | Possible | NULL | Sterkenburg E, et al. 2015. New Phytologist 207:1145-1158 ((https://doi.org/10.1111/nph.13426)) |
| Otu32769 | 1 | k__Fungi(100);p__Ascomycota(100);c__Saccharomycetes(100);o__Saccharomycetales(100);f__Metschnikowiaceae(100);g__Metschnikowia(100);s__Metschnikowia_pulcherrima(100); | Metschnikowia | 13 | Saprotroph | \|Undefined Saprotroph\| | Yeast | NULL | Probable | As nectar-tap_saprotroph (Põlme et al. 2020); Decay sugar-rich substrates (Põlme et al. 2020) | Põlme S, et al. 2020. Fungal Diversity 105:1-16 ((https://doi.org/10.1007/s13225-020-00466-2)) |
| Otu32771 | 1 | k__Fungi(100);p__Ascomycota(100);c__Saccharomycetes(100);o__Saccharomycetales(100);f__Metschnikowiaceae(100);g__Metschnikowia(100);s__Metschnikowia_pulcherrima(100); | Metschnikowia | 13 | Saprotroph | \|Undefined Saprotroph\| | Yeast | NULL | Probable | As nectar-tap_saprotroph (Põlme et al. 2020); Decay sugar-rich substrates (Põlme et al. 2020) | Põlme S, et al. 2020. Fungal Diversity 105:1-16 ((https://doi.org/10.1007/s13225-020-00466-2)) |
| Otu32772 | 1 | k__Fungi(100);p__Ascomycota(100);c__Saccharomycetes(100);o__Saccharomycetales(100);f__Saccharomycetales_family_Incertae_sedis(100);f__Saccharomycetales_family_Incertae_sedis_unclassified(100);f__Saccharomycetales_family_Incertae_sedis_unclassified(100); | Saccharomycetales | 7 | Saprotroph | Undefined Saprotroph | Yeast | NULL | Possible | NULL | Sterkenburg E, et al. 2015. New Phytologist 207:1145-1158 ((https://doi.org/10.1111/nph.13426)) |
| Otu32776 | 1 | k__Fungi(100);p__Ascomycota(100);c__Saccharomycetes(100);o__Saccharomycetales(100);f__Metschnikowiaceae(100);g__Metschnikowia(100);g__Metschnikowia_unclassified(100); | Metschnikowia | 13 | Saprotroph | \|Undefined Saprotroph\| | Yeast | NULL | Probable | As nectar-tap_saprotroph (Põlme et al. 2020); Decay sugar-rich substrates (Põlme et al. 2020) | Põlme S, et al. 2020. Fungal Diversity 105:1-16 ((https://doi.org/10.1007/s13225-020-00466-2)) |
| Otu32777 | 1 | k__Fungi(100);p__Ascomycota(100);c__Saccharomycetes(100);o__Saccharomycetales(100);f__Metschnikowiaceae(100);g__Metschnikowia(100);s__Metschnikowia_pulcherrima(100); | Metschnikowia | 13 | Saprotroph | \|Undefined Saprotroph\| | Yeast | NULL | Probable | As nectar-tap_saprotroph (Põlme et al. 2020); Decay sugar-rich substrates (Põlme et al. 2020) | Põlme S, et al. 2020. Fungal Diversity 105:1-16 ((https://doi.org/10.1007/s13225-020-00466-2)) |
| Otu32779 | 1 | k__Fungi(100);p__Ascomycota(100);c__Saccharomycetes(100);o__Saccharomycetales(100);f__Saccharomycetales_family_Incertae_sedis(100);g__Candida(100);g__Candida_unclassified(100); | Candida | 13 | Pathotroph-Saprotroph-Symbiotroph | Animal Pathogen-Endophyte-Endosymbiont-Epiphyte-\|Undefined Saprotroph\| | Dimorphic Yeast-Yeast | NULL | Probable | As nectar-tap_saprotroph (Põlme et al. 2020); Decay sugar-rich substrates (Põlme et al. 2020); This asexual genus is polyphyletic and contain species that function very differently depend on what they are. Caution should be taken when interpreting guilds for Candida species. | Manolakaki D, et al. 2010. Virulence. 1: 367-375 ((https://doi.org/10.4161/viru.1.5.12796)); Mekbib SB, et al. 2011. Tropical Plant Pathology 36:233-240 ((https://doi.org/10.1590/S1982-56762011000400004)); Põlme S, et al. 2020. Fungal Diversity 105:1-16 ((https://doi.org/10.1007/s13225-020-00466-2)) |
| Otu32783 | 1 | k__Fungi(100);p__Ascomycota(100);c__Saccharomycetes(100);o__Saccharomycetales(100);f__Metschnikowiaceae(100);g__Metschnikowia(100);g__Metschnikowia_unclassified(100); | Metschnikowia | 13 | Saprotroph | \|Undefined Saprotroph\| | Yeast | NULL | Probable | As nectar-tap_saprotroph (Põlme et al. 2020); Decay sugar-rich substrates (Põlme et al. 2020) | Põlme S, et al. 2020. Fungal Diversity 105:1-16 ((https://doi.org/10.1007/s13225-020-00466-2)) |
| Otu32784 | 1 | k__Fungi(100);p__Ascomycota(100);c__Saccharomycetes(100);o__Saccharomycetales(100);f__Metschnikowiaceae(100);g__Metschnikowia(100);s__Metschnikowia_pulcherrima(100); | Metschnikowia | 13 | Saprotroph | \|Undefined Saprotroph\| | Yeast | NULL | Probable | As nectar-tap_saprotroph (Põlme et al. 2020); Decay sugar-rich substrates (Põlme et al. 2020) | Põlme S, et al. 2020. Fungal Diversity 105:1-16 ((https://doi.org/10.1007/s13225-020-00466-2)) |
| Otu32788 | 1 | k__Fungi(100);p__Ascomycota(100);c__Saccharomycetes(100);o__Saccharomycetales(100);f__Metschnikowiaceae(100);g__Metschnikowia(100);s__Metschnikowia_pulcherrima(100); | Metschnikowia | 13 | Saprotroph | \|Undefined Saprotroph\| | Yeast | NULL | Probable | As nectar-tap_saprotroph (Põlme et al. 2020); Decay sugar-rich substrates (Põlme et al. 2020) | Põlme S, et al. 2020. Fungal Diversity 105:1-16 ((https://doi.org/10.1007/s13225-020-00466-2)) |
| Otu32789 | 1 | k__Fungi(100);p__Ascomycota(100);c__Saccharomycetes(100);o__Saccharomycetales(100);f__Metschnikowiaceae(100);g__Metschnikowia(100);s__Metschnikowia_pulcherrima(100); | Metschnikowia | 13 | Saprotroph | \|Undefined Saprotroph\| | Yeast | NULL | Probable | As nectar-tap_saprotroph (Põlme et al. 2020); Decay sugar-rich substrates (Põlme et al. 2020) | Põlme S, et al. 2020. Fungal Diversity 105:1-16 ((https://doi.org/10.1007/s13225-020-00466-2)) |
| Otu32790 | 1 | k__Fungi(100);p__Ascomycota(100);c__Saccharomycetes(100);o__Saccharomycetales(100);f__Metschnikowiaceae(100);g__Metschnikowia(100);s__Metschnikowia_pulcherrima(100); | Metschnikowia | 13 | Saprotroph | \|Undefined Saprotroph\| | Yeast | NULL | Probable | As nectar-tap_saprotroph (Põlme et al. 2020); Decay sugar-rich substrates (Põlme et al. 2020) | Põlme S, et al. 2020. Fungal Diversity 105:1-16 ((https://doi.org/10.1007/s13225-020-00466-2)) |
| Otu32791 | 1 | k__Fungi(100);p__Ascomycota(100);c__Saccharomycetes(100);o__Saccharomycetales(100);f__Saccharomycetales_family_Incertae_sedis(100);g__Candida(100);s__Candida_cretensis(100); | Candida | 13 | Pathotroph-Saprotroph-Symbiotroph | Animal Pathogen-Endophyte-Endosymbiont-Epiphyte-\|Undefined Saprotroph\| | Dimorphic Yeast-Yeast | NULL | Probable | As nectar-tap_saprotroph (Põlme et al. 2020); Decay sugar-rich substrates (Põlme et al. 2020); This asexual genus is polyphyletic and contain species that function very differently depend on what they are. Caution should be taken when interpreting guilds for Candida species. | Manolakaki D, et al. 2010. Virulence. 1: 367-375 ((https://doi.org/10.4161/viru.1.5.12796)); Mekbib SB, et al. 2011. Tropical Plant Pathology 36:233-240 ((https://doi.org/10.1590/S1982-56762011000400004)); Põlme S, et al. 2020. Fungal Diversity 105:1-16 ((https://doi.org/10.1007/s13225-020-00466-2)) |
| Otu32792 | 1 | k__Fungi(100);p__Ascomycota(100);c__Saccharomycetes(100);o__Saccharomycetales(100);f__Metschnikowiaceae(100);g__Metschnikowia(100);s__Metschnikowia_pulcherrima(100); | Metschnikowia | 13 | Saprotroph | \|Undefined Saprotroph\| | Yeast | NULL | Probable | As nectar-tap_saprotroph (Põlme et al. 2020); Decay sugar-rich substrates (Põlme et al. 2020) | Põlme S, et al. 2020. Fungal Diversity 105:1-16 ((https://doi.org/10.1007/s13225-020-00466-2)) |
| Otu32794 | 1 | k__Fungi(100);p__Ascomycota(100);c__Saccharomycetes(100);o__Saccharomycetales(100);f__Metschnikowiaceae(100);g__Metschnikowia(100);s__Metschnikowia_pulcherrima(100); | Metschnikowia | 13 | Saprotroph | \|Undefined Saprotroph\| | Yeast | NULL | Probable | As nectar-tap_saprotroph (Põlme et al. 2020); Decay sugar-rich substrates (Põlme et al. 2020) | Põlme S, et al. 2020. Fungal Diversity 105:1-16 ((https://doi.org/10.1007/s13225-020-00466-2)) |
| Otu32796 | 1 | k__Fungi(100);p__Ascomycota(100);c__Saccharomycetes(100);o__Saccharomycetales(100);f__Metschnikowiaceae(100);g__Metschnikowia(100);g__Metschnikowia_unclassified(100); | Metschnikowia | 13 | Saprotroph | \|Undefined Saprotroph\| | Yeast | NULL | Probable | As nectar-tap_saprotroph (Põlme et al. 2020); Decay sugar-rich substrates (Põlme et al. 2020) | Põlme S, et al. 2020. Fungal Diversity 105:1-16 ((https://doi.org/10.1007/s13225-020-00466-2)) |
| Otu32797 | 1 | k__Fungi(100);p__Ascomycota(100);c__Saccharomycetes(100);o__Saccharomycetales(100);f__Metschnikowiaceae(100);g__Metschnikowia(100);g__Metschnikowia_unclassified(100); | Metschnikowia | 13 | Saprotroph | \|Undefined Saprotroph\| | Yeast | NULL | Probable | As nectar-tap_saprotroph (Põlme et al. 2020); Decay sugar-rich substrates (Põlme et al. 2020) | Põlme S, et al. 2020. Fungal Diversity 105:1-16 ((https://doi.org/10.1007/s13225-020-00466-2)) |
| Otu32798 | 1 | k__Fungi(100);p__Ascomycota(100);c__Saccharomycetes(100);o__Saccharomycetales(100);f__Metschnikowiaceae(100);g__Metschnikowia(100);s__Metschnikowia_pulcherrima(100); | Metschnikowia | 13 | Saprotroph | \|Undefined Saprotroph\| | Yeast | NULL | Probable | As nectar-tap_saprotroph (Põlme et al. 2020); Decay sugar-rich substrates (Põlme et al. 2020) | Põlme S, et al. 2020. Fungal Diversity 105:1-16 ((https://doi.org/10.1007/s13225-020-00466-2)) |
| Otu32799 | 1 | k__Fungi(100);p__Ascomycota(100);c__Saccharomycetes(100);o__Saccharomycetales(100);f__Metschnikowiaceae(100);g__Metschnikowia(100);s__Metschnikowia_pulcherrima(100); | Metschnikowia | 13 | Saprotroph | \|Undefined Saprotroph\| | Yeast | NULL | Probable | As nectar-tap_saprotroph (Põlme et al. 2020); Decay sugar-rich substrates (Põlme et al. 2020) | Põlme S, et al. 2020. Fungal Diversity 105:1-16 ((https://doi.org/10.1007/s13225-020-00466-2)) |
| Otu32801 | 1 | k__Fungi(100);p__Ascomycota(100);c__Saccharomycetes(100);o__Saccharomycetales(100);f__Metschnikowiaceae(100);g__Metschnikowia(100);g__Metschnikowia_unclassified(100); | Metschnikowia | 13 | Saprotroph | \|Undefined Saprotroph\| | Yeast | NULL | Probable | As nectar-tap_saprotroph (Põlme et al. 2020); Decay sugar-rich substrates (Põlme et al. 2020) | Põlme S, et al. 2020. Fungal Diversity 105:1-16 ((https://doi.org/10.1007/s13225-020-00466-2)) |
| Otu32803 | 1 | k__Fungi(100);p__Ascomycota(100);c__Saccharomycetes(100);o__Saccharomycetales(100);f__Metschnikowiaceae(100);g__Metschnikowia(100);s__Metschnikowia_pulcherrima(100); | Metschnikowia | 13 | Saprotroph | \|Undefined Saprotroph\| | Yeast | NULL | Probable | As nectar-tap_saprotroph (Põlme et al. 2020); Decay sugar-rich substrates (Põlme et al. 2020) | Põlme S, et al. 2020. Fungal Diversity 105:1-16 ((https://doi.org/10.1007/s13225-020-00466-2)) |
| Otu32804 | 1 | k__Fungi(100);p__Ascomycota(100);c__Saccharomycetes(100);o__Saccharomycetales(100);f__Metschnikowiaceae(100);g__Metschnikowia(100);s__Metschnikowia_pulcherrima(100); | Metschnikowia | 13 | Saprotroph | \|Undefined Saprotroph\| | Yeast | NULL | Probable | As nectar-tap_saprotroph (Põlme et al. 2020); Decay sugar-rich substrates (Põlme et al. 2020) | Põlme S, et al. 2020. Fungal Diversity 105:1-16 ((https://doi.org/10.1007/s13225-020-00466-2)) |
| Otu32805 | 1 | k__Fungi(100);p__Ascomycota(100);c__Saccharomycetes(100);o__Saccharomycetales(100);f__Metschnikowiaceae(100);g__Metschnikowia(100);s__Metschnikowia_pulcherrima(100); | Metschnikowia | 13 | Saprotroph | \|Undefined Saprotroph\| | Yeast | NULL | Probable | As nectar-tap_saprotroph (Põlme et al. 2020); Decay sugar-rich substrates (Põlme et al. 2020) | Põlme S, et al. 2020. Fungal Diversity 105:1-16 ((https://doi.org/10.1007/s13225-020-00466-2)) |
| Otu32806 | 1 | k__Fungi(100);p__Ascomycota(100);c__Saccharomycetes(100);o__Saccharomycetales(100);f__Metschnikowiaceae(100);g__Metschnikowia(100);g__Metschnikowia_unclassified(100); | Metschnikowia | 13 | Saprotroph | \|Undefined Saprotroph\| | Yeast | NULL | Probable | As nectar-tap_saprotroph (Põlme et al. 2020); Decay sugar-rich substrates (Põlme et al. 2020) | Põlme S, et al. 2020. Fungal Diversity 105:1-16 ((https://doi.org/10.1007/s13225-020-00466-2)) |
| Otu32807 | 1 | k__Fungi(100);p__Ascomycota(100);c__Saccharomycetes(100);o__Saccharomycetales(100);f__Metschnikowiaceae(100);g__Metschnikowia(100);s__Metschnikowia_pulcherrima(100); | Metschnikowia | 13 | Saprotroph | \|Undefined Saprotroph\| | Yeast | NULL | Probable | As nectar-tap_saprotroph (Põlme et al. 2020); Decay sugar-rich substrates (Põlme et al. 2020) | Põlme S, et al. 2020. Fungal Diversity 105:1-16 ((https://doi.org/10.1007/s13225-020-00466-2)) |
| Otu32808 | 1 | k__Fungi(100);p__Ascomycota(100);c__Saccharomycetes(100);o__Saccharomycetales(100);f__Metschnikowiaceae(100);g__Metschnikowia(100);s__Metschnikowia_pulcherrima(100); | Metschnikowia | 13 | Saprotroph | \|Undefined Saprotroph\| | Yeast | NULL | Probable | As nectar-tap_saprotroph (Põlme et al. 2020); Decay sugar-rich substrates (Põlme et al. 2020) | Põlme S, et al. 2020. Fungal Diversity 105:1-16 ((https://doi.org/10.1007/s13225-020-00466-2)) |
| Otu32809 | 1 | k__Fungi(100);p__Ascomycota(100);c__Saccharomycetes(100);o__Saccharomycetales(100);f__Metschnikowiaceae(100);g__Metschnikowia(100);s__Metschnikowia_pulcherrima(100); | Metschnikowia | 13 | Saprotroph | \|Undefined Saprotroph\| | Yeast | NULL | Probable | As nectar-tap_saprotroph (Põlme et al. 2020); Decay sugar-rich substrates (Põlme et al. 2020) | Põlme S, et al. 2020. Fungal Diversity 105:1-16 ((https://doi.org/10.1007/s13225-020-00466-2)) |
| Otu32810 | 1 | k__Fungi(100);p__Ascomycota(100);c__Saccharomycetes(100);o__Saccharomycetales(100);f__Metschnikowiaceae(100);g__Metschnikowia(100);s__Metschnikowia_pulcherrima(100); | Metschnikowia | 13 | Saprotroph | \|Undefined Saprotroph\| | Yeast | NULL | Probable | As nectar-tap_saprotroph (Põlme et al. 2020); Decay sugar-rich substrates (Põlme et al. 2020) | Põlme S, et al. 2020. Fungal Diversity 105:1-16 ((https://doi.org/10.1007/s13225-020-00466-2)) |
| Otu32811 | 1 | k__Fungi(100);p__Ascomycota(100);c__Saccharomycetes(100);o__Saccharomycetales(100);f__Metschnikowiaceae(100);g__Metschnikowia(100);s__Metschnikowia_pulcherrima(100); | Metschnikowia | 13 | Saprotroph | \|Undefined Saprotroph\| | Yeast | NULL | Probable | As nectar-tap_saprotroph (Põlme et al. 2020); Decay sugar-rich substrates (Põlme et al. 2020) | Põlme S, et al. 2020. Fungal Diversity 105:1-16 ((https://doi.org/10.1007/s13225-020-00466-2)) |
| Otu32812 | 1 | k__Fungi(100);p__Ascomycota(100);c__Saccharomycetes(100);o__Saccharomycetales(100);f__Metschnikowiaceae(100);g__Metschnikowia(100);s__Metschnikowia_pulcherrima(100); | Metschnikowia | 13 | Saprotroph | \|Undefined Saprotroph\| | Yeast | NULL | Probable | As nectar-tap_saprotroph (Põlme et al. 2020); Decay sugar-rich substrates (Põlme et al. 2020) | Põlme S, et al. 2020. Fungal Diversity 105:1-16 ((https://doi.org/10.1007/s13225-020-00466-2)) |
| Otu32813 | 1 | k__Fungi(100);p__Ascomycota(100);c__Saccharomycetes(100);o__Saccharomycetales(100);f__Metschnikowiaceae(100);g__Metschnikowia(100);g__Metschnikowia_unclassified(100); | Metschnikowia | 13 | Saprotroph | \|Undefined Saprotroph\| | Yeast | NULL | Probable | As nectar-tap_saprotroph (Põlme et al. 2020); Decay sugar-rich substrates (Põlme et al. 2020) | Põlme S, et al. 2020. Fungal Diversity 105:1-16 ((https://doi.org/10.1007/s13225-020-00466-2)) |
| Otu32815 | 1 | k__Fungi(100);p__Ascomycota(100);c__Saccharomycetes(100);o__Saccharomycetales(100);f__Metschnikowiaceae(100);g__Metschnikowia(100);s__Metschnikowia_pulcherrima(100); | Metschnikowia | 13 | Saprotroph | \|Undefined Saprotroph\| | Yeast | NULL | Probable | As nectar-tap_saprotroph (Põlme et al. 2020); Decay sugar-rich substrates (Põlme et al. 2020) | Põlme S, et al. 2020. Fungal Diversity 105:1-16 ((https://doi.org/10.1007/s13225-020-00466-2)) |
| Otu32817 | 1 | k__Fungi(100);p__Ascomycota(100);c__Saccharomycetes(100);o__Saccharomycetales(100);f__Metschnikowiaceae(100);g__Metschnikowia(100);s__Metschnikowia_pulcherrima(100); | Metschnikowia | 13 | Saprotroph | \|Undefined Saprotroph\| | Yeast | NULL | Probable | As nectar-tap_saprotroph (Põlme et al. 2020); Decay sugar-rich substrates (Põlme et al. 2020) | Põlme S, et al. 2020. Fungal Diversity 105:1-16 ((https://doi.org/10.1007/s13225-020-00466-2)) |
| Otu32818 | 1 | k__Fungi(100);p__Ascomycota(100);c__Saccharomycetes(100);o__Saccharomycetales(100);f__Metschnikowiaceae(100);g__Metschnikowia(100);s__Metschnikowia_pulcherrima(100); | Metschnikowia | 13 | Saprotroph | \|Undefined Saprotroph\| | Yeast | NULL | Probable | As nectar-tap_saprotroph (Põlme et al. 2020); Decay sugar-rich substrates (Põlme et al. 2020) | Põlme S, et al. 2020. Fungal Diversity 105:1-16 ((https://doi.org/10.1007/s13225-020-00466-2)) |
| Otu32819 | 1 | k__Fungi(100);p__Ascomycota(100);c__Saccharomycetes(100);o__Saccharomycetales(100);f__Metschnikowiaceae(100);g__Metschnikowia(100);g__Metschnikowia_unclassified(100); | Metschnikowia | 13 | Saprotroph | \|Undefined Saprotroph\| | Yeast | NULL | Probable | As nectar-tap_saprotroph (Põlme et al. 2020); Decay sugar-rich substrates (Põlme et al. 2020) | Põlme S, et al. 2020. Fungal Diversity 105:1-16 ((https://doi.org/10.1007/s13225-020-00466-2)) |
| Otu32820 | 1 | k__Fungi(100);p__Ascomycota(100);c__Saccharomycetes(100);o__Saccharomycetales(100);f__Metschnikowiaceae(100);g__Metschnikowia(100);g__Metschnikowia_unclassified(100); | Metschnikowia | 13 | Saprotroph | \|Undefined Saprotroph\| | Yeast | NULL | Probable | As nectar-tap_saprotroph (Põlme et al. 2020); Decay sugar-rich substrates (Põlme et al. 2020) | Põlme S, et al. 2020. Fungal Diversity 105:1-16 ((https://doi.org/10.1007/s13225-020-00466-2)) |
| Otu32821 | 1 | k__Fungi(100);p__Ascomycota(100);c__Saccharomycetes(100);o__Saccharomycetales(100);f__Metschnikowiaceae(100);g__Metschnikowia(100);s__Metschnikowia_pulcherrima(100); | Metschnikowia | 13 | Saprotroph | \|Undefined Saprotroph\| | Yeast | NULL | Probable | As nectar-tap_saprotroph (Põlme et al. 2020); Decay sugar-rich substrates (Põlme et al. 2020) | Põlme S, et al. 2020. Fungal Diversity 105:1-16 ((https://doi.org/10.1007/s13225-020-00466-2)) |
| Otu32822 | 1 | k__Fungi(100);p__Ascomycota(100);c__Saccharomycetes(100);o__Saccharomycetales(100);f__Metschnikowiaceae(100);g__Metschnikowia(100);s__Metschnikowia_pulcherrima(100); | Metschnikowia | 13 | Saprotroph | \|Undefined Saprotroph\| | Yeast | NULL | Probable | As nectar-tap_saprotroph (Põlme et al. 2020); Decay sugar-rich substrates (Põlme et al. 2020) | Põlme S, et al. 2020. Fungal Diversity 105:1-16 ((https://doi.org/10.1007/s13225-020-00466-2)) |
| Otu32823 | 1 | k__Fungi(100);p__Ascomycota(100);c__Saccharomycetes(100);o__Saccharomycetales(100);f__Metschnikowiaceae(100);g__Metschnikowia(100);s__Metschnikowia_pulcherrima(100); | Metschnikowia | 13 | Saprotroph | \|Undefined Saprotroph\| | Yeast | NULL | Probable | As nectar-tap_saprotroph (Põlme et al. 2020); Decay sugar-rich substrates (Põlme et al. 2020) | Põlme S, et al. 2020. Fungal Diversity 105:1-16 ((https://doi.org/10.1007/s13225-020-00466-2)) |
| Otu32824 | 1 | k__Fungi(100);p__Ascomycota(100);c__Saccharomycetes(100);o__Saccharomycetales(100);f__Metschnikowiaceae(100);g__Metschnikowia(100);s__Metschnikowia_pulcherrima(100); | Metschnikowia | 13 | Saprotroph | \|Undefined Saprotroph\| | Yeast | NULL | Probable | As nectar-tap_saprotroph (Põlme et al. 2020); Decay sugar-rich substrates (Põlme et al. 2020) | Põlme S, et al. 2020. Fungal Diversity 105:1-16 ((https://doi.org/10.1007/s13225-020-00466-2)) |
| Otu32825 | 1 | k__Fungi(100);p__Ascomycota(100);c__Saccharomycetes(100);o__Saccharomycetales(100);f__Metschnikowiaceae(100);g__Metschnikowia(100);s__Metschnikowia_pulcherrima(100); | Metschnikowia | 13 | Saprotroph | \|Undefined Saprotroph\| | Yeast | NULL | Probable | As nectar-tap_saprotroph (Põlme et al. 2020); Decay sugar-rich substrates (Põlme et al. 2020) | Põlme S, et al. 2020. Fungal Diversity 105:1-16 ((https://doi.org/10.1007/s13225-020-00466-2)) |
| Otu32826 | 1 | k__Fungi(100);p__Ascomycota(100);c__Saccharomycetes(100);o__Saccharomycetales(100);f__Metschnikowiaceae(100);g__Metschnikowia(100);s__Metschnikowia_pulcherrima(100); | Metschnikowia | 13 | Saprotroph | \|Undefined Saprotroph\| | Yeast | NULL | Probable | As nectar-tap_saprotroph (Põlme et al. 2020); Decay sugar-rich substrates (Põlme et al. 2020) | Põlme S, et al. 2020. Fungal Diversity 105:1-16 ((https://doi.org/10.1007/s13225-020-00466-2)) |
| Otu32827 | 1 | k__Fungi(100);p__Ascomycota(100);c__Saccharomycetes(100);o__Saccharomycetales(100);f__Metschnikowiaceae(100);g__Metschnikowia(100);s__Metschnikowia_pulcherrima(100); | Metschnikowia | 13 | Saprotroph | \|Undefined Saprotroph\| | Yeast | NULL | Probable | As nectar-tap_saprotroph (Põlme et al. 2020); Decay sugar-rich substrates (Põlme et al. 2020) | Põlme S, et al. 2020. Fungal Diversity 105:1-16 ((https://doi.org/10.1007/s13225-020-00466-2)) |
| Otu32830 | 1 | k__Fungi(100);p__Ascomycota(100);c__Saccharomycetes(100);o__Saccharomycetales(100);f__Metschnikowiaceae(100);g__Metschnikowia(100);s__Metschnikowia_pulcherrima(100); | Metschnikowia | 13 | Saprotroph | \|Undefined Saprotroph\| | Yeast | NULL | Probable | As nectar-tap_saprotroph (Põlme et al. 2020); Decay sugar-rich substrates (Põlme et al. 2020) | Põlme S, et al. 2020. Fungal Diversity 105:1-16 ((https://doi.org/10.1007/s13225-020-00466-2)) |
| Otu32831 | 1 | k__Fungi(100);p__Ascomycota(100);c__Saccharomycetes(100);o__Saccharomycetales(100);f__Metschnikowiaceae(100);g__Metschnikowia(100);s__Metschnikowia_pulcherrima(100); | Metschnikowia | 13 | Saprotroph | \|Undefined Saprotroph\| | Yeast | NULL | Probable | As nectar-tap_saprotroph (Põlme et al. 2020); Decay sugar-rich substrates (Põlme et al. 2020) | Põlme S, et al. 2020. Fungal Diversity 105:1-16 ((https://doi.org/10.1007/s13225-020-00466-2)) |
| Otu32832 | 1 | k__Fungi(100);p__Ascomycota(100);c__Saccharomycetes(100);o__Saccharomycetales(100);f__Metschnikowiaceae(100);g__Metschnikowia(100);g__Metschnikowia_unclassified(100); | Metschnikowia | 13 | Saprotroph | \|Undefined Saprotroph\| | Yeast | NULL | Probable | As nectar-tap_saprotroph (Põlme et al. 2020); Decay sugar-rich substrates (Põlme et al. 2020) | Põlme S, et al. 2020. Fungal Diversity 105:1-16 ((https://doi.org/10.1007/s13225-020-00466-2)) |
| Otu32833 | 1 | k__Fungi(100);p__Ascomycota(100);c__Saccharomycetes(100);o__Saccharomycetales(100);f__Saccharomycetales_family_Incertae_sedis(100);g__Candida(100);s__Candida_cretensis(100); | Candida | 13 | Pathotroph-Saprotroph-Symbiotroph | Animal Pathogen-Endophyte-Endosymbiont-Epiphyte-\|Undefined Saprotroph\| | Dimorphic Yeast-Yeast | NULL | Probable | As nectar-tap_saprotroph (Põlme et al. 2020); Decay sugar-rich substrates (Põlme et al. 2020); This asexual genus is polyphyletic and contain species that function very differently depend on what they are. Caution should be taken when interpreting guilds for Candida species. | Manolakaki D, et al. 2010. Virulence. 1: 367-375 ((https://doi.org/10.4161/viru.1.5.12796)); Mekbib SB, et al. 2011. Tropical Plant Pathology 36:233-240 ((https://doi.org/10.1590/S1982-56762011000400004)); Põlme S, et al. 2020. Fungal Diversity 105:1-16 ((https://doi.org/10.1007/s13225-020-00466-2)) |
| Otu32834 | 1 | k__Fungi(100);p__Ascomycota(100);c__Saccharomycetes(100);o__Saccharomycetales(100);f__Metschnikowiaceae(100);g__Metschnikowia(100);s__Metschnikowia_pulcherrima(100); | Metschnikowia | 13 | Saprotroph | \|Undefined Saprotroph\| | Yeast | NULL | Probable | As nectar-tap_saprotroph (Põlme et al. 2020); Decay sugar-rich substrates (Põlme et al. 2020) | Põlme S, et al. 2020. Fungal Diversity 105:1-16 ((https://doi.org/10.1007/s13225-020-00466-2)) |
| Otu32835 | 1 | k__Fungi(100);p__Ascomycota(100);c__Saccharomycetes(100);o__Saccharomycetales(100);f__Metschnikowiaceae(100);g__Metschnikowia(100);s__Metschnikowia_pulcherrima(100); | Metschnikowia | 13 | Saprotroph | \|Undefined Saprotroph\| | Yeast | NULL | Probable | As nectar-tap_saprotroph (Põlme et al. 2020); Decay sugar-rich substrates (Põlme et al. 2020) | Põlme S, et al. 2020. Fungal Diversity 105:1-16 ((https://doi.org/10.1007/s13225-020-00466-2)) |
| Otu32838 | 1 | k__Fungi(100);p__Ascomycota(100);c__Saccharomycetes(100);o__Saccharomycetales(100);f__Metschnikowiaceae(100);g__Metschnikowia(100);s__Metschnikowia_pulcherrima(100); | Metschnikowia | 13 | Saprotroph | \|Undefined Saprotroph\| | Yeast | NULL | Probable | As nectar-tap_saprotroph (Põlme et al. 2020); Decay sugar-rich substrates (Põlme et al. 2020) | Põlme S, et al. 2020. Fungal Diversity 105:1-16 ((https://doi.org/10.1007/s13225-020-00466-2)) |
| Otu32840 | 1 | k__Fungi(100);p__Ascomycota(100);c__Saccharomycetes(100);o__Saccharomycetales(100);f__Metschnikowiaceae(100);g__Metschnikowia(100);g__Metschnikowia_unclassified(100); | Metschnikowia | 13 | Saprotroph | \|Undefined Saprotroph\| | Yeast | NULL | Probable | As nectar-tap_saprotroph (Põlme et al. 2020); Decay sugar-rich substrates (Põlme et al. 2020) | Põlme S, et al. 2020. Fungal Diversity 105:1-16 ((https://doi.org/10.1007/s13225-020-00466-2)) |
| Otu32842 | 1 | k__Fungi(100);p__Ascomycota(100);c__Saccharomycetes(100);o__Saccharomycetales(100);f__Metschnikowiaceae(100);g__Metschnikowia(100);s__Metschnikowia_pulcherrima(100); | Metschnikowia | 13 | Saprotroph | \|Undefined Saprotroph\| | Yeast | NULL | Probable | As nectar-tap_saprotroph (Põlme et al. 2020); Decay sugar-rich substrates (Põlme et al. 2020) | Põlme S, et al. 2020. Fungal Diversity 105:1-16 ((https://doi.org/10.1007/s13225-020-00466-2)) |
| Otu32844 | 1 | k__Fungi(100);p__Ascomycota(100);c__Saccharomycetes(100);o__Saccharomycetales(100);f__Metschnikowiaceae(100);g__Metschnikowia(100);s__Metschnikowia_pulcherrima(100); | Metschnikowia | 13 | Saprotroph | \|Undefined Saprotroph\| | Yeast | NULL | Probable | As nectar-tap_saprotroph (Põlme et al. 2020); Decay sugar-rich substrates (Põlme et al. 2020) | Põlme S, et al. 2020. Fungal Diversity 105:1-16 ((https://doi.org/10.1007/s13225-020-00466-2)) |
| Otu32845 | 1 | k__Fungi(100);p__Ascomycota(100);c__Saccharomycetes(100);o__Saccharomycetales(100);f__Metschnikowiaceae(100);g__Metschnikowia(100);g__Metschnikowia_unclassified(100); | Metschnikowia | 13 | Saprotroph | \|Undefined Saprotroph\| | Yeast | NULL | Probable | As nectar-tap_saprotroph (Põlme et al. 2020); Decay sugar-rich substrates (Põlme et al. 2020) | Põlme S, et al. 2020. Fungal Diversity 105:1-16 ((https://doi.org/10.1007/s13225-020-00466-2)) |
| Otu32846 | 1 | k__Fungi(100);p__Ascomycota(100);c__Saccharomycetes(100);o__Saccharomycetales(100);f__Metschnikowiaceae(100);g__Metschnikowia(100);g__Metschnikowia_unclassified(100); | Metschnikowia | 13 | Saprotroph | \|Undefined Saprotroph\| | Yeast | NULL | Probable | As nectar-tap_saprotroph (Põlme et al. 2020); Decay sugar-rich substrates (Põlme et al. 2020) | Põlme S, et al. 2020. Fungal Diversity 105:1-16 ((https://doi.org/10.1007/s13225-020-00466-2)) |
| Otu32847 | 1 | k__Fungi(100);p__Ascomycota(100);c__Saccharomycetes(100);o__Saccharomycetales(100);f__Metschnikowiaceae(100);g__Metschnikowia(100);g__Metschnikowia_unclassified(100); | Metschnikowia | 13 | Saprotroph | \|Undefined Saprotroph\| | Yeast | NULL | Probable | As nectar-tap_saprotroph (Põlme et al. 2020); Decay sugar-rich substrates (Põlme et al. 2020) | Põlme S, et al. 2020. Fungal Diversity 105:1-16 ((https://doi.org/10.1007/s13225-020-00466-2)) |
| Otu32848 | 1 | k__Fungi(100);p__Ascomycota(100);c__Saccharomycetes(100);o__Saccharomycetales(100);f__Metschnikowiaceae(100);g__Metschnikowia(100);s__Metschnikowia_pulcherrima(100); | Metschnikowia | 13 | Saprotroph | \|Undefined Saprotroph\| | Yeast | NULL | Probable | As nectar-tap_saprotroph (Põlme et al. 2020); Decay sugar-rich substrates (Põlme et al. 2020) | Põlme S, et al. 2020. Fungal Diversity 105:1-16 ((https://doi.org/10.1007/s13225-020-00466-2)) |
| Otu32849 | 1 | k__Fungi(100);p__Ascomycota(100);c__Saccharomycetes(100);o__Saccharomycetales(100);f__Metschnikowiaceae(100);g__Metschnikowia(100);s__Metschnikowia_pulcherrima(100); | Metschnikowia | 13 | Saprotroph | \|Undefined Saprotroph\| | Yeast | NULL | Probable | As nectar-tap_saprotroph (Põlme et al. 2020); Decay sugar-rich substrates (Põlme et al. 2020) | Põlme S, et al. 2020. Fungal Diversity 105:1-16 ((https://doi.org/10.1007/s13225-020-00466-2)) |
| Otu32850 | 1 | k__Fungi(100);p__Ascomycota(100);c__Saccharomycetes(100);o__Saccharomycetales(100);f__Metschnikowiaceae(100);g__Metschnikowia(100);s__Metschnikowia_pulcherrima(100); | Metschnikowia | 13 | Saprotroph | \|Undefined Saprotroph\| | Yeast | NULL | Probable | As nectar-tap_saprotroph (Põlme et al. 2020); Decay sugar-rich substrates (Põlme et al. 2020) | Põlme S, et al. 2020. Fungal Diversity 105:1-16 ((https://doi.org/10.1007/s13225-020-00466-2)) |
| Otu32851 | 1 | k__Fungi(100);p__Ascomycota(100);c__Saccharomycetes(100);o__Saccharomycetales(100);f__Metschnikowiaceae(100);g__Metschnikowia(100);s__Metschnikowia_pulcherrima(100); | Metschnikowia | 13 | Saprotroph | \|Undefined Saprotroph\| | Yeast | NULL | Probable | As nectar-tap_saprotroph (Põlme et al. 2020); Decay sugar-rich substrates (Põlme et al. 2020) | Põlme S, et al. 2020. Fungal Diversity 105:1-16 ((https://doi.org/10.1007/s13225-020-00466-2)) |
| Otu32852 | 1 | k__Fungi(100);p__Ascomycota(100);c__Saccharomycetes(100);o__Saccharomycetales(100);f__Metschnikowiaceae(100);g__Metschnikowia(100);g__Metschnikowia_unclassified(100); | Metschnikowia | 13 | Saprotroph | \|Undefined Saprotroph\| | Yeast | NULL | Probable | As nectar-tap_saprotroph (Põlme et al. 2020); Decay sugar-rich substrates (Põlme et al. 2020) | Põlme S, et al. 2020. Fungal Diversity 105:1-16 ((https://doi.org/10.1007/s13225-020-00466-2)) |
| Otu32853 | 1 | k__Fungi(100);p__Ascomycota(100);c__Saccharomycetes(100);o__Saccharomycetales(100);f__Metschnikowiaceae(100);g__Metschnikowia(100);s__Metschnikowia_pulcherrima(100); | Metschnikowia | 13 | Saprotroph | \|Undefined Saprotroph\| | Yeast | NULL | Probable | As nectar-tap_saprotroph (Põlme et al. 2020); Decay sugar-rich substrates (Põlme et al. 2020) | Põlme S, et al. 2020. Fungal Diversity 105:1-16 ((https://doi.org/10.1007/s13225-020-00466-2)) |
| Otu32854 | 1 | k__Fungi(100);p__Ascomycota(100);c__Saccharomycetes(100);o__Saccharomycetales(100);f__Metschnikowiaceae(100);g__Metschnikowia(100);s__Metschnikowia_pulcherrima(100); | Metschnikowia | 13 | Saprotroph | \|Undefined Saprotroph\| | Yeast | NULL | Probable | As nectar-tap_saprotroph (Põlme et al. 2020); Decay sugar-rich substrates (Põlme et al. 2020) | Põlme S, et al. 2020. Fungal Diversity 105:1-16 ((https://doi.org/10.1007/s13225-020-00466-2)) |
| Otu32856 | 1 | k__Fungi(100);p__Ascomycota(100);c__Saccharomycetes(100);o__Saccharomycetales(100);f__Metschnikowiaceae(100);g__Metschnikowia(100);s__Metschnikowia_pulcherrima(100); | Metschnikowia | 13 | Saprotroph | \|Undefined Saprotroph\| | Yeast | NULL | Probable | As nectar-tap_saprotroph (Põlme et al. 2020); Decay sugar-rich substrates (Põlme et al. 2020) | Põlme S, et al. 2020. Fungal Diversity 105:1-16 ((https://doi.org/10.1007/s13225-020-00466-2)) |
| Otu32857 | 1 | k__Fungi(100);p__Ascomycota(100);c__Saccharomycetes(100);o__Saccharomycetales(100);f__Metschnikowiaceae(100);g__Metschnikowia(100);g__Metschnikowia_unclassified(100); | Metschnikowia | 13 | Saprotroph | \|Undefined Saprotroph\| | Yeast | NULL | Probable | As nectar-tap_saprotroph (Põlme et al. 2020); Decay sugar-rich substrates (Põlme et al. 2020) | Põlme S, et al. 2020. Fungal Diversity 105:1-16 ((https://doi.org/10.1007/s13225-020-00466-2)) |
| Otu32858 | 1 | k__Fungi(100);p__Ascomycota(100);c__Saccharomycetes(100);o__Saccharomycetales(100);f__Metschnikowiaceae(100);g__Metschnikowia(100);g__Metschnikowia_unclassified(100); | Metschnikowia | 13 | Saprotroph | \|Undefined Saprotroph\| | Yeast | NULL | Probable | As nectar-tap_saprotroph (Põlme et al. 2020); Decay sugar-rich substrates (Põlme et al. 2020) | Põlme S, et al. 2020. Fungal Diversity 105:1-16 ((https://doi.org/10.1007/s13225-020-00466-2)) |
| Otu32859 | 1 | k__Fungi(100);p__Ascomycota(100);c__Saccharomycetes(100);o__Saccharomycetales(100);f__Metschnikowiaceae(100);g__Metschnikowia(100);s__Metschnikowia_pulcherrima(100); | Metschnikowia | 13 | Saprotroph | \|Undefined Saprotroph\| | Yeast | NULL | Probable | As nectar-tap_saprotroph (Põlme et al. 2020); Decay sugar-rich substrates (Põlme et al. 2020) | Põlme S, et al. 2020. Fungal Diversity 105:1-16 ((https://doi.org/10.1007/s13225-020-00466-2)) |
| Otu32860 | 1 | k__Fungi(100);p__Ascomycota(100);c__Saccharomycetes(100);o__Saccharomycetales(100);f__Metschnikowiaceae(100);g__Metschnikowia(100);s__Metschnikowia_pulcherrima(100); | Metschnikowia | 13 | Saprotroph | \|Undefined Saprotroph\| | Yeast | NULL | Probable | As nectar-tap_saprotroph (Põlme et al. 2020); Decay sugar-rich substrates (Põlme et al. 2020) | Põlme S, et al. 2020. Fungal Diversity 105:1-16 ((https://doi.org/10.1007/s13225-020-00466-2)) |
| Otu32861 | 1 | k__Fungi(100);p__Ascomycota(100);c__Saccharomycetes(100);o__Saccharomycetales(100);f__Metschnikowiaceae(100);g__Metschnikowia(100);s__Metschnikowia_pulcherrima(100); | Metschnikowia | 13 | Saprotroph | \|Undefined Saprotroph\| | Yeast | NULL | Probable | As nectar-tap_saprotroph (Põlme et al. 2020); Decay sugar-rich substrates (Põlme et al. 2020) | Põlme S, et al. 2020. Fungal Diversity 105:1-16 ((https://doi.org/10.1007/s13225-020-00466-2)) |
| Otu32862 | 1 | k__Fungi(100);p__Ascomycota(100);c__Saccharomycetes(100);o__Saccharomycetales(100);f__Metschnikowiaceae(100);g__Metschnikowia(100);s__Metschnikowia_pulcherrima(100); | Metschnikowia | 13 | Saprotroph | \|Undefined Saprotroph\| | Yeast | NULL | Probable | As nectar-tap_saprotroph (Põlme et al. 2020); Decay sugar-rich substrates (Põlme et al. 2020) | Põlme S, et al. 2020. Fungal Diversity 105:1-16 ((https://doi.org/10.1007/s13225-020-00466-2)) |
| Otu32863 | 1 | k__Fungi(100);p__Ascomycota(100);c__Saccharomycetes(100);o__Saccharomycetales(100);f__Metschnikowiaceae(100);g__Metschnikowia(100);s__Metschnikowia_pulcherrima(100); | Metschnikowia | 13 | Saprotroph | \|Undefined Saprotroph\| | Yeast | NULL | Probable | As nectar-tap_saprotroph (Põlme et al. 2020); Decay sugar-rich substrates (Põlme et al. 2020) | Põlme S, et al. 2020. Fungal Diversity 105:1-16 ((https://doi.org/10.1007/s13225-020-00466-2)) |
| Otu32864 | 1 | k__Fungi(100);p__Ascomycota(100);c__Saccharomycetes(100);o__Saccharomycetales(100);f__Metschnikowiaceae(100);g__Metschnikowia(100);g__Metschnikowia_unclassified(100); | Metschnikowia | 13 | Saprotroph | \|Undefined Saprotroph\| | Yeast | NULL | Probable | As nectar-tap_saprotroph (Põlme et al. 2020); Decay sugar-rich substrates (Põlme et al. 2020) | Põlme S, et al. 2020. Fungal Diversity 105:1-16 ((https://doi.org/10.1007/s13225-020-00466-2)) |
| Otu32866 | 1 | k__Fungi(100);p__Ascomycota(100);c__Saccharomycetes(100);o__Saccharomycetales(100);f__Saccharomycetales_family_Incertae_sedis(100);f__Saccharomycetales_family_Incertae_sedis_unclassified(100);f__Saccharomycetales_family_Incertae_sedis_unclassified(100); | Saccharomycetales | 7 | Saprotroph | Undefined Saprotroph | Yeast | NULL | Possible | NULL | Sterkenburg E, et al. 2015. New Phytologist 207:1145-1158 ((https://doi.org/10.1111/nph.13426)) |
| Otu32867 | 1 | k__Fungi(100);p__Ascomycota(100);c__Saccharomycetes(100);o__Saccharomycetales(100);f__Metschnikowiaceae(100);g__Metschnikowia(100);g__Metschnikowia_unclassified(100); | Metschnikowia | 13 | Saprotroph | \|Undefined Saprotroph\| | Yeast | NULL | Probable | As nectar-tap_saprotroph (Põlme et al. 2020); Decay sugar-rich substrates (Põlme et al. 2020) | Põlme S, et al. 2020. Fungal Diversity 105:1-16 ((https://doi.org/10.1007/s13225-020-00466-2)) |
| Otu32868 | 1 | k__Fungi(100);p__Ascomycota(100);c__Saccharomycetes(100);o__Saccharomycetales(100);f__Metschnikowiaceae(100);g__Metschnikowia(100);s__Metschnikowia_pulcherrima(100); | Metschnikowia | 13 | Saprotroph | \|Undefined Saprotroph\| | Yeast | NULL | Probable | As nectar-tap_saprotroph (Põlme et al. 2020); Decay sugar-rich substrates (Põlme et al. 2020) | Põlme S, et al. 2020. Fungal Diversity 105:1-16 ((https://doi.org/10.1007/s13225-020-00466-2)) |
| Otu32869 | 1 | k__Fungi(100);p__Ascomycota(100);c__Saccharomycetes(100);o__Saccharomycetales(100);f__Metschnikowiaceae(100);g__Metschnikowia(100);g__Metschnikowia_unclassified(100); | Metschnikowia | 13 | Saprotroph | \|Undefined Saprotroph\| | Yeast | NULL | Probable | As nectar-tap_saprotroph (Põlme et al. 2020); Decay sugar-rich substrates (Põlme et al. 2020) | Põlme S, et al. 2020. Fungal Diversity 105:1-16 ((https://doi.org/10.1007/s13225-020-00466-2)) |
| Otu32870 | 1 | k__Fungi(100);p__Ascomycota(100);c__Saccharomycetes(100);o__Saccharomycetales(100);f__Metschnikowiaceae(100);g__Metschnikowia(100);s__Metschnikowia_pulcherrima(100); | Metschnikowia | 13 | Saprotroph | \|Undefined Saprotroph\| | Yeast | NULL | Probable | As nectar-tap_saprotroph (Põlme et al. 2020); Decay sugar-rich substrates (Põlme et al. 2020) | Põlme S, et al. 2020. Fungal Diversity 105:1-16 ((https://doi.org/10.1007/s13225-020-00466-2)) |
| Otu32872 | 1 | k__Fungi(100);p__Ascomycota(100);c__Saccharomycetes(100);o__Saccharomycetales(100);f__Metschnikowiaceae(100);g__Metschnikowia(100);s__Metschnikowia_pulcherrima(100); | Metschnikowia | 13 | Saprotroph | \|Undefined Saprotroph\| | Yeast | NULL | Probable | As nectar-tap_saprotroph (Põlme et al. 2020); Decay sugar-rich substrates (Põlme et al. 2020) | Põlme S, et al. 2020. Fungal Diversity 105:1-16 ((https://doi.org/10.1007/s13225-020-00466-2)) |
| Otu32873 | 1 | k__Fungi(100);p__Ascomycota(100);c__Saccharomycetes(100);o__Saccharomycetales(100);f__Metschnikowiaceae(100);g__Metschnikowia(100);s__Metschnikowia_pulcherrima(100); | Metschnikowia | 13 | Saprotroph | \|Undefined Saprotroph\| | Yeast | NULL | Probable | As nectar-tap_saprotroph (Põlme et al. 2020); Decay sugar-rich substrates (Põlme et al. 2020) | Põlme S, et al. 2020. Fungal Diversity 105:1-16 ((https://doi.org/10.1007/s13225-020-00466-2)) |
| Otu32874 | 1 | k__Fungi(100);p__Ascomycota(100);c__Saccharomycetes(100);o__Saccharomycetales(100);f__Metschnikowiaceae(100);g__Metschnikowia(100);s__Metschnikowia_pulcherrima(100); | Metschnikowia | 13 | Saprotroph | \|Undefined Saprotroph\| | Yeast | NULL | Probable | As nectar-tap_saprotroph (Põlme et al. 2020); Decay sugar-rich substrates (Põlme et al. 2020) | Põlme S, et al. 2020. Fungal Diversity 105:1-16 ((https://doi.org/10.1007/s13225-020-00466-2)) |
| Otu32875 | 1 | k__Fungi(100);p__Ascomycota(100);c__Saccharomycetes(100);o__Saccharomycetales(100);f__Metschnikowiaceae(100);g__Metschnikowia(100);g__Metschnikowia_unclassified(100); | Metschnikowia | 13 | Saprotroph | \|Undefined Saprotroph\| | Yeast | NULL | Probable | As nectar-tap_saprotroph (Põlme et al. 2020); Decay sugar-rich substrates (Põlme et al. 2020) | Põlme S, et al. 2020. Fungal Diversity 105:1-16 ((https://doi.org/10.1007/s13225-020-00466-2)) |
| Otu32876 | 1 | k__Fungi(100);p__Ascomycota(100);c__Saccharomycetes(100);o__Saccharomycetales(100);f__Metschnikowiaceae(100);g__Metschnikowia(100);s__Metschnikowia_pulcherrima(100); | Metschnikowia | 13 | Saprotroph | \|Undefined Saprotroph\| | Yeast | NULL | Probable | As nectar-tap_saprotroph (Põlme et al. 2020); Decay sugar-rich substrates (Põlme et al. 2020) | Põlme S, et al. 2020. Fungal Diversity 105:1-16 ((https://doi.org/10.1007/s13225-020-00466-2)) |
| Otu32877 | 1 | k__Fungi(100);p__Ascomycota(100);c__Saccharomycetes(100);o__Saccharomycetales(100);f__Metschnikowiaceae(100);g__Metschnikowia(100);g__Metschnikowia_unclassified(100); | Metschnikowia | 13 | Saprotroph | \|Undefined Saprotroph\| | Yeast | NULL | Probable | As nectar-tap_saprotroph (Põlme et al. 2020); Decay sugar-rich substrates (Põlme et al. 2020) | Põlme S, et al. 2020. Fungal Diversity 105:1-16 ((https://doi.org/10.1007/s13225-020-00466-2)) |
| Otu32879 | 1 | k__Fungi(100);p__Ascomycota(100);c__Saccharomycetes(100);o__Saccharomycetales(100);f__Metschnikowiaceae(100);g__Metschnikowia(100);g__Metschnikowia_unclassified(100); | Metschnikowia | 13 | Saprotroph | \|Undefined Saprotroph\| | Yeast | NULL | Probable | As nectar-tap_saprotroph (Põlme et al. 2020); Decay sugar-rich substrates (Põlme et al. 2020) | Põlme S, et al. 2020. Fungal Diversity 105:1-16 ((https://doi.org/10.1007/s13225-020-00466-2)) |
| Otu32880 | 1 | k__Fungi(100);p__Ascomycota(100);c__Saccharomycetes(100);o__Saccharomycetales(100);f__Metschnikowiaceae(100);g__Metschnikowia(100);s__Metschnikowia_pulcherrima(100); | Metschnikowia | 13 | Saprotroph | \|Undefined Saprotroph\| | Yeast | NULL | Probable | As nectar-tap_saprotroph (Põlme et al. 2020); Decay sugar-rich substrates (Põlme et al. 2020) | Põlme S, et al. 2020. Fungal Diversity 105:1-16 ((https://doi.org/10.1007/s13225-020-00466-2)) |
| Otu32881 | 1 | k__Fungi(100);p__Ascomycota(100);c__Saccharomycetes(100);o__Saccharomycetales(100);f__Metschnikowiaceae(100);g__Metschnikowia(100);s__Metschnikowia_pulcherrima(100); | Metschnikowia | 13 | Saprotroph | \|Undefined Saprotroph\| | Yeast | NULL | Probable | As nectar-tap_saprotroph (Põlme et al. 2020); Decay sugar-rich substrates (Põlme et al. 2020) | Põlme S, et al. 2020. Fungal Diversity 105:1-16 ((https://doi.org/10.1007/s13225-020-00466-2)) |
| Otu32883 | 1 | k__Fungi(100);p__Ascomycota(100);c__Saccharomycetes(100);o__Saccharomycetales(100);f__Metschnikowiaceae(100);g__Metschnikowia(100);s__Metschnikowia_pulcherrima(100); | Metschnikowia | 13 | Saprotroph | \|Undefined Saprotroph\| | Yeast | NULL | Probable | As nectar-tap_saprotroph (Põlme et al. 2020); Decay sugar-rich substrates (Põlme et al. 2020) | Põlme S, et al. 2020. Fungal Diversity 105:1-16 ((https://doi.org/10.1007/s13225-020-00466-2)) |

| | | | | | | | | | | | |
|---|---|---|---|---|---|---|---|---|---|---|---|
| Otu32884 | 1 | k__Fungi(100);p__Ascomycota(100);c__Saccharomycetes(100);o__Saccharomycetales(100);f__Metschnikowiaceae(100);g__Metschnikowia(100);s__Metschnikowia_pulcherrima(100); | Metschnikowia | 13 | Saprotroph | \|Undefined Saprotroph\| | Yeast | NULL | Probable | As nectar-tap_saprotroph (Põlme et al. 2020); Decay sugar-rich substrates (Põlme et al. 2020) | Põlme S, et al. 2020. Fungal Diversity 105:1-16 ((https://doi.org/10.1007/s13225-020-00466-2)) |
| Otu32885 | 1 | k__Fungi(100);p__Ascomycota(100);c__Saccharomycetes(100);o__Saccharomycetales(100);f__Saccharomycetales_family_Incertae_sedis(100);g__Candida(100);s__Candida_sp_AS23486(100); | Candida | 13 | Pathotroph-Saprotroph-Symbiotroph | Animal Pathogen-Endophyte-Endosymbiont-Epiphyte-\|Undefined Saprotroph\| | Dimorphic Yeast-Yeast | NULL | Probable | As nectar-tap_saprotroph (Põlme et al. 2020); Decay sugar-rich substrates (Põlme et al. 2020); This asexual genus is polyphyletic and contain species that function very differently depend on what they are. Caution should be taken when interpreting guilds for Candida species. | Manolakaki D, et al. 2010. Virulence. 1: 367-375 ((https://doi.org/10.4161/viru.1.5.12796)); Mekbib SB, et al. 2011. Tropical Plant Pathology 36:233-240 ((https://doi.org/10.1590/S1982-56762011000400004)); Põlme S, et al. 2020. Fungal Diversity 105:1-16 ((https://doi.org/10.1007/s13225-020-00466-2)) |
| Otu32886 | 1 | k__Fungi(100);p__Ascomycota(100);c__Saccharomycetes(100);o__Saccharomycetales(100);f__Metschnikowiaceae(100);g__Metschnikowia(100);s__Metschnikowia_pulcherrima(100); | Metschnikowia | 13 | Saprotroph | \|Undefined Saprotroph\| | Yeast | NULL | Probable | As nectar-tap_saprotroph (Põlme et al. 2020); Decay sugar-rich substrates (Põlme et al. 2020) | Põlme S, et al. 2020. Fungal Diversity 105:1-16 ((https://doi.org/10.1007/s13225-020-00466-2)) |
| Otu32887 | 1 | k__Fungi(100);p__Ascomycota(100);c__Saccharomycetes(100);o__Saccharomycetales(100);f__Metschnikowiaceae(100);g__Metschnikowia(100);s__Metschnikowia_pulcherrima(100); | Metschnikowia | 13 | Saprotroph | \|Undefined Saprotroph\| | Yeast | NULL | Probable | As nectar-tap_saprotroph (Põlme et al. 2020); Decay sugar-rich substrates (Põlme et al. 2020) | Põlme S, et al. 2020. Fungal Diversity 105:1-16 ((https://doi.org/10.1007/s13225-020-00466-2)) |
| Otu32888 | 1 | k__Fungi(100);p__Ascomycota(100);c__Saccharomycetes(100);o__Saccharomycetales(100);f__Metschnikowiaceae(100);g__Metschnikowia(100);g__Metschnikowia_unclassified(100); | Metschnikowia | 13 | Saprotroph | \|Undefined Saprotroph\| | Yeast | NULL | Probable | As nectar-tap_saprotroph (Põlme et al. 2020); Decay sugar-rich substrates (Põlme et al. 2020) | Põlme S, et al. 2020. Fungal Diversity 105:1-16 ((https://doi.org/10.1007/s13225-020-00466-2)) |
| Otu32889 | 1 | k__Fungi(100);p__Ascomycota(100);c__Saccharomycetes(100);o__Saccharomycetales(100);f__Metschnikowiaceae(100);g__Metschnikowia(100);g__Metschnikowia_unclassified(100); | Metschnikowia | 13 | Saprotroph | \|Undefined Saprotroph\| | Yeast | NULL | Probable | As nectar-tap_saprotroph (Põlme et al. 2020); Decay sugar-rich substrates (Põlme et al. 2020) | Põlme S, et al. 2020. Fungal Diversity 105:1-16 ((https://doi.org/10.1007/s13225-020-00466-2)) |
| Otu32890 | 1 | k__Fungi(100);p__Ascomycota(100);c__Saccharomycetes(100);o__Saccharomycetales(100);f__Metschnikowiaceae(100);g__Metschnikowia(100);g__Metschnikowia_unclassified(100); | Metschnikowia | 13 | Saprotroph | \|Undefined Saprotroph\| | Yeast | NULL | Probable | As nectar-tap_saprotroph (Põlme et al. 2020); Decay sugar-rich substrates (Põlme et al. 2020) | Põlme S, et al. 2020. Fungal Diversity 105:1-16 ((https://doi.org/10.1007/s13225-020-00466-2)) |
| Otu32891 | 1 | k__Fungi(100);p__Ascomycota(100);c__Saccharomycetes(100);o__Saccharomycetales(100);f__Metschnikowiaceae(100);g__Metschnikowia(100);g__Metschnikowia_unclassified(100); | Metschnikowia | 13 | Saprotroph | \|Undefined Saprotroph\| | Yeast | NULL | Probable | As nectar-tap_saprotroph (Põlme et al. 2020); Decay sugar-rich substrates (Põlme et al. 2020) | Põlme S, et al. 2020. Fungal Diversity 105:1-16 ((https://doi.org/10.1007/s13225-020-00466-2)) |
| Otu32892 | 1 | k__Fungi(100);p__Ascomycota(100);c__Saccharomycetes(100);o__Saccharomycetales(100);f__Metschnikowiaceae(100);g__Metschnikowia(100);s__Metschnikowia_pulcherrima(100); | Metschnikowia | 13 | Saprotroph | \|Undefined Saprotroph\| | Yeast | NULL | Probable | As nectar-tap_saprotroph (Põlme et al. 2020); Decay sugar-rich substrates (Põlme et al. 2020) | Põlme S, et al. 2020. Fungal Diversity 105:1-16 ((https://doi.org/10.1007/s13225-020-00466-2)) |
| Otu32893 | 1 | k__Fungi(100);p__Ascomycota(100);c__Saccharomycetes(100);o__Saccharomycetales(100);f__Metschnikowiaceae(100);g__Metschnikowia(100);g__Metschnikowia_unclassified(100); | Metschnikowia | 13 | Saprotroph | \|Undefined Saprotroph\| | Yeast | NULL | Probable | As nectar-tap_saprotroph (Põlme et al. 2020); Decay sugar-rich substrates (Põlme et al. 2020) | Põlme S, et al. 2020. Fungal Diversity 105:1-16 ((https://doi.org/10.1007/s13225-020-00466-2)) |
| Otu32894 | 1 | k__Fungi(100);p__Ascomycota(100);c__Saccharomycetes(100);o__Saccharomycetales(100);f__Metschnikowiaceae(100);g__Metschnikowia(100);s__Metschnikowia_pulcherrima(100); | Metschnikowia | 13 | Saprotroph | \|Undefined Saprotroph\| | Yeast | NULL | Probable | As nectar-tap_saprotroph (Põlme et al. 2020); Decay sugar-rich substrates (Põlme et al. 2020) | Põlme S, et al. 2020. Fungal Diversity 105:1-16 ((https://doi.org/10.1007/s13225-020-00466-2)) |
| Otu32895 | 1 | k__Fungi(100);p__Ascomycota(100);c__Saccharomycetes(100);o__Saccharomycetales(100);f__Metschnikowiaceae(100);g__Metschnikowia(100);s__Metschnikowia_pulcherrima(100); | Metschnikowia | 13 | Saprotroph | \|Undefined Saprotroph\| | Yeast | NULL | Probable | As nectar-tap_saprotroph (Põlme et al. 2020); Decay sugar-rich substrates (Põlme et al. 2020) | Põlme S, et al. 2020. Fungal Diversity 105:1-16 ((https://doi.org/10.1007/s13225-020-00466-2)) |
| Otu32896 | 1 | k__Fungi(100);p__Ascomycota(100);c__Saccharomycetes(100);o__Saccharomycetales(100);f__Metschnikowiaceae(100);g__Metschnikowia(100);g__Metschnikowia_unclassified(100); | Metschnikowia | 13 | Saprotroph | \|Undefined Saprotroph\| | Yeast | NULL | Probable | As nectar-tap_saprotroph (Põlme et al. 2020); Decay sugar-rich substrates (Põlme et al. 2020) | Põlme S, et al. 2020. Fungal Diversity 105:1-16 ((https://doi.org/10.1007/s13225-020-00466-2)) |
| Otu32899 | 1 | k__Fungi(100);p__Ascomycota(100);c__Saccharomycetes(100);o__Saccharomycetales(100);f__Metschnikowiaceae(100);g__Metschnikowia(100);s__Metschnikowia_pulcherrima(100); | Metschnikowia | 13 | Saprotroph | \|Undefined Saprotroph\| | Yeast | NULL | Probable | As nectar-tap_saprotroph (Põlme et al. 2020); Decay sugar-rich substrates (Põlme et al. 2020) | Põlme S, et al. 2020. Fungal Diversity 105:1-16 ((https://doi.org/10.1007/s13225-020-00466-2)) |
| Otu32901 | 1 | k__Fungi(100);p__Ascomycota(100);c__Saccharomycetes(100);o__Saccharomycetales(100);f__Saccharomycetales_family_Incertae_sedis(100);g__Candida(100);g__Candida_unclassified(100); | Candida | 13 | Pathotroph-Saprotroph-Symbiotroph | Animal Pathogen-Endophyte-Endosymbiont-Epiphyte-\|Undefined Saprotroph\| | Dimorphic Yeast-Yeast | NULL | Probable | As nectar-tap_saprotroph (Põlme et al. 2020); Decay sugar-rich substrates (Põlme et al. 2020); This asexual genus is polyphyletic and contain species that function very differently depend on what they are. Caution should be taken when interpreting guilds for Candida species. | Manolakaki D, et al. 2010. Virulence. 1: 367-375 ((https://doi.org/10.4161/viru.1.5.12796)); Mekbib SB, et al. 2011. Tropical Plant Pathology 36:233-240 ((https://doi.org/10.1590/S1982-56762011000400004)); Põlme S, et al. 2020. Fungal Diversity 105:1-16 ((https://doi.org/10.1007/s13225-020-00466-2)) |
| Otu32902 | 1 | k__Fungi(100);p__Ascomycota(100);c__Saccharomycetes(100);o__Saccharomycetales(100);f__Saccharomycetales_family_Incertae_sedis(100);g__Candida(100);g__Candida_unclassified(100); | Candida | 13 | Pathotroph-Saprotroph-Symbiotroph | Animal Pathogen-Endophyte-Endosymbiont-Epiphyte-\|Undefined Saprotroph\| | Dimorphic Yeast-Yeast | NULL | Probable | As nectar-tap_saprotroph (Põlme et al. 2020); Decay sugar-rich substrates (Põlme et al. 2020); This asexual genus is polyphyletic and contain species that function very differently depend on what they are. Caution should be taken when interpreting guilds for Candida species. | Manolakaki D, et al. 2010. Virulence. 1: 367-375 ((https://doi.org/10.4161/viru.1.5.12796)); Mekbib SB, et al. 2011. Tropical Plant Pathology 36:233-240 ((https://doi.org/10.1590/S1982-56762011000400004)); Põlme S, et al. 2020. Fungal Diversity 105:1-16 ((https://doi.org/10.1007/s13225-020-00466-2)) |
| Otu32910 | 1 | k__Fungi(100);p__Ascomycota(100);c__Saccharomycetes(100);o__Saccharomycetales(100);f__Metschnikowiaceae(100);g__Metschnikowia(100);s__Metschnikowia_pulcherrima(100); | Metschnikowia | 13 | Saprotroph | \|Undefined Saprotroph\| | Yeast | NULL | Probable | As nectar-tap_saprotroph (Põlme et al. 2020); Decay sugar-rich substrates (Põlme et al. 2020) | Põlme S, et al. 2020. Fungal Diversity 105:1-16 ((https://doi.org/10.1007/s13225-020-00466-2)) |
| Otu32912 | 1 | k__Fungi(100);p__Ascomycota(100);c__Saccharomycetes(100);o__Saccharomycetales(100);f__Metschnikowiaceae(100);g__Metschnikowia(100);g__Metschnikowia_unclassified(100); | Metschnikowia | 13 | Saprotroph | \|Undefined Saprotroph\| | Yeast | NULL | Probable | As nectar-tap_saprotroph (Põlme et al. 2020); Decay sugar-rich substrates (Põlme et al. 2020) | Põlme S, et al. 2020. Fungal Diversity 105:1-16 ((https://doi.org/10.1007/s13225-020-00466-2)) |
| Otu32913 | 1 | k__Fungi(100);p__Ascomycota(100);c__Saccharomycetes(100);o__Saccharomycetales(100);f__Metschnikowiaceae(100);g__Metschnikowia(100);s__Metschnikowia_pulcherrima(100); | Metschnikowia | 13 | Saprotroph | \|Undefined Saprotroph\| | Yeast | NULL | Probable | As nectar-tap_saprotroph (Põlme et al. 2020); Decay sugar-rich substrates (Põlme et al. 2020) | Põlme S, et al. 2020. Fungal Diversity 105:1-16 ((https://doi.org/10.1007/s13225-020-00466-2)) |
| Otu32914 | 1 | k__Fungi(100);p__Ascomycota(100);c__Saccharomycetes(100);o__Saccharomycetales(100);f__Metschnikowiaceae(100);g__Metschnikowia(100);s__Metschnikowia_pulcherrima(100); | Metschnikowia | 13 | Saprotroph | \|Undefined Saprotroph\| | Yeast | NULL | Probable | As nectar-tap_saprotroph (Põlme et al. 2020); Decay sugar-rich substrates (Põlme et al. 2020) | Põlme S, et al. 2020. Fungal Diversity 105:1-16 ((https://doi.org/10.1007/s13225-020-00466-2)) |
| Otu32915 | 1 | k__Fungi(100);p__Ascomycota(100);c__Saccharomycetes(100);o__Saccharomycetales(100);f__Metschnikowiaceae(100);g__Metschnikowia(100);s__Metschnikowia_pulcherrima(100); | Metschnikowia | 13 | Saprotroph | \|Undefined Saprotroph\| | Yeast | NULL | Probable | As nectar-tap_saprotroph (Põlme et al. 2020); Decay sugar-rich substrates (Põlme et al. 2020) | Põlme S, et al. 2020. Fungal Diversity 105:1-16 ((https://doi.org/10.1007/s13225-020-00466-2)) |
| Otu32916 | 1 | k__Fungi(100);p__Ascomycota(100);c__Saccharomycetes(100);o__Saccharomycetales(100);f__Metschnikowiaceae(100);g__Metschnikowia(100);s__Metschnikowia_pulcherrima(100); | Metschnikowia | 13 | Saprotroph | \|Undefined Saprotroph\| | Yeast | NULL | Probable | As nectar-tap_saprotroph (Põlme et al. 2020); Decay sugar-rich substrates (Põlme et al. 2020) | Põlme S, et al. 2020. Fungal Diversity 105:1-16 ((https://doi.org/10.1007/s13225-020-00466-2)) |
| Otu32919 | 1 | k__Fungi(100);p__Ascomycota(100);c__Saccharomycetes(100);o__Saccharomycetales(100);f__Metschnikowiaceae(100);g__Metschnikowia(100);g__Metschnikowia_unclassified(100); | Metschnikowia | 13 | Saprotroph | \|Undefined Saprotroph\| | Yeast | NULL | Probable | As nectar-tap_saprotroph (Põlme et al. 2020); Decay sugar-rich substrates (Põlme et al. 2020) | Põlme S, et al. 2020. Fungal Diversity 105:1-16 ((https://doi.org/10.1007/s13225-020-00466-2)) |
| Otu32920 | 1 | k__Fungi(100);p__Ascomycota(100);c__Saccharomycetes(100);o__Saccharomycetales(100);f__Metschnikowiaceae(100);g__Metschnikowia(100);g__Metschnikowia_unclassified(100); | Metschnikowia | 13 | Saprotroph | \|Undefined Saprotroph\| | Yeast | NULL | Probable | As nectar-tap_saprotroph (Põlme et al. 2020); Decay sugar-rich substrates (Põlme et al. 2020) | Põlme S, et al. 2020. Fungal Diversity 105:1-16 ((https://doi.org/10.1007/s13225-020-00466-2)) |
| Otu32921 | 1 | k__Fungi(100);p__Ascomycota(100);c__Saccharomycetes(100);o__Saccharomycetales(100);f__Metschnikowiaceae(100);g__Metschnikowia(100);s__Metschnikowia_pulcherrima(100); | Metschnikowia | 13 | Saprotroph | \|Undefined Saprotroph\| | Yeast | NULL | Probable | As nectar-tap_saprotroph (Põlme et al. 2020); Decay sugar-rich substrates (Põlme et al. 2020) | Põlme S, et al. 2020. Fungal Diversity 105:1-16 ((https://doi.org/10.1007/s13225-020-00466-2)) |
| Otu32922 | 1 | k__Fungi(100);p__Ascomycota(100);c__Saccharomycetes(100);o__Saccharomycetales(100);f__Metschnikowiaceae(100);g__Metschnikowia(100);s__Metschnikowia_pulcherrima(100); | Metschnikowia | 13 | Saprotroph | \|Undefined Saprotroph\| | Yeast | NULL | Probable | As nectar-tap_saprotroph (Põlme et al. 2020); Decay sugar-rich substrates (Põlme et al. 2020) | Põlme S, et al. 2020. Fungal Diversity 105:1-16 ((https://doi.org/10.1007/s13225-020-00466-2)) |
| Otu32925 | 1 | k__Fungi(100);p__Ascomycota(100);c__Saccharomycetes(100);o__Saccharomycetales(100);f__Metschnikowiaceae(100);g__Metschnikowia(100);g__Metschnikowia_unclassified(100); | Metschnikowia | 13 | Saprotroph | \|Undefined Saprotroph\| | Yeast | NULL | Probable | As nectar-tap_saprotroph (Põlme et al. 2020); Decay sugar-rich substrates (Põlme et al. 2020) | Põlme S, et al. 2020. Fungal Diversity 105:1-16 ((https://doi.org/10.1007/s13225-020-00466-2)) |
| Otu32926 | 1 | k__Fungi(100);p__Ascomycota(100);c__Saccharomycetes(100);o__Saccharomycetales(100);f__Metschnikowiaceae(100);g__Metschnikowia(100);s__Metschnikowia_pulcherrima(100); | Metschnikowia | 13 | Saprotroph | \|Undefined Saprotroph\| | Yeast | NULL | Probable | As nectar-tap_saprotroph (Põlme et al. 2020); Decay sugar-rich substrates (Põlme et al. 2020) | Põlme S, et al. 2020. Fungal Diversity 105:1-16 ((https://doi.org/10.1007/s13225-020-00466-2)) |
| Otu32930 | 1 | k__Fungi(100);p__Ascomycota(100);c__Saccharomycetes(100);o__Saccharomycetales(100);f__Metschnikowiaceae(100);g__Metschnikowia(100);g__Metschnikowia_unclassified(100); | Metschnikowia | 13 | Saprotroph | \|Undefined Saprotroph\| | Yeast | NULL | Probable | As nectar-tap_saprotroph (Põlme et al. 2020); Decay sugar-rich substrates (Põlme et al. 2020) | Põlme S, et al. 2020. Fungal Diversity 105:1-16 ((https://doi.org/10.1007/s13225-020-00466-2)) |
| Otu32933 | 1 | k__Fungi(100);p__Ascomycota(100);c__Saccharomycetes(100);o__Saccharomycetales(100);f__Metschnikowiaceae(100);g__Metschnikowia(100);s__Metschnikowia_pulcherrima(100); | Metschnikowia | 13 | Saprotroph | \|Undefined Saprotroph\| | Yeast | NULL | Probable | As nectar-tap_saprotroph (Põlme et al. 2020); Decay sugar-rich substrates (Põlme et al. 2020) | Põlme S, et al. 2020. Fungal Diversity 105:1-16 ((https://doi.org/10.1007/s13225-020-00466-2)) |
| Otu32934 | 1 | k__Fungi(100);p__Ascomycota(100);c__Saccharomycetes(100);o__Saccharomycetales(100);f__Metschnikowiaceae(100);g__Metschnikowia(100);g__Metschnikowia_unclassified(100); | Metschnikowia | 13 | Saprotroph | \|Undefined Saprotroph\| | Yeast | NULL | Probable | As nectar-tap_saprotroph (Põlme et al. 2020); Decay sugar-rich substrates (Põlme et al. 2020) | Põlme S, et al. 2020. Fungal Diversity 105:1-16 ((https://doi.org/10.1007/s13225-020-00466-2)) |
| Otu32936 | 1 | k__Fungi(100);p__Ascomycota(100);c__Saccharomycetes(100);o__Saccharomycetales(100);f__Metschnikowiaceae(100);g__Metschnikowia(100);s__Metschnikowia_pulcherrima(100); | Metschnikowia | 13 | Saprotroph | \|Undefined Saprotroph\| | Yeast | NULL | Probable | As nectar-tap_saprotroph (Põlme et al. 2020); Decay sugar-rich substrates (Põlme et al. 2020) | Põlme S, et al. 2020. Fungal Diversity 105:1-16 ((https://doi.org/10.1007/s13225-020-00466-2)) |
| Otu32949 | 1 | k__Fungi(100);p__Ascomycota(100);c__Saccharomycetes(100);o__Saccharomycetales(100);f__Metschnikowiaceae(100);g__Metschnikowia(100);g__Metschnikowia_unclassified(100); | Metschnikowia | 13 | Saprotroph | \|Undefined Saprotroph\| | Yeast | NULL | Probable | As nectar-tap_saprotroph (Põlme et al. 2020); Decay sugar-rich substrates (Põlme et al. 2020) | Põlme S, et al. 2020. Fungal Diversity 105:1-16 ((https://doi.org/10.1007/s13225-020-00466-2)) |
| Otu32951 | 1 | k__Fungi(100);p__Ascomycota(100);c__Saccharomycetes(100);o__Saccharomycetales(100);f__Metschnikowiaceae(100);g__Metschnikowia(100);s__Metschnikowia_pulcherrima(100); | Metschnikowia | 13 | Saprotroph | \|Undefined Saprotroph\| | Yeast | NULL | Probable | As nectar-tap_saprotroph (Põlme et al. 2020); Decay sugar-rich substrates (Põlme et al. 2020) | Põlme S, et al. 2020. Fungal Diversity 105:1-16 ((https://doi.org/10.1007/s13225-020-00466-2)) |
| Otu32952 | 1 | k__Fungi(100);p__Ascomycota(100);c__Saccharomycetes(100);o__Saccharomycetales(100);f__Metschnikowiaceae(100);g__Metschnikowia(100);s__Metschnikowia_pulcherrima(100); | Metschnikowia | 13 | Saprotroph | \|Undefined Saprotroph\| | Yeast | NULL | Probable | As nectar-tap_saprotroph (Põlme et al. 2020); Decay sugar-rich substrates (Põlme et al. 2020) | Põlme S, et al. 2020. Fungal Diversity 105:1-16 ((https://doi.org/10.1007/s13225-020-00466-2)) |
| Otu32953 | 1 | k__Fungi(100);p__Ascomycota(100);c__Saccharomycetes(100);o__Saccharomycetales(100);f__Metschnikowiaceae(100);g__Metschnikowia(100);g__Metschnikowia_unclassified(100); | Metschnikowia | 13 | Saprotroph | \|Undefined Saprotroph\| | Yeast | NULL | Probable | As nectar-tap_saprotroph (Põlme et al. 2020); Decay sugar-rich substrates (Põlme et al. 2020) | Põlme S, et al. 2020. Fungal Diversity 105:1-16 ((https://doi.org/10.1007/s13225-020-00466-2)) |
| Otu32954 | 1 | k__Fungi(100);p__Ascomycota(100);c__Saccharomycetes(100);o__Saccharomycetales(100);f__Metschnikowiaceae(100);g__Metschnikowia(100);s__Metschnikowia_pulcherrima(100); | Metschnikowia | 13 | Saprotroph | \|Undefined Saprotroph\| | Yeast | NULL | Probable | As nectar-tap_saprotroph (Põlme et al. 2020); Decay sugar-rich substrates (Põlme et al. 2020) | Põlme S, et al. 2020. Fungal Diversity 105:1-16 ((https://doi.org/10.1007/s13225-020-00466-2)) |
| Otu32955 | 1 | k__Fungi(100);p__Ascomycota(100);c__Saccharomycetes(100);o__Saccharomycetales(100);f__Metschnikowiaceae(100);g__Metschnikowia(100);s__Metschnikowia_pulcherrima(100); | Metschnikowia | 13 | Saprotroph | \|Undefined Saprotroph\| | Yeast | NULL | Probable | As nectar-tap_saprotroph (Põlme et al. 2020); Decay sugar-rich substrates (Põlme et al. 2020) | Põlme S, et al. 2020. Fungal Diversity 105:1-16 ((https://doi.org/10.1007/s13225-020-00466-2)) |
| Otu32956 | 1 | k__Fungi(100);p__Ascomycota(100);c__Saccharomycetes(100);o__Saccharomycetales(100);f__Metschnikowiaceae(100);g__Metschnikowia(100);s__Metschnikowia_pulcherrima(100); | Metschnikowia | 13 | Saprotroph | \|Undefined Saprotroph\| | Yeast | NULL | Probable | As nectar-tap_saprotroph (Põlme et al. 2020); Decay sugar-rich substrates (Põlme et al. 2020) | Põlme S, et al. 2020. Fungal Diversity 105:1-16 ((https://doi.org/10.1007/s13225-020-00466-2)) |
| Otu32957 | 1 | k__Fungi(100);p__Ascomycota(100);c__Saccharomycetes(100);o__Saccharomycetales(100);f__Metschnikowiaceae(100);g__Metschnikowia(100);s__Metschnikowia_pulcherrima(100); | Metschnikowia | 13 | Saprotroph | \|Undefined Saprotroph\| | Yeast | NULL | Probable | As nectar-tap_saprotroph (Põlme et al. 2020); Decay sugar-rich substrates (Põlme et al. 2020) | Põlme S, et al. 2020. Fungal Diversity 105:1-16 ((https://doi.org/10.1007/s13225-020-00466-2)) |
| Otu32959 | 1 | k__Fungi(100);p__Ascomycota(100);c__Saccharomycetes(100);o__Saccharomycetales(100);f__Metschnikowiaceae(100);g__Metschnikowia(100);s__Metschnikowia_pulcherrima(100); | Metschnikowia | 13 | Saprotroph | \|Undefined Saprotroph\| | Yeast | NULL | Probable | As nectar-tap_saprotroph (Põlme et al. 2020); Decay sugar-rich substrates (Põlme et al. 2020) | Põlme S, et al. 2020. Fungal Diversity 105:1-16 ((https://doi.org/10.1007/s13225-020-00466-2)) |
| Otu32961 | 1 | k__Fungi(100);p__Ascomycota(100);c__Saccharomycetes(100);o__Saccharomycetales(100);f__Metschnikowiaceae(100);g__Metschnikowia(100);g__Metschnikowia_unclassified(100); | Metschnikowia | 13 | Saprotroph | \|Undefined Saprotroph\| | Yeast | NULL | Probable | As nectar-tap_saprotroph (Põlme et al. 2020); Decay sugar-rich substrates (Põlme et al. 2020) | Põlme S, et al. 2020. Fungal Diversity 105:1-16 ((https://doi.org/10.1007/s13225-020-00466-2)) |
| Otu32962 | 1 | k__Fungi(100);p__Ascomycota(100);c__Saccharomycetes(100);o__Saccharomycetales(100);f__Metschnikowiaceae(100);g__Metschnikowia(100);s__Metschnikowia_pulcherrima(100); | Metschnikowia | 13 | Saprotroph | \|Undefined Saprotroph\| | Yeast | NULL | Probable | As nectar-tap_saprotroph (Põlme et al. 2020); Decay sugar-rich substrates (Põlme et al. 2020) | Põlme S, et al. 2020. Fungal Diversity 105:1-16 ((https://doi.org/10.1007/s13225-020-00466-2)) |
| Otu32964 | 1 | k__Fungi(100);p__Ascomycota(100);c__Saccharomycetes(100);o__Saccharomycetales(100);o__Saccharomycetales_unclassified(100);o__Saccharomycetales_unclassified(100);o__Saccharomycetales_unclassified(100); | Saccharomycetales | 7 | Saprotroph | Undefined Saprotroph | Yeast | NULL | Possible | NULL | Sterkenburg E, et al. 2015. New Phytologist 207:1145-1158 ((https://doi.org/10.1111/nph.13426)) |
| Otu32966 | 1 | k__Fungi(100);p__Ascomycota(100);c__Saccharomycetes(100);o__Saccharomycetales(100);f__Metschnikowiaceae(100);g__Metschnikowia(100);s__Metschnikowia_pulcherrima(100); | Metschnikowia | 13 | Saprotroph | \|Undefined Saprotroph\| | Yeast | NULL | Probable | As nectar-tap_saprotroph (Põlme et al. 2020); Decay sugar-rich substrates (Põlme et al. 2020) | Põlme S, et al. 2020. Fungal Diversity 105:1-16 ((https://doi.org/10.1007/s13225-020-00466-2)) |
| Otu32969 | 1 | k__Fungi(100);p__Ascomycota(100);c__Saccharomycetes(100);o__Saccharomycetales(100);f__Metschnikowiaceae(100);g__Metschnikowia(100);s__Metschnikowia_pulcherrima(100); | Metschnikowia | 13 | Saprotroph | \|Undefined Saprotroph\| | Yeast | NULL | Probable | As nectar-tap_saprotroph (Põlme et al. 2020); Decay sugar-rich substrates (Põlme et al. 2020) | Põlme S, et al. 2020. Fungal Diversity 105:1-16 ((https://doi.org/10.1007/s13225-020-00466-2)) |
| Otu32976 | 1 | k__Fungi(100);p__Ascomycota(100);c__Saccharomycetes(100);o__Saccharomycetales(100);f__Metschnikowiaceae(100);g__Metschnikowia(100);s__Metschnikowia_pulcherrima(100); | Metschnikowia | 13 | Saprotroph | \|Undefined Saprotroph\| | Yeast | NULL | Probable | As nectar-tap_saprotroph (Põlme et al. 2020); Decay sugar-rich substrates (Põlme et al. 2020) | Põlme S, et al. 2020. Fungal Diversity 105:1-16 ((https://doi.org/10.1007/s13225-020-00466-2)) |
| Otu32977 | 1 | k__Fungi(100);p__Ascomycota(100);c__Saccharomycetes(100);o__Saccharomycetales(100);f__Metschnikowiaceae(100);g__Metschnikowia(100);g__Metschnikowia_unclassified(100); | Metschnikowia | 13 | Saprotroph | \|Undefined Saprotroph\| | Yeast | NULL | Probable | As nectar-tap_saprotroph (Põlme et al. 2020); Decay sugar-rich substrates (Põlme et al. 2020) | Põlme S, et al. 2020. Fungal Diversity 105:1-16 ((https://doi.org/10.1007/s13225-020-00466-2)) |
| Otu32978 | 1 | k__Fungi(100);p__Ascomycota(100);c__Saccharomycetes(100);o__Saccharomycetales(100);f__Metschnikowiaceae(100);g__Metschnikowia(100);s__Metschnikowia_pulcherrima(100); | Metschnikowia | 13 | Saprotroph | \|Undefined Saprotroph\| | Yeast | NULL | Probable | As nectar-tap_saprotroph (Põlme et al. 2020); Decay sugar-rich substrates (Põlme et al. 2020) | Põlme S, et al. 2020. Fungal Diversity 105:1-16 ((https://doi.org/10.1007/s13225-020-00466-2)) |
| Otu32979 | 1 | k__Fungi(100);p__Ascomycota(100);c__Saccharomycetes(100);o__Saccharomycetales(100);f__Metschnikowiaceae(100);g__Metschnikowia(100);s__Metschnikowia_pulcherrima(100); | Metschnikowia | 13 | Saprotroph | \|Undefined Saprotroph\| | Yeast | NULL | Probable | As nectar-tap_saprotroph (Põlme et al. 2020); Decay sugar-rich substrates (Põlme et al. 2020) | Põlme S, et al. 2020. Fungal Diversity 105:1-16 ((https://doi.org/10.1007/s13225-020-00466-2)) |
| Otu32980 | 1 | k__Fungi(100);p__Ascomycota(100);c__Saccharomycetes(100);o__Saccharomycetales(100);f__Metschnikowiaceae(100);g__Metschnikowia(100);g__Metschnikowia_unclassified(100); | Metschnikowia | 13 | Saprotroph | \|Undefined Saprotroph\| | Yeast | NULL | Probable | As nectar-tap_saprotroph (Põlme et al. 2020); Decay sugar-rich substrates (Põlme et al. 2020) | Põlme S, et al. 2020. Fungal Diversity 105:1-16 ((https://doi.org/10.1007/s13225-020-00466-2)) |
| Otu32981 | 1 | k__Fungi(100);p__Ascomycota(100);c__Saccharomycetes(100);o__Saccharomycetales(100);f__Metschnikowiaceae(100);g__Metschnikowia(100);g__Metschnikowia_unclassified(100); | Metschnikowia | 13 | Saprotroph | \|Undefined Saprotroph\| | Yeast | NULL | Probable | As nectar-tap_saprotroph (Põlme et al. 2020); Decay sugar-rich substrates (Põlme et al. 2020) | Põlme S, et al. 2020. Fungal Diversity 105:1-16 ((https://doi.org/10.1007/s13225-020-00466-2)) |
| Otu32982 | 1 | k__Fungi(100);p__Ascomycota(100);c__Saccharomycetes(100);o__Saccharomycetales(100);f__Metschnikowiaceae(100);g__Metschnikowia(100);g__Metschnikowia_unclassified(100); | Metschnikowia | 13 | Saprotroph | \|Undefined Saprotroph\| | Yeast | NULL | Probable | As nectar-tap_saprotroph (Põlme et al. 2020); Decay sugar-rich substrates (Põlme et al. 2020) | Põlme S, et al. 2020. Fungal Diversity 105:1-16 ((https://doi.org/10.1007/s13225-020-00466-2)) |
| Otu32983 | 1 | k__Fungi(100);p__Ascomycota(100);c__Saccharomycetes(100);o__Saccharomycetales(100);f__Metschnikowiaceae(100);g__Metschnikowia(100);s__Metschnikowia_pulcherrima(100); | Metschnikowia | 13 | Saprotroph | \|Undefined Saprotroph\| | Yeast | NULL | Probable | As nectar-tap_saprotroph (Põlme et al. 2020); Decay sugar-rich substrates (Põlme et al. 2020) | Põlme S, et al. 2020. Fungal Diversity 105:1-16 ((https://doi.org/10.1007/s13225-020-00466-2)) |
| Otu32984 | 1 | k__Fungi(100);p__Ascomycota(100);c__Saccharomycetes(100);o__Saccharomycetales(100);f__Metschnikowiaceae(100);g__Metschnikowia(100);s__Metschnikowia_pulcherrima(100); | Metschnikowia | 13 | Saprotroph | \|Undefined Saprotroph\| | Yeast | NULL | Probable | As nectar-tap_saprotroph (Põlme et al. 2020); Decay sugar-rich substrates (Põlme et al. 2020) | Põlme S, et al. 2020. Fungal Diversity 105:1-16 ((https://doi.org/10.1007/s13225-020-00466-2)) |
| Otu32987 | 1 | k__Fungi(100);p__Ascomycota(100);c__Saccharomycetes(100);o__Saccharomycetales(100);f__Metschnikowiaceae(100);g__Metschnikowia(100);g__Metschnikowia_unclassified(100); | Metschnikowia | 13 | Saprotroph | \|Undefined Saprotroph\| | Yeast | NULL | Probable | As nectar-tap_saprotroph (Põlme et al. 2020); Decay sugar-rich substrates (Põlme et al. 2020) | Põlme S, et al. 2020. Fungal Diversity 105:1-16 ((https://doi.org/10.1007/s13225-020-00466-2)) |
| Otu32988 | 1 | k__Fungi(100);p__Ascomycota(100);c__Saccharomycetes(100);o__Saccharomycetales(100);f__Metschnikowiaceae(100);g__Metschnikowia(100);s__Metschnikowia_pulcherrima(100); | Metschnikowia | 13 | Saprotroph | \|Undefined Saprotroph\| | Yeast | NULL | Probable | As nectar-tap_saprotroph (Põlme et al. 2020); Decay sugar-rich substrates (Põlme et al. 2020) | Põlme S, et al. 2020. Fungal Diversity 105:1-16 ((https://doi.org/10.1007/s13225-020-00466-2)) |
| Otu32989 | 1 | k__Fungi(100);p__Ascomycota(100);c__Saccharomycetes(100);o__Saccharomycetales(100);f__Metschnikowiaceae(100);g__Metschnikowia(100);s__Metschnikowia_pulcherrima(100); | Metschnikowia | 13 | Saprotroph | \|Undefined Saprotroph\| | Yeast | NULL | Probable | As nectar-tap_saprotroph (Põlme et al. 2020); Decay sugar-rich substrates (Põlme et al. 2020) | Põlme S, et al. 2020. Fungal Diversity 105:1-16 ((https://doi.org/10.1007/s13225-020-00466-2)) |
| Otu32990 | 1 | k__Fungi(100);p__Ascomycota(100);c__Saccharomycetes(100);o__Saccharomycetales(100);f__Metschnikowiaceae(100);g__Metschnikowia(100);g__Metschnikowia_unclassified(100); | Metschnikowia | 13 | Saprotroph | \|Undefined Saprotroph\| | Yeast | NULL | Probable | As nectar-tap_saprotroph (Põlme et al. 2020); Decay sugar-rich substrates (Põlme et al. 2020) | Põlme S, et al. 2020. Fungal Diversity 105:1-16 ((https://doi.org/10.1007/s13225-020-00466-2)) |
| Otu32991 | 1 | k__Fungi(100);p__Ascomycota(100);c__Saccharomycetes(100);o__Saccharomycetales(100);f__Metschnikowiaceae(100);g__Metschnikowia(100);s__Metschnikowia_pulcherrima(100); | Metschnikowia | 13 | Saprotroph | \|Undefined Saprotroph\| | Yeast | NULL | Probable | As nectar-tap_saprotroph (Põlme et al. 2020); Decay sugar-rich substrates (Põlme et al. 2020) | Põlme S, et al. 2020. Fungal Diversity 105:1-16 ((https://doi.org/10.1007/s13225-020-00466-2)) |
| Otu32992 | 1 | k__Fungi(100);p__Ascomycota(100);c__Saccharomycetes(100);o__Saccharomycetales(100);f__Metschnikowiaceae(100);g__Metschnikowia(100);s__Metschnikowia_pulcherrima(100); | Metschnikowia | 13 | Saprotroph | \|Undefined Saprotroph\| | Yeast | NULL | Probable | As nectar-tap_saprotroph (Põlme et al. 2020); Decay sugar-rich substrates (Põlme et al. 2020) | Põlme S, et al. 2020. Fungal Diversity 105:1-16 ((https://doi.org/10.1007/s13225-020-00466-2)) |
| Otu32993 | 1 | k__Fungi(100);p__Ascomycota(100);c__Saccharomycetes(100);o__Saccharomycetales(100);f__Metschnikowiaceae(100);g__Metschnikowia(100);g__Metschnikowia_unclassified(100); | Metschnikowia | 13 | Saprotroph | \|Undefined Saprotroph\| | Yeast | NULL | Probable | As nectar-tap_saprotroph (Põlme et al. 2020); Decay sugar-rich substrates (Põlme et al. 2020) | Põlme S, et al. 2020. Fungal Diversity 105:1-16 ((https://doi.org/10.1007/s13225-020-00466-2)) |
| Otu32994 | 1 | k__Fungi(100);p__Ascomycota(100);c__Saccharomycetes(100);o__Saccharomycetales(100);f__Metschnikowiaceae(100);g__Metschnikowia(100);s__Metschnikowia_pulcherrima(100); | Metschnikowia | 13 | Saprotroph | \|Undefined Saprotroph\| | Yeast | NULL | Probable | As nectar-tap_saprotroph (Põlme et al. 2020); Decay sugar-rich substrates (Põlme et al. 2020) | Põlme S, et al. 2020. Fungal Diversity 105:1-16 ((https://doi.org/10.1007/s13225-020-00466-2)) |
| Otu32995 | 1 | k__Fungi(100);p__Ascomycota(100);c__Saccharomycetes(100);o__Saccharomycetales(100);f__Metschnikowiaceae(100);g__Metschnikowia(100);s__Metschnikowia_pulcherrima(100); | Metschnikowia | 13 | Saprotroph | \|Undefined Saprotroph\| | Yeast | NULL | Probable | As nectar-tap_saprotroph (Põlme et al. 2020); Decay sugar-rich substrates (Põlme et al. 2020) | Põlme S, et al. 2020. Fungal Diversity 105:1-16 ((https://doi.org/10.1007/s13225-020-00466-2)) |
| Otu32997 | 1 | k__Fungi(100);p__Ascomycota(100);c__Saccharomycetes(100);o__Saccharomycetales(100);f__Metschnikowiaceae(100);g__Metschnikowia(100);s__Metschnikowia_pulcherrima(100); | Metschnikowia | 13 | Saprotroph | \|Undefined Saprotroph\| | Yeast | NULL | Probable | As nectar-tap_saprotroph (Põlme et al. 2020); Decay sugar-rich substrates (Põlme et al. 2020) | Põlme S, et al. 2020. Fungal Diversity 105:1-16 ((https://doi.org/10.1007/s13225-020-00466-2)) |
| Otu33001 | 1 | k__Fungi(100);p__Ascomycota(100);c__Saccharomycetes(100);o__Saccharomycetales(100);f__Metschnikowiaceae(100);g__Metschnikowia(100);s__Metschnikowia_pulcherrima(100); | Metschnikowia | 13 | Saprotroph | \|Undefined Saprotroph\| | Yeast | NULL | Probable | As nectar-tap_saprotroph (Põlme et al. 2020); Decay sugar-rich substrates (Põlme et al. 2020) | Põlme S, et al. 2020. Fungal Diversity 105:1-16 ((https://doi.org/10.1007/s13225-020-00466-2)) |
| Otu33002 | 1 | k__Fungi(100);p__Ascomycota(100);c__Saccharomycetes(100);o__Saccharomycetales(100);f__Metschnikowiaceae(100);g__Metschnikowia(100);g__Metschnikowia_unclassified(100); | Metschnikowia | 13 | Saprotroph | \|Undefined Saprotroph\| | Yeast | NULL | Probable | As nectar-tap_saprotroph (Põlme et al. 2020); Decay sugar-rich substrates (Põlme et al. 2020) | Põlme S, et al. 2020. Fungal Diversity 105:1-16 ((https://doi.org/10.1007/s13225-020-00466-2)) |
| Otu33003 | 1 | k__Fungi(100);p__Ascomycota(100);c__Saccharomycetes(100);o__Saccharomycetales(100);f__Metschnikowiaceae(100);g__Metschnikowia(100);s__Metschnikowia_pulcherrima(100); | Metschnikowia | 13 | Saprotroph | \|Undefined Saprotroph\| | Yeast | NULL | Probable | As nectar-tap_saprotroph (Põlme et al. 2020); Decay sugar-rich substrates (Põlme et al. 2020) | Põlme S, et al. 2020. Fungal Diversity 105:1-16 ((https://doi.org/10.1007/s13225-020-00466-2)) |
| Otu33004 | 1 | k__Fungi(100);p__Ascomycota(100);c__Saccharomycetes(100);o__Saccharomycetales(100);f__Metschnikowiaceae(100);g__Metschnikowia(100);s__Metschnikowia_pulcherrima(100); | Metschnikowia | 13 | Saprotroph | \|Undefined Saprotroph\| | Yeast | NULL | Probable | As nectar-tap_saprotroph (Põlme et al. 2020); Decay sugar-rich substrates (Põlme et al. 2020) | Põlme S, et al. 2020. Fungal Diversity 105:1-16 ((https://doi.org/10.1007/s13225-020-00466-2)) |
| Otu33005 | 1 | k__Fungi(100);p__Ascomycota(100);c__Saccharomycetes(100);o__Saccharomycetales(100);f__Metschnikowiaceae(100);g__Metschnikowia(100);g__Metschnikowia_unclassified(100); | Metschnikowia | 13 | Saprotroph | \|Undefined Saprotroph\| | Yeast | NULL | Probable | As nectar-tap_saprotroph (Põlme et al. 2020); Decay sugar-rich substrates (Põlme et al. 2020) | Põlme S, et al. 2020. Fungal Diversity 105:1-16 ((https://doi.org/10.1007/s13225-020-00466-2)) |
| Otu33006 | 1 | k__Fungi(100);p__Ascomycota(100);c__Saccharomycetes(100);o__Saccharomycetales(100);f__Metschnikowiaceae(100);g__Metschnikowia(100);s__Metschnikowia_pulcherrima(100); | Metschnikowia | 13 | Saprotroph | \|Undefined Saprotroph\| | Yeast | NULL | Probable | As nectar-tap_saprotroph (Põlme et al. 2020); Decay sugar-rich substrates (Põlme et al. 2020) | Põlme S, et al. 2020. Fungal Diversity 105:1-16 ((https://doi.org/10.1007/s13225-020-00466-2)) |
| Otu33007 | 1 | k__Fungi(100);p__Ascomycota(100);c__Saccharomycetes(100);o__Saccharomycetales(100);f__Metschnikowiaceae(100);g__Metschnikowia(100);s__Metschnikowia_pulcherrima(100); | Metschnikowia | 13 | Saprotroph | \|Undefined Saprotroph\| | Yeast | NULL | Probable | As nectar-tap_saprotroph (Põlme et al. 2020); Decay sugar-rich substrates (Põlme et al. 2020) | Põlme S, et al. 2020. Fungal Diversity 105:1-16 ((https://doi.org/10.1007/s13225-020-00466-2)) |
| Otu33008 | 1 | k__Fungi(100);p__Ascomycota(100);c__Saccharomycetes(100);o__Saccharomycetales(100);f__Metschnikowiaceae(100);g__Metschnikowia(100);g__Metschnikowia_unclassified(100); | Metschnikowia | 13 | Saprotroph | \|Undefined Saprotroph\| | Yeast | NULL | Probable | As nectar-tap_saprotroph (Põlme et al. 2020); Decay sugar-rich substrates (Põlme et al. 2020) | Põlme S, et al. 2020. Fungal Diversity 105:1-16 ((https://doi.org/10.1007/s13225-020-00466-2)) |
| Otu33009 | 1 | k__Fungi(100);p__Ascomycota(100);c__Saccharomycetes(100);o__Saccharomycetales(100);f__Metschnikowiaceae(100);g__Metschnikowia(100);s__Metschnikowia_pulcherrima(100); | Metschnikowia | 13 | Saprotroph | \|Undefined Saprotroph\| | Yeast | NULL | Probable | As nectar-tap_saprotroph (Põlme et al. 2020); Decay sugar-rich substrates (Põlme et al. 2020) | Põlme S, et al. 2020. Fungal Diversity 105:1-16 ((https://doi.org/10.1007/s13225-020-00466-2)) |
| Otu33010 | 1 | k__Fungi(100);p__Ascomycota(100);c__Saccharomycetes(100);o__Saccharomycetales(100);f__Metschnikowiaceae(100);g__Metschnikowia(100);g__Metschnikowia_unclassified(100); | Metschnikowia | 13 | Saprotroph | \|Undefined Saprotroph\| | Yeast | NULL | Probable | As nectar-tap_saprotroph (Põlme et al. 2020); Decay sugar-rich substrates (Põlme et al. 2020) | Põlme S, et al. 2020. Fungal Diversity 105:1-16 ((https://doi.org/10.1007/s13225-020-00466-2)) |
| Otu33011 | 1 | k__Fungi(100);p__Ascomycota(100);c__Saccharomycetes(100);o__Saccharomycetales(100);f__Metschnikowiaceae(100);g__Metschnikowia(100);s__Metschnikowia_pulcherrima(100); | Metschnikowia | 13 | Saprotroph | \|Undefined Saprotroph\| | Yeast | NULL | Probable | As nectar-tap_saprotroph (Põlme et al. 2020); Decay sugar-rich substrates (Põlme et al. 2020) | Põlme S, et al. 2020. Fungal Diversity 105:1-16 ((https://doi.org/10.1007/s13225-020-00466-2)) |
| Otu33012 | 1 | k__Fungi(100);p__Ascomycota(100);c__Saccharomycetes(100);o__Saccharomycetales(100);f__Metschnikowiaceae(100);g__Metschnikowia(100);s__Metschnikowia_pulcherrima(100); | Metschnikowia | 13 | Saprotroph | \|Undefined Saprotroph\| | Yeast | NULL | Probable | As nectar-tap_saprotroph (Põlme et al. 2020); Decay sugar-rich substrates (Põlme et al. 2020) | Põlme S, et al. 2020. Fungal Diversity 105:1-16 ((https://doi.org/10.1007/s13225-020-00466-2)) |
| Otu33013 | 1 | k__Fungi(100);p__Ascomycota(100);c__Saccharomycetes(100);o__Saccharomycetales(100);f__Metschnikowiaceae(100);g__Metschnikowia(100);s__Metschnikowia_pulcherrima(100); | Metschnikowia | 13 | Saprotroph | \|Undefined Saprotroph\| | Yeast | NULL | Probable | As nectar-tap_saprotroph (Põlme et al. 2020); Decay sugar-rich substrates (Põlme et al. 2020) | Põlme S, et al. 2020. Fungal Diversity 105:1-16 ((https://doi.org/10.1007/s13225-020-00466-2)) |
| Otu33014 | 1 | k__Fungi(100);p__Ascomycota(100);c__Saccharomycetes(100);o__Saccharomycetales(100);f__Metschnikowiaceae(100);g__Metschnikowia(100);g__Metschnikowia_unclassified(100); | Metschnikowia | 13 | Saprotroph | \|Undefined Saprotroph\| | Yeast | NULL | Probable | As nectar-tap_saprotroph (Põlme et al. 2020); Decay sugar-rich substrates (Põlme et al. 2020) | Põlme S, et al. 2020. Fungal Diversity 105:1-16 ((https://doi.org/10.1007/s13225-020-00466-2)) |
| Otu33015 | 1 | k__Fungi(100);p__Ascomycota(100);c__Saccharomycetes(100);o__Saccharomycetales(100);f__Metschnikowiaceae(100);g__Metschnikowia(100);s__Metschnikowia_pulcherrima(100); | Metschnikowia | 13 | Saprotroph | \|Undefined Saprotroph\| | Yeast | NULL | Probable | As nectar-tap_saprotroph (Põlme et al. 2020); Decay sugar-rich substrates (Põlme et al. 2020) | Põlme S, et al. 2020. Fungal Diversity 105:1-16 ((https://doi.org/10.1007/s13225-020-00466-2)) |
| Otu33016 | 1 | k__Fungi(100);p__Ascomycota(100);c__Saccharomycetes(100);o__Saccharomycetales(100);f__Metschnikowiaceae(100);g__Metschnikowia(100);s__Metschnikowia_pulcherrima(100); | Metschnikowia | 13 | Saprotroph | \|Undefined Saprotroph\| | Yeast | NULL | Probable | As nectar-tap_saprotroph (Põlme et al. 2020); Decay sugar-rich substrates (Põlme et al. 2020) | Põlme S, et al. 2020. Fungal Diversity 105:1-16 ((https://doi.org/10.1007/s13225-020-00466-2)) |
| Otu33020 | 1 | k__Fungi(100);p__Ascomycota(100);c__Saccharomycetes(100);o__Saccharomycetales(100);f__Metschnikowiaceae(100);g__Metschnikowia(100);g__Metschnikowia_unclassified(100); | Metschnikowia | 13 | Saprotroph | \|Undefined Saprotroph\| | Yeast | NULL | Probable | As nectar-tap_saprotroph (Põlme et al. 2020); Decay sugar-rich substrates (Põlme et al. 2020) | Põlme S, et al. 2020. Fungal Diversity 105:1-16 ((https://doi.org/10.1007/s13225-020-00466-2)) |
| Otu33021 | 1 | k__Fungi(100);p__Ascomycota(100);c__Saccharomycetes(100);o__Saccharomycetales(100);f__Metschnikowiaceae(100);g__Metschnikowia(100);g__Metschnikowia_unclassified(100); | Metschnikowia | 13 | Saprotroph | \|Undefined Saprotroph\| | Yeast | NULL | Probable | As nectar-tap_saprotroph (Põlme et al. 2020); Decay sugar-rich substrates (Põlme et al. 2020) | Põlme S, et al. 2020. Fungal Diversity 105:1-16 ((https://doi.org/10.1007/s13225-020-00466-2)) |
| Otu33022 | 1 | k__Fungi(100);p__Ascomycota(100);c__Saccharomycetes(100);o__Saccharomycetales(100);f__Metschnikowiaceae(100);g__Metschnikowia(100);g__Metschnikowia_unclassified(100); | Metschnikowia | 13 | Saprotroph | \|Undefined Saprotroph\| | Yeast | NULL | Probable | As nectar-tap_saprotroph (Põlme et al. 2020); Decay sugar-rich substrates (Põlme et al. 2020) | Põlme S, et al. 2020. Fungal Diversity 105:1-16 ((https://doi.org/10.1007/s13225-020-00466-2)) |
| Otu33023 | 1 | k__Fungi(100);p__Ascomycota(100);c__Saccharomycetes(100);o__Saccharomycetales(100);f__Metschnikowiaceae(100);g__Metschnikowia(100);s__Metschnikowia_pulcherrima(100); | Metschnikowia | 13 | Saprotroph | \|Undefined Saprotroph\| | Yeast | NULL | Probable | As nectar-tap_saprotroph (Põlme et al. 2020); Decay sugar-rich substrates (Põlme et al. 2020) | Põlme S, et al. 2020. Fungal Diversity 105:1-16 ((https://doi.org/10.1007/s13225-020-00466-2)) |
| Otu33024 | 1 | k__Fungi(100);p__Ascomycota(100);c__Saccharomycetes(100);o__Saccharomycetales(100);f__Metschnikowiaceae(100);g__Metschnikowia(100);s__Metschnikowia_pulcherrima(100); | Metschnikowia | 13 | Saprotroph | \|Undefined Saprotroph\| | Yeast | NULL | Probable | As nectar-tap_saprotroph (Põlme et al. 2020); Decay sugar-rich substrates (Põlme et al. 2020) | Põlme S, et al. 2020. Fungal Diversity 105:1-16 ((https://doi.org/10.1007/s13225-020-00466-2)) |
| Otu33025 | 1 | k__Fungi(100);p__Ascomycota(100);c__Dothideomycetes(100);o__Capnodiales(100);f__Mycosphaerellaceae(100);f__Mycosphaerellaceae_unclassified(100);f__Mycosphaerellaceae_unclassified(100); | Mycosphaerellaceae | 9 | Pathotroph-Saprotroph | Plant Pathogen-Undefined Saprotroph | Microfungus | NULL | Probable | Biotrophic, necrotrophic or saprobic on various plant tissues (Cannon & Kirk 2007) | Cannon PF, Kirk PM. 2007. Fungal Families of the World. CAB International, Cambridge (ISBN: 978-0851998275) |
| Otu33026 | 1 | k__Fungi(100);p__Ascomycota(100);c__Saccharomycetes(100);o__Saccharomycetales(100);f__Metschnikowiaceae(100);g__Metschnikowia(100);s__Metschnikowia_pulcherrima(100); | Metschnikowia | 13 | Saprotroph | \|Undefined Saprotroph\| | Yeast | NULL | Probable | As nectar-tap_saprotroph (Põlme et al. 2020); Decay sugar-rich substrates (Põlme et al. 2020) | Põlme S, et al. 2020. Fungal Diversity 105:1-16 ((https://doi.org/10.1007/s13225-020-00466-2)) |
| Otu33027 | 1 | k__Fungi(100);p__Ascomycota(100);c__Saccharomycetes(100);o__Saccharomycetales(100);f__Metschnikowiaceae(100);g__Metschnikowia(100);s__Metschnikowia_pulcherrima(100); | Metschnikowia | 13 | Saprotroph | \|Undefined Saprotroph\| | Yeast | NULL | Probable | As nectar-tap_saprotroph (Põlme et al. 2020); Decay sugar-rich substrates (Põlme et al. 2020) | Põlme S, et al. 2020. Fungal Diversity 105:1-16 ((https://doi.org/10.1007/s13225-020-00466-2)) |
| Otu33028 | 1 | k__Fungi(100);p__Ascomycota(100);c__Saccharomycetes(100);o__Saccharomycetales(100);f__Metschnikowiaceae(100);g__Metschnikowia(100);s__Metschnikowia_pulcherrima(100); | Metschnikowia | 13 | Saprotroph | \|Undefined Saprotroph\| | Yeast | NULL | Probable | As nectar-tap_saprotroph (Põlme et al. 2020); Decay sugar-rich substrates (Põlme et al. 2020) | Põlme S, et al. 2020. Fungal Diversity 105:1-16 ((https://doi.org/10.1007/s13225-020-00466-2)) |
| Otu33029 | 1 | k__Fungi(100);p__Ascomycota(100);c__Saccharomycetes(100);o__Saccharomycetales(100);f__Metschnikowiaceae(100);g__Metschnikowia(100);g__Metschnikowia_unclassified(100); | Metschnikowia | 13 | Saprotroph | \|Undefined Saprotroph\| | Yeast | NULL | Probable | As nectar-tap_saprotroph (Põlme et al. 2020); Decay sugar-rich substrates (Põlme et al. 2020) | Põlme S, et al. 2020. Fungal Diversity 105:1-16 ((https://doi.org/10.1007/s13225-020-00466-2)) |
| Otu33031 | 1 | k__Fungi(100);p__Ascomycota(100);c__Saccharomycetes(100);o__Saccharomycetales(100);f__Metschnikowiaceae(100);g__Metschnikowia(100);s__Metschnikowia_pulcherrima(100); | Metschnikowia | 13 | Saprotroph | \|Undefined Saprotroph\| | Yeast | NULL | Probable | As nectar-tap_saprotroph (Põlme et al. 2020); Decay sugar-rich substrates (Põlme et al. 2020) | Põlme S, et al. 2020. Fungal Diversity 105:1-16 ((https://doi.org/10.1007/s13225-020-00466-2)) |
| Otu33032 | 1 | k__Fungi(100);p__Ascomycota(100);c__Saccharomycetes(100);o__Saccharomycetales(100);f__Saccharomycetales_family_Incertae_sedis(100);g__Candida(100);g__Candida_unclassified(100); | Candida | 13 | Pathotroph-Saprotroph-Symbiotroph | Animal Pathogen-Endophyte-Endosymbiont-Epiphyte-\|Undefined Saprotroph\| | Dimorphic Yeast-Yeast | NULL | Probable | As nectar-tap_saprotroph (Põlme et al. 2020); Decay sugar-rich substrates (Põlme et al. 2020); This asexual genus is polyphyletic and contain species that function very differently depend on what they are. Caution should be taken when interpreting guilds for Candida species. | Manolakaki D, et al. 2010. Virulence. 1: 367-375 ((https://doi.org/10.4161/viru.1.5.12796)); Mekbib SB, et al. 2011. Tropical Plant Pathology 36:233-240 ((https://doi.org/10.1590/S1982-56762011000400004)); Põlme S, et al. 2020. Fungal Diversity 105:1-16 ((https://doi.org/10.1007/s13225-020-00466-2)) |
| Otu33034 | 1 | k__Fungi(100);p__Ascomycota(100);c__Saccharomycetes(100);o__Saccharomycetales(100);f__Metschnikowiaceae(100);g__Metschnikowia(100);s__Metschnikowia_pulcherrima(100); | Metschnikowia | 13 | Saprotroph | \|Undefined Saprotroph\| | Yeast | NULL | Probable | As nectar-tap_saprotroph (Põlme et al. 2020); Decay sugar-rich substrates (Põlme et al. 2020) | Põlme S, et al. 2020. Fungal Diversity 105:1-16 ((https://doi.org/10.1007/s13225-020-00466-2)) |
| Otu33035 | 1 | k__Fungi(100);p__Ascomycota(100);c__Saccharomycetes(100);o__Saccharomycetales(100);f__Metschnikowiaceae(100);g__Metschnikowia(100);s__Metschnikowia_pulcherrima(100); | Metschnikowia | 13 | Saprotroph | \|Undefined Saprotroph\| | Yeast | NULL | Probable | As nectar-tap_saprotroph (Põlme et al. 2020); Decay sugar-rich substrates (Põlme et al. 2020) | Põlme S, et al. 2020. Fungal Diversity 105:1-16 ((https://doi.org/10.1007/s13225-020-00466-2)) |
| Otu33036 | 1 | k__Fungi(100);p__Ascomycota(100);c__Saccharomycetes(100);o__Saccharomycetales(100);f__Metschnikowiaceae(100);g__Metschnikowia(100);s__Metschnikowia_pulcherrima(100); | Metschnikowia | 13 | Saprotroph | \|Undefined Saprotroph\| | Yeast | NULL | Probable | As nectar-tap_saprotroph (Põlme et al. 2020); Decay sugar-rich substrates (Põlme et al. 2020) | Põlme S, et al. 2020. Fungal Diversity 105:1-16 ((https://doi.org/10.1007/s13225-020-00466-2)) |
| Otu33038 | 1 | k__Fungi(100);p__Ascomycota(100);c__Saccharomycetes(100);o__Saccharomycetales(100);f__Metschnikowiaceae(100);g__Metschnikowia(100);g__Metschnikowia_unclassified(100); | Metschnikowia | 13 | Saprotroph | \|Undefined Saprotroph\| | Yeast | NULL | Probable | As nectar-tap_saprotroph (Põlme et al. 2020); Decay sugar-rich substrates (Põlme et al. 2020) | Põlme S, et al. 2020. Fungal Diversity 105:1-16 ((https://doi.org/10.1007/s13225-020-00466-2)) |
| Otu33039 | 1 | k__Fungi(100);p__Ascomycota(100);c__Saccharomycetes(100);o__Saccharomycetales(100);f__Metschnikowiaceae(100);g__Metschnikowia(100);g__Metschnikowia_unclassified(100); | Metschnikowia | 13 | Saprotroph | \|Undefined Saprotroph\| | Yeast | NULL | Probable | As nectar-tap_saprotroph (Põlme et al. 2020); Decay sugar-rich substrates (Põlme et al. 2020) | Põlme S, et al. 2020. Fungal Diversity 105:1-16 ((https://doi.org/10.1007/s13225-020-00466-2)) |
| Otu33040 | 1 | k__Fungi(100);p__Ascomycota(100);c__Saccharomycetes(100);o__Saccharomycetales(100);f__Metschnikowiaceae(100);g__Metschnikowia(100);g__Metschnikowia_unclassified(100); | Metschnikowia | 13 | Saprotroph | \|Undefined Saprotroph\| | Yeast | NULL | Probable | As nectar-tap_saprotroph (Põlme et al. 2020); Decay sugar-rich substrates (Põlme et al. 2020) | Põlme S, et al. 2020. Fungal Diversity 105:1-16 ((https://doi.org/10.1007/s13225-020-00466-2)) |
| Otu33041 | 1 | k__Fungi(100);p__Ascomycota(100);c__Saccharomycetes(100);o__Saccharomycetales(100);f__Metschnikowiaceae(100);g__Metschnikowia(100);g__Metschnikowia_unclassified(100); | Metschnikowia | 13 | Saprotroph | \|Undefined Saprotroph\| | Yeast | NULL | Probable | As nectar-tap_saprotroph (Põlme et al. 2020); Decay sugar-rich substrates (Põlme et al. 2020) | Põlme S, et al. 2020. Fungal Diversity 105:1-16 ((https://doi.org/10.1007/s13225-020-00466-2)) |
| Otu33042 | 1 | k__Fungi(100);p__Ascomycota(100);c__Saccharomycetes(100);o__Saccharomycetales(100);f__Metschnikowiaceae(100);g__Metschnikowia(100);s__Metschnikowia_pulcherrima(100); | Metschnikowia | 13 | Saprotroph | \|Undefined Saprotroph\| | Yeast | NULL | Probable | As nectar-tap_saprotroph (Põlme et al. 2020); Decay sugar-rich substrates (Põlme et al. 2020) | Põlme S, et al. 2020. Fungal Diversity 105:1-16 ((https://doi.org/10.1007/s13225-020-00466-2)) |
| Otu33043 | 1 | k__Fungi(100);p__Ascomycota(100);c__Saccharomycetes(100);o__Saccharomycetales(100);f__Metschnikowiaceae(100);g__Metschnikowia(100);g__Metschnikowia_unclassified(100); | Metschnikowia | 13 | Saprotroph | \|Undefined Saprotroph\| | Yeast | NULL | Probable | As nectar-tap_saprotroph (Põlme et al. 2020); Decay sugar-rich substrates (Põlme et al. 2020) | Põlme S, et al. 2020. Fungal Diversity 105:1-16 ((https://doi.org/10.1007/s13225-020-00466-2)) |
| Otu33044 | 1 | k__Fungi(100);p__Ascomycota(100);c__Saccharomycetes(100);o__Saccharomycetales(100);f__Metschnikowiaceae(100);g__Metschnikowia(100);g__Metschnikowia_unclassified(100); | Metschnikowia | 13 | Saprotroph | \|Undefined Saprotroph\| | Yeast | NULL | Probable | As nectar-tap_saprotroph (Põlme et al. 2020); Decay sugar-rich substrates (Põlme et al. 2020) | Põlme S, et al. 2020. Fungal Diversity 105:1-16 ((https://doi.org/10.1007/s13225-020-00466-2)) |
| Otu33045 | 1 | k__Fungi(100);p__Ascomycota(100);c__Saccharomycetes(100);o__Saccharomycetales(100);f__Metschnikowiaceae(100);g__Metschnikowia(100);s__Metschnikowia_pulcherrima(100); | Metschnikowia | 13 | Saprotroph | \|Undefined Saprotroph\| | Yeast | NULL | Probable | As nectar-tap_saprotroph (Põlme et al. 2020); Decay sugar-rich substrates (Põlme et al. 2020) | Põlme S, et al. 2020. Fungal Diversity 105:1-16 ((https://doi.org/10.1007/s13225-020-00466-2)) |
| Otu33046 | 1 | k__Fungi(100);p__Ascomycota(100);c__Saccharomycetes(100);o__Saccharomycetales(100);f__Metschnikowiaceae(100);g__Metschnikowia(100);s__Metschnikowia_pulcherrima(100); | Metschnikowia | 13 | Saprotroph | \|Undefined Saprotroph\| | Yeast | NULL | Probable | As nectar-tap_saprotroph (Põlme et al. 2020); Decay sugar-rich substrates (Põlme et al. 2020) | Põlme S, et al. 2020. Fungal Diversity 105:1-16 ((https://doi.org/10.1007/s13225-020-00466-2)) |
| Otu33047 | 1 | k__Fungi(100);p__Ascomycota(100);c__Saccharomycetes(100);o__Saccharomycetales(100);f__Metschnikowiaceae(100);g__Metschnikowia(100);s__Metschnikowia_pulcherrima(100); | Metschnikowia | 13 | Saprotroph | \|Undefined Saprotroph\| | Yeast | NULL | Probable | As nectar-tap_saprotroph (Põlme et al. 2020); Decay sugar-rich substrates (Põlme et al. 2020) | Põlme S, et al. 2020. Fungal Diversity 105:1-16 ((https://doi.org/10.1007/s13225-020-00466-2)) |
| Otu33048 | 1 | k__Fungi(100);p__Ascomycota(100);c__Saccharomycetes(100);o__Saccharomycetales(100);f__Metschnikowiaceae(100);g__Metschnikowia(100);s__Metschnikowia_pulcherrima(100); | Metschnikowia | 13 | Saprotroph | \|Undefined Saprotroph\| | Yeast | NULL | Probable | As nectar-tap_saprotroph (Põlme et al. 2020); Decay sugar-rich substrates (Põlme et al. 2020) | Põlme S, et al. 2020. Fungal Diversity 105:1-16 ((https://doi.org/10.1007/s13225-020-00466-2)) |
| Otu33049 | 1 | k__Fungi(100);p__Ascomycota(100);c__Saccharomycetes(100);o__Saccharomycetales(100);f__Metschnikowiaceae(100);g__Metschnikowia(100);s__Metschnikowia_pulcherrima(100); | Metschnikowia | 13 | Saprotroph | \|Undefined Saprotroph\| | Yeast | NULL | Probable | As nectar-tap_saprotroph (Põlme et al. 2020); Decay sugar-rich substrates (Põlme et al. 2020) | Põlme S, et al. 2020. Fungal Diversity 105:1-16 ((https://doi.org/10.1007/s13225-020-00466-2)) |
| Otu33050 | 1 | k__Fungi(100);p__Ascomycota(100);c__Saccharomycetes(100);o__Saccharomycetales(100);f__Metschnikowiaceae(100);g__Metschnikowia(100);s__Metschnikowia_pulcherrima(100); | Metschnikowia | 13 | Saprotroph | \|Undefined Saprotroph\| | Yeast | NULL | Probable | As nectar-tap_saprotroph (Põlme et al. 2020); Decay sugar-rich substrates (Põlme et al. 2020) | Põlme S, et al. 2020. Fungal Diversity 105:1-16 ((https://doi.org/10.1007/s13225-020-00466-2)) |
| Otu33051 | 1 | k__Fungi(100);p__Ascomycota(100);c__Saccharomycetes(100);o__Saccharomycetales(100);f__Metschnikowiaceae(100);g__Metschnikowia(100);g__Metschnikowia_unclassified(100); | Metschnikowia | 13 | Saprotroph | \|Undefined Saprotroph\| | Yeast | NULL | Probable | As nectar-tap_saprotroph (Põlme et al. 2020); Decay sugar-rich substrates (Põlme et al. 2020) | Põlme S, et al. 2020. Fungal Diversity 105:1-16 ((https://doi.org/10.1007/s13225-020-00466-2)) |
| Otu33052 | 1 | k__Fungi(100);p__Ascomycota(100);c__Saccharomycetes(100);o__Saccharomycetales(100);f__Metschnikowiaceae(100);g__Metschnikowia(100);s__Metschnikowia_pulcherrima(100); | Metschnikowia | 13 | Saprotroph | \|Undefined Saprotroph\| | Yeast | NULL | Probable | As nectar-tap_saprotroph (Põlme et al. 2020); Decay sugar-rich substrates (Põlme et al. 2020) | Põlme S, et al. 2020. Fungal Diversity 105:1-16 ((https://doi.org/10.1007/s13225-020-00466-2)) |
| Otu33053 | 1 | k__Fungi(100);p__Ascomycota(100);c__Saccharomycetes(100);o__Saccharomycetales(100);f__Metschnikowiaceae(100);g__Metschnikowia(100);g__Metschnikowia_unclassified(100); | Metschnikowia | 13 | Saprotroph | \|Undefined Saprotroph\| | Yeast | NULL | Probable | As nectar-tap_saprotroph (Põlme et al. 2020); Decay sugar-rich substrates (Põlme et al. 2020) | Põlme S, et al. 2020. Fungal Diversity 105:1-16 ((https://doi.org/10.1007/s13225-020-00466-2)) |
| Otu33054 | 1 | k__Fungi(100);p__Ascomycota(100);c__Saccharomycetes(100);o__Saccharomycetales(100);f__Metschnikowiaceae(100);g__Metschnikowia(100);s__Metschnikowia_pulcherrima(100); | Metschnikowia | 13 | Saprotroph | \|Undefined Saprotroph\| | Yeast | NULL | Probable | As nectar-tap_saprotroph (Põlme et al. 2020); Decay sugar-rich substrates (Põlme et al. 2020) | Põlme S, et al. 2020. Fungal Diversity 105:1-16 ((https://doi.org/10.1007/s13225-020-00466-2)) |
| Otu33055 | 1 | k__Fungi(100);p__Ascomycota(100);c__Saccharomycetes(100);o__Saccharomycetales(100);f__Metschnikowiaceae(100);g__Metschnikowia(100);s__Metschnikowia_pulcherrima(100); | Metschnikowia | 13 | Saprotroph | \|Undefined Saprotroph\| | Yeast | NULL | Probable | As nectar-tap_saprotroph (Põlme et al. 2020); Decay sugar-rich substrates (Põlme et al. 2020) | Põlme S, et al. 2020. Fungal Diversity 105:1-16 ((https://doi.org/10.1007/s13225-020-00466-2)) |
| Otu33056 | 1 | k__Fungi(100);p__Ascomycota(100);c__Saccharomycetes(100);o__Saccharomycetales(100);f__Metschnikowiaceae(100);g__Metschnikowia(100);s__Metschnikowia_pulcherrima(100); | Metschnikowia | 13 | Saprotroph | \|Undefined Saprotroph\| | Yeast | NULL | Probable | As nectar-tap_saprotroph (Põlme et al. 2020); Decay sugar-rich substrates (Põlme et al. 2020) | Põlme S, et al. 2020. Fungal Diversity 105:1-16 ((https://doi.org/10.1007/s13225-020-00466-2)) |
| Otu33058 | 1 | k__Fungi(100);p__Ascomycota(100);c__Saccharomycetes(100);o__Saccharomycetales(100);f__Metschnikowiaceae(100);g__Metschnikowia(100);s__Metschnikowia_pulcherrima(100); | Metschnikowia | 13 | Saprotroph | \|Undefined Saprotroph\| | Yeast | NULL | Probable | As nectar-tap_saprotroph (Põlme et al. 2020); Decay sugar-rich substrates (Põlme et al. 2020) | Põlme S, et al. 2020. Fungal Diversity 105:1-16 ((https://doi.org/10.1007/s13225-020-00466-2)) |
| Otu33059 | 1 | k__Fungi(100);p__Ascomycota(100);c__Saccharomycetes(100);o__Saccharomycetales(100);f__Metschnikowiaceae(100);g__Metschnikowia(100);s__Metschnikowia_pulcherrima(100); | Metschnikowia | 13 | Saprotroph | \|Undefined Saprotroph\| | Yeast | NULL | Probable | As nectar-tap_saprotroph (Põlme et al. 2020); Decay sugar-rich substrates (Põlme et al. 2020) | Põlme S, et al. 2020. Fungal Diversity 105:1-16 ((https://doi.org/10.1007/s13225-020-00466-2)) |
| Otu33060 | 1 | k__Fungi(100);p__Ascomycota(100);c__Saccharomycetes(100);o__Saccharomycetales(100);f__Metschnikowiaceae(100);g__Metschnikowia(100);s__Metschnikowia_pulcherrima(100); | Metschnikowia | 13 | Saprotroph | \|Undefined Saprotroph\| | Yeast | NULL | Probable | As nectar-tap_saprotroph (Põlme et al. 2020); Decay sugar-rich substrates (Põlme et al. 2020) | Põlme S, et al. 2020. Fungal Diversity 105:1-16 ((https://doi.org/10.1007/s13225-020-00466-2)) |
| Otu33061 | 1 | k__Fungi(100);p__Ascomycota(100);c__Saccharomycetes(100);o__Saccharomycetales(100);f__Metschnikowiaceae(100);g__Metschnikowia(100);s__Metschnikowia_pulcherrima(100); | Metschnikowia | 13 | Saprotroph | \|Undefined Saprotroph\| | Yeast | NULL | Probable | As nectar-tap_saprotroph (Põlme et al. 2020); Decay sugar-rich substrates (Põlme et al. 2020) | Põlme S, et al. 2020. Fungal Diversity 105:1-16 ((https://doi.org/10.1007/s13225-020-00466-2)) |
| Otu33062 | 1 | k__Fungi(100);p__Ascomycota(100);c__Saccharomycetes(100);o__Saccharomycetales(100);f__Metschnikowiaceae(100);g__Metschnikowia(100);g__Metschnikowia_unclassified(100); | Metschnikowia | 13 | Saprotroph | \|Undefined Saprotroph\| | Yeast | NULL | Probable | As nectar-tap_saprotroph (Põlme et al. 2020); Decay sugar-rich substrates (Põlme et al. 2020) | Põlme S, et al. 2020. Fungal Diversity 105:1-16 ((https://doi.org/10.1007/s13225-020-00466-2)) |
| Otu33063 | 1 | k__Fungi(100);p__Ascomycota(100);c__Saccharomycetes(100);o__Saccharomycetales(100);f__Metschnikowiaceae(100);g__Metschnikowia(100);g__Metschnikowia_unclassified(100); | Metschnikowia | 13 | Saprotroph | \|Undefined Saprotroph\| | Yeast | NULL | Probable | As nectar-tap_saprotroph (Põlme et al. 2020); Decay sugar-rich substrates (Põlme et al. 2020) | Põlme S, et al. 2020. Fungal Diversity 105:1-16 ((https://doi.org/10.1007/s13225-020-00466-2)) |
| Otu33064 | 1 | k__Fungi(100);p__Ascomycota(100);c__Saccharomycetes(100);o__Saccharomycetales(100);f__Metschnikowiaceae(100);g__Metschnikowia(100);s__Metschnikowia_pulcherrima(100); | Metschnikowia | 13 | Saprotroph | \|Undefined Saprotroph\| | Yeast | NULL | Probable | As nectar-tap_saprotroph (Põlme et al. 2020); Decay sugar-rich substrates (Põlme et al. 2020) | Põlme S, et al. 2020. Fungal Diversity 105:1-16 ((https://doi.org/10.1007/s13225-020-00466-2)) |
| Otu33065 | 1 | k__Fungi(100);p__Ascomycota(100);c__Saccharomycetes(100);o__Saccharomycetales(100);f__Metschnikowiaceae(100);g__Metschnikowia(100);g__Metschnikowia_unclassified(100); | Metschnikowia | 13 | Saprotroph | \|Undefined Saprotroph\| | Yeast | NULL | Probable | As nectar-tap_saprotroph (Põlme et al. 2020); Decay sugar-rich substrates (Põlme et al. 2020) | Põlme S, et al. 2020. Fungal Diversity 105:1-16 ((https://doi.org/10.1007/s13225-020-00466-2)) |
| Otu33066 | 1 | k__Fungi(100);p__Ascomycota(100);c__Saccharomycetes(100);o__Saccharomycetales(100);f__Metschnikowiaceae(100);g__Metschnikowia(100);s__Metschnikowia_pulcherrima(100); | Metschnikowia | 13 | Saprotroph | \|Undefined Saprotroph\| | Yeast | NULL | Probable | As nectar-tap_saprotroph (Põlme et al. 2020); Decay sugar-rich substrates (Põlme et al. 2020) | Põlme S, et al. 2020. Fungal Diversity 105:1-16 ((https://doi.org/10.1007/s13225-020-00466-2)) |
| Otu33067 | 1 | k__Fungi(100);p__Ascomycota(100);c__Saccharomycetes(100);o__Saccharomycetales(100);f__Metschnikowiaceae(100);g__Metschnikowia(100);s__Metschnikowia_pulcherrima(100); | Metschnikowia | 13 | Saprotroph | \|Undefined Saprotroph\| | Yeast | NULL | Probable | As nectar-tap_saprotroph (Põlme et al. 2020); Decay sugar-rich substrates (Põlme et al. 2020) | Põlme S, et al. 2020. Fungal Diversity 105:1-16 ((https://doi.org/10.1007/s13225-020-00466-2)) |
| Otu33068 | 1 | k__Fungi(100);p__Ascomycota(100);c__Saccharomycetes(100);o__Saccharomycetales(100);f__Metschnikowiaceae(100);g__Metschnikowia(100);s__Metschnikowia_pulcherrima(100); | Metschnikowia | 13 | Saprotroph | \|Undefined Saprotroph\| | Yeast | NULL | Probable | As nectar-tap_saprotroph (Põlme et al. 2020); Decay sugar-rich substrates (Põlme et al. 2020) | Põlme S, et al. 2020. Fungal Diversity 105:1-16 ((https://doi.org/10.1007/s13225-020-00466-2)) |
| Otu33069 | 1 | k__Fungi(100);p__Ascomycota(100);c__Saccharomycetes(100);o__Saccharomycetales(100);f__Metschnikowiaceae(100);g__Metschnikowia(100);s__Metschnikowia_pulcherrima(100); | Metschnikowia | 13 | Saprotroph | \|Undefined Saprotroph\| | Yeast | NULL | Probable | As nectar-tap_saprotroph (Põlme et al. 2020); Decay sugar-rich substrates (Põlme et al. 2020) | Põlme S, et al. 2020. Fungal Diversity 105:1-16 ((https://doi.org/10.1007/s13225-020-00466-2)) |
| Otu33070 | 1 | k__Fungi(100);p__Ascomycota(100);c__Saccharomycetes(100);o__Saccharomycetales(100);f__Metschnikowiaceae(100);g__Metschnikowia(100);s__Metschnikowia_pulcherrima(100); | Metschnikowia | 13 | Saprotroph | \|Undefined Saprotroph\| | Yeast | NULL | Probable | As nectar-tap_saprotroph (Põlme et al. 2020); Decay sugar-rich substrates (Põlme et al. 2020) | Põlme S, et al. 2020. Fungal Diversity 105:1-16 ((https://doi.org/10.1007/s13225-020-00466-2)) |
| Otu33071 | 1 | k__Fungi(100);p__Ascomycota(100);c__Saccharomycetes(100);o__Saccharomycetales(100);f__Metschnikowiaceae(100);g__Metschnikowia(100);s__Metschnikowia_pulcherrima(100); | Metschnikowia | 13 | Saprotroph | \|Undefined Saprotroph\| | Yeast | NULL | Probable | As nectar-tap_saprotroph (Põlme et al. 2020); Decay sugar-rich substrates (Põlme et al. 2020) | Põlme S, et al. 2020. Fungal Diversity 105:1-16 ((https://doi.org/10.1007/s13225-020-00466-2)) |
| Otu33072 | 1 | k__Fungi(100);p__Ascomycota(100);c__Saccharomycetes(100);o__Saccharomycetales(100);f__Metschnikowiaceae(100);g__Metschnikowia(100);s__Metschnikowia_pulcherrima(100); | Metschnikowia | 13 | Saprotroph | \|Undefined Saprotroph\| | Yeast | NULL | Probable | As nectar-tap_saprotroph (Põlme et al. 2020); Decay sugar-rich substrates (Põlme et al. 2020) | Põlme S, et al. 2020. Fungal Diversity 105:1-16 ((https://doi.org/10.1007/s13225-020-00466-2)) |
| Otu33073 | 1 | k__Fungi(100);p__Ascomycota(100);c__Saccharomycetes(100);o__Saccharomycetales(100);f__Metschnikowiaceae(100);g__Metschnikowia(100);s__Metschnikowia_pulcherrima(100); | Metschnikowia | 13 | Saprotroph | \|Undefined Saprotroph\| | Yeast | NULL | Probable | As nectar-tap_saprotroph (Põlme et al. 2020); Decay sugar-rich substrates (Põlme et al. 2020) | Põlme S, et al. 2020. Fungal Diversity 105:1-16 ((https://doi.org/10.1007/s13225-020-00466-2)) |
| Otu33074 | 1 | k__Fungi(100);p__Ascomycota(100);c__Saccharomycetes(100);o__Saccharomycetales(100);f__Metschnikowiaceae(100);g__Metschnikowia(100);s__Metschnikowia_pulcherrima(100); | Metschnikowia | 13 | Saprotroph | \|Undefined Saprotroph\| | Yeast | NULL | Probable | As nectar-tap_saprotroph (Põlme et al. 2020); Decay sugar-rich substrates (Põlme et al. 2020) | Põlme S, et al. 2020. Fungal Diversity 105:1-16 ((https://doi.org/10.1007/s13225-020-00466-2)) |
| Otu33076 | 1 | k__Fungi(100);p__Ascomycota(100);c__Saccharomycetes(100);o__Saccharomycetales(100);f__Metschnikowiaceae(100);g__Metschnikowia(100);s__Metschnikowia_pulcherrima(100); | Metschnikowia | 13 | Saprotroph | \|Undefined Saprotroph\| | Yeast | NULL | Probable | As nectar-tap_saprotroph (Põlme et al. 2020); Decay sugar-rich substrates (Põlme et al. 2020) | Põlme S, et al. 2020. Fungal Diversity 105:1-16 ((https://doi.org/10.1007/s13225-020-00466-2)) |
| Otu33077 | 1 | k__Fungi(100);p__Ascomycota(100);c__Saccharomycetes(100);o__Saccharomycetales(100);f__Metschnikowiaceae(100);g__Metschnikowia(100);s__Metschnikowia_pulcherrima(100); | Metschnikowia | 13 | Saprotroph | \|Undefined Saprotroph\| | Yeast | NULL | Probable | As nectar-tap_saprotroph (Põlme et al. 2020); Decay sugar-rich substrates (Põlme et al. 2020) | Põlme S, et al. 2020. Fungal Diversity 105:1-16 ((https://doi.org/10.1007/s13225-020-00466-2)) |
| Otu33078 | 1 | k__Fungi(100);p__Ascomycota(100);c__Saccharomycetes(100);o__Saccharomycetales(100);f__Metschnikowiaceae(100);g__Metschnikowia(100);s__Metschnikowia_pulcherrima(100); | Metschnikowia | 13 | Saprotroph | \|Undefined Saprotroph\| | Yeast | NULL | Probable | As nectar-tap_saprotroph (Põlme et al. 2020); Decay sugar-rich substrates (Põlme et al. 2020) | Põlme S, et al. 2020. Fungal Diversity 105:1-16 ((https://doi.org/10.1007/s13225-020-00466-2)) |
| Otu33079 | 1 | k__Fungi(100);p__Ascomycota(100);c__Saccharomycetes(100);o__Saccharomycetales(100);f__Metschnikowiaceae(100);g__Metschnikowia(100);s__Metschnikowia_pulcherrima(100); | Metschnikowia | 13 | Saprotroph | \|Undefined Saprotroph\| | Yeast | NULL | Probable | As nectar-tap_saprotroph (Põlme et al. 2020); Decay sugar-rich substrates (Põlme et al. 2020) | Põlme S, et al. 2020. Fungal Diversity 105:1-16 ((https://doi.org/10.1007/s13225-020-00466-2)) |
| Otu33080 | 1 | k__Fungi(100);p__Ascomycota(100);c__Saccharomycetes(100);o__Saccharomycetales(100);f__Metschnikowiaceae(100);g__Metschnikowia(100);s__Metschnikowia_pulcherrima(100); | Metschnikowia | 13 | Saprotroph | \|Undefined Saprotroph\| | Yeast | NULL | Probable | As nectar-tap_saprotroph (Põlme et al. 2020); Decay sugar-rich substrates (Põlme et al. 2020) | Põlme S, et al. 2020. Fungal Diversity 105:1-16 ((https://doi.org/10.1007/s13225-020-00466-2)) |
| Otu33081 | 1 | k__Fungi(100);p__Ascomycota(100);c__Saccharomycetes(100);o__Saccharomycetales(100);f__Metschnikowiaceae(100);g__Metschnikowia(100);s__Metschnikowia_pulcherrima(100); | Metschnikowia | 13 | Saprotroph | \|Undefined Saprotroph\| | Yeast | NULL | Probable | As nectar-tap_saprotroph (Põlme et al. 2020); Decay sugar-rich substrates (Põlme et al. 2020) | Põlme S, et al. 2020. Fungal Diversity 105:1-16 ((https://doi.org/10.1007/s13225-020-00466-2)) |
| Otu33082 | 1 | k__Fungi(100);p__Ascomycota(100);c__Saccharomycetes(100);o__Saccharomycetales(100);f__Metschnikowiaceae(100);g__Metschnikowia(100);s__Metschnikowia_pulcherrima(100); | Metschnikowia | 13 | Saprotroph | \|Undefined Saprotroph\| | Yeast | NULL | Probable | As nectar-tap_saprotroph (Põlme et al. 2020); Decay sugar-rich substrates (Põlme et al. 2020) | Põlme S, et al. 2020. Fungal Diversity 105:1-16 ((https://doi.org/10.1007/s13225-020-00466-2)) |
| Otu33083 | 1 | k__Fungi(100);p__Ascomycota(100);c__Saccharomycetes(100);o__Saccharomycetales(100);f__Metschnikowiaceae(100);g__Metschnikowia(100);s__Metschnikowia_pulcherrima(100); | Metschnikowia | 13 | Saprotroph | \|Undefined Saprotroph\| | Yeast | NULL | Probable | As nectar-tap_saprotroph (Põlme et al. 2020); Decay sugar-rich substrates (Põlme et al. 2020) | Põlme S, et al. 2020. Fungal Diversity 105:1-16 ((https://doi.org/10.1007/s13225-020-00466-2)) |
| Otu33084 | 1 | k__Fungi(100);p__Ascomycota(100);c__Saccharomycetes(100);o__Saccharomycetales(100);f__Metschnikowiaceae(100);g__Metschnikowia(100);s__Metschnikowia_pulcherrima(100); | Metschnikowia | 13 | Saprotroph | \|Undefined Saprotroph\| | Yeast | NULL | Probable | As nectar-tap_saprotroph (Põlme et al. 2020); Decay sugar-rich substrates (Põlme et al. 2020) | Põlme S, et al. 2020. Fungal Diversity 105:1-16 ((https://doi.org/10.1007/s13225-020-00466-2)) |
| Otu33085 | 1 | k__Fungi(100);p__Ascomycota(100);c__Saccharomycetes(100);o__Saccharomycetales(100);f__Metschnikowiaceae(100);g__Metschnikowia(100);s__Metschnikowia_pulcherrima(100); | Metschnikowia | 13 | Saprotroph | \|Undefined Saprotroph\| | Yeast | NULL | Probable | As nectar-tap_saprotroph (Põlme et al. 2020); Decay sugar-rich substrates (Põlme et al. 2020) | Põlme S, et al. 2020. Fungal Diversity 105:1-16 ((https://doi.org/10.1007/s13225-020-00466-2)) |
| Otu33086 | 1 | k__Fungi(100);p__Ascomycota(100);c__Saccharomycetes(100);o__Saccharomycetales(100);f__Metschnikowiaceae(100);g__Metschnikowia(100);s__Metschnikowia_pulcherrima(100); | Metschnikowia | 13 | Saprotroph | \|Undefined Saprotroph\| | Yeast | NULL | Probable | As nectar-tap_saprotroph (Põlme et al. 2020); Decay sugar-rich substrates (Põlme et al. 2020) | Põlme S, et al. 2020. Fungal Diversity 105:1-16 ((https://doi.org/10.1007/s13225-020-00466-2)) |
| Otu33087 | 1 | k__Fungi(100);p__Ascomycota(100);c__Saccharomycetes(100);o__Saccharomycetales(100);f__Metschnikowiaceae(100);g__Metschnikowia(100);s__Metschnikowia_pulcherrima(100); | Metschnikowia | 13 | Saprotroph | \|Undefined Saprotroph\| | Yeast | NULL | Probable | As nectar-tap_saprotroph (Põlme et al. 2020); Decay sugar-rich substrates (Põlme et al. 2020) | Põlme S, et al. 2020. Fungal Diversity 105:1-16 ((https://doi.org/10.1007/s13225-020-00466-2)) |
| Otu33088 | 1 | k__Fungi(100);p__Ascomycota(100);c__Saccharomycetes(100);o__Saccharomycetales(100);f__Metschnikowiaceae(100);g__Metschnikowia(100);s__Metschnikowia_pulcherrima(100); | Metschnikowia | 13 | Saprotroph | \|Undefined Saprotroph\| | Yeast | NULL | Probable | As nectar-tap_saprotroph (Põlme et al. 2020); Decay sugar-rich substrates (Põlme et al. 2020) | Põlme S, et al. 2020. Fungal Diversity 105:1-16 ((https://doi.org/10.1007/s13225-020-00466-2)) |
| Otu33089 | 1 | k__Fungi(100);p__Ascomycota(100);c__Saccharomycetes(100);o__Saccharomycetales(100);f__Metschnikowiaceae(100);g__Metschnikowia(100);s__Metschnikowia_pulcherrima(100); | Metschnikowia | 13 | Saprotroph | \|Undefined Saprotroph\| | Yeast | NULL | Probable | As nectar-tap_saprotroph (Põlme et al. 2020); Decay sugar-rich substrates (Põlme et al. 2020) | Põlme S, et al. 2020. Fungal Diversity 105:1-16 ((https://doi.org/10.1007/s13225-020-00466-2)) |
| Otu33090 | 1 | k__Fungi(100);p__Ascomycota(100);c__Saccharomycetes(100);o__Saccharomycetales(100);f__Metschnikowiaceae(100);g__Metschnikowia(100);s__Metschnikowia_pulcherrima(100); | Metschnikowia | 13 | Saprotroph | \|Undefined Saprotroph\| | Yeast | NULL | Probable | As nectar-tap_saprotroph (Põlme et al. 2020); Decay sugar-rich substrates (Põlme et al. 2020) | Põlme S, et al. 2020. Fungal Diversity 105:1-16 ((https://doi.org/10.1007/s13225-020-00466-2)) |
| Otu33091 | 1 | k__Fungi(100);p__Ascomycota(100);c__Saccharomycetes(100);o__Saccharomycetales(100);f__Metschnikowiaceae(100);g__Metschnikowia(100);s__Metschnikowia_pulcherrima(100); | Metschnikowia | 13 | Saprotroph | \|Undefined Saprotroph\| | Yeast | NULL | Probable | As nectar-tap_saprotroph (Põlme et al. 2020); Decay sugar-rich substrates (Põlme et al. 2020) | Põlme S, et al. 2020. Fungal Diversity 105:1-16 ((https://doi.org/10.1007/s13225-020-00466-2)) |
| Otu33092 | 1 | k__Fungi(100);p__Ascomycota(100);c__Saccharomycetes(100);o__Saccharomycetales(100);f__Metschnikowiaceae(100);g__Metschnikowia(100);s__Metschnikowia_pulcherrima(100); | Metschnikowia | 13 | Saprotroph | \|Undefined Saprotroph\| | Yeast | NULL | Probable | As nectar-tap_saprotroph (Põlme et al. 2020); Decay sugar-rich substrates (Põlme et al. 2020) | Põlme S, et al. 2020. Fungal Diversity 105:1-16 ((https://doi.org/10.1007/s13225-020-00466-2)) |
| Otu33097 | 1 | k__Fungi(100);p__Ascomycota(100);c__Saccharomycetes(100);o__Saccharomycetales(100);f__Metschnikowiaceae(100);g__Metschnikowia(100);s__Metschnikowia_pulcherrima(100); | Metschnikowia | 13 | Saprotroph | \|Undefined Saprotroph\| | Yeast | NULL | Probable | As nectar-tap_saprotroph (Põlme et al. 2020); Decay sugar-rich substrates (Põlme et al. 2020) | Põlme S, et al. 2020. Fungal Diversity 105:1-16 ((https://doi.org/10.1007/s13225-020-00466-2)) |
| Otu33098 | 1 | k__Fungi(100);p__Ascomycota(100);c__Saccharomycetes(100);o__Saccharomycetales(100);f__Metschnikowiaceae(100);g__Metschnikowia(100);s__Metschnikowia_pulcherrima(100); | Metschnikowia | 13 | Saprotroph | \|Undefined Saprotroph\| | Yeast | NULL | Probable | As nectar-tap_saprotroph (Põlme et al. 2020); Decay sugar-rich substrates (Põlme et al. 2020) | Põlme S, et al. 2020. Fungal Diversity 105:1-16 ((https://doi.org/10.1007/s13225-020-00466-2)) |
| Otu33099 | 1 | k__Fungi(100);p__Ascomycota(100);c__Saccharomycetes(100);o__Saccharomycetales(100);f__Metschnikowiaceae(100);g__Metschnikowia(100);s__Metschnikowia_pulcherrima(100); | Metschnikowia | 13 | Saprotroph | \|Undefined Saprotroph\| | Yeast | NULL | Probable | As nectar-tap_saprotroph (Põlme et al. 2020); Decay sugar-rich substrates (Põlme et al. 2020) | Põlme S, et al. 2020. Fungal Diversity 105:1-16 ((https://doi.org/10.1007/s13225-020-00466-2)) |
| Otu33102 | 1 | k__Fungi(100);p__Ascomycota(100);c__Saccharomycetes(100);o__Saccharomycetales(100);f__Metschnikowiaceae(100);g__Metschnikowia(100);g__Metschnikowia_unclassified(100); | Metschnikowia | 13 | Saprotroph | \|Undefined Saprotroph\| | Yeast | NULL | Probable | As nectar-tap_saprotroph (Põlme et al. 2020); Decay sugar-rich substrates (Põlme et al. 2020) | Põlme S, et al. 2020. Fungal Diversity 105:1-16 ((https://doi.org/10.1007/s13225-020-00466-2)) |
| Otu33103 | 1 | k__Fungi(100);p__Ascomycota(100);c__Saccharomycetes(100);o__Saccharomycetales(100);f__Metschnikowiaceae(100);g__Metschnikowia(100);s__Metschnikowia_pulcherrima(100); | Metschnikowia | 13 | Saprotroph | \|Undefined Saprotroph\| | Yeast | NULL | Probable | As nectar-tap_saprotroph (Põlme et al. 2020); Decay sugar-rich substrates (Põlme et al. 2020) | Põlme S, et al. 2020. Fungal Diversity 105:1-16 ((https://doi.org/10.1007/s13225-020-00466-2)) |
| Otu33104 | 1 | k__Fungi(100);p__Ascomycota(100);c__Saccharomycetes(100);o__Saccharomycetales(100);f__Metschnikowiaceae(100);g__Metschnikowia(100);s__Metschnikowia_pulcherrima(100); | Metschnikowia | 13 | Saprotroph | \|Undefined Saprotroph\| | Yeast | NULL | Probable | As nectar-tap_saprotroph (Põlme et al. 2020); Decay sugar-rich substrates (Põlme et al. 2020) | Põlme S, et al. 2020. Fungal Diversity 105:1-16 ((https://doi.org/10.1007/s13225-020-00466-2)) |
| Otu33105 | 1 | k__Fungi(100);p__Ascomycota(100);c__Saccharomycetes(100);o__Saccharomycetales(100);f__Metschnikowiaceae(100);g__Metschnikowia(100);s__Metschnikowia_pulcherrima(100); | Metschnikowia | 13 | Saprotroph | \|Undefined Saprotroph\| | Yeast | NULL | Probable | As nectar-tap_saprotroph (Põlme et al. 2020); Decay sugar-rich substrates (Põlme et al. 2020) | Põlme S, et al. 2020. Fungal Diversity 105:1-16 ((https://doi.org/10.1007/s13225-020-00466-2)) |
| Otu33106 | 1 | k__Fungi(100);p__Ascomycota(100);c__Saccharomycetes(100);o__Saccharomycetales(100);f__Metschnikowiaceae(100);g__Metschnikowia(100);s__Metschnikowia_pulcherrima(100); | Metschnikowia | 13 | Saprotroph | \|Undefined Saprotroph\| | Yeast | NULL | Probable | As nectar-tap_saprotroph (Põlme et al. 2020); Decay sugar-rich substrates (Põlme et al. 2020) | Põlme S, et al. 2020. Fungal Diversity 105:1-16 ((https://doi.org/10.1007/s13225-020-00466-2)) |
| Otu33107 | 1 | k__Fungi(100);p__Ascomycota(100);c__Saccharomycetes(100);o__Saccharomycetales(100);f__Metschnikowiaceae(100);g__Metschnikowia(100);s__Metschnikowia_pulcherrima(100); | Metschnikowia | 13 | Saprotroph | \|Undefined Saprotroph\| | Yeast | NULL | Probable | As nectar-tap_saprotroph (Põlme et al. 2020); Decay sugar-rich substrates (Põlme et al. 2020) | Põlme S, et al. 2020. Fungal Diversity 105:1-16 ((https://doi.org/10.1007/s13225-020-00466-2)) |
| Otu33117 | 1 | k__Fungi(100);p__Ascomycota(100);c__Saccharomycetes(100);o__Saccharomycetales(100);f__Saccharomycetales_family_Incertae_sedis(100);f__Saccharomycetales_family_Incertae_sedis_unclassified(100);f__Saccharomycetales_family_Incertae_sedis_unclassified(100); | Saccharomycetales | 7 | Saprotroph | Undefined Saprotroph | Yeast | NULL | Possible | NULL | Sterkenburg E, et al. 2015. New Phytologist 207:1145-1158 ((https://doi.org/10.1111/nph.13426)) |
| Otu33122 | 1 | k__Fungi(100);p__Ascomycota(100);c__Saccharomycetes(100);o__Saccharomycetales(100);f__Metschnikowiaceae(100);g__Metschnikowia(100);s__Metschnikowia_pulcherrima(100); | Metschnikowia | 13 | Saprotroph | \|Undefined Saprotroph\| | Yeast | NULL | Probable | As nectar-tap_saprotroph (Põlme et al. 2020); Decay sugar-rich substrates (Põlme et al. 2020) | Põlme S, et al. 2020. Fungal Diversity 105:1-16 ((https://doi.org/10.1007/s13225-020-00466-2)) |
| Otu33124 | 1 | k__Fungi(100);p__Ascomycota(100);c__Saccharomycetes(100);o__Saccharomycetales(100);f__Metschnikowiaceae(100);g__Metschnikowia(100);s__Metschnikowia_pulcherrima(100); | Metschnikowia | 13 | Saprotroph | \|Undefined Saprotroph\| | Yeast | NULL | Probable | As nectar-tap_saprotroph (Põlme et al. 2020); Decay sugar-rich substrates (Põlme et al. 2020) | Põlme S, et al. 2020. Fungal Diversity 105:1-16 ((https://doi.org/10.1007/s13225-020-00466-2)) |
| Otu33126 | 1 | k__Fungi(100);p__Ascomycota(100);c__Saccharomycetes(100);o__Saccharomycetales(100);f__Metschnikowiaceae(100);g__Metschnikowia(100);g__Metschnikowia_unclassified(100); | Metschnikowia | 13 | Saprotroph | \|Undefined Saprotroph\| | Yeast | NULL | Probable | As nectar-tap_saprotroph (Põlme et al. 2020); Decay sugar-rich substrates (Põlme et al. 2020) | Põlme S, et al. 2020. Fungal Diversity 105:1-16 ((https://doi.org/10.1007/s13225-020-00466-2)) |
| Otu33127 | 1 | k__Fungi(100);p__Ascomycota(100);c__Saccharomycetes(100);o__Saccharomycetales(100);f__Metschnikowiaceae(100);g__Metschnikowia(100);g__Metschnikowia_unclassified(100); | Metschnikowia | 13 | Saprotroph | \|Undefined Saprotroph\| | Yeast | NULL | Probable | As nectar-tap_saprotroph (Põlme et al. 2020); Decay sugar-rich substrates (Põlme et al. 2020) | Põlme S, et al. 2020. Fungal Diversity 105:1-16 ((https://doi.org/10.1007/s13225-020-00466-2)) |
| Otu33128 | 1 | k__Fungi(100);p__Ascomycota(100);c__Saccharomycetes(100);o__Saccharomycetales(100);f__Metschnikowiaceae(100);g__Metschnikowia(100);g__Metschnikowia_unclassified(100); | Metschnikowia | 13 | Saprotroph | \|Undefined Saprotroph\| | Yeast | NULL | Probable | As nectar-tap_saprotroph (Põlme et al. 2020); Decay sugar-rich substrates (Põlme et al. 2020) | Põlme S, et al. 2020. Fungal Diversity 105:1-16 ((https://doi.org/10.1007/s13225-020-00466-2)) |
| Otu33129 | 1 | k__Fungi(100);p__Ascomycota(100);c__Saccharomycetes(100);o__Saccharomycetales(100);f__Metschnikowiaceae(100);g__Metschnikowia(100);g__Metschnikowia_unclassified(100); | Metschnikowia | 13 | Saprotroph | \|Undefined Saprotroph\| | Yeast | NULL | Probable | As nectar-tap_saprotroph (Põlme et al. 2020); Decay sugar-rich substrates (Põlme et al. 2020) | Põlme S, et al. 2020. Fungal Diversity 105:1-16 ((https://doi.org/10.1007/s13225-020-00466-2)) |
| Otu33130 | 1 | k__Fungi(100);p__Ascomycota(100);c__Saccharomycetes(100);o__Saccharomycetales(100);f__Metschnikowiaceae(100);g__Metschnikowia(100);g__Metschnikowia_unclassified(100); | Metschnikowia | 13 | Saprotroph | \|Undefined Saprotroph\| | Yeast | NULL | Probable | As nectar-tap_saprotroph (Põlme et al. 2020); Decay sugar-rich substrates (Põlme et al. 2020) | Põlme S, et al. 2020. Fungal Diversity 105:1-16 ((https://doi.org/10.1007/s13225-020-00466-2)) |
| Otu33131 | 1 | k__Fungi(100);p__Ascomycota(100);c__Saccharomycetes(100);o__Saccharomycetales(100);f__Metschnikowiaceae(100);g__Metschnikowia(100);g__Metschnikowia_unclassified(100); | Metschnikowia | 13 | Saprotroph | \|Undefined Saprotroph\| | Yeast | NULL | Probable | As nectar-tap_saprotroph (Põlme et al. 2020); Decay sugar-rich substrates (Põlme et al. 2020) | Põlme S, et al. 2020. Fungal Diversity 105:1-16 ((https://doi.org/10.1007/s13225-020-00466-2)) |
| Otu33132 | 1 | k__Fungi(100);p__Ascomycota(100);c__Saccharomycetes(100);o__Saccharomycetales(100);f__Metschnikowiaceae(100);g__Metschnikowia(100);g__Metschnikowia_unclassified(100); | Metschnikowia | 13 | Saprotroph | \|Undefined Saprotroph\| | Yeast | NULL | Probable | As nectar-tap_saprotroph (Põlme et al. 2020); Decay sugar-rich substrates (Põlme et al. 2020) | Põlme S, et al. 2020. Fungal Diversity 105:1-16 ((https://doi.org/10.1007/s13225-020-00466-2)) |
| Otu33133 | 1 | k__Fungi(100);p__Ascomycota(100);c__Saccharomycetes(100);o__Saccharomycetales(100);f__Metschnikowiaceae(100);g__Metschnikowia(100);s__Metschnikowia_pulcherrima(100); | Metschnikowia | 13 | Saprotroph | \|Undefined Saprotroph\| | Yeast | NULL | Probable | As nectar-tap_saprotroph (Põlme et al. 2020); Decay sugar-rich substrates (Põlme et al. 2020) | Põlme S, et al. 2020. Fungal Diversity 105:1-16 ((https://doi.org/10.1007/s13225-020-00466-2)) |
| Otu33136 | 1 | k__Fungi(100);p__Ascomycota(100);c__Saccharomycetes(100);o__Saccharomycetales(100);f__Metschnikowiaceae(100);g__Metschnikowia(100);g__Metschnikowia_unclassified(100); | Metschnikowia | 13 | Saprotroph | \|Undefined Saprotroph\| | Yeast | NULL | Probable | As nectar-tap_saprotroph (Põlme et al. 2020); Decay sugar-rich substrates (Põlme et al. 2020) | Põlme S, et al. 2020. Fungal Diversity 105:1-16 ((https://doi.org/10.1007/s13225-020-00466-2)) |
| Otu33137 | 1 | k__Fungi(100);p__Ascomycota(100);c__Saccharomycetes(100);o__Saccharomycetales(100);f__Metschnikowiaceae(100);g__Metschnikowia(100);s__Metschnikowia_pulcherrima(100); | Metschnikowia | 13 | Saprotroph | \|Undefined Saprotroph\| | Yeast | NULL | Probable | As nectar-tap_saprotroph (Põlme et al. 2020); Decay sugar-rich substrates (Põlme et al. 2020) | Põlme S, et al. 2020. Fungal Diversity 105:1-16 ((https://doi.org/10.1007/s13225-020-00466-2)) |
| Otu33138 | 1 | k__Fungi(100);p__Ascomycota(100);c__Saccharomycetes(100);o__Saccharomycetales(100);f__Metschnikowiaceae(100);g__Metschnikowia(100);g__Metschnikowia_unclassified(100); | Metschnikowia | 13 | Saprotroph | \|Undefined Saprotroph\| | Yeast | NULL | Probable | As nectar-tap_saprotroph (Põlme et al. 2020); Decay sugar-rich substrates (Põlme et al. 2020) | Põlme S, et al. 2020. Fungal Diversity 105:1-16 ((https://doi.org/10.1007/s13225-020-00466-2)) |
| Otu33139 | 1 | k__Fungi(100);p__Ascomycota(100);c__Saccharomycetes(100);o__Saccharomycetales(100);f__Metschnikowiaceae(100);g__Metschnikowia(100);s__Metschnikowia_pulcherrima(100); | Metschnikowia | 13 | Saprotroph | \|Undefined Saprotroph\| | Yeast | NULL | Probable | As nectar-tap_saprotroph (Põlme et al. 2020); Decay sugar-rich substrates (Põlme et al. 2020) | Põlme S, et al. 2020. Fungal Diversity 105:1-16 ((https://doi.org/10.1007/s13225-020-00466-2)) |
| Otu33140 | 1 | k__Fungi(100);p__Ascomycota(100);c__Saccharomycetes(100);o__Saccharomycetales(100);f__Metschnikowiaceae(100);g__Metschnikowia(100);s__Metschnikowia_pulcherrima(100); | Metschnikowia | 13 | Saprotroph | \|Undefined Saprotroph\| | Yeast | NULL | Probable | As nectar-tap_saprotroph (Põlme et al. 2020); Decay sugar-rich substrates (Põlme et al. 2020) | Põlme S, et al. 2020. Fungal Diversity 105:1-16 ((https://doi.org/10.1007/s13225-020-00466-2)) |
| Otu33141 | 1 | k__Fungi(100);p__Ascomycota(100);c__Saccharomycetes(100);o__Saccharomycetales(100);f__Metschnikowiaceae(100);g__Metschnikowia(100);s__Metschnikowia_pulcherrima(100); | Metschnikowia | 13 | Saprotroph | \|Undefined Saprotroph\| | Yeast | NULL | Probable | As nectar-tap_saprotroph (Põlme et al. 2020); Decay sugar-rich substrates (Põlme et al. 2020) | Põlme S, et al. 2020. Fungal Diversity 105:1-16 ((https://doi.org/10.1007/s13225-020-00466-2)) |
| Otu33142 | 1 | k__Fungi(100);p__Ascomycota(100);c__Saccharomycetes(100);o__Saccharomycetales(100);f__Metschnikowiaceae(100);g__Metschnikowia(100);s__Metschnikowia_pulcherrima(100); | Metschnikowia | 13 | Saprotroph | \|Undefined Saprotroph\| | Yeast | NULL | Probable | As nectar-tap_saprotroph (Põlme et al. 2020); Decay sugar-rich substrates (Põlme et al. 2020) | Põlme S, et al. 2020. Fungal Diversity 105:1-16 ((https://doi.org/10.1007/s13225-020-00466-2)) |
| Otu33143 | 1 | k__Fungi(100);p__Ascomycota(100);c__Saccharomycetes(100);o__Saccharomycetales(100);f__Metschnikowiaceae(100);g__Metschnikowia(100);s__Metschnikowia_pulcherrima(100); | Metschnikowia | 13 | Saprotroph | \|Undefined Saprotroph\| | Yeast | NULL | Probable | As nectar-tap_saprotroph (Põlme et al. 2020); Decay sugar-rich substrates (Põlme et al. 2020) | Põlme S, et al. 2020. Fungal Diversity 105:1-16 ((https://doi.org/10.1007/s13225-020-00466-2)) |
| Otu33144 | 1 | k__Fungi(100);p__Ascomycota(100);c__Saccharomycetes(100);o__Saccharomycetales(100);f__Metschnikowiaceae(100);g__Metschnikowia(100);s__Metschnikowia_pulcherrima(100); | Metschnikowia | 13 | Saprotroph | \|Undefined Saprotroph\| | Yeast | NULL | Probable | As nectar-tap_saprotroph (Põlme et al. 2020); Decay sugar-rich substrates (Põlme et al. 2020) | Põlme S, et al. 2020. Fungal Diversity 105:1-16 ((https://doi.org/10.1007/s13225-020-00466-2)) |
| Otu33145 | 1 | k__Fungi(100);p__Ascomycota(100);c__Saccharomycetes(100);o__Saccharomycetales(100);f__Metschnikowiaceae(100);g__Metschnikowia(100);s__Metschnikowia_pulcherrima(100); | Metschnikowia | 13 | Saprotroph | \|Undefined Saprotroph\| | Yeast | NULL | Probable | As nectar-tap_saprotroph (Põlme et al. 2020); Decay sugar-rich substrates (Põlme et al. 2020) | Põlme S, et al. 2020. Fungal Diversity 105:1-16 ((https://doi.org/10.1007/s13225-020-00466-2)) |
| Otu33146 | 1 | k__Fungi(100);p__Ascomycota(100);c__Saccharomycetes(100);o__Saccharomycetales(100);f__Metschnikowiaceae(100);g__Metschnikowia(100);s__Metschnikowia_pulcherrima(100); | Metschnikowia | 13 | Saprotroph | \|Undefined Saprotroph\| | Yeast | NULL | Probable | As nectar-tap_saprotroph (Põlme et al. 2020); Decay sugar-rich substrates (Põlme et al. 2020) | Põlme S, et al. 2020. Fungal Diversity 105:1-16 ((https://doi.org/10.1007/s13225-020-00466-2)) |
| Otu33148 | 1 | k__Fungi(100);p__Ascomycota(100);c__Saccharomycetes(100);o__Saccharomycetales(100);f__Metschnikowiaceae(100);g__Metschnikowia(100);g__Metschnikowia_unclassified(100); | Metschnikowia | 13 | Saprotroph | \|Undefined Saprotroph\| | Yeast | NULL | Probable | As nectar-tap_saprotroph (Põlme et al. 2020); Decay sugar-rich substrates (Põlme et al. 2020) | Põlme S, et al. 2020. Fungal Diversity 105:1-16 ((https://doi.org/10.1007/s13225-020-00466-2)) |
| Otu33149 | 1 | k__Fungi(100);p__Ascomycota(100);c__Saccharomycetes(100);o__Saccharomycetales(100);f__Metschnikowiaceae(100);g__Metschnikowia(100);s__Metschnikowia_pulcherrima(100); | Metschnikowia | 13 | Saprotroph | \|Undefined Saprotroph\| | Yeast | NULL | Probable | As nectar-tap_saprotroph (Põlme et al. 2020); Decay sugar-rich substrates (Põlme et al. 2020) | Põlme S, et al. 2020. Fungal Diversity 105:1-16 ((https://doi.org/10.1007/s13225-020-00466-2)) |
| Otu33150 | 1 | k__Fungi(100);p__Ascomycota(100);c__Saccharomycetes(100);o__Saccharomycetales(100);f__Metschnikowiaceae(100);g__Metschnikowia(100);g__Metschnikowia_unclassified(100); | Metschnikowia | 13 | Saprotroph | \|Undefined Saprotroph\| | Yeast | NULL | Probable | As nectar-tap_saprotroph (Põlme et al. 2020); Decay sugar-rich substrates (Põlme et al. 2020) | Põlme S, et al. 2020. Fungal Diversity 105:1-16 ((https://doi.org/10.1007/s13225-020-00466-2)) |
| Otu33151 | 1 | k__Fungi(100);p__Ascomycota(100);c__Saccharomycetes(100);o__Saccharomycetales(100);f__Metschnikowiaceae(100);g__Metschnikowia(100);s__Metschnikowia_pulcherrima(100); | Metschnikowia | 13 | Saprotroph | \|Undefined Saprotroph\| | Yeast | NULL | Probable | As nectar-tap_saprotroph (Põlme et al. 2020); Decay sugar-rich substrates (Põlme et al. 2020) | Põlme S, et al. 2020. Fungal Diversity 105:1-16 ((https://doi.org/10.1007/s13225-020-00466-2)) |
| Otu33152 | 1 | k__Fungi(100);p__Ascomycota(100);c__Saccharomycetes(100);o__Saccharomycetales(100);f__Metschnikowiaceae(100);g__Metschnikowia(100);s__Metschnikowia_pulcherrima(100); | Metschnikowia | 13 | Saprotroph | \|Undefined Saprotroph\| | Yeast | NULL | Probable | As nectar-tap_saprotroph (Põlme et al. 2020); Decay sugar-rich substrates (Põlme et al. 2020) | Põlme S, et al. 2020. Fungal Diversity 105:1-16 ((https://doi.org/10.1007/s13225-020-00466-2)) |
| Otu33153 | 1 | k__Fungi(100);p__Ascomycota(100);c__Saccharomycetes(100);o__Saccharomycetales(100);f__Metschnikowiaceae(100);g__Metschnikowia(100);g__Metschnikowia_unclassified(100); | Metschnikowia | 13 | Saprotroph | \|Undefined Saprotroph\| | Yeast | NULL | Probable | As nectar-tap_saprotroph (Põlme et al. 2020); Decay sugar-rich substrates (Põlme et al. 2020) | Põlme S, et al. 2020. Fungal Diversity 105:1-16 ((https://doi.org/10.1007/s13225-020-00466-2)) |
| Otu33154 | 1 | k__Fungi(100);p__Ascomycota(100);c__Saccharomycetes(100);o__Saccharomycetales(100);f__Metschnikowiaceae(100);g__Metschnikowia(100);g__Metschnikowia_unclassified(100); | Metschnikowia | 13 | Saprotroph | \|Undefined Saprotroph\| | Yeast | NULL | Probable | As nectar-tap_saprotroph (Põlme et al. 2020); Decay sugar-rich substrates (Põlme et al. 2020) | Põlme S, et al. 2020. Fungal Diversity 105:1-16 ((https://doi.org/10.1007/s13225-020-00466-2)) |
| Otu33155 | 1 | k__Fungi(100);p__Ascomycota(100);c__Saccharomycetes(100);o__Saccharomycetales(100);f__Metschnikowiaceae(100);g__Metschnikowia(100);s__Metschnikowia_pulcherrima(100); | Metschnikowia | 13 | Saprotroph | \|Undefined Saprotroph\| | Yeast | NULL | Probable | As nectar-tap_saprotroph (Põlme et al. 2020); Decay sugar-rich substrates (Põlme et al. 2020) | Põlme S, et al. 2020. Fungal Diversity 105:1-16 ((https://doi.org/10.1007/s13225-020-00466-2)) |
| Otu33156 | 1 | k__Fungi(100);p__Ascomycota(100);c__Saccharomycetes(100);o__Saccharomycetales(100);f__Metschnikowiaceae(100);g__Metschnikowia(100);s__Metschnikowia_pulcherrima(100); | Metschnikowia | 13 | Saprotroph | \|Undefined Saprotroph\| | Yeast | NULL | Probable | As nectar-tap_saprotroph (Põlme et al. 2020); Decay sugar-rich substrates (Põlme et al. 2020) | Põlme S, et al. 2020. Fungal Diversity 105:1-16 ((https://doi.org/10.1007/s13225-020-00466-2)) |
| Otu33157 | 1 | k__Fungi(100);p__Ascomycota(100);c__Saccharomycetes(100);o__Saccharomycetales(100);f__Metschnikowiaceae(100);g__Metschnikowia(100);s__Metschnikowia_pulcherrima(100); | Metschnikowia | 13 | Saprotroph | \|Undefined Saprotroph\| | Yeast | NULL | Probable | As nectar-tap_saprotroph (Põlme et al. 2020); Decay sugar-rich substrates (Põlme et al. 2020) | Põlme S, et al. 2020. Fungal Diversity 105:1-16 ((https://doi.org/10.1007/s13225-020-00466-2)) |
| Otu33158 | 1 | k__Fungi(100);p__Ascomycota(100);c__Saccharomycetes(100);o__Saccharomycetales(100);f__Metschnikowiaceae(100);g__Metschnikowia(100);s__Metschnikowia_pulcherrima(100); | Metschnikowia | 13 | Saprotroph | \|Undefined Saprotroph\| | Yeast | NULL | Probable | As nectar-tap_saprotroph (Põlme et al. 2020); Decay sugar-rich substrates (Põlme et al. 2020) | Põlme S, et al. 2020. Fungal Diversity 105:1-16 ((https://doi.org/10.1007/s13225-020-00466-2)) |
| Otu33159 | 1 | k__Fungi(100);p__Ascomycota(100);c__Saccharomycetes(100);o__Saccharomycetales(100);f__Saccharomycetales_family_Incertae_sedis(100);g__Candida(100);g__Candida_unclassified(100); | Candida | 13 | Pathotroph-Saprotroph-Symbiotroph | Animal Pathogen-Endophyte-Endosymbiont-Epiphyte-\|Undefined Saprotroph\| | Dimorphic Yeast-Yeast | NULL | Probable | As nectar-tap_saprotroph (Põlme et al. 2020); Decay sugar-rich substrates (Põlme et al. 2020); This asexual genus is polyphyletic and contain species that function very differently depend on what they are. Caution should be taken when interpreting guilds for Candida species. | Manolakaki D, et al. 2010. Virulence. 1: 367-375 ((https://doi.org/10.4161/viru.1.5.12796)); Mekbib SB, et al. 2011. Tropical Plant Pathology 36:233-240 ((https://doi.org/10.1590/S1982-56762011000400004)); Põlme S, et al. 2020. Fungal Diversity 105:1-16 ((https://doi.org/10.1007/s13225-020-00466-2)) |
| Otu33161 | 1 | k__Fungi(100);p__Ascomycota(100);c__Saccharomycetes(100);o__Saccharomycetales(100);f__Metschnikowiaceae(100);g__Metschnikowia(100);s__Metschnikowia_pulcherrima(100); | Metschnikowia | 13 | Saprotroph | \|Undefined Saprotroph\| | Yeast | NULL | Probable | As nectar-tap_saprotroph (Põlme et al. 2020); Decay sugar-rich substrates (Põlme et al. 2020) | Põlme S, et al. 2020. Fungal Diversity 105:1-16 ((https://doi.org/10.1007/s13225-020-00466-2)) |
| Otu33164 | 1 | k__Fungi(100);p__Ascomycota(100);c__Saccharomycetes(100);o__Saccharomycetales(100);f__Metschnikowiaceae(100);g__Metschnikowia(100);s__Metschnikowia_pulcherrima(100); | Metschnikowia | 13 | Saprotroph | \|Undefined Saprotroph\| | Yeast | NULL | Probable | As nectar-tap_saprotroph (Põlme et al. 2020); Decay sugar-rich substrates (Põlme et al. 2020) | Põlme S, et al. 2020. Fungal Diversity 105:1-16 ((https://doi.org/10.1007/s13225-020-00466-2)) |
| Otu33165 | 1 | k__Fungi(100);p__Ascomycota(100);c__Saccharomycetes(100);o__Saccharomycetales(100);f__Metschnikowiaceae(100);g__Metschnikowia(100);s__Metschnikowia_pulcherrima(100); | Metschnikowia | 13 | Saprotroph | \|Undefined Saprotroph\| | Yeast | NULL | Probable | As nectar-tap_saprotroph (Põlme et al. 2020); Decay sugar-rich substrates (Põlme et al. 2020) | Põlme S, et al. 2020. Fungal Diversity 105:1-16 ((https://doi.org/10.1007/s13225-020-00466-2)) |
| Otu33167 | 1 | k__Fungi(100);p__Ascomycota(100);c__Saccharomycetes(100);o__Saccharomycetales(100);f__Saccharomycetales_family_Incertae_sedis(100);f__Saccharomycetales_family_Incertae_sedis_unclassified(100);f__Saccharomycetales_family_Incertae_sedis_unclassified(100); | Saccharomycetales | 7 | Saprotroph | Undefined Saprotroph | Yeast | NULL | Possible | NULL | Sterkenburg E, et al. 2015. New Phytologist 207:1145-1158 ((https://doi.org/10.1111/nph.13426)) |
| Otu33169 | 1 | k__Fungi(100);p__Ascomycota(100);c__Saccharomycetes(100);o__Saccharomycetales(100);f__Metschnikowiaceae(100);g__Metschnikowia(100);s__Metschnikowia_pulcherrima(100); | Metschnikowia | 13 | Saprotroph | \|Undefined Saprotroph\| | Yeast | NULL | Probable | As nectar-tap_saprotroph (Põlme et al. 2020); Decay sugar-rich substrates (Põlme et al. 2020) | Põlme S, et al. 2020. Fungal Diversity 105:1-16 ((https://doi.org/10.1007/s13225-020-00466-2)) |
| Otu33170 | 1 | k__Fungi(100);p__Ascomycota(100);c__Saccharomycetes(100);o__Saccharomycetales(100);f__Metschnikowiaceae(100);g__Metschnikowia(100);s__Metschnikowia_pulcherrima(100); | Metschnikowia | 13 | Saprotroph | \|Undefined Saprotroph\| | Yeast | NULL | Probable | As nectar-tap_saprotroph (Põlme et al. 2020); Decay sugar-rich substrates (Põlme et al. 2020) | Põlme S, et al. 2020. Fungal Diversity 105:1-16 ((https://doi.org/10.1007/s13225-020-00466-2)) |
| Otu33171 | 1 | k__Fungi(100);p__Ascomycota(100);c__Saccharomycetes(100);o__Saccharomycetales(100);f__Metschnikowiaceae(100);g__Metschnikowia(100);g__Metschnikowia_unclassified(100); | Metschnikowia | 13 | Saprotroph | \|Undefined Saprotroph\| | Yeast | NULL | Probable | As nectar-tap_saprotroph (Põlme et al. 2020); Decay sugar-rich substrates (Põlme et al. 2020) | Põlme S, et al. 2020. Fungal Diversity 105:1-16 ((https://doi.org/10.1007/s13225-020-00466-2)) |
| Otu33172 | 1 | k__Fungi(100);p__Ascomycota(100);c__Saccharomycetes(100);o__Saccharomycetales(100);f__Metschnikowiaceae(100);g__Metschnikowia(100);s__Metschnikowia_pulcherrima(100); | Metschnikowia | 13 | Saprotroph | \|Undefined Saprotroph\| | Yeast | NULL | Probable | As nectar-tap_saprotroph (Põlme et al. 2020); Decay sugar-rich substrates (Põlme et al. 2020) | Põlme S, et al. 2020. Fungal Diversity 105:1-16 ((https://doi.org/10.1007/s13225-020-00466-2)) |
| Otu33173 | 1 | k__Fungi(100);p__Ascomycota(100);c__Saccharomycetes(100);o__Saccharomycetales(100);f__Metschnikowiaceae(100);g__Metschnikowia(100);s__Metschnikowia_pulcherrima(100); | Metschnikowia | 13 | Saprotroph | \|Undefined Saprotroph\| | Yeast | NULL | Probable | As nectar-tap_saprotroph (Põlme et al. 2020); Decay sugar-rich substrates (Põlme et al. 2020) | Põlme S, et al. 2020. Fungal Diversity 105:1-16 ((https://doi.org/10.1007/s13225-020-00466-2)) |
| Otu33174 | 1 | k__Fungi(100);p__Ascomycota(100);c__Saccharomycetes(100);o__Saccharomycetales(100);f__Metschnikowiaceae(100);g__Metschnikowia(100);g__Metschnikowia_unclassified(100); | Metschnikowia | 13 | Saprotroph | \|Undefined Saprotroph\| | Yeast | NULL | Probable | As nectar-tap_saprotroph (Põlme et al. 2020); Decay sugar-rich substrates (Põlme et al. 2020) | Põlme S, et al. 2020. Fungal Diversity 105:1-16 ((https://doi.org/10.1007/s13225-020-00466-2)) |
| Otu33176 | 1 | k__Fungi(100);p__Ascomycota(100);c__Saccharomycetes(100);o__Saccharomycetales(100);f__Metschnikowiaceae(100);g__Metschnikowia(100);s__Metschnikowia_pulcherrima(100); | Metschnikowia | 13 | Saprotroph | \|Undefined Saprotroph\| | Yeast | NULL | Probable | As nectar-tap_saprotroph (Põlme et al. 2020); Decay sugar-rich substrates (Põlme et al. 2020) | Põlme S, et al. 2020. Fungal Diversity 105:1-16 ((https://doi.org/10.1007/s13225-020-00466-2)) |
| Otu33177 | 1 | k__Fungi(100);p__Ascomycota(100);c__Saccharomycetes(100);o__Saccharomycetales(100);f__Metschnikowiaceae(100);g__Metschnikowia(100);s__Metschnikowia_pulcherrima(100); | Metschnikowia | 13 | Saprotroph | \|Undefined Saprotroph\| | Yeast | NULL | Probable | As nectar-tap_saprotroph (Põlme et al. 2020); Decay sugar-rich substrates (Põlme et al. 2020) | Põlme S, et al. 2020. Fungal Diversity 105:1-16 ((https://doi.org/10.1007/s13225-020-00466-2)) |
| Otu33178 | 1 | k__Fungi(100);p__Ascomycota(100);c__Saccharomycetes(100);o__Saccharomycetales(100);f__Metschnikowiaceae(100);g__Metschnikowia(100);s__Metschnikowia_pulcherrima(100); | Metschnikowia | 13 | Saprotroph | \|Undefined Saprotroph\| | Yeast | NULL | Probable | As nectar-tap_saprotroph (Põlme et al. 2020); Decay sugar-rich substrates (Põlme et al. 2020) | Põlme S, et al. 2020. Fungal Diversity 105:1-16 ((https://doi.org/10.1007/s13225-020-00466-2)) |
| Otu33179 | 1 | k__Fungi(100);p__Ascomycota(100);c__Saccharomycetes(100);o__Saccharomycetales(100);f__Metschnikowiaceae(100);g__Metschnikowia(100);s__Metschnikowia_pulcherrima(100); | Metschnikowia | 13 | Saprotroph | \|Undefined Saprotroph\| | Yeast | NULL | Probable | As nectar-tap_saprotroph (Põlme et al. 2020); Decay sugar-rich substrates (Põlme et al. 2020) | Põlme S, et al. 2020. Fungal Diversity 105:1-16 ((https://doi.org/10.1007/s13225-020-00466-2)) |
| Otu33180 | 1 | k__Fungi(100);p__Ascomycota(100);c__Saccharomycetes(100);o__Saccharomycetales(100);f__Metschnikowiaceae(100);g__Metschnikowia(100);s__Metschnikowia_pulcherrima(100); | Metschnikowia | 13 | Saprotroph | \|Undefined Saprotroph\| | Yeast | NULL | Probable | As nectar-tap_saprotroph (Põlme et al. 2020); Decay sugar-rich substrates (Põlme et al. 2020) | Põlme S, et al. 2020. Fungal Diversity 105:1-16 ((https://doi.org/10.1007/s13225-020-00466-2)) |
| Otu33186 | 1 | k__Fungi(100);p__Ascomycota(100);c__Saccharomycetes(100);o__Saccharomycetales(100);f__Metschnikowiaceae(100);g__Metschnikowia(100);s__Metschnikowia_pulcherrima(100); | Metschnikowia | 13 | Saprotroph | \|Undefined Saprotroph\| | Yeast | NULL | Probable | As nectar-tap_saprotroph (Põlme et al. 2020); Decay sugar-rich substrates (Põlme et al. 2020) | Põlme S, et al. 2020. Fungal Diversity 105:1-16 ((https://doi.org/10.1007/s13225-020-00466-2)) |
| Otu33187 | 1 | k__Fungi(100);p__Ascomycota(100);c__Saccharomycetes(100);o__Saccharomycetales(100);f__Saccharomycetales_family_Incertae_sedis(100);f__Saccharomycetales_family_Incertae_sedis_unclassified(100);f__Saccharomycetales_family_Incertae_sedis_unclassified(100); | Saccharomycetales | 7 | Saprotroph | Undefined Saprotroph | Yeast | NULL | Possible | NULL | Sterkenburg E, et al. 2015. New Phytologist 207:1145-1158 ((https://doi.org/10.1111/nph.13426)) |

| | | | | | | | | | | | |
|---|---|---|---|---|---|---|---|---|---|---|---|
| Otu33188 | 1 | k__Fungi(100);p__Ascomycota(100);c__Saccharomycetes(100);o__Saccharomycetales(100);f__Metschnikowiaceae(100);g__Metschnikowia(100);g__Metschnikowia_unclassified(100); | Metschnikowia | 13 | Saprotroph | \|Undefined Saprotroph\| | Yeast | NULL | Probable | As nectar-tap_saprotroph (Põlme et al. 2020); Decay sugar-rich substrates (Põlme et al. 2020) | Põlme S, et al. 2020. Fungal Diversity 105:1-16 ((https://doi.org/10.1007/s13225-020-00466-2)) |
| Otu33189 | 1 | k__Fungi(100);p__Ascomycota(100);c__Saccharomycetes(100);o__Saccharomycetales(100);f__Saccharomycetales_family_Incertae_sedis(100);g__Candida(100);g__Candida_unclassified(100); | Candida | 13 | Pathotroph-Saprotroph-Symbiotroph | Animal Pathogen-Endophyte-Endosymbiont-Epiphyte-\|Undefined Saprotroph\| | Dimorphic Yeast-Yeast | NULL | Probable | As nectar-tap_saprotroph (Põlme et al. 2020); Decay sugar-rich substrates (Põlme et al. 2020); This asexual genus is polyphyletic and contain species that function very differently depend on what they are. Caution should be taken when interpreting guilds for Candida species. | Manolakaki D, et al. 2010. Virulence. 1: 367-375 ((https://doi.org/10.4161/viru.1.5.12796)); Mekbib SB, et al. 2011. Tropical Plant Pathology 36:233-240 ((https://doi.org/10.1590/S1982-56762011000400004)); Põlme S, et al. 2020. Fungal Diversity 105:1-16 ((https://doi.org/10.1007/s13225-020-00466-2)) |
| Otu33190 | 1 | k__Fungi(100);p__Ascomycota(100);c__Saccharomycetes(100);o__Saccharomycetales(100);f__Metschnikowiaceae(100);g__Metschnikowia(100);s__Metschnikowia_pulcherrima(100); | Metschnikowia | 13 | Saprotroph | \|Undefined Saprotroph\| | Yeast | NULL | Probable | As nectar-tap_saprotroph (Põlme et al. 2020); Decay sugar-rich substrates (Põlme et al. 2020) | Põlme S, et al. 2020. Fungal Diversity 105:1-16 ((https://doi.org/10.1007/s13225-020-00466-2)) |
| Otu33191 | 1 | k__Fungi(100);p__Ascomycota(100);c__Saccharomycetes(100);o__Saccharomycetales(100);f__Saccharomycetales_family_Incertae_sedis(100);g__Candida(100);s__Candida_cretensis(100); | Candida | 13 | Pathotroph-Saprotroph-Symbiotroph | Animal Pathogen-Endophyte-Endosymbiont-Epiphyte-\|Undefined Saprotroph\| | Dimorphic Yeast-Yeast | NULL | Probable | As nectar-tap_saprotroph (Põlme et al. 2020); Decay sugar-rich substrates (Põlme et al. 2020); This asexual genus is polyphyletic and contain species that function very differently depend on what they are. Caution should be taken when interpreting guilds for Candida species. | Manolakaki D, et al. 2010. Virulence. 1: 367-375 ((https://doi.org/10.4161/viru.1.5.12796)); Mekbib SB, et al. 2011. Tropical Plant Pathology 36:233-240 ((https://doi.org/10.1590/S1982-56762011000400004)); Põlme S, et al. 2020. Fungal Diversity 105:1-16 ((https://doi.org/10.1007/s13225-020-00466-2)) |
| Otu33192 | 1 | k__Fungi(100);p__Ascomycota(100);c__Saccharomycetes(100);o__Saccharomycetales(100);f__Metschnikowiaceae(100);g__Metschnikowia(100);s__Metschnikowia_pulcherrima(100); | Metschnikowia | 13 | Saprotroph | \|Undefined Saprotroph\| | Yeast | NULL | Probable | As nectar-tap_saprotroph (Põlme et al. 2020); Decay sugar-rich substrates (Põlme et al. 2020) | Põlme S, et al. 2020. Fungal Diversity 105:1-16 ((https://doi.org/10.1007/s13225-020-00466-2)) |
| Otu33193 | 1 | k__Fungi(100);p__Ascomycota(100);c__Saccharomycetes(100);o__Saccharomycetales(100);o__Saccharomycetales_unclassified(100);o__Saccharomycetales_unclassified(100);o__Saccharomycetales_unclassified(100); | Saccharomycetales | 7 | Saprotroph | Undefined Saprotroph | Yeast | NULL | Possible | NULL | Sterkenburg E, et al. 2015. New Phytologist 207:1145-1158 ((https://doi.org/10.1111/nph.13426)) |
| Otu33194 | 1 | k__Fungi(100);p__Ascomycota(100);c__Saccharomycetes(100);o__Saccharomycetales(100);f__Metschnikowiaceae(100);g__Metschnikowia(100);s__Metschnikowia_pulcherrima(100); | Metschnikowia | 13 | Saprotroph | \|Undefined Saprotroph\| | Yeast | NULL | Probable | As nectar-tap_saprotroph (Põlme et al. 2020); Decay sugar-rich substrates (Põlme et al. 2020) | Põlme S, et al. 2020. Fungal Diversity 105:1-16 ((https://doi.org/10.1007/s13225-020-00466-2)) |
| Otu33195 | 1 | k__Fungi(100);p__Ascomycota(100);c__Saccharomycetes(100);o__Saccharomycetales(100);f__Metschnikowiaceae(100);g__Metschnikowia(100);s__Metschnikowia_pulcherrima(100); | Metschnikowia | 13 | Saprotroph | \|Undefined Saprotroph\| | Yeast | NULL | Probable | As nectar-tap_saprotroph (Põlme et al. 2020); Decay sugar-rich substrates (Põlme et al. 2020) | Põlme S, et al. 2020. Fungal Diversity 105:1-16 ((https://doi.org/10.1007/s13225-020-00466-2)) |
| Otu33198 | 1 | k__Fungi(100);p__Ascomycota(100);c__Saccharomycetes(100);o__Saccharomycetales(100);f__Metschnikowiaceae(100);g__Metschnikowia(100);s__Metschnikowia_pulcherrima(100); | Metschnikowia | 13 | Saprotroph | \|Undefined Saprotroph\| | Yeast | NULL | Probable | As nectar-tap_saprotroph (Põlme et al. 2020); Decay sugar-rich substrates (Põlme et al. 2020) | Põlme S, et al. 2020. Fungal Diversity 105:1-16 ((https://doi.org/10.1007/s13225-020-00466-2)) |
| Otu33199 | 1 | k__Fungi(100);p__Ascomycota(100);c__Saccharomycetes(100);o__Saccharomycetales(100);f__Saccharomycetales_family_Incertae_sedis(100);g__Candida(100);g__Candida_unclassified(100); | Candida | 13 | Pathotroph-Saprotroph-Symbiotroph | Animal Pathogen-Endophyte-Endosymbiont-Epiphyte-\|Undefined Saprotroph\| | Dimorphic Yeast-Yeast | NULL | Probable | As nectar-tap_saprotroph (Põlme et al. 2020); Decay sugar-rich substrates (Põlme et al. 2020); This asexual genus is polyphyletic and contain species that function very differently depend on what they are. Caution should be taken when interpreting guilds for Candida species. | Manolakaki D, et al. 2010. Virulence. 1: 367-375 ((https://doi.org/10.4161/viru.1.5.12796)); Mekbib SB, et al. 2011. Tropical Plant Pathology 36:233-240 ((https://doi.org/10.1590/S1982-56762011000400004)); Põlme S, et al. 2020. Fungal Diversity 105:1-16 ((https://doi.org/10.1007/s13225-020-00466-2)) |
| Otu33200 | 1 | k__Fungi(100);p__Ascomycota(100);c__Saccharomycetes(100);o__Saccharomycetales(100);f__Metschnikowiaceae(100);g__Metschnikowia(100);g__Metschnikowia_unclassified(100); | Metschnikowia | 13 | Saprotroph | \|Undefined Saprotroph\| | Yeast | NULL | Probable | As nectar-tap_saprotroph (Põlme et al. 2020); Decay sugar-rich substrates (Põlme et al. 2020) | Põlme S, et al. 2020. Fungal Diversity 105:1-16 ((https://doi.org/10.1007/s13225-020-00466-2)) |
| Otu33201 | 1 | k__Fungi(100);p__Ascomycota(100);c__Saccharomycetes(100);o__Saccharomycetales(100);f__Metschnikowiaceae(100);g__Metschnikowia(100);s__Metschnikowia_pulcherrima(100); | Metschnikowia | 13 | Saprotroph | \|Undefined Saprotroph\| | Yeast | NULL | Probable | As nectar-tap_saprotroph (Põlme et al. 2020); Decay sugar-rich substrates (Põlme et al. 2020) | Põlme S, et al. 2020. Fungal Diversity 105:1-16 ((https://doi.org/10.1007/s13225-020-00466-2)) |
| Otu33206 | 1 | k__Fungi(100);p__Ascomycota(100);c__Saccharomycetes(100);o__Saccharomycetales(100);f__Saccharomycetales_family_Incertae_sedis(100);f__Saccharomycetales_family_Incertae_sedis_unclassified(100);f__Saccharomycetales_family_Incertae_sedis_unclassified(100); | Saccharomycetales | 7 | Saprotroph | Undefined Saprotroph | Yeast | NULL | Possible | NULL | Sterkenburg E, et al. 2015. New Phytologist 207:1145-1158 ((https://doi.org/10.1111/nph.13426)) |
| Otu33210 | 1 | k__Fungi(100);p__Ascomycota(100);c__Saccharomycetes(100);o__Saccharomycetales(100);f__Saccharomycetales_family_Incertae_sedis(100);f__Saccharomycetales_family_Incertae_sedis_unclassified(100);f__Saccharomycetales_family_Incertae_sedis_unclassified(100); | Saccharomycetales | 7 | Saprotroph | Undefined Saprotroph | Yeast | NULL | Possible | NULL | Sterkenburg E, et al. 2015. New Phytologist 207:1145-1158 ((https://doi.org/10.1111/nph.13426)) |
| Otu33213 | 1 | k__Fungi(100);p__Ascomycota(100);c__Saccharomycetes(100);o__Saccharomycetales(100);f__Metschnikowiaceae(100);g__Metschnikowia(100);g__Metschnikowia_unclassified(100); | Metschnikowia | 13 | Saprotroph | \|Undefined Saprotroph\| | Yeast | NULL | Probable | As nectar-tap_saprotroph (Põlme et al. 2020); Decay sugar-rich substrates (Põlme et al. 2020) | Põlme S, et al. 2020. Fungal Diversity 105:1-16 ((https://doi.org/10.1007/s13225-020-00466-2)) |
| Otu33215 | 1 | k__Fungi(100);p__Ascomycota(100);c__Saccharomycetes(100);o__Saccharomycetales(100);f__Saccharomycetales_family_Incertae_sedis(100);g__Candida(100);g__Candida_unclassified(100); | Candida | 13 | Pathotroph-Saprotroph-Symbiotroph | Animal Pathogen-Endophyte-Endosymbiont-Epiphyte-\|Undefined Saprotroph\| | Dimorphic Yeast-Yeast | NULL | Probable | As nectar-tap_saprotroph (Põlme et al. 2020); Decay sugar-rich substrates (Põlme et al. 2020); This asexual genus is polyphyletic and contain species that function very differently depend on what they are. Caution should be taken when interpreting guilds for Candida species. | Manolakaki D, et al. 2010. Virulence. 1: 367-375 ((https://doi.org/10.4161/viru.1.5.12796)); Mekbib SB, et al. 2011. Tropical Plant Pathology 36:233-240 ((https://doi.org/10.1590/S1982-56762011000400004)); Põlme S, et al. 2020. Fungal Diversity 105:1-16 ((https://doi.org/10.1007/s13225-020-00466-2)) |
| Otu33217 | 1 | k__Fungi(100);p__Ascomycota(100);c__Saccharomycetes(100);o__Saccharomycetales(100);f__Metschnikowiaceae(100);g__Metschnikowia(100);s__Metschnikowia_pulcherrima(100); | Metschnikowia | 13 | Saprotroph | \|Undefined Saprotroph\| | Yeast | NULL | Probable | As nectar-tap_saprotroph (Põlme et al. 2020); Decay sugar-rich substrates (Põlme et al. 2020) | Põlme S, et al. 2020. Fungal Diversity 105:1-16 ((https://doi.org/10.1007/s13225-020-00466-2)) |
| Otu33226 | 1 | k__Fungi(100);p__Ascomycota(100);c__Saccharomycetes(100);o__Saccharomycetales(100);f__Saccharomycetales_family_Incertae_sedis(100);g__Candida(100);g__Candida_unclassified(100); | Candida | 13 | Pathotroph-Saprotroph-Symbiotroph | Animal Pathogen-Endophyte-Endosymbiont-Epiphyte-\|Undefined Saprotroph\| | Dimorphic Yeast-Yeast | NULL | Probable | As nectar-tap_saprotroph (Põlme et al. 2020); Decay sugar-rich substrates (Põlme et al. 2020); This asexual genus is polyphyletic and contain species that function very differently depend on what they are. Caution should be taken when interpreting guilds for Candida species. | Manolakaki D, et al. 2010. Virulence. 1: 367-375 ((https://doi.org/10.4161/viru.1.5.12796)); Mekbib SB, et al. 2011. Tropical Plant Pathology 36:233-240 ((https://doi.org/10.1590/S1982-56762011000400004)); Põlme S, et al. 2020. Fungal Diversity 105:1-16 ((https://doi.org/10.1007/s13225-020-00466-2)) |
| Otu33227 | 1 | k__Fungi(100);p__Ascomycota(100);c__Saccharomycetes(100);o__Saccharomycetales(100);o__Saccharomycetales_unclassified(100);o__Saccharomycetales_unclassified(100);o__Saccharomycetales_unclassified(100); | Saccharomycetales | 7 | Saprotroph | Undefined Saprotroph | Yeast | NULL | Possible | NULL | Sterkenburg E, et al. 2015. New Phytologist 207:1145-1158 ((https://doi.org/10.1111/nph.13426)) |
| Otu33231 | 1 | k__Fungi(100);p__Ascomycota(100);c__Saccharomycetes(100);o__Saccharomycetales(100);f__Saccharomycetales_family_Incertae_sedis(100);g__Candida(100);g__Candida_unclassified(100); | Candida | 13 | Pathotroph-Saprotroph-Symbiotroph | Animal Pathogen-Endophyte-Endosymbiont-Epiphyte-\|Undefined Saprotroph\| | Dimorphic Yeast-Yeast | NULL | Probable | As nectar-tap_saprotroph (Põlme et al. 2020); Decay sugar-rich substrates (Põlme et al. 2020); This asexual genus is polyphyletic and contain species that function very differently depend on what they are. Caution should be taken when interpreting guilds for Candida species. | Manolakaki D, et al. 2010. Virulence. 1: 367-375 ((https://doi.org/10.4161/viru.1.5.12796)); Mekbib SB, et al. 2011. Tropical Plant Pathology 36:233-240 ((https://doi.org/10.1590/S1982-56762011000400004)); Põlme S, et al. 2020. Fungal Diversity 105:1-16 ((https://doi.org/10.1007/s13225-020-00466-2)) |
| Otu33234 | 1 | k__Fungi(100);p__Ascomycota(100);c__Saccharomycetes(100);o__Saccharomycetales(100);f__Saccharomycetales_family_Incertae_sedis(100);g__Candida(100);g__Candida_unclassified(100); | Candida | 13 | Pathotroph-Saprotroph-Symbiotroph | Animal Pathogen-Endophyte-Endosymbiont-Epiphyte-\|Undefined Saprotroph\| | Dimorphic Yeast-Yeast | NULL | Probable | As nectar-tap_saprotroph (Põlme et al. 2020); Decay sugar-rich substrates (Põlme et al. 2020); This asexual genus is polyphyletic and contain species that function very differently depend on what they are. Caution should be taken when interpreting guilds for Candida species. | Manolakaki D, et al. 2010. Virulence. 1: 367-375 ((https://doi.org/10.4161/viru.1.5.12796)); Mekbib SB, et al. 2011. Tropical Plant Pathology 36:233-240 ((https://doi.org/10.1590/S1982-56762011000400004)); Põlme S, et al. 2020. Fungal Diversity 105:1-16 ((https://doi.org/10.1007/s13225-020-00466-2)) |
| Otu33236 | 1 | k__Fungi(100);p__Ascomycota(100);c__Saccharomycetes(100);o__Saccharomycetales(100);f__Saccharomycetales_family_Incertae_sedis(100);g__Candida(100);g__Candida_unclassified(100); | Candida | 13 | Pathotroph-Saprotroph-Symbiotroph | Animal Pathogen-Endophyte-Endosymbiont-Epiphyte-\|Undefined Saprotroph\| | Dimorphic Yeast-Yeast | NULL | Probable | As nectar-tap_saprotroph (Põlme et al. 2020); Decay sugar-rich substrates (Põlme et al. 2020); This asexual genus is polyphyletic and contain species that function very differently depend on what they are. Caution should be taken when interpreting guilds for Candida species. | Manolakaki D, et al. 2010. Virulence. 1: 367-375 ((https://doi.org/10.4161/viru.1.5.12796)); Mekbib SB, et al. 2011. Tropical Plant Pathology 36:233-240 ((https://doi.org/10.1590/S1982-56762011000400004)); Põlme S, et al. 2020. Fungal Diversity 105:1-16 ((https://doi.org/10.1007/s13225-020-00466-2)) |
| Otu33237 | 1 | k__Fungi(100);p__Ascomycota(100);c__Saccharomycetes(100);o__Saccharomycetales(100);f__Metschnikowiaceae(100);g__Metschnikowia(100);g__Metschnikowia_unclassified(100); | Metschnikowia | 13 | Saprotroph | \|Undefined Saprotroph\| | Yeast | NULL | Probable | As nectar-tap_saprotroph (Põlme et al. 2020); Decay sugar-rich substrates (Põlme et al. 2020) | Põlme S, et al. 2020. Fungal Diversity 105:1-16 ((https://doi.org/10.1007/s13225-020-00466-2)) |
| Otu33250 | 1 | k__Fungi(100);p__Ascomycota(100);c__Saccharomycetes(100);o__Saccharomycetales(100);f__Saccharomycetales_family_Incertae_sedis(100);f__Saccharomycetales_family_Incertae_sedis_unclassified(100);f__Saccharomycetales_family_Incertae_sedis_unclassified(100); | Saccharomycetales | 7 | Saprotroph | Undefined Saprotroph | Yeast | NULL | Possible | NULL | Sterkenburg E, et al. 2015. New Phytologist 207:1145-1158 ((https://doi.org/10.1111/nph.13426)) |
| Otu33255 | 1 | k__Fungi(100);p__Ascomycota(100);c__Saccharomycetes(100);o__Saccharomycetales(100);f__Metschnikowiaceae(100);g__Metschnikowia(100);s__Metschnikowia_pulcherrima(100); | Metschnikowia | 13 | Saprotroph | \|Undefined Saprotroph\| | Yeast | NULL | Probable | As nectar-tap_saprotroph (Põlme et al. 2020); Decay sugar-rich substrates (Põlme et al. 2020) | Põlme S, et al. 2020. Fungal Diversity 105:1-16 ((https://doi.org/10.1007/s13225-020-00466-2)) |
| Otu33256 | 1 | k__Fungi(100);p__Ascomycota(100);c__Saccharomycetes(100);o__Saccharomycetales(100);f__Metschnikowiaceae(100);g__Metschnikowia(100);s__Metschnikowia_pulcherrima(100); | Metschnikowia | 13 | Saprotroph | \|Undefined Saprotroph\| | Yeast | NULL | Probable | As nectar-tap_saprotroph (Põlme et al. 2020); Decay sugar-rich substrates (Põlme et al. 2020) | Põlme S, et al. 2020. Fungal Diversity 105:1-16 ((https://doi.org/10.1007/s13225-020-00466-2)) |
| Otu33257 | 1 | k__Fungi(100);p__Ascomycota(100);c__Saccharomycetes(100);o__Saccharomycetales(100);f__Metschnikowiaceae(100);g__Metschnikowia(100);s__Metschnikowia_pulcherrima(100); | Metschnikowia | 13 | Saprotroph | \|Undefined Saprotroph\| | Yeast | NULL | Probable | As nectar-tap_saprotroph (Põlme et al. 2020); Decay sugar-rich substrates (Põlme et al. 2020) | Põlme S, et al. 2020. Fungal Diversity 105:1-16 ((https://doi.org/10.1007/s13225-020-00466-2)) |
| Otu33258 | 1 | k__Fungi(100);p__Ascomycota(100);c__Saccharomycetes(100);o__Saccharomycetales(100);f__Metschnikowiaceae(100);g__Metschnikowia(100);s__Metschnikowia_pulcherrima(100); | Metschnikowia | 13 | Saprotroph | \|Undefined Saprotroph\| | Yeast | NULL | Probable | As nectar-tap_saprotroph (Põlme et al. 2020); Decay sugar-rich substrates (Põlme et al. 2020) | Põlme S, et al. 2020. Fungal Diversity 105:1-16 ((https://doi.org/10.1007/s13225-020-00466-2)) |
| Otu33259 | 1 | k__Fungi(100);p__Ascomycota(100);c__Saccharomycetes(100);o__Saccharomycetales(100);f__Metschnikowiaceae(100);g__Metschnikowia(100);s__Metschnikowia_pulcherrima(100); | Metschnikowia | 13 | Saprotroph | \|Undefined Saprotroph\| | Yeast | NULL | Probable | As nectar-tap_saprotroph (Põlme et al. 2020); Decay sugar-rich substrates (Põlme et al. 2020) | Põlme S, et al. 2020. Fungal Diversity 105:1-16 ((https://doi.org/10.1007/s13225-020-00466-2)) |
| Otu33260 | 1 | k__Fungi(100);p__Ascomycota(100);c__Saccharomycetes(100);o__Saccharomycetales(100);f__Metschnikowiaceae(100);g__Metschnikowia(100);s__Metschnikowia_pulcherrima(100); | Metschnikowia | 13 | Saprotroph | \|Undefined Saprotroph\| | Yeast | NULL | Probable | As nectar-tap_saprotroph (Põlme et al. 2020); Decay sugar-rich substrates (Põlme et al. 2020) | Põlme S, et al. 2020. Fungal Diversity 105:1-16 ((https://doi.org/10.1007/s13225-020-00466-2)) |
| Otu33262 | 1 | k__Fungi(100);p__Ascomycota(100);c__Saccharomycetes(100);o__Saccharomycetales(100);f__Saccharomycetales_family_Incertae_sedis(100);g__Candida(100);s__Candida_cretensis(100); | Candida | 13 | Pathotroph-Saprotroph-Symbiotroph | Animal Pathogen-Endophyte-Endosymbiont-Epiphyte-\|Undefined Saprotroph\| | Dimorphic Yeast-Yeast | NULL | Probable | As nectar-tap_saprotroph (Põlme et al. 2020); Decay sugar-rich substrates (Põlme et al. 2020); This asexual genus is polyphyletic and contain species that function very differently depend on what they are. Caution should be taken when interpreting guilds for Candida species. | Manolakaki D, et al. 2010. Virulence. 1: 367-375 ((https://doi.org/10.4161/viru.1.5.12796)); Mekbib SB, et al. 2011. Tropical Plant Pathology 36:233-240 ((https://doi.org/10.1590/S1982-56762011000400004)); Põlme S, et al. 2020. Fungal Diversity 105:1-16 ((https://doi.org/10.1007/s13225-020-00466-2)) |
| Otu33263 | 1 | k__Fungi(100);p__Ascomycota(100);c__Saccharomycetes(100);o__Saccharomycetales(100);f__Metschnikowiaceae(100);g__Metschnikowia(100);s__Metschnikowia_pulcherrima(100); | Metschnikowia | 13 | Saprotroph | \|Undefined Saprotroph\| | Yeast | NULL | Probable | As nectar-tap_saprotroph (Põlme et al. 2020); Decay sugar-rich substrates (Põlme et al. 2020) | Põlme S, et al. 2020. Fungal Diversity 105:1-16 ((https://doi.org/10.1007/s13225-020-00466-2)) |
| Otu33264 | 1 | k__Fungi(100);p__Ascomycota(100);c__Saccharomycetes(100);o__Saccharomycetales(100);f__Metschnikowiaceae(100);g__Metschnikowia(100);s__Metschnikowia_pulcherrima(100); | Metschnikowia | 13 | Saprotroph | \|Undefined Saprotroph\| | Yeast | NULL | Probable | As nectar-tap_saprotroph (Põlme et al. 2020); Decay sugar-rich substrates (Põlme et al. 2020) | Põlme S, et al. 2020. Fungal Diversity 105:1-16 ((https://doi.org/10.1007/s13225-020-00466-2)) |
| Otu33265 | 1 | k__Fungi(100);p__Ascomycota(100);c__Saccharomycetes(100);o__Saccharomycetales(100);f__Metschnikowiaceae(100);g__Metschnikowia(100);s__Metschnikowia_pulcherrima(100); | Metschnikowia | 13 | Saprotroph | \|Undefined Saprotroph\| | Yeast | NULL | Probable | As nectar-tap_saprotroph (Põlme et al. 2020); Decay sugar-rich substrates (Põlme et al. 2020) | Põlme S, et al. 2020. Fungal Diversity 105:1-16 ((https://doi.org/10.1007/s13225-020-00466-2)) |
| Otu33266 | 1 | k__Fungi(100);p__Ascomycota(100);c__Saccharomycetes(100);o__Saccharomycetales(100);f__Saccharomycetales_family_Incertae_sedis(100);f__Saccharomycetales_family_Incertae_sedis_unclassified(100);f__Saccharomycetales_family_Incertae_sedis_unclassified(100); | Saccharomycetales | 7 | Saprotroph | Undefined Saprotroph | Yeast | NULL | Possible | NULL | Sterkenburg E, et al. 2015. New Phytologist 207:1145-1158 ((https://doi.org/10.1111/nph.13426)) |
| Otu33270 | 1 | k__Fungi(100);p__Ascomycota(100);c__Saccharomycetes(100);o__Saccharomycetales(100);f__Saccharomycetales_family_Incertae_sedis(100);g__Candida(100);g__Candida_unclassified(100); | Candida | 13 | Pathotroph-Saprotroph-Symbiotroph | Animal Pathogen-Endophyte-Endosymbiont-Epiphyte-\|Undefined Saprotroph\| | Dimorphic Yeast-Yeast | NULL | Probable | As nectar-tap_saprotroph (Põlme et al. 2020); Decay sugar-rich substrates (Põlme et al. 2020); This asexual genus is polyphyletic and contain species that function very differently depend on what they are. Caution should be taken when interpreting guilds for Candida species. | Manolakaki D, et al. 2010. Virulence. 1: 367-375 ((https://doi.org/10.4161/viru.1.5.12796)); Mekbib SB, et al. 2011. Tropical Plant Pathology 36:233-240 ((https://doi.org/10.1590/S1982-56762011000400004)); Põlme S, et al. 2020. Fungal Diversity 105:1-16 ((https://doi.org/10.1007/s13225-020-00466-2)) |
| Otu33271 | 1 | k__Fungi(100);p__Ascomycota(100);c__Saccharomycetes(100);o__Saccharomycetales(100);f__Metschnikowiaceae(100);g__Metschnikowia(100);s__Metschnikowia_pulcherrima(100); | Metschnikowia | 13 | Saprotroph | \|Undefined Saprotroph\| | Yeast | NULL | Probable | As nectar-tap_saprotroph (Põlme et al. 2020); Decay sugar-rich substrates (Põlme et al. 2020) | Põlme S, et al. 2020. Fungal Diversity 105:1-16 ((https://doi.org/10.1007/s13225-020-00466-2)) |
| Otu33272 | 1 | k__Fungi(100);p__Ascomycota(100);c__Saccharomycetes(100);o__Saccharomycetales(100);f__Metschnikowiaceae(100);g__Metschnikowia(100);s__Metschnikowia_pulcherrima(100); | Metschnikowia | 13 | Saprotroph | \|Undefined Saprotroph\| | Yeast | NULL | Probable | As nectar-tap_saprotroph (Põlme et al. 2020); Decay sugar-rich substrates (Põlme et al. 2020) | Põlme S, et al. 2020. Fungal Diversity 105:1-16 ((https://doi.org/10.1007/s13225-020-00466-2)) |
| Otu33276 | 1 | k__Fungi(100);p__Ascomycota(100);c__Saccharomycetes(100);o__Saccharomycetales(100);f__Metschnikowiaceae(100);g__Metschnikowia(100);g__Metschnikowia_unclassified(100); | Metschnikowia | 13 | Saprotroph | \|Undefined Saprotroph\| | Yeast | NULL | Probable | As nectar-tap_saprotroph (Põlme et al. 2020); Decay sugar-rich substrates (Põlme et al. 2020) | Põlme S, et al. 2020. Fungal Diversity 105:1-16 ((https://doi.org/10.1007/s13225-020-00466-2)) |
| Otu33277 | 1 | k__Fungi(100);p__Ascomycota(100);c__Saccharomycetes(100);o__Saccharomycetales(100);f__Metschnikowiaceae(100);g__Metschnikowia(100);s__Metschnikowia_pulcherrima(100); | Metschnikowia | 13 | Saprotroph | \|Undefined Saprotroph\| | Yeast | NULL | Probable | As nectar-tap_saprotroph (Põlme et al. 2020); Decay sugar-rich substrates (Põlme et al. 2020) | Põlme S, et al. 2020. Fungal Diversity 105:1-16 ((https://doi.org/10.1007/s13225-020-00466-2)) |
| Otu33279 | 1 | k__Fungi(100);p__Ascomycota(100);c__Saccharomycetes(100);o__Saccharomycetales(100);f__Metschnikowiaceae(100);g__Metschnikowia(100);g__Metschnikowia_unclassified(100); | Metschnikowia | 13 | Saprotroph | \|Undefined Saprotroph\| | Yeast | NULL | Probable | As nectar-tap_saprotroph (Põlme et al. 2020); Decay sugar-rich substrates (Põlme et al. 2020) | Põlme S, et al. 2020. Fungal Diversity 105:1-16 ((https://doi.org/10.1007/s13225-020-00466-2)) |
| Otu33280 | 1 | k__Fungi(100);p__Ascomycota(100);c__Saccharomycetes(100);o__Saccharomycetales(100);f__Metschnikowiaceae(100);g__Metschnikowia(100);s__Metschnikowia_pulcherrima(100); | Metschnikowia | 13 | Saprotroph | \|Undefined Saprotroph\| | Yeast | NULL | Probable | As nectar-tap_saprotroph (Põlme et al. 2020); Decay sugar-rich substrates (Põlme et al. 2020) | Põlme S, et al. 2020. Fungal Diversity 105:1-16 ((https://doi.org/10.1007/s13225-020-00466-2)) |
| Otu33282 | 1 | k__Fungi(100);p__Ascomycota(100);c__Saccharomycetes(100);o__Saccharomycetales(100);f__Metschnikowiaceae(100);g__Metschnikowia(100);s__Metschnikowia_pulcherrima(100); | Metschnikowia | 13 | Saprotroph | \|Undefined Saprotroph\| | Yeast | NULL | Probable | As nectar-tap_saprotroph (Põlme et al. 2020); Decay sugar-rich substrates (Põlme et al. 2020) | Põlme S, et al. 2020. Fungal Diversity 105:1-16 ((https://doi.org/10.1007/s13225-020-00466-2)) |
| Otu33283 | 1 | k__Fungi(100);p__Ascomycota(100);c__Saccharomycetes(100);o__Saccharomycetales(100);f__Metschnikowiaceae(100);g__Metschnikowia(100);s__Metschnikowia_pulcherrima(100); | Metschnikowia | 13 | Saprotroph | \|Undefined Saprotroph\| | Yeast | NULL | Probable | As nectar-tap_saprotroph (Põlme et al. 2020); Decay sugar-rich substrates (Põlme et al. 2020) | Põlme S, et al. 2020. Fungal Diversity 105:1-16 ((https://doi.org/10.1007/s13225-020-00466-2)) |
| Otu33285 | 1 | k__Fungi(100);p__Ascomycota(100);c__Saccharomycetes(100);o__Saccharomycetales(100);f__Metschnikowiaceae(100);g__Metschnikowia(100);s__Metschnikowia_pulcherrima(100); | Metschnikowia | 13 | Saprotroph | \|Undefined Saprotroph\| | Yeast | NULL | Probable | As nectar-tap_saprotroph (Põlme et al. 2020); Decay sugar-rich substrates (Põlme et al. 2020) | Põlme S, et al. 2020. Fungal Diversity 105:1-16 ((https://doi.org/10.1007/s13225-020-00466-2)) |
| Otu33286 | 1 | k__Fungi(100);p__Ascomycota(100);c__Saccharomycetes(100);o__Saccharomycetales(100);f__Metschnikowiaceae(100);g__Metschnikowia(100);g__Metschnikowia_unclassified(100); | Metschnikowia | 13 | Saprotroph | \|Undefined Saprotroph\| | Yeast | NULL | Probable | As nectar-tap_saprotroph (Põlme et al. 2020); Decay sugar-rich substrates (Põlme et al. 2020) | Põlme S, et al. 2020. Fungal Diversity 105:1-16 ((https://doi.org/10.1007/s13225-020-00466-2)) |
| Otu33287 | 1 | k__Fungi(100);p__Ascomycota(100);c__Saccharomycetes(100);o__Saccharomycetales(100);f__Metschnikowiaceae(100);g__Metschnikowia(100);g__Metschnikowia_unclassified(100); | Metschnikowia | 13 | Saprotroph | \|Undefined Saprotroph\| | Yeast | NULL | Probable | As nectar-tap_saprotroph (Põlme et al. 2020); Decay sugar-rich substrates (Põlme et al. 2020) | Põlme S, et al. 2020. Fungal Diversity 105:1-16 ((https://doi.org/10.1007/s13225-020-00466-2)) |
| Otu33288 | 1 | k__Fungi(100);p__Ascomycota(100);c__Saccharomycetes(100);o__Saccharomycetales(100);f__Metschnikowiaceae(100);g__Metschnikowia(100);s__Metschnikowia_pulcherrima(100); | Metschnikowia | 13 | Saprotroph | \|Undefined Saprotroph\| | Yeast | NULL | Probable | As nectar-tap_saprotroph (Põlme et al. 2020); Decay sugar-rich substrates (Põlme et al. 2020) | Põlme S, et al. 2020. Fungal Diversity 105:1-16 ((https://doi.org/10.1007/s13225-020-00466-2)) |
| Otu33289 | 1 | k__Fungi(100);p__Ascomycota(100);c__Saccharomycetes(100);o__Saccharomycetales(100);f__Metschnikowiaceae(100);g__Metschnikowia(100);s__Metschnikowia_pulcherrima(100); | Metschnikowia | 13 | Saprotroph | \|Undefined Saprotroph\| | Yeast | NULL | Probable | As nectar-tap_saprotroph (Põlme et al. 2020); Decay sugar-rich substrates (Põlme et al. 2020) | Põlme S, et al. 2020. Fungal Diversity 105:1-16 ((https://doi.org/10.1007/s13225-020-00466-2)) |
| Otu33290 | 1 | k__Fungi(100);p__Ascomycota(100);c__Saccharomycetes(100);o__Saccharomycetales(100);f__Metschnikowiaceae(100);g__Metschnikowia(100);s__Metschnikowia_pulcherrima(100); | Metschnikowia | 13 | Saprotroph | \|Undefined Saprotroph\| | Yeast | NULL | Probable | As nectar-tap_saprotroph (Põlme et al. 2020); Decay sugar-rich substrates (Põlme et al. 2020) | Põlme S, et al. 2020. Fungal Diversity 105:1-16 ((https://doi.org/10.1007/s13225-020-00466-2)) |
| Otu33291 | 1 | k__Fungi(100);p__Ascomycota(100);c__Saccharomycetes(100);o__Saccharomycetales(100);f__Metschnikowiaceae(100);g__Metschnikowia(100);g__Metschnikowia_unclassified(100); | Metschnikowia | 13 | Saprotroph | \|Undefined Saprotroph\| | Yeast | NULL | Probable | As nectar-tap_saprotroph (Põlme et al. 2020); Decay sugar-rich substrates (Põlme et al. 2020) | Põlme S, et al. 2020. Fungal Diversity 105:1-16 ((https://doi.org/10.1007/s13225-020-00466-2)) |
| Otu33292 | 1 | k__Fungi(100);p__Ascomycota(100);c__Saccharomycetes(100);o__Saccharomycetales(100);f__Metschnikowiaceae(100);g__Metschnikowia(100);s__Metschnikowia_pulcherrima(100); | Metschnikowia | 13 | Saprotroph | \|Undefined Saprotroph\| | Yeast | NULL | Probable | As nectar-tap_saprotroph (Põlme et al. 2020); Decay sugar-rich substrates (Põlme et al. 2020) | Põlme S, et al. 2020. Fungal Diversity 105:1-16 ((https://doi.org/10.1007/s13225-020-00466-2)) |
| Otu33293 | 1 | k__Fungi(100);p__Ascomycota(100);c__Saccharomycetes(100);o__Saccharomycetales(100);f__Metschnikowiaceae(100);g__Metschnikowia(100);s__Metschnikowia_pulcherrima(100); | Metschnikowia | 13 | Saprotroph | \|Undefined Saprotroph\| | Yeast | NULL | Probable | As nectar-tap_saprotroph (Põlme et al. 2020); Decay sugar-rich substrates (Põlme et al. 2020) | Põlme S, et al. 2020. Fungal Diversity 105:1-16 ((https://doi.org/10.1007/s13225-020-00466-2)) |
| Otu33294 | 1 | k__Fungi(100);p__Ascomycota(100);c__Saccharomycetes(100);o__Saccharomycetales(100);f__Metschnikowiaceae(100);g__Metschnikowia(100);s__Metschnikowia_pulcherrima(100); | Metschnikowia | 13 | Saprotroph | \|Undefined Saprotroph\| | Yeast | NULL | Probable | As nectar-tap_saprotroph (Põlme et al. 2020); Decay sugar-rich substrates (Põlme et al. 2020) | Põlme S, et al. 2020. Fungal Diversity 105:1-16 ((https://doi.org/10.1007/s13225-020-00466-2)) |
| Otu33297 | 1 | k__Fungi(100);p__Ascomycota(100);c__Saccharomycetes(100);o__Saccharomycetales(100);f__Metschnikowiaceae(100);g__Metschnikowia(100);s__Metschnikowia_pulcherrima(100); | Metschnikowia | 13 | Saprotroph | \|Undefined Saprotroph\| | Yeast | NULL | Probable | As nectar-tap_saprotroph (Põlme et al. 2020); Decay sugar-rich substrates (Põlme et al. 2020) | Põlme S, et al. 2020. Fungal Diversity 105:1-16 ((https://doi.org/10.1007/s13225-020-00466-2)) |
| Otu33300 | 1 | k__Fungi(100);p__Ascomycota(100);c__Saccharomycetes(100);o__Saccharomycetales(100);f__Metschnikowiaceae(100);g__Metschnikowia(100);s__Metschnikowia_pulcherrima(100); | Metschnikowia | 13 | Saprotroph | \|Undefined Saprotroph\| | Yeast | NULL | Probable | As nectar-tap_saprotroph (Põlme et al. 2020); Decay sugar-rich substrates (Põlme et al. 2020) | Põlme S, et al. 2020. Fungal Diversity 105:1-16 ((https://doi.org/10.1007/s13225-020-00466-2)) |
| Otu33302 | 1 | k__Fungi(100);p__Ascomycota(100);c__Saccharomycetes(100);o__Saccharomycetales(100);f__Saccharomycetales_family_Incertae_sedis(100);g__Candida(100);g__Candida_unclassified(100); | Candida | 13 | Pathotroph-Saprotroph-Symbiotroph | Animal Pathogen-Endophyte-Endosymbiont-Epiphyte-\|Undefined Saprotroph\| | Dimorphic Yeast-Yeast | NULL | Probable | As nectar-tap_saprotroph (Põlme et al. 2020); Decay sugar-rich substrates (Põlme et al. 2020); This asexual genus is polyphyletic and contain species that function very differently depend on what they are. Caution should be taken when interpreting guilds for Candida species. | Manolakaki D, et al. 2010. Virulence. 1: 367-375 ((https://doi.org/10.4161/viru.1.5.12796)); Mekbib SB, et al. 2011. Tropical Plant Pathology 36:233-240 ((https://doi.org/10.1590/S1982-56762011000400004)); Põlme S, et al. 2020. Fungal Diversity 105:1-16 ((https://doi.org/10.1007/s13225-020-00466-2)) |
| Otu33303 | 1 | k__Fungi(100);p__Ascomycota(100);c__Saccharomycetes(100);o__Saccharomycetales(100);f__Metschnikowiaceae(100);g__Metschnikowia(100);g__Metschnikowia_unclassified(100); | Metschnikowia | 13 | Saprotroph | \|Undefined Saprotroph\| | Yeast | NULL | Probable | As nectar-tap_saprotroph (Põlme et al. 2020); Decay sugar-rich substrates (Põlme et al. 2020) | Põlme S, et al. 2020. Fungal Diversity 105:1-16 ((https://doi.org/10.1007/s13225-020-00466-2)) |
| Otu33304 | 1 | k__Fungi(100);p__Ascomycota(100);c__Saccharomycetes(100);o__Saccharomycetales(100);f__Metschnikowiaceae(100);g__Metschnikowia(100);s__Metschnikowia_pulcherrima(100); | Metschnikowia | 13 | Saprotroph | \|Undefined Saprotroph\| | Yeast | NULL | Probable | As nectar-tap_saprotroph (Põlme et al. 2020); Decay sugar-rich substrates (Põlme et al. 2020) | Põlme S, et al. 2020. Fungal Diversity 105:1-16 ((https://doi.org/10.1007/s13225-020-00466-2)) |
| Otu33305 | 1 | k__Fungi(100);p__Ascomycota(100);c__Saccharomycetes(100);o__Saccharomycetales(100);f__Metschnikowiaceae(100);g__Metschnikowia(100);s__Metschnikowia_pulcherrima(100); | Metschnikowia | 13 | Saprotroph | \|Undefined Saprotroph\| | Yeast | NULL | Probable | As nectar-tap_saprotroph (Põlme et al. 2020); Decay sugar-rich substrates (Põlme et al. 2020) | Põlme S, et al. 2020. Fungal Diversity 105:1-16 ((https://doi.org/10.1007/s13225-020-00466-2)) |
| Otu33307 | 1 | k__Fungi(100);p__Ascomycota(100);c__Saccharomycetes(100);o__Saccharomycetales(100);f__Metschnikowiaceae(100);g__Metschnikowia(100);g__Metschnikowia_unclassified(100); | Metschnikowia | 13 | Saprotroph | \|Undefined Saprotroph\| | Yeast | NULL | Probable | As nectar-tap_saprotroph (Põlme et al. 2020); Decay sugar-rich substrates (Põlme et al. 2020) | Põlme S, et al. 2020. Fungal Diversity 105:1-16 ((https://doi.org/10.1007/s13225-020-00466-2)) |
| Otu33308 | 1 | k__Fungi(100);p__Ascomycota(100);c__Saccharomycetes(100);o__Saccharomycetales(100);f__Metschnikowiaceae(100);g__Metschnikowia(100);s__Metschnikowia_pulcherrima(100); | Metschnikowia | 13 | Saprotroph | \|Undefined Saprotroph\| | Yeast | NULL | Probable | As nectar-tap_saprotroph (Põlme et al. 2020); Decay sugar-rich substrates (Põlme et al. 2020) | Põlme S, et al. 2020. Fungal Diversity 105:1-16 ((https://doi.org/10.1007/s13225-020-00466-2)) |
| Otu33309 | 1 | k__Fungi(100);p__Ascomycota(100);c__Saccharomycetes(100);o__Saccharomycetales(100);f__Metschnikowiaceae(100);g__Metschnikowia(100);s__Metschnikowia_pulcherrima(100); | Metschnikowia | 13 | Saprotroph | \|Undefined Saprotroph\| | Yeast | NULL | Probable | As nectar-tap_saprotroph (Põlme et al. 2020); Decay sugar-rich substrates (Põlme et al. 2020) | Põlme S, et al. 2020. Fungal Diversity 105:1-16 ((https://doi.org/10.1007/s13225-020-00466-2)) |
| Otu33310 | 1 | k__Fungi(100);p__Ascomycota(100);c__Saccharomycetes(100);o__Saccharomycetales(100);f__Metschnikowiaceae(100);g__Metschnikowia(100);g__Metschnikowia_unclassified(100); | Metschnikowia | 13 | Saprotroph | \|Undefined Saprotroph\| | Yeast | NULL | Probable | As nectar-tap_saprotroph (Põlme et al. 2020); Decay sugar-rich substrates (Põlme et al. 2020) | Põlme S, et al. 2020. Fungal Diversity 105:1-16 ((https://doi.org/10.1007/s13225-020-00466-2)) |
| Otu33312 | 1 | k__Fungi(100);p__Ascomycota(100);c__Saccharomycetes(100);o__Saccharomycetales(100);f__Metschnikowiaceae(100);g__Metschnikowia(100);s__Metschnikowia_pulcherrima(100); | Metschnikowia | 13 | Saprotroph | \|Undefined Saprotroph\| | Yeast | NULL | Probable | As nectar-tap_saprotroph (Põlme et al. 2020); Decay sugar-rich substrates (Põlme et al. 2020) | Põlme S, et al. 2020. Fungal Diversity 105:1-16 ((https://doi.org/10.1007/s13225-020-00466-2)) |
| Otu33313 | 1 | k__Fungi(100);p__Ascomycota(100);c__Saccharomycetes(100);o__Saccharomycetales(100);f__Metschnikowiaceae(100);g__Metschnikowia(100);s__Metschnikowia_pulcherrima(100); | Metschnikowia | 13 | Saprotroph | \|Undefined Saprotroph\| | Yeast | NULL | Probable | As nectar-tap_saprotroph (Põlme et al. 2020); Decay sugar-rich substrates (Põlme et al. 2020) | Põlme S, et al. 2020. Fungal Diversity 105:1-16 ((https://doi.org/10.1007/s13225-020-00466-2)) |
| Otu33314 | 1 | k__Fungi(100);p__Ascomycota(100);c__Saccharomycetes(100);o__Saccharomycetales(100);f__Saccharomycetales_family_Incertae_sedis(100);f__Saccharomycetales_family_Incertae_sedis_unclassified(100);f__Saccharomycetales_family_Incertae_sedis_unclassified(100); | Saccharomycetales | 7 | Saprotroph | Undefined Saprotroph | Yeast | NULL | Possible | NULL | Sterkenburg E, et al. 2015. New Phytologist 207:1145-1158 ((https://doi.org/10.1111/nph.13426)) |
| Otu33315 | 1 | k__Fungi(100);p__Ascomycota(100);c__Saccharomycetes(100);o__Saccharomycetales(100);f__Metschnikowiaceae(100);g__Metschnikowia(100);s__Metschnikowia_pulcherrima(100); | Metschnikowia | 13 | Saprotroph | \|Undefined Saprotroph\| | Yeast | NULL | Probable | As nectar-tap_saprotroph (Põlme et al. 2020); Decay sugar-rich substrates (Põlme et al. 2020) | Põlme S, et al. 2020. Fungal Diversity 105:1-16 ((https://doi.org/10.1007/s13225-020-00466-2)) |
| Otu33316 | 1 | k__Fungi(100);p__Ascomycota(100);c__Saccharomycetes(100);o__Saccharomycetales(100);f__Metschnikowiaceae(100);g__Metschnikowia(100);g__Metschnikowia_unclassified(100); | Metschnikowia | 13 | Saprotroph | \|Undefined Saprotroph\| | Yeast | NULL | Probable | As nectar-tap_saprotroph (Põlme et al. 2020); Decay sugar-rich substrates (Põlme et al. 2020) | Põlme S, et al. 2020. Fungal Diversity 105:1-16 ((https://doi.org/10.1007/s13225-020-00466-2)) |
| Otu33317 | 1 | k__Fungi(100);p__Ascomycota(100);c__Saccharomycetes(100);o__Saccharomycetales(100);f__Saccharomycetales_family_Incertae_sedis(100);g__Candida(100);g__Candida_unclassified(100); | Candida | 13 | Pathotroph-Saprotroph-Symbiotroph | Animal Pathogen-Endophyte-Endosymbiont-Epiphyte-\|Undefined Saprotroph\| | Dimorphic Yeast-Yeast | NULL | Probable | As nectar-tap_saprotroph (Põlme et al. 2020); Decay sugar-rich substrates (Põlme et al. 2020); This asexual genus is polyphyletic and contain species that function very differently depend on what they are. Caution should be taken when interpreting guilds for Candida species. | Manolakaki D, et al. 2010. Virulence. 1: 367-375 ((https://doi.org/10.4161/viru.1.5.12796)); Mekbib SB, et al. 2011. Tropical Plant Pathology 36:233-240 ((https://doi.org/10.1590/S1982-56762011000400004)); Põlme S, et al. 2020. Fungal Diversity 105:1-16 ((https://doi.org/10.1007/s13225-020-00466-2)) |
| Otu33323 | 1 | k__Fungi(100);p__Ascomycota(100);c__Saccharomycetes(100);o__Saccharomycetales(100);f__Metschnikowiaceae(100);g__Metschnikowia(100);g__Metschnikowia_unclassified(100); | Metschnikowia | 13 | Saprotroph | \|Undefined Saprotroph\| | Yeast | NULL | Probable | As nectar-tap_saprotroph (Põlme et al. 2020); Decay sugar-rich substrates (Põlme et al. 2020) | Põlme S, et al. 2020. Fungal Diversity 105:1-16 ((https://doi.org/10.1007/s13225-020-00466-2)) |
| Otu33324 | 1 | k__Fungi(100);p__Ascomycota(100);c__Saccharomycetes(100);o__Saccharomycetales(100);f__Saccharomycetales_family_Incertae_sedis(100);g__Candida(100);g__Candida_unclassified(100); | Candida | 13 | Pathotroph-Saprotroph-Symbiotroph | Animal Pathogen-Endophyte-Endosymbiont-Epiphyte-\|Undefined Saprotroph\| | Dimorphic Yeast-Yeast | NULL | Probable | As nectar-tap_saprotroph (Põlme et al. 2020); Decay sugar-rich substrates (Põlme et al. 2020); This asexual genus is polyphyletic and contain species that function very differently depend on what they are. Caution should be taken when interpreting guilds for Candida species. | Manolakaki D, et al. 2010. Virulence. 1: 367-375 ((https://doi.org/10.4161/viru.1.5.12796)); Mekbib SB, et al. 2011. Tropical Plant Pathology 36:233-240 ((https://doi.org/10.1590/S1982-56762011000400004)); Põlme S, et al. 2020. Fungal Diversity 105:1-16 ((https://doi.org/10.1007/s13225-020-00466-2)) |
| Otu33326 | 1 | k__Fungi(100);p__Ascomycota(100);c__Saccharomycetes(100);o__Saccharomycetales(100);f__Saccharomycetales_family_Incertae_sedis(100);g__Candida(100);g__Candida_unclassified(100); | Candida | 13 | Pathotroph-Saprotroph-Symbiotroph | Animal Pathogen-Endophyte-Endosymbiont-Epiphyte-\|Undefined Saprotroph\| | Dimorphic Yeast-Yeast | NULL | Probable | As nectar-tap_saprotroph (Põlme et al. 2020); Decay sugar-rich substrates (Põlme et al. 2020); This asexual genus is polyphyletic and contain species that function very differently depend on what they are. Caution should be taken when interpreting guilds for Candida species. | Manolakaki D, et al. 2010. Virulence. 1: 367-375 ((https://doi.org/10.4161/viru.1.5.12796)); Mekbib SB, et al. 2011. Tropical Plant Pathology 36:233-240 ((https://doi.org/10.1590/S1982-56762011000400004)); Põlme S, et al. 2020. Fungal Diversity 105:1-16 ((https://doi.org/10.1007/s13225-020-00466-2)) |
| Otu33328 | 1 | k__Fungi(100);p__Ascomycota(100);c__Saccharomycetes(100);o__Saccharomycetales(100);f__Saccharomycetales_family_Incertae_sedis(100);g__Candida(100);g__Candida_unclassified(100); | Candida | 13 | Pathotroph-Saprotroph-Symbiotroph | Animal Pathogen-Endophyte-Endosymbiont-Epiphyte-\|Undefined Saprotroph\| | Dimorphic Yeast-Yeast | NULL | Probable | As nectar-tap_saprotroph (Põlme et al. 2020); Decay sugar-rich substrates (Põlme et al. 2020); This asexual genus is polyphyletic and contain species that function very differently depend on what they are. Caution should be taken when interpreting guilds for Candida species. | Manolakaki D, et al. 2010. Virulence. 1: 367-375 ((https://doi.org/10.4161/viru.1.5.12796)); Mekbib SB, et al. 2011. Tropical Plant Pathology 36:233-240 ((https://doi.org/10.1590/S1982-56762011000400004)); Põlme S, et al. 2020. Fungal Diversity 105:1-16 ((https://doi.org/10.1007/s13225-020-00466-2)) |
| Otu33332 | 1 | k__Fungi(100);p__Ascomycota(100);c__Saccharomycetes(100);o__Saccharomycetales(100);f__Saccharomycetales_family_Incertae_sedis(100);f__Saccharomycetales_family_Incertae_sedis_unclassified(100);f__Saccharomycetales_family_Incertae_sedis_unclassified(100); | Saccharomycetales | 7 | Saprotroph | Undefined Saprotroph | Yeast | NULL | Possible | NULL | Sterkenburg E, et al. 2015. New Phytologist 207:1145-1158 ((https://doi.org/10.1111/nph.13426)) |
| Otu33338 | 1 | k__Fungi(100);p__Ascomycota(100);c__Saccharomycetes(100);o__Saccharomycetales(100);f__Saccharomycetales_family_Incertae_sedis(100);g__Candida(100);g__Candida_unclassified(100); | Candida | 13 | Pathotroph-Saprotroph-Symbiotroph | Animal Pathogen-Endophyte-Endosymbiont-Epiphyte-\|Undefined Saprotroph\| | Dimorphic Yeast-Yeast | NULL | Probable | As nectar-tap_saprotroph (Põlme et al. 2020); Decay sugar-rich substrates (Põlme et al. 2020); This asexual genus is polyphyletic and contain species that function very differently depend on what they are. Caution should be taken when interpreting guilds for Candida species. | Manolakaki D, et al. 2010. Virulence. 1: 367-375 ((https://doi.org/10.4161/viru.1.5.12796)); Mekbib SB, et al. 2011. Tropical Plant Pathology 36:233-240 ((https://doi.org/10.1590/S1982-56762011000400004)); Põlme S, et al. 2020. Fungal Diversity 105:1-16 ((https://doi.org/10.1007/s13225-020-00466-2)) |
| Otu33342 | 1 | k__Fungi(100);p__Ascomycota(100);c__Saccharomycetes(100);o__Saccharomycetales(100);f__Metschnikowiaceae(100);g__Metschnikowia(100);s__Metschnikowia_pulcherrima(100); | Metschnikowia | 13 | Saprotroph | \|Undefined Saprotroph\| | Yeast | NULL | Probable | As nectar-tap_saprotroph (Põlme et al. 2020); Decay sugar-rich substrates (Põlme et al. 2020) | Põlme S, et al. 2020. Fungal Diversity 105:1-16 ((https://doi.org/10.1007/s13225-020-00466-2)) |
| Otu33343 | 1 | k__Fungi(100);p__Ascomycota(100);c__Saccharomycetes(100);o__Saccharomycetales(100);f__Saccharomycetales_family_Incertae_sedis(100);g__Candida(100);g__Candida_unclassified(100); | Candida | 13 | Pathotroph-Saprotroph-Symbiotroph | Animal Pathogen-Endophyte-Endosymbiont-Epiphyte-\|Undefined Saprotroph\| | Dimorphic Yeast-Yeast | NULL | Probable | As nectar-tap_saprotroph (Põlme et al. 2020); Decay sugar-rich substrates (Põlme et al. 2020); This asexual genus is polyphyletic and contain species that function very differently depend on what they are. Caution should be taken when interpreting guilds for Candida species. | Manolakaki D, et al. 2010. Virulence. 1: 367-375 ((https://doi.org/10.4161/viru.1.5.12796)); Mekbib SB, et al. 2011. Tropical Plant Pathology 36:233-240 ((https://doi.org/10.1590/S1982-56762011000400004)); Põlme S, et al. 2020. Fungal Diversity 105:1-16 ((https://doi.org/10.1007/s13225-020-00466-2)) |
| Otu33350 | 1 | k__Fungi(100);p__Ascomycota(100);c__Saccharomycetes(100);o__Saccharomycetales(100);f__Metschnikowiaceae(100);g__Metschnikowia(100);s__Metschnikowia_pulcherrima(100); | Metschnikowia | 13 | Saprotroph | \|Undefined Saprotroph\| | Yeast | NULL | Probable | As nectar-tap_saprotroph (Põlme et al. 2020); Decay sugar-rich substrates (Põlme et al. 2020) | Põlme S, et al. 2020. Fungal Diversity 105:1-16 ((https://doi.org/10.1007/s13225-020-00466-2)) |
| Otu33361 | 1 | k__Fungi(100);p__Ascomycota(100);c__Saccharomycetes(100);o__Saccharomycetales(100);f__Metschnikowiaceae(100);g__Metschnikowia(100);g__Metschnikowia_unclassified(100); | Metschnikowia | 13 | Saprotroph | \|Undefined Saprotroph\| | Yeast | NULL | Probable | As nectar-tap_saprotroph (Põlme et al. 2020); Decay sugar-rich substrates (Põlme et al. 2020) | Põlme S, et al. 2020. Fungal Diversity 105:1-16 ((https://doi.org/10.1007/s13225-020-00466-2)) |
| Otu33365 | 1 | k__Fungi(100);p__Ascomycota(100);c__Saccharomycetes(100);o__Saccharomycetales(100);f__Metschnikowiaceae(100);g__Metschnikowia(100);g__Metschnikowia_unclassified(100); | Metschnikowia | 13 | Saprotroph | \|Undefined Saprotroph\| | Yeast | NULL | Probable | As nectar-tap_saprotroph (Põlme et al. 2020); Decay sugar-rich substrates (Põlme et al. 2020) | Põlme S, et al. 2020. Fungal Diversity 105:1-16 ((https://doi.org/10.1007/s13225-020-00466-2)) |
| Otu33368 | 1 | k__Fungi(100);p__Ascomycota(100);c__Saccharomycetes(100);o__Saccharomycetales(100);o__Saccharomycetales_unclassified(100);o__Saccharomycetales_unclassified(100);o__Saccharomycetales_unclassified(100); | Saccharomycetales | 7 | Saprotroph | Undefined Saprotroph | Yeast | NULL | Possible | NULL | Sterkenburg E, et al. 2015. New Phytologist 207:1145-1158 ((https://doi.org/10.1111/nph.13426)) |
| Otu33373 | 1 | k__Fungi(100);p__Ascomycota(100);c__Saccharomycetes(100);o__Saccharomycetales(100);f__Metschnikowiaceae(100);g__Metschnikowia(100);s__Metschnikowia_pulcherrima(100); | Metschnikowia | 13 | Saprotroph | \|Undefined Saprotroph\| | Yeast | NULL | Probable | As nectar-tap_saprotroph (Põlme et al. 2020); Decay sugar-rich substrates (Põlme et al. 2020) | Põlme S, et al. 2020. Fungal Diversity 105:1-16 ((https://doi.org/10.1007/s13225-020-00466-2)) |
| Otu33374 | 1 | k__Fungi(100);p__Ascomycota(100);c__Saccharomycetes(100);o__Saccharomycetales(100);f__Metschnikowiaceae(100);g__Metschnikowia(100);s__Metschnikowia_pulcherrima(100); | Metschnikowia | 13 | Saprotroph | \|Undefined Saprotroph\| | Yeast | NULL | Probable | As nectar-tap_saprotroph (Põlme et al. 2020); Decay sugar-rich substrates (Põlme et al. 2020) | Põlme S, et al. 2020. Fungal Diversity 105:1-16 ((https://doi.org/10.1007/s13225-020-00466-2)) |
| Otu33375 | 1 | k__Fungi(100);p__Ascomycota(100);c__Saccharomycetes(100);o__Saccharomycetales(100);f__Metschnikowiaceae(100);g__Metschnikowia(100);s__Metschnikowia_pulcherrima(100); | Metschnikowia | 13 | Saprotroph | \|Undefined Saprotroph\| | Yeast | NULL | Probable | As nectar-tap_saprotroph (Põlme et al. 2020); Decay sugar-rich substrates (Põlme et al. 2020) | Põlme S, et al. 2020. Fungal Diversity 105:1-16 ((https://doi.org/10.1007/s13225-020-00466-2)) |
| Otu33376 | 1 | k__Fungi(100);p__Ascomycota(100);c__Saccharomycetes(100);o__Saccharomycetales(100);f__Metschnikowiaceae(100);g__Metschnikowia(100);s__Metschnikowia_pulcherrima(100); | Metschnikowia | 13 | Saprotroph | \|Undefined Saprotroph\| | Yeast | NULL | Probable | As nectar-tap_saprotroph (Põlme et al. 2020); Decay sugar-rich substrates (Põlme et al. 2020) | Põlme S, et al. 2020. Fungal Diversity 105:1-16 ((https://doi.org/10.1007/s13225-020-00466-2)) |
| Otu33377 | 1 | k__Fungi(100);p__Ascomycota(100);c__Saccharomycetes(100);o__Saccharomycetales(100);f__Metschnikowiaceae(100);g__Metschnikowia(100);g__Metschnikowia_unclassified(100); | Metschnikowia | 13 | Saprotroph | \|Undefined Saprotroph\| | Yeast | NULL | Probable | As nectar-tap_saprotroph (Põlme et al. 2020); Decay sugar-rich substrates (Põlme et al. 2020) | Põlme S, et al. 2020. Fungal Diversity 105:1-16 ((https://doi.org/10.1007/s13225-020-00466-2)) |
| Otu33378 | 1 | k__Fungi(100);p__Ascomycota(100);c__Saccharomycetes(100);o__Saccharomycetales(100);f__Metschnikowiaceae(100);g__Metschnikowia(100);g__Metschnikowia_unclassified(100); | Metschnikowia | 13 | Saprotroph | \|Undefined Saprotroph\| | Yeast | NULL | Probable | As nectar-tap_saprotroph (Põlme et al. 2020); Decay sugar-rich substrates (Põlme et al. 2020) | Põlme S, et al. 2020. Fungal Diversity 105:1-16 ((https://doi.org/10.1007/s13225-020-00466-2)) |
| Otu33379 | 1 | k__Fungi(100);p__Ascomycota(100);c__Saccharomycetes(100);o__Saccharomycetales(100);f__Metschnikowiaceae(100);g__Metschnikowia(100);g__Metschnikowia_unclassified(100); | Metschnikowia | 13 | Saprotroph | \|Undefined Saprotroph\| | Yeast | NULL | Probable | As nectar-tap_saprotroph (Põlme et al. 2020); Decay sugar-rich substrates (Põlme et al. 2020) | Põlme S, et al. 2020. Fungal Diversity 105:1-16 ((https://doi.org/10.1007/s13225-020-00466-2)) |
| Otu33381 | 1 | k__Fungi(100);p__Ascomycota(100);c__Saccharomycetes(100);o__Saccharomycetales(100);f__Metschnikowiaceae(100);g__Metschnikowia(100);s__Metschnikowia_pulcherrima(100); | Metschnikowia | 13 | Saprotroph | \|Undefined Saprotroph\| | Yeast | NULL | Probable | As nectar-tap_saprotroph (Põlme et al. 2020); Decay sugar-rich substrates (Põlme et al. 2020) | Põlme S, et al. 2020. Fungal Diversity 105:1-16 ((https://doi.org/10.1007/s13225-020-00466-2)) |
| Otu33384 | 1 | k__Fungi(100);p__Ascomycota(100);c__Saccharomycetes(100);o__Saccharomycetales(100);f__Metschnikowiaceae(100);g__Metschnikowia(100);g__Metschnikowia_unclassified(100); | Metschnikowia | 13 | Saprotroph | \|Undefined Saprotroph\| | Yeast | NULL | Probable | As nectar-tap_saprotroph (Põlme et al. 2020); Decay sugar-rich substrates (Põlme et al. 2020) | Põlme S, et al. 2020. Fungal Diversity 105:1-16 ((https://doi.org/10.1007/s13225-020-00466-2)) |
| Otu33385 | 1 | k__Fungi(100);p__Ascomycota(100);c__Saccharomycetes(100);o__Saccharomycetales(100);f__Metschnikowiaceae(100);g__Metschnikowia(100);g__Metschnikowia_unclassified(100); | Metschnikowia | 13 | Saprotroph | \|Undefined Saprotroph\| | Yeast | NULL | Probable | As nectar-tap_saprotroph (Põlme et al. 2020); Decay sugar-rich substrates (Põlme et al. 2020) | Põlme S, et al. 2020. Fungal Diversity 105:1-16 ((https://doi.org/10.1007/s13225-020-00466-2)) |
| Otu33386 | 1 | k__Fungi(100);p__Ascomycota(100);c__Saccharomycetes(100);o__Saccharomycetales(100);f__Metschnikowiaceae(100);g__Metschnikowia(100);g__Metschnikowia_unclassified(100); | Metschnikowia | 13 | Saprotroph | \|Undefined Saprotroph\| | Yeast | NULL | Probable | As nectar-tap_saprotroph (Põlme et al. 2020); Decay sugar-rich substrates (Põlme et al. 2020) | Põlme S, et al. 2020. Fungal Diversity 105:1-16 ((https://doi.org/10.1007/s13225-020-00466-2)) |
| Otu33387 | 1 | k__Fungi(100);p__Ascomycota(100);c__Saccharomycetes(100);o__Saccharomycetales(100);f__Metschnikowiaceae(100);g__Metschnikowia(100);g__Metschnikowia_unclassified(100); | Metschnikowia | 13 | Saprotroph | \|Undefined Saprotroph\| | Yeast | NULL | Probable | As nectar-tap_saprotroph (Põlme et al. 2020); Decay sugar-rich substrates (Põlme et al. 2020) | Põlme S, et al. 2020. Fungal Diversity 105:1-16 ((https://doi.org/10.1007/s13225-020-00466-2)) |
| Otu33388 | 1 | k__Fungi(100);p__Ascomycota(100);c__Saccharomycetes(100);o__Saccharomycetales(100);f__Metschnikowiaceae(100);g__Metschnikowia(100);s__Metschnikowia_pulcherrima(100); | Metschnikowia | 13 | Saprotroph | \|Undefined Saprotroph\| | Yeast | NULL | Probable | As nectar-tap_saprotroph (Põlme et al. 2020); Decay sugar-rich substrates (Põlme et al. 2020) | Põlme S, et al. 2020. Fungal Diversity 105:1-16 ((https://doi.org/10.1007/s13225-020-00466-2)) |
| Otu33389 | 1 | k__Fungi(100);p__Ascomycota(100);c__Saccharomycetes(100);o__Saccharomycetales(100);f__Metschnikowiaceae(100);g__Metschnikowia(100);s__Metschnikowia_pulcherrima(100); | Metschnikowia | 13 | Saprotroph | \|Undefined Saprotroph\| | Yeast | NULL | Probable | As nectar-tap_saprotroph (Põlme et al. 2020); Decay sugar-rich substrates (Põlme et al. 2020) | Põlme S, et al. 2020. Fungal Diversity 105:1-16 ((https://doi.org/10.1007/s13225-020-00466-2)) |
| Otu33390 | 1 | k__Fungi(100);p__Ascomycota(100);c__Saccharomycetes(100);o__Saccharomycetales(100);f__Metschnikowiaceae(100);g__Metschnikowia(100);s__Metschnikowia_pulcherrima(100); | Metschnikowia | 13 | Saprotroph | \|Undefined Saprotroph\| | Yeast | NULL | Probable | As nectar-tap_saprotroph (Põlme et al. 2020); Decay sugar-rich substrates (Põlme et al. 2020) | Põlme S, et al. 2020. Fungal Diversity 105:1-16 ((https://doi.org/10.1007/s13225-020-00466-2)) |
| Otu33391 | 1 | k__Fungi(100);p__Ascomycota(100);c__Saccharomycetes(100);o__Saccharomycetales(100);f__Metschnikowiaceae(100);g__Metschnikowia(100);s__Metschnikowia_pulcherrima(100); | Metschnikowia | 13 | Saprotroph | \|Undefined Saprotroph\| | Yeast | NULL | Probable | As nectar-tap_saprotroph (Põlme et al. 2020); Decay sugar-rich substrates (Põlme et al. 2020) | Põlme S, et al. 2020. Fungal Diversity 105:1-16 ((https://doi.org/10.1007/s13225-020-00466-2)) |
| Otu33392 | 1 | k__Fungi(100);p__Ascomycota(100);c__Saccharomycetes(100);o__Saccharomycetales(100);f__Metschnikowiaceae(100);g__Metschnikowia(100);g__Metschnikowia_unclassified(100); | Metschnikowia | 13 | Saprotroph | \|Undefined Saprotroph\| | Yeast | NULL | Probable | As nectar-tap_saprotroph (Põlme et al. 2020); Decay sugar-rich substrates (Põlme et al. 2020) | Põlme S, et al. 2020. Fungal Diversity 105:1-16 ((https://doi.org/10.1007/s13225-020-00466-2)) |
| Otu33393 | 1 | k__Fungi(100);p__Ascomycota(100);c__Saccharomycetes(100);o__Saccharomycetales(100);f__Metschnikowiaceae(100);g__Metschnikowia(100);s__Metschnikowia_pulcherrima(100); | Metschnikowia | 13 | Saprotroph | \|Undefined Saprotroph\| | Yeast | NULL | Probable | As nectar-tap_saprotroph (Põlme et al. 2020); Decay sugar-rich substrates (Põlme et al. 2020) | Põlme S, et al. 2020. Fungal Diversity 105:1-16 ((https://doi.org/10.1007/s13225-020-00466-2)) |
| Otu33394 | 1 | k__Fungi(100);p__Ascomycota(100);c__Saccharomycetes(100);o__Saccharomycetales(100);f__Metschnikowiaceae(100);g__Metschnikowia(100);s__Metschnikowia_pulcherrima(100); | Metschnikowia | 13 | Saprotroph | \|Undefined Saprotroph\| | Yeast | NULL | Probable | As nectar-tap_saprotroph (Põlme et al. 2020); Decay sugar-rich substrates (Põlme et al. 2020) | Põlme S, et al. 2020. Fungal Diversity 105:1-16 ((https://doi.org/10.1007/s13225-020-00466-2)) |
| Otu33396 | 1 | k__Fungi(100);p__Ascomycota(100);c__Saccharomycetes(100);o__Saccharomycetales(100);f__Metschnikowiaceae(100);g__Metschnikowia(100);s__Metschnikowia_pulcherrima(100); | Metschnikowia | 13 | Saprotroph | \|Undefined Saprotroph\| | Yeast | NULL | Probable | As nectar-tap_saprotroph (Põlme et al. 2020); Decay sugar-rich substrates (Põlme et al. 2020) | Põlme S, et al. 2020. Fungal Diversity 105:1-16 ((https://doi.org/10.1007/s13225-020-00466-2)) |
| Otu33397 | 1 | k__Fungi(100);p__Ascomycota(100);c__Saccharomycetes(100);o__Saccharomycetales(100);f__Metschnikowiaceae(100);g__Metschnikowia(100);s__Metschnikowia_pulcherrima(100); | Metschnikowia | 13 | Saprotroph | \|Undefined Saprotroph\| | Yeast | NULL | Probable | As nectar-tap_saprotroph (Põlme et al. 2020); Decay sugar-rich substrates (Põlme et al. 2020) | Põlme S, et al. 2020. Fungal Diversity 105:1-16 ((https://doi.org/10.1007/s13225-020-00466-2)) |
| Otu33398 | 1 | k__Fungi(100);p__Ascomycota(100);c__Saccharomycetes(100);o__Saccharomycetales(100);f__Metschnikowiaceae(100);g__Metschnikowia(100);g__Metschnikowia_unclassified(100); | Metschnikowia | 13 | Saprotroph | \|Undefined Saprotroph\| | Yeast | NULL | Probable | As nectar-tap_saprotroph (Põlme et al. 2020); Decay sugar-rich substrates (Põlme et al. 2020) | Põlme S, et al. 2020. Fungal Diversity 105:1-16 ((https://doi.org/10.1007/s13225-020-00466-2)) |
| Otu33399 | 1 | k__Fungi(100);p__Ascomycota(100);c__Saccharomycetes(100);o__Saccharomycetales(100);f__Metschnikowiaceae(100);g__Metschnikowia(100);s__Metschnikowia_pulcherrima(100); | Metschnikowia | 13 | Saprotroph | \|Undefined Saprotroph\| | Yeast | NULL | Probable | As nectar-tap_saprotroph (Põlme et al. 2020); Decay sugar-rich substrates (Põlme et al. 2020) | Põlme S, et al. 2020. Fungal Diversity 105:1-16 ((https://doi.org/10.1007/s13225-020-00466-2)) |
| Otu33401 | 1 | k__Fungi(100);p__Ascomycota(100);c__Saccharomycetes(100);o__Saccharomycetales(100);f__Metschnikowiaceae(100);g__Metschnikowia(100);g__Metschnikowia_unclassified(100); | Metschnikowia | 13 | Saprotroph | \|Undefined Saprotroph\| | Yeast | NULL | Probable | As nectar-tap_saprotroph (Põlme et al. 2020); Decay sugar-rich substrates (Põlme et al. 2020) | Põlme S, et al. 2020. Fungal Diversity 105:1-16 ((https://doi.org/10.1007/s13225-020-00466-2)) |
| Otu33402 | 1 | k__Fungi(100);p__Ascomycota(100);c__Saccharomycetes(100);o__Saccharomycetales(100);f__Metschnikowiaceae(100);g__Metschnikowia(100);s__Metschnikowia_pulcherrima(100); | Metschnikowia | 13 | Saprotroph | \|Undefined Saprotroph\| | Yeast | NULL | Probable | As nectar-tap_saprotroph (Põlme et al. 2020); Decay sugar-rich substrates (Põlme et al. 2020) | Põlme S, et al. 2020. Fungal Diversity 105:1-16 ((https://doi.org/10.1007/s13225-020-00466-2)) |
| Otu33407 | 1 | k__Fungi(100);p__Ascomycota(100);c__Saccharomycetes(100);o__Saccharomycetales(100);f__Metschnikowiaceae(100);g__Metschnikowia(100);s__Metschnikowia_pulcherrima(100); | Metschnikowia | 13 | Saprotroph | \|Undefined Saprotroph\| | Yeast | NULL | Probable | As nectar-tap_saprotroph (Põlme et al. 2020); Decay sugar-rich substrates (Põlme et al. 2020) | Põlme S, et al. 2020. Fungal Diversity 105:1-16 ((https://doi.org/10.1007/s13225-020-00466-2)) |
| Otu33410 | 1 | k__Fungi(100);p__Ascomycota(100);c__Saccharomycetes(100);o__Saccharomycetales(100);f__Saccharomycetales_family_Incertae_sedis(100);g__Candida(100);g__Candida_unclassified(100); | Candida | 13 | Pathotroph-Saprotroph-Symbiotroph | Animal Pathogen-Endophyte-Endosymbiont-Epiphyte-\|Undefined Saprotroph\| | Dimorphic Yeast-Yeast | NULL | Probable | As nectar-tap_saprotroph (Põlme et al. 2020); Decay sugar-rich substrates (Põlme et al. 2020); This asexual genus is polyphyletic and contain species that function very differently depend on what they are. Caution should be taken when interpreting guilds for Candida species. | Manolakaki D, et al. 2010. Virulence. 1: 367-375 ((https://doi.org/10.4161/viru.1.5.12796)); Mekbib SB, et al. 2011. Tropical Plant Pathology 36:233-240 ((https://doi.org/10.1590/S1982-56762011000400004)); Põlme S, et al. 2020. Fungal Diversity 105:1-16 ((https://doi.org/10.1007/s13225-020-00466-2)) |
| Otu33412 | 1 | k__Fungi(100);p__Ascomycota(100);c__Saccharomycetes(100);o__Saccharomycetales(100);f__Metschnikowiaceae(100);g__Metschnikowia(100);g__Metschnikowia_unclassified(100); | Metschnikowia | 13 | Saprotroph | \|Undefined Saprotroph\| | Yeast | NULL | Probable | As nectar-tap_saprotroph (Põlme et al. 2020); Decay sugar-rich substrates (Põlme et al. 2020) | Põlme S, et al. 2020. Fungal Diversity 105:1-16 ((https://doi.org/10.1007/s13225-020-00466-2)) |
| Otu33413 | 1 | k__Fungi(100);p__Ascomycota(100);c__Saccharomycetes(100);o__Saccharomycetales(100);f__Metschnikowiaceae(100);g__Metschnikowia(100);s__Metschnikowia_pulcherrima(100); | Metschnikowia | 13 | Saprotroph | \|Undefined Saprotroph\| | Yeast | NULL | Probable | As nectar-tap_saprotroph (Põlme et al. 2020); Decay sugar-rich substrates (Põlme et al. 2020) | Põlme S, et al. 2020. Fungal Diversity 105:1-16 ((https://doi.org/10.1007/s13225-020-00466-2)) |
| Otu33415 | 1 | k__Fungi(100);p__Basidiomycota(100);c__Microbotryomycetes(100);o__Sporidiobolales(100);f__Sporidiobolales_family_Incertae_sedis(100);g__Sporobolomyces(100);s__Sporobolomyces_lactosus(100); | Sporobolomyces | 13 | Pathotroph-Saprotroph | \|Fungal Parasite\|-Plant Saprotroph | Yeast | NULL | Probable | As litter_saprotroph (Põlme et al. 2020); As mycoparasite (Põlme et al. 2020); Decay fungal material (Põlme et al. 2020) | Tedersoo L, et al. 2014. Science 346:e1256688 ((https://doi.org/10.1126/science.1256688)); Purahong W, et al. 2016. Molecular Ecology 25:4059-4074 ((https://doi.org/10.1111/mec.13739)); Põlme S, et al. 2020. Fungal Diversity 105:1-16 ((https://doi.org/10.1007/s13225-020-00466-2)) |
| Otu33416 | 1 | k__Fungi(100);p__Ascomycota(100);c__Saccharomycetes(100);o__Saccharomycetales(100);f__Metschnikowiaceae(100);g__Metschnikowia(100);s__Metschnikowia_pulcherrima(100); | Metschnikowia | 13 | Saprotroph | \|Undefined Saprotroph\| | Yeast | NULL | Probable | As nectar-tap_saprotroph (Põlme et al. 2020); Decay sugar-rich substrates (Põlme et al. 2020) | Põlme S, et al. 2020. Fungal Diversity 105:1-16 ((https://doi.org/10.1007/s13225-020-00466-2)) |
| Otu33417 | 1 | k__Fungi(100);p__Ascomycota(100);c__Saccharomycetes(100);o__Saccharomycetales(100);f__Metschnikowiaceae(100);g__Metschnikowia(100);s__Metschnikowia_pulcherrima(100); | Metschnikowia | 13 | Saprotroph | \|Undefined Saprotroph\| | Yeast | NULL | Probable | As nectar-tap_saprotroph (Põlme et al. 2020); Decay sugar-rich substrates (Põlme et al. 2020) | Põlme S, et al. 2020. Fungal Diversity 105:1-16 ((https://doi.org/10.1007/s13225-020-00466-2)) |
| Otu33418 | 1 | k__Fungi(100);p__Ascomycota(100);c__Saccharomycetes(100);o__Saccharomycetales(100);f__Metschnikowiaceae(100);g__Metschnikowia(100);s__Metschnikowia_pulcherrima(100); | Metschnikowia | 13 | Saprotroph | \|Undefined Saprotroph\| | Yeast | NULL | Probable | As nectar-tap_saprotroph (Põlme et al. 2020); Decay sugar-rich substrates (Põlme et al. 2020) | Põlme S, et al. 2020. Fungal Diversity 105:1-16 ((https://doi.org/10.1007/s13225-020-00466-2)) |
| Otu33419 | 1 | k__Fungi(100);p__Ascomycota(100);c__Saccharomycetes(100);o__Saccharomycetales(100);f__Metschnikowiaceae(100);g__Metschnikowia(100);s__Metschnikowia_pulcherrima(100); | Metschnikowia | 13 | Saprotroph | \|Undefined Saprotroph\| | Yeast | NULL | Probable | As nectar-tap_saprotroph (Põlme et al. 2020); Decay sugar-rich substrates (Põlme et al. 2020) | Põlme S, et al. 2020. Fungal Diversity 105:1-16 ((https://doi.org/10.1007/s13225-020-00466-2)) |
| Otu33420 | 1 | k__Fungi(100);p__Ascomycota(100);c__Saccharomycetes(100);o__Saccharomycetales(100);f__Metschnikowiaceae(100);g__Metschnikowia(100);s__Metschnikowia_pulcherrima(100); | Metschnikowia | 13 | Saprotroph | \|Undefined Saprotroph\| | Yeast | NULL | Probable | As nectar-tap_saprotroph (Põlme et al. 2020); Decay sugar-rich substrates (Põlme et al. 2020) | Põlme S, et al. 2020. Fungal Diversity 105:1-16 ((https://doi.org/10.1007/s13225-020-00466-2)) |
| Otu33421 | 1 | k__Fungi(100);p__Ascomycota(100);c__Saccharomycetes(100);o__Saccharomycetales(100);f__Metschnikowiaceae(100);g__Metschnikowia(100);s__Metschnikowia_pulcherrima(100); | Metschnikowia | 13 | Saprotroph | \|Undefined Saprotroph\| | Yeast | NULL | Probable | As nectar-tap_saprotroph (Põlme et al. 2020); Decay sugar-rich substrates (Põlme et al. 2020) | Põlme S, et al. 2020. Fungal Diversity 105:1-16 ((https://doi.org/10.1007/s13225-020-00466-2)) |
| Otu33422 | 1 | k__Fungi(100);p__Ascomycota(100);c__Saccharomycetes(100);o__Saccharomycetales(100);f__Metschnikowiaceae(100);g__Metschnikowia(100);s__Metschnikowia_pulcherrima(100); | Metschnikowia | 13 | Saprotroph | \|Undefined Saprotroph\| | Yeast | NULL | Probable | As nectar-tap_saprotroph (Põlme et al. 2020); Decay sugar-rich substrates (Põlme et al. 2020) | Põlme S, et al. 2020. Fungal Diversity 105:1-16 ((https://doi.org/10.1007/s13225-020-00466-2)) |
| Otu33423 | 1 | k__Fungi(100);p__Ascomycota(100);c__Saccharomycetes(100);o__Saccharomycetales(100);f__Metschnikowiaceae(100);g__Metschnikowia(100);s__Metschnikowia_pulcherrima(100); | Metschnikowia | 13 | Saprotroph | \|Undefined Saprotroph\| | Yeast | NULL | Probable | As nectar-tap_saprotroph (Põlme et al. 2020); Decay sugar-rich substrates (Põlme et al. 2020) | Põlme S, et al. 2020. Fungal Diversity 105:1-16 ((https://doi.org/10.1007/s13225-020-00466-2)) |
| Otu33424 | 1 | k__Fungi(100);p__Ascomycota(100);c__Saccharomycetes(100);o__Saccharomycetales(100);f__Metschnikowiaceae(100);g__Metschnikowia(100);s__Metschnikowia_pulcherrima(100); | Metschnikowia | 13 | Saprotroph | \|Undefined Saprotroph\| | Yeast | NULL | Probable | As nectar-tap_saprotroph (Põlme et al. 2020); Decay sugar-rich substrates (Põlme et al. 2020) | Põlme S, et al. 2020. Fungal Diversity 105:1-16 ((https://doi.org/10.1007/s13225-020-00466-2)) |
| Otu33425 | 1 | k__Fungi(100);p__Ascomycota(100);c__Saccharomycetes(100);o__Saccharomycetales(100);f__Metschnikowiaceae(100);g__Metschnikowia(100);s__Metschnikowia_pulcherrima(100); | Metschnikowia | 13 | Saprotroph | \|Undefined Saprotroph\| | Yeast | NULL | Probable | As nectar-tap_saprotroph (Põlme et al. 2020); Decay sugar-rich substrates (Põlme et al. 2020) | Põlme S, et al. 2020. Fungal Diversity 105:1-16 ((https://doi.org/10.1007/s13225-020-00466-2)) |
| Otu33426 | 1 | k__Fungi(100);p__Ascomycota(100);c__Saccharomycetes(100);o__Saccharomycetales(100);f__Metschnikowiaceae(100);g__Metschnikowia(100);s__Metschnikowia_pulcherrima(100); | Metschnikowia | 13 | Saprotroph | \|Undefined Saprotroph\| | Yeast | NULL | Probable | As nectar-tap_saprotroph (Põlme et al. 2020); Decay sugar-rich substrates (Põlme et al. 2020) | Põlme S, et al. 2020. Fungal Diversity 105:1-16 ((https://doi.org/10.1007/s13225-020-00466-2)) |
| Otu33428 | 1 | k__Fungi(100);p__Ascomycota(100);c__Saccharomycetes(100);o__Saccharomycetales(100);f__Saccharomycetales_family_Incertae_sedis(100);g__Candida(100);g__Candida_unclassified(100); | Candida | 13 | Pathotroph-Saprotroph-Symbiotroph | Animal Pathogen-Endophyte-Endosymbiont-Epiphyte-\|Undefined Saprotroph\| | Dimorphic Yeast-Yeast | NULL | Probable | As nectar-tap_saprotroph (Põlme et al. 2020); Decay sugar-rich substrates (Põlme et al. 2020); This asexual genus is polyphyletic and contain species that function very differently depend on what they are. Caution should be taken when interpreting guilds for Candida species. | Manolakaki D, et al. 2010. Virulence. 1: 367-375 ((https://doi.org/10.4161/viru.1.5.12796)); Mekbib SB, et al. 2011. Tropical Plant Pathology 36:233-240 ((https://doi.org/10.1590/S1982-56762011000400004)); Põlme S, et al. 2020. Fungal Diversity 105:1-16 ((https://doi.org/10.1007/s13225-020-00466-2)) |
| Otu33430 | 1 | k__Fungi(100);p__Ascomycota(100);c__Saccharomycetes(100);o__Saccharomycetales(100);f__Metschnikowiaceae(100);g__Metschnikowia(100);g__Metschnikowia_unclassified(100); | Metschnikowia | 13 | Saprotroph | \|Undefined Saprotroph\| | Yeast | NULL | Probable | As nectar-tap_saprotroph (Põlme et al. 2020); Decay sugar-rich substrates (Põlme et al. 2020) | Põlme S, et al. 2020. Fungal Diversity 105:1-16 ((https://doi.org/10.1007/s13225-020-00466-2)) |
| Otu33431 | 1 | k__Fungi(100);p__Ascomycota(100);c__Saccharomycetes(100);o__Saccharomycetales(100);f__Metschnikowiaceae(100);g__Metschnikowia(100);s__Metschnikowia_pulcherrima(100); | Metschnikowia | 13 | Saprotroph | \|Undefined Saprotroph\| | Yeast | NULL | Probable | As nectar-tap_saprotroph (Põlme et al. 2020); Decay sugar-rich substrates (Põlme et al. 2020) | Põlme S, et al. 2020. Fungal Diversity 105:1-16 ((https://doi.org/10.1007/s13225-020-00466-2)) |
| Otu33432 | 1 | k__Fungi(100);p__Ascomycota(100);c__Dothideomycetes(100);o__Dothideomycetes_order_Incertae_sedis(100);f__Pseudeurotiaceae(100);g__Pseudeurotium(100);g__Pseudeurotium_unclassified(100); | Pseudeurotium | 13 | Saprotroph | \|Dung Saprotroph\|-Undefined Saprotroph | Microfungus | NULL | Probable | As soil_saprotroph (Põlme et al. 2020); Decay soil, dung (Põlme et al. 2020) | Tedersoo L, et al. 2014. Science 346:e1256688 ((https://doi.org/10.1126/science.1256688)); Põlme S, et al. 2020. Fungal Diversity 105:1-16 ((https://doi.org/10.1007/s13225-020-00466-2)) |
| Otu33433 | 1 | k__Fungi(100);p__Ascomycota(100);c__Saccharomycetes(100);o__Saccharomycetales(100);f__Metschnikowiaceae(100);g__Metschnikowia(100);s__Metschnikowia_pulcherrima(100); | Metschnikowia | 13 | Saprotroph | \|Undefined Saprotroph\| | Yeast | NULL | Probable | As nectar-tap_saprotroph (Põlme et al. 2020); Decay sugar-rich substrates (Põlme et al. 2020) | Põlme S, et al. 2020. Fungal Diversity 105:1-16 ((https://doi.org/10.1007/s13225-020-00466-2)) |
| Otu33434 | 1 | k__Fungi(100);p__Ascomycota(100);c__Saccharomycetes(100);o__Saccharomycetales(100);f__Metschnikowiaceae(100);g__Metschnikowia(100);s__Metschnikowia_pulcherrima(100); | Metschnikowia | 13 | Saprotroph | \|Undefined Saprotroph\| | Yeast | NULL | Probable | As nectar-tap_saprotroph (Põlme et al. 2020); Decay sugar-rich substrates (Põlme et al. 2020) | Põlme S, et al. 2020. Fungal Diversity 105:1-16 ((https://doi.org/10.1007/s13225-020-00466-2)) |
| Otu33436 | 1 | k__Fungi(100);p__Ascomycota(100);c__Saccharomycetes(100);o__Saccharomycetales(100);f__Saccharomycetales_family_Incertae_sedis(100);g__Candida(100);g__Candida_unclassified(100); | Candida | 13 | Pathotroph-Saprotroph-Symbiotroph | Animal Pathogen-Endophyte-Endosymbiont-Epiphyte-\|Undefined Saprotroph\| | Dimorphic Yeast-Yeast | NULL | Probable | As nectar-tap_saprotroph (Põlme et al. 2020); Decay sugar-rich substrates (Põlme et al. 2020); This asexual genus is polyphyletic and contain species that function very differently depend on what they are. Caution should be taken when interpreting guilds for Candida species. | Manolakaki D, et al. 2010. Virulence. 1: 367-375 ((https://doi.org/10.4161/viru.1.5.12796)); Mekbib SB, et al. 2011. Tropical Plant Pathology 36:233-240 ((https://doi.org/10.1590/S1982-56762011000400004)); Põlme S, et al. 2020. Fungal Diversity 105:1-16 ((https://doi.org/10.1007/s13225-020-00466-2)) |
| Otu33437 | 1 | k__Fungi(100);p__Ascomycota(100);c__Saccharomycetes(100);o__Saccharomycetales(100);f__Metschnikowiaceae(100);g__Metschnikowia(100);s__Metschnikowia_pulcherrima(100); | Metschnikowia | 13 | Saprotroph | \|Undefined Saprotroph\| | Yeast | NULL | Probable | As nectar-tap_saprotroph (Põlme et al. 2020); Decay sugar-rich substrates (Põlme et al. 2020) | Põlme S, et al. 2020. Fungal Diversity 105:1-16 ((https://doi.org/10.1007/s13225-020-00466-2)) |
| Otu33438 | 1 | k__Fungi(100);p__Ascomycota(100);c__Saccharomycetes(100);o__Saccharomycetales(100);f__Metschnikowiaceae(100);g__Metschnikowia(100);g__Metschnikowia_unclassified(100); | Metschnikowia | 13 | Saprotroph | \|Undefined Saprotroph\| | Yeast | NULL | Probable | As nectar-tap_saprotroph (Põlme et al. 2020); Decay sugar-rich substrates (Põlme et al. 2020) | Põlme S, et al. 2020. Fungal Diversity 105:1-16 ((https://doi.org/10.1007/s13225-020-00466-2)) |
| Otu33439 | 1 | k__Fungi(100);p__Ascomycota(100);c__Saccharomycetes(100);o__Saccharomycetales(100);f__Metschnikowiaceae(100);g__Metschnikowia(100);g__Metschnikowia_unclassified(100); | Metschnikowia | 13 | Saprotroph | \|Undefined Saprotroph\| | Yeast | NULL | Probable | As nectar-tap_saprotroph (Põlme et al. 2020); Decay sugar-rich substrates (Põlme et al. 2020) | Põlme S, et al. 2020. Fungal Diversity 105:1-16 ((https://doi.org/10.1007/s13225-020-00466-2)) |
| Otu33440 | 1 | k__Fungi(100);p__Ascomycota(100);c__Saccharomycetes(100);o__Saccharomycetales(100);f__Metschnikowiaceae(100);g__Metschnikowia(100);s__Metschnikowia_pulcherrima(100); | Metschnikowia | 13 | Saprotroph | \|Undefined Saprotroph\| | Yeast | NULL | Probable | As nectar-tap_saprotroph (Põlme et al. 2020); Decay sugar-rich substrates (Põlme et al. 2020) | Põlme S, et al. 2020. Fungal Diversity 105:1-16 ((https://doi.org/10.1007/s13225-020-00466-2)) |
| Otu33441 | 1 | k__Fungi(100);p__Ascomycota(100);c__Saccharomycetes(100);o__Saccharomycetales(100);f__Metschnikowiaceae(100);g__Metschnikowia(100);g__Metschnikowia_unclassified(100); | Metschnikowia | 13 | Saprotroph | \|Undefined Saprotroph\| | Yeast | NULL | Probable | As nectar-tap_saprotroph (Põlme et al. 2020); Decay sugar-rich substrates (Põlme et al. 2020) | Põlme S, et al. 2020. Fungal Diversity 105:1-16 ((https://doi.org/10.1007/s13225-020-00466-2)) |
| Otu33442 | 1 | k__Fungi(100);p__Ascomycota(100);c__Saccharomycetes(100);o__Saccharomycetales(100);f__Metschnikowiaceae(100);g__Metschnikowia(100);s__Metschnikowia_pulcherrima(100); | Metschnikowia | 13 | Saprotroph | \|Undefined Saprotroph\| | Yeast | NULL | Probable | As nectar-tap_saprotroph (Põlme et al. 2020); Decay sugar-rich substrates (Põlme et al. 2020) | Põlme S, et al. 2020. Fungal Diversity 105:1-16 ((https://doi.org/10.1007/s13225-020-00466-2)) |
| Otu33443 | 1 | k__Fungi(100);p__Ascomycota(100);c__Saccharomycetes(100);o__Saccharomycetales(100);f__Metschnikowiaceae(100);g__Metschnikowia(100);g__Metschnikowia_unclassified(100); | Metschnikowia | 13 | Saprotroph | \|Undefined Saprotroph\| | Yeast | NULL | Probable | As nectar-tap_saprotroph (Põlme et al. 2020); Decay sugar-rich substrates (Põlme et al. 2020) | Põlme S, et al. 2020. Fungal Diversity 105:1-16 ((https://doi.org/10.1007/s13225-020-00466-2)) |
| Otu33446 | 1 | k__Fungi(100);p__Ascomycota(100);c__Saccharomycetes(100);o__Saccharomycetales(100);f__Metschnikowiaceae(100);g__Metschnikowia(100);s__Metschnikowia_pulcherrima(100); | Metschnikowia | 13 | Saprotroph | \|Undefined Saprotroph\| | Yeast | NULL | Probable | As nectar-tap_saprotroph (Põlme et al. 2020); Decay sugar-rich substrates (Põlme et al. 2020) | Põlme S, et al. 2020. Fungal Diversity 105:1-16 ((https://doi.org/10.1007/s13225-020-00466-2)) |
| Otu33447 | 1 | k__Fungi(100);p__Ascomycota(100);c__Saccharomycetes(100);o__Saccharomycetales(100);f__Metschnikowiaceae(100);g__Metschnikowia(100);s__Metschnikowia_pulcherrima(100); | Metschnikowia | 13 | Saprotroph | \|Undefined Saprotroph\| | Yeast | NULL | Probable | As nectar-tap_saprotroph (Põlme et al. 2020); Decay sugar-rich substrates (Põlme et al. 2020) | Põlme S, et al. 2020. Fungal Diversity 105:1-16 ((https://doi.org/10.1007/s13225-020-00466-2)) |
| Otu33448 | 1 | k__Fungi(100);p__Ascomycota(100);c__Saccharomycetes(100);o__Saccharomycetales(100);f__Metschnikowiaceae(100);g__Metschnikowia(100);s__Metschnikowia_pulcherrima(100); | Metschnikowia | 13 | Saprotroph | \|Undefined Saprotroph\| | Yeast | NULL | Probable | As nectar-tap_saprotroph (Põlme et al. 2020); Decay sugar-rich substrates (Põlme et al. 2020) | Põlme S, et al. 2020. Fungal Diversity 105:1-16 ((https://doi.org/10.1007/s13225-020-00466-2)) |
| Otu33449 | 1 | k__Fungi(100);p__Ascomycota(100);c__Saccharomycetes(100);o__Saccharomycetales(100);f__Metschnikowiaceae(100);g__Metschnikowia(100);s__Metschnikowia_pulcherrima(100); | Metschnikowia | 13 | Saprotroph | \|Undefined Saprotroph\| | Yeast | NULL | Probable | As nectar-tap_saprotroph (Põlme et al. 2020); Decay sugar-rich substrates (Põlme et al. 2020) | Põlme S, et al. 2020. Fungal Diversity 105:1-16 ((https://doi.org/10.1007/s13225-020-00466-2)) |
| Otu33454 | 1 | k__Fungi(100);p__Basidiomycota(100);c__Microbotryomycetes(100);o__Sporidiobolales(100);f__Sporidiobolales_family_Incertae_sedis(100);g__Sporobolomyces(100);s__Sporobolomyces_lactosus(100); | Sporobolomyces | 13 | Pathotroph-Saprotroph | \|Fungal Parasite\|-Plant Saprotroph | Yeast | NULL | Probable | As litter_saprotroph (Põlme et al. 2020); As mycoparasite (Põlme et al. 2020); Decay fungal material (Põlme et al. 2020) | Tedersoo L, et al. 2014. Science 346:e1256688 ((https://doi.org/10.1126/science.1256688)); Purahong W, et al. 2016. Molecular Ecology 25:4059-4074 ((https://doi.org/10.1111/mec.13739)); Põlme S, et al. 2020. Fungal Diversity 105:1-16 ((https://doi.org/10.1007/s13225-020-00466-2)) |
| Otu33459 | 1 | k__Fungi(100);p__Ascomycota(100);c__Saccharomycetes(100);o__Saccharomycetales(100);f__Saccharomycetales_family_Incertae_sedis(100);g__Candida(100);g__Candida_unclassified(100); | Candida | 13 | Pathotroph-Saprotroph-Symbiotroph | Animal Pathogen-Endophyte-Endosymbiont-Epiphyte-\|Undefined Saprotroph\| | Dimorphic Yeast-Yeast | NULL | Probable | As nectar-tap_saprotroph (Põlme et al. 2020); Decay sugar-rich substrates (Põlme et al. 2020); This asexual genus is polyphyletic and contain species that function very differently depend on what they are. Caution should be taken when interpreting guilds for Candida species. | Manolakaki D, et al. 2010. Virulence. 1: 367-375 ((https://doi.org/10.4161/viru.1.5.12796)); Mekbib SB, et al. 2011. Tropical Plant Pathology 36:233-240 ((https://doi.org/10.1590/S1982-56762011000400004)); Põlme S, et al. 2020. Fungal Diversity 105:1-16 ((https://doi.org/10.1007/s13225-020-00466-2)) |
| Otu33462 | 1 | k__Fungi(100);p__Ascomycota(100);c__Saccharomycetes(100);o__Saccharomycetales(100);f__Saccharomycetales_family_Incertae_sedis(100);f__Saccharomycetales_family_Incertae_sedis_unclassified(100);f__Saccharomycetales_family_Incertae_sedis_unclassified(100); | Saccharomycetales | 7 | Saprotroph | Undefined Saprotroph | Yeast | NULL | Possible | NULL | Sterkenburg E, et al. 2015. New Phytologist 207:1145-1158 ((https://doi.org/10.1111/nph.13426)) |
| Otu33463 | 1 | k__Fungi(100);p__Basidiomycota(100);c__Tremellomycetes(100);o__Tremellales(100);f__Tremellales_family_Incertae_sedis(100);g__Cryptococcus(100);s__Cryptococcus_sp_JCM_11356(100); | Cryptococcus | 13 | Pathotroph-Saprotroph-Symbiotroph | Animal Pathogen-Endophyte-Epiphyte-\|Undefined Saprotroph\| | Dimorphic Yeast-Yeast | NULL | Possible | As foliar_endophyte (Põlme et al. 2020); Decay leaf, fruit, seed, soil, dung, animal material (Põlme et al. 2020) | Andrews JH, et al. 1983. Phytopathology 73:228-234 ((https://doi.org/10.1094/Phyto-73-228)); Kurtzman CP, et al. (eds.) 2011. The Yeasts, a Taxonomic Study. Fifth Edition. Vols 1-3. Elsevier, San Diego (ISBN: 9780444521491); Põlme S, et al. 2020. Fungal Diversity 105:1-16 ((https://doi.org/10.1007/s13225-020-00466-2)) |
| Otu33464 | 1 | k__Fungi(100);p__Ascomycota(100);c__Saccharomycetes(100);o__Saccharomycetales(100);f__Saccharomycetales_family_Incertae_sedis(100);g__Candida(100);g__Candida_unclassified(100); | Candida | 13 | Pathotroph-Saprotroph-Symbiotroph | Animal Pathogen-Endophyte-Endosymbiont-Epiphyte-\|Undefined Saprotroph\| | Dimorphic Yeast-Yeast | NULL | Probable | As nectar-tap_saprotroph (Põlme et al. 2020); Decay sugar-rich substrates (Põlme et al. 2020); This asexual genus is polyphyletic and contain species that function very differently depend on what they are. Caution should be taken when interpreting guilds for Candida species. | Manolakaki D, et al. 2010. Virulence. 1: 367-375 ((https://doi.org/10.4161/viru.1.5.12796)); Mekbib SB, et al. 2011. Tropical Plant Pathology 36:233-240 ((https://doi.org/10.1590/S1982-56762011000400004)); Põlme S, et al. 2020. Fungal Diversity 105:1-16 ((https://doi.org/10.1007/s13225-020-00466-2)) |
| Otu33466 | 1 | k__Fungi(100);p__Ascomycota(100);c__Saccharomycetes(100);o__Saccharomycetales(100);f__Metschnikowiaceae(100);g__Metschnikowia(100);g__Metschnikowia_unclassified(100); | Metschnikowia | 13 | Saprotroph | \|Undefined Saprotroph\| | Yeast | NULL | Probable | As nectar-tap_saprotroph (Põlme et al. 2020); Decay sugar-rich substrates (Põlme et al. 2020) | Põlme S, et al. 2020. Fungal Diversity 105:1-16 ((https://doi.org/10.1007/s13225-020-00466-2)) |
| Otu33468 | 1 | k__Fungi(100);p__Ascomycota(100);c__Saccharomycetes(100);o__Saccharomycetales(100);f__Metschnikowiaceae(100);g__Metschnikowia(100);g__Metschnikowia_unclassified(100); | Metschnikowia | 13 | Saprotroph | \|Undefined Saprotroph\| | Yeast | NULL | Probable | As nectar-tap_saprotroph (Põlme et al. 2020); Decay sugar-rich substrates (Põlme et al. 2020) | Põlme S, et al. 2020. Fungal Diversity 105:1-16 ((https://doi.org/10.1007/s13225-020-00466-2)) |
| Otu33470 | 1 | k__Fungi(100);p__Ascomycota(100);c__Saccharomycetes(100);o__Saccharomycetales(100);f__Metschnikowiaceae(100);g__Metschnikowia(100);s__Metschnikowia_pulcherrima(100); | Metschnikowia | 13 | Saprotroph | \|Undefined Saprotroph\| | Yeast | NULL | Probable | As nectar-tap_saprotroph (Põlme et al. 2020); Decay sugar-rich substrates (Põlme et al. 2020) | Põlme S, et al. 2020. Fungal Diversity 105:1-16 ((https://doi.org/10.1007/s13225-020-00466-2)) |
| Otu33471 | 1 | k__Fungi(100);p__Ascomycota(100);c__Saccharomycetes(100);o__Saccharomycetales(100);f__Metschnikowiaceae(100);g__Metschnikowia(100);g__Metschnikowia_unclassified(100); | Metschnikowia | 13 | Saprotroph | \|Undefined Saprotroph\| | Yeast | NULL | Probable | As nectar-tap_saprotroph (Põlme et al. 2020); Decay sugar-rich substrates (Põlme et al. 2020) | Põlme S, et al. 2020. Fungal Diversity 105:1-16 ((https://doi.org/10.1007/s13225-020-00466-2)) |
| Otu33472 | 1 | k__Fungi(100);p__Ascomycota(100);c__Saccharomycetes(100);o__Saccharomycetales(100);f__Metschnikowiaceae(100);g__Metschnikowia(100);s__Metschnikowia_pulcherrima(100); | Metschnikowia | 13 | Saprotroph | \|Undefined Saprotroph\| | Yeast | NULL | Probable | As nectar-tap_saprotroph (Põlme et al. 2020); Decay sugar-rich substrates (Põlme et al. 2020) | Põlme S, et al. 2020. Fungal Diversity 105:1-16 ((https://doi.org/10.1007/s13225-020-00466-2)) |
| Otu33473 | 1 | k__Fungi(100);p__Ascomycota(100);c__Saccharomycetes(100);o__Saccharomycetales(100);f__Metschnikowiaceae(100);g__Metschnikowia(100);s__Metschnikowia_pulcherrima(100); | Metschnikowia | 13 | Saprotroph | \|Undefined Saprotroph\| | Yeast | NULL | Probable | As nectar-tap_saprotroph (Põlme et al. 2020); Decay sugar-rich substrates (Põlme et al. 2020) | Põlme S, et al. 2020. Fungal Diversity 105:1-16 ((https://doi.org/10.1007/s13225-020-00466-2)) |
| Otu33474 | 1 | k__Fungi(100);p__Ascomycota(100);c__Saccharomycetes(100);o__Saccharomycetales(100);f__Metschnikowiaceae(100);g__Metschnikowia(100);g__Metschnikowia_unclassified(100); | Metschnikowia | 13 | Saprotroph | \|Undefined Saprotroph\| | Yeast | NULL | Probable | As nectar-tap_saprotroph (Põlme et al. 2020); Decay sugar-rich substrates (Põlme et al. 2020) | Põlme S, et al. 2020. Fungal Diversity 105:1-16 ((https://doi.org/10.1007/s13225-020-00466-2)) |
| Otu33475 | 1 | k__Fungi(100);p__Basidiomycota(100);c__Microbotryomycetes(100);o__Sporidiobolales(100);f__Sporidiobolales_family_Incertae_sedis(100);g__Sporobolomyces(100);s__Sporobolomyces_lactosus(100); | Sporobolomyces | 13 | Pathotroph-Saprotroph | \|Fungal Parasite\|-Plant Saprotroph | Yeast | NULL | Probable | As litter_saprotroph (Põlme et al. 2020); As mycoparasite (Põlme et al. 2020); Decay fungal material (Põlme et al. 2020) | Tedersoo L, et al. 2014. Science 346:e1256688 ((https://doi.org/10.1126/science.1256688)); Purahong W, et al. 2016. Molecular Ecology 25:4059-4074 ((https://doi.org/10.1111/mec.13739)); Põlme S, et al. 2020. Fungal Diversity 105:1-16 ((https://doi.org/10.1007/s13225-020-00466-2)) |
| Otu33476 | 1 | k__Fungi(100);p__Ascomycota(100);c__Leotiomycetes(100);o__Thelebolales(100);f__Thelebolaceae(100);g__Antarctomyces(100);s__Antarctomyces_psychrotrophicus(100); | Antarctomyces | 13 | Saprotroph | \|Undefined Saprotroph\| | Microfungus | NULL | Probable | As soil_saprotroph (Põlme et al. 2020); Decay soil (Põlme et al. 2020) | Tedersoo L, et al. 2014. Science 346:e1256688 ((https://doi.org/10.1126/science.1256688)); Põlme S, et al. 2020. Fungal Diversity 105:1-16 ((https://doi.org/10.1007/s13225-020-00466-2)) |
| Otu33477 | 1 | k__Fungi(100);p__Basidiomycota(100);c__Microbotryomycetes(100);o__Sporidiobolales(100);f__Sporidiobolales_family_Incertae_sedis(100);g__Sporobolomyces(100);s__Sporobolomyces_lactosus(100); | Sporobolomyces | 13 | Pathotroph-Saprotroph | \|Fungal Parasite\|-Plant Saprotroph | Yeast | NULL | Probable | As litter_saprotroph (Põlme et al. 2020); As mycoparasite (Põlme et al. 2020); Decay fungal material (Põlme et al. 2020) | Tedersoo L, et al. 2014. Science 346:e1256688 ((https://doi.org/10.1126/science.1256688)); Purahong W, et al. 2016. Molecular Ecology 25:4059-4074 ((https://doi.org/10.1111/mec.13739)); Põlme S, et al. 2020. Fungal Diversity 105:1-16 ((https://doi.org/10.1007/s13225-020-00466-2)) |
| Otu33479 | 1 | k__Fungi(100);p__Basidiomycota(100);c__Microbotryomycetes(100);o__Sporidiobolales(100);f__Sporidiobolales_family_Incertae_sedis(100);g__Sporobolomyces(100);s__Sporobolomyces_lactosus(100); | Sporobolomyces | 13 | Pathotroph-Saprotroph | \|Fungal Parasite\|-Plant Saprotroph | Yeast | NULL | Probable | As litter_saprotroph (Põlme et al. 2020); As mycoparasite (Põlme et al. 2020); Decay fungal material (Põlme et al. 2020) | Tedersoo L, et al. 2014. Science 346:e1256688 ((https://doi.org/10.1126/science.1256688)); Purahong W, et al. 2016. Molecular Ecology 25:4059-4074 ((https://doi.org/10.1111/mec.13739)); Põlme S, et al. 2020. Fungal Diversity 105:1-16 ((https://doi.org/10.1007/s13225-020-00466-2)) |
| Otu33483 | 1 | k__Fungi(100);p__Ascomycota(100);c__Saccharomycetes(100);o__Saccharomycetales(100);f__Metschnikowiaceae(100);g__Metschnikowia(100);s__Metschnikowia_pulcherrima(100); | Metschnikowia | 13 | Saprotroph | \|Undefined Saprotroph\| | Yeast | NULL | Probable | As nectar-tap_saprotroph (Põlme et al. 2020); Decay sugar-rich substrates (Põlme et al. 2020) | Põlme S, et al. 2020. Fungal Diversity 105:1-16 ((https://doi.org/10.1007/s13225-020-00466-2)) |
| Otu33484 | 1 | k__Fungi(100);p__Ascomycota(100);c__Saccharomycetes(100);o__Saccharomycetales(100);f__Metschnikowiaceae(100);g__Metschnikowia(100);g__Metschnikowia_unclassified(100); | Metschnikowia | 13 | Saprotroph | \|Undefined Saprotroph\| | Yeast | NULL | Probable | As nectar-tap_saprotroph (Põlme et al. 2020); Decay sugar-rich substrates (Põlme et al. 2020) | Põlme S, et al. 2020. Fungal Diversity 105:1-16 ((https://doi.org/10.1007/s13225-020-00466-2)) |
| Otu33485 | 1 | k__Fungi(100);p__Ascomycota(100);c__Saccharomycetes(100);o__Saccharomycetales(100);f__Metschnikowiaceae(100);g__Metschnikowia(100);g__Metschnikowia_unclassified(100); | Metschnikowia | 13 | Saprotroph | \|Undefined Saprotroph\| | Yeast | NULL | Probable | As nectar-tap_saprotroph (Põlme et al. 2020); Decay sugar-rich substrates (Põlme et al. 2020) | Põlme S, et al. 2020. Fungal Diversity 105:1-16 ((https://doi.org/10.1007/s13225-020-00466-2)) |
| Otu33486 | 1 | k__Fungi(100);p__Ascomycota(100);c__Saccharomycetes(100);o__Saccharomycetales(100);f__Metschnikowiaceae(100);g__Metschnikowia(100);g__Metschnikowia_unclassified(100); | Metschnikowia | 13 | Saprotroph | \|Undefined Saprotroph\| | Yeast | NULL | Probable | As nectar-tap_saprotroph (Põlme et al. 2020); Decay sugar-rich substrates (Põlme et al. 2020) | Põlme S, et al. 2020. Fungal Diversity 105:1-16 ((https://doi.org/10.1007/s13225-020-00466-2)) |
| Otu33489 | 1 | k__Fungi(100);p__Ascomycota(100);c__Saccharomycetes(100);o__Saccharomycetales(100);f__Metschnikowiaceae(100);g__Metschnikowia(100);s__Metschnikowia_pulcherrima(100); | Metschnikowia | 13 | Saprotroph | \|Undefined Saprotroph\| | Yeast | NULL | Probable | As nectar-tap_saprotroph (Põlme et al. 2020); Decay sugar-rich substrates (Põlme et al. 2020) | Põlme S, et al. 2020. Fungal Diversity 105:1-16 ((https://doi.org/10.1007/s13225-020-00466-2)) |
| Otu33494 | 1 | k__Fungi(100);p__Ascomycota(100);c__Saccharomycetes(100);o__Saccharomycetales(100);f__Metschnikowiaceae(100);g__Metschnikowia(100);g__Metschnikowia_unclassified(100); | Metschnikowia | 13 | Saprotroph | \|Undefined Saprotroph\| | Yeast | NULL | Probable | As nectar-tap_saprotroph (Põlme et al. 2020); Decay sugar-rich substrates (Põlme et al. 2020) | Põlme S, et al. 2020. Fungal Diversity 105:1-16 ((https://doi.org/10.1007/s13225-020-00466-2)) |
| Otu33498 | 1 | k__Fungi(100);p__Ascomycota(100);c__Saccharomycetes(100);o__Saccharomycetales(100);f__Metschnikowiaceae(100);g__Metschnikowia(100);g__Metschnikowia_unclassified(100); | Metschnikowia | 13 | Saprotroph | \|Undefined Saprotroph\| | Yeast | NULL | Probable | As nectar-tap_saprotroph (Põlme et al. 2020); Decay sugar-rich substrates (Põlme et al. 2020) | Põlme S, et al. 2020. Fungal Diversity 105:1-16 ((https://doi.org/10.1007/s13225-020-00466-2)) |
| Otu33511 | 1 | k__Fungi(100);p__Ascomycota(100);c__Saccharomycetes(100);o__Saccharomycetales(100);f__Metschnikowiaceae(100);g__Metschnikowia(100);s__Metschnikowia_pulcherrima(100); | Metschnikowia | 13 | Saprotroph | \|Undefined Saprotroph\| | Yeast | NULL | Probable | As nectar-tap_saprotroph (Põlme et al. 2020); Decay sugar-rich substrates (Põlme et al. 2020) | Põlme S, et al. 2020. Fungal Diversity 105:1-16 ((https://doi.org/10.1007/s13225-020-00466-2)) |
| Otu33513 | 1 | k__Fungi(100);p__Ascomycota(100);c__Saccharomycetes(100);o__Saccharomycetales(100);f__Metschnikowiaceae(100);g__Metschnikowia(100);g__Metschnikowia_unclassified(100); | Metschnikowia | 13 | Saprotroph | \|Undefined Saprotroph\| | Yeast | NULL | Probable | As nectar-tap_saprotroph (Põlme et al. 2020); Decay sugar-rich substrates (Põlme et al. 2020) | Põlme S, et al. 2020. Fungal Diversity 105:1-16 ((https://doi.org/10.1007/s13225-020-00466-2)) |
| Otu33515 | 1 | k__Fungi(100);p__Ascomycota(100);c__Saccharomycetes(100);o__Saccharomycetales(100);f__Metschnikowiaceae(100);g__Metschnikowia(100);s__Metschnikowia_pulcherrima(100); | Metschnikowia | 13 | Saprotroph | \|Undefined Saprotroph\| | Yeast | NULL | Probable | As nectar-tap_saprotroph (Põlme et al. 2020); Decay sugar-rich substrates (Põlme et al. 2020) | Põlme S, et al. 2020. Fungal Diversity 105:1-16 ((https://doi.org/10.1007/s13225-020-00466-2)) |
| Otu33517 | 1 | k__Fungi(100);p__Ascomycota(100);c__Saccharomycetes(100);o__Saccharomycetales(100);f__Saccharomycetales_family_Incertae_sedis(100);g__Candida(100);g__Candida_unclassified(100); | Candida | 13 | Pathotroph-Saprotroph-Symbiotroph | Animal Pathogen-Endophyte-Endosymbiont-Epiphyte-\|Undefined Saprotroph\| | Dimorphic Yeast-Yeast | NULL | Probable | As nectar-tap_saprotroph (Põlme et al. 2020); Decay sugar-rich substrates (Põlme et al. 2020); This asexual genus is polyphyletic and contain species that function very differently depend on what they are. Caution should be taken when interpreting guilds for Candida species. | Manolakaki D, et al. 2010. Virulence. 1: 367-375 ((https://doi.org/10.4161/viru.1.5.12796)); Mekbib SB, et al. 2011. Tropical Plant Pathology 36:233-240 ((https://doi.org/10.1590/S1982-56762011000400004)); Põlme S, et al. 2020. Fungal Diversity 105:1-16 ((https://doi.org/10.1007/s13225-020-00466-2)) |
| Otu33518 | 1 | k__Fungi(100);p__Ascomycota(100);c__Saccharomycetes(100);o__Saccharomycetales(100);f__Metschnikowiaceae(100);g__Metschnikowia(100);s__Metschnikowia_pulcherrima(100); | Metschnikowia | 13 | Saprotroph | \|Undefined Saprotroph\| | Yeast | NULL | Probable | As nectar-tap_saprotroph (Põlme et al. 2020); Decay sugar-rich substrates (Põlme et al. 2020) | Põlme S, et al. 2020. Fungal Diversity 105:1-16 ((https://doi.org/10.1007/s13225-020-00466-2)) |
| Otu33522 | 1 | k__Fungi(100);p__Ascomycota(100);c__Saccharomycetes(100);o__Saccharomycetales(100);f__Metschnikowiaceae(100);g__Metschnikowia(100);s__Metschnikowia_pulcherrima(100); | Metschnikowia | 13 | Saprotroph | \|Undefined Saprotroph\| | Yeast | NULL | Probable | As nectar-tap_saprotroph (Põlme et al. 2020); Decay sugar-rich substrates (Põlme et al. 2020) | Põlme S, et al. 2020. Fungal Diversity 105:1-16 ((https://doi.org/10.1007/s13225-020-00466-2)) |
| Otu33523 | 1 | k__Fungi(100);p__Ascomycota(100);c__Saccharomycetes(100);o__Saccharomycetales(100);f__Metschnikowiaceae(100);g__Metschnikowia(100);s__Metschnikowia_pulcherrima(100); | Metschnikowia | 13 | Saprotroph | \|Undefined Saprotroph\| | Yeast | NULL | Probable | As nectar-tap_saprotroph (Põlme et al. 2020); Decay sugar-rich substrates (Põlme et al. 2020) | Põlme S, et al. 2020. Fungal Diversity 105:1-16 ((https://doi.org/10.1007/s13225-020-00466-2)) |
| Otu33524 | 1 | k__Fungi(100);p__Ascomycota(100);c__Saccharomycetes(100);o__Saccharomycetales(100);f__Metschnikowiaceae(100);g__Metschnikowia(100);s__Metschnikowia_pulcherrima(100); | Metschnikowia | 13 | Saprotroph | \|Undefined Saprotroph\| | Yeast | NULL | Probable | As nectar-tap_saprotroph (Põlme et al. 2020); Decay sugar-rich substrates (Põlme et al. 2020) | Põlme S, et al. 2020. Fungal Diversity 105:1-16 ((https://doi.org/10.1007/s13225-020-00466-2)) |
| Otu33527 | 1 | k__Fungi(100);p__Ascomycota(100);c__Saccharomycetes(100);o__Saccharomycetales(100);f__Metschnikowiaceae(100);g__Metschnikowia(100);g__Metschnikowia_unclassified(100); | Metschnikowia | 13 | Saprotroph | \|Undefined Saprotroph\| | Yeast | NULL | Probable | As nectar-tap_saprotroph (Põlme et al. 2020); Decay sugar-rich substrates (Põlme et al. 2020) | Põlme S, et al. 2020. Fungal Diversity 105:1-16 ((https://doi.org/10.1007/s13225-020-00466-2)) |
| Otu33528 | 1 | k__Fungi(100);p__Ascomycota(100);c__Saccharomycetes(100);o__Saccharomycetales(100);o__Saccharomycetales_unclassified(100);o__Saccharomycetales_unclassified(100);o__Saccharomycetales_unclassified(100); | Saccharomycetales | 7 | Saprotroph | Undefined Saprotroph | Yeast | NULL | Possible | NULL | Sterkenburg E, et al. 2015. New Phytologist 207:1145-1158 ((https://doi.org/10.1111/nph.13426)) |
| Otu33529 | 1 | k__Fungi(100);p__Ascomycota(100);c__Saccharomycetes(100);o__Saccharomycetales(100);f__Metschnikowiaceae(100);g__Metschnikowia(100);g__Metschnikowia_unclassified(100); | Metschnikowia | 13 | Saprotroph | \|Undefined Saprotroph\| | Yeast | NULL | Probable | As nectar-tap_saprotroph (Põlme et al. 2020); Decay sugar-rich substrates (Põlme et al. 2020) | Põlme S, et al. 2020. Fungal Diversity 105:1-16 ((https://doi.org/10.1007/s13225-020-00466-2)) |

| | | | | | | | | | | | |
|---|---|---|---|---|---|---|---|---|---|---|---|
| Otu33531 | 1 | k__Fungi(100);p__Ascomycota(100);c__Saccharomycetes(100);o__Saccharomycetales(100);f__Metschnikowiaceae(100);g__Metschnikowia(100);s__Metschnikowia_pulcherrima(100); | Metschnikowia | 13 | Saprotroph | \|Undefined Saprotroph\| | Yeast | NULL | Probable | As nectar-tap_saprotroph (Põlme et al. 2020); Decay sugar-rich substrates (Põlme et al. 2020) | Põlme S, et al. 2020. Fungal Diversity 105:1-16 ((https://doi.org/10.1007/s13225-020-00466-2)) |
| Otu33533 | 1 | k__Fungi(100);p__Ascomycota(100);c__Saccharomycetes(100);o__Saccharomycetales(100);f__Metschnikowiaceae(100);g__Metschnikowia(100);s__Metschnikowia_pulcherrima(100); | Metschnikowia | 13 | Saprotroph | \|Undefined Saprotroph\| | Yeast | NULL | Probable | As nectar-tap_saprotroph (Põlme et al. 2020); Decay sugar-rich substrates (Põlme et al. 2020) | Põlme S, et al. 2020. Fungal Diversity 105:1-16 ((https://doi.org/10.1007/s13225-020-00466-2)) |
| Otu33535 | 1 | k__Fungi(100);p__Ascomycota(100);c__Saccharomycetes(100);o__Saccharomycetales(100);f__Metschnikowiaceae(100);g__Metschnikowia(100);s__Metschnikowia_pulcherrima(100); | Metschnikowia | 13 | Saprotroph | \|Undefined Saprotroph\| | Yeast | NULL | Probable | As nectar-tap_saprotroph (Põlme et al. 2020); Decay sugar-rich substrates (Põlme et al. 2020) | Põlme S, et al. 2020. Fungal Diversity 105:1-16 ((https://doi.org/10.1007/s13225-020-00466-2)) |
| Otu33536 | 1 | k__Fungi(100);p__Ascomycota(100);c__Saccharomycetes(100);o__Saccharomycetales(100);f__Metschnikowiaceae(100);g__Metschnikowia(100);g__Metschnikowia_unclassified(100); | Metschnikowia | 13 | Saprotroph | \|Undefined Saprotroph\| | Yeast | NULL | Probable | As nectar-tap_saprotroph (Põlme et al. 2020); Decay sugar-rich substrates (Põlme et al. 2020) | Põlme S, et al. 2020. Fungal Diversity 105:1-16 ((https://doi.org/10.1007/s13225-020-00466-2)) |
| Otu33537 | 1 | k__Fungi(100);p__Ascomycota(100);c__Saccharomycetes(100);o__Saccharomycetales(100);f__Metschnikowiaceae(100);g__Metschnikowia(100);s__Metschnikowia_pulcherrima(100); | Metschnikowia | 13 | Saprotroph | \|Undefined Saprotroph\| | Yeast | NULL | Probable | As nectar-tap_saprotroph (Põlme et al. 2020); Decay sugar-rich substrates (Põlme et al. 2020) | Põlme S, et al. 2020. Fungal Diversity 105:1-16 ((https://doi.org/10.1007/s13225-020-00466-2)) |
| Otu33538 | 1 | k__Fungi(100);p__Ascomycota(100);c__Saccharomycetes(100);o__Saccharomycetales(100);f__Metschnikowiaceae(100);g__Metschnikowia(100);s__Metschnikowia_pulcherrima(100); | Metschnikowia | 13 | Saprotroph | \|Undefined Saprotroph\| | Yeast | NULL | Probable | As nectar-tap_saprotroph (Põlme et al. 2020); Decay sugar-rich substrates (Põlme et al. 2020) | Põlme S, et al. 2020. Fungal Diversity 105:1-16 ((https://doi.org/10.1007/s13225-020-00466-2)) |
| Otu33539 | 1 | k__Fungi(100);p__Ascomycota(100);c__Saccharomycetes(100);o__Saccharomycetales(100);f__Metschnikowiaceae(100);g__Metschnikowia(100);s__Metschnikowia_pulcherrima(100); | Metschnikowia | 13 | Saprotroph | \|Undefined Saprotroph\| | Yeast | NULL | Probable | As nectar-tap_saprotroph (Põlme et al. 2020); Decay sugar-rich substrates (Põlme et al. 2020) | Põlme S, et al. 2020. Fungal Diversity 105:1-16 ((https://doi.org/10.1007/s13225-020-00466-2)) |
| Otu33540 | 1 | k__Fungi(100);p__Ascomycota(100);c__Saccharomycetes(100);o__Saccharomycetales(100);f__Metschnikowiaceae(100);g__Metschnikowia(100);s__Metschnikowia_pulcherrima(100); | Metschnikowia | 13 | Saprotroph | \|Undefined Saprotroph\| | Yeast | NULL | Probable | As nectar-tap_saprotroph (Põlme et al. 2020); Decay sugar-rich substrates (Põlme et al. 2020) | Põlme S, et al. 2020. Fungal Diversity 105:1-16 ((https://doi.org/10.1007/s13225-020-00466-2)) |
| Otu33541 | 1 | k__Fungi(100);p__Ascomycota(100);c__Saccharomycetes(100);o__Saccharomycetales(100);f__Metschnikowiaceae(100);g__Metschnikowia(100);s__Metschnikowia_pulcherrima(100); | Metschnikowia | 13 | Saprotroph | \|Undefined Saprotroph\| | Yeast | NULL | Probable | As nectar-tap_saprotroph (Põlme et al. 2020); Decay sugar-rich substrates (Põlme et al. 2020) | Põlme S, et al. 2020. Fungal Diversity 105:1-16 ((https://doi.org/10.1007/s13225-020-00466-2)) |
| Otu33542 | 1 | k__Fungi(100);p__Ascomycota(100);c__Saccharomycetes(100);o__Saccharomycetales(100);f__Metschnikowiaceae(100);g__Metschnikowia(100);s__Metschnikowia_pulcherrima(100); | Metschnikowia | 13 | Saprotroph | \|Undefined Saprotroph\| | Yeast | NULL | Probable | As nectar-tap_saprotroph (Põlme et al. 2020); Decay sugar-rich substrates (Põlme et al. 2020) | Põlme S, et al. 2020. Fungal Diversity 105:1-16 ((https://doi.org/10.1007/s13225-020-00466-2)) |
| Otu33543 | 1 | k__Fungi(100);p__Ascomycota(100);c__Saccharomycetes(100);o__Saccharomycetales(100);f__Metschnikowiaceae(100);g__Metschnikowia(100);g__Metschnikowia_unclassified(100); | Metschnikowia | 13 | Saprotroph | \|Undefined Saprotroph\| | Yeast | NULL | Probable | As nectar-tap_saprotroph (Põlme et al. 2020); Decay sugar-rich substrates (Põlme et al. 2020) | Põlme S, et al. 2020. Fungal Diversity 105:1-16 ((https://doi.org/10.1007/s13225-020-00466-2)) |
| Otu33544 | 1 | k__Fungi(100);p__Ascomycota(100);c__Saccharomycetes(100);o__Saccharomycetales(100);f__Metschnikowiaceae(100);g__Metschnikowia(100);g__Metschnikowia_unclassified(100); | Metschnikowia | 13 | Saprotroph | \|Undefined Saprotroph\| | Yeast | NULL | Probable | As nectar-tap_saprotroph (Põlme et al. 2020); Decay sugar-rich substrates (Põlme et al. 2020) | Põlme S, et al. 2020. Fungal Diversity 105:1-16 ((https://doi.org/10.1007/s13225-020-00466-2)) |
| Otu33548 | 1 | k__Fungi(100);p__Ascomycota(100);c__Saccharomycetes(100);o__Saccharomycetales(100);f__Metschnikowiaceae(100);g__Metschnikowia(100);s__Metschnikowia_pulcherrima(100); | Metschnikowia | 13 | Saprotroph | \|Undefined Saprotroph\| | Yeast | NULL | Probable | As nectar-tap_saprotroph (Põlme et al. 2020); Decay sugar-rich substrates (Põlme et al. 2020) | Põlme S, et al. 2020. Fungal Diversity 105:1-16 ((https://doi.org/10.1007/s13225-020-00466-2)) |
| Otu33549 | 1 | k__Fungi(100);p__Ascomycota(100);c__Saccharomycetes(100);o__Saccharomycetales(100);f__Metschnikowiaceae(100);g__Metschnikowia(100);s__Metschnikowia_pulcherrima(100); | Metschnikowia | 13 | Saprotroph | \|Undefined Saprotroph\| | Yeast | NULL | Probable | As nectar-tap_saprotroph (Põlme et al. 2020); Decay sugar-rich substrates (Põlme et al. 2020) | Põlme S, et al. 2020. Fungal Diversity 105:1-16 ((https://doi.org/10.1007/s13225-020-00466-2)) |
| Otu33553 | 1 | k__Fungi(100);p__Ascomycota(100);c__Saccharomycetes(100);o__Saccharomycetales(100);f__Metschnikowiaceae(100);g__Metschnikowia(100);g__Metschnikowia_unclassified(100); | Metschnikowia | 13 | Saprotroph | \|Undefined Saprotroph\| | Yeast | NULL | Probable | As nectar-tap_saprotroph (Põlme et al. 2020); Decay sugar-rich substrates (Põlme et al. 2020) | Põlme S, et al. 2020. Fungal Diversity 105:1-16 ((https://doi.org/10.1007/s13225-020-00466-2)) |
| Otu33554 | 1 | k__Fungi(100);p__Ascomycota(100);c__Saccharomycetes(100);o__Saccharomycetales(100);f__Metschnikowiaceae(100);g__Metschnikowia(100);s__Metschnikowia_pulcherrima(100); | Metschnikowia | 13 | Saprotroph | \|Undefined Saprotroph\| | Yeast | NULL | Probable | As nectar-tap_saprotroph (Põlme et al. 2020); Decay sugar-rich substrates (Põlme et al. 2020) | Põlme S, et al. 2020. Fungal Diversity 105:1-16 ((https://doi.org/10.1007/s13225-020-00466-2)) |
| Otu33556 | 1 | k__Fungi(100);p__Ascomycota(100);c__Saccharomycetes(100);o__Saccharomycetales(100);f__Metschnikowiaceae(100);g__Metschnikowia(100);g__Metschnikowia_unclassified(100); | Metschnikowia | 13 | Saprotroph | \|Undefined Saprotroph\| | Yeast | NULL | Probable | As nectar-tap_saprotroph (Põlme et al. 2020); Decay sugar-rich substrates (Põlme et al. 2020) | Põlme S, et al. 2020. Fungal Diversity 105:1-16 ((https://doi.org/10.1007/s13225-020-00466-2)) |
| Otu33557 | 1 | k__Fungi(100);p__Ascomycota(100);c__Saccharomycetes(100);o__Saccharomycetales(100);f__Metschnikowiaceae(100);g__Metschnikowia(100);s__Metschnikowia_pulcherrima(100); | Metschnikowia | 13 | Saprotroph | \|Undefined Saprotroph\| | Yeast | NULL | Probable | As nectar-tap_saprotroph (Põlme et al. 2020); Decay sugar-rich substrates (Põlme et al. 2020) | Põlme S, et al. 2020. Fungal Diversity 105:1-16 ((https://doi.org/10.1007/s13225-020-00466-2)) |
| Otu33558 | 1 | k__Fungi(100);p__Ascomycota(100);c__Saccharomycetes(100);o__Saccharomycetales(100);f__Metschnikowiaceae(100);g__Metschnikowia(100);s__Metschnikowia_pulcherrima(100); | Metschnikowia | 13 | Saprotroph | \|Undefined Saprotroph\| | Yeast | NULL | Probable | As nectar-tap_saprotroph (Põlme et al. 2020); Decay sugar-rich substrates (Põlme et al. 2020) | Põlme S, et al. 2020. Fungal Diversity 105:1-16 ((https://doi.org/10.1007/s13225-020-00466-2)) |
| Otu33559 | 1 | k__Fungi(100);p__Ascomycota(100);c__Saccharomycetes(100);o__Saccharomycetales(100);f__Metschnikowiaceae(100);g__Metschnikowia(100);g__Metschnikowia_unclassified(100); | Metschnikowia | 13 | Saprotroph | \|Undefined Saprotroph\| | Yeast | NULL | Probable | As nectar-tap_saprotroph (Põlme et al. 2020); Decay sugar-rich substrates (Põlme et al. 2020) | Põlme S, et al. 2020. Fungal Diversity 105:1-16 ((https://doi.org/10.1007/s13225-020-00466-2)) |
| Otu33560 | 1 | k__Fungi(100);p__Ascomycota(100);c__Saccharomycetes(100);o__Saccharomycetales(100);f__Metschnikowiaceae(100);g__Metschnikowia(100);s__Metschnikowia_pulcherrima(100); | Metschnikowia | 13 | Saprotroph | \|Undefined Saprotroph\| | Yeast | NULL | Probable | As nectar-tap_saprotroph (Põlme et al. 2020); Decay sugar-rich substrates (Põlme et al. 2020) | Põlme S, et al. 2020. Fungal Diversity 105:1-16 ((https://doi.org/10.1007/s13225-020-00466-2)) |
| Otu33561 | 1 | k__Fungi(100);p__Ascomycota(100);c__Saccharomycetes(100);o__Saccharomycetales(100);f__Metschnikowiaceae(100);g__Metschnikowia(100);s__Metschnikowia_pulcherrima(100); | Metschnikowia | 13 | Saprotroph | \|Undefined Saprotroph\| | Yeast | NULL | Probable | As nectar-tap_saprotroph (Põlme et al. 2020); Decay sugar-rich substrates (Põlme et al. 2020) | Põlme S, et al. 2020. Fungal Diversity 105:1-16 ((https://doi.org/10.1007/s13225-020-00466-2)) |
| Otu33562 | 1 | k__Fungi(100);p__Ascomycota(100);c__Saccharomycetes(100);o__Saccharomycetales(100);f__Metschnikowiaceae(100);g__Metschnikowia(100);s__Metschnikowia_pulcherrima(100); | Metschnikowia | 13 | Saprotroph | \|Undefined Saprotroph\| | Yeast | NULL | Probable | As nectar-tap_saprotroph (Põlme et al. 2020); Decay sugar-rich substrates (Põlme et al. 2020) | Põlme S, et al. 2020. Fungal Diversity 105:1-16 ((https://doi.org/10.1007/s13225-020-00466-2)) |
| Otu33564 | 1 | k__Fungi(100);p__Ascomycota(100);c__Saccharomycetes(100);o__Saccharomycetales(100);f__Metschnikowiaceae(100);g__Metschnikowia(100);s__Metschnikowia_pulcherrima(100); | Metschnikowia | 13 | Saprotroph | \|Undefined Saprotroph\| | Yeast | NULL | Probable | As nectar-tap_saprotroph (Põlme et al. 2020); Decay sugar-rich substrates (Põlme et al. 2020) | Põlme S, et al. 2020. Fungal Diversity 105:1-16 ((https://doi.org/10.1007/s13225-020-00466-2)) |
| Otu33565 | 1 | k__Fungi(100);p__Ascomycota(100);c__Saccharomycetes(100);o__Saccharomycetales(100);f__Metschnikowiaceae(100);g__Metschnikowia(100);g__Metschnikowia_unclassified(100); | Metschnikowia | 13 | Saprotroph | \|Undefined Saprotroph\| | Yeast | NULL | Probable | As nectar-tap_saprotroph (Põlme et al. 2020); Decay sugar-rich substrates (Põlme et al. 2020) | Põlme S, et al. 2020. Fungal Diversity 105:1-16 ((https://doi.org/10.1007/s13225-020-00466-2)) |
| Otu33566 | 1 | k__Fungi(100);p__Ascomycota(100);c__Saccharomycetes(100);o__Saccharomycetales(100);f__Metschnikowiaceae(100);g__Metschnikowia(100);s__Metschnikowia_pulcherrima(100); | Metschnikowia | 13 | Saprotroph | \|Undefined Saprotroph\| | Yeast | NULL | Probable | As nectar-tap_saprotroph (Põlme et al. 2020); Decay sugar-rich substrates (Põlme et al. 2020) | Põlme S, et al. 2020. Fungal Diversity 105:1-16 ((https://doi.org/10.1007/s13225-020-00466-2)) |
| Otu33567 | 1 | k__Fungi(100);p__Ascomycota(100);c__Saccharomycetes(100);o__Saccharomycetales(100);f__Metschnikowiaceae(100);g__Metschnikowia(100);s__Metschnikowia_pulcherrima(100); | Metschnikowia | 13 | Saprotroph | \|Undefined Saprotroph\| | Yeast | NULL | Probable | As nectar-tap_saprotroph (Põlme et al. 2020); Decay sugar-rich substrates (Põlme et al. 2020) | Põlme S, et al. 2020. Fungal Diversity 105:1-16 ((https://doi.org/10.1007/s13225-020-00466-2)) |
| Otu33568 | 1 | k__Fungi(100);p__Ascomycota(100);c__Saccharomycetes(100);o__Saccharomycetales(100);f__Metschnikowiaceae(100);g__Metschnikowia(100);g__Metschnikowia_unclassified(100); | Metschnikowia | 13 | Saprotroph | \|Undefined Saprotroph\| | Yeast | NULL | Probable | As nectar-tap_saprotroph (Põlme et al. 2020); Decay sugar-rich substrates (Põlme et al. 2020) | Põlme S, et al. 2020. Fungal Diversity 105:1-16 ((https://doi.org/10.1007/s13225-020-00466-2)) |
| Otu33569 | 1 | k__Fungi(100);p__Ascomycota(100);c__Saccharomycetes(100);o__Saccharomycetales(100);f__Metschnikowiaceae(100);g__Metschnikowia(100);s__Metschnikowia_pulcherrima(100); | Metschnikowia | 13 | Saprotroph | \|Undefined Saprotroph\| | Yeast | NULL | Probable | As nectar-tap_saprotroph (Põlme et al. 2020); Decay sugar-rich substrates (Põlme et al. 2020) | Põlme S, et al. 2020. Fungal Diversity 105:1-16 ((https://doi.org/10.1007/s13225-020-00466-2)) |
| Otu33570 | 1 | k__Fungi(100);p__Ascomycota(100);c__Saccharomycetes(100);o__Saccharomycetales(100);f__Metschnikowiaceae(100);g__Metschnikowia(100);s__Metschnikowia_pulcherrima(100); | Metschnikowia | 13 | Saprotroph | \|Undefined Saprotroph\| | Yeast | NULL | Probable | As nectar-tap_saprotroph (Põlme et al. 2020); Decay sugar-rich substrates (Põlme et al. 2020) | Põlme S, et al. 2020. Fungal Diversity 105:1-16 ((https://doi.org/10.1007/s13225-020-00466-2)) |
| Otu33571 | 1 | k__Fungi(100);p__Ascomycota(100);c__Saccharomycetes(100);o__Saccharomycetales(100);f__Metschnikowiaceae(100);g__Metschnikowia(100);s__Metschnikowia_pulcherrima(100); | Metschnikowia | 13 | Saprotroph | \|Undefined Saprotroph\| | Yeast | NULL | Probable | As nectar-tap_saprotroph (Põlme et al. 2020); Decay sugar-rich substrates (Põlme et al. 2020) | Põlme S, et al. 2020. Fungal Diversity 105:1-16 ((https://doi.org/10.1007/s13225-020-00466-2)) |
| Otu33573 | 1 | k__Fungi(100);p__Ascomycota(100);c__Saccharomycetes(100);o__Saccharomycetales(100);f__Metschnikowiaceae(100);g__Metschnikowia(100);s__Metschnikowia_pulcherrima(100); | Metschnikowia | 13 | Saprotroph | \|Undefined Saprotroph\| | Yeast | NULL | Probable | As nectar-tap_saprotroph (Põlme et al. 2020); Decay sugar-rich substrates (Põlme et al. 2020) | Põlme S, et al. 2020. Fungal Diversity 105:1-16 ((https://doi.org/10.1007/s13225-020-00466-2)) |
| Otu33576 | 1 | k__Fungi(100);p__Ascomycota(100);c__Saccharomycetes(100);o__Saccharomycetales(100);f__Metschnikowiaceae(100);g__Metschnikowia(100);s__Metschnikowia_pulcherrima(100); | Metschnikowia | 13 | Saprotroph | \|Undefined Saprotroph\| | Yeast | NULL | Probable | As nectar-tap_saprotroph (Põlme et al. 2020); Decay sugar-rich substrates (Põlme et al. 2020) | Põlme S, et al. 2020. Fungal Diversity 105:1-16 ((https://doi.org/10.1007/s13225-020-00466-2)) |
| Otu33579 | 1 | k__Fungi(100);p__Ascomycota(100);c__Saccharomycetes(100);o__Saccharomycetales(100);f__Metschnikowiaceae(100);g__Metschnikowia(100);s__Metschnikowia_pulcherrima(100); | Metschnikowia | 13 | Saprotroph | \|Undefined Saprotroph\| | Yeast | NULL | Probable | As nectar-tap_saprotroph (Põlme et al. 2020); Decay sugar-rich substrates (Põlme et al. 2020) | Põlme S, et al. 2020. Fungal Diversity 105:1-16 ((https://doi.org/10.1007/s13225-020-00466-2)) |
| Otu33581 | 1 | k__Fungi(100);p__Ascomycota(100);c__Saccharomycetes(100);o__Saccharomycetales(100);f__Metschnikowiaceae(100);g__Metschnikowia(100);s__Metschnikowia_pulcherrima(100); | Metschnikowia | 13 | Saprotroph | \|Undefined Saprotroph\| | Yeast | NULL | Probable | As nectar-tap_saprotroph (Põlme et al. 2020); Decay sugar-rich substrates (Põlme et al. 2020) | Põlme S, et al. 2020. Fungal Diversity 105:1-16 ((https://doi.org/10.1007/s13225-020-00466-2)) |
| Otu33582 | 1 | k__Fungi(100);p__Ascomycota(100);c__Saccharomycetes(100);o__Saccharomycetales(100);f__Metschnikowiaceae(100);g__Metschnikowia(100);s__Metschnikowia_pulcherrima(100); | Metschnikowia | 13 | Saprotroph | \|Undefined Saprotroph\| | Yeast | NULL | Probable | As nectar-tap_saprotroph (Põlme et al. 2020); Decay sugar-rich substrates (Põlme et al. 2020) | Põlme S, et al. 2020. Fungal Diversity 105:1-16 ((https://doi.org/10.1007/s13225-020-00466-2)) |
| Otu33583 | 1 | k__Fungi(100);p__Ascomycota(100);c__Saccharomycetes(100);o__Saccharomycetales(100);f__Metschnikowiaceae(100);g__Metschnikowia(100);s__Metschnikowia_pulcherrima(100); | Metschnikowia | 13 | Saprotroph | \|Undefined Saprotroph\| | Yeast | NULL | Probable | As nectar-tap_saprotroph (Põlme et al. 2020); Decay sugar-rich substrates (Põlme et al. 2020) | Põlme S, et al. 2020. Fungal Diversity 105:1-16 ((https://doi.org/10.1007/s13225-020-00466-2)) |
| Otu33586 | 1 | k__Fungi(100);p__Ascomycota(100);c__Saccharomycetes(100);o__Saccharomycetales(100);f__Metschnikowiaceae(100);g__Metschnikowia(100);g__Metschnikowia_unclassified(100); | Metschnikowia | 13 | Saprotroph | \|Undefined Saprotroph\| | Yeast | NULL | Probable | As nectar-tap_saprotroph (Põlme et al. 2020); Decay sugar-rich substrates (Põlme et al. 2020) | Põlme S, et al. 2020. Fungal Diversity 105:1-16 ((https://doi.org/10.1007/s13225-020-00466-2)) |
| Otu33587 | 1 | k__Fungi(100);p__Ascomycota(100);c__Saccharomycetes(100);o__Saccharomycetales(100);f__Metschnikowiaceae(100);g__Metschnikowia(100);s__Metschnikowia_pulcherrima(100); | Metschnikowia | 13 | Saprotroph | \|Undefined Saprotroph\| | Yeast | NULL | Probable | As nectar-tap_saprotroph (Põlme et al. 2020); Decay sugar-rich substrates (Põlme et al. 2020) | Põlme S, et al. 2020. Fungal Diversity 105:1-16 ((https://doi.org/10.1007/s13225-020-00466-2)) |
| Otu33588 | 1 | k__Fungi(100);p__Ascomycota(100);c__Saccharomycetes(100);o__Saccharomycetales(100);f__Metschnikowiaceae(100);g__Metschnikowia(100);s__Metschnikowia_pulcherrima(100); | Metschnikowia | 13 | Saprotroph | \|Undefined Saprotroph\| | Yeast | NULL | Probable | As nectar-tap_saprotroph (Põlme et al. 2020); Decay sugar-rich substrates (Põlme et al. 2020) | Põlme S, et al. 2020. Fungal Diversity 105:1-16 ((https://doi.org/10.1007/s13225-020-00466-2)) |
| Otu33589 | 1 | k__Fungi(100);p__Ascomycota(100);c__Saccharomycetes(100);o__Saccharomycetales(100);f__Metschnikowiaceae(100);g__Metschnikowia(100);s__Metschnikowia_pulcherrima(100); | Metschnikowia | 13 | Saprotroph | \|Undefined Saprotroph\| | Yeast | NULL | Probable | As nectar-tap_saprotroph (Põlme et al. 2020); Decay sugar-rich substrates (Põlme et al. 2020) | Põlme S, et al. 2020. Fungal Diversity 105:1-16 ((https://doi.org/10.1007/s13225-020-00466-2)) |
| Otu33592 | 1 | k__Fungi(100);p__Ascomycota(100);c__Saccharomycetes(100);o__Saccharomycetales(100);f__Metschnikowiaceae(100);g__Metschnikowia(100);s__Metschnikowia_pulcherrima(100); | Metschnikowia | 13 | Saprotroph | \|Undefined Saprotroph\| | Yeast | NULL | Probable | As nectar-tap_saprotroph (Põlme et al. 2020); Decay sugar-rich substrates (Põlme et al. 2020) | Põlme S, et al. 2020. Fungal Diversity 105:1-16 ((https://doi.org/10.1007/s13225-020-00466-2)) |
| Otu33594 | 1 | k__Fungi(100);p__Ascomycota(100);c__Saccharomycetes(100);o__Saccharomycetales(100);f__Metschnikowiaceae(100);g__Metschnikowia(100);s__Metschnikowia_pulcherrima(100); | Metschnikowia | 13 | Saprotroph | \|Undefined Saprotroph\| | Yeast | NULL | Probable | As nectar-tap_saprotroph (Põlme et al. 2020); Decay sugar-rich substrates (Põlme et al. 2020) | Põlme S, et al. 2020. Fungal Diversity 105:1-16 ((https://doi.org/10.1007/s13225-020-00466-2)) |
| Otu33595 | 1 | k__Fungi(100);p__Ascomycota(100);c__Saccharomycetes(100);o__Saccharomycetales(100);f__Metschnikowiaceae(100);g__Metschnikowia(100);s__Metschnikowia_pulcherrima(100); | Metschnikowia | 13 | Saprotroph | \|Undefined Saprotroph\| | Yeast | NULL | Probable | As nectar-tap_saprotroph (Põlme et al. 2020); Decay sugar-rich substrates (Põlme et al. 2020) | Põlme S, et al. 2020. Fungal Diversity 105:1-16 ((https://doi.org/10.1007/s13225-020-00466-2)) |
| Otu33597 | 1 | k__Fungi(100);p__Ascomycota(100);c__Saccharomycetes(100);o__Saccharomycetales(100);f__Metschnikowiaceae(100);g__Metschnikowia(100);s__Metschnikowia_pulcherrima(100); | Metschnikowia | 13 | Saprotroph | \|Undefined Saprotroph\| | Yeast | NULL | Probable | As nectar-tap_saprotroph (Põlme et al. 2020); Decay sugar-rich substrates (Põlme et al. 2020) | Põlme S, et al. 2020. Fungal Diversity 105:1-16 ((https://doi.org/10.1007/s13225-020-00466-2)) |
| Otu33598 | 1 | k__Fungi(100);p__Ascomycota(100);c__Saccharomycetes(100);o__Saccharomycetales(100);f__Metschnikowiaceae(100);g__Metschnikowia(100);g__Metschnikowia_unclassified(100); | Metschnikowia | 13 | Saprotroph | \|Undefined Saprotroph\| | Yeast | NULL | Probable | As nectar-tap_saprotroph (Põlme et al. 2020); Decay sugar-rich substrates (Põlme et al. 2020) | Põlme S, et al. 2020. Fungal Diversity 105:1-16 ((https://doi.org/10.1007/s13225-020-00466-2)) |
| Otu33599 | 1 | k__Fungi(100);p__Ascomycota(100);c__Saccharomycetes(100);o__Saccharomycetales(100);f__Metschnikowiaceae(100);g__Metschnikowia(100);g__Metschnikowia_unclassified(100); | Metschnikowia | 13 | Saprotroph | \|Undefined Saprotroph\| | Yeast | NULL | Probable | As nectar-tap_saprotroph (Põlme et al. 2020); Decay sugar-rich substrates (Põlme et al. 2020) | Põlme S, et al. 2020. Fungal Diversity 105:1-16 ((https://doi.org/10.1007/s13225-020-00466-2)) |
| Otu33600 | 1 | k__Fungi(100);p__Ascomycota(100);c__Saccharomycetes(100);o__Saccharomycetales(100);f__Metschnikowiaceae(100);g__Metschnikowia(100);s__Metschnikowia_pulcherrima(100); | Metschnikowia | 13 | Saprotroph | \|Undefined Saprotroph\| | Yeast | NULL | Probable | As nectar-tap_saprotroph (Põlme et al. 2020); Decay sugar-rich substrates (Põlme et al. 2020) | Põlme S, et al. 2020. Fungal Diversity 105:1-16 ((https://doi.org/10.1007/s13225-020-00466-2)) |
| Otu33601 | 1 | k__Fungi(100);p__Ascomycota(100);c__Saccharomycetes(100);o__Saccharomycetales(100);f__Metschnikowiaceae(100);g__Metschnikowia(100);g__Metschnikowia_unclassified(100); | Metschnikowia | 13 | Saprotroph | \|Undefined Saprotroph\| | Yeast | NULL | Probable | As nectar-tap_saprotroph (Põlme et al. 2020); Decay sugar-rich substrates (Põlme et al. 2020) | Põlme S, et al. 2020. Fungal Diversity 105:1-16 ((https://doi.org/10.1007/s13225-020-00466-2)) |
| Otu33602 | 1 | k__Fungi(100);p__Ascomycota(100);c__Saccharomycetes(100);o__Saccharomycetales(100);f__Metschnikowiaceae(100);g__Metschnikowia(100);g__Metschnikowia_unclassified(100); | Metschnikowia | 13 | Saprotroph | \|Undefined Saprotroph\| | Yeast | NULL | Probable | As nectar-tap_saprotroph (Põlme et al. 2020); Decay sugar-rich substrates (Põlme et al. 2020) | Põlme S, et al. 2020. Fungal Diversity 105:1-16 ((https://doi.org/10.1007/s13225-020-00466-2)) |
| Otu33603 | 1 | k__Fungi(100);p__Ascomycota(100);c__Saccharomycetes(100);o__Saccharomycetales(100);f__Metschnikowiaceae(100);g__Metschnikowia(100);s__Metschnikowia_pulcherrima(100); | Metschnikowia | 13 | Saprotroph | \|Undefined Saprotroph\| | Yeast | NULL | Probable | As nectar-tap_saprotroph (Põlme et al. 2020); Decay sugar-rich substrates (Põlme et al. 2020) | Põlme S, et al. 2020. Fungal Diversity 105:1-16 ((https://doi.org/10.1007/s13225-020-00466-2)) |
| Otu33604 | 1 | k__Fungi(100);p__Ascomycota(100);c__Saccharomycetes(100);o__Saccharomycetales(100);f__Metschnikowiaceae(100);g__Metschnikowia(100);s__Metschnikowia_pulcherrima(100); | Metschnikowia | 13 | Saprotroph | \|Undefined Saprotroph\| | Yeast | NULL | Probable | As nectar-tap_saprotroph (Põlme et al. 2020); Decay sugar-rich substrates (Põlme et al. 2020) | Põlme S, et al. 2020. Fungal Diversity 105:1-16 ((https://doi.org/10.1007/s13225-020-00466-2)) |
| Otu33605 | 1 | k__Fungi(100);p__Ascomycota(100);c__Saccharomycetes(100);o__Saccharomycetales(100);f__Metschnikowiaceae(100);g__Metschnikowia(100);s__Metschnikowia_pulcherrima(100); | Metschnikowia | 13 | Saprotroph | \|Undefined Saprotroph\| | Yeast | NULL | Probable | As nectar-tap_saprotroph (Põlme et al. 2020); Decay sugar-rich substrates (Põlme et al. 2020) | Põlme S, et al. 2020. Fungal Diversity 105:1-16 ((https://doi.org/10.1007/s13225-020-00466-2)) |
| Otu33606 | 1 | k__Fungi(100);p__Ascomycota(100);c__Saccharomycetes(100);o__Saccharomycetales(100);f__Metschnikowiaceae(100);g__Metschnikowia(100);s__Metschnikowia_pulcherrima(100); | Metschnikowia | 13 | Saprotroph | \|Undefined Saprotroph\| | Yeast | NULL | Probable | As nectar-tap_saprotroph (Põlme et al. 2020); Decay sugar-rich substrates (Põlme et al. 2020) | Põlme S, et al. 2020. Fungal Diversity 105:1-16 ((https://doi.org/10.1007/s13225-020-00466-2)) |
| Otu33607 | 1 | k__Fungi(100);p__Ascomycota(100);c__Saccharomycetes(100);o__Saccharomycetales(100);f__Metschnikowiaceae(100);g__Metschnikowia(100);s__Metschnikowia_pulcherrima(100); | Metschnikowia | 13 | Saprotroph | \|Undefined Saprotroph\| | Yeast | NULL | Probable | As nectar-tap_saprotroph (Põlme et al. 2020); Decay sugar-rich substrates (Põlme et al. 2020) | Põlme S, et al. 2020. Fungal Diversity 105:1-16 ((https://doi.org/10.1007/s13225-020-00466-2)) |
| Otu33608 | 1 | k__Fungi(100);p__Ascomycota(100);c__Saccharomycetes(100);o__Saccharomycetales(100);f__Metschnikowiaceae(100);g__Metschnikowia(100);s__Metschnikowia_pulcherrima(100); | Metschnikowia | 13 | Saprotroph | \|Undefined Saprotroph\| | Yeast | NULL | Probable | As nectar-tap_saprotroph (Põlme et al. 2020); Decay sugar-rich substrates (Põlme et al. 2020) | Põlme S, et al. 2020. Fungal Diversity 105:1-16 ((https://doi.org/10.1007/s13225-020-00466-2)) |
| Otu33609 | 1 | k__Fungi(100);p__Ascomycota(100);c__Saccharomycetes(100);o__Saccharomycetales(100);f__Metschnikowiaceae(100);g__Metschnikowia(100);s__Metschnikowia_pulcherrima(100); | Metschnikowia | 13 | Saprotroph | \|Undefined Saprotroph\| | Yeast | NULL | Probable | As nectar-tap_saprotroph (Põlme et al. 2020); Decay sugar-rich substrates (Põlme et al. 2020) | Põlme S, et al. 2020. Fungal Diversity 105:1-16 ((https://doi.org/10.1007/s13225-020-00466-2)) |
| Otu33610 | 1 | k__Fungi(100);p__Ascomycota(100);c__Saccharomycetes(100);o__Saccharomycetales(100);f__Metschnikowiaceae(100);g__Metschnikowia(100);s__Metschnikowia_pulcherrima(100); | Metschnikowia | 13 | Saprotroph | \|Undefined Saprotroph\| | Yeast | NULL | Probable | As nectar-tap_saprotroph (Põlme et al. 2020); Decay sugar-rich substrates (Põlme et al. 2020) | Põlme S, et al. 2020. Fungal Diversity 105:1-16 ((https://doi.org/10.1007/s13225-020-00466-2)) |
| Otu33611 | 1 | k__Fungi(100);p__Ascomycota(100);c__Saccharomycetes(100);o__Saccharomycetales(100);f__Metschnikowiaceae(100);g__Metschnikowia(100);s__Metschnikowia_pulcherrima(100); | Metschnikowia | 13 | Saprotroph | \|Undefined Saprotroph\| | Yeast | NULL | Probable | As nectar-tap_saprotroph (Põlme et al. 2020); Decay sugar-rich substrates (Põlme et al. 2020) | Põlme S, et al. 2020. Fungal Diversity 105:1-16 ((https://doi.org/10.1007/s13225-020-00466-2)) |
| Otu33612 | 1 | k__Fungi(100);p__Ascomycota(100);c__Saccharomycetes(100);o__Saccharomycetales(100);f__Metschnikowiaceae(100);g__Metschnikowia(100);s__Metschnikowia_pulcherrima(100); | Metschnikowia | 13 | Saprotroph | \|Undefined Saprotroph\| | Yeast | NULL | Probable | As nectar-tap_saprotroph (Põlme et al. 2020); Decay sugar-rich substrates (Põlme et al. 2020) | Põlme S, et al. 2020. Fungal Diversity 105:1-16 ((https://doi.org/10.1007/s13225-020-00466-2)) |
| Otu33614 | 1 | k__Fungi(100);p__Ascomycota(100);c__Saccharomycetes(100);o__Saccharomycetales(100);f__Metschnikowiaceae(100);g__Metschnikowia(100);g__Metschnikowia_unclassified(100); | Metschnikowia | 13 | Saprotroph | \|Undefined Saprotroph\| | Yeast | NULL | Probable | As nectar-tap_saprotroph (Põlme et al. 2020); Decay sugar-rich substrates (Põlme et al. 2020) | Põlme S, et al. 2020. Fungal Diversity 105:1-16 ((https://doi.org/10.1007/s13225-020-00466-2)) |
| Otu33615 | 1 | k__Fungi(100);p__Ascomycota(100);c__Saccharomycetes(100);o__Saccharomycetales(100);f__Metschnikowiaceae(100);g__Metschnikowia(100);s__Metschnikowia_pulcherrima(100); | Metschnikowia | 13 | Saprotroph | \|Undefined Saprotroph\| | Yeast | NULL | Probable | As nectar-tap_saprotroph (Põlme et al. 2020); Decay sugar-rich substrates (Põlme et al. 2020) | Põlme S, et al. 2020. Fungal Diversity 105:1-16 ((https://doi.org/10.1007/s13225-020-00466-2)) |
| Otu33616 | 1 | k__Fungi(100);p__Ascomycota(100);c__Saccharomycetes(100);o__Saccharomycetales(100);f__Metschnikowiaceae(100);g__Metschnikowia(100);s__Metschnikowia_pulcherrima(100); | Metschnikowia | 13 | Saprotroph | \|Undefined Saprotroph\| | Yeast | NULL | Probable | As nectar-tap_saprotroph (Põlme et al. 2020); Decay sugar-rich substrates (Põlme et al. 2020) | Põlme S, et al. 2020. Fungal Diversity 105:1-16 ((https://doi.org/10.1007/s13225-020-00466-2)) |
| Otu33617 | 1 | k__Fungi(100);p__Ascomycota(100);c__Saccharomycetes(100);o__Saccharomycetales(100);f__Metschnikowiaceae(100);g__Metschnikowia(100);s__Metschnikowia_pulcherrima(100); | Metschnikowia | 13 | Saprotroph | \|Undefined Saprotroph\| | Yeast | NULL | Probable | As nectar-tap_saprotroph (Põlme et al. 2020); Decay sugar-rich substrates (Põlme et al. 2020) | Põlme S, et al. 2020. Fungal Diversity 105:1-16 ((https://doi.org/10.1007/s13225-020-00466-2)) |
| Otu33618 | 1 | k__Fungi(100);p__Ascomycota(100);c__Saccharomycetes(100);o__Saccharomycetales(100);f__Metschnikowiaceae(100);g__Metschnikowia(100);g__Metschnikowia_unclassified(100); | Metschnikowia | 13 | Saprotroph | \|Undefined Saprotroph\| | Yeast | NULL | Probable | As nectar-tap_saprotroph (Põlme et al. 2020); Decay sugar-rich substrates (Põlme et al. 2020) | Põlme S, et al. 2020. Fungal Diversity 105:1-16 ((https://doi.org/10.1007/s13225-020-00466-2)) |
| Otu33620 | 1 | k__Fungi(100);p__Ascomycota(100);c__Saccharomycetes(100);o__Saccharomycetales(100);f__Metschnikowiaceae(100);g__Metschnikowia(100);s__Metschnikowia_pulcherrima(100); | Metschnikowia | 13 | Saprotroph | \|Undefined Saprotroph\| | Yeast | NULL | Probable | As nectar-tap_saprotroph (Põlme et al. 2020); Decay sugar-rich substrates (Põlme et al. 2020) | Põlme S, et al. 2020. Fungal Diversity 105:1-16 ((https://doi.org/10.1007/s13225-020-00466-2)) |
| Otu33621 | 1 | k__Fungi(100);p__Ascomycota(100);c__Saccharomycetes(100);o__Saccharomycetales(100);f__Metschnikowiaceae(100);g__Metschnikowia(100);s__Metschnikowia_pulcherrima(100); | Metschnikowia | 13 | Saprotroph | \|Undefined Saprotroph\| | Yeast | NULL | Probable | As nectar-tap_saprotroph (Põlme et al. 2020); Decay sugar-rich substrates (Põlme et al. 2020) | Põlme S, et al. 2020. Fungal Diversity 105:1-16 ((https://doi.org/10.1007/s13225-020-00466-2)) |
| Otu33622 | 1 | k__Fungi(100);p__Ascomycota(100);c__Saccharomycetes(100);o__Saccharomycetales(100);f__Metschnikowiaceae(100);g__Metschnikowia(100);s__Metschnikowia_pulcherrima(100); | Metschnikowia | 13 | Saprotroph | \|Undefined Saprotroph\| | Yeast | NULL | Probable | As nectar-tap_saprotroph (Põlme et al. 2020); Decay sugar-rich substrates (Põlme et al. 2020) | Põlme S, et al. 2020. Fungal Diversity 105:1-16 ((https://doi.org/10.1007/s13225-020-00466-2)) |
| Otu33623 | 1 | k__Fungi(100);p__Ascomycota(100);c__Saccharomycetes(100);o__Saccharomycetales(100);f__Metschnikowiaceae(100);g__Metschnikowia(100);s__Metschnikowia_pulcherrima(100); | Metschnikowia | 13 | Saprotroph | \|Undefined Saprotroph\| | Yeast | NULL | Probable | As nectar-tap_saprotroph (Põlme et al. 2020); Decay sugar-rich substrates (Põlme et al. 2020) | Põlme S, et al. 2020. Fungal Diversity 105:1-16 ((https://doi.org/10.1007/s13225-020-00466-2)) |
| Otu33624 | 1 | k__Fungi(100);p__Ascomycota(100);c__Saccharomycetes(100);o__Saccharomycetales(100);f__Metschnikowiaceae(100);g__Metschnikowia(100);s__Metschnikowia_pulcherrima(100); | Metschnikowia | 13 | Saprotroph | \|Undefined Saprotroph\| | Yeast | NULL | Probable | As nectar-tap_saprotroph (Põlme et al. 2020); Decay sugar-rich substrates (Põlme et al. 2020) | Põlme S, et al. 2020. Fungal Diversity 105:1-16 ((https://doi.org/10.1007/s13225-020-00466-2)) |
| Otu33625 | 1 | k__Fungi(100);p__Ascomycota(100);c__Saccharomycetes(100);o__Saccharomycetales(100);f__Metschnikowiaceae(100);g__Metschnikowia(100);s__Metschnikowia_pulcherrima(100); | Metschnikowia | 13 | Saprotroph | \|Undefined Saprotroph\| | Yeast | NULL | Probable | As nectar-tap_saprotroph (Põlme et al. 2020); Decay sugar-rich substrates (Põlme et al. 2020) | Põlme S, et al. 2020. Fungal Diversity 105:1-16 ((https://doi.org/10.1007/s13225-020-00466-2)) |
| Otu33626 | 1 | k__Fungi(100);p__Ascomycota(100);c__Saccharomycetes(100);o__Saccharomycetales(100);f__Metschnikowiaceae(100);g__Metschnikowia(100);s__Metschnikowia_pulcherrima(100); | Metschnikowia | 13 | Saprotroph | \|Undefined Saprotroph\| | Yeast | NULL | Probable | As nectar-tap_saprotroph (Põlme et al. 2020); Decay sugar-rich substrates (Põlme et al. 2020) | Põlme S, et al. 2020. Fungal Diversity 105:1-16 ((https://doi.org/10.1007/s13225-020-00466-2)) |
| Otu33627 | 1 | k__Fungi(100);p__Ascomycota(100);c__Saccharomycetes(100);o__Saccharomycetales(100);f__Metschnikowiaceae(100);g__Metschnikowia(100);s__Metschnikowia_pulcherrima(100); | Metschnikowia | 13 | Saprotroph | \|Undefined Saprotroph\| | Yeast | NULL | Probable | As nectar-tap_saprotroph (Põlme et al. 2020); Decay sugar-rich substrates (Põlme et al. 2020) | Põlme S, et al. 2020. Fungal Diversity 105:1-16 ((https://doi.org/10.1007/s13225-020-00466-2)) |
| Otu33629 | 1 | k__Fungi(100);p__Ascomycota(100);c__Saccharomycetes(100);o__Saccharomycetales(100);f__Metschnikowiaceae(100);g__Metschnikowia(100);s__Metschnikowia_pulcherrima(100); | Metschnikowia | 13 | Saprotroph | \|Undefined Saprotroph\| | Yeast | NULL | Probable | As nectar-tap_saprotroph (Põlme et al. 2020); Decay sugar-rich substrates (Põlme et al. 2020) | Põlme S, et al. 2020. Fungal Diversity 105:1-16 ((https://doi.org/10.1007/s13225-020-00466-2)) |
| Otu33633 | 1 | k__Fungi(100);p__Ascomycota(100);c__Saccharomycetes(100);o__Saccharomycetales(100);f__Metschnikowiaceae(100);g__Metschnikowia(100);s__Metschnikowia_pulcherrima(100); | Metschnikowia | 13 | Saprotroph | \|Undefined Saprotroph\| | Yeast | NULL | Probable | As nectar-tap_saprotroph (Põlme et al. 2020); Decay sugar-rich substrates (Põlme et al. 2020) | Põlme S, et al. 2020. Fungal Diversity 105:1-16 ((https://doi.org/10.1007/s13225-020-00466-2)) |
| Otu33634 | 1 | k__Fungi(100);p__Ascomycota(100);c__Saccharomycetes(100);o__Saccharomycetales(100);f__Metschnikowiaceae(100);g__Metschnikowia(100);s__Metschnikowia_pulcherrima(100); | Metschnikowia | 13 | Saprotroph | \|Undefined Saprotroph\| | Yeast | NULL | Probable | As nectar-tap_saprotroph (Põlme et al. 2020); Decay sugar-rich substrates (Põlme et al. 2020) | Põlme S, et al. 2020. Fungal Diversity 105:1-16 ((https://doi.org/10.1007/s13225-020-00466-2)) |
| Otu33635 | 1 | k__Fungi(100);p__Ascomycota(100);c__Saccharomycetes(100);o__Saccharomycetales(100);f__Metschnikowiaceae(100);g__Metschnikowia(100);s__Metschnikowia_pulcherrima(100); | Metschnikowia | 13 | Saprotroph | \|Undefined Saprotroph\| | Yeast | NULL | Probable | As nectar-tap_saprotroph (Põlme et al. 2020); Decay sugar-rich substrates (Põlme et al. 2020) | Põlme S, et al. 2020. Fungal Diversity 105:1-16 ((https://doi.org/10.1007/s13225-020-00466-2)) |
| Otu33640 | 1 | k__Fungi(100);p__Ascomycota(100);c__Saccharomycetes(100);o__Saccharomycetales(100);f__Metschnikowiaceae(100);g__Metschnikowia(100);g__Metschnikowia_unclassified(100); | Metschnikowia | 13 | Saprotroph | \|Undefined Saprotroph\| | Yeast | NULL | Probable | As nectar-tap_saprotroph (Põlme et al. 2020); Decay sugar-rich substrates (Põlme et al. 2020) | Põlme S, et al. 2020. Fungal Diversity 105:1-16 ((https://doi.org/10.1007/s13225-020-00466-2)) |
| Otu33641 | 1 | k__Fungi(100);p__Ascomycota(100);c__Saccharomycetes(100);o__Saccharomycetales(100);f__Metschnikowiaceae(100);g__Metschnikowia(100);s__Metschnikowia_pulcherrima(100); | Metschnikowia | 13 | Saprotroph | \|Undefined Saprotroph\| | Yeast | NULL | Probable | As nectar-tap_saprotroph (Põlme et al. 2020); Decay sugar-rich substrates (Põlme et al. 2020) | Põlme S, et al. 2020. Fungal Diversity 105:1-16 ((https://doi.org/10.1007/s13225-020-00466-2)) |
| Otu33642 | 1 | k__Fungi(100);p__Ascomycota(100);c__Saccharomycetes(100);o__Saccharomycetales(100);f__Metschnikowiaceae(100);g__Metschnikowia(100);g__Metschnikowia_unclassified(100); | Metschnikowia | 13 | Saprotroph | \|Undefined Saprotroph\| | Yeast | NULL | Probable | As nectar-tap_saprotroph (Põlme et al. 2020); Decay sugar-rich substrates (Põlme et al. 2020) | Põlme S, et al. 2020. Fungal Diversity 105:1-16 ((https://doi.org/10.1007/s13225-020-00466-2)) |
| Otu33643 | 1 | k__Fungi(100);p__Ascomycota(100);c__Saccharomycetes(100);o__Saccharomycetales(100);f__Metschnikowiaceae(100);g__Metschnikowia(100);s__Metschnikowia_pulcherrima(100); | Metschnikowia | 13 | Saprotroph | \|Undefined Saprotroph\| | Yeast | NULL | Probable | As nectar-tap_saprotroph (Põlme et al. 2020); Decay sugar-rich substrates (Põlme et al. 2020) | Põlme S, et al. 2020. Fungal Diversity 105:1-16 ((https://doi.org/10.1007/s13225-020-00466-2)) |
| Otu33644 | 1 | k__Fungi(100);p__Ascomycota(100);c__Saccharomycetes(100);o__Saccharomycetales(100);f__Metschnikowiaceae(100);g__Metschnikowia(100);s__Metschnikowia_pulcherrima(100); | Metschnikowia | 13 | Saprotroph | \|Undefined Saprotroph\| | Yeast | NULL | Probable | As nectar-tap_saprotroph (Põlme et al. 2020); Decay sugar-rich substrates (Põlme et al. 2020) | Põlme S, et al. 2020. Fungal Diversity 105:1-16 ((https://doi.org/10.1007/s13225-020-00466-2)) |
| Otu33645 | 1 | k__Fungi(100);p__Ascomycota(100);c__Saccharomycetes(100);o__Saccharomycetales(100);f__Metschnikowiaceae(100);g__Metschnikowia(100);s__Metschnikowia_pulcherrima(100); | Metschnikowia | 13 | Saprotroph | \|Undefined Saprotroph\| | Yeast | NULL | Probable | As nectar-tap_saprotroph (Põlme et al. 2020); Decay sugar-rich substrates (Põlme et al. 2020) | Põlme S, et al. 2020. Fungal Diversity 105:1-16 ((https://doi.org/10.1007/s13225-020-00466-2)) |
| Otu33646 | 1 | k__Fungi(100);p__Ascomycota(100);c__Saccharomycetes(100);o__Saccharomycetales(100);f__Metschnikowiaceae(100);g__Metschnikowia(100);g__Metschnikowia_unclassified(100); | Metschnikowia | 13 | Saprotroph | \|Undefined Saprotroph\| | Yeast | NULL | Probable | As nectar-tap_saprotroph (Põlme et al. 2020); Decay sugar-rich substrates (Põlme et al. 2020) | Põlme S, et al. 2020. Fungal Diversity 105:1-16 ((https://doi.org/10.1007/s13225-020-00466-2)) |
| Otu33647 | 1 | k__Fungi(100);p__Ascomycota(100);c__Saccharomycetes(100);o__Saccharomycetales(100);f__Metschnikowiaceae(100);g__Metschnikowia(100);s__Metschnikowia_pulcherrima(100); | Metschnikowia | 13 | Saprotroph | \|Undefined Saprotroph\| | Yeast | NULL | Probable | As nectar-tap_saprotroph (Põlme et al. 2020); Decay sugar-rich substrates (Põlme et al. 2020) | Põlme S, et al. 2020. Fungal Diversity 105:1-16 ((https://doi.org/10.1007/s13225-020-00466-2)) |
| Otu33648 | 1 | k__Fungi(100);p__Ascomycota(100);c__Saccharomycetes(100);o__Saccharomycetales(100);f__Metschnikowiaceae(100);g__Metschnikowia(100);s__Metschnikowia_pulcherrima(100); | Metschnikowia | 13 | Saprotroph | \|Undefined Saprotroph\| | Yeast | NULL | Probable | As nectar-tap_saprotroph (Põlme et al. 2020); Decay sugar-rich substrates (Põlme et al. 2020) | Põlme S, et al. 2020. Fungal Diversity 105:1-16 ((https://doi.org/10.1007/s13225-020-00466-2)) |
| Otu33649 | 1 | k__Fungi(100);p__Ascomycota(100);c__Saccharomycetes(100);o__Saccharomycetales(100);f__Metschnikowiaceae(100);g__Metschnikowia(100);s__Metschnikowia_pulcherrima(100); | Metschnikowia | 13 | Saprotroph | \|Undefined Saprotroph\| | Yeast | NULL | Probable | As nectar-tap_saprotroph (Põlme et al. 2020); Decay sugar-rich substrates (Põlme et al. 2020) | Põlme S, et al. 2020. Fungal Diversity 105:1-16 ((https://doi.org/10.1007/s13225-020-00466-2)) |
| Otu33650 | 1 | k__Fungi(100);p__Ascomycota(100);c__Saccharomycetes(100);o__Saccharomycetales(100);f__Metschnikowiaceae(100);g__Metschnikowia(100);s__Metschnikowia_pulcherrima(100); | Metschnikowia | 13 | Saprotroph | \|Undefined Saprotroph\| | Yeast | NULL | Probable | As nectar-tap_saprotroph (Põlme et al. 2020); Decay sugar-rich substrates (Põlme et al. 2020) | Põlme S, et al. 2020. Fungal Diversity 105:1-16 ((https://doi.org/10.1007/s13225-020-00466-2)) |
| Otu33651 | 1 | k__Fungi(100);p__Ascomycota(100);c__Saccharomycetes(100);o__Saccharomycetales(100);f__Metschnikowiaceae(100);g__Metschnikowia(100);s__Metschnikowia_pulcherrima(100); | Metschnikowia | 13 | Saprotroph | \|Undefined Saprotroph\| | Yeast | NULL | Probable | As nectar-tap_saprotroph (Põlme et al. 2020); Decay sugar-rich substrates (Põlme et al. 2020) | Põlme S, et al. 2020. Fungal Diversity 105:1-16 ((https://doi.org/10.1007/s13225-020-00466-2)) |
| Otu33652 | 1 | k__Fungi(100);p__Ascomycota(100);c__Saccharomycetes(100);o__Saccharomycetales(100);f__Metschnikowiaceae(100);g__Metschnikowia(100);s__Metschnikowia_pulcherrima(100); | Metschnikowia | 13 | Saprotroph | \|Undefined Saprotroph\| | Yeast | NULL | Probable | As nectar-tap_saprotroph (Põlme et al. 2020); Decay sugar-rich substrates (Põlme et al. 2020) | Põlme S, et al. 2020. Fungal Diversity 105:1-16 ((https://doi.org/10.1007/s13225-020-00466-2)) |
| Otu33653 | 1 | k__Fungi(100);p__Ascomycota(100);c__Saccharomycetes(100);o__Saccharomycetales(100);f__Metschnikowiaceae(100);g__Metschnikowia(100);s__Metschnikowia_pulcherrima(100); | Metschnikowia | 13 | Saprotroph | \|Undefined Saprotroph\| | Yeast | NULL | Probable | As nectar-tap_saprotroph (Põlme et al. 2020); Decay sugar-rich substrates (Põlme et al. 2020) | Põlme S, et al. 2020. Fungal Diversity 105:1-16 ((https://doi.org/10.1007/s13225-020-00466-2)) |
| Otu33654 | 1 | k__Fungi(100);p__Ascomycota(100);c__Saccharomycetes(100);o__Saccharomycetales(100);f__Metschnikowiaceae(100);g__Metschnikowia(100);s__Metschnikowia_pulcherrima(100); | Metschnikowia | 13 | Saprotroph | \|Undefined Saprotroph\| | Yeast | NULL | Probable | As nectar-tap_saprotroph (Põlme et al. 2020); Decay sugar-rich substrates (Põlme et al. 2020) | Põlme S, et al. 2020. Fungal Diversity 105:1-16 ((https://doi.org/10.1007/s13225-020-00466-2)) |
| Otu33655 | 1 | k__Fungi(100);p__Ascomycota(100);c__Saccharomycetes(100);o__Saccharomycetales(100);f__Metschnikowiaceae(100);g__Metschnikowia(100);s__Metschnikowia_pulcherrima(100); | Metschnikowia | 13 | Saprotroph | \|Undefined Saprotroph\| | Yeast | NULL | Probable | As nectar-tap_saprotroph (Põlme et al. 2020); Decay sugar-rich substrates (Põlme et al. 2020) | Põlme S, et al. 2020. Fungal Diversity 105:1-16 ((https://doi.org/10.1007/s13225-020-00466-2)) |
| Otu33656 | 1 | k__Fungi(100);p__Ascomycota(100);c__Saccharomycetes(100);o__Saccharomycetales(100);f__Metschnikowiaceae(100);g__Metschnikowia(100);s__Metschnikowia_pulcherrima(100); | Metschnikowia | 13 | Saprotroph | \|Undefined Saprotroph\| | Yeast | NULL | Probable | As nectar-tap_saprotroph (Põlme et al. 2020); Decay sugar-rich substrates (Põlme et al. 2020) | Põlme S, et al. 2020. Fungal Diversity 105:1-16 ((https://doi.org/10.1007/s13225-020-00466-2)) |
| Otu33657 | 1 | k__Fungi(100);p__Ascomycota(100);c__Saccharomycetes(100);o__Saccharomycetales(100);f__Metschnikowiaceae(100);g__Metschnikowia(100);s__Metschnikowia_pulcherrima(100); | Metschnikowia | 13 | Saprotroph | \|Undefined Saprotroph\| | Yeast | NULL | Probable | As nectar-tap_saprotroph (Põlme et al. 2020); Decay sugar-rich substrates (Põlme et al. 2020) | Põlme S, et al. 2020. Fungal Diversity 105:1-16 ((https://doi.org/10.1007/s13225-020-00466-2)) |
| Otu33658 | 1 | k__Fungi(100);p__Ascomycota(100);c__Saccharomycetes(100);o__Saccharomycetales(100);f__Metschnikowiaceae(100);g__Metschnikowia(100);s__Metschnikowia_pulcherrima(100); | Metschnikowia | 13 | Saprotroph | \|Undefined Saprotroph\| | Yeast | NULL | Probable | As nectar-tap_saprotroph (Põlme et al. 2020); Decay sugar-rich substrates (Põlme et al. 2020) | Põlme S, et al. 2020. Fungal Diversity 105:1-16 ((https://doi.org/10.1007/s13225-020-00466-2)) |
| Otu33660 | 1 | k__Fungi(100);p__Ascomycota(100);c__Saccharomycetes(100);o__Saccharomycetales(100);f__Metschnikowiaceae(100);g__Metschnikowia(100);s__Metschnikowia_pulcherrima(100); | Metschnikowia | 13 | Saprotroph | \|Undefined Saprotroph\| | Yeast | NULL | Probable | As nectar-tap_saprotroph (Põlme et al. 2020); Decay sugar-rich substrates (Põlme et al. 2020) | Põlme S, et al. 2020. Fungal Diversity 105:1-16 ((https://doi.org/10.1007/s13225-020-00466-2)) |
| Otu33661 | 1 | k__Fungi(100);p__Ascomycota(100);c__Saccharomycetes(100);o__Saccharomycetales(100);f__Metschnikowiaceae(100);g__Metschnikowia(100);s__Metschnikowia_pulcherrima(100); | Metschnikowia | 13 | Saprotroph | \|Undefined Saprotroph\| | Yeast | NULL | Probable | As nectar-tap_saprotroph (Põlme et al. 2020); Decay sugar-rich substrates (Põlme et al. 2020) | Põlme S, et al. 2020. Fungal Diversity 105:1-16 ((https://doi.org/10.1007/s13225-020-00466-2)) |
| Otu33662 | 1 | k__Fungi(100);p__Ascomycota(100);c__Saccharomycetes(100);o__Saccharomycetales(100);f__Metschnikowiaceae(100);g__Metschnikowia(100);g__Metschnikowia_unclassified(100); | Metschnikowia | 13 | Saprotroph | \|Undefined Saprotroph\| | Yeast | NULL | Probable | As nectar-tap_saprotroph (Põlme et al. 2020); Decay sugar-rich substrates (Põlme et al. 2020) | Põlme S, et al. 2020. Fungal Diversity 105:1-16 ((https://doi.org/10.1007/s13225-020-00466-2)) |
| Otu33664 | 1 | k__Fungi(100);p__Ascomycota(100);c__Saccharomycetes(100);o__Saccharomycetales(100);f__Metschnikowiaceae(100);g__Metschnikowia(100);s__Metschnikowia_pulcherrima(100); | Metschnikowia | 13 | Saprotroph | \|Undefined Saprotroph\| | Yeast | NULL | Probable | As nectar-tap_saprotroph (Põlme et al. 2020); Decay sugar-rich substrates (Põlme et al. 2020) | Põlme S, et al. 2020. Fungal Diversity 105:1-16 ((https://doi.org/10.1007/s13225-020-00466-2)) |
| Otu33665 | 1 | k__Fungi(100);p__Ascomycota(100);c__Saccharomycetes(100);o__Saccharomycetales(100);f__Metschnikowiaceae(100);g__Metschnikowia(100);s__Metschnikowia_pulcherrima(100); | Metschnikowia | 13 | Saprotroph | \|Undefined Saprotroph\| | Yeast | NULL | Probable | As nectar-tap_saprotroph (Põlme et al. 2020); Decay sugar-rich substrates (Põlme et al. 2020) | Põlme S, et al. 2020. Fungal Diversity 105:1-16 ((https://doi.org/10.1007/s13225-020-00466-2)) |
| Otu33670 | 1 | k__Fungi(100);p__Ascomycota(100);c__Saccharomycetes(100);o__Saccharomycetales(100);f__Metschnikowiaceae(100);g__Metschnikowia(100);s__Metschnikowia_pulcherrima(100); | Metschnikowia | 13 | Saprotroph | \|Undefined Saprotroph\| | Yeast | NULL | Probable | As nectar-tap_saprotroph (Põlme et al. 2020); Decay sugar-rich substrates (Põlme et al. 2020) | Põlme S, et al. 2020. Fungal Diversity 105:1-16 ((https://doi.org/10.1007/s13225-020-00466-2)) |
| Otu33671 | 1 | k__Fungi(100);p__Ascomycota(100);c__Saccharomycetes(100);o__Saccharomycetales(100);f__Metschnikowiaceae(100);g__Metschnikowia(100);s__Metschnikowia_pulcherrima(100); | Metschnikowia | 13 | Saprotroph | \|Undefined Saprotroph\| | Yeast | NULL | Probable | As nectar-tap_saprotroph (Põlme et al. 2020); Decay sugar-rich substrates (Põlme et al. 2020) | Põlme S, et al. 2020. Fungal Diversity 105:1-16 ((https://doi.org/10.1007/s13225-020-00466-2)) |
| Otu33673 | 1 | k__Fungi(100);p__Ascomycota(100);c__Saccharomycetes(100);o__Saccharomycetales(100);f__Metschnikowiaceae(100);g__Metschnikowia(100);s__Metschnikowia_pulcherrima(100); | Metschnikowia | 13 | Saprotroph | \|Undefined Saprotroph\| | Yeast | NULL | Probable | As nectar-tap_saprotroph (Põlme et al. 2020); Decay sugar-rich substrates (Põlme et al. 2020) | Põlme S, et al. 2020. Fungal Diversity 105:1-16 ((https://doi.org/10.1007/s13225-020-00466-2)) |
| Otu33674 | 1 | k__Fungi(100);p__Ascomycota(100);c__Saccharomycetes(100);o__Saccharomycetales(100);f__Metschnikowiaceae(100);g__Metschnikowia(100);g__Metschnikowia_unclassified(100); | Metschnikowia | 13 | Saprotroph | \|Undefined Saprotroph\| | Yeast | NULL | Probable | As nectar-tap_saprotroph (Põlme et al. 2020); Decay sugar-rich substrates (Põlme et al. 2020) | Põlme S, et al. 2020. Fungal Diversity 105:1-16 ((https://doi.org/10.1007/s13225-020-00466-2)) |
| Otu33675 | 1 | k__Fungi(100);p__Ascomycota(100);c__Saccharomycetes(100);o__Saccharomycetales(100);f__Metschnikowiaceae(100);g__Metschnikowia(100);g__Metschnikowia_unclassified(100); | Metschnikowia | 13 | Saprotroph | \|Undefined Saprotroph\| | Yeast | NULL | Probable | As nectar-tap_saprotroph (Põlme et al. 2020); Decay sugar-rich substrates (Põlme et al. 2020) | Põlme S, et al. 2020. Fungal Diversity 105:1-16 ((https://doi.org/10.1007/s13225-020-00466-2)) |
| Otu33679 | 1 | k__Fungi(100);p__Ascomycota(100);c__Saccharomycetes(100);o__Saccharomycetales(100);f__Metschnikowiaceae(100);g__Metschnikowia(100);s__Metschnikowia_pulcherrima(100); | Metschnikowia | 13 | Saprotroph | \|Undefined Saprotroph\| | Yeast | NULL | Probable | As nectar-tap_saprotroph (Põlme et al. 2020); Decay sugar-rich substrates (Põlme et al. 2020) | Põlme S, et al. 2020. Fungal Diversity 105:1-16 ((https://doi.org/10.1007/s13225-020-00466-2)) |
| Otu33680 | 1 | k__Fungi(100);p__Ascomycota(100);c__Saccharomycetes(100);o__Saccharomycetales(100);f__Metschnikowiaceae(100);g__Metschnikowia(100);s__Metschnikowia_pulcherrima(100); | Metschnikowia | 13 | Saprotroph | \|Undefined Saprotroph\| | Yeast | NULL | Probable | As nectar-tap_saprotroph (Põlme et al. 2020); Decay sugar-rich substrates (Põlme et al. 2020) | Põlme S, et al. 2020. Fungal Diversity 105:1-16 ((https://doi.org/10.1007/s13225-020-00466-2)) |
| Otu33681 | 1 | k__Fungi(100);p__Ascomycota(100);c__Saccharomycetes(100);o__Saccharomycetales(100);f__Metschnikowiaceae(100);g__Metschnikowia(100);s__Metschnikowia_pulcherrima(100); | Metschnikowia | 13 | Saprotroph | \|Undefined Saprotroph\| | Yeast | NULL | Probable | As nectar-tap_saprotroph (Põlme et al. 2020); Decay sugar-rich substrates (Põlme et al. 2020) | Põlme S, et al. 2020. Fungal Diversity 105:1-16 ((https://doi.org/10.1007/s13225-020-00466-2)) |
| Otu33683 | 1 | k__Fungi(100);p__Ascomycota(100);c__Saccharomycetes(100);o__Saccharomycetales(100);f__Metschnikowiaceae(100);g__Metschnikowia(100);s__Metschnikowia_pulcherrima(100); | Metschnikowia | 13 | Saprotroph | \|Undefined Saprotroph\| | Yeast | NULL | Probable | As nectar-tap_saprotroph (Põlme et al. 2020); Decay sugar-rich substrates (Põlme et al. 2020) | Põlme S, et al. 2020. Fungal Diversity 105:1-16 ((https://doi.org/10.1007/s13225-020-00466-2)) |
| Otu33684 | 1 | k__Fungi(100);p__Ascomycota(100);c__Saccharomycetes(100);o__Saccharomycetales(100);f__Metschnikowiaceae(100);g__Metschnikowia(100);s__Metschnikowia_pulcherrima(100); | Metschnikowia | 13 | Saprotroph | \|Undefined Saprotroph\| | Yeast | NULL | Probable | As nectar-tap_saprotroph (Põlme et al. 2020); Decay sugar-rich substrates (Põlme et al. 2020) | Põlme S, et al. 2020. Fungal Diversity 105:1-16 ((https://doi.org/10.1007/s13225-020-00466-2)) |
| Otu33685 | 1 | k__Fungi(100);p__Ascomycota(100);c__Saccharomycetes(100);o__Saccharomycetales(100);f__Metschnikowiaceae(100);g__Metschnikowia(100);s__Metschnikowia_pulcherrima(100); | Metschnikowia | 13 | Saprotroph | \|Undefined Saprotroph\| | Yeast | NULL | Probable | As nectar-tap_saprotroph (Põlme et al. 2020); Decay sugar-rich substrates (Põlme et al. 2020) | Põlme S, et al. 2020. Fungal Diversity 105:1-16 ((https://doi.org/10.1007/s13225-020-00466-2)) |
| Otu33686 | 1 | k__Fungi(100);p__Ascomycota(100);c__Saccharomycetes(100);o__Saccharomycetales(100);f__Metschnikowiaceae(100);g__Metschnikowia(100);s__Metschnikowia_pulcherrima(100); | Metschnikowia | 13 | Saprotroph | \|Undefined Saprotroph\| | Yeast | NULL | Probable | As nectar-tap_saprotroph (Põlme et al. 2020); Decay sugar-rich substrates (Põlme et al. 2020) | Põlme S, et al. 2020. Fungal Diversity 105:1-16 ((https://doi.org/10.1007/s13225-020-00466-2)) |
| Otu33688 | 1 | k__Fungi(100);p__Ascomycota(100);c__Saccharomycetes(100);o__Saccharomycetales(100);f__Metschnikowiaceae(100);g__Metschnikowia(100);s__Metschnikowia_pulcherrima(100); | Metschnikowia | 13 | Saprotroph | \|Undefined Saprotroph\| | Yeast | NULL | Probable | As nectar-tap_saprotroph (Põlme et al. 2020); Decay sugar-rich substrates (Põlme et al. 2020) | Põlme S, et al. 2020. Fungal Diversity 105:1-16 ((https://doi.org/10.1007/s13225-020-00466-2)) |
| Otu33689 | 1 | k__Fungi(100);p__Ascomycota(100);c__Saccharomycetes(100);o__Saccharomycetales(100);f__Metschnikowiaceae(100);g__Metschnikowia(100);s__Metschnikowia_pulcherrima(100); | Metschnikowia | 13 | Saprotroph | \|Undefined Saprotroph\| | Yeast | NULL | Probable | As nectar-tap_saprotroph (Põlme et al. 2020); Decay sugar-rich substrates (Põlme et al. 2020) | Põlme S, et al. 2020. Fungal Diversity 105:1-16 ((https://doi.org/10.1007/s13225-020-00466-2)) |
| Otu33690 | 1 | k__Fungi(100);p__Ascomycota(100);c__Saccharomycetes(100);o__Saccharomycetales(100);f__Metschnikowiaceae(100);g__Metschnikowia(100);g__Metschnikowia_unclassified(100); | Metschnikowia | 13 | Saprotroph | \|Undefined Saprotroph\| | Yeast | NULL | Probable | As nectar-tap_saprotroph (Põlme et al. 2020); Decay sugar-rich substrates (Põlme et al. 2020) | Põlme S, et al. 2020. Fungal Diversity 105:1-16 ((https://doi.org/10.1007/s13225-020-00466-2)) |
| Otu33691 | 1 | k__Fungi(100);p__Ascomycota(100);c__Saccharomycetes(100);o__Saccharomycetales(100);f__Metschnikowiaceae(100);g__Metschnikowia(100);s__Metschnikowia_pulcherrima(100); | Metschnikowia | 13 | Saprotroph | \|Undefined Saprotroph\| | Yeast | NULL | Probable | As nectar-tap_saprotroph (Põlme et al. 2020); Decay sugar-rich substrates (Põlme et al. 2020) | Põlme S, et al. 2020. Fungal Diversity 105:1-16 ((https://doi.org/10.1007/s13225-020-00466-2)) |
| Otu33692 | 1 | k__Fungi(100);p__Ascomycota(100);c__Saccharomycetes(100);o__Saccharomycetales(100);f__Metschnikowiaceae(100);g__Metschnikowia(100);s__Metschnikowia_pulcherrima(100); | Metschnikowia | 13 | Saprotroph | \|Undefined Saprotroph\| | Yeast | NULL | Probable | As nectar-tap_saprotroph (Põlme et al. 2020); Decay sugar-rich substrates (Põlme et al. 2020) | Põlme S, et al. 2020. Fungal Diversity 105:1-16 ((https://doi.org/10.1007/s13225-020-00466-2)) |
| Otu33693 | 1 | k__Fungi(100);p__Ascomycota(100);c__Saccharomycetes(100);o__Saccharomycetales(100);f__Metschnikowiaceae(100);g__Metschnikowia(100);s__Metschnikowia_pulcherrima(100); | Metschnikowia | 13 | Saprotroph | \|Undefined Saprotroph\| | Yeast | NULL | Probable | As nectar-tap_saprotroph (Põlme et al. 2020); Decay sugar-rich substrates (Põlme et al. 2020) | Põlme S, et al. 2020. Fungal Diversity 105:1-16 ((https://doi.org/10.1007/s13225-020-00466-2)) |
| Otu33694 | 1 | k__Fungi(100);p__Ascomycota(100);c__Saccharomycetes(100);o__Saccharomycetales(100);f__Metschnikowiaceae(100);g__Metschnikowia(100);g__Metschnikowia_unclassified(100); | Metschnikowia | 13 | Saprotroph | \|Undefined Saprotroph\| | Yeast | NULL | Probable | As nectar-tap_saprotroph (Põlme et al. 2020); Decay sugar-rich substrates (Põlme et al. 2020) | Põlme S, et al. 2020. Fungal Diversity 105:1-16 ((https://doi.org/10.1007/s13225-020-00466-2)) |
| Otu33695 | 1 | k__Fungi(100);p__Ascomycota(100);c__Saccharomycetes(100);o__Saccharomycetales(100);f__Metschnikowiaceae(100);g__Metschnikowia(100);s__Metschnikowia_pulcherrima(100); | Metschnikowia | 13 | Saprotroph | \|Undefined Saprotroph\| | Yeast | NULL | Probable | As nectar-tap_saprotroph (Põlme et al. 2020); Decay sugar-rich substrates (Põlme et al. 2020) | Põlme S, et al. 2020. Fungal Diversity 105:1-16 ((https://doi.org/10.1007/s13225-020-00466-2)) |
| Otu33696 | 1 | k__Fungi(100);p__Ascomycota(100);c__Saccharomycetes(100);o__Saccharomycetales(100);f__Metschnikowiaceae(100);g__Metschnikowia(100);g__Metschnikowia_unclassified(100); | Metschnikowia | 13 | Saprotroph | \|Undefined Saprotroph\| | Yeast | NULL | Probable | As nectar-tap_saprotroph (Põlme et al. 2020); Decay sugar-rich substrates (Põlme et al. 2020) | Põlme S, et al. 2020. Fungal Diversity 105:1-16 ((https://doi.org/10.1007/s13225-020-00466-2)) |
| Otu33697 | 1 | k__Fungi(100);p__Ascomycota(100);c__Saccharomycetes(100);o__Saccharomycetales(100);f__Metschnikowiaceae(100);g__Metschnikowia(100);s__Metschnikowia_pulcherrima(100); | Metschnikowia | 13 | Saprotroph | \|Undefined Saprotroph\| | Yeast | NULL | Probable | As nectar-tap_saprotroph (Põlme et al. 2020); Decay sugar-rich substrates (Põlme et al. 2020) | Põlme S, et al. 2020. Fungal Diversity 105:1-16 ((https://doi.org/10.1007/s13225-020-00466-2)) |
| Otu33698 | 1 | k__Fungi(100);p__Ascomycota(100);c__Saccharomycetes(100);o__Saccharomycetales(100);f__Metschnikowiaceae(100);g__Metschnikowia(100);g__Metschnikowia_unclassified(100); | Metschnikowia | 13 | Saprotroph | \|Undefined Saprotroph\| | Yeast | NULL | Probable | As nectar-tap_saprotroph (Põlme et al. 2020); Decay sugar-rich substrates (Põlme et al. 2020) | Põlme S, et al. 2020. Fungal Diversity 105:1-16 ((https://doi.org/10.1007/s13225-020-00466-2)) |
| Otu33699 | 1 | k__Fungi(100);p__Ascomycota(100);c__Saccharomycetes(100);o__Saccharomycetales(100);f__Metschnikowiaceae(100);g__Metschnikowia(100);g__Metschnikowia_unclassified(100); | Metschnikowia | 13 | Saprotroph | \|Undefined Saprotroph\| | Yeast | NULL | Probable | As nectar-tap_saprotroph (Põlme et al. 2020); Decay sugar-rich substrates (Põlme et al. 2020) | Põlme S, et al. 2020. Fungal Diversity 105:1-16 ((https://doi.org/10.1007/s13225-020-00466-2)) |
| Otu33700 | 1 | k__Fungi(100);p__Ascomycota(100);c__Saccharomycetes(100);o__Saccharomycetales(100);f__Metschnikowiaceae(100);g__Metschnikowia(100);g__Metschnikowia_unclassified(100); | Metschnikowia | 13 | Saprotroph | \|Undefined Saprotroph\| | Yeast | NULL | Probable | As nectar-tap_saprotroph (Põlme et al. 2020); Decay sugar-rich substrates (Põlme et al. 2020) | Põlme S, et al. 2020. Fungal Diversity 105:1-16 ((https://doi.org/10.1007/s13225-020-00466-2)) |
| Otu33701 | 1 | k__Fungi(100);p__Ascomycota(100);c__Saccharomycetes(100);o__Saccharomycetales(100);f__Metschnikowiaceae(100);g__Metschnikowia(100);s__Metschnikowia_pulcherrima(100); | Metschnikowia | 13 | Saprotroph | \|Undefined Saprotroph\| | Yeast | NULL | Probable | As nectar-tap_saprotroph (Põlme et al. 2020); Decay sugar-rich substrates (Põlme et al. 2020) | Põlme S, et al. 2020. Fungal Diversity 105:1-16 ((https://doi.org/10.1007/s13225-020-00466-2)) |
| Otu33702 | 1 | k__Fungi(100);p__Ascomycota(100);c__Saccharomycetes(100);o__Saccharomycetales(100);f__Metschnikowiaceae(100);g__Metschnikowia(100);g__Metschnikowia_unclassified(100); | Metschnikowia | 13 | Saprotroph | \|Undefined Saprotroph\| | Yeast | NULL | Probable | As nectar-tap_saprotroph (Põlme et al. 2020); Decay sugar-rich substrates (Põlme et al. 2020) | Põlme S, et al. 2020. Fungal Diversity 105:1-16 ((https://doi.org/10.1007/s13225-020-00466-2)) |
| Otu33703 | 1 | k__Fungi(100);p__Ascomycota(100);c__Saccharomycetes(100);o__Saccharomycetales(100);f__Metschnikowiaceae(100);g__Metschnikowia(100);s__Metschnikowia_pulcherrima(100); | Metschnikowia | 13 | Saprotroph | \|Undefined Saprotroph\| | Yeast | NULL | Probable | As nectar-tap_saprotroph (Põlme et al. 2020); Decay sugar-rich substrates (Põlme et al. 2020) | Põlme S, et al. 2020. Fungal Diversity 105:1-16 ((https://doi.org/10.1007/s13225-020-00466-2)) |
| Otu33704 | 1 | k__Fungi(100);p__Ascomycota(100);c__Saccharomycetes(100);o__Saccharomycetales(100);f__Metschnikowiaceae(100);g__Metschnikowia(100);g__Metschnikowia_unclassified(100); | Metschnikowia | 13 | Saprotroph | \|Undefined Saprotroph\| | Yeast | NULL | Probable | As nectar-tap_saprotroph (Põlme et al. 2020); Decay sugar-rich substrates (Põlme et al. 2020) | Põlme S, et al. 2020. Fungal Diversity 105:1-16 ((https://doi.org/10.1007/s13225-020-00466-2)) |
| Otu33706 | 1 | k__Fungi(100);p__Ascomycota(100);c__Saccharomycetes(100);o__Saccharomycetales(100);f__Metschnikowiaceae(100);g__Metschnikowia(100);g__Metschnikowia_unclassified(100); | Metschnikowia | 13 | Saprotroph | \|Undefined Saprotroph\| | Yeast | NULL | Probable | As nectar-tap_saprotroph (Põlme et al. 2020); Decay sugar-rich substrates (Põlme et al. 2020) | Põlme S, et al. 2020. Fungal Diversity 105:1-16 ((https://doi.org/10.1007/s13225-020-00466-2)) |
| Otu33709 | 1 | k__Fungi(100);p__Ascomycota(100);c__Saccharomycetes(100);o__Saccharomycetales(100);f__Metschnikowiaceae(100);g__Metschnikowia(100);s__Metschnikowia_pulcherrima(100); | Metschnikowia | 13 | Saprotroph | \|Undefined Saprotroph\| | Yeast | NULL | Probable | As nectar-tap_saprotroph (Põlme et al. 2020); Decay sugar-rich substrates (Põlme et al. 2020) | Põlme S, et al. 2020. Fungal Diversity 105:1-16 ((https://doi.org/10.1007/s13225-020-00466-2)) |
| Otu33710 | 1 | k__Fungi(100);p__Ascomycota(100);c__Saccharomycetes(100);o__Saccharomycetales(100);f__Metschnikowiaceae(100);g__Metschnikowia(100);s__Metschnikowia_pulcherrima(100); | Metschnikowia | 13 | Saprotroph | \|Undefined Saprotroph\| | Yeast | NULL | Probable | As nectar-tap_saprotroph (Põlme et al. 2020); Decay sugar-rich substrates (Põlme et al. 2020) | Põlme S, et al. 2020. Fungal Diversity 105:1-16 ((https://doi.org/10.1007/s13225-020-00466-2)) |
| Otu33713 | 1 | k__Fungi(100);p__Ascomycota(100);c__Saccharomycetes(100);o__Saccharomycetales(100);f__Metschnikowiaceae(100);g__Metschnikowia(100);s__Metschnikowia_pulcherrima(100); | Metschnikowia | 13 | Saprotroph | \|Undefined Saprotroph\| | Yeast | NULL | Probable | As nectar-tap_saprotroph (Põlme et al. 2020); Decay sugar-rich substrates (Põlme et al. 2020) | Põlme S, et al. 2020. Fungal Diversity 105:1-16 ((https://doi.org/10.1007/s13225-020-00466-2)) |
| Otu33715 | 1 | k__Fungi(100);p__Ascomycota(100);c__Saccharomycetes(100);o__Saccharomycetales(100);f__Metschnikowiaceae(100);g__Metschnikowia(100);s__Metschnikowia_pulcherrima(100); | Metschnikowia | 13 | Saprotroph | \|Undefined Saprotroph\| | Yeast | NULL | Probable | As nectar-tap_saprotroph (Põlme et al. 2020); Decay sugar-rich substrates (Põlme et al. 2020) | Põlme S, et al. 2020. Fungal Diversity 105:1-16 ((https://doi.org/10.1007/s13225-020-00466-2)) |
| Otu33716 | 1 | k__Fungi(100);p__Ascomycota(100);c__Saccharomycetes(100);o__Saccharomycetales(100);f__Metschnikowiaceae(100);g__Metschnikowia(100);s__Metschnikowia_pulcherrima(100); | Metschnikowia | 13 | Saprotroph | \|Undefined Saprotroph\| | Yeast | NULL | Probable | As nectar-tap_saprotroph (Põlme et al. 2020); Decay sugar-rich substrates (Põlme et al. 2020) | Põlme S, et al. 2020. Fungal Diversity 105:1-16 ((https://doi.org/10.1007/s13225-020-00466-2)) |
| Otu33717 | 1 | k__Fungi(100);p__Ascomycota(100);c__Saccharomycetes(100);o__Saccharomycetales(100);f__Metschnikowiaceae(100);g__Metschnikowia(100);g__Metschnikowia_unclassified(100); | Metschnikowia | 13 | Saprotroph | \|Undefined Saprotroph\| | Yeast | NULL | Probable | As nectar-tap_saprotroph (Põlme et al. 2020); Decay sugar-rich substrates (Põlme et al. 2020) | Põlme S, et al. 2020. Fungal Diversity 105:1-16 ((https://doi.org/10.1007/s13225-020-00466-2)) |
| Otu33718 | 1 | k__Fungi(100);p__Ascomycota(100);c__Saccharomycetes(100);o__Saccharomycetales(100);f__Metschnikowiaceae(100);g__Metschnikowia(100);s__Metschnikowia_pulcherrima(100); | Metschnikowia | 13 | Saprotroph | \|Undefined Saprotroph\| | Yeast | NULL | Probable | As nectar-tap_saprotroph (Põlme et al. 2020); Decay sugar-rich substrates (Põlme et al. 2020) | Põlme S, et al. 2020. Fungal Diversity 105:1-16 ((https://doi.org/10.1007/s13225-020-00466-2)) |
| Otu33719 | 1 | k__Fungi(100);p__Ascomycota(100);c__Saccharomycetes(100);o__Saccharomycetales(100);f__Metschnikowiaceae(100);g__Metschnikowia(100);s__Metschnikowia_pulcherrima(100); | Metschnikowia | 13 | Saprotroph | \|Undefined Saprotroph\| | Yeast | NULL | Probable | As nectar-tap_saprotroph (Põlme et al. 2020); Decay sugar-rich substrates (Põlme et al. 2020) | Põlme S, et al. 2020. Fungal Diversity 105:1-16 ((https://doi.org/10.1007/s13225-020-00466-2)) |
| Otu33720 | 1 | k__Fungi(100);p__Ascomycota(100);c__Saccharomycetes(100);o__Saccharomycetales(100);f__Metschnikowiaceae(100);g__Metschnikowia(100);s__Metschnikowia_pulcherrima(100); | Metschnikowia | 13 | Saprotroph | \|Undefined Saprotroph\| | Yeast | NULL | Probable | As nectar-tap_saprotroph (Põlme et al. 2020); Decay sugar-rich substrates (Põlme et al. 2020) | Põlme S, et al. 2020. Fungal Diversity 105:1-16 ((https://doi.org/10.1007/s13225-020-00466-2)) |
| Otu33721 | 1 | k__Fungi(100);p__Ascomycota(100);c__Saccharomycetes(100);o__Saccharomycetales(100);f__Metschnikowiaceae(100);g__Metschnikowia(100);g__Metschnikowia_unclassified(100); | Metschnikowia | 13 | Saprotroph | \|Undefined Saprotroph\| | Yeast | NULL | Probable | As nectar-tap_saprotroph (Põlme et al. 2020); Decay sugar-rich substrates (Põlme et al. 2020) | Põlme S, et al. 2020. Fungal Diversity 105:1-16 ((https://doi.org/10.1007/s13225-020-00466-2)) |
| Otu33722 | 1 | k__Fungi(100);p__Ascomycota(100);c__Saccharomycetes(100);o__Saccharomycetales(100);f__Metschnikowiaceae(100);g__Metschnikowia(100);g__Metschnikowia_unclassified(100); | Metschnikowia | 13 | Saprotroph | \|Undefined Saprotroph\| | Yeast | NULL | Probable | As nectar-tap_saprotroph (Põlme et al. 2020); Decay sugar-rich substrates (Põlme et al. 2020) | Põlme S, et al. 2020. Fungal Diversity 105:1-16 ((https://doi.org/10.1007/s13225-020-00466-2)) |
| Otu33723 | 1 | k__Fungi(100);p__Ascomycota(100);c__Saccharomycetes(100);o__Saccharomycetales(100);f__Metschnikowiaceae(100);g__Metschnikowia(100);s__Metschnikowia_pulcherrima(100); | Metschnikowia | 13 | Saprotroph | \|Undefined Saprotroph\| | Yeast | NULL | Probable | As nectar-tap_saprotroph (Põlme et al. 2020); Decay sugar-rich substrates (Põlme et al. 2020) | Põlme S, et al. 2020. Fungal Diversity 105:1-16 ((https://doi.org/10.1007/s13225-020-00466-2)) |
| Otu33724 | 1 | k__Fungi(100);p__Ascomycota(100);c__Saccharomycetes(100);o__Saccharomycetales(100);f__Metschnikowiaceae(100);g__Metschnikowia(100);g__Metschnikowia_unclassified(100); | Metschnikowia | 13 | Saprotroph | \|Undefined Saprotroph\| | Yeast | NULL | Probable | As nectar-tap_saprotroph (Põlme et al. 2020); Decay sugar-rich substrates (Põlme et al. 2020) | Põlme S, et al. 2020. Fungal Diversity 105:1-16 ((https://doi.org/10.1007/s13225-020-00466-2)) |
| Otu33725 | 1 | k__Fungi(100);p__Ascomycota(100);c__Saccharomycetes(100);o__Saccharomycetales(100);f__Metschnikowiaceae(100);g__Metschnikowia(100);s__Metschnikowia_pulcherrima(100); | Metschnikowia | 13 | Saprotroph | \|Undefined Saprotroph\| | Yeast | NULL | Probable | As nectar-tap_saprotroph (Põlme et al. 2020); Decay sugar-rich substrates (Põlme et al. 2020) | Põlme S, et al. 2020. Fungal Diversity 105:1-16 ((https://doi.org/10.1007/s13225-020-00466-2)) |
| Otu33726 | 1 | k__Fungi(100);p__Ascomycota(100);c__Saccharomycetes(100);o__Saccharomycetales(100);f__Metschnikowiaceae(100);g__Metschnikowia(100);s__Metschnikowia_pulcherrima(100); | Metschnikowia | 13 | Saprotroph | \|Undefined Saprotroph\| | Yeast | NULL | Probable | As nectar-tap_saprotroph (Põlme et al. 2020); Decay sugar-rich substrates (Põlme et al. 2020) | Põlme S, et al. 2020. Fungal Diversity 105:1-16 ((https://doi.org/10.1007/s13225-020-00466-2)) |
| Otu33727 | 1 | k__Fungi(100);p__Ascomycota(100);c__Saccharomycetes(100);o__Saccharomycetales(100);f__Metschnikowiaceae(100);g__Metschnikowia(100);s__Metschnikowia_pulcherrima(100); | Metschnikowia | 13 | Saprotroph | \|Undefined Saprotroph\| | Yeast | NULL | Probable | As nectar-tap_saprotroph (Põlme et al. 2020); Decay sugar-rich substrates (Põlme et al. 2020) | Põlme S, et al. 2020. Fungal Diversity 105:1-16 ((https://doi.org/10.1007/s13225-020-00466-2)) |
| Otu33728 | 1 | k__Fungi(100);p__Ascomycota(100);c__Saccharomycetes(100);o__Saccharomycetales(100);f__Metschnikowiaceae(100);g__Metschnikowia(100);s__Metschnikowia_pulcherrima(100); | Metschnikowia | 13 | Saprotroph | \|Undefined Saprotroph\| | Yeast | NULL | Probable | As nectar-tap_saprotroph (Põlme et al. 2020); Decay sugar-rich substrates (Põlme et al. 2020) | Põlme S, et al. 2020. Fungal Diversity 105:1-16 ((https://doi.org/10.1007/s13225-020-00466-2)) |
| Otu33729 | 1 | k__Fungi(100);p__Ascomycota(100);c__Saccharomycetes(100);o__Saccharomycetales(100);f__Metschnikowiaceae(100);g__Metschnikowia(100);g__Metschnikowia_unclassified(100); | Metschnikowia | 13 | Saprotroph | \|Undefined Saprotroph\| | Yeast | NULL | Probable | As nectar-tap_saprotroph (Põlme et al. 2020); Decay sugar-rich substrates (Põlme et al. 2020) | Põlme S, et al. 2020. Fungal Diversity 105:1-16 ((https://doi.org/10.1007/s13225-020-00466-2)) |
| Otu33730 | 1 | k__Fungi(100);p__Ascomycota(100);c__Saccharomycetes(100);o__Saccharomycetales(100);f__Metschnikowiaceae(100);g__Metschnikowia(100);s__Metschnikowia_pulcherrima(100); | Metschnikowia | 13 | Saprotroph | \|Undefined Saprotroph\| | Yeast | NULL | Probable | As nectar-tap_saprotroph (Põlme et al. 2020); Decay sugar-rich substrates (Põlme et al. 2020) | Põlme S, et al. 2020. Fungal Diversity 105:1-16 ((https://doi.org/10.1007/s13225-020-00466-2)) |
| Otu33731 | 1 | k__Fungi(100);p__Ascomycota(100);c__Saccharomycetes(100);o__Saccharomycetales(100);f__Metschnikowiaceae(100);g__Metschnikowia(100);g__Metschnikowia_unclassified(100); | Metschnikowia | 13 | Saprotroph | \|Undefined Saprotroph\| | Yeast | NULL | Probable | As nectar-tap_saprotroph (Põlme et al. 2020); Decay sugar-rich substrates (Põlme et al. 2020) | Põlme S, et al. 2020. Fungal Diversity 105:1-16 ((https://doi.org/10.1007/s13225-020-00466-2)) |
| Otu33733 | 1 | k__Fungi(100);p__Ascomycota(100);c__Saccharomycetes(100);o__Saccharomycetales(100);f__Metschnikowiaceae(100);g__Metschnikowia(100);s__Metschnikowia_pulcherrima(100); | Metschnikowia | 13 | Saprotroph | \|Undefined Saprotroph\| | Yeast | NULL | Probable | As nectar-tap_saprotroph (Põlme et al. 2020); Decay sugar-rich substrates (Põlme et al. 2020) | Põlme S, et al. 2020. Fungal Diversity 105:1-16 ((https://doi.org/10.1007/s13225-020-00466-2)) |
| Otu33734 | 1 | k__Fungi(100);p__Ascomycota(100);c__Saccharomycetes(100);o__Saccharomycetales(100);f__Metschnikowiaceae(100);g__Metschnikowia(100);s__Metschnikowia_pulcherrima(100); | Metschnikowia | 13 | Saprotroph | \|Undefined Saprotroph\| | Yeast | NULL | Probable | As nectar-tap_saprotroph (Põlme et al. 2020); Decay sugar-rich substrates (Põlme et al. 2020) | Põlme S, et al. 2020. Fungal Diversity 105:1-16 ((https://doi.org/10.1007/s13225-020-00466-2)) |
| Otu33735 | 1 | k__Fungi(100);p__Ascomycota(100);c__Saccharomycetes(100);o__Saccharomycetales(100);f__Metschnikowiaceae(100);g__Metschnikowia(100);s__Metschnikowia_pulcherrima(100); | Metschnikowia | 13 | Saprotroph | \|Undefined Saprotroph\| | Yeast | NULL | Probable | As nectar-tap_saprotroph (Põlme et al. 2020); Decay sugar-rich substrates (Põlme et al. 2020) | Põlme S, et al. 2020. Fungal Diversity 105:1-16 ((https://doi.org/10.1007/s13225-020-00466-2)) |
| Otu33736 | 1 | k__Fungi(100);p__Ascomycota(100);c__Saccharomycetes(100);o__Saccharomycetales(100);f__Metschnikowiaceae(100);g__Metschnikowia(100);s__Metschnikowia_pulcherrima(100); | Metschnikowia | 13 | Saprotroph | \|Undefined Saprotroph\| | Yeast | NULL | Probable | As nectar-tap_saprotroph (Põlme et al. 2020); Decay sugar-rich substrates (Põlme et al. 2020) | Põlme S, et al. 2020. Fungal Diversity 105:1-16 ((https://doi.org/10.1007/s13225-020-00466-2)) |
| Otu33737 | 1 | k__Fungi(100);p__Ascomycota(100);c__Saccharomycetes(100);o__Saccharomycetales(100);f__Metschnikowiaceae(100);g__Metschnikowia(100);s__Metschnikowia_pulcherrima(100); | Metschnikowia | 13 | Saprotroph | \|Undefined Saprotroph\| | Yeast | NULL | Probable | As nectar-tap_saprotroph (Põlme et al. 2020); Decay sugar-rich substrates (Põlme et al. 2020) | Põlme S, et al. 2020. Fungal Diversity 105:1-16 ((https://doi.org/10.1007/s13225-020-00466-2)) |
| Otu33738 | 1 | k__Fungi(100);p__Ascomycota(100);c__Saccharomycetes(100);o__Saccharomycetales(100);f__Metschnikowiaceae(100);g__Metschnikowia(100);s__Metschnikowia_pulcherrima(100); | Metschnikowia | 13 | Saprotroph | \|Undefined Saprotroph\| | Yeast | NULL | Probable | As nectar-tap_saprotroph (Põlme et al. 2020); Decay sugar-rich substrates (Põlme et al. 2020) | Põlme S, et al. 2020. Fungal Diversity 105:1-16 ((https://doi.org/10.1007/s13225-020-00466-2)) |
| Otu33739 | 1 | k__Fungi(100);p__Ascomycota(100);c__Saccharomycetes(100);o__Saccharomycetales(100);f__Metschnikowiaceae(100);g__Metschnikowia(100);s__Metschnikowia_pulcherrima(100); | Metschnikowia | 13 | Saprotroph | \|Undefined Saprotroph\| | Yeast | NULL | Probable | As nectar-tap_saprotroph (Põlme et al. 2020); Decay sugar-rich substrates (Põlme et al. 2020) | Põlme S, et al. 2020. Fungal Diversity 105:1-16 ((https://doi.org/10.1007/s13225-020-00466-2)) |
| Otu33740 | 1 | k__Fungi(100);p__Ascomycota(100);c__Saccharomycetes(100);o__Saccharomycetales(100);f__Metschnikowiaceae(100);g__Metschnikowia(100);s__Metschnikowia_pulcherrima(100); | Metschnikowia | 13 | Saprotroph | \|Undefined Saprotroph\| | Yeast | NULL | Probable | As nectar-tap_saprotroph (Põlme et al. 2020); Decay sugar-rich substrates (Põlme et al. 2020) | Põlme S, et al. 2020. Fungal Diversity 105:1-16 ((https://doi.org/10.1007/s13225-020-00466-2)) |
| Otu33741 | 1 | k__Fungi(100);p__Ascomycota(100);c__Saccharomycetes(100);o__Saccharomycetales(100);f__Metschnikowiaceae(100);g__Metschnikowia(100);s__Metschnikowia_pulcherrima(100); | Metschnikowia | 13 | Saprotroph | \|Undefined Saprotroph\| | Yeast | NULL | Probable | As nectar-tap_saprotroph (Põlme et al. 2020); Decay sugar-rich substrates (Põlme et al. 2020) | Põlme S, et al. 2020. Fungal Diversity 105:1-16 ((https://doi.org/10.1007/s13225-020-00466-2)) |
| Otu33742 | 1 | k__Fungi(100);p__Ascomycota(100);c__Saccharomycetes(100);o__Saccharomycetales(100);f__Metschnikowiaceae(100);g__Metschnikowia(100);s__Metschnikowia_pulcherrima(100); | Metschnikowia | 13 | Saprotroph | \|Undefined Saprotroph\| | Yeast | NULL | Probable | As nectar-tap_saprotroph (Põlme et al. 2020); Decay sugar-rich substrates (Põlme et al. 2020) | Põlme S, et al. 2020. Fungal Diversity 105:1-16 ((https://doi.org/10.1007/s13225-020-00466-2)) |
| Otu33743 | 1 | k__Fungi(100);p__Ascomycota(100);c__Saccharomycetes(100);o__Saccharomycetales(100);f__Metschnikowiaceae(100);g__Metschnikowia(100);s__Metschnikowia_pulcherrima(100); | Metschnikowia | 13 | Saprotroph | \|Undefined Saprotroph\| | Yeast | NULL | Probable | As nectar-tap_saprotroph (Põlme et al. 2020); Decay sugar-rich substrates (Põlme et al. 2020) | Põlme S, et al. 2020. Fungal Diversity 105:1-16 ((https://doi.org/10.1007/s13225-020-00466-2)) |
| Otu33748 | 1 | k__Fungi(100);p__Ascomycota(100);c__Saccharomycetes(100);o__Saccharomycetales(100);f__Saccharomycetales_family_Incertae_sedis(100);g__Candida(100);g__Candida_unclassified(100); | Candida | 13 | Pathotroph-Saprotroph-Symbiotroph | Animal Pathogen-Endophyte-Endosymbiont-Epiphyte-\|Undefined Saprotroph\| | Dimorphic Yeast-Yeast | NULL | Probable | As nectar-tap_saprotroph (Põlme et al. 2020); Decay sugar-rich substrates (Põlme et al. 2020); This asexual genus is polyphyletic and contain species that function very differently depend on what they are. Caution should be taken when interpreting guilds for Candida species. | Manolakaki D, et al. 2010. Virulence. 1: 367-375 ((https://doi.org/10.4161/viru.1.5.12796)); Mekbib SB, et al. 2011. Tropical Plant Pathology 36:233-240 ((https://doi.org/10.1590/S1982-56762011000400004)); Põlme S, et al. 2020. Fungal Diversity 105:1-16 ((https://doi.org/10.1007/s13225-020-00466-2)) |
| Otu33750 | 1 | k__Fungi(100);p__Ascomycota(100);c__Saccharomycetes(100);o__Saccharomycetales(100);f__Metschnikowiaceae(100);g__Metschnikowia(100);s__Metschnikowia_pulcherrima(100); | Metschnikowia | 13 | Saprotroph | \|Undefined Saprotroph\| | Yeast | NULL | Probable | As nectar-tap_saprotroph (Põlme et al. 2020); Decay sugar-rich substrates (Põlme et al. 2020) | Põlme S, et al. 2020. Fungal Diversity 105:1-16 ((https://doi.org/10.1007/s13225-020-00466-2)) |
| Otu33751 | 1 | k__Fungi(100);p__Ascomycota(100);c__Saccharomycetes(100);o__Saccharomycetales(100);f__Metschnikowiaceae(100);g__Metschnikowia(100);s__Metschnikowia_pulcherrima(100); | Metschnikowia | 13 | Saprotroph | \|Undefined Saprotroph\| | Yeast | NULL | Probable | As nectar-tap_saprotroph (Põlme et al. 2020); Decay sugar-rich substrates (Põlme et al. 2020) | Põlme S, et al. 2020. Fungal Diversity 105:1-16 ((https://doi.org/10.1007/s13225-020-00466-2)) |
| Otu33755 | 1 | k__Fungi(100);p__Ascomycota(100);c__Saccharomycetes(100);o__Saccharomycetales(100);f__Metschnikowiaceae(100);g__Metschnikowia(100);g__Metschnikowia_unclassified(100); | Metschnikowia | 13 | Saprotroph | \|Undefined Saprotroph\| | Yeast | NULL | Probable | As nectar-tap_saprotroph (Põlme et al. 2020); Decay sugar-rich substrates (Põlme et al. 2020) | Põlme S, et al. 2020. Fungal Diversity 105:1-16 ((https://doi.org/10.1007/s13225-020-00466-2)) |
| Otu33756 | 1 | k__Fungi(100);p__Ascomycota(100);c__Saccharomycetes(100);o__Saccharomycetales(100);f__Metschnikowiaceae(100);g__Metschnikowia(100);s__Metschnikowia_pulcherrima(100); | Metschnikowia | 13 | Saprotroph | \|Undefined Saprotroph\| | Yeast | NULL | Probable | As nectar-tap_saprotroph (Põlme et al. 2020); Decay sugar-rich substrates (Põlme et al. 2020) | Põlme S, et al. 2020. Fungal Diversity 105:1-16 ((https://doi.org/10.1007/s13225-020-00466-2)) |
| Otu33758 | 1 | k__Fungi(100);p__Ascomycota(100);c__Saccharomycetes(100);o__Saccharomycetales(100);f__Saccharomycetales_family_Incertae_sedis(100);g__Candida(100);g__Candida_unclassified(100); | Candida | 13 | Pathotroph-Saprotroph-Symbiotroph | Animal Pathogen-Endophyte-Endosymbiont-Epiphyte-\|Undefined Saprotroph\| | Dimorphic Yeast-Yeast | NULL | Probable | As nectar-tap_saprotroph (Põlme et al. 2020); Decay sugar-rich substrates (Põlme et al. 2020); This asexual genus is polyphyletic and contain species that function very differently depend on what they are. Caution should be taken when interpreting guilds for Candida species. | Manolakaki D, et al. 2010. Virulence. 1: 367-375 ((https://doi.org/10.4161/viru.1.5.12796)); Mekbib SB, et al. 2011. Tropical Plant Pathology 36:233-240 ((https://doi.org/10.1590/S1982-56762011000400004)); Põlme S, et al. 2020. Fungal Diversity 105:1-16 ((https://doi.org/10.1007/s13225-020-00466-2)) |
| Otu33760 | 1 | k__Fungi(100);p__Ascomycota(100);c__Saccharomycetes(100);o__Saccharomycetales(100);f__Metschnikowiaceae(100);g__Metschnikowia(100);s__Metschnikowia_pulcherrima(100); | Metschnikowia | 13 | Saprotroph | \|Undefined Saprotroph\| | Yeast | NULL | Probable | As nectar-tap_saprotroph (Põlme et al. 2020); Decay sugar-rich substrates (Põlme et al. 2020) | Põlme S, et al. 2020. Fungal Diversity 105:1-16 ((https://doi.org/10.1007/s13225-020-00466-2)) |
| Otu33761 | 1 | k__Fungi(100);p__Ascomycota(100);c__Saccharomycetes(100);o__Saccharomycetales(100);f__Metschnikowiaceae(100);g__Metschnikowia(100);g__Metschnikowia_unclassified(100); | Metschnikowia | 13 | Saprotroph | \|Undefined Saprotroph\| | Yeast | NULL | Probable | As nectar-tap_saprotroph (Põlme et al. 2020); Decay sugar-rich substrates (Põlme et al. 2020) | Põlme S, et al. 2020. Fungal Diversity 105:1-16 ((https://doi.org/10.1007/s13225-020-00466-2)) |
| Otu33762 | 1 | k__Fungi(100);p__Basidiomycota(100);c__Microbotryomycetes(100);o__Sporidiobolales(100);f__Sporidiobolales_family_Incertae_sedis(100);g__Sporobolomyces(100);s__Sporobolomyces_lactosus(100); | Sporobolomyces | 13 | Pathotroph-Saprotroph | \|Fungal Parasite\|-Plant Saprotroph | Yeast | NULL | Probable | As litter_saprotroph (Põlme et al. 2020); As mycoparasite (Põlme et al. 2020); Decay fungal material (Põlme et al. 2020) | Tedersoo L, et al. 2014. Science 346:e1256688 ((https://doi.org/10.1126/science.1256688)); Purahong W, et al. 2016. Molecular Ecology 25:4059-4074 ((https://doi.org/10.1111/mec.13739)); Põlme S, et al. 2020. Fungal Diversity 105:1-16 ((https://doi.org/10.1007/s13225-020-00466-2)) |
| Otu33763 | 1 | k__Fungi(100);p__Ascomycota(100);c__Saccharomycetes(100);o__Saccharomycetales(100);f__Metschnikowiaceae(100);g__Metschnikowia(100);s__Metschnikowia_pulcherrima(100); | Metschnikowia | 13 | Saprotroph | \|Undefined Saprotroph\| | Yeast | NULL | Probable | As nectar-tap_saprotroph (Põlme et al. 2020); Decay sugar-rich substrates (Põlme et al. 2020) | Põlme S, et al. 2020. Fungal Diversity 105:1-16 ((https://doi.org/10.1007/s13225-020-00466-2)) |
| Otu33764 | 1 | k__Fungi(100);p__Ascomycota(100);c__Saccharomycetes(100);o__Saccharomycetales(100);f__Metschnikowiaceae(100);g__Metschnikowia(100);s__Metschnikowia_pulcherrima(100); | Metschnikowia | 13 | Saprotroph | \|Undefined Saprotroph\| | Yeast | NULL | Probable | As nectar-tap_saprotroph (Põlme et al. 2020); Decay sugar-rich substrates (Põlme et al. 2020) | Põlme S, et al. 2020. Fungal Diversity 105:1-16 ((https://doi.org/10.1007/s13225-020-00466-2)) |
| Otu33767 | 1 | k__Fungi(100);p__Ascomycota(100);c__Saccharomycetes(100);o__Saccharomycetales(100);f__Metschnikowiaceae(100);g__Metschnikowia(100);s__Metschnikowia_pulcherrima(100); | Metschnikowia | 13 | Saprotroph | \|Undefined Saprotroph\| | Yeast | NULL | Probable | As nectar-tap_saprotroph (Põlme et al. 2020); Decay sugar-rich substrates (Põlme et al. 2020) | Põlme S, et al. 2020. Fungal Diversity 105:1-16 ((https://doi.org/10.1007/s13225-020-00466-2)) |
| Otu33768 | 1 | k__Fungi(100);p__Ascomycota(100);c__Saccharomycetes(100);o__Saccharomycetales(100);f__Metschnikowiaceae(100);g__Metschnikowia(100);s__Metschnikowia_pulcherrima(100); | Metschnikowia | 13 | Saprotroph | \|Undefined Saprotroph\| | Yeast | NULL | Probable | As nectar-tap_saprotroph (Põlme et al. 2020); Decay sugar-rich substrates (Põlme et al. 2020) | Põlme S, et al. 2020. Fungal Diversity 105:1-16 ((https://doi.org/10.1007/s13225-020-00466-2)) |
| Otu33769 | 1 | k__Fungi(100);p__Ascomycota(100);c__Saccharomycetes(100);o__Saccharomycetales(100);f__Metschnikowiaceae(100);g__Metschnikowia(100);s__Metschnikowia_pulcherrima(100); | Metschnikowia | 13 | Saprotroph | \|Undefined Saprotroph\| | Yeast | NULL | Probable | As nectar-tap_saprotroph (Põlme et al. 2020); Decay sugar-rich substrates (Põlme et al. 2020) | Põlme S, et al. 2020. Fungal Diversity 105:1-16 ((https://doi.org/10.1007/s13225-020-00466-2)) |
| Otu33770 | 1 | k__Fungi(100);p__Ascomycota(100);c__Saccharomycetes(100);o__Saccharomycetales(100);f__Metschnikowiaceae(100);g__Metschnikowia(100);s__Metschnikowia_pulcherrima(100); | Metschnikowia | 13 | Saprotroph | \|Undefined Saprotroph\| | Yeast | NULL | Probable | As nectar-tap_saprotroph (Põlme et al. 2020); Decay sugar-rich substrates (Põlme et al. 2020) | Põlme S, et al. 2020. Fungal Diversity 105:1-16 ((https://doi.org/10.1007/s13225-020-00466-2)) |
| Otu33771 | 1 | k__Fungi(100);p__Ascomycota(100);c__Saccharomycetes(100);o__Saccharomycetales(100);f__Metschnikowiaceae(100);g__Metschnikowia(100);s__Metschnikowia_pulcherrima(100); | Metschnikowia | 13 | Saprotroph | \|Undefined Saprotroph\| | Yeast | NULL | Probable | As nectar-tap_saprotroph (Põlme et al. 2020); Decay sugar-rich substrates (Põlme et al. 2020) | Põlme S, et al. 2020. Fungal Diversity 105:1-16 ((https://doi.org/10.1007/s13225-020-00466-2)) |
| Otu33772 | 1 | k__Fungi(100);p__Ascomycota(100);c__Saccharomycetes(100);o__Saccharomycetales(100);f__Metschnikowiaceae(100);g__Metschnikowia(100);g__Metschnikowia_unclassified(100); | Metschnikowia | 13 | Saprotroph | \|Undefined Saprotroph\| | Yeast | NULL | Probable | As nectar-tap_saprotroph (Põlme et al. 2020); Decay sugar-rich substrates (Põlme et al. 2020) | Põlme S, et al. 2020. Fungal Diversity 105:1-16 ((https://doi.org/10.1007/s13225-020-00466-2)) |
| Otu33775 | 1 | k__Fungi(100);p__Ascomycota(100);c__Saccharomycetes(100);o__Saccharomycetales(100);f__Saccharomycetales_family_Incertae_sedis(100);g__Candida(100);g__Candida_unclassified(100); | Candida | 13 | Pathotroph-Saprotroph-Symbiotroph | Animal Pathogen-Endophyte-Endosymbiont-Epiphyte-\|Undefined Saprotroph\| | Dimorphic Yeast-Yeast | NULL | Probable | As nectar-tap_saprotroph (Põlme et al. 2020); Decay sugar-rich substrates (Põlme et al. 2020); This asexual genus is polyphyletic and contain species that function very differently depend on what they are. Caution should be taken when interpreting guilds for Candida species. | Manolakaki D, et al. 2010. Virulence. 1: 367-375 ((https://doi.org/10.4161/viru.1.5.12796)); Mekbib SB, et al. 2011. Tropical Plant Pathology 36:233-240 ((https://doi.org/10.1590/S1982-56762011000400004)); Põlme S, et al. 2020. Fungal Diversity 105:1-16 ((https://doi.org/10.1007/s13225-020-00466-2)) |
| Otu33777 | 1 | k__Fungi(100);p__Ascomycota(100);c__Saccharomycetes(100);o__Saccharomycetales(100);f__Metschnikowiaceae(100);g__Metschnikowia(100);s__Metschnikowia_pulcherrima(100); | Metschnikowia | 13 | Saprotroph | \|Undefined Saprotroph\| | Yeast | NULL | Probable | As nectar-tap_saprotroph (Põlme et al. 2020); Decay sugar-rich substrates (Põlme et al. 2020) | Põlme S, et al. 2020. Fungal Diversity 105:1-16 ((https://doi.org/10.1007/s13225-020-00466-2)) |
| Otu33778 | 1 | k__Fungi(100);p__Ascomycota(100);c__Saccharomycetes(100);o__Saccharomycetales(100);f__Metschnikowiaceae(100);g__Metschnikowia(100);s__Metschnikowia_pulcherrima(100); | Metschnikowia | 13 | Saprotroph | \|Undefined Saprotroph\| | Yeast | NULL | Probable | As nectar-tap_saprotroph (Põlme et al. 2020); Decay sugar-rich substrates (Põlme et al. 2020) | Põlme S, et al. 2020. Fungal Diversity 105:1-16 ((https://doi.org/10.1007/s13225-020-00466-2)) |
| Otu33779 | 1 | k__Fungi(100);p__Ascomycota(100);c__Saccharomycetes(100);o__Saccharomycetales(100);f__Metschnikowiaceae(100);g__Metschnikowia(100);g__Metschnikowia_unclassified(100); | Metschnikowia | 13 | Saprotroph | \|Undefined Saprotroph\| | Yeast | NULL | Probable | As nectar-tap_saprotroph (Põlme et al. 2020); Decay sugar-rich substrates (Põlme et al. 2020) | Põlme S, et al. 2020. Fungal Diversity 105:1-16 ((https://doi.org/10.1007/s13225-020-00466-2)) |
| Otu33780 | 1 | k__Fungi(100);p__Ascomycota(100);c__Saccharomycetes(100);o__Saccharomycetales(100);f__Metschnikowiaceae(100);g__Metschnikowia(100);s__Metschnikowia_pulcherrima(100); | Metschnikowia | 13 | Saprotroph | \|Undefined Saprotroph\| | Yeast | NULL | Probable | As nectar-tap_saprotroph (Põlme et al. 2020); Decay sugar-rich substrates (Põlme et al. 2020) | Põlme S, et al. 2020. Fungal Diversity 105:1-16 ((https://doi.org/10.1007/s13225-020-00466-2)) |
| Otu33781 | 1 | k__Fungi(100);p__Ascomycota(100);c__Saccharomycetes(100);o__Saccharomycetales(100);f__Metschnikowiaceae(100);g__Metschnikowia(100);s__Metschnikowia_pulcherrima(100); | Metschnikowia | 13 | Saprotroph | \|Undefined Saprotroph\| | Yeast | NULL | Probable | As nectar-tap_saprotroph (Põlme et al. 2020); Decay sugar-rich substrates (Põlme et al. 2020) | Põlme S, et al. 2020. Fungal Diversity 105:1-16 ((https://doi.org/10.1007/s13225-020-00466-2)) |
| Otu33783 | 1 | k__Fungi(100);p__Ascomycota(100);c__Saccharomycetes(100);o__Saccharomycetales(100);f__Metschnikowiaceae(100);g__Metschnikowia(100);s__Metschnikowia_pulcherrima(100); | Metschnikowia | 13 | Saprotroph | \|Undefined Saprotroph\| | Yeast | NULL | Probable | As nectar-tap_saprotroph (Põlme et al. 2020); Decay sugar-rich substrates (Põlme et al. 2020) | Põlme S, et al. 2020. Fungal Diversity 105:1-16 ((https://doi.org/10.1007/s13225-020-00466-2)) |
| Otu33784 | 1 | k__Fungi(100);p__Ascomycota(100);c__Saccharomycetes(100);o__Saccharomycetales(100);f__Metschnikowiaceae(100);g__Metschnikowia(100);g__Metschnikowia_unclassified(100); | Metschnikowia | 13 | Saprotroph | \|Undefined Saprotroph\| | Yeast | NULL | Probable | As nectar-tap_saprotroph (Põlme et al. 2020); Decay sugar-rich substrates (Põlme et al. 2020) | Põlme S, et al. 2020. Fungal Diversity 105:1-16 ((https://doi.org/10.1007/s13225-020-00466-2)) |
| Otu33785 | 1 | k__Fungi(100);p__Ascomycota(100);c__Saccharomycetes(100);o__Saccharomycetales(100);f__Metschnikowiaceae(100);g__Metschnikowia(100);s__Metschnikowia_pulcherrima(100); | Metschnikowia | 13 | Saprotroph | \|Undefined Saprotroph\| | Yeast | NULL | Probable | As nectar-tap_saprotroph (Põlme et al. 2020); Decay sugar-rich substrates (Põlme et al. 2020) | Põlme S, et al. 2020. Fungal Diversity 105:1-16 ((https://doi.org/10.1007/s13225-020-00466-2)) |
| Otu33786 | 1 | k__Fungi(100);p__Ascomycota(100);c__Saccharomycetes(100);o__Saccharomycetales(100);f__Metschnikowiaceae(100);g__Metschnikowia(100);s__Metschnikowia_pulcherrima(100); | Metschnikowia | 13 | Saprotroph | \|Undefined Saprotroph\| | Yeast | NULL | Probable | As nectar-tap_saprotroph (Põlme et al. 2020); Decay sugar-rich substrates (Põlme et al. 2020) | Põlme S, et al. 2020. Fungal Diversity 105:1-16 ((https://doi.org/10.1007/s13225-020-00466-2)) |
| Otu33788 | 1 | k__Fungi(100);p__Ascomycota(100);c__Saccharomycetes(100);o__Saccharomycetales(100);f__Metschnikowiaceae(100);g__Metschnikowia(100);s__Metschnikowia_pulcherrima(100); | Metschnikowia | 13 | Saprotroph | \|Undefined Saprotroph\| | Yeast | NULL | Probable | As nectar-tap_saprotroph (Põlme et al. 2020); Decay sugar-rich substrates (Põlme et al. 2020) | Põlme S, et al. 2020. Fungal Diversity 105:1-16 ((https://doi.org/10.1007/s13225-020-00466-2)) |
| Otu33793 | 1 | k__Fungi(100);p__Ascomycota(100);c__Saccharomycetes(100);o__Saccharomycetales(100);f__Metschnikowiaceae(100);g__Metschnikowia(100);s__Metschnikowia_pulcherrima(100); | Metschnikowia | 13 | Saprotroph | \|Undefined Saprotroph\| | Yeast | NULL | Probable | As nectar-tap_saprotroph (Põlme et al. 2020); Decay sugar-rich substrates (Põlme et al. 2020) | Põlme S, et al. 2020. Fungal Diversity 105:1-16 ((https://doi.org/10.1007/s13225-020-00466-2)) |
| Otu33794 | 1 | k__Fungi(100);p__Ascomycota(100);c__Saccharomycetes(100);o__Saccharomycetales(100);f__Metschnikowiaceae(100);g__Metschnikowia(100);s__Metschnikowia_pulcherrima(100); | Metschnikowia | 13 | Saprotroph | \|Undefined Saprotroph\| | Yeast | NULL | Probable | As nectar-tap_saprotroph (Põlme et al. 2020); Decay sugar-rich substrates (Põlme et al. 2020) | Põlme S, et al. 2020. Fungal Diversity 105:1-16 ((https://doi.org/10.1007/s13225-020-00466-2)) |
| Otu33795 | 1 | k__Fungi(100);p__Ascomycota(100);c__Saccharomycetes(100);o__Saccharomycetales(100);f__Metschnikowiaceae(100);g__Metschnikowia(100);s__Metschnikowia_pulcherrima(100); | Metschnikowia | 13 | Saprotroph | \|Undefined Saprotroph\| | Yeast | NULL | Probable | As nectar-tap_saprotroph (Põlme et al. 2020); Decay sugar-rich substrates (Põlme et al. 2020) | Põlme S, et al. 2020. Fungal Diversity 105:1-16 ((https://doi.org/10.1007/s13225-020-00466-2)) |
| Otu33800 | 1 | k__Fungi(100);p__Basidiomycota(100);c__Microbotryomycetes(100);o__Sporidiobolales(100);f__Sporidiobolales_family_Incertae_sedis(100);g__Sporobolomyces(100);s__Sporobolomyces_lactosus(100); | Sporobolomyces | 13 | Pathotroph-Saprotroph | \|Fungal Parasite\|-Plant Saprotroph | Yeast | NULL | Probable | As litter_saprotroph (Põlme et al. 2020); As mycoparasite (Põlme et al. 2020); Decay fungal material (Põlme et al. 2020) | Tedersoo L, et al. 2014. Science 346:e1256688 ((https://doi.org/10.1126/science.1256688)); Purahong W, et al. 2016. Molecular Ecology 25:4059-4074 ((https://doi.org/10.1111/mec.13739)); Põlme S, et al. 2020. Fungal Diversity 105:1-16 ((https://doi.org/10.1007/s13225-020-00466-2)) |
| Otu33801 | 1 | k__Fungi(100);p__Ascomycota(100);c__Saccharomycetes(100);o__Saccharomycetales(100);f__Metschnikowiaceae(100);g__Metschnikowia(100);s__Metschnikowia_pulcherrima(100); | Metschnikowia | 13 | Saprotroph | \|Undefined Saprotroph\| | Yeast | NULL | Probable | As nectar-tap_saprotroph (Põlme et al. 2020); Decay sugar-rich substrates (Põlme et al. 2020) | Põlme S, et al. 2020. Fungal Diversity 105:1-16 ((https://doi.org/10.1007/s13225-020-00466-2)) |
| Otu33803 | 1 | k__Fungi(100);p__Ascomycota(100);c__Saccharomycetes(100);o__Saccharomycetales(100);f__Metschnikowiaceae(100);g__Metschnikowia(100);g__Metschnikowia_unclassified(100); | Metschnikowia | 13 | Saprotroph | \|Undefined Saprotroph\| | Yeast | NULL | Probable | As nectar-tap_saprotroph (Põlme et al. 2020); Decay sugar-rich substrates (Põlme et al. 2020) | Põlme S, et al. 2020. Fungal Diversity 105:1-16 ((https://doi.org/10.1007/s13225-020-00466-2)) |
| Otu33804 | 1 | k__Fungi(100);p__Ascomycota(100);c__Saccharomycetes(100);o__Saccharomycetales(100);f__Metschnikowiaceae(100);g__Metschnikowia(100);s__Metschnikowia_pulcherrima(100); | Metschnikowia | 13 | Saprotroph | \|Undefined Saprotroph\| | Yeast | NULL | Probable | As nectar-tap_saprotroph (Põlme et al. 2020); Decay sugar-rich substrates (Põlme et al. 2020) | Põlme S, et al. 2020. Fungal Diversity 105:1-16 ((https://doi.org/10.1007/s13225-020-00466-2)) |
| Otu33806 | 1 | k__Fungi(100);p__Ascomycota(100);c__Saccharomycetes(100);o__Saccharomycetales(100);f__Metschnikowiaceae(100);g__Metschnikowia(100);s__Metschnikowia_pulcherrima(100); | Metschnikowia | 13 | Saprotroph | \|Undefined Saprotroph\| | Yeast | NULL | Probable | As nectar-tap_saprotroph (Põlme et al. 2020); Decay sugar-rich substrates (Põlme et al. 2020) | Põlme S, et al. 2020. Fungal Diversity 105:1-16 ((https://doi.org/10.1007/s13225-020-00466-2)) |
| Otu33819 | 1 | k__Fungi(100);p__Ascomycota(100);c__Saccharomycetes(100);o__Saccharomycetales(100);f__Metschnikowiaceae(100);g__Metschnikowia(100);s__Metschnikowia_pulcherrima(100); | Metschnikowia | 13 | Saprotroph | \|Undefined Saprotroph\| | Yeast | NULL | Probable | As nectar-tap_saprotroph (Põlme et al. 2020); Decay sugar-rich substrates (Põlme et al. 2020) | Põlme S, et al. 2020. Fungal Diversity 105:1-16 ((https://doi.org/10.1007/s13225-020-00466-2)) |
| Otu33823 | 1 | k__Fungi(100);p__Ascomycota(100);c__Saccharomycetes(100);o__Saccharomycetales(100);f__Metschnikowiaceae(100);g__Metschnikowia(100);s__Metschnikowia_pulcherrima(100); | Metschnikowia | 13 | Saprotroph | \|Undefined Saprotroph\| | Yeast | NULL | Probable | As nectar-tap_saprotroph (Põlme et al. 2020); Decay sugar-rich substrates (Põlme et al. 2020) | Põlme S, et al. 2020. Fungal Diversity 105:1-16 ((https://doi.org/10.1007/s13225-020-00466-2)) |
| Otu33827 | 1 | k__Fungi(100);p__Ascomycota(100);c__Saccharomycetes(100);o__Saccharomycetales(100);f__Metschnikowiaceae(100);g__Metschnikowia(100);g__Metschnikowia_unclassified(100); | Metschnikowia | 13 | Saprotroph | \|Undefined Saprotroph\| | Yeast | NULL | Probable | As nectar-tap_saprotroph (Põlme et al. 2020); Decay sugar-rich substrates (Põlme et al. 2020) | Põlme S, et al. 2020. Fungal Diversity 105:1-16 ((https://doi.org/10.1007/s13225-020-00466-2)) |
| Otu33828 | 1 | k__Fungi(100);p__Ascomycota(100);c__Saccharomycetes(100);o__Saccharomycetales(100);f__Metschnikowiaceae(100);g__Metschnikowia(100);s__Metschnikowia_pulcherrima(100); | Metschnikowia | 13 | Saprotroph | \|Undefined Saprotroph\| | Yeast | NULL | Probable | As nectar-tap_saprotroph (Põlme et al. 2020); Decay sugar-rich substrates (Põlme et al. 2020) | Põlme S, et al. 2020. Fungal Diversity 105:1-16 ((https://doi.org/10.1007/s13225-020-00466-2)) |
| Otu33829 | 1 | k__Fungi(100);p__Ascomycota(100);c__Saccharomycetes(100);o__Saccharomycetales(100);f__Metschnikowiaceae(100);g__Metschnikowia(100);s__Metschnikowia_pulcherrima(100); | Metschnikowia | 13 | Saprotroph | \|Undefined Saprotroph\| | Yeast | NULL | Probable | As nectar-tap_saprotroph (Põlme et al. 2020); Decay sugar-rich substrates (Põlme et al. 2020) | Põlme S, et al. 2020. Fungal Diversity 105:1-16 ((https://doi.org/10.1007/s13225-020-00466-2)) |
| Otu33830 | 1 | k__Fungi(100);p__Ascomycota(100);c__Saccharomycetes(100);o__Saccharomycetales(100);f__Metschnikowiaceae(100);g__Metschnikowia(100);g__Metschnikowia_unclassified(100); | Metschnikowia | 13 | Saprotroph | \|Undefined Saprotroph\| | Yeast | NULL | Probable | As nectar-tap_saprotroph (Põlme et al. 2020); Decay sugar-rich substrates (Põlme et al. 2020) | Põlme S, et al. 2020. Fungal Diversity 105:1-16 ((https://doi.org/10.1007/s13225-020-00466-2)) |
| Otu33834 | 1 | k__Fungi(100);p__Ascomycota(100);c__Saccharomycetes(100);o__Saccharomycetales(100);f__Metschnikowiaceae(100);g__Metschnikowia(100);s__Metschnikowia_pulcherrima(100); | Metschnikowia | 13 | Saprotroph | \|Undefined Saprotroph\| | Yeast | NULL | Probable | As nectar-tap_saprotroph (Põlme et al. 2020); Decay sugar-rich substrates (Põlme et al. 2020) | Põlme S, et al. 2020. Fungal Diversity 105:1-16 ((https://doi.org/10.1007/s13225-020-00466-2)) |
| Otu33838 | 1 | k__Fungi(100);p__Ascomycota(100);c__Saccharomycetes(100);o__Saccharomycetales(100);f__Metschnikowiaceae(100);g__Metschnikowia(100);s__Metschnikowia_pulcherrima(100); | Metschnikowia | 13 | Saprotroph | \|Undefined Saprotroph\| | Yeast | NULL | Probable | As nectar-tap_saprotroph (Põlme et al. 2020); Decay sugar-rich substrates (Põlme et al. 2020) | Põlme S, et al. 2020. Fungal Diversity 105:1-16 ((https://doi.org/10.1007/s13225-020-00466-2)) |
| Otu33842 | 1 | k__Fungi(100);p__Ascomycota(100);c__Saccharomycetes(100);o__Saccharomycetales(100);f__Metschnikowiaceae(100);g__Metschnikowia(100);g__Metschnikowia_unclassified(100); | Metschnikowia | 13 | Saprotroph | \|Undefined Saprotroph\| | Yeast | NULL | Probable | As nectar-tap_saprotroph (Põlme et al. 2020); Decay sugar-rich substrates (Põlme et al. 2020) | Põlme S, et al. 2020. Fungal Diversity 105:1-16 ((https://doi.org/10.1007/s13225-020-00466-2)) |
| Otu33844 | 1 | k__Fungi(100);p__Ascomycota(100);c__Saccharomycetes(100);o__Saccharomycetales(100);f__Metschnikowiaceae(100);g__Metschnikowia(100);s__Metschnikowia_pulcherrima(100); | Metschnikowia | 13 | Saprotroph | \|Undefined Saprotroph\| | Yeast | NULL | Probable | As nectar-tap_saprotroph (Põlme et al. 2020); Decay sugar-rich substrates (Põlme et al. 2020) | Põlme S, et al. 2020. Fungal Diversity 105:1-16 ((https://doi.org/10.1007/s13225-020-00466-2)) |
| Otu33845 | 1 | k__Fungi(100);p__Ascomycota(100);c__Saccharomycetes(100);o__Saccharomycetales(100);f__Metschnikowiaceae(100);g__Metschnikowia(100);s__Metschnikowia_pulcherrima(100); | Metschnikowia | 13 | Saprotroph | \|Undefined Saprotroph\| | Yeast | NULL | Probable | As nectar-tap_saprotroph (Põlme et al. 2020); Decay sugar-rich substrates (Põlme et al. 2020) | Põlme S, et al. 2020. Fungal Diversity 105:1-16 ((https://doi.org/10.1007/s13225-020-00466-2)) |

| | | | | | | | | | | | |
|---|---|---|---|---|---|---|---|---|---|---|---|
| Otu33847 | 1 | k__Fungi(100);p__Ascomycota(100);c__Saccharomycetes(100);o__Saccharomycetales(100);f__Metschnikowiaceae(100);g__Metschnikowia(100);g__Metschnikowia_unclassified(100); | Metschnikowia | 13 | Saprotroph | \|Undefined Saprotroph\| | Yeast | NULL | Probable | As nectar-tap_saprotroph (Põlme et al. 2020); Decay sugar-rich substrates (Põlme et al. 2020) | Põlme S, et al. 2020. Fungal Diversity 105:1-16 ((https://doi.org/10.1007/s13225-020-00466-2)) |
| Otu33848 | 1 | k__Fungi(100);p__Ascomycota(100);c__Saccharomycetes(100);o__Saccharomycetales(100);f__Metschnikowiaceae(100);g__Metschnikowia(100);s__Metschnikowia_pulcherrima(100); | Metschnikowia | 13 | Saprotroph | \|Undefined Saprotroph\| | Yeast | NULL | Probable | As nectar-tap_saprotroph (Põlme et al. 2020); Decay sugar-rich substrates (Põlme et al. 2020) | Põlme S, et al. 2020. Fungal Diversity 105:1-16 ((https://doi.org/10.1007/s13225-020-00466-2)) |
| Otu33850 | 1 | k__Fungi(100);p__Ascomycota(100);c__Saccharomycetes(100);o__Saccharomycetales(100);f__Metschnikowiaceae(100);g__Metschnikowia(100);s__Metschnikowia_pulcherrima(100); | Metschnikowia | 13 | Saprotroph | \|Undefined Saprotroph\| | Yeast | NULL | Probable | As nectar-tap_saprotroph (Põlme et al. 2020); Decay sugar-rich substrates (Põlme et al. 2020) | Põlme S, et al. 2020. Fungal Diversity 105:1-16 ((https://doi.org/10.1007/s13225-020-00466-2)) |
| Otu33851 | 1 | k__Fungi(100);p__Ascomycota(100);c__Saccharomycetes(100);o__Saccharomycetales(100);f__Metschnikowiaceae(100);g__Metschnikowia(100);s__Metschnikowia_pulcherrima(100); | Metschnikowia | 13 | Saprotroph | \|Undefined Saprotroph\| | Yeast | NULL | Probable | As nectar-tap_saprotroph (Põlme et al. 2020); Decay sugar-rich substrates (Põlme et al. 2020) | Põlme S, et al. 2020. Fungal Diversity 105:1-16 ((https://doi.org/10.1007/s13225-020-00466-2)) |
| Otu33853 | 1 | k__Fungi(100);p__Ascomycota(100);c__Saccharomycetes(100);o__Saccharomycetales(100);f__Saccharomycetales_family_Incertae_sedis(100);g__Candida(100);s__Candida_cretensis(100); | Candida | 13 | Pathotroph-Saprotroph-Symbiotroph | Animal Pathogen-Endophyte-Endosymbiont-Epiphyte-\|Undefined Saprotroph\| | Dimorphic Yeast-Yeast | NULL | Probable | As nectar-tap_saprotroph (Põlme et al. 2020); Decay sugar-rich substrates (Põlme et al. 2020); This asexual genus is polyphyletic and contain species that function very differently depend on what they are. Caution should be taken when interpreting guilds for Candida species. | Manolakaki D, et al. 2010. Virulence. 1: 367-375 ((https://doi.org/10.4161/viru.1.5.12796)); Mekbib SB, et al. 2011. Tropical Plant Pathology 36:233-240 ((https://doi.org/10.1590/S1982-56762011000400004)); Põlme S, et al. 2020. Fungal Diversity 105:1-16 ((https://doi.org/10.1007/s13225-020-00466-2)) |
| Otu33854 | 1 | k__Fungi(100);p__Ascomycota(100);c__Saccharomycetes(100);o__Saccharomycetales(100);f__Metschnikowiaceae(100);g__Metschnikowia(100);s__Metschnikowia_pulcherrima(100); | Metschnikowia | 13 | Saprotroph | \|Undefined Saprotroph\| | Yeast | NULL | Probable | As nectar-tap_saprotroph (Põlme et al. 2020); Decay sugar-rich substrates (Põlme et al. 2020) | Põlme S, et al. 2020. Fungal Diversity 105:1-16 ((https://doi.org/10.1007/s13225-020-00466-2)) |
| Otu33856 | 1 | k__Fungi(100);p__Ascomycota(100);c__Saccharomycetes(100);o__Saccharomycetales(100);f__Metschnikowiaceae(100);g__Metschnikowia(100);g__Metschnikowia_unclassified(100); | Metschnikowia | 13 | Saprotroph | \|Undefined Saprotroph\| | Yeast | NULL | Probable | As nectar-tap_saprotroph (Põlme et al. 2020); Decay sugar-rich substrates (Põlme et al. 2020) | Põlme S, et al. 2020. Fungal Diversity 105:1-16 ((https://doi.org/10.1007/s13225-020-00466-2)) |
| Otu33861 | 1 | k__Fungi(100);p__Ascomycota(100);c__Dothideomycetes(100);o__Dothideomycetes_order_Incertae_sedis(100);f__Myxotrichaceae(100);g__Oidiodendron(100);s__Oidiodendron_maius(100); | Oidiodendron | 13 | Pathotroph-Saprotroph-Symbiotroph | Endophyte-Ericoid Mycorrhizal-\|Undefined Saprotroph\| | Microfungus | NULL | Probable | As root_endophyte (Põlme et al. 2020); As soil_saprotroph (Põlme et al. 2020); Dark Septate Endophyte; Decay soil (Põlme et al. 2020) | Newsham KK. 2011. New Phytologist 190:783-793 ((https://doi.org/10.1111/j.1469-8137.2010.03611.x)) (as dark septate root endophytes); Põlme S, et al. 2020. Fungal Diversity 105:1-16 ((https://doi.org/10.1007/s13225-020-00466-2)) |
| Otu33865 | 1 | k__Fungi(100);p__Ascomycota(100);c__Saccharomycetes(100);o__Saccharomycetales(100);f__Metschnikowiaceae(100);g__Metschnikowia(100);s__Metschnikowia_pulcherrima(100); | Metschnikowia | 13 | Saprotroph | \|Undefined Saprotroph\| | Yeast | NULL | Probable | As nectar-tap_saprotroph (Põlme et al. 2020); Decay sugar-rich substrates (Põlme et al. 2020) | Põlme S, et al. 2020. Fungal Diversity 105:1-16 ((https://doi.org/10.1007/s13225-020-00466-2)) |
| Otu33866 | 1 | k__Fungi(100);p__Ascomycota(100);c__Saccharomycetes(100);o__Saccharomycetales(100);f__Metschnikowiaceae(100);g__Metschnikowia(100);g__Metschnikowia_unclassified(100); | Metschnikowia | 13 | Saprotroph | \|Undefined Saprotroph\| | Yeast | NULL | Probable | As nectar-tap_saprotroph (Põlme et al. 2020); Decay sugar-rich substrates (Põlme et al. 2020) | Põlme S, et al. 2020. Fungal Diversity 105:1-16 ((https://doi.org/10.1007/s13225-020-00466-2)) |
| Otu33867 | 1 | k__Fungi(100);p__Ascomycota(100);c__Saccharomycetes(100);o__Saccharomycetales(100);f__Metschnikowiaceae(100);g__Metschnikowia(100);s__Metschnikowia_pulcherrima(100); | Metschnikowia | 13 | Saprotroph | \|Undefined Saprotroph\| | Yeast | NULL | Probable | As nectar-tap_saprotroph (Põlme et al. 2020); Decay sugar-rich substrates (Põlme et al. 2020) | Põlme S, et al. 2020. Fungal Diversity 105:1-16 ((https://doi.org/10.1007/s13225-020-00466-2)) |
| Otu33872 | 1 | k__Fungi(100);p__Ascomycota(100);c__Saccharomycetes(100);o__Saccharomycetales(100);f__Metschnikowiaceae(100);g__Metschnikowia(100);g__Metschnikowia_unclassified(100); | Metschnikowia | 13 | Saprotroph | \|Undefined Saprotroph\| | Yeast | NULL | Probable | As nectar-tap_saprotroph (Põlme et al. 2020); Decay sugar-rich substrates (Põlme et al. 2020) | Põlme S, et al. 2020. Fungal Diversity 105:1-16 ((https://doi.org/10.1007/s13225-020-00466-2)) |
| Otu33873 | 1 | k__Fungi(100);p__Ascomycota(100);c__Saccharomycetes(100);o__Saccharomycetales(100);f__Metschnikowiaceae(100);g__Metschnikowia(100);s__Metschnikowia_pulcherrima(100); | Metschnikowia | 13 | Saprotroph | \|Undefined Saprotroph\| | Yeast | NULL | Probable | As nectar-tap_saprotroph (Põlme et al. 2020); Decay sugar-rich substrates (Põlme et al. 2020) | Põlme S, et al. 2020. Fungal Diversity 105:1-16 ((https://doi.org/10.1007/s13225-020-00466-2)) |
| Otu33874 | 1 | k__Fungi(100);p__Ascomycota(100);c__Saccharomycetes(100);o__Saccharomycetales(100);f__Metschnikowiaceae(100);g__Metschnikowia(100);s__Metschnikowia_pulcherrima(100); | Metschnikowia | 13 | Saprotroph | \|Undefined Saprotroph\| | Yeast | NULL | Probable | As nectar-tap_saprotroph (Põlme et al. 2020); Decay sugar-rich substrates (Põlme et al. 2020) | Põlme S, et al. 2020. Fungal Diversity 105:1-16 ((https://doi.org/10.1007/s13225-020-00466-2)) |
| Otu33875 | 1 | k__Fungi(100);p__Ascomycota(100);c__Saccharomycetes(100);o__Saccharomycetales(100);f__Metschnikowiaceae(100);g__Metschnikowia(100);g__Metschnikowia_unclassified(100); | Metschnikowia | 13 | Saprotroph | \|Undefined Saprotroph\| | Yeast | NULL | Probable | As nectar-tap_saprotroph (Põlme et al. 2020); Decay sugar-rich substrates (Põlme et al. 2020) | Põlme S, et al. 2020. Fungal Diversity 105:1-16 ((https://doi.org/10.1007/s13225-020-00466-2)) |
| Otu33876 | 1 | k__Fungi(100);p__Ascomycota(100);c__Saccharomycetes(100);o__Saccharomycetales(100);f__Metschnikowiaceae(100);g__Metschnikowia(100);s__Metschnikowia_pulcherrima(100); | Metschnikowia | 13 | Saprotroph | \|Undefined Saprotroph\| | Yeast | NULL | Probable | As nectar-tap_saprotroph (Põlme et al. 2020); Decay sugar-rich substrates (Põlme et al. 2020) | Põlme S, et al. 2020. Fungal Diversity 105:1-16 ((https://doi.org/10.1007/s13225-020-00466-2)) |
| Otu33877 | 1 | k__Fungi(100);p__Ascomycota(100);c__Saccharomycetes(100);o__Saccharomycetales(100);f__Metschnikowiaceae(100);g__Metschnikowia(100);s__Metschnikowia_pulcherrima(100); | Metschnikowia | 13 | Saprotroph | \|Undefined Saprotroph\| | Yeast | NULL | Probable | As nectar-tap_saprotroph (Põlme et al. 2020); Decay sugar-rich substrates (Põlme et al. 2020) | Põlme S, et al. 2020. Fungal Diversity 105:1-16 ((https://doi.org/10.1007/s13225-020-00466-2)) |
| Otu33881 | 1 | k__Fungi(100);p__Ascomycota(100);c__Saccharomycetes(100);o__Saccharomycetales(100);f__Metschnikowiaceae(100);g__Metschnikowia(100);s__Metschnikowia_pulcherrima(100); | Metschnikowia | 13 | Saprotroph | \|Undefined Saprotroph\| | Yeast | NULL | Probable | As nectar-tap_saprotroph (Põlme et al. 2020); Decay sugar-rich substrates (Põlme et al. 2020) | Põlme S, et al. 2020. Fungal Diversity 105:1-16 ((https://doi.org/10.1007/s13225-020-00466-2)) |
| Otu33882 | 1 | k__Fungi(100);p__Ascomycota(100);c__Saccharomycetes(100);o__Saccharomycetales(100);f__Metschnikowiaceae(100);g__Metschnikowia(100);s__Metschnikowia_pulcherrima(100); | Metschnikowia | 13 | Saprotroph | \|Undefined Saprotroph\| | Yeast | NULL | Probable | As nectar-tap_saprotroph (Põlme et al. 2020); Decay sugar-rich substrates (Põlme et al. 2020) | Põlme S, et al. 2020. Fungal Diversity 105:1-16 ((https://doi.org/10.1007/s13225-020-00466-2)) |
| Otu33889 | 1 | k__Fungi(100);p__Ascomycota(100);c__Saccharomycetes(100);o__Saccharomycetales(100);f__Metschnikowiaceae(100);g__Metschnikowia(100);g__Metschnikowia_unclassified(100); | Metschnikowia | 13 | Saprotroph | \|Undefined Saprotroph\| | Yeast | NULL | Probable | As nectar-tap_saprotroph (Põlme et al. 2020); Decay sugar-rich substrates (Põlme et al. 2020) | Põlme S, et al. 2020. Fungal Diversity 105:1-16 ((https://doi.org/10.1007/s13225-020-00466-2)) |
| Otu33892 | 1 | k__Fungi(100);p__Ascomycota(100);c__Saccharomycetes(100);o__Saccharomycetales(100);f__Saccharomycetales_family_Incertae_sedis(100);g__Candida(100);g__Candida_unclassified(100); | Candida | 13 | Pathotroph-Saprotroph-Symbiotroph | Animal Pathogen-Endophyte-Endosymbiont-Epiphyte-\|Undefined Saprotroph\| | Dimorphic Yeast-Yeast | NULL | Probable | As nectar-tap_saprotroph (Põlme et al. 2020); Decay sugar-rich substrates (Põlme et al. 2020); This asexual genus is polyphyletic and contain species that function very differently depend on what they are. Caution should be taken when interpreting guilds for Candida species. | Manolakaki D, et al. 2010. Virulence. 1: 367-375 ((https://doi.org/10.4161/viru.1.5.12796)); Mekbib SB, et al. 2011. Tropical Plant Pathology 36:233-240 ((https://doi.org/10.1590/S1982-56762011000400004)); Põlme S, et al. 2020. Fungal Diversity 105:1-16 ((https://doi.org/10.1007/s13225-020-00466-2)) |
| Otu33893 | 1 | k__Fungi(100);p__Ascomycota(100);c__Saccharomycetes(100);o__Saccharomycetales(100);f__Saccharomycetales_family_Incertae_sedis(100);g__Candida(100);g__Candida_unclassified(100); | Candida | 13 | Pathotroph-Saprotroph-Symbiotroph | Animal Pathogen-Endophyte-Endosymbiont-Epiphyte-\|Undefined Saprotroph\| | Dimorphic Yeast-Yeast | NULL | Probable | As nectar-tap_saprotroph (Põlme et al. 2020); Decay sugar-rich substrates (Põlme et al. 2020); This asexual genus is polyphyletic and contain species that function very differently depend on what they are. Caution should be taken when interpreting guilds for Candida species. | Manolakaki D, et al. 2010. Virulence. 1: 367-375 ((https://doi.org/10.4161/viru.1.5.12796)); Mekbib SB, et al. 2011. Tropical Plant Pathology 36:233-240 ((https://doi.org/10.1590/S1982-56762011000400004)); Põlme S, et al. 2020. Fungal Diversity 105:1-16 ((https://doi.org/10.1007/s13225-020-00466-2)) |
| Otu33895 | 1 | k__Fungi(100);p__Ascomycota(100);c__Saccharomycetes(100);o__Saccharomycetales(100);f__Metschnikowiaceae(100);g__Metschnikowia(100);g__Metschnikowia_unclassified(100); | Metschnikowia | 13 | Saprotroph | \|Undefined Saprotroph\| | Yeast | NULL | Probable | As nectar-tap_saprotroph (Põlme et al. 2020); Decay sugar-rich substrates (Põlme et al. 2020) | Põlme S, et al. 2020. Fungal Diversity 105:1-16 ((https://doi.org/10.1007/s13225-020-00466-2)) |
| Otu33896 | 1 | k__Fungi(100);p__Ascomycota(100);c__Saccharomycetes(100);o__Saccharomycetales(100);f__Metschnikowiaceae(100);g__Metschnikowia(100);s__Metschnikowia_pulcherrima(100); | Metschnikowia | 13 | Saprotroph | \|Undefined Saprotroph\| | Yeast | NULL | Probable | As nectar-tap_saprotroph (Põlme et al. 2020); Decay sugar-rich substrates (Põlme et al. 2020) | Põlme S, et al. 2020. Fungal Diversity 105:1-16 ((https://doi.org/10.1007/s13225-020-00466-2)) |
| Otu33898 | 1 | k__Fungi(100);p__Ascomycota(100);c__Saccharomycetes(100);o__Saccharomycetales(100);f__Metschnikowiaceae(100);g__Metschnikowia(100);s__Metschnikowia_pulcherrima(100); | Metschnikowia | 13 | Saprotroph | \|Undefined Saprotroph\| | Yeast | NULL | Probable | As nectar-tap_saprotroph (Põlme et al. 2020); Decay sugar-rich substrates (Põlme et al. 2020) | Põlme S, et al. 2020. Fungal Diversity 105:1-16 ((https://doi.org/10.1007/s13225-020-00466-2)) |
| Otu33903 | 1 | k__Fungi(100);p__Ascomycota(100);c__Saccharomycetes(100);o__Saccharomycetales(100);f__Metschnikowiaceae(100);g__Metschnikowia(100);g__Metschnikowia_unclassified(100); | Metschnikowia | 13 | Saprotroph | \|Undefined Saprotroph\| | Yeast | NULL | Probable | As nectar-tap_saprotroph (Põlme et al. 2020); Decay sugar-rich substrates (Põlme et al. 2020) | Põlme S, et al. 2020. Fungal Diversity 105:1-16 ((https://doi.org/10.1007/s13225-020-00466-2)) |
| Otu33904 | 1 | k__Fungi(100);p__Ascomycota(100);c__Saccharomycetes(100);o__Saccharomycetales(100);f__Metschnikowiaceae(100);g__Metschnikowia(100);s__Metschnikowia_pulcherrima(100); | Metschnikowia | 13 | Saprotroph | \|Undefined Saprotroph\| | Yeast | NULL | Probable | As nectar-tap_saprotroph (Põlme et al. 2020); Decay sugar-rich substrates (Põlme et al. 2020) | Põlme S, et al. 2020. Fungal Diversity 105:1-16 ((https://doi.org/10.1007/s13225-020-00466-2)) |
| Otu33905 | 1 | k__Fungi(100);p__Ascomycota(100);c__Saccharomycetes(100);o__Saccharomycetales(100);f__Metschnikowiaceae(100);g__Metschnikowia(100);s__Metschnikowia_pulcherrima(100); | Metschnikowia | 13 | Saprotroph | \|Undefined Saprotroph\| | Yeast | NULL | Probable | As nectar-tap_saprotroph (Põlme et al. 2020); Decay sugar-rich substrates (Põlme et al. 2020) | Põlme S, et al. 2020. Fungal Diversity 105:1-16 ((https://doi.org/10.1007/s13225-020-00466-2)) |
| Otu33906 | 1 | k__Fungi(100);p__Ascomycota(100);c__Saccharomycetes(100);o__Saccharomycetales(100);f__Metschnikowiaceae(100);g__Metschnikowia(100);s__Metschnikowia_pulcherrima(100); | Metschnikowia | 13 | Saprotroph | \|Undefined Saprotroph\| | Yeast | NULL | Probable | As nectar-tap_saprotroph (Põlme et al. 2020); Decay sugar-rich substrates (Põlme et al. 2020) | Põlme S, et al. 2020. Fungal Diversity 105:1-16 ((https://doi.org/10.1007/s13225-020-00466-2)) |
| Otu33907 | 1 | k__Fungi(100);p__Ascomycota(100);c__Saccharomycetes(100);o__Saccharomycetales(100);f__Metschnikowiaceae(100);g__Metschnikowia(100);s__Metschnikowia_pulcherrima(100); | Metschnikowia | 13 | Saprotroph | \|Undefined Saprotroph\| | Yeast | NULL | Probable | As nectar-tap_saprotroph (Põlme et al. 2020); Decay sugar-rich substrates (Põlme et al. 2020) | Põlme S, et al. 2020. Fungal Diversity 105:1-16 ((https://doi.org/10.1007/s13225-020-00466-2)) |
| Otu33908 | 1 | k__Fungi(100);p__Ascomycota(100);c__Saccharomycetes(100);o__Saccharomycetales(100);f__Metschnikowiaceae(100);g__Metschnikowia(100);g__Metschnikowia_unclassified(100); | Metschnikowia | 13 | Saprotroph | \|Undefined Saprotroph\| | Yeast | NULL | Probable | As nectar-tap_saprotroph (Põlme et al. 2020); Decay sugar-rich substrates (Põlme et al. 2020) | Põlme S, et al. 2020. Fungal Diversity 105:1-16 ((https://doi.org/10.1007/s13225-020-00466-2)) |
| Otu33909 | 1 | k__Fungi(100);p__Ascomycota(100);c__Saccharomycetes(100);o__Saccharomycetales(100);f__Metschnikowiaceae(100);g__Metschnikowia(100);s__Metschnikowia_pulcherrima(100); | Metschnikowia | 13 | Saprotroph | \|Undefined Saprotroph\| | Yeast | NULL | Probable | As nectar-tap_saprotroph (Põlme et al. 2020); Decay sugar-rich substrates (Põlme et al. 2020) | Põlme S, et al. 2020. Fungal Diversity 105:1-16 ((https://doi.org/10.1007/s13225-020-00466-2)) |
| Otu33910 | 1 | k__Fungi(100);p__Ascomycota(100);c__Saccharomycetes(100);o__Saccharomycetales(100);f__Metschnikowiaceae(100);g__Metschnikowia(100);s__Metschnikowia_pulcherrima(100); | Metschnikowia | 13 | Saprotroph | \|Undefined Saprotroph\| | Yeast | NULL | Probable | As nectar-tap_saprotroph (Põlme et al. 2020); Decay sugar-rich substrates (Põlme et al. 2020) | Põlme S, et al. 2020. Fungal Diversity 105:1-16 ((https://doi.org/10.1007/s13225-020-00466-2)) |
| Otu33911 | 1 | k__Fungi(100);p__Ascomycota(100);c__Saccharomycetes(100);o__Saccharomycetales(100);f__Metschnikowiaceae(100);g__Metschnikowia(100);s__Metschnikowia_pulcherrima(100); | Metschnikowia | 13 | Saprotroph | \|Undefined Saprotroph\| | Yeast | NULL | Probable | As nectar-tap_saprotroph (Põlme et al. 2020); Decay sugar-rich substrates (Põlme et al. 2020) | Põlme S, et al. 2020. Fungal Diversity 105:1-16 ((https://doi.org/10.1007/s13225-020-00466-2)) |
| Otu33912 | 1 | k__Fungi(100);p__Ascomycota(100);c__Saccharomycetes(100);o__Saccharomycetales(100);f__Metschnikowiaceae(100);g__Metschnikowia(100);g__Metschnikowia_unclassified(100); | Metschnikowia | 13 | Saprotroph | \|Undefined Saprotroph\| | Yeast | NULL | Probable | As nectar-tap_saprotroph (Põlme et al. 2020); Decay sugar-rich substrates (Põlme et al. 2020) | Põlme S, et al. 2020. Fungal Diversity 105:1-16 ((https://doi.org/10.1007/s13225-020-00466-2)) |
| Otu33913 | 1 | k__Fungi(100);p__Ascomycota(100);c__Saccharomycetes(100);o__Saccharomycetales(100);f__Metschnikowiaceae(100);g__Metschnikowia(100);s__Metschnikowia_pulcherrima(100); | Metschnikowia | 13 | Saprotroph | \|Undefined Saprotroph\| | Yeast | NULL | Probable | As nectar-tap_saprotroph (Põlme et al. 2020); Decay sugar-rich substrates (Põlme et al. 2020) | Põlme S, et al. 2020. Fungal Diversity 105:1-16 ((https://doi.org/10.1007/s13225-020-00466-2)) |
| Otu33915 | 1 | k__Fungi(100);p__Ascomycota(100);c__Saccharomycetes(100);o__Saccharomycetales(100);f__Metschnikowiaceae(100);g__Metschnikowia(100);s__Metschnikowia_pulcherrima(100); | Metschnikowia | 13 | Saprotroph | \|Undefined Saprotroph\| | Yeast | NULL | Probable | As nectar-tap_saprotroph (Põlme et al. 2020); Decay sugar-rich substrates (Põlme et al. 2020) | Põlme S, et al. 2020. Fungal Diversity 105:1-16 ((https://doi.org/10.1007/s13225-020-00466-2)) |
| Otu33916 | 1 | k__Fungi(100);p__Ascomycota(100);c__Saccharomycetes(100);o__Saccharomycetales(100);f__Metschnikowiaceae(100);g__Metschnikowia(100);s__Metschnikowia_pulcherrima(100); | Metschnikowia | 13 | Saprotroph | \|Undefined Saprotroph\| | Yeast | NULL | Probable | As nectar-tap_saprotroph (Põlme et al. 2020); Decay sugar-rich substrates (Põlme et al. 2020) | Põlme S, et al. 2020. Fungal Diversity 105:1-16 ((https://doi.org/10.1007/s13225-020-00466-2)) |
| Otu33917 | 1 | k__Fungi(100);p__Ascomycota(100);c__Saccharomycetes(100);o__Saccharomycetales(100);f__Metschnikowiaceae(100);g__Metschnikowia(100);s__Metschnikowia_pulcherrima(100); | Metschnikowia | 13 | Saprotroph | \|Undefined Saprotroph\| | Yeast | NULL | Probable | As nectar-tap_saprotroph (Põlme et al. 2020); Decay sugar-rich substrates (Põlme et al. 2020) | Põlme S, et al. 2020. Fungal Diversity 105:1-16 ((https://doi.org/10.1007/s13225-020-00466-2)) |
| Otu33918 | 1 | k__Fungi(100);p__Ascomycota(100);c__Saccharomycetes(100);o__Saccharomycetales(100);f__Metschnikowiaceae(100);g__Metschnikowia(100);s__Metschnikowia_pulcherrima(100); | Metschnikowia | 13 | Saprotroph | \|Undefined Saprotroph\| | Yeast | NULL | Probable | As nectar-tap_saprotroph (Põlme et al. 2020); Decay sugar-rich substrates (Põlme et al. 2020) | Põlme S, et al. 2020. Fungal Diversity 105:1-16 ((https://doi.org/10.1007/s13225-020-00466-2)) |
| Otu33919 | 1 | k__Fungi(100);p__Ascomycota(100);c__Saccharomycetes(100);o__Saccharomycetales(100);f__Metschnikowiaceae(100);g__Metschnikowia(100);g__Metschnikowia_unclassified(100); | Metschnikowia | 13 | Saprotroph | \|Undefined Saprotroph\| | Yeast | NULL | Probable | As nectar-tap_saprotroph (Põlme et al. 2020); Decay sugar-rich substrates (Põlme et al. 2020) | Põlme S, et al. 2020. Fungal Diversity 105:1-16 ((https://doi.org/10.1007/s13225-020-00466-2)) |
| Otu33920 | 1 | k__Fungi(100);p__Ascomycota(100);c__Saccharomycetes(100);o__Saccharomycetales(100);f__Metschnikowiaceae(100);g__Metschnikowia(100);s__Metschnikowia_pulcherrima(100); | Metschnikowia | 13 | Saprotroph | \|Undefined Saprotroph\| | Yeast | NULL | Probable | As nectar-tap_saprotroph (Põlme et al. 2020); Decay sugar-rich substrates (Põlme et al. 2020) | Põlme S, et al. 2020. Fungal Diversity 105:1-16 ((https://doi.org/10.1007/s13225-020-00466-2)) |
| Otu33921 | 1 | k__Fungi(100);p__Ascomycota(100);c__Saccharomycetes(100);o__Saccharomycetales(100);f__Metschnikowiaceae(100);g__Metschnikowia(100);s__Metschnikowia_pulcherrima(100); | Metschnikowia | 13 | Saprotroph | \|Undefined Saprotroph\| | Yeast | NULL | Probable | As nectar-tap_saprotroph (Põlme et al. 2020); Decay sugar-rich substrates (Põlme et al. 2020) | Põlme S, et al. 2020. Fungal Diversity 105:1-16 ((https://doi.org/10.1007/s13225-020-00466-2)) |
| Otu33922 | 1 | k__Fungi(100);p__Basidiomycota(100);c__Microbotryomycetes(100);o__Sporidiobolales(100);f__Sporidiobolales_family_Incertae_sedis(100);g__Sporobolomyces(100);s__Sporobolomyces_lactosus(100); | Sporobolomyces | 13 | Pathotroph-Saprotroph | \|Fungal Parasite\|-Plant Saprotroph | Yeast | NULL | Probable | As litter_saprotroph (Põlme et al. 2020); As mycoparasite (Põlme et al. 2020); Decay fungal material (Põlme et al. 2020) | Tedersoo L, et al. 2014. Science 346:e1256688 ((https://doi.org/10.1126/science.1256688)); Purahong W, et al. 2016. Molecular Ecology 25:4059-4074 ((https://doi.org/10.1111/mec.13739)); Põlme S, et al. 2020. Fungal Diversity 105:1-16 ((https://doi.org/10.1007/s13225-020-00466-2)) |
| Otu33923 | 1 | k__Fungi(100);p__Ascomycota(100);c__Saccharomycetes(100);o__Saccharomycetales(100);f__Metschnikowiaceae(100);g__Metschnikowia(100);s__Metschnikowia_pulcherrima(100); | Metschnikowia | 13 | Saprotroph | \|Undefined Saprotroph\| | Yeast | NULL | Probable | As nectar-tap_saprotroph (Põlme et al. 2020); Decay sugar-rich substrates (Põlme et al. 2020) | Põlme S, et al. 2020. Fungal Diversity 105:1-16 ((https://doi.org/10.1007/s13225-020-00466-2)) |
| Otu33924 | 1 | k__Fungi(100);p__Ascomycota(100);c__Saccharomycetes(100);o__Saccharomycetales(100);f__Metschnikowiaceae(100);g__Metschnikowia(100);s__Metschnikowia_pulcherrima(100); | Metschnikowia | 13 | Saprotroph | \|Undefined Saprotroph\| | Yeast | NULL | Probable | As nectar-tap_saprotroph (Põlme et al. 2020); Decay sugar-rich substrates (Põlme et al. 2020) | Põlme S, et al. 2020. Fungal Diversity 105:1-16 ((https://doi.org/10.1007/s13225-020-00466-2)) |
| Otu33925 | 1 | k__Fungi(100);p__Ascomycota(100);c__Saccharomycetes(100);o__Saccharomycetales(100);f__Metschnikowiaceae(100);g__Metschnikowia(100);s__Metschnikowia_pulcherrima(100); | Metschnikowia | 13 | Saprotroph | \|Undefined Saprotroph\| | Yeast | NULL | Probable | As nectar-tap_saprotroph (Põlme et al. 2020); Decay sugar-rich substrates (Põlme et al. 2020) | Põlme S, et al. 2020. Fungal Diversity 105:1-16 ((https://doi.org/10.1007/s13225-020-00466-2)) |
| Otu33930 | 1 | k__Fungi(100);p__Ascomycota(100);c__Saccharomycetes(100);o__Saccharomycetales(100);f__Metschnikowiaceae(100);g__Metschnikowia(100);s__Metschnikowia_pulcherrima(100); | Metschnikowia | 13 | Saprotroph | \|Undefined Saprotroph\| | Yeast | NULL | Probable | As nectar-tap_saprotroph (Põlme et al. 2020); Decay sugar-rich substrates (Põlme et al. 2020) | Põlme S, et al. 2020. Fungal Diversity 105:1-16 ((https://doi.org/10.1007/s13225-020-00466-2)) |
| Otu33931 | 1 | k__Fungi(100);p__Ascomycota(100);c__Saccharomycetes(100);o__Saccharomycetales(100);f__Metschnikowiaceae(100);g__Metschnikowia(100);s__Metschnikowia_pulcherrima(100); | Metschnikowia | 13 | Saprotroph | \|Undefined Saprotroph\| | Yeast | NULL | Probable | As nectar-tap_saprotroph (Põlme et al. 2020); Decay sugar-rich substrates (Põlme et al. 2020) | Põlme S, et al. 2020. Fungal Diversity 105:1-16 ((https://doi.org/10.1007/s13225-020-00466-2)) |
| Otu33934 | 1 | k__Fungi(100);p__Ascomycota(100);c__Saccharomycetes(100);o__Saccharomycetales(100);f__Metschnikowiaceae(100);g__Metschnikowia(100);s__Metschnikowia_pulcherrima(100); | Metschnikowia | 13 | Saprotroph | \|Undefined Saprotroph\| | Yeast | NULL | Probable | As nectar-tap_saprotroph (Põlme et al. 2020); Decay sugar-rich substrates (Põlme et al. 2020) | Põlme S, et al. 2020. Fungal Diversity 105:1-16 ((https://doi.org/10.1007/s13225-020-00466-2)) |
| Otu33937 | 1 | k__Fungi(100);p__Ascomycota(100);c__Saccharomycetes(100);o__Saccharomycetales(100);f__Metschnikowiaceae(100);g__Metschnikowia(100);g__Metschnikowia_unclassified(100); | Metschnikowia | 13 | Saprotroph | \|Undefined Saprotroph\| | Yeast | NULL | Probable | As nectar-tap_saprotroph (Põlme et al. 2020); Decay sugar-rich substrates (Põlme et al. 2020) | Põlme S, et al. 2020. Fungal Diversity 105:1-16 ((https://doi.org/10.1007/s13225-020-00466-2)) |
| Otu33938 | 1 | k__Fungi(100);p__Ascomycota(100);c__Saccharomycetes(100);o__Saccharomycetales(100);f__Metschnikowiaceae(100);g__Metschnikowia(100);g__Metschnikowia_unclassified(100); | Metschnikowia | 13 | Saprotroph | \|Undefined Saprotroph\| | Yeast | NULL | Probable | As nectar-tap_saprotroph (Põlme et al. 2020); Decay sugar-rich substrates (Põlme et al. 2020) | Põlme S, et al. 2020. Fungal Diversity 105:1-16 ((https://doi.org/10.1007/s13225-020-00466-2)) |
| Otu33940 | 1 | k__Fungi(100);p__Ascomycota(100);c__Saccharomycetes(100);o__Saccharomycetales(100);f__Metschnikowiaceae(100);g__Metschnikowia(100);s__Metschnikowia_pulcherrima(100); | Metschnikowia | 13 | Saprotroph | \|Undefined Saprotroph\| | Yeast | NULL | Probable | As nectar-tap_saprotroph (Põlme et al. 2020); Decay sugar-rich substrates (Põlme et al. 2020) | Põlme S, et al. 2020. Fungal Diversity 105:1-16 ((https://doi.org/10.1007/s13225-020-00466-2)) |
| Otu33941 | 1 | k__Fungi(100);p__Ascomycota(100);c__Saccharomycetes(100);o__Saccharomycetales(100);f__Metschnikowiaceae(100);g__Metschnikowia(100);s__Metschnikowia_pulcherrima(100); | Metschnikowia | 13 | Saprotroph | \|Undefined Saprotroph\| | Yeast | NULL | Probable | As nectar-tap_saprotroph (Põlme et al. 2020); Decay sugar-rich substrates (Põlme et al. 2020) | Põlme S, et al. 2020. Fungal Diversity 105:1-16 ((https://doi.org/10.1007/s13225-020-00466-2)) |
| Otu33943 | 1 | k__Fungi(100);p__Ascomycota(100);c__Saccharomycetes(100);o__Saccharomycetales(100);f__Metschnikowiaceae(100);g__Metschnikowia(100);s__Metschnikowia_pulcherrima(100); | Metschnikowia | 13 | Saprotroph | \|Undefined Saprotroph\| | Yeast | NULL | Probable | As nectar-tap_saprotroph (Põlme et al. 2020); Decay sugar-rich substrates (Põlme et al. 2020) | Põlme S, et al. 2020. Fungal Diversity 105:1-16 ((https://doi.org/10.1007/s13225-020-00466-2)) |
| Otu33944 | 1 | k__Fungi(100);p__Ascomycota(100);c__Saccharomycetes(100);o__Saccharomycetales(100);f__Metschnikowiaceae(100);g__Metschnikowia(100);s__Metschnikowia_pulcherrima(100); | Metschnikowia | 13 | Saprotroph | \|Undefined Saprotroph\| | Yeast | NULL | Probable | As nectar-tap_saprotroph (Põlme et al. 2020); Decay sugar-rich substrates (Põlme et al. 2020) | Põlme S, et al. 2020. Fungal Diversity 105:1-16 ((https://doi.org/10.1007/s13225-020-00466-2)) |
| Otu33945 | 1 | k__Fungi(100);p__Ascomycota(100);c__Saccharomycetes(100);o__Saccharomycetales(100);f__Metschnikowiaceae(100);g__Metschnikowia(100);s__Metschnikowia_pulcherrima(100); | Metschnikowia | 13 | Saprotroph | \|Undefined Saprotroph\| | Yeast | NULL | Probable | As nectar-tap_saprotroph (Põlme et al. 2020); Decay sugar-rich substrates (Põlme et al. 2020) | Põlme S, et al. 2020. Fungal Diversity 105:1-16 ((https://doi.org/10.1007/s13225-020-00466-2)) |
| Otu33952 | 1 | k__Fungi(100);p__Ascomycota(100);c__Saccharomycetes(100);o__Saccharomycetales(100);f__Metschnikowiaceae(100);g__Metschnikowia(100);s__Metschnikowia_pulcherrima(100); | Metschnikowia | 13 | Saprotroph | \|Undefined Saprotroph\| | Yeast | NULL | Probable | As nectar-tap_saprotroph (Põlme et al. 2020); Decay sugar-rich substrates (Põlme et al. 2020) | Põlme S, et al. 2020. Fungal Diversity 105:1-16 ((https://doi.org/10.1007/s13225-020-00466-2)) |
| Otu33955 | 1 | k__Fungi(100);p__Ascomycota(100);c__Saccharomycetes(100);o__Saccharomycetales(100);f__Metschnikowiaceae(100);g__Metschnikowia(100);s__Metschnikowia_pulcherrima(100); | Metschnikowia | 13 | Saprotroph | \|Undefined Saprotroph\| | Yeast | NULL | Probable | As nectar-tap_saprotroph (Põlme et al. 2020); Decay sugar-rich substrates (Põlme et al. 2020) | Põlme S, et al. 2020. Fungal Diversity 105:1-16 ((https://doi.org/10.1007/s13225-020-00466-2)) |
| Otu33958 | 1 | k__Fungi(100);p__Ascomycota(100);c__Saccharomycetes(100);o__Saccharomycetales(100);f__Saccharomycetales_family_Incertae_sedis(100);g__Candida(100);s__Candida_cretensis(100); | Candida | 13 | Pathotroph-Saprotroph-Symbiotroph | Animal Pathogen-Endophyte-Endosymbiont-Epiphyte-\|Undefined Saprotroph\| | Dimorphic Yeast-Yeast | NULL | Probable | As nectar-tap_saprotroph (Põlme et al. 2020); Decay sugar-rich substrates (Põlme et al. 2020); This asexual genus is polyphyletic and contain species that function very differently depend on what they are. Caution should be taken when interpreting guilds for Candida species. | Manolakaki D, et al. 2010. Virulence. 1: 367-375 ((https://doi.org/10.4161/viru.1.5.12796)); Mekbib SB, et al. 2011. Tropical Plant Pathology 36:233-240 ((https://doi.org/10.1590/S1982-56762011000400004)); Põlme S, et al. 2020. Fungal Diversity 105:1-16 ((https://doi.org/10.1007/s13225-020-00466-2)) |
| Otu33961 | 1 | k__Fungi(100);p__Ascomycota(100);c__Saccharomycetes(100);o__Saccharomycetales(100);f__Saccharomycetales_family_Incertae_sedis(100);f__Saccharomycetales_family_Incertae_sedis_unclassified(100);f__Saccharomycetales_family_Incertae_sedis_unclassified(100); | Saccharomycetales | 7 | Saprotroph | Undefined Saprotroph | Yeast | NULL | Possible | NULL | Sterkenburg E, et al. 2015. New Phytologist 207:1145-1158 ((https://doi.org/10.1111/nph.13426)) |
| Otu33962 | 1 | k__Fungi(100);p__Ascomycota(100);c__Saccharomycetes(100);o__Saccharomycetales(100);f__Saccharomycetales_family_Incertae_sedis(100);g__Candida(100);s__Candida_cretensis(100); | Candida | 13 | Pathotroph-Saprotroph-Symbiotroph | Animal Pathogen-Endophyte-Endosymbiont-Epiphyte-\|Undefined Saprotroph\| | Dimorphic Yeast-Yeast | NULL | Probable | As nectar-tap_saprotroph (Põlme et al. 2020); Decay sugar-rich substrates (Põlme et al. 2020); This asexual genus is polyphyletic and contain species that function very differently depend on what they are. Caution should be taken when interpreting guilds for Candida species. | Manolakaki D, et al. 2010. Virulence. 1: 367-375 ((https://doi.org/10.4161/viru.1.5.12796)); Mekbib SB, et al. 2011. Tropical Plant Pathology 36:233-240 ((https://doi.org/10.1590/S1982-56762011000400004)); Põlme S, et al. 2020. Fungal Diversity 105:1-16 ((https://doi.org/10.1007/s13225-020-00466-2)) |
| Otu33963 | 1 | k__Fungi(100);p__Ascomycota(100);c__Saccharomycetes(100);o__Saccharomycetales(100);o__Saccharomycetales_unclassified(100);o__Saccharomycetales_unclassified(100);o__Saccharomycetales_unclassified(100); | Saccharomycetales | 7 | Saprotroph | Undefined Saprotroph | Yeast | NULL | Possible | NULL | Sterkenburg E, et al. 2015. New Phytologist 207:1145-1158 ((https://doi.org/10.1111/nph.13426)) |
| Otu33967 | 1 | k__Fungi(100);p__Ascomycota(100);c__Saccharomycetes(100);o__Saccharomycetales(100);f__Saccharomycetales_family_Incertae_sedis(100);g__Candida(100);g__Candida_unclassified(100); | Candida | 13 | Pathotroph-Saprotroph-Symbiotroph | Animal Pathogen-Endophyte-Endosymbiont-Epiphyte-\|Undefined Saprotroph\| | Dimorphic Yeast-Yeast | NULL | Probable | As nectar-tap_saprotroph (Põlme et al. 2020); Decay sugar-rich substrates (Põlme et al. 2020); This asexual genus is polyphyletic and contain species that function very differently depend on what they are. Caution should be taken when interpreting guilds for Candida species. | Manolakaki D, et al. 2010. Virulence. 1: 367-375 ((https://doi.org/10.4161/viru.1.5.12796)); Mekbib SB, et al. 2011. Tropical Plant Pathology 36:233-240 ((https://doi.org/10.1590/S1982-56762011000400004)); Põlme S, et al. 2020. Fungal Diversity 105:1-16 ((https://doi.org/10.1007/s13225-020-00466-2)) |
| Otu33969 | 1 | k__Fungi(100);p__Ascomycota(100);c__Saccharomycetes(100);o__Saccharomycetales(100);f__Saccharomycetales_family_Incertae_sedis(100);f__Saccharomycetales_family_Incertae_sedis_unclassified(100);f__Saccharomycetales_family_Incertae_sedis_unclassified(100); | Saccharomycetales | 7 | Saprotroph | Undefined Saprotroph | Yeast | NULL | Possible | NULL | Sterkenburg E, et al. 2015. New Phytologist 207:1145-1158 ((https://doi.org/10.1111/nph.13426)) |
| Otu33973 | 1 | k__Fungi(100);p__Ascomycota(100);c__Saccharomycetes(100);o__Saccharomycetales(100);f__Saccharomycetales_family_Incertae_sedis(100);g__Candida(100);g__Candida_unclassified(100); | Candida | 13 | Pathotroph-Saprotroph-Symbiotroph | Animal Pathogen-Endophyte-Endosymbiont-Epiphyte-\|Undefined Saprotroph\| | Dimorphic Yeast-Yeast | NULL | Probable | As nectar-tap_saprotroph (Põlme et al. 2020); Decay sugar-rich substrates (Põlme et al. 2020); This asexual genus is polyphyletic and contain species that function very differently depend on what they are. Caution should be taken when interpreting guilds for Candida species. | Manolakaki D, et al. 2010. Virulence. 1: 367-375 ((https://doi.org/10.4161/viru.1.5.12796)); Mekbib SB, et al. 2011. Tropical Plant Pathology 36:233-240 ((https://doi.org/10.1590/S1982-56762011000400004)); Põlme S, et al. 2020. Fungal Diversity 105:1-16 ((https://doi.org/10.1007/s13225-020-00466-2)) |
| Otu33975 | 1 | k__Fungi(100);p__Ascomycota(100);c__Saccharomycetes(100);o__Saccharomycetales(100);o__Saccharomycetales_unclassified(100);o__Saccharomycetales_unclassified(100);o__Saccharomycetales_unclassified(100); | Saccharomycetales | 7 | Saprotroph | Undefined Saprotroph | Yeast | NULL | Possible | NULL | Sterkenburg E, et al. 2015. New Phytologist 207:1145-1158 ((https://doi.org/10.1111/nph.13426)) |
| Otu33977 | 1 | k__Fungi(100);p__Ascomycota(100);c__Saccharomycetes(100);o__Saccharomycetales(100);f__Metschnikowiaceae(100);g__Metschnikowia(100);g__Metschnikowia_unclassified(100); | Metschnikowia | 13 | Saprotroph | \|Undefined Saprotroph\| | Yeast | NULL | Probable | As nectar-tap_saprotroph (Põlme et al. 2020); Decay sugar-rich substrates (Põlme et al. 2020) | Põlme S, et al. 2020. Fungal Diversity 105:1-16 ((https://doi.org/10.1007/s13225-020-00466-2)) |
| Otu33978 | 1 | k__Fungi(100);p__Ascomycota(100);c__Saccharomycetes(100);o__Saccharomycetales(100);f__Metschnikowiaceae(100);g__Metschnikowia(100);s__Metschnikowia_pulcherrima(100); | Metschnikowia | 13 | Saprotroph | \|Undefined Saprotroph\| | Yeast | NULL | Probable | As nectar-tap_saprotroph (Põlme et al. 2020); Decay sugar-rich substrates (Põlme et al. 2020) | Põlme S, et al. 2020. Fungal Diversity 105:1-16 ((https://doi.org/10.1007/s13225-020-00466-2)) |
| Otu33981 | 1 | k__Fungi(100);p__Ascomycota(100);c__Saccharomycetes(100);o__Saccharomycetales(100);f__Saccharomycetales_family_Incertae_sedis(100);g__Candida(100);g__Candida_unclassified(100); | Candida | 13 | Pathotroph-Saprotroph-Symbiotroph | Animal Pathogen-Endophyte-Endosymbiont-Epiphyte-\|Undefined Saprotroph\| | Dimorphic Yeast-Yeast | NULL | Probable | As nectar-tap_saprotroph (Põlme et al. 2020); Decay sugar-rich substrates (Põlme et al. 2020); This asexual genus is polyphyletic and contain species that function very differently depend on what they are. Caution should be taken when interpreting guilds for Candida species. | Manolakaki D, et al. 2010. Virulence. 1: 367-375 ((https://doi.org/10.4161/viru.1.5.12796)); Mekbib SB, et al. 2011. Tropical Plant Pathology 36:233-240 ((https://doi.org/10.1590/S1982-56762011000400004)); Põlme S, et al. 2020. Fungal Diversity 105:1-16 ((https://doi.org/10.1007/s13225-020-00466-2)) |
| Otu33982 | 1 | k__Fungi(100);p__Ascomycota(100);c__Saccharomycetes(100);o__Saccharomycetales(100);o__Saccharomycetales_unclassified(100);o__Saccharomycetales_unclassified(100);o__Saccharomycetales_unclassified(100); | Saccharomycetales | 7 | Saprotroph | Undefined Saprotroph | Yeast | NULL | Possible | NULL | Sterkenburg E, et al. 2015. New Phytologist 207:1145-1158 ((https://doi.org/10.1111/nph.13426)) |
| Otu33983 | 1 | k__Fungi(100);p__Ascomycota(100);c__Saccharomycetes(100);o__Saccharomycetales(100);f__Saccharomycetales_family_Incertae_sedis(100);g__Candida(100);g__Candida_unclassified(100); | Candida | 13 | Pathotroph-Saprotroph-Symbiotroph | Animal Pathogen-Endophyte-Endosymbiont-Epiphyte-\|Undefined Saprotroph\| | Dimorphic Yeast-Yeast | NULL | Probable | As nectar-tap_saprotroph (Põlme et al. 2020); Decay sugar-rich substrates (Põlme et al. 2020); This asexual genus is polyphyletic and contain species that function very differently depend on what they are. Caution should be taken when interpreting guilds for Candida species. | Manolakaki D, et al. 2010. Virulence. 1: 367-375 ((https://doi.org/10.4161/viru.1.5.12796)); Mekbib SB, et al. 2011. Tropical Plant Pathology 36:233-240 ((https://doi.org/10.1590/S1982-56762011000400004)); Põlme S, et al. 2020. Fungal Diversity 105:1-16 ((https://doi.org/10.1007/s13225-020-00466-2)) |
| Otu33985 | 1 | k__Fungi(100);p__Ascomycota(100);c__Saccharomycetes(100);o__Saccharomycetales(100);f__Metschnikowiaceae(100);g__Metschnikowia(100);s__Metschnikowia_pulcherrima(100); | Metschnikowia | 13 | Saprotroph | \|Undefined Saprotroph\| | Yeast | NULL | Probable | As nectar-tap_saprotroph (Põlme et al. 2020); Decay sugar-rich substrates (Põlme et al. 2020) | Põlme S, et al. 2020. Fungal Diversity 105:1-16 ((https://doi.org/10.1007/s13225-020-00466-2)) |
| Otu33991 | 1 | k__Fungi(100);p__Ascomycota(100);c__Saccharomycetes(100);o__Saccharomycetales(100);f__Metschnikowiaceae(100);g__Metschnikowia(100);g__Metschnikowia_unclassified(100); | Metschnikowia | 13 | Saprotroph | \|Undefined Saprotroph\| | Yeast | NULL | Probable | As nectar-tap_saprotroph (Põlme et al. 2020); Decay sugar-rich substrates (Põlme et al. 2020) | Põlme S, et al. 2020. Fungal Diversity 105:1-16 ((https://doi.org/10.1007/s13225-020-00466-2)) |
| Otu33992 | 1 | k__Fungi(100);p__Ascomycota(100);c__Saccharomycetes(100);o__Saccharomycetales(100);f__Metschnikowiaceae(100);g__Metschnikowia(100);s__Metschnikowia_pulcherrima(100); | Metschnikowia | 13 | Saprotroph | \|Undefined Saprotroph\| | Yeast | NULL | Probable | As nectar-tap_saprotroph (Põlme et al. 2020); Decay sugar-rich substrates (Põlme et al. 2020) | Põlme S, et al. 2020. Fungal Diversity 105:1-16 ((https://doi.org/10.1007/s13225-020-00466-2)) |
| Otu33993 | 1 | k__Fungi(100);p__Ascomycota(100);c__Saccharomycetes(100);o__Saccharomycetales(100);f__Metschnikowiaceae(100);g__Metschnikowia(100);s__Metschnikowia_pulcherrima(100); | Metschnikowia | 13 | Saprotroph | \|Undefined Saprotroph\| | Yeast | NULL | Probable | As nectar-tap_saprotroph (Põlme et al. 2020); Decay sugar-rich substrates (Põlme et al. 2020) | Põlme S, et al. 2020. Fungal Diversity 105:1-16 ((https://doi.org/10.1007/s13225-020-00466-2)) |
| Otu33994 | 1 | k__Fungi(100);p__Ascomycota(100);c__Saccharomycetes(100);o__Saccharomycetales(100);f__Metschnikowiaceae(100);g__Metschnikowia(100);s__Metschnikowia_pulcherrima(100); | Metschnikowia | 13 | Saprotroph | \|Undefined Saprotroph\| | Yeast | NULL | Probable | As nectar-tap_saprotroph (Põlme et al. 2020); Decay sugar-rich substrates (Põlme et al. 2020) | Põlme S, et al. 2020. Fungal Diversity 105:1-16 ((https://doi.org/10.1007/s13225-020-00466-2)) |
| Otu33995 | 1 | k__Fungi(100);p__Ascomycota(100);c__Saccharomycetes(100);o__Saccharomycetales(100);o__Saccharomycetales_unclassified(100);o__Saccharomycetales_unclassified(100);o__Saccharomycetales_unclassified(100); | Saccharomycetales | 7 | Saprotroph | Undefined Saprotroph | Yeast | NULL | Possible | NULL | Sterkenburg E, et al. 2015. New Phytologist 207:1145-1158 ((https://doi.org/10.1111/nph.13426)) |
| Otu33998 | 1 | k__Fungi(100);p__Ascomycota(100);c__Saccharomycetes(100);o__Saccharomycetales(100);f__Saccharomycetales_family_Incertae_sedis(100);g__Candida(100);s__Candida_cretensis(100); | Candida | 13 | Pathotroph-Saprotroph-Symbiotroph | Animal Pathogen-Endophyte-Endosymbiont-Epiphyte-\|Undefined Saprotroph\| | Dimorphic Yeast-Yeast | NULL | Probable | As nectar-tap_saprotroph (Põlme et al. 2020); Decay sugar-rich substrates (Põlme et al. 2020); This asexual genus is polyphyletic and contain species that function very differently depend on what they are. Caution should be taken when interpreting guilds for Candida species. | Manolakaki D, et al. 2010. Virulence. 1: 367-375 ((https://doi.org/10.4161/viru.1.5.12796)); Mekbib SB, et al. 2011. Tropical Plant Pathology 36:233-240 ((https://doi.org/10.1590/S1982-56762011000400004)); Põlme S, et al. 2020. Fungal Diversity 105:1-16 ((https://doi.org/10.1007/s13225-020-00466-2)) |
| Otu34000 | 1 | k__Fungi(100);p__Ascomycota(100);c__Saccharomycetes(100);o__Saccharomycetales(100);f__Metschnikowiaceae(100);g__Metschnikowia(100);g__Metschnikowia_unclassified(100); | Metschnikowia | 13 | Saprotroph | \|Undefined Saprotroph\| | Yeast | NULL | Probable | As nectar-tap_saprotroph (Põlme et al. 2020); Decay sugar-rich substrates (Põlme et al. 2020) | Põlme S, et al. 2020. Fungal Diversity 105:1-16 ((https://doi.org/10.1007/s13225-020-00466-2)) |
| Otu34001 | 1 | k__Fungi(100);p__Ascomycota(100);c__Saccharomycetes(100);o__Saccharomycetales(100);f__Metschnikowiaceae(100);g__Metschnikowia(100);s__Metschnikowia_pulcherrima(100); | Metschnikowia | 13 | Saprotroph | \|Undefined Saprotroph\| | Yeast | NULL | Probable | As nectar-tap_saprotroph (Põlme et al. 2020); Decay sugar-rich substrates (Põlme et al. 2020) | Põlme S, et al. 2020. Fungal Diversity 105:1-16 ((https://doi.org/10.1007/s13225-020-00466-2)) |
| Otu34002 | 1 | k__Fungi(100);p__Ascomycota(100);c__Saccharomycetes(100);o__Saccharomycetales(100);f__Metschnikowiaceae(100);g__Metschnikowia(100);s__Metschnikowia_pulcherrima(100); | Metschnikowia | 13 | Saprotroph | \|Undefined Saprotroph\| | Yeast | NULL | Probable | As nectar-tap_saprotroph (Põlme et al. 2020); Decay sugar-rich substrates (Põlme et al. 2020) | Põlme S, et al. 2020. Fungal Diversity 105:1-16 ((https://doi.org/10.1007/s13225-020-00466-2)) |
| Otu34011 | 1 | k__Fungi(100);p__Ascomycota(100);c__Saccharomycetes(100);o__Saccharomycetales(100);f__Metschnikowiaceae(100);g__Metschnikowia(100);s__Metschnikowia_pulcherrima(100); | Metschnikowia | 13 | Saprotroph | \|Undefined Saprotroph\| | Yeast | NULL | Probable | As nectar-tap_saprotroph (Põlme et al. 2020); Decay sugar-rich substrates (Põlme et al. 2020) | Põlme S, et al. 2020. Fungal Diversity 105:1-16 ((https://doi.org/10.1007/s13225-020-00466-2)) |
| Otu34012 | 1 | k__Fungi(100);p__Ascomycota(100);c__Saccharomycetes(100);o__Saccharomycetales(100);f__Metschnikowiaceae(100);g__Metschnikowia(100);s__Metschnikowia_pulcherrima(100); | Metschnikowia | 13 | Saprotroph | \|Undefined Saprotroph\| | Yeast | NULL | Probable | As nectar-tap_saprotroph (Põlme et al. 2020); Decay sugar-rich substrates (Põlme et al. 2020) | Põlme S, et al. 2020. Fungal Diversity 105:1-16 ((https://doi.org/10.1007/s13225-020-00466-2)) |
| Otu34013 | 1 | k__Fungi(100);p__Ascomycota(100);c__Saccharomycetes(100);o__Saccharomycetales(100);f__Saccharomycetales_family_Incertae_sedis(100);f__Saccharomycetales_family_Incertae_sedis_unclassified(100);f__Saccharomycetales_family_Incertae_sedis_unclassified(100); | Saccharomycetales | 7 | Saprotroph | Undefined Saprotroph | Yeast | NULL | Possible | NULL | Sterkenburg E, et al. 2015. New Phytologist 207:1145-1158 ((https://doi.org/10.1111/nph.13426)) |
| Otu34017 | 1 | k__Fungi(100);p__Ascomycota(100);c__Saccharomycetes(100);o__Saccharomycetales(100);f__Metschnikowiaceae(100);g__Metschnikowia(100);s__Metschnikowia_pulcherrima(100); | Metschnikowia | 13 | Saprotroph | \|Undefined Saprotroph\| | Yeast | NULL | Probable | As nectar-tap_saprotroph (Põlme et al. 2020); Decay sugar-rich substrates (Põlme et al. 2020) | Põlme S, et al. 2020. Fungal Diversity 105:1-16 ((https://doi.org/10.1007/s13225-020-00466-2)) |
| Otu34025 | 1 | k__Fungi(100);p__Ascomycota(100);c__Saccharomycetes(100);o__Saccharomycetales(100);f__Metschnikowiaceae(100);g__Metschnikowia(100);g__Metschnikowia_unclassified(100); | Metschnikowia | 13 | Saprotroph | \|Undefined Saprotroph\| | Yeast | NULL | Probable | As nectar-tap_saprotroph (Põlme et al. 2020); Decay sugar-rich substrates (Põlme et al. 2020) | Põlme S, et al. 2020. Fungal Diversity 105:1-16 ((https://doi.org/10.1007/s13225-020-00466-2)) |
| Otu34030 | 1 | k__Fungi(100);p__Ascomycota(100);c__Saccharomycetes(100);o__Saccharomycetales(100);f__Metschnikowiaceae(100);g__Metschnikowia(100);s__Metschnikowia_pulcherrima(100); | Metschnikowia | 13 | Saprotroph | \|Undefined Saprotroph\| | Yeast | NULL | Probable | As nectar-tap_saprotroph (Põlme et al. 2020); Decay sugar-rich substrates (Põlme et al. 2020) | Põlme S, et al. 2020. Fungal Diversity 105:1-16 ((https://doi.org/10.1007/s13225-020-00466-2)) |
| Otu34032 | 1 | k__Fungi(100);p__Ascomycota(100);c__Saccharomycetes(100);o__Saccharomycetales(100);o__Saccharomycetales_unclassified(100);o__Saccharomycetales_unclassified(100);o__Saccharomycetales_unclassified(100); | Saccharomycetales | 7 | Saprotroph | Undefined Saprotroph | Yeast | NULL | Possible | NULL | Sterkenburg E, et al. 2015. New Phytologist 207:1145-1158 ((https://doi.org/10.1111/nph.13426)) |
| Otu34033 | 1 | k__Fungi(100);p__Ascomycota(100);c__Saccharomycetes(100);o__Saccharomycetales(100);f__Saccharomycetales_family_Incertae_sedis(100);g__Candida(100);s__Candida_cretensis(100); | Candida | 13 | Pathotroph-Saprotroph-Symbiotroph | Animal Pathogen-Endophyte-Endosymbiont-Epiphyte-\|Undefined Saprotroph\| | Dimorphic Yeast-Yeast | NULL | Probable | As nectar-tap_saprotroph (Põlme et al. 2020); Decay sugar-rich substrates (Põlme et al. 2020); This asexual genus is polyphyletic and contain species that function very differently depend on what they are. Caution should be taken when interpreting guilds for Candida species. | Manolakaki D, et al. 2010. Virulence. 1: 367-375 ((https://doi.org/10.4161/viru.1.5.12796)); Mekbib SB, et al. 2011. Tropical Plant Pathology 36:233-240 ((https://doi.org/10.1590/S1982-56762011000400004)); Põlme S, et al. 2020. Fungal Diversity 105:1-16 ((https://doi.org/10.1007/s13225-020-00466-2)) |
| Otu34035 | 1 | k__Fungi(100);p__Ascomycota(100);c__Saccharomycetes(100);o__Saccharomycetales(100);f__Metschnikowiaceae(100);g__Metschnikowia(100);s__Metschnikowia_pulcherrima(100); | Metschnikowia | 13 | Saprotroph | \|Undefined Saprotroph\| | Yeast | NULL | Probable | As nectar-tap_saprotroph (Põlme et al. 2020); Decay sugar-rich substrates (Põlme et al. 2020) | Põlme S, et al. 2020. Fungal Diversity 105:1-16 ((https://doi.org/10.1007/s13225-020-00466-2)) |
| Otu34037 | 1 | k__Fungi(100);p__Ascomycota(100);c__Saccharomycetes(100);o__Saccharomycetales(100);f__Metschnikowiaceae(100);g__Metschnikowia(100);g__Metschnikowia_unclassified(100); | Metschnikowia | 13 | Saprotroph | \|Undefined Saprotroph\| | Yeast | NULL | Probable | As nectar-tap_saprotroph (Põlme et al. 2020); Decay sugar-rich substrates (Põlme et al. 2020) | Põlme S, et al. 2020. Fungal Diversity 105:1-16 ((https://doi.org/10.1007/s13225-020-00466-2)) |
| Otu34038 | 1 | k__Fungi(100);p__Ascomycota(100);c__Saccharomycetes(100);o__Saccharomycetales(100);f__Metschnikowiaceae(100);g__Metschnikowia(100);g__Metschnikowia_unclassified(100); | Metschnikowia | 13 | Saprotroph | \|Undefined Saprotroph\| | Yeast | NULL | Probable | As nectar-tap_saprotroph (Põlme et al. 2020); Decay sugar-rich substrates (Põlme et al. 2020) | Põlme S, et al. 2020. Fungal Diversity 105:1-16 ((https://doi.org/10.1007/s13225-020-00466-2)) |
| Otu34039 | 1 | k__Fungi(100);p__Ascomycota(100);c__Saccharomycetes(100);o__Saccharomycetales(100);f__Metschnikowiaceae(100);g__Metschnikowia(100);s__Metschnikowia_pulcherrima(100); | Metschnikowia | 13 | Saprotroph | \|Undefined Saprotroph\| | Yeast | NULL | Probable | As nectar-tap_saprotroph (Põlme et al. 2020); Decay sugar-rich substrates (Põlme et al. 2020) | Põlme S, et al. 2020. Fungal Diversity 105:1-16 ((https://doi.org/10.1007/s13225-020-00466-2)) |
| Otu34040 | 1 | k__Fungi(100);p__Ascomycota(100);c__Saccharomycetes(100);o__Saccharomycetales(100);f__Metschnikowiaceae(100);g__Metschnikowia(100);g__Metschnikowia_unclassified(100); | Metschnikowia | 13 | Saprotroph | \|Undefined Saprotroph\| | Yeast | NULL | Probable | As nectar-tap_saprotroph (Põlme et al. 2020); Decay sugar-rich substrates (Põlme et al. 2020) | Põlme S, et al. 2020. Fungal Diversity 105:1-16 ((https://doi.org/10.1007/s13225-020-00466-2)) |
| Otu34042 | 1 | k__Fungi(100);p__Ascomycota(100);c__Saccharomycetes(100);o__Saccharomycetales(100);f__Saccharomycetales_family_Incertae_sedis(100);g__Candida(100);g__Candida_unclassified(100); | Candida | 13 | Pathotroph-Saprotroph-Symbiotroph | Animal Pathogen-Endophyte-Endosymbiont-Epiphyte-\|Undefined Saprotroph\| | Dimorphic Yeast-Yeast | NULL | Probable | As nectar-tap_saprotroph (Põlme et al. 2020); Decay sugar-rich substrates (Põlme et al. 2020); This asexual genus is polyphyletic and contain species that function very differently depend on what they are. Caution should be taken when interpreting guilds for Candida species. | Manolakaki D, et al. 2010. Virulence. 1: 367-375 ((https://doi.org/10.4161/viru.1.5.12796)); Mekbib SB, et al. 2011. Tropical Plant Pathology 36:233-240 ((https://doi.org/10.1590/S1982-56762011000400004)); Põlme S, et al. 2020. Fungal Diversity 105:1-16 ((https://doi.org/10.1007/s13225-020-00466-2)) |
| Otu34044 | 1 | k__Fungi(100);p__Ascomycota(100);c__Saccharomycetes(100);o__Saccharomycetales(100);f__Saccharomycetales_family_Incertae_sedis(100);g__Candida(100);s__Candida_cretensis(100); | Candida | 13 | Pathotroph-Saprotroph-Symbiotroph | Animal Pathogen-Endophyte-Endosymbiont-Epiphyte-\|Undefined Saprotroph\| | Dimorphic Yeast-Yeast | NULL | Probable | As nectar-tap_saprotroph (Põlme et al. 2020); Decay sugar-rich substrates (Põlme et al. 2020); This asexual genus is polyphyletic and contain species that function very differently depend on what they are. Caution should be taken when interpreting guilds for Candida species. | Manolakaki D, et al. 2010. Virulence. 1: 367-375 ((https://doi.org/10.4161/viru.1.5.12796)); Mekbib SB, et al. 2011. Tropical Plant Pathology 36:233-240 ((https://doi.org/10.1590/S1982-56762011000400004)); Põlme S, et al. 2020. Fungal Diversity 105:1-16 ((https://doi.org/10.1007/s13225-020-00466-2)) |
| Otu34045 | 1 | k__Fungi(100);p__Ascomycota(100);c__Saccharomycetes(100);o__Saccharomycetales(100);f__Saccharomycetales_family_Incertae_sedis(100);g__Candida(100);g__Candida_unclassified(100); | Candida | 13 | Pathotroph-Saprotroph-Symbiotroph | Animal Pathogen-Endophyte-Endosymbiont-Epiphyte-\|Undefined Saprotroph\| | Dimorphic Yeast-Yeast | NULL | Probable | As nectar-tap_saprotroph (Põlme et al. 2020); Decay sugar-rich substrates (Põlme et al. 2020); This asexual genus is polyphyletic and contain species that function very differently depend on what they are. Caution should be taken when interpreting guilds for Candida species. | Manolakaki D, et al. 2010. Virulence. 1: 367-375 ((https://doi.org/10.4161/viru.1.5.12796)); Mekbib SB, et al. 2011. Tropical Plant Pathology 36:233-240 ((https://doi.org/10.1590/S1982-56762011000400004)); Põlme S, et al. 2020. Fungal Diversity 105:1-16 ((https://doi.org/10.1007/s13225-020-00466-2)) |
| Otu34047 | 1 | k__Fungi(100);p__Ascomycota(100);c__Saccharomycetes(100);o__Saccharomycetales(100);f__Saccharomycetales_family_Incertae_sedis(100);g__Candida(100);g__Candida_unclassified(100); | Candida | 13 | Pathotroph-Saprotroph-Symbiotroph | Animal Pathogen-Endophyte-Endosymbiont-Epiphyte-\|Undefined Saprotroph\| | Dimorphic Yeast-Yeast | NULL | Probable | As nectar-tap_saprotroph (Põlme et al. 2020); Decay sugar-rich substrates (Põlme et al. 2020); This asexual genus is polyphyletic and contain species that function very differently depend on what they are. Caution should be taken when interpreting guilds for Candida species. | Manolakaki D, et al. 2010. Virulence. 1: 367-375 ((https://doi.org/10.4161/viru.1.5.12796)); Mekbib SB, et al. 2011. Tropical Plant Pathology 36:233-240 ((https://doi.org/10.1590/S1982-56762011000400004)); Põlme S, et al. 2020. Fungal Diversity 105:1-16 ((https://doi.org/10.1007/s13225-020-00466-2)) |
| Otu34048 | 1 | k__Fungi(100);p__Ascomycota(100);c__Saccharomycetes(100);o__Saccharomycetales(100);f__Metschnikowiaceae(100);g__Metschnikowia(100);s__Metschnikowia_pulcherrima(100); | Metschnikowia | 13 | Saprotroph | \|Undefined Saprotroph\| | Yeast | NULL | Probable | As nectar-tap_saprotroph (Põlme et al. 2020); Decay sugar-rich substrates (Põlme et al. 2020) | Põlme S, et al. 2020. Fungal Diversity 105:1-16 ((https://doi.org/10.1007/s13225-020-00466-2)) |
| Otu34049 | 1 | k__Fungi(100);p__Ascomycota(100);c__Saccharomycetes(100);o__Saccharomycetales(100);f__Metschnikowiaceae(100);g__Metschnikowia(100);s__Metschnikowia_pulcherrima(100); | Metschnikowia | 13 | Saprotroph | \|Undefined Saprotroph\| | Yeast | NULL | Probable | As nectar-tap_saprotroph (Põlme et al. 2020); Decay sugar-rich substrates (Põlme et al. 2020) | Põlme S, et al. 2020. Fungal Diversity 105:1-16 ((https://doi.org/10.1007/s13225-020-00466-2)) |
| Otu34050 | 1 | k__Fungi(100);p__Basidiomycota(100);c__Microbotryomycetes(100);o__Sporidiobolales(100);f__Sporidiobolales_family_Incertae_sedis(100);g__Sporobolomyces(100);s__Sporobolomyces_lactosus(100); | Sporobolomyces | 13 | Pathotroph-Saprotroph | \|Fungal Parasite\|-Plant Saprotroph | Yeast | NULL | Probable | As litter_saprotroph (Põlme et al. 2020); As mycoparasite (Põlme et al. 2020); Decay fungal material (Põlme et al. 2020) | Tedersoo L, et al. 2014. Science 346:e1256688 ((https://doi.org/10.1126/science.1256688)); Purahong W, et al. 2016. Molecular Ecology 25:4059-4074 ((https://doi.org/10.1111/mec.13739)); Põlme S, et al. 2020. Fungal Diversity 105:1-16 ((https://doi.org/10.1007/s13225-020-00466-2)) |
| Otu34052 | 1 | k__Fungi(100);p__Basidiomycota(100);c__Microbotryomycetes(100);o__Sporidiobolales(100);f__Sporidiobolales_family_Incertae_sedis(100);g__Sporobolomyces(100);s__Sporobolomyces_lactosus(100); | Sporobolomyces | 13 | Pathotroph-Saprotroph | \|Fungal Parasite\|-Plant Saprotroph | Yeast | NULL | Probable | As litter_saprotroph (Põlme et al. 2020); As mycoparasite (Põlme et al. 2020); Decay fungal material (Põlme et al. 2020) | Tedersoo L, et al. 2014. Science 346:e1256688 ((https://doi.org/10.1126/science.1256688)); Purahong W, et al. 2016. Molecular Ecology 25:4059-4074 ((https://doi.org/10.1111/mec.13739)); Põlme S, et al. 2020. Fungal Diversity 105:1-16 ((https://doi.org/10.1007/s13225-020-00466-2)) |
| Otu34053 | 1 | k__Fungi(100);p__Ascomycota(100);c__Saccharomycetes(100);o__Saccharomycetales(100);f__Metschnikowiaceae(100);g__Metschnikowia(100);s__Metschnikowia_pulcherrima(100); | Metschnikowia | 13 | Saprotroph | \|Undefined Saprotroph\| | Yeast | NULL | Probable | As nectar-tap_saprotroph (Põlme et al. 2020); Decay sugar-rich substrates (Põlme et al. 2020) | Põlme S, et al. 2020. Fungal Diversity 105:1-16 ((https://doi.org/10.1007/s13225-020-00466-2)) |
| Otu34054 | 1 | k__Fungi(100);p__Ascomycota(100);c__Saccharomycetes(100);o__Saccharomycetales(100);o__Saccharomycetales_unclassified(100);o__Saccharomycetales_unclassified(100);o__Saccharomycetales_unclassified(100); | Saccharomycetales | 7 | Saprotroph | Undefined Saprotroph | Yeast | NULL | Possible | NULL | Sterkenburg E, et al. 2015. New Phytologist 207:1145-1158 ((https://doi.org/10.1111/nph.13426)) |
| Otu34055 | 1 | k__Fungi(100);p__Ascomycota(100);c__Saccharomycetes(100);o__Saccharomycetales(100);f__Metschnikowiaceae(100);g__Metschnikowia(100);s__Metschnikowia_pulcherrima(100); | Metschnikowia | 13 | Saprotroph | \|Undefined Saprotroph\| | Yeast | NULL | Probable | As nectar-tap_saprotroph (Põlme et al. 2020); Decay sugar-rich substrates (Põlme et al. 2020) | Põlme S, et al. 2020. Fungal Diversity 105:1-16 ((https://doi.org/10.1007/s13225-020-00466-2)) |
| Otu34056 | 1 | k__Fungi(100);p__Ascomycota(100);c__Saccharomycetes(100);o__Saccharomycetales(100);f__Metschnikowiaceae(100);g__Metschnikowia(100);s__Metschnikowia_pulcherrima(100); | Metschnikowia | 13 | Saprotroph | \|Undefined Saprotroph\| | Yeast | NULL | Probable | As nectar-tap_saprotroph (Põlme et al. 2020); Decay sugar-rich substrates (Põlme et al. 2020) | Põlme S, et al. 2020. Fungal Diversity 105:1-16 ((https://doi.org/10.1007/s13225-020-00466-2)) |
| Otu34058 | 1 | k__Fungi(100);p__Ascomycota(100);c__Saccharomycetes(100);o__Saccharomycetales(100);f__Metschnikowiaceae(100);g__Metschnikowia(100);s__Metschnikowia_pulcherrima(100); | Metschnikowia | 13 | Saprotroph | \|Undefined Saprotroph\| | Yeast | NULL | Probable | As nectar-tap_saprotroph (Põlme et al. 2020); Decay sugar-rich substrates (Põlme et al. 2020) | Põlme S, et al. 2020. Fungal Diversity 105:1-16 ((https://doi.org/10.1007/s13225-020-00466-2)) |
| Otu34061 | 1 | k__Fungi(100);p__Ascomycota(100);c__Saccharomycetes(100);o__Saccharomycetales(100);f__Metschnikowiaceae(100);g__Metschnikowia(100);s__Metschnikowia_pulcherrima(100); | Metschnikowia | 13 | Saprotroph | \|Undefined Saprotroph\| | Yeast | NULL | Probable | As nectar-tap_saprotroph (Põlme et al. 2020); Decay sugar-rich substrates (Põlme et al. 2020) | Põlme S, et al. 2020. Fungal Diversity 105:1-16 ((https://doi.org/10.1007/s13225-020-00466-2)) |
| Otu34065 | 1 | k__Fungi(100);p__Ascomycota(100);c__Saccharomycetes(100);o__Saccharomycetales(100);f__Metschnikowiaceae(100);g__Metschnikowia(100);s__Metschnikowia_pulcherrima(100); | Metschnikowia | 13 | Saprotroph | \|Undefined Saprotroph\| | Yeast | NULL | Probable | As nectar-tap_saprotroph (Põlme et al. 2020); Decay sugar-rich substrates (Põlme et al. 2020) | Põlme S, et al. 2020. Fungal Diversity 105:1-16 ((https://doi.org/10.1007/s13225-020-00466-2)) |
| Otu34066 | 1 | k__Fungi(100);p__Ascomycota(100);c__Saccharomycetes(100);o__Saccharomycetales(100);f__Metschnikowiaceae(100);g__Metschnikowia(100);g__Metschnikowia_unclassified(100); | Metschnikowia | 13 | Saprotroph | \|Undefined Saprotroph\| | Yeast | NULL | Probable | As nectar-tap_saprotroph (Põlme et al. 2020); Decay sugar-rich substrates (Põlme et al. 2020) | Põlme S, et al. 2020. Fungal Diversity 105:1-16 ((https://doi.org/10.1007/s13225-020-00466-2)) |
| Otu34067 | 1 | k__Fungi(100);p__Ascomycota(100);c__Saccharomycetes(100);o__Saccharomycetales(100);f__Metschnikowiaceae(100);g__Metschnikowia(100);s__Metschnikowia_pulcherrima(100); | Metschnikowia | 13 | Saprotroph | \|Undefined Saprotroph\| | Yeast | NULL | Probable | As nectar-tap_saprotroph (Põlme et al. 2020); Decay sugar-rich substrates (Põlme et al. 2020) | Põlme S, et al. 2020. Fungal Diversity 105:1-16 ((https://doi.org/10.1007/s13225-020-00466-2)) |
| Otu34068 | 1 | k__Fungi(100);p__Ascomycota(100);c__Saccharomycetes(100);o__Saccharomycetales(100);f__Metschnikowiaceae(100);g__Metschnikowia(100);g__Metschnikowia_unclassified(100); | Metschnikowia | 13 | Saprotroph | \|Undefined Saprotroph\| | Yeast | NULL | Probable | As nectar-tap_saprotroph (Põlme et al. 2020); Decay sugar-rich substrates (Põlme et al. 2020) | Põlme S, et al. 2020. Fungal Diversity 105:1-16 ((https://doi.org/10.1007/s13225-020-00466-2)) |
| Otu34069 | 1 | k__Fungi(100);p__Ascomycota(100);c__Saccharomycetes(100);o__Saccharomycetales(100);f__Metschnikowiaceae(100);g__Metschnikowia(100);s__Metschnikowia_pulcherrima(100); | Metschnikowia | 13 | Saprotroph | \|Undefined Saprotroph\| | Yeast | NULL | Probable | As nectar-tap_saprotroph (Põlme et al. 2020); Decay sugar-rich substrates (Põlme et al. 2020) | Põlme S, et al. 2020. Fungal Diversity 105:1-16 ((https://doi.org/10.1007/s13225-020-00466-2)) |
| Otu34075 | 1 | k__Fungi(100);p__Ascomycota(100);c__Saccharomycetes(100);o__Saccharomycetales(100);f__Metschnikowiaceae(100);g__Metschnikowia(100);s__Metschnikowia_pulcherrima(100); | Metschnikowia | 13 | Saprotroph | \|Undefined Saprotroph\| | Yeast | NULL | Probable | As nectar-tap_saprotroph (Põlme et al. 2020); Decay sugar-rich substrates (Põlme et al. 2020) | Põlme S, et al. 2020. Fungal Diversity 105:1-16 ((https://doi.org/10.1007/s13225-020-00466-2)) |
| Otu34078 | 1 | k__Fungi(100);p__Ascomycota(100);c__Saccharomycetes(100);o__Saccharomycetales(100);f__Metschnikowiaceae(100);g__Metschnikowia(100);s__Metschnikowia_pulcherrima(100); | Metschnikowia | 13 | Saprotroph | \|Undefined Saprotroph\| | Yeast | NULL | Probable | As nectar-tap_saprotroph (Põlme et al. 2020); Decay sugar-rich substrates (Põlme et al. 2020) | Põlme S, et al. 2020. Fungal Diversity 105:1-16 ((https://doi.org/10.1007/s13225-020-00466-2)) |
| Otu34079 | 1 | k__Fungi(100);p__Ascomycota(100);c__Saccharomycetes(100);o__Saccharomycetales(100);f__Metschnikowiaceae(100);g__Metschnikowia(100);s__Metschnikowia_pulcherrima(100); | Metschnikowia | 13 | Saprotroph | \|Undefined Saprotroph\| | Yeast | NULL | Probable | As nectar-tap_saprotroph (Põlme et al. 2020); Decay sugar-rich substrates (Põlme et al. 2020) | Põlme S, et al. 2020. Fungal Diversity 105:1-16 ((https://doi.org/10.1007/s13225-020-00466-2)) |
| Otu34080 | 1 | k__Fungi(100);p__Ascomycota(100);c__Saccharomycetes(100);o__Saccharomycetales(100);f__Metschnikowiaceae(100);g__Metschnikowia(100);s__Metschnikowia_pulcherrima(100); | Metschnikowia | 13 | Saprotroph | \|Undefined Saprotroph\| | Yeast | NULL | Probable | As nectar-tap_saprotroph (Põlme et al. 2020); Decay sugar-rich substrates (Põlme et al. 2020) | Põlme S, et al. 2020. Fungal Diversity 105:1-16 ((https://doi.org/10.1007/s13225-020-00466-2)) |
| Otu34083 | 1 | k__Fungi(100);p__Ascomycota(100);c__Saccharomycetes(100);o__Saccharomycetales(100);f__Metschnikowiaceae(100);g__Metschnikowia(100);g__Metschnikowia_unclassified(100); | Metschnikowia | 13 | Saprotroph | \|Undefined Saprotroph\| | Yeast | NULL | Probable | As nectar-tap_saprotroph (Põlme et al. 2020); Decay sugar-rich substrates (Põlme et al. 2020) | Põlme S, et al. 2020. Fungal Diversity 105:1-16 ((https://doi.org/10.1007/s13225-020-00466-2)) |
| Otu34084 | 1 | k__Fungi(100);p__Ascomycota(100);c__Saccharomycetes(100);o__Saccharomycetales(100);f__Metschnikowiaceae(100);g__Metschnikowia(100);g__Metschnikowia_unclassified(100); | Metschnikowia | 13 | Saprotroph | \|Undefined Saprotroph\| | Yeast | NULL | Probable | As nectar-tap_saprotroph (Põlme et al. 2020); Decay sugar-rich substrates (Põlme et al. 2020) | Põlme S, et al. 2020. Fungal Diversity 105:1-16 ((https://doi.org/10.1007/s13225-020-00466-2)) |
| Otu34085 | 1 | k__Fungi(100);p__Ascomycota(100);c__Saccharomycetes(100);o__Saccharomycetales(100);o__Saccharomycetales_unclassified(100);o__Saccharomycetales_unclassified(100);o__Saccharomycetales_unclassified(100); | Saccharomycetales | 7 | Saprotroph | Undefined Saprotroph | Yeast | NULL | Possible | NULL | Sterkenburg E, et al. 2015. New Phytologist 207:1145-1158 ((https://doi.org/10.1111/nph.13426)) |
| Otu34087 | 1 | k__Fungi(100);p__Ascomycota(100);c__Saccharomycetes(100);o__Saccharomycetales(100);f__Metschnikowiaceae(100);g__Metschnikowia(100);g__Metschnikowia_unclassified(100); | Metschnikowia | 13 | Saprotroph | \|Undefined Saprotroph\| | Yeast | NULL | Probable | As nectar-tap_saprotroph (Põlme et al. 2020); Decay sugar-rich substrates (Põlme et al. 2020) | Põlme S, et al. 2020. Fungal Diversity 105:1-16 ((https://doi.org/10.1007/s13225-020-00466-2)) |
| Otu34088 | 1 | k__Fungi(100);p__Ascomycota(100);c__Saccharomycetes(100);o__Saccharomycetales(100);f__Metschnikowiaceae(100);g__Metschnikowia(100);g__Metschnikowia_unclassified(100); | Metschnikowia | 13 | Saprotroph | \|Undefined Saprotroph\| | Yeast | NULL | Probable | As nectar-tap_saprotroph (Põlme et al. 2020); Decay sugar-rich substrates (Põlme et al. 2020) | Põlme S, et al. 2020. Fungal Diversity 105:1-16 ((https://doi.org/10.1007/s13225-020-00466-2)) |
| Otu34089 | 1 | k__Fungi(100);p__Ascomycota(100);c__Saccharomycetes(100);o__Saccharomycetales(100);f__Metschnikowiaceae(100);g__Metschnikowia(100);s__Metschnikowia_pulcherrima(100); | Metschnikowia | 13 | Saprotroph | \|Undefined Saprotroph\| | Yeast | NULL | Probable | As nectar-tap_saprotroph (Põlme et al. 2020); Decay sugar-rich substrates (Põlme et al. 2020) | Põlme S, et al. 2020. Fungal Diversity 105:1-16 ((https://doi.org/10.1007/s13225-020-00466-2)) |
| Otu34090 | 1 | k__Fungi(100);p__Ascomycota(100);c__Saccharomycetes(100);o__Saccharomycetales(100);f__Metschnikowiaceae(100);g__Metschnikowia(100);s__Metschnikowia_pulcherrima(100); | Metschnikowia | 13 | Saprotroph | \|Undefined Saprotroph\| | Yeast | NULL | Probable | As nectar-tap_saprotroph (Põlme et al. 2020); Decay sugar-rich substrates (Põlme et al. 2020) | Põlme S, et al. 2020. Fungal Diversity 105:1-16 ((https://doi.org/10.1007/s13225-020-00466-2)) |
| Otu34097 | 1 | k__Fungi(100);p__Ascomycota(100);c__Saccharomycetes(100);o__Saccharomycetales(100);f__Saccharomycetales_family_Incertae_sedis(100);g__Candida(100);g__Candida_unclassified(100); | Candida | 13 | Pathotroph-Saprotroph-Symbiotroph | Animal Pathogen-Endophyte-Endosymbiont-Epiphyte-\|Undefined Saprotroph\| | Dimorphic Yeast-Yeast | NULL | Probable | As nectar-tap_saprotroph (Põlme et al. 2020); Decay sugar-rich substrates (Põlme et al. 2020); This asexual genus is polyphyletic and contain species that function very differently depend on what they are. Caution should be taken when interpreting guilds for Candida species. | Manolakaki D, et al. 2010. Virulence. 1: 367-375 ((https://doi.org/10.4161/viru.1.5.12796)); Mekbib SB, et al. 2011. Tropical Plant Pathology 36:233-240 ((https://doi.org/10.1590/S1982-56762011000400004)); Põlme S, et al. 2020. Fungal Diversity 105:1-16 ((https://doi.org/10.1007/s13225-020-00466-2)) |
| Otu34098 | 1 | k__Fungi(100);p__Ascomycota(100);c__Saccharomycetes(100);o__Saccharomycetales(100);f__Metschnikowiaceae(100);g__Metschnikowia(100);s__Metschnikowia_pulcherrima(100); | Metschnikowia | 13 | Saprotroph | \|Undefined Saprotroph\| | Yeast | NULL | Probable | As nectar-tap_saprotroph (Põlme et al. 2020); Decay sugar-rich substrates (Põlme et al. 2020) | Põlme S, et al. 2020. Fungal Diversity 105:1-16 ((https://doi.org/10.1007/s13225-020-00466-2)) |
| Otu34099 | 1 | k__Fungi(100);p__Ascomycota(100);c__Saccharomycetes(100);o__Saccharomycetales(100);f__Saccharomycetales_family_Incertae_sedis(100);g__Candida(100);g__Candida_unclassified(100); | Candida | 13 | Pathotroph-Saprotroph-Symbiotroph | Animal Pathogen-Endophyte-Endosymbiont-Epiphyte-\|Undefined Saprotroph\| | Dimorphic Yeast-Yeast | NULL | Probable | As nectar-tap_saprotroph (Põlme et al. 2020); Decay sugar-rich substrates (Põlme et al. 2020); This asexual genus is polyphyletic and contain species that function very differently depend on what they are. Caution should be taken when interpreting guilds for Candida species. | Manolakaki D, et al. 2010. Virulence. 1: 367-375 ((https://doi.org/10.4161/viru.1.5.12796)); Mekbib SB, et al. 2011. Tropical Plant Pathology 36:233-240 ((https://doi.org/10.1590/S1982-56762011000400004)); Põlme S, et al. 2020. Fungal Diversity 105:1-16 ((https://doi.org/10.1007/s13225-020-00466-2)) |
| Otu34102 | 1 | k__Fungi(100);p__Ascomycota(100);c__Saccharomycetes(100);o__Saccharomycetales(100);f__Saccharomycetales_family_Incertae_sedis(100);g__Candida(100);s__Candida_cretensis(100); | Candida | 13 | Pathotroph-Saprotroph-Symbiotroph | Animal Pathogen-Endophyte-Endosymbiont-Epiphyte-\|Undefined Saprotroph\| | Dimorphic Yeast-Yeast | NULL | Probable | As nectar-tap_saprotroph (Põlme et al. 2020); Decay sugar-rich substrates (Põlme et al. 2020); This asexual genus is polyphyletic and contain species that function very differently depend on what they are. Caution should be taken when interpreting guilds for Candida species. | Manolakaki D, et al. 2010. Virulence. 1: 367-375 ((https://doi.org/10.4161/viru.1.5.12796)); Mekbib SB, et al. 2011. Tropical Plant Pathology 36:233-240 ((https://doi.org/10.1590/S1982-56762011000400004)); Põlme S, et al. 2020. Fungal Diversity 105:1-16 ((https://doi.org/10.1007/s13225-020-00466-2)) |
| Otu34104 | 1 | k__Fungi(100);p__Ascomycota(100);c__Saccharomycetes(100);o__Saccharomycetales(100);f__Metschnikowiaceae(100);g__Metschnikowia(100);g__Metschnikowia_unclassified(100); | Metschnikowia | 13 | Saprotroph | \|Undefined Saprotroph\| | Yeast | NULL | Probable | As nectar-tap_saprotroph (Põlme et al. 2020); Decay sugar-rich substrates (Põlme et al. 2020) | Põlme S, et al. 2020. Fungal Diversity 105:1-16 ((https://doi.org/10.1007/s13225-020-00466-2)) |
| Otu34105 | 1 | k__Fungi(100);p__Ascomycota(100);c__Saccharomycetes(100);o__Saccharomycetales(100);o__Saccharomycetales_unclassified(100);o__Saccharomycetales_unclassified(100);o__Saccharomycetales_unclassified(100); | Saccharomycetales | 7 | Saprotroph | Undefined Saprotroph | Yeast | NULL | Possible | NULL | Sterkenburg E, et al. 2015. New Phytologist 207:1145-1158 ((https://doi.org/10.1111/nph.13426)) |
| Otu34115 | 1 | k__Fungi(100);p__Ascomycota(100);c__Saccharomycetes(100);o__Saccharomycetales(100);f__Metschnikowiaceae(100);g__Metschnikowia(100);s__Metschnikowia_pulcherrima(100); | Metschnikowia | 13 | Saprotroph | \|Undefined Saprotroph\| | Yeast | NULL | Probable | As nectar-tap_saprotroph (Põlme et al. 2020); Decay sugar-rich substrates (Põlme et al. 2020) | Põlme S, et al. 2020. Fungal Diversity 105:1-16 ((https://doi.org/10.1007/s13225-020-00466-2)) |
| Otu34122 | 1 | k__Fungi(100);p__Ascomycota(100);c__Saccharomycetes(100);o__Saccharomycetales(100);o__Saccharomycetales_unclassified(100);o__Saccharomycetales_unclassified(100);o__Saccharomycetales_unclassified(100); | Saccharomycetales | 7 | Saprotroph | Undefined Saprotroph | Yeast | NULL | Possible | NULL | Sterkenburg E, et al. 2015. New Phytologist 207:1145-1158 ((https://doi.org/10.1111/nph.13426)) |
| Otu34130 | 1 | k__Fungi(100);p__Ascomycota(100);c__Saccharomycetes(100);o__Saccharomycetales(100);f__Metschnikowiaceae(100);g__Metschnikowia(100);s__Metschnikowia_pulcherrima(100); | Metschnikowia | 13 | Saprotroph | \|Undefined Saprotroph\| | Yeast | NULL | Probable | As nectar-tap_saprotroph (Põlme et al. 2020); Decay sugar-rich substrates (Põlme et al. 2020) | Põlme S, et al. 2020. Fungal Diversity 105:1-16 ((https://doi.org/10.1007/s13225-020-00466-2)) |
| Otu34134 | 1 | k__Fungi(100);p__Ascomycota(100);c__Saccharomycetes(100);o__Saccharomycetales(100);f__Metschnikowiaceae(100);g__Metschnikowia(100);g__Metschnikowia_unclassified(100); | Metschnikowia | 13 | Saprotroph | \|Undefined Saprotroph\| | Yeast | NULL | Probable | As nectar-tap_saprotroph (Põlme et al. 2020); Decay sugar-rich substrates (Põlme et al. 2020) | Põlme S, et al. 2020. Fungal Diversity 105:1-16 ((https://doi.org/10.1007/s13225-020-00466-2)) |
| Otu34137 | 1 | k__Fungi(100);p__Ascomycota(100);c__Saccharomycetes(100);o__Saccharomycetales(100);o__Saccharomycetales_unclassified(100);o__Saccharomycetales_unclassified(100);o__Saccharomycetales_unclassified(100); | Saccharomycetales | 7 | Saprotroph | Undefined Saprotroph | Yeast | NULL | Possible | NULL | Sterkenburg E, et al. 2015. New Phytologist 207:1145-1158 ((https://doi.org/10.1111/nph.13426)) |
| Otu34144 | 1 | k__Fungi(100);p__Ascomycota(100);c__Saccharomycetes(100);o__Saccharomycetales(100);f__Metschnikowiaceae(100);g__Metschnikowia(100);s__Metschnikowia_pulcherrima(100); | Metschnikowia | 13 | Saprotroph | \|Undefined Saprotroph\| | Yeast | NULL | Probable | As nectar-tap_saprotroph (Põlme et al. 2020); Decay sugar-rich substrates (Põlme et al. 2020) | Põlme S, et al. 2020. Fungal Diversity 105:1-16 ((https://doi.org/10.1007/s13225-020-00466-2)) |
| Otu34145 | 1 | k__Fungi(100);p__Ascomycota(100);c__Saccharomycetes(100);o__Saccharomycetales(100);f__Metschnikowiaceae(100);g__Metschnikowia(100);s__Metschnikowia_pulcherrima(100); | Metschnikowia | 13 | Saprotroph | \|Undefined Saprotroph\| | Yeast | NULL | Probable | As nectar-tap_saprotroph (Põlme et al. 2020); Decay sugar-rich substrates (Põlme et al. 2020) | Põlme S, et al. 2020. Fungal Diversity 105:1-16 ((https://doi.org/10.1007/s13225-020-00466-2)) |
| Otu34146 | 1 | k__Fungi(100);p__Ascomycota(100);c__Saccharomycetes(100);o__Saccharomycetales(100);f__Metschnikowiaceae(100);g__Metschnikowia(100);s__Metschnikowia_pulcherrima(100); | Metschnikowia | 13 | Saprotroph | \|Undefined Saprotroph\| | Yeast | NULL | Probable | As nectar-tap_saprotroph (Põlme et al. 2020); Decay sugar-rich substrates (Põlme et al. 2020) | Põlme S, et al. 2020. Fungal Diversity 105:1-16 ((https://doi.org/10.1007/s13225-020-00466-2)) |
| Otu34147 | 1 | k__Fungi(100);p__Ascomycota(100);c__Saccharomycetes(100);o__Saccharomycetales(100);f__Metschnikowiaceae(100);g__Metschnikowia(100);s__Metschnikowia_pulcherrima(100); | Metschnikowia | 13 | Saprotroph | \|Undefined Saprotroph\| | Yeast | NULL | Probable | As nectar-tap_saprotroph (Põlme et al. 2020); Decay sugar-rich substrates (Põlme et al. 2020) | Põlme S, et al. 2020. Fungal Diversity 105:1-16 ((https://doi.org/10.1007/s13225-020-00466-2)) |
| Otu34148 | 1 | k__Fungi(100);p__Ascomycota(100);c__Saccharomycetes(100);o__Saccharomycetales(100);f__Metschnikowiaceae(100);g__Metschnikowia(100);s__Metschnikowia_pulcherrima(100); | Metschnikowia | 13 | Saprotroph | \|Undefined Saprotroph\| | Yeast | NULL | Probable | As nectar-tap_saprotroph (Põlme et al. 2020); Decay sugar-rich substrates (Põlme et al. 2020) | Põlme S, et al. 2020. Fungal Diversity 105:1-16 ((https://doi.org/10.1007/s13225-020-00466-2)) |
| Otu34150 | 1 | k__Fungi(100);p__Ascomycota(100);c__Saccharomycetes(100);o__Saccharomycetales(100);f__Metschnikowiaceae(100);g__Metschnikowia(100);s__Metschnikowia_pulcherrima(100); | Metschnikowia | 13 | Saprotroph | \|Undefined Saprotroph\| | Yeast | NULL | Probable | As nectar-tap_saprotroph (Põlme et al. 2020); Decay sugar-rich substrates (Põlme et al. 2020) | Põlme S, et al. 2020. Fungal Diversity 105:1-16 ((https://doi.org/10.1007/s13225-020-00466-2)) |
| Otu34153 | 1 | k__Fungi(100);p__Basidiomycota(100);c__Microbotryomycetes(100);o__Sporidiobolales(100);f__Sporidiobolales_family_Incertae_sedis(100);g__Sporobolomyces(100);s__Sporobolomyces_lactosus(100); | Sporobolomyces | 13 | Pathotroph-Saprotroph | \|Fungal Parasite\|-Plant Saprotroph | Yeast | NULL | Probable | As litter_saprotroph (Põlme et al. 2020); As mycoparasite (Põlme et al. 2020); Decay fungal material (Põlme et al. 2020) | Tedersoo L, et al. 2014. Science 346:e1256688 ((https://doi.org/10.1126/science.1256688)); Purahong W, et al. 2016. Molecular Ecology 25:4059-4074 ((https://doi.org/10.1111/mec.13739)); Põlme S, et al. 2020. Fungal Diversity 105:1-16 ((https://doi.org/10.1007/s13225-020-00466-2)) |
| Otu34154 | 1 | k__Fungi(100);p__Ascomycota(100);c__Saccharomycetes(100);o__Saccharomycetales(100);f__Metschnikowiaceae(100);g__Metschnikowia(100);g__Metschnikowia_unclassified(100); | Metschnikowia | 13 | Saprotroph | \|Undefined Saprotroph\| | Yeast | NULL | Probable | As nectar-tap_saprotroph (Põlme et al. 2020); Decay sugar-rich substrates (Põlme et al. 2020) | Põlme S, et al. 2020. Fungal Diversity 105:1-16 ((https://doi.org/10.1007/s13225-020-00466-2)) |
| Otu34155 | 1 | k__Fungi(100);p__Ascomycota(100);c__Saccharomycetes(100);o__Saccharomycetales(100);f__Metschnikowiaceae(100);g__Metschnikowia(100);g__Metschnikowia_unclassified(100); | Metschnikowia | 13 | Saprotroph | \|Undefined Saprotroph\| | Yeast | NULL | Probable | As nectar-tap_saprotroph (Põlme et al. 2020); Decay sugar-rich substrates (Põlme et al. 2020) | Põlme S, et al. 2020. Fungal Diversity 105:1-16 ((https://doi.org/10.1007/s13225-020-00466-2)) |
| Otu34156 | 1 | k__Fungi(100);p__Ascomycota(100);c__Saccharomycetes(100);o__Saccharomycetales(100);f__Saccharomycetales_family_Incertae_sedis(100);g__Candida(100);g__Candida_unclassified(100); | Candida | 13 | Pathotroph-Saprotroph-Symbiotroph | Animal Pathogen-Endophyte-Endosymbiont-Epiphyte-\|Undefined Saprotroph\| | Dimorphic Yeast-Yeast | NULL | Probable | As nectar-tap_saprotroph (Põlme et al. 2020); Decay sugar-rich substrates (Põlme et al. 2020); This asexual genus is polyphyletic and contain species that function very differently depend on what they are. Caution should be taken when interpreting guilds for Candida species. | Manolakaki D, et al. 2010. Virulence. 1: 367-375 ((https://doi.org/10.4161/viru.1.5.12796)); Mekbib SB, et al. 2011. Tropical Plant Pathology 36:233-240 ((https://doi.org/10.1590/S1982-56762011000400004)); Põlme S, et al. 2020. Fungal Diversity 105:1-16 ((https://doi.org/10.1007/s13225-020-00466-2)) |
| Otu34157 | 1 | k__Fungi(100);p__Ascomycota(100);c__Saccharomycetes(100);o__Saccharomycetales(100);f__Metschnikowiaceae(100);g__Metschnikowia(100);s__Metschnikowia_pulcherrima(100); | Metschnikowia | 13 | Saprotroph | \|Undefined Saprotroph\| | Yeast | NULL | Probable | As nectar-tap_saprotroph (Põlme et al. 2020); Decay sugar-rich substrates (Põlme et al. 2020) | Põlme S, et al. 2020. Fungal Diversity 105:1-16 ((https://doi.org/10.1007/s13225-020-00466-2)) |
| Otu34159 | 1 | k__Fungi(100);p__Ascomycota(100);c__Saccharomycetes(100);o__Saccharomycetales(100);f__Metschnikowiaceae(100);g__Metschnikowia(100);s__Metschnikowia_pulcherrima(100); | Metschnikowia | 13 | Saprotroph | \|Undefined Saprotroph\| | Yeast | NULL | Probable | As nectar-tap_saprotroph (Põlme et al. 2020); Decay sugar-rich substrates (Põlme et al. 2020) | Põlme S, et al. 2020. Fungal Diversity 105:1-16 ((https://doi.org/10.1007/s13225-020-00466-2)) |
| Otu34160 | 1 | k__Fungi(100);p__Ascomycota(100);c__Saccharomycetes(100);o__Saccharomycetales(100);f__Metschnikowiaceae(100);g__Metschnikowia(100);s__Metschnikowia_pulcherrima(100); | Metschnikowia | 13 | Saprotroph | \|Undefined Saprotroph\| | Yeast | NULL | Probable | As nectar-tap_saprotroph (Põlme et al. 2020); Decay sugar-rich substrates (Põlme et al. 2020) | Põlme S, et al. 2020. Fungal Diversity 105:1-16 ((https://doi.org/10.1007/s13225-020-00466-2)) |
| Otu34162 | 1 | k__Fungi(100);p__Ascomycota(100);c__Saccharomycetes(100);o__Saccharomycetales(100);f__Metschnikowiaceae(100);g__Metschnikowia(100);s__Metschnikowia_pulcherrima(100); | Metschnikowia | 13 | Saprotroph | \|Undefined Saprotroph\| | Yeast | NULL | Probable | As nectar-tap_saprotroph (Põlme et al. 2020); Decay sugar-rich substrates (Põlme et al. 2020) | Põlme S, et al. 2020. Fungal Diversity 105:1-16 ((https://doi.org/10.1007/s13225-020-00466-2)) |
| Otu34163 | 1 | k__Fungi(100);p__Ascomycota(100);c__Saccharomycetes(100);o__Saccharomycetales(100);f__Metschnikowiaceae(100);g__Metschnikowia(100);s__Metschnikowia_pulcherrima(100); | Metschnikowia | 13 | Saprotroph | \|Undefined Saprotroph\| | Yeast | NULL | Probable | As nectar-tap_saprotroph (Põlme et al. 2020); Decay sugar-rich substrates (Põlme et al. 2020) | Põlme S, et al. 2020. Fungal Diversity 105:1-16 ((https://doi.org/10.1007/s13225-020-00466-2)) |
| Otu34165 | 1 | k__Fungi(100);p__Ascomycota(100);c__Saccharomycetes(100);o__Saccharomycetales(100);f__Metschnikowiaceae(100);g__Metschnikowia(100);g__Metschnikowia_unclassified(100); | Metschnikowia | 13 | Saprotroph | \|Undefined Saprotroph\| | Yeast | NULL | Probable | As nectar-tap_saprotroph (Põlme et al. 2020); Decay sugar-rich substrates (Põlme et al. 2020) | Põlme S, et al. 2020. Fungal Diversity 105:1-16 ((https://doi.org/10.1007/s13225-020-00466-2)) |
| Otu34166 | 1 | k__Fungi(100);p__Ascomycota(100);c__Saccharomycetes(100);o__Saccharomycetales(100);f__Metschnikowiaceae(100);g__Metschnikowia(100);s__Metschnikowia_pulcherrima(100); | Metschnikowia | 13 | Saprotroph | \|Undefined Saprotroph\| | Yeast | NULL | Probable | As nectar-tap_saprotroph (Põlme et al. 2020); Decay sugar-rich substrates (Põlme et al. 2020) | Põlme S, et al. 2020. Fungal Diversity 105:1-16 ((https://doi.org/10.1007/s13225-020-00466-2)) |
| Otu34168 | 1 | k__Fungi(100);p__Ascomycota(100);c__Saccharomycetes(100);o__Saccharomycetales(100);f__Metschnikowiaceae(100);g__Metschnikowia(100);s__Metschnikowia_pulcherrima(100); | Metschnikowia | 13 | Saprotroph | \|Undefined Saprotroph\| | Yeast | NULL | Probable | As nectar-tap_saprotroph (Põlme et al. 2020); Decay sugar-rich substrates (Põlme et al. 2020) | Põlme S, et al. 2020. Fungal Diversity 105:1-16 ((https://doi.org/10.1007/s13225-020-00466-2)) |
| Otu34169 | 1 | k__Fungi(100);p__Ascomycota(100);c__Saccharomycetes(100);o__Saccharomycetales(100);f__Metschnikowiaceae(100);g__Metschnikowia(100);g__Metschnikowia_unclassified(100); | Metschnikowia | 13 | Saprotroph | \|Undefined Saprotroph\| | Yeast | NULL | Probable | As nectar-tap_saprotroph (Põlme et al. 2020); Decay sugar-rich substrates (Põlme et al. 2020) | Põlme S, et al. 2020. Fungal Diversity 105:1-16 ((https://doi.org/10.1007/s13225-020-00466-2)) |
| Otu34170 | 1 | k__Fungi(100);p__Ascomycota(100);c__Saccharomycetes(100);o__Saccharomycetales(100);f__Metschnikowiaceae(100);g__Metschnikowia(100);g__Metschnikowia_unclassified(100); | Metschnikowia | 13 | Saprotroph | \|Undefined Saprotroph\| | Yeast | NULL | Probable | As nectar-tap_saprotroph (Põlme et al. 2020); Decay sugar-rich substrates (Põlme et al. 2020) | Põlme S, et al. 2020. Fungal Diversity 105:1-16 ((https://doi.org/10.1007/s13225-020-00466-2)) |
| Otu34173 | 1 | k__Fungi(100);p__Ascomycota(100);c__Saccharomycetes(100);o__Saccharomycetales(100);f__Metschnikowiaceae(100);g__Metschnikowia(100);s__Metschnikowia_pulcherrima(100); | Metschnikowia | 13 | Saprotroph | \|Undefined Saprotroph\| | Yeast | NULL | Probable | As nectar-tap_saprotroph (Põlme et al. 2020); Decay sugar-rich substrates (Põlme et al. 2020) | Põlme S, et al. 2020. Fungal Diversity 105:1-16 ((https://doi.org/10.1007/s13225-020-00466-2)) |
| Otu34177 | 1 | k__Fungi(100);p__Ascomycota(100);c__Saccharomycetes(100);o__Saccharomycetales(100);f__Metschnikowiaceae(100);g__Metschnikowia(100);s__Metschnikowia_pulcherrima(100); | Metschnikowia | 13 | Saprotroph | \|Undefined Saprotroph\| | Yeast | NULL | Probable | As nectar-tap_saprotroph (Põlme et al. 2020); Decay sugar-rich substrates (Põlme et al. 2020) | Põlme S, et al. 2020. Fungal Diversity 105:1-16 ((https://doi.org/10.1007/s13225-020-00466-2)) |
| Otu34178 | 1 | k__Fungi(100);p__Ascomycota(100);c__Saccharomycetes(100);o__Saccharomycetales(100);f__Metschnikowiaceae(100);g__Metschnikowia(100);s__Metschnikowia_pulcherrima(100); | Metschnikowia | 13 | Saprotroph | \|Undefined Saprotroph\| | Yeast | NULL | Probable | As nectar-tap_saprotroph (Põlme et al. 2020); Decay sugar-rich substrates (Põlme et al. 2020) | Põlme S, et al. 2020. Fungal Diversity 105:1-16 ((https://doi.org/10.1007/s13225-020-00466-2)) |
| Otu34180 | 1 | k__Fungi(100);p__Ascomycota(100);c__Saccharomycetes(100);o__Saccharomycetales(100);f__Metschnikowiaceae(100);g__Metschnikowia(100);g__Metschnikowia_unclassified(100); | Metschnikowia | 13 | Saprotroph | \|Undefined Saprotroph\| | Yeast | NULL | Probable | As nectar-tap_saprotroph (Põlme et al. 2020); Decay sugar-rich substrates (Põlme et al. 2020) | Põlme S, et al. 2020. Fungal Diversity 105:1-16 ((https://doi.org/10.1007/s13225-020-00466-2)) |
| Otu34181 | 1 | k__Fungi(100);p__Ascomycota(100);c__Saccharomycetes(100);o__Saccharomycetales(100);f__Metschnikowiaceae(100);g__Metschnikowia(100);s__Metschnikowia_pulcherrima(100); | Metschnikowia | 13 | Saprotroph | \|Undefined Saprotroph\| | Yeast | NULL | Probable | As nectar-tap_saprotroph (Põlme et al. 2020); Decay sugar-rich substrates (Põlme et al. 2020) | Põlme S, et al. 2020. Fungal Diversity 105:1-16 ((https://doi.org/10.1007/s13225-020-00466-2)) |
| Otu34182 | 1 | k__Fungi(100);p__Ascomycota(100);c__Saccharomycetes(100);o__Saccharomycetales(100);f__Metschnikowiaceae(100);g__Metschnikowia(100);s__Metschnikowia_pulcherrima(100); | Metschnikowia | 13 | Saprotroph | \|Undefined Saprotroph\| | Yeast | NULL | Probable | As nectar-tap_saprotroph (Põlme et al. 2020); Decay sugar-rich substrates (Põlme et al. 2020) | Põlme S, et al. 2020. Fungal Diversity 105:1-16 ((https://doi.org/10.1007/s13225-020-00466-2)) |
| Otu34185 | 1 | k__Fungi(100);p__Ascomycota(100);c__Saccharomycetes(100);o__Saccharomycetales(100);f__Metschnikowiaceae(100);g__Metschnikowia(100);s__Metschnikowia_pulcherrima(100); | Metschnikowia | 13 | Saprotroph | \|Undefined Saprotroph\| | Yeast | NULL | Probable | As nectar-tap_saprotroph (Põlme et al. 2020); Decay sugar-rich substrates (Põlme et al. 2020) | Põlme S, et al. 2020. Fungal Diversity 105:1-16 ((https://doi.org/10.1007/s13225-020-00466-2)) |
| Otu34186 | 1 | k__Fungi(100);p__Ascomycota(100);c__Saccharomycetes(100);o__Saccharomycetales(100);f__Metschnikowiaceae(100);g__Metschnikowia(100);s__Metschnikowia_pulcherrima(100); | Metschnikowia | 13 | Saprotroph | \|Undefined Saprotroph\| | Yeast | NULL | Probable | As nectar-tap_saprotroph (Põlme et al. 2020); Decay sugar-rich substrates (Põlme et al. 2020) | Põlme S, et al. 2020. Fungal Diversity 105:1-16 ((https://doi.org/10.1007/s13225-020-00466-2)) |
| Otu34188 | 1 | k__Fungi(100);p__Ascomycota(100);c__Saccharomycetes(100);o__Saccharomycetales(100);f__Metschnikowiaceae(100);g__Metschnikowia(100);s__Metschnikowia_pulcherrima(100); | Metschnikowia | 13 | Saprotroph | \|Undefined Saprotroph\| | Yeast | NULL | Probable | As nectar-tap_saprotroph (Põlme et al. 2020); Decay sugar-rich substrates (Põlme et al. 2020) | Põlme S, et al. 2020. Fungal Diversity 105:1-16 ((https://doi.org/10.1007/s13225-020-00466-2)) |
| Otu34195 | 1 | k__Fungi(100);p__Ascomycota(100);c__Saccharomycetes(100);o__Saccharomycetales(100);f__Metschnikowiaceae(100);g__Metschnikowia(100);s__Metschnikowia_pulcherrima(100); | Metschnikowia | 13 | Saprotroph | \|Undefined Saprotroph\| | Yeast | NULL | Probable | As nectar-tap_saprotroph (Põlme et al. 2020); Decay sugar-rich substrates (Põlme et al. 2020) | Põlme S, et al. 2020. Fungal Diversity 105:1-16 ((https://doi.org/10.1007/s13225-020-00466-2)) |
| Otu34196 | 1 | k__Fungi(100);p__Ascomycota(100);c__Saccharomycetes(100);o__Saccharomycetales(100);f__Metschnikowiaceae(100);g__Metschnikowia(100);s__Metschnikowia_pulcherrima(100); | Metschnikowia | 13 | Saprotroph | \|Undefined Saprotroph\| | Yeast | NULL | Probable | As nectar-tap_saprotroph (Põlme et al. 2020); Decay sugar-rich substrates (Põlme et al. 2020) | Põlme S, et al. 2020. Fungal Diversity 105:1-16 ((https://doi.org/10.1007/s13225-020-00466-2)) |
| Otu34198 | 1 | k__Fungi(100);p__Ascomycota(100);c__Saccharomycetes(100);o__Saccharomycetales(100);f__Metschnikowiaceae(100);g__Metschnikowia(100);s__Metschnikowia_pulcherrima(100); | Metschnikowia | 13 | Saprotroph | \|Undefined Saprotroph\| | Yeast | NULL | Probable | As nectar-tap_saprotroph (Põlme et al. 2020); Decay sugar-rich substrates (Põlme et al. 2020) | Põlme S, et al. 2020. Fungal Diversity 105:1-16 ((https://doi.org/10.1007/s13225-020-00466-2)) |
| Otu34199 | 1 | k__Fungi(100);p__Ascomycota(100);c__Saccharomycetes(100);o__Saccharomycetales(100);f__Metschnikowiaceae(100);g__Metschnikowia(100);s__Metschnikowia_pulcherrima(100); | Metschnikowia | 13 | Saprotroph | \|Undefined Saprotroph\| | Yeast | NULL | Probable | As nectar-tap_saprotroph (Põlme et al. 2020); Decay sugar-rich substrates (Põlme et al. 2020) | Põlme S, et al. 2020. Fungal Diversity 105:1-16 ((https://doi.org/10.1007/s13225-020-00466-2)) |
| Otu34200 | 1 | k__Fungi(100);p__Ascomycota(100);c__Saccharomycetes(100);o__Saccharomycetales(100);f__Metschnikowiaceae(100);g__Metschnikowia(100);g__Metschnikowia_unclassified(100); | Metschnikowia | 13 | Saprotroph | \|Undefined Saprotroph\| | Yeast | NULL | Probable | As nectar-tap_saprotroph (Põlme et al. 2020); Decay sugar-rich substrates (Põlme et al. 2020) | Põlme S, et al. 2020. Fungal Diversity 105:1-16 ((https://doi.org/10.1007/s13225-020-00466-2)) |
| Otu34201 | 1 | k__Fungi(100);p__Ascomycota(100);c__Saccharomycetes(100);o__Saccharomycetales(100);f__Metschnikowiaceae(100);g__Metschnikowia(100);g__Metschnikowia_unclassified(100); | Metschnikowia | 13 | Saprotroph | \|Undefined Saprotroph\| | Yeast | NULL | Probable | As nectar-tap_saprotroph (Põlme et al. 2020); Decay sugar-rich substrates (Põlme et al. 2020) | Põlme S, et al. 2020. Fungal Diversity 105:1-16 ((https://doi.org/10.1007/s13225-020-00466-2)) |
| Otu34202 | 1 | k__Fungi(100);p__Ascomycota(100);c__Saccharomycetes(100);o__Saccharomycetales(100);f__Metschnikowiaceae(100);g__Metschnikowia(100);g__Metschnikowia_unclassified(100); | Metschnikowia | 13 | Saprotroph | \|Undefined Saprotroph\| | Yeast | NULL | Probable | As nectar-tap_saprotroph (Põlme et al. 2020); Decay sugar-rich substrates (Põlme et al. 2020) | Põlme S, et al. 2020. Fungal Diversity 105:1-16 ((https://doi.org/10.1007/s13225-020-00466-2)) |
| Otu34203 | 1 | k__Fungi(100);p__Ascomycota(100);c__Saccharomycetes(100);o__Saccharomycetales(100);f__Metschnikowiaceae(100);g__Metschnikowia(100);s__Metschnikowia_pulcherrima(100); | Metschnikowia | 13 | Saprotroph | \|Undefined Saprotroph\| | Yeast | NULL | Probable | As nectar-tap_saprotroph (Põlme et al. 2020); Decay sugar-rich substrates (Põlme et al. 2020) | Põlme S, et al. 2020. Fungal Diversity 105:1-16 ((https://doi.org/10.1007/s13225-020-00466-2)) |
| Otu34205 | 1 | k__Fungi(100);p__Ascomycota(100);c__Saccharomycetes(100);o__Saccharomycetales(100);f__Metschnikowiaceae(100);g__Metschnikowia(100);s__Metschnikowia_pulcherrima(100); | Metschnikowia | 13 | Saprotroph | \|Undefined Saprotroph\| | Yeast | NULL | Probable | As nectar-tap_saprotroph (Põlme et al. 2020); Decay sugar-rich substrates (Põlme et al. 2020) | Põlme S, et al. 2020. Fungal Diversity 105:1-16 ((https://doi.org/10.1007/s13225-020-00466-2)) |
| Otu34206 | 1 | k__Fungi(100);p__Ascomycota(100);c__Saccharomycetes(100);o__Saccharomycetales(100);f__Metschnikowiaceae(100);g__Metschnikowia(100);s__Metschnikowia_pulcherrima(100); | Metschnikowia | 13 | Saprotroph | \|Undefined Saprotroph\| | Yeast | NULL | Probable | As nectar-tap_saprotroph (Põlme et al. 2020); Decay sugar-rich substrates (Põlme et al. 2020) | Põlme S, et al. 2020. Fungal Diversity 105:1-16 ((https://doi.org/10.1007/s13225-020-00466-2)) |
| Otu34207 | 1 | k__Fungi(100);p__Ascomycota(100);c__Saccharomycetes(100);o__Saccharomycetales(100);f__Metschnikowiaceae(100);g__Metschnikowia(100);s__Metschnikowia_pulcherrima(100); | Metschnikowia | 13 | Saprotroph | \|Undefined Saprotroph\| | Yeast | NULL | Probable | As nectar-tap_saprotroph (Põlme et al. 2020); Decay sugar-rich substrates (Põlme et al. 2020) | Põlme S, et al. 2020. Fungal Diversity 105:1-16 ((https://doi.org/10.1007/s13225-020-00466-2)) |
| Otu34208 | 1 | k__Fungi(100);p__Ascomycota(100);c__Saccharomycetes(100);o__Saccharomycetales(100);f__Metschnikowiaceae(100);g__Metschnikowia(100);s__Metschnikowia_pulcherrima(100); | Metschnikowia | 13 | Saprotroph | \|Undefined Saprotroph\| | Yeast | NULL | Probable | As nectar-tap_saprotroph (Põlme et al. 2020); Decay sugar-rich substrates (Põlme et al. 2020) | Põlme S, et al. 2020. Fungal Diversity 105:1-16 ((https://doi.org/10.1007/s13225-020-00466-2)) |
| Otu34210 | 1 | k__Fungi(100);p__Ascomycota(100);c__Saccharomycetes(100);o__Saccharomycetales(100);f__Metschnikowiaceae(100);g__Metschnikowia(100);s__Metschnikowia_pulcherrima(100); | Metschnikowia | 13 | Saprotroph | \|Undefined Saprotroph\| | Yeast | NULL | Probable | As nectar-tap_saprotroph (Põlme et al. 2020); Decay sugar-rich substrates (Põlme et al. 2020) | Põlme S, et al. 2020. Fungal Diversity 105:1-16 ((https://doi.org/10.1007/s13225-020-00466-2)) |
| Otu34211 | 1 | k__Fungi(100);p__Ascomycota(100);c__Saccharomycetes(100);o__Saccharomycetales(100);f__Metschnikowiaceae(100);g__Metschnikowia(100);s__Metschnikowia_pulcherrima(100); | Metschnikowia | 13 | Saprotroph | \|Undefined Saprotroph\| | Yeast | NULL | Probable | As nectar-tap_saprotroph (Põlme et al. 2020); Decay sugar-rich substrates (Põlme et al. 2020) | Põlme S, et al. 2020. Fungal Diversity 105:1-16 ((https://doi.org/10.1007/s13225-020-00466-2)) |
| Otu34212 | 1 | k__Fungi(100);p__Ascomycota(100);c__Saccharomycetes(100);o__Saccharomycetales(100);f__Metschnikowiaceae(100);g__Metschnikowia(100);s__Metschnikowia_pulcherrima(100); | Metschnikowia | 13 | Saprotroph | \|Undefined Saprotroph\| | Yeast | NULL | Probable | As nectar-tap_saprotroph (Põlme et al. 2020); Decay sugar-rich substrates (Põlme et al. 2020) | Põlme S, et al. 2020. Fungal Diversity 105:1-16 ((https://doi.org/10.1007/s13225-020-00466-2)) |
| Otu34213 | 1 | k__Fungi(100);p__Ascomycota(100);c__Saccharomycetes(100);o__Saccharomycetales(100);f__Metschnikowiaceae(100);g__Metschnikowia(100);g__Metschnikowia_unclassified(100); | Metschnikowia | 13 | Saprotroph | \|Undefined Saprotroph\| | Yeast | NULL | Probable | As nectar-tap_saprotroph (Põlme et al. 2020); Decay sugar-rich substrates (Põlme et al. 2020) | Põlme S, et al. 2020. Fungal Diversity 105:1-16 ((https://doi.org/10.1007/s13225-020-00466-2)) |
| Otu34216 | 1 | k__Fungi(100);p__Ascomycota(100);c__Saccharomycetes(100);o__Saccharomycetales(100);f__Metschnikowiaceae(100);g__Metschnikowia(100);s__Metschnikowia_pulcherrima(100); | Metschnikowia | 13 | Saprotroph | \|Undefined Saprotroph\| | Yeast | NULL | Probable | As nectar-tap_saprotroph (Põlme et al. 2020); Decay sugar-rich substrates (Põlme et al. 2020) | Põlme S, et al. 2020. Fungal Diversity 105:1-16 ((https://doi.org/10.1007/s13225-020-00466-2)) |
| Otu34220 | 1 | k__Fungi(100);p__Ascomycota(100);c__Saccharomycetes(100);o__Saccharomycetales(100);f__Metschnikowiaceae(100);g__Metschnikowia(100);s__Metschnikowia_pulcherrima(100); | Metschnikowia | 13 | Saprotroph | \|Undefined Saprotroph\| | Yeast | NULL | Probable | As nectar-tap_saprotroph (Põlme et al. 2020); Decay sugar-rich substrates (Põlme et al. 2020) | Põlme S, et al. 2020. Fungal Diversity 105:1-16 ((https://doi.org/10.1007/s13225-020-00466-2)) |
| Otu34222 | 1 | k__Fungi(100);p__Ascomycota(100);c__Saccharomycetes(100);o__Saccharomycetales(100);f__Metschnikowiaceae(100);g__Metschnikowia(100);s__Metschnikowia_pulcherrima(100); | Metschnikowia | 13 | Saprotroph | \|Undefined Saprotroph\| | Yeast | NULL | Probable | As nectar-tap_saprotroph (Põlme et al. 2020); Decay sugar-rich substrates (Põlme et al. 2020) | Põlme S, et al. 2020. Fungal Diversity 105:1-16 ((https://doi.org/10.1007/s13225-020-00466-2)) |

| | | | | | | | | | | | |
|---|---|---|---|---|---|---|---|---|---|---|---|
| Otu34223 | 1 | k__Fungi(100);p__Ascomycota(100);c__Saccharomycetes(100);o__Saccharomycetales(100);f__Metschnikowiaceae(100);g__Metschnikowia(100);s__Metschnikowia_pulcherrima(100); | Metschnikowia | 13 | Saprotroph | \|Undefined Saprotroph\| | Yeast | NULL | Probable | As nectar-tap_saprotroph (Põlme et al. 2020); Decay sugar-rich substrates (Põlme et al. 2020) | Põlme S, et al. 2020. Fungal Diversity 105:1-16 ((https://doi.org/10.1007/s13225-020-00466-2)) |
| Otu34224 | 1 | k__Fungi(100);p__Ascomycota(100);c__Saccharomycetes(100);o__Saccharomycetales(100);f__Metschnikowiaceae(100);g__Metschnikowia(100);s__Metschnikowia_pulcherrima(100); | Metschnikowia | 13 | Saprotroph | \|Undefined Saprotroph\| | Yeast | NULL | Probable | As nectar-tap_saprotroph (Põlme et al. 2020); Decay sugar-rich substrates (Põlme et al. 2020) | Põlme S, et al. 2020. Fungal Diversity 105:1-16 ((https://doi.org/10.1007/s13225-020-00466-2)) |
| Otu34225 | 1 | k__Fungi(100);p__Ascomycota(100);c__Saccharomycetes(100);o__Saccharomycetales(100);f__Metschnikowiaceae(100);g__Metschnikowia(100);s__Metschnikowia_pulcherrima(100); | Metschnikowia | 13 | Saprotroph | \|Undefined Saprotroph\| | Yeast | NULL | Probable | As nectar-tap_saprotroph (Põlme et al. 2020); Decay sugar-rich substrates (Põlme et al. 2020) | Põlme S, et al. 2020. Fungal Diversity 105:1-16 ((https://doi.org/10.1007/s13225-020-00466-2)) |
| Otu34226 | 1 | k__Fungi(100);p__Ascomycota(100);c__Saccharomycetes(100);o__Saccharomycetales(100);f__Metschnikowiaceae(100);g__Metschnikowia(100);s__Metschnikowia_pulcherrima(100); | Metschnikowia | 13 | Saprotroph | \|Undefined Saprotroph\| | Yeast | NULL | Probable | As nectar-tap_saprotroph (Põlme et al. 2020); Decay sugar-rich substrates (Põlme et al. 2020) | Põlme S, et al. 2020. Fungal Diversity 105:1-16 ((https://doi.org/10.1007/s13225-020-00466-2)) |
| Otu34227 | 1 | k__Fungi(100);p__Ascomycota(100);c__Saccharomycetes(100);o__Saccharomycetales(100);f__Metschnikowiaceae(100);g__Metschnikowia(100);s__Metschnikowia_pulcherrima(100); | Metschnikowia | 13 | Saprotroph | \|Undefined Saprotroph\| | Yeast | NULL | Probable | As nectar-tap_saprotroph (Põlme et al. 2020); Decay sugar-rich substrates (Põlme et al. 2020) | Põlme S, et al. 2020. Fungal Diversity 105:1-16 ((https://doi.org/10.1007/s13225-020-00466-2)) |
| Otu34228 | 1 | k__Fungi(100);p__Ascomycota(100);c__Saccharomycetes(100);o__Saccharomycetales(100);f__Metschnikowiaceae(100);g__Metschnikowia(100);s__Metschnikowia_pulcherrima(100); | Metschnikowia | 13 | Saprotroph | \|Undefined Saprotroph\| | Yeast | NULL | Probable | As nectar-tap_saprotroph (Põlme et al. 2020); Decay sugar-rich substrates (Põlme et al. 2020) | Põlme S, et al. 2020. Fungal Diversity 105:1-16 ((https://doi.org/10.1007/s13225-020-00466-2)) |
| Otu34229 | 1 | k__Fungi(100);p__Ascomycota(100);c__Saccharomycetes(100);o__Saccharomycetales(100);f__Metschnikowiaceae(100);g__Metschnikowia(100);g__Metschnikowia_unclassified(100); | Metschnikowia | 13 | Saprotroph | \|Undefined Saprotroph\| | Yeast | NULL | Probable | As nectar-tap_saprotroph (Põlme et al. 2020); Decay sugar-rich substrates (Põlme et al. 2020) | Põlme S, et al. 2020. Fungal Diversity 105:1-16 ((https://doi.org/10.1007/s13225-020-00466-2)) |
| Otu34230 | 1 | k__Fungi(100);p__Ascomycota(100);c__Saccharomycetes(100);o__Saccharomycetales(100);f__Metschnikowiaceae(100);g__Metschnikowia(100);s__Metschnikowia_pulcherrima(100); | Metschnikowia | 13 | Saprotroph | \|Undefined Saprotroph\| | Yeast | NULL | Probable | As nectar-tap_saprotroph (Põlme et al. 2020); Decay sugar-rich substrates (Põlme et al. 2020) | Põlme S, et al. 2020. Fungal Diversity 105:1-16 ((https://doi.org/10.1007/s13225-020-00466-2)) |
| Otu34231 | 1 | k__Fungi(100);p__Ascomycota(100);c__Saccharomycetes(100);o__Saccharomycetales(100);f__Metschnikowiaceae(100);g__Metschnikowia(100);s__Metschnikowia_pulcherrima(100); | Metschnikowia | 13 | Saprotroph | \|Undefined Saprotroph\| | Yeast | NULL | Probable | As nectar-tap_saprotroph (Põlme et al. 2020); Decay sugar-rich substrates (Põlme et al. 2020) | Põlme S, et al. 2020. Fungal Diversity 105:1-16 ((https://doi.org/10.1007/s13225-020-00466-2)) |
| Otu34232 | 1 | k__Fungi(100);p__Ascomycota(100);c__Saccharomycetes(100);o__Saccharomycetales(100);f__Metschnikowiaceae(100);g__Metschnikowia(100);s__Metschnikowia_pulcherrima(100); | Metschnikowia | 13 | Saprotroph | \|Undefined Saprotroph\| | Yeast | NULL | Probable | As nectar-tap_saprotroph (Põlme et al. 2020); Decay sugar-rich substrates (Põlme et al. 2020) | Põlme S, et al. 2020. Fungal Diversity 105:1-16 ((https://doi.org/10.1007/s13225-020-00466-2)) |
| Otu34233 | 1 | k__Fungi(100);p__Ascomycota(100);c__Saccharomycetes(100);o__Saccharomycetales(100);f__Metschnikowiaceae(100);g__Metschnikowia(100);s__Metschnikowia_pulcherrima(100); | Metschnikowia | 13 | Saprotroph | \|Undefined Saprotroph\| | Yeast | NULL | Probable | As nectar-tap_saprotroph (Põlme et al. 2020); Decay sugar-rich substrates (Põlme et al. 2020) | Põlme S, et al. 2020. Fungal Diversity 105:1-16 ((https://doi.org/10.1007/s13225-020-00466-2)) |
| Otu34234 | 1 | k__Fungi(100);p__Ascomycota(100);c__Saccharomycetes(100);o__Saccharomycetales(100);f__Metschnikowiaceae(100);g__Metschnikowia(100);s__Metschnikowia_pulcherrima(100); | Metschnikowia | 13 | Saprotroph | \|Undefined Saprotroph\| | Yeast | NULL | Probable | As nectar-tap_saprotroph (Põlme et al. 2020); Decay sugar-rich substrates (Põlme et al. 2020) | Põlme S, et al. 2020. Fungal Diversity 105:1-16 ((https://doi.org/10.1007/s13225-020-00466-2)) |
| Otu34235 | 1 | k__Fungi(100);p__Ascomycota(100);c__Saccharomycetes(100);o__Saccharomycetales(100);f__Metschnikowiaceae(100);g__Metschnikowia(100);s__Metschnikowia_pulcherrima(100); | Metschnikowia | 13 | Saprotroph | \|Undefined Saprotroph\| | Yeast | NULL | Probable | As nectar-tap_saprotroph (Põlme et al. 2020); Decay sugar-rich substrates (Põlme et al. 2020) | Põlme S, et al. 2020. Fungal Diversity 105:1-16 ((https://doi.org/10.1007/s13225-020-00466-2)) |
| Otu34236 | 1 | k__Fungi(100);p__Ascomycota(100);c__Saccharomycetes(100);o__Saccharomycetales(100);f__Metschnikowiaceae(100);g__Metschnikowia(100);s__Metschnikowia_pulcherrima(100); | Metschnikowia | 13 | Saprotroph | \|Undefined Saprotroph\| | Yeast | NULL | Probable | As nectar-tap_saprotroph (Põlme et al. 2020); Decay sugar-rich substrates (Põlme et al. 2020) | Põlme S, et al. 2020. Fungal Diversity 105:1-16 ((https://doi.org/10.1007/s13225-020-00466-2)) |
| Otu34237 | 1 | k__Fungi(100);p__Ascomycota(100);c__Saccharomycetes(100);o__Saccharomycetales(100);f__Metschnikowiaceae(100);g__Metschnikowia(100);s__Metschnikowia_pulcherrima(100); | Metschnikowia | 13 | Saprotroph | \|Undefined Saprotroph\| | Yeast | NULL | Probable | As nectar-tap_saprotroph (Põlme et al. 2020); Decay sugar-rich substrates (Põlme et al. 2020) | Põlme S, et al. 2020. Fungal Diversity 105:1-16 ((https://doi.org/10.1007/s13225-020-00466-2)) |
| Otu34238 | 1 | k__Fungi(100);p__Ascomycota(100);c__Saccharomycetes(100);o__Saccharomycetales(100);f__Metschnikowiaceae(100);g__Metschnikowia(100);g__Metschnikowia_unclassified(100); | Metschnikowia | 13 | Saprotroph | \|Undefined Saprotroph\| | Yeast | NULL | Probable | As nectar-tap_saprotroph (Põlme et al. 2020); Decay sugar-rich substrates (Põlme et al. 2020) | Põlme S, et al. 2020. Fungal Diversity 105:1-16 ((https://doi.org/10.1007/s13225-020-00466-2)) |
| Otu34239 | 1 | k__Fungi(100);p__Ascomycota(100);c__Saccharomycetes(100);o__Saccharomycetales(100);f__Metschnikowiaceae(100);g__Metschnikowia(100);s__Metschnikowia_pulcherrima(100); | Metschnikowia | 13 | Saprotroph | \|Undefined Saprotroph\| | Yeast | NULL | Probable | As nectar-tap_saprotroph (Põlme et al. 2020); Decay sugar-rich substrates (Põlme et al. 2020) | Põlme S, et al. 2020. Fungal Diversity 105:1-16 ((https://doi.org/10.1007/s13225-020-00466-2)) |
| Otu34240 | 1 | k__Fungi(100);p__Ascomycota(100);c__Saccharomycetes(100);o__Saccharomycetales(100);f__Metschnikowiaceae(100);g__Metschnikowia(100);g__Metschnikowia_unclassified(100); | Metschnikowia | 13 | Saprotroph | \|Undefined Saprotroph\| | Yeast | NULL | Probable | As nectar-tap_saprotroph (Põlme et al. 2020); Decay sugar-rich substrates (Põlme et al. 2020) | Põlme S, et al. 2020. Fungal Diversity 105:1-16 ((https://doi.org/10.1007/s13225-020-00466-2)) |
| Otu34241 | 1 | k__Fungi(100);p__Ascomycota(100);c__Saccharomycetes(100);o__Saccharomycetales(100);f__Metschnikowiaceae(100);g__Metschnikowia(100);s__Metschnikowia_pulcherrima(100); | Metschnikowia | 13 | Saprotroph | \|Undefined Saprotroph\| | Yeast | NULL | Probable | As nectar-tap_saprotroph (Põlme et al. 2020); Decay sugar-rich substrates (Põlme et al. 2020) | Põlme S, et al. 2020. Fungal Diversity 105:1-16 ((https://doi.org/10.1007/s13225-020-00466-2)) |
| Otu34242 | 1 | k__Fungi(100);p__Ascomycota(100);c__Saccharomycetes(100);o__Saccharomycetales(100);f__Metschnikowiaceae(100);g__Metschnikowia(100);g__Metschnikowia_unclassified(100); | Metschnikowia | 13 | Saprotroph | \|Undefined Saprotroph\| | Yeast | NULL | Probable | As nectar-tap_saprotroph (Põlme et al. 2020); Decay sugar-rich substrates (Põlme et al. 2020) | Põlme S, et al. 2020. Fungal Diversity 105:1-16 ((https://doi.org/10.1007/s13225-020-00466-2)) |
| Otu34244 | 1 | k__Fungi(100);p__Ascomycota(100);c__Saccharomycetes(100);o__Saccharomycetales(100);f__Metschnikowiaceae(100);g__Metschnikowia(100);s__Metschnikowia_pulcherrima(100); | Metschnikowia | 13 | Saprotroph | \|Undefined Saprotroph\| | Yeast | NULL | Probable | As nectar-tap_saprotroph (Põlme et al. 2020); Decay sugar-rich substrates (Põlme et al. 2020) | Põlme S, et al. 2020. Fungal Diversity 105:1-16 ((https://doi.org/10.1007/s13225-020-00466-2)) |
| Otu34245 | 1 | k__Fungi(100);p__Ascomycota(100);c__Saccharomycetes(100);o__Saccharomycetales(100);f__Saccharomycetales_family_Incertae_sedis(100);f__Saccharomycetales_family_Incertae_sedis_unclassified(100);f__Saccharomycetales_family_Incertae_sedis_unclassified(100); | Saccharomycetales | 7 | Saprotroph | Undefined Saprotroph | Yeast | NULL | Probable | NULL | Sterkenburg E, et al. 2015. New Phytologist 207:1145-1158 (https://doi.org/10.1111/nph.13405) |
| Otu34246 | 1 | k__Fungi(100);p__Ascomycota(100);c__Saccharomycetes(100);o__Saccharomycetales(100);f__Metschnikowiaceae(100);g__Metschnikowia(100);s__Metschnikowia_pulcherrima(100); | Metschnikowia | 13 | Saprotroph | \|Undefined Saprotroph\| | Yeast | NULL | Probable | As nectar-tap_saprotroph (Põlme et al. 2020); Decay sugar-rich substrates (Põlme et al. 2020) | Põlme S, et al. 2020. Fungal Diversity 105:1-16 ((https://doi.org/10.1007/s13225-020-00466-2)) |
| Otu34247 | 1 | k__Fungi(100);p__Ascomycota(100);c__Saccharomycetes(100);o__Saccharomycetales(100);f__Metschnikowiaceae(100);g__Metschnikowia(100);s__Metschnikowia_pulcherrima(100); | Metschnikowia | 13 | Saprotroph | \|Undefined Saprotroph\| | Yeast | NULL | Probable | As nectar-tap_saprotroph (Põlme et al. 2020); Decay sugar-rich substrates (Põlme et al. 2020) | Põlme S, et al. 2020. Fungal Diversity 105:1-16 ((https://doi.org/10.1007/s13225-020-00466-2)) |
| Otu34248 | 1 | k__Fungi(100);p__Ascomycota(100);c__Saccharomycetes(100);o__Saccharomycetales(100);f__Metschnikowiaceae(100);g__Metschnikowia(100);s__Metschnikowia_pulcherrima(100); | Metschnikowia | 13 | Saprotroph | \|Undefined Saprotroph\| | Yeast | NULL | Probable | As nectar-tap_saprotroph (Põlme et al. 2020); Decay sugar-rich substrates (Põlme et al. 2020) | Põlme S, et al. 2020. Fungal Diversity 105:1-16 ((https://doi.org/10.1007/s13225-020-00466-2)) |
| Otu34251 | 1 | k__Fungi(100);p__Ascomycota(100);c__Saccharomycetes(100);o__Saccharomycetales(100);f__Metschnikowiaceae(100);g__Metschnikowia(100);s__Metschnikowia_pulcherrima(100); | Metschnikowia | 13 | Saprotroph | \|Undefined Saprotroph\| | Yeast | NULL | Probable | As nectar-tap_saprotroph (Põlme et al. 2020); Decay sugar-rich substrates (Põlme et al. 2020) | Põlme S, et al. 2020. Fungal Diversity 105:1-16 ((https://doi.org/10.1007/s13225-020-00466-2)) |
| Otu34252 | 1 | k__Fungi(100);p__Ascomycota(100);c__Saccharomycetes(100);o__Saccharomycetales(100);f__Metschnikowiaceae(100);g__Metschnikowia(100);s__Metschnikowia_pulcherrima(100); | Metschnikowia | 13 | Saprotroph | \|Undefined Saprotroph\| | Yeast | NULL | Probable | As nectar-tap_saprotroph (Põlme et al. 2020); Decay sugar-rich substrates (Põlme et al. 2020) | Põlme S, et al. 2020. Fungal Diversity 105:1-16 ((https://doi.org/10.1007/s13225-020-00466-2)) |
| Otu34254 | 1 | k__Fungi(100);p__Ascomycota(100);c__Saccharomycetes(100);o__Saccharomycetales(100);f__Metschnikowiaceae(100);g__Metschnikowia(100);s__Metschnikowia_pulcherrima(100); | Metschnikowia | 13 | Saprotroph | \|Undefined Saprotroph\| | Yeast | NULL | Probable | As nectar-tap_saprotroph (Põlme et al. 2020); Decay sugar-rich substrates (Põlme et al. 2020) | Põlme S, et al. 2020. Fungal Diversity 105:1-16 ((https://doi.org/10.1007/s13225-020-00466-2)) |
| Otu34256 | 1 | k__Fungi(100);p__Ascomycota(100);c__Saccharomycetes(100);o__Saccharomycetales(100);f__Metschnikowiaceae(100);g__Metschnikowia(100);g__Metschnikowia_unclassified(100); | Metschnikowia | 13 | Saprotroph | \|Undefined Saprotroph\| | Yeast | NULL | Probable | As nectar-tap_saprotroph (Põlme et al. 2020); Decay sugar-rich substrates (Põlme et al. 2020) | Põlme S, et al. 2020. Fungal Diversity 105:1-16 ((https://doi.org/10.1007/s13225-020-00466-2)) |
| Otu34257 | 1 | k__Fungi(100);p__Ascomycota(100);c__Saccharomycetes(100);o__Saccharomycetales(100);f__Metschnikowiaceae(100);g__Metschnikowia(100);s__Metschnikowia_pulcherrima(100); | Metschnikowia | 13 | Saprotroph | \|Undefined Saprotroph\| | Yeast | NULL | Probable | As nectar-tap_saprotroph (Põlme et al. 2020); Decay sugar-rich substrates (Põlme et al. 2020) | Põlme S, et al. 2020. Fungal Diversity 105:1-16 ((https://doi.org/10.1007/s13225-020-00466-2)) |
| Otu34259 | 1 | k__Fungi(100);p__Ascomycota(100);c__Saccharomycetes(100);o__Saccharomycetales(100);f__Metschnikowiaceae(100);g__Metschnikowia(100);s__Metschnikowia_pulcherrima(100); | Metschnikowia | 13 | Saprotroph | \|Undefined Saprotroph\| | Yeast | NULL | Probable | As nectar-tap_saprotroph (Põlme et al. 2020); Decay sugar-rich substrates (Põlme et al. 2020) | Põlme S, et al. 2020. Fungal Diversity 105:1-16 ((https://doi.org/10.1007/s13225-020-00466-2)) |
| Otu34261 | 1 | k__Fungi(100);p__Ascomycota(100);c__Saccharomycetes(100);o__Saccharomycetales(100);f__Metschnikowiaceae(100);g__Metschnikowia(100);g__Metschnikowia_unclassified(100); | Metschnikowia | 13 | Saprotroph | \|Undefined Saprotroph\| | Yeast | NULL | Probable | As nectar-tap_saprotroph (Põlme et al. 2020); Decay sugar-rich substrates (Põlme et al. 2020) | Põlme S, et al. 2020. Fungal Diversity 105:1-16 ((https://doi.org/10.1007/s13225-020-00466-2)) |
| Otu34262 | 1 | k__Fungi(100);p__Ascomycota(100);c__Saccharomycetes(100);o__Saccharomycetales(100);f__Metschnikowiaceae(100);g__Metschnikowia(100);g__Metschnikowia_unclassified(100); | Metschnikowia | 13 | Saprotroph | \|Undefined Saprotroph\| | Yeast | NULL | Probable | As nectar-tap_saprotroph (Põlme et al. 2020); Decay sugar-rich substrates (Põlme et al. 2020) | Põlme S, et al. 2020. Fungal Diversity 105:1-16 ((https://doi.org/10.1007/s13225-020-00466-2)) |
| Otu34263 | 1 | k__Fungi(100);p__Ascomycota(100);c__Saccharomycetes(100);o__Saccharomycetales(100);f__Metschnikowiaceae(100);g__Metschnikowia(100);s__Metschnikowia_pulcherrima(100); | Metschnikowia | 13 | Saprotroph | \|Undefined Saprotroph\| | Yeast | NULL | Probable | As nectar-tap_saprotroph (Põlme et al. 2020); Decay sugar-rich substrates (Põlme et al. 2020) | Põlme S, et al. 2020. Fungal Diversity 105:1-16 ((https://doi.org/10.1007/s13225-020-00466-2)) |
| Otu34264 | 1 | k__Fungi(100);p__Ascomycota(100);c__Saccharomycetes(100);o__Saccharomycetales(100);f__Metschnikowiaceae(100);g__Metschnikowia(100);s__Metschnikowia_pulcherrima(100); | Metschnikowia | 13 | Saprotroph | \|Undefined Saprotroph\| | Yeast | NULL | Probable | As nectar-tap_saprotroph (Põlme et al. 2020); Decay sugar-rich substrates (Põlme et al. 2020) | Põlme S, et al. 2020. Fungal Diversity 105:1-16 ((https://doi.org/10.1007/s13225-020-00466-2)) |
| Otu34265 | 1 | k__Fungi(100);p__Ascomycota(100);c__Saccharomycetes(100);o__Saccharomycetales(100);f__Metschnikowiaceae(100);g__Metschnikowia(100);s__Metschnikowia_pulcherrima(100); | Metschnikowia | 13 | Saprotroph | \|Undefined Saprotroph\| | Yeast | NULL | Probable | As nectar-tap_saprotroph (Põlme et al. 2020); Decay sugar-rich substrates (Põlme et al. 2020) | Põlme S, et al. 2020. Fungal Diversity 105:1-16 ((https://doi.org/10.1007/s13225-020-00466-2)) |
| Otu34266 | 1 | k__Fungi(100);p__Ascomycota(100);c__Saccharomycetes(100);o__Saccharomycetales(100);f__Metschnikowiaceae(100);g__Metschnikowia(100);s__Metschnikowia_pulcherrima(100); | Metschnikowia | 13 | Saprotroph | \|Undefined Saprotroph\| | Yeast | NULL | Probable | As nectar-tap_saprotroph (Põlme et al. 2020); Decay sugar-rich substrates (Põlme et al. 2020) | Põlme S, et al. 2020. Fungal Diversity 105:1-16 ((https://doi.org/10.1007/s13225-020-00466-2)) |
| Otu34267 | 1 | k__Fungi(100);p__Ascomycota(100);c__Saccharomycetes(100);o__Saccharomycetales(100);f__Metschnikowiaceae(100);g__Metschnikowia(100);g__Metschnikowia_unclassified(100); | Metschnikowia | 13 | Saprotroph | \|Undefined Saprotroph\| | Yeast | NULL | Probable | As nectar-tap_saprotroph (Põlme et al. 2020); Decay sugar-rich substrates (Põlme et al. 2020) | Põlme S, et al. 2020. Fungal Diversity 105:1-16 ((https://doi.org/10.1007/s13225-020-00466-2)) |
| Otu34268 | 1 | k__Fungi(100);p__Ascomycota(100);c__Saccharomycetes(100);o__Saccharomycetales(100);f__Metschnikowiaceae(100);g__Metschnikowia(100);s__Metschnikowia_pulcherrima(100); | Metschnikowia | 13 | Saprotroph | \|Undefined Saprotroph\| | Yeast | NULL | Probable | As nectar-tap_saprotroph (Põlme et al. 2020); Decay sugar-rich substrates (Põlme et al. 2020) | Põlme S, et al. 2020. Fungal Diversity 105:1-16 ((https://doi.org/10.1007/s13225-020-00466-2)) |
| Otu34269 | 1 | k__Fungi(100);p__Ascomycota(100);c__Saccharomycetes(100);o__Saccharomycetales(100);f__Metschnikowiaceae(100);g__Metschnikowia(100);s__Metschnikowia_pulcherrima(100); | Metschnikowia | 13 | Saprotroph | \|Undefined Saprotroph\| | Yeast | NULL | Probable | As nectar-tap_saprotroph (Põlme et al. 2020); Decay sugar-rich substrates (Põlme et al. 2020) | Põlme S, et al. 2020. Fungal Diversity 105:1-16 ((https://doi.org/10.1007/s13225-020-00466-2)) |
| Otu34270 | 1 | k__Fungi(100);p__Ascomycota(100);c__Saccharomycetes(100);o__Saccharomycetales(100);f__Metschnikowiaceae(100);g__Metschnikowia(100);s__Metschnikowia_pulcherrima(100); | Metschnikowia | 13 | Saprotroph | \|Undefined Saprotroph\| | Yeast | NULL | Probable | As nectar-tap_saprotroph (Põlme et al. 2020); Decay sugar-rich substrates (Põlme et al. 2020) | Põlme S, et al. 2020. Fungal Diversity 105:1-16 ((https://doi.org/10.1007/s13225-020-00466-2)) |
| Otu34271 | 1 | k__Fungi(100);p__Ascomycota(100);c__Saccharomycetes(100);o__Saccharomycetales(100);f__Metschnikowiaceae(100);g__Metschnikowia(100);s__Metschnikowia_pulcherrima(100); | Metschnikowia | 13 | Saprotroph | \|Undefined Saprotroph\| | Yeast | NULL | Probable | As nectar-tap_saprotroph (Põlme et al. 2020); Decay sugar-rich substrates (Põlme et al. 2020) | Põlme S, et al. 2020. Fungal Diversity 105:1-16 ((https://doi.org/10.1007/s13225-020-00466-2)) |
| Otu34272 | 1 | k__Fungi(100);p__Ascomycota(100);c__Saccharomycetes(100);o__Saccharomycetales(100);f__Metschnikowiaceae(100);g__Metschnikowia(100);s__Metschnikowia_pulcherrima(100); | Metschnikowia | 13 | Saprotroph | \|Undefined Saprotroph\| | Yeast | NULL | Probable | As nectar-tap_saprotroph (Põlme et al. 2020); Decay sugar-rich substrates (Põlme et al. 2020) | Põlme S, et al. 2020. Fungal Diversity 105:1-16 ((https://doi.org/10.1007/s13225-020-00466-2)) |
| Otu34273 | 1 | k__Fungi(100);p__Ascomycota(100);c__Saccharomycetes(100);o__Saccharomycetales(100);f__Metschnikowiaceae(100);g__Metschnikowia(100);s__Metschnikowia_pulcherrima(100); | Metschnikowia | 13 | Saprotroph | \|Undefined Saprotroph\| | Yeast | NULL | Probable | As nectar-tap_saprotroph (Põlme et al. 2020); Decay sugar-rich substrates (Põlme et al. 2020) | Põlme S, et al. 2020. Fungal Diversity 105:1-16 ((https://doi.org/10.1007/s13225-020-00466-2)) |
| Otu34274 | 1 | k__Fungi(100);p__Ascomycota(100);c__Saccharomycetes(100);o__Saccharomycetales(100);f__Metschnikowiaceae(100);g__Metschnikowia(100);s__Metschnikowia_pulcherrima(100); | Metschnikowia | 13 | Saprotroph | \|Undefined Saprotroph\| | Yeast | NULL | Probable | As nectar-tap_saprotroph (Põlme et al. 2020); Decay sugar-rich substrates (Põlme et al. 2020) | Põlme S, et al. 2020. Fungal Diversity 105:1-16 ((https://doi.org/10.1007/s13225-020-00466-2)) |
| Otu34275 | 1 | k__Fungi(100);p__Ascomycota(100);c__Saccharomycetes(100);o__Saccharomycetales(100);f__Metschnikowiaceae(100);g__Metschnikowia(100);s__Metschnikowia_pulcherrima(100); | Metschnikowia | 13 | Saprotroph | \|Undefined Saprotroph\| | Yeast | NULL | Probable | As nectar-tap_saprotroph (Põlme et al. 2020); Decay sugar-rich substrates (Põlme et al. 2020) | Põlme S, et al. 2020. Fungal Diversity 105:1-16 ((https://doi.org/10.1007/s13225-020-00466-2)) |
| Otu34276 | 1 | k__Fungi(100);p__Ascomycota(100);c__Saccharomycetes(100);o__Saccharomycetales(100);f__Metschnikowiaceae(100);g__Metschnikowia(100);s__Metschnikowia_pulcherrima(100); | Metschnikowia | 13 | Saprotroph | \|Undefined Saprotroph\| | Yeast | NULL | Probable | As nectar-tap_saprotroph (Põlme et al. 2020); Decay sugar-rich substrates (Põlme et al. 2020) | Põlme S, et al. 2020. Fungal Diversity 105:1-16 ((https://doi.org/10.1007/s13225-020-00466-2)) |
| Otu34278 | 1 | k__Fungi(100);p__Ascomycota(100);c__Saccharomycetes(100);o__Saccharomycetales(100);f__Metschnikowiaceae(100);g__Metschnikowia(100);s__Metschnikowia_pulcherrima(100); | Metschnikowia | 13 | Saprotroph | \|Undefined Saprotroph\| | Yeast | NULL | Probable | As nectar-tap_saprotroph (Põlme et al. 2020); Decay sugar-rich substrates (Põlme et al. 2020) | Põlme S, et al. 2020. Fungal Diversity 105:1-16 ((https://doi.org/10.1007/s13225-020-00466-2)) |
| Otu34279 | 1 | k__Fungi(100);p__Ascomycota(100);c__Saccharomycetes(100);o__Saccharomycetales(100);f__Metschnikowiaceae(100);g__Metschnikowia(100);s__Metschnikowia_pulcherrima(100); | Metschnikowia | 13 | Saprotroph | \|Undefined Saprotroph\| | Yeast | NULL | Probable | As nectar-tap_saprotroph (Põlme et al. 2020); Decay sugar-rich substrates (Põlme et al. 2020) | Põlme S, et al. 2020. Fungal Diversity 105:1-16 ((https://doi.org/10.1007/s13225-020-00466-2)) |
| Otu34280 | 1 | k__Fungi(100);p__Ascomycota(100);c__Saccharomycetes(100);o__Saccharomycetales(100);f__Metschnikowiaceae(100);g__Metschnikowia(100);s__Metschnikowia_pulcherrima(100); | Metschnikowia | 13 | Saprotroph | \|Undefined Saprotroph\| | Yeast | NULL | Probable | As nectar-tap_saprotroph (Põlme et al. 2020); Decay sugar-rich substrates (Põlme et al. 2020) | Põlme S, et al. 2020. Fungal Diversity 105:1-16 ((https://doi.org/10.1007/s13225-020-00466-2)) |
| Otu34281 | 1 | k__Fungi(100);p__Ascomycota(100);c__Saccharomycetes(100);o__Saccharomycetales(100);f__Metschnikowiaceae(100);g__Metschnikowia(100);g__Metschnikowia_unclassified(100); | Metschnikowia | 13 | Saprotroph | \|Undefined Saprotroph\| | Yeast | NULL | Probable | As nectar-tap_saprotroph (Põlme et al. 2020); Decay sugar-rich substrates (Põlme et al. 2020) | Põlme S, et al. 2020. Fungal Diversity 105:1-16 ((https://doi.org/10.1007/s13225-020-00466-2)) |
| Otu34282 | 1 | k__Fungi(100);p__Ascomycota(100);c__Saccharomycetes(100);o__Saccharomycetales(100);f__Metschnikowiaceae(100);g__Metschnikowia(100);s__Metschnikowia_pulcherrima(100); | Metschnikowia | 13 | Saprotroph | \|Undefined Saprotroph\| | Yeast | NULL | Probable | As nectar-tap_saprotroph (Põlme et al. 2020); Decay sugar-rich substrates (Põlme et al. 2020) | Põlme S, et al. 2020. Fungal Diversity 105:1-16 ((https://doi.org/10.1007/s13225-020-00466-2)) |
| Otu34284 | 1 | k__Fungi(100);p__Ascomycota(100);c__Saccharomycetes(100);o__Saccharomycetales(100);f__Metschnikowiaceae(100);g__Metschnikowia(100);g__Metschnikowia_unclassified(100); | Metschnikowia | 13 | Saprotroph | \|Undefined Saprotroph\| | Yeast | NULL | Probable | As nectar-tap_saprotroph (Põlme et al. 2020); Decay sugar-rich substrates (Põlme et al. 2020) | Põlme S, et al. 2020. Fungal Diversity 105:1-16 ((https://doi.org/10.1007/s13225-020-00466-2)) |
| Otu34285 | 1 | k__Fungi(100);p__Ascomycota(100);c__Saccharomycetes(100);o__Saccharomycetales(100);f__Metschnikowiaceae(100);g__Metschnikowia(100);s__Metschnikowia_pulcherrima(100); | Metschnikowia | 13 | Saprotroph | \|Undefined Saprotroph\| | Yeast | NULL | Probable | As nectar-tap_saprotroph (Põlme et al. 2020); Decay sugar-rich substrates (Põlme et al. 2020) | Põlme S, et al. 2020. Fungal Diversity 105:1-16 ((https://doi.org/10.1007/s13225-020-00466-2)) |
| Otu34286 | 1 | k__Fungi(100);p__Ascomycota(100);c__Saccharomycetes(100);o__Saccharomycetales(100);f__Metschnikowiaceae(100);g__Metschnikowia(100);g__Metschnikowia_unclassified(100); | Metschnikowia | 13 | Saprotroph | \|Undefined Saprotroph\| | Yeast | NULL | Probable | As nectar-tap_saprotroph (Põlme et al. 2020); Decay sugar-rich substrates (Põlme et al. 2020) | Põlme S, et al. 2020. Fungal Diversity 105:1-16 ((https://doi.org/10.1007/s13225-020-00466-2)) |
| Otu34287 | 1 | k__Fungi(100);p__Ascomycota(100);c__Saccharomycetes(100);o__Saccharomycetales(100);f__Metschnikowiaceae(100);g__Metschnikowia(100);g__Metschnikowia_unclassified(100); | Metschnikowia | 13 | Saprotroph | \|Undefined Saprotroph\| | Yeast | NULL | Probable | As nectar-tap_saprotroph (Põlme et al. 2020); Decay sugar-rich substrates (Põlme et al. 2020) | Põlme S, et al. 2020. Fungal Diversity 105:1-16 ((https://doi.org/10.1007/s13225-020-00466-2)) |
| Otu34288 | 1 | k__Fungi(100);p__Ascomycota(100);c__Saccharomycetes(100);o__Saccharomycetales(100);f__Metschnikowiaceae(100);g__Metschnikowia(100);s__Metschnikowia_pulcherrima(100); | Metschnikowia | 13 | Saprotroph | \|Undefined Saprotroph\| | Yeast | NULL | Probable | As nectar-tap_saprotroph (Põlme et al. 2020); Decay sugar-rich substrates (Põlme et al. 2020) | Põlme S, et al. 2020. Fungal Diversity 105:1-16 ((https://doi.org/10.1007/s13225-020-00466-2)) |
| Otu34289 | 1 | k__Fungi(100);p__Ascomycota(100);c__Saccharomycetes(100);o__Saccharomycetales(100);f__Metschnikowiaceae(100);g__Metschnikowia(100);s__Metschnikowia_pulcherrima(100); | Metschnikowia | 13 | Saprotroph | \|Undefined Saprotroph\| | Yeast | NULL | Probable | As nectar-tap_saprotroph (Põlme et al. 2020); Decay sugar-rich substrates (Põlme et al. 2020) | Põlme S, et al. 2020. Fungal Diversity 105:1-16 ((https://doi.org/10.1007/s13225-020-00466-2)) |
| Otu34290 | 1 | k__Fungi(100);p__Ascomycota(100);c__Saccharomycetes(100);o__Saccharomycetales(100);f__Metschnikowiaceae(100);g__Metschnikowia(100);g__Metschnikowia_unclassified(100); | Metschnikowia | 13 | Saprotroph | \|Undefined Saprotroph\| | Yeast | NULL | Probable | As nectar-tap_saprotroph (Põlme et al. 2020); Decay sugar-rich substrates (Põlme et al. 2020) | Põlme S, et al. 2020. Fungal Diversity 105:1-16 ((https://doi.org/10.1007/s13225-020-00466-2)) |
| Otu34291 | 1 | k__Fungi(100);p__Ascomycota(100);c__Saccharomycetes(100);o__Saccharomycetales(100);f__Metschnikowiaceae(100);g__Metschnikowia(100);g__Metschnikowia_unclassified(100); | Metschnikowia | 13 | Saprotroph | \|Undefined Saprotroph\| | Yeast | NULL | Probable | As nectar-tap_saprotroph (Põlme et al. 2020); Decay sugar-rich substrates (Põlme et al. 2020) | Põlme S, et al. 2020. Fungal Diversity 105:1-16 ((https://doi.org/10.1007/s13225-020-00466-2)) |
| Otu34306 | 1 | k__Fungi(100);p__Ascomycota(100);c__Saccharomycetes(100);o__Saccharomycetales(100);f__Metschnikowiaceae(100);g__Metschnikowia(100);s__Metschnikowia_pulcherrima(100); | Metschnikowia | 13 | Saprotroph | \|Undefined Saprotroph\| | Yeast | NULL | Probable | As nectar-tap_saprotroph (Põlme et al. 2020); Decay sugar-rich substrates (Põlme et al. 2020) | Põlme S, et al. 2020. Fungal Diversity 105:1-16 ((https://doi.org/10.1007/s13225-020-00466-2)) |
| Otu34308 | 1 | k__Fungi(100);p__Ascomycota(100);c__Saccharomycetes(100);o__Saccharomycetales(100);f__Metschnikowiaceae(100);g__Metschnikowia(100);s__Metschnikowia_pulcherrima(100); | Metschnikowia | 13 | Saprotroph | \|Undefined Saprotroph\| | Yeast | NULL | Probable | As nectar-tap_saprotroph (Põlme et al. 2020); Decay sugar-rich substrates (Põlme et al. 2020) | Põlme S, et al. 2020. Fungal Diversity 105:1-16 ((https://doi.org/10.1007/s13225-020-00466-2)) |
| Otu34309 | 1 | k__Fungi(100);p__Ascomycota(100);c__Saccharomycetes(100);o__Saccharomycetales(100);f__Metschnikowiaceae(100);g__Metschnikowia(100);g__Metschnikowia_unclassified(100); | Metschnikowia | 13 | Saprotroph | \|Undefined Saprotroph\| | Yeast | NULL | Probable | As nectar-tap_saprotroph (Põlme et al. 2020); Decay sugar-rich substrates (Põlme et al. 2020) | Põlme S, et al. 2020. Fungal Diversity 105:1-16 ((https://doi.org/10.1007/s13225-020-00466-2)) |
| Otu34310 | 1 | k__Fungi(100);p__Ascomycota(100);c__Saccharomycetes(100);o__Saccharomycetales(100);f__Metschnikowiaceae(100);g__Metschnikowia(100);s__Metschnikowia_pulcherrima(100); | Metschnikowia | 13 | Saprotroph | \|Undefined Saprotroph\| | Yeast | NULL | Probable | As nectar-tap_saprotroph (Põlme et al. 2020); Decay sugar-rich substrates (Põlme et al. 2020) | Põlme S, et al. 2020. Fungal Diversity 105:1-16 ((https://doi.org/10.1007/s13225-020-00466-2)) |
| Otu34313 | 1 | k__Fungi(100);p__Ascomycota(100);c__Saccharomycetes(100);o__Saccharomycetales(100);f__Metschnikowiaceae(100);g__Metschnikowia(100);s__Metschnikowia_pulcherrima(100); | Metschnikowia | 13 | Saprotroph | \|Undefined Saprotroph\| | Yeast | NULL | Probable | As nectar-tap_saprotroph (Põlme et al. 2020); Decay sugar-rich substrates (Põlme et al. 2020) | Põlme S, et al. 2020. Fungal Diversity 105:1-16 ((https://doi.org/10.1007/s13225-020-00466-2)) |
| Otu34315 | 1 | k__Fungi(100);p__Ascomycota(100);c__Saccharomycetes(100);o__Saccharomycetales(100);f__Metschnikowiaceae(100);g__Metschnikowia(100);g__Metschnikowia_unclassified(100); | Metschnikowia | 13 | Saprotroph | \|Undefined Saprotroph\| | Yeast | NULL | Probable | As nectar-tap_saprotroph (Põlme et al. 2020); Decay sugar-rich substrates (Põlme et al. 2020) | Põlme S, et al. 2020. Fungal Diversity 105:1-16 ((https://doi.org/10.1007/s13225-020-00466-2)) |
| Otu34316 | 1 | k__Fungi(100);p__Ascomycota(100);c__Saccharomycetes(100);o__Saccharomycetales(100);f__Metschnikowiaceae(100);g__Metschnikowia(100);s__Metschnikowia_pulcherrima(100); | Metschnikowia | 13 | Saprotroph | \|Undefined Saprotroph\| | Yeast | NULL | Probable | As nectar-tap_saprotroph (Põlme et al. 2020); Decay sugar-rich substrates (Põlme et al. 2020) | Põlme S, et al. 2020. Fungal Diversity 105:1-16 ((https://doi.org/10.1007/s13225-020-00466-2)) |
| Otu34317 | 1 | k__Fungi(100);p__Ascomycota(100);c__Saccharomycetes(100);o__Saccharomycetales(100);f__Metschnikowiaceae(100);g__Metschnikowia(100);g__Metschnikowia_unclassified(100); | Metschnikowia | 13 | Saprotroph | \|Undefined Saprotroph\| | Yeast | NULL | Probable | As nectar-tap_saprotroph (Põlme et al. 2020); Decay sugar-rich substrates (Põlme et al. 2020) | Põlme S, et al. 2020. Fungal Diversity 105:1-16 ((https://doi.org/10.1007/s13225-020-00466-2)) |
| Otu34324 | 1 | k__Fungi(100);p__Ascomycota(100);c__Saccharomycetes(100);o__Saccharomycetales(100);f__Metschnikowiaceae(100);g__Metschnikowia(100);s__Metschnikowia_pulcherrima(100); | Metschnikowia | 13 | Saprotroph | \|Undefined Saprotroph\| | Yeast | NULL | Probable | As nectar-tap_saprotroph (Põlme et al. 2020); Decay sugar-rich substrates (Põlme et al. 2020) | Põlme S, et al. 2020. Fungal Diversity 105:1-16 ((https://doi.org/10.1007/s13225-020-00466-2)) |
| Otu34325 | 1 | k__Fungi(100);p__Ascomycota(100);c__Saccharomycetes(100);o__Saccharomycetales(100);f__Metschnikowiaceae(100);g__Metschnikowia(100);s__Metschnikowia_pulcherrima(100); | Metschnikowia | 13 | Saprotroph | \|Undefined Saprotroph\| | Yeast | NULL | Probable | As nectar-tap_saprotroph (Põlme et al. 2020); Decay sugar-rich substrates (Põlme et al. 2020) | Põlme S, et al. 2020. Fungal Diversity 105:1-16 ((https://doi.org/10.1007/s13225-020-00466-2)) |
| Otu34326 | 1 | k__Fungi(100);p__Ascomycota(100);c__Saccharomycetes(100);o__Saccharomycetales(100);f__Metschnikowiaceae(100);g__Metschnikowia(100);s__Metschnikowia_pulcherrima(100); | Metschnikowia | 13 | Saprotroph | \|Undefined Saprotroph\| | Yeast | NULL | Probable | As nectar-tap_saprotroph (Põlme et al. 2020); Decay sugar-rich substrates (Põlme et al. 2020) | Põlme S, et al. 2020. Fungal Diversity 105:1-16 ((https://doi.org/10.1007/s13225-020-00466-2)) |
| Otu34327 | 1 | k__Fungi(100);p__Ascomycota(100);c__Saccharomycetes(100);o__Saccharomycetales(100);f__Metschnikowiaceae(100);g__Metschnikowia(100);s__Metschnikowia_pulcherrima(100); | Metschnikowia | 13 | Saprotroph | \|Undefined Saprotroph\| | Yeast | NULL | Probable | As nectar-tap_saprotroph (Põlme et al. 2020); Decay sugar-rich substrates (Põlme et al. 2020) | Põlme S, et al. 2020. Fungal Diversity 105:1-16 ((https://doi.org/10.1007/s13225-020-00466-2)) |
| Otu34328 | 1 | k__Fungi(100);p__Ascomycota(100);c__Saccharomycetes(100);o__Saccharomycetales(100);f__Metschnikowiaceae(100);g__Metschnikowia(100);s__Metschnikowia_pulcherrima(100); | Metschnikowia | 13 | Saprotroph | \|Undefined Saprotroph\| | Yeast | NULL | Probable | As nectar-tap_saprotroph (Põlme et al. 2020); Decay sugar-rich substrates (Põlme et al. 2020) | Põlme S, et al. 2020. Fungal Diversity 105:1-16 ((https://doi.org/10.1007/s13225-020-00466-2)) |
| Otu34329 | 1 | k__Fungi(100);p__Ascomycota(100);c__Saccharomycetes(100);o__Saccharomycetales(100);f__Metschnikowiaceae(100);g__Metschnikowia(100);s__Metschnikowia_pulcherrima(100); | Metschnikowia | 13 | Saprotroph | \|Undefined Saprotroph\| | Yeast | NULL | Probable | As nectar-tap_saprotroph (Põlme et al. 2020); Decay sugar-rich substrates (Põlme et al. 2020) | Põlme S, et al. 2020. Fungal Diversity 105:1-16 ((https://doi.org/10.1007/s13225-020-00466-2)) |
| Otu34330 | 1 | k__Fungi(100);p__Ascomycota(100);c__Saccharomycetes(100);o__Saccharomycetales(100);f__Metschnikowiaceae(100);g__Metschnikowia(100);s__Metschnikowia_pulcherrima(100); | Metschnikowia | 13 | Saprotroph | \|Undefined Saprotroph\| | Yeast | NULL | Probable | As nectar-tap_saprotroph (Põlme et al. 2020); Decay sugar-rich substrates (Põlme et al. 2020) | Põlme S, et al. 2020. Fungal Diversity 105:1-16 ((https://doi.org/10.1007/s13225-020-00466-2)) |
| Otu34334 | 1 | k__Fungi(100);p__Ascomycota(100);c__Saccharomycetes(100);o__Saccharomycetales(100);f__Metschnikowiaceae(100);g__Metschnikowia(100);s__Metschnikowia_pulcherrima(100); | Metschnikowia | 13 | Saprotroph | \|Undefined Saprotroph\| | Yeast | NULL | Probable | As nectar-tap_saprotroph (Põlme et al. 2020); Decay sugar-rich substrates (Põlme et al. 2020) | Põlme S, et al. 2020. Fungal Diversity 105:1-16 ((https://doi.org/10.1007/s13225-020-00466-2)) |
| Otu34338 | 1 | k__Fungi(100);p__Ascomycota(100);c__Saccharomycetes(100);o__Saccharomycetales(100);f__Metschnikowiaceae(100);g__Metschnikowia(100);s__Metschnikowia_pulcherrima(100); | Metschnikowia | 13 | Saprotroph | \|Undefined Saprotroph\| | Yeast | NULL | Probable | As nectar-tap_saprotroph (Põlme et al. 2020); Decay sugar-rich substrates (Põlme et al. 2020) | Põlme S, et al. 2020. Fungal Diversity 105:1-16 ((https://doi.org/10.1007/s13225-020-00466-2)) |
| Otu34340 | 1 | k__Fungi(100);p__Ascomycota(100);c__Saccharomycetes(100);o__Saccharomycetales(100);f__Metschnikowiaceae(100);g__Metschnikowia(100);s__Metschnikowia_pulcherrima(100); | Metschnikowia | 13 | Saprotroph | \|Undefined Saprotroph\| | Yeast | NULL | Probable | As nectar-tap_saprotroph (Põlme et al. 2020); Decay sugar-rich substrates (Põlme et al. 2020) | Põlme S, et al. 2020. Fungal Diversity 105:1-16 ((https://doi.org/10.1007/s13225-020-00466-2)) |
| Otu34341 | 1 | k__Fungi(100);p__Ascomycota(100);c__Saccharomycetes(100);o__Saccharomycetales(100);f__Metschnikowiaceae(100);g__Metschnikowia(100);s__Metschnikowia_pulcherrima(100); | Metschnikowia | 13 | Saprotroph | \|Undefined Saprotroph\| | Yeast | NULL | Probable | As nectar-tap_saprotroph (Põlme et al. 2020); Decay sugar-rich substrates (Põlme et al. 2020) | Põlme S, et al. 2020. Fungal Diversity 105:1-16 ((https://doi.org/10.1007/s13225-020-00466-2)) |
| Otu34342 | 1 | k__Fungi(100);p__Ascomycota(100);c__Saccharomycetes(100);o__Saccharomycetales(100);f__Metschnikowiaceae(100);g__Metschnikowia(100);s__Metschnikowia_pulcherrima(100); | Metschnikowia | 13 | Saprotroph | \|Undefined Saprotroph\| | Yeast | NULL | Probable | As nectar-tap_saprotroph (Põlme et al. 2020); Decay sugar-rich substrates (Põlme et al. 2020) | Põlme S, et al. 2020. Fungal Diversity 105:1-16 ((https://doi.org/10.1007/s13225-020-00466-2)) |
| Otu34344 | 1 | k__Fungi(100);p__Ascomycota(100);c__Saccharomycetes(100);o__Saccharomycetales(100);f__Metschnikowiaceae(100);g__Metschnikowia(100);g__Metschnikowia_unclassified(100); | Metschnikowia | 13 | Saprotroph | \|Undefined Saprotroph\| | Yeast | NULL | Probable | As nectar-tap_saprotroph (Põlme et al. 2020); Decay sugar-rich substrates (Põlme et al. 2020) | Põlme S, et al. 2020. Fungal Diversity 105:1-16 ((https://doi.org/10.1007/s13225-020-00466-2)) |
| Otu34345 | 1 | k__Fungi(100);p__Ascomycota(100);c__Saccharomycetes(100);o__Saccharomycetales(100);f__Metschnikowiaceae(100);g__Metschnikowia(100);s__Metschnikowia_pulcherrima(100); | Metschnikowia | 13 | Saprotroph | \|Undefined Saprotroph\| | Yeast | NULL | Probable | As nectar-tap_saprotroph (Põlme et al. 2020); Decay sugar-rich substrates (Põlme et al. 2020) | Põlme S, et al. 2020. Fungal Diversity 105:1-16 ((https://doi.org/10.1007/s13225-020-00466-2)) |
| Otu34346 | 1 | k__Fungi(100);p__Ascomycota(100);c__Saccharomycetes(100);o__Saccharomycetales(100);f__Metschnikowiaceae(100);g__Metschnikowia(100);s__Metschnikowia_pulcherrima(100); | Metschnikowia | 13 | Saprotroph | \|Undefined Saprotroph\| | Yeast | NULL | Probable | As nectar-tap_saprotroph (Põlme et al. 2020); Decay sugar-rich substrates (Põlme et al. 2020) | Põlme S, et al. 2020. Fungal Diversity 105:1-16 ((https://doi.org/10.1007/s13225-020-00466-2)) |
| Otu34347 | 1 | k__Fungi(100);p__Ascomycota(100);c__Saccharomycetes(100);o__Saccharomycetales(100);f__Metschnikowiaceae(100);g__Metschnikowia(100);s__Metschnikowia_pulcherrima(100); | Metschnikowia | 13 | Saprotroph | \|Undefined Saprotroph\| | Yeast | NULL | Probable | As nectar-tap_saprotroph (Põlme et al. 2020); Decay sugar-rich substrates (Põlme et al. 2020) | Põlme S, et al. 2020. Fungal Diversity 105:1-16 ((https://doi.org/10.1007/s13225-020-00466-2)) |
| Otu34349 | 1 | k__Fungi(100);p__Ascomycota(100);c__Saccharomycetes(100);o__Saccharomycetales(100);f__Metschnikowiaceae(100);g__Metschnikowia(100);s__Metschnikowia_pulcherrima(100); | Metschnikowia | 13 | Saprotroph | \|Undefined Saprotroph\| | Yeast | NULL | Probable | As nectar-tap_saprotroph (Põlme et al. 2020); Decay sugar-rich substrates (Põlme et al. 2020) | Põlme S, et al. 2020. Fungal Diversity 105:1-16 ((https://doi.org/10.1007/s13225-020-00466-2)) |
| Otu34351 | 1 | k__Fungi(100);p__Ascomycota(100);c__Saccharomycetes(100);o__Saccharomycetales(100);f__Metschnikowiaceae(100);g__Metschnikowia(100);g__Metschnikowia_unclassified(100); | Metschnikowia | 13 | Saprotroph | \|Undefined Saprotroph\| | Yeast | NULL | Probable | As nectar-tap_saprotroph (Põlme et al. 2020); Decay sugar-rich substrates (Põlme et al. 2020) | Põlme S, et al. 2020. Fungal Diversity 105:1-16 ((https://doi.org/10.1007/s13225-020-00466-2)) |
| Otu34352 | 1 | k__Fungi(100);p__Ascomycota(100);c__Saccharomycetes(100);o__Saccharomycetales(100);f__Metschnikowiaceae(100);g__Metschnikowia(100);g__Metschnikowia_unclassified(100); | Metschnikowia | 13 | Saprotroph | \|Undefined Saprotroph\| | Yeast | NULL | Probable | As nectar-tap_saprotroph (Põlme et al. 2020); Decay sugar-rich substrates (Põlme et al. 2020) | Põlme S, et al. 2020. Fungal Diversity 105:1-16 ((https://doi.org/10.1007/s13225-020-00466-2)) |
| Otu34353 | 1 | k__Fungi(100);p__Ascomycota(100);c__Saccharomycetes(100);o__Saccharomycetales(100);f__Metschnikowiaceae(100);g__Metschnikowia(100);s__Metschnikowia_pulcherrima(100); | Metschnikowia | 13 | Saprotroph | \|Undefined Saprotroph\| | Yeast | NULL | Probable | As nectar-tap_saprotroph (Põlme et al. 2020); Decay sugar-rich substrates (Põlme et al. 2020) | Põlme S, et al. 2020. Fungal Diversity 105:1-16 ((https://doi.org/10.1007/s13225-020-00466-2)) |
| Otu34354 | 1 | k__Fungi(100);p__Ascomycota(100);c__Saccharomycetes(100);o__Saccharomycetales(100);f__Metschnikowiaceae(100);g__Metschnikowia(100);s__Metschnikowia_pulcherrima(100); | Metschnikowia | 13 | Saprotroph | \|Undefined Saprotroph\| | Yeast | NULL | Probable | As nectar-tap_saprotroph (Põlme et al. 2020); Decay sugar-rich substrates (Põlme et al. 2020) | Põlme S, et al. 2020. Fungal Diversity 105:1-16 ((https://doi.org/10.1007/s13225-020-00466-2)) |
| Otu34355 | 1 | k__Fungi(100);p__Ascomycota(100);c__Saccharomycetes(100);o__Saccharomycetales(100);f__Metschnikowiaceae(100);g__Metschnikowia(100);g__Metschnikowia_unclassified(100); | Metschnikowia | 13 | Saprotroph | \|Undefined Saprotroph\| | Yeast | NULL | Probable | As nectar-tap_saprotroph (Põlme et al. 2020); Decay sugar-rich substrates (Põlme et al. 2020) | Põlme S, et al. 2020. Fungal Diversity 105:1-16 ((https://doi.org/10.1007/s13225-020-00466-2)) |
| Otu34356 | 1 | k__Fungi(100);p__Ascomycota(100);c__Saccharomycetes(100);o__Saccharomycetales(100);f__Metschnikowiaceae(100);g__Metschnikowia(100);g__Metschnikowia_unclassified(100); | Metschnikowia | 13 | Saprotroph | \|Undefined Saprotroph\| | Yeast | NULL | Probable | As nectar-tap_saprotroph (Põlme et al. 2020); Decay sugar-rich substrates (Põlme et al. 2020) | Põlme S, et al. 2020. Fungal Diversity 105:1-16 ((https://doi.org/10.1007/s13225-020-00466-2)) |
| Otu34358 | 1 | k__Fungi(100);p__Ascomycota(100);c__Saccharomycetes(100);o__Saccharomycetales(100);f__Metschnikowiaceae(100);g__Metschnikowia(100);s__Metschnikowia_pulcherrima(100); | Metschnikowia | 13 | Saprotroph | \|Undefined Saprotroph\| | Yeast | NULL | Probable | As nectar-tap_saprotroph (Põlme et al. 2020); Decay sugar-rich substrates (Põlme et al. 2020) | Põlme S, et al. 2020. Fungal Diversity 105:1-16 ((https://doi.org/10.1007/s13225-020-00466-2)) |
| Otu34359 | 1 | k__Fungi(100);p__Ascomycota(100);c__Saccharomycetes(100);o__Saccharomycetales(100);f__Metschnikowiaceae(100);g__Metschnikowia(100);g__Metschnikowia_unclassified(100); | Metschnikowia | 13 | Saprotroph | \|Undefined Saprotroph\| | Yeast | NULL | Probable | As nectar-tap_saprotroph (Põlme et al. 2020); Decay sugar-rich substrates (Põlme et al. 2020) | Põlme S, et al. 2020. Fungal Diversity 105:1-16 ((https://doi.org/10.1007/s13225-020-00466-2)) |
| Otu34360 | 1 | k__Fungi(100);p__Ascomycota(100);c__Saccharomycetes(100);o__Saccharomycetales(100);f__Metschnikowiaceae(100);g__Metschnikowia(100);s__Metschnikowia_pulcherrima(100); | Metschnikowia | 13 | Saprotroph | \|Undefined Saprotroph\| | Yeast | NULL | Probable | As nectar-tap_saprotroph (Põlme et al. 2020); Decay sugar-rich substrates (Põlme et al. 2020) | Põlme S, et al. 2020. Fungal Diversity 105:1-16 ((https://doi.org/10.1007/s13225-020-00466-2)) |
| Otu34361 | 1 | k__Fungi(100);p__Ascomycota(100);c__Saccharomycetes(100);o__Saccharomycetales(100);f__Metschnikowiaceae(100);g__Metschnikowia(100);g__Metschnikowia_unclassified(100); | Metschnikowia | 13 | Saprotroph | \|Undefined Saprotroph\| | Yeast | NULL | Probable | As nectar-tap_saprotroph (Põlme et al. 2020); Decay sugar-rich substrates (Põlme et al. 2020) | Põlme S, et al. 2020. Fungal Diversity 105:1-16 ((https://doi.org/10.1007/s13225-020-00466-2)) |
| Otu34363 | 1 | k__Fungi(100);p__Ascomycota(100);c__Saccharomycetes(100);o__Saccharomycetales(100);f__Metschnikowiaceae(100);g__Metschnikowia(100);g__Metschnikowia_unclassified(100); | Metschnikowia | 13 | Saprotroph | \|Undefined Saprotroph\| | Yeast | NULL | Probable | As nectar-tap_saprotroph (Põlme et al. 2020); Decay sugar-rich substrates (Põlme et al. 2020) | Põlme S, et al. 2020. Fungal Diversity 105:1-16 ((https://doi.org/10.1007/s13225-020-00466-2)) |
| Otu34365 | 1 | k__Fungi(100);p__Ascomycota(100);c__Saccharomycetes(100);o__Saccharomycetales(100);f__Metschnikowiaceae(100);g__Metschnikowia(100);s__Metschnikowia_pulcherrima(100); | Metschnikowia | 13 | Saprotroph | \|Undefined Saprotroph\| | Yeast | NULL | Probable | As nectar-tap_saprotroph (Põlme et al. 2020); Decay sugar-rich substrates (Põlme et al. 2020) | Põlme S, et al. 2020. Fungal Diversity 105:1-16 ((https://doi.org/10.1007/s13225-020-00466-2)) |
| Otu34366 | 1 | k__Fungi(100);p__Ascomycota(100);c__Saccharomycetes(100);o__Saccharomycetales(100);f__Metschnikowiaceae(100);g__Metschnikowia(100);s__Metschnikowia_pulcherrima(100); | Metschnikowia | 13 | Saprotroph | \|Undefined Saprotroph\| | Yeast | NULL | Probable | As nectar-tap_saprotroph (Põlme et al. 2020); Decay sugar-rich substrates (Põlme et al. 2020) | Põlme S, et al. 2020. Fungal Diversity 105:1-16 ((https://doi.org/10.1007/s13225-020-00466-2)) |
| Otu34367 | 1 | k__Fungi(100);p__Ascomycota(100);c__Saccharomycetes(100);o__Saccharomycetales(100);f__Metschnikowiaceae(100);g__Metschnikowia(100);g__Metschnikowia_unclassified(100); | Metschnikowia | 13 | Saprotroph | \|Undefined Saprotroph\| | Yeast | NULL | Probable | As nectar-tap_saprotroph (Põlme et al. 2020); Decay sugar-rich substrates (Põlme et al. 2020) | Põlme S, et al. 2020. Fungal Diversity 105:1-16 ((https://doi.org/10.1007/s13225-020-00466-2)) |
| Otu34368 | 1 | k__Fungi(100);p__Ascomycota(100);c__Saccharomycetes(100);o__Saccharomycetales(100);f__Metschnikowiaceae(100);g__Metschnikowia(100);s__Metschnikowia_pulcherrima(100); | Metschnikowia | 13 | Saprotroph | \|Undefined Saprotroph\| | Yeast | NULL | Probable | As nectar-tap_saprotroph (Põlme et al. 2020); Decay sugar-rich substrates (Põlme et al. 2020) | Põlme S, et al. 2020. Fungal Diversity 105:1-16 ((https://doi.org/10.1007/s13225-020-00466-2)) |
| Otu34369 | 1 | k__Fungi(100);p__Ascomycota(100);c__Saccharomycetes(100);o__Saccharomycetales(100);f__Metschnikowiaceae(100);g__Metschnikowia(100);s__Metschnikowia_pulcherrima(100); | Metschnikowia | 13 | Saprotroph | \|Undefined Saprotroph\| | Yeast | NULL | Probable | As nectar-tap_saprotroph (Põlme et al. 2020); Decay sugar-rich substrates (Põlme et al. 2020) | Põlme S, et al. 2020. Fungal Diversity 105:1-16 ((https://doi.org/10.1007/s13225-020-00466-2)) |
| Otu34371 | 1 | k__Fungi(100);p__Ascomycota(100);c__Saccharomycetes(100);o__Saccharomycetales(100);f__Metschnikowiaceae(100);g__Metschnikowia(100);s__Metschnikowia_pulcherrima(100); | Metschnikowia | 13 | Saprotroph | \|Undefined Saprotroph\| | Yeast | NULL | Probable | As nectar-tap_saprotroph (Põlme et al. 2020); Decay sugar-rich substrates (Põlme et al. 2020) | Põlme S, et al. 2020. Fungal Diversity 105:1-16 ((https://doi.org/10.1007/s13225-020-00466-2)) |
| Otu34373 | 1 | k__Fungi(100);p__Ascomycota(100);c__Saccharomycetes(100);o__Saccharomycetales(100);f__Metschnikowiaceae(100);g__Metschnikowia(100);g__Metschnikowia_unclassified(100); | Metschnikowia | 13 | Saprotroph | \|Undefined Saprotroph\| | Yeast | NULL | Probable | As nectar-tap_saprotroph (Põlme et al. 2020); Decay sugar-rich substrates (Põlme et al. 2020) | Põlme S, et al. 2020. Fungal Diversity 105:1-16 ((https://doi.org/10.1007/s13225-020-00466-2)) |
| Otu34374 | 1 | k__Fungi(100);p__Ascomycota(100);c__Saccharomycetes(100);o__Saccharomycetales(100);f__Metschnikowiaceae(100);g__Metschnikowia(100);s__Metschnikowia_pulcherrima(100); | Metschnikowia | 13 | Saprotroph | \|Undefined Saprotroph\| | Yeast | NULL | Probable | As nectar-tap_saprotroph (Põlme et al. 2020); Decay sugar-rich substrates (Põlme et al. 2020) | Põlme S, et al. 2020. Fungal Diversity 105:1-16 ((https://doi.org/10.1007/s13225-020-00466-2)) |
| Otu34375 | 1 | k__Fungi(100);p__Ascomycota(100);c__Saccharomycetes(100);o__Saccharomycetales(100);f__Metschnikowiaceae(100);g__Metschnikowia(100);s__Metschnikowia_pulcherrima(100); | Metschnikowia | 13 | Saprotroph | \|Undefined Saprotroph\| | Yeast | NULL | Probable | As nectar-tap_saprotroph (Põlme et al. 2020); Decay sugar-rich substrates (Põlme et al. 2020) | Põlme S, et al. 2020. Fungal Diversity 105:1-16 ((https://doi.org/10.1007/s13225-020-00466-2)) |
| Otu34376 | 1 | k__Fungi(100);p__Ascomycota(100);c__Saccharomycetes(100);o__Saccharomycetales(100);f__Metschnikowiaceae(100);g__Metschnikowia(100);s__Metschnikowia_pulcherrima(100); | Metschnikowia | 13 | Saprotroph | \|Undefined Saprotroph\| | Yeast | NULL | Probable | As nectar-tap_saprotroph (Põlme et al. 2020); Decay sugar-rich substrates (Põlme et al. 2020) | Põlme S, et al. 2020. Fungal Diversity 105:1-16 ((https://doi.org/10.1007/s13225-020-00466-2)) |
| Otu34377 | 1 | k__Fungi(100);p__Ascomycota(100);c__Saccharomycetes(100);o__Saccharomycetales(100);f__Metschnikowiaceae(100);g__Metschnikowia(100);g__Metschnikowia_unclassified(100); | Metschnikowia | 13 | Saprotroph | \|Undefined Saprotroph\| | Yeast | NULL | Probable | As nectar-tap_saprotroph (Põlme et al. 2020); Decay sugar-rich substrates (Põlme et al. 2020) | Põlme S, et al. 2020. Fungal Diversity 105:1-16 ((https://doi.org/10.1007/s13225-020-00466-2)) |
| Otu34378 | 1 | k__Fungi(100);p__Ascomycota(100);c__Saccharomycetes(100);o__Saccharomycetales(100);f__Metschnikowiaceae(100);g__Metschnikowia(100);s__Metschnikowia_pulcherrima(100); | Metschnikowia | 13 | Saprotroph | \|Undefined Saprotroph\| | Yeast | NULL | Probable | As nectar-tap_saprotroph (Põlme et al. 2020); Decay sugar-rich substrates (Põlme et al. 2020) | Põlme S, et al. 2020. Fungal Diversity 105:1-16 ((https://doi.org/10.1007/s13225-020-00466-2)) |
| Otu34379 | 1 | k__Fungi(100);p__Ascomycota(100);c__Saccharomycetes(100);o__Saccharomycetales(100);f__Metschnikowiaceae(100);g__Metschnikowia(100);g__Metschnikowia_unclassified(100); | Metschnikowia | 13 | Saprotroph | \|Undefined Saprotroph\| | Yeast | NULL | Probable | As nectar-tap_saprotroph (Põlme et al. 2020); Decay sugar-rich substrates (Põlme et al. 2020) | Põlme S, et al. 2020. Fungal Diversity 105:1-16 ((https://doi.org/10.1007/s13225-020-00466-2)) |
| Otu34380 | 1 | k__Fungi(100);p__Ascomycota(100);c__Saccharomycetes(100);o__Saccharomycetales(100);f__Metschnikowiaceae(100);g__Metschnikowia(100);s__Metschnikowia_pulcherrima(100); | Metschnikowia | 13 | Saprotroph | \|Undefined Saprotroph\| | Yeast | NULL | Probable | As nectar-tap_saprotroph (Põlme et al. 2020); Decay sugar-rich substrates (Põlme et al. 2020) | Põlme S, et al. 2020. Fungal Diversity 105:1-16 ((https://doi.org/10.1007/s13225-020-00466-2)) |
| Otu34382 | 1 | k__Fungi(100);p__Ascomycota(100);c__Saccharomycetes(100);o__Saccharomycetales(100);f__Metschnikowiaceae(100);g__Metschnikowia(100);g__Metschnikowia_unclassified(100); | Metschnikowia | 13 | Saprotroph | \|Undefined Saprotroph\| | Yeast | NULL | Probable | As nectar-tap_saprotroph (Põlme et al. 2020); Decay sugar-rich substrates (Põlme et al. 2020) | Põlme S, et al. 2020. Fungal Diversity 105:1-16 ((https://doi.org/10.1007/s13225-020-00466-2)) |
| Otu34383 | 1 | k__Fungi(100);p__Ascomycota(100);c__Saccharomycetes(100);o__Saccharomycetales(100);f__Metschnikowiaceae(100);g__Metschnikowia(100);s__Metschnikowia_pulcherrima(100); | Metschnikowia | 13 | Saprotroph | \|Undefined Saprotroph\| | Yeast | NULL | Probable | As nectar-tap_saprotroph (Põlme et al. 2020); Decay sugar-rich substrates (Põlme et al. 2020) | Põlme S, et al. 2020. Fungal Diversity 105:1-16 ((https://doi.org/10.1007/s13225-020-00466-2)) |
| Otu34384 | 1 | k__Fungi(100);p__Ascomycota(100);c__Saccharomycetes(100);o__Saccharomycetales(100);f__Metschnikowiaceae(100);g__Metschnikowia(100);g__Metschnikowia_unclassified(100); | Metschnikowia | 13 | Saprotroph | \|Undefined Saprotroph\| | Yeast | NULL | Probable | As nectar-tap_saprotroph (Põlme et al. 2020); Decay sugar-rich substrates (Põlme et al. 2020) | Põlme S, et al. 2020. Fungal Diversity 105:1-16 ((https://doi.org/10.1007/s13225-020-00466-2)) |
| Otu34385 | 1 | k__Fungi(100);p__Ascomycota(100);c__Saccharomycetes(100);o__Saccharomycetales(100);f__Metschnikowiaceae(100);g__Metschnikowia(100);s__Metschnikowia_pulcherrima(100); | Metschnikowia | 13 | Saprotroph | \|Undefined Saprotroph\| | Yeast | NULL | Probable | As nectar-tap_saprotroph (Põlme et al. 2020); Decay sugar-rich substrates (Põlme et al. 2020) | Põlme S, et al. 2020. Fungal Diversity 105:1-16 ((https://doi.org/10.1007/s13225-020-00466-2)) |
| Otu34386 | 1 | k__Fungi(100);p__Ascomycota(100);c__Saccharomycetes(100);o__Saccharomycetales(100);f__Metschnikowiaceae(100);g__Metschnikowia(100);s__Metschnikowia_pulcherrima(100); | Metschnikowia | 13 | Saprotroph | \|Undefined Saprotroph\| | Yeast | NULL | Probable | As nectar-tap_saprotroph (Põlme et al. 2020); Decay sugar-rich substrates (Põlme et al. 2020) | Põlme S, et al. 2020. Fungal Diversity 105:1-16 ((https://doi.org/10.1007/s13225-020-00466-2)) |
| Otu34387 | 1 | k__Fungi(100);p__Ascomycota(100);c__Saccharomycetes(100);o__Saccharomycetales(100);f__Metschnikowiaceae(100);g__Metschnikowia(100);s__Metschnikowia_pulcherrima(100); | Metschnikowia | 13 | Saprotroph | \|Undefined Saprotroph\| | Yeast | NULL | Probable | As nectar-tap_saprotroph (Põlme et al. 2020); Decay sugar-rich substrates (Põlme et al. 2020) | Põlme S, et al. 2020. Fungal Diversity 105:1-16 ((https://doi.org/10.1007/s13225-020-00466-2)) |
| Otu34389 | 1 | k__Fungi(100);p__Ascomycota(100);c__Saccharomycetes(100);o__Saccharomycetales(100);f__Metschnikowiaceae(100);g__Metschnikowia(100);s__Metschnikowia_pulcherrima(100); | Metschnikowia | 13 | Saprotroph | \|Undefined Saprotroph\| | Yeast | NULL | Probable | As nectar-tap_saprotroph (Põlme et al. 2020); Decay sugar-rich substrates (Põlme et al. 2020) | Põlme S, et al. 2020. Fungal Diversity 105:1-16 ((https://doi.org/10.1007/s13225-020-00466-2)) |
| Otu34390 | 1 | k__Fungi(100);p__Ascomycota(100);c__Saccharomycetes(100);o__Saccharomycetales(100);f__Metschnikowiaceae(100);g__Metschnikowia(100);s__Metschnikowia_pulcherrima(100); | Metschnikowia | 13 | Saprotroph | \|Undefined Saprotroph\| | Yeast | NULL | Probable | As nectar-tap_saprotroph (Põlme et al. 2020); Decay sugar-rich substrates (Põlme et al. 2020) | Põlme S, et al. 2020. Fungal Diversity 105:1-16 ((https://doi.org/10.1007/s13225-020-00466-2)) |
| Otu34391 | 1 | k__Fungi(100);p__Ascomycota(100);c__Saccharomycetes(100);o__Saccharomycetales(100);f__Metschnikowiaceae(100);g__Metschnikowia(100);s__Metschnikowia_pulcherrima(100); | Metschnikowia | 13 | Saprotroph | \|Undefined Saprotroph\| | Yeast | NULL | Probable | As nectar-tap_saprotroph (Põlme et al. 2020); Decay sugar-rich substrates (Põlme et al. 2020) | Põlme S, et al. 2020. Fungal Diversity 105:1-16 ((https://doi.org/10.1007/s13225-020-00466-2)) |
| Otu34392 | 1 | k__Fungi(100);p__Ascomycota(100);c__Saccharomycetes(100);o__Saccharomycetales(100);f__Metschnikowiaceae(100);g__Metschnikowia(100);s__Metschnikowia_pulcherrima(100); | Metschnikowia | 13 | Saprotroph | \|Undefined Saprotroph\| | Yeast | NULL | Probable | As nectar-tap_saprotroph (Põlme et al. 2020); Decay sugar-rich substrates (Põlme et al. 2020) | Põlme S, et al. 2020. Fungal Diversity 105:1-16 ((https://doi.org/10.1007/s13225-020-00466-2)) |
| Otu34393 | 1 | k__Fungi(100);p__Ascomycota(100);c__Saccharomycetes(100);o__Saccharomycetales(100);f__Metschnikowiaceae(100);g__Metschnikowia(100);s__Metschnikowia_pulcherrima(100); | Metschnikowia | 13 | Saprotroph | \|Undefined Saprotroph\| | Yeast | NULL | Probable | As nectar-tap_saprotroph (Põlme et al. 2020); Decay sugar-rich substrates (Põlme et al. 2020) | Põlme S, et al. 2020. Fungal Diversity 105:1-16 ((https://doi.org/10.1007/s13225-020-00466-2)) |
| Otu34394 | 1 | k__Fungi(100);p__Ascomycota(100);c__Saccharomycetes(100);o__Saccharomycetales(100);f__Metschnikowiaceae(100);g__Metschnikowia(100);s__Metschnikowia_pulcherrima(100); | Metschnikowia | 13 | Saprotroph | \|Undefined Saprotroph\| | Yeast | NULL | Probable | As nectar-tap_saprotroph (Põlme et al. 2020); Decay sugar-rich substrates (Põlme et al. 2020) | Põlme S, et al. 2020. Fungal Diversity 105:1-16 ((https://doi.org/10.1007/s13225-020-00466-2)) |
| Otu34395 | 1 | k__Fungi(100);p__Ascomycota(100);c__Dothideomycetes(100);o__Dothideomycetes_order_Incertae_sedis(100);f__Pseudeurotiaceae(100);g__Pseudeurotium(100);g__Pseudeurotium_unclassified(100); | Pseudeurotium | 13 | Saprotroph | \|Dung Saprotroph\|-Undefined Saprotroph | Microfungus | NULL | Probable | As soil_saprotroph (Põlme et al. 2020); Decay soil, dung (Põlme et al. 2020) | Tedersoo L, et al. 2014. Science 346:e1256688 ((https://doi.org/10.1126/science.1256688)); Põlme S, et al. 2020. Fungal Diversity 105:1-16 ((https://doi.org/10.1007/s13225-020-00466-2)) |
| Otu34396 | 1 | k__Fungi(100);p__Ascomycota(100);c__Saccharomycetes(100);o__Saccharomycetales(100);f__Metschnikowiaceae(100);g__Metschnikowia(100);g__Metschnikowia_unclassified(100); | Metschnikowia | 13 | Saprotroph | \|Undefined Saprotroph\| | Yeast | NULL | Probable | As nectar-tap_saprotroph (Põlme et al. 2020); Decay sugar-rich substrates (Põlme et al. 2020) | Põlme S, et al. 2020. Fungal Diversity 105:1-16 ((https://doi.org/10.1007/s13225-020-00466-2)) |
| Otu34397 | 1 | k__Fungi(100);p__Ascomycota(100);c__Saccharomycetes(100);o__Saccharomycetales(100);f__Metschnikowiaceae(100);g__Metschnikowia(100);s__Metschnikowia_pulcherrima(100); | Metschnikowia | 13 | Saprotroph | \|Undefined Saprotroph\| | Yeast | NULL | Probable | As nectar-tap_saprotroph (Põlme et al. 2020); Decay sugar-rich substrates (Põlme et al. 2020) | Põlme S, et al. 2020. Fungal Diversity 105:1-16 ((https://doi.org/10.1007/s13225-020-00466-2)) |
| Otu34399 | 1 | k__Fungi(100);p__Ascomycota(100);c__Saccharomycetes(100);o__Saccharomycetales(100);f__Saccharomycetales_family_Incertae_sedis(100);g__Candida(100);g__Candida_unclassified(100); | Candida | 13 | Pathotroph-Saprotroph-Symbiotroph | Animal Pathogen-Endophyte-Endosymbiont-Epiphyte-\|Undefined Saprotroph\| | Dimorphic Yeast-Yeast | NULL | Probable | As nectar-tap_saprotroph (Põlme et al. 2020); Decay sugar-rich substrates (Põlme et al. 2020); This asexual genus is polyphyletic and contain species that function very differently depend on what they are. Caution should be taken when interpreting guilds for Candida species. | Manolakaki D, et al. 2010. Virulence. 1: 367-375 ((https://doi.org/10.4161/viru.1.5.12796)); Mekbib SB, et al. 2011. Tropical Plant Pathology 36:233-240 ((https://doi.org/10.1590/S1982-56762011000400004)); Põlme S, et al. 2020. Fungal Diversity 105:1-16 ((https://doi.org/10.1007/s13225-020-00466-2)) |
| Otu34400 | 1 | k__Fungi(100);p__Ascomycota(100);c__Saccharomycetes(100);o__Saccharomycetales(100);f__Metschnikowiaceae(100);g__Metschnikowia(100);s__Metschnikowia_pulcherrima(100); | Metschnikowia | 13 | Saprotroph | \|Undefined Saprotroph\| | Yeast | NULL | Probable | As nectar-tap_saprotroph (Põlme et al. 2020); Decay sugar-rich substrates (Põlme et al. 2020) | Põlme S, et al. 2020. Fungal Diversity 105:1-16 ((https://doi.org/10.1007/s13225-020-00466-2)) |
| Otu34401 | 1 | k__Fungi(100);p__Ascomycota(100);c__Saccharomycetes(100);o__Saccharomycetales(100);f__Metschnikowiaceae(100);g__Metschnikowia(100);s__Metschnikowia_pulcherrima(100); | Metschnikowia | 13 | Saprotroph | \|Undefined Saprotroph\| | Yeast | NULL | Probable | As nectar-tap_saprotroph (Põlme et al. 2020); Decay sugar-rich substrates (Põlme et al. 2020) | Põlme S, et al. 2020. Fungal Diversity 105:1-16 ((https://doi.org/10.1007/s13225-020-00466-2)) |
| Otu34402 | 1 | k__Fungi(100);p__Ascomycota(100);c__Saccharomycetes(100);o__Saccharomycetales(100);f__Metschnikowiaceae(100);g__Metschnikowia(100);s__Metschnikowia_pulcherrima(100); | Metschnikowia | 13 | Saprotroph | \|Undefined Saprotroph\| | Yeast | NULL | Probable | As nectar-tap_saprotroph (Põlme et al. 2020); Decay sugar-rich substrates (Põlme et al. 2020) | Põlme S, et al. 2020. Fungal Diversity 105:1-16 ((https://doi.org/10.1007/s13225-020-00466-2)) |
| Otu34403 | 1 | k__Fungi(100);p__Ascomycota(100);c__Saccharomycetes(100);o__Saccharomycetales(100);f__Metschnikowiaceae(100);g__Metschnikowia(100);s__Metschnikowia_pulcherrima(100); | Metschnikowia | 13 | Saprotroph | \|Undefined Saprotroph\| | Yeast | NULL | Probable | As nectar-tap_saprotroph (Põlme et al. 2020); Decay sugar-rich substrates (Põlme et al. 2020) | Põlme S, et al. 2020. Fungal Diversity 105:1-16 ((https://doi.org/10.1007/s13225-020-00466-2)) |
| Otu34404 | 1 | k__Fungi(100);p__Ascomycota(100);c__Saccharomycetes(100);o__Saccharomycetales(100);f__Metschnikowiaceae(100);g__Metschnikowia(100);s__Metschnikowia_pulcherrima(100); | Metschnikowia | 13 | Saprotroph | \|Undefined Saprotroph\| | Yeast | NULL | Probable | As nectar-tap_saprotroph (Põlme et al. 2020); Decay sugar-rich substrates (Põlme et al. 2020) | Põlme S, et al. 2020. Fungal Diversity 105:1-16 ((https://doi.org/10.1007/s13225-020-00466-2)) |
| Otu34406 | 1 | k__Fungi(100);p__Ascomycota(100);c__Saccharomycetes(100);o__Saccharomycetales(100);f__Metschnikowiaceae(100);g__Metschnikowia(100);s__Metschnikowia_pulcherrima(100); | Metschnikowia | 13 | Saprotroph | \|Undefined Saprotroph\| | Yeast | NULL | Probable | As nectar-tap_saprotroph (Põlme et al. 2020); Decay sugar-rich substrates (Põlme et al. 2020) | Põlme S, et al. 2020. Fungal Diversity 105:1-16 ((https://doi.org/10.1007/s13225-020-00466-2)) |
| Otu34407 | 1 | k__Fungi(100);p__Ascomycota(100);c__Saccharomycetes(100);o__Saccharomycetales(100);f__Metschnikowiaceae(100);g__Metschnikowia(100);g__Metschnikowia_unclassified(100); | Metschnikowia | 13 | Saprotroph | \|Undefined Saprotroph\| | Yeast | NULL | Probable | As nectar-tap_saprotroph (Põlme et al. 2020); Decay sugar-rich substrates (Põlme et al. 2020) | Põlme S, et al. 2020. Fungal Diversity 105:1-16 ((https://doi.org/10.1007/s13225-020-00466-2)) |
| Otu34408 | 1 | k__Fungi(100);p__Ascomycota(100);c__Saccharomycetes(100);o__Saccharomycetales(100);f__Metschnikowiaceae(100);g__Metschnikowia(100);s__Metschnikowia_pulcherrima(100); | Metschnikowia | 13 | Saprotroph | \|Undefined Saprotroph\| | Yeast | NULL | Probable | As nectar-tap_saprotroph (Põlme et al. 2020); Decay sugar-rich substrates (Põlme et al. 2020) | Põlme S, et al. 2020. Fungal Diversity 105:1-16 ((https://doi.org/10.1007/s13225-020-00466-2)) |
| Otu34409 | 1 | k__Fungi(100);p__Ascomycota(100);c__Saccharomycetes(100);o__Saccharomycetales(100);f__Metschnikowiaceae(100);g__Metschnikowia(100);s__Metschnikowia_pulcherrima(100); | Metschnikowia | 13 | Saprotroph | \|Undefined Saprotroph\| | Yeast | NULL | Probable | As nectar-tap_saprotroph (Põlme et al. 2020); Decay sugar-rich substrates (Põlme et al. 2020) | Põlme S, et al. 2020. Fungal Diversity 105:1-16 ((https://doi.org/10.1007/s13225-020-00466-2)) |
| Otu34410 | 1 | k__Fungi(100);p__Ascomycota(100);c__Saccharomycetes(100);o__Saccharomycetales(100);f__Metschnikowiaceae(100);g__Metschnikowia(100);g__Metschnikowia_unclassified(100); | Metschnikowia | 13 | Saprotroph | \|Undefined Saprotroph\| | Yeast | NULL | Probable | As nectar-tap_saprotroph (Põlme et al. 2020); Decay sugar-rich substrates (Põlme et al. 2020) | Põlme S, et al. 2020. Fungal Diversity 105:1-16 ((https://doi.org/10.1007/s13225-020-00466-2)) |
| Otu34413 | 1 | k__Fungi(100);p__Ascomycota(100);c__Saccharomycetes(100);o__Saccharomycetales(100);f__Metschnikowiaceae(100);g__Metschnikowia(100);s__Metschnikowia_pulcherrima(100); | Metschnikowia | 13 | Saprotroph | \|Undefined Saprotroph\| | Yeast | NULL | Probable | As nectar-tap_saprotroph (Põlme et al. 2020); Decay sugar-rich substrates (Põlme et al. 2020) | Põlme S, et al. 2020. Fungal Diversity 105:1-16 ((https://doi.org/10.1007/s13225-020-00466-2)) |
| Otu34414 | 1 | k__Fungi(100);p__Ascomycota(100);c__Saccharomycetes(100);o__Saccharomycetales(100);f__Metschnikowiaceae(100);g__Metschnikowia(100);g__Metschnikowia_unclassified(100); | Metschnikowia | 13 | Saprotroph | \|Undefined Saprotroph\| | Yeast | NULL | Probable | As nectar-tap_saprotroph (Põlme et al. 2020); Decay sugar-rich substrates (Põlme et al. 2020) | Põlme S, et al. 2020. Fungal Diversity 105:1-16 ((https://doi.org/10.1007/s13225-020-00466-2)) |
| Otu34415 | 1 | k__Fungi(100);p__Ascomycota(100);c__Saccharomycetes(100);o__Saccharomycetales(100);f__Metschnikowiaceae(100);g__Metschnikowia(100);s__Metschnikowia_pulcherrima(100); | Metschnikowia | 13 | Saprotroph | \|Undefined Saprotroph\| | Yeast | NULL | Probable | As nectar-tap_saprotroph (Põlme et al. 2020); Decay sugar-rich substrates (Põlme et al. 2020) | Põlme S, et al. 2020. Fungal Diversity 105:1-16 ((https://doi.org/10.1007/s13225-020-00466-2)) |
| Otu34419 | 1 | k__Fungi(100);p__Ascomycota(100);c__Saccharomycetes(100);o__Saccharomycetales(100);f__Saccharomycetales_family_Incertae_sedis(100);g__Candida(100);g__Candida_unclassified(100); | Candida | 13 | Pathotroph-Saprotroph-Symbiotroph | Animal Pathogen-Endophyte-Endosymbiont-Epiphyte-\|Undefined Saprotroph\| | Dimorphic Yeast-Yeast | NULL | Probable | As nectar-tap_saprotroph (Põlme et al. 2020); Decay sugar-rich substrates (Põlme et al. 2020); This asexual genus is polyphyletic and contain species that function very differently depend on what they are. Caution should be taken when interpreting guilds for Candida species. | Manolakaki D, et al. 2010. Virulence. 1: 367-375 ((https://doi.org/10.4161/viru.1.5.12796)); Mekbib SB, et al. 2011. Tropical Plant Pathology 36:233-240 ((https://doi.org/10.1590/S1982-56762011000400004)); Põlme S, et al. 2020. Fungal Diversity 105:1-16 ((https://doi.org/10.1007/s13225-020-00466-2)) |
| Otu34420 | 1 | k__Fungi(100);p__Ascomycota(100);c__Saccharomycetes(100);o__Saccharomycetales(100);f__Metschnikowiaceae(100);g__Metschnikowia(100);s__Metschnikowia_pulcherrima(100); | Metschnikowia | 13 | Saprotroph | \|Undefined Saprotroph\| | Yeast | NULL | Probable | As nectar-tap_saprotroph (Põlme et al. 2020); Decay sugar-rich substrates (Põlme et al. 2020) | Põlme S, et al. 2020. Fungal Diversity 105:1-16 ((https://doi.org/10.1007/s13225-020-00466-2)) |
| Otu34421 | 1 | k__Fungi(100);p__Ascomycota(100);c__Saccharomycetes(100);o__Saccharomycetales(100);f__Metschnikowiaceae(100);g__Metschnikowia(100);g__Metschnikowia_unclassified(100); | Metschnikowia | 13 | Saprotroph | \|Undefined Saprotroph\| | Yeast | NULL | Probable | As nectar-tap_saprotroph (Põlme et al. 2020); Decay sugar-rich substrates (Põlme et al. 2020) | Põlme S, et al. 2020. Fungal Diversity 105:1-16 ((https://doi.org/10.1007/s13225-020-00466-2)) |
| Otu34423 | 1 | k__Fungi(100);p__Ascomycota(100);c__Saccharomycetes(100);o__Saccharomycetales(100);f__Metschnikowiaceae(100);g__Metschnikowia(100);s__Metschnikowia_pulcherrima(100); | Metschnikowia | 13 | Saprotroph | \|Undefined Saprotroph\| | Yeast | NULL | Probable | As nectar-tap_saprotroph (Põlme et al. 2020); Decay sugar-rich substrates (Põlme et al. 2020) | Põlme S, et al. 2020. Fungal Diversity 105:1-16 ((https://doi.org/10.1007/s13225-020-00466-2)) |
| Otu34427 | 1 | k__Fungi(100);p__Ascomycota(100);c__Saccharomycetes(100);o__Saccharomycetales(100);f__Metschnikowiaceae(100);g__Metschnikowia(100);s__Metschnikowia_pulcherrima(100); | Metschnikowia | 13 | Saprotroph | \|Undefined Saprotroph\| | Yeast | NULL | Probable | As nectar-tap_saprotroph (Põlme et al. 2020); Decay sugar-rich substrates (Põlme et al. 2020) | Põlme S, et al. 2020. Fungal Diversity 105:1-16 ((https://doi.org/10.1007/s13225-020-00466-2)) |
| Otu34428 | 1 | k__Fungi(100);p__Ascomycota(100);c__Saccharomycetes(100);o__Saccharomycetales(100);f__Metschnikowiaceae(100);g__Metschnikowia(100);s__Metschnikowia_pulcherrima(100); | Metschnikowia | 13 | Saprotroph | \|Undefined Saprotroph\| | Yeast | NULL | Probable | As nectar-tap_saprotroph (Põlme et al. 2020); Decay sugar-rich substrates (Põlme et al. 2020) | Põlme S, et al. 2020. Fungal Diversity 105:1-16 ((https://doi.org/10.1007/s13225-020-00466-2)) |
| Otu34429 | 1 | k__Fungi(100);p__Ascomycota(100);c__Saccharomycetes(100);o__Saccharomycetales(100);f__Metschnikowiaceae(100);g__Metschnikowia(100);s__Metschnikowia_pulcherrima(100); | Metschnikowia | 13 | Saprotroph | \|Undefined Saprotroph\| | Yeast | NULL | Probable | As nectar-tap_saprotroph (Põlme et al. 2020); Decay sugar-rich substrates (Põlme et al. 2020) | Põlme S, et al. 2020. Fungal Diversity 105:1-16 ((https://doi.org/10.1007/s13225-020-00466-2)) |
| Otu34432 | 1 | k__Fungi(100);p__Ascomycota(100);c__Saccharomycetes(100);o__Saccharomycetales(100);f__Metschnikowiaceae(100);g__Metschnikowia(100);s__Metschnikowia_pulcherrima(100); | Metschnikowia | 13 | Saprotroph | \|Undefined Saprotroph\| | Yeast | NULL | Probable | As nectar-tap_saprotroph (Põlme et al. 2020); Decay sugar-rich substrates (Põlme et al. 2020) | Põlme S, et al. 2020. Fungal Diversity 105:1-16 ((https://doi.org/10.1007/s13225-020-00466-2)) |
| Otu34433 | 1 | k__Fungi(100);p__Ascomycota(100);c__Saccharomycetes(100);o__Saccharomycetales(100);f__Metschnikowiaceae(100);g__Metschnikowia(100);s__Metschnikowia_pulcherrima(100); | Metschnikowia | 13 | Saprotroph | \|Undefined Saprotroph\| | Yeast | NULL | Probable | As nectar-tap_saprotroph (Põlme et al. 2020); Decay sugar-rich substrates (Põlme et al. 2020) | Põlme S, et al. 2020. Fungal Diversity 105:1-16 ((https://doi.org/10.1007/s13225-020-00466-2)) |
| Otu34434 | 1 | k__Fungi(100);p__Ascomycota(100);c__Saccharomycetes(100);o__Saccharomycetales(100);f__Metschnikowiaceae(100);g__Metschnikowia(100);g__Metschnikowia_unclassified(100); | Metschnikowia | 13 | Saprotroph | \|Undefined Saprotroph\| | Yeast | NULL | Probable | As nectar-tap_saprotroph (Põlme et al. 2020); Decay sugar-rich substrates (Põlme et al. 2020) | Põlme S, et al. 2020. Fungal Diversity 105:1-16 ((https://doi.org/10.1007/s13225-020-00466-2)) |
| Otu34435 | 1 | k__Fungi(100);p__Ascomycota(100);c__Saccharomycetes(100);o__Saccharomycetales(100);f__Metschnikowiaceae(100);g__Metschnikowia(100);g__Metschnikowia_unclassified(100); | Metschnikowia | 13 | Saprotroph | \|Undefined Saprotroph\| | Yeast | NULL | Probable | As nectar-tap_saprotroph (Põlme et al. 2020); Decay sugar-rich substrates (Põlme et al. 2020) | Põlme S, et al. 2020. Fungal Diversity 105:1-16 ((https://doi.org/10.1007/s13225-020-00466-2)) |
| Otu34439 | 1 | k__Fungi(100);p__Ascomycota(100);c__Saccharomycetes(100);o__Saccharomycetales(100);f__Metschnikowiaceae(100);g__Metschnikowia(100);s__Metschnikowia_pulcherrima(100); | Metschnikowia | 13 | Saprotroph | \|Undefined Saprotroph\| | Yeast | NULL | Probable | As nectar-tap_saprotroph (Põlme et al. 2020); Decay sugar-rich substrates (Põlme et al. 2020) | Põlme S, et al. 2020. Fungal Diversity 105:1-16 ((https://doi.org/10.1007/s13225-020-00466-2)) |
| Otu34440 | 1 | k__Fungi(100);p__Ascomycota(100);c__Saccharomycetes(100);o__Saccharomycetales(100);f__Metschnikowiaceae(100);g__Metschnikowia(100);s__Metschnikowia_pulcherrima(100); | Metschnikowia | 13 | Saprotroph | \|Undefined Saprotroph\| | Yeast | NULL | Probable | As nectar-tap_saprotroph (Põlme et al. 2020); Decay sugar-rich substrates (Põlme et al. 2020) | Põlme S, et al. 2020. Fungal Diversity 105:1-16 ((https://doi.org/10.1007/s13225-020-00466-2)) |
| Otu34441 | 1 | k__Fungi(100);p__Ascomycota(100);c__Saccharomycetes(100);o__Saccharomycetales(100);f__Metschnikowiaceae(100);g__Metschnikowia(100);s__Metschnikowia_pulcherrima(100); | Metschnikowia | 13 | Saprotroph | \|Undefined Saprotroph\| | Yeast | NULL | Probable | As nectar-tap_saprotroph (Põlme et al. 2020); Decay sugar-rich substrates (Põlme et al. 2020) | Põlme S, et al. 2020. Fungal Diversity 105:1-16 ((https://doi.org/10.1007/s13225-020-00466-2)) |
| Otu34443 | 1 | k__Fungi(100);p__Ascomycota(100);c__Saccharomycetes(100);o__Saccharomycetales(100);f__Metschnikowiaceae(100);g__Metschnikowia(100);s__Metschnikowia_pulcherrima(100); | Metschnikowia | 13 | Saprotroph | \|Undefined Saprotroph\| | Yeast | NULL | Probable | As nectar-tap_saprotroph (Põlme et al. 2020); Decay sugar-rich substrates (Põlme et al. 2020) | Põlme S, et al. 2020. Fungal Diversity 105:1-16 ((https://doi.org/10.1007/s13225-020-00466-2)) |
| Otu34444 | 1 | k__Fungi(100);p__Ascomycota(100);c__Saccharomycetes(100);o__Saccharomycetales(100);f__Metschnikowiaceae(100);g__Metschnikowia(100);g__Metschnikowia_unclassified(100); | Metschnikowia | 13 | Saprotroph | \|Undefined Saprotroph\| | Yeast | NULL | Probable | As nectar-tap_saprotroph (Põlme et al. 2020); Decay sugar-rich substrates (Põlme et al. 2020) | Põlme S, et al. 2020. Fungal Diversity 105:1-16 ((https://doi.org/10.1007/s13225-020-00466-2)) |
| Otu34445 | 1 | k__Fungi(100);p__Ascomycota(100);c__Saccharomycetes(100);o__Saccharomycetales(100);f__Metschnikowiaceae(100);g__Metschnikowia(100);g__Metschnikowia_unclassified(100); | Metschnikowia | 13 | Saprotroph | \|Undefined Saprotroph\| | Yeast | NULL | Probable | As nectar-tap_saprotroph (Põlme et al. 2020); Decay sugar-rich substrates (Põlme et al. 2020) | Põlme S, et al. 2020. Fungal Diversity 105:1-16 ((https://doi.org/10.1007/s13225-020-00466-2)) |
| Otu34446 | 1 | k__Fungi(100);p__Ascomycota(100);c__Saccharomycetes(100);o__Saccharomycetales(100);f__Metschnikowiaceae(100);g__Metschnikowia(100);s__Metschnikowia_pulcherrima(100); | Metschnikowia | 13 | Saprotroph | \|Undefined Saprotroph\| | Yeast | NULL | Probable | As nectar-tap_saprotroph (Põlme et al. 2020); Decay sugar-rich substrates (Põlme et al. 2020) | Põlme S, et al. 2020. Fungal Diversity 105:1-16 ((https://doi.org/10.1007/s13225-020-00466-2)) |
| Otu34454 | 1 | k__Fungi(100);p__Ascomycota(100);c__Saccharomycetes(100);o__Saccharomycetales(100);f__Metschnikowiaceae(100);g__Metschnikowia(100);s__Metschnikowia_pulcherrima(100); | Metschnikowia | 13 | Saprotroph | \|Undefined Saprotroph\| | Yeast | NULL | Probable | As nectar-tap_saprotroph (Põlme et al. 2020); Decay sugar-rich substrates (Põlme et al. 2020) | Põlme S, et al. 2020. Fungal Diversity 105:1-16 ((https://doi.org/10.1007/s13225-020-00466-2)) |
| Otu34459 | 1 | k__Fungi(100);p__Ascomycota(100);c__Saccharomycetes(100);o__Saccharomycetales(100);f__Metschnikowiaceae(100);g__Metschnikowia(100);g__Metschnikowia_unclassified(100); | Metschnikowia | 13 | Saprotroph | \|Undefined Saprotroph\| | Yeast | NULL | Probable | As nectar-tap_saprotroph (Põlme et al. 2020); Decay sugar-rich substrates (Põlme et al. 2020) | Põlme S, et al. 2020. Fungal Diversity 105:1-16 ((https://doi.org/10.1007/s13225-020-00466-2)) |
| Otu34460 | 1 | k__Fungi(100);p__Ascomycota(100);c__Saccharomycetes(100);o__Saccharomycetales(100);f__Metschnikowiaceae(100);g__Metschnikowia(100);g__Metschnikowia_unclassified(100); | Metschnikowia | 13 | Saprotroph | \|Undefined Saprotroph\| | Yeast | NULL | Probable | As nectar-tap_saprotroph (Põlme et al. 2020); Decay sugar-rich substrates (Põlme et al. 2020) | Põlme S, et al. 2020. Fungal Diversity 105:1-16 ((https://doi.org/10.1007/s13225-020-00466-2)) |
| Otu34461 | 1 | k__Fungi(100);p__Ascomycota(100);c__Saccharomycetes(100);o__Saccharomycetales(100);f__Metschnikowiaceae(100);g__Metschnikowia(100);s__Metschnikowia_pulcherrima(100); | Metschnikowia | 13 | Saprotroph | \|Undefined Saprotroph\| | Yeast | NULL | Probable | As nectar-tap_saprotroph (Põlme et al. 2020); Decay sugar-rich substrates (Põlme et al. 2020) | Põlme S, et al. 2020. Fungal Diversity 105:1-16 ((https://doi.org/10.1007/s13225-020-00466-2)) |
| Otu34464 | 1 | k__Fungi(100);p__Ascomycota(100);c__Saccharomycetes(100);o__Saccharomycetales(100);f__Metschnikowiaceae(100);g__Metschnikowia(100);s__Metschnikowia_pulcherrima(100); | Metschnikowia | 13 | Saprotroph | \|Undefined Saprotroph\| | Yeast | NULL | Probable | As nectar-tap_saprotroph (Põlme et al. 2020); Decay sugar-rich substrates (Põlme et al. 2020) | Põlme S, et al. 2020. Fungal Diversity 105:1-16 ((https://doi.org/10.1007/s13225-020-00466-2)) |
| Otu34465 | 1 | k__Fungi(100);p__Ascomycota(100);c__Saccharomycetes(100);o__Saccharomycetales(100);f__Metschnikowiaceae(100);g__Metschnikowia(100);s__Metschnikowia_pulcherrima(100); | Metschnikowia | 13 | Saprotroph | \|Undefined Saprotroph\| | Yeast | NULL | Probable | As nectar-tap_saprotroph (Põlme et al. 2020); Decay sugar-rich substrates (Põlme et al. 2020) | Põlme S, et al. 2020. Fungal Diversity 105:1-16 ((https://doi.org/10.1007/s13225-020-00466-2)) |
| Otu34466 | 1 | k__Fungi(100);p__Ascomycota(100);c__Saccharomycetes(100);o__Saccharomycetales(100);f__Metschnikowiaceae(100);g__Metschnikowia(100);s__Metschnikowia_pulcherrima(100); | Metschnikowia | 13 | Saprotroph | \|Undefined Saprotroph\| | Yeast | NULL | Probable | As nectar-tap_saprotroph (Põlme et al. 2020); Decay sugar-rich substrates (Põlme et al. 2020) | Põlme S, et al. 2020. Fungal Diversity 105:1-16 ((https://doi.org/10.1007/s13225-020-00466-2)) |
| Otu34467 | 1 | k__Fungi(100);p__Ascomycota(100);c__Saccharomycetes(100);o__Saccharomycetales(100);f__Metschnikowiaceae(100);g__Metschnikowia(100);g__Metschnikowia_unclassified(100); | Metschnikowia | 13 | Saprotroph | \|Undefined Saprotroph\| | Yeast | NULL | Probable | As nectar-tap_saprotroph (Põlme et al. 2020); Decay sugar-rich substrates (Põlme et al. 2020) | Põlme S, et al. 2020. Fungal Diversity 105:1-16 ((https://doi.org/10.1007/s13225-020-00466-2)) |
| Otu34468 | 1 | k__Fungi(100);p__Ascomycota(100);c__Saccharomycetes(100);o__Saccharomycetales(100);f__Metschnikowiaceae(100);g__Metschnikowia(100);s__Metschnikowia_pulcherrima(100); | Metschnikowia | 13 | Saprotroph | \|Undefined Saprotroph\| | Yeast | NULL | Probable | As nectar-tap_saprotroph (Põlme et al. 2020); Decay sugar-rich substrates (Põlme et al. 2020) | Põlme S, et al. 2020. Fungal Diversity 105:1-16 ((https://doi.org/10.1007/s13225-020-00466-2)) |
| Otu34474 | 1 | k__Fungi(100);p__Ascomycota(100);c__Saccharomycetes(100);o__Saccharomycetales(100);f__Metschnikowiaceae(100);g__Metschnikowia(100);s__Metschnikowia_pulcherrima(100); | Metschnikowia | 13 | Saprotroph | \|Undefined Saprotroph\| | Yeast | NULL | Probable | As nectar-tap_saprotroph (Põlme et al. 2020); Decay sugar-rich substrates (Põlme et al. 2020) | Põlme S, et al. 2020. Fungal Diversity 105:1-16 ((https://doi.org/10.1007/s13225-020-00466-2)) |
| Otu34475 | 1 | k__Fungi(100);p__Ascomycota(100);c__Saccharomycetes(100);o__Saccharomycetales(100);f__Metschnikowiaceae(100);g__Metschnikowia(100);g__Metschnikowia_unclassified(100); | Metschnikowia | 13 | Saprotroph | \|Undefined Saprotroph\| | Yeast | NULL | Probable | As nectar-tap_saprotroph (Põlme et al. 2020); Decay sugar-rich substrates (Põlme et al. 2020) | Põlme S, et al. 2020. Fungal Diversity 105:1-16 ((https://doi.org/10.1007/s13225-020-00466-2)) |
| Otu34477 | 1 | k__Fungi(100);p__Ascomycota(100);c__Saccharomycetes(100);o__Saccharomycetales(100);f__Metschnikowiaceae(100);g__Metschnikowia(100);s__Metschnikowia_pulcherrima(100); | Metschnikowia | 13 | Saprotroph | \|Undefined Saprotroph\| | Yeast | NULL | Probable | As nectar-tap_saprotroph (Põlme et al. 2020); Decay sugar-rich substrates (Põlme et al. 2020) | Põlme S, et al. 2020. Fungal Diversity 105:1-16 ((https://doi.org/10.1007/s13225-020-00466-2)) |
| Otu34483 | 1 | k__Fungi(100);p__Ascomycota(100);c__Saccharomycetes(100);o__Saccharomycetales(100);f__Metschnikowiaceae(100);g__Metschnikowia(100);s__Metschnikowia_pulcherrima(100); | Metschnikowia | 13 | Saprotroph | \|Undefined Saprotroph\| | Yeast | NULL | Probable | As nectar-tap_saprotroph (Põlme et al. 2020); Decay sugar-rich substrates (Põlme et al. 2020) | Põlme S, et al. 2020. Fungal Diversity 105:1-16 ((https://doi.org/10.1007/s13225-020-00466-2)) |
| Otu34484 | 1 | k__Fungi(100);p__Ascomycota(100);c__Saccharomycetes(100);o__Saccharomycetales(100);f__Metschnikowiaceae(100);g__Metschnikowia(100);g__Metschnikowia_unclassified(100); | Metschnikowia | 13 | Saprotroph | \|Undefined Saprotroph\| | Yeast | NULL | Probable | As nectar-tap_saprotroph (Põlme et al. 2020); Decay sugar-rich substrates (Põlme et al. 2020) | Põlme S, et al. 2020. Fungal Diversity 105:1-16 ((https://doi.org/10.1007/s13225-020-00466-2)) |
| Otu34485 | 1 | k__Fungi(100);p__Ascomycota(100);c__Saccharomycetes(100);o__Saccharomycetales(100);f__Metschnikowiaceae(100);g__Metschnikowia(100);s__Metschnikowia_pulcherrima(100); | Metschnikowia | 13 | Saprotroph | \|Undefined Saprotroph\| | Yeast | NULL | Probable | As nectar-tap_saprotroph (Põlme et al. 2020); Decay sugar-rich substrates (Põlme et al. 2020) | Põlme S, et al. 2020. Fungal Diversity 105:1-16 ((https://doi.org/10.1007/s13225-020-00466-2)) |
| Otu34486 | 1 | k__Fungi(100);p__Ascomycota(100);c__Saccharomycetes(100);o__Saccharomycetales(100);f__Metschnikowiaceae(100);g__Metschnikowia(100);g__Metschnikowia_unclassified(100); | Metschnikowia | 13 | Saprotroph | \|Undefined Saprotroph\| | Yeast | NULL | Probable | As nectar-tap_saprotroph (Põlme et al. 2020); Decay sugar-rich substrates (Põlme et al. 2020) | Põlme S, et al. 2020. Fungal Diversity 105:1-16 ((https://doi.org/10.1007/s13225-020-00466-2)) |
| Otu34488 | 1 | k__Fungi(100);p__Ascomycota(100);c__Saccharomycetes(100);o__Saccharomycetales(100);f__Metschnikowiaceae(100);g__Metschnikowia(100);s__Metschnikowia_pulcherrima(100); | Metschnikowia | 13 | Saprotroph | \|Undefined Saprotroph\| | Yeast | NULL | Probable | As nectar-tap_saprotroph (Põlme et al. 2020); Decay sugar-rich substrates (Põlme et al. 2020) | Põlme S, et al. 2020. Fungal Diversity 105:1-16 ((https://doi.org/10.1007/s13225-020-00466-2)) |
| Otu34497 | 1 | k__Fungi(100);p__Ascomycota(100);c__Saccharomycetes(100);o__Saccharomycetales(100);f__Metschnikowiaceae(100);g__Metschnikowia(100);s__Metschnikowia_pulcherrima(100); | Metschnikowia | 13 | Saprotroph | \|Undefined Saprotroph\| | Yeast | NULL | Probable | As nectar-tap_saprotroph (Põlme et al. 2020); Decay sugar-rich substrates (Põlme et al. 2020) | Põlme S, et al. 2020. Fungal Diversity 105:1-16 ((https://doi.org/10.1007/s13225-020-00466-2)) |
| Otu34500 | 1 | k__Fungi(100);p__Ascomycota(100);c__Saccharomycetes(100);o__Saccharomycetales(100);f__Metschnikowiaceae(100);g__Metschnikowia(100);s__Metschnikowia_pulcherrima(100); | Metschnikowia | 13 | Saprotroph | \|Undefined Saprotroph\| | Yeast | NULL | Probable | As nectar-tap_saprotroph (Põlme et al. 2020); Decay sugar-rich substrates (Põlme et al. 2020) | Põlme S, et al. 2020. Fungal Diversity 105:1-16 ((https://doi.org/10.1007/s13225-020-00466-2)) |
| Otu34501 | 1 | k__Fungi(100);p__Ascomycota(100);c__Saccharomycetes(100);o__Saccharomycetales(100);f__Metschnikowiaceae(100);g__Metschnikowia(100);g__Metschnikowia_unclassified(100); | Metschnikowia | 13 | Saprotroph | \|Undefined Saprotroph\| | Yeast | NULL | Probable | As nectar-tap_saprotroph (Põlme et al. 2020); Decay sugar-rich substrates (Põlme et al. 2020) | Põlme S, et al. 2020. Fungal Diversity 105:1-16 ((https://doi.org/10.1007/s13225-020-00466-2)) |
| Otu34502 | 1 | k__Fungi(100);p__Ascomycota(100);c__Saccharomycetes(100);o__Saccharomycetales(100);f__Metschnikowiaceae(100);g__Metschnikowia(100);g__Metschnikowia_unclassified(100); | Metschnikowia | 13 | Saprotroph | \|Undefined Saprotroph\| | Yeast | NULL | Probable | As nectar-tap_saprotroph (Põlme et al. 2020); Decay sugar-rich substrates (Põlme et al. 2020) | Põlme S, et al. 2020. Fungal Diversity 105:1-16 ((https://doi.org/10.1007/s13225-020-00466-2)) |
| Otu34504 | 1 | k__Fungi(100);p__Ascomycota(100);c__Saccharomycetes(100);o__Saccharomycetales(100);f__Metschnikowiaceae(100);g__Metschnikowia(100);g__Metschnikowia_unclassified(100); | Metschnikowia | 13 | Saprotroph | \|Undefined Saprotroph\| | Yeast | NULL | Probable | As nectar-tap_saprotroph (Põlme et al. 2020); Decay sugar-rich substrates (Põlme et al. 2020) | Põlme S, et al. 2020. Fungal Diversity 105:1-16 ((https://doi.org/10.1007/s13225-020-00466-2)) |
| Otu34507 | 1 | k__Fungi(100);p__Ascomycota(100);c__Saccharomycetes(100);o__Saccharomycetales(100);f__Metschnikowiaceae(100);g__Metschnikowia(100);s__Metschnikowia_pulcherrima(100); | Metschnikowia | 13 | Saprotroph | \|Undefined Saprotroph\| | Yeast | NULL | Probable | As nectar-tap_saprotroph (Põlme et al. 2020); Decay sugar-rich substrates (Põlme et al. 2020) | Põlme S, et al. 2020. Fungal Diversity 105:1-16 ((https://doi.org/10.1007/s13225-020-00466-2)) |
| Otu34508 | 1 | k__Fungi(100);p__Ascomycota(100);c__Saccharomycetes(100);o__Saccharomycetales(100);f__Metschnikowiaceae(100);g__Metschnikowia(100);s__Metschnikowia_pulcherrima(100); | Metschnikowia | 13 | Saprotroph | \|Undefined Saprotroph\| | Yeast | NULL | Probable | As nectar-tap_saprotroph (Põlme et al. 2020); Decay sugar-rich substrates (Põlme et al. 2020) | Põlme S, et al. 2020. Fungal Diversity 105:1-16 ((https://doi.org/10.1007/s13225-020-00466-2)) |
| Otu34509 | 1 | k__Fungi(100);p__Ascomycota(100);c__Saccharomycetes(100);o__Saccharomycetales(100);f__Metschnikowiaceae(100);g__Metschnikowia(100);s__Metschnikowia_pulcherrima(100); | Metschnikowia | 13 | Saprotroph | \|Undefined Saprotroph\| | Yeast | NULL | Probable | As nectar-tap_saprotroph (Põlme et al. 2020); Decay sugar-rich substrates (Põlme et al. 2020) | Põlme S, et al. 2020. Fungal Diversity 105:1-16 ((https://doi.org/10.1007/s13225-020-00466-2)) |
| Otu34510 | 1 | k__Fungi(100);p__Ascomycota(100);c__Saccharomycetes(100);o__Saccharomycetales(100);f__Metschnikowiaceae(100);g__Metschnikowia(100);s__Metschnikowia_pulcherrima(100); | Metschnikowia | 13 | Saprotroph | \|Undefined Saprotroph\| | Yeast | NULL | Probable | As nectar-tap_saprotroph (Põlme et al. 2020); Decay sugar-rich substrates (Põlme et al. 2020) | Põlme S, et al. 2020. Fungal Diversity 105:1-16 ((https://doi.org/10.1007/s13225-020-00466-2)) |
| Otu34511 | 1 | k__Fungi(100);p__Ascomycota(100);c__Saccharomycetes(100);o__Saccharomycetales(100);f__Metschnikowiaceae(100);g__Metschnikowia(100);s__Metschnikowia_pulcherrima(100); | Metschnikowia | 13 | Saprotroph | \|Undefined Saprotroph\| | Yeast | NULL | Probable | As nectar-tap_saprotroph (Põlme et al. 2020); Decay sugar-rich substrates (Põlme et al. 2020) | Põlme S, et al. 2020. Fungal Diversity 105:1-16 ((https://doi.org/10.1007/s13225-020-00466-2)) |
| Otu34512 | 1 | k__Fungi(100);p__Ascomycota(100);c__Saccharomycetes(100);o__Saccharomycetales(100);f__Metschnikowiaceae(100);g__Metschnikowia(100);s__Metschnikowia_pulcherrima(100); | Metschnikowia | 13 | Saprotroph | \|Undefined Saprotroph\| | Yeast | NULL | Probable | As nectar-tap_saprotroph (Põlme et al. 2020); Decay sugar-rich substrates (Põlme et al. 2020) | Põlme S, et al. 2020. Fungal Diversity 105:1-16 ((https://doi.org/10.1007/s13225-020-00466-2)) |
| Otu34513 | 1 | k__Fungi(100);p__Ascomycota(100);c__Saccharomycetes(100);o__Saccharomycetales(100);f__Metschnikowiaceae(100);g__Metschnikowia(100);s__Metschnikowia_pulcherrima(100); | Metschnikowia | 13 | Saprotroph | \|Undefined Saprotroph\| | Yeast | NULL | Probable | As nectar-tap_saprotroph (Põlme et al. 2020); Decay sugar-rich substrates (Põlme et al. 2020) | Põlme S, et al. 2020. Fungal Diversity 105:1-16 ((https://doi.org/10.1007/s13225-020-00466-2)) |
| Otu34514 | 1 | k__Fungi(100);p__Ascomycota(100);c__Saccharomycetes(100);o__Saccharomycetales(100);f__Metschnikowiaceae(100);g__Metschnikowia(100);g__Metschnikowia_unclassified(100); | Metschnikowia | 13 | Saprotroph | \|Undefined Saprotroph\| | Yeast | NULL | Probable | As nectar-tap_saprotroph (Põlme et al. 2020); Decay sugar-rich substrates (Põlme et al. 2020) | Põlme S, et al. 2020. Fungal Diversity 105:1-16 ((https://doi.org/10.1007/s13225-020-00466-2)) |
| Otu34515 | 1 | k__Fungi(100);p__Ascomycota(100);c__Saccharomycetes(100);o__Saccharomycetales(100);f__Metschnikowiaceae(100);g__Metschnikowia(100);s__Metschnikowia_pulcherrima(100); | Metschnikowia | 13 | Saprotroph | \|Undefined Saprotroph\| | Yeast | NULL | Probable | As nectar-tap_saprotroph (Põlme et al. 2020); Decay sugar-rich substrates (Põlme et al. 2020) | Põlme S, et al. 2020. Fungal Diversity 105:1-16 ((https://doi.org/10.1007/s13225-020-00466-2)) |
| Otu34516 | 1 | k__Fungi(100);p__Ascomycota(100);c__Saccharomycetes(100);o__Saccharomycetales(100);f__Metschnikowiaceae(100);g__Metschnikowia(100);s__Metschnikowia_pulcherrima(100); | Metschnikowia | 13 | Saprotroph | \|Undefined Saprotroph\| | Yeast | NULL | Probable | As nectar-tap_saprotroph (Põlme et al. 2020); Decay sugar-rich substrates (Põlme et al. 2020) | Põlme S, et al. 2020. Fungal Diversity 105:1-16 ((https://doi.org/10.1007/s13225-020-00466-2)) |
| Otu34518 | 1 | k__Fungi(100);p__Ascomycota(100);c__Saccharomycetes(100);o__Saccharomycetales(100);f__Metschnikowiaceae(100);g__Metschnikowia(100);s__Metschnikowia_pulcherrima(100); | Metschnikowia | 13 | Saprotroph | \|Undefined Saprotroph\| | Yeast | NULL | Probable | As nectar-tap_saprotroph (Põlme et al. 2020); Decay sugar-rich substrates (Põlme et al. 2020) | Põlme S, et al. 2020. Fungal Diversity 105:1-16 ((https://doi.org/10.1007/s13225-020-00466-2)) |
| Otu34519 | 1 | k__Fungi(100);p__Ascomycota(100);c__Saccharomycetes(100);o__Saccharomycetales(100);f__Metschnikowiaceae(100);g__Metschnikowia(100);s__Metschnikowia_pulcherrima(100); | Metschnikowia | 13 | Saprotroph | \|Undefined Saprotroph\| | Yeast | NULL | Probable | As nectar-tap_saprotroph (Põlme et al. 2020); Decay sugar-rich substrates (Põlme et al. 2020) | Põlme S, et al. 2020. Fungal Diversity 105:1-16 ((https://doi.org/10.1007/s13225-020-00466-2)) |
| Otu34520 | 1 | k__Fungi(100);p__Ascomycota(100);c__Saccharomycetes(100);o__Saccharomycetales(100);f__Metschnikowiaceae(100);g__Metschnikowia(100);s__Metschnikowia_pulcherrima(100); | Metschnikowia | 13 | Saprotroph | \|Undefined Saprotroph\| | Yeast | NULL | Probable | As nectar-tap_saprotroph (Põlme et al. 2020); Decay sugar-rich substrates (Põlme et al. 2020) | Põlme S, et al. 2020. Fungal Diversity 105:1-16 ((https://doi.org/10.1007/s13225-020-00466-2)) |
| Otu34521 | 1 | k__Fungi(100);p__Ascomycota(100);c__Saccharomycetes(100);o__Saccharomycetales(100);f__Metschnikowiaceae(100);g__Metschnikowia(100);s__Metschnikowia_pulcherrima(100); | Metschnikowia | 13 | Saprotroph | \|Undefined Saprotroph\| | Yeast | NULL | Probable | As nectar-tap_saprotroph (Põlme et al. 2020); Decay sugar-rich substrates (Põlme et al. 2020) | Põlme S, et al. 2020. Fungal Diversity 105:1-16 ((https://doi.org/10.1007/s13225-020-00466-2)) |
| Otu34522 | 1 | k__Fungi(100);p__Ascomycota(100);c__Saccharomycetes(100);o__Saccharomycetales(100);f__Metschnikowiaceae(100);g__Metschnikowia(100);s__Metschnikowia_pulcherrima(100); | Metschnikowia | 13 | Saprotroph | \|Undefined Saprotroph\| | Yeast | NULL | Probable | As nectar-tap_saprotroph (Põlme et al. 2020); Decay sugar-rich substrates (Põlme et al. 2020) | Põlme S, et al. 2020. Fungal Diversity 105:1-16 ((https://doi.org/10.1007/s13225-020-00466-2)) |
| Otu34523 | 1 | k__Fungi(100);p__Ascomycota(100);c__Saccharomycetes(100);o__Saccharomycetales(100);f__Metschnikowiaceae(100);g__Metschnikowia(100);s__Metschnikowia_pulcherrima(100); | Metschnikowia | 13 | Saprotroph | \|Undefined Saprotroph\| | Yeast | NULL | Probable | As nectar-tap_saprotroph (Põlme et al. 2020); Decay sugar-rich substrates (Põlme et al. 2020) | Põlme S, et al. 2020. Fungal Diversity 105:1-16 ((https://doi.org/10.1007/s13225-020-00466-2)) |
| Otu34524 | 1 | k__Fungi(100);p__Ascomycota(100);c__Saccharomycetes(100);o__Saccharomycetales(100);f__Metschnikowiaceae(100);g__Metschnikowia(100);g__Metschnikowia_unclassified(100); | Metschnikowia | 13 | Saprotroph | \|Undefined Saprotroph\| | Yeast | NULL | Probable | As nectar-tap_saprotroph (Põlme et al. 2020); Decay sugar-rich substrates (Põlme et al. 2020) | Põlme S, et al. 2020. Fungal Diversity 105:1-16 ((https://doi.org/10.1007/s13225-020-00466-2)) |
| Otu34525 | 1 | k__Fungi(100);p__Ascomycota(100);c__Saccharomycetes(100);o__Saccharomycetales(100);f__Metschnikowiaceae(100);g__Metschnikowia(100);s__Metschnikowia_pulcherrima(100); | Metschnikowia | 13 | Saprotroph | \|Undefined Saprotroph\| | Yeast | NULL | Probable | As nectar-tap_saprotroph (Põlme et al. 2020); Decay sugar-rich substrates (Põlme et al. 2020) | Põlme S, et al. 2020. Fungal Diversity 105:1-16 ((https://doi.org/10.1007/s13225-020-00466-2)) |
| Otu34526 | 1 | k__Fungi(100);p__Ascomycota(100);c__Saccharomycetes(100);o__Saccharomycetales(100);f__Metschnikowiaceae(100);g__Metschnikowia(100);s__Metschnikowia_pulcherrima(100); | Metschnikowia | 13 | Saprotroph | \|Undefined Saprotroph\| | Yeast | NULL | Probable | As nectar-tap_saprotroph (Põlme et al. 2020); Decay sugar-rich substrates (Põlme et al. 2020) | Põlme S, et al. 2020. Fungal Diversity 105:1-16 ((https://doi.org/10.1007/s13225-020-00466-2)) |
| Otu34527 | 1 | k__Fungi(100);p__Ascomycota(100);c__Saccharomycetes(100);o__Saccharomycetales(100);f__Metschnikowiaceae(100);g__Metschnikowia(100);g__Metschnikowia_unclassified(100); | Metschnikowia | 13 | Saprotroph | \|Undefined Saprotroph\| | Yeast | NULL | Probable | As nectar-tap_saprotroph (Põlme et al. 2020); Decay sugar-rich substrates (Põlme et al. 2020) | Põlme S, et al. 2020. Fungal Diversity 105:1-16 ((https://doi.org/10.1007/s13225-020-00466-2)) |
| Otu34528 | 1 | k__Fungi(100);p__Ascomycota(100);c__Saccharomycetes(100);o__Saccharomycetales(100);f__Metschnikowiaceae(100);g__Metschnikowia(100);g__Metschnikowia_unclassified(100); | Metschnikowia | 13 | Saprotroph | \|Undefined Saprotroph\| | Yeast | NULL | Probable | As nectar-tap_saprotroph (Põlme et al. 2020); Decay sugar-rich substrates (Põlme et al. 2020) | Põlme S, et al. 2020. Fungal Diversity 105:1-16 ((https://doi.org/10.1007/s13225-020-00466-2)) |
| Otu34529 | 1 | k__Fungi(100);p__Ascomycota(100);c__Saccharomycetes(100);o__Saccharomycetales(100);f__Metschnikowiaceae(100);g__Metschnikowia(100);s__Metschnikowia_pulcherrima(100); | Metschnikowia | 13 | Saprotroph | \|Undefined Saprotroph\| | Yeast | NULL | Probable | As nectar-tap_saprotroph (Põlme et al. 2020); Decay sugar-rich substrates (Põlme et al. 2020) | Põlme S, et al. 2020. Fungal Diversity 105:1-16 ((https://doi.org/10.1007/s13225-020-00466-2)) |
| Otu34530 | 1 | k__Fungi(100);p__Ascomycota(100);c__Saccharomycetes(100);o__Saccharomycetales(100);f__Metschnikowiaceae(100);g__Metschnikowia(100);s__Metschnikowia_pulcherrima(100); | Metschnikowia | 13 | Saprotroph | \|Undefined Saprotroph\| | Yeast | NULL | Probable | As nectar-tap_saprotroph (Põlme et al. 2020); Decay sugar-rich substrates (Põlme et al. 2020) | Põlme S, et al. 2020. Fungal Diversity 105:1-16 ((https://doi.org/10.1007/s13225-020-00466-2)) |
| Otu34532 | 1 | k__Fungi(100);p__Ascomycota(100);c__Saccharomycetes(100);o__Saccharomycetales(100);f__Metschnikowiaceae(100);g__Metschnikowia(100);s__Metschnikowia_pulcherrima(100); | Metschnikowia | 13 | Saprotroph | \|Undefined Saprotroph\| | Yeast | NULL | Probable | As nectar-tap_saprotroph (Põlme et al. 2020); Decay sugar-rich substrates (Põlme et al. 2020) | Põlme S, et al. 2020. Fungal Diversity 105:1-16 ((https://doi.org/10.1007/s13225-020-00466-2)) |
| Otu34533 | 1 | k__Fungi(100);p__Ascomycota(100);c__Saccharomycetes(100);o__Saccharomycetales(100);f__Metschnikowiaceae(100);g__Metschnikowia(100);g__Metschnikowia_unclassified(100); | Metschnikowia | 13 | Saprotroph | \|Undefined Saprotroph\| | Yeast | NULL | Probable | As nectar-tap_saprotroph (Põlme et al. 2020); Decay sugar-rich substrates (Põlme et al. 2020) | Põlme S, et al. 2020. Fungal Diversity 105:1-16 ((https://doi.org/10.1007/s13225-020-00466-2)) |
| Otu34534 | 1 | k__Fungi(100);p__Ascomycota(100);c__Saccharomycetes(100);o__Saccharomycetales(100);f__Metschnikowiaceae(100);g__Metschnikowia(100);s__Metschnikowia_pulcherrima(100); | Metschnikowia | 13 | Saprotroph | \|Undefined Saprotroph\| | Yeast | NULL | Probable | As nectar-tap_saprotroph (Põlme et al. 2020); Decay sugar-rich substrates (Põlme et al. 2020) | Põlme S, et al. 2020. Fungal Diversity 105:1-16 ((https://doi.org/10.1007/s13225-020-00466-2)) |
| Otu34535 | 1 | k__Fungi(100);p__Ascomycota(100);c__Saccharomycetes(100);o__Saccharomycetales(100);f__Metschnikowiaceae(100);g__Metschnikowia(100);s__Metschnikowia_pulcherrima(100); | Metschnikowia | 13 | Saprotroph | \|Undefined Saprotroph\| | Yeast | NULL | Probable | As nectar-tap_saprotroph (Põlme et al. 2020); Decay sugar-rich substrates (Põlme et al. 2020) | Põlme S, et al. 2020. Fungal Diversity 105:1-16 ((https://doi.org/10.1007/s13225-020-00466-2)) |
| Otu34538 | 1 | k__Fungi(100);p__Ascomycota(100);c__Saccharomycetes(100);o__Saccharomycetales(100);f__Metschnikowiaceae(100);g__Metschnikowia(100);s__Metschnikowia_pulcherrima(100); | Metschnikowia | 13 | Saprotroph | \|Undefined Saprotroph\| | Yeast | NULL | Probable | As nectar-tap_saprotroph (Põlme et al. 2020); Decay sugar-rich substrates (Põlme et al. 2020) | Põlme S, et al. 2020. Fungal Diversity 105:1-16 ((https://doi.org/10.1007/s13225-020-00466-2)) |
| Otu34539 | 1 | k__Fungi(100);p__Ascomycota(100);c__Saccharomycetes(100);o__Saccharomycetales(100);f__Saccharomycetales_family_Incertae_sedis(100);g__Candida(100);s__Candida_stellata(100); | Candida | 13 | Pathotroph-Saprotroph-Symbiotroph | Animal Pathogen-Endophyte-Endosymbiont-Epiphyte-\|Undefined Saprotroph\| | Dimorphic Yeast-Yeast | NULL | Probable | As nectar-tap_saprotroph (Põlme et al. 2020); Decay sugar-rich substrates (Põlme et al. 2020); This asexual genus is polyphyletic and contain species that function very differently depend on what they are. Caution should be taken when interpreting guilds for Candida species. | Manolakaki D, et al. 2010. Virulence. 1: 367-375 ((https://doi.org/10.4161/viru.1.5.12796)); Mekbib SB, et al. 2011. Tropical Plant Pathology 36:233-240 ((https://doi.org/10.1590/S1982-56762011000400004)); Põlme S, et al. 2020. Fungal Diversity 105:1-16 ((https://doi.org/10.1007/s13225-020-00466-2)) |
| Otu34543 | 1 | k__Fungi(100);p__Ascomycota(100);c__Saccharomycetes(100);o__Saccharomycetales(100);f__Metschnikowiaceae(100);g__Metschnikowia(100);g__Metschnikowia_unclassified(100); | Metschnikowia | 13 | Saprotroph | \|Undefined Saprotroph\| | Yeast | NULL | Probable | As nectar-tap_saprotroph (Põlme et al. 2020); Decay sugar-rich substrates (Põlme et al. 2020) | Põlme S, et al. 2020. Fungal Diversity 105:1-16 ((https://doi.org/10.1007/s13225-020-00466-2)) |
| Otu34554 | 1 | k__Fungi(100);p__Ascomycota(100);c__Saccharomycetes(100);o__Saccharomycetales(100);f__Metschnikowiaceae(100);g__Metschnikowia(100);g__Metschnikowia_unclassified(100); | Metschnikowia | 13 | Saprotroph | \|Undefined Saprotroph\| | Yeast | NULL | Probable | As nectar-tap_saprotroph (Põlme et al. 2020); Decay sugar-rich substrates (Põlme et al. 2020) | Põlme S, et al. 2020. Fungal Diversity 105:1-16 ((https://doi.org/10.1007/s13225-020-00466-2)) |
| Otu34555 | 1 | k__Fungi(100);p__Ascomycota(100);c__Saccharomycetes(100);o__Saccharomycetales(100);f__Metschnikowiaceae(100);g__Metschnikowia(100);s__Metschnikowia_pulcherrima(100); | Metschnikowia | 13 | Saprotroph | \|Undefined Saprotroph\| | Yeast | NULL | Probable | As nectar-tap_saprotroph (Põlme et al. 2020); Decay sugar-rich substrates (Põlme et al. 2020) | Põlme S, et al. 2020. Fungal Diversity 105:1-16 ((https://doi.org/10.1007/s13225-020-00466-2)) |
| Otu34556 | 1 | k__Fungi(100);p__Ascomycota(100);c__Saccharomycetes(100);o__Saccharomycetales(100);f__Metschnikowiaceae(100);g__Metschnikowia(100);g__Metschnikowia_unclassified(100); | Metschnikowia | 13 | Saprotroph | \|Undefined Saprotroph\| | Yeast | NULL | Probable | As nectar-tap_saprotroph (Põlme et al. 2020); Decay sugar-rich substrates (Põlme et al. 2020) | Põlme S, et al. 2020. Fungal Diversity 105:1-16 ((https://doi.org/10.1007/s13225-020-00466-2)) |

| | | | | | | | | | | | |
|---|---|---|---|---|---|---|---|---|---|---|---|
| Otu34558 | 1 | k__Fungi(100);p__Ascomycota(100);c__Saccharomycetes(100);o__Saccharomycetales(100);f__Saccharomycetales_family_Incertae_sedis(100);g__Candida(100);g__Candida_unclassified(100); | Candida | 13 | Pathotroph-Saprotroph-Symbiotroph | Animal Pathogen-Endophyte-Endosymbiont-Epiphyte-\|Undefined Saprotroph\| | Dimorphic Yeast-Yeast | NULL | Probable | As nectar-tap_saprotroph (Põlme et al. 2020); Decay sugar-rich substrates (Põlme et al. 2020); This asexual genus is polyphyletic and contain species that function very differently depend on what they are. Caution should be taken when interpreting guilds for Candida species. | Manolakaki D, et al. 2010. Virulence. 1: 367-375 ((https://doi.org/10.4161/viru.1.5.12796)); Mekbib SB, et al. 2011. Tropical Plant Pathology 36:233-240 ((https://doi.org/10.1590/S1982-56762011000400004)); Põlme S, et al. 2020. Fungal Diversity 105:1-16 ((https://doi.org/10.1007/s13225-020-00466-2)) |
| Otu34561 | 1 | k__Fungi(100);p__Ascomycota(100);c__Saccharomycetes(100);o__Saccharomycetales(100);f__Metschnikowiaceae(100);g__Metschnikowia(100);g__Metschnikowia_unclassified(100); | Metschnikowia | 13 | Saprotroph | \|Undefined Saprotroph\| | Yeast | NULL | Probable | As nectar-tap_saprotroph (Põlme et al. 2020); Decay sugar-rich substrates (Põlme et al. 2020) | Põlme S, et al. 2020. Fungal Diversity 105:1-16 ((https://doi.org/10.1007/s13225-020-00466-2)) |
| Otu34562 | 1 | k__Fungi(100);p__Ascomycota(100);c__Saccharomycetes(100);o__Saccharomycetales(100);f__Metschnikowiaceae(100);g__Metschnikowia(100);s__Metschnikowia_pulcherrima(100); | Metschnikowia | 13 | Saprotroph | \|Undefined Saprotroph\| | Yeast | NULL | Probable | As nectar-tap_saprotroph (Põlme et al. 2020); Decay sugar-rich substrates (Põlme et al. 2020) | Põlme S, et al. 2020. Fungal Diversity 105:1-16 ((https://doi.org/10.1007/s13225-020-00466-2)) |
| Otu34563 | 1 | k__Fungi(100);p__Ascomycota(100);c__Saccharomycetes(100);o__Saccharomycetales(100);f__Metschnikowiaceae(100);g__Metschnikowia(100);s__Metschnikowia_pulcherrima(100); | Metschnikowia | 13 | Saprotroph | \|Undefined Saprotroph\| | Yeast | NULL | Probable | As nectar-tap_saprotroph (Põlme et al. 2020); Decay sugar-rich substrates (Põlme et al. 2020) | Põlme S, et al. 2020. Fungal Diversity 105:1-16 ((https://doi.org/10.1007/s13225-020-00466-2)) |
| Otu34564 | 1 | k__Fungi(100);p__Ascomycota(100);c__Saccharomycetes(100);o__Saccharomycetales(100);f__Metschnikowiaceae(100);g__Metschnikowia(100);s__Metschnikowia_pulcherrima(100); | Metschnikowia | 13 | Saprotroph | \|Undefined Saprotroph\| | Yeast | NULL | Probable | As nectar-tap_saprotroph (Põlme et al. 2020); Decay sugar-rich substrates (Põlme et al. 2020) | Põlme S, et al. 2020. Fungal Diversity 105:1-16 ((https://doi.org/10.1007/s13225-020-00466-2)) |
| Otu34565 | 1 | k__Fungi(100);p__Ascomycota(100);c__Saccharomycetes(100);o__Saccharomycetales(100);f__Metschnikowiaceae(100);g__Metschnikowia(100);s__Metschnikowia_pulcherrima(100); | Metschnikowia | 13 | Saprotroph | \|Undefined Saprotroph\| | Yeast | NULL | Probable | As nectar-tap_saprotroph (Põlme et al. 2020); Decay sugar-rich substrates (Põlme et al. 2020) | Põlme S, et al. 2020. Fungal Diversity 105:1-16 ((https://doi.org/10.1007/s13225-020-00466-2)) |
| Otu34566 | 1 | k__Fungi(100);p__Ascomycota(100);c__Saccharomycetes(100);o__Saccharomycetales(100);f__Metschnikowiaceae(100);g__Metschnikowia(100);s__Metschnikowia_pulcherrima(100); | Metschnikowia | 13 | Saprotroph | \|Undefined Saprotroph\| | Yeast | NULL | Probable | As nectar-tap_saprotroph (Põlme et al. 2020); Decay sugar-rich substrates (Põlme et al. 2020) | Põlme S, et al. 2020. Fungal Diversity 105:1-16 ((https://doi.org/10.1007/s13225-020-00466-2)) |
| Otu34570 | 1 | k__Fungi(100);p__Ascomycota(100);c__Saccharomycetes(100);o__Saccharomycetales(100);f__Metschnikowiaceae(100);g__Metschnikowia(100);s__Metschnikowia_pulcherrima(100); | Metschnikowia | 13 | Saprotroph | \|Undefined Saprotroph\| | Yeast | NULL | Probable | As nectar-tap_saprotroph (Põlme et al. 2020); Decay sugar-rich substrates (Põlme et al. 2020) | Põlme S, et al. 2020. Fungal Diversity 105:1-16 ((https://doi.org/10.1007/s13225-020-00466-2)) |
| Otu34571 | 1 | k__Fungi(100);p__Ascomycota(100);c__Saccharomycetes(100);o__Saccharomycetales(100);f__Metschnikowiaceae(100);g__Metschnikowia(100);s__Metschnikowia_pulcherrima(100); | Metschnikowia | 13 | Saprotroph | \|Undefined Saprotroph\| | Yeast | NULL | Probable | As nectar-tap_saprotroph (Põlme et al. 2020); Decay sugar-rich substrates (Põlme et al. 2020) | Põlme S, et al. 2020. Fungal Diversity 105:1-16 ((https://doi.org/10.1007/s13225-020-00466-2)) |
| Otu34573 | 1 | k__Fungi(100);p__Ascomycota(100);c__Saccharomycetes(100);o__Saccharomycetales(100);o__Saccharomycetales_unclassified(100);o__Saccharomycetales_unclassified(100);o__Saccharomycetales_unclassified(100); | Saccharomycetales | 7 | Saprotroph | Undefined Saprotroph | Yeast | NULL | Possible | NULL | Sterkenburg E, et al. 2015. New Phytologist 207:1145-1158 ((https://doi.org/10.1111/nph.13426)) |
| Otu34574 | 1 | k__Fungi(100);p__Ascomycota(100);c__Saccharomycetes(100);o__Saccharomycetales(100);f__Metschnikowiaceae(100);g__Metschnikowia(100);s__Metschnikowia_pulcherrima(100); | Metschnikowia | 13 | Saprotroph | \|Undefined Saprotroph\| | Yeast | NULL | Probable | As nectar-tap_saprotroph (Põlme et al. 2020); Decay sugar-rich substrates (Põlme et al. 2020) | Põlme S, et al. 2020. Fungal Diversity 105:1-16 ((https://doi.org/10.1007/s13225-020-00466-2)) |
| Otu34575 | 1 | k__Fungi(100);p__Ascomycota(100);c__Saccharomycetes(100);o__Saccharomycetales(100);f__Metschnikowiaceae(100);g__Metschnikowia(100);g__Metschnikowia_unclassified(100); | Metschnikowia | 13 | Saprotroph | \|Undefined Saprotroph\| | Yeast | NULL | Probable | As nectar-tap_saprotroph (Põlme et al. 2020); Decay sugar-rich substrates (Põlme et al. 2020) | Põlme S, et al. 2020. Fungal Diversity 105:1-16 ((https://doi.org/10.1007/s13225-020-00466-2)) |
| Otu34576 | 1 | k__Fungi(100);p__Ascomycota(100);c__Saccharomycetes(100);o__Saccharomycetales(100);f__Metschnikowiaceae(100);g__Metschnikowia(100);g__Metschnikowia_unclassified(100); | Metschnikowia | 13 | Saprotroph | \|Undefined Saprotroph\| | Yeast | NULL | Probable | As nectar-tap_saprotroph (Põlme et al. 2020); Decay sugar-rich substrates (Põlme et al. 2020) | Põlme S, et al. 2020. Fungal Diversity 105:1-16 ((https://doi.org/10.1007/s13225-020-00466-2)) |
| Otu34577 | 1 | k__Fungi(100);p__Ascomycota(100);c__Saccharomycetes(100);o__Saccharomycetales(100);f__Saccharomycetales_family_Incertae_sedis(100);g__Candida(100);g__Candida_unclassified(100); | Candida | 13 | Pathotroph-Saprotroph-Symbiotroph | Animal Pathogen-Endophyte-Endosymbiont-Epiphyte-\|Undefined Saprotroph\| | Dimorphic Yeast-Yeast | NULL | Probable | As nectar-tap_saprotroph (Põlme et al. 2020); Decay sugar-rich substrates (Põlme et al. 2020); This asexual genus is polyphyletic and contain species that function very differently depend on what they are. Caution should be taken when interpreting guilds for Candida species. | Manolakaki D, et al. 2010. Virulence. 1: 367-375 ((https://doi.org/10.4161/viru.1.5.12796)); Mekbib SB, et al. 2011. Tropical Plant Pathology 36:233-240 ((https://doi.org/10.1590/S1982-56762011000400004)); Põlme S, et al. 2020. Fungal Diversity 105:1-16 ((https://doi.org/10.1007/s13225-020-00466-2)) |
| Otu34578 | 1 | k__Fungi(100);p__Ascomycota(100);c__Saccharomycetes(100);o__Saccharomycetales(100);f__Metschnikowiaceae(100);g__Metschnikowia(100);s__Metschnikowia_pulcherrima(100); | Metschnikowia | 13 | Saprotroph | \|Undefined Saprotroph\| | Yeast | NULL | Probable | As nectar-tap_saprotroph (Põlme et al. 2020); Decay sugar-rich substrates (Põlme et al. 2020) | Põlme S, et al. 2020. Fungal Diversity 105:1-16 ((https://doi.org/10.1007/s13225-020-00466-2)) |
| Otu34579 | 1 | k__Fungi(100);p__Ascomycota(100);c__Saccharomycetes(100);o__Saccharomycetales(100);f__Saccharomycetales_family_Incertae_sedis(100);g__Candida(100);g__Candida_unclassified(100); | Candida | 13 | Pathotroph-Saprotroph-Symbiotroph | Animal Pathogen-Endophyte-Endosymbiont-Epiphyte-\|Undefined Saprotroph\| | Dimorphic Yeast-Yeast | NULL | Probable | As nectar-tap_saprotroph (Põlme et al. 2020); Decay sugar-rich substrates (Põlme et al. 2020); This asexual genus is polyphyletic and contain species that function very differently depend on what they are. Caution should be taken when interpreting guilds for Candida species. | Manolakaki D, et al. 2010. Virulence. 1: 367-375 ((https://doi.org/10.4161/viru.1.5.12796)); Mekbib SB, et al. 2011. Tropical Plant Pathology 36:233-240 ((https://doi.org/10.1590/S1982-56762011000400004)); Põlme S, et al. 2020. Fungal Diversity 105:1-16 ((https://doi.org/10.1007/s13225-020-00466-2)) |
| Otu34582 | 1 | k__Fungi(100);p__Ascomycota(100);c__Saccharomycetes(100);o__Saccharomycetales(100);f__Metschnikowiaceae(100);g__Metschnikowia(100);g__Metschnikowia_unclassified(100); | Metschnikowia | 13 | Saprotroph | \|Undefined Saprotroph\| | Yeast | NULL | Probable | As nectar-tap_saprotroph (Põlme et al. 2020); Decay sugar-rich substrates (Põlme et al. 2020) | Põlme S, et al. 2020. Fungal Diversity 105:1-16 ((https://doi.org/10.1007/s13225-020-00466-2)) |
| Otu34583 | 1 | k__Fungi(100);p__Ascomycota(100);c__Saccharomycetes(100);o__Saccharomycetales(100);f__Metschnikowiaceae(100);g__Metschnikowia(100);s__Metschnikowia_pulcherrima(100); | Metschnikowia | 13 | Saprotroph | \|Undefined Saprotroph\| | Yeast | NULL | Probable | As nectar-tap_saprotroph (Põlme et al. 2020); Decay sugar-rich substrates (Põlme et al. 2020) | Põlme S, et al. 2020. Fungal Diversity 105:1-16 ((https://doi.org/10.1007/s13225-020-00466-2)) |
| Otu34588 | 1 | k__Fungi(100);p__Ascomycota(100);c__Saccharomycetes(100);o__Saccharomycetales(100);f__Metschnikowiaceae(100);g__Metschnikowia(100);g__Metschnikowia_unclassified(100); | Metschnikowia | 13 | Saprotroph | \|Undefined Saprotroph\| | Yeast | NULL | Probable | As nectar-tap_saprotroph (Põlme et al. 2020); Decay sugar-rich substrates (Põlme et al. 2020) | Põlme S, et al. 2020. Fungal Diversity 105:1-16 ((https://doi.org/10.1007/s13225-020-00466-2)) |
| Otu34589 | 1 | k__Fungi(100);p__Ascomycota(100);c__Saccharomycetes(100);o__Saccharomycetales(100);f__Metschnikowiaceae(100);g__Metschnikowia(100);s__Metschnikowia_pulcherrima(100); | Metschnikowia | 13 | Saprotroph | \|Undefined Saprotroph\| | Yeast | NULL | Probable | As nectar-tap_saprotroph (Põlme et al. 2020); Decay sugar-rich substrates (Põlme et al. 2020) | Põlme S, et al. 2020. Fungal Diversity 105:1-16 ((https://doi.org/10.1007/s13225-020-00466-2)) |
| Otu34590 | 1 | k__Fungi(100);p__Ascomycota(100);c__Saccharomycetes(100);o__Saccharomycetales(100);f__Metschnikowiaceae(100);g__Metschnikowia(100);s__Metschnikowia_pulcherrima(100); | Metschnikowia | 13 | Saprotroph | \|Undefined Saprotroph\| | Yeast | NULL | Probable | As nectar-tap_saprotroph (Põlme et al. 2020); Decay sugar-rich substrates (Põlme et al. 2020) | Põlme S, et al. 2020. Fungal Diversity 105:1-16 ((https://doi.org/10.1007/s13225-020-00466-2)) |
| Otu34593 | 1 | k__Fungi(100);p__Ascomycota(100);c__Saccharomycetes(100);o__Saccharomycetales(100);f__Metschnikowiaceae(100);g__Metschnikowia(100);g__Metschnikowia_unclassified(100); | Metschnikowia | 13 | Saprotroph | \|Undefined Saprotroph\| | Yeast | NULL | Probable | As nectar-tap_saprotroph (Põlme et al. 2020); Decay sugar-rich substrates (Põlme et al. 2020) | Põlme S, et al. 2020. Fungal Diversity 105:1-16 ((https://doi.org/10.1007/s13225-020-00466-2)) |
| Otu34594 | 1 | k__Fungi(100);p__Ascomycota(100);c__Saccharomycetes(100);o__Saccharomycetales(100);f__Metschnikowiaceae(100);g__Metschnikowia(100);s__Metschnikowia_pulcherrima(100); | Metschnikowia | 13 | Saprotroph | \|Undefined Saprotroph\| | Yeast | NULL | Probable | As nectar-tap_saprotroph (Põlme et al. 2020); Decay sugar-rich substrates (Põlme et al. 2020) | Põlme S, et al. 2020. Fungal Diversity 105:1-16 ((https://doi.org/10.1007/s13225-020-00466-2)) |
| Otu34595 | 1 | k__Fungi(100);p__Ascomycota(100);c__Saccharomycetes(100);o__Saccharomycetales(100);f__Metschnikowiaceae(100);g__Metschnikowia(100);s__Metschnikowia_pulcherrima(100); | Metschnikowia | 13 | Saprotroph | \|Undefined Saprotroph\| | Yeast | NULL | Probable | As nectar-tap_saprotroph (Põlme et al. 2020); Decay sugar-rich substrates (Põlme et al. 2020) | Põlme S, et al. 2020. Fungal Diversity 105:1-16 ((https://doi.org/10.1007/s13225-020-00466-2)) |
| Otu34596 | 1 | k__Fungi(100);p__Ascomycota(100);c__Saccharomycetes(100);o__Saccharomycetales(100);f__Metschnikowiaceae(100);g__Metschnikowia(100);s__Metschnikowia_pulcherrima(100); | Metschnikowia | 13 | Saprotroph | \|Undefined Saprotroph\| | Yeast | NULL | Probable | As nectar-tap_saprotroph (Põlme et al. 2020); Decay sugar-rich substrates (Põlme et al. 2020) | Põlme S, et al. 2020. Fungal Diversity 105:1-16 ((https://doi.org/10.1007/s13225-020-00466-2)) |
| Otu34597 | 1 | k__Fungi(100);p__Ascomycota(100);c__Saccharomycetes(100);o__Saccharomycetales(100);f__Metschnikowiaceae(100);g__Metschnikowia(100);s__Metschnikowia_pulcherrima(100); | Metschnikowia | 13 | Saprotroph | \|Undefined Saprotroph\| | Yeast | NULL | Probable | As nectar-tap_saprotroph (Põlme et al. 2020); Decay sugar-rich substrates (Põlme et al. 2020) | Põlme S, et al. 2020. Fungal Diversity 105:1-16 ((https://doi.org/10.1007/s13225-020-00466-2)) |
| Otu34598 | 1 | k__Fungi(100);p__Ascomycota(100);c__Saccharomycetes(100);o__Saccharomycetales(100);f__Metschnikowiaceae(100);g__Metschnikowia(100);s__Metschnikowia_pulcherrima(100); | Metschnikowia | 13 | Saprotroph | \|Undefined Saprotroph\| | Yeast | NULL | Probable | As nectar-tap_saprotroph (Põlme et al. 2020); Decay sugar-rich substrates (Põlme et al. 2020) | Põlme S, et al. 2020. Fungal Diversity 105:1-16 ((https://doi.org/10.1007/s13225-020-00466-2)) |
| Otu34599 | 1 | k__Fungi(100);p__Ascomycota(100);c__Saccharomycetes(100);o__Saccharomycetales(100);f__Metschnikowiaceae(100);g__Metschnikowia(100);s__Metschnikowia_pulcherrima(100); | Metschnikowia | 13 | Saprotroph | \|Undefined Saprotroph\| | Yeast | NULL | Probable | As nectar-tap_saprotroph (Põlme et al. 2020); Decay sugar-rich substrates (Põlme et al. 2020) | Põlme S, et al. 2020. Fungal Diversity 105:1-16 ((https://doi.org/10.1007/s13225-020-00466-2)) |
| Otu34600 | 1 | k__Fungi(100);p__Ascomycota(100);c__Saccharomycetes(100);o__Saccharomycetales(100);f__Metschnikowiaceae(100);g__Metschnikowia(100);s__Metschnikowia_pulcherrima(100); | Metschnikowia | 13 | Saprotroph | \|Undefined Saprotroph\| | Yeast | NULL | Probable | As nectar-tap_saprotroph (Põlme et al. 2020); Decay sugar-rich substrates (Põlme et al. 2020) | Põlme S, et al. 2020. Fungal Diversity 105:1-16 ((https://doi.org/10.1007/s13225-020-00466-2)) |
| Otu34601 | 1 | k__Fungi(100);p__Ascomycota(100);c__Saccharomycetes(100);o__Saccharomycetales(100);f__Metschnikowiaceae(100);g__Metschnikowia(100);s__Metschnikowia_pulcherrima(100); | Metschnikowia | 13 | Saprotroph | \|Undefined Saprotroph\| | Yeast | NULL | Probable | As nectar-tap_saprotroph (Põlme et al. 2020); Decay sugar-rich substrates (Põlme et al. 2020) | Põlme S, et al. 2020. Fungal Diversity 105:1-16 ((https://doi.org/10.1007/s13225-020-00466-2)) |
| Otu34602 | 1 | k__Fungi(100);p__Ascomycota(100);c__Saccharomycetes(100);o__Saccharomycetales(100);f__Metschnikowiaceae(100);g__Metschnikowia(100);s__Metschnikowia_pulcherrima(100); | Metschnikowia | 13 | Saprotroph | \|Undefined Saprotroph\| | Yeast | NULL | Probable | As nectar-tap_saprotroph (Põlme et al. 2020); Decay sugar-rich substrates (Põlme et al. 2020) | Põlme S, et al. 2020. Fungal Diversity 105:1-16 ((https://doi.org/10.1007/s13225-020-00466-2)) |
| Otu34603 | 1 | k__Fungi(100);p__Ascomycota(100);c__Saccharomycetes(100);o__Saccharomycetales(100);f__Metschnikowiaceae(100);g__Metschnikowia(100);s__Metschnikowia_pulcherrima(100); | Metschnikowia | 13 | Saprotroph | \|Undefined Saprotroph\| | Yeast | NULL | Probable | As nectar-tap_saprotroph (Põlme et al. 2020); Decay sugar-rich substrates (Põlme et al. 2020) | Põlme S, et al. 2020. Fungal Diversity 105:1-16 ((https://doi.org/10.1007/s13225-020-00466-2)) |
| Otu34604 | 1 | k__Fungi(100);p__Ascomycota(100);c__Saccharomycetes(100);o__Saccharomycetales(100);f__Metschnikowiaceae(100);g__Metschnikowia(100);s__Metschnikowia_pulcherrima(100); | Metschnikowia | 13 | Saprotroph | \|Undefined Saprotroph\| | Yeast | NULL | Probable | As nectar-tap_saprotroph (Põlme et al. 2020); Decay sugar-rich substrates (Põlme et al. 2020) | Põlme S, et al. 2020. Fungal Diversity 105:1-16 ((https://doi.org/10.1007/s13225-020-00466-2)) |
| Otu34605 | 1 | k__Fungi(100);p__Ascomycota(100);c__Saccharomycetes(100);o__Saccharomycetales(100);f__Metschnikowiaceae(100);g__Metschnikowia(100);s__Metschnikowia_pulcherrima(100); | Metschnikowia | 13 | Saprotroph | \|Undefined Saprotroph\| | Yeast | NULL | Probable | As nectar-tap_saprotroph (Põlme et al. 2020); Decay sugar-rich substrates (Põlme et al. 2020) | Põlme S, et al. 2020. Fungal Diversity 105:1-16 ((https://doi.org/10.1007/s13225-020-00466-2)) |
| Otu34606 | 1 | k__Fungi(100);p__Ascomycota(100);c__Saccharomycetes(100);o__Saccharomycetales(100);f__Metschnikowiaceae(100);g__Metschnikowia(100);s__Metschnikowia_pulcherrima(100); | Metschnikowia | 13 | Saprotroph | \|Undefined Saprotroph\| | Yeast | NULL | Probable | As nectar-tap_saprotroph (Põlme et al. 2020); Decay sugar-rich substrates (Põlme et al. 2020) | Põlme S, et al. 2020. Fungal Diversity 105:1-16 ((https://doi.org/10.1007/s13225-020-00466-2)) |
| Otu34607 | 1 | k__Fungi(100);p__Ascomycota(100);c__Saccharomycetes(100);o__Saccharomycetales(100);f__Metschnikowiaceae(100);g__Metschnikowia(100);g__Metschnikowia_unclassified(100); | Metschnikowia | 13 | Saprotroph | \|Undefined Saprotroph\| | Yeast | NULL | Probable | As nectar-tap_saprotroph (Põlme et al. 2020); Decay sugar-rich substrates (Põlme et al. 2020) | Põlme S, et al. 2020. Fungal Diversity 105:1-16 ((https://doi.org/10.1007/s13225-020-00466-2)) |
| Otu34608 | 1 | k__Fungi(100);p__Ascomycota(100);c__Saccharomycetes(100);o__Saccharomycetales(100);f__Metschnikowiaceae(100);g__Metschnikowia(100);g__Metschnikowia_unclassified(100); | Metschnikowia | 13 | Saprotroph | \|Undefined Saprotroph\| | Yeast | NULL | Probable | As nectar-tap_saprotroph (Põlme et al. 2020); Decay sugar-rich substrates (Põlme et al. 2020) | Põlme S, et al. 2020. Fungal Diversity 105:1-16 ((https://doi.org/10.1007/s13225-020-00466-2)) |
| Otu34609 | 1 | k__Fungi(100);p__Ascomycota(100);c__Saccharomycetes(100);o__Saccharomycetales(100);f__Metschnikowiaceae(100);g__Metschnikowia(100);g__Metschnikowia_unclassified(100); | Metschnikowia | 13 | Saprotroph | \|Undefined Saprotroph\| | Yeast | NULL | Probable | As nectar-tap_saprotroph (Põlme et al. 2020); Decay sugar-rich substrates (Põlme et al. 2020) | Põlme S, et al. 2020. Fungal Diversity 105:1-16 ((https://doi.org/10.1007/s13225-020-00466-2)) |
| Otu34610 | 1 | k__Fungi(100);p__Ascomycota(100);c__Saccharomycetes(100);o__Saccharomycetales(100);f__Metschnikowiaceae(100);g__Metschnikowia(100);g__Metschnikowia_unclassified(100); | Metschnikowia | 13 | Saprotroph | \|Undefined Saprotroph\| | Yeast | NULL | Probable | As nectar-tap_saprotroph (Põlme et al. 2020); Decay sugar-rich substrates (Põlme et al. 2020) | Põlme S, et al. 2020. Fungal Diversity 105:1-16 ((https://doi.org/10.1007/s13225-020-00466-2)) |
| Otu34611 | 1 | k__Fungi(100);p__Ascomycota(100);c__Saccharomycetes(100);o__Saccharomycetales(100);f__Metschnikowiaceae(100);g__Metschnikowia(100);s__Metschnikowia_pulcherrima(100); | Metschnikowia | 13 | Saprotroph | \|Undefined Saprotroph\| | Yeast | NULL | Probable | As nectar-tap_saprotroph (Põlme et al. 2020); Decay sugar-rich substrates (Põlme et al. 2020) | Põlme S, et al. 2020. Fungal Diversity 105:1-16 ((https://doi.org/10.1007/s13225-020-00466-2)) |
| Otu34613 | 1 | k__Fungi(100);p__Ascomycota(100);c__Saccharomycetes(100);o__Saccharomycetales(100);f__Metschnikowiaceae(100);g__Metschnikowia(100);g__Metschnikowia_unclassified(100); | Metschnikowia | 13 | Saprotroph | \|Undefined Saprotroph\| | Yeast | NULL | Probable | As nectar-tap_saprotroph (Põlme et al. 2020); Decay sugar-rich substrates (Põlme et al. 2020) | Põlme S, et al. 2020. Fungal Diversity 105:1-16 ((https://doi.org/10.1007/s13225-020-00466-2)) |
| Otu34614 | 1 | k__Fungi(100);p__Ascomycota(100);c__Saccharomycetes(100);o__Saccharomycetales(100);f__Metschnikowiaceae(100);g__Metschnikowia(100);s__Metschnikowia_pulcherrima(100); | Metschnikowia | 13 | Saprotroph | \|Undefined Saprotroph\| | Yeast | NULL | Probable | As nectar-tap_saprotroph (Põlme et al. 2020); Decay sugar-rich substrates (Põlme et al. 2020) | Põlme S, et al. 2020. Fungal Diversity 105:1-16 ((https://doi.org/10.1007/s13225-020-00466-2)) |
| Otu34615 | 1 | k__Fungi(100);p__Ascomycota(100);c__Saccharomycetes(100);o__Saccharomycetales(100);f__Metschnikowiaceae(100);g__Metschnikowia(100);g__Metschnikowia_unclassified(100); | Metschnikowia | 13 | Saprotroph | \|Undefined Saprotroph\| | Yeast | NULL | Probable | As nectar-tap_saprotroph (Põlme et al. 2020); Decay sugar-rich substrates (Põlme et al. 2020) | Põlme S, et al. 2020. Fungal Diversity 105:1-16 ((https://doi.org/10.1007/s13225-020-00466-2)) |
| Otu34616 | 1 | k__Fungi(100);p__Ascomycota(100);c__Saccharomycetes(100);o__Saccharomycetales(100);f__Metschnikowiaceae(100);g__Metschnikowia(100);s__Metschnikowia_pulcherrima(100); | Metschnikowia | 13 | Saprotroph | \|Undefined Saprotroph\| | Yeast | NULL | Probable | As nectar-tap_saprotroph (Põlme et al. 2020); Decay sugar-rich substrates (Põlme et al. 2020) | Põlme S, et al. 2020. Fungal Diversity 105:1-16 ((https://doi.org/10.1007/s13225-020-00466-2)) |
| Otu34620 | 1 | k__Fungi(100);p__Ascomycota(100);c__Saccharomycetes(100);o__Saccharomycetales(100);f__Metschnikowiaceae(100);g__Metschnikowia(100);s__Metschnikowia_pulcherrima(100); | Metschnikowia | 13 | Saprotroph | \|Undefined Saprotroph\| | Yeast | NULL | Probable | As nectar-tap_saprotroph (Põlme et al. 2020); Decay sugar-rich substrates (Põlme et al. 2020) | Põlme S, et al. 2020. Fungal Diversity 105:1-16 ((https://doi.org/10.1007/s13225-020-00466-2)) |
| Otu34621 | 1 | k__Fungi(100);p__Ascomycota(100);c__Saccharomycetes(100);o__Saccharomycetales(100);f__Metschnikowiaceae(100);g__Metschnikowia(100);s__Metschnikowia_pulcherrima(100); | Metschnikowia | 13 | Saprotroph | \|Undefined Saprotroph\| | Yeast | NULL | Probable | As nectar-tap_saprotroph (Põlme et al. 2020); Decay sugar-rich substrates (Põlme et al. 2020) | Põlme S, et al. 2020. Fungal Diversity 105:1-16 ((https://doi.org/10.1007/s13225-020-00466-2)) |
| Otu34623 | 1 | k__Fungi(100);p__Ascomycota(100);c__Saccharomycetes(100);o__Saccharomycetales(100);f__Metschnikowiaceae(100);g__Metschnikowia(100);g__Metschnikowia_unclassified(100); | Metschnikowia | 13 | Saprotroph | \|Undefined Saprotroph\| | Yeast | NULL | Probable | As nectar-tap_saprotroph (Põlme et al. 2020); Decay sugar-rich substrates (Põlme et al. 2020) | Põlme S, et al. 2020. Fungal Diversity 105:1-16 ((https://doi.org/10.1007/s13225-020-00466-2)) |
| Otu34624 | 1 | k__Fungi(100);p__Ascomycota(100);c__Saccharomycetes(100);o__Saccharomycetales(100);f__Metschnikowiaceae(100);g__Metschnikowia(100);s__Metschnikowia_pulcherrima(100); | Metschnikowia | 13 | Saprotroph | \|Undefined Saprotroph\| | Yeast | NULL | Probable | As nectar-tap_saprotroph (Põlme et al. 2020); Decay sugar-rich substrates (Põlme et al. 2020) | Põlme S, et al. 2020. Fungal Diversity 105:1-16 ((https://doi.org/10.1007/s13225-020-00466-2)) |
| Otu34627 | 1 | k__Fungi(100);p__Ascomycota(100);c__Saccharomycetes(100);o__Saccharomycetales(100);f__Metschnikowiaceae(100);g__Metschnikowia(100);s__Metschnikowia_pulcherrima(100); | Metschnikowia | 13 | Saprotroph | \|Undefined Saprotroph\| | Yeast | NULL | Probable | As nectar-tap_saprotroph (Põlme et al. 2020); Decay sugar-rich substrates (Põlme et al. 2020) | Põlme S, et al. 2020. Fungal Diversity 105:1-16 ((https://doi.org/10.1007/s13225-020-00466-2)) |
| Otu34628 | 1 | k__Fungi(100);p__Ascomycota(100);c__Saccharomycetes(100);o__Saccharomycetales(100);f__Metschnikowiaceae(100);g__Metschnikowia(100);s__Metschnikowia_pulcherrima(100); | Metschnikowia | 13 | Saprotroph | \|Undefined Saprotroph\| | Yeast | NULL | Probable | As nectar-tap_saprotroph (Põlme et al. 2020); Decay sugar-rich substrates (Põlme et al. 2020) | Põlme S, et al. 2020. Fungal Diversity 105:1-16 ((https://doi.org/10.1007/s13225-020-00466-2)) |
| Otu34629 | 1 | k__Fungi(100);p__Ascomycota(100);c__Saccharomycetes(100);o__Saccharomycetales(100);f__Metschnikowiaceae(100);g__Metschnikowia(100);s__Metschnikowia_pulcherrima(100); | Metschnikowia | 13 | Saprotroph | \|Undefined Saprotroph\| | Yeast | NULL | Probable | As nectar-tap_saprotroph (Põlme et al. 2020); Decay sugar-rich substrates (Põlme et al. 2020) | Põlme S, et al. 2020. Fungal Diversity 105:1-16 ((https://doi.org/10.1007/s13225-020-00466-2)) |
| Otu34630 | 1 | k__Fungi(100);p__Ascomycota(100);c__Saccharomycetes(100);o__Saccharomycetales(100);f__Metschnikowiaceae(100);g__Metschnikowia(100);s__Metschnikowia_pulcherrima(100); | Metschnikowia | 13 | Saprotroph | \|Undefined Saprotroph\| | Yeast | NULL | Probable | As nectar-tap_saprotroph (Põlme et al. 2020); Decay sugar-rich substrates (Põlme et al. 2020) | Põlme S, et al. 2020. Fungal Diversity 105:1-16 ((https://doi.org/10.1007/s13225-020-00466-2)) |
| Otu34631 | 1 | k__Fungi(100);p__Ascomycota(100);c__Saccharomycetes(100);o__Saccharomycetales(100);f__Metschnikowiaceae(100);g__Metschnikowia(100);s__Metschnikowia_pulcherrima(100); | Metschnikowia | 13 | Saprotroph | \|Undefined Saprotroph\| | Yeast | NULL | Probable | As nectar-tap_saprotroph (Põlme et al. 2020); Decay sugar-rich substrates (Põlme et al. 2020) | Põlme S, et al. 2020. Fungal Diversity 105:1-16 ((https://doi.org/10.1007/s13225-020-00466-2)) |
| Otu34632 | 1 | k__Fungi(100);p__Ascomycota(100);c__Saccharomycetes(100);o__Saccharomycetales(100);f__Metschnikowiaceae(100);g__Metschnikowia(100);s__Metschnikowia_pulcherrima(100); | Metschnikowia | 13 | Saprotroph | \|Undefined Saprotroph\| | Yeast | NULL | Probable | As nectar-tap_saprotroph (Põlme et al. 2020); Decay sugar-rich substrates (Põlme et al. 2020) | Põlme S, et al. 2020. Fungal Diversity 105:1-16 ((https://doi.org/10.1007/s13225-020-00466-2)) |
| Otu34634 | 1 | k__Fungi(100);p__Ascomycota(100);c__Saccharomycetes(100);o__Saccharomycetales(100);f__Metschnikowiaceae(100);g__Metschnikowia(100);s__Metschnikowia_pulcherrima(100); | Metschnikowia | 13 | Saprotroph | \|Undefined Saprotroph\| | Yeast | NULL | Probable | As nectar-tap_saprotroph (Põlme et al. 2020); Decay sugar-rich substrates (Põlme et al. 2020) | Põlme S, et al. 2020. Fungal Diversity 105:1-16 ((https://doi.org/10.1007/s13225-020-00466-2)) |
| Otu34635 | 1 | k__Fungi(100);p__Ascomycota(100);c__Saccharomycetes(100);o__Saccharomycetales(100);f__Metschnikowiaceae(100);g__Metschnikowia(100);s__Metschnikowia_pulcherrima(100); | Metschnikowia | 13 | Saprotroph | \|Undefined Saprotroph\| | Yeast | NULL | Probable | As nectar-tap_saprotroph (Põlme et al. 2020); Decay sugar-rich substrates (Põlme et al. 2020) | Põlme S, et al. 2020. Fungal Diversity 105:1-16 ((https://doi.org/10.1007/s13225-020-00466-2)) |
| Otu34638 | 1 | k__Fungi(100);p__Ascomycota(100);c__Saccharomycetes(100);o__Saccharomycetales(100);f__Metschnikowiaceae(100);g__Metschnikowia(100);s__Metschnikowia_pulcherrima(100); | Metschnikowia | 13 | Saprotroph | \|Undefined Saprotroph\| | Yeast | NULL | Probable | As nectar-tap_saprotroph (Põlme et al. 2020); Decay sugar-rich substrates (Põlme et al. 2020) | Põlme S, et al. 2020. Fungal Diversity 105:1-16 ((https://doi.org/10.1007/s13225-020-00466-2)) |
| Otu34639 | 1 | k__Fungi(100);p__Ascomycota(100);c__Saccharomycetes(100);o__Saccharomycetales(100);f__Metschnikowiaceae(100);g__Metschnikowia(100);s__Metschnikowia_pulcherrima(100); | Metschnikowia | 13 | Saprotroph | \|Undefined Saprotroph\| | Yeast | NULL | Probable | As nectar-tap_saprotroph (Põlme et al. 2020); Decay sugar-rich substrates (Põlme et al. 2020) | Põlme S, et al. 2020. Fungal Diversity 105:1-16 ((https://doi.org/10.1007/s13225-020-00466-2)) |
| Otu34641 | 1 | k__Fungi(100);p__Ascomycota(100);c__Saccharomycetes(100);o__Saccharomycetales(100);f__Metschnikowiaceae(100);g__Metschnikowia(100);s__Metschnikowia_pulcherrima(100); | Metschnikowia | 13 | Saprotroph | \|Undefined Saprotroph\| | Yeast | NULL | Probable | As nectar-tap_saprotroph (Põlme et al. 2020); Decay sugar-rich substrates (Põlme et al. 2020) | Põlme S, et al. 2020. Fungal Diversity 105:1-16 ((https://doi.org/10.1007/s13225-020-00466-2)) |
| Otu34642 | 1 | k__Fungi(100);p__Ascomycota(100);c__Saccharomycetes(100);o__Saccharomycetales(100);f__Metschnikowiaceae(100);g__Metschnikowia(100);g__Metschnikowia_unclassified(100); | Metschnikowia | 13 | Saprotroph | \|Undefined Saprotroph\| | Yeast | NULL | Probable | As nectar-tap_saprotroph (Põlme et al. 2020); Decay sugar-rich substrates (Põlme et al. 2020) | Põlme S, et al. 2020. Fungal Diversity 105:1-16 ((https://doi.org/10.1007/s13225-020-00466-2)) |
| Otu34643 | 1 | k__Fungi(100);p__Ascomycota(100);c__Saccharomycetes(100);o__Saccharomycetales(100);f__Metschnikowiaceae(100);g__Metschnikowia(100);g__Metschnikowia_unclassified(100); | Metschnikowia | 13 | Saprotroph | \|Undefined Saprotroph\| | Yeast | NULL | Probable | As nectar-tap_saprotroph (Põlme et al. 2020); Decay sugar-rich substrates (Põlme et al. 2020) | Põlme S, et al. 2020. Fungal Diversity 105:1-16 ((https://doi.org/10.1007/s13225-020-00466-2)) |
| Otu34644 | 1 | k__Fungi(100);p__Ascomycota(100);c__Saccharomycetes(100);o__Saccharomycetales(100);f__Metschnikowiaceae(100);g__Metschnikowia(100);g__Metschnikowia_unclassified(100); | Metschnikowia | 13 | Saprotroph | \|Undefined Saprotroph\| | Yeast | NULL | Probable | As nectar-tap_saprotroph (Põlme et al. 2020); Decay sugar-rich substrates (Põlme et al. 2020) | Põlme S, et al. 2020. Fungal Diversity 105:1-16 ((https://doi.org/10.1007/s13225-020-00466-2)) |
| Otu34645 | 1 | k__Fungi(100);p__Ascomycota(100);c__Saccharomycetes(100);o__Saccharomycetales(100);f__Metschnikowiaceae(100);g__Metschnikowia(100);s__Metschnikowia_pulcherrima(100); | Metschnikowia | 13 | Saprotroph | \|Undefined Saprotroph\| | Yeast | NULL | Probable | As nectar-tap_saprotroph (Põlme et al. 2020); Decay sugar-rich substrates (Põlme et al. 2020) | Põlme S, et al. 2020. Fungal Diversity 105:1-16 ((https://doi.org/10.1007/s13225-020-00466-2)) |
| Otu34646 | 1 | k__Fungi(100);p__Ascomycota(100);c__Saccharomycetes(100);o__Saccharomycetales(100);f__Metschnikowiaceae(100);g__Metschnikowia(100);s__Metschnikowia_pulcherrima(100); | Metschnikowia | 13 | Saprotroph | \|Undefined Saprotroph\| | Yeast | NULL | Probable | As nectar-tap_saprotroph (Põlme et al. 2020); Decay sugar-rich substrates (Põlme et al. 2020) | Põlme S, et al. 2020. Fungal Diversity 105:1-16 ((https://doi.org/10.1007/s13225-020-00466-2)) |
| Otu34647 | 1 | k__Fungi(100);p__Ascomycota(100);c__Saccharomycetes(100);o__Saccharomycetales(100);f__Metschnikowiaceae(100);g__Metschnikowia(100);g__Metschnikowia_unclassified(100); | Metschnikowia | 13 | Saprotroph | \|Undefined Saprotroph\| | Yeast | NULL | Probable | As nectar-tap_saprotroph (Põlme et al. 2020); Decay sugar-rich substrates (Põlme et al. 2020) | Põlme S, et al. 2020. Fungal Diversity 105:1-16 ((https://doi.org/10.1007/s13225-020-00466-2)) |
| Otu34648 | 1 | k__Fungi(100);p__Ascomycota(100);c__Saccharomycetes(100);o__Saccharomycetales(100);f__Metschnikowiaceae(100);g__Metschnikowia(100);s__Metschnikowia_pulcherrima(100); | Metschnikowia | 13 | Saprotroph | \|Undefined Saprotroph\| | Yeast | NULL | Probable | As nectar-tap_saprotroph (Põlme et al. 2020); Decay sugar-rich substrates (Põlme et al. 2020) | Põlme S, et al. 2020. Fungal Diversity 105:1-16 ((https://doi.org/10.1007/s13225-020-00466-2)) |
| Otu34649 | 1 | k__Fungi(100);p__Ascomycota(100);c__Saccharomycetes(100);o__Saccharomycetales(100);f__Metschnikowiaceae(100);g__Metschnikowia(100);s__Metschnikowia_pulcherrima(100); | Metschnikowia | 13 | Saprotroph | \|Undefined Saprotroph\| | Yeast | NULL | Probable | As nectar-tap_saprotroph (Põlme et al. 2020); Decay sugar-rich substrates (Põlme et al. 2020) | Põlme S, et al. 2020. Fungal Diversity 105:1-16 ((https://doi.org/10.1007/s13225-020-00466-2)) |
| Otu34650 | 1 | k__Fungi(100);p__Ascomycota(100);c__Saccharomycetes(100);o__Saccharomycetales(100);f__Metschnikowiaceae(100);g__Metschnikowia(100);s__Metschnikowia_pulcherrima(100); | Metschnikowia | 13 | Saprotroph | \|Undefined Saprotroph\| | Yeast | NULL | Probable | As nectar-tap_saprotroph (Põlme et al. 2020); Decay sugar-rich substrates (Põlme et al. 2020) | Põlme S, et al. 2020. Fungal Diversity 105:1-16 ((https://doi.org/10.1007/s13225-020-00466-2)) |
| Otu34652 | 1 | k__Fungi(100);p__Ascomycota(100);c__Saccharomycetes(100);o__Saccharomycetales(100);f__Metschnikowiaceae(100);g__Metschnikowia(100);s__Metschnikowia_pulcherrima(100); | Metschnikowia | 13 | Saprotroph | \|Undefined Saprotroph\| | Yeast | NULL | Probable | As nectar-tap_saprotroph (Põlme et al. 2020); Decay sugar-rich substrates (Põlme et al. 2020) | Põlme S, et al. 2020. Fungal Diversity 105:1-16 ((https://doi.org/10.1007/s13225-020-00466-2)) |
| Otu34653 | 1 | k__Fungi(100);p__Ascomycota(100);c__Saccharomycetes(100);o__Saccharomycetales(100);f__Metschnikowiaceae(100);g__Metschnikowia(100);s__Metschnikowia_pulcherrima(100); | Metschnikowia | 13 | Saprotroph | \|Undefined Saprotroph\| | Yeast | NULL | Probable | As nectar-tap_saprotroph (Põlme et al. 2020); Decay sugar-rich substrates (Põlme et al. 2020) | Põlme S, et al. 2020. Fungal Diversity 105:1-16 ((https://doi.org/10.1007/s13225-020-00466-2)) |
| Otu34655 | 1 | k__Fungi(100);p__Ascomycota(100);c__Saccharomycetes(100);o__Saccharomycetales(100);f__Metschnikowiaceae(100);g__Metschnikowia(100);s__Metschnikowia_pulcherrima(100); | Metschnikowia | 13 | Saprotroph | \|Undefined Saprotroph\| | Yeast | NULL | Probable | As nectar-tap_saprotroph (Põlme et al. 2020); Decay sugar-rich substrates (Põlme et al. 2020) | Põlme S, et al. 2020. Fungal Diversity 105:1-16 ((https://doi.org/10.1007/s13225-020-00466-2)) |
| Otu34656 | 1 | k__Fungi(100);p__Ascomycota(100);c__Saccharomycetes(100);o__Saccharomycetales(100);f__Metschnikowiaceae(100);g__Metschnikowia(100);s__Metschnikowia_pulcherrima(100); | Metschnikowia | 13 | Saprotroph | \|Undefined Saprotroph\| | Yeast | NULL | Probable | As nectar-tap_saprotroph (Põlme et al. 2020); Decay sugar-rich substrates (Põlme et al. 2020) | Põlme S, et al. 2020. Fungal Diversity 105:1-16 ((https://doi.org/10.1007/s13225-020-00466-2)) |
| Otu34657 | 1 | k__Fungi(100);p__Ascomycota(100);c__Saccharomycetes(100);o__Saccharomycetales(100);f__Metschnikowiaceae(100);g__Metschnikowia(100);s__Metschnikowia_pulcherrima(100); | Metschnikowia | 13 | Saprotroph | \|Undefined Saprotroph\| | Yeast | NULL | Probable | As nectar-tap_saprotroph (Põlme et al. 2020); Decay sugar-rich substrates (Põlme et al. 2020) | Põlme S, et al. 2020. Fungal Diversity 105:1-16 ((https://doi.org/10.1007/s13225-020-00466-2)) |
| Otu34658 | 1 | k__Fungi(100);p__Ascomycota(100);c__Saccharomycetes(100);o__Saccharomycetales(100);f__Metschnikowiaceae(100);g__Metschnikowia(100);s__Metschnikowia_pulcherrima(100); | Metschnikowia | 13 | Saprotroph | \|Undefined Saprotroph\| | Yeast | NULL | Probable | As nectar-tap_saprotroph (Põlme et al. 2020); Decay sugar-rich substrates (Põlme et al. 2020) | Põlme S, et al. 2020. Fungal Diversity 105:1-16 ((https://doi.org/10.1007/s13225-020-00466-2)) |
| Otu34659 | 1 | k__Fungi(100);p__Ascomycota(100);c__Saccharomycetes(100);o__Saccharomycetales(100);f__Metschnikowiaceae(100);g__Metschnikowia(100);g__Metschnikowia_unclassified(100); | Metschnikowia | 13 | Saprotroph | \|Undefined Saprotroph\| | Yeast | NULL | Probable | As nectar-tap_saprotroph (Põlme et al. 2020); Decay sugar-rich substrates (Põlme et al. 2020) | Põlme S, et al. 2020. Fungal Diversity 105:1-16 ((https://doi.org/10.1007/s13225-020-00466-2)) |
| Otu34661 | 1 | k__Fungi(100);p__Ascomycota(100);c__Saccharomycetes(100);o__Saccharomycetales(100);f__Metschnikowiaceae(100);g__Metschnikowia(100);g__Metschnikowia_unclassified(100); | Metschnikowia | 13 | Saprotroph | \|Undefined Saprotroph\| | Yeast | NULL | Probable | As nectar-tap_saprotroph (Põlme et al. 2020); Decay sugar-rich substrates (Põlme et al. 2020) | Põlme S, et al. 2020. Fungal Diversity 105:1-16 ((https://doi.org/10.1007/s13225-020-00466-2)) |
| Otu34663 | 1 | k__Fungi(100);p__Ascomycota(100);c__Saccharomycetes(100);o__Saccharomycetales(100);f__Metschnikowiaceae(100);g__Metschnikowia(100);s__Metschnikowia_pulcherrima(100); | Metschnikowia | 13 | Saprotroph | \|Undefined Saprotroph\| | Yeast | NULL | Probable | As nectar-tap_saprotroph (Põlme et al. 2020); Decay sugar-rich substrates (Põlme et al. 2020) | Põlme S, et al. 2020. Fungal Diversity 105:1-16 ((https://doi.org/10.1007/s13225-020-00466-2)) |
| Otu34664 | 1 | k__Fungi(100);p__Ascomycota(100);c__Saccharomycetes(100);o__Saccharomycetales(100);f__Metschnikowiaceae(100);g__Metschnikowia(100);s__Metschnikowia_pulcherrima(100); | Metschnikowia | 13 | Saprotroph | \|Undefined Saprotroph\| | Yeast | NULL | Probable | As nectar-tap_saprotroph (Põlme et al. 2020); Decay sugar-rich substrates (Põlme et al. 2020) | Põlme S, et al. 2020. Fungal Diversity 105:1-16 ((https://doi.org/10.1007/s13225-020-00466-2)) |
| Otu34665 | 1 | k__Fungi(100);p__Ascomycota(100);c__Saccharomycetes(100);o__Saccharomycetales(100);o__Saccharomycetales_unclassified(100);o__Saccharomycetales_unclassified(100);o__Saccharomycetales_unclassified(100); | Saccharomycetales | 7 | Saprotroph | Undefined Saprotroph | Yeast | NULL | Possible | NULL | Sterkenburg E, et al. 2015. New Phytologist 207:1145-1158 ((https://doi.org/10.1111/nph.13426)) |
| Otu34667 | 1 | k__Fungi(100);p__Ascomycota(100);c__Saccharomycetes(100);o__Saccharomycetales(100);f__Metschnikowiaceae(100);g__Metschnikowia(100);s__Metschnikowia_pulcherrima(100); | Metschnikowia | 13 | Saprotroph | \|Undefined Saprotroph\| | Yeast | NULL | Probable | As nectar-tap_saprotroph (Põlme et al. 2020); Decay sugar-rich substrates (Põlme et al. 2020) | Põlme S, et al. 2020. Fungal Diversity 105:1-16 ((https://doi.org/10.1007/s13225-020-00466-2)) |
| Otu34668 | 1 | k__Fungi(100);p__Ascomycota(100);c__Saccharomycetes(100);o__Saccharomycetales(100);f__Metschnikowiaceae(100);g__Metschnikowia(100);s__Metschnikowia_pulcherrima(100); | Metschnikowia | 13 | Saprotroph | \|Undefined Saprotroph\| | Yeast | NULL | Probable | As nectar-tap_saprotroph (Põlme et al. 2020); Decay sugar-rich substrates (Põlme et al. 2020) | Põlme S, et al. 2020. Fungal Diversity 105:1-16 ((https://doi.org/10.1007/s13225-020-00466-2)) |
| Otu34669 | 1 | k__Fungi(100);p__Ascomycota(100);c__Saccharomycetes(100);o__Saccharomycetales(100);f__Metschnikowiaceae(100);g__Metschnikowia(100);s__Metschnikowia_pulcherrima(100); | Metschnikowia | 13 | Saprotroph | \|Undefined Saprotroph\| | Yeast | NULL | Probable | As nectar-tap_saprotroph (Põlme et al. 2020); Decay sugar-rich substrates (Põlme et al. 2020) | Põlme S, et al. 2020. Fungal Diversity 105:1-16 ((https://doi.org/10.1007/s13225-020-00466-2)) |
| Otu34670 | 1 | k__Fungi(100);p__Ascomycota(100);c__Saccharomycetes(100);o__Saccharomycetales(100);f__Metschnikowiaceae(100);g__Metschnikowia(100);s__Metschnikowia_pulcherrima(100); | Metschnikowia | 13 | Saprotroph | \|Undefined Saprotroph\| | Yeast | NULL | Probable | As nectar-tap_saprotroph (Põlme et al. 2020); Decay sugar-rich substrates (Põlme et al. 2020) | Põlme S, et al. 2020. Fungal Diversity 105:1-16 ((https://doi.org/10.1007/s13225-020-00466-2)) |
| Otu34671 | 1 | k__Fungi(100);p__Ascomycota(100);c__Saccharomycetes(100);o__Saccharomycetales(100);f__Metschnikowiaceae(100);g__Metschnikowia(100);s__Metschnikowia_pulcherrima(100); | Metschnikowia | 13 | Saprotroph | \|Undefined Saprotroph\| | Yeast | NULL | Probable | As nectar-tap_saprotroph (Põlme et al. 2020); Decay sugar-rich substrates (Põlme et al. 2020) | Põlme S, et al. 2020. Fungal Diversity 105:1-16 ((https://doi.org/10.1007/s13225-020-00466-2)) |
| Otu34672 | 1 | k__Fungi(100);p__Ascomycota(100);c__Saccharomycetes(100);o__Saccharomycetales(100);f__Metschnikowiaceae(100);g__Metschnikowia(100);s__Metschnikowia_pulcherrima(100); | Metschnikowia | 13 | Saprotroph | \|Undefined Saprotroph\| | Yeast | NULL | Probable | As nectar-tap_saprotroph (Põlme et al. 2020); Decay sugar-rich substrates (Põlme et al. 2020) | Põlme S, et al. 2020. Fungal Diversity 105:1-16 ((https://doi.org/10.1007/s13225-020-00466-2)) |
| Otu34673 | 1 | k__Fungi(100);p__Ascomycota(100);c__Saccharomycetes(100);o__Saccharomycetales(100);f__Metschnikowiaceae(100);g__Metschnikowia(100);s__Metschnikowia_pulcherrima(100); | Metschnikowia | 13 | Saprotroph | \|Undefined Saprotroph\| | Yeast | NULL | Probable | As nectar-tap_saprotroph (Põlme et al. 2020); Decay sugar-rich substrates (Põlme et al. 2020) | Põlme S, et al. 2020. Fungal Diversity 105:1-16 ((https://doi.org/10.1007/s13225-020-00466-2)) |
| Otu34674 | 1 | k__Fungi(100);p__Ascomycota(100);c__Saccharomycetes(100);o__Saccharomycetales(100);f__Metschnikowiaceae(100);g__Metschnikowia(100);s__Metschnikowia_pulcherrima(100); | Metschnikowia | 13 | Saprotroph | \|Undefined Saprotroph\| | Yeast | NULL | Probable | As nectar-tap_saprotroph (Põlme et al. 2020); Decay sugar-rich substrates (Põlme et al. 2020) | Põlme S, et al. 2020. Fungal Diversity 105:1-16 ((https://doi.org/10.1007/s13225-020-00466-2)) |
| Otu34675 | 1 | k__Fungi(100);p__Ascomycota(100);c__Saccharomycetes(100);o__Saccharomycetales(100);f__Metschnikowiaceae(100);g__Metschnikowia(100);s__Metschnikowia_pulcherrima(100); | Metschnikowia | 13 | Saprotroph | \|Undefined Saprotroph\| | Yeast | NULL | Probable | As nectar-tap_saprotroph (Põlme et al. 2020); Decay sugar-rich substrates (Põlme et al. 2020) | Põlme S, et al. 2020. Fungal Diversity 105:1-16 ((https://doi.org/10.1007/s13225-020-00466-2)) |
| Otu34676 | 1 | k__Fungi(100);p__Ascomycota(100);c__Saccharomycetes(100);o__Saccharomycetales(100);f__Metschnikowiaceae(100);g__Metschnikowia(100);s__Metschnikowia_pulcherrima(100); | Metschnikowia | 13 | Saprotroph | \|Undefined Saprotroph\| | Yeast | NULL | Probable | As nectar-tap_saprotroph (Põlme et al. 2020); Decay sugar-rich substrates (Põlme et al. 2020) | Põlme S, et al. 2020. Fungal Diversity 105:1-16 ((https://doi.org/10.1007/s13225-020-00466-2)) |
| Otu34677 | 1 | k__Fungi(100);p__Ascomycota(100);c__Saccharomycetes(100);o__Saccharomycetales(100);f__Metschnikowiaceae(100);g__Metschnikowia(100);s__Metschnikowia_pulcherrima(100); | Metschnikowia | 13 | Saprotroph | \|Undefined Saprotroph\| | Yeast | NULL | Probable | As nectar-tap_saprotroph (Põlme et al. 2020); Decay sugar-rich substrates (Põlme et al. 2020) | Põlme S, et al. 2020. Fungal Diversity 105:1-16 ((https://doi.org/10.1007/s13225-020-00466-2)) |
| Otu34680 | 1 | k__Fungi(100);p__Ascomycota(100);c__Saccharomycetes(100);o__Saccharomycetales(100);f__Metschnikowiaceae(100);g__Metschnikowia(100);s__Metschnikowia_pulcherrima(100); | Metschnikowia | 13 | Saprotroph | \|Undefined Saprotroph\| | Yeast | NULL | Probable | As nectar-tap_saprotroph (Põlme et al. 2020); Decay sugar-rich substrates (Põlme et al. 2020) | Põlme S, et al. 2020. Fungal Diversity 105:1-16 ((https://doi.org/10.1007/s13225-020-00466-2)) |
| Otu34682 | 1 | k__Fungi(100);p__Ascomycota(100);c__Saccharomycetes(100);o__Saccharomycetales(100);f__Metschnikowiaceae(100);g__Metschnikowia(100);s__Metschnikowia_pulcherrima(100); | Metschnikowia | 13 | Saprotroph | \|Undefined Saprotroph\| | Yeast | NULL | Probable | As nectar-tap_saprotroph (Põlme et al. 2020); Decay sugar-rich substrates (Põlme et al. 2020) | Põlme S, et al. 2020. Fungal Diversity 105:1-16 ((https://doi.org/10.1007/s13225-020-00466-2)) |
| Otu34684 | 1 | k__Fungi(100);p__Ascomycota(100);c__Saccharomycetes(100);o__Saccharomycetales(100);f__Metschnikowiaceae(100);g__Metschnikowia(100);s__Metschnikowia_pulcherrima(100); | Metschnikowia | 13 | Saprotroph | \|Undefined Saprotroph\| | Yeast | NULL | Probable | As nectar-tap_saprotroph (Põlme et al. 2020); Decay sugar-rich substrates (Põlme et al. 2020) | Põlme S, et al. 2020. Fungal Diversity 105:1-16 ((https://doi.org/10.1007/s13225-020-00466-2)) |
| Otu34685 | 1 | k__Fungi(100);p__Ascomycota(100);c__Saccharomycetes(100);o__Saccharomycetales(100);f__Metschnikowiaceae(100);g__Metschnikowia(100);s__Metschnikowia_pulcherrima(100); | Metschnikowia | 13 | Saprotroph | \|Undefined Saprotroph\| | Yeast | NULL | Probable | As nectar-tap_saprotroph (Põlme et al. 2020); Decay sugar-rich substrates (Põlme et al. 2020) | Põlme S, et al. 2020. Fungal Diversity 105:1-16 ((https://doi.org/10.1007/s13225-020-00466-2)) |
| Otu34686 | 1 | k__Fungi(100);p__Ascomycota(100);c__Saccharomycetes(100);o__Saccharomycetales(100);f__Metschnikowiaceae(100);g__Metschnikowia(100);s__Metschnikowia_pulcherrima(100); | Metschnikowia | 13 | Saprotroph | \|Undefined Saprotroph\| | Yeast | NULL | Probable | As nectar-tap_saprotroph (Põlme et al. 2020); Decay sugar-rich substrates (Põlme et al. 2020) | Põlme S, et al. 2020. Fungal Diversity 105:1-16 ((https://doi.org/10.1007/s13225-020-00466-2)) |
| Otu34687 | 1 | k__Fungi(100);p__Ascomycota(100);c__Saccharomycetes(100);o__Saccharomycetales(100);f__Metschnikowiaceae(100);g__Metschnikowia(100);s__Metschnikowia_pulcherrima(100); | Metschnikowia | 13 | Saprotroph | \|Undefined Saprotroph\| | Yeast | NULL | Probable | As nectar-tap_saprotroph (Põlme et al. 2020); Decay sugar-rich substrates (Põlme et al. 2020) | Põlme S, et al. 2020. Fungal Diversity 105:1-16 ((https://doi.org/10.1007/s13225-020-00466-2)) |
| Otu34688 | 1 | k__Fungi(100);p__Ascomycota(100);c__Saccharomycetes(100);o__Saccharomycetales(100);f__Metschnikowiaceae(100);g__Metschnikowia(100);s__Metschnikowia_pulcherrima(100); | Metschnikowia | 13 | Saprotroph | \|Undefined Saprotroph\| | Yeast | NULL | Probable | As nectar-tap_saprotroph (Põlme et al. 2020); Decay sugar-rich substrates (Põlme et al. 2020) | Põlme S, et al. 2020. Fungal Diversity 105:1-16 ((https://doi.org/10.1007/s13225-020-00466-2)) |
| Otu34690 | 1 | k__Fungi(100);p__Ascomycota(100);c__Saccharomycetes(100);o__Saccharomycetales(100);f__Metschnikowiaceae(100);g__Metschnikowia(100);s__Metschnikowia_pulcherrima(100); | Metschnikowia | 13 | Saprotroph | \|Undefined Saprotroph\| | Yeast | NULL | Probable | As nectar-tap_saprotroph (Põlme et al. 2020); Decay sugar-rich substrates (Põlme et al. 2020) | Põlme S, et al. 2020. Fungal Diversity 105:1-16 ((https://doi.org/10.1007/s13225-020-00466-2)) |
| Otu34691 | 1 | k__Fungi(100);p__Ascomycota(100);c__Saccharomycetes(100);o__Saccharomycetales(100);f__Metschnikowiaceae(100);g__Metschnikowia(100);s__Metschnikowia_pulcherrima(100); | Metschnikowia | 13 | Saprotroph | \|Undefined Saprotroph\| | Yeast | NULL | Probable | As nectar-tap_saprotroph (Põlme et al. 2020); Decay sugar-rich substrates (Põlme et al. 2020) | Põlme S, et al. 2020. Fungal Diversity 105:1-16 ((https://doi.org/10.1007/s13225-020-00466-2)) |
| Otu34693 | 1 | k__Fungi(100);p__Ascomycota(100);c__Saccharomycetes(100);o__Saccharomycetales(100);f__Metschnikowiaceae(100);g__Metschnikowia(100);g__Metschnikowia_unclassified(100); | Metschnikowia | 13 | Saprotroph | \|Undefined Saprotroph\| | Yeast | NULL | Probable | As nectar-tap_saprotroph (Põlme et al. 2020); Decay sugar-rich substrates (Põlme et al. 2020) | Põlme S, et al. 2020. Fungal Diversity 105:1-16 ((https://doi.org/10.1007/s13225-020-00466-2)) |
| Otu34694 | 1 | k__Fungi(100);p__Ascomycota(100);c__Saccharomycetes(100);o__Saccharomycetales(100);f__Metschnikowiaceae(100);g__Metschnikowia(100);g__Metschnikowia_unclassified(100); | Metschnikowia | 13 | Saprotroph | \|Undefined Saprotroph\| | Yeast | NULL | Probable | As nectar-tap_saprotroph (Põlme et al. 2020); Decay sugar-rich substrates (Põlme et al. 2020) | Põlme S, et al. 2020. Fungal Diversity 105:1-16 ((https://doi.org/10.1007/s13225-020-00466-2)) |
| Otu34695 | 1 | k__Fungi(100);p__Ascomycota(100);c__Saccharomycetes(100);o__Saccharomycetales(100);f__Metschnikowiaceae(100);g__Metschnikowia(100);s__Metschnikowia_pulcherrima(100); | Metschnikowia | 13 | Saprotroph | \|Undefined Saprotroph\| | Yeast | NULL | Probable | As nectar-tap_saprotroph (Põlme et al. 2020); Decay sugar-rich substrates (Põlme et al. 2020) | Põlme S, et al. 2020. Fungal Diversity 105:1-16 ((https://doi.org/10.1007/s13225-020-00466-2)) |
| Otu34696 | 1 | k__Fungi(100);p__Ascomycota(100);c__Saccharomycetes(100);o__Saccharomycetales(100);f__Metschnikowiaceae(100);g__Metschnikowia(100);g__Metschnikowia_unclassified(100); | Metschnikowia | 13 | Saprotroph | \|Undefined Saprotroph\| | Yeast | NULL | Probable | As nectar-tap_saprotroph (Põlme et al. 2020); Decay sugar-rich substrates (Põlme et al. 2020) | Põlme S, et al. 2020. Fungal Diversity 105:1-16 ((https://doi.org/10.1007/s13225-020-00466-2)) |
| Otu34699 | 1 | k__Fungi(100);p__Ascomycota(100);c__Saccharomycetes(100);o__Saccharomycetales(100);f__Metschnikowiaceae(100);g__Metschnikowia(100);s__Metschnikowia_pulcherrima(100); | Metschnikowia | 13 | Saprotroph | \|Undefined Saprotroph\| | Yeast | NULL | Probable | As nectar-tap_saprotroph (Põlme et al. 2020); Decay sugar-rich substrates (Põlme et al. 2020) | Põlme S, et al. 2020. Fungal Diversity 105:1-16 ((https://doi.org/10.1007/s13225-020-00466-2)) |
| Otu34700 | 1 | k__Fungi(100);p__Ascomycota(100);c__Saccharomycetes(100);o__Saccharomycetales(100);f__Metschnikowiaceae(100);g__Metschnikowia(100);s__Metschnikowia_pulcherrima(100); | Metschnikowia | 13 | Saprotroph | \|Undefined Saprotroph\| | Yeast | NULL | Probable | As nectar-tap_saprotroph (Põlme et al. 2020); Decay sugar-rich substrates (Põlme et al. 2020) | Põlme S, et al. 2020. Fungal Diversity 105:1-16 ((https://doi.org/10.1007/s13225-020-00466-2)) |
| Otu34702 | 1 | k__Fungi(100);p__Ascomycota(100);c__Saccharomycetes(100);o__Saccharomycetales(100);f__Metschnikowiaceae(100);g__Metschnikowia(100);s__Metschnikowia_pulcherrima(100); | Metschnikowia | 13 | Saprotroph | \|Undefined Saprotroph\| | Yeast | NULL | Probable | As nectar-tap_saprotroph (Põlme et al. 2020); Decay sugar-rich substrates (Põlme et al. 2020) | Põlme S, et al. 2020. Fungal Diversity 105:1-16 ((https://doi.org/10.1007/s13225-020-00466-2)) |
| Otu34703 | 1 | k__Fungi(100);p__Ascomycota(100);c__Saccharomycetes(100);o__Saccharomycetales(100);f__Metschnikowiaceae(100);g__Metschnikowia(100);g__Metschnikowia_unclassified(100); | Metschnikowia | 13 | Saprotroph | \|Undefined Saprotroph\| | Yeast | NULL | Probable | As nectar-tap_saprotroph (Põlme et al. 2020); Decay sugar-rich substrates (Põlme et al. 2020) | Põlme S, et al. 2020. Fungal Diversity 105:1-16 ((https://doi.org/10.1007/s13225-020-00466-2)) |
| Otu34704 | 1 | k__Fungi(100);p__Ascomycota(100);c__Saccharomycetes(100);o__Saccharomycetales(100);f__Metschnikowiaceae(100);g__Metschnikowia(100);s__Metschnikowia_pulcherrima(100); | Metschnikowia | 13 | Saprotroph | \|Undefined Saprotroph\| | Yeast | NULL | Probable | As nectar-tap_saprotroph (Põlme et al. 2020); Decay sugar-rich substrates (Põlme et al. 2020) | Põlme S, et al. 2020. Fungal Diversity 105:1-16 ((https://doi.org/10.1007/s13225-020-00466-2)) |
| Otu34705 | 1 | k__Fungi(100);p__Ascomycota(100);c__Saccharomycetes(100);o__Saccharomycetales(100);f__Metschnikowiaceae(100);g__Metschnikowia(100);s__Metschnikowia_pulcherrima(100); | Metschnikowia | 13 | Saprotroph | \|Undefined Saprotroph\| | Yeast | NULL | Probable | As nectar-tap_saprotroph (Põlme et al. 2020); Decay sugar-rich substrates (Põlme et al. 2020) | Põlme S, et al. 2020. Fungal Diversity 105:1-16 ((https://doi.org/10.1007/s13225-020-00466-2)) |
| Otu34706 | 1 | k__Fungi(100);p__Ascomycota(100);c__Saccharomycetes(100);o__Saccharomycetales(100);f__Metschnikowiaceae(100);g__Metschnikowia(100);s__Metschnikowia_pulcherrima(100); | Metschnikowia | 13 | Saprotroph | \|Undefined Saprotroph\| | Yeast | NULL | Probable | As nectar-tap_saprotroph (Põlme et al. 2020); Decay sugar-rich substrates (Põlme et al. 2020) | Põlme S, et al. 2020. Fungal Diversity 105:1-16 ((https://doi.org/10.1007/s13225-020-00466-2)) |
| Otu34707 | 1 | k__Fungi(100);p__Ascomycota(100);c__Saccharomycetes(100);o__Saccharomycetales(100);f__Metschnikowiaceae(100);g__Metschnikowia(100);g__Metschnikowia_unclassified(100); | Metschnikowia | 13 | Saprotroph | \|Undefined Saprotroph\| | Yeast | NULL | Probable | As nectar-tap_saprotroph (Põlme et al. 2020); Decay sugar-rich substrates (Põlme et al. 2020) | Põlme S, et al. 2020. Fungal Diversity 105:1-16 ((https://doi.org/10.1007/s13225-020-00466-2)) |
| Otu34708 | 1 | k__Fungi(100);p__Ascomycota(100);c__Saccharomycetes(100);o__Saccharomycetales(100);f__Metschnikowiaceae(100);g__Metschnikowia(100);s__Metschnikowia_pulcherrima(100); | Metschnikowia | 13 | Saprotroph | \|Undefined Saprotroph\| | Yeast | NULL | Probable | As nectar-tap_saprotroph (Põlme et al. 2020); Decay sugar-rich substrates (Põlme et al. 2020) | Põlme S, et al. 2020. Fungal Diversity 105:1-16 ((https://doi.org/10.1007/s13225-020-00466-2)) |
| Otu34709 | 1 | k__Fungi(100);p__Ascomycota(100);c__Saccharomycetes(100);o__Saccharomycetales(100);f__Metschnikowiaceae(100);g__Metschnikowia(100);s__Metschnikowia_pulcherrima(100); | Metschnikowia | 13 | Saprotroph | \|Undefined Saprotroph\| | Yeast | NULL | Probable | As nectar-tap_saprotroph (Põlme et al. 2020); Decay sugar-rich substrates (Põlme et al. 2020) | Põlme S, et al. 2020. Fungal Diversity 105:1-16 ((https://doi.org/10.1007/s13225-020-00466-2)) |
| Otu34710 | 1 | k__Fungi(100);p__Ascomycota(100);c__Saccharomycetes(100);o__Saccharomycetales(100);f__Metschnikowiaceae(100);g__Metschnikowia(100);s__Metschnikowia_pulcherrima(100); | Metschnikowia | 13 | Saprotroph | \|Undefined Saprotroph\| | Yeast | NULL | Probable | As nectar-tap_saprotroph (Põlme et al. 2020); Decay sugar-rich substrates (Põlme et al. 2020) | Põlme S, et al. 2020. Fungal Diversity 105:1-16 ((https://doi.org/10.1007/s13225-020-00466-2)) |
| Otu34711 | 1 | k__Fungi(100);p__Ascomycota(100);c__Saccharomycetes(100);o__Saccharomycetales(100);f__Metschnikowiaceae(100);g__Metschnikowia(100);g__Metschnikowia_unclassified(100); | Metschnikowia | 13 | Saprotroph | \|Undefined Saprotroph\| | Yeast | NULL | Probable | As nectar-tap_saprotroph (Põlme et al. 2020); Decay sugar-rich substrates (Põlme et al. 2020) | Põlme S, et al. 2020. Fungal Diversity 105:1-16 ((https://doi.org/10.1007/s13225-020-00466-2)) |
| Otu34712 | 1 | k__Fungi(100);p__Ascomycota(100);c__Saccharomycetes(100);o__Saccharomycetales(100);f__Metschnikowiaceae(100);g__Metschnikowia(100);s__Metschnikowia_pulcherrima(100); | Metschnikowia | 13 | Saprotroph | \|Undefined Saprotroph\| | Yeast | NULL | Probable | As nectar-tap_saprotroph (Põlme et al. 2020); Decay sugar-rich substrates (Põlme et al. 2020) | Põlme S, et al. 2020. Fungal Diversity 105:1-16 ((https://doi.org/10.1007/s13225-020-00466-2)) |
| Otu34713 | 1 | k__Fungi(100);p__Ascomycota(100);c__Saccharomycetes(100);o__Saccharomycetales(100);f__Metschnikowiaceae(100);g__Metschnikowia(100);s__Metschnikowia_pulcherrima(100); | Metschnikowia | 13 | Saprotroph | \|Undefined Saprotroph\| | Yeast | NULL | Probable | As nectar-tap_saprotroph (Põlme et al. 2020); Decay sugar-rich substrates (Põlme et al. 2020) | Põlme S, et al. 2020. Fungal Diversity 105:1-16 ((https://doi.org/10.1007/s13225-020-00466-2)) |
| Otu34714 | 1 | k__Fungi(100);p__Ascomycota(100);c__Saccharomycetes(100);o__Saccharomycetales(100);f__Metschnikowiaceae(100);g__Metschnikowia(100);s__Metschnikowia_pulcherrima(100); | Metschnikowia | 13 | Saprotroph | \|Undefined Saprotroph\| | Yeast | NULL | Probable | As nectar-tap_saprotroph (Põlme et al. 2020); Decay sugar-rich substrates (Põlme et al. 2020) | Põlme S, et al. 2020. Fungal Diversity 105:1-16 ((https://doi.org/10.1007/s13225-020-00466-2)) |
| Otu34715 | 1 | k__Fungi(100);p__Ascomycota(100);c__Saccharomycetes(100);o__Saccharomycetales(100);f__Metschnikowiaceae(100);g__Metschnikowia(100);g__Metschnikowia_unclassified(100); | Metschnikowia | 13 | Saprotroph | \|Undefined Saprotroph\| | Yeast | NULL | Probable | As nectar-tap_saprotroph (Põlme et al. 2020); Decay sugar-rich substrates (Põlme et al. 2020) | Põlme S, et al. 2020. Fungal Diversity 105:1-16 ((https://doi.org/10.1007/s13225-020-00466-2)) |
| Otu34716 | 1 | k__Fungi(100);p__Ascomycota(100);c__Saccharomycetes(100);o__Saccharomycetales(100);f__Metschnikowiaceae(100);g__Metschnikowia(100);s__Metschnikowia_pulcherrima(100); | Metschnikowia | 13 | Saprotroph | \|Undefined Saprotroph\| | Yeast | NULL | Probable | As nectar-tap_saprotroph (Põlme et al. 2020); Decay sugar-rich substrates (Põlme et al. 2020) | Põlme S, et al. 2020. Fungal Diversity 105:1-16 ((https://doi.org/10.1007/s13225-020-00466-2)) |
| Otu34717 | 1 | k__Fungi(100);p__Ascomycota(100);c__Saccharomycetes(100);o__Saccharomycetales(100);f__Metschnikowiaceae(100);g__Metschnikowia(100);s__Metschnikowia_pulcherrima(100); | Metschnikowia | 13 | Saprotroph | \|Undefined Saprotroph\| | Yeast | NULL | Probable | As nectar-tap_saprotroph (Põlme et al. 2020); Decay sugar-rich substrates (Põlme et al. 2020) | Põlme S, et al. 2020. Fungal Diversity 105:1-16 ((https://doi.org/10.1007/s13225-020-00466-2)) |
| Otu34718 | 1 | k__Fungi(100);p__Ascomycota(100);c__Saccharomycetes(100);o__Saccharomycetales(100);f__Metschnikowiaceae(100);g__Metschnikowia(100);g__Metschnikowia_unclassified(100); | Metschnikowia | 13 | Saprotroph | \|Undefined Saprotroph\| | Yeast | NULL | Probable | As nectar-tap_saprotroph (Põlme et al. 2020); Decay sugar-rich substrates (Põlme et al. 2020) | Põlme S, et al. 2020. Fungal Diversity 105:1-16 ((https://doi.org/10.1007/s13225-020-00466-2)) |
| Otu34719 | 1 | k__Fungi(100);p__Ascomycota(100);c__Saccharomycetes(100);o__Saccharomycetales(100);f__Metschnikowiaceae(100);g__Metschnikowia(100);s__Metschnikowia_pulcherrima(100); | Metschnikowia | 13 | Saprotroph | \|Undefined Saprotroph\| | Yeast | NULL | Probable | As nectar-tap_saprotroph (Põlme et al. 2020); Decay sugar-rich substrates (Põlme et al. 2020) | Põlme S, et al. 2020. Fungal Diversity 105:1-16 ((https://doi.org/10.1007/s13225-020-00466-2)) |
| Otu34720 | 1 | k__Fungi(100);p__Ascomycota(100);c__Saccharomycetes(100);o__Saccharomycetales(100);f__Metschnikowiaceae(100);g__Metschnikowia(100);s__Metschnikowia_pulcherrima(100); | Metschnikowia | 13 | Saprotroph | \|Undefined Saprotroph\| | Yeast | NULL | Probable | As nectar-tap_saprotroph (Põlme et al. 2020); Decay sugar-rich substrates (Põlme et al. 2020) | Põlme S, et al. 2020. Fungal Diversity 105:1-16 ((https://doi.org/10.1007/s13225-020-00466-2)) |
| Otu34721 | 1 | k__Fungi(100);p__Ascomycota(100);c__Saccharomycetes(100);o__Saccharomycetales(100);f__Metschnikowiaceae(100);g__Metschnikowia(100);g__Metschnikowia_unclassified(100); | Metschnikowia | 13 | Saprotroph | \|Undefined Saprotroph\| | Yeast | NULL | Probable | As nectar-tap_saprotroph (Põlme et al. 2020); Decay sugar-rich substrates (Põlme et al. 2020) | Põlme S, et al. 2020. Fungal Diversity 105:1-16 ((https://doi.org/10.1007/s13225-020-00466-2)) |
| Otu34722 | 1 | k__Fungi(100);p__Ascomycota(100);c__Saccharomycetes(100);o__Saccharomycetales(100);f__Metschnikowiaceae(100);g__Metschnikowia(100);g__Metschnikowia_unclassified(100); | Metschnikowia | 13 | Saprotroph | \|Undefined Saprotroph\| | Yeast | NULL | Probable | As nectar-tap_saprotroph (Põlme et al. 2020); Decay sugar-rich substrates (Põlme et al. 2020) | Põlme S, et al. 2020. Fungal Diversity 105:1-16 ((https://doi.org/10.1007/s13225-020-00466-2)) |
| Otu34723 | 1 | k__Fungi(100);p__Ascomycota(100);c__Saccharomycetes(100);o__Saccharomycetales(100);f__Metschnikowiaceae(100);g__Metschnikowia(100);s__Metschnikowia_pulcherrima(100); | Metschnikowia | 13 | Saprotroph | \|Undefined Saprotroph\| | Yeast | NULL | Probable | As nectar-tap_saprotroph (Põlme et al. 2020); Decay sugar-rich substrates (Põlme et al. 2020) | Põlme S, et al. 2020. Fungal Diversity 105:1-16 ((https://doi.org/10.1007/s13225-020-00466-2)) |
| Otu34724 | 1 | k__Fungi(100);p__Ascomycota(100);c__Saccharomycetes(100);o__Saccharomycetales(100);f__Metschnikowiaceae(100);g__Metschnikowia(100);s__Metschnikowia_pulcherrima(100); | Metschnikowia | 13 | Saprotroph | \|Undefined Saprotroph\| | Yeast | NULL | Probable | As nectar-tap_saprotroph (Põlme et al. 2020); Decay sugar-rich substrates (Põlme et al. 2020) | Põlme S, et al. 2020. Fungal Diversity 105:1-16 ((https://doi.org/10.1007/s13225-020-00466-2)) |
| Otu34725 | 1 | k__Fungi(100);p__Ascomycota(100);c__Saccharomycetes(100);o__Saccharomycetales(100);f__Metschnikowiaceae(100);g__Metschnikowia(100);s__Metschnikowia_pulcherrima(100); | Metschnikowia | 13 | Saprotroph | \|Undefined Saprotroph\| | Yeast | NULL | Probable | As nectar-tap_saprotroph (Põlme et al. 2020); Decay sugar-rich substrates (Põlme et al. 2020) | Põlme S, et al. 2020. Fungal Diversity 105:1-16 ((https://doi.org/10.1007/s13225-020-00466-2)) |
| Otu34726 | 1 | k__Fungi(100);p__Ascomycota(100);c__Saccharomycetes(100);o__Saccharomycetales(100);f__Metschnikowiaceae(100);g__Metschnikowia(100);g__Metschnikowia_unclassified(100); | Metschnikowia | 13 | Saprotroph | \|Undefined Saprotroph\| | Yeast | NULL | Probable | As nectar-tap_saprotroph (Põlme et al. 2020); Decay sugar-rich substrates (Põlme et al. 2020) | Põlme S, et al. 2020. Fungal Diversity 105:1-16 ((https://doi.org/10.1007/s13225-020-00466-2)) |
| Otu34727 | 1 | k__Fungi(100);p__Ascomycota(100);c__Saccharomycetes(100);o__Saccharomycetales(100);f__Metschnikowiaceae(100);g__Metschnikowia(100);s__Metschnikowia_pulcherrima(100); | Metschnikowia | 13 | Saprotroph | \|Undefined Saprotroph\| | Yeast | NULL | Probable | As nectar-tap_saprotroph (Põlme et al. 2020); Decay sugar-rich substrates (Põlme et al. 2020) | Põlme S, et al. 2020. Fungal Diversity 105:1-16 ((https://doi.org/10.1007/s13225-020-00466-2)) |
| Otu34728 | 1 | k__Fungi(100);p__Ascomycota(100);c__Saccharomycetes(100);o__Saccharomycetales(100);f__Metschnikowiaceae(100);g__Metschnikowia(100);s__Metschnikowia_pulcherrima(100); | Metschnikowia | 13 | Saprotroph | \|Undefined Saprotroph\| | Yeast | NULL | Probable | As nectar-tap_saprotroph (Põlme et al. 2020); Decay sugar-rich substrates (Põlme et al. 2020) | Põlme S, et al. 2020. Fungal Diversity 105:1-16 ((https://doi.org/10.1007/s13225-020-00466-2)) |
| Otu34729 | 1 | k__Fungi(100);p__Ascomycota(100);c__Saccharomycetes(100);o__Saccharomycetales(100);f__Metschnikowiaceae(100);g__Metschnikowia(100);s__Metschnikowia_pulcherrima(100); | Metschnikowia | 13 | Saprotroph | \|Undefined Saprotroph\| | Yeast | NULL | Probable | As nectar-tap_saprotroph (Põlme et al. 2020); Decay sugar-rich substrates (Põlme et al. 2020) | Põlme S, et al. 2020. Fungal Diversity 105:1-16 ((https://doi.org/10.1007/s13225-020-00466-2)) |
| Otu34730 | 1 | k__Fungi(100);p__Ascomycota(100);c__Dothideomycetes(100);o__Capnodiales(100);f__Davidiellaceae(100);g__Cladosporium(100);s__Cladosporium_sphaerospermum(100); | Cladosporium | 13 | Pathotroph-Saprotroph-Symbiotroph | Animal Pathogen-Endophyte-Lichen Parasite-Plant Pathogen-\|Plant Saprotroph\|-Wood Saprotroph | Microfungus | NULL | Possible | As foliar_endophyte (Põlme et al. 2020); As litter_saprotroph (Põlme et al. 2020); Decay wood, leaf, fruit, seed, soil, dung, animal material (Põlme et al. 2020) | Seehann G, et al. 1975. List of Fungi in Soft-Rot Tissue Tests. Institute fur Holzbiologie und Holzschutz, Bundesforschungsanstalt fur Forst - und Holzwirtschaft, Hamburg - Lohbrugge, Germany; Lawrey JD, Diederich P. 2016. http://www.lichenicolous.net (accessed 24 May 2017) ((http://www.lichenicolous.net)); Põlme S, et al. 2020. Fungal Diversity 105:1-16 ((https://doi.org/10.1007/s13225-020-00466-2)); The American Phytopathological Society Common Names of Plant Diseases ((http://www.apsnet.org/publications/commonnames/Pages/default.aspx)) |
| Otu34731 | 1 | k__Fungi(100);p__Ascomycota(100);c__Saccharomycetes(100);o__Saccharomycetales(100);f__Metschnikowiaceae(100);g__Metschnikowia(100);s__Metschnikowia_pulcherrima(100); | Metschnikowia | 13 | Saprotroph | \|Undefined Saprotroph\| | Yeast | NULL | Probable | As nectar-tap_saprotroph (Põlme et al. 2020); Decay sugar-rich substrates (Põlme et al. 2020) | Põlme S, et al. 2020. Fungal Diversity 105:1-16 ((https://doi.org/10.1007/s13225-020-00466-2)) |
| Otu34732 | 1 | k__Fungi(100);p__Ascomycota(100);c__Saccharomycetes(100);o__Saccharomycetales(100);f__Metschnikowiaceae(100);g__Metschnikowia(100);g__Metschnikowia_unclassified(100); | Metschnikowia | 13 | Saprotroph | \|Undefined Saprotroph\| | Yeast | NULL | Probable | As nectar-tap_saprotroph (Põlme et al. 2020); Decay sugar-rich substrates (Põlme et al. 2020) | Põlme S, et al. 2020. Fungal Diversity 105:1-16 ((https://doi.org/10.1007/s13225-020-00466-2)) |
| Otu34733 | 1 | k__Fungi(100);p__Ascomycota(100);c__Saccharomycetes(100);o__Saccharomycetales(100);f__Metschnikowiaceae(100);g__Metschnikowia(100);s__Metschnikowia_pulcherrima(100); | Metschnikowia | 13 | Saprotroph | \|Undefined Saprotroph\| | Yeast | NULL | Probable | As nectar-tap_saprotroph (Põlme et al. 2020); Decay sugar-rich substrates (Põlme et al. 2020) | Põlme S, et al. 2020. Fungal Diversity 105:1-16 ((https://doi.org/10.1007/s13225-020-00466-2)) |
| Otu34734 | 1 | k__Fungi(100);p__Ascomycota(100);c__Saccharomycetes(100);o__Saccharomycetales(100);f__Metschnikowiaceae(100);g__Metschnikowia(100);s__Metschnikowia_pulcherrima(100); | Metschnikowia | 13 | Saprotroph | \|Undefined Saprotroph\| | Yeast | NULL | Probable | As nectar-tap_saprotroph (Põlme et al. 2020); Decay sugar-rich substrates (Põlme et al. 2020) | Põlme S, et al. 2020. Fungal Diversity 105:1-16 ((https://doi.org/10.1007/s13225-020-00466-2)) |
| Otu34735 | 1 | k__Fungi(100);p__Ascomycota(100);c__Saccharomycetes(100);o__Saccharomycetales(100);f__Metschnikowiaceae(100);g__Metschnikowia(100);s__Metschnikowia_pulcherrima(100); | Metschnikowia | 13 | Saprotroph | \|Undefined Saprotroph\| | Yeast | NULL | Probable | As nectar-tap_saprotroph (Põlme et al. 2020); Decay sugar-rich substrates (Põlme et al. 2020) | Põlme S, et al. 2020. Fungal Diversity 105:1-16 ((https://doi.org/10.1007/s13225-020-00466-2)) |
| Otu34736 | 1 | k__Fungi(100);p__Ascomycota(100);c__Saccharomycetes(100);o__Saccharomycetales(100);f__Metschnikowiaceae(100);g__Metschnikowia(100);s__Metschnikowia_pulcherrima(100); | Metschnikowia | 13 | Saprotroph | \|Undefined Saprotroph\| | Yeast | NULL | Probable | As nectar-tap_saprotroph (Põlme et al. 2020); Decay sugar-rich substrates (Põlme et al. 2020) | Põlme S, et al. 2020. Fungal Diversity 105:1-16 ((https://doi.org/10.1007/s13225-020-00466-2)) |
| Otu34737 | 1 | k__Fungi(100);p__Ascomycota(100);c__Saccharomycetes(100);o__Saccharomycetales(100);f__Metschnikowiaceae(100);g__Metschnikowia(100);s__Metschnikowia_pulcherrima(100); | Metschnikowia | 13 | Saprotroph | \|Undefined Saprotroph\| | Yeast | NULL | Probable | As nectar-tap_saprotroph (Põlme et al. 2020); Decay sugar-rich substrates (Põlme et al. 2020) | Põlme S, et al. 2020. Fungal Diversity 105:1-16 ((https://doi.org/10.1007/s13225-020-00466-2)) |
| Otu34738 | 1 | k__Fungi(100);p__Ascomycota(100);c__Saccharomycetes(100);o__Saccharomycetales(100);f__Metschnikowiaceae(100);g__Metschnikowia(100);g__Metschnikowia_unclassified(100); | Metschnikowia | 13 | Saprotroph | \|Undefined Saprotroph\| | Yeast | NULL | Probable | As nectar-tap_saprotroph (Põlme et al. 2020); Decay sugar-rich substrates (Põlme et al. 2020) | Põlme S, et al. 2020. Fungal Diversity 105:1-16 ((https://doi.org/10.1007/s13225-020-00466-2)) |
| Otu34739 | 1 | k__Fungi(100);p__Ascomycota(100);c__Saccharomycetes(100);o__Saccharomycetales(100);f__Metschnikowiaceae(100);g__Metschnikowia(100);g__Metschnikowia_unclassified(100); | Metschnikowia | 13 | Saprotroph | \|Undefined Saprotroph\| | Yeast | NULL | Probable | As nectar-tap_saprotroph (Põlme et al. 2020); Decay sugar-rich substrates (Põlme et al. 2020) | Põlme S, et al. 2020. Fungal Diversity 105:1-16 ((https://doi.org/10.1007/s13225-020-00466-2)) |
| Otu34740 | 1 | k__Fungi(100);p__Ascomycota(100);c__Saccharomycetes(100);o__Saccharomycetales(100);f__Metschnikowiaceae(100);g__Metschnikowia(100);g__Metschnikowia_unclassified(100); | Metschnikowia | 13 | Saprotroph | \|Undefined Saprotroph\| | Yeast | NULL | Probable | As nectar-tap_saprotroph (Põlme et al. 2020); Decay sugar-rich substrates (Põlme et al. 2020) | Põlme S, et al. 2020. Fungal Diversity 105:1-16 ((https://doi.org/10.1007/s13225-020-00466-2)) |
| Otu34741 | 1 | k__Fungi(100);p__Ascomycota(100);c__Saccharomycetes(100);o__Saccharomycetales(100);f__Metschnikowiaceae(100);g__Metschnikowia(100);s__Metschnikowia_pulcherrima(100); | Metschnikowia | 13 | Saprotroph | \|Undefined Saprotroph\| | Yeast | NULL | Probable | As nectar-tap_saprotroph (Põlme et al. 2020); Decay sugar-rich substrates (Põlme et al. 2020) | Põlme S, et al. 2020. Fungal Diversity 105:1-16 ((https://doi.org/10.1007/s13225-020-00466-2)) |
| Otu34742 | 1 | k__Fungi(100);p__Ascomycota(100);c__Saccharomycetes(100);o__Saccharomycetales(100);f__Metschnikowiaceae(100);g__Metschnikowia(100);s__Metschnikowia_pulcherrima(100); | Metschnikowia | 13 | Saprotroph | \|Undefined Saprotroph\| | Yeast | NULL | Probable | As nectar-tap_saprotroph (Põlme et al. 2020); Decay sugar-rich substrates (Põlme et al. 2020) | Põlme S, et al. 2020. Fungal Diversity 105:1-16 ((https://doi.org/10.1007/s13225-020-00466-2)) |
| Otu34743 | 1 | k__Fungi(100);p__Ascomycota(100);c__Saccharomycetes(100);o__Saccharomycetales(100);f__Metschnikowiaceae(100);g__Metschnikowia(100);g__Metschnikowia_unclassified(100); | Metschnikowia | 13 | Saprotroph | \|Undefined Saprotroph\| | Yeast | NULL | Probable | As nectar-tap_saprotroph (Põlme et al. 2020); Decay sugar-rich substrates (Põlme et al. 2020) | Põlme S, et al. 2020. Fungal Diversity 105:1-16 ((https://doi.org/10.1007/s13225-020-00466-2)) |
| Otu34744 | 1 | k__Fungi(100);p__Ascomycota(100);c__Saccharomycetes(100);o__Saccharomycetales(100);f__Metschnikowiaceae(100);g__Metschnikowia(100);s__Metschnikowia_pulcherrima(100); | Metschnikowia | 13 | Saprotroph | \|Undefined Saprotroph\| | Yeast | NULL | Probable | As nectar-tap_saprotroph (Põlme et al. 2020); Decay sugar-rich substrates (Põlme et al. 2020) | Põlme S, et al. 2020. Fungal Diversity 105:1-16 ((https://doi.org/10.1007/s13225-020-00466-2)) |
| Otu34745 | 1 | k__Fungi(100);p__Ascomycota(100);c__Saccharomycetes(100);o__Saccharomycetales(100);f__Metschnikowiaceae(100);g__Metschnikowia(100);g__Metschnikowia_unclassified(100); | Metschnikowia | 13 | Saprotroph | \|Undefined Saprotroph\| | Yeast | NULL | Probable | As nectar-tap_saprotroph (Põlme et al. 2020); Decay sugar-rich substrates (Põlme et al. 2020) | Põlme S, et al. 2020. Fungal Diversity 105:1-16 ((https://doi.org/10.1007/s13225-020-00466-2)) |
| Otu34746 | 1 | k__Fungi(100);p__Ascomycota(100);c__Saccharomycetes(100);o__Saccharomycetales(100);f__Metschnikowiaceae(100);g__Metschnikowia(100);g__Metschnikowia_unclassified(100); | Metschnikowia | 13 | Saprotroph | \|Undefined Saprotroph\| | Yeast | NULL | Probable | As nectar-tap_saprotroph (Põlme et al. 2020); Decay sugar-rich substrates (Põlme et al. 2020) | Põlme S, et al. 2020. Fungal Diversity 105:1-16 ((https://doi.org/10.1007/s13225-020-00466-2)) |
| Otu34747 | 1 | k__Fungi(100);p__Ascomycota(100);c__Saccharomycetes(100);o__Saccharomycetales(100);f__Metschnikowiaceae(100);g__Metschnikowia(100);s__Metschnikowia_pulcherrima(100); | Metschnikowia | 13 | Saprotroph | \|Undefined Saprotroph\| | Yeast | NULL | Probable | As nectar-tap_saprotroph (Põlme et al. 2020); Decay sugar-rich substrates (Põlme et al. 2020) | Põlme S, et al. 2020. Fungal Diversity 105:1-16 ((https://doi.org/10.1007/s13225-020-00466-2)) |
| Otu34748 | 1 | k__Fungi(100);p__Ascomycota(100);c__Saccharomycetes(100);o__Saccharomycetales(100);f__Metschnikowiaceae(100);g__Metschnikowia(100);g__Metschnikowia_unclassified(100); | Metschnikowia | 13 | Saprotroph | \|Undefined Saprotroph\| | Yeast | NULL | Probable | As nectar-tap_saprotroph (Põlme et al. 2020); Decay sugar-rich substrates (Põlme et al. 2020) | Põlme S, et al. 2020. Fungal Diversity 105:1-16 ((https://doi.org/10.1007/s13225-020-00466-2)) |
| Otu34749 | 1 | k__Fungi(100);p__Ascomycota(100);c__Saccharomycetes(100);o__Saccharomycetales(100);f__Metschnikowiaceae(100);g__Metschnikowia(100);g__Metschnikowia_unclassified(100); | Metschnikowia | 13 | Saprotroph | \|Undefined Saprotroph\| | Yeast | NULL | Probable | As nectar-tap_saprotroph (Põlme et al. 2020); Decay sugar-rich substrates (Põlme et al. 2020) | Põlme S, et al. 2020. Fungal Diversity 105:1-16 ((https://doi.org/10.1007/s13225-020-00466-2)) |
| Otu34750 | 1 | k__Fungi(100);p__Ascomycota(100);c__Saccharomycetes(100);o__Saccharomycetales(100);f__Metschnikowiaceae(100);g__Metschnikowia(100);g__Metschnikowia_unclassified(100); | Metschnikowia | 13 | Saprotroph | \|Undefined Saprotroph\| | Yeast | NULL | Probable | As nectar-tap_saprotroph (Põlme et al. 2020); Decay sugar-rich substrates (Põlme et al. 2020) | Põlme S, et al. 2020. Fungal Diversity 105:1-16 ((https://doi.org/10.1007/s13225-020-00466-2)) |
| Otu34751 | 1 | k__Fungi(100);p__Ascomycota(100);c__Saccharomycetes(100);o__Saccharomycetales(100);f__Metschnikowiaceae(100);g__Metschnikowia(100);s__Metschnikowia_pulcherrima(100); | Metschnikowia | 13 | Saprotroph | \|Undefined Saprotroph\| | Yeast | NULL | Probable | As nectar-tap_saprotroph (Põlme et al. 2020); Decay sugar-rich substrates (Põlme et al. 2020) | Põlme S, et al. 2020. Fungal Diversity 105:1-16 ((https://doi.org/10.1007/s13225-020-00466-2)) |
| Otu34752 | 1 | k__Fungi(100);p__Ascomycota(100);c__Saccharomycetes(100);o__Saccharomycetales(100);f__Metschnikowiaceae(100);g__Metschnikowia(100);s__Metschnikowia_pulcherrima(100); | Metschnikowia | 13 | Saprotroph | \|Undefined Saprotroph\| | Yeast | NULL | Probable | As nectar-tap_saprotroph (Põlme et al. 2020); Decay sugar-rich substrates (Põlme et al. 2020) | Põlme S, et al. 2020. Fungal Diversity 105:1-16 ((https://doi.org/10.1007/s13225-020-00466-2)) |
| Otu34753 | 1 | k__Fungi(100);p__Ascomycota(100);c__Saccharomycetes(100);o__Saccharomycetales(100);f__Metschnikowiaceae(100);g__Metschnikowia(100);g__Metschnikowia_unclassified(100); | Metschnikowia | 13 | Saprotroph | \|Undefined Saprotroph\| | Yeast | NULL | Probable | As nectar-tap_saprotroph (Põlme et al. 2020); Decay sugar-rich substrates (Põlme et al. 2020) | Põlme S, et al. 2020. Fungal Diversity 105:1-16 ((https://doi.org/10.1007/s13225-020-00466-2)) |
| Otu34754 | 1 | k__Fungi(100);p__Ascomycota(100);c__Saccharomycetes(100);o__Saccharomycetales(100);f__Metschnikowiaceae(100);g__Metschnikowia(100);s__Metschnikowia_pulcherrima(100); | Metschnikowia | 13 | Saprotroph | \|Undefined Saprotroph\| | Yeast | NULL | Probable | As nectar-tap_saprotroph (Põlme et al. 2020); Decay sugar-rich substrates (Põlme et al. 2020) | Põlme S, et al. 2020. Fungal Diversity 105:1-16 ((https://doi.org/10.1007/s13225-020-00466-2)) |
| Otu34757 | 1 | k__Fungi(100);p__Ascomycota(100);c__Saccharomycetes(100);o__Saccharomycetales(100);f__Metschnikowiaceae(100);g__Metschnikowia(100);s__Metschnikowia_pulcherrima(100); | Metschnikowia | 13 | Saprotroph | \|Undefined Saprotroph\| | Yeast | NULL | Probable | As nectar-tap_saprotroph (Põlme et al. 2020); Decay sugar-rich substrates (Põlme et al. 2020) | Põlme S, et al. 2020. Fungal Diversity 105:1-16 ((https://doi.org/10.1007/s13225-020-00466-2)) |
| Otu34759 | 1 | k__Fungi(100);p__Ascomycota(100);c__Saccharomycetes(100);o__Saccharomycetales(100);f__Metschnikowiaceae(100);g__Metschnikowia(100);s__Metschnikowia_pulcherrima(100); | Metschnikowia | 13 | Saprotroph | \|Undefined Saprotroph\| | Yeast | NULL | Probable | As nectar-tap_saprotroph (Põlme et al. 2020); Decay sugar-rich substrates (Põlme et al. 2020) | Põlme S, et al. 2020. Fungal Diversity 105:1-16 ((https://doi.org/10.1007/s13225-020-00466-2)) |
| Otu34760 | 1 | k__Fungi(100);p__Ascomycota(100);c__Saccharomycetes(100);o__Saccharomycetales(100);f__Metschnikowiaceae(100);g__Metschnikowia(100);s__Metschnikowia_pulcherrima(100); | Metschnikowia | 13 | Saprotroph | \|Undefined Saprotroph\| | Yeast | NULL | Probable | As nectar-tap_saprotroph (Põlme et al. 2020); Decay sugar-rich substrates (Põlme et al. 2020) | Põlme S, et al. 2020. Fungal Diversity 105:1-16 ((https://doi.org/10.1007/s13225-020-00466-2)) |
| Otu34761 | 1 | k__Fungi(100);p__Ascomycota(100);c__Saccharomycetes(100);o__Saccharomycetales(100);f__Metschnikowiaceae(100);g__Metschnikowia(100);g__Metschnikowia_unclassified(100); | Metschnikowia | 13 | Saprotroph | \|Undefined Saprotroph\| | Yeast | NULL | Probable | As nectar-tap_saprotroph (Põlme et al. 2020); Decay sugar-rich substrates (Põlme et al. 2020) | Põlme S, et al. 2020. Fungal Diversity 105:1-16 ((https://doi.org/10.1007/s13225-020-00466-2)) |
| Otu34762 | 1 | k__Fungi(100);p__Ascomycota(100);c__Saccharomycetes(100);o__Saccharomycetales(100);f__Metschnikowiaceae(100);g__Metschnikowia(100);s__Metschnikowia_pulcherrima(100); | Metschnikowia | 13 | Saprotroph | \|Undefined Saprotroph\| | Yeast | NULL | Probable | As nectar-tap_saprotroph (Põlme et al. 2020); Decay sugar-rich substrates (Põlme et al. 2020) | Põlme S, et al. 2020. Fungal Diversity 105:1-16 ((https://doi.org/10.1007/s13225-020-00466-2)) |
| Otu34769 | 1 | k__Fungi(100);p__Ascomycota(100);c__Saccharomycetes(100);o__Saccharomycetales(100);f__Metschnikowiaceae(100);g__Metschnikowia(100);g__Metschnikowia_unclassified(100); | Metschnikowia | 13 | Saprotroph | \|Undefined Saprotroph\| | Yeast | NULL | Probable | As nectar-tap_saprotroph (Põlme et al. 2020); Decay sugar-rich substrates (Põlme et al. 2020) | Põlme S, et al. 2020. Fungal Diversity 105:1-16 ((https://doi.org/10.1007/s13225-020-00466-2)) |
| Otu34774 | 1 | k__Fungi(100);p__Ascomycota(100);c__Saccharomycetes(100);o__Saccharomycetales(100);f__Metschnikowiaceae(100);g__Metschnikowia(100);s__Metschnikowia_pulcherrima(100); | Metschnikowia | 13 | Saprotroph | \|Undefined Saprotroph\| | Yeast | NULL | Probable | As nectar-tap_saprotroph (Põlme et al. 2020); Decay sugar-rich substrates (Põlme et al. 2020) | Põlme S, et al. 2020. Fungal Diversity 105:1-16 ((https://doi.org/10.1007/s13225-020-00466-2)) |
| Otu34775 | 1 | k__Fungi(100);p__Ascomycota(100);c__Saccharomycetes(100);o__Saccharomycetales(100);f__Metschnikowiaceae(100);g__Metschnikowia(100);s__Metschnikowia_pulcherrima(100); | Metschnikowia | 13 | Saprotroph | \|Undefined Saprotroph\| | Yeast | NULL | Probable | As nectar-tap_saprotroph (Põlme et al. 2020); Decay sugar-rich substrates (Põlme et al. 2020) | Põlme S, et al. 2020. Fungal Diversity 105:1-16 ((https://doi.org/10.1007/s13225-020-00466-2)) |
| Otu34776 | 1 | k__Fungi(100);p__Ascomycota(100);c__Saccharomycetes(100);o__Saccharomycetales(100);f__Metschnikowiaceae(100);g__Metschnikowia(100);g__Metschnikowia_unclassified(100); | Metschnikowia | 13 | Saprotroph | \|Undefined Saprotroph\| | Yeast | NULL | Probable | As nectar-tap_saprotroph (Põlme et al. 2020); Decay sugar-rich substrates (Põlme et al. 2020) | Põlme S, et al. 2020. Fungal Diversity 105:1-16 ((https://doi.org/10.1007/s13225-020-00466-2)) |
| Otu34784 | 1 | k__Fungi(100);p__Ascomycota(100);c__Saccharomycetes(100);o__Saccharomycetales(100);f__Metschnikowiaceae(100);g__Metschnikowia(100);g__Metschnikowia_unclassified(100); | Metschnikowia | 13 | Saprotroph | \|Undefined Saprotroph\| | Yeast | NULL | Probable | As nectar-tap_saprotroph (Põlme et al. 2020); Decay sugar-rich substrates (Põlme et al. 2020) | Põlme S, et al. 2020. Fungal Diversity 105:1-16 ((https://doi.org/10.1007/s13225-020-00466-2)) |
| Otu34785 | 1 | k__Fungi(100);p__Ascomycota(100);c__Saccharomycetes(100);o__Saccharomycetales(100);f__Metschnikowiaceae(100);g__Metschnikowia(100);s__Metschnikowia_pulcherrima(100); | Metschnikowia | 13 | Saprotroph | \|Undefined Saprotroph\| | Yeast | NULL | Probable | As nectar-tap_saprotroph (Põlme et al. 2020); Decay sugar-rich substrates (Põlme et al. 2020) | Põlme S, et al. 2020. Fungal Diversity 105:1-16 ((https://doi.org/10.1007/s13225-020-00466-2)) |
| Otu34786 | 1 | k__Fungi(100);p__Ascomycota(100);c__Saccharomycetes(100);o__Saccharomycetales(100);f__Metschnikowiaceae(100);g__Metschnikowia(100);s__Metschnikowia_pulcherrima(100); | Metschnikowia | 13 | Saprotroph | \|Undefined Saprotroph\| | Yeast | NULL | Probable | As nectar-tap_saprotroph (Põlme et al. 2020); Decay sugar-rich substrates (Põlme et al. 2020) | Põlme S, et al. 2020. Fungal Diversity 105:1-16 ((https://doi.org/10.1007/s13225-020-00466-2)) |
| Otu34788 | 1 | k__Fungi(100);p__Ascomycota(100);c__Saccharomycetes(100);o__Saccharomycetales(100);f__Metschnikowiaceae(100);g__Metschnikowia(100);g__Metschnikowia_unclassified(100); | Metschnikowia | 13 | Saprotroph | \|Undefined Saprotroph\| | Yeast | NULL | Probable | As nectar-tap_saprotroph (Põlme et al. 2020); Decay sugar-rich substrates (Põlme et al. 2020) | Põlme S, et al. 2020. Fungal Diversity 105:1-16 ((https://doi.org/10.1007/s13225-020-00466-2)) |
| Otu34791 | 1 | k__Fungi(100);p__Ascomycota(100);c__Saccharomycetes(100);o__Saccharomycetales(100);f__Metschnikowiaceae(100);g__Metschnikowia(100);s__Metschnikowia_pulcherrima(100); | Metschnikowia | 13 | Saprotroph | \|Undefined Saprotroph\| | Yeast | NULL | Probable | As nectar-tap_saprotroph (Põlme et al. 2020); Decay sugar-rich substrates (Põlme et al. 2020) | Põlme S, et al. 2020. Fungal Diversity 105:1-16 ((https://doi.org/10.1007/s13225-020-00466-2)) |
| Otu34795 | 1 | k__Fungi(100);p__Ascomycota(100);c__Saccharomycetes(100);o__Saccharomycetales(100);f__Metschnikowiaceae(100);g__Metschnikowia(100);s__Metschnikowia_pulcherrima(100); | Metschnikowia | 13 | Saprotroph | \|Undefined Saprotroph\| | Yeast | NULL | Probable | As nectar-tap_saprotroph (Põlme et al. 2020); Decay sugar-rich substrates (Põlme et al. 2020) | Põlme S, et al. 2020. Fungal Diversity 105:1-16 ((https://doi.org/10.1007/s13225-020-00466-2)) |
| Otu34801 | 1 | k__Fungi(100);p__Ascomycota(100);c__Saccharomycetes(100);o__Saccharomycetales(100);f__Metschnikowiaceae(100);g__Metschnikowia(100);g__Metschnikowia_unclassified(100); | Metschnikowia | 13 | Saprotroph | \|Undefined Saprotroph\| | Yeast | NULL | Probable | As nectar-tap_saprotroph (Põlme et al. 2020); Decay sugar-rich substrates (Põlme et al. 2020) | Põlme S, et al. 2020. Fungal Diversity 105:1-16 ((https://doi.org/10.1007/s13225-020-00466-2)) |
| Otu34803 | 1 | k__Fungi(100);p__Ascomycota(100);c__Saccharomycetes(100);o__Saccharomycetales(100);f__Metschnikowiaceae(100);g__Metschnikowia(100);g__Metschnikowia_unclassified(100); | Metschnikowia | 13 | Saprotroph | \|Undefined Saprotroph\| | Yeast | NULL | Probable | As nectar-tap_saprotroph (Põlme et al. 2020); Decay sugar-rich substrates (Põlme et al. 2020) | Põlme S, et al. 2020. Fungal Diversity 105:1-16 ((https://doi.org/10.1007/s13225-020-00466-2)) |
| Otu34805 | 1 | k__Fungi(100);p__Ascomycota(100);c__Saccharomycetes(100);o__Saccharomycetales(100);f__Metschnikowiaceae(100);g__Metschnikowia(100);s__Metschnikowia_pulcherrima(100); | Metschnikowia | 13 | Saprotroph | \|Undefined Saprotroph\| | Yeast | NULL | Probable | As nectar-tap_saprotroph (Põlme et al. 2020); Decay sugar-rich substrates (Põlme et al. 2020) | Põlme S, et al. 2020. Fungal Diversity 105:1-16 ((https://doi.org/10.1007/s13225-020-00466-2)) |
| Otu34806 | 1 | k__Fungi(100);p__Ascomycota(100);c__Saccharomycetes(100);o__Saccharomycetales(100);f__Metschnikowiaceae(100);g__Metschnikowia(100);s__Metschnikowia_pulcherrima(100); | Metschnikowia | 13 | Saprotroph | \|Undefined Saprotroph\| | Yeast | NULL | Probable | As nectar-tap_saprotroph (Põlme et al. 2020); Decay sugar-rich substrates (Põlme et al. 2020) | Põlme S, et al. 2020. Fungal Diversity 105:1-16 ((https://doi.org/10.1007/s13225-020-00466-2)) |
| Otu34807 | 1 | k__Fungi(100);p__Ascomycota(100);c__Saccharomycetes(100);o__Saccharomycetales(100);f__Metschnikowiaceae(100);g__Metschnikowia(100);g__Metschnikowia_unclassified(100); | Metschnikowia | 13 | Saprotroph | \|Undefined Saprotroph\| | Yeast | NULL | Probable | As nectar-tap_saprotroph (Põlme et al. 2020); Decay sugar-rich substrates (Põlme et al. 2020) | Põlme S, et al. 2020. Fungal Diversity 105:1-16 ((https://doi.org/10.1007/s13225-020-00466-2)) |
| Otu34809 | 1 | k__Fungi(100);p__Ascomycota(100);c__Saccharomycetes(100);o__Saccharomycetales(100);f__Metschnikowiaceae(100);g__Metschnikowia(100);s__Metschnikowia_pulcherrima(100); | Metschnikowia | 13 | Saprotroph | \|Undefined Saprotroph\| | Yeast | NULL | Probable | As nectar-tap_saprotroph (Põlme et al. 2020); Decay sugar-rich substrates (Põlme et al. 2020) | Põlme S, et al. 2020. Fungal Diversity 105:1-16 ((https://doi.org/10.1007/s13225-020-00466-2)) |
| Otu34811 | 1 | k__Fungi(100);p__Ascomycota(100);c__Saccharomycetes(100);o__Saccharomycetales(100);f__Metschnikowiaceae(100);g__Metschnikowia(100);s__Metschnikowia_pulcherrima(100); | Metschnikowia | 13 | Saprotroph | \|Undefined Saprotroph\| | Yeast | NULL | Probable | As nectar-tap_saprotroph (Põlme et al. 2020); Decay sugar-rich substrates (Põlme et al. 2020) | Põlme S, et al. 2020. Fungal Diversity 105:1-16 ((https://doi.org/10.1007/s13225-020-00466-2)) |
| Otu34812 | 1 | k__Fungi(100);p__Ascomycota(100);c__Saccharomycetes(100);o__Saccharomycetales(100);f__Metschnikowiaceae(100);g__Metschnikowia(100);s__Metschnikowia_pulcherrima(100); | Metschnikowia | 13 | Saprotroph | \|Undefined Saprotroph\| | Yeast | NULL | Probable | As nectar-tap_saprotroph (Põlme et al. 2020); Decay sugar-rich substrates (Põlme et al. 2020) | Põlme S, et al. 2020. Fungal Diversity 105:1-16 ((https://doi.org/10.1007/s13225-020-00466-2)) |
| Otu34813 | 1 | k__Fungi(100);p__Ascomycota(100);c__Saccharomycetes(100);o__Saccharomycetales(100);f__Metschnikowiaceae(100);g__Metschnikowia(100);s__Metschnikowia_pulcherrima(100); | Metschnikowia | 13 | Saprotroph | \|Undefined Saprotroph\| | Yeast | NULL | Probable | As nectar-tap_saprotroph (Põlme et al. 2020); Decay sugar-rich substrates (Põlme et al. 2020) | Põlme S, et al. 2020. Fungal Diversity 105:1-16 ((https://doi.org/10.1007/s13225-020-00466-2)) |
| Otu34814 | 1 | k__Fungi(100);p__Ascomycota(100);c__Saccharomycetes(100);o__Saccharomycetales(100);f__Metschnikowiaceae(100);g__Metschnikowia(100);s__Metschnikowia_pulcherrima(100); | Metschnikowia | 13 | Saprotroph | \|Undefined Saprotroph\| | Yeast | NULL | Probable | As nectar-tap_saprotroph (Põlme et al. 2020); Decay sugar-rich substrates (Põlme et al. 2020) | Põlme S, et al. 2020. Fungal Diversity 105:1-16 ((https://doi.org/10.1007/s13225-020-00466-2)) |
| Otu34815 | 1 | k__Fungi(100);p__Ascomycota(100);c__Saccharomycetes(100);o__Saccharomycetales(100);f__Metschnikowiaceae(100);g__Metschnikowia(100);s__Metschnikowia_pulcherrima(100); | Metschnikowia | 13 | Saprotroph | \|Undefined Saprotroph\| | Yeast | NULL | Probable | As nectar-tap_saprotroph (Põlme et al. 2020); Decay sugar-rich substrates (Põlme et al. 2020) | Põlme S, et al. 2020. Fungal Diversity 105:1-16 ((https://doi.org/10.1007/s13225-020-00466-2)) |
| Otu34816 | 1 | k__Fungi(100);p__Ascomycota(100);c__Saccharomycetes(100);o__Saccharomycetales(100);f__Metschnikowiaceae(100);g__Metschnikowia(100);g__Metschnikowia_unclassified(100); | Metschnikowia | 13 | Saprotroph | \|Undefined Saprotroph\| | Yeast | NULL | Probable | As nectar-tap_saprotroph (Põlme et al. 2020); Decay sugar-rich substrates (Põlme et al. 2020) | Põlme S, et al. 2020. Fungal Diversity 105:1-16 ((https://doi.org/10.1007/s13225-020-00466-2)) |
| Otu34818 | 1 | k__Fungi(100);p__Ascomycota(100);c__Saccharomycetes(100);o__Saccharomycetales(100);f__Metschnikowiaceae(100);g__Metschnikowia(100);s__Metschnikowia_pulcherrima(100); | Metschnikowia | 13 | Saprotroph | \|Undefined Saprotroph\| | Yeast | NULL | Probable | As nectar-tap_saprotroph (Põlme et al. 2020); Decay sugar-rich substrates (Põlme et al. 2020) | Põlme S, et al. 2020. Fungal Diversity 105:1-16 ((https://doi.org/10.1007/s13225-020-00466-2)) |
| Otu34819 | 1 | k__Fungi(100);p__Ascomycota(100);c__Saccharomycetes(100);o__Saccharomycetales(100);f__Metschnikowiaceae(100);g__Metschnikowia(100);s__Metschnikowia_pulcherrima(100); | Metschnikowia | 13 | Saprotroph | \|Undefined Saprotroph\| | Yeast | NULL | Probable | As nectar-tap_saprotroph (Põlme et al. 2020); Decay sugar-rich substrates (Põlme et al. 2020) | Põlme S, et al. 2020. Fungal Diversity 105:1-16 ((https://doi.org/10.1007/s13225-020-00466-2)) |
| Otu34821 | 1 | k__Fungi(100);p__Ascomycota(100);c__Saccharomycetes(100);o__Saccharomycetales(100);f__Metschnikowiaceae(100);g__Metschnikowia(100);s__Metschnikowia_pulcherrima(100); | Metschnikowia | 13 | Saprotroph | \|Undefined Saprotroph\| | Yeast | NULL | Probable | As nectar-tap_saprotroph (Põlme et al. 2020); Decay sugar-rich substrates (Põlme et al. 2020) | Põlme S, et al. 2020. Fungal Diversity 105:1-16 ((https://doi.org/10.1007/s13225-020-00466-2)) |
| Otu34822 | 1 | k__Fungi(100);p__Ascomycota(100);c__Saccharomycetes(100);o__Saccharomycetales(100);f__Metschnikowiaceae(100);g__Metschnikowia(100);s__Metschnikowia_pulcherrima(100); | Metschnikowia | 13 | Saprotroph | \|Undefined Saprotroph\| | Yeast | NULL | Probable | As nectar-tap_saprotroph (Põlme et al. 2020); Decay sugar-rich substrates (Põlme et al. 2020) | Põlme S, et al. 2020. Fungal Diversity 105:1-16 ((https://doi.org/10.1007/s13225-020-00466-2)) |
| Otu34829 | 1 | k__Fungi(100);p__Ascomycota(100);c__Saccharomycetes(100);o__Saccharomycetales(100);f__Metschnikowiaceae(100);g__Metschnikowia(100);s__Metschnikowia_pulcherrima(100); | Metschnikowia | 13 | Saprotroph | \|Undefined Saprotroph\| | Yeast | NULL | Probable | As nectar-tap_saprotroph (Põlme et al. 2020); Decay sugar-rich substrates (Põlme et al. 2020) | Põlme S, et al. 2020. Fungal Diversity 105:1-16 ((https://doi.org/10.1007/s13225-020-00466-2)) |
| Otu34835 | 1 | k__Fungi(100);p__Ascomycota(100);c__Saccharomycetes(100);o__Saccharomycetales(100);f__Metschnikowiaceae(100);g__Metschnikowia(100);s__Metschnikowia_pulcherrima(100); | Metschnikowia | 13 | Saprotroph | \|Undefined Saprotroph\| | Yeast | NULL | Probable | As nectar-tap_saprotroph (Põlme et al. 2020); Decay sugar-rich substrates (Põlme et al. 2020) | Põlme S, et al. 2020. Fungal Diversity 105:1-16 ((https://doi.org/10.1007/s13225-020-00466-2)) |
| Otu34836 | 1 | k__Fungi(100);p__Ascomycota(100);c__Saccharomycetes(100);o__Saccharomycetales(100);f__Metschnikowiaceae(100);g__Metschnikowia(100);s__Metschnikowia_pulcherrima(100); | Metschnikowia | 13 | Saprotroph | \|Undefined Saprotroph\| | Yeast | NULL | Probable | As nectar-tap_saprotroph (Põlme et al. 2020); Decay sugar-rich substrates (Põlme et al. 2020) | Põlme S, et al. 2020. Fungal Diversity 105:1-16 ((https://doi.org/10.1007/s13225-020-00466-2)) |
| Otu34837 | 1 | k__Fungi(100);p__Ascomycota(100);c__Saccharomycetes(100);o__Saccharomycetales(100);f__Metschnikowiaceae(100);g__Metschnikowia(100);g__Metschnikowia_unclassified(100); | Metschnikowia | 13 | Saprotroph | \|Undefined Saprotroph\| | Yeast | NULL | Probable | As nectar-tap_saprotroph (Põlme et al. 2020); Decay sugar-rich substrates (Põlme et al. 2020) | Põlme S, et al. 2020. Fungal Diversity 105:1-16 ((https://doi.org/10.1007/s13225-020-00466-2)) |
| Otu34838 | 1 | k__Fungi(100);p__Ascomycota(100);c__Saccharomycetes(100);o__Saccharomycetales(100);f__Metschnikowiaceae(100);g__Metschnikowia(100);g__Metschnikowia_unclassified(100); | Metschnikowia | 13 | Saprotroph | \|Undefined Saprotroph\| | Yeast | NULL | Probable | As nectar-tap_saprotroph (Põlme et al. 2020); Decay sugar-rich substrates (Põlme et al. 2020) | Põlme S, et al. 2020. Fungal Diversity 105:1-16 ((https://doi.org/10.1007/s13225-020-00466-2)) |
| Otu34839 | 1 | k__Fungi(100);p__Ascomycota(100);c__Saccharomycetes(100);o__Saccharomycetales(100);f__Metschnikowiaceae(100);g__Metschnikowia(100);s__Metschnikowia_pulcherrima(100); | Metschnikowia | 13 | Saprotroph | \|Undefined Saprotroph\| | Yeast | NULL | Probable | As nectar-tap_saprotroph (Põlme et al. 2020); Decay sugar-rich substrates (Põlme et al. 2020) | Põlme S, et al. 2020. Fungal Diversity 105:1-16 ((https://doi.org/10.1007/s13225-020-00466-2)) |
| Otu34840 | 1 | k__Fungi(100);p__Ascomycota(100);c__Saccharomycetes(100);o__Saccharomycetales(100);f__Metschnikowiaceae(100);g__Metschnikowia(100);g__Metschnikowia_unclassified(100); | Metschnikowia | 13 | Saprotroph | \|Undefined Saprotroph\| | Yeast | NULL | Probable | As nectar-tap_saprotroph (Põlme et al. 2020); Decay sugar-rich substrates (Põlme et al. 2020) | Põlme S, et al. 2020. Fungal Diversity 105:1-16 ((https://doi.org/10.1007/s13225-020-00466-2)) |
| Otu34841 | 1 | k__Fungi(100);p__Ascomycota(100);c__Saccharomycetes(100);o__Saccharomycetales(100);f__Metschnikowiaceae(100);g__Metschnikowia(100);s__Metschnikowia_pulcherrima(100); | Metschnikowia | 13 | Saprotroph | \|Undefined Saprotroph\| | Yeast | NULL | Probable | As nectar-tap_saprotroph (Põlme et al. 2020); Decay sugar-rich substrates (Põlme et al. 2020) | Põlme S, et al. 2020. Fungal Diversity 105:1-16 ((https://doi.org/10.1007/s13225-020-00466-2)) |
| Otu34842 | 1 | k__Fungi(100);p__Ascomycota(100);c__Saccharomycetes(100);o__Saccharomycetales(100);f__Metschnikowiaceae(100);g__Metschnikowia(100);g__Metschnikowia_unclassified(100); | Metschnikowia | 13 | Saprotroph | \|Undefined Saprotroph\| | Yeast | NULL | Probable | As nectar-tap_saprotroph (Põlme et al. 2020); Decay sugar-rich substrates (Põlme et al. 2020) | Põlme S, et al. 2020. Fungal Diversity 105:1-16 ((https://doi.org/10.1007/s13225-020-00466-2)) |
| Otu34843 | 1 | k__Fungi(100);p__Ascomycota(100);c__Saccharomycetes(100);o__Saccharomycetales(100);f__Metschnikowiaceae(100);g__Metschnikowia(100);s__Metschnikowia_pulcherrima(100); | Metschnikowia | 13 | Saprotroph | \|Undefined Saprotroph\| | Yeast | NULL | Probable | As nectar-tap_saprotroph (Põlme et al. 2020); Decay sugar-rich substrates (Põlme et al. 2020) | Põlme S, et al. 2020. Fungal Diversity 105:1-16 ((https://doi.org/10.1007/s13225-020-00466-2)) |
| Otu34844 | 1 | k__Fungi(100);p__Ascomycota(100);c__Saccharomycetes(100);o__Saccharomycetales(100);f__Metschnikowiaceae(100);g__Metschnikowia(100);s__Metschnikowia_pulcherrima(100); | Metschnikowia | 13 | Saprotroph | \|Undefined Saprotroph\| | Yeast | NULL | Probable | As nectar-tap_saprotroph (Põlme et al. 2020); Decay sugar-rich substrates (Põlme et al. 2020) | Põlme S, et al. 2020. Fungal Diversity 105:1-16 ((https://doi.org/10.1007/s13225-020-00466-2)) |
| Otu34845 | 1 | k__Fungi(100);p__Ascomycota(100);c__Saccharomycetes(100);o__Saccharomycetales(100);f__Metschnikowiaceae(100);g__Metschnikowia(100);s__Metschnikowia_pulcherrima(100); | Metschnikowia | 13 | Saprotroph | \|Undefined Saprotroph\| | Yeast | NULL | Probable | As nectar-tap_saprotroph (Põlme et al. 2020); Decay sugar-rich substrates (Põlme et al. 2020) | Põlme S, et al. 2020. Fungal Diversity 105:1-16 ((https://doi.org/10.1007/s13225-020-00466-2)) |
| Otu34846 | 1 | k__Fungi(100);p__Ascomycota(100);c__Saccharomycetes(100);o__Saccharomycetales(100);f__Metschnikowiaceae(100);g__Metschnikowia(100);s__Metschnikowia_pulcherrima(100); | Metschnikowia | 13 | Saprotroph | \|Undefined Saprotroph\| | Yeast | NULL | Probable | As nectar-tap_saprotroph (Põlme et al. 2020); Decay sugar-rich substrates (Põlme et al. 2020) | Põlme S, et al. 2020. Fungal Diversity 105:1-16 ((https://doi.org/10.1007/s13225-020-00466-2)) |
| Otu34847 | 1 | k__Fungi(100);p__Ascomycota(100);c__Saccharomycetes(100);o__Saccharomycetales(100);f__Metschnikowiaceae(100);g__Metschnikowia(100);s__Metschnikowia_pulcherrima(100); | Metschnikowia | 13 | Saprotroph | \|Undefined Saprotroph\| | Yeast | NULL | Probable | As nectar-tap_saprotroph (Põlme et al. 2020); Decay sugar-rich substrates (Põlme et al. 2020) | Põlme S, et al. 2020. Fungal Diversity 105:1-16 ((https://doi.org/10.1007/s13225-020-00466-2)) |
| Otu34849 | 1 | k__Fungi(100);p__Ascomycota(100);c__Saccharomycetes(100);o__Saccharomycetales(100);f__Metschnikowiaceae(100);g__Metschnikowia(100);g__Metschnikowia_unclassified(100); | Metschnikowia | 13 | Saprotroph | \|Undefined Saprotroph\| | Yeast | NULL | Probable | As nectar-tap_saprotroph (Põlme et al. 2020); Decay sugar-rich substrates (Põlme et al. 2020) | Põlme S, et al. 2020. Fungal Diversity 105:1-16 ((https://doi.org/10.1007/s13225-020-00466-2)) |
| Otu34850 | 1 | k__Fungi(100);p__Ascomycota(100);c__Saccharomycetes(100);o__Saccharomycetales(100);f__Metschnikowiaceae(100);g__Metschnikowia(100);s__Metschnikowia_pulcherrima(100); | Metschnikowia | 13 | Saprotroph | \|Undefined Saprotroph\| | Yeast | NULL | Probable | As nectar-tap_saprotroph (Põlme et al. 2020); Decay sugar-rich substrates (Põlme et al. 2020) | Põlme S, et al. 2020. Fungal Diversity 105:1-16 ((https://doi.org/10.1007/s13225-020-00466-2)) |
| Otu34851 | 1 | k__Fungi(100);p__Ascomycota(100);c__Saccharomycetes(100);o__Saccharomycetales(100);f__Metschnikowiaceae(100);g__Metschnikowia(100);s__Metschnikowia_pulcherrima(100); | Metschnikowia | 13 | Saprotroph | \|Undefined Saprotroph\| | Yeast | NULL | Probable | As nectar-tap_saprotroph (Põlme et al. 2020); Decay sugar-rich substrates (Põlme et al. 2020) | Põlme S, et al. 2020. Fungal Diversity 105:1-16 ((https://doi.org/10.1007/s13225-020-00466-2)) |
| Otu34852 | 1 | k__Fungi(100);p__Ascomycota(100);c__Saccharomycetes(100);o__Saccharomycetales(100);f__Metschnikowiaceae(100);g__Metschnikowia(100);s__Metschnikowia_pulcherrima(100); | Metschnikowia | 13 | Saprotroph | \|Undefined Saprotroph\| | Yeast | NULL | Probable | As nectar-tap_saprotroph (Põlme et al. 2020); Decay sugar-rich substrates (Põlme et al. 2020) | Põlme S, et al. 2020. Fungal Diversity 105:1-16 ((https://doi.org/10.1007/s13225-020-00466-2)) |
| Otu34854 | 1 | k__Fungi(100);p__Ascomycota(100);c__Saccharomycetes(100);o__Saccharomycetales(100);f__Metschnikowiaceae(100);g__Metschnikowia(100);s__Metschnikowia_pulcherrima(100); | Metschnikowia | 13 | Saprotroph | \|Undefined Saprotroph\| | Yeast | NULL | Probable | As nectar-tap_saprotroph (Põlme et al. 2020); Decay sugar-rich substrates (Põlme et al. 2020) | Põlme S, et al. 2020. Fungal Diversity 105:1-16 ((https://doi.org/10.1007/s13225-020-00466-2)) |
| Otu34855 | 1 | k__Fungi(100);p__Ascomycota(100);c__Saccharomycetes(100);o__Saccharomycetales(100);f__Metschnikowiaceae(100);g__Metschnikowia(100);g__Metschnikowia_unclassified(100); | Metschnikowia | 13 | Saprotroph | \|Undefined Saprotroph\| | Yeast | NULL | Probable | As nectar-tap_saprotroph (Põlme et al. 2020); Decay sugar-rich substrates (Põlme et al. 2020) | Põlme S, et al. 2020. Fungal Diversity 105:1-16 ((https://doi.org/10.1007/s13225-020-00466-2)) |

| | | | | | | | | | | | |
|---|---|---|---|---|---|---|---|---|---|---|---|
| Otu34856 | 1 | k__Fungi(100);p__Ascomycota(100);c__Saccharomycetes(100);o__Saccharomycetales(100);f__Metschnikowiaceae(100);g__Metschnikowia(100);g__Metschnikowia_unclassified(100); | Metschnikowia | 13 | Saprotroph | \|Undefined Saprotroph\| | Yeast | NULL | Probable | As nectar-tap_saprotroph (Põlme et al. 2020); Decay sugar-rich substrates (Põlme et al. 2020) | Põlme S, et al. 2020. Fungal Diversity 105:1-16 ((https://doi.org/10.1007/s13225-020-00466-2)) |
| Otu34858 | 1 | k__Fungi(100);p__Ascomycota(100);c__Saccharomycetes(100);o__Saccharomycetales(100);f__Metschnikowiaceae(100);g__Metschnikowia(100);g__Metschnikowia_unclassified(100); | Metschnikowia | 13 | Saprotroph | \|Undefined Saprotroph\| | Yeast | NULL | Probable | As nectar-tap_saprotroph (Põlme et al. 2020); Decay sugar-rich substrates (Põlme et al. 2020) | Põlme S, et al. 2020. Fungal Diversity 105:1-16 ((https://doi.org/10.1007/s13225-020-00466-2)) |
| Otu34859 | 1 | k__Fungi(100);p__Ascomycota(100);c__Saccharomycetes(100);o__Saccharomycetales(100);f__Metschnikowiaceae(100);g__Metschnikowia(100);g__Metschnikowia_unclassified(100); | Metschnikowia | 13 | Saprotroph | \|Undefined Saprotroph\| | Yeast | NULL | Probable | As nectar-tap_saprotroph (Põlme et al. 2020); Decay sugar-rich substrates (Põlme et al. 2020) | Põlme S, et al. 2020. Fungal Diversity 105:1-16 ((https://doi.org/10.1007/s13225-020-00466-2)) |
| Otu34860 | 1 | k__Fungi(100);p__Ascomycota(100);c__Saccharomycetes(100);o__Saccharomycetales(100);f__Metschnikowiaceae(100);g__Metschnikowia(100);g__Metschnikowia_unclassified(100); | Metschnikowia | 13 | Saprotroph | \|Undefined Saprotroph\| | Yeast | NULL | Probable | As nectar-tap_saprotroph (Põlme et al. 2020); Decay sugar-rich substrates (Põlme et al. 2020) | Põlme S, et al. 2020. Fungal Diversity 105:1-16 ((https://doi.org/10.1007/s13225-020-00466-2)) |
| Otu34861 | 1 | k__Fungi(100);p__Ascomycota(100);c__Saccharomycetes(100);o__Saccharomycetales(100);f__Metschnikowiaceae(100);g__Metschnikowia(100);s__Metschnikowia_pulcherrima(100); | Metschnikowia | 13 | Saprotroph | \|Undefined Saprotroph\| | Yeast | NULL | Probable | As nectar-tap_saprotroph (Põlme et al. 2020); Decay sugar-rich substrates (Põlme et al. 2020) | Põlme S, et al. 2020. Fungal Diversity 105:1-16 ((https://doi.org/10.1007/s13225-020-00466-2)) |
| Otu34868 | 1 | k__Fungi(100);p__Ascomycota(100);c__Saccharomycetes(100);o__Saccharomycetales(100);f__Metschnikowiaceae(100);g__Metschnikowia(100);s__Metschnikowia_pulcherrima(100); | Metschnikowia | 13 | Saprotroph | \|Undefined Saprotroph\| | Yeast | NULL | Probable | As nectar-tap_saprotroph (Põlme et al. 2020); Decay sugar-rich substrates (Põlme et al. 2020) | Põlme S, et al. 2020. Fungal Diversity 105:1-16 ((https://doi.org/10.1007/s13225-020-00466-2)) |
| Otu34869 | 1 | k__Fungi(100);p__Ascomycota(100);c__Saccharomycetes(100);o__Saccharomycetales(100);f__Metschnikowiaceae(100);g__Metschnikowia(100);g__Metschnikowia_unclassified(100); | Metschnikowia | 13 | Saprotroph | \|Undefined Saprotroph\| | Yeast | NULL | Probable | As nectar-tap_saprotroph (Põlme et al. 2020); Decay sugar-rich substrates (Põlme et al. 2020) | Põlme S, et al. 2020. Fungal Diversity 105:1-16 ((https://doi.org/10.1007/s13225-020-00466-2)) |
| Otu34870 | 1 | k__Fungi(100);p__Basidiomycota(100);c__Microbotryomycetes(100);o__Sporidiobolales(100);f__Sporidiobolales_family_Incertae_sedis(100);g__Rhodotorula(100);s__Rhodotorula_nothofagi(100); | Rhodotorula | 13 | Pathotroph-Saprotroph-Symbiotroph | Animal Pathogen-Animal Symbiotroph-Endophyte-Plant Pathogen-\|Undefined Saprotroph\| | Yeast | NULL | Probable | As foliar_endophyte (Põlme et al. 2020); Opportunistic human pathogen | Põlme S, et al. 2020. Fungal Diversity 105:1-16 ((https://doi.org/10.1007/s13225-020-00466-2)); https://www.cdc.gov ((https://www.cdc.gov)) |
| Otu34871 | 1 | k__Fungi(100);p__Ascomycota(100);c__Saccharomycetes(100);o__Saccharomycetales(100);f__Metschnikowiaceae(100);g__Metschnikowia(100);s__Metschnikowia_pulcherrima(100); | Metschnikowia | 13 | Saprotroph | \|Undefined Saprotroph\| | Yeast | NULL | Probable | As nectar-tap_saprotroph (Põlme et al. 2020); Decay sugar-rich substrates (Põlme et al. 2020) | Põlme S, et al. 2020. Fungal Diversity 105:1-16 ((https://doi.org/10.1007/s13225-020-00466-2)) |
| Otu34872 | 1 | k__Fungi(100);p__Ascomycota(100);c__Saccharomycetes(100);o__Saccharomycetales(100);f__Metschnikowiaceae(100);g__Metschnikowia(100);s__Metschnikowia_pulcherrima(100); | Metschnikowia | 13 | Saprotroph | \|Undefined Saprotroph\| | Yeast | NULL | Probable | As nectar-tap_saprotroph (Põlme et al. 2020); Decay sugar-rich substrates (Põlme et al. 2020) | Põlme S, et al. 2020. Fungal Diversity 105:1-16 ((https://doi.org/10.1007/s13225-020-00466-2)) |
| Otu34873 | 1 | k__Fungi(100);p__Ascomycota(100);c__Saccharomycetes(100);o__Saccharomycetales(100);f__Metschnikowiaceae(100);g__Metschnikowia(100);s__Metschnikowia_pulcherrima(100); | Metschnikowia | 13 | Saprotroph | \|Undefined Saprotroph\| | Yeast | NULL | Probable | As nectar-tap_saprotroph (Põlme et al. 2020); Decay sugar-rich substrates (Põlme et al. 2020) | Põlme S, et al. 2020. Fungal Diversity 105:1-16 ((https://doi.org/10.1007/s13225-020-00466-2)) |
| Otu34876 | 1 | k__Fungi(100);p__Ascomycota(100);c__Saccharomycetes(100);o__Saccharomycetales(100);f__Metschnikowiaceae(100);g__Metschnikowia(100);s__Metschnikowia_pulcherrima(100); | Metschnikowia | 13 | Saprotroph | \|Undefined Saprotroph\| | Yeast | NULL | Probable | As nectar-tap_saprotroph (Põlme et al. 2020); Decay sugar-rich substrates (Põlme et al. 2020) | Põlme S, et al. 2020. Fungal Diversity 105:1-16 ((https://doi.org/10.1007/s13225-020-00466-2)) |
| Otu34878 | 1 | k__Fungi(100);p__Ascomycota(100);c__Saccharomycetes(100);o__Saccharomycetales(100);f__Metschnikowiaceae(100);g__Metschnikowia(100);s__Metschnikowia_pulcherrima(100); | Metschnikowia | 13 | Saprotroph | \|Undefined Saprotroph\| | Yeast | NULL | Probable | As nectar-tap_saprotroph (Põlme et al. 2020); Decay sugar-rich substrates (Põlme et al. 2020) | Põlme S, et al. 2020. Fungal Diversity 105:1-16 ((https://doi.org/10.1007/s13225-020-00466-2)) |
| Otu34880 | 1 | k__Fungi(100);p__Ascomycota(100);c__Saccharomycetes(100);o__Saccharomycetales(100);f__Metschnikowiaceae(100);g__Metschnikowia(100);s__Metschnikowia_pulcherrima(100); | Metschnikowia | 13 | Saprotroph | \|Undefined Saprotroph\| | Yeast | NULL | Probable | As nectar-tap_saprotroph (Põlme et al. 2020); Decay sugar-rich substrates (Põlme et al. 2020) | Põlme S, et al. 2020. Fungal Diversity 105:1-16 ((https://doi.org/10.1007/s13225-020-00466-2)) |
| Otu34881 | 1 | k__Fungi(100);p__Ascomycota(100);c__Saccharomycetes(100);o__Saccharomycetales(100);f__Metschnikowiaceae(100);g__Metschnikowia(100);g__Metschnikowia_unclassified(100); | Metschnikowia | 13 | Saprotroph | \|Undefined Saprotroph\| | Yeast | NULL | Probable | As nectar-tap_saprotroph (Põlme et al. 2020); Decay sugar-rich substrates (Põlme et al. 2020) | Põlme S, et al. 2020. Fungal Diversity 105:1-16 ((https://doi.org/10.1007/s13225-020-00466-2)) |
| Otu34887 | 1 | k__Fungi(100);p__Ascomycota(100);c__Saccharomycetes(100);o__Saccharomycetales(100);f__Metschnikowiaceae(100);g__Metschnikowia(100);g__Metschnikowia_unclassified(100); | Metschnikowia | 13 | Saprotroph | \|Undefined Saprotroph\| | Yeast | NULL | Probable | As nectar-tap_saprotroph (Põlme et al. 2020); Decay sugar-rich substrates (Põlme et al. 2020) | Põlme S, et al. 2020. Fungal Diversity 105:1-16 ((https://doi.org/10.1007/s13225-020-00466-2)) |
| Otu34914 | 1 | k__Fungi(100);p__Ascomycota(100);c__Saccharomycetes(100);o__Saccharomycetales(100);f__Metschnikowiaceae(100);g__Metschnikowia(100);s__Metschnikowia_pulcherrima(100); | Metschnikowia | 13 | Saprotroph | \|Undefined Saprotroph\| | Yeast | NULL | Probable | As nectar-tap_saprotroph (Põlme et al. 2020); Decay sugar-rich substrates (Põlme et al. 2020) | Põlme S, et al. 2020. Fungal Diversity 105:1-16 ((https://doi.org/10.1007/s13225-020-00466-2)) |
| Otu34915 | 1 | k__Fungi(100);p__Ascomycota(100);c__Saccharomycetes(100);o__Saccharomycetales(100);f__Metschnikowiaceae(100);g__Metschnikowia(100);s__Metschnikowia_pulcherrima(100); | Metschnikowia | 13 | Saprotroph | \|Undefined Saprotroph\| | Yeast | NULL | Probable | As nectar-tap_saprotroph (Põlme et al. 2020); Decay sugar-rich substrates (Põlme et al. 2020) | Põlme S, et al. 2020. Fungal Diversity 105:1-16 ((https://doi.org/10.1007/s13225-020-00466-2)) |
| Otu34916 | 1 | k__Fungi(100);p__Ascomycota(100);c__Saccharomycetes(100);o__Saccharomycetales(100);f__Metschnikowiaceae(100);g__Metschnikowia(100);s__Metschnikowia_pulcherrima(100); | Metschnikowia | 13 | Saprotroph | \|Undefined Saprotroph\| | Yeast | NULL | Probable | As nectar-tap_saprotroph (Põlme et al. 2020); Decay sugar-rich substrates (Põlme et al. 2020) | Põlme S, et al. 2020. Fungal Diversity 105:1-16 ((https://doi.org/10.1007/s13225-020-00466-2)) |
| Otu34917 | 1 | k__Fungi(100);p__Ascomycota(100);c__Saccharomycetes(100);o__Saccharomycetales(100);f__Metschnikowiaceae(100);g__Metschnikowia(100);g__Metschnikowia_unclassified(100); | Metschnikowia | 13 | Saprotroph | \|Undefined Saprotroph\| | Yeast | NULL | Probable | As nectar-tap_saprotroph (Põlme et al. 2020); Decay sugar-rich substrates (Põlme et al. 2020) | Põlme S, et al. 2020. Fungal Diversity 105:1-16 ((https://doi.org/10.1007/s13225-020-00466-2)) |
| Otu34918 | 1 | k__Fungi(100);p__Ascomycota(100);c__Saccharomycetes(100);o__Saccharomycetales(100);f__Metschnikowiaceae(100);g__Metschnikowia(100);s__Metschnikowia_sp(100); | Metschnikowia | 13 | Saprotroph | \|Undefined Saprotroph\| | Yeast | NULL | Probable | As nectar-tap_saprotroph (Põlme et al. 2020); Decay sugar-rich substrates (Põlme et al. 2020) | Põlme S, et al. 2020. Fungal Diversity 105:1-16 ((https://doi.org/10.1007/s13225-020-00466-2)) |
| Otu34919 | 1 | k__Fungi(100);p__Ascomycota(100);c__Saccharomycetes(100);o__Saccharomycetales(100);f__Metschnikowiaceae(100);g__Metschnikowia(100);s__Metschnikowia_pulcherrima(100); | Metschnikowia | 13 | Saprotroph | \|Undefined Saprotroph\| | Yeast | NULL | Probable | As nectar-tap_saprotroph (Põlme et al. 2020); Decay sugar-rich substrates (Põlme et al. 2020) | Põlme S, et al. 2020. Fungal Diversity 105:1-16 ((https://doi.org/10.1007/s13225-020-00466-2)) |
| Otu34920 | 1 | k__Fungi(100);p__Ascomycota(100);c__Saccharomycetes(100);o__Saccharomycetales(100);f__Metschnikowiaceae(100);g__Metschnikowia(100);s__Metschnikowia_pulcherrima(100); | Metschnikowia | 13 | Saprotroph | \|Undefined Saprotroph\| | Yeast | NULL | Probable | As nectar-tap_saprotroph (Põlme et al. 2020); Decay sugar-rich substrates (Põlme et al. 2020) | Põlme S, et al. 2020. Fungal Diversity 105:1-16 ((https://doi.org/10.1007/s13225-020-00466-2)) |
| Otu34921 | 1 | k__Fungi(100);p__Ascomycota(100);c__Saccharomycetes(100);o__Saccharomycetales(100);f__Metschnikowiaceae(100);g__Metschnikowia(100);s__Metschnikowia_pulcherrima(100); | Metschnikowia | 13 | Saprotroph | \|Undefined Saprotroph\| | Yeast | NULL | Probable | As nectar-tap_saprotroph (Põlme et al. 2020); Decay sugar-rich substrates (Põlme et al. 2020) | Põlme S, et al. 2020. Fungal Diversity 105:1-16 ((https://doi.org/10.1007/s13225-020-00466-2)) |
| Otu34922 | 1 | k__Fungi(100);p__Ascomycota(100);c__Saccharomycetes(100);o__Saccharomycetales(100);f__Metschnikowiaceae(100);g__Metschnikowia(100);s__Metschnikowia_pulcherrima(100); | Metschnikowia | 13 | Saprotroph | \|Undefined Saprotroph\| | Yeast | NULL | Probable | As nectar-tap_saprotroph (Põlme et al. 2020); Decay sugar-rich substrates (Põlme et al. 2020) | Põlme S, et al. 2020. Fungal Diversity 105:1-16 ((https://doi.org/10.1007/s13225-020-00466-2)) |
| Otu34923 | 1 | k__Fungi(100);p__Ascomycota(100);c__Saccharomycetes(100);o__Saccharomycetales(100);f__Metschnikowiaceae(100);g__Metschnikowia(100);s__Metschnikowia_pulcherrima(100); | Metschnikowia | 13 | Saprotroph | \|Undefined Saprotroph\| | Yeast | NULL | Probable | As nectar-tap_saprotroph (Põlme et al. 2020); Decay sugar-rich substrates (Põlme et al. 2020) | Põlme S, et al. 2020. Fungal Diversity 105:1-16 ((https://doi.org/10.1007/s13225-020-00466-2)) |
| Otu34924 | 1 | k__Fungi(100);p__Ascomycota(100);c__Saccharomycetes(100);o__Saccharomycetales(100);f__Metschnikowiaceae(100);g__Metschnikowia(100);g__Metschnikowia_unclassified(100); | Metschnikowia | 13 | Saprotroph | \|Undefined Saprotroph\| | Yeast | NULL | Probable | As nectar-tap_saprotroph (Põlme et al. 2020); Decay sugar-rich substrates (Põlme et al. 2020) | Põlme S, et al. 2020. Fungal Diversity 105:1-16 ((https://doi.org/10.1007/s13225-020-00466-2)) |
| Otu34925 | 1 | k__Fungi(100);p__Ascomycota(100);c__Saccharomycetes(100);o__Saccharomycetales(100);f__Metschnikowiaceae(100);g__Metschnikowia(100);s__Metschnikowia_pulcherrima(100); | Metschnikowia | 13 | Saprotroph | \|Undefined Saprotroph\| | Yeast | NULL | Probable | As nectar-tap_saprotroph (Põlme et al. 2020); Decay sugar-rich substrates (Põlme et al. 2020) | Põlme S, et al. 2020. Fungal Diversity 105:1-16 ((https://doi.org/10.1007/s13225-020-00466-2)) |
| Otu34926 | 1 | k__Fungi(100);p__Ascomycota(100);c__Saccharomycetes(100);o__Saccharomycetales(100);f__Metschnikowiaceae(100);g__Metschnikowia(100);s__Metschnikowia_pulcherrima(100); | Metschnikowia | 13 | Saprotroph | \|Undefined Saprotroph\| | Yeast | NULL | Probable | As nectar-tap_saprotroph (Põlme et al. 2020); Decay sugar-rich substrates (Põlme et al. 2020) | Põlme S, et al. 2020. Fungal Diversity 105:1-16 ((https://doi.org/10.1007/s13225-020-00466-2)) |
| Otu34927 | 1 | k__Fungi(100);p__Ascomycota(100);c__Saccharomycetes(100);o__Saccharomycetales(100);f__Metschnikowiaceae(100);g__Metschnikowia(100);s__Metschnikowia_pulcherrima(100); | Metschnikowia | 13 | Saprotroph | \|Undefined Saprotroph\| | Yeast | NULL | Probable | As nectar-tap_saprotroph (Põlme et al. 2020); Decay sugar-rich substrates (Põlme et al. 2020) | Põlme S, et al. 2020. Fungal Diversity 105:1-16 ((https://doi.org/10.1007/s13225-020-00466-2)) |
| Otu34928 | 1 | k__Fungi(100);p__Ascomycota(100);c__Saccharomycetes(100);o__Saccharomycetales(100);f__Metschnikowiaceae(100);g__Metschnikowia(100);s__Metschnikowia_pulcherrima(100); | Metschnikowia | 13 | Saprotroph | \|Undefined Saprotroph\| | Yeast | NULL | Probable | As nectar-tap_saprotroph (Põlme et al. 2020); Decay sugar-rich substrates (Põlme et al. 2020) | Põlme S, et al. 2020. Fungal Diversity 105:1-16 ((https://doi.org/10.1007/s13225-020-00466-2)) |
| Otu34929 | 1 | k__Fungi(100);p__Ascomycota(100);c__Saccharomycetes(100);o__Saccharomycetales(100);f__Metschnikowiaceae(100);g__Metschnikowia(100);s__Metschnikowia_pulcherrima(100); | Metschnikowia | 13 | Saprotroph | \|Undefined Saprotroph\| | Yeast | NULL | Probable | As nectar-tap_saprotroph (Põlme et al. 2020); Decay sugar-rich substrates (Põlme et al. 2020) | Põlme S, et al. 2020. Fungal Diversity 105:1-16 ((https://doi.org/10.1007/s13225-020-00466-2)) |
| Otu34930 | 1 | k__Fungi(100);p__Ascomycota(100);c__Saccharomycetes(100);o__Saccharomycetales(100);f__Metschnikowiaceae(100);g__Metschnikowia(100);s__Metschnikowia_pulcherrima(100); | Metschnikowia | 13 | Saprotroph | \|Undefined Saprotroph\| | Yeast | NULL | Probable | As nectar-tap_saprotroph (Põlme et al. 2020); Decay sugar-rich substrates (Põlme et al. 2020) | Põlme S, et al. 2020. Fungal Diversity 105:1-16 ((https://doi.org/10.1007/s13225-020-00466-2)) |
| Otu34931 | 1 | k__Fungi(100);p__Ascomycota(100);c__Saccharomycetes(100);o__Saccharomycetales(100);f__Metschnikowiaceae(100);g__Metschnikowia(100);s__Metschnikowia_pulcherrima(100); | Metschnikowia | 13 | Saprotroph | \|Undefined Saprotroph\| | Yeast | NULL | Probable | As nectar-tap_saprotroph (Põlme et al. 2020); Decay sugar-rich substrates (Põlme et al. 2020) | Põlme S, et al. 2020. Fungal Diversity 105:1-16 ((https://doi.org/10.1007/s13225-020-00466-2)) |
| Otu34932 | 1 | k__Fungi(100);p__Ascomycota(100);c__Saccharomycetes(100);o__Saccharomycetales(100);f__Metschnikowiaceae(100);g__Metschnikowia(100);g__Metschnikowia_unclassified(100); | Metschnikowia | 13 | Saprotroph | \|Undefined Saprotroph\| | Yeast | NULL | Probable | As nectar-tap_saprotroph (Põlme et al. 2020); Decay sugar-rich substrates (Põlme et al. 2020) | Põlme S, et al. 2020. Fungal Diversity 105:1-16 ((https://doi.org/10.1007/s13225-020-00466-2)) |
| Otu34933 | 1 | k__Fungi(100);p__Ascomycota(100);c__Saccharomycetes(100);o__Saccharomycetales(100);f__Metschnikowiaceae(100);g__Metschnikowia(100);g__Metschnikowia_unclassified(100); | Metschnikowia | 13 | Saprotroph | \|Undefined Saprotroph\| | Yeast | NULL | Probable | As nectar-tap_saprotroph (Põlme et al. 2020); Decay sugar-rich substrates (Põlme et al. 2020) | Põlme S, et al. 2020. Fungal Diversity 105:1-16 ((https://doi.org/10.1007/s13225-020-00466-2)) |
| Otu34934 | 1 | k__Fungi(100);p__Ascomycota(100);c__Saccharomycetes(100);o__Saccharomycetales(100);f__Metschnikowiaceae(100);g__Metschnikowia(100);s__Metschnikowia_pulcherrima(100); | Metschnikowia | 13 | Saprotroph | \|Undefined Saprotroph\| | Yeast | NULL | Probable | As nectar-tap_saprotroph (Põlme et al. 2020); Decay sugar-rich substrates (Põlme et al. 2020) | Põlme S, et al. 2020. Fungal Diversity 105:1-16 ((https://doi.org/10.1007/s13225-020-00466-2)) |
| Otu34935 | 1 | k__Fungi(100);p__Ascomycota(100);c__Saccharomycetes(100);o__Saccharomycetales(100);f__Metschnikowiaceae(100);g__Metschnikowia(100);s__Metschnikowia_pulcherrima(100); | Metschnikowia | 13 | Saprotroph | \|Undefined Saprotroph\| | Yeast | NULL | Probable | As nectar-tap_saprotroph (Põlme et al. 2020); Decay sugar-rich substrates (Põlme et al. 2020) | Põlme S, et al. 2020. Fungal Diversity 105:1-16 ((https://doi.org/10.1007/s13225-020-00466-2)) |
| Otu34936 | 1 | k__Fungi(100);p__Ascomycota(100);c__Saccharomycetes(100);o__Saccharomycetales(100);f__Metschnikowiaceae(100);g__Metschnikowia(100);g__Metschnikowia_unclassified(100); | Metschnikowia | 13 | Saprotroph | \|Undefined Saprotroph\| | Yeast | NULL | Probable | As nectar-tap_saprotroph (Põlme et al. 2020); Decay sugar-rich substrates (Põlme et al. 2020) | Põlme S, et al. 2020. Fungal Diversity 105:1-16 ((https://doi.org/10.1007/s13225-020-00466-2)) |
| Otu34937 | 1 | k__Fungi(100);p__Ascomycota(100);c__Saccharomycetes(100);o__Saccharomycetales(100);f__Metschnikowiaceae(100);g__Metschnikowia(100);s__Metschnikowia_pulcherrima(100); | Metschnikowia | 13 | Saprotroph | \|Undefined Saprotroph\| | Yeast | NULL | Probable | As nectar-tap_saprotroph (Põlme et al. 2020); Decay sugar-rich substrates (Põlme et al. 2020) | Põlme S, et al. 2020. Fungal Diversity 105:1-16 ((https://doi.org/10.1007/s13225-020-00466-2)) |
| Otu34938 | 1 | k__Fungi(100);p__Ascomycota(100);c__Saccharomycetes(100);o__Saccharomycetales(100);f__Metschnikowiaceae(100);g__Metschnikowia(100);s__Metschnikowia_pulcherrima(100); | Metschnikowia | 13 | Saprotroph | \|Undefined Saprotroph\| | Yeast | NULL | Probable | As nectar-tap_saprotroph (Põlme et al. 2020); Decay sugar-rich substrates (Põlme et al. 2020) | Põlme S, et al. 2020. Fungal Diversity 105:1-16 ((https://doi.org/10.1007/s13225-020-00466-2)) |
| Otu34939 | 1 | k__Fungi(100);p__Ascomycota(100);c__Saccharomycetes(100);o__Saccharomycetales(100);f__Metschnikowiaceae(100);g__Metschnikowia(100);g__Metschnikowia_unclassified(100); | Metschnikowia | 13 | Saprotroph | \|Undefined Saprotroph\| | Yeast | NULL | Probable | As nectar-tap_saprotroph (Põlme et al. 2020); Decay sugar-rich substrates (Põlme et al. 2020) | Põlme S, et al. 2020. Fungal Diversity 105:1-16 ((https://doi.org/10.1007/s13225-020-00466-2)) |
| Otu34940 | 1 | k__Fungi(100);p__Ascomycota(100);c__Saccharomycetes(100);o__Saccharomycetales(100);f__Metschnikowiaceae(100);g__Metschnikowia(100);g__Metschnikowia_unclassified(100); | Metschnikowia | 13 | Saprotroph | \|Undefined Saprotroph\| | Yeast | NULL | Probable | As nectar-tap_saprotroph (Põlme et al. 2020); Decay sugar-rich substrates (Põlme et al. 2020) | Põlme S, et al. 2020. Fungal Diversity 105:1-16 ((https://doi.org/10.1007/s13225-020-00466-2)) |
| Otu34941 | 1 | k__Fungi(100);p__Ascomycota(100);c__Saccharomycetes(100);o__Saccharomycetales(100);f__Metschnikowiaceae(100);g__Metschnikowia(100);s__Metschnikowia_pulcherrima(100); | Metschnikowia | 13 | Saprotroph | \|Undefined Saprotroph\| | Yeast | NULL | Probable | As nectar-tap_saprotroph (Põlme et al. 2020); Decay sugar-rich substrates (Põlme et al. 2020) | Põlme S, et al. 2020. Fungal Diversity 105:1-16 ((https://doi.org/10.1007/s13225-020-00466-2)) |
| Otu34942 | 1 | k__Fungi(100);p__Ascomycota(100);c__Saccharomycetes(100);o__Saccharomycetales(100);f__Metschnikowiaceae(100);g__Metschnikowia(100);g__Metschnikowia_unclassified(100); | Metschnikowia | 13 | Saprotroph | \|Undefined Saprotroph\| | Yeast | NULL | Probable | As nectar-tap_saprotroph (Põlme et al. 2020); Decay sugar-rich substrates (Põlme et al. 2020) | Põlme S, et al. 2020. Fungal Diversity 105:1-16 ((https://doi.org/10.1007/s13225-020-00466-2)) |
| Otu34943 | 1 | k__Fungi(100);p__Ascomycota(100);c__Saccharomycetes(100);o__Saccharomycetales(100);f__Metschnikowiaceae(100);g__Metschnikowia(100);s__Metschnikowia_pulcherrima(100); | Metschnikowia | 13 | Saprotroph | \|Undefined Saprotroph\| | Yeast | NULL | Probable | As nectar-tap_saprotroph (Põlme et al. 2020); Decay sugar-rich substrates (Põlme et al. 2020) | Põlme S, et al. 2020. Fungal Diversity 105:1-16 ((https://doi.org/10.1007/s13225-020-00466-2)) |
| Otu34944 | 1 | k__Fungi(100);p__Ascomycota(100);c__Saccharomycetes(100);o__Saccharomycetales(100);f__Metschnikowiaceae(100);g__Metschnikowia(100);s__Metschnikowia_pulcherrima(100); | Metschnikowia | 13 | Saprotroph | \|Undefined Saprotroph\| | Yeast | NULL | Probable | As nectar-tap_saprotroph (Põlme et al. 2020); Decay sugar-rich substrates (Põlme et al. 2020) | Põlme S, et al. 2020. Fungal Diversity 105:1-16 ((https://doi.org/10.1007/s13225-020-00466-2)) |
| Otu34945 | 1 | k__Fungi(100);p__Ascomycota(100);c__Saccharomycetes(100);o__Saccharomycetales(100);f__Metschnikowiaceae(100);g__Metschnikowia(100);g__Metschnikowia_unclassified(100); | Metschnikowia | 13 | Saprotroph | \|Undefined Saprotroph\| | Yeast | NULL | Probable | As nectar-tap_saprotroph (Põlme et al. 2020); Decay sugar-rich substrates (Põlme et al. 2020) | Põlme S, et al. 2020. Fungal Diversity 105:1-16 ((https://doi.org/10.1007/s13225-020-00466-2)) |
| Otu34946 | 1 | k__Fungi(100);p__Ascomycota(100);c__Saccharomycetes(100);o__Saccharomycetales(100);f__Metschnikowiaceae(100);g__Metschnikowia(100);g__Metschnikowia_unclassified(100); | Metschnikowia | 13 | Saprotroph | \|Undefined Saprotroph\| | Yeast | NULL | Probable | As nectar-tap_saprotroph (Põlme et al. 2020); Decay sugar-rich substrates (Põlme et al. 2020) | Põlme S, et al. 2020. Fungal Diversity 105:1-16 ((https://doi.org/10.1007/s13225-020-00466-2)) |
| Otu34947 | 1 | k__Fungi(100);p__Ascomycota(100);c__Saccharomycetes(100);o__Saccharomycetales(100);f__Metschnikowiaceae(100);g__Metschnikowia(100);s__Metschnikowia_pulcherrima(100); | Metschnikowia | 13 | Saprotroph | \|Undefined Saprotroph\| | Yeast | NULL | Probable | As nectar-tap_saprotroph (Põlme et al. 2020); Decay sugar-rich substrates (Põlme et al. 2020) | Põlme S, et al. 2020. Fungal Diversity 105:1-16 ((https://doi.org/10.1007/s13225-020-00466-2)) |
| Otu34948 | 1 | k__Fungi(100);p__Ascomycota(100);c__Saccharomycetes(100);o__Saccharomycetales(100);f__Metschnikowiaceae(100);g__Metschnikowia(100);s__Metschnikowia_pulcherrima(100); | Metschnikowia | 13 | Saprotroph | \|Undefined Saprotroph\| | Yeast | NULL | Probable | As nectar-tap_saprotroph (Põlme et al. 2020); Decay sugar-rich substrates (Põlme et al. 2020) | Põlme S, et al. 2020. Fungal Diversity 105:1-16 ((https://doi.org/10.1007/s13225-020-00466-2)) |
| Otu34949 | 1 | k__Fungi(100);p__Ascomycota(100);c__Saccharomycetes(100);o__Saccharomycetales(100);f__Metschnikowiaceae(100);g__Metschnikowia(100);s__Metschnikowia_pulcherrima(100); | Metschnikowia | 13 | Saprotroph | \|Undefined Saprotroph\| | Yeast | NULL | Probable | As nectar-tap_saprotroph (Põlme et al. 2020); Decay sugar-rich substrates (Põlme et al. 2020) | Põlme S, et al. 2020. Fungal Diversity 105:1-16 ((https://doi.org/10.1007/s13225-020-00466-2)) |
| Otu34950 | 1 | k__Fungi(100);p__Ascomycota(100);c__Saccharomycetes(100);o__Saccharomycetales(100);f__Metschnikowiaceae(100);g__Metschnikowia(100);g__Metschnikowia_unclassified(100); | Metschnikowia | 13 | Saprotroph | \|Undefined Saprotroph\| | Yeast | NULL | Probable | As nectar-tap_saprotroph (Põlme et al. 2020); Decay sugar-rich substrates (Põlme et al. 2020) | Põlme S, et al. 2020. Fungal Diversity 105:1-16 ((https://doi.org/10.1007/s13225-020-00466-2)) |
| Otu34951 | 1 | k__Fungi(100);p__Ascomycota(100);c__Saccharomycetes(100);o__Saccharomycetales(100);f__Metschnikowiaceae(100);g__Metschnikowia(100);s__Metschnikowia_pulcherrima(100); | Metschnikowia | 13 | Saprotroph | \|Undefined Saprotroph\| | Yeast | NULL | Probable | As nectar-tap_saprotroph (Põlme et al. 2020); Decay sugar-rich substrates (Põlme et al. 2020) | Põlme S, et al. 2020. Fungal Diversity 105:1-16 ((https://doi.org/10.1007/s13225-020-00466-2)) |
| Otu34952 | 1 | k__Fungi(100);p__Ascomycota(100);c__Saccharomycetes(100);o__Saccharomycetales(100);f__Metschnikowiaceae(100);g__Metschnikowia(100);g__Metschnikowia_unclassified(100); | Metschnikowia | 13 | Saprotroph | \|Undefined Saprotroph\| | Yeast | NULL | Probable | As nectar-tap_saprotroph (Põlme et al. 2020); Decay sugar-rich substrates (Põlme et al. 2020) | Põlme S, et al. 2020. Fungal Diversity 105:1-16 ((https://doi.org/10.1007/s13225-020-00466-2)) |
| Otu34953 | 1 | k__Fungi(100);p__Ascomycota(100);c__Saccharomycetes(100);o__Saccharomycetales(100);f__Metschnikowiaceae(100);g__Metschnikowia(100);s__Metschnikowia_pulcherrima(100); | Metschnikowia | 13 | Saprotroph | \|Undefined Saprotroph\| | Yeast | NULL | Probable | As nectar-tap_saprotroph (Põlme et al. 2020); Decay sugar-rich substrates (Põlme et al. 2020) | Põlme S, et al. 2020. Fungal Diversity 105:1-16 ((https://doi.org/10.1007/s13225-020-00466-2)) |
| Otu34954 | 1 | k__Fungi(100);p__Ascomycota(100);c__Saccharomycetes(100);o__Saccharomycetales(100);f__Metschnikowiaceae(100);g__Metschnikowia(100);s__Metschnikowia_pulcherrima(100); | Metschnikowia | 13 | Saprotroph | \|Undefined Saprotroph\| | Yeast | NULL | Probable | As nectar-tap_saprotroph (Põlme et al. 2020); Decay sugar-rich substrates (Põlme et al. 2020) | Põlme S, et al. 2020. Fungal Diversity 105:1-16 ((https://doi.org/10.1007/s13225-020-00466-2)) |
| Otu34955 | 1 | k__Fungi(100);p__Ascomycota(100);c__Saccharomycetes(100);o__Saccharomycetales(100);f__Metschnikowiaceae(100);g__Metschnikowia(100);s__Metschnikowia_pulcherrima(100); | Metschnikowia | 13 | Saprotroph | \|Undefined Saprotroph\| | Yeast | NULL | Probable | As nectar-tap_saprotroph (Põlme et al. 2020); Decay sugar-rich substrates (Põlme et al. 2020) | Põlme S, et al. 2020. Fungal Diversity 105:1-16 ((https://doi.org/10.1007/s13225-020-00466-2)) |
| Otu34956 | 1 | k__Fungi(100);p__Ascomycota(100);c__Saccharomycetes(100);o__Saccharomycetales(100);f__Metschnikowiaceae(100);g__Metschnikowia(100);s__Metschnikowia_pulcherrima(100); | Metschnikowia | 13 | Saprotroph | \|Undefined Saprotroph\| | Yeast | NULL | Probable | As nectar-tap_saprotroph (Põlme et al. 2020); Decay sugar-rich substrates (Põlme et al. 2020) | Põlme S, et al. 2020. Fungal Diversity 105:1-16 ((https://doi.org/10.1007/s13225-020-00466-2)) |
| Otu34957 | 1 | k__Fungi(100);p__Ascomycota(100);c__Saccharomycetes(100);o__Saccharomycetales(100);f__Metschnikowiaceae(100);g__Metschnikowia(100);s__Metschnikowia_pulcherrima(100); | Metschnikowia | 13 | Saprotroph | \|Undefined Saprotroph\| | Yeast | NULL | Probable | As nectar-tap_saprotroph (Põlme et al. 2020); Decay sugar-rich substrates (Põlme et al. 2020) | Põlme S, et al. 2020. Fungal Diversity 105:1-16 ((https://doi.org/10.1007/s13225-020-00466-2)) |
| Otu34958 | 1 | k__Fungi(100);p__Ascomycota(100);c__Saccharomycetes(100);o__Saccharomycetales(100);f__Metschnikowiaceae(100);g__Metschnikowia(100);g__Metschnikowia_unclassified(100); | Metschnikowia | 13 | Saprotroph | \|Undefined Saprotroph\| | Yeast | NULL | Probable | As nectar-tap_saprotroph (Põlme et al. 2020); Decay sugar-rich substrates (Põlme et al. 2020) | Põlme S, et al. 2020. Fungal Diversity 105:1-16 ((https://doi.org/10.1007/s13225-020-00466-2)) |
| Otu34959 | 1 | k__Fungi(100);p__Ascomycota(100);c__Saccharomycetes(100);o__Saccharomycetales(100);f__Metschnikowiaceae(100);g__Metschnikowia(100);s__Metschnikowia_pulcherrima(100); | Metschnikowia | 13 | Saprotroph | \|Undefined Saprotroph\| | Yeast | NULL | Probable | As nectar-tap_saprotroph (Põlme et al. 2020); Decay sugar-rich substrates (Põlme et al. 2020) | Põlme S, et al. 2020. Fungal Diversity 105:1-16 ((https://doi.org/10.1007/s13225-020-00466-2)) |
| Otu34961 | 1 | k__Fungi(100);p__Ascomycota(100);c__Saccharomycetes(100);o__Saccharomycetales(100);f__Metschnikowiaceae(100);g__Metschnikowia(100);s__Metschnikowia_pulcherrima(100); | Metschnikowia | 13 | Saprotroph | \|Undefined Saprotroph\| | Yeast | NULL | Probable | As nectar-tap_saprotroph (Põlme et al. 2020); Decay sugar-rich substrates (Põlme et al. 2020) | Põlme S, et al. 2020. Fungal Diversity 105:1-16 ((https://doi.org/10.1007/s13225-020-00466-2)) |
| Otu34962 | 1 | k__Fungi(100);p__Ascomycota(100);c__Saccharomycetes(100);o__Saccharomycetales(100);f__Metschnikowiaceae(100);g__Metschnikowia(100);s__Metschnikowia_pulcherrima(100); | Metschnikowia | 13 | Saprotroph | \|Undefined Saprotroph\| | Yeast | NULL | Probable | As nectar-tap_saprotroph (Põlme et al. 2020); Decay sugar-rich substrates (Põlme et al. 2020) | Põlme S, et al. 2020. Fungal Diversity 105:1-16 ((https://doi.org/10.1007/s13225-020-00466-2)) |
| Otu34963 | 1 | k__Fungi(100);p__Ascomycota(100);c__Saccharomycetes(100);o__Saccharomycetales(100);f__Metschnikowiaceae(100);g__Metschnikowia(100);s__Metschnikowia_pulcherrima(100); | Metschnikowia | 13 | Saprotroph | \|Undefined Saprotroph\| | Yeast | NULL | Probable | As nectar-tap_saprotroph (Põlme et al. 2020); Decay sugar-rich substrates (Põlme et al. 2020) | Põlme S, et al. 2020. Fungal Diversity 105:1-16 ((https://doi.org/10.1007/s13225-020-00466-2)) |
| Otu34964 | 1 | k__Fungi(100);p__Ascomycota(100);c__Saccharomycetes(100);o__Saccharomycetales(100);f__Metschnikowiaceae(100);g__Metschnikowia(100);s__Metschnikowia_pulcherrima(100); | Metschnikowia | 13 | Saprotroph | \|Undefined Saprotroph\| | Yeast | NULL | Probable | As nectar-tap_saprotroph (Põlme et al. 2020); Decay sugar-rich substrates (Põlme et al. 2020) | Põlme S, et al. 2020. Fungal Diversity 105:1-16 ((https://doi.org/10.1007/s13225-020-00466-2)) |
| Otu34965 | 1 | k__Fungi(100);p__Ascomycota(100);c__Saccharomycetes(100);o__Saccharomycetales(100);f__Metschnikowiaceae(100);g__Metschnikowia(100);g__Metschnikowia_unclassified(100); | Metschnikowia | 13 | Saprotroph | \|Undefined Saprotroph\| | Yeast | NULL | Probable | As nectar-tap_saprotroph (Põlme et al. 2020); Decay sugar-rich substrates (Põlme et al. 2020) | Põlme S, et al. 2020. Fungal Diversity 105:1-16 ((https://doi.org/10.1007/s13225-020-00466-2)) |
| Otu34966 | 1 | k__Fungi(100);p__Ascomycota(100);c__Saccharomycetes(100);o__Saccharomycetales(100);f__Metschnikowiaceae(100);g__Metschnikowia(100);g__Metschnikowia_unclassified(100); | Metschnikowia | 13 | Saprotroph | \|Undefined Saprotroph\| | Yeast | NULL | Probable | As nectar-tap_saprotroph (Põlme et al. 2020); Decay sugar-rich substrates (Põlme et al. 2020) | Põlme S, et al. 2020. Fungal Diversity 105:1-16 ((https://doi.org/10.1007/s13225-020-00466-2)) |
| Otu34967 | 1 | k__Fungi(100);p__Ascomycota(100);c__Saccharomycetes(100);o__Saccharomycetales(100);f__Metschnikowiaceae(100);g__Metschnikowia(100);g__Metschnikowia_unclassified(100); | Metschnikowia | 13 | Saprotroph | \|Undefined Saprotroph\| | Yeast | NULL | Probable | As nectar-tap_saprotroph (Põlme et al. 2020); Decay sugar-rich substrates (Põlme et al. 2020) | Põlme S, et al. 2020. Fungal Diversity 105:1-16 ((https://doi.org/10.1007/s13225-020-00466-2)) |
| Otu34973 | 1 | k__Fungi(100);p__Ascomycota(100);c__Saccharomycetes(100);o__Saccharomycetales(100);f__Metschnikowiaceae(100);g__Metschnikowia(100);s__Metschnikowia_pulcherrima(100); | Metschnikowia | 13 | Saprotroph | \|Undefined Saprotroph\| | Yeast | NULL | Probable | As nectar-tap_saprotroph (Põlme et al. 2020); Decay sugar-rich substrates (Põlme et al. 2020) | Põlme S, et al. 2020. Fungal Diversity 105:1-16 ((https://doi.org/10.1007/s13225-020-00466-2)) |
| Otu34974 | 1 | k__Fungi(100);p__Ascomycota(100);c__Saccharomycetes(100);o__Saccharomycetales(100);f__Metschnikowiaceae(100);g__Metschnikowia(100);s__Metschnikowia_pulcherrima(100); | Metschnikowia | 13 | Saprotroph | \|Undefined Saprotroph\| | Yeast | NULL | Probable | As nectar-tap_saprotroph (Põlme et al. 2020); Decay sugar-rich substrates (Põlme et al. 2020) | Põlme S, et al. 2020. Fungal Diversity 105:1-16 ((https://doi.org/10.1007/s13225-020-00466-2)) |
| Otu34975 | 1 | k__Fungi(100);p__Ascomycota(100);c__Saccharomycetes(100);o__Saccharomycetales(100);f__Metschnikowiaceae(100);g__Metschnikowia(100);s__Metschnikowia_pulcherrima(100); | Metschnikowia | 13 | Saprotroph | \|Undefined Saprotroph\| | Yeast | NULL | Probable | As nectar-tap_saprotroph (Põlme et al. 2020); Decay sugar-rich substrates (Põlme et al. 2020) | Põlme S, et al. 2020. Fungal Diversity 105:1-16 ((https://doi.org/10.1007/s13225-020-00466-2)) |
| Otu34976 | 1 | k__Fungi(100);p__Ascomycota(100);c__Dothideomycetes(100);o__Pleosporales(100);f__Lophiostomataceae(100);g__Lophiostoma(100);g__Lophiostoma_unclassified(100); | Lophiostoma | 13 | Saprotroph | Plant Saprotroph-Undefined Saprotroph-\|Wood Saprotroph\| | Microfungus | NULL | Probable | As litter_saprotroph (Põlme et al. 2020); Decay wood, leaf, fruit, seed (Põlme et al. 2020) | Tedersoo L, et al. 2014. Science 346:e1256688 ((https://doi.org/10.1126/science.1256688)); Põlme S, et al. 2020. Fungal Diversity 105:1-16 ((https://doi.org/10.1007/s13225-020-00466-2)) |
| Otu34978 | 1 | k__Fungi(100);p__Ascomycota(100);c__Saccharomycetes(100);o__Saccharomycetales(100);f__Metschnikowiaceae(100);g__Metschnikowia(100);s__Metschnikowia_pulcherrima(100); | Metschnikowia | 13 | Saprotroph | \|Undefined Saprotroph\| | Yeast | NULL | Probable | As nectar-tap_saprotroph (Põlme et al. 2020); Decay sugar-rich substrates (Põlme et al. 2020) | Põlme S, et al. 2020. Fungal Diversity 105:1-16 ((https://doi.org/10.1007/s13225-020-00466-2)) |
| Otu34979 | 1 | k__Fungi(100);p__Ascomycota(100);c__Saccharomycetes(100);o__Saccharomycetales(100);f__Metschnikowiaceae(100);g__Metschnikowia(100);s__Metschnikowia_pulcherrima(100); | Metschnikowia | 13 | Saprotroph | \|Undefined Saprotroph\| | Yeast | NULL | Probable | As nectar-tap_saprotroph (Põlme et al. 2020); Decay sugar-rich substrates (Põlme et al. 2020) | Põlme S, et al. 2020. Fungal Diversity 105:1-16 ((https://doi.org/10.1007/s13225-020-00466-2)) |
| Otu34981 | 1 | k__Fungi(100);p__Ascomycota(100);c__Saccharomycetes(100);o__Saccharomycetales(100);f__Metschnikowiaceae(100);g__Metschnikowia(100);s__Metschnikowia_pulcherrima(100); | Metschnikowia | 13 | Saprotroph | \|Undefined Saprotroph\| | Yeast | NULL | Probable | As nectar-tap_saprotroph (Põlme et al. 2020); Decay sugar-rich substrates (Põlme et al. 2020) | Põlme S, et al. 2020. Fungal Diversity 105:1-16 ((https://doi.org/10.1007/s13225-020-00466-2)) |
| Otu34984 | 1 | k__Fungi(100);p__Ascomycota(100);c__Saccharomycetes(100);o__Saccharomycetales(100);f__Metschnikowiaceae(100);g__Metschnikowia(100);s__Metschnikowia_pulcherrima(100); | Metschnikowia | 13 | Saprotroph | \|Undefined Saprotroph\| | Yeast | NULL | Probable | As nectar-tap_saprotroph (Põlme et al. 2020); Decay sugar-rich substrates (Põlme et al. 2020) | Põlme S, et al. 2020. Fungal Diversity 105:1-16 ((https://doi.org/10.1007/s13225-020-00466-2)) |
| Otu34985 | 1 | k__Fungi(100);p__Ascomycota(100);c__Saccharomycetes(100);o__Saccharomycetales(100);f__Metschnikowiaceae(100);g__Metschnikowia(100);g__Metschnikowia_unclassified(100); | Metschnikowia | 13 | Saprotroph | \|Undefined Saprotroph\| | Yeast | NULL | Probable | As nectar-tap_saprotroph (Põlme et al. 2020); Decay sugar-rich substrates (Põlme et al. 2020) | Põlme S, et al. 2020. Fungal Diversity 105:1-16 ((https://doi.org/10.1007/s13225-020-00466-2)) |
| Otu34986 | 1 | k__Fungi(100);p__Ascomycota(100);c__Saccharomycetes(100);o__Saccharomycetales(100);f__Metschnikowiaceae(100);g__Metschnikowia(100);g__Metschnikowia_unclassified(100); | Metschnikowia | 13 | Saprotroph | \|Undefined Saprotroph\| | Yeast | NULL | Probable | As nectar-tap_saprotroph (Põlme et al. 2020); Decay sugar-rich substrates (Põlme et al. 2020) | Põlme S, et al. 2020. Fungal Diversity 105:1-16 ((https://doi.org/10.1007/s13225-020-00466-2)) |
| Otu34988 | 1 | k__Fungi(100);p__Ascomycota(100);c__Saccharomycetes(100);o__Saccharomycetales(100);f__Metschnikowiaceae(100);g__Metschnikowia(100);s__Metschnikowia_pulcherrima(100); | Metschnikowia | 13 | Saprotroph | \|Undefined Saprotroph\| | Yeast | NULL | Probable | As nectar-tap_saprotroph (Põlme et al. 2020); Decay sugar-rich substrates (Põlme et al. 2020) | Põlme S, et al. 2020. Fungal Diversity 105:1-16 ((https://doi.org/10.1007/s13225-020-00466-2)) |
| Otu34989 | 1 | k__Fungi(100);p__Ascomycota(100);c__Saccharomycetes(100);o__Saccharomycetales(100);f__Metschnikowiaceae(100);g__Metschnikowia(100);s__Metschnikowia_pulcherrima(100); | Metschnikowia | 13 | Saprotroph | \|Undefined Saprotroph\| | Yeast | NULL | Probable | As nectar-tap_saprotroph (Põlme et al. 2020); Decay sugar-rich substrates (Põlme et al. 2020) | Põlme S, et al. 2020. Fungal Diversity 105:1-16 ((https://doi.org/10.1007/s13225-020-00466-2)) |
| Otu34990 | 1 | k__Fungi(100);p__Ascomycota(100);c__Saccharomycetes(100);o__Saccharomycetales(100);f__Metschnikowiaceae(100);g__Metschnikowia(100);s__Metschnikowia_pulcherrima(100); | Metschnikowia | 13 | Saprotroph | \|Undefined Saprotroph\| | Yeast | NULL | Probable | As nectar-tap_saprotroph (Põlme et al. 2020); Decay sugar-rich substrates (Põlme et al. 2020) | Põlme S, et al. 2020. Fungal Diversity 105:1-16 ((https://doi.org/10.1007/s13225-020-00466-2)) |
| Otu34991 | 1 | k__Fungi(100);p__Ascomycota(100);c__Saccharomycetes(100);o__Saccharomycetales(100);f__Metschnikowiaceae(100);g__Metschnikowia(100);s__Metschnikowia_pulcherrima(100); | Metschnikowia | 13 | Saprotroph | \|Undefined Saprotroph\| | Yeast | NULL | Probable | As nectar-tap_saprotroph (Põlme et al. 2020); Decay sugar-rich substrates (Põlme et al. 2020) | Põlme S, et al. 2020. Fungal Diversity 105:1-16 ((https://doi.org/10.1007/s13225-020-00466-2)) |
| Otu34993 | 1 | k__Fungi(100);p__Ascomycota(100);c__Saccharomycetes(100);o__Saccharomycetales(100);f__Metschnikowiaceae(100);g__Metschnikowia(100);g__Metschnikowia_unclassified(100); | Metschnikowia | 13 | Saprotroph | \|Undefined Saprotroph\| | Yeast | NULL | Probable | As nectar-tap_saprotroph (Põlme et al. 2020); Decay sugar-rich substrates (Põlme et al. 2020) | Põlme S, et al. 2020. Fungal Diversity 105:1-16 ((https://doi.org/10.1007/s13225-020-00466-2)) |
| Otu34994 | 1 | k__Fungi(100);p__Ascomycota(100);c__Saccharomycetes(100);o__Saccharomycetales(100);f__Metschnikowiaceae(100);g__Metschnikowia(100);s__Metschnikowia_pulcherrima(100); | Metschnikowia | 13 | Saprotroph | \|Undefined Saprotroph\| | Yeast | NULL | Probable | As nectar-tap_saprotroph (Põlme et al. 2020); Decay sugar-rich substrates (Põlme et al. 2020) | Põlme S, et al. 2020. Fungal Diversity 105:1-16 ((https://doi.org/10.1007/s13225-020-00466-2)) |
| Otu34995 | 1 | k__Fungi(100);p__Ascomycota(100);c__Saccharomycetes(100);o__Saccharomycetales(100);f__Metschnikowiaceae(100);g__Metschnikowia(100);g__Metschnikowia_unclassified(100); | Metschnikowia | 13 | Saprotroph | \|Undefined Saprotroph\| | Yeast | NULL | Probable | As nectar-tap_saprotroph (Põlme et al. 2020); Decay sugar-rich substrates (Põlme et al. 2020) | Põlme S, et al. 2020. Fungal Diversity 105:1-16 ((https://doi.org/10.1007/s13225-020-00466-2)) |
| Otu34996 | 1 | k__Fungi(100);p__Ascomycota(100);c__Saccharomycetes(100);o__Saccharomycetales(100);f__Metschnikowiaceae(100);g__Metschnikowia(100);s__Metschnikowia_pulcherrima(100); | Metschnikowia | 13 | Saprotroph | \|Undefined Saprotroph\| | Yeast | NULL | Probable | As nectar-tap_saprotroph (Põlme et al. 2020); Decay sugar-rich substrates (Põlme et al. 2020) | Põlme S, et al. 2020. Fungal Diversity 105:1-16 ((https://doi.org/10.1007/s13225-020-00466-2)) |
| Otu34997 | 1 | k__Fungi(100);p__Ascomycota(100);c__Saccharomycetes(100);o__Saccharomycetales(100);f__Metschnikowiaceae(100);g__Metschnikowia(100);s__Metschnikowia_pulcherrima(100); | Metschnikowia | 13 | Saprotroph | \|Undefined Saprotroph\| | Yeast | NULL | Probable | As nectar-tap_saprotroph (Põlme et al. 2020); Decay sugar-rich substrates (Põlme et al. 2020) | Põlme S, et al. 2020. Fungal Diversity 105:1-16 ((https://doi.org/10.1007/s13225-020-00466-2)) |
| Otu34998 | 1 | k__Fungi(100);p__Ascomycota(100);c__Saccharomycetes(100);o__Saccharomycetales(100);f__Metschnikowiaceae(100);g__Metschnikowia(100);s__Metschnikowia_pulcherrima(100); | Metschnikowia | 13 | Saprotroph | \|Undefined Saprotroph\| | Yeast | NULL | Probable | As nectar-tap_saprotroph (Põlme et al. 2020); Decay sugar-rich substrates (Põlme et al. 2020) | Põlme S, et al. 2020. Fungal Diversity 105:1-16 ((https://doi.org/10.1007/s13225-020-00466-2)) |
| Otu34999 | 1 | k__Fungi(100);p__Ascomycota(100);c__Saccharomycetes(100);o__Saccharomycetales(100);f__Metschnikowiaceae(100);g__Metschnikowia(100);g__Metschnikowia_unclassified(100); | Metschnikowia | 13 | Saprotroph | \|Undefined Saprotroph\| | Yeast | NULL | Probable | As nectar-tap_saprotroph (Põlme et al. 2020); Decay sugar-rich substrates (Põlme et al. 2020) | Põlme S, et al. 2020. Fungal Diversity 105:1-16 ((https://doi.org/10.1007/s13225-020-00466-2)) |
| Otu35000 | 1 | k__Fungi(100);p__Ascomycota(100);c__Saccharomycetes(100);o__Saccharomycetales(100);f__Metschnikowiaceae(100);g__Metschnikowia(100);s__Metschnikowia_pulcherrima(100); | Metschnikowia | 13 | Saprotroph | \|Undefined Saprotroph\| | Yeast | NULL | Probable | As nectar-tap_saprotroph (Põlme et al. 2020); Decay sugar-rich substrates (Põlme et al. 2020) | Põlme S, et al. 2020. Fungal Diversity 105:1-16 ((https://doi.org/10.1007/s13225-020-00466-2)) |
| Otu35001 | 1 | k__Fungi(100);p__Ascomycota(100);c__Saccharomycetes(100);o__Saccharomycetales(100);f__Metschnikowiaceae(100);g__Metschnikowia(100);s__Metschnikowia_pulcherrima(100); | Metschnikowia | 13 | Saprotroph | \|Undefined Saprotroph\| | Yeast | NULL | Probable | As nectar-tap_saprotroph (Põlme et al. 2020); Decay sugar-rich substrates (Põlme et al. 2020) | Põlme S, et al. 2020. Fungal Diversity 105:1-16 ((https://doi.org/10.1007/s13225-020-00466-2)) |
| Otu35002 | 1 | k__Fungi(100);p__Ascomycota(100);c__Saccharomycetes(100);o__Saccharomycetales(100);f__Metschnikowiaceae(100);g__Metschnikowia(100);s__Metschnikowia_pulcherrima(100); | Metschnikowia | 13 | Saprotroph | \|Undefined Saprotroph\| | Yeast | NULL | Probable | As nectar-tap_saprotroph (Põlme et al. 2020); Decay sugar-rich substrates (Põlme et al. 2020) | Põlme S, et al. 2020. Fungal Diversity 105:1-16 ((https://doi.org/10.1007/s13225-020-00466-2)) |
| Otu35003 | 1 | k__Fungi(100);p__Ascomycota(100);c__Saccharomycetes(100);o__Saccharomycetales(100);f__Metschnikowiaceae(100);g__Metschnikowia(100);s__Metschnikowia_pulcherrima(100); | Metschnikowia | 13 | Saprotroph | \|Undefined Saprotroph\| | Yeast | NULL | Probable | As nectar-tap_saprotroph (Põlme et al. 2020); Decay sugar-rich substrates (Põlme et al. 2020) | Põlme S, et al. 2020. Fungal Diversity 105:1-16 ((https://doi.org/10.1007/s13225-020-00466-2)) |
| Otu35004 | 1 | k__Fungi(100);p__Ascomycota(100);c__Saccharomycetes(100);o__Saccharomycetales(100);f__Metschnikowiaceae(100);g__Metschnikowia(100);g__Metschnikowia_unclassified(100); | Metschnikowia | 13 | Saprotroph | \|Undefined Saprotroph\| | Yeast | NULL | Probable | As nectar-tap_saprotroph (Põlme et al. 2020); Decay sugar-rich substrates (Põlme et al. 2020) | Põlme S, et al. 2020. Fungal Diversity 105:1-16 ((https://doi.org/10.1007/s13225-020-00466-2)) |
| Otu35005 | 1 | k__Fungi(100);p__Ascomycota(100);c__Saccharomycetes(100);o__Saccharomycetales(100);f__Metschnikowiaceae(100);g__Metschnikowia(100);s__Metschnikowia_pulcherrima(100); | Metschnikowia | 13 | Saprotroph | \|Undefined Saprotroph\| | Yeast | NULL | Probable | As nectar-tap_saprotroph (Põlme et al. 2020); Decay sugar-rich substrates (Põlme et al. 2020) | Põlme S, et al. 2020. Fungal Diversity 105:1-16 ((https://doi.org/10.1007/s13225-020-00466-2)) |
| Otu35006 | 1 | k__Fungi(100);p__Ascomycota(100);c__Saccharomycetes(100);o__Saccharomycetales(100);f__Metschnikowiaceae(100);g__Metschnikowia(100);s__Metschnikowia_pulcherrima(100); | Metschnikowia | 13 | Saprotroph | \|Undefined Saprotroph\| | Yeast | NULL | Probable | As nectar-tap_saprotroph (Põlme et al. 2020); Decay sugar-rich substrates (Põlme et al. 2020) | Põlme S, et al. 2020. Fungal Diversity 105:1-16 ((https://doi.org/10.1007/s13225-020-00466-2)) |
| Otu35008 | 1 | k__Fungi(100);p__Ascomycota(100);c__Saccharomycetes(100);o__Saccharomycetales(100);f__Metschnikowiaceae(100);g__Metschnikowia(100);s__Metschnikowia_pulcherrima(100); | Metschnikowia | 13 | Saprotroph | \|Undefined Saprotroph\| | Yeast | NULL | Probable | As nectar-tap_saprotroph (Põlme et al. 2020); Decay sugar-rich substrates (Põlme et al. 2020) | Põlme S, et al. 2020. Fungal Diversity 105:1-16 ((https://doi.org/10.1007/s13225-020-00466-2)) |
| Otu35009 | 1 | k__Fungi(100);p__Ascomycota(100);c__Saccharomycetes(100);o__Saccharomycetales(100);f__Metschnikowiaceae(100);g__Metschnikowia(100);s__Metschnikowia_pulcherrima(100); | Metschnikowia | 13 | Saprotroph | \|Undefined Saprotroph\| | Yeast | NULL | Probable | As nectar-tap_saprotroph (Põlme et al. 2020); Decay sugar-rich substrates (Põlme et al. 2020) | Põlme S, et al. 2020. Fungal Diversity 105:1-16 ((https://doi.org/10.1007/s13225-020-00466-2)) |
| Otu35010 | 1 | k__Fungi(100);p__Ascomycota(100);c__Saccharomycetes(100);o__Saccharomycetales(100);f__Metschnikowiaceae(100);g__Metschnikowia(100);s__Metschnikowia_pulcherrima(100); | Metschnikowia | 13 | Saprotroph | \|Undefined Saprotroph\| | Yeast | NULL | Probable | As nectar-tap_saprotroph (Põlme et al. 2020); Decay sugar-rich substrates (Põlme et al. 2020) | Põlme S, et al. 2020. Fungal Diversity 105:1-16 ((https://doi.org/10.1007/s13225-020-00466-2)) |
| Otu35011 | 1 | k__Fungi(100);p__Ascomycota(100);c__Saccharomycetes(100);o__Saccharomycetales(100);f__Metschnikowiaceae(100);g__Metschnikowia(100);s__Metschnikowia_pulcherrima(100); | Metschnikowia | 13 | Saprotroph | \|Undefined Saprotroph\| | Yeast | NULL | Probable | As nectar-tap_saprotroph (Põlme et al. 2020); Decay sugar-rich substrates (Põlme et al. 2020) | Põlme S, et al. 2020. Fungal Diversity 105:1-16 ((https://doi.org/10.1007/s13225-020-00466-2)) |
| Otu35012 | 1 | k__Fungi(100);p__Ascomycota(100);c__Saccharomycetes(100);o__Saccharomycetales(100);f__Metschnikowiaceae(100);g__Metschnikowia(100);g__Metschnikowia_unclassified(100); | Metschnikowia | 13 | Saprotroph | \|Undefined Saprotroph\| | Yeast | NULL | Probable | As nectar-tap_saprotroph (Põlme et al. 2020); Decay sugar-rich substrates (Põlme et al. 2020) | Põlme S, et al. 2020. Fungal Diversity 105:1-16 ((https://doi.org/10.1007/s13225-020-00466-2)) |
| Otu35013 | 1 | k__Fungi(100);p__Ascomycota(100);c__Saccharomycetes(100);o__Saccharomycetales(100);f__Metschnikowiaceae(100);g__Metschnikowia(100);s__Metschnikowia_pulcherrima(100); | Metschnikowia | 13 | Saprotroph | \|Undefined Saprotroph\| | Yeast | NULL | Probable | As nectar-tap_saprotroph (Põlme et al. 2020); Decay sugar-rich substrates (Põlme et al. 2020) | Põlme S, et al. 2020. Fungal Diversity 105:1-16 ((https://doi.org/10.1007/s13225-020-00466-2)) |
| Otu35014 | 1 | k__Fungi(100);p__Ascomycota(100);c__Saccharomycetes(100);o__Saccharomycetales(100);f__Metschnikowiaceae(100);g__Metschnikowia(100);g__Metschnikowia_unclassified(100); | Metschnikowia | 13 | Saprotroph | \|Undefined Saprotroph\| | Yeast | NULL | Probable | As nectar-tap_saprotroph (Põlme et al. 2020); Decay sugar-rich substrates (Põlme et al. 2020) | Põlme S, et al. 2020. Fungal Diversity 105:1-16 ((https://doi.org/10.1007/s13225-020-00466-2)) |
| Otu35015 | 1 | k__Fungi(100);p__Ascomycota(100);c__Saccharomycetes(100);o__Saccharomycetales(100);f__Metschnikowiaceae(100);g__Metschnikowia(100);s__Metschnikowia_pulcherrima(100); | Metschnikowia | 13 | Saprotroph | \|Undefined Saprotroph\| | Yeast | NULL | Probable | As nectar-tap_saprotroph (Põlme et al. 2020); Decay sugar-rich substrates (Põlme et al. 2020) | Põlme S, et al. 2020. Fungal Diversity 105:1-16 ((https://doi.org/10.1007/s13225-020-00466-2)) |
| Otu35016 | 1 | k__Fungi(100);p__Ascomycota(100);c__Saccharomycetes(100);o__Saccharomycetales(100);f__Metschnikowiaceae(100);g__Metschnikowia(100);s__Metschnikowia_pulcherrima(100); | Metschnikowia | 13 | Saprotroph | \|Undefined Saprotroph\| | Yeast | NULL | Probable | As nectar-tap_saprotroph (Põlme et al. 2020); Decay sugar-rich substrates (Põlme et al. 2020) | Põlme S, et al. 2020. Fungal Diversity 105:1-16 ((https://doi.org/10.1007/s13225-020-00466-2)) |
| Otu35018 | 1 | k__Fungi(100);p__Ascomycota(100);c__Saccharomycetes(100);o__Saccharomycetales(100);f__Metschnikowiaceae(100);g__Metschnikowia(100);g__Metschnikowia_unclassified(100); | Metschnikowia | 13 | Saprotroph | \|Undefined Saprotroph\| | Yeast | NULL | Probable | As nectar-tap_saprotroph (Põlme et al. 2020); Decay sugar-rich substrates (Põlme et al. 2020) | Põlme S, et al. 2020. Fungal Diversity 105:1-16 ((https://doi.org/10.1007/s13225-020-00466-2)) |
| Otu35037 | 1 | k__Fungi(100);p__Ascomycota(100);c__Saccharomycetes(100);o__Saccharomycetales(100);f__Metschnikowiaceae(100);g__Metschnikowia(100);s__Metschnikowia_pulcherrima(100); | Metschnikowia | 13 | Saprotroph | \|Undefined Saprotroph\| | Yeast | NULL | Probable | As nectar-tap_saprotroph (Põlme et al. 2020); Decay sugar-rich substrates (Põlme et al. 2020) | Põlme S, et al. 2020. Fungal Diversity 105:1-16 ((https://doi.org/10.1007/s13225-020-00466-2)) |
| Otu35038 | 1 | k__Fungi(100);p__Ascomycota(100);c__Saccharomycetes(100);o__Saccharomycetales(100);f__Metschnikowiaceae(100);g__Metschnikowia(100);s__Metschnikowia_pulcherrima(100); | Metschnikowia | 13 | Saprotroph | \|Undefined Saprotroph\| | Yeast | NULL | Probable | As nectar-tap_saprotroph (Põlme et al. 2020); Decay sugar-rich substrates (Põlme et al. 2020) | Põlme S, et al. 2020. Fungal Diversity 105:1-16 ((https://doi.org/10.1007/s13225-020-00466-2)) |
| Otu35040 | 1 | k__Fungi(100);p__Ascomycota(100);c__Saccharomycetes(100);o__Saccharomycetales(100);o__Saccharomycetales_unclassified(100);o__Saccharomycetales_unclassified(100);o__Saccharomycetales_unclassified(100); | Saccharomycetales | 7 | Saprotroph | Undefined Saprotroph | Yeast | NULL | Possible | NULL | Sterkenburg E, et al. 2015. New Phytologist 207:1145-1158 ((https://doi.org/10.1111/nph.13426)) |
| Otu35045 | 1 | k__Fungi(100);p__Ascomycota(100);c__Saccharomycetes(100);o__Saccharomycetales(100);f__Metschnikowiaceae(100);g__Metschnikowia(100);g__Metschnikowia_unclassified(100); | Metschnikowia | 13 | Saprotroph | \|Undefined Saprotroph\| | Yeast | NULL | Probable | As nectar-tap_saprotroph (Põlme et al. 2020); Decay sugar-rich substrates (Põlme et al. 2020) | Põlme S, et al. 2020. Fungal Diversity 105:1-16 ((https://doi.org/10.1007/s13225-020-00466-2)) |
| Otu35048 | 1 | k__Fungi(100);p__Ascomycota(100);c__Saccharomycetes(100);o__Saccharomycetales(100);f__Metschnikowiaceae(100);g__Metschnikowia(100);s__Metschnikowia_pulcherrima(100); | Metschnikowia | 13 | Saprotroph | \|Undefined Saprotroph\| | Yeast | NULL | Probable | As nectar-tap_saprotroph (Põlme et al. 2020); Decay sugar-rich substrates (Põlme et al. 2020) | Põlme S, et al. 2020. Fungal Diversity 105:1-16 ((https://doi.org/10.1007/s13225-020-00466-2)) |
| Otu35052 | 1 | k__Fungi(100);p__Ascomycota(100);c__Saccharomycetes(100);o__Saccharomycetales(100);f__Metschnikowiaceae(100);g__Metschnikowia(100);s__Metschnikowia_pulcherrima(100); | Metschnikowia | 13 | Saprotroph | \|Undefined Saprotroph\| | Yeast | NULL | Probable | As nectar-tap_saprotroph (Põlme et al. 2020); Decay sugar-rich substrates (Põlme et al. 2020) | Põlme S, et al. 2020. Fungal Diversity 105:1-16 ((https://doi.org/10.1007/s13225-020-00466-2)) |
| Otu35056 | 1 | k__Fungi(100);p__Ascomycota(100);c__Saccharomycetes(100);o__Saccharomycetales(100);f__Metschnikowiaceae(100);g__Metschnikowia(100);s__Metschnikowia_pulcherrima(100); | Metschnikowia | 13 | Saprotroph | \|Undefined Saprotroph\| | Yeast | NULL | Probable | As nectar-tap_saprotroph (Põlme et al. 2020); Decay sugar-rich substrates (Põlme et al. 2020) | Põlme S, et al. 2020. Fungal Diversity 105:1-16 ((https://doi.org/10.1007/s13225-020-00466-2)) |
| Otu35060 | 1 | k__Fungi(100);p__Ascomycota(100);c__Saccharomycetes(100);o__Saccharomycetales(100);f__Metschnikowiaceae(100);g__Metschnikowia(100);s__Metschnikowia_pulcherrima(100); | Metschnikowia | 13 | Saprotroph | \|Undefined Saprotroph\| | Yeast | NULL | Probable | As nectar-tap_saprotroph (Põlme et al. 2020); Decay sugar-rich substrates (Põlme et al. 2020) | Põlme S, et al. 2020. Fungal Diversity 105:1-16 ((https://doi.org/10.1007/s13225-020-00466-2)) |
| Otu35076 | 1 | k__Fungi(100);p__Ascomycota(100);c__Saccharomycetes(100);o__Saccharomycetales(100);f__Metschnikowiaceae(100);g__Metschnikowia(100);g__Metschnikowia_unclassified(100); | Metschnikowia | 13 | Saprotroph | \|Undefined Saprotroph\| | Yeast | NULL | Probable | As nectar-tap_saprotroph (Põlme et al. 2020); Decay sugar-rich substrates (Põlme et al. 2020) | Põlme S, et al. 2020. Fungal Diversity 105:1-16 ((https://doi.org/10.1007/s13225-020-00466-2)) |
| Otu35077 | 1 | k__Fungi(100);p__Ascomycota(100);c__Saccharomycetes(100);o__Saccharomycetales(100);f__Metschnikowiaceae(100);g__Metschnikowia(100);s__Metschnikowia_pulcherrima(100); | Metschnikowia | 13 | Saprotroph | \|Undefined Saprotroph\| | Yeast | NULL | Probable | As nectar-tap_saprotroph (Põlme et al. 2020); Decay sugar-rich substrates (Põlme et al. 2020) | Põlme S, et al. 2020. Fungal Diversity 105:1-16 ((https://doi.org/10.1007/s13225-020-00466-2)) |
| Otu35078 | 1 | k__Fungi(100);p__Ascomycota(100);c__Saccharomycetes(100);o__Saccharomycetales(100);f__Metschnikowiaceae(100);g__Metschnikowia(100);g__Metschnikowia_unclassified(100); | Metschnikowia | 13 | Saprotroph | \|Undefined Saprotroph\| | Yeast | NULL | Probable | As nectar-tap_saprotroph (Põlme et al. 2020); Decay sugar-rich substrates (Põlme et al. 2020) | Põlme S, et al. 2020. Fungal Diversity 105:1-16 ((https://doi.org/10.1007/s13225-020-00466-2)) |
| Otu35079 | 1 | k__Fungi(100);p__Ascomycota(100);c__Saccharomycetes(100);o__Saccharomycetales(100);f__Metschnikowiaceae(100);g__Metschnikowia(100);s__Metschnikowia_pulcherrima(100); | Metschnikowia | 13 | Saprotroph | \|Undefined Saprotroph\| | Yeast | NULL | Probable | As nectar-tap_saprotroph (Põlme et al. 2020); Decay sugar-rich substrates (Põlme et al. 2020) | Põlme S, et al. 2020. Fungal Diversity 105:1-16 ((https://doi.org/10.1007/s13225-020-00466-2)) |
| Otu35080 | 1 | k__Fungi(100);p__Ascomycota(100);c__Saccharomycetes(100);o__Saccharomycetales(100);f__Metschnikowiaceae(100);g__Metschnikowia(100);s__Metschnikowia_pulcherrima(100); | Metschnikowia | 13 | Saprotroph | \|Undefined Saprotroph\| | Yeast | NULL | Probable | As nectar-tap_saprotroph (Põlme et al. 2020); Decay sugar-rich substrates (Põlme et al. 2020) | Põlme S, et al. 2020. Fungal Diversity 105:1-16 ((https://doi.org/10.1007/s13225-020-00466-2)) |
| Otu35083 | 1 | k__Fungi(100);p__Ascomycota(100);c__Saccharomycetes(100);o__Saccharomycetales(100);f__Metschnikowiaceae(100);g__Metschnikowia(100);s__Metschnikowia_pulcherrima(100); | Metschnikowia | 13 | Saprotroph | \|Undefined Saprotroph\| | Yeast | NULL | Probable | As nectar-tap_saprotroph (Põlme et al. 2020); Decay sugar-rich substrates (Põlme et al. 2020) | Põlme S, et al. 2020. Fungal Diversity 105:1-16 ((https://doi.org/10.1007/s13225-020-00466-2)) |
| Otu35084 | 1 | k__Fungi(100);p__Ascomycota(100);c__Saccharomycetes(100);o__Saccharomycetales(100);f__Metschnikowiaceae(100);g__Metschnikowia(100);g__Metschnikowia_unclassified(100); | Metschnikowia | 13 | Saprotroph | \|Undefined Saprotroph\| | Yeast | NULL | Probable | As nectar-tap_saprotroph (Põlme et al. 2020); Decay sugar-rich substrates (Põlme et al. 2020) | Põlme S, et al. 2020. Fungal Diversity 105:1-16 ((https://doi.org/10.1007/s13225-020-00466-2)) |
| Otu35085 | 1 | k__Fungi(100);p__Ascomycota(100);c__Saccharomycetes(100);o__Saccharomycetales(100);f__Metschnikowiaceae(100);g__Metschnikowia(100);s__Metschnikowia_pulcherrima(100); | Metschnikowia | 13 | Saprotroph | \|Undefined Saprotroph\| | Yeast | NULL | Probable | As nectar-tap_saprotroph (Põlme et al. 2020); Decay sugar-rich substrates (Põlme et al. 2020) | Põlme S, et al. 2020. Fungal Diversity 105:1-16 ((https://doi.org/10.1007/s13225-020-00466-2)) |
| Otu35086 | 1 | k__Fungi(100);p__Ascomycota(100);c__Saccharomycetes(100);o__Saccharomycetales(100);f__Metschnikowiaceae(100);g__Metschnikowia(100);g__Metschnikowia_unclassified(100); | Metschnikowia | 13 | Saprotroph | \|Undefined Saprotroph\| | Yeast | NULL | Probable | As nectar-tap_saprotroph (Põlme et al. 2020); Decay sugar-rich substrates (Põlme et al. 2020) | Põlme S, et al. 2020. Fungal Diversity 105:1-16 ((https://doi.org/10.1007/s13225-020-00466-2)) |
| Otu35087 | 1 | k__Fungi(100);p__Ascomycota(100);c__Saccharomycetes(100);o__Saccharomycetales(100);f__Metschnikowiaceae(100);g__Metschnikowia(100);s__Metschnikowia_pulcherrima(100); | Metschnikowia | 13 | Saprotroph | \|Undefined Saprotroph\| | Yeast | NULL | Probable | As nectar-tap_saprotroph (Põlme et al. 2020); Decay sugar-rich substrates (Põlme et al. 2020) | Põlme S, et al. 2020. Fungal Diversity 105:1-16 ((https://doi.org/10.1007/s13225-020-00466-2)) |
| Otu35088 | 1 | k__Fungi(100);p__Ascomycota(100);c__Saccharomycetes(100);o__Saccharomycetales(100);f__Metschnikowiaceae(100);g__Metschnikowia(100);s__Metschnikowia_pulcherrima(100); | Metschnikowia | 13 | Saprotroph | \|Undefined Saprotroph\| | Yeast | NULL | Probable | As nectar-tap_saprotroph (Põlme et al. 2020); Decay sugar-rich substrates (Põlme et al. 2020) | Põlme S, et al. 2020. Fungal Diversity 105:1-16 ((https://doi.org/10.1007/s13225-020-00466-2)) |
| Otu35109 | 1 | k__Fungi(100);p__Ascomycota(100);c__Saccharomycetes(100);o__Saccharomycetales(100);f__Metschnikowiaceae(100);g__Metschnikowia(100);s__Metschnikowia_pulcherrima(100); | Metschnikowia | 13 | Saprotroph | \|Undefined Saprotroph\| | Yeast | NULL | Probable | As nectar-tap_saprotroph (Põlme et al. 2020); Decay sugar-rich substrates (Põlme et al. 2020) | Põlme S, et al. 2020. Fungal Diversity 105:1-16 ((https://doi.org/10.1007/s13225-020-00466-2)) |
| Otu35111 | 1 | k__Fungi(100);p__Ascomycota(100);c__Saccharomycetes(100);o__Saccharomycetales(100);f__Metschnikowiaceae(100);g__Metschnikowia(100);s__Metschnikowia_pulcherrima(100); | Metschnikowia | 13 | Saprotroph | \|Undefined Saprotroph\| | Yeast | NULL | Probable | As nectar-tap_saprotroph (Põlme et al. 2020); Decay sugar-rich substrates (Põlme et al. 2020) | Põlme S, et al. 2020. Fungal Diversity 105:1-16 ((https://doi.org/10.1007/s13225-020-00466-2)) |
| Otu35113 | 1 | k__Fungi(100);p__Ascomycota(100);c__Saccharomycetes(100);o__Saccharomycetales(100);f__Metschnikowiaceae(100);g__Metschnikowia(100);s__Metschnikowia_pulcherrima(100); | Metschnikowia | 13 | Saprotroph | \|Undefined Saprotroph\| | Yeast | NULL | Probable | As nectar-tap_saprotroph (Põlme et al. 2020); Decay sugar-rich substrates (Põlme et al. 2020) | Põlme S, et al. 2020. Fungal Diversity 105:1-16 ((https://doi.org/10.1007/s13225-020-00466-2)) |
| Otu35114 | 1 | k__Fungi(100);p__Ascomycota(100);c__Saccharomycetes(100);o__Saccharomycetales(100);f__Metschnikowiaceae(100);g__Metschnikowia(100);s__Metschnikowia_pulcherrima(100); | Metschnikowia | 13 | Saprotroph | \|Undefined Saprotroph\| | Yeast | NULL | Probable | As nectar-tap_saprotroph (Põlme et al. 2020); Decay sugar-rich substrates (Põlme et al. 2020) | Põlme S, et al. 2020. Fungal Diversity 105:1-16 ((https://doi.org/10.1007/s13225-020-00466-2)) |
| Otu35116 | 1 | k__Fungi(100);p__Ascomycota(100);c__Saccharomycetes(100);o__Saccharomycetales(100);f__Metschnikowiaceae(100);g__Metschnikowia(100);s__Metschnikowia_pulcherrima(100); | Metschnikowia | 13 | Saprotroph | \|Undefined Saprotroph\| | Yeast | NULL | Probable | As nectar-tap_saprotroph (Põlme et al. 2020); Decay sugar-rich substrates (Põlme et al. 2020) | Põlme S, et al. 2020. Fungal Diversity 105:1-16 ((https://doi.org/10.1007/s13225-020-00466-2)) |
| Otu35119 | 1 | k__Fungi(100);p__Ascomycota(100);c__Saccharomycetes(100);o__Saccharomycetales(100);f__Metschnikowiaceae(100);g__Metschnikowia(100);g__Metschnikowia_unclassified(100); | Metschnikowia | 13 | Saprotroph | \|Undefined Saprotroph\| | Yeast | NULL | Probable | As nectar-tap_saprotroph (Põlme et al. 2020); Decay sugar-rich substrates (Põlme et al. 2020) | Põlme S, et al. 2020. Fungal Diversity 105:1-16 ((https://doi.org/10.1007/s13225-020-00466-2)) |
| Otu35125 | 1 | k__Fungi(100);p__Ascomycota(100);c__Saccharomycetes(100);o__Saccharomycetales(100);f__Metschnikowiaceae(100);g__Metschnikowia(100);s__Metschnikowia_pulcherrima(100); | Metschnikowia | 13 | Saprotroph | \|Undefined Saprotroph\| | Yeast | NULL | Probable | As nectar-tap_saprotroph (Põlme et al. 2020); Decay sugar-rich substrates (Põlme et al. 2020) | Põlme S, et al. 2020. Fungal Diversity 105:1-16 ((https://doi.org/10.1007/s13225-020-00466-2)) |
| Otu35127 | 1 | k__Fungi(100);p__Ascomycota(100);c__Saccharomycetes(100);o__Saccharomycetales(100);f__Metschnikowiaceae(100);g__Metschnikowia(100);s__Metschnikowia_pulcherrima(100); | Metschnikowia | 13 | Saprotroph | \|Undefined Saprotroph\| | Yeast | NULL | Probable | As nectar-tap_saprotroph (Põlme et al. 2020); Decay sugar-rich substrates (Põlme et al. 2020) | Põlme S, et al. 2020. Fungal Diversity 105:1-16 ((https://doi.org/10.1007/s13225-020-00466-2)) |
| Otu35130 | 1 | k__Fungi(100);p__Ascomycota(100);c__Saccharomycetes(100);o__Saccharomycetales(100);f__Metschnikowiaceae(100);g__Metschnikowia(100);s__Metschnikowia_pulcherrima(100); | Metschnikowia | 13 | Saprotroph | \|Undefined Saprotroph\| | Yeast | NULL | Probable | As nectar-tap_saprotroph (Põlme et al. 2020); Decay sugar-rich substrates (Põlme et al. 2020) | Põlme S, et al. 2020. Fungal Diversity 105:1-16 ((https://doi.org/10.1007/s13225-020-00466-2)) |
| Otu35132 | 1 | k__Fungi(100);p__Ascomycota(100);c__Saccharomycetes(100);o__Saccharomycetales(100);f__Metschnikowiaceae(100);g__Metschnikowia(100);s__Metschnikowia_pulcherrima(100); | Metschnikowia | 13 | Saprotroph | \|Undefined Saprotroph\| | Yeast | NULL | Probable | As nectar-tap_saprotroph (Põlme et al. 2020); Decay sugar-rich substrates (Põlme et al. 2020) | Põlme S, et al. 2020. Fungal Diversity 105:1-16 ((https://doi.org/10.1007/s13225-020-00466-2)) |
| Otu35133 | 1 | k__Fungi(100);p__Ascomycota(100);c__Saccharomycetes(100);o__Saccharomycetales(100);f__Metschnikowiaceae(100);g__Metschnikowia(100);s__Metschnikowia_pulcherrima(100); | Metschnikowia | 13 | Saprotroph | \|Undefined Saprotroph\| | Yeast | NULL | Probable | As nectar-tap_saprotroph (Põlme et al. 2020); Decay sugar-rich substrates (Põlme et al. 2020) | Põlme S, et al. 2020. Fungal Diversity 105:1-16 ((https://doi.org/10.1007/s13225-020-00466-2)) |
| Otu35134 | 1 | k__Fungi(100);p__Ascomycota(100);c__Saccharomycetes(100);o__Saccharomycetales(100);f__Metschnikowiaceae(100);g__Metschnikowia(100);g__Metschnikowia_unclassified(100); | Metschnikowia | 13 | Saprotroph | \|Undefined Saprotroph\| | Yeast | NULL | Probable | As nectar-tap_saprotroph (Põlme et al. 2020); Decay sugar-rich substrates (Põlme et al. 2020) | Põlme S, et al. 2020. Fungal Diversity 105:1-16 ((https://doi.org/10.1007/s13225-020-00466-2)) |
| Otu35135 | 1 | k__Fungi(100);p__Ascomycota(100);c__Saccharomycetes(100);o__Saccharomycetales(100);f__Metschnikowiaceae(100);g__Metschnikowia(100);s__Metschnikowia_pulcherrima(100); | Metschnikowia | 13 | Saprotroph | \|Undefined Saprotroph\| | Yeast | NULL | Probable | As nectar-tap_saprotroph (Põlme et al. 2020); Decay sugar-rich substrates (Põlme et al. 2020) | Põlme S, et al. 2020. Fungal Diversity 105:1-16 ((https://doi.org/10.1007/s13225-020-00466-2)) |
| Otu35136 | 1 | k__Fungi(100);p__Ascomycota(100);c__Saccharomycetes(100);o__Saccharomycetales(100);f__Metschnikowiaceae(100);g__Metschnikowia(100);s__Metschnikowia_pulcherrima(100); | Metschnikowia | 13 | Saprotroph | \|Undefined Saprotroph\| | Yeast | NULL | Probable | As nectar-tap_saprotroph (Põlme et al. 2020); Decay sugar-rich substrates (Põlme et al. 2020) | Põlme S, et al. 2020. Fungal Diversity 105:1-16 ((https://doi.org/10.1007/s13225-020-00466-2)) |
| Otu35137 | 1 | k__Fungi(100);p__Ascomycota(100);c__Saccharomycetes(100);o__Saccharomycetales(100);f__Metschnikowiaceae(100);g__Metschnikowia(100);s__Metschnikowia_pulcherrima(100); | Metschnikowia | 13 | Saprotroph | \|Undefined Saprotroph\| | Yeast | NULL | Probable | As nectar-tap_saprotroph (Põlme et al. 2020); Decay sugar-rich substrates (Põlme et al. 2020) | Põlme S, et al. 2020. Fungal Diversity 105:1-16 ((https://doi.org/10.1007/s13225-020-00466-2)) |
| Otu35138 | 1 | k__Fungi(100);p__Ascomycota(100);c__Saccharomycetes(100);o__Saccharomycetales(100);f__Metschnikowiaceae(100);g__Metschnikowia(100);g__Metschnikowia_unclassified(100); | Metschnikowia | 13 | Saprotroph | \|Undefined Saprotroph\| | Yeast | NULL | Probable | As nectar-tap_saprotroph (Põlme et al. 2020); Decay sugar-rich substrates (Põlme et al. 2020) | Põlme S, et al. 2020. Fungal Diversity 105:1-16 ((https://doi.org/10.1007/s13225-020-00466-2)) |
| Otu35139 | 1 | k__Fungi(100);p__Ascomycota(100);c__Saccharomycetes(100);o__Saccharomycetales(100);f__Metschnikowiaceae(100);g__Metschnikowia(100);s__Metschnikowia_pulcherrima(100); | Metschnikowia | 13 | Saprotroph | \|Undefined Saprotroph\| | Yeast | NULL | Probable | As nectar-tap_saprotroph (Põlme et al. 2020); Decay sugar-rich substrates (Põlme et al. 2020) | Põlme S, et al. 2020. Fungal Diversity 105:1-16 ((https://doi.org/10.1007/s13225-020-00466-2)) |
| Otu35140 | 1 | k__Fungi(100);p__Ascomycota(100);c__Saccharomycetes(100);o__Saccharomycetales(100);f__Metschnikowiaceae(100);g__Metschnikowia(100);s__Metschnikowia_pulcherrima(100); | Metschnikowia | 13 | Saprotroph | \|Undefined Saprotroph\| | Yeast | NULL | Probable | As nectar-tap_saprotroph (Põlme et al. 2020); Decay sugar-rich substrates (Põlme et al. 2020) | Põlme S, et al. 2020. Fungal Diversity 105:1-16 ((https://doi.org/10.1007/s13225-020-00466-2)) |
| Otu35142 | 1 | k__Fungi(100);p__Ascomycota(100);c__Saccharomycetes(100);o__Saccharomycetales(100);f__Metschnikowiaceae(100);g__Metschnikowia(100);s__Metschnikowia_pulcherrima(100); | Metschnikowia | 13 | Saprotroph | \|Undefined Saprotroph\| | Yeast | NULL | Probable | As nectar-tap_saprotroph (Põlme et al. 2020); Decay sugar-rich substrates (Põlme et al. 2020) | Põlme S, et al. 2020. Fungal Diversity 105:1-16 ((https://doi.org/10.1007/s13225-020-00466-2)) |
| Otu35143 | 1 | k__Fungi(100);p__Ascomycota(100);c__Saccharomycetes(100);o__Saccharomycetales(100);f__Metschnikowiaceae(100);g__Metschnikowia(100);s__Metschnikowia_pulcherrima(100); | Metschnikowia | 13 | Saprotroph | \|Undefined Saprotroph\| | Yeast | NULL | Probable | As nectar-tap_saprotroph (Põlme et al. 2020); Decay sugar-rich substrates (Põlme et al. 2020) | Põlme S, et al. 2020. Fungal Diversity 105:1-16 ((https://doi.org/10.1007/s13225-020-00466-2)) |
| Otu35144 | 1 | k__Fungi(100);p__Ascomycota(100);c__Saccharomycetes(100);o__Saccharomycetales(100);f__Metschnikowiaceae(100);g__Metschnikowia(100);g__Metschnikowia_unclassified(100); | Metschnikowia | 13 | Saprotroph | \|Undefined Saprotroph\| | Yeast | NULL | Probable | As nectar-tap_saprotroph (Põlme et al. 2020); Decay sugar-rich substrates (Põlme et al. 2020) | Põlme S, et al. 2020. Fungal Diversity 105:1-16 ((https://doi.org/10.1007/s13225-020-00466-2)) |
| Otu35145 | 1 | k__Fungi(100);p__Ascomycota(100);c__Saccharomycetes(100);o__Saccharomycetales(100);f__Metschnikowiaceae(100);g__Metschnikowia(100);s__Metschnikowia_pulcherrima(100); | Metschnikowia | 13 | Saprotroph | \|Undefined Saprotroph\| | Yeast | NULL | Probable | As nectar-tap_saprotroph (Põlme et al. 2020); Decay sugar-rich substrates (Põlme et al. 2020) | Põlme S, et al. 2020. Fungal Diversity 105:1-16 ((https://doi.org/10.1007/s13225-020-00466-2)) |
| Otu35146 | 1 | k__Fungi(100);p__Ascomycota(100);c__Saccharomycetes(100);o__Saccharomycetales(100);f__Metschnikowiaceae(100);g__Metschnikowia(100);s__Metschnikowia_pulcherrima(100); | Metschnikowia | 13 | Saprotroph | \|Undefined Saprotroph\| | Yeast | NULL | Probable | As nectar-tap_saprotroph (Põlme et al. 2020); Decay sugar-rich substrates (Põlme et al. 2020) | Põlme S, et al. 2020. Fungal Diversity 105:1-16 ((https://doi.org/10.1007/s13225-020-00466-2)) |
| Otu35147 | 1 | k__Fungi(100);p__Ascomycota(100);c__Saccharomycetes(100);o__Saccharomycetales(100);f__Metschnikowiaceae(100);g__Metschnikowia(100);s__Metschnikowia_pulcherrima(100); | Metschnikowia | 13 | Saprotroph | \|Undefined Saprotroph\| | Yeast | NULL | Probable | As nectar-tap_saprotroph (Põlme et al. 2020); Decay sugar-rich substrates (Põlme et al. 2020) | Põlme S, et al. 2020. Fungal Diversity 105:1-16 ((https://doi.org/10.1007/s13225-020-00466-2)) |
| Otu35148 | 1 | k__Fungi(100);p__Ascomycota(100);c__Saccharomycetes(100);o__Saccharomycetales(100);f__Metschnikowiaceae(100);g__Metschnikowia(100);s__Metschnikowia_pulcherrima(100); | Metschnikowia | 13 | Saprotroph | \|Undefined Saprotroph\| | Yeast | NULL | Probable | As nectar-tap_saprotroph (Põlme et al. 2020); Decay sugar-rich substrates (Põlme et al. 2020) | Põlme S, et al. 2020. Fungal Diversity 105:1-16 ((https://doi.org/10.1007/s13225-020-00466-2)) |
| Otu35149 | 1 | k__Fungi(100);p__Ascomycota(100);c__Saccharomycetes(100);o__Saccharomycetales(100);f__Metschnikowiaceae(100);g__Metschnikowia(100);s__Metschnikowia_pulcherrima(100); | Metschnikowia | 13 | Saprotroph | \|Undefined Saprotroph\| | Yeast | NULL | Probable | As nectar-tap_saprotroph (Põlme et al. 2020); Decay sugar-rich substrates (Põlme et al. 2020) | Põlme S, et al. 2020. Fungal Diversity 105:1-16 ((https://doi.org/10.1007/s13225-020-00466-2)) |
| Otu35150 | 1 | k__Fungi(100);p__Ascomycota(100);c__Saccharomycetes(100);o__Saccharomycetales(100);f__Metschnikowiaceae(100);g__Metschnikowia(100);s__Metschnikowia_pulcherrima(100); | Metschnikowia | 13 | Saprotroph | \|Undefined Saprotroph\| | Yeast | NULL | Probable | As nectar-tap_saprotroph (Põlme et al. 2020); Decay sugar-rich substrates (Põlme et al. 2020) | Põlme S, et al. 2020. Fungal Diversity 105:1-16 ((https://doi.org/10.1007/s13225-020-00466-2)) |
| Otu35151 | 1 | k__Fungi(100);p__Ascomycota(100);c__Saccharomycetes(100);o__Saccharomycetales(100);f__Metschnikowiaceae(100);g__Metschnikowia(100);s__Metschnikowia_pulcherrima(100); | Metschnikowia | 13 | Saprotroph | \|Undefined Saprotroph\| | Yeast | NULL | Probable | As nectar-tap_saprotroph (Põlme et al. 2020); Decay sugar-rich substrates (Põlme et al. 2020) | Põlme S, et al. 2020. Fungal Diversity 105:1-16 ((https://doi.org/10.1007/s13225-020-00466-2)) |
| Otu35152 | 1 | k__Fungi(100);p__Ascomycota(100);c__Saccharomycetes(100);o__Saccharomycetales(100);f__Metschnikowiaceae(100);g__Metschnikowia(100);s__Metschnikowia_pulcherrima(100); | Metschnikowia | 13 | Saprotroph | \|Undefined Saprotroph\| | Yeast | NULL | Probable | As nectar-tap_saprotroph (Põlme et al. 2020); Decay sugar-rich substrates (Põlme et al. 2020) | Põlme S, et al. 2020. Fungal Diversity 105:1-16 ((https://doi.org/10.1007/s13225-020-00466-2)) |
| Otu35153 | 1 | k__Fungi(100);p__Ascomycota(100);c__Saccharomycetes(100);o__Saccharomycetales(100);f__Metschnikowiaceae(100);g__Metschnikowia(100);s__Metschnikowia_pulcherrima(100); | Metschnikowia | 13 | Saprotroph | \|Undefined Saprotroph\| | Yeast | NULL | Probable | As nectar-tap_saprotroph (Põlme et al. 2020); Decay sugar-rich substrates (Põlme et al. 2020) | Põlme S, et al. 2020. Fungal Diversity 105:1-16 ((https://doi.org/10.1007/s13225-020-00466-2)) |
| Otu35154 | 1 | k__Fungi(100);p__Ascomycota(100);c__Saccharomycetes(100);o__Saccharomycetales(100);f__Metschnikowiaceae(100);g__Metschnikowia(100);s__Metschnikowia_pulcherrima(100); | Metschnikowia | 13 | Saprotroph | \|Undefined Saprotroph\| | Yeast | NULL | Probable | As nectar-tap_saprotroph (Põlme et al. 2020); Decay sugar-rich substrates (Põlme et al. 2020) | Põlme S, et al. 2020. Fungal Diversity 105:1-16 ((https://doi.org/10.1007/s13225-020-00466-2)) |
| Otu35156 | 1 | k__Fungi(100);p__Ascomycota(100);c__Saccharomycetes(100);o__Saccharomycetales(100);f__Metschnikowiaceae(100);g__Metschnikowia(100);s__Metschnikowia_pulcherrima(100); | Metschnikowia | 13 | Saprotroph | \|Undefined Saprotroph\| | Yeast | NULL | Probable | As nectar-tap_saprotroph (Põlme et al. 2020); Decay sugar-rich substrates (Põlme et al. 2020) | Põlme S, et al. 2020. Fungal Diversity 105:1-16 ((https://doi.org/10.1007/s13225-020-00466-2)) |
| Otu35158 | 1 | k__Fungi(100);p__Ascomycota(100);c__Saccharomycetes(100);o__Saccharomycetales(100);f__Metschnikowiaceae(100);g__Metschnikowia(100);s__Metschnikowia_pulcherrima(100); | Metschnikowia | 13 | Saprotroph | \|Undefined Saprotroph\| | Yeast | NULL | Probable | As nectar-tap_saprotroph (Põlme et al. 2020); Decay sugar-rich substrates (Põlme et al. 2020) | Põlme S, et al. 2020. Fungal Diversity 105:1-16 ((https://doi.org/10.1007/s13225-020-00466-2)) |
| Otu35159 | 1 | k__Fungi(100);p__Ascomycota(100);c__Saccharomycetes(100);o__Saccharomycetales(100);f__Metschnikowiaceae(100);g__Metschnikowia(100);s__Metschnikowia_pulcherrima(100); | Metschnikowia | 13 | Saprotroph | \|Undefined Saprotroph\| | Yeast | NULL | Probable | As nectar-tap_saprotroph (Põlme et al. 2020); Decay sugar-rich substrates (Põlme et al. 2020) | Põlme S, et al. 2020. Fungal Diversity 105:1-16 ((https://doi.org/10.1007/s13225-020-00466-2)) |
| Otu35160 | 1 | k__Fungi(100);p__Ascomycota(100);c__Saccharomycetes(100);o__Saccharomycetales(100);f__Metschnikowiaceae(100);g__Metschnikowia(100);g__Metschnikowia_unclassified(100); | Metschnikowia | 13 | Saprotroph | \|Undefined Saprotroph\| | Yeast | NULL | Probable | As nectar-tap_saprotroph (Põlme et al. 2020); Decay sugar-rich substrates (Põlme et al. 2020) | Põlme S, et al. 2020. Fungal Diversity 105:1-16 ((https://doi.org/10.1007/s13225-020-00466-2)) |
| Otu35162 | 1 | k__Fungi(100);p__Ascomycota(100);c__Saccharomycetes(100);o__Saccharomycetales(100);f__Metschnikowiaceae(100);g__Metschnikowia(100);g__Metschnikowia_unclassified(100); | Metschnikowia | 13 | Saprotroph | \|Undefined Saprotroph\| | Yeast | NULL | Probable | As nectar-tap_saprotroph (Põlme et al. 2020); Decay sugar-rich substrates (Põlme et al. 2020) | Põlme S, et al. 2020. Fungal Diversity 105:1-16 ((https://doi.org/10.1007/s13225-020-00466-2)) |
| Otu35163 | 1 | k__Fungi(100);p__Ascomycota(100);c__Saccharomycetes(100);o__Saccharomycetales(100);f__Metschnikowiaceae(100);g__Metschnikowia(100);s__Metschnikowia_pulcherrima(100); | Metschnikowia | 13 | Saprotroph | \|Undefined Saprotroph\| | Yeast | NULL | Probable | As nectar-tap_saprotroph (Põlme et al. 2020); Decay sugar-rich substrates (Põlme et al. 2020) | Põlme S, et al. 2020. Fungal Diversity 105:1-16 ((https://doi.org/10.1007/s13225-020-00466-2)) |
| Otu35164 | 1 | k__Fungi(100);p__Ascomycota(100);c__Saccharomycetes(100);o__Saccharomycetales(100);f__Metschnikowiaceae(100);g__Metschnikowia(100);s__Metschnikowia_pulcherrima(100); | Metschnikowia | 13 | Saprotroph | \|Undefined Saprotroph\| | Yeast | NULL | Probable | As nectar-tap_saprotroph (Põlme et al. 2020); Decay sugar-rich substrates (Põlme et al. 2020) | Põlme S, et al. 2020. Fungal Diversity 105:1-16 ((https://doi.org/10.1007/s13225-020-00466-2)) |
| Otu35165 | 1 | k__Fungi(100);p__Ascomycota(100);c__Saccharomycetes(100);o__Saccharomycetales(100);f__Metschnikowiaceae(100);g__Metschnikowia(100);s__Metschnikowia_pulcherrima(100); | Metschnikowia | 13 | Saprotroph | \|Undefined Saprotroph\| | Yeast | NULL | Probable | As nectar-tap_saprotroph (Põlme et al. 2020); Decay sugar-rich substrates (Põlme et al. 2020) | Põlme S, et al. 2020. Fungal Diversity 105:1-16 ((https://doi.org/10.1007/s13225-020-00466-2)) |
| Otu35166 | 1 | k__Fungi(100);p__Ascomycota(100);c__Saccharomycetes(100);o__Saccharomycetales(100);f__Metschnikowiaceae(100);g__Metschnikowia(100);s__Metschnikowia_pulcherrima(100); | Metschnikowia | 13 | Saprotroph | \|Undefined Saprotroph\| | Yeast | NULL | Probable | As nectar-tap_saprotroph (Põlme et al. 2020); Decay sugar-rich substrates (Põlme et al. 2020) | Põlme S, et al. 2020. Fungal Diversity 105:1-16 ((https://doi.org/10.1007/s13225-020-00466-2)) |
| Otu35167 | 1 | k__Fungi(100);p__Ascomycota(100);c__Saccharomycetes(100);o__Saccharomycetales(100);f__Metschnikowiaceae(100);g__Metschnikowia(100);s__Metschnikowia_pulcherrima(100); | Metschnikowia | 13 | Saprotroph | \|Undefined Saprotroph\| | Yeast | NULL | Probable | As nectar-tap_saprotroph (Põlme et al. 2020); Decay sugar-rich substrates (Põlme et al. 2020) | Põlme S, et al. 2020. Fungal Diversity 105:1-16 ((https://doi.org/10.1007/s13225-020-00466-2)) |
| Otu35168 | 1 | k__Fungi(100);p__Ascomycota(100);c__Saccharomycetes(100);o__Saccharomycetales(100);f__Metschnikowiaceae(100);g__Metschnikowia(100);g__Metschnikowia_unclassified(100); | Metschnikowia | 13 | Saprotroph | \|Undefined Saprotroph\| | Yeast | NULL | Probable | As nectar-tap_saprotroph (Põlme et al. 2020); Decay sugar-rich substrates (Põlme et al. 2020) | Põlme S, et al. 2020. Fungal Diversity 105:1-16 ((https://doi.org/10.1007/s13225-020-00466-2)) |
| Otu35169 | 1 | k__Fungi(100);p__Ascomycota(100);c__Saccharomycetes(100);o__Saccharomycetales(100);f__Metschnikowiaceae(100);g__Metschnikowia(100);s__Metschnikowia_pulcherrima(100); | Metschnikowia | 13 | Saprotroph | \|Undefined Saprotroph\| | Yeast | NULL | Probable | As nectar-tap_saprotroph (Põlme et al. 2020); Decay sugar-rich substrates (Põlme et al. 2020) | Põlme S, et al. 2020. Fungal Diversity 105:1-16 ((https://doi.org/10.1007/s13225-020-00466-2)) |
| Otu35170 | 1 | k__Fungi(100);p__Ascomycota(100);c__Saccharomycetes(100);o__Saccharomycetales(100);f__Metschnikowiaceae(100);g__Metschnikowia(100);s__Metschnikowia_pulcherrima(100); | Metschnikowia | 13 | Saprotroph | \|Undefined Saprotroph\| | Yeast | NULL | Probable | As nectar-tap_saprotroph (Põlme et al. 2020); Decay sugar-rich substrates (Põlme et al. 2020) | Põlme S, et al. 2020. Fungal Diversity 105:1-16 ((https://doi.org/10.1007/s13225-020-00466-2)) |
| Otu35172 | 1 | k__Fungi(100);p__Ascomycota(100);c__Saccharomycetes(100);o__Saccharomycetales(100);f__Metschnikowiaceae(100);g__Metschnikowia(100);s__Metschnikowia_pulcherrima(100); | Metschnikowia | 13 | Saprotroph | \|Undefined Saprotroph\| | Yeast | NULL | Probable | As nectar-tap_saprotroph (Põlme et al. 2020); Decay sugar-rich substrates (Põlme et al. 2020) | Põlme S, et al. 2020. Fungal Diversity 105:1-16 ((https://doi.org/10.1007/s13225-020-00466-2)) |
| Otu35178 | 1 | k__Fungi(100);p__Ascomycota(100);c__Saccharomycetes(100);o__Saccharomycetales(100);f__Metschnikowiaceae(100);g__Metschnikowia(100);s__Metschnikowia_pulcherrima(100); | Metschnikowia | 13 | Saprotroph | \|Undefined Saprotroph\| | Yeast | NULL | Probable | As nectar-tap_saprotroph (Põlme et al. 2020); Decay sugar-rich substrates (Põlme et al. 2020) | Põlme S, et al. 2020. Fungal Diversity 105:1-16 ((https://doi.org/10.1007/s13225-020-00466-2)) |
| Otu35179 | 1 | k__Fungi(100);p__Ascomycota(100);c__Saccharomycetes(100);o__Saccharomycetales(100);f__Metschnikowiaceae(100);g__Metschnikowia(100);s__Metschnikowia_pulcherrima(100); | Metschnikowia | 13 | Saprotroph | \|Undefined Saprotroph\| | Yeast | NULL | Probable | As nectar-tap_saprotroph (Põlme et al. 2020); Decay sugar-rich substrates (Põlme et al. 2020) | Põlme S, et al. 2020. Fungal Diversity 105:1-16 ((https://doi.org/10.1007/s13225-020-00466-2)) |
| Otu35180 | 1 | k__Fungi(100);p__Ascomycota(100);c__Saccharomycetes(100);o__Saccharomycetales(100);f__Metschnikowiaceae(100);g__Metschnikowia(100);s__Metschnikowia_pulcherrima(100); | Metschnikowia | 13 | Saprotroph | \|Undefined Saprotroph\| | Yeast | NULL | Probable | As nectar-tap_saprotroph (Põlme et al. 2020); Decay sugar-rich substrates (Põlme et al. 2020) | Põlme S, et al. 2020. Fungal Diversity 105:1-16 ((https://doi.org/10.1007/s13225-020-00466-2)) |
| Otu35181 | 1 | k__Fungi(100);p__Ascomycota(100);c__Saccharomycetes(100);o__Saccharomycetales(100);f__Metschnikowiaceae(100);g__Metschnikowia(100);s__Metschnikowia_pulcherrima(100); | Metschnikowia | 13 | Saprotroph | \|Undefined Saprotroph\| | Yeast | NULL | Probable | As nectar-tap_saprotroph (Põlme et al. 2020); Decay sugar-rich substrates (Põlme et al. 2020) | Põlme S, et al. 2020. Fungal Diversity 105:1-16 ((https://doi.org/10.1007/s13225-020-00466-2)) |
| Otu35183 | 1 | k__Fungi(100);p__Ascomycota(100);c__Saccharomycetes(100);o__Saccharomycetales(100);f__Metschnikowiaceae(100);g__Metschnikowia(100);s__Metschnikowia_pulcherrima(100); | Metschnikowia | 13 | Saprotroph | \|Undefined Saprotroph\| | Yeast | NULL | Probable | As nectar-tap_saprotroph (Põlme et al. 2020); Decay sugar-rich substrates (Põlme et al. 2020) | Põlme S, et al. 2020. Fungal Diversity 105:1-16 ((https://doi.org/10.1007/s13225-020-00466-2)) |
| Otu35184 | 1 | k__Fungi(100);p__Ascomycota(100);c__Saccharomycetes(100);o__Saccharomycetales(100);f__Metschnikowiaceae(100);g__Metschnikowia(100);s__Metschnikowia_pulcherrima(100); | Metschnikowia | 13 | Saprotroph | \|Undefined Saprotroph\| | Yeast | NULL | Probable | As nectar-tap_saprotroph (Põlme et al. 2020); Decay sugar-rich substrates (Põlme et al. 2020) | Põlme S, et al. 2020. Fungal Diversity 105:1-16 ((https://doi.org/10.1007/s13225-020-00466-2)) |
| Otu35185 | 1 | k__Fungi(100);p__Ascomycota(100);c__Saccharomycetes(100);o__Saccharomycetales(100);f__Metschnikowiaceae(100);g__Metschnikowia(100);g__Metschnikowia_unclassified(100); | Metschnikowia | 13 | Saprotroph | \|Undefined Saprotroph\| | Yeast | NULL | Probable | As nectar-tap_saprotroph (Põlme et al. 2020); Decay sugar-rich substrates (Põlme et al. 2020) | Põlme S, et al. 2020. Fungal Diversity 105:1-16 ((https://doi.org/10.1007/s13225-020-00466-2)) |
| Otu35186 | 1 | k__Fungi(100);p__Ascomycota(100);c__Saccharomycetes(100);o__Saccharomycetales(100);f__Metschnikowiaceae(100);g__Metschnikowia(100);s__Metschnikowia_pulcherrima(100); | Metschnikowia | 13 | Saprotroph | \|Undefined Saprotroph\| | Yeast | NULL | Probable | As nectar-tap_saprotroph (Põlme et al. 2020); Decay sugar-rich substrates (Põlme et al. 2020) | Põlme S, et al. 2020. Fungal Diversity 105:1-16 ((https://doi.org/10.1007/s13225-020-00466-2)) |
| Otu35187 | 1 | k__Fungi(100);p__Ascomycota(100);c__Saccharomycetes(100);o__Saccharomycetales(100);f__Metschnikowiaceae(100);g__Metschnikowia(100);s__Metschnikowia_pulcherrima(100); | Metschnikowia | 13 | Saprotroph | \|Undefined Saprotroph\| | Yeast | NULL | Probable | As nectar-tap_saprotroph (Põlme et al. 2020); Decay sugar-rich substrates (Põlme et al. 2020) | Põlme S, et al. 2020. Fungal Diversity 105:1-16 ((https://doi.org/10.1007/s13225-020-00466-2)) |
| Otu35188 | 1 | k__Fungi(100);p__Ascomycota(100);c__Saccharomycetes(100);o__Saccharomycetales(100);f__Metschnikowiaceae(100);g__Metschnikowia(100);s__Metschnikowia_pulcherrima(100); | Metschnikowia | 13 | Saprotroph | \|Undefined Saprotroph\| | Yeast | NULL | Probable | As nectar-tap_saprotroph (Põlme et al. 2020); Decay sugar-rich substrates (Põlme et al. 2020) | Põlme S, et al. 2020. Fungal Diversity 105:1-16 ((https://doi.org/10.1007/s13225-020-00466-2)) |
| Otu35189 | 1 | k__Fungi(100);p__Ascomycota(100);c__Saccharomycetes(100);o__Saccharomycetales(100);f__Metschnikowiaceae(100);g__Metschnikowia(100);g__Metschnikowia_unclassified(100); | Metschnikowia | 13 | Saprotroph | \|Undefined Saprotroph\| | Yeast | NULL | Probable | As nectar-tap_saprotroph (Põlme et al. 2020); Decay sugar-rich substrates (Põlme et al. 2020) | Põlme S, et al. 2020. Fungal Diversity 105:1-16 ((https://doi.org/10.1007/s13225-020-00466-2)) |
| Otu35190 | 1 | k__Fungi(100);p__Ascomycota(100);c__Saccharomycetes(100);o__Saccharomycetales(100);f__Metschnikowiaceae(100);g__Metschnikowia(100);s__Metschnikowia_pulcherrima(100); | Metschnikowia | 13 | Saprotroph | \|Undefined Saprotroph\| | Yeast | NULL | Probable | As nectar-tap_saprotroph (Põlme et al. 2020); Decay sugar-rich substrates (Põlme et al. 2020) | Põlme S, et al. 2020. Fungal Diversity 105:1-16 ((https://doi.org/10.1007/s13225-020-00466-2)) |
| Otu35191 | 1 | k__Fungi(100);p__Ascomycota(100);c__Saccharomycetes(100);o__Saccharomycetales(100);f__Metschnikowiaceae(100);g__Metschnikowia(100);s__Metschnikowia_pulcherrima(100); | Metschnikowia | 13 | Saprotroph | \|Undefined Saprotroph\| | Yeast | NULL | Probable | As nectar-tap_saprotroph (Põlme et al. 2020); Decay sugar-rich substrates (Põlme et al. 2020) | Põlme S, et al. 2020. Fungal Diversity 105:1-16 ((https://doi.org/10.1007/s13225-020-00466-2)) |
| Otu35192 | 1 | k__Fungi(100);p__Ascomycota(100);c__Saccharomycetes(100);o__Saccharomycetales(100);f__Metschnikowiaceae(100);g__Metschnikowia(100);s__Metschnikowia_pulcherrima(100); | Metschnikowia | 13 | Saprotroph | \|Undefined Saprotroph\| | Yeast | NULL | Probable | As nectar-tap_saprotroph (Põlme et al. 2020); Decay sugar-rich substrates (Põlme et al. 2020) | Põlme S, et al. 2020. Fungal Diversity 105:1-16 ((https://doi.org/10.1007/s13225-020-00466-2)) |
| Otu35193 | 1 | k__Fungi(100);p__Ascomycota(100);c__Saccharomycetes(100);o__Saccharomycetales(100);f__Metschnikowiaceae(100);g__Metschnikowia(100);g__Metschnikowia_unclassified(100); | Metschnikowia | 13 | Saprotroph | \|Undefined Saprotroph\| | Yeast | NULL | Probable | As nectar-tap_saprotroph (Põlme et al. 2020); Decay sugar-rich substrates (Põlme et al. 2020) | Põlme S, et al. 2020. Fungal Diversity 105:1-16 ((https://doi.org/10.1007/s13225-020-00466-2)) |
| Otu35194 | 1 | k__Fungi(100);p__Ascomycota(100);c__Saccharomycetes(100);o__Saccharomycetales(100);f__Metschnikowiaceae(100);g__Metschnikowia(100);s__Metschnikowia_pulcherrima(100); | Metschnikowia | 13 | Saprotroph | \|Undefined Saprotroph\| | Yeast | NULL | Probable | As nectar-tap_saprotroph (Põlme et al. 2020); Decay sugar-rich substrates (Põlme et al. 2020) | Põlme S, et al. 2020. Fungal Diversity 105:1-16 ((https://doi.org/10.1007/s13225-020-00466-2)) |
| Otu35195 | 1 | k__Fungi(100);p__Ascomycota(100);c__Saccharomycetes(100);o__Saccharomycetales(100);f__Metschnikowiaceae(100);g__Metschnikowia(100);g__Metschnikowia_unclassified(100); | Metschnikowia | 13 | Saprotroph | \|Undefined Saprotroph\| | Yeast | NULL | Probable | As nectar-tap_saprotroph (Põlme et al. 2020); Decay sugar-rich substrates (Põlme et al. 2020) | Põlme S, et al. 2020. Fungal Diversity 105:1-16 ((https://doi.org/10.1007/s13225-020-00466-2)) |
| Otu35196 | 1 | k__Fungi(100);p__Ascomycota(100);c__Saccharomycetes(100);o__Saccharomycetales(100);f__Metschnikowiaceae(100);g__Metschnikowia(100);s__Metschnikowia_pulcherrima(100); | Metschnikowia | 13 | Saprotroph | \|Undefined Saprotroph\| | Yeast | NULL | Probable | As nectar-tap_saprotroph (Põlme et al. 2020); Decay sugar-rich substrates (Põlme et al. 2020) | Põlme S, et al. 2020. Fungal Diversity 105:1-16 ((https://doi.org/10.1007/s13225-020-00466-2)) |
| Otu35197 | 1 | k__Fungi(100);p__Ascomycota(100);c__Saccharomycetes(100);o__Saccharomycetales(100);f__Metschnikowiaceae(100);g__Metschnikowia(100);s__Metschnikowia_pulcherrima(100); | Metschnikowia | 13 | Saprotroph | \|Undefined Saprotroph\| | Yeast | NULL | Probable | As nectar-tap_saprotroph (Põlme et al. 2020); Decay sugar-rich substrates (Põlme et al. 2020) | Põlme S, et al. 2020. Fungal Diversity 105:1-16 ((https://doi.org/10.1007/s13225-020-00466-2)) |
| Otu35199 | 1 | k__Fungi(100);p__Ascomycota(100);c__Saccharomycetes(100);o__Saccharomycetales(100);f__Metschnikowiaceae(100);g__Metschnikowia(100);g__Metschnikowia_unclassified(100); | Metschnikowia | 13 | Saprotroph | \|Undefined Saprotroph\| | Yeast | NULL | Probable | As nectar-tap_saprotroph (Põlme et al. 2020); Decay sugar-rich substrates (Põlme et al. 2020) | Põlme S, et al. 2020. Fungal Diversity 105:1-16 ((https://doi.org/10.1007/s13225-020-00466-2)) |
| Otu35200 | 1 | k__Fungi(100);p__Ascomycota(100);c__Saccharomycetes(100);o__Saccharomycetales(100);f__Metschnikowiaceae(100);g__Metschnikowia(100);s__Metschnikowia_pulcherrima(100); | Metschnikowia | 13 | Saprotroph | \|Undefined Saprotroph\| | Yeast | NULL | Probable | As nectar-tap_saprotroph (Põlme et al. 2020); Decay sugar-rich substrates (Põlme et al. 2020) | Põlme S, et al. 2020. Fungal Diversity 105:1-16 ((https://doi.org/10.1007/s13225-020-00466-2)) |
| Otu35201 | 1 | k__Fungi(100);p__Ascomycota(100);c__Saccharomycetes(100);o__Saccharomycetales(100);f__Metschnikowiaceae(100);g__Metschnikowia(100);g__Metschnikowia_unclassified(100); | Metschnikowia | 13 | Saprotroph | \|Undefined Saprotroph\| | Yeast | NULL | Probable | As nectar-tap_saprotroph (Põlme et al. 2020); Decay sugar-rich substrates (Põlme et al. 2020) | Põlme S, et al. 2020. Fungal Diversity 105:1-16 ((https://doi.org/10.1007/s13225-020-00466-2)) |
| Otu35203 | 1 | k__Fungi(100);p__Ascomycota(100);c__Saccharomycetes(100);o__Saccharomycetales(100);f__Metschnikowiaceae(100);g__Metschnikowia(100);g__Metschnikowia_unclassified(100); | Metschnikowia | 13 | Saprotroph | \|Undefined Saprotroph\| | Yeast | NULL | Probable | As nectar-tap_saprotroph (Põlme et al. 2020); Decay sugar-rich substrates (Põlme et al. 2020) | Põlme S, et al. 2020. Fungal Diversity 105:1-16 ((https://doi.org/10.1007/s13225-020-00466-2)) |
| Otu35204 | 1 | k__Fungi(100);p__Ascomycota(100);c__Saccharomycetes(100);o__Saccharomycetales(100);f__Metschnikowiaceae(100);g__Metschnikowia(100);s__Metschnikowia_pulcherrima(100); | Metschnikowia | 13 | Saprotroph | \|Undefined Saprotroph\| | Yeast | NULL | Probable | As nectar-tap_saprotroph (Põlme et al. 2020); Decay sugar-rich substrates (Põlme et al. 2020) | Põlme S, et al. 2020. Fungal Diversity 105:1-16 ((https://doi.org/10.1007/s13225-020-00466-2)) |
| Otu35205 | 1 | k__Fungi(100);p__Ascomycota(100);c__Saccharomycetes(100);o__Saccharomycetales(100);f__Metschnikowiaceae(100);g__Metschnikowia(100);s__Metschnikowia_pulcherrima(100); | Metschnikowia | 13 | Saprotroph | \|Undefined Saprotroph\| | Yeast | NULL | Probable | As nectar-tap_saprotroph (Põlme et al. 2020); Decay sugar-rich substrates (Põlme et al. 2020) | Põlme S, et al. 2020. Fungal Diversity 105:1-16 ((https://doi.org/10.1007/s13225-020-00466-2)) |
| Otu35206 | 1 | k__Fungi(100);p__Ascomycota(100);c__Saccharomycetes(100);o__Saccharomycetales(100);f__Metschnikowiaceae(100);g__Metschnikowia(100);s__Metschnikowia_pulcherrima(100); | Metschnikowia | 13 | Saprotroph | \|Undefined Saprotroph\| | Yeast | NULL | Probable | As nectar-tap_saprotroph (Põlme et al. 2020); Decay sugar-rich substrates (Põlme et al. 2020) | Põlme S, et al. 2020. Fungal Diversity 105:1-16 ((https://doi.org/10.1007/s13225-020-00466-2)) |
| Otu35207 | 1 | k__Fungi(100);p__Ascomycota(100);c__Saccharomycetes(100);o__Saccharomycetales(100);f__Metschnikowiaceae(100);g__Metschnikowia(100);s__Metschnikowia_pulcherrima(100); | Metschnikowia | 13 | Saprotroph | \|Undefined Saprotroph\| | Yeast | NULL | Probable | As nectar-tap_saprotroph (Põlme et al. 2020); Decay sugar-rich substrates (Põlme et al. 2020) | Põlme S, et al. 2020. Fungal Diversity 105:1-16 ((https://doi.org/10.1007/s13225-020-00466-2)) |
| Otu35208 | 1 | k__Fungi(100);p__Ascomycota(100);c__Saccharomycetes(100);o__Saccharomycetales(100);f__Metschnikowiaceae(100);g__Metschnikowia(100);s__Metschnikowia_pulcherrima(100); | Metschnikowia | 13 | Saprotroph | \|Undefined Saprotroph\| | Yeast | NULL | Probable | As nectar-tap_saprotroph (Põlme et al. 2020); Decay sugar-rich substrates (Põlme et al. 2020) | Põlme S, et al. 2020. Fungal Diversity 105:1-16 ((https://doi.org/10.1007/s13225-020-00466-2)) |
| Otu35209 | 1 | k__Fungi(100);p__Ascomycota(100);c__Saccharomycetes(100);o__Saccharomycetales(100);f__Metschnikowiaceae(100);g__Metschnikowia(100);s__Metschnikowia_pulcherrima(100); | Metschnikowia | 13 | Saprotroph | \|Undefined Saprotroph\| | Yeast | NULL | Probable | As nectar-tap_saprotroph (Põlme et al. 2020); Decay sugar-rich substrates (Põlme et al. 2020) | Põlme S, et al. 2020. Fungal Diversity 105:1-16 ((https://doi.org/10.1007/s13225-020-00466-2)) |
| Otu35210 | 1 | k__Fungi(100);p__Ascomycota(100);c__Saccharomycetes(100);o__Saccharomycetales(100);f__Metschnikowiaceae(100);g__Metschnikowia(100);s__Metschnikowia_pulcherrima(100); | Metschnikowia | 13 | Saprotroph | \|Undefined Saprotroph\| | Yeast | NULL | Probable | As nectar-tap_saprotroph (Põlme et al. 2020); Decay sugar-rich substrates (Põlme et al. 2020) | Põlme S, et al. 2020. Fungal Diversity 105:1-16 ((https://doi.org/10.1007/s13225-020-00466-2)) |
| Otu35214 | 1 | k__Fungi(100);p__Ascomycota(100);c__Saccharomycetes(100);o__Saccharomycetales(100);f__Metschnikowiaceae(100);g__Metschnikowia(100);s__Metschnikowia_pulcherrima(100); | Metschnikowia | 13 | Saprotroph | \|Undefined Saprotroph\| | Yeast | NULL | Probable | As nectar-tap_saprotroph (Põlme et al. 2020); Decay sugar-rich substrates (Põlme et al. 2020) | Põlme S, et al. 2020. Fungal Diversity 105:1-16 ((https://doi.org/10.1007/s13225-020-00466-2)) |
| Otu35215 | 1 | k__Fungi(100);p__Ascomycota(100);c__Saccharomycetes(100);o__Saccharomycetales(100);f__Metschnikowiaceae(100);g__Metschnikowia(100);s__Metschnikowia_pulcherrima(100); | Metschnikowia | 13 | Saprotroph | \|Undefined Saprotroph\| | Yeast | NULL | Probable | As nectar-tap_saprotroph (Põlme et al. 2020); Decay sugar-rich substrates (Põlme et al. 2020) | Põlme S, et al. 2020. Fungal Diversity 105:1-16 ((https://doi.org/10.1007/s13225-020-00466-2)) |
| Otu35216 | 1 | k__Fungi(100);p__Ascomycota(100);c__Saccharomycetes(100);o__Saccharomycetales(100);f__Metschnikowiaceae(100);g__Metschnikowia(100);s__Metschnikowia_pulcherrima(100); | Metschnikowia | 13 | Saprotroph | \|Undefined Saprotroph\| | Yeast | NULL | Probable | As nectar-tap_saprotroph (Põlme et al. 2020); Decay sugar-rich substrates (Põlme et al. 2020) | Põlme S, et al. 2020. Fungal Diversity 105:1-16 ((https://doi.org/10.1007/s13225-020-00466-2)) |
| Otu35217 | 1 | k__Fungi(100);p__Ascomycota(100);c__Saccharomycetes(100);o__Saccharomycetales(100);f__Metschnikowiaceae(100);g__Metschnikowia(100);s__Metschnikowia_pulcherrima(100); | Metschnikowia | 13 | Saprotroph | \|Undefined Saprotroph\| | Yeast | NULL | Probable | As nectar-tap_saprotroph (Põlme et al. 2020); Decay sugar-rich substrates (Põlme et al. 2020) | Põlme S, et al. 2020. Fungal Diversity 105:1-16 ((https://doi.org/10.1007/s13225-020-00466-2)) |
| Otu35218 | 1 | k__Fungi(100);p__Ascomycota(100);c__Saccharomycetes(100);o__Saccharomycetales(100);f__Metschnikowiaceae(100);g__Metschnikowia(100);s__Metschnikowia_pulcherrima(100); | Metschnikowia | 13 | Saprotroph | \|Undefined Saprotroph\| | Yeast | NULL | Probable | As nectar-tap_saprotroph (Põlme et al. 2020); Decay sugar-rich substrates (Põlme et al. 2020) | Põlme S, et al. 2020. Fungal Diversity 105:1-16 ((https://doi.org/10.1007/s13225-020-00466-2)) |
| Otu35219 | 1 | k__Fungi(100);p__Ascomycota(100);c__Saccharomycetes(100);o__Saccharomycetales(100);f__Metschnikowiaceae(100);g__Metschnikowia(100);s__Metschnikowia_pulcherrima(100); | Metschnikowia | 13 | Saprotroph | \|Undefined Saprotroph\| | Yeast | NULL | Probable | As nectar-tap_saprotroph (Põlme et al. 2020); Decay sugar-rich substrates (Põlme et al. 2020) | Põlme S, et al. 2020. Fungal Diversity 105:1-16 ((https://doi.org/10.1007/s13225-020-00466-2)) |
| Otu35221 | 1 | k__Fungi(100);p__Ascomycota(100);c__Saccharomycetes(100);o__Saccharomycetales(100);f__Metschnikowiaceae(100);g__Metschnikowia(100);s__Metschnikowia_pulcherrima(100); | Metschnikowia | 13 | Saprotroph | \|Undefined Saprotroph\| | Yeast | NULL | Probable | As nectar-tap_saprotroph (Põlme et al. 2020); Decay sugar-rich substrates (Põlme et al. 2020) | Põlme S, et al. 2020. Fungal Diversity 105:1-16 ((https://doi.org/10.1007/s13225-020-00466-2)) |
| Otu35222 | 1 | k__Fungi(100);p__Ascomycota(100);c__Saccharomycetes(100);o__Saccharomycetales(100);f__Metschnikowiaceae(100);g__Metschnikowia(100);s__Metschnikowia_pulcherrima(100); | Metschnikowia | 13 | Saprotroph | \|Undefined Saprotroph\| | Yeast | NULL | Probable | As nectar-tap_saprotroph (Põlme et al. 2020); Decay sugar-rich substrates (Põlme et al. 2020) | Põlme S, et al. 2020. Fungal Diversity 105:1-16 ((https://doi.org/10.1007/s13225-020-00466-2)) |
| Otu35223 | 1 | k__Fungi(100);p__Ascomycota(100);c__Saccharomycetes(100);o__Saccharomycetales(100);f__Metschnikowiaceae(100);g__Metschnikowia(100);s__Metschnikowia_pulcherrima(100); | Metschnikowia | 13 | Saprotroph | \|Undefined Saprotroph\| | Yeast | NULL | Probable | As nectar-tap_saprotroph (Põlme et al. 2020); Decay sugar-rich substrates (Põlme et al. 2020) | Põlme S, et al. 2020. Fungal Diversity 105:1-16 ((https://doi.org/10.1007/s13225-020-00466-2)) |
| Otu35224 | 1 | k__Fungi(100);p__Ascomycota(100);c__Saccharomycetes(100);o__Saccharomycetales(100);f__Metschnikowiaceae(100);g__Metschnikowia(100);s__Metschnikowia_pulcherrima(100); | Metschnikowia | 13 | Saprotroph | \|Undefined Saprotroph\| | Yeast | NULL | Probable | As nectar-tap_saprotroph (Põlme et al. 2020); Decay sugar-rich substrates (Põlme et al. 2020) | Põlme S, et al. 2020. Fungal Diversity 105:1-16 ((https://doi.org/10.1007/s13225-020-00466-2)) |
| Otu35225 | 1 | k__Fungi(100);p__Ascomycota(100);c__Saccharomycetes(100);o__Saccharomycetales(100);f__Metschnikowiaceae(100);g__Metschnikowia(100);s__Metschnikowia_pulcherrima(100); | Metschnikowia | 13 | Saprotroph | \|Undefined Saprotroph\| | Yeast | NULL | Probable | As nectar-tap_saprotroph (Põlme et al. 2020); Decay sugar-rich substrates (Põlme et al. 2020) | Põlme S, et al. 2020. Fungal Diversity 105:1-16 ((https://doi.org/10.1007/s13225-020-00466-2)) |
| Otu35226 | 1 | k__Fungi(100);p__Ascomycota(100);c__Saccharomycetes(100);o__Saccharomycetales(100);f__Metschnikowiaceae(100);g__Metschnikowia(100);s__Metschnikowia_pulcherrima(100); | Metschnikowia | 13 | Saprotroph | \|Undefined Saprotroph\| | Yeast | NULL | Probable | As nectar-tap_saprotroph (Põlme et al. 2020); Decay sugar-rich substrates (Põlme et al. 2020) | Põlme S, et al. 2020. Fungal Diversity 105:1-16 ((https://doi.org/10.1007/s13225-020-00466-2)) |
| Otu35229 | 1 | k__Fungi(100);p__Ascomycota(100);c__Saccharomycetes(100);o__Saccharomycetales(100);f__Metschnikowiaceae(100);g__Metschnikowia(100);g__Metschnikowia_unclassified(100); | Metschnikowia | 13 | Saprotroph | \|Undefined Saprotroph\| | Yeast | NULL | Probable | As nectar-tap_saprotroph (Põlme et al. 2020); Decay sugar-rich substrates (Põlme et al. 2020) | Põlme S, et al. 2020. Fungal Diversity 105:1-16 ((https://doi.org/10.1007/s13225-020-00466-2)) |
| Otu35230 | 1 | k__Fungi(100);p__Ascomycota(100);c__Saccharomycetes(100);o__Saccharomycetales(100);f__Metschnikowiaceae(100);g__Metschnikowia(100);g__Metschnikowia_unclassified(100); | Metschnikowia | 13 | Saprotroph | \|Undefined Saprotroph\| | Yeast | NULL | Probable | As nectar-tap_saprotroph (Põlme et al. 2020); Decay sugar-rich substrates (Põlme et al. 2020) | Põlme S, et al. 2020. Fungal Diversity 105:1-16 ((https://doi.org/10.1007/s13225-020-00466-2)) |

| | | | | | | | | | | | |
|---|---|---|---|---|---|---|---|---|---|---|---|
| Otu35231 | 1 | k__Fungi(100);p__Ascomycota(100);c__Saccharomycetes(100);o__Saccharomycetales(100);f__Metschnikowiaceae(100);g__Metschnikowia(100);s__Metschnikowia_pulcherrima(100); | Metschnikowia | 13 | Saprotroph | \|Undefined Saprotroph\| | Yeast | NULL | Probable | As nectar-tap_saprotroph (Põlme et al. 2020); Decay sugar-rich substrates (Põlme et al. 2020) | Põlme S, et al. 2020. Fungal Diversity 105:1-16 ((https://doi.org/10.1007/s13225-020-00466-2)) |
| Otu35232 | 1 | k__Fungi(100);p__Ascomycota(100);c__Saccharomycetes(100);o__Saccharomycetales(100);f__Metschnikowiaceae(100);g__Metschnikowia(100);s__Metschnikowia_pulcherrima(100); | Metschnikowia | 13 | Saprotroph | \|Undefined Saprotroph\| | Yeast | NULL | Probable | As nectar-tap_saprotroph (Põlme et al. 2020); Decay sugar-rich substrates (Põlme et al. 2020) | Põlme S, et al. 2020. Fungal Diversity 105:1-16 ((https://doi.org/10.1007/s13225-020-00466-2)) |
| Otu35233 | 1 | k__Fungi(100);p__Ascomycota(100);c__Saccharomycetes(100);o__Saccharomycetales(100);f__Metschnikowiaceae(100);g__Metschnikowia(100);g__Metschnikowia_unclassified(100); | Metschnikowia | 13 | Saprotroph | \|Undefined Saprotroph\| | Yeast | NULL | Probable | As nectar-tap_saprotroph (Põlme et al. 2020); Decay sugar-rich substrates (Põlme et al. 2020) | Põlme S, et al. 2020. Fungal Diversity 105:1-16 ((https://doi.org/10.1007/s13225-020-00466-2)) |
| Otu35234 | 1 | k__Fungi(100);p__Ascomycota(100);c__Saccharomycetes(100);o__Saccharomycetales(100);f__Metschnikowiaceae(100);g__Metschnikowia(100);s__Metschnikowia_pulcherrima(100); | Metschnikowia | 13 | Saprotroph | \|Undefined Saprotroph\| | Yeast | NULL | Probable | As nectar-tap_saprotroph (Põlme et al. 2020); Decay sugar-rich substrates (Põlme et al. 2020) | Põlme S, et al. 2020. Fungal Diversity 105:1-16 ((https://doi.org/10.1007/s13225-020-00466-2)) |
| Otu35235 | 1 | k__Fungi(100);p__Ascomycota(100);c__Saccharomycetes(100);o__Saccharomycetales(100);f__Metschnikowiaceae(100);g__Metschnikowia(100);s__Metschnikowia_pulcherrima(100); | Metschnikowia | 13 | Saprotroph | \|Undefined Saprotroph\| | Yeast | NULL | Probable | As nectar-tap_saprotroph (Põlme et al. 2020); Decay sugar-rich substrates (Põlme et al. 2020) | Põlme S, et al. 2020. Fungal Diversity 105:1-16 ((https://doi.org/10.1007/s13225-020-00466-2)) |
| Otu35237 | 1 | k__Fungi(100);p__Ascomycota(100);c__Saccharomycetes(100);o__Saccharomycetales(100);f__Metschnikowiaceae(100);g__Metschnikowia(100);s__Metschnikowia_pulcherrima(100); | Metschnikowia | 13 | Saprotroph | \|Undefined Saprotroph\| | Yeast | NULL | Probable | As nectar-tap_saprotroph (Põlme et al. 2020); Decay sugar-rich substrates (Põlme et al. 2020) | Põlme S, et al. 2020. Fungal Diversity 105:1-16 ((https://doi.org/10.1007/s13225-020-00466-2)) |
| Otu35238 | 1 | k__Fungi(100);p__Ascomycota(100);c__Saccharomycetes(100);o__Saccharomycetales(100);f__Metschnikowiaceae(100);g__Metschnikowia(100);g__Metschnikowia_unclassified(100); | Metschnikowia | 13 | Saprotroph | \|Undefined Saprotroph\| | Yeast | NULL | Probable | As nectar-tap_saprotroph (Põlme et al. 2020); Decay sugar-rich substrates (Põlme et al. 2020) | Põlme S, et al. 2020. Fungal Diversity 105:1-16 ((https://doi.org/10.1007/s13225-020-00466-2)) |
| Otu35241 | 1 | k__Fungi(100);p__Ascomycota(100);c__Saccharomycetes(100);o__Saccharomycetales(100);f__Metschnikowiaceae(100);g__Metschnikowia(100);s__Metschnikowia_pulcherrima(100); | Metschnikowia | 13 | Saprotroph | \|Undefined Saprotroph\| | Yeast | NULL | Probable | As nectar-tap_saprotroph (Põlme et al. 2020); Decay sugar-rich substrates (Põlme et al. 2020) | Põlme S, et al. 2020. Fungal Diversity 105:1-16 ((https://doi.org/10.1007/s13225-020-00466-2)) |
| Otu35246 | 1 | k__Fungi(100);p__Ascomycota(100);c__Saccharomycetes(100);o__Saccharomycetales(100);f__Metschnikowiaceae(100);g__Metschnikowia(100);s__Metschnikowia_pulcherrima(100); | Metschnikowia | 13 | Saprotroph | \|Undefined Saprotroph\| | Yeast | NULL | Probable | As nectar-tap_saprotroph (Põlme et al. 2020); Decay sugar-rich substrates (Põlme et al. 2020) | Põlme S, et al. 2020. Fungal Diversity 105:1-16 ((https://doi.org/10.1007/s13225-020-00466-2)) |
| Otu35247 | 1 | k__Fungi(100);p__Ascomycota(100);c__Saccharomycetes(100);o__Saccharomycetales(100);f__Metschnikowiaceae(100);g__Metschnikowia(100);g__Metschnikowia_unclassified(100); | Metschnikowia | 13 | Saprotroph | \|Undefined Saprotroph\| | Yeast | NULL | Probable | As nectar-tap_saprotroph (Põlme et al. 2020); Decay sugar-rich substrates (Põlme et al. 2020) | Põlme S, et al. 2020. Fungal Diversity 105:1-16 ((https://doi.org/10.1007/s13225-020-00466-2)) |
| Otu35248 | 1 | k__Fungi(100);p__Ascomycota(100);c__Saccharomycetes(100);o__Saccharomycetales(100);f__Metschnikowiaceae(100);g__Metschnikowia(100);g__Metschnikowia_unclassified(100); | Metschnikowia | 13 | Saprotroph | \|Undefined Saprotroph\| | Yeast | NULL | Probable | As nectar-tap_saprotroph (Põlme et al. 2020); Decay sugar-rich substrates (Põlme et al. 2020) | Põlme S, et al. 2020. Fungal Diversity 105:1-16 ((https://doi.org/10.1007/s13225-020-00466-2)) |
| Otu35249 | 1 | k__Fungi(100);p__Ascomycota(100);c__Saccharomycetes(100);o__Saccharomycetales(100);f__Metschnikowiaceae(100);g__Metschnikowia(100);s__Metschnikowia_pulcherrima(100); | Metschnikowia | 13 | Saprotroph | \|Undefined Saprotroph\| | Yeast | NULL | Probable | As nectar-tap_saprotroph (Põlme et al. 2020); Decay sugar-rich substrates (Põlme et al. 2020) | Põlme S, et al. 2020. Fungal Diversity 105:1-16 ((https://doi.org/10.1007/s13225-020-00466-2)) |
| Otu35250 | 1 | k__Fungi(100);p__Ascomycota(100);c__Saccharomycetes(100);o__Saccharomycetales(100);f__Metschnikowiaceae(100);g__Metschnikowia(100);g__Metschnikowia_unclassified(100); | Metschnikowia | 13 | Saprotroph | \|Undefined Saprotroph\| | Yeast | NULL | Probable | As nectar-tap_saprotroph (Põlme et al. 2020); Decay sugar-rich substrates (Põlme et al. 2020) | Põlme S, et al. 2020. Fungal Diversity 105:1-16 ((https://doi.org/10.1007/s13225-020-00466-2)) |
| Otu35251 | 1 | k__Fungi(100);p__Ascomycota(100);c__Saccharomycetes(100);o__Saccharomycetales(100);f__Metschnikowiaceae(100);g__Metschnikowia(100);s__Metschnikowia_pulcherrima(100); | Metschnikowia | 13 | Saprotroph | \|Undefined Saprotroph\| | Yeast | NULL | Probable | As nectar-tap_saprotroph (Põlme et al. 2020); Decay sugar-rich substrates (Põlme et al. 2020) | Põlme S, et al. 2020. Fungal Diversity 105:1-16 ((https://doi.org/10.1007/s13225-020-00466-2)) |
| Otu35252 | 1 | k__Fungi(100);p__Ascomycota(100);c__Saccharomycetes(100);o__Saccharomycetales(100);f__Metschnikowiaceae(100);g__Metschnikowia(100);g__Metschnikowia_unclassified(100); | Metschnikowia | 13 | Saprotroph | \|Undefined Saprotroph\| | Yeast | NULL | Probable | As nectar-tap_saprotroph (Põlme et al. 2020); Decay sugar-rich substrates (Põlme et al. 2020) | Põlme S, et al. 2020. Fungal Diversity 105:1-16 ((https://doi.org/10.1007/s13225-020-00466-2)) |
| Otu35253 | 1 | k__Fungi(100);p__Ascomycota(100);c__Saccharomycetes(100);o__Saccharomycetales(100);f__Metschnikowiaceae(100);g__Metschnikowia(100);s__Metschnikowia_pulcherrima(100); | Metschnikowia | 13 | Saprotroph | \|Undefined Saprotroph\| | Yeast | NULL | Probable | As nectar-tap_saprotroph (Põlme et al. 2020); Decay sugar-rich substrates (Põlme et al. 2020) | Põlme S, et al. 2020. Fungal Diversity 105:1-16 ((https://doi.org/10.1007/s13225-020-00466-2)) |
| Otu35254 | 1 | k__Fungi(100);p__Ascomycota(100);c__Saccharomycetes(100);o__Saccharomycetales(100);f__Metschnikowiaceae(100);g__Metschnikowia(100);s__Metschnikowia_pulcherrima(100); | Metschnikowia | 13 | Saprotroph | \|Undefined Saprotroph\| | Yeast | NULL | Probable | As nectar-tap_saprotroph (Põlme et al. 2020); Decay sugar-rich substrates (Põlme et al. 2020) | Põlme S, et al. 2020. Fungal Diversity 105:1-16 ((https://doi.org/10.1007/s13225-020-00466-2)) |
| Otu35255 | 1 | k__Fungi(100);p__Ascomycota(100);c__Saccharomycetes(100);o__Saccharomycetales(100);f__Metschnikowiaceae(100);g__Metschnikowia(100);s__Metschnikowia_pulcherrima(100); | Metschnikowia | 13 | Saprotroph | \|Undefined Saprotroph\| | Yeast | NULL | Probable | As nectar-tap_saprotroph (Põlme et al. 2020); Decay sugar-rich substrates (Põlme et al. 2020) | Põlme S, et al. 2020. Fungal Diversity 105:1-16 ((https://doi.org/10.1007/s13225-020-00466-2)) |
| Otu35256 | 1 | k__Fungi(100);p__Ascomycota(100);c__Saccharomycetes(100);o__Saccharomycetales(100);f__Metschnikowiaceae(100);g__Metschnikowia(100);s__Metschnikowia_pulcherrima(100); | Metschnikowia | 13 | Saprotroph | \|Undefined Saprotroph\| | Yeast | NULL | Probable | As nectar-tap_saprotroph (Põlme et al. 2020); Decay sugar-rich substrates (Põlme et al. 2020) | Põlme S, et al. 2020. Fungal Diversity 105:1-16 ((https://doi.org/10.1007/s13225-020-00466-2)) |
| Otu35257 | 1 | k__Fungi(100);p__Ascomycota(100);c__Saccharomycetes(100);o__Saccharomycetales(100);f__Metschnikowiaceae(100);g__Metschnikowia(100);g__Metschnikowia_unclassified(100); | Metschnikowia | 13 | Saprotroph | \|Undefined Saprotroph\| | Yeast | NULL | Probable | As nectar-tap_saprotroph (Põlme et al. 2020); Decay sugar-rich substrates (Põlme et al. 2020) | Põlme S, et al. 2020. Fungal Diversity 105:1-16 ((https://doi.org/10.1007/s13225-020-00466-2)) |
| Otu35258 | 1 | k__Fungi(100);p__Ascomycota(100);c__Saccharomycetes(100);o__Saccharomycetales(100);f__Metschnikowiaceae(100);g__Metschnikowia(100);s__Metschnikowia_pulcherrima(100); | Metschnikowia | 13 | Saprotroph | \|Undefined Saprotroph\| | Yeast | NULL | Probable | As nectar-tap_saprotroph (Põlme et al. 2020); Decay sugar-rich substrates (Põlme et al. 2020) | Põlme S, et al. 2020. Fungal Diversity 105:1-16 ((https://doi.org/10.1007/s13225-020-00466-2)) |
| Otu35259 | 1 | k__Fungi(100);p__Ascomycota(100);c__Saccharomycetes(100);o__Saccharomycetales(100);f__Metschnikowiaceae(100);g__Metschnikowia(100);s__Metschnikowia_pulcherrima(100); | Metschnikowia | 13 | Saprotroph | \|Undefined Saprotroph\| | Yeast | NULL | Probable | As nectar-tap_saprotroph (Põlme et al. 2020); Decay sugar-rich substrates (Põlme et al. 2020) | Põlme S, et al. 2020. Fungal Diversity 105:1-16 ((https://doi.org/10.1007/s13225-020-00466-2)) |
| Otu35263 | 1 | k__Fungi(100);p__Ascomycota(100);c__Saccharomycetes(100);o__Saccharomycetales(100);f__Metschnikowiaceae(100);g__Metschnikowia(100);s__Metschnikowia_pulcherrima(100); | Metschnikowia | 13 | Saprotroph | \|Undefined Saprotroph\| | Yeast | NULL | Probable | As nectar-tap_saprotroph (Põlme et al. 2020); Decay sugar-rich substrates (Põlme et al. 2020) | Põlme S, et al. 2020. Fungal Diversity 105:1-16 ((https://doi.org/10.1007/s13225-020-00466-2)) |
| Otu35264 | 1 | k__Fungi(100);p__Ascomycota(100);c__Saccharomycetes(100);o__Saccharomycetales(100);f__Metschnikowiaceae(100);g__Metschnikowia(100);s__Metschnikowia_pulcherrima(100); | Metschnikowia | 13 | Saprotroph | \|Undefined Saprotroph\| | Yeast | NULL | Probable | As nectar-tap_saprotroph (Põlme et al. 2020); Decay sugar-rich substrates (Põlme et al. 2020) | Põlme S, et al. 2020. Fungal Diversity 105:1-16 ((https://doi.org/10.1007/s13225-020-00466-2)) |
| Otu35267 | 1 | k__Fungi(100);p__Ascomycota(100);c__Saccharomycetes(100);o__Saccharomycetales(100);f__Metschnikowiaceae(100);g__Metschnikowia(100);g__Metschnikowia_unclassified(100); | Metschnikowia | 13 | Saprotroph | \|Undefined Saprotroph\| | Yeast | NULL | Probable | As nectar-tap_saprotroph (Põlme et al. 2020); Decay sugar-rich substrates (Põlme et al. 2020) | Põlme S, et al. 2020. Fungal Diversity 105:1-16 ((https://doi.org/10.1007/s13225-020-00466-2)) |
| Otu35268 | 1 | k__Fungi(100);p__Ascomycota(100);c__Saccharomycetes(100);o__Saccharomycetales(100);f__Metschnikowiaceae(100);g__Metschnikowia(100);s__Metschnikowia_pulcherrima(100); | Metschnikowia | 13 | Saprotroph | \|Undefined Saprotroph\| | Yeast | NULL | Probable | As nectar-tap_saprotroph (Põlme et al. 2020); Decay sugar-rich substrates (Põlme et al. 2020) | Põlme S, et al. 2020. Fungal Diversity 105:1-16 ((https://doi.org/10.1007/s13225-020-00466-2)) |
| Otu35272 | 1 | k__Fungi(100);p__Ascomycota(100);c__Saccharomycetes(100);o__Saccharomycetales(100);f__Metschnikowiaceae(100);g__Metschnikowia(100);g__Metschnikowia_unclassified(100); | Metschnikowia | 13 | Saprotroph | \|Undefined Saprotroph\| | Yeast | NULL | Probable | As nectar-tap_saprotroph (Põlme et al. 2020); Decay sugar-rich substrates (Põlme et al. 2020) | Põlme S, et al. 2020. Fungal Diversity 105:1-16 ((https://doi.org/10.1007/s13225-020-00466-2)) |
| Otu35273 | 1 | k__Fungi(100);p__Ascomycota(100);c__Saccharomycetes(100);o__Saccharomycetales(100);f__Metschnikowiaceae(100);g__Metschnikowia(100);s__Metschnikowia_pulcherrima(100); | Metschnikowia | 13 | Saprotroph | \|Undefined Saprotroph\| | Yeast | NULL | Probable | As nectar-tap_saprotroph (Põlme et al. 2020); Decay sugar-rich substrates (Põlme et al. 2020) | Põlme S, et al. 2020. Fungal Diversity 105:1-16 ((https://doi.org/10.1007/s13225-020-00466-2)) |
| Otu35275 | 1 | k__Fungi(100);p__Ascomycota(100);c__Saccharomycetes(100);o__Saccharomycetales(100);f__Metschnikowiaceae(100);g__Metschnikowia(100);s__Metschnikowia_pulcherrima(100); | Metschnikowia | 13 | Saprotroph | \|Undefined Saprotroph\| | Yeast | NULL | Probable | As nectar-tap_saprotroph (Põlme et al. 2020); Decay sugar-rich substrates (Põlme et al. 2020) | Põlme S, et al. 2020. Fungal Diversity 105:1-16 ((https://doi.org/10.1007/s13225-020-00466-2)) |
| Otu35276 | 1 | k__Fungi(100);p__Ascomycota(100);c__Saccharomycetes(100);o__Saccharomycetales(100);f__Metschnikowiaceae(100);g__Metschnikowia(100);s__Metschnikowia_pulcherrima(100); | Metschnikowia | 13 | Saprotroph | \|Undefined Saprotroph\| | Yeast | NULL | Probable | As nectar-tap_saprotroph (Põlme et al. 2020); Decay sugar-rich substrates (Põlme et al. 2020) | Põlme S, et al. 2020. Fungal Diversity 105:1-16 ((https://doi.org/10.1007/s13225-020-00466-2)) |
| Otu35277 | 1 | k__Fungi(100);p__Ascomycota(100);c__Saccharomycetes(100);o__Saccharomycetales(100);f__Metschnikowiaceae(100);g__Metschnikowia(100);g__Metschnikowia_unclassified(100); | Metschnikowia | 13 | Saprotroph | \|Undefined Saprotroph\| | Yeast | NULL | Probable | As nectar-tap_saprotroph (Põlme et al. 2020); Decay sugar-rich substrates (Põlme et al. 2020) | Põlme S, et al. 2020. Fungal Diversity 105:1-16 ((https://doi.org/10.1007/s13225-020-00466-2)) |
| Otu35279 | 1 | k__Fungi(100);p__Ascomycota(100);c__Saccharomycetes(100);o__Saccharomycetales(100);f__Metschnikowiaceae(100);g__Metschnikowia(100);g__Metschnikowia_unclassified(100); | Metschnikowia | 13 | Saprotroph | \|Undefined Saprotroph\| | Yeast | NULL | Probable | As nectar-tap_saprotroph (Põlme et al. 2020); Decay sugar-rich substrates (Põlme et al. 2020) | Põlme S, et al. 2020. Fungal Diversity 105:1-16 ((https://doi.org/10.1007/s13225-020-00466-2)) |
| Otu35283 | 1 | k__Fungi(100);p__Ascomycota(100);c__Saccharomycetes(100);o__Saccharomycetales(100);f__Metschnikowiaceae(100);g__Metschnikowia(100);s__Metschnikowia_pulcherrima(100); | Metschnikowia | 13 | Saprotroph | \|Undefined Saprotroph\| | Yeast | NULL | Probable | As nectar-tap_saprotroph (Põlme et al. 2020); Decay sugar-rich substrates (Põlme et al. 2020) | Põlme S, et al. 2020. Fungal Diversity 105:1-16 ((https://doi.org/10.1007/s13225-020-00466-2)) |
| Otu35284 | 1 | k__Fungi(100);p__Ascomycota(100);c__Saccharomycetes(100);o__Saccharomycetales(100);f__Metschnikowiaceae(100);g__Metschnikowia(100);s__Metschnikowia_pulcherrima(100); | Metschnikowia | 13 | Saprotroph | \|Undefined Saprotroph\| | Yeast | NULL | Probable | As nectar-tap_saprotroph (Põlme et al. 2020); Decay sugar-rich substrates (Põlme et al. 2020) | Põlme S, et al. 2020. Fungal Diversity 105:1-16 ((https://doi.org/10.1007/s13225-020-00466-2)) |
| Otu35285 | 1 | k__Fungi(100);p__Ascomycota(100);c__Saccharomycetes(100);o__Saccharomycetales(100);f__Metschnikowiaceae(100);g__Metschnikowia(100);s__Metschnikowia_pulcherrima(100); | Metschnikowia | 13 | Saprotroph | \|Undefined Saprotroph\| | Yeast | NULL | Probable | As nectar-tap_saprotroph (Põlme et al. 2020); Decay sugar-rich substrates (Põlme et al. 2020) | Põlme S, et al. 2020. Fungal Diversity 105:1-16 ((https://doi.org/10.1007/s13225-020-00466-2)) |
| Otu35286 | 1 | k__Fungi(100);p__Ascomycota(100);c__Saccharomycetes(100);o__Saccharomycetales(100);f__Metschnikowiaceae(100);g__Metschnikowia(100);s__Metschnikowia_pulcherrima(100); | Metschnikowia | 13 | Saprotroph | \|Undefined Saprotroph\| | Yeast | NULL | Probable | As nectar-tap_saprotroph (Põlme et al. 2020); Decay sugar-rich substrates (Põlme et al. 2020) | Põlme S, et al. 2020. Fungal Diversity 105:1-16 ((https://doi.org/10.1007/s13225-020-00466-2)) |
| Otu35298 | 1 | k__Fungi(100);p__Ascomycota(100);c__Saccharomycetes(100);o__Saccharomycetales(100);f__Metschnikowiaceae(100);g__Metschnikowia(100);g__Metschnikowia_unclassified(100); | Metschnikowia | 13 | Saprotroph | \|Undefined Saprotroph\| | Yeast | NULL | Probable | As nectar-tap_saprotroph (Põlme et al. 2020); Decay sugar-rich substrates (Põlme et al. 2020) | Põlme S, et al. 2020. Fungal Diversity 105:1-16 ((https://doi.org/10.1007/s13225-020-00466-2)) |
| Otu35302 | 1 | k__Fungi(100);p__Ascomycota(100);c__Saccharomycetes(100);o__Saccharomycetales(100);f__Metschnikowiaceae(100);g__Metschnikowia(100);s__Metschnikowia_pulcherrima(100); | Metschnikowia | 13 | Saprotroph | \|Undefined Saprotroph\| | Yeast | NULL | Probable | As nectar-tap_saprotroph (Põlme et al. 2020); Decay sugar-rich substrates (Põlme et al. 2020) | Põlme S, et al. 2020. Fungal Diversity 105:1-16 ((https://doi.org/10.1007/s13225-020-00466-2)) |
| Otu35306 | 1 | k__Fungi(100);p__Ascomycota(100);c__Saccharomycetes(100);o__Saccharomycetales(100);f__Metschnikowiaceae(100);g__Metschnikowia(100);s__Metschnikowia_pulcherrima(100); | Metschnikowia | 13 | Saprotroph | \|Undefined Saprotroph\| | Yeast | NULL | Probable | As nectar-tap_saprotroph (Põlme et al. 2020); Decay sugar-rich substrates (Põlme et al. 2020) | Põlme S, et al. 2020. Fungal Diversity 105:1-16 ((https://doi.org/10.1007/s13225-020-00466-2)) |
| Otu35309 | 1 | k__Fungi(100);p__Ascomycota(100);c__Saccharomycetes(100);o__Saccharomycetales(100);f__Metschnikowiaceae(100);g__Metschnikowia(100);s__Metschnikowia_pulcherrima(100); | Metschnikowia | 13 | Saprotroph | \|Undefined Saprotroph\| | Yeast | NULL | Probable | As nectar-tap_saprotroph (Põlme et al. 2020); Decay sugar-rich substrates (Põlme et al. 2020) | Põlme S, et al. 2020. Fungal Diversity 105:1-16 ((https://doi.org/10.1007/s13225-020-00466-2)) |
| Otu35310 | 1 | k__Fungi(100);p__Ascomycota(100);c__Saccharomycetes(100);o__Saccharomycetales(100);f__Metschnikowiaceae(100);g__Metschnikowia(100);s__Metschnikowia_pulcherrima(100); | Metschnikowia | 13 | Saprotroph | \|Undefined Saprotroph\| | Yeast | NULL | Probable | As nectar-tap_saprotroph (Põlme et al. 2020); Decay sugar-rich substrates (Põlme et al. 2020) | Põlme S, et al. 2020. Fungal Diversity 105:1-16 ((https://doi.org/10.1007/s13225-020-00466-2)) |
| Otu35312 | 1 | k__Fungi(100);p__Ascomycota(100);c__Saccharomycetes(100);o__Saccharomycetales(100);f__Metschnikowiaceae(100);g__Metschnikowia(100);g__Metschnikowia_unclassified(100); | Metschnikowia | 13 | Saprotroph | \|Undefined Saprotroph\| | Yeast | NULL | Probable | As nectar-tap_saprotroph (Põlme et al. 2020); Decay sugar-rich substrates (Põlme et al. 2020) | Põlme S, et al. 2020. Fungal Diversity 105:1-16 ((https://doi.org/10.1007/s13225-020-00466-2)) |
| Otu35313 | 1 | k__Fungi(100);p__Ascomycota(100);c__Saccharomycetes(100);o__Saccharomycetales(100);f__Metschnikowiaceae(100);g__Metschnikowia(100);s__Metschnikowia_pulcherrima(100); | Metschnikowia | 13 | Saprotroph | \|Undefined Saprotroph\| | Yeast | NULL | Probable | As nectar-tap_saprotroph (Põlme et al. 2020); Decay sugar-rich substrates (Põlme et al. 2020) | Põlme S, et al. 2020. Fungal Diversity 105:1-16 ((https://doi.org/10.1007/s13225-020-00466-2)) |
| Otu35315 | 1 | k__Fungi(100);p__Ascomycota(100);c__Saccharomycetes(100);o__Saccharomycetales(100);f__Metschnikowiaceae(100);g__Metschnikowia(100);s__Metschnikowia_pulcherrima(100); | Metschnikowia | 13 | Saprotroph | \|Undefined Saprotroph\| | Yeast | NULL | Probable | As nectar-tap_saprotroph (Põlme et al. 2020); Decay sugar-rich substrates (Põlme et al. 2020) | Põlme S, et al. 2020. Fungal Diversity 105:1-16 ((https://doi.org/10.1007/s13225-020-00466-2)) |
| Otu35317 | 1 | k__Fungi(100);p__Ascomycota(100);c__Saccharomycetes(100);o__Saccharomycetales(100);f__Metschnikowiaceae(100);g__Metschnikowia(100);s__Metschnikowia_pulcherrima(100); | Metschnikowia | 13 | Saprotroph | \|Undefined Saprotroph\| | Yeast | NULL | Probable | As nectar-tap_saprotroph (Põlme et al. 2020); Decay sugar-rich substrates (Põlme et al. 2020) | Põlme S, et al. 2020. Fungal Diversity 105:1-16 ((https://doi.org/10.1007/s13225-020-00466-2)) |
| Otu35319 | 1 | k__Fungi(100);p__Ascomycota(100);c__Saccharomycetes(100);o__Saccharomycetales(100);f__Metschnikowiaceae(100);g__Metschnikowia(100);g__Metschnikowia_unclassified(100); | Metschnikowia | 13 | Saprotroph | \|Undefined Saprotroph\| | Yeast | NULL | Probable | As nectar-tap_saprotroph (Põlme et al. 2020); Decay sugar-rich substrates (Põlme et al. 2020) | Põlme S, et al. 2020. Fungal Diversity 105:1-16 ((https://doi.org/10.1007/s13225-020-00466-2)) |
| Otu35320 | 1 | k__Fungi(100);p__Ascomycota(100);c__Saccharomycetes(100);o__Saccharomycetales(100);f__Metschnikowiaceae(100);g__Metschnikowia(100);s__Metschnikowia_pulcherrima(100); | Metschnikowia | 13 | Saprotroph | \|Undefined Saprotroph\| | Yeast | NULL | Probable | As nectar-tap_saprotroph (Põlme et al. 2020); Decay sugar-rich substrates (Põlme et al. 2020) | Põlme S, et al. 2020. Fungal Diversity 105:1-16 ((https://doi.org/10.1007/s13225-020-00466-2)) |
| Otu35321 | 1 | k__Fungi(100);p__Ascomycota(100);c__Saccharomycetes(100);o__Saccharomycetales(100);f__Metschnikowiaceae(100);g__Metschnikowia(100);s__Metschnikowia_pulcherrima(100); | Metschnikowia | 13 | Saprotroph | \|Undefined Saprotroph\| | Yeast | NULL | Probable | As nectar-tap_saprotroph (Põlme et al. 2020); Decay sugar-rich substrates (Põlme et al. 2020) | Põlme S, et al. 2020. Fungal Diversity 105:1-16 ((https://doi.org/10.1007/s13225-020-00466-2)) |
| Otu35322 | 1 | k__Fungi(100);p__Ascomycota(100);c__Saccharomycetes(100);o__Saccharomycetales(100);f__Metschnikowiaceae(100);g__Metschnikowia(100);s__Metschnikowia_pulcherrima(100); | Metschnikowia | 13 | Saprotroph | \|Undefined Saprotroph\| | Yeast | NULL | Probable | As nectar-tap_saprotroph (Põlme et al. 2020); Decay sugar-rich substrates (Põlme et al. 2020) | Põlme S, et al. 2020. Fungal Diversity 105:1-16 ((https://doi.org/10.1007/s13225-020-00466-2)) |
| Otu35323 | 1 | k__Fungi(100);p__Ascomycota(100);c__Saccharomycetes(100);o__Saccharomycetales(100);f__Metschnikowiaceae(100);g__Metschnikowia(100);g__Metschnikowia_unclassified(100); | Metschnikowia | 13 | Saprotroph | \|Undefined Saprotroph\| | Yeast | NULL | Probable | As nectar-tap_saprotroph (Põlme et al. 2020); Decay sugar-rich substrates (Põlme et al. 2020) | Põlme S, et al. 2020. Fungal Diversity 105:1-16 ((https://doi.org/10.1007/s13225-020-00466-2)) |
| Otu35324 | 1 | k__Fungi(100);p__Ascomycota(100);c__Saccharomycetes(100);o__Saccharomycetales(100);f__Metschnikowiaceae(100);g__Metschnikowia(100);s__Metschnikowia_pulcherrima(100); | Metschnikowia | 13 | Saprotroph | \|Undefined Saprotroph\| | Yeast | NULL | Probable | As nectar-tap_saprotroph (Põlme et al. 2020); Decay sugar-rich substrates (Põlme et al. 2020) | Põlme S, et al. 2020. Fungal Diversity 105:1-16 ((https://doi.org/10.1007/s13225-020-00466-2)) |
| Otu35325 | 1 | k__Fungi(100);p__Basidiomycota(100);c__Microbotryomycetes(100);o__Sporidiobolales(100);f__Sporidiobolales_family_Incertae_sedis(100);g__Sporobolomyces(100);s__Sporobolomyces_lactosus(100); | Sporobolomyces | 13 | Pathotroph-Saprotroph | \|Fungal Parasite\|-Plant Saprotroph | Yeast | NULL | Probable | As litter_saprotroph (Põlme et al. 2020); As mycoparasite (Põlme et al. 2020); Decay fungal material (Põlme et al. 2020) | Tedersoo L, et al. 2014. Science 346:e1256688 ((https://doi.org/10.1126/science.1256688)); Purahong W, et al. 2016. Molecular Ecology 25:4059-4074 ((https://doi.org/10.1111/mec.13739)); Põlme S, et al. 2020. Fungal Diversity 105:1-16 ((https://doi.org/10.1007/s13225-020-00466-2)) |
| Otu35326 | 1 | k__Fungi(100);p__Ascomycota(100);c__Saccharomycetes(100);o__Saccharomycetales(100);f__Metschnikowiaceae(100);g__Metschnikowia(100);s__Metschnikowia_pulcherrima(100); | Metschnikowia | 13 | Saprotroph | \|Undefined Saprotroph\| | Yeast | NULL | Probable | As nectar-tap_saprotroph (Põlme et al. 2020); Decay sugar-rich substrates (Põlme et al. 2020) | Põlme S, et al. 2020. Fungal Diversity 105:1-16 ((https://doi.org/10.1007/s13225-020-00466-2)) |
| Otu35327 | 1 | k__Fungi(100);p__Ascomycota(100);c__Saccharomycetes(100);o__Saccharomycetales(100);f__Metschnikowiaceae(100);g__Metschnikowia(100);g__Metschnikowia_unclassified(100); | Metschnikowia | 13 | Saprotroph | \|Undefined Saprotroph\| | Yeast | NULL | Probable | As nectar-tap_saprotroph (Põlme et al. 2020); Decay sugar-rich substrates (Põlme et al. 2020) | Põlme S, et al. 2020. Fungal Diversity 105:1-16 ((https://doi.org/10.1007/s13225-020-00466-2)) |
| Otu35328 | 1 | k__Fungi(100);p__Ascomycota(100);c__Saccharomycetes(100);o__Saccharomycetales(100);f__Metschnikowiaceae(100);g__Metschnikowia(100);s__Metschnikowia_pulcherrima(100); | Metschnikowia | 13 | Saprotroph | \|Undefined Saprotroph\| | Yeast | NULL | Probable | As nectar-tap_saprotroph (Põlme et al. 2020); Decay sugar-rich substrates (Põlme et al. 2020) | Põlme S, et al. 2020. Fungal Diversity 105:1-16 ((https://doi.org/10.1007/s13225-020-00466-2)) |
| Otu35329 | 1 | k__Fungi(100);p__Ascomycota(100);c__Saccharomycetes(100);o__Saccharomycetales(100);f__Metschnikowiaceae(100);g__Metschnikowia(100);s__Metschnikowia_pulcherrima(100); | Metschnikowia | 13 | Saprotroph | \|Undefined Saprotroph\| | Yeast | NULL | Probable | As nectar-tap_saprotroph (Põlme et al. 2020); Decay sugar-rich substrates (Põlme et al. 2020) | Põlme S, et al. 2020. Fungal Diversity 105:1-16 ((https://doi.org/10.1007/s13225-020-00466-2)) |
| Otu35330 | 1 | k__Fungi(100);p__Ascomycota(100);c__Saccharomycetes(100);o__Saccharomycetales(100);f__Metschnikowiaceae(100);g__Metschnikowia(100);s__Metschnikowia_pulcherrima(100); | Metschnikowia | 13 | Saprotroph | \|Undefined Saprotroph\| | Yeast | NULL | Probable | As nectar-tap_saprotroph (Põlme et al. 2020); Decay sugar-rich substrates (Põlme et al. 2020) | Põlme S, et al. 2020. Fungal Diversity 105:1-16 ((https://doi.org/10.1007/s13225-020-00466-2)) |
| Otu35331 | 1 | k__Fungi(100);p__Ascomycota(100);c__Saccharomycetes(100);o__Saccharomycetales(100);f__Metschnikowiaceae(100);g__Metschnikowia(100);s__Metschnikowia_pulcherrima(100); | Metschnikowia | 13 | Saprotroph | \|Undefined Saprotroph\| | Yeast | NULL | Probable | As nectar-tap_saprotroph (Põlme et al. 2020); Decay sugar-rich substrates (Põlme et al. 2020) | Põlme S, et al. 2020. Fungal Diversity 105:1-16 ((https://doi.org/10.1007/s13225-020-00466-2)) |
| Otu35332 | 1 | k__Fungi(100);p__Ascomycota(100);c__Saccharomycetes(100);o__Saccharomycetales(100);f__Metschnikowiaceae(100);g__Metschnikowia(100);s__Metschnikowia_pulcherrima(100); | Metschnikowia | 13 | Saprotroph | \|Undefined Saprotroph\| | Yeast | NULL | Probable | As nectar-tap_saprotroph (Põlme et al. 2020); Decay sugar-rich substrates (Põlme et al. 2020) | Põlme S, et al. 2020. Fungal Diversity 105:1-16 ((https://doi.org/10.1007/s13225-020-00466-2)) |
| Otu35335 | 1 | k__Fungi(100);p__Ascomycota(100);c__Saccharomycetes(100);o__Saccharomycetales(100);f__Metschnikowiaceae(100);g__Metschnikowia(100);g__Metschnikowia_unclassified(100); | Metschnikowia | 13 | Saprotroph | \|Undefined Saprotroph\| | Yeast | NULL | Probable | As nectar-tap_saprotroph (Põlme et al. 2020); Decay sugar-rich substrates (Põlme et al. 2020) | Põlme S, et al. 2020. Fungal Diversity 105:1-16 ((https://doi.org/10.1007/s13225-020-00466-2)) |
| Otu35336 | 1 | k__Fungi(100);p__Ascomycota(100);c__Saccharomycetes(100);o__Saccharomycetales(100);f__Metschnikowiaceae(100);g__Metschnikowia(100);s__Metschnikowia_pulcherrima(100); | Metschnikowia | 13 | Saprotroph | \|Undefined Saprotroph\| | Yeast | NULL | Probable | As nectar-tap_saprotroph (Põlme et al. 2020); Decay sugar-rich substrates (Põlme et al. 2020) | Põlme S, et al. 2020. Fungal Diversity 105:1-16 ((https://doi.org/10.1007/s13225-020-00466-2)) |
| Otu35339 | 1 | k__Fungi(100);p__Ascomycota(100);c__Saccharomycetes(100);o__Saccharomycetales(100);f__Metschnikowiaceae(100);g__Metschnikowia(100);s__Metschnikowia_pulcherrima(100); | Metschnikowia | 13 | Saprotroph | \|Undefined Saprotroph\| | Yeast | NULL | Probable | As nectar-tap_saprotroph (Põlme et al. 2020); Decay sugar-rich substrates (Põlme et al. 2020) | Põlme S, et al. 2020. Fungal Diversity 105:1-16 ((https://doi.org/10.1007/s13225-020-00466-2)) |
| Otu35343 | 1 | k__Fungi(100);p__Ascomycota(100);c__Saccharomycetes(100);o__Saccharomycetales(100);f__Metschnikowiaceae(100);g__Metschnikowia(100);s__Metschnikowia_pulcherrima(100); | Metschnikowia | 13 | Saprotroph | \|Undefined Saprotroph\| | Yeast | NULL | Probable | As nectar-tap_saprotroph (Põlme et al. 2020); Decay sugar-rich substrates (Põlme et al. 2020) | Põlme S, et al. 2020. Fungal Diversity 105:1-16 ((https://doi.org/10.1007/s13225-020-00466-2)) |
| Otu35344 | 1 | k__Fungi(100);p__Ascomycota(100);c__Saccharomycetes(100);o__Saccharomycetales(100);f__Metschnikowiaceae(100);g__Metschnikowia(100);s__Metschnikowia_pulcherrima(100); | Metschnikowia | 13 | Saprotroph | \|Undefined Saprotroph\| | Yeast | NULL | Probable | As nectar-tap_saprotroph (Põlme et al. 2020); Decay sugar-rich substrates (Põlme et al. 2020) | Põlme S, et al. 2020. Fungal Diversity 105:1-16 ((https://doi.org/10.1007/s13225-020-00466-2)) |
| Otu35345 | 1 | k__Fungi(100);p__Ascomycota(100);c__Saccharomycetes(100);o__Saccharomycetales(100);f__Metschnikowiaceae(100);g__Metschnikowia(100);g__Metschnikowia_unclassified(100); | Metschnikowia | 13 | Saprotroph | \|Undefined Saprotroph\| | Yeast | NULL | Probable | As nectar-tap_saprotroph (Põlme et al. 2020); Decay sugar-rich substrates (Põlme et al. 2020) | Põlme S, et al. 2020. Fungal Diversity 105:1-16 ((https://doi.org/10.1007/s13225-020-00466-2)) |
| Otu35349 | 1 | k__Fungi(100);p__Ascomycota(100);c__Saccharomycetes(100);o__Saccharomycetales(100);f__Saccharomycetales_family_Incertae_sedis(100);f__Saccharomycetales_family_Incertae_sedis_unclassified(100);f__Saccharomycetales_family_Incertae_sedis_unclassified(100); | Saccharomycetales | 7 | Saprotroph | Undefined Saprotroph | Yeast | NULL | Possible | NULL | Sterkenburg E, et al. 2015. New Phytologist 207:1145-1158 ((https://doi.org/10.1111/nph.13426)) |
| Otu35351 | 1 | k__Fungi(100);p__Ascomycota(100);c__Saccharomycetes(100);o__Saccharomycetales(100);f__Metschnikowiaceae(100);g__Metschnikowia(100);s__Metschnikowia_pulcherrima(100); | Metschnikowia | 13 | Saprotroph | \|Undefined Saprotroph\| | Yeast | NULL | Probable | As nectar-tap_saprotroph (Põlme et al. 2020); Decay sugar-rich substrates (Põlme et al. 2020) | Põlme S, et al. 2020. Fungal Diversity 105:1-16 ((https://doi.org/10.1007/s13225-020-00466-2)) |
| Otu35356 | 1 | k__Fungi(100);p__Ascomycota(100);c__Saccharomycetes(100);o__Saccharomycetales(100);f__Metschnikowiaceae(100);g__Metschnikowia(100);g__Metschnikowia_unclassified(100); | Metschnikowia | 13 | Saprotroph | \|Undefined Saprotroph\| | Yeast | NULL | Probable | As nectar-tap_saprotroph (Põlme et al. 2020); Decay sugar-rich substrates (Põlme et al. 2020) | Põlme S, et al. 2020. Fungal Diversity 105:1-16 ((https://doi.org/10.1007/s13225-020-00466-2)) |
| Otu35357 | 1 | k__Fungi(100);p__Ascomycota(100);c__Saccharomycetes(100);o__Saccharomycetales(100);f__Metschnikowiaceae(100);g__Metschnikowia(100);g__Metschnikowia_unclassified(100); | Metschnikowia | 13 | Saprotroph | \|Undefined Saprotroph\| | Yeast | NULL | Probable | As nectar-tap_saprotroph (Põlme et al. 2020); Decay sugar-rich substrates (Põlme et al. 2020) | Põlme S, et al. 2020. Fungal Diversity 105:1-16 ((https://doi.org/10.1007/s13225-020-00466-2)) |
| Otu35358 | 1 | k__Fungi(100);p__Ascomycota(100);c__Saccharomycetes(100);o__Saccharomycetales(100);f__Metschnikowiaceae(100);g__Metschnikowia(100);s__Metschnikowia_pulcherrima(100); | Metschnikowia | 13 | Saprotroph | \|Undefined Saprotroph\| | Yeast | NULL | Probable | As nectar-tap_saprotroph (Põlme et al. 2020); Decay sugar-rich substrates (Põlme et al. 2020) | Põlme S, et al. 2020. Fungal Diversity 105:1-16 ((https://doi.org/10.1007/s13225-020-00466-2)) |
| Otu35359 | 1 | k__Fungi(100);p__Ascomycota(100);c__Saccharomycetes(100);o__Saccharomycetales(100);f__Metschnikowiaceae(100);g__Metschnikowia(100);s__Metschnikowia_pulcherrima(100); | Metschnikowia | 13 | Saprotroph | \|Undefined Saprotroph\| | Yeast | NULL | Probable | As nectar-tap_saprotroph (Põlme et al. 2020); Decay sugar-rich substrates (Põlme et al. 2020) | Põlme S, et al. 2020. Fungal Diversity 105:1-16 ((https://doi.org/10.1007/s13225-020-00466-2)) |
| Otu35360 | 1 | k__Fungi(100);p__Ascomycota(100);c__Saccharomycetes(100);o__Saccharomycetales(100);f__Metschnikowiaceae(100);g__Metschnikowia(100);s__Metschnikowia_pulcherrima(100); | Metschnikowia | 13 | Saprotroph | \|Undefined Saprotroph\| | Yeast | NULL | Probable | As nectar-tap_saprotroph (Põlme et al. 2020); Decay sugar-rich substrates (Põlme et al. 2020) | Põlme S, et al. 2020. Fungal Diversity 105:1-16 ((https://doi.org/10.1007/s13225-020-00466-2)) |
| Otu35361 | 1 | k__Fungi(100);p__Ascomycota(100);c__Saccharomycetes(100);o__Saccharomycetales(100);f__Metschnikowiaceae(100);g__Metschnikowia(100);s__Metschnikowia_pulcherrima(100); | Metschnikowia | 13 | Saprotroph | \|Undefined Saprotroph\| | Yeast | NULL | Probable | As nectar-tap_saprotroph (Põlme et al. 2020); Decay sugar-rich substrates (Põlme et al. 2020) | Põlme S, et al. 2020. Fungal Diversity 105:1-16 ((https://doi.org/10.1007/s13225-020-00466-2)) |
| Otu35362 | 1 | k__Fungi(100);p__Ascomycota(100);c__Saccharomycetes(100);o__Saccharomycetales(100);f__Metschnikowiaceae(100);g__Metschnikowia(100);s__Metschnikowia_pulcherrima(100); | Metschnikowia | 13 | Saprotroph | \|Undefined Saprotroph\| | Yeast | NULL | Probable | As nectar-tap_saprotroph (Põlme et al. 2020); Decay sugar-rich substrates (Põlme et al. 2020) | Põlme S, et al. 2020. Fungal Diversity 105:1-16 ((https://doi.org/10.1007/s13225-020-00466-2)) |
| Otu35363 | 1 | k__Fungi(100);p__Ascomycota(100);c__Saccharomycetes(100);o__Saccharomycetales(100);f__Metschnikowiaceae(100);g__Metschnikowia(100);g__Metschnikowia_unclassified(100); | Metschnikowia | 13 | Saprotroph | \|Undefined Saprotroph\| | Yeast | NULL | Probable | As nectar-tap_saprotroph (Põlme et al. 2020); Decay sugar-rich substrates (Põlme et al. 2020) | Põlme S, et al. 2020. Fungal Diversity 105:1-16 ((https://doi.org/10.1007/s13225-020-00466-2)) |
| Otu35364 | 1 | k__Fungi(100);p__Ascomycota(100);c__Saccharomycetes(100);o__Saccharomycetales(100);f__Metschnikowiaceae(100);g__Metschnikowia(100);s__Metschnikowia_pulcherrima(100); | Metschnikowia | 13 | Saprotroph | \|Undefined Saprotroph\| | Yeast | NULL | Probable | As nectar-tap_saprotroph (Põlme et al. 2020); Decay sugar-rich substrates (Põlme et al. 2020) | Põlme S, et al. 2020. Fungal Diversity 105:1-16 ((https://doi.org/10.1007/s13225-020-00466-2)) |
| Otu35365 | 1 | k__Fungi(100);p__Ascomycota(100);c__Saccharomycetes(100);o__Saccharomycetales(100);f__Metschnikowiaceae(100);g__Metschnikowia(100);s__Metschnikowia_pulcherrima(100); | Metschnikowia | 13 | Saprotroph | \|Undefined Saprotroph\| | Yeast | NULL | Probable | As nectar-tap_saprotroph (Põlme et al. 2020); Decay sugar-rich substrates (Põlme et al. 2020) | Põlme S, et al. 2020. Fungal Diversity 105:1-16 ((https://doi.org/10.1007/s13225-020-00466-2)) |
| Otu35370 | 1 | k__Fungi(100);p__Ascomycota(100);c__Saccharomycetes(100);o__Saccharomycetales(100);f__Metschnikowiaceae(100);g__Metschnikowia(100);g__Metschnikowia_unclassified(100); | Metschnikowia | 13 | Saprotroph | \|Undefined Saprotroph\| | Yeast | NULL | Probable | As nectar-tap_saprotroph (Põlme et al. 2020); Decay sugar-rich substrates (Põlme et al. 2020) | Põlme S, et al. 2020. Fungal Diversity 105:1-16 ((https://doi.org/10.1007/s13225-020-00466-2)) |
| Otu35373 | 1 | k__Fungi(100);p__Ascomycota(100);c__Saccharomycetes(100);o__Saccharomycetales(100);f__Metschnikowiaceae(100);g__Metschnikowia(100);g__Metschnikowia_unclassified(100); | Metschnikowia | 13 | Saprotroph | \|Undefined Saprotroph\| | Yeast | NULL | Probable | As nectar-tap_saprotroph (Põlme et al. 2020); Decay sugar-rich substrates (Põlme et al. 2020) | Põlme S, et al. 2020. Fungal Diversity 105:1-16 ((https://doi.org/10.1007/s13225-020-00466-2)) |
| Otu35377 | 1 | k__Fungi(100);p__Ascomycota(100);c__Saccharomycetes(100);o__Saccharomycetales(100);f__Metschnikowiaceae(100);g__Metschnikowia(100);s__Metschnikowia_pulcherrima(100); | Metschnikowia | 13 | Saprotroph | \|Undefined Saprotroph\| | Yeast | NULL | Probable | As nectar-tap_saprotroph (Põlme et al. 2020); Decay sugar-rich substrates (Põlme et al. 2020) | Põlme S, et al. 2020. Fungal Diversity 105:1-16 ((https://doi.org/10.1007/s13225-020-00466-2)) |
| Otu35379 | 1 | k__Fungi(100);p__Ascomycota(100);c__Saccharomycetes(100);o__Saccharomycetales(100);f__Metschnikowiaceae(100);g__Metschnikowia(100);g__Metschnikowia_unclassified(100); | Metschnikowia | 13 | Saprotroph | \|Undefined Saprotroph\| | Yeast | NULL | Probable | As nectar-tap_saprotroph (Põlme et al. 2020); Decay sugar-rich substrates (Põlme et al. 2020) | Põlme S, et al. 2020. Fungal Diversity 105:1-16 ((https://doi.org/10.1007/s13225-020-00466-2)) |
| Otu35381 | 1 | k__Fungi(100);p__Ascomycota(100);c__Saccharomycetes(100);o__Saccharomycetales(100);o__Saccharomycetales_unclassified(100);o__Saccharomycetales_unclassified(100);o__Saccharomycetales_unclassified(100); | Saccharomycetales | 7 | Saprotroph | Undefined Saprotroph | Yeast | NULL | Possible | NULL | Sterkenburg E, et al. 2015. New Phytologist 207:1145-1158 ((https://doi.org/10.1111/nph.13426)) |
| Otu35385 | 1 | k__Fungi(100);p__Ascomycota(100);c__Saccharomycetes(100);o__Saccharomycetales(100);f__Metschnikowiaceae(100);g__Metschnikowia(100);g__Metschnikowia_unclassified(100); | Metschnikowia | 13 | Saprotroph | \|Undefined Saprotroph\| | Yeast | NULL | Probable | As nectar-tap_saprotroph (Põlme et al. 2020); Decay sugar-rich substrates (Põlme et al. 2020) | Põlme S, et al. 2020. Fungal Diversity 105:1-16 ((https://doi.org/10.1007/s13225-020-00466-2)) |
| Otu35388 | 1 | k__Fungi(100);p__Ascomycota(100);c__Saccharomycetes(100);o__Saccharomycetales(100);f__Metschnikowiaceae(100);g__Metschnikowia(100);g__Metschnikowia_unclassified(100); | Metschnikowia | 13 | Saprotroph | \|Undefined Saprotroph\| | Yeast | NULL | Probable | As nectar-tap_saprotroph (Põlme et al. 2020); Decay sugar-rich substrates (Põlme et al. 2020) | Põlme S, et al. 2020. Fungal Diversity 105:1-16 ((https://doi.org/10.1007/s13225-020-00466-2)) |
| Otu35390 | 1 | k__Fungi(100);p__Ascomycota(100);c__Saccharomycetes(100);o__Saccharomycetales(100);f__Metschnikowiaceae(100);g__Metschnikowia(100);s__Metschnikowia_pulcherrima(100); | Metschnikowia | 13 | Saprotroph | \|Undefined Saprotroph\| | Yeast | NULL | Probable | As nectar-tap_saprotroph (Põlme et al. 2020); Decay sugar-rich substrates (Põlme et al. 2020) | Põlme S, et al. 2020. Fungal Diversity 105:1-16 ((https://doi.org/10.1007/s13225-020-00466-2)) |
| Otu35395 | 1 | k__Fungi(100);p__Ascomycota(100);c__Saccharomycetes(100);o__Saccharomycetales(100);f__Metschnikowiaceae(100);g__Metschnikowia(100);s__Metschnikowia_pulcherrima(100); | Metschnikowia | 13 | Saprotroph | \|Undefined Saprotroph\| | Yeast | NULL | Probable | As nectar-tap_saprotroph (Põlme et al. 2020); Decay sugar-rich substrates (Põlme et al. 2020) | Põlme S, et al. 2020. Fungal Diversity 105:1-16 ((https://doi.org/10.1007/s13225-020-00466-2)) |
| Otu35396 | 1 | k__Fungi(100);p__Ascomycota(100);c__Saccharomycetes(100);o__Saccharomycetales(100);f__Metschnikowiaceae(100);g__Metschnikowia(100);s__Metschnikowia_pulcherrima(100); | Metschnikowia | 13 | Saprotroph | \|Undefined Saprotroph\| | Yeast | NULL | Probable | As nectar-tap_saprotroph (Põlme et al. 2020); Decay sugar-rich substrates (Põlme et al. 2020) | Põlme S, et al. 2020. Fungal Diversity 105:1-16 ((https://doi.org/10.1007/s13225-020-00466-2)) |
| Otu35411 | 1 | k__Fungi(100);p__Ascomycota(100);c__Saccharomycetes(100);o__Saccharomycetales(100);f__Metschnikowiaceae(100);g__Metschnikowia(100);g__Metschnikowia_unclassified(100); | Metschnikowia | 13 | Saprotroph | \|Undefined Saprotroph\| | Yeast | NULL | Probable | As nectar-tap_saprotroph (Põlme et al. 2020); Decay sugar-rich substrates (Põlme et al. 2020) | Põlme S, et al. 2020. Fungal Diversity 105:1-16 ((https://doi.org/10.1007/s13225-020-00466-2)) |
| Otu35412 | 1 | k__Fungi(100);p__Ascomycota(100);c__Saccharomycetes(100);o__Saccharomycetales(100);f__Metschnikowiaceae(100);g__Metschnikowia(100);s__Metschnikowia_pulcherrima(100); | Metschnikowia | 13 | Saprotroph | \|Undefined Saprotroph\| | Yeast | NULL | Probable | As nectar-tap_saprotroph (Põlme et al. 2020); Decay sugar-rich substrates (Põlme et al. 2020) | Põlme S, et al. 2020. Fungal Diversity 105:1-16 ((https://doi.org/10.1007/s13225-020-00466-2)) |
| Otu35417 | 1 | k__Fungi(100);p__Ascomycota(100);c__Saccharomycetes(100);o__Saccharomycetales(100);f__Metschnikowiaceae(100);g__Metschnikowia(100);s__Metschnikowia_pulcherrima(100); | Metschnikowia | 13 | Saprotroph | \|Undefined Saprotroph\| | Yeast | NULL | Probable | As nectar-tap_saprotroph (Põlme et al. 2020); Decay sugar-rich substrates (Põlme et al. 2020) | Põlme S, et al. 2020. Fungal Diversity 105:1-16 ((https://doi.org/10.1007/s13225-020-00466-2)) |
| Otu35418 | 1 | k__Fungi(100);p__Ascomycota(100);c__Saccharomycetes(100);o__Saccharomycetales(100);f__Metschnikowiaceae(100);g__Metschnikowia(100);s__Metschnikowia_pulcherrima(100); | Metschnikowia | 13 | Saprotroph | \|Undefined Saprotroph\| | Yeast | NULL | Probable | As nectar-tap_saprotroph (Põlme et al. 2020); Decay sugar-rich substrates (Põlme et al. 2020) | Põlme S, et al. 2020. Fungal Diversity 105:1-16 ((https://doi.org/10.1007/s13225-020-00466-2)) |
| Otu35419 | 1 | k__Fungi(100);p__Ascomycota(100);c__Saccharomycetes(100);o__Saccharomycetales(100);f__Metschnikowiaceae(100);g__Metschnikowia(100);s__Metschnikowia_pulcherrima(100); | Metschnikowia | 13 | Saprotroph | \|Undefined Saprotroph\| | Yeast | NULL | Probable | As nectar-tap_saprotroph (Põlme et al. 2020); Decay sugar-rich substrates (Põlme et al. 2020) | Põlme S, et al. 2020. Fungal Diversity 105:1-16 ((https://doi.org/10.1007/s13225-020-00466-2)) |
| Otu35420 | 1 | k__Fungi(100);p__Ascomycota(100);c__Saccharomycetes(100);o__Saccharomycetales(100);f__Metschnikowiaceae(100);g__Metschnikowia(100);s__Metschnikowia_pulcherrima(100); | Metschnikowia | 13 | Saprotroph | \|Undefined Saprotroph\| | Yeast | NULL | Probable | As nectar-tap_saprotroph (Põlme et al. 2020); Decay sugar-rich substrates (Põlme et al. 2020) | Põlme S, et al. 2020. Fungal Diversity 105:1-16 ((https://doi.org/10.1007/s13225-020-00466-2)) |
| Otu35421 | 1 | k__Fungi(100);p__Ascomycota(100);c__Saccharomycetes(100);o__Saccharomycetales(100);f__Metschnikowiaceae(100);g__Metschnikowia(100);s__Metschnikowia_pulcherrima(100); | Metschnikowia | 13 | Saprotroph | \|Undefined Saprotroph\| | Yeast | NULL | Probable | As nectar-tap_saprotroph (Põlme et al. 2020); Decay sugar-rich substrates (Põlme et al. 2020) | Põlme S, et al. 2020. Fungal Diversity 105:1-16 ((https://doi.org/10.1007/s13225-020-00466-2)) |
| Otu35423 | 1 | k__Fungi(100);p__Ascomycota(100);c__Saccharomycetes(100);o__Saccharomycetales(100);f__Metschnikowiaceae(100);g__Metschnikowia(100);s__Metschnikowia_pulcherrima(100); | Metschnikowia | 13 | Saprotroph | \|Undefined Saprotroph\| | Yeast | NULL | Probable | As nectar-tap_saprotroph (Põlme et al. 2020); Decay sugar-rich substrates (Põlme et al. 2020) | Põlme S, et al. 2020. Fungal Diversity 105:1-16 ((https://doi.org/10.1007/s13225-020-00466-2)) |
| Otu35424 | 1 | k__Fungi(100);p__Ascomycota(100);c__Saccharomycetes(100);o__Saccharomycetales(100);f__Metschnikowiaceae(100);g__Metschnikowia(100);s__Metschnikowia_pulcherrima(100); | Metschnikowia | 13 | Saprotroph | \|Undefined Saprotroph\| | Yeast | NULL | Probable | As nectar-tap_saprotroph (Põlme et al. 2020); Decay sugar-rich substrates (Põlme et al. 2020) | Põlme S, et al. 2020. Fungal Diversity 105:1-16 ((https://doi.org/10.1007/s13225-020-00466-2)) |
| Otu35425 | 1 | k__Fungi(100);p__Ascomycota(100);c__Saccharomycetes(100);o__Saccharomycetales(100);f__Metschnikowiaceae(100);g__Metschnikowia(100);s__Metschnikowia_pulcherrima(100); | Metschnikowia | 13 | Saprotroph | \|Undefined Saprotroph\| | Yeast | NULL | Probable | As nectar-tap_saprotroph (Põlme et al. 2020); Decay sugar-rich substrates (Põlme et al. 2020) | Põlme S, et al. 2020. Fungal Diversity 105:1-16 ((https://doi.org/10.1007/s13225-020-00466-2)) |
| Otu35426 | 1 | k__Fungi(100);p__Ascomycota(100);c__Saccharomycetes(100);o__Saccharomycetales(100);f__Metschnikowiaceae(100);g__Metschnikowia(100);s__Metschnikowia_pulcherrima(100); | Metschnikowia | 13 | Saprotroph | \|Undefined Saprotroph\| | Yeast | NULL | Probable | As nectar-tap_saprotroph (Põlme et al. 2020); Decay sugar-rich substrates (Põlme et al. 2020) | Põlme S, et al. 2020. Fungal Diversity 105:1-16 ((https://doi.org/10.1007/s13225-020-00466-2)) |
| Otu35427 | 1 | k__Fungi(100);p__Ascomycota(100);c__Saccharomycetes(100);o__Saccharomycetales(100);f__Metschnikowiaceae(100);g__Metschnikowia(100);s__Metschnikowia_pulcherrima(100); | Metschnikowia | 13 | Saprotroph | \|Undefined Saprotroph\| | Yeast | NULL | Probable | As nectar-tap_saprotroph (Põlme et al. 2020); Decay sugar-rich substrates (Põlme et al. 2020) | Põlme S, et al. 2020. Fungal Diversity 105:1-16 ((https://doi.org/10.1007/s13225-020-00466-2)) |
| Otu35429 | 1 | k__Fungi(100);p__Ascomycota(100);c__Saccharomycetes(100);o__Saccharomycetales(100);f__Metschnikowiaceae(100);g__Metschnikowia(100);s__Metschnikowia_pulcherrima(100); | Metschnikowia | 13 | Saprotroph | \|Undefined Saprotroph\| | Yeast | NULL | Probable | As nectar-tap_saprotroph (Põlme et al. 2020); Decay sugar-rich substrates (Põlme et al. 2020) | Põlme S, et al. 2020. Fungal Diversity 105:1-16 ((https://doi.org/10.1007/s13225-020-00466-2)) |
| Otu35430 | 1 | k__Fungi(100);p__Ascomycota(100);c__Saccharomycetes(100);o__Saccharomycetales(100);f__Metschnikowiaceae(100);g__Metschnikowia(100);s__Metschnikowia_pulcherrima(100); | Metschnikowia | 13 | Saprotroph | \|Undefined Saprotroph\| | Yeast | NULL | Probable | As nectar-tap_saprotroph (Põlme et al. 2020); Decay sugar-rich substrates (Põlme et al. 2020) | Põlme S, et al. 2020. Fungal Diversity 105:1-16 ((https://doi.org/10.1007/s13225-020-00466-2)) |
| Otu35431 | 1 | k__Fungi(100);p__Ascomycota(100);c__Saccharomycetes(100);o__Saccharomycetales(100);f__Metschnikowiaceae(100);g__Metschnikowia(100);s__Metschnikowia_pulcherrima(100); | Metschnikowia | 13 | Saprotroph | \|Undefined Saprotroph\| | Yeast | NULL | Probable | As nectar-tap_saprotroph (Põlme et al. 2020); Decay sugar-rich substrates (Põlme et al. 2020) | Põlme S, et al. 2020. Fungal Diversity 105:1-16 ((https://doi.org/10.1007/s13225-020-00466-2)) |
| Otu35432 | 1 | k__Fungi(100);p__Ascomycota(100);c__Saccharomycetes(100);o__Saccharomycetales(100);f__Metschnikowiaceae(100);g__Metschnikowia(100);s__Metschnikowia_pulcherrima(100); | Metschnikowia | 13 | Saprotroph | \|Undefined Saprotroph\| | Yeast | NULL | Probable | As nectar-tap_saprotroph (Põlme et al. 2020); Decay sugar-rich substrates (Põlme et al. 2020) | Põlme S, et al. 2020. Fungal Diversity 105:1-16 ((https://doi.org/10.1007/s13225-020-00466-2)) |
| Otu35433 | 1 | k__Fungi(100);p__Ascomycota(100);c__Saccharomycetes(100);o__Saccharomycetales(100);f__Metschnikowiaceae(100);g__Metschnikowia(100);g__Metschnikowia_unclassified(100); | Metschnikowia | 13 | Saprotroph | \|Undefined Saprotroph\| | Yeast | NULL | Probable | As nectar-tap_saprotroph (Põlme et al. 2020); Decay sugar-rich substrates (Põlme et al. 2020) | Põlme S, et al. 2020. Fungal Diversity 105:1-16 ((https://doi.org/10.1007/s13225-020-00466-2)) |
| Otu35434 | 1 | k__Fungi(100);p__Ascomycota(100);c__Saccharomycetes(100);o__Saccharomycetales(100);f__Metschnikowiaceae(100);g__Metschnikowia(100);s__Metschnikowia_pulcherrima(100); | Metschnikowia | 13 | Saprotroph | \|Undefined Saprotroph\| | Yeast | NULL | Probable | As nectar-tap_saprotroph (Põlme et al. 2020); Decay sugar-rich substrates (Põlme et al. 2020) | Põlme S, et al. 2020. Fungal Diversity 105:1-16 ((https://doi.org/10.1007/s13225-020-00466-2)) |
| Otu35435 | 1 | k__Fungi(100);p__Ascomycota(100);c__Saccharomycetes(100);o__Saccharomycetales(100);f__Metschnikowiaceae(100);g__Metschnikowia(100);s__Metschnikowia_pulcherrima(100); | Metschnikowia | 13 | Saprotroph | \|Undefined Saprotroph\| | Yeast | NULL | Probable | As nectar-tap_saprotroph (Põlme et al. 2020); Decay sugar-rich substrates (Põlme et al. 2020) | Põlme S, et al. 2020. Fungal Diversity 105:1-16 ((https://doi.org/10.1007/s13225-020-00466-2)) |
| Otu35436 | 1 | k__Fungi(100);p__Ascomycota(100);c__Saccharomycetes(100);o__Saccharomycetales(100);f__Metschnikowiaceae(100);g__Metschnikowia(100);g__Metschnikowia_unclassified(100); | Metschnikowia | 13 | Saprotroph | \|Undefined Saprotroph\| | Yeast | NULL | Probable | As nectar-tap_saprotroph (Põlme et al. 2020); Decay sugar-rich substrates (Põlme et al. 2020) | Põlme S, et al. 2020. Fungal Diversity 105:1-16 ((https://doi.org/10.1007/s13225-020-00466-2)) |
| Otu35437 | 1 | k__Fungi(100);p__Ascomycota(100);c__Saccharomycetes(100);o__Saccharomycetales(100);f__Metschnikowiaceae(100);g__Metschnikowia(100);s__Metschnikowia_pulcherrima(100); | Metschnikowia | 13 | Saprotroph | \|Undefined Saprotroph\| | Yeast | NULL | Probable | As nectar-tap_saprotroph (Põlme et al. 2020); Decay sugar-rich substrates (Põlme et al. 2020) | Põlme S, et al. 2020. Fungal Diversity 105:1-16 ((https://doi.org/10.1007/s13225-020-00466-2)) |
| Otu35438 | 1 | k__Fungi(100);p__Ascomycota(100);c__Saccharomycetes(100);o__Saccharomycetales(100);f__Metschnikowiaceae(100);g__Metschnikowia(100);s__Metschnikowia_pulcherrima(100); | Metschnikowia | 13 | Saprotroph | \|Undefined Saprotroph\| | Yeast | NULL | Probable | As nectar-tap_saprotroph (Põlme et al. 2020); Decay sugar-rich substrates (Põlme et al. 2020) | Põlme S, et al. 2020. Fungal Diversity 105:1-16 ((https://doi.org/10.1007/s13225-020-00466-2)) |
| Otu35439 | 1 | k__Fungi(100);p__Ascomycota(100);c__Saccharomycetes(100);o__Saccharomycetales(100);f__Metschnikowiaceae(100);g__Metschnikowia(100);s__Metschnikowia_pulcherrima(100); | Metschnikowia | 13 | Saprotroph | \|Undefined Saprotroph\| | Yeast | NULL | Probable | As nectar-tap_saprotroph (Põlme et al. 2020); Decay sugar-rich substrates (Põlme et al. 2020) | Põlme S, et al. 2020. Fungal Diversity 105:1-16 ((https://doi.org/10.1007/s13225-020-00466-2)) |
| Otu35441 | 1 | k__Fungi(100);p__Ascomycota(100);c__Saccharomycetes(100);o__Saccharomycetales(100);f__Metschnikowiaceae(100);g__Metschnikowia(100);s__Metschnikowia_pulcherrima(100); | Metschnikowia | 13 | Saprotroph | \|Undefined Saprotroph\| | Yeast | NULL | Probable | As nectar-tap_saprotroph (Põlme et al. 2020); Decay sugar-rich substrates (Põlme et al. 2020) | Põlme S, et al. 2020. Fungal Diversity 105:1-16 ((https://doi.org/10.1007/s13225-020-00466-2)) |
| Otu35442 | 1 | k__Fungi(100);p__Ascomycota(100);c__Saccharomycetes(100);o__Saccharomycetales(100);f__Metschnikowiaceae(100);g__Metschnikowia(100);s__Metschnikowia_pulcherrima(100); | Metschnikowia | 13 | Saprotroph | \|Undefined Saprotroph\| | Yeast | NULL | Probable | As nectar-tap_saprotroph (Põlme et al. 2020); Decay sugar-rich substrates (Põlme et al. 2020) | Põlme S, et al. 2020. Fungal Diversity 105:1-16 ((https://doi.org/10.1007/s13225-020-00466-2)) |
| Otu35443 | 1 | k__Fungi(100);p__Ascomycota(100);c__Saccharomycetes(100);o__Saccharomycetales(100);f__Metschnikowiaceae(100);g__Metschnikowia(100);s__Metschnikowia_pulcherrima(100); | Metschnikowia | 13 | Saprotroph | \|Undefined Saprotroph\| | Yeast | NULL | Probable | As nectar-tap_saprotroph (Põlme et al. 2020); Decay sugar-rich substrates (Põlme et al. 2020) | Põlme S, et al. 2020. Fungal Diversity 105:1-16 ((https://doi.org/10.1007/s13225-020-00466-2)) |
| Otu35445 | 1 | k__Fungi(100);p__Ascomycota(100);c__Saccharomycetes(100);o__Saccharomycetales(100);f__Metschnikowiaceae(100);g__Metschnikowia(100);s__Metschnikowia_pulcherrima(100); | Metschnikowia | 13 | Saprotroph | \|Undefined Saprotroph\| | Yeast | NULL | Probable | As nectar-tap_saprotroph (Põlme et al. 2020); Decay sugar-rich substrates (Põlme et al. 2020) | Põlme S, et al. 2020. Fungal Diversity 105:1-16 ((https://doi.org/10.1007/s13225-020-00466-2)) |
| Otu35446 | 1 | k__Fungi(100);p__Ascomycota(100);c__Saccharomycetes(100);o__Saccharomycetales(100);f__Metschnikowiaceae(100);g__Metschnikowia(100);g__Metschnikowia_unclassified(100); | Metschnikowia | 13 | Saprotroph | \|Undefined Saprotroph\| | Yeast | NULL | Probable | As nectar-tap_saprotroph (Põlme et al. 2020); Decay sugar-rich substrates (Põlme et al. 2020) | Põlme S, et al. 2020. Fungal Diversity 105:1-16 ((https://doi.org/10.1007/s13225-020-00466-2)) |
| Otu35448 | 1 | k__Fungi(100);p__Ascomycota(100);c__Saccharomycetes(100);o__Saccharomycetales(100);f__Metschnikowiaceae(100);g__Metschnikowia(100);s__Metschnikowia_pulcherrima(100); | Metschnikowia | 13 | Saprotroph | \|Undefined Saprotroph\| | Yeast | NULL | Probable | As nectar-tap_saprotroph (Põlme et al. 2020); Decay sugar-rich substrates (Põlme et al. 2020) | Põlme S, et al. 2020. Fungal Diversity 105:1-16 ((https://doi.org/10.1007/s13225-020-00466-2)) |
| Otu35449 | 1 | k__Fungi(100);p__Ascomycota(100);c__Saccharomycetes(100);o__Saccharomycetales(100);f__Metschnikowiaceae(100);g__Metschnikowia(100);s__Metschnikowia_pulcherrima(100); | Metschnikowia | 13 | Saprotroph | \|Undefined Saprotroph\| | Yeast | NULL | Probable | As nectar-tap_saprotroph (Põlme et al. 2020); Decay sugar-rich substrates (Põlme et al. 2020) | Põlme S, et al. 2020. Fungal Diversity 105:1-16 ((https://doi.org/10.1007/s13225-020-00466-2)) |
| Otu35450 | 1 | k__Fungi(100);p__Ascomycota(100);c__Saccharomycetes(100);o__Saccharomycetales(100);f__Metschnikowiaceae(100);g__Metschnikowia(100);g__Metschnikowia_unclassified(100); | Metschnikowia | 13 | Saprotroph | \|Undefined Saprotroph\| | Yeast | NULL | Probable | As nectar-tap_saprotroph (Põlme et al. 2020); Decay sugar-rich substrates (Põlme et al. 2020) | Põlme S, et al. 2020. Fungal Diversity 105:1-16 ((https://doi.org/10.1007/s13225-020-00466-2)) |
| Otu35451 | 1 | k__Fungi(100);p__Ascomycota(100);c__Saccharomycetes(100);o__Saccharomycetales(100);f__Metschnikowiaceae(100);g__Metschnikowia(100);s__Metschnikowia_pulcherrima(100); | Metschnikowia | 13 | Saprotroph | \|Undefined Saprotroph\| | Yeast | NULL | Probable | As nectar-tap_saprotroph (Põlme et al. 2020); Decay sugar-rich substrates (Põlme et al. 2020) | Põlme S, et al. 2020. Fungal Diversity 105:1-16 ((https://doi.org/10.1007/s13225-020-00466-2)) |
| Otu35452 | 1 | k__Fungi(100);p__Ascomycota(100);c__Saccharomycetes(100);o__Saccharomycetales(100);f__Metschnikowiaceae(100);g__Metschnikowia(100);s__Metschnikowia_pulcherrima(100); | Metschnikowia | 13 | Saprotroph | \|Undefined Saprotroph\| | Yeast | NULL | Probable | As nectar-tap_saprotroph (Põlme et al. 2020); Decay sugar-rich substrates (Põlme et al. 2020) | Põlme S, et al. 2020. Fungal Diversity 105:1-16 ((https://doi.org/10.1007/s13225-020-00466-2)) |
| Otu35453 | 1 | k__Fungi(100);p__Ascomycota(100);c__Saccharomycetes(100);o__Saccharomycetales(100);f__Metschnikowiaceae(100);g__Metschnikowia(100);s__Metschnikowia_pulcherrima(100); | Metschnikowia | 13 | Saprotroph | \|Undefined Saprotroph\| | Yeast | NULL | Probable | As nectar-tap_saprotroph (Põlme et al. 2020); Decay sugar-rich substrates (Põlme et al. 2020) | Põlme S, et al. 2020. Fungal Diversity 105:1-16 ((https://doi.org/10.1007/s13225-020-00466-2)) |
| Otu35455 | 1 | k__Fungi(100);p__Ascomycota(100);c__Saccharomycetes(100);o__Saccharomycetales(100);f__Metschnikowiaceae(100);g__Metschnikowia(100);g__Metschnikowia_unclassified(100); | Metschnikowia | 13 | Saprotroph | \|Undefined Saprotroph\| | Yeast | NULL | Probable | As nectar-tap_saprotroph (Põlme et al. 2020); Decay sugar-rich substrates (Põlme et al. 2020) | Põlme S, et al. 2020. Fungal Diversity 105:1-16 ((https://doi.org/10.1007/s13225-020-00466-2)) |
| Otu35456 | 1 | k__Fungi(100);p__Ascomycota(100);c__Saccharomycetes(100);o__Saccharomycetales(100);f__Metschnikowiaceae(100);g__Metschnikowia(100);s__Metschnikowia_pulcherrima(100); | Metschnikowia | 13 | Saprotroph | \|Undefined Saprotroph\| | Yeast | NULL | Probable | As nectar-tap_saprotroph (Põlme et al. 2020); Decay sugar-rich substrates (Põlme et al. 2020) | Põlme S, et al. 2020. Fungal Diversity 105:1-16 ((https://doi.org/10.1007/s13225-020-00466-2)) |
| Otu35458 | 1 | k__Fungi(100);p__Ascomycota(100);c__Saccharomycetes(100);o__Saccharomycetales(100);f__Metschnikowiaceae(100);g__Metschnikowia(100);s__Metschnikowia_pulcherrima(100); | Metschnikowia | 13 | Saprotroph | \|Undefined Saprotroph\| | Yeast | NULL | Probable | As nectar-tap_saprotroph (Põlme et al. 2020); Decay sugar-rich substrates (Põlme et al. 2020) | Põlme S, et al. 2020. Fungal Diversity 105:1-16 ((https://doi.org/10.1007/s13225-020-00466-2)) |
| Otu35459 | 1 | k__Fungi(100);p__Ascomycota(100);c__Saccharomycetes(100);o__Saccharomycetales(100);f__Metschnikowiaceae(100);g__Metschnikowia(100);s__Metschnikowia_pulcherrima(100); | Metschnikowia | 13 | Saprotroph | \|Undefined Saprotroph\| | Yeast | NULL | Probable | As nectar-tap_saprotroph (Põlme et al. 2020); Decay sugar-rich substrates (Põlme et al. 2020) | Põlme S, et al. 2020. Fungal Diversity 105:1-16 ((https://doi.org/10.1007/s13225-020-00466-2)) |
| Otu35460 | 1 | k__Fungi(100);p__Ascomycota(100);c__Saccharomycetes(100);o__Saccharomycetales(100);f__Metschnikowiaceae(100);g__Metschnikowia(100);s__Metschnikowia_pulcherrima(100); | Metschnikowia | 13 | Saprotroph | \|Undefined Saprotroph\| | Yeast | NULL | Probable | As nectar-tap_saprotroph (Põlme et al. 2020); Decay sugar-rich substrates (Põlme et al. 2020) | Põlme S, et al. 2020. Fungal Diversity 105:1-16 ((https://doi.org/10.1007/s13225-020-00466-2)) |
| Otu35463 | 1 | k__Fungi(100);p__Ascomycota(100);c__Saccharomycetes(100);o__Saccharomycetales(100);f__Metschnikowiaceae(100);g__Metschnikowia(100);s__Metschnikowia_pulcherrima(100); | Metschnikowia | 13 | Saprotroph | \|Undefined Saprotroph\| | Yeast | NULL | Probable | As nectar-tap_saprotroph (Põlme et al. 2020); Decay sugar-rich substrates (Põlme et al. 2020) | Põlme S, et al. 2020. Fungal Diversity 105:1-16 ((https://doi.org/10.1007/s13225-020-00466-2)) |
| Otu35464 | 1 | k__Fungi(100);p__Ascomycota(100);c__Saccharomycetes(100);o__Saccharomycetales(100);f__Metschnikowiaceae(100);g__Metschnikowia(100);s__Metschnikowia_pulcherrima(100); | Metschnikowia | 13 | Saprotroph | \|Undefined Saprotroph\| | Yeast | NULL | Probable | As nectar-tap_saprotroph (Põlme et al. 2020); Decay sugar-rich substrates (Põlme et al. 2020) | Põlme S, et al. 2020. Fungal Diversity 105:1-16 ((https://doi.org/10.1007/s13225-020-00466-2)) |
| Otu35467 | 1 | k__Fungi(100);p__Ascomycota(100);c__Saccharomycetes(100);o__Saccharomycetales(100);f__Metschnikowiaceae(100);g__Metschnikowia(100);s__Metschnikowia_pulcherrima(100); | Metschnikowia | 13 | Saprotroph | \|Undefined Saprotroph\| | Yeast | NULL | Probable | As nectar-tap_saprotroph (Põlme et al. 2020); Decay sugar-rich substrates (Põlme et al. 2020) | Põlme S, et al. 2020. Fungal Diversity 105:1-16 ((https://doi.org/10.1007/s13225-020-00466-2)) |
| Otu35468 | 1 | k__Fungi(100);p__Ascomycota(100);c__Saccharomycetes(100);o__Saccharomycetales(100);f__Metschnikowiaceae(100);g__Metschnikowia(100);g__Metschnikowia_unclassified(100); | Metschnikowia | 13 | Saprotroph | \|Undefined Saprotroph\| | Yeast | NULL | Probable | As nectar-tap_saprotroph (Põlme et al. 2020); Decay sugar-rich substrates (Põlme et al. 2020) | Põlme S, et al. 2020. Fungal Diversity 105:1-16 ((https://doi.org/10.1007/s13225-020-00466-2)) |
| Otu35471 | 1 | k__Fungi(100);p__Ascomycota(100);c__Saccharomycetes(100);o__Saccharomycetales(100);f__Metschnikowiaceae(100);g__Metschnikowia(100);g__Metschnikowia_unclassified(100); | Metschnikowia | 13 | Saprotroph | \|Undefined Saprotroph\| | Yeast | NULL | Probable | As nectar-tap_saprotroph (Põlme et al. 2020); Decay sugar-rich substrates (Põlme et al. 2020) | Põlme S, et al. 2020. Fungal Diversity 105:1-16 ((https://doi.org/10.1007/s13225-020-00466-2)) |
| Otu35472 | 1 | k__Fungi(100);p__Ascomycota(100);c__Saccharomycetes(100);o__Saccharomycetales(100);f__Metschnikowiaceae(100);g__Metschnikowia(100);g__Metschnikowia_unclassified(100); | Metschnikowia | 13 | Saprotroph | \|Undefined Saprotroph\| | Yeast | NULL | Probable | As nectar-tap_saprotroph (Põlme et al. 2020); Decay sugar-rich substrates (Põlme et al. 2020) | Põlme S, et al. 2020. Fungal Diversity 105:1-16 ((https://doi.org/10.1007/s13225-020-00466-2)) |
| Otu35475 | 1 | k__Fungi(100);p__Ascomycota(100);c__Saccharomycetes(100);o__Saccharomycetales(100);f__Metschnikowiaceae(100);g__Metschnikowia(100);s__Metschnikowia_pulcherrima(100); | Metschnikowia | 13 | Saprotroph | \|Undefined Saprotroph\| | Yeast | NULL | Probable | As nectar-tap_saprotroph (Põlme et al. 2020); Decay sugar-rich substrates (Põlme et al. 2020) | Põlme S, et al. 2020. Fungal Diversity 105:1-16 ((https://doi.org/10.1007/s13225-020-00466-2)) |
| Otu35476 | 1 | k__Fungi(100);p__Ascomycota(100);c__Saccharomycetes(100);o__Saccharomycetales(100);f__Metschnikowiaceae(100);g__Metschnikowia(100);s__Metschnikowia_pulcherrima(100); | Metschnikowia | 13 | Saprotroph | \|Undefined Saprotroph\| | Yeast | NULL | Probable | As nectar-tap_saprotroph (Põlme et al. 2020); Decay sugar-rich substrates (Põlme et al. 2020) | Põlme S, et al. 2020. Fungal Diversity 105:1-16 ((https://doi.org/10.1007/s13225-020-00466-2)) |
| Otu35477 | 1 | k__Fungi(100);p__Ascomycota(100);c__Saccharomycetes(100);o__Saccharomycetales(100);f__Metschnikowiaceae(100);g__Metschnikowia(100);s__Metschnikowia_pulcherrima(100); | Metschnikowia | 13 | Saprotroph | \|Undefined Saprotroph\| | Yeast | NULL | Probable | As nectar-tap_saprotroph (Põlme et al. 2020); Decay sugar-rich substrates (Põlme et al. 2020) | Põlme S, et al. 2020. Fungal Diversity 105:1-16 ((https://doi.org/10.1007/s13225-020-00466-2)) |
| Otu35478 | 1 | k__Fungi(100);p__Ascomycota(100);c__Saccharomycetes(100);o__Saccharomycetales(100);f__Metschnikowiaceae(100);g__Metschnikowia(100);g__Metschnikowia_unclassified(100); | Metschnikowia | 13 | Saprotroph | \|Undefined Saprotroph\| | Yeast | NULL | Probable | As nectar-tap_saprotroph (Põlme et al. 2020); Decay sugar-rich substrates (Põlme et al. 2020) | Põlme S, et al. 2020. Fungal Diversity 105:1-16 ((https://doi.org/10.1007/s13225-020-00466-2)) |
| Otu35480 | 1 | k__Fungi(100);p__Ascomycota(100);c__Saccharomycetes(100);o__Saccharomycetales(100);f__Metschnikowiaceae(100);g__Metschnikowia(100);g__Metschnikowia_unclassified(100); | Metschnikowia | 13 | Saprotroph | \|Undefined Saprotroph\| | Yeast | NULL | Probable | As nectar-tap_saprotroph (Põlme et al. 2020); Decay sugar-rich substrates (Põlme et al. 2020) | Põlme S, et al. 2020. Fungal Diversity 105:1-16 ((https://doi.org/10.1007/s13225-020-00466-2)) |
| Otu35481 | 1 | k__Fungi(100);p__Ascomycota(100);c__Saccharomycetes(100);o__Saccharomycetales(100);f__Metschnikowiaceae(100);g__Metschnikowia(100);s__Metschnikowia_pulcherrima(100); | Metschnikowia | 13 | Saprotroph | \|Undefined Saprotroph\| | Yeast | NULL | Probable | As nectar-tap_saprotroph (Põlme et al. 2020); Decay sugar-rich substrates (Põlme et al. 2020) | Põlme S, et al. 2020. Fungal Diversity 105:1-16 ((https://doi.org/10.1007/s13225-020-00466-2)) |
| Otu35482 | 1 | k__Fungi(100);p__Ascomycota(100);c__Saccharomycetes(100);o__Saccharomycetales(100);f__Metschnikowiaceae(100);g__Metschnikowia(100);s__Metschnikowia_pulcherrima(100); | Metschnikowia | 13 | Saprotroph | \|Undefined Saprotroph\| | Yeast | NULL | Probable | As nectar-tap_saprotroph (Põlme et al. 2020); Decay sugar-rich substrates (Põlme et al. 2020) | Põlme S, et al. 2020. Fungal Diversity 105:1-16 ((https://doi.org/10.1007/s13225-020-00466-2)) |
| Otu35484 | 1 | k__Fungi(100);p__Ascomycota(100);c__Saccharomycetes(100);o__Saccharomycetales(100);f__Metschnikowiaceae(100);g__Metschnikowia(100);s__Metschnikowia_pulcherrima(100); | Metschnikowia | 13 | Saprotroph | \|Undefined Saprotroph\| | Yeast | NULL | Probable | As nectar-tap_saprotroph (Põlme et al. 2020); Decay sugar-rich substrates (Põlme et al. 2020) | Põlme S, et al. 2020. Fungal Diversity 105:1-16 ((https://doi.org/10.1007/s13225-020-00466-2)) |
| Otu35485 | 1 | k__Fungi(100);p__Ascomycota(100);c__Saccharomycetes(100);o__Saccharomycetales(100);f__Metschnikowiaceae(100);g__Metschnikowia(100);s__Metschnikowia_pulcherrima(100); | Metschnikowia | 13 | Saprotroph | \|Undefined Saprotroph\| | Yeast | NULL | Probable | As nectar-tap_saprotroph (Põlme et al. 2020); Decay sugar-rich substrates (Põlme et al. 2020) | Põlme S, et al. 2020. Fungal Diversity 105:1-16 ((https://doi.org/10.1007/s13225-020-00466-2)) |
| Otu35486 | 1 | k__Fungi(100);p__Ascomycota(100);c__Saccharomycetes(100);o__Saccharomycetales(100);f__Metschnikowiaceae(100);g__Metschnikowia(100);s__Metschnikowia_pulcherrima(100); | Metschnikowia | 13 | Saprotroph | \|Undefined Saprotroph\| | Yeast | NULL | Probable | As nectar-tap_saprotroph (Põlme et al. 2020); Decay sugar-rich substrates (Põlme et al. 2020) | Põlme S, et al. 2020. Fungal Diversity 105:1-16 ((https://doi.org/10.1007/s13225-020-00466-2)) |
| Otu35488 | 1 | k__Fungi(100);p__Ascomycota(100);c__Saccharomycetes(100);o__Saccharomycetales(100);f__Metschnikowiaceae(100);g__Metschnikowia(100);s__Metschnikowia_pulcherrima(100); | Metschnikowia | 13 | Saprotroph | \|Undefined Saprotroph\| | Yeast | NULL | Probable | As nectar-tap_saprotroph (Põlme et al. 2020); Decay sugar-rich substrates (Põlme et al. 2020) | Põlme S, et al. 2020. Fungal Diversity 105:1-16 ((https://doi.org/10.1007/s13225-020-00466-2)) |
| Otu35489 | 1 | k__Fungi(100);p__Ascomycota(100);c__Saccharomycetes(100);o__Saccharomycetales(100);f__Metschnikowiaceae(100);g__Metschnikowia(100);g__Metschnikowia_unclassified(100); | Metschnikowia | 13 | Saprotroph | \|Undefined Saprotroph\| | Yeast | NULL | Probable | As nectar-tap_saprotroph (Põlme et al. 2020); Decay sugar-rich substrates (Põlme et al. 2020) | Põlme S, et al. 2020. Fungal Diversity 105:1-16 ((https://doi.org/10.1007/s13225-020-00466-2)) |
| Otu35490 | 1 | k__Fungi(100);p__Ascomycota(100);c__Saccharomycetes(100);o__Saccharomycetales(100);f__Metschnikowiaceae(100);g__Metschnikowia(100);s__Metschnikowia_pulcherrima(100); | Metschnikowia | 13 | Saprotroph | \|Undefined Saprotroph\| | Yeast | NULL | Probable | As nectar-tap_saprotroph (Põlme et al. 2020); Decay sugar-rich substrates (Põlme et al. 2020) | Põlme S, et al. 2020. Fungal Diversity 105:1-16 ((https://doi.org/10.1007/s13225-020-00466-2)) |
| Otu35491 | 1 | k__Fungi(100);p__Ascomycota(100);c__Saccharomycetes(100);o__Saccharomycetales(100);f__Metschnikowiaceae(100);g__Metschnikowia(100);s__Metschnikowia_pulcherrima(100); | Metschnikowia | 13 | Saprotroph | \|Undefined Saprotroph\| | Yeast | NULL | Probable | As nectar-tap_saprotroph (Põlme et al. 2020); Decay sugar-rich substrates (Põlme et al. 2020) | Põlme S, et al. 2020. Fungal Diversity 105:1-16 ((https://doi.org/10.1007/s13225-020-00466-2)) |
| Otu35495 | 1 | k__Fungi(100);p__Ascomycota(100);c__Saccharomycetes(100);o__Saccharomycetales(100);f__Metschnikowiaceae(100);g__Metschnikowia(100);s__Metschnikowia_pulcherrima(100); | Metschnikowia | 13 | Saprotroph | \|Undefined Saprotroph\| | Yeast | NULL | Probable | As nectar-tap_saprotroph (Põlme et al. 2020); Decay sugar-rich substrates (Põlme et al. 2020) | Põlme S, et al. 2020. Fungal Diversity 105:1-16 ((https://doi.org/10.1007/s13225-020-00466-2)) |
| Otu35496 | 1 | k__Fungi(100);p__Ascomycota(100);c__Saccharomycetes(100);o__Saccharomycetales(100);f__Metschnikowiaceae(100);g__Metschnikowia(100);s__Metschnikowia_pulcherrima(100); | Metschnikowia | 13 | Saprotroph | \|Undefined Saprotroph\| | Yeast | NULL | Probable | As nectar-tap_saprotroph (Põlme et al. 2020); Decay sugar-rich substrates (Põlme et al. 2020) | Põlme S, et al. 2020. Fungal Diversity 105:1-16 ((https://doi.org/10.1007/s13225-020-00466-2)) |
| Otu35497 | 1 | k__Fungi(100);p__Ascomycota(100);c__Saccharomycetes(100);o__Saccharomycetales(100);f__Metschnikowiaceae(100);g__Metschnikowia(100);g__Metschnikowia_unclassified(100); | Metschnikowia | 13 | Saprotroph | \|Undefined Saprotroph\| | Yeast | NULL | Probable | As nectar-tap_saprotroph (Põlme et al. 2020); Decay sugar-rich substrates (Põlme et al. 2020) | Põlme S, et al. 2020. Fungal Diversity 105:1-16 ((https://doi.org/10.1007/s13225-020-00466-2)) |
| Otu35498 | 1 | k__Fungi(100);p__Ascomycota(100);c__Saccharomycetes(100);o__Saccharomycetales(100);f__Metschnikowiaceae(100);g__Metschnikowia(100);s__Metschnikowia_pulcherrima(100); | Metschnikowia | 13 | Saprotroph | \|Undefined Saprotroph\| | Yeast | NULL | Probable | As nectar-tap_saprotroph (Põlme et al. 2020); Decay sugar-rich substrates (Põlme et al. 2020) | Põlme S, et al. 2020. Fungal Diversity 105:1-16 ((https://doi.org/10.1007/s13225-020-00466-2)) |
| Otu35499 | 1 | k__Fungi(100);p__Ascomycota(100);c__Saccharomycetes(100);o__Saccharomycetales(100);f__Metschnikowiaceae(100);g__Metschnikowia(100);s__Metschnikowia_pulcherrima(100); | Metschnikowia | 13 | Saprotroph | \|Undefined Saprotroph\| | Yeast | NULL | Probable | As nectar-tap_saprotroph (Põlme et al. 2020); Decay sugar-rich substrates (Põlme et al. 2020) | Põlme S, et al. 2020. Fungal Diversity 105:1-16 ((https://doi.org/10.1007/s13225-020-00466-2)) |
| Otu35500 | 1 | k__Fungi(100);p__Ascomycota(100);c__Saccharomycetes(100);o__Saccharomycetales(100);f__Metschnikowiaceae(100);g__Metschnikowia(100);s__Metschnikowia_pulcherrima(100); | Metschnikowia | 13 | Saprotroph | \|Undefined Saprotroph\| | Yeast | NULL | Probable | As nectar-tap_saprotroph (Põlme et al. 2020); Decay sugar-rich substrates (Põlme et al. 2020) | Põlme S, et al. 2020. Fungal Diversity 105:1-16 ((https://doi.org/10.1007/s13225-020-00466-2)) |
| Otu35501 | 1 | k__Fungi(100);p__Ascomycota(100);c__Saccharomycetes(100);o__Saccharomycetales(100);f__Metschnikowiaceae(100);g__Metschnikowia(100);s__Metschnikowia_pulcherrima(100); | Metschnikowia | 13 | Saprotroph | \|Undefined Saprotroph\| | Yeast | NULL | Probable | As nectar-tap_saprotroph (Põlme et al. 2020); Decay sugar-rich substrates (Põlme et al. 2020) | Põlme S, et al. 2020. Fungal Diversity 105:1-16 ((https://doi.org/10.1007/s13225-020-00466-2)) |
| Otu35502 | 1 | k__Fungi(100);p__Ascomycota(100);c__Saccharomycetes(100);o__Saccharomycetales(100);f__Metschnikowiaceae(100);g__Metschnikowia(100);s__Metschnikowia_pulcherrima(100); | Metschnikowia | 13 | Saprotroph | \|Undefined Saprotroph\| | Yeast | NULL | Probable | As nectar-tap_saprotroph (Põlme et al. 2020); Decay sugar-rich substrates (Põlme et al. 2020) | Põlme S, et al. 2020. Fungal Diversity 105:1-16 ((https://doi.org/10.1007/s13225-020-00466-2)) |
| Otu35503 | 1 | k__Fungi(100);p__Ascomycota(100);c__Saccharomycetes(100);o__Saccharomycetales(100);f__Metschnikowiaceae(100);g__Metschnikowia(100);s__Metschnikowia_pulcherrima(100); | Metschnikowia | 13 | Saprotroph | \|Undefined Saprotroph\| | Yeast | NULL | Probable | As nectar-tap_saprotroph (Põlme et al. 2020); Decay sugar-rich substrates (Põlme et al. 2020) | Põlme S, et al. 2020. Fungal Diversity 105:1-16 ((https://doi.org/10.1007/s13225-020-00466-2)) |
| Otu35504 | 1 | k__Fungi(100);p__Ascomycota(100);c__Saccharomycetes(100);o__Saccharomycetales(100);f__Metschnikowiaceae(100);g__Metschnikowia(100);g__Metschnikowia_unclassified(100); | Metschnikowia | 13 | Saprotroph | \|Undefined Saprotroph\| | Yeast | NULL | Probable | As nectar-tap_saprotroph (Põlme et al. 2020); Decay sugar-rich substrates (Põlme et al. 2020) | Põlme S, et al. 2020. Fungal Diversity 105:1-16 ((https://doi.org/10.1007/s13225-020-00466-2)) |
| Otu35505 | 1 | k__Fungi(100);p__Ascomycota(100);c__Saccharomycetes(100);o__Saccharomycetales(100);f__Metschnikowiaceae(100);g__Metschnikowia(100);s__Metschnikowia_pulcherrima(100); | Metschnikowia | 13 | Saprotroph | \|Undefined Saprotroph\| | Yeast | NULL | Probable | As nectar-tap_saprotroph (Põlme et al. 2020); Decay sugar-rich substrates (Põlme et al. 2020) | Põlme S, et al. 2020. Fungal Diversity 105:1-16 ((https://doi.org/10.1007/s13225-020-00466-2)) |
| Otu35506 | 1 | k__Fungi(100);p__Ascomycota(100);c__Saccharomycetes(100);o__Saccharomycetales(100);f__Metschnikowiaceae(100);g__Metschnikowia(100);s__Metschnikowia_pulcherrima(100); | Metschnikowia | 13 | Saprotroph | \|Undefined Saprotroph\| | Yeast | NULL | Probable | As nectar-tap_saprotroph (Põlme et al. 2020); Decay sugar-rich substrates (Põlme et al. 2020) | Põlme S, et al. 2020. Fungal Diversity 105:1-16 ((https://doi.org/10.1007/s13225-020-00466-2)) |
| Otu35507 | 1 | k__Fungi(100);p__Ascomycota(100);c__Saccharomycetes(100);o__Saccharomycetales(100);f__Metschnikowiaceae(100);g__Metschnikowia(100);s__Metschnikowia_pulcherrima(100); | Metschnikowia | 13 | Saprotroph | \|Undefined Saprotroph\| | Yeast | NULL | Probable | As nectar-tap_saprotroph (Põlme et al. 2020); Decay sugar-rich substrates (Põlme et al. 2020) | Põlme S, et al. 2020. Fungal Diversity 105:1-16 ((https://doi.org/10.1007/s13225-020-00466-2)) |
| Otu35508 | 1 | k__Fungi(100);p__Ascomycota(100);c__Saccharomycetes(100);o__Saccharomycetales(100);f__Metschnikowiaceae(100);g__Metschnikowia(100);s__Metschnikowia_pulcherrima(100); | Metschnikowia | 13 | Saprotroph | \|Undefined Saprotroph\| | Yeast | NULL | Probable | As nectar-tap_saprotroph (Põlme et al. 2020); Decay sugar-rich substrates (Põlme et al. 2020) | Põlme S, et al. 2020. Fungal Diversity 105:1-16 ((https://doi.org/10.1007/s13225-020-00466-2)) |
| Otu35509 | 1 | k__Fungi(100);p__Ascomycota(100);c__Saccharomycetes(100);o__Saccharomycetales(100);f__Metschnikowiaceae(100);g__Metschnikowia(100);s__Metschnikowia_pulcherrima(100); | Metschnikowia | 13 | Saprotroph | \|Undefined Saprotroph\| | Yeast | NULL | Probable | As nectar-tap_saprotroph (Põlme et al. 2020); Decay sugar-rich substrates (Põlme et al. 2020) | Põlme S, et al. 2020. Fungal Diversity 105:1-16 ((https://doi.org/10.1007/s13225-020-00466-2)) |
| Otu35510 | 1 | k__Fungi(100);p__Ascomycota(100);c__Saccharomycetes(100);o__Saccharomycetales(100);f__Metschnikowiaceae(100);g__Metschnikowia(100);s__Metschnikowia_pulcherrima(100); | Metschnikowia | 13 | Saprotroph | \|Undefined Saprotroph\| | Yeast | NULL | Probable | As nectar-tap_saprotroph (Põlme et al. 2020); Decay sugar-rich substrates (Põlme et al. 2020) | Põlme S, et al. 2020. Fungal Diversity 105:1-16 ((https://doi.org/10.1007/s13225-020-00466-2)) |
| Otu35511 | 1 | k__Fungi(100);p__Ascomycota(100);c__Saccharomycetes(100);o__Saccharomycetales(100);f__Metschnikowiaceae(100);g__Metschnikowia(100);s__Metschnikowia_pulcherrima(100); | Metschnikowia | 13 | Saprotroph | \|Undefined Saprotroph\| | Yeast | NULL | Probable | As nectar-tap_saprotroph (Põlme et al. 2020); Decay sugar-rich substrates (Põlme et al. 2020) | Põlme S, et al. 2020. Fungal Diversity 105:1-16 ((https://doi.org/10.1007/s13225-020-00466-2)) |
| Otu35512 | 1 | k__Fungi(100);p__Ascomycota(100);c__Saccharomycetes(100);o__Saccharomycetales(100);f__Metschnikowiaceae(100);g__Metschnikowia(100);s__Metschnikowia_pulcherrima(100); | Metschnikowia | 13 | Saprotroph | \|Undefined Saprotroph\| | Yeast | NULL | Probable | As nectar-tap_saprotroph (Põlme et al. 2020); Decay sugar-rich substrates (Põlme et al. 2020) | Põlme S, et al. 2020. Fungal Diversity 105:1-16 ((https://doi.org/10.1007/s13225-020-00466-2)) |
| Otu35513 | 1 | k__Fungi(100);p__Ascomycota(100);c__Saccharomycetes(100);o__Saccharomycetales(100);f__Metschnikowiaceae(100);g__Metschnikowia(100);s__Metschnikowia_pulcherrima(100); | Metschnikowia | 13 | Saprotroph | \|Undefined Saprotroph\| | Yeast | NULL | Probable | As nectar-tap_saprotroph (Põlme et al. 2020); Decay sugar-rich substrates (Põlme et al. 2020) | Põlme S, et al. 2020. Fungal Diversity 105:1-16 ((https://doi.org/10.1007/s13225-020-00466-2)) |
| Otu35514 | 1 | k__Fungi(100);p__Ascomycota(100);c__Saccharomycetes(100);o__Saccharomycetales(100);f__Metschnikowiaceae(100);g__Metschnikowia(100);s__Metschnikowia_pulcherrima(100); | Metschnikowia | 13 | Saprotroph | \|Undefined Saprotroph\| | Yeast | NULL | Probable | As nectar-tap_saprotroph (Põlme et al. 2020); Decay sugar-rich substrates (Põlme et al. 2020) | Põlme S, et al. 2020. Fungal Diversity 105:1-16 ((https://doi.org/10.1007/s13225-020-00466-2)) |
| Otu35515 | 1 | k__Fungi(100);p__Ascomycota(100);c__Saccharomycetes(100);o__Saccharomycetales(100);f__Metschnikowiaceae(100);g__Metschnikowia(100);s__Metschnikowia_pulcherrima(100); | Metschnikowia | 13 | Saprotroph | \|Undefined Saprotroph\| | Yeast | NULL | Probable | As nectar-tap_saprotroph (Põlme et al. 2020); Decay sugar-rich substrates (Põlme et al. 2020) | Põlme S, et al. 2020. Fungal Diversity 105:1-16 ((https://doi.org/10.1007/s13225-020-00466-2)) |
| Otu35516 | 1 | k__Fungi(100);p__Ascomycota(100);c__Saccharomycetes(100);o__Saccharomycetales(100);f__Metschnikowiaceae(100);g__Metschnikowia(100);s__Metschnikowia_pulcherrima(100); | Metschnikowia | 13 | Saprotroph | \|Undefined Saprotroph\| | Yeast | NULL | Probable | As nectar-tap_saprotroph (Põlme et al. 2020); Decay sugar-rich substrates (Põlme et al. 2020) | Põlme S, et al. 2020. Fungal Diversity 105:1-16 ((https://doi.org/10.1007/s13225-020-00466-2)) |
| Otu35517 | 1 | k__Fungi(100);p__Ascomycota(100);c__Saccharomycetes(100);o__Saccharomycetales(100);f__Metschnikowiaceae(100);g__Metschnikowia(100);s__Metschnikowia_pulcherrima(100); | Metschnikowia | 13 | Saprotroph | \|Undefined Saprotroph\| | Yeast | NULL | Probable | As nectar-tap_saprotroph (Põlme et al. 2020); Decay sugar-rich substrates (Põlme et al. 2020) | Põlme S, et al. 2020. Fungal Diversity 105:1-16 ((https://doi.org/10.1007/s13225-020-00466-2)) |
| Otu35518 | 1 | k__Fungi(100);p__Ascomycota(100);c__Saccharomycetes(100);o__Saccharomycetales(100);f__Metschnikowiaceae(100);g__Metschnikowia(100);s__Metschnikowia_pulcherrima(100); | Metschnikowia | 13 | Saprotroph | \|Undefined Saprotroph\| | Yeast | NULL | Probable | As nectar-tap_saprotroph (Põlme et al. 2020); Decay sugar-rich substrates (Põlme et al. 2020) | Põlme S, et al. 2020. Fungal Diversity 105:1-16 ((https://doi.org/10.1007/s13225-020-00466-2)) |
| Otu35519 | 1 | k__Fungi(100);p__Ascomycota(100);c__Saccharomycetes(100);o__Saccharomycetales(100);f__Metschnikowiaceae(100);g__Metschnikowia(100);s__Metschnikowia_pulcherrima(100); | Metschnikowia | 13 | Saprotroph | \|Undefined Saprotroph\| | Yeast | NULL | Probable | As nectar-tap_saprotroph (Põlme et al. 2020); Decay sugar-rich substrates (Põlme et al. 2020) | Põlme S, et al. 2020. Fungal Diversity 105:1-16 ((https://doi.org/10.1007/s13225-020-00466-2)) |
| Otu35520 | 1 | k__Fungi(100);p__Ascomycota(100);c__Saccharomycetes(100);o__Saccharomycetales(100);f__Metschnikowiaceae(100);g__Metschnikowia(100);s__Metschnikowia_pulcherrima(100); | Metschnikowia | 13 | Saprotroph | \|Undefined Saprotroph\| | Yeast | NULL | Probable | As nectar-tap_saprotroph (Põlme et al. 2020); Decay sugar-rich substrates (Põlme et al. 2020) | Põlme S, et al. 2020. Fungal Diversity 105:1-16 ((https://doi.org/10.1007/s13225-020-00466-2)) |
| Otu35521 | 1 | k__Fungi(100);p__Ascomycota(100);c__Saccharomycetes(100);o__Saccharomycetales(100);f__Metschnikowiaceae(100);g__Metschnikowia(100);s__Metschnikowia_pulcherrima(100); | Metschnikowia | 13 | Saprotroph | \|Undefined Saprotroph\| | Yeast | NULL | Probable | As nectar-tap_saprotroph (Põlme et al. 2020); Decay sugar-rich substrates (Põlme et al. 2020) | Põlme S, et al. 2020. Fungal Diversity 105:1-16 ((https://doi.org/10.1007/s13225-020-00466-2)) |
| Otu35522 | 1 | k__Fungi(100);p__Ascomycota(100);c__Saccharomycetes(100);o__Saccharomycetales(100);f__Metschnikowiaceae(100);g__Metschnikowia(100);s__Metschnikowia_pulcherrima(100); | Metschnikowia | 13 | Saprotroph | \|Undefined Saprotroph\| | Yeast | NULL | Probable | As nectar-tap_saprotroph (Põlme et al. 2020); Decay sugar-rich substrates (Põlme et al. 2020) | Põlme S, et al. 2020. Fungal Diversity 105:1-16 ((https://doi.org/10.1007/s13225-020-00466-2)) |
| Otu35524 | 1 | k__Fungi(100);p__Ascomycota(100);c__Saccharomycetes(100);o__Saccharomycetales(100);f__Metschnikowiaceae(100);g__Metschnikowia(100);s__Metschnikowia_pulcherrima(100); | Metschnikowia | 13 | Saprotroph | \|Undefined Saprotroph\| | Yeast | NULL | Probable | As nectar-tap_saprotroph (Põlme et al. 2020); Decay sugar-rich substrates (Põlme et al. 2020) | Põlme S, et al. 2020. Fungal Diversity 105:1-16 ((https://doi.org/10.1007/s13225-020-00466-2)) |
| Otu35525 | 1 | k__Fungi(100);p__Ascomycota(100);c__Saccharomycetes(100);o__Saccharomycetales(100);f__Metschnikowiaceae(100);g__Metschnikowia(100);s__Metschnikowia_pulcherrima(100); | Metschnikowia | 13 | Saprotroph | \|Undefined Saprotroph\| | Yeast | NULL | Probable | As nectar-tap_saprotroph (Põlme et al. 2020); Decay sugar-rich substrates (Põlme et al. 2020) | Põlme S, et al. 2020. Fungal Diversity 105:1-16 ((https://doi.org/10.1007/s13225-020-00466-2)) |
| Otu35526 | 1 | k__Fungi(100);p__Ascomycota(100);c__Saccharomycetes(100);o__Saccharomycetales(100);f__Metschnikowiaceae(100);g__Metschnikowia(100);s__Metschnikowia_pulcherrima(100); | Metschnikowia | 13 | Saprotroph | \|Undefined Saprotroph\| | Yeast | NULL | Probable | As nectar-tap_saprotroph (Põlme et al. 2020); Decay sugar-rich substrates (Põlme et al. 2020) | Põlme S, et al. 2020. Fungal Diversity 105:1-16 ((https://doi.org/10.1007/s13225-020-00466-2)) |
| Otu35527 | 1 | k__Fungi(100);p__Ascomycota(100);c__Saccharomycetes(100);o__Saccharomycetales(100);f__Metschnikowiaceae(100);g__Metschnikowia(100);s__Metschnikowia_pulcherrima(100); | Metschnikowia | 13 | Saprotroph | \|Undefined Saprotroph\| | Yeast | NULL | Probable | As nectar-tap_saprotroph (Põlme et al. 2020); Decay sugar-rich substrates (Põlme et al. 2020) | Põlme S, et al. 2020. Fungal Diversity 105:1-16 ((https://doi.org/10.1007/s13225-020-00466-2)) |
| Otu35528 | 1 | k__Fungi(100);p__Ascomycota(100);c__Saccharomycetes(100);o__Saccharomycetales(100);f__Metschnikowiaceae(100);g__Metschnikowia(100);g__Metschnikowia_unclassified(100); | Metschnikowia | 13 | Saprotroph | \|Undefined Saprotroph\| | Yeast | NULL | Probable | As nectar-tap_saprotroph (Põlme et al. 2020); Decay sugar-rich substrates (Põlme et al. 2020) | Põlme S, et al. 2020. Fungal Diversity 105:1-16 ((https://doi.org/10.1007/s13225-020-00466-2)) |
| Otu35530 | 1 | k__Fungi(100);p__Ascomycota(100);c__Saccharomycetes(100);o__Saccharomycetales(100);f__Metschnikowiaceae(100);g__Metschnikowia(100);s__Metschnikowia_pulcherrima(100); | Metschnikowia | 13 | Saprotroph | \|Undefined Saprotroph\| | Yeast | NULL | Probable | As nectar-tap_saprotroph (Põlme et al. 2020); Decay sugar-rich substrates (Põlme et al. 2020) | Põlme S, et al. 2020. Fungal Diversity 105:1-16 ((https://doi.org/10.1007/s13225-020-00466-2)) |
| Otu35531 | 1 | k__Fungi(100);p__Ascomycota(100);c__Saccharomycetes(100);o__Saccharomycetales(100);f__Metschnikowiaceae(100);g__Metschnikowia(100);s__Metschnikowia_pulcherrima(100); | Metschnikowia | 13 | Saprotroph | \|Undefined Saprotroph\| | Yeast | NULL | Probable | As nectar-tap_saprotroph (Põlme et al. 2020); Decay sugar-rich substrates (Põlme et al. 2020) | Põlme S, et al. 2020. Fungal Diversity 105:1-16 ((https://doi.org/10.1007/s13225-020-00466-2)) |
| Otu35532 | 1 | k__Fungi(100);p__Ascomycota(100);c__Saccharomycetes(100);o__Saccharomycetales(100);f__Metschnikowiaceae(100);g__Metschnikowia(100);s__Metschnikowia_pulcherrima(100); | Metschnikowia | 13 | Saprotroph | \|Undefined Saprotroph\| | Yeast | NULL | Probable | As nectar-tap_saprotroph (Põlme et al. 2020); Decay sugar-rich substrates (Põlme et al. 2020) | Põlme S, et al. 2020. Fungal Diversity 105:1-16 ((https://doi.org/10.1007/s13225-020-00466-2)) |
| Otu35533 | 1 | k__Fungi(100);p__Ascomycota(100);c__Saccharomycetes(100);o__Saccharomycetales(100);f__Metschnikowiaceae(100);g__Metschnikowia(100);g__Metschnikowia_unclassified(100); | Metschnikowia | 13 | Saprotroph | \|Undefined Saprotroph\| | Yeast | NULL | Probable | As nectar-tap_saprotroph (Põlme et al. 2020); Decay sugar-rich substrates (Põlme et al. 2020) | Põlme S, et al. 2020. Fungal Diversity 105:1-16 ((https://doi.org/10.1007/s13225-020-00466-2)) |
| Otu35534 | 1 | k__Fungi(100);p__Ascomycota(100);c__Saccharomycetes(100);o__Saccharomycetales(100);f__Metschnikowiaceae(100);g__Metschnikowia(100);g__Metschnikowia_unclassified(100); | Metschnikowia | 13 | Saprotroph | \|Undefined Saprotroph\| | Yeast | NULL | Probable | As nectar-tap_saprotroph (Põlme et al. 2020); Decay sugar-rich substrates (Põlme et al. 2020) | Põlme S, et al. 2020. Fungal Diversity 105:1-16 ((https://doi.org/10.1007/s13225-020-00466-2)) |
| Otu35535 | 1 | k__Fungi(100);p__Ascomycota(100);c__Saccharomycetes(100);o__Saccharomycetales(100);f__Metschnikowiaceae(100);g__Metschnikowia(100);s__Metschnikowia_pulcherrima(100); | Metschnikowia | 13 | Saprotroph | \|Undefined Saprotroph\| | Yeast | NULL | Probable | As nectar-tap_saprotroph (Põlme et al. 2020); Decay sugar-rich substrates (Põlme et al. 2020) | Põlme S, et al. 2020. Fungal Diversity 105:1-16 ((https://doi.org/10.1007/s13225-020-00466-2)) |
| Otu35536 | 1 | k__Fungi(100);p__Ascomycota(100);c__Saccharomycetes(100);o__Saccharomycetales(100);f__Metschnikowiaceae(100);g__Metschnikowia(100);s__Metschnikowia_pulcherrima(100); | Metschnikowia | 13 | Saprotroph | \|Undefined Saprotroph\| | Yeast | NULL | Probable | As nectar-tap_saprotroph (Põlme et al. 2020); Decay sugar-rich substrates (Põlme et al. 2020) | Põlme S, et al. 2020. Fungal Diversity 105:1-16 ((https://doi.org/10.1007/s13225-020-00466-2)) |
| Otu35537 | 1 | k__Fungi(100);p__Ascomycota(100);c__Saccharomycetes(100);o__Saccharomycetales(100);f__Metschnikowiaceae(100);g__Metschnikowia(100);s__Metschnikowia_pulcherrima(100); | Metschnikowia | 13 | Saprotroph | \|Undefined Saprotroph\| | Yeast | NULL | Probable | As nectar-tap_saprotroph (Põlme et al. 2020); Decay sugar-rich substrates (Põlme et al. 2020) | Põlme S, et al. 2020. Fungal Diversity 105:1-16 ((https://doi.org/10.1007/s13225-020-00466-2)) |
| Otu35538 | 1 | k__Fungi(100);p__Ascomycota(100);c__Saccharomycetes(100);o__Saccharomycetales(100);f__Metschnikowiaceae(100);g__Metschnikowia(100);s__Metschnikowia_pulcherrima(100); | Metschnikowia | 13 | Saprotroph | \|Undefined Saprotroph\| | Yeast | NULL | Probable | As nectar-tap_saprotroph (Põlme et al. 2020); Decay sugar-rich substrates (Põlme et al. 2020) | Põlme S, et al. 2020. Fungal Diversity 105:1-16 ((https://doi.org/10.1007/s13225-020-00466-2)) |
| Otu35540 | 1 | k__Fungi(100);p__Ascomycota(100);c__Saccharomycetes(100);o__Saccharomycetales(100);f__Metschnikowiaceae(100);g__Metschnikowia(100);s__Metschnikowia_pulcherrima(100); | Metschnikowia | 13 | Saprotroph | \|Undefined Saprotroph\| | Yeast | NULL | Probable | As nectar-tap_saprotroph (Põlme et al. 2020); Decay sugar-rich substrates (Põlme et al. 2020) | Põlme S, et al. 2020. Fungal Diversity 105:1-16 ((https://doi.org/10.1007/s13225-020-00466-2)) |
| Otu35541 | 1 | k__Fungi(100);p__Ascomycota(100);c__Saccharomycetes(100);o__Saccharomycetales(100);f__Metschnikowiaceae(100);g__Metschnikowia(100);g__Metschnikowia_unclassified(100); | Metschnikowia | 13 | Saprotroph | \|Undefined Saprotroph\| | Yeast | NULL | Probable | As nectar-tap_saprotroph (Põlme et al. 2020); Decay sugar-rich substrates (Põlme et al. 2020) | Põlme S, et al. 2020. Fungal Diversity 105:1-16 ((https://doi.org/10.1007/s13225-020-00466-2)) |
| Otu35542 | 1 | k__Fungi(100);p__Ascomycota(100);c__Saccharomycetes(100);o__Saccharomycetales(100);f__Metschnikowiaceae(100);g__Metschnikowia(100);s__Metschnikowia_pulcherrima(100); | Metschnikowia | 13 | Saprotroph | \|Undefined Saprotroph\| | Yeast | NULL | Probable | As nectar-tap_saprotroph (Põlme et al. 2020); Decay sugar-rich substrates (Põlme et al. 2020) | Põlme S, et al. 2020. Fungal Diversity 105:1-16 ((https://doi.org/10.1007/s13225-020-00466-2)) |
| Otu35543 | 1 | k__Fungi(100);p__Ascomycota(100);c__Saccharomycetes(100);o__Saccharomycetales(100);f__Metschnikowiaceae(100);g__Metschnikowia(100);s__Metschnikowia_pulcherrima(100); | Metschnikowia | 13 | Saprotroph | \|Undefined Saprotroph\| | Yeast | NULL | Probable | As nectar-tap_saprotroph (Põlme et al. 2020); Decay sugar-rich substrates (Põlme et al. 2020) | Põlme S, et al. 2020. Fungal Diversity 105:1-16 ((https://doi.org/10.1007/s13225-020-00466-2)) |
| Otu35544 | 1 | k__Fungi(100);p__Ascomycota(100);c__Saccharomycetes(100);o__Saccharomycetales(100);f__Metschnikowiaceae(100);g__Metschnikowia(100);g__Metschnikowia_unclassified(100); | Metschnikowia | 13 | Saprotroph | \|Undefined Saprotroph\| | Yeast | NULL | Probable | As nectar-tap_saprotroph (Põlme et al. 2020); Decay sugar-rich substrates (Põlme et al. 2020) | Põlme S, et al. 2020. Fungal Diversity 105:1-16 ((https://doi.org/10.1007/s13225-020-00466-2)) |
| Otu35545 | 1 | k__Fungi(100);p__Ascomycota(100);c__Saccharomycetes(100);o__Saccharomycetales(100);f__Metschnikowiaceae(100);g__Metschnikowia(100);s__Metschnikowia_pulcherrima(100); | Metschnikowia | 13 | Saprotroph | \|Undefined Saprotroph\| | Yeast | NULL | Probable | As nectar-tap_saprotroph (Põlme et al. 2020); Decay sugar-rich substrates (Põlme et al. 2020) | Põlme S, et al. 2020. Fungal Diversity 105:1-16 ((https://doi.org/10.1007/s13225-020-00466-2)) |
| Otu35547 | 1 | k__Fungi(100);p__Ascomycota(100);c__Saccharomycetes(100);o__Saccharomycetales(100);f__Metschnikowiaceae(100);g__Metschnikowia(100);g__Metschnikowia_unclassified(100); | Metschnikowia | 13 | Saprotroph | \|Undefined Saprotroph\| | Yeast | NULL | Probable | As nectar-tap_saprotroph (Põlme et al. 2020); Decay sugar-rich substrates (Põlme et al. 2020) | Põlme S, et al. 2020. Fungal Diversity 105:1-16 ((https://doi.org/10.1007/s13225-020-00466-2)) |
| Otu35548 | 1 | k__Fungi(100);p__Ascomycota(100);c__Saccharomycetes(100);o__Saccharomycetales(100);f__Metschnikowiaceae(100);g__Metschnikowia(100);s__Metschnikowia_pulcherrima(100); | Metschnikowia | 13 | Saprotroph | \|Undefined Saprotroph\| | Yeast | NULL | Probable | As nectar-tap_saprotroph (Põlme et al. 2020); Decay sugar-rich substrates (Põlme et al. 2020) | Põlme S, et al. 2020. Fungal Diversity 105:1-16 ((https://doi.org/10.1007/s13225-020-00466-2)) |
| Otu35549 | 1 | k__Fungi(100);p__Ascomycota(100);c__Saccharomycetes(100);o__Saccharomycetales(100);f__Metschnikowiaceae(100);g__Metschnikowia(100);s__Metschnikowia_pulcherrima(100); | Metschnikowia | 13 | Saprotroph | \|Undefined Saprotroph\| | Yeast | NULL | Probable | As nectar-tap_saprotroph (Põlme et al. 2020); Decay sugar-rich substrates (Põlme et al. 2020) | Põlme S, et al. 2020. Fungal Diversity 105:1-16 ((https://doi.org/10.1007/s13225-020-00466-2)) |
| Otu35550 | 1 | k__Fungi(100);p__Ascomycota(100);c__Saccharomycetes(100);o__Saccharomycetales(100);f__Metschnikowiaceae(100);g__Metschnikowia(100);s__Metschnikowia_pulcherrima(100); | Metschnikowia | 13 | Saprotroph | \|Undefined Saprotroph\| | Yeast | NULL | Probable | As nectar-tap_saprotroph (Põlme et al. 2020); Decay sugar-rich substrates (Põlme et al. 2020) | Põlme S, et al. 2020. Fungal Diversity 105:1-16 ((https://doi.org/10.1007/s13225-020-00466-2)) |
| Otu35552 | 1 | k__Fungi(100);p__Ascomycota(100);c__Saccharomycetes(100);o__Saccharomycetales(100);f__Metschnikowiaceae(100);g__Metschnikowia(100);s__Metschnikowia_pulcherrima(100); | Metschnikowia | 13 | Saprotroph | \|Undefined Saprotroph\| | Yeast | NULL | Probable | As nectar-tap_saprotroph (Põlme et al. 2020); Decay sugar-rich substrates (Põlme et al. 2020) | Põlme S, et al. 2020. Fungal Diversity 105:1-16 ((https://doi.org/10.1007/s13225-020-00466-2)) |
| Otu35553 | 1 | k__Fungi(100);p__Ascomycota(100);c__Saccharomycetes(100);o__Saccharomycetales(100);f__Metschnikowiaceae(100);g__Metschnikowia(100);s__Metschnikowia_pulcherrima(100); | Metschnikowia | 13 | Saprotroph | \|Undefined Saprotroph\| | Yeast | NULL | Probable | As nectar-tap_saprotroph (Põlme et al. 2020); Decay sugar-rich substrates (Põlme et al. 2020) | Põlme S, et al. 2020. Fungal Diversity 105:1-16 ((https://doi.org/10.1007/s13225-020-00466-2)) |
| Otu35554 | 1 | k__Fungi(100);p__Ascomycota(100);c__Saccharomycetes(100);o__Saccharomycetales(100);f__Metschnikowiaceae(100);g__Metschnikowia(100);s__Metschnikowia_pulcherrima(100); | Metschnikowia | 13 | Saprotroph | \|Undefined Saprotroph\| | Yeast | NULL | Probable | As nectar-tap_saprotroph (Põlme et al. 2020); Decay sugar-rich substrates (Põlme et al. 2020) | Põlme S, et al. 2020. Fungal Diversity 105:1-16 ((https://doi.org/10.1007/s13225-020-00466-2)) |
| Otu35555 | 1 | k__Fungi(100);p__Ascomycota(100);c__Saccharomycetes(100);o__Saccharomycetales(100);f__Metschnikowiaceae(100);g__Metschnikowia(100);s__Metschnikowia_pulcherrima(100); | Metschnikowia | 13 | Saprotroph | \|Undefined Saprotroph\| | Yeast | NULL | Probable | As nectar-tap_saprotroph (Põlme et al. 2020); Decay sugar-rich substrates (Põlme et al. 2020) | Põlme S, et al. 2020. Fungal Diversity 105:1-16 ((https://doi.org/10.1007/s13225-020-00466-2)) |
| Otu35556 | 1 | k__Fungi(100);p__Ascomycota(100);c__Saccharomycetes(100);o__Saccharomycetales(100);f__Metschnikowiaceae(100);g__Metschnikowia(100);s__Metschnikowia_pulcherrima(100); | Metschnikowia | 13 | Saprotroph | \|Undefined Saprotroph\| | Yeast | NULL | Probable | As nectar-tap_saprotroph (Põlme et al. 2020); Decay sugar-rich substrates (Põlme et al. 2020) | Põlme S, et al. 2020. Fungal Diversity 105:1-16 ((https://doi.org/10.1007/s13225-020-00466-2)) |
| Otu35557 | 1 | k__Fungi(100);p__Ascomycota(100);c__Saccharomycetes(100);o__Saccharomycetales(100);f__Metschnikowiaceae(100);g__Metschnikowia(100);g__Metschnikowia_unclassified(100); | Metschnikowia | 13 | Saprotroph | \|Undefined Saprotroph\| | Yeast | NULL | Probable | As nectar-tap_saprotroph (Põlme et al. 2020); Decay sugar-rich substrates (Põlme et al. 2020) | Põlme S, et al. 2020. Fungal Diversity 105:1-16 ((https://doi.org/10.1007/s13225-020-00466-2)) |
| Otu35558 | 1 | k__Fungi(100);p__Ascomycota(100);c__Saccharomycetes(100);o__Saccharomycetales(100);f__Metschnikowiaceae(100);g__Metschnikowia(100);s__Metschnikowia_pulcherrima(100); | Metschnikowia | 13 | Saprotroph | \|Undefined Saprotroph\| | Yeast | NULL | Probable | As nectar-tap_saprotroph (Põlme et al. 2020); Decay sugar-rich substrates (Põlme et al. 2020) | Põlme S, et al. 2020. Fungal Diversity 105:1-16 ((https://doi.org/10.1007/s13225-020-00466-2)) |
| Otu35559 | 1 | k__Fungi(100);p__Ascomycota(100);c__Saccharomycetes(100);o__Saccharomycetales(100);f__Metschnikowiaceae(100);g__Metschnikowia(100);g__Metschnikowia_unclassified(100); | Metschnikowia | 13 | Saprotroph | \|Undefined Saprotroph\| | Yeast | NULL | Probable | As nectar-tap_saprotroph (Põlme et al. 2020); Decay sugar-rich substrates (Põlme et al. 2020) | Põlme S, et al. 2020. Fungal Diversity 105:1-16 ((https://doi.org/10.1007/s13225-020-00466-2)) |
| Otu35561 | 1 | k__Fungi(100);p__Ascomycota(100);c__Saccharomycetes(100);o__Saccharomycetales(100);f__Metschnikowiaceae(100);g__Metschnikowia(100);s__Metschnikowia_pulcherrima(100); | Metschnikowia | 13 | Saprotroph | \|Undefined Saprotroph\| | Yeast | NULL | Probable | As nectar-tap_saprotroph (Põlme et al. 2020); Decay sugar-rich substrates (Põlme et al. 2020) | Põlme S, et al. 2020. Fungal Diversity 105:1-16 ((https://doi.org/10.1007/s13225-020-00466-2)) |
| Otu35562 | 1 | k__Fungi(100);p__Ascomycota(100);c__Saccharomycetes(100);o__Saccharomycetales(100);f__Metschnikowiaceae(100);g__Metschnikowia(100);s__Metschnikowia_pulcherrima(100); | Metschnikowia | 13 | Saprotroph | \|Undefined Saprotroph\| | Yeast | NULL | Probable | As nectar-tap_saprotroph (Põlme et al. 2020); Decay sugar-rich substrates (Põlme et al. 2020) | Põlme S, et al. 2020. Fungal Diversity 105:1-16 ((https://doi.org/10.1007/s13225-020-00466-2)) |
| Otu35563 | 1 | k__Fungi(100);p__Ascomycota(100);c__Saccharomycetes(100);o__Saccharomycetales(100);f__Metschnikowiaceae(100);g__Metschnikowia(100);s__Metschnikowia_pulcherrima(100); | Metschnikowia | 13 | Saprotroph | \|Undefined Saprotroph\| | Yeast | NULL | Probable | As nectar-tap_saprotroph (Põlme et al. 2020); Decay sugar-rich substrates (Põlme et al. 2020) | Põlme S, et al. 2020. Fungal Diversity 105:1-16 ((https://doi.org/10.1007/s13225-020-00466-2)) |
| Otu35564 | 1 | k__Fungi(100);p__Ascomycota(100);c__Saccharomycetes(100);o__Saccharomycetales(100);f__Metschnikowiaceae(100);g__Metschnikowia(100);g__Metschnikowia_unclassified(100); | Metschnikowia | 13 | Saprotroph | \|Undefined Saprotroph\| | Yeast | NULL | Probable | As nectar-tap_saprotroph (Põlme et al. 2020); Decay sugar-rich substrates (Põlme et al. 2020) | Põlme S, et al. 2020. Fungal Diversity 105:1-16 ((https://doi.org/10.1007/s13225-020-00466-2)) |
| Otu35565 | 1 | k__Fungi(100);p__Ascomycota(100);c__Saccharomycetes(100);o__Saccharomycetales(100);f__Metschnikowiaceae(100);g__Metschnikowia(100);s__Metschnikowia_pulcherrima(100); | Metschnikowia | 13 | Saprotroph | \|Undefined Saprotroph\| | Yeast | NULL | Probable | As nectar-tap_saprotroph (Põlme et al. 2020); Decay sugar-rich substrates (Põlme et al. 2020) | Põlme S, et al. 2020. Fungal Diversity 105:1-16 ((https://doi.org/10.1007/s13225-020-00466-2)) |
| Otu35566 | 1 | k__Fungi(100);p__Ascomycota(100);c__Saccharomycetes(100);o__Saccharomycetales(100);f__Metschnikowiaceae(100);g__Metschnikowia(100);g__Metschnikowia_unclassified(100); | Metschnikowia | 13 | Saprotroph | \|Undefined Saprotroph\| | Yeast | NULL | Probable | As nectar-tap_saprotroph (Põlme et al. 2020); Decay sugar-rich substrates (Põlme et al. 2020) | Põlme S, et al. 2020. Fungal Diversity 105:1-16 ((https://doi.org/10.1007/s13225-020-00466-2)) |
| Otu35567 | 1 | k__Fungi(100);p__Ascomycota(100);c__Saccharomycetes(100);o__Saccharomycetales(100);f__Metschnikowiaceae(100);g__Metschnikowia(100);s__Metschnikowia_pulcherrima(100); | Metschnikowia | 13 | Saprotroph | \|Undefined Saprotroph\| | Yeast | NULL | Probable | As nectar-tap_saprotroph (Põlme et al. 2020); Decay sugar-rich substrates (Põlme et al. 2020) | Põlme S, et al. 2020. Fungal Diversity 105:1-16 ((https://doi.org/10.1007/s13225-020-00466-2)) |
| Otu35571 | 1 | k__Fungi(100);p__Ascomycota(100);c__Saccharomycetes(100);o__Saccharomycetales(100);f__Metschnikowiaceae(100);g__Metschnikowia(100);s__Metschnikowia_pulcherrima(100); | Metschnikowia | 13 | Saprotroph | \|Undefined Saprotroph\| | Yeast | NULL | Probable | As nectar-tap_saprotroph (Põlme et al. 2020); Decay sugar-rich substrates (Põlme et al. 2020) | Põlme S, et al. 2020. Fungal Diversity 105:1-16 ((https://doi.org/10.1007/s13225-020-00466-2)) |
| Otu35574 | 1 | k__Fungi(100);p__Ascomycota(100);c__Saccharomycetes(100);o__Saccharomycetales(100);f__Metschnikowiaceae(100);g__Metschnikowia(100);s__Metschnikowia_pulcherrima(100); | Metschnikowia | 13 | Saprotroph | \|Undefined Saprotroph\| | Yeast | NULL | Probable | As nectar-tap_saprotroph (Põlme et al. 2020); Decay sugar-rich substrates (Põlme et al. 2020) | Põlme S, et al. 2020. Fungal Diversity 105:1-16 ((https://doi.org/10.1007/s13225-020-00466-2)) |

| | | | | | | | | | | | |
|---|---|---|---|---|---|---|---|---|---|---|---|
| Otu35575 | 1 | k__Fungi(100);p__Ascomycota(100);c__Sordariomycetes(100);o__Sordariales(100);f__Sordariales_family_Incertae_sedis(100);g__Ramophialophora(100);s__Ramophialophora_sp_FMR_9523(100); | Ramophialophora | 13 | Saprotroph | \|Undefined Saprotroph\| | Microfungus | NULL | Probable | As soil_saprotroph (Põlme et al. 2020); Decay soil (Põlme et al. 2020) | Põlme S, et al. 2020. Fungal Diversity 105:1-16 ((https://doi.org/10.1007/s13225-020-00466-2)) |
| Otu35577 | 1 | k__Fungi(100);p__Ascomycota(100);c__Saccharomycetes(100);o__Saccharomycetales(100);f__Metschnikowiaceae(100);g__Metschnikowia(100);g__Metschnikowia_unclassified(100); | Metschnikowia | 13 | Saprotroph | \|Undefined Saprotroph\| | Yeast | NULL | Probable | As nectar-tap_saprotroph (Põlme et al. 2020); Decay sugar-rich substrates (Põlme et al. 2020) | Põlme S, et al. 2020. Fungal Diversity 105:1-16 ((https://doi.org/10.1007/s13225-020-00466-2)) |
| Otu35579 | 1 | k__Fungi(100);p__Ascomycota(100);c__Saccharomycetes(100);o__Saccharomycetales(100);f__Metschnikowiaceae(100);g__Metschnikowia(100);g__Metschnikowia_unclassified(100); | Metschnikowia | 13 | Saprotroph | \|Undefined Saprotroph\| | Yeast | NULL | Probable | As nectar-tap_saprotroph (Põlme et al. 2020); Decay sugar-rich substrates (Põlme et al. 2020) | Põlme S, et al. 2020. Fungal Diversity 105:1-16 ((https://doi.org/10.1007/s13225-020-00466-2)) |
| Otu35580 | 1 | k__Fungi(100);p__Ascomycota(100);c__Saccharomycetes(100);o__Saccharomycetales(100);f__Metschnikowiaceae(100);g__Metschnikowia(100);s__Metschnikowia_pulcherrima(100); | Metschnikowia | 13 | Saprotroph | \|Undefined Saprotroph\| | Yeast | NULL | Probable | As nectar-tap_saprotroph (Põlme et al. 2020); Decay sugar-rich substrates (Põlme et al. 2020) | Põlme S, et al. 2020. Fungal Diversity 105:1-16 ((https://doi.org/10.1007/s13225-020-00466-2)) |
| Otu35581 | 1 | k__Fungi(100);p__Ascomycota(100);c__Saccharomycetes(100);o__Saccharomycetales(100);f__Metschnikowiaceae(100);g__Metschnikowia(100);s__Metschnikowia_pulcherrima(100); | Metschnikowia | 13 | Saprotroph | \|Undefined Saprotroph\| | Yeast | NULL | Probable | As nectar-tap_saprotroph (Põlme et al. 2020); Decay sugar-rich substrates (Põlme et al. 2020) | Põlme S, et al. 2020. Fungal Diversity 105:1-16 ((https://doi.org/10.1007/s13225-020-00466-2)) |
| Otu35582 | 1 | k__Fungi(100);p__Ascomycota(100);c__Saccharomycetes(100);o__Saccharomycetales(100);f__Metschnikowiaceae(100);g__Metschnikowia(100);s__Metschnikowia_pulcherrima(100); | Metschnikowia | 13 | Saprotroph | \|Undefined Saprotroph\| | Yeast | NULL | Probable | As nectar-tap_saprotroph (Põlme et al. 2020); Decay sugar-rich substrates (Põlme et al. 2020) | Põlme S, et al. 2020. Fungal Diversity 105:1-16 ((https://doi.org/10.1007/s13225-020-00466-2)) |
| Otu35583 | 1 | k__Fungi(100);p__Ascomycota(100);c__Saccharomycetes(100);o__Saccharomycetales(100);f__Metschnikowiaceae(100);g__Metschnikowia(100);s__Metschnikowia_pulcherrima(100); | Metschnikowia | 13 | Saprotroph | \|Undefined Saprotroph\| | Yeast | NULL | Probable | As nectar-tap_saprotroph (Põlme et al. 2020); Decay sugar-rich substrates (Põlme et al. 2020) | Põlme S, et al. 2020. Fungal Diversity 105:1-16 ((https://doi.org/10.1007/s13225-020-00466-2)) |
| Otu35584 | 1 | k__Fungi(100);p__Ascomycota(100);c__Saccharomycetes(100);o__Saccharomycetales(100);f__Metschnikowiaceae(100);g__Metschnikowia(100);s__Metschnikowia_pulcherrima(100); | Metschnikowia | 13 | Saprotroph | \|Undefined Saprotroph\| | Yeast | NULL | Probable | As nectar-tap_saprotroph (Põlme et al. 2020); Decay sugar-rich substrates (Põlme et al. 2020) | Põlme S, et al. 2020. Fungal Diversity 105:1-16 ((https://doi.org/10.1007/s13225-020-00466-2)) |
| Otu35586 | 1 | k__Fungi(100);p__Ascomycota(100);c__Saccharomycetes(100);o__Saccharomycetales(100);f__Metschnikowiaceae(100);g__Metschnikowia(100);s__Metschnikowia_pulcherrima(100); | Metschnikowia | 13 | Saprotroph | \|Undefined Saprotroph\| | Yeast | NULL | Probable | As nectar-tap_saprotroph (Põlme et al. 2020); Decay sugar-rich substrates (Põlme et al. 2020) | Põlme S, et al. 2020. Fungal Diversity 105:1-16 ((https://doi.org/10.1007/s13225-020-00466-2)) |
| Otu35587 | 1 | k__Fungi(100);p__Ascomycota(100);c__Saccharomycetes(100);o__Saccharomycetales(100);f__Metschnikowiaceae(100);g__Metschnikowia(100);s__Metschnikowia_pulcherrima(100); | Metschnikowia | 13 | Saprotroph | \|Undefined Saprotroph\| | Yeast | NULL | Probable | As nectar-tap_saprotroph (Põlme et al. 2020); Decay sugar-rich substrates (Põlme et al. 2020) | Põlme S, et al. 2020. Fungal Diversity 105:1-16 ((https://doi.org/10.1007/s13225-020-00466-2)) |
| Otu35588 | 1 | k__Fungi(100);p__Ascomycota(100);c__Saccharomycetes(100);o__Saccharomycetales(100);f__Metschnikowiaceae(100);g__Metschnikowia(100);s__Metschnikowia_pulcherrima(100); | Metschnikowia | 13 | Saprotroph | \|Undefined Saprotroph\| | Yeast | NULL | Probable | As nectar-tap_saprotroph (Põlme et al. 2020); Decay sugar-rich substrates (Põlme et al. 2020) | Põlme S, et al. 2020. Fungal Diversity 105:1-16 ((https://doi.org/10.1007/s13225-020-00466-2)) |
| Otu35589 | 1 | k__Fungi(100);p__Ascomycota(100);c__Saccharomycetes(100);o__Saccharomycetales(100);f__Metschnikowiaceae(100);g__Metschnikowia(100);s__Metschnikowia_pulcherrima(100); | Metschnikowia | 13 | Saprotroph | \|Undefined Saprotroph\| | Yeast | NULL | Probable | As nectar-tap_saprotroph (Põlme et al. 2020); Decay sugar-rich substrates (Põlme et al. 2020) | Põlme S, et al. 2020. Fungal Diversity 105:1-16 ((https://doi.org/10.1007/s13225-020-00466-2)) |
| Otu35590 | 1 | k__Fungi(100);p__Ascomycota(100);c__Saccharomycetes(100);o__Saccharomycetales(100);f__Metschnikowiaceae(100);g__Metschnikowia(100);s__Metschnikowia_pulcherrima(100); | Metschnikowia | 13 | Saprotroph | \|Undefined Saprotroph\| | Yeast | NULL | Probable | As nectar-tap_saprotroph (Põlme et al. 2020); Decay sugar-rich substrates (Põlme et al. 2020) | Põlme S, et al. 2020. Fungal Diversity 105:1-16 ((https://doi.org/10.1007/s13225-020-00466-2)) |
| Otu35591 | 1 | k__Fungi(100);p__Ascomycota(100);c__Saccharomycetes(100);o__Saccharomycetales(100);f__Metschnikowiaceae(100);g__Metschnikowia(100);s__Metschnikowia_pulcherrima(100); | Metschnikowia | 13 | Saprotroph | \|Undefined Saprotroph\| | Yeast | NULL | Probable | As nectar-tap_saprotroph (Põlme et al. 2020); Decay sugar-rich substrates (Põlme et al. 2020) | Põlme S, et al. 2020. Fungal Diversity 105:1-16 ((https://doi.org/10.1007/s13225-020-00466-2)) |
| Otu35592 | 1 | k__Fungi(100);p__Ascomycota(100);c__Saccharomycetes(100);o__Saccharomycetales(100);f__Metschnikowiaceae(100);g__Metschnikowia(100);g__Metschnikowia_unclassified(100); | Metschnikowia | 13 | Saprotroph | \|Undefined Saprotroph\| | Yeast | NULL | Probable | As nectar-tap_saprotroph (Põlme et al. 2020); Decay sugar-rich substrates (Põlme et al. 2020) | Põlme S, et al. 2020. Fungal Diversity 105:1-16 ((https://doi.org/10.1007/s13225-020-00466-2)) |
| Otu35593 | 1 | k__Fungi(100);p__Ascomycota(100);c__Saccharomycetes(100);o__Saccharomycetales(100);f__Metschnikowiaceae(100);g__Metschnikowia(100);s__Metschnikowia_pulcherrima(100); | Metschnikowia | 13 | Saprotroph | \|Undefined Saprotroph\| | Yeast | NULL | Probable | As nectar-tap_saprotroph (Põlme et al. 2020); Decay sugar-rich substrates (Põlme et al. 2020) | Põlme S, et al. 2020. Fungal Diversity 105:1-16 ((https://doi.org/10.1007/s13225-020-00466-2)) |
| Otu35595 | 1 | k__Fungi(100);p__Ascomycota(100);c__Saccharomycetes(100);o__Saccharomycetales(100);f__Metschnikowiaceae(100);g__Metschnikowia(100);s__Metschnikowia_pulcherrima(100); | Metschnikowia | 13 | Saprotroph | \|Undefined Saprotroph\| | Yeast | NULL | Probable | As nectar-tap_saprotroph (Põlme et al. 2020); Decay sugar-rich substrates (Põlme et al. 2020) | Põlme S, et al. 2020. Fungal Diversity 105:1-16 ((https://doi.org/10.1007/s13225-020-00466-2)) |
| Otu35597 | 1 | k__Fungi(100);p__Ascomycota(100);c__Saccharomycetes(100);o__Saccharomycetales(100);f__Metschnikowiaceae(100);g__Metschnikowia(100);s__Metschnikowia_pulcherrima(100); | Metschnikowia | 13 | Saprotroph | \|Undefined Saprotroph\| | Yeast | NULL | Probable | As nectar-tap_saprotroph (Põlme et al. 2020); Decay sugar-rich substrates (Põlme et al. 2020) | Põlme S, et al. 2020. Fungal Diversity 105:1-16 ((https://doi.org/10.1007/s13225-020-00466-2)) |
| Otu35598 | 1 | k__Fungi(100);p__Ascomycota(100);c__Saccharomycetes(100);o__Saccharomycetales(100);f__Metschnikowiaceae(100);g__Metschnikowia(100);s__Metschnikowia_pulcherrima(100); | Metschnikowia | 13 | Saprotroph | \|Undefined Saprotroph\| | Yeast | NULL | Probable | As nectar-tap_saprotroph (Põlme et al. 2020); Decay sugar-rich substrates (Põlme et al. 2020) | Põlme S, et al. 2020. Fungal Diversity 105:1-16 ((https://doi.org/10.1007/s13225-020-00466-2)) |
| Otu35599 | 1 | k__Fungi(100);p__Ascomycota(100);c__Saccharomycetes(100);o__Saccharomycetales(100);f__Metschnikowiaceae(100);g__Metschnikowia(100);s__Metschnikowia_pulcherrima(100); | Metschnikowia | 13 | Saprotroph | \|Undefined Saprotroph\| | Yeast | NULL | Probable | As nectar-tap_saprotroph (Põlme et al. 2020); Decay sugar-rich substrates (Põlme et al. 2020) | Põlme S, et al. 2020. Fungal Diversity 105:1-16 ((https://doi.org/10.1007/s13225-020-00466-2)) |
| Otu35600 | 1 | k__Fungi(100);p__Ascomycota(100);c__Saccharomycetes(100);o__Saccharomycetales(100);f__Metschnikowiaceae(100);g__Metschnikowia(100);s__Metschnikowia_pulcherrima(100); | Metschnikowia | 13 | Saprotroph | \|Undefined Saprotroph\| | Yeast | NULL | Probable | As nectar-tap_saprotroph (Põlme et al. 2020); Decay sugar-rich substrates (Põlme et al. 2020) | Põlme S, et al. 2020. Fungal Diversity 105:1-16 ((https://doi.org/10.1007/s13225-020-00466-2)) |
| Otu35601 | 1 | k__Fungi(100);p__Ascomycota(100);c__Saccharomycetes(100);o__Saccharomycetales(100);f__Metschnikowiaceae(100);g__Metschnikowia(100);s__Metschnikowia_pulcherrima(100); | Metschnikowia | 13 | Saprotroph | \|Undefined Saprotroph\| | Yeast | NULL | Probable | As nectar-tap_saprotroph (Põlme et al. 2020); Decay sugar-rich substrates (Põlme et al. 2020) | Põlme S, et al. 2020. Fungal Diversity 105:1-16 ((https://doi.org/10.1007/s13225-020-00466-2)) |
| Otu35602 | 1 | k__Fungi(100);p__Ascomycota(100);c__Saccharomycetes(100);o__Saccharomycetales(100);f__Metschnikowiaceae(100);g__Metschnikowia(100);s__Metschnikowia_pulcherrima(100); | Metschnikowia | 13 | Saprotroph | \|Undefined Saprotroph\| | Yeast | NULL | Probable | As nectar-tap_saprotroph (Põlme et al. 2020); Decay sugar-rich substrates (Põlme et al. 2020) | Põlme S, et al. 2020. Fungal Diversity 105:1-16 ((https://doi.org/10.1007/s13225-020-00466-2)) |
| Otu35603 | 1 | k__Fungi(100);p__Ascomycota(100);c__Saccharomycetes(100);o__Saccharomycetales(100);f__Metschnikowiaceae(100);g__Metschnikowia(100);s__Metschnikowia_pulcherrima(100); | Metschnikowia | 13 | Saprotroph | \|Undefined Saprotroph\| | Yeast | NULL | Probable | As nectar-tap_saprotroph (Põlme et al. 2020); Decay sugar-rich substrates (Põlme et al. 2020) | Põlme S, et al. 2020. Fungal Diversity 105:1-16 ((https://doi.org/10.1007/s13225-020-00466-2)) |
| Otu35604 | 1 | k__Fungi(100);p__Ascomycota(100);c__Saccharomycetes(100);o__Saccharomycetales(100);f__Metschnikowiaceae(100);g__Metschnikowia(100);g__Metschnikowia_unclassified(100); | Metschnikowia | 13 | Saprotroph | \|Undefined Saprotroph\| | Yeast | NULL | Probable | As nectar-tap_saprotroph (Põlme et al. 2020); Decay sugar-rich substrates (Põlme et al. 2020) | Põlme S, et al. 2020. Fungal Diversity 105:1-16 ((https://doi.org/10.1007/s13225-020-00466-2)) |
| Otu35606 | 1 | k__Fungi(100);p__Ascomycota(100);c__Saccharomycetes(100);o__Saccharomycetales(100);f__Metschnikowiaceae(100);g__Metschnikowia(100);s__Metschnikowia_pulcherrima(100); | Metschnikowia | 13 | Saprotroph | \|Undefined Saprotroph\| | Yeast | NULL | Probable | As nectar-tap_saprotroph (Põlme et al. 2020); Decay sugar-rich substrates (Põlme et al. 2020) | Põlme S, et al. 2020. Fungal Diversity 105:1-16 ((https://doi.org/10.1007/s13225-020-00466-2)) |
| Otu35610 | 1 | k__Fungi(100);p__Ascomycota(100);c__Saccharomycetes(100);o__Saccharomycetales(100);f__Metschnikowiaceae(100);g__Metschnikowia(100);s__Metschnikowia_pulcherrima(100); | Metschnikowia | 13 | Saprotroph | \|Undefined Saprotroph\| | Yeast | NULL | Probable | As nectar-tap_saprotroph (Põlme et al. 2020); Decay sugar-rich substrates (Põlme et al. 2020) | Põlme S, et al. 2020. Fungal Diversity 105:1-16 ((https://doi.org/10.1007/s13225-020-00466-2)) |
| Otu35612 | 1 | k__Fungi(100);p__Ascomycota(100);c__Saccharomycetes(100);o__Saccharomycetales(100);f__Metschnikowiaceae(100);g__Metschnikowia(100);s__Metschnikowia_pulcherrima(100); | Metschnikowia | 13 | Saprotroph | \|Undefined Saprotroph\| | Yeast | NULL | Probable | As nectar-tap_saprotroph (Põlme et al. 2020); Decay sugar-rich substrates (Põlme et al. 2020) | Põlme S, et al. 2020. Fungal Diversity 105:1-16 ((https://doi.org/10.1007/s13225-020-00466-2)) |
| Otu35619 | 1 | k__Fungi(100);p__Ascomycota(100);c__Saccharomycetes(100);o__Saccharomycetales(100);f__Metschnikowiaceae(100);g__Metschnikowia(100);g__Metschnikowia_unclassified(100); | Metschnikowia | 13 | Saprotroph | \|Undefined Saprotroph\| | Yeast | NULL | Probable | As nectar-tap_saprotroph (Põlme et al. 2020); Decay sugar-rich substrates (Põlme et al. 2020) | Põlme S, et al. 2020. Fungal Diversity 105:1-16 ((https://doi.org/10.1007/s13225-020-00466-2)) |
| Otu35620 | 1 | k__Fungi(100);p__Ascomycota(100);c__Saccharomycetes(100);o__Saccharomycetales(100);f__Metschnikowiaceae(100);g__Metschnikowia(100);s__Metschnikowia_pulcherrima(100); | Metschnikowia | 13 | Saprotroph | \|Undefined Saprotroph\| | Yeast | NULL | Probable | As nectar-tap_saprotroph (Põlme et al. 2020); Decay sugar-rich substrates (Põlme et al. 2020) | Põlme S, et al. 2020. Fungal Diversity 105:1-16 ((https://doi.org/10.1007/s13225-020-00466-2)) |
| Otu35624 | 1 | k__Fungi(100);p__Ascomycota(100);c__Saccharomycetes(100);o__Saccharomycetales(100);f__Metschnikowiaceae(100);g__Metschnikowia(100);s__Metschnikowia_pulcherrima(100); | Metschnikowia | 13 | Saprotroph | \|Undefined Saprotroph\| | Yeast | NULL | Probable | As nectar-tap_saprotroph (Põlme et al. 2020); Decay sugar-rich substrates (Põlme et al. 2020) | Põlme S, et al. 2020. Fungal Diversity 105:1-16 ((https://doi.org/10.1007/s13225-020-00466-2)) |
| Otu35625 | 1 | k__Fungi(100);p__Ascomycota(100);c__Saccharomycetes(100);o__Saccharomycetales(100);f__Metschnikowiaceae(100);g__Metschnikowia(100);s__Metschnikowia_pulcherrima(100); | Metschnikowia | 13 | Saprotroph | \|Undefined Saprotroph\| | Yeast | NULL | Probable | As nectar-tap_saprotroph (Põlme et al. 2020); Decay sugar-rich substrates (Põlme et al. 2020) | Põlme S, et al. 2020. Fungal Diversity 105:1-16 ((https://doi.org/10.1007/s13225-020-00466-2)) |
| Otu35626 | 1 | k__Fungi(100);p__Ascomycota(100);c__Saccharomycetes(100);o__Saccharomycetales(100);f__Metschnikowiaceae(100);g__Metschnikowia(100);g__Metschnikowia_unclassified(100); | Metschnikowia | 13 | Saprotroph | \|Undefined Saprotroph\| | Yeast | NULL | Probable | As nectar-tap_saprotroph (Põlme et al. 2020); Decay sugar-rich substrates (Põlme et al. 2020) | Põlme S, et al. 2020. Fungal Diversity 105:1-16 ((https://doi.org/10.1007/s13225-020-00466-2)) |
| Otu35631 | 1 | k__Fungi(100);p__Ascomycota(100);c__Saccharomycetes(100);o__Saccharomycetales(100);f__Saccharomycetales_family_Incertae_sedis(100);g__Candida(100);g__Candida_unclassified(100); | Candida | 13 | Pathotroph-Saprotroph-Symbiotroph | Animal Pathogen-Endophyte-Endosymbiont-Epiphyte-\|Undefined Saprotroph\| | Dimorphic Yeast-Yeast | NULL | Probable | As nectar-tap_saprotroph (Põlme et al. 2020); Decay sugar-rich substrates (Põlme et al. 2020); This asexual genus is polyphyletic and contain species that function very differently depend on what they are. Caution should be taken when interpreting guilds for Candida species. | Manolakaki D, et al. 2010. Virulence. 1: 367-375 ((https://doi.org/10.4161/viru.1.5.12796)); Mekbib SB, et al. 2011. Tropical Plant Pathology 36:233-240 ((https://doi.org/10.1590/S1982-56762011000400004)); Põlme S, et al. 2020. Fungal Diversity 105:1-16 ((https://doi.org/10.1007/s13225-020-00466-2)) |
| Otu35632 | 1 | k__Fungi(100);p__Ascomycota(100);c__Saccharomycetes(100);o__Saccharomycetales(100);f__Metschnikowiaceae(100);g__Metschnikowia(100);s__Metschnikowia_pulcherrima(100); | Metschnikowia | 13 | Saprotroph | \|Undefined Saprotroph\| | Yeast | NULL | Probable | As nectar-tap_saprotroph (Põlme et al. 2020); Decay sugar-rich substrates (Põlme et al. 2020) | Põlme S, et al. 2020. Fungal Diversity 105:1-16 ((https://doi.org/10.1007/s13225-020-00466-2)) |
| Otu35635 | 1 | k__Fungi(100);p__Ascomycota(100);c__Saccharomycetes(100);o__Saccharomycetales(100);f__Metschnikowiaceae(100);g__Metschnikowia(100);s__Metschnikowia_pulcherrima(100); | Metschnikowia | 13 | Saprotroph | \|Undefined Saprotroph\| | Yeast | NULL | Probable | As nectar-tap_saprotroph (Põlme et al. 2020); Decay sugar-rich substrates (Põlme et al. 2020) | Põlme S, et al. 2020. Fungal Diversity 105:1-16 ((https://doi.org/10.1007/s13225-020-00466-2)) |
| Otu35636 | 1 | k__Fungi(100);p__Ascomycota(100);c__Saccharomycetes(100);o__Saccharomycetales(100);f__Metschnikowiaceae(100);g__Metschnikowia(100);s__Metschnikowia_pulcherrima(100); | Metschnikowia | 13 | Saprotroph | \|Undefined Saprotroph\| | Yeast | NULL | Probable | As nectar-tap_saprotroph (Põlme et al. 2020); Decay sugar-rich substrates (Põlme et al. 2020) | Põlme S, et al. 2020. Fungal Diversity 105:1-16 ((https://doi.org/10.1007/s13225-020-00466-2)) |
| Otu35637 | 1 | k__Fungi(100);p__Ascomycota(100);c__Saccharomycetes(100);o__Saccharomycetales(100);f__Metschnikowiaceae(100);g__Metschnikowia(100);s__Metschnikowia_pulcherrima(100); | Metschnikowia | 13 | Saprotroph | \|Undefined Saprotroph\| | Yeast | NULL | Probable | As nectar-tap_saprotroph (Põlme et al. 2020); Decay sugar-rich substrates (Põlme et al. 2020) | Põlme S, et al. 2020. Fungal Diversity 105:1-16 ((https://doi.org/10.1007/s13225-020-00466-2)) |
| Otu35638 | 1 | k__Fungi(100);p__Ascomycota(100);c__Saccharomycetes(100);o__Saccharomycetales(100);f__Metschnikowiaceae(100);g__Metschnikowia(100);s__Metschnikowia_pulcherrima(100); | Metschnikowia | 13 | Saprotroph | \|Undefined Saprotroph\| | Yeast | NULL | Probable | As nectar-tap_saprotroph (Põlme et al. 2020); Decay sugar-rich substrates (Põlme et al. 2020) | Põlme S, et al. 2020. Fungal Diversity 105:1-16 ((https://doi.org/10.1007/s13225-020-00466-2)) |
| Otu35639 | 1 | k__Fungi(100);p__Ascomycota(100);c__Saccharomycetes(100);o__Saccharomycetales(100);f__Metschnikowiaceae(100);g__Metschnikowia(100);s__Metschnikowia_pulcherrima(100); | Metschnikowia | 13 | Saprotroph | \|Undefined Saprotroph\| | Yeast | NULL | Probable | As nectar-tap_saprotroph (Põlme et al. 2020); Decay sugar-rich substrates (Põlme et al. 2020) | Põlme S, et al. 2020. Fungal Diversity 105:1-16 ((https://doi.org/10.1007/s13225-020-00466-2)) |
| Otu35642 | 1 | k__Fungi(100);p__Ascomycota(100);c__Saccharomycetes(100);o__Saccharomycetales(100);f__Metschnikowiaceae(100);g__Metschnikowia(100);s__Metschnikowia_pulcherrima(100); | Metschnikowia | 13 | Saprotroph | \|Undefined Saprotroph\| | Yeast | NULL | Probable | As nectar-tap_saprotroph (Põlme et al. 2020); Decay sugar-rich substrates (Põlme et al. 2020) | Põlme S, et al. 2020. Fungal Diversity 105:1-16 ((https://doi.org/10.1007/s13225-020-00466-2)) |
| Otu35643 | 1 | k__Fungi(100);p__Ascomycota(100);c__Saccharomycetes(100);o__Saccharomycetales(100);f__Metschnikowiaceae(100);g__Metschnikowia(100);g__Metschnikowia_unclassified(100); | Metschnikowia | 13 | Saprotroph | \|Undefined Saprotroph\| | Yeast | NULL | Probable | As nectar-tap_saprotroph (Põlme et al. 2020); Decay sugar-rich substrates (Põlme et al. 2020) | Põlme S, et al. 2020. Fungal Diversity 105:1-16 ((https://doi.org/10.1007/s13225-020-00466-2)) |
| Otu35645 | 1 | k__Fungi(100);p__Ascomycota(100);c__Saccharomycetes(100);o__Saccharomycetales(100);f__Metschnikowiaceae(100);g__Metschnikowia(100);s__Metschnikowia_pulcherrima(100); | Metschnikowia | 13 | Saprotroph | \|Undefined Saprotroph\| | Yeast | NULL | Probable | As nectar-tap_saprotroph (Põlme et al. 2020); Decay sugar-rich substrates (Põlme et al. 2020) | Põlme S, et al. 2020. Fungal Diversity 105:1-16 ((https://doi.org/10.1007/s13225-020-00466-2)) |
| Otu35646 | 1 | k__Fungi(100);p__Ascomycota(100);c__Saccharomycetes(100);o__Saccharomycetales(100);f__Metschnikowiaceae(100);g__Metschnikowia(100);s__Metschnikowia_pulcherrima(100); | Metschnikowia | 13 | Saprotroph | \|Undefined Saprotroph\| | Yeast | NULL | Probable | As nectar-tap_saprotroph (Põlme et al. 2020); Decay sugar-rich substrates (Põlme et al. 2020) | Põlme S, et al. 2020. Fungal Diversity 105:1-16 ((https://doi.org/10.1007/s13225-020-00466-2)) |
| Otu35647 | 1 | k__Fungi(100);p__Ascomycota(100);c__Saccharomycetes(100);o__Saccharomycetales(100);f__Metschnikowiaceae(100);g__Metschnikowia(100);g__Metschnikowia_unclassified(100); | Metschnikowia | 13 | Saprotroph | \|Undefined Saprotroph\| | Yeast | NULL | Probable | As nectar-tap_saprotroph (Põlme et al. 2020); Decay sugar-rich substrates (Põlme et al. 2020) | Põlme S, et al. 2020. Fungal Diversity 105:1-16 ((https://doi.org/10.1007/s13225-020-00466-2)) |
| Otu35648 | 1 | k__Fungi(100);p__Ascomycota(100);c__Saccharomycetes(100);o__Saccharomycetales(100);f__Metschnikowiaceae(100);g__Metschnikowia(100);g__Metschnikowia_unclassified(100); | Metschnikowia | 13 | Saprotroph | \|Undefined Saprotroph\| | Yeast | NULL | Probable | As nectar-tap_saprotroph (Põlme et al. 2020); Decay sugar-rich substrates (Põlme et al. 2020) | Põlme S, et al. 2020. Fungal Diversity 105:1-16 ((https://doi.org/10.1007/s13225-020-00466-2)) |
| Otu35649 | 1 | k__Fungi(100);p__Ascomycota(100);c__Saccharomycetes(100);o__Saccharomycetales(100);f__Metschnikowiaceae(100);g__Metschnikowia(100);g__Metschnikowia_unclassified(100); | Metschnikowia | 13 | Saprotroph | \|Undefined Saprotroph\| | Yeast | NULL | Probable | As nectar-tap_saprotroph (Põlme et al. 2020); Decay sugar-rich substrates (Põlme et al. 2020) | Põlme S, et al. 2020. Fungal Diversity 105:1-16 ((https://doi.org/10.1007/s13225-020-00466-2)) |
| Otu35650 | 1 | k__Fungi(100);p__Ascomycota(100);c__Saccharomycetes(100);o__Saccharomycetales(100);f__Metschnikowiaceae(100);g__Metschnikowia(100);s__Metschnikowia_pulcherrima(100); | Metschnikowia | 13 | Saprotroph | \|Undefined Saprotroph\| | Yeast | NULL | Probable | As nectar-tap_saprotroph (Põlme et al. 2020); Decay sugar-rich substrates (Põlme et al. 2020) | Põlme S, et al. 2020. Fungal Diversity 105:1-16 ((https://doi.org/10.1007/s13225-020-00466-2)) |
| Otu35651 | 1 | k__Fungi(100);p__Ascomycota(100);c__Dothideomycetes(100);o__Dothideomycetes_order_Incertae_sedis(100);f__Myxotrichaceae(100);g__Oidiodendron(100);g__Oidiodendron_unclassified(100); | Oidiodendron | 13 | Pathotroph-Saprotroph-Symbiotroph | Endophyte-Ericoid Mycorrhizal-\|Undefined Saprotroph\| | Microfungus | NULL | Probable | As root_endophyte (Põlme et al. 2020); As soil_saprotroph (Põlme et al. 2020); Dark Septate Endophyte; Decay soil (Põlme et al. 2020) | Newsham KK. 2011. New Phytologist 190:783-793 ((https://doi.org/10.1111/j.1469-8137.2010.03611.x)) (as dark septate root endophytes); Põlme S, et al. 2020. Fungal Diversity 105:1-16 ((https://doi.org/10.1007/s13225-020-00466-2)) |
| Otu35652 | 1 | k__Fungi(100);p__Ascomycota(100);c__Saccharomycetes(100);o__Saccharomycetales(100);f__Metschnikowiaceae(100);g__Metschnikowia(100);s__Metschnikowia_pulcherrima(100); | Metschnikowia | 13 | Saprotroph | \|Undefined Saprotroph\| | Yeast | NULL | Probable | As nectar-tap_saprotroph (Põlme et al. 2020); Decay sugar-rich substrates (Põlme et al. 2020) | Põlme S, et al. 2020. Fungal Diversity 105:1-16 ((https://doi.org/10.1007/s13225-020-00466-2)) |
| Otu35653 | 1 | k__Fungi(100);p__Ascomycota(100);c__Saccharomycetes(100);o__Saccharomycetales(100);f__Metschnikowiaceae(100);g__Metschnikowia(100);s__Metschnikowia_pulcherrima(100); | Metschnikowia | 13 | Saprotroph | \|Undefined Saprotroph\| | Yeast | NULL | Probable | As nectar-tap_saprotroph (Põlme et al. 2020); Decay sugar-rich substrates (Põlme et al. 2020) | Põlme S, et al. 2020. Fungal Diversity 105:1-16 ((https://doi.org/10.1007/s13225-020-00466-2)) |
| Otu35654 | 1 | k__Fungi(100);p__Ascomycota(100);c__Saccharomycetes(100);o__Saccharomycetales(100);f__Metschnikowiaceae(100);g__Metschnikowia(100);s__Metschnikowia_pulcherrima(100); | Metschnikowia | 13 | Saprotroph | \|Undefined Saprotroph\| | Yeast | NULL | Probable | As nectar-tap_saprotroph (Põlme et al. 2020); Decay sugar-rich substrates (Põlme et al. 2020) | Põlme S, et al. 2020. Fungal Diversity 105:1-16 ((https://doi.org/10.1007/s13225-020-00466-2)) |
| Otu35655 | 1 | k__Fungi(100);p__Ascomycota(100);c__Saccharomycetes(100);o__Saccharomycetales(100);f__Metschnikowiaceae(100);g__Metschnikowia(100);s__Metschnikowia_pulcherrima(100); | Metschnikowia | 13 | Saprotroph | \|Undefined Saprotroph\| | Yeast | NULL | Probable | As nectar-tap_saprotroph (Põlme et al. 2020); Decay sugar-rich substrates (Põlme et al. 2020) | Põlme S, et al. 2020. Fungal Diversity 105:1-16 ((https://doi.org/10.1007/s13225-020-00466-2)) |
| Otu35656 | 1 | k__Fungi(100);p__Ascomycota(100);c__Saccharomycetes(100);o__Saccharomycetales(100);f__Metschnikowiaceae(100);g__Metschnikowia(100);g__Metschnikowia_unclassified(100); | Metschnikowia | 13 | Saprotroph | \|Undefined Saprotroph\| | Yeast | NULL | Probable | As nectar-tap_saprotroph (Põlme et al. 2020); Decay sugar-rich substrates (Põlme et al. 2020) | Põlme S, et al. 2020. Fungal Diversity 105:1-16 ((https://doi.org/10.1007/s13225-020-00466-2)) |
| Otu35657 | 1 | k__Fungi(100);p__Ascomycota(100);c__Saccharomycetes(100);o__Saccharomycetales(100);f__Metschnikowiaceae(100);g__Metschnikowia(100);s__Metschnikowia_pulcherrima(100); | Metschnikowia | 13 | Saprotroph | \|Undefined Saprotroph\| | Yeast | NULL | Probable | As nectar-tap_saprotroph (Põlme et al. 2020); Decay sugar-rich substrates (Põlme et al. 2020) | Põlme S, et al. 2020. Fungal Diversity 105:1-16 ((https://doi.org/10.1007/s13225-020-00466-2)) |
| Otu35658 | 1 | k__Fungi(100);p__Ascomycota(100);c__Saccharomycetes(100);o__Saccharomycetales(100);f__Metschnikowiaceae(100);g__Metschnikowia(100);g__Metschnikowia_unclassified(100); | Metschnikowia | 13 | Saprotroph | \|Undefined Saprotroph\| | Yeast | NULL | Probable | As nectar-tap_saprotroph (Põlme et al. 2020); Decay sugar-rich substrates (Põlme et al. 2020) | Põlme S, et al. 2020. Fungal Diversity 105:1-16 ((https://doi.org/10.1007/s13225-020-00466-2)) |
| Otu35659 | 1 | k__Fungi(100);p__Ascomycota(100);c__Saccharomycetes(100);o__Saccharomycetales(100);f__Saccharomycetales_family_Incertae_sedis(100);g__Candida(100);g__Candida_unclassified(100); | Candida | 13 | Pathotroph-Saprotroph-Symbiotroph | Animal Pathogen-Endophyte-Endosymbiont-Epiphyte-\|Undefined Saprotroph\| | Dimorphic Yeast-Yeast | NULL | Probable | As nectar-tap_saprotroph (Põlme et al. 2020); Decay sugar-rich substrates (Põlme et al. 2020); This asexual genus is polyphyletic and contain species that function very differently depend on what they are. Caution should be taken when interpreting guilds for Candida species. | Manolakaki D, et al. 2010. Virulence. 1: 367-375 ((https://doi.org/10.4161/viru.1.5.12796)); Mekbib SB, et al. 2011. Tropical Plant Pathology 36:233-240 ((https://doi.org/10.1590/S1982-56762011000400004)); Põlme S, et al. 2020. Fungal Diversity 105:1-16 ((https://doi.org/10.1007/s13225-020-00466-2)) |
| Otu35660 | 1 | k__Fungi(100);p__Ascomycota(100);c__Saccharomycetes(100);o__Saccharomycetales(100);f__Metschnikowiaceae(100);g__Metschnikowia(100);s__Metschnikowia_pulcherrima(100); | Metschnikowia | 13 | Saprotroph | \|Undefined Saprotroph\| | Yeast | NULL | Probable | As nectar-tap_saprotroph (Põlme et al. 2020); Decay sugar-rich substrates (Põlme et al. 2020) | Põlme S, et al. 2020. Fungal Diversity 105:1-16 ((https://doi.org/10.1007/s13225-020-00466-2)) |
| Otu35661 | 1 | k__Fungi(100);p__Ascomycota(100);c__Saccharomycetes(100);o__Saccharomycetales(100);f__Metschnikowiaceae(100);g__Metschnikowia(100);s__Metschnikowia_pulcherrima(100); | Metschnikowia | 13 | Saprotroph | \|Undefined Saprotroph\| | Yeast | NULL | Probable | As nectar-tap_saprotroph (Põlme et al. 2020); Decay sugar-rich substrates (Põlme et al. 2020) | Põlme S, et al. 2020. Fungal Diversity 105:1-16 ((https://doi.org/10.1007/s13225-020-00466-2)) |
| Otu35662 | 1 | k__Fungi(100);p__Ascomycota(100);c__Saccharomycetes(100);o__Saccharomycetales(100);f__Metschnikowiaceae(100);g__Metschnikowia(100);s__Metschnikowia_pulcherrima(100); | Metschnikowia | 13 | Saprotroph | \|Undefined Saprotroph\| | Yeast | NULL | Probable | As nectar-tap_saprotroph (Põlme et al. 2020); Decay sugar-rich substrates (Põlme et al. 2020) | Põlme S, et al. 2020. Fungal Diversity 105:1-16 ((https://doi.org/10.1007/s13225-020-00466-2)) |
| Otu35665 | 1 | k__Fungi(100);p__Ascomycota(100);c__Saccharomycetes(100);o__Saccharomycetales(100);f__Metschnikowiaceae(100);g__Metschnikowia(100);s__Metschnikowia_pulcherrima(100); | Metschnikowia | 13 | Saprotroph | \|Undefined Saprotroph\| | Yeast | NULL | Probable | As nectar-tap_saprotroph (Põlme et al. 2020); Decay sugar-rich substrates (Põlme et al. 2020) | Põlme S, et al. 2020. Fungal Diversity 105:1-16 ((https://doi.org/10.1007/s13225-020-00466-2)) |
| Otu35666 | 1 | k__Fungi(100);p__Ascomycota(100);c__Saccharomycetes(100);o__Saccharomycetales(100);f__Metschnikowiaceae(100);g__Metschnikowia(100);s__Metschnikowia_pulcherrima(100); | Metschnikowia | 13 | Saprotroph | \|Undefined Saprotroph\| | Yeast | NULL | Probable | As nectar-tap_saprotroph (Põlme et al. 2020); Decay sugar-rich substrates (Põlme et al. 2020) | Põlme S, et al. 2020. Fungal Diversity 105:1-16 ((https://doi.org/10.1007/s13225-020-00466-2)) |
| Otu35667 | 1 | k__Fungi(100);p__Ascomycota(100);c__Saccharomycetes(100);o__Saccharomycetales(100);f__Metschnikowiaceae(100);g__Metschnikowia(100);g__Metschnikowia_unclassified(100); | Metschnikowia | 13 | Saprotroph | \|Undefined Saprotroph\| | Yeast | NULL | Probable | As nectar-tap_saprotroph (Põlme et al. 2020); Decay sugar-rich substrates (Põlme et al. 2020) | Põlme S, et al. 2020. Fungal Diversity 105:1-16 ((https://doi.org/10.1007/s13225-020-00466-2)) |
| Otu35668 | 1 | k__Fungi(100);p__Ascomycota(100);c__Saccharomycetes(100);o__Saccharomycetales(100);f__Saccharomycetales_family_Incertae_sedis(100);g__Candida(100);s__Candida_cretensis(100); | Candida | 13 | Pathotroph-Saprotroph-Symbiotroph | Animal Pathogen-Endophyte-Endosymbiont-Epiphyte-\|Undefined Saprotroph\| | Dimorphic Yeast-Yeast | NULL | Probable | As nectar-tap_saprotroph (Põlme et al. 2020); Decay sugar-rich substrates (Põlme et al. 2020); This asexual genus is polyphyletic and contain species that function very differently depend on what they are. Caution should be taken when interpreting guilds for Candida species. | Manolakaki D, et al. 2010. Virulence. 1: 367-375 ((https://doi.org/10.4161/viru.1.5.12796)); Mekbib SB, et al. 2011. Tropical Plant Pathology 36:233-240 ((https://doi.org/10.1590/S1982-56762011000400004)); Põlme S, et al. 2020. Fungal Diversity 105:1-16 ((https://doi.org/10.1007/s13225-020-00466-2)) |
| Otu35669 | 1 | k__Fungi(100);p__Ascomycota(100);c__Saccharomycetes(100);o__Saccharomycetales(100);f__Metschnikowiaceae(100);g__Metschnikowia(100);s__Metschnikowia_pulcherrima(100); | Metschnikowia | 13 | Saprotroph | \|Undefined Saprotroph\| | Yeast | NULL | Probable | As nectar-tap_saprotroph (Põlme et al. 2020); Decay sugar-rich substrates (Põlme et al. 2020) | Põlme S, et al. 2020. Fungal Diversity 105:1-16 ((https://doi.org/10.1007/s13225-020-00466-2)) |
| Otu35670 | 1 | k__Fungi(100);p__Ascomycota(100);c__Saccharomycetes(100);o__Saccharomycetales(100);f__Metschnikowiaceae(100);g__Metschnikowia(100);s__Metschnikowia_pulcherrima(100); | Metschnikowia | 13 | Saprotroph | \|Undefined Saprotroph\| | Yeast | NULL | Probable | As nectar-tap_saprotroph (Põlme et al. 2020); Decay sugar-rich substrates (Põlme et al. 2020) | Põlme S, et al. 2020. Fungal Diversity 105:1-16 ((https://doi.org/10.1007/s13225-020-00466-2)) |
| Otu35671 | 1 | k__Fungi(100);p__Ascomycota(100);c__Saccharomycetes(100);o__Saccharomycetales(100);f__Metschnikowiaceae(100);g__Metschnikowia(100);s__Metschnikowia_pulcherrima(100); | Metschnikowia | 13 | Saprotroph | \|Undefined Saprotroph\| | Yeast | NULL | Probable | As nectar-tap_saprotroph (Põlme et al. 2020); Decay sugar-rich substrates (Põlme et al. 2020) | Põlme S, et al. 2020. Fungal Diversity 105:1-16 ((https://doi.org/10.1007/s13225-020-00466-2)) |
| Otu35672 | 1 | k__Fungi(100);p__Ascomycota(100);c__Saccharomycetes(100);o__Saccharomycetales(100);f__Metschnikowiaceae(100);g__Metschnikowia(100);s__Metschnikowia_pulcherrima(100); | Metschnikowia | 13 | Saprotroph | \|Undefined Saprotroph\| | Yeast | NULL | Probable | As nectar-tap_saprotroph (Põlme et al. 2020); Decay sugar-rich substrates (Põlme et al. 2020) | Põlme S, et al. 2020. Fungal Diversity 105:1-16 ((https://doi.org/10.1007/s13225-020-00466-2)) |
| Otu35673 | 1 | k__Fungi(100);p__Ascomycota(100);c__Saccharomycetes(100);o__Saccharomycetales(100);f__Metschnikowiaceae(100);g__Metschnikowia(100);g__Metschnikowia_unclassified(100); | Metschnikowia | 13 | Saprotroph | \|Undefined Saprotroph\| | Yeast | NULL | Probable | As nectar-tap_saprotroph (Põlme et al. 2020); Decay sugar-rich substrates (Põlme et al. 2020) | Põlme S, et al. 2020. Fungal Diversity 105:1-16 ((https://doi.org/10.1007/s13225-020-00466-2)) |
| Otu35674 | 1 | k__Fungi(100);p__Ascomycota(100);c__Saccharomycetes(100);o__Saccharomycetales(100);f__Metschnikowiaceae(100);g__Metschnikowia(100);g__Metschnikowia_unclassified(100); | Metschnikowia | 13 | Saprotroph | \|Undefined Saprotroph\| | Yeast | NULL | Probable | As nectar-tap_saprotroph (Põlme et al. 2020); Decay sugar-rich substrates (Põlme et al. 2020) | Põlme S, et al. 2020. Fungal Diversity 105:1-16 ((https://doi.org/10.1007/s13225-020-00466-2)) |
| Otu35675 | 1 | k__Fungi(100);p__Ascomycota(100);c__Saccharomycetes(100);o__Saccharomycetales(100);f__Metschnikowiaceae(100);g__Metschnikowia(100);s__Metschnikowia_pulcherrima(100); | Metschnikowia | 13 | Saprotroph | \|Undefined Saprotroph\| | Yeast | NULL | Probable | As nectar-tap_saprotroph (Põlme et al. 2020); Decay sugar-rich substrates (Põlme et al. 2020) | Põlme S, et al. 2020. Fungal Diversity 105:1-16 ((https://doi.org/10.1007/s13225-020-00466-2)) |
| Otu35676 | 1 | k__Fungi(100);p__Ascomycota(100);c__Saccharomycetes(100);o__Saccharomycetales(100);f__Metschnikowiaceae(100);g__Metschnikowia(100);s__Metschnikowia_pulcherrima(100); | Metschnikowia | 13 | Saprotroph | \|Undefined Saprotroph\| | Yeast | NULL | Probable | As nectar-tap_saprotroph (Põlme et al. 2020); Decay sugar-rich substrates (Põlme et al. 2020) | Põlme S, et al. 2020. Fungal Diversity 105:1-16 ((https://doi.org/10.1007/s13225-020-00466-2)) |
| Otu35677 | 1 | k__Fungi(100);p__Ascomycota(100);c__Saccharomycetes(100);o__Saccharomycetales(100);f__Metschnikowiaceae(100);g__Metschnikowia(100);g__Metschnikowia_unclassified(100); | Metschnikowia | 13 | Saprotroph | \|Undefined Saprotroph\| | Yeast | NULL | Probable | As nectar-tap_saprotroph (Põlme et al. 2020); Decay sugar-rich substrates (Põlme et al. 2020) | Põlme S, et al. 2020. Fungal Diversity 105:1-16 ((https://doi.org/10.1007/s13225-020-00466-2)) |
| Otu35678 | 1 | k__Fungi(100);p__Ascomycota(100);c__Saccharomycetes(100);o__Saccharomycetales(100);f__Metschnikowiaceae(100);g__Metschnikowia(100);s__Metschnikowia_pulcherrima(100); | Metschnikowia | 13 | Saprotroph | \|Undefined Saprotroph\| | Yeast | NULL | Probable | As nectar-tap_saprotroph (Põlme et al. 2020); Decay sugar-rich substrates (Põlme et al. 2020) | Põlme S, et al. 2020. Fungal Diversity 105:1-16 ((https://doi.org/10.1007/s13225-020-00466-2)) |
| Otu35679 | 1 | k__Fungi(100);p__Ascomycota(100);c__Saccharomycetes(100);o__Saccharomycetales(100);f__Metschnikowiaceae(100);g__Metschnikowia(100);s__Metschnikowia_pulcherrima(100); | Metschnikowia | 13 | Saprotroph | \|Undefined Saprotroph\| | Yeast | NULL | Probable | As nectar-tap_saprotroph (Põlme et al. 2020); Decay sugar-rich substrates (Põlme et al. 2020) | Põlme S, et al. 2020. Fungal Diversity 105:1-16 ((https://doi.org/10.1007/s13225-020-00466-2)) |
| Otu35680 | 1 | k__Fungi(100);p__Ascomycota(100);c__Saccharomycetes(100);o__Saccharomycetales(100);f__Metschnikowiaceae(100);g__Metschnikowia(100);s__Metschnikowia_pulcherrima(100); | Metschnikowia | 13 | Saprotroph | \|Undefined Saprotroph\| | Yeast | NULL | Probable | As nectar-tap_saprotroph (Põlme et al. 2020); Decay sugar-rich substrates (Põlme et al. 2020) | Põlme S, et al. 2020. Fungal Diversity 105:1-16 ((https://doi.org/10.1007/s13225-020-00466-2)) |
| Otu35681 | 1 | k__Fungi(100);p__Ascomycota(100);c__Saccharomycetes(100);o__Saccharomycetales(100);f__Metschnikowiaceae(100);g__Metschnikowia(100);g__Metschnikowia_unclassified(100); | Metschnikowia | 13 | Saprotroph | \|Undefined Saprotroph\| | Yeast | NULL | Probable | As nectar-tap_saprotroph (Põlme et al. 2020); Decay sugar-rich substrates (Põlme et al. 2020) | Põlme S, et al. 2020. Fungal Diversity 105:1-16 ((https://doi.org/10.1007/s13225-020-00466-2)) |
| Otu35682 | 1 | k__Fungi(100);p__Ascomycota(100);c__Saccharomycetes(100);o__Saccharomycetales(100);f__Metschnikowiaceae(100);g__Metschnikowia(100);g__Metschnikowia_unclassified(100); | Metschnikowia | 13 | Saprotroph | \|Undefined Saprotroph\| | Yeast | NULL | Probable | As nectar-tap_saprotroph (Põlme et al. 2020); Decay sugar-rich substrates (Põlme et al. 2020) | Põlme S, et al. 2020. Fungal Diversity 105:1-16 ((https://doi.org/10.1007/s13225-020-00466-2)) |
| Otu35683 | 1 | k__Fungi(100);p__Ascomycota(100);c__Saccharomycetes(100);o__Saccharomycetales(100);f__Metschnikowiaceae(100);g__Metschnikowia(100);g__Metschnikowia_unclassified(100); | Metschnikowia | 13 | Saprotroph | \|Undefined Saprotroph\| | Yeast | NULL | Probable | As nectar-tap_saprotroph (Põlme et al. 2020); Decay sugar-rich substrates (Põlme et al. 2020) | Põlme S, et al. 2020. Fungal Diversity 105:1-16 ((https://doi.org/10.1007/s13225-020-00466-2)) |
| Otu35684 | 1 | k__Fungi(100);p__Ascomycota(100);c__Saccharomycetes(100);o__Saccharomycetales(100);f__Metschnikowiaceae(100);g__Metschnikowia(100);s__Metschnikowia_pulcherrima(100); | Metschnikowia | 13 | Saprotroph | \|Undefined Saprotroph\| | Yeast | NULL | Probable | As nectar-tap_saprotroph (Põlme et al. 2020); Decay sugar-rich substrates (Põlme et al. 2020) | Põlme S, et al. 2020. Fungal Diversity 105:1-16 ((https://doi.org/10.1007/s13225-020-00466-2)) |
| Otu35685 | 1 | k__Fungi(100);p__Ascomycota(100);c__Saccharomycetes(100);o__Saccharomycetales(100);f__Metschnikowiaceae(100);g__Metschnikowia(100);s__Metschnikowia_pulcherrima(100); | Metschnikowia | 13 | Saprotroph | \|Undefined Saprotroph\| | Yeast | NULL | Probable | As nectar-tap_saprotroph (Põlme et al. 2020); Decay sugar-rich substrates (Põlme et al. 2020) | Põlme S, et al. 2020. Fungal Diversity 105:1-16 ((https://doi.org/10.1007/s13225-020-00466-2)) |
| Otu35686 | 1 | k__Fungi(100);p__Ascomycota(100);c__Saccharomycetes(100);o__Saccharomycetales(100);f__Metschnikowiaceae(100);g__Metschnikowia(100);s__Metschnikowia_pulcherrima(100); | Metschnikowia | 13 | Saprotroph | \|Undefined Saprotroph\| | Yeast | NULL | Probable | As nectar-tap_saprotroph (Põlme et al. 2020); Decay sugar-rich substrates (Põlme et al. 2020) | Põlme S, et al. 2020. Fungal Diversity 105:1-16 ((https://doi.org/10.1007/s13225-020-00466-2)) |
| Otu35689 | 1 | k__Fungi(100);p__Ascomycota(100);c__Saccharomycetes(100);o__Saccharomycetales(100);f__Metschnikowiaceae(100);g__Metschnikowia(100);g__Metschnikowia_unclassified(100); | Metschnikowia | 13 | Saprotroph | \|Undefined Saprotroph\| | Yeast | NULL | Probable | As nectar-tap_saprotroph (Põlme et al. 2020); Decay sugar-rich substrates (Põlme et al. 2020) | Põlme S, et al. 2020. Fungal Diversity 105:1-16 ((https://doi.org/10.1007/s13225-020-00466-2)) |
| Otu35691 | 1 | k__Fungi(100);p__Ascomycota(100);c__Saccharomycetes(100);o__Saccharomycetales(100);f__Metschnikowiaceae(100);g__Metschnikowia(100);s__Metschnikowia_pulcherrima(100); | Metschnikowia | 13 | Saprotroph | \|Undefined Saprotroph\| | Yeast | NULL | Probable | As nectar-tap_saprotroph (Põlme et al. 2020); Decay sugar-rich substrates (Põlme et al. 2020) | Põlme S, et al. 2020. Fungal Diversity 105:1-16 ((https://doi.org/10.1007/s13225-020-00466-2)) |
| Otu35692 | 1 | k__Fungi(100);p__Ascomycota(100);c__Saccharomycetes(100);o__Saccharomycetales(100);f__Metschnikowiaceae(100);g__Metschnikowia(100);s__Metschnikowia_pulcherrima(100); | Metschnikowia | 13 | Saprotroph | \|Undefined Saprotroph\| | Yeast | NULL | Probable | As nectar-tap_saprotroph (Põlme et al. 2020); Decay sugar-rich substrates (Põlme et al. 2020) | Põlme S, et al. 2020. Fungal Diversity 105:1-16 ((https://doi.org/10.1007/s13225-020-00466-2)) |
| Otu35693 | 1 | k__Fungi(100);p__Ascomycota(100);c__Saccharomycetes(100);o__Saccharomycetales(100);f__Metschnikowiaceae(100);g__Metschnikowia(100);s__Metschnikowia_pulcherrima(100); | Metschnikowia | 13 | Saprotroph | \|Undefined Saprotroph\| | Yeast | NULL | Probable | As nectar-tap_saprotroph (Põlme et al. 2020); Decay sugar-rich substrates (Põlme et al. 2020) | Põlme S, et al. 2020. Fungal Diversity 105:1-16 ((https://doi.org/10.1007/s13225-020-00466-2)) |
| Otu35694 | 1 | k__Fungi(100);p__Ascomycota(100);c__Saccharomycetes(100);o__Saccharomycetales(100);f__Metschnikowiaceae(100);g__Metschnikowia(100);g__Metschnikowia_unclassified(100); | Metschnikowia | 13 | Saprotroph | \|Undefined Saprotroph\| | Yeast | NULL | Probable | As nectar-tap_saprotroph (Põlme et al. 2020); Decay sugar-rich substrates (Põlme et al. 2020) | Põlme S, et al. 2020. Fungal Diversity 105:1-16 ((https://doi.org/10.1007/s13225-020-00466-2)) |
| Otu35696 | 1 | k__Fungi(100);p__Ascomycota(100);c__Saccharomycetes(100);o__Saccharomycetales(100);f__Metschnikowiaceae(100);g__Metschnikowia(100);g__Metschnikowia_unclassified(100); | Metschnikowia | 13 | Saprotroph | \|Undefined Saprotroph\| | Yeast | NULL | Probable | As nectar-tap_saprotroph (Põlme et al. 2020); Decay sugar-rich substrates (Põlme et al. 2020) | Põlme S, et al. 2020. Fungal Diversity 105:1-16 ((https://doi.org/10.1007/s13225-020-00466-2)) |
| Otu35697 | 1 | k__Fungi(100);p__Ascomycota(100);c__Saccharomycetes(100);o__Saccharomycetales(100);f__Metschnikowiaceae(100);g__Metschnikowia(100);s__Metschnikowia_pulcherrima(100); | Metschnikowia | 13 | Saprotroph | \|Undefined Saprotroph\| | Yeast | NULL | Probable | As nectar-tap_saprotroph (Põlme et al. 2020); Decay sugar-rich substrates (Põlme et al. 2020) | Põlme S, et al. 2020. Fungal Diversity 105:1-16 ((https://doi.org/10.1007/s13225-020-00466-2)) |
| Otu35698 | 1 | k__Fungi(100);p__Ascomycota(100);c__Saccharomycetes(100);o__Saccharomycetales(100);f__Metschnikowiaceae(100);g__Metschnikowia(100);g__Metschnikowia_unclassified(100); | Metschnikowia | 13 | Saprotroph | \|Undefined Saprotroph\| | Yeast | NULL | Probable | As nectar-tap_saprotroph (Põlme et al. 2020); Decay sugar-rich substrates (Põlme et al. 2020) | Põlme S, et al. 2020. Fungal Diversity 105:1-16 ((https://doi.org/10.1007/s13225-020-00466-2)) |
| Otu35699 | 1 | k__Fungi(100);p__Ascomycota(100);c__Saccharomycetes(100);o__Saccharomycetales(100);f__Metschnikowiaceae(100);g__Metschnikowia(100);g__Metschnikowia_unclassified(100); | Metschnikowia | 13 | Saprotroph | \|Undefined Saprotroph\| | Yeast | NULL | Probable | As nectar-tap_saprotroph (Põlme et al. 2020); Decay sugar-rich substrates (Põlme et al. 2020) | Põlme S, et al. 2020. Fungal Diversity 105:1-16 ((https://doi.org/10.1007/s13225-020-00466-2)) |
| Otu35700 | 1 | k__Fungi(100);p__Ascomycota(100);c__Saccharomycetes(100);o__Saccharomycetales(100);f__Metschnikowiaceae(100);g__Metschnikowia(100);g__Metschnikowia_unclassified(100); | Metschnikowia | 13 | Saprotroph | \|Undefined Saprotroph\| | Yeast | NULL | Probable | As nectar-tap_saprotroph (Põlme et al. 2020); Decay sugar-rich substrates (Põlme et al. 2020) | Põlme S, et al. 2020. Fungal Diversity 105:1-16 ((https://doi.org/10.1007/s13225-020-00466-2)) |
| Otu35701 | 1 | k__Fungi(100);p__Ascomycota(100);c__Saccharomycetes(100);o__Saccharomycetales(100);f__Metschnikowiaceae(100);g__Metschnikowia(100);s__Metschnikowia_pulcherrima(100); | Metschnikowia | 13 | Saprotroph | \|Undefined Saprotroph\| | Yeast | NULL | Probable | As nectar-tap_saprotroph (Põlme et al. 2020); Decay sugar-rich substrates (Põlme et al. 2020) | Põlme S, et al. 2020. Fungal Diversity 105:1-16 ((https://doi.org/10.1007/s13225-020-00466-2)) |
| Otu35702 | 1 | k__Fungi(100);p__Ascomycota(100);c__Saccharomycetes(100);o__Saccharomycetales(100);f__Metschnikowiaceae(100);g__Metschnikowia(100);g__Metschnikowia_unclassified(100); | Metschnikowia | 13 | Saprotroph | \|Undefined Saprotroph\| | Yeast | NULL | Probable | As nectar-tap_saprotroph (Põlme et al. 2020); Decay sugar-rich substrates (Põlme et al. 2020) | Põlme S, et al. 2020. Fungal Diversity 105:1-16 ((https://doi.org/10.1007/s13225-020-00466-2)) |
| Otu35704 | 1 | k__Fungi(100);p__Ascomycota(100);c__Saccharomycetes(100);o__Saccharomycetales(100);f__Metschnikowiaceae(100);g__Metschnikowia(100);g__Metschnikowia_unclassified(100); | Metschnikowia | 13 | Saprotroph | \|Undefined Saprotroph\| | Yeast | NULL | Probable | As nectar-tap_saprotroph (Põlme et al. 2020); Decay sugar-rich substrates (Põlme et al. 2020) | Põlme S, et al. 2020. Fungal Diversity 105:1-16 ((https://doi.org/10.1007/s13225-020-00466-2)) |
| Otu35705 | 1 | k__Fungi(100);p__Ascomycota(100);c__Saccharomycetes(100);o__Saccharomycetales(100);f__Metschnikowiaceae(100);g__Metschnikowia(100);s__Metschnikowia_pulcherrima(100); | Metschnikowia | 13 | Saprotroph | \|Undefined Saprotroph\| | Yeast | NULL | Probable | As nectar-tap_saprotroph (Põlme et al. 2020); Decay sugar-rich substrates (Põlme et al. 2020) | Põlme S, et al. 2020. Fungal Diversity 105:1-16 ((https://doi.org/10.1007/s13225-020-00466-2)) |
| Otu35706 | 1 | k__Fungi(100);p__Ascomycota(100);c__Saccharomycetes(100);o__Saccharomycetales(100);f__Metschnikowiaceae(100);g__Metschnikowia(100);s__Metschnikowia_pulcherrima(100); | Metschnikowia | 13 | Saprotroph | \|Undefined Saprotroph\| | Yeast | NULL | Probable | As nectar-tap_saprotroph (Põlme et al. 2020); Decay sugar-rich substrates (Põlme et al. 2020) | Põlme S, et al. 2020. Fungal Diversity 105:1-16 ((https://doi.org/10.1007/s13225-020-00466-2)) |
| Otu35707 | 1 | k__Fungi(100);p__Ascomycota(100);c__Saccharomycetes(100);o__Saccharomycetales(100);f__Metschnikowiaceae(100);g__Metschnikowia(100);s__Metschnikowia_pulcherrima(100); | Metschnikowia | 13 | Saprotroph | \|Undefined Saprotroph\| | Yeast | NULL | Probable | As nectar-tap_saprotroph (Põlme et al. 2020); Decay sugar-rich substrates (Põlme et al. 2020) | Põlme S, et al. 2020. Fungal Diversity 105:1-16 ((https://doi.org/10.1007/s13225-020-00466-2)) |
| Otu35708 | 1 | k__Fungi(100);p__Ascomycota(100);c__Saccharomycetes(100);o__Saccharomycetales(100);f__Metschnikowiaceae(100);g__Metschnikowia(100);g__Metschnikowia_unclassified(100); | Metschnikowia | 13 | Saprotroph | \|Undefined Saprotroph\| | Yeast | NULL | Probable | As nectar-tap_saprotroph (Põlme et al. 2020); Decay sugar-rich substrates (Põlme et al. 2020) | Põlme S, et al. 2020. Fungal Diversity 105:1-16 ((https://doi.org/10.1007/s13225-020-00466-2)) |
| Otu35709 | 1 | k__Fungi(100);p__Ascomycota(100);c__Saccharomycetes(100);o__Saccharomycetales(100);f__Metschnikowiaceae(100);g__Metschnikowia(100);s__Metschnikowia_pulcherrima(100); | Metschnikowia | 13 | Saprotroph | \|Undefined Saprotroph\| | Yeast | NULL | Probable | As nectar-tap_saprotroph (Põlme et al. 2020); Decay sugar-rich substrates (Põlme et al. 2020) | Põlme S, et al. 2020. Fungal Diversity 105:1-16 ((https://doi.org/10.1007/s13225-020-00466-2)) |
| Otu35710 | 1 | k__Fungi(100);p__Ascomycota(100);c__Saccharomycetes(100);o__Saccharomycetales(100);f__Metschnikowiaceae(100);g__Metschnikowia(100);s__Metschnikowia_pulcherrima(100); | Metschnikowia | 13 | Saprotroph | \|Undefined Saprotroph\| | Yeast | NULL | Probable | As nectar-tap_saprotroph (Põlme et al. 2020); Decay sugar-rich substrates (Põlme et al. 2020) | Põlme S, et al. 2020. Fungal Diversity 105:1-16 ((https://doi.org/10.1007/s13225-020-00466-2)) |
| Otu35717 | 1 | k__Fungi(100);p__Ascomycota(100);c__Saccharomycetes(100);o__Saccharomycetales(100);f__Metschnikowiaceae(100);g__Metschnikowia(100);s__Metschnikowia_pulcherrima(100); | Metschnikowia | 13 | Saprotroph | \|Undefined Saprotroph\| | Yeast | NULL | Probable | As nectar-tap_saprotroph (Põlme et al. 2020); Decay sugar-rich substrates (Põlme et al. 2020) | Põlme S, et al. 2020. Fungal Diversity 105:1-16 ((https://doi.org/10.1007/s13225-020-00466-2)) |
| Otu35718 | 1 | k__Fungi(100);p__Ascomycota(100);c__Saccharomycetes(100);o__Saccharomycetales(100);f__Metschnikowiaceae(100);g__Metschnikowia(100);s__Metschnikowia_pulcherrima(100); | Metschnikowia | 13 | Saprotroph | \|Undefined Saprotroph\| | Yeast | NULL | Probable | As nectar-tap_saprotroph (Põlme et al. 2020); Decay sugar-rich substrates (Põlme et al. 2020) | Põlme S, et al. 2020. Fungal Diversity 105:1-16 ((https://doi.org/10.1007/s13225-020-00466-2)) |
| Otu35719 | 1 | k__Fungi(100);p__Ascomycota(100);c__Saccharomycetes(100);o__Saccharomycetales(100);f__Metschnikowiaceae(100);g__Metschnikowia(100);g__Metschnikowia_unclassified(100); | Metschnikowia | 13 | Saprotroph | \|Undefined Saprotroph\| | Yeast | NULL | Probable | As nectar-tap_saprotroph (Põlme et al. 2020); Decay sugar-rich substrates (Põlme et al. 2020) | Põlme S, et al. 2020. Fungal Diversity 105:1-16 ((https://doi.org/10.1007/s13225-020-00466-2)) |
| Otu35721 | 1 | k__Fungi(100);p__Ascomycota(100);c__Saccharomycetes(100);o__Saccharomycetales(100);f__Metschnikowiaceae(100);g__Metschnikowia(100);s__Metschnikowia_pulcherrima(100); | Metschnikowia | 13 | Saprotroph | \|Undefined Saprotroph\| | Yeast | NULL | Probable | As nectar-tap_saprotroph (Põlme et al. 2020); Decay sugar-rich substrates (Põlme et al. 2020) | Põlme S, et al. 2020. Fungal Diversity 105:1-16 ((https://doi.org/10.1007/s13225-020-00466-2)) |
| Otu35722 | 1 | k__Fungi(100);p__Ascomycota(100);c__Saccharomycetes(100);o__Saccharomycetales(100);f__Metschnikowiaceae(100);g__Metschnikowia(100);g__Metschnikowia_unclassified(100); | Metschnikowia | 13 | Saprotroph | \|Undefined Saprotroph\| | Yeast | NULL | Probable | As nectar-tap_saprotroph (Põlme et al. 2020); Decay sugar-rich substrates (Põlme et al. 2020) | Põlme S, et al. 2020. Fungal Diversity 105:1-16 ((https://doi.org/10.1007/s13225-020-00466-2)) |
| Otu35723 | 1 | k__Fungi(100);p__Ascomycota(100);c__Saccharomycetes(100);o__Saccharomycetales(100);f__Metschnikowiaceae(100);g__Metschnikowia(100);g__Metschnikowia_unclassified(100); | Metschnikowia | 13 | Saprotroph | \|Undefined Saprotroph\| | Yeast | NULL | Probable | As nectar-tap_saprotroph (Põlme et al. 2020); Decay sugar-rich substrates (Põlme et al. 2020) | Põlme S, et al. 2020. Fungal Diversity 105:1-16 ((https://doi.org/10.1007/s13225-020-00466-2)) |
| Otu35724 | 1 | k__Fungi(100);p__Ascomycota(100);c__Saccharomycetes(100);o__Saccharomycetales(100);f__Metschnikowiaceae(100);g__Metschnikowia(100);s__Metschnikowia_pulcherrima(100); | Metschnikowia | 13 | Saprotroph | \|Undefined Saprotroph\| | Yeast | NULL | Probable | As nectar-tap_saprotroph (Põlme et al. 2020); Decay sugar-rich substrates (Põlme et al. 2020) | Põlme S, et al. 2020. Fungal Diversity 105:1-16 ((https://doi.org/10.1007/s13225-020-00466-2)) |
| Otu35726 | 1 | k__Fungi(100);p__Ascomycota(100);c__Saccharomycetes(100);o__Saccharomycetales(100);f__Metschnikowiaceae(100);g__Metschnikowia(100);s__Metschnikowia_pulcherrima(100); | Metschnikowia | 13 | Saprotroph | \|Undefined Saprotroph\| | Yeast | NULL | Probable | As nectar-tap_saprotroph (Põlme et al. 2020); Decay sugar-rich substrates (Põlme et al. 2020) | Põlme S, et al. 2020. Fungal Diversity 105:1-16 ((https://doi.org/10.1007/s13225-020-00466-2)) |
| Otu35729 | 1 | k__Fungi(100);p__Ascomycota(100);c__Saccharomycetes(100);o__Saccharomycetales(100);f__Metschnikowiaceae(100);g__Metschnikowia(100);g__Metschnikowia_unclassified(100); | Metschnikowia | 13 | Saprotroph | \|Undefined Saprotroph\| | Yeast | NULL | Probable | As nectar-tap_saprotroph (Põlme et al. 2020); Decay sugar-rich substrates (Põlme et al. 2020) | Põlme S, et al. 2020. Fungal Diversity 105:1-16 ((https://doi.org/10.1007/s13225-020-00466-2)) |
| Otu35732 | 1 | k__Fungi(100);p__Ascomycota(100);c__Saccharomycetes(100);o__Saccharomycetales(100);f__Metschnikowiaceae(100);g__Metschnikowia(100);g__Metschnikowia_unclassified(100); | Metschnikowia | 13 | Saprotroph | \|Undefined Saprotroph\| | Yeast | NULL | Probable | As nectar-tap_saprotroph (Põlme et al. 2020); Decay sugar-rich substrates (Põlme et al. 2020) | Põlme S, et al. 2020. Fungal Diversity 105:1-16 ((https://doi.org/10.1007/s13225-020-00466-2)) |
| Otu35733 | 1 | k__Fungi(100);p__Ascomycota(100);c__Saccharomycetes(100);o__Saccharomycetales(100);f__Metschnikowiaceae(100);g__Metschnikowia(100);g__Metschnikowia_unclassified(100); | Metschnikowia | 13 | Saprotroph | \|Undefined Saprotroph\| | Yeast | NULL | Probable | As nectar-tap_saprotroph (Põlme et al. 2020); Decay sugar-rich substrates (Põlme et al. 2020) | Põlme S, et al. 2020. Fungal Diversity 105:1-16 ((https://doi.org/10.1007/s13225-020-00466-2)) |
| Otu35734 | 1 | k__Fungi(100);p__Ascomycota(100);c__Saccharomycetes(100);o__Saccharomycetales(100);f__Metschnikowiaceae(100);g__Metschnikowia(100);s__Metschnikowia_pulcherrima(100); | Metschnikowia | 13 | Saprotroph | \|Undefined Saprotroph\| | Yeast | NULL | Probable | As nectar-tap_saprotroph (Põlme et al. 2020); Decay sugar-rich substrates (Põlme et al. 2020) | Põlme S, et al. 2020. Fungal Diversity 105:1-16 ((https://doi.org/10.1007/s13225-020-00466-2)) |
| Otu35736 | 1 | k__Fungi(100);p__Ascomycota(100);c__Saccharomycetes(100);o__Saccharomycetales(100);f__Metschnikowiaceae(100);g__Metschnikowia(100);s__Metschnikowia_pulcherrima(100); | Metschnikowia | 13 | Saprotroph | \|Undefined Saprotroph\| | Yeast | NULL | Probable | As nectar-tap_saprotroph (Põlme et al. 2020); Decay sugar-rich substrates (Põlme et al. 2020) | Põlme S, et al. 2020. Fungal Diversity 105:1-16 ((https://doi.org/10.1007/s13225-020-00466-2)) |
| Otu35737 | 1 | k__Fungi(100);p__Ascomycota(100);c__Saccharomycetes(100);o__Saccharomycetales(100);f__Metschnikowiaceae(100);g__Metschnikowia(100);s__Metschnikowia_pulcherrima(100); | Metschnikowia | 13 | Saprotroph | \|Undefined Saprotroph\| | Yeast | NULL | Probable | As nectar-tap_saprotroph (Põlme et al. 2020); Decay sugar-rich substrates (Põlme et al. 2020) | Põlme S, et al. 2020. Fungal Diversity 105:1-16 ((https://doi.org/10.1007/s13225-020-00466-2)) |
| Otu35738 | 1 | k__Fungi(100);p__Ascomycota(100);c__Saccharomycetes(100);o__Saccharomycetales(100);f__Metschnikowiaceae(100);g__Metschnikowia(100);s__Metschnikowia_pulcherrima(100); | Metschnikowia | 13 | Saprotroph | \|Undefined Saprotroph\| | Yeast | NULL | Probable | As nectar-tap_saprotroph (Põlme et al. 2020); Decay sugar-rich substrates (Põlme et al. 2020) | Põlme S, et al. 2020. Fungal Diversity 105:1-16 ((https://doi.org/10.1007/s13225-020-00466-2)) |
| Otu35740 | 1 | k__Fungi(100);p__Ascomycota(100);c__Saccharomycetes(100);o__Saccharomycetales(100);f__Metschnikowiaceae(100);g__Metschnikowia(100);s__Metschnikowia_pulcherrima(100); | Metschnikowia | 13 | Saprotroph | \|Undefined Saprotroph\| | Yeast | NULL | Probable | As nectar-tap_saprotroph (Põlme et al. 2020); Decay sugar-rich substrates (Põlme et al. 2020) | Põlme S, et al. 2020. Fungal Diversity 105:1-16 ((https://doi.org/10.1007/s13225-020-00466-2)) |
| Otu35741 | 1 | k__Fungi(100);p__Ascomycota(100);c__Saccharomycetes(100);o__Saccharomycetales(100);f__Metschnikowiaceae(100);g__Metschnikowia(100);s__Metschnikowia_pulcherrima(100); | Metschnikowia | 13 | Saprotroph | \|Undefined Saprotroph\| | Yeast | NULL | Probable | As nectar-tap_saprotroph (Põlme et al. 2020); Decay sugar-rich substrates (Põlme et al. 2020) | Põlme S, et al. 2020. Fungal Diversity 105:1-16 ((https://doi.org/10.1007/s13225-020-00466-2)) |
| Otu35743 | 1 | k__Fungi(100);p__Ascomycota(100);c__Saccharomycetes(100);o__Saccharomycetales(100);f__Metschnikowiaceae(100);g__Metschnikowia(100);g__Metschnikowia_unclassified(100); | Metschnikowia | 13 | Saprotroph | \|Undefined Saprotroph\| | Yeast | NULL | Probable | As nectar-tap_saprotroph (Põlme et al. 2020); Decay sugar-rich substrates (Põlme et al. 2020) | Põlme S, et al. 2020. Fungal Diversity 105:1-16 ((https://doi.org/10.1007/s13225-020-00466-2)) |
| Otu35744 | 1 | k__Fungi(100);p__Ascomycota(100);c__Saccharomycetes(100);o__Saccharomycetales(100);f__Metschnikowiaceae(100);g__Metschnikowia(100);s__Metschnikowia_pulcherrima(100); | Metschnikowia | 13 | Saprotroph | \|Undefined Saprotroph\| | Yeast | NULL | Probable | As nectar-tap_saprotroph (Põlme et al. 2020); Decay sugar-rich substrates (Põlme et al. 2020) | Põlme S, et al. 2020. Fungal Diversity 105:1-16 ((https://doi.org/10.1007/s13225-020-00466-2)) |
| Otu35745 | 1 | k__Fungi(100);p__Ascomycota(100);c__Saccharomycetes(100);o__Saccharomycetales(100);f__Metschnikowiaceae(100);g__Metschnikowia(100);s__Metschnikowia_pulcherrima(100); | Metschnikowia | 13 | Saprotroph | \|Undefined Saprotroph\| | Yeast | NULL | Probable | As nectar-tap_saprotroph (Põlme et al. 2020); Decay sugar-rich substrates (Põlme et al. 2020) | Põlme S, et al. 2020. Fungal Diversity 105:1-16 ((https://doi.org/10.1007/s13225-020-00466-2)) |
| Otu35746 | 1 | k__Fungi(100);p__Ascomycota(100);c__Saccharomycetes(100);o__Saccharomycetales(100);f__Metschnikowiaceae(100);g__Metschnikowia(100);s__Metschnikowia_pulcherrima(100); | Metschnikowia | 13 | Saprotroph | \|Undefined Saprotroph\| | Yeast | NULL | Probable | As nectar-tap_saprotroph (Põlme et al. 2020); Decay sugar-rich substrates (Põlme et al. 2020) | Põlme S, et al. 2020. Fungal Diversity 105:1-16 ((https://doi.org/10.1007/s13225-020-00466-2)) |
| Otu35747 | 1 | k__Fungi(100);p__Ascomycota(100);c__Saccharomycetes(100);o__Saccharomycetales(100);f__Metschnikowiaceae(100);g__Metschnikowia(100);s__Metschnikowia_pulcherrima(100); | Metschnikowia | 13 | Saprotroph | \|Undefined Saprotroph\| | Yeast | NULL | Probable | As nectar-tap_saprotroph (Põlme et al. 2020); Decay sugar-rich substrates (Põlme et al. 2020) | Põlme S, et al. 2020. Fungal Diversity 105:1-16 ((https://doi.org/10.1007/s13225-020-00466-2)) |
| Otu35748 | 1 | k__Fungi(100);p__Ascomycota(100);c__Saccharomycetes(100);o__Saccharomycetales(100);f__Metschnikowiaceae(100);g__Metschnikowia(100);s__Metschnikowia_pulcherrima(100); | Metschnikowia | 13 | Saprotroph | \|Undefined Saprotroph\| | Yeast | NULL | Probable | As nectar-tap_saprotroph (Põlme et al. 2020); Decay sugar-rich substrates (Põlme et al. 2020) | Põlme S, et al. 2020. Fungal Diversity 105:1-16 ((https://doi.org/10.1007/s13225-020-00466-2)) |
| Otu35749 | 1 | k__Fungi(100);p__Ascomycota(100);c__Saccharomycetes(100);o__Saccharomycetales(100);f__Metschnikowiaceae(100);g__Metschnikowia(100);s__Metschnikowia_pulcherrima(100); | Metschnikowia | 13 | Saprotroph | \|Undefined Saprotroph\| | Yeast | NULL | Probable | As nectar-tap_saprotroph (Põlme et al. 2020); Decay sugar-rich substrates (Põlme et al. 2020) | Põlme S, et al. 2020. Fungal Diversity 105:1-16 ((https://doi.org/10.1007/s13225-020-00466-2)) |
| Otu35750 | 1 | k__Fungi(100);p__Ascomycota(100);c__Saccharomycetes(100);o__Saccharomycetales(100);f__Metschnikowiaceae(100);g__Metschnikowia(100);s__Metschnikowia_pulcherrima(100); | Metschnikowia | 13 | Saprotroph | \|Undefined Saprotroph\| | Yeast | NULL | Probable | As nectar-tap_saprotroph (Põlme et al. 2020); Decay sugar-rich substrates (Põlme et al. 2020) | Põlme S, et al. 2020. Fungal Diversity 105:1-16 ((https://doi.org/10.1007/s13225-020-00466-2)) |
| Otu35751 | 1 | k__Fungi(100);p__Ascomycota(100);c__Saccharomycetes(100);o__Saccharomycetales(100);f__Metschnikowiaceae(100);g__Metschnikowia(100);s__Metschnikowia_pulcherrima(100); | Metschnikowia | 13 | Saprotroph | \|Undefined Saprotroph\| | Yeast | NULL | Probable | As nectar-tap_saprotroph (Põlme et al. 2020); Decay sugar-rich substrates (Põlme et al. 2020) | Põlme S, et al. 2020. Fungal Diversity 105:1-16 ((https://doi.org/10.1007/s13225-020-00466-2)) |
| Otu35752 | 1 | k__Fungi(100);p__Ascomycota(100);c__Saccharomycetes(100);o__Saccharomycetales(100);f__Metschnikowiaceae(100);g__Metschnikowia(100);s__Metschnikowia_pulcherrima(100); | Metschnikowia | 13 | Saprotroph | \|Undefined Saprotroph\| | Yeast | NULL | Probable | As nectar-tap_saprotroph (Põlme et al. 2020); Decay sugar-rich substrates (Põlme et al. 2020) | Põlme S, et al. 2020. Fungal Diversity 105:1-16 ((https://doi.org/10.1007/s13225-020-00466-2)) |
| Otu35753 | 1 | k__Fungi(100);p__Ascomycota(100);c__Saccharomycetes(100);o__Saccharomycetales(100);f__Metschnikowiaceae(100);g__Metschnikowia(100);s__Metschnikowia_pulcherrima(100); | Metschnikowia | 13 | Saprotroph | \|Undefined Saprotroph\| | Yeast | NULL | Probable | As nectar-tap_saprotroph (Põlme et al. 2020); Decay sugar-rich substrates (Põlme et al. 2020) | Põlme S, et al. 2020. Fungal Diversity 105:1-16 ((https://doi.org/10.1007/s13225-020-00466-2)) |
| Otu35754 | 1 | k__Fungi(100);p__Ascomycota(100);c__Saccharomycetes(100);o__Saccharomycetales(100);f__Metschnikowiaceae(100);g__Metschnikowia(100);s__Metschnikowia_pulcherrima(100); | Metschnikowia | 13 | Saprotroph | \|Undefined Saprotroph\| | Yeast | NULL | Probable | As nectar-tap_saprotroph (Põlme et al. 2020); Decay sugar-rich substrates (Põlme et al. 2020) | Põlme S, et al. 2020. Fungal Diversity 105:1-16 ((https://doi.org/10.1007/s13225-020-00466-2)) |
| Otu35755 | 1 | k__Fungi(100);p__Ascomycota(100);c__Saccharomycetes(100);o__Saccharomycetales(100);f__Metschnikowiaceae(100);g__Metschnikowia(100);s__Metschnikowia_pulcherrima(100); | Metschnikowia | 13 | Saprotroph | \|Undefined Saprotroph\| | Yeast | NULL | Probable | As nectar-tap_saprotroph (Põlme et al. 2020); Decay sugar-rich substrates (Põlme et al. 2020) | Põlme S, et al. 2020. Fungal Diversity 105:1-16 ((https://doi.org/10.1007/s13225-020-00466-2)) |
| Otu35756 | 1 | k__Fungi(100);p__Ascomycota(100);c__Saccharomycetes(100);o__Saccharomycetales(100);f__Metschnikowiaceae(100);g__Metschnikowia(100);s__Metschnikowia_pulcherrima(100); | Metschnikowia | 13 | Saprotroph | \|Undefined Saprotroph\| | Yeast | NULL | Probable | As nectar-tap_saprotroph (Põlme et al. 2020); Decay sugar-rich substrates (Põlme et al. 2020) | Põlme S, et al. 2020. Fungal Diversity 105:1-16 ((https://doi.org/10.1007/s13225-020-00466-2)) |
| Otu35757 | 1 | k__Fungi(100);p__Ascomycota(100);c__Saccharomycetes(100);o__Saccharomycetales(100);f__Metschnikowiaceae(100);g__Metschnikowia(100);g__Metschnikowia_unclassified(100); | Metschnikowia | 13 | Saprotroph | \|Undefined Saprotroph\| | Yeast | NULL | Probable | As nectar-tap_saprotroph (Põlme et al. 2020); Decay sugar-rich substrates (Põlme et al. 2020) | Põlme S, et al. 2020. Fungal Diversity 105:1-16 ((https://doi.org/10.1007/s13225-020-00466-2)) |
| Otu35758 | 1 | k__Fungi(100);p__Ascomycota(100);c__Saccharomycetes(100);o__Saccharomycetales(100);f__Metschnikowiaceae(100);g__Metschnikowia(100);g__Metschnikowia_unclassified(100); | Metschnikowia | 13 | Saprotroph | \|Undefined Saprotroph\| | Yeast | NULL | Probable | As nectar-tap_saprotroph (Põlme et al. 2020); Decay sugar-rich substrates (Põlme et al. 2020) | Põlme S, et al. 2020. Fungal Diversity 105:1-16 ((https://doi.org/10.1007/s13225-020-00466-2)) |
| Otu35765 | 1 | k__Fungi(100);p__Ascomycota(100);c__Saccharomycetes(100);o__Saccharomycetales(100);f__Metschnikowiaceae(100);g__Metschnikowia(100);s__Metschnikowia_pulcherrima(100); | Metschnikowia | 13 | Saprotroph | \|Undefined Saprotroph\| | Yeast | NULL | Probable | As nectar-tap_saprotroph (Põlme et al. 2020); Decay sugar-rich substrates (Põlme et al. 2020) | Põlme S, et al. 2020. Fungal Diversity 105:1-16 ((https://doi.org/10.1007/s13225-020-00466-2)) |
| Otu35766 | 1 | k__Fungi(100);p__Ascomycota(100);c__Saccharomycetes(100);o__Saccharomycetales(100);f__Metschnikowiaceae(100);g__Metschnikowia(100);s__Metschnikowia_pulcherrima(100); | Metschnikowia | 13 | Saprotroph | \|Undefined Saprotroph\| | Yeast | NULL | Probable | As nectar-tap_saprotroph (Põlme et al. 2020); Decay sugar-rich substrates (Põlme et al. 2020) | Põlme S, et al. 2020. Fungal Diversity 105:1-16 ((https://doi.org/10.1007/s13225-020-00466-2)) |
| Otu35767 | 1 | k__Fungi(100);p__Ascomycota(100);c__Saccharomycetes(100);o__Saccharomycetales(100);f__Metschnikowiaceae(100);g__Metschnikowia(100);s__Metschnikowia_pulcherrima(100); | Metschnikowia | 13 | Saprotroph | \|Undefined Saprotroph\| | Yeast | NULL | Probable | As nectar-tap_saprotroph (Põlme et al. 2020); Decay sugar-rich substrates (Põlme et al. 2020) | Põlme S, et al. 2020. Fungal Diversity 105:1-16 ((https://doi.org/10.1007/s13225-020-00466-2)) |
| Otu35769 | 1 | k__Fungi(100);p__Ascomycota(100);c__Saccharomycetes(100);o__Saccharomycetales(100);f__Metschnikowiaceae(100);g__Metschnikowia(100);s__Metschnikowia_pulcherrima(100); | Metschnikowia | 13 | Saprotroph | \|Undefined Saprotroph\| | Yeast | NULL | Probable | As nectar-tap_saprotroph (Põlme et al. 2020); Decay sugar-rich substrates (Põlme et al. 2020) | Põlme S, et al. 2020. Fungal Diversity 105:1-16 ((https://doi.org/10.1007/s13225-020-00466-2)) |
| Otu35770 | 1 | k__Fungi(100);p__Ascomycota(100);c__Saccharomycetes(100);o__Saccharomycetales(100);f__Metschnikowiaceae(100);g__Metschnikowia(100);s__Metschnikowia_pulcherrima(100); | Metschnikowia | 13 | Saprotroph | \|Undefined Saprotroph\| | Yeast | NULL | Probable | As nectar-tap_saprotroph (Põlme et al. 2020); Decay sugar-rich substrates (Põlme et al. 2020) | Põlme S, et al. 2020. Fungal Diversity 105:1-16 ((https://doi.org/10.1007/s13225-020-00466-2)) |
| Otu35771 | 1 | k__Fungi(100);p__Ascomycota(100);c__Saccharomycetes(100);o__Saccharomycetales(100);f__Metschnikowiaceae(100);g__Metschnikowia(100);s__Metschnikowia_pulcherrima(100); | Metschnikowia | 13 | Saprotroph | \|Undefined Saprotroph\| | Yeast | NULL | Probable | As nectar-tap_saprotroph (Põlme et al. 2020); Decay sugar-rich substrates (Põlme et al. 2020) | Põlme S, et al. 2020. Fungal Diversity 105:1-16 ((https://doi.org/10.1007/s13225-020-00466-2)) |
| Otu35772 | 1 | k__Fungi(100);p__Ascomycota(100);c__Saccharomycetes(100);o__Saccharomycetales(100);f__Metschnikowiaceae(100);g__Metschnikowia(100);s__Metschnikowia_pulcherrima(100); | Metschnikowia | 13 | Saprotroph | \|Undefined Saprotroph\| | Yeast | NULL | Probable | As nectar-tap_saprotroph (Põlme et al. 2020); Decay sugar-rich substrates (Põlme et al. 2020) | Põlme S, et al. 2020. Fungal Diversity 105:1-16 ((https://doi.org/10.1007/s13225-020-00466-2)) |
| Otu35773 | 1 | k__Fungi(100);p__Ascomycota(100);c__Saccharomycetes(100);o__Saccharomycetales(100);f__Metschnikowiaceae(100);g__Metschnikowia(100);g__Metschnikowia_unclassified(100); | Metschnikowia | 13 | Saprotroph | \|Undefined Saprotroph\| | Yeast | NULL | Probable | As nectar-tap_saprotroph (Põlme et al. 2020); Decay sugar-rich substrates (Põlme et al. 2020) | Põlme S, et al. 2020. Fungal Diversity 105:1-16 ((https://doi.org/10.1007/s13225-020-00466-2)) |
| Otu35774 | 1 | k__Fungi(100);p__Ascomycota(100);c__Saccharomycetes(100);o__Saccharomycetales(100);f__Metschnikowiaceae(100);g__Metschnikowia(100);g__Metschnikowia_unclassified(100); | Metschnikowia | 13 | Saprotroph | \|Undefined Saprotroph\| | Yeast | NULL | Probable | As nectar-tap_saprotroph (Põlme et al. 2020); Decay sugar-rich substrates (Põlme et al. 2020) | Põlme S, et al. 2020. Fungal Diversity 105:1-16 ((https://doi.org/10.1007/s13225-020-00466-2)) |
| Otu35775 | 1 | k__Fungi(100);p__Ascomycota(100);c__Saccharomycetes(100);o__Saccharomycetales(100);f__Metschnikowiaceae(100);g__Metschnikowia(100);s__Metschnikowia_pulcherrima(100); | Metschnikowia | 13 | Saprotroph | \|Undefined Saprotroph\| | Yeast | NULL | Probable | As nectar-tap_saprotroph (Põlme et al. 2020); Decay sugar-rich substrates (Põlme et al. 2020) | Põlme S, et al. 2020. Fungal Diversity 105:1-16 ((https://doi.org/10.1007/s13225-020-00466-2)) |
| Otu35777 | 1 | k__Fungi(100);p__Ascomycota(100);c__Saccharomycetes(100);o__Saccharomycetales(100);f__Metschnikowiaceae(100);g__Metschnikowia(100);s__Metschnikowia_pulcherrima(100); | Metschnikowia | 13 | Saprotroph | \|Undefined Saprotroph\| | Yeast | NULL | Probable | As nectar-tap_saprotroph (Põlme et al. 2020); Decay sugar-rich substrates (Põlme et al. 2020) | Põlme S, et al. 2020. Fungal Diversity 105:1-16 ((https://doi.org/10.1007/s13225-020-00466-2)) |
| Otu35778 | 1 | k__Fungi(100);p__Ascomycota(100);c__Saccharomycetes(100);o__Saccharomycetales(100);f__Metschnikowiaceae(100);g__Metschnikowia(100);s__Metschnikowia_pulcherrima(100); | Metschnikowia | 13 | Saprotroph | \|Undefined Saprotroph\| | Yeast | NULL | Probable | As nectar-tap_saprotroph (Põlme et al. 2020); Decay sugar-rich substrates (Põlme et al. 2020) | Põlme S, et al. 2020. Fungal Diversity 105:1-16 ((https://doi.org/10.1007/s13225-020-00466-2)) |
| Otu35779 | 1 | k__Fungi(100);p__Ascomycota(100);c__Saccharomycetes(100);o__Saccharomycetales(100);f__Metschnikowiaceae(100);g__Metschnikowia(100);g__Metschnikowia_unclassified(100); | Metschnikowia | 13 | Saprotroph | \|Undefined Saprotroph\| | Yeast | NULL | Probable | As nectar-tap_saprotroph (Põlme et al. 2020); Decay sugar-rich substrates (Põlme et al. 2020) | Põlme S, et al. 2020. Fungal Diversity 105:1-16 ((https://doi.org/10.1007/s13225-020-00466-2)) |
| Otu35780 | 1 | k__Fungi(100);p__Ascomycota(100);c__Saccharomycetes(100);o__Saccharomycetales(100);f__Metschnikowiaceae(100);g__Metschnikowia(100);s__Metschnikowia_pulcherrima(100); | Metschnikowia | 13 | Saprotroph | \|Undefined Saprotroph\| | Yeast | NULL | Probable | As nectar-tap_saprotroph (Põlme et al. 2020); Decay sugar-rich substrates (Põlme et al. 2020) | Põlme S, et al. 2020. Fungal Diversity 105:1-16 ((https://doi.org/10.1007/s13225-020-00466-2)) |
| Otu35782 | 1 | k__Fungi(100);p__Ascomycota(100);c__Saccharomycetes(100);o__Saccharomycetales(100);f__Metschnikowiaceae(100);g__Metschnikowia(100);g__Metschnikowia_unclassified(100); | Metschnikowia | 13 | Saprotroph | \|Undefined Saprotroph\| | Yeast | NULL | Probable | As nectar-tap_saprotroph (Põlme et al. 2020); Decay sugar-rich substrates (Põlme et al. 2020) | Põlme S, et al. 2020. Fungal Diversity 105:1-16 ((https://doi.org/10.1007/s13225-020-00466-2)) |
| Otu35783 | 1 | k__Fungi(100);p__Ascomycota(100);c__Saccharomycetes(100);o__Saccharomycetales(100);f__Metschnikowiaceae(100);g__Metschnikowia(100);s__Metschnikowia_pulcherrima(100); | Metschnikowia | 13 | Saprotroph | \|Undefined Saprotroph\| | Yeast | NULL | Probable | As nectar-tap_saprotroph (Põlme et al. 2020); Decay sugar-rich substrates (Põlme et al. 2020) | Põlme S, et al. 2020. Fungal Diversity 105:1-16 ((https://doi.org/10.1007/s13225-020-00466-2)) |
| Otu35785 | 1 | k__Fungi(100);p__Ascomycota(100);c__Saccharomycetes(100);o__Saccharomycetales(100);f__Metschnikowiaceae(100);g__Metschnikowia(100);s__Metschnikowia_pulcherrima(100); | Metschnikowia | 13 | Saprotroph | \|Undefined Saprotroph\| | Yeast | NULL | Probable | As nectar-tap_saprotroph (Põlme et al. 2020); Decay sugar-rich substrates (Põlme et al. 2020) | Põlme S, et al. 2020. Fungal Diversity 105:1-16 ((https://doi.org/10.1007/s13225-020-00466-2)) |
| Otu35787 | 1 | k__Fungi(100);p__Ascomycota(100);c__Saccharomycetes(100);o__Saccharomycetales(100);f__Metschnikowiaceae(100);g__Metschnikowia(100);s__Metschnikowia_pulcherrima(100); | Metschnikowia | 13 | Saprotroph | \|Undefined Saprotroph\| | Yeast | NULL | Probable | As nectar-tap_saprotroph (Põlme et al. 2020); Decay sugar-rich substrates (Põlme et al. 2020) | Põlme S, et al. 2020. Fungal Diversity 105:1-16 ((https://doi.org/10.1007/s13225-020-00466-2)) |
| Otu35788 | 1 | k__Fungi(100);p__Ascomycota(100);c__Saccharomycetes(100);o__Saccharomycetales(100);f__Metschnikowiaceae(100);g__Metschnikowia(100);s__Metschnikowia_pulcherrima(100); | Metschnikowia | 13 | Saprotroph | \|Undefined Saprotroph\| | Yeast | NULL | Probable | As nectar-tap_saprotroph (Põlme et al. 2020); Decay sugar-rich substrates (Põlme et al. 2020) | Põlme S, et al. 2020. Fungal Diversity 105:1-16 ((https://doi.org/10.1007/s13225-020-00466-2)) |
| Otu35789 | 1 | k__Fungi(100);p__Ascomycota(100);c__Saccharomycetes(100);o__Saccharomycetales(100);f__Metschnikowiaceae(100);g__Metschnikowia(100);g__Metschnikowia_unclassified(100); | Metschnikowia | 13 | Saprotroph | \|Undefined Saprotroph\| | Yeast | NULL | Probable | As nectar-tap_saprotroph (Põlme et al. 2020); Decay sugar-rich substrates (Põlme et al. 2020) | Põlme S, et al. 2020. Fungal Diversity 105:1-16 ((https://doi.org/10.1007/s13225-020-00466-2)) |
| Otu35790 | 1 | k__Fungi(100);p__Ascomycota(100);c__Saccharomycetes(100);o__Saccharomycetales(100);f__Metschnikowiaceae(100);g__Metschnikowia(100);s__Metschnikowia_pulcherrima(100); | Metschnikowia | 13 | Saprotroph | \|Undefined Saprotroph\| | Yeast | NULL | Probable | As nectar-tap_saprotroph (Põlme et al. 2020); Decay sugar-rich substrates (Põlme et al. 2020) | Põlme S, et al. 2020. Fungal Diversity 105:1-16 ((https://doi.org/10.1007/s13225-020-00466-2)) |
| Otu35791 | 1 | k__Fungi(100);p__Ascomycota(100);c__Saccharomycetes(100);o__Saccharomycetales(100);f__Metschnikowiaceae(100);g__Metschnikowia(100);s__Metschnikowia_pulcherrima(100); | Metschnikowia | 13 | Saprotroph | \|Undefined Saprotroph\| | Yeast | NULL | Probable | As nectar-tap_saprotroph (Põlme et al. 2020); Decay sugar-rich substrates (Põlme et al. 2020) | Põlme S, et al. 2020. Fungal Diversity 105:1-16 ((https://doi.org/10.1007/s13225-020-00466-2)) |
| Otu35792 | 1 | k__Fungi(100);p__Ascomycota(100);c__Saccharomycetes(100);o__Saccharomycetales(100);f__Metschnikowiaceae(100);g__Metschnikowia(100);s__Metschnikowia_pulcherrima(100); | Metschnikowia | 13 | Saprotroph | \|Undefined Saprotroph\| | Yeast | NULL | Probable | As nectar-tap_saprotroph (Põlme et al. 2020); Decay sugar-rich substrates (Põlme et al. 2020) | Põlme S, et al. 2020. Fungal Diversity 105:1-16 ((https://doi.org/10.1007/s13225-020-00466-2)) |
| Otu35793 | 1 | k__Fungi(100);p__Ascomycota(100);c__Saccharomycetes(100);o__Saccharomycetales(100);f__Metschnikowiaceae(100);g__Metschnikowia(100);s__Metschnikowia_pulcherrima(100); | Metschnikowia | 13 | Saprotroph | \|Undefined Saprotroph\| | Yeast | NULL | Probable | As nectar-tap_saprotroph (Põlme et al. 2020); Decay sugar-rich substrates (Põlme et al. 2020) | Põlme S, et al. 2020. Fungal Diversity 105:1-16 ((https://doi.org/10.1007/s13225-020-00466-2)) |
| Otu35796 | 1 | k__Fungi(100);p__Ascomycota(100);c__Saccharomycetes(100);o__Saccharomycetales(100);f__Metschnikowiaceae(100);g__Metschnikowia(100);s__Metschnikowia_pulcherrima(100); | Metschnikowia | 13 | Saprotroph | \|Undefined Saprotroph\| | Yeast | NULL | Probable | As nectar-tap_saprotroph (Põlme et al. 2020); Decay sugar-rich substrates (Põlme et al. 2020) | Põlme S, et al. 2020. Fungal Diversity 105:1-16 ((https://doi.org/10.1007/s13225-020-00466-2)) |
| Otu35797 | 1 | k__Fungi(100);p__Ascomycota(100);c__Saccharomycetes(100);o__Saccharomycetales(100);f__Metschnikowiaceae(100);g__Metschnikowia(100);s__Metschnikowia_pulcherrima(100); | Metschnikowia | 13 | Saprotroph | \|Undefined Saprotroph\| | Yeast | NULL | Probable | As nectar-tap_saprotroph (Põlme et al. 2020); Decay sugar-rich substrates (Põlme et al. 2020) | Põlme S, et al. 2020. Fungal Diversity 105:1-16 ((https://doi.org/10.1007/s13225-020-00466-2)) |
| Otu35799 | 1 | k__Fungi(100);p__Ascomycota(100);c__Saccharomycetes(100);o__Saccharomycetales(100);f__Metschnikowiaceae(100);g__Metschnikowia(100);g__Metschnikowia_unclassified(100); | Metschnikowia | 13 | Saprotroph | \|Undefined Saprotroph\| | Yeast | NULL | Probable | As nectar-tap_saprotroph (Põlme et al. 2020); Decay sugar-rich substrates (Põlme et al. 2020) | Põlme S, et al. 2020. Fungal Diversity 105:1-16 ((https://doi.org/10.1007/s13225-020-00466-2)) |
| Otu35800 | 1 | k__Fungi(100);p__Ascomycota(100);c__Saccharomycetes(100);o__Saccharomycetales(100);f__Metschnikowiaceae(100);g__Metschnikowia(100);g__Metschnikowia_unclassified(100); | Metschnikowia | 13 | Saprotroph | \|Undefined Saprotroph\| | Yeast | NULL | Probable | As nectar-tap_saprotroph (Põlme et al. 2020); Decay sugar-rich substrates (Põlme et al. 2020) | Põlme S, et al. 2020. Fungal Diversity 105:1-16 ((https://doi.org/10.1007/s13225-020-00466-2)) |
| Otu35802 | 1 | k__Fungi(100);p__Ascomycota(100);c__Saccharomycetes(100);o__Saccharomycetales(100);f__Metschnikowiaceae(100);g__Metschnikowia(100);s__Metschnikowia_pulcherrima(100); | Metschnikowia | 13 | Saprotroph | \|Undefined Saprotroph\| | Yeast | NULL | Probable | As nectar-tap_saprotroph (Põlme et al. 2020); Decay sugar-rich substrates (Põlme et al. 2020) | Põlme S, et al. 2020. Fungal Diversity 105:1-16 ((https://doi.org/10.1007/s13225-020-00466-2)) |
| Otu35803 | 1 | k__Fungi(100);p__Ascomycota(100);c__Saccharomycetes(100);o__Saccharomycetales(100);f__Metschnikowiaceae(100);g__Metschnikowia(100);s__Metschnikowia_pulcherrima(100); | Metschnikowia | 13 | Saprotroph | \|Undefined Saprotroph\| | Yeast | NULL | Probable | As nectar-tap_saprotroph (Põlme et al. 2020); Decay sugar-rich substrates (Põlme et al. 2020) | Põlme S, et al. 2020. Fungal Diversity 105:1-16 ((https://doi.org/10.1007/s13225-020-00466-2)) |
| Otu35804 | 1 | k__Fungi(100);p__Ascomycota(100);c__Saccharomycetes(100);o__Saccharomycetales(100);f__Metschnikowiaceae(100);g__Metschnikowia(100);s__Metschnikowia_pulcherrima(100); | Metschnikowia | 13 | Saprotroph | \|Undefined Saprotroph\| | Yeast | NULL | Probable | As nectar-tap_saprotroph (Põlme et al. 2020); Decay sugar-rich substrates (Põlme et al. 2020) | Põlme S, et al. 2020. Fungal Diversity 105:1-16 ((https://doi.org/10.1007/s13225-020-00466-2)) |
| Otu35806 | 1 | k__Fungi(100);p__Ascomycota(100);c__Saccharomycetes(100);o__Saccharomycetales(100);f__Metschnikowiaceae(100);g__Metschnikowia(100);s__Metschnikowia_pulcherrima(100); | Metschnikowia | 13 | Saprotroph | \|Undefined Saprotroph\| | Yeast | NULL | Probable | As nectar-tap_saprotroph (Põlme et al. 2020); Decay sugar-rich substrates (Põlme et al. 2020) | Põlme S, et al. 2020. Fungal Diversity 105:1-16 ((https://doi.org/10.1007/s13225-020-00466-2)) |
| Otu35807 | 1 | k__Fungi(100);p__Ascomycota(100);c__Saccharomycetes(100);o__Saccharomycetales(100);f__Metschnikowiaceae(100);g__Metschnikowia(100);s__Metschnikowia_pulcherrima(100); | Metschnikowia | 13 | Saprotroph | \|Undefined Saprotroph\| | Yeast | NULL | Probable | As nectar-tap_saprotroph (Põlme et al. 2020); Decay sugar-rich substrates (Põlme et al. 2020) | Põlme S, et al. 2020. Fungal Diversity 105:1-16 ((https://doi.org/10.1007/s13225-020-00466-2)) |
| Otu35808 | 1 | k__Fungi(100);p__Ascomycota(100);c__Saccharomycetes(100);o__Saccharomycetales(100);f__Metschnikowiaceae(100);g__Metschnikowia(100);g__Metschnikowia_unclassified(100); | Metschnikowia | 13 | Saprotroph | \|Undefined Saprotroph\| | Yeast | NULL | Probable | As nectar-tap_saprotroph (Põlme et al. 2020); Decay sugar-rich substrates (Põlme et al. 2020) | Põlme S, et al. 2020. Fungal Diversity 105:1-16 ((https://doi.org/10.1007/s13225-020-00466-2)) |
| Otu35809 | 1 | k__Fungi(100);p__Ascomycota(100);c__Saccharomycetes(100);o__Saccharomycetales(100);f__Metschnikowiaceae(100);g__Metschnikowia(100);g__Metschnikowia_unclassified(100); | Metschnikowia | 13 | Saprotroph | \|Undefined Saprotroph\| | Yeast | NULL | Probable | As nectar-tap_saprotroph (Põlme et al. 2020); Decay sugar-rich substrates (Põlme et al. 2020) | Põlme S, et al. 2020. Fungal Diversity 105:1-16 ((https://doi.org/10.1007/s13225-020-00466-2)) |
| Otu35810 | 1 | k__Fungi(100);p__Ascomycota(100);c__Saccharomycetes(100);o__Saccharomycetales(100);f__Metschnikowiaceae(100);g__Metschnikowia(100);g__Metschnikowia_unclassified(100); | Metschnikowia | 13 | Saprotroph | \|Undefined Saprotroph\| | Yeast | NULL | Probable | As nectar-tap_saprotroph (Põlme et al. 2020); Decay sugar-rich substrates (Põlme et al. 2020) | Põlme S, et al. 2020. Fungal Diversity 105:1-16 ((https://doi.org/10.1007/s13225-020-00466-2)) |
| Otu35811 | 1 | k__Fungi(100);p__Ascomycota(100);c__Saccharomycetes(100);o__Saccharomycetales(100);f__Metschnikowiaceae(100);g__Metschnikowia(100);g__Metschnikowia_unclassified(100); | Metschnikowia | 13 | Saprotroph | \|Undefined Saprotroph\| | Yeast | NULL | Probable | As nectar-tap_saprotroph (Põlme et al. 2020); Decay sugar-rich substrates (Põlme et al. 2020) | Põlme S, et al. 2020. Fungal Diversity 105:1-16 ((https://doi.org/10.1007/s13225-020-00466-2)) |
| Otu35812 | 1 | k__Fungi(100);p__Ascomycota(100);c__Saccharomycetes(100);o__Saccharomycetales(100);f__Metschnikowiaceae(100);g__Metschnikowia(100);s__Metschnikowia_pulcherrima(100); | Metschnikowia | 13 | Saprotroph | \|Undefined Saprotroph\| | Yeast | NULL | Probable | As nectar-tap_saprotroph (Põlme et al. 2020); Decay sugar-rich substrates (Põlme et al. 2020) | Põlme S, et al. 2020. Fungal Diversity 105:1-16 ((https://doi.org/10.1007/s13225-020-00466-2)) |
| Otu35813 | 1 | k__Fungi(100);p__Ascomycota(100);c__Saccharomycetes(100);o__Saccharomycetales(100);f__Metschnikowiaceae(100);g__Metschnikowia(100);s__Metschnikowia_pulcherrima(100); | Metschnikowia | 13 | Saprotroph | \|Undefined Saprotroph\| | Yeast | NULL | Probable | As nectar-tap_saprotroph (Põlme et al. 2020); Decay sugar-rich substrates (Põlme et al. 2020) | Põlme S, et al. 2020. Fungal Diversity 105:1-16 ((https://doi.org/10.1007/s13225-020-00466-2)) |
| Otu35814 | 1 | k__Fungi(100);p__Ascomycota(100);c__Saccharomycetes(100);o__Saccharomycetales(100);f__Metschnikowiaceae(100);g__Metschnikowia(100);s__Metschnikowia_pulcherrima(100); | Metschnikowia | 13 | Saprotroph | \|Undefined Saprotroph\| | Yeast | NULL | Probable | As nectar-tap_saprotroph (Põlme et al. 2020); Decay sugar-rich substrates (Põlme et al. 2020) | Põlme S, et al. 2020. Fungal Diversity 105:1-16 ((https://doi.org/10.1007/s13225-020-00466-2)) |
| Otu35815 | 1 | k__Fungi(100);p__Ascomycota(100);c__Saccharomycetes(100);o__Saccharomycetales(100);f__Metschnikowiaceae(100);g__Metschnikowia(100);s__Metschnikowia_pulcherrima(100); | Metschnikowia | 13 | Saprotroph | \|Undefined Saprotroph\| | Yeast | NULL | Probable | As nectar-tap_saprotroph (Põlme et al. 2020); Decay sugar-rich substrates (Põlme et al. 2020) | Põlme S, et al. 2020. Fungal Diversity 105:1-16 ((https://doi.org/10.1007/s13225-020-00466-2)) |
| Otu35816 | 1 | k__Fungi(100);p__Ascomycota(100);c__Saccharomycetes(100);o__Saccharomycetales(100);f__Metschnikowiaceae(100);g__Metschnikowia(100);s__Metschnikowia_pulcherrima(100); | Metschnikowia | 13 | Saprotroph | \|Undefined Saprotroph\| | Yeast | NULL | Probable | As nectar-tap_saprotroph (Põlme et al. 2020); Decay sugar-rich substrates (Põlme et al. 2020) | Põlme S, et al. 2020. Fungal Diversity 105:1-16 ((https://doi.org/10.1007/s13225-020-00466-2)) |
| Otu35817 | 1 | k__Fungi(100);p__Ascomycota(100);c__Saccharomycetes(100);o__Saccharomycetales(100);f__Metschnikowiaceae(100);g__Metschnikowia(100);g__Metschnikowia_unclassified(100); | Metschnikowia | 13 | Saprotroph | \|Undefined Saprotroph\| | Yeast | NULL | Probable | As nectar-tap_saprotroph (Põlme et al. 2020); Decay sugar-rich substrates (Põlme et al. 2020) | Põlme S, et al. 2020. Fungal Diversity 105:1-16 ((https://doi.org/10.1007/s13225-020-00466-2)) |
| Otu35818 | 1 | k__Fungi(100);p__Ascomycota(100);c__Saccharomycetes(100);o__Saccharomycetales(100);f__Metschnikowiaceae(100);g__Metschnikowia(100);s__Metschnikowia_pulcherrima(100); | Metschnikowia | 13 | Saprotroph | \|Undefined Saprotroph\| | Yeast | NULL | Probable | As nectar-tap_saprotroph (Põlme et al. 2020); Decay sugar-rich substrates (Põlme et al. 2020) | Põlme S, et al. 2020. Fungal Diversity 105:1-16 ((https://doi.org/10.1007/s13225-020-00466-2)) |
| Otu35819 | 1 | k__Fungi(100);p__Ascomycota(100);c__Saccharomycetes(100);o__Saccharomycetales(100);f__Metschnikowiaceae(100);g__Metschnikowia(100);g__Metschnikowia_unclassified(100); | Metschnikowia | 13 | Saprotroph | \|Undefined Saprotroph\| | Yeast | NULL | Probable | As nectar-tap_saprotroph (Põlme et al. 2020); Decay sugar-rich substrates (Põlme et al. 2020) | Põlme S, et al. 2020. Fungal Diversity 105:1-16 ((https://doi.org/10.1007/s13225-020-00466-2)) |
| Otu35820 | 1 | k__Fungi(100);p__Ascomycota(100);c__Saccharomycetes(100);o__Saccharomycetales(100);f__Metschnikowiaceae(100);g__Metschnikowia(100);s__Metschnikowia_pulcherrima(100); | Metschnikowia | 13 | Saprotroph | \|Undefined Saprotroph\| | Yeast | NULL | Probable | As nectar-tap_saprotroph (Põlme et al. 2020); Decay sugar-rich substrates (Põlme et al. 2020) | Põlme S, et al. 2020. Fungal Diversity 105:1-16 ((https://doi.org/10.1007/s13225-020-00466-2)) |
| Otu35822 | 1 | k__Fungi(100);p__Ascomycota(100);c__Saccharomycetes(100);o__Saccharomycetales(100);f__Metschnikowiaceae(100);g__Metschnikowia(100);s__Metschnikowia_pulcherrima(100); | Metschnikowia | 13 | Saprotroph | \|Undefined Saprotroph\| | Yeast | NULL | Probable | As nectar-tap_saprotroph (Põlme et al. 2020); Decay sugar-rich substrates (Põlme et al. 2020) | Põlme S, et al. 2020. Fungal Diversity 105:1-16 ((https://doi.org/10.1007/s13225-020-00466-2)) |
| Otu35823 | 1 | k__Fungi(100);p__Ascomycota(100);c__Saccharomycetes(100);o__Saccharomycetales(100);f__Metschnikowiaceae(100);g__Metschnikowia(100);s__Metschnikowia_pulcherrima(100); | Metschnikowia | 13 | Saprotroph | \|Undefined Saprotroph\| | Yeast | NULL | Probable | As nectar-tap_saprotroph (Põlme et al. 2020); Decay sugar-rich substrates (Põlme et al. 2020) | Põlme S, et al. 2020. Fungal Diversity 105:1-16 ((https://doi.org/10.1007/s13225-020-00466-2)) |
| Otu35824 | 1 | k__Fungi(100);p__Ascomycota(100);c__Saccharomycetes(100);o__Saccharomycetales(100);f__Metschnikowiaceae(100);g__Metschnikowia(100);s__Metschnikowia_pulcherrima(100); | Metschnikowia | 13 | Saprotroph | \|Undefined Saprotroph\| | Yeast | NULL | Probable | As nectar-tap_saprotroph (Põlme et al. 2020); Decay sugar-rich substrates (Põlme et al. 2020) | Põlme S, et al. 2020. Fungal Diversity 105:1-16 ((https://doi.org/10.1007/s13225-020-00466-2)) |
| Otu35825 | 1 | k__Fungi(100);p__Ascomycota(100);c__Saccharomycetes(100);o__Saccharomycetales(100);f__Metschnikowiaceae(100);g__Metschnikowia(100);s__Metschnikowia_pulcherrima(100); | Metschnikowia | 13 | Saprotroph | \|Undefined Saprotroph\| | Yeast | NULL | Probable | As nectar-tap_saprotroph (Põlme et al. 2020); Decay sugar-rich substrates (Põlme et al. 2020) | Põlme S, et al. 2020. Fungal Diversity 105:1-16 ((https://doi.org/10.1007/s13225-020-00466-2)) |
| Otu35826 | 1 | k__Fungi(100);p__Ascomycota(100);c__Saccharomycetes(100);o__Saccharomycetales(100);f__Metschnikowiaceae(100);g__Metschnikowia(100);s__Metschnikowia_pulcherrima(100); | Metschnikowia | 13 | Saprotroph | \|Undefined Saprotroph\| | Yeast | NULL | Probable | As nectar-tap_saprotroph (Põlme et al. 2020); Decay sugar-rich substrates (Põlme et al. 2020) | Põlme S, et al. 2020. Fungal Diversity 105:1-16 ((https://doi.org/10.1007/s13225-020-00466-2)) |
| Otu35827 | 1 | k__Fungi(100);p__Ascomycota(100);c__Saccharomycetes(100);o__Saccharomycetales(100);f__Metschnikowiaceae(100);g__Metschnikowia(100);s__Metschnikowia_pulcherrima(100); | Metschnikowia | 13 | Saprotroph | \|Undefined Saprotroph\| | Yeast | NULL | Probable | As nectar-tap_saprotroph (Põlme et al. 2020); Decay sugar-rich substrates (Põlme et al. 2020) | Põlme S, et al. 2020. Fungal Diversity 105:1-16 ((https://doi.org/10.1007/s13225-020-00466-2)) |
| Otu35829 | 1 | k__Fungi(100);p__Ascomycota(100);c__Saccharomycetes(100);o__Saccharomycetales(100);f__Metschnikowiaceae(100);g__Metschnikowia(100);s__Metschnikowia_pulcherrima(100); | Metschnikowia | 13 | Saprotroph | \|Undefined Saprotroph\| | Yeast | NULL | Probable | As nectar-tap_saprotroph (Põlme et al. 2020); Decay sugar-rich substrates (Põlme et al. 2020) | Põlme S, et al. 2020. Fungal Diversity 105:1-16 ((https://doi.org/10.1007/s13225-020-00466-2)) |
| Otu35830 | 1 | k__Fungi(100);p__Ascomycota(100);c__Saccharomycetes(100);o__Saccharomycetales(100);f__Metschnikowiaceae(100);g__Metschnikowia(100);s__Metschnikowia_pulcherrima(100); | Metschnikowia | 13 | Saprotroph | \|Undefined Saprotroph\| | Yeast | NULL | Probable | As nectar-tap_saprotroph (Põlme et al. 2020); Decay sugar-rich substrates (Põlme et al. 2020) | Põlme S, et al. 2020. Fungal Diversity 105:1-16 ((https://doi.org/10.1007/s13225-020-00466-2)) |
| Otu35831 | 1 | k__Fungi(100);p__Ascomycota(100);c__Saccharomycetes(100);o__Saccharomycetales(100);f__Metschnikowiaceae(100);g__Metschnikowia(100);s__Metschnikowia_pulcherrima(100); | Metschnikowia | 13 | Saprotroph | \|Undefined Saprotroph\| | Yeast | NULL | Probable | As nectar-tap_saprotroph (Põlme et al. 2020); Decay sugar-rich substrates (Põlme et al. 2020) | Põlme S, et al. 2020. Fungal Diversity 105:1-16 ((https://doi.org/10.1007/s13225-020-00466-2)) |
| Otu35832 | 1 | k__Fungi(100);p__Ascomycota(100);c__Saccharomycetes(100);o__Saccharomycetales(100);f__Metschnikowiaceae(100);g__Metschnikowia(100);g__Metschnikowia_unclassified(100); | Metschnikowia | 13 | Saprotroph | \|Undefined Saprotroph\| | Yeast | NULL | Probable | As nectar-tap_saprotroph (Põlme et al. 2020); Decay sugar-rich substrates (Põlme et al. 2020) | Põlme S, et al. 2020. Fungal Diversity 105:1-16 ((https://doi.org/10.1007/s13225-020-00466-2)) |
| Otu35833 | 1 | k__Fungi(100);p__Ascomycota(100);c__Saccharomycetes(100);o__Saccharomycetales(100);f__Metschnikowiaceae(100);g__Metschnikowia(100);s__Metschnikowia_pulcherrima(100); | Metschnikowia | 13 | Saprotroph | \|Undefined Saprotroph\| | Yeast | NULL | Probable | As nectar-tap_saprotroph (Põlme et al. 2020); Decay sugar-rich substrates (Põlme et al. 2020) | Põlme S, et al. 2020. Fungal Diversity 105:1-16 ((https://doi.org/10.1007/s13225-020-00466-2)) |
| Otu35834 | 1 | k__Fungi(100);p__Ascomycota(100);c__Saccharomycetes(100);o__Saccharomycetales(100);f__Metschnikowiaceae(100);g__Metschnikowia(100);s__Metschnikowia_pulcherrima(100); | Metschnikowia | 13 | Saprotroph | \|Undefined Saprotroph\| | Yeast | NULL | Probable | As nectar-tap_saprotroph (Põlme et al. 2020); Decay sugar-rich substrates (Põlme et al. 2020) | Põlme S, et al. 2020. Fungal Diversity 105:1-16 ((https://doi.org/10.1007/s13225-020-00466-2)) |
| Otu35835 | 1 | k__Fungi(100);p__Ascomycota(100);c__Saccharomycetes(100);o__Saccharomycetales(100);f__Metschnikowiaceae(100);g__Metschnikowia(100);s__Metschnikowia_pulcherrima(100); | Metschnikowia | 13 | Saprotroph | \|Undefined Saprotroph\| | Yeast | NULL | Probable | As nectar-tap_saprotroph (Põlme et al. 2020); Decay sugar-rich substrates (Põlme et al. 2020) | Põlme S, et al. 2020. Fungal Diversity 105:1-16 ((https://doi.org/10.1007/s13225-020-00466-2)) |
| Otu35836 | 1 | k__Fungi(100);p__Ascomycota(100);c__Saccharomycetes(100);o__Saccharomycetales(100);f__Metschnikowiaceae(100);g__Metschnikowia(100);g__Metschnikowia_unclassified(100); | Metschnikowia | 13 | Saprotroph | \|Undefined Saprotroph\| | Yeast | NULL | Probable | As nectar-tap_saprotroph (Põlme et al. 2020); Decay sugar-rich substrates (Põlme et al. 2020) | Põlme S, et al. 2020. Fungal Diversity 105:1-16 ((https://doi.org/10.1007/s13225-020-00466-2)) |
| Otu35837 | 1 | k__Fungi(100);p__Ascomycota(100);c__Saccharomycetes(100);o__Saccharomycetales(100);f__Metschnikowiaceae(100);g__Metschnikowia(100);s__Metschnikowia_pulcherrima(100); | Metschnikowia | 13 | Saprotroph | \|Undefined Saprotroph\| | Yeast | NULL | Probable | As nectar-tap_saprotroph (Põlme et al. 2020); Decay sugar-rich substrates (Põlme et al. 2020) | Põlme S, et al. 2020. Fungal Diversity 105:1-16 ((https://doi.org/10.1007/s13225-020-00466-2)) |
| Otu35838 | 1 | k__Fungi(100);p__Ascomycota(100);c__Saccharomycetes(100);o__Saccharomycetales(100);f__Metschnikowiaceae(100);g__Metschnikowia(100);s__Metschnikowia_pulcherrima(100); | Metschnikowia | 13 | Saprotroph | \|Undefined Saprotroph\| | Yeast | NULL | Probable | As nectar-tap_saprotroph (Põlme et al. 2020); Decay sugar-rich substrates (Põlme et al. 2020) | Põlme S, et al. 2020. Fungal Diversity 105:1-16 ((https://doi.org/10.1007/s13225-020-00466-2)) |
| Otu35839 | 1 | k__Fungi(100);p__Ascomycota(100);c__Saccharomycetes(100);o__Saccharomycetales(100);f__Metschnikowiaceae(100);g__Metschnikowia(100);s__Metschnikowia_pulcherrima(100); | Metschnikowia | 13 | Saprotroph | \|Undefined Saprotroph\| | Yeast | NULL | Probable | As nectar-tap_saprotroph (Põlme et al. 2020); Decay sugar-rich substrates (Põlme et al. 2020) | Põlme S, et al. 2020. Fungal Diversity 105:1-16 ((https://doi.org/10.1007/s13225-020-00466-2)) |
| Otu35840 | 1 | k__Fungi(100);p__Ascomycota(100);c__Saccharomycetes(100);o__Saccharomycetales(100);f__Metschnikowiaceae(100);g__Metschnikowia(100);g__Metschnikowia_unclassified(100); | Metschnikowia | 13 | Saprotroph | \|Undefined Saprotroph\| | Yeast | NULL | Probable | As nectar-tap_saprotroph (Põlme et al. 2020); Decay sugar-rich substrates (Põlme et al. 2020) | Põlme S, et al. 2020. Fungal Diversity 105:1-16 ((https://doi.org/10.1007/s13225-020-00466-2)) |
| Otu35841 | 1 | k__Fungi(100);p__Ascomycota(100);c__Saccharomycetes(100);o__Saccharomycetales(100);f__Metschnikowiaceae(100);g__Metschnikowia(100);s__Metschnikowia_pulcherrima(100); | Metschnikowia | 13 | Saprotroph | \|Undefined Saprotroph\| | Yeast | NULL | Probable | As nectar-tap_saprotroph (Põlme et al. 2020); Decay sugar-rich substrates (Põlme et al. 2020) | Põlme S, et al. 2020. Fungal Diversity 105:1-16 ((https://doi.org/10.1007/s13225-020-00466-2)) |
| Otu35842 | 1 | k__Fungi(100);p__Ascomycota(100);c__Saccharomycetes(100);o__Saccharomycetales(100);f__Metschnikowiaceae(100);g__Metschnikowia(100);s__Metschnikowia_pulcherrima(100); | Metschnikowia | 13 | Saprotroph | \|Undefined Saprotroph\| | Yeast | NULL | Probable | As nectar-tap_saprotroph (Põlme et al. 2020); Decay sugar-rich substrates (Põlme et al. 2020) | Põlme S, et al. 2020. Fungal Diversity 105:1-16 ((https://doi.org/10.1007/s13225-020-00466-2)) |
| Otu35843 | 1 | k__Fungi(100);p__Ascomycota(100);c__Saccharomycetes(100);o__Saccharomycetales(100);f__Metschnikowiaceae(100);g__Metschnikowia(100);g__Metschnikowia_unclassified(100); | Metschnikowia | 13 | Saprotroph | \|Undefined Saprotroph\| | Yeast | NULL | Probable | As nectar-tap_saprotroph (Põlme et al. 2020); Decay sugar-rich substrates (Põlme et al. 2020) | Põlme S, et al. 2020. Fungal Diversity 105:1-16 ((https://doi.org/10.1007/s13225-020-00466-2)) |
| Otu35844 | 1 | k__Fungi(100);p__Ascomycota(100);c__Saccharomycetes(100);o__Saccharomycetales(100);f__Metschnikowiaceae(100);g__Metschnikowia(100);s__Metschnikowia_pulcherrima(100); | Metschnikowia | 13 | Saprotroph | \|Undefined Saprotroph\| | Yeast | NULL | Probable | As nectar-tap_saprotroph (Põlme et al. 2020); Decay sugar-rich substrates (Põlme et al. 2020) | Põlme S, et al. 2020. Fungal Diversity 105:1-16 ((https://doi.org/10.1007/s13225-020-00466-2)) |
| Otu35845 | 1 | k__Fungi(100);p__Ascomycota(100);c__Saccharomycetes(100);o__Saccharomycetales(100);f__Metschnikowiaceae(100);g__Metschnikowia(100);s__Metschnikowia_pulcherrima(100); | Metschnikowia | 13 | Saprotroph | \|Undefined Saprotroph\| | Yeast | NULL | Probable | As nectar-tap_saprotroph (Põlme et al. 2020); Decay sugar-rich substrates (Põlme et al. 2020) | Põlme S, et al. 2020. Fungal Diversity 105:1-16 ((https://doi.org/10.1007/s13225-020-00466-2)) |
| Otu35846 | 1 | k__Fungi(100);p__Ascomycota(100);c__Saccharomycetes(100);o__Saccharomycetales(100);f__Metschnikowiaceae(100);g__Metschnikowia(100);s__Metschnikowia_pulcherrima(100); | Metschnikowia | 13 | Saprotroph | \|Undefined Saprotroph\| | Yeast | NULL | Probable | As nectar-tap_saprotroph (Põlme et al. 2020); Decay sugar-rich substrates (Põlme et al. 2020) | Põlme S, et al. 2020. Fungal Diversity 105:1-16 ((https://doi.org/10.1007/s13225-020-00466-2)) |
| Otu35847 | 1 | k__Fungi(100);p__Ascomycota(100);c__Saccharomycetes(100);o__Saccharomycetales(100);f__Metschnikowiaceae(100);g__Metschnikowia(100);s__Metschnikowia_pulcherrima(100); | Metschnikowia | 13 | Saprotroph | \|Undefined Saprotroph\| | Yeast | NULL | Probable | As nectar-tap_saprotroph (Põlme et al. 2020); Decay sugar-rich substrates (Põlme et al. 2020) | Põlme S, et al. 2020. Fungal Diversity 105:1-16 ((https://doi.org/10.1007/s13225-020-00466-2)) |
| Otu35848 | 1 | k__Fungi(100);p__Ascomycota(100);c__Saccharomycetes(100);o__Saccharomycetales(100);f__Metschnikowiaceae(100);g__Metschnikowia(100);s__Metschnikowia_pulcherrima(100); | Metschnikowia | 13 | Saprotroph | \|Undefined Saprotroph\| | Yeast | NULL | Probable | As nectar-tap_saprotroph (Põlme et al. 2020); Decay sugar-rich substrates (Põlme et al. 2020) | Põlme S, et al. 2020. Fungal Diversity 105:1-16 ((https://doi.org/10.1007/s13225-020-00466-2)) |
| Otu35849 | 1 | k__Fungi(100);p__Ascomycota(100);c__Saccharomycetes(100);o__Saccharomycetales(100);f__Metschnikowiaceae(100);g__Metschnikowia(100);g__Metschnikowia_unclassified(100); | Metschnikowia | 13 | Saprotroph | \|Undefined Saprotroph\| | Yeast | NULL | Probable | As nectar-tap_saprotroph (Põlme et al. 2020); Decay sugar-rich substrates (Põlme et al. 2020) | Põlme S, et al. 2020. Fungal Diversity 105:1-16 ((https://doi.org/10.1007/s13225-020-00466-2)) |
| Otu35851 | 1 | k__Fungi(100);p__Ascomycota(100);c__Saccharomycetes(100);o__Saccharomycetales(100);f__Metschnikowiaceae(100);g__Metschnikowia(100);g__Metschnikowia_unclassified(100); | Metschnikowia | 13 | Saprotroph | \|Undefined Saprotroph\| | Yeast | NULL | Probable | As nectar-tap_saprotroph (Põlme et al. 2020); Decay sugar-rich substrates (Põlme et al. 2020) | Põlme S, et al. 2020. Fungal Diversity 105:1-16 ((https://doi.org/10.1007/s13225-020-00466-2)) |
| Otu35852 | 1 | k__Fungi(100);p__Ascomycota(100);c__Saccharomycetes(100);o__Saccharomycetales(100);f__Metschnikowiaceae(100);g__Metschnikowia(100);g__Metschnikowia_unclassified(100); | Metschnikowia | 13 | Saprotroph | \|Undefined Saprotroph\| | Yeast | NULL | Probable | As nectar-tap_saprotroph (Põlme et al. 2020); Decay sugar-rich substrates (Põlme et al. 2020) | Põlme S, et al. 2020. Fungal Diversity 105:1-16 ((https://doi.org/10.1007/s13225-020-00466-2)) |
| Otu35855 | 1 | k__Fungi(100);p__Ascomycota(100);c__Saccharomycetes(100);o__Saccharomycetales(100);f__Metschnikowiaceae(100);g__Metschnikowia(100);g__Metschnikowia_unclassified(100); | Metschnikowia | 13 | Saprotroph | \|Undefined Saprotroph\| | Yeast | NULL | Probable | As nectar-tap_saprotroph (Põlme et al. 2020); Decay sugar-rich substrates (Põlme et al. 2020) | Põlme S, et al. 2020. Fungal Diversity 105:1-16 ((https://doi.org/10.1007/s13225-020-00466-2)) |
| Otu35856 | 1 | k__Fungi(100);p__Ascomycota(100);c__Saccharomycetes(100);o__Saccharomycetales(100);f__Metschnikowiaceae(100);g__Metschnikowia(100);s__Metschnikowia_pulcherrima(100); | Metschnikowia | 13 | Saprotroph | \|Undefined Saprotroph\| | Yeast | NULL | Probable | As nectar-tap_saprotroph (Põlme et al. 2020); Decay sugar-rich substrates (Põlme et al. 2020) | Põlme S, et al. 2020. Fungal Diversity 105:1-16 ((https://doi.org/10.1007/s13225-020-00466-2)) |
| Otu35857 | 1 | k__Fungi(100);p__Ascomycota(100);c__Saccharomycetes(100);o__Saccharomycetales(100);f__Metschnikowiaceae(100);g__Metschnikowia(100);s__Metschnikowia_pulcherrima(100); | Metschnikowia | 13 | Saprotroph | \|Undefined Saprotroph\| | Yeast | NULL | Probable | As nectar-tap_saprotroph (Põlme et al. 2020); Decay sugar-rich substrates (Põlme et al. 2020) | Põlme S, et al. 2020. Fungal Diversity 105:1-16 ((https://doi.org/10.1007/s13225-020-00466-2)) |

| | | | | | | | | | | | |
|---|---|---|---|---|---|---|---|---|---|---|---|
| Otu35858 | 1 | k__Fungi(100);p__Ascomycota(100);c__Saccharomycetes(100);o__Saccharomycetales(100);f__Metschnikowiaceae(100);g__Metschnikowia(100);s__Metschnikowia_pulcherrima(100); | Metschnikowia | 13 | Saprotroph | \|Undefined Saprotroph\| | Yeast | NULL | Probable | As nectar-tap_saprotroph (Põlme et al. 2020); Decay sugar-rich substrates (Põlme et al. 2020) | Põlme S, et al. 2020. Fungal Diversity 105:1-16 ((https://doi.org/10.1007/s13225-020-00466-2)) |
| Otu35859 | 1 | k__Fungi(100);p__Ascomycota(100);c__Saccharomycetes(100);o__Saccharomycetales(100);f__Metschnikowiaceae(100);g__Metschnikowia(100);s__Metschnikowia_pulcherrima(100); | Metschnikowia | 13 | Saprotroph | \|Undefined Saprotroph\| | Yeast | NULL | Probable | As nectar-tap_saprotroph (Põlme et al. 2020); Decay sugar-rich substrates (Põlme et al. 2020) | Põlme S, et al. 2020. Fungal Diversity 105:1-16 ((https://doi.org/10.1007/s13225-020-00466-2)) |
| Otu35860 | 1 | k__Fungi(100);p__Ascomycota(100);c__Saccharomycetes(100);o__Saccharomycetales(100);f__Metschnikowiaceae(100);g__Metschnikowia(100);s__Metschnikowia_pulcherrima(100); | Metschnikowia | 13 | Saprotroph | \|Undefined Saprotroph\| | Yeast | NULL | Probable | As nectar-tap_saprotroph (Põlme et al. 2020); Decay sugar-rich substrates (Põlme et al. 2020) | Põlme S, et al. 2020. Fungal Diversity 105:1-16 ((https://doi.org/10.1007/s13225-020-00466-2)) |
| Otu35861 | 1 | k__Fungi(100);p__Ascomycota(100);c__Saccharomycetes(100);o__Saccharomycetales(100);f__Metschnikowiaceae(100);g__Metschnikowia(100);s__Metschnikowia_pulcherrima(100); | Metschnikowia | 13 | Saprotroph | \|Undefined Saprotroph\| | Yeast | NULL | Probable | As nectar-tap_saprotroph (Põlme et al. 2020); Decay sugar-rich substrates (Põlme et al. 2020) | Põlme S, et al. 2020. Fungal Diversity 105:1-16 ((https://doi.org/10.1007/s13225-020-00466-2)) |
| Otu35862 | 1 | k__Fungi(100);p__Ascomycota(100);c__Saccharomycetes(100);o__Saccharomycetales(100);f__Metschnikowiaceae(100);g__Metschnikowia(100);s__Metschnikowia_pulcherrima(100); | Metschnikowia | 13 | Saprotroph | \|Undefined Saprotroph\| | Yeast | NULL | Probable | As nectar-tap_saprotroph (Põlme et al. 2020); Decay sugar-rich substrates (Põlme et al. 2020) | Põlme S, et al. 2020. Fungal Diversity 105:1-16 ((https://doi.org/10.1007/s13225-020-00466-2)) |
| Otu35863 | 1 | k__Fungi(100);p__Ascomycota(100);c__Saccharomycetes(100);o__Saccharomycetales(100);f__Metschnikowiaceae(100);g__Metschnikowia(100);s__Metschnikowia_pulcherrima(100); | Metschnikowia | 13 | Saprotroph | \|Undefined Saprotroph\| | Yeast | NULL | Probable | As nectar-tap_saprotroph (Põlme et al. 2020); Decay sugar-rich substrates (Põlme et al. 2020) | Põlme S, et al. 2020. Fungal Diversity 105:1-16 ((https://doi.org/10.1007/s13225-020-00466-2)) |
| Otu35864 | 1 | k__Fungi(100);p__Ascomycota(100);c__Saccharomycetes(100);o__Saccharomycetales(100);f__Metschnikowiaceae(100);g__Metschnikowia(100);g__Metschnikowia_unclassified(100); | Metschnikowia | 13 | Saprotroph | \|Undefined Saprotroph\| | Yeast | NULL | Probable | As nectar-tap_saprotroph (Põlme et al. 2020); Decay sugar-rich substrates (Põlme et al. 2020) | Põlme S, et al. 2020. Fungal Diversity 105:1-16 ((https://doi.org/10.1007/s13225-020-00466-2)) |
| Otu35865 | 1 | k__Fungi(100);p__Ascomycota(100);c__Saccharomycetes(100);o__Saccharomycetales(100);f__Metschnikowiaceae(100);g__Metschnikowia(100);g__Metschnikowia_unclassified(100); | Metschnikowia | 13 | Saprotroph | \|Undefined Saprotroph\| | Yeast | NULL | Probable | As nectar-tap_saprotroph (Põlme et al. 2020); Decay sugar-rich substrates (Põlme et al. 2020) | Põlme S, et al. 2020. Fungal Diversity 105:1-16 ((https://doi.org/10.1007/s13225-020-00466-2)) |
| Otu35866 | 1 | k__Fungi(100);p__Ascomycota(100);c__Saccharomycetes(100);o__Saccharomycetales(100);f__Metschnikowiaceae(100);g__Metschnikowia(100);s__Metschnikowia_pulcherrima(100); | Metschnikowia | 13 | Saprotroph | \|Undefined Saprotroph\| | Yeast | NULL | Probable | As nectar-tap_saprotroph (Põlme et al. 2020); Decay sugar-rich substrates (Põlme et al. 2020) | Põlme S, et al. 2020. Fungal Diversity 105:1-16 ((https://doi.org/10.1007/s13225-020-00466-2)) |
| Otu35867 | 1 | k__Fungi(100);p__Ascomycota(100);c__Saccharomycetes(100);o__Saccharomycetales(100);f__Metschnikowiaceae(100);g__Metschnikowia(100);g__Metschnikowia_unclassified(100); | Metschnikowia | 13 | Saprotroph | \|Undefined Saprotroph\| | Yeast | NULL | Probable | As nectar-tap_saprotroph (Põlme et al. 2020); Decay sugar-rich substrates (Põlme et al. 2020) | Põlme S, et al. 2020. Fungal Diversity 105:1-16 ((https://doi.org/10.1007/s13225-020-00466-2)) |
| Otu35868 | 1 | k__Fungi(100);p__Ascomycota(100);c__Saccharomycetes(100);o__Saccharomycetales(100);f__Metschnikowiaceae(100);g__Metschnikowia(100);s__Metschnikowia_pulcherrima(100); | Metschnikowia | 13 | Saprotroph | \|Undefined Saprotroph\| | Yeast | NULL | Probable | As nectar-tap_saprotroph (Põlme et al. 2020); Decay sugar-rich substrates (Põlme et al. 2020) | Põlme S, et al. 2020. Fungal Diversity 105:1-16 ((https://doi.org/10.1007/s13225-020-00466-2)) |
| Otu35869 | 1 | k__Fungi(100);p__Ascomycota(100);c__Saccharomycetes(100);o__Saccharomycetales(100);f__Metschnikowiaceae(100);g__Metschnikowia(100);s__Metschnikowia_pulcherrima(100); | Metschnikowia | 13 | Saprotroph | \|Undefined Saprotroph\| | Yeast | NULL | Probable | As nectar-tap_saprotroph (Põlme et al. 2020); Decay sugar-rich substrates (Põlme et al. 2020) | Põlme S, et al. 2020. Fungal Diversity 105:1-16 ((https://doi.org/10.1007/s13225-020-00466-2)) |
| Otu35872 | 1 | k__Fungi(100);p__Ascomycota(100);c__Saccharomycetes(100);o__Saccharomycetales(100);f__Metschnikowiaceae(100);g__Metschnikowia(100);s__Metschnikowia_pulcherrima(100); | Metschnikowia | 13 | Saprotroph | \|Undefined Saprotroph\| | Yeast | NULL | Probable | As nectar-tap_saprotroph (Põlme et al. 2020); Decay sugar-rich substrates (Põlme et al. 2020) | Põlme S, et al. 2020. Fungal Diversity 105:1-16 ((https://doi.org/10.1007/s13225-020-00466-2)) |
| Otu35873 | 1 | k__Fungi(100);p__Ascomycota(100);c__Saccharomycetes(100);o__Saccharomycetales(100);f__Metschnikowiaceae(100);g__Metschnikowia(100);s__Metschnikowia_pulcherrima(100); | Metschnikowia | 13 | Saprotroph | \|Undefined Saprotroph\| | Yeast | NULL | Probable | As nectar-tap_saprotroph (Põlme et al. 2020); Decay sugar-rich substrates (Põlme et al. 2020) | Põlme S, et al. 2020. Fungal Diversity 105:1-16 ((https://doi.org/10.1007/s13225-020-00466-2)) |
| Otu35874 | 1 | k__Fungi(100);p__Ascomycota(100);c__Saccharomycetes(100);o__Saccharomycetales(100);f__Metschnikowiaceae(100);g__Metschnikowia(100);s__Metschnikowia_pulcherrima(100); | Metschnikowia | 13 | Saprotroph | \|Undefined Saprotroph\| | Yeast | NULL | Probable | As nectar-tap_saprotroph (Põlme et al. 2020); Decay sugar-rich substrates (Põlme et al. 2020) | Põlme S, et al. 2020. Fungal Diversity 105:1-16 ((https://doi.org/10.1007/s13225-020-00466-2)) |
| Otu35878 | 1 | k__Fungi(100);p__Ascomycota(100);c__Saccharomycetes(100);o__Saccharomycetales(100);f__Metschnikowiaceae(100);g__Metschnikowia(100);s__Metschnikowia_pulcherrima(100); | Metschnikowia | 13 | Saprotroph | \|Undefined Saprotroph\| | Yeast | NULL | Probable | As nectar-tap_saprotroph (Põlme et al. 2020); Decay sugar-rich substrates (Põlme et al. 2020) | Põlme S, et al. 2020. Fungal Diversity 105:1-16 ((https://doi.org/10.1007/s13225-020-00466-2)) |
| Otu35879 | 1 | k__Fungi(100);p__Ascomycota(100);c__Saccharomycetes(100);o__Saccharomycetales(100);f__Metschnikowiaceae(100);g__Metschnikowia(100);s__Metschnikowia_pulcherrima(100); | Metschnikowia | 13 | Saprotroph | \|Undefined Saprotroph\| | Yeast | NULL | Probable | As nectar-tap_saprotroph (Põlme et al. 2020); Decay sugar-rich substrates (Põlme et al. 2020) | Põlme S, et al. 2020. Fungal Diversity 105:1-16 ((https://doi.org/10.1007/s13225-020-00466-2)) |
| Otu35881 | 1 | k__Fungi(100);p__Ascomycota(100);c__Saccharomycetes(100);o__Saccharomycetales(100);f__Metschnikowiaceae(100);g__Metschnikowia(100);g__Metschnikowia_unclassified(100); | Metschnikowia | 13 | Saprotroph | \|Undefined Saprotroph\| | Yeast | NULL | Probable | As nectar-tap_saprotroph (Põlme et al. 2020); Decay sugar-rich substrates (Põlme et al. 2020) | Põlme S, et al. 2020. Fungal Diversity 105:1-16 ((https://doi.org/10.1007/s13225-020-00466-2)) |
| Otu35882 | 1 | k__Fungi(100);p__Ascomycota(100);c__Saccharomycetes(100);o__Saccharomycetales(100);f__Metschnikowiaceae(100);g__Metschnikowia(100);s__Metschnikowia_pulcherrima(100); | Metschnikowia | 13 | Saprotroph | \|Undefined Saprotroph\| | Yeast | NULL | Probable | As nectar-tap_saprotroph (Põlme et al. 2020); Decay sugar-rich substrates (Põlme et al. 2020) | Põlme S, et al. 2020. Fungal Diversity 105:1-16 ((https://doi.org/10.1007/s13225-020-00466-2)) |
| Otu35883 | 1 | k__Fungi(100);p__Ascomycota(100);c__Saccharomycetes(100);o__Saccharomycetales(100);f__Metschnikowiaceae(100);g__Metschnikowia(100);s__Metschnikowia_pulcherrima(100); | Metschnikowia | 13 | Saprotroph | \|Undefined Saprotroph\| | Yeast | NULL | Probable | As nectar-tap_saprotroph (Põlme et al. 2020); Decay sugar-rich substrates (Põlme et al. 2020) | Põlme S, et al. 2020. Fungal Diversity 105:1-16 ((https://doi.org/10.1007/s13225-020-00466-2)) |
| Otu35884 | 1 | k__Fungi(100);p__Ascomycota(100);c__Saccharomycetes(100);o__Saccharomycetales(100);f__Metschnikowiaceae(100);g__Metschnikowia(100);s__Metschnikowia_pulcherrima(100); | Metschnikowia | 13 | Saprotroph | \|Undefined Saprotroph\| | Yeast | NULL | Probable | As nectar-tap_saprotroph (Põlme et al. 2020); Decay sugar-rich substrates (Põlme et al. 2020) | Põlme S, et al. 2020. Fungal Diversity 105:1-16 ((https://doi.org/10.1007/s13225-020-00466-2)) |
| Otu35885 | 1 | k__Fungi(100);p__Ascomycota(100);c__Saccharomycetes(100);o__Saccharomycetales(100);f__Metschnikowiaceae(100);g__Metschnikowia(100);g__Metschnikowia_unclassified(100); | Metschnikowia | 13 | Saprotroph | \|Undefined Saprotroph\| | Yeast | NULL | Probable | As nectar-tap_saprotroph (Põlme et al. 2020); Decay sugar-rich substrates (Põlme et al. 2020) | Põlme S, et al. 2020. Fungal Diversity 105:1-16 ((https://doi.org/10.1007/s13225-020-00466-2)) |
| Otu35887 | 1 | k__Fungi(100);p__Ascomycota(100);c__Saccharomycetes(100);o__Saccharomycetales(100);f__Metschnikowiaceae(100);g__Metschnikowia(100);s__Metschnikowia_pulcherrima(100); | Metschnikowia | 13 | Saprotroph | \|Undefined Saprotroph\| | Yeast | NULL | Probable | As nectar-tap_saprotroph (Põlme et al. 2020); Decay sugar-rich substrates (Põlme et al. 2020) | Põlme S, et al. 2020. Fungal Diversity 105:1-16 ((https://doi.org/10.1007/s13225-020-00466-2)) |
| Otu35889 | 1 | k__Fungi(100);p__Ascomycota(100);c__Saccharomycetes(100);o__Saccharomycetales(100);f__Metschnikowiaceae(100);g__Metschnikowia(100);s__Metschnikowia_pulcherrima(100); | Metschnikowia | 13 | Saprotroph | \|Undefined Saprotroph\| | Yeast | NULL | Probable | As nectar-tap_saprotroph (Põlme et al. 2020); Decay sugar-rich substrates (Põlme et al. 2020) | Põlme S, et al. 2020. Fungal Diversity 105:1-16 ((https://doi.org/10.1007/s13225-020-00466-2)) |
| Otu35897 | 1 | k__Fungi(100);p__Ascomycota(100);c__Saccharomycetes(100);o__Saccharomycetales(100);f__Metschnikowiaceae(100);g__Metschnikowia(100);s__Metschnikowia_pulcherrima(100); | Metschnikowia | 13 | Saprotroph | \|Undefined Saprotroph\| | Yeast | NULL | Probable | As nectar-tap_saprotroph (Põlme et al. 2020); Decay sugar-rich substrates (Põlme et al. 2020) | Põlme S, et al. 2020. Fungal Diversity 105:1-16 ((https://doi.org/10.1007/s13225-020-00466-2)) |
| Otu35898 | 1 | k__Fungi(100);p__Ascomycota(100);c__Saccharomycetes(100);o__Saccharomycetales(100);f__Metschnikowiaceae(100);g__Metschnikowia(100);g__Metschnikowia_unclassified(100); | Metschnikowia | 13 | Saprotroph | \|Undefined Saprotroph\| | Yeast | NULL | Probable | As nectar-tap_saprotroph (Põlme et al. 2020); Decay sugar-rich substrates (Põlme et al. 2020) | Põlme S, et al. 2020. Fungal Diversity 105:1-16 ((https://doi.org/10.1007/s13225-020-00466-2)) |
| Otu35901 | 1 | k__Fungi(100);p__Ascomycota(100);c__Saccharomycetes(100);o__Saccharomycetales(100);f__Metschnikowiaceae(100);g__Metschnikowia(100);g__Metschnikowia_unclassified(100); | Metschnikowia | 13 | Saprotroph | \|Undefined Saprotroph\| | Yeast | NULL | Probable | As nectar-tap_saprotroph (Põlme et al. 2020); Decay sugar-rich substrates (Põlme et al. 2020) | Põlme S, et al. 2020. Fungal Diversity 105:1-16 ((https://doi.org/10.1007/s13225-020-00466-2)) |
| Otu35902 | 1 | k__Fungi(100);p__Ascomycota(100);c__Saccharomycetes(100);o__Saccharomycetales(100);f__Metschnikowiaceae(100);g__Metschnikowia(100);s__Metschnikowia_pulcherrima(100); | Metschnikowia | 13 | Saprotroph | \|Undefined Saprotroph\| | Yeast | NULL | Probable | As nectar-tap_saprotroph (Põlme et al. 2020); Decay sugar-rich substrates (Põlme et al. 2020) | Põlme S, et al. 2020. Fungal Diversity 105:1-16 ((https://doi.org/10.1007/s13225-020-00466-2)) |
| Otu35903 | 1 | k__Fungi(100);p__Ascomycota(100);c__Saccharomycetes(100);o__Saccharomycetales(100);f__Metschnikowiaceae(100);g__Metschnikowia(100);s__Metschnikowia_pulcherrima(100); | Metschnikowia | 13 | Saprotroph | \|Undefined Saprotroph\| | Yeast | NULL | Probable | As nectar-tap_saprotroph (Põlme et al. 2020); Decay sugar-rich substrates (Põlme et al. 2020) | Põlme S, et al. 2020. Fungal Diversity 105:1-16 ((https://doi.org/10.1007/s13225-020-00466-2)) |
| Otu35906 | 1 | k__Fungi(100);p__Ascomycota(100);c__Saccharomycetes(100);o__Saccharomycetales(100);f__Metschnikowiaceae(100);g__Metschnikowia(100);s__Metschnikowia_pulcherrima(100); | Metschnikowia | 13 | Saprotroph | \|Undefined Saprotroph\| | Yeast | NULL | Probable | As nectar-tap_saprotroph (Põlme et al. 2020); Decay sugar-rich substrates (Põlme et al. 2020) | Põlme S, et al. 2020. Fungal Diversity 105:1-16 ((https://doi.org/10.1007/s13225-020-00466-2)) |
| Otu35907 | 1 | k__Fungi(100);p__Ascomycota(100);c__Saccharomycetes(100);o__Saccharomycetales(100);f__Metschnikowiaceae(100);g__Metschnikowia(100);s__Metschnikowia_pulcherrima(100); | Metschnikowia | 13 | Saprotroph | \|Undefined Saprotroph\| | Yeast | NULL | Probable | As nectar-tap_saprotroph (Põlme et al. 2020); Decay sugar-rich substrates (Põlme et al. 2020) | Põlme S, et al. 2020. Fungal Diversity 105:1-16 ((https://doi.org/10.1007/s13225-020-00466-2)) |
| Otu35908 | 1 | k__Fungi(100);p__Ascomycota(100);c__Saccharomycetes(100);o__Saccharomycetales(100);f__Metschnikowiaceae(100);g__Metschnikowia(100);s__Metschnikowia_pulcherrima(100); | Metschnikowia | 13 | Saprotroph | \|Undefined Saprotroph\| | Yeast | NULL | Probable | As nectar-tap_saprotroph (Põlme et al. 2020); Decay sugar-rich substrates (Põlme et al. 2020) | Põlme S, et al. 2020. Fungal Diversity 105:1-16 ((https://doi.org/10.1007/s13225-020-00466-2)) |
| Otu35910 | 1 | k__Fungi(100);p__Ascomycota(100);c__Saccharomycetes(100);o__Saccharomycetales(100);f__Metschnikowiaceae(100);g__Metschnikowia(100);g__Metschnikowia_unclassified(100); | Metschnikowia | 13 | Saprotroph | \|Undefined Saprotroph\| | Yeast | NULL | Probable | As nectar-tap_saprotroph (Põlme et al. 2020); Decay sugar-rich substrates (Põlme et al. 2020) | Põlme S, et al. 2020. Fungal Diversity 105:1-16 ((https://doi.org/10.1007/s13225-020-00466-2)) |
| Otu35911 | 1 | k__Fungi(100);p__Ascomycota(100);c__Saccharomycetes(100);o__Saccharomycetales(100);f__Metschnikowiaceae(100);g__Metschnikowia(100);s__Metschnikowia_pulcherrima(100); | Metschnikowia | 13 | Saprotroph | \|Undefined Saprotroph\| | Yeast | NULL | Probable | As nectar-tap_saprotroph (Põlme et al. 2020); Decay sugar-rich substrates (Põlme et al. 2020) | Põlme S, et al. 2020. Fungal Diversity 105:1-16 ((https://doi.org/10.1007/s13225-020-00466-2)) |
| Otu35912 | 1 | k__Fungi(100);p__Ascomycota(100);c__Saccharomycetes(100);o__Saccharomycetales(100);f__Metschnikowiaceae(100);g__Metschnikowia(100);s__Metschnikowia_pulcherrima(100); | Metschnikowia | 13 | Saprotroph | \|Undefined Saprotroph\| | Yeast | NULL | Probable | As nectar-tap_saprotroph (Põlme et al. 2020); Decay sugar-rich substrates (Põlme et al. 2020) | Põlme S, et al. 2020. Fungal Diversity 105:1-16 ((https://doi.org/10.1007/s13225-020-00466-2)) |
| Otu35913 | 1 | k__Fungi(100);p__Ascomycota(100);c__Saccharomycetes(100);o__Saccharomycetales(100);f__Metschnikowiaceae(100);g__Metschnikowia(100);g__Metschnikowia_unclassified(100); | Metschnikowia | 13 | Saprotroph | \|Undefined Saprotroph\| | Yeast | NULL | Probable | As nectar-tap_saprotroph (Põlme et al. 2020); Decay sugar-rich substrates (Põlme et al. 2020) | Põlme S, et al. 2020. Fungal Diversity 105:1-16 ((https://doi.org/10.1007/s13225-020-00466-2)) |
| Otu35915 | 1 | k__Fungi(100);p__Ascomycota(100);c__Saccharomycetes(100);o__Saccharomycetales(100);f__Metschnikowiaceae(100);g__Metschnikowia(100);g__Metschnikowia_unclassified(100); | Metschnikowia | 13 | Saprotroph | \|Undefined Saprotroph\| | Yeast | NULL | Probable | As nectar-tap_saprotroph (Põlme et al. 2020); Decay sugar-rich substrates (Põlme et al. 2020) | Põlme S, et al. 2020. Fungal Diversity 105:1-16 ((https://doi.org/10.1007/s13225-020-00466-2)) |
| Otu35916 | 1 | k__Fungi(100);p__Ascomycota(100);c__Saccharomycetes(100);o__Saccharomycetales(100);f__Metschnikowiaceae(100);g__Metschnikowia(100);g__Metschnikowia_unclassified(100); | Metschnikowia | 13 | Saprotroph | \|Undefined Saprotroph\| | Yeast | NULL | Probable | As nectar-tap_saprotroph (Põlme et al. 2020); Decay sugar-rich substrates (Põlme et al. 2020) | Põlme S, et al. 2020. Fungal Diversity 105:1-16 ((https://doi.org/10.1007/s13225-020-00466-2)) |
| Otu35917 | 1 | k__Fungi(100);p__Ascomycota(100);c__Saccharomycetes(100);o__Saccharomycetales(100);f__Metschnikowiaceae(100);g__Metschnikowia(100);s__Metschnikowia_pulcherrima(100); | Metschnikowia | 13 | Saprotroph | \|Undefined Saprotroph\| | Yeast | NULL | Probable | As nectar-tap_saprotroph (Põlme et al. 2020); Decay sugar-rich substrates (Põlme et al. 2020) | Põlme S, et al. 2020. Fungal Diversity 105:1-16 ((https://doi.org/10.1007/s13225-020-00466-2)) |
| Otu35918 | 1 | k__Fungi(100);p__Ascomycota(100);c__Saccharomycetes(100);o__Saccharomycetales(100);f__Metschnikowiaceae(100);g__Metschnikowia(100);s__Metschnikowia_pulcherrima(100); | Metschnikowia | 13 | Saprotroph | \|Undefined Saprotroph\| | Yeast | NULL | Probable | As nectar-tap_saprotroph (Põlme et al. 2020); Decay sugar-rich substrates (Põlme et al. 2020) | Põlme S, et al. 2020. Fungal Diversity 105:1-16 ((https://doi.org/10.1007/s13225-020-00466-2)) |
| Otu35919 | 1 | k__Fungi(100);p__Ascomycota(100);c__Saccharomycetes(100);o__Saccharomycetales(100);f__Metschnikowiaceae(100);g__Metschnikowia(100);g__Metschnikowia_unclassified(100); | Metschnikowia | 13 | Saprotroph | \|Undefined Saprotroph\| | Yeast | NULL | Probable | As nectar-tap_saprotroph (Põlme et al. 2020); Decay sugar-rich substrates (Põlme et al. 2020) | Põlme S, et al. 2020. Fungal Diversity 105:1-16 ((https://doi.org/10.1007/s13225-020-00466-2)) |
| Otu35920 | 1 | k__Fungi(100);p__Ascomycota(100);c__Saccharomycetes(100);o__Saccharomycetales(100);f__Metschnikowiaceae(100);g__Metschnikowia(100);s__Metschnikowia_pulcherrima(100); | Metschnikowia | 13 | Saprotroph | \|Undefined Saprotroph\| | Yeast | NULL | Probable | As nectar-tap_saprotroph (Põlme et al. 2020); Decay sugar-rich substrates (Põlme et al. 2020) | Põlme S, et al. 2020. Fungal Diversity 105:1-16 ((https://doi.org/10.1007/s13225-020-00466-2)) |
| Otu35921 | 1 | k__Fungi(100);p__Ascomycota(100);c__Saccharomycetes(100);o__Saccharomycetales(100);f__Saccharomycetales_family_Incertae_sedis(100);g__Candida(100);g__Candida_unclassified(100); | Candida | 13 | Pathotroph-Saprotroph-Symbiotroph | Animal Pathogen-Endophyte-Endosymbiont-Epiphyte-\|Undefined Saprotroph\| | Dimorphic Yeast-Yeast | NULL | Probable | As nectar-tap_saprotroph (Põlme et al. 2020); Decay sugar-rich substrates (Põlme et al. 2020); This asexual genus is polyphyletic and contain species that function very differently depend on what they are. Caution should be taken when interpreting guilds for Candida species. | Manolakaki D, et al. 2010. Virulence. 1: 367-375 ((https://doi.org/10.4161/viru.1.5.12796)); Mekbib SB, et al. 2011. Tropical Plant Pathology 36:233-240 ((https://doi.org/10.1590/S1982-56762011000400004)); Põlme S, et al. 2020. Fungal Diversity 105:1-16 ((https://doi.org/10.1007/s13225-020-00466-2)) |
| Otu35922 | 1 | k__Fungi(100);p__Ascomycota(100);c__Saccharomycetes(100);o__Saccharomycetales(100);f__Metschnikowiaceae(100);g__Metschnikowia(100);s__Metschnikowia_pulcherrima(100); | Metschnikowia | 13 | Saprotroph | \|Undefined Saprotroph\| | Yeast | NULL | Probable | As nectar-tap_saprotroph (Põlme et al. 2020); Decay sugar-rich substrates (Põlme et al. 2020) | Põlme S, et al. 2020. Fungal Diversity 105:1-16 ((https://doi.org/10.1007/s13225-020-00466-2)) |
| Otu35923 | 1 | k__Fungi(100);p__Ascomycota(100);c__Saccharomycetes(100);o__Saccharomycetales(100);f__Metschnikowiaceae(100);g__Metschnikowia(100);s__Metschnikowia_pulcherrima(100); | Metschnikowia | 13 | Saprotroph | \|Undefined Saprotroph\| | Yeast | NULL | Probable | As nectar-tap_saprotroph (Põlme et al. 2020); Decay sugar-rich substrates (Põlme et al. 2020) | Põlme S, et al. 2020. Fungal Diversity 105:1-16 ((https://doi.org/10.1007/s13225-020-00466-2)) |
| Otu35925 | 1 | k__Fungi(100);p__Ascomycota(100);c__Saccharomycetes(100);o__Saccharomycetales(100);f__Metschnikowiaceae(100);g__Metschnikowia(100);s__Metschnikowia_pulcherrima(100); | Metschnikowia | 13 | Saprotroph | \|Undefined Saprotroph\| | Yeast | NULL | Probable | As nectar-tap_saprotroph (Põlme et al. 2020); Decay sugar-rich substrates (Põlme et al. 2020) | Põlme S, et al. 2020. Fungal Diversity 105:1-16 ((https://doi.org/10.1007/s13225-020-00466-2)) |
| Otu35926 | 1 | k__Fungi(100);p__Ascomycota(100);c__Saccharomycetes(100);o__Saccharomycetales(100);f__Metschnikowiaceae(100);g__Metschnikowia(100);s__Metschnikowia_pulcherrima(100); | Metschnikowia | 13 | Saprotroph | \|Undefined Saprotroph\| | Yeast | NULL | Probable | As nectar-tap_saprotroph (Põlme et al. 2020); Decay sugar-rich substrates (Põlme et al. 2020) | Põlme S, et al. 2020. Fungal Diversity 105:1-16 ((https://doi.org/10.1007/s13225-020-00466-2)) |
| Otu35927 | 1 | k__Fungi(100);p__Ascomycota(100);c__Saccharomycetes(100);o__Saccharomycetales(100);f__Metschnikowiaceae(100);g__Metschnikowia(100);s__Metschnikowia_pulcherrima(100); | Metschnikowia | 13 | Saprotroph | \|Undefined Saprotroph\| | Yeast | NULL | Probable | As nectar-tap_saprotroph (Põlme et al. 2020); Decay sugar-rich substrates (Põlme et al. 2020) | Põlme S, et al. 2020. Fungal Diversity 105:1-16 ((https://doi.org/10.1007/s13225-020-00466-2)) |
| Otu35932 | 1 | k__Fungi(100);p__Ascomycota(100);c__Saccharomycetes(100);o__Saccharomycetales(100);f__Metschnikowiaceae(100);g__Metschnikowia(100);s__Metschnikowia_pulcherrima(100); | Metschnikowia | 13 | Saprotroph | \|Undefined Saprotroph\| | Yeast | NULL | Probable | As nectar-tap_saprotroph (Põlme et al. 2020); Decay sugar-rich substrates (Põlme et al. 2020) | Põlme S, et al. 2020. Fungal Diversity 105:1-16 ((https://doi.org/10.1007/s13225-020-00466-2)) |
| Otu35933 | 1 | k__Fungi(100);p__Ascomycota(100);c__Saccharomycetes(100);o__Saccharomycetales(100);f__Metschnikowiaceae(100);g__Metschnikowia(100);s__Metschnikowia_pulcherrima(100); | Metschnikowia | 13 | Saprotroph | \|Undefined Saprotroph\| | Yeast | NULL | Probable | As nectar-tap_saprotroph (Põlme et al. 2020); Decay sugar-rich substrates (Põlme et al. 2020) | Põlme S, et al. 2020. Fungal Diversity 105:1-16 ((https://doi.org/10.1007/s13225-020-00466-2)) |
| Otu35935 | 1 | k__Fungi(100);p__Ascomycota(100);c__Saccharomycetes(100);o__Saccharomycetales(100);f__Metschnikowiaceae(100);g__Metschnikowia(100);s__Metschnikowia_pulcherrima(100); | Metschnikowia | 13 | Saprotroph | \|Undefined Saprotroph\| | Yeast | NULL | Probable | As nectar-tap_saprotroph (Põlme et al. 2020); Decay sugar-rich substrates (Põlme et al. 2020) | Põlme S, et al. 2020. Fungal Diversity 105:1-16 ((https://doi.org/10.1007/s13225-020-00466-2)) |
| Otu35936 | 1 | k__Fungi(100);p__Ascomycota(100);c__Saccharomycetes(100);o__Saccharomycetales(100);f__Metschnikowiaceae(100);g__Metschnikowia(100);s__Metschnikowia_pulcherrima(100); | Metschnikowia | 13 | Saprotroph | \|Undefined Saprotroph\| | Yeast | NULL | Probable | As nectar-tap_saprotroph (Põlme et al. 2020); Decay sugar-rich substrates (Põlme et al. 2020) | Põlme S, et al. 2020. Fungal Diversity 105:1-16 ((https://doi.org/10.1007/s13225-020-00466-2)) |
| Otu35938 | 1 | k__Fungi(100);p__Ascomycota(100);c__Saccharomycetes(100);o__Saccharomycetales(100);f__Metschnikowiaceae(100);g__Metschnikowia(100);s__Metschnikowia_pulcherrima(100); | Metschnikowia | 13 | Saprotroph | \|Undefined Saprotroph\| | Yeast | NULL | Probable | As nectar-tap_saprotroph (Põlme et al. 2020); Decay sugar-rich substrates (Põlme et al. 2020) | Põlme S, et al. 2020. Fungal Diversity 105:1-16 ((https://doi.org/10.1007/s13225-020-00466-2)) |
| Otu35943 | 1 | k__Fungi(100);p__Ascomycota(100);c__Saccharomycetes(100);o__Saccharomycetales(100);f__Metschnikowiaceae(100);g__Metschnikowia(100);s__Metschnikowia_pulcherrima(100); | Metschnikowia | 13 | Saprotroph | \|Undefined Saprotroph\| | Yeast | NULL | Probable | As nectar-tap_saprotroph (Põlme et al. 2020); Decay sugar-rich substrates (Põlme et al. 2020) | Põlme S, et al. 2020. Fungal Diversity 105:1-16 ((https://doi.org/10.1007/s13225-020-00466-2)) |
| Otu35947 | 1 | k__Fungi(100);p__Ascomycota(100);c__Saccharomycetes(100);o__Saccharomycetales(100);f__Metschnikowiaceae(100);g__Metschnikowia(100);s__Metschnikowia_pulcherrima(100); | Metschnikowia | 13 | Saprotroph | \|Undefined Saprotroph\| | Yeast | NULL | Probable | As nectar-tap_saprotroph (Põlme et al. 2020); Decay sugar-rich substrates (Põlme et al. 2020) | Põlme S, et al. 2020. Fungal Diversity 105:1-16 ((https://doi.org/10.1007/s13225-020-00466-2)) |
| Otu35948 | 1 | k__Fungi(100);p__Ascomycota(100);c__Saccharomycetes(100);o__Saccharomycetales(100);f__Metschnikowiaceae(100);g__Metschnikowia(100);s__Metschnikowia_pulcherrima(100); | Metschnikowia | 13 | Saprotroph | \|Undefined Saprotroph\| | Yeast | NULL | Probable | As nectar-tap_saprotroph (Põlme et al. 2020); Decay sugar-rich substrates (Põlme et al. 2020) | Põlme S, et al. 2020. Fungal Diversity 105:1-16 ((https://doi.org/10.1007/s13225-020-00466-2)) |
| Otu35949 | 1 | k__Fungi(100);p__Ascomycota(100);c__Saccharomycetes(100);o__Saccharomycetales(100);f__Metschnikowiaceae(100);g__Metschnikowia(100);s__Metschnikowia_pulcherrima(100); | Metschnikowia | 13 | Saprotroph | \|Undefined Saprotroph\| | Yeast | NULL | Probable | As nectar-tap_saprotroph (Põlme et al. 2020); Decay sugar-rich substrates (Põlme et al. 2020) | Põlme S, et al. 2020. Fungal Diversity 105:1-16 ((https://doi.org/10.1007/s13225-020-00466-2)) |
| Otu35950 | 1 | k__Fungi(100);p__Ascomycota(100);c__Saccharomycetes(100);o__Saccharomycetales(100);f__Metschnikowiaceae(100);g__Metschnikowia(100);s__Metschnikowia_pulcherrima(100); | Metschnikowia | 13 | Saprotroph | \|Undefined Saprotroph\| | Yeast | NULL | Probable | As nectar-tap_saprotroph (Põlme et al. 2020); Decay sugar-rich substrates (Põlme et al. 2020) | Põlme S, et al. 2020. Fungal Diversity 105:1-16 ((https://doi.org/10.1007/s13225-020-00466-2)) |
| Otu35951 | 1 | k__Fungi(100);p__Ascomycota(100);c__Saccharomycetes(100);o__Saccharomycetales(100);f__Metschnikowiaceae(100);g__Metschnikowia(100);s__Metschnikowia_pulcherrima(100); | Metschnikowia | 13 | Saprotroph | \|Undefined Saprotroph\| | Yeast | NULL | Probable | As nectar-tap_saprotroph (Põlme et al. 2020); Decay sugar-rich substrates (Põlme et al. 2020) | Põlme S, et al. 2020. Fungal Diversity 105:1-16 ((https://doi.org/10.1007/s13225-020-00466-2)) |
| Otu35952 | 1 | k__Fungi(100);p__Ascomycota(100);c__Saccharomycetes(100);o__Saccharomycetales(100);f__Metschnikowiaceae(100);g__Metschnikowia(100);s__Metschnikowia_pulcherrima(100); | Metschnikowia | 13 | Saprotroph | \|Undefined Saprotroph\| | Yeast | NULL | Probable | As nectar-tap_saprotroph (Põlme et al. 2020); Decay sugar-rich substrates (Põlme et al. 2020) | Põlme S, et al. 2020. Fungal Diversity 105:1-16 ((https://doi.org/10.1007/s13225-020-00466-2)) |
| Otu35953 | 1 | k__Fungi(100);p__Ascomycota(100);c__Saccharomycetes(100);o__Saccharomycetales(100);f__Metschnikowiaceae(100);g__Metschnikowia(100);s__Metschnikowia_pulcherrima(100); | Metschnikowia | 13 | Saprotroph | \|Undefined Saprotroph\| | Yeast | NULL | Probable | As nectar-tap_saprotroph (Põlme et al. 2020); Decay sugar-rich substrates (Põlme et al. 2020) | Põlme S, et al. 2020. Fungal Diversity 105:1-16 ((https://doi.org/10.1007/s13225-020-00466-2)) |
| Otu35956 | 1 | k__Fungi(100);p__Ascomycota(100);c__Saccharomycetes(100);o__Saccharomycetales(100);f__Metschnikowiaceae(100);g__Metschnikowia(100);s__Metschnikowia_pulcherrima(100); | Metschnikowia | 13 | Saprotroph | \|Undefined Saprotroph\| | Yeast | NULL | Probable | As nectar-tap_saprotroph (Põlme et al. 2020); Decay sugar-rich substrates (Põlme et al. 2020) | Põlme S, et al. 2020. Fungal Diversity 105:1-16 ((https://doi.org/10.1007/s13225-020-00466-2)) |
| Otu35959 | 1 | k__Fungi(100);p__Ascomycota(100);c__Saccharomycetes(100);o__Saccharomycetales(100);f__Metschnikowiaceae(100);g__Metschnikowia(100);s__Metschnikowia_pulcherrima(100); | Metschnikowia | 13 | Saprotroph | \|Undefined Saprotroph\| | Yeast | NULL | Probable | As nectar-tap_saprotroph (Põlme et al. 2020); Decay sugar-rich substrates (Põlme et al. 2020) | Põlme S, et al. 2020. Fungal Diversity 105:1-16 ((https://doi.org/10.1007/s13225-020-00466-2)) |
| Otu35962 | 1 | k__Fungi(100);p__Ascomycota(100);c__Saccharomycetes(100);o__Saccharomycetales(100);f__Metschnikowiaceae(100);g__Metschnikowia(100);g__Metschnikowia_unclassified(100); | Metschnikowia | 13 | Saprotroph | \|Undefined Saprotroph\| | Yeast | NULL | Probable | As nectar-tap_saprotroph (Põlme et al. 2020); Decay sugar-rich substrates (Põlme et al. 2020) | Põlme S, et al. 2020. Fungal Diversity 105:1-16 ((https://doi.org/10.1007/s13225-020-00466-2)) |
| Otu35963 | 1 | k__Fungi(100);p__Ascomycota(100);c__Saccharomycetes(100);o__Saccharomycetales(100);f__Metschnikowiaceae(100);g__Metschnikowia(100);s__Metschnikowia_pulcherrima(100); | Metschnikowia | 13 | Saprotroph | \|Undefined Saprotroph\| | Yeast | NULL | Probable | As nectar-tap_saprotroph (Põlme et al. 2020); Decay sugar-rich substrates (Põlme et al. 2020) | Põlme S, et al. 2020. Fungal Diversity 105:1-16 ((https://doi.org/10.1007/s13225-020-00466-2)) |
| Otu35964 | 1 | k__Fungi(100);p__Ascomycota(100);c__Saccharomycetes(100);o__Saccharomycetales(100);f__Metschnikowiaceae(100);g__Metschnikowia(100);s__Metschnikowia_pulcherrima(100); | Metschnikowia | 13 | Saprotroph | \|Undefined Saprotroph\| | Yeast | NULL | Probable | As nectar-tap_saprotroph (Põlme et al. 2020); Decay sugar-rich substrates (Põlme et al. 2020) | Põlme S, et al. 2020. Fungal Diversity 105:1-16 ((https://doi.org/10.1007/s13225-020-00466-2)) |
| Otu35965 | 1 | k__Fungi(100);p__Ascomycota(100);c__Saccharomycetes(100);o__Saccharomycetales(100);f__Metschnikowiaceae(100);g__Metschnikowia(100);s__Metschnikowia_pulcherrima(100); | Metschnikowia | 13 | Saprotroph | \|Undefined Saprotroph\| | Yeast | NULL | Probable | As nectar-tap_saprotroph (Põlme et al. 2020); Decay sugar-rich substrates (Põlme et al. 2020) | Põlme S, et al. 2020. Fungal Diversity 105:1-16 ((https://doi.org/10.1007/s13225-020-00466-2)) |
| Otu35968 | 1 | k__Fungi(100);p__Ascomycota(100);c__Saccharomycetes(100);o__Saccharomycetales(100);f__Metschnikowiaceae(100);g__Metschnikowia(100);g__Metschnikowia_unclassified(100); | Metschnikowia | 13 | Saprotroph | \|Undefined Saprotroph\| | Yeast | NULL | Probable | As nectar-tap_saprotroph (Põlme et al. 2020); Decay sugar-rich substrates (Põlme et al. 2020) | Põlme S, et al. 2020. Fungal Diversity 105:1-16 ((https://doi.org/10.1007/s13225-020-00466-2)) |
| Otu35969 | 1 | k__Fungi(100);p__Ascomycota(100);c__Saccharomycetes(100);o__Saccharomycetales(100);f__Metschnikowiaceae(100);g__Metschnikowia(100);g__Metschnikowia_unclassified(100); | Metschnikowia | 13 | Saprotroph | \|Undefined Saprotroph\| | Yeast | NULL | Probable | As nectar-tap_saprotroph (Põlme et al. 2020); Decay sugar-rich substrates (Põlme et al. 2020) | Põlme S, et al. 2020. Fungal Diversity 105:1-16 ((https://doi.org/10.1007/s13225-020-00466-2)) |
| Otu35970 | 1 | k__Fungi(100);p__Ascomycota(100);c__Saccharomycetes(100);o__Saccharomycetales(100);f__Metschnikowiaceae(100);g__Metschnikowia(100);s__Metschnikowia_pulcherrima(100); | Metschnikowia | 13 | Saprotroph | \|Undefined Saprotroph\| | Yeast | NULL | Probable | As nectar-tap_saprotroph (Põlme et al. 2020); Decay sugar-rich substrates (Põlme et al. 2020) | Põlme S, et al. 2020. Fungal Diversity 105:1-16 ((https://doi.org/10.1007/s13225-020-00466-2)) |
| Otu35971 | 1 | k__Fungi(100);p__Ascomycota(100);c__Saccharomycetes(100);o__Saccharomycetales(100);f__Metschnikowiaceae(100);g__Metschnikowia(100);s__Metschnikowia_pulcherrima(100); | Metschnikowia | 13 | Saprotroph | \|Undefined Saprotroph\| | Yeast | NULL | Probable | As nectar-tap_saprotroph (Põlme et al. 2020); Decay sugar-rich substrates (Põlme et al. 2020) | Põlme S, et al. 2020. Fungal Diversity 105:1-16 ((https://doi.org/10.1007/s13225-020-00466-2)) |
| Otu35972 | 1 | k__Fungi(100);p__Ascomycota(100);c__Saccharomycetes(100);o__Saccharomycetales(100);f__Metschnikowiaceae(100);g__Metschnikowia(100);s__Metschnikowia_pulcherrima(100); | Metschnikowia | 13 | Saprotroph | \|Undefined Saprotroph\| | Yeast | NULL | Probable | As nectar-tap_saprotroph (Põlme et al. 2020); Decay sugar-rich substrates (Põlme et al. 2020) | Põlme S, et al. 2020. Fungal Diversity 105:1-16 ((https://doi.org/10.1007/s13225-020-00466-2)) |
| Otu35974 | 1 | k__Fungi(100);p__Ascomycota(100);c__Saccharomycetes(100);o__Saccharomycetales(100);f__Metschnikowiaceae(100);g__Metschnikowia(100);s__Metschnikowia_pulcherrima(100); | Metschnikowia | 13 | Saprotroph | \|Undefined Saprotroph\| | Yeast | NULL | Probable | As nectar-tap_saprotroph (Põlme et al. 2020); Decay sugar-rich substrates (Põlme et al. 2020) | Põlme S, et al. 2020. Fungal Diversity 105:1-16 ((https://doi.org/10.1007/s13225-020-00466-2)) |
| Otu35975 | 1 | k__Fungi(100);p__Ascomycota(100);c__Saccharomycetes(100);o__Saccharomycetales(100);f__Metschnikowiaceae(100);g__Metschnikowia(100);s__Metschnikowia_pulcherrima(100); | Metschnikowia | 13 | Saprotroph | \|Undefined Saprotroph\| | Yeast | NULL | Probable | As nectar-tap_saprotroph (Põlme et al. 2020); Decay sugar-rich substrates (Põlme et al. 2020) | Põlme S, et al. 2020. Fungal Diversity 105:1-16 ((https://doi.org/10.1007/s13225-020-00466-2)) |
| Otu35976 | 1 | k__Fungi(100);p__Ascomycota(100);c__Saccharomycetes(100);o__Saccharomycetales(100);f__Metschnikowiaceae(100);g__Metschnikowia(100);g__Metschnikowia_unclassified(100); | Metschnikowia | 13 | Saprotroph | \|Undefined Saprotroph\| | Yeast | NULL | Probable | As nectar-tap_saprotroph (Põlme et al. 2020); Decay sugar-rich substrates (Põlme et al. 2020) | Põlme S, et al. 2020. Fungal Diversity 105:1-16 ((https://doi.org/10.1007/s13225-020-00466-2)) |
| Otu35990 | 1 | k__Fungi(100);p__Ascomycota(100);c__Saccharomycetes(100);o__Saccharomycetales(100);f__Saccharomycetales_family_Incertae_sedis(100);f__Saccharomycetales_family_Incertae_sedis_unclassified(100);f__Saccharomycetales_family_Incertae_sedis_unclassified(100); | Saccharomycetales | 7 | Saprotroph | Undefined Saprotroph | Yeast | NULL | Possible | NULL | Sterkenburg E, et al. 2015. New Phytologist 207:1145-1158 ((https://doi.org/10.1111/nph.13426)) |
| Otu35995 | 1 | k__Fungi(100);p__Ascomycota(100);c__Saccharomycetes(100);o__Saccharomycetales(100);f__Metschnikowiaceae(100);g__Metschnikowia(100);s__Metschnikowia_pulcherrima(100); | Metschnikowia | 13 | Saprotroph | \|Undefined Saprotroph\| | Yeast | NULL | Probable | As nectar-tap_saprotroph (Põlme et al. 2020); Decay sugar-rich substrates (Põlme et al. 2020) | Põlme S, et al. 2020. Fungal Diversity 105:1-16 ((https://doi.org/10.1007/s13225-020-00466-2)) |
| Otu36001 | 1 | k__Fungi(100);p__Ascomycota(100);c__Saccharomycetes(100);o__Saccharomycetales(100);f__Metschnikowiaceae(100);g__Metschnikowia(100);s__Metschnikowia_pulcherrima(100); | Metschnikowia | 13 | Saprotroph | \|Undefined Saprotroph\| | Yeast | NULL | Probable | As nectar-tap_saprotroph (Põlme et al. 2020); Decay sugar-rich substrates (Põlme et al. 2020) | Põlme S, et al. 2020. Fungal Diversity 105:1-16 ((https://doi.org/10.1007/s13225-020-00466-2)) |
| Otu36002 | 1 | k__Fungi(100);p__Ascomycota(100);c__Saccharomycetes(100);o__Saccharomycetales(100);f__Metschnikowiaceae(100);g__Metschnikowia(100);s__Metschnikowia_pulcherrima(100); | Metschnikowia | 13 | Saprotroph | \|Undefined Saprotroph\| | Yeast | NULL | Probable | As nectar-tap_saprotroph (Põlme et al. 2020); Decay sugar-rich substrates (Põlme et al. 2020) | Põlme S, et al. 2020. Fungal Diversity 105:1-16 ((https://doi.org/10.1007/s13225-020-00466-2)) |
| Otu36003 | 1 | k__Fungi(100);p__Ascomycota(100);c__Saccharomycetes(100);o__Saccharomycetales(100);f__Metschnikowiaceae(100);g__Metschnikowia(100);g__Metschnikowia_unclassified(100); | Metschnikowia | 13 | Saprotroph | \|Undefined Saprotroph\| | Yeast | NULL | Probable | As nectar-tap_saprotroph (Põlme et al. 2020); Decay sugar-rich substrates (Põlme et al. 2020) | Põlme S, et al. 2020. Fungal Diversity 105:1-16 ((https://doi.org/10.1007/s13225-020-00466-2)) |
| Otu36005 | 1 | k__Fungi(100);p__Ascomycota(100);c__Saccharomycetes(100);o__Saccharomycetales(100);f__Metschnikowiaceae(100);g__Metschnikowia(100);s__Metschnikowia_pulcherrima(100); | Metschnikowia | 13 | Saprotroph | \|Undefined Saprotroph\| | Yeast | NULL | Probable | As nectar-tap_saprotroph (Põlme et al. 2020); Decay sugar-rich substrates (Põlme et al. 2020) | Põlme S, et al. 2020. Fungal Diversity 105:1-16 ((https://doi.org/10.1007/s13225-020-00466-2)) |
| Otu36006 | 1 | k__Fungi(100);p__Ascomycota(100);c__Saccharomycetes(100);o__Saccharomycetales(100);f__Metschnikowiaceae(100);g__Metschnikowia(100);g__Metschnikowia_unclassified(100); | Metschnikowia | 13 | Saprotroph | \|Undefined Saprotroph\| | Yeast | NULL | Probable | As nectar-tap_saprotroph (Põlme et al. 2020); Decay sugar-rich substrates (Põlme et al. 2020) | Põlme S, et al. 2020. Fungal Diversity 105:1-16 ((https://doi.org/10.1007/s13225-020-00466-2)) |
| Otu36008 | 1 | k__Fungi(100);p__Ascomycota(100);c__Saccharomycetes(100);o__Saccharomycetales(100);f__Metschnikowiaceae(100);g__Metschnikowia(100);g__Metschnikowia_unclassified(100); | Metschnikowia | 13 | Saprotroph | \|Undefined Saprotroph\| | Yeast | NULL | Probable | As nectar-tap_saprotroph (Põlme et al. 2020); Decay sugar-rich substrates (Põlme et al. 2020) | Põlme S, et al. 2020. Fungal Diversity 105:1-16 ((https://doi.org/10.1007/s13225-020-00466-2)) |
| Otu36009 | 1 | k__Fungi(100);p__Ascomycota(100);c__Saccharomycetes(100);o__Saccharomycetales(100);f__Metschnikowiaceae(100);g__Metschnikowia(100);s__Metschnikowia_pulcherrima(100); | Metschnikowia | 13 | Saprotroph | \|Undefined Saprotroph\| | Yeast | NULL | Probable | As nectar-tap_saprotroph (Põlme et al. 2020); Decay sugar-rich substrates (Põlme et al. 2020) | Põlme S, et al. 2020. Fungal Diversity 105:1-16 ((https://doi.org/10.1007/s13225-020-00466-2)) |
| Otu36010 | 1 | k__Fungi(100);p__Ascomycota(100);c__Saccharomycetes(100);o__Saccharomycetales(100);f__Metschnikowiaceae(100);g__Metschnikowia(100);s__Metschnikowia_pulcherrima(100); | Metschnikowia | 13 | Saprotroph | \|Undefined Saprotroph\| | Yeast | NULL | Probable | As nectar-tap_saprotroph (Põlme et al. 2020); Decay sugar-rich substrates (Põlme et al. 2020) | Põlme S, et al. 2020. Fungal Diversity 105:1-16 ((https://doi.org/10.1007/s13225-020-00466-2)) |
| Otu36012 | 1 | k__Fungi(100);p__Ascomycota(100);c__Saccharomycetes(100);o__Saccharomycetales(100);f__Metschnikowiaceae(100);g__Metschnikowia(100);s__Metschnikowia_pulcherrima(100); | Metschnikowia | 13 | Saprotroph | \|Undefined Saprotroph\| | Yeast | NULL | Probable | As nectar-tap_saprotroph (Põlme et al. 2020); Decay sugar-rich substrates (Põlme et al. 2020) | Põlme S, et al. 2020. Fungal Diversity 105:1-16 ((https://doi.org/10.1007/s13225-020-00466-2)) |
| Otu36013 | 1 | k__Fungi(100);p__Ascomycota(100);c__Saccharomycetes(100);o__Saccharomycetales(100);f__Metschnikowiaceae(100);g__Metschnikowia(100);s__Metschnikowia_pulcherrima(100); | Metschnikowia | 13 | Saprotroph | \|Undefined Saprotroph\| | Yeast | NULL | Probable | As nectar-tap_saprotroph (Põlme et al. 2020); Decay sugar-rich substrates (Põlme et al. 2020) | Põlme S, et al. 2020. Fungal Diversity 105:1-16 ((https://doi.org/10.1007/s13225-020-00466-2)) |
| Otu36015 | 1 | k__Fungi(100);p__Ascomycota(100);c__Saccharomycetes(100);o__Saccharomycetales(100);f__Metschnikowiaceae(100);g__Metschnikowia(100);s__Metschnikowia_pulcherrima(100); | Metschnikowia | 13 | Saprotroph | \|Undefined Saprotroph\| | Yeast | NULL | Probable | As nectar-tap_saprotroph (Põlme et al. 2020); Decay sugar-rich substrates (Põlme et al. 2020) | Põlme S, et al. 2020. Fungal Diversity 105:1-16 ((https://doi.org/10.1007/s13225-020-00466-2)) |
| Otu36016 | 1 | k__Fungi(100);p__Ascomycota(100);c__Saccharomycetes(100);o__Saccharomycetales(100);f__Metschnikowiaceae(100);g__Metschnikowia(100);s__Metschnikowia_pulcherrima(100); | Metschnikowia | 13 | Saprotroph | \|Undefined Saprotroph\| | Yeast | NULL | Probable | As nectar-tap_saprotroph (Põlme et al. 2020); Decay sugar-rich substrates (Põlme et al. 2020) | Põlme S, et al. 2020. Fungal Diversity 105:1-16 ((https://doi.org/10.1007/s13225-020-00466-2)) |
| Otu36019 | 1 | k__Fungi(100);p__Ascomycota(100);c__Saccharomycetes(100);o__Saccharomycetales(100);f__Metschnikowiaceae(100);g__Metschnikowia(100);g__Metschnikowia_unclassified(100); | Metschnikowia | 13 | Saprotroph | \|Undefined Saprotroph\| | Yeast | NULL | Probable | As nectar-tap_saprotroph (Põlme et al. 2020); Decay sugar-rich substrates (Põlme et al. 2020) | Põlme S, et al. 2020. Fungal Diversity 105:1-16 ((https://doi.org/10.1007/s13225-020-00466-2)) |
| Otu36030 | 1 | k__Fungi(100);p__Ascomycota(100);c__Saccharomycetes(100);o__Saccharomycetales(100);f__Metschnikowiaceae(100);g__Metschnikowia(100);s__Metschnikowia_pulcherrima(100); | Metschnikowia | 13 | Saprotroph | \|Undefined Saprotroph\| | Yeast | NULL | Probable | As nectar-tap_saprotroph (Põlme et al. 2020); Decay sugar-rich substrates (Põlme et al. 2020) | Põlme S, et al. 2020. Fungal Diversity 105:1-16 ((https://doi.org/10.1007/s13225-020-00466-2)) |
| Otu36031 | 1 | k__Fungi(100);p__Ascomycota(100);c__Saccharomycetes(100);o__Saccharomycetales(100);f__Saccharomycetales_family_Incertae_sedis(100);g__Candida(100);g__Candida_unclassified(100); | Candida | 13 | Pathotroph-Saprotroph-Symbiotroph | Animal Pathogen-Endophyte-Endosymbiont-Epiphyte-\|Undefined Saprotroph\| | Dimorphic Yeast-Yeast | NULL | Probable | As nectar-tap_saprotroph (Põlme et al. 2020); Decay sugar-rich substrates (Põlme et al. 2020); This asexual genus is polyphyletic and contain species that function very differently depend on what they are. Caution should be taken when interpreting guilds for Candida species. | Manolakaki D, et al. 2010. Virulence. 1: 367-375 ((https://doi.org/10.4161/viru.1.5.12796)); Mekbib SB, et al. 2011. Tropical Plant Pathology 36:233-240 ((https://doi.org/10.1590/S1982-56762011000400004)); Põlme S, et al. 2020. Fungal Diversity 105:1-16 ((https://doi.org/10.1007/s13225-020-00466-2)) |
| Otu36032 | 1 | k__Fungi(100);p__Ascomycota(100);c__Saccharomycetes(100);o__Saccharomycetales(100);f__Metschnikowiaceae(100);g__Metschnikowia(100);g__Metschnikowia_unclassified(100); | Metschnikowia | 13 | Saprotroph | \|Undefined Saprotroph\| | Yeast | NULL | Probable | As nectar-tap_saprotroph (Põlme et al. 2020); Decay sugar-rich substrates (Põlme et al. 2020) | Põlme S, et al. 2020. Fungal Diversity 105:1-16 ((https://doi.org/10.1007/s13225-020-00466-2)) |
| Otu36033 | 1 | k__Fungi(100);p__Ascomycota(100);c__Saccharomycetes(100);o__Saccharomycetales(100);o__Saccharomycetales_unclassified(100);o__Saccharomycetales_unclassified(100);o__Saccharomycetales_unclassified(100); | Saccharomycetales | 7 | Saprotroph | Undefined Saprotroph | Yeast | NULL | Possible | NULL | Sterkenburg E, et al. 2015. New Phytologist 207:1145-1158 ((https://doi.org/10.1111/nph.13426)) |
| Otu36034 | 1 | k__Fungi(100);p__Ascomycota(100);c__Saccharomycetes(100);o__Saccharomycetales(100);f__Saccharomycetales_family_Incertae_sedis(100);g__Candida(100);g__Candida_unclassified(100); | Candida | 13 | Pathotroph-Saprotroph-Symbiotroph | Animal Pathogen-Endophyte-Endosymbiont-Epiphyte-\|Undefined Saprotroph\| | Dimorphic Yeast-Yeast | NULL | Probable | As nectar-tap_saprotroph (Põlme et al. 2020); Decay sugar-rich substrates (Põlme et al. 2020); This asexual genus is polyphyletic and contain species that function very differently depend on what they are. Caution should be taken when interpreting guilds for Candida species. | Manolakaki D, et al. 2010. Virulence. 1: 367-375 ((https://doi.org/10.4161/viru.1.5.12796)); Mekbib SB, et al. 2011. Tropical Plant Pathology 36:233-240 ((https://doi.org/10.1590/S1982-56762011000400004)); Põlme S, et al. 2020. Fungal Diversity 105:1-16 ((https://doi.org/10.1007/s13225-020-00466-2)) |
| Otu36037 | 1 | k__Fungi(100);p__Ascomycota(100);c__Saccharomycetes(100);o__Saccharomycetales(100);f__Metschnikowiaceae(100);g__Metschnikowia(100);s__Metschnikowia_pulcherrima(100); | Metschnikowia | 13 | Saprotroph | \|Undefined Saprotroph\| | Yeast | NULL | Probable | As nectar-tap_saprotroph (Põlme et al. 2020); Decay sugar-rich substrates (Põlme et al. 2020) | Põlme S, et al. 2020. Fungal Diversity 105:1-16 ((https://doi.org/10.1007/s13225-020-00466-2)) |
| Otu36038 | 1 | k__Fungi(100);p__Ascomycota(100);c__Saccharomycetes(100);o__Saccharomycetales(100);f__Metschnikowiaceae(100);g__Metschnikowia(100);s__Metschnikowia_pulcherrima(100); | Metschnikowia | 13 | Saprotroph | \|Undefined Saprotroph\| | Yeast | NULL | Probable | As nectar-tap_saprotroph (Põlme et al. 2020); Decay sugar-rich substrates (Põlme et al. 2020) | Põlme S, et al. 2020. Fungal Diversity 105:1-16 ((https://doi.org/10.1007/s13225-020-00466-2)) |
| Otu36039 | 1 | k__Fungi(100);p__Ascomycota(100);c__Saccharomycetes(100);o__Saccharomycetales(100);f__Metschnikowiaceae(100);g__Metschnikowia(100);s__Metschnikowia_pulcherrima(100); | Metschnikowia | 13 | Saprotroph | \|Undefined Saprotroph\| | Yeast | NULL | Probable | As nectar-tap_saprotroph (Põlme et al. 2020); Decay sugar-rich substrates (Põlme et al. 2020) | Põlme S, et al. 2020. Fungal Diversity 105:1-16 ((https://doi.org/10.1007/s13225-020-00466-2)) |
| Otu36040 | 1 | k__Fungi(100);p__Ascomycota(100);c__Saccharomycetes(100);o__Saccharomycetales(100);f__Metschnikowiaceae(100);g__Metschnikowia(100);s__Metschnikowia_pulcherrima(100); | Metschnikowia | 13 | Saprotroph | \|Undefined Saprotroph\| | Yeast | NULL | Probable | As nectar-tap_saprotroph (Põlme et al. 2020); Decay sugar-rich substrates (Põlme et al. 2020) | Põlme S, et al. 2020. Fungal Diversity 105:1-16 ((https://doi.org/10.1007/s13225-020-00466-2)) |
| Otu36041 | 1 | k__Fungi(100);p__Ascomycota(100);c__Saccharomycetes(100);o__Saccharomycetales(100);f__Saccharomycetales_family_Incertae_sedis(100);g__Candida(100);s__Candida_cretensis(100); | Candida | 13 | Pathotroph-Saprotroph-Symbiotroph | Animal Pathogen-Endophyte-Endosymbiont-Epiphyte-\|Undefined Saprotroph\| | Dimorphic Yeast-Yeast | NULL | Probable | As nectar-tap_saprotroph (Põlme et al. 2020); Decay sugar-rich substrates (Põlme et al. 2020); This asexual genus is polyphyletic and contain species that function very differently depend on what they are. Caution should be taken when interpreting guilds for Candida species. | Manolakaki D, et al. 2010. Virulence. 1: 367-375 ((https://doi.org/10.4161/viru.1.5.12796)); Mekbib SB, et al. 2011. Tropical Plant Pathology 36:233-240 ((https://doi.org/10.1590/S1982-56762011000400004)); Põlme S, et al. 2020. Fungal Diversity 105:1-16 ((https://doi.org/10.1007/s13225-020-00466-2)) |
| Otu36043 | 1 | k__Fungi(100);p__Ascomycota(100);c__Saccharomycetes(100);o__Saccharomycetales(100);f__Metschnikowiaceae(100);g__Metschnikowia(100);s__Metschnikowia_pulcherrima(100); | Metschnikowia | 13 | Saprotroph | \|Undefined Saprotroph\| | Yeast | NULL | Probable | As nectar-tap_saprotroph (Põlme et al. 2020); Decay sugar-rich substrates (Põlme et al. 2020) | Põlme S, et al. 2020. Fungal Diversity 105:1-16 ((https://doi.org/10.1007/s13225-020-00466-2)) |
| Otu36051 | 1 | k__Fungi(100);p__Ascomycota(100);c__Saccharomycetes(100);o__Saccharomycetales(100);f__Metschnikowiaceae(100);g__Metschnikowia(100);g__Metschnikowia_unclassified(100); | Metschnikowia | 13 | Saprotroph | \|Undefined Saprotroph\| | Yeast | NULL | Probable | As nectar-tap_saprotroph (Põlme et al. 2020); Decay sugar-rich substrates (Põlme et al. 2020) | Põlme S, et al. 2020. Fungal Diversity 105:1-16 ((https://doi.org/10.1007/s13225-020-00466-2)) |
| Otu36053 | 1 | k__Fungi(100);p__Ascomycota(100);c__Saccharomycetes(100);o__Saccharomycetales(100);f__Metschnikowiaceae(100);g__Metschnikowia(100);s__Metschnikowia_pulcherrima(100); | Metschnikowia | 13 | Saprotroph | \|Undefined Saprotroph\| | Yeast | NULL | Probable | As nectar-tap_saprotroph (Põlme et al. 2020); Decay sugar-rich substrates (Põlme et al. 2020) | Põlme S, et al. 2020. Fungal Diversity 105:1-16 ((https://doi.org/10.1007/s13225-020-00466-2)) |
| Otu36054 | 1 | k__Fungi(100);p__Ascomycota(100);c__Saccharomycetes(100);o__Saccharomycetales(100);f__Metschnikowiaceae(100);g__Metschnikowia(100);s__Metschnikowia_pulcherrima(100); | Metschnikowia | 13 | Saprotroph | \|Undefined Saprotroph\| | Yeast | NULL | Probable | As nectar-tap_saprotroph (Põlme et al. 2020); Decay sugar-rich substrates (Põlme et al. 2020) | Põlme S, et al. 2020. Fungal Diversity 105:1-16 ((https://doi.org/10.1007/s13225-020-00466-2)) |
| Otu36057 | 1 | k__Fungi(100);p__Ascomycota(100);c__Saccharomycetes(100);o__Saccharomycetales(100);f__Metschnikowiaceae(100);g__Metschnikowia(100);s__Metschnikowia_pulcherrima(100); | Metschnikowia | 13 | Saprotroph | \|Undefined Saprotroph\| | Yeast | NULL | Probable | As nectar-tap_saprotroph (Põlme et al. 2020); Decay sugar-rich substrates (Põlme et al. 2020) | Põlme S, et al. 2020. Fungal Diversity 105:1-16 ((https://doi.org/10.1007/s13225-020-00466-2)) |
| Otu36062 | 1 | k__Fungi(100);p__Ascomycota(100);c__Saccharomycetes(100);o__Saccharomycetales(100);f__Metschnikowiaceae(100);g__Metschnikowia(100);s__Metschnikowia_pulcherrima(100); | Metschnikowia | 13 | Saprotroph | \|Undefined Saprotroph\| | Yeast | NULL | Probable | As nectar-tap_saprotroph (Põlme et al. 2020); Decay sugar-rich substrates (Põlme et al. 2020) | Põlme S, et al. 2020. Fungal Diversity 105:1-16 ((https://doi.org/10.1007/s13225-020-00466-2)) |
| Otu36063 | 1 | k__Fungi(100);p__Ascomycota(100);c__Saccharomycetes(100);o__Saccharomycetales(100);f__Saccharomycetales_family_Incertae_sedis(100);g__Candida(100);g__Candida_unclassified(100); | Candida | 13 | Pathotroph-Saprotroph-Symbiotroph | Animal Pathogen-Endophyte-Endosymbiont-Epiphyte-\|Undefined Saprotroph\| | Dimorphic Yeast-Yeast | NULL | Probable | As nectar-tap_saprotroph (Põlme et al. 2020); Decay sugar-rich substrates (Põlme et al. 2020); This asexual genus is polyphyletic and contain species that function very differently depend on what they are. Caution should be taken when interpreting guilds for Candida species. | Manolakaki D, et al. 2010. Virulence. 1: 367-375 ((https://doi.org/10.4161/viru.1.5.12796)); Mekbib SB, et al. 2011. Tropical Plant Pathology 36:233-240 ((https://doi.org/10.1590/S1982-56762011000400004)); Põlme S, et al. 2020. Fungal Diversity 105:1-16 ((https://doi.org/10.1007/s13225-020-00466-2)) |
| Otu36064 | 1 | k__Fungi(100);p__Ascomycota(100);c__Saccharomycetes(100);o__Saccharomycetales(100);f__Metschnikowiaceae(100);g__Metschnikowia(100);s__Metschnikowia_pulcherrima(100); | Metschnikowia | 13 | Saprotroph | \|Undefined Saprotroph\| | Yeast | NULL | Probable | As nectar-tap_saprotroph (Põlme et al. 2020); Decay sugar-rich substrates (Põlme et al. 2020) | Põlme S, et al. 2020. Fungal Diversity 105:1-16 ((https://doi.org/10.1007/s13225-020-00466-2)) |
| Otu36065 | 1 | k__Fungi(100);p__Ascomycota(100);c__Saccharomycetes(100);o__Saccharomycetales(100);f__Metschnikowiaceae(100);g__Metschnikowia(100);s__Metschnikowia_pulcherrima(100); | Metschnikowia | 13 | Saprotroph | \|Undefined Saprotroph\| | Yeast | NULL | Probable | As nectar-tap_saprotroph (Põlme et al. 2020); Decay sugar-rich substrates (Põlme et al. 2020) | Põlme S, et al. 2020. Fungal Diversity 105:1-16 ((https://doi.org/10.1007/s13225-020-00466-2)) |
| Otu36066 | 1 | k__Fungi(100);p__Ascomycota(100);c__Saccharomycetes(100);o__Saccharomycetales(100);f__Metschnikowiaceae(100);g__Metschnikowia(100);g__Metschnikowia_unclassified(100); | Metschnikowia | 13 | Saprotroph | \|Undefined Saprotroph\| | Yeast | NULL | Probable | As nectar-tap_saprotroph (Põlme et al. 2020); Decay sugar-rich substrates (Põlme et al. 2020) | Põlme S, et al. 2020. Fungal Diversity 105:1-16 ((https://doi.org/10.1007/s13225-020-00466-2)) |
| Otu36069 | 1 | k__Fungi(100);p__Ascomycota(100);c__Saccharomycetes(100);o__Saccharomycetales(100);f__Saccharomycetales_family_Incertae_sedis(100);g__Candida(100);g__Candida_unclassified(100); | Candida | 13 | Pathotroph-Saprotroph-Symbiotroph | Animal Pathogen-Endophyte-Endosymbiont-Epiphyte-\|Undefined Saprotroph\| | Dimorphic Yeast-Yeast | NULL | Probable | As nectar-tap_saprotroph (Põlme et al. 2020); Decay sugar-rich substrates (Põlme et al. 2020); This asexual genus is polyphyletic and contain species that function very differently depend on what they are. Caution should be taken when interpreting guilds for Candida species. | Manolakaki D, et al. 2010. Virulence. 1: 367-375 ((https://doi.org/10.4161/viru.1.5.12796)); Mekbib SB, et al. 2011. Tropical Plant Pathology 36:233-240 ((https://doi.org/10.1590/S1982-56762011000400004)); Põlme S, et al. 2020. Fungal Diversity 105:1-16 ((https://doi.org/10.1007/s13225-020-00466-2)) |
| Otu36072 | 1 | k__Fungi(100);p__Ascomycota(100);c__Saccharomycetes(100);o__Saccharomycetales(100);f__Metschnikowiaceae(100);g__Metschnikowia(100);s__Metschnikowia_pulcherrima(100); | Metschnikowia | 13 | Saprotroph | \|Undefined Saprotroph\| | Yeast | NULL | Probable | As nectar-tap_saprotroph (Põlme et al. 2020); Decay sugar-rich substrates (Põlme et al. 2020) | Põlme S, et al. 2020. Fungal Diversity 105:1-16 ((https://doi.org/10.1007/s13225-020-00466-2)) |
| Otu36073 | 1 | k__Fungi(100);p__Ascomycota(100);c__Saccharomycetes(100);o__Saccharomycetales(100);f__Metschnikowiaceae(100);g__Metschnikowia(100);s__Metschnikowia_pulcherrima(100); | Metschnikowia | 13 | Saprotroph | \|Undefined Saprotroph\| | Yeast | NULL | Probable | As nectar-tap_saprotroph (Põlme et al. 2020); Decay sugar-rich substrates (Põlme et al. 2020) | Põlme S, et al. 2020. Fungal Diversity 105:1-16 ((https://doi.org/10.1007/s13225-020-00466-2)) |
| Otu36074 | 1 | k__Fungi(100);p__Ascomycota(100);c__Saccharomycetes(100);o__Saccharomycetales(100);f__Metschnikowiaceae(100);g__Metschnikowia(100);g__Metschnikowia_unclassified(100); | Metschnikowia | 13 | Saprotroph | \|Undefined Saprotroph\| | Yeast | NULL | Probable | As nectar-tap_saprotroph (Põlme et al. 2020); Decay sugar-rich substrates (Põlme et al. 2020) | Põlme S, et al. 2020. Fungal Diversity 105:1-16 ((https://doi.org/10.1007/s13225-020-00466-2)) |
| Otu36077 | 1 | k__Fungi(100);p__Ascomycota(100);c__Saccharomycetes(100);o__Saccharomycetales(100);f__Metschnikowiaceae(100);g__Metschnikowia(100);g__Metschnikowia_unclassified(100); | Metschnikowia | 13 | Saprotroph | \|Undefined Saprotroph\| | Yeast | NULL | Probable | As nectar-tap_saprotroph (Põlme et al. 2020); Decay sugar-rich substrates (Põlme et al. 2020) | Põlme S, et al. 2020. Fungal Diversity 105:1-16 ((https://doi.org/10.1007/s13225-020-00466-2)) |
| Otu36080 | 1 | k__Fungi(100);p__Ascomycota(100);c__Saccharomycetes(100);o__Saccharomycetales(100);f__Saccharomycetales_family_Incertae_sedis(100);f__Saccharomycetales_family_Incertae_sedis_unclassified(100);f__Saccharomycetales_family_Incertae_sedis_unclassified(100); | Saccharomycetales | 7 | Saprotroph | Undefined Saprotroph | Yeast | NULL | Possible | NULL | Sterkenburg E, et al. 2015. New Phytologist 207:1145-1158 ((https://doi.org/10.1111/nph.13426)) |
| Otu36084 | 1 | k__Fungi(100);p__Ascomycota(100);c__Saccharomycetes(100);o__Saccharomycetales(100);f__Metschnikowiaceae(100);g__Metschnikowia(100);s__Metschnikowia_pulcherrima(100); | Metschnikowia | 13 | Saprotroph | \|Undefined Saprotroph\| | Yeast | NULL | Probable | As nectar-tap_saprotroph (Põlme et al. 2020); Decay sugar-rich substrates (Põlme et al. 2020) | Põlme S, et al. 2020. Fungal Diversity 105:1-16 ((https://doi.org/10.1007/s13225-020-00466-2)) |
| Otu36085 | 1 | k__Fungi(100);p__Ascomycota(100);c__Saccharomycetes(100);o__Saccharomycetales(100);f__Metschnikowiaceae(100);g__Metschnikowia(100);s__Metschnikowia_pulcherrima(100); | Metschnikowia | 13 | Saprotroph | \|Undefined Saprotroph\| | Yeast | NULL | Probable | As nectar-tap_saprotroph (Põlme et al. 2020); Decay sugar-rich substrates (Põlme et al. 2020) | Põlme S, et al. 2020. Fungal Diversity 105:1-16 ((https://doi.org/10.1007/s13225-020-00466-2)) |
| Otu36086 | 1 | k__Fungi(100);p__Ascomycota(100);c__Saccharomycetes(100);o__Saccharomycetales(100);f__Metschnikowiaceae(100);g__Metschnikowia(100);s__Metschnikowia_pulcherrima(100); | Metschnikowia | 13 | Saprotroph | \|Undefined Saprotroph\| | Yeast | NULL | Probable | As nectar-tap_saprotroph (Põlme et al. 2020); Decay sugar-rich substrates (Põlme et al. 2020) | Põlme S, et al. 2020. Fungal Diversity 105:1-16 ((https://doi.org/10.1007/s13225-020-00466-2)) |
| Otu36092 | 1 | k__Fungi(100);p__Ascomycota(100);c__Saccharomycetes(100);o__Saccharomycetales(100);f__Metschnikowiaceae(100);g__Metschnikowia(100);g__Metschnikowia_unclassified(100); | Metschnikowia | 13 | Saprotroph | \|Undefined Saprotroph\| | Yeast | NULL | Probable | As nectar-tap_saprotroph (Põlme et al. 2020); Decay sugar-rich substrates (Põlme et al. 2020) | Põlme S, et al. 2020. Fungal Diversity 105:1-16 ((https://doi.org/10.1007/s13225-020-00466-2)) |
| Otu36097 | 1 | k__Fungi(100);p__Ascomycota(100);c__Saccharomycetes(100);o__Saccharomycetales(100);f__Metschnikowiaceae(100);g__Metschnikowia(100);s__Metschnikowia_pulcherrima(100); | Metschnikowia | 13 | Saprotroph | \|Undefined Saprotroph\| | Yeast | NULL | Probable | As nectar-tap_saprotroph (Põlme et al. 2020); Decay sugar-rich substrates (Põlme et al. 2020) | Põlme S, et al. 2020. Fungal Diversity 105:1-16 ((https://doi.org/10.1007/s13225-020-00466-2)) |
| Otu36104 | 1 | k__Fungi(100);p__Ascomycota(100);c__Saccharomycetes(100);o__Saccharomycetales(100);f__Metschnikowiaceae(100);g__Metschnikowia(100);g__Metschnikowia_unclassified(100); | Metschnikowia | 13 | Saprotroph | \|Undefined Saprotroph\| | Yeast | NULL | Probable | As nectar-tap_saprotroph (Põlme et al. 2020); Decay sugar-rich substrates (Põlme et al. 2020) | Põlme S, et al. 2020. Fungal Diversity 105:1-16 ((https://doi.org/10.1007/s13225-020-00466-2)) |
| Otu36106 | 1 | k__Fungi(100);p__Ascomycota(100);c__Saccharomycetes(100);o__Saccharomycetales(100);f__Saccharomycetales_family_Incertae_sedis(100);g__Candida(100);s__Candida_cretensis(100); | Candida | 13 | Pathotroph-Saprotroph-Symbiotroph | Animal Pathogen-Endophyte-Endosymbiont-Epiphyte-\|Undefined Saprotroph\| | Dimorphic Yeast-Yeast | NULL | Probable | As nectar-tap_saprotroph (Põlme et al. 2020); Decay sugar-rich substrates (Põlme et al. 2020); This asexual genus is polyphyletic and contain species that function very differently depend on what they are. Caution should be taken when interpreting guilds for Candida species. | Manolakaki D, et al. 2010. Virulence. 1: 367-375 ((https://doi.org/10.4161/viru.1.5.12796)); Mekbib SB, et al. 2011. Tropical Plant Pathology 36:233-240 ((https://doi.org/10.1590/S1982-56762011000400004)); Põlme S, et al. 2020. Fungal Diversity 105:1-16 ((https://doi.org/10.1007/s13225-020-00466-2)) |
| Otu36107 | 1 | k__Fungi(100);p__Ascomycota(100);c__Saccharomycetes(100);o__Saccharomycetales(100);f__Saccharomycetales_family_Incertae_sedis(100);f__Saccharomycetales_family_Incertae_sedis_unclassified(100);f__Saccharomycetales_family_Incertae_sedis_unclassified(100); | Saccharomycetales | 7 | Saprotroph | Undefined Saprotroph | Yeast | NULL | Possible | NULL | Sterkenburg E, et al. 2015. New Phytologist 207:1145-1158 ((https://doi.org/10.1111/nph.13426)) |
| Otu36108 | 1 | k__Fungi(100);p__Ascomycota(100);c__Saccharomycetes(100);o__Saccharomycetales(100);f__Saccharomycetales_family_Incertae_sedis(100);g__Candida(100);g__Candida_unclassified(100); | Candida | 13 | Pathotroph-Saprotroph-Symbiotroph | Animal Pathogen-Endophyte-Endosymbiont-Epiphyte-\|Undefined Saprotroph\| | Dimorphic Yeast-Yeast | NULL | Probable | As nectar-tap_saprotroph (Põlme et al. 2020); Decay sugar-rich substrates (Põlme et al. 2020); This asexual genus is polyphyletic and contain species that function very differently depend on what they are. Caution should be taken when interpreting guilds for Candida species. | Manolakaki D, et al. 2010. Virulence. 1: 367-375 ((https://doi.org/10.4161/viru.1.5.12796)); Mekbib SB, et al. 2011. Tropical Plant Pathology 36:233-240 ((https://doi.org/10.1590/S1982-56762011000400004)); Põlme S, et al. 2020. Fungal Diversity 105:1-16 ((https://doi.org/10.1007/s13225-020-00466-2)) |
| Otu36109 | 1 | k__Fungi(100);p__Ascomycota(100);c__Saccharomycetes(100);o__Saccharomycetales(100);f__Saccharomycetales_family_Incertae_sedis(100);g__Candida(100);s__Candida_cretensis(100); | Candida | 13 | Pathotroph-Saprotroph-Symbiotroph | Animal Pathogen-Endophyte-Endosymbiont-Epiphyte-\|Undefined Saprotroph\| | Dimorphic Yeast-Yeast | NULL | Probable | As nectar-tap_saprotroph (Põlme et al. 2020); Decay sugar-rich substrates (Põlme et al. 2020); This asexual genus is polyphyletic and contain species that function very differently depend on what they are. Caution should be taken when interpreting guilds for Candida species. | Manolakaki D, et al. 2010. Virulence. 1: 367-375 ((https://doi.org/10.4161/viru.1.5.12796)); Mekbib SB, et al. 2011. Tropical Plant Pathology 36:233-240 ((https://doi.org/10.1590/S1982-56762011000400004)); Põlme S, et al. 2020. Fungal Diversity 105:1-16 ((https://doi.org/10.1007/s13225-020-00466-2)) |
| Otu36110 | 1 | k__Fungi(100);p__Ascomycota(100);c__Saccharomycetes(100);o__Saccharomycetales(100);f__Metschnikowiaceae(100);g__Metschnikowia(100);g__Metschnikowia_unclassified(100); | Metschnikowia | 13 | Saprotroph | \|Undefined Saprotroph\| | Yeast | NULL | Probable | As nectar-tap_saprotroph (Põlme et al. 2020); Decay sugar-rich substrates (Põlme et al. 2020) | Põlme S, et al. 2020. Fungal Diversity 105:1-16 ((https://doi.org/10.1007/s13225-020-00466-2)) |
| Otu36113 | 1 | k__Fungi(100);p__Ascomycota(100);c__Saccharomycetes(100);o__Saccharomycetales(100);f__Metschnikowiaceae(100);g__Metschnikowia(100);s__Metschnikowia_pulcherrima(100); | Metschnikowia | 13 | Saprotroph | \|Undefined Saprotroph\| | Yeast | NULL | Probable | As nectar-tap_saprotroph (Põlme et al. 2020); Decay sugar-rich substrates (Põlme et al. 2020) | Põlme S, et al. 2020. Fungal Diversity 105:1-16 ((https://doi.org/10.1007/s13225-020-00466-2)) |
| Otu36116 | 1 | k__Fungi(100);p__Ascomycota(100);c__Saccharomycetes(100);o__Saccharomycetales(100);f__Metschnikowiaceae(100);g__Metschnikowia(100);s__Metschnikowia_pulcherrima(100); | Metschnikowia | 13 | Saprotroph | \|Undefined Saprotroph\| | Yeast | NULL | Probable | As nectar-tap_saprotroph (Põlme et al. 2020); Decay sugar-rich substrates (Põlme et al. 2020) | Põlme S, et al. 2020. Fungal Diversity 105:1-16 ((https://doi.org/10.1007/s13225-020-00466-2)) |
| Otu36117 | 1 | k__Fungi(100);p__Ascomycota(100);c__Saccharomycetes(100);o__Saccharomycetales(100);f__Metschnikowiaceae(100);g__Metschnikowia(100);s__Metschnikowia_pulcherrima(100); | Metschnikowia | 13 | Saprotroph | \|Undefined Saprotroph\| | Yeast | NULL | Probable | As nectar-tap_saprotroph (Põlme et al. 2020); Decay sugar-rich substrates (Põlme et al. 2020) | Põlme S, et al. 2020. Fungal Diversity 105:1-16 ((https://doi.org/10.1007/s13225-020-00466-2)) |
| Otu36118 | 1 | k__Fungi(100);p__Ascomycota(100);c__Saccharomycetes(100);o__Saccharomycetales(100);f__Metschnikowiaceae(100);g__Metschnikowia(100);s__Metschnikowia_pulcherrima(100); | Metschnikowia | 13 | Saprotroph | \|Undefined Saprotroph\| | Yeast | NULL | Probable | As nectar-tap_saprotroph (Põlme et al. 2020); Decay sugar-rich substrates (Põlme et al. 2020) | Põlme S, et al. 2020. Fungal Diversity 105:1-16 ((https://doi.org/10.1007/s13225-020-00466-2)) |
| Otu36120 | 1 | k__Fungi(100);p__Ascomycota(100);c__Saccharomycetes(100);o__Saccharomycetales(100);f__Metschnikowiaceae(100);g__Metschnikowia(100);s__Metschnikowia_pulcherrima(100); | Metschnikowia | 13 | Saprotroph | \|Undefined Saprotroph\| | Yeast | NULL | Probable | As nectar-tap_saprotroph (Põlme et al. 2020); Decay sugar-rich substrates (Põlme et al. 2020) | Põlme S, et al. 2020. Fungal Diversity 105:1-16 ((https://doi.org/10.1007/s13225-020-00466-2)) |
| Otu36126 | 1 | k__Fungi(100);p__Ascomycota(100);c__Saccharomycetes(100);o__Saccharomycetales(100);f__Metschnikowiaceae(100);g__Metschnikowia(100);s__Metschnikowia_pulcherrima(100); | Metschnikowia | 13 | Saprotroph | \|Undefined Saprotroph\| | Yeast | NULL | Probable | As nectar-tap_saprotroph (Põlme et al. 2020); Decay sugar-rich substrates (Põlme et al. 2020) | Põlme S, et al. 2020. Fungal Diversity 105:1-16 ((https://doi.org/10.1007/s13225-020-00466-2)) |
| Otu36134 | 1 | k__Fungi(100);p__Ascomycota(100);c__Saccharomycetes(100);o__Saccharomycetales(100);f__Metschnikowiaceae(100);g__Metschnikowia(100);s__Metschnikowia_pulcherrima(100); | Metschnikowia | 13 | Saprotroph | \|Undefined Saprotroph\| | Yeast | NULL | Probable | As nectar-tap_saprotroph (Põlme et al. 2020); Decay sugar-rich substrates (Põlme et al. 2020) | Põlme S, et al. 2020. Fungal Diversity 105:1-16 ((https://doi.org/10.1007/s13225-020-00466-2)) |
| Otu36135 | 1 | k__Fungi(100);p__Ascomycota(100);c__Saccharomycetes(100);o__Saccharomycetales(100);f__Metschnikowiaceae(100);g__Metschnikowia(100);s__Metschnikowia_pulcherrima(100); | Metschnikowia | 13 | Saprotroph | \|Undefined Saprotroph\| | Yeast | NULL | Probable | As nectar-tap_saprotroph (Põlme et al. 2020); Decay sugar-rich substrates (Põlme et al. 2020) | Põlme S, et al. 2020. Fungal Diversity 105:1-16 ((https://doi.org/10.1007/s13225-020-00466-2)) |
| Otu36136 | 1 | k__Fungi(100);p__Ascomycota(100);c__Saccharomycetes(100);o__Saccharomycetales(100);f__Metschnikowiaceae(100);g__Metschnikowia(100);s__Metschnikowia_pulcherrima(100); | Metschnikowia | 13 | Saprotroph | \|Undefined Saprotroph\| | Yeast | NULL | Probable | As nectar-tap_saprotroph (Põlme et al. 2020); Decay sugar-rich substrates (Põlme et al. 2020) | Põlme S, et al. 2020. Fungal Diversity 105:1-16 ((https://doi.org/10.1007/s13225-020-00466-2)) |
| Otu36137 | 1 | k__Fungi(100);p__Ascomycota(100);c__Saccharomycetes(100);o__Saccharomycetales(100);f__Metschnikowiaceae(100);g__Metschnikowia(100);s__Metschnikowia_pulcherrima(100); | Metschnikowia | 13 | Saprotroph | \|Undefined Saprotroph\| | Yeast | NULL | Probable | As nectar-tap_saprotroph (Põlme et al. 2020); Decay sugar-rich substrates (Põlme et al. 2020) | Põlme S, et al. 2020. Fungal Diversity 105:1-16 ((https://doi.org/10.1007/s13225-020-00466-2)) |
| Otu36138 | 1 | k__Fungi(100);p__Ascomycota(100);c__Saccharomycetes(100);o__Saccharomycetales(100);f__Metschnikowiaceae(100);g__Metschnikowia(100);s__Metschnikowia_pulcherrima(100); | Metschnikowia | 13 | Saprotroph | \|Undefined Saprotroph\| | Yeast | NULL | Probable | As nectar-tap_saprotroph (Põlme et al. 2020); Decay sugar-rich substrates (Põlme et al. 2020) | Põlme S, et al. 2020. Fungal Diversity 105:1-16 ((https://doi.org/10.1007/s13225-020-00466-2)) |
| Otu36139 | 1 | k__Fungi(100);p__Ascomycota(100);c__Saccharomycetes(100);o__Saccharomycetales(100);f__Metschnikowiaceae(100);g__Metschnikowia(100);s__Metschnikowia_pulcherrima(100); | Metschnikowia | 13 | Saprotroph | \|Undefined Saprotroph\| | Yeast | NULL | Probable | As nectar-tap_saprotroph (Põlme et al. 2020); Decay sugar-rich substrates (Põlme et al. 2020) | Põlme S, et al. 2020. Fungal Diversity 105:1-16 ((https://doi.org/10.1007/s13225-020-00466-2)) |
| Otu36140 | 1 | k__Fungi(100);p__Ascomycota(100);c__Saccharomycetes(100);o__Saccharomycetales(100);f__Metschnikowiaceae(100);g__Metschnikowia(100);s__Metschnikowia_pulcherrima(100); | Metschnikowia | 13 | Saprotroph | \|Undefined Saprotroph\| | Yeast | NULL | Probable | As nectar-tap_saprotroph (Põlme et al. 2020); Decay sugar-rich substrates (Põlme et al. 2020) | Põlme S, et al. 2020. Fungal Diversity 105:1-16 ((https://doi.org/10.1007/s13225-020-00466-2)) |
| Otu36141 | 1 | k__Fungi(100);p__Ascomycota(100);c__Saccharomycetes(100);o__Saccharomycetales(100);f__Metschnikowiaceae(100);g__Metschnikowia(100);s__Metschnikowia_pulcherrima(100); | Metschnikowia | 13 | Saprotroph | \|Undefined Saprotroph\| | Yeast | NULL | Probable | As nectar-tap_saprotroph (Põlme et al. 2020); Decay sugar-rich substrates (Põlme et al. 2020) | Põlme S, et al. 2020. Fungal Diversity 105:1-16 ((https://doi.org/10.1007/s13225-020-00466-2)) |
| Otu36142 | 1 | k__Fungi(100);p__Ascomycota(100);c__Saccharomycetes(100);o__Saccharomycetales(100);f__Metschnikowiaceae(100);g__Metschnikowia(100);s__Metschnikowia_pulcherrima(100); | Metschnikowia | 13 | Saprotroph | \|Undefined Saprotroph\| | Yeast | NULL | Probable | As nectar-tap_saprotroph (Põlme et al. 2020); Decay sugar-rich substrates (Põlme et al. 2020) | Põlme S, et al. 2020. Fungal Diversity 105:1-16 ((https://doi.org/10.1007/s13225-020-00466-2)) |
| Otu36143 | 1 | k__Fungi(100);p__Basidiomycota(100);c__Microbotryomycetes(100);o__Sporidiobolales(100);f__Sporidiobolales_family_Incertae_sedis(100);g__Sporobolomyces(100);s__Sporobolomyces_lactosus(100); | Sporobolomyces | 13 | Pathotroph-Saprotroph | \|Fungal Parasite\|-Plant Saprotroph | Yeast | NULL | Probable | As litter_saprotroph (Põlme et al. 2020); As mycoparasite (Põlme et al. 2020); Decay fungal material (Põlme et al. 2020) | Tedersoo L, et al. 2014. Science 346:e1256688 ((https://doi.org/10.1126/science.1256688)); Purahong W, et al. 2016. Molecular Ecology 25:4059-4074 ((https://doi.org/10.1111/mec.13739)); Põlme S, et al. 2020. Fungal Diversity 105:1-16 ((https://doi.org/10.1007/s13225-020-00466-2)) |
| Otu36144 | 1 | k__Fungi(100);p__Ascomycota(100);c__Saccharomycetes(100);o__Saccharomycetales(100);f__Metschnikowiaceae(100);g__Metschnikowia(100);g__Metschnikowia_unclassified(100); | Metschnikowia | 13 | Saprotroph | \|Undefined Saprotroph\| | Yeast | NULL | Probable | As nectar-tap_saprotroph (Põlme et al. 2020); Decay sugar-rich substrates (Põlme et al. 2020) | Põlme S, et al. 2020. Fungal Diversity 105:1-16 ((https://doi.org/10.1007/s13225-020-00466-2)) |
| Otu36145 | 1 | k__Fungi(100);p__Ascomycota(100);c__Saccharomycetes(100);o__Saccharomycetales(100);f__Metschnikowiaceae(100);g__Metschnikowia(100);s__Metschnikowia_pulcherrima(100); | Metschnikowia | 13 | Saprotroph | \|Undefined Saprotroph\| | Yeast | NULL | Probable | As nectar-tap_saprotroph (Põlme et al. 2020); Decay sugar-rich substrates (Põlme et al. 2020) | Põlme S, et al. 2020. Fungal Diversity 105:1-16 ((https://doi.org/10.1007/s13225-020-00466-2)) |
| Otu36146 | 1 | k__Fungi(100);p__Ascomycota(100);c__Saccharomycetes(100);o__Saccharomycetales(100);f__Saccharomycetales_family_Incertae_sedis(100);f__Saccharomycetales_family_Incertae_sedis_unclassified(100);f__Saccharomycetales_family_Incertae_sedis_unclassified(100); | Saccharomycetales | 7 | Saprotroph | Undefined Saprotroph | Yeast | NULL | Possible | NULL | Sterkenburg E, et al. 2015. New Phytologist 207:1145-1158 ((https://doi.org/10.1111/nph.13426)) |
| Otu36147 | 1 | k__Fungi(100);p__Ascomycota(100);c__Saccharomycetes(100);o__Saccharomycetales(100);f__Metschnikowiaceae(100);g__Metschnikowia(100);g__Metschnikowia_unclassified(100); | Metschnikowia | 13 | Saprotroph | \|Undefined Saprotroph\| | Yeast | NULL | Probable | As nectar-tap_saprotroph (Põlme et al. 2020); Decay sugar-rich substrates (Põlme et al. 2020) | Põlme S, et al. 2020. Fungal Diversity 105:1-16 ((https://doi.org/10.1007/s13225-020-00466-2)) |
| Otu36148 | 1 | k__Fungi(100);p__Ascomycota(100);c__Saccharomycetes(100);o__Saccharomycetales(100);f__Metschnikowiaceae(100);g__Metschnikowia(100);s__Metschnikowia_pulcherrima(100); | Metschnikowia | 13 | Saprotroph | \|Undefined Saprotroph\| | Yeast | NULL | Probable | As nectar-tap_saprotroph (Põlme et al. 2020); Decay sugar-rich substrates (Põlme et al. 2020) | Põlme S, et al. 2020. Fungal Diversity 105:1-16 ((https://doi.org/10.1007/s13225-020-00466-2)) |
| Otu36149 | 1 | k__Fungi(100);p__Ascomycota(100);c__Saccharomycetes(100);o__Saccharomycetales(100);f__Metschnikowiaceae(100);g__Metschnikowia(100);s__Metschnikowia_pulcherrima(100); | Metschnikowia | 13 | Saprotroph | \|Undefined Saprotroph\| | Yeast | NULL | Probable | As nectar-tap_saprotroph (Põlme et al. 2020); Decay sugar-rich substrates (Põlme et al. 2020) | Põlme S, et al. 2020. Fungal Diversity 105:1-16 ((https://doi.org/10.1007/s13225-020-00466-2)) |
| Otu36150 | 1 | k__Fungi(100);p__Ascomycota(100);c__Saccharomycetes(100);o__Saccharomycetales(100);f__Metschnikowiaceae(100);g__Metschnikowia(100);s__Metschnikowia_pulcherrima(100); | Metschnikowia | 13 | Saprotroph | \|Undefined Saprotroph\| | Yeast | NULL | Probable | As nectar-tap_saprotroph (Põlme et al. 2020); Decay sugar-rich substrates (Põlme et al. 2020) | Põlme S, et al. 2020. Fungal Diversity 105:1-16 ((https://doi.org/10.1007/s13225-020-00466-2)) |
| Otu36151 | 1 | k__Fungi(100);p__Ascomycota(100);c__Saccharomycetes(100);o__Saccharomycetales(100);f__Metschnikowiaceae(100);g__Metschnikowia(100);s__Metschnikowia_pulcherrima(100); | Metschnikowia | 13 | Saprotroph | \|Undefined Saprotroph\| | Yeast | NULL | Probable | As nectar-tap_saprotroph (Põlme et al. 2020); Decay sugar-rich substrates (Põlme et al. 2020) | Põlme S, et al. 2020. Fungal Diversity 105:1-16 ((https://doi.org/10.1007/s13225-020-00466-2)) |
| Otu36152 | 1 | k__Fungi(100);p__Ascomycota(100);c__Saccharomycetes(100);o__Saccharomycetales(100);f__Metschnikowiaceae(100);g__Metschnikowia(100);g__Metschnikowia_unclassified(100); | Metschnikowia | 13 | Saprotroph | \|Undefined Saprotroph\| | Yeast | NULL | Probable | As nectar-tap_saprotroph (Põlme et al. 2020); Decay sugar-rich substrates (Põlme et al. 2020) | Põlme S, et al. 2020. Fungal Diversity 105:1-16 ((https://doi.org/10.1007/s13225-020-00466-2)) |
| Otu36156 | 1 | k__Fungi(100);p__Ascomycota(100);c__Saccharomycetes(100);o__Saccharomycetales(100);f__Metschnikowiaceae(100);g__Metschnikowia(100);s__Metschnikowia_pulcherrima(100); | Metschnikowia | 13 | Saprotroph | \|Undefined Saprotroph\| | Yeast | NULL | Probable | As nectar-tap_saprotroph (Põlme et al. 2020); Decay sugar-rich substrates (Põlme et al. 2020) | Põlme S, et al. 2020. Fungal Diversity 105:1-16 ((https://doi.org/10.1007/s13225-020-00466-2)) |
| Otu36158 | 1 | k__Fungi(100);p__Ascomycota(100);c__Saccharomycetes(100);o__Saccharomycetales(100);f__Metschnikowiaceae(100);g__Metschnikowia(100);s__Metschnikowia_pulcherrima(100); | Metschnikowia | 13 | Saprotroph | \|Undefined Saprotroph\| | Yeast | NULL | Probable | As nectar-tap_saprotroph (Põlme et al. 2020); Decay sugar-rich substrates (Põlme et al. 2020) | Põlme S, et al. 2020. Fungal Diversity 105:1-16 ((https://doi.org/10.1007/s13225-020-00466-2)) |
| Otu36165 | 1 | k__Fungi(100);p__Ascomycota(100);c__Saccharomycetes(100);o__Saccharomycetales(100);f__Metschnikowiaceae(100);g__Metschnikowia(100);s__Metschnikowia_pulcherrima(100); | Metschnikowia | 13 | Saprotroph | \|Undefined Saprotroph\| | Yeast | NULL | Probable | As nectar-tap_saprotroph (Põlme et al. 2020); Decay sugar-rich substrates (Põlme et al. 2020) | Põlme S, et al. 2020. Fungal Diversity 105:1-16 ((https://doi.org/10.1007/s13225-020-00466-2)) |
| Otu36166 | 1 | k__Fungi(100);p__Ascomycota(100);c__Saccharomycetes(100);o__Saccharomycetales(100);f__Saccharomycetales_family_Incertae_sedis(100);g__Candida(100);s__Candida_cretensis(100); | Candida | 13 | Pathotroph-Saprotroph-Symbiotroph | Animal Pathogen-Endophyte-Endosymbiont-Epiphyte-\|Undefined Saprotroph\| | Dimorphic Yeast-Yeast | NULL | Probable | As nectar-tap_saprotroph (Põlme et al. 2020); Decay sugar-rich substrates (Põlme et al. 2020); This asexual genus is polyphyletic and contain species that function very differently depend on what they are. Caution should be taken when interpreting guilds for Candida species. | Manolakaki D, et al. 2010. Virulence. 1: 367-375 ((https://doi.org/10.4161/viru.1.5.12796)); Mekbib SB, et al. 2011. Tropical Plant Pathology 36:233-240 ((https://doi.org/10.1590/S1982-56762011000400004)); Põlme S, et al. 2020. Fungal Diversity 105:1-16 ((https://doi.org/10.1007/s13225-020-00466-2)) |
| Otu36167 | 1 | k__Fungi(100);p__Ascomycota(100);c__Saccharomycetes(100);o__Saccharomycetales(100);f__Metschnikowiaceae(100);g__Metschnikowia(100);g__Metschnikowia_unclassified(100); | Metschnikowia | 13 | Saprotroph | \|Undefined Saprotroph\| | Yeast | NULL | Probable | As nectar-tap_saprotroph (Põlme et al. 2020); Decay sugar-rich substrates (Põlme et al. 2020) | Põlme S, et al. 2020. Fungal Diversity 105:1-16 ((https://doi.org/10.1007/s13225-020-00466-2)) |
| Otu36170 | 1 | k__Fungi(100);p__Ascomycota(100);c__Saccharomycetes(100);o__Saccharomycetales(100);f__Metschnikowiaceae(100);g__Metschnikowia(100);s__Metschnikowia_pulcherrima(100); | Metschnikowia | 13 | Saprotroph | \|Undefined Saprotroph\| | Yeast | NULL | Probable | As nectar-tap_saprotroph (Põlme et al. 2020); Decay sugar-rich substrates (Põlme et al. 2020) | Põlme S, et al. 2020. Fungal Diversity 105:1-16 ((https://doi.org/10.1007/s13225-020-00466-2)) |
| Otu36171 | 1 | k__Fungi(100);p__Ascomycota(100);c__Saccharomycetes(100);o__Saccharomycetales(100);f__Metschnikowiaceae(100);g__Metschnikowia(100);s__Metschnikowia_pulcherrima(100); | Metschnikowia | 13 | Saprotroph | \|Undefined Saprotroph\| | Yeast | NULL | Probable | As nectar-tap_saprotroph (Põlme et al. 2020); Decay sugar-rich substrates (Põlme et al. 2020) | Põlme S, et al. 2020. Fungal Diversity 105:1-16 ((https://doi.org/10.1007/s13225-020-00466-2)) |
| Otu36172 | 1 | k__Fungi(100);p__Ascomycota(100);c__Saccharomycetes(100);o__Saccharomycetales(100);f__Metschnikowiaceae(100);g__Metschnikowia(100);s__Metschnikowia_pulcherrima(100); | Metschnikowia | 13 | Saprotroph | \|Undefined Saprotroph\| | Yeast | NULL | Probable | As nectar-tap_saprotroph (Põlme et al. 2020); Decay sugar-rich substrates (Põlme et al. 2020) | Põlme S, et al. 2020. Fungal Diversity 105:1-16 ((https://doi.org/10.1007/s13225-020-00466-2)) |
| Otu36173 | 1 | k__Fungi(100);p__Ascomycota(100);c__Saccharomycetes(100);o__Saccharomycetales(100);f__Metschnikowiaceae(100);g__Metschnikowia(100);s__Metschnikowia_pulcherrima(100); | Metschnikowia | 13 | Saprotroph | \|Undefined Saprotroph\| | Yeast | NULL | Probable | As nectar-tap_saprotroph (Põlme et al. 2020); Decay sugar-rich substrates (Põlme et al. 2020) | Põlme S, et al. 2020. Fungal Diversity 105:1-16 ((https://doi.org/10.1007/s13225-020-00466-2)) |
| Otu36175 | 1 | k__Fungi(100);p__Ascomycota(100);c__Saccharomycetes(100);o__Saccharomycetales(100);f__Metschnikowiaceae(100);g__Metschnikowia(100);g__Metschnikowia_unclassified(100); | Metschnikowia | 13 | Saprotroph | \|Undefined Saprotroph\| | Yeast | NULL | Probable | As nectar-tap_saprotroph (Põlme et al. 2020); Decay sugar-rich substrates (Põlme et al. 2020) | Põlme S, et al. 2020. Fungal Diversity 105:1-16 ((https://doi.org/10.1007/s13225-020-00466-2)) |
| Otu36176 | 1 | k__Fungi(100);p__Ascomycota(100);c__Saccharomycetes(100);o__Saccharomycetales(100);f__Saccharomycetales_family_Incertae_sedis(100);g__Candida(100);g__Candida_unclassified(100); | Candida | 13 | Pathotroph-Saprotroph-Symbiotroph | Animal Pathogen-Endophyte-Endosymbiont-Epiphyte-\|Undefined Saprotroph\| | Dimorphic Yeast-Yeast | NULL | Probable | As nectar-tap_saprotroph (Põlme et al. 2020); Decay sugar-rich substrates (Põlme et al. 2020); This asexual genus is polyphyletic and contain species that function very differently depend on what they are. Caution should be taken when interpreting guilds for Candida species. | Manolakaki D, et al. 2010. Virulence. 1: 367-375 ((https://doi.org/10.4161/viru.1.5.12796)); Mekbib SB, et al. 2011. Tropical Plant Pathology 36:233-240 ((https://doi.org/10.1590/S1982-56762011000400004)); Põlme S, et al. 2020. Fungal Diversity 105:1-16 ((https://doi.org/10.1007/s13225-020-00466-2)) |
| Otu36177 | 1 | k__Fungi(100);p__Ascomycota(100);c__Saccharomycetes(100);o__Saccharomycetales(100);f__Metschnikowiaceae(100);g__Metschnikowia(100);s__Metschnikowia_pulcherrima(100); | Metschnikowia | 13 | Saprotroph | \|Undefined Saprotroph\| | Yeast | NULL | Probable | As nectar-tap_saprotroph (Põlme et al. 2020); Decay sugar-rich substrates (Põlme et al. 2020) | Põlme S, et al. 2020. Fungal Diversity 105:1-16 ((https://doi.org/10.1007/s13225-020-00466-2)) |
| Otu36178 | 1 | k__Fungi(100);p__Ascomycota(100);c__Saccharomycetes(100);o__Saccharomycetales(100);f__Metschnikowiaceae(100);g__Metschnikowia(100);s__Metschnikowia_pulcherrima(100); | Metschnikowia | 13 | Saprotroph | \|Undefined Saprotroph\| | Yeast | NULL | Probable | As nectar-tap_saprotroph (Põlme et al. 2020); Decay sugar-rich substrates (Põlme et al. 2020) | Põlme S, et al. 2020. Fungal Diversity 105:1-16 ((https://doi.org/10.1007/s13225-020-00466-2)) |
| Otu36179 | 1 | k__Fungi(100);p__Ascomycota(100);c__Saccharomycetes(100);o__Saccharomycetales(100);f__Metschnikowiaceae(100);g__Metschnikowia(100);s__Metschnikowia_pulcherrima(100); | Metschnikowia | 13 | Saprotroph | \|Undefined Saprotroph\| | Yeast | NULL | Probable | As nectar-tap_saprotroph (Põlme et al. 2020); Decay sugar-rich substrates (Põlme et al. 2020) | Põlme S, et al. 2020. Fungal Diversity 105:1-16 ((https://doi.org/10.1007/s13225-020-00466-2)) |
| Otu36180 | 1 | k__Fungi(100);p__Ascomycota(100);c__Saccharomycetes(100);o__Saccharomycetales(100);f__Metschnikowiaceae(100);g__Metschnikowia(100);g__Metschnikowia_unclassified(100); | Metschnikowia | 13 | Saprotroph | \|Undefined Saprotroph\| | Yeast | NULL | Probable | As nectar-tap_saprotroph (Põlme et al. 2020); Decay sugar-rich substrates (Põlme et al. 2020) | Põlme S, et al. 2020. Fungal Diversity 105:1-16 ((https://doi.org/10.1007/s13225-020-00466-2)) |
| Otu36181 | 1 | k__Fungi(100);p__Ascomycota(100);c__Saccharomycetes(100);o__Saccharomycetales(100);f__Metschnikowiaceae(100);g__Metschnikowia(100);s__Metschnikowia_pulcherrima(100); | Metschnikowia | 13 | Saprotroph | \|Undefined Saprotroph\| | Yeast | NULL | Probable | As nectar-tap_saprotroph (Põlme et al. 2020); Decay sugar-rich substrates (Põlme et al. 2020) | Põlme S, et al. 2020. Fungal Diversity 105:1-16 ((https://doi.org/10.1007/s13225-020-00466-2)) |
| Otu36182 | 1 | k__Fungi(100);p__Ascomycota(100);c__Saccharomycetes(100);o__Saccharomycetales(100);f__Metschnikowiaceae(100);g__Metschnikowia(100);s__Metschnikowia_pulcherrima(100); | Metschnikowia | 13 | Saprotroph | \|Undefined Saprotroph\| | Yeast | NULL | Probable | As nectar-tap_saprotroph (Põlme et al. 2020); Decay sugar-rich substrates (Põlme et al. 2020) | Põlme S, et al. 2020. Fungal Diversity 105:1-16 ((https://doi.org/10.1007/s13225-020-00466-2)) |
| Otu36183 | 1 | k__Fungi(100);p__Ascomycota(100);c__Saccharomycetes(100);o__Saccharomycetales(100);f__Metschnikowiaceae(100);g__Metschnikowia(100);g__Metschnikowia_unclassified(100); | Metschnikowia | 13 | Saprotroph | \|Undefined Saprotroph\| | Yeast | NULL | Probable | As nectar-tap_saprotroph (Põlme et al. 2020); Decay sugar-rich substrates (Põlme et al. 2020) | Põlme S, et al. 2020. Fungal Diversity 105:1-16 ((https://doi.org/10.1007/s13225-020-00466-2)) |
| Otu36184 | 1 | k__Fungi(100);p__Ascomycota(100);c__Saccharomycetes(100);o__Saccharomycetales(100);f__Metschnikowiaceae(100);g__Metschnikowia(100);s__Metschnikowia_pulcherrima(100); | Metschnikowia | 13 | Saprotroph | \|Undefined Saprotroph\| | Yeast | NULL | Probable | As nectar-tap_saprotroph (Põlme et al. 2020); Decay sugar-rich substrates (Põlme et al. 2020) | Põlme S, et al. 2020. Fungal Diversity 105:1-16 ((https://doi.org/10.1007/s13225-020-00466-2)) |
| Otu36185 | 1 | k__Fungi(100);p__Ascomycota(100);c__Saccharomycetes(100);o__Saccharomycetales(100);f__Metschnikowiaceae(100);g__Metschnikowia(100);s__Metschnikowia_pulcherrima(100); | Metschnikowia | 13 | Saprotroph | \|Undefined Saprotroph\| | Yeast | NULL | Probable | As nectar-tap_saprotroph (Põlme et al. 2020); Decay sugar-rich substrates (Põlme et al. 2020) | Põlme S, et al. 2020. Fungal Diversity 105:1-16 ((https://doi.org/10.1007/s13225-020-00466-2)) |
| Otu36186 | 1 | k__Fungi(100);p__Ascomycota(100);c__Saccharomycetes(100);o__Saccharomycetales(100);f__Metschnikowiaceae(100);g__Metschnikowia(100);s__Metschnikowia_pulcherrima(100); | Metschnikowia | 13 | Saprotroph | \|Undefined Saprotroph\| | Yeast | NULL | Probable | As nectar-tap_saprotroph (Põlme et al. 2020); Decay sugar-rich substrates (Põlme et al. 2020) | Põlme S, et al. 2020. Fungal Diversity 105:1-16 ((https://doi.org/10.1007/s13225-020-00466-2)) |
| Otu36187 | 1 | k__Fungi(100);p__Ascomycota(100);c__Saccharomycetes(100);o__Saccharomycetales(100);f__Metschnikowiaceae(100);g__Metschnikowia(100);g__Metschnikowia_unclassified(100); | Metschnikowia | 13 | Saprotroph | \|Undefined Saprotroph\| | Yeast | NULL | Probable | As nectar-tap_saprotroph (Põlme et al. 2020); Decay sugar-rich substrates (Põlme et al. 2020) | Põlme S, et al. 2020. Fungal Diversity 105:1-16 ((https://doi.org/10.1007/s13225-020-00466-2)) |
| Otu36188 | 1 | k__Fungi(100);p__Ascomycota(100);c__Saccharomycetes(100);o__Saccharomycetales(100);f__Metschnikowiaceae(100);g__Metschnikowia(100);s__Metschnikowia_pulcherrima(100); | Metschnikowia | 13 | Saprotroph | \|Undefined Saprotroph\| | Yeast | NULL | Probable | As nectar-tap_saprotroph (Põlme et al. 2020); Decay sugar-rich substrates (Põlme et al. 2020) | Põlme S, et al. 2020. Fungal Diversity 105:1-16 ((https://doi.org/10.1007/s13225-020-00466-2)) |
| Otu36189 | 1 | k__Fungi(100);p__Ascomycota(100);c__Saccharomycetes(100);o__Saccharomycetales(100);f__Metschnikowiaceae(100);g__Metschnikowia(100);s__Metschnikowia_pulcherrima(100); | Metschnikowia | 13 | Saprotroph | \|Undefined Saprotroph\| | Yeast | NULL | Probable | As nectar-tap_saprotroph (Põlme et al. 2020); Decay sugar-rich substrates (Põlme et al. 2020) | Põlme S, et al. 2020. Fungal Diversity 105:1-16 ((https://doi.org/10.1007/s13225-020-00466-2)) |
| Otu36191 | 1 | k__Fungi(100);p__Ascomycota(100);c__Saccharomycetes(100);o__Saccharomycetales(100);f__Metschnikowiaceae(100);g__Metschnikowia(100);s__Metschnikowia_pulcherrima(100); | Metschnikowia | 13 | Saprotroph | \|Undefined Saprotroph\| | Yeast | NULL | Probable | As nectar-tap_saprotroph (Põlme et al. 2020); Decay sugar-rich substrates (Põlme et al. 2020) | Põlme S, et al. 2020. Fungal Diversity 105:1-16 ((https://doi.org/10.1007/s13225-020-00466-2)) |
| Otu36195 | 1 | k__Fungi(100);p__Ascomycota(100);c__Saccharomycetes(100);o__Saccharomycetales(100);f__Metschnikowiaceae(100);g__Metschnikowia(100);g__Metschnikowia_unclassified(100); | Metschnikowia | 13 | Saprotroph | \|Undefined Saprotroph\| | Yeast | NULL | Probable | As nectar-tap_saprotroph (Põlme et al. 2020); Decay sugar-rich substrates (Põlme et al. 2020) | Põlme S, et al. 2020. Fungal Diversity 105:1-16 ((https://doi.org/10.1007/s13225-020-00466-2)) |
| Otu36197 | 1 | k__Fungi(100);p__Ascomycota(100);c__Saccharomycetes(100);o__Saccharomycetales(100);f__Metschnikowiaceae(100);g__Metschnikowia(100);g__Metschnikowia_unclassified(100); | Metschnikowia | 13 | Saprotroph | \|Undefined Saprotroph\| | Yeast | NULL | Probable | As nectar-tap_saprotroph (Põlme et al. 2020); Decay sugar-rich substrates (Põlme et al. 2020) | Põlme S, et al. 2020. Fungal Diversity 105:1-16 ((https://doi.org/10.1007/s13225-020-00466-2)) |
| Otu36200 | 1 | k__Fungi(100);p__Ascomycota(100);c__Saccharomycetes(100);o__Saccharomycetales(100);f__Metschnikowiaceae(100);g__Metschnikowia(100);s__Metschnikowia_pulcherrima(100); | Metschnikowia | 13 | Saprotroph | \|Undefined Saprotroph\| | Yeast | NULL | Probable | As nectar-tap_saprotroph (Põlme et al. 2020); Decay sugar-rich substrates (Põlme et al. 2020) | Põlme S, et al. 2020. Fungal Diversity 105:1-16 ((https://doi.org/10.1007/s13225-020-00466-2)) |
| Otu36201 | 1 | k__Fungi(100);p__Ascomycota(100);c__Saccharomycetes(100);o__Saccharomycetales(100);f__Metschnikowiaceae(100);g__Metschnikowia(100);s__Metschnikowia_pulcherrima(100); | Metschnikowia | 13 | Saprotroph | \|Undefined Saprotroph\| | Yeast | NULL | Probable | As nectar-tap_saprotroph (Põlme et al. 2020); Decay sugar-rich substrates (Põlme et al. 2020) | Põlme S, et al. 2020. Fungal Diversity 105:1-16 ((https://doi.org/10.1007/s13225-020-00466-2)) |
| Otu36203 | 1 | k__Fungi(100);p__Ascomycota(100);c__Saccharomycetes(100);o__Saccharomycetales(100);f__Metschnikowiaceae(100);g__Metschnikowia(100);s__Metschnikowia_pulcherrima(100); | Metschnikowia | 13 | Saprotroph | \|Undefined Saprotroph\| | Yeast | NULL | Probable | As nectar-tap_saprotroph (Põlme et al. 2020); Decay sugar-rich substrates (Põlme et al. 2020) | Põlme S, et al. 2020. Fungal Diversity 105:1-16 ((https://doi.org/10.1007/s13225-020-00466-2)) |
| Otu36204 | 1 | k__Fungi(100);p__Ascomycota(100);c__Saccharomycetes(100);o__Saccharomycetales(100);f__Metschnikowiaceae(100);g__Metschnikowia(100);s__Metschnikowia_pulcherrima(100); | Metschnikowia | 13 | Saprotroph | \|Undefined Saprotroph\| | Yeast | NULL | Probable | As nectar-tap_saprotroph (Põlme et al. 2020); Decay sugar-rich substrates (Põlme et al. 2020) | Põlme S, et al. 2020. Fungal Diversity 105:1-16 ((https://doi.org/10.1007/s13225-020-00466-2)) |
| Otu36205 | 1 | k__Fungi(100);p__Ascomycota(100);c__Saccharomycetes(100);o__Saccharomycetales(100);f__Metschnikowiaceae(100);g__Metschnikowia(100);s__Metschnikowia_pulcherrima(100); | Metschnikowia | 13 | Saprotroph | \|Undefined Saprotroph\| | Yeast | NULL | Probable | As nectar-tap_saprotroph (Põlme et al. 2020); Decay sugar-rich substrates (Põlme et al. 2020) | Põlme S, et al. 2020. Fungal Diversity 105:1-16 ((https://doi.org/10.1007/s13225-020-00466-2)) |
| Otu36206 | 1 | k__Fungi(100);p__Ascomycota(100);c__Saccharomycetes(100);o__Saccharomycetales(100);f__Metschnikowiaceae(100);g__Metschnikowia(100);s__Metschnikowia_pulcherrima(100); | Metschnikowia | 13 | Saprotroph | \|Undefined Saprotroph\| | Yeast | NULL | Probable | As nectar-tap_saprotroph (Põlme et al. 2020); Decay sugar-rich substrates (Põlme et al. 2020) | Põlme S, et al. 2020. Fungal Diversity 105:1-16 ((https://doi.org/10.1007/s13225-020-00466-2)) |
| Otu36207 | 1 | k__Fungi(100);p__Ascomycota(100);c__Saccharomycetes(100);o__Saccharomycetales(100);f__Metschnikowiaceae(100);g__Metschnikowia(100);s__Metschnikowia_pulcherrima(100); | Metschnikowia | 13 | Saprotroph | \|Undefined Saprotroph\| | Yeast | NULL | Probable | As nectar-tap_saprotroph (Põlme et al. 2020); Decay sugar-rich substrates (Põlme et al. 2020) | Põlme S, et al. 2020. Fungal Diversity 105:1-16 ((https://doi.org/10.1007/s13225-020-00466-2)) |
| Otu36210 | 1 | k__Fungi(100);p__Ascomycota(100);c__Saccharomycetes(100);o__Saccharomycetales(100);f__Metschnikowiaceae(100);g__Metschnikowia(100);g__Metschnikowia_unclassified(100); | Metschnikowia | 13 | Saprotroph | \|Undefined Saprotroph\| | Yeast | NULL | Probable | As nectar-tap_saprotroph (Põlme et al. 2020); Decay sugar-rich substrates (Põlme et al. 2020) | Põlme S, et al. 2020. Fungal Diversity 105:1-16 ((https://doi.org/10.1007/s13225-020-00466-2)) |
| Otu36211 | 1 | k__Fungi(100);p__Ascomycota(100);c__Saccharomycetes(100);o__Saccharomycetales(100);f__Metschnikowiaceae(100);g__Metschnikowia(100);s__Metschnikowia_pulcherrima(100); | Metschnikowia | 13 | Saprotroph | \|Undefined Saprotroph\| | Yeast | NULL | Probable | As nectar-tap_saprotroph (Põlme et al. 2020); Decay sugar-rich substrates (Põlme et al. 2020) | Põlme S, et al. 2020. Fungal Diversity 105:1-16 ((https://doi.org/10.1007/s13225-020-00466-2)) |
| Otu36212 | 1 | k__Fungi(100);p__Ascomycota(100);c__Saccharomycetes(100);o__Saccharomycetales(100);f__Metschnikowiaceae(100);g__Metschnikowia(100);s__Metschnikowia_pulcherrima(100); | Metschnikowia | 13 | Saprotroph | \|Undefined Saprotroph\| | Yeast | NULL | Probable | As nectar-tap_saprotroph (Põlme et al. 2020); Decay sugar-rich substrates (Põlme et al. 2020) | Põlme S, et al. 2020. Fungal Diversity 105:1-16 ((https://doi.org/10.1007/s13225-020-00466-2)) |
| Otu36214 | 1 | k__Fungi(100);p__Ascomycota(100);c__Saccharomycetes(100);o__Saccharomycetales(100);f__Metschnikowiaceae(100);g__Metschnikowia(100);s__Metschnikowia_pulcherrima(100); | Metschnikowia | 13 | Saprotroph | \|Undefined Saprotroph\| | Yeast | NULL | Probable | As nectar-tap_saprotroph (Põlme et al. 2020); Decay sugar-rich substrates (Põlme et al. 2020) | Põlme S, et al. 2020. Fungal Diversity 105:1-16 ((https://doi.org/10.1007/s13225-020-00466-2)) |
| Otu36215 | 1 | k__Fungi(100);p__Ascomycota(100);c__Saccharomycetes(100);o__Saccharomycetales(100);f__Metschnikowiaceae(100);g__Metschnikowia(100);s__Metschnikowia_pulcherrima(100); | Metschnikowia | 13 | Saprotroph | \|Undefined Saprotroph\| | Yeast | NULL | Probable | As nectar-tap_saprotroph (Põlme et al. 2020); Decay sugar-rich substrates (Põlme et al. 2020) | Põlme S, et al. 2020. Fungal Diversity 105:1-16 ((https://doi.org/10.1007/s13225-020-00466-2)) |
| Otu36216 | 1 | k__Fungi(100);p__Ascomycota(100);c__Saccharomycetes(100);o__Saccharomycetales(100);f__Metschnikowiaceae(100);g__Metschnikowia(100);s__Metschnikowia_pulcherrima(100); | Metschnikowia | 13 | Saprotroph | \|Undefined Saprotroph\| | Yeast | NULL | Probable | As nectar-tap_saprotroph (Põlme et al. 2020); Decay sugar-rich substrates (Põlme et al. 2020) | Põlme S, et al. 2020. Fungal Diversity 105:1-16 ((https://doi.org/10.1007/s13225-020-00466-2)) |
| Otu36219 | 1 | k__Fungi(100);p__Ascomycota(100);c__Saccharomycetes(100);o__Saccharomycetales(100);f__Metschnikowiaceae(100);g__Metschnikowia(100);s__Metschnikowia_pulcherrima(100); | Metschnikowia | 13 | Saprotroph | \|Undefined Saprotroph\| | Yeast | NULL | Probable | As nectar-tap_saprotroph (Põlme et al. 2020); Decay sugar-rich substrates (Põlme et al. 2020) | Põlme S, et al. 2020. Fungal Diversity 105:1-16 ((https://doi.org/10.1007/s13225-020-00466-2)) |
| Otu36225 | 1 | k__Fungi(100);p__Ascomycota(100);c__Saccharomycetes(100);o__Saccharomycetales(100);f__Metschnikowiaceae(100);g__Metschnikowia(100);g__Metschnikowia_unclassified(100); | Metschnikowia | 13 | Saprotroph | \|Undefined Saprotroph\| | Yeast | NULL | Probable | As nectar-tap_saprotroph (Põlme et al. 2020); Decay sugar-rich substrates (Põlme et al. 2020) | Põlme S, et al. 2020. Fungal Diversity 105:1-16 ((https://doi.org/10.1007/s13225-020-00466-2)) |
| Otu36227 | 1 | k__Fungi(100);p__Ascomycota(100);c__Saccharomycetes(100);o__Saccharomycetales(100);f__Metschnikowiaceae(100);g__Metschnikowia(100);s__Metschnikowia_pulcherrima(100); | Metschnikowia | 13 | Saprotroph | \|Undefined Saprotroph\| | Yeast | NULL | Probable | As nectar-tap_saprotroph (Põlme et al. 2020); Decay sugar-rich substrates (Põlme et al. 2020) | Põlme S, et al. 2020. Fungal Diversity 105:1-16 ((https://doi.org/10.1007/s13225-020-00466-2)) |
| Otu36228 | 1 | k__Fungi(100);p__Ascomycota(100);c__Saccharomycetes(100);o__Saccharomycetales(100);f__Metschnikowiaceae(100);g__Metschnikowia(100);s__Metschnikowia_pulcherrima(100); | Metschnikowia | 13 | Saprotroph | \|Undefined Saprotroph\| | Yeast | NULL | Probable | As nectar-tap_saprotroph (Põlme et al. 2020); Decay sugar-rich substrates (Põlme et al. 2020) | Põlme S, et al. 2020. Fungal Diversity 105:1-16 ((https://doi.org/10.1007/s13225-020-00466-2)) |
| Otu36229 | 1 | k__Fungi(100);p__Ascomycota(100);c__Saccharomycetes(100);o__Saccharomycetales(100);f__Metschnikowiaceae(100);g__Metschnikowia(100);s__Metschnikowia_pulcherrima(100); | Metschnikowia | 13 | Saprotroph | \|Undefined Saprotroph\| | Yeast | NULL | Probable | As nectar-tap_saprotroph (Põlme et al. 2020); Decay sugar-rich substrates (Põlme et al. 2020) | Põlme S, et al. 2020. Fungal Diversity 105:1-16 ((https://doi.org/10.1007/s13225-020-00466-2)) |
| Otu36232 | 1 | k__Fungi(100);p__Ascomycota(100);c__Saccharomycetes(100);o__Saccharomycetales(100);f__Metschnikowiaceae(100);g__Metschnikowia(100);s__Metschnikowia_pulcherrima(100); | Metschnikowia | 13 | Saprotroph | \|Undefined Saprotroph\| | Yeast | NULL | Probable | As nectar-tap_saprotroph (Põlme et al. 2020); Decay sugar-rich substrates (Põlme et al. 2020) | Põlme S, et al. 2020. Fungal Diversity 105:1-16 ((https://doi.org/10.1007/s13225-020-00466-2)) |
| Otu36233 | 1 | k__Fungi(100);p__Ascomycota(100);c__Saccharomycetes(100);o__Saccharomycetales(100);f__Saccharomycetales_family_Incertae_sedis(100);f__Saccharomycetales_family_Incertae_sedis_unclassified(100);f__Saccharomycetales_family_Incertae_sedis_unclassified(100); | Saccharomycetales | 7 | Saprotroph | Undefined Saprotroph | Yeast | NULL | Possible | NULL | Sterkenburg E, et al. 2015. New Phytologist 207:1145-1158 ((https://doi.org/10.1111/nph.13426)) |
| Otu36234 | 1 | k__Fungi(100);p__Ascomycota(100);c__Saccharomycetes(100);o__Saccharomycetales(100);f__Metschnikowiaceae(100);g__Metschnikowia(100);g__Metschnikowia_unclassified(100); | Metschnikowia | 13 | Saprotroph | \|Undefined Saprotroph\| | Yeast | NULL | Probable | As nectar-tap_saprotroph (Põlme et al. 2020); Decay sugar-rich substrates (Põlme et al. 2020) | Põlme S, et al. 2020. Fungal Diversity 105:1-16 ((https://doi.org/10.1007/s13225-020-00466-2)) |

| | | | | | | | | | | | |
|---|---|---|---|---|---|---|---|---|---|---|---|
| Otu36235 | 1 | k__Fungi(100);p__Ascomycota(100);c__Saccharomycetes(100);o__Saccharomycetales(100);f__Metschnikowiaceae(100);g__Metschnikowia(100);s__Metschnikowia_pulcherrima(100); | Metschnikowia | 13 | Saprotroph | \|Undefined Saprotroph\| | Yeast | NULL | Probable | As nectar-tap_saprotroph (Põlme et al. 2020); Decay sugar-rich substrates (Põlme et al. 2020) | Põlme S, et al. 2020. Fungal Diversity 105:1-16 ((https://doi.org/10.1007/s13225-020-00466-2)) |
| Otu36236 | 1 | k__Fungi(100);p__Ascomycota(100);c__Saccharomycetes(100);o__Saccharomycetales(100);f__Metschnikowiaceae(100);g__Metschnikowia(100);g__Metschnikowia_unclassified(100); | Metschnikowia | 13 | Saprotroph | \|Undefined Saprotroph\| | Yeast | NULL | Probable | As nectar-tap_saprotroph (Põlme et al. 2020); Decay sugar-rich substrates (Põlme et al. 2020) | Põlme S, et al. 2020. Fungal Diversity 105:1-16 ((https://doi.org/10.1007/s13225-020-00466-2)) |
| Otu36237 | 1 | k__Fungi(100);p__Ascomycota(100);c__Dothideomycetes(100);o__Capnodiales(100);f__Davidiellaceae(100);g__Cladosporium(100);g__Cladosporium_unclassified(100); | Cladosporium | 13 | Pathotroph-Saprotroph-Symbiotroph | Animal Pathogen-Endophyte-Lichen Parasite-Plant Pathogen-\|Plant Saprotroph\|-Wood Saprotroph | Microfungus | NULL | Possible | As foliar_endophyte (Põlme et al. 2020); As litter_saprotroph (Põlme et al. 2020); Decay wood, leaf, fruit, seed, soil, dung, animal material (Põlme et al. 2020) | Seehann G, et al. 1975. List of Fungi in Soft-Rot Tissue Tests. Institute fur Holzbiologie und Holzschutz, Bundesforschungsanstalt fur Forst - und Holzwirtshaft, Hamburg - Lohbrugge, Germany; Lawrey JD, Diederich P. 2016. http://www.lichenicolous.net (accessed 24 May 2017) ((http://www.lichenicolous.net)); Põlme S, et al. 2020. Fungal Diversity 105:1-16 ((https://doi.org/10.1007/s13225-020-00466-2)); The American Phytopathological Society Common Names of Plant Diseases ((http://www.apsnet.org/publications/commonnames/Pages/default.aspx)) |
| Otu36238 | 1 | k__Fungi(100);p__Ascomycota(100);c__Saccharomycetes(100);o__Saccharomycetales(100);f__Metschnikowiaceae(100);g__Metschnikowia(100);g__Metschnikowia_unclassified(100); | Metschnikowia | 13 | Saprotroph | \|Undefined Saprotroph\| | Yeast | NULL | Probable | As nectar-tap_saprotroph (Põlme et al. 2020); Decay sugar-rich substrates (Põlme et al. 2020) | Põlme S, et al. 2020. Fungal Diversity 105:1-16 ((https://doi.org/10.1007/s13225-020-00466-2)) |
| Otu36239 | 1 | k__Fungi(100);p__Ascomycota(100);c__Dothideomycetes(100);o__Capnodiales(100);f__Davidiellaceae(100);g__Cladosporium(100);g__Cladosporium_unclassified(100); | Cladosporium | 13 | Pathotroph-Saprotroph-Symbiotroph | Animal Pathogen-Endophyte-Lichen Parasite-Plant Pathogen-\|Plant Saprotroph\|-Wood Saprotroph | Microfungus | NULL | Possible | As foliar_endophyte (Põlme et al. 2020); As litter_saprotroph (Põlme et al. 2020); Decay wood, leaf, fruit, seed, soil, dung, animal material (Põlme et al. 2020) | Seehann G, et al. 1975. List of Fungi in Soft-Rot Tissue Tests. Institute fur Holzbiologie und Holzschutz, Bundesforschungsanstalt fur Forst - und Holzwirtshaft, Hamburg - Lohbrugge, Germany; Lawrey JD, Diederich P. 2016. http://www.lichenicolous.net (accessed 24 May 2017) ((http://www.lichenicolous.net)); Põlme S, et al. 2020. Fungal Diversity 105:1-16 ((https://doi.org/10.1007/s13225-020-00466-2)); The American Phytopathological Society Common Names of Plant Diseases ((http://www.apsnet.org/publications/commonnames/Pages/default.aspx)) |
| Otu36257 | 1 | k__Fungi(100);p__Ascomycota(100);c__Saccharomycetes(100);o__Saccharomycetales(100);f__Metschnikowiaceae(100);g__Metschnikowia(100);s__Metschnikowia_pulcherrima(100); | Metschnikowia | 13 | Saprotroph | \|Undefined Saprotroph\| | Yeast | NULL | Probable | As nectar-tap_saprotroph (Põlme et al. 2020); Decay sugar-rich substrates (Põlme et al. 2020) | Põlme S, et al. 2020. Fungal Diversity 105:1-16 ((https://doi.org/10.1007/s13225-020-00466-2)) |
| Otu36259 | 1 | k__Fungi(100);p__Zygomycota(100);c__Zygomycota_class_Incertae_sedis(100);o__Mucorales(100);f__Umbelopsidaceae(100);g__Umbelopsis(100);s__Umbelopsis_ramanniana(100); | Umbelopsis | 13 | Saprotroph | \|Undefined Saprotroph\| | Microfungus | NULL | Probable | As soil_saprotroph (Põlme et al. 2020); Decay roots, soil (Põlme et al. 2020) | Tedersoo L, et al. 2014. Science 346:e1256688 ((https://doi.org/10.1126/science.1256688)); Põlme S, et al. 2020. Fungal Diversity 105:1-16 ((https://doi.org/10.1007/s13225-020-00466-2)) |
| Otu36261 | 1 | k__Fungi(100);p__Ascomycota(100);c__Saccharomycetes(100);o__Saccharomycetales(100);f__Metschnikowiaceae(100);g__Metschnikowia(100);g__Metschnikowia_unclassified(100); | Metschnikowia | 13 | Saprotroph | \|Undefined Saprotroph\| | Yeast | NULL | Probable | As nectar-tap_saprotroph (Põlme et al. 2020); Decay sugar-rich substrates (Põlme et al. 2020) | Põlme S, et al. 2020. Fungal Diversity 105:1-16 ((https://doi.org/10.1007/s13225-020-00466-2)) |
| Otu36262 | 1 | k__Fungi(100);p__Ascomycota(100);c__Saccharomycetes(100);o__Saccharomycetales(100);f__Metschnikowiaceae(100);g__Metschnikowia(100);g__Metschnikowia_unclassified(100); | Metschnikowia | 13 | Saprotroph | \|Undefined Saprotroph\| | Yeast | NULL | Probable | As nectar-tap_saprotroph (Põlme et al. 2020); Decay sugar-rich substrates (Põlme et al. 2020) | Põlme S, et al. 2020. Fungal Diversity 105:1-16 ((https://doi.org/10.1007/s13225-020-00466-2)) |
| Otu36263 | 1 | k__Fungi(100);p__Ascomycota(100);c__Saccharomycetes(100);o__Saccharomycetales(100);f__Metschnikowiaceae(100);g__Metschnikowia(100);g__Metschnikowia_unclassified(100); | Metschnikowia | 13 | Saprotroph | \|Undefined Saprotroph\| | Yeast | NULL | Probable | As nectar-tap_saprotroph (Põlme et al. 2020); Decay sugar-rich substrates (Põlme et al. 2020) | Põlme S, et al. 2020. Fungal Diversity 105:1-16 ((https://doi.org/10.1007/s13225-020-00466-2)) |
| Otu36264 | 1 | k__Fungi(100);p__Ascomycota(100);c__Saccharomycetes(100);o__Saccharomycetales(100);f__Metschnikowiaceae(100);g__Metschnikowia(100);g__Metschnikowia_unclassified(100); | Metschnikowia | 13 | Saprotroph | \|Undefined Saprotroph\| | Yeast | NULL | Probable | As nectar-tap_saprotroph (Põlme et al. 2020); Decay sugar-rich substrates (Põlme et al. 2020) | Põlme S, et al. 2020. Fungal Diversity 105:1-16 ((https://doi.org/10.1007/s13225-020-00466-2)) |
| Otu36267 | 1 | k__Fungi(100);p__Basidiomycota(100);c__Microbotryomycetes(100);o__Sporidiobolales(100);f__Sporidiobolales_family_Incertae_sedis(100);g__Rhodotorula(100);s__Rhodotorula_graminis(100); | Rhodotorula | 13 | Pathotroph-Saprotroph-Symbiotroph | Animal Pathogen-Animal Symbiotroph-Endophyte-Plant Pathogen-\|Undefined Saprotroph\| | Yeast | NULL | Probable | As foliar_endophyte (Põlme et al. 2020); Opportunistic human pathogen | Põlme S, et al. 2020. Fungal Diversity 105:1-16 ((https://doi.org/10.1007/s13225-020-00466-2)); https://www.cdc.gov ((https://www.cdc.gov)) |
| Otu36268 | 1 | k__Fungi(100);p__Basidiomycota(100);c__Agaricomycetes(100);o__Polyporales(100);f__Meripilaceae(100);g__Rigidoporus(100);s__Rigidoporus_sp_JV0511_23(100); | Rigidoporus | 13 | Saprotroph | \|Wood Saprotroph\| | Polyporoid | White Rot | Highly Probable | As white_rot (Põlme et al. 2020); Decay wood (Põlme et al. 2020) | Gilbertson RL, Ryvarden L. 1987-1987. North American Polypores. Fungiflora, Oslo (ISBN: 978-0945345060); Põlme S, et al. 2020. Fungal Diversity 105:1-16 ((https://doi.org/10.1007/s13225-020-00466-2)) |
| Otu36269 | 1 | k__Fungi(100);p__Ascomycota(100);c__Saccharomycetes(100);o__Saccharomycetales(100);f__Metschnikowiaceae(100);g__Metschnikowia(100);s__Metschnikowia_pulcherrima(100); | Metschnikowia | 13 | Saprotroph | \|Undefined Saprotroph\| | Yeast | NULL | Probable | As nectar-tap_saprotroph (Põlme et al. 2020); Decay sugar-rich substrates (Põlme et al. 2020) | Põlme S, et al. 2020. Fungal Diversity 105:1-16 ((https://doi.org/10.1007/s13225-020-00466-2)) |
| Otu36270 | 1 | k__Fungi(100);p__Ascomycota(100);c__Saccharomycetes(100);o__Saccharomycetales(100);f__Metschnikowiaceae(100);g__Metschnikowia(100);s__Metschnikowia_pulcherrima(100); | Metschnikowia | 13 | Saprotroph | \|Undefined Saprotroph\| | Yeast | NULL | Probable | As nectar-tap_saprotroph (Põlme et al. 2020); Decay sugar-rich substrates (Põlme et al. 2020) | Põlme S, et al. 2020. Fungal Diversity 105:1-16 ((https://doi.org/10.1007/s13225-020-00466-2)) |
| Otu36272 | 1 | k__Fungi(100);p__Ascomycota(100);c__Saccharomycetes(100);o__Saccharomycetales(100);f__Metschnikowiaceae(100);g__Metschnikowia(100);s__Metschnikowia_pulcherrima(100); | Metschnikowia | 13 | Saprotroph | \|Undefined Saprotroph\| | Yeast | NULL | Probable | As nectar-tap_saprotroph (Põlme et al. 2020); Decay sugar-rich substrates (Põlme et al. 2020) | Põlme S, et al. 2020. Fungal Diversity 105:1-16 ((https://doi.org/10.1007/s13225-020-00466-2)) |
| Otu36273 | 1 | k__Fungi(100);p__Ascomycota(100);c__Saccharomycetes(100);o__Saccharomycetales(100);f__Metschnikowiaceae(100);g__Metschnikowia(100);s__Metschnikowia_pulcherrima(100); | Metschnikowia | 13 | Saprotroph | \|Undefined Saprotroph\| | Yeast | NULL | Probable | As nectar-tap_saprotroph (Põlme et al. 2020); Decay sugar-rich substrates (Põlme et al. 2020) | Põlme S, et al. 2020. Fungal Diversity 105:1-16 ((https://doi.org/10.1007/s13225-020-00466-2)) |
| Otu36274 | 1 | k__Fungi(100);p__Ascomycota(100);c__Saccharomycetes(100);o__Saccharomycetales(100);f__Metschnikowiaceae(100);g__Metschnikowia(100);g__Metschnikowia_unclassified(100); | Metschnikowia | 13 | Saprotroph | \|Undefined Saprotroph\| | Yeast | NULL | Probable | As nectar-tap_saprotroph (Põlme et al. 2020); Decay sugar-rich substrates (Põlme et al. 2020) | Põlme S, et al. 2020. Fungal Diversity 105:1-16 ((https://doi.org/10.1007/s13225-020-00466-2)) |
| Otu36275 | 1 | k__Fungi(100);p__Ascomycota(100);c__Saccharomycetes(100);o__Saccharomycetales(100);f__Metschnikowiaceae(100);g__Metschnikowia(100);s__Metschnikowia_pulcherrima(100); | Metschnikowia | 13 | Saprotroph | \|Undefined Saprotroph\| | Yeast | NULL | Probable | As nectar-tap_saprotroph (Põlme et al. 2020); Decay sugar-rich substrates (Põlme et al. 2020) | Põlme S, et al. 2020. Fungal Diversity 105:1-16 ((https://doi.org/10.1007/s13225-020-00466-2)) |
| Otu36276 | 1 | k__Fungi(100);p__Ascomycota(100);c__Saccharomycetes(100);o__Saccharomycetales(100);f__Metschnikowiaceae(100);g__Metschnikowia(100);s__Metschnikowia_pulcherrima(100); | Metschnikowia | 13 | Saprotroph | \|Undefined Saprotroph\| | Yeast | NULL | Probable | As nectar-tap_saprotroph (Põlme et al. 2020); Decay sugar-rich substrates (Põlme et al. 2020) | Põlme S, et al. 2020. Fungal Diversity 105:1-16 ((https://doi.org/10.1007/s13225-020-00466-2)) |
| Otu36277 | 1 | k__Fungi(100);p__Ascomycota(100);c__Saccharomycetes(100);o__Saccharomycetales(100);f__Metschnikowiaceae(100);g__Metschnikowia(100);s__Metschnikowia_pulcherrima(100); | Metschnikowia | 13 | Saprotroph | \|Undefined Saprotroph\| | Yeast | NULL | Probable | As nectar-tap_saprotroph (Põlme et al. 2020); Decay sugar-rich substrates (Põlme et al. 2020) | Põlme S, et al. 2020. Fungal Diversity 105:1-16 ((https://doi.org/10.1007/s13225-020-00466-2)) |
| Otu36281 | 1 | k__Fungi(100);p__Ascomycota(100);c__Saccharomycetes(100);o__Saccharomycetales(100);f__Metschnikowiaceae(100);g__Metschnikowia(100);s__Metschnikowia_pulcherrima(100); | Metschnikowia | 13 | Saprotroph | \|Undefined Saprotroph\| | Yeast | NULL | Probable | As nectar-tap_saprotroph (Põlme et al. 2020); Decay sugar-rich substrates (Põlme et al. 2020) | Põlme S, et al. 2020. Fungal Diversity 105:1-16 ((https://doi.org/10.1007/s13225-020-00466-2)) |
| Otu36282 | 1 | k__Fungi(100);p__Ascomycota(100);c__Saccharomycetes(100);o__Saccharomycetales(100);f__Metschnikowiaceae(100);g__Metschnikowia(100);g__Metschnikowia_unclassified(100); | Metschnikowia | 13 | Saprotroph | \|Undefined Saprotroph\| | Yeast | NULL | Probable | As nectar-tap_saprotroph (Põlme et al. 2020); Decay sugar-rich substrates (Põlme et al. 2020) | Põlme S, et al. 2020. Fungal Diversity 105:1-16 ((https://doi.org/10.1007/s13225-020-00466-2)) |
| Otu36283 | 1 | k__Fungi(100);p__Ascomycota(100);c__Saccharomycetes(100);o__Saccharomycetales(100);f__Metschnikowiaceae(100);g__Metschnikowia(100);s__Metschnikowia_pulcherrima(100); | Metschnikowia | 13 | Saprotroph | \|Undefined Saprotroph\| | Yeast | NULL | Probable | As nectar-tap_saprotroph (Põlme et al. 2020); Decay sugar-rich substrates (Põlme et al. 2020) | Põlme S, et al. 2020. Fungal Diversity 105:1-16 ((https://doi.org/10.1007/s13225-020-00466-2)) |
| Otu36284 | 1 | k__Fungi(100);p__Ascomycota(100);c__Saccharomycetes(100);o__Saccharomycetales(100);f__Metschnikowiaceae(100);g__Metschnikowia(100);s__Metschnikowia_pulcherrima(100); | Metschnikowia | 13 | Saprotroph | \|Undefined Saprotroph\| | Yeast | NULL | Probable | As nectar-tap_saprotroph (Põlme et al. 2020); Decay sugar-rich substrates (Põlme et al. 2020) | Põlme S, et al. 2020. Fungal Diversity 105:1-16 ((https://doi.org/10.1007/s13225-020-00466-2)) |
| Otu36285 | 1 | k__Fungi(100);p__Ascomycota(100);c__Dothideomycetes(100);o__Capnodiales(100);f__Davidiellaceae(100);g__Cladosporium(100);s__Cladosporium_sphaerospermum(100); | Cladosporium | 13 | Pathotroph-Saprotroph-Symbiotroph | Animal Pathogen-Endophyte-Lichen Parasite-Plant Pathogen-\|Plant Saprotroph\|-Wood Saprotroph | Microfungus | NULL | Possible | As foliar_endophyte (Põlme et al. 2020); As litter_saprotroph (Põlme et al. 2020); Decay wood, leaf, fruit, seed, soil, dung, animal material (Põlme et al. 2020) | Seehann G, et al. 1975. List of Fungi in Soft-Rot Tissue Tests. Institute fur Holzbiologie und Holzschutz, Bundesforschungsanstalt fur Forst - und Holzwirtshaft, Hamburg - Lohbrugge, Germany; Lawrey JD, Diederich P. 2016. http://www.lichenicolous.net (accessed 24 May 2017) ((http://www.lichenicolous.net)); Põlme S, et al. 2020. Fungal Diversity 105:1-16 ((https://doi.org/10.1007/s13225-020-00466-2)); The American Phytopathological Society Common Names of Plant Diseases ((http://www.apsnet.org/publications/commonnames/Pages/default.aspx)) |
| Otu36286 | 1 | k__Fungi(100);p__Ascomycota(100);c__Saccharomycetes(100);o__Saccharomycetales(100);f__Metschnikowiaceae(100);g__Metschnikowia(100);s__Metschnikowia_pulcherrima(100); | Metschnikowia | 13 | Saprotroph | \|Undefined Saprotroph\| | Yeast | NULL | Probable | As nectar-tap_saprotroph (Põlme et al. 2020); Decay sugar-rich substrates (Põlme et al. 2020) | Põlme S, et al. 2020. Fungal Diversity 105:1-16 ((https://doi.org/10.1007/s13225-020-00466-2)) |
| Otu36287 | 1 | k__Fungi(100);p__Ascomycota(100);c__Saccharomycetes(100);o__Saccharomycetales(100);f__Metschnikowiaceae(100);g__Metschnikowia(100);s__Metschnikowia_pulcherrima(100); | Metschnikowia | 13 | Saprotroph | \|Undefined Saprotroph\| | Yeast | NULL | Probable | As nectar-tap_saprotroph (Põlme et al. 2020); Decay sugar-rich substrates (Põlme et al. 2020) | Põlme S, et al. 2020. Fungal Diversity 105:1-16 ((https://doi.org/10.1007/s13225-020-00466-2)) |
| Otu36288 | 1 | k__Fungi(100);p__Ascomycota(100);c__Saccharomycetes(100);o__Saccharomycetales(100);f__Metschnikowiaceae(100);g__Metschnikowia(100);s__Metschnikowia_pulcherrima(100); | Metschnikowia | 13 | Saprotroph | \|Undefined Saprotroph\| | Yeast | NULL | Probable | As nectar-tap_saprotroph (Põlme et al. 2020); Decay sugar-rich substrates (Põlme et al. 2020) | Põlme S, et al. 2020. Fungal Diversity 105:1-16 ((https://doi.org/10.1007/s13225-020-00466-2)) |
| Otu36289 | 1 | k__Fungi(100);p__Ascomycota(100);c__Saccharomycetes(100);o__Saccharomycetales(100);f__Metschnikowiaceae(100);g__Metschnikowia(100);s__Metschnikowia_pulcherrima(100); | Metschnikowia | 13 | Saprotroph | \|Undefined Saprotroph\| | Yeast | NULL | Probable | As nectar-tap_saprotroph (Põlme et al. 2020); Decay sugar-rich substrates (Põlme et al. 2020) | Põlme S, et al. 2020. Fungal Diversity 105:1-16 ((https://doi.org/10.1007/s13225-020-00466-2)) |
| Otu36290 | 1 | k__Fungi(100);p__Ascomycota(100);c__Saccharomycetes(100);o__Saccharomycetales(100);f__Metschnikowiaceae(100);g__Metschnikowia(100);s__Metschnikowia_pulcherrima(100); | Metschnikowia | 13 | Saprotroph | \|Undefined Saprotroph\| | Yeast | NULL | Probable | As nectar-tap_saprotroph (Põlme et al. 2020); Decay sugar-rich substrates (Põlme et al. 2020) | Põlme S, et al. 2020. Fungal Diversity 105:1-16 ((https://doi.org/10.1007/s13225-020-00466-2)) |
| Otu36292 | 1 | k__Fungi(100);p__Ascomycota(100);c__Saccharomycetes(100);o__Saccharomycetales(100);f__Metschnikowiaceae(100);g__Metschnikowia(100);g__Metschnikowia_unclassified(100); | Metschnikowia | 13 | Saprotroph | \|Undefined Saprotroph\| | Yeast | NULL | Probable | As nectar-tap_saprotroph (Põlme et al. 2020); Decay sugar-rich substrates (Põlme et al. 2020) | Põlme S, et al. 2020. Fungal Diversity 105:1-16 ((https://doi.org/10.1007/s13225-020-00466-2)) |
| Otu36295 | 1 | k__Fungi(100);p__Ascomycota(100);c__Saccharomycetes(100);o__Saccharomycetales(100);f__Metschnikowiaceae(100);g__Metschnikowia(100);s__Metschnikowia_pulcherrima(100); | Metschnikowia | 13 | Saprotroph | \|Undefined Saprotroph\| | Yeast | NULL | Probable | As nectar-tap_saprotroph (Põlme et al. 2020); Decay sugar-rich substrates (Põlme et al. 2020) | Põlme S, et al. 2020. Fungal Diversity 105:1-16 ((https://doi.org/10.1007/s13225-020-00466-2)) |
| Otu36296 | 1 | k__Fungi(100);p__Ascomycota(100);c__Saccharomycetes(100);o__Saccharomycetales(100);f__Metschnikowiaceae(100);g__Metschnikowia(100);s__Metschnikowia_pulcherrima(100); | Metschnikowia | 13 | Saprotroph | \|Undefined Saprotroph\| | Yeast | NULL | Probable | As nectar-tap_saprotroph (Põlme et al. 2020); Decay sugar-rich substrates (Põlme et al. 2020) | Põlme S, et al. 2020. Fungal Diversity 105:1-16 ((https://doi.org/10.1007/s13225-020-00466-2)) |
| Otu36297 | 1 | k__Fungi(100);p__Ascomycota(100);c__Saccharomycetes(100);o__Saccharomycetales(100);f__Metschnikowiaceae(100);g__Metschnikowia(100);s__Metschnikowia_pulcherrima(100); | Metschnikowia | 13 | Saprotroph | \|Undefined Saprotroph\| | Yeast | NULL | Probable | As nectar-tap_saprotroph (Põlme et al. 2020); Decay sugar-rich substrates (Põlme et al. 2020) | Põlme S, et al. 2020. Fungal Diversity 105:1-16 ((https://doi.org/10.1007/s13225-020-00466-2)) |
| Otu36298 | 1 | k__Fungi(100);p__Ascomycota(100);c__Saccharomycetes(100);o__Saccharomycetales(100);f__Metschnikowiaceae(100);g__Metschnikowia(100);s__Metschnikowia_pulcherrima(100); | Metschnikowia | 13 | Saprotroph | \|Undefined Saprotroph\| | Yeast | NULL | Probable | As nectar-tap_saprotroph (Põlme et al. 2020); Decay sugar-rich substrates (Põlme et al. 2020) | Põlme S, et al. 2020. Fungal Diversity 105:1-16 ((https://doi.org/10.1007/s13225-020-00466-2)) |
| Otu36299 | 1 | k__Fungi(100);p__Ascomycota(100);c__Saccharomycetes(100);o__Saccharomycetales(100);f__Metschnikowiaceae(100);g__Metschnikowia(100);s__Metschnikowia_pulcherrima(100); | Metschnikowia | 13 | Saprotroph | \|Undefined Saprotroph\| | Yeast | NULL | Probable | As nectar-tap_saprotroph (Põlme et al. 2020); Decay sugar-rich substrates (Põlme et al. 2020) | Põlme S, et al. 2020. Fungal Diversity 105:1-16 ((https://doi.org/10.1007/s13225-020-00466-2)) |
| Otu36300 | 1 | k__Fungi(100);p__Ascomycota(100);c__Saccharomycetes(100);o__Saccharomycetales(100);f__Metschnikowiaceae(100);g__Metschnikowia(100);g__Metschnikowia_unclassified(100); | Metschnikowia | 13 | Saprotroph | \|Undefined Saprotroph\| | Yeast | NULL | Probable | As nectar-tap_saprotroph (Põlme et al. 2020); Decay sugar-rich substrates (Põlme et al. 2020) | Põlme S, et al. 2020. Fungal Diversity 105:1-16 ((https://doi.org/10.1007/s13225-020-00466-2)) |
| Otu36301 | 1 | k__Fungi(100);p__Ascomycota(100);c__Saccharomycetes(100);o__Saccharomycetales(100);f__Metschnikowiaceae(100);g__Metschnikowia(100);g__Metschnikowia_unclassified(100); | Metschnikowia | 13 | Saprotroph | \|Undefined Saprotroph\| | Yeast | NULL | Probable | As nectar-tap_saprotroph (Põlme et al. 2020); Decay sugar-rich substrates (Põlme et al. 2020) | Põlme S, et al. 2020. Fungal Diversity 105:1-16 ((https://doi.org/10.1007/s13225-020-00466-2)) |
| Otu36302 | 1 | k__Fungi(100);p__Ascomycota(100);c__Saccharomycetes(100);o__Saccharomycetales(100);f__Metschnikowiaceae(100);g__Metschnikowia(100);s__Metschnikowia_pulcherrima(100); | Metschnikowia | 13 | Saprotroph | \|Undefined Saprotroph\| | Yeast | NULL | Probable | As nectar-tap_saprotroph (Põlme et al. 2020); Decay sugar-rich substrates (Põlme et al. 2020) | Põlme S, et al. 2020. Fungal Diversity 105:1-16 ((https://doi.org/10.1007/s13225-020-00466-2)) |
| Otu36303 | 1 | k__Fungi(100);p__Ascomycota(100);c__Saccharomycetes(100);o__Saccharomycetales(100);f__Metschnikowiaceae(100);g__Metschnikowia(100);s__Metschnikowia_pulcherrima(100); | Metschnikowia | 13 | Saprotroph | \|Undefined Saprotroph\| | Yeast | NULL | Probable | As nectar-tap_saprotroph (Põlme et al. 2020); Decay sugar-rich substrates (Põlme et al. 2020) | Põlme S, et al. 2020. Fungal Diversity 105:1-16 ((https://doi.org/10.1007/s13225-020-00466-2)) |
| Otu36304 | 1 | k__Fungi(100);p__Ascomycota(100);c__Saccharomycetes(100);o__Saccharomycetales(100);f__Metschnikowiaceae(100);g__Metschnikowia(100);s__Metschnikowia_pulcherrima(100); | Metschnikowia | 13 | Saprotroph | \|Undefined Saprotroph\| | Yeast | NULL | Probable | As nectar-tap_saprotroph (Põlme et al. 2020); Decay sugar-rich substrates (Põlme et al. 2020) | Põlme S, et al. 2020. Fungal Diversity 105:1-16 ((https://doi.org/10.1007/s13225-020-00466-2)) |
| Otu36305 | 1 | k__Fungi(100);p__Ascomycota(100);c__Saccharomycetes(100);o__Saccharomycetales(100);f__Metschnikowiaceae(100);g__Metschnikowia(100);s__Metschnikowia_pulcherrima(100); | Metschnikowia | 13 | Saprotroph | \|Undefined Saprotroph\| | Yeast | NULL | Probable | As nectar-tap_saprotroph (Põlme et al. 2020); Decay sugar-rich substrates (Põlme et al. 2020) | Põlme S, et al. 2020. Fungal Diversity 105:1-16 ((https://doi.org/10.1007/s13225-020-00466-2)) |
| Otu36306 | 1 | k__Fungi(100);p__Ascomycota(100);c__Saccharomycetes(100);o__Saccharomycetales(100);f__Metschnikowiaceae(100);g__Metschnikowia(100);s__Metschnikowia_pulcherrima(100); | Metschnikowia | 13 | Saprotroph | \|Undefined Saprotroph\| | Yeast | NULL | Probable | As nectar-tap_saprotroph (Põlme et al. 2020); Decay sugar-rich substrates (Põlme et al. 2020) | Põlme S, et al. 2020. Fungal Diversity 105:1-16 ((https://doi.org/10.1007/s13225-020-00466-2)) |
| Otu36307 | 1 | k__Fungi(100);p__Ascomycota(100);c__Saccharomycetes(100);o__Saccharomycetales(100);f__Metschnikowiaceae(100);g__Metschnikowia(100);s__Metschnikowia_pulcherrima(100); | Metschnikowia | 13 | Saprotroph | \|Undefined Saprotroph\| | Yeast | NULL | Probable | As nectar-tap_saprotroph (Põlme et al. 2020); Decay sugar-rich substrates (Põlme et al. 2020) | Põlme S, et al. 2020. Fungal Diversity 105:1-16 ((https://doi.org/10.1007/s13225-020-00466-2)) |
| Otu36308 | 1 | k__Fungi(100);p__Ascomycota(100);c__Saccharomycetes(100);o__Saccharomycetales(100);f__Metschnikowiaceae(100);g__Metschnikowia(100);g__Metschnikowia_unclassified(100); | Metschnikowia | 13 | Saprotroph | \|Undefined Saprotroph\| | Yeast | NULL | Probable | As nectar-tap_saprotroph (Põlme et al. 2020); Decay sugar-rich substrates (Põlme et al. 2020) | Põlme S, et al. 2020. Fungal Diversity 105:1-16 ((https://doi.org/10.1007/s13225-020-00466-2)) |
| Otu36309 | 1 | k__Fungi(100);p__Ascomycota(100);c__Saccharomycetes(100);o__Saccharomycetales(100);f__Metschnikowiaceae(100);g__Metschnikowia(100);s__Metschnikowia_pulcherrima(100); | Metschnikowia | 13 | Saprotroph | \|Undefined Saprotroph\| | Yeast | NULL | Probable | As nectar-tap_saprotroph (Põlme et al. 2020); Decay sugar-rich substrates (Põlme et al. 2020) | Põlme S, et al. 2020. Fungal Diversity 105:1-16 ((https://doi.org/10.1007/s13225-020-00466-2)) |
| Otu36310 | 1 | k__Fungi(100);p__Ascomycota(100);c__Saccharomycetes(100);o__Saccharomycetales(100);f__Metschnikowiaceae(100);g__Metschnikowia(100);s__Metschnikowia_pulcherrima(100); | Metschnikowia | 13 | Saprotroph | \|Undefined Saprotroph\| | Yeast | NULL | Probable | As nectar-tap_saprotroph (Põlme et al. 2020); Decay sugar-rich substrates (Põlme et al. 2020) | Põlme S, et al. 2020. Fungal Diversity 105:1-16 ((https://doi.org/10.1007/s13225-020-00466-2)) |
| Otu36314 | 1 | k__Fungi(100);p__Ascomycota(100);c__Saccharomycetes(100);o__Saccharomycetales(100);f__Metschnikowiaceae(100);g__Metschnikowia(100);s__Metschnikowia_pulcherrima(100); | Metschnikowia | 13 | Saprotroph | \|Undefined Saprotroph\| | Yeast | NULL | Probable | As nectar-tap_saprotroph (Põlme et al. 2020); Decay sugar-rich substrates (Põlme et al. 2020) | Põlme S, et al. 2020. Fungal Diversity 105:1-16 ((https://doi.org/10.1007/s13225-020-00466-2)) |
| Otu36316 | 1 | k__Fungi(100);p__Ascomycota(100);c__Saccharomycetes(100);o__Saccharomycetales(100);f__Metschnikowiaceae(100);g__Metschnikowia(100);s__Metschnikowia_pulcherrima(100); | Metschnikowia | 13 | Saprotroph | \|Undefined Saprotroph\| | Yeast | NULL | Probable | As nectar-tap_saprotroph (Põlme et al. 2020); Decay sugar-rich substrates (Põlme et al. 2020) | Põlme S, et al. 2020. Fungal Diversity 105:1-16 ((https://doi.org/10.1007/s13225-020-00466-2)) |
| Otu36322 | 1 | k__Fungi(100);p__Ascomycota(100);c__Saccharomycetes(100);o__Saccharomycetales(100);f__Metschnikowiaceae(100);g__Metschnikowia(100);s__Metschnikowia_pulcherrima(100); | Metschnikowia | 13 | Saprotroph | \|Undefined Saprotroph\| | Yeast | NULL | Probable | As nectar-tap_saprotroph (Põlme et al. 2020); Decay sugar-rich substrates (Põlme et al. 2020) | Põlme S, et al. 2020. Fungal Diversity 105:1-16 ((https://doi.org/10.1007/s13225-020-00466-2)) |
| Otu36323 | 1 | k__Fungi(100);p__Ascomycota(100);c__Saccharomycetes(100);o__Saccharomycetales(100);f__Metschnikowiaceae(100);g__Metschnikowia(100);s__Metschnikowia_pulcherrima(100); | Metschnikowia | 13 | Saprotroph | \|Undefined Saprotroph\| | Yeast | NULL | Probable | As nectar-tap_saprotroph (Põlme et al. 2020); Decay sugar-rich substrates (Põlme et al. 2020) | Põlme S, et al. 2020. Fungal Diversity 105:1-16 ((https://doi.org/10.1007/s13225-020-00466-2)) |
| Otu36324 | 1 | k__Fungi(100);p__Ascomycota(100);c__Saccharomycetes(100);o__Saccharomycetales(100);f__Metschnikowiaceae(100);g__Metschnikowia(100);s__Metschnikowia_pulcherrima(100); | Metschnikowia | 13 | Saprotroph | \|Undefined Saprotroph\| | Yeast | NULL | Probable | As nectar-tap_saprotroph (Põlme et al. 2020); Decay sugar-rich substrates (Põlme et al. 2020) | Põlme S, et al. 2020. Fungal Diversity 105:1-16 ((https://doi.org/10.1007/s13225-020-00466-2)) |
| Otu36325 | 1 | k__Fungi(100);p__Ascomycota(100);c__Saccharomycetes(100);o__Saccharomycetales(100);f__Metschnikowiaceae(100);g__Metschnikowia(100);s__Metschnikowia_pulcherrima(100); | Metschnikowia | 13 | Saprotroph | \|Undefined Saprotroph\| | Yeast | NULL | Probable | As nectar-tap_saprotroph (Põlme et al. 2020); Decay sugar-rich substrates (Põlme et al. 2020) | Põlme S, et al. 2020. Fungal Diversity 105:1-16 ((https://doi.org/10.1007/s13225-020-00466-2)) |
| Otu36326 | 1 | k__Fungi(100);p__Ascomycota(100);c__Saccharomycetes(100);o__Saccharomycetales(100);f__Metschnikowiaceae(100);g__Metschnikowia(100);s__Metschnikowia_pulcherrima(100); | Metschnikowia | 13 | Saprotroph | \|Undefined Saprotroph\| | Yeast | NULL | Probable | As nectar-tap_saprotroph (Põlme et al. 2020); Decay sugar-rich substrates (Põlme et al. 2020) | Põlme S, et al. 2020. Fungal Diversity 105:1-16 ((https://doi.org/10.1007/s13225-020-00466-2)) |
| Otu36327 | 1 | k__Fungi(100);p__Ascomycota(100);c__Saccharomycetes(100);o__Saccharomycetales(100);f__Metschnikowiaceae(100);g__Metschnikowia(100);s__Metschnikowia_pulcherrima(100); | Metschnikowia | 13 | Saprotroph | \|Undefined Saprotroph\| | Yeast | NULL | Probable | As nectar-tap_saprotroph (Põlme et al. 2020); Decay sugar-rich substrates (Põlme et al. 2020) | Põlme S, et al. 2020. Fungal Diversity 105:1-16 ((https://doi.org/10.1007/s13225-020-00466-2)) |
| Otu36328 | 1 | k__Fungi(100);p__Ascomycota(100);c__Saccharomycetes(100);o__Saccharomycetales(100);f__Metschnikowiaceae(100);g__Metschnikowia(100);s__Metschnikowia_pulcherrima(100); | Metschnikowia | 13 | Saprotroph | \|Undefined Saprotroph\| | Yeast | NULL | Probable | As nectar-tap_saprotroph (Põlme et al. 2020); Decay sugar-rich substrates (Põlme et al. 2020) | Põlme S, et al. 2020. Fungal Diversity 105:1-16 ((https://doi.org/10.1007/s13225-020-00466-2)) |
| Otu36330 | 1 | k__Fungi(100);p__Ascomycota(100);c__Saccharomycetes(100);o__Saccharomycetales(100);f__Metschnikowiaceae(100);g__Metschnikowia(100);s__Metschnikowia_pulcherrima(100); | Metschnikowia | 13 | Saprotroph | \|Undefined Saprotroph\| | Yeast | NULL | Probable | As nectar-tap_saprotroph (Põlme et al. 2020); Decay sugar-rich substrates (Põlme et al. 2020) | Põlme S, et al. 2020. Fungal Diversity 105:1-16 ((https://doi.org/10.1007/s13225-020-00466-2)) |
| Otu36331 | 1 | k__Fungi(100);p__Ascomycota(100);c__Saccharomycetes(100);o__Saccharomycetales(100);f__Metschnikowiaceae(100);g__Metschnikowia(100);s__Metschnikowia_pulcherrima(100); | Metschnikowia | 13 | Saprotroph | \|Undefined Saprotroph\| | Yeast | NULL | Probable | As nectar-tap_saprotroph (Põlme et al. 2020); Decay sugar-rich substrates (Põlme et al. 2020) | Põlme S, et al. 2020. Fungal Diversity 105:1-16 ((https://doi.org/10.1007/s13225-020-00466-2)) |
| Otu36333 | 1 | k__Fungi(100);p__Ascomycota(100);c__Saccharomycetes(100);o__Saccharomycetales(100);f__Metschnikowiaceae(100);g__Metschnikowia(100);s__Metschnikowia_pulcherrima(100); | Metschnikowia | 13 | Saprotroph | \|Undefined Saprotroph\| | Yeast | NULL | Probable | As nectar-tap_saprotroph (Põlme et al. 2020); Decay sugar-rich substrates (Põlme et al. 2020) | Põlme S, et al. 2020. Fungal Diversity 105:1-16 ((https://doi.org/10.1007/s13225-020-00466-2)) |
| Otu36334 | 1 | k__Fungi(100);p__Ascomycota(100);c__Saccharomycetes(100);o__Saccharomycetales(100);f__Metschnikowiaceae(100);g__Metschnikowia(100);s__Metschnikowia_pulcherrima(100); | Metschnikowia | 13 | Saprotroph | \|Undefined Saprotroph\| | Yeast | NULL | Probable | As nectar-tap_saprotroph (Põlme et al. 2020); Decay sugar-rich substrates (Põlme et al. 2020) | Põlme S, et al. 2020. Fungal Diversity 105:1-16 ((https://doi.org/10.1007/s13225-020-00466-2)) |
| Otu36336 | 1 | k__Fungi(100);p__Ascomycota(100);c__Saccharomycetes(100);o__Saccharomycetales(100);f__Metschnikowiaceae(100);g__Metschnikowia(100);s__Metschnikowia_pulcherrima(100); | Metschnikowia | 13 | Saprotroph | \|Undefined Saprotroph\| | Yeast | NULL | Probable | As nectar-tap_saprotroph (Põlme et al. 2020); Decay sugar-rich substrates (Põlme et al. 2020) | Põlme S, et al. 2020. Fungal Diversity 105:1-16 ((https://doi.org/10.1007/s13225-020-00466-2)) |
| Otu36337 | 1 | k__Fungi(100);p__Ascomycota(100);c__Saccharomycetes(100);o__Saccharomycetales(100);f__Metschnikowiaceae(100);g__Metschnikowia(100);s__Metschnikowia_pulcherrima(100); | Metschnikowia | 13 | Saprotroph | \|Undefined Saprotroph\| | Yeast | NULL | Probable | As nectar-tap_saprotroph (Põlme et al. 2020); Decay sugar-rich substrates (Põlme et al. 2020) | Põlme S, et al. 2020. Fungal Diversity 105:1-16 ((https://doi.org/10.1007/s13225-020-00466-2)) |
| Otu36338 | 1 | k__Fungi(100);p__Ascomycota(100);c__Saccharomycetes(100);o__Saccharomycetales(100);f__Metschnikowiaceae(100);g__Metschnikowia(100);g__Metschnikowia_unclassified(100); | Metschnikowia | 13 | Saprotroph | \|Undefined Saprotroph\| | Yeast | NULL | Probable | As nectar-tap_saprotroph (Põlme et al. 2020); Decay sugar-rich substrates (Põlme et al. 2020) | Põlme S, et al. 2020. Fungal Diversity 105:1-16 ((https://doi.org/10.1007/s13225-020-00466-2)) |
| Otu36339 | 1 | k__Fungi(100);p__Ascomycota(100);c__Saccharomycetes(100);o__Saccharomycetales(100);f__Metschnikowiaceae(100);g__Metschnikowia(100);s__Metschnikowia_pulcherrima(100); | Metschnikowia | 13 | Saprotroph | \|Undefined Saprotroph\| | Yeast | NULL | Probable | As nectar-tap_saprotroph (Põlme et al. 2020); Decay sugar-rich substrates (Põlme et al. 2020) | Põlme S, et al. 2020. Fungal Diversity 105:1-16 ((https://doi.org/10.1007/s13225-020-00466-2)) |
| Otu36340 | 1 | k__Fungi(100);p__Ascomycota(100);c__Saccharomycetes(100);o__Saccharomycetales(100);f__Metschnikowiaceae(100);g__Metschnikowia(100);s__Metschnikowia_pulcherrima(100); | Metschnikowia | 13 | Saprotroph | \|Undefined Saprotroph\| | Yeast | NULL | Probable | As nectar-tap_saprotroph (Põlme et al. 2020); Decay sugar-rich substrates (Põlme et al. 2020) | Põlme S, et al. 2020. Fungal Diversity 105:1-16 ((https://doi.org/10.1007/s13225-020-00466-2)) |
| Otu36341 | 1 | k__Fungi(100);p__Ascomycota(100);c__Saccharomycetes(100);o__Saccharomycetales(100);f__Metschnikowiaceae(100);g__Metschnikowia(100);s__Metschnikowia_pulcherrima(100); | Metschnikowia | 13 | Saprotroph | \|Undefined Saprotroph\| | Yeast | NULL | Probable | As nectar-tap_saprotroph (Põlme et al. 2020); Decay sugar-rich substrates (Põlme et al. 2020) | Põlme S, et al. 2020. Fungal Diversity 105:1-16 ((https://doi.org/10.1007/s13225-020-00466-2)) |
| Otu36342 | 1 | k__Fungi(100);p__Ascomycota(100);c__Saccharomycetes(100);o__Saccharomycetales(100);f__Metschnikowiaceae(100);g__Metschnikowia(100);s__Metschnikowia_pulcherrima(100); | Metschnikowia | 13 | Saprotroph | \|Undefined Saprotroph\| | Yeast | NULL | Probable | As nectar-tap_saprotroph (Põlme et al. 2020); Decay sugar-rich substrates (Põlme et al. 2020) | Põlme S, et al. 2020. Fungal Diversity 105:1-16 ((https://doi.org/10.1007/s13225-020-00466-2)) |
| Otu36343 | 1 | k__Fungi(100);p__Ascomycota(100);c__Saccharomycetes(100);o__Saccharomycetales(100);f__Metschnikowiaceae(100);g__Metschnikowia(100);s__Metschnikowia_pulcherrima(100); | Metschnikowia | 13 | Saprotroph | \|Undefined Saprotroph\| | Yeast | NULL | Probable | As nectar-tap_saprotroph (Põlme et al. 2020); Decay sugar-rich substrates (Põlme et al. 2020) | Põlme S, et al. 2020. Fungal Diversity 105:1-16 ((https://doi.org/10.1007/s13225-020-00466-2)) |
| Otu36344 | 1 | k__Fungi(100);p__Ascomycota(100);c__Saccharomycetes(100);o__Saccharomycetales(100);f__Metschnikowiaceae(100);g__Metschnikowia(100);s__Metschnikowia_pulcherrima(100); | Metschnikowia | 13 | Saprotroph | \|Undefined Saprotroph\| | Yeast | NULL | Probable | As nectar-tap_saprotroph (Põlme et al. 2020); Decay sugar-rich substrates (Põlme et al. 2020) | Põlme S, et al. 2020. Fungal Diversity 105:1-16 ((https://doi.org/10.1007/s13225-020-00466-2)) |
| Otu36345 | 1 | k__Fungi(100);p__Ascomycota(100);c__Saccharomycetes(100);o__Saccharomycetales(100);f__Metschnikowiaceae(100);g__Metschnikowia(100);s__Metschnikowia_pulcherrima(100); | Metschnikowia | 13 | Saprotroph | \|Undefined Saprotroph\| | Yeast | NULL | Probable | As nectar-tap_saprotroph (Põlme et al. 2020); Decay sugar-rich substrates (Põlme et al. 2020) | Põlme S, et al. 2020. Fungal Diversity 105:1-16 ((https://doi.org/10.1007/s13225-020-00466-2)) |
| Otu36346 | 1 | k__Fungi(100);p__Ascomycota(100);c__Saccharomycetes(100);o__Saccharomycetales(100);f__Metschnikowiaceae(100);g__Metschnikowia(100);s__Metschnikowia_pulcherrima(100); | Metschnikowia | 13 | Saprotroph | \|Undefined Saprotroph\| | Yeast | NULL | Probable | As nectar-tap_saprotroph (Põlme et al. 2020); Decay sugar-rich substrates (Põlme et al. 2020) | Põlme S, et al. 2020. Fungal Diversity 105:1-16 ((https://doi.org/10.1007/s13225-020-00466-2)) |
| Otu36347 | 1 | k__Fungi(100);p__Ascomycota(100);c__Saccharomycetes(100);o__Saccharomycetales(100);f__Metschnikowiaceae(100);g__Metschnikowia(100);s__Metschnikowia_pulcherrima(100); | Metschnikowia | 13 | Saprotroph | \|Undefined Saprotroph\| | Yeast | NULL | Probable | As nectar-tap_saprotroph (Põlme et al. 2020); Decay sugar-rich substrates (Põlme et al. 2020) | Põlme S, et al. 2020. Fungal Diversity 105:1-16 ((https://doi.org/10.1007/s13225-020-00466-2)) |
| Otu36348 | 1 | k__Fungi(100);p__Ascomycota(100);c__Saccharomycetes(100);o__Saccharomycetales(100);f__Metschnikowiaceae(100);g__Metschnikowia(100);s__Metschnikowia_pulcherrima(100); | Metschnikowia | 13 | Saprotroph | \|Undefined Saprotroph\| | Yeast | NULL | Probable | As nectar-tap_saprotroph (Põlme et al. 2020); Decay sugar-rich substrates (Põlme et al. 2020) | Põlme S, et al. 2020. Fungal Diversity 105:1-16 ((https://doi.org/10.1007/s13225-020-00466-2)) |
| Otu36349 | 1 | k__Fungi(100);p__Ascomycota(100);c__Saccharomycetes(100);o__Saccharomycetales(100);f__Metschnikowiaceae(100);g__Metschnikowia(100);s__Metschnikowia_pulcherrima(100); | Metschnikowia | 13 | Saprotroph | \|Undefined Saprotroph\| | Yeast | NULL | Probable | As nectar-tap_saprotroph (Põlme et al. 2020); Decay sugar-rich substrates (Põlme et al. 2020) | Põlme S, et al. 2020. Fungal Diversity 105:1-16 ((https://doi.org/10.1007/s13225-020-00466-2)) |
| Otu36350 | 1 | k__Fungi(100);p__Ascomycota(100);c__Saccharomycetes(100);o__Saccharomycetales(100);f__Metschnikowiaceae(100);g__Metschnikowia(100);s__Metschnikowia_pulcherrima(100); | Metschnikowia | 13 | Saprotroph | \|Undefined Saprotroph\| | Yeast | NULL | Probable | As nectar-tap_saprotroph (Põlme et al. 2020); Decay sugar-rich substrates (Põlme et al. 2020) | Põlme S, et al. 2020. Fungal Diversity 105:1-16 ((https://doi.org/10.1007/s13225-020-00466-2)) |
| Otu36351 | 1 | k__Fungi(100);p__Ascomycota(100);c__Saccharomycetes(100);o__Saccharomycetales(100);f__Metschnikowiaceae(100);g__Metschnikowia(100);s__Metschnikowia_pulcherrima(100); | Metschnikowia | 13 | Saprotroph | \|Undefined Saprotroph\| | Yeast | NULL | Probable | As nectar-tap_saprotroph (Põlme et al. 2020); Decay sugar-rich substrates (Põlme et al. 2020) | Põlme S, et al. 2020. Fungal Diversity 105:1-16 ((https://doi.org/10.1007/s13225-020-00466-2)) |
| Otu36354 | 1 | k__Fungi(100);p__Ascomycota(100);c__Saccharomycetes(100);o__Saccharomycetales(100);f__Metschnikowiaceae(100);g__Metschnikowia(100);s__Metschnikowia_pulcherrima(100); | Metschnikowia | 13 | Saprotroph | \|Undefined Saprotroph\| | Yeast | NULL | Probable | As nectar-tap_saprotroph (Põlme et al. 2020); Decay sugar-rich substrates (Põlme et al. 2020) | Põlme S, et al. 2020. Fungal Diversity 105:1-16 ((https://doi.org/10.1007/s13225-020-00466-2)) |
| Otu36355 | 1 | k__Fungi(100);p__Ascomycota(100);c__Saccharomycetes(100);o__Saccharomycetales(100);f__Metschnikowiaceae(100);g__Metschnikowia(100);g__Metschnikowia_unclassified(100); | Metschnikowia | 13 | Saprotroph | \|Undefined Saprotroph\| | Yeast | NULL | Probable | As nectar-tap_saprotroph (Põlme et al. 2020); Decay sugar-rich substrates (Põlme et al. 2020) | Põlme S, et al. 2020. Fungal Diversity 105:1-16 ((https://doi.org/10.1007/s13225-020-00466-2)) |
| Otu36356 | 1 | k__Fungi(100);p__Ascomycota(100);c__Saccharomycetes(100);o__Saccharomycetales(100);f__Metschnikowiaceae(100);g__Metschnikowia(100);s__Metschnikowia_pulcherrima(100); | Metschnikowia | 13 | Saprotroph | \|Undefined Saprotroph\| | Yeast | NULL | Probable | As nectar-tap_saprotroph (Põlme et al. 2020); Decay sugar-rich substrates (Põlme et al. 2020) | Põlme S, et al. 2020. Fungal Diversity 105:1-16 ((https://doi.org/10.1007/s13225-020-00466-2)) |
| Otu36357 | 1 | k__Fungi(100);p__Ascomycota(100);c__Saccharomycetes(100);o__Saccharomycetales(100);f__Metschnikowiaceae(100);g__Metschnikowia(100);s__Metschnikowia_pulcherrima(100); | Metschnikowia | 13 | Saprotroph | \|Undefined Saprotroph\| | Yeast | NULL | Probable | As nectar-tap_saprotroph (Põlme et al. 2020); Decay sugar-rich substrates (Põlme et al. 2020) | Põlme S, et al. 2020. Fungal Diversity 105:1-16 ((https://doi.org/10.1007/s13225-020-00466-2)) |
| Otu36358 | 1 | k__Fungi(100);p__Ascomycota(100);c__Saccharomycetes(100);o__Saccharomycetales(100);f__Metschnikowiaceae(100);g__Metschnikowia(100);s__Metschnikowia_pulcherrima(100); | Metschnikowia | 13 | Saprotroph | \|Undefined Saprotroph\| | Yeast | NULL | Probable | As nectar-tap_saprotroph (Põlme et al. 2020); Decay sugar-rich substrates (Põlme et al. 2020) | Põlme S, et al. 2020. Fungal Diversity 105:1-16 ((https://doi.org/10.1007/s13225-020-00466-2)) |
| Otu36359 | 1 | k__Fungi(100);p__Ascomycota(100);c__Saccharomycetes(100);o__Saccharomycetales(100);f__Metschnikowiaceae(100);g__Metschnikowia(100);s__Metschnikowia_pulcherrima(100); | Metschnikowia | 13 | Saprotroph | \|Undefined Saprotroph\| | Yeast | NULL | Probable | As nectar-tap_saprotroph (Põlme et al. 2020); Decay sugar-rich substrates (Põlme et al. 2020) | Põlme S, et al. 2020. Fungal Diversity 105:1-16 ((https://doi.org/10.1007/s13225-020-00466-2)) |
| Otu36360 | 1 | k__Fungi(100);p__Ascomycota(100);c__Saccharomycetes(100);o__Saccharomycetales(100);f__Metschnikowiaceae(100);g__Metschnikowia(100);s__Metschnikowia_pulcherrima(100); | Metschnikowia | 13 | Saprotroph | \|Undefined Saprotroph\| | Yeast | NULL | Probable | As nectar-tap_saprotroph (Põlme et al. 2020); Decay sugar-rich substrates (Põlme et al. 2020) | Põlme S, et al. 2020. Fungal Diversity 105:1-16 ((https://doi.org/10.1007/s13225-020-00466-2)) |
| Otu36361 | 1 | k__Fungi(100);p__Ascomycota(100);c__Saccharomycetes(100);o__Saccharomycetales(100);f__Metschnikowiaceae(100);g__Metschnikowia(100);s__Metschnikowia_pulcherrima(100); | Metschnikowia | 13 | Saprotroph | \|Undefined Saprotroph\| | Yeast | NULL | Probable | As nectar-tap_saprotroph (Põlme et al. 2020); Decay sugar-rich substrates (Põlme et al. 2020) | Põlme S, et al. 2020. Fungal Diversity 105:1-16 ((https://doi.org/10.1007/s13225-020-00466-2)) |
| Otu36362 | 1 | k__Fungi(100);p__Ascomycota(100);c__Saccharomycetes(100);o__Saccharomycetales(100);f__Metschnikowiaceae(100);g__Metschnikowia(100);s__Metschnikowia_pulcherrima(100); | Metschnikowia | 13 | Saprotroph | \|Undefined Saprotroph\| | Yeast | NULL | Probable | As nectar-tap_saprotroph (Põlme et al. 2020); Decay sugar-rich substrates (Põlme et al. 2020) | Põlme S, et al. 2020. Fungal Diversity 105:1-16 ((https://doi.org/10.1007/s13225-020-00466-2)) |
| Otu36363 | 1 | k__Fungi(100);p__Ascomycota(100);c__Saccharomycetes(100);o__Saccharomycetales(100);f__Metschnikowiaceae(100);g__Metschnikowia(100);s__Metschnikowia_pulcherrima(100); | Metschnikowia | 13 | Saprotroph | \|Undefined Saprotroph\| | Yeast | NULL | Probable | As nectar-tap_saprotroph (Põlme et al. 2020); Decay sugar-rich substrates (Põlme et al. 2020) | Põlme S, et al. 2020. Fungal Diversity 105:1-16 ((https://doi.org/10.1007/s13225-020-00466-2)) |
| Otu36364 | 1 | k__Fungi(100);p__Ascomycota(100);c__Saccharomycetes(100);o__Saccharomycetales(100);f__Metschnikowiaceae(100);g__Metschnikowia(100);s__Metschnikowia_pulcherrima(100); | Metschnikowia | 13 | Saprotroph | \|Undefined Saprotroph\| | Yeast | NULL | Probable | As nectar-tap_saprotroph (Põlme et al. 2020); Decay sugar-rich substrates (Põlme et al. 2020) | Põlme S, et al. 2020. Fungal Diversity 105:1-16 ((https://doi.org/10.1007/s13225-020-00466-2)) |
| Otu36365 | 1 | k__Fungi(100);p__Ascomycota(100);c__Saccharomycetes(100);o__Saccharomycetales(100);f__Metschnikowiaceae(100);g__Metschnikowia(100);g__Metschnikowia_unclassified(100); | Metschnikowia | 13 | Saprotroph | \|Undefined Saprotroph\| | Yeast | NULL | Probable | As nectar-tap_saprotroph (Põlme et al. 2020); Decay sugar-rich substrates (Põlme et al. 2020) | Põlme S, et al. 2020. Fungal Diversity 105:1-16 ((https://doi.org/10.1007/s13225-020-00466-2)) |
| Otu36366 | 1 | k__Fungi(100);p__Ascomycota(100);c__Saccharomycetes(100);o__Saccharomycetales(100);f__Metschnikowiaceae(100);g__Metschnikowia(100);g__Metschnikowia_unclassified(100); | Metschnikowia | 13 | Saprotroph | \|Undefined Saprotroph\| | Yeast | NULL | Probable | As nectar-tap_saprotroph (Põlme et al. 2020); Decay sugar-rich substrates (Põlme et al. 2020) | Põlme S, et al. 2020. Fungal Diversity 105:1-16 ((https://doi.org/10.1007/s13225-020-00466-2)) |
| Otu36367 | 1 | k__Fungi(100);p__Ascomycota(100);c__Saccharomycetes(100);o__Saccharomycetales(100);f__Metschnikowiaceae(100);g__Metschnikowia(100);s__Metschnikowia_pulcherrima(100); | Metschnikowia | 13 | Saprotroph | \|Undefined Saprotroph\| | Yeast | NULL | Probable | As nectar-tap_saprotroph (Põlme et al. 2020); Decay sugar-rich substrates (Põlme et al. 2020) | Põlme S, et al. 2020. Fungal Diversity 105:1-16 ((https://doi.org/10.1007/s13225-020-00466-2)) |
| Otu36369 | 1 | k__Fungi(100);p__Ascomycota(100);c__Saccharomycetes(100);o__Saccharomycetales(100);f__Metschnikowiaceae(100);g__Metschnikowia(100);s__Metschnikowia_pulcherrima(100); | Metschnikowia | 13 | Saprotroph | \|Undefined Saprotroph\| | Yeast | NULL | Probable | As nectar-tap_saprotroph (Põlme et al. 2020); Decay sugar-rich substrates (Põlme et al. 2020) | Põlme S, et al. 2020. Fungal Diversity 105:1-16 ((https://doi.org/10.1007/s13225-020-00466-2)) |
| Otu36370 | 1 | k__Fungi(100);p__Ascomycota(100);c__Saccharomycetes(100);o__Saccharomycetales(100);f__Metschnikowiaceae(100);g__Metschnikowia(100);s__Metschnikowia_pulcherrima(100); | Metschnikowia | 13 | Saprotroph | \|Undefined Saprotroph\| | Yeast | NULL | Probable | As nectar-tap_saprotroph (Põlme et al. 2020); Decay sugar-rich substrates (Põlme et al. 2020) | Põlme S, et al. 2020. Fungal Diversity 105:1-16 ((https://doi.org/10.1007/s13225-020-00466-2)) |
| Otu36371 | 1 | k__Fungi(100);p__Ascomycota(100);c__Saccharomycetes(100);o__Saccharomycetales(100);f__Metschnikowiaceae(100);g__Metschnikowia(100);g__Metschnikowia_unclassified(100); | Metschnikowia | 13 | Saprotroph | \|Undefined Saprotroph\| | Yeast | NULL | Probable | As nectar-tap_saprotroph (Põlme et al. 2020); Decay sugar-rich substrates (Põlme et al. 2020) | Põlme S, et al. 2020. Fungal Diversity 105:1-16 ((https://doi.org/10.1007/s13225-020-00466-2)) |
| Otu36372 | 1 | k__Fungi(100);p__Ascomycota(100);c__Saccharomycetes(100);o__Saccharomycetales(100);f__Metschnikowiaceae(100);g__Metschnikowia(100);s__Metschnikowia_pulcherrima(100); | Metschnikowia | 13 | Saprotroph | \|Undefined Saprotroph\| | Yeast | NULL | Probable | As nectar-tap_saprotroph (Põlme et al. 2020); Decay sugar-rich substrates (Põlme et al. 2020) | Põlme S, et al. 2020. Fungal Diversity 105:1-16 ((https://doi.org/10.1007/s13225-020-00466-2)) |
| Otu36373 | 1 | k__Fungi(100);p__Ascomycota(100);c__Saccharomycetes(100);o__Saccharomycetales(100);f__Metschnikowiaceae(100);g__Metschnikowia(100);s__Metschnikowia_pulcherrima(100); | Metschnikowia | 13 | Saprotroph | \|Undefined Saprotroph\| | Yeast | NULL | Probable | As nectar-tap_saprotroph (Põlme et al. 2020); Decay sugar-rich substrates (Põlme et al. 2020) | Põlme S, et al. 2020. Fungal Diversity 105:1-16 ((https://doi.org/10.1007/s13225-020-00466-2)) |
| Otu36374 | 1 | k__Fungi(100);p__Ascomycota(100);c__Saccharomycetes(100);o__Saccharomycetales(100);f__Metschnikowiaceae(100);g__Metschnikowia(100);s__Metschnikowia_pulcherrima(100); | Metschnikowia | 13 | Saprotroph | \|Undefined Saprotroph\| | Yeast | NULL | Probable | As nectar-tap_saprotroph (Põlme et al. 2020); Decay sugar-rich substrates (Põlme et al. 2020) | Põlme S, et al. 2020. Fungal Diversity 105:1-16 ((https://doi.org/10.1007/s13225-020-00466-2)) |
| Otu36375 | 1 | k__Fungi(100);p__Ascomycota(100);c__Saccharomycetes(100);o__Saccharomycetales(100);f__Metschnikowiaceae(100);g__Metschnikowia(100);s__Metschnikowia_pulcherrima(100); | Metschnikowia | 13 | Saprotroph | \|Undefined Saprotroph\| | Yeast | NULL | Probable | As nectar-tap_saprotroph (Põlme et al. 2020); Decay sugar-rich substrates (Põlme et al. 2020) | Põlme S, et al. 2020. Fungal Diversity 105:1-16 ((https://doi.org/10.1007/s13225-020-00466-2)) |
| Otu36379 | 1 | k__Fungi(100);p__Ascomycota(100);c__Saccharomycetes(100);o__Saccharomycetales(100);f__Metschnikowiaceae(100);g__Metschnikowia(100);s__Metschnikowia_pulcherrima(100); | Metschnikowia | 13 | Saprotroph | \|Undefined Saprotroph\| | Yeast | NULL | Probable | As nectar-tap_saprotroph (Põlme et al. 2020); Decay sugar-rich substrates (Põlme et al. 2020) | Põlme S, et al. 2020. Fungal Diversity 105:1-16 ((https://doi.org/10.1007/s13225-020-00466-2)) |
| Otu36381 | 1 | k__Fungi(100);p__Ascomycota(100);c__Saccharomycetes(100);o__Saccharomycetales(100);f__Metschnikowiaceae(100);g__Metschnikowia(100);g__Metschnikowia_unclassified(100); | Metschnikowia | 13 | Saprotroph | \|Undefined Saprotroph\| | Yeast | NULL | Probable | As nectar-tap_saprotroph (Põlme et al. 2020); Decay sugar-rich substrates (Põlme et al. 2020) | Põlme S, et al. 2020. Fungal Diversity 105:1-16 ((https://doi.org/10.1007/s13225-020-00466-2)) |
| Otu36382 | 1 | k__Fungi(100);p__Ascomycota(100);c__Saccharomycetes(100);o__Saccharomycetales(100);f__Metschnikowiaceae(100);g__Metschnikowia(100);s__Metschnikowia_pulcherrima(100); | Metschnikowia | 13 | Saprotroph | \|Undefined Saprotroph\| | Yeast | NULL | Probable | As nectar-tap_saprotroph (Põlme et al. 2020); Decay sugar-rich substrates (Põlme et al. 2020) | Põlme S, et al. 2020. Fungal Diversity 105:1-16 ((https://doi.org/10.1007/s13225-020-00466-2)) |
| Otu36383 | 1 | k__Fungi(100);p__Ascomycota(100);c__Saccharomycetes(100);o__Saccharomycetales(100);f__Metschnikowiaceae(100);g__Metschnikowia(100);g__Metschnikowia_unclassified(100); | Metschnikowia | 13 | Saprotroph | \|Undefined Saprotroph\| | Yeast | NULL | Probable | As nectar-tap_saprotroph (Põlme et al. 2020); Decay sugar-rich substrates (Põlme et al. 2020) | Põlme S, et al. 2020. Fungal Diversity 105:1-16 ((https://doi.org/10.1007/s13225-020-00466-2)) |
| Otu36385 | 1 | k__Fungi(100);p__Ascomycota(100);c__Saccharomycetes(100);o__Saccharomycetales(100);f__Metschnikowiaceae(100);g__Metschnikowia(100);s__Metschnikowia_pulcherrima(100); | Metschnikowia | 13 | Saprotroph | \|Undefined Saprotroph\| | Yeast | NULL | Probable | As nectar-tap_saprotroph (Põlme et al. 2020); Decay sugar-rich substrates (Põlme et al. 2020) | Põlme S, et al. 2020. Fungal Diversity 105:1-16 ((https://doi.org/10.1007/s13225-020-00466-2)) |
| Otu36386 | 1 | k__Fungi(100);p__Ascomycota(100);c__Saccharomycetes(100);o__Saccharomycetales(100);f__Metschnikowiaceae(100);g__Metschnikowia(100);s__Metschnikowia_pulcherrima(100); | Metschnikowia | 13 | Saprotroph | \|Undefined Saprotroph\| | Yeast | NULL | Probable | As nectar-tap_saprotroph (Põlme et al. 2020); Decay sugar-rich substrates (Põlme et al. 2020) | Põlme S, et al. 2020. Fungal Diversity 105:1-16 ((https://doi.org/10.1007/s13225-020-00466-2)) |
| Otu36387 | 1 | k__Fungi(100);p__Ascomycota(100);c__Saccharomycetes(100);o__Saccharomycetales(100);f__Metschnikowiaceae(100);g__Metschnikowia(100);g__Metschnikowia_unclassified(100); | Metschnikowia | 13 | Saprotroph | \|Undefined Saprotroph\| | Yeast | NULL | Probable | As nectar-tap_saprotroph (Põlme et al. 2020); Decay sugar-rich substrates (Põlme et al. 2020) | Põlme S, et al. 2020. Fungal Diversity 105:1-16 ((https://doi.org/10.1007/s13225-020-00466-2)) |
| Otu36388 | 1 | k__Fungi(100);p__Ascomycota(100);c__Saccharomycetes(100);o__Saccharomycetales(100);f__Metschnikowiaceae(100);g__Metschnikowia(100);s__Metschnikowia_pulcherrima(100); | Metschnikowia | 13 | Saprotroph | \|Undefined Saprotroph\| | Yeast | NULL | Probable | As nectar-tap_saprotroph (Põlme et al. 2020); Decay sugar-rich substrates (Põlme et al. 2020) | Põlme S, et al. 2020. Fungal Diversity 105:1-16 ((https://doi.org/10.1007/s13225-020-00466-2)) |
| Otu36389 | 1 | k__Fungi(100);p__Ascomycota(100);c__Saccharomycetes(100);o__Saccharomycetales(100);f__Metschnikowiaceae(100);g__Metschnikowia(100);s__Metschnikowia_pulcherrima(100); | Metschnikowia | 13 | Saprotroph | \|Undefined Saprotroph\| | Yeast | NULL | Probable | As nectar-tap_saprotroph (Põlme et al. 2020); Decay sugar-rich substrates (Põlme et al. 2020) | Põlme S, et al. 2020. Fungal Diversity 105:1-16 ((https://doi.org/10.1007/s13225-020-00466-2)) |
| Otu36390 | 1 | k__Fungi(100);p__Ascomycota(100);c__Saccharomycetes(100);o__Saccharomycetales(100);f__Metschnikowiaceae(100);g__Metschnikowia(100);s__Metschnikowia_pulcherrima(100); | Metschnikowia | 13 | Saprotroph | \|Undefined Saprotroph\| | Yeast | NULL | Probable | As nectar-tap_saprotroph (Põlme et al. 2020); Decay sugar-rich substrates (Põlme et al. 2020) | Põlme S, et al. 2020. Fungal Diversity 105:1-16 ((https://doi.org/10.1007/s13225-020-00466-2)) |
| Otu36391 | 1 | k__Fungi(100);p__Ascomycota(100);c__Saccharomycetes(100);o__Saccharomycetales(100);f__Metschnikowiaceae(100);g__Metschnikowia(100);g__Metschnikowia_unclassified(100); | Metschnikowia | 13 | Saprotroph | \|Undefined Saprotroph\| | Yeast | NULL | Probable | As nectar-tap_saprotroph (Põlme et al. 2020); Decay sugar-rich substrates (Põlme et al. 2020) | Põlme S, et al. 2020. Fungal Diversity 105:1-16 ((https://doi.org/10.1007/s13225-020-00466-2)) |
| Otu36392 | 1 | k__Fungi(100);p__Ascomycota(100);c__Saccharomycetes(100);o__Saccharomycetales(100);f__Metschnikowiaceae(100);g__Metschnikowia(100);s__Metschnikowia_pulcherrima(100); | Metschnikowia | 13 | Saprotroph | \|Undefined Saprotroph\| | Yeast | NULL | Probable | As nectar-tap_saprotroph (Põlme et al. 2020); Decay sugar-rich substrates (Põlme et al. 2020) | Põlme S, et al. 2020. Fungal Diversity 105:1-16 ((https://doi.org/10.1007/s13225-020-00466-2)) |
| Otu36393 | 1 | k__Fungi(100);p__Ascomycota(100);c__Saccharomycetes(100);o__Saccharomycetales(100);f__Metschnikowiaceae(100);g__Metschnikowia(100);s__Metschnikowia_pulcherrima(100); | Metschnikowia | 13 | Saprotroph | \|Undefined Saprotroph\| | Yeast | NULL | Probable | As nectar-tap_saprotroph (Põlme et al. 2020); Decay sugar-rich substrates (Põlme et al. 2020) | Põlme S, et al. 2020. Fungal Diversity 105:1-16 ((https://doi.org/10.1007/s13225-020-00466-2)) |
| Otu36395 | 1 | k__Fungi(100);p__Ascomycota(100);c__Saccharomycetes(100);o__Saccharomycetales(100);f__Metschnikowiaceae(100);g__Metschnikowia(100);s__Metschnikowia_pulcherrima(100); | Metschnikowia | 13 | Saprotroph | \|Undefined Saprotroph\| | Yeast | NULL | Probable | As nectar-tap_saprotroph (Põlme et al. 2020); Decay sugar-rich substrates (Põlme et al. 2020) | Põlme S, et al. 2020. Fungal Diversity 105:1-16 ((https://doi.org/10.1007/s13225-020-00466-2)) |
| Otu36396 | 1 | k__Fungi(100);p__Ascomycota(100);c__Saccharomycetes(100);o__Saccharomycetales(100);f__Metschnikowiaceae(100);g__Metschnikowia(100);g__Metschnikowia_unclassified(100); | Metschnikowia | 13 | Saprotroph | \|Undefined Saprotroph\| | Yeast | NULL | Probable | As nectar-tap_saprotroph (Põlme et al. 2020); Decay sugar-rich substrates (Põlme et al. 2020) | Põlme S, et al. 2020. Fungal Diversity 105:1-16 ((https://doi.org/10.1007/s13225-020-00466-2)) |
| Otu36398 | 1 | k__Fungi(100);p__Ascomycota(100);c__Saccharomycetes(100);o__Saccharomycetales(100);f__Metschnikowiaceae(100);g__Metschnikowia(100);s__Metschnikowia_pulcherrima(100); | Metschnikowia | 13 | Saprotroph | \|Undefined Saprotroph\| | Yeast | NULL | Probable | As nectar-tap_saprotroph (Põlme et al. 2020); Decay sugar-rich substrates (Põlme et al. 2020) | Põlme S, et al. 2020. Fungal Diversity 105:1-16 ((https://doi.org/10.1007/s13225-020-00466-2)) |
| Otu36400 | 1 | k__Fungi(100);p__Ascomycota(100);c__Saccharomycetes(100);o__Saccharomycetales(100);f__Metschnikowiaceae(100);g__Metschnikowia(100);s__Metschnikowia_pulcherrima(100); | Metschnikowia | 13 | Saprotroph | \|Undefined Saprotroph\| | Yeast | NULL | Probable | As nectar-tap_saprotroph (Põlme et al. 2020); Decay sugar-rich substrates (Põlme et al. 2020) | Põlme S, et al. 2020. Fungal Diversity 105:1-16 ((https://doi.org/10.1007/s13225-020-00466-2)) |
| Otu36401 | 1 | k__Fungi(100);p__Ascomycota(100);c__Saccharomycetes(100);o__Saccharomycetales(100);f__Metschnikowiaceae(100);g__Metschnikowia(100);s__Metschnikowia_pulcherrima(100); | Metschnikowia | 13 | Saprotroph | \|Undefined Saprotroph\| | Yeast | NULL | Probable | As nectar-tap_saprotroph (Põlme et al. 2020); Decay sugar-rich substrates (Põlme et al. 2020) | Põlme S, et al. 2020. Fungal Diversity 105:1-16 ((https://doi.org/10.1007/s13225-020-00466-2)) |
| Otu36403 | 1 | k__Fungi(100);p__Ascomycota(100);c__Saccharomycetes(100);o__Saccharomycetales(100);f__Metschnikowiaceae(100);g__Metschnikowia(100);s__Metschnikowia_pulcherrima(100); | Metschnikowia | 13 | Saprotroph | \|Undefined Saprotroph\| | Yeast | NULL | Probable | As nectar-tap_saprotroph (Põlme et al. 2020); Decay sugar-rich substrates (Põlme et al. 2020) | Põlme S, et al. 2020. Fungal Diversity 105:1-16 ((https://doi.org/10.1007/s13225-020-00466-2)) |
| Otu36404 | 1 | k__Fungi(100);p__Ascomycota(100);c__Saccharomycetes(100);o__Saccharomycetales(100);f__Metschnikowiaceae(100);g__Metschnikowia(100);g__Metschnikowia_unclassified(100); | Metschnikowia | 13 | Saprotroph | \|Undefined Saprotroph\| | Yeast | NULL | Probable | As nectar-tap_saprotroph (Põlme et al. 2020); Decay sugar-rich substrates (Põlme et al. 2020) | Põlme S, et al. 2020. Fungal Diversity 105:1-16 ((https://doi.org/10.1007/s13225-020-00466-2)) |
| Otu36405 | 1 | k__Fungi(100);p__Ascomycota(100);c__Saccharomycetes(100);o__Saccharomycetales(100);f__Metschnikowiaceae(100);g__Metschnikowia(100);s__Metschnikowia_pulcherrima(100); | Metschnikowia | 13 | Saprotroph | \|Undefined Saprotroph\| | Yeast | NULL | Probable | As nectar-tap_saprotroph (Põlme et al. 2020); Decay sugar-rich substrates (Põlme et al. 2020) | Põlme S, et al. 2020. Fungal Diversity 105:1-16 ((https://doi.org/10.1007/s13225-020-00466-2)) |
| Otu36408 | 1 | k__Fungi(100);p__Ascomycota(100);c__Saccharomycetes(100);o__Saccharomycetales(100);f__Metschnikowiaceae(100);g__Metschnikowia(100);s__Metschnikowia_pulcherrima(100); | Metschnikowia | 13 | Saprotroph | \|Undefined Saprotroph\| | Yeast | NULL | Probable | As nectar-tap_saprotroph (Põlme et al. 2020); Decay sugar-rich substrates (Põlme et al. 2020) | Põlme S, et al. 2020. Fungal Diversity 105:1-16 ((https://doi.org/10.1007/s13225-020-00466-2)) |
| Otu36409 | 1 | k__Fungi(100);p__Ascomycota(100);c__Saccharomycetes(100);o__Saccharomycetales(100);f__Metschnikowiaceae(100);g__Metschnikowia(100);s__Metschnikowia_pulcherrima(100); | Metschnikowia | 13 | Saprotroph | \|Undefined Saprotroph\| | Yeast | NULL | Probable | As nectar-tap_saprotroph (Põlme et al. 2020); Decay sugar-rich substrates (Põlme et al. 2020) | Põlme S, et al. 2020. Fungal Diversity 105:1-16 ((https://doi.org/10.1007/s13225-020-00466-2)) |
| Otu36410 | 1 | k__Fungi(100);p__Ascomycota(100);c__Saccharomycetes(100);o__Saccharomycetales(100);f__Metschnikowiaceae(100);g__Metschnikowia(100);s__Metschnikowia_pulcherrima(100); | Metschnikowia | 13 | Saprotroph | \|Undefined Saprotroph\| | Yeast | NULL | Probable | As nectar-tap_saprotroph (Põlme et al. 2020); Decay sugar-rich substrates (Põlme et al. 2020) | Põlme S, et al. 2020. Fungal Diversity 105:1-16 ((https://doi.org/10.1007/s13225-020-00466-2)) |
| Otu36411 | 1 | k__Fungi(100);p__Ascomycota(100);c__Saccharomycetes(100);o__Saccharomycetales(100);f__Metschnikowiaceae(100);g__Metschnikowia(100);g__Metschnikowia_unclassified(100); | Metschnikowia | 13 | Saprotroph | \|Undefined Saprotroph\| | Yeast | NULL | Probable | As nectar-tap_saprotroph (Põlme et al. 2020); Decay sugar-rich substrates (Põlme et al. 2020) | Põlme S, et al. 2020. Fungal Diversity 105:1-16 ((https://doi.org/10.1007/s13225-020-00466-2)) |
| Otu36413 | 1 | k__Fungi(100);p__Ascomycota(100);c__Saccharomycetes(100);o__Saccharomycetales(100);f__Metschnikowiaceae(100);g__Metschnikowia(100);s__Metschnikowia_pulcherrima(100); | Metschnikowia | 13 | Saprotroph | \|Undefined Saprotroph\| | Yeast | NULL | Probable | As nectar-tap_saprotroph (Põlme et al. 2020); Decay sugar-rich substrates (Põlme et al. 2020) | Põlme S, et al. 2020. Fungal Diversity 105:1-16 ((https://doi.org/10.1007/s13225-020-00466-2)) |
| Otu36416 | 1 | k__Fungi(100);p__Ascomycota(100);c__Saccharomycetes(100);o__Saccharomycetales(100);f__Metschnikowiaceae(100);g__Metschnikowia(100);s__Metschnikowia_pulcherrima(100); | Metschnikowia | 13 | Saprotroph | \|Undefined Saprotroph\| | Yeast | NULL | Probable | As nectar-tap_saprotroph (Põlme et al. 2020); Decay sugar-rich substrates (Põlme et al. 2020) | Põlme S, et al. 2020. Fungal Diversity 105:1-16 ((https://doi.org/10.1007/s13225-020-00466-2)) |
| Otu36423 | 1 | k__Fungi(100);p__Ascomycota(100);c__Saccharomycetes(100);o__Saccharomycetales(100);f__Metschnikowiaceae(100);g__Metschnikowia(100);g__Metschnikowia_unclassified(100); | Metschnikowia | 13 | Saprotroph | \|Undefined Saprotroph\| | Yeast | NULL | Probable | As nectar-tap_saprotroph (Põlme et al. 2020); Decay sugar-rich substrates (Põlme et al. 2020) | Põlme S, et al. 2020. Fungal Diversity 105:1-16 ((https://doi.org/10.1007/s13225-020-00466-2)) |
| Otu36424 | 1 | k__Fungi(100);p__Ascomycota(100);c__Saccharomycetes(100);o__Saccharomycetales(100);f__Metschnikowiaceae(100);g__Metschnikowia(100);g__Metschnikowia_unclassified(100); | Metschnikowia | 13 | Saprotroph | \|Undefined Saprotroph\| | Yeast | NULL | Probable | As nectar-tap_saprotroph (Põlme et al. 2020); Decay sugar-rich substrates (Põlme et al. 2020) | Põlme S, et al. 2020. Fungal Diversity 105:1-16 ((https://doi.org/10.1007/s13225-020-00466-2)) |
| Otu36429 | 1 | k__Fungi(100);p__Ascomycota(100);c__Saccharomycetes(100);o__Saccharomycetales(100);f__Metschnikowiaceae(100);g__Metschnikowia(100);s__Metschnikowia_pulcherrima(100); | Metschnikowia | 13 | Saprotroph | \|Undefined Saprotroph\| | Yeast | NULL | Probable | As nectar-tap_saprotroph (Põlme et al. 2020); Decay sugar-rich substrates (Põlme et al. 2020) | Põlme S, et al. 2020. Fungal Diversity 105:1-16 ((https://doi.org/10.1007/s13225-020-00466-2)) |
| Otu36430 | 1 | k__Fungi(100);p__Ascomycota(100);c__Saccharomycetes(100);o__Saccharomycetales(100);f__Metschnikowiaceae(100);g__Metschnikowia(100);s__Metschnikowia_pulcherrima(100); | Metschnikowia | 13 | Saprotroph | \|Undefined Saprotroph\| | Yeast | NULL | Probable | As nectar-tap_saprotroph (Põlme et al. 2020); Decay sugar-rich substrates (Põlme et al. 2020) | Põlme S, et al. 2020. Fungal Diversity 105:1-16 ((https://doi.org/10.1007/s13225-020-00466-2)) |
| Otu36431 | 1 | k__Fungi(100);p__Ascomycota(100);c__Saccharomycetes(100);o__Saccharomycetales(100);f__Metschnikowiaceae(100);g__Metschnikowia(100);s__Metschnikowia_pulcherrima(100); | Metschnikowia | 13 | Saprotroph | \|Undefined Saprotroph\| | Yeast | NULL | Probable | As nectar-tap_saprotroph (Põlme et al. 2020); Decay sugar-rich substrates (Põlme et al. 2020) | Põlme S, et al. 2020. Fungal Diversity 105:1-16 ((https://doi.org/10.1007/s13225-020-00466-2)) |
| Otu36432 | 1 | k__Fungi(100);p__Ascomycota(100);c__Saccharomycetes(100);o__Saccharomycetales(100);f__Metschnikowiaceae(100);g__Metschnikowia(100);s__Metschnikowia_pulcherrima(100); | Metschnikowia | 13 | Saprotroph | \|Undefined Saprotroph\| | Yeast | NULL | Probable | As nectar-tap_saprotroph (Põlme et al. 2020); Decay sugar-rich substrates (Põlme et al. 2020) | Põlme S, et al. 2020. Fungal Diversity 105:1-16 ((https://doi.org/10.1007/s13225-020-00466-2)) |
| Otu36433 | 1 | k__Fungi(100);p__Ascomycota(100);c__Saccharomycetes(100);o__Saccharomycetales(100);f__Metschnikowiaceae(100);g__Metschnikowia(100);s__Metschnikowia_pulcherrima(100); | Metschnikowia | 13 | Saprotroph | \|Undefined Saprotroph\| | Yeast | NULL | Probable | As nectar-tap_saprotroph (Põlme et al. 2020); Decay sugar-rich substrates (Põlme et al. 2020) | Põlme S, et al. 2020. Fungal Diversity 105:1-16 ((https://doi.org/10.1007/s13225-020-00466-2)) |
| Otu36434 | 1 | k__Fungi(100);p__Ascomycota(100);c__Saccharomycetes(100);o__Saccharomycetales(100);f__Metschnikowiaceae(100);g__Metschnikowia(100);s__Metschnikowia_pulcherrima(100); | Metschnikowia | 13 | Saprotroph | \|Undefined Saprotroph\| | Yeast | NULL | Probable | As nectar-tap_saprotroph (Põlme et al. 2020); Decay sugar-rich substrates (Põlme et al. 2020) | Põlme S, et al. 2020. Fungal Diversity 105:1-16 ((https://doi.org/10.1007/s13225-020-00466-2)) |
| Otu36435 | 1 | k__Fungi(100);p__Ascomycota(100);c__Saccharomycetes(100);o__Saccharomycetales(100);f__Metschnikowiaceae(100);g__Metschnikowia(100);s__Metschnikowia_pulcherrima(100); | Metschnikowia | 13 | Saprotroph | \|Undefined Saprotroph\| | Yeast | NULL | Probable | As nectar-tap_saprotroph (Põlme et al. 2020); Decay sugar-rich substrates (Põlme et al. 2020) | Põlme S, et al. 2020. Fungal Diversity 105:1-16 ((https://doi.org/10.1007/s13225-020-00466-2)) |
| Otu36436 | 1 | k__Fungi(100);p__Ascomycota(100);c__Saccharomycetes(100);o__Saccharomycetales(100);f__Metschnikowiaceae(100);g__Metschnikowia(100);s__Metschnikowia_pulcherrima(100); | Metschnikowia | 13 | Saprotroph | \|Undefined Saprotroph\| | Yeast | NULL | Probable | As nectar-tap_saprotroph (Põlme et al. 2020); Decay sugar-rich substrates (Põlme et al. 2020) | Põlme S, et al. 2020. Fungal Diversity 105:1-16 ((https://doi.org/10.1007/s13225-020-00466-2)) |
| Otu36437 | 1 | k__Fungi(100);p__Ascomycota(100);c__Saccharomycetes(100);o__Saccharomycetales(100);f__Metschnikowiaceae(100);g__Metschnikowia(100);s__Metschnikowia_pulcherrima(100); | Metschnikowia | 13 | Saprotroph | \|Undefined Saprotroph\| | Yeast | NULL | Probable | As nectar-tap_saprotroph (Põlme et al. 2020); Decay sugar-rich substrates (Põlme et al. 2020) | Põlme S, et al. 2020. Fungal Diversity 105:1-16 ((https://doi.org/10.1007/s13225-020-00466-2)) |
| Otu36441 | 1 | k__Fungi(100);p__Ascomycota(100);c__Saccharomycetes(100);o__Saccharomycetales(100);f__Metschnikowiaceae(100);g__Metschnikowia(100);g__Metschnikowia_unclassified(100); | Metschnikowia | 13 | Saprotroph | \|Undefined Saprotroph\| | Yeast | NULL | Probable | As nectar-tap_saprotroph (Põlme et al. 2020); Decay sugar-rich substrates (Põlme et al. 2020) | Põlme S, et al. 2020. Fungal Diversity 105:1-16 ((https://doi.org/10.1007/s13225-020-00466-2)) |
| Otu36442 | 1 | k__Fungi(100);p__Ascomycota(100);c__Saccharomycetes(100);o__Saccharomycetales(100);f__Metschnikowiaceae(100);g__Metschnikowia(100);s__Metschnikowia_pulcherrima(100); | Metschnikowia | 13 | Saprotroph | \|Undefined Saprotroph\| | Yeast | NULL | Probable | As nectar-tap_saprotroph (Põlme et al. 2020); Decay sugar-rich substrates (Põlme et al. 2020) | Põlme S, et al. 2020. Fungal Diversity 105:1-16 ((https://doi.org/10.1007/s13225-020-00466-2)) |
| Otu36449 | 1 | k__Fungi(100);p__Ascomycota(100);c__Saccharomycetes(100);o__Saccharomycetales(100);f__Metschnikowiaceae(100);g__Metschnikowia(100);s__Metschnikowia_pulcherrima(100); | Metschnikowia | 13 | Saprotroph | \|Undefined Saprotroph\| | Yeast | NULL | Probable | As nectar-tap_saprotroph (Põlme et al. 2020); Decay sugar-rich substrates (Põlme et al. 2020) | Põlme S, et al. 2020. Fungal Diversity 105:1-16 ((https://doi.org/10.1007/s13225-020-00466-2)) |
| Otu36452 | 1 | k__Fungi(100);p__Ascomycota(100);c__Saccharomycetes(100);o__Saccharomycetales(100);f__Metschnikowiaceae(100);g__Metschnikowia(100);s__Metschnikowia_pulcherrima(100); | Metschnikowia | 13 | Saprotroph | \|Undefined Saprotroph\| | Yeast | NULL | Probable | As nectar-tap_saprotroph (Põlme et al. 2020); Decay sugar-rich substrates (Põlme et al. 2020) | Põlme S, et al. 2020. Fungal Diversity 105:1-16 ((https://doi.org/10.1007/s13225-020-00466-2)) |
| Otu36453 | 1 | k__Fungi(100);p__Ascomycota(100);c__Saccharomycetes(100);o__Saccharomycetales(100);f__Metschnikowiaceae(100);g__Metschnikowia(100);g__Metschnikowia_unclassified(100); | Metschnikowia | 13 | Saprotroph | \|Undefined Saprotroph\| | Yeast | NULL | Probable | As nectar-tap_saprotroph (Põlme et al. 2020); Decay sugar-rich substrates (Põlme et al. 2020) | Põlme S, et al. 2020. Fungal Diversity 105:1-16 ((https://doi.org/10.1007/s13225-020-00466-2)) |
| Otu36454 | 1 | k__Fungi(100);p__Ascomycota(100);c__Saccharomycetes(100);o__Saccharomycetales(100);f__Metschnikowiaceae(100);g__Metschnikowia(100);s__Metschnikowia_pulcherrima(100); | Metschnikowia | 13 | Saprotroph | \|Undefined Saprotroph\| | Yeast | NULL | Probable | As nectar-tap_saprotroph (Põlme et al. 2020); Decay sugar-rich substrates (Põlme et al. 2020) | Põlme S, et al. 2020. Fungal Diversity 105:1-16 ((https://doi.org/10.1007/s13225-020-00466-2)) |
| Otu36455 | 1 | k__Fungi(100);p__Ascomycota(100);c__Saccharomycetes(100);o__Saccharomycetales(100);f__Metschnikowiaceae(100);g__Metschnikowia(100);s__Metschnikowia_pulcherrima(100); | Metschnikowia | 13 | Saprotroph | \|Undefined Saprotroph\| | Yeast | NULL | Probable | As nectar-tap_saprotroph (Põlme et al. 2020); Decay sugar-rich substrates (Põlme et al. 2020) | Põlme S, et al. 2020. Fungal Diversity 105:1-16 ((https://doi.org/10.1007/s13225-020-00466-2)) |
| Otu36456 | 1 | k__Fungi(100);p__Ascomycota(100);c__Saccharomycetes(100);o__Saccharomycetales(100);f__Metschnikowiaceae(100);g__Metschnikowia(100);g__Metschnikowia_unclassified(100); | Metschnikowia | 13 | Saprotroph | \|Undefined Saprotroph\| | Yeast | NULL | Probable | As nectar-tap_saprotroph (Põlme et al. 2020); Decay sugar-rich substrates (Põlme et al. 2020) | Põlme S, et al. 2020. Fungal Diversity 105:1-16 ((https://doi.org/10.1007/s13225-020-00466-2)) |
| Otu36457 | 1 | k__Fungi(100);p__Ascomycota(100);c__Saccharomycetes(100);o__Saccharomycetales(100);f__Metschnikowiaceae(100);g__Metschnikowia(100);s__Metschnikowia_pulcherrima(100); | Metschnikowia | 13 | Saprotroph | \|Undefined Saprotroph\| | Yeast | NULL | Probable | As nectar-tap_saprotroph (Põlme et al. 2020); Decay sugar-rich substrates (Põlme et al. 2020) | Põlme S, et al. 2020. Fungal Diversity 105:1-16 ((https://doi.org/10.1007/s13225-020-00466-2)) |
| Otu36458 | 1 | k__Fungi(100);p__Ascomycota(100);c__Saccharomycetes(100);o__Saccharomycetales(100);f__Metschnikowiaceae(100);g__Metschnikowia(100);s__Metschnikowia_pulcherrima(100); | Metschnikowia | 13 | Saprotroph | \|Undefined Saprotroph\| | Yeast | NULL | Probable | As nectar-tap_saprotroph (Põlme et al. 2020); Decay sugar-rich substrates (Põlme et al. 2020) | Põlme S, et al. 2020. Fungal Diversity 105:1-16 ((https://doi.org/10.1007/s13225-020-00466-2)) |
| Otu36459 | 1 | k__Fungi(100);p__Ascomycota(100);c__Saccharomycetes(100);o__Saccharomycetales(100);f__Metschnikowiaceae(100);g__Metschnikowia(100);s__Metschnikowia_pulcherrima(100); | Metschnikowia | 13 | Saprotroph | \|Undefined Saprotroph\| | Yeast | NULL | Probable | As nectar-tap_saprotroph (Põlme et al. 2020); Decay sugar-rich substrates (Põlme et al. 2020) | Põlme S, et al. 2020. Fungal Diversity 105:1-16 ((https://doi.org/10.1007/s13225-020-00466-2)) |
| Otu36460 | 1 | k__Fungi(100);p__Ascomycota(100);c__Saccharomycetes(100);o__Saccharomycetales(100);f__Metschnikowiaceae(100);g__Metschnikowia(100);s__Metschnikowia_pulcherrima(100); | Metschnikowia | 13 | Saprotroph | \|Undefined Saprotroph\| | Yeast | NULL | Probable | As nectar-tap_saprotroph (Põlme et al. 2020); Decay sugar-rich substrates (Põlme et al. 2020) | Põlme S, et al. 2020. Fungal Diversity 105:1-16 ((https://doi.org/10.1007/s13225-020-00466-2)) |
| Otu36461 | 1 | k__Fungi(100);p__Ascomycota(100);c__Saccharomycetes(100);o__Saccharomycetales(100);f__Metschnikowiaceae(100);g__Metschnikowia(100);s__Metschnikowia_pulcherrima(100); | Metschnikowia | 13 | Saprotroph | \|Undefined Saprotroph\| | Yeast | NULL | Probable | As nectar-tap_saprotroph (Põlme et al. 2020); Decay sugar-rich substrates (Põlme et al. 2020) | Põlme S, et al. 2020. Fungal Diversity 105:1-16 ((https://doi.org/10.1007/s13225-020-00466-2)) |
| Otu36462 | 1 | k__Fungi(100);p__Ascomycota(100);c__Saccharomycetes(100);o__Saccharomycetales(100);f__Metschnikowiaceae(100);g__Metschnikowia(100);s__Metschnikowia_pulcherrima(100); | Metschnikowia | 13 | Saprotroph | \|Undefined Saprotroph\| | Yeast | NULL | Probable | As nectar-tap_saprotroph (Põlme et al. 2020); Decay sugar-rich substrates (Põlme et al. 2020) | Põlme S, et al. 2020. Fungal Diversity 105:1-16 ((https://doi.org/10.1007/s13225-020-00466-2)) |
| Otu36463 | 1 | k__Fungi(100);p__Ascomycota(100);c__Saccharomycetes(100);o__Saccharomycetales(100);f__Metschnikowiaceae(100);g__Metschnikowia(100);s__Metschnikowia_pulcherrima(100); | Metschnikowia | 13 | Saprotroph | \|Undefined Saprotroph\| | Yeast | NULL | Probable | As nectar-tap_saprotroph (Põlme et al. 2020); Decay sugar-rich substrates (Põlme et al. 2020) | Põlme S, et al. 2020. Fungal Diversity 105:1-16 ((https://doi.org/10.1007/s13225-020-00466-2)) |
| Otu36464 | 1 | k__Fungi(100);p__Ascomycota(100);c__Saccharomycetes(100);o__Saccharomycetales(100);f__Metschnikowiaceae(100);g__Metschnikowia(100);s__Metschnikowia_pulcherrima(100); | Metschnikowia | 13 | Saprotroph | \|Undefined Saprotroph\| | Yeast | NULL | Probable | As nectar-tap_saprotroph (Põlme et al. 2020); Decay sugar-rich substrates (Põlme et al. 2020) | Põlme S, et al. 2020. Fungal Diversity 105:1-16 ((https://doi.org/10.1007/s13225-020-00466-2)) |
| Otu36465 | 1 | k__Fungi(100);p__Ascomycota(100);c__Saccharomycetes(100);o__Saccharomycetales(100);f__Metschnikowiaceae(100);g__Metschnikowia(100);s__Metschnikowia_pulcherrima(100); | Metschnikowia | 13 | Saprotroph | \|Undefined Saprotroph\| | Yeast | NULL | Probable | As nectar-tap_saprotroph (Põlme et al. 2020); Decay sugar-rich substrates (Põlme et al. 2020) | Põlme S, et al. 2020. Fungal Diversity 105:1-16 ((https://doi.org/10.1007/s13225-020-00466-2)) |
| Otu36466 | 1 | k__Fungi(100);p__Ascomycota(100);c__Saccharomycetes(100);o__Saccharomycetales(100);f__Metschnikowiaceae(100);g__Metschnikowia(100);s__Metschnikowia_pulcherrima(100); | Metschnikowia | 13 | Saprotroph | \|Undefined Saprotroph\| | Yeast | NULL | Probable | As nectar-tap_saprotroph (Põlme et al. 2020); Decay sugar-rich substrates (Põlme et al. 2020) | Põlme S, et al. 2020. Fungal Diversity 105:1-16 ((https://doi.org/10.1007/s13225-020-00466-2)) |
| Otu36467 | 1 | k__Fungi(100);p__Ascomycota(100);c__Saccharomycetes(100);o__Saccharomycetales(100);f__Metschnikowiaceae(100);g__Metschnikowia(100);s__Metschnikowia_pulcherrima(100); | Metschnikowia | 13 | Saprotroph | \|Undefined Saprotroph\| | Yeast | NULL | Probable | As nectar-tap_saprotroph (Põlme et al. 2020); Decay sugar-rich substrates (Põlme et al. 2020) | Põlme S, et al. 2020. Fungal Diversity 105:1-16 ((https://doi.org/10.1007/s13225-020-00466-2)) |
| Otu36468 | 1 | k__Fungi(100);p__Ascomycota(100);c__Saccharomycetes(100);o__Saccharomycetales(100);f__Metschnikowiaceae(100);g__Metschnikowia(100);s__Metschnikowia_pulcherrima(100); | Metschnikowia | 13 | Saprotroph | \|Undefined Saprotroph\| | Yeast | NULL | Probable | As nectar-tap_saprotroph (Põlme et al. 2020); Decay sugar-rich substrates (Põlme et al. 2020) | Põlme S, et al. 2020. Fungal Diversity 105:1-16 ((https://doi.org/10.1007/s13225-020-00466-2)) |
| Otu36469 | 1 | k__Fungi(100);p__Ascomycota(100);c__Saccharomycetes(100);o__Saccharomycetales(100);f__Metschnikowiaceae(100);g__Metschnikowia(100);g__Metschnikowia_unclassified(100); | Metschnikowia | 13 | Saprotroph | \|Undefined Saprotroph\| | Yeast | NULL | Probable | As nectar-tap_saprotroph (Põlme et al. 2020); Decay sugar-rich substrates (Põlme et al. 2020) | Põlme S, et al. 2020. Fungal Diversity 105:1-16 ((https://doi.org/10.1007/s13225-020-00466-2)) |
| Otu36470 | 1 | k__Fungi(100);p__Ascomycota(100);c__Saccharomycetes(100);o__Saccharomycetales(100);f__Metschnikowiaceae(100);g__Metschnikowia(100);s__Metschnikowia_pulcherrima(100); | Metschnikowia | 13 | Saprotroph | \|Undefined Saprotroph\| | Yeast | NULL | Probable | As nectar-tap_saprotroph (Põlme et al. 2020); Decay sugar-rich substrates (Põlme et al. 2020) | Põlme S, et al. 2020. Fungal Diversity 105:1-16 ((https://doi.org/10.1007/s13225-020-00466-2)) |
| Otu36471 | 1 | k__Fungi(100);p__Ascomycota(100);c__Saccharomycetes(100);o__Saccharomycetales(100);f__Metschnikowiaceae(100);g__Metschnikowia(100);s__Metschnikowia_pulcherrima(100); | Metschnikowia | 13 | Saprotroph | \|Undefined Saprotroph\| | Yeast | NULL | Probable | As nectar-tap_saprotroph (Põlme et al. 2020); Decay sugar-rich substrates (Põlme et al. 2020) | Põlme S, et al. 2020. Fungal Diversity 105:1-16 ((https://doi.org/10.1007/s13225-020-00466-2)) |
| Otu36476 | 1 | k__Fungi(100);p__Ascomycota(100);c__Saccharomycetes(100);o__Saccharomycetales(100);f__Metschnikowiaceae(100);g__Metschnikowia(100);g__Metschnikowia_unclassified(100); | Metschnikowia | 13 | Saprotroph | \|Undefined Saprotroph\| | Yeast | NULL | Probable | As nectar-tap_saprotroph (Põlme et al. 2020); Decay sugar-rich substrates (Põlme et al. 2020) | Põlme S, et al. 2020. Fungal Diversity 105:1-16 ((https://doi.org/10.1007/s13225-020-00466-2)) |
| Otu36477 | 1 | k__Fungi(100);p__Ascomycota(100);c__Saccharomycetes(100);o__Saccharomycetales(100);f__Metschnikowiaceae(100);g__Metschnikowia(100);s__Metschnikowia_pulcherrima(100); | Metschnikowia | 13 | Saprotroph | \|Undefined Saprotroph\| | Yeast | NULL | Probable | As nectar-tap_saprotroph (Põlme et al. 2020); Decay sugar-rich substrates (Põlme et al. 2020) | Põlme S, et al. 2020. Fungal Diversity 105:1-16 ((https://doi.org/10.1007/s13225-020-00466-2)) |
| Otu36478 | 1 | k__Fungi(100);p__Ascomycota(100);c__Saccharomycetes(100);o__Saccharomycetales(100);f__Metschnikowiaceae(100);g__Metschnikowia(100);s__Metschnikowia_pulcherrima(100); | Metschnikowia | 13 | Saprotroph | \|Undefined Saprotroph\| | Yeast | NULL | Probable | As nectar-tap_saprotroph (Põlme et al. 2020); Decay sugar-rich substrates (Põlme et al. 2020) | Põlme S, et al. 2020. Fungal Diversity 105:1-16 ((https://doi.org/10.1007/s13225-020-00466-2)) |
| Otu36480 | 1 | k__Fungi(100);p__Ascomycota(100);c__Saccharomycetes(100);o__Saccharomycetales(100);f__Metschnikowiaceae(100);g__Metschnikowia(100);g__Metschnikowia_unclassified(100); | Metschnikowia | 13 | Saprotroph | \|Undefined Saprotroph\| | Yeast | NULL | Probable | As nectar-tap_saprotroph (Põlme et al. 2020); Decay sugar-rich substrates (Põlme et al. 2020) | Põlme S, et al. 2020. Fungal Diversity 105:1-16 ((https://doi.org/10.1007/s13225-020-00466-2)) |
| Otu36481 | 1 | k__Fungi(100);p__Ascomycota(100);c__Saccharomycetes(100);o__Saccharomycetales(100);f__Metschnikowiaceae(100);g__Metschnikowia(100);s__Metschnikowia_pulcherrima(100); | Metschnikowia | 13 | Saprotroph | \|Undefined Saprotroph\| | Yeast | NULL | Probable | As nectar-tap_saprotroph (Põlme et al. 2020); Decay sugar-rich substrates (Põlme et al. 2020) | Põlme S, et al. 2020. Fungal Diversity 105:1-16 ((https://doi.org/10.1007/s13225-020-00466-2)) |
| Otu36482 | 1 | k__Fungi(100);p__Ascomycota(100);c__Saccharomycetes(100);o__Saccharomycetales(100);f__Metschnikowiaceae(100);g__Metschnikowia(100);s__Metschnikowia_pulcherrima(100); | Metschnikowia | 13 | Saprotroph | \|Undefined Saprotroph\| | Yeast | NULL | Probable | As nectar-tap_saprotroph (Põlme et al. 2020); Decay sugar-rich substrates (Põlme et al. 2020) | Põlme S, et al. 2020. Fungal Diversity 105:1-16 ((https://doi.org/10.1007/s13225-020-00466-2)) |
| Otu36483 | 1 | k__Fungi(100);p__Ascomycota(100);c__Saccharomycetes(100);o__Saccharomycetales(100);f__Metschnikowiaceae(100);g__Metschnikowia(100);s__Metschnikowia_pulcherrima(100); | Metschnikowia | 13 | Saprotroph | \|Undefined Saprotroph\| | Yeast | NULL | Probable | As nectar-tap_saprotroph (Põlme et al. 2020); Decay sugar-rich substrates (Põlme et al. 2020) | Põlme S, et al. 2020. Fungal Diversity 105:1-16 ((https://doi.org/10.1007/s13225-020-00466-2)) |
| Otu36484 | 1 | k__Fungi(100);p__Ascomycota(100);c__Saccharomycetes(100);o__Saccharomycetales(100);f__Metschnikowiaceae(100);g__Metschnikowia(100);s__Metschnikowia_pulcherrima(100); | Metschnikowia | 13 | Saprotroph | \|Undefined Saprotroph\| | Yeast | NULL | Probable | As nectar-tap_saprotroph (Põlme et al. 2020); Decay sugar-rich substrates (Põlme et al. 2020) | Põlme S, et al. 2020. Fungal Diversity 105:1-16 ((https://doi.org/10.1007/s13225-020-00466-2)) |
| Otu36485 | 1 | k__Fungi(100);p__Ascomycota(100);c__Saccharomycetes(100);o__Saccharomycetales(100);f__Metschnikowiaceae(100);g__Metschnikowia(100);s__Metschnikowia_pulcherrima(100); | Metschnikowia | 13 | Saprotroph | \|Undefined Saprotroph\| | Yeast | NULL | Probable | As nectar-tap_saprotroph (Põlme et al. 2020); Decay sugar-rich substrates (Põlme et al. 2020) | Põlme S, et al. 2020. Fungal Diversity 105:1-16 ((https://doi.org/10.1007/s13225-020-00466-2)) |
| Otu36488 | 1 | k__Fungi(100);p__Ascomycota(100);c__Saccharomycetes(100);o__Saccharomycetales(100);f__Metschnikowiaceae(100);g__Metschnikowia(100);s__Metschnikowia_pulcherrima(100); | Metschnikowia | 13 | Saprotroph | \|Undefined Saprotroph\| | Yeast | NULL | Probable | As nectar-tap_saprotroph (Põlme et al. 2020); Decay sugar-rich substrates (Põlme et al. 2020) | Põlme S, et al. 2020. Fungal Diversity 105:1-16 ((https://doi.org/10.1007/s13225-020-00466-2)) |
| Otu36491 | 1 | k__Fungi(100);p__Ascomycota(100);c__Saccharomycetes(100);o__Saccharomycetales(100);f__Metschnikowiaceae(100);g__Metschnikowia(100);s__Metschnikowia_pulcherrima(100); | Metschnikowia | 13 | Saprotroph | \|Undefined Saprotroph\| | Yeast | NULL | Probable | As nectar-tap_saprotroph (Põlme et al. 2020); Decay sugar-rich substrates (Põlme et al. 2020) | Põlme S, et al. 2020. Fungal Diversity 105:1-16 ((https://doi.org/10.1007/s13225-020-00466-2)) |
| Otu36493 | 1 | k__Fungi(100);p__Ascomycota(100);c__Saccharomycetes(100);o__Saccharomycetales(100);f__Metschnikowiaceae(100);g__Metschnikowia(100);g__Metschnikowia_unclassified(100); | Metschnikowia | 13 | Saprotroph | \|Undefined Saprotroph\| | Yeast | NULL | Probable | As nectar-tap_saprotroph (Põlme et al. 2020); Decay sugar-rich substrates (Põlme et al. 2020) | Põlme S, et al. 2020. Fungal Diversity 105:1-16 ((https://doi.org/10.1007/s13225-020-00466-2)) |
| Otu36494 | 1 | k__Fungi(100);p__Ascomycota(100);c__Sordariomycetes(100);o__Coniochaetales(100);f__Coniochaetaceae(100);g__Coniochaeta(100);g__Coniochaeta_unclassified(100); | Coniochaeta | 13 | Pathotroph-Saprotroph-Symbiotroph | Animal Pathogen-Dung Saprotroph-Endophyte-Lichen Parasite-Plant Pathogen-\|Undefined Saprotroph\| | Microfungus | NULL | Possible | As foliar_endophyte (Põlme et al. 2020); Decay leaf, fruit, seed, soil, dung, animal material (Põlme et al. 2020) | Bell A. 1983. Dung Fungi: An Illustrated Guide to Coprophilous Fungi in New Zealand. Victoria University Press, Wellington (ISBN: 978-0864730015); Damm U, et al. 2010. Persoonia 24:60-80 ((http://doi.org/10.3767/003158510X500705)); Costa IPMW, et al. 2012. Mycotaxon 119:e26 ((http://mycotaxon.com/resources/checklists/costa_v119_checklist.pdf)); Tedersoo L, et al. 2014. Science 346:1256688 ((https://doi.org/10.1126/science.1256688)); Irinyi L, et al. 2015. Medical Mycology 53:313-337 ((https://doi.org/10.1093/mmy/myv008)); Lawrey JD, Diederich P. 2016. http://www.lichenicolous.net (accessed 24 May 2017) ((http://www.lichenicolous.net)); Põlme S, et al. 2020. Fungal Diversity 105:1-16 ((https://doi.org/10.1007/s13225-020-00466-2)) |
| Otu36495 | 1 | k__Fungi(100);p__Ascomycota(100);c__Saccharomycetes(100);o__Saccharomycetales(100);f__Metschnikowiaceae(100);g__Metschnikowia(100);g__Metschnikowia_unclassified(100); | Metschnikowia | 13 | Saprotroph | \|Undefined Saprotroph\| | Yeast | NULL | Probable | As nectar-tap_saprotroph (Põlme et al. 2020); Decay sugar-rich substrates (Põlme et al. 2020) | Põlme S, et al. 2020. Fungal Diversity 105:1-16 ((https://doi.org/10.1007/s13225-020-00466-2)) |
| Otu36496 | 1 | k__Fungi(100);p__Ascomycota(100);c__Saccharomycetes(100);o__Saccharomycetales(100);f__Metschnikowiaceae(100);g__Metschnikowia(100);g__Metschnikowia_unclassified(100); | Metschnikowia | 13 | Saprotroph | \|Undefined Saprotroph\| | Yeast | NULL | Probable | As nectar-tap_saprotroph (Põlme et al. 2020); Decay sugar-rich substrates (Põlme et al. 2020) | Põlme S, et al. 2020. Fungal Diversity 105:1-16 ((https://doi.org/10.1007/s13225-020-00466-2)) |
| Otu36497 | 1 | k__Fungi(100);p__Ascomycota(100);c__Saccharomycetes(100);o__Saccharomycetales(100);f__Metschnikowiaceae(100);g__Metschnikowia(100);g__Metschnikowia_unclassified(100); | Metschnikowia | 13 | Saprotroph | \|Undefined Saprotroph\| | Yeast | NULL | Probable | As nectar-tap_saprotroph (Põlme et al. 2020); Decay sugar-rich substrates (Põlme et al. 2020) | Põlme S, et al. 2020. Fungal Diversity 105:1-16 ((https://doi.org/10.1007/s13225-020-00466-2)) |
| Otu36498 | 1 | k__Fungi(100);p__Ascomycota(100);c__Saccharomycetes(100);o__Saccharomycetales(100);f__Metschnikowiaceae(100);g__Metschnikowia(100);g__Metschnikowia_unclassified(100); | Metschnikowia | 13 | Saprotroph | \|Undefined Saprotroph\| | Yeast | NULL | Probable | As nectar-tap_saprotroph (Põlme et al. 2020); Decay sugar-rich substrates (Põlme et al. 2020) | Põlme S, et al. 2020. Fungal Diversity 105:1-16 ((https://doi.org/10.1007/s13225-020-00466-2)) |
| Otu36499 | 1 | k__Fungi(100);p__Ascomycota(100);c__Saccharomycetes(100);o__Saccharomycetales(100);f__Metschnikowiaceae(100);g__Metschnikowia(100);s__Metschnikowia_pulcherrima(100); | Metschnikowia | 13 | Saprotroph | \|Undefined Saprotroph\| | Yeast | NULL | Probable | As nectar-tap_saprotroph (Põlme et al. 2020); Decay sugar-rich substrates (Põlme et al. 2020) | Põlme S, et al. 2020. Fungal Diversity 105:1-16 ((https://doi.org/10.1007/s13225-020-00466-2)) |
| Otu36500 | 1 | k__Fungi(100);p__Ascomycota(100);c__Saccharomycetes(100);o__Saccharomycetales(100);f__Metschnikowiaceae(100);g__Metschnikowia(100);s__Metschnikowia_pulcherrima(100); | Metschnikowia | 13 | Saprotroph | \|Undefined Saprotroph\| | Yeast | NULL | Probable | As nectar-tap_saprotroph (Põlme et al. 2020); Decay sugar-rich substrates (Põlme et al. 2020) | Põlme S, et al. 2020. Fungal Diversity 105:1-16 ((https://doi.org/10.1007/s13225-020-00466-2)) |
| Otu36501 | 1 | k__Fungi(100);p__Ascomycota(100);c__Saccharomycetes(100);o__Saccharomycetales(100);f__Metschnikowiaceae(100);g__Metschnikowia(100);s__Metschnikowia_pulcherrima(100); | Metschnikowia | 13 | Saprotroph | \|Undefined Saprotroph\| | Yeast | NULL | Probable | As nectar-tap_saprotroph (Põlme et al. 2020); Decay sugar-rich substrates (Põlme et al. 2020) | Põlme S, et al. 2020. Fungal Diversity 105:1-16 ((https://doi.org/10.1007/s13225-020-00466-2)) |
| Otu36502 | 1 | k__Fungi(100);p__Ascomycota(100);c__Saccharomycetes(100);o__Saccharomycetales(100);f__Metschnikowiaceae(100);g__Metschnikowia(100);s__Metschnikowia_pulcherrima(100); | Metschnikowia | 13 | Saprotroph | \|Undefined Saprotroph\| | Yeast | NULL | Probable | As nectar-tap_saprotroph (Põlme et al. 2020); Decay sugar-rich substrates (Põlme et al. 2020) | Põlme S, et al. 2020. Fungal Diversity 105:1-16 ((https://doi.org/10.1007/s13225-020-00466-2)) |
| Otu36503 | 1 | k__Fungi(100);p__Ascomycota(100);c__Saccharomycetes(100);o__Saccharomycetales(100);f__Metschnikowiaceae(100);g__Metschnikowia(100);s__Metschnikowia_pulcherrima(100); | Metschnikowia | 13 | Saprotroph | \|Undefined Saprotroph\| | Yeast | NULL | Probable | As nectar-tap_saprotroph (Põlme et al. 2020); Decay sugar-rich substrates (Põlme et al. 2020) | Põlme S, et al. 2020. Fungal Diversity 105:1-16 ((https://doi.org/10.1007/s13225-020-00466-2)) |
| Otu36504 | 1 | k__Fungi(100);p__Ascomycota(100);c__Saccharomycetes(100);o__Saccharomycetales(100);f__Metschnikowiaceae(100);g__Metschnikowia(100);g__Metschnikowia_unclassified(100); | Metschnikowia | 13 | Saprotroph | \|Undefined Saprotroph\| | Yeast | NULL | Probable | As nectar-tap_saprotroph (Põlme et al. 2020); Decay sugar-rich substrates (Põlme et al. 2020) | Põlme S, et al. 2020. Fungal Diversity 105:1-16 ((https://doi.org/10.1007/s13225-020-00466-2)) |
| Otu36505 | 1 | k__Fungi(100);p__Ascomycota(100);c__Saccharomycetes(100);o__Saccharomycetales(100);f__Metschnikowiaceae(100);g__Metschnikowia(100);s__Metschnikowia_pulcherrima(100); | Metschnikowia | 13 | Saprotroph | \|Undefined Saprotroph\| | Yeast | NULL | Probable | As nectar-tap_saprotroph (Põlme et al. 2020); Decay sugar-rich substrates (Põlme et al. 2020) | Põlme S, et al. 2020. Fungal Diversity 105:1-16 ((https://doi.org/10.1007/s13225-020-00466-2)) |
| Otu36507 | 1 | k__Fungi(100);p__Ascomycota(100);c__Saccharomycetes(100);o__Saccharomycetales(100);f__Metschnikowiaceae(100);g__Metschnikowia(100);g__Metschnikowia_unclassified(100); | Metschnikowia | 13 | Saprotroph | \|Undefined Saprotroph\| | Yeast | NULL | Probable | As nectar-tap_saprotroph (Põlme et al. 2020); Decay sugar-rich substrates (Põlme et al. 2020) | Põlme S, et al. 2020. Fungal Diversity 105:1-16 ((https://doi.org/10.1007/s13225-020-00466-2)) |
| Otu36508 | 1 | k__Fungi(100);p__Ascomycota(100);c__Saccharomycetes(100);o__Saccharomycetales(100);f__Metschnikowiaceae(100);g__Metschnikowia(100);s__Metschnikowia_pulcherrima(100); | Metschnikowia | 13 | Saprotroph | \|Undefined Saprotroph\| | Yeast | NULL | Probable | As nectar-tap_saprotroph (Põlme et al. 2020); Decay sugar-rich substrates (Põlme et al. 2020) | Põlme S, et al. 2020. Fungal Diversity 105:1-16 ((https://doi.org/10.1007/s13225-020-00466-2)) |
| Otu36509 | 1 | k__Fungi(100);p__Ascomycota(100);c__Saccharomycetes(100);o__Saccharomycetales(100);f__Metschnikowiaceae(100);g__Metschnikowia(100);g__Metschnikowia_unclassified(100); | Metschnikowia | 13 | Saprotroph | \|Undefined Saprotroph\| | Yeast | NULL | Probable | As nectar-tap_saprotroph (Põlme et al. 2020); Decay sugar-rich substrates (Põlme et al. 2020) | Põlme S, et al. 2020. Fungal Diversity 105:1-16 ((https://doi.org/10.1007/s13225-020-00466-2)) |
| Otu36510 | 1 | k__Fungi(100);p__Ascomycota(100);c__Saccharomycetes(100);o__Saccharomycetales(100);f__Metschnikowiaceae(100);g__Metschnikowia(100);g__Metschnikowia_unclassified(100); | Metschnikowia | 13 | Saprotroph | \|Undefined Saprotroph\| | Yeast | NULL | Probable | As nectar-tap_saprotroph (Põlme et al. 2020); Decay sugar-rich substrates (Põlme et al. 2020) | Põlme S, et al. 2020. Fungal Diversity 105:1-16 ((https://doi.org/10.1007/s13225-020-00466-2)) |
| Otu36511 | 1 | k__Fungi(100);p__Ascomycota(100);c__Saccharomycetes(100);o__Saccharomycetales(100);f__Metschnikowiaceae(100);g__Metschnikowia(100);s__Metschnikowia_pulcherrima(100); | Metschnikowia | 13 | Saprotroph | \|Undefined Saprotroph\| | Yeast | NULL | Probable | As nectar-tap_saprotroph (Põlme et al. 2020); Decay sugar-rich substrates (Põlme et al. 2020) | Põlme S, et al. 2020. Fungal Diversity 105:1-16 ((https://doi.org/10.1007/s13225-020-00466-2)) |
| Otu36512 | 1 | k__Fungi(100);p__Ascomycota(100);c__Saccharomycetes(100);o__Saccharomycetales(100);f__Metschnikowiaceae(100);g__Metschnikowia(100);g__Metschnikowia_unclassified(100); | Metschnikowia | 13 | Saprotroph | \|Undefined Saprotroph\| | Yeast | NULL | Probable | As nectar-tap_saprotroph (Põlme et al. 2020); Decay sugar-rich substrates (Põlme et al. 2020) | Põlme S, et al. 2020. Fungal Diversity 105:1-16 ((https://doi.org/10.1007/s13225-020-00466-2)) |
| Otu36513 | 1 | k__Fungi(100);p__Ascomycota(100);c__Saccharomycetes(100);o__Saccharomycetales(100);f__Metschnikowiaceae(100);g__Metschnikowia(100);s__Metschnikowia_pulcherrima(100); | Metschnikowia | 13 | Saprotroph | \|Undefined Saprotroph\| | Yeast | NULL | Probable | As nectar-tap_saprotroph (Põlme et al. 2020); Decay sugar-rich substrates (Põlme et al. 2020) | Põlme S, et al. 2020. Fungal Diversity 105:1-16 ((https://doi.org/10.1007/s13225-020-00466-2)) |
| Otu36514 | 1 | k__Fungi(100);p__Ascomycota(100);c__Saccharomycetes(100);o__Saccharomycetales(100);f__Metschnikowiaceae(100);g__Metschnikowia(100);s__Metschnikowia_pulcherrima(100); | Metschnikowia | 13 | Saprotroph | \|Undefined Saprotroph\| | Yeast | NULL | Probable | As nectar-tap_saprotroph (Põlme et al. 2020); Decay sugar-rich substrates (Põlme et al. 2020) | Põlme S, et al. 2020. Fungal Diversity 105:1-16 ((https://doi.org/10.1007/s13225-020-00466-2)) |
| Otu36515 | 1 | k__Fungi(100);p__Ascomycota(100);c__Saccharomycetes(100);o__Saccharomycetales(100);f__Metschnikowiaceae(100);g__Metschnikowia(100);s__Metschnikowia_pulcherrima(100); | Metschnikowia | 13 | Saprotroph | \|Undefined Saprotroph\| | Yeast | NULL | Probable | As nectar-tap_saprotroph (Põlme et al. 2020); Decay sugar-rich substrates (Põlme et al. 2020) | Põlme S, et al. 2020. Fungal Diversity 105:1-16 ((https://doi.org/10.1007/s13225-020-00466-2)) |
| Otu36516 | 1 | k__Fungi(100);p__Ascomycota(100);c__Saccharomycetes(100);o__Saccharomycetales(100);f__Metschnikowiaceae(100);g__Metschnikowia(100);g__Metschnikowia_unclassified(100); | Metschnikowia | 13 | Saprotroph | \|Undefined Saprotroph\| | Yeast | NULL | Probable | As nectar-tap_saprotroph (Põlme et al. 2020); Decay sugar-rich substrates (Põlme et al. 2020) | Põlme S, et al. 2020. Fungal Diversity 105:1-16 ((https://doi.org/10.1007/s13225-020-00466-2)) |
| Otu36517 | 1 | k__Fungi(100);p__Ascomycota(100);c__Saccharomycetes(100);o__Saccharomycetales(100);f__Metschnikowiaceae(100);g__Metschnikowia(100);g__Metschnikowia_unclassified(100); | Metschnikowia | 13 | Saprotroph | \|Undefined Saprotroph\| | Yeast | NULL | Probable | As nectar-tap_saprotroph (Põlme et al. 2020); Decay sugar-rich substrates (Põlme et al. 2020) | Põlme S, et al. 2020. Fungal Diversity 105:1-16 ((https://doi.org/10.1007/s13225-020-00466-2)) |
| Otu36518 | 1 | k__Fungi(100);p__Ascomycota(100);c__Saccharomycetes(100);o__Saccharomycetales(100);f__Metschnikowiaceae(100);g__Metschnikowia(100);s__Metschnikowia_pulcherrima(100); | Metschnikowia | 13 | Saprotroph | \|Undefined Saprotroph\| | Yeast | NULL | Probable | As nectar-tap_saprotroph (Põlme et al. 2020); Decay sugar-rich substrates (Põlme et al. 2020) | Põlme S, et al. 2020. Fungal Diversity 105:1-16 ((https://doi.org/10.1007/s13225-020-00466-2)) |
| Otu36519 | 1 | k__Fungi(100);p__Ascomycota(100);c__Saccharomycetes(100);o__Saccharomycetales(100);f__Metschnikowiaceae(100);g__Metschnikowia(100);s__Metschnikowia_pulcherrima(100); | Metschnikowia | 13 | Saprotroph | \|Undefined Saprotroph\| | Yeast | NULL | Probable | As nectar-tap_saprotroph (Põlme et al. 2020); Decay sugar-rich substrates (Põlme et al. 2020) | Põlme S, et al. 2020. Fungal Diversity 105:1-16 ((https://doi.org/10.1007/s13225-020-00466-2)) |
| Otu36520 | 1 | k__Fungi(100);p__Ascomycota(100);c__Saccharomycetes(100);o__Saccharomycetales(100);f__Metschnikowiaceae(100);g__Metschnikowia(100);s__Metschnikowia_pulcherrima(100); | Metschnikowia | 13 | Saprotroph | \|Undefined Saprotroph\| | Yeast | NULL | Probable | As nectar-tap_saprotroph (Põlme et al. 2020); Decay sugar-rich substrates (Põlme et al. 2020) | Põlme S, et al. 2020. Fungal Diversity 105:1-16 ((https://doi.org/10.1007/s13225-020-00466-2)) |
| Otu36521 | 1 | k__Fungi(100);p__Ascomycota(100);c__Saccharomycetes(100);o__Saccharomycetales(100);f__Metschnikowiaceae(100);g__Metschnikowia(100);s__Metschnikowia_pulcherrima(100); | Metschnikowia | 13 | Saprotroph | \|Undefined Saprotroph\| | Yeast | NULL | Probable | As nectar-tap_saprotroph (Põlme et al. 2020); Decay sugar-rich substrates (Põlme et al. 2020) | Põlme S, et al. 2020. Fungal Diversity 105:1-16 ((https://doi.org/10.1007/s13225-020-00466-2)) |
| Otu36522 | 1 | k__Fungi(100);p__Ascomycota(100);c__Saccharomycetes(100);o__Saccharomycetales(100);f__Metschnikowiaceae(100);g__Metschnikowia(100);s__Metschnikowia_pulcherrima(100); | Metschnikowia | 13 | Saprotroph | \|Undefined Saprotroph\| | Yeast | NULL | Probable | As nectar-tap_saprotroph (Põlme et al. 2020); Decay sugar-rich substrates (Põlme et al. 2020) | Põlme S, et al. 2020. Fungal Diversity 105:1-16 ((https://doi.org/10.1007/s13225-020-00466-2)) |
| Otu36523 | 1 | k__Fungi(100);p__Ascomycota(100);c__Saccharomycetes(100);o__Saccharomycetales(100);f__Metschnikowiaceae(100);g__Metschnikowia(100);s__Metschnikowia_pulcherrima(100); | Metschnikowia | 13 | Saprotroph | \|Undefined Saprotroph\| | Yeast | NULL | Probable | As nectar-tap_saprotroph (Põlme et al. 2020); Decay sugar-rich substrates (Põlme et al. 2020) | Põlme S, et al. 2020. Fungal Diversity 105:1-16 ((https://doi.org/10.1007/s13225-020-00466-2)) |
| Otu36524 | 1 | k__Fungi(100);p__Ascomycota(100);c__Saccharomycetes(100);o__Saccharomycetales(100);f__Metschnikowiaceae(100);g__Metschnikowia(100);s__Metschnikowia_pulcherrima(100); | Metschnikowia | 13 | Saprotroph | \|Undefined Saprotroph\| | Yeast | NULL | Probable | As nectar-tap_saprotroph (Põlme et al. 2020); Decay sugar-rich substrates (Põlme et al. 2020) | Põlme S, et al. 2020. Fungal Diversity 105:1-16 ((https://doi.org/10.1007/s13225-020-00466-2)) |
| Otu36525 | 1 | k__Fungi(100);p__Ascomycota(100);c__Saccharomycetes(100);o__Saccharomycetales(100);f__Metschnikowiaceae(100);g__Metschnikowia(100);s__Metschnikowia_pulcherrima(100); | Metschnikowia | 13 | Saprotroph | \|Undefined Saprotroph\| | Yeast | NULL | Probable | As nectar-tap_saprotroph (Põlme et al. 2020); Decay sugar-rich substrates (Põlme et al. 2020) | Põlme S, et al. 2020. Fungal Diversity 105:1-16 ((https://doi.org/10.1007/s13225-020-00466-2)) |

| | | | | | | | | | | | |
|---|---|---|---|---|---|---|---|---|---|---|---|
| Otu36526 | 1 | k__Fungi(100);p__Ascomycota(100);c__Saccharomycetes(100);o__Saccharomycetales(100);f__Metschnikowiaceae(100);g__Metschnikowia(100);s__Metschnikowia_pulcherrima(100); | Metschnikowia | 13 | Saprotroph | \|Undefined Saprotroph\| | Yeast | NULL | Probable | As nectar-tap_saprotroph (Põlme et al. 2020); Decay sugar-rich substrates (Põlme et al. 2020) | Põlme S, et al. 2020. Fungal Diversity 105:1-16 ((https://doi.org/10.1007/s13225-020-00466-2)) |
| Otu36527 | 1 | k__Fungi(100);p__Ascomycota(100);c__Saccharomycetes(100);o__Saccharomycetales(100);f__Metschnikowiaceae(100);g__Metschnikowia(100);s__Metschnikowia_pulcherrima(100); | Metschnikowia | 13 | Saprotroph | \|Undefined Saprotroph\| | Yeast | NULL | Probable | As nectar-tap_saprotroph (Põlme et al. 2020); Decay sugar-rich substrates (Põlme et al. 2020) | Põlme S, et al. 2020. Fungal Diversity 105:1-16 ((https://doi.org/10.1007/s13225-020-00466-2)) |
| Otu36529 | 1 | k__Fungi(100);p__Ascomycota(100);c__Saccharomycetes(100);o__Saccharomycetales(100);f__Metschnikowiaceae(100);g__Metschnikowia(100);s__Metschnikowia_pulcherrima(100); | Metschnikowia | 13 | Saprotroph | \|Undefined Saprotroph\| | Yeast | NULL | Probable | As nectar-tap_saprotroph (Põlme et al. 2020); Decay sugar-rich substrates (Põlme et al. 2020) | Põlme S, et al. 2020. Fungal Diversity 105:1-16 ((https://doi.org/10.1007/s13225-020-00466-2)) |
| Otu36531 | 1 | k__Fungi(100);p__Ascomycota(100);c__Saccharomycetes(100);o__Saccharomycetales(100);f__Metschnikowiaceae(100);g__Metschnikowia(100);s__Metschnikowia_pulcherrima(100); | Metschnikowia | 13 | Saprotroph | \|Undefined Saprotroph\| | Yeast | NULL | Probable | As nectar-tap_saprotroph (Põlme et al. 2020); Decay sugar-rich substrates (Põlme et al. 2020) | Põlme S, et al. 2020. Fungal Diversity 105:1-16 ((https://doi.org/10.1007/s13225-020-00466-2)) |
| Otu36532 | 1 | k__Fungi(100);p__Ascomycota(100);c__Saccharomycetes(100);o__Saccharomycetales(100);f__Metschnikowiaceae(100);g__Metschnikowia(100);s__Metschnikowia_pulcherrima(100); | Metschnikowia | 13 | Saprotroph | \|Undefined Saprotroph\| | Yeast | NULL | Probable | As nectar-tap_saprotroph (Põlme et al. 2020); Decay sugar-rich substrates (Põlme et al. 2020) | Põlme S, et al. 2020. Fungal Diversity 105:1-16 ((https://doi.org/10.1007/s13225-020-00466-2)) |
| Otu36533 | 1 | k__Fungi(100);p__Ascomycota(100);c__Saccharomycetes(100);o__Saccharomycetales(100);f__Metschnikowiaceae(100);g__Metschnikowia(100);s__Metschnikowia_pulcherrima(100); | Metschnikowia | 13 | Saprotroph | \|Undefined Saprotroph\| | Yeast | NULL | Probable | As nectar-tap_saprotroph (Põlme et al. 2020); Decay sugar-rich substrates (Põlme et al. 2020) | Põlme S, et al. 2020. Fungal Diversity 105:1-16 ((https://doi.org/10.1007/s13225-020-00466-2)) |
| Otu36534 | 1 | k__Fungi(100);p__Ascomycota(100);c__Saccharomycetes(100);o__Saccharomycetales(100);f__Metschnikowiaceae(100);g__Metschnikowia(100);g__Metschnikowia_unclassified(100); | Metschnikowia | 13 | Saprotroph | \|Undefined Saprotroph\| | Yeast | NULL | Probable | As nectar-tap_saprotroph (Põlme et al. 2020); Decay sugar-rich substrates (Põlme et al. 2020) | Põlme S, et al. 2020. Fungal Diversity 105:1-16 ((https://doi.org/10.1007/s13225-020-00466-2)) |
| Otu36535 | 1 | k__Fungi(100);p__Ascomycota(100);c__Saccharomycetes(100);o__Saccharomycetales(100);f__Metschnikowiaceae(100);g__Metschnikowia(100);s__Metschnikowia_pulcherrima(100); | Metschnikowia | 13 | Saprotroph | \|Undefined Saprotroph\| | Yeast | NULL | Probable | As nectar-tap_saprotroph (Põlme et al. 2020); Decay sugar-rich substrates (Põlme et al. 2020) | Põlme S, et al. 2020. Fungal Diversity 105:1-16 ((https://doi.org/10.1007/s13225-020-00466-2)) |
| Otu36536 | 1 | k__Fungi(100);p__Ascomycota(100);c__Saccharomycetes(100);o__Saccharomycetales(100);f__Metschnikowiaceae(100);g__Metschnikowia(100);g__Metschnikowia_unclassified(100); | Metschnikowia | 13 | Saprotroph | \|Undefined Saprotroph\| | Yeast | NULL | Probable | As nectar-tap_saprotroph (Põlme et al. 2020); Decay sugar-rich substrates (Põlme et al. 2020) | Põlme S, et al. 2020. Fungal Diversity 105:1-16 ((https://doi.org/10.1007/s13225-020-00466-2)) |
| Otu36537 | 1 | k__Fungi(100);p__Ascomycota(100);c__Saccharomycetes(100);o__Saccharomycetales(100);f__Metschnikowiaceae(100);g__Metschnikowia(100);s__Metschnikowia_pulcherrima(100); | Metschnikowia | 13 | Saprotroph | \|Undefined Saprotroph\| | Yeast | NULL | Probable | As nectar-tap_saprotroph (Põlme et al. 2020); Decay sugar-rich substrates (Põlme et al. 2020) | Põlme S, et al. 2020. Fungal Diversity 105:1-16 ((https://doi.org/10.1007/s13225-020-00466-2)) |
| Otu36538 | 1 | k__Fungi(100);p__Ascomycota(100);c__Saccharomycetes(100);o__Saccharomycetales(100);f__Metschnikowiaceae(100);g__Metschnikowia(100);s__Metschnikowia_pulcherrima(100); | Metschnikowia | 13 | Saprotroph | \|Undefined Saprotroph\| | Yeast | NULL | Probable | As nectar-tap_saprotroph (Põlme et al. 2020); Decay sugar-rich substrates (Põlme et al. 2020) | Põlme S, et al. 2020. Fungal Diversity 105:1-16 ((https://doi.org/10.1007/s13225-020-00466-2)) |
| Otu36539 | 1 | k__Fungi(100);p__Ascomycota(100);c__Saccharomycetes(100);o__Saccharomycetales(100);f__Metschnikowiaceae(100);g__Metschnikowia(100);g__Metschnikowia_unclassified(100); | Metschnikowia | 13 | Saprotroph | \|Undefined Saprotroph\| | Yeast | NULL | Probable | As nectar-tap_saprotroph (Põlme et al. 2020); Decay sugar-rich substrates (Põlme et al. 2020) | Põlme S, et al. 2020. Fungal Diversity 105:1-16 ((https://doi.org/10.1007/s13225-020-00466-2)) |
| Otu36540 | 1 | k__Fungi(100);p__Ascomycota(100);c__Saccharomycetes(100);o__Saccharomycetales(100);f__Metschnikowiaceae(100);g__Metschnikowia(100);s__Metschnikowia_pulcherrima(100); | Metschnikowia | 13 | Saprotroph | \|Undefined Saprotroph\| | Yeast | NULL | Probable | As nectar-tap_saprotroph (Põlme et al. 2020); Decay sugar-rich substrates (Põlme et al. 2020) | Põlme S, et al. 2020. Fungal Diversity 105:1-16 ((https://doi.org/10.1007/s13225-020-00466-2)) |
| Otu36541 | 1 | k__Fungi(100);p__Ascomycota(100);c__Saccharomycetes(100);o__Saccharomycetales(100);f__Metschnikowiaceae(100);g__Metschnikowia(100);s__Metschnikowia_pulcherrima(100); | Metschnikowia | 13 | Saprotroph | \|Undefined Saprotroph\| | Yeast | NULL | Probable | As nectar-tap_saprotroph (Põlme et al. 2020); Decay sugar-rich substrates (Põlme et al. 2020) | Põlme S, et al. 2020. Fungal Diversity 105:1-16 ((https://doi.org/10.1007/s13225-020-00466-2)) |
| Otu36542 | 1 | k__Fungi(100);p__Ascomycota(100);c__Saccharomycetes(100);o__Saccharomycetales(100);f__Metschnikowiaceae(100);g__Metschnikowia(100);s__Metschnikowia_pulcherrima(100); | Metschnikowia | 13 | Saprotroph | \|Undefined Saprotroph\| | Yeast | NULL | Probable | As nectar-tap_saprotroph (Põlme et al. 2020); Decay sugar-rich substrates (Põlme et al. 2020) | Põlme S, et al. 2020. Fungal Diversity 105:1-16 ((https://doi.org/10.1007/s13225-020-00466-2)) |
| Otu36543 | 1 | k__Fungi(100);p__Ascomycota(100);c__Saccharomycetes(100);o__Saccharomycetales(100);f__Metschnikowiaceae(100);g__Metschnikowia(100);s__Metschnikowia_pulcherrima(100); | Metschnikowia | 13 | Saprotroph | \|Undefined Saprotroph\| | Yeast | NULL | Probable | As nectar-tap_saprotroph (Põlme et al. 2020); Decay sugar-rich substrates (Põlme et al. 2020) | Põlme S, et al. 2020. Fungal Diversity 105:1-16 ((https://doi.org/10.1007/s13225-020-00466-2)) |
| Otu36544 | 1 | k__Fungi(100);p__Ascomycota(100);c__Saccharomycetes(100);o__Saccharomycetales(100);f__Metschnikowiaceae(100);g__Metschnikowia(100);s__Metschnikowia_pulcherrima(100); | Metschnikowia | 13 | Saprotroph | \|Undefined Saprotroph\| | Yeast | NULL | Probable | As nectar-tap_saprotroph (Põlme et al. 2020); Decay sugar-rich substrates (Põlme et al. 2020) | Põlme S, et al. 2020. Fungal Diversity 105:1-16 ((https://doi.org/10.1007/s13225-020-00466-2)) |
| Otu36545 | 1 | k__Fungi(100);p__Ascomycota(100);c__Saccharomycetes(100);o__Saccharomycetales(100);f__Metschnikowiaceae(100);g__Metschnikowia(100);s__Metschnikowia_pulcherrima(100); | Metschnikowia | 13 | Saprotroph | \|Undefined Saprotroph\| | Yeast | NULL | Probable | As nectar-tap_saprotroph (Põlme et al. 2020); Decay sugar-rich substrates (Põlme et al. 2020) | Põlme S, et al. 2020. Fungal Diversity 105:1-16 ((https://doi.org/10.1007/s13225-020-00466-2)) |
| Otu36546 | 1 | k__Fungi(100);p__Ascomycota(100);c__Saccharomycetes(100);o__Saccharomycetales(100);f__Metschnikowiaceae(100);g__Metschnikowia(100);g__Metschnikowia_unclassified(100); | Metschnikowia | 13 | Saprotroph | \|Undefined Saprotroph\| | Yeast | NULL | Probable | As nectar-tap_saprotroph (Põlme et al. 2020); Decay sugar-rich substrates (Põlme et al. 2020) | Põlme S, et al. 2020. Fungal Diversity 105:1-16 ((https://doi.org/10.1007/s13225-020-00466-2)) |
| Otu36547 | 1 | k__Fungi(100);p__Ascomycota(100);c__Saccharomycetes(100);o__Saccharomycetales(100);f__Metschnikowiaceae(100);g__Metschnikowia(100);s__Metschnikowia_pulcherrima(100); | Metschnikowia | 13 | Saprotroph | \|Undefined Saprotroph\| | Yeast | NULL | Probable | As nectar-tap_saprotroph (Põlme et al. 2020); Decay sugar-rich substrates (Põlme et al. 2020) | Põlme S, et al. 2020. Fungal Diversity 105:1-16 ((https://doi.org/10.1007/s13225-020-00466-2)) |
| Otu36548 | 1 | k__Fungi(100);p__Ascomycota(100);c__Saccharomycetes(100);o__Saccharomycetales(100);f__Metschnikowiaceae(100);g__Metschnikowia(100);s__Metschnikowia_pulcherrima(100); | Metschnikowia | 13 | Saprotroph | \|Undefined Saprotroph\| | Yeast | NULL | Probable | As nectar-tap_saprotroph (Põlme et al. 2020); Decay sugar-rich substrates (Põlme et al. 2020) | Põlme S, et al. 2020. Fungal Diversity 105:1-16 ((https://doi.org/10.1007/s13225-020-00466-2)) |
| Otu36549 | 1 | k__Fungi(100);p__Ascomycota(100);c__Saccharomycetes(100);o__Saccharomycetales(100);f__Metschnikowiaceae(100);g__Metschnikowia(100);s__Metschnikowia_pulcherrima(100); | Metschnikowia | 13 | Saprotroph | \|Undefined Saprotroph\| | Yeast | NULL | Probable | As nectar-tap_saprotroph (Põlme et al. 2020); Decay sugar-rich substrates (Põlme et al. 2020) | Põlme S, et al. 2020. Fungal Diversity 105:1-16 ((https://doi.org/10.1007/s13225-020-00466-2)) |
| Otu36550 | 1 | k__Fungi(100);p__Ascomycota(100);c__Saccharomycetes(100);o__Saccharomycetales(100);f__Metschnikowiaceae(100);g__Metschnikowia(100);g__Metschnikowia_unclassified(100); | Metschnikowia | 13 | Saprotroph | \|Undefined Saprotroph\| | Yeast | NULL | Probable | As nectar-tap_saprotroph (Põlme et al. 2020); Decay sugar-rich substrates (Põlme et al. 2020) | Põlme S, et al. 2020. Fungal Diversity 105:1-16 ((https://doi.org/10.1007/s13225-020-00466-2)) |
| Otu36551 | 1 | k__Fungi(100);p__Ascomycota(100);c__Saccharomycetes(100);o__Saccharomycetales(100);f__Metschnikowiaceae(100);g__Metschnikowia(100);s__Metschnikowia_pulcherrima(100); | Metschnikowia | 13 | Saprotroph | \|Undefined Saprotroph\| | Yeast | NULL | Probable | As nectar-tap_saprotroph (Põlme et al. 2020); Decay sugar-rich substrates (Põlme et al. 2020) | Põlme S, et al. 2020. Fungal Diversity 105:1-16 ((https://doi.org/10.1007/s13225-020-00466-2)) |
| Otu36552 | 1 | k__Fungi(100);p__Ascomycota(100);c__Saccharomycetes(100);o__Saccharomycetales(100);f__Metschnikowiaceae(100);g__Metschnikowia(100);s__Metschnikowia_pulcherrima(100); | Metschnikowia | 13 | Saprotroph | \|Undefined Saprotroph\| | Yeast | NULL | Probable | As nectar-tap_saprotroph (Põlme et al. 2020); Decay sugar-rich substrates (Põlme et al. 2020) | Põlme S, et al. 2020. Fungal Diversity 105:1-16 ((https://doi.org/10.1007/s13225-020-00466-2)) |
| Otu36553 | 1 | k__Fungi(100);p__Ascomycota(100);c__Saccharomycetes(100);o__Saccharomycetales(100);f__Metschnikowiaceae(100);g__Metschnikowia(100);s__Metschnikowia_pulcherrima(100); | Metschnikowia | 13 | Saprotroph | \|Undefined Saprotroph\| | Yeast | NULL | Probable | As nectar-tap_saprotroph (Põlme et al. 2020); Decay sugar-rich substrates (Põlme et al. 2020) | Põlme S, et al. 2020. Fungal Diversity 105:1-16 ((https://doi.org/10.1007/s13225-020-00466-2)) |
| Otu36554 | 1 | k__Fungi(100);p__Ascomycota(100);c__Saccharomycetes(100);o__Saccharomycetales(100);f__Metschnikowiaceae(100);g__Metschnikowia(100);g__Metschnikowia_unclassified(100); | Metschnikowia | 13 | Saprotroph | \|Undefined Saprotroph\| | Yeast | NULL | Probable | As nectar-tap_saprotroph (Põlme et al. 2020); Decay sugar-rich substrates (Põlme et al. 2020) | Põlme S, et al. 2020. Fungal Diversity 105:1-16 ((https://doi.org/10.1007/s13225-020-00466-2)) |
| Otu36555 | 1 | k__Fungi(100);p__Ascomycota(100);c__Saccharomycetes(100);o__Saccharomycetales(100);f__Metschnikowiaceae(100);g__Metschnikowia(100);s__Metschnikowia_pulcherrima(100); | Metschnikowia | 13 | Saprotroph | \|Undefined Saprotroph\| | Yeast | NULL | Probable | As nectar-tap_saprotroph (Põlme et al. 2020); Decay sugar-rich substrates (Põlme et al. 2020) | Põlme S, et al. 2020. Fungal Diversity 105:1-16 ((https://doi.org/10.1007/s13225-020-00466-2)) |
| Otu36556 | 1 | k__Fungi(100);p__Ascomycota(100);c__Saccharomycetes(100);o__Saccharomycetales(100);f__Metschnikowiaceae(100);g__Metschnikowia(100);s__Metschnikowia_pulcherrima(100); | Metschnikowia | 13 | Saprotroph | \|Undefined Saprotroph\| | Yeast | NULL | Probable | As nectar-tap_saprotroph (Põlme et al. 2020); Decay sugar-rich substrates (Põlme et al. 2020) | Põlme S, et al. 2020. Fungal Diversity 105:1-16 ((https://doi.org/10.1007/s13225-020-00466-2)) |
| Otu36557 | 1 | k__Fungi(100);p__Ascomycota(100);c__Saccharomycetes(100);o__Saccharomycetales(100);f__Metschnikowiaceae(100);g__Metschnikowia(100);s__Metschnikowia_pulcherrima(100); | Metschnikowia | 13 | Saprotroph | \|Undefined Saprotroph\| | Yeast | NULL | Probable | As nectar-tap_saprotroph (Põlme et al. 2020); Decay sugar-rich substrates (Põlme et al. 2020) | Põlme S, et al. 2020. Fungal Diversity 105:1-16 ((https://doi.org/10.1007/s13225-020-00466-2)) |
| Otu36558 | 1 | k__Fungi(100);p__Ascomycota(100);c__Saccharomycetes(100);o__Saccharomycetales(100);f__Metschnikowiaceae(100);g__Metschnikowia(100);s__Metschnikowia_pulcherrima(100); | Metschnikowia | 13 | Saprotroph | \|Undefined Saprotroph\| | Yeast | NULL | Probable | As nectar-tap_saprotroph (Põlme et al. 2020); Decay sugar-rich substrates (Põlme et al. 2020) | Põlme S, et al. 2020. Fungal Diversity 105:1-16 ((https://doi.org/10.1007/s13225-020-00466-2)) |
| Otu36559 | 1 | k__Fungi(100);p__Ascomycota(100);c__Saccharomycetes(100);o__Saccharomycetales(100);f__Metschnikowiaceae(100);g__Metschnikowia(100);s__Metschnikowia_pulcherrima(100); | Metschnikowia | 13 | Saprotroph | \|Undefined Saprotroph\| | Yeast | NULL | Probable | As nectar-tap_saprotroph (Põlme et al. 2020); Decay sugar-rich substrates (Põlme et al. 2020) | Põlme S, et al. 2020. Fungal Diversity 105:1-16 ((https://doi.org/10.1007/s13225-020-00466-2)) |
| Otu36560 | 1 | k__Fungi(100);p__Ascomycota(100);c__Saccharomycetes(100);o__Saccharomycetales(100);f__Metschnikowiaceae(100);g__Metschnikowia(100);s__Metschnikowia_pulcherrima(100); | Metschnikowia | 13 | Saprotroph | \|Undefined Saprotroph\| | Yeast | NULL | Probable | As nectar-tap_saprotroph (Põlme et al. 2020); Decay sugar-rich substrates (Põlme et al. 2020) | Põlme S, et al. 2020. Fungal Diversity 105:1-16 ((https://doi.org/10.1007/s13225-020-00466-2)) |
| Otu36561 | 1 | k__Fungi(100);p__Ascomycota(100);c__Saccharomycetes(100);o__Saccharomycetales(100);f__Metschnikowiaceae(100);g__Metschnikowia(100);s__Metschnikowia_pulcherrima(100); | Metschnikowia | 13 | Saprotroph | \|Undefined Saprotroph\| | Yeast | NULL | Probable | As nectar-tap_saprotroph (Põlme et al. 2020); Decay sugar-rich substrates (Põlme et al. 2020) | Põlme S, et al. 2020. Fungal Diversity 105:1-16 ((https://doi.org/10.1007/s13225-020-00466-2)) |
| Otu36563 | 1 | k__Fungi(100);p__Ascomycota(100);c__Saccharomycetes(100);o__Saccharomycetales(100);f__Metschnikowiaceae(100);g__Metschnikowia(100);g__Metschnikowia_unclassified(100); | Metschnikowia | 13 | Saprotroph | \|Undefined Saprotroph\| | Yeast | NULL | Probable | As nectar-tap_saprotroph (Põlme et al. 2020); Decay sugar-rich substrates (Põlme et al. 2020) | Põlme S, et al. 2020. Fungal Diversity 105:1-16 ((https://doi.org/10.1007/s13225-020-00466-2)) |
| Otu36564 | 1 | k__Fungi(100);p__Ascomycota(100);c__Saccharomycetes(100);o__Saccharomycetales(100);f__Metschnikowiaceae(100);g__Metschnikowia(100);s__Metschnikowia_pulcherrima(100); | Metschnikowia | 13 | Saprotroph | \|Undefined Saprotroph\| | Yeast | NULL | Probable | As nectar-tap_saprotroph (Põlme et al. 2020); Decay sugar-rich substrates (Põlme et al. 2020) | Põlme S, et al. 2020. Fungal Diversity 105:1-16 ((https://doi.org/10.1007/s13225-020-00466-2)) |
| Otu36565 | 1 | k__Fungi(100);p__Ascomycota(100);c__Saccharomycetes(100);o__Saccharomycetales(100);f__Metschnikowiaceae(100);g__Metschnikowia(100);g__Metschnikowia_unclassified(100); | Metschnikowia | 13 | Saprotroph | \|Undefined Saprotroph\| | Yeast | NULL | Probable | As nectar-tap_saprotroph (Põlme et al. 2020); Decay sugar-rich substrates (Põlme et al. 2020) | Põlme S, et al. 2020. Fungal Diversity 105:1-16 ((https://doi.org/10.1007/s13225-020-00466-2)) |
| Otu36566 | 1 | k__Fungi(100);p__Ascomycota(100);c__Saccharomycetes(100);o__Saccharomycetales(100);f__Metschnikowiaceae(100);g__Metschnikowia(100);s__Metschnikowia_pulcherrima(100); | Metschnikowia | 13 | Saprotroph | \|Undefined Saprotroph\| | Yeast | NULL | Probable | As nectar-tap_saprotroph (Põlme et al. 2020); Decay sugar-rich substrates (Põlme et al. 2020) | Põlme S, et al. 2020. Fungal Diversity 105:1-16 ((https://doi.org/10.1007/s13225-020-00466-2)) |
| Otu36567 | 1 | k__Fungi(100);p__Ascomycota(100);c__Saccharomycetes(100);o__Saccharomycetales(100);f__Metschnikowiaceae(100);g__Metschnikowia(100);s__Metschnikowia_pulcherrima(100); | Metschnikowia | 13 | Saprotroph | \|Undefined Saprotroph\| | Yeast | NULL | Probable | As nectar-tap_saprotroph (Põlme et al. 2020); Decay sugar-rich substrates (Põlme et al. 2020) | Põlme S, et al. 2020. Fungal Diversity 105:1-16 ((https://doi.org/10.1007/s13225-020-00466-2)) |
| Otu36568 | 1 | k__Fungi(100);p__Ascomycota(100);c__Saccharomycetes(100);o__Saccharomycetales(100);f__Metschnikowiaceae(100);g__Metschnikowia(100);s__Metschnikowia_pulcherrima(100); | Metschnikowia | 13 | Saprotroph | \|Undefined Saprotroph\| | Yeast | NULL | Probable | As nectar-tap_saprotroph (Põlme et al. 2020); Decay sugar-rich substrates (Põlme et al. 2020) | Põlme S, et al. 2020. Fungal Diversity 105:1-16 ((https://doi.org/10.1007/s13225-020-00466-2)) |
| Otu36572 | 1 | k__Fungi(100);p__Ascomycota(100);c__Saccharomycetes(100);o__Saccharomycetales(100);f__Metschnikowiaceae(100);g__Metschnikowia(100);s__Metschnikowia_pulcherrima(100); | Metschnikowia | 13 | Saprotroph | \|Undefined Saprotroph\| | Yeast | NULL | Probable | As nectar-tap_saprotroph (Põlme et al. 2020); Decay sugar-rich substrates (Põlme et al. 2020) | Põlme S, et al. 2020. Fungal Diversity 105:1-16 ((https://doi.org/10.1007/s13225-020-00466-2)) |
| Otu36573 | 1 | k__Fungi(100);p__Ascomycota(100);c__Saccharomycetes(100);o__Saccharomycetales(100);f__Metschnikowiaceae(100);g__Metschnikowia(100);s__Metschnikowia_pulcherrima(100); | Metschnikowia | 13 | Saprotroph | \|Undefined Saprotroph\| | Yeast | NULL | Probable | As nectar-tap_saprotroph (Põlme et al. 2020); Decay sugar-rich substrates (Põlme et al. 2020) | Põlme S, et al. 2020. Fungal Diversity 105:1-16 ((https://doi.org/10.1007/s13225-020-00466-2)) |
| Otu36574 | 1 | k__Fungi(100);p__Ascomycota(100);c__Saccharomycetes(100);o__Saccharomycetales(100);f__Metschnikowiaceae(100);g__Metschnikowia(100);s__Metschnikowia_pulcherrima(100); | Metschnikowia | 13 | Saprotroph | \|Undefined Saprotroph\| | Yeast | NULL | Probable | As nectar-tap_saprotroph (Põlme et al. 2020); Decay sugar-rich substrates (Põlme et al. 2020) | Põlme S, et al. 2020. Fungal Diversity 105:1-16 ((https://doi.org/10.1007/s13225-020-00466-2)) |
| Otu36575 | 1 | k__Fungi(100);p__Ascomycota(100);c__Saccharomycetes(100);o__Saccharomycetales(100);f__Metschnikowiaceae(100);g__Metschnikowia(100);s__Metschnikowia_pulcherrima(100); | Metschnikowia | 13 | Saprotroph | \|Undefined Saprotroph\| | Yeast | NULL | Probable | As nectar-tap_saprotroph (Põlme et al. 2020); Decay sugar-rich substrates (Põlme et al. 2020) | Põlme S, et al. 2020. Fungal Diversity 105:1-16 ((https://doi.org/10.1007/s13225-020-00466-2)) |
| Otu36579 | 1 | k__Fungi(100);p__Ascomycota(100);c__Saccharomycetes(100);o__Saccharomycetales(100);f__Metschnikowiaceae(100);g__Metschnikowia(100);g__Metschnikowia_unclassified(100); | Metschnikowia | 13 | Saprotroph | \|Undefined Saprotroph\| | Yeast | NULL | Probable | As nectar-tap_saprotroph (Põlme et al. 2020); Decay sugar-rich substrates (Põlme et al. 2020) | Põlme S, et al. 2020. Fungal Diversity 105:1-16 ((https://doi.org/10.1007/s13225-020-00466-2)) |
| Otu36580 | 1 | k__Fungi(100);p__Ascomycota(100);c__Saccharomycetes(100);o__Saccharomycetales(100);f__Metschnikowiaceae(100);g__Metschnikowia(100);s__Metschnikowia_pulcherrima(100); | Metschnikowia | 13 | Saprotroph | \|Undefined Saprotroph\| | Yeast | NULL | Probable | As nectar-tap_saprotroph (Põlme et al. 2020); Decay sugar-rich substrates (Põlme et al. 2020) | Põlme S, et al. 2020. Fungal Diversity 105:1-16 ((https://doi.org/10.1007/s13225-020-00466-2)) |
| Otu36583 | 1 | k__Fungi(100);p__Ascomycota(100);c__Saccharomycetes(100);o__Saccharomycetales(100);f__Metschnikowiaceae(100);g__Metschnikowia(100);g__Metschnikowia_unclassified(100); | Metschnikowia | 13 | Saprotroph | \|Undefined Saprotroph\| | Yeast | NULL | Probable | As nectar-tap_saprotroph (Põlme et al. 2020); Decay sugar-rich substrates (Põlme et al. 2020) | Põlme S, et al. 2020. Fungal Diversity 105:1-16 ((https://doi.org/10.1007/s13225-020-00466-2)) |
| Otu36584 | 1 | k__Fungi(100);p__Ascomycota(100);c__Saccharomycetes(100);o__Saccharomycetales(100);f__Metschnikowiaceae(100);g__Metschnikowia(100);s__Metschnikowia_pulcherrima(100); | Metschnikowia | 13 | Saprotroph | \|Undefined Saprotroph\| | Yeast | NULL | Probable | As nectar-tap_saprotroph (Põlme et al. 2020); Decay sugar-rich substrates (Põlme et al. 2020) | Põlme S, et al. 2020. Fungal Diversity 105:1-16 ((https://doi.org/10.1007/s13225-020-00466-2)) |
| Otu36586 | 1 | k__Fungi(100);p__Ascomycota(100);c__Saccharomycetes(100);o__Saccharomycetales(100);f__Metschnikowiaceae(100);g__Metschnikowia(100);s__Metschnikowia_pulcherrima(100); | Metschnikowia | 13 | Saprotroph | \|Undefined Saprotroph\| | Yeast | NULL | Probable | As nectar-tap_saprotroph (Põlme et al. 2020); Decay sugar-rich substrates (Põlme et al. 2020) | Põlme S, et al. 2020. Fungal Diversity 105:1-16 ((https://doi.org/10.1007/s13225-020-00466-2)) |
| Otu36587 | 1 | k__Fungi(100);p__Ascomycota(100);c__Saccharomycetes(100);o__Saccharomycetales(100);f__Metschnikowiaceae(100);g__Metschnikowia(100);s__Metschnikowia_pulcherrima(100); | Metschnikowia | 13 | Saprotroph | \|Undefined Saprotroph\| | Yeast | NULL | Probable | As nectar-tap_saprotroph (Põlme et al. 2020); Decay sugar-rich substrates (Põlme et al. 2020) | Põlme S, et al. 2020. Fungal Diversity 105:1-16 ((https://doi.org/10.1007/s13225-020-00466-2)) |
| Otu36589 | 1 | k__Fungi(100);p__Ascomycota(100);c__Saccharomycetes(100);o__Saccharomycetales(100);f__Metschnikowiaceae(100);g__Metschnikowia(100);s__Metschnikowia_pulcherrima(100); | Metschnikowia | 13 | Saprotroph | \|Undefined Saprotroph\| | Yeast | NULL | Probable | As nectar-tap_saprotroph (Põlme et al. 2020); Decay sugar-rich substrates (Põlme et al. 2020) | Põlme S, et al. 2020. Fungal Diversity 105:1-16 ((https://doi.org/10.1007/s13225-020-00466-2)) |
| Otu36590 | 1 | k__Fungi(100);p__Ascomycota(100);c__Saccharomycetes(100);o__Saccharomycetales(100);f__Metschnikowiaceae(100);g__Metschnikowia(100);s__Metschnikowia_pulcherrima(100); | Metschnikowia | 13 | Saprotroph | \|Undefined Saprotroph\| | Yeast | NULL | Probable | As nectar-tap_saprotroph (Põlme et al. 2020); Decay sugar-rich substrates (Põlme et al. 2020) | Põlme S, et al. 2020. Fungal Diversity 105:1-16 ((https://doi.org/10.1007/s13225-020-00466-2)) |
| Otu36591 | 1 | k__Fungi(100);p__Ascomycota(100);c__Saccharomycetes(100);o__Saccharomycetales(100);f__Metschnikowiaceae(100);g__Metschnikowia(100);s__Metschnikowia_pulcherrima(100); | Metschnikowia | 13 | Saprotroph | \|Undefined Saprotroph\| | Yeast | NULL | Probable | As nectar-tap_saprotroph (Põlme et al. 2020); Decay sugar-rich substrates (Põlme et al. 2020) | Põlme S, et al. 2020. Fungal Diversity 105:1-16 ((https://doi.org/10.1007/s13225-020-00466-2)) |
| Otu36593 | 1 | k__Fungi(100);p__Ascomycota(100);c__Saccharomycetes(100);o__Saccharomycetales(100);f__Metschnikowiaceae(100);g__Metschnikowia(100);s__Metschnikowia_pulcherrima(100); | Metschnikowia | 13 | Saprotroph | \|Undefined Saprotroph\| | Yeast | NULL | Probable | As nectar-tap_saprotroph (Põlme et al. 2020); Decay sugar-rich substrates (Põlme et al. 2020) | Põlme S, et al. 2020. Fungal Diversity 105:1-16 ((https://doi.org/10.1007/s13225-020-00466-2)) |
| Otu36594 | 1 | k__Fungi(100);p__Ascomycota(100);c__Saccharomycetes(100);o__Saccharomycetales(100);f__Metschnikowiaceae(100);g__Metschnikowia(100);s__Metschnikowia_pulcherrima(100); | Metschnikowia | 13 | Saprotroph | \|Undefined Saprotroph\| | Yeast | NULL | Probable | As nectar-tap_saprotroph (Põlme et al. 2020); Decay sugar-rich substrates (Põlme et al. 2020) | Põlme S, et al. 2020. Fungal Diversity 105:1-16 ((https://doi.org/10.1007/s13225-020-00466-2)) |
| Otu36597 | 1 | k__Fungi(100);p__Ascomycota(100);c__Saccharomycetes(100);o__Saccharomycetales(100);f__Metschnikowiaceae(100);g__Metschnikowia(100);s__Metschnikowia_pulcherrima(100); | Metschnikowia | 13 | Saprotroph | \|Undefined Saprotroph\| | Yeast | NULL | Probable | As nectar-tap_saprotroph (Põlme et al. 2020); Decay sugar-rich substrates (Põlme et al. 2020) | Põlme S, et al. 2020. Fungal Diversity 105:1-16 ((https://doi.org/10.1007/s13225-020-00466-2)) |
| Otu36598 | 1 | k__Fungi(100);p__Ascomycota(100);c__Saccharomycetes(100);o__Saccharomycetales(100);f__Metschnikowiaceae(100);g__Metschnikowia(100);s__Metschnikowia_pulcherrima(100); | Metschnikowia | 13 | Saprotroph | \|Undefined Saprotroph\| | Yeast | NULL | Probable | As nectar-tap_saprotroph (Põlme et al. 2020); Decay sugar-rich substrates (Põlme et al. 2020) | Põlme S, et al. 2020. Fungal Diversity 105:1-16 ((https://doi.org/10.1007/s13225-020-00466-2)) |
| Otu36599 | 1 | k__Fungi(100);p__Ascomycota(100);c__Saccharomycetes(100);o__Saccharomycetales(100);f__Metschnikowiaceae(100);g__Metschnikowia(100);s__Metschnikowia_pulcherrima(100); | Metschnikowia | 13 | Saprotroph | \|Undefined Saprotroph\| | Yeast | NULL | Probable | As nectar-tap_saprotroph (Põlme et al. 2020); Decay sugar-rich substrates (Põlme et al. 2020) | Põlme S, et al. 2020. Fungal Diversity 105:1-16 ((https://doi.org/10.1007/s13225-020-00466-2)) |
| Otu36600 | 1 | k__Fungi(100);p__Ascomycota(100);c__Saccharomycetes(100);o__Saccharomycetales(100);f__Metschnikowiaceae(100);g__Metschnikowia(100);s__Metschnikowia_pulcherrima(100); | Metschnikowia | 13 | Saprotroph | \|Undefined Saprotroph\| | Yeast | NULL | Probable | As nectar-tap_saprotroph (Põlme et al. 2020); Decay sugar-rich substrates (Põlme et al. 2020) | Põlme S, et al. 2020. Fungal Diversity 105:1-16 ((https://doi.org/10.1007/s13225-020-00466-2)) |
| Otu36609 | 1 | k__Fungi(100);p__Ascomycota(100);c__Saccharomycetes(100);o__Saccharomycetales(100);f__Metschnikowiaceae(100);g__Metschnikowia(100);s__Metschnikowia_pulcherrima(100); | Metschnikowia | 13 | Saprotroph | \|Undefined Saprotroph\| | Yeast | NULL | Probable | As nectar-tap_saprotroph (Põlme et al. 2020); Decay sugar-rich substrates (Põlme et al. 2020) | Põlme S, et al. 2020. Fungal Diversity 105:1-16 ((https://doi.org/10.1007/s13225-020-00466-2)) |
| Otu36610 | 1 | k__Fungi(100);p__Ascomycota(100);c__Saccharomycetes(100);o__Saccharomycetales(100);f__Metschnikowiaceae(100);g__Metschnikowia(100);s__Metschnikowia_pulcherrima(100); | Metschnikowia | 13 | Saprotroph | \|Undefined Saprotroph\| | Yeast | NULL | Probable | As nectar-tap_saprotroph (Põlme et al. 2020); Decay sugar-rich substrates (Põlme et al. 2020) | Põlme S, et al. 2020. Fungal Diversity 105:1-16 ((https://doi.org/10.1007/s13225-020-00466-2)) |
| Otu36614 | 1 | k__Fungi(100);p__Ascomycota(100);c__Saccharomycetes(100);o__Saccharomycetales(100);f__Metschnikowiaceae(100);g__Metschnikowia(100);s__Metschnikowia_pulcherrima(100); | Metschnikowia | 13 | Saprotroph | \|Undefined Saprotroph\| | Yeast | NULL | Probable | As nectar-tap_saprotroph (Põlme et al. 2020); Decay sugar-rich substrates (Põlme et al. 2020) | Põlme S, et al. 2020. Fungal Diversity 105:1-16 ((https://doi.org/10.1007/s13225-020-00466-2)) |
| Otu36615 | 1 | k__Fungi(100);p__Ascomycota(100);c__Saccharomycetes(100);o__Saccharomycetales(100);f__Metschnikowiaceae(100);g__Metschnikowia(100);s__Metschnikowia_pulcherrima(100); | Metschnikowia | 13 | Saprotroph | \|Undefined Saprotroph\| | Yeast | NULL | Probable | As nectar-tap_saprotroph (Põlme et al. 2020); Decay sugar-rich substrates (Põlme et al. 2020) | Põlme S, et al. 2020. Fungal Diversity 105:1-16 ((https://doi.org/10.1007/s13225-020-00466-2)) |
| Otu36616 | 1 | k__Fungi(100);p__Ascomycota(100);c__Saccharomycetes(100);o__Saccharomycetales(100);f__Metschnikowiaceae(100);g__Metschnikowia(100);g__Metschnikowia_unclassified(100); | Metschnikowia | 13 | Saprotroph | \|Undefined Saprotroph\| | Yeast | NULL | Probable | As nectar-tap_saprotroph (Põlme et al. 2020); Decay sugar-rich substrates (Põlme et al. 2020) | Põlme S, et al. 2020. Fungal Diversity 105:1-16 ((https://doi.org/10.1007/s13225-020-00466-2)) |
| Otu36617 | 1 | k__Fungi(100);p__Ascomycota(100);c__Saccharomycetes(100);o__Saccharomycetales(100);f__Metschnikowiaceae(100);g__Metschnikowia(100);s__Metschnikowia_pulcherrima(100); | Metschnikowia | 13 | Saprotroph | \|Undefined Saprotroph\| | Yeast | NULL | Probable | As nectar-tap_saprotroph (Põlme et al. 2020); Decay sugar-rich substrates (Põlme et al. 2020) | Põlme S, et al. 2020. Fungal Diversity 105:1-16 ((https://doi.org/10.1007/s13225-020-00466-2)) |
| Otu36618 | 1 | k__Fungi(100);p__Ascomycota(100);c__Saccharomycetes(100);o__Saccharomycetales(100);f__Metschnikowiaceae(100);g__Metschnikowia(100);s__Metschnikowia_pulcherrima(100); | Metschnikowia | 13 | Saprotroph | \|Undefined Saprotroph\| | Yeast | NULL | Probable | As nectar-tap_saprotroph (Põlme et al. 2020); Decay sugar-rich substrates (Põlme et al. 2020) | Põlme S, et al. 2020. Fungal Diversity 105:1-16 ((https://doi.org/10.1007/s13225-020-00466-2)) |
| Otu36622 | 1 | k__Fungi(100);p__Ascomycota(100);c__Saccharomycetes(100);o__Saccharomycetales(100);f__Metschnikowiaceae(100);g__Metschnikowia(100);g__Metschnikowia_unclassified(100); | Metschnikowia | 13 | Saprotroph | \|Undefined Saprotroph\| | Yeast | NULL | Probable | As nectar-tap_saprotroph (Põlme et al. 2020); Decay sugar-rich substrates (Põlme et al. 2020) | Põlme S, et al. 2020. Fungal Diversity 105:1-16 ((https://doi.org/10.1007/s13225-020-00466-2)) |
| Otu36623 | 1 | k__Fungi(100);p__Ascomycota(100);c__Saccharomycetes(100);o__Saccharomycetales(100);f__Metschnikowiaceae(100);g__Metschnikowia(100);g__Metschnikowia_unclassified(100); | Metschnikowia | 13 | Saprotroph | \|Undefined Saprotroph\| | Yeast | NULL | Probable | As nectar-tap_saprotroph (Põlme et al. 2020); Decay sugar-rich substrates (Põlme et al. 2020) | Põlme S, et al. 2020. Fungal Diversity 105:1-16 ((https://doi.org/10.1007/s13225-020-00466-2)) |
| Otu36630 | 1 | k__Fungi(100);p__Ascomycota(100);c__Saccharomycetes(100);o__Saccharomycetales(100);f__Saccharomycetales_family_Incertae_sedis(100);g__Candida(100);g__Candida_unclassified(100); | Candida | 13 | Pathotroph-Saprotroph-Symbiotroph | Animal Pathogen-Endophyte-Endosymbiont-Epiphyte-\|Undefined Saprotroph\| | Dimorphic Yeast-Yeast | NULL | Probable | As nectar-tap_saprotroph (Põlme et al. 2020); Decay sugar-rich substrates (Põlme et al. 2020); This asexual genus is polyphyletic and contain species that function very differently depend on what they are. Caution should be taken when interpreting guilds for Candida species. | Manolakaki D, et al. 2010. Virulence. 1: 367-375 ((https://doi.org/10.4161/viru.1.5.12796)); Mekbib SB, et al. 2011. Tropical Plant Pathology 36:233-240 ((https://doi.org/10.1590/S1982-56762011000400004)); Põlme S, et al. 2020. Fungal Diversity 105:1-16 ((https://doi.org/10.1007/s13225-020-00466-2)) |
| Otu36635 | 1 | k__Fungi(100);p__Ascomycota(100);c__Saccharomycetes(100);o__Saccharomycetales(100);f__Metschnikowiaceae(100);g__Metschnikowia(100);g__Metschnikowia_unclassified(100); | Metschnikowia | 13 | Saprotroph | \|Undefined Saprotroph\| | Yeast | NULL | Probable | As nectar-tap_saprotroph (Põlme et al. 2020); Decay sugar-rich substrates (Põlme et al. 2020) | Põlme S, et al. 2020. Fungal Diversity 105:1-16 ((https://doi.org/10.1007/s13225-020-00466-2)) |
| Otu36636 | 1 | k__Fungi(100);p__Ascomycota(100);c__Saccharomycetes(100);o__Saccharomycetales(100);f__Metschnikowiaceae(100);g__Metschnikowia(100);s__Metschnikowia_pulcherrima(100); | Metschnikowia | 13 | Saprotroph | \|Undefined Saprotroph\| | Yeast | NULL | Probable | As nectar-tap_saprotroph (Põlme et al. 2020); Decay sugar-rich substrates (Põlme et al. 2020) | Põlme S, et al. 2020. Fungal Diversity 105:1-16 ((https://doi.org/10.1007/s13225-020-00466-2)) |
| Otu36639 | 1 | k__Fungi(100);p__Ascomycota(100);c__Saccharomycetes(100);o__Saccharomycetales(100);f__Saccharomycetales_family_Incertae_sedis(100);g__Candida(100);g__Candida_unclassified(100); | Candida | 13 | Pathotroph-Saprotroph-Symbiotroph | Animal Pathogen-Endophyte-Endosymbiont-Epiphyte-\|Undefined Saprotroph\| | Dimorphic Yeast-Yeast | NULL | Probable | As nectar-tap_saprotroph (Põlme et al. 2020); Decay sugar-rich substrates (Põlme et al. 2020); This asexual genus is polyphyletic and contain species that function very differently depend on what they are. Caution should be taken when interpreting guilds for Candida species. | Manolakaki D, et al. 2010. Virulence. 1: 367-375 ((https://doi.org/10.4161/viru.1.5.12796)); Mekbib SB, et al. 2011. Tropical Plant Pathology 36:233-240 ((https://doi.org/10.1590/S1982-56762011000400004)); Põlme S, et al. 2020. Fungal Diversity 105:1-16 ((https://doi.org/10.1007/s13225-020-00466-2)) |
| Otu36641 | 1 | k__Fungi(100);p__Ascomycota(100);c__Saccharomycetes(100);o__Saccharomycetales(100);f__Metschnikowiaceae(100);g__Metschnikowia(100);g__Metschnikowia_unclassified(100); | Metschnikowia | 13 | Saprotroph | \|Undefined Saprotroph\| | Yeast | NULL | Probable | As nectar-tap_saprotroph (Põlme et al. 2020); Decay sugar-rich substrates (Põlme et al. 2020) | Põlme S, et al. 2020. Fungal Diversity 105:1-16 ((https://doi.org/10.1007/s13225-020-00466-2)) |
| Otu36642 | 1 | k__Fungi(100);p__Ascomycota(100);c__Saccharomycetes(100);o__Saccharomycetales(100);f__Metschnikowiaceae(100);g__Metschnikowia(100);s__Metschnikowia_pulcherrima(100); | Metschnikowia | 13 | Saprotroph | \|Undefined Saprotroph\| | Yeast | NULL | Probable | As nectar-tap_saprotroph (Põlme et al. 2020); Decay sugar-rich substrates (Põlme et al. 2020) | Põlme S, et al. 2020. Fungal Diversity 105:1-16 ((https://doi.org/10.1007/s13225-020-00466-2)) |
| Otu36643 | 1 | k__Fungi(100);p__Ascomycota(100);c__Saccharomycetes(100);o__Saccharomycetales(100);f__Metschnikowiaceae(100);g__Metschnikowia(100);s__Metschnikowia_pulcherrima(100); | Metschnikowia | 13 | Saprotroph | \|Undefined Saprotroph\| | Yeast | NULL | Probable | As nectar-tap_saprotroph (Põlme et al. 2020); Decay sugar-rich substrates (Põlme et al. 2020) | Põlme S, et al. 2020. Fungal Diversity 105:1-16 ((https://doi.org/10.1007/s13225-020-00466-2)) |
| Otu36644 | 1 | k__Fungi(100);p__Ascomycota(100);c__Saccharomycetes(100);o__Saccharomycetales(100);f__Metschnikowiaceae(100);g__Metschnikowia(100);g__Metschnikowia_unclassified(100); | Metschnikowia | 13 | Saprotroph | \|Undefined Saprotroph\| | Yeast | NULL | Probable | As nectar-tap_saprotroph (Põlme et al. 2020); Decay sugar-rich substrates (Põlme et al. 2020) | Põlme S, et al. 2020. Fungal Diversity 105:1-16 ((https://doi.org/10.1007/s13225-020-00466-2)) |
| Otu36645 | 1 | k__Fungi(100);p__Ascomycota(100);c__Saccharomycetes(100);o__Saccharomycetales(100);f__Metschnikowiaceae(100);g__Metschnikowia(100);s__Metschnikowia_pulcherrima(100); | Metschnikowia | 13 | Saprotroph | \|Undefined Saprotroph\| | Yeast | NULL | Probable | As nectar-tap_saprotroph (Põlme et al. 2020); Decay sugar-rich substrates (Põlme et al. 2020) | Põlme S, et al. 2020. Fungal Diversity 105:1-16 ((https://doi.org/10.1007/s13225-020-00466-2)) |
| Otu36646 | 1 | k__Fungi(100);p__Ascomycota(100);c__Saccharomycetes(100);o__Saccharomycetales(100);f__Metschnikowiaceae(100);g__Metschnikowia(100);s__Metschnikowia_pulcherrima(100); | Metschnikowia | 13 | Saprotroph | \|Undefined Saprotroph\| | Yeast | NULL | Probable | As nectar-tap_saprotroph (Põlme et al. 2020); Decay sugar-rich substrates (Põlme et al. 2020) | Põlme S, et al. 2020. Fungal Diversity 105:1-16 ((https://doi.org/10.1007/s13225-020-00466-2)) |
| Otu36647 | 1 | k__Fungi(100);p__Ascomycota(100);c__Saccharomycetes(100);o__Saccharomycetales(100);f__Metschnikowiaceae(100);g__Metschnikowia(100);s__Metschnikowia_pulcherrima(100); | Metschnikowia | 13 | Saprotroph | \|Undefined Saprotroph\| | Yeast | NULL | Probable | As nectar-tap_saprotroph (Põlme et al. 2020); Decay sugar-rich substrates (Põlme et al. 2020) | Põlme S, et al. 2020. Fungal Diversity 105:1-16 ((https://doi.org/10.1007/s13225-020-00466-2)) |
| Otu36649 | 1 | k__Fungi(100);p__Ascomycota(100);c__Saccharomycetes(100);o__Saccharomycetales(100);f__Metschnikowiaceae(100);g__Metschnikowia(100);g__Metschnikowia_unclassified(100); | Metschnikowia | 13 | Saprotroph | \|Undefined Saprotroph\| | Yeast | NULL | Probable | As nectar-tap_saprotroph (Põlme et al. 2020); Decay sugar-rich substrates (Põlme et al. 2020) | Põlme S, et al. 2020. Fungal Diversity 105:1-16 ((https://doi.org/10.1007/s13225-020-00466-2)) |
| Otu36650 | 1 | k__Fungi(100);p__Ascomycota(100);c__Saccharomycetes(100);o__Saccharomycetales(100);f__Metschnikowiaceae(100);g__Metschnikowia(100);s__Metschnikowia_pulcherrima(100); | Metschnikowia | 13 | Saprotroph | \|Undefined Saprotroph\| | Yeast | NULL | Probable | As nectar-tap_saprotroph (Põlme et al. 2020); Decay sugar-rich substrates (Põlme et al. 2020) | Põlme S, et al. 2020. Fungal Diversity 105:1-16 ((https://doi.org/10.1007/s13225-020-00466-2)) |
| Otu36652 | 1 | k__Fungi(100);p__Ascomycota(100);c__Saccharomycetes(100);o__Saccharomycetales(100);f__Metschnikowiaceae(100);g__Metschnikowia(100);s__Metschnikowia_pulcherrima(100); | Metschnikowia | 13 | Saprotroph | \|Undefined Saprotroph\| | Yeast | NULL | Probable | As nectar-tap_saprotroph (Põlme et al. 2020); Decay sugar-rich substrates (Põlme et al. 2020) | Põlme S, et al. 2020. Fungal Diversity 105:1-16 ((https://doi.org/10.1007/s13225-020-00466-2)) |
| Otu36654 | 1 | k__Fungi(100);p__Ascomycota(100);c__Saccharomycetes(100);o__Saccharomycetales(100);f__Metschnikowiaceae(100);g__Metschnikowia(100);s__Metschnikowia_pulcherrima(100); | Metschnikowia | 13 | Saprotroph | \|Undefined Saprotroph\| | Yeast | NULL | Probable | As nectar-tap_saprotroph (Põlme et al. 2020); Decay sugar-rich substrates (Põlme et al. 2020) | Põlme S, et al. 2020. Fungal Diversity 105:1-16 ((https://doi.org/10.1007/s13225-020-00466-2)) |
| Otu36656 | 1 | k__Fungi(100);p__Ascomycota(100);c__Saccharomycetes(100);o__Saccharomycetales(100);f__Metschnikowiaceae(100);g__Metschnikowia(100);s__Metschnikowia_pulcherrima(100); | Metschnikowia | 13 | Saprotroph | \|Undefined Saprotroph\| | Yeast | NULL | Probable | As nectar-tap_saprotroph (Põlme et al. 2020); Decay sugar-rich substrates (Põlme et al. 2020) | Põlme S, et al. 2020. Fungal Diversity 105:1-16 ((https://doi.org/10.1007/s13225-020-00466-2)) |
| Otu36657 | 1 | k__Fungi(100);p__Ascomycota(100);c__Saccharomycetes(100);o__Saccharomycetales(100);f__Metschnikowiaceae(100);g__Metschnikowia(100);s__Metschnikowia_pulcherrima(100); | Metschnikowia | 13 | Saprotroph | \|Undefined Saprotroph\| | Yeast | NULL | Probable | As nectar-tap_saprotroph (Põlme et al. 2020); Decay sugar-rich substrates (Põlme et al. 2020) | Põlme S, et al. 2020. Fungal Diversity 105:1-16 ((https://doi.org/10.1007/s13225-020-00466-2)) |
| Otu36658 | 1 | k__Fungi(100);p__Ascomycota(100);c__Saccharomycetes(100);o__Saccharomycetales(100);f__Metschnikowiaceae(100);g__Metschnikowia(100);s__Metschnikowia_pulcherrima(100); | Metschnikowia | 13 | Saprotroph | \|Undefined Saprotroph\| | Yeast | NULL | Probable | As nectar-tap_saprotroph (Põlme et al. 2020); Decay sugar-rich substrates (Põlme et al. 2020) | Põlme S, et al. 2020. Fungal Diversity 105:1-16 ((https://doi.org/10.1007/s13225-020-00466-2)) |
| Otu36659 | 1 | k__Fungi(100);p__Ascomycota(100);c__Saccharomycetes(100);o__Saccharomycetales(100);f__Saccharomycetales_family_Incertae_sedis(100);g__Candida(100);g__Candida_unclassified(100); | Candida | 13 | Pathotroph-Saprotroph-Symbiotroph | Animal Pathogen-Endophyte-Endosymbiont-Epiphyte-\|Undefined Saprotroph\| | Dimorphic Yeast-Yeast | NULL | Probable | As nectar-tap_saprotroph (Põlme et al. 2020); Decay sugar-rich substrates (Põlme et al. 2020); This asexual genus is polyphyletic and contain species that function very differently depend on what they are. Caution should be taken when interpreting guilds for Candida species. | Manolakaki D, et al. 2010. Virulence. 1: 367-375 ((https://doi.org/10.4161/viru.1.5.12796)); Mekbib SB, et al. 2011. Tropical Plant Pathology 36:233-240 ((https://doi.org/10.1590/S1982-56762011000400004)); Põlme S, et al. 2020. Fungal Diversity 105:1-16 ((https://doi.org/10.1007/s13225-020-00466-2)) |
| Otu36661 | 1 | k__Fungi(100);p__Ascomycota(100);c__Saccharomycetes(100);o__Saccharomycetales(100);f__Metschnikowiaceae(100);g__Metschnikowia(100);s__Metschnikowia_pulcherrima(100); | Metschnikowia | 13 | Saprotroph | \|Undefined Saprotroph\| | Yeast | NULL | Probable | As nectar-tap_saprotroph (Põlme et al. 2020); Decay sugar-rich substrates (Põlme et al. 2020) | Põlme S, et al. 2020. Fungal Diversity 105:1-16 ((https://doi.org/10.1007/s13225-020-00466-2)) |
| Otu36666 | 1 | k__Fungi(100);p__Ascomycota(100);c__Saccharomycetes(100);o__Saccharomycetales(100);f__Metschnikowiaceae(100);g__Metschnikowia(100);s__Metschnikowia_pulcherrima(100); | Metschnikowia | 13 | Saprotroph | \|Undefined Saprotroph\| | Yeast | NULL | Probable | As nectar-tap_saprotroph (Põlme et al. 2020); Decay sugar-rich substrates (Põlme et al. 2020) | Põlme S, et al. 2020. Fungal Diversity 105:1-16 ((https://doi.org/10.1007/s13225-020-00466-2)) |
| Otu36667 | 1 | k__Fungi(100);p__Ascomycota(100);c__Saccharomycetes(100);o__Saccharomycetales(100);f__Metschnikowiaceae(100);g__Metschnikowia(100);s__Metschnikowia_pulcherrima(100); | Metschnikowia | 13 | Saprotroph | \|Undefined Saprotroph\| | Yeast | NULL | Probable | As nectar-tap_saprotroph (Põlme et al. 2020); Decay sugar-rich substrates (Põlme et al. 2020) | Põlme S, et al. 2020. Fungal Diversity 105:1-16 ((https://doi.org/10.1007/s13225-020-00466-2)) |
| Otu36668 | 1 | k__Fungi(100);p__Ascomycota(100);c__Saccharomycetes(100);o__Saccharomycetales(100);f__Saccharomycetales_family_Incertae_sedis(100);g__Candida(100);g__Candida_unclassified(100); | Candida | 13 | Pathotroph-Saprotroph-Symbiotroph | Animal Pathogen-Endophyte-Endosymbiont-Epiphyte-\|Undefined Saprotroph\| | Dimorphic Yeast-Yeast | NULL | Probable | As nectar-tap_saprotroph (Põlme et al. 2020); Decay sugar-rich substrates (Põlme et al. 2020); This asexual genus is polyphyletic and contain species that function very differently depend on what they are. Caution should be taken when interpreting guilds for Candida species. | Manolakaki D, et al. 2010. Virulence. 1: 367-375 ((https://doi.org/10.4161/viru.1.5.12796)); Mekbib SB, et al. 2011. Tropical Plant Pathology 36:233-240 ((https://doi.org/10.1590/S1982-56762011000400004)); Põlme S, et al. 2020. Fungal Diversity 105:1-16 ((https://doi.org/10.1007/s13225-020-00466-2)) |
| Otu36670 | 1 | k__Fungi(100);p__Ascomycota(100);c__Saccharomycetes(100);o__Saccharomycetales(100);f__Metschnikowiaceae(100);g__Metschnikowia(100);s__Metschnikowia_pulcherrima(100); | Metschnikowia | 13 | Saprotroph | \|Undefined Saprotroph\| | Yeast | NULL | Probable | As nectar-tap_saprotroph (Põlme et al. 2020); Decay sugar-rich substrates (Põlme et al. 2020) | Põlme S, et al. 2020. Fungal Diversity 105:1-16 ((https://doi.org/10.1007/s13225-020-00466-2)) |
| Otu36671 | 1 | k__Fungi(100);p__Ascomycota(100);c__Saccharomycetes(100);o__Saccharomycetales(100);f__Metschnikowiaceae(100);g__Metschnikowia(100);s__Metschnikowia_pulcherrima(100); | Metschnikowia | 13 | Saprotroph | \|Undefined Saprotroph\| | Yeast | NULL | Probable | As nectar-tap_saprotroph (Põlme et al. 2020); Decay sugar-rich substrates (Põlme et al. 2020) | Põlme S, et al. 2020. Fungal Diversity 105:1-16 ((https://doi.org/10.1007/s13225-020-00466-2)) |
| Otu36672 | 1 | k__Fungi(100);p__Ascomycota(100);c__Saccharomycetes(100);o__Saccharomycetales(100);f__Metschnikowiaceae(100);g__Metschnikowia(100);s__Metschnikowia_pulcherrima(100); | Metschnikowia | 13 | Saprotroph | \|Undefined Saprotroph\| | Yeast | NULL | Probable | As nectar-tap_saprotroph (Põlme et al. 2020); Decay sugar-rich substrates (Põlme et al. 2020) | Põlme S, et al. 2020. Fungal Diversity 105:1-16 ((https://doi.org/10.1007/s13225-020-00466-2)) |
| Otu36673 | 1 | k__Fungi(100);p__Ascomycota(100);c__Saccharomycetes(100);o__Saccharomycetales(100);f__Metschnikowiaceae(100);g__Metschnikowia(100);g__Metschnikowia_unclassified(100); | Metschnikowia | 13 | Saprotroph | \|Undefined Saprotroph\| | Yeast | NULL | Probable | As nectar-tap_saprotroph (Põlme et al. 2020); Decay sugar-rich substrates (Põlme et al. 2020) | Põlme S, et al. 2020. Fungal Diversity 105:1-16 ((https://doi.org/10.1007/s13225-020-00466-2)) |
| Otu36674 | 1 | k__Fungi(100);p__Ascomycota(100);c__Saccharomycetes(100);o__Saccharomycetales(100);f__Metschnikowiaceae(100);g__Metschnikowia(100);s__Metschnikowia_pulcherrima(100); | Metschnikowia | 13 | Saprotroph | \|Undefined Saprotroph\| | Yeast | NULL | Probable | As nectar-tap_saprotroph (Põlme et al. 2020); Decay sugar-rich substrates (Põlme et al. 2020) | Põlme S, et al. 2020. Fungal Diversity 105:1-16 ((https://doi.org/10.1007/s13225-020-00466-2)) |
| Otu36675 | 1 | k__Fungi(100);p__Ascomycota(100);c__Saccharomycetes(100);o__Saccharomycetales(100);f__Metschnikowiaceae(100);g__Metschnikowia(100);g__Metschnikowia_unclassified(100); | Metschnikowia | 13 | Saprotroph | \|Undefined Saprotroph\| | Yeast | NULL | Probable | As nectar-tap_saprotroph (Põlme et al. 2020); Decay sugar-rich substrates (Põlme et al. 2020) | Põlme S, et al. 2020. Fungal Diversity 105:1-16 ((https://doi.org/10.1007/s13225-020-00466-2)) |
| Otu36676 | 1 | k__Fungi(100);p__Ascomycota(100);c__Saccharomycetes(100);o__Saccharomycetales(100);f__Metschnikowiaceae(100);g__Metschnikowia(100);s__Metschnikowia_pulcherrima(100); | Metschnikowia | 13 | Saprotroph | \|Undefined Saprotroph\| | Yeast | NULL | Probable | As nectar-tap_saprotroph (Põlme et al. 2020); Decay sugar-rich substrates (Põlme et al. 2020) | Põlme S, et al. 2020. Fungal Diversity 105:1-16 ((https://doi.org/10.1007/s13225-020-00466-2)) |
| Otu36677 | 1 | k__Fungi(100);p__Ascomycota(100);c__Saccharomycetes(100);o__Saccharomycetales(100);f__Metschnikowiaceae(100);g__Metschnikowia(100);s__Metschnikowia_pulcherrima(100); | Metschnikowia | 13 | Saprotroph | \|Undefined Saprotroph\| | Yeast | NULL | Probable | As nectar-tap_saprotroph (Põlme et al. 2020); Decay sugar-rich substrates (Põlme et al. 2020) | Põlme S, et al. 2020. Fungal Diversity 105:1-16 ((https://doi.org/10.1007/s13225-020-00466-2)) |
| Otu36678 | 1 | k__Fungi(100);p__Ascomycota(100);c__Saccharomycetes(100);o__Saccharomycetales(100);f__Metschnikowiaceae(100);g__Metschnikowia(100);s__Metschnikowia_pulcherrima(100); | Metschnikowia | 13 | Saprotroph | \|Undefined Saprotroph\| | Yeast | NULL | Probable | As nectar-tap_saprotroph (Põlme et al. 2020); Decay sugar-rich substrates (Põlme et al. 2020) | Põlme S, et al. 2020. Fungal Diversity 105:1-16 ((https://doi.org/10.1007/s13225-020-00466-2)) |
| Otu36679 | 1 | k__Fungi(100);p__Ascomycota(100);c__Saccharomycetes(100);o__Saccharomycetales(100);f__Metschnikowiaceae(100);g__Metschnikowia(100);g__Metschnikowia_unclassified(100); | Metschnikowia | 13 | Saprotroph | \|Undefined Saprotroph\| | Yeast | NULL | Probable | As nectar-tap_saprotroph (Põlme et al. 2020); Decay sugar-rich substrates (Põlme et al. 2020) | Põlme S, et al. 2020. Fungal Diversity 105:1-16 ((https://doi.org/10.1007/s13225-020-00466-2)) |
| Otu36680 | 1 | k__Fungi(100);p__Ascomycota(100);c__Saccharomycetes(100);o__Saccharomycetales(100);f__Metschnikowiaceae(100);g__Metschnikowia(100);s__Metschnikowia_pulcherrima(100); | Metschnikowia | 13 | Saprotroph | \|Undefined Saprotroph\| | Yeast | NULL | Probable | As nectar-tap_saprotroph (Põlme et al. 2020); Decay sugar-rich substrates (Põlme et al. 2020) | Põlme S, et al. 2020. Fungal Diversity 105:1-16 ((https://doi.org/10.1007/s13225-020-00466-2)) |
| Otu36681 | 1 | k__Fungi(100);p__Ascomycota(100);c__Saccharomycetes(100);o__Saccharomycetales(100);f__Metschnikowiaceae(100);g__Metschnikowia(100);s__Metschnikowia_pulcherrima(100); | Metschnikowia | 13 | Saprotroph | \|Undefined Saprotroph\| | Yeast | NULL | Probable | As nectar-tap_saprotroph (Põlme et al. 2020); Decay sugar-rich substrates (Põlme et al. 2020) | Põlme S, et al. 2020. Fungal Diversity 105:1-16 ((https://doi.org/10.1007/s13225-020-00466-2)) |
| Otu36682 | 1 | k__Fungi(100);p__Ascomycota(100);c__Saccharomycetes(100);o__Saccharomycetales(100);f__Metschnikowiaceae(100);g__Metschnikowia(100);s__Metschnikowia_pulcherrima(100); | Metschnikowia | 13 | Saprotroph | \|Undefined Saprotroph\| | Yeast | NULL | Probable | As nectar-tap_saprotroph (Põlme et al. 2020); Decay sugar-rich substrates (Põlme et al. 2020) | Põlme S, et al. 2020. Fungal Diversity 105:1-16 ((https://doi.org/10.1007/s13225-020-00466-2)) |
| Otu36683 | 1 | k__Fungi(100);p__Ascomycota(100);c__Saccharomycetes(100);o__Saccharomycetales(100);f__Metschnikowiaceae(100);g__Metschnikowia(100);s__Metschnikowia_pulcherrima(100); | Metschnikowia | 13 | Saprotroph | \|Undefined Saprotroph\| | Yeast | NULL | Probable | As nectar-tap_saprotroph (Põlme et al. 2020); Decay sugar-rich substrates (Põlme et al. 2020) | Põlme S, et al. 2020. Fungal Diversity 105:1-16 ((https://doi.org/10.1007/s13225-020-00466-2)) |
| Otu36684 | 1 | k__Fungi(100);p__Ascomycota(100);c__Saccharomycetes(100);o__Saccharomycetales(100);f__Metschnikowiaceae(100);g__Metschnikowia(100);s__Metschnikowia_pulcherrima(100); | Metschnikowia | 13 | Saprotroph | \|Undefined Saprotroph\| | Yeast | NULL | Probable | As nectar-tap_saprotroph (Põlme et al. 2020); Decay sugar-rich substrates (Põlme et al. 2020) | Põlme S, et al. 2020. Fungal Diversity 105:1-16 ((https://doi.org/10.1007/s13225-020-00466-2)) |
| Otu36685 | 1 | k__Fungi(100);p__Ascomycota(100);c__Saccharomycetes(100);o__Saccharomycetales(100);f__Metschnikowiaceae(100);g__Metschnikowia(100);s__Metschnikowia_pulcherrima(100); | Metschnikowia | 13 | Saprotroph | \|Undefined Saprotroph\| | Yeast | NULL | Probable | As nectar-tap_saprotroph (Põlme et al. 2020); Decay sugar-rich substrates (Põlme et al. 2020) | Põlme S, et al. 2020. Fungal Diversity 105:1-16 ((https://doi.org/10.1007/s13225-020-00466-2)) |
| Otu36686 | 1 | k__Fungi(100);p__Ascomycota(100);c__Saccharomycetes(100);o__Saccharomycetales(100);f__Metschnikowiaceae(100);g__Metschnikowia(100);s__Metschnikowia_pulcherrima(100); | Metschnikowia | 13 | Saprotroph | \|Undefined Saprotroph\| | Yeast | NULL | Probable | As nectar-tap_saprotroph (Põlme et al. 2020); Decay sugar-rich substrates (Põlme et al. 2020) | Põlme S, et al. 2020. Fungal Diversity 105:1-16 ((https://doi.org/10.1007/s13225-020-00466-2)) |
| Otu36687 | 1 | k__Fungi(100);p__Ascomycota(100);c__Saccharomycetes(100);o__Saccharomycetales(100);f__Metschnikowiaceae(100);g__Metschnikowia(100);s__Metschnikowia_pulcherrima(100); | Metschnikowia | 13 | Saprotroph | \|Undefined Saprotroph\| | Yeast | NULL | Probable | As nectar-tap_saprotroph (Põlme et al. 2020); Decay sugar-rich substrates (Põlme et al. 2020) | Põlme S, et al. 2020. Fungal Diversity 105:1-16 ((https://doi.org/10.1007/s13225-020-00466-2)) |
| Otu36688 | 1 | k__Fungi(100);p__Ascomycota(100);c__Saccharomycetes(100);o__Saccharomycetales(100);f__Metschnikowiaceae(100);g__Metschnikowia(100);s__Metschnikowia_pulcherrima(100); | Metschnikowia | 13 | Saprotroph | \|Undefined Saprotroph\| | Yeast | NULL | Probable | As nectar-tap_saprotroph (Põlme et al. 2020); Decay sugar-rich substrates (Põlme et al. 2020) | Põlme S, et al. 2020. Fungal Diversity 105:1-16 ((https://doi.org/10.1007/s13225-020-00466-2)) |
| Otu36689 | 1 | k__Fungi(100);p__Ascomycota(100);c__Saccharomycetes(100);o__Saccharomycetales(100);f__Metschnikowiaceae(100);g__Metschnikowia(100);s__Metschnikowia_pulcherrima(100); | Metschnikowia | 13 | Saprotroph | \|Undefined Saprotroph\| | Yeast | NULL | Probable | As nectar-tap_saprotroph (Põlme et al. 2020); Decay sugar-rich substrates (Põlme et al. 2020) | Põlme S, et al. 2020. Fungal Diversity 105:1-16 ((https://doi.org/10.1007/s13225-020-00466-2)) |
| Otu36690 | 1 | k__Fungi(100);p__Ascomycota(100);c__Saccharomycetes(100);o__Saccharomycetales(100);f__Metschnikowiaceae(100);g__Metschnikowia(100);s__Metschnikowia_pulcherrima(100); | Metschnikowia | 13 | Saprotroph | \|Undefined Saprotroph\| | Yeast | NULL | Probable | As nectar-tap_saprotroph (Põlme et al. 2020); Decay sugar-rich substrates (Põlme et al. 2020) | Põlme S, et al. 2020. Fungal Diversity 105:1-16 ((https://doi.org/10.1007/s13225-020-00466-2)) |
| Otu36692 | 1 | k__Fungi(100);p__Ascomycota(100);c__Saccharomycetes(100);o__Saccharomycetales(100);f__Metschnikowiaceae(100);g__Metschnikowia(100);s__Metschnikowia_pulcherrima(100); | Metschnikowia | 13 | Saprotroph | \|Undefined Saprotroph\| | Yeast | NULL | Probable | As nectar-tap_saprotroph (Põlme et al. 2020); Decay sugar-rich substrates (Põlme et al. 2020) | Põlme S, et al. 2020. Fungal Diversity 105:1-16 ((https://doi.org/10.1007/s13225-020-00466-2)) |
| Otu36693 | 1 | k__Fungi(100);p__Ascomycota(100);c__Saccharomycetes(100);o__Saccharomycetales(100);f__Metschnikowiaceae(100);g__Metschnikowia(100);s__Metschnikowia_pulcherrima(100); | Metschnikowia | 13 | Saprotroph | \|Undefined Saprotroph\| | Yeast | NULL | Probable | As nectar-tap_saprotroph (Põlme et al. 2020); Decay sugar-rich substrates (Põlme et al. 2020) | Põlme S, et al. 2020. Fungal Diversity 105:1-16 ((https://doi.org/10.1007/s13225-020-00466-2)) |
| Otu36694 | 1 | k__Fungi(100);p__Ascomycota(100);c__Saccharomycetes(100);o__Saccharomycetales(100);f__Metschnikowiaceae(100);g__Metschnikowia(100);g__Metschnikowia_unclassified(100); | Metschnikowia | 13 | Saprotroph | \|Undefined Saprotroph\| | Yeast | NULL | Probable | As nectar-tap_saprotroph (Põlme et al. 2020); Decay sugar-rich substrates (Põlme et al. 2020) | Põlme S, et al. 2020. Fungal Diversity 105:1-16 ((https://doi.org/10.1007/s13225-020-00466-2)) |
| Otu36695 | 1 | k__Fungi(100);p__Ascomycota(100);c__Dothideomycetes(100);o__Dothideomycetes_order_Incertae_sedis(100);f__Myxotrichaceae(100);g__Oidiodendron(100);s__Oidiodendron_cereale(100); | Oidiodendron | 13 | Pathotroph-Saprotroph-Symbiotroph | Endophyte-Ericoid Mycorrhizal-\|Undefined Saprotroph\| | Microfungus | NULL | Probable | As root_endophyte (Põlme et al. 2020); As soil_saprotroph (Põlme et al. 2020); Dark Septate Endophyte; Decay soil (Põlme et al. 2020) | Newsham KK. 2011. New Phytologist 190:783-793 ((https://doi.org/10.1111/j.1469-8137.2010.03611.x)) (as dark septate root endophytes); Põlme S, et al. 2020. Fungal Diversity 105:1-16 ((https://doi.org/10.1007/s13225-020-00466-2)) |
| Otu36696 | 1 | k__Fungi(100);p__Ascomycota(100);c__Saccharomycetes(100);o__Saccharomycetales(100);f__Metschnikowiaceae(100);g__Metschnikowia(100);s__Metschnikowia_pulcherrima(100); | Metschnikowia | 13 | Saprotroph | \|Undefined Saprotroph\| | Yeast | NULL | Probable | As nectar-tap_saprotroph (Põlme et al. 2020); Decay sugar-rich substrates (Põlme et al. 2020) | Põlme S, et al. 2020. Fungal Diversity 105:1-16 ((https://doi.org/10.1007/s13225-020-00466-2)) |
| Otu36698 | 1 | k__Fungi(100);p__Ascomycota(100);c__Saccharomycetes(100);o__Saccharomycetales(100);f__Metschnikowiaceae(100);g__Metschnikowia(100);s__Metschnikowia_pulcherrima(100); | Metschnikowia | 13 | Saprotroph | \|Undefined Saprotroph\| | Yeast | NULL | Probable | As nectar-tap_saprotroph (Põlme et al. 2020); Decay sugar-rich substrates (Põlme et al. 2020) | Põlme S, et al. 2020. Fungal Diversity 105:1-16 ((https://doi.org/10.1007/s13225-020-00466-2)) |
| Otu36699 | 1 | k__Fungi(100);p__Ascomycota(100);c__Saccharomycetes(100);o__Saccharomycetales(100);f__Metschnikowiaceae(100);g__Metschnikowia(100);s__Metschnikowia_pulcherrima(100); | Metschnikowia | 13 | Saprotroph | \|Undefined Saprotroph\| | Yeast | NULL | Probable | As nectar-tap_saprotroph (Põlme et al. 2020); Decay sugar-rich substrates (Põlme et al. 2020) | Põlme S, et al. 2020. Fungal Diversity 105:1-16 ((https://doi.org/10.1007/s13225-020-00466-2)) |
| Otu36700 | 1 | k__Fungi(100);p__Ascomycota(100);c__Saccharomycetes(100);o__Saccharomycetales(100);f__Metschnikowiaceae(100);g__Metschnikowia(100);g__Metschnikowia_unclassified(100); | Metschnikowia | 13 | Saprotroph | \|Undefined Saprotroph\| | Yeast | NULL | Probable | As nectar-tap_saprotroph (Põlme et al. 2020); Decay sugar-rich substrates (Põlme et al. 2020) | Põlme S, et al. 2020. Fungal Diversity 105:1-16 ((https://doi.org/10.1007/s13225-020-00466-2)) |
| Otu36702 | 1 | k__Fungi(100);p__Ascomycota(100);c__Saccharomycetes(100);o__Saccharomycetales(100);f__Metschnikowiaceae(100);g__Metschnikowia(100);s__Metschnikowia_pulcherrima(100); | Metschnikowia | 13 | Saprotroph | \|Undefined Saprotroph\| | Yeast | NULL | Probable | As nectar-tap_saprotroph (Põlme et al. 2020); Decay sugar-rich substrates (Põlme et al. 2020) | Põlme S, et al. 2020. Fungal Diversity 105:1-16 ((https://doi.org/10.1007/s13225-020-00466-2)) |
| Otu36706 | 1 | k__Fungi(100);p__Ascomycota(100);c__Saccharomycetes(100);o__Saccharomycetales(100);f__Metschnikowiaceae(100);g__Metschnikowia(100);s__Metschnikowia_pulcherrima(100); | Metschnikowia | 13 | Saprotroph | \|Undefined Saprotroph\| | Yeast | NULL | Probable | As nectar-tap_saprotroph (Põlme et al. 2020); Decay sugar-rich substrates (Põlme et al. 2020) | Põlme S, et al. 2020. Fungal Diversity 105:1-16 ((https://doi.org/10.1007/s13225-020-00466-2)) |
| Otu36707 | 1 | k__Fungi(100);p__Ascomycota(100);c__Saccharomycetes(100);o__Saccharomycetales(100);f__Metschnikowiaceae(100);g__Metschnikowia(100);s__Metschnikowia_pulcherrima(100); | Metschnikowia | 13 | Saprotroph | \|Undefined Saprotroph\| | Yeast | NULL | Probable | As nectar-tap_saprotroph (Põlme et al. 2020); Decay sugar-rich substrates (Põlme et al. 2020) | Põlme S, et al. 2020. Fungal Diversity 105:1-16 ((https://doi.org/10.1007/s13225-020-00466-2)) |
| Otu36708 | 1 | k__Fungi(100);p__Ascomycota(100);c__Saccharomycetes(100);o__Saccharomycetales(100);f__Metschnikowiaceae(100);g__Metschnikowia(100);g__Metschnikowia_unclassified(100); | Metschnikowia | 13 | Saprotroph | \|Undefined Saprotroph\| | Yeast | NULL | Probable | As nectar-tap_saprotroph (Põlme et al. 2020); Decay sugar-rich substrates (Põlme et al. 2020) | Põlme S, et al. 2020. Fungal Diversity 105:1-16 ((https://doi.org/10.1007/s13225-020-00466-2)) |
| Otu36710 | 1 | k__Fungi(100);p__Ascomycota(100);c__Saccharomycetes(100);o__Saccharomycetales(100);f__Metschnikowiaceae(100);g__Metschnikowia(100);s__Metschnikowia_pulcherrima(100); | Metschnikowia | 13 | Saprotroph | \|Undefined Saprotroph\| | Yeast | NULL | Probable | As nectar-tap_saprotroph (Põlme et al. 2020); Decay sugar-rich substrates (Põlme et al. 2020) | Põlme S, et al. 2020. Fungal Diversity 105:1-16 ((https://doi.org/10.1007/s13225-020-00466-2)) |
| Otu36712 | 1 | k__Fungi(100);p__Ascomycota(100);c__Saccharomycetes(100);o__Saccharomycetales(100);f__Metschnikowiaceae(100);g__Metschnikowia(100);s__Metschnikowia_pulcherrima(100); | Metschnikowia | 13 | Saprotroph | \|Undefined Saprotroph\| | Yeast | NULL | Probable | As nectar-tap_saprotroph (Põlme et al. 2020); Decay sugar-rich substrates (Põlme et al. 2020) | Põlme S, et al. 2020. Fungal Diversity 105:1-16 ((https://doi.org/10.1007/s13225-020-00466-2)) |
| Otu36713 | 1 | k__Fungi(100);p__Ascomycota(100);c__Saccharomycetes(100);o__Saccharomycetales(100);f__Metschnikowiaceae(100);g__Metschnikowia(100);s__Metschnikowia_pulcherrima(100); | Metschnikowia | 13 | Saprotroph | \|Undefined Saprotroph\| | Yeast | NULL | Probable | As nectar-tap_saprotroph (Põlme et al. 2020); Decay sugar-rich substrates (Põlme et al. 2020) | Põlme S, et al. 2020. Fungal Diversity 105:1-16 ((https://doi.org/10.1007/s13225-020-00466-2)) |
| Otu36714 | 1 | k__Fungi(100);p__Ascomycota(100);c__Saccharomycetes(100);o__Saccharomycetales(100);f__Metschnikowiaceae(100);g__Metschnikowia(100);s__Metschnikowia_pulcherrima(100); | Metschnikowia | 13 | Saprotroph | \|Undefined Saprotroph\| | Yeast | NULL | Probable | As nectar-tap_saprotroph (Põlme et al. 2020); Decay sugar-rich substrates (Põlme et al. 2020) | Põlme S, et al. 2020. Fungal Diversity 105:1-16 ((https://doi.org/10.1007/s13225-020-00466-2)) |
| Otu36715 | 1 | k__Fungi(100);p__Ascomycota(100);c__Saccharomycetes(100);o__Saccharomycetales(100);f__Metschnikowiaceae(100);g__Metschnikowia(100);s__Metschnikowia_pulcherrima(100); | Metschnikowia | 13 | Saprotroph | \|Undefined Saprotroph\| | Yeast | NULL | Probable | As nectar-tap_saprotroph (Põlme et al. 2020); Decay sugar-rich substrates (Põlme et al. 2020) | Põlme S, et al. 2020. Fungal Diversity 105:1-16 ((https://doi.org/10.1007/s13225-020-00466-2)) |
| Otu36717 | 1 | k__Fungi(100);p__Ascomycota(100);c__Saccharomycetes(100);o__Saccharomycetales(100);f__Saccharomycetales_family_Incertae_sedis(100);g__Candida(100);g__Candida_unclassified(100); | Candida | 13 | Pathotroph-Saprotroph-Symbiotroph | Animal Pathogen-Endophyte-Endosymbiont-Epiphyte-\|Undefined Saprotroph\| | Dimorphic Yeast-Yeast | NULL | Probable | As nectar-tap_saprotroph (Põlme et al. 2020); Decay sugar-rich substrates (Põlme et al. 2020); This asexual genus is polyphyletic and contain species that function very differently depend on what they are. Caution should be taken when interpreting guilds for Candida species. | Manolakaki D, et al. 2010. Virulence. 1: 367-375 ((https://doi.org/10.4161/viru.1.5.12796)); Mekbib SB, et al. 2011. Tropical Plant Pathology 36:233-240 ((https://doi.org/10.1590/S1982-56762011000400004)); Põlme S, et al. 2020. Fungal Diversity 105:1-16 ((https://doi.org/10.1007/s13225-020-00466-2)) |
| Otu36719 | 1 | k__Fungi(100);p__Ascomycota(100);c__Saccharomycetes(100);o__Saccharomycetales(100);f__Metschnikowiaceae(100);g__Metschnikowia(100);s__Metschnikowia_pulcherrima(100); | Metschnikowia | 13 | Saprotroph | \|Undefined Saprotroph\| | Yeast | NULL | Probable | As nectar-tap_saprotroph (Põlme et al. 2020); Decay sugar-rich substrates (Põlme et al. 2020) | Põlme S, et al. 2020. Fungal Diversity 105:1-16 ((https://doi.org/10.1007/s13225-020-00466-2)) |
| Otu36720 | 1 | k__Fungi(100);p__Ascomycota(100);c__Saccharomycetes(100);o__Saccharomycetales(100);f__Metschnikowiaceae(100);g__Metschnikowia(100);s__Metschnikowia_pulcherrima(100); | Metschnikowia | 13 | Saprotroph | \|Undefined Saprotroph\| | Yeast | NULL | Probable | As nectar-tap_saprotroph (Põlme et al. 2020); Decay sugar-rich substrates (Põlme et al. 2020) | Põlme S, et al. 2020. Fungal Diversity 105:1-16 ((https://doi.org/10.1007/s13225-020-00466-2)) |
| Otu36721 | 1 | k__Fungi(100);p__Ascomycota(100);c__Saccharomycetes(100);o__Saccharomycetales(100);f__Metschnikowiaceae(100);g__Metschnikowia(100);s__Metschnikowia_pulcherrima(100); | Metschnikowia | 13 | Saprotroph | \|Undefined Saprotroph\| | Yeast | NULL | Probable | As nectar-tap_saprotroph (Põlme et al. 2020); Decay sugar-rich substrates (Põlme et al. 2020) | Põlme S, et al. 2020. Fungal Diversity 105:1-16 ((https://doi.org/10.1007/s13225-020-00466-2)) |
| Otu36722 | 1 | k__Fungi(100);p__Ascomycota(100);c__Saccharomycetes(100);o__Saccharomycetales(100);f__Metschnikowiaceae(100);g__Metschnikowia(100);g__Metschnikowia_unclassified(100); | Metschnikowia | 13 | Saprotroph | \|Undefined Saprotroph\| | Yeast | NULL | Probable | As nectar-tap_saprotroph (Põlme et al. 2020); Decay sugar-rich substrates (Põlme et al. 2020) | Põlme S, et al. 2020. Fungal Diversity 105:1-16 ((https://doi.org/10.1007/s13225-020-00466-2)) |
| Otu36723 | 1 | k__Fungi(100);p__Ascomycota(100);c__Saccharomycetes(100);o__Saccharomycetales(100);f__Metschnikowiaceae(100);g__Metschnikowia(100);s__Metschnikowia_pulcherrima(100); | Metschnikowia | 13 | Saprotroph | \|Undefined Saprotroph\| | Yeast | NULL | Probable | As nectar-tap_saprotroph (Põlme et al. 2020); Decay sugar-rich substrates (Põlme et al. 2020) | Põlme S, et al. 2020. Fungal Diversity 105:1-16 ((https://doi.org/10.1007/s13225-020-00466-2)) |
| Otu36724 | 1 | k__Fungi(100);p__Ascomycota(100);c__Saccharomycetes(100);o__Saccharomycetales(100);f__Metschnikowiaceae(100);g__Metschnikowia(100);s__Metschnikowia_pulcherrima(100); | Metschnikowia | 13 | Saprotroph | \|Undefined Saprotroph\| | Yeast | NULL | Probable | As nectar-tap_saprotroph (Põlme et al. 2020); Decay sugar-rich substrates (Põlme et al. 2020) | Põlme S, et al. 2020. Fungal Diversity 105:1-16 ((https://doi.org/10.1007/s13225-020-00466-2)) |
| Otu36725 | 1 | k__Fungi(100);p__Ascomycota(100);c__Saccharomycetes(100);o__Saccharomycetales(100);f__Metschnikowiaceae(100);g__Metschnikowia(100);s__Metschnikowia_pulcherrima(100); | Metschnikowia | 13 | Saprotroph | \|Undefined Saprotroph\| | Yeast | NULL | Probable | As nectar-tap_saprotroph (Põlme et al. 2020); Decay sugar-rich substrates (Põlme et al. 2020) | Põlme S, et al. 2020. Fungal Diversity 105:1-16 ((https://doi.org/10.1007/s13225-020-00466-2)) |
| Otu36726 | 1 | k__Fungi(100);p__Ascomycota(100);c__Saccharomycetes(100);o__Saccharomycetales(100);f__Metschnikowiaceae(100);g__Metschnikowia(100);s__Metschnikowia_pulcherrima(100); | Metschnikowia | 13 | Saprotroph | \|Undefined Saprotroph\| | Yeast | NULL | Probable | As nectar-tap_saprotroph (Põlme et al. 2020); Decay sugar-rich substrates (Põlme et al. 2020) | Põlme S, et al. 2020. Fungal Diversity 105:1-16 ((https://doi.org/10.1007/s13225-020-00466-2)) |
| Otu36727 | 1 | k__Fungi(100);p__Ascomycota(100);c__Saccharomycetes(100);o__Saccharomycetales(100);f__Metschnikowiaceae(100);g__Metschnikowia(100);s__Metschnikowia_pulcherrima(100); | Metschnikowia | 13 | Saprotroph | \|Undefined Saprotroph\| | Yeast | NULL | Probable | As nectar-tap_saprotroph (Põlme et al. 2020); Decay sugar-rich substrates (Põlme et al. 2020) | Põlme S, et al. 2020. Fungal Diversity 105:1-16 ((https://doi.org/10.1007/s13225-020-00466-2)) |
| Otu36728 | 1 | k__Fungi(100);p__Ascomycota(100);c__Saccharomycetes(100);o__Saccharomycetales(100);f__Metschnikowiaceae(100);g__Metschnikowia(100);g__Metschnikowia_unclassified(100); | Metschnikowia | 13 | Saprotroph | \|Undefined Saprotroph\| | Yeast | NULL | Probable | As nectar-tap_saprotroph (Põlme et al. 2020); Decay sugar-rich substrates (Põlme et al. 2020) | Põlme S, et al. 2020. Fungal Diversity 105:1-16 ((https://doi.org/10.1007/s13225-020-00466-2)) |
| Otu36729 | 1 | k__Fungi(100);p__Ascomycota(100);c__Saccharomycetes(100);o__Saccharomycetales(100);f__Metschnikowiaceae(100);g__Metschnikowia(100);g__Metschnikowia_unclassified(100); | Metschnikowia | 13 | Saprotroph | \|Undefined Saprotroph\| | Yeast | NULL | Probable | As nectar-tap_saprotroph (Põlme et al. 2020); Decay sugar-rich substrates (Põlme et al. 2020) | Põlme S, et al. 2020. Fungal Diversity 105:1-16 ((https://doi.org/10.1007/s13225-020-00466-2)) |
| Otu36730 | 1 | k__Fungi(100);p__Ascomycota(100);c__Saccharomycetes(100);o__Saccharomycetales(100);f__Metschnikowiaceae(100);g__Metschnikowia(100);s__Metschnikowia_pulcherrima(100); | Metschnikowia | 13 | Saprotroph | \|Undefined Saprotroph\| | Yeast | NULL | Probable | As nectar-tap_saprotroph (Põlme et al. 2020); Decay sugar-rich substrates (Põlme et al. 2020) | Põlme S, et al. 2020. Fungal Diversity 105:1-16 ((https://doi.org/10.1007/s13225-020-00466-2)) |
| Otu36731 | 1 | k__Fungi(100);p__Ascomycota(100);c__Saccharomycetes(100);o__Saccharomycetales(100);f__Metschnikowiaceae(100);g__Metschnikowia(100);s__Metschnikowia_pulcherrima(100); | Metschnikowia | 13 | Saprotroph | \|Undefined Saprotroph\| | Yeast | NULL | Probable | As nectar-tap_saprotroph (Põlme et al. 2020); Decay sugar-rich substrates (Põlme et al. 2020) | Põlme S, et al. 2020. Fungal Diversity 105:1-16 ((https://doi.org/10.1007/s13225-020-00466-2)) |
| Otu36732 | 1 | k__Fungi(100);p__Ascomycota(100);c__Saccharomycetes(100);o__Saccharomycetales(100);f__Metschnikowiaceae(100);g__Metschnikowia(100);s__Metschnikowia_pulcherrima(100); | Metschnikowia | 13 | Saprotroph | \|Undefined Saprotroph\| | Yeast | NULL | Probable | As nectar-tap_saprotroph (Põlme et al. 2020); Decay sugar-rich substrates (Põlme et al. 2020) | Põlme S, et al. 2020. Fungal Diversity 105:1-16 ((https://doi.org/10.1007/s13225-020-00466-2)) |
| Otu36733 | 1 | k__Fungi(100);p__Ascomycota(100);c__Saccharomycetes(100);o__Saccharomycetales(100);f__Metschnikowiaceae(100);g__Metschnikowia(100);s__Metschnikowia_pulcherrima(100); | Metschnikowia | 13 | Saprotroph | \|Undefined Saprotroph\| | Yeast | NULL | Probable | As nectar-tap_saprotroph (Põlme et al. 2020); Decay sugar-rich substrates (Põlme et al. 2020) | Põlme S, et al. 2020. Fungal Diversity 105:1-16 ((https://doi.org/10.1007/s13225-020-00466-2)) |
| Otu36734 | 1 | k__Fungi(100);p__Ascomycota(100);c__Saccharomycetes(100);o__Saccharomycetales(100);f__Metschnikowiaceae(100);g__Metschnikowia(100);g__Metschnikowia_unclassified(100); | Metschnikowia | 13 | Saprotroph | \|Undefined Saprotroph\| | Yeast | NULL | Probable | As nectar-tap_saprotroph (Põlme et al. 2020); Decay sugar-rich substrates (Põlme et al. 2020) | Põlme S, et al. 2020. Fungal Diversity 105:1-16 ((https://doi.org/10.1007/s13225-020-00466-2)) |
| Otu36735 | 1 | k__Fungi(100);p__Ascomycota(100);c__Saccharomycetes(100);o__Saccharomycetales(100);f__Metschnikowiaceae(100);g__Metschnikowia(100);s__Metschnikowia_pulcherrima(100); | Metschnikowia | 13 | Saprotroph | \|Undefined Saprotroph\| | Yeast | NULL | Probable | As nectar-tap_saprotroph (Põlme et al. 2020); Decay sugar-rich substrates (Põlme et al. 2020) | Põlme S, et al. 2020. Fungal Diversity 105:1-16 ((https://doi.org/10.1007/s13225-020-00466-2)) |
| Otu36736 | 1 | k__Fungi(100);p__Ascomycota(100);c__Saccharomycetes(100);o__Saccharomycetales(100);f__Metschnikowiaceae(100);g__Metschnikowia(100);g__Metschnikowia_unclassified(100); | Metschnikowia | 13 | Saprotroph | \|Undefined Saprotroph\| | Yeast | NULL | Probable | As nectar-tap_saprotroph (Põlme et al. 2020); Decay sugar-rich substrates (Põlme et al. 2020) | Põlme S, et al. 2020. Fungal Diversity 105:1-16 ((https://doi.org/10.1007/s13225-020-00466-2)) |
| Otu36737 | 1 | k__Fungi(100);p__Ascomycota(100);c__Saccharomycetes(100);o__Saccharomycetales(100);f__Metschnikowiaceae(100);g__Metschnikowia(100);g__Metschnikowia_unclassified(100); | Metschnikowia | 13 | Saprotroph | \|Undefined Saprotroph\| | Yeast | NULL | Probable | As nectar-tap_saprotroph (Põlme et al. 2020); Decay sugar-rich substrates (Põlme et al. 2020) | Põlme S, et al. 2020. Fungal Diversity 105:1-16 ((https://doi.org/10.1007/s13225-020-00466-2)) |
| Otu36738 | 1 | k__Fungi(100);p__Ascomycota(100);c__Saccharomycetes(100);o__Saccharomycetales(100);f__Metschnikowiaceae(100);g__Metschnikowia(100);s__Metschnikowia_pulcherrima(100); | Metschnikowia | 13 | Saprotroph | \|Undefined Saprotroph\| | Yeast | NULL | Probable | As nectar-tap_saprotroph (Põlme et al. 2020); Decay sugar-rich substrates (Põlme et al. 2020) | Põlme S, et al. 2020. Fungal Diversity 105:1-16 ((https://doi.org/10.1007/s13225-020-00466-2)) |
| Otu36739 | 1 | k__Fungi(100);p__Ascomycota(100);c__Saccharomycetes(100);o__Saccharomycetales(100);f__Metschnikowiaceae(100);g__Metschnikowia(100);s__Metschnikowia_pulcherrima(100); | Metschnikowia | 13 | Saprotroph | \|Undefined Saprotroph\| | Yeast | NULL | Probable | As nectar-tap_saprotroph (Põlme et al. 2020); Decay sugar-rich substrates (Põlme et al. 2020) | Põlme S, et al. 2020. Fungal Diversity 105:1-16 ((https://doi.org/10.1007/s13225-020-00466-2)) |
| Otu36740 | 1 | k__Fungi(100);p__Ascomycota(100);c__Saccharomycetes(100);o__Saccharomycetales(100);f__Metschnikowiaceae(100);g__Metschnikowia(100);g__Metschnikowia_unclassified(100); | Metschnikowia | 13 | Saprotroph | \|Undefined Saprotroph\| | Yeast | NULL | Probable | As nectar-tap_saprotroph (Põlme et al. 2020); Decay sugar-rich substrates (Põlme et al. 2020) | Põlme S, et al. 2020. Fungal Diversity 105:1-16 ((https://doi.org/10.1007/s13225-020-00466-2)) |
| Otu36741 | 1 | k__Fungi(100);p__Ascomycota(100);c__Saccharomycetes(100);o__Saccharomycetales(100);f__Metschnikowiaceae(100);g__Metschnikowia(100);s__Metschnikowia_pulcherrima(100); | Metschnikowia | 13 | Saprotroph | \|Undefined Saprotroph\| | Yeast | NULL | Probable | As nectar-tap_saprotroph (Põlme et al. 2020); Decay sugar-rich substrates (Põlme et al. 2020) | Põlme S, et al. 2020. Fungal Diversity 105:1-16 ((https://doi.org/10.1007/s13225-020-00466-2)) |
| Otu36742 | 1 | k__Fungi(100);p__Ascomycota(100);c__Saccharomycetes(100);o__Saccharomycetales(100);f__Metschnikowiaceae(100);g__Metschnikowia(100);g__Metschnikowia_unclassified(100); | Metschnikowia | 13 | Saprotroph | \|Undefined Saprotroph\| | Yeast | NULL | Probable | As nectar-tap_saprotroph (Põlme et al. 2020); Decay sugar-rich substrates (Põlme et al. 2020) | Põlme S, et al. 2020. Fungal Diversity 105:1-16 ((https://doi.org/10.1007/s13225-020-00466-2)) |
| Otu36743 | 1 | k__Fungi(100);p__Ascomycota(100);c__Saccharomycetes(100);o__Saccharomycetales(100);f__Metschnikowiaceae(100);g__Metschnikowia(100);s__Metschnikowia_pulcherrima(100); | Metschnikowia | 13 | Saprotroph | \|Undefined Saprotroph\| | Yeast | NULL | Probable | As nectar-tap_saprotroph (Põlme et al. 2020); Decay sugar-rich substrates (Põlme et al. 2020) | Põlme S, et al. 2020. Fungal Diversity 105:1-16 ((https://doi.org/10.1007/s13225-020-00466-2)) |
| Otu36744 | 1 | k__Fungi(100);p__Ascomycota(100);c__Saccharomycetes(100);o__Saccharomycetales(100);f__Metschnikowiaceae(100);g__Metschnikowia(100);s__Metschnikowia_pulcherrima(100); | Metschnikowia | 13 | Saprotroph | \|Undefined Saprotroph\| | Yeast | NULL | Probable | As nectar-tap_saprotroph (Põlme et al. 2020); Decay sugar-rich substrates (Põlme et al. 2020) | Põlme S, et al. 2020. Fungal Diversity 105:1-16 ((https://doi.org/10.1007/s13225-020-00466-2)) |
| Otu36745 | 1 | k__Fungi(100);p__Ascomycota(100);c__Saccharomycetes(100);o__Saccharomycetales(100);f__Metschnikowiaceae(100);g__Metschnikowia(100);s__Metschnikowia_viticola(100); | Metschnikowia | 13 | Saprotroph | \|Undefined Saprotroph\| | Yeast | NULL | Probable | As nectar-tap_saprotroph (Põlme et al. 2020); Decay sugar-rich substrates (Põlme et al. 2020) | Põlme S, et al. 2020. Fungal Diversity 105:1-16 ((https://doi.org/10.1007/s13225-020-00466-2)) |
| Otu36746 | 1 | k__Fungi(100);p__Ascomycota(100);c__Saccharomycetes(100);o__Saccharomycetales(100);f__Metschnikowiaceae(100);g__Metschnikowia(100);s__Metschnikowia_pulcherrima(100); | Metschnikowia | 13 | Saprotroph | \|Undefined Saprotroph\| | Yeast | NULL | Probable | As nectar-tap_saprotroph (Põlme et al. 2020); Decay sugar-rich substrates (Põlme et al. 2020) | Põlme S, et al. 2020. Fungal Diversity 105:1-16 ((https://doi.org/10.1007/s13225-020-00466-2)) |
| Otu36747 | 1 | k__Fungi(100);p__Ascomycota(100);c__Saccharomycetes(100);o__Saccharomycetales(100);f__Metschnikowiaceae(100);g__Metschnikowia(100);s__Metschnikowia_pulcherrima(100); | Metschnikowia | 13 | Saprotroph | \|Undefined Saprotroph\| | Yeast | NULL | Probable | As nectar-tap_saprotroph (Põlme et al. 2020); Decay sugar-rich substrates (Põlme et al. 2020) | Põlme S, et al. 2020. Fungal Diversity 105:1-16 ((https://doi.org/10.1007/s13225-020-00466-2)) |
| Otu36748 | 1 | k__Fungi(100);p__Ascomycota(100);c__Saccharomycetes(100);o__Saccharomycetales(100);f__Metschnikowiaceae(100);g__Metschnikowia(100);g__Metschnikowia_unclassified(100); | Metschnikowia | 13 | Saprotroph | \|Undefined Saprotroph\| | Yeast | NULL | Probable | As nectar-tap_saprotroph (Põlme et al. 2020); Decay sugar-rich substrates (Põlme et al. 2020) | Põlme S, et al. 2020. Fungal Diversity 105:1-16 ((https://doi.org/10.1007/s13225-020-00466-2)) |
| Otu36749 | 1 | k__Fungi(100);p__Ascomycota(100);c__Saccharomycetes(100);o__Saccharomycetales(100);f__Metschnikowiaceae(100);g__Metschnikowia(100);s__Metschnikowia_pulcherrima(100); | Metschnikowia | 13 | Saprotroph | \|Undefined Saprotroph\| | Yeast | NULL | Probable | As nectar-tap_saprotroph (Põlme et al. 2020); Decay sugar-rich substrates (Põlme et al. 2020) | Põlme S, et al. 2020. Fungal Diversity 105:1-16 ((https://doi.org/10.1007/s13225-020-00466-2)) |
| Otu36750 | 1 | k__Fungi(100);p__Ascomycota(100);c__Saccharomycetes(100);o__Saccharomycetales(100);f__Metschnikowiaceae(100);g__Metschnikowia(100);s__Metschnikowia_pulcherrima(100); | Metschnikowia | 13 | Saprotroph | \|Undefined Saprotroph\| | Yeast | NULL | Probable | As nectar-tap_saprotroph (Põlme et al. 2020); Decay sugar-rich substrates (Põlme et al. 2020) | Põlme S, et al. 2020. Fungal Diversity 105:1-16 ((https://doi.org/10.1007/s13225-020-00466-2)) |
| Otu36751 | 1 | k__Fungi(100);p__Ascomycota(100);c__Saccharomycetes(100);o__Saccharomycetales(100);f__Metschnikowiaceae(100);g__Metschnikowia(100);s__Metschnikowia_pulcherrima(100); | Metschnikowia | 13 | Saprotroph | \|Undefined Saprotroph\| | Yeast | NULL | Probable | As nectar-tap_saprotroph (Põlme et al. 2020); Decay sugar-rich substrates (Põlme et al. 2020) | Põlme S, et al. 2020. Fungal Diversity 105:1-16 ((https://doi.org/10.1007/s13225-020-00466-2)) |
| Otu36752 | 1 | k__Fungi(100);p__Ascomycota(100);c__Saccharomycetes(100);o__Saccharomycetales(100);f__Metschnikowiaceae(100);g__Metschnikowia(100);s__Metschnikowia_pulcherrima(100); | Metschnikowia | 13 | Saprotroph | \|Undefined Saprotroph\| | Yeast | NULL | Probable | As nectar-tap_saprotroph (Põlme et al. 2020); Decay sugar-rich substrates (Põlme et al. 2020) | Põlme S, et al. 2020. Fungal Diversity 105:1-16 ((https://doi.org/10.1007/s13225-020-00466-2)) |
| Otu36753 | 1 | k__Fungi(100);p__Ascomycota(100);c__Saccharomycetes(100);o__Saccharomycetales(100);f__Metschnikowiaceae(100);g__Metschnikowia(100);s__Metschnikowia_pulcherrima(100); | Metschnikowia | 13 | Saprotroph | \|Undefined Saprotroph\| | Yeast | NULL | Probable | As nectar-tap_saprotroph (Põlme et al. 2020); Decay sugar-rich substrates (Põlme et al. 2020) | Põlme S, et al. 2020. Fungal Diversity 105:1-16 ((https://doi.org/10.1007/s13225-020-00466-2)) |
| Otu36754 | 1 | k__Fungi(100);p__Ascomycota(100);c__Saccharomycetes(100);o__Saccharomycetales(100);f__Metschnikowiaceae(100);g__Metschnikowia(100);s__Metschnikowia_pulcherrima(100); | Metschnikowia | 13 | Saprotroph | \|Undefined Saprotroph\| | Yeast | NULL | Probable | As nectar-tap_saprotroph (Põlme et al. 2020); Decay sugar-rich substrates (Põlme et al. 2020) | Põlme S, et al. 2020. Fungal Diversity 105:1-16 ((https://doi.org/10.1007/s13225-020-00466-2)) |
| Otu36755 | 1 | k__Fungi(100);p__Ascomycota(100);c__Saccharomycetes(100);o__Saccharomycetales(100);f__Metschnikowiaceae(100);g__Metschnikowia(100);s__Metschnikowia_pulcherrima(100); | Metschnikowia | 13 | Saprotroph | \|Undefined Saprotroph\| | Yeast | NULL | Probable | As nectar-tap_saprotroph (Põlme et al. 2020); Decay sugar-rich substrates (Põlme et al. 2020) | Põlme S, et al. 2020. Fungal Diversity 105:1-16 ((https://doi.org/10.1007/s13225-020-00466-2)) |
| Otu36756 | 1 | k__Fungi(100);p__Ascomycota(100);c__Saccharomycetes(100);o__Saccharomycetales(100);f__Metschnikowiaceae(100);g__Metschnikowia(100);s__Metschnikowia_pulcherrima(100); | Metschnikowia | 13 | Saprotroph | \|Undefined Saprotroph\| | Yeast | NULL | Probable | As nectar-tap_saprotroph (Põlme et al. 2020); Decay sugar-rich substrates (Põlme et al. 2020) | Põlme S, et al. 2020. Fungal Diversity 105:1-16 ((https://doi.org/10.1007/s13225-020-00466-2)) |
| Otu36757 | 1 | k__Fungi(100);p__Ascomycota(100);c__Saccharomycetes(100);o__Saccharomycetales(100);f__Metschnikowiaceae(100);g__Metschnikowia(100);s__Metschnikowia_pulcherrima(100); | Metschnikowia | 13 | Saprotroph | \|Undefined Saprotroph\| | Yeast | NULL | Probable | As nectar-tap_saprotroph (Põlme et al. 2020); Decay sugar-rich substrates (Põlme et al. 2020) | Põlme S, et al. 2020. Fungal Diversity 105:1-16 ((https://doi.org/10.1007/s13225-020-00466-2)) |
| Otu36758 | 1 | k__Fungi(100);p__Ascomycota(100);c__Saccharomycetes(100);o__Saccharomycetales(100);f__Metschnikowiaceae(100);g__Metschnikowia(100);s__Metschnikowia_pulcherrima(100); | Metschnikowia | 13 | Saprotroph | \|Undefined Saprotroph\| | Yeast | NULL | Probable | As nectar-tap_saprotroph (Põlme et al. 2020); Decay sugar-rich substrates (Põlme et al. 2020) | Põlme S, et al. 2020. Fungal Diversity 105:1-16 ((https://doi.org/10.1007/s13225-020-00466-2)) |
| Otu36759 | 1 | k__Fungi(100);p__Ascomycota(100);c__Saccharomycetes(100);o__Saccharomycetales(100);f__Metschnikowiaceae(100);g__Metschnikowia(100);s__Metschnikowia_pulcherrima(100); | Metschnikowia | 13 | Saprotroph | \|Undefined Saprotroph\| | Yeast | NULL | Probable | As nectar-tap_saprotroph (Põlme et al. 2020); Decay sugar-rich substrates (Põlme et al. 2020) | Põlme S, et al. 2020. Fungal Diversity 105:1-16 ((https://doi.org/10.1007/s13225-020-00466-2)) |
| Otu36760 | 1 | k__Fungi(100);p__Ascomycota(100);c__Saccharomycetes(100);o__Saccharomycetales(100);f__Metschnikowiaceae(100);g__Metschnikowia(100);s__Metschnikowia_pulcherrima(100); | Metschnikowia | 13 | Saprotroph | \|Undefined Saprotroph\| | Yeast | NULL | Probable | As nectar-tap_saprotroph (Põlme et al. 2020); Decay sugar-rich substrates (Põlme et al. 2020) | Põlme S, et al. 2020. Fungal Diversity 105:1-16 ((https://doi.org/10.1007/s13225-020-00466-2)) |
| Otu36761 | 1 | k__Fungi(100);p__Ascomycota(100);c__Saccharomycetes(100);o__Saccharomycetales(100);f__Metschnikowiaceae(100);g__Metschnikowia(100);s__Metschnikowia_pulcherrima(100); | Metschnikowia | 13 | Saprotroph | \|Undefined Saprotroph\| | Yeast | NULL | Probable | As nectar-tap_saprotroph (Põlme et al. 2020); Decay sugar-rich substrates (Põlme et al. 2020) | Põlme S, et al. 2020. Fungal Diversity 105:1-16 ((https://doi.org/10.1007/s13225-020-00466-2)) |
| Otu36762 | 1 | k__Fungi(100);p__Ascomycota(100);c__Saccharomycetes(100);o__Saccharomycetales(100);f__Metschnikowiaceae(100);g__Metschnikowia(100);s__Metschnikowia_pulcherrima(100); | Metschnikowia | 13 | Saprotroph | \|Undefined Saprotroph\| | Yeast | NULL | Probable | As nectar-tap_saprotroph (Põlme et al. 2020); Decay sugar-rich substrates (Põlme et al. 2020) | Põlme S, et al. 2020. Fungal Diversity 105:1-16 ((https://doi.org/10.1007/s13225-020-00466-2)) |
| Otu36763 | 1 | k__Fungi(100);p__Ascomycota(100);c__Saccharomycetes(100);o__Saccharomycetales(100);f__Metschnikowiaceae(100);g__Metschnikowia(100);s__Metschnikowia_pulcherrima(100); | Metschnikowia | 13 | Saprotroph | \|Undefined Saprotroph\| | Yeast | NULL | Probable | As nectar-tap_saprotroph (Põlme et al. 2020); Decay sugar-rich substrates (Põlme et al. 2020) | Põlme S, et al. 2020. Fungal Diversity 105:1-16 ((https://doi.org/10.1007/s13225-020-00466-2)) |
| Otu36764 | 1 | k__Fungi(100);p__Ascomycota(100);c__Saccharomycetes(100);o__Saccharomycetales(100);f__Metschnikowiaceae(100);g__Metschnikowia(100);s__Metschnikowia_pulcherrima(100); | Metschnikowia | 13 | Saprotroph | \|Undefined Saprotroph\| | Yeast | NULL | Probable | As nectar-tap_saprotroph (Põlme et al. 2020); Decay sugar-rich substrates (Põlme et al. 2020) | Põlme S, et al. 2020. Fungal Diversity 105:1-16 ((https://doi.org/10.1007/s13225-020-00466-2)) |
| Otu36765 | 1 | k__Fungi(100);p__Ascomycota(100);c__Saccharomycetes(100);o__Saccharomycetales(100);f__Metschnikowiaceae(100);g__Metschnikowia(100);s__Metschnikowia_pulcherrima(100); | Metschnikowia | 13 | Saprotroph | \|Undefined Saprotroph\| | Yeast | NULL | Probable | As nectar-tap_saprotroph (Põlme et al. 2020); Decay sugar-rich substrates (Põlme et al. 2020) | Põlme S, et al. 2020. Fungal Diversity 105:1-16 ((https://doi.org/10.1007/s13225-020-00466-2)) |
| Otu36766 | 1 | k__Fungi(100);p__Ascomycota(100);c__Saccharomycetes(100);o__Saccharomycetales(100);f__Metschnikowiaceae(100);g__Metschnikowia(100);s__Metschnikowia_pulcherrima(100); | Metschnikowia | 13 | Saprotroph | \|Undefined Saprotroph\| | Yeast | NULL | Probable | As nectar-tap_saprotroph (Põlme et al. 2020); Decay sugar-rich substrates (Põlme et al. 2020) | Põlme S, et al. 2020. Fungal Diversity 105:1-16 ((https://doi.org/10.1007/s13225-020-00466-2)) |
| Otu36767 | 1 | k__Fungi(100);p__Ascomycota(100);c__Saccharomycetes(100);o__Saccharomycetales(100);f__Metschnikowiaceae(100);g__Metschnikowia(100);s__Metschnikowia_pulcherrima(100); | Metschnikowia | 13 | Saprotroph | \|Undefined Saprotroph\| | Yeast | NULL | Probable | As nectar-tap_saprotroph (Põlme et al. 2020); Decay sugar-rich substrates (Põlme et al. 2020) | Põlme S, et al. 2020. Fungal Diversity 105:1-16 ((https://doi.org/10.1007/s13225-020-00466-2)) |
| Otu36768 | 1 | k__Fungi(100);p__Ascomycota(100);c__Saccharomycetes(100);o__Saccharomycetales(100);f__Metschnikowiaceae(100);g__Metschnikowia(100);s__Metschnikowia_pulcherrima(100); | Metschnikowia | 13 | Saprotroph | \|Undefined Saprotroph\| | Yeast | NULL | Probable | As nectar-tap_saprotroph (Põlme et al. 2020); Decay sugar-rich substrates (Põlme et al. 2020) | Põlme S, et al. 2020. Fungal Diversity 105:1-16 ((https://doi.org/10.1007/s13225-020-00466-2)) |
| Otu36769 | 1 | k__Fungi(100);p__Ascomycota(100);c__Saccharomycetes(100);o__Saccharomycetales(100);f__Metschnikowiaceae(100);g__Metschnikowia(100);s__Metschnikowia_pulcherrima(100); | Metschnikowia | 13 | Saprotroph | \|Undefined Saprotroph\| | Yeast | NULL | Probable | As nectar-tap_saprotroph (Põlme et al. 2020); Decay sugar-rich substrates (Põlme et al. 2020) | Põlme S, et al. 2020. Fungal Diversity 105:1-16 ((https://doi.org/10.1007/s13225-020-00466-2)) |
| Otu36770 | 1 | k__Fungi(100);p__Ascomycota(100);c__Saccharomycetes(100);o__Saccharomycetales(100);f__Metschnikowiaceae(100);g__Metschnikowia(100);g__Metschnikowia_unclassified(100); | Metschnikowia | 13 | Saprotroph | \|Undefined Saprotroph\| | Yeast | NULL | Probable | As nectar-tap_saprotroph (Põlme et al. 2020); Decay sugar-rich substrates (Põlme et al. 2020) | Põlme S, et al. 2020. Fungal Diversity 105:1-16 ((https://doi.org/10.1007/s13225-020-00466-2)) |
| Otu36771 | 1 | k__Fungi(100);p__Ascomycota(100);c__Saccharomycetes(100);o__Saccharomycetales(100);f__Metschnikowiaceae(100);g__Metschnikowia(100);g__Metschnikowia_unclassified(100); | Metschnikowia | 13 | Saprotroph | \|Undefined Saprotroph\| | Yeast | NULL | Probable | As nectar-tap_saprotroph (Põlme et al. 2020); Decay sugar-rich substrates (Põlme et al. 2020) | Põlme S, et al. 2020. Fungal Diversity 105:1-16 ((https://doi.org/10.1007/s13225-020-00466-2)) |
| Otu36772 | 1 | k__Fungi(100);p__Ascomycota(100);c__Saccharomycetes(100);o__Saccharomycetales(100);f__Metschnikowiaceae(100);g__Metschnikowia(100);g__Metschnikowia_unclassified(100); | Metschnikowia | 13 | Saprotroph | \|Undefined Saprotroph\| | Yeast | NULL | Probable | As nectar-tap_saprotroph (Põlme et al. 2020); Decay sugar-rich substrates (Põlme et al. 2020) | Põlme S, et al. 2020. Fungal Diversity 105:1-16 ((https://doi.org/10.1007/s13225-020-00466-2)) |
| Otu36773 | 1 | k__Fungi(100);p__Ascomycota(100);c__Saccharomycetes(100);o__Saccharomycetales(100);f__Metschnikowiaceae(100);g__Metschnikowia(100);s__Metschnikowia_pulcherrima(100); | Metschnikowia | 13 | Saprotroph | \|Undefined Saprotroph\| | Yeast | NULL | Probable | As nectar-tap_saprotroph (Põlme et al. 2020); Decay sugar-rich substrates (Põlme et al. 2020) | Põlme S, et al. 2020. Fungal Diversity 105:1-16 ((https://doi.org/10.1007/s13225-020-00466-2)) |
| Otu36774 | 1 | k__Fungi(100);p__Ascomycota(100);c__Saccharomycetes(100);o__Saccharomycetales(100);f__Metschnikowiaceae(100);g__Metschnikowia(100);g__Metschnikowia_unclassified(100); | Metschnikowia | 13 | Saprotroph | \|Undefined Saprotroph\| | Yeast | NULL | Probable | As nectar-tap_saprotroph (Põlme et al. 2020); Decay sugar-rich substrates (Põlme et al. 2020) | Põlme S, et al. 2020. Fungal Diversity 105:1-16 ((https://doi.org/10.1007/s13225-020-00466-2)) |
| Otu36775 | 1 | k__Fungi(100);p__Ascomycota(100);c__Saccharomycetes(100);o__Saccharomycetales(100);f__Metschnikowiaceae(100);g__Metschnikowia(100);s__Metschnikowia_pulcherrima(100); | Metschnikowia | 13 | Saprotroph | \|Undefined Saprotroph\| | Yeast | NULL | Probable | As nectar-tap_saprotroph (Põlme et al. 2020); Decay sugar-rich substrates (Põlme et al. 2020) | Põlme S, et al. 2020. Fungal Diversity 105:1-16 ((https://doi.org/10.1007/s13225-020-00466-2)) |
| Otu36776 | 1 | k__Fungi(100);p__Ascomycota(100);c__Saccharomycetes(100);o__Saccharomycetales(100);f__Metschnikowiaceae(100);g__Metschnikowia(100);g__Metschnikowia_unclassified(100); | Metschnikowia | 13 | Saprotroph | \|Undefined Saprotroph\| | Yeast | NULL | Probable | As nectar-tap_saprotroph (Põlme et al. 2020); Decay sugar-rich substrates (Põlme et al. 2020) | Põlme S, et al. 2020. Fungal Diversity 105:1-16 ((https://doi.org/10.1007/s13225-020-00466-2)) |
| Otu36777 | 1 | k__Fungi(100);p__Ascomycota(100);c__Saccharomycetes(100);o__Saccharomycetales(100);f__Metschnikowiaceae(100);g__Metschnikowia(100);s__Metschnikowia_pulcherrima(100); | Metschnikowia | 13 | Saprotroph | \|Undefined Saprotroph\| | Yeast | NULL | Probable | As nectar-tap_saprotroph (Põlme et al. 2020); Decay sugar-rich substrates (Põlme et al. 2020) | Põlme S, et al. 2020. Fungal Diversity 105:1-16 ((https://doi.org/10.1007/s13225-020-00466-2)) |
| Otu36778 | 1 | k__Fungi(100);p__Ascomycota(100);c__Saccharomycetes(100);o__Saccharomycetales(100);f__Metschnikowiaceae(100);g__Metschnikowia(100);g__Metschnikowia_unclassified(100); | Metschnikowia | 13 | Saprotroph | \|Undefined Saprotroph\| | Yeast | NULL | Probable | As nectar-tap_saprotroph (Põlme et al. 2020); Decay sugar-rich substrates (Põlme et al. 2020) | Põlme S, et al. 2020. Fungal Diversity 105:1-16 ((https://doi.org/10.1007/s13225-020-00466-2)) |
| Otu36779 | 1 | k__Fungi(100);p__Ascomycota(100);c__Saccharomycetes(100);o__Saccharomycetales(100);f__Metschnikowiaceae(100);g__Metschnikowia(100);s__Metschnikowia_pulcherrima(100); | Metschnikowia | 13 | Saprotroph | \|Undefined Saprotroph\| | Yeast | NULL | Probable | As nectar-tap_saprotroph (Põlme et al. 2020); Decay sugar-rich substrates (Põlme et al. 2020) | Põlme S, et al. 2020. Fungal Diversity 105:1-16 ((https://doi.org/10.1007/s13225-020-00466-2)) |
| Otu36780 | 1 | k__Fungi(100);p__Ascomycota(100);c__Saccharomycetes(100);o__Saccharomycetales(100);f__Metschnikowiaceae(100);g__Metschnikowia(100);s__Metschnikowia_pulcherrima(100); | Metschnikowia | 13 | Saprotroph | \|Undefined Saprotroph\| | Yeast | NULL | Probable | As nectar-tap_saprotroph (Põlme et al. 2020); Decay sugar-rich substrates (Põlme et al. 2020) | Põlme S, et al. 2020. Fungal Diversity 105:1-16 ((https://doi.org/10.1007/s13225-020-00466-2)) |
| Otu36781 | 1 | k__Fungi(100);p__Ascomycota(100);c__Saccharomycetes(100);o__Saccharomycetales(100);f__Metschnikowiaceae(100);g__Metschnikowia(100);s__Metschnikowia_pulcherrima(100); | Metschnikowia | 13 | Saprotroph | \|Undefined Saprotroph\| | Yeast | NULL | Probable | As nectar-tap_saprotroph (Põlme et al. 2020); Decay sugar-rich substrates (Põlme et al. 2020) | Põlme S, et al. 2020. Fungal Diversity 105:1-16 ((https://doi.org/10.1007/s13225-020-00466-2)) |
| Otu36782 | 1 | k__Fungi(100);p__Ascomycota(100);c__Saccharomycetes(100);o__Saccharomycetales(100);f__Metschnikowiaceae(100);g__Metschnikowia(100);g__Metschnikowia_unclassified(100); | Metschnikowia | 13 | Saprotroph | \|Undefined Saprotroph\| | Yeast | NULL | Probable | As nectar-tap_saprotroph (Põlme et al. 2020); Decay sugar-rich substrates (Põlme et al. 2020) | Põlme S, et al. 2020. Fungal Diversity 105:1-16 ((https://doi.org/10.1007/s13225-020-00466-2)) |
| Otu36783 | 1 | k__Fungi(100);p__Ascomycota(100);c__Saccharomycetes(100);o__Saccharomycetales(100);f__Metschnikowiaceae(100);g__Metschnikowia(100);g__Metschnikowia_unclassified(100); | Metschnikowia | 13 | Saprotroph | \|Undefined Saprotroph\| | Yeast | NULL | Probable | As nectar-tap_saprotroph (Põlme et al. 2020); Decay sugar-rich substrates (Põlme et al. 2020) | Põlme S, et al. 2020. Fungal Diversity 105:1-16 ((https://doi.org/10.1007/s13225-020-00466-2)) |
| Otu36784 | 1 | k__Fungi(100);p__Ascomycota(100);c__Saccharomycetes(100);o__Saccharomycetales(100);f__Metschnikowiaceae(100);g__Metschnikowia(100);s__Metschnikowia_pulcherrima(100); | Metschnikowia | 13 | Saprotroph | \|Undefined Saprotroph\| | Yeast | NULL | Probable | As nectar-tap_saprotroph (Põlme et al. 2020); Decay sugar-rich substrates (Põlme et al. 2020) | Põlme S, et al. 2020. Fungal Diversity 105:1-16 ((https://doi.org/10.1007/s13225-020-00466-2)) |
| Otu36785 | 1 | k__Fungi(100);p__Ascomycota(100);c__Saccharomycetes(100);o__Saccharomycetales(100);f__Metschnikowiaceae(100);g__Metschnikowia(100);g__Metschnikowia_unclassified(100); | Metschnikowia | 13 | Saprotroph | \|Undefined Saprotroph\| | Yeast | NULL | Probable | As nectar-tap_saprotroph (Põlme et al. 2020); Decay sugar-rich substrates (Põlme et al. 2020) | Põlme S, et al. 2020. Fungal Diversity 105:1-16 ((https://doi.org/10.1007/s13225-020-00466-2)) |
| Otu36786 | 1 | k__Fungi(100);p__Ascomycota(100);c__Saccharomycetes(100);o__Saccharomycetales(100);f__Metschnikowiaceae(100);g__Metschnikowia(100);s__Metschnikowia_pulcherrima(100); | Metschnikowia | 13 | Saprotroph | \|Undefined Saprotroph\| | Yeast | NULL | Probable | As nectar-tap_saprotroph (Põlme et al. 2020); Decay sugar-rich substrates (Põlme et al. 2020) | Põlme S, et al. 2020. Fungal Diversity 105:1-16 ((https://doi.org/10.1007/s13225-020-00466-2)) |
| Otu36787 | 1 | k__Fungi(100);p__Ascomycota(100);c__Saccharomycetes(100);o__Saccharomycetales(100);f__Metschnikowiaceae(100);g__Metschnikowia(100);s__Metschnikowia_pulcherrima(100); | Metschnikowia | 13 | Saprotroph | \|Undefined Saprotroph\| | Yeast | NULL | Probable | As nectar-tap_saprotroph (Põlme et al. 2020); Decay sugar-rich substrates (Põlme et al. 2020) | Põlme S, et al. 2020. Fungal Diversity 105:1-16 ((https://doi.org/10.1007/s13225-020-00466-2)) |
| Otu36788 | 1 | k__Fungi(100);p__Ascomycota(100);c__Saccharomycetes(100);o__Saccharomycetales(100);f__Metschnikowiaceae(100);g__Metschnikowia(100);s__Metschnikowia_pulcherrima(100); | Metschnikowia | 13 | Saprotroph | \|Undefined Saprotroph\| | Yeast | NULL | Probable | As nectar-tap_saprotroph (Põlme et al. 2020); Decay sugar-rich substrates (Põlme et al. 2020) | Põlme S, et al. 2020. Fungal Diversity 105:1-16 ((https://doi.org/10.1007/s13225-020-00466-2)) |
| Otu36789 | 1 | k__Fungi(100);p__Ascomycota(100);c__Saccharomycetes(100);o__Saccharomycetales(100);f__Metschnikowiaceae(100);g__Metschnikowia(100);g__Metschnikowia_unclassified(100); | Metschnikowia | 13 | Saprotroph | \|Undefined Saprotroph\| | Yeast | NULL | Probable | As nectar-tap_saprotroph (Põlme et al. 2020); Decay sugar-rich substrates (Põlme et al. 2020) | Põlme S, et al. 2020. Fungal Diversity 105:1-16 ((https://doi.org/10.1007/s13225-020-00466-2)) |
| Otu36790 | 1 | k__Fungi(100);p__Ascomycota(100);c__Saccharomycetes(100);o__Saccharomycetales(100);f__Metschnikowiaceae(100);g__Metschnikowia(100);s__Metschnikowia_pulcherrima(100); | Metschnikowia | 13 | Saprotroph | \|Undefined Saprotroph\| | Yeast | NULL | Probable | As nectar-tap_saprotroph (Põlme et al. 2020); Decay sugar-rich substrates (Põlme et al. 2020) | Põlme S, et al. 2020. Fungal Diversity 105:1-16 ((https://doi.org/10.1007/s13225-020-00466-2)) |
| Otu36791 | 1 | k__Fungi(100);p__Ascomycota(100);c__Saccharomycetes(100);o__Saccharomycetales(100);f__Metschnikowiaceae(100);g__Metschnikowia(100);s__Metschnikowia_pulcherrima(100); | Metschnikowia | 13 | Saprotroph | \|Undefined Saprotroph\| | Yeast | NULL | Probable | As nectar-tap_saprotroph (Põlme et al. 2020); Decay sugar-rich substrates (Põlme et al. 2020) | Põlme S, et al. 2020. Fungal Diversity 105:1-16 ((https://doi.org/10.1007/s13225-020-00466-2)) |
| Otu36792 | 1 | k__Fungi(100);p__Ascomycota(100);c__Saccharomycetes(100);o__Saccharomycetales(100);f__Metschnikowiaceae(100);g__Metschnikowia(100);s__Metschnikowia_pulcherrima(100); | Metschnikowia | 13 | Saprotroph | \|Undefined Saprotroph\| | Yeast | NULL | Probable | As nectar-tap_saprotroph (Põlme et al. 2020); Decay sugar-rich substrates (Põlme et al. 2020) | Põlme S, et al. 2020. Fungal Diversity 105:1-16 ((https://doi.org/10.1007/s13225-020-00466-2)) |
| Otu36793 | 1 | k__Fungi(100);p__Ascomycota(100);c__Saccharomycetes(100);o__Saccharomycetales(100);f__Metschnikowiaceae(100);g__Metschnikowia(100);s__Metschnikowia_pulcherrima(100); | Metschnikowia | 13 | Saprotroph | \|Undefined Saprotroph\| | Yeast | NULL | Probable | As nectar-tap_saprotroph (Põlme et al. 2020); Decay sugar-rich substrates (Põlme et al. 2020) | Põlme S, et al. 2020. Fungal Diversity 105:1-16 ((https://doi.org/10.1007/s13225-020-00466-2)) |
| Otu36794 | 1 | k__Fungi(100);p__Ascomycota(100);c__Saccharomycetes(100);o__Saccharomycetales(100);f__Metschnikowiaceae(100);g__Metschnikowia(100);s__Metschnikowia_pulcherrima(100); | Metschnikowia | 13 | Saprotroph | \|Undefined Saprotroph\| | Yeast | NULL | Probable | As nectar-tap_saprotroph (Põlme et al. 2020); Decay sugar-rich substrates (Põlme et al. 2020) | Põlme S, et al. 2020. Fungal Diversity 105:1-16 ((https://doi.org/10.1007/s13225-020-00466-2)) |
| Otu36795 | 1 | k__Fungi(100);p__Ascomycota(100);c__Saccharomycetes(100);o__Saccharomycetales(100);f__Metschnikowiaceae(100);g__Metschnikowia(100);s__Metschnikowia_pulcherrima(100); | Metschnikowia | 13 | Saprotroph | \|Undefined Saprotroph\| | Yeast | NULL | Probable | As nectar-tap_saprotroph (Põlme et al. 2020); Decay sugar-rich substrates (Põlme et al. 2020) | Põlme S, et al. 2020. Fungal Diversity 105:1-16 ((https://doi.org/10.1007/s13225-020-00466-2)) |
| Otu36796 | 1 | k__Fungi(100);p__Ascomycota(100);c__Saccharomycetes(100);o__Saccharomycetales(100);f__Metschnikowiaceae(100);g__Metschnikowia(100);g__Metschnikowia_unclassified(100); | Metschnikowia | 13 | Saprotroph | \|Undefined Saprotroph\| | Yeast | NULL | Probable | As nectar-tap_saprotroph (Põlme et al. 2020); Decay sugar-rich substrates (Põlme et al. 2020) | Põlme S, et al. 2020. Fungal Diversity 105:1-16 ((https://doi.org/10.1007/s13225-020-00466-2)) |
| Otu36797 | 1 | k__Fungi(100);p__Ascomycota(100);c__Saccharomycetes(100);o__Saccharomycetales(100);f__Metschnikowiaceae(100);g__Metschnikowia(100);g__Metschnikowia_unclassified(100); | Metschnikowia | 13 | Saprotroph | \|Undefined Saprotroph\| | Yeast | NULL | Probable | As nectar-tap_saprotroph (Põlme et al. 2020); Decay sugar-rich substrates (Põlme et al. 2020) | Põlme S, et al. 2020. Fungal Diversity 105:1-16 ((https://doi.org/10.1007/s13225-020-00466-2)) |

| | | | | | | | | | | | |
|---|---|---|---|---|---|---|---|---|---|---|---|
| Otu36798 | 1 | k__Fungi(100);p__Ascomycota(100);c__Saccharomycetes(100);o__Saccharomycetales(100);f__Metschnikowiaceae(100);g__Metschnikowia(100);g__Metschnikowia_unclassified(100); | Metschnikowia | 13 | Saprotroph | \|Undefined Saprotroph\| | Yeast | NULL | Probable | As nectar-tap_saprotroph (Põlme et al. 2020); Decay sugar-rich substrates (Põlme et al. 2020) | Põlme S, et al. 2020. Fungal Diversity 105:1-16 ((https://doi.org/10.1007/s13225-020-00466-2)) |
| Otu36799 | 1 | k__Fungi(100);p__Ascomycota(100);c__Saccharomycetes(100);o__Saccharomycetales(100);f__Metschnikowiaceae(100);g__Metschnikowia(100);s__Metschnikowia_pulcherrima(100); | Metschnikowia | 13 | Saprotroph | \|Undefined Saprotroph\| | Yeast | NULL | Probable | As nectar-tap_saprotroph (Põlme et al. 2020); Decay sugar-rich substrates (Põlme et al. 2020) | Põlme S, et al. 2020. Fungal Diversity 105:1-16 ((https://doi.org/10.1007/s13225-020-00466-2)) |
| Otu36800 | 1 | k__Fungi(100);p__Ascomycota(100);c__Saccharomycetes(100);o__Saccharomycetales(100);f__Metschnikowiaceae(100);g__Metschnikowia(100);s__Metschnikowia_pulcherrima(100); | Metschnikowia | 13 | Saprotroph | \|Undefined Saprotroph\| | Yeast | NULL | Probable | As nectar-tap_saprotroph (Põlme et al. 2020); Decay sugar-rich substrates (Põlme et al. 2020) | Põlme S, et al. 2020. Fungal Diversity 105:1-16 ((https://doi.org/10.1007/s13225-020-00466-2)) |
| Otu36801 | 1 | k__Fungi(100);p__Ascomycota(100);c__Saccharomycetes(100);o__Saccharomycetales(100);f__Metschnikowiaceae(100);g__Metschnikowia(100);g__Metschnikowia_unclassified(100); | Metschnikowia | 13 | Saprotroph | \|Undefined Saprotroph\| | Yeast | NULL | Probable | As nectar-tap_saprotroph (Põlme et al. 2020); Decay sugar-rich substrates (Põlme et al. 2020) | Põlme S, et al. 2020. Fungal Diversity 105:1-16 ((https://doi.org/10.1007/s13225-020-00466-2)) |
| Otu36802 | 1 | k__Fungi(100);p__Ascomycota(100);c__Saccharomycetes(100);o__Saccharomycetales(100);f__Metschnikowiaceae(100);g__Metschnikowia(100);s__Metschnikowia_pulcherrima(100); | Metschnikowia | 13 | Saprotroph | \|Undefined Saprotroph\| | Yeast | NULL | Probable | As nectar-tap_saprotroph (Põlme et al. 2020); Decay sugar-rich substrates (Põlme et al. 2020) | Põlme S, et al. 2020. Fungal Diversity 105:1-16 ((https://doi.org/10.1007/s13225-020-00466-2)) |
| Otu36803 | 1 | k__Fungi(100);p__Ascomycota(100);c__Saccharomycetes(100);o__Saccharomycetales(100);f__Metschnikowiaceae(100);g__Metschnikowia(100);g__Metschnikowia_unclassified(100); | Metschnikowia | 13 | Saprotroph | \|Undefined Saprotroph\| | Yeast | NULL | Probable | As nectar-tap_saprotroph (Põlme et al. 2020); Decay sugar-rich substrates (Põlme et al. 2020) | Põlme S, et al. 2020. Fungal Diversity 105:1-16 ((https://doi.org/10.1007/s13225-020-00466-2)) |
| Otu36804 | 1 | k__Fungi(100);p__Ascomycota(100);c__Saccharomycetes(100);o__Saccharomycetales(100);f__Metschnikowiaceae(100);g__Metschnikowia(100);s__Metschnikowia_pulcherrima(100); | Metschnikowia | 13 | Saprotroph | \|Undefined Saprotroph\| | Yeast | NULL | Probable | As nectar-tap_saprotroph (Põlme et al. 2020); Decay sugar-rich substrates (Põlme et al. 2020) | Põlme S, et al. 2020. Fungal Diversity 105:1-16 ((https://doi.org/10.1007/s13225-020-00466-2)) |
| Otu36805 | 1 | k__Fungi(100);p__Ascomycota(100);c__Saccharomycetes(100);o__Saccharomycetales(100);f__Metschnikowiaceae(100);g__Metschnikowia(100);s__Metschnikowia_pulcherrima(100); | Metschnikowia | 13 | Saprotroph | \|Undefined Saprotroph\| | Yeast | NULL | Probable | As nectar-tap_saprotroph (Põlme et al. 2020); Decay sugar-rich substrates (Põlme et al. 2020) | Põlme S, et al. 2020. Fungal Diversity 105:1-16 ((https://doi.org/10.1007/s13225-020-00466-2)) |
| Otu36806 | 1 | k__Fungi(100);p__Ascomycota(100);c__Saccharomycetes(100);o__Saccharomycetales(100);f__Metschnikowiaceae(100);g__Metschnikowia(100);s__Metschnikowia_pulcherrima(100); | Metschnikowia | 13 | Saprotroph | \|Undefined Saprotroph\| | Yeast | NULL | Probable | As nectar-tap_saprotroph (Põlme et al. 2020); Decay sugar-rich substrates (Põlme et al. 2020) | Põlme S, et al. 2020. Fungal Diversity 105:1-16 ((https://doi.org/10.1007/s13225-020-00466-2)) |
| Otu36807 | 1 | k__Fungi(100);p__Ascomycota(100);c__Saccharomycetes(100);o__Saccharomycetales(100);f__Metschnikowiaceae(100);g__Metschnikowia(100);g__Metschnikowia_unclassified(100); | Metschnikowia | 13 | Saprotroph | \|Undefined Saprotroph\| | Yeast | NULL | Probable | As nectar-tap_saprotroph (Põlme et al. 2020); Decay sugar-rich substrates (Põlme et al. 2020) | Põlme S, et al. 2020. Fungal Diversity 105:1-16 ((https://doi.org/10.1007/s13225-020-00466-2)) |
| Otu36808 | 1 | k__Fungi(100);p__Ascomycota(100);c__Saccharomycetes(100);o__Saccharomycetales(100);f__Metschnikowiaceae(100);g__Metschnikowia(100);g__Metschnikowia_unclassified(100); | Metschnikowia | 13 | Saprotroph | \|Undefined Saprotroph\| | Yeast | NULL | Probable | As nectar-tap_saprotroph (Põlme et al. 2020); Decay sugar-rich substrates (Põlme et al. 2020) | Põlme S, et al. 2020. Fungal Diversity 105:1-16 ((https://doi.org/10.1007/s13225-020-00466-2)) |
| Otu36809 | 1 | k__Fungi(100);p__Ascomycota(100);c__Saccharomycetes(100);o__Saccharomycetales(100);f__Metschnikowiaceae(100);g__Metschnikowia(100);s__Metschnikowia_pulcherrima(100); | Metschnikowia | 13 | Saprotroph | \|Undefined Saprotroph\| | Yeast | NULL | Probable | As nectar-tap_saprotroph (Põlme et al. 2020); Decay sugar-rich substrates (Põlme et al. 2020) | Põlme S, et al. 2020. Fungal Diversity 105:1-16 ((https://doi.org/10.1007/s13225-020-00466-2)) |
| Otu36810 | 1 | k__Fungi(100);p__Ascomycota(100);c__Saccharomycetes(100);o__Saccharomycetales(100);f__Metschnikowiaceae(100);g__Metschnikowia(100);g__Metschnikowia_unclassified(100); | Metschnikowia | 13 | Saprotroph | \|Undefined Saprotroph\| | Yeast | NULL | Probable | As nectar-tap_saprotroph (Põlme et al. 2020); Decay sugar-rich substrates (Põlme et al. 2020) | Põlme S, et al. 2020. Fungal Diversity 105:1-16 ((https://doi.org/10.1007/s13225-020-00466-2)) |
| Otu36811 | 1 | k__Fungi(100);p__Ascomycota(100);c__Saccharomycetes(100);o__Saccharomycetales(100);f__Metschnikowiaceae(100);g__Metschnikowia(100);s__Metschnikowia_pulcherrima(100); | Metschnikowia | 13 | Saprotroph | \|Undefined Saprotroph\| | Yeast | NULL | Probable | As nectar-tap_saprotroph (Põlme et al. 2020); Decay sugar-rich substrates (Põlme et al. 2020) | Põlme S, et al. 2020. Fungal Diversity 105:1-16 ((https://doi.org/10.1007/s13225-020-00466-2)) |
| Otu36812 | 1 | k__Fungi(100);p__Ascomycota(100);c__Saccharomycetes(100);o__Saccharomycetales(100);f__Metschnikowiaceae(100);g__Metschnikowia(100);s__Metschnikowia_pulcherrima(100); | Metschnikowia | 13 | Saprotroph | \|Undefined Saprotroph\| | Yeast | NULL | Probable | As nectar-tap_saprotroph (Põlme et al. 2020); Decay sugar-rich substrates (Põlme et al. 2020) | Põlme S, et al. 2020. Fungal Diversity 105:1-16 ((https://doi.org/10.1007/s13225-020-00466-2)) |
| Otu36814 | 1 | k__Fungi(100);p__Ascomycota(100);c__Saccharomycetes(100);o__Saccharomycetales(100);f__Metschnikowiaceae(100);g__Metschnikowia(100);s__Metschnikowia_pulcherrima(100); | Metschnikowia | 13 | Saprotroph | \|Undefined Saprotroph\| | Yeast | NULL | Probable | As nectar-tap_saprotroph (Põlme et al. 2020); Decay sugar-rich substrates (Põlme et al. 2020) | Põlme S, et al. 2020. Fungal Diversity 105:1-16 ((https://doi.org/10.1007/s13225-020-00466-2)) |
| Otu36816 | 1 | k__Fungi(100);p__Ascomycota(100);c__Saccharomycetes(100);o__Saccharomycetales(100);f__Metschnikowiaceae(100);g__Metschnikowia(100);s__Metschnikowia_pulcherrima(100); | Metschnikowia | 13 | Saprotroph | \|Undefined Saprotroph\| | Yeast | NULL | Probable | As nectar-tap_saprotroph (Põlme et al. 2020); Decay sugar-rich substrates (Põlme et al. 2020) | Põlme S, et al. 2020. Fungal Diversity 105:1-16 ((https://doi.org/10.1007/s13225-020-00466-2)) |
| Otu36817 | 1 | k__Fungi(100);p__Ascomycota(100);c__Saccharomycetes(100);o__Saccharomycetales(100);f__Metschnikowiaceae(100);g__Metschnikowia(100);g__Metschnikowia_unclassified(100); | Metschnikowia | 13 | Saprotroph | \|Undefined Saprotroph\| | Yeast | NULL | Probable | As nectar-tap_saprotroph (Põlme et al. 2020); Decay sugar-rich substrates (Põlme et al. 2020) | Põlme S, et al. 2020. Fungal Diversity 105:1-16 ((https://doi.org/10.1007/s13225-020-00466-2)) |
| Otu36818 | 1 | k__Fungi(100);p__Ascomycota(100);c__Saccharomycetes(100);o__Saccharomycetales(100);f__Metschnikowiaceae(100);g__Metschnikowia(100);g__Metschnikowia_unclassified(100); | Metschnikowia | 13 | Saprotroph | \|Undefined Saprotroph\| | Yeast | NULL | Probable | As nectar-tap_saprotroph (Põlme et al. 2020); Decay sugar-rich substrates (Põlme et al. 2020) | Põlme S, et al. 2020. Fungal Diversity 105:1-16 ((https://doi.org/10.1007/s13225-020-00466-2)) |
| Otu36819 | 1 | k__Fungi(100);p__Ascomycota(100);c__Saccharomycetes(100);o__Saccharomycetales(100);f__Metschnikowiaceae(100);g__Metschnikowia(100);s__Metschnikowia_pulcherrima(100); | Metschnikowia | 13 | Saprotroph | \|Undefined Saprotroph\| | Yeast | NULL | Probable | As nectar-tap_saprotroph (Põlme et al. 2020); Decay sugar-rich substrates (Põlme et al. 2020) | Põlme S, et al. 2020. Fungal Diversity 105:1-16 ((https://doi.org/10.1007/s13225-020-00466-2)) |
| Otu36820 | 1 | k__Fungi(100);p__Ascomycota(100);c__Saccharomycetes(100);o__Saccharomycetales(100);f__Metschnikowiaceae(100);g__Metschnikowia(100);s__Metschnikowia_pulcherrima(100); | Metschnikowia | 13 | Saprotroph | \|Undefined Saprotroph\| | Yeast | NULL | Probable | As nectar-tap_saprotroph (Põlme et al. 2020); Decay sugar-rich substrates (Põlme et al. 2020) | Põlme S, et al. 2020. Fungal Diversity 105:1-16 ((https://doi.org/10.1007/s13225-020-00466-2)) |
| Otu36821 | 1 | k__Fungi(100);p__Ascomycota(100);c__Saccharomycetes(100);o__Saccharomycetales(100);f__Metschnikowiaceae(100);g__Metschnikowia(100);s__Metschnikowia_pulcherrima(100); | Metschnikowia | 13 | Saprotroph | \|Undefined Saprotroph\| | Yeast | NULL | Probable | As nectar-tap_saprotroph (Põlme et al. 2020); Decay sugar-rich substrates (Põlme et al. 2020) | Põlme S, et al. 2020. Fungal Diversity 105:1-16 ((https://doi.org/10.1007/s13225-020-00466-2)) |
| Otu36822 | 1 | k__Fungi(100);p__Ascomycota(100);c__Saccharomycetes(100);o__Saccharomycetales(100);f__Metschnikowiaceae(100);g__Metschnikowia(100);s__Metschnikowia_pulcherrima(100); | Metschnikowia | 13 | Saprotroph | \|Undefined Saprotroph\| | Yeast | NULL | Probable | As nectar-tap_saprotroph (Põlme et al. 2020); Decay sugar-rich substrates (Põlme et al. 2020) | Põlme S, et al. 2020. Fungal Diversity 105:1-16 ((https://doi.org/10.1007/s13225-020-00466-2)) |
| Otu36823 | 1 | k__Fungi(100);p__Ascomycota(100);c__Saccharomycetes(100);o__Saccharomycetales(100);f__Metschnikowiaceae(100);g__Metschnikowia(100);g__Metschnikowia_unclassified(100); | Metschnikowia | 13 | Saprotroph | \|Undefined Saprotroph\| | Yeast | NULL | Probable | As nectar-tap_saprotroph (Põlme et al. 2020); Decay sugar-rich substrates (Põlme et al. 2020) | Põlme S, et al. 2020. Fungal Diversity 105:1-16 ((https://doi.org/10.1007/s13225-020-00466-2)) |
| Otu36824 | 1 | k__Fungi(100);p__Ascomycota(100);c__Saccharomycetes(100);o__Saccharomycetales(100);f__Metschnikowiaceae(100);g__Metschnikowia(100);s__Metschnikowia_pulcherrima(100); | Metschnikowia | 13 | Saprotroph | \|Undefined Saprotroph\| | Yeast | NULL | Probable | As nectar-tap_saprotroph (Põlme et al. 2020); Decay sugar-rich substrates (Põlme et al. 2020) | Põlme S, et al. 2020. Fungal Diversity 105:1-16 ((https://doi.org/10.1007/s13225-020-00466-2)) |
| Otu36825 | 1 | k__Fungi(100);p__Ascomycota(100);c__Saccharomycetes(100);o__Saccharomycetales(100);f__Metschnikowiaceae(100);g__Metschnikowia(100);g__Metschnikowia_unclassified(100); | Metschnikowia | 13 | Saprotroph | \|Undefined Saprotroph\| | Yeast | NULL | Probable | As nectar-tap_saprotroph (Põlme et al. 2020); Decay sugar-rich substrates (Põlme et al. 2020) | Põlme S, et al. 2020. Fungal Diversity 105:1-16 ((https://doi.org/10.1007/s13225-020-00466-2)) |
| Otu36826 | 1 | k__Fungi(100);p__Ascomycota(100);c__Saccharomycetes(100);o__Saccharomycetales(100);f__Metschnikowiaceae(100);g__Metschnikowia(100);s__Metschnikowia_pulcherrima(100); | Metschnikowia | 13 | Saprotroph | \|Undefined Saprotroph\| | Yeast | NULL | Probable | As nectar-tap_saprotroph (Põlme et al. 2020); Decay sugar-rich substrates (Põlme et al. 2020) | Põlme S, et al. 2020. Fungal Diversity 105:1-16 ((https://doi.org/10.1007/s13225-020-00466-2)) |
| Otu36827 | 1 | k__Fungi(100);p__Ascomycota(100);c__Saccharomycetes(100);o__Saccharomycetales(100);f__Metschnikowiaceae(100);g__Metschnikowia(100);g__Metschnikowia_unclassified(100); | Metschnikowia | 13 | Saprotroph | \|Undefined Saprotroph\| | Yeast | NULL | Probable | As nectar-tap_saprotroph (Põlme et al. 2020); Decay sugar-rich substrates (Põlme et al. 2020) | Põlme S, et al. 2020. Fungal Diversity 105:1-16 ((https://doi.org/10.1007/s13225-020-00466-2)) |
| Otu36828 | 1 | k__Fungi(100);p__Ascomycota(100);c__Saccharomycetes(100);o__Saccharomycetales(100);f__Metschnikowiaceae(100);g__Metschnikowia(100);s__Metschnikowia_pulcherrima(100); | Metschnikowia | 13 | Saprotroph | \|Undefined Saprotroph\| | Yeast | NULL | Probable | As nectar-tap_saprotroph (Põlme et al. 2020); Decay sugar-rich substrates (Põlme et al. 2020) | Põlme S, et al. 2020. Fungal Diversity 105:1-16 ((https://doi.org/10.1007/s13225-020-00466-2)) |
| Otu36829 | 1 | k__Fungi(100);p__Ascomycota(100);c__Saccharomycetes(100);o__Saccharomycetales(100);f__Metschnikowiaceae(100);g__Metschnikowia(100);g__Metschnikowia_unclassified(100); | Metschnikowia | 13 | Saprotroph | \|Undefined Saprotroph\| | Yeast | NULL | Probable | As nectar-tap_saprotroph (Põlme et al. 2020); Decay sugar-rich substrates (Põlme et al. 2020) | Põlme S, et al. 2020. Fungal Diversity 105:1-16 ((https://doi.org/10.1007/s13225-020-00466-2)) |
| Otu36830 | 1 | k__Fungi(100);p__Ascomycota(100);c__Saccharomycetes(100);o__Saccharomycetales(100);f__Metschnikowiaceae(100);g__Metschnikowia(100);s__Metschnikowia_pulcherrima(100); | Metschnikowia | 13 | Saprotroph | \|Undefined Saprotroph\| | Yeast | NULL | Probable | As nectar-tap_saprotroph (Põlme et al. 2020); Decay sugar-rich substrates (Põlme et al. 2020) | Põlme S, et al. 2020. Fungal Diversity 105:1-16 ((https://doi.org/10.1007/s13225-020-00466-2)) |
| Otu36831 | 1 | k__Fungi(100);p__Ascomycota(100);c__Saccharomycetes(100);o__Saccharomycetales(100);f__Metschnikowiaceae(100);g__Metschnikowia(100);g__Metschnikowia_unclassified(100); | Metschnikowia | 13 | Saprotroph | \|Undefined Saprotroph\| | Yeast | NULL | Probable | As nectar-tap_saprotroph (Põlme et al. 2020); Decay sugar-rich substrates (Põlme et al. 2020) | Põlme S, et al. 2020. Fungal Diversity 105:1-16 ((https://doi.org/10.1007/s13225-020-00466-2)) |
| Otu36832 | 1 | k__Fungi(100);p__Ascomycota(100);c__Saccharomycetes(100);o__Saccharomycetales(100);f__Metschnikowiaceae(100);g__Metschnikowia(100);s__Metschnikowia_pulcherrima(100); | Metschnikowia | 13 | Saprotroph | \|Undefined Saprotroph\| | Yeast | NULL | Probable | As nectar-tap_saprotroph (Põlme et al. 2020); Decay sugar-rich substrates (Põlme et al. 2020) | Põlme S, et al. 2020. Fungal Diversity 105:1-16 ((https://doi.org/10.1007/s13225-020-00466-2)) |
| Otu36833 | 1 | k__Fungi(100);p__Ascomycota(100);c__Saccharomycetes(100);o__Saccharomycetales(100);f__Metschnikowiaceae(100);g__Metschnikowia(100);s__Metschnikowia_pulcherrima(100); | Metschnikowia | 13 | Saprotroph | \|Undefined Saprotroph\| | Yeast | NULL | Probable | As nectar-tap_saprotroph (Põlme et al. 2020); Decay sugar-rich substrates (Põlme et al. 2020) | Põlme S, et al. 2020. Fungal Diversity 105:1-16 ((https://doi.org/10.1007/s13225-020-00466-2)) |
| Otu36834 | 1 | k__Fungi(100);p__Ascomycota(100);c__Saccharomycetes(100);o__Saccharomycetales(100);f__Metschnikowiaceae(100);g__Metschnikowia(100);s__Metschnikowia_pulcherrima(100); | Metschnikowia | 13 | Saprotroph | \|Undefined Saprotroph\| | Yeast | NULL | Probable | As nectar-tap_saprotroph (Põlme et al. 2020); Decay sugar-rich substrates (Põlme et al. 2020) | Põlme S, et al. 2020. Fungal Diversity 105:1-16 ((https://doi.org/10.1007/s13225-020-00466-2)) |
| Otu36835 | 1 | k__Fungi(100);p__Ascomycota(100);c__Saccharomycetes(100);o__Saccharomycetales(100);f__Metschnikowiaceae(100);g__Metschnikowia(100);s__Metschnikowia_pulcherrima(100); | Metschnikowia | 13 | Saprotroph | \|Undefined Saprotroph\| | Yeast | NULL | Probable | As nectar-tap_saprotroph (Põlme et al. 2020); Decay sugar-rich substrates (Põlme et al. 2020) | Põlme S, et al. 2020. Fungal Diversity 105:1-16 ((https://doi.org/10.1007/s13225-020-00466-2)) |
| Otu36836 | 1 | k__Fungi(100);p__Ascomycota(100);c__Saccharomycetes(100);o__Saccharomycetales(100);f__Metschnikowiaceae(100);g__Metschnikowia(100);s__Metschnikowia_pulcherrima(100); | Metschnikowia | 13 | Saprotroph | \|Undefined Saprotroph\| | Yeast | NULL | Probable | As nectar-tap_saprotroph (Põlme et al. 2020); Decay sugar-rich substrates (Põlme et al. 2020) | Põlme S, et al. 2020. Fungal Diversity 105:1-16 ((https://doi.org/10.1007/s13225-020-00466-2)) |
| Otu36837 | 1 | k__Fungi(100);p__Ascomycota(100);c__Saccharomycetes(100);o__Saccharomycetales(100);f__Metschnikowiaceae(100);g__Metschnikowia(100);s__Metschnikowia_pulcherrima(100); | Metschnikowia | 13 | Saprotroph | \|Undefined Saprotroph\| | Yeast | NULL | Probable | As nectar-tap_saprotroph (Põlme et al. 2020); Decay sugar-rich substrates (Põlme et al. 2020) | Põlme S, et al. 2020. Fungal Diversity 105:1-16 ((https://doi.org/10.1007/s13225-020-00466-2)) |
| Otu36838 | 1 | k__Fungi(100);p__Ascomycota(100);c__Saccharomycetes(100);o__Saccharomycetales(100);f__Metschnikowiaceae(100);g__Metschnikowia(100);s__Metschnikowia_pulcherrima(100); | Metschnikowia | 13 | Saprotroph | \|Undefined Saprotroph\| | Yeast | NULL | Probable | As nectar-tap_saprotroph (Põlme et al. 2020); Decay sugar-rich substrates (Põlme et al. 2020) | Põlme S, et al. 2020. Fungal Diversity 105:1-16 ((https://doi.org/10.1007/s13225-020-00466-2)) |
| Otu36839 | 1 | k__Fungi(100);p__Ascomycota(100);c__Saccharomycetes(100);o__Saccharomycetales(100);f__Metschnikowiaceae(100);g__Metschnikowia(100);s__Metschnikowia_pulcherrima(100); | Metschnikowia | 13 | Saprotroph | \|Undefined Saprotroph\| | Yeast | NULL | Probable | As nectar-tap_saprotroph (Põlme et al. 2020); Decay sugar-rich substrates (Põlme et al. 2020) | Põlme S, et al. 2020. Fungal Diversity 105:1-16 ((https://doi.org/10.1007/s13225-020-00466-2)) |
| Otu36840 | 1 | k__Fungi(100);p__Ascomycota(100);c__Saccharomycetes(100);o__Saccharomycetales(100);f__Metschnikowiaceae(100);g__Metschnikowia(100);s__Metschnikowia_pulcherrima(100); | Metschnikowia | 13 | Saprotroph | \|Undefined Saprotroph\| | Yeast | NULL | Probable | As nectar-tap_saprotroph (Põlme et al. 2020); Decay sugar-rich substrates (Põlme et al. 2020) | Põlme S, et al. 2020. Fungal Diversity 105:1-16 ((https://doi.org/10.1007/s13225-020-00466-2)) |
| Otu36841 | 1 | k__Fungi(100);p__Ascomycota(100);c__Saccharomycetes(100);o__Saccharomycetales(100);f__Metschnikowiaceae(100);g__Metschnikowia(100);s__Metschnikowia_pulcherrima(100); | Metschnikowia | 13 | Saprotroph | \|Undefined Saprotroph\| | Yeast | NULL | Probable | As nectar-tap_saprotroph (Põlme et al. 2020); Decay sugar-rich substrates (Põlme et al. 2020) | Põlme S, et al. 2020. Fungal Diversity 105:1-16 ((https://doi.org/10.1007/s13225-020-00466-2)) |
| Otu36842 | 1 | k__Fungi(100);p__Ascomycota(100);c__Saccharomycetes(100);o__Saccharomycetales(100);f__Metschnikowiaceae(100);g__Metschnikowia(100);s__Metschnikowia_pulcherrima(100); | Metschnikowia | 13 | Saprotroph | \|Undefined Saprotroph\| | Yeast | NULL | Probable | As nectar-tap_saprotroph (Põlme et al. 2020); Decay sugar-rich substrates (Põlme et al. 2020) | Põlme S, et al. 2020. Fungal Diversity 105:1-16 ((https://doi.org/10.1007/s13225-020-00466-2)) |
| Otu36843 | 1 | k__Fungi(100);p__Ascomycota(100);c__Saccharomycetes(100);o__Saccharomycetales(100);f__Metschnikowiaceae(100);g__Metschnikowia(100);s__Metschnikowia_pulcherrima(100); | Metschnikowia | 13 | Saprotroph | \|Undefined Saprotroph\| | Yeast | NULL | Probable | As nectar-tap_saprotroph (Põlme et al. 2020); Decay sugar-rich substrates (Põlme et al. 2020) | Põlme S, et al. 2020. Fungal Diversity 105:1-16 ((https://doi.org/10.1007/s13225-020-00466-2)) |
| Otu36844 | 1 | k__Fungi(100);p__Ascomycota(100);c__Saccharomycetes(100);o__Saccharomycetales(100);f__Metschnikowiaceae(100);g__Metschnikowia(100);g__Metschnikowia_unclassified(100); | Metschnikowia | 13 | Saprotroph | \|Undefined Saprotroph\| | Yeast | NULL | Probable | As nectar-tap_saprotroph (Põlme et al. 2020); Decay sugar-rich substrates (Põlme et al. 2020) | Põlme S, et al. 2020. Fungal Diversity 105:1-16 ((https://doi.org/10.1007/s13225-020-00466-2)) |
| Otu36845 | 1 | k__Fungi(100);p__Ascomycota(100);c__Saccharomycetes(100);o__Saccharomycetales(100);f__Metschnikowiaceae(100);g__Metschnikowia(100);s__Metschnikowia_pulcherrima(100); | Metschnikowia | 13 | Saprotroph | \|Undefined Saprotroph\| | Yeast | NULL | Probable | As nectar-tap_saprotroph (Põlme et al. 2020); Decay sugar-rich substrates (Põlme et al. 2020) | Põlme S, et al. 2020. Fungal Diversity 105:1-16 ((https://doi.org/10.1007/s13225-020-00466-2)) |
| Otu36846 | 1 | k__Fungi(100);p__Ascomycota(100);c__Saccharomycetes(100);o__Saccharomycetales(100);f__Metschnikowiaceae(100);g__Metschnikowia(100);s__Metschnikowia_pulcherrima(100); | Metschnikowia | 13 | Saprotroph | \|Undefined Saprotroph\| | Yeast | NULL | Probable | As nectar-tap_saprotroph (Põlme et al. 2020); Decay sugar-rich substrates (Põlme et al. 2020) | Põlme S, et al. 2020. Fungal Diversity 105:1-16 ((https://doi.org/10.1007/s13225-020-00466-2)) |
| Otu36847 | 1 | k__Fungi(100);p__Ascomycota(100);c__Saccharomycetes(100);o__Saccharomycetales(100);f__Metschnikowiaceae(100);g__Metschnikowia(100);s__Metschnikowia_pulcherrima(100); | Metschnikowia | 13 | Saprotroph | \|Undefined Saprotroph\| | Yeast | NULL | Probable | As nectar-tap_saprotroph (Põlme et al. 2020); Decay sugar-rich substrates (Põlme et al. 2020) | Põlme S, et al. 2020. Fungal Diversity 105:1-16 ((https://doi.org/10.1007/s13225-020-00466-2)) |
| Otu36848 | 1 | k__Fungi(100);p__Ascomycota(100);c__Saccharomycetes(100);o__Saccharomycetales(100);f__Metschnikowiaceae(100);g__Metschnikowia(100);g__Metschnikowia_unclassified(100); | Metschnikowia | 13 | Saprotroph | \|Undefined Saprotroph\| | Yeast | NULL | Probable | As nectar-tap_saprotroph (Põlme et al. 2020); Decay sugar-rich substrates (Põlme et al. 2020) | Põlme S, et al. 2020. Fungal Diversity 105:1-16 ((https://doi.org/10.1007/s13225-020-00466-2)) |
| Otu36849 | 1 | k__Fungi(100);p__Ascomycota(100);c__Saccharomycetes(100);o__Saccharomycetales(100);f__Metschnikowiaceae(100);g__Metschnikowia(100);s__Metschnikowia_pulcherrima(100); | Metschnikowia | 13 | Saprotroph | \|Undefined Saprotroph\| | Yeast | NULL | Probable | As nectar-tap_saprotroph (Põlme et al. 2020); Decay sugar-rich substrates (Põlme et al. 2020) | Põlme S, et al. 2020. Fungal Diversity 105:1-16 ((https://doi.org/10.1007/s13225-020-00466-2)) |
| Otu36850 | 1 | k__Fungi(100);p__Ascomycota(100);c__Saccharomycetes(100);o__Saccharomycetales(100);f__Metschnikowiaceae(100);g__Metschnikowia(100);s__Metschnikowia_pulcherrima(100); | Metschnikowia | 13 | Saprotroph | \|Undefined Saprotroph\| | Yeast | NULL | Probable | As nectar-tap_saprotroph (Põlme et al. 2020); Decay sugar-rich substrates (Põlme et al. 2020) | Põlme S, et al. 2020. Fungal Diversity 105:1-16 ((https://doi.org/10.1007/s13225-020-00466-2)) |
| Otu36851 | 1 | k__Fungi(100);p__Ascomycota(100);c__Saccharomycetes(100);o__Saccharomycetales(100);f__Metschnikowiaceae(100);g__Metschnikowia(100);s__Metschnikowia_pulcherrima(100); | Metschnikowia | 13 | Saprotroph | \|Undefined Saprotroph\| | Yeast | NULL | Probable | As nectar-tap_saprotroph (Põlme et al. 2020); Decay sugar-rich substrates (Põlme et al. 2020) | Põlme S, et al. 2020. Fungal Diversity 105:1-16 ((https://doi.org/10.1007/s13225-020-00466-2)) |
| Otu36852 | 1 | k__Fungi(100);p__Ascomycota(100);c__Saccharomycetes(100);o__Saccharomycetales(100);f__Metschnikowiaceae(100);g__Metschnikowia(100);s__Metschnikowia_pulcherrima(100); | Metschnikowia | 13 | Saprotroph | \|Undefined Saprotroph\| | Yeast | NULL | Probable | As nectar-tap_saprotroph (Põlme et al. 2020); Decay sugar-rich substrates (Põlme et al. 2020) | Põlme S, et al. 2020. Fungal Diversity 105:1-16 ((https://doi.org/10.1007/s13225-020-00466-2)) |
| Otu36853 | 1 | k__Fungi(100);p__Ascomycota(100);c__Saccharomycetes(100);o__Saccharomycetales(100);f__Metschnikowiaceae(100);g__Metschnikowia(100);s__Metschnikowia_pulcherrima(100); | Metschnikowia | 13 | Saprotroph | \|Undefined Saprotroph\| | Yeast | NULL | Probable | As nectar-tap_saprotroph (Põlme et al. 2020); Decay sugar-rich substrates (Põlme et al. 2020) | Põlme S, et al. 2020. Fungal Diversity 105:1-16 ((https://doi.org/10.1007/s13225-020-00466-2)) |
| Otu36854 | 1 | k__Fungi(100);p__Ascomycota(100);c__Saccharomycetes(100);o__Saccharomycetales(100);f__Metschnikowiaceae(100);g__Metschnikowia(100);s__Metschnikowia_pulcherrima(100); | Metschnikowia | 13 | Saprotroph | \|Undefined Saprotroph\| | Yeast | NULL | Probable | As nectar-tap_saprotroph (Põlme et al. 2020); Decay sugar-rich substrates (Põlme et al. 2020) | Põlme S, et al. 2020. Fungal Diversity 105:1-16 ((https://doi.org/10.1007/s13225-020-00466-2)) |
| Otu36855 | 1 | k__Fungi(100);p__Ascomycota(100);c__Saccharomycetes(100);o__Saccharomycetales(100);f__Metschnikowiaceae(100);g__Metschnikowia(100);s__Metschnikowia_pulcherrima(100); | Metschnikowia | 13 | Saprotroph | \|Undefined Saprotroph\| | Yeast | NULL | Probable | As nectar-tap_saprotroph (Põlme et al. 2020); Decay sugar-rich substrates (Põlme et al. 2020) | Põlme S, et al. 2020. Fungal Diversity 105:1-16 ((https://doi.org/10.1007/s13225-020-00466-2)) |
| Otu36856 | 1 | k__Fungi(100);p__Ascomycota(100);c__Saccharomycetes(100);o__Saccharomycetales(100);f__Metschnikowiaceae(100);g__Metschnikowia(100);s__Metschnikowia_pulcherrima(100); | Metschnikowia | 13 | Saprotroph | \|Undefined Saprotroph\| | Yeast | NULL | Probable | As nectar-tap_saprotroph (Põlme et al. 2020); Decay sugar-rich substrates (Põlme et al. 2020) | Põlme S, et al. 2020. Fungal Diversity 105:1-16 ((https://doi.org/10.1007/s13225-020-00466-2)) |
| Otu36857 | 1 | k__Fungi(100);p__Ascomycota(100);c__Saccharomycetes(100);o__Saccharomycetales(100);f__Metschnikowiaceae(100);g__Metschnikowia(100);s__Metschnikowia_pulcherrima(100); | Metschnikowia | 13 | Saprotroph | \|Undefined Saprotroph\| | Yeast | NULL | Probable | As nectar-tap_saprotroph (Põlme et al. 2020); Decay sugar-rich substrates (Põlme et al. 2020) | Põlme S, et al. 2020. Fungal Diversity 105:1-16 ((https://doi.org/10.1007/s13225-020-00466-2)) |
| Otu36859 | 1 | k__Fungi(100);p__Ascomycota(100);c__Saccharomycetes(100);o__Saccharomycetales(100);f__Metschnikowiaceae(100);g__Metschnikowia(100);s__Metschnikowia_pulcherrima(100); | Metschnikowia | 13 | Saprotroph | \|Undefined Saprotroph\| | Yeast | NULL | Probable | As nectar-tap_saprotroph (Põlme et al. 2020); Decay sugar-rich substrates (Põlme et al. 2020) | Põlme S, et al. 2020. Fungal Diversity 105:1-16 ((https://doi.org/10.1007/s13225-020-00466-2)) |
| Otu36860 | 1 | k__Fungi(100);p__Ascomycota(100);c__Saccharomycetes(100);o__Saccharomycetales(100);f__Metschnikowiaceae(100);g__Metschnikowia(100);s__Metschnikowia_pulcherrima(100); | Metschnikowia | 13 | Saprotroph | \|Undefined Saprotroph\| | Yeast | NULL | Probable | As nectar-tap_saprotroph (Põlme et al. 2020); Decay sugar-rich substrates (Põlme et al. 2020) | Põlme S, et al. 2020. Fungal Diversity 105:1-16 ((https://doi.org/10.1007/s13225-020-00466-2)) |
| Otu36861 | 1 | k__Fungi(100);p__Ascomycota(100);c__Saccharomycetes(100);o__Saccharomycetales(100);f__Metschnikowiaceae(100);g__Metschnikowia(100);s__Metschnikowia_pulcherrima(100); | Metschnikowia | 13 | Saprotroph | \|Undefined Saprotroph\| | Yeast | NULL | Probable | As nectar-tap_saprotroph (Põlme et al. 2020); Decay sugar-rich substrates (Põlme et al. 2020) | Põlme S, et al. 2020. Fungal Diversity 105:1-16 ((https://doi.org/10.1007/s13225-020-00466-2)) |
| Otu36862 | 1 | k__Fungi(100);p__Ascomycota(100);c__Saccharomycetes(100);o__Saccharomycetales(100);f__Metschnikowiaceae(100);g__Metschnikowia(100);s__Metschnikowia_pulcherrima(100); | Metschnikowia | 13 | Saprotroph | \|Undefined Saprotroph\| | Yeast | NULL | Probable | As nectar-tap_saprotroph (Põlme et al. 2020); Decay sugar-rich substrates (Põlme et al. 2020) | Põlme S, et al. 2020. Fungal Diversity 105:1-16 ((https://doi.org/10.1007/s13225-020-00466-2)) |
| Otu36863 | 1 | k__Fungi(100);p__Ascomycota(100);c__Saccharomycetes(100);o__Saccharomycetales(100);f__Metschnikowiaceae(100);g__Metschnikowia(100);s__Metschnikowia_pulcherrima(100); | Metschnikowia | 13 | Saprotroph | \|Undefined Saprotroph\| | Yeast | NULL | Probable | As nectar-tap_saprotroph (Põlme et al. 2020); Decay sugar-rich substrates (Põlme et al. 2020) | Põlme S, et al. 2020. Fungal Diversity 105:1-16 ((https://doi.org/10.1007/s13225-020-00466-2)) |
| Otu36864 | 1 | k__Fungi(100);p__Ascomycota(100);c__Saccharomycetes(100);o__Saccharomycetales(100);f__Metschnikowiaceae(100);g__Metschnikowia(100);s__Metschnikowia_pulcherrima(100); | Metschnikowia | 13 | Saprotroph | \|Undefined Saprotroph\| | Yeast | NULL | Probable | As nectar-tap_saprotroph (Põlme et al. 2020); Decay sugar-rich substrates (Põlme et al. 2020) | Põlme S, et al. 2020. Fungal Diversity 105:1-16 ((https://doi.org/10.1007/s13225-020-00466-2)) |
| Otu36865 | 1 | k__Fungi(100);p__Ascomycota(100);c__Saccharomycetes(100);o__Saccharomycetales(100);f__Metschnikowiaceae(100);g__Metschnikowia(100);g__Metschnikowia_unclassified(100); | Metschnikowia | 13 | Saprotroph | \|Undefined Saprotroph\| | Yeast | NULL | Probable | As nectar-tap_saprotroph (Põlme et al. 2020); Decay sugar-rich substrates (Põlme et al. 2020) | Põlme S, et al. 2020. Fungal Diversity 105:1-16 ((https://doi.org/10.1007/s13225-020-00466-2)) |
| Otu36866 | 1 | k__Fungi(100);p__Ascomycota(100);c__Saccharomycetes(100);o__Saccharomycetales(100);f__Metschnikowiaceae(100);g__Metschnikowia(100);g__Metschnikowia_unclassified(100); | Metschnikowia | 13 | Saprotroph | \|Undefined Saprotroph\| | Yeast | NULL | Probable | As nectar-tap_saprotroph (Põlme et al. 2020); Decay sugar-rich substrates (Põlme et al. 2020) | Põlme S, et al. 2020. Fungal Diversity 105:1-16 ((https://doi.org/10.1007/s13225-020-00466-2)) |
| Otu36867 | 1 | k__Fungi(100);p__Ascomycota(100);c__Saccharomycetes(100);o__Saccharomycetales(100);f__Metschnikowiaceae(100);g__Metschnikowia(100);s__Metschnikowia_pulcherrima(100); | Metschnikowia | 13 | Saprotroph | \|Undefined Saprotroph\| | Yeast | NULL | Probable | As nectar-tap_saprotroph (Põlme et al. 2020); Decay sugar-rich substrates (Põlme et al. 2020) | Põlme S, et al. 2020. Fungal Diversity 105:1-16 ((https://doi.org/10.1007/s13225-020-00466-2)) |
| Otu36868 | 1 | k__Fungi(100);p__Ascomycota(100);c__Saccharomycetes(100);o__Saccharomycetales(100);f__Metschnikowiaceae(100);g__Metschnikowia(100);s__Metschnikowia_pulcherrima(100); | Metschnikowia | 13 | Saprotroph | \|Undefined Saprotroph\| | Yeast | NULL | Probable | As nectar-tap_saprotroph (Põlme et al. 2020); Decay sugar-rich substrates (Põlme et al. 2020) | Põlme S, et al. 2020. Fungal Diversity 105:1-16 ((https://doi.org/10.1007/s13225-020-00466-2)) |
| Otu36869 | 1 | k__Fungi(100);p__Ascomycota(100);c__Saccharomycetes(100);o__Saccharomycetales(100);f__Metschnikowiaceae(100);g__Metschnikowia(100);s__Metschnikowia_pulcherrima(100); | Metschnikowia | 13 | Saprotroph | \|Undefined Saprotroph\| | Yeast | NULL | Probable | As nectar-tap_saprotroph (Põlme et al. 2020); Decay sugar-rich substrates (Põlme et al. 2020) | Põlme S, et al. 2020. Fungal Diversity 105:1-16 ((https://doi.org/10.1007/s13225-020-00466-2)) |
| Otu36870 | 1 | k__Fungi(100);p__Ascomycota(100);c__Saccharomycetes(100);o__Saccharomycetales(100);f__Metschnikowiaceae(100);g__Metschnikowia(100);s__Metschnikowia_pulcherrima(100); | Metschnikowia | 13 | Saprotroph | \|Undefined Saprotroph\| | Yeast | NULL | Probable | As nectar-tap_saprotroph (Põlme et al. 2020); Decay sugar-rich substrates (Põlme et al. 2020) | Põlme S, et al. 2020. Fungal Diversity 105:1-16 ((https://doi.org/10.1007/s13225-020-00466-2)) |
| Otu36871 | 1 | k__Fungi(100);p__Ascomycota(100);c__Saccharomycetes(100);o__Saccharomycetales(100);f__Metschnikowiaceae(100);g__Metschnikowia(100);s__Metschnikowia_pulcherrima(100); | Metschnikowia | 13 | Saprotroph | \|Undefined Saprotroph\| | Yeast | NULL | Probable | As nectar-tap_saprotroph (Põlme et al. 2020); Decay sugar-rich substrates (Põlme et al. 2020) | Põlme S, et al. 2020. Fungal Diversity 105:1-16 ((https://doi.org/10.1007/s13225-020-00466-2)) |
| Otu36872 | 1 | k__Fungi(100);p__Ascomycota(100);c__Saccharomycetes(100);o__Saccharomycetales(100);f__Metschnikowiaceae(100);g__Metschnikowia(100);s__Metschnikowia_pulcherrima(100); | Metschnikowia | 13 | Saprotroph | \|Undefined Saprotroph\| | Yeast | NULL | Probable | As nectar-tap_saprotroph (Põlme et al. 2020); Decay sugar-rich substrates (Põlme et al. 2020) | Põlme S, et al. 2020. Fungal Diversity 105:1-16 ((https://doi.org/10.1007/s13225-020-00466-2)) |
| Otu36873 | 1 | k__Fungi(100);p__Ascomycota(100);c__Saccharomycetes(100);o__Saccharomycetales(100);f__Metschnikowiaceae(100);g__Metschnikowia(100);s__Metschnikowia_pulcherrima(100); | Metschnikowia | 13 | Saprotroph | \|Undefined Saprotroph\| | Yeast | NULL | Probable | As nectar-tap_saprotroph (Põlme et al. 2020); Decay sugar-rich substrates (Põlme et al. 2020) | Põlme S, et al. 2020. Fungal Diversity 105:1-16 ((https://doi.org/10.1007/s13225-020-00466-2)) |
| Otu36874 | 1 | k__Fungi(100);p__Ascomycota(100);c__Saccharomycetes(100);o__Saccharomycetales(100);f__Metschnikowiaceae(100);g__Metschnikowia(100);s__Metschnikowia_pulcherrima(100); | Metschnikowia | 13 | Saprotroph | \|Undefined Saprotroph\| | Yeast | NULL | Probable | As nectar-tap_saprotroph (Põlme et al. 2020); Decay sugar-rich substrates (Põlme et al. 2020) | Põlme S, et al. 2020. Fungal Diversity 105:1-16 ((https://doi.org/10.1007/s13225-020-00466-2)) |
| Otu36875 | 1 | k__Fungi(100);p__Ascomycota(100);c__Saccharomycetes(100);o__Saccharomycetales(100);f__Metschnikowiaceae(100);g__Metschnikowia(100);s__Metschnikowia_pulcherrima(100); | Metschnikowia | 13 | Saprotroph | \|Undefined Saprotroph\| | Yeast | NULL | Probable | As nectar-tap_saprotroph (Põlme et al. 2020); Decay sugar-rich substrates (Põlme et al. 2020) | Põlme S, et al. 2020. Fungal Diversity 105:1-16 ((https://doi.org/10.1007/s13225-020-00466-2)) |
| Otu36876 | 1 | k__Fungi(100);p__Ascomycota(100);c__Saccharomycetes(100);o__Saccharomycetales(100);f__Metschnikowiaceae(100);g__Metschnikowia(100);g__Metschnikowia_unclassified(100); | Metschnikowia | 13 | Saprotroph | \|Undefined Saprotroph\| | Yeast | NULL | Probable | As nectar-tap_saprotroph (Põlme et al. 2020); Decay sugar-rich substrates (Põlme et al. 2020) | Põlme S, et al. 2020. Fungal Diversity 105:1-16 ((https://doi.org/10.1007/s13225-020-00466-2)) |
| Otu36877 | 1 | k__Fungi(100);p__Ascomycota(100);c__Saccharomycetes(100);o__Saccharomycetales(100);f__Metschnikowiaceae(100);g__Metschnikowia(100);s__Metschnikowia_pulcherrima(100); | Metschnikowia | 13 | Saprotroph | \|Undefined Saprotroph\| | Yeast | NULL | Probable | As nectar-tap_saprotroph (Põlme et al. 2020); Decay sugar-rich substrates (Põlme et al. 2020) | Põlme S, et al. 2020. Fungal Diversity 105:1-16 ((https://doi.org/10.1007/s13225-020-00466-2)) |
| Otu36878 | 1 | k__Fungi(100);p__Ascomycota(100);c__Saccharomycetes(100);o__Saccharomycetales(100);f__Metschnikowiaceae(100);g__Metschnikowia(100);s__Metschnikowia_pulcherrima(100); | Metschnikowia | 13 | Saprotroph | \|Undefined Saprotroph\| | Yeast | NULL | Probable | As nectar-tap_saprotroph (Põlme et al. 2020); Decay sugar-rich substrates (Põlme et al. 2020) | Põlme S, et al. 2020. Fungal Diversity 105:1-16 ((https://doi.org/10.1007/s13225-020-00466-2)) |
| Otu36880 | 1 | k__Fungi(100);p__Ascomycota(100);c__Saccharomycetes(100);o__Saccharomycetales(100);f__Metschnikowiaceae(100);g__Metschnikowia(100);s__Metschnikowia_pulcherrima(100); | Metschnikowia | 13 | Saprotroph | \|Undefined Saprotroph\| | Yeast | NULL | Probable | As nectar-tap_saprotroph (Põlme et al. 2020); Decay sugar-rich substrates (Põlme et al. 2020) | Põlme S, et al. 2020. Fungal Diversity 105:1-16 ((https://doi.org/10.1007/s13225-020-00466-2)) |
| Otu36881 | 1 | k__Fungi(100);p__Ascomycota(100);c__Saccharomycetes(100);o__Saccharomycetales(100);f__Metschnikowiaceae(100);g__Metschnikowia(100);s__Metschnikowia_pulcherrima(100); | Metschnikowia | 13 | Saprotroph | \|Undefined Saprotroph\| | Yeast | NULL | Probable | As nectar-tap_saprotroph (Põlme et al. 2020); Decay sugar-rich substrates (Põlme et al. 2020) | Põlme S, et al. 2020. Fungal Diversity 105:1-16 ((https://doi.org/10.1007/s13225-020-00466-2)) |
| Otu36882 | 1 | k__Fungi(100);p__Ascomycota(100);c__Saccharomycetes(100);o__Saccharomycetales(100);f__Metschnikowiaceae(100);g__Metschnikowia(100);s__Metschnikowia_pulcherrima(100); | Metschnikowia | 13 | Saprotroph | \|Undefined Saprotroph\| | Yeast | NULL | Probable | As nectar-tap_saprotroph (Põlme et al. 2020); Decay sugar-rich substrates (Põlme et al. 2020) | Põlme S, et al. 2020. Fungal Diversity 105:1-16 ((https://doi.org/10.1007/s13225-020-00466-2)) |
| Otu36883 | 1 | k__Fungi(100);p__Ascomycota(100);c__Saccharomycetes(100);o__Saccharomycetales(100);f__Metschnikowiaceae(100);g__Metschnikowia(100);s__Metschnikowia_pulcherrima(100); | Metschnikowia | 13 | Saprotroph | \|Undefined Saprotroph\| | Yeast | NULL | Probable | As nectar-tap_saprotroph (Põlme et al. 2020); Decay sugar-rich substrates (Põlme et al. 2020) | Põlme S, et al. 2020. Fungal Diversity 105:1-16 ((https://doi.org/10.1007/s13225-020-00466-2)) |
| Otu36884 | 1 | k__Fungi(100);p__Ascomycota(100);c__Saccharomycetes(100);o__Saccharomycetales(100);f__Metschnikowiaceae(100);g__Metschnikowia(100);s__Metschnikowia_pulcherrima(100); | Metschnikowia | 13 | Saprotroph | \|Undefined Saprotroph\| | Yeast | NULL | Probable | As nectar-tap_saprotroph (Põlme et al. 2020); Decay sugar-rich substrates (Põlme et al. 2020) | Põlme S, et al. 2020. Fungal Diversity 105:1-16 ((https://doi.org/10.1007/s13225-020-00466-2)) |
| Otu36885 | 1 | k__Fungi(100);p__Ascomycota(100);c__Saccharomycetes(100);o__Saccharomycetales(100);f__Metschnikowiaceae(100);g__Metschnikowia(100);s__Metschnikowia_pulcherrima(100); | Metschnikowia | 13 | Saprotroph | \|Undefined Saprotroph\| | Yeast | NULL | Probable | As nectar-tap_saprotroph (Põlme et al. 2020); Decay sugar-rich substrates (Põlme et al. 2020) | Põlme S, et al. 2020. Fungal Diversity 105:1-16 ((https://doi.org/10.1007/s13225-020-00466-2)) |
| Otu36886 | 1 | k__Fungi(100);p__Ascomycota(100);c__Saccharomycetes(100);o__Saccharomycetales(100);f__Metschnikowiaceae(100);g__Metschnikowia(100);s__Metschnikowia_pulcherrima(100); | Metschnikowia | 13 | Saprotroph | \|Undefined Saprotroph\| | Yeast | NULL | Probable | As nectar-tap_saprotroph (Põlme et al. 2020); Decay sugar-rich substrates (Põlme et al. 2020) | Põlme S, et al. 2020. Fungal Diversity 105:1-16 ((https://doi.org/10.1007/s13225-020-00466-2)) |
| Otu36887 | 1 | k__Fungi(100);p__Ascomycota(100);c__Saccharomycetes(100);o__Saccharomycetales(100);f__Metschnikowiaceae(100);g__Metschnikowia(100);s__Metschnikowia_pulcherrima(100); | Metschnikowia | 13 | Saprotroph | \|Undefined Saprotroph\| | Yeast | NULL | Probable | As nectar-tap_saprotroph (Põlme et al. 2020); Decay sugar-rich substrates (Põlme et al. 2020) | Põlme S, et al. 2020. Fungal Diversity 105:1-16 ((https://doi.org/10.1007/s13225-020-00466-2)) |
| Otu36888 | 1 | k__Fungi(100);p__Ascomycota(100);c__Saccharomycetes(100);o__Saccharomycetales(100);f__Metschnikowiaceae(100);g__Metschnikowia(100);g__Metschnikowia_unclassified(100); | Metschnikowia | 13 | Saprotroph | \|Undefined Saprotroph\| | Yeast | NULL | Probable | As nectar-tap_saprotroph (Põlme et al. 2020); Decay sugar-rich substrates (Põlme et al. 2020) | Põlme S, et al. 2020. Fungal Diversity 105:1-16 ((https://doi.org/10.1007/s13225-020-00466-2)) |
| Otu36889 | 1 | k__Fungi(100);p__Ascomycota(100);c__Saccharomycetes(100);o__Saccharomycetales(100);f__Metschnikowiaceae(100);g__Metschnikowia(100);s__Metschnikowia_pulcherrima(100); | Metschnikowia | 13 | Saprotroph | \|Undefined Saprotroph\| | Yeast | NULL | Probable | As nectar-tap_saprotroph (Põlme et al. 2020); Decay sugar-rich substrates (Põlme et al. 2020) | Põlme S, et al. 2020. Fungal Diversity 105:1-16 ((https://doi.org/10.1007/s13225-020-00466-2)) |
| Otu36890 | 1 | k__Fungi(100);p__Ascomycota(100);c__Saccharomycetes(100);o__Saccharomycetales(100);f__Metschnikowiaceae(100);g__Metschnikowia(100);s__Metschnikowia_pulcherrima(100); | Metschnikowia | 13 | Saprotroph | \|Undefined Saprotroph\| | Yeast | NULL | Probable | As nectar-tap_saprotroph (Põlme et al. 2020); Decay sugar-rich substrates (Põlme et al. 2020) | Põlme S, et al. 2020. Fungal Diversity 105:1-16 ((https://doi.org/10.1007/s13225-020-00466-2)) |
| Otu36891 | 1 | k__Fungi(100);p__Ascomycota(100);c__Saccharomycetes(100);o__Saccharomycetales(100);f__Metschnikowiaceae(100);g__Metschnikowia(100);g__Metschnikowia_unclassified(100); | Metschnikowia | 13 | Saprotroph | \|Undefined Saprotroph\| | Yeast | NULL | Probable | As nectar-tap_saprotroph (Põlme et al. 2020); Decay sugar-rich substrates (Põlme et al. 2020) | Põlme S, et al. 2020. Fungal Diversity 105:1-16 ((https://doi.org/10.1007/s13225-020-00466-2)) |
| Otu36892 | 1 | k__Fungi(100);p__Ascomycota(100);c__Saccharomycetes(100);o__Saccharomycetales(100);f__Metschnikowiaceae(100);g__Metschnikowia(100);s__Metschnikowia_pulcherrima(100); | Metschnikowia | 13 | Saprotroph | \|Undefined Saprotroph\| | Yeast | NULL | Probable | As nectar-tap_saprotroph (Põlme et al. 2020); Decay sugar-rich substrates (Põlme et al. 2020) | Põlme S, et al. 2020. Fungal Diversity 105:1-16 ((https://doi.org/10.1007/s13225-020-00466-2)) |
| Otu36893 | 1 | k__Fungi(100);p__Ascomycota(100);c__Saccharomycetes(100);o__Saccharomycetales(100);f__Metschnikowiaceae(100);g__Metschnikowia(100);s__Metschnikowia_pulcherrima(100); | Metschnikowia | 13 | Saprotroph | \|Undefined Saprotroph\| | Yeast | NULL | Probable | As nectar-tap_saprotroph (Põlme et al. 2020); Decay sugar-rich substrates (Põlme et al. 2020) | Põlme S, et al. 2020. Fungal Diversity 105:1-16 ((https://doi.org/10.1007/s13225-020-00466-2)) |
| Otu36894 | 1 | k__Fungi(100);p__Ascomycota(100);c__Saccharomycetes(100);o__Saccharomycetales(100);f__Metschnikowiaceae(100);g__Metschnikowia(100);s__Metschnikowia_pulcherrima(100); | Metschnikowia | 13 | Saprotroph | \|Undefined Saprotroph\| | Yeast | NULL | Probable | As nectar-tap_saprotroph (Põlme et al. 2020); Decay sugar-rich substrates (Põlme et al. 2020) | Põlme S, et al. 2020. Fungal Diversity 105:1-16 ((https://doi.org/10.1007/s13225-020-00466-2)) |
| Otu36895 | 1 | k__Fungi(100);p__Ascomycota(100);c__Saccharomycetes(100);o__Saccharomycetales(100);f__Metschnikowiaceae(100);g__Metschnikowia(100);s__Metschnikowia_pulcherrima(100); | Metschnikowia | 13 | Saprotroph | \|Undefined Saprotroph\| | Yeast | NULL | Probable | As nectar-tap_saprotroph (Põlme et al. 2020); Decay sugar-rich substrates (Põlme et al. 2020) | Põlme S, et al. 2020. Fungal Diversity 105:1-16 ((https://doi.org/10.1007/s13225-020-00466-2)) |
| Otu36896 | 1 | k__Fungi(100);p__Ascomycota(100);c__Saccharomycetes(100);o__Saccharomycetales(100);f__Metschnikowiaceae(100);g__Metschnikowia(100);g__Metschnikowia_unclassified(100); | Metschnikowia | 13 | Saprotroph | \|Undefined Saprotroph\| | Yeast | NULL | Probable | As nectar-tap_saprotroph (Põlme et al. 2020); Decay sugar-rich substrates (Põlme et al. 2020) | Põlme S, et al. 2020. Fungal Diversity 105:1-16 ((https://doi.org/10.1007/s13225-020-00466-2)) |
| Otu36897 | 1 | k__Fungi(100);p__Ascomycota(100);c__Saccharomycetes(100);o__Saccharomycetales(100);f__Metschnikowiaceae(100);g__Metschnikowia(100);g__Metschnikowia_unclassified(100); | Metschnikowia | 13 | Saprotroph | \|Undefined Saprotroph\| | Yeast | NULL | Probable | As nectar-tap_saprotroph (Põlme et al. 2020); Decay sugar-rich substrates (Põlme et al. 2020) | Põlme S, et al. 2020. Fungal Diversity 105:1-16 ((https://doi.org/10.1007/s13225-020-00466-2)) |
| Otu36898 | 1 | k__Fungi(100);p__Ascomycota(100);c__Saccharomycetes(100);o__Saccharomycetales(100);f__Metschnikowiaceae(100);g__Metschnikowia(100);s__Metschnikowia_pulcherrima(100); | Metschnikowia | 13 | Saprotroph | \|Undefined Saprotroph\| | Yeast | NULL | Probable | As nectar-tap_saprotroph (Põlme et al. 2020); Decay sugar-rich substrates (Põlme et al. 2020) | Põlme S, et al. 2020. Fungal Diversity 105:1-16 ((https://doi.org/10.1007/s13225-020-00466-2)) |
| Otu36899 | 1 | k__Fungi(100);p__Ascomycota(100);c__Saccharomycetes(100);o__Saccharomycetales(100);f__Metschnikowiaceae(100);g__Metschnikowia(100);s__Metschnikowia_pulcherrima(100); | Metschnikowia | 13 | Saprotroph | \|Undefined Saprotroph\| | Yeast | NULL | Probable | As nectar-tap_saprotroph (Põlme et al. 2020); Decay sugar-rich substrates (Põlme et al. 2020) | Põlme S, et al. 2020. Fungal Diversity 105:1-16 ((https://doi.org/10.1007/s13225-020-00466-2)) |
| Otu36900 | 1 | k__Fungi(100);p__Ascomycota(100);c__Saccharomycetes(100);o__Saccharomycetales(100);f__Metschnikowiaceae(100);g__Metschnikowia(100);s__Metschnikowia_pulcherrima(100); | Metschnikowia | 13 | Saprotroph | \|Undefined Saprotroph\| | Yeast | NULL | Probable | As nectar-tap_saprotroph (Põlme et al. 2020); Decay sugar-rich substrates (Põlme et al. 2020) | Põlme S, et al. 2020. Fungal Diversity 105:1-16 ((https://doi.org/10.1007/s13225-020-00466-2)) |
| Otu36901 | 1 | k__Fungi(100);p__Ascomycota(100);c__Saccharomycetes(100);o__Saccharomycetales(100);f__Metschnikowiaceae(100);g__Metschnikowia(100);g__Metschnikowia_unclassified(100); | Metschnikowia | 13 | Saprotroph | \|Undefined Saprotroph\| | Yeast | NULL | Probable | As nectar-tap_saprotroph (Põlme et al. 2020); Decay sugar-rich substrates (Põlme et al. 2020) | Põlme S, et al. 2020. Fungal Diversity 105:1-16 ((https://doi.org/10.1007/s13225-020-00466-2)) |
| Otu36902 | 1 | k__Fungi(100);p__Ascomycota(100);c__Saccharomycetes(100);o__Saccharomycetales(100);f__Metschnikowiaceae(100);g__Metschnikowia(100);s__Metschnikowia_pulcherrima(100); | Metschnikowia | 13 | Saprotroph | \|Undefined Saprotroph\| | Yeast | NULL | Probable | As nectar-tap_saprotroph (Põlme et al. 2020); Decay sugar-rich substrates (Põlme et al. 2020) | Põlme S, et al. 2020. Fungal Diversity 105:1-16 ((https://doi.org/10.1007/s13225-020-00466-2)) |
| Otu36903 | 1 | k__Fungi(100);p__Ascomycota(100);c__Saccharomycetes(100);o__Saccharomycetales(100);f__Metschnikowiaceae(100);g__Metschnikowia(100);s__Metschnikowia_pulcherrima(100); | Metschnikowia | 13 | Saprotroph | \|Undefined Saprotroph\| | Yeast | NULL | Probable | As nectar-tap_saprotroph (Põlme et al. 2020); Decay sugar-rich substrates (Põlme et al. 2020) | Põlme S, et al. 2020. Fungal Diversity 105:1-16 ((https://doi.org/10.1007/s13225-020-00466-2)) |
| Otu36904 | 1 | k__Fungi(100);p__Ascomycota(100);c__Saccharomycetes(100);o__Saccharomycetales(100);f__Metschnikowiaceae(100);g__Metschnikowia(100);s__Metschnikowia_pulcherrima(100); | Metschnikowia | 13 | Saprotroph | \|Undefined Saprotroph\| | Yeast | NULL | Probable | As nectar-tap_saprotroph (Põlme et al. 2020); Decay sugar-rich substrates (Põlme et al. 2020) | Põlme S, et al. 2020. Fungal Diversity 105:1-16 ((https://doi.org/10.1007/s13225-020-00466-2)) |
| Otu36905 | 1 | k__Fungi(100);p__Ascomycota(100);c__Saccharomycetes(100);o__Saccharomycetales(100);f__Metschnikowiaceae(100);g__Metschnikowia(100);g__Metschnikowia_unclassified(100); | Metschnikowia | 13 | Saprotroph | \|Undefined Saprotroph\| | Yeast | NULL | Probable | As nectar-tap_saprotroph (Põlme et al. 2020); Decay sugar-rich substrates (Põlme et al. 2020) | Põlme S, et al. 2020. Fungal Diversity 105:1-16 ((https://doi.org/10.1007/s13225-020-00466-2)) |
| Otu36906 | 1 | k__Fungi(100);p__Ascomycota(100);c__Saccharomycetes(100);o__Saccharomycetales(100);f__Metschnikowiaceae(100);g__Metschnikowia(100);g__Metschnikowia_unclassified(100); | Metschnikowia | 13 | Saprotroph | \|Undefined Saprotroph\| | Yeast | NULL | Probable | As nectar-tap_saprotroph (Põlme et al. 2020); Decay sugar-rich substrates (Põlme et al. 2020) | Põlme S, et al. 2020. Fungal Diversity 105:1-16 ((https://doi.org/10.1007/s13225-020-00466-2)) |
| Otu36907 | 1 | k__Fungi(100);p__Ascomycota(100);c__Saccharomycetes(100);o__Saccharomycetales(100);f__Metschnikowiaceae(100);g__Metschnikowia(100);s__Metschnikowia_pulcherrima(100); | Metschnikowia | 13 | Saprotroph | \|Undefined Saprotroph\| | Yeast | NULL | Probable | As nectar-tap_saprotroph (Põlme et al. 2020); Decay sugar-rich substrates (Põlme et al. 2020) | Põlme S, et al. 2020. Fungal Diversity 105:1-16 ((https://doi.org/10.1007/s13225-020-00466-2)) |
| Otu36908 | 1 | k__Fungi(100);p__Ascomycota(100);c__Saccharomycetes(100);o__Saccharomycetales(100);f__Metschnikowiaceae(100);g__Metschnikowia(100);g__Metschnikowia_unclassified(100); | Metschnikowia | 13 | Saprotroph | \|Undefined Saprotroph\| | Yeast | NULL | Probable | As nectar-tap_saprotroph (Põlme et al. 2020); Decay sugar-rich substrates (Põlme et al. 2020) | Põlme S, et al. 2020. Fungal Diversity 105:1-16 ((https://doi.org/10.1007/s13225-020-00466-2)) |
| Otu36909 | 1 | k__Fungi(100);p__Ascomycota(100);c__Saccharomycetes(100);o__Saccharomycetales(100);f__Metschnikowiaceae(100);g__Metschnikowia(100);s__Metschnikowia_pulcherrima(100); | Metschnikowia | 13 | Saprotroph | \|Undefined Saprotroph\| | Yeast | NULL | Probable | As nectar-tap_saprotroph (Põlme et al. 2020); Decay sugar-rich substrates (Põlme et al. 2020) | Põlme S, et al. 2020. Fungal Diversity 105:1-16 ((https://doi.org/10.1007/s13225-020-00466-2)) |
| Otu36910 | 1 | k__Fungi(100);p__Ascomycota(100);c__Saccharomycetes(100);o__Saccharomycetales(100);f__Metschnikowiaceae(100);g__Metschnikowia(100);s__Metschnikowia_pulcherrima(100); | Metschnikowia | 13 | Saprotroph | \|Undefined Saprotroph\| | Yeast | NULL | Probable | As nectar-tap_saprotroph (Põlme et al. 2020); Decay sugar-rich substrates (Põlme et al. 2020) | Põlme S, et al. 2020. Fungal Diversity 105:1-16 ((https://doi.org/10.1007/s13225-020-00466-2)) |
| Otu36911 | 1 | k__Fungi(100);p__Ascomycota(100);c__Saccharomycetes(100);o__Saccharomycetales(100);f__Metschnikowiaceae(100);g__Metschnikowia(100);s__Metschnikowia_pulcherrima(100); | Metschnikowia | 13 | Saprotroph | \|Undefined Saprotroph\| | Yeast | NULL | Probable | As nectar-tap_saprotroph (Põlme et al. 2020); Decay sugar-rich substrates (Põlme et al. 2020) | Põlme S, et al. 2020. Fungal Diversity 105:1-16 ((https://doi.org/10.1007/s13225-020-00466-2)) |
| Otu36912 | 1 | k__Fungi(100);p__Ascomycota(100);c__Saccharomycetes(100);o__Saccharomycetales(100);f__Metschnikowiaceae(100);g__Metschnikowia(100);s__Metschnikowia_pulcherrima(100); | Metschnikowia | 13 | Saprotroph | \|Undefined Saprotroph\| | Yeast | NULL | Probable | As nectar-tap_saprotroph (Põlme et al. 2020); Decay sugar-rich substrates (Põlme et al. 2020) | Põlme S, et al. 2020. Fungal Diversity 105:1-16 ((https://doi.org/10.1007/s13225-020-00466-2)) |
| Otu36913 | 1 | k__Fungi(100);p__Ascomycota(100);c__Saccharomycetes(100);o__Saccharomycetales(100);f__Metschnikowiaceae(100);g__Metschnikowia(100);g__Metschnikowia_unclassified(100); | Metschnikowia | 13 | Saprotroph | \|Undefined Saprotroph\| | Yeast | NULL | Probable | As nectar-tap_saprotroph (Põlme et al. 2020); Decay sugar-rich substrates (Põlme et al. 2020) | Põlme S, et al. 2020. Fungal Diversity 105:1-16 ((https://doi.org/10.1007/s13225-020-00466-2)) |
| Otu36914 | 1 | k__Fungi(100);p__Ascomycota(100);c__Saccharomycetes(100);o__Saccharomycetales(100);f__Metschnikowiaceae(100);g__Metschnikowia(100);g__Metschnikowia_unclassified(100); | Metschnikowia | 13 | Saprotroph | \|Undefined Saprotroph\| | Yeast | NULL | Probable | As nectar-tap_saprotroph (Põlme et al. 2020); Decay sugar-rich substrates (Põlme et al. 2020) | Põlme S, et al. 2020. Fungal Diversity 105:1-16 ((https://doi.org/10.1007/s13225-020-00466-2)) |
| Otu36915 | 1 | k__Fungi(100);p__Ascomycota(100);c__Saccharomycetes(100);o__Saccharomycetales(100);f__Metschnikowiaceae(100);g__Metschnikowia(100);s__Metschnikowia_pulcherrima(100); | Metschnikowia | 13 | Saprotroph | \|Undefined Saprotroph\| | Yeast | NULL | Probable | As nectar-tap_saprotroph (Põlme et al. 2020); Decay sugar-rich substrates (Põlme et al. 2020) | Põlme S, et al. 2020. Fungal Diversity 105:1-16 ((https://doi.org/10.1007/s13225-020-00466-2)) |
| Otu36916 | 1 | k__Fungi(100);p__Ascomycota(100);c__Saccharomycetes(100);o__Saccharomycetales(100);f__Metschnikowiaceae(100);g__Metschnikowia(100);s__Metschnikowia_pulcherrima(100); | Metschnikowia | 13 | Saprotroph | \|Undefined Saprotroph\| | Yeast | NULL | Probable | As nectar-tap_saprotroph (Põlme et al. 2020); Decay sugar-rich substrates (Põlme et al. 2020) | Põlme S, et al. 2020. Fungal Diversity 105:1-16 ((https://doi.org/10.1007/s13225-020-00466-2)) |
| Otu36917 | 1 | k__Fungi(100);p__Ascomycota(100);c__Saccharomycetes(100);o__Saccharomycetales(100);f__Metschnikowiaceae(100);g__Metschnikowia(100);s__Metschnikowia_pulcherrima(100); | Metschnikowia | 13 | Saprotroph | \|Undefined Saprotroph\| | Yeast | NULL | Probable | As nectar-tap_saprotroph (Põlme et al. 2020); Decay sugar-rich substrates (Põlme et al. 2020) | Põlme S, et al. 2020. Fungal Diversity 105:1-16 ((https://doi.org/10.1007/s13225-020-00466-2)) |
| Otu36918 | 1 | k__Fungi(100);p__Ascomycota(100);c__Saccharomycetes(100);o__Saccharomycetales(100);f__Metschnikowiaceae(100);g__Metschnikowia(100);s__Metschnikowia_pulcherrima(100); | Metschnikowia | 13 | Saprotroph | \|Undefined Saprotroph\| | Yeast | NULL | Probable | As nectar-tap_saprotroph (Põlme et al. 2020); Decay sugar-rich substrates (Põlme et al. 2020) | Põlme S, et al. 2020. Fungal Diversity 105:1-16 ((https://doi.org/10.1007/s13225-020-00466-2)) |
| Otu36919 | 1 | k__Fungi(100);p__Ascomycota(100);c__Saccharomycetes(100);o__Saccharomycetales(100);f__Metschnikowiaceae(100);g__Metschnikowia(100);g__Metschnikowia_unclassified(100); | Metschnikowia | 13 | Saprotroph | \|Undefined Saprotroph\| | Yeast | NULL | Probable | As nectar-tap_saprotroph (Põlme et al. 2020); Decay sugar-rich substrates (Põlme et al. 2020) | Põlme S, et al. 2020. Fungal Diversity 105:1-16 ((https://doi.org/10.1007/s13225-020-00466-2)) |
| Otu36920 | 1 | k__Fungi(100);p__Ascomycota(100);c__Saccharomycetes(100);o__Saccharomycetales(100);f__Metschnikowiaceae(100);g__Metschnikowia(100);s__Metschnikowia_pulcherrima(100); | Metschnikowia | 13 | Saprotroph | \|Undefined Saprotroph\| | Yeast | NULL | Probable | As nectar-tap_saprotroph (Põlme et al. 2020); Decay sugar-rich substrates (Põlme et al. 2020) | Põlme S, et al. 2020. Fungal Diversity 105:1-16 ((https://doi.org/10.1007/s13225-020-00466-2)) |
| Otu36921 | 1 | k__Fungi(100);p__Ascomycota(100);c__Saccharomycetes(100);o__Saccharomycetales(100);f__Metschnikowiaceae(100);g__Metschnikowia(100);g__Metschnikowia_unclassified(100); | Metschnikowia | 13 | Saprotroph | \|Undefined Saprotroph\| | Yeast | NULL | Probable | As nectar-tap_saprotroph (Põlme et al. 2020); Decay sugar-rich substrates (Põlme et al. 2020) | Põlme S, et al. 2020. Fungal Diversity 105:1-16 ((https://doi.org/10.1007/s13225-020-00466-2)) |
| Otu36922 | 1 | k__Fungi(100);p__Ascomycota(100);c__Saccharomycetes(100);o__Saccharomycetales(100);f__Metschnikowiaceae(100);g__Metschnikowia(100);g__Metschnikowia_unclassified(100); | Metschnikowia | 13 | Saprotroph | \|Undefined Saprotroph\| | Yeast | NULL | Probable | As nectar-tap_saprotroph (Põlme et al. 2020); Decay sugar-rich substrates (Põlme et al. 2020) | Põlme S, et al. 2020. Fungal Diversity 105:1-16 ((https://doi.org/10.1007/s13225-020-00466-2)) |
| Otu36923 | 1 | k__Fungi(100);p__Ascomycota(100);c__Saccharomycetes(100);o__Saccharomycetales(100);f__Metschnikowiaceae(100);g__Metschnikowia(100);g__Metschnikowia_unclassified(100); | Metschnikowia | 13 | Saprotroph | \|Undefined Saprotroph\| | Yeast | NULL | Probable | As nectar-tap_saprotroph (Põlme et al. 2020); Decay sugar-rich substrates (Põlme et al. 2020) | Põlme S, et al. 2020. Fungal Diversity 105:1-16 ((https://doi.org/10.1007/s13225-020-00466-2)) |
| Otu36924 | 1 | k__Fungi(100);p__Ascomycota(100);c__Saccharomycetes(100);o__Saccharomycetales(100);f__Metschnikowiaceae(100);g__Metschnikowia(100);s__Metschnikowia_pulcherrima(100); | Metschnikowia | 13 | Saprotroph | \|Undefined Saprotroph\| | Yeast | NULL | Probable | As nectar-tap_saprotroph (Põlme et al. 2020); Decay sugar-rich substrates (Põlme et al. 2020) | Põlme S, et al. 2020. Fungal Diversity 105:1-16 ((https://doi.org/10.1007/s13225-020-00466-2)) |
| Otu36925 | 1 | k__Fungi(100);p__Ascomycota(100);c__Saccharomycetes(100);o__Saccharomycetales(100);f__Metschnikowiaceae(100);g__Metschnikowia(100);s__Metschnikowia_pulcherrima(100); | Metschnikowia | 13 | Saprotroph | \|Undefined Saprotroph\| | Yeast | NULL | Probable | As nectar-tap_saprotroph (Põlme et al. 2020); Decay sugar-rich substrates (Põlme et al. 2020) | Põlme S, et al. 2020. Fungal Diversity 105:1-16 ((https://doi.org/10.1007/s13225-020-00466-2)) |
| Otu36926 | 1 | k__Fungi(100);p__Ascomycota(100);c__Saccharomycetes(100);o__Saccharomycetales(100);f__Saccharomycetales_family_Incertae_sedis(100);g__Candida(100);g__Candida_unclassified(100); | Candida | 13 | Pathotroph-Saprotroph-Symbiotroph | Animal Pathogen-Endophyte-Endosymbiont-Epiphyte-\|Undefined Saprotroph\| | Dimorphic Yeast-Yeast | NULL | Probable | As nectar-tap_saprotroph (Põlme et al. 2020); Decay sugar-rich substrates (Põlme et al. 2020); This asexual genus is polyphyletic and contain species that function very differently depend on what they are. Caution should be taken when interpreting guilds for Candida species. | Manolakaki D, et al. 2010. Virulence. 1: 367-375 ((https://doi.org/10.4161/viru.1.5.12796)); Mekbib SB, et al. 2011. Tropical Plant Pathology 36:233-240 ((https://doi.org/10.1590/S1982-56762011000400004)); Põlme S, et al. 2020. Fungal Diversity 105:1-16 ((https://doi.org/10.1007/s13225-020-00466-2)) |
| Otu36927 | 1 | k__Fungi(100);p__Ascomycota(100);c__Saccharomycetes(100);o__Saccharomycetales(100);f__Metschnikowiaceae(100);g__Metschnikowia(100);g__Metschnikowia_unclassified(100); | Metschnikowia | 13 | Saprotroph | \|Undefined Saprotroph\| | Yeast | NULL | Probable | As nectar-tap_saprotroph (Põlme et al. 2020); Decay sugar-rich substrates (Põlme et al. 2020) | Põlme S, et al. 2020. Fungal Diversity 105:1-16 ((https://doi.org/10.1007/s13225-020-00466-2)) |
| Otu36928 | 1 | k__Fungi(100);p__Ascomycota(100);c__Saccharomycetes(100);o__Saccharomycetales(100);f__Metschnikowiaceae(100);g__Metschnikowia(100);s__Metschnikowia_pulcherrima(100); | Metschnikowia | 13 | Saprotroph | \|Undefined Saprotroph\| | Yeast | NULL | Probable | As nectar-tap_saprotroph (Põlme et al. 2020); Decay sugar-rich substrates (Põlme et al. 2020) | Põlme S, et al. 2020. Fungal Diversity 105:1-16 ((https://doi.org/10.1007/s13225-020-00466-2)) |
| Otu36929 | 1 | k__Fungi(100);p__Ascomycota(100);c__Saccharomycetes(100);o__Saccharomycetales(100);f__Metschnikowiaceae(100);g__Metschnikowia(100);s__Metschnikowia_pulcherrima(100); | Metschnikowia | 13 | Saprotroph | \|Undefined Saprotroph\| | Yeast | NULL | Probable | As nectar-tap_saprotroph (Põlme et al. 2020); Decay sugar-rich substrates (Põlme et al. 2020) | Põlme S, et al. 2020. Fungal Diversity 105:1-16 ((https://doi.org/10.1007/s13225-020-00466-2)) |
| Otu36930 | 1 | k__Fungi(100);p__Ascomycota(100);c__Saccharomycetes(100);o__Saccharomycetales(100);f__Metschnikowiaceae(100);g__Metschnikowia(100);g__Metschnikowia_unclassified(100); | Metschnikowia | 13 | Saprotroph | \|Undefined Saprotroph\| | Yeast | NULL | Probable | As nectar-tap_saprotroph (Põlme et al. 2020); Decay sugar-rich substrates (Põlme et al. 2020) | Põlme S, et al. 2020. Fungal Diversity 105:1-16 ((https://doi.org/10.1007/s13225-020-00466-2)) |
| Otu36931 | 1 | k__Fungi(100);p__Ascomycota(100);c__Saccharomycetes(100);o__Saccharomycetales(100);f__Metschnikowiaceae(100);g__Metschnikowia(100);s__Metschnikowia_pulcherrima(100); | Metschnikowia | 13 | Saprotroph | \|Undefined Saprotroph\| | Yeast | NULL | Probable | As nectar-tap_saprotroph (Põlme et al. 2020); Decay sugar-rich substrates (Põlme et al. 2020) | Põlme S, et al. 2020. Fungal Diversity 105:1-16 ((https://doi.org/10.1007/s13225-020-00466-2)) |
| Otu36932 | 1 | k__Fungi(100);p__Ascomycota(100);c__Saccharomycetes(100);o__Saccharomycetales(100);f__Metschnikowiaceae(100);g__Metschnikowia(100);s__Metschnikowia_pulcherrima(100); | Metschnikowia | 13 | Saprotroph | \|Undefined Saprotroph\| | Yeast | NULL | Probable | As nectar-tap_saprotroph (Põlme et al. 2020); Decay sugar-rich substrates (Põlme et al. 2020) | Põlme S, et al. 2020. Fungal Diversity 105:1-16 ((https://doi.org/10.1007/s13225-020-00466-2)) |
| Otu36933 | 1 | k__Fungi(100);p__Ascomycota(100);c__Saccharomycetes(100);o__Saccharomycetales(100);f__Metschnikowiaceae(100);g__Metschnikowia(100);g__Metschnikowia_unclassified(100); | Metschnikowia | 13 | Saprotroph | \|Undefined Saprotroph\| | Yeast | NULL | Probable | As nectar-tap_saprotroph (Põlme et al. 2020); Decay sugar-rich substrates (Põlme et al. 2020) | Põlme S, et al. 2020. Fungal Diversity 105:1-16 ((https://doi.org/10.1007/s13225-020-00466-2)) |
| Otu36934 | 1 | k__Fungi(100);p__Ascomycota(100);c__Saccharomycetes(100);o__Saccharomycetales(100);f__Metschnikowiaceae(100);g__Metschnikowia(100);g__Metschnikowia_unclassified(100); | Metschnikowia | 13 | Saprotroph | \|Undefined Saprotroph\| | Yeast | NULL | Probable | As nectar-tap_saprotroph (Põlme et al. 2020); Decay sugar-rich substrates (Põlme et al. 2020) | Põlme S, et al. 2020. Fungal Diversity 105:1-16 ((https://doi.org/10.1007/s13225-020-00466-2)) |
| Otu36935 | 1 | k__Fungi(100);p__Ascomycota(100);c__Saccharomycetes(100);o__Saccharomycetales(100);f__Metschnikowiaceae(100);g__Metschnikowia(100);g__Metschnikowia_unclassified(100); | Metschnikowia | 13 | Saprotroph | \|Undefined Saprotroph\| | Yeast | NULL | Probable | As nectar-tap_saprotroph (Põlme et al. 2020); Decay sugar-rich substrates (Põlme et al. 2020) | Põlme S, et al. 2020. Fungal Diversity 105:1-16 ((https://doi.org/10.1007/s13225-020-00466-2)) |
| Otu36936 | 1 | k__Fungi(100);p__Ascomycota(100);c__Sordariomycetes(100);o__Sordariomycetes_order_Incertae_sedis(100);f__Sordariomycetes_family_Incertae_sedis(100);g__Myrmecridium(100);s__Myrmecridium_sp(100); | Myrmecridium | 13 | Saprotroph | \|Undefined Saprotroph\| | Microfungus | NULL | Probable | As soil_saprotroph (Põlme et al. 2020); Decay soil (Põlme et al. 2020) | Tedersoo L, et al. 2014. Science 346:e1256688 ((https://doi.org/10.1126/science.1256688)); Põlme S, et al. 2020. Fungal Diversity 105:1-16 ((https://doi.org/10.1007/s13225-020-00466-2)) |
| Otu36937 | 1 | k__Fungi(100);p__Ascomycota(100);c__Saccharomycetes(100);o__Saccharomycetales(100);f__Metschnikowiaceae(100);g__Metschnikowia(100);g__Metschnikowia_unclassified(100); | Metschnikowia | 13 | Saprotroph | \|Undefined Saprotroph\| | Yeast | NULL | Probable | As nectar-tap_saprotroph (Põlme et al. 2020); Decay sugar-rich substrates (Põlme et al. 2020) | Põlme S, et al. 2020. Fungal Diversity 105:1-16 ((https://doi.org/10.1007/s13225-020-00466-2)) |
| Otu36938 | 1 | k__Fungi(100);p__Ascomycota(100);c__Saccharomycetes(100);o__Saccharomycetales(100);f__Metschnikowiaceae(100);g__Metschnikowia(100);s__Metschnikowia_pulcherrima(100); | Metschnikowia | 13 | Saprotroph | \|Undefined Saprotroph\| | Yeast | NULL | Probable | As nectar-tap_saprotroph (Põlme et al. 2020); Decay sugar-rich substrates (Põlme et al. 2020) | Põlme S, et al. 2020. Fungal Diversity 105:1-16 ((https://doi.org/10.1007/s13225-020-00466-2)) |
| Otu36939 | 1 | k__Fungi(100);p__Ascomycota(100);c__Saccharomycetes(100);o__Saccharomycetales(100);f__Metschnikowiaceae(100);g__Metschnikowia(100);s__Metschnikowia_pulcherrima(100); | Metschnikowia | 13 | Saprotroph | \|Undefined Saprotroph\| | Yeast | NULL | Probable | As nectar-tap_saprotroph (Põlme et al. 2020); Decay sugar-rich substrates (Põlme et al. 2020) | Põlme S, et al. 2020. Fungal Diversity 105:1-16 ((https://doi.org/10.1007/s13225-020-00466-2)) |
| Otu36940 | 1 | k__Fungi(100);p__Ascomycota(100);c__Saccharomycetes(100);o__Saccharomycetales(100);f__Metschnikowiaceae(100);g__Metschnikowia(100);g__Metschnikowia_unclassified(100); | Metschnikowia | 13 | Saprotroph | \|Undefined Saprotroph\| | Yeast | NULL | Probable | As nectar-tap_saprotroph (Põlme et al. 2020); Decay sugar-rich substrates (Põlme et al. 2020) | Põlme S, et al. 2020. Fungal Diversity 105:1-16 ((https://doi.org/10.1007/s13225-020-00466-2)) |
| Otu36941 | 1 | k__Fungi(100);p__Ascomycota(100);c__Saccharomycetes(100);o__Saccharomycetales(100);f__Metschnikowiaceae(100);g__Metschnikowia(100);s__Metschnikowia_pulcherrima(100); | Metschnikowia | 13 | Saprotroph | \|Undefined Saprotroph\| | Yeast | NULL | Probable | As nectar-tap_saprotroph (Põlme et al. 2020); Decay sugar-rich substrates (Põlme et al. 2020) | Põlme S, et al. 2020. Fungal Diversity 105:1-16 ((https://doi.org/10.1007/s13225-020-00466-2)) |
| Otu36942 | 1 | k__Fungi(100);p__Ascomycota(100);c__Saccharomycetes(100);o__Saccharomycetales(100);f__Metschnikowiaceae(100);g__Metschnikowia(100);s__Metschnikowia_pulcherrima(100); | Metschnikowia | 13 | Saprotroph | \|Undefined Saprotroph\| | Yeast | NULL | Probable | As nectar-tap_saprotroph (Põlme et al. 2020); Decay sugar-rich substrates (Põlme et al. 2020) | Põlme S, et al. 2020. Fungal Diversity 105:1-16 ((https://doi.org/10.1007/s13225-020-00466-2)) |
| Otu36943 | 1 | k__Fungi(100);p__Ascomycota(100);c__Saccharomycetes(100);o__Saccharomycetales(100);f__Metschnikowiaceae(100);g__Metschnikowia(100);g__Metschnikowia_unclassified(100); | Metschnikowia | 13 | Saprotroph | \|Undefined Saprotroph\| | Yeast | NULL | Probable | As nectar-tap_saprotroph (Põlme et al. 2020); Decay sugar-rich substrates (Põlme et al. 2020) | Põlme S, et al. 2020. Fungal Diversity 105:1-16 ((https://doi.org/10.1007/s13225-020-00466-2)) |
| Otu36944 | 1 | k__Fungi(100);p__Ascomycota(100);c__Saccharomycetes(100);o__Saccharomycetales(100);f__Metschnikowiaceae(100);g__Metschnikowia(100);s__Metschnikowia_pulcherrima(100); | Metschnikowia | 13 | Saprotroph | \|Undefined Saprotroph\| | Yeast | NULL | Probable | As nectar-tap_saprotroph (Põlme et al. 2020); Decay sugar-rich substrates (Põlme et al. 2020) | Põlme S, et al. 2020. Fungal Diversity 105:1-16 ((https://doi.org/10.1007/s13225-020-00466-2)) |
| Otu36945 | 1 | k__Fungi(100);p__Ascomycota(100);c__Saccharomycetes(100);o__Saccharomycetales(100);f__Metschnikowiaceae(100);g__Metschnikowia(100);s__Metschnikowia_pulcherrima(100); | Metschnikowia | 13 | Saprotroph | \|Undefined Saprotroph\| | Yeast | NULL | Probable | As nectar-tap_saprotroph (Põlme et al. 2020); Decay sugar-rich substrates (Põlme et al. 2020) | Põlme S, et al. 2020. Fungal Diversity 105:1-16 ((https://doi.org/10.1007/s13225-020-00466-2)) |
| Otu36946 | 1 | k__Fungi(100);p__Ascomycota(100);c__Saccharomycetes(100);o__Saccharomycetales(100);f__Metschnikowiaceae(100);g__Metschnikowia(100);s__Metschnikowia_pulcherrima(100); | Metschnikowia | 13 | Saprotroph | \|Undefined Saprotroph\| | Yeast | NULL | Probable | As nectar-tap_saprotroph (Põlme et al. 2020); Decay sugar-rich substrates (Põlme et al. 2020) | Põlme S, et al. 2020. Fungal Diversity 105:1-16 ((https://doi.org/10.1007/s13225-020-00466-2)) |
| Otu36947 | 1 | k__Fungi(100);p__Ascomycota(100);c__Saccharomycetes(100);o__Saccharomycetales(100);f__Metschnikowiaceae(100);g__Metschnikowia(100);g__Metschnikowia_unclassified(100); | Metschnikowia | 13 | Saprotroph | \|Undefined Saprotroph\| | Yeast | NULL | Probable | As nectar-tap_saprotroph (Põlme et al. 2020); Decay sugar-rich substrates (Põlme et al. 2020) | Põlme S, et al. 2020. Fungal Diversity 105:1-16 ((https://doi.org/10.1007/s13225-020-00466-2)) |
| Otu36948 | 1 | k__Fungi(100);p__Ascomycota(100);c__Saccharomycetes(100);o__Saccharomycetales(100);f__Metschnikowiaceae(100);g__Metschnikowia(100);s__Metschnikowia_pulcherrima(100); | Metschnikowia | 13 | Saprotroph | \|Undefined Saprotroph\| | Yeast | NULL | Probable | As nectar-tap_saprotroph (Põlme et al. 2020); Decay sugar-rich substrates (Põlme et al. 2020) | Põlme S, et al. 2020. Fungal Diversity 105:1-16 ((https://doi.org/10.1007/s13225-020-00466-2)) |
| Otu36949 | 1 | k__Fungi(100);p__Ascomycota(100);c__Saccharomycetes(100);o__Saccharomycetales(100);f__Metschnikowiaceae(100);g__Metschnikowia(100);s__Metschnikowia_pulcherrima(100); | Metschnikowia | 13 | Saprotroph | \|Undefined Saprotroph\| | Yeast | NULL | Probable | As nectar-tap_saprotroph (Põlme et al. 2020); Decay sugar-rich substrates (Põlme et al. 2020) | Põlme S, et al. 2020. Fungal Diversity 105:1-16 ((https://doi.org/10.1007/s13225-020-00466-2)) |
| Otu36950 | 1 | k__Fungi(100);p__Ascomycota(100);c__Saccharomycetes(100);o__Saccharomycetales(100);f__Metschnikowiaceae(100);g__Metschnikowia(100);s__Metschnikowia_pulcherrima(100); | Metschnikowia | 13 | Saprotroph | \|Undefined Saprotroph\| | Yeast | NULL | Probable | As nectar-tap_saprotroph (Põlme et al. 2020); Decay sugar-rich substrates (Põlme et al. 2020) | Põlme S, et al. 2020. Fungal Diversity 105:1-16 ((https://doi.org/10.1007/s13225-020-00466-2)) |
| Otu36951 | 1 | k__Fungi(100);p__Ascomycota(100);c__Saccharomycetes(100);o__Saccharomycetales(100);f__Metschnikowiaceae(100);g__Metschnikowia(100);s__Metschnikowia_pulcherrima(100); | Metschnikowia | 13 | Saprotroph | \|Undefined Saprotroph\| | Yeast | NULL | Probable | As nectar-tap_saprotroph (Põlme et al. 2020); Decay sugar-rich substrates (Põlme et al. 2020) | Põlme S, et al. 2020. Fungal Diversity 105:1-16 ((https://doi.org/10.1007/s13225-020-00466-2)) |
| Otu36952 | 1 | k__Fungi(100);p__Ascomycota(100);c__Saccharomycetes(100);o__Saccharomycetales(100);f__Metschnikowiaceae(100);g__Metschnikowia(100);s__Metschnikowia_pulcherrima(100); | Metschnikowia | 13 | Saprotroph | \|Undefined Saprotroph\| | Yeast | NULL | Probable | As nectar-tap_saprotroph (Põlme et al. 2020); Decay sugar-rich substrates (Põlme et al. 2020) | Põlme S, et al. 2020. Fungal Diversity 105:1-16 ((https://doi.org/10.1007/s13225-020-00466-2)) |
| Otu36953 | 1 | k__Fungi(100);p__Ascomycota(100);c__Saccharomycetes(100);o__Saccharomycetales(100);f__Metschnikowiaceae(100);g__Metschnikowia(100);g__Metschnikowia_unclassified(100); | Metschnikowia | 13 | Saprotroph | \|Undefined Saprotroph\| | Yeast | NULL | Probable | As nectar-tap_saprotroph (Põlme et al. 2020); Decay sugar-rich substrates (Põlme et al. 2020) | Põlme S, et al. 2020. Fungal Diversity 105:1-16 ((https://doi.org/10.1007/s13225-020-00466-2)) |
| Otu36954 | 1 | k__Fungi(100);p__Ascomycota(100);c__Saccharomycetes(100);o__Saccharomycetales(100);f__Metschnikowiaceae(100);g__Metschnikowia(100);s__Metschnikowia_pulcherrima(100); | Metschnikowia | 13 | Saprotroph | \|Undefined Saprotroph\| | Yeast | NULL | Probable | As nectar-tap_saprotroph (Põlme et al. 2020); Decay sugar-rich substrates (Põlme et al. 2020) | Põlme S, et al. 2020. Fungal Diversity 105:1-16 ((https://doi.org/10.1007/s13225-020-00466-2)) |
| Otu36956 | 1 | k__Fungi(100);p__Ascomycota(100);c__Saccharomycetes(100);o__Saccharomycetales(100);f__Metschnikowiaceae(100);g__Metschnikowia(100);g__Metschnikowia_unclassified(100); | Metschnikowia | 13 | Saprotroph | \|Undefined Saprotroph\| | Yeast | NULL | Probable | As nectar-tap_saprotroph (Põlme et al. 2020); Decay sugar-rich substrates (Põlme et al. 2020) | Põlme S, et al. 2020. Fungal Diversity 105:1-16 ((https://doi.org/10.1007/s13225-020-00466-2)) |
| Otu36957 | 1 | k__Fungi(100);p__Ascomycota(100);c__Saccharomycetes(100);o__Saccharomycetales(100);f__Metschnikowiaceae(100);g__Metschnikowia(100);s__Metschnikowia_pulcherrima(100); | Metschnikowia | 13 | Saprotroph | \|Undefined Saprotroph\| | Yeast | NULL | Probable | As nectar-tap_saprotroph (Põlme et al. 2020); Decay sugar-rich substrates (Põlme et al. 2020) | Põlme S, et al. 2020. Fungal Diversity 105:1-16 ((https://doi.org/10.1007/s13225-020-00466-2)) |
| Otu36958 | 1 | k__Fungi(100);p__Ascomycota(100);c__Saccharomycetes(100);o__Saccharomycetales(100);f__Metschnikowiaceae(100);g__Metschnikowia(100);g__Metschnikowia_unclassified(100); | Metschnikowia | 13 | Saprotroph | \|Undefined Saprotroph\| | Yeast | NULL | Probable | As nectar-tap_saprotroph (Põlme et al. 2020); Decay sugar-rich substrates (Põlme et al. 2020) | Põlme S, et al. 2020. Fungal Diversity 105:1-16 ((https://doi.org/10.1007/s13225-020-00466-2)) |
| Otu36960 | 1 | k__Fungi(100);p__Ascomycota(100);c__Saccharomycetes(100);o__Saccharomycetales(100);f__Metschnikowiaceae(100);g__Metschnikowia(100);s__Metschnikowia_pulcherrima(100); | Metschnikowia | 13 | Saprotroph | \|Undefined Saprotroph\| | Yeast | NULL | Probable | As nectar-tap_saprotroph (Põlme et al. 2020); Decay sugar-rich substrates (Põlme et al. 2020) | Põlme S, et al. 2020. Fungal Diversity 105:1-16 ((https://doi.org/10.1007/s13225-020-00466-2)) |
| Otu36961 | 1 | k__Fungi(100);p__Ascomycota(100);c__Saccharomycetes(100);o__Saccharomycetales(100);f__Metschnikowiaceae(100);g__Metschnikowia(100);s__Metschnikowia_pulcherrima(100); | Metschnikowia | 13 | Saprotroph | \|Undefined Saprotroph\| | Yeast | NULL | Probable | As nectar-tap_saprotroph (Põlme et al. 2020); Decay sugar-rich substrates (Põlme et al. 2020) | Põlme S, et al. 2020. Fungal Diversity 105:1-16 ((https://doi.org/10.1007/s13225-020-00466-2)) |
| Otu36963 | 1 | k__Fungi(100);p__Ascomycota(100);c__Saccharomycetes(100);o__Saccharomycetales(100);f__Metschnikowiaceae(100);g__Metschnikowia(100);s__Metschnikowia_pulcherrima(100); | Metschnikowia | 13 | Saprotroph | \|Undefined Saprotroph\| | Yeast | NULL | Probable | As nectar-tap_saprotroph (Põlme et al. 2020); Decay sugar-rich substrates (Põlme et al. 2020) | Põlme S, et al. 2020. Fungal Diversity 105:1-16 ((https://doi.org/10.1007/s13225-020-00466-2)) |
| Otu36964 | 1 | k__Fungi(100);p__Ascomycota(100);c__Saccharomycetes(100);o__Saccharomycetales(100);f__Metschnikowiaceae(100);g__Metschnikowia(100);s__Metschnikowia_pulcherrima(100); | Metschnikowia | 13 | Saprotroph | \|Undefined Saprotroph\| | Yeast | NULL | Probable | As nectar-tap_saprotroph (Põlme et al. 2020); Decay sugar-rich substrates (Põlme et al. 2020) | Põlme S, et al. 2020. Fungal Diversity 105:1-16 ((https://doi.org/10.1007/s13225-020-00466-2)) |
| Otu36965 | 1 | k__Fungi(100);p__Ascomycota(100);c__Saccharomycetes(100);o__Saccharomycetales(100);f__Metschnikowiaceae(100);g__Metschnikowia(100);s__Metschnikowia_pulcherrima(100); | Metschnikowia | 13 | Saprotroph | \|Undefined Saprotroph\| | Yeast | NULL | Probable | As nectar-tap_saprotroph (Põlme et al. 2020); Decay sugar-rich substrates (Põlme et al. 2020) | Põlme S, et al. 2020. Fungal Diversity 105:1-16 ((https://doi.org/10.1007/s13225-020-00466-2)) |
| Otu36966 | 1 | k__Fungi(100);p__Ascomycota(100);c__Saccharomycetes(100);o__Saccharomycetales(100);f__Metschnikowiaceae(100);g__Metschnikowia(100);g__Metschnikowia_unclassified(100); | Metschnikowia | 13 | Saprotroph | \|Undefined Saprotroph\| | Yeast | NULL | Probable | As nectar-tap_saprotroph (Põlme et al. 2020); Decay sugar-rich substrates (Põlme et al. 2020) | Põlme S, et al. 2020. Fungal Diversity 105:1-16 ((https://doi.org/10.1007/s13225-020-00466-2)) |
| Otu36967 | 1 | k__Fungi(100);p__Ascomycota(100);c__Saccharomycetes(100);o__Saccharomycetales(100);f__Metschnikowiaceae(100);g__Metschnikowia(100);s__Metschnikowia_pulcherrima(100); | Metschnikowia | 13 | Saprotroph | \|Undefined Saprotroph\| | Yeast | NULL | Probable | As nectar-tap_saprotroph (Põlme et al. 2020); Decay sugar-rich substrates (Põlme et al. 2020) | Põlme S, et al. 2020. Fungal Diversity 105:1-16 ((https://doi.org/10.1007/s13225-020-00466-2)) |
| Otu36970 | 1 | k__Fungi(100);p__Ascomycota(100);c__Saccharomycetes(100);o__Saccharomycetales(100);f__Metschnikowiaceae(100);g__Metschnikowia(100);s__Metschnikowia_pulcherrima(100); | Metschnikowia | 13 | Saprotroph | \|Undefined Saprotroph\| | Yeast | NULL | Probable | As nectar-tap_saprotroph (Põlme et al. 2020); Decay sugar-rich substrates (Põlme et al. 2020) | Põlme S, et al. 2020. Fungal Diversity 105:1-16 ((https://doi.org/10.1007/s13225-020-00466-2)) |
| Otu36971 | 1 | k__Fungi(100);p__Ascomycota(100);c__Saccharomycetes(100);o__Saccharomycetales(100);f__Metschnikowiaceae(100);g__Metschnikowia(100);s__Metschnikowia_pulcherrima(100); | Metschnikowia | 13 | Saprotroph | \|Undefined Saprotroph\| | Yeast | NULL | Probable | As nectar-tap_saprotroph (Põlme et al. 2020); Decay sugar-rich substrates (Põlme et al. 2020) | Põlme S, et al. 2020. Fungal Diversity 105:1-16 ((https://doi.org/10.1007/s13225-020-00466-2)) |
| Otu36972 | 1 | k__Fungi(100);p__Ascomycota(100);c__Saccharomycetes(100);o__Saccharomycetales(100);f__Metschnikowiaceae(100);g__Metschnikowia(100);s__Metschnikowia_pulcherrima(100); | Metschnikowia | 13 | Saprotroph | \|Undefined Saprotroph\| | Yeast | NULL | Probable | As nectar-tap_saprotroph (Põlme et al. 2020); Decay sugar-rich substrates (Põlme et al. 2020) | Põlme S, et al. 2020. Fungal Diversity 105:1-16 ((https://doi.org/10.1007/s13225-020-00466-2)) |
| Otu36973 | 1 | k__Fungi(100);p__Ascomycota(100);c__Saccharomycetes(100);o__Saccharomycetales(100);f__Metschnikowiaceae(100);g__Metschnikowia(100);s__Metschnikowia_pulcherrima(100); | Metschnikowia | 13 | Saprotroph | \|Undefined Saprotroph\| | Yeast | NULL | Probable | As nectar-tap_saprotroph (Põlme et al. 2020); Decay sugar-rich substrates (Põlme et al. 2020) | Põlme S, et al. 2020. Fungal Diversity 105:1-16 ((https://doi.org/10.1007/s13225-020-00466-2)) |
| Otu36974 | 1 | k__Fungi(100);p__Ascomycota(100);c__Saccharomycetes(100);o__Saccharomycetales(100);f__Metschnikowiaceae(100);g__Metschnikowia(100);s__Metschnikowia_pulcherrima(100); | Metschnikowia | 13 | Saprotroph | \|Undefined Saprotroph\| | Yeast | NULL | Probable | As nectar-tap_saprotroph (Põlme et al. 2020); Decay sugar-rich substrates (Põlme et al. 2020) | Põlme S, et al. 2020. Fungal Diversity 105:1-16 ((https://doi.org/10.1007/s13225-020-00466-2)) |
| Otu36975 | 1 | k__Fungi(100);p__Ascomycota(100);c__Saccharomycetes(100);o__Saccharomycetales(100);f__Metschnikowiaceae(100);g__Metschnikowia(100);g__Metschnikowia_unclassified(100); | Metschnikowia | 13 | Saprotroph | \|Undefined Saprotroph\| | Yeast | NULL | Probable | As nectar-tap_saprotroph (Põlme et al. 2020); Decay sugar-rich substrates (Põlme et al. 2020) | Põlme S, et al. 2020. Fungal Diversity 105:1-16 ((https://doi.org/10.1007/s13225-020-00466-2)) |
| Otu36976 | 1 | k__Fungi(100);p__Ascomycota(100);c__Saccharomycetes(100);o__Saccharomycetales(100);f__Metschnikowiaceae(100);g__Metschnikowia(100);s__Metschnikowia_pulcherrima(100); | Metschnikowia | 13 | Saprotroph | \|Undefined Saprotroph\| | Yeast | NULL | Probable | As nectar-tap_saprotroph (Põlme et al. 2020); Decay sugar-rich substrates (Põlme et al. 2020) | Põlme S, et al. 2020. Fungal Diversity 105:1-16 ((https://doi.org/10.1007/s13225-020-00466-2)) |
| Otu36977 | 1 | k__Fungi(100);p__Ascomycota(100);c__Saccharomycetes(100);o__Saccharomycetales(100);f__Metschnikowiaceae(100);g__Metschnikowia(100);s__Metschnikowia_pulcherrima(100); | Metschnikowia | 13 | Saprotroph | \|Undefined Saprotroph\| | Yeast | NULL | Probable | As nectar-tap_saprotroph (Põlme et al. 2020); Decay sugar-rich substrates (Põlme et al. 2020) | Põlme S, et al. 2020. Fungal Diversity 105:1-16 ((https://doi.org/10.1007/s13225-020-00466-2)) |
| Otu36978 | 1 | k__Fungi(100);p__Ascomycota(100);c__Saccharomycetes(100);o__Saccharomycetales(100);f__Metschnikowiaceae(100);g__Metschnikowia(100);g__Metschnikowia_unclassified(100); | Metschnikowia | 13 | Saprotroph | \|Undefined Saprotroph\| | Yeast | NULL | Probable | As nectar-tap_saprotroph (Põlme et al. 2020); Decay sugar-rich substrates (Põlme et al. 2020) | Põlme S, et al. 2020. Fungal Diversity 105:1-16 ((https://doi.org/10.1007/s13225-020-00466-2)) |
| Otu36979 | 1 | k__Fungi(100);p__Ascomycota(100);c__Saccharomycetes(100);o__Saccharomycetales(100);f__Metschnikowiaceae(100);g__Metschnikowia(100);g__Metschnikowia_unclassified(100); | Metschnikowia | 13 | Saprotroph | \|Undefined Saprotroph\| | Yeast | NULL | Probable | As nectar-tap_saprotroph (Põlme et al. 2020); Decay sugar-rich substrates (Põlme et al. 2020) | Põlme S, et al. 2020. Fungal Diversity 105:1-16 ((https://doi.org/10.1007/s13225-020-00466-2)) |
| Otu36980 | 1 | k__Fungi(100);p__Ascomycota(100);c__Saccharomycetes(100);o__Saccharomycetales(100);f__Metschnikowiaceae(100);g__Metschnikowia(100);s__Metschnikowia_pulcherrima(100); | Metschnikowia | 13 | Saprotroph | \|Undefined Saprotroph\| | Yeast | NULL | Probable | As nectar-tap_saprotroph (Põlme et al. 2020); Decay sugar-rich substrates (Põlme et al. 2020) | Põlme S, et al. 2020. Fungal Diversity 105:1-16 ((https://doi.org/10.1007/s13225-020-00466-2)) |
| Otu36981 | 1 | k__Fungi(100);p__Ascomycota(100);c__Saccharomycetes(100);o__Saccharomycetales(100);f__Metschnikowiaceae(100);g__Metschnikowia(100);g__Metschnikowia_unclassified(100); | Metschnikowia | 13 | Saprotroph | \|Undefined Saprotroph\| | Yeast | NULL | Probable | As nectar-tap_saprotroph (Põlme et al. 2020); Decay sugar-rich substrates (Põlme et al. 2020) | Põlme S, et al. 2020. Fungal Diversity 105:1-16 ((https://doi.org/10.1007/s13225-020-00466-2)) |
| Otu36982 | 1 | k__Fungi(100);p__Ascomycota(100);c__Saccharomycetes(100);o__Saccharomycetales(100);f__Metschnikowiaceae(100);g__Metschnikowia(100);s__Metschnikowia_pulcherrima(100); | Metschnikowia | 13 | Saprotroph | \|Undefined Saprotroph\| | Yeast | NULL | Probable | As nectar-tap_saprotroph (Põlme et al. 2020); Decay sugar-rich substrates (Põlme et al. 2020) | Põlme S, et al. 2020. Fungal Diversity 105:1-16 ((https://doi.org/10.1007/s13225-020-00466-2)) |
| Otu36983 | 1 | k__Fungi(100);p__Ascomycota(100);c__Saccharomycetes(100);o__Saccharomycetales(100);f__Metschnikowiaceae(100);g__Metschnikowia(100);s__Metschnikowia_pulcherrima(100); | Metschnikowia | 13 | Saprotroph | \|Undefined Saprotroph\| | Yeast | NULL | Probable | As nectar-tap_saprotroph (Põlme et al. 2020); Decay sugar-rich substrates (Põlme et al. 2020) | Põlme S, et al. 2020. Fungal Diversity 105:1-16 ((https://doi.org/10.1007/s13225-020-00466-2)) |
| Otu36984 | 1 | k__Fungi(100);p__Ascomycota(100);c__Saccharomycetes(100);o__Saccharomycetales(100);f__Metschnikowiaceae(100);g__Metschnikowia(100);s__Metschnikowia_pulcherrima(100); | Metschnikowia | 13 | Saprotroph | \|Undefined Saprotroph\| | Yeast | NULL | Probable | As nectar-tap_saprotroph (Põlme et al. 2020); Decay sugar-rich substrates (Põlme et al. 2020) | Põlme S, et al. 2020. Fungal Diversity 105:1-16 ((https://doi.org/10.1007/s13225-020-00466-2)) |
| Otu36985 | 1 | k__Fungi(100);p__Ascomycota(100);c__Saccharomycetes(100);o__Saccharomycetales(100);f__Metschnikowiaceae(100);g__Metschnikowia(100);s__Metschnikowia_pulcherrima(100); | Metschnikowia | 13 | Saprotroph | \|Undefined Saprotroph\| | Yeast | NULL | Probable | As nectar-tap_saprotroph (Põlme et al. 2020); Decay sugar-rich substrates (Põlme et al. 2020) | Põlme S, et al. 2020. Fungal Diversity 105:1-16 ((https://doi.org/10.1007/s13225-020-00466-2)) |
| Otu36986 | 1 | k__Fungi(100);p__Ascomycota(100);c__Saccharomycetes(100);o__Saccharomycetales(100);f__Metschnikowiaceae(100);g__Metschnikowia(100);s__Metschnikowia_pulcherrima(100); | Metschnikowia | 13 | Saprotroph | \|Undefined Saprotroph\| | Yeast | NULL | Probable | As nectar-tap_saprotroph (Põlme et al. 2020); Decay sugar-rich substrates (Põlme et al. 2020) | Põlme S, et al. 2020. Fungal Diversity 105:1-16 ((https://doi.org/10.1007/s13225-020-00466-2)) |
| Otu36987 | 1 | k__Fungi(100);p__Ascomycota(100);c__Saccharomycetes(100);o__Saccharomycetales(100);f__Metschnikowiaceae(100);g__Metschnikowia(100);g__Metschnikowia_unclassified(100); | Metschnikowia | 13 | Saprotroph | \|Undefined Saprotroph\| | Yeast | NULL | Probable | As nectar-tap_saprotroph (Põlme et al. 2020); Decay sugar-rich substrates (Põlme et al. 2020) | Põlme S, et al. 2020. Fungal Diversity 105:1-16 ((https://doi.org/10.1007/s13225-020-00466-2)) |
| Otu36988 | 1 | k__Fungi(100);p__Ascomycota(100);c__Saccharomycetes(100);o__Saccharomycetales(100);f__Metschnikowiaceae(100);g__Metschnikowia(100);s__Metschnikowia_pulcherrima(100); | Metschnikowia | 13 | Saprotroph | \|Undefined Saprotroph\| | Yeast | NULL | Probable | As nectar-tap_saprotroph (Põlme et al. 2020); Decay sugar-rich substrates (Põlme et al. 2020) | Põlme S, et al. 2020. Fungal Diversity 105:1-16 ((https://doi.org/10.1007/s13225-020-00466-2)) |
| Otu36989 | 1 | k__Fungi(100);p__Ascomycota(100);c__Saccharomycetes(100);o__Saccharomycetales(100);f__Metschnikowiaceae(100);g__Metschnikowia(100);s__Metschnikowia_pulcherrima(100); | Metschnikowia | 13 | Saprotroph | \|Undefined Saprotroph\| | Yeast | NULL | Probable | As nectar-tap_saprotroph (Põlme et al. 2020); Decay sugar-rich substrates (Põlme et al. 2020) | Põlme S, et al. 2020. Fungal Diversity 105:1-16 ((https://doi.org/10.1007/s13225-020-00466-2)) |
| Otu36990 | 1 | k__Fungi(100);p__Ascomycota(100);c__Saccharomycetes(100);o__Saccharomycetales(100);f__Metschnikowiaceae(100);g__Metschnikowia(100);s__Metschnikowia_pulcherrima(100); | Metschnikowia | 13 | Saprotroph | \|Undefined Saprotroph\| | Yeast | NULL | Probable | As nectar-tap_saprotroph (Põlme et al. 2020); Decay sugar-rich substrates (Põlme et al. 2020) | Põlme S, et al. 2020. Fungal Diversity 105:1-16 ((https://doi.org/10.1007/s13225-020-00466-2)) |
| Otu36991 | 1 | k__Fungi(100);p__Ascomycota(100);c__Saccharomycetes(100);o__Saccharomycetales(100);f__Metschnikowiaceae(100);g__Metschnikowia(100);s__Metschnikowia_pulcherrima(100); | Metschnikowia | 13 | Saprotroph | \|Undefined Saprotroph\| | Yeast | NULL | Probable | As nectar-tap_saprotroph (Põlme et al. 2020); Decay sugar-rich substrates (Põlme et al. 2020) | Põlme S, et al. 2020. Fungal Diversity 105:1-16 ((https://doi.org/10.1007/s13225-020-00466-2)) |
| Otu36992 | 1 | k__Fungi(100);p__Ascomycota(100);c__Saccharomycetes(100);o__Saccharomycetales(100);f__Metschnikowiaceae(100);g__Metschnikowia(100);s__Metschnikowia_pulcherrima(100); | Metschnikowia | 13 | Saprotroph | \|Undefined Saprotroph\| | Yeast | NULL | Probable | As nectar-tap_saprotroph (Põlme et al. 2020); Decay sugar-rich substrates (Põlme et al. 2020) | Põlme S, et al. 2020. Fungal Diversity 105:1-16 ((https://doi.org/10.1007/s13225-020-00466-2)) |
| Otu36993 | 1 | k__Fungi(100);p__Ascomycota(100);c__Saccharomycetes(100);o__Saccharomycetales(100);f__Metschnikowiaceae(100);g__Metschnikowia(100);s__Metschnikowia_pulcherrima(100); | Metschnikowia | 13 | Saprotroph | \|Undefined Saprotroph\| | Yeast | NULL | Probable | As nectar-tap_saprotroph (Põlme et al. 2020); Decay sugar-rich substrates (Põlme et al. 2020) | Põlme S, et al. 2020. Fungal Diversity 105:1-16 ((https://doi.org/10.1007/s13225-020-00466-2)) |
| Otu36994 | 1 | k__Fungi(100);p__Ascomycota(100);c__Saccharomycetes(100);o__Saccharomycetales(100);f__Metschnikowiaceae(100);g__Metschnikowia(100);g__Metschnikowia_unclassified(100); | Metschnikowia | 13 | Saprotroph | \|Undefined Saprotroph\| | Yeast | NULL | Probable | As nectar-tap_saprotroph (Põlme et al. 2020); Decay sugar-rich substrates (Põlme et al. 2020) | Põlme S, et al. 2020. Fungal Diversity 105:1-16 ((https://doi.org/10.1007/s13225-020-00466-2)) |
| Otu36995 | 1 | k__Fungi(100);p__Ascomycota(100);c__Saccharomycetes(100);o__Saccharomycetales(100);f__Metschnikowiaceae(100);g__Metschnikowia(100);g__Metschnikowia_unclassified(100); | Metschnikowia | 13 | Saprotroph | \|Undefined Saprotroph\| | Yeast | NULL | Probable | As nectar-tap_saprotroph (Põlme et al. 2020); Decay sugar-rich substrates (Põlme et al. 2020) | Põlme S, et al. 2020. Fungal Diversity 105:1-16 ((https://doi.org/10.1007/s13225-020-00466-2)) |
| Otu36996 | 1 | k__Fungi(100);p__Ascomycota(100);c__Saccharomycetes(100);o__Saccharomycetales(100);f__Metschnikowiaceae(100);g__Metschnikowia(100);s__Metschnikowia_pulcherrima(100); | Metschnikowia | 13 | Saprotroph | \|Undefined Saprotroph\| | Yeast | NULL | Probable | As nectar-tap_saprotroph (Põlme et al. 2020); Decay sugar-rich substrates (Põlme et al. 2020) | Põlme S, et al. 2020. Fungal Diversity 105:1-16 ((https://doi.org/10.1007/s13225-020-00466-2)) |
| Otu36997 | 1 | k__Fungi(100);p__Ascomycota(100);c__Saccharomycetes(100);o__Saccharomycetales(100);f__Metschnikowiaceae(100);g__Metschnikowia(100);s__Metschnikowia_pulcherrima(100); | Metschnikowia | 13 | Saprotroph | \|Undefined Saprotroph\| | Yeast | NULL | Probable | As nectar-tap_saprotroph (Põlme et al. 2020); Decay sugar-rich substrates (Põlme et al. 2020) | Põlme S, et al. 2020. Fungal Diversity 105:1-16 ((https://doi.org/10.1007/s13225-020-00466-2)) |
| Otu36998 | 1 | k__Fungi(100);p__Ascomycota(100);c__Saccharomycetes(100);o__Saccharomycetales(100);f__Metschnikowiaceae(100);g__Metschnikowia(100);g__Metschnikowia_unclassified(100); | Metschnikowia | 13 | Saprotroph | \|Undefined Saprotroph\| | Yeast | NULL | Probable | As nectar-tap_saprotroph (Põlme et al. 2020); Decay sugar-rich substrates (Põlme et al. 2020) | Põlme S, et al. 2020. Fungal Diversity 105:1-16 ((https://doi.org/10.1007/s13225-020-00466-2)) |
| Otu36999 | 1 | k__Fungi(100);p__Ascomycota(100);c__Saccharomycetes(100);o__Saccharomycetales(100);f__Metschnikowiaceae(100);g__Metschnikowia(100);s__Metschnikowia_pulcherrima(100); | Metschnikowia | 13 | Saprotroph | \|Undefined Saprotroph\| | Yeast | NULL | Probable | As nectar-tap_saprotroph (Põlme et al. 2020); Decay sugar-rich substrates (Põlme et al. 2020) | Põlme S, et al. 2020. Fungal Diversity 105:1-16 ((https://doi.org/10.1007/s13225-020-00466-2)) |
| Otu37001 | 1 | k__Fungi(100);p__Ascomycota(100);c__Saccharomycetes(100);o__Saccharomycetales(100);f__Metschnikowiaceae(100);g__Metschnikowia(100);s__Metschnikowia_pulcherrima(100); | Metschnikowia | 13 | Saprotroph | \|Undefined Saprotroph\| | Yeast | NULL | Probable | As nectar-tap_saprotroph (Põlme et al. 2020); Decay sugar-rich substrates (Põlme et al. 2020) | Põlme S, et al. 2020. Fungal Diversity 105:1-16 ((https://doi.org/10.1007/s13225-020-00466-2)) |
| Otu37002 | 1 | k__Fungi(100);p__Ascomycota(100);c__Saccharomycetes(100);o__Saccharomycetales(100);f__Metschnikowiaceae(100);g__Metschnikowia(100);s__Metschnikowia_pulcherrima(100); | Metschnikowia | 13 | Saprotroph | \|Undefined Saprotroph\| | Yeast | NULL | Probable | As nectar-tap_saprotroph (Põlme et al. 2020); Decay sugar-rich substrates (Põlme et al. 2020) | Põlme S, et al. 2020. Fungal Diversity 105:1-16 ((https://doi.org/10.1007/s13225-020-00466-2)) |
| Otu37003 | 1 | k__Fungi(100);p__Ascomycota(100);c__Saccharomycetes(100);o__Saccharomycetales(100);f__Metschnikowiaceae(100);g__Metschnikowia(100);s__Metschnikowia_pulcherrima(100); | Metschnikowia | 13 | Saprotroph | \|Undefined Saprotroph\| | Yeast | NULL | Probable | As nectar-tap_saprotroph (Põlme et al. 2020); Decay sugar-rich substrates (Põlme et al. 2020) | Põlme S, et al. 2020. Fungal Diversity 105:1-16 ((https://doi.org/10.1007/s13225-020-00466-2)) |
| Otu37004 | 1 | k__Fungi(100);p__Ascomycota(100);c__Saccharomycetes(100);o__Saccharomycetales(100);f__Metschnikowiaceae(100);g__Metschnikowia(100);s__Metschnikowia_pulcherrima(100); | Metschnikowia | 13 | Saprotroph | \|Undefined Saprotroph\| | Yeast | NULL | Probable | As nectar-tap_saprotroph (Põlme et al. 2020); Decay sugar-rich substrates (Põlme et al. 2020) | Põlme S, et al. 2020. Fungal Diversity 105:1-16 ((https://doi.org/10.1007/s13225-020-00466-2)) |
| Otu37005 | 1 | k__Fungi(100);p__Ascomycota(100);c__Saccharomycetes(100);o__Saccharomycetales(100);f__Metschnikowiaceae(100);g__Metschnikowia(100);g__Metschnikowia_unclassified(100); | Metschnikowia | 13 | Saprotroph | \|Undefined Saprotroph\| | Yeast | NULL | Probable | As nectar-tap_saprotroph (Põlme et al. 2020); Decay sugar-rich substrates (Põlme et al. 2020) | Põlme S, et al. 2020. Fungal Diversity 105:1-16 ((https://doi.org/10.1007/s13225-020-00466-2)) |
| Otu37006 | 1 | k__Fungi(100);p__Ascomycota(100);c__Saccharomycetes(100);o__Saccharomycetales(100);f__Metschnikowiaceae(100);g__Metschnikowia(100);s__Metschnikowia_pulcherrima(100); | Metschnikowia | 13 | Saprotroph | \|Undefined Saprotroph\| | Yeast | NULL | Probable | As nectar-tap_saprotroph (Põlme et al. 2020); Decay sugar-rich substrates (Põlme et al. 2020) | Põlme S, et al. 2020. Fungal Diversity 105:1-16 ((https://doi.org/10.1007/s13225-020-00466-2)) |
| Otu37008 | 1 | k__Fungi(100);p__Ascomycota(100);c__Saccharomycetes(100);o__Saccharomycetales(100);f__Metschnikowiaceae(100);g__Metschnikowia(100);s__Metschnikowia_pulcherrima(100); | Metschnikowia | 13 | Saprotroph | \|Undefined Saprotroph\| | Yeast | NULL | Probable | As nectar-tap_saprotroph (Põlme et al. 2020); Decay sugar-rich substrates (Põlme et al. 2020) | Põlme S, et al. 2020. Fungal Diversity 105:1-16 ((https://doi.org/10.1007/s13225-020-00466-2)) |
| Otu37009 | 1 | k__Fungi(100);p__Ascomycota(100);c__Saccharomycetes(100);o__Saccharomycetales(100);f__Metschnikowiaceae(100);g__Metschnikowia(100);s__Metschnikowia_pulcherrima(100); | Metschnikowia | 13 | Saprotroph | \|Undefined Saprotroph\| | Yeast | NULL | Probable | As nectar-tap_saprotroph (Põlme et al. 2020); Decay sugar-rich substrates (Põlme et al. 2020) | Põlme S, et al. 2020. Fungal Diversity 105:1-16 ((https://doi.org/10.1007/s13225-020-00466-2)) |
| Otu37010 | 1 | k__Fungi(100);p__Ascomycota(100);c__Saccharomycetes(100);o__Saccharomycetales(100);f__Metschnikowiaceae(100);g__Metschnikowia(100);s__Metschnikowia_pulcherrima(100); | Metschnikowia | 13 | Saprotroph | \|Undefined Saprotroph\| | Yeast | NULL | Probable | As nectar-tap_saprotroph (Põlme et al. 2020); Decay sugar-rich substrates (Põlme et al. 2020) | Põlme S, et al. 2020. Fungal Diversity 105:1-16 ((https://doi.org/10.1007/s13225-020-00466-2)) |
| Otu37011 | 1 | k__Fungi(100);p__Ascomycota(100);c__Saccharomycetes(100);o__Saccharomycetales(100);f__Metschnikowiaceae(100);g__Metschnikowia(100);s__Metschnikowia_pulcherrima(100); | Metschnikowia | 13 | Saprotroph | \|Undefined Saprotroph\| | Yeast | NULL | Probable | As nectar-tap_saprotroph (Põlme et al. 2020); Decay sugar-rich substrates (Põlme et al. 2020) | Põlme S, et al. 2020. Fungal Diversity 105:1-16 ((https://doi.org/10.1007/s13225-020-00466-2)) |
| Otu37012 | 1 | k__Fungi(100);p__Ascomycota(100);c__Saccharomycetes(100);o__Saccharomycetales(100);f__Metschnikowiaceae(100);g__Metschnikowia(100);s__Metschnikowia_pulcherrima(100); | Metschnikowia | 13 | Saprotroph | \|Undefined Saprotroph\| | Yeast | NULL | Probable | As nectar-tap_saprotroph (Põlme et al. 2020); Decay sugar-rich substrates (Põlme et al. 2020) | Põlme S, et al. 2020. Fungal Diversity 105:1-16 ((https://doi.org/10.1007/s13225-020-00466-2)) |
| Otu37013 | 1 | k__Fungi(100);p__Ascomycota(100);c__Saccharomycetes(100);o__Saccharomycetales(100);f__Metschnikowiaceae(100);g__Metschnikowia(100);s__Metschnikowia_pulcherrima(100); | Metschnikowia | 13 | Saprotroph | \|Undefined Saprotroph\| | Yeast | NULL | Probable | As nectar-tap_saprotroph (Põlme et al. 2020); Decay sugar-rich substrates (Põlme et al. 2020) | Põlme S, et al. 2020. Fungal Diversity 105:1-16 ((https://doi.org/10.1007/s13225-020-00466-2)) |
| Otu37014 | 1 | k__Fungi(100);p__Ascomycota(100);c__Saccharomycetes(100);o__Saccharomycetales(100);f__Metschnikowiaceae(100);g__Metschnikowia(100);g__Metschnikowia_unclassified(100); | Metschnikowia | 13 | Saprotroph | \|Undefined Saprotroph\| | Yeast | NULL | Probable | As nectar-tap_saprotroph (Põlme et al. 2020); Decay sugar-rich substrates (Põlme et al. 2020) | Põlme S, et al. 2020. Fungal Diversity 105:1-16 ((https://doi.org/10.1007/s13225-020-00466-2)) |
| Otu37015 | 1 | k__Fungi(100);p__Ascomycota(100);c__Saccharomycetes(100);o__Saccharomycetales(100);f__Metschnikowiaceae(100);g__Metschnikowia(100);s__Metschnikowia_pulcherrima(100); | Metschnikowia | 13 | Saprotroph | \|Undefined Saprotroph\| | Yeast | NULL | Probable | As nectar-tap_saprotroph (Põlme et al. 2020); Decay sugar-rich substrates (Põlme et al. 2020) | Põlme S, et al. 2020. Fungal Diversity 105:1-16 ((https://doi.org/10.1007/s13225-020-00466-2)) |
| Otu37016 | 1 | k__Fungi(100);p__Ascomycota(100);c__Saccharomycetes(100);o__Saccharomycetales(100);f__Metschnikowiaceae(100);g__Metschnikowia(100);s__Metschnikowia_pulcherrima(100); | Metschnikowia | 13 | Saprotroph | \|Undefined Saprotroph\| | Yeast | NULL | Probable | As nectar-tap_saprotroph (Põlme et al. 2020); Decay sugar-rich substrates (Põlme et al. 2020) | Põlme S, et al. 2020. Fungal Diversity 105:1-16 ((https://doi.org/10.1007/s13225-020-00466-2)) |
| Otu37017 | 1 | k__Fungi(100);p__Ascomycota(100);c__Saccharomycetes(100);o__Saccharomycetales(100);f__Metschnikowiaceae(100);g__Metschnikowia(100);s__Metschnikowia_pulcherrima(100); | Metschnikowia | 13 | Saprotroph | \|Undefined Saprotroph\| | Yeast | NULL | Probable | As nectar-tap_saprotroph (Põlme et al. 2020); Decay sugar-rich substrates (Põlme et al. 2020) | Põlme S, et al. 2020. Fungal Diversity 105:1-16 ((https://doi.org/10.1007/s13225-020-00466-2)) |
| Otu37018 | 1 | k__Fungi(100);p__Ascomycota(100);c__Saccharomycetes(100);o__Saccharomycetales(100);f__Metschnikowiaceae(100);g__Metschnikowia(100);s__Metschnikowia_pulcherrima(100); | Metschnikowia | 13 | Saprotroph | \|Undefined Saprotroph\| | Yeast | NULL | Probable | As nectar-tap_saprotroph (Põlme et al. 2020); Decay sugar-rich substrates (Põlme et al. 2020) | Põlme S, et al. 2020. Fungal Diversity 105:1-16 ((https://doi.org/10.1007/s13225-020-00466-2)) |
| Otu37019 | 1 | k__Fungi(100);p__Ascomycota(100);c__Saccharomycetes(100);o__Saccharomycetales(100);f__Metschnikowiaceae(100);g__Metschnikowia(100);s__Metschnikowia_pulcherrima(100); | Metschnikowia | 13 | Saprotroph | \|Undefined Saprotroph\| | Yeast | NULL | Probable | As nectar-tap_saprotroph (Põlme et al. 2020); Decay sugar-rich substrates (Põlme et al. 2020) | Põlme S, et al. 2020. Fungal Diversity 105:1-16 ((https://doi.org/10.1007/s13225-020-00466-2)) |
| Otu37020 | 1 | k__Fungi(100);p__Ascomycota(100);c__Saccharomycetes(100);o__Saccharomycetales(100);f__Metschnikowiaceae(100);g__Metschnikowia(100);g__Metschnikowia_unclassified(100); | Metschnikowia | 13 | Saprotroph | \|Undefined Saprotroph\| | Yeast | NULL | Probable | As nectar-tap_saprotroph (Põlme et al. 2020); Decay sugar-rich substrates (Põlme et al. 2020) | Põlme S, et al. 2020. Fungal Diversity 105:1-16 ((https://doi.org/10.1007/s13225-020-00466-2)) |
| Otu37021 | 1 | k__Fungi(100);p__Ascomycota(100);c__Saccharomycetes(100);o__Saccharomycetales(100);f__Metschnikowiaceae(100);g__Metschnikowia(100);s__Metschnikowia_pulcherrima(100); | Metschnikowia | 13 | Saprotroph | \|Undefined Saprotroph\| | Yeast | NULL | Probable | As nectar-tap_saprotroph (Põlme et al. 2020); Decay sugar-rich substrates (Põlme et al. 2020) | Põlme S, et al. 2020. Fungal Diversity 105:1-16 ((https://doi.org/10.1007/s13225-020-00466-2)) |
| Otu37022 | 1 | k__Fungi(100);p__Ascomycota(100);c__Saccharomycetes(100);o__Saccharomycetales(100);f__Metschnikowiaceae(100);g__Metschnikowia(100);g__Metschnikowia_unclassified(100); | Metschnikowia | 13 | Saprotroph | \|Undefined Saprotroph\| | Yeast | NULL | Probable | As nectar-tap_saprotroph (Põlme et al. 2020); Decay sugar-rich substrates (Põlme et al. 2020) | Põlme S, et al. 2020. Fungal Diversity 105:1-16 ((https://doi.org/10.1007/s13225-020-00466-2)) |

| | | | | | | | | | | | |
|---|---|---|---|---|---|---|---|---|---|---|---|
| Otu37023 | 1 | k__Fungi(100);p__Ascomycota(100);c__Saccharomycetes(100);o__Saccharomycetales(100);f__Metschnikowiaceae(100);g__Metschnikowia(100);g__Metschnikowia_unclassified(100); | Metschnikowia | 13 | Saprotroph | \|Undefined Saprotroph\| | Yeast | NULL | Probable | As nectar-tap_saprotroph (Põlme et al. 2020); Decay sugar-rich substrates (Põlme et al. 2020) | Põlme S, et al. 2020. Fungal Diversity 105:1-16 ((https://doi.org/10.1007/s13225-020-00466-2)) |
| Otu37024 | 1 | k__Fungi(100);p__Ascomycota(100);c__Saccharomycetes(100);o__Saccharomycetales(100);f__Metschnikowiaceae(100);g__Metschnikowia(100);g__Metschnikowia_unclassified(100); | Metschnikowia | 13 | Saprotroph | \|Undefined Saprotroph\| | Yeast | NULL | Probable | As nectar-tap_saprotroph (Põlme et al. 2020); Decay sugar-rich substrates (Põlme et al. 2020) | Põlme S, et al. 2020. Fungal Diversity 105:1-16 ((https://doi.org/10.1007/s13225-020-00466-2)) |
| Otu37025 | 1 | k__Fungi(100);p__Ascomycota(100);c__Saccharomycetes(100);o__Saccharomycetales(100);f__Metschnikowiaceae(100);g__Metschnikowia(100);g__Metschnikowia_unclassified(100); | Metschnikowia | 13 | Saprotroph | \|Undefined Saprotroph\| | Yeast | NULL | Probable | As nectar-tap_saprotroph (Põlme et al. 2020); Decay sugar-rich substrates (Põlme et al. 2020) | Põlme S, et al. 2020. Fungal Diversity 105:1-16 ((https://doi.org/10.1007/s13225-020-00466-2)) |
| Otu37026 | 1 | k__Fungi(100);p__Ascomycota(100);c__Saccharomycetes(100);o__Saccharomycetales(100);f__Metschnikowiaceae(100);g__Metschnikowia(100);g__Metschnikowia_unclassified(100); | Metschnikowia | 13 | Saprotroph | \|Undefined Saprotroph\| | Yeast | NULL | Probable | As nectar-tap_saprotroph (Põlme et al. 2020); Decay sugar-rich substrates (Põlme et al. 2020) | Põlme S, et al. 2020. Fungal Diversity 105:1-16 ((https://doi.org/10.1007/s13225-020-00466-2)) |
| Otu37027 | 1 | k__Fungi(100);p__Ascomycota(100);c__Saccharomycetes(100);o__Saccharomycetales(100);f__Metschnikowiaceae(100);g__Metschnikowia(100);g__Metschnikowia_unclassified(100); | Metschnikowia | 13 | Saprotroph | \|Undefined Saprotroph\| | Yeast | NULL | Probable | As nectar-tap_saprotroph (Põlme et al. 2020); Decay sugar-rich substrates (Põlme et al. 2020) | Põlme S, et al. 2020. Fungal Diversity 105:1-16 ((https://doi.org/10.1007/s13225-020-00466-2)) |
| Otu37028 | 1 | k__Fungi(100);p__Ascomycota(100);c__Saccharomycetes(100);o__Saccharomycetales(100);f__Metschnikowiaceae(100);g__Metschnikowia(100);s__Metschnikowia_pulcherrima(100); | Metschnikowia | 13 | Saprotroph | \|Undefined Saprotroph\| | Yeast | NULL | Probable | As nectar-tap_saprotroph (Põlme et al. 2020); Decay sugar-rich substrates (Põlme et al. 2020) | Põlme S, et al. 2020. Fungal Diversity 105:1-16 ((https://doi.org/10.1007/s13225-020-00466-2)) |
| Otu37029 | 1 | k__Fungi(100);p__Ascomycota(100);c__Saccharomycetes(100);o__Saccharomycetales(100);f__Metschnikowiaceae(100);g__Metschnikowia(100);s__Metschnikowia_pulcherrima(100); | Metschnikowia | 13 | Saprotroph | \|Undefined Saprotroph\| | Yeast | NULL | Probable | As nectar-tap_saprotroph (Põlme et al. 2020); Decay sugar-rich substrates (Põlme et al. 2020) | Põlme S, et al. 2020. Fungal Diversity 105:1-16 ((https://doi.org/10.1007/s13225-020-00466-2)) |
| Otu37030 | 1 | k__Fungi(100);p__Ascomycota(100);c__Saccharomycetes(100);o__Saccharomycetales(100);f__Metschnikowiaceae(100);g__Metschnikowia(100);s__Metschnikowia_pulcherrima(100); | Metschnikowia | 13 | Saprotroph | \|Undefined Saprotroph\| | Yeast | NULL | Probable | As nectar-tap_saprotroph (Põlme et al. 2020); Decay sugar-rich substrates (Põlme et al. 2020) | Põlme S, et al. 2020. Fungal Diversity 105:1-16 ((https://doi.org/10.1007/s13225-020-00466-2)) |
| Otu37031 | 1 | k__Fungi(100);p__Ascomycota(100);c__Saccharomycetes(100);o__Saccharomycetales(100);f__Metschnikowiaceae(100);g__Metschnikowia(100);s__Metschnikowia_pulcherrima(100); | Metschnikowia | 13 | Saprotroph | \|Undefined Saprotroph\| | Yeast | NULL | Probable | As nectar-tap_saprotroph (Põlme et al. 2020); Decay sugar-rich substrates (Põlme et al. 2020) | Põlme S, et al. 2020. Fungal Diversity 105:1-16 ((https://doi.org/10.1007/s13225-020-00466-2)) |
| Otu37035 | 1 | k__Fungi(100);p__Ascomycota(100);c__Saccharomycetes(100);o__Saccharomycetales(100);f__Metschnikowiaceae(100);g__Metschnikowia(100);s__Metschnikowia_pulcherrima(100); | Metschnikowia | 13 | Saprotroph | \|Undefined Saprotroph\| | Yeast | NULL | Probable | As nectar-tap_saprotroph (Põlme et al. 2020); Decay sugar-rich substrates (Põlme et al. 2020) | Põlme S, et al. 2020. Fungal Diversity 105:1-16 ((https://doi.org/10.1007/s13225-020-00466-2)) |
| Otu37036 | 1 | k__Fungi(100);p__Ascomycota(100);c__Saccharomycetes(100);o__Saccharomycetales(100);f__Metschnikowiaceae(100);g__Metschnikowia(100);g__Metschnikowia_unclassified(100); | Metschnikowia | 13 | Saprotroph | \|Undefined Saprotroph\| | Yeast | NULL | Probable | As nectar-tap_saprotroph (Põlme et al. 2020); Decay sugar-rich substrates (Põlme et al. 2020) | Põlme S, et al. 2020. Fungal Diversity 105:1-16 ((https://doi.org/10.1007/s13225-020-00466-2)) |
| Otu37037 | 1 | k__Fungi(100);p__Ascomycota(100);c__Saccharomycetes(100);o__Saccharomycetales(100);f__Metschnikowiaceae(100);g__Metschnikowia(100);g__Metschnikowia_unclassified(100); | Metschnikowia | 13 | Saprotroph | \|Undefined Saprotroph\| | Yeast | NULL | Probable | As nectar-tap_saprotroph (Põlme et al. 2020); Decay sugar-rich substrates (Põlme et al. 2020) | Põlme S, et al. 2020. Fungal Diversity 105:1-16 ((https://doi.org/10.1007/s13225-020-00466-2)) |
| Otu37038 | 1 | k__Fungi(100);p__Ascomycota(100);c__Saccharomycetes(100);o__Saccharomycetales(100);f__Metschnikowiaceae(100);g__Metschnikowia(100);s__Metschnikowia_pulcherrima(100); | Metschnikowia | 13 | Saprotroph | \|Undefined Saprotroph\| | Yeast | NULL | Probable | As nectar-tap_saprotroph (Põlme et al. 2020); Decay sugar-rich substrates (Põlme et al. 2020) | Põlme S, et al. 2020. Fungal Diversity 105:1-16 ((https://doi.org/10.1007/s13225-020-00466-2)) |
| Otu37039 | 1 | k__Fungi(100);p__Ascomycota(100);c__Saccharomycetes(100);o__Saccharomycetales(100);f__Metschnikowiaceae(100);g__Metschnikowia(100);s__Metschnikowia_pulcherrima(100); | Metschnikowia | 13 | Saprotroph | \|Undefined Saprotroph\| | Yeast | NULL | Probable | As nectar-tap_saprotroph (Põlme et al. 2020); Decay sugar-rich substrates (Põlme et al. 2020) | Põlme S, et al. 2020. Fungal Diversity 105:1-16 ((https://doi.org/10.1007/s13225-020-00466-2)) |
| Otu37040 | 1 | k__Fungi(100);p__Ascomycota(100);c__Saccharomycetes(100);o__Saccharomycetales(100);f__Metschnikowiaceae(100);g__Metschnikowia(100);s__Metschnikowia_pulcherrima(100); | Metschnikowia | 13 | Saprotroph | \|Undefined Saprotroph\| | Yeast | NULL | Probable | As nectar-tap_saprotroph (Põlme et al. 2020); Decay sugar-rich substrates (Põlme et al. 2020) | Põlme S, et al. 2020. Fungal Diversity 105:1-16 ((https://doi.org/10.1007/s13225-020-00466-2)) |
| Otu37041 | 1 | k__Fungi(100);p__Ascomycota(100);c__Saccharomycetes(100);o__Saccharomycetales(100);f__Metschnikowiaceae(100);g__Metschnikowia(100);s__Metschnikowia_pulcherrima(100); | Metschnikowia | 13 | Saprotroph | \|Undefined Saprotroph\| | Yeast | NULL | Probable | As nectar-tap_saprotroph (Põlme et al. 2020); Decay sugar-rich substrates (Põlme et al. 2020) | Põlme S, et al. 2020. Fungal Diversity 105:1-16 ((https://doi.org/10.1007/s13225-020-00466-2)) |
| Otu37042 | 1 | k__Fungi(100);p__Ascomycota(100);c__Saccharomycetes(100);o__Saccharomycetales(100);f__Metschnikowiaceae(100);g__Metschnikowia(100);s__Metschnikowia_pulcherrima(100); | Metschnikowia | 13 | Saprotroph | \|Undefined Saprotroph\| | Yeast | NULL | Probable | As nectar-tap_saprotroph (Põlme et al. 2020); Decay sugar-rich substrates (Põlme et al. 2020) | Põlme S, et al. 2020. Fungal Diversity 105:1-16 ((https://doi.org/10.1007/s13225-020-00466-2)) |
| Otu37043 | 1 | k__Fungi(100);p__Ascomycota(100);c__Saccharomycetes(100);o__Saccharomycetales(100);f__Metschnikowiaceae(100);g__Metschnikowia(100);s__Metschnikowia_pulcherrima(100); | Metschnikowia | 13 | Saprotroph | \|Undefined Saprotroph\| | Yeast | NULL | Probable | As nectar-tap_saprotroph (Põlme et al. 2020); Decay sugar-rich substrates (Põlme et al. 2020) | Põlme S, et al. 2020. Fungal Diversity 105:1-16 ((https://doi.org/10.1007/s13225-020-00466-2)) |
| Otu37044 | 1 | k__Fungi(100);p__Ascomycota(100);c__Saccharomycetes(100);o__Saccharomycetales(100);f__Metschnikowiaceae(100);g__Metschnikowia(100);s__Metschnikowia_pulcherrima(100); | Metschnikowia | 13 | Saprotroph | \|Undefined Saprotroph\| | Yeast | NULL | Probable | As nectar-tap_saprotroph (Põlme et al. 2020); Decay sugar-rich substrates (Põlme et al. 2020) | Põlme S, et al. 2020. Fungal Diversity 105:1-16 ((https://doi.org/10.1007/s13225-020-00466-2)) |
| Otu37045 | 1 | k__Fungi(100);p__Ascomycota(100);c__Saccharomycetes(100);o__Saccharomycetales(100);f__Metschnikowiaceae(100);g__Metschnikowia(100);g__Metschnikowia_unclassified(100); | Metschnikowia | 13 | Saprotroph | \|Undefined Saprotroph\| | Yeast | NULL | Probable | As nectar-tap_saprotroph (Põlme et al. 2020); Decay sugar-rich substrates (Põlme et al. 2020) | Põlme S, et al. 2020. Fungal Diversity 105:1-16 ((https://doi.org/10.1007/s13225-020-00466-2)) |
| Otu37046 | 1 | k__Fungi(100);p__Ascomycota(100);c__Saccharomycetes(100);o__Saccharomycetales(100);f__Metschnikowiaceae(100);g__Metschnikowia(100);s__Metschnikowia_pulcherrima(100); | Metschnikowia | 13 | Saprotroph | \|Undefined Saprotroph\| | Yeast | NULL | Probable | As nectar-tap_saprotroph (Põlme et al. 2020); Decay sugar-rich substrates (Põlme et al. 2020) | Põlme S, et al. 2020. Fungal Diversity 105:1-16 ((https://doi.org/10.1007/s13225-020-00466-2)) |
| Otu37047 | 1 | k__Fungi(100);p__Ascomycota(100);c__Saccharomycetes(100);o__Saccharomycetales(100);f__Metschnikowiaceae(100);g__Metschnikowia(100);s__Metschnikowia_pulcherrima(100); | Metschnikowia | 13 | Saprotroph | \|Undefined Saprotroph\| | Yeast | NULL | Probable | As nectar-tap_saprotroph (Põlme et al. 2020); Decay sugar-rich substrates (Põlme et al. 2020) | Põlme S, et al. 2020. Fungal Diversity 105:1-16 ((https://doi.org/10.1007/s13225-020-00466-2)) |
| Otu37048 | 1 | k__Fungi(100);p__Ascomycota(100);c__Saccharomycetes(100);o__Saccharomycetales(100);f__Metschnikowiaceae(100);g__Metschnikowia(100);s__Metschnikowia_pulcherrima(100); | Metschnikowia | 13 | Saprotroph | \|Undefined Saprotroph\| | Yeast | NULL | Probable | As nectar-tap_saprotroph (Põlme et al. 2020); Decay sugar-rich substrates (Põlme et al. 2020) | Põlme S, et al. 2020. Fungal Diversity 105:1-16 ((https://doi.org/10.1007/s13225-020-00466-2)) |
| Otu37049 | 1 | k__Fungi(100);p__Ascomycota(100);c__Saccharomycetes(100);o__Saccharomycetales(100);f__Metschnikowiaceae(100);g__Metschnikowia(100);s__Metschnikowia_pulcherrima(100); | Metschnikowia | 13 | Saprotroph | \|Undefined Saprotroph\| | Yeast | NULL | Probable | As nectar-tap_saprotroph (Põlme et al. 2020); Decay sugar-rich substrates (Põlme et al. 2020) | Põlme S, et al. 2020. Fungal Diversity 105:1-16 ((https://doi.org/10.1007/s13225-020-00466-2)) |
| Otu37050 | 1 | k__Fungi(100);p__Ascomycota(100);c__Saccharomycetes(100);o__Saccharomycetales(100);f__Metschnikowiaceae(100);g__Metschnikowia(100);s__Metschnikowia_pulcherrima(100); | Metschnikowia | 13 | Saprotroph | \|Undefined Saprotroph\| | Yeast | NULL | Probable | As nectar-tap_saprotroph (Põlme et al. 2020); Decay sugar-rich substrates (Põlme et al. 2020) | Põlme S, et al. 2020. Fungal Diversity 105:1-16 ((https://doi.org/10.1007/s13225-020-00466-2)) |
| Otu37051 | 1 | k__Fungi(100);p__Ascomycota(100);c__Saccharomycetes(100);o__Saccharomycetales(100);f__Metschnikowiaceae(100);g__Metschnikowia(100);s__Metschnikowia_pulcherrima(100); | Metschnikowia | 13 | Saprotroph | \|Undefined Saprotroph\| | Yeast | NULL | Probable | As nectar-tap_saprotroph (Põlme et al. 2020); Decay sugar-rich substrates (Põlme et al. 2020) | Põlme S, et al. 2020. Fungal Diversity 105:1-16 ((https://doi.org/10.1007/s13225-020-00466-2)) |
| Otu37052 | 1 | k__Fungi(100);p__Ascomycota(100);c__Saccharomycetes(100);o__Saccharomycetales(100);f__Metschnikowiaceae(100);g__Metschnikowia(100);s__Metschnikowia_pulcherrima(100); | Metschnikowia | 13 | Saprotroph | \|Undefined Saprotroph\| | Yeast | NULL | Probable | As nectar-tap_saprotroph (Põlme et al. 2020); Decay sugar-rich substrates (Põlme et al. 2020) | Põlme S, et al. 2020. Fungal Diversity 105:1-16 ((https://doi.org/10.1007/s13225-020-00466-2)) |
| Otu37053 | 1 | k__Fungi(100);p__Ascomycota(100);c__Saccharomycetes(100);o__Saccharomycetales(100);f__Metschnikowiaceae(100);g__Metschnikowia(100);s__Metschnikowia_pulcherrima(100); | Metschnikowia | 13 | Saprotroph | \|Undefined Saprotroph\| | Yeast | NULL | Probable | As nectar-tap_saprotroph (Põlme et al. 2020); Decay sugar-rich substrates (Põlme et al. 2020) | Põlme S, et al. 2020. Fungal Diversity 105:1-16 ((https://doi.org/10.1007/s13225-020-00466-2)) |
| Otu37058 | 1 | k__Fungi(100);p__Ascomycota(100);c__Saccharomycetes(100);o__Saccharomycetales(100);f__Metschnikowiaceae(100);g__Metschnikowia(100);s__Metschnikowia_pulcherrima(100); | Metschnikowia | 13 | Saprotroph | \|Undefined Saprotroph\| | Yeast | NULL | Probable | As nectar-tap_saprotroph (Põlme et al. 2020); Decay sugar-rich substrates (Põlme et al. 2020) | Põlme S, et al. 2020. Fungal Diversity 105:1-16 ((https://doi.org/10.1007/s13225-020-00466-2)) |
| Otu37059 | 1 | k__Fungi(100);p__Ascomycota(100);c__Saccharomycetes(100);o__Saccharomycetales(100);f__Metschnikowiaceae(100);g__Metschnikowia(100);s__Metschnikowia_pulcherrima(100); | Metschnikowia | 13 | Saprotroph | \|Undefined Saprotroph\| | Yeast | NULL | Probable | As nectar-tap_saprotroph (Põlme et al. 2020); Decay sugar-rich substrates (Põlme et al. 2020) | Põlme S, et al. 2020. Fungal Diversity 105:1-16 ((https://doi.org/10.1007/s13225-020-00466-2)) |
| Otu37060 | 1 | k__Fungi(100);p__Ascomycota(100);c__Saccharomycetes(100);o__Saccharomycetales(100);f__Metschnikowiaceae(100);g__Metschnikowia(100);s__Metschnikowia_pulcherrima(100); | Metschnikowia | 13 | Saprotroph | \|Undefined Saprotroph\| | Yeast | NULL | Probable | As nectar-tap_saprotroph (Põlme et al. 2020); Decay sugar-rich substrates (Põlme et al. 2020) | Põlme S, et al. 2020. Fungal Diversity 105:1-16 ((https://doi.org/10.1007/s13225-020-00466-2)) |
| Otu37061 | 1 | k__Fungi(100);p__Ascomycota(100);c__Saccharomycetes(100);o__Saccharomycetales(100);f__Metschnikowiaceae(100);g__Metschnikowia(100);s__Metschnikowia_pulcherrima(100); | Metschnikowia | 13 | Saprotroph | \|Undefined Saprotroph\| | Yeast | NULL | Probable | As nectar-tap_saprotroph (Põlme et al. 2020); Decay sugar-rich substrates (Põlme et al. 2020) | Põlme S, et al. 2020. Fungal Diversity 105:1-16 ((https://doi.org/10.1007/s13225-020-00466-2)) |
| Otu37062 | 1 | k__Fungi(100);p__Ascomycota(100);c__Saccharomycetes(100);o__Saccharomycetales(100);f__Metschnikowiaceae(100);g__Metschnikowia(100);s__Metschnikowia_pulcherrima(100); | Metschnikowia | 13 | Saprotroph | \|Undefined Saprotroph\| | Yeast | NULL | Probable | As nectar-tap_saprotroph (Põlme et al. 2020); Decay sugar-rich substrates (Põlme et al. 2020) | Põlme S, et al. 2020. Fungal Diversity 105:1-16 ((https://doi.org/10.1007/s13225-020-00466-2)) |
| Otu37063 | 1 | k__Fungi(100);p__Ascomycota(100);c__Saccharomycetes(100);o__Saccharomycetales(100);f__Metschnikowiaceae(100);g__Metschnikowia(100);s__Metschnikowia_pulcherrima(100); | Metschnikowia | 13 | Saprotroph | \|Undefined Saprotroph\| | Yeast | NULL | Probable | As nectar-tap_saprotroph (Põlme et al. 2020); Decay sugar-rich substrates (Põlme et al. 2020) | Põlme S, et al. 2020. Fungal Diversity 105:1-16 ((https://doi.org/10.1007/s13225-020-00466-2)) |
| Otu37077 | 1 | k__Fungi(100);p__Ascomycota(100);c__Saccharomycetes(100);o__Saccharomycetales(100);f__Metschnikowiaceae(100);g__Metschnikowia(100);s__Metschnikowia_pulcherrima(100); | Metschnikowia | 13 | Saprotroph | \|Undefined Saprotroph\| | Yeast | NULL | Probable | As nectar-tap_saprotroph (Põlme et al. 2020); Decay sugar-rich substrates (Põlme et al. 2020) | Põlme S, et al. 2020. Fungal Diversity 105:1-16 ((https://doi.org/10.1007/s13225-020-00466-2)) |
| Otu37080 | 1 | k__Fungi(100);p__Ascomycota(100);c__Saccharomycetes(100);o__Saccharomycetales(100);f__Metschnikowiaceae(100);g__Metschnikowia(100);g__Metschnikowia_unclassified(100); | Metschnikowia | 13 | Saprotroph | \|Undefined Saprotroph\| | Yeast | NULL | Probable | As nectar-tap_saprotroph (Põlme et al. 2020); Decay sugar-rich substrates (Põlme et al. 2020) | Põlme S, et al. 2020. Fungal Diversity 105:1-16 ((https://doi.org/10.1007/s13225-020-00466-2)) |
| Otu37081 | 1 | k__Fungi(100);p__Ascomycota(100);c__Saccharomycetes(100);o__Saccharomycetales(100);f__Metschnikowiaceae(100);g__Metschnikowia(100);g__Metschnikowia_unclassified(100); | Metschnikowia | 13 | Saprotroph | \|Undefined Saprotroph\| | Yeast | NULL | Probable | As nectar-tap_saprotroph (Põlme et al. 2020); Decay sugar-rich substrates (Põlme et al. 2020) | Põlme S, et al. 2020. Fungal Diversity 105:1-16 ((https://doi.org/10.1007/s13225-020-00466-2)) |
| Otu37082 | 1 | k__Fungi(100);p__Ascomycota(100);c__Saccharomycetes(100);o__Saccharomycetales(100);f__Metschnikowiaceae(100);g__Metschnikowia(100);s__Metschnikowia_pulcherrima(100); | Metschnikowia | 13 | Saprotroph | \|Undefined Saprotroph\| | Yeast | NULL | Probable | As nectar-tap_saprotroph (Põlme et al. 2020); Decay sugar-rich substrates (Põlme et al. 2020) | Põlme S, et al. 2020. Fungal Diversity 105:1-16 ((https://doi.org/10.1007/s13225-020-00466-2)) |
| Otu37083 | 1 | k__Fungi(100);p__Ascomycota(100);c__Saccharomycetes(100);o__Saccharomycetales(100);f__Metschnikowiaceae(100);g__Metschnikowia(100);g__Metschnikowia_unclassified(100); | Metschnikowia | 13 | Saprotroph | \|Undefined Saprotroph\| | Yeast | NULL | Probable | As nectar-tap_saprotroph (Põlme et al. 2020); Decay sugar-rich substrates (Põlme et al. 2020) | Põlme S, et al. 2020. Fungal Diversity 105:1-16 ((https://doi.org/10.1007/s13225-020-00466-2)) |
| Otu37084 | 1 | k__Fungi(100);p__Ascomycota(100);c__Saccharomycetes(100);o__Saccharomycetales(100);f__Metschnikowiaceae(100);g__Metschnikowia(100);s__Metschnikowia_pulcherrima(100); | Metschnikowia | 13 | Saprotroph | \|Undefined Saprotroph\| | Yeast | NULL | Probable | As nectar-tap_saprotroph (Põlme et al. 2020); Decay sugar-rich substrates (Põlme et al. 2020) | Põlme S, et al. 2020. Fungal Diversity 105:1-16 ((https://doi.org/10.1007/s13225-020-00466-2)) |
| Otu37085 | 1 | k__Fungi(100);p__Ascomycota(100);c__Saccharomycetes(100);o__Saccharomycetales(100);f__Metschnikowiaceae(100);g__Metschnikowia(100);s__Metschnikowia_pulcherrima(100); | Metschnikowia | 13 | Saprotroph | \|Undefined Saprotroph\| | Yeast | NULL | Probable | As nectar-tap_saprotroph (Põlme et al. 2020); Decay sugar-rich substrates (Põlme et al. 2020) | Põlme S, et al. 2020. Fungal Diversity 105:1-16 ((https://doi.org/10.1007/s13225-020-00466-2)) |
| Otu37086 | 1 | k__Fungi(100);p__Ascomycota(100);c__Saccharomycetes(100);o__Saccharomycetales(100);f__Metschnikowiaceae(100);g__Metschnikowia(100);s__Metschnikowia_pulcherrima(100); | Metschnikowia | 13 | Saprotroph | \|Undefined Saprotroph\| | Yeast | NULL | Probable | As nectar-tap_saprotroph (Põlme et al. 2020); Decay sugar-rich substrates (Põlme et al. 2020) | Põlme S, et al. 2020. Fungal Diversity 105:1-16 ((https://doi.org/10.1007/s13225-020-00466-2)) |
| Otu37087 | 1 | k__Fungi(100);p__Ascomycota(100);c__Saccharomycetes(100);o__Saccharomycetales(100);f__Metschnikowiaceae(100);g__Metschnikowia(100);s__Metschnikowia_pulcherrima(100); | Metschnikowia | 13 | Saprotroph | \|Undefined Saprotroph\| | Yeast | NULL | Probable | As nectar-tap_saprotroph (Põlme et al. 2020); Decay sugar-rich substrates (Põlme et al. 2020) | Põlme S, et al. 2020. Fungal Diversity 105:1-16 ((https://doi.org/10.1007/s13225-020-00466-2)) |
| Otu37088 | 1 | k__Fungi(100);p__Ascomycota(100);c__Saccharomycetes(100);o__Saccharomycetales(100);f__Metschnikowiaceae(100);g__Metschnikowia(100);s__Metschnikowia_pulcherrima(100); | Metschnikowia | 13 | Saprotroph | \|Undefined Saprotroph\| | Yeast | NULL | Probable | As nectar-tap_saprotroph (Põlme et al. 2020); Decay sugar-rich substrates (Põlme et al. 2020) | Põlme S, et al. 2020. Fungal Diversity 105:1-16 ((https://doi.org/10.1007/s13225-020-00466-2)) |
| Otu37089 | 1 | k__Fungi(100);p__Ascomycota(100);c__Saccharomycetes(100);o__Saccharomycetales(100);f__Metschnikowiaceae(100);g__Metschnikowia(100);s__Metschnikowia_pulcherrima(100); | Metschnikowia | 13 | Saprotroph | \|Undefined Saprotroph\| | Yeast | NULL | Probable | As nectar-tap_saprotroph (Põlme et al. 2020); Decay sugar-rich substrates (Põlme et al. 2020) | Põlme S, et al. 2020. Fungal Diversity 105:1-16 ((https://doi.org/10.1007/s13225-020-00466-2)) |
| Otu37090 | 1 | k__Fungi(100);p__Ascomycota(100);c__Saccharomycetes(100);o__Saccharomycetales(100);f__Metschnikowiaceae(100);g__Metschnikowia(100);s__Metschnikowia_pulcherrima(100); | Metschnikowia | 13 | Saprotroph | \|Undefined Saprotroph\| | Yeast | NULL | Probable | As nectar-tap_saprotroph (Põlme et al. 2020); Decay sugar-rich substrates (Põlme et al. 2020) | Põlme S, et al. 2020. Fungal Diversity 105:1-16 ((https://doi.org/10.1007/s13225-020-00466-2)) |
| Otu37091 | 1 | k__Fungi(100);p__Ascomycota(100);c__Saccharomycetes(100);o__Saccharomycetales(100);f__Metschnikowiaceae(100);g__Metschnikowia(100);s__Metschnikowia_pulcherrima(100); | Metschnikowia | 13 | Saprotroph | \|Undefined Saprotroph\| | Yeast | NULL | Probable | As nectar-tap_saprotroph (Põlme et al. 2020); Decay sugar-rich substrates (Põlme et al. 2020) | Põlme S, et al. 2020. Fungal Diversity 105:1-16 ((https://doi.org/10.1007/s13225-020-00466-2)) |
| Otu37092 | 1 | k__Fungi(100);p__Ascomycota(100);c__Saccharomycetes(100);o__Saccharomycetales(100);f__Metschnikowiaceae(100);g__Metschnikowia(100);s__Metschnikowia_pulcherrima(100); | Metschnikowia | 13 | Saprotroph | \|Undefined Saprotroph\| | Yeast | NULL | Probable | As nectar-tap_saprotroph (Põlme et al. 2020); Decay sugar-rich substrates (Põlme et al. 2020) | Põlme S, et al. 2020. Fungal Diversity 105:1-16 ((https://doi.org/10.1007/s13225-020-00466-2)) |
| Otu37093 | 1 | k__Fungi(100);p__Ascomycota(100);c__Saccharomycetes(100);o__Saccharomycetales(100);f__Metschnikowiaceae(100);g__Metschnikowia(100);g__Metschnikowia_unclassified(100); | Metschnikowia | 13 | Saprotroph | \|Undefined Saprotroph\| | Yeast | NULL | Probable | As nectar-tap_saprotroph (Põlme et al. 2020); Decay sugar-rich substrates (Põlme et al. 2020) | Põlme S, et al. 2020. Fungal Diversity 105:1-16 ((https://doi.org/10.1007/s13225-020-00466-2)) |
| Otu37094 | 1 | k__Fungi(100);p__Ascomycota(100);c__Saccharomycetes(100);o__Saccharomycetales(100);f__Metschnikowiaceae(100);g__Metschnikowia(100);s__Metschnikowia_pulcherrima(100); | Metschnikowia | 13 | Saprotroph | \|Undefined Saprotroph\| | Yeast | NULL | Probable | As nectar-tap_saprotroph (Põlme et al. 2020); Decay sugar-rich substrates (Põlme et al. 2020) | Põlme S, et al. 2020. Fungal Diversity 105:1-16 ((https://doi.org/10.1007/s13225-020-00466-2)) |
| Otu37095 | 1 | k__Fungi(100);p__Ascomycota(100);c__Saccharomycetes(100);o__Saccharomycetales(100);f__Metschnikowiaceae(100);g__Metschnikowia(100);s__Metschnikowia_pulcherrima(100); | Metschnikowia | 13 | Saprotroph | \|Undefined Saprotroph\| | Yeast | NULL | Probable | As nectar-tap_saprotroph (Põlme et al. 2020); Decay sugar-rich substrates (Põlme et al. 2020) | Põlme S, et al. 2020. Fungal Diversity 105:1-16 ((https://doi.org/10.1007/s13225-020-00466-2)) |
| Otu37096 | 1 | k__Fungi(100);p__Ascomycota(100);c__Saccharomycetes(100);o__Saccharomycetales(100);f__Metschnikowiaceae(100);g__Metschnikowia(100);g__Metschnikowia_unclassified(100); | Metschnikowia | 13 | Saprotroph | \|Undefined Saprotroph\| | Yeast | NULL | Probable | As nectar-tap_saprotroph (Põlme et al. 2020); Decay sugar-rich substrates (Põlme et al. 2020) | Põlme S, et al. 2020. Fungal Diversity 105:1-16 ((https://doi.org/10.1007/s13225-020-00466-2)) |
| Otu37097 | 1 | k__Fungi(100);p__Ascomycota(100);c__Saccharomycetes(100);o__Saccharomycetales(100);f__Metschnikowiaceae(100);g__Metschnikowia(100);g__Metschnikowia_unclassified(100); | Metschnikowia | 13 | Saprotroph | \|Undefined Saprotroph\| | Yeast | NULL | Probable | As nectar-tap_saprotroph (Põlme et al. 2020); Decay sugar-rich substrates (Põlme et al. 2020) | Põlme S, et al. 2020. Fungal Diversity 105:1-16 ((https://doi.org/10.1007/s13225-020-00466-2)) |
| Otu37098 | 1 | k__Fungi(100);p__Ascomycota(100);c__Saccharomycetes(100);o__Saccharomycetales(100);f__Metschnikowiaceae(100);g__Metschnikowia(100);s__Metschnikowia_pulcherrima(100); | Metschnikowia | 13 | Saprotroph | \|Undefined Saprotroph\| | Yeast | NULL | Probable | As nectar-tap_saprotroph (Põlme et al. 2020); Decay sugar-rich substrates (Põlme et al. 2020) | Põlme S, et al. 2020. Fungal Diversity 105:1-16 ((https://doi.org/10.1007/s13225-020-00466-2)) |
| Otu37099 | 1 | k__Fungi(100);p__Ascomycota(100);c__Saccharomycetes(100);o__Saccharomycetales(100);f__Metschnikowiaceae(100);g__Metschnikowia(100);s__Metschnikowia_pulcherrima(100); | Metschnikowia | 13 | Saprotroph | \|Undefined Saprotroph\| | Yeast | NULL | Probable | As nectar-tap_saprotroph (Põlme et al. 2020); Decay sugar-rich substrates (Põlme et al. 2020) | Põlme S, et al. 2020. Fungal Diversity 105:1-16 ((https://doi.org/10.1007/s13225-020-00466-2)) |
| Otu37100 | 1 | k__Fungi(100);p__Ascomycota(100);c__Saccharomycetes(100);o__Saccharomycetales(100);f__Metschnikowiaceae(100);g__Metschnikowia(100);s__Metschnikowia_pulcherrima(100); | Metschnikowia | 13 | Saprotroph | \|Undefined Saprotroph\| | Yeast | NULL | Probable | As nectar-tap_saprotroph (Põlme et al. 2020); Decay sugar-rich substrates (Põlme et al. 2020) | Põlme S, et al. 2020. Fungal Diversity 105:1-16 ((https://doi.org/10.1007/s13225-020-00466-2)) |
| Otu37101 | 1 | k__Fungi(100);p__Ascomycota(100);c__Saccharomycetes(100);o__Saccharomycetales(100);f__Metschnikowiaceae(100);g__Metschnikowia(100);s__Metschnikowia_pulcherrima(100); | Metschnikowia | 13 | Saprotroph | \|Undefined Saprotroph\| | Yeast | NULL | Probable | As nectar-tap_saprotroph (Põlme et al. 2020); Decay sugar-rich substrates (Põlme et al. 2020) | Põlme S, et al. 2020. Fungal Diversity 105:1-16 ((https://doi.org/10.1007/s13225-020-00466-2)) |
| Otu37102 | 1 | k__Fungi(100);p__Ascomycota(100);c__Saccharomycetes(100);o__Saccharomycetales(100);f__Metschnikowiaceae(100);g__Metschnikowia(100);s__Metschnikowia_pulcherrima(100); | Metschnikowia | 13 | Saprotroph | \|Undefined Saprotroph\| | Yeast | NULL | Probable | As nectar-tap_saprotroph (Põlme et al. 2020); Decay sugar-rich substrates (Põlme et al. 2020) | Põlme S, et al. 2020. Fungal Diversity 105:1-16 ((https://doi.org/10.1007/s13225-020-00466-2)) |
| Otu37103 | 1 | k__Fungi(100);p__Ascomycota(100);c__Saccharomycetes(100);o__Saccharomycetales(100);f__Metschnikowiaceae(100);g__Metschnikowia(100);g__Metschnikowia_unclassified(100); | Metschnikowia | 13 | Saprotroph | \|Undefined Saprotroph\| | Yeast | NULL | Probable | As nectar-tap_saprotroph (Põlme et al. 2020); Decay sugar-rich substrates (Põlme et al. 2020) | Põlme S, et al. 2020. Fungal Diversity 105:1-16 ((https://doi.org/10.1007/s13225-020-00466-2)) |
| Otu37104 | 1 | k__Fungi(100);p__Ascomycota(100);c__Saccharomycetes(100);o__Saccharomycetales(100);f__Metschnikowiaceae(100);g__Metschnikowia(100);g__Metschnikowia_unclassified(100); | Metschnikowia | 13 | Saprotroph | \|Undefined Saprotroph\| | Yeast | NULL | Probable | As nectar-tap_saprotroph (Põlme et al. 2020); Decay sugar-rich substrates (Põlme et al. 2020) | Põlme S, et al. 2020. Fungal Diversity 105:1-16 ((https://doi.org/10.1007/s13225-020-00466-2)) |
| Otu37105 | 1 | k__Fungi(100);p__Ascomycota(100);c__Saccharomycetes(100);o__Saccharomycetales(100);f__Metschnikowiaceae(100);g__Metschnikowia(100);s__Metschnikowia_pulcherrima(100); | Metschnikowia | 13 | Saprotroph | \|Undefined Saprotroph\| | Yeast | NULL | Probable | As nectar-tap_saprotroph (Põlme et al. 2020); Decay sugar-rich substrates (Põlme et al. 2020) | Põlme S, et al. 2020. Fungal Diversity 105:1-16 ((https://doi.org/10.1007/s13225-020-00466-2)) |
| Otu37106 | 1 | k__Fungi(100);p__Ascomycota(100);c__Saccharomycetes(100);o__Saccharomycetales(100);f__Metschnikowiaceae(100);g__Metschnikowia(100);g__Metschnikowia_unclassified(100); | Metschnikowia | 13 | Saprotroph | \|Undefined Saprotroph\| | Yeast | NULL | Probable | As nectar-tap_saprotroph (Põlme et al. 2020); Decay sugar-rich substrates (Põlme et al. 2020) | Põlme S, et al. 2020. Fungal Diversity 105:1-16 ((https://doi.org/10.1007/s13225-020-00466-2)) |
| Otu37107 | 1 | k__Fungi(100);p__Ascomycota(100);c__Saccharomycetes(100);o__Saccharomycetales(100);f__Metschnikowiaceae(100);g__Metschnikowia(100);s__Metschnikowia_pulcherrima(100); | Metschnikowia | 13 | Saprotroph | \|Undefined Saprotroph\| | Yeast | NULL | Probable | As nectar-tap_saprotroph (Põlme et al. 2020); Decay sugar-rich substrates (Põlme et al. 2020) | Põlme S, et al. 2020. Fungal Diversity 105:1-16 ((https://doi.org/10.1007/s13225-020-00466-2)) |
| Otu37109 | 1 | k__Fungi(100);p__Ascomycota(100);c__Saccharomycetes(100);o__Saccharomycetales(100);f__Metschnikowiaceae(100);g__Metschnikowia(100);s__Metschnikowia_pulcherrima(100); | Metschnikowia | 13 | Saprotroph | \|Undefined Saprotroph\| | Yeast | NULL | Probable | As nectar-tap_saprotroph (Põlme et al. 2020); Decay sugar-rich substrates (Põlme et al. 2020) | Põlme S, et al. 2020. Fungal Diversity 105:1-16 ((https://doi.org/10.1007/s13225-020-00466-2)) |
| Otu37110 | 1 | k__Fungi(100);p__Ascomycota(100);c__Saccharomycetes(100);o__Saccharomycetales(100);f__Metschnikowiaceae(100);g__Metschnikowia(100);s__Metschnikowia_pulcherrima(100); | Metschnikowia | 13 | Saprotroph | \|Undefined Saprotroph\| | Yeast | NULL | Probable | As nectar-tap_saprotroph (Põlme et al. 2020); Decay sugar-rich substrates (Põlme et al. 2020) | Põlme S, et al. 2020. Fungal Diversity 105:1-16 ((https://doi.org/10.1007/s13225-020-00466-2)) |
| Otu37111 | 1 | k__Fungi(100);p__Ascomycota(100);c__Saccharomycetes(100);o__Saccharomycetales(100);f__Metschnikowiaceae(100);g__Metschnikowia(100);s__Metschnikowia_pulcherrima(100); | Metschnikowia | 13 | Saprotroph | \|Undefined Saprotroph\| | Yeast | NULL | Probable | As nectar-tap_saprotroph (Põlme et al. 2020); Decay sugar-rich substrates (Põlme et al. 2020) | Põlme S, et al. 2020. Fungal Diversity 105:1-16 ((https://doi.org/10.1007/s13225-020-00466-2)) |
| Otu37112 | 1 | k__Fungi(100);p__Ascomycota(100);c__Saccharomycetes(100);o__Saccharomycetales(100);f__Metschnikowiaceae(100);g__Metschnikowia(100);s__Metschnikowia_pulcherrima(100); | Metschnikowia | 13 | Saprotroph | \|Undefined Saprotroph\| | Yeast | NULL | Probable | As nectar-tap_saprotroph (Põlme et al. 2020); Decay sugar-rich substrates (Põlme et al. 2020) | Põlme S, et al. 2020. Fungal Diversity 105:1-16 ((https://doi.org/10.1007/s13225-020-00466-2)) |
| Otu37115 | 1 | k__Fungi(100);p__Ascomycota(100);c__Saccharomycetes(100);o__Saccharomycetales(100);f__Metschnikowiaceae(100);g__Metschnikowia(100);s__Metschnikowia_pulcherrima(100); | Metschnikowia | 13 | Saprotroph | \|Undefined Saprotroph\| | Yeast | NULL | Probable | As nectar-tap_saprotroph (Põlme et al. 2020); Decay sugar-rich substrates (Põlme et al. 2020) | Põlme S, et al. 2020. Fungal Diversity 105:1-16 ((https://doi.org/10.1007/s13225-020-00466-2)) |
| Otu37117 | 1 | k__Fungi(100);p__Ascomycota(100);c__Saccharomycetes(100);o__Saccharomycetales(100);f__Metschnikowiaceae(100);g__Metschnikowia(100);g__Metschnikowia_unclassified(100); | Metschnikowia | 13 | Saprotroph | \|Undefined Saprotroph\| | Yeast | NULL | Probable | As nectar-tap_saprotroph (Põlme et al. 2020); Decay sugar-rich substrates (Põlme et al. 2020) | Põlme S, et al. 2020. Fungal Diversity 105:1-16 ((https://doi.org/10.1007/s13225-020-00466-2)) |
| Otu37118 | 1 | k__Fungi(100);p__Ascomycota(100);c__Saccharomycetes(100);o__Saccharomycetales(100);f__Metschnikowiaceae(100);g__Metschnikowia(100);g__Metschnikowia_unclassified(100); | Metschnikowia | 13 | Saprotroph | \|Undefined Saprotroph\| | Yeast | NULL | Probable | As nectar-tap_saprotroph (Põlme et al. 2020); Decay sugar-rich substrates (Põlme et al. 2020) | Põlme S, et al. 2020. Fungal Diversity 105:1-16 ((https://doi.org/10.1007/s13225-020-00466-2)) |
| Otu37119 | 1 | k__Fungi(100);p__Ascomycota(100);c__Saccharomycetes(100);o__Saccharomycetales(100);f__Metschnikowiaceae(100);g__Metschnikowia(100);s__Metschnikowia_pulcherrima(100); | Metschnikowia | 13 | Saprotroph | \|Undefined Saprotroph\| | Yeast | NULL | Probable | As nectar-tap_saprotroph (Põlme et al. 2020); Decay sugar-rich substrates (Põlme et al. 2020) | Põlme S, et al. 2020. Fungal Diversity 105:1-16 ((https://doi.org/10.1007/s13225-020-00466-2)) |
| Otu37120 | 1 | k__Fungi(100);p__Ascomycota(100);c__Saccharomycetes(100);o__Saccharomycetales(100);f__Metschnikowiaceae(100);g__Metschnikowia(100);s__Metschnikowia_pulcherrima(100); | Metschnikowia | 13 | Saprotroph | \|Undefined Saprotroph\| | Yeast | NULL | Probable | As nectar-tap_saprotroph (Põlme et al. 2020); Decay sugar-rich substrates (Põlme et al. 2020) | Põlme S, et al. 2020. Fungal Diversity 105:1-16 ((https://doi.org/10.1007/s13225-020-00466-2)) |
| Otu37121 | 1 | k__Fungi(100);p__Ascomycota(100);c__Saccharomycetes(100);o__Saccharomycetales(100);f__Metschnikowiaceae(100);g__Metschnikowia(100);s__Metschnikowia_pulcherrima(100); | Metschnikowia | 13 | Saprotroph | \|Undefined Saprotroph\| | Yeast | NULL | Probable | As nectar-tap_saprotroph (Põlme et al. 2020); Decay sugar-rich substrates (Põlme et al. 2020) | Põlme S, et al. 2020. Fungal Diversity 105:1-16 ((https://doi.org/10.1007/s13225-020-00466-2)) |
| Otu37122 | 1 | k__Fungi(100);p__Ascomycota(100);c__Saccharomycetes(100);o__Saccharomycetales(100);f__Metschnikowiaceae(100);g__Metschnikowia(100);s__Metschnikowia_pulcherrima(100); | Metschnikowia | 13 | Saprotroph | \|Undefined Saprotroph\| | Yeast | NULL | Probable | As nectar-tap_saprotroph (Põlme et al. 2020); Decay sugar-rich substrates (Põlme et al. 2020) | Põlme S, et al. 2020. Fungal Diversity 105:1-16 ((https://doi.org/10.1007/s13225-020-00466-2)) |
| Otu37123 | 1 | k__Fungi(100);p__Ascomycota(100);c__Saccharomycetes(100);o__Saccharomycetales(100);f__Metschnikowiaceae(100);g__Metschnikowia(100);s__Metschnikowia_pulcherrima(100); | Metschnikowia | 13 | Saprotroph | \|Undefined Saprotroph\| | Yeast | NULL | Probable | As nectar-tap_saprotroph (Põlme et al. 2020); Decay sugar-rich substrates (Põlme et al. 2020) | Põlme S, et al. 2020. Fungal Diversity 105:1-16 ((https://doi.org/10.1007/s13225-020-00466-2)) |
| Otu37124 | 1 | k__Fungi(100);p__Ascomycota(100);c__Saccharomycetes(100);o__Saccharomycetales(100);f__Metschnikowiaceae(100);g__Metschnikowia(100);s__Metschnikowia_pulcherrima(100); | Metschnikowia | 13 | Saprotroph | \|Undefined Saprotroph\| | Yeast | NULL | Probable | As nectar-tap_saprotroph (Põlme et al. 2020); Decay sugar-rich substrates (Põlme et al. 2020) | Põlme S, et al. 2020. Fungal Diversity 105:1-16 ((https://doi.org/10.1007/s13225-020-00466-2)) |
| Otu37127 | 1 | k__Fungi(100);p__Ascomycota(100);c__Saccharomycetes(100);o__Saccharomycetales(100);f__Metschnikowiaceae(100);g__Metschnikowia(100);s__Metschnikowia_pulcherrima(100); | Metschnikowia | 13 | Saprotroph | \|Undefined Saprotroph\| | Yeast | NULL | Probable | As nectar-tap_saprotroph (Põlme et al. 2020); Decay sugar-rich substrates (Põlme et al. 2020) | Põlme S, et al. 2020. Fungal Diversity 105:1-16 ((https://doi.org/10.1007/s13225-020-00466-2)) |
| Otu37128 | 1 | k__Fungi(100);p__Ascomycota(100);c__Saccharomycetes(100);o__Saccharomycetales(100);f__Metschnikowiaceae(100);g__Metschnikowia(100);s__Metschnikowia_pulcherrima(100); | Metschnikowia | 13 | Saprotroph | \|Undefined Saprotroph\| | Yeast | NULL | Probable | As nectar-tap_saprotroph (Põlme et al. 2020); Decay sugar-rich substrates (Põlme et al. 2020) | Põlme S, et al. 2020. Fungal Diversity 105:1-16 ((https://doi.org/10.1007/s13225-020-00466-2)) |
| Otu37130 | 1 | k__Fungi(100);p__Ascomycota(100);c__Saccharomycetes(100);o__Saccharomycetales(100);f__Metschnikowiaceae(100);g__Metschnikowia(100);s__Metschnikowia_pulcherrima(100); | Metschnikowia | 13 | Saprotroph | \|Undefined Saprotroph\| | Yeast | NULL | Probable | As nectar-tap_saprotroph (Põlme et al. 2020); Decay sugar-rich substrates (Põlme et al. 2020) | Põlme S, et al. 2020. Fungal Diversity 105:1-16 ((https://doi.org/10.1007/s13225-020-00466-2)) |
| Otu37131 | 1 | k__Fungi(100);p__Ascomycota(100);c__Saccharomycetes(100);o__Saccharomycetales(100);f__Metschnikowiaceae(100);g__Metschnikowia(100);s__Metschnikowia_pulcherrima(100); | Metschnikowia | 13 | Saprotroph | \|Undefined Saprotroph\| | Yeast | NULL | Probable | As nectar-tap_saprotroph (Põlme et al. 2020); Decay sugar-rich substrates (Põlme et al. 2020) | Põlme S, et al. 2020. Fungal Diversity 105:1-16 ((https://doi.org/10.1007/s13225-020-00466-2)) |
| Otu37133 | 1 | k__Fungi(100);p__Ascomycota(100);c__Saccharomycetes(100);o__Saccharomycetales(100);f__Metschnikowiaceae(100);g__Metschnikowia(100);g__Metschnikowia_unclassified(100); | Metschnikowia | 13 | Saprotroph | \|Undefined Saprotroph\| | Yeast | NULL | Probable | As nectar-tap_saprotroph (Põlme et al. 2020); Decay sugar-rich substrates (Põlme et al. 2020) | Põlme S, et al. 2020. Fungal Diversity 105:1-16 ((https://doi.org/10.1007/s13225-020-00466-2)) |
| Otu37134 | 1 | k__Fungi(100);p__Ascomycota(100);c__Saccharomycetes(100);o__Saccharomycetales(100);f__Metschnikowiaceae(100);g__Metschnikowia(100);s__Metschnikowia_pulcherrima(100); | Metschnikowia | 13 | Saprotroph | \|Undefined Saprotroph\| | Yeast | NULL | Probable | As nectar-tap_saprotroph (Põlme et al. 2020); Decay sugar-rich substrates (Põlme et al. 2020) | Põlme S, et al. 2020. Fungal Diversity 105:1-16 ((https://doi.org/10.1007/s13225-020-00466-2)) |
| Otu37135 | 1 | k__Fungi(100);p__Ascomycota(100);c__Saccharomycetes(100);o__Saccharomycetales(100);f__Metschnikowiaceae(100);g__Metschnikowia(100);s__Metschnikowia_pulcherrima(100); | Metschnikowia | 13 | Saprotroph | \|Undefined Saprotroph\| | Yeast | NULL | Probable | As nectar-tap_saprotroph (Põlme et al. 2020); Decay sugar-rich substrates (Põlme et al. 2020) | Põlme S, et al. 2020. Fungal Diversity 105:1-16 ((https://doi.org/10.1007/s13225-020-00466-2)) |
| Otu37136 | 1 | k__Fungi(100);p__Ascomycota(100);c__Saccharomycetes(100);o__Saccharomycetales(100);f__Metschnikowiaceae(100);g__Metschnikowia(100);s__Metschnikowia_pulcherrima(100); | Metschnikowia | 13 | Saprotroph | \|Undefined Saprotroph\| | Yeast | NULL | Probable | As nectar-tap_saprotroph (Põlme et al. 2020); Decay sugar-rich substrates (Põlme et al. 2020) | Põlme S, et al. 2020. Fungal Diversity 105:1-16 ((https://doi.org/10.1007/s13225-020-00466-2)) |
| Otu37137 | 1 | k__Fungi(100);p__Ascomycota(100);c__Saccharomycetes(100);o__Saccharomycetales(100);f__Metschnikowiaceae(100);g__Metschnikowia(100);s__Metschnikowia_pulcherrima(100); | Metschnikowia | 13 | Saprotroph | \|Undefined Saprotroph\| | Yeast | NULL | Probable | As nectar-tap_saprotroph (Põlme et al. 2020); Decay sugar-rich substrates (Põlme et al. 2020) | Põlme S, et al. 2020. Fungal Diversity 105:1-16 ((https://doi.org/10.1007/s13225-020-00466-2)) |
| Otu37138 | 1 | k__Fungi(100);p__Ascomycota(100);c__Saccharomycetes(100);o__Saccharomycetales(100);f__Metschnikowiaceae(100);g__Metschnikowia(100);g__Metschnikowia_unclassified(100); | Metschnikowia | 13 | Saprotroph | \|Undefined Saprotroph\| | Yeast | NULL | Probable | As nectar-tap_saprotroph (Põlme et al. 2020); Decay sugar-rich substrates (Põlme et al. 2020) | Põlme S, et al. 2020. Fungal Diversity 105:1-16 ((https://doi.org/10.1007/s13225-020-00466-2)) |
| Otu37139 | 1 | k__Fungi(100);p__Ascomycota(100);c__Saccharomycetes(100);o__Saccharomycetales(100);f__Metschnikowiaceae(100);g__Metschnikowia(100);s__Metschnikowia_pulcherrima(100); | Metschnikowia | 13 | Saprotroph | \|Undefined Saprotroph\| | Yeast | NULL | Probable | As nectar-tap_saprotroph (Põlme et al. 2020); Decay sugar-rich substrates (Põlme et al. 2020) | Põlme S, et al. 2020. Fungal Diversity 105:1-16 ((https://doi.org/10.1007/s13225-020-00466-2)) |
| Otu37140 | 1 | k__Fungi(100);p__Ascomycota(100);c__Saccharomycetes(100);o__Saccharomycetales(100);f__Metschnikowiaceae(100);g__Metschnikowia(100);g__Metschnikowia_unclassified(100); | Metschnikowia | 13 | Saprotroph | \|Undefined Saprotroph\| | Yeast | NULL | Probable | As nectar-tap_saprotroph (Põlme et al. 2020); Decay sugar-rich substrates (Põlme et al. 2020) | Põlme S, et al. 2020. Fungal Diversity 105:1-16 ((https://doi.org/10.1007/s13225-020-00466-2)) |
| Otu37141 | 1 | k__Fungi(100);p__Ascomycota(100);c__Saccharomycetes(100);o__Saccharomycetales(100);f__Metschnikowiaceae(100);g__Metschnikowia(100);s__Metschnikowia_pulcherrima(100); | Metschnikowia | 13 | Saprotroph | \|Undefined Saprotroph\| | Yeast | NULL | Probable | As nectar-tap_saprotroph (Põlme et al. 2020); Decay sugar-rich substrates (Põlme et al. 2020) | Põlme S, et al. 2020. Fungal Diversity 105:1-16 ((https://doi.org/10.1007/s13225-020-00466-2)) |
| Otu37142 | 1 | k__Fungi(100);p__Ascomycota(100);c__Saccharomycetes(100);o__Saccharomycetales(100);f__Metschnikowiaceae(100);g__Metschnikowia(100);g__Metschnikowia_unclassified(100); | Metschnikowia | 13 | Saprotroph | \|Undefined Saprotroph\| | Yeast | NULL | Probable | As nectar-tap_saprotroph (Põlme et al. 2020); Decay sugar-rich substrates (Põlme et al. 2020) | Põlme S, et al. 2020. Fungal Diversity 105:1-16 ((https://doi.org/10.1007/s13225-020-00466-2)) |
| Otu37143 | 1 | k__Fungi(100);p__Ascomycota(100);c__Saccharomycetes(100);o__Saccharomycetales(100);f__Metschnikowiaceae(100);g__Metschnikowia(100);s__Metschnikowia_pulcherrima(100); | Metschnikowia | 13 | Saprotroph | \|Undefined Saprotroph\| | Yeast | NULL | Probable | As nectar-tap_saprotroph (Põlme et al. 2020); Decay sugar-rich substrates (Põlme et al. 2020) | Põlme S, et al. 2020. Fungal Diversity 105:1-16 ((https://doi.org/10.1007/s13225-020-00466-2)) |
| Otu37144 | 1 | k__Fungi(100);p__Ascomycota(100);c__Saccharomycetes(100);o__Saccharomycetales(100);f__Metschnikowiaceae(100);g__Metschnikowia(100);g__Metschnikowia_unclassified(100); | Metschnikowia | 13 | Saprotroph | \|Undefined Saprotroph\| | Yeast | NULL | Probable | As nectar-tap_saprotroph (Põlme et al. 2020); Decay sugar-rich substrates (Põlme et al. 2020) | Põlme S, et al. 2020. Fungal Diversity 105:1-16 ((https://doi.org/10.1007/s13225-020-00466-2)) |
| Otu37145 | 1 | k__Fungi(100);p__Ascomycota(100);c__Saccharomycetes(100);o__Saccharomycetales(100);f__Metschnikowiaceae(100);g__Metschnikowia(100);g__Metschnikowia_unclassified(100); | Metschnikowia | 13 | Saprotroph | \|Undefined Saprotroph\| | Yeast | NULL | Probable | As nectar-tap_saprotroph (Põlme et al. 2020); Decay sugar-rich substrates (Põlme et al. 2020) | Põlme S, et al. 2020. Fungal Diversity 105:1-16 ((https://doi.org/10.1007/s13225-020-00466-2)) |
| Otu37146 | 1 | k__Fungi(100);p__Ascomycota(100);c__Saccharomycetes(100);o__Saccharomycetales(100);f__Metschnikowiaceae(100);g__Metschnikowia(100);g__Metschnikowia_unclassified(100); | Metschnikowia | 13 | Saprotroph | \|Undefined Saprotroph\| | Yeast | NULL | Probable | As nectar-tap_saprotroph (Põlme et al. 2020); Decay sugar-rich substrates (Põlme et al. 2020) | Põlme S, et al. 2020. Fungal Diversity 105:1-16 ((https://doi.org/10.1007/s13225-020-00466-2)) |
| Otu37147 | 1 | k__Fungi(100);p__Ascomycota(100);c__Saccharomycetes(100);o__Saccharomycetales(100);f__Metschnikowiaceae(100);g__Metschnikowia(100);s__Metschnikowia_pulcherrima(100); | Metschnikowia | 13 | Saprotroph | \|Undefined Saprotroph\| | Yeast | NULL | Probable | As nectar-tap_saprotroph (Põlme et al. 2020); Decay sugar-rich substrates (Põlme et al. 2020) | Põlme S, et al. 2020. Fungal Diversity 105:1-16 ((https://doi.org/10.1007/s13225-020-00466-2)) |
| Otu37148 | 1 | k__Fungi(100);p__Ascomycota(100);c__Saccharomycetes(100);o__Saccharomycetales(100);f__Metschnikowiaceae(100);g__Metschnikowia(100);s__Metschnikowia_pulcherrima(100); | Metschnikowia | 13 | Saprotroph | \|Undefined Saprotroph\| | Yeast | NULL | Probable | As nectar-tap_saprotroph (Põlme et al. 2020); Decay sugar-rich substrates (Põlme et al. 2020) | Põlme S, et al. 2020. Fungal Diversity 105:1-16 ((https://doi.org/10.1007/s13225-020-00466-2)) |
| Otu37149 | 1 | k__Fungi(100);p__Ascomycota(100);c__Saccharomycetes(100);o__Saccharomycetales(100);f__Metschnikowiaceae(100);g__Metschnikowia(100);s__Metschnikowia_pulcherrima(100); | Metschnikowia | 13 | Saprotroph | \|Undefined Saprotroph\| | Yeast | NULL | Probable | As nectar-tap_saprotroph (Põlme et al. 2020); Decay sugar-rich substrates (Põlme et al. 2020) | Põlme S, et al. 2020. Fungal Diversity 105:1-16 ((https://doi.org/10.1007/s13225-020-00466-2)) |
| Otu37150 | 1 | k__Fungi(100);p__Ascomycota(100);c__Saccharomycetes(100);o__Saccharomycetales(100);f__Metschnikowiaceae(100);g__Metschnikowia(100);s__Metschnikowia_pulcherrima(100); | Metschnikowia | 13 | Saprotroph | \|Undefined Saprotroph\| | Yeast | NULL | Probable | As nectar-tap_saprotroph (Põlme et al. 2020); Decay sugar-rich substrates (Põlme et al. 2020) | Põlme S, et al. 2020. Fungal Diversity 105:1-16 ((https://doi.org/10.1007/s13225-020-00466-2)) |
| Otu37151 | 1 | k__Fungi(100);p__Ascomycota(100);c__Saccharomycetes(100);o__Saccharomycetales(100);f__Metschnikowiaceae(100);g__Metschnikowia(100);g__Metschnikowia_unclassified(100); | Metschnikowia | 13 | Saprotroph | \|Undefined Saprotroph\| | Yeast | NULL | Probable | As nectar-tap_saprotroph (Põlme et al. 2020); Decay sugar-rich substrates (Põlme et al. 2020) | Põlme S, et al. 2020. Fungal Diversity 105:1-16 ((https://doi.org/10.1007/s13225-020-00466-2)) |
| Otu37152 | 1 | k__Fungi(100);p__Ascomycota(100);c__Saccharomycetes(100);o__Saccharomycetales(100);f__Metschnikowiaceae(100);g__Metschnikowia(100);g__Metschnikowia_unclassified(100); | Metschnikowia | 13 | Saprotroph | \|Undefined Saprotroph\| | Yeast | NULL | Probable | As nectar-tap_saprotroph (Põlme et al. 2020); Decay sugar-rich substrates (Põlme et al. 2020) | Põlme S, et al. 2020. Fungal Diversity 105:1-16 ((https://doi.org/10.1007/s13225-020-00466-2)) |
| Otu37153 | 1 | k__Fungi(100);p__Ascomycota(100);c__Saccharomycetes(100);o__Saccharomycetales(100);f__Metschnikowiaceae(100);g__Metschnikowia(100);s__Metschnikowia_pulcherrima(100); | Metschnikowia | 13 | Saprotroph | \|Undefined Saprotroph\| | Yeast | NULL | Probable | As nectar-tap_saprotroph (Põlme et al. 2020); Decay sugar-rich substrates (Põlme et al. 2020) | Põlme S, et al. 2020. Fungal Diversity 105:1-16 ((https://doi.org/10.1007/s13225-020-00466-2)) |
| Otu37154 | 1 | k__Fungi(100);p__Ascomycota(100);c__Saccharomycetes(100);o__Saccharomycetales(100);f__Metschnikowiaceae(100);g__Metschnikowia(100);s__Metschnikowia_pulcherrima(100); | Metschnikowia | 13 | Saprotroph | \|Undefined Saprotroph\| | Yeast | NULL | Probable | As nectar-tap_saprotroph (Põlme et al. 2020); Decay sugar-rich substrates (Põlme et al. 2020) | Põlme S, et al. 2020. Fungal Diversity 105:1-16 ((https://doi.org/10.1007/s13225-020-00466-2)) |
| Otu37155 | 1 | k__Fungi(100);p__Ascomycota(100);c__Saccharomycetes(100);o__Saccharomycetales(100);f__Metschnikowiaceae(100);g__Metschnikowia(100);s__Metschnikowia_pulcherrima(100); | Metschnikowia | 13 | Saprotroph | \|Undefined Saprotroph\| | Yeast | NULL | Probable | As nectar-tap_saprotroph (Põlme et al. 2020); Decay sugar-rich substrates (Põlme et al. 2020) | Põlme S, et al. 2020. Fungal Diversity 105:1-16 ((https://doi.org/10.1007/s13225-020-00466-2)) |
| Otu37156 | 1 | k__Fungi(100);p__Ascomycota(100);c__Saccharomycetes(100);o__Saccharomycetales(100);f__Metschnikowiaceae(100);g__Metschnikowia(100);s__Metschnikowia_pulcherrima(100); | Metschnikowia | 13 | Saprotroph | \|Undefined Saprotroph\| | Yeast | NULL | Probable | As nectar-tap_saprotroph (Põlme et al. 2020); Decay sugar-rich substrates (Põlme et al. 2020) | Põlme S, et al. 2020. Fungal Diversity 105:1-16 ((https://doi.org/10.1007/s13225-020-00466-2)) |
| Otu37157 | 1 | k__Fungi(100);p__Ascomycota(100);c__Saccharomycetes(100);o__Saccharomycetales(100);f__Metschnikowiaceae(100);g__Metschnikowia(100);g__Metschnikowia_unclassified(100); | Metschnikowia | 13 | Saprotroph | \|Undefined Saprotroph\| | Yeast | NULL | Probable | As nectar-tap_saprotroph (Põlme et al. 2020); Decay sugar-rich substrates (Põlme et al. 2020) | Põlme S, et al. 2020. Fungal Diversity 105:1-16 ((https://doi.org/10.1007/s13225-020-00466-2)) |
| Otu37158 | 1 | k__Fungi(100);p__Ascomycota(100);c__Saccharomycetes(100);o__Saccharomycetales(100);f__Metschnikowiaceae(100);g__Metschnikowia(100);g__Metschnikowia_unclassified(100); | Metschnikowia | 13 | Saprotroph | \|Undefined Saprotroph\| | Yeast | NULL | Probable | As nectar-tap_saprotroph (Põlme et al. 2020); Decay sugar-rich substrates (Põlme et al. 2020) | Põlme S, et al. 2020. Fungal Diversity 105:1-16 ((https://doi.org/10.1007/s13225-020-00466-2)) |
| Otu37159 | 1 | k__Fungi(100);p__Ascomycota(100);c__Saccharomycetes(100);o__Saccharomycetales(100);f__Metschnikowiaceae(100);g__Metschnikowia(100);g__Metschnikowia_unclassified(100); | Metschnikowia | 13 | Saprotroph | \|Undefined Saprotroph\| | Yeast | NULL | Probable | As nectar-tap_saprotroph (Põlme et al. 2020); Decay sugar-rich substrates (Põlme et al. 2020) | Põlme S, et al. 2020. Fungal Diversity 105:1-16 ((https://doi.org/10.1007/s13225-020-00466-2)) |
| Otu37160 | 1 | k__Fungi(100);p__Ascomycota(100);c__Saccharomycetes(100);o__Saccharomycetales(100);f__Metschnikowiaceae(100);g__Metschnikowia(100);s__Metschnikowia_pulcherrima(100); | Metschnikowia | 13 | Saprotroph | \|Undefined Saprotroph\| | Yeast | NULL | Probable | As nectar-tap_saprotroph (Põlme et al. 2020); Decay sugar-rich substrates (Põlme et al. 2020) | Põlme S, et al. 2020. Fungal Diversity 105:1-16 ((https://doi.org/10.1007/s13225-020-00466-2)) |
| Otu37161 | 1 | k__Fungi(100);p__Ascomycota(100);c__Saccharomycetes(100);o__Saccharomycetales(100);f__Metschnikowiaceae(100);g__Metschnikowia(100);s__Metschnikowia_pulcherrima(100); | Metschnikowia | 13 | Saprotroph | \|Undefined Saprotroph\| | Yeast | NULL | Probable | As nectar-tap_saprotroph (Põlme et al. 2020); Decay sugar-rich substrates (Põlme et al. 2020) | Põlme S, et al. 2020. Fungal Diversity 105:1-16 ((https://doi.org/10.1007/s13225-020-00466-2)) |
| Otu37162 | 1 | k__Fungi(100);p__Ascomycota(100);c__Saccharomycetes(100);o__Saccharomycetales(100);f__Metschnikowiaceae(100);g__Metschnikowia(100);s__Metschnikowia_pulcherrima(100); | Metschnikowia | 13 | Saprotroph | \|Undefined Saprotroph\| | Yeast | NULL | Probable | As nectar-tap_saprotroph (Põlme et al. 2020); Decay sugar-rich substrates (Põlme et al. 2020) | Põlme S, et al. 2020. Fungal Diversity 105:1-16 ((https://doi.org/10.1007/s13225-020-00466-2)) |
| Otu37163 | 1 | k__Fungi(100);p__Ascomycota(100);c__Saccharomycetes(100);o__Saccharomycetales(100);f__Metschnikowiaceae(100);g__Metschnikowia(100);s__Metschnikowia_pulcherrima(100); | Metschnikowia | 13 | Saprotroph | \|Undefined Saprotroph\| | Yeast | NULL | Probable | As nectar-tap_saprotroph (Põlme et al. 2020); Decay sugar-rich substrates (Põlme et al. 2020) | Põlme S, et al. 2020. Fungal Diversity 105:1-16 ((https://doi.org/10.1007/s13225-020-00466-2)) |
| Otu37164 | 1 | k__Fungi(100);p__Ascomycota(100);c__Saccharomycetes(100);o__Saccharomycetales(100);f__Metschnikowiaceae(100);g__Metschnikowia(100);g__Metschnikowia_unclassified(100); | Metschnikowia | 13 | Saprotroph | \|Undefined Saprotroph\| | Yeast | NULL | Probable | As nectar-tap_saprotroph (Põlme et al. 2020); Decay sugar-rich substrates (Põlme et al. 2020) | Põlme S, et al. 2020. Fungal Diversity 105:1-16 ((https://doi.org/10.1007/s13225-020-00466-2)) |
| Otu37165 | 1 | k__Fungi(100);p__Ascomycota(100);c__Saccharomycetes(100);o__Saccharomycetales(100);f__Metschnikowiaceae(100);g__Metschnikowia(100);g__Metschnikowia_unclassified(100); | Metschnikowia | 13 | Saprotroph | \|Undefined Saprotroph\| | Yeast | NULL | Probable | As nectar-tap_saprotroph (Põlme et al. 2020); Decay sugar-rich substrates (Põlme et al. 2020) | Põlme S, et al. 2020. Fungal Diversity 105:1-16 ((https://doi.org/10.1007/s13225-020-00466-2)) |
| Otu37166 | 1 | k__Fungi(100);p__Ascomycota(100);c__Saccharomycetes(100);o__Saccharomycetales(100);f__Metschnikowiaceae(100);g__Metschnikowia(100);s__Metschnikowia_pulcherrima(100); | Metschnikowia | 13 | Saprotroph | \|Undefined Saprotroph\| | Yeast | NULL | Probable | As nectar-tap_saprotroph (Põlme et al. 2020); Decay sugar-rich substrates (Põlme et al. 2020) | Põlme S, et al. 2020. Fungal Diversity 105:1-16 ((https://doi.org/10.1007/s13225-020-00466-2)) |
| Otu37167 | 1 | k__Fungi(100);p__Ascomycota(100);c__Saccharomycetes(100);o__Saccharomycetales(100);f__Metschnikowiaceae(100);g__Metschnikowia(100);s__Metschnikowia_pulcherrima(100); | Metschnikowia | 13 | Saprotroph | \|Undefined Saprotroph\| | Yeast | NULL | Probable | As nectar-tap_saprotroph (Põlme et al. 2020); Decay sugar-rich substrates (Põlme et al. 2020) | Põlme S, et al. 2020. Fungal Diversity 105:1-16 ((https://doi.org/10.1007/s13225-020-00466-2)) |
| Otu37168 | 1 | k__Fungi(100);p__Ascomycota(100);c__Saccharomycetes(100);o__Saccharomycetales(100);f__Metschnikowiaceae(100);g__Metschnikowia(100);s__Metschnikowia_pulcherrima(100); | Metschnikowia | 13 | Saprotroph | \|Undefined Saprotroph\| | Yeast | NULL | Probable | As nectar-tap_saprotroph (Põlme et al. 2020); Decay sugar-rich substrates (Põlme et al. 2020) | Põlme S, et al. 2020. Fungal Diversity 105:1-16 ((https://doi.org/10.1007/s13225-020-00466-2)) |
| Otu37169 | 1 | k__Fungi(100);p__Ascomycota(100);c__Saccharomycetes(100);o__Saccharomycetales(100);f__Metschnikowiaceae(100);g__Metschnikowia(100);g__Metschnikowia_unclassified(100); | Metschnikowia | 13 | Saprotroph | \|Undefined Saprotroph\| | Yeast | NULL | Probable | As nectar-tap_saprotroph (Põlme et al. 2020); Decay sugar-rich substrates (Põlme et al. 2020) | Põlme S, et al. 2020. Fungal Diversity 105:1-16 ((https://doi.org/10.1007/s13225-020-00466-2)) |
| Otu37171 | 1 | k__Fungi(100);p__Ascomycota(100);c__Saccharomycetes(100);o__Saccharomycetales(100);f__Metschnikowiaceae(100);g__Metschnikowia(100);s__Metschnikowia_pulcherrima(100); | Metschnikowia | 13 | Saprotroph | \|Undefined Saprotroph\| | Yeast | NULL | Probable | As nectar-tap_saprotroph (Põlme et al. 2020); Decay sugar-rich substrates (Põlme et al. 2020) | Põlme S, et al. 2020. Fungal Diversity 105:1-16 ((https://doi.org/10.1007/s13225-020-00466-2)) |
| Otu37172 | 1 | k__Fungi(100);p__Ascomycota(100);c__Saccharomycetes(100);o__Saccharomycetales(100);f__Metschnikowiaceae(100);g__Metschnikowia(100);s__Metschnikowia_pulcherrima(100); | Metschnikowia | 13 | Saprotroph | \|Undefined Saprotroph\| | Yeast | NULL | Probable | As nectar-tap_saprotroph (Põlme et al. 2020); Decay sugar-rich substrates (Põlme et al. 2020) | Põlme S, et al. 2020. Fungal Diversity 105:1-16 ((https://doi.org/10.1007/s13225-020-00466-2)) |
| Otu37173 | 1 | k__Fungi(100);p__Ascomycota(100);c__Saccharomycetes(100);o__Saccharomycetales(100);f__Metschnikowiaceae(100);g__Metschnikowia(100);s__Metschnikowia_pulcherrima(100); | Metschnikowia | 13 | Saprotroph | \|Undefined Saprotroph\| | Yeast | NULL | Probable | As nectar-tap_saprotroph (Põlme et al. 2020); Decay sugar-rich substrates (Põlme et al. 2020) | Põlme S, et al. 2020. Fungal Diversity 105:1-16 ((https://doi.org/10.1007/s13225-020-00466-2)) |
| Otu37174 | 1 | k__Fungi(100);p__Ascomycota(100);c__Saccharomycetes(100);o__Saccharomycetales(100);f__Metschnikowiaceae(100);g__Metschnikowia(100);s__Metschnikowia_pulcherrima(100); | Metschnikowia | 13 | Saprotroph | \|Undefined Saprotroph\| | Yeast | NULL | Probable | As nectar-tap_saprotroph (Põlme et al. 2020); Decay sugar-rich substrates (Põlme et al. 2020) | Põlme S, et al. 2020. Fungal Diversity 105:1-16 ((https://doi.org/10.1007/s13225-020-00466-2)) |
| Otu37175 | 1 | k__Fungi(100);p__Ascomycota(100);c__Saccharomycetes(100);o__Saccharomycetales(100);f__Metschnikowiaceae(100);g__Metschnikowia(100);s__Metschnikowia_pulcherrima(100); | Metschnikowia | 13 | Saprotroph | \|Undefined Saprotroph\| | Yeast | NULL | Probable | As nectar-tap_saprotroph (Põlme et al. 2020); Decay sugar-rich substrates (Põlme et al. 2020) | Põlme S, et al. 2020. Fungal Diversity 105:1-16 ((https://doi.org/10.1007/s13225-020-00466-2)) |
| Otu37176 | 1 | k__Fungi(100);p__Ascomycota(100);c__Saccharomycetes(100);o__Saccharomycetales(100);f__Metschnikowiaceae(100);g__Metschnikowia(100);g__Metschnikowia_unclassified(100); | Metschnikowia | 13 | Saprotroph | \|Undefined Saprotroph\| | Yeast | NULL | Probable | As nectar-tap_saprotroph (Põlme et al. 2020); Decay sugar-rich substrates (Põlme et al. 2020) | Põlme S, et al. 2020. Fungal Diversity 105:1-16 ((https://doi.org/10.1007/s13225-020-00466-2)) |
| Otu37177 | 1 | k__Fungi(100);p__Ascomycota(100);c__Saccharomycetes(100);o__Saccharomycetales(100);f__Metschnikowiaceae(100);g__Metschnikowia(100);s__Metschnikowia_pulcherrima(100); | Metschnikowia | 13 | Saprotroph | \|Undefined Saprotroph\| | Yeast | NULL | Probable | As nectar-tap_saprotroph (Põlme et al. 2020); Decay sugar-rich substrates (Põlme et al. 2020) | Põlme S, et al. 2020. Fungal Diversity 105:1-16 ((https://doi.org/10.1007/s13225-020-00466-2)) |
| Otu37178 | 1 | k__Fungi(100);p__Ascomycota(100);c__Saccharomycetes(100);o__Saccharomycetales(100);f__Metschnikowiaceae(100);g__Metschnikowia(100);s__Metschnikowia_pulcherrima(100); | Metschnikowia | 13 | Saprotroph | \|Undefined Saprotroph\| | Yeast | NULL | Probable | As nectar-tap_saprotroph (Põlme et al. 2020); Decay sugar-rich substrates (Põlme et al. 2020) | Põlme S, et al. 2020. Fungal Diversity 105:1-16 ((https://doi.org/10.1007/s13225-020-00466-2)) |
| Otu37179 | 1 | k__Fungi(100);p__Ascomycota(100);c__Saccharomycetes(100);o__Saccharomycetales(100);f__Metschnikowiaceae(100);g__Metschnikowia(100);s__Metschnikowia_pulcherrima(100); | Metschnikowia | 13 | Saprotroph | \|Undefined Saprotroph\| | Yeast | NULL | Probable | As nectar-tap_saprotroph (Põlme et al. 2020); Decay sugar-rich substrates (Põlme et al. 2020) | Põlme S, et al. 2020. Fungal Diversity 105:1-16 ((https://doi.org/10.1007/s13225-020-00466-2)) |
| Otu37180 | 1 | k__Fungi(100);p__Ascomycota(100);c__Saccharomycetes(100);o__Saccharomycetales(100);f__Metschnikowiaceae(100);g__Metschnikowia(100);s__Metschnikowia_pulcherrima(100); | Metschnikowia | 13 | Saprotroph | \|Undefined Saprotroph\| | Yeast | NULL | Probable | As nectar-tap_saprotroph (Põlme et al. 2020); Decay sugar-rich substrates (Põlme et al. 2020) | Põlme S, et al. 2020. Fungal Diversity 105:1-16 ((https://doi.org/10.1007/s13225-020-00466-2)) |
| Otu37181 | 1 | k__Fungi(100);p__Ascomycota(100);c__Saccharomycetes(100);o__Saccharomycetales(100);f__Metschnikowiaceae(100);g__Metschnikowia(100);g__Metschnikowia_unclassified(100); | Metschnikowia | 13 | Saprotroph | \|Undefined Saprotroph\| | Yeast | NULL | Probable | As nectar-tap_saprotroph (Põlme et al. 2020); Decay sugar-rich substrates (Põlme et al. 2020) | Põlme S, et al. 2020. Fungal Diversity 105:1-16 ((https://doi.org/10.1007/s13225-020-00466-2)) |
| Otu37182 | 1 | k__Fungi(100);p__Ascomycota(100);c__Saccharomycetes(100);o__Saccharomycetales(100);f__Metschnikowiaceae(100);g__Metschnikowia(100);g__Metschnikowia_unclassified(100); | Metschnikowia | 13 | Saprotroph | \|Undefined Saprotroph\| | Yeast | NULL | Probable | As nectar-tap_saprotroph (Põlme et al. 2020); Decay sugar-rich substrates (Põlme et al. 2020) | Põlme S, et al. 2020. Fungal Diversity 105:1-16 ((https://doi.org/10.1007/s13225-020-00466-2)) |
| Otu37183 | 1 | k__Fungi(100);p__Ascomycota(100);c__Saccharomycetes(100);o__Saccharomycetales(100);f__Metschnikowiaceae(100);g__Metschnikowia(100);g__Metschnikowia_unclassified(100); | Metschnikowia | 13 | Saprotroph | \|Undefined Saprotroph\| | Yeast | NULL | Probable | As nectar-tap_saprotroph (Põlme et al. 2020); Decay sugar-rich substrates (Põlme et al. 2020) | Põlme S, et al. 2020. Fungal Diversity 105:1-16 ((https://doi.org/10.1007/s13225-020-00466-2)) |
| Otu37184 | 1 | k__Fungi(100);p__Ascomycota(100);c__Saccharomycetes(100);o__Saccharomycetales(100);f__Metschnikowiaceae(100);g__Metschnikowia(100);s__Metschnikowia_pulcherrima(100); | Metschnikowia | 13 | Saprotroph | \|Undefined Saprotroph\| | Yeast | NULL | Probable | As nectar-tap_saprotroph (Põlme et al. 2020); Decay sugar-rich substrates (Põlme et al. 2020) | Põlme S, et al. 2020. Fungal Diversity 105:1-16 ((https://doi.org/10.1007/s13225-020-00466-2)) |
| Otu37185 | 1 | k__Fungi(100);p__Ascomycota(100);c__Saccharomycetes(100);o__Saccharomycetales(100);f__Metschnikowiaceae(100);g__Metschnikowia(100);g__Metschnikowia_unclassified(100); | Metschnikowia | 13 | Saprotroph | \|Undefined Saprotroph\| | Yeast | NULL | Probable | As nectar-tap_saprotroph (Põlme et al. 2020); Decay sugar-rich substrates (Põlme et al. 2020) | Põlme S, et al. 2020. Fungal Diversity 105:1-16 ((https://doi.org/10.1007/s13225-020-00466-2)) |
| Otu37186 | 1 | k__Fungi(100);p__Ascomycota(100);c__Saccharomycetes(100);o__Saccharomycetales(100);f__Metschnikowiaceae(100);g__Metschnikowia(100);s__Metschnikowia_pulcherrima(100); | Metschnikowia | 13 | Saprotroph | \|Undefined Saprotroph\| | Yeast | NULL | Probable | As nectar-tap_saprotroph (Põlme et al. 2020); Decay sugar-rich substrates (Põlme et al. 2020) | Põlme S, et al. 2020. Fungal Diversity 105:1-16 ((https://doi.org/10.1007/s13225-020-00466-2)) |
| Otu37187 | 1 | k__Fungi(100);p__Ascomycota(100);c__Saccharomycetes(100);o__Saccharomycetales(100);f__Metschnikowiaceae(100);g__Metschnikowia(100);s__Metschnikowia_pulcherrima(100); | Metschnikowia | 13 | Saprotroph | \|Undefined Saprotroph\| | Yeast | NULL | Probable | As nectar-tap_saprotroph (Põlme et al. 2020); Decay sugar-rich substrates (Põlme et al. 2020) | Põlme S, et al. 2020. Fungal Diversity 105:1-16 ((https://doi.org/10.1007/s13225-020-00466-2)) |
| Otu37188 | 1 | k__Fungi(100);p__Ascomycota(100);c__Saccharomycetes(100);o__Saccharomycetales(100);f__Metschnikowiaceae(100);g__Metschnikowia(100);s__Metschnikowia_pulcherrima(100); | Metschnikowia | 13 | Saprotroph | \|Undefined Saprotroph\| | Yeast | NULL | Probable | As nectar-tap_saprotroph (Põlme et al. 2020); Decay sugar-rich substrates (Põlme et al. 2020) | Põlme S, et al. 2020. Fungal Diversity 105:1-16 ((https://doi.org/10.1007/s13225-020-00466-2)) |
| Otu37189 | 1 | k__Fungi(100);p__Ascomycota(100);c__Saccharomycetes(100);o__Saccharomycetales(100);f__Metschnikowiaceae(100);g__Metschnikowia(100);s__Metschnikowia_pulcherrima(100); | Metschnikowia | 13 | Saprotroph | \|Undefined Saprotroph\| | Yeast | NULL | Probable | As nectar-tap_saprotroph (Põlme et al. 2020); Decay sugar-rich substrates (Põlme et al. 2020) | Põlme S, et al. 2020. Fungal Diversity 105:1-16 ((https://doi.org/10.1007/s13225-020-00466-2)) |
| Otu37190 | 1 | k__Fungi(100);p__Ascomycota(100);c__Saccharomycetes(100);o__Saccharomycetales(100);f__Metschnikowiaceae(100);g__Metschnikowia(100);g__Metschnikowia_unclassified(100); | Metschnikowia | 13 | Saprotroph | \|Undefined Saprotroph\| | Yeast | NULL | Probable | As nectar-tap_saprotroph (Põlme et al. 2020); Decay sugar-rich substrates (Põlme et al. 2020) | Põlme S, et al. 2020. Fungal Diversity 105:1-16 ((https://doi.org/10.1007/s13225-020-00466-2)) |
| Otu37191 | 1 | k__Fungi(100);p__Ascomycota(100);c__Saccharomycetes(100);o__Saccharomycetales(100);f__Metschnikowiaceae(100);g__Metschnikowia(100);g__Metschnikowia_unclassified(100); | Metschnikowia | 13 | Saprotroph | \|Undefined Saprotroph\| | Yeast | NULL | Probable | As nectar-tap_saprotroph (Põlme et al. 2020); Decay sugar-rich substrates (Põlme et al. 2020) | Põlme S, et al. 2020. Fungal Diversity 105:1-16 ((https://doi.org/10.1007/s13225-020-00466-2)) |
| Otu37192 | 1 | k__Fungi(100);p__Ascomycota(100);c__Saccharomycetes(100);o__Saccharomycetales(100);f__Metschnikowiaceae(100);g__Metschnikowia(100);s__Metschnikowia_pulcherrima(100); | Metschnikowia | 13 | Saprotroph | \|Undefined Saprotroph\| | Yeast | NULL | Probable | As nectar-tap_saprotroph (Põlme et al. 2020); Decay sugar-rich substrates (Põlme et al. 2020) | Põlme S, et al. 2020. Fungal Diversity 105:1-16 ((https://doi.org/10.1007/s13225-020-00466-2)) |
| Otu37193 | 1 | k__Fungi(100);p__Ascomycota(100);c__Saccharomycetes(100);o__Saccharomycetales(100);f__Metschnikowiaceae(100);g__Metschnikowia(100);s__Metschnikowia_pulcherrima(100); | Metschnikowia | 13 | Saprotroph | \|Undefined Saprotroph\| | Yeast | NULL | Probable | As nectar-tap_saprotroph (Põlme et al. 2020); Decay sugar-rich substrates (Põlme et al. 2020) | Põlme S, et al. 2020. Fungal Diversity 105:1-16 ((https://doi.org/10.1007/s13225-020-00466-2)) |
| Otu37194 | 1 | k__Fungi(100);p__Ascomycota(100);c__Saccharomycetes(100);o__Saccharomycetales(100);f__Metschnikowiaceae(100);g__Metschnikowia(100);s__Metschnikowia_pulcherrima(100); | Metschnikowia | 13 | Saprotroph | \|Undefined Saprotroph\| | Yeast | NULL | Probable | As nectar-tap_saprotroph (Põlme et al. 2020); Decay sugar-rich substrates (Põlme et al. 2020) | Põlme S, et al. 2020. Fungal Diversity 105:1-16 ((https://doi.org/10.1007/s13225-020-00466-2)) |
| Otu37195 | 1 | k__Fungi(100);p__Ascomycota(100);c__Saccharomycetes(100);o__Saccharomycetales(100);f__Metschnikowiaceae(100);g__Metschnikowia(100);g__Metschnikowia_unclassified(100); | Metschnikowia | 13 | Saprotroph | \|Undefined Saprotroph\| | Yeast | NULL | Probable | As nectar-tap_saprotroph (Põlme et al. 2020); Decay sugar-rich substrates (Põlme et al. 2020) | Põlme S, et al. 2020. Fungal Diversity 105:1-16 ((https://doi.org/10.1007/s13225-020-00466-2)) |
| Otu37196 | 1 | k__Fungi(100);p__Ascomycota(100);c__Saccharomycetes(100);o__Saccharomycetales(100);f__Metschnikowiaceae(100);g__Metschnikowia(100);s__Metschnikowia_pulcherrima(100); | Metschnikowia | 13 | Saprotroph | \|Undefined Saprotroph\| | Yeast | NULL | Probable | As nectar-tap_saprotroph (Põlme et al. 2020); Decay sugar-rich substrates (Põlme et al. 2020) | Põlme S, et al. 2020. Fungal Diversity 105:1-16 ((https://doi.org/10.1007/s13225-020-00466-2)) |
| Otu37197 | 1 | k__Fungi(100);p__Ascomycota(100);c__Saccharomycetes(100);o__Saccharomycetales(100);f__Metschnikowiaceae(100);g__Metschnikowia(100);s__Metschnikowia_pulcherrima(100); | Metschnikowia | 13 | Saprotroph | \|Undefined Saprotroph\| | Yeast | NULL | Probable | As nectar-tap_saprotroph (Põlme et al. 2020); Decay sugar-rich substrates (Põlme et al. 2020) | Põlme S, et al. 2020. Fungal Diversity 105:1-16 ((https://doi.org/10.1007/s13225-020-00466-2)) |
| Otu37198 | 1 | k__Fungi(100);p__Ascomycota(100);c__Saccharomycetes(100);o__Saccharomycetales(100);f__Metschnikowiaceae(100);g__Metschnikowia(100);s__Metschnikowia_pulcherrima(100); | Metschnikowia | 13 | Saprotroph | \|Undefined Saprotroph\| | Yeast | NULL | Probable | As nectar-tap_saprotroph (Põlme et al. 2020); Decay sugar-rich substrates (Põlme et al. 2020) | Põlme S, et al. 2020. Fungal Diversity 105:1-16 ((https://doi.org/10.1007/s13225-020-00466-2)) |
| Otu37200 | 1 | k__Fungi(100);p__Ascomycota(100);c__Saccharomycetes(100);o__Saccharomycetales(100);f__Metschnikowiaceae(100);g__Metschnikowia(100);s__Metschnikowia_pulcherrima(100); | Metschnikowia | 13 | Saprotroph | \|Undefined Saprotroph\| | Yeast | NULL | Probable | As nectar-tap_saprotroph (Põlme et al. 2020); Decay sugar-rich substrates (Põlme et al. 2020) | Põlme S, et al. 2020. Fungal Diversity 105:1-16 ((https://doi.org/10.1007/s13225-020-00466-2)) |
| Otu37201 | 1 | k__Fungi(100);p__Ascomycota(100);c__Saccharomycetes(100);o__Saccharomycetales(100);f__Metschnikowiaceae(100);g__Metschnikowia(100);s__Metschnikowia_pulcherrima(100); | Metschnikowia | 13 | Saprotroph | \|Undefined Saprotroph\| | Yeast | NULL | Probable | As nectar-tap_saprotroph (Põlme et al. 2020); Decay sugar-rich substrates (Põlme et al. 2020) | Põlme S, et al. 2020. Fungal Diversity 105:1-16 ((https://doi.org/10.1007/s13225-020-00466-2)) |
| Otu37202 | 1 | k__Fungi(100);p__Ascomycota(100);c__Saccharomycetes(100);o__Saccharomycetales(100);f__Metschnikowiaceae(100);g__Metschnikowia(100);s__Metschnikowia_pulcherrima(100); | Metschnikowia | 13 | Saprotroph | \|Undefined Saprotroph\| | Yeast | NULL | Probable | As nectar-tap_saprotroph (Põlme et al. 2020); Decay sugar-rich substrates (Põlme et al. 2020) | Põlme S, et al. 2020. Fungal Diversity 105:1-16 ((https://doi.org/10.1007/s13225-020-00466-2)) |
| Otu37203 | 1 | k__Fungi(100);p__Ascomycota(100);c__Saccharomycetes(100);o__Saccharomycetales(100);f__Metschnikowiaceae(100);g__Metschnikowia(100);s__Metschnikowia_pulcherrima(100); | Metschnikowia | 13 | Saprotroph | \|Undefined Saprotroph\| | Yeast | NULL | Probable | As nectar-tap_saprotroph (Põlme et al. 2020); Decay sugar-rich substrates (Põlme et al. 2020) | Põlme S, et al. 2020. Fungal Diversity 105:1-16 ((https://doi.org/10.1007/s13225-020-00466-2)) |
| Otu37204 | 1 | k__Fungi(100);p__Ascomycota(100);c__Saccharomycetes(100);o__Saccharomycetales(100);f__Metschnikowiaceae(100);g__Metschnikowia(100);s__Metschnikowia_pulcherrima(100); | Metschnikowia | 13 | Saprotroph | \|Undefined Saprotroph\| | Yeast | NULL | Probable | As nectar-tap_saprotroph (Põlme et al. 2020); Decay sugar-rich substrates (Põlme et al. 2020) | Põlme S, et al. 2020. Fungal Diversity 105:1-16 ((https://doi.org/10.1007/s13225-020-00466-2)) |
| Otu37205 | 1 | k__Fungi(100);p__Ascomycota(100);c__Saccharomycetes(100);o__Saccharomycetales(100);f__Metschnikowiaceae(100);g__Metschnikowia(100);s__Metschnikowia_pulcherrima(100); | Metschnikowia | 13 | Saprotroph | \|Undefined Saprotroph\| | Yeast | NULL | Probable | As nectar-tap_saprotroph (Põlme et al. 2020); Decay sugar-rich substrates (Põlme et al. 2020) | Põlme S, et al. 2020. Fungal Diversity 105:1-16 ((https://doi.org/10.1007/s13225-020-00466-2)) |
| Otu37206 | 1 | k__Fungi(100);p__Ascomycota(100);c__Saccharomycetes(100);o__Saccharomycetales(100);f__Metschnikowiaceae(100);g__Metschnikowia(100);s__Metschnikowia_pulcherrima(100); | Metschnikowia | 13 | Saprotroph | \|Undefined Saprotroph\| | Yeast | NULL | Probable | As nectar-tap_saprotroph (Põlme et al. 2020); Decay sugar-rich substrates (Põlme et al. 2020) | Põlme S, et al. 2020. Fungal Diversity 105:1-16 ((https://doi.org/10.1007/s13225-020-00466-2)) |
| Otu37207 | 1 | k__Fungi(100);p__Ascomycota(100);c__Saccharomycetes(100);o__Saccharomycetales(100);f__Metschnikowiaceae(100);g__Metschnikowia(100);g__Metschnikowia_unclassified(100); | Metschnikowia | 13 | Saprotroph | \|Undefined Saprotroph\| | Yeast | NULL | Probable | As nectar-tap_saprotroph (Põlme et al. 2020); Decay sugar-rich substrates (Põlme et al. 2020) | Põlme S, et al. 2020. Fungal Diversity 105:1-16 ((https://doi.org/10.1007/s13225-020-00466-2)) |
| Otu37208 | 1 | k__Fungi(100);p__Ascomycota(100);c__Saccharomycetes(100);o__Saccharomycetales(100);f__Metschnikowiaceae(100);g__Metschnikowia(100);s__Metschnikowia_pulcherrima(100); | Metschnikowia | 13 | Saprotroph | \|Undefined Saprotroph\| | Yeast | NULL | Probable | As nectar-tap_saprotroph (Põlme et al. 2020); Decay sugar-rich substrates (Põlme et al. 2020) | Põlme S, et al. 2020. Fungal Diversity 105:1-16 ((https://doi.org/10.1007/s13225-020-00466-2)) |
| Otu37209 | 1 | k__Fungi(100);p__Ascomycota(100);c__Saccharomycetes(100);o__Saccharomycetales(100);f__Metschnikowiaceae(100);g__Metschnikowia(100);g__Metschnikowia_unclassified(100); | Metschnikowia | 13 | Saprotroph | \|Undefined Saprotroph\| | Yeast | NULL | Probable | As nectar-tap_saprotroph (Põlme et al. 2020); Decay sugar-rich substrates (Põlme et al. 2020) | Põlme S, et al. 2020. Fungal Diversity 105:1-16 ((https://doi.org/10.1007/s13225-020-00466-2)) |
| Otu37210 | 1 | k__Fungi(100);p__Ascomycota(100);c__Saccharomycetes(100);o__Saccharomycetales(100);f__Metschnikowiaceae(100);g__Metschnikowia(100);s__Metschnikowia_pulcherrima(100); | Metschnikowia | 13 | Saprotroph | \|Undefined Saprotroph\| | Yeast | NULL | Probable | As nectar-tap_saprotroph (Põlme et al. 2020); Decay sugar-rich substrates (Põlme et al. 2020) | Põlme S, et al. 2020. Fungal Diversity 105:1-16 ((https://doi.org/10.1007/s13225-020-00466-2)) |
| Otu37211 | 1 | k__Fungi(100);p__Ascomycota(100);c__Saccharomycetes(100);o__Saccharomycetales(100);f__Metschnikowiaceae(100);g__Metschnikowia(100);s__Metschnikowia_pulcherrima(100); | Metschnikowia | 13 | Saprotroph | \|Undefined Saprotroph\| | Yeast | NULL | Probable | As nectar-tap_saprotroph (Põlme et al. 2020); Decay sugar-rich substrates (Põlme et al. 2020) | Põlme S, et al. 2020. Fungal Diversity 105:1-16 ((https://doi.org/10.1007/s13225-020-00466-2)) |
| Otu37212 | 1 | k__Fungi(100);p__Ascomycota(100);c__Saccharomycetes(100);o__Saccharomycetales(100);f__Metschnikowiaceae(100);g__Metschnikowia(100);s__Metschnikowia_pulcherrima(100); | Metschnikowia | 13 | Saprotroph | \|Undefined Saprotroph\| | Yeast | NULL | Probable | As nectar-tap_saprotroph (Põlme et al. 2020); Decay sugar-rich substrates (Põlme et al. 2020) | Põlme S, et al. 2020. Fungal Diversity 105:1-16 ((https://doi.org/10.1007/s13225-020-00466-2)) |
| Otu37213 | 1 | k__Fungi(100);p__Ascomycota(100);c__Saccharomycetes(100);o__Saccharomycetales(100);f__Metschnikowiaceae(100);g__Metschnikowia(100);s__Metschnikowia_pulcherrima(100); | Metschnikowia | 13 | Saprotroph | \|Undefined Saprotroph\| | Yeast | NULL | Probable | As nectar-tap_saprotroph (Põlme et al. 2020); Decay sugar-rich substrates (Põlme et al. 2020) | Põlme S, et al. 2020. Fungal Diversity 105:1-16 ((https://doi.org/10.1007/s13225-020-00466-2)) |
| Otu37214 | 1 | k__Fungi(100);p__Ascomycota(100);c__Saccharomycetes(100);o__Saccharomycetales(100);f__Metschnikowiaceae(100);g__Metschnikowia(100);s__Metschnikowia_pulcherrima(100); | Metschnikowia | 13 | Saprotroph | \|Undefined Saprotroph\| | Yeast | NULL | Probable | As nectar-tap_saprotroph (Põlme et al. 2020); Decay sugar-rich substrates (Põlme et al. 2020) | Põlme S, et al. 2020. Fungal Diversity 105:1-16 ((https://doi.org/10.1007/s13225-020-00466-2)) |
| Otu37215 | 1 | k__Fungi(100);p__Ascomycota(100);c__Saccharomycetes(100);o__Saccharomycetales(100);f__Metschnikowiaceae(100);g__Metschnikowia(100);s__Metschnikowia_pulcherrima(100); | Metschnikowia | 13 | Saprotroph | \|Undefined Saprotroph\| | Yeast | NULL | Probable | As nectar-tap_saprotroph (Põlme et al. 2020); Decay sugar-rich substrates (Põlme et al. 2020) | Põlme S, et al. 2020. Fungal Diversity 105:1-16 ((https://doi.org/10.1007/s13225-020-00466-2)) |
| Otu37216 | 1 | k__Fungi(100);p__Ascomycota(100);c__Saccharomycetes(100);o__Saccharomycetales(100);f__Metschnikowiaceae(100);g__Metschnikowia(100);g__Metschnikowia_unclassified(100); | Metschnikowia | 13 | Saprotroph | \|Undefined Saprotroph\| | Yeast | NULL | Probable | As nectar-tap_saprotroph (Põlme et al. 2020); Decay sugar-rich substrates (Põlme et al. 2020) | Põlme S, et al. 2020. Fungal Diversity 105:1-16 ((https://doi.org/10.1007/s13225-020-00466-2)) |
| Otu37217 | 1 | k__Fungi(100);p__Ascomycota(100);c__Saccharomycetes(100);o__Saccharomycetales(100);f__Metschnikowiaceae(100);g__Metschnikowia(100);s__Metschnikowia_pulcherrima(100); | Metschnikowia | 13 | Saprotroph | \|Undefined Saprotroph\| | Yeast | NULL | Probable | As nectar-tap_saprotroph (Põlme et al. 2020); Decay sugar-rich substrates (Põlme et al. 2020) | Põlme S, et al. 2020. Fungal Diversity 105:1-16 ((https://doi.org/10.1007/s13225-020-00466-2)) |
| Otu37218 | 1 | k__Fungi(100);p__Ascomycota(100);c__Saccharomycetes(100);o__Saccharomycetales(100);f__Metschnikowiaceae(100);g__Metschnikowia(100);s__Metschnikowia_pulcherrima(100); | Metschnikowia | 13 | Saprotroph | \|Undefined Saprotroph\| | Yeast | NULL | Probable | As nectar-tap_saprotroph (Põlme et al. 2020); Decay sugar-rich substrates (Põlme et al. 2020) | Põlme S, et al. 2020. Fungal Diversity 105:1-16 ((https://doi.org/10.1007/s13225-020-00466-2)) |
| Otu37219 | 1 | k__Fungi(100);p__Ascomycota(100);c__Saccharomycetes(100);o__Saccharomycetales(100);f__Metschnikowiaceae(100);g__Metschnikowia(100);s__Metschnikowia_pulcherrima(100); | Metschnikowia | 13 | Saprotroph | \|Undefined Saprotroph\| | Yeast | NULL | Probable | As nectar-tap_saprotroph (Põlme et al. 2020); Decay sugar-rich substrates (Põlme et al. 2020) | Põlme S, et al. 2020. Fungal Diversity 105:1-16 ((https://doi.org/10.1007/s13225-020-00466-2)) |
| Otu37220 | 1 | k__Fungi(100);p__Ascomycota(100);c__Saccharomycetes(100);o__Saccharomycetales(100);f__Metschnikowiaceae(100);g__Metschnikowia(100);s__Metschnikowia_pulcherrima(100); | Metschnikowia | 13 | Saprotroph | \|Undefined Saprotroph\| | Yeast | NULL | Probable | As nectar-tap_saprotroph (Põlme et al. 2020); Decay sugar-rich substrates (Põlme et al. 2020) | Põlme S, et al. 2020. Fungal Diversity 105:1-16 ((https://doi.org/10.1007/s13225-020-00466-2)) |
| Otu37221 | 1 | k__Fungi(100);p__Ascomycota(100);c__Saccharomycetes(100);o__Saccharomycetales(100);f__Metschnikowiaceae(100);g__Metschnikowia(100);g__Metschnikowia_unclassified(100); | Metschnikowia | 13 | Saprotroph | \|Undefined Saprotroph\| | Yeast | NULL | Probable | As nectar-tap_saprotroph (Põlme et al. 2020); Decay sugar-rich substrates (Põlme et al. 2020) | Põlme S, et al. 2020. Fungal Diversity 105:1-16 ((https://doi.org/10.1007/s13225-020-00466-2)) |
| Otu37222 | 1 | k__Fungi(100);p__Ascomycota(100);c__Saccharomycetes(100);o__Saccharomycetales(100);f__Metschnikowiaceae(100);g__Metschnikowia(100);s__Metschnikowia_pulcherrima(100); | Metschnikowia | 13 | Saprotroph | \|Undefined Saprotroph\| | Yeast | NULL | Probable | As nectar-tap_saprotroph (Põlme et al. 2020); Decay sugar-rich substrates (Põlme et al. 2020) | Põlme S, et al. 2020. Fungal Diversity 105:1-16 ((https://doi.org/10.1007/s13225-020-00466-2)) |
| Otu37223 | 1 | k__Fungi(100);p__Ascomycota(100);c__Saccharomycetes(100);o__Saccharomycetales(100);f__Metschnikowiaceae(100);g__Metschnikowia(100);s__Metschnikowia_pulcherrima(100); | Metschnikowia | 13 | Saprotroph | \|Undefined Saprotroph\| | Yeast | NULL | Probable | As nectar-tap_saprotroph (Põlme et al. 2020); Decay sugar-rich substrates (Põlme et al. 2020) | Põlme S, et al. 2020. Fungal Diversity 105:1-16 ((https://doi.org/10.1007/s13225-020-00466-2)) |
| Otu37224 | 1 | k__Fungi(100);p__Ascomycota(100);c__Saccharomycetes(100);o__Saccharomycetales(100);f__Metschnikowiaceae(100);g__Metschnikowia(100);g__Metschnikowia_unclassified(100); | Metschnikowia | 13 | Saprotroph | \|Undefined Saprotroph\| | Yeast | NULL | Probable | As nectar-tap_saprotroph (Põlme et al. 2020); Decay sugar-rich substrates (Põlme et al. 2020) | Põlme S, et al. 2020. Fungal Diversity 105:1-16 ((https://doi.org/10.1007/s13225-020-00466-2)) |
| Otu37225 | 1 | k__Fungi(100);p__Ascomycota(100);c__Saccharomycetes(100);o__Saccharomycetales(100);f__Metschnikowiaceae(100);g__Metschnikowia(100);s__Metschnikowia_pulcherrima(100); | Metschnikowia | 13 | Saprotroph | \|Undefined Saprotroph\| | Yeast | NULL | Probable | As nectar-tap_saprotroph (Põlme et al. 2020); Decay sugar-rich substrates (Põlme et al. 2020) | Põlme S, et al. 2020. Fungal Diversity 105:1-16 ((https://doi.org/10.1007/s13225-020-00466-2)) |
| Otu37226 | 1 | k__Fungi(100);p__Ascomycota(100);c__Saccharomycetes(100);o__Saccharomycetales(100);f__Metschnikowiaceae(100);g__Metschnikowia(100);g__Metschnikowia_unclassified(100); | Metschnikowia | 13 | Saprotroph | \|Undefined Saprotroph\| | Yeast | NULL | Probable | As nectar-tap_saprotroph (Põlme et al. 2020); Decay sugar-rich substrates (Põlme et al. 2020) | Põlme S, et al. 2020. Fungal Diversity 105:1-16 ((https://doi.org/10.1007/s13225-020-00466-2)) |
| Otu37227 | 1 | k__Fungi(100);p__Ascomycota(100);c__Saccharomycetes(100);o__Saccharomycetales(100);f__Metschnikowiaceae(100);g__Metschnikowia(100);g__Metschnikowia_unclassified(100); | Metschnikowia | 13 | Saprotroph | \|Undefined Saprotroph\| | Yeast | NULL | Probable | As nectar-tap_saprotroph (Põlme et al. 2020); Decay sugar-rich substrates (Põlme et al. 2020) | Põlme S, et al. 2020. Fungal Diversity 105:1-16 ((https://doi.org/10.1007/s13225-020-00466-2)) |
| Otu37228 | 1 | k__Fungi(100);p__Ascomycota(100);c__Saccharomycetes(100);o__Saccharomycetales(100);f__Metschnikowiaceae(100);g__Metschnikowia(100);s__Metschnikowia_pulcherrima(100); | Metschnikowia | 13 | Saprotroph | \|Undefined Saprotroph\| | Yeast | NULL | Probable | As nectar-tap_saprotroph (Põlme et al. 2020); Decay sugar-rich substrates (Põlme et al. 2020) | Põlme S, et al. 2020. Fungal Diversity 105:1-16 ((https://doi.org/10.1007/s13225-020-00466-2)) |
| Otu37229 | 1 | k__Fungi(100);p__Ascomycota(100);c__Saccharomycetes(100);o__Saccharomycetales(100);f__Metschnikowiaceae(100);g__Metschnikowia(100);g__Metschnikowia_unclassified(100); | Metschnikowia | 13 | Saprotroph | \|Undefined Saprotroph\| | Yeast | NULL | Probable | As nectar-tap_saprotroph (Põlme et al. 2020); Decay sugar-rich substrates (Põlme et al. 2020) | Põlme S, et al. 2020. Fungal Diversity 105:1-16 ((https://doi.org/10.1007/s13225-020-00466-2)) |
| Otu37230 | 1 | k__Fungi(100);p__Ascomycota(100);c__Saccharomycetes(100);o__Saccharomycetales(100);f__Metschnikowiaceae(100);g__Metschnikowia(100);g__Metschnikowia_unclassified(100); | Metschnikowia | 13 | Saprotroph | \|Undefined Saprotroph\| | Yeast | NULL | Probable | As nectar-tap_saprotroph (Põlme et al. 2020); Decay sugar-rich substrates (Põlme et al. 2020) | Põlme S, et al. 2020. Fungal Diversity 105:1-16 ((https://doi.org/10.1007/s13225-020-00466-2)) |
| Otu37231 | 1 | k__Fungi(100);p__Ascomycota(100);c__Saccharomycetes(100);o__Saccharomycetales(100);f__Metschnikowiaceae(100);g__Metschnikowia(100);s__Metschnikowia_pulcherrima(100); | Metschnikowia | 13 | Saprotroph | \|Undefined Saprotroph\| | Yeast | NULL | Probable | As nectar-tap_saprotroph (Põlme et al. 2020); Decay sugar-rich substrates (Põlme et al. 2020) | Põlme S, et al. 2020. Fungal Diversity 105:1-16 ((https://doi.org/10.1007/s13225-020-00466-2)) |
| Otu37232 | 1 | k__Fungi(100);p__Ascomycota(100);c__Saccharomycetes(100);o__Saccharomycetales(100);f__Metschnikowiaceae(100);g__Metschnikowia(100);s__Metschnikowia_pulcherrima(100); | Metschnikowia | 13 | Saprotroph | \|Undefined Saprotroph\| | Yeast | NULL | Probable | As nectar-tap_saprotroph (Põlme et al. 2020); Decay sugar-rich substrates (Põlme et al. 2020) | Põlme S, et al. 2020. Fungal Diversity 105:1-16 ((https://doi.org/10.1007/s13225-020-00466-2)) |
| Otu37233 | 1 | k__Fungi(100);p__Ascomycota(100);c__Saccharomycetes(100);o__Saccharomycetales(100);f__Metschnikowiaceae(100);g__Metschnikowia(100);g__Metschnikowia_unclassified(100); | Metschnikowia | 13 | Saprotroph | \|Undefined Saprotroph\| | Yeast | NULL | Probable | As nectar-tap_saprotroph (Põlme et al. 2020); Decay sugar-rich substrates (Põlme et al. 2020) | Põlme S, et al. 2020. Fungal Diversity 105:1-16 ((https://doi.org/10.1007/s13225-020-00466-2)) |
| Otu37234 | 1 | k__Fungi(100);p__Ascomycota(100);c__Saccharomycetes(100);o__Saccharomycetales(100);f__Metschnikowiaceae(100);g__Metschnikowia(100);s__Metschnikowia_pulcherrima(100); | Metschnikowia | 13 | Saprotroph | \|Undefined Saprotroph\| | Yeast | NULL | Probable | As nectar-tap_saprotroph (Põlme et al. 2020); Decay sugar-rich substrates (Põlme et al. 2020) | Põlme S, et al. 2020. Fungal Diversity 105:1-16 ((https://doi.org/10.1007/s13225-020-00466-2)) |
| Otu37235 | 1 | k__Fungi(100);p__Ascomycota(100);c__Saccharomycetes(100);o__Saccharomycetales(100);f__Metschnikowiaceae(100);g__Metschnikowia(100);g__Metschnikowia_unclassified(100); | Metschnikowia | 13 | Saprotroph | \|Undefined Saprotroph\| | Yeast | NULL | Probable | As nectar-tap_saprotroph (Põlme et al. 2020); Decay sugar-rich substrates (Põlme et al. 2020) | Põlme S, et al. 2020. Fungal Diversity 105:1-16 ((https://doi.org/10.1007/s13225-020-00466-2)) |
| Otu37236 | 1 | k__Fungi(100);p__Ascomycota(100);c__Saccharomycetes(100);o__Saccharomycetales(100);f__Metschnikowiaceae(100);g__Metschnikowia(100);s__Metschnikowia_pulcherrima(100); | Metschnikowia | 13 | Saprotroph | \|Undefined Saprotroph\| | Yeast | NULL | Probable | As nectar-tap_saprotroph (Põlme et al. 2020); Decay sugar-rich substrates (Põlme et al. 2020) | Põlme S, et al. 2020. Fungal Diversity 105:1-16 ((https://doi.org/10.1007/s13225-020-00466-2)) |
| Otu37237 | 1 | k__Fungi(100);p__Ascomycota(100);c__Saccharomycetes(100);o__Saccharomycetales(100);f__Metschnikowiaceae(100);g__Metschnikowia(100);s__Metschnikowia_pulcherrima(100); | Metschnikowia | 13 | Saprotroph | \|Undefined Saprotroph\| | Yeast | NULL | Probable | As nectar-tap_saprotroph (Põlme et al. 2020); Decay sugar-rich substrates (Põlme et al. 2020) | Põlme S, et al. 2020. Fungal Diversity 105:1-16 ((https://doi.org/10.1007/s13225-020-00466-2)) |
| Otu37238 | 1 | k__Fungi(100);p__Ascomycota(100);c__Saccharomycetes(100);o__Saccharomycetales(100);f__Metschnikowiaceae(100);g__Metschnikowia(100);s__Metschnikowia_pulcherrima(100); | Metschnikowia | 13 | Saprotroph | \|Undefined Saprotroph\| | Yeast | NULL | Probable | As nectar-tap_saprotroph (Põlme et al. 2020); Decay sugar-rich substrates (Põlme et al. 2020) | Põlme S, et al. 2020. Fungal Diversity 105:1-16 ((https://doi.org/10.1007/s13225-020-00466-2)) |
| Otu37239 | 1 | k__Fungi(100);p__Ascomycota(100);c__Saccharomycetes(100);o__Saccharomycetales(100);f__Metschnikowiaceae(100);g__Metschnikowia(100);s__Metschnikowia_pulcherrima(100); | Metschnikowia | 13 | Saprotroph | \|Undefined Saprotroph\| | Yeast | NULL | Probable | As nectar-tap_saprotroph (Põlme et al. 2020); Decay sugar-rich substrates (Põlme et al. 2020) | Põlme S, et al. 2020. Fungal Diversity 105:1-16 ((https://doi.org/10.1007/s13225-020-00466-2)) |
| Otu37240 | 1 | k__Fungi(100);p__Ascomycota(100);c__Saccharomycetes(100);o__Saccharomycetales(100);f__Metschnikowiaceae(100);g__Metschnikowia(100);s__Metschnikowia_pulcherrima(100); | Metschnikowia | 13 | Saprotroph | \|Undefined Saprotroph\| | Yeast | NULL | Probable | As nectar-tap_saprotroph (Põlme et al. 2020); Decay sugar-rich substrates (Põlme et al. 2020) | Põlme S, et al. 2020. Fungal Diversity 105:1-16 ((https://doi.org/10.1007/s13225-020-00466-2)) |
| Otu37241 | 1 | k__Fungi(100);p__Ascomycota(100);c__Saccharomycetes(100);o__Saccharomycetales(100);f__Metschnikowiaceae(100);g__Metschnikowia(100);s__Metschnikowia_pulcherrima(100); | Metschnikowia | 13 | Saprotroph | \|Undefined Saprotroph\| | Yeast | NULL | Probable | As nectar-tap_saprotroph (Põlme et al. 2020); Decay sugar-rich substrates (Põlme et al. 2020) | Põlme S, et al. 2020. Fungal Diversity 105:1-16 ((https://doi.org/10.1007/s13225-020-00466-2)) |
| Otu37242 | 1 | k__Fungi(100);p__Ascomycota(100);c__Saccharomycetes(100);o__Saccharomycetales(100);f__Metschnikowiaceae(100);g__Metschnikowia(100);g__Metschnikowia_unclassified(100); | Metschnikowia | 13 | Saprotroph | \|Undefined Saprotroph\| | Yeast | NULL | Probable | As nectar-tap_saprotroph (Põlme et al. 2020); Decay sugar-rich substrates (Põlme et al. 2020) | Põlme S, et al. 2020. Fungal Diversity 105:1-16 ((https://doi.org/10.1007/s13225-020-00466-2)) |
| Otu37243 | 1 | k__Fungi(100);p__Ascomycota(100);c__Saccharomycetes(100);o__Saccharomycetales(100);f__Metschnikowiaceae(100);g__Metschnikowia(100);g__Metschnikowia_unclassified(100); | Metschnikowia | 13 | Saprotroph | \|Undefined Saprotroph\| | Yeast | NULL | Probable | As nectar-tap_saprotroph (Põlme et al. 2020); Decay sugar-rich substrates (Põlme et al. 2020) | Põlme S, et al. 2020. Fungal Diversity 105:1-16 ((https://doi.org/10.1007/s13225-020-00466-2)) |
| Otu37244 | 1 | k__Fungi(100);p__Ascomycota(100);c__Saccharomycetes(100);o__Saccharomycetales(100);f__Metschnikowiaceae(100);g__Metschnikowia(100);g__Metschnikowia_unclassified(100); | Metschnikowia | 13 | Saprotroph | \|Undefined Saprotroph\| | Yeast | NULL | Probable | As nectar-tap_saprotroph (Põlme et al. 2020); Decay sugar-rich substrates (Põlme et al. 2020) | Põlme S, et al. 2020. Fungal Diversity 105:1-16 ((https://doi.org/10.1007/s13225-020-00466-2)) |
| Otu37245 | 1 | k__Fungi(100);p__Ascomycota(100);c__Saccharomycetes(100);o__Saccharomycetales(100);f__Metschnikowiaceae(100);g__Metschnikowia(100);g__Metschnikowia_unclassified(100); | Metschnikowia | 13 | Saprotroph | \|Undefined Saprotroph\| | Yeast | NULL | Probable | As nectar-tap_saprotroph (Põlme et al. 2020); Decay sugar-rich substrates (Põlme et al. 2020) | Põlme S, et al. 2020. Fungal Diversity 105:1-16 ((https://doi.org/10.1007/s13225-020-00466-2)) |
| Otu37246 | 1 | k__Fungi(100);p__Ascomycota(100);c__Saccharomycetes(100);o__Saccharomycetales(100);f__Metschnikowiaceae(100);g__Metschnikowia(100);s__Metschnikowia_pulcherrima(100); | Metschnikowia | 13 | Saprotroph | \|Undefined Saprotroph\| | Yeast | NULL | Probable | As nectar-tap_saprotroph (Põlme et al. 2020); Decay sugar-rich substrates (Põlme et al. 2020) | Põlme S, et al. 2020. Fungal Diversity 105:1-16 ((https://doi.org/10.1007/s13225-020-00466-2)) |
| Otu37247 | 1 | k__Fungi(100);p__Ascomycota(100);c__Saccharomycetes(100);o__Saccharomycetales(100);f__Metschnikowiaceae(100);g__Metschnikowia(100);g__Metschnikowia_unclassified(100); | Metschnikowia | 13 | Saprotroph | \|Undefined Saprotroph\| | Yeast | NULL | Probable | As nectar-tap_saprotroph (Põlme et al. 2020); Decay sugar-rich substrates (Põlme et al. 2020) | Põlme S, et al. 2020. Fungal Diversity 105:1-16 ((https://doi.org/10.1007/s13225-020-00466-2)) |
| Otu37248 | 1 | k__Fungi(100);p__Ascomycota(100);c__Saccharomycetes(100);o__Saccharomycetales(100);f__Metschnikowiaceae(100);g__Metschnikowia(100);g__Metschnikowia_unclassified(100); | Metschnikowia | 13 | Saprotroph | \|Undefined Saprotroph\| | Yeast | NULL | Probable | As nectar-tap_saprotroph (Põlme et al. 2020); Decay sugar-rich substrates (Põlme et al. 2020) | Põlme S, et al. 2020. Fungal Diversity 105:1-16 ((https://doi.org/10.1007/s13225-020-00466-2)) |
| Otu37249 | 1 | k__Fungi(100);p__Ascomycota(100);c__Saccharomycetes(100);o__Saccharomycetales(100);f__Metschnikowiaceae(100);g__Metschnikowia(100);s__Metschnikowia_pulcherrima(100); | Metschnikowia | 13 | Saprotroph | \|Undefined Saprotroph\| | Yeast | NULL | Probable | As nectar-tap_saprotroph (Põlme et al. 2020); Decay sugar-rich substrates (Põlme et al. 2020) | Põlme S, et al. 2020. Fungal Diversity 105:1-16 ((https://doi.org/10.1007/s13225-020-00466-2)) |
| Otu37250 | 1 | k__Fungi(100);p__Ascomycota(100);c__Saccharomycetes(100);o__Saccharomycetales(100);f__Metschnikowiaceae(100);g__Metschnikowia(100);s__Metschnikowia_pulcherrima(100); | Metschnikowia | 13 | Saprotroph | \|Undefined Saprotroph\| | Yeast | NULL | Probable | As nectar-tap_saprotroph (Põlme et al. 2020); Decay sugar-rich substrates (Põlme et al. 2020) | Põlme S, et al. 2020. Fungal Diversity 105:1-16 ((https://doi.org/10.1007/s13225-020-00466-2)) |
| Otu37251 | 1 | k__Fungi(100);p__Ascomycota(100);c__Saccharomycetes(100);o__Saccharomycetales(100);f__Metschnikowiaceae(100);g__Metschnikowia(100);s__Metschnikowia_pulcherrima(100); | Metschnikowia | 13 | Saprotroph | \|Undefined Saprotroph\| | Yeast | NULL | Probable | As nectar-tap_saprotroph (Põlme et al. 2020); Decay sugar-rich substrates (Põlme et al. 2020) | Põlme S, et al. 2020. Fungal Diversity 105:1-16 ((https://doi.org/10.1007/s13225-020-00466-2)) |
| Otu37252 | 1 | k__Fungi(100);p__Ascomycota(100);c__Saccharomycetes(100);o__Saccharomycetales(100);f__Metschnikowiaceae(100);g__Metschnikowia(100);s__Metschnikowia_pulcherrima(100); | Metschnikowia | 13 | Saprotroph | \|Undefined Saprotroph\| | Yeast | NULL | Probable | As nectar-tap_saprotroph (Põlme et al. 2020); Decay sugar-rich substrates (Põlme et al. 2020) | Põlme S, et al. 2020. Fungal Diversity 105:1-16 ((https://doi.org/10.1007/s13225-020-00466-2)) |
| Otu37253 | 1 | k__Fungi(100);p__Ascomycota(100);c__Saccharomycetes(100);o__Saccharomycetales(100);f__Metschnikowiaceae(100);g__Metschnikowia(100);s__Metschnikowia_pulcherrima(100); | Metschnikowia | 13 | Saprotroph | \|Undefined Saprotroph\| | Yeast | NULL | Probable | As nectar-tap_saprotroph (Põlme et al. 2020); Decay sugar-rich substrates (Põlme et al. 2020) | Põlme S, et al. 2020. Fungal Diversity 105:1-16 ((https://doi.org/10.1007/s13225-020-00466-2)) |
| Otu37254 | 1 | k__Fungi(100);p__Ascomycota(100);c__Saccharomycetes(100);o__Saccharomycetales(100);f__Metschnikowiaceae(100);g__Metschnikowia(100);s__Metschnikowia_pulcherrima(100); | Metschnikowia | 13 | Saprotroph | \|Undefined Saprotroph\| | Yeast | NULL | Probable | As nectar-tap_saprotroph (Põlme et al. 2020); Decay sugar-rich substrates (Põlme et al. 2020) | Põlme S, et al. 2020. Fungal Diversity 105:1-16 ((https://doi.org/10.1007/s13225-020-00466-2)) |
| Otu37255 | 1 | k__Fungi(100);p__Ascomycota(100);c__Saccharomycetes(100);o__Saccharomycetales(100);f__Metschnikowiaceae(100);g__Metschnikowia(100);s__Metschnikowia_pulcherrima(100); | Metschnikowia | 13 | Saprotroph | \|Undefined Saprotroph\| | Yeast | NULL | Probable | As nectar-tap_saprotroph (Põlme et al. 2020); Decay sugar-rich substrates (Põlme et al. 2020) | Põlme S, et al. 2020. Fungal Diversity 105:1-16 ((https://doi.org/10.1007/s13225-020-00466-2)) |
| Otu37256 | 1 | k__Fungi(100);p__Ascomycota(100);c__Saccharomycetes(100);o__Saccharomycetales(100);f__Metschnikowiaceae(100);g__Metschnikowia(100);g__Metschnikowia_unclassified(100); | Metschnikowia | 13 | Saprotroph | \|Undefined Saprotroph\| | Yeast | NULL | Probable | As nectar-tap_saprotroph (Põlme et al. 2020); Decay sugar-rich substrates (Põlme et al. 2020) | Põlme S, et al. 2020. Fungal Diversity 105:1-16 ((https://doi.org/10.1007/s13225-020-00466-2)) |
| Otu37257 | 1 | k__Fungi(100);p__Ascomycota(100);c__Saccharomycetes(100);o__Saccharomycetales(100);f__Metschnikowiaceae(100);g__Metschnikowia(100);g__Metschnikowia_unclassified(100); | Metschnikowia | 13 | Saprotroph | \|Undefined Saprotroph\| | Yeast | NULL | Probable | As nectar-tap_saprotroph (Põlme et al. 2020); Decay sugar-rich substrates (Põlme et al. 2020) | Põlme S, et al. 2020. Fungal Diversity 105:1-16 ((https://doi.org/10.1007/s13225-020-00466-2)) |
| Otu37258 | 1 | k__Fungi(100);p__Ascomycota(100);c__Saccharomycetes(100);o__Saccharomycetales(100);f__Metschnikowiaceae(100);g__Metschnikowia(100);s__Metschnikowia_pulcherrima(100); | Metschnikowia | 13 | Saprotroph | \|Undefined Saprotroph\| | Yeast | NULL | Probable | As nectar-tap_saprotroph (Põlme et al. 2020); Decay sugar-rich substrates (Põlme et al. 2020) | Põlme S, et al. 2020. Fungal Diversity 105:1-16 ((https://doi.org/10.1007/s13225-020-00466-2)) |
| Otu37259 | 1 | k__Fungi(100);p__Ascomycota(100);c__Saccharomycetes(100);o__Saccharomycetales(100);f__Metschnikowiaceae(100);g__Metschnikowia(100);s__Metschnikowia_pulcherrima(100); | Metschnikowia | 13 | Saprotroph | \|Undefined Saprotroph\| | Yeast | NULL | Probable | As nectar-tap_saprotroph (Põlme et al. 2020); Decay sugar-rich substrates (Põlme et al. 2020) | Põlme S, et al. 2020. Fungal Diversity 105:1-16 ((https://doi.org/10.1007/s13225-020-00466-2)) |
| Otu37260 | 1 | k__Fungi(100);p__Ascomycota(100);c__Saccharomycetes(100);o__Saccharomycetales(100);f__Metschnikowiaceae(100);g__Metschnikowia(100);s__Metschnikowia_pulcherrima(100); | Metschnikowia | 13 | Saprotroph | \|Undefined Saprotroph\| | Yeast | NULL | Probable | As nectar-tap_saprotroph (Põlme et al. 2020); Decay sugar-rich substrates (Põlme et al. 2020) | Põlme S, et al. 2020. Fungal Diversity 105:1-16 ((https://doi.org/10.1007/s13225-020-00466-2)) |
| Otu37261 | 1 | k__Fungi(100);p__Ascomycota(100);c__Saccharomycetes(100);o__Saccharomycetales(100);f__Metschnikowiaceae(100);g__Metschnikowia(100);s__Metschnikowia_pulcherrima(100); | Metschnikowia | 13 | Saprotroph | \|Undefined Saprotroph\| | Yeast | NULL | Probable | As nectar-tap_saprotroph (Põlme et al. 2020); Decay sugar-rich substrates (Põlme et al. 2020) | Põlme S, et al. 2020. Fungal Diversity 105:1-16 ((https://doi.org/10.1007/s13225-020-00466-2)) |
| Otu37263 | 1 | k__Fungi(100);p__Ascomycota(100);c__Saccharomycetes(100);o__Saccharomycetales(100);f__Metschnikowiaceae(100);g__Metschnikowia(100);s__Metschnikowia_pulcherrima(100); | Metschnikowia | 13 | Saprotroph | \|Undefined Saprotroph\| | Yeast | NULL | Probable | As nectar-tap_saprotroph (Põlme et al. 2020); Decay sugar-rich substrates (Põlme et al. 2020) | Põlme S, et al. 2020. Fungal Diversity 105:1-16 ((https://doi.org/10.1007/s13225-020-00466-2)) |
| Otu37265 | 1 | k__Fungi(100);p__Ascomycota(100);c__Saccharomycetes(100);o__Saccharomycetales(100);f__Metschnikowiaceae(100);g__Metschnikowia(100);s__Metschnikowia_pulcherrima(100); | Metschnikowia | 13 | Saprotroph | \|Undefined Saprotroph\| | Yeast | NULL | Probable | As nectar-tap_saprotroph (Põlme et al. 2020); Decay sugar-rich substrates (Põlme et al. 2020) | Põlme S, et al. 2020. Fungal Diversity 105:1-16 ((https://doi.org/10.1007/s13225-020-00466-2)) |
| Otu37266 | 1 | k__Fungi(100);p__Ascomycota(100);c__Saccharomycetes(100);o__Saccharomycetales(100);f__Metschnikowiaceae(100);g__Metschnikowia(100);s__Metschnikowia_pulcherrima(100); | Metschnikowia | 13 | Saprotroph | \|Undefined Saprotroph\| | Yeast | NULL | Probable | As nectar-tap_saprotroph (Põlme et al. 2020); Decay sugar-rich substrates (Põlme et al. 2020) | Põlme S, et al. 2020. Fungal Diversity 105:1-16 ((https://doi.org/10.1007/s13225-020-00466-2)) |
| Otu37267 | 1 | k__Fungi(100);p__Ascomycota(100);c__Saccharomycetes(100);o__Saccharomycetales(100);f__Metschnikowiaceae(100);g__Metschnikowia(100);s__Metschnikowia_pulcherrima(100); | Metschnikowia | 13 | Saprotroph | \|Undefined Saprotroph\| | Yeast | NULL | Probable | As nectar-tap_saprotroph (Põlme et al. 2020); Decay sugar-rich substrates (Põlme et al. 2020) | Põlme S, et al. 2020. Fungal Diversity 105:1-16 ((https://doi.org/10.1007/s13225-020-00466-2)) |
| Otu37268 | 1 | k__Fungi(100);p__Ascomycota(100);c__Saccharomycetes(100);o__Saccharomycetales(100);f__Metschnikowiaceae(100);g__Metschnikowia(100);s__Metschnikowia_pulcherrima(100); | Metschnikowia | 13 | Saprotroph | \|Undefined Saprotroph\| | Yeast | NULL | Probable | As nectar-tap_saprotroph (Põlme et al. 2020); Decay sugar-rich substrates (Põlme et al. 2020) | Põlme S, et al. 2020. Fungal Diversity 105:1-16 ((https://doi.org/10.1007/s13225-020-00466-2)) |
| Otu37269 | 1 | k__Fungi(100);p__Ascomycota(100);c__Saccharomycetes(100);o__Saccharomycetales(100);f__Metschnikowiaceae(100);g__Metschnikowia(100);g__Metschnikowia_unclassified(100); | Metschnikowia | 13 | Saprotroph | \|Undefined Saprotroph\| | Yeast | NULL | Probable | As nectar-tap_saprotroph (Põlme et al. 2020); Decay sugar-rich substrates (Põlme et al. 2020) | Põlme S, et al. 2020. Fungal Diversity 105:1-16 ((https://doi.org/10.1007/s13225-020-00466-2)) |
| Otu37270 | 1 | k__Fungi(100);p__Ascomycota(100);c__Saccharomycetes(100);o__Saccharomycetales(100);f__Metschnikowiaceae(100);g__Metschnikowia(100);g__Metschnikowia_unclassified(100); | Metschnikowia | 13 | Saprotroph | \|Undefined Saprotroph\| | Yeast | NULL | Probable | As nectar-tap_saprotroph (Põlme et al. 2020); Decay sugar-rich substrates (Põlme et al. 2020) | Põlme S, et al. 2020. Fungal Diversity 105:1-16 ((https://doi.org/10.1007/s13225-020-00466-2)) |
| Otu37271 | 1 | k__Fungi(100);p__Ascomycota(100);c__Saccharomycetes(100);o__Saccharomycetales(100);f__Metschnikowiaceae(100);g__Metschnikowia(100);g__Metschnikowia_unclassified(100); | Metschnikowia | 13 | Saprotroph | \|Undefined Saprotroph\| | Yeast | NULL | Probable | As nectar-tap_saprotroph (Põlme et al. 2020); Decay sugar-rich substrates (Põlme et al. 2020) | Põlme S, et al. 2020. Fungal Diversity 105:1-16 ((https://doi.org/10.1007/s13225-020-00466-2)) |
| Otu37272 | 1 | k__Fungi(100);p__Ascomycota(100);c__Saccharomycetes(100);o__Saccharomycetales(100);f__Metschnikowiaceae(100);g__Metschnikowia(100);s__Metschnikowia_pulcherrima(100); | Metschnikowia | 13 | Saprotroph | \|Undefined Saprotroph\| | Yeast | NULL | Probable | As nectar-tap_saprotroph (Põlme et al. 2020); Decay sugar-rich substrates (Põlme et al. 2020) | Põlme S, et al. 2020. Fungal Diversity 105:1-16 ((https://doi.org/10.1007/s13225-020-00466-2)) |

| | | | | | | | | | | | |
|---|---|---|---|---|---|---|---|---|---|---|---|
| Otu37274 | 1 | k__Fungi(100);p__Ascomycota(100);c__Saccharomycetes(100);o__Saccharomycetales(100);f__Metschnikowiaceae(100);g__Metschnikowia(100);s__Metschnikowia_pulcherrima(100); | Metschnikowia | 13 | Saprotroph | \|Undefined Saprotroph\| | Yeast | NULL | Probable | As nectar-tap_saprotroph (Põlme et al. 2020); Decay sugar-rich substrates (Põlme et al. 2020) | Põlme S, et al. 2020. Fungal Diversity 105:1-16 ((https://doi.org/10.1007/s13225-020-00466-2)) |
| Otu37275 | 1 | k__Fungi(100);p__Ascomycota(100);c__Saccharomycetes(100);o__Saccharomycetales(100);f__Metschnikowiaceae(100);g__Metschnikowia(100);g__Metschnikowia_unclassified(100); | Metschnikowia | 13 | Saprotroph | \|Undefined Saprotroph\| | Yeast | NULL | Probable | As nectar-tap_saprotroph (Põlme et al. 2020); Decay sugar-rich substrates (Põlme et al. 2020) | Põlme S, et al. 2020. Fungal Diversity 105:1-16 ((https://doi.org/10.1007/s13225-020-00466-2)) |
| Otu37276 | 1 | k__Fungi(100);p__Ascomycota(100);c__Saccharomycetes(100);o__Saccharomycetales(100);f__Metschnikowiaceae(100);g__Metschnikowia(100);s__Metschnikowia_pulcherrima(100); | Metschnikowia | 13 | Saprotroph | \|Undefined Saprotroph\| | Yeast | NULL | Probable | As nectar-tap_saprotroph (Põlme et al. 2020); Decay sugar-rich substrates (Põlme et al. 2020) | Põlme S, et al. 2020. Fungal Diversity 105:1-16 ((https://doi.org/10.1007/s13225-020-00466-2)) |
| Otu37277 | 1 | k__Fungi(100);p__Ascomycota(100);c__Saccharomycetes(100);o__Saccharomycetales(100);f__Metschnikowiaceae(100);g__Metschnikowia(100);s__Metschnikowia_pulcherrima(100); | Metschnikowia | 13 | Saprotroph | \|Undefined Saprotroph\| | Yeast | NULL | Probable | As nectar-tap_saprotroph (Põlme et al. 2020); Decay sugar-rich substrates (Põlme et al. 2020) | Põlme S, et al. 2020. Fungal Diversity 105:1-16 ((https://doi.org/10.1007/s13225-020-00466-2)) |
| Otu37281 | 1 | k__Fungi(100);p__Ascomycota(100);c__Saccharomycetes(100);o__Saccharomycetales(100);f__Metschnikowiaceae(100);g__Metschnikowia(100);g__Metschnikowia_unclassified(100); | Metschnikowia | 13 | Saprotroph | \|Undefined Saprotroph\| | Yeast | NULL | Probable | As nectar-tap_saprotroph (Põlme et al. 2020); Decay sugar-rich substrates (Põlme et al. 2020) | Põlme S, et al. 2020. Fungal Diversity 105:1-16 ((https://doi.org/10.1007/s13225-020-00466-2)) |
| Otu37282 | 1 | k__Fungi(100);p__Ascomycota(100);c__Saccharomycetes(100);o__Saccharomycetales(100);f__Metschnikowiaceae(100);g__Metschnikowia(100);s__Metschnikowia_pulcherrima(100); | Metschnikowia | 13 | Saprotroph | \|Undefined Saprotroph\| | Yeast | NULL | Probable | As nectar-tap_saprotroph (Põlme et al. 2020); Decay sugar-rich substrates (Põlme et al. 2020) | Põlme S, et al. 2020. Fungal Diversity 105:1-16 ((https://doi.org/10.1007/s13225-020-00466-2)) |
| Otu37283 | 1 | k__Fungi(100);p__Ascomycota(100);c__Saccharomycetes(100);o__Saccharomycetales(100);f__Metschnikowiaceae(100);g__Metschnikowia(100);s__Metschnikowia_pulcherrima(100); | Metschnikowia | 13 | Saprotroph | \|Undefined Saprotroph\| | Yeast | NULL | Probable | As nectar-tap_saprotroph (Põlme et al. 2020); Decay sugar-rich substrates (Põlme et al. 2020) | Põlme S, et al. 2020. Fungal Diversity 105:1-16 ((https://doi.org/10.1007/s13225-020-00466-2)) |
| Otu37284 | 1 | k__Fungi(100);p__Ascomycota(100);c__Saccharomycetes(100);o__Saccharomycetales(100);f__Metschnikowiaceae(100);g__Metschnikowia(100);s__Metschnikowia_pulcherrima(100); | Metschnikowia | 13 | Saprotroph | \|Undefined Saprotroph\| | Yeast | NULL | Probable | As nectar-tap_saprotroph (Põlme et al. 2020); Decay sugar-rich substrates (Põlme et al. 2020) | Põlme S, et al. 2020. Fungal Diversity 105:1-16 ((https://doi.org/10.1007/s13225-020-00466-2)) |
| Otu37286 | 1 | k__Fungi(100);p__Ascomycota(100);c__Saccharomycetes(100);o__Saccharomycetales(100);f__Metschnikowiaceae(100);g__Metschnikowia(100);s__Metschnikowia_pulcherrima(100); | Metschnikowia | 13 | Saprotroph | \|Undefined Saprotroph\| | Yeast | NULL | Probable | As nectar-tap_saprotroph (Põlme et al. 2020); Decay sugar-rich substrates (Põlme et al. 2020) | Põlme S, et al. 2020. Fungal Diversity 105:1-16 ((https://doi.org/10.1007/s13225-020-00466-2)) |
| Otu37287 | 1 | k__Fungi(100);p__Ascomycota(100);c__Saccharomycetes(100);o__Saccharomycetales(100);f__Metschnikowiaceae(100);g__Metschnikowia(100);s__Metschnikowia_pulcherrima(100); | Metschnikowia | 13 | Saprotroph | \|Undefined Saprotroph\| | Yeast | NULL | Probable | As nectar-tap_saprotroph (Põlme et al. 2020); Decay sugar-rich substrates (Põlme et al. 2020) | Põlme S, et al. 2020. Fungal Diversity 105:1-16 ((https://doi.org/10.1007/s13225-020-00466-2)) |
| Otu37288 | 1 | k__Fungi(100);p__Ascomycota(100);c__Saccharomycetes(100);o__Saccharomycetales(100);f__Metschnikowiaceae(100);g__Metschnikowia(100);s__Metschnikowia_pulcherrima(100); | Metschnikowia | 13 | Saprotroph | \|Undefined Saprotroph\| | Yeast | NULL | Probable | As nectar-tap_saprotroph (Põlme et al. 2020); Decay sugar-rich substrates (Põlme et al. 2020) | Põlme S, et al. 2020. Fungal Diversity 105:1-16 ((https://doi.org/10.1007/s13225-020-00466-2)) |
| Otu37289 | 1 | k__Fungi(100);p__Ascomycota(100);c__Saccharomycetes(100);o__Saccharomycetales(100);f__Metschnikowiaceae(100);g__Metschnikowia(100);s__Metschnikowia_pulcherrima(100); | Metschnikowia | 13 | Saprotroph | \|Undefined Saprotroph\| | Yeast | NULL | Probable | As nectar-tap_saprotroph (Põlme et al. 2020); Decay sugar-rich substrates (Põlme et al. 2020) | Põlme S, et al. 2020. Fungal Diversity 105:1-16 ((https://doi.org/10.1007/s13225-020-00466-2)) |
| Otu37290 | 1 | k__Fungi(100);p__Ascomycota(100);c__Saccharomycetes(100);o__Saccharomycetales(100);f__Metschnikowiaceae(100);g__Metschnikowia(100);s__Metschnikowia_pulcherrima(100); | Metschnikowia | 13 | Saprotroph | \|Undefined Saprotroph\| | Yeast | NULL | Probable | As nectar-tap_saprotroph (Põlme et al. 2020); Decay sugar-rich substrates (Põlme et al. 2020) | Põlme S, et al. 2020. Fungal Diversity 105:1-16 ((https://doi.org/10.1007/s13225-020-00466-2)) |
| Otu37291 | 1 | k__Fungi(100);p__Ascomycota(100);c__Saccharomycetes(100);o__Saccharomycetales(100);f__Metschnikowiaceae(100);g__Metschnikowia(100);s__Metschnikowia_pulcherrima(100); | Metschnikowia | 13 | Saprotroph | \|Undefined Saprotroph\| | Yeast | NULL | Probable | As nectar-tap_saprotroph (Põlme et al. 2020); Decay sugar-rich substrates (Põlme et al. 2020) | Põlme S, et al. 2020. Fungal Diversity 105:1-16 ((https://doi.org/10.1007/s13225-020-00466-2)) |
| Otu37294 | 1 | k__Fungi(100);p__Ascomycota(100);c__Saccharomycetes(100);o__Saccharomycetales(100);f__Metschnikowiaceae(100);g__Metschnikowia(100);s__Metschnikowia_pulcherrima(100); | Metschnikowia | 13 | Saprotroph | \|Undefined Saprotroph\| | Yeast | NULL | Probable | As nectar-tap_saprotroph (Põlme et al. 2020); Decay sugar-rich substrates (Põlme et al. 2020) | Põlme S, et al. 2020. Fungal Diversity 105:1-16 ((https://doi.org/10.1007/s13225-020-00466-2)) |
| Otu37295 | 1 | k__Fungi(100);p__Ascomycota(100);c__Saccharomycetes(100);o__Saccharomycetales(100);f__Metschnikowiaceae(100);g__Metschnikowia(100);g__Metschnikowia_unclassified(100); | Metschnikowia | 13 | Saprotroph | \|Undefined Saprotroph\| | Yeast | NULL | Probable | As nectar-tap_saprotroph (Põlme et al. 2020); Decay sugar-rich substrates (Põlme et al. 2020) | Põlme S, et al. 2020. Fungal Diversity 105:1-16 ((https://doi.org/10.1007/s13225-020-00466-2)) |
| Otu37296 | 1 | k__Fungi(100);p__Ascomycota(100);c__Saccharomycetes(100);o__Saccharomycetales(100);f__Metschnikowiaceae(100);g__Metschnikowia(100);s__Metschnikowia_pulcherrima(100); | Metschnikowia | 13 | Saprotroph | \|Undefined Saprotroph\| | Yeast | NULL | Probable | As nectar-tap_saprotroph (Põlme et al. 2020); Decay sugar-rich substrates (Põlme et al. 2020) | Põlme S, et al. 2020. Fungal Diversity 105:1-16 ((https://doi.org/10.1007/s13225-020-00466-2)) |
| Otu37297 | 1 | k__Fungi(100);p__Ascomycota(100);c__Saccharomycetes(100);o__Saccharomycetales(100);f__Metschnikowiaceae(100);g__Metschnikowia(100);s__Metschnikowia_pulcherrima(100); | Metschnikowia | 13 | Saprotroph | \|Undefined Saprotroph\| | Yeast | NULL | Probable | As nectar-tap_saprotroph (Põlme et al. 2020); Decay sugar-rich substrates (Põlme et al. 2020) | Põlme S, et al. 2020. Fungal Diversity 105:1-16 ((https://doi.org/10.1007/s13225-020-00466-2)) |
| Otu37298 | 1 | k__Fungi(100);p__Ascomycota(100);c__Saccharomycetes(100);o__Saccharomycetales(100);f__Metschnikowiaceae(100);g__Metschnikowia(100);g__Metschnikowia_unclassified(100); | Metschnikowia | 13 | Saprotroph | \|Undefined Saprotroph\| | Yeast | NULL | Probable | As nectar-tap_saprotroph (Põlme et al. 2020); Decay sugar-rich substrates (Põlme et al. 2020) | Põlme S, et al. 2020. Fungal Diversity 105:1-16 ((https://doi.org/10.1007/s13225-020-00466-2)) |
| Otu37299 | 1 | k__Fungi(100);p__Ascomycota(100);c__Saccharomycetes(100);o__Saccharomycetales(100);f__Metschnikowiaceae(100);g__Metschnikowia(100);s__Metschnikowia_pulcherrima(100); | Metschnikowia | 13 | Saprotroph | \|Undefined Saprotroph\| | Yeast | NULL | Probable | As nectar-tap_saprotroph (Põlme et al. 2020); Decay sugar-rich substrates (Põlme et al. 2020) | Põlme S, et al. 2020. Fungal Diversity 105:1-16 ((https://doi.org/10.1007/s13225-020-00466-2)) |
| Otu37304 | 1 | k__Fungi(100);p__Ascomycota(100);c__Saccharomycetes(100);o__Saccharomycetales(100);f__Metschnikowiaceae(100);g__Metschnikowia(100);s__Metschnikowia_pulcherrima(100); | Metschnikowia | 13 | Saprotroph | \|Undefined Saprotroph\| | Yeast | NULL | Probable | As nectar-tap_saprotroph (Põlme et al. 2020); Decay sugar-rich substrates (Põlme et al. 2020) | Põlme S, et al. 2020. Fungal Diversity 105:1-16 ((https://doi.org/10.1007/s13225-020-00466-2)) |
| Otu37305 | 1 | k__Fungi(100);p__Ascomycota(100);c__Saccharomycetes(100);o__Saccharomycetales(100);f__Metschnikowiaceae(100);g__Metschnikowia(100);s__Metschnikowia_pulcherrima(100); | Metschnikowia | 13 | Saprotroph | \|Undefined Saprotroph\| | Yeast | NULL | Probable | As nectar-tap_saprotroph (Põlme et al. 2020); Decay sugar-rich substrates (Põlme et al. 2020) | Põlme S, et al. 2020. Fungal Diversity 105:1-16 ((https://doi.org/10.1007/s13225-020-00466-2)) |
| Otu37306 | 1 | k__Fungi(100);p__Ascomycota(100);c__Saccharomycetes(100);o__Saccharomycetales(100);f__Metschnikowiaceae(100);g__Metschnikowia(100);s__Metschnikowia_pulcherrima(100); | Metschnikowia | 13 | Saprotroph | \|Undefined Saprotroph\| | Yeast | NULL | Probable | As nectar-tap_saprotroph (Põlme et al. 2020); Decay sugar-rich substrates (Põlme et al. 2020) | Põlme S, et al. 2020. Fungal Diversity 105:1-16 ((https://doi.org/10.1007/s13225-020-00466-2)) |
| Otu37307 | 1 | k__Fungi(100);p__Ascomycota(100);c__Saccharomycetes(100);o__Saccharomycetales(100);f__Metschnikowiaceae(100);g__Metschnikowia(100);g__Metschnikowia_unclassified(100); | Metschnikowia | 13 | Saprotroph | \|Undefined Saprotroph\| | Yeast | NULL | Probable | As nectar-tap_saprotroph (Põlme et al. 2020); Decay sugar-rich substrates (Põlme et al. 2020) | Põlme S, et al. 2020. Fungal Diversity 105:1-16 ((https://doi.org/10.1007/s13225-020-00466-2)) |
| Otu37308 | 1 | k__Fungi(100);p__Ascomycota(100);c__Saccharomycetes(100);o__Saccharomycetales(100);f__Metschnikowiaceae(100);g__Metschnikowia(100);s__Metschnikowia_pulcherrima(100); | Metschnikowia | 13 | Saprotroph | \|Undefined Saprotroph\| | Yeast | NULL | Probable | As nectar-tap_saprotroph (Põlme et al. 2020); Decay sugar-rich substrates (Põlme et al. 2020) | Põlme S, et al. 2020. Fungal Diversity 105:1-16 ((https://doi.org/10.1007/s13225-020-00466-2)) |
| Otu37309 | 1 | k__Fungi(100);p__Ascomycota(100);c__Saccharomycetes(100);o__Saccharomycetales(100);f__Metschnikowiaceae(100);g__Metschnikowia(100);g__Metschnikowia_unclassified(100); | Metschnikowia | 13 | Saprotroph | \|Undefined Saprotroph\| | Yeast | NULL | Probable | As nectar-tap_saprotroph (Põlme et al. 2020); Decay sugar-rich substrates (Põlme et al. 2020) | Põlme S, et al. 2020. Fungal Diversity 105:1-16 ((https://doi.org/10.1007/s13225-020-00466-2)) |
| Otu37310 | 1 | k__Fungi(100);p__Ascomycota(100);c__Saccharomycetes(100);o__Saccharomycetales(100);f__Metschnikowiaceae(100);g__Metschnikowia(100);s__Metschnikowia_pulcherrima(100); | Metschnikowia | 13 | Saprotroph | \|Undefined Saprotroph\| | Yeast | NULL | Probable | As nectar-tap_saprotroph (Põlme et al. 2020); Decay sugar-rich substrates (Põlme et al. 2020) | Põlme S, et al. 2020. Fungal Diversity 105:1-16 ((https://doi.org/10.1007/s13225-020-00466-2)) |
| Otu37312 | 1 | k__Fungi(100);p__Ascomycota(100);c__Saccharomycetes(100);o__Saccharomycetales(100);f__Metschnikowiaceae(100);g__Metschnikowia(100);s__Metschnikowia_pulcherrima(100); | Metschnikowia | 13 | Saprotroph | \|Undefined Saprotroph\| | Yeast | NULL | Probable | As nectar-tap_saprotroph (Põlme et al. 2020); Decay sugar-rich substrates (Põlme et al. 2020) | Põlme S, et al. 2020. Fungal Diversity 105:1-16 ((https://doi.org/10.1007/s13225-020-00466-2)) |
| Otu37314 | 1 | k__Fungi(100);p__Ascomycota(100);c__Saccharomycetes(100);o__Saccharomycetales(100);f__Metschnikowiaceae(100);g__Metschnikowia(100);s__Metschnikowia_pulcherrima(100); | Metschnikowia | 13 | Saprotroph | \|Undefined Saprotroph\| | Yeast | NULL | Probable | As nectar-tap_saprotroph (Põlme et al. 2020); Decay sugar-rich substrates (Põlme et al. 2020) | Põlme S, et al. 2020. Fungal Diversity 105:1-16 ((https://doi.org/10.1007/s13225-020-00466-2)) |
| Otu37315 | 1 | k__Fungi(100);p__Ascomycota(100);c__Saccharomycetes(100);o__Saccharomycetales(100);f__Metschnikowiaceae(100);g__Metschnikowia(100);s__Metschnikowia_pulcherrima(100); | Metschnikowia | 13 | Saprotroph | \|Undefined Saprotroph\| | Yeast | NULL | Probable | As nectar-tap_saprotroph (Põlme et al. 2020); Decay sugar-rich substrates (Põlme et al. 2020) | Põlme S, et al. 2020. Fungal Diversity 105:1-16 ((https://doi.org/10.1007/s13225-020-00466-2)) |
| Otu37316 | 1 | k__Fungi(100);p__Ascomycota(100);c__Saccharomycetes(100);o__Saccharomycetales(100);f__Metschnikowiaceae(100);g__Metschnikowia(100);g__Metschnikowia_unclassified(100); | Metschnikowia | 13 | Saprotroph | \|Undefined Saprotroph\| | Yeast | NULL | Probable | As nectar-tap_saprotroph (Põlme et al. 2020); Decay sugar-rich substrates (Põlme et al. 2020) | Põlme S, et al. 2020. Fungal Diversity 105:1-16 ((https://doi.org/10.1007/s13225-020-00466-2)) |
| Otu37317 | 1 | k__Fungi(100);p__Ascomycota(100);c__Saccharomycetes(100);o__Saccharomycetales(100);f__Metschnikowiaceae(100);g__Metschnikowia(100);g__Metschnikowia_unclassified(100); | Metschnikowia | 13 | Saprotroph | \|Undefined Saprotroph\| | Yeast | NULL | Probable | As nectar-tap_saprotroph (Põlme et al. 2020); Decay sugar-rich substrates (Põlme et al. 2020) | Põlme S, et al. 2020. Fungal Diversity 105:1-16 ((https://doi.org/10.1007/s13225-020-00466-2)) |
| Otu37318 | 1 | k__Fungi(100);p__Ascomycota(100);c__Saccharomycetes(100);o__Saccharomycetales(100);f__Metschnikowiaceae(100);g__Metschnikowia(100);s__Metschnikowia_pulcherrima(100); | Metschnikowia | 13 | Saprotroph | \|Undefined Saprotroph\| | Yeast | NULL | Probable | As nectar-tap_saprotroph (Põlme et al. 2020); Decay sugar-rich substrates (Põlme et al. 2020) | Põlme S, et al. 2020. Fungal Diversity 105:1-16 ((https://doi.org/10.1007/s13225-020-00466-2)) |
| Otu37319 | 1 | k__Fungi(100);p__Ascomycota(100);c__Saccharomycetes(100);o__Saccharomycetales(100);f__Metschnikowiaceae(100);g__Metschnikowia(100);s__Metschnikowia_pulcherrima(100); | Metschnikowia | 13 | Saprotroph | \|Undefined Saprotroph\| | Yeast | NULL | Probable | As nectar-tap_saprotroph (Põlme et al. 2020); Decay sugar-rich substrates (Põlme et al. 2020) | Põlme S, et al. 2020. Fungal Diversity 105:1-16 ((https://doi.org/10.1007/s13225-020-00466-2)) |
| Otu37320 | 1 | k__Fungi(100);p__Ascomycota(100);c__Saccharomycetes(100);o__Saccharomycetales(100);f__Metschnikowiaceae(100);g__Metschnikowia(100);s__Metschnikowia_pulcherrima(100); | Metschnikowia | 13 | Saprotroph | \|Undefined Saprotroph\| | Yeast | NULL | Probable | As nectar-tap_saprotroph (Põlme et al. 2020); Decay sugar-rich substrates (Põlme et al. 2020) | Põlme S, et al. 2020. Fungal Diversity 105:1-16 ((https://doi.org/10.1007/s13225-020-00466-2)) |
| Otu37321 | 1 | k__Fungi(100);p__Ascomycota(100);c__Saccharomycetes(100);o__Saccharomycetales(100);f__Metschnikowiaceae(100);g__Metschnikowia(100);s__Metschnikowia_pulcherrima(100); | Metschnikowia | 13 | Saprotroph | \|Undefined Saprotroph\| | Yeast | NULL | Probable | As nectar-tap_saprotroph (Põlme et al. 2020); Decay sugar-rich substrates (Põlme et al. 2020) | Põlme S, et al. 2020. Fungal Diversity 105:1-16 ((https://doi.org/10.1007/s13225-020-00466-2)) |
| Otu37322 | 1 | k__Fungi(100);p__Ascomycota(100);c__Saccharomycetes(100);o__Saccharomycetales(100);f__Metschnikowiaceae(100);g__Metschnikowia(100);s__Metschnikowia_pulcherrima(100); | Metschnikowia | 13 | Saprotroph | \|Undefined Saprotroph\| | Yeast | NULL | Probable | As nectar-tap_saprotroph (Põlme et al. 2020); Decay sugar-rich substrates (Põlme et al. 2020) | Põlme S, et al. 2020. Fungal Diversity 105:1-16 ((https://doi.org/10.1007/s13225-020-00466-2)) |
| Otu37323 | 1 | k__Fungi(100);p__Ascomycota(100);c__Saccharomycetes(100);o__Saccharomycetales(100);f__Metschnikowiaceae(100);g__Metschnikowia(100);g__Metschnikowia_unclassified(100); | Metschnikowia | 13 | Saprotroph | \|Undefined Saprotroph\| | Yeast | NULL | Probable | As nectar-tap_saprotroph (Põlme et al. 2020); Decay sugar-rich substrates (Põlme et al. 2020) | Põlme S, et al. 2020. Fungal Diversity 105:1-16 ((https://doi.org/10.1007/s13225-020-00466-2)) |
| Otu37324 | 1 | k__Fungi(100);p__Ascomycota(100);c__Saccharomycetes(100);o__Saccharomycetales(100);f__Metschnikowiaceae(100);g__Metschnikowia(100);s__Metschnikowia_pulcherrima(100); | Metschnikowia | 13 | Saprotroph | \|Undefined Saprotroph\| | Yeast | NULL | Probable | As nectar-tap_saprotroph (Põlme et al. 2020); Decay sugar-rich substrates (Põlme et al. 2020) | Põlme S, et al. 2020. Fungal Diversity 105:1-16 ((https://doi.org/10.1007/s13225-020-00466-2)) |
| Otu37325 | 1 | k__Fungi(100);p__Ascomycota(100);c__Saccharomycetes(100);o__Saccharomycetales(100);f__Metschnikowiaceae(100);g__Metschnikowia(100);s__Metschnikowia_pulcherrima(100); | Metschnikowia | 13 | Saprotroph | \|Undefined Saprotroph\| | Yeast | NULL | Probable | As nectar-tap_saprotroph (Põlme et al. 2020); Decay sugar-rich substrates (Põlme et al. 2020) | Põlme S, et al. 2020. Fungal Diversity 105:1-16 ((https://doi.org/10.1007/s13225-020-00466-2)) |
| Otu37326 | 1 | k__Fungi(100);p__Ascomycota(100);c__Saccharomycetes(100);o__Saccharomycetales(100);f__Metschnikowiaceae(100);g__Metschnikowia(100);s__Metschnikowia_pulcherrima(100); | Metschnikowia | 13 | Saprotroph | \|Undefined Saprotroph\| | Yeast | NULL | Probable | As nectar-tap_saprotroph (Põlme et al. 2020); Decay sugar-rich substrates (Põlme et al. 2020) | Põlme S, et al. 2020. Fungal Diversity 105:1-16 ((https://doi.org/10.1007/s13225-020-00466-2)) |
| Otu37327 | 1 | k__Fungi(100);p__Ascomycota(100);c__Saccharomycetes(100);o__Saccharomycetales(100);f__Metschnikowiaceae(100);g__Metschnikowia(100);s__Metschnikowia_pulcherrima(100); | Metschnikowia | 13 | Saprotroph | \|Undefined Saprotroph\| | Yeast | NULL | Probable | As nectar-tap_saprotroph (Põlme et al. 2020); Decay sugar-rich substrates (Põlme et al. 2020) | Põlme S, et al. 2020. Fungal Diversity 105:1-16 ((https://doi.org/10.1007/s13225-020-00466-2)) |
| Otu37328 | 1 | k__Fungi(100);p__Ascomycota(100);c__Saccharomycetes(100);o__Saccharomycetales(100);f__Metschnikowiaceae(100);g__Metschnikowia(100);s__Metschnikowia_pulcherrima(100); | Metschnikowia | 13 | Saprotroph | \|Undefined Saprotroph\| | Yeast | NULL | Probable | As nectar-tap_saprotroph (Põlme et al. 2020); Decay sugar-rich substrates (Põlme et al. 2020) | Põlme S, et al. 2020. Fungal Diversity 105:1-16 ((https://doi.org/10.1007/s13225-020-00466-2)) |
| Otu37329 | 1 | k__Fungi(100);p__Ascomycota(100);c__Saccharomycetes(100);o__Saccharomycetales(100);f__Metschnikowiaceae(100);g__Metschnikowia(100);s__Metschnikowia_pulcherrima(100); | Metschnikowia | 13 | Saprotroph | \|Undefined Saprotroph\| | Yeast | NULL | Probable | As nectar-tap_saprotroph (Põlme et al. 2020); Decay sugar-rich substrates (Põlme et al. 2020) | Põlme S, et al. 2020. Fungal Diversity 105:1-16 ((https://doi.org/10.1007/s13225-020-00466-2)) |
| Otu37330 | 1 | k__Fungi(100);p__Ascomycota(100);c__Saccharomycetes(100);o__Saccharomycetales(100);f__Metschnikowiaceae(100);g__Metschnikowia(100);s__Metschnikowia_pulcherrima(100); | Metschnikowia | 13 | Saprotroph | \|Undefined Saprotroph\| | Yeast | NULL | Probable | As nectar-tap_saprotroph (Põlme et al. 2020); Decay sugar-rich substrates (Põlme et al. 2020) | Põlme S, et al. 2020. Fungal Diversity 105:1-16 ((https://doi.org/10.1007/s13225-020-00466-2)) |
| Otu37331 | 1 | k__Fungi(100);p__Ascomycota(100);c__Saccharomycetes(100);o__Saccharomycetales(100);f__Metschnikowiaceae(100);g__Metschnikowia(100);s__Metschnikowia_pulcherrima(100); | Metschnikowia | 13 | Saprotroph | \|Undefined Saprotroph\| | Yeast | NULL | Probable | As nectar-tap_saprotroph (Põlme et al. 2020); Decay sugar-rich substrates (Põlme et al. 2020) | Põlme S, et al. 2020. Fungal Diversity 105:1-16 ((https://doi.org/10.1007/s13225-020-00466-2)) |
| Otu37332 | 1 | k__Fungi(100);p__Ascomycota(100);c__Saccharomycetes(100);o__Saccharomycetales(100);f__Metschnikowiaceae(100);g__Metschnikowia(100);s__Metschnikowia_pulcherrima(100); | Metschnikowia | 13 | Saprotroph | \|Undefined Saprotroph\| | Yeast | NULL | Probable | As nectar-tap_saprotroph (Põlme et al. 2020); Decay sugar-rich substrates (Põlme et al. 2020) | Põlme S, et al. 2020. Fungal Diversity 105:1-16 ((https://doi.org/10.1007/s13225-020-00466-2)) |
| Otu37333 | 1 | k__Fungi(100);p__Ascomycota(100);c__Saccharomycetes(100);o__Saccharomycetales(100);f__Metschnikowiaceae(100);g__Metschnikowia(100);s__Metschnikowia_pulcherrima(100); | Metschnikowia | 13 | Saprotroph | \|Undefined Saprotroph\| | Yeast | NULL | Probable | As nectar-tap_saprotroph (Põlme et al. 2020); Decay sugar-rich substrates (Põlme et al. 2020) | Põlme S, et al. 2020. Fungal Diversity 105:1-16 ((https://doi.org/10.1007/s13225-020-00466-2)) |
| Otu37334 | 1 | k__Fungi(100);p__Ascomycota(100);c__Saccharomycetes(100);o__Saccharomycetales(100);f__Metschnikowiaceae(100);g__Metschnikowia(100);s__Metschnikowia_pulcherrima(100); | Metschnikowia | 13 | Saprotroph | \|Undefined Saprotroph\| | Yeast | NULL | Probable | As nectar-tap_saprotroph (Põlme et al. 2020); Decay sugar-rich substrates (Põlme et al. 2020) | Põlme S, et al. 2020. Fungal Diversity 105:1-16 ((https://doi.org/10.1007/s13225-020-00466-2)) |
| Otu37335 | 1 | k__Fungi(100);p__Ascomycota(100);c__Saccharomycetes(100);o__Saccharomycetales(100);f__Metschnikowiaceae(100);g__Metschnikowia(100);g__Metschnikowia_unclassified(100); | Metschnikowia | 13 | Saprotroph | \|Undefined Saprotroph\| | Yeast | NULL | Probable | As nectar-tap_saprotroph (Põlme et al. 2020); Decay sugar-rich substrates (Põlme et al. 2020) | Põlme S, et al. 2020. Fungal Diversity 105:1-16 ((https://doi.org/10.1007/s13225-020-00466-2)) |
| Otu37336 | 1 | k__Fungi(100);p__Ascomycota(100);c__Saccharomycetes(100);o__Saccharomycetales(100);f__Metschnikowiaceae(100);g__Metschnikowia(100);s__Metschnikowia_pulcherrima(100); | Metschnikowia | 13 | Saprotroph | \|Undefined Saprotroph\| | Yeast | NULL | Probable | As nectar-tap_saprotroph (Põlme et al. 2020); Decay sugar-rich substrates (Põlme et al. 2020) | Põlme S, et al. 2020. Fungal Diversity 105:1-16 ((https://doi.org/10.1007/s13225-020-00466-2)) |
| Otu37342 | 1 | k__Fungi(100);p__Ascomycota(100);c__Saccharomycetes(100);o__Saccharomycetales(100);f__Metschnikowiaceae(100);g__Metschnikowia(100);s__Metschnikowia_pulcherrima(100); | Metschnikowia | 13 | Saprotroph | \|Undefined Saprotroph\| | Yeast | NULL | Probable | As nectar-tap_saprotroph (Põlme et al. 2020); Decay sugar-rich substrates (Põlme et al. 2020) | Põlme S, et al. 2020. Fungal Diversity 105:1-16 ((https://doi.org/10.1007/s13225-020-00466-2)) |
| Otu37343 | 1 | k__Fungi(100);p__Ascomycota(100);c__Saccharomycetes(100);o__Saccharomycetales(100);f__Metschnikowiaceae(100);g__Metschnikowia(100);s__Metschnikowia_pulcherrima(100); | Metschnikowia | 13 | Saprotroph | \|Undefined Saprotroph\| | Yeast | NULL | Probable | As nectar-tap_saprotroph (Põlme et al. 2020); Decay sugar-rich substrates (Põlme et al. 2020) | Põlme S, et al. 2020. Fungal Diversity 105:1-16 ((https://doi.org/10.1007/s13225-020-00466-2)) |
| Otu37344 | 1 | k__Fungi(100);p__Ascomycota(100);c__Saccharomycetes(100);o__Saccharomycetales(100);f__Metschnikowiaceae(100);g__Metschnikowia(100);s__Metschnikowia_pulcherrima(100); | Metschnikowia | 13 | Saprotroph | \|Undefined Saprotroph\| | Yeast | NULL | Probable | As nectar-tap_saprotroph (Põlme et al. 2020); Decay sugar-rich substrates (Põlme et al. 2020) | Põlme S, et al. 2020. Fungal Diversity 105:1-16 ((https://doi.org/10.1007/s13225-020-00466-2)) |
| Otu37345 | 1 | k__Fungi(100);p__Ascomycota(100);c__Saccharomycetes(100);o__Saccharomycetales(100);f__Metschnikowiaceae(100);g__Metschnikowia(100);s__Metschnikowia_pulcherrima(100); | Metschnikowia | 13 | Saprotroph | \|Undefined Saprotroph\| | Yeast | NULL | Probable | As nectar-tap_saprotroph (Põlme et al. 2020); Decay sugar-rich substrates (Põlme et al. 2020) | Põlme S, et al. 2020. Fungal Diversity 105:1-16 ((https://doi.org/10.1007/s13225-020-00466-2)) |
| Otu37346 | 1 | k__Fungi(100);p__Ascomycota(100);c__Saccharomycetes(100);o__Saccharomycetales(100);f__Metschnikowiaceae(100);g__Metschnikowia(100);s__Metschnikowia_pulcherrima(100); | Metschnikowia | 13 | Saprotroph | \|Undefined Saprotroph\| | Yeast | NULL | Probable | As nectar-tap_saprotroph (Põlme et al. 2020); Decay sugar-rich substrates (Põlme et al. 2020) | Põlme S, et al. 2020. Fungal Diversity 105:1-16 ((https://doi.org/10.1007/s13225-020-00466-2)) |
| Otu37347 | 1 | k__Fungi(100);p__Ascomycota(100);c__Saccharomycetes(100);o__Saccharomycetales(100);f__Metschnikowiaceae(100);g__Metschnikowia(100);g__Metschnikowia_unclassified(100); | Metschnikowia | 13 | Saprotroph | \|Undefined Saprotroph\| | Yeast | NULL | Probable | As nectar-tap_saprotroph (Põlme et al. 2020); Decay sugar-rich substrates (Põlme et al. 2020) | Põlme S, et al. 2020. Fungal Diversity 105:1-16 ((https://doi.org/10.1007/s13225-020-00466-2)) |
| Otu37348 | 1 | k__Fungi(100);p__Ascomycota(100);c__Saccharomycetes(100);o__Saccharomycetales(100);f__Metschnikowiaceae(100);g__Metschnikowia(100);s__Metschnikowia_pulcherrima(100); | Metschnikowia | 13 | Saprotroph | \|Undefined Saprotroph\| | Yeast | NULL | Probable | As nectar-tap_saprotroph (Põlme et al. 2020); Decay sugar-rich substrates (Põlme et al. 2020) | Põlme S, et al. 2020. Fungal Diversity 105:1-16 ((https://doi.org/10.1007/s13225-020-00466-2)) |
| Otu37349 | 1 | k__Fungi(100);p__Ascomycota(100);c__Saccharomycetes(100);o__Saccharomycetales(100);f__Metschnikowiaceae(100);g__Metschnikowia(100);s__Metschnikowia_pulcherrima(100); | Metschnikowia | 13 | Saprotroph | \|Undefined Saprotroph\| | Yeast | NULL | Probable | As nectar-tap_saprotroph (Põlme et al. 2020); Decay sugar-rich substrates (Põlme et al. 2020) | Põlme S, et al. 2020. Fungal Diversity 105:1-16 ((https://doi.org/10.1007/s13225-020-00466-2)) |
| Otu37350 | 1 | k__Fungi(100);p__Ascomycota(100);c__Saccharomycetes(100);o__Saccharomycetales(100);f__Metschnikowiaceae(100);g__Metschnikowia(100);g__Metschnikowia_unclassified(100); | Metschnikowia | 13 | Saprotroph | \|Undefined Saprotroph\| | Yeast | NULL | Probable | As nectar-tap_saprotroph (Põlme et al. 2020); Decay sugar-rich substrates (Põlme et al. 2020) | Põlme S, et al. 2020. Fungal Diversity 105:1-16 ((https://doi.org/10.1007/s13225-020-00466-2)) |
| Otu37351 | 1 | k__Fungi(100);p__Ascomycota(100);c__Saccharomycetes(100);o__Saccharomycetales(100);f__Metschnikowiaceae(100);g__Metschnikowia(100);g__Metschnikowia_unclassified(100); | Metschnikowia | 13 | Saprotroph | \|Undefined Saprotroph\| | Yeast | NULL | Probable | As nectar-tap_saprotroph (Põlme et al. 2020); Decay sugar-rich substrates (Põlme et al. 2020) | Põlme S, et al. 2020. Fungal Diversity 105:1-16 ((https://doi.org/10.1007/s13225-020-00466-2)) |
| Otu37352 | 1 | k__Fungi(100);p__Ascomycota(100);c__Saccharomycetes(100);o__Saccharomycetales(100);f__Metschnikowiaceae(100);g__Metschnikowia(100);s__Metschnikowia_pulcherrima(100); | Metschnikowia | 13 | Saprotroph | \|Undefined Saprotroph\| | Yeast | NULL | Probable | As nectar-tap_saprotroph (Põlme et al. 2020); Decay sugar-rich substrates (Põlme et al. 2020) | Põlme S, et al. 2020. Fungal Diversity 105:1-16 ((https://doi.org/10.1007/s13225-020-00466-2)) |
| Otu37353 | 1 | k__Fungi(100);p__Ascomycota(100);c__Saccharomycetes(100);o__Saccharomycetales(100);f__Metschnikowiaceae(100);g__Metschnikowia(100);s__Metschnikowia_pulcherrima(100); | Metschnikowia | 13 | Saprotroph | \|Undefined Saprotroph\| | Yeast | NULL | Probable | As nectar-tap_saprotroph (Põlme et al. 2020); Decay sugar-rich substrates (Põlme et al. 2020) | Põlme S, et al. 2020. Fungal Diversity 105:1-16 ((https://doi.org/10.1007/s13225-020-00466-2)) |
| Otu37354 | 1 | k__Fungi(100);p__Ascomycota(100);c__Saccharomycetes(100);o__Saccharomycetales(100);f__Metschnikowiaceae(100);g__Metschnikowia(100);g__Metschnikowia_unclassified(100); | Metschnikowia | 13 | Saprotroph | \|Undefined Saprotroph\| | Yeast | NULL | Probable | As nectar-tap_saprotroph (Põlme et al. 2020); Decay sugar-rich substrates (Põlme et al. 2020) | Põlme S, et al. 2020. Fungal Diversity 105:1-16 ((https://doi.org/10.1007/s13225-020-00466-2)) |
| Otu37355 | 1 | k__Fungi(100);p__Ascomycota(100);c__Saccharomycetes(100);o__Saccharomycetales(100);f__Metschnikowiaceae(100);g__Metschnikowia(100);g__Metschnikowia_unclassified(100); | Metschnikowia | 13 | Saprotroph | \|Undefined Saprotroph\| | Yeast | NULL | Probable | As nectar-tap_saprotroph (Põlme et al. 2020); Decay sugar-rich substrates (Põlme et al. 2020) | Põlme S, et al. 2020. Fungal Diversity 105:1-16 ((https://doi.org/10.1007/s13225-020-00466-2)) |
| Otu37356 | 1 | k__Fungi(100);p__Ascomycota(100);c__Saccharomycetes(100);o__Saccharomycetales(100);f__Metschnikowiaceae(100);g__Metschnikowia(100);s__Metschnikowia_pulcherrima(100); | Metschnikowia | 13 | Saprotroph | \|Undefined Saprotroph\| | Yeast | NULL | Probable | As nectar-tap_saprotroph (Põlme et al. 2020); Decay sugar-rich substrates (Põlme et al. 2020) | Põlme S, et al. 2020. Fungal Diversity 105:1-16 ((https://doi.org/10.1007/s13225-020-00466-2)) |
| Otu37357 | 1 | k__Fungi(100);p__Ascomycota(100);c__Saccharomycetes(100);o__Saccharomycetales(100);f__Metschnikowiaceae(100);g__Metschnikowia(100);s__Metschnikowia_pulcherrima(100); | Metschnikowia | 13 | Saprotroph | \|Undefined Saprotroph\| | Yeast | NULL | Probable | As nectar-tap_saprotroph (Põlme et al. 2020); Decay sugar-rich substrates (Põlme et al. 2020) | Põlme S, et al. 2020. Fungal Diversity 105:1-16 ((https://doi.org/10.1007/s13225-020-00466-2)) |
| Otu37358 | 1 | k__Fungi(100);p__Ascomycota(100);c__Saccharomycetes(100);o__Saccharomycetales(100);f__Metschnikowiaceae(100);g__Metschnikowia(100);s__Metschnikowia_pulcherrima(100); | Metschnikowia | 13 | Saprotroph | \|Undefined Saprotroph\| | Yeast | NULL | Probable | As nectar-tap_saprotroph (Põlme et al. 2020); Decay sugar-rich substrates (Põlme et al. 2020) | Põlme S, et al. 2020. Fungal Diversity 105:1-16 ((https://doi.org/10.1007/s13225-020-00466-2)) |
| Otu37359 | 1 | k__Fungi(100);p__Ascomycota(100);c__Saccharomycetes(100);o__Saccharomycetales(100);f__Metschnikowiaceae(100);g__Metschnikowia(100);s__Metschnikowia_pulcherrima(100); | Metschnikowia | 13 | Saprotroph | \|Undefined Saprotroph\| | Yeast | NULL | Probable | As nectar-tap_saprotroph (Põlme et al. 2020); Decay sugar-rich substrates (Põlme et al. 2020) | Põlme S, et al. 2020. Fungal Diversity 105:1-16 ((https://doi.org/10.1007/s13225-020-00466-2)) |
| Otu37360 | 1 | k__Fungi(100);p__Ascomycota(100);c__Saccharomycetes(100);o__Saccharomycetales(100);f__Metschnikowiaceae(100);g__Metschnikowia(100);s__Metschnikowia_pulcherrima(100); | Metschnikowia | 13 | Saprotroph | \|Undefined Saprotroph\| | Yeast | NULL | Probable | As nectar-tap_saprotroph (Põlme et al. 2020); Decay sugar-rich substrates (Põlme et al. 2020) | Põlme S, et al. 2020. Fungal Diversity 105:1-16 ((https://doi.org/10.1007/s13225-020-00466-2)) |
| Otu37361 | 1 | k__Fungi(100);p__Ascomycota(100);c__Saccharomycetes(100);o__Saccharomycetales(100);f__Metschnikowiaceae(100);g__Metschnikowia(100);s__Metschnikowia_pulcherrima(100); | Metschnikowia | 13 | Saprotroph | \|Undefined Saprotroph\| | Yeast | NULL | Probable | As nectar-tap_saprotroph (Põlme et al. 2020); Decay sugar-rich substrates (Põlme et al. 2020) | Põlme S, et al. 2020. Fungal Diversity 105:1-16 ((https://doi.org/10.1007/s13225-020-00466-2)) |
| Otu37362 | 1 | k__Fungi(100);p__Ascomycota(100);c__Saccharomycetes(100);o__Saccharomycetales(100);f__Metschnikowiaceae(100);g__Metschnikowia(100);s__Metschnikowia_pulcherrima(100); | Metschnikowia | 13 | Saprotroph | \|Undefined Saprotroph\| | Yeast | NULL | Probable | As nectar-tap_saprotroph (Põlme et al. 2020); Decay sugar-rich substrates (Põlme et al. 2020) | Põlme S, et al. 2020. Fungal Diversity 105:1-16 ((https://doi.org/10.1007/s13225-020-00466-2)) |
| Otu37363 | 1 | k__Fungi(100);p__Ascomycota(100);c__Saccharomycetes(100);o__Saccharomycetales(100);f__Metschnikowiaceae(100);g__Metschnikowia(100);s__Metschnikowia_pulcherrima(100); | Metschnikowia | 13 | Saprotroph | \|Undefined Saprotroph\| | Yeast | NULL | Probable | As nectar-tap_saprotroph (Põlme et al. 2020); Decay sugar-rich substrates (Põlme et al. 2020) | Põlme S, et al. 2020. Fungal Diversity 105:1-16 ((https://doi.org/10.1007/s13225-020-00466-2)) |
| Otu37364 | 1 | k__Fungi(100);p__Ascomycota(100);c__Saccharomycetes(100);o__Saccharomycetales(100);f__Metschnikowiaceae(100);g__Metschnikowia(100);s__Metschnikowia_pulcherrima(100); | Metschnikowia | 13 | Saprotroph | \|Undefined Saprotroph\| | Yeast | NULL | Probable | As nectar-tap_saprotroph (Põlme et al. 2020); Decay sugar-rich substrates (Põlme et al. 2020) | Põlme S, et al. 2020. Fungal Diversity 105:1-16 ((https://doi.org/10.1007/s13225-020-00466-2)) |
| Otu37365 | 1 | k__Fungi(100);p__Ascomycota(100);c__Saccharomycetes(100);o__Saccharomycetales(100);f__Metschnikowiaceae(100);g__Metschnikowia(100);s__Metschnikowia_pulcherrima(100); | Metschnikowia | 13 | Saprotroph | \|Undefined Saprotroph\| | Yeast | NULL | Probable | As nectar-tap_saprotroph (Põlme et al. 2020); Decay sugar-rich substrates (Põlme et al. 2020) | Põlme S, et al. 2020. Fungal Diversity 105:1-16 ((https://doi.org/10.1007/s13225-020-00466-2)) |
| Otu37366 | 1 | k__Fungi(100);p__Ascomycota(100);c__Saccharomycetes(100);o__Saccharomycetales(100);f__Metschnikowiaceae(100);g__Metschnikowia(100);s__Metschnikowia_pulcherrima(100); | Metschnikowia | 13 | Saprotroph | \|Undefined Saprotroph\| | Yeast | NULL | Probable | As nectar-tap_saprotroph (Põlme et al. 2020); Decay sugar-rich substrates (Põlme et al. 2020) | Põlme S, et al. 2020. Fungal Diversity 105:1-16 ((https://doi.org/10.1007/s13225-020-00466-2)) |
| Otu37367 | 1 | k__Fungi(100);p__Ascomycota(100);c__Saccharomycetes(100);o__Saccharomycetales(100);f__Metschnikowiaceae(100);g__Metschnikowia(100);s__Metschnikowia_pulcherrima(100); | Metschnikowia | 13 | Saprotroph | \|Undefined Saprotroph\| | Yeast | NULL | Probable | As nectar-tap_saprotroph (Põlme et al. 2020); Decay sugar-rich substrates (Põlme et al. 2020) | Põlme S, et al. 2020. Fungal Diversity 105:1-16 ((https://doi.org/10.1007/s13225-020-00466-2)) |
| Otu37368 | 1 | k__Fungi(100);p__Ascomycota(100);c__Saccharomycetes(100);o__Saccharomycetales(100);f__Metschnikowiaceae(100);g__Metschnikowia(100);g__Metschnikowia_unclassified(100); | Metschnikowia | 13 | Saprotroph | \|Undefined Saprotroph\| | Yeast | NULL | Probable | As nectar-tap_saprotroph (Põlme et al. 2020); Decay sugar-rich substrates (Põlme et al. 2020) | Põlme S, et al. 2020. Fungal Diversity 105:1-16 ((https://doi.org/10.1007/s13225-020-00466-2)) |
| Otu37369 | 1 | k__Fungi(100);p__Ascomycota(100);c__Saccharomycetes(100);o__Saccharomycetales(100);f__Metschnikowiaceae(100);g__Metschnikowia(100);s__Metschnikowia_pulcherrima(100); | Metschnikowia | 13 | Saprotroph | \|Undefined Saprotroph\| | Yeast | NULL | Probable | As nectar-tap_saprotroph (Põlme et al. 2020); Decay sugar-rich substrates (Põlme et al. 2020) | Põlme S, et al. 2020. Fungal Diversity 105:1-16 ((https://doi.org/10.1007/s13225-020-00466-2)) |
| Otu37370 | 1 | k__Fungi(100);p__Ascomycota(100);c__Saccharomycetes(100);o__Saccharomycetales(100);f__Metschnikowiaceae(100);g__Metschnikowia(100);g__Metschnikowia_unclassified(100); | Metschnikowia | 13 | Saprotroph | \|Undefined Saprotroph\| | Yeast | NULL | Probable | As nectar-tap_saprotroph (Põlme et al. 2020); Decay sugar-rich substrates (Põlme et al. 2020) | Põlme S, et al. 2020. Fungal Diversity 105:1-16 ((https://doi.org/10.1007/s13225-020-00466-2)) |
| Otu37371 | 1 | k__Fungi(100);p__Ascomycota(100);c__Saccharomycetes(100);o__Saccharomycetales(100);f__Metschnikowiaceae(100);g__Metschnikowia(100);g__Metschnikowia_unclassified(100); | Metschnikowia | 13 | Saprotroph | \|Undefined Saprotroph\| | Yeast | NULL | Probable | As nectar-tap_saprotroph (Põlme et al. 2020); Decay sugar-rich substrates (Põlme et al. 2020) | Põlme S, et al. 2020. Fungal Diversity 105:1-16 ((https://doi.org/10.1007/s13225-020-00466-2)) |
| Otu37372 | 1 | k__Fungi(100);p__Ascomycota(100);c__Saccharomycetes(100);o__Saccharomycetales(100);f__Metschnikowiaceae(100);g__Metschnikowia(100);s__Metschnikowia_pulcherrima(100); | Metschnikowia | 13 | Saprotroph | \|Undefined Saprotroph\| | Yeast | NULL | Probable | As nectar-tap_saprotroph (Põlme et al. 2020); Decay sugar-rich substrates (Põlme et al. 2020) | Põlme S, et al. 2020. Fungal Diversity 105:1-16 ((https://doi.org/10.1007/s13225-020-00466-2)) |
| Otu37373 | 1 | k__Fungi(100);p__Ascomycota(100);c__Saccharomycetes(100);o__Saccharomycetales(100);f__Metschnikowiaceae(100);g__Metschnikowia(100);s__Metschnikowia_pulcherrima(100); | Metschnikowia | 13 | Saprotroph | \|Undefined Saprotroph\| | Yeast | NULL | Probable | As nectar-tap_saprotroph (Põlme et al. 2020); Decay sugar-rich substrates (Põlme et al. 2020) | Põlme S, et al. 2020. Fungal Diversity 105:1-16 ((https://doi.org/10.1007/s13225-020-00466-2)) |
| Otu37374 | 1 | k__Fungi(100);p__Ascomycota(100);c__Saccharomycetes(100);o__Saccharomycetales(100);f__Metschnikowiaceae(100);g__Metschnikowia(100);s__Metschnikowia_pulcherrima(100); | Metschnikowia | 13 | Saprotroph | \|Undefined Saprotroph\| | Yeast | NULL | Probable | As nectar-tap_saprotroph (Põlme et al. 2020); Decay sugar-rich substrates (Põlme et al. 2020) | Põlme S, et al. 2020. Fungal Diversity 105:1-16 ((https://doi.org/10.1007/s13225-020-00466-2)) |
| Otu37375 | 1 | k__Fungi(100);p__Ascomycota(100);c__Saccharomycetes(100);o__Saccharomycetales(100);f__Metschnikowiaceae(100);g__Metschnikowia(100);s__Metschnikowia_pulcherrima(100); | Metschnikowia | 13 | Saprotroph | \|Undefined Saprotroph\| | Yeast | NULL | Probable | As nectar-tap_saprotroph (Põlme et al. 2020); Decay sugar-rich substrates (Põlme et al. 2020) | Põlme S, et al. 2020. Fungal Diversity 105:1-16 ((https://doi.org/10.1007/s13225-020-00466-2)) |
| Otu37376 | 1 | k__Fungi(100);p__Ascomycota(100);c__Saccharomycetes(100);o__Saccharomycetales(100);f__Metschnikowiaceae(100);g__Metschnikowia(100);s__Metschnikowia_pulcherrima(100); | Metschnikowia | 13 | Saprotroph | \|Undefined Saprotroph\| | Yeast | NULL | Probable | As nectar-tap_saprotroph (Põlme et al. 2020); Decay sugar-rich substrates (Põlme et al. 2020) | Põlme S, et al. 2020. Fungal Diversity 105:1-16 ((https://doi.org/10.1007/s13225-020-00466-2)) |
| Otu37377 | 1 | k__Fungi(100);p__Ascomycota(100);c__Saccharomycetes(100);o__Saccharomycetales(100);f__Metschnikowiaceae(100);g__Metschnikowia(100);s__Metschnikowia_pulcherrima(100); | Metschnikowia | 13 | Saprotroph | \|Undefined Saprotroph\| | Yeast | NULL | Probable | As nectar-tap_saprotroph (Põlme et al. 2020); Decay sugar-rich substrates (Põlme et al. 2020) | Põlme S, et al. 2020. Fungal Diversity 105:1-16 ((https://doi.org/10.1007/s13225-020-00466-2)) |
| Otu37378 | 1 | k__Fungi(100);p__Ascomycota(100);c__Saccharomycetes(100);o__Saccharomycetales(100);f__Metschnikowiaceae(100);g__Metschnikowia(100);s__Metschnikowia_pulcherrima(100); | Metschnikowia | 13 | Saprotroph | \|Undefined Saprotroph\| | Yeast | NULL | Probable | As nectar-tap_saprotroph (Põlme et al. 2020); Decay sugar-rich substrates (Põlme et al. 2020) | Põlme S, et al. 2020. Fungal Diversity 105:1-16 ((https://doi.org/10.1007/s13225-020-00466-2)) |
| Otu37379 | 1 | k__Fungi(100);p__Ascomycota(100);c__Saccharomycetes(100);o__Saccharomycetales(100);f__Metschnikowiaceae(100);g__Metschnikowia(100);s__Metschnikowia_pulcherrima(100); | Metschnikowia | 13 | Saprotroph | \|Undefined Saprotroph\| | Yeast | NULL | Probable | As nectar-tap_saprotroph (Põlme et al. 2020); Decay sugar-rich substrates (Põlme et al. 2020) | Põlme S, et al. 2020. Fungal Diversity 105:1-16 ((https://doi.org/10.1007/s13225-020-00466-2)) |
| Otu37380 | 1 | k__Fungi(100);p__Ascomycota(100);c__Saccharomycetes(100);o__Saccharomycetales(100);f__Metschnikowiaceae(100);g__Metschnikowia(100);s__Metschnikowia_pulcherrima(100); | Metschnikowia | 13 | Saprotroph | \|Undefined Saprotroph\| | Yeast | NULL | Probable | As nectar-tap_saprotroph (Põlme et al. 2020); Decay sugar-rich substrates (Põlme et al. 2020) | Põlme S, et al. 2020. Fungal Diversity 105:1-16 ((https://doi.org/10.1007/s13225-020-00466-2)) |
| Otu37381 | 1 | k__Fungi(100);p__Ascomycota(100);c__Saccharomycetes(100);o__Saccharomycetales(100);f__Metschnikowiaceae(100);g__Metschnikowia(100);g__Metschnikowia_unclassified(100); | Metschnikowia | 13 | Saprotroph | \|Undefined Saprotroph\| | Yeast | NULL | Probable | As nectar-tap_saprotroph (Põlme et al. 2020); Decay sugar-rich substrates (Põlme et al. 2020) | Põlme S, et al. 2020. Fungal Diversity 105:1-16 ((https://doi.org/10.1007/s13225-020-00466-2)) |
| Otu37382 | 1 | k__Fungi(100);p__Ascomycota(100);c__Saccharomycetes(100);o__Saccharomycetales(100);f__Metschnikowiaceae(100);g__Metschnikowia(100);s__Metschnikowia_pulcherrima(100); | Metschnikowia | 13 | Saprotroph | \|Undefined Saprotroph\| | Yeast | NULL | Probable | As nectar-tap_saprotroph (Põlme et al. 2020); Decay sugar-rich substrates (Põlme et al. 2020) | Põlme S, et al. 2020. Fungal Diversity 105:1-16 ((https://doi.org/10.1007/s13225-020-00466-2)) |
| Otu37383 | 1 | k__Fungi(100);p__Ascomycota(100);c__Saccharomycetes(100);o__Saccharomycetales(100);f__Metschnikowiaceae(100);g__Metschnikowia(100);s__Metschnikowia_pulcherrima(100); | Metschnikowia | 13 | Saprotroph | \|Undefined Saprotroph\| | Yeast | NULL | Probable | As nectar-tap_saprotroph (Põlme et al. 2020); Decay sugar-rich substrates (Põlme et al. 2020) | Põlme S, et al. 2020. Fungal Diversity 105:1-16 ((https://doi.org/10.1007/s13225-020-00466-2)) |
| Otu37384 | 1 | k__Fungi(100);p__Ascomycota(100);c__Saccharomycetes(100);o__Saccharomycetales(100);f__Metschnikowiaceae(100);g__Metschnikowia(100);s__Metschnikowia_pulcherrima(100); | Metschnikowia | 13 | Saprotroph | \|Undefined Saprotroph\| | Yeast | NULL | Probable | As nectar-tap_saprotroph (Põlme et al. 2020); Decay sugar-rich substrates (Põlme et al. 2020) | Põlme S, et al. 2020. Fungal Diversity 105:1-16 ((https://doi.org/10.1007/s13225-020-00466-2)) |
| Otu37385 | 1 | k__Fungi(100);p__Ascomycota(100);c__Saccharomycetes(100);o__Saccharomycetales(100);f__Metschnikowiaceae(100);g__Metschnikowia(100);g__Metschnikowia_unclassified(100); | Metschnikowia | 13 | Saprotroph | \|Undefined Saprotroph\| | Yeast | NULL | Probable | As nectar-tap_saprotroph (Põlme et al. 2020); Decay sugar-rich substrates (Põlme et al. 2020) | Põlme S, et al. 2020. Fungal Diversity 105:1-16 ((https://doi.org/10.1007/s13225-020-00466-2)) |
| Otu37386 | 1 | k__Fungi(100);p__Ascomycota(100);c__Saccharomycetes(100);o__Saccharomycetales(100);f__Metschnikowiaceae(100);g__Metschnikowia(100);s__Metschnikowia_pulcherrima(100); | Metschnikowia | 13 | Saprotroph | \|Undefined Saprotroph\| | Yeast | NULL | Probable | As nectar-tap_saprotroph (Põlme et al. 2020); Decay sugar-rich substrates (Põlme et al. 2020) | Põlme S, et al. 2020. Fungal Diversity 105:1-16 ((https://doi.org/10.1007/s13225-020-00466-2)) |
| Otu37387 | 1 | k__Fungi(100);p__Ascomycota(100);c__Saccharomycetes(100);o__Saccharomycetales(100);f__Metschnikowiaceae(100);g__Metschnikowia(100);s__Metschnikowia_pulcherrima(100); | Metschnikowia | 13 | Saprotroph | \|Undefined Saprotroph\| | Yeast | NULL | Probable | As nectar-tap_saprotroph (Põlme et al. 2020); Decay sugar-rich substrates (Põlme et al. 2020) | Põlme S, et al. 2020. Fungal Diversity 105:1-16 ((https://doi.org/10.1007/s13225-020-00466-2)) |
| Otu37388 | 1 | k__Fungi(100);p__Ascomycota(100);c__Saccharomycetes(100);o__Saccharomycetales(100);f__Metschnikowiaceae(100);g__Metschnikowia(100);s__Metschnikowia_pulcherrima(100); | Metschnikowia | 13 | Saprotroph | \|Undefined Saprotroph\| | Yeast | NULL | Probable | As nectar-tap_saprotroph (Põlme et al. 2020); Decay sugar-rich substrates (Põlme et al. 2020) | Põlme S, et al. 2020. Fungal Diversity 105:1-16 ((https://doi.org/10.1007/s13225-020-00466-2)) |
| Otu37389 | 1 | k__Fungi(100);p__Ascomycota(100);c__Saccharomycetes(100);o__Saccharomycetales(100);f__Metschnikowiaceae(100);g__Metschnikowia(100);s__Metschnikowia_pulcherrima(100); | Metschnikowia | 13 | Saprotroph | \|Undefined Saprotroph\| | Yeast | NULL | Probable | As nectar-tap_saprotroph (Põlme et al. 2020); Decay sugar-rich substrates (Põlme et al. 2020) | Põlme S, et al. 2020. Fungal Diversity 105:1-16 ((https://doi.org/10.1007/s13225-020-00466-2)) |
| Otu37390 | 1 | k__Fungi(100);p__Ascomycota(100);c__Saccharomycetes(100);o__Saccharomycetales(100);f__Metschnikowiaceae(100);g__Metschnikowia(100);g__Metschnikowia_unclassified(100); | Metschnikowia | 13 | Saprotroph | \|Undefined Saprotroph\| | Yeast | NULL | Probable | As nectar-tap_saprotroph (Põlme et al. 2020); Decay sugar-rich substrates (Põlme et al. 2020) | Põlme S, et al. 2020. Fungal Diversity 105:1-16 ((https://doi.org/10.1007/s13225-020-00466-2)) |
| Otu37391 | 1 | k__Fungi(100);p__Ascomycota(100);c__Saccharomycetes(100);o__Saccharomycetales(100);f__Metschnikowiaceae(100);g__Metschnikowia(100);g__Metschnikowia_unclassified(100); | Metschnikowia | 13 | Saprotroph | \|Undefined Saprotroph\| | Yeast | NULL | Probable | As nectar-tap_saprotroph (Põlme et al. 2020); Decay sugar-rich substrates (Põlme et al. 2020) | Põlme S, et al. 2020. Fungal Diversity 105:1-16 ((https://doi.org/10.1007/s13225-020-00466-2)) |
| Otu37392 | 1 | k__Fungi(100);p__Ascomycota(100);c__Saccharomycetes(100);o__Saccharomycetales(100);f__Metschnikowiaceae(100);g__Metschnikowia(100);s__Metschnikowia_pulcherrima(100); | Metschnikowia | 13 | Saprotroph | \|Undefined Saprotroph\| | Yeast | NULL | Probable | As nectar-tap_saprotroph (Põlme et al. 2020); Decay sugar-rich substrates (Põlme et al. 2020) | Põlme S, et al. 2020. Fungal Diversity 105:1-16 ((https://doi.org/10.1007/s13225-020-00466-2)) |
| Otu37393 | 1 | k__Fungi(100);p__Ascomycota(100);c__Saccharomycetes(100);o__Saccharomycetales(100);f__Metschnikowiaceae(100);g__Metschnikowia(100);s__Metschnikowia_pulcherrima(100); | Metschnikowia | 13 | Saprotroph | \|Undefined Saprotroph\| | Yeast | NULL | Probable | As nectar-tap_saprotroph (Põlme et al. 2020); Decay sugar-rich substrates (Põlme et al. 2020) | Põlme S, et al. 2020. Fungal Diversity 105:1-16 ((https://doi.org/10.1007/s13225-020-00466-2)) |
| Otu37394 | 1 | k__Fungi(100);p__Ascomycota(100);c__Saccharomycetes(100);o__Saccharomycetales(100);f__Metschnikowiaceae(100);g__Metschnikowia(100);g__Metschnikowia_unclassified(100); | Metschnikowia | 13 | Saprotroph | \|Undefined Saprotroph\| | Yeast | NULL | Probable | As nectar-tap_saprotroph (Põlme et al. 2020); Decay sugar-rich substrates (Põlme et al. 2020) | Põlme S, et al. 2020. Fungal Diversity 105:1-16 ((https://doi.org/10.1007/s13225-020-00466-2)) |
| Otu37395 | 1 | k__Fungi(100);p__Ascomycota(100);c__Saccharomycetes(100);o__Saccharomycetales(100);f__Metschnikowiaceae(100);g__Metschnikowia(100);g__Metschnikowia_unclassified(100); | Metschnikowia | 13 | Saprotroph | \|Undefined Saprotroph\| | Yeast | NULL | Probable | As nectar-tap_saprotroph (Põlme et al. 2020); Decay sugar-rich substrates (Põlme et al. 2020) | Põlme S, et al. 2020. Fungal Diversity 105:1-16 ((https://doi.org/10.1007/s13225-020-00466-2)) |
| Otu37396 | 1 | k__Fungi(100);p__Ascomycota(100);c__Saccharomycetes(100);o__Saccharomycetales(100);f__Metschnikowiaceae(100);g__Metschnikowia(100);s__Metschnikowia_pulcherrima(100); | Metschnikowia | 13 | Saprotroph | \|Undefined Saprotroph\| | Yeast | NULL | Probable | As nectar-tap_saprotroph (Põlme et al. 2020); Decay sugar-rich substrates (Põlme et al. 2020) | Põlme S, et al. 2020. Fungal Diversity 105:1-16 ((https://doi.org/10.1007/s13225-020-00466-2)) |
| Otu37397 | 1 | k__Fungi(100);p__Ascomycota(100);c__Saccharomycetes(100);o__Saccharomycetales(100);f__Metschnikowiaceae(100);g__Metschnikowia(100);s__Metschnikowia_pulcherrima(100); | Metschnikowia | 13 | Saprotroph | \|Undefined Saprotroph\| | Yeast | NULL | Probable | As nectar-tap_saprotroph (Põlme et al. 2020); Decay sugar-rich substrates (Põlme et al. 2020) | Põlme S, et al. 2020. Fungal Diversity 105:1-16 ((https://doi.org/10.1007/s13225-020-00466-2)) |
| Otu37398 | 1 | k__Fungi(100);p__Ascomycota(100);c__Saccharomycetes(100);o__Saccharomycetales(100);f__Metschnikowiaceae(100);g__Metschnikowia(100);s__Metschnikowia_pulcherrima(100); | Metschnikowia | 13 | Saprotroph | \|Undefined Saprotroph\| | Yeast | NULL | Probable | As nectar-tap_saprotroph (Põlme et al. 2020); Decay sugar-rich substrates (Põlme et al. 2020) | Põlme S, et al. 2020. Fungal Diversity 105:1-16 ((https://doi.org/10.1007/s13225-020-00466-2)) |
| Otu37399 | 1 | k__Fungi(100);p__Ascomycota(100);c__Saccharomycetes(100);o__Saccharomycetales(100);f__Metschnikowiaceae(100);g__Metschnikowia(100);s__Metschnikowia_pulcherrima(100); | Metschnikowia | 13 | Saprotroph | \|Undefined Saprotroph\| | Yeast | NULL | Probable | As nectar-tap_saprotroph (Põlme et al. 2020); Decay sugar-rich substrates (Põlme et al. 2020) | Põlme S, et al. 2020. Fungal Diversity 105:1-16 ((https://doi.org/10.1007/s13225-020-00466-2)) |
| Otu37400 | 1 | k__Fungi(100);p__Ascomycota(100);c__Saccharomycetes(100);o__Saccharomycetales(100);f__Metschnikowiaceae(100);g__Metschnikowia(100);s__Metschnikowia_pulcherrima(100); | Metschnikowia | 13 | Saprotroph | \|Undefined Saprotroph\| | Yeast | NULL | Probable | As nectar-tap_saprotroph (Põlme et al. 2020); Decay sugar-rich substrates (Põlme et al. 2020) | Põlme S, et al. 2020. Fungal Diversity 105:1-16 ((https://doi.org/10.1007/s13225-020-00466-2)) |
| Otu37401 | 1 | k__Fungi(100);p__Ascomycota(100);c__Saccharomycetes(100);o__Saccharomycetales(100);f__Metschnikowiaceae(100);g__Metschnikowia(100);s__Metschnikowia_pulcherrima(100); | Metschnikowia | 13 | Saprotroph | \|Undefined Saprotroph\| | Yeast | NULL | Probable | As nectar-tap_saprotroph (Põlme et al. 2020); Decay sugar-rich substrates (Põlme et al. 2020) | Põlme S, et al. 2020. Fungal Diversity 105:1-16 ((https://doi.org/10.1007/s13225-020-00466-2)) |
| Otu37402 | 1 | k__Fungi(100);p__Ascomycota(100);c__Saccharomycetes(100);o__Saccharomycetales(100);f__Metschnikowiaceae(100);g__Metschnikowia(100);s__Metschnikowia_pulcherrima(100); | Metschnikowia | 13 | Saprotroph | \|Undefined Saprotroph\| | Yeast | NULL | Probable | As nectar-tap_saprotroph (Põlme et al. 2020); Decay sugar-rich substrates (Põlme et al. 2020) | Põlme S, et al. 2020. Fungal Diversity 105:1-16 ((https://doi.org/10.1007/s13225-020-00466-2)) |
| Otu37403 | 1 | k__Fungi(100);p__Ascomycota(100);c__Saccharomycetes(100);o__Saccharomycetales(100);f__Metschnikowiaceae(100);g__Metschnikowia(100);s__Metschnikowia_pulcherrima(100); | Metschnikowia | 13 | Saprotroph | \|Undefined Saprotroph\| | Yeast | NULL | Probable | As nectar-tap_saprotroph (Põlme et al. 2020); Decay sugar-rich substrates (Põlme et al. 2020) | Põlme S, et al. 2020. Fungal Diversity 105:1-16 ((https://doi.org/10.1007/s13225-020-00466-2)) |
| Otu37404 | 1 | k__Fungi(100);p__Ascomycota(100);c__Saccharomycetes(100);o__Saccharomycetales(100);f__Metschnikowiaceae(100);g__Metschnikowia(100);g__Metschnikowia_unclassified(100); | Metschnikowia | 13 | Saprotroph | \|Undefined Saprotroph\| | Yeast | NULL | Probable | As nectar-tap_saprotroph (Põlme et al. 2020); Decay sugar-rich substrates (Põlme et al. 2020) | Põlme S, et al. 2020. Fungal Diversity 105:1-16 ((https://doi.org/10.1007/s13225-020-00466-2)) |
| Otu37405 | 1 | k__Fungi(100);p__Ascomycota(100);c__Saccharomycetes(100);o__Saccharomycetales(100);f__Metschnikowiaceae(100);g__Metschnikowia(100);s__Metschnikowia_pulcherrima(100); | Metschnikowia | 13 | Saprotroph | \|Undefined Saprotroph\| | Yeast | NULL | Probable | As nectar-tap_saprotroph (Põlme et al. 2020); Decay sugar-rich substrates (Põlme et al. 2020) | Põlme S, et al. 2020. Fungal Diversity 105:1-16 ((https://doi.org/10.1007/s13225-020-00466-2)) |
| Otu37406 | 1 | k__Fungi(100);p__Ascomycota(100);c__Saccharomycetes(100);o__Saccharomycetales(100);f__Metschnikowiaceae(100);g__Metschnikowia(100);g__Metschnikowia_unclassified(100); | Metschnikowia | 13 | Saprotroph | \|Undefined Saprotroph\| | Yeast | NULL | Probable | As nectar-tap_saprotroph (Põlme et al. 2020); Decay sugar-rich substrates (Põlme et al. 2020) | Põlme S, et al. 2020. Fungal Diversity 105:1-16 ((https://doi.org/10.1007/s13225-020-00466-2)) |
| Otu37407 | 1 | k__Fungi(100);p__Ascomycota(100);c__Saccharomycetes(100);o__Saccharomycetales(100);f__Metschnikowiaceae(100);g__Metschnikowia(100);s__Metschnikowia_pulcherrima(100); | Metschnikowia | 13 | Saprotroph | \|Undefined Saprotroph\| | Yeast | NULL | Probable | As nectar-tap_saprotroph (Põlme et al. 2020); Decay sugar-rich substrates (Põlme et al. 2020) | Põlme S, et al. 2020. Fungal Diversity 105:1-16 ((https://doi.org/10.1007/s13225-020-00466-2)) |
| Otu37408 | 1 | k__Fungi(100);p__Ascomycota(100);c__Saccharomycetes(100);o__Saccharomycetales(100);f__Metschnikowiaceae(100);g__Metschnikowia(100);g__Metschnikowia_unclassified(100); | Metschnikowia | 13 | Saprotroph | \|Undefined Saprotroph\| | Yeast | NULL | Probable | As nectar-tap_saprotroph (Põlme et al. 2020); Decay sugar-rich substrates (Põlme et al. 2020) | Põlme S, et al. 2020. Fungal Diversity 105:1-16 ((https://doi.org/10.1007/s13225-020-00466-2)) |
| Otu37409 | 1 | k__Fungi(100);p__Ascomycota(100);c__Saccharomycetes(100);o__Saccharomycetales(100);f__Metschnikowiaceae(100);g__Metschnikowia(100);s__Metschnikowia_pulcherrima(100); | Metschnikowia | 13 | Saprotroph | \|Undefined Saprotroph\| | Yeast | NULL | Probable | As nectar-tap_saprotroph (Põlme et al. 2020); Decay sugar-rich substrates (Põlme et al. 2020) | Põlme S, et al. 2020. Fungal Diversity 105:1-16 ((https://doi.org/10.1007/s13225-020-00466-2)) |
| Otu37410 | 1 | k__Fungi(100);p__Ascomycota(100);c__Saccharomycetes(100);o__Saccharomycetales(100);f__Metschnikowiaceae(100);g__Metschnikowia(100);s__Metschnikowia_pulcherrima(100); | Metschnikowia | 13 | Saprotroph | \|Undefined Saprotroph\| | Yeast | NULL | Probable | As nectar-tap_saprotroph (Põlme et al. 2020); Decay sugar-rich substrates (Põlme et al. 2020) | Põlme S, et al. 2020. Fungal Diversity 105:1-16 ((https://doi.org/10.1007/s13225-020-00466-2)) |
| Otu37411 | 1 | k__Fungi(100);p__Ascomycota(100);c__Saccharomycetes(100);o__Saccharomycetales(100);f__Metschnikowiaceae(100);g__Metschnikowia(100);s__Metschnikowia_pulcherrima(100); | Metschnikowia | 13 | Saprotroph | \|Undefined Saprotroph\| | Yeast | NULL | Probable | As nectar-tap_saprotroph (Põlme et al. 2020); Decay sugar-rich substrates (Põlme et al. 2020) | Põlme S, et al. 2020. Fungal Diversity 105:1-16 ((https://doi.org/10.1007/s13225-020-00466-2)) |
| Otu37412 | 1 | k__Fungi(100);p__Ascomycota(100);c__Saccharomycetes(100);o__Saccharomycetales(100);f__Metschnikowiaceae(100);g__Metschnikowia(100);s__Metschnikowia_pulcherrima(100); | Metschnikowia | 13 | Saprotroph | \|Undefined Saprotroph\| | Yeast | NULL | Probable | As nectar-tap_saprotroph (Põlme et al. 2020); Decay sugar-rich substrates (Põlme et al. 2020) | Põlme S, et al. 2020. Fungal Diversity 105:1-16 ((https://doi.org/10.1007/s13225-020-00466-2)) |
| Otu37413 | 1 | k__Fungi(100);p__Ascomycota(100);c__Saccharomycetes(100);o__Saccharomycetales(100);f__Metschnikowiaceae(100);g__Metschnikowia(100);g__Metschnikowia_unclassified(100); | Metschnikowia | 13 | Saprotroph | \|Undefined Saprotroph\| | Yeast | NULL | Probable | As nectar-tap_saprotroph (Põlme et al. 2020); Decay sugar-rich substrates (Põlme et al. 2020) | Põlme S, et al. 2020. Fungal Diversity 105:1-16 ((https://doi.org/10.1007/s13225-020-00466-2)) |
| Otu37414 | 1 | k__Fungi(100);p__Ascomycota(100);c__Saccharomycetes(100);o__Saccharomycetales(100);f__Metschnikowiaceae(100);g__Metschnikowia(100);s__Metschnikowia_pulcherrima(100); | Metschnikowia | 13 | Saprotroph | \|Undefined Saprotroph\| | Yeast | NULL | Probable | As nectar-tap_saprotroph (Põlme et al. 2020); Decay sugar-rich substrates (Põlme et al. 2020) | Põlme S, et al. 2020. Fungal Diversity 105:1-16 ((https://doi.org/10.1007/s13225-020-00466-2)) |
| Otu37415 | 1 | k__Fungi(100);p__Ascomycota(100);c__Saccharomycetes(100);o__Saccharomycetales(100);f__Metschnikowiaceae(100);g__Metschnikowia(100);s__Metschnikowia_pulcherrima(100); | Metschnikowia | 13 | Saprotroph | \|Undefined Saprotroph\| | Yeast | NULL | Probable | As nectar-tap_saprotroph (Põlme et al. 2020); Decay sugar-rich substrates (Põlme et al. 2020) | Põlme S, et al. 2020. Fungal Diversity 105:1-16 ((https://doi.org/10.1007/s13225-020-00466-2)) |
| Otu37416 | 1 | k__Fungi(100);p__Ascomycota(100);c__Saccharomycetes(100);o__Saccharomycetales(100);f__Metschnikowiaceae(100);g__Metschnikowia(100);s__Metschnikowia_pulcherrima(100); | Metschnikowia | 13 | Saprotroph | \|Undefined Saprotroph\| | Yeast | NULL | Probable | As nectar-tap_saprotroph (Põlme et al. 2020); Decay sugar-rich substrates (Põlme et al. 2020) | Põlme S, et al. 2020. Fungal Diversity 105:1-16 ((https://doi.org/10.1007/s13225-020-00466-2)) |
| Otu37417 | 1 | k__Fungi(100);p__Ascomycota(100);c__Saccharomycetes(100);o__Saccharomycetales(100);f__Metschnikowiaceae(100);g__Metschnikowia(100);s__Metschnikowia_pulcherrima(100); | Metschnikowia | 13 | Saprotroph | \|Undefined Saprotroph\| | Yeast | NULL | Probable | As nectar-tap_saprotroph (Põlme et al. 2020); Decay sugar-rich substrates (Põlme et al. 2020) | Põlme S, et al. 2020. Fungal Diversity 105:1-16 ((https://doi.org/10.1007/s13225-020-00466-2)) |
| Otu37418 | 1 | k__Fungi(100);p__Ascomycota(100);c__Saccharomycetes(100);o__Saccharomycetales(100);f__Metschnikowiaceae(100);g__Metschnikowia(100);s__Metschnikowia_pulcherrima(100); | Metschnikowia | 13 | Saprotroph | \|Undefined Saprotroph\| | Yeast | NULL | Probable | As nectar-tap_saprotroph (Põlme et al. 2020); Decay sugar-rich substrates (Põlme et al. 2020) | Põlme S, et al. 2020. Fungal Diversity 105:1-16 ((https://doi.org/10.1007/s13225-020-00466-2)) |
| Otu37419 | 1 | k__Fungi(100);p__Ascomycota(100);c__Saccharomycetes(100);o__Saccharomycetales(100);f__Metschnikowiaceae(100);g__Metschnikowia(100);g__Metschnikowia_unclassified(100); | Metschnikowia | 13 | Saprotroph | \|Undefined Saprotroph\| | Yeast | NULL | Probable | As nectar-tap_saprotroph (Põlme et al. 2020); Decay sugar-rich substrates (Põlme et al. 2020) | Põlme S, et al. 2020. Fungal Diversity 105:1-16 ((https://doi.org/10.1007/s13225-020-00466-2)) |
| Otu37420 | 1 | k__Fungi(100);p__Ascomycota(100);c__Saccharomycetes(100);o__Saccharomycetales(100);f__Metschnikowiaceae(100);g__Metschnikowia(100);g__Metschnikowia_unclassified(100); | Metschnikowia | 13 | Saprotroph | \|Undefined Saprotroph\| | Yeast | NULL | Probable | As nectar-tap_saprotroph (Põlme et al. 2020); Decay sugar-rich substrates (Põlme et al. 2020) | Põlme S, et al. 2020. Fungal Diversity 105:1-16 ((https://doi.org/10.1007/s13225-020-00466-2)) |
| Otu37421 | 1 | k__Fungi(100);p__Ascomycota(100);c__Saccharomycetes(100);o__Saccharomycetales(100);f__Metschnikowiaceae(100);g__Metschnikowia(100);s__Metschnikowia_pulcherrima(100); | Metschnikowia | 13 | Saprotroph | \|Undefined Saprotroph\| | Yeast | NULL | Probable | As nectar-tap_saprotroph (Põlme et al. 2020); Decay sugar-rich substrates (Põlme et al. 2020) | Põlme S, et al. 2020. Fungal Diversity 105:1-16 ((https://doi.org/10.1007/s13225-020-00466-2)) |
| Otu37422 | 1 | k__Fungi(100);p__Ascomycota(100);c__Saccharomycetes(100);o__Saccharomycetales(100);f__Metschnikowiaceae(100);g__Metschnikowia(100);s__Metschnikowia_pulcherrima(100); | Metschnikowia | 13 | Saprotroph | \|Undefined Saprotroph\| | Yeast | NULL | Probable | As nectar-tap_saprotroph (Põlme et al. 2020); Decay sugar-rich substrates (Põlme et al. 2020) | Põlme S, et al. 2020. Fungal Diversity 105:1-16 ((https://doi.org/10.1007/s13225-020-00466-2)) |
| Otu37423 | 1 | k__Fungi(100);p__Ascomycota(100);c__Saccharomycetes(100);o__Saccharomycetales(100);f__Metschnikowiaceae(100);g__Metschnikowia(100);g__Metschnikowia_unclassified(100); | Metschnikowia | 13 | Saprotroph | \|Undefined Saprotroph\| | Yeast | NULL | Probable | As nectar-tap_saprotroph (Põlme et al. 2020); Decay sugar-rich substrates (Põlme et al. 2020) | Põlme S, et al. 2020. Fungal Diversity 105:1-16 ((https://doi.org/10.1007/s13225-020-00466-2)) |
| Otu37424 | 1 | k__Fungi(100);p__Ascomycota(100);c__Saccharomycetes(100);o__Saccharomycetales(100);f__Metschnikowiaceae(100);g__Metschnikowia(100);g__Metschnikowia_unclassified(100); | Metschnikowia | 13 | Saprotroph | \|Undefined Saprotroph\| | Yeast | NULL | Probable | As nectar-tap_saprotroph (Põlme et al. 2020); Decay sugar-rich substrates (Põlme et al. 2020) | Põlme S, et al. 2020. Fungal Diversity 105:1-16 ((https://doi.org/10.1007/s13225-020-00466-2)) |
| Otu37425 | 1 | k__Fungi(100);p__Ascomycota(100);c__Saccharomycetes(100);o__Saccharomycetales(100);f__Metschnikowiaceae(100);g__Metschnikowia(100);s__Metschnikowia_pulcherrima(100); | Metschnikowia | 13 | Saprotroph | \|Undefined Saprotroph\| | Yeast | NULL | Probable | As nectar-tap_saprotroph (Põlme et al. 2020); Decay sugar-rich substrates (Põlme et al. 2020) | Põlme S, et al. 2020. Fungal Diversity 105:1-16 ((https://doi.org/10.1007/s13225-020-00466-2)) |
| Otu37426 | 1 | k__Fungi(100);p__Ascomycota(100);c__Saccharomycetes(100);o__Saccharomycetales(100);f__Metschnikowiaceae(100);g__Metschnikowia(100);g__Metschnikowia_unclassified(100); | Metschnikowia | 13 | Saprotroph | \|Undefined Saprotroph\| | Yeast | NULL | Probable | As nectar-tap_saprotroph (Põlme et al. 2020); Decay sugar-rich substrates (Põlme et al. 2020) | Põlme S, et al. 2020. Fungal Diversity 105:1-16 ((https://doi.org/10.1007/s13225-020-00466-2)) |
| Otu37427 | 1 | k__Fungi(100);p__Ascomycota(100);c__Saccharomycetes(100);o__Saccharomycetales(100);f__Metschnikowiaceae(100);g__Metschnikowia(100);g__Metschnikowia_unclassified(100); | Metschnikowia | 13 | Saprotroph | \|Undefined Saprotroph\| | Yeast | NULL | Probable | As nectar-tap_saprotroph (Põlme et al. 2020); Decay sugar-rich substrates (Põlme et al. 2020) | Põlme S, et al. 2020. Fungal Diversity 105:1-16 ((https://doi.org/10.1007/s13225-020-00466-2)) |
| Otu37428 | 1 | k__Fungi(100);p__Ascomycota(100);c__Saccharomycetes(100);o__Saccharomycetales(100);f__Metschnikowiaceae(100);g__Metschnikowia(100);s__Metschnikowia_pulcherrima(100); | Metschnikowia | 13 | Saprotroph | \|Undefined Saprotroph\| | Yeast | NULL | Probable | As nectar-tap_saprotroph (Põlme et al. 2020); Decay sugar-rich substrates (Põlme et al. 2020) | Põlme S, et al. 2020. Fungal Diversity 105:1-16 ((https://doi.org/10.1007/s13225-020-00466-2)) |
| Otu37429 | 1 | k__Fungi(100);p__Ascomycota(100);c__Saccharomycetes(100);o__Saccharomycetales(100);f__Metschnikowiaceae(100);g__Metschnikowia(100);s__Metschnikowia_pulcherrima(100); | Metschnikowia | 13 | Saprotroph | \|Undefined Saprotroph\| | Yeast | NULL | Probable | As nectar-tap_saprotroph (Põlme et al. 2020); Decay sugar-rich substrates (Põlme et al. 2020) | Põlme S, et al. 2020. Fungal Diversity 105:1-16 ((https://doi.org/10.1007/s13225-020-00466-2)) |
| Otu37430 | 1 | k__Fungi(100);p__Ascomycota(100);c__Saccharomycetes(100);o__Saccharomycetales(100);f__Metschnikowiaceae(100);g__Metschnikowia(100);g__Metschnikowia_unclassified(100); | Metschnikowia | 13 | Saprotroph | \|Undefined Saprotroph\| | Yeast | NULL | Probable | As nectar-tap_saprotroph (Põlme et al. 2020); Decay sugar-rich substrates (Põlme et al. 2020) | Põlme S, et al. 2020. Fungal Diversity 105:1-16 ((https://doi.org/10.1007/s13225-020-00466-2)) |
| Otu37431 | 1 | k__Fungi(100);p__Ascomycota(100);c__Saccharomycetes(100);o__Saccharomycetales(100);f__Metschnikowiaceae(100);g__Metschnikowia(100);s__Metschnikowia_pulcherrima(100); | Metschnikowia | 13 | Saprotroph | \|Undefined Saprotroph\| | Yeast | NULL | Probable | As nectar-tap_saprotroph (Põlme et al. 2020); Decay sugar-rich substrates (Põlme et al. 2020) | Põlme S, et al. 2020. Fungal Diversity 105:1-16 ((https://doi.org/10.1007/s13225-020-00466-2)) |
| Otu37432 | 1 | k__Fungi(100);p__Ascomycota(100);c__Saccharomycetes(100);o__Saccharomycetales(100);f__Metschnikowiaceae(100);g__Metschnikowia(100);s__Metschnikowia_pulcherrima(100); | Metschnikowia | 13 | Saprotroph | \|Undefined Saprotroph\| | Yeast | NULL | Probable | As nectar-tap_saprotroph (Põlme et al. 2020); Decay sugar-rich substrates (Põlme et al. 2020) | Põlme S, et al. 2020. Fungal Diversity 105:1-16 ((https://doi.org/10.1007/s13225-020-00466-2)) |
| Otu37433 | 1 | k__Fungi(100);p__Ascomycota(100);c__Saccharomycetes(100);o__Saccharomycetales(100);f__Metschnikowiaceae(100);g__Metschnikowia(100);s__Metschnikowia_pulcherrima(100); | Metschnikowia | 13 | Saprotroph | \|Undefined Saprotroph\| | Yeast | NULL | Probable | As nectar-tap_saprotroph (Põlme et al. 2020); Decay sugar-rich substrates (Põlme et al. 2020) | Põlme S, et al. 2020. Fungal Diversity 105:1-16 ((https://doi.org/10.1007/s13225-020-00466-2)) |
| Otu37434 | 1 | k__Fungi(100);p__Ascomycota(100);c__Saccharomycetes(100);o__Saccharomycetales(100);f__Metschnikowiaceae(100);g__Metschnikowia(100);g__Metschnikowia_unclassified(100); | Metschnikowia | 13 | Saprotroph | \|Undefined Saprotroph\| | Yeast | NULL | Probable | As nectar-tap_saprotroph (Põlme et al. 2020); Decay sugar-rich substrates (Põlme et al. 2020) | Põlme S, et al. 2020. Fungal Diversity 105:1-16 ((https://doi.org/10.1007/s13225-020-00466-2)) |
| Otu37435 | 1 | k__Fungi(100);p__Ascomycota(100);c__Saccharomycetes(100);o__Saccharomycetales(100);f__Metschnikowiaceae(100);g__Metschnikowia(100);s__Metschnikowia_pulcherrima(100); | Metschnikowia | 13 | Saprotroph | \|Undefined Saprotroph\| | Yeast | NULL | Probable | As nectar-tap_saprotroph (Põlme et al. 2020); Decay sugar-rich substrates (Põlme et al. 2020) | Põlme S, et al. 2020. Fungal Diversity 105:1-16 ((https://doi.org/10.1007/s13225-020-00466-2)) |
| Otu37436 | 1 | k__Fungi(100);p__Ascomycota(100);c__Saccharomycetes(100);o__Saccharomycetales(100);f__Metschnikowiaceae(100);g__Metschnikowia(100);s__Metschnikowia_pulcherrima(100); | Metschnikowia | 13 | Saprotroph | \|Undefined Saprotroph\| | Yeast | NULL | Probable | As nectar-tap_saprotroph (Põlme et al. 2020); Decay sugar-rich substrates (Põlme et al. 2020) | Põlme S, et al. 2020. Fungal Diversity 105:1-16 ((https://doi.org/10.1007/s13225-020-00466-2)) |
| Otu37437 | 1 | k__Fungi(100);p__Ascomycota(100);c__Saccharomycetes(100);o__Saccharomycetales(100);f__Metschnikowiaceae(100);g__Metschnikowia(100);g__Metschnikowia_unclassified(100); | Metschnikowia | 13 | Saprotroph | \|Undefined Saprotroph\| | Yeast | NULL | Probable | As nectar-tap_saprotroph (Põlme et al. 2020); Decay sugar-rich substrates (Põlme et al. 2020) | Põlme S, et al. 2020. Fungal Diversity 105:1-16 ((https://doi.org/10.1007/s13225-020-00466-2)) |
| Otu37438 | 1 | k__Fungi(100);p__Ascomycota(100);c__Saccharomycetes(100);o__Saccharomycetales(100);f__Metschnikowiaceae(100);g__Metschnikowia(100);g__Metschnikowia_unclassified(100); | Metschnikowia | 13 | Saprotroph | \|Undefined Saprotroph\| | Yeast | NULL | Probable | As nectar-tap_saprotroph (Põlme et al. 2020); Decay sugar-rich substrates (Põlme et al. 2020) | Põlme S, et al. 2020. Fungal Diversity 105:1-16 ((https://doi.org/10.1007/s13225-020-00466-2)) |
| Otu37439 | 1 | k__Fungi(100);p__Ascomycota(100);c__Saccharomycetes(100);o__Saccharomycetales(100);f__Metschnikowiaceae(100);g__Metschnikowia(100);s__Metschnikowia_pulcherrima(100); | Metschnikowia | 13 | Saprotroph | \|Undefined Saprotroph\| | Yeast | NULL | Probable | As nectar-tap_saprotroph (Põlme et al. 2020); Decay sugar-rich substrates (Põlme et al. 2020) | Põlme S, et al. 2020. Fungal Diversity 105:1-16 ((https://doi.org/10.1007/s13225-020-00466-2)) |
| Otu37440 | 1 | k__Fungi(100);p__Ascomycota(100);c__Saccharomycetes(100);o__Saccharomycetales(100);f__Metschnikowiaceae(100);g__Metschnikowia(100);s__Metschnikowia_pulcherrima(100); | Metschnikowia | 13 | Saprotroph | \|Undefined Saprotroph\| | Yeast | NULL | Probable | As nectar-tap_saprotroph (Põlme et al. 2020); Decay sugar-rich substrates (Põlme et al. 2020) | Põlme S, et al. 2020. Fungal Diversity 105:1-16 ((https://doi.org/10.1007/s13225-020-00466-2)) |
| Otu37441 | 1 | k__Fungi(100);p__Ascomycota(100);c__Saccharomycetes(100);o__Saccharomycetales(100);f__Metschnikowiaceae(100);g__Metschnikowia(100);s__Metschnikowia_pulcherrima(100); | Metschnikowia | 13 | Saprotroph | \|Undefined Saprotroph\| | Yeast | NULL | Probable | As nectar-tap_saprotroph (Põlme et al. 2020); Decay sugar-rich substrates (Põlme et al. 2020) | Põlme S, et al. 2020. Fungal Diversity 105:1-16 ((https://doi.org/10.1007/s13225-020-00466-2)) |
| Otu37442 | 1 | k__Fungi(100);p__Ascomycota(100);c__Saccharomycetes(100);o__Saccharomycetales(100);f__Metschnikowiaceae(100);g__Metschnikowia(100);s__Metschnikowia_pulcherrima(100); | Metschnikowia | 13 | Saprotroph | \|Undefined Saprotroph\| | Yeast | NULL | Probable | As nectar-tap_saprotroph (Põlme et al. 2020); Decay sugar-rich substrates (Põlme et al. 2020) | Põlme S, et al. 2020. Fungal Diversity 105:1-16 ((https://doi.org/10.1007/s13225-020-00466-2)) |
| Otu37443 | 1 | k__Fungi(100);p__Ascomycota(100);c__Saccharomycetes(100);o__Saccharomycetales(100);f__Metschnikowiaceae(100);g__Metschnikowia(100);g__Metschnikowia_unclassified(100); | Metschnikowia | 13 | Saprotroph | \|Undefined Saprotroph\| | Yeast | NULL | Probable | As nectar-tap_saprotroph (Põlme et al. 2020); Decay sugar-rich substrates (Põlme et al. 2020) | Põlme S, et al. 2020. Fungal Diversity 105:1-16 ((https://doi.org/10.1007/s13225-020-00466-2)) |
| Otu37444 | 1 | k__Fungi(100);p__Ascomycota(100);c__Saccharomycetes(100);o__Saccharomycetales(100);f__Metschnikowiaceae(100);g__Metschnikowia(100);g__Metschnikowia_unclassified(100); | Metschnikowia | 13 | Saprotroph | \|Undefined Saprotroph\| | Yeast | NULL | Probable | As nectar-tap_saprotroph (Põlme et al. 2020); Decay sugar-rich substrates (Põlme et al. 2020) | Põlme S, et al. 2020. Fungal Diversity 105:1-16 ((https://doi.org/10.1007/s13225-020-00466-2)) |
| Otu37445 | 1 | k__Fungi(100);p__Ascomycota(100);c__Saccharomycetes(100);o__Saccharomycetales(100);f__Metschnikowiaceae(100);g__Metschnikowia(100);s__Metschnikowia_pulcherrima(100); | Metschnikowia | 13 | Saprotroph | \|Undefined Saprotroph\| | Yeast | NULL | Probable | As nectar-tap_saprotroph (Põlme et al. 2020); Decay sugar-rich substrates (Põlme et al. 2020) | Põlme S, et al. 2020. Fungal Diversity 105:1-16 ((https://doi.org/10.1007/s13225-020-00466-2)) |
| Otu37446 | 1 | k__Fungi(100);p__Ascomycota(100);c__Saccharomycetes(100);o__Saccharomycetales(100);f__Metschnikowiaceae(100);g__Metschnikowia(100);s__Metschnikowia_pulcherrima(100); | Metschnikowia | 13 | Saprotroph | \|Undefined Saprotroph\| | Yeast | NULL | Probable | As nectar-tap_saprotroph (Põlme et al. 2020); Decay sugar-rich substrates (Põlme et al. 2020) | Põlme S, et al. 2020. Fungal Diversity 105:1-16 ((https://doi.org/10.1007/s13225-020-00466-2)) |
| Otu37447 | 1 | k__Fungi(100);p__Ascomycota(100);c__Saccharomycetes(100);o__Saccharomycetales(100);f__Metschnikowiaceae(100);g__Metschnikowia(100);s__Metschnikowia_pulcherrima(100); | Metschnikowia | 13 | Saprotroph | \|Undefined Saprotroph\| | Yeast | NULL | Probable | As nectar-tap_saprotroph (Põlme et al. 2020); Decay sugar-rich substrates (Põlme et al. 2020) | Põlme S, et al. 2020. Fungal Diversity 105:1-16 ((https://doi.org/10.1007/s13225-020-00466-2)) |
| Otu37448 | 1 | k__Fungi(100);p__Ascomycota(100);c__Saccharomycetes(100);o__Saccharomycetales(100);f__Metschnikowiaceae(100);g__Metschnikowia(100);s__Metschnikowia_pulcherrima(100); | Metschnikowia | 13 | Saprotroph | \|Undefined Saprotroph\| | Yeast | NULL | Probable | As nectar-tap_saprotroph (Põlme et al. 2020); Decay sugar-rich substrates (Põlme et al. 2020) | Põlme S, et al. 2020. Fungal Diversity 105:1-16 ((https://doi.org/10.1007/s13225-020-00466-2)) |
| Otu37449 | 1 | k__Fungi(100);p__Ascomycota(100);c__Saccharomycetes(100);o__Saccharomycetales(100);f__Metschnikowiaceae(100);g__Metschnikowia(100);s__Metschnikowia_pulcherrima(100); | Metschnikowia | 13 | Saprotroph | \|Undefined Saprotroph\| | Yeast | NULL | Probable | As nectar-tap_saprotroph (Põlme et al. 2020); Decay sugar-rich substrates (Põlme et al. 2020) | Põlme S, et al. 2020. Fungal Diversity 105:1-16 ((https://doi.org/10.1007/s13225-020-00466-2)) |
| Otu37450 | 1 | k__Fungi(100);p__Ascomycota(100);c__Saccharomycetes(100);o__Saccharomycetales(100);f__Metschnikowiaceae(100);g__Metschnikowia(100);s__Metschnikowia_pulcherrima(100); | Metschnikowia | 13 | Saprotroph | \|Undefined Saprotroph\| | Yeast | NULL | Probable | As nectar-tap_saprotroph (Põlme et al. 2020); Decay sugar-rich substrates (Põlme et al. 2020) | Põlme S, et al. 2020. Fungal Diversity 105:1-16 ((https://doi.org/10.1007/s13225-020-00466-2)) |
| Otu37451 | 1 | k__Fungi(100);p__Ascomycota(100);c__Saccharomycetes(100);o__Saccharomycetales(100);f__Metschnikowiaceae(100);g__Metschnikowia(100);s__Metschnikowia_pulcherrima(100); | Metschnikowia | 13 | Saprotroph | \|Undefined Saprotroph\| | Yeast | NULL | Probable | As nectar-tap_saprotroph (Põlme et al. 2020); Decay sugar-rich substrates (Põlme et al. 2020) | Põlme S, et al. 2020. Fungal Diversity 105:1-16 ((https://doi.org/10.1007/s13225-020-00466-2)) |
| Otu37452 | 1 | k__Fungi(100);p__Ascomycota(100);c__Saccharomycetes(100);o__Saccharomycetales(100);f__Metschnikowiaceae(100);g__Metschnikowia(100);g__Metschnikowia_unclassified(100); | Metschnikowia | 13 | Saprotroph | \|Undefined Saprotroph\| | Yeast | NULL | Probable | As nectar-tap_saprotroph (Põlme et al. 2020); Decay sugar-rich substrates (Põlme et al. 2020) | Põlme S, et al. 2020. Fungal Diversity 105:1-16 ((https://doi.org/10.1007/s13225-020-00466-2)) |
| Otu37453 | 1 | k__Fungi(100);p__Ascomycota(100);c__Saccharomycetes(100);o__Saccharomycetales(100);f__Metschnikowiaceae(100);g__Metschnikowia(100);g__Metschnikowia_unclassified(100); | Metschnikowia | 13 | Saprotroph | \|Undefined Saprotroph\| | Yeast | NULL | Probable | As nectar-tap_saprotroph (Põlme et al. 2020); Decay sugar-rich substrates (Põlme et al. 2020) | Põlme S, et al. 2020. Fungal Diversity 105:1-16 ((https://doi.org/10.1007/s13225-020-00466-2)) |
| Otu37454 | 1 | k__Fungi(100);p__Ascomycota(100);c__Saccharomycetes(100);o__Saccharomycetales(100);f__Metschnikowiaceae(100);g__Metschnikowia(100);g__Metschnikowia_unclassified(100); | Metschnikowia | 13 | Saprotroph | \|Undefined Saprotroph\| | Yeast | NULL | Probable | As nectar-tap_saprotroph (Põlme et al. 2020); Decay sugar-rich substrates (Põlme et al. 2020) | Põlme S, et al. 2020. Fungal Diversity 105:1-16 ((https://doi.org/10.1007/s13225-020-00466-2)) |
| Otu37455 | 1 | k__Fungi(100);p__Ascomycota(100);c__Saccharomycetes(100);o__Saccharomycetales(100);f__Metschnikowiaceae(100);g__Metschnikowia(100);g__Metschnikowia_unclassified(100); | Metschnikowia | 13 | Saprotroph | \|Undefined Saprotroph\| | Yeast | NULL | Probable | As nectar-tap_saprotroph (Põlme et al. 2020); Decay sugar-rich substrates (Põlme et al. 2020) | Põlme S, et al. 2020. Fungal Diversity 105:1-16 ((https://doi.org/10.1007/s13225-020-00466-2)) |
| Otu37456 | 1 | k__Fungi(100);p__Ascomycota(100);c__Saccharomycetes(100);o__Saccharomycetales(100);f__Metschnikowiaceae(100);g__Metschnikowia(100);s__Metschnikowia_pulcherrima(100); | Metschnikowia | 13 | Saprotroph | \|Undefined Saprotroph\| | Yeast | NULL | Probable | As nectar-tap_saprotroph (Põlme et al. 2020); Decay sugar-rich substrates (Põlme et al. 2020) | Põlme S, et al. 2020. Fungal Diversity 105:1-16 ((https://doi.org/10.1007/s13225-020-00466-2)) |
| Otu37457 | 1 | k__Fungi(100);p__Ascomycota(100);c__Saccharomycetes(100);o__Saccharomycetales(100);f__Metschnikowiaceae(100);g__Metschnikowia(100);s__Metschnikowia_pulcherrima(100); | Metschnikowia | 13 | Saprotroph | \|Undefined Saprotroph\| | Yeast | NULL | Probable | As nectar-tap_saprotroph (Põlme et al. 2020); Decay sugar-rich substrates (Põlme et al. 2020) | Põlme S, et al. 2020. Fungal Diversity 105:1-16 ((https://doi.org/10.1007/s13225-020-00466-2)) |
| Otu37458 | 1 | k__Fungi(100);p__Ascomycota(100);c__Saccharomycetes(100);o__Saccharomycetales(100);f__Metschnikowiaceae(100);g__Metschnikowia(100);g__Metschnikowia_unclassified(100); | Metschnikowia | 13 | Saprotroph | \|Undefined Saprotroph\| | Yeast | NULL | Probable | As nectar-tap_saprotroph (Põlme et al. 2020); Decay sugar-rich substrates (Põlme et al. 2020) | Põlme S, et al. 2020. Fungal Diversity 105:1-16 ((https://doi.org/10.1007/s13225-020-00466-2)) |
| Otu37459 | 1 | k__Fungi(100);p__Ascomycota(100);c__Saccharomycetes(100);o__Saccharomycetales(100);f__Metschnikowiaceae(100);g__Metschnikowia(100);g__Metschnikowia_unclassified(100); | Metschnikowia | 13 | Saprotroph | \|Undefined Saprotroph\| | Yeast | NULL | Probable | As nectar-tap_saprotroph (Põlme et al. 2020); Decay sugar-rich substrates (Põlme et al. 2020) | Põlme S, et al. 2020. Fungal Diversity 105:1-16 ((https://doi.org/10.1007/s13225-020-00466-2)) |
| Otu37460 | 1 | k__Fungi(100);p__Ascomycota(100);c__Saccharomycetes(100);o__Saccharomycetales(100);f__Metschnikowiaceae(100);g__Metschnikowia(100);s__Metschnikowia_pulcherrima(100); | Metschnikowia | 13 | Saprotroph | \|Undefined Saprotroph\| | Yeast | NULL | Probable | As nectar-tap_saprotroph (Põlme et al. 2020); Decay sugar-rich substrates (Põlme et al. 2020) | Põlme S, et al. 2020. Fungal Diversity 105:1-16 ((https://doi.org/10.1007/s13225-020-00466-2)) |
| Otu37461 | 1 | k__Fungi(100);p__Ascomycota(100);c__Saccharomycetes(100);o__Saccharomycetales(100);f__Metschnikowiaceae(100);g__Metschnikowia(100);s__Metschnikowia_pulcherrima(100); | Metschnikowia | 13 | Saprotroph | \|Undefined Saprotroph\| | Yeast | NULL | Probable | As nectar-tap_saprotroph (Põlme et al. 2020); Decay sugar-rich substrates (Põlme et al. 2020) | Põlme S, et al. 2020. Fungal Diversity 105:1-16 ((https://doi.org/10.1007/s13225-020-00466-2)) |
| Otu37462 | 1 | k__Fungi(100);p__Ascomycota(100);c__Saccharomycetes(100);o__Saccharomycetales(100);f__Metschnikowiaceae(100);g__Metschnikowia(100);s__Metschnikowia_pulcherrima(100); | Metschnikowia | 13 | Saprotroph | \|Undefined Saprotroph\| | Yeast | NULL | Probable | As nectar-tap_saprotroph (Põlme et al. 2020); Decay sugar-rich substrates (Põlme et al. 2020) | Põlme S, et al. 2020. Fungal Diversity 105:1-16 ((https://doi.org/10.1007/s13225-020-00466-2)) |
| Otu37463 | 1 | k__Fungi(100);p__Ascomycota(100);c__Saccharomycetes(100);o__Saccharomycetales(100);f__Metschnikowiaceae(100);g__Metschnikowia(100);s__Metschnikowia_pulcherrima(100); | Metschnikowia | 13 | Saprotroph | \|Undefined Saprotroph\| | Yeast | NULL | Probable | As nectar-tap_saprotroph (Põlme et al. 2020); Decay sugar-rich substrates (Põlme et al. 2020) | Põlme S, et al. 2020. Fungal Diversity 105:1-16 ((https://doi.org/10.1007/s13225-020-00466-2)) |
| Otu37464 | 1 | k__Fungi(100);p__Ascomycota(100);c__Saccharomycetes(100);o__Saccharomycetales(100);f__Metschnikowiaceae(100);g__Metschnikowia(100);s__Metschnikowia_pulcherrima(100); | Metschnikowia | 13 | Saprotroph | \|Undefined Saprotroph\| | Yeast | NULL | Probable | As nectar-tap_saprotroph (Põlme et al. 2020); Decay sugar-rich substrates (Põlme et al. 2020) | Põlme S, et al. 2020. Fungal Diversity 105:1-16 ((https://doi.org/10.1007/s13225-020-00466-2)) |
| Otu37465 | 1 | k__Fungi(100);p__Ascomycota(100);c__Saccharomycetes(100);o__Saccharomycetales(100);f__Metschnikowiaceae(100);g__Metschnikowia(100);s__Metschnikowia_pulcherrima(100); | Metschnikowia | 13 | Saprotroph | \|Undefined Saprotroph\| | Yeast | NULL | Probable | As nectar-tap_saprotroph (Põlme et al. 2020); Decay sugar-rich substrates (Põlme et al. 2020) | Põlme S, et al. 2020. Fungal Diversity 105:1-16 ((https://doi.org/10.1007/s13225-020-00466-2)) |
| Otu37466 | 1 | k__Fungi(100);p__Ascomycota(100);c__Saccharomycetes(100);o__Saccharomycetales(100);f__Metschnikowiaceae(100);g__Metschnikowia(100);s__Metschnikowia_pulcherrima(100); | Metschnikowia | 13 | Saprotroph | \|Undefined Saprotroph\| | Yeast | NULL | Probable | As nectar-tap_saprotroph (Põlme et al. 2020); Decay sugar-rich substrates (Põlme et al. 2020) | Põlme S, et al. 2020. Fungal Diversity 105:1-16 ((https://doi.org/10.1007/s13225-020-00466-2)) |
| Otu37467 | 1 | k__Fungi(100);p__Ascomycota(100);c__Saccharomycetes(100);o__Saccharomycetales(100);f__Metschnikowiaceae(100);g__Metschnikowia(100);g__Metschnikowia_unclassified(100); | Metschnikowia | 13 | Saprotroph | \|Undefined Saprotroph\| | Yeast | NULL | Probable | As nectar-tap_saprotroph (Põlme et al. 2020); Decay sugar-rich substrates (Põlme et al. 2020) | Põlme S, et al. 2020. Fungal Diversity 105:1-16 ((https://doi.org/10.1007/s13225-020-00466-2)) |
| Otu37468 | 1 | k__Fungi(100);p__Ascomycota(100);c__Saccharomycetes(100);o__Saccharomycetales(100);f__Metschnikowiaceae(100);g__Metschnikowia(100);g__Metschnikowia_unclassified(100); | Metschnikowia | 13 | Saprotroph | \|Undefined Saprotroph\| | Yeast | NULL | Probable | As nectar-tap_saprotroph (Põlme et al. 2020); Decay sugar-rich substrates (Põlme et al. 2020) | Põlme S, et al. 2020. Fungal Diversity 105:1-16 ((https://doi.org/10.1007/s13225-020-00466-2)) |
| Otu37469 | 1 | k__Fungi(100);p__Ascomycota(100);c__Saccharomycetes(100);o__Saccharomycetales(100);f__Metschnikowiaceae(100);g__Metschnikowia(100);g__Metschnikowia_unclassified(100); | Metschnikowia | 13 | Saprotroph | \|Undefined Saprotroph\| | Yeast | NULL | Probable | As nectar-tap_saprotroph (Põlme et al. 2020); Decay sugar-rich substrates (Põlme et al. 2020) | Põlme S, et al. 2020. Fungal Diversity 105:1-16 ((https://doi.org/10.1007/s13225-020-00466-2)) |
| Otu37470 | 1 | k__Fungi(100);p__Ascomycota(100);c__Saccharomycetes(100);o__Saccharomycetales(100);f__Metschnikowiaceae(100);g__Metschnikowia(100);g__Metschnikowia_unclassified(100); | Metschnikowia | 13 | Saprotroph | \|Undefined Saprotroph\| | Yeast | NULL | Probable | As nectar-tap_saprotroph (Põlme et al. 2020); Decay sugar-rich substrates (Põlme et al. 2020) | Põlme S, et al. 2020. Fungal Diversity 105:1-16 ((https://doi.org/10.1007/s13225-020-00466-2)) |
| Otu37471 | 1 | k__Fungi(100);p__Ascomycota(100);c__Saccharomycetes(100);o__Saccharomycetales(100);f__Metschnikowiaceae(100);g__Metschnikowia(100);g__Metschnikowia_unclassified(100); | Metschnikowia | 13 | Saprotroph | \|Undefined Saprotroph\| | Yeast | NULL | Probable | As nectar-tap_saprotroph (Põlme et al. 2020); Decay sugar-rich substrates (Põlme et al. 2020) | Põlme S, et al. 2020. Fungal Diversity 105:1-16 ((https://doi.org/10.1007/s13225-020-00466-2)) |
| Otu37472 | 1 | k__Fungi(100);p__Ascomycota(100);c__Saccharomycetes(100);o__Saccharomycetales(100);f__Metschnikowiaceae(100);g__Metschnikowia(100);g__Metschnikowia_unclassified(100); | Metschnikowia | 13 | Saprotroph | \|Undefined Saprotroph\| | Yeast | NULL | Probable | As nectar-tap_saprotroph (Põlme et al. 2020); Decay sugar-rich substrates (Põlme et al. 2020) | Põlme S, et al. 2020. Fungal Diversity 105:1-16 ((https://doi.org/10.1007/s13225-020-00466-2)) |
| Otu37473 | 1 | k__Fungi(100);p__Ascomycota(100);c__Saccharomycetes(100);o__Saccharomycetales(100);f__Metschnikowiaceae(100);g__Metschnikowia(100);s__Metschnikowia_pulcherrima(100); | Metschnikowia | 13 | Saprotroph | \|Undefined Saprotroph\| | Yeast | NULL | Probable | As nectar-tap_saprotroph (Põlme et al. 2020); Decay sugar-rich substrates (Põlme et al. 2020) | Põlme S, et al. 2020. Fungal Diversity 105:1-16 ((https://doi.org/10.1007/s13225-020-00466-2)) |
| Otu37474 | 1 | k__Fungi(100);p__Ascomycota(100);c__Saccharomycetes(100);o__Saccharomycetales(100);f__Metschnikowiaceae(100);g__Metschnikowia(100);s__Metschnikowia_pulcherrima(100); | Metschnikowia | 13 | Saprotroph | \|Undefined Saprotroph\| | Yeast | NULL | Probable | As nectar-tap_saprotroph (Põlme et al. 2020); Decay sugar-rich substrates (Põlme et al. 2020) | Põlme S, et al. 2020. Fungal Diversity 105:1-16 ((https://doi.org/10.1007/s13225-020-00466-2)) |
| Otu37475 | 1 | k__Fungi(100);p__Ascomycota(100);c__Saccharomycetes(100);o__Saccharomycetales(100);f__Metschnikowiaceae(100);g__Metschnikowia(100);g__Metschnikowia_unclassified(100); | Metschnikowia | 13 | Saprotroph | \|Undefined Saprotroph\| | Yeast | NULL | Probable | As nectar-tap_saprotroph (Põlme et al. 2020); Decay sugar-rich substrates (Põlme et al. 2020) | Põlme S, et al. 2020. Fungal Diversity 105:1-16 ((https://doi.org/10.1007/s13225-020-00466-2)) |
| Otu37476 | 1 | k__Fungi(100);p__Ascomycota(100);c__Saccharomycetes(100);o__Saccharomycetales(100);f__Metschnikowiaceae(100);g__Metschnikowia(100);s__Metschnikowia_pulcherrima(100); | Metschnikowia | 13 | Saprotroph | \|Undefined Saprotroph\| | Yeast | NULL | Probable | As nectar-tap_saprotroph (Põlme et al. 2020); Decay sugar-rich substrates (Põlme et al. 2020) | Põlme S, et al. 2020. Fungal Diversity 105:1-16 ((https://doi.org/10.1007/s13225-020-00466-2)) |
| Otu37477 | 1 | k__Fungi(100);p__Ascomycota(100);c__Saccharomycetes(100);o__Saccharomycetales(100);f__Metschnikowiaceae(100);g__Metschnikowia(100);s__Metschnikowia_pulcherrima(100); | Metschnikowia | 13 | Saprotroph | \|Undefined Saprotroph\| | Yeast | NULL | Probable | As nectar-tap_saprotroph (Põlme et al. 2020); Decay sugar-rich substrates (Põlme et al. 2020) | Põlme S, et al. 2020. Fungal Diversity 105:1-16 ((https://doi.org/10.1007/s13225-020-00466-2)) |
| Otu37478 | 1 | k__Fungi(100);p__Ascomycota(100);c__Saccharomycetes(100);o__Saccharomycetales(100);f__Metschnikowiaceae(100);g__Metschnikowia(100);s__Metschnikowia_pulcherrima(100); | Metschnikowia | 13 | Saprotroph | \|Undefined Saprotroph\| | Yeast | NULL | Probable | As nectar-tap_saprotroph (Põlme et al. 2020); Decay sugar-rich substrates (Põlme et al. 2020) | Põlme S, et al. 2020. Fungal Diversity 105:1-16 ((https://doi.org/10.1007/s13225-020-00466-2)) |
| Otu37479 | 1 | k__Fungi(100);p__Ascomycota(100);c__Saccharomycetes(100);o__Saccharomycetales(100);f__Metschnikowiaceae(100);g__Metschnikowia(100);s__Metschnikowia_pulcherrima(100); | Metschnikowia | 13 | Saprotroph | \|Undefined Saprotroph\| | Yeast | NULL | Probable | As nectar-tap_saprotroph (Põlme et al. 2020); Decay sugar-rich substrates (Põlme et al. 2020) | Põlme S, et al. 2020. Fungal Diversity 105:1-16 ((https://doi.org/10.1007/s13225-020-00466-2)) |
| Otu37481 | 1 | k__Fungi(100);p__Ascomycota(100);c__Saccharomycetes(100);o__Saccharomycetales(100);f__Metschnikowiaceae(100);g__Metschnikowia(100);s__Metschnikowia_pulcherrima(100); | Metschnikowia | 13 | Saprotroph | \|Undefined Saprotroph\| | Yeast | NULL | Probable | As nectar-tap_saprotroph (Põlme et al. 2020); Decay sugar-rich substrates (Põlme et al. 2020) | Põlme S, et al. 2020. Fungal Diversity 105:1-16 ((https://doi.org/10.1007/s13225-020-00466-2)) |
| Otu37482 | 1 | k__Fungi(100);p__Ascomycota(100);c__Saccharomycetes(100);o__Saccharomycetales(100);f__Metschnikowiaceae(100);g__Metschnikowia(100);g__Metschnikowia_unclassified(100); | Metschnikowia | 13 | Saprotroph | \|Undefined Saprotroph\| | Yeast | NULL | Probable | As nectar-tap_saprotroph (Põlme et al. 2020); Decay sugar-rich substrates (Põlme et al. 2020) | Põlme S, et al. 2020. Fungal Diversity 105:1-16 ((https://doi.org/10.1007/s13225-020-00466-2)) |
| Otu37483 | 1 | k__Fungi(100);p__Ascomycota(100);c__Saccharomycetes(100);o__Saccharomycetales(100);f__Metschnikowiaceae(100);g__Metschnikowia(100);s__Metschnikowia_pulcherrima(100); | Metschnikowia | 13 | Saprotroph | \|Undefined Saprotroph\| | Yeast | NULL | Probable | As nectar-tap_saprotroph (Põlme et al. 2020); Decay sugar-rich substrates (Põlme et al. 2020) | Põlme S, et al. 2020. Fungal Diversity 105:1-16 ((https://doi.org/10.1007/s13225-020-00466-2)) |
| Otu37484 | 1 | k__Fungi(100);p__Ascomycota(100);c__Saccharomycetes(100);o__Saccharomycetales(100);f__Metschnikowiaceae(100);g__Metschnikowia(100);s__Metschnikowia_pulcherrima(100); | Metschnikowia | 13 | Saprotroph | \|Undefined Saprotroph\| | Yeast | NULL | Probable | As nectar-tap_saprotroph (Põlme et al. 2020); Decay sugar-rich substrates (Põlme et al. 2020) | Põlme S, et al. 2020. Fungal Diversity 105:1-16 ((https://doi.org/10.1007/s13225-020-00466-2)) |
| Otu37485 | 1 | k__Fungi(100);p__Ascomycota(100);c__Saccharomycetes(100);o__Saccharomycetales(100);f__Metschnikowiaceae(100);g__Metschnikowia(100);g__Metschnikowia_unclassified(100); | Metschnikowia | 13 | Saprotroph | \|Undefined Saprotroph\| | Yeast | NULL | Probable | As nectar-tap_saprotroph (Põlme et al. 2020); Decay sugar-rich substrates (Põlme et al. 2020) | Põlme S, et al. 2020. Fungal Diversity 105:1-16 ((https://doi.org/10.1007/s13225-020-00466-2)) |
| Otu37486 | 1 | k__Fungi(100);p__Ascomycota(100);c__Saccharomycetes(100);o__Saccharomycetales(100);f__Metschnikowiaceae(100);g__Metschnikowia(100);s__Metschnikowia_pulcherrima(100); | Metschnikowia | 13 | Saprotroph | \|Undefined Saprotroph\| | Yeast | NULL | Probable | As nectar-tap_saprotroph (Põlme et al. 2020); Decay sugar-rich substrates (Põlme et al. 2020) | Põlme S, et al. 2020. Fungal Diversity 105:1-16 ((https://doi.org/10.1007/s13225-020-00466-2)) |
| Otu37487 | 1 | k__Fungi(100);p__Ascomycota(100);c__Saccharomycetes(100);o__Saccharomycetales(100);f__Metschnikowiaceae(100);g__Metschnikowia(100);s__Metschnikowia_pulcherrima(100); | Metschnikowia | 13 | Saprotroph | \|Undefined Saprotroph\| | Yeast | NULL | Probable | As nectar-tap_saprotroph (Põlme et al. 2020); Decay sugar-rich substrates (Põlme et al. 2020) | Põlme S, et al. 2020. Fungal Diversity 105:1-16 ((https://doi.org/10.1007/s13225-020-00466-2)) |
| Otu37488 | 1 | k__Fungi(100);p__Ascomycota(100);c__Saccharomycetes(100);o__Saccharomycetales(100);f__Metschnikowiaceae(100);g__Metschnikowia(100);s__Metschnikowia_pulcherrima(100); | Metschnikowia | 13 | Saprotroph | \|Undefined Saprotroph\| | Yeast | NULL | Probable | As nectar-tap_saprotroph (Põlme et al. 2020); Decay sugar-rich substrates (Põlme et al. 2020) | Põlme S, et al. 2020. Fungal Diversity 105:1-16 ((https://doi.org/10.1007/s13225-020-00466-2)) |
| Otu37489 | 1 | k__Fungi(100);p__Ascomycota(100);c__Saccharomycetes(100);o__Saccharomycetales(100);f__Metschnikowiaceae(100);g__Metschnikowia(100);s__Metschnikowia_pulcherrima(100); | Metschnikowia | 13 | Saprotroph | \|Undefined Saprotroph\| | Yeast | NULL | Probable | As nectar-tap_saprotroph (Põlme et al. 2020); Decay sugar-rich substrates (Põlme et al. 2020) | Põlme S, et al. 2020. Fungal Diversity 105:1-16 ((https://doi.org/10.1007/s13225-020-00466-2)) |
| Otu37490 | 1 | k__Fungi(100);p__Ascomycota(100);c__Saccharomycetes(100);o__Saccharomycetales(100);f__Metschnikowiaceae(100);g__Metschnikowia(100);s__Metschnikowia_pulcherrima(100); | Metschnikowia | 13 | Saprotroph | \|Undefined Saprotroph\| | Yeast | NULL | Probable | As nectar-tap_saprotroph (Põlme et al. 2020); Decay sugar-rich substrates (Põlme et al. 2020) | Põlme S, et al. 2020. Fungal Diversity 105:1-16 ((https://doi.org/10.1007/s13225-020-00466-2)) |
| Otu37491 | 1 | k__Fungi(100);p__Ascomycota(100);c__Saccharomycetes(100);o__Saccharomycetales(100);f__Metschnikowiaceae(100);g__Metschnikowia(100);g__Metschnikowia_unclassified(100); | Metschnikowia | 13 | Saprotroph | \|Undefined Saprotroph\| | Yeast | NULL | Probable | As nectar-tap_saprotroph (Põlme et al. 2020); Decay sugar-rich substrates (Põlme et al. 2020) | Põlme S, et al. 2020. Fungal Diversity 105:1-16 ((https://doi.org/10.1007/s13225-020-00466-2)) |
| Otu37492 | 1 | k__Fungi(100);p__Ascomycota(100);c__Saccharomycetes(100);o__Saccharomycetales(100);f__Metschnikowiaceae(100);g__Metschnikowia(100);s__Metschnikowia_pulcherrima(100); | Metschnikowia | 13 | Saprotroph | \|Undefined Saprotroph\| | Yeast | NULL | Probable | As nectar-tap_saprotroph (Põlme et al. 2020); Decay sugar-rich substrates (Põlme et al. 2020) | Põlme S, et al. 2020. Fungal Diversity 105:1-16 ((https://doi.org/10.1007/s13225-020-00466-2)) |
| Otu37493 | 1 | k__Fungi(100);p__Ascomycota(100);c__Saccharomycetes(100);o__Saccharomycetales(100);f__Metschnikowiaceae(100);g__Metschnikowia(100);s__Metschnikowia_pulcherrima(100); | Metschnikowia | 13 | Saprotroph | \|Undefined Saprotroph\| | Yeast | NULL | Probable | As nectar-tap_saprotroph (Põlme et al. 2020); Decay sugar-rich substrates (Põlme et al. 2020) | Põlme S, et al. 2020. Fungal Diversity 105:1-16 ((https://doi.org/10.1007/s13225-020-00466-2)) |
| Otu37494 | 1 | k__Fungi(100);p__Ascomycota(100);c__Saccharomycetes(100);o__Saccharomycetales(100);f__Metschnikowiaceae(100);g__Metschnikowia(100);g__Metschnikowia_unclassified(100); | Metschnikowia | 13 | Saprotroph | \|Undefined Saprotroph\| | Yeast | NULL | Probable | As nectar-tap_saprotroph (Põlme et al. 2020); Decay sugar-rich substrates (Põlme et al. 2020) | Põlme S, et al. 2020. Fungal Diversity 105:1-16 ((https://doi.org/10.1007/s13225-020-00466-2)) |
| Otu37495 | 1 | k__Fungi(100);p__Ascomycota(100);c__Saccharomycetes(100);o__Saccharomycetales(100);f__Metschnikowiaceae(100);g__Metschnikowia(100);s__Metschnikowia_pulcherrima(100); | Metschnikowia | 13 | Saprotroph | \|Undefined Saprotroph\| | Yeast | NULL | Probable | As nectar-tap_saprotroph (Põlme et al. 2020); Decay sugar-rich substrates (Põlme et al. 2020) | Põlme S, et al. 2020. Fungal Diversity 105:1-16 ((https://doi.org/10.1007/s13225-020-00466-2)) |
| Otu37496 | 1 | k__Fungi(100);p__Ascomycota(100);c__Saccharomycetes(100);o__Saccharomycetales(100);f__Metschnikowiaceae(100);g__Metschnikowia(100);s__Metschnikowia_pulcherrima(100); | Metschnikowia | 13 | Saprotroph | \|Undefined Saprotroph\| | Yeast | NULL | Probable | As nectar-tap_saprotroph (Põlme et al. 2020); Decay sugar-rich substrates (Põlme et al. 2020) | Põlme S, et al. 2020. Fungal Diversity 105:1-16 ((https://doi.org/10.1007/s13225-020-00466-2)) |
| Otu37497 | 1 | k__Fungi(100);p__Ascomycota(100);c__Saccharomycetes(100);o__Saccharomycetales(100);f__Metschnikowiaceae(100);g__Metschnikowia(100);s__Metschnikowia_pulcherrima(100); | Metschnikowia | 13 | Saprotroph | \|Undefined Saprotroph\| | Yeast | NULL | Probable | As nectar-tap_saprotroph (Põlme et al. 2020); Decay sugar-rich substrates (Põlme et al. 2020) | Põlme S, et al. 2020. Fungal Diversity 105:1-16 ((https://doi.org/10.1007/s13225-020-00466-2)) |
| Otu37498 | 1 | k__Fungi(100);p__Ascomycota(100);c__Saccharomycetes(100);o__Saccharomycetales(100);f__Metschnikowiaceae(100);g__Metschnikowia(100);s__Metschnikowia_pulcherrima(100); | Metschnikowia | 13 | Saprotroph | \|Undefined Saprotroph\| | Yeast | NULL | Probable | As nectar-tap_saprotroph (Põlme et al. 2020); Decay sugar-rich substrates (Põlme et al. 2020) | Põlme S, et al. 2020. Fungal Diversity 105:1-16 ((https://doi.org/10.1007/s13225-020-00466-2)) |
| Otu37499 | 1 | k__Fungi(100);p__Ascomycota(100);c__Saccharomycetes(100);o__Saccharomycetales(100);f__Metschnikowiaceae(100);g__Metschnikowia(100);s__Metschnikowia_pulcherrima(100); | Metschnikowia | 13 | Saprotroph | \|Undefined Saprotroph\| | Yeast | NULL | Probable | As nectar-tap_saprotroph (Põlme et al. 2020); Decay sugar-rich substrates (Põlme et al. 2020) | Põlme S, et al. 2020. Fungal Diversity 105:1-16 ((https://doi.org/10.1007/s13225-020-00466-2)) |
| Otu37500 | 1 | k__Fungi(100);p__Ascomycota(100);c__Saccharomycetes(100);o__Saccharomycetales(100);f__Metschnikowiaceae(100);g__Metschnikowia(100);s__Metschnikowia_pulcherrima(100); | Metschnikowia | 13 | Saprotroph | \|Undefined Saprotroph\| | Yeast | NULL | Probable | As nectar-tap_saprotroph (Põlme et al. 2020); Decay sugar-rich substrates (Põlme et al. 2020) | Põlme S, et al. 2020. Fungal Diversity 105:1-16 ((https://doi.org/10.1007/s13225-020-00466-2)) |
| Otu37501 | 1 | k__Fungi(100);p__Ascomycota(100);c__Saccharomycetes(100);o__Saccharomycetales(100);f__Metschnikowiaceae(100);g__Metschnikowia(100);s__Metschnikowia_pulcherrima(100); | Metschnikowia | 13 | Saprotroph | \|Undefined Saprotroph\| | Yeast | NULL | Probable | As nectar-tap_saprotroph (Põlme et al. 2020); Decay sugar-rich substrates (Põlme et al. 2020) | Põlme S, et al. 2020. Fungal Diversity 105:1-16 ((https://doi.org/10.1007/s13225-020-00466-2)) |
| Otu37502 | 1 | k__Fungi(100);p__Ascomycota(100);c__Saccharomycetes(100);o__Saccharomycetales(100);f__Metschnikowiaceae(100);g__Metschnikowia(100);s__Metschnikowia_pulcherrima(100); | Metschnikowia | 13 | Saprotroph | \|Undefined Saprotroph\| | Yeast | NULL | Probable | As nectar-tap_saprotroph (Põlme et al. 2020); Decay sugar-rich substrates (Põlme et al. 2020) | Põlme S, et al. 2020. Fungal Diversity 105:1-16 ((https://doi.org/10.1007/s13225-020-00466-2)) |
| Otu37503 | 1 | k__Fungi(100);p__Ascomycota(100);c__Saccharomycetes(100);o__Saccharomycetales(100);f__Metschnikowiaceae(100);g__Metschnikowia(100);s__Metschnikowia_pulcherrima(100); | Metschnikowia | 13 | Saprotroph | \|Undefined Saprotroph\| | Yeast | NULL | Probable | As nectar-tap_saprotroph (Põlme et al. 2020); Decay sugar-rich substrates (Põlme et al. 2020) | Põlme S, et al. 2020. Fungal Diversity 105:1-16 ((https://doi.org/10.1007/s13225-020-00466-2)) |
| Otu37504 | 1 | k__Fungi(100);p__Ascomycota(100);c__Saccharomycetes(100);o__Saccharomycetales(100);f__Metschnikowiaceae(100);g__Metschnikowia(100);s__Metschnikowia_pulcherrima(100); | Metschnikowia | 13 | Saprotroph | \|Undefined Saprotroph\| | Yeast | NULL | Probable | As nectar-tap_saprotroph (Põlme et al. 2020); Decay sugar-rich substrates (Põlme et al. 2020) | Põlme S, et al. 2020. Fungal Diversity 105:1-16 ((https://doi.org/10.1007/s13225-020-00466-2)) |
| Otu37505 | 1 | k__Fungi(100);p__Ascomycota(100);c__Saccharomycetes(100);o__Saccharomycetales(100);f__Metschnikowiaceae(100);g__Metschnikowia(100);s__Metschnikowia_pulcherrima(100); | Metschnikowia | 13 | Saprotroph | \|Undefined Saprotroph\| | Yeast | NULL | Probable | As nectar-tap_saprotroph (Põlme et al. 2020); Decay sugar-rich substrates (Põlme et al. 2020) | Põlme S, et al. 2020. Fungal Diversity 105:1-16 ((https://doi.org/10.1007/s13225-020-00466-2)) |
| Otu37506 | 1 | k__Fungi(100);p__Ascomycota(100);c__Saccharomycetes(100);o__Saccharomycetales(100);f__Metschnikowiaceae(100);g__Metschnikowia(100);s__Metschnikowia_pulcherrima(100); | Metschnikowia | 13 | Saprotroph | \|Undefined Saprotroph\| | Yeast | NULL | Probable | As nectar-tap_saprotroph (Põlme et al. 2020); Decay sugar-rich substrates (Põlme et al. 2020) | Põlme S, et al. 2020. Fungal Diversity 105:1-16 ((https://doi.org/10.1007/s13225-020-00466-2)) |
| Otu37507 | 1 | k__Fungi(100);p__Ascomycota(100);c__Saccharomycetes(100);o__Saccharomycetales(100);f__Metschnikowiaceae(100);g__Metschnikowia(100);s__Metschnikowia_pulcherrima(100); | Metschnikowia | 13 | Saprotroph | \|Undefined Saprotroph\| | Yeast | NULL | Probable | As nectar-tap_saprotroph (Põlme et al. 2020); Decay sugar-rich substrates (Põlme et al. 2020) | Põlme S, et al. 2020. Fungal Diversity 105:1-16 ((https://doi.org/10.1007/s13225-020-00466-2)) |

| | | | | | | | | | | | |
|---|---|---|---|---|---|---|---|---|---|---|---|
| Otu37508 | 1 | k__Fungi(100);p__Ascomycota(100);c__Saccharomycetes(100);o__Saccharomycetales(100);f__Metschnikowiaceae(100);g__Metschnikowia(100);s__Metschnikowia_pulcherrima(100); | Metschnikowia | 13 | Saprotroph | \|Undefined Saprotroph\| | Yeast | NULL | Probable | As nectar-tap_saprotroph (Põlme et al. 2020); Decay sugar-rich substrates (Põlme et al. 2020) | Põlme S, et al. 2020. Fungal Diversity 105:1-16 ((https://doi.org/10.1007/s13225-020-00466-2)) |
| Otu37509 | 1 | k__Fungi(100);p__Ascomycota(100);c__Saccharomycetes(100);o__Saccharomycetales(100);f__Metschnikowiaceae(100);g__Metschnikowia(100);s__Metschnikowia_pulcherrima(100); | Metschnikowia | 13 | Saprotroph | \|Undefined Saprotroph\| | Yeast | NULL | Probable | As nectar-tap_saprotroph (Põlme et al. 2020); Decay sugar-rich substrates (Põlme et al. 2020) | Põlme S, et al. 2020. Fungal Diversity 105:1-16 ((https://doi.org/10.1007/s13225-020-00466-2)) |
| Otu37510 | 1 | k__Fungi(100);p__Ascomycota(100);c__Saccharomycetes(100);o__Saccharomycetales(100);f__Metschnikowiaceae(100);g__Metschnikowia(100);g__Metschnikowia_unclassified(100); | Metschnikowia | 13 | Saprotroph | \|Undefined Saprotroph\| | Yeast | NULL | Probable | As nectar-tap_saprotroph (Põlme et al. 2020); Decay sugar-rich substrates (Põlme et al. 2020) | Põlme S, et al. 2020. Fungal Diversity 105:1-16 ((https://doi.org/10.1007/s13225-020-00466-2)) |
| Otu37511 | 1 | k__Fungi(100);p__Ascomycota(100);c__Saccharomycetes(100);o__Saccharomycetales(100);f__Metschnikowiaceae(100);g__Metschnikowia(100);s__Metschnikowia_pulcherrima(100); | Metschnikowia | 13 | Saprotroph | \|Undefined Saprotroph\| | Yeast | NULL | Probable | As nectar-tap_saprotroph (Põlme et al. 2020); Decay sugar-rich substrates (Põlme et al. 2020) | Põlme S, et al. 2020. Fungal Diversity 105:1-16 ((https://doi.org/10.1007/s13225-020-00466-2)) |
| Otu37512 | 1 | k__Fungi(100);p__Ascomycota(100);c__Saccharomycetes(100);o__Saccharomycetales(100);f__Metschnikowiaceae(100);g__Metschnikowia(100);s__Metschnikowia_pulcherrima(100); | Metschnikowia | 13 | Saprotroph | \|Undefined Saprotroph\| | Yeast | NULL | Probable | As nectar-tap_saprotroph (Põlme et al. 2020); Decay sugar-rich substrates (Põlme et al. 2020) | Põlme S, et al. 2020. Fungal Diversity 105:1-16 ((https://doi.org/10.1007/s13225-020-00466-2)) |
| Otu37513 | 1 | k__Fungi(100);p__Ascomycota(100);c__Saccharomycetes(100);o__Saccharomycetales(100);f__Metschnikowiaceae(100);g__Metschnikowia(100);s__Metschnikowia_pulcherrima(100); | Metschnikowia | 13 | Saprotroph | \|Undefined Saprotroph\| | Yeast | NULL | Probable | As nectar-tap_saprotroph (Põlme et al. 2020); Decay sugar-rich substrates (Põlme et al. 2020) | Põlme S, et al. 2020. Fungal Diversity 105:1-16 ((https://doi.org/10.1007/s13225-020-00466-2)) |
| Otu37514 | 1 | k__Fungi(100);p__Ascomycota(100);c__Saccharomycetes(100);o__Saccharomycetales(100);f__Metschnikowiaceae(100);g__Metschnikowia(100);g__Metschnikowia_unclassified(100); | Metschnikowia | 13 | Saprotroph | \|Undefined Saprotroph\| | Yeast | NULL | Probable | As nectar-tap_saprotroph (Põlme et al. 2020); Decay sugar-rich substrates (Põlme et al. 2020) | Põlme S, et al. 2020. Fungal Diversity 105:1-16 ((https://doi.org/10.1007/s13225-020-00466-2)) |
| Otu37515 | 1 | k__Fungi(100);p__Ascomycota(100);c__Saccharomycetes(100);o__Saccharomycetales(100);f__Metschnikowiaceae(100);g__Metschnikowia(100);s__Metschnikowia_pulcherrima(100); | Metschnikowia | 13 | Saprotroph | \|Undefined Saprotroph\| | Yeast | NULL | Probable | As nectar-tap_saprotroph (Põlme et al. 2020); Decay sugar-rich substrates (Põlme et al. 2020) | Põlme S, et al. 2020. Fungal Diversity 105:1-16 ((https://doi.org/10.1007/s13225-020-00466-2)) |
| Otu37518 | 1 | k__Fungi(100);p__Ascomycota(100);c__Saccharomycetes(100);o__Saccharomycetales(100);f__Metschnikowiaceae(100);g__Metschnikowia(100);s__Metschnikowia_pulcherrima(100); | Metschnikowia | 13 | Saprotroph | \|Undefined Saprotroph\| | Yeast | NULL | Probable | As nectar-tap_saprotroph (Põlme et al. 2020); Decay sugar-rich substrates (Põlme et al. 2020) | Põlme S, et al. 2020. Fungal Diversity 105:1-16 ((https://doi.org/10.1007/s13225-020-00466-2)) |
| Otu37519 | 1 | k__Fungi(100);p__Ascomycota(100);c__Saccharomycetes(100);o__Saccharomycetales(100);f__Metschnikowiaceae(100);g__Metschnikowia(100);s__Metschnikowia_pulcherrima(100); | Metschnikowia | 13 | Saprotroph | \|Undefined Saprotroph\| | Yeast | NULL | Probable | As nectar-tap_saprotroph (Põlme et al. 2020); Decay sugar-rich substrates (Põlme et al. 2020) | Põlme S, et al. 2020. Fungal Diversity 105:1-16 ((https://doi.org/10.1007/s13225-020-00466-2)) |
| Otu37520 | 1 | k__Fungi(100);p__Ascomycota(100);c__Saccharomycetes(100);o__Saccharomycetales(100);f__Metschnikowiaceae(100);g__Metschnikowia(100);s__Metschnikowia_pulcherrima(100); | Metschnikowia | 13 | Saprotroph | \|Undefined Saprotroph\| | Yeast | NULL | Probable | As nectar-tap_saprotroph (Põlme et al. 2020); Decay sugar-rich substrates (Põlme et al. 2020) | Põlme S, et al. 2020. Fungal Diversity 105:1-16 ((https://doi.org/10.1007/s13225-020-00466-2)) |
| Otu37521 | 1 | k__Fungi(100);p__Ascomycota(100);c__Saccharomycetes(100);o__Saccharomycetales(100);f__Metschnikowiaceae(100);g__Metschnikowia(100);s__Metschnikowia_pulcherrima(100); | Metschnikowia | 13 | Saprotroph | \|Undefined Saprotroph\| | Yeast | NULL | Probable | As nectar-tap_saprotroph (Põlme et al. 2020); Decay sugar-rich substrates (Põlme et al. 2020) | Põlme S, et al. 2020. Fungal Diversity 105:1-16 ((https://doi.org/10.1007/s13225-020-00466-2)) |
| Otu37522 | 1 | k__Fungi(100);p__Ascomycota(100);c__Saccharomycetes(100);o__Saccharomycetales(100);f__Metschnikowiaceae(100);g__Metschnikowia(100);s__Metschnikowia_pulcherrima(100); | Metschnikowia | 13 | Saprotroph | \|Undefined Saprotroph\| | Yeast | NULL | Probable | As nectar-tap_saprotroph (Põlme et al. 2020); Decay sugar-rich substrates (Põlme et al. 2020) | Põlme S, et al. 2020. Fungal Diversity 105:1-16 ((https://doi.org/10.1007/s13225-020-00466-2)) |
| Otu37523 | 1 | k__Fungi(100);p__Ascomycota(100);c__Saccharomycetes(100);o__Saccharomycetales(100);f__Metschnikowiaceae(100);g__Metschnikowia(100);s__Metschnikowia_pulcherrima(100); | Metschnikowia | 13 | Saprotroph | \|Undefined Saprotroph\| | Yeast | NULL | Probable | As nectar-tap_saprotroph (Põlme et al. 2020); Decay sugar-rich substrates (Põlme et al. 2020) | Põlme S, et al. 2020. Fungal Diversity 105:1-16 ((https://doi.org/10.1007/s13225-020-00466-2)) |
| Otu37524 | 1 | k__Fungi(100);p__Ascomycota(100);c__Saccharomycetes(100);o__Saccharomycetales(100);f__Metschnikowiaceae(100);g__Metschnikowia(100);s__Metschnikowia_pulcherrima(100); | Metschnikowia | 13 | Saprotroph | \|Undefined Saprotroph\| | Yeast | NULL | Probable | As nectar-tap_saprotroph (Põlme et al. 2020); Decay sugar-rich substrates (Põlme et al. 2020) | Põlme S, et al. 2020. Fungal Diversity 105:1-16 ((https://doi.org/10.1007/s13225-020-00466-2)) |
| Otu37525 | 1 | k__Fungi(100);p__Ascomycota(100);c__Saccharomycetes(100);o__Saccharomycetales(100);f__Metschnikowiaceae(100);g__Metschnikowia(100);g__Metschnikowia_unclassified(100); | Metschnikowia | 13 | Saprotroph | \|Undefined Saprotroph\| | Yeast | NULL | Probable | As nectar-tap_saprotroph (Põlme et al. 2020); Decay sugar-rich substrates (Põlme et al. 2020) | Põlme S, et al. 2020. Fungal Diversity 105:1-16 ((https://doi.org/10.1007/s13225-020-00466-2)) |
| Otu37526 | 1 | k__Fungi(100);p__Ascomycota(100);c__Saccharomycetes(100);o__Saccharomycetales(100);f__Metschnikowiaceae(100);g__Metschnikowia(100);g__Metschnikowia_unclassified(100); | Metschnikowia | 13 | Saprotroph | \|Undefined Saprotroph\| | Yeast | NULL | Probable | As nectar-tap_saprotroph (Põlme et al. 2020); Decay sugar-rich substrates (Põlme et al. 2020) | Põlme S, et al. 2020. Fungal Diversity 105:1-16 ((https://doi.org/10.1007/s13225-020-00466-2)) |
| Otu37527 | 1 | k__Fungi(100);p__Ascomycota(100);c__Saccharomycetes(100);o__Saccharomycetales(100);f__Metschnikowiaceae(100);g__Metschnikowia(100);g__Metschnikowia_unclassified(100); | Metschnikowia | 13 | Saprotroph | \|Undefined Saprotroph\| | Yeast | NULL | Probable | As nectar-tap_saprotroph (Põlme et al. 2020); Decay sugar-rich substrates (Põlme et al. 2020) | Põlme S, et al. 2020. Fungal Diversity 105:1-16 ((https://doi.org/10.1007/s13225-020-00466-2)) |
| Otu37528 | 1 | k__Fungi(100);p__Ascomycota(100);c__Saccharomycetes(100);o__Saccharomycetales(100);f__Metschnikowiaceae(100);g__Metschnikowia(100);g__Metschnikowia_unclassified(100); | Metschnikowia | 13 | Saprotroph | \|Undefined Saprotroph\| | Yeast | NULL | Probable | As nectar-tap_saprotroph (Põlme et al. 2020); Decay sugar-rich substrates (Põlme et al. 2020) | Põlme S, et al. 2020. Fungal Diversity 105:1-16 ((https://doi.org/10.1007/s13225-020-00466-2)) |
| Otu37529 | 1 | k__Fungi(100);p__Ascomycota(100);c__Saccharomycetes(100);o__Saccharomycetales(100);f__Metschnikowiaceae(100);g__Metschnikowia(100);g__Metschnikowia_unclassified(100); | Metschnikowia | 13 | Saprotroph | \|Undefined Saprotroph\| | Yeast | NULL | Probable | As nectar-tap_saprotroph (Põlme et al. 2020); Decay sugar-rich substrates (Põlme et al. 2020) | Põlme S, et al. 2020. Fungal Diversity 105:1-16 ((https://doi.org/10.1007/s13225-020-00466-2)) |
| Otu37530 | 1 | k__Fungi(100);p__Ascomycota(100);c__Saccharomycetes(100);o__Saccharomycetales(100);f__Metschnikowiaceae(100);g__Metschnikowia(100);g__Metschnikowia_unclassified(100); | Metschnikowia | 13 | Saprotroph | \|Undefined Saprotroph\| | Yeast | NULL | Probable | As nectar-tap_saprotroph (Põlme et al. 2020); Decay sugar-rich substrates (Põlme et al. 2020) | Põlme S, et al. 2020. Fungal Diversity 105:1-16 ((https://doi.org/10.1007/s13225-020-00466-2)) |
| Otu37531 | 1 | k__Fungi(100);p__Ascomycota(100);c__Saccharomycetes(100);o__Saccharomycetales(100);f__Metschnikowiaceae(100);g__Metschnikowia(100);g__Metschnikowia_unclassified(100); | Metschnikowia | 13 | Saprotroph | \|Undefined Saprotroph\| | Yeast | NULL | Probable | As nectar-tap_saprotroph (Põlme et al. 2020); Decay sugar-rich substrates (Põlme et al. 2020) | Põlme S, et al. 2020. Fungal Diversity 105:1-16 ((https://doi.org/10.1007/s13225-020-00466-2)) |
| Otu37532 | 1 | k__Fungi(100);p__Ascomycota(100);c__Saccharomycetes(100);o__Saccharomycetales(100);f__Metschnikowiaceae(100);g__Metschnikowia(100);g__Metschnikowia_unclassified(100); | Metschnikowia | 13 | Saprotroph | \|Undefined Saprotroph\| | Yeast | NULL | Probable | As nectar-tap_saprotroph (Põlme et al. 2020); Decay sugar-rich substrates (Põlme et al. 2020) | Põlme S, et al. 2020. Fungal Diversity 105:1-16 ((https://doi.org/10.1007/s13225-020-00466-2)) |
| Otu37533 | 1 | k__Fungi(100);p__Ascomycota(100);c__Saccharomycetes(100);o__Saccharomycetales(100);f__Metschnikowiaceae(100);g__Metschnikowia(100);s__Metschnikowia_pulcherrima(100); | Metschnikowia | 13 | Saprotroph | \|Undefined Saprotroph\| | Yeast | NULL | Probable | As nectar-tap_saprotroph (Põlme et al. 2020); Decay sugar-rich substrates (Põlme et al. 2020) | Põlme S, et al. 2020. Fungal Diversity 105:1-16 ((https://doi.org/10.1007/s13225-020-00466-2)) |
| Otu37534 | 1 | k__Fungi(100);p__Ascomycota(100);c__Saccharomycetes(100);o__Saccharomycetales(100);f__Metschnikowiaceae(100);g__Metschnikowia(100);s__Metschnikowia_pulcherrima(100); | Metschnikowia | 13 | Saprotroph | \|Undefined Saprotroph\| | Yeast | NULL | Probable | As nectar-tap_saprotroph (Põlme et al. 2020); Decay sugar-rich substrates (Põlme et al. 2020) | Põlme S, et al. 2020. Fungal Diversity 105:1-16 ((https://doi.org/10.1007/s13225-020-00466-2)) |
| Otu37535 | 1 | k__Fungi(100);p__Ascomycota(100);c__Saccharomycetes(100);o__Saccharomycetales(100);f__Metschnikowiaceae(100);g__Metschnikowia(100);s__Metschnikowia_pulcherrima(100); | Metschnikowia | 13 | Saprotroph | \|Undefined Saprotroph\| | Yeast | NULL | Probable | As nectar-tap_saprotroph (Põlme et al. 2020); Decay sugar-rich substrates (Põlme et al. 2020) | Põlme S, et al. 2020. Fungal Diversity 105:1-16 ((https://doi.org/10.1007/s13225-020-00466-2)) |
| Otu37536 | 1 | k__Fungi(100);p__Ascomycota(100);c__Saccharomycetes(100);o__Saccharomycetales(100);f__Metschnikowiaceae(100);g__Metschnikowia(100);s__Metschnikowia_pulcherrima(100); | Metschnikowia | 13 | Saprotroph | \|Undefined Saprotroph\| | Yeast | NULL | Probable | As nectar-tap_saprotroph (Põlme et al. 2020); Decay sugar-rich substrates (Põlme et al. 2020) | Põlme S, et al. 2020. Fungal Diversity 105:1-16 ((https://doi.org/10.1007/s13225-020-00466-2)) |
| Otu37537 | 1 | k__Fungi(100);p__Ascomycota(100);c__Saccharomycetes(100);o__Saccharomycetales(100);f__Metschnikowiaceae(100);g__Metschnikowia(100);g__Metschnikowia_unclassified(100); | Metschnikowia | 13 | Saprotroph | \|Undefined Saprotroph\| | Yeast | NULL | Probable | As nectar-tap_saprotroph (Põlme et al. 2020); Decay sugar-rich substrates (Põlme et al. 2020) | Põlme S, et al. 2020. Fungal Diversity 105:1-16 ((https://doi.org/10.1007/s13225-020-00466-2)) |
| Otu37538 | 1 | k__Fungi(100);p__Ascomycota(100);c__Saccharomycetes(100);o__Saccharomycetales(100);f__Metschnikowiaceae(100);g__Metschnikowia(100);g__Metschnikowia_unclassified(100); | Metschnikowia | 13 | Saprotroph | \|Undefined Saprotroph\| | Yeast | NULL | Probable | As nectar-tap_saprotroph (Põlme et al. 2020); Decay sugar-rich substrates (Põlme et al. 2020) | Põlme S, et al. 2020. Fungal Diversity 105:1-16 ((https://doi.org/10.1007/s13225-020-00466-2)) |
| Otu37539 | 1 | k__Fungi(100);p__Ascomycota(100);c__Saccharomycetes(100);o__Saccharomycetales(100);f__Metschnikowiaceae(100);g__Metschnikowia(100);s__Metschnikowia_pulcherrima(100); | Metschnikowia | 13 | Saprotroph | \|Undefined Saprotroph\| | Yeast | NULL | Probable | As nectar-tap_saprotroph (Põlme et al. 2020); Decay sugar-rich substrates (Põlme et al. 2020) | Põlme S, et al. 2020. Fungal Diversity 105:1-16 ((https://doi.org/10.1007/s13225-020-00466-2)) |
| Otu37540 | 1 | k__Fungi(100);p__Ascomycota(100);c__Saccharomycetes(100);o__Saccharomycetales(100);f__Metschnikowiaceae(100);g__Metschnikowia(100);s__Metschnikowia_pulcherrima(100); | Metschnikowia | 13 | Saprotroph | \|Undefined Saprotroph\| | Yeast | NULL | Probable | As nectar-tap_saprotroph (Põlme et al. 2020); Decay sugar-rich substrates (Põlme et al. 2020) | Põlme S, et al. 2020. Fungal Diversity 105:1-16 ((https://doi.org/10.1007/s13225-020-00466-2)) |
| Otu37541 | 1 | k__Fungi(100);p__Ascomycota(100);c__Saccharomycetes(100);o__Saccharomycetales(100);f__Metschnikowiaceae(100);g__Metschnikowia(100);s__Metschnikowia_pulcherrima(100); | Metschnikowia | 13 | Saprotroph | \|Undefined Saprotroph\| | Yeast | NULL | Probable | As nectar-tap_saprotroph (Põlme et al. 2020); Decay sugar-rich substrates (Põlme et al. 2020) | Põlme S, et al. 2020. Fungal Diversity 105:1-16 ((https://doi.org/10.1007/s13225-020-00466-2)) |
| Otu37542 | 1 | k__Fungi(100);p__Ascomycota(100);c__Saccharomycetes(100);o__Saccharomycetales(100);f__Metschnikowiaceae(100);g__Metschnikowia(100);s__Metschnikowia_pulcherrima(100); | Metschnikowia | 13 | Saprotroph | \|Undefined Saprotroph\| | Yeast | NULL | Probable | As nectar-tap_saprotroph (Põlme et al. 2020); Decay sugar-rich substrates (Põlme et al. 2020) | Põlme S, et al. 2020. Fungal Diversity 105:1-16 ((https://doi.org/10.1007/s13225-020-00466-2)) |
| Otu37543 | 1 | k__Fungi(100);p__Ascomycota(100);c__Saccharomycetes(100);o__Saccharomycetales(100);f__Metschnikowiaceae(100);g__Metschnikowia(100);g__Metschnikowia_unclassified(100); | Metschnikowia | 13 | Saprotroph | \|Undefined Saprotroph\| | Yeast | NULL | Probable | As nectar-tap_saprotroph (Põlme et al. 2020); Decay sugar-rich substrates (Põlme et al. 2020) | Põlme S, et al. 2020. Fungal Diversity 105:1-16 ((https://doi.org/10.1007/s13225-020-00466-2)) |
| Otu37544 | 1 | k__Fungi(100);p__Ascomycota(100);c__Saccharomycetes(100);o__Saccharomycetales(100);f__Metschnikowiaceae(100);g__Metschnikowia(100);g__Metschnikowia_unclassified(100); | Metschnikowia | 13 | Saprotroph | \|Undefined Saprotroph\| | Yeast | NULL | Probable | As nectar-tap_saprotroph (Põlme et al. 2020); Decay sugar-rich substrates (Põlme et al. 2020) | Põlme S, et al. 2020. Fungal Diversity 105:1-16 ((https://doi.org/10.1007/s13225-020-00466-2)) |
| Otu37545 | 1 | k__Fungi(100);p__Ascomycota(100);c__Saccharomycetes(100);o__Saccharomycetales(100);f__Metschnikowiaceae(100);g__Metschnikowia(100);g__Metschnikowia_unclassified(100); | Metschnikowia | 13 | Saprotroph | \|Undefined Saprotroph\| | Yeast | NULL | Probable | As nectar-tap_saprotroph (Põlme et al. 2020); Decay sugar-rich substrates (Põlme et al. 2020) | Põlme S, et al. 2020. Fungal Diversity 105:1-16 ((https://doi.org/10.1007/s13225-020-00466-2)) |
| Otu37546 | 1 | k__Fungi(100);p__Ascomycota(100);c__Saccharomycetes(100);o__Saccharomycetales(100);f__Metschnikowiaceae(100);g__Metschnikowia(100);s__Metschnikowia_pulcherrima(100); | Metschnikowia | 13 | Saprotroph | \|Undefined Saprotroph\| | Yeast | NULL | Probable | As nectar-tap_saprotroph (Põlme et al. 2020); Decay sugar-rich substrates (Põlme et al. 2020) | Põlme S, et al. 2020. Fungal Diversity 105:1-16 ((https://doi.org/10.1007/s13225-020-00466-2)) |
| Otu37547 | 1 | k__Fungi(100);p__Ascomycota(100);c__Saccharomycetes(100);o__Saccharomycetales(100);f__Metschnikowiaceae(100);g__Metschnikowia(100);s__Metschnikowia_pulcherrima(100); | Metschnikowia | 13 | Saprotroph | \|Undefined Saprotroph\| | Yeast | NULL | Probable | As nectar-tap_saprotroph (Põlme et al. 2020); Decay sugar-rich substrates (Põlme et al. 2020) | Põlme S, et al. 2020. Fungal Diversity 105:1-16 ((https://doi.org/10.1007/s13225-020-00466-2)) |
| Otu37548 | 1 | k__Fungi(100);p__Ascomycota(100);c__Saccharomycetes(100);o__Saccharomycetales(100);f__Metschnikowiaceae(100);g__Metschnikowia(100);g__Metschnikowia_unclassified(100); | Metschnikowia | 13 | Saprotroph | \|Undefined Saprotroph\| | Yeast | NULL | Probable | As nectar-tap_saprotroph (Põlme et al. 2020); Decay sugar-rich substrates (Põlme et al. 2020) | Põlme S, et al. 2020. Fungal Diversity 105:1-16 ((https://doi.org/10.1007/s13225-020-00466-2)) |
| Otu37549 | 1 | k__Fungi(100);p__Ascomycota(100);c__Saccharomycetes(100);o__Saccharomycetales(100);f__Metschnikowiaceae(100);g__Metschnikowia(100);s__Metschnikowia_pulcherrima(100); | Metschnikowia | 13 | Saprotroph | \|Undefined Saprotroph\| | Yeast | NULL | Probable | As nectar-tap_saprotroph (Põlme et al. 2020); Decay sugar-rich substrates (Põlme et al. 2020) | Põlme S, et al. 2020. Fungal Diversity 105:1-16 ((https://doi.org/10.1007/s13225-020-00466-2)) |
| Otu37550 | 1 | k__Fungi(100);p__Ascomycota(100);c__Saccharomycetes(100);o__Saccharomycetales(100);f__Metschnikowiaceae(100);g__Metschnikowia(100);s__Metschnikowia_pulcherrima(100); | Metschnikowia | 13 | Saprotroph | \|Undefined Saprotroph\| | Yeast | NULL | Probable | As nectar-tap_saprotroph (Põlme et al. 2020); Decay sugar-rich substrates (Põlme et al. 2020) | Põlme S, et al. 2020. Fungal Diversity 105:1-16 ((https://doi.org/10.1007/s13225-020-00466-2)) |
| Otu37551 | 1 | k__Fungi(100);p__Ascomycota(100);c__Saccharomycetes(100);o__Saccharomycetales(100);f__Metschnikowiaceae(100);g__Metschnikowia(100);s__Metschnikowia_pulcherrima(100); | Metschnikowia | 13 | Saprotroph | \|Undefined Saprotroph\| | Yeast | NULL | Probable | As nectar-tap_saprotroph (Põlme et al. 2020); Decay sugar-rich substrates (Põlme et al. 2020) | Põlme S, et al. 2020. Fungal Diversity 105:1-16 ((https://doi.org/10.1007/s13225-020-00466-2)) |
| Otu37552 | 1 | k__Fungi(100);p__Ascomycota(100);c__Saccharomycetes(100);o__Saccharomycetales(100);f__Metschnikowiaceae(100);g__Metschnikowia(100);g__Metschnikowia_unclassified(100); | Metschnikowia | 13 | Saprotroph | \|Undefined Saprotroph\| | Yeast | NULL | Probable | As nectar-tap_saprotroph (Põlme et al. 2020); Decay sugar-rich substrates (Põlme et al. 2020) | Põlme S, et al. 2020. Fungal Diversity 105:1-16 ((https://doi.org/10.1007/s13225-020-00466-2)) |
| Otu37553 | 1 | k__Fungi(100);p__Ascomycota(100);c__Saccharomycetes(100);o__Saccharomycetales(100);f__Metschnikowiaceae(100);g__Metschnikowia(100);s__Metschnikowia_pulcherrima(100); | Metschnikowia | 13 | Saprotroph | \|Undefined Saprotroph\| | Yeast | NULL | Probable | As nectar-tap_saprotroph (Põlme et al. 2020); Decay sugar-rich substrates (Põlme et al. 2020) | Põlme S, et al. 2020. Fungal Diversity 105:1-16 ((https://doi.org/10.1007/s13225-020-00466-2)) |
| Otu37554 | 1 | k__Fungi(100);p__Ascomycota(100);c__Saccharomycetes(100);o__Saccharomycetales(100);f__Metschnikowiaceae(100);g__Metschnikowia(100);s__Metschnikowia_pulcherrima(100); | Metschnikowia | 13 | Saprotroph | \|Undefined Saprotroph\| | Yeast | NULL | Probable | As nectar-tap_saprotroph (Põlme et al. 2020); Decay sugar-rich substrates (Põlme et al. 2020) | Põlme S, et al. 2020. Fungal Diversity 105:1-16 ((https://doi.org/10.1007/s13225-020-00466-2)) |
| Otu37555 | 1 | k__Fungi(100);p__Ascomycota(100);c__Saccharomycetes(100);o__Saccharomycetales(100);f__Metschnikowiaceae(100);g__Metschnikowia(100);g__Metschnikowia_unclassified(100); | Metschnikowia | 13 | Saprotroph | \|Undefined Saprotroph\| | Yeast | NULL | Probable | As nectar-tap_saprotroph (Põlme et al. 2020); Decay sugar-rich substrates (Põlme et al. 2020) | Põlme S, et al. 2020. Fungal Diversity 105:1-16 ((https://doi.org/10.1007/s13225-020-00466-2)) |
| Otu37557 | 1 | k__Fungi(100);p__Ascomycota(100);c__Saccharomycetes(100);o__Saccharomycetales(100);f__Metschnikowiaceae(100);g__Metschnikowia(100);g__Metschnikowia_unclassified(100); | Metschnikowia | 13 | Saprotroph | \|Undefined Saprotroph\| | Yeast | NULL | Probable | As nectar-tap_saprotroph (Põlme et al. 2020); Decay sugar-rich substrates (Põlme et al. 2020) | Põlme S, et al. 2020. Fungal Diversity 105:1-16 ((https://doi.org/10.1007/s13225-020-00466-2)) |
| Otu37558 | 1 | k__Fungi(100);p__Ascomycota(100);c__Saccharomycetes(100);o__Saccharomycetales(100);f__Metschnikowiaceae(100);g__Metschnikowia(100);g__Metschnikowia_unclassified(100); | Metschnikowia | 13 | Saprotroph | \|Undefined Saprotroph\| | Yeast | NULL | Probable | As nectar-tap_saprotroph (Põlme et al. 2020); Decay sugar-rich substrates (Põlme et al. 2020) | Põlme S, et al. 2020. Fungal Diversity 105:1-16 ((https://doi.org/10.1007/s13225-020-00466-2)) |
| Otu37559 | 1 | k__Fungi(100);p__Ascomycota(100);c__Saccharomycetes(100);o__Saccharomycetales(100);f__Metschnikowiaceae(100);g__Metschnikowia(100);s__Metschnikowia_pulcherrima(100); | Metschnikowia | 13 | Saprotroph | \|Undefined Saprotroph\| | Yeast | NULL | Probable | As nectar-tap_saprotroph (Põlme et al. 2020); Decay sugar-rich substrates (Põlme et al. 2020) | Põlme S, et al. 2020. Fungal Diversity 105:1-16 ((https://doi.org/10.1007/s13225-020-00466-2)) |
| Otu37560 | 1 | k__Fungi(100);p__Ascomycota(100);c__Saccharomycetes(100);o__Saccharomycetales(100);f__Metschnikowiaceae(100);g__Metschnikowia(100);s__Metschnikowia_pulcherrima(100); | Metschnikowia | 13 | Saprotroph | \|Undefined Saprotroph\| | Yeast | NULL | Probable | As nectar-tap_saprotroph (Põlme et al. 2020); Decay sugar-rich substrates (Põlme et al. 2020) | Põlme S, et al. 2020. Fungal Diversity 105:1-16 ((https://doi.org/10.1007/s13225-020-00466-2)) |
| Otu37561 | 1 | k__Fungi(100);p__Ascomycota(100);c__Saccharomycetes(100);o__Saccharomycetales(100);f__Metschnikowiaceae(100);g__Metschnikowia(100);s__Metschnikowia_pulcherrima(100); | Metschnikowia | 13 | Saprotroph | \|Undefined Saprotroph\| | Yeast | NULL | Probable | As nectar-tap_saprotroph (Põlme et al. 2020); Decay sugar-rich substrates (Põlme et al. 2020) | Põlme S, et al. 2020. Fungal Diversity 105:1-16 ((https://doi.org/10.1007/s13225-020-00466-2)) |
| Otu37562 | 1 | k__Fungi(100);p__Ascomycota(100);c__Saccharomycetes(100);o__Saccharomycetales(100);f__Metschnikowiaceae(100);g__Metschnikowia(100);g__Metschnikowia_unclassified(100); | Metschnikowia | 13 | Saprotroph | \|Undefined Saprotroph\| | Yeast | NULL | Probable | As nectar-tap_saprotroph (Põlme et al. 2020); Decay sugar-rich substrates (Põlme et al. 2020) | Põlme S, et al. 2020. Fungal Diversity 105:1-16 ((https://doi.org/10.1007/s13225-020-00466-2)) |
| Otu37563 | 1 | k__Fungi(100);p__Ascomycota(100);c__Saccharomycetes(100);o__Saccharomycetales(100);f__Metschnikowiaceae(100);g__Metschnikowia(100);s__Metschnikowia_pulcherrima(100); | Metschnikowia | 13 | Saprotroph | \|Undefined Saprotroph\| | Yeast | NULL | Probable | As nectar-tap_saprotroph (Põlme et al. 2020); Decay sugar-rich substrates (Põlme et al. 2020) | Põlme S, et al. 2020. Fungal Diversity 105:1-16 ((https://doi.org/10.1007/s13225-020-00466-2)) |
| Otu37564 | 1 | k__Fungi(100);p__Ascomycota(100);c__Saccharomycetes(100);o__Saccharomycetales(100);f__Metschnikowiaceae(100);g__Metschnikowia(100);g__Metschnikowia_unclassified(100); | Metschnikowia | 13 | Saprotroph | \|Undefined Saprotroph\| | Yeast | NULL | Probable | As nectar-tap_saprotroph (Põlme et al. 2020); Decay sugar-rich substrates (Põlme et al. 2020) | Põlme S, et al. 2020. Fungal Diversity 105:1-16 ((https://doi.org/10.1007/s13225-020-00466-2)) |
| Otu37565 | 1 | k__Fungi(100);p__Ascomycota(100);c__Saccharomycetes(100);o__Saccharomycetales(100);f__Metschnikowiaceae(100);g__Metschnikowia(100);s__Metschnikowia_pulcherrima(100); | Metschnikowia | 13 | Saprotroph | \|Undefined Saprotroph\| | Yeast | NULL | Probable | As nectar-tap_saprotroph (Põlme et al. 2020); Decay sugar-rich substrates (Põlme et al. 2020) | Põlme S, et al. 2020. Fungal Diversity 105:1-16 ((https://doi.org/10.1007/s13225-020-00466-2)) |
| Otu37566 | 1 | k__Fungi(100);p__Ascomycota(100);c__Saccharomycetes(100);o__Saccharomycetales(100);f__Metschnikowiaceae(100);g__Metschnikowia(100);s__Metschnikowia_pulcherrima(100); | Metschnikowia | 13 | Saprotroph | \|Undefined Saprotroph\| | Yeast | NULL | Probable | As nectar-tap_saprotroph (Põlme et al. 2020); Decay sugar-rich substrates (Põlme et al. 2020) | Põlme S, et al. 2020. Fungal Diversity 105:1-16 ((https://doi.org/10.1007/s13225-020-00466-2)) |
| Otu37567 | 1 | k__Fungi(100);p__Ascomycota(100);c__Saccharomycetes(100);o__Saccharomycetales(100);f__Metschnikowiaceae(100);g__Metschnikowia(100);s__Metschnikowia_pulcherrima(100); | Metschnikowia | 13 | Saprotroph | \|Undefined Saprotroph\| | Yeast | NULL | Probable | As nectar-tap_saprotroph (Põlme et al. 2020); Decay sugar-rich substrates (Põlme et al. 2020) | Põlme S, et al. 2020. Fungal Diversity 105:1-16 ((https://doi.org/10.1007/s13225-020-00466-2)) |
| Otu37571 | 1 | k__Fungi(100);p__Ascomycota(100);c__Saccharomycetes(100);o__Saccharomycetales(100);f__Metschnikowiaceae(100);g__Metschnikowia(100);s__Metschnikowia_pulcherrima(100); | Metschnikowia | 13 | Saprotroph | \|Undefined Saprotroph\| | Yeast | NULL | Probable | As nectar-tap_saprotroph (Põlme et al. 2020); Decay sugar-rich substrates (Põlme et al. 2020) | Põlme S, et al. 2020. Fungal Diversity 105:1-16 ((https://doi.org/10.1007/s13225-020-00466-2)) |
| Otu37572 | 1 | k__Fungi(100);p__Ascomycota(100);c__Saccharomycetes(100);o__Saccharomycetales(100);f__Metschnikowiaceae(100);g__Metschnikowia(100);s__Metschnikowia_pulcherrima(100); | Metschnikowia | 13 | Saprotroph | \|Undefined Saprotroph\| | Yeast | NULL | Probable | As nectar-tap_saprotroph (Põlme et al. 2020); Decay sugar-rich substrates (Põlme et al. 2020) | Põlme S, et al. 2020. Fungal Diversity 105:1-16 ((https://doi.org/10.1007/s13225-020-00466-2)) |
| Otu37573 | 1 | k__Fungi(100);p__Ascomycota(100);c__Saccharomycetes(100);o__Saccharomycetales(100);f__Metschnikowiaceae(100);g__Metschnikowia(100);g__Metschnikowia_unclassified(100); | Metschnikowia | 13 | Saprotroph | \|Undefined Saprotroph\| | Yeast | NULL | Probable | As nectar-tap_saprotroph (Põlme et al. 2020); Decay sugar-rich substrates (Põlme et al. 2020) | Põlme S, et al. 2020. Fungal Diversity 105:1-16 ((https://doi.org/10.1007/s13225-020-00466-2)) |
| Otu37574 | 1 | k__Fungi(100);p__Ascomycota(100);c__Saccharomycetes(100);o__Saccharomycetales(100);f__Metschnikowiaceae(100);g__Metschnikowia(100);s__Metschnikowia_pulcherrima(100); | Metschnikowia | 13 | Saprotroph | \|Undefined Saprotroph\| | Yeast | NULL | Probable | As nectar-tap_saprotroph (Põlme et al. 2020); Decay sugar-rich substrates (Põlme et al. 2020) | Põlme S, et al. 2020. Fungal Diversity 105:1-16 ((https://doi.org/10.1007/s13225-020-00466-2)) |
| Otu37575 | 1 | k__Fungi(100);p__Ascomycota(100);c__Saccharomycetes(100);o__Saccharomycetales(100);f__Metschnikowiaceae(100);g__Metschnikowia(100);s__Metschnikowia_pulcherrima(100); | Metschnikowia | 13 | Saprotroph | \|Undefined Saprotroph\| | Yeast | NULL | Probable | As nectar-tap_saprotroph (Põlme et al. 2020); Decay sugar-rich substrates (Põlme et al. 2020) | Põlme S, et al. 2020. Fungal Diversity 105:1-16 ((https://doi.org/10.1007/s13225-020-00466-2)) |
| Otu37576 | 1 | k__Fungi(100);p__Ascomycota(100);c__Saccharomycetes(100);o__Saccharomycetales(100);f__Metschnikowiaceae(100);g__Metschnikowia(100);g__Metschnikowia_unclassified(100); | Metschnikowia | 13 | Saprotroph | \|Undefined Saprotroph\| | Yeast | NULL | Probable | As nectar-tap_saprotroph (Põlme et al. 2020); Decay sugar-rich substrates (Põlme et al. 2020) | Põlme S, et al. 2020. Fungal Diversity 105:1-16 ((https://doi.org/10.1007/s13225-020-00466-2)) |
| Otu37577 | 1 | k__Fungi(100);p__Ascomycota(100);c__Saccharomycetes(100);o__Saccharomycetales(100);f__Metschnikowiaceae(100);g__Metschnikowia(100);s__Metschnikowia_pulcherrima(100); | Metschnikowia | 13 | Saprotroph | \|Undefined Saprotroph\| | Yeast | NULL | Probable | As nectar-tap_saprotroph (Põlme et al. 2020); Decay sugar-rich substrates (Põlme et al. 2020) | Põlme S, et al. 2020. Fungal Diversity 105:1-16 ((https://doi.org/10.1007/s13225-020-00466-2)) |
| Otu37578 | 1 | k__Fungi(100);p__Ascomycota(100);c__Saccharomycetes(100);o__Saccharomycetales(100);f__Metschnikowiaceae(100);g__Metschnikowia(100);s__Metschnikowia_pulcherrima(100); | Metschnikowia | 13 | Saprotroph | \|Undefined Saprotroph\| | Yeast | NULL | Probable | As nectar-tap_saprotroph (Põlme et al. 2020); Decay sugar-rich substrates (Põlme et al. 2020) | Põlme S, et al. 2020. Fungal Diversity 105:1-16 ((https://doi.org/10.1007/s13225-020-00466-2)) |
| Otu37579 | 1 | k__Fungi(100);p__Ascomycota(100);c__Saccharomycetes(100);o__Saccharomycetales(100);f__Metschnikowiaceae(100);g__Metschnikowia(100);s__Metschnikowia_pulcherrima(100); | Metschnikowia | 13 | Saprotroph | \|Undefined Saprotroph\| | Yeast | NULL | Probable | As nectar-tap_saprotroph (Põlme et al. 2020); Decay sugar-rich substrates (Põlme et al. 2020) | Põlme S, et al. 2020. Fungal Diversity 105:1-16 ((https://doi.org/10.1007/s13225-020-00466-2)) |
| Otu37580 | 1 | k__Fungi(100);p__Ascomycota(100);c__Saccharomycetes(100);o__Saccharomycetales(100);f__Metschnikowiaceae(100);g__Metschnikowia(100);s__Metschnikowia_pulcherrima(100); | Metschnikowia | 13 | Saprotroph | \|Undefined Saprotroph\| | Yeast | NULL | Probable | As nectar-tap_saprotroph (Põlme et al. 2020); Decay sugar-rich substrates (Põlme et al. 2020) | Põlme S, et al. 2020. Fungal Diversity 105:1-16 ((https://doi.org/10.1007/s13225-020-00466-2)) |
| Otu37581 | 1 | k__Fungi(100);p__Ascomycota(100);c__Saccharomycetes(100);o__Saccharomycetales(100);f__Metschnikowiaceae(100);g__Metschnikowia(100);g__Metschnikowia_unclassified(100); | Metschnikowia | 13 | Saprotroph | \|Undefined Saprotroph\| | Yeast | NULL | Probable | As nectar-tap_saprotroph (Põlme et al. 2020); Decay sugar-rich substrates (Põlme et al. 2020) | Põlme S, et al. 2020. Fungal Diversity 105:1-16 ((https://doi.org/10.1007/s13225-020-00466-2)) |
| Otu37582 | 1 | k__Fungi(100);p__Ascomycota(100);c__Saccharomycetes(100);o__Saccharomycetales(100);f__Metschnikowiaceae(100);g__Metschnikowia(100);s__Metschnikowia_pulcherrima(100); | Metschnikowia | 13 | Saprotroph | \|Undefined Saprotroph\| | Yeast | NULL | Probable | As nectar-tap_saprotroph (Põlme et al. 2020); Decay sugar-rich substrates (Põlme et al. 2020) | Põlme S, et al. 2020. Fungal Diversity 105:1-16 ((https://doi.org/10.1007/s13225-020-00466-2)) |
| Otu37584 | 1 | k__Fungi(100);p__Ascomycota(100);c__Saccharomycetes(100);o__Saccharomycetales(100);f__Metschnikowiaceae(100);g__Metschnikowia(100);s__Metschnikowia_pulcherrima(100); | Metschnikowia | 13 | Saprotroph | \|Undefined Saprotroph\| | Yeast | NULL | Probable | As nectar-tap_saprotroph (Põlme et al. 2020); Decay sugar-rich substrates (Põlme et al. 2020) | Põlme S, et al. 2020. Fungal Diversity 105:1-16 ((https://doi.org/10.1007/s13225-020-00466-2)) |
| Otu37585 | 1 | k__Fungi(100);p__Ascomycota(100);c__Saccharomycetes(100);o__Saccharomycetales(100);f__Metschnikowiaceae(100);g__Metschnikowia(100);g__Metschnikowia_unclassified(100); | Metschnikowia | 13 | Saprotroph | \|Undefined Saprotroph\| | Yeast | NULL | Probable | As nectar-tap_saprotroph (Põlme et al. 2020); Decay sugar-rich substrates (Põlme et al. 2020) | Põlme S, et al. 2020. Fungal Diversity 105:1-16 ((https://doi.org/10.1007/s13225-020-00466-2)) |
| Otu37586 | 1 | k__Fungi(100);p__Ascomycota(100);c__Saccharomycetes(100);o__Saccharomycetales(100);f__Metschnikowiaceae(100);g__Metschnikowia(100);s__Metschnikowia_pulcherrima(100); | Metschnikowia | 13 | Saprotroph | \|Undefined Saprotroph\| | Yeast | NULL | Probable | As nectar-tap_saprotroph (Põlme et al. 2020); Decay sugar-rich substrates (Põlme et al. 2020) | Põlme S, et al. 2020. Fungal Diversity 105:1-16 ((https://doi.org/10.1007/s13225-020-00466-2)) |
| Otu37587 | 1 | k__Fungi(100);p__Ascomycota(100);c__Saccharomycetes(100);o__Saccharomycetales(100);f__Metschnikowiaceae(100);g__Metschnikowia(100);s__Metschnikowia_pulcherrima(100); | Metschnikowia | 13 | Saprotroph | \|Undefined Saprotroph\| | Yeast | NULL | Probable | As nectar-tap_saprotroph (Põlme et al. 2020); Decay sugar-rich substrates (Põlme et al. 2020) | Põlme S, et al. 2020. Fungal Diversity 105:1-16 ((https://doi.org/10.1007/s13225-020-00466-2)) |
| Otu37588 | 1 | k__Fungi(100);p__Ascomycota(100);c__Saccharomycetes(100);o__Saccharomycetales(100);f__Metschnikowiaceae(100);g__Metschnikowia(100);s__Metschnikowia_pulcherrima(100); | Metschnikowia | 13 | Saprotroph | \|Undefined Saprotroph\| | Yeast | NULL | Probable | As nectar-tap_saprotroph (Põlme et al. 2020); Decay sugar-rich substrates (Põlme et al. 2020) | Põlme S, et al. 2020. Fungal Diversity 105:1-16 ((https://doi.org/10.1007/s13225-020-00466-2)) |
| Otu37589 | 1 | k__Fungi(100);p__Ascomycota(100);c__Saccharomycetes(100);o__Saccharomycetales(100);f__Metschnikowiaceae(100);g__Metschnikowia(100);g__Metschnikowia_unclassified(100); | Metschnikowia | 13 | Saprotroph | \|Undefined Saprotroph\| | Yeast | NULL | Probable | As nectar-tap_saprotroph (Põlme et al. 2020); Decay sugar-rich substrates (Põlme et al. 2020) | Põlme S, et al. 2020. Fungal Diversity 105:1-16 ((https://doi.org/10.1007/s13225-020-00466-2)) |
| Otu37590 | 1 | k__Fungi(100);p__Ascomycota(100);c__Saccharomycetes(100);o__Saccharomycetales(100);f__Metschnikowiaceae(100);g__Metschnikowia(100);s__Metschnikowia_pulcherrima(100); | Metschnikowia | 13 | Saprotroph | \|Undefined Saprotroph\| | Yeast | NULL | Probable | As nectar-tap_saprotroph (Põlme et al. 2020); Decay sugar-rich substrates (Põlme et al. 2020) | Põlme S, et al. 2020. Fungal Diversity 105:1-16 ((https://doi.org/10.1007/s13225-020-00466-2)) |
| Otu37591 | 1 | k__Fungi(100);p__Ascomycota(100);c__Saccharomycetes(100);o__Saccharomycetales(100);f__Metschnikowiaceae(100);g__Metschnikowia(100);s__Metschnikowia_pulcherrima(100); | Metschnikowia | 13 | Saprotroph | \|Undefined Saprotroph\| | Yeast | NULL | Probable | As nectar-tap_saprotroph (Põlme et al. 2020); Decay sugar-rich substrates (Põlme et al. 2020) | Põlme S, et al. 2020. Fungal Diversity 105:1-16 ((https://doi.org/10.1007/s13225-020-00466-2)) |
| Otu37592 | 1 | k__Fungi(100);p__Ascomycota(100);c__Saccharomycetes(100);o__Saccharomycetales(100);f__Metschnikowiaceae(100);g__Metschnikowia(100);s__Metschnikowia_pulcherrima(100); | Metschnikowia | 13 | Saprotroph | \|Undefined Saprotroph\| | Yeast | NULL | Probable | As nectar-tap_saprotroph (Põlme et al. 2020); Decay sugar-rich substrates (Põlme et al. 2020) | Põlme S, et al. 2020. Fungal Diversity 105:1-16 ((https://doi.org/10.1007/s13225-020-00466-2)) |
| Otu37593 | 1 | k__Fungi(100);p__Ascomycota(100);c__Saccharomycetes(100);o__Saccharomycetales(100);f__Metschnikowiaceae(100);g__Metschnikowia(100);s__Metschnikowia_pulcherrima(100); | Metschnikowia | 13 | Saprotroph | \|Undefined Saprotroph\| | Yeast | NULL | Probable | As nectar-tap_saprotroph (Põlme et al. 2020); Decay sugar-rich substrates (Põlme et al. 2020) | Põlme S, et al. 2020. Fungal Diversity 105:1-16 ((https://doi.org/10.1007/s13225-020-00466-2)) |
| Otu37594 | 1 | k__Fungi(100);p__Ascomycota(100);c__Saccharomycetes(100);o__Saccharomycetales(100);f__Metschnikowiaceae(100);g__Metschnikowia(100);s__Metschnikowia_pulcherrima(100); | Metschnikowia | 13 | Saprotroph | \|Undefined Saprotroph\| | Yeast | NULL | Probable | As nectar-tap_saprotroph (Põlme et al. 2020); Decay sugar-rich substrates (Põlme et al. 2020) | Põlme S, et al. 2020. Fungal Diversity 105:1-16 ((https://doi.org/10.1007/s13225-020-00466-2)) |
| Otu37595 | 1 | k__Fungi(100);p__Ascomycota(100);c__Saccharomycetes(100);o__Saccharomycetales(100);f__Metschnikowiaceae(100);g__Metschnikowia(100);s__Metschnikowia_pulcherrima(100); | Metschnikowia | 13 | Saprotroph | \|Undefined Saprotroph\| | Yeast | NULL | Probable | As nectar-tap_saprotroph (Põlme et al. 2020); Decay sugar-rich substrates (Põlme et al. 2020) | Põlme S, et al. 2020. Fungal Diversity 105:1-16 ((https://doi.org/10.1007/s13225-020-00466-2)) |
| Otu37596 | 1 | k__Fungi(100);p__Ascomycota(100);c__Saccharomycetes(100);o__Saccharomycetales(100);f__Metschnikowiaceae(100);g__Metschnikowia(100);s__Metschnikowia_pulcherrima(100); | Metschnikowia | 13 | Saprotroph | \|Undefined Saprotroph\| | Yeast | NULL | Probable | As nectar-tap_saprotroph (Põlme et al. 2020); Decay sugar-rich substrates (Põlme et al. 2020) | Põlme S, et al. 2020. Fungal Diversity 105:1-16 ((https://doi.org/10.1007/s13225-020-00466-2)) |
| Otu37597 | 1 | k__Fungi(100);p__Ascomycota(100);c__Saccharomycetes(100);o__Saccharomycetales(100);f__Metschnikowiaceae(100);g__Metschnikowia(100);g__Metschnikowia_unclassified(100); | Metschnikowia | 13 | Saprotroph | \|Undefined Saprotroph\| | Yeast | NULL | Probable | As nectar-tap_saprotroph (Põlme et al. 2020); Decay sugar-rich substrates (Põlme et al. 2020) | Põlme S, et al. 2020. Fungal Diversity 105:1-16 ((https://doi.org/10.1007/s13225-020-00466-2)) |
| Otu37598 | 1 | k__Fungi(100);p__Ascomycota(100);c__Saccharomycetes(100);o__Saccharomycetales(100);f__Metschnikowiaceae(100);g__Metschnikowia(100);g__Metschnikowia_unclassified(100); | Metschnikowia | 13 | Saprotroph | \|Undefined Saprotroph\| | Yeast | NULL | Probable | As nectar-tap_saprotroph (Põlme et al. 2020); Decay sugar-rich substrates (Põlme et al. 2020) | Põlme S, et al. 2020. Fungal Diversity 105:1-16 ((https://doi.org/10.1007/s13225-020-00466-2)) |
| Otu37599 | 1 | k__Fungi(100);p__Ascomycota(100);c__Saccharomycetes(100);o__Saccharomycetales(100);f__Metschnikowiaceae(100);g__Metschnikowia(100);g__Metschnikowia_unclassified(100); | Metschnikowia | 13 | Saprotroph | \|Undefined Saprotroph\| | Yeast | NULL | Probable | As nectar-tap_saprotroph (Põlme et al. 2020); Decay sugar-rich substrates (Põlme et al. 2020) | Põlme S, et al. 2020. Fungal Diversity 105:1-16 ((https://doi.org/10.1007/s13225-020-00466-2)) |
| Otu37600 | 1 | k__Fungi(100);p__Ascomycota(100);c__Saccharomycetes(100);o__Saccharomycetales(100);f__Metschnikowiaceae(100);g__Metschnikowia(100);g__Metschnikowia_unclassified(100); | Metschnikowia | 13 | Saprotroph | \|Undefined Saprotroph\| | Yeast | NULL | Probable | As nectar-tap_saprotroph (Põlme et al. 2020); Decay sugar-rich substrates (Põlme et al. 2020) | Põlme S, et al. 2020. Fungal Diversity 105:1-16 ((https://doi.org/10.1007/s13225-020-00466-2)) |
| Otu37601 | 1 | k__Fungi(100);p__Ascomycota(100);c__Saccharomycetes(100);o__Saccharomycetales(100);f__Metschnikowiaceae(100);g__Metschnikowia(100);s__Metschnikowia_pulcherrima(100); | Metschnikowia | 13 | Saprotroph | \|Undefined Saprotroph\| | Yeast | NULL | Probable | As nectar-tap_saprotroph (Põlme et al. 2020); Decay sugar-rich substrates (Põlme et al. 2020) | Põlme S, et al. 2020. Fungal Diversity 105:1-16 ((https://doi.org/10.1007/s13225-020-00466-2)) |
| Otu37602 | 1 | k__Fungi(100);p__Ascomycota(100);c__Saccharomycetes(100);o__Saccharomycetales(100);f__Metschnikowiaceae(100);g__Metschnikowia(100);s__Metschnikowia_pulcherrima(100); | Metschnikowia | 13 | Saprotroph | \|Undefined Saprotroph\| | Yeast | NULL | Probable | As nectar-tap_saprotroph (Põlme et al. 2020); Decay sugar-rich substrates (Põlme et al. 2020) | Põlme S, et al. 2020. Fungal Diversity 105:1-16 ((https://doi.org/10.1007/s13225-020-00466-2)) |
| Otu37603 | 1 | k__Fungi(100);p__Ascomycota(100);c__Saccharomycetes(100);o__Saccharomycetales(100);f__Metschnikowiaceae(100);g__Metschnikowia(100);s__Metschnikowia_pulcherrima(100); | Metschnikowia | 13 | Saprotroph | \|Undefined Saprotroph\| | Yeast | NULL | Probable | As nectar-tap_saprotroph (Põlme et al. 2020); Decay sugar-rich substrates (Põlme et al. 2020) | Põlme S, et al. 2020. Fungal Diversity 105:1-16 ((https://doi.org/10.1007/s13225-020-00466-2)) |
| Otu37604 | 1 | k__Fungi(100);p__Ascomycota(100);c__Saccharomycetes(100);o__Saccharomycetales(100);f__Metschnikowiaceae(100);g__Metschnikowia(100);s__Metschnikowia_pulcherrima(100); | Metschnikowia | 13 | Saprotroph | \|Undefined Saprotroph\| | Yeast | NULL | Probable | As nectar-tap_saprotroph (Põlme et al. 2020); Decay sugar-rich substrates (Põlme et al. 2020) | Põlme S, et al. 2020. Fungal Diversity 105:1-16 ((https://doi.org/10.1007/s13225-020-00466-2)) |
| Otu37605 | 1 | k__Fungi(100);p__Ascomycota(100);c__Saccharomycetes(100);o__Saccharomycetales(100);f__Metschnikowiaceae(100);g__Metschnikowia(100);s__Metschnikowia_pulcherrima(100); | Metschnikowia | 13 | Saprotroph | \|Undefined Saprotroph\| | Yeast | NULL | Probable | As nectar-tap_saprotroph (Põlme et al. 2020); Decay sugar-rich substrates (Põlme et al. 2020) | Põlme S, et al. 2020. Fungal Diversity 105:1-16 ((https://doi.org/10.1007/s13225-020-00466-2)) |
| Otu37606 | 1 | k__Fungi(100);p__Ascomycota(100);c__Saccharomycetes(100);o__Saccharomycetales(100);f__Metschnikowiaceae(100);g__Metschnikowia(100);g__Metschnikowia_unclassified(100); | Metschnikowia | 13 | Saprotroph | \|Undefined Saprotroph\| | Yeast | NULL | Probable | As nectar-tap_saprotroph (Põlme et al. 2020); Decay sugar-rich substrates (Põlme et al. 2020) | Põlme S, et al. 2020. Fungal Diversity 105:1-16 ((https://doi.org/10.1007/s13225-020-00466-2)) |
| Otu37607 | 1 | k__Fungi(100);p__Ascomycota(100);c__Saccharomycetes(100);o__Saccharomycetales(100);f__Metschnikowiaceae(100);g__Metschnikowia(100);s__Metschnikowia_pulcherrima(100); | Metschnikowia | 13 | Saprotroph | \|Undefined Saprotroph\| | Yeast | NULL | Probable | As nectar-tap_saprotroph (Põlme et al. 2020); Decay sugar-rich substrates (Põlme et al. 2020) | Põlme S, et al. 2020. Fungal Diversity 105:1-16 ((https://doi.org/10.1007/s13225-020-00466-2)) |
| Otu37608 | 1 | k__Fungi(100);p__Ascomycota(100);c__Saccharomycetes(100);o__Saccharomycetales(100);f__Metschnikowiaceae(100);g__Metschnikowia(100);s__Metschnikowia_pulcherrima(100); | Metschnikowia | 13 | Saprotroph | \|Undefined Saprotroph\| | Yeast | NULL | Probable | As nectar-tap_saprotroph (Põlme et al. 2020); Decay sugar-rich substrates (Põlme et al. 2020) | Põlme S, et al. 2020. Fungal Diversity 105:1-16 ((https://doi.org/10.1007/s13225-020-00466-2)) |
| Otu37609 | 1 | k__Fungi(100);p__Ascomycota(100);c__Saccharomycetes(100);o__Saccharomycetales(100);f__Metschnikowiaceae(100);g__Metschnikowia(100);g__Metschnikowia_unclassified(100); | Metschnikowia | 13 | Saprotroph | \|Undefined Saprotroph\| | Yeast | NULL | Probable | As nectar-tap_saprotroph (Põlme et al. 2020); Decay sugar-rich substrates (Põlme et al. 2020) | Põlme S, et al. 2020. Fungal Diversity 105:1-16 ((https://doi.org/10.1007/s13225-020-00466-2)) |
| Otu37610 | 1 | k__Fungi(100);p__Ascomycota(100);c__Saccharomycetes(100);o__Saccharomycetales(100);f__Metschnikowiaceae(100);g__Metschnikowia(100);g__Metschnikowia_unclassified(100); | Metschnikowia | 13 | Saprotroph | \|Undefined Saprotroph\| | Yeast | NULL | Probable | As nectar-tap_saprotroph (Põlme et al. 2020); Decay sugar-rich substrates (Põlme et al. 2020) | Põlme S, et al. 2020. Fungal Diversity 105:1-16 ((https://doi.org/10.1007/s13225-020-00466-2)) |
| Otu37611 | 1 | k__Fungi(100);p__Ascomycota(100);c__Saccharomycetes(100);o__Saccharomycetales(100);f__Metschnikowiaceae(100);g__Metschnikowia(100);s__Metschnikowia_pulcherrima(100); | Metschnikowia | 13 | Saprotroph | \|Undefined Saprotroph\| | Yeast | NULL | Probable | As nectar-tap_saprotroph (Põlme et al. 2020); Decay sugar-rich substrates (Põlme et al. 2020) | Põlme S, et al. 2020. Fungal Diversity 105:1-16 ((https://doi.org/10.1007/s13225-020-00466-2)) |
| Otu37612 | 1 | k__Fungi(100);p__Ascomycota(100);c__Saccharomycetes(100);o__Saccharomycetales(100);f__Metschnikowiaceae(100);g__Metschnikowia(100);s__Metschnikowia_pulcherrima(100); | Metschnikowia | 13 | Saprotroph | \|Undefined Saprotroph\| | Yeast | NULL | Probable | As nectar-tap_saprotroph (Põlme et al. 2020); Decay sugar-rich substrates (Põlme et al. 2020) | Põlme S, et al. 2020. Fungal Diversity 105:1-16 ((https://doi.org/10.1007/s13225-020-00466-2)) |
| Otu37613 | 1 | k__Fungi(100);p__Ascomycota(100);c__Saccharomycetes(100);o__Saccharomycetales(100);f__Metschnikowiaceae(100);g__Metschnikowia(100);s__Metschnikowia_pulcherrima(100); | Metschnikowia | 13 | Saprotroph | \|Undefined Saprotroph\| | Yeast | NULL | Probable | As nectar-tap_saprotroph (Põlme et al. 2020); Decay sugar-rich substrates (Põlme et al. 2020) | Põlme S, et al. 2020. Fungal Diversity 105:1-16 ((https://doi.org/10.1007/s13225-020-00466-2)) |
| Otu37614 | 1 | k__Fungi(100);p__Ascomycota(100);c__Saccharomycetes(100);o__Saccharomycetales(100);f__Metschnikowiaceae(100);g__Metschnikowia(100);s__Metschnikowia_pulcherrima(100); | Metschnikowia | 13 | Saprotroph | \|Undefined Saprotroph\| | Yeast | NULL | Probable | As nectar-tap_saprotroph (Põlme et al. 2020); Decay sugar-rich substrates (Põlme et al. 2020) | Põlme S, et al. 2020. Fungal Diversity 105:1-16 ((https://doi.org/10.1007/s13225-020-00466-2)) |
| Otu37615 | 1 | k__Fungi(100);p__Ascomycota(100);c__Saccharomycetes(100);o__Saccharomycetales(100);f__Metschnikowiaceae(100);g__Metschnikowia(100);s__Metschnikowia_pulcherrima(100); | Metschnikowia | 13 | Saprotroph | \|Undefined Saprotroph\| | Yeast | NULL | Probable | As nectar-tap_saprotroph (Põlme et al. 2020); Decay sugar-rich substrates (Põlme et al. 2020) | Põlme S, et al. 2020. Fungal Diversity 105:1-16 ((https://doi.org/10.1007/s13225-020-00466-2)) |
| Otu37616 | 1 | k__Fungi(100);p__Ascomycota(100);c__Saccharomycetes(100);o__Saccharomycetales(100);f__Metschnikowiaceae(100);g__Metschnikowia(100);s__Metschnikowia_pulcherrima(100); | Metschnikowia | 13 | Saprotroph | \|Undefined Saprotroph\| | Yeast | NULL | Probable | As nectar-tap_saprotroph (Põlme et al. 2020); Decay sugar-rich substrates (Põlme et al. 2020) | Põlme S, et al. 2020. Fungal Diversity 105:1-16 ((https://doi.org/10.1007/s13225-020-00466-2)) |
| Otu37617 | 1 | k__Fungi(100);p__Ascomycota(100);c__Saccharomycetes(100);o__Saccharomycetales(100);f__Metschnikowiaceae(100);g__Metschnikowia(100);s__Metschnikowia_pulcherrima(100); | Metschnikowia | 13 | Saprotroph | \|Undefined Saprotroph\| | Yeast | NULL | Probable | As nectar-tap_saprotroph (Põlme et al. 2020); Decay sugar-rich substrates (Põlme et al. 2020) | Põlme S, et al. 2020. Fungal Diversity 105:1-16 ((https://doi.org/10.1007/s13225-020-00466-2)) |
| Otu37618 | 1 | k__Fungi(100);p__Ascomycota(100);c__Saccharomycetes(100);o__Saccharomycetales(100);f__Metschnikowiaceae(100);g__Metschnikowia(100);s__Metschnikowia_pulcherrima(100); | Metschnikowia | 13 | Saprotroph | \|Undefined Saprotroph\| | Yeast | NULL | Probable | As nectar-tap_saprotroph (Põlme et al. 2020); Decay sugar-rich substrates (Põlme et al. 2020) | Põlme S, et al. 2020. Fungal Diversity 105:1-16 ((https://doi.org/10.1007/s13225-020-00466-2)) |
| Otu37619 | 1 | k__Fungi(100);p__Ascomycota(100);c__Saccharomycetes(100);o__Saccharomycetales(100);f__Metschnikowiaceae(100);g__Metschnikowia(100);s__Metschnikowia_pulcherrima(100); | Metschnikowia | 13 | Saprotroph | \|Undefined Saprotroph\| | Yeast | NULL | Probable | As nectar-tap_saprotroph (Põlme et al. 2020); Decay sugar-rich substrates (Põlme et al. 2020) | Põlme S, et al. 2020. Fungal Diversity 105:1-16 ((https://doi.org/10.1007/s13225-020-00466-2)) |
| Otu37620 | 1 | k__Fungi(100);p__Ascomycota(100);c__Saccharomycetes(100);o__Saccharomycetales(100);f__Metschnikowiaceae(100);g__Metschnikowia(100);g__Metschnikowia_unclassified(100); | Metschnikowia | 13 | Saprotroph | \|Undefined Saprotroph\| | Yeast | NULL | Probable | As nectar-tap_saprotroph (Põlme et al. 2020); Decay sugar-rich substrates (Põlme et al. 2020) | Põlme S, et al. 2020. Fungal Diversity 105:1-16 ((https://doi.org/10.1007/s13225-020-00466-2)) |
| Otu37621 | 1 | k__Fungi(100);p__Ascomycota(100);c__Saccharomycetes(100);o__Saccharomycetales(100);f__Metschnikowiaceae(100);g__Metschnikowia(100);g__Metschnikowia_unclassified(100); | Metschnikowia | 13 | Saprotroph | \|Undefined Saprotroph\| | Yeast | NULL | Probable | As nectar-tap_saprotroph (Põlme et al. 2020); Decay sugar-rich substrates (Põlme et al. 2020) | Põlme S, et al. 2020. Fungal Diversity 105:1-16 ((https://doi.org/10.1007/s13225-020-00466-2)) |
| Otu37622 | 1 | k__Fungi(100);p__Ascomycota(100);c__Saccharomycetes(100);o__Saccharomycetales(100);f__Metschnikowiaceae(100);g__Metschnikowia(100);g__Metschnikowia_unclassified(100); | Metschnikowia | 13 | Saprotroph | \|Undefined Saprotroph\| | Yeast | NULL | Probable | As nectar-tap_saprotroph (Põlme et al. 2020); Decay sugar-rich substrates (Põlme et al. 2020) | Põlme S, et al. 2020. Fungal Diversity 105:1-16 ((https://doi.org/10.1007/s13225-020-00466-2)) |
| Otu37623 | 1 | k__Fungi(100);p__Ascomycota(100);c__Saccharomycetes(100);o__Saccharomycetales(100);f__Metschnikowiaceae(100);g__Metschnikowia(100);s__Metschnikowia_pulcherrima(100); | Metschnikowia | 13 | Saprotroph | \|Undefined Saprotroph\| | Yeast | NULL | Probable | As nectar-tap_saprotroph (Põlme et al. 2020); Decay sugar-rich substrates (Põlme et al. 2020) | Põlme S, et al. 2020. Fungal Diversity 105:1-16 ((https://doi.org/10.1007/s13225-020-00466-2)) |
| Otu37624 | 1 | k__Fungi(100);p__Ascomycota(100);c__Saccharomycetes(100);o__Saccharomycetales(100);f__Metschnikowiaceae(100);g__Metschnikowia(100);s__Metschnikowia_pulcherrima(100); | Metschnikowia | 13 | Saprotroph | \|Undefined Saprotroph\| | Yeast | NULL | Probable | As nectar-tap_saprotroph (Põlme et al. 2020); Decay sugar-rich substrates (Põlme et al. 2020) | Põlme S, et al. 2020. Fungal Diversity 105:1-16 ((https://doi.org/10.1007/s13225-020-00466-2)) |
| Otu37625 | 1 | k__Fungi(100);p__Ascomycota(100);c__Saccharomycetes(100);o__Saccharomycetales(100);f__Metschnikowiaceae(100);g__Metschnikowia(100);g__Metschnikowia_unclassified(100); | Metschnikowia | 13 | Saprotroph | \|Undefined Saprotroph\| | Yeast | NULL | Probable | As nectar-tap_saprotroph (Põlme et al. 2020); Decay sugar-rich substrates (Põlme et al. 2020) | Põlme S, et al. 2020. Fungal Diversity 105:1-16 ((https://doi.org/10.1007/s13225-020-00466-2)) |
| Otu37626 | 1 | k__Fungi(100);p__Ascomycota(100);c__Saccharomycetes(100);o__Saccharomycetales(100);f__Metschnikowiaceae(100);g__Metschnikowia(100);g__Metschnikowia_unclassified(100); | Metschnikowia | 13 | Saprotroph | \|Undefined Saprotroph\| | Yeast | NULL | Probable | As nectar-tap_saprotroph (Põlme et al. 2020); Decay sugar-rich substrates (Põlme et al. 2020) | Põlme S, et al. 2020. Fungal Diversity 105:1-16 ((https://doi.org/10.1007/s13225-020-00466-2)) |
| Otu37627 | 1 | k__Fungi(100);p__Ascomycota(100);c__Saccharomycetes(100);o__Saccharomycetales(100);f__Metschnikowiaceae(100);g__Metschnikowia(100);g__Metschnikowia_unclassified(100); | Metschnikowia | 13 | Saprotroph | \|Undefined Saprotroph\| | Yeast | NULL | Probable | As nectar-tap_saprotroph (Põlme et al. 2020); Decay sugar-rich substrates (Põlme et al. 2020) | Põlme S, et al. 2020. Fungal Diversity 105:1-16 ((https://doi.org/10.1007/s13225-020-00466-2)) |
| Otu37630 | 1 | k__Fungi(100);p__Ascomycota(100);c__Saccharomycetes(100);o__Saccharomycetales(100);f__Metschnikowiaceae(100);g__Metschnikowia(100);g__Metschnikowia_unclassified(100); | Metschnikowia | 13 | Saprotroph | \|Undefined Saprotroph\| | Yeast | NULL | Probable | As nectar-tap_saprotroph (Põlme et al. 2020); Decay sugar-rich substrates (Põlme et al. 2020) | Põlme S, et al. 2020. Fungal Diversity 105:1-16 ((https://doi.org/10.1007/s13225-020-00466-2)) |
| Otu37631 | 1 | k__Fungi(100);p__Ascomycota(100);c__Saccharomycetes(100);o__Saccharomycetales(100);f__Metschnikowiaceae(100);g__Metschnikowia(100);s__Metschnikowia_pulcherrima(100); | Metschnikowia | 13 | Saprotroph | \|Undefined Saprotroph\| | Yeast | NULL | Probable | As nectar-tap_saprotroph (Põlme et al. 2020); Decay sugar-rich substrates (Põlme et al. 2020) | Põlme S, et al. 2020. Fungal Diversity 105:1-16 ((https://doi.org/10.1007/s13225-020-00466-2)) |
| Otu37633 | 1 | k__Fungi(100);p__Ascomycota(100);c__Saccharomycetes(100);o__Saccharomycetales(100);f__Metschnikowiaceae(100);g__Metschnikowia(100);s__Metschnikowia_pulcherrima(100); | Metschnikowia | 13 | Saprotroph | \|Undefined Saprotroph\| | Yeast | NULL | Probable | As nectar-tap_saprotroph (Põlme et al. 2020); Decay sugar-rich substrates (Põlme et al. 2020) | Põlme S, et al. 2020. Fungal Diversity 105:1-16 ((https://doi.org/10.1007/s13225-020-00466-2)) |
| Otu37634 | 1 | k__Fungi(100);p__Ascomycota(100);c__Saccharomycetes(100);o__Saccharomycetales(100);f__Metschnikowiaceae(100);g__Metschnikowia(100);s__Metschnikowia_pulcherrima(100); | Metschnikowia | 13 | Saprotroph | \|Undefined Saprotroph\| | Yeast | NULL | Probable | As nectar-tap_saprotroph (Põlme et al. 2020); Decay sugar-rich substrates (Põlme et al. 2020) | Põlme S, et al. 2020. Fungal Diversity 105:1-16 ((https://doi.org/10.1007/s13225-020-00466-2)) |
| Otu37636 | 1 | k__Fungi(100);p__Ascomycota(100);c__Saccharomycetes(100);o__Saccharomycetales(100);f__Metschnikowiaceae(100);g__Metschnikowia(100);g__Metschnikowia_unclassified(100); | Metschnikowia | 13 | Saprotroph | \|Undefined Saprotroph\| | Yeast | NULL | Probable | As nectar-tap_saprotroph (Põlme et al. 2020); Decay sugar-rich substrates (Põlme et al. 2020) | Põlme S, et al. 2020. Fungal Diversity 105:1-16 ((https://doi.org/10.1007/s13225-020-00466-2)) |
| Otu37637 | 1 | k__Fungi(100);p__Ascomycota(100);c__Saccharomycetes(100);o__Saccharomycetales(100);f__Metschnikowiaceae(100);g__Metschnikowia(100);g__Metschnikowia_unclassified(100); | Metschnikowia | 13 | Saprotroph | \|Undefined Saprotroph\| | Yeast | NULL | Probable | As nectar-tap_saprotroph (Põlme et al. 2020); Decay sugar-rich substrates (Põlme et al. 2020) | Põlme S, et al. 2020. Fungal Diversity 105:1-16 ((https://doi.org/10.1007/s13225-020-00466-2)) |
| Otu37639 | 1 | k__Fungi(100);p__Ascomycota(100);c__Saccharomycetes(100);o__Saccharomycetales(100);f__Metschnikowiaceae(100);g__Metschnikowia(100);g__Metschnikowia_unclassified(100); | Metschnikowia | 13 | Saprotroph | \|Undefined Saprotroph\| | Yeast | NULL | Probable | As nectar-tap_saprotroph (Põlme et al. 2020); Decay sugar-rich substrates (Põlme et al. 2020) | Põlme S, et al. 2020. Fungal Diversity 105:1-16 ((https://doi.org/10.1007/s13225-020-00466-2)) |
| Otu37640 | 1 | k__Fungi(100);p__Ascomycota(100);c__Saccharomycetes(100);o__Saccharomycetales(100);f__Metschnikowiaceae(100);g__Metschnikowia(100);g__Metschnikowia_unclassified(100); | Metschnikowia | 13 | Saprotroph | \|Undefined Saprotroph\| | Yeast | NULL | Probable | As nectar-tap_saprotroph (Põlme et al. 2020); Decay sugar-rich substrates (Põlme et al. 2020) | Põlme S, et al. 2020. Fungal Diversity 105:1-16 ((https://doi.org/10.1007/s13225-020-00466-2)) |
| Otu37641 | 1 | k__Fungi(100);p__Ascomycota(100);c__Saccharomycetes(100);o__Saccharomycetales(100);f__Metschnikowiaceae(100);g__Metschnikowia(100);s__Metschnikowia_pulcherrima(100); | Metschnikowia | 13 | Saprotroph | \|Undefined Saprotroph\| | Yeast | NULL | Probable | As nectar-tap_saprotroph (Põlme et al. 2020); Decay sugar-rich substrates (Põlme et al. 2020) | Põlme S, et al. 2020. Fungal Diversity 105:1-16 ((https://doi.org/10.1007/s13225-020-00466-2)) |
| Otu37642 | 1 | k__Fungi(100);p__Ascomycota(100);c__Saccharomycetes(100);o__Saccharomycetales(100);f__Metschnikowiaceae(100);g__Metschnikowia(100);s__Metschnikowia_pulcherrima(100); | Metschnikowia | 13 | Saprotroph | \|Undefined Saprotroph\| | Yeast | NULL | Probable | As nectar-tap_saprotroph (Põlme et al. 2020); Decay sugar-rich substrates (Põlme et al. 2020) | Põlme S, et al. 2020. Fungal Diversity 105:1-16 ((https://doi.org/10.1007/s13225-020-00466-2)) |
| Otu37643 | 1 | k__Fungi(100);p__Ascomycota(100);c__Saccharomycetes(100);o__Saccharomycetales(100);f__Metschnikowiaceae(100);g__Metschnikowia(100);s__Metschnikowia_pulcherrima(100); | Metschnikowia | 13 | Saprotroph | \|Undefined Saprotroph\| | Yeast | NULL | Probable | As nectar-tap_saprotroph (Põlme et al. 2020); Decay sugar-rich substrates (Põlme et al. 2020) | Põlme S, et al. 2020. Fungal Diversity 105:1-16 ((https://doi.org/10.1007/s13225-020-00466-2)) |
| Otu37644 | 1 | k__Fungi(100);p__Ascomycota(100);c__Saccharomycetes(100);o__Saccharomycetales(100);f__Metschnikowiaceae(100);g__Metschnikowia(100);s__Metschnikowia_pulcherrima(100); | Metschnikowia | 13 | Saprotroph | \|Undefined Saprotroph\| | Yeast | NULL | Probable | As nectar-tap_saprotroph (Põlme et al. 2020); Decay sugar-rich substrates (Põlme et al. 2020) | Põlme S, et al. 2020. Fungal Diversity 105:1-16 ((https://doi.org/10.1007/s13225-020-00466-2)) |
| Otu37645 | 1 | k__Fungi(100);p__Ascomycota(100);c__Saccharomycetes(100);o__Saccharomycetales(100);f__Metschnikowiaceae(100);g__Metschnikowia(100);s__Metschnikowia_pulcherrima(100); | Metschnikowia | 13 | Saprotroph | \|Undefined Saprotroph\| | Yeast | NULL | Probable | As nectar-tap_saprotroph (Põlme et al. 2020); Decay sugar-rich substrates (Põlme et al. 2020) | Põlme S, et al. 2020. Fungal Diversity 105:1-16 ((https://doi.org/10.1007/s13225-020-00466-2)) |
| Otu37646 | 1 | k__Fungi(100);p__Ascomycota(100);c__Saccharomycetes(100);o__Saccharomycetales(100);f__Metschnikowiaceae(100);g__Metschnikowia(100);s__Metschnikowia_pulcherrima(100); | Metschnikowia | 13 | Saprotroph | \|Undefined Saprotroph\| | Yeast | NULL | Probable | As nectar-tap_saprotroph (Põlme et al. 2020); Decay sugar-rich substrates (Põlme et al. 2020) | Põlme S, et al. 2020. Fungal Diversity 105:1-16 ((https://doi.org/10.1007/s13225-020-00466-2)) |
| Otu37647 | 1 | k__Fungi(100);p__Ascomycota(100);c__Saccharomycetes(100);o__Saccharomycetales(100);f__Metschnikowiaceae(100);g__Metschnikowia(100);g__Metschnikowia_unclassified(100); | Metschnikowia | 13 | Saprotroph | \|Undefined Saprotroph\| | Yeast | NULL | Probable | As nectar-tap_saprotroph (Põlme et al. 2020); Decay sugar-rich substrates (Põlme et al. 2020) | Põlme S, et al. 2020. Fungal Diversity 105:1-16 ((https://doi.org/10.1007/s13225-020-00466-2)) |
| Otu37648 | 1 | k__Fungi(100);p__Ascomycota(100);c__Saccharomycetes(100);o__Saccharomycetales(100);f__Metschnikowiaceae(100);g__Metschnikowia(100);s__Metschnikowia_pulcherrima(100); | Metschnikowia | 13 | Saprotroph | \|Undefined Saprotroph\| | Yeast | NULL | Probable | As nectar-tap_saprotroph (Põlme et al. 2020); Decay sugar-rich substrates (Põlme et al. 2020) | Põlme S, et al. 2020. Fungal Diversity 105:1-16 ((https://doi.org/10.1007/s13225-020-00466-2)) |
| Otu37649 | 1 | k__Fungi(100);p__Ascomycota(100);c__Saccharomycetes(100);o__Saccharomycetales(100);f__Metschnikowiaceae(100);g__Metschnikowia(100);s__Metschnikowia_pulcherrima(100); | Metschnikowia | 13 | Saprotroph | \|Undefined Saprotroph\| | Yeast | NULL | Probable | As nectar-tap_saprotroph (Põlme et al. 2020); Decay sugar-rich substrates (Põlme et al. 2020) | Põlme S, et al. 2020. Fungal Diversity 105:1-16 ((https://doi.org/10.1007/s13225-020-00466-2)) |
| Otu37650 | 1 | k__Fungi(100);p__Ascomycota(100);c__Saccharomycetes(100);o__Saccharomycetales(100);f__Metschnikowiaceae(100);g__Metschnikowia(100);s__Metschnikowia_pulcherrima(100); | Metschnikowia | 13 | Saprotroph | \|Undefined Saprotroph\| | Yeast | NULL | Probable | As nectar-tap_saprotroph (Põlme et al. 2020); Decay sugar-rich substrates (Põlme et al. 2020) | Põlme S, et al. 2020. Fungal Diversity 105:1-16 ((https://doi.org/10.1007/s13225-020-00466-2)) |
| Otu37651 | 1 | k__Fungi(100);p__Ascomycota(100);c__Saccharomycetes(100);o__Saccharomycetales(100);f__Metschnikowiaceae(100);g__Metschnikowia(100);s__Metschnikowia_pulcherrima(100); | Metschnikowia | 13 | Saprotroph | \|Undefined Saprotroph\| | Yeast | NULL | Probable | As nectar-tap_saprotroph (Põlme et al. 2020); Decay sugar-rich substrates (Põlme et al. 2020) | Põlme S, et al. 2020. Fungal Diversity 105:1-16 ((https://doi.org/10.1007/s13225-020-00466-2)) |
| Otu37652 | 1 | k__Fungi(100);p__Ascomycota(100);c__Saccharomycetes(100);o__Saccharomycetales(100);f__Metschnikowiaceae(100);g__Metschnikowia(100);s__Metschnikowia_pulcherrima(100); | Metschnikowia | 13 | Saprotroph | \|Undefined Saprotroph\| | Yeast | NULL | Probable | As nectar-tap_saprotroph (Põlme et al. 2020); Decay sugar-rich substrates (Põlme et al. 2020) | Põlme S, et al. 2020. Fungal Diversity 105:1-16 ((https://doi.org/10.1007/s13225-020-00466-2)) |
| Otu37653 | 1 | k__Fungi(100);p__Ascomycota(100);c__Saccharomycetes(100);o__Saccharomycetales(100);f__Metschnikowiaceae(100);g__Metschnikowia(100);s__Metschnikowia_pulcherrima(100); | Metschnikowia | 13 | Saprotroph | \|Undefined Saprotroph\| | Yeast | NULL | Probable | As nectar-tap_saprotroph (Põlme et al. 2020); Decay sugar-rich substrates (Põlme et al. 2020) | Põlme S, et al. 2020. Fungal Diversity 105:1-16 ((https://doi.org/10.1007/s13225-020-00466-2)) |
| Otu37654 | 1 | k__Fungi(100);p__Ascomycota(100);c__Saccharomycetes(100);o__Saccharomycetales(100);f__Metschnikowiaceae(100);g__Metschnikowia(100);s__Metschnikowia_pulcherrima(100); | Metschnikowia | 13 | Saprotroph | \|Undefined Saprotroph\| | Yeast | NULL | Probable | As nectar-tap_saprotroph (Põlme et al. 2020); Decay sugar-rich substrates (Põlme et al. 2020) | Põlme S, et al. 2020. Fungal Diversity 105:1-16 ((https://doi.org/10.1007/s13225-020-00466-2)) |
| Otu37656 | 1 | k__Fungi(100);p__Ascomycota(100);c__Saccharomycetes(100);o__Saccharomycetales(100);f__Metschnikowiaceae(100);g__Metschnikowia(100);s__Metschnikowia_pulcherrima(100); | Metschnikowia | 13 | Saprotroph | \|Undefined Saprotroph\| | Yeast | NULL | Probable | As nectar-tap_saprotroph (Põlme et al. 2020); Decay sugar-rich substrates (Põlme et al. 2020) | Põlme S, et al. 2020. Fungal Diversity 105:1-16 ((https://doi.org/10.1007/s13225-020-00466-2)) |
| Otu37657 | 1 | k__Fungi(100);p__Ascomycota(100);c__Saccharomycetes(100);o__Saccharomycetales(100);f__Metschnikowiaceae(100);g__Metschnikowia(100);s__Metschnikowia_pulcherrima(100); | Metschnikowia | 13 | Saprotroph | \|Undefined Saprotroph\| | Yeast | NULL | Probable | As nectar-tap_saprotroph (Põlme et al. 2020); Decay sugar-rich substrates (Põlme et al. 2020) | Põlme S, et al. 2020. Fungal Diversity 105:1-16 ((https://doi.org/10.1007/s13225-020-00466-2)) |
| Otu37658 | 1 | k__Fungi(100);p__Ascomycota(100);c__Saccharomycetes(100);o__Saccharomycetales(100);f__Metschnikowiaceae(100);g__Metschnikowia(100);s__Metschnikowia_pulcherrima(100); | Metschnikowia | 13 | Saprotroph | \|Undefined Saprotroph\| | Yeast | NULL | Probable | As nectar-tap_saprotroph (Põlme et al. 2020); Decay sugar-rich substrates (Põlme et al. 2020) | Põlme S, et al. 2020. Fungal Diversity 105:1-16 ((https://doi.org/10.1007/s13225-020-00466-2)) |
| Otu37659 | 1 | k__Fungi(100);p__Ascomycota(100);c__Saccharomycetes(100);o__Saccharomycetales(100);f__Metschnikowiaceae(100);g__Metschnikowia(100);s__Metschnikowia_pulcherrima(100); | Metschnikowia | 13 | Saprotroph | \|Undefined Saprotroph\| | Yeast | NULL | Probable | As nectar-tap_saprotroph (Põlme et al. 2020); Decay sugar-rich substrates (Põlme et al. 2020) | Põlme S, et al. 2020. Fungal Diversity 105:1-16 ((https://doi.org/10.1007/s13225-020-00466-2)) |
| Otu37661 | 1 | k__Fungi(100);p__Ascomycota(100);c__Saccharomycetes(100);o__Saccharomycetales(100);f__Metschnikowiaceae(100);g__Metschnikowia(100);g__Metschnikowia_unclassified(100); | Metschnikowia | 13 | Saprotroph | \|Undefined Saprotroph\| | Yeast | NULL | Probable | As nectar-tap_saprotroph (Põlme et al. 2020); Decay sugar-rich substrates (Põlme et al. 2020) | Põlme S, et al. 2020. Fungal Diversity 105:1-16 ((https://doi.org/10.1007/s13225-020-00466-2)) |
| Otu37662 | 1 | k__Fungi(100);p__Ascomycota(100);c__Saccharomycetes(100);o__Saccharomycetales(100);f__Metschnikowiaceae(100);g__Metschnikowia(100);s__Metschnikowia_pulcherrima(100); | Metschnikowia | 13 | Saprotroph | \|Undefined Saprotroph\| | Yeast | NULL | Probable | As nectar-tap_saprotroph (Põlme et al. 2020); Decay sugar-rich substrates (Põlme et al. 2020) | Põlme S, et al. 2020. Fungal Diversity 105:1-16 ((https://doi.org/10.1007/s13225-020-00466-2)) |
| Otu37663 | 1 | k__Fungi(100);p__Ascomycota(100);c__Saccharomycetes(100);o__Saccharomycetales(100);f__Metschnikowiaceae(100);g__Metschnikowia(100);s__Metschnikowia_pulcherrima(100); | Metschnikowia | 13 | Saprotroph | \|Undefined Saprotroph\| | Yeast | NULL | Probable | As nectar-tap_saprotroph (Põlme et al. 2020); Decay sugar-rich substrates (Põlme et al. 2020) | Põlme S, et al. 2020. Fungal Diversity 105:1-16 ((https://doi.org/10.1007/s13225-020-00466-2)) |
| Otu37664 | 1 | k__Fungi(100);p__Ascomycota(100);c__Saccharomycetes(100);o__Saccharomycetales(100);f__Metschnikowiaceae(100);g__Metschnikowia(100);s__Metschnikowia_pulcherrima(100); | Metschnikowia | 13 | Saprotroph | \|Undefined Saprotroph\| | Yeast | NULL | Probable | As nectar-tap_saprotroph (Põlme et al. 2020); Decay sugar-rich substrates (Põlme et al. 2020) | Põlme S, et al. 2020. Fungal Diversity 105:1-16 ((https://doi.org/10.1007/s13225-020-00466-2)) |
| Otu37665 | 1 | k__Fungi(100);p__Ascomycota(100);c__Saccharomycetes(100);o__Saccharomycetales(100);f__Metschnikowiaceae(100);g__Metschnikowia(100);s__Metschnikowia_pulcherrima(100); | Metschnikowia | 13 | Saprotroph | \|Undefined Saprotroph\| | Yeast | NULL | Probable | As nectar-tap_saprotroph (Põlme et al. 2020); Decay sugar-rich substrates (Põlme et al. 2020) | Põlme S, et al. 2020. Fungal Diversity 105:1-16 ((https://doi.org/10.1007/s13225-020-00466-2)) |
| Otu37666 | 1 | k__Fungi(100);p__Ascomycota(100);c__Saccharomycetes(100);o__Saccharomycetales(100);f__Metschnikowiaceae(100);g__Metschnikowia(100);g__Metschnikowia_unclassified(100); | Metschnikowia | 13 | Saprotroph | \|Undefined Saprotroph\| | Yeast | NULL | Probable | As nectar-tap_saprotroph (Põlme et al. 2020); Decay sugar-rich substrates (Põlme et al. 2020) | Põlme S, et al. 2020. Fungal Diversity 105:1-16 ((https://doi.org/10.1007/s13225-020-00466-2)) |
| Otu37667 | 1 | k__Fungi(100);p__Ascomycota(100);c__Saccharomycetes(100);o__Saccharomycetales(100);f__Metschnikowiaceae(100);g__Metschnikowia(100);s__Metschnikowia_pulcherrima(100); | Metschnikowia | 13 | Saprotroph | \|Undefined Saprotroph\| | Yeast | NULL | Probable | As nectar-tap_saprotroph (Põlme et al. 2020); Decay sugar-rich substrates (Põlme et al. 2020) | Põlme S, et al. 2020. Fungal Diversity 105:1-16 ((https://doi.org/10.1007/s13225-020-00466-2)) |
| Otu37668 | 1 | k__Fungi(100);p__Ascomycota(100);c__Saccharomycetes(100);o__Saccharomycetales(100);f__Metschnikowiaceae(100);g__Metschnikowia(100);s__Metschnikowia_pulcherrima(100); | Metschnikowia | 13 | Saprotroph | \|Undefined Saprotroph\| | Yeast | NULL | Probable | As nectar-tap_saprotroph (Põlme et al. 2020); Decay sugar-rich substrates (Põlme et al. 2020) | Põlme S, et al. 2020. Fungal Diversity 105:1-16 ((https://doi.org/10.1007/s13225-020-00466-2)) |
| Otu37669 | 1 | k__Fungi(100);p__Basidiomycota(100);c__Dacrymycetes(100);o__Dacrymycetales(100);f__Dacrymycetaceae(100);g__Cerinosterus(100);s__Cerinosterus_sp(100); | Cerinosterus | 13 | Saprotroph | \|Undefined Saprotroph\| | Microfungus | Brown Rot | Probable | NULL | Tedersoo L, et al. 2014. Science 346:e1256688 ((https://doi.org/10.1126/science.1256688)); Põlme S, et al. 2020. Fungal Diversity 105:1-16 ((https://doi.org/10.1007/s13225-020-00466-2)) |
| Otu37671 | 1 | k__Fungi(100);p__Ascomycota(100);c__Leotiomycetes(100);o__Leotiomycetes_order_Incertae_sedis(100);f__Leotiomycetes_family_Incertae_sedis(100);g__Meliniomyces(100);g__Meliniomyces_unclassified(100); | Meliniomyces | 13 | Saprotroph-Symbiotroph | Ectomycorrhizal-\|Endophyte\|-Ericoid Mycorrhizal-Orchid Mycorrhizal-Plant Saprotroph-Undefined Saprotroph | Microfungus | NULL | Probable | As root_endophyte (Põlme et al. 2020); As soil_saprotroph (Põlme et al. 2020); Dark Septate Endophyte; Decay soil (Põlme et al. 2020) | Tedersoo L, et al. 2014. Science 346:e1256688 ((https://doi.org/10.1126/science.1256688)); Põlme S, et al. 2020. Fungal Diversity 105:1-16 ((https://doi.org/10.1007/s13225-020-00466-2)) |
| Otu37676 | 1 | k__Fungi(100);p__Ascomycota(100);c__Saccharomycetes(100);o__Saccharomycetales(100);f__Metschnikowiaceae(100);g__Metschnikowia(100);s__Metschnikowia_pulcherrima(100); | Metschnikowia | 13 | Saprotroph | \|Undefined Saprotroph\| | Yeast | NULL | Probable | As nectar-tap_saprotroph (Põlme et al. 2020); Decay sugar-rich substrates (Põlme et al. 2020) | Põlme S, et al. 2020. Fungal Diversity 105:1-16 ((https://doi.org/10.1007/s13225-020-00466-2)) |
| Otu37677 | 1 | k__Fungi(100);p__Basidiomycota(100);c__Microbotryomycetes(100);o__Sporidiobolales(100);f__Sporidiobolales_family_Incertae_sedis(100);g__Sporobolomyces(100);s__Sporobolomyces_lactosus(100); | Sporobolomyces | 13 | Pathotroph-Saprotroph | \|Fungal Parasite\|-Plant Saprotroph | Yeast | NULL | Probable | As litter_saprotroph (Põlme et al. 2020); As mycoparasite (Põlme et al. 2020); Decay fungal material (Põlme et al. 2020) | Tedersoo L, et al. 2014. Science 346:e1256688 ((https://doi.org/10.1126/science.1256688)); Purahong W, et al. 2016. Molecular Ecology 25:4059-4074 ((https://doi.org/10.1111/mec.13739)); Põlme S, et al. 2020. Fungal Diversity 105:1-16 ((https://doi.org/10.1007/s13225-020-00466-2)) |
| Otu37678 | 1 | k__Fungi(100);p__Ascomycota(100);c__Saccharomycetes(100);o__Saccharomycetales(100);f__Metschnikowiaceae(100);g__Metschnikowia(100);s__Metschnikowia_pulcherrima(100); | Metschnikowia | 13 | Saprotroph | \|Undefined Saprotroph\| | Yeast | NULL | Probable | As nectar-tap_saprotroph (Põlme et al. 2020); Decay sugar-rich substrates (Põlme et al. 2020) | Põlme S, et al. 2020. Fungal Diversity 105:1-16 ((https://doi.org/10.1007/s13225-020-00466-2)) |
| Otu37679 | 1 | k__Fungi(100);p__Ascomycota(100);c__Saccharomycetes(100);o__Saccharomycetales(100);f__Metschnikowiaceae(100);g__Metschnikowia(100);s__Metschnikowia_pulcherrima(100); | Metschnikowia | 13 | Saprotroph | \|Undefined Saprotroph\| | Yeast | NULL | Probable | As nectar-tap_saprotroph (Põlme et al. 2020); Decay sugar-rich substrates (Põlme et al. 2020) | Põlme S, et al. 2020. Fungal Diversity 105:1-16 ((https://doi.org/10.1007/s13225-020-00466-2)) |
| Otu37680 | 1 | k__Fungi(100);p__Ascomycota(100);c__Saccharomycetes(100);o__Saccharomycetales(100);f__Metschnikowiaceae(100);g__Metschnikowia(100);g__Metschnikowia_unclassified(100); | Metschnikowia | 13 | Saprotroph | \|Undefined Saprotroph\| | Yeast | NULL | Probable | As nectar-tap_saprotroph (Põlme et al. 2020); Decay sugar-rich substrates (Põlme et al. 2020) | Põlme S, et al. 2020. Fungal Diversity 105:1-16 ((https://doi.org/10.1007/s13225-020-00466-2)) |
| Otu37681 | 1 | k__Fungi(100);p__Ascomycota(100);c__Saccharomycetes(100);o__Saccharomycetales(100);f__Metschnikowiaceae(100);g__Metschnikowia(100);s__Metschnikowia_pulcherrima(100); | Metschnikowia | 13 | Saprotroph | \|Undefined Saprotroph\| | Yeast | NULL | Probable | As nectar-tap_saprotroph (Põlme et al. 2020); Decay sugar-rich substrates (Põlme et al. 2020) | Põlme S, et al. 2020. Fungal Diversity 105:1-16 ((https://doi.org/10.1007/s13225-020-00466-2)) |
| Otu37682 | 1 | k__Fungi(100);p__Ascomycota(100);c__Saccharomycetes(100);o__Saccharomycetales(100);f__Metschnikowiaceae(100);g__Metschnikowia(100);s__Metschnikowia_pulcherrima(100); | Metschnikowia | 13 | Saprotroph | \|Undefined Saprotroph\| | Yeast | NULL | Probable | As nectar-tap_saprotroph (Põlme et al. 2020); Decay sugar-rich substrates (Põlme et al. 2020) | Põlme S, et al. 2020. Fungal Diversity 105:1-16 ((https://doi.org/10.1007/s13225-020-00466-2)) |
| Otu37683 | 1 | k__Fungi(100);p__Ascomycota(100);c__Saccharomycetes(100);o__Saccharomycetales(100);f__Metschnikowiaceae(100);g__Metschnikowia(100);g__Metschnikowia_unclassified(100); | Metschnikowia | 13 | Saprotroph | \|Undefined Saprotroph\| | Yeast | NULL | Probable | As nectar-tap_saprotroph (Põlme et al. 2020); Decay sugar-rich substrates (Põlme et al. 2020) | Põlme S, et al. 2020. Fungal Diversity 105:1-16 ((https://doi.org/10.1007/s13225-020-00466-2)) |
| Otu37684 | 1 | k__Fungi(100);p__Ascomycota(100);c__Saccharomycetes(100);o__Saccharomycetales(100);f__Metschnikowiaceae(100);g__Metschnikowia(100);s__Metschnikowia_pulcherrima(100); | Metschnikowia | 13 | Saprotroph | \|Undefined Saprotroph\| | Yeast | NULL | Probable | As nectar-tap_saprotroph (Põlme et al. 2020); Decay sugar-rich substrates (Põlme et al. 2020) | Põlme S, et al. 2020. Fungal Diversity 105:1-16 ((https://doi.org/10.1007/s13225-020-00466-2)) |
| Otu37685 | 1 | k__Fungi(100);p__Ascomycota(100);c__Saccharomycetes(100);o__Saccharomycetales(100);f__Metschnikowiaceae(100);g__Metschnikowia(100);s__Metschnikowia_pulcherrima(100); | Metschnikowia | 13 | Saprotroph | \|Undefined Saprotroph\| | Yeast | NULL | Probable | As nectar-tap_saprotroph (Põlme et al. 2020); Decay sugar-rich substrates (Põlme et al. 2020) | Põlme S, et al. 2020. Fungal Diversity 105:1-16 ((https://doi.org/10.1007/s13225-020-00466-2)) |
| Otu37686 | 1 | k__Fungi(100);p__Ascomycota(100);c__Saccharomycetes(100);o__Saccharomycetales(100);f__Metschnikowiaceae(100);g__Metschnikowia(100);s__Metschnikowia_pulcherrima(100); | Metschnikowia | 13 | Saprotroph | \|Undefined Saprotroph\| | Yeast | NULL | Probable | As nectar-tap_saprotroph (Põlme et al. 2020); Decay sugar-rich substrates (Põlme et al. 2020) | Põlme S, et al. 2020. Fungal Diversity 105:1-16 ((https://doi.org/10.1007/s13225-020-00466-2)) |
| Otu37687 | 1 | k__Fungi(100);p__Ascomycota(100);c__Saccharomycetes(100);o__Saccharomycetales(100);f__Metschnikowiaceae(100);g__Metschnikowia(100);s__Metschnikowia_pulcherrima(100); | Metschnikowia | 13 | Saprotroph | \|Undefined Saprotroph\| | Yeast | NULL | Probable | As nectar-tap_saprotroph (Põlme et al. 2020); Decay sugar-rich substrates (Põlme et al. 2020) | Põlme S, et al. 2020. Fungal Diversity 105:1-16 ((https://doi.org/10.1007/s13225-020-00466-2)) |
| Otu37688 | 1 | k__Fungi(100);p__Ascomycota(100);c__Saccharomycetes(100);o__Saccharomycetales(100);f__Metschnikowiaceae(100);g__Metschnikowia(100);s__Metschnikowia_pulcherrima(100); | Metschnikowia | 13 | Saprotroph | \|Undefined Saprotroph\| | Yeast | NULL | Probable | As nectar-tap_saprotroph (Põlme et al. 2020); Decay sugar-rich substrates (Põlme et al. 2020) | Põlme S, et al. 2020. Fungal Diversity 105:1-16 ((https://doi.org/10.1007/s13225-020-00466-2)) |
| Otu37689 | 1 | k__Fungi(100);p__Ascomycota(100);c__Saccharomycetes(100);o__Saccharomycetales(100);f__Metschnikowiaceae(100);g__Metschnikowia(100);s__Metschnikowia_pulcherrima(100); | Metschnikowia | 13 | Saprotroph | \|Undefined Saprotroph\| | Yeast | NULL | Probable | As nectar-tap_saprotroph (Põlme et al. 2020); Decay sugar-rich substrates (Põlme et al. 2020) | Põlme S, et al. 2020. Fungal Diversity 105:1-16 ((https://doi.org/10.1007/s13225-020-00466-2)) |
| Otu37690 | 1 | k__Fungi(100);p__Ascomycota(100);c__Saccharomycetes(100);o__Saccharomycetales(100);f__Metschnikowiaceae(100);g__Metschnikowia(100);s__Metschnikowia_pulcherrima(100); | Metschnikowia | 13 | Saprotroph | \|Undefined Saprotroph\| | Yeast | NULL | Probable | As nectar-tap_saprotroph (Põlme et al. 2020); Decay sugar-rich substrates (Põlme et al. 2020) | Põlme S, et al. 2020. Fungal Diversity 105:1-16 ((https://doi.org/10.1007/s13225-020-00466-2)) |
| Otu37696 | 1 | k__Fungi(100);p__Ascomycota(100);c__Saccharomycetes(100);o__Saccharomycetales(100);f__Metschnikowiaceae(100);g__Metschnikowia(100);s__Metschnikowia_pulcherrima(100); | Metschnikowia | 13 | Saprotroph | \|Undefined Saprotroph\| | Yeast | NULL | Probable | As nectar-tap_saprotroph (Põlme et al. 2020); Decay sugar-rich substrates (Põlme et al. 2020) | Põlme S, et al. 2020. Fungal Diversity 105:1-16 ((https://doi.org/10.1007/s13225-020-00466-2)) |
| Otu37698 | 1 | k__Fungi(100);p__Ascomycota(100);c__Saccharomycetes(100);o__Saccharomycetales(100);f__Metschnikowiaceae(100);g__Metschnikowia(100);s__Metschnikowia_pulcherrima(100); | Metschnikowia | 13 | Saprotroph | \|Undefined Saprotroph\| | Yeast | NULL | Probable | As nectar-tap_saprotroph (Põlme et al. 2020); Decay sugar-rich substrates (Põlme et al. 2020) | Põlme S, et al. 2020. Fungal Diversity 105:1-16 ((https://doi.org/10.1007/s13225-020-00466-2)) |
| Otu37699 | 1 | k__Fungi(100);p__Ascomycota(100);c__Saccharomycetes(100);o__Saccharomycetales(100);f__Metschnikowiaceae(100);g__Metschnikowia(100);g__Metschnikowia_unclassified(100); | Metschnikowia | 13 | Saprotroph | \|Undefined Saprotroph\| | Yeast | NULL | Probable | As nectar-tap_saprotroph (Põlme et al. 2020); Decay sugar-rich substrates (Põlme et al. 2020) | Põlme S, et al. 2020. Fungal Diversity 105:1-16 ((https://doi.org/10.1007/s13225-020-00466-2)) |
| Otu37700 | 1 | k__Fungi(100);p__Ascomycota(100);c__Saccharomycetes(100);o__Saccharomycetales(100);f__Metschnikowiaceae(100);g__Metschnikowia(100);s__Metschnikowia_pulcherrima(100); | Metschnikowia | 13 | Saprotroph | \|Undefined Saprotroph\| | Yeast | NULL | Probable | As nectar-tap_saprotroph (Põlme et al. 2020); Decay sugar-rich substrates (Põlme et al. 2020) | Põlme S, et al. 2020. Fungal Diversity 105:1-16 ((https://doi.org/10.1007/s13225-020-00466-2)) |
| Otu37701 | 1 | k__Fungi(100);p__Ascomycota(100);c__Saccharomycetes(100);o__Saccharomycetales(100);f__Metschnikowiaceae(100);g__Metschnikowia(100);s__Metschnikowia_pulcherrima(100); | Metschnikowia | 13 | Saprotroph | \|Undefined Saprotroph\| | Yeast | NULL | Probable | As nectar-tap_saprotroph (Põlme et al. 2020); Decay sugar-rich substrates (Põlme et al. 2020) | Põlme S, et al. 2020. Fungal Diversity 105:1-16 ((https://doi.org/10.1007/s13225-020-00466-2)) |
| Otu37703 | 1 | k__Fungi(100);p__Ascomycota(100);c__Saccharomycetes(100);o__Saccharomycetales(100);f__Metschnikowiaceae(100);g__Metschnikowia(100);g__Metschnikowia_unclassified(100); | Metschnikowia | 13 | Saprotroph | \|Undefined Saprotroph\| | Yeast | NULL | Probable | As nectar-tap_saprotroph (Põlme et al. 2020); Decay sugar-rich substrates (Põlme et al. 2020) | Põlme S, et al. 2020. Fungal Diversity 105:1-16 ((https://doi.org/10.1007/s13225-020-00466-2)) |
| Otu37704 | 1 | k__Fungi(100);p__Ascomycota(100);c__Saccharomycetes(100);o__Saccharomycetales(100);f__Metschnikowiaceae(100);g__Metschnikowia(100);s__Metschnikowia_pulcherrima(100); | Metschnikowia | 13 | Saprotroph | \|Undefined Saprotroph\| | Yeast | NULL | Probable | As nectar-tap_saprotroph (Põlme et al. 2020); Decay sugar-rich substrates (Põlme et al. 2020) | Põlme S, et al. 2020. Fungal Diversity 105:1-16 ((https://doi.org/10.1007/s13225-020-00466-2)) |
| Otu37705 | 1 | k__Fungi(100);p__Ascomycota(100);c__Saccharomycetes(100);o__Saccharomycetales(100);f__Metschnikowiaceae(100);g__Metschnikowia(100);s__Metschnikowia_pulcherrima(100); | Metschnikowia | 13 | Saprotroph | \|Undefined Saprotroph\| | Yeast | NULL | Probable | As nectar-tap_saprotroph (Põlme et al. 2020); Decay sugar-rich substrates (Põlme et al. 2020) | Põlme S, et al. 2020. Fungal Diversity 105:1-16 ((https://doi.org/10.1007/s13225-020-00466-2)) |
| Otu37706 | 1 | k__Fungi(100);p__Ascomycota(100);c__Saccharomycetes(100);o__Saccharomycetales(100);f__Metschnikowiaceae(100);g__Metschnikowia(100);s__Metschnikowia_pulcherrima(100); | Metschnikowia | 13 | Saprotroph | \|Undefined Saprotroph\| | Yeast | NULL | Probable | As nectar-tap_saprotroph (Põlme et al. 2020); Decay sugar-rich substrates (Põlme et al. 2020) | Põlme S, et al. 2020. Fungal Diversity 105:1-16 ((https://doi.org/10.1007/s13225-020-00466-2)) |
| Otu37707 | 1 | k__Fungi(100);p__Ascomycota(100);c__Saccharomycetes(100);o__Saccharomycetales(100);f__Metschnikowiaceae(100);g__Metschnikowia(100);s__Metschnikowia_pulcherrima(100); | Metschnikowia | 13 | Saprotroph | \|Undefined Saprotroph\| | Yeast | NULL | Probable | As nectar-tap_saprotroph (Põlme et al. 2020); Decay sugar-rich substrates (Põlme et al. 2020) | Põlme S, et al. 2020. Fungal Diversity 105:1-16 ((https://doi.org/10.1007/s13225-020-00466-2)) |
| Otu37708 | 1 | k__Fungi(100);p__Ascomycota(100);c__Saccharomycetes(100);o__Saccharomycetales(100);f__Metschnikowiaceae(100);g__Metschnikowia(100);g__Metschnikowia_unclassified(100); | Metschnikowia | 13 | Saprotroph | \|Undefined Saprotroph\| | Yeast | NULL | Probable | As nectar-tap_saprotroph (Põlme et al. 2020); Decay sugar-rich substrates (Põlme et al. 2020) | Põlme S, et al. 2020. Fungal Diversity 105:1-16 ((https://doi.org/10.1007/s13225-020-00466-2)) |
| Otu37709 | 1 | k__Fungi(100);p__Ascomycota(100);c__Saccharomycetes(100);o__Saccharomycetales(100);f__Metschnikowiaceae(100);g__Metschnikowia(100);s__Metschnikowia_pulcherrima(100); | Metschnikowia | 13 | Saprotroph | \|Undefined Saprotroph\| | Yeast | NULL | Probable | As nectar-tap_saprotroph (Põlme et al. 2020); Decay sugar-rich substrates (Põlme et al. 2020) | Põlme S, et al. 2020. Fungal Diversity 105:1-16 ((https://doi.org/10.1007/s13225-020-00466-2)) |
| Otu37710 | 1 | k__Fungi(100);p__Ascomycota(100);c__Saccharomycetes(100);o__Saccharomycetales(100);f__Metschnikowiaceae(100);g__Metschnikowia(100);s__Metschnikowia_pulcherrima(100); | Metschnikowia | 13 | Saprotroph | \|Undefined Saprotroph\| | Yeast | NULL | Probable | As nectar-tap_saprotroph (Põlme et al. 2020); Decay sugar-rich substrates (Põlme et al. 2020) | Põlme S, et al. 2020. Fungal Diversity 105:1-16 ((https://doi.org/10.1007/s13225-020-00466-2)) |
| Otu37711 | 1 | k__Fungi(100);p__Ascomycota(100);c__Saccharomycetes(100);o__Saccharomycetales(100);f__Metschnikowiaceae(100);g__Metschnikowia(100);s__Metschnikowia_pulcherrima(100); | Metschnikowia | 13 | Saprotroph | \|Undefined Saprotroph\| | Yeast | NULL | Probable | As nectar-tap_saprotroph (Põlme et al. 2020); Decay sugar-rich substrates (Põlme et al. 2020) | Põlme S, et al. 2020. Fungal Diversity 105:1-16 ((https://doi.org/10.1007/s13225-020-00466-2)) |
| Otu37712 | 1 | k__Fungi(100);p__Ascomycota(100);c__Saccharomycetes(100);o__Saccharomycetales(100);f__Metschnikowiaceae(100);g__Metschnikowia(100);g__Metschnikowia_unclassified(100); | Metschnikowia | 13 | Saprotroph | \|Undefined Saprotroph\| | Yeast | NULL | Probable | As nectar-tap_saprotroph (Põlme et al. 2020); Decay sugar-rich substrates (Põlme et al. 2020) | Põlme S, et al. 2020. Fungal Diversity 105:1-16 ((https://doi.org/10.1007/s13225-020-00466-2)) |
| Otu37713 | 1 | k__Fungi(100);p__Ascomycota(100);c__Saccharomycetes(100);o__Saccharomycetales(100);f__Metschnikowiaceae(100);g__Metschnikowia(100);s__Metschnikowia_pulcherrima(100); | Metschnikowia | 13 | Saprotroph | \|Undefined Saprotroph\| | Yeast | NULL | Probable | As nectar-tap_saprotroph (Põlme et al. 2020); Decay sugar-rich substrates (Põlme et al. 2020) | Põlme S, et al. 2020. Fungal Diversity 105:1-16 ((https://doi.org/10.1007/s13225-020-00466-2)) |
| Otu37714 | 1 | k__Fungi(100);p__Ascomycota(100);c__Saccharomycetes(100);o__Saccharomycetales(100);f__Metschnikowiaceae(100);g__Metschnikowia(100);s__Metschnikowia_pulcherrima(100); | Metschnikowia | 13 | Saprotroph | \|Undefined Saprotroph\| | Yeast | NULL | Probable | As nectar-tap_saprotroph (Põlme et al. 2020); Decay sugar-rich substrates (Põlme et al. 2020) | Põlme S, et al. 2020. Fungal Diversity 105:1-16 ((https://doi.org/10.1007/s13225-020-00466-2)) |
| Otu37715 | 1 | k__Fungi(100);p__Ascomycota(100);c__Saccharomycetes(100);o__Saccharomycetales(100);f__Metschnikowiaceae(100);g__Metschnikowia(100);s__Metschnikowia_pulcherrima(100); | Metschnikowia | 13 | Saprotroph | \|Undefined Saprotroph\| | Yeast | NULL | Probable | As nectar-tap_saprotroph (Põlme et al. 2020); Decay sugar-rich substrates (Põlme et al. 2020) | Põlme S, et al. 2020. Fungal Diversity 105:1-16 ((https://doi.org/10.1007/s13225-020-00466-2)) |
| Otu37716 | 1 | k__Fungi(100);p__Ascomycota(100);c__Saccharomycetes(100);o__Saccharomycetales(100);f__Metschnikowiaceae(100);g__Metschnikowia(100);s__Metschnikowia_pulcherrima(100); | Metschnikowia | 13 | Saprotroph | \|Undefined Saprotroph\| | Yeast | NULL | Probable | As nectar-tap_saprotroph (Põlme et al. 2020); Decay sugar-rich substrates (Põlme et al. 2020) | Põlme S, et al. 2020. Fungal Diversity 105:1-16 ((https://doi.org/10.1007/s13225-020-00466-2)) |
| Otu37717 | 1 | k__Fungi(100);p__Ascomycota(100);c__Saccharomycetes(100);o__Saccharomycetales(100);f__Metschnikowiaceae(100);g__Metschnikowia(100);s__Metschnikowia_pulcherrima(100); | Metschnikowia | 13 | Saprotroph | \|Undefined Saprotroph\| | Yeast | NULL | Probable | As nectar-tap_saprotroph (Põlme et al. 2020); Decay sugar-rich substrates (Põlme et al. 2020) | Põlme S, et al. 2020. Fungal Diversity 105:1-16 ((https://doi.org/10.1007/s13225-020-00466-2)) |
| Otu37718 | 1 | k__Fungi(100);p__Ascomycota(100);c__Saccharomycetes(100);o__Saccharomycetales(100);f__Metschnikowiaceae(100);g__Metschnikowia(100);s__Metschnikowia_pulcherrima(100); | Metschnikowia | 13 | Saprotroph | \|Undefined Saprotroph\| | Yeast | NULL | Probable | As nectar-tap_saprotroph (Põlme et al. 2020); Decay sugar-rich substrates (Põlme et al. 2020) | Põlme S, et al. 2020. Fungal Diversity 105:1-16 ((https://doi.org/10.1007/s13225-020-00466-2)) |
| Otu37719 | 1 | k__Fungi(100);p__Ascomycota(100);c__Saccharomycetes(100);o__Saccharomycetales(100);f__Metschnikowiaceae(100);g__Metschnikowia(100);s__Metschnikowia_pulcherrima(100); | Metschnikowia | 13 | Saprotroph | \|Undefined Saprotroph\| | Yeast | NULL | Probable | As nectar-tap_saprotroph (Põlme et al. 2020); Decay sugar-rich substrates (Põlme et al. 2020) | Põlme S, et al. 2020. Fungal Diversity 105:1-16 ((https://doi.org/10.1007/s13225-020-00466-2)) |
| Otu37721 | 1 | k__Fungi(100);p__Ascomycota(100);c__Saccharomycetes(100);o__Saccharomycetales(100);f__Metschnikowiaceae(100);g__Metschnikowia(100);s__Metschnikowia_pulcherrima(100); | Metschnikowia | 13 | Saprotroph | \|Undefined Saprotroph\| | Yeast | NULL | Probable | As nectar-tap_saprotroph (Põlme et al. 2020); Decay sugar-rich substrates (Põlme et al. 2020) | Põlme S, et al. 2020. Fungal Diversity 105:1-16 ((https://doi.org/10.1007/s13225-020-00466-2)) |
| Otu37722 | 1 | k__Fungi(100);p__Ascomycota(100);c__Saccharomycetes(100);o__Saccharomycetales(100);f__Metschnikowiaceae(100);g__Metschnikowia(100);s__Metschnikowia_pulcherrima(100); | Metschnikowia | 13 | Saprotroph | \|Undefined Saprotroph\| | Yeast | NULL | Probable | As nectar-tap_saprotroph (Põlme et al. 2020); Decay sugar-rich substrates (Põlme et al. 2020) | Põlme S, et al. 2020. Fungal Diversity 105:1-16 ((https://doi.org/10.1007/s13225-020-00466-2)) |
| Otu37723 | 1 | k__Fungi(100);p__Ascomycota(100);c__Saccharomycetes(100);o__Saccharomycetales(100);f__Metschnikowiaceae(100);g__Metschnikowia(100);s__Metschnikowia_pulcherrima(100); | Metschnikowia | 13 | Saprotroph | \|Undefined Saprotroph\| | Yeast | NULL | Probable | As nectar-tap_saprotroph (Põlme et al. 2020); Decay sugar-rich substrates (Põlme et al. 2020) | Põlme S, et al. 2020. Fungal Diversity 105:1-16 ((https://doi.org/10.1007/s13225-020-00466-2)) |
| Otu37724 | 1 | k__Fungi(100);p__Ascomycota(100);c__Saccharomycetes(100);o__Saccharomycetales(100);f__Metschnikowiaceae(100);g__Metschnikowia(100);s__Metschnikowia_pulcherrima(100); | Metschnikowia | 13 | Saprotroph | \|Undefined Saprotroph\| | Yeast | NULL | Probable | As nectar-tap_saprotroph (Põlme et al. 2020); Decay sugar-rich substrates (Põlme et al. 2020) | Põlme S, et al. 2020. Fungal Diversity 105:1-16 ((https://doi.org/10.1007/s13225-020-00466-2)) |
| Otu37725 | 1 | k__Fungi(100);p__Ascomycota(100);c__Saccharomycetes(100);o__Saccharomycetales(100);f__Metschnikowiaceae(100);g__Metschnikowia(100);g__Metschnikowia_unclassified(100); | Metschnikowia | 13 | Saprotroph | \|Undefined Saprotroph\| | Yeast | NULL | Probable | As nectar-tap_saprotroph (Põlme et al. 2020); Decay sugar-rich substrates (Põlme et al. 2020) | Põlme S, et al. 2020. Fungal Diversity 105:1-16 ((https://doi.org/10.1007/s13225-020-00466-2)) |
| Otu37726 | 1 | k__Fungi(100);p__Ascomycota(100);c__Saccharomycetes(100);o__Saccharomycetales(100);f__Metschnikowiaceae(100);g__Metschnikowia(100);g__Metschnikowia_unclassified(100); | Metschnikowia | 13 | Saprotroph | \|Undefined Saprotroph\| | Yeast | NULL | Probable | As nectar-tap_saprotroph (Põlme et al. 2020); Decay sugar-rich substrates (Põlme et al. 2020) | Põlme S, et al. 2020. Fungal Diversity 105:1-16 ((https://doi.org/10.1007/s13225-020-00466-2)) |
| Otu37727 | 1 | k__Fungi(100);p__Ascomycota(100);c__Saccharomycetes(100);o__Saccharomycetales(100);f__Metschnikowiaceae(100);g__Metschnikowia(100);s__Metschnikowia_pulcherrima(100); | Metschnikowia | 13 | Saprotroph | \|Undefined Saprotroph\| | Yeast | NULL | Probable | As nectar-tap_saprotroph (Põlme et al. 2020); Decay sugar-rich substrates (Põlme et al. 2020) | Põlme S, et al. 2020. Fungal Diversity 105:1-16 ((https://doi.org/10.1007/s13225-020-00466-2)) |
| Otu37728 | 1 | k__Fungi(100);p__Ascomycota(100);c__Saccharomycetes(100);o__Saccharomycetales(100);f__Metschnikowiaceae(100);g__Metschnikowia(100);g__Metschnikowia_unclassified(100); | Metschnikowia | 13 | Saprotroph | \|Undefined Saprotroph\| | Yeast | NULL | Probable | As nectar-tap_saprotroph (Põlme et al. 2020); Decay sugar-rich substrates (Põlme et al. 2020) | Põlme S, et al. 2020. Fungal Diversity 105:1-16 ((https://doi.org/10.1007/s13225-020-00466-2)) |
| Otu37729 | 1 | k__Fungi(100);p__Ascomycota(100);c__Saccharomycetes(100);o__Saccharomycetales(100);f__Metschnikowiaceae(100);g__Metschnikowia(100);g__Metschnikowia_unclassified(100); | Metschnikowia | 13 | Saprotroph | \|Undefined Saprotroph\| | Yeast | NULL | Probable | As nectar-tap_saprotroph (Põlme et al. 2020); Decay sugar-rich substrates (Põlme et al. 2020) | Põlme S, et al. 2020. Fungal Diversity 105:1-16 ((https://doi.org/10.1007/s13225-020-00466-2)) |
| Otu37730 | 1 | k__Fungi(100);p__Ascomycota(100);c__Saccharomycetes(100);o__Saccharomycetales(100);f__Metschnikowiaceae(100);g__Metschnikowia(100);s__Metschnikowia_pulcherrima(100); | Metschnikowia | 13 | Saprotroph | \|Undefined Saprotroph\| | Yeast | NULL | Probable | As nectar-tap_saprotroph (Põlme et al. 2020); Decay sugar-rich substrates (Põlme et al. 2020) | Põlme S, et al. 2020. Fungal Diversity 105:1-16 ((https://doi.org/10.1007/s13225-020-00466-2)) |
| Otu37732 | 1 | k__Fungi(100);p__Ascomycota(100);c__Saccharomycetes(100);o__Saccharomycetales(100);f__Metschnikowiaceae(100);g__Metschnikowia(100);s__Metschnikowia_pulcherrima(100); | Metschnikowia | 13 | Saprotroph | \|Undefined Saprotroph\| | Yeast | NULL | Probable | As nectar-tap_saprotroph (Põlme et al. 2020); Decay sugar-rich substrates (Põlme et al. 2020) | Põlme S, et al. 2020. Fungal Diversity 105:1-16 ((https://doi.org/10.1007/s13225-020-00466-2)) |
| Otu37733 | 1 | k__Fungi(100);p__Ascomycota(100);c__Saccharomycetes(100);o__Saccharomycetales(100);f__Metschnikowiaceae(100);g__Metschnikowia(100);g__Metschnikowia_unclassified(100); | Metschnikowia | 13 | Saprotroph | \|Undefined Saprotroph\| | Yeast | NULL | Probable | As nectar-tap_saprotroph (Põlme et al. 2020); Decay sugar-rich substrates (Põlme et al. 2020) | Põlme S, et al. 2020. Fungal Diversity 105:1-16 ((https://doi.org/10.1007/s13225-020-00466-2)) |
| Otu37734 | 1 | k__Fungi(100);p__Ascomycota(100);c__Saccharomycetes(100);o__Saccharomycetales(100);f__Metschnikowiaceae(100);g__Metschnikowia(100);s__Metschnikowia_sp(100); | Metschnikowia | 13 | Saprotroph | \|Undefined Saprotroph\| | Yeast | NULL | Probable | As nectar-tap_saprotroph (Põlme et al. 2020); Decay sugar-rich substrates (Põlme et al. 2020) | Põlme S, et al. 2020. Fungal Diversity 105:1-16 ((https://doi.org/10.1007/s13225-020-00466-2)) |
| Otu37735 | 1 | k__Fungi(100);p__Ascomycota(100);c__Sordariomycetes(100);o__Chaetosphaeriales(100);f__Chaetosphaeriaceae(100);f__Chaetosphaeriaceae_unclassified(100);f__Chaetosphaeriaceae_unclassified(100); | Chaetosphaeriaceae | 9 | Saprotroph | Plant Saprotroph-Wood Saprotroph | Microfungus | NULL | Probable | Saprobes associated with decaying woody and herbaceous plant material, including timber and timber products in damp buildings (Cannon & Kirk 2007) | Cannon PF, Kirk PM. 2007. Fungal Families of the World. CAB International, Cambridge (ISBN: 978-0851998275) |
| Otu37736 | 1 | k__Fungi(100);p__Ascomycota(100);c__Saccharomycetes(100);o__Saccharomycetales(100);f__Metschnikowiaceae(100);g__Metschnikowia(100);s__Metschnikowia_pulcherrima(100); | Metschnikowia | 13 | Saprotroph | \|Undefined Saprotroph\| | Yeast | NULL | Probable | As nectar-tap_saprotroph (Põlme et al. 2020); Decay sugar-rich substrates (Põlme et al. 2020) | Põlme S, et al. 2020. Fungal Diversity 105:1-16 ((https://doi.org/10.1007/s13225-020-00466-2)) |
| Otu37737 | 1 | k__Fungi(100);p__Ascomycota(100);c__Saccharomycetes(100);o__Saccharomycetales(100);f__Metschnikowiaceae(100);g__Metschnikowia(100);g__Metschnikowia_unclassified(100); | Metschnikowia | 13 | Saprotroph | \|Undefined Saprotroph\| | Yeast | NULL | Probable | As nectar-tap_saprotroph (Põlme et al. 2020); Decay sugar-rich substrates (Põlme et al. 2020) | Põlme S, et al. 2020. Fungal Diversity 105:1-16 ((https://doi.org/10.1007/s13225-020-00466-2)) |
| Otu37738 | 1 | k__Fungi(100);p__Ascomycota(100);c__Saccharomycetes(100);o__Saccharomycetales(100);f__Metschnikowiaceae(100);g__Metschnikowia(100);s__Metschnikowia_pulcherrima(100); | Metschnikowia | 13 | Saprotroph | \|Undefined Saprotroph\| | Yeast | NULL | Probable | As nectar-tap_saprotroph (Põlme et al. 2020); Decay sugar-rich substrates (Põlme et al. 2020) | Põlme S, et al. 2020. Fungal Diversity 105:1-16 ((https://doi.org/10.1007/s13225-020-00466-2)) |
| Otu37739 | 1 | k__Fungi(100);p__Ascomycota(100);c__Saccharomycetes(100);o__Saccharomycetales(100);f__Metschnikowiaceae(100);g__Metschnikowia(100);g__Metschnikowia_unclassified(100); | Metschnikowia | 13 | Saprotroph | \|Undefined Saprotroph\| | Yeast | NULL | Probable | As nectar-tap_saprotroph (Põlme et al. 2020); Decay sugar-rich substrates (Põlme et al. 2020) | Põlme S, et al. 2020. Fungal Diversity 105:1-16 ((https://doi.org/10.1007/s13225-020-00466-2)) |
| Otu37740 | 1 | k__Fungi(100);p__Ascomycota(100);c__Saccharomycetes(100);o__Saccharomycetales(100);f__Metschnikowiaceae(100);g__Metschnikowia(100);g__Metschnikowia_unclassified(100); | Metschnikowia | 13 | Saprotroph | \|Undefined Saprotroph\| | Yeast | NULL | Probable | As nectar-tap_saprotroph (Põlme et al. 2020); Decay sugar-rich substrates (Põlme et al. 2020) | Põlme S, et al. 2020. Fungal Diversity 105:1-16 ((https://doi.org/10.1007/s13225-020-00466-2)) |
| Otu37741 | 1 | k__Fungi(100);p__Ascomycota(100);c__Saccharomycetes(100);o__Saccharomycetales(100);f__Metschnikowiaceae(100);g__Metschnikowia(100);g__Metschnikowia_unclassified(100); | Metschnikowia | 13 | Saprotroph | \|Undefined Saprotroph\| | Yeast | NULL | Probable | As nectar-tap_saprotroph (Põlme et al. 2020); Decay sugar-rich substrates (Põlme et al. 2020) | Põlme S, et al. 2020. Fungal Diversity 105:1-16 ((https://doi.org/10.1007/s13225-020-00466-2)) |
| Otu37742 | 1 | k__Fungi(100);p__Ascomycota(100);c__Saccharomycetes(100);o__Saccharomycetales(100);f__Metschnikowiaceae(100);g__Metschnikowia(100);g__Metschnikowia_unclassified(100); | Metschnikowia | 13 | Saprotroph | \|Undefined Saprotroph\| | Yeast | NULL | Probable | As nectar-tap_saprotroph (Põlme et al. 2020); Decay sugar-rich substrates (Põlme et al. 2020) | Põlme S, et al. 2020. Fungal Diversity 105:1-16 ((https://doi.org/10.1007/s13225-020-00466-2)) |
| Otu37743 | 1 | k__Fungi(100);p__Ascomycota(100);c__Saccharomycetes(100);o__Saccharomycetales(100);f__Metschnikowiaceae(100);g__Metschnikowia(100);g__Metschnikowia_unclassified(100); | Metschnikowia | 13 | Saprotroph | \|Undefined Saprotroph\| | Yeast | NULL | Probable | As nectar-tap_saprotroph (Põlme et al. 2020); Decay sugar-rich substrates (Põlme et al. 2020) | Põlme S, et al. 2020. Fungal Diversity 105:1-16 ((https://doi.org/10.1007/s13225-020-00466-2)) |
| Otu37744 | 1 | k__Fungi(100);p__Ascomycota(100);c__Saccharomycetes(100);o__Saccharomycetales(100);f__Metschnikowiaceae(100);g__Metschnikowia(100);s__Metschnikowia_pulcherrima(100); | Metschnikowia | 13 | Saprotroph | \|Undefined Saprotroph\| | Yeast | NULL | Probable | As nectar-tap_saprotroph (Põlme et al. 2020); Decay sugar-rich substrates (Põlme et al. 2020) | Põlme S, et al. 2020. Fungal Diversity 105:1-16 ((https://doi.org/10.1007/s13225-020-00466-2)) |
| Otu37745 | 1 | k__Fungi(100);p__Ascomycota(100);c__Saccharomycetes(100);o__Saccharomycetales(100);f__Metschnikowiaceae(100);g__Metschnikowia(100);s__Metschnikowia_pulcherrima(100); | Metschnikowia | 13 | Saprotroph | \|Undefined Saprotroph\| | Yeast | NULL | Probable | As nectar-tap_saprotroph (Põlme et al. 2020); Decay sugar-rich substrates (Põlme et al. 2020) | Põlme S, et al. 2020. Fungal Diversity 105:1-16 ((https://doi.org/10.1007/s13225-020-00466-2)) |
| Otu37746 | 1 | k__Fungi(100);p__Ascomycota(100);c__Saccharomycetes(100);o__Saccharomycetales(100);f__Metschnikowiaceae(100);g__Metschnikowia(100);s__Metschnikowia_pulcherrima(100); | Metschnikowia | 13 | Saprotroph | \|Undefined Saprotroph\| | Yeast | NULL | Probable | As nectar-tap_saprotroph (Põlme et al. 2020); Decay sugar-rich substrates (Põlme et al. 2020) | Põlme S, et al. 2020. Fungal Diversity 105:1-16 ((https://doi.org/10.1007/s13225-020-00466-2)) |
| Otu37747 | 1 | k__Fungi(100);p__Ascomycota(100);c__Saccharomycetes(100);o__Saccharomycetales(100);f__Metschnikowiaceae(100);g__Metschnikowia(100);s__Metschnikowia_pulcherrima(100); | Metschnikowia | 13 | Saprotroph | \|Undefined Saprotroph\| | Yeast | NULL | Probable | As nectar-tap_saprotroph (Põlme et al. 2020); Decay sugar-rich substrates (Põlme et al. 2020) | Põlme S, et al. 2020. Fungal Diversity 105:1-16 ((https://doi.org/10.1007/s13225-020-00466-2)) |
| Otu37748 | 1 | k__Fungi(100);p__Ascomycota(100);c__Saccharomycetes(100);o__Saccharomycetales(100);f__Metschnikowiaceae(100);g__Metschnikowia(100);s__Metschnikowia_pulcherrima(100); | Metschnikowia | 13 | Saprotroph | \|Undefined Saprotroph\| | Yeast | NULL | Probable | As nectar-tap_saprotroph (Põlme et al. 2020); Decay sugar-rich substrates (Põlme et al. 2020) | Põlme S, et al. 2020. Fungal Diversity 105:1-16 ((https://doi.org/10.1007/s13225-020-00466-2)) |
| Otu37749 | 1 | k__Fungi(100);p__Ascomycota(100);c__Saccharomycetes(100);o__Saccharomycetales(100);f__Metschnikowiaceae(100);g__Metschnikowia(100);g__Metschnikowia_unclassified(100); | Metschnikowia | 13 | Saprotroph | \|Undefined Saprotroph\| | Yeast | NULL | Probable | As nectar-tap_saprotroph (Põlme et al. 2020); Decay sugar-rich substrates (Põlme et al. 2020) | Põlme S, et al. 2020. Fungal Diversity 105:1-16 ((https://doi.org/10.1007/s13225-020-00466-2)) |

| | | | | | | | | | | | |
|---|---|---|---|---|---|---|---|---|---|---|---|
| Otu37750 | 1 | k__Fungi(100);p__Ascomycota(100);c__Saccharomycetes(100);o__Saccharomycetales(100);f__Metschnikowiaceae(100);g__Metschnikowia(100);g__Metschnikowia_unclassified(100); | Metschnikowia | 13 | Saprotroph | \|Undefined Saprotroph\| | Yeast | NULL | Probable | As nectar-tap_saprotroph (Põlme et al. 2020); Decay sugar-rich substrates (Põlme et al. 2020) | Põlme S, et al. 2020. Fungal Diversity 105:1-16 ((https://doi.org/10.1007/s13225-020-00466-2)) |
| Otu37751 | 1 | k__Fungi(100);p__Ascomycota(100);c__Saccharomycetes(100);o__Saccharomycetales(100);f__Metschnikowiaceae(100);g__Metschnikowia(100);s__Metschnikowia_pulcherrima(100); | Metschnikowia | 13 | Saprotroph | \|Undefined Saprotroph\| | Yeast | NULL | Probable | As nectar-tap_saprotroph (Põlme et al. 2020); Decay sugar-rich substrates (Põlme et al. 2020) | Põlme S, et al. 2020. Fungal Diversity 105:1-16 ((https://doi.org/10.1007/s13225-020-00466-2)) |
| Otu37752 | 1 | k__Fungi(100);p__Ascomycota(100);c__Saccharomycetes(100);o__Saccharomycetales(100);f__Metschnikowiaceae(100);g__Metschnikowia(100);s__Metschnikowia_pulcherrima(100); | Metschnikowia | 13 | Saprotroph | \|Undefined Saprotroph\| | Yeast | NULL | Probable | As nectar-tap_saprotroph (Põlme et al. 2020); Decay sugar-rich substrates (Põlme et al. 2020) | Põlme S, et al. 2020. Fungal Diversity 105:1-16 ((https://doi.org/10.1007/s13225-020-00466-2)) |
| Otu37753 | 1 | k__Fungi(100);p__Ascomycota(100);c__Saccharomycetes(100);o__Saccharomycetales(100);f__Metschnikowiaceae(100);g__Metschnikowia(100);g__Metschnikowia_unclassified(100); | Metschnikowia | 13 | Saprotroph | \|Undefined Saprotroph\| | Yeast | NULL | Probable | As nectar-tap_saprotroph (Põlme et al. 2020); Decay sugar-rich substrates (Põlme et al. 2020) | Põlme S, et al. 2020. Fungal Diversity 105:1-16 ((https://doi.org/10.1007/s13225-020-00466-2)) |
| Otu37754 | 1 | k__Fungi(100);p__Ascomycota(100);c__Saccharomycetes(100);o__Saccharomycetales(100);f__Metschnikowiaceae(100);g__Metschnikowia(100);s__Metschnikowia_pulcherrima(100); | Metschnikowia | 13 | Saprotroph | \|Undefined Saprotroph\| | Yeast | NULL | Probable | As nectar-tap_saprotroph (Põlme et al. 2020); Decay sugar-rich substrates (Põlme et al. 2020) | Põlme S, et al. 2020. Fungal Diversity 105:1-16 ((https://doi.org/10.1007/s13225-020-00466-2)) |
| Otu37755 | 1 | k__Fungi(100);p__Ascomycota(100);c__Saccharomycetes(100);o__Saccharomycetales(100);f__Metschnikowiaceae(100);g__Metschnikowia(100);s__Metschnikowia_pulcherrima(100); | Metschnikowia | 13 | Saprotroph | \|Undefined Saprotroph\| | Yeast | NULL | Probable | As nectar-tap_saprotroph (Põlme et al. 2020); Decay sugar-rich substrates (Põlme et al. 2020) | Põlme S, et al. 2020. Fungal Diversity 105:1-16 ((https://doi.org/10.1007/s13225-020-00466-2)) |
| Otu37756 | 1 | k__Fungi(100);p__Ascomycota(100);c__Saccharomycetes(100);o__Saccharomycetales(100);f__Metschnikowiaceae(100);g__Metschnikowia(100);s__Metschnikowia_pulcherrima(100); | Metschnikowia | 13 | Saprotroph | \|Undefined Saprotroph\| | Yeast | NULL | Probable | As nectar-tap_saprotroph (Põlme et al. 2020); Decay sugar-rich substrates (Põlme et al. 2020) | Põlme S, et al. 2020. Fungal Diversity 105:1-16 ((https://doi.org/10.1007/s13225-020-00466-2)) |
| Otu37757 | 1 | k__Fungi(100);p__Ascomycota(100);c__Saccharomycetes(100);o__Saccharomycetales(100);f__Metschnikowiaceae(100);g__Metschnikowia(100);s__Metschnikowia_pulcherrima(100); | Metschnikowia | 13 | Saprotroph | \|Undefined Saprotroph\| | Yeast | NULL | Probable | As nectar-tap_saprotroph (Põlme et al. 2020); Decay sugar-rich substrates (Põlme et al. 2020) | Põlme S, et al. 2020. Fungal Diversity 105:1-16 ((https://doi.org/10.1007/s13225-020-00466-2)) |
| Otu37758 | 1 | k__Fungi(100);p__Ascomycota(100);c__Saccharomycetes(100);o__Saccharomycetales(100);f__Metschnikowiaceae(100);g__Metschnikowia(100);g__Metschnikowia_unclassified(100); | Metschnikowia | 13 | Saprotroph | \|Undefined Saprotroph\| | Yeast | NULL | Probable | As nectar-tap_saprotroph (Põlme et al. 2020); Decay sugar-rich substrates (Põlme et al. 2020) | Põlme S, et al. 2020. Fungal Diversity 105:1-16 ((https://doi.org/10.1007/s13225-020-00466-2)) |
| Otu37759 | 1 | k__Fungi(100);p__Ascomycota(100);c__Saccharomycetes(100);o__Saccharomycetales(100);f__Metschnikowiaceae(100);g__Metschnikowia(100);s__Metschnikowia_pulcherrima(100); | Metschnikowia | 13 | Saprotroph | \|Undefined Saprotroph\| | Yeast | NULL | Probable | As nectar-tap_saprotroph (Põlme et al. 2020); Decay sugar-rich substrates (Põlme et al. 2020) | Põlme S, et al. 2020. Fungal Diversity 105:1-16 ((https://doi.org/10.1007/s13225-020-00466-2)) |
| Otu37760 | 1 | k__Fungi(100);p__Ascomycota(100);c__Saccharomycetes(100);o__Saccharomycetales(100);f__Metschnikowiaceae(100);g__Metschnikowia(100);g__Metschnikowia_unclassified(100); | Metschnikowia | 13 | Saprotroph | \|Undefined Saprotroph\| | Yeast | NULL | Probable | As nectar-tap_saprotroph (Põlme et al. 2020); Decay sugar-rich substrates (Põlme et al. 2020) | Põlme S, et al. 2020. Fungal Diversity 105:1-16 ((https://doi.org/10.1007/s13225-020-00466-2)) |
| Otu37761 | 1 | k__Fungi(100);p__Ascomycota(100);c__Saccharomycetes(100);o__Saccharomycetales(100);f__Metschnikowiaceae(100);g__Metschnikowia(100);g__Metschnikowia_unclassified(100); | Metschnikowia | 13 | Saprotroph | \|Undefined Saprotroph\| | Yeast | NULL | Probable | As nectar-tap_saprotroph (Põlme et al. 2020); Decay sugar-rich substrates (Põlme et al. 2020) | Põlme S, et al. 2020. Fungal Diversity 105:1-16 ((https://doi.org/10.1007/s13225-020-00466-2)) |
| Otu37762 | 1 | k__Fungi(100);p__Ascomycota(100);c__Saccharomycetes(100);o__Saccharomycetales(100);f__Metschnikowiaceae(100);g__Metschnikowia(100);s__Metschnikowia_pulcherrima(100); | Metschnikowia | 13 | Saprotroph | \|Undefined Saprotroph\| | Yeast | NULL | Probable | As nectar-tap_saprotroph (Põlme et al. 2020); Decay sugar-rich substrates (Põlme et al. 2020) | Põlme S, et al. 2020. Fungal Diversity 105:1-16 ((https://doi.org/10.1007/s13225-020-00466-2)) |
| Otu37763 | 1 | k__Fungi(100);p__Ascomycota(100);c__Saccharomycetes(100);o__Saccharomycetales(100);f__Metschnikowiaceae(100);g__Metschnikowia(100);s__Metschnikowia_pulcherrima(100); | Metschnikowia | 13 | Saprotroph | \|Undefined Saprotroph\| | Yeast | NULL | Probable | As nectar-tap_saprotroph (Põlme et al. 2020); Decay sugar-rich substrates (Põlme et al. 2020) | Põlme S, et al. 2020. Fungal Diversity 105:1-16 ((https://doi.org/10.1007/s13225-020-00466-2)) |
| Otu37764 | 1 | k__Fungi(100);p__Ascomycota(100);c__Saccharomycetes(100);o__Saccharomycetales(100);f__Metschnikowiaceae(100);g__Metschnikowia(100);s__Metschnikowia_pulcherrima(100); | Metschnikowia | 13 | Saprotroph | \|Undefined Saprotroph\| | Yeast | NULL | Probable | As nectar-tap_saprotroph (Põlme et al. 2020); Decay sugar-rich substrates (Põlme et al. 2020) | Põlme S, et al. 2020. Fungal Diversity 105:1-16 ((https://doi.org/10.1007/s13225-020-00466-2)) |
| Otu37765 | 1 | k__Fungi(100);p__Ascomycota(100);c__Saccharomycetes(100);o__Saccharomycetales(100);f__Metschnikowiaceae(100);g__Metschnikowia(100);s__Metschnikowia_pulcherrima(100); | Metschnikowia | 13 | Saprotroph | \|Undefined Saprotroph\| | Yeast | NULL | Probable | As nectar-tap_saprotroph (Põlme et al. 2020); Decay sugar-rich substrates (Põlme et al. 2020) | Põlme S, et al. 2020. Fungal Diversity 105:1-16 ((https://doi.org/10.1007/s13225-020-00466-2)) |
| Otu37766 | 1 | k__Fungi(100);p__Ascomycota(100);c__Saccharomycetes(100);o__Saccharomycetales(100);f__Metschnikowiaceae(100);g__Metschnikowia(100);s__Metschnikowia_pulcherrima(100); | Metschnikowia | 13 | Saprotroph | \|Undefined Saprotroph\| | Yeast | NULL | Probable | As nectar-tap_saprotroph (Põlme et al. 2020); Decay sugar-rich substrates (Põlme et al. 2020) | Põlme S, et al. 2020. Fungal Diversity 105:1-16 ((https://doi.org/10.1007/s13225-020-00466-2)) |
| Otu37767 | 1 | k__Fungi(100);p__Ascomycota(100);c__Saccharomycetes(100);o__Saccharomycetales(100);f__Metschnikowiaceae(100);g__Metschnikowia(100);s__Metschnikowia_pulcherrima(100); | Metschnikowia | 13 | Saprotroph | \|Undefined Saprotroph\| | Yeast | NULL | Probable | As nectar-tap_saprotroph (Põlme et al. 2020); Decay sugar-rich substrates (Põlme et al. 2020) | Põlme S, et al. 2020. Fungal Diversity 105:1-16 ((https://doi.org/10.1007/s13225-020-00466-2)) |
| Otu37768 | 1 | k__Fungi(100);p__Ascomycota(100);c__Saccharomycetes(100);o__Saccharomycetales(100);f__Metschnikowiaceae(100);g__Metschnikowia(100);s__Metschnikowia_pulcherrima(100); | Metschnikowia | 13 | Saprotroph | \|Undefined Saprotroph\| | Yeast | NULL | Probable | As nectar-tap_saprotroph (Põlme et al. 2020); Decay sugar-rich substrates (Põlme et al. 2020) | Põlme S, et al. 2020. Fungal Diversity 105:1-16 ((https://doi.org/10.1007/s13225-020-00466-2)) |
| Otu37769 | 1 | k__Fungi(100);p__Ascomycota(100);c__Saccharomycetes(100);o__Saccharomycetales(100);f__Metschnikowiaceae(100);g__Metschnikowia(100);s__Metschnikowia_pulcherrima(100); | Metschnikowia | 13 | Saprotroph | \|Undefined Saprotroph\| | Yeast | NULL | Probable | As nectar-tap_saprotroph (Põlme et al. 2020); Decay sugar-rich substrates (Põlme et al. 2020) | Põlme S, et al. 2020. Fungal Diversity 105:1-16 ((https://doi.org/10.1007/s13225-020-00466-2)) |
| Otu37770 | 1 | k__Fungi(100);p__Ascomycota(100);c__Saccharomycetes(100);o__Saccharomycetales(100);f__Metschnikowiaceae(100);g__Metschnikowia(100);s__Metschnikowia_pulcherrima(100); | Metschnikowia | 13 | Saprotroph | \|Undefined Saprotroph\| | Yeast | NULL | Probable | As nectar-tap_saprotroph (Põlme et al. 2020); Decay sugar-rich substrates (Põlme et al. 2020) | Põlme S, et al. 2020. Fungal Diversity 105:1-16 ((https://doi.org/10.1007/s13225-020-00466-2)) |
| Otu37771 | 1 | k__Fungi(100);p__Ascomycota(100);c__Saccharomycetes(100);o__Saccharomycetales(100);f__Metschnikowiaceae(100);g__Metschnikowia(100);g__Metschnikowia_unclassified(100); | Metschnikowia | 13 | Saprotroph | \|Undefined Saprotroph\| | Yeast | NULL | Probable | As nectar-tap_saprotroph (Põlme et al. 2020); Decay sugar-rich substrates (Põlme et al. 2020) | Põlme S, et al. 2020. Fungal Diversity 105:1-16 ((https://doi.org/10.1007/s13225-020-00466-2)) |
| Otu37772 | 1 | k__Fungi(100);p__Ascomycota(100);c__Saccharomycetes(100);o__Saccharomycetales(100);f__Metschnikowiaceae(100);g__Metschnikowia(100);g__Metschnikowia_unclassified(100); | Metschnikowia | 13 | Saprotroph | \|Undefined Saprotroph\| | Yeast | NULL | Probable | As nectar-tap_saprotroph (Põlme et al. 2020); Decay sugar-rich substrates (Põlme et al. 2020) | Põlme S, et al. 2020. Fungal Diversity 105:1-16 ((https://doi.org/10.1007/s13225-020-00466-2)) |
| Otu37773 | 1 | k__Fungi(100);p__Ascomycota(100);c__Saccharomycetes(100);o__Saccharomycetales(100);f__Metschnikowiaceae(100);g__Metschnikowia(100);s__Metschnikowia_pulcherrima(100); | Metschnikowia | 13 | Saprotroph | \|Undefined Saprotroph\| | Yeast | NULL | Probable | As nectar-tap_saprotroph (Põlme et al. 2020); Decay sugar-rich substrates (Põlme et al. 2020) | Põlme S, et al. 2020. Fungal Diversity 105:1-16 ((https://doi.org/10.1007/s13225-020-00466-2)) |
| Otu37774 | 1 | k__Fungi(100);p__Ascomycota(100);c__Saccharomycetes(100);o__Saccharomycetales(100);f__Metschnikowiaceae(100);g__Metschnikowia(100);s__Metschnikowia_pulcherrima(100); | Metschnikowia | 13 | Saprotroph | \|Undefined Saprotroph\| | Yeast | NULL | Probable | As nectar-tap_saprotroph (Põlme et al. 2020); Decay sugar-rich substrates (Põlme et al. 2020) | Põlme S, et al. 2020. Fungal Diversity 105:1-16 ((https://doi.org/10.1007/s13225-020-00466-2)) |
| Otu37775 | 1 | k__Fungi(100);p__Ascomycota(100);c__Saccharomycetes(100);o__Saccharomycetales(100);f__Metschnikowiaceae(100);g__Metschnikowia(100);s__Metschnikowia_pulcherrima(100); | Metschnikowia | 13 | Saprotroph | \|Undefined Saprotroph\| | Yeast | NULL | Probable | As nectar-tap_saprotroph (Põlme et al. 2020); Decay sugar-rich substrates (Põlme et al. 2020) | Põlme S, et al. 2020. Fungal Diversity 105:1-16 ((https://doi.org/10.1007/s13225-020-00466-2)) |
| Otu37776 | 1 | k__Fungi(100);p__Ascomycota(100);c__Saccharomycetes(100);o__Saccharomycetales(100);f__Metschnikowiaceae(100);g__Metschnikowia(100);g__Metschnikowia_unclassified(100); | Metschnikowia | 13 | Saprotroph | \|Undefined Saprotroph\| | Yeast | NULL | Probable | As nectar-tap_saprotroph (Põlme et al. 2020); Decay sugar-rich substrates (Põlme et al. 2020) | Põlme S, et al. 2020. Fungal Diversity 105:1-16 ((https://doi.org/10.1007/s13225-020-00466-2)) |
| Otu37777 | 1 | k__Fungi(100);p__Ascomycota(100);c__Saccharomycetes(100);o__Saccharomycetales(100);f__Metschnikowiaceae(100);g__Metschnikowia(100);s__Metschnikowia_pulcherrima(100); | Metschnikowia | 13 | Saprotroph | \|Undefined Saprotroph\| | Yeast | NULL | Probable | As nectar-tap_saprotroph (Põlme et al. 2020); Decay sugar-rich substrates (Põlme et al. 2020) | Põlme S, et al. 2020. Fungal Diversity 105:1-16 ((https://doi.org/10.1007/s13225-020-00466-2)) |
| Otu37778 | 1 | k__Fungi(100);p__Ascomycota(100);c__Saccharomycetes(100);o__Saccharomycetales(100);f__Metschnikowiaceae(100);g__Metschnikowia(100);s__Metschnikowia_pulcherrima(100); | Metschnikowia | 13 | Saprotroph | \|Undefined Saprotroph\| | Yeast | NULL | Probable | As nectar-tap_saprotroph (Põlme et al. 2020); Decay sugar-rich substrates (Põlme et al. 2020) | Põlme S, et al. 2020. Fungal Diversity 105:1-16 ((https://doi.org/10.1007/s13225-020-00466-2)) |
| Otu37779 | 1 | k__Fungi(100);p__Ascomycota(100);c__Saccharomycetes(100);o__Saccharomycetales(100);f__Metschnikowiaceae(100);g__Metschnikowia(100);s__Metschnikowia_pulcherrima(100); | Metschnikowia | 13 | Saprotroph | \|Undefined Saprotroph\| | Yeast | NULL | Probable | As nectar-tap_saprotroph (Põlme et al. 2020); Decay sugar-rich substrates (Põlme et al. 2020) | Põlme S, et al. 2020. Fungal Diversity 105:1-16 ((https://doi.org/10.1007/s13225-020-00466-2)) |
| Otu37780 | 1 | k__Fungi(100);p__Ascomycota(100);c__Saccharomycetes(100);o__Saccharomycetales(100);f__Metschnikowiaceae(100);g__Metschnikowia(100);s__Metschnikowia_pulcherrima(100); | Metschnikowia | 13 | Saprotroph | \|Undefined Saprotroph\| | Yeast | NULL | Probable | As nectar-tap_saprotroph (Põlme et al. 2020); Decay sugar-rich substrates (Põlme et al. 2020) | Põlme S, et al. 2020. Fungal Diversity 105:1-16 ((https://doi.org/10.1007/s13225-020-00466-2)) |
| Otu37781 | 1 | k__Fungi(100);p__Ascomycota(100);c__Saccharomycetes(100);o__Saccharomycetales(100);f__Metschnikowiaceae(100);g__Metschnikowia(100);s__Metschnikowia_pulcherrima(100); | Metschnikowia | 13 | Saprotroph | \|Undefined Saprotroph\| | Yeast | NULL | Probable | As nectar-tap_saprotroph (Põlme et al. 2020); Decay sugar-rich substrates (Põlme et al. 2020) | Põlme S, et al. 2020. Fungal Diversity 105:1-16 ((https://doi.org/10.1007/s13225-020-00466-2)) |
| Otu37782 | 1 | k__Fungi(100);p__Ascomycota(100);c__Saccharomycetes(100);o__Saccharomycetales(100);f__Metschnikowiaceae(100);g__Metschnikowia(100);s__Metschnikowia_pulcherrima(100); | Metschnikowia | 13 | Saprotroph | \|Undefined Saprotroph\| | Yeast | NULL | Probable | As nectar-tap_saprotroph (Põlme et al. 2020); Decay sugar-rich substrates (Põlme et al. 2020) | Põlme S, et al. 2020. Fungal Diversity 105:1-16 ((https://doi.org/10.1007/s13225-020-00466-2)) |
| Otu37783 | 1 | k__Fungi(100);p__Ascomycota(100);c__Saccharomycetes(100);o__Saccharomycetales(100);f__Metschnikowiaceae(100);g__Metschnikowia(100);g__Metschnikowia_unclassified(100); | Metschnikowia | 13 | Saprotroph | \|Undefined Saprotroph\| | Yeast | NULL | Probable | As nectar-tap_saprotroph (Põlme et al. 2020); Decay sugar-rich substrates (Põlme et al. 2020) | Põlme S, et al. 2020. Fungal Diversity 105:1-16 ((https://doi.org/10.1007/s13225-020-00466-2)) |
| Otu37784 | 1 | k__Fungi(100);p__Ascomycota(100);c__Saccharomycetes(100);o__Saccharomycetales(100);f__Metschnikowiaceae(100);g__Metschnikowia(100);s__Metschnikowia_pulcherrima(100); | Metschnikowia | 13 | Saprotroph | \|Undefined Saprotroph\| | Yeast | NULL | Probable | As nectar-tap_saprotroph (Põlme et al. 2020); Decay sugar-rich substrates (Põlme et al. 2020) | Põlme S, et al. 2020. Fungal Diversity 105:1-16 ((https://doi.org/10.1007/s13225-020-00466-2)) |
| Otu37785 | 1 | k__Fungi(100);p__Ascomycota(100);c__Saccharomycetes(100);o__Saccharomycetales(100);f__Metschnikowiaceae(100);g__Metschnikowia(100);g__Metschnikowia_unclassified(100); | Metschnikowia | 13 | Saprotroph | \|Undefined Saprotroph\| | Yeast | NULL | Probable | As nectar-tap_saprotroph (Põlme et al. 2020); Decay sugar-rich substrates (Põlme et al. 2020) | Põlme S, et al. 2020. Fungal Diversity 105:1-16 ((https://doi.org/10.1007/s13225-020-00466-2)) |
| Otu37786 | 1 | k__Fungi(100);p__Ascomycota(100);c__Saccharomycetes(100);o__Saccharomycetales(100);f__Metschnikowiaceae(100);g__Metschnikowia(100);g__Metschnikowia_unclassified(100); | Metschnikowia | 13 | Saprotroph | \|Undefined Saprotroph\| | Yeast | NULL | Probable | As nectar-tap_saprotroph (Põlme et al. 2020); Decay sugar-rich substrates (Põlme et al. 2020) | Põlme S, et al. 2020. Fungal Diversity 105:1-16 ((https://doi.org/10.1007/s13225-020-00466-2)) |
| Otu37787 | 1 | k__Fungi(100);p__Ascomycota(100);c__Saccharomycetes(100);o__Saccharomycetales(100);f__Metschnikowiaceae(100);g__Metschnikowia(100);g__Metschnikowia_unclassified(100); | Metschnikowia | 13 | Saprotroph | \|Undefined Saprotroph\| | Yeast | NULL | Probable | As nectar-tap_saprotroph (Põlme et al. 2020); Decay sugar-rich substrates (Põlme et al. 2020) | Põlme S, et al. 2020. Fungal Diversity 105:1-16 ((https://doi.org/10.1007/s13225-020-00466-2)) |
| Otu37788 | 1 | k__Fungi(100);p__Ascomycota(100);c__Saccharomycetes(100);o__Saccharomycetales(100);f__Metschnikowiaceae(100);g__Metschnikowia(100);g__Metschnikowia_unclassified(100); | Metschnikowia | 13 | Saprotroph | \|Undefined Saprotroph\| | Yeast | NULL | Probable | As nectar-tap_saprotroph (Põlme et al. 2020); Decay sugar-rich substrates (Põlme et al. 2020) | Põlme S, et al. 2020. Fungal Diversity 105:1-16 ((https://doi.org/10.1007/s13225-020-00466-2)) |
| Otu37789 | 1 | k__Fungi(100);p__Ascomycota(100);c__Saccharomycetes(100);o__Saccharomycetales(100);f__Metschnikowiaceae(100);g__Metschnikowia(100);g__Metschnikowia_unclassified(100); | Metschnikowia | 13 | Saprotroph | \|Undefined Saprotroph\| | Yeast | NULL | Probable | As nectar-tap_saprotroph (Põlme et al. 2020); Decay sugar-rich substrates (Põlme et al. 2020) | Põlme S, et al. 2020. Fungal Diversity 105:1-16 ((https://doi.org/10.1007/s13225-020-00466-2)) |
| Otu37790 | 1 | k__Fungi(100);p__Ascomycota(100);c__Saccharomycetes(100);o__Saccharomycetales(100);f__Metschnikowiaceae(100);g__Metschnikowia(100);s__Metschnikowia_pulcherrima(100); | Metschnikowia | 13 | Saprotroph | \|Undefined Saprotroph\| | Yeast | NULL | Probable | As nectar-tap_saprotroph (Põlme et al. 2020); Decay sugar-rich substrates (Põlme et al. 2020) | Põlme S, et al. 2020. Fungal Diversity 105:1-16 ((https://doi.org/10.1007/s13225-020-00466-2)) |
| Otu37791 | 1 | k__Fungi(100);p__Ascomycota(100);c__Saccharomycetes(100);o__Saccharomycetales(100);f__Metschnikowiaceae(100);g__Metschnikowia(100);g__Metschnikowia_unclassified(100); | Metschnikowia | 13 | Saprotroph | \|Undefined Saprotroph\| | Yeast | NULL | Probable | As nectar-tap_saprotroph (Põlme et al. 2020); Decay sugar-rich substrates (Põlme et al. 2020) | Põlme S, et al. 2020. Fungal Diversity 105:1-16 ((https://doi.org/10.1007/s13225-020-00466-2)) |
| Otu37792 | 1 | k__Fungi(100);p__Ascomycota(100);c__Saccharomycetes(100);o__Saccharomycetales(100);f__Metschnikowiaceae(100);g__Metschnikowia(100);s__Metschnikowia_pulcherrima(100); | Metschnikowia | 13 | Saprotroph | \|Undefined Saprotroph\| | Yeast | NULL | Probable | As nectar-tap_saprotroph (Põlme et al. 2020); Decay sugar-rich substrates (Põlme et al. 2020) | Põlme S, et al. 2020. Fungal Diversity 105:1-16 ((https://doi.org/10.1007/s13225-020-00466-2)) |
| Otu37793 | 1 | k__Fungi(100);p__Ascomycota(100);c__Saccharomycetes(100);o__Saccharomycetales(100);f__Metschnikowiaceae(100);g__Metschnikowia(100);g__Metschnikowia_unclassified(100); | Metschnikowia | 13 | Saprotroph | \|Undefined Saprotroph\| | Yeast | NULL | Probable | As nectar-tap_saprotroph (Põlme et al. 2020); Decay sugar-rich substrates (Põlme et al. 2020) | Põlme S, et al. 2020. Fungal Diversity 105:1-16 ((https://doi.org/10.1007/s13225-020-00466-2)) |
| Otu37794 | 1 | k__Fungi(100);p__Ascomycota(100);c__Saccharomycetes(100);o__Saccharomycetales(100);f__Metschnikowiaceae(100);g__Metschnikowia(100);g__Metschnikowia_unclassified(100); | Metschnikowia | 13 | Saprotroph | \|Undefined Saprotroph\| | Yeast | NULL | Probable | As nectar-tap_saprotroph (Põlme et al. 2020); Decay sugar-rich substrates (Põlme et al. 2020) | Põlme S, et al. 2020. Fungal Diversity 105:1-16 ((https://doi.org/10.1007/s13225-020-00466-2)) |
| Otu37795 | 1 | k__Fungi(100);p__Ascomycota(100);c__Saccharomycetes(100);o__Saccharomycetales(100);f__Metschnikowiaceae(100);g__Metschnikowia(100);s__Metschnikowia_pulcherrima(100); | Metschnikowia | 13 | Saprotroph | \|Undefined Saprotroph\| | Yeast | NULL | Probable | As nectar-tap_saprotroph (Põlme et al. 2020); Decay sugar-rich substrates (Põlme et al. 2020) | Põlme S, et al. 2020. Fungal Diversity 105:1-16 ((https://doi.org/10.1007/s13225-020-00466-2)) |
| Otu37796 | 1 | k__Fungi(100);p__Ascomycota(100);c__Saccharomycetes(100);o__Saccharomycetales(100);f__Metschnikowiaceae(100);g__Metschnikowia(100);s__Metschnikowia_pulcherrima(100); | Metschnikowia | 13 | Saprotroph | \|Undefined Saprotroph\| | Yeast | NULL | Probable | As nectar-tap_saprotroph (Põlme et al. 2020); Decay sugar-rich substrates (Põlme et al. 2020) | Põlme S, et al. 2020. Fungal Diversity 105:1-16 ((https://doi.org/10.1007/s13225-020-00466-2)) |
| Otu37797 | 1 | k__Fungi(100);p__Ascomycota(100);c__Saccharomycetes(100);o__Saccharomycetales(100);f__Metschnikowiaceae(100);g__Metschnikowia(100);s__Metschnikowia_pulcherrima(100); | Metschnikowia | 13 | Saprotroph | \|Undefined Saprotroph\| | Yeast | NULL | Probable | As nectar-tap_saprotroph (Põlme et al. 2020); Decay sugar-rich substrates (Põlme et al. 2020) | Põlme S, et al. 2020. Fungal Diversity 105:1-16 ((https://doi.org/10.1007/s13225-020-00466-2)) |
| Otu37798 | 1 | k__Fungi(100);p__Ascomycota(100);c__Saccharomycetes(100);o__Saccharomycetales(100);f__Metschnikowiaceae(100);g__Metschnikowia(100);s__Metschnikowia_pulcherrima(100); | Metschnikowia | 13 | Saprotroph | \|Undefined Saprotroph\| | Yeast | NULL | Probable | As nectar-tap_saprotroph (Põlme et al. 2020); Decay sugar-rich substrates (Põlme et al. 2020) | Põlme S, et al. 2020. Fungal Diversity 105:1-16 ((https://doi.org/10.1007/s13225-020-00466-2)) |
| Otu37799 | 1 | k__Fungi(100);p__Ascomycota(100);c__Saccharomycetes(100);o__Saccharomycetales(100);f__Metschnikowiaceae(100);g__Metschnikowia(100);s__Metschnikowia_pulcherrima(100); | Metschnikowia | 13 | Saprotroph | \|Undefined Saprotroph\| | Yeast | NULL | Probable | As nectar-tap_saprotroph (Põlme et al. 2020); Decay sugar-rich substrates (Põlme et al. 2020) | Põlme S, et al. 2020. Fungal Diversity 105:1-16 ((https://doi.org/10.1007/s13225-020-00466-2)) |
| Otu37800 | 1 | k__Fungi(100);p__Ascomycota(100);c__Saccharomycetes(100);o__Saccharomycetales(100);f__Metschnikowiaceae(100);g__Metschnikowia(100);g__Metschnikowia_unclassified(100); | Metschnikowia | 13 | Saprotroph | \|Undefined Saprotroph\| | Yeast | NULL | Probable | As nectar-tap_saprotroph (Põlme et al. 2020); Decay sugar-rich substrates (Põlme et al. 2020) | Põlme S, et al. 2020. Fungal Diversity 105:1-16 ((https://doi.org/10.1007/s13225-020-00466-2)) |
| Otu37801 | 1 | k__Fungi(100);p__Ascomycota(100);c__Saccharomycetes(100);o__Saccharomycetales(100);f__Metschnikowiaceae(100);g__Metschnikowia(100);s__Metschnikowia_pulcherrima(100); | Metschnikowia | 13 | Saprotroph | \|Undefined Saprotroph\| | Yeast | NULL | Probable | As nectar-tap_saprotroph (Põlme et al. 2020); Decay sugar-rich substrates (Põlme et al. 2020) | Põlme S, et al. 2020. Fungal Diversity 105:1-16 ((https://doi.org/10.1007/s13225-020-00466-2)) |
| Otu37804 | 1 | k__Fungi(100);p__Ascomycota(100);c__Saccharomycetes(100);o__Saccharomycetales(100);f__Metschnikowiaceae(100);g__Metschnikowia(100);s__Metschnikowia_pulcherrima(100); | Metschnikowia | 13 | Saprotroph | \|Undefined Saprotroph\| | Yeast | NULL | Probable | As nectar-tap_saprotroph (Põlme et al. 2020); Decay sugar-rich substrates (Põlme et al. 2020) | Põlme S, et al. 2020. Fungal Diversity 105:1-16 ((https://doi.org/10.1007/s13225-020-00466-2)) |
| Otu37806 | 1 | k__Fungi(100);p__Ascomycota(100);c__Saccharomycetes(100);o__Saccharomycetales(100);f__Metschnikowiaceae(100);g__Metschnikowia(100);s__Metschnikowia_pulcherrima(100); | Metschnikowia | 13 | Saprotroph | \|Undefined Saprotroph\| | Yeast | NULL | Probable | As nectar-tap_saprotroph (Põlme et al. 2020); Decay sugar-rich substrates (Põlme et al. 2020) | Põlme S, et al. 2020. Fungal Diversity 105:1-16 ((https://doi.org/10.1007/s13225-020-00466-2)) |
| Otu37807 | 1 | k__Fungi(100);p__Ascomycota(100);c__Saccharomycetes(100);o__Saccharomycetales(100);f__Metschnikowiaceae(100);g__Metschnikowia(100);s__Metschnikowia_pulcherrima(100); | Metschnikowia | 13 | Saprotroph | \|Undefined Saprotroph\| | Yeast | NULL | Probable | As nectar-tap_saprotroph (Põlme et al. 2020); Decay sugar-rich substrates (Põlme et al. 2020) | Põlme S, et al. 2020. Fungal Diversity 105:1-16 ((https://doi.org/10.1007/s13225-020-00466-2)) |
| Otu37808 | 1 | k__Fungi(100);p__Ascomycota(100);c__Saccharomycetes(100);o__Saccharomycetales(100);f__Metschnikowiaceae(100);g__Metschnikowia(100);s__Metschnikowia_pulcherrima(100); | Metschnikowia | 13 | Saprotroph | \|Undefined Saprotroph\| | Yeast | NULL | Probable | As nectar-tap_saprotroph (Põlme et al. 2020); Decay sugar-rich substrates (Põlme et al. 2020) | Põlme S, et al. 2020. Fungal Diversity 105:1-16 ((https://doi.org/10.1007/s13225-020-00466-2)) |
| Otu37809 | 1 | k__Fungi(100);p__Ascomycota(100);c__Saccharomycetes(100);o__Saccharomycetales(100);f__Metschnikowiaceae(100);g__Metschnikowia(100);s__Metschnikowia_pulcherrima(100); | Metschnikowia | 13 | Saprotroph | \|Undefined Saprotroph\| | Yeast | NULL | Probable | As nectar-tap_saprotroph (Põlme et al. 2020); Decay sugar-rich substrates (Põlme et al. 2020) | Põlme S, et al. 2020. Fungal Diversity 105:1-16 ((https://doi.org/10.1007/s13225-020-00466-2)) |
| Otu37812 | 1 | k__Fungi(100);p__Ascomycota(100);c__Saccharomycetes(100);o__Saccharomycetales(100);f__Metschnikowiaceae(100);g__Metschnikowia(100);g__Metschnikowia_unclassified(100); | Metschnikowia | 13 | Saprotroph | \|Undefined Saprotroph\| | Yeast | NULL | Probable | As nectar-tap_saprotroph (Põlme et al. 2020); Decay sugar-rich substrates (Põlme et al. 2020) | Põlme S, et al. 2020. Fungal Diversity 105:1-16 ((https://doi.org/10.1007/s13225-020-00466-2)) |
| Otu37819 | 1 | k__Fungi(100);p__Ascomycota(100);c__Saccharomycetes(100);o__Saccharomycetales(100);f__Metschnikowiaceae(100);g__Metschnikowia(100);s__Metschnikowia_pulcherrima(100); | Metschnikowia | 13 | Saprotroph | \|Undefined Saprotroph\| | Yeast | NULL | Probable | As nectar-tap_saprotroph (Põlme et al. 2020); Decay sugar-rich substrates (Põlme et al. 2020) | Põlme S, et al. 2020. Fungal Diversity 105:1-16 ((https://doi.org/10.1007/s13225-020-00466-2)) |
| Otu37820 | 1 | k__Fungi(100);p__Ascomycota(100);c__Saccharomycetes(100);o__Saccharomycetales(100);f__Metschnikowiaceae(100);g__Metschnikowia(100);s__Metschnikowia_pulcherrima(100); | Metschnikowia | 13 | Saprotroph | \|Undefined Saprotroph\| | Yeast | NULL | Probable | As nectar-tap_saprotroph (Põlme et al. 2020); Decay sugar-rich substrates (Põlme et al. 2020) | Põlme S, et al. 2020. Fungal Diversity 105:1-16 ((https://doi.org/10.1007/s13225-020-00466-2)) |
| Otu37823 | 1 | k__Fungi(100);p__Ascomycota(100);c__Saccharomycetes(100);o__Saccharomycetales(100);f__Metschnikowiaceae(100);g__Metschnikowia(100);s__Metschnikowia_pulcherrima(100); | Metschnikowia | 13 | Saprotroph | \|Undefined Saprotroph\| | Yeast | NULL | Probable | As nectar-tap_saprotroph (Põlme et al. 2020); Decay sugar-rich substrates (Põlme et al. 2020) | Põlme S, et al. 2020. Fungal Diversity 105:1-16 ((https://doi.org/10.1007/s13225-020-00466-2)) |
| Otu37824 | 1 | k__Fungi(100);p__Ascomycota(100);c__Saccharomycetes(100);o__Saccharomycetales(100);f__Metschnikowiaceae(100);g__Metschnikowia(100);s__Metschnikowia_pulcherrima(100); | Metschnikowia | 13 | Saprotroph | \|Undefined Saprotroph\| | Yeast | NULL | Probable | As nectar-tap_saprotroph (Põlme et al. 2020); Decay sugar-rich substrates (Põlme et al. 2020) | Põlme S, et al. 2020. Fungal Diversity 105:1-16 ((https://doi.org/10.1007/s13225-020-00466-2)) |
| Otu37825 | 1 | k__Fungi(100);p__Ascomycota(100);c__Saccharomycetes(100);o__Saccharomycetales(100);f__Metschnikowiaceae(100);g__Metschnikowia(100);s__Metschnikowia_pulcherrima(100); | Metschnikowia | 13 | Saprotroph | \|Undefined Saprotroph\| | Yeast | NULL | Probable | As nectar-tap_saprotroph (Põlme et al. 2020); Decay sugar-rich substrates (Põlme et al. 2020) | Põlme S, et al. 2020. Fungal Diversity 105:1-16 ((https://doi.org/10.1007/s13225-020-00466-2)) |
| Otu37826 | 1 | k__Fungi(100);p__Ascomycota(100);c__Saccharomycetes(100);o__Saccharomycetales(100);f__Metschnikowiaceae(100);g__Metschnikowia(100);s__Metschnikowia_pulcherrima(100); | Metschnikowia | 13 | Saprotroph | \|Undefined Saprotroph\| | Yeast | NULL | Probable | As nectar-tap_saprotroph (Põlme et al. 2020); Decay sugar-rich substrates (Põlme et al. 2020) | Põlme S, et al. 2020. Fungal Diversity 105:1-16 ((https://doi.org/10.1007/s13225-020-00466-2)) |
| Otu37827 | 1 | k__Fungi(100);p__Ascomycota(100);c__Saccharomycetes(100);o__Saccharomycetales(100);f__Metschnikowiaceae(100);g__Metschnikowia(100);g__Metschnikowia_unclassified(100); | Metschnikowia | 13 | Saprotroph | \|Undefined Saprotroph\| | Yeast | NULL | Probable | As nectar-tap_saprotroph (Põlme et al. 2020); Decay sugar-rich substrates (Põlme et al. 2020) | Põlme S, et al. 2020. Fungal Diversity 105:1-16 ((https://doi.org/10.1007/s13225-020-00466-2)) |
| Otu37829 | 1 | k__Fungi(100);p__Ascomycota(100);c__Saccharomycetes(100);o__Saccharomycetales(100);f__Metschnikowiaceae(100);g__Metschnikowia(100);g__Metschnikowia_unclassified(100); | Metschnikowia | 13 | Saprotroph | \|Undefined Saprotroph\| | Yeast | NULL | Probable | As nectar-tap_saprotroph (Põlme et al. 2020); Decay sugar-rich substrates (Põlme et al. 2020) | Põlme S, et al. 2020. Fungal Diversity 105:1-16 ((https://doi.org/10.1007/s13225-020-00466-2)) |
| Otu37830 | 1 | k__Fungi(100);p__Ascomycota(100);c__Saccharomycetes(100);o__Saccharomycetales(100);f__Metschnikowiaceae(100);g__Metschnikowia(100);g__Metschnikowia_unclassified(100); | Metschnikowia | 13 | Saprotroph | \|Undefined Saprotroph\| | Yeast | NULL | Probable | As nectar-tap_saprotroph (Põlme et al. 2020); Decay sugar-rich substrates (Põlme et al. 2020) | Põlme S, et al. 2020. Fungal Diversity 105:1-16 ((https://doi.org/10.1007/s13225-020-00466-2)) |
| Otu37831 | 1 | k__Fungi(100);p__Ascomycota(100);c__Saccharomycetes(100);o__Saccharomycetales(100);f__Metschnikowiaceae(100);g__Metschnikowia(100);g__Metschnikowia_unclassified(100); | Metschnikowia | 13 | Saprotroph | \|Undefined Saprotroph\| | Yeast | NULL | Probable | As nectar-tap_saprotroph (Põlme et al. 2020); Decay sugar-rich substrates (Põlme et al. 2020) | Põlme S, et al. 2020. Fungal Diversity 105:1-16 ((https://doi.org/10.1007/s13225-020-00466-2)) |
| Otu37834 | 1 | k__Fungi(100);p__Ascomycota(100);c__Saccharomycetes(100);o__Saccharomycetales(100);f__Metschnikowiaceae(100);g__Metschnikowia(100);s__Metschnikowia_pulcherrima(100); | Metschnikowia | 13 | Saprotroph | \|Undefined Saprotroph\| | Yeast | NULL | Probable | As nectar-tap_saprotroph (Põlme et al. 2020); Decay sugar-rich substrates (Põlme et al. 2020) | Põlme S, et al. 2020. Fungal Diversity 105:1-16 ((https://doi.org/10.1007/s13225-020-00466-2)) |
| Otu37836 | 1 | k__Fungi(100);p__Ascomycota(100);c__Saccharomycetes(100);o__Saccharomycetales(100);f__Metschnikowiaceae(100);g__Metschnikowia(100);s__Metschnikowia_pulcherrima(100); | Metschnikowia | 13 | Saprotroph | \|Undefined Saprotroph\| | Yeast | NULL | Probable | As nectar-tap_saprotroph (Põlme et al. 2020); Decay sugar-rich substrates (Põlme et al. 2020) | Põlme S, et al. 2020. Fungal Diversity 105:1-16 ((https://doi.org/10.1007/s13225-020-00466-2)) |
| Otu37837 | 1 | k__Fungi(100);p__Ascomycota(100);c__Saccharomycetes(100);o__Saccharomycetales(100);f__Metschnikowiaceae(100);g__Metschnikowia(100);s__Metschnikowia_pulcherrima(100); | Metschnikowia | 13 | Saprotroph | \|Undefined Saprotroph\| | Yeast | NULL | Probable | As nectar-tap_saprotroph (Põlme et al. 2020); Decay sugar-rich substrates (Põlme et al. 2020) | Põlme S, et al. 2020. Fungal Diversity 105:1-16 ((https://doi.org/10.1007/s13225-020-00466-2)) |
| Otu37842 | 1 | k__Fungi(100);p__Ascomycota(100);c__Saccharomycetes(100);o__Saccharomycetales(100);f__Metschnikowiaceae(100);g__Metschnikowia(100);s__Metschnikowia_pulcherrima(100); | Metschnikowia | 13 | Saprotroph | \|Undefined Saprotroph\| | Yeast | NULL | Probable | As nectar-tap_saprotroph (Põlme et al. 2020); Decay sugar-rich substrates (Põlme et al. 2020) | Põlme S, et al. 2020. Fungal Diversity 105:1-16 ((https://doi.org/10.1007/s13225-020-00466-2)) |
| Otu37844 | 1 | k__Fungi(100);p__Ascomycota(100);c__Saccharomycetes(100);o__Saccharomycetales(100);f__Metschnikowiaceae(100);g__Metschnikowia(100);s__Metschnikowia_pulcherrima(100); | Metschnikowia | 13 | Saprotroph | \|Undefined Saprotroph\| | Yeast | NULL | Probable | As nectar-tap_saprotroph (Põlme et al. 2020); Decay sugar-rich substrates (Põlme et al. 2020) | Põlme S, et al. 2020. Fungal Diversity 105:1-16 ((https://doi.org/10.1007/s13225-020-00466-2)) |
| Otu37845 | 1 | k__Fungi(100);p__Ascomycota(100);c__Saccharomycetes(100);o__Saccharomycetales(100);f__Metschnikowiaceae(100);g__Metschnikowia(100);s__Metschnikowia_pulcherrima(100); | Metschnikowia | 13 | Saprotroph | \|Undefined Saprotroph\| | Yeast | NULL | Probable | As nectar-tap_saprotroph (Põlme et al. 2020); Decay sugar-rich substrates (Põlme et al. 2020) | Põlme S, et al. 2020. Fungal Diversity 105:1-16 ((https://doi.org/10.1007/s13225-020-00466-2)) |
| Otu37846 | 1 | k__Fungi(100);p__Ascomycota(100);c__Saccharomycetes(100);o__Saccharomycetales(100);f__Metschnikowiaceae(100);g__Metschnikowia(100);s__Metschnikowia_pulcherrima(100); | Metschnikowia | 13 | Saprotroph | \|Undefined Saprotroph\| | Yeast | NULL | Probable | As nectar-tap_saprotroph (Põlme et al. 2020); Decay sugar-rich substrates (Põlme et al. 2020) | Põlme S, et al. 2020. Fungal Diversity 105:1-16 ((https://doi.org/10.1007/s13225-020-00466-2)) |
| Otu37847 | 1 | k__Fungi(100);p__Ascomycota(100);c__Saccharomycetes(100);o__Saccharomycetales(100);f__Metschnikowiaceae(100);g__Metschnikowia(100);g__Metschnikowia_unclassified(100); | Metschnikowia | 13 | Saprotroph | \|Undefined Saprotroph\| | Yeast | NULL | Probable | As nectar-tap_saprotroph (Põlme et al. 2020); Decay sugar-rich substrates (Põlme et al. 2020) | Põlme S, et al. 2020. Fungal Diversity 105:1-16 ((https://doi.org/10.1007/s13225-020-00466-2)) |
| Otu37848 | 1 | k__Fungi(100);p__Ascomycota(100);c__Saccharomycetes(100);o__Saccharomycetales(100);f__Metschnikowiaceae(100);g__Metschnikowia(100);s__Metschnikowia_pulcherrima(100); | Metschnikowia | 13 | Saprotroph | \|Undefined Saprotroph\| | Yeast | NULL | Probable | As nectar-tap_saprotroph (Põlme et al. 2020); Decay sugar-rich substrates (Põlme et al. 2020) | Põlme S, et al. 2020. Fungal Diversity 105:1-16 ((https://doi.org/10.1007/s13225-020-00466-2)) |
| Otu37849 | 1 | k__Fungi(100);p__Ascomycota(100);c__Saccharomycetes(100);o__Saccharomycetales(100);f__Metschnikowiaceae(100);g__Metschnikowia(100);g__Metschnikowia_unclassified(100); | Metschnikowia | 13 | Saprotroph | \|Undefined Saprotroph\| | Yeast | NULL | Probable | As nectar-tap_saprotroph (Põlme et al. 2020); Decay sugar-rich substrates (Põlme et al. 2020) | Põlme S, et al. 2020. Fungal Diversity 105:1-16 ((https://doi.org/10.1007/s13225-020-00466-2)) |
| Otu37850 | 1 | k__Fungi(100);p__Ascomycota(100);c__Saccharomycetes(100);o__Saccharomycetales(100);f__Metschnikowiaceae(100);g__Metschnikowia(100);s__Metschnikowia_pulcherrima(100); | Metschnikowia | 13 | Saprotroph | \|Undefined Saprotroph\| | Yeast | NULL | Probable | As nectar-tap_saprotroph (Põlme et al. 2020); Decay sugar-rich substrates (Põlme et al. 2020) | Põlme S, et al. 2020. Fungal Diversity 105:1-16 ((https://doi.org/10.1007/s13225-020-00466-2)) |
| Otu37851 | 1 | k__Fungi(100);p__Ascomycota(100);c__Saccharomycetes(100);o__Saccharomycetales(100);f__Metschnikowiaceae(100);g__Metschnikowia(100);g__Metschnikowia_unclassified(100); | Metschnikowia | 13 | Saprotroph | \|Undefined Saprotroph\| | Yeast | NULL | Probable | As nectar-tap_saprotroph (Põlme et al. 2020); Decay sugar-rich substrates (Põlme et al. 2020) | Põlme S, et al. 2020. Fungal Diversity 105:1-16 ((https://doi.org/10.1007/s13225-020-00466-2)) |
| Otu37852 | 1 | k__Fungi(100);p__Ascomycota(100);c__Saccharomycetes(100);o__Saccharomycetales(100);f__Metschnikowiaceae(100);g__Metschnikowia(100);s__Metschnikowia_pulcherrima(100); | Metschnikowia | 13 | Saprotroph | \|Undefined Saprotroph\| | Yeast | NULL | Probable | As nectar-tap_saprotroph (Põlme et al. 2020); Decay sugar-rich substrates (Põlme et al. 2020) | Põlme S, et al. 2020. Fungal Diversity 105:1-16 ((https://doi.org/10.1007/s13225-020-00466-2)) |
| Otu37853 | 1 | k__Fungi(100);p__Ascomycota(100);c__Saccharomycetes(100);o__Saccharomycetales(100);f__Metschnikowiaceae(100);g__Metschnikowia(100);s__Metschnikowia_pulcherrima(100); | Metschnikowia | 13 | Saprotroph | \|Undefined Saprotroph\| | Yeast | NULL | Probable | As nectar-tap_saprotroph (Põlme et al. 2020); Decay sugar-rich substrates (Põlme et al. 2020) | Põlme S, et al. 2020. Fungal Diversity 105:1-16 ((https://doi.org/10.1007/s13225-020-00466-2)) |
| Otu37854 | 1 | k__Fungi(100);p__Ascomycota(100);c__Saccharomycetes(100);o__Saccharomycetales(100);f__Metschnikowiaceae(100);g__Metschnikowia(100);s__Metschnikowia_pulcherrima(100); | Metschnikowia | 13 | Saprotroph | \|Undefined Saprotroph\| | Yeast | NULL | Probable | As nectar-tap_saprotroph (Põlme et al. 2020); Decay sugar-rich substrates (Põlme et al. 2020) | Põlme S, et al. 2020. Fungal Diversity 105:1-16 ((https://doi.org/10.1007/s13225-020-00466-2)) |
| Otu37856 | 1 | k__Fungi(100);p__Ascomycota(100);c__Saccharomycetes(100);o__Saccharomycetales(100);f__Metschnikowiaceae(100);g__Metschnikowia(100);s__Metschnikowia_pulcherrima(100); | Metschnikowia | 13 | Saprotroph | \|Undefined Saprotroph\| | Yeast | NULL | Probable | As nectar-tap_saprotroph (Põlme et al. 2020); Decay sugar-rich substrates (Põlme et al. 2020) | Põlme S, et al. 2020. Fungal Diversity 105:1-16 ((https://doi.org/10.1007/s13225-020-00466-2)) |
| Otu37862 | 1 | k__Fungi(100);p__Ascomycota(100);c__Saccharomycetes(100);o__Saccharomycetales(100);f__Metschnikowiaceae(100);g__Metschnikowia(100);s__Metschnikowia_pulcherrima(100); | Metschnikowia | 13 | Saprotroph | \|Undefined Saprotroph\| | Yeast | NULL | Probable | As nectar-tap_saprotroph (Põlme et al. 2020); Decay sugar-rich substrates (Põlme et al. 2020) | Põlme S, et al. 2020. Fungal Diversity 105:1-16 ((https://doi.org/10.1007/s13225-020-00466-2)) |
| Otu37865 | 1 | k__Fungi(100);p__Ascomycota(100);c__Saccharomycetes(100);o__Saccharomycetales(100);f__Metschnikowiaceae(100);g__Metschnikowia(100);s__Metschnikowia_pulcherrima(100); | Metschnikowia | 13 | Saprotroph | \|Undefined Saprotroph\| | Yeast | NULL | Probable | As nectar-tap_saprotroph (Põlme et al. 2020); Decay sugar-rich substrates (Põlme et al. 2020) | Põlme S, et al. 2020. Fungal Diversity 105:1-16 ((https://doi.org/10.1007/s13225-020-00466-2)) |
| Otu37867 | 1 | k__Fungi(100);p__Ascomycota(100);c__Saccharomycetes(100);o__Saccharomycetales(100);f__Metschnikowiaceae(100);g__Metschnikowia(100);s__Metschnikowia_pulcherrima(100); | Metschnikowia | 13 | Saprotroph | \|Undefined Saprotroph\| | Yeast | NULL | Probable | As nectar-tap_saprotroph (Põlme et al. 2020); Decay sugar-rich substrates (Põlme et al. 2020) | Põlme S, et al. 2020. Fungal Diversity 105:1-16 ((https://doi.org/10.1007/s13225-020-00466-2)) |
| Otu37868 | 1 | k__Fungi(100);p__Ascomycota(100);c__Saccharomycetes(100);o__Saccharomycetales(100);f__Metschnikowiaceae(100);g__Metschnikowia(100);s__Metschnikowia_pulcherrima(100); | Metschnikowia | 13 | Saprotroph | \|Undefined Saprotroph\| | Yeast | NULL | Probable | As nectar-tap_saprotroph (Põlme et al. 2020); Decay sugar-rich substrates (Põlme et al. 2020) | Põlme S, et al. 2020. Fungal Diversity 105:1-16 ((https://doi.org/10.1007/s13225-020-00466-2)) |
| Otu37869 | 1 | k__Fungi(100);p__Ascomycota(100);c__Saccharomycetes(100);o__Saccharomycetales(100);f__Metschnikowiaceae(100);g__Metschnikowia(100);s__Metschnikowia_pulcherrima(100); | Metschnikowia | 13 | Saprotroph | \|Undefined Saprotroph\| | Yeast | NULL | Probable | As nectar-tap_saprotroph (Põlme et al. 2020); Decay sugar-rich substrates (Põlme et al. 2020) | Põlme S, et al. 2020. Fungal Diversity 105:1-16 ((https://doi.org/10.1007/s13225-020-00466-2)) |
| Otu37870 | 1 | k__Fungi(100);p__Ascomycota(100);c__Saccharomycetes(100);o__Saccharomycetales(100);f__Metschnikowiaceae(100);g__Metschnikowia(100);g__Metschnikowia_unclassified(100); | Metschnikowia | 13 | Saprotroph | \|Undefined Saprotroph\| | Yeast | NULL | Probable | As nectar-tap_saprotroph (Põlme et al. 2020); Decay sugar-rich substrates (Põlme et al. 2020) | Põlme S, et al. 2020. Fungal Diversity 105:1-16 ((https://doi.org/10.1007/s13225-020-00466-2)) |
| Otu37871 | 1 | k__Fungi(100);p__Ascomycota(100);c__Saccharomycetes(100);o__Saccharomycetales(100);f__Metschnikowiaceae(100);g__Metschnikowia(100);s__Metschnikowia_pulcherrima(100); | Metschnikowia | 13 | Saprotroph | \|Undefined Saprotroph\| | Yeast | NULL | Probable | As nectar-tap_saprotroph (Põlme et al. 2020); Decay sugar-rich substrates (Põlme et al. 2020) | Põlme S, et al. 2020. Fungal Diversity 105:1-16 ((https://doi.org/10.1007/s13225-020-00466-2)) |
| Otu37872 | 1 | k__Fungi(100);p__Ascomycota(100);c__Saccharomycetes(100);o__Saccharomycetales(100);f__Metschnikowiaceae(100);g__Metschnikowia(100);s__Metschnikowia_pulcherrima(100); | Metschnikowia | 13 | Saprotroph | \|Undefined Saprotroph\| | Yeast | NULL | Probable | As nectar-tap_saprotroph (Põlme et al. 2020); Decay sugar-rich substrates (Põlme et al. 2020) | Põlme S, et al. 2020. Fungal Diversity 105:1-16 ((https://doi.org/10.1007/s13225-020-00466-2)) |
| Otu37873 | 1 | k__Fungi(100);p__Ascomycota(100);c__Saccharomycetes(100);o__Saccharomycetales(100);f__Metschnikowiaceae(100);g__Metschnikowia(100);s__Metschnikowia_pulcherrima(100); | Metschnikowia | 13 | Saprotroph | \|Undefined Saprotroph\| | Yeast | NULL | Probable | As nectar-tap_saprotroph (Põlme et al. 2020); Decay sugar-rich substrates (Põlme et al. 2020) | Põlme S, et al. 2020. Fungal Diversity 105:1-16 ((https://doi.org/10.1007/s13225-020-00466-2)) |
| Otu37874 | 1 | k__Fungi(100);p__Ascomycota(100);c__Saccharomycetes(100);o__Saccharomycetales(100);f__Metschnikowiaceae(100);g__Metschnikowia(100);s__Metschnikowia_pulcherrima(100); | Metschnikowia | 13 | Saprotroph | \|Undefined Saprotroph\| | Yeast | NULL | Probable | As nectar-tap_saprotroph (Põlme et al. 2020); Decay sugar-rich substrates (Põlme et al. 2020) | Põlme S, et al. 2020. Fungal Diversity 105:1-16 ((https://doi.org/10.1007/s13225-020-00466-2)) |
| Otu37875 | 1 | k__Fungi(100);p__Ascomycota(100);c__Saccharomycetes(100);o__Saccharomycetales(100);f__Metschnikowiaceae(100);g__Metschnikowia(100);g__Metschnikowia_unclassified(100); | Metschnikowia | 13 | Saprotroph | \|Undefined Saprotroph\| | Yeast | NULL | Probable | As nectar-tap_saprotroph (Põlme et al. 2020); Decay sugar-rich substrates (Põlme et al. 2020) | Põlme S, et al. 2020. Fungal Diversity 105:1-16 ((https://doi.org/10.1007/s13225-020-00466-2)) |
| Otu37876 | 1 | k__Fungi(100);p__Ascomycota(100);c__Saccharomycetes(100);o__Saccharomycetales(100);f__Metschnikowiaceae(100);g__Metschnikowia(100);s__Metschnikowia_pulcherrima(100); | Metschnikowia | 13 | Saprotroph | \|Undefined Saprotroph\| | Yeast | NULL | Probable | As nectar-tap_saprotroph (Põlme et al. 2020); Decay sugar-rich substrates (Põlme et al. 2020) | Põlme S, et al. 2020. Fungal Diversity 105:1-16 ((https://doi.org/10.1007/s13225-020-00466-2)) |
| Otu37877 | 1 | k__Fungi(100);p__Ascomycota(100);c__Saccharomycetes(100);o__Saccharomycetales(100);f__Metschnikowiaceae(100);g__Metschnikowia(100);s__Metschnikowia_pulcherrima(100); | Metschnikowia | 13 | Saprotroph | \|Undefined Saprotroph\| | Yeast | NULL | Probable | As nectar-tap_saprotroph (Põlme et al. 2020); Decay sugar-rich substrates (Põlme et al. 2020) | Põlme S, et al. 2020. Fungal Diversity 105:1-16 ((https://doi.org/10.1007/s13225-020-00466-2)) |
| Otu37878 | 1 | k__Fungi(100);p__Ascomycota(100);c__Saccharomycetes(100);o__Saccharomycetales(100);f__Metschnikowiaceae(100);g__Metschnikowia(100);s__Metschnikowia_pulcherrima(100); | Metschnikowia | 13 | Saprotroph | \|Undefined Saprotroph\| | Yeast | NULL | Probable | As nectar-tap_saprotroph (Põlme et al. 2020); Decay sugar-rich substrates (Põlme et al. 2020) | Põlme S, et al. 2020. Fungal Diversity 105:1-16 ((https://doi.org/10.1007/s13225-020-00466-2)) |
| Otu37879 | 1 | k__Fungi(100);p__Ascomycota(100);c__Saccharomycetes(100);o__Saccharomycetales(100);f__Metschnikowiaceae(100);g__Metschnikowia(100);g__Metschnikowia_unclassified(100); | Metschnikowia | 13 | Saprotroph | \|Undefined Saprotroph\| | Yeast | NULL | Probable | As nectar-tap_saprotroph (Põlme et al. 2020); Decay sugar-rich substrates (Põlme et al. 2020) | Põlme S, et al. 2020. Fungal Diversity 105:1-16 ((https://doi.org/10.1007/s13225-020-00466-2)) |
| Otu37880 | 1 | k__Fungi(100);p__Ascomycota(100);c__Saccharomycetes(100);o__Saccharomycetales(100);f__Metschnikowiaceae(100);g__Metschnikowia(100);s__Metschnikowia_pulcherrima(100); | Metschnikowia | 13 | Saprotroph | \|Undefined Saprotroph\| | Yeast | NULL | Probable | As nectar-tap_saprotroph (Põlme et al. 2020); Decay sugar-rich substrates (Põlme et al. 2020) | Põlme S, et al. 2020. Fungal Diversity 105:1-16 ((https://doi.org/10.1007/s13225-020-00466-2)) |
| Otu37881 | 1 | k__Fungi(100);p__Ascomycota(100);c__Saccharomycetes(100);o__Saccharomycetales(100);f__Metschnikowiaceae(100);g__Metschnikowia(100);g__Metschnikowia_unclassified(100); | Metschnikowia | 13 | Saprotroph | \|Undefined Saprotroph\| | Yeast | NULL | Probable | As nectar-tap_saprotroph (Põlme et al. 2020); Decay sugar-rich substrates (Põlme et al. 2020) | Põlme S, et al. 2020. Fungal Diversity 105:1-16 ((https://doi.org/10.1007/s13225-020-00466-2)) |
| Otu37882 | 1 | k__Fungi(100);p__Ascomycota(100);c__Saccharomycetes(100);o__Saccharomycetales(100);f__Metschnikowiaceae(100);g__Metschnikowia(100);s__Metschnikowia_pulcherrima(100); | Metschnikowia | 13 | Saprotroph | \|Undefined Saprotroph\| | Yeast | NULL | Probable | As nectar-tap_saprotroph (Põlme et al. 2020); Decay sugar-rich substrates (Põlme et al. 2020) | Põlme S, et al. 2020. Fungal Diversity 105:1-16 ((https://doi.org/10.1007/s13225-020-00466-2)) |
| Otu37883 | 1 | k__Fungi(100);p__Ascomycota(100);c__Saccharomycetes(100);o__Saccharomycetales(100);f__Metschnikowiaceae(100);g__Metschnikowia(100);s__Metschnikowia_pulcherrima(100); | Metschnikowia | 13 | Saprotroph | \|Undefined Saprotroph\| | Yeast | NULL | Probable | As nectar-tap_saprotroph (Põlme et al. 2020); Decay sugar-rich substrates (Põlme et al. 2020) | Põlme S, et al. 2020. Fungal Diversity 105:1-16 ((https://doi.org/10.1007/s13225-020-00466-2)) |
| Otu37884 | 1 | k__Fungi(100);p__Ascomycota(100);c__Saccharomycetes(100);o__Saccharomycetales(100);f__Metschnikowiaceae(100);g__Metschnikowia(100);g__Metschnikowia_unclassified(100); | Metschnikowia | 13 | Saprotroph | \|Undefined Saprotroph\| | Yeast | NULL | Probable | As nectar-tap_saprotroph (Põlme et al. 2020); Decay sugar-rich substrates (Põlme et al. 2020) | Põlme S, et al. 2020. Fungal Diversity 105:1-16 ((https://doi.org/10.1007/s13225-020-00466-2)) |
| Otu37885 | 1 | k__Fungi(100);p__Ascomycota(100);c__Saccharomycetes(100);o__Saccharomycetales(100);f__Metschnikowiaceae(100);g__Metschnikowia(100);s__Metschnikowia_pulcherrima(100); | Metschnikowia | 13 | Saprotroph | \|Undefined Saprotroph\| | Yeast | NULL | Probable | As nectar-tap_saprotroph (Põlme et al. 2020); Decay sugar-rich substrates (Põlme et al. 2020) | Põlme S, et al. 2020. Fungal Diversity 105:1-16 ((https://doi.org/10.1007/s13225-020-00466-2)) |
| Otu37886 | 1 | k__Fungi(100);p__Ascomycota(100);c__Saccharomycetes(100);o__Saccharomycetales(100);f__Metschnikowiaceae(100);g__Metschnikowia(100);s__Metschnikowia_pulcherrima(100); | Metschnikowia | 13 | Saprotroph | \|Undefined Saprotroph\| | Yeast | NULL | Probable | As nectar-tap_saprotroph (Põlme et al. 2020); Decay sugar-rich substrates (Põlme et al. 2020) | Põlme S, et al. 2020. Fungal Diversity 105:1-16 ((https://doi.org/10.1007/s13225-020-00466-2)) |
| Otu37887 | 1 | k__Fungi(100);p__Ascomycota(100);c__Saccharomycetes(100);o__Saccharomycetales(100);f__Metschnikowiaceae(100);g__Metschnikowia(100);s__Metschnikowia_pulcherrima(100); | Metschnikowia | 13 | Saprotroph | \|Undefined Saprotroph\| | Yeast | NULL | Probable | As nectar-tap_saprotroph (Põlme et al. 2020); Decay sugar-rich substrates (Põlme et al. 2020) | Põlme S, et al. 2020. Fungal Diversity 105:1-16 ((https://doi.org/10.1007/s13225-020-00466-2)) |
| Otu37888 | 1 | k__Fungi(100);p__Ascomycota(100);c__Saccharomycetes(100);o__Saccharomycetales(100);f__Metschnikowiaceae(100);g__Metschnikowia(100);s__Metschnikowia_pulcherrima(100); | Metschnikowia | 13 | Saprotroph | \|Undefined Saprotroph\| | Yeast | NULL | Probable | As nectar-tap_saprotroph (Põlme et al. 2020); Decay sugar-rich substrates (Põlme et al. 2020) | Põlme S, et al. 2020. Fungal Diversity 105:1-16 ((https://doi.org/10.1007/s13225-020-00466-2)) |
| Otu37889 | 1 | k__Fungi(100);p__Ascomycota(100);c__Saccharomycetes(100);o__Saccharomycetales(100);f__Metschnikowiaceae(100);g__Metschnikowia(100);s__Metschnikowia_pulcherrima(100); | Metschnikowia | 13 | Saprotroph | \|Undefined Saprotroph\| | Yeast | NULL | Probable | As nectar-tap_saprotroph (Põlme et al. 2020); Decay sugar-rich substrates (Põlme et al. 2020) | Põlme S, et al. 2020. Fungal Diversity 105:1-16 ((https://doi.org/10.1007/s13225-020-00466-2)) |
| Otu37890 | 1 | k__Fungi(100);p__Ascomycota(100);c__Saccharomycetes(100);o__Saccharomycetales(100);f__Metschnikowiaceae(100);g__Metschnikowia(100);s__Metschnikowia_pulcherrima(100); | Metschnikowia | 13 | Saprotroph | \|Undefined Saprotroph\| | Yeast | NULL | Probable | As nectar-tap_saprotroph (Põlme et al. 2020); Decay sugar-rich substrates (Põlme et al. 2020) | Põlme S, et al. 2020. Fungal Diversity 105:1-16 ((https://doi.org/10.1007/s13225-020-00466-2)) |
| Otu37891 | 1 | k__Fungi(100);p__Ascomycota(100);c__Saccharomycetes(100);o__Saccharomycetales(100);f__Metschnikowiaceae(100);g__Metschnikowia(100);s__Metschnikowia_pulcherrima(100); | Metschnikowia | 13 | Saprotroph | \|Undefined Saprotroph\| | Yeast | NULL | Probable | As nectar-tap_saprotroph (Põlme et al. 2020); Decay sugar-rich substrates (Põlme et al. 2020) | Põlme S, et al. 2020. Fungal Diversity 105:1-16 ((https://doi.org/10.1007/s13225-020-00466-2)) |
| Otu37893 | 1 | k__Fungi(100);p__Ascomycota(100);c__Saccharomycetes(100);o__Saccharomycetales(100);f__Metschnikowiaceae(100);g__Metschnikowia(100);s__Metschnikowia_pulcherrima(100); | Metschnikowia | 13 | Saprotroph | \|Undefined Saprotroph\| | Yeast | NULL | Probable | As nectar-tap_saprotroph (Põlme et al. 2020); Decay sugar-rich substrates (Põlme et al. 2020) | Põlme S, et al. 2020. Fungal Diversity 105:1-16 ((https://doi.org/10.1007/s13225-020-00466-2)) |
| Otu37894 | 1 | k__Fungi(100);p__Ascomycota(100);c__Saccharomycetes(100);o__Saccharomycetales(100);f__Metschnikowiaceae(100);g__Metschnikowia(100);g__Metschnikowia_unclassified(100); | Metschnikowia | 13 | Saprotroph | \|Undefined Saprotroph\| | Yeast | NULL | Probable | As nectar-tap_saprotroph (Põlme et al. 2020); Decay sugar-rich substrates (Põlme et al. 2020) | Põlme S, et al. 2020. Fungal Diversity 105:1-16 ((https://doi.org/10.1007/s13225-020-00466-2)) |
| Otu37895 | 1 | k__Fungi(100);p__Ascomycota(100);c__Saccharomycetes(100);o__Saccharomycetales(100);f__Metschnikowiaceae(100);g__Metschnikowia(100);g__Metschnikowia_unclassified(100); | Metschnikowia | 13 | Saprotroph | \|Undefined Saprotroph\| | Yeast | NULL | Probable | As nectar-tap_saprotroph (Põlme et al. 2020); Decay sugar-rich substrates (Põlme et al. 2020) | Põlme S, et al. 2020. Fungal Diversity 105:1-16 ((https://doi.org/10.1007/s13225-020-00466-2)) |
| Otu37896 | 1 | k__Fungi(100);p__Ascomycota(100);c__Saccharomycetes(100);o__Saccharomycetales(100);f__Metschnikowiaceae(100);g__Metschnikowia(100);s__Metschnikowia_pulcherrima(100); | Metschnikowia | 13 | Saprotroph | \|Undefined Saprotroph\| | Yeast | NULL | Probable | As nectar-tap_saprotroph (Põlme et al. 2020); Decay sugar-rich substrates (Põlme et al. 2020) | Põlme S, et al. 2020. Fungal Diversity 105:1-16 ((https://doi.org/10.1007/s13225-020-00466-2)) |
| Otu37900 | 1 | k__Fungi(100);p__Ascomycota(100);c__Saccharomycetes(100);o__Saccharomycetales(100);f__Metschnikowiaceae(100);g__Metschnikowia(100);s__Metschnikowia_pulcherrima(100); | Metschnikowia | 13 | Saprotroph | \|Undefined Saprotroph\| | Yeast | NULL | Probable | As nectar-tap_saprotroph (Põlme et al. 2020); Decay sugar-rich substrates (Põlme et al. 2020) | Põlme S, et al. 2020. Fungal Diversity 105:1-16 ((https://doi.org/10.1007/s13225-020-00466-2)) |
| Otu37901 | 1 | k__Fungi(100);p__Ascomycota(100);c__Saccharomycetes(100);o__Saccharomycetales(100);f__Metschnikowiaceae(100);g__Metschnikowia(100);g__Metschnikowia_unclassified(100); | Metschnikowia | 13 | Saprotroph | \|Undefined Saprotroph\| | Yeast | NULL | Probable | As nectar-tap_saprotroph (Põlme et al. 2020); Decay sugar-rich substrates (Põlme et al. 2020) | Põlme S, et al. 2020. Fungal Diversity 105:1-16 ((https://doi.org/10.1007/s13225-020-00466-2)) |
| Otu37902 | 1 | k__Fungi(100);p__Ascomycota(100);c__Saccharomycetes(100);o__Saccharomycetales(100);f__Metschnikowiaceae(100);g__Metschnikowia(100);g__Metschnikowia_unclassified(100); | Metschnikowia | 13 | Saprotroph | \|Undefined Saprotroph\| | Yeast | NULL | Probable | As nectar-tap_saprotroph (Põlme et al. 2020); Decay sugar-rich substrates (Põlme et al. 2020) | Põlme S, et al. 2020. Fungal Diversity 105:1-16 ((https://doi.org/10.1007/s13225-020-00466-2)) |
| Otu37903 | 1 | k__Fungi(100);p__Ascomycota(100);c__Saccharomycetes(100);o__Saccharomycetales(100);f__Metschnikowiaceae(100);g__Metschnikowia(100);s__Metschnikowia_pulcherrima(100); | Metschnikowia | 13 | Saprotroph | \|Undefined Saprotroph\| | Yeast | NULL | Probable | As nectar-tap_saprotroph (Põlme et al. 2020); Decay sugar-rich substrates (Põlme et al. 2020) | Põlme S, et al. 2020. Fungal Diversity 105:1-16 ((https://doi.org/10.1007/s13225-020-00466-2)) |
| Otu37904 | 1 | k__Fungi(100);p__Ascomycota(100);c__Saccharomycetes(100);o__Saccharomycetales(100);f__Metschnikowiaceae(100);g__Metschnikowia(100);g__Metschnikowia_unclassified(100); | Metschnikowia | 13 | Saprotroph | \|Undefined Saprotroph\| | Yeast | NULL | Probable | As nectar-tap_saprotroph (Põlme et al. 2020); Decay sugar-rich substrates (Põlme et al. 2020) | Põlme S, et al. 2020. Fungal Diversity 105:1-16 ((https://doi.org/10.1007/s13225-020-00466-2)) |
| Otu37905 | 1 | k__Fungi(100);p__Ascomycota(100);c__Saccharomycetes(100);o__Saccharomycetales(100);f__Metschnikowiaceae(100);g__Metschnikowia(100);g__Metschnikowia_unclassified(100); | Metschnikowia | 13 | Saprotroph | \|Undefined Saprotroph\| | Yeast | NULL | Probable | As nectar-tap_saprotroph (Põlme et al. 2020); Decay sugar-rich substrates (Põlme et al. 2020) | Põlme S, et al. 2020. Fungal Diversity 105:1-16 ((https://doi.org/10.1007/s13225-020-00466-2)) |
| Otu37906 | 1 | k__Fungi(100);p__Ascomycota(100);c__Saccharomycetes(100);o__Saccharomycetales(100);f__Metschnikowiaceae(100);g__Metschnikowia(100);g__Metschnikowia_unclassified(100); | Metschnikowia | 13 | Saprotroph | \|Undefined Saprotroph\| | Yeast | NULL | Probable | As nectar-tap_saprotroph (Põlme et al. 2020); Decay sugar-rich substrates (Põlme et al. 2020) | Põlme S, et al. 2020. Fungal Diversity 105:1-16 ((https://doi.org/10.1007/s13225-020-00466-2)) |
| Otu37907 | 1 | k__Fungi(100);p__Ascomycota(100);c__Saccharomycetes(100);o__Saccharomycetales(100);f__Metschnikowiaceae(100);g__Metschnikowia(100);s__Metschnikowia_pulcherrima(100); | Metschnikowia | 13 | Saprotroph | \|Undefined Saprotroph\| | Yeast | NULL | Probable | As nectar-tap_saprotroph (Põlme et al. 2020); Decay sugar-rich substrates (Põlme et al. 2020) | Põlme S, et al. 2020. Fungal Diversity 105:1-16 ((https://doi.org/10.1007/s13225-020-00466-2)) |
| Otu37908 | 1 | k__Fungi(100);p__Ascomycota(100);c__Saccharomycetes(100);o__Saccharomycetales(100);f__Metschnikowiaceae(100);g__Metschnikowia(100);s__Metschnikowia_pulcherrima(100); | Metschnikowia | 13 | Saprotroph | \|Undefined Saprotroph\| | Yeast | NULL | Probable | As nectar-tap_saprotroph (Põlme et al. 2020); Decay sugar-rich substrates (Põlme et al. 2020) | Põlme S, et al. 2020. Fungal Diversity 105:1-16 ((https://doi.org/10.1007/s13225-020-00466-2)) |
| Otu37910 | 1 | k__Fungi(100);p__Ascomycota(100);c__Saccharomycetes(100);o__Saccharomycetales(100);f__Metschnikowiaceae(100);g__Metschnikowia(100);g__Metschnikowia_unclassified(100); | Metschnikowia | 13 | Saprotroph | \|Undefined Saprotroph\| | Yeast | NULL | Probable | As nectar-tap_saprotroph (Põlme et al. 2020); Decay sugar-rich substrates (Põlme et al. 2020) | Põlme S, et al. 2020. Fungal Diversity 105:1-16 ((https://doi.org/10.1007/s13225-020-00466-2)) |
| Otu37911 | 1 | k__Fungi(100);p__Ascomycota(100);c__Saccharomycetes(100);o__Saccharomycetales(100);f__Metschnikowiaceae(100);g__Metschnikowia(100);s__Metschnikowia_pulcherrima(100); | Metschnikowia | 13 | Saprotroph | \|Undefined Saprotroph\| | Yeast | NULL | Probable | As nectar-tap_saprotroph (Põlme et al. 2020); Decay sugar-rich substrates (Põlme et al. 2020) | Põlme S, et al. 2020. Fungal Diversity 105:1-16 ((https://doi.org/10.1007/s13225-020-00466-2)) |
| Otu37912 | 1 | k__Fungi(100);p__Ascomycota(100);c__Saccharomycetes(100);o__Saccharomycetales(100);f__Metschnikowiaceae(100);g__Metschnikowia(100);s__Metschnikowia_pulcherrima(100); | Metschnikowia | 13 | Saprotroph | \|Undefined Saprotroph\| | Yeast | NULL | Probable | As nectar-tap_saprotroph (Põlme et al. 2020); Decay sugar-rich substrates (Põlme et al. 2020) | Põlme S, et al. 2020. Fungal Diversity 105:1-16 ((https://doi.org/10.1007/s13225-020-00466-2)) |
| Otu37913 | 1 | k__Fungi(100);p__Ascomycota(100);c__Saccharomycetes(100);o__Saccharomycetales(100);f__Metschnikowiaceae(100);g__Metschnikowia(100);g__Metschnikowia_unclassified(100); | Metschnikowia | 13 | Saprotroph | \|Undefined Saprotroph\| | Yeast | NULL | Probable | As nectar-tap_saprotroph (Põlme et al. 2020); Decay sugar-rich substrates (Põlme et al. 2020) | Põlme S, et al. 2020. Fungal Diversity 105:1-16 ((https://doi.org/10.1007/s13225-020-00466-2)) |
| Otu37914 | 1 | k__Fungi(100);p__Ascomycota(100);c__Saccharomycetes(100);o__Saccharomycetales(100);f__Metschnikowiaceae(100);g__Metschnikowia(100);s__Metschnikowia_pulcherrima(100); | Metschnikowia | 13 | Saprotroph | \|Undefined Saprotroph\| | Yeast | NULL | Probable | As nectar-tap_saprotroph (Põlme et al. 2020); Decay sugar-rich substrates (Põlme et al. 2020) | Põlme S, et al. 2020. Fungal Diversity 105:1-16 ((https://doi.org/10.1007/s13225-020-00466-2)) |
| Otu37915 | 1 | k__Fungi(100);p__Ascomycota(100);c__Saccharomycetes(100);o__Saccharomycetales(100);f__Metschnikowiaceae(100);g__Metschnikowia(100);g__Metschnikowia_unclassified(100); | Metschnikowia | 13 | Saprotroph | \|Undefined Saprotroph\| | Yeast | NULL | Probable | As nectar-tap_saprotroph (Põlme et al. 2020); Decay sugar-rich substrates (Põlme et al. 2020) | Põlme S, et al. 2020. Fungal Diversity 105:1-16 ((https://doi.org/10.1007/s13225-020-00466-2)) |
| Otu37916 | 1 | k__Fungi(100);p__Ascomycota(100);c__Saccharomycetes(100);o__Saccharomycetales(100);f__Metschnikowiaceae(100);g__Metschnikowia(100);s__Metschnikowia_pulcherrima(100); | Metschnikowia | 13 | Saprotroph | \|Undefined Saprotroph\| | Yeast | NULL | Probable | As nectar-tap_saprotroph (Põlme et al. 2020); Decay sugar-rich substrates (Põlme et al. 2020) | Põlme S, et al. 2020. Fungal Diversity 105:1-16 ((https://doi.org/10.1007/s13225-020-00466-2)) |
| Otu37917 | 1 | k__Fungi(100);p__Ascomycota(100);c__Saccharomycetes(100);o__Saccharomycetales(100);f__Metschnikowiaceae(100);g__Metschnikowia(100);s__Metschnikowia_pulcherrima(100); | Metschnikowia | 13 | Saprotroph | \|Undefined Saprotroph\| | Yeast | NULL | Probable | As nectar-tap_saprotroph (Põlme et al. 2020); Decay sugar-rich substrates (Põlme et al. 2020) | Põlme S, et al. 2020. Fungal Diversity 105:1-16 ((https://doi.org/10.1007/s13225-020-00466-2)) |
| Otu37918 | 1 | k__Fungi(100);p__Ascomycota(100);c__Saccharomycetes(100);o__Saccharomycetales(100);f__Metschnikowiaceae(100);g__Metschnikowia(100);s__Metschnikowia_pulcherrima(100); | Metschnikowia | 13 | Saprotroph | \|Undefined Saprotroph\| | Yeast | NULL | Probable | As nectar-tap_saprotroph (Põlme et al. 2020); Decay sugar-rich substrates (Põlme et al. 2020) | Põlme S, et al. 2020. Fungal Diversity 105:1-16 ((https://doi.org/10.1007/s13225-020-00466-2)) |
| Otu37919 | 1 | k__Fungi(100);p__Ascomycota(100);c__Saccharomycetes(100);o__Saccharomycetales(100);f__Metschnikowiaceae(100);g__Metschnikowia(100);s__Metschnikowia_pulcherrima(100); | Metschnikowia | 13 | Saprotroph | \|Undefined Saprotroph\| | Yeast | NULL | Probable | As nectar-tap_saprotroph (Põlme et al. 2020); Decay sugar-rich substrates (Põlme et al. 2020) | Põlme S, et al. 2020. Fungal Diversity 105:1-16 ((https://doi.org/10.1007/s13225-020-00466-2)) |
| Otu37920 | 1 | k__Fungi(100);p__Ascomycota(100);c__Saccharomycetes(100);o__Saccharomycetales(100);f__Metschnikowiaceae(100);g__Metschnikowia(100);s__Metschnikowia_pulcherrima(100); | Metschnikowia | 13 | Saprotroph | \|Undefined Saprotroph\| | Yeast | NULL | Probable | As nectar-tap_saprotroph (Põlme et al. 2020); Decay sugar-rich substrates (Põlme et al. 2020) | Põlme S, et al. 2020. Fungal Diversity 105:1-16 ((https://doi.org/10.1007/s13225-020-00466-2)) |
| Otu37921 | 1 | k__Fungi(100);p__Ascomycota(100);c__Saccharomycetes(100);o__Saccharomycetales(100);f__Metschnikowiaceae(100);g__Metschnikowia(100);s__Metschnikowia_pulcherrima(100); | Metschnikowia | 13 | Saprotroph | \|Undefined Saprotroph\| | Yeast | NULL | Probable | As nectar-tap_saprotroph (Põlme et al. 2020); Decay sugar-rich substrates (Põlme et al. 2020) | Põlme S, et al. 2020. Fungal Diversity 105:1-16 ((https://doi.org/10.1007/s13225-020-00466-2)) |
| Otu37922 | 1 | k__Fungi(100);p__Ascomycota(100);c__Saccharomycetes(100);o__Saccharomycetales(100);f__Metschnikowiaceae(100);g__Metschnikowia(100);g__Metschnikowia_unclassified(100); | Metschnikowia | 13 | Saprotroph | \|Undefined Saprotroph\| | Yeast | NULL | Probable | As nectar-tap_saprotroph (Põlme et al. 2020); Decay sugar-rich substrates (Põlme et al. 2020) | Põlme S, et al. 2020. Fungal Diversity 105:1-16 ((https://doi.org/10.1007/s13225-020-00466-2)) |
| Otu37923 | 1 | k__Fungi(100);p__Ascomycota(100);c__Saccharomycetes(100);o__Saccharomycetales(100);f__Metschnikowiaceae(100);g__Metschnikowia(100);s__Metschnikowia_pulcherrima(100); | Metschnikowia | 13 | Saprotroph | \|Undefined Saprotroph\| | Yeast | NULL | Probable | As nectar-tap_saprotroph (Põlme et al. 2020); Decay sugar-rich substrates (Põlme et al. 2020) | Põlme S, et al. 2020. Fungal Diversity 105:1-16 ((https://doi.org/10.1007/s13225-020-00466-2)) |
| Otu37924 | 1 | k__Fungi(100);p__Ascomycota(100);c__Saccharomycetes(100);o__Saccharomycetales(100);f__Metschnikowiaceae(100);g__Metschnikowia(100);s__Metschnikowia_pulcherrima(100); | Metschnikowia | 13 | Saprotroph | \|Undefined Saprotroph\| | Yeast | NULL | Probable | As nectar-tap_saprotroph (Põlme et al. 2020); Decay sugar-rich substrates (Põlme et al. 2020) | Põlme S, et al. 2020. Fungal Diversity 105:1-16 ((https://doi.org/10.1007/s13225-020-00466-2)) |
| Otu37925 | 1 | k__Fungi(100);p__Ascomycota(100);c__Saccharomycetes(100);o__Saccharomycetales(100);f__Metschnikowiaceae(100);g__Metschnikowia(100);s__Metschnikowia_pulcherrima(100); | Metschnikowia | 13 | Saprotroph | \|Undefined Saprotroph\| | Yeast | NULL | Probable | As nectar-tap_saprotroph (Põlme et al. 2020); Decay sugar-rich substrates (Põlme et al. 2020) | Põlme S, et al. 2020. Fungal Diversity 105:1-16 ((https://doi.org/10.1007/s13225-020-00466-2)) |
| Otu37926 | 1 | k__Fungi(100);p__Ascomycota(100);c__Saccharomycetes(100);o__Saccharomycetales(100);f__Metschnikowiaceae(100);g__Metschnikowia(100);g__Metschnikowia_unclassified(100); | Metschnikowia | 13 | Saprotroph | \|Undefined Saprotroph\| | Yeast | NULL | Probable | As nectar-tap_saprotroph (Põlme et al. 2020); Decay sugar-rich substrates (Põlme et al. 2020) | Põlme S, et al. 2020. Fungal Diversity 105:1-16 ((https://doi.org/10.1007/s13225-020-00466-2)) |
| Otu37927 | 1 | k__Fungi(100);p__Ascomycota(100);c__Saccharomycetes(100);o__Saccharomycetales(100);f__Metschnikowiaceae(100);g__Metschnikowia(100);s__Metschnikowia_pulcherrima(100); | Metschnikowia | 13 | Saprotroph | \|Undefined Saprotroph\| | Yeast | NULL | Probable | As nectar-tap_saprotroph (Põlme et al. 2020); Decay sugar-rich substrates (Põlme et al. 2020) | Põlme S, et al. 2020. Fungal Diversity 105:1-16 ((https://doi.org/10.1007/s13225-020-00466-2)) |
| Otu37928 | 1 | k__Fungi(100);p__Ascomycota(100);c__Saccharomycetes(100);o__Saccharomycetales(100);f__Metschnikowiaceae(100);g__Metschnikowia(100);s__Metschnikowia_pulcherrima(100); | Metschnikowia | 13 | Saprotroph | \|Undefined Saprotroph\| | Yeast | NULL | Probable | As nectar-tap_saprotroph (Põlme et al. 2020); Decay sugar-rich substrates (Põlme et al. 2020) | Põlme S, et al. 2020. Fungal Diversity 105:1-16 ((https://doi.org/10.1007/s13225-020-00466-2)) |
| Otu37929 | 1 | k__Fungi(100);p__Ascomycota(100);c__Saccharomycetes(100);o__Saccharomycetales(100);f__Metschnikowiaceae(100);g__Metschnikowia(100);s__Metschnikowia_pulcherrima(100); | Metschnikowia | 13 | Saprotroph | \|Undefined Saprotroph\| | Yeast | NULL | Probable | As nectar-tap_saprotroph (Põlme et al. 2020); Decay sugar-rich substrates (Põlme et al. 2020) | Põlme S, et al. 2020. Fungal Diversity 105:1-16 ((https://doi.org/10.1007/s13225-020-00466-2)) |
| Otu37930 | 1 | k__Fungi(100);p__Ascomycota(100);c__Saccharomycetes(100);o__Saccharomycetales(100);f__Metschnikowiaceae(100);g__Metschnikowia(100);s__Metschnikowia_pulcherrima(100); | Metschnikowia | 13 | Saprotroph | \|Undefined Saprotroph\| | Yeast | NULL | Probable | As nectar-tap_saprotroph (Põlme et al. 2020); Decay sugar-rich substrates (Põlme et al. 2020) | Põlme S, et al. 2020. Fungal Diversity 105:1-16 ((https://doi.org/10.1007/s13225-020-00466-2)) |
| Otu37931 | 1 | k__Fungi(100);p__Ascomycota(100);c__Saccharomycetes(100);o__Saccharomycetales(100);f__Metschnikowiaceae(100);g__Metschnikowia(100);s__Metschnikowia_pulcherrima(100); | Metschnikowia | 13 | Saprotroph | \|Undefined Saprotroph\| | Yeast | NULL | Probable | As nectar-tap_saprotroph (Põlme et al. 2020); Decay sugar-rich substrates (Põlme et al. 2020) | Põlme S, et al. 2020. Fungal Diversity 105:1-16 ((https://doi.org/10.1007/s13225-020-00466-2)) |
| Otu37932 | 1 | k__Fungi(100);p__Ascomycota(100);c__Saccharomycetes(100);o__Saccharomycetales(100);f__Metschnikowiaceae(100);g__Metschnikowia(100);s__Metschnikowia_pulcherrima(100); | Metschnikowia | 13 | Saprotroph | \|Undefined Saprotroph\| | Yeast | NULL | Probable | As nectar-tap_saprotroph (Põlme et al. 2020); Decay sugar-rich substrates (Põlme et al. 2020) | Põlme S, et al. 2020. Fungal Diversity 105:1-16 ((https://doi.org/10.1007/s13225-020-00466-2)) |
| Otu37933 | 1 | k__Fungi(100);p__Ascomycota(100);c__Saccharomycetes(100);o__Saccharomycetales(100);f__Metschnikowiaceae(100);g__Metschnikowia(100);s__Metschnikowia_pulcherrima(100); | Metschnikowia | 13 | Saprotroph | \|Undefined Saprotroph\| | Yeast | NULL | Probable | As nectar-tap_saprotroph (Põlme et al. 2020); Decay sugar-rich substrates (Põlme et al. 2020) | Põlme S, et al. 2020. Fungal Diversity 105:1-16 ((https://doi.org/10.1007/s13225-020-00466-2)) |
| Otu37934 | 1 | k__Fungi(100);p__Ascomycota(100);c__Saccharomycetes(100);o__Saccharomycetales(100);f__Metschnikowiaceae(100);g__Metschnikowia(100);s__Metschnikowia_pulcherrima(100); | Metschnikowia | 13 | Saprotroph | \|Undefined Saprotroph\| | Yeast | NULL | Probable | As nectar-tap_saprotroph (Põlme et al. 2020); Decay sugar-rich substrates (Põlme et al. 2020) | Põlme S, et al. 2020. Fungal Diversity 105:1-16 ((https://doi.org/10.1007/s13225-020-00466-2)) |
| Otu37935 | 1 | k__Fungi(100);p__Ascomycota(100);c__Saccharomycetes(100);o__Saccharomycetales(100);f__Metschnikowiaceae(100);g__Metschnikowia(100);s__Metschnikowia_pulcherrima(100); | Metschnikowia | 13 | Saprotroph | \|Undefined Saprotroph\| | Yeast | NULL | Probable | As nectar-tap_saprotroph (Põlme et al. 2020); Decay sugar-rich substrates (Põlme et al. 2020) | Põlme S, et al. 2020. Fungal Diversity 105:1-16 ((https://doi.org/10.1007/s13225-020-00466-2)) |
| Otu37936 | 1 | k__Fungi(100);p__Ascomycota(100);c__Saccharomycetes(100);o__Saccharomycetales(100);f__Metschnikowiaceae(100);g__Metschnikowia(100);s__Metschnikowia_pulcherrima(100); | Metschnikowia | 13 | Saprotroph | \|Undefined Saprotroph\| | Yeast | NULL | Probable | As nectar-tap_saprotroph (Põlme et al. 2020); Decay sugar-rich substrates (Põlme et al. 2020) | Põlme S, et al. 2020. Fungal Diversity 105:1-16 ((https://doi.org/10.1007/s13225-020-00466-2)) |
| Otu37937 | 1 | k__Fungi(100);p__Ascomycota(100);c__Saccharomycetes(100);o__Saccharomycetales(100);f__Metschnikowiaceae(100);g__Metschnikowia(100);s__Metschnikowia_pulcherrima(100); | Metschnikowia | 13 | Saprotroph | \|Undefined Saprotroph\| | Yeast | NULL | Probable | As nectar-tap_saprotroph (Põlme et al. 2020); Decay sugar-rich substrates (Põlme et al. 2020) | Põlme S, et al. 2020. Fungal Diversity 105:1-16 ((https://doi.org/10.1007/s13225-020-00466-2)) |
| Otu37938 | 1 | k__Fungi(100);p__Ascomycota(100);c__Saccharomycetes(100);o__Saccharomycetales(100);f__Metschnikowiaceae(100);g__Metschnikowia(100);g__Metschnikowia_unclassified(100); | Metschnikowia | 13 | Saprotroph | \|Undefined Saprotroph\| | Yeast | NULL | Probable | As nectar-tap_saprotroph (Põlme et al. 2020); Decay sugar-rich substrates (Põlme et al. 2020) | Põlme S, et al. 2020. Fungal Diversity 105:1-16 ((https://doi.org/10.1007/s13225-020-00466-2)) |
| Otu37939 | 1 | k__Fungi(100);p__Ascomycota(100);c__Saccharomycetes(100);o__Saccharomycetales(100);f__Metschnikowiaceae(100);g__Metschnikowia(100);s__Metschnikowia_pulcherrima(100); | Metschnikowia | 13 | Saprotroph | \|Undefined Saprotroph\| | Yeast | NULL | Probable | As nectar-tap_saprotroph (Põlme et al. 2020); Decay sugar-rich substrates (Põlme et al. 2020) | Põlme S, et al. 2020. Fungal Diversity 105:1-16 ((https://doi.org/10.1007/s13225-020-00466-2)) |
| Otu37940 | 1 | k__Fungi(100);p__Ascomycota(100);c__Saccharomycetes(100);o__Saccharomycetales(100);f__Metschnikowiaceae(100);g__Metschnikowia(100);s__Metschnikowia_pulcherrima(100); | Metschnikowia | 13 | Saprotroph | \|Undefined Saprotroph\| | Yeast | NULL | Probable | As nectar-tap_saprotroph (Põlme et al. 2020); Decay sugar-rich substrates (Põlme et al. 2020) | Põlme S, et al. 2020. Fungal Diversity 105:1-16 ((https://doi.org/10.1007/s13225-020-00466-2)) |
| Otu37941 | 1 | k__Fungi(100);p__Ascomycota(100);c__Saccharomycetes(100);o__Saccharomycetales(100);f__Metschnikowiaceae(100);g__Metschnikowia(100);s__Metschnikowia_pulcherrima(100); | Metschnikowia | 13 | Saprotroph | \|Undefined Saprotroph\| | Yeast | NULL | Probable | As nectar-tap_saprotroph (Põlme et al. 2020); Decay sugar-rich substrates (Põlme et al. 2020) | Põlme S, et al. 2020. Fungal Diversity 105:1-16 ((https://doi.org/10.1007/s13225-020-00466-2)) |
| Otu37942 | 1 | k__Fungi(100);p__Ascomycota(100);c__Saccharomycetes(100);o__Saccharomycetales(100);f__Metschnikowiaceae(100);g__Metschnikowia(100);s__Metschnikowia_pulcherrima(100); | Metschnikowia | 13 | Saprotroph | \|Undefined Saprotroph\| | Yeast | NULL | Probable | As nectar-tap_saprotroph (Põlme et al. 2020); Decay sugar-rich substrates (Põlme et al. 2020) | Põlme S, et al. 2020. Fungal Diversity 105:1-16 ((https://doi.org/10.1007/s13225-020-00466-2)) |
| Otu37943 | 1 | k__Fungi(100);p__Ascomycota(100);c__Saccharomycetes(100);o__Saccharomycetales(100);f__Metschnikowiaceae(100);g__Metschnikowia(100);s__Metschnikowia_pulcherrima(100); | Metschnikowia | 13 | Saprotroph | \|Undefined Saprotroph\| | Yeast | NULL | Probable | As nectar-tap_saprotroph (Põlme et al. 2020); Decay sugar-rich substrates (Põlme et al. 2020) | Põlme S, et al. 2020. Fungal Diversity 105:1-16 ((https://doi.org/10.1007/s13225-020-00466-2)) |
| Otu37944 | 1 | k__Fungi(100);p__Ascomycota(100);c__Saccharomycetes(100);o__Saccharomycetales(100);f__Metschnikowiaceae(100);g__Metschnikowia(100);s__Metschnikowia_pulcherrima(100); | Metschnikowia | 13 | Saprotroph | \|Undefined Saprotroph\| | Yeast | NULL | Probable | As nectar-tap_saprotroph (Põlme et al. 2020); Decay sugar-rich substrates (Põlme et al. 2020) | Põlme S, et al. 2020. Fungal Diversity 105:1-16 ((https://doi.org/10.1007/s13225-020-00466-2)) |
| Otu37945 | 1 | k__Fungi(100);p__Ascomycota(100);c__Saccharomycetes(100);o__Saccharomycetales(100);f__Metschnikowiaceae(100);g__Metschnikowia(100);s__Metschnikowia_pulcherrima(100); | Metschnikowia | 13 | Saprotroph | \|Undefined Saprotroph\| | Yeast | NULL | Probable | As nectar-tap_saprotroph (Põlme et al. 2020); Decay sugar-rich substrates (Põlme et al. 2020) | Põlme S, et al. 2020. Fungal Diversity 105:1-16 ((https://doi.org/10.1007/s13225-020-00466-2)) |
| Otu37946 | 1 | k__Fungi(100);p__Ascomycota(100);c__Saccharomycetes(100);o__Saccharomycetales(100);f__Metschnikowiaceae(100);g__Metschnikowia(100);g__Metschnikowia_unclassified(100); | Metschnikowia | 13 | Saprotroph | \|Undefined Saprotroph\| | Yeast | NULL | Probable | As nectar-tap_saprotroph (Põlme et al. 2020); Decay sugar-rich substrates (Põlme et al. 2020) | Põlme S, et al. 2020. Fungal Diversity 105:1-16 ((https://doi.org/10.1007/s13225-020-00466-2)) |
| Otu37947 | 1 | k__Fungi(100);p__Ascomycota(100);c__Saccharomycetes(100);o__Saccharomycetales(100);f__Metschnikowiaceae(100);g__Metschnikowia(100);s__Metschnikowia_pulcherrima(100); | Metschnikowia | 13 | Saprotroph | \|Undefined Saprotroph\| | Yeast | NULL | Probable | As nectar-tap_saprotroph (Põlme et al. 2020); Decay sugar-rich substrates (Põlme et al. 2020) | Põlme S, et al. 2020. Fungal Diversity 105:1-16 ((https://doi.org/10.1007/s13225-020-00466-2)) |
| Otu37948 | 1 | k__Fungi(100);p__Ascomycota(100);c__Saccharomycetes(100);o__Saccharomycetales(100);f__Metschnikowiaceae(100);g__Metschnikowia(100);s__Metschnikowia_pulcherrima(100); | Metschnikowia | 13 | Saprotroph | \|Undefined Saprotroph\| | Yeast | NULL | Probable | As nectar-tap_saprotroph (Põlme et al. 2020); Decay sugar-rich substrates (Põlme et al. 2020) | Põlme S, et al. 2020. Fungal Diversity 105:1-16 ((https://doi.org/10.1007/s13225-020-00466-2)) |
| Otu37949 | 1 | k__Fungi(100);p__Ascomycota(100);c__Saccharomycetes(100);o__Saccharomycetales(100);f__Metschnikowiaceae(100);g__Metschnikowia(100);s__Metschnikowia_pulcherrima(100); | Metschnikowia | 13 | Saprotroph | \|Undefined Saprotroph\| | Yeast | NULL | Probable | As nectar-tap_saprotroph (Põlme et al. 2020); Decay sugar-rich substrates (Põlme et al. 2020) | Põlme S, et al. 2020. Fungal Diversity 105:1-16 ((https://doi.org/10.1007/s13225-020-00466-2)) |
| Otu37950 | 1 | k__Fungi(100);p__Ascomycota(100);c__Saccharomycetes(100);o__Saccharomycetales(100);f__Metschnikowiaceae(100);g__Metschnikowia(100);g__Metschnikowia_unclassified(100); | Metschnikowia | 13 | Saprotroph | \|Undefined Saprotroph\| | Yeast | NULL | Probable | As nectar-tap_saprotroph (Põlme et al. 2020); Decay sugar-rich substrates (Põlme et al. 2020) | Põlme S, et al. 2020. Fungal Diversity 105:1-16 ((https://doi.org/10.1007/s13225-020-00466-2)) |
| Otu37952 | 1 | k__Fungi(100);p__Ascomycota(100);c__Saccharomycetes(100);o__Saccharomycetales(100);f__Metschnikowiaceae(100);g__Metschnikowia(100);s__Metschnikowia_pulcherrima(100); | Metschnikowia | 13 | Saprotroph | \|Undefined Saprotroph\| | Yeast | NULL | Probable | As nectar-tap_saprotroph (Põlme et al. 2020); Decay sugar-rich substrates (Põlme et al. 2020) | Põlme S, et al. 2020. Fungal Diversity 105:1-16 ((https://doi.org/10.1007/s13225-020-00466-2)) |
| Otu37954 | 1 | k__Fungi(100);p__Ascomycota(100);c__Saccharomycetes(100);o__Saccharomycetales(100);f__Metschnikowiaceae(100);g__Metschnikowia(100);s__Metschnikowia_pulcherrima(100); | Metschnikowia | 13 | Saprotroph | \|Undefined Saprotroph\| | Yeast | NULL | Probable | As nectar-tap_saprotroph (Põlme et al. 2020); Decay sugar-rich substrates (Põlme et al. 2020) | Põlme S, et al. 2020. Fungal Diversity 105:1-16 ((https://doi.org/10.1007/s13225-020-00466-2)) |
| Otu37955 | 1 | k__Fungi(100);p__Ascomycota(100);c__Saccharomycetes(100);o__Saccharomycetales(100);f__Metschnikowiaceae(100);g__Metschnikowia(100);g__Metschnikowia_unclassified(100); | Metschnikowia | 13 | Saprotroph | \|Undefined Saprotroph\| | Yeast | NULL | Probable | As nectar-tap_saprotroph (Põlme et al. 2020); Decay sugar-rich substrates (Põlme et al. 2020) | Põlme S, et al. 2020. Fungal Diversity 105:1-16 ((https://doi.org/10.1007/s13225-020-00466-2)) |
| Otu37956 | 1 | k__Fungi(100);p__Ascomycota(100);c__Saccharomycetes(100);o__Saccharomycetales(100);f__Metschnikowiaceae(100);g__Metschnikowia(100);g__Metschnikowia_unclassified(100); | Metschnikowia | 13 | Saprotroph | \|Undefined Saprotroph\| | Yeast | NULL | Probable | As nectar-tap_saprotroph (Põlme et al. 2020); Decay sugar-rich substrates (Põlme et al. 2020) | Põlme S, et al. 2020. Fungal Diversity 105:1-16 ((https://doi.org/10.1007/s13225-020-00466-2)) |
| Otu37957 | 1 | k__Fungi(100);p__Ascomycota(100);c__Saccharomycetes(100);o__Saccharomycetales(100);f__Metschnikowiaceae(100);g__Metschnikowia(100);s__Metschnikowia_pulcherrima(100); | Metschnikowia | 13 | Saprotroph | \|Undefined Saprotroph\| | Yeast | NULL | Probable | As nectar-tap_saprotroph (Põlme et al. 2020); Decay sugar-rich substrates (Põlme et al. 2020) | Põlme S, et al. 2020. Fungal Diversity 105:1-16 ((https://doi.org/10.1007/s13225-020-00466-2)) |
| Otu37958 | 1 | k__Fungi(100);p__Ascomycota(100);c__Saccharomycetes(100);o__Saccharomycetales(100);f__Metschnikowiaceae(100);g__Metschnikowia(100);s__Metschnikowia_pulcherrima(100); | Metschnikowia | 13 | Saprotroph | \|Undefined Saprotroph\| | Yeast | NULL | Probable | As nectar-tap_saprotroph (Põlme et al. 2020); Decay sugar-rich substrates (Põlme et al. 2020) | Põlme S, et al. 2020. Fungal Diversity 105:1-16 ((https://doi.org/10.1007/s13225-020-00466-2)) |
| Otu37959 | 1 | k__Fungi(100);p__Ascomycota(100);c__Saccharomycetes(100);o__Saccharomycetales(100);f__Metschnikowiaceae(100);g__Metschnikowia(100);s__Metschnikowia_pulcherrima(100); | Metschnikowia | 13 | Saprotroph | \|Undefined Saprotroph\| | Yeast | NULL | Probable | As nectar-tap_saprotroph (Põlme et al. 2020); Decay sugar-rich substrates (Põlme et al. 2020) | Põlme S, et al. 2020. Fungal Diversity 105:1-16 ((https://doi.org/10.1007/s13225-020-00466-2)) |
| Otu37960 | 1 | k__Fungi(100);p__Ascomycota(100);c__Saccharomycetes(100);o__Saccharomycetales(100);f__Metschnikowiaceae(100);g__Metschnikowia(100);s__Metschnikowia_pulcherrima(100); | Metschnikowia | 13 | Saprotroph | \|Undefined Saprotroph\| | Yeast | NULL | Probable | As nectar-tap_saprotroph (Põlme et al. 2020); Decay sugar-rich substrates (Põlme et al. 2020) | Põlme S, et al. 2020. Fungal Diversity 105:1-16 ((https://doi.org/10.1007/s13225-020-00466-2)) |
| Otu37961 | 1 | k__Fungi(100);p__Ascomycota(100);c__Saccharomycetes(100);o__Saccharomycetales(100);f__Metschnikowiaceae(100);g__Metschnikowia(100);s__Metschnikowia_pulcherrima(100); | Metschnikowia | 13 | Saprotroph | \|Undefined Saprotroph\| | Yeast | NULL | Probable | As nectar-tap_saprotroph (Põlme et al. 2020); Decay sugar-rich substrates (Põlme et al. 2020) | Põlme S, et al. 2020. Fungal Diversity 105:1-16 ((https://doi.org/10.1007/s13225-020-00466-2)) |
| Otu37962 | 1 | k__Fungi(100);p__Ascomycota(100);c__Saccharomycetes(100);o__Saccharomycetales(100);f__Metschnikowiaceae(100);g__Metschnikowia(100);s__Metschnikowia_pulcherrima(100); | Metschnikowia | 13 | Saprotroph | \|Undefined Saprotroph\| | Yeast | NULL | Probable | As nectar-tap_saprotroph (Põlme et al. 2020); Decay sugar-rich substrates (Põlme et al. 2020) | Põlme S, et al. 2020. Fungal Diversity 105:1-16 ((https://doi.org/10.1007/s13225-020-00466-2)) |
| Otu37963 | 1 | k__Fungi(100);p__Ascomycota(100);c__Saccharomycetes(100);o__Saccharomycetales(100);f__Metschnikowiaceae(100);g__Metschnikowia(100);s__Metschnikowia_pulcherrima(100); | Metschnikowia | 13 | Saprotroph | \|Undefined Saprotroph\| | Yeast | NULL | Probable | As nectar-tap_saprotroph (Põlme et al. 2020); Decay sugar-rich substrates (Põlme et al. 2020) | Põlme S, et al. 2020. Fungal Diversity 105:1-16 ((https://doi.org/10.1007/s13225-020-00466-2)) |
| Otu37964 | 1 | k__Fungi(100);p__Ascomycota(100);c__Saccharomycetes(100);o__Saccharomycetales(100);f__Metschnikowiaceae(100);g__Metschnikowia(100);g__Metschnikowia_unclassified(100); | Metschnikowia | 13 | Saprotroph | \|Undefined Saprotroph\| | Yeast | NULL | Probable | As nectar-tap_saprotroph (Põlme et al. 2020); Decay sugar-rich substrates (Põlme et al. 2020) | Põlme S, et al. 2020. Fungal Diversity 105:1-16 ((https://doi.org/10.1007/s13225-020-00466-2)) |
| Otu37965 | 1 | k__Fungi(100);p__Ascomycota(100);c__Saccharomycetes(100);o__Saccharomycetales(100);f__Metschnikowiaceae(100);g__Metschnikowia(100);s__Metschnikowia_pulcherrima(100); | Metschnikowia | 13 | Saprotroph | \|Undefined Saprotroph\| | Yeast | NULL | Probable | As nectar-tap_saprotroph (Põlme et al. 2020); Decay sugar-rich substrates (Põlme et al. 2020) | Põlme S, et al. 2020. Fungal Diversity 105:1-16 ((https://doi.org/10.1007/s13225-020-00466-2)) |
| Otu37966 | 1 | k__Fungi(100);p__Ascomycota(100);c__Saccharomycetes(100);o__Saccharomycetales(100);f__Metschnikowiaceae(100);g__Metschnikowia(100);s__Metschnikowia_pulcherrima(100); | Metschnikowia | 13 | Saprotroph | \|Undefined Saprotroph\| | Yeast | NULL | Probable | As nectar-tap_saprotroph (Põlme et al. 2020); Decay sugar-rich substrates (Põlme et al. 2020) | Põlme S, et al. 2020. Fungal Diversity 105:1-16 ((https://doi.org/10.1007/s13225-020-00466-2)) |
| Otu37967 | 1 | k__Fungi(100);p__Ascomycota(100);c__Saccharomycetes(100);o__Saccharomycetales(100);f__Metschnikowiaceae(100);g__Metschnikowia(100);g__Metschnikowia_unclassified(100); | Metschnikowia | 13 | Saprotroph | \|Undefined Saprotroph\| | Yeast | NULL | Probable | As nectar-tap_saprotroph (Põlme et al. 2020); Decay sugar-rich substrates (Põlme et al. 2020) | Põlme S, et al. 2020. Fungal Diversity 105:1-16 ((https://doi.org/10.1007/s13225-020-00466-2)) |
| Otu37968 | 1 | k__Fungi(100);p__Ascomycota(100);c__Saccharomycetes(100);o__Saccharomycetales(100);f__Metschnikowiaceae(100);g__Metschnikowia(100);s__Metschnikowia_pulcherrima(100); | Metschnikowia | 13 | Saprotroph | \|Undefined Saprotroph\| | Yeast | NULL | Probable | As nectar-tap_saprotroph (Põlme et al. 2020); Decay sugar-rich substrates (Põlme et al. 2020) | Põlme S, et al. 2020. Fungal Diversity 105:1-16 ((https://doi.org/10.1007/s13225-020-00466-2)) |
| Otu37969 | 1 | k__Fungi(100);p__Ascomycota(100);c__Saccharomycetes(100);o__Saccharomycetales(100);f__Metschnikowiaceae(100);g__Metschnikowia(100);s__Metschnikowia_pulcherrima(100); | Metschnikowia | 13 | Saprotroph | \|Undefined Saprotroph\| | Yeast | NULL | Probable | As nectar-tap_saprotroph (Põlme et al. 2020); Decay sugar-rich substrates (Põlme et al. 2020) | Põlme S, et al. 2020. Fungal Diversity 105:1-16 ((https://doi.org/10.1007/s13225-020-00466-2)) |
| Otu37970 | 1 | k__Fungi(100);p__Ascomycota(100);c__Saccharomycetes(100);o__Saccharomycetales(100);f__Metschnikowiaceae(100);g__Metschnikowia(100);s__Metschnikowia_pulcherrima(100); | Metschnikowia | 13 | Saprotroph | \|Undefined Saprotroph\| | Yeast | NULL | Probable | As nectar-tap_saprotroph (Põlme et al. 2020); Decay sugar-rich substrates (Põlme et al. 2020) | Põlme S, et al. 2020. Fungal Diversity 105:1-16 ((https://doi.org/10.1007/s13225-020-00466-2)) |
| Otu37971 | 1 | k__Fungi(100);p__Ascomycota(100);c__Saccharomycetes(100);o__Saccharomycetales(100);f__Metschnikowiaceae(100);g__Metschnikowia(100);s__Metschnikowia_pulcherrima(100); | Metschnikowia | 13 | Saprotroph | \|Undefined Saprotroph\| | Yeast | NULL | Probable | As nectar-tap_saprotroph (Põlme et al. 2020); Decay sugar-rich substrates (Põlme et al. 2020) | Põlme S, et al. 2020. Fungal Diversity 105:1-16 ((https://doi.org/10.1007/s13225-020-00466-2)) |
| Otu37972 | 1 | k__Fungi(100);p__Ascomycota(100);c__Saccharomycetes(100);o__Saccharomycetales(100);f__Metschnikowiaceae(100);g__Metschnikowia(100);s__Metschnikowia_pulcherrima(100); | Metschnikowia | 13 | Saprotroph | \|Undefined Saprotroph\| | Yeast | NULL | Probable | As nectar-tap_saprotroph (Põlme et al. 2020); Decay sugar-rich substrates (Põlme et al. 2020) | Põlme S, et al. 2020. Fungal Diversity 105:1-16 ((https://doi.org/10.1007/s13225-020-00466-2)) |
| Otu37973 | 1 | k__Fungi(100);p__Ascomycota(100);c__Saccharomycetes(100);o__Saccharomycetales(100);f__Metschnikowiaceae(100);g__Metschnikowia(100);g__Metschnikowia_unclassified(100); | Metschnikowia | 13 | Saprotroph | \|Undefined Saprotroph\| | Yeast | NULL | Probable | As nectar-tap_saprotroph (Põlme et al. 2020); Decay sugar-rich substrates (Põlme et al. 2020) | Põlme S, et al. 2020. Fungal Diversity 105:1-16 ((https://doi.org/10.1007/s13225-020-00466-2)) |
| Otu37974 | 1 | k__Fungi(100);p__Ascomycota(100);c__Saccharomycetes(100);o__Saccharomycetales(100);f__Metschnikowiaceae(100);g__Metschnikowia(100);s__Metschnikowia_pulcherrima(100); | Metschnikowia | 13 | Saprotroph | \|Undefined Saprotroph\| | Yeast | NULL | Probable | As nectar-tap_saprotroph (Põlme et al. 2020); Decay sugar-rich substrates (Põlme et al. 2020) | Põlme S, et al. 2020. Fungal Diversity 105:1-16 ((https://doi.org/10.1007/s13225-020-00466-2)) |
| Otu37975 | 1 | k__Fungi(100);p__Ascomycota(100);c__Saccharomycetes(100);o__Saccharomycetales(100);f__Metschnikowiaceae(100);g__Metschnikowia(100);s__Metschnikowia_pulcherrima(100); | Metschnikowia | 13 | Saprotroph | \|Undefined Saprotroph\| | Yeast | NULL | Probable | As nectar-tap_saprotroph (Põlme et al. 2020); Decay sugar-rich substrates (Põlme et al. 2020) | Põlme S, et al. 2020. Fungal Diversity 105:1-16 ((https://doi.org/10.1007/s13225-020-00466-2)) |
| Otu37976 | 1 | k__Fungi(100);p__Ascomycota(100);c__Saccharomycetes(100);o__Saccharomycetales(100);f__Metschnikowiaceae(100);g__Metschnikowia(100);g__Metschnikowia_unclassified(100); | Metschnikowia | 13 | Saprotroph | \|Undefined Saprotroph\| | Yeast | NULL | Probable | As nectar-tap_saprotroph (Põlme et al. 2020); Decay sugar-rich substrates (Põlme et al. 2020) | Põlme S, et al. 2020. Fungal Diversity 105:1-16 ((https://doi.org/10.1007/s13225-020-00466-2)) |
| Otu37977 | 1 | k__Fungi(100);p__Ascomycota(100);c__Saccharomycetes(100);o__Saccharomycetales(100);f__Metschnikowiaceae(100);g__Metschnikowia(100);g__Metschnikowia_unclassified(100); | Metschnikowia | 13 | Saprotroph | \|Undefined Saprotroph\| | Yeast | NULL | Probable | As nectar-tap_saprotroph (Põlme et al. 2020); Decay sugar-rich substrates (Põlme et al. 2020) | Põlme S, et al. 2020. Fungal Diversity 105:1-16 ((https://doi.org/10.1007/s13225-020-00466-2)) |
| Otu37978 | 1 | k__Fungi(100);p__Ascomycota(100);c__Saccharomycetes(100);o__Saccharomycetales(100);f__Metschnikowiaceae(100);g__Metschnikowia(100);g__Metschnikowia_unclassified(100); | Metschnikowia | 13 | Saprotroph | \|Undefined Saprotroph\| | Yeast | NULL | Probable | As nectar-tap_saprotroph (Põlme et al. 2020); Decay sugar-rich substrates (Põlme et al. 2020) | Põlme S, et al. 2020. Fungal Diversity 105:1-16 ((https://doi.org/10.1007/s13225-020-00466-2)) |
| Otu37979 | 1 | k__Fungi(100);p__Ascomycota(100);c__Saccharomycetes(100);o__Saccharomycetales(100);f__Metschnikowiaceae(100);g__Metschnikowia(100);s__Metschnikowia_pulcherrima(100); | Metschnikowia | 13 | Saprotroph | \|Undefined Saprotroph\| | Yeast | NULL | Probable | As nectar-tap_saprotroph (Põlme et al. 2020); Decay sugar-rich substrates (Põlme et al. 2020) | Põlme S, et al. 2020. Fungal Diversity 105:1-16 ((https://doi.org/10.1007/s13225-020-00466-2)) |
| Otu37980 | 1 | k__Fungi(100);p__Ascomycota(100);c__Saccharomycetes(100);o__Saccharomycetales(100);f__Metschnikowiaceae(100);g__Metschnikowia(100);g__Metschnikowia_unclassified(100); | Metschnikowia | 13 | Saprotroph | \|Undefined Saprotroph\| | Yeast | NULL | Probable | As nectar-tap_saprotroph (Põlme et al. 2020); Decay sugar-rich substrates (Põlme et al. 2020) | Põlme S, et al. 2020. Fungal Diversity 105:1-16 ((https://doi.org/10.1007/s13225-020-00466-2)) |
| Otu37981 | 1 | k__Fungi(100);p__Ascomycota(100);c__Saccharomycetes(100);o__Saccharomycetales(100);f__Metschnikowiaceae(100);g__Metschnikowia(100);s__Metschnikowia_pulcherrima(100); | Metschnikowia | 13 | Saprotroph | \|Undefined Saprotroph\| | Yeast | NULL | Probable | As nectar-tap_saprotroph (Põlme et al. 2020); Decay sugar-rich substrates (Põlme et al. 2020) | Põlme S, et al. 2020. Fungal Diversity 105:1-16 ((https://doi.org/10.1007/s13225-020-00466-2)) |
| Otu37982 | 1 | k__Fungi(100);p__Ascomycota(100);c__Saccharomycetes(100);o__Saccharomycetales(100);f__Metschnikowiaceae(100);g__Metschnikowia(100);s__Metschnikowia_pulcherrima(100); | Metschnikowia | 13 | Saprotroph | \|Undefined Saprotroph\| | Yeast | NULL | Probable | As nectar-tap_saprotroph (Põlme et al. 2020); Decay sugar-rich substrates (Põlme et al. 2020) | Põlme S, et al. 2020. Fungal Diversity 105:1-16 ((https://doi.org/10.1007/s13225-020-00466-2)) |
| Otu37983 | 1 | k__Fungi(100);p__Ascomycota(100);c__Saccharomycetes(100);o__Saccharomycetales(100);f__Metschnikowiaceae(100);g__Metschnikowia(100);s__Metschnikowia_pulcherrima(100); | Metschnikowia | 13 | Saprotroph | \|Undefined Saprotroph\| | Yeast | NULL | Probable | As nectar-tap_saprotroph (Põlme et al. 2020); Decay sugar-rich substrates (Põlme et al. 2020) | Põlme S, et al. 2020. Fungal Diversity 105:1-16 ((https://doi.org/10.1007/s13225-020-00466-2)) |
| Otu37984 | 1 | k__Fungi(100);p__Ascomycota(100);c__Saccharomycetes(100);o__Saccharomycetales(100);f__Metschnikowiaceae(100);g__Metschnikowia(100);s__Metschnikowia_pulcherrima(100); | Metschnikowia | 13 | Saprotroph | \|Undefined Saprotroph\| | Yeast | NULL | Probable | As nectar-tap_saprotroph (Põlme et al. 2020); Decay sugar-rich substrates (Põlme et al. 2020) | Põlme S, et al. 2020. Fungal Diversity 105:1-16 ((https://doi.org/10.1007/s13225-020-00466-2)) |
| Otu37985 | 1 | k__Fungi(100);p__Ascomycota(100);c__Saccharomycetes(100);o__Saccharomycetales(100);f__Metschnikowiaceae(100);g__Metschnikowia(100);s__Metschnikowia_pulcherrima(100); | Metschnikowia | 13 | Saprotroph | \|Undefined Saprotroph\| | Yeast | NULL | Probable | As nectar-tap_saprotroph (Põlme et al. 2020); Decay sugar-rich substrates (Põlme et al. 2020) | Põlme S, et al. 2020. Fungal Diversity 105:1-16 ((https://doi.org/10.1007/s13225-020-00466-2)) |
| Otu37986 | 1 | k__Fungi(100);p__Ascomycota(100);c__Saccharomycetes(100);o__Saccharomycetales(100);f__Metschnikowiaceae(100);g__Metschnikowia(100);s__Metschnikowia_pulcherrima(100); | Metschnikowia | 13 | Saprotroph | \|Undefined Saprotroph\| | Yeast | NULL | Probable | As nectar-tap_saprotroph (Põlme et al. 2020); Decay sugar-rich substrates (Põlme et al. 2020) | Põlme S, et al. 2020. Fungal Diversity 105:1-16 ((https://doi.org/10.1007/s13225-020-00466-2)) |
| Otu37987 | 1 | k__Fungi(100);p__Ascomycota(100);c__Saccharomycetes(100);o__Saccharomycetales(100);f__Metschnikowiaceae(100);g__Metschnikowia(100);g__Metschnikowia_unclassified(100); | Metschnikowia | 13 | Saprotroph | \|Undefined Saprotroph\| | Yeast | NULL | Probable | As nectar-tap_saprotroph (Põlme et al. 2020); Decay sugar-rich substrates (Põlme et al. 2020) | Põlme S, et al. 2020. Fungal Diversity 105:1-16 ((https://doi.org/10.1007/s13225-020-00466-2)) |
| Otu37988 | 1 | k__Fungi(100);p__Ascomycota(100);c__Saccharomycetes(100);o__Saccharomycetales(100);f__Metschnikowiaceae(100);g__Metschnikowia(100);s__Metschnikowia_pulcherrima(100); | Metschnikowia | 13 | Saprotroph | \|Undefined Saprotroph\| | Yeast | NULL | Probable | As nectar-tap_saprotroph (Põlme et al. 2020); Decay sugar-rich substrates (Põlme et al. 2020) | Põlme S, et al. 2020. Fungal Diversity 105:1-16 ((https://doi.org/10.1007/s13225-020-00466-2)) |
| Otu37989 | 1 | k__Fungi(100);p__Ascomycota(100);c__Saccharomycetes(100);o__Saccharomycetales(100);f__Metschnikowiaceae(100);g__Metschnikowia(100);s__Metschnikowia_pulcherrima(100); | Metschnikowia | 13 | Saprotroph | \|Undefined Saprotroph\| | Yeast | NULL | Probable | As nectar-tap_saprotroph (Põlme et al. 2020); Decay sugar-rich substrates (Põlme et al. 2020) | Põlme S, et al. 2020. Fungal Diversity 105:1-16 ((https://doi.org/10.1007/s13225-020-00466-2)) |
| Otu37990 | 1 | k__Fungi(100);p__Ascomycota(100);c__Saccharomycetes(100);o__Saccharomycetales(100);f__Metschnikowiaceae(100);g__Metschnikowia(100);g__Metschnikowia_unclassified(100); | Metschnikowia | 13 | Saprotroph | \|Undefined Saprotroph\| | Yeast | NULL | Probable | As nectar-tap_saprotroph (Põlme et al. 2020); Decay sugar-rich substrates (Põlme et al. 2020) | Põlme S, et al. 2020. Fungal Diversity 105:1-16 ((https://doi.org/10.1007/s13225-020-00466-2)) |
| Otu37991 | 1 | k__Fungi(100);p__Ascomycota(100);c__Saccharomycetes(100);o__Saccharomycetales(100);f__Metschnikowiaceae(100);g__Metschnikowia(100);s__Metschnikowia_pulcherrima(100); | Metschnikowia | 13 | Saprotroph | \|Undefined Saprotroph\| | Yeast | NULL | Probable | As nectar-tap_saprotroph (Põlme et al. 2020); Decay sugar-rich substrates (Põlme et al. 2020) | Põlme S, et al. 2020. Fungal Diversity 105:1-16 ((https://doi.org/10.1007/s13225-020-00466-2)) |
| Otu37992 | 1 | k__Fungi(100);p__Ascomycota(100);c__Saccharomycetes(100);o__Saccharomycetales(100);f__Metschnikowiaceae(100);g__Metschnikowia(100);s__Metschnikowia_pulcherrima(100); | Metschnikowia | 13 | Saprotroph | \|Undefined Saprotroph\| | Yeast | NULL | Probable | As nectar-tap_saprotroph (Põlme et al. 2020); Decay sugar-rich substrates (Põlme et al. 2020) | Põlme S, et al. 2020. Fungal Diversity 105:1-16 ((https://doi.org/10.1007/s13225-020-00466-2)) |
| Otu37993 | 1 | k__Fungi(100);p__Ascomycota(100);c__Saccharomycetes(100);o__Saccharomycetales(100);f__Metschnikowiaceae(100);g__Metschnikowia(100);s__Metschnikowia_pulcherrima(100); | Metschnikowia | 13 | Saprotroph | \|Undefined Saprotroph\| | Yeast | NULL | Probable | As nectar-tap_saprotroph (Põlme et al. 2020); Decay sugar-rich substrates (Põlme et al. 2020) | Põlme S, et al. 2020. Fungal Diversity 105:1-16 ((https://doi.org/10.1007/s13225-020-00466-2)) |
| Otu37995 | 1 | k__Fungi(100);p__Ascomycota(100);c__Saccharomycetes(100);o__Saccharomycetales(100);f__Metschnikowiaceae(100);g__Metschnikowia(100);s__Metschnikowia_pulcherrima(100); | Metschnikowia | 13 | Saprotroph | \|Undefined Saprotroph\| | Yeast | NULL | Probable | As nectar-tap_saprotroph (Põlme et al. 2020); Decay sugar-rich substrates (Põlme et al. 2020) | Põlme S, et al. 2020. Fungal Diversity 105:1-16 ((https://doi.org/10.1007/s13225-020-00466-2)) |
| Otu37996 | 1 | k__Fungi(100);p__Ascomycota(100);c__Saccharomycetes(100);o__Saccharomycetales(100);f__Metschnikowiaceae(100);g__Metschnikowia(100);s__Metschnikowia_pulcherrima(100); | Metschnikowia | 13 | Saprotroph | \|Undefined Saprotroph\| | Yeast | NULL | Probable | As nectar-tap_saprotroph (Põlme et al. 2020); Decay sugar-rich substrates (Põlme et al. 2020) | Põlme S, et al. 2020. Fungal Diversity 105:1-16 ((https://doi.org/10.1007/s13225-020-00466-2)) |
| Otu37997 | 1 | k__Fungi(100);p__Ascomycota(100);c__Saccharomycetes(100);o__Saccharomycetales(100);f__Metschnikowiaceae(100);g__Metschnikowia(100);g__Metschnikowia_unclassified(100); | Metschnikowia | 13 | Saprotroph | \|Undefined Saprotroph\| | Yeast | NULL | Probable | As nectar-tap_saprotroph (Põlme et al. 2020); Decay sugar-rich substrates (Põlme et al. 2020) | Põlme S, et al. 2020. Fungal Diversity 105:1-16 ((https://doi.org/10.1007/s13225-020-00466-2)) |
| Otu37998 | 1 | k__Fungi(100);p__Ascomycota(100);c__Saccharomycetes(100);o__Saccharomycetales(100);f__Metschnikowiaceae(100);g__Metschnikowia(100);s__Metschnikowia_pulcherrima(100); | Metschnikowia | 13 | Saprotroph | \|Undefined Saprotroph\| | Yeast | NULL | Probable | As nectar-tap_saprotroph (Põlme et al. 2020); Decay sugar-rich substrates (Põlme et al. 2020) | Põlme S, et al. 2020. Fungal Diversity 105:1-16 ((https://doi.org/10.1007/s13225-020-00466-2)) |
| Otu37999 | 1 | k__Fungi(100);p__Ascomycota(100);c__Saccharomycetes(100);o__Saccharomycetales(100);f__Metschnikowiaceae(100);g__Metschnikowia(100);s__Metschnikowia_pulcherrima(100); | Metschnikowia | 13 | Saprotroph | \|Undefined Saprotroph\| | Yeast | NULL | Probable | As nectar-tap_saprotroph (Põlme et al. 2020); Decay sugar-rich substrates (Põlme et al. 2020) | Põlme S, et al. 2020. Fungal Diversity 105:1-16 ((https://doi.org/10.1007/s13225-020-00466-2)) |
| Otu38000 | 1 | k__Fungi(100);p__Ascomycota(100);c__Saccharomycetes(100);o__Saccharomycetales(100);f__Metschnikowiaceae(100);g__Metschnikowia(100);g__Metschnikowia_unclassified(100); | Metschnikowia | 13 | Saprotroph | \|Undefined Saprotroph\| | Yeast | NULL | Probable | As nectar-tap_saprotroph (Põlme et al. 2020); Decay sugar-rich substrates (Põlme et al. 2020) | Põlme S, et al. 2020. Fungal Diversity 105:1-16 ((https://doi.org/10.1007/s13225-020-00466-2)) |
| Otu38001 | 1 | k__Fungi(100);p__Ascomycota(100);c__Saccharomycetes(100);o__Saccharomycetales(100);f__Metschnikowiaceae(100);g__Metschnikowia(100);g__Metschnikowia_unclassified(100); | Metschnikowia | 13 | Saprotroph | \|Undefined Saprotroph\| | Yeast | NULL | Probable | As nectar-tap_saprotroph (Põlme et al. 2020); Decay sugar-rich substrates (Põlme et al. 2020) | Põlme S, et al. 2020. Fungal Diversity 105:1-16 ((https://doi.org/10.1007/s13225-020-00466-2)) |
| Otu38002 | 1 | k__Fungi(100);p__Ascomycota(100);c__Saccharomycetes(100);o__Saccharomycetales(100);f__Metschnikowiaceae(100);g__Metschnikowia(100);s__Metschnikowia_pulcherrima(100); | Metschnikowia | 13 | Saprotroph | \|Undefined Saprotroph\| | Yeast | NULL | Probable | As nectar-tap_saprotroph (Põlme et al. 2020); Decay sugar-rich substrates (Põlme et al. 2020) | Põlme S, et al. 2020. Fungal Diversity 105:1-16 ((https://doi.org/10.1007/s13225-020-00466-2)) |
| Otu38003 | 1 | k__Fungi(100);p__Ascomycota(100);c__Saccharomycetes(100);o__Saccharomycetales(100);f__Metschnikowiaceae(100);g__Metschnikowia(100);s__Metschnikowia_pulcherrima(100); | Metschnikowia | 13 | Saprotroph | \|Undefined Saprotroph\| | Yeast | NULL | Probable | As nectar-tap_saprotroph (Põlme et al. 2020); Decay sugar-rich substrates (Põlme et al. 2020) | Põlme S, et al. 2020. Fungal Diversity 105:1-16 ((https://doi.org/10.1007/s13225-020-00466-2)) |
| Otu38004 | 1 | k__Fungi(100);p__Ascomycota(100);c__Saccharomycetes(100);o__Saccharomycetales(100);f__Metschnikowiaceae(100);g__Metschnikowia(100);s__Metschnikowia_pulcherrima(100); | Metschnikowia | 13 | Saprotroph | \|Undefined Saprotroph\| | Yeast | NULL | Probable | As nectar-tap_saprotroph (Põlme et al. 2020); Decay sugar-rich substrates (Põlme et al. 2020) | Põlme S, et al. 2020. Fungal Diversity 105:1-16 ((https://doi.org/10.1007/s13225-020-00466-2)) |

| | | | | | | | | | | | |
|---|---|---|---|---|---|---|---|---|---|---|---|
| Otu38005 | 1 | k__Fungi(100);p__Ascomycota(100);c__Saccharomycetes(100);o__Saccharomycetales(100);f__Metschnikowiaceae(100);g__Metschnikowia(100);s__Metschnikowia_pulcherrima(100); | Metschnikowia | 13 | Saprotroph | \|Undefined Saprotroph\| | Yeast | NULL | Probable | As nectar-tap_saprotroph (Põlme et al. 2020); Decay sugar-rich substrates (Põlme et al. 2020) | Põlme S, et al. 2020. Fungal Diversity 105:1-16 ((https://doi.org/10.1007/s13225-020-00466-2)) |
| Otu38006 | 1 | k__Fungi(100);p__Ascomycota(100);c__Saccharomycetes(100);o__Saccharomycetales(100);f__Metschnikowiaceae(100);g__Metschnikowia(100);s__Metschnikowia_pulcherrima(100); | Metschnikowia | 13 | Saprotroph | \|Undefined Saprotroph\| | Yeast | NULL | Probable | As nectar-tap_saprotroph (Põlme et al. 2020); Decay sugar-rich substrates (Põlme et al. 2020) | Põlme S, et al. 2020. Fungal Diversity 105:1-16 ((https://doi.org/10.1007/s13225-020-00466-2)) |
| Otu38007 | 1 | k__Fungi(100);p__Ascomycota(100);c__Saccharomycetes(100);o__Saccharomycetales(100);f__Metschnikowiaceae(100);g__Metschnikowia(100);s__Metschnikowia_pulcherrima(100); | Metschnikowia | 13 | Saprotroph | \|Undefined Saprotroph\| | Yeast | NULL | Probable | As nectar-tap_saprotroph (Põlme et al. 2020); Decay sugar-rich substrates (Põlme et al. 2020) | Põlme S, et al. 2020. Fungal Diversity 105:1-16 ((https://doi.org/10.1007/s13225-020-00466-2)) |
| Otu38008 | 1 | k__Fungi(100);p__Ascomycota(100);c__Saccharomycetes(100);o__Saccharomycetales(100);f__Metschnikowiaceae(100);g__Metschnikowia(100);s__Metschnikowia_pulcherrima(100); | Metschnikowia | 13 | Saprotroph | \|Undefined Saprotroph\| | Yeast | NULL | Probable | As nectar-tap_saprotroph (Põlme et al. 2020); Decay sugar-rich substrates (Põlme et al. 2020) | Põlme S, et al. 2020. Fungal Diversity 105:1-16 ((https://doi.org/10.1007/s13225-020-00466-2)) |
| Otu38009 | 1 | k__Fungi(100);p__Ascomycota(100);c__Saccharomycetes(100);o__Saccharomycetales(100);f__Metschnikowiaceae(100);g__Metschnikowia(100);s__Metschnikowia_pulcherrima(100); | Metschnikowia | 13 | Saprotroph | \|Undefined Saprotroph\| | Yeast | NULL | Probable | As nectar-tap_saprotroph (Põlme et al. 2020); Decay sugar-rich substrates (Põlme et al. 2020) | Põlme S, et al. 2020. Fungal Diversity 105:1-16 ((https://doi.org/10.1007/s13225-020-00466-2)) |
| Otu38010 | 1 | k__Fungi(100);p__Ascomycota(100);c__Saccharomycetes(100);o__Saccharomycetales(100);f__Metschnikowiaceae(100);g__Metschnikowia(100);s__Metschnikowia_pulcherrima(100); | Metschnikowia | 13 | Saprotroph | \|Undefined Saprotroph\| | Yeast | NULL | Probable | As nectar-tap_saprotroph (Põlme et al. 2020); Decay sugar-rich substrates (Põlme et al. 2020) | Põlme S, et al. 2020. Fungal Diversity 105:1-16 ((https://doi.org/10.1007/s13225-020-00466-2)) |
| Otu38011 | 1 | k__Fungi(100);p__Ascomycota(100);c__Saccharomycetes(100);o__Saccharomycetales(100);f__Metschnikowiaceae(100);g__Metschnikowia(100);s__Metschnikowia_pulcherrima(100); | Metschnikowia | 13 | Saprotroph | \|Undefined Saprotroph\| | Yeast | NULL | Probable | As nectar-tap_saprotroph (Põlme et al. 2020); Decay sugar-rich substrates (Põlme et al. 2020) | Põlme S, et al. 2020. Fungal Diversity 105:1-16 ((https://doi.org/10.1007/s13225-020-00466-2)) |
| Otu38012 | 1 | k__Fungi(100);p__Ascomycota(100);c__Saccharomycetes(100);o__Saccharomycetales(100);f__Metschnikowiaceae(100);g__Metschnikowia(100);s__Metschnikowia_pulcherrima(100); | Metschnikowia | 13 | Saprotroph | \|Undefined Saprotroph\| | Yeast | NULL | Probable | As nectar-tap_saprotroph (Põlme et al. 2020); Decay sugar-rich substrates (Põlme et al. 2020) | Põlme S, et al. 2020. Fungal Diversity 105:1-16 ((https://doi.org/10.1007/s13225-020-00466-2)) |
| Otu38013 | 1 | k__Fungi(100);p__Ascomycota(100);c__Saccharomycetes(100);o__Saccharomycetales(100);f__Metschnikowiaceae(100);g__Metschnikowia(100);s__Metschnikowia_pulcherrima(100); | Metschnikowia | 13 | Saprotroph | \|Undefined Saprotroph\| | Yeast | NULL | Probable | As nectar-tap_saprotroph (Põlme et al. 2020); Decay sugar-rich substrates (Põlme et al. 2020) | Põlme S, et al. 2020. Fungal Diversity 105:1-16 ((https://doi.org/10.1007/s13225-020-00466-2)) |
| Otu38014 | 1 | k__Fungi(100);p__Ascomycota(100);c__Saccharomycetes(100);o__Saccharomycetales(100);f__Metschnikowiaceae(100);g__Metschnikowia(100);s__Metschnikowia_pulcherrima(100); | Metschnikowia | 13 | Saprotroph | \|Undefined Saprotroph\| | Yeast | NULL | Probable | As nectar-tap_saprotroph (Põlme et al. 2020); Decay sugar-rich substrates (Põlme et al. 2020) | Põlme S, et al. 2020. Fungal Diversity 105:1-16 ((https://doi.org/10.1007/s13225-020-00466-2)) |
| Otu38015 | 1 | k__Fungi(100);p__Ascomycota(100);c__Saccharomycetes(100);o__Saccharomycetales(100);f__Metschnikowiaceae(100);g__Metschnikowia(100);s__Metschnikowia_pulcherrima(100); | Metschnikowia | 13 | Saprotroph | \|Undefined Saprotroph\| | Yeast | NULL | Probable | As nectar-tap_saprotroph (Põlme et al. 2020); Decay sugar-rich substrates (Põlme et al. 2020) | Põlme S, et al. 2020. Fungal Diversity 105:1-16 ((https://doi.org/10.1007/s13225-020-00466-2)) |
| Otu38017 | 1 | k__Fungi(100);p__Ascomycota(100);c__Saccharomycetes(100);o__Saccharomycetales(100);f__Metschnikowiaceae(100);g__Metschnikowia(100);s__Metschnikowia_pulcherrima(100); | Metschnikowia | 13 | Saprotroph | \|Undefined Saprotroph\| | Yeast | NULL | Probable | As nectar-tap_saprotroph (Põlme et al. 2020); Decay sugar-rich substrates (Põlme et al. 2020) | Põlme S, et al. 2020. Fungal Diversity 105:1-16 ((https://doi.org/10.1007/s13225-020-00466-2)) |
| Otu38018 | 1 | k__Fungi(100);p__Ascomycota(100);c__Saccharomycetes(100);o__Saccharomycetales(100);f__Metschnikowiaceae(100);g__Metschnikowia(100);s__Metschnikowia_pulcherrima(100); | Metschnikowia | 13 | Saprotroph | \|Undefined Saprotroph\| | Yeast | NULL | Probable | As nectar-tap_saprotroph (Põlme et al. 2020); Decay sugar-rich substrates (Põlme et al. 2020) | Põlme S, et al. 2020. Fungal Diversity 105:1-16 ((https://doi.org/10.1007/s13225-020-00466-2)) |
| Otu38019 | 1 | k__Fungi(100);p__Ascomycota(100);c__Saccharomycetes(100);o__Saccharomycetales(100);f__Metschnikowiaceae(100);g__Metschnikowia(100);s__Metschnikowia_pulcherrima(100); | Metschnikowia | 13 | Saprotroph | \|Undefined Saprotroph\| | Yeast | NULL | Probable | As nectar-tap_saprotroph (Põlme et al. 2020); Decay sugar-rich substrates (Põlme et al. 2020) | Põlme S, et al. 2020. Fungal Diversity 105:1-16 ((https://doi.org/10.1007/s13225-020-00466-2)) |
| Otu38020 | 1 | k__Fungi(100);p__Ascomycota(100);c__Saccharomycetes(100);o__Saccharomycetales(100);f__Metschnikowiaceae(100);g__Metschnikowia(100);s__Metschnikowia_pulcherrima(100); | Metschnikowia | 13 | Saprotroph | \|Undefined Saprotroph\| | Yeast | NULL | Probable | As nectar-tap_saprotroph (Põlme et al. 2020); Decay sugar-rich substrates (Põlme et al. 2020) | Põlme S, et al. 2020. Fungal Diversity 105:1-16 ((https://doi.org/10.1007/s13225-020-00466-2)) |
| Otu38021 | 1 | k__Fungi(100);p__Ascomycota(100);c__Saccharomycetes(100);o__Saccharomycetales(100);f__Metschnikowiaceae(100);g__Metschnikowia(100);g__Metschnikowia_unclassified(100); | Metschnikowia | 13 | Saprotroph | \|Undefined Saprotroph\| | Yeast | NULL | Probable | As nectar-tap_saprotroph (Põlme et al. 2020); Decay sugar-rich substrates (Põlme et al. 2020) | Põlme S, et al. 2020. Fungal Diversity 105:1-16 ((https://doi.org/10.1007/s13225-020-00466-2)) |
| Otu38022 | 1 | k__Fungi(100);p__Ascomycota(100);c__Saccharomycetes(100);o__Saccharomycetales(100);f__Metschnikowiaceae(100);g__Metschnikowia(100);g__Metschnikowia_unclassified(100); | Metschnikowia | 13 | Saprotroph | \|Undefined Saprotroph\| | Yeast | NULL | Probable | As nectar-tap_saprotroph (Põlme et al. 2020); Decay sugar-rich substrates (Põlme et al. 2020) | Põlme S, et al. 2020. Fungal Diversity 105:1-16 ((https://doi.org/10.1007/s13225-020-00466-2)) |
| Otu38023 | 1 | k__Fungi(100);p__Ascomycota(100);c__Saccharomycetes(100);o__Saccharomycetales(100);f__Metschnikowiaceae(100);g__Metschnikowia(100);s__Metschnikowia_pulcherrima(100); | Metschnikowia | 13 | Saprotroph | \|Undefined Saprotroph\| | Yeast | NULL | Probable | As nectar-tap_saprotroph (Põlme et al. 2020); Decay sugar-rich substrates (Põlme et al. 2020) | Põlme S, et al. 2020. Fungal Diversity 105:1-16 ((https://doi.org/10.1007/s13225-020-00466-2)) |
| Otu38024 | 1 | k__Fungi(100);p__Ascomycota(100);c__Saccharomycetes(100);o__Saccharomycetales(100);f__Metschnikowiaceae(100);g__Metschnikowia(100);g__Metschnikowia_unclassified(100); | Metschnikowia | 13 | Saprotroph | \|Undefined Saprotroph\| | Yeast | NULL | Probable | As nectar-tap_saprotroph (Põlme et al. 2020); Decay sugar-rich substrates (Põlme et al. 2020) | Põlme S, et al. 2020. Fungal Diversity 105:1-16 ((https://doi.org/10.1007/s13225-020-00466-2)) |
| Otu38026 | 1 | k__Fungi(100);p__Ascomycota(100);c__Saccharomycetes(100);o__Saccharomycetales(100);f__Metschnikowiaceae(100);g__Metschnikowia(100);g__Metschnikowia_unclassified(100); | Metschnikowia | 13 | Saprotroph | \|Undefined Saprotroph\| | Yeast | NULL | Probable | As nectar-tap_saprotroph (Põlme et al. 2020); Decay sugar-rich substrates (Põlme et al. 2020) | Põlme S, et al. 2020. Fungal Diversity 105:1-16 ((https://doi.org/10.1007/s13225-020-00466-2)) |
| Otu38027 | 1 | k__Fungi(100);p__Ascomycota(100);c__Saccharomycetes(100);o__Saccharomycetales(100);f__Metschnikowiaceae(100);g__Metschnikowia(100);s__Metschnikowia_pulcherrima(100); | Metschnikowia | 13 | Saprotroph | \|Undefined Saprotroph\| | Yeast | NULL | Probable | As nectar-tap_saprotroph (Põlme et al. 2020); Decay sugar-rich substrates (Põlme et al. 2020) | Põlme S, et al. 2020. Fungal Diversity 105:1-16 ((https://doi.org/10.1007/s13225-020-00466-2)) |
| Otu38028 | 1 | k__Fungi(100);p__Ascomycota(100);c__Saccharomycetes(100);o__Saccharomycetales(100);f__Metschnikowiaceae(100);g__Metschnikowia(100);s__Metschnikowia_pulcherrima(100); | Metschnikowia | 13 | Saprotroph | \|Undefined Saprotroph\| | Yeast | NULL | Probable | As nectar-tap_saprotroph (Põlme et al. 2020); Decay sugar-rich substrates (Põlme et al. 2020) | Põlme S, et al. 2020. Fungal Diversity 105:1-16 ((https://doi.org/10.1007/s13225-020-00466-2)) |
| Otu38029 | 1 | k__Fungi(100);p__Ascomycota(100);c__Saccharomycetes(100);o__Saccharomycetales(100);f__Metschnikowiaceae(100);g__Metschnikowia(100);s__Metschnikowia_pulcherrima(100); | Metschnikowia | 13 | Saprotroph | \|Undefined Saprotroph\| | Yeast | NULL | Probable | As nectar-tap_saprotroph (Põlme et al. 2020); Decay sugar-rich substrates (Põlme et al. 2020) | Põlme S, et al. 2020. Fungal Diversity 105:1-16 ((https://doi.org/10.1007/s13225-020-00466-2)) |
| Otu38030 | 1 | k__Fungi(100);p__Ascomycota(100);c__Saccharomycetes(100);o__Saccharomycetales(100);f__Metschnikowiaceae(100);g__Metschnikowia(100);s__Metschnikowia_pulcherrima(100); | Metschnikowia | 13 | Saprotroph | \|Undefined Saprotroph\| | Yeast | NULL | Probable | As nectar-tap_saprotroph (Põlme et al. 2020); Decay sugar-rich substrates (Põlme et al. 2020) | Põlme S, et al. 2020. Fungal Diversity 105:1-16 ((https://doi.org/10.1007/s13225-020-00466-2)) |
| Otu38031 | 1 | k__Fungi(100);p__Ascomycota(100);c__Saccharomycetes(100);o__Saccharomycetales(100);f__Metschnikowiaceae(100);g__Metschnikowia(100);s__Metschnikowia_pulcherrima(100); | Metschnikowia | 13 | Saprotroph | \|Undefined Saprotroph\| | Yeast | NULL | Probable | As nectar-tap_saprotroph (Põlme et al. 2020); Decay sugar-rich substrates (Põlme et al. 2020) | Põlme S, et al. 2020. Fungal Diversity 105:1-16 ((https://doi.org/10.1007/s13225-020-00466-2)) |
| Otu38032 | 1 | k__Fungi(100);p__Ascomycota(100);c__Saccharomycetes(100);o__Saccharomycetales(100);f__Metschnikowiaceae(100);g__Metschnikowia(100);s__Metschnikowia_pulcherrima(100); | Metschnikowia | 13 | Saprotroph | \|Undefined Saprotroph\| | Yeast | NULL | Probable | As nectar-tap_saprotroph (Põlme et al. 2020); Decay sugar-rich substrates (Põlme et al. 2020) | Põlme S, et al. 2020. Fungal Diversity 105:1-16 ((https://doi.org/10.1007/s13225-020-00466-2)) |
| Otu38033 | 1 | k__Fungi(100);p__Ascomycota(100);c__Saccharomycetes(100);o__Saccharomycetales(100);f__Metschnikowiaceae(100);g__Metschnikowia(100);s__Metschnikowia_pulcherrima(100); | Metschnikowia | 13 | Saprotroph | \|Undefined Saprotroph\| | Yeast | NULL | Probable | As nectar-tap_saprotroph (Põlme et al. 2020); Decay sugar-rich substrates (Põlme et al. 2020) | Põlme S, et al. 2020. Fungal Diversity 105:1-16 ((https://doi.org/10.1007/s13225-020-00466-2)) |
| Otu38034 | 1 | k__Fungi(100);p__Ascomycota(100);c__Saccharomycetes(100);o__Saccharomycetales(100);f__Metschnikowiaceae(100);g__Metschnikowia(100);s__Metschnikowia_pulcherrima(100); | Metschnikowia | 13 | Saprotroph | \|Undefined Saprotroph\| | Yeast | NULL | Probable | As nectar-tap_saprotroph (Põlme et al. 2020); Decay sugar-rich substrates (Põlme et al. 2020) | Põlme S, et al. 2020. Fungal Diversity 105:1-16 ((https://doi.org/10.1007/s13225-020-00466-2)) |
| Otu38035 | 1 | k__Fungi(100);p__Ascomycota(100);c__Saccharomycetes(100);o__Saccharomycetales(100);f__Metschnikowiaceae(100);g__Metschnikowia(100);s__Metschnikowia_pulcherrima(100); | Metschnikowia | 13 | Saprotroph | \|Undefined Saprotroph\| | Yeast | NULL | Probable | As nectar-tap_saprotroph (Põlme et al. 2020); Decay sugar-rich substrates (Põlme et al. 2020) | Põlme S, et al. 2020. Fungal Diversity 105:1-16 ((https://doi.org/10.1007/s13225-020-00466-2)) |
| Otu38037 | 1 | k__Fungi(100);p__Ascomycota(100);c__Saccharomycetes(100);o__Saccharomycetales(100);f__Metschnikowiaceae(100);g__Metschnikowia(100);g__Metschnikowia_unclassified(100); | Metschnikowia | 13 | Saprotroph | \|Undefined Saprotroph\| | Yeast | NULL | Probable | As nectar-tap_saprotroph (Põlme et al. 2020); Decay sugar-rich substrates (Põlme et al. 2020) | Põlme S, et al. 2020. Fungal Diversity 105:1-16 ((https://doi.org/10.1007/s13225-020-00466-2)) |
| Otu38038 | 1 | k__Fungi(100);p__Ascomycota(100);c__Saccharomycetes(100);o__Saccharomycetales(100);f__Metschnikowiaceae(100);g__Metschnikowia(100);s__Metschnikowia_pulcherrima(100); | Metschnikowia | 13 | Saprotroph | \|Undefined Saprotroph\| | Yeast | NULL | Probable | As nectar-tap_saprotroph (Põlme et al. 2020); Decay sugar-rich substrates (Põlme et al. 2020) | Põlme S, et al. 2020. Fungal Diversity 105:1-16 ((https://doi.org/10.1007/s13225-020-00466-2)) |
| Otu38039 | 1 | k__Fungi(100);p__Ascomycota(100);c__Saccharomycetes(100);o__Saccharomycetales(100);f__Metschnikowiaceae(100);g__Metschnikowia(100);s__Metschnikowia_pulcherrima(100); | Metschnikowia | 13 | Saprotroph | \|Undefined Saprotroph\| | Yeast | NULL | Probable | As nectar-tap_saprotroph (Põlme et al. 2020); Decay sugar-rich substrates (Põlme et al. 2020) | Põlme S, et al. 2020. Fungal Diversity 105:1-16 ((https://doi.org/10.1007/s13225-020-00466-2)) |
| Otu38040 | 1 | k__Fungi(100);p__Ascomycota(100);c__Leotiomycetes(100);o__Leotiomycetes_order_Incertae_sedis(100);f__Leotiomycetes_family_Incertae_sedis(100);g__Geomyces(100);g__Geomyces_unclassified(100); | Geomyces | 13 | Saprotroph | \|Undefined Saprotroph\| | Microfungus | NULL | Probable | As soil_saprotroph (Põlme et al. 2020); Decay soil (Põlme et al. 2020); Nose Bat Syndrome causing Geomyces destructans, are now placed in the genus Pseudogymnoascus (see Minnis & Lindner 2013); This genus is now restricted to one species Geomyces auratus; however, it is important to note that the majority of species that once comprised this genus, including the White | Minnis AM, Lindner DL. 2013. Fungal Biology 117:638-649 ((https://doi.org/10.1016/j.funbio.2013.07.001)); Tedersoo L, et al. 2014. Science 346:e1256688 ((https://doi.org/10.1126/science.1256688)); Põlme S, et al. 2020. Fungal Diversity 105:1-16 ((https://doi.org/10.1007/s13225-020-00466-2)) |
| Otu38041 | 1 | k__Fungi(100);p__Ascomycota(100);c__Dothideomycetes(100);o__Dothideomycetes_order_Incertae_sedis(100);f__Pseudeurotiaceae(100);f__Pseudeurotiaceae_unclassified(100);f__Pseudeurotiaceae_unclassified(100); | Pseudeurotiaceae | 9 | Saprotroph | Plant Saprotroph-Wood Saprotroph | Microfungus | NULL | Probable | Saprobic on woody tissue and rotting vegetation, often isolated from soil (Cannon & Kirk 2007) | Cannon PF, Kirk PM. 2007. Fungal Families of the World. CAB International, Cambridge (ISBN: 978-0851998275) |
| Otu38042 | 1 | k__Fungi(100);p__Ascomycota(100);c__Saccharomycetes(100);o__Saccharomycetales(100);f__Metschnikowiaceae(100);g__Metschnikowia(100);s__Metschnikowia_pulcherrima(100); | Metschnikowia | 13 | Saprotroph | \|Undefined Saprotroph\| | Yeast | NULL | Probable | As nectar-tap_saprotroph (Põlme et al. 2020); Decay sugar-rich substrates (Põlme et al. 2020) | Põlme S, et al. 2020. Fungal Diversity 105:1-16 ((https://doi.org/10.1007/s13225-020-00466-2)) |
| Otu38043 | 1 | k__Fungi(100);p__Ascomycota(100);c__Saccharomycetes(100);o__Saccharomycetales(100);f__Metschnikowiaceae(100);g__Metschnikowia(100);g__Metschnikowia_unclassified(100); | Metschnikowia | 13 | Saprotroph | \|Undefined Saprotroph\| | Yeast | NULL | Probable | As nectar-tap_saprotroph (Põlme et al. 2020); Decay sugar-rich substrates (Põlme et al. 2020) | Põlme S, et al. 2020. Fungal Diversity 105:1-16 ((https://doi.org/10.1007/s13225-020-00466-2)) |
| Otu38044 | 1 | k__Fungi(100);p__Ascomycota(100);c__Saccharomycetes(100);o__Saccharomycetales(100);f__Metschnikowiaceae(100);g__Metschnikowia(100);s__Metschnikowia_pulcherrima(100); | Metschnikowia | 13 | Saprotroph | \|Undefined Saprotroph\| | Yeast | NULL | Probable | As nectar-tap_saprotroph (Põlme et al. 2020); Decay sugar-rich substrates (Põlme et al. 2020) | Põlme S, et al. 2020. Fungal Diversity 105:1-16 ((https://doi.org/10.1007/s13225-020-00466-2)) |
| Otu38045 | 1 | k__Fungi(100);p__Ascomycota(100);c__Saccharomycetes(100);o__Saccharomycetales(100);f__Metschnikowiaceae(100);g__Metschnikowia(100);g__Metschnikowia_unclassified(100); | Metschnikowia | 13 | Saprotroph | \|Undefined Saprotroph\| | Yeast | NULL | Probable | As nectar-tap_saprotroph (Põlme et al. 2020); Decay sugar-rich substrates (Põlme et al. 2020) | Põlme S, et al. 2020. Fungal Diversity 105:1-16 ((https://doi.org/10.1007/s13225-020-00466-2)) |
| Otu38046 | 1 | k__Fungi(100);p__Ascomycota(100);c__Saccharomycetes(100);o__Saccharomycetales(100);f__Metschnikowiaceae(100);g__Metschnikowia(100);s__Metschnikowia_pulcherrima(100); | Metschnikowia | 13 | Saprotroph | \|Undefined Saprotroph\| | Yeast | NULL | Probable | As nectar-tap_saprotroph (Põlme et al. 2020); Decay sugar-rich substrates (Põlme et al. 2020) | Põlme S, et al. 2020. Fungal Diversity 105:1-16 ((https://doi.org/10.1007/s13225-020-00466-2)) |
| Otu38047 | 1 | k__Fungi(100);p__Ascomycota(100);c__Saccharomycetes(100);o__Saccharomycetales(100);f__Metschnikowiaceae(100);g__Metschnikowia(100);s__Metschnikowia_pulcherrima(100); | Metschnikowia | 13 | Saprotroph | \|Undefined Saprotroph\| | Yeast | NULL | Probable | As nectar-tap_saprotroph (Põlme et al. 2020); Decay sugar-rich substrates (Põlme et al. 2020) | Põlme S, et al. 2020. Fungal Diversity 105:1-16 ((https://doi.org/10.1007/s13225-020-00466-2)) |
| Otu38048 | 1 | k__Fungi(100);p__Ascomycota(100);c__Saccharomycetes(100);o__Saccharomycetales(100);f__Metschnikowiaceae(100);g__Metschnikowia(100);g__Metschnikowia_unclassified(100); | Metschnikowia | 13 | Saprotroph | \|Undefined Saprotroph\| | Yeast | NULL | Probable | As nectar-tap_saprotroph (Põlme et al. 2020); Decay sugar-rich substrates (Põlme et al. 2020) | Põlme S, et al. 2020. Fungal Diversity 105:1-16 ((https://doi.org/10.1007/s13225-020-00466-2)) |
| Otu38049 | 1 | k__Fungi(100);p__Ascomycota(100);c__Saccharomycetes(100);o__Saccharomycetales(100);f__Metschnikowiaceae(100);g__Metschnikowia(100);s__Metschnikowia_pulcherrima(100); | Metschnikowia | 13 | Saprotroph | \|Undefined Saprotroph\| | Yeast | NULL | Probable | As nectar-tap_saprotroph (Põlme et al. 2020); Decay sugar-rich substrates (Põlme et al. 2020) | Põlme S, et al. 2020. Fungal Diversity 105:1-16 ((https://doi.org/10.1007/s13225-020-00466-2)) |
| Otu38050 | 1 | k__Fungi(100);p__Ascomycota(100);c__Saccharomycetes(100);o__Saccharomycetales(100);f__Metschnikowiaceae(100);g__Metschnikowia(100);g__Metschnikowia_unclassified(100); | Metschnikowia | 13 | Saprotroph | \|Undefined Saprotroph\| | Yeast | NULL | Probable | As nectar-tap_saprotroph (Põlme et al. 2020); Decay sugar-rich substrates (Põlme et al. 2020) | Põlme S, et al. 2020. Fungal Diversity 105:1-16 ((https://doi.org/10.1007/s13225-020-00466-2)) |
| Otu38051 | 1 | k__Fungi(100);p__Ascomycota(100);c__Saccharomycetes(100);o__Saccharomycetales(100);f__Metschnikowiaceae(100);g__Metschnikowia(100);g__Metschnikowia_unclassified(100); | Metschnikowia | 13 | Saprotroph | \|Undefined Saprotroph\| | Yeast | NULL | Probable | As nectar-tap_saprotroph (Põlme et al. 2020); Decay sugar-rich substrates (Põlme et al. 2020) | Põlme S, et al. 2020. Fungal Diversity 105:1-16 ((https://doi.org/10.1007/s13225-020-00466-2)) |
| Otu38052 | 1 | k__Fungi(100);p__Ascomycota(100);c__Saccharomycetes(100);o__Saccharomycetales(100);f__Metschnikowiaceae(100);g__Metschnikowia(100);g__Metschnikowia_unclassified(100); | Metschnikowia | 13 | Saprotroph | \|Undefined Saprotroph\| | Yeast | NULL | Probable | As nectar-tap_saprotroph (Põlme et al. 2020); Decay sugar-rich substrates (Põlme et al. 2020) | Põlme S, et al. 2020. Fungal Diversity 105:1-16 ((https://doi.org/10.1007/s13225-020-00466-2)) |
| Otu38053 | 1 | k__Fungi(100);p__Ascomycota(100);c__Saccharomycetes(100);o__Saccharomycetales(100);f__Metschnikowiaceae(100);g__Metschnikowia(100);s__Metschnikowia_pulcherrima(100); | Metschnikowia | 13 | Saprotroph | \|Undefined Saprotroph\| | Yeast | NULL | Probable | As nectar-tap_saprotroph (Põlme et al. 2020); Decay sugar-rich substrates (Põlme et al. 2020) | Põlme S, et al. 2020. Fungal Diversity 105:1-16 ((https://doi.org/10.1007/s13225-020-00466-2)) |
| Otu38054 | 1 | k__Fungi(100);p__Ascomycota(100);c__Saccharomycetes(100);o__Saccharomycetales(100);f__Metschnikowiaceae(100);g__Metschnikowia(100);s__Metschnikowia_pulcherrima(100); | Metschnikowia | 13 | Saprotroph | \|Undefined Saprotroph\| | Yeast | NULL | Probable | As nectar-tap_saprotroph (Põlme et al. 2020); Decay sugar-rich substrates (Põlme et al. 2020) | Põlme S, et al. 2020. Fungal Diversity 105:1-16 ((https://doi.org/10.1007/s13225-020-00466-2)) |
| Otu38055 | 1 | k__Fungi(100);p__Ascomycota(100);c__Saccharomycetes(100);o__Saccharomycetales(100);f__Metschnikowiaceae(100);g__Metschnikowia(100);s__Metschnikowia_pulcherrima(100); | Metschnikowia | 13 | Saprotroph | \|Undefined Saprotroph\| | Yeast | NULL | Probable | As nectar-tap_saprotroph (Põlme et al. 2020); Decay sugar-rich substrates (Põlme et al. 2020) | Põlme S, et al. 2020. Fungal Diversity 105:1-16 ((https://doi.org/10.1007/s13225-020-00466-2)) |
| Otu38056 | 1 | k__Fungi(100);p__Ascomycota(100);c__Saccharomycetes(100);o__Saccharomycetales(100);f__Metschnikowiaceae(100);g__Metschnikowia(100);g__Metschnikowia_unclassified(100); | Metschnikowia | 13 | Saprotroph | \|Undefined Saprotroph\| | Yeast | NULL | Probable | As nectar-tap_saprotroph (Põlme et al. 2020); Decay sugar-rich substrates (Põlme et al. 2020) | Põlme S, et al. 2020. Fungal Diversity 105:1-16 ((https://doi.org/10.1007/s13225-020-00466-2)) |
| Otu38057 | 1 | k__Fungi(100);p__Ascomycota(100);c__Saccharomycetes(100);o__Saccharomycetales(100);f__Metschnikowiaceae(100);g__Metschnikowia(100);s__Metschnikowia_pulcherrima(100); | Metschnikowia | 13 | Saprotroph | \|Undefined Saprotroph\| | Yeast | NULL | Probable | As nectar-tap_saprotroph (Põlme et al. 2020); Decay sugar-rich substrates (Põlme et al. 2020) | Põlme S, et al. 2020. Fungal Diversity 105:1-16 ((https://doi.org/10.1007/s13225-020-00466-2)) |
| Otu38058 | 1 | k__Fungi(100);p__Ascomycota(100);c__Saccharomycetes(100);o__Saccharomycetales(100);f__Metschnikowiaceae(100);g__Metschnikowia(100);s__Metschnikowia_pulcherrima(100); | Metschnikowia | 13 | Saprotroph | \|Undefined Saprotroph\| | Yeast | NULL | Probable | As nectar-tap_saprotroph (Põlme et al. 2020); Decay sugar-rich substrates (Põlme et al. 2020) | Põlme S, et al. 2020. Fungal Diversity 105:1-16 ((https://doi.org/10.1007/s13225-020-00466-2)) |
| Otu38059 | 1 | k__Fungi(100);p__Ascomycota(100);c__Saccharomycetes(100);o__Saccharomycetales(100);f__Metschnikowiaceae(100);g__Metschnikowia(100);g__Metschnikowia_unclassified(100); | Metschnikowia | 13 | Saprotroph | \|Undefined Saprotroph\| | Yeast | NULL | Probable | As nectar-tap_saprotroph (Põlme et al. 2020); Decay sugar-rich substrates (Põlme et al. 2020) | Põlme S, et al. 2020. Fungal Diversity 105:1-16 ((https://doi.org/10.1007/s13225-020-00466-2)) |
| Otu38060 | 1 | k__Fungi(100);p__Ascomycota(100);c__Saccharomycetes(100);o__Saccharomycetales(100);f__Metschnikowiaceae(100);g__Metschnikowia(100);s__Metschnikowia_pulcherrima(100); | Metschnikowia | 13 | Saprotroph | \|Undefined Saprotroph\| | Yeast | NULL | Probable | As nectar-tap_saprotroph (Põlme et al. 2020); Decay sugar-rich substrates (Põlme et al. 2020) | Põlme S, et al. 2020. Fungal Diversity 105:1-16 ((https://doi.org/10.1007/s13225-020-00466-2)) |
| Otu38061 | 1 | k__Fungi(100);p__Ascomycota(100);c__Saccharomycetes(100);o__Saccharomycetales(100);f__Metschnikowiaceae(100);g__Metschnikowia(100);s__Metschnikowia_pulcherrima(100); | Metschnikowia | 13 | Saprotroph | \|Undefined Saprotroph\| | Yeast | NULL | Probable | As nectar-tap_saprotroph (Põlme et al. 2020); Decay sugar-rich substrates (Põlme et al. 2020) | Põlme S, et al. 2020. Fungal Diversity 105:1-16 ((https://doi.org/10.1007/s13225-020-00466-2)) |
| Otu38062 | 1 | k__Fungi(100);p__Ascomycota(100);c__Saccharomycetes(100);o__Saccharomycetales(100);f__Metschnikowiaceae(100);g__Metschnikowia(100);s__Metschnikowia_pulcherrima(100); | Metschnikowia | 13 | Saprotroph | \|Undefined Saprotroph\| | Yeast | NULL | Probable | As nectar-tap_saprotroph (Põlme et al. 2020); Decay sugar-rich substrates (Põlme et al. 2020) | Põlme S, et al. 2020. Fungal Diversity 105:1-16 ((https://doi.org/10.1007/s13225-020-00466-2)) |
| Otu38063 | 1 | k__Fungi(100);p__Ascomycota(100);c__Saccharomycetes(100);o__Saccharomycetales(100);f__Metschnikowiaceae(100);g__Metschnikowia(100);s__Metschnikowia_pulcherrima(100); | Metschnikowia | 13 | Saprotroph | \|Undefined Saprotroph\| | Yeast | NULL | Probable | As nectar-tap_saprotroph (Põlme et al. 2020); Decay sugar-rich substrates (Põlme et al. 2020) | Põlme S, et al. 2020. Fungal Diversity 105:1-16 ((https://doi.org/10.1007/s13225-020-00466-2)) |
| Otu38064 | 1 | k__Fungi(100);p__Ascomycota(100);c__Saccharomycetes(100);o__Saccharomycetales(100);f__Metschnikowiaceae(100);g__Metschnikowia(100);s__Metschnikowia_pulcherrima(100); | Metschnikowia | 13 | Saprotroph | \|Undefined Saprotroph\| | Yeast | NULL | Probable | As nectar-tap_saprotroph (Põlme et al. 2020); Decay sugar-rich substrates (Põlme et al. 2020) | Põlme S, et al. 2020. Fungal Diversity 105:1-16 ((https://doi.org/10.1007/s13225-020-00466-2)) |
| Otu38065 | 1 | k__Fungi(100);p__Ascomycota(100);c__Saccharomycetes(100);o__Saccharomycetales(100);f__Metschnikowiaceae(100);g__Metschnikowia(100);s__Metschnikowia_pulcherrima(100); | Metschnikowia | 13 | Saprotroph | \|Undefined Saprotroph\| | Yeast | NULL | Probable | As nectar-tap_saprotroph (Põlme et al. 2020); Decay sugar-rich substrates (Põlme et al. 2020) | Põlme S, et al. 2020. Fungal Diversity 105:1-16 ((https://doi.org/10.1007/s13225-020-00466-2)) |
| Otu38066 | 1 | k__Fungi(100);p__Ascomycota(100);c__Saccharomycetes(100);o__Saccharomycetales(100);f__Metschnikowiaceae(100);g__Metschnikowia(100);g__Metschnikowia_unclassified(100); | Metschnikowia | 13 | Saprotroph | \|Undefined Saprotroph\| | Yeast | NULL | Probable | As nectar-tap_saprotroph (Põlme et al. 2020); Decay sugar-rich substrates (Põlme et al. 2020) | Põlme S, et al. 2020. Fungal Diversity 105:1-16 ((https://doi.org/10.1007/s13225-020-00466-2)) |
| Otu38067 | 1 | k__Fungi(100);p__Ascomycota(100);c__Saccharomycetes(100);o__Saccharomycetales(100);f__Metschnikowiaceae(100);g__Metschnikowia(100);s__Metschnikowia_pulcherrima(100); | Metschnikowia | 13 | Saprotroph | \|Undefined Saprotroph\| | Yeast | NULL | Probable | As nectar-tap_saprotroph (Põlme et al. 2020); Decay sugar-rich substrates (Põlme et al. 2020) | Põlme S, et al. 2020. Fungal Diversity 105:1-16 ((https://doi.org/10.1007/s13225-020-00466-2)) |
| Otu38068 | 1 | k__Fungi(100);p__Ascomycota(100);c__Saccharomycetes(100);o__Saccharomycetales(100);f__Metschnikowiaceae(100);g__Metschnikowia(100);g__Metschnikowia_unclassified(100); | Metschnikowia | 13 | Saprotroph | \|Undefined Saprotroph\| | Yeast | NULL | Probable | As nectar-tap_saprotroph (Põlme et al. 2020); Decay sugar-rich substrates (Põlme et al. 2020) | Põlme S, et al. 2020. Fungal Diversity 105:1-16 ((https://doi.org/10.1007/s13225-020-00466-2)) |
| Otu38069 | 1 | k__Fungi(100);p__Ascomycota(100);c__Saccharomycetes(100);o__Saccharomycetales(100);f__Metschnikowiaceae(100);g__Metschnikowia(100);s__Metschnikowia_pulcherrima(100); | Metschnikowia | 13 | Saprotroph | \|Undefined Saprotroph\| | Yeast | NULL | Probable | As nectar-tap_saprotroph (Põlme et al. 2020); Decay sugar-rich substrates (Põlme et al. 2020) | Põlme S, et al. 2020. Fungal Diversity 105:1-16 ((https://doi.org/10.1007/s13225-020-00466-2)) |
| Otu38070 | 1 | k__Fungi(100);p__Ascomycota(100);c__Saccharomycetes(100);o__Saccharomycetales(100);f__Metschnikowiaceae(100);g__Metschnikowia(100);s__Metschnikowia_pulcherrima(100); | Metschnikowia | 13 | Saprotroph | \|Undefined Saprotroph\| | Yeast | NULL | Probable | As nectar-tap_saprotroph (Põlme et al. 2020); Decay sugar-rich substrates (Põlme et al. 2020) | Põlme S, et al. 2020. Fungal Diversity 105:1-16 ((https://doi.org/10.1007/s13225-020-00466-2)) |
| Otu38071 | 1 | k__Fungi(100);p__Ascomycota(100);c__Saccharomycetes(100);o__Saccharomycetales(100);f__Metschnikowiaceae(100);g__Metschnikowia(100);s__Metschnikowia_pulcherrima(100); | Metschnikowia | 13 | Saprotroph | \|Undefined Saprotroph\| | Yeast | NULL | Probable | As nectar-tap_saprotroph (Põlme et al. 2020); Decay sugar-rich substrates (Põlme et al. 2020) | Põlme S, et al. 2020. Fungal Diversity 105:1-16 ((https://doi.org/10.1007/s13225-020-00466-2)) |
| Otu38072 | 1 | k__Fungi(100);p__Ascomycota(100);c__Saccharomycetes(100);o__Saccharomycetales(100);f__Metschnikowiaceae(100);g__Metschnikowia(100);s__Metschnikowia_pulcherrima(100); | Metschnikowia | 13 | Saprotroph | \|Undefined Saprotroph\| | Yeast | NULL | Probable | As nectar-tap_saprotroph (Põlme et al. 2020); Decay sugar-rich substrates (Põlme et al. 2020) | Põlme S, et al. 2020. Fungal Diversity 105:1-16 ((https://doi.org/10.1007/s13225-020-00466-2)) |
| Otu38073 | 1 | k__Fungi(100);p__Ascomycota(100);c__Saccharomycetes(100);o__Saccharomycetales(100);f__Metschnikowiaceae(100);g__Metschnikowia(100);s__Metschnikowia_pulcherrima(100); | Metschnikowia | 13 | Saprotroph | \|Undefined Saprotroph\| | Yeast | NULL | Probable | As nectar-tap_saprotroph (Põlme et al. 2020); Decay sugar-rich substrates (Põlme et al. 2020) | Põlme S, et al. 2020. Fungal Diversity 105:1-16 ((https://doi.org/10.1007/s13225-020-00466-2)) |
| Otu38074 | 1 | k__Fungi(100);p__Ascomycota(100);c__Saccharomycetes(100);o__Saccharomycetales(100);f__Metschnikowiaceae(100);g__Metschnikowia(100);s__Metschnikowia_pulcherrima(100); | Metschnikowia | 13 | Saprotroph | \|Undefined Saprotroph\| | Yeast | NULL | Probable | As nectar-tap_saprotroph (Põlme et al. 2020); Decay sugar-rich substrates (Põlme et al. 2020) | Põlme S, et al. 2020. Fungal Diversity 105:1-16 ((https://doi.org/10.1007/s13225-020-00466-2)) |
| Otu38075 | 1 | k__Fungi(100);p__Ascomycota(100);c__Saccharomycetes(100);o__Saccharomycetales(100);f__Metschnikowiaceae(100);g__Metschnikowia(100);g__Metschnikowia_unclassified(100); | Metschnikowia | 13 | Saprotroph | \|Undefined Saprotroph\| | Yeast | NULL | Probable | As nectar-tap_saprotroph (Põlme et al. 2020); Decay sugar-rich substrates (Põlme et al. 2020) | Põlme S, et al. 2020. Fungal Diversity 105:1-16 ((https://doi.org/10.1007/s13225-020-00466-2)) |
| Otu38076 | 1 | k__Fungi(100);p__Ascomycota(100);c__Saccharomycetes(100);o__Saccharomycetales(100);f__Metschnikowiaceae(100);g__Metschnikowia(100);s__Metschnikowia_pulcherrima(100); | Metschnikowia | 13 | Saprotroph | \|Undefined Saprotroph\| | Yeast | NULL | Probable | As nectar-tap_saprotroph (Põlme et al. 2020); Decay sugar-rich substrates (Põlme et al. 2020) | Põlme S, et al. 2020. Fungal Diversity 105:1-16 ((https://doi.org/10.1007/s13225-020-00466-2)) |
| Otu38077 | 1 | k__Fungi(100);p__Ascomycota(100);c__Saccharomycetes(100);o__Saccharomycetales(100);f__Metschnikowiaceae(100);g__Metschnikowia(100);s__Metschnikowia_pulcherrima(100); | Metschnikowia | 13 | Saprotroph | \|Undefined Saprotroph\| | Yeast | NULL | Probable | As nectar-tap_saprotroph (Põlme et al. 2020); Decay sugar-rich substrates (Põlme et al. 2020) | Põlme S, et al. 2020. Fungal Diversity 105:1-16 ((https://doi.org/10.1007/s13225-020-00466-2)) |
| Otu38078 | 1 | k__Fungi(100);p__Ascomycota(100);c__Saccharomycetes(100);o__Saccharomycetales(100);f__Metschnikowiaceae(100);g__Metschnikowia(100);s__Metschnikowia_pulcherrima(100); | Metschnikowia | 13 | Saprotroph | \|Undefined Saprotroph\| | Yeast | NULL | Probable | As nectar-tap_saprotroph (Põlme et al. 2020); Decay sugar-rich substrates (Põlme et al. 2020) | Põlme S, et al. 2020. Fungal Diversity 105:1-16 ((https://doi.org/10.1007/s13225-020-00466-2)) |
| Otu38079 | 1 | k__Fungi(100);p__Ascomycota(100);c__Saccharomycetes(100);o__Saccharomycetales(100);f__Metschnikowiaceae(100);g__Metschnikowia(100);s__Metschnikowia_pulcherrima(100); | Metschnikowia | 13 | Saprotroph | \|Undefined Saprotroph\| | Yeast | NULL | Probable | As nectar-tap_saprotroph (Põlme et al. 2020); Decay sugar-rich substrates (Põlme et al. 2020) | Põlme S, et al. 2020. Fungal Diversity 105:1-16 ((https://doi.org/10.1007/s13225-020-00466-2)) |
| Otu38080 | 1 | k__Fungi(100);p__Ascomycota(100);c__Saccharomycetes(100);o__Saccharomycetales(100);f__Metschnikowiaceae(100);g__Metschnikowia(100);s__Metschnikowia_pulcherrima(100); | Metschnikowia | 13 | Saprotroph | \|Undefined Saprotroph\| | Yeast | NULL | Probable | As nectar-tap_saprotroph (Põlme et al. 2020); Decay sugar-rich substrates (Põlme et al. 2020) | Põlme S, et al. 2020. Fungal Diversity 105:1-16 ((https://doi.org/10.1007/s13225-020-00466-2)) |
| Otu38081 | 1 | k__Fungi(100);p__Ascomycota(100);c__Saccharomycetes(100);o__Saccharomycetales(100);f__Metschnikowiaceae(100);g__Metschnikowia(100);g__Metschnikowia_unclassified(100); | Metschnikowia | 13 | Saprotroph | \|Undefined Saprotroph\| | Yeast | NULL | Probable | As nectar-tap_saprotroph (Põlme et al. 2020); Decay sugar-rich substrates (Põlme et al. 2020) | Põlme S, et al. 2020. Fungal Diversity 105:1-16 ((https://doi.org/10.1007/s13225-020-00466-2)) |
| Otu38082 | 1 | k__Fungi(100);p__Ascomycota(100);c__Saccharomycetes(100);o__Saccharomycetales(100);f__Metschnikowiaceae(100);g__Metschnikowia(100);s__Metschnikowia_pulcherrima(100); | Metschnikowia | 13 | Saprotroph | \|Undefined Saprotroph\| | Yeast | NULL | Probable | As nectar-tap_saprotroph (Põlme et al. 2020); Decay sugar-rich substrates (Põlme et al. 2020) | Põlme S, et al. 2020. Fungal Diversity 105:1-16 ((https://doi.org/10.1007/s13225-020-00466-2)) |
| Otu38083 | 1 | k__Fungi(100);p__Ascomycota(100);c__Saccharomycetes(100);o__Saccharomycetales(100);f__Metschnikowiaceae(100);g__Metschnikowia(100);s__Metschnikowia_pulcherrima(100); | Metschnikowia | 13 | Saprotroph | \|Undefined Saprotroph\| | Yeast | NULL | Probable | As nectar-tap_saprotroph (Põlme et al. 2020); Decay sugar-rich substrates (Põlme et al. 2020) | Põlme S, et al. 2020. Fungal Diversity 105:1-16 ((https://doi.org/10.1007/s13225-020-00466-2)) |
| Otu38084 | 1 | k__Fungi(100);p__Ascomycota(100);c__Saccharomycetes(100);o__Saccharomycetales(100);f__Metschnikowiaceae(100);g__Metschnikowia(100);s__Metschnikowia_pulcherrima(100); | Metschnikowia | 13 | Saprotroph | \|Undefined Saprotroph\| | Yeast | NULL | Probable | As nectar-tap_saprotroph (Põlme et al. 2020); Decay sugar-rich substrates (Põlme et al. 2020) | Põlme S, et al. 2020. Fungal Diversity 105:1-16 ((https://doi.org/10.1007/s13225-020-00466-2)) |
| Otu38085 | 1 | k__Fungi(100);p__Ascomycota(100);c__Saccharomycetes(100);o__Saccharomycetales(100);f__Metschnikowiaceae(100);g__Metschnikowia(100);s__Metschnikowia_pulcherrima(100); | Metschnikowia | 13 | Saprotroph | \|Undefined Saprotroph\| | Yeast | NULL | Probable | As nectar-tap_saprotroph (Põlme et al. 2020); Decay sugar-rich substrates (Põlme et al. 2020) | Põlme S, et al. 2020. Fungal Diversity 105:1-16 ((https://doi.org/10.1007/s13225-020-00466-2)) |
| Otu38086 | 1 | k__Fungi(100);p__Ascomycota(100);c__Saccharomycetes(100);o__Saccharomycetales(100);f__Metschnikowiaceae(100);g__Metschnikowia(100);g__Metschnikowia_unclassified(100); | Metschnikowia | 13 | Saprotroph | \|Undefined Saprotroph\| | Yeast | NULL | Probable | As nectar-tap_saprotroph (Põlme et al. 2020); Decay sugar-rich substrates (Põlme et al. 2020) | Põlme S, et al. 2020. Fungal Diversity 105:1-16 ((https://doi.org/10.1007/s13225-020-00466-2)) |
| Otu38087 | 1 | k__Fungi(100);p__Ascomycota(100);c__Saccharomycetes(100);o__Saccharomycetales(100);f__Metschnikowiaceae(100);g__Metschnikowia(100);s__Metschnikowia_pulcherrima(100); | Metschnikowia | 13 | Saprotroph | \|Undefined Saprotroph\| | Yeast | NULL | Probable | As nectar-tap_saprotroph (Põlme et al. 2020); Decay sugar-rich substrates (Põlme et al. 2020) | Põlme S, et al. 2020. Fungal Diversity 105:1-16 ((https://doi.org/10.1007/s13225-020-00466-2)) |
| Otu38088 | 1 | k__Fungi(100);p__Ascomycota(100);c__Saccharomycetes(100);o__Saccharomycetales(100);f__Metschnikowiaceae(100);g__Metschnikowia(100);g__Metschnikowia_unclassified(100); | Metschnikowia | 13 | Saprotroph | \|Undefined Saprotroph\| | Yeast | NULL | Probable | As nectar-tap_saprotroph (Põlme et al. 2020); Decay sugar-rich substrates (Põlme et al. 2020) | Põlme S, et al. 2020. Fungal Diversity 105:1-16 ((https://doi.org/10.1007/s13225-020-00466-2)) |
| Otu38090 | 1 | k__Fungi(100);p__Ascomycota(100);c__Saccharomycetes(100);o__Saccharomycetales(100);f__Metschnikowiaceae(100);g__Metschnikowia(100);g__Metschnikowia_unclassified(100); | Metschnikowia | 13 | Saprotroph | \|Undefined Saprotroph\| | Yeast | NULL | Probable | As nectar-tap_saprotroph (Põlme et al. 2020); Decay sugar-rich substrates (Põlme et al. 2020) | Põlme S, et al. 2020. Fungal Diversity 105:1-16 ((https://doi.org/10.1007/s13225-020-00466-2)) |
| Otu38091 | 1 | k__Fungi(100);p__Ascomycota(100);c__Saccharomycetes(100);o__Saccharomycetales(100);f__Metschnikowiaceae(100);g__Metschnikowia(100);s__Metschnikowia_pulcherrima(100); | Metschnikowia | 13 | Saprotroph | \|Undefined Saprotroph\| | Yeast | NULL | Probable | As nectar-tap_saprotroph (Põlme et al. 2020); Decay sugar-rich substrates (Põlme et al. 2020) | Põlme S, et al. 2020. Fungal Diversity 105:1-16 ((https://doi.org/10.1007/s13225-020-00466-2)) |
| Otu38092 | 1 | k__Fungi(100);p__Ascomycota(100);c__Saccharomycetes(100);o__Saccharomycetales(100);f__Metschnikowiaceae(100);g__Metschnikowia(100);s__Metschnikowia_pulcherrima(100); | Metschnikowia | 13 | Saprotroph | \|Undefined Saprotroph\| | Yeast | NULL | Probable | As nectar-tap_saprotroph (Põlme et al. 2020); Decay sugar-rich substrates (Põlme et al. 2020) | Põlme S, et al. 2020. Fungal Diversity 105:1-16 ((https://doi.org/10.1007/s13225-020-00466-2)) |
| Otu38093 | 1 | k__Fungi(100);p__Ascomycota(100);c__Saccharomycetes(100);o__Saccharomycetales(100);f__Metschnikowiaceae(100);g__Metschnikowia(100);s__Metschnikowia_pulcherrima(100); | Metschnikowia | 13 | Saprotroph | \|Undefined Saprotroph\| | Yeast | NULL | Probable | As nectar-tap_saprotroph (Põlme et al. 2020); Decay sugar-rich substrates (Põlme et al. 2020) | Põlme S, et al. 2020. Fungal Diversity 105:1-16 ((https://doi.org/10.1007/s13225-020-00466-2)) |
| Otu38094 | 1 | k__Fungi(100);p__Ascomycota(100);c__Saccharomycetes(100);o__Saccharomycetales(100);f__Metschnikowiaceae(100);g__Metschnikowia(100);s__Metschnikowia_pulcherrima(100); | Metschnikowia | 13 | Saprotroph | \|Undefined Saprotroph\| | Yeast | NULL | Probable | As nectar-tap_saprotroph (Põlme et al. 2020); Decay sugar-rich substrates (Põlme et al. 2020) | Põlme S, et al. 2020. Fungal Diversity 105:1-16 ((https://doi.org/10.1007/s13225-020-00466-2)) |
| Otu38095 | 1 | k__Fungi(100);p__Ascomycota(100);c__Saccharomycetes(100);o__Saccharomycetales(100);f__Metschnikowiaceae(100);g__Metschnikowia(100);s__Metschnikowia_pulcherrima(100); | Metschnikowia | 13 | Saprotroph | \|Undefined Saprotroph\| | Yeast | NULL | Probable | As nectar-tap_saprotroph (Põlme et al. 2020); Decay sugar-rich substrates (Põlme et al. 2020) | Põlme S, et al. 2020. Fungal Diversity 105:1-16 ((https://doi.org/10.1007/s13225-020-00466-2)) |
| Otu38096 | 1 | k__Fungi(100);p__Ascomycota(100);c__Saccharomycetes(100);o__Saccharomycetales(100);f__Metschnikowiaceae(100);g__Metschnikowia(100);s__Metschnikowia_pulcherrima(100); | Metschnikowia | 13 | Saprotroph | \|Undefined Saprotroph\| | Yeast | NULL | Probable | As nectar-tap_saprotroph (Põlme et al. 2020); Decay sugar-rich substrates (Põlme et al. 2020) | Põlme S, et al. 2020. Fungal Diversity 105:1-16 ((https://doi.org/10.1007/s13225-020-00466-2)) |
| Otu38097 | 1 | k__Fungi(100);p__Ascomycota(100);c__Saccharomycetes(100);o__Saccharomycetales(100);f__Metschnikowiaceae(100);g__Metschnikowia(100);g__Metschnikowia_unclassified(100); | Metschnikowia | 13 | Saprotroph | \|Undefined Saprotroph\| | Yeast | NULL | Probable | As nectar-tap_saprotroph (Põlme et al. 2020); Decay sugar-rich substrates (Põlme et al. 2020) | Põlme S, et al. 2020. Fungal Diversity 105:1-16 ((https://doi.org/10.1007/s13225-020-00466-2)) |
| Otu38098 | 1 | k__Fungi(100);p__Ascomycota(100);c__Saccharomycetes(100);o__Saccharomycetales(100);f__Metschnikowiaceae(100);g__Metschnikowia(100);g__Metschnikowia_unclassified(100); | Metschnikowia | 13 | Saprotroph | \|Undefined Saprotroph\| | Yeast | NULL | Probable | As nectar-tap_saprotroph (Põlme et al. 2020); Decay sugar-rich substrates (Põlme et al. 2020) | Põlme S, et al. 2020. Fungal Diversity 105:1-16 ((https://doi.org/10.1007/s13225-020-00466-2)) |
| Otu38099 | 1 | k__Fungi(100);p__Ascomycota(100);c__Saccharomycetes(100);o__Saccharomycetales(100);f__Metschnikowiaceae(100);g__Metschnikowia(100);g__Metschnikowia_unclassified(100); | Metschnikowia | 13 | Saprotroph | \|Undefined Saprotroph\| | Yeast | NULL | Probable | As nectar-tap_saprotroph (Põlme et al. 2020); Decay sugar-rich substrates (Põlme et al. 2020) | Põlme S, et al. 2020. Fungal Diversity 105:1-16 ((https://doi.org/10.1007/s13225-020-00466-2)) |
| Otu38100 | 1 | k__Fungi(100);p__Ascomycota(100);c__Saccharomycetes(100);o__Saccharomycetales(100);f__Metschnikowiaceae(100);g__Metschnikowia(100);s__Metschnikowia_pulcherrima(100); | Metschnikowia | 13 | Saprotroph | \|Undefined Saprotroph\| | Yeast | NULL | Probable | As nectar-tap_saprotroph (Põlme et al. 2020); Decay sugar-rich substrates (Põlme et al. 2020) | Põlme S, et al. 2020. Fungal Diversity 105:1-16 ((https://doi.org/10.1007/s13225-020-00466-2)) |
| Otu38101 | 1 | k__Fungi(100);p__Ascomycota(100);c__Saccharomycetes(100);o__Saccharomycetales(100);f__Metschnikowiaceae(100);g__Metschnikowia(100);s__Metschnikowia_pulcherrima(100); | Metschnikowia | 13 | Saprotroph | \|Undefined Saprotroph\| | Yeast | NULL | Probable | As nectar-tap_saprotroph (Põlme et al. 2020); Decay sugar-rich substrates (Põlme et al. 2020) | Põlme S, et al. 2020. Fungal Diversity 105:1-16 ((https://doi.org/10.1007/s13225-020-00466-2)) |
| Otu38102 | 1 | k__Fungi(100);p__Ascomycota(100);c__Saccharomycetes(100);o__Saccharomycetales(100);f__Metschnikowiaceae(100);g__Metschnikowia(100);s__Metschnikowia_pulcherrima(100); | Metschnikowia | 13 | Saprotroph | \|Undefined Saprotroph\| | Yeast | NULL | Probable | As nectar-tap_saprotroph (Põlme et al. 2020); Decay sugar-rich substrates (Põlme et al. 2020) | Põlme S, et al. 2020. Fungal Diversity 105:1-16 ((https://doi.org/10.1007/s13225-020-00466-2)) |
| Otu38103 | 1 | k__Fungi(100);p__Ascomycota(100);c__Saccharomycetes(100);o__Saccharomycetales(100);f__Metschnikowiaceae(100);g__Metschnikowia(100);g__Metschnikowia_unclassified(100); | Metschnikowia | 13 | Saprotroph | \|Undefined Saprotroph\| | Yeast | NULL | Probable | As nectar-tap_saprotroph (Põlme et al. 2020); Decay sugar-rich substrates (Põlme et al. 2020) | Põlme S, et al. 2020. Fungal Diversity 105:1-16 ((https://doi.org/10.1007/s13225-020-00466-2)) |
| Otu38104 | 1 | k__Fungi(100);p__Ascomycota(100);c__Saccharomycetes(100);o__Saccharomycetales(100);f__Metschnikowiaceae(100);g__Metschnikowia(100);s__Metschnikowia_pulcherrima(100); | Metschnikowia | 13 | Saprotroph | \|Undefined Saprotroph\| | Yeast | NULL | Probable | As nectar-tap_saprotroph (Põlme et al. 2020); Decay sugar-rich substrates (Põlme et al. 2020) | Põlme S, et al. 2020. Fungal Diversity 105:1-16 ((https://doi.org/10.1007/s13225-020-00466-2)) |
| Otu38105 | 1 | k__Fungi(100);p__Ascomycota(100);c__Saccharomycetes(100);o__Saccharomycetales(100);f__Metschnikowiaceae(100);g__Metschnikowia(100);s__Metschnikowia_pulcherrima(100); | Metschnikowia | 13 | Saprotroph | \|Undefined Saprotroph\| | Yeast | NULL | Probable | As nectar-tap_saprotroph (Põlme et al. 2020); Decay sugar-rich substrates (Põlme et al. 2020) | Põlme S, et al. 2020. Fungal Diversity 105:1-16 ((https://doi.org/10.1007/s13225-020-00466-2)) |
| Otu38106 | 1 | k__Fungi(100);p__Ascomycota(100);c__Saccharomycetes(100);o__Saccharomycetales(100);f__Metschnikowiaceae(100);g__Metschnikowia(100);s__Metschnikowia_pulcherrima(100); | Metschnikowia | 13 | Saprotroph | \|Undefined Saprotroph\| | Yeast | NULL | Probable | As nectar-tap_saprotroph (Põlme et al. 2020); Decay sugar-rich substrates (Põlme et al. 2020) | Põlme S, et al. 2020. Fungal Diversity 105:1-16 ((https://doi.org/10.1007/s13225-020-00466-2)) |
| Otu38107 | 1 | k__Fungi(100);p__Ascomycota(100);c__Saccharomycetes(100);o__Saccharomycetales(100);f__Metschnikowiaceae(100);g__Metschnikowia(100);s__Metschnikowia_pulcherrima(100); | Metschnikowia | 13 | Saprotroph | \|Undefined Saprotroph\| | Yeast | NULL | Probable | As nectar-tap_saprotroph (Põlme et al. 2020); Decay sugar-rich substrates (Põlme et al. 2020) | Põlme S, et al. 2020. Fungal Diversity 105:1-16 ((https://doi.org/10.1007/s13225-020-00466-2)) |
| Otu38108 | 1 | k__Fungi(100);p__Ascomycota(100);c__Saccharomycetes(100);o__Saccharomycetales(100);f__Metschnikowiaceae(100);g__Metschnikowia(100);s__Metschnikowia_pulcherrima(100); | Metschnikowia | 13 | Saprotroph | \|Undefined Saprotroph\| | Yeast | NULL | Probable | As nectar-tap_saprotroph (Põlme et al. 2020); Decay sugar-rich substrates (Põlme et al. 2020) | Põlme S, et al. 2020. Fungal Diversity 105:1-16 ((https://doi.org/10.1007/s13225-020-00466-2)) |
| Otu38109 | 1 | k__Fungi(100);p__Ascomycota(100);c__Saccharomycetes(100);o__Saccharomycetales(100);f__Metschnikowiaceae(100);g__Metschnikowia(100);s__Metschnikowia_pulcherrima(100); | Metschnikowia | 13 | Saprotroph | \|Undefined Saprotroph\| | Yeast | NULL | Probable | As nectar-tap_saprotroph (Põlme et al. 2020); Decay sugar-rich substrates (Põlme et al. 2020) | Põlme S, et al. 2020. Fungal Diversity 105:1-16 ((https://doi.org/10.1007/s13225-020-00466-2)) |
| Otu38110 | 1 | k__Fungi(100);p__Ascomycota(100);c__Saccharomycetes(100);o__Saccharomycetales(100);f__Metschnikowiaceae(100);g__Metschnikowia(100);s__Metschnikowia_pulcherrima(100); | Metschnikowia | 13 | Saprotroph | \|Undefined Saprotroph\| | Yeast | NULL | Probable | As nectar-tap_saprotroph (Põlme et al. 2020); Decay sugar-rich substrates (Põlme et al. 2020) | Põlme S, et al. 2020. Fungal Diversity 105:1-16 ((https://doi.org/10.1007/s13225-020-00466-2)) |
| Otu38111 | 1 | k__Fungi(100);p__Ascomycota(100);c__Saccharomycetes(100);o__Saccharomycetales(100);f__Metschnikowiaceae(100);g__Metschnikowia(100);g__Metschnikowia_unclassified(100); | Metschnikowia | 13 | Saprotroph | \|Undefined Saprotroph\| | Yeast | NULL | Probable | As nectar-tap_saprotroph (Põlme et al. 2020); Decay sugar-rich substrates (Põlme et al. 2020) | Põlme S, et al. 2020. Fungal Diversity 105:1-16 ((https://doi.org/10.1007/s13225-020-00466-2)) |
| Otu38112 | 1 | k__Fungi(100);p__Ascomycota(100);c__Saccharomycetes(100);o__Saccharomycetales(100);f__Metschnikowiaceae(100);g__Metschnikowia(100);s__Metschnikowia_pulcherrima(100); | Metschnikowia | 13 | Saprotroph | \|Undefined Saprotroph\| | Yeast | NULL | Probable | As nectar-tap_saprotroph (Põlme et al. 2020); Decay sugar-rich substrates (Põlme et al. 2020) | Põlme S, et al. 2020. Fungal Diversity 105:1-16 ((https://doi.org/10.1007/s13225-020-00466-2)) |
| Otu38113 | 1 | k__Fungi(100);p__Ascomycota(100);c__Saccharomycetes(100);o__Saccharomycetales(100);f__Metschnikowiaceae(100);g__Metschnikowia(100);g__Metschnikowia_unclassified(100); | Metschnikowia | 13 | Saprotroph | \|Undefined Saprotroph\| | Yeast | NULL | Probable | As nectar-tap_saprotroph (Põlme et al. 2020); Decay sugar-rich substrates (Põlme et al. 2020) | Põlme S, et al. 2020. Fungal Diversity 105:1-16 ((https://doi.org/10.1007/s13225-020-00466-2)) |
| Otu38114 | 1 | k__Fungi(100);p__Ascomycota(100);c__Saccharomycetes(100);o__Saccharomycetales(100);f__Metschnikowiaceae(100);g__Metschnikowia(100);s__Metschnikowia_pulcherrima(100); | Metschnikowia | 13 | Saprotroph | \|Undefined Saprotroph\| | Yeast | NULL | Probable | As nectar-tap_saprotroph (Põlme et al. 2020); Decay sugar-rich substrates (Põlme et al. 2020) | Põlme S, et al. 2020. Fungal Diversity 105:1-16 ((https://doi.org/10.1007/s13225-020-00466-2)) |
| Otu38115 | 1 | k__Fungi(100);p__Ascomycota(100);c__Saccharomycetes(100);o__Saccharomycetales(100);f__Metschnikowiaceae(100);g__Metschnikowia(100);g__Metschnikowia_unclassified(100); | Metschnikowia | 13 | Saprotroph | \|Undefined Saprotroph\| | Yeast | NULL | Probable | As nectar-tap_saprotroph (Põlme et al. 2020); Decay sugar-rich substrates (Põlme et al. 2020) | Põlme S, et al. 2020. Fungal Diversity 105:1-16 ((https://doi.org/10.1007/s13225-020-00466-2)) |
| Otu38116 | 1 | k__Fungi(100);p__Ascomycota(100);c__Saccharomycetes(100);o__Saccharomycetales(100);f__Metschnikowiaceae(100);g__Metschnikowia(100);s__Metschnikowia_pulcherrima(100); | Metschnikowia | 13 | Saprotroph | \|Undefined Saprotroph\| | Yeast | NULL | Probable | As nectar-tap_saprotroph (Põlme et al. 2020); Decay sugar-rich substrates (Põlme et al. 2020) | Põlme S, et al. 2020. Fungal Diversity 105:1-16 ((https://doi.org/10.1007/s13225-020-00466-2)) |
| Otu38118 | 1 | k__Fungi(100);p__Ascomycota(100);c__Saccharomycetes(100);o__Saccharomycetales(100);f__Metschnikowiaceae(100);g__Metschnikowia(100);s__Metschnikowia_pulcherrima(100); | Metschnikowia | 13 | Saprotroph | \|Undefined Saprotroph\| | Yeast | NULL | Probable | As nectar-tap_saprotroph (Põlme et al. 2020); Decay sugar-rich substrates (Põlme et al. 2020) | Põlme S, et al. 2020. Fungal Diversity 105:1-16 ((https://doi.org/10.1007/s13225-020-00466-2)) |
| Otu38119 | 1 | k__Fungi(100);p__Ascomycota(100);c__Saccharomycetes(100);o__Saccharomycetales(100);f__Metschnikowiaceae(100);g__Metschnikowia(100);s__Metschnikowia_pulcherrima(100); | Metschnikowia | 13 | Saprotroph | \|Undefined Saprotroph\| | Yeast | NULL | Probable | As nectar-tap_saprotroph (Põlme et al. 2020); Decay sugar-rich substrates (Põlme et al. 2020) | Põlme S, et al. 2020. Fungal Diversity 105:1-16 ((https://doi.org/10.1007/s13225-020-00466-2)) |
| Otu38120 | 1 | k__Fungi(100);p__Ascomycota(100);c__Saccharomycetes(100);o__Saccharomycetales(100);f__Metschnikowiaceae(100);g__Metschnikowia(100);s__Metschnikowia_pulcherrima(100); | Metschnikowia | 13 | Saprotroph | \|Undefined Saprotroph\| | Yeast | NULL | Probable | As nectar-tap_saprotroph (Põlme et al. 2020); Decay sugar-rich substrates (Põlme et al. 2020) | Põlme S, et al. 2020. Fungal Diversity 105:1-16 ((https://doi.org/10.1007/s13225-020-00466-2)) |
| Otu38121 | 1 | k__Fungi(100);p__Ascomycota(100);c__Saccharomycetes(100);o__Saccharomycetales(100);f__Metschnikowiaceae(100);g__Metschnikowia(100);s__Metschnikowia_pulcherrima(100); | Metschnikowia | 13 | Saprotroph | \|Undefined Saprotroph\| | Yeast | NULL | Probable | As nectar-tap_saprotroph (Põlme et al. 2020); Decay sugar-rich substrates (Põlme et al. 2020) | Põlme S, et al. 2020. Fungal Diversity 105:1-16 ((https://doi.org/10.1007/s13225-020-00466-2)) |
| Otu38122 | 1 | k__Fungi(100);p__Ascomycota(100);c__Saccharomycetes(100);o__Saccharomycetales(100);f__Metschnikowiaceae(100);g__Metschnikowia(100);s__Metschnikowia_pulcherrima(100); | Metschnikowia | 13 | Saprotroph | \|Undefined Saprotroph\| | Yeast | NULL | Probable | As nectar-tap_saprotroph (Põlme et al. 2020); Decay sugar-rich substrates (Põlme et al. 2020) | Põlme S, et al. 2020. Fungal Diversity 105:1-16 ((https://doi.org/10.1007/s13225-020-00466-2)) |
| Otu38123 | 1 | k__Fungi(100);p__Ascomycota(100);c__Saccharomycetes(100);o__Saccharomycetales(100);f__Metschnikowiaceae(100);g__Metschnikowia(100);s__Metschnikowia_pulcherrima(100); | Metschnikowia | 13 | Saprotroph | \|Undefined Saprotroph\| | Yeast | NULL | Probable | As nectar-tap_saprotroph (Põlme et al. 2020); Decay sugar-rich substrates (Põlme et al. 2020) | Põlme S, et al. 2020. Fungal Diversity 105:1-16 ((https://doi.org/10.1007/s13225-020-00466-2)) |
| Otu38124 | 1 | k__Fungi(100);p__Ascomycota(100);c__Saccharomycetes(100);o__Saccharomycetales(100);f__Metschnikowiaceae(100);g__Metschnikowia(100);s__Metschnikowia_pulcherrima(100); | Metschnikowia | 13 | Saprotroph | \|Undefined Saprotroph\| | Yeast | NULL | Probable | As nectar-tap_saprotroph (Põlme et al. 2020); Decay sugar-rich substrates (Põlme et al. 2020) | Põlme S, et al. 2020. Fungal Diversity 105:1-16 ((https://doi.org/10.1007/s13225-020-00466-2)) |
| Otu38125 | 1 | k__Fungi(100);p__Ascomycota(100);c__Saccharomycetes(100);o__Saccharomycetales(100);f__Metschnikowiaceae(100);g__Metschnikowia(100);s__Metschnikowia_pulcherrima(100); | Metschnikowia | 13 | Saprotroph | \|Undefined Saprotroph\| | Yeast | NULL | Probable | As nectar-tap_saprotroph (Põlme et al. 2020); Decay sugar-rich substrates (Põlme et al. 2020) | Põlme S, et al. 2020. Fungal Diversity 105:1-16 ((https://doi.org/10.1007/s13225-020-00466-2)) |
| Otu38126 | 1 | k__Fungi(100);p__Ascomycota(100);c__Saccharomycetes(100);o__Saccharomycetales(100);f__Metschnikowiaceae(100);g__Metschnikowia(100);s__Metschnikowia_pulcherrima(100); | Metschnikowia | 13 | Saprotroph | \|Undefined Saprotroph\| | Yeast | NULL | Probable | As nectar-tap_saprotroph (Põlme et al. 2020); Decay sugar-rich substrates (Põlme et al. 2020) | Põlme S, et al. 2020. Fungal Diversity 105:1-16 ((https://doi.org/10.1007/s13225-020-00466-2)) |
| Otu38127 | 1 | k__Fungi(100);p__Ascomycota(100);c__Saccharomycetes(100);o__Saccharomycetales(100);f__Metschnikowiaceae(100);g__Metschnikowia(100);s__Metschnikowia_pulcherrima(100); | Metschnikowia | 13 | Saprotroph | \|Undefined Saprotroph\| | Yeast | NULL | Probable | As nectar-tap_saprotroph (Põlme et al. 2020); Decay sugar-rich substrates (Põlme et al. 2020) | Põlme S, et al. 2020. Fungal Diversity 105:1-16 ((https://doi.org/10.1007/s13225-020-00466-2)) |
| Otu38128 | 1 | k__Fungi(100);p__Ascomycota(100);c__Saccharomycetes(100);o__Saccharomycetales(100);f__Metschnikowiaceae(100);g__Metschnikowia(100);s__Metschnikowia_pulcherrima(100); | Metschnikowia | 13 | Saprotroph | \|Undefined Saprotroph\| | Yeast | NULL | Probable | As nectar-tap_saprotroph (Põlme et al. 2020); Decay sugar-rich substrates (Põlme et al. 2020) | Põlme S, et al. 2020. Fungal Diversity 105:1-16 ((https://doi.org/10.1007/s13225-020-00466-2)) |
| Otu38129 | 1 | k__Fungi(100);p__Ascomycota(100);c__Saccharomycetes(100);o__Saccharomycetales(100);f__Metschnikowiaceae(100);g__Metschnikowia(100);s__Metschnikowia_pulcherrima(100); | Metschnikowia | 13 | Saprotroph | \|Undefined Saprotroph\| | Yeast | NULL | Probable | As nectar-tap_saprotroph (Põlme et al. 2020); Decay sugar-rich substrates (Põlme et al. 2020) | Põlme S, et al. 2020. Fungal Diversity 105:1-16 ((https://doi.org/10.1007/s13225-020-00466-2)) |
| Otu38130 | 1 | k__Fungi(100);p__Ascomycota(100);c__Saccharomycetes(100);o__Saccharomycetales(100);f__Metschnikowiaceae(100);g__Metschnikowia(100);s__Metschnikowia_pulcherrima(100); | Metschnikowia | 13 | Saprotroph | \|Undefined Saprotroph\| | Yeast | NULL | Probable | As nectar-tap_saprotroph (Põlme et al. 2020); Decay sugar-rich substrates (Põlme et al. 2020) | Põlme S, et al. 2020. Fungal Diversity 105:1-16 ((https://doi.org/10.1007/s13225-020-00466-2)) |
| Otu38131 | 1 | k__Fungi(100);p__Ascomycota(100);c__Saccharomycetes(100);o__Saccharomycetales(100);f__Metschnikowiaceae(100);g__Metschnikowia(100);s__Metschnikowia_pulcherrima(100); | Metschnikowia | 13 | Saprotroph | \|Undefined Saprotroph\| | Yeast | NULL | Probable | As nectar-tap_saprotroph (Põlme et al. 2020); Decay sugar-rich substrates (Põlme et al. 2020) | Põlme S, et al. 2020. Fungal Diversity 105:1-16 ((https://doi.org/10.1007/s13225-020-00466-2)) |
| Otu38132 | 1 | k__Fungi(100);p__Ascomycota(100);c__Saccharomycetes(100);o__Saccharomycetales(100);f__Metschnikowiaceae(100);g__Metschnikowia(100);s__Metschnikowia_pulcherrima(100); | Metschnikowia | 13 | Saprotroph | \|Undefined Saprotroph\| | Yeast | NULL | Probable | As nectar-tap_saprotroph (Põlme et al. 2020); Decay sugar-rich substrates (Põlme et al. 2020) | Põlme S, et al. 2020. Fungal Diversity 105:1-16 ((https://doi.org/10.1007/s13225-020-00466-2)) |
| Otu38133 | 1 | k__Fungi(100);p__Ascomycota(100);c__Saccharomycetes(100);o__Saccharomycetales(100);f__Metschnikowiaceae(100);g__Metschnikowia(100);g__Metschnikowia_unclassified(100); | Metschnikowia | 13 | Saprotroph | \|Undefined Saprotroph\| | Yeast | NULL | Probable | As nectar-tap_saprotroph (Põlme et al. 2020); Decay sugar-rich substrates (Põlme et al. 2020) | Põlme S, et al. 2020. Fungal Diversity 105:1-16 ((https://doi.org/10.1007/s13225-020-00466-2)) |
| Otu38134 | 1 | k__Fungi(100);p__Ascomycota(100);c__Saccharomycetes(100);o__Saccharomycetales(100);f__Metschnikowiaceae(100);g__Metschnikowia(100);g__Metschnikowia_unclassified(100); | Metschnikowia | 13 | Saprotroph | \|Undefined Saprotroph\| | Yeast | NULL | Probable | As nectar-tap_saprotroph (Põlme et al. 2020); Decay sugar-rich substrates (Põlme et al. 2020) | Põlme S, et al. 2020. Fungal Diversity 105:1-16 ((https://doi.org/10.1007/s13225-020-00466-2)) |
| Otu38135 | 1 | k__Fungi(100);p__Ascomycota(100);c__Saccharomycetes(100);o__Saccharomycetales(100);f__Metschnikowiaceae(100);g__Metschnikowia(100);g__Metschnikowia_unclassified(100); | Metschnikowia | 13 | Saprotroph | \|Undefined Saprotroph\| | Yeast | NULL | Probable | As nectar-tap_saprotroph (Põlme et al. 2020); Decay sugar-rich substrates (Põlme et al. 2020) | Põlme S, et al. 2020. Fungal Diversity 105:1-16 ((https://doi.org/10.1007/s13225-020-00466-2)) |
| Otu38136 | 1 | k__Fungi(100);p__Ascomycota(100);c__Saccharomycetes(100);o__Saccharomycetales(100);f__Metschnikowiaceae(100);g__Metschnikowia(100);g__Metschnikowia_unclassified(100); | Metschnikowia | 13 | Saprotroph | \|Undefined Saprotroph\| | Yeast | NULL | Probable | As nectar-tap_saprotroph (Põlme et al. 2020); Decay sugar-rich substrates (Põlme et al. 2020) | Põlme S, et al. 2020. Fungal Diversity 105:1-16 ((https://doi.org/10.1007/s13225-020-00466-2)) |
| Otu38137 | 1 | k__Fungi(100);p__Ascomycota(100);c__Saccharomycetes(100);o__Saccharomycetales(100);f__Metschnikowiaceae(100);g__Metschnikowia(100);g__Metschnikowia_unclassified(100); | Metschnikowia | 13 | Saprotroph | \|Undefined Saprotroph\| | Yeast | NULL | Probable | As nectar-tap_saprotroph (Põlme et al. 2020); Decay sugar-rich substrates (Põlme et al. 2020) | Põlme S, et al. 2020. Fungal Diversity 105:1-16 ((https://doi.org/10.1007/s13225-020-00466-2)) |
| Otu38138 | 1 | k__Fungi(100);p__Ascomycota(100);c__Saccharomycetes(100);o__Saccharomycetales(100);f__Metschnikowiaceae(100);g__Metschnikowia(100);s__Metschnikowia_pulcherrima(100); | Metschnikowia | 13 | Saprotroph | \|Undefined Saprotroph\| | Yeast | NULL | Probable | As nectar-tap_saprotroph (Põlme et al. 2020); Decay sugar-rich substrates (Põlme et al. 2020) | Põlme S, et al. 2020. Fungal Diversity 105:1-16 ((https://doi.org/10.1007/s13225-020-00466-2)) |
| Otu38139 | 1 | k__Fungi(100);p__Ascomycota(100);c__Saccharomycetes(100);o__Saccharomycetales(100);f__Metschnikowiaceae(100);g__Metschnikowia(100);s__Metschnikowia_pulcherrima(100); | Metschnikowia | 13 | Saprotroph | \|Undefined Saprotroph\| | Yeast | NULL | Probable | As nectar-tap_saprotroph (Põlme et al. 2020); Decay sugar-rich substrates (Põlme et al. 2020) | Põlme S, et al. 2020. Fungal Diversity 105:1-16 ((https://doi.org/10.1007/s13225-020-00466-2)) |
| Otu38140 | 1 | k__Fungi(100);p__Ascomycota(100);c__Saccharomycetes(100);o__Saccharomycetales(100);f__Metschnikowiaceae(100);g__Metschnikowia(100);s__Metschnikowia_pulcherrima(100); | Metschnikowia | 13 | Saprotroph | \|Undefined Saprotroph\| | Yeast | NULL | Probable | As nectar-tap_saprotroph (Põlme et al. 2020); Decay sugar-rich substrates (Põlme et al. 2020) | Põlme S, et al. 2020. Fungal Diversity 105:1-16 ((https://doi.org/10.1007/s13225-020-00466-2)) |
| Otu38141 | 1 | k__Fungi(100);p__Ascomycota(100);c__Saccharomycetes(100);o__Saccharomycetales(100);f__Metschnikowiaceae(100);g__Metschnikowia(100);s__Metschnikowia_pulcherrima(100); | Metschnikowia | 13 | Saprotroph | \|Undefined Saprotroph\| | Yeast | NULL | Probable | As nectar-tap_saprotroph (Põlme et al. 2020); Decay sugar-rich substrates (Põlme et al. 2020) | Põlme S, et al. 2020. Fungal Diversity 105:1-16 ((https://doi.org/10.1007/s13225-020-00466-2)) |
| Otu38142 | 1 | k__Fungi(100);p__Ascomycota(100);c__Saccharomycetes(100);o__Saccharomycetales(100);f__Metschnikowiaceae(100);g__Metschnikowia(100);g__Metschnikowia_unclassified(100); | Metschnikowia | 13 | Saprotroph | \|Undefined Saprotroph\| | Yeast | NULL | Probable | As nectar-tap_saprotroph (Põlme et al. 2020); Decay sugar-rich substrates (Põlme et al. 2020) | Põlme S, et al. 2020. Fungal Diversity 105:1-16 ((https://doi.org/10.1007/s13225-020-00466-2)) |
| Otu38143 | 1 | k__Fungi(100);p__Ascomycota(100);c__Saccharomycetes(100);o__Saccharomycetales(100);f__Metschnikowiaceae(100);g__Metschnikowia(100);s__Metschnikowia_pulcherrima(100); | Metschnikowia | 13 | Saprotroph | \|Undefined Saprotroph\| | Yeast | NULL | Probable | As nectar-tap_saprotroph (Põlme et al. 2020); Decay sugar-rich substrates (Põlme et al. 2020) | Põlme S, et al. 2020. Fungal Diversity 105:1-16 ((https://doi.org/10.1007/s13225-020-00466-2)) |
| Otu38144 | 1 | k__Fungi(100);p__Ascomycota(100);c__Saccharomycetes(100);o__Saccharomycetales(100);f__Metschnikowiaceae(100);g__Metschnikowia(100);g__Metschnikowia_unclassified(100); | Metschnikowia | 13 | Saprotroph | \|Undefined Saprotroph\| | Yeast | NULL | Probable | As nectar-tap_saprotroph (Põlme et al. 2020); Decay sugar-rich substrates (Põlme et al. 2020) | Põlme S, et al. 2020. Fungal Diversity 105:1-16 ((https://doi.org/10.1007/s13225-020-00466-2)) |
| Otu38145 | 1 | k__Fungi(100);p__Ascomycota(100);c__Saccharomycetes(100);o__Saccharomycetales(100);f__Metschnikowiaceae(100);g__Metschnikowia(100);g__Metschnikowia_unclassified(100); | Metschnikowia | 13 | Saprotroph | \|Undefined Saprotroph\| | Yeast | NULL | Probable | As nectar-tap_saprotroph (Põlme et al. 2020); Decay sugar-rich substrates (Põlme et al. 2020) | Põlme S, et al. 2020. Fungal Diversity 105:1-16 ((https://doi.org/10.1007/s13225-020-00466-2)) |
| Otu38146 | 1 | k__Fungi(100);p__Ascomycota(100);c__Saccharomycetes(100);o__Saccharomycetales(100);f__Metschnikowiaceae(100);g__Metschnikowia(100);s__Metschnikowia_pulcherrima(100); | Metschnikowia | 13 | Saprotroph | \|Undefined Saprotroph\| | Yeast | NULL | Probable | As nectar-tap_saprotroph (Põlme et al. 2020); Decay sugar-rich substrates (Põlme et al. 2020) | Põlme S, et al. 2020. Fungal Diversity 105:1-16 ((https://doi.org/10.1007/s13225-020-00466-2)) |
| Otu38147 | 1 | k__Fungi(100);p__Ascomycota(100);c__Saccharomycetes(100);o__Saccharomycetales(100);f__Metschnikowiaceae(100);g__Metschnikowia(100);g__Metschnikowia_unclassified(100); | Metschnikowia | 13 | Saprotroph | \|Undefined Saprotroph\| | Yeast | NULL | Probable | As nectar-tap_saprotroph (Põlme et al. 2020); Decay sugar-rich substrates (Põlme et al. 2020) | Põlme S, et al. 2020. Fungal Diversity 105:1-16 ((https://doi.org/10.1007/s13225-020-00466-2)) |
| Otu38148 | 1 | k__Fungi(100);p__Ascomycota(100);c__Saccharomycetes(100);o__Saccharomycetales(100);f__Metschnikowiaceae(100);g__Metschnikowia(100);s__Metschnikowia_pulcherrima(100); | Metschnikowia | 13 | Saprotroph | \|Undefined Saprotroph\| | Yeast | NULL | Probable | As nectar-tap_saprotroph (Põlme et al. 2020); Decay sugar-rich substrates (Põlme et al. 2020) | Põlme S, et al. 2020. Fungal Diversity 105:1-16 ((https://doi.org/10.1007/s13225-020-00466-2)) |
| Otu38149 | 1 | k__Fungi(100);p__Ascomycota(100);c__Saccharomycetes(100);o__Saccharomycetales(100);f__Metschnikowiaceae(100);g__Metschnikowia(100);s__Metschnikowia_pulcherrima(100); | Metschnikowia | 13 | Saprotroph | \|Undefined Saprotroph\| | Yeast | NULL | Probable | As nectar-tap_saprotroph (Põlme et al. 2020); Decay sugar-rich substrates (Põlme et al. 2020) | Põlme S, et al. 2020. Fungal Diversity 105:1-16 ((https://doi.org/10.1007/s13225-020-00466-2)) |
| Otu38150 | 1 | k__Fungi(100);p__Ascomycota(100);c__Saccharomycetes(100);o__Saccharomycetales(100);f__Metschnikowiaceae(100);g__Metschnikowia(100);g__Metschnikowia_unclassified(100); | Metschnikowia | 13 | Saprotroph | \|Undefined Saprotroph\| | Yeast | NULL | Probable | As nectar-tap_saprotroph (Põlme et al. 2020); Decay sugar-rich substrates (Põlme et al. 2020) | Põlme S, et al. 2020. Fungal Diversity 105:1-16 ((https://doi.org/10.1007/s13225-020-00466-2)) |
| Otu38151 | 1 | k__Fungi(100);p__Ascomycota(100);c__Saccharomycetes(100);o__Saccharomycetales(100);f__Metschnikowiaceae(100);g__Metschnikowia(100);s__Metschnikowia_pulcherrima(100); | Metschnikowia | 13 | Saprotroph | \|Undefined Saprotroph\| | Yeast | NULL | Probable | As nectar-tap_saprotroph (Põlme et al. 2020); Decay sugar-rich substrates (Põlme et al. 2020) | Põlme S, et al. 2020. Fungal Diversity 105:1-16 ((https://doi.org/10.1007/s13225-020-00466-2)) |
| Otu38152 | 1 | k__Fungi(100);p__Ascomycota(100);c__Saccharomycetes(100);o__Saccharomycetales(100);f__Metschnikowiaceae(100);g__Metschnikowia(100);g__Metschnikowia_unclassified(100); | Metschnikowia | 13 | Saprotroph | \|Undefined Saprotroph\| | Yeast | NULL | Probable | As nectar-tap_saprotroph (Põlme et al. 2020); Decay sugar-rich substrates (Põlme et al. 2020) | Põlme S, et al. 2020. Fungal Diversity 105:1-16 ((https://doi.org/10.1007/s13225-020-00466-2)) |
| Otu38153 | 1 | k__Fungi(100);p__Ascomycota(100);c__Saccharomycetes(100);o__Saccharomycetales(100);f__Metschnikowiaceae(100);g__Metschnikowia(100);s__Metschnikowia_pulcherrima(100); | Metschnikowia | 13 | Saprotroph | \|Undefined Saprotroph\| | Yeast | NULL | Probable | As nectar-tap_saprotroph (Põlme et al. 2020); Decay sugar-rich substrates (Põlme et al. 2020) | Põlme S, et al. 2020. Fungal Diversity 105:1-16 ((https://doi.org/10.1007/s13225-020-00466-2)) |
| Otu38154 | 1 | k__Fungi(100);p__Ascomycota(100);c__Saccharomycetes(100);o__Saccharomycetales(100);f__Metschnikowiaceae(100);g__Metschnikowia(100);g__Metschnikowia_unclassified(100); | Metschnikowia | 13 | Saprotroph | \|Undefined Saprotroph\| | Yeast | NULL | Probable | As nectar-tap_saprotroph (Põlme et al. 2020); Decay sugar-rich substrates (Põlme et al. 2020) | Põlme S, et al. 2020. Fungal Diversity 105:1-16 ((https://doi.org/10.1007/s13225-020-00466-2)) |
| Otu38155 | 1 | k__Fungi(100);p__Ascomycota(100);c__Saccharomycetes(100);o__Saccharomycetales(100);f__Metschnikowiaceae(100);g__Metschnikowia(100);g__Metschnikowia_unclassified(100); | Metschnikowia | 13 | Saprotroph | \|Undefined Saprotroph\| | Yeast | NULL | Probable | As nectar-tap_saprotroph (Põlme et al. 2020); Decay sugar-rich substrates (Põlme et al. 2020) | Põlme S, et al. 2020. Fungal Diversity 105:1-16 ((https://doi.org/10.1007/s13225-020-00466-2)) |
| Otu38156 | 1 | k__Fungi(100);p__Ascomycota(100);c__Saccharomycetes(100);o__Saccharomycetales(100);f__Metschnikowiaceae(100);g__Metschnikowia(100);s__Metschnikowia_pulcherrima(100); | Metschnikowia | 13 | Saprotroph | \|Undefined Saprotroph\| | Yeast | NULL | Probable | As nectar-tap_saprotroph (Põlme et al. 2020); Decay sugar-rich substrates (Põlme et al. 2020) | Põlme S, et al. 2020. Fungal Diversity 105:1-16 ((https://doi.org/10.1007/s13225-020-00466-2)) |
| Otu38157 | 1 | k__Fungi(100);p__Ascomycota(100);c__Saccharomycetes(100);o__Saccharomycetales(100);f__Metschnikowiaceae(100);g__Metschnikowia(100);s__Metschnikowia_pulcherrima(100); | Metschnikowia | 13 | Saprotroph | \|Undefined Saprotroph\| | Yeast | NULL | Probable | As nectar-tap_saprotroph (Põlme et al. 2020); Decay sugar-rich substrates (Põlme et al. 2020) | Põlme S, et al. 2020. Fungal Diversity 105:1-16 ((https://doi.org/10.1007/s13225-020-00466-2)) |
| Otu38158 | 1 | k__Fungi(100);p__Ascomycota(100);c__Saccharomycetes(100);o__Saccharomycetales(100);f__Metschnikowiaceae(100);g__Metschnikowia(100);s__Metschnikowia_pulcherrima(100); | Metschnikowia | 13 | Saprotroph | \|Undefined Saprotroph\| | Yeast | NULL | Probable | As nectar-tap_saprotroph (Põlme et al. 2020); Decay sugar-rich substrates (Põlme et al. 2020) | Põlme S, et al. 2020. Fungal Diversity 105:1-16 ((https://doi.org/10.1007/s13225-020-00466-2)) |
| Otu38159 | 1 | k__Fungi(100);p__Ascomycota(100);c__Saccharomycetes(100);o__Saccharomycetales(100);f__Metschnikowiaceae(100);g__Metschnikowia(100);s__Metschnikowia_pulcherrima(100); | Metschnikowia | 13 | Saprotroph | \|Undefined Saprotroph\| | Yeast | NULL | Probable | As nectar-tap_saprotroph (Põlme et al. 2020); Decay sugar-rich substrates (Põlme et al. 2020) | Põlme S, et al. 2020. Fungal Diversity 105:1-16 ((https://doi.org/10.1007/s13225-020-00466-2)) |
| Otu38160 | 1 | k__Fungi(100);p__Ascomycota(100);c__Saccharomycetes(100);o__Saccharomycetales(100);f__Metschnikowiaceae(100);g__Metschnikowia(100);g__Metschnikowia_unclassified(100); | Metschnikowia | 13 | Saprotroph | \|Undefined Saprotroph\| | Yeast | NULL | Probable | As nectar-tap_saprotroph (Põlme et al. 2020); Decay sugar-rich substrates (Põlme et al. 2020) | Põlme S, et al. 2020. Fungal Diversity 105:1-16 ((https://doi.org/10.1007/s13225-020-00466-2)) |
| Otu38161 | 1 | k__Fungi(100);p__Ascomycota(100);c__Saccharomycetes(100);o__Saccharomycetales(100);f__Metschnikowiaceae(100);g__Metschnikowia(100);s__Metschnikowia_pulcherrima(100); | Metschnikowia | 13 | Saprotroph | \|Undefined Saprotroph\| | Yeast | NULL | Probable | As nectar-tap_saprotroph (Põlme et al. 2020); Decay sugar-rich substrates (Põlme et al. 2020) | Põlme S, et al. 2020. Fungal Diversity 105:1-16 ((https://doi.org/10.1007/s13225-020-00466-2)) |
| Otu38162 | 1 | k__Fungi(100);p__Ascomycota(100);c__Saccharomycetes(100);o__Saccharomycetales(100);f__Metschnikowiaceae(100);g__Metschnikowia(100);g__Metschnikowia_unclassified(100); | Metschnikowia | 13 | Saprotroph | \|Undefined Saprotroph\| | Yeast | NULL | Probable | As nectar-tap_saprotroph (Põlme et al. 2020); Decay sugar-rich substrates (Põlme et al. 2020) | Põlme S, et al. 2020. Fungal Diversity 105:1-16 ((https://doi.org/10.1007/s13225-020-00466-2)) |
| Otu38163 | 1 | k__Fungi(100);p__Ascomycota(100);c__Saccharomycetes(100);o__Saccharomycetales(100);f__Metschnikowiaceae(100);g__Metschnikowia(100);s__Metschnikowia_pulcherrima(100); | Metschnikowia | 13 | Saprotroph | \|Undefined Saprotroph\| | Yeast | NULL | Probable | As nectar-tap_saprotroph (Põlme et al. 2020); Decay sugar-rich substrates (Põlme et al. 2020) | Põlme S, et al. 2020. Fungal Diversity 105:1-16 ((https://doi.org/10.1007/s13225-020-00466-2)) |
| Otu38164 | 1 | k__Fungi(100);p__Ascomycota(100);c__Saccharomycetes(100);o__Saccharomycetales(100);f__Metschnikowiaceae(100);g__Metschnikowia(100);s__Metschnikowia_pulcherrima(100); | Metschnikowia | 13 | Saprotroph | \|Undefined Saprotroph\| | Yeast | NULL | Probable | As nectar-tap_saprotroph (Põlme et al. 2020); Decay sugar-rich substrates (Põlme et al. 2020) | Põlme S, et al. 2020. Fungal Diversity 105:1-16 ((https://doi.org/10.1007/s13225-020-00466-2)) |
| Otu38165 | 1 | k__Fungi(100);p__Ascomycota(100);c__Saccharomycetes(100);o__Saccharomycetales(100);f__Metschnikowiaceae(100);g__Metschnikowia(100);g__Metschnikowia_unclassified(100); | Metschnikowia | 13 | Saprotroph | \|Undefined Saprotroph\| | Yeast | NULL | Probable | As nectar-tap_saprotroph (Põlme et al. 2020); Decay sugar-rich substrates (Põlme et al. 2020) | Põlme S, et al. 2020. Fungal Diversity 105:1-16 ((https://doi.org/10.1007/s13225-020-00466-2)) |
| Otu38166 | 1 | k__Fungi(100);p__Ascomycota(100);c__Saccharomycetes(100);o__Saccharomycetales(100);f__Metschnikowiaceae(100);g__Metschnikowia(100);s__Metschnikowia_pulcherrima(100); | Metschnikowia | 13 | Saprotroph | \|Undefined Saprotroph\| | Yeast | NULL | Probable | As nectar-tap_saprotroph (Põlme et al. 2020); Decay sugar-rich substrates (Põlme et al. 2020) | Põlme S, et al. 2020. Fungal Diversity 105:1-16 ((https://doi.org/10.1007/s13225-020-00466-2)) |
| Otu38167 | 1 | k__Fungi(100);p__Ascomycota(100);c__Saccharomycetes(100);o__Saccharomycetales(100);f__Metschnikowiaceae(100);g__Metschnikowia(100);s__Metschnikowia_pulcherrima(100); | Metschnikowia | 13 | Saprotroph | \|Undefined Saprotroph\| | Yeast | NULL | Probable | As nectar-tap_saprotroph (Põlme et al. 2020); Decay sugar-rich substrates (Põlme et al. 2020) | Põlme S, et al. 2020. Fungal Diversity 105:1-16 ((https://doi.org/10.1007/s13225-020-00466-2)) |
| Otu38168 | 1 | k__Fungi(100);p__Ascomycota(100);c__Saccharomycetes(100);o__Saccharomycetales(100);f__Metschnikowiaceae(100);g__Metschnikowia(100);g__Metschnikowia_unclassified(100); | Metschnikowia | 13 | Saprotroph | \|Undefined Saprotroph\| | Yeast | NULL | Probable | As nectar-tap_saprotroph (Põlme et al. 2020); Decay sugar-rich substrates (Põlme et al. 2020) | Põlme S, et al. 2020. Fungal Diversity 105:1-16 ((https://doi.org/10.1007/s13225-020-00466-2)) |
| Otu38169 | 1 | k__Fungi(100);p__Ascomycota(100);c__Saccharomycetes(100);o__Saccharomycetales(100);f__Metschnikowiaceae(100);g__Metschnikowia(100);s__Metschnikowia_pulcherrima(100); | Metschnikowia | 13 | Saprotroph | \|Undefined Saprotroph\| | Yeast | NULL | Probable | As nectar-tap_saprotroph (Põlme et al. 2020); Decay sugar-rich substrates (Põlme et al. 2020) | Põlme S, et al. 2020. Fungal Diversity 105:1-16 ((https://doi.org/10.1007/s13225-020-00466-2)) |
| Otu38170 | 1 | k__Fungi(100);p__Ascomycota(100);c__Saccharomycetes(100);o__Saccharomycetales(100);f__Metschnikowiaceae(100);g__Metschnikowia(100);s__Metschnikowia_pulcherrima(100); | Metschnikowia | 13 | Saprotroph | \|Undefined Saprotroph\| | Yeast | NULL | Probable | As nectar-tap_saprotroph (Põlme et al. 2020); Decay sugar-rich substrates (Põlme et al. 2020) | Põlme S, et al. 2020. Fungal Diversity 105:1-16 ((https://doi.org/10.1007/s13225-020-00466-2)) |
| Otu38171 | 1 | k__Fungi(100);p__Ascomycota(100);c__Saccharomycetes(100);o__Saccharomycetales(100);f__Metschnikowiaceae(100);g__Metschnikowia(100);s__Metschnikowia_pulcherrima(100); | Metschnikowia | 13 | Saprotroph | \|Undefined Saprotroph\| | Yeast | NULL | Probable | As nectar-tap_saprotroph (Põlme et al. 2020); Decay sugar-rich substrates (Põlme et al. 2020) | Põlme S, et al. 2020. Fungal Diversity 105:1-16 ((https://doi.org/10.1007/s13225-020-00466-2)) |
| Otu38172 | 1 | k__Fungi(100);p__Ascomycota(100);c__Saccharomycetes(100);o__Saccharomycetales(100);f__Metschnikowiaceae(100);g__Metschnikowia(100);s__Metschnikowia_pulcherrima(100); | Metschnikowia | 13 | Saprotroph | \|Undefined Saprotroph\| | Yeast | NULL | Probable | As nectar-tap_saprotroph (Põlme et al. 2020); Decay sugar-rich substrates (Põlme et al. 2020) | Põlme S, et al. 2020. Fungal Diversity 105:1-16 ((https://doi.org/10.1007/s13225-020-00466-2)) |
| Otu38173 | 1 | k__Fungi(100);p__Ascomycota(100);c__Saccharomycetes(100);o__Saccharomycetales(100);f__Metschnikowiaceae(100);g__Metschnikowia(100);g__Metschnikowia_unclassified(100); | Metschnikowia | 13 | Saprotroph | \|Undefined Saprotroph\| | Yeast | NULL | Probable | As nectar-tap_saprotroph (Põlme et al. 2020); Decay sugar-rich substrates (Põlme et al. 2020) | Põlme S, et al. 2020. Fungal Diversity 105:1-16 ((https://doi.org/10.1007/s13225-020-00466-2)) |
| Otu38174 | 1 | k__Fungi(100);p__Ascomycota(100);c__Saccharomycetes(100);o__Saccharomycetales(100);f__Metschnikowiaceae(100);g__Metschnikowia(100);s__Metschnikowia_pulcherrima(100); | Metschnikowia | 13 | Saprotroph | \|Undefined Saprotroph\| | Yeast | NULL | Probable | As nectar-tap_saprotroph (Põlme et al. 2020); Decay sugar-rich substrates (Põlme et al. 2020) | Põlme S, et al. 2020. Fungal Diversity 105:1-16 ((https://doi.org/10.1007/s13225-020-00466-2)) |
| Otu38175 | 1 | k__Fungi(100);p__Ascomycota(100);c__Saccharomycetes(100);o__Saccharomycetales(100);f__Metschnikowiaceae(100);g__Metschnikowia(100);s__Metschnikowia_pulcherrima(100); | Metschnikowia | 13 | Saprotroph | \|Undefined Saprotroph\| | Yeast | NULL | Probable | As nectar-tap_saprotroph (Põlme et al. 2020); Decay sugar-rich substrates (Põlme et al. 2020) | Põlme S, et al. 2020. Fungal Diversity 105:1-16 ((https://doi.org/10.1007/s13225-020-00466-2)) |
| Otu38176 | 1 | k__Fungi(100);p__Ascomycota(100);c__Saccharomycetes(100);o__Saccharomycetales(100);f__Metschnikowiaceae(100);g__Metschnikowia(100);s__Metschnikowia_pulcherrima(100); | Metschnikowia | 13 | Saprotroph | \|Undefined Saprotroph\| | Yeast | NULL | Probable | As nectar-tap_saprotroph (Põlme et al. 2020); Decay sugar-rich substrates (Põlme et al. 2020) | Põlme S, et al. 2020. Fungal Diversity 105:1-16 ((https://doi.org/10.1007/s13225-020-00466-2)) |
| Otu38177 | 1 | k__Fungi(100);p__Ascomycota(100);c__Saccharomycetes(100);o__Saccharomycetales(100);f__Metschnikowiaceae(100);g__Metschnikowia(100);s__Metschnikowia_pulcherrima(100); | Metschnikowia | 13 | Saprotroph | \|Undefined Saprotroph\| | Yeast | NULL | Probable | As nectar-tap_saprotroph (Põlme et al. 2020); Decay sugar-rich substrates (Põlme et al. 2020) | Põlme S, et al. 2020. Fungal Diversity 105:1-16 ((https://doi.org/10.1007/s13225-020-00466-2)) |
| Otu38178 | 1 | k__Fungi(100);p__Ascomycota(100);c__Saccharomycetes(100);o__Saccharomycetales(100);f__Metschnikowiaceae(100);g__Metschnikowia(100);s__Metschnikowia_pulcherrima(100); | Metschnikowia | 13 | Saprotroph | \|Undefined Saprotroph\| | Yeast | NULL | Probable | As nectar-tap_saprotroph (Põlme et al. 2020); Decay sugar-rich substrates (Põlme et al. 2020) | Põlme S, et al. 2020. Fungal Diversity 105:1-16 ((https://doi.org/10.1007/s13225-020-00466-2)) |
| Otu38179 | 1 | k__Fungi(100);p__Ascomycota(100);c__Saccharomycetes(100);o__Saccharomycetales(100);f__Metschnikowiaceae(100);g__Metschnikowia(100);s__Metschnikowia_pulcherrima(100); | Metschnikowia | 13 | Saprotroph | \|Undefined Saprotroph\| | Yeast | NULL | Probable | As nectar-tap_saprotroph (Põlme et al. 2020); Decay sugar-rich substrates (Põlme et al. 2020) | Põlme S, et al. 2020. Fungal Diversity 105:1-16 ((https://doi.org/10.1007/s13225-020-00466-2)) |
| Otu38180 | 1 | k__Fungi(100);p__Ascomycota(100);c__Saccharomycetes(100);o__Saccharomycetales(100);f__Metschnikowiaceae(100);g__Metschnikowia(100);s__Metschnikowia_pulcherrima(100); | Metschnikowia | 13 | Saprotroph | \|Undefined Saprotroph\| | Yeast | NULL | Probable | As nectar-tap_saprotroph (Põlme et al. 2020); Decay sugar-rich substrates (Põlme et al. 2020) | Põlme S, et al. 2020. Fungal Diversity 105:1-16 ((https://doi.org/10.1007/s13225-020-00466-2)) |
| Otu38181 | 1 | k__Fungi(100);p__Ascomycota(100);c__Saccharomycetes(100);o__Saccharomycetales(100);f__Metschnikowiaceae(100);g__Metschnikowia(100);s__Metschnikowia_pulcherrima(100); | Metschnikowia | 13 | Saprotroph | \|Undefined Saprotroph\| | Yeast | NULL | Probable | As nectar-tap_saprotroph (Põlme et al. 2020); Decay sugar-rich substrates (Põlme et al. 2020) | Põlme S, et al. 2020. Fungal Diversity 105:1-16 ((https://doi.org/10.1007/s13225-020-00466-2)) |
| Otu38182 | 1 | k__Fungi(100);p__Ascomycota(100);c__Saccharomycetes(100);o__Saccharomycetales(100);f__Metschnikowiaceae(100);g__Metschnikowia(100);g__Metschnikowia_unclassified(100); | Metschnikowia | 13 | Saprotroph | \|Undefined Saprotroph\| | Yeast | NULL | Probable | As nectar-tap_saprotroph (Põlme et al. 2020); Decay sugar-rich substrates (Põlme et al. 2020) | Põlme S, et al. 2020. Fungal Diversity 105:1-16 ((https://doi.org/10.1007/s13225-020-00466-2)) |
| Otu38183 | 1 | k__Fungi(100);p__Ascomycota(100);c__Saccharomycetes(100);o__Saccharomycetales(100);f__Metschnikowiaceae(100);g__Metschnikowia(100);s__Metschnikowia_pulcherrima(100); | Metschnikowia | 13 | Saprotroph | \|Undefined Saprotroph\| | Yeast | NULL | Probable | As nectar-tap_saprotroph (Põlme et al. 2020); Decay sugar-rich substrates (Põlme et al. 2020) | Põlme S, et al. 2020. Fungal Diversity 105:1-16 ((https://doi.org/10.1007/s13225-020-00466-2)) |
| Otu38184 | 1 | k__Fungi(100);p__Ascomycota(100);c__Saccharomycetes(100);o__Saccharomycetales(100);f__Metschnikowiaceae(100);g__Metschnikowia(100);g__Metschnikowia_unclassified(100); | Metschnikowia | 13 | Saprotroph | \|Undefined Saprotroph\| | Yeast | NULL | Probable | As nectar-tap_saprotroph (Põlme et al. 2020); Decay sugar-rich substrates (Põlme et al. 2020) | Põlme S, et al. 2020. Fungal Diversity 105:1-16 ((https://doi.org/10.1007/s13225-020-00466-2)) |
| Otu38185 | 1 | k__Fungi(100);p__Ascomycota(100);c__Saccharomycetes(100);o__Saccharomycetales(100);f__Metschnikowiaceae(100);g__Metschnikowia(100);s__Metschnikowia_pulcherrima(100); | Metschnikowia | 13 | Saprotroph | \|Undefined Saprotroph\| | Yeast | NULL | Probable | As nectar-tap_saprotroph (Põlme et al. 2020); Decay sugar-rich substrates (Põlme et al. 2020) | Põlme S, et al. 2020. Fungal Diversity 105:1-16 ((https://doi.org/10.1007/s13225-020-00466-2)) |
| Otu38186 | 1 | k__Fungi(100);p__Ascomycota(100);c__Saccharomycetes(100);o__Saccharomycetales(100);f__Metschnikowiaceae(100);g__Metschnikowia(100);s__Metschnikowia_pulcherrima(100); | Metschnikowia | 13 | Saprotroph | \|Undefined Saprotroph\| | Yeast | NULL | Probable | As nectar-tap_saprotroph (Põlme et al. 2020); Decay sugar-rich substrates (Põlme et al. 2020) | Põlme S, et al. 2020. Fungal Diversity 105:1-16 ((https://doi.org/10.1007/s13225-020-00466-2)) |
| Otu38187 | 1 | k__Fungi(100);p__Ascomycota(100);c__Saccharomycetes(100);o__Saccharomycetales(100);f__Metschnikowiaceae(100);g__Metschnikowia(100);s__Metschnikowia_pulcherrima(100); | Metschnikowia | 13 | Saprotroph | \|Undefined Saprotroph\| | Yeast | NULL | Probable | As nectar-tap_saprotroph (Põlme et al. 2020); Decay sugar-rich substrates (Põlme et al. 2020) | Põlme S, et al. 2020. Fungal Diversity 105:1-16 ((https://doi.org/10.1007/s13225-020-00466-2)) |
| Otu38188 | 1 | k__Fungi(100);p__Ascomycota(100);c__Saccharomycetes(100);o__Saccharomycetales(100);f__Metschnikowiaceae(100);g__Metschnikowia(100);s__Metschnikowia_pulcherrima(100); | Metschnikowia | 13 | Saprotroph | \|Undefined Saprotroph\| | Yeast | NULL | Probable | As nectar-tap_saprotroph (Põlme et al. 2020); Decay sugar-rich substrates (Põlme et al. 2020) | Põlme S, et al. 2020. Fungal Diversity 105:1-16 ((https://doi.org/10.1007/s13225-020-00466-2)) |
| Otu38189 | 1 | k__Fungi(100);p__Ascomycota(100);c__Saccharomycetes(100);o__Saccharomycetales(100);f__Metschnikowiaceae(100);g__Metschnikowia(100);g__Metschnikowia_unclassified(100); | Metschnikowia | 13 | Saprotroph | \|Undefined Saprotroph\| | Yeast | NULL | Probable | As nectar-tap_saprotroph (Põlme et al. 2020); Decay sugar-rich substrates (Põlme et al. 2020) | Põlme S, et al. 2020. Fungal Diversity 105:1-16 ((https://doi.org/10.1007/s13225-020-00466-2)) |
| Otu38190 | 1 | k__Fungi(100);p__Ascomycota(100);c__Saccharomycetes(100);o__Saccharomycetales(100);f__Metschnikowiaceae(100);g__Metschnikowia(100);s__Metschnikowia_pulcherrima(100); | Metschnikowia | 13 | Saprotroph | \|Undefined Saprotroph\| | Yeast | NULL | Probable | As nectar-tap_saprotroph (Põlme et al. 2020); Decay sugar-rich substrates (Põlme et al. 2020) | Põlme S, et al. 2020. Fungal Diversity 105:1-16 ((https://doi.org/10.1007/s13225-020-00466-2)) |
| Otu38191 | 1 | k__Fungi(100);p__Ascomycota(100);c__Saccharomycetes(100);o__Saccharomycetales(100);f__Metschnikowiaceae(100);g__Metschnikowia(100);g__Metschnikowia_unclassified(100); | Metschnikowia | 13 | Saprotroph | \|Undefined Saprotroph\| | Yeast | NULL | Probable | As nectar-tap_saprotroph (Põlme et al. 2020); Decay sugar-rich substrates (Põlme et al. 2020) | Põlme S, et al. 2020. Fungal Diversity 105:1-16 ((https://doi.org/10.1007/s13225-020-00466-2)) |
| Otu38192 | 1 | k__Fungi(100);p__Ascomycota(100);c__Saccharomycetes(100);o__Saccharomycetales(100);f__Metschnikowiaceae(100);g__Metschnikowia(100);s__Metschnikowia_pulcherrima(100); | Metschnikowia | 13 | Saprotroph | \|Undefined Saprotroph\| | Yeast | NULL | Probable | As nectar-tap_saprotroph (Põlme et al. 2020); Decay sugar-rich substrates (Põlme et al. 2020) | Põlme S, et al. 2020. Fungal Diversity 105:1-16 ((https://doi.org/10.1007/s13225-020-00466-2)) |
| Otu38193 | 1 | k__Fungi(100);p__Ascomycota(100);c__Saccharomycetes(100);o__Saccharomycetales(100);f__Metschnikowiaceae(100);g__Metschnikowia(100);s__Metschnikowia_pulcherrima(100); | Metschnikowia | 13 | Saprotroph | \|Undefined Saprotroph\| | Yeast | NULL | Probable | As nectar-tap_saprotroph (Põlme et al. 2020); Decay sugar-rich substrates (Põlme et al. 2020) | Põlme S, et al. 2020. Fungal Diversity 105:1-16 ((https://doi.org/10.1007/s13225-020-00466-2)) |
| Otu38195 | 1 | k__Fungi(100);p__Ascomycota(100);c__Saccharomycetes(100);o__Saccharomycetales(100);f__Metschnikowiaceae(100);g__Metschnikowia(100);s__Metschnikowia_pulcherrima(100); | Metschnikowia | 13 | Saprotroph | \|Undefined Saprotroph\| | Yeast | NULL | Probable | As nectar-tap_saprotroph (Põlme et al. 2020); Decay sugar-rich substrates (Põlme et al. 2020) | Põlme S, et al. 2020. Fungal Diversity 105:1-16 ((https://doi.org/10.1007/s13225-020-00466-2)) |
| Otu38196 | 1 | k__Fungi(100);p__Ascomycota(100);c__Saccharomycetes(100);o__Saccharomycetales(100);f__Metschnikowiaceae(100);g__Metschnikowia(100);g__Metschnikowia_unclassified(100); | Metschnikowia | 13 | Saprotroph | \|Undefined Saprotroph\| | Yeast | NULL | Probable | As nectar-tap_saprotroph (Põlme et al. 2020); Decay sugar-rich substrates (Põlme et al. 2020) | Põlme S, et al. 2020. Fungal Diversity 105:1-16 ((https://doi.org/10.1007/s13225-020-00466-2)) |
| Otu38197 | 1 | k__Fungi(100);p__Ascomycota(100);c__Saccharomycetes(100);o__Saccharomycetales(100);f__Metschnikowiaceae(100);g__Metschnikowia(100);s__Metschnikowia_pulcherrima(100); | Metschnikowia | 13 | Saprotroph | \|Undefined Saprotroph\| | Yeast | NULL | Probable | As nectar-tap_saprotroph (Põlme et al. 2020); Decay sugar-rich substrates (Põlme et al. 2020) | Põlme S, et al. 2020. Fungal Diversity 105:1-16 ((https://doi.org/10.1007/s13225-020-00466-2)) |
| Otu38198 | 1 | k__Fungi(100);p__Ascomycota(100);c__Saccharomycetes(100);o__Saccharomycetales(100);f__Metschnikowiaceae(100);g__Metschnikowia(100);s__Metschnikowia_pulcherrima(100); | Metschnikowia | 13 | Saprotroph | \|Undefined Saprotroph\| | Yeast | NULL | Probable | As nectar-tap_saprotroph (Põlme et al. 2020); Decay sugar-rich substrates (Põlme et al. 2020) | Põlme S, et al. 2020. Fungal Diversity 105:1-16 ((https://doi.org/10.1007/s13225-020-00466-2)) |
| Otu38199 | 1 | k__Fungi(100);p__Ascomycota(100);c__Saccharomycetes(100);o__Saccharomycetales(100);f__Metschnikowiaceae(100);g__Metschnikowia(100);s__Metschnikowia_pulcherrima(100); | Metschnikowia | 13 | Saprotroph | \|Undefined Saprotroph\| | Yeast | NULL | Probable | As nectar-tap_saprotroph (Põlme et al. 2020); Decay sugar-rich substrates (Põlme et al. 2020) | Põlme S, et al. 2020. Fungal Diversity 105:1-16 ((https://doi.org/10.1007/s13225-020-00466-2)) |
| Otu38200 | 1 | k__Fungi(100);p__Ascomycota(100);c__Saccharomycetes(100);o__Saccharomycetales(100);f__Metschnikowiaceae(100);g__Metschnikowia(100);s__Metschnikowia_pulcherrima(100); | Metschnikowia | 13 | Saprotroph | \|Undefined Saprotroph\| | Yeast | NULL | Probable | As nectar-tap_saprotroph (Põlme et al. 2020); Decay sugar-rich substrates (Põlme et al. 2020) | Põlme S, et al. 2020. Fungal Diversity 105:1-16 ((https://doi.org/10.1007/s13225-020-00466-2)) |
| Otu38201 | 1 | k__Fungi(100);p__Ascomycota(100);c__Saccharomycetes(100);o__Saccharomycetales(100);f__Metschnikowiaceae(100);g__Metschnikowia(100);s__Metschnikowia_pulcherrima(100); | Metschnikowia | 13 | Saprotroph | \|Undefined Saprotroph\| | Yeast | NULL | Probable | As nectar-tap_saprotroph (Põlme et al. 2020); Decay sugar-rich substrates (Põlme et al. 2020) | Põlme S, et al. 2020. Fungal Diversity 105:1-16 ((https://doi.org/10.1007/s13225-020-00466-2)) |
| Otu38202 | 1 | k__Fungi(100);p__Ascomycota(100);c__Saccharomycetes(100);o__Saccharomycetales(100);f__Metschnikowiaceae(100);g__Metschnikowia(100);s__Metschnikowia_pulcherrima(100); | Metschnikowia | 13 | Saprotroph | \|Undefined Saprotroph\| | Yeast | NULL | Probable | As nectar-tap_saprotroph (Põlme et al. 2020); Decay sugar-rich substrates (Põlme et al. 2020) | Põlme S, et al. 2020. Fungal Diversity 105:1-16 ((https://doi.org/10.1007/s13225-020-00466-2)) |
| Otu38203 | 1 | k__Fungi(100);p__Ascomycota(100);c__Saccharomycetes(100);o__Saccharomycetales(100);f__Metschnikowiaceae(100);g__Metschnikowia(100);s__Metschnikowia_pulcherrima(100); | Metschnikowia | 13 | Saprotroph | \|Undefined Saprotroph\| | Yeast | NULL | Probable | As nectar-tap_saprotroph (Põlme et al. 2020); Decay sugar-rich substrates (Põlme et al. 2020) | Põlme S, et al. 2020. Fungal Diversity 105:1-16 ((https://doi.org/10.1007/s13225-020-00466-2)) |
| Otu38204 | 1 | k__Fungi(100);p__Ascomycota(100);c__Saccharomycetes(100);o__Saccharomycetales(100);f__Metschnikowiaceae(100);g__Metschnikowia(100);s__Metschnikowia_pulcherrima(100); | Metschnikowia | 13 | Saprotroph | \|Undefined Saprotroph\| | Yeast | NULL | Probable | As nectar-tap_saprotroph (Põlme et al. 2020); Decay sugar-rich substrates (Põlme et al. 2020) | Põlme S, et al. 2020. Fungal Diversity 105:1-16 ((https://doi.org/10.1007/s13225-020-00466-2)) |
| Otu38205 | 1 | k__Fungi(100);p__Ascomycota(100);c__Saccharomycetes(100);o__Saccharomycetales(100);f__Metschnikowiaceae(100);g__Metschnikowia(100);g__Metschnikowia_unclassified(100); | Metschnikowia | 13 | Saprotroph | \|Undefined Saprotroph\| | Yeast | NULL | Probable | As nectar-tap_saprotroph (Põlme et al. 2020); Decay sugar-rich substrates (Põlme et al. 2020) | Põlme S, et al. 2020. Fungal Diversity 105:1-16 ((https://doi.org/10.1007/s13225-020-00466-2)) |
| Otu38206 | 1 | k__Fungi(100);p__Ascomycota(100);c__Saccharomycetes(100);o__Saccharomycetales(100);f__Metschnikowiaceae(100);g__Metschnikowia(100);s__Metschnikowia_pulcherrima(100); | Metschnikowia | 13 | Saprotroph | \|Undefined Saprotroph\| | Yeast | NULL | Probable | As nectar-tap_saprotroph (Põlme et al. 2020); Decay sugar-rich substrates (Põlme et al. 2020) | Põlme S, et al. 2020. Fungal Diversity 105:1-16 ((https://doi.org/10.1007/s13225-020-00466-2)) |
| Otu38207 | 1 | k__Fungi(100);p__Ascomycota(100);c__Saccharomycetes(100);o__Saccharomycetales(100);f__Metschnikowiaceae(100);g__Metschnikowia(100);s__Metschnikowia_pulcherrima(100); | Metschnikowia | 13 | Saprotroph | \|Undefined Saprotroph\| | Yeast | NULL | Probable | As nectar-tap_saprotroph (Põlme et al. 2020); Decay sugar-rich substrates (Põlme et al. 2020) | Põlme S, et al. 2020. Fungal Diversity 105:1-16 ((https://doi.org/10.1007/s13225-020-00466-2)) |
| Otu38208 | 1 | k__Fungi(100);p__Ascomycota(100);c__Saccharomycetes(100);o__Saccharomycetales(100);f__Metschnikowiaceae(100);g__Metschnikowia(100);s__Metschnikowia_pulcherrima(100); | Metschnikowia | 13 | Saprotroph | \|Undefined Saprotroph\| | Yeast | NULL | Probable | As nectar-tap_saprotroph (Põlme et al. 2020); Decay sugar-rich substrates (Põlme et al. 2020) | Põlme S, et al. 2020. Fungal Diversity 105:1-16 ((https://doi.org/10.1007/s13225-020-00466-2)) |
| Otu38209 | 1 | k__Fungi(100);p__Ascomycota(100);c__Saccharomycetes(100);o__Saccharomycetales(100);f__Metschnikowiaceae(100);g__Metschnikowia(100);s__Metschnikowia_pulcherrima(100); | Metschnikowia | 13 | Saprotroph | \|Undefined Saprotroph\| | Yeast | NULL | Probable | As nectar-tap_saprotroph (Põlme et al. 2020); Decay sugar-rich substrates (Põlme et al. 2020) | Põlme S, et al. 2020. Fungal Diversity 105:1-16 ((https://doi.org/10.1007/s13225-020-00466-2)) |
| Otu38210 | 1 | k__Fungi(100);p__Ascomycota(100);c__Saccharomycetes(100);o__Saccharomycetales(100);f__Metschnikowiaceae(100);g__Metschnikowia(100);s__Metschnikowia_pulcherrima(100); | Metschnikowia | 13 | Saprotroph | \|Undefined Saprotroph\| | Yeast | NULL | Probable | As nectar-tap_saprotroph (Põlme et al. 2020); Decay sugar-rich substrates (Põlme et al. 2020) | Põlme S, et al. 2020. Fungal Diversity 105:1-16 ((https://doi.org/10.1007/s13225-020-00466-2)) |
| Otu38212 | 1 | k__Fungi(100);p__Ascomycota(100);c__Saccharomycetes(100);o__Saccharomycetales(100);f__Metschnikowiaceae(100);g__Metschnikowia(100);s__Metschnikowia_pulcherrima(100); | Metschnikowia | 13 | Saprotroph | \|Undefined Saprotroph\| | Yeast | NULL | Probable | As nectar-tap_saprotroph (Põlme et al. 2020); Decay sugar-rich substrates (Põlme et al. 2020) | Põlme S, et al. 2020. Fungal Diversity 105:1-16 ((https://doi.org/10.1007/s13225-020-00466-2)) |
| Otu38213 | 1 | k__Fungi(100);p__Ascomycota(100);c__Saccharomycetes(100);o__Saccharomycetales(100);f__Metschnikowiaceae(100);g__Metschnikowia(100);s__Metschnikowia_pulcherrima(100); | Metschnikowia | 13 | Saprotroph | \|Undefined Saprotroph\| | Yeast | NULL | Probable | As nectar-tap_saprotroph (Põlme et al. 2020); Decay sugar-rich substrates (Põlme et al. 2020) | Põlme S, et al. 2020. Fungal Diversity 105:1-16 ((https://doi.org/10.1007/s13225-020-00466-2)) |
| Otu38214 | 1 | k__Fungi(100);p__Ascomycota(100);c__Saccharomycetes(100);o__Saccharomycetales(100);f__Metschnikowiaceae(100);g__Metschnikowia(100);s__Metschnikowia_pulcherrima(100); | Metschnikowia | 13 | Saprotroph | \|Undefined Saprotroph\| | Yeast | NULL | Probable | As nectar-tap_saprotroph (Põlme et al. 2020); Decay sugar-rich substrates (Põlme et al. 2020) | Põlme S, et al. 2020. Fungal Diversity 105:1-16 ((https://doi.org/10.1007/s13225-020-00466-2)) |
| Otu38216 | 1 | k__Fungi(100);p__Ascomycota(100);c__Saccharomycetes(100);o__Saccharomycetales(100);f__Metschnikowiaceae(100);g__Metschnikowia(100);g__Metschnikowia_unclassified(100); | Metschnikowia | 13 | Saprotroph | \|Undefined Saprotroph\| | Yeast | NULL | Probable | As nectar-tap_saprotroph (Põlme et al. 2020); Decay sugar-rich substrates (Põlme et al. 2020) | Põlme S, et al. 2020. Fungal Diversity 105:1-16 ((https://doi.org/10.1007/s13225-020-00466-2)) |
| Otu38217 | 1 | k__Fungi(100);p__Ascomycota(100);c__Saccharomycetes(100);o__Saccharomycetales(100);f__Metschnikowiaceae(100);g__Metschnikowia(100);s__Metschnikowia_pulcherrima(100); | Metschnikowia | 13 | Saprotroph | \|Undefined Saprotroph\| | Yeast | NULL | Probable | As nectar-tap_saprotroph (Põlme et al. 2020); Decay sugar-rich substrates (Põlme et al. 2020) | Põlme S, et al. 2020. Fungal Diversity 105:1-16 ((https://doi.org/10.1007/s13225-020-00466-2)) |
| Otu38218 | 1 | k__Fungi(100);p__Ascomycota(100);c__Saccharomycetes(100);o__Saccharomycetales(100);f__Metschnikowiaceae(100);g__Metschnikowia(100);g__Metschnikowia_unclassified(100); | Metschnikowia | 13 | Saprotroph | \|Undefined Saprotroph\| | Yeast | NULL | Probable | As nectar-tap_saprotroph (Põlme et al. 2020); Decay sugar-rich substrates (Põlme et al. 2020) | Põlme S, et al. 2020. Fungal Diversity 105:1-16 ((https://doi.org/10.1007/s13225-020-00466-2)) |
| Otu38219 | 1 | k__Fungi(100);p__Ascomycota(100);c__Saccharomycetes(100);o__Saccharomycetales(100);f__Metschnikowiaceae(100);g__Metschnikowia(100);s__Metschnikowia_pulcherrima(100); | Metschnikowia | 13 | Saprotroph | \|Undefined Saprotroph\| | Yeast | NULL | Probable | As nectar-tap_saprotroph (Põlme et al. 2020); Decay sugar-rich substrates (Põlme et al. 2020) | Põlme S, et al. 2020. Fungal Diversity 105:1-16 ((https://doi.org/10.1007/s13225-020-00466-2)) |
| Otu38220 | 1 | k__Fungi(100);p__Ascomycota(100);c__Saccharomycetes(100);o__Saccharomycetales(100);f__Metschnikowiaceae(100);g__Metschnikowia(100);s__Metschnikowia_pulcherrima(100); | Metschnikowia | 13 | Saprotroph | \|Undefined Saprotroph\| | Yeast | NULL | Probable | As nectar-tap_saprotroph (Põlme et al. 2020); Decay sugar-rich substrates (Põlme et al. 2020) | Põlme S, et al. 2020. Fungal Diversity 105:1-16 ((https://doi.org/10.1007/s13225-020-00466-2)) |
| Otu38221 | 1 | k__Fungi(100);p__Ascomycota(100);c__Saccharomycetes(100);o__Saccharomycetales(100);f__Metschnikowiaceae(100);g__Metschnikowia(100);s__Metschnikowia_pulcherrima(100); | Metschnikowia | 13 | Saprotroph | \|Undefined Saprotroph\| | Yeast | NULL | Probable | As nectar-tap_saprotroph (Põlme et al. 2020); Decay sugar-rich substrates (Põlme et al. 2020) | Põlme S, et al. 2020. Fungal Diversity 105:1-16 ((https://doi.org/10.1007/s13225-020-00466-2)) |
| Otu38222 | 1 | k__Fungi(100);p__Ascomycota(100);c__Saccharomycetes(100);o__Saccharomycetales(100);f__Metschnikowiaceae(100);g__Metschnikowia(100);s__Metschnikowia_pulcherrima(100); | Metschnikowia | 13 | Saprotroph | \|Undefined Saprotroph\| | Yeast | NULL | Probable | As nectar-tap_saprotroph (Põlme et al. 2020); Decay sugar-rich substrates (Põlme et al. 2020) | Põlme S, et al. 2020. Fungal Diversity 105:1-16 ((https://doi.org/10.1007/s13225-020-00466-2)) |
| Otu38223 | 1 | k__Fungi(100);p__Ascomycota(100);c__Saccharomycetes(100);o__Saccharomycetales(100);f__Metschnikowiaceae(100);g__Metschnikowia(100);s__Metschnikowia_pulcherrima(100); | Metschnikowia | 13 | Saprotroph | \|Undefined Saprotroph\| | Yeast | NULL | Probable | As nectar-tap_saprotroph (Põlme et al. 2020); Decay sugar-rich substrates (Põlme et al. 2020) | Põlme S, et al. 2020. Fungal Diversity 105:1-16 ((https://doi.org/10.1007/s13225-020-00466-2)) |
| Otu38224 | 1 | k__Fungi(100);p__Ascomycota(100);c__Saccharomycetes(100);o__Saccharomycetales(100);f__Metschnikowiaceae(100);g__Metschnikowia(100);s__Metschnikowia_pulcherrima(100); | Metschnikowia | 13 | Saprotroph | \|Undefined Saprotroph\| | Yeast | NULL | Probable | As nectar-tap_saprotroph (Põlme et al. 2020); Decay sugar-rich substrates (Põlme et al. 2020) | Põlme S, et al. 2020. Fungal Diversity 105:1-16 ((https://doi.org/10.1007/s13225-020-00466-2)) |
| Otu38225 | 1 | k__Fungi(100);p__Ascomycota(100);c__Saccharomycetes(100);o__Saccharomycetales(100);f__Metschnikowiaceae(100);g__Metschnikowia(100);g__Metschnikowia_unclassified(100); | Metschnikowia | 13 | Saprotroph | \|Undefined Saprotroph\| | Yeast | NULL | Probable | As nectar-tap_saprotroph (Põlme et al. 2020); Decay sugar-rich substrates (Põlme et al. 2020) | Põlme S, et al. 2020. Fungal Diversity 105:1-16 ((https://doi.org/10.1007/s13225-020-00466-2)) |
| Otu38226 | 1 | k__Fungi(100);p__Ascomycota(100);c__Saccharomycetes(100);o__Saccharomycetales(100);f__Metschnikowiaceae(100);g__Metschnikowia(100);s__Metschnikowia_pulcherrima(100); | Metschnikowia | 13 | Saprotroph | \|Undefined Saprotroph\| | Yeast | NULL | Probable | As nectar-tap_saprotroph (Põlme et al. 2020); Decay sugar-rich substrates (Põlme et al. 2020) | Põlme S, et al. 2020. Fungal Diversity 105:1-16 ((https://doi.org/10.1007/s13225-020-00466-2)) |
| Otu38227 | 1 | k__Fungi(100);p__Ascomycota(100);c__Saccharomycetes(100);o__Saccharomycetales(100);f__Metschnikowiaceae(100);g__Metschnikowia(100);s__Metschnikowia_pulcherrima(100); | Metschnikowia | 13 | Saprotroph | \|Undefined Saprotroph\| | Yeast | NULL | Probable | As nectar-tap_saprotroph (Põlme et al. 2020); Decay sugar-rich substrates (Põlme et al. 2020) | Põlme S, et al. 2020. Fungal Diversity 105:1-16 ((https://doi.org/10.1007/s13225-020-00466-2)) |

| | | | | | | | | | | | |
|---|---|---|---|---|---|---|---|---|---|---|---|
| Otu38228 | 1 | k__Fungi(100);p__Ascomycota(100);c__Saccharomycetes(100);o__Saccharomycetales(100);f__Metschnikowiaceae(100);g__Metschnikowia(100);s__Metschnikowia_pulcherrima(100); | Metschnikowia | 13 | Saprotroph | \|Undefined Saprotroph\| | Yeast | NULL | Probable | As nectar-tap_saprotroph (Põlme et al. 2020); Decay sugar-rich substrates (Põlme et al. 2020) | Põlme S, et al. 2020. Fungal Diversity 105:1-16 ((https://doi.org/10.1007/s13225-020-00466-2)) |
| Otu38229 | 1 | k__Fungi(100);p__Ascomycota(100);c__Saccharomycetes(100);o__Saccharomycetales(100);f__Metschnikowiaceae(100);g__Metschnikowia(100);s__Metschnikowia_pulcherrima(100); | Metschnikowia | 13 | Saprotroph | \|Undefined Saprotroph\| | Yeast | NULL | Probable | As nectar-tap_saprotroph (Põlme et al. 2020); Decay sugar-rich substrates (Põlme et al. 2020) | Põlme S, et al. 2020. Fungal Diversity 105:1-16 ((https://doi.org/10.1007/s13225-020-00466-2)) |
| Otu38230 | 1 | k__Fungi(100);p__Ascomycota(100);c__Saccharomycetes(100);o__Saccharomycetales(100);f__Metschnikowiaceae(100);g__Metschnikowia(100);s__Metschnikowia_pulcherrima(100); | Metschnikowia | 13 | Saprotroph | \|Undefined Saprotroph\| | Yeast | NULL | Probable | As nectar-tap_saprotroph (Põlme et al. 2020); Decay sugar-rich substrates (Põlme et al. 2020) | Põlme S, et al. 2020. Fungal Diversity 105:1-16 ((https://doi.org/10.1007/s13225-020-00466-2)) |
| Otu38231 | 1 | k__Fungi(100);p__Ascomycota(100);c__Saccharomycetes(100);o__Saccharomycetales(100);f__Metschnikowiaceae(100);g__Metschnikowia(100);s__Metschnikowia_pulcherrima(100); | Metschnikowia | 13 | Saprotroph | \|Undefined Saprotroph\| | Yeast | NULL | Probable | As nectar-tap_saprotroph (Põlme et al. 2020); Decay sugar-rich substrates (Põlme et al. 2020) | Põlme S, et al. 2020. Fungal Diversity 105:1-16 ((https://doi.org/10.1007/s13225-020-00466-2)) |
| Otu38233 | 1 | k__Fungi(100);p__Ascomycota(100);c__Saccharomycetes(100);o__Saccharomycetales(100);f__Metschnikowiaceae(100);g__Metschnikowia(100);g__Metschnikowia_unclassified(100); | Metschnikowia | 13 | Saprotroph | \|Undefined Saprotroph\| | Yeast | NULL | Probable | As nectar-tap_saprotroph (Põlme et al. 2020); Decay sugar-rich substrates (Põlme et al. 2020) | Põlme S, et al. 2020. Fungal Diversity 105:1-16 ((https://doi.org/10.1007/s13225-020-00466-2)) |
| Otu38234 | 1 | k__Fungi(100);p__Ascomycota(100);c__Saccharomycetes(100);o__Saccharomycetales(100);f__Metschnikowiaceae(100);g__Metschnikowia(100);s__Metschnikowia_pulcherrima(100); | Metschnikowia | 13 | Saprotroph | \|Undefined Saprotroph\| | Yeast | NULL | Probable | As nectar-tap_saprotroph (Põlme et al. 2020); Decay sugar-rich substrates (Põlme et al. 2020) | Põlme S, et al. 2020. Fungal Diversity 105:1-16 ((https://doi.org/10.1007/s13225-020-00466-2)) |
| Otu38235 | 1 | k__Fungi(100);p__Ascomycota(100);c__Saccharomycetes(100);o__Saccharomycetales(100);f__Metschnikowiaceae(100);g__Metschnikowia(100);s__Metschnikowia_pulcherrima(100); | Metschnikowia | 13 | Saprotroph | \|Undefined Saprotroph\| | Yeast | NULL | Probable | As nectar-tap_saprotroph (Põlme et al. 2020); Decay sugar-rich substrates (Põlme et al. 2020) | Põlme S, et al. 2020. Fungal Diversity 105:1-16 ((https://doi.org/10.1007/s13225-020-00466-2)) |
| Otu38236 | 1 | k__Fungi(100);p__Ascomycota(100);c__Saccharomycetes(100);o__Saccharomycetales(100);f__Metschnikowiaceae(100);g__Metschnikowia(100);s__Metschnikowia_pulcherrima(100); | Metschnikowia | 13 | Saprotroph | \|Undefined Saprotroph\| | Yeast | NULL | Probable | As nectar-tap_saprotroph (Põlme et al. 2020); Decay sugar-rich substrates (Põlme et al. 2020) | Põlme S, et al. 2020. Fungal Diversity 105:1-16 ((https://doi.org/10.1007/s13225-020-00466-2)) |
| Otu38237 | 1 | k__Fungi(100);p__Ascomycota(100);c__Saccharomycetes(100);o__Saccharomycetales(100);f__Metschnikowiaceae(100);g__Metschnikowia(100);s__Metschnikowia_pulcherrima(100); | Metschnikowia | 13 | Saprotroph | \|Undefined Saprotroph\| | Yeast | NULL | Probable | As nectar-tap_saprotroph (Põlme et al. 2020); Decay sugar-rich substrates (Põlme et al. 2020) | Põlme S, et al. 2020. Fungal Diversity 105:1-16 ((https://doi.org/10.1007/s13225-020-00466-2)) |
| Otu38238 | 1 | k__Fungi(100);p__Ascomycota(100);c__Saccharomycetes(100);o__Saccharomycetales(100);f__Metschnikowiaceae(100);g__Metschnikowia(100);g__Metschnikowia_unclassified(100); | Metschnikowia | 13 | Saprotroph | \|Undefined Saprotroph\| | Yeast | NULL | Probable | As nectar-tap_saprotroph (Põlme et al. 2020); Decay sugar-rich substrates (Põlme et al. 2020) | Põlme S, et al. 2020. Fungal Diversity 105:1-16 ((https://doi.org/10.1007/s13225-020-00466-2)) |
| Otu38239 | 1 | k__Fungi(100);p__Ascomycota(100);c__Saccharomycetes(100);o__Saccharomycetales(100);f__Metschnikowiaceae(100);g__Metschnikowia(100);s__Metschnikowia_pulcherrima(100); | Metschnikowia | 13 | Saprotroph | \|Undefined Saprotroph\| | Yeast | NULL | Probable | As nectar-tap_saprotroph (Põlme et al. 2020); Decay sugar-rich substrates (Põlme et al. 2020) | Põlme S, et al. 2020. Fungal Diversity 105:1-16 ((https://doi.org/10.1007/s13225-020-00466-2)) |
| Otu38241 | 1 | k__Fungi(100);p__Ascomycota(100);c__Saccharomycetes(100);o__Saccharomycetales(100);f__Metschnikowiaceae(100);g__Metschnikowia(100);s__Metschnikowia_pulcherrima(100); | Metschnikowia | 13 | Saprotroph | \|Undefined Saprotroph\| | Yeast | NULL | Probable | As nectar-tap_saprotroph (Põlme et al. 2020); Decay sugar-rich substrates (Põlme et al. 2020) | Põlme S, et al. 2020. Fungal Diversity 105:1-16 ((https://doi.org/10.1007/s13225-020-00466-2)) |
| Otu38242 | 1 | k__Fungi(100);p__Ascomycota(100);c__Saccharomycetes(100);o__Saccharomycetales(100);f__Metschnikowiaceae(100);g__Metschnikowia(100);s__Metschnikowia_pulcherrima(100); | Metschnikowia | 13 | Saprotroph | \|Undefined Saprotroph\| | Yeast | NULL | Probable | As nectar-tap_saprotroph (Põlme et al. 2020); Decay sugar-rich substrates (Põlme et al. 2020) | Põlme S, et al. 2020. Fungal Diversity 105:1-16 ((https://doi.org/10.1007/s13225-020-00466-2)) |
| Otu38243 | 1 | k__Fungi(100);p__Ascomycota(100);c__Saccharomycetes(100);o__Saccharomycetales(100);f__Metschnikowiaceae(100);g__Metschnikowia(100);s__Metschnikowia_pulcherrima(100); | Metschnikowia | 13 | Saprotroph | \|Undefined Saprotroph\| | Yeast | NULL | Probable | As nectar-tap_saprotroph (Põlme et al. 2020); Decay sugar-rich substrates (Põlme et al. 2020) | Põlme S, et al. 2020. Fungal Diversity 105:1-16 ((https://doi.org/10.1007/s13225-020-00466-2)) |
| Otu38244 | 1 | k__Fungi(100);p__Ascomycota(100);c__Saccharomycetes(100);o__Saccharomycetales(100);f__Metschnikowiaceae(100);g__Metschnikowia(100);s__Metschnikowia_pulcherrima(100); | Metschnikowia | 13 | Saprotroph | \|Undefined Saprotroph\| | Yeast | NULL | Probable | As nectar-tap_saprotroph (Põlme et al. 2020); Decay sugar-rich substrates (Põlme et al. 2020) | Põlme S, et al. 2020. Fungal Diversity 105:1-16 ((https://doi.org/10.1007/s13225-020-00466-2)) |
| Otu38245 | 1 | k__Fungi(100);p__Ascomycota(100);c__Saccharomycetes(100);o__Saccharomycetales(100);f__Metschnikowiaceae(100);g__Metschnikowia(100);s__Metschnikowia_pulcherrima(100); | Metschnikowia | 13 | Saprotroph | \|Undefined Saprotroph\| | Yeast | NULL | Probable | As nectar-tap_saprotroph (Põlme et al. 2020); Decay sugar-rich substrates (Põlme et al. 2020) | Põlme S, et al. 2020. Fungal Diversity 105:1-16 ((https://doi.org/10.1007/s13225-020-00466-2)) |
| Otu38246 | 1 | k__Fungi(100);p__Ascomycota(100);c__Saccharomycetes(100);o__Saccharomycetales(100);f__Metschnikowiaceae(100);g__Metschnikowia(100);s__Metschnikowia_pulcherrima(100); | Metschnikowia | 13 | Saprotroph | \|Undefined Saprotroph\| | Yeast | NULL | Probable | As nectar-tap_saprotroph (Põlme et al. 2020); Decay sugar-rich substrates (Põlme et al. 2020) | Põlme S, et al. 2020. Fungal Diversity 105:1-16 ((https://doi.org/10.1007/s13225-020-00466-2)) |
| Otu38247 | 1 | k__Fungi(100);p__Ascomycota(100);c__Saccharomycetes(100);o__Saccharomycetales(100);f__Metschnikowiaceae(100);g__Metschnikowia(100);g__Metschnikowia_unclassified(100); | Metschnikowia | 13 | Saprotroph | \|Undefined Saprotroph\| | Yeast | NULL | Probable | As nectar-tap_saprotroph (Põlme et al. 2020); Decay sugar-rich substrates (Põlme et al. 2020) | Põlme S, et al. 2020. Fungal Diversity 105:1-16 ((https://doi.org/10.1007/s13225-020-00466-2)) |
| Otu38248 | 1 | k__Fungi(100);p__Ascomycota(100);c__Saccharomycetes(100);o__Saccharomycetales(100);f__Metschnikowiaceae(100);g__Metschnikowia(100);s__Metschnikowia_pulcherrima(100); | Metschnikowia | 13 | Saprotroph | \|Undefined Saprotroph\| | Yeast | NULL | Probable | As nectar-tap_saprotroph (Põlme et al. 2020); Decay sugar-rich substrates (Põlme et al. 2020) | Põlme S, et al. 2020. Fungal Diversity 105:1-16 ((https://doi.org/10.1007/s13225-020-00466-2)) |
| Otu38249 | 1 | k__Fungi(100);p__Ascomycota(100);c__Leotiomycetes(100);o__Helotiales(100);f__Helotiaceae(100);g__Neobulgaria(100);s__Neobulgaria_sp(100); | Neobulgaria | 13 | Saprotroph | Undefined Saprotroph-\|Wood Saprotroph\| | Microfungus | NULL | Probable | Decay wood (Põlme et al. 2020) | Tedersoo L, et al. 2014. Science 346:e1256688 ((https://doi.org/10.1126/science.1256688)); Põlme S, et al. 2020. Fungal Diversity 105:1-16 ((https://doi.org/10.1007/s13225-020-00466-2)) |
| Otu38250 | 1 | k__Fungi(100);p__Ascomycota(100);c__Saccharomycetes(100);o__Saccharomycetales(100);f__Metschnikowiaceae(100);g__Metschnikowia(100);s__Metschnikowia_pulcherrima(100); | Metschnikowia | 13 | Saprotroph | \|Undefined Saprotroph\| | Yeast | NULL | Probable | As nectar-tap_saprotroph (Põlme et al. 2020); Decay sugar-rich substrates (Põlme et al. 2020) | Põlme S, et al. 2020. Fungal Diversity 105:1-16 ((https://doi.org/10.1007/s13225-020-00466-2)) |
| Otu38251 | 1 | k__Fungi(100);p__Ascomycota(100);c__Saccharomycetes(100);o__Saccharomycetales(100);f__Metschnikowiaceae(100);g__Metschnikowia(100);s__Metschnikowia_pulcherrima(100); | Metschnikowia | 13 | Saprotroph | \|Undefined Saprotroph\| | Yeast | NULL | Probable | As nectar-tap_saprotroph (Põlme et al. 2020); Decay sugar-rich substrates (Põlme et al. 2020) | Põlme S, et al. 2020. Fungal Diversity 105:1-16 ((https://doi.org/10.1007/s13225-020-00466-2)) |
| Otu38252 | 1 | k__Fungi(100);p__Ascomycota(100);c__Saccharomycetes(100);o__Saccharomycetales(100);f__Metschnikowiaceae(100);g__Metschnikowia(100);s__Metschnikowia_pulcherrima(100); | Metschnikowia | 13 | Saprotroph | \|Undefined Saprotroph\| | Yeast | NULL | Probable | As nectar-tap_saprotroph (Põlme et al. 2020); Decay sugar-rich substrates (Põlme et al. 2020) | Põlme S, et al. 2020. Fungal Diversity 105:1-16 ((https://doi.org/10.1007/s13225-020-00466-2)) |
| Otu38253 | 1 | k__Fungi(100);p__Ascomycota(100);c__Saccharomycetes(100);o__Saccharomycetales(100);f__Metschnikowiaceae(100);g__Metschnikowia(100);s__Metschnikowia_pulcherrima(100); | Metschnikowia | 13 | Saprotroph | \|Undefined Saprotroph\| | Yeast | NULL | Probable | As nectar-tap_saprotroph (Põlme et al. 2020); Decay sugar-rich substrates (Põlme et al. 2020) | Põlme S, et al. 2020. Fungal Diversity 105:1-16 ((https://doi.org/10.1007/s13225-020-00466-2)) |
| Otu38254 | 1 | k__Fungi(100);p__Ascomycota(100);c__Saccharomycetes(100);o__Saccharomycetales(100);f__Metschnikowiaceae(100);g__Metschnikowia(100);s__Metschnikowia_pulcherrima(100); | Metschnikowia | 13 | Saprotroph | \|Undefined Saprotroph\| | Yeast | NULL | Probable | As nectar-tap_saprotroph (Põlme et al. 2020); Decay sugar-rich substrates (Põlme et al. 2020) | Põlme S, et al. 2020. Fungal Diversity 105:1-16 ((https://doi.org/10.1007/s13225-020-00466-2)) |
| Otu38255 | 1 | k__Fungi(100);p__Ascomycota(100);c__Saccharomycetes(100);o__Saccharomycetales(100);f__Metschnikowiaceae(100);g__Metschnikowia(100);s__Metschnikowia_pulcherrima(100); | Metschnikowia | 13 | Saprotroph | \|Undefined Saprotroph\| | Yeast | NULL | Probable | As nectar-tap_saprotroph (Põlme et al. 2020); Decay sugar-rich substrates (Põlme et al. 2020) | Põlme S, et al. 2020. Fungal Diversity 105:1-16 ((https://doi.org/10.1007/s13225-020-00466-2)) |
| Otu38256 | 1 | k__Fungi(100);p__Ascomycota(100);c__Saccharomycetes(100);o__Saccharomycetales(100);f__Metschnikowiaceae(100);g__Metschnikowia(100);s__Metschnikowia_pulcherrima(100); | Metschnikowia | 13 | Saprotroph | \|Undefined Saprotroph\| | Yeast | NULL | Probable | As nectar-tap_saprotroph (Põlme et al. 2020); Decay sugar-rich substrates (Põlme et al. 2020) | Põlme S, et al. 2020. Fungal Diversity 105:1-16 ((https://doi.org/10.1007/s13225-020-00466-2)) |
| Otu38257 | 1 | k__Fungi(100);p__Ascomycota(100);c__Saccharomycetes(100);o__Saccharomycetales(100);f__Metschnikowiaceae(100);g__Metschnikowia(100);g__Metschnikowia_unclassified(100); | Metschnikowia | 13 | Saprotroph | \|Undefined Saprotroph\| | Yeast | NULL | Probable | As nectar-tap_saprotroph (Põlme et al. 2020); Decay sugar-rich substrates (Põlme et al. 2020) | Põlme S, et al. 2020. Fungal Diversity 105:1-16 ((https://doi.org/10.1007/s13225-020-00466-2)) |
| Otu38258 | 1 | k__Fungi(100);p__Ascomycota(100);c__Saccharomycetes(100);o__Saccharomycetales(100);f__Metschnikowiaceae(100);g__Metschnikowia(100);g__Metschnikowia_unclassified(100); | Metschnikowia | 13 | Saprotroph | \|Undefined Saprotroph\| | Yeast | NULL | Probable | As nectar-tap_saprotroph (Põlme et al. 2020); Decay sugar-rich substrates (Põlme et al. 2020) | Põlme S, et al. 2020. Fungal Diversity 105:1-16 ((https://doi.org/10.1007/s13225-020-00466-2)) |
| Otu38259 | 1 | k__Fungi(100);p__Ascomycota(100);c__Saccharomycetes(100);o__Saccharomycetales(100);f__Metschnikowiaceae(100);g__Metschnikowia(100);s__Metschnikowia_pulcherrima(100); | Metschnikowia | 13 | Saprotroph | \|Undefined Saprotroph\| | Yeast | NULL | Probable | As nectar-tap_saprotroph (Põlme et al. 2020); Decay sugar-rich substrates (Põlme et al. 2020) | Põlme S, et al. 2020. Fungal Diversity 105:1-16 ((https://doi.org/10.1007/s13225-020-00466-2)) |
| Otu38260 | 1 | k__Fungi(100);p__Ascomycota(100);c__Saccharomycetes(100);o__Saccharomycetales(100);f__Metschnikowiaceae(100);g__Metschnikowia(100);s__Metschnikowia_pulcherrima(100); | Metschnikowia | 13 | Saprotroph | \|Undefined Saprotroph\| | Yeast | NULL | Probable | As nectar-tap_saprotroph (Põlme et al. 2020); Decay sugar-rich substrates (Põlme et al. 2020) | Põlme S, et al. 2020. Fungal Diversity 105:1-16 ((https://doi.org/10.1007/s13225-020-00466-2)) |
| Otu38261 | 1 | k__Fungi(100);p__Ascomycota(100);c__Saccharomycetes(100);o__Saccharomycetales(100);f__Metschnikowiaceae(100);g__Metschnikowia(100);s__Metschnikowia_pulcherrima(100); | Metschnikowia | 13 | Saprotroph | \|Undefined Saprotroph\| | Yeast | NULL | Probable | As nectar-tap_saprotroph (Põlme et al. 2020); Decay sugar-rich substrates (Põlme et al. 2020) | Põlme S, et al. 2020. Fungal Diversity 105:1-16 ((https://doi.org/10.1007/s13225-020-00466-2)) |
| Otu38262 | 1 | k__Fungi(100);p__Ascomycota(100);c__Saccharomycetes(100);o__Saccharomycetales(100);f__Metschnikowiaceae(100);g__Metschnikowia(100);s__Metschnikowia_pulcherrima(100); | Metschnikowia | 13 | Saprotroph | \|Undefined Saprotroph\| | Yeast | NULL | Probable | As nectar-tap_saprotroph (Põlme et al. 2020); Decay sugar-rich substrates (Põlme et al. 2020) | Põlme S, et al. 2020. Fungal Diversity 105:1-16 ((https://doi.org/10.1007/s13225-020-00466-2)) |
| Otu38263 | 1 | k__Fungi(100);p__Ascomycota(100);c__Saccharomycetes(100);o__Saccharomycetales(100);f__Metschnikowiaceae(100);g__Metschnikowia(100);s__Metschnikowia_pulcherrima(100); | Metschnikowia | 13 | Saprotroph | \|Undefined Saprotroph\| | Yeast | NULL | Probable | As nectar-tap_saprotroph (Põlme et al. 2020); Decay sugar-rich substrates (Põlme et al. 2020) | Põlme S, et al. 2020. Fungal Diversity 105:1-16 ((https://doi.org/10.1007/s13225-020-00466-2)) |
| Otu38264 | 1 | k__Fungi(100);p__Ascomycota(100);c__Saccharomycetes(100);o__Saccharomycetales(100);f__Metschnikowiaceae(100);g__Metschnikowia(100);s__Metschnikowia_pulcherrima(100); | Metschnikowia | 13 | Saprotroph | \|Undefined Saprotroph\| | Yeast | NULL | Probable | As nectar-tap_saprotroph (Põlme et al. 2020); Decay sugar-rich substrates (Põlme et al. 2020) | Põlme S, et al. 2020. Fungal Diversity 105:1-16 ((https://doi.org/10.1007/s13225-020-00466-2)) |
| Otu38265 | 1 | k__Fungi(100);p__Ascomycota(100);c__Saccharomycetes(100);o__Saccharomycetales(100);f__Metschnikowiaceae(100);g__Metschnikowia(100);g__Metschnikowia_unclassified(100); | Metschnikowia | 13 | Saprotroph | \|Undefined Saprotroph\| | Yeast | NULL | Probable | As nectar-tap_saprotroph (Põlme et al. 2020); Decay sugar-rich substrates (Põlme et al. 2020) | Põlme S, et al. 2020. Fungal Diversity 105:1-16 ((https://doi.org/10.1007/s13225-020-00466-2)) |
| Otu38266 | 1 | k__Fungi(100);p__Ascomycota(100);c__Saccharomycetes(100);o__Saccharomycetales(100);f__Metschnikowiaceae(100);g__Metschnikowia(100);s__Metschnikowia_pulcherrima(100); | Metschnikowia | 13 | Saprotroph | \|Undefined Saprotroph\| | Yeast | NULL | Probable | As nectar-tap_saprotroph (Põlme et al. 2020); Decay sugar-rich substrates (Põlme et al. 2020) | Põlme S, et al. 2020. Fungal Diversity 105:1-16 ((https://doi.org/10.1007/s13225-020-00466-2)) |
| Otu38267 | 1 | k__Fungi(100);p__Ascomycota(100);c__Saccharomycetes(100);o__Saccharomycetales(100);f__Metschnikowiaceae(100);g__Metschnikowia(100);s__Metschnikowia_pulcherrima(100); | Metschnikowia | 13 | Saprotroph | \|Undefined Saprotroph\| | Yeast | NULL | Probable | As nectar-tap_saprotroph (Põlme et al. 2020); Decay sugar-rich substrates (Põlme et al. 2020) | Põlme S, et al. 2020. Fungal Diversity 105:1-16 ((https://doi.org/10.1007/s13225-020-00466-2)) |
| Otu38268 | 1 | k__Fungi(100);p__Ascomycota(100);c__Saccharomycetes(100);o__Saccharomycetales(100);f__Metschnikowiaceae(100);g__Metschnikowia(100);s__Metschnikowia_pulcherrima(100); | Metschnikowia | 13 | Saprotroph | \|Undefined Saprotroph\| | Yeast | NULL | Probable | As nectar-tap_saprotroph (Põlme et al. 2020); Decay sugar-rich substrates (Põlme et al. 2020) | Põlme S, et al. 2020. Fungal Diversity 105:1-16 ((https://doi.org/10.1007/s13225-020-00466-2)) |
| Otu38269 | 1 | k__Fungi(100);p__Ascomycota(100);c__Saccharomycetes(100);o__Saccharomycetales(100);f__Metschnikowiaceae(100);g__Metschnikowia(100);s__Metschnikowia_pulcherrima(100); | Metschnikowia | 13 | Saprotroph | \|Undefined Saprotroph\| | Yeast | NULL | Probable | As nectar-tap_saprotroph (Põlme et al. 2020); Decay sugar-rich substrates (Põlme et al. 2020) | Põlme S, et al. 2020. Fungal Diversity 105:1-16 ((https://doi.org/10.1007/s13225-020-00466-2)) |
| Otu38270 | 1 | k__Fungi(100);p__Ascomycota(100);c__Saccharomycetes(100);o__Saccharomycetales(100);f__Metschnikowiaceae(100);g__Metschnikowia(100);s__Metschnikowia_pulcherrima(100); | Metschnikowia | 13 | Saprotroph | \|Undefined Saprotroph\| | Yeast | NULL | Probable | As nectar-tap_saprotroph (Põlme et al. 2020); Decay sugar-rich substrates (Põlme et al. 2020) | Põlme S, et al. 2020. Fungal Diversity 105:1-16 ((https://doi.org/10.1007/s13225-020-00466-2)) |
| Otu38271 | 1 | k__Fungi(100);p__Ascomycota(100);c__Saccharomycetes(100);o__Saccharomycetales(100);f__Metschnikowiaceae(100);g__Metschnikowia(100);s__Metschnikowia_pulcherrima(100); | Metschnikowia | 13 | Saprotroph | \|Undefined Saprotroph\| | Yeast | NULL | Probable | As nectar-tap_saprotroph (Põlme et al. 2020); Decay sugar-rich substrates (Põlme et al. 2020) | Põlme S, et al. 2020. Fungal Diversity 105:1-16 ((https://doi.org/10.1007/s13225-020-00466-2)) |
| Otu38272 | 1 | k__Fungi(100);p__Ascomycota(100);c__Saccharomycetes(100);o__Saccharomycetales(100);f__Metschnikowiaceae(100);g__Metschnikowia(100);g__Metschnikowia_unclassified(100); | Metschnikowia | 13 | Saprotroph | \|Undefined Saprotroph\| | Yeast | NULL | Probable | As nectar-tap_saprotroph (Põlme et al. 2020); Decay sugar-rich substrates (Põlme et al. 2020) | Põlme S, et al. 2020. Fungal Diversity 105:1-16 ((https://doi.org/10.1007/s13225-020-00466-2)) |
| Otu38274 | 1 | k__Fungi(100);p__Ascomycota(100);c__Saccharomycetes(100);o__Saccharomycetales(100);f__Metschnikowiaceae(100);g__Metschnikowia(100);g__Metschnikowia_unclassified(100); | Metschnikowia | 13 | Saprotroph | \|Undefined Saprotroph\| | Yeast | NULL | Probable | As nectar-tap_saprotroph (Põlme et al. 2020); Decay sugar-rich substrates (Põlme et al. 2020) | Põlme S, et al. 2020. Fungal Diversity 105:1-16 ((https://doi.org/10.1007/s13225-020-00466-2)) |
| Otu38275 | 1 | k__Fungi(100);p__Ascomycota(100);c__Saccharomycetes(100);o__Saccharomycetales(100);f__Metschnikowiaceae(100);g__Metschnikowia(100);s__Metschnikowia_pulcherrima(100); | Metschnikowia | 13 | Saprotroph | \|Undefined Saprotroph\| | Yeast | NULL | Probable | As nectar-tap_saprotroph (Põlme et al. 2020); Decay sugar-rich substrates (Põlme et al. 2020) | Põlme S, et al. 2020. Fungal Diversity 105:1-16 ((https://doi.org/10.1007/s13225-020-00466-2)) |
| Otu38276 | 1 | k__Fungi(100);p__Ascomycota(100);c__Saccharomycetes(100);o__Saccharomycetales(100);f__Metschnikowiaceae(100);g__Metschnikowia(100);g__Metschnikowia_unclassified(100); | Metschnikowia | 13 | Saprotroph | \|Undefined Saprotroph\| | Yeast | NULL | Probable | As nectar-tap_saprotroph (Põlme et al. 2020); Decay sugar-rich substrates (Põlme et al. 2020) | Põlme S, et al. 2020. Fungal Diversity 105:1-16 ((https://doi.org/10.1007/s13225-020-00466-2)) |
| Otu38277 | 1 | k__Fungi(100);p__Ascomycota(100);c__Saccharomycetes(100);o__Saccharomycetales(100);f__Metschnikowiaceae(100);g__Metschnikowia(100);s__Metschnikowia_pulcherrima(100); | Metschnikowia | 13 | Saprotroph | \|Undefined Saprotroph\| | Yeast | NULL | Probable | As nectar-tap_saprotroph (Põlme et al. 2020); Decay sugar-rich substrates (Põlme et al. 2020) | Põlme S, et al. 2020. Fungal Diversity 105:1-16 ((https://doi.org/10.1007/s13225-020-00466-2)) |
| Otu38278 | 1 | k__Fungi(100);p__Ascomycota(100);c__Saccharomycetes(100);o__Saccharomycetales(100);f__Metschnikowiaceae(100);g__Metschnikowia(100);s__Metschnikowia_pulcherrima(100); | Metschnikowia | 13 | Saprotroph | \|Undefined Saprotroph\| | Yeast | NULL | Probable | As nectar-tap_saprotroph (Põlme et al. 2020); Decay sugar-rich substrates (Põlme et al. 2020) | Põlme S, et al. 2020. Fungal Diversity 105:1-16 ((https://doi.org/10.1007/s13225-020-00466-2)) |
| Otu38279 | 1 | k__Fungi(100);p__Ascomycota(100);c__Saccharomycetes(100);o__Saccharomycetales(100);f__Metschnikowiaceae(100);g__Metschnikowia(100);s__Metschnikowia_pulcherrima(100); | Metschnikowia | 13 | Saprotroph | \|Undefined Saprotroph\| | Yeast | NULL | Probable | As nectar-tap_saprotroph (Põlme et al. 2020); Decay sugar-rich substrates (Põlme et al. 2020) | Põlme S, et al. 2020. Fungal Diversity 105:1-16 ((https://doi.org/10.1007/s13225-020-00466-2)) |
| Otu38280 | 1 | k__Fungi(100);p__Ascomycota(100);c__Saccharomycetes(100);o__Saccharomycetales(100);f__Metschnikowiaceae(100);g__Metschnikowia(100);s__Metschnikowia_pulcherrima(100); | Metschnikowia | 13 | Saprotroph | \|Undefined Saprotroph\| | Yeast | NULL | Probable | As nectar-tap_saprotroph (Põlme et al. 2020); Decay sugar-rich substrates (Põlme et al. 2020) | Põlme S, et al. 2020. Fungal Diversity 105:1-16 ((https://doi.org/10.1007/s13225-020-00466-2)) |
| Otu38281 | 1 | k__Fungi(100);p__Ascomycota(100);c__Saccharomycetes(100);o__Saccharomycetales(100);f__Metschnikowiaceae(100);g__Metschnikowia(100);s__Metschnikowia_pulcherrima(100); | Metschnikowia | 13 | Saprotroph | \|Undefined Saprotroph\| | Yeast | NULL | Probable | As nectar-tap_saprotroph (Põlme et al. 2020); Decay sugar-rich substrates (Põlme et al. 2020) | Põlme S, et al. 2020. Fungal Diversity 105:1-16 ((https://doi.org/10.1007/s13225-020-00466-2)) |
| Otu38282 | 1 | k__Fungi(100);p__Ascomycota(100);c__Saccharomycetes(100);o__Saccharomycetales(100);f__Metschnikowiaceae(100);g__Metschnikowia(100);s__Metschnikowia_pulcherrima(100); | Metschnikowia | 13 | Saprotroph | \|Undefined Saprotroph\| | Yeast | NULL | Probable | As nectar-tap_saprotroph (Põlme et al. 2020); Decay sugar-rich substrates (Põlme et al. 2020) | Põlme S, et al. 2020. Fungal Diversity 105:1-16 ((https://doi.org/10.1007/s13225-020-00466-2)) |
| Otu38283 | 1 | k__Fungi(100);p__Ascomycota(100);c__Saccharomycetes(100);o__Saccharomycetales(100);f__Metschnikowiaceae(100);g__Metschnikowia(100);s__Metschnikowia_pulcherrima(100); | Metschnikowia | 13 | Saprotroph | \|Undefined Saprotroph\| | Yeast | NULL | Probable | As nectar-tap_saprotroph (Põlme et al. 2020); Decay sugar-rich substrates (Põlme et al. 2020) | Põlme S, et al. 2020. Fungal Diversity 105:1-16 ((https://doi.org/10.1007/s13225-020-00466-2)) |
| Otu38284 | 1 | k__Fungi(100);p__Ascomycota(100);c__Saccharomycetes(100);o__Saccharomycetales(100);f__Metschnikowiaceae(100);g__Metschnikowia(100);g__Metschnikowia_unclassified(100); | Metschnikowia | 13 | Saprotroph | \|Undefined Saprotroph\| | Yeast | NULL | Probable | As nectar-tap_saprotroph (Põlme et al. 2020); Decay sugar-rich substrates (Põlme et al. 2020) | Põlme S, et al. 2020. Fungal Diversity 105:1-16 ((https://doi.org/10.1007/s13225-020-00466-2)) |
| Otu38285 | 1 | k__Fungi(100);p__Ascomycota(100);c__Saccharomycetes(100);o__Saccharomycetales(100);f__Metschnikowiaceae(100);g__Metschnikowia(100);s__Metschnikowia_pulcherrima(100); | Metschnikowia | 13 | Saprotroph | \|Undefined Saprotroph\| | Yeast | NULL | Probable | As nectar-tap_saprotroph (Põlme et al. 2020); Decay sugar-rich substrates (Põlme et al. 2020) | Põlme S, et al. 2020. Fungal Diversity 105:1-16 ((https://doi.org/10.1007/s13225-020-00466-2)) |
| Otu38286 | 1 | k__Fungi(100);p__Ascomycota(100);c__Saccharomycetes(100);o__Saccharomycetales(100);f__Metschnikowiaceae(100);g__Metschnikowia(100);s__Metschnikowia_pulcherrima(100); | Metschnikowia | 13 | Saprotroph | \|Undefined Saprotroph\| | Yeast | NULL | Probable | As nectar-tap_saprotroph (Põlme et al. 2020); Decay sugar-rich substrates (Põlme et al. 2020) | Põlme S, et al. 2020. Fungal Diversity 105:1-16 ((https://doi.org/10.1007/s13225-020-00466-2)) |
| Otu38287 | 1 | k__Fungi(100);p__Ascomycota(100);c__Saccharomycetes(100);o__Saccharomycetales(100);f__Metschnikowiaceae(100);g__Metschnikowia(100);s__Metschnikowia_pulcherrima(100); | Metschnikowia | 13 | Saprotroph | \|Undefined Saprotroph\| | Yeast | NULL | Probable | As nectar-tap_saprotroph (Põlme et al. 2020); Decay sugar-rich substrates (Põlme et al. 2020) | Põlme S, et al. 2020. Fungal Diversity 105:1-16 ((https://doi.org/10.1007/s13225-020-00466-2)) |
| Otu38288 | 1 | k__Fungi(100);p__Ascomycota(100);c__Saccharomycetes(100);o__Saccharomycetales(100);f__Metschnikowiaceae(100);g__Metschnikowia(100);s__Metschnikowia_pulcherrima(100); | Metschnikowia | 13 | Saprotroph | \|Undefined Saprotroph\| | Yeast | NULL | Probable | As nectar-tap_saprotroph (Põlme et al. 2020); Decay sugar-rich substrates (Põlme et al. 2020) | Põlme S, et al. 2020. Fungal Diversity 105:1-16 ((https://doi.org/10.1007/s13225-020-00466-2)) |
| Otu38289 | 1 | k__Fungi(100);p__Ascomycota(100);c__Saccharomycetes(100);o__Saccharomycetales(100);f__Metschnikowiaceae(100);g__Metschnikowia(100);s__Metschnikowia_pulcherrima(100); | Metschnikowia | 13 | Saprotroph | \|Undefined Saprotroph\| | Yeast | NULL | Probable | As nectar-tap_saprotroph (Põlme et al. 2020); Decay sugar-rich substrates (Põlme et al. 2020) | Põlme S, et al. 2020. Fungal Diversity 105:1-16 ((https://doi.org/10.1007/s13225-020-00466-2)) |
| Otu38290 | 1 | k__Fungi(100);p__Ascomycota(100);c__Saccharomycetes(100);o__Saccharomycetales(100);f__Metschnikowiaceae(100);g__Metschnikowia(100);s__Metschnikowia_pulcherrima(100); | Metschnikowia | 13 | Saprotroph | \|Undefined Saprotroph\| | Yeast | NULL | Probable | As nectar-tap_saprotroph (Põlme et al. 2020); Decay sugar-rich substrates (Põlme et al. 2020) | Põlme S, et al. 2020. Fungal Diversity 105:1-16 ((https://doi.org/10.1007/s13225-020-00466-2)) |
| Otu38291 | 1 | k__Fungi(100);p__Ascomycota(100);c__Saccharomycetes(100);o__Saccharomycetales(100);f__Metschnikowiaceae(100);g__Metschnikowia(100);s__Metschnikowia_pulcherrima(100); | Metschnikowia | 13 | Saprotroph | \|Undefined Saprotroph\| | Yeast | NULL | Probable | As nectar-tap_saprotroph (Põlme et al. 2020); Decay sugar-rich substrates (Põlme et al. 2020) | Põlme S, et al. 2020. Fungal Diversity 105:1-16 ((https://doi.org/10.1007/s13225-020-00466-2)) |
| Otu38292 | 1 | k__Fungi(100);p__Ascomycota(100);c__Saccharomycetes(100);o__Saccharomycetales(100);f__Metschnikowiaceae(100);g__Metschnikowia(100);g__Metschnikowia_unclassified(100); | Metschnikowia | 13 | Saprotroph | \|Undefined Saprotroph\| | Yeast | NULL | Probable | As nectar-tap_saprotroph (Põlme et al. 2020); Decay sugar-rich substrates (Põlme et al. 2020) | Põlme S, et al. 2020. Fungal Diversity 105:1-16 ((https://doi.org/10.1007/s13225-020-00466-2)) |
| Otu38293 | 1 | k__Fungi(100);p__Ascomycota(100);c__Saccharomycetes(100);o__Saccharomycetales(100);f__Metschnikowiaceae(100);g__Metschnikowia(100);g__Metschnikowia_unclassified(100); | Metschnikowia | 13 | Saprotroph | \|Undefined Saprotroph\| | Yeast | NULL | Probable | As nectar-tap_saprotroph (Põlme et al. 2020); Decay sugar-rich substrates (Põlme et al. 2020) | Põlme S, et al. 2020. Fungal Diversity 105:1-16 ((https://doi.org/10.1007/s13225-020-00466-2)) |
| Otu38294 | 1 | k__Fungi(100);p__Ascomycota(100);c__Saccharomycetes(100);o__Saccharomycetales(100);f__Metschnikowiaceae(100);g__Metschnikowia(100);s__Metschnikowia_pulcherrima(100); | Metschnikowia | 13 | Saprotroph | \|Undefined Saprotroph\| | Yeast | NULL | Probable | As nectar-tap_saprotroph (Põlme et al. 2020); Decay sugar-rich substrates (Põlme et al. 2020) | Põlme S, et al. 2020. Fungal Diversity 105:1-16 ((https://doi.org/10.1007/s13225-020-00466-2)) |
| Otu38295 | 1 | k__Fungi(100);p__Ascomycota(100);c__Saccharomycetes(100);o__Saccharomycetales(100);f__Metschnikowiaceae(100);g__Metschnikowia(100);s__Metschnikowia_pulcherrima(100); | Metschnikowia | 13 | Saprotroph | \|Undefined Saprotroph\| | Yeast | NULL | Probable | As nectar-tap_saprotroph (Põlme et al. 2020); Decay sugar-rich substrates (Põlme et al. 2020) | Põlme S, et al. 2020. Fungal Diversity 105:1-16 ((https://doi.org/10.1007/s13225-020-00466-2)) |
| Otu38296 | 1 | k__Fungi(100);p__Ascomycota(100);c__Saccharomycetes(100);o__Saccharomycetales(100);f__Metschnikowiaceae(100);g__Metschnikowia(100);s__Metschnikowia_pulcherrima(100); | Metschnikowia | 13 | Saprotroph | \|Undefined Saprotroph\| | Yeast | NULL | Probable | As nectar-tap_saprotroph (Põlme et al. 2020); Decay sugar-rich substrates (Põlme et al. 2020) | Põlme S, et al. 2020. Fungal Diversity 105:1-16 ((https://doi.org/10.1007/s13225-020-00466-2)) |
| Otu38297 | 1 | k__Fungi(100);p__Ascomycota(100);c__Saccharomycetes(100);o__Saccharomycetales(100);f__Metschnikowiaceae(100);g__Metschnikowia(100);s__Metschnikowia_pulcherrima(100); | Metschnikowia | 13 | Saprotroph | \|Undefined Saprotroph\| | Yeast | NULL | Probable | As nectar-tap_saprotroph (Põlme et al. 2020); Decay sugar-rich substrates (Põlme et al. 2020) | Põlme S, et al. 2020. Fungal Diversity 105:1-16 ((https://doi.org/10.1007/s13225-020-00466-2)) |
| Otu38298 | 1 | k__Fungi(100);p__Ascomycota(100);c__Saccharomycetes(100);o__Saccharomycetales(100);f__Metschnikowiaceae(100);g__Metschnikowia(100);g__Metschnikowia_unclassified(100); | Metschnikowia | 13 | Saprotroph | \|Undefined Saprotroph\| | Yeast | NULL | Probable | As nectar-tap_saprotroph (Põlme et al. 2020); Decay sugar-rich substrates (Põlme et al. 2020) | Põlme S, et al. 2020. Fungal Diversity 105:1-16 ((https://doi.org/10.1007/s13225-020-00466-2)) |
| Otu38299 | 1 | k__Fungi(100);p__Ascomycota(100);c__Saccharomycetes(100);o__Saccharomycetales(100);f__Metschnikowiaceae(100);g__Metschnikowia(100);s__Metschnikowia_pulcherrima(100); | Metschnikowia | 13 | Saprotroph | \|Undefined Saprotroph\| | Yeast | NULL | Probable | As nectar-tap_saprotroph (Põlme et al. 2020); Decay sugar-rich substrates (Põlme et al. 2020) | Põlme S, et al. 2020. Fungal Diversity 105:1-16 ((https://doi.org/10.1007/s13225-020-00466-2)) |
| Otu38300 | 1 | k__Fungi(100);p__Ascomycota(100);c__Saccharomycetes(100);o__Saccharomycetales(100);f__Metschnikowiaceae(100);g__Metschnikowia(100);s__Metschnikowia_pulcherrima(100); | Metschnikowia | 13 | Saprotroph | \|Undefined Saprotroph\| | Yeast | NULL | Probable | As nectar-tap_saprotroph (Põlme et al. 2020); Decay sugar-rich substrates (Põlme et al. 2020) | Põlme S, et al. 2020. Fungal Diversity 105:1-16 ((https://doi.org/10.1007/s13225-020-00466-2)) |
| Otu38301 | 1 | k__Fungi(100);p__Ascomycota(100);c__Saccharomycetes(100);o__Saccharomycetales(100);f__Metschnikowiaceae(100);g__Metschnikowia(100);s__Metschnikowia_pulcherrima(100); | Metschnikowia | 13 | Saprotroph | \|Undefined Saprotroph\| | Yeast | NULL | Probable | As nectar-tap_saprotroph (Põlme et al. 2020); Decay sugar-rich substrates (Põlme et al. 2020) | Põlme S, et al. 2020. Fungal Diversity 105:1-16 ((https://doi.org/10.1007/s13225-020-00466-2)) |
| Otu38302 | 1 | k__Fungi(100);p__Ascomycota(100);c__Saccharomycetes(100);o__Saccharomycetales(100);f__Metschnikowiaceae(100);g__Metschnikowia(100);g__Metschnikowia_unclassified(100); | Metschnikowia | 13 | Saprotroph | \|Undefined Saprotroph\| | Yeast | NULL | Probable | As nectar-tap_saprotroph (Põlme et al. 2020); Decay sugar-rich substrates (Põlme et al. 2020) | Põlme S, et al. 2020. Fungal Diversity 105:1-16 ((https://doi.org/10.1007/s13225-020-00466-2)) |
| Otu38305 | 1 | k__Fungi(100);p__Ascomycota(100);c__Saccharomycetes(100);o__Saccharomycetales(100);f__Metschnikowiaceae(100);g__Metschnikowia(100);s__Metschnikowia_pulcherrima(100); | Metschnikowia | 13 | Saprotroph | \|Undefined Saprotroph\| | Yeast | NULL | Probable | As nectar-tap_saprotroph (Põlme et al. 2020); Decay sugar-rich substrates (Põlme et al. 2020) | Põlme S, et al. 2020. Fungal Diversity 105:1-16 ((https://doi.org/10.1007/s13225-020-00466-2)) |
| Otu38306 | 1 | k__Fungi(100);p__Ascomycota(100);c__Saccharomycetes(100);o__Saccharomycetales(100);f__Metschnikowiaceae(100);g__Metschnikowia(100);g__Metschnikowia_unclassified(100); | Metschnikowia | 13 | Saprotroph | \|Undefined Saprotroph\| | Yeast | NULL | Probable | As nectar-tap_saprotroph (Põlme et al. 2020); Decay sugar-rich substrates (Põlme et al. 2020) | Põlme S, et al. 2020. Fungal Diversity 105:1-16 ((https://doi.org/10.1007/s13225-020-00466-2)) |
| Otu38307 | 1 | k__Fungi(100);p__Ascomycota(100);c__Saccharomycetes(100);o__Saccharomycetales(100);f__Metschnikowiaceae(100);g__Metschnikowia(100);g__Metschnikowia_unclassified(100); | Metschnikowia | 13 | Saprotroph | \|Undefined Saprotroph\| | Yeast | NULL | Probable | As nectar-tap_saprotroph (Põlme et al. 2020); Decay sugar-rich substrates (Põlme et al. 2020) | Põlme S, et al. 2020. Fungal Diversity 105:1-16 ((https://doi.org/10.1007/s13225-020-00466-2)) |
| Otu38308 | 1 | k__Fungi(100);p__Ascomycota(100);c__Saccharomycetes(100);o__Saccharomycetales(100);f__Metschnikowiaceae(100);g__Metschnikowia(100);g__Metschnikowia_unclassified(100); | Metschnikowia | 13 | Saprotroph | \|Undefined Saprotroph\| | Yeast | NULL | Probable | As nectar-tap_saprotroph (Põlme et al. 2020); Decay sugar-rich substrates (Põlme et al. 2020) | Põlme S, et al. 2020. Fungal Diversity 105:1-16 ((https://doi.org/10.1007/s13225-020-00466-2)) |
| Otu38309 | 1 | k__Fungi(100);p__Ascomycota(100);c__Saccharomycetes(100);o__Saccharomycetales(100);f__Metschnikowiaceae(100);g__Metschnikowia(100);s__Metschnikowia_pulcherrima(100); | Metschnikowia | 13 | Saprotroph | \|Undefined Saprotroph\| | Yeast | NULL | Probable | As nectar-tap_saprotroph (Põlme et al. 2020); Decay sugar-rich substrates (Põlme et al. 2020) | Põlme S, et al. 2020. Fungal Diversity 105:1-16 ((https://doi.org/10.1007/s13225-020-00466-2)) |
| Otu38310 | 1 | k__Fungi(100);p__Ascomycota(100);c__Saccharomycetes(100);o__Saccharomycetales(100);f__Metschnikowiaceae(100);g__Metschnikowia(100);g__Metschnikowia_unclassified(100); | Metschnikowia | 13 | Saprotroph | \|Undefined Saprotroph\| | Yeast | NULL | Probable | As nectar-tap_saprotroph (Põlme et al. 2020); Decay sugar-rich substrates (Põlme et al. 2020) | Põlme S, et al. 2020. Fungal Diversity 105:1-16 ((https://doi.org/10.1007/s13225-020-00466-2)) |
| Otu38311 | 1 | k__Fungi(100);p__Ascomycota(100);c__Saccharomycetes(100);o__Saccharomycetales(100);f__Metschnikowiaceae(100);g__Metschnikowia(100);s__Metschnikowia_pulcherrima(100); | Metschnikowia | 13 | Saprotroph | \|Undefined Saprotroph\| | Yeast | NULL | Probable | As nectar-tap_saprotroph (Põlme et al. 2020); Decay sugar-rich substrates (Põlme et al. 2020) | Põlme S, et al. 2020. Fungal Diversity 105:1-16 ((https://doi.org/10.1007/s13225-020-00466-2)) |
| Otu38312 | 1 | k__Fungi(100);p__Ascomycota(100);c__Saccharomycetes(100);o__Saccharomycetales(100);f__Metschnikowiaceae(100);g__Metschnikowia(100);g__Metschnikowia_unclassified(100); | Metschnikowia | 13 | Saprotroph | \|Undefined Saprotroph\| | Yeast | NULL | Probable | As nectar-tap_saprotroph (Põlme et al. 2020); Decay sugar-rich substrates (Põlme et al. 2020) | Põlme S, et al. 2020. Fungal Diversity 105:1-16 ((https://doi.org/10.1007/s13225-020-00466-2)) |
| Otu38313 | 1 | k__Fungi(100);p__Ascomycota(100);c__Saccharomycetes(100);o__Saccharomycetales(100);f__Metschnikowiaceae(100);g__Metschnikowia(100);s__Metschnikowia_pulcherrima(100); | Metschnikowia | 13 | Saprotroph | \|Undefined Saprotroph\| | Yeast | NULL | Probable | As nectar-tap_saprotroph (Põlme et al. 2020); Decay sugar-rich substrates (Põlme et al. 2020) | Põlme S, et al. 2020. Fungal Diversity 105:1-16 ((https://doi.org/10.1007/s13225-020-00466-2)) |
| Otu38314 | 1 | k__Fungi(100);p__Ascomycota(100);c__Saccharomycetes(100);o__Saccharomycetales(100);f__Metschnikowiaceae(100);g__Metschnikowia(100);s__Metschnikowia_pulcherrima(100); | Metschnikowia | 13 | Saprotroph | \|Undefined Saprotroph\| | Yeast | NULL | Probable | As nectar-tap_saprotroph (Põlme et al. 2020); Decay sugar-rich substrates (Põlme et al. 2020) | Põlme S, et al. 2020. Fungal Diversity 105:1-16 ((https://doi.org/10.1007/s13225-020-00466-2)) |
| Otu38316 | 1 | k__Fungi(100);p__Ascomycota(100);c__Saccharomycetes(100);o__Saccharomycetales(100);f__Metschnikowiaceae(100);g__Metschnikowia(100);s__Metschnikowia_pulcherrima(100); | Metschnikowia | 13 | Saprotroph | \|Undefined Saprotroph\| | Yeast | NULL | Probable | As nectar-tap_saprotroph (Põlme et al. 2020); Decay sugar-rich substrates (Põlme et al. 2020) | Põlme S, et al. 2020. Fungal Diversity 105:1-16 ((https://doi.org/10.1007/s13225-020-00466-2)) |
| Otu38317 | 1 | k__Fungi(100);p__Ascomycota(100);c__Saccharomycetes(100);o__Saccharomycetales(100);f__Metschnikowiaceae(100);g__Metschnikowia(100);g__Metschnikowia_unclassified(100); | Metschnikowia | 13 | Saprotroph | \|Undefined Saprotroph\| | Yeast | NULL | Probable | As nectar-tap_saprotroph (Põlme et al. 2020); Decay sugar-rich substrates (Põlme et al. 2020) | Põlme S, et al. 2020. Fungal Diversity 105:1-16 ((https://doi.org/10.1007/s13225-020-00466-2)) |
| Otu38318 | 1 | k__Fungi(100);p__Ascomycota(100);c__Saccharomycetes(100);o__Saccharomycetales(100);f__Metschnikowiaceae(100);g__Metschnikowia(100);g__Metschnikowia_unclassified(100); | Metschnikowia | 13 | Saprotroph | \|Undefined Saprotroph\| | Yeast | NULL | Probable | As nectar-tap_saprotroph (Põlme et al. 2020); Decay sugar-rich substrates (Põlme et al. 2020) | Põlme S, et al. 2020. Fungal Diversity 105:1-16 ((https://doi.org/10.1007/s13225-020-00466-2)) |
| Otu38319 | 1 | k__Fungi(100);p__Ascomycota(100);c__Saccharomycetes(100);o__Saccharomycetales(100);f__Metschnikowiaceae(100);g__Metschnikowia(100);g__Metschnikowia_unclassified(100); | Metschnikowia | 13 | Saprotroph | \|Undefined Saprotroph\| | Yeast | NULL | Probable | As nectar-tap_saprotroph (Põlme et al. 2020); Decay sugar-rich substrates (Põlme et al. 2020) | Põlme S, et al. 2020. Fungal Diversity 105:1-16 ((https://doi.org/10.1007/s13225-020-00466-2)) |
| Otu38320 | 1 | k__Fungi(100);p__Ascomycota(100);c__Saccharomycetes(100);o__Saccharomycetales(100);f__Metschnikowiaceae(100);g__Metschnikowia(100);s__Metschnikowia_pulcherrima(100); | Metschnikowia | 13 | Saprotroph | \|Undefined Saprotroph\| | Yeast | NULL | Probable | As nectar-tap_saprotroph (Põlme et al. 2020); Decay sugar-rich substrates (Põlme et al. 2020) | Põlme S, et al. 2020. Fungal Diversity 105:1-16 ((https://doi.org/10.1007/s13225-020-00466-2)) |
| Otu38321 | 1 | k__Fungi(100);p__Ascomycota(100);c__Saccharomycetes(100);o__Saccharomycetales(100);f__Metschnikowiaceae(100);g__Metschnikowia(100);s__Metschnikowia_pulcherrima(100); | Metschnikowia | 13 | Saprotroph | \|Undefined Saprotroph\| | Yeast | NULL | Probable | As nectar-tap_saprotroph (Põlme et al. 2020); Decay sugar-rich substrates (Põlme et al. 2020) | Põlme S, et al. 2020. Fungal Diversity 105:1-16 ((https://doi.org/10.1007/s13225-020-00466-2)) |
| Otu38322 | 1 | k__Fungi(100);p__Ascomycota(100);c__Saccharomycetes(100);o__Saccharomycetales(100);f__Metschnikowiaceae(100);g__Metschnikowia(100);s__Metschnikowia_pulcherrima(100); | Metschnikowia | 13 | Saprotroph | \|Undefined Saprotroph\| | Yeast | NULL | Probable | As nectar-tap_saprotroph (Põlme et al. 2020); Decay sugar-rich substrates (Põlme et al. 2020) | Põlme S, et al. 2020. Fungal Diversity 105:1-16 ((https://doi.org/10.1007/s13225-020-00466-2)) |
| Otu38323 | 1 | k__Fungi(100);p__Ascomycota(100);c__Saccharomycetes(100);o__Saccharomycetales(100);f__Metschnikowiaceae(100);g__Metschnikowia(100);s__Metschnikowia_pulcherrima(100); | Metschnikowia | 13 | Saprotroph | \|Undefined Saprotroph\| | Yeast | NULL | Probable | As nectar-tap_saprotroph (Põlme et al. 2020); Decay sugar-rich substrates (Põlme et al. 2020) | Põlme S, et al. 2020. Fungal Diversity 105:1-16 ((https://doi.org/10.1007/s13225-020-00466-2)) |
| Otu38324 | 1 | k__Fungi(100);p__Ascomycota(100);c__Saccharomycetes(100);o__Saccharomycetales(100);f__Metschnikowiaceae(100);g__Metschnikowia(100);s__Metschnikowia_pulcherrima(100); | Metschnikowia | 13 | Saprotroph | \|Undefined Saprotroph\| | Yeast | NULL | Probable | As nectar-tap_saprotroph (Põlme et al. 2020); Decay sugar-rich substrates (Põlme et al. 2020) | Põlme S, et al. 2020. Fungal Diversity 105:1-16 ((https://doi.org/10.1007/s13225-020-00466-2)) |
| Otu38325 | 1 | k__Fungi(100);p__Ascomycota(100);c__Saccharomycetes(100);o__Saccharomycetales(100);f__Metschnikowiaceae(100);g__Metschnikowia(100);g__Metschnikowia_unclassified(100); | Metschnikowia | 13 | Saprotroph | \|Undefined Saprotroph\| | Yeast | NULL | Probable | As nectar-tap_saprotroph (Põlme et al. 2020); Decay sugar-rich substrates (Põlme et al. 2020) | Põlme S, et al. 2020. Fungal Diversity 105:1-16 ((https://doi.org/10.1007/s13225-020-00466-2)) |
| Otu38326 | 1 | k__Fungi(100);p__Ascomycota(100);c__Saccharomycetes(100);o__Saccharomycetales(100);f__Metschnikowiaceae(100);g__Metschnikowia(100);g__Metschnikowia_unclassified(100); | Metschnikowia | 13 | Saprotroph | \|Undefined Saprotroph\| | Yeast | NULL | Probable | As nectar-tap_saprotroph (Põlme et al. 2020); Decay sugar-rich substrates (Põlme et al. 2020) | Põlme S, et al. 2020. Fungal Diversity 105:1-16 ((https://doi.org/10.1007/s13225-020-00466-2)) |
| Otu38327 | 1 | k__Fungi(100);p__Ascomycota(100);c__Saccharomycetes(100);o__Saccharomycetales(100);f__Metschnikowiaceae(100);g__Metschnikowia(100);s__Metschnikowia_pulcherrima(100); | Metschnikowia | 13 | Saprotroph | \|Undefined Saprotroph\| | Yeast | NULL | Probable | As nectar-tap_saprotroph (Põlme et al. 2020); Decay sugar-rich substrates (Põlme et al. 2020) | Põlme S, et al. 2020. Fungal Diversity 105:1-16 ((https://doi.org/10.1007/s13225-020-00466-2)) |
| Otu38328 | 1 | k__Fungi(100);p__Ascomycota(100);c__Saccharomycetes(100);o__Saccharomycetales(100);f__Metschnikowiaceae(100);g__Metschnikowia(100);s__Metschnikowia_pulcherrima(100); | Metschnikowia | 13 | Saprotroph | \|Undefined Saprotroph\| | Yeast | NULL | Probable | As nectar-tap_saprotroph (Põlme et al. 2020); Decay sugar-rich substrates (Põlme et al. 2020) | Põlme S, et al. 2020. Fungal Diversity 105:1-16 ((https://doi.org/10.1007/s13225-020-00466-2)) |
| Otu38329 | 1 | k__Fungi(100);p__Ascomycota(100);c__Saccharomycetes(100);o__Saccharomycetales(100);f__Metschnikowiaceae(100);g__Metschnikowia(100);s__Metschnikowia_pulcherrima(100); | Metschnikowia | 13 | Saprotroph | \|Undefined Saprotroph\| | Yeast | NULL | Probable | As nectar-tap_saprotroph (Põlme et al. 2020); Decay sugar-rich substrates (Põlme et al. 2020) | Põlme S, et al. 2020. Fungal Diversity 105:1-16 ((https://doi.org/10.1007/s13225-020-00466-2)) |
| Otu38330 | 1 | k__Fungi(100);p__Ascomycota(100);c__Saccharomycetes(100);o__Saccharomycetales(100);f__Metschnikowiaceae(100);g__Metschnikowia(100);s__Metschnikowia_pulcherrima(100); | Metschnikowia | 13 | Saprotroph | \|Undefined Saprotroph\| | Yeast | NULL | Probable | As nectar-tap_saprotroph (Põlme et al. 2020); Decay sugar-rich substrates (Põlme et al. 2020) | Põlme S, et al. 2020. Fungal Diversity 105:1-16 ((https://doi.org/10.1007/s13225-020-00466-2)) |
| Otu38331 | 1 | k__Fungi(100);p__Ascomycota(100);c__Saccharomycetes(100);o__Saccharomycetales(100);f__Metschnikowiaceae(100);g__Metschnikowia(100);g__Metschnikowia_unclassified(100); | Metschnikowia | 13 | Saprotroph | \|Undefined Saprotroph\| | Yeast | NULL | Probable | As nectar-tap_saprotroph (Põlme et al. 2020); Decay sugar-rich substrates (Põlme et al. 2020) | Põlme S, et al. 2020. Fungal Diversity 105:1-16 ((https://doi.org/10.1007/s13225-020-00466-2)) |
| Otu38332 | 1 | k__Fungi(100);p__Ascomycota(100);c__Saccharomycetes(100);o__Saccharomycetales(100);f__Metschnikowiaceae(100);g__Metschnikowia(100);s__Metschnikowia_pulcherrima(100); | Metschnikowia | 13 | Saprotroph | \|Undefined Saprotroph\| | Yeast | NULL | Probable | As nectar-tap_saprotroph (Põlme et al. 2020); Decay sugar-rich substrates (Põlme et al. 2020) | Põlme S, et al. 2020. Fungal Diversity 105:1-16 ((https://doi.org/10.1007/s13225-020-00466-2)) |
| Otu38333 | 1 | k__Fungi(100);p__Ascomycota(100);c__Saccharomycetes(100);o__Saccharomycetales(100);f__Metschnikowiaceae(100);g__Metschnikowia(100);s__Metschnikowia_pulcherrima(100); | Metschnikowia | 13 | Saprotroph | \|Undefined Saprotroph\| | Yeast | NULL | Probable | As nectar-tap_saprotroph (Põlme et al. 2020); Decay sugar-rich substrates (Põlme et al. 2020) | Põlme S, et al. 2020. Fungal Diversity 105:1-16 ((https://doi.org/10.1007/s13225-020-00466-2)) |
| Otu38334 | 1 | k__Fungi(100);p__Ascomycota(100);c__Saccharomycetes(100);o__Saccharomycetales(100);f__Metschnikowiaceae(100);g__Metschnikowia(100);s__Metschnikowia_pulcherrima(100); | Metschnikowia | 13 | Saprotroph | \|Undefined Saprotroph\| | Yeast | NULL | Probable | As nectar-tap_saprotroph (Põlme et al. 2020); Decay sugar-rich substrates (Põlme et al. 2020) | Põlme S, et al. 2020. Fungal Diversity 105:1-16 ((https://doi.org/10.1007/s13225-020-00466-2)) |
| Otu38335 | 1 | k__Fungi(100);p__Ascomycota(100);c__Saccharomycetes(100);o__Saccharomycetales(100);f__Metschnikowiaceae(100);g__Metschnikowia(100);s__Metschnikowia_pulcherrima(100); | Metschnikowia | 13 | Saprotroph | \|Undefined Saprotroph\| | Yeast | NULL | Probable | As nectar-tap_saprotroph (Põlme et al. 2020); Decay sugar-rich substrates (Põlme et al. 2020) | Põlme S, et al. 2020. Fungal Diversity 105:1-16 ((https://doi.org/10.1007/s13225-020-00466-2)) |
| Otu38336 | 1 | k__Fungi(100);p__Ascomycota(100);c__Saccharomycetes(100);o__Saccharomycetales(100);f__Metschnikowiaceae(100);g__Metschnikowia(100);g__Metschnikowia_unclassified(100); | Metschnikowia | 13 | Saprotroph | \|Undefined Saprotroph\| | Yeast | NULL | Probable | As nectar-tap_saprotroph (Põlme et al. 2020); Decay sugar-rich substrates (Põlme et al. 2020) | Põlme S, et al. 2020. Fungal Diversity 105:1-16 ((https://doi.org/10.1007/s13225-020-00466-2)) |
| Otu38337 | 1 | k__Fungi(100);p__Ascomycota(100);c__Saccharomycetes(100);o__Saccharomycetales(100);f__Metschnikowiaceae(100);g__Metschnikowia(100);g__Metschnikowia_unclassified(100); | Metschnikowia | 13 | Saprotroph | \|Undefined Saprotroph\| | Yeast | NULL | Probable | As nectar-tap_saprotroph (Põlme et al. 2020); Decay sugar-rich substrates (Põlme et al. 2020) | Põlme S, et al. 2020. Fungal Diversity 105:1-16 ((https://doi.org/10.1007/s13225-020-00466-2)) |
| Otu38338 | 1 | k__Fungi(100);p__Ascomycota(100);c__Saccharomycetes(100);o__Saccharomycetales(100);f__Metschnikowiaceae(100);g__Metschnikowia(100);s__Metschnikowia_pulcherrima(100); | Metschnikowia | 13 | Saprotroph | \|Undefined Saprotroph\| | Yeast | NULL | Probable | As nectar-tap_saprotroph (Põlme et al. 2020); Decay sugar-rich substrates (Põlme et al. 2020) | Põlme S, et al. 2020. Fungal Diversity 105:1-16 ((https://doi.org/10.1007/s13225-020-00466-2)) |
| Otu38339 | 1 | k__Fungi(100);p__Ascomycota(100);c__Saccharomycetes(100);o__Saccharomycetales(100);f__Metschnikowiaceae(100);g__Metschnikowia(100);s__Metschnikowia_pulcherrima(100); | Metschnikowia | 13 | Saprotroph | \|Undefined Saprotroph\| | Yeast | NULL | Probable | As nectar-tap_saprotroph (Põlme et al. 2020); Decay sugar-rich substrates (Põlme et al. 2020) | Põlme S, et al. 2020. Fungal Diversity 105:1-16 ((https://doi.org/10.1007/s13225-020-00466-2)) |
| Otu38340 | 1 | k__Fungi(100);p__Ascomycota(100);c__Saccharomycetes(100);o__Saccharomycetales(100);f__Metschnikowiaceae(100);g__Metschnikowia(100);s__Metschnikowia_pulcherrima(100); | Metschnikowia | 13 | Saprotroph | \|Undefined Saprotroph\| | Yeast | NULL | Probable | As nectar-tap_saprotroph (Põlme et al. 2020); Decay sugar-rich substrates (Põlme et al. 2020) | Põlme S, et al. 2020. Fungal Diversity 105:1-16 ((https://doi.org/10.1007/s13225-020-00466-2)) |
| Otu38341 | 1 | k__Fungi(100);p__Ascomycota(100);c__Saccharomycetes(100);o__Saccharomycetales(100);f__Metschnikowiaceae(100);g__Metschnikowia(100);s__Metschnikowia_pulcherrima(100); | Metschnikowia | 13 | Saprotroph | \|Undefined Saprotroph\| | Yeast | NULL | Probable | As nectar-tap_saprotroph (Põlme et al. 2020); Decay sugar-rich substrates (Põlme et al. 2020) | Põlme S, et al. 2020. Fungal Diversity 105:1-16 ((https://doi.org/10.1007/s13225-020-00466-2)) |
| Otu38342 | 1 | k__Fungi(100);p__Ascomycota(100);c__Saccharomycetes(100);o__Saccharomycetales(100);f__Metschnikowiaceae(100);g__Metschnikowia(100);s__Metschnikowia_pulcherrima(100); | Metschnikowia | 13 | Saprotroph | \|Undefined Saprotroph\| | Yeast | NULL | Probable | As nectar-tap_saprotroph (Põlme et al. 2020); Decay sugar-rich substrates (Põlme et al. 2020) | Põlme S, et al. 2020. Fungal Diversity 105:1-16 ((https://doi.org/10.1007/s13225-020-00466-2)) |
| Otu38343 | 1 | k__Fungi(100);p__Ascomycota(100);c__Saccharomycetes(100);o__Saccharomycetales(100);f__Metschnikowiaceae(100);g__Metschnikowia(100);s__Metschnikowia_pulcherrima(100); | Metschnikowia | 13 | Saprotroph | \|Undefined Saprotroph\| | Yeast | NULL | Probable | As nectar-tap_saprotroph (Põlme et al. 2020); Decay sugar-rich substrates (Põlme et al. 2020) | Põlme S, et al. 2020. Fungal Diversity 105:1-16 ((https://doi.org/10.1007/s13225-020-00466-2)) |
| Otu38345 | 1 | k__Fungi(100);p__Ascomycota(100);c__Saccharomycetes(100);o__Saccharomycetales(100);f__Metschnikowiaceae(100);g__Metschnikowia(100);s__Metschnikowia_pulcherrima(100); | Metschnikowia | 13 | Saprotroph | \|Undefined Saprotroph\| | Yeast | NULL | Probable | As nectar-tap_saprotroph (Põlme et al. 2020); Decay sugar-rich substrates (Põlme et al. 2020) | Põlme S, et al. 2020. Fungal Diversity 105:1-16 ((https://doi.org/10.1007/s13225-020-00466-2)) |
| Otu38346 | 1 | k__Fungi(100);p__Ascomycota(100);c__Saccharomycetes(100);o__Saccharomycetales(100);f__Metschnikowiaceae(100);g__Metschnikowia(100);g__Metschnikowia_unclassified(100); | Metschnikowia | 13 | Saprotroph | \|Undefined Saprotroph\| | Yeast | NULL | Probable | As nectar-tap_saprotroph (Põlme et al. 2020); Decay sugar-rich substrates (Põlme et al. 2020) | Põlme S, et al. 2020. Fungal Diversity 105:1-16 ((https://doi.org/10.1007/s13225-020-00466-2)) |
| Otu38347 | 1 | k__Fungi(100);p__Ascomycota(100);c__Saccharomycetes(100);o__Saccharomycetales(100);f__Metschnikowiaceae(100);g__Metschnikowia(100);s__Metschnikowia_pulcherrima(100); | Metschnikowia | 13 | Saprotroph | \|Undefined Saprotroph\| | Yeast | NULL | Probable | As nectar-tap_saprotroph (Põlme et al. 2020); Decay sugar-rich substrates (Põlme et al. 2020) | Põlme S, et al. 2020. Fungal Diversity 105:1-16 ((https://doi.org/10.1007/s13225-020-00466-2)) |
| Otu38348 | 1 | k__Fungi(100);p__Ascomycota(100);c__Saccharomycetes(100);o__Saccharomycetales(100);f__Metschnikowiaceae(100);g__Metschnikowia(100);g__Metschnikowia_unclassified(100); | Metschnikowia | 13 | Saprotroph | \|Undefined Saprotroph\| | Yeast | NULL | Probable | As nectar-tap_saprotroph (Põlme et al. 2020); Decay sugar-rich substrates (Põlme et al. 2020) | Põlme S, et al. 2020. Fungal Diversity 105:1-16 ((https://doi.org/10.1007/s13225-020-00466-2)) |
| Otu38349 | 1 | k__Fungi(100);p__Ascomycota(100);c__Saccharomycetes(100);o__Saccharomycetales(100);f__Metschnikowiaceae(100);g__Metschnikowia(100);g__Metschnikowia_unclassified(100); | Metschnikowia | 13 | Saprotroph | \|Undefined Saprotroph\| | Yeast | NULL | Probable | As nectar-tap_saprotroph (Põlme et al. 2020); Decay sugar-rich substrates (Põlme et al. 2020) | Põlme S, et al. 2020. Fungal Diversity 105:1-16 ((https://doi.org/10.1007/s13225-020-00466-2)) |
| Otu38350 | 1 | k__Fungi(100);p__Ascomycota(100);c__Saccharomycetes(100);o__Saccharomycetales(100);f__Metschnikowiaceae(100);g__Metschnikowia(100);g__Metschnikowia_unclassified(100); | Metschnikowia | 13 | Saprotroph | \|Undefined Saprotroph\| | Yeast | NULL | Probable | As nectar-tap_saprotroph (Põlme et al. 2020); Decay sugar-rich substrates (Põlme et al. 2020) | Põlme S, et al. 2020. Fungal Diversity 105:1-16 ((https://doi.org/10.1007/s13225-020-00466-2)) |
| Otu38351 | 1 | k__Fungi(100);p__Ascomycota(100);c__Saccharomycetes(100);o__Saccharomycetales(100);f__Metschnikowiaceae(100);g__Metschnikowia(100);s__Metschnikowia_pulcherrima(100); | Metschnikowia | 13 | Saprotroph | \|Undefined Saprotroph\| | Yeast | NULL | Probable | As nectar-tap_saprotroph (Põlme et al. 2020); Decay sugar-rich substrates (Põlme et al. 2020) | Põlme S, et al. 2020. Fungal Diversity 105:1-16 ((https://doi.org/10.1007/s13225-020-00466-2)) |
| Otu38352 | 1 | k__Fungi(100);p__Ascomycota(100);c__Saccharomycetes(100);o__Saccharomycetales(100);f__Metschnikowiaceae(100);g__Metschnikowia(100);s__Metschnikowia_pulcherrima(100); | Metschnikowia | 13 | Saprotroph | \|Undefined Saprotroph\| | Yeast | NULL | Probable | As nectar-tap_saprotroph (Põlme et al. 2020); Decay sugar-rich substrates (Põlme et al. 2020) | Põlme S, et al. 2020. Fungal Diversity 105:1-16 ((https://doi.org/10.1007/s13225-020-00466-2)) |
| Otu38353 | 1 | k__Fungi(100);p__Ascomycota(100);c__Saccharomycetes(100);o__Saccharomycetales(100);f__Metschnikowiaceae(100);g__Metschnikowia(100);s__Metschnikowia_pulcherrima(100); | Metschnikowia | 13 | Saprotroph | \|Undefined Saprotroph\| | Yeast | NULL | Probable | As nectar-tap_saprotroph (Põlme et al. 2020); Decay sugar-rich substrates (Põlme et al. 2020) | Põlme S, et al. 2020. Fungal Diversity 105:1-16 ((https://doi.org/10.1007/s13225-020-00466-2)) |
| Otu38354 | 1 | k__Fungi(100);p__Ascomycota(100);c__Saccharomycetes(100);o__Saccharomycetales(100);f__Metschnikowiaceae(100);g__Metschnikowia(100);s__Metschnikowia_pulcherrima(100); | Metschnikowia | 13 | Saprotroph | \|Undefined Saprotroph\| | Yeast | NULL | Probable | As nectar-tap_saprotroph (Põlme et al. 2020); Decay sugar-rich substrates (Põlme et al. 2020) | Põlme S, et al. 2020. Fungal Diversity 105:1-16 ((https://doi.org/10.1007/s13225-020-00466-2)) |
| Otu38355 | 1 | k__Fungi(100);p__Ascomycota(100);c__Saccharomycetes(100);o__Saccharomycetales(100);f__Metschnikowiaceae(100);g__Metschnikowia(100);g__Metschnikowia_unclassified(100); | Metschnikowia | 13 | Saprotroph | \|Undefined Saprotroph\| | Yeast | NULL | Probable | As nectar-tap_saprotroph (Põlme et al. 2020); Decay sugar-rich substrates (Põlme et al. 2020) | Põlme S, et al. 2020. Fungal Diversity 105:1-16 ((https://doi.org/10.1007/s13225-020-00466-2)) |
| Otu38356 | 1 | k__Fungi(100);p__Ascomycota(100);c__Saccharomycetes(100);o__Saccharomycetales(100);f__Metschnikowiaceae(100);g__Metschnikowia(100);s__Metschnikowia_pulcherrima(100); | Metschnikowia | 13 | Saprotroph | \|Undefined Saprotroph\| | Yeast | NULL | Probable | As nectar-tap_saprotroph (Põlme et al. 2020); Decay sugar-rich substrates (Põlme et al. 2020) | Põlme S, et al. 2020. Fungal Diversity 105:1-16 ((https://doi.org/10.1007/s13225-020-00466-2)) |
| Otu38357 | 1 | k__Fungi(100);p__Ascomycota(100);c__Saccharomycetes(100);o__Saccharomycetales(100);f__Metschnikowiaceae(100);g__Metschnikowia(100);g__Metschnikowia_unclassified(100); | Metschnikowia | 13 | Saprotroph | \|Undefined Saprotroph\| | Yeast | NULL | Probable | As nectar-tap_saprotroph (Põlme et al. 2020); Decay sugar-rich substrates (Põlme et al. 2020) | Põlme S, et al. 2020. Fungal Diversity 105:1-16 ((https://doi.org/10.1007/s13225-020-00466-2)) |
| Otu38358 | 1 | k__Fungi(100);p__Ascomycota(100);c__Saccharomycetes(100);o__Saccharomycetales(100);f__Metschnikowiaceae(100);g__Metschnikowia(100);s__Metschnikowia_pulcherrima(100); | Metschnikowia | 13 | Saprotroph | \|Undefined Saprotroph\| | Yeast | NULL | Probable | As nectar-tap_saprotroph (Põlme et al. 2020); Decay sugar-rich substrates (Põlme et al. 2020) | Põlme S, et al. 2020. Fungal Diversity 105:1-16 ((https://doi.org/10.1007/s13225-020-00466-2)) |
| Otu38359 | 1 | k__Fungi(100);p__Ascomycota(100);c__Saccharomycetes(100);o__Saccharomycetales(100);f__Metschnikowiaceae(100);g__Metschnikowia(100);s__Metschnikowia_pulcherrima(100); | Metschnikowia | 13 | Saprotroph | \|Undefined Saprotroph\| | Yeast | NULL | Probable | As nectar-tap_saprotroph (Põlme et al. 2020); Decay sugar-rich substrates (Põlme et al. 2020) | Põlme S, et al. 2020. Fungal Diversity 105:1-16 ((https://doi.org/10.1007/s13225-020-00466-2)) |
| Otu38360 | 1 | k__Fungi(100);p__Ascomycota(100);c__Saccharomycetes(100);o__Saccharomycetales(100);f__Metschnikowiaceae(100);g__Metschnikowia(100);s__Metschnikowia_pulcherrima(100); | Metschnikowia | 13 | Saprotroph | \|Undefined Saprotroph\| | Yeast | NULL | Probable | As nectar-tap_saprotroph (Põlme et al. 2020); Decay sugar-rich substrates (Põlme et al. 2020) | Põlme S, et al. 2020. Fungal Diversity 105:1-16 ((https://doi.org/10.1007/s13225-020-00466-2)) |
| Otu38361 | 1 | k__Fungi(100);p__Ascomycota(100);c__Saccharomycetes(100);o__Saccharomycetales(100);f__Metschnikowiaceae(100);g__Metschnikowia(100);s__Metschnikowia_pulcherrima(100); | Metschnikowia | 13 | Saprotroph | \|Undefined Saprotroph\| | Yeast | NULL | Probable | As nectar-tap_saprotroph (Põlme et al. 2020); Decay sugar-rich substrates (Põlme et al. 2020) | Põlme S, et al. 2020. Fungal Diversity 105:1-16 ((https://doi.org/10.1007/s13225-020-00466-2)) |
| Otu38362 | 1 | k__Fungi(100);p__Ascomycota(100);c__Saccharomycetes(100);o__Saccharomycetales(100);f__Metschnikowiaceae(100);g__Metschnikowia(100);g__Metschnikowia_unclassified(100); | Metschnikowia | 13 | Saprotroph | \|Undefined Saprotroph\| | Yeast | NULL | Probable | As nectar-tap_saprotroph (Põlme et al. 2020); Decay sugar-rich substrates (Põlme et al. 2020) | Põlme S, et al. 2020. Fungal Diversity 105:1-16 ((https://doi.org/10.1007/s13225-020-00466-2)) |
| Otu38363 | 1 | k__Fungi(100);p__Ascomycota(100);c__Saccharomycetes(100);o__Saccharomycetales(100);f__Metschnikowiaceae(100);g__Metschnikowia(100);g__Metschnikowia_unclassified(100); | Metschnikowia | 13 | Saprotroph | \|Undefined Saprotroph\| | Yeast | NULL | Probable | As nectar-tap_saprotroph (Põlme et al. 2020); Decay sugar-rich substrates (Põlme et al. 2020) | Põlme S, et al. 2020. Fungal Diversity 105:1-16 ((https://doi.org/10.1007/s13225-020-00466-2)) |
| Otu38364 | 1 | k__Fungi(100);p__Ascomycota(100);c__Saccharomycetes(100);o__Saccharomycetales(100);f__Metschnikowiaceae(100);g__Metschnikowia(100);s__Metschnikowia_pulcherrima(100); | Metschnikowia | 13 | Saprotroph | \|Undefined Saprotroph\| | Yeast | NULL | Probable | As nectar-tap_saprotroph (Põlme et al. 2020); Decay sugar-rich substrates (Põlme et al. 2020) | Põlme S, et al. 2020. Fungal Diversity 105:1-16 ((https://doi.org/10.1007/s13225-020-00466-2)) |
| Otu38365 | 1 | k__Fungi(100);p__Ascomycota(100);c__Saccharomycetes(100);o__Saccharomycetales(100);f__Metschnikowiaceae(100);g__Metschnikowia(100);s__Metschnikowia_pulcherrima(100); | Metschnikowia | 13 | Saprotroph | \|Undefined Saprotroph\| | Yeast | NULL | Probable | As nectar-tap_saprotroph (Põlme et al. 2020); Decay sugar-rich substrates (Põlme et al. 2020) | Põlme S, et al. 2020. Fungal Diversity 105:1-16 ((https://doi.org/10.1007/s13225-020-00466-2)) |
| Otu38371 | 1 | k__Fungi(100);p__Ascomycota(100);c__Saccharomycetes(100);o__Saccharomycetales(100);f__Metschnikowiaceae(100);g__Metschnikowia(100);g__Metschnikowia_unclassified(100); | Metschnikowia | 13 | Saprotroph | \|Undefined Saprotroph\| | Yeast | NULL | Probable | As nectar-tap_saprotroph (Põlme et al. 2020); Decay sugar-rich substrates (Põlme et al. 2020) | Põlme S, et al. 2020. Fungal Diversity 105:1-16 ((https://doi.org/10.1007/s13225-020-00466-2)) |
| Otu38372 | 1 | k__Fungi(100);p__Ascomycota(100);c__Saccharomycetes(100);o__Saccharomycetales(100);f__Metschnikowiaceae(100);g__Metschnikowia(100);s__Metschnikowia_pulcherrima(100); | Metschnikowia | 13 | Saprotroph | \|Undefined Saprotroph\| | Yeast | NULL | Probable | As nectar-tap_saprotroph (Põlme et al. 2020); Decay sugar-rich substrates (Põlme et al. 2020) | Põlme S, et al. 2020. Fungal Diversity 105:1-16 ((https://doi.org/10.1007/s13225-020-00466-2)) |
| Otu38373 | 1 | k__Fungi(100);p__Ascomycota(100);c__Saccharomycetes(100);o__Saccharomycetales(100);f__Metschnikowiaceae(100);g__Metschnikowia(100);s__Metschnikowia_pulcherrima(100); | Metschnikowia | 13 | Saprotroph | \|Undefined Saprotroph\| | Yeast | NULL | Probable | As nectar-tap_saprotroph (Põlme et al. 2020); Decay sugar-rich substrates (Põlme et al. 2020) | Põlme S, et al. 2020. Fungal Diversity 105:1-16 ((https://doi.org/10.1007/s13225-020-00466-2)) |
| Otu38374 | 1 | k__Fungi(100);p__Ascomycota(100);c__Saccharomycetes(100);o__Saccharomycetales(100);f__Metschnikowiaceae(100);g__Metschnikowia(100);s__Metschnikowia_pulcherrima(100); | Metschnikowia | 13 | Saprotroph | \|Undefined Saprotroph\| | Yeast | NULL | Probable | As nectar-tap_saprotroph (Põlme et al. 2020); Decay sugar-rich substrates (Põlme et al. 2020) | Põlme S, et al. 2020. Fungal Diversity 105:1-16 ((https://doi.org/10.1007/s13225-020-00466-2)) |
| Otu38375 | 1 | k__Fungi(100);p__Ascomycota(100);c__Saccharomycetes(100);o__Saccharomycetales(100);f__Metschnikowiaceae(100);g__Metschnikowia(100);s__Metschnikowia_pulcherrima(100); | Metschnikowia | 13 | Saprotroph | \|Undefined Saprotroph\| | Yeast | NULL | Probable | As nectar-tap_saprotroph (Põlme et al. 2020); Decay sugar-rich substrates (Põlme et al. 2020) | Põlme S, et al. 2020. Fungal Diversity 105:1-16 ((https://doi.org/10.1007/s13225-020-00466-2)) |
| Otu38376 | 1 | k__Fungi(100);p__Ascomycota(100);c__Saccharomycetes(100);o__Saccharomycetales(100);f__Metschnikowiaceae(100);g__Metschnikowia(100);g__Metschnikowia_unclassified(100); | Metschnikowia | 13 | Saprotroph | \|Undefined Saprotroph\| | Yeast | NULL | Probable | As nectar-tap_saprotroph (Põlme et al. 2020); Decay sugar-rich substrates (Põlme et al. 2020) | Põlme S, et al. 2020. Fungal Diversity 105:1-16 ((https://doi.org/10.1007/s13225-020-00466-2)) |
| Otu38377 | 1 | k__Fungi(100);p__Ascomycota(100);c__Saccharomycetes(100);o__Saccharomycetales(100);f__Metschnikowiaceae(100);g__Metschnikowia(100);s__Metschnikowia_pulcherrima(100); | Metschnikowia | 13 | Saprotroph | \|Undefined Saprotroph\| | Yeast | NULL | Probable | As nectar-tap_saprotroph (Põlme et al. 2020); Decay sugar-rich substrates (Põlme et al. 2020) | Põlme S, et al. 2020. Fungal Diversity 105:1-16 ((https://doi.org/10.1007/s13225-020-00466-2)) |
| Otu38378 | 1 | k__Fungi(100);p__Ascomycota(100);c__Saccharomycetes(100);o__Saccharomycetales(100);f__Metschnikowiaceae(100);g__Metschnikowia(100);s__Metschnikowia_pulcherrima(100); | Metschnikowia | 13 | Saprotroph | \|Undefined Saprotroph\| | Yeast | NULL | Probable | As nectar-tap_saprotroph (Põlme et al. 2020); Decay sugar-rich substrates (Põlme et al. 2020) | Põlme S, et al. 2020. Fungal Diversity 105:1-16 ((https://doi.org/10.1007/s13225-020-00466-2)) |
| Otu38380 | 1 | k__Fungi(100);p__Ascomycota(100);c__Saccharomycetes(100);o__Saccharomycetales(100);f__Metschnikowiaceae(100);g__Metschnikowia(100);g__Metschnikowia_unclassified(100); | Metschnikowia | 13 | Saprotroph | \|Undefined Saprotroph\| | Yeast | NULL | Probable | As nectar-tap_saprotroph (Põlme et al. 2020); Decay sugar-rich substrates (Põlme et al. 2020) | Põlme S, et al. 2020. Fungal Diversity 105:1-16 ((https://doi.org/10.1007/s13225-020-00466-2)) |
| Otu38382 | 1 | k__Fungi(100);p__Ascomycota(100);c__Saccharomycetes(100);o__Saccharomycetales(100);f__Metschnikowiaceae(100);g__Metschnikowia(100);s__Metschnikowia_pulcherrima(100); | Metschnikowia | 13 | Saprotroph | \|Undefined Saprotroph\| | Yeast | NULL | Probable | As nectar-tap_saprotroph (Põlme et al. 2020); Decay sugar-rich substrates (Põlme et al. 2020) | Põlme S, et al. 2020. Fungal Diversity 105:1-16 ((https://doi.org/10.1007/s13225-020-00466-2)) |
| Otu38383 | 1 | k__Fungi(100);p__Ascomycota(100);c__Saccharomycetes(100);o__Saccharomycetales(100);f__Metschnikowiaceae(100);g__Metschnikowia(100);s__Metschnikowia_pulcherrima(100); | Metschnikowia | 13 | Saprotroph | \|Undefined Saprotroph\| | Yeast | NULL | Probable | As nectar-tap_saprotroph (Põlme et al. 2020); Decay sugar-rich substrates (Põlme et al. 2020) | Põlme S, et al. 2020. Fungal Diversity 105:1-16 ((https://doi.org/10.1007/s13225-020-00466-2)) |
| Otu38384 | 1 | k__Fungi(100);p__Ascomycota(100);c__Saccharomycetes(100);o__Saccharomycetales(100);f__Metschnikowiaceae(100);g__Metschnikowia(100);s__Metschnikowia_pulcherrima(100); | Metschnikowia | 13 | Saprotroph | \|Undefined Saprotroph\| | Yeast | NULL | Probable | As nectar-tap_saprotroph (Põlme et al. 2020); Decay sugar-rich substrates (Põlme et al. 2020) | Põlme S, et al. 2020. Fungal Diversity 105:1-16 ((https://doi.org/10.1007/s13225-020-00466-2)) |
| Otu38385 | 1 | k__Fungi(100);p__Ascomycota(100);c__Saccharomycetes(100);o__Saccharomycetales(100);f__Metschnikowiaceae(100);g__Metschnikowia(100);s__Metschnikowia_pulcherrima(100); | Metschnikowia | 13 | Saprotroph | \|Undefined Saprotroph\| | Yeast | NULL | Probable | As nectar-tap_saprotroph (Põlme et al. 2020); Decay sugar-rich substrates (Põlme et al. 2020) | Põlme S, et al. 2020. Fungal Diversity 105:1-16 ((https://doi.org/10.1007/s13225-020-00466-2)) |
| Otu38388 | 1 | k__Fungi(100);p__Ascomycota(100);c__Saccharomycetes(100);o__Saccharomycetales(100);f__Metschnikowiaceae(100);g__Metschnikowia(100);s__Metschnikowia_pulcherrima(100); | Metschnikowia | 13 | Saprotroph | \|Undefined Saprotroph\| | Yeast | NULL | Probable | As nectar-tap_saprotroph (Põlme et al. 2020); Decay sugar-rich substrates (Põlme et al. 2020) | Põlme S, et al. 2020. Fungal Diversity 105:1-16 ((https://doi.org/10.1007/s13225-020-00466-2)) |
| Otu38389 | 1 | k__Fungi(100);p__Ascomycota(100);c__Saccharomycetes(100);o__Saccharomycetales(100);f__Metschnikowiaceae(100);g__Metschnikowia(100);s__Metschnikowia_pulcherrima(100); | Metschnikowia | 13 | Saprotroph | \|Undefined Saprotroph\| | Yeast | NULL | Probable | As nectar-tap_saprotroph (Põlme et al. 2020); Decay sugar-rich substrates (Põlme et al. 2020) | Põlme S, et al. 2020. Fungal Diversity 105:1-16 ((https://doi.org/10.1007/s13225-020-00466-2)) |
| Otu38390 | 1 | k__Fungi(100);p__Ascomycota(100);c__Saccharomycetes(100);o__Saccharomycetales(100);f__Metschnikowiaceae(100);g__Metschnikowia(100);s__Metschnikowia_pulcherrima(100); | Metschnikowia | 13 | Saprotroph | \|Undefined Saprotroph\| | Yeast | NULL | Probable | As nectar-tap_saprotroph (Põlme et al. 2020); Decay sugar-rich substrates (Põlme et al. 2020) | Põlme S, et al. 2020. Fungal Diversity 105:1-16 ((https://doi.org/10.1007/s13225-020-00466-2)) |
| Otu38391 | 1 | k__Fungi(100);p__Ascomycota(100);c__Saccharomycetes(100);o__Saccharomycetales(100);f__Metschnikowiaceae(100);g__Metschnikowia(100);s__Metschnikowia_pulcherrima(100); | Metschnikowia | 13 | Saprotroph | \|Undefined Saprotroph\| | Yeast | NULL | Probable | As nectar-tap_saprotroph (Põlme et al. 2020); Decay sugar-rich substrates (Põlme et al. 2020) | Põlme S, et al. 2020. Fungal Diversity 105:1-16 ((https://doi.org/10.1007/s13225-020-00466-2)) |
| Otu38392 | 1 | k__Fungi(100);p__Ascomycota(100);c__Saccharomycetes(100);o__Saccharomycetales(100);f__Metschnikowiaceae(100);g__Metschnikowia(100);g__Metschnikowia_unclassified(100); | Metschnikowia | 13 | Saprotroph | \|Undefined Saprotroph\| | Yeast | NULL | Probable | As nectar-tap_saprotroph (Põlme et al. 2020); Decay sugar-rich substrates (Põlme et al. 2020) | Põlme S, et al. 2020. Fungal Diversity 105:1-16 ((https://doi.org/10.1007/s13225-020-00466-2)) |
| Otu38393 | 1 | k__Fungi(100);p__Ascomycota(100);c__Saccharomycetes(100);o__Saccharomycetales(100);f__Metschnikowiaceae(100);g__Metschnikowia(100);s__Metschnikowia_pulcherrima(100); | Metschnikowia | 13 | Saprotroph | \|Undefined Saprotroph\| | Yeast | NULL | Probable | As nectar-tap_saprotroph (Põlme et al. 2020); Decay sugar-rich substrates (Põlme et al. 2020) | Põlme S, et al. 2020. Fungal Diversity 105:1-16 ((https://doi.org/10.1007/s13225-020-00466-2)) |
| Otu38394 | 1 | k__Fungi(100);p__Ascomycota(100);c__Saccharomycetes(100);o__Saccharomycetales(100);f__Metschnikowiaceae(100);g__Metschnikowia(100);s__Metschnikowia_pulcherrima(100); | Metschnikowia | 13 | Saprotroph | \|Undefined Saprotroph\| | Yeast | NULL | Probable | As nectar-tap_saprotroph (Põlme et al. 2020); Decay sugar-rich substrates (Põlme et al. 2020) | Põlme S, et al. 2020. Fungal Diversity 105:1-16 ((https://doi.org/10.1007/s13225-020-00466-2)) |
| Otu38395 | 1 | k__Fungi(100);p__Ascomycota(100);c__Saccharomycetes(100);o__Saccharomycetales(100);f__Metschnikowiaceae(100);g__Metschnikowia(100);s__Metschnikowia_pulcherrima(100); | Metschnikowia | 13 | Saprotroph | \|Undefined Saprotroph\| | Yeast | NULL | Probable | As nectar-tap_saprotroph (Põlme et al. 2020); Decay sugar-rich substrates (Põlme et al. 2020) | Põlme S, et al. 2020. Fungal Diversity 105:1-16 ((https://doi.org/10.1007/s13225-020-00466-2)) |
| Otu38396 | 1 | k__Fungi(100);p__Ascomycota(100);c__Saccharomycetes(100);o__Saccharomycetales(100);f__Metschnikowiaceae(100);g__Metschnikowia(100);g__Metschnikowia_unclassified(100); | Metschnikowia | 13 | Saprotroph | \|Undefined Saprotroph\| | Yeast | NULL | Probable | As nectar-tap_saprotroph (Põlme et al. 2020); Decay sugar-rich substrates (Põlme et al. 2020) | Põlme S, et al. 2020. Fungal Diversity 105:1-16 ((https://doi.org/10.1007/s13225-020-00466-2)) |
| Otu38397 | 1 | k__Fungi(100);p__Ascomycota(100);c__Saccharomycetes(100);o__Saccharomycetales(100);f__Metschnikowiaceae(100);g__Metschnikowia(100);s__Metschnikowia_pulcherrima(100); | Metschnikowia | 13 | Saprotroph | \|Undefined Saprotroph\| | Yeast | NULL | Probable | As nectar-tap_saprotroph (Põlme et al. 2020); Decay sugar-rich substrates (Põlme et al. 2020) | Põlme S, et al. 2020. Fungal Diversity 105:1-16 ((https://doi.org/10.1007/s13225-020-00466-2)) |
| Otu38398 | 1 | k__Fungi(100);p__Ascomycota(100);c__Saccharomycetes(100);o__Saccharomycetales(100);f__Metschnikowiaceae(100);g__Metschnikowia(100);s__Metschnikowia_pulcherrima(100); | Metschnikowia | 13 | Saprotroph | \|Undefined Saprotroph\| | Yeast | NULL | Probable | As nectar-tap_saprotroph (Põlme et al. 2020); Decay sugar-rich substrates (Põlme et al. 2020) | Põlme S, et al. 2020. Fungal Diversity 105:1-16 ((https://doi.org/10.1007/s13225-020-00466-2)) |
| Otu38399 | 1 | k__Fungi(100);p__Ascomycota(100);c__Saccharomycetes(100);o__Saccharomycetales(100);f__Metschnikowiaceae(100);g__Metschnikowia(100);s__Metschnikowia_pulcherrima(100); | Metschnikowia | 13 | Saprotroph | \|Undefined Saprotroph\| | Yeast | NULL | Probable | As nectar-tap_saprotroph (Põlme et al. 2020); Decay sugar-rich substrates (Põlme et al. 2020) | Põlme S, et al. 2020. Fungal Diversity 105:1-16 ((https://doi.org/10.1007/s13225-020-00466-2)) |
| Otu38400 | 1 | k__Fungi(100);p__Ascomycota(100);c__Saccharomycetes(100);o__Saccharomycetales(100);f__Metschnikowiaceae(100);g__Metschnikowia(100);g__Metschnikowia_unclassified(100); | Metschnikowia | 13 | Saprotroph | \|Undefined Saprotroph\| | Yeast | NULL | Probable | As nectar-tap_saprotroph (Põlme et al. 2020); Decay sugar-rich substrates (Põlme et al. 2020) | Põlme S, et al. 2020. Fungal Diversity 105:1-16 ((https://doi.org/10.1007/s13225-020-00466-2)) |
| Otu38401 | 1 | k__Fungi(100);p__Basidiomycota(100);c__Microbotryomycetes(100);o__Sporidiobolales(100);f__Sporidiobolales_family_Incertae_sedis(100);g__Sporobolomyces(100);s__Sporobolomyces_lactosus(100); | Sporobolomyces | 13 | Pathotroph-Saprotroph | \|Fungal Parasite\|-Plant Saprotroph | Yeast | NULL | Probable | As litter_saprotroph (Põlme et al. 2020); As mycoparasite (Põlme et al. 2020); Decay fungal material (Põlme et al. 2020) | Tedersoo L, et al. 2014. Science 346:e1256688 ((https://doi.org/10.1126/science.1256688)); Purahong W, et al. 2016. Molecular Ecology 25:4059-4074 ((https://doi.org/10.1111/mec.13739)); Põlme S, et al. 2020. Fungal Diversity 105:1-16 ((https://doi.org/10.1007/s13225-020-00466-2)) |
| Otu38402 | 1 | k__Fungi(100);p__Ascomycota(100);c__Saccharomycetes(100);o__Saccharomycetales(100);f__Metschnikowiaceae(100);g__Metschnikowia(100);s__Metschnikowia_pulcherrima(100); | Metschnikowia | 13 | Saprotroph | \|Undefined Saprotroph\| | Yeast | NULL | Probable | As nectar-tap_saprotroph (Põlme et al. 2020); Decay sugar-rich substrates (Põlme et al. 2020) | Põlme S, et al. 2020. Fungal Diversity 105:1-16 ((https://doi.org/10.1007/s13225-020-00466-2)) |
| Otu38403 | 1 | k__Fungi(100);p__Ascomycota(100);c__Saccharomycetes(100);o__Saccharomycetales(100);f__Metschnikowiaceae(100);g__Metschnikowia(100);s__Metschnikowia_pulcherrima(100); | Metschnikowia | 13 | Saprotroph | \|Undefined Saprotroph\| | Yeast | NULL | Probable | As nectar-tap_saprotroph (Põlme et al. 2020); Decay sugar-rich substrates (Põlme et al. 2020) | Põlme S, et al. 2020. Fungal Diversity 105:1-16 ((https://doi.org/10.1007/s13225-020-00466-2)) |
| Otu38404 | 1 | k__Fungi(100);p__Ascomycota(100);c__Saccharomycetes(100);o__Saccharomycetales(100);f__Metschnikowiaceae(100);g__Metschnikowia(100);s__Metschnikowia_pulcherrima(100); | Metschnikowia | 13 | Saprotroph | \|Undefined Saprotroph\| | Yeast | NULL | Probable | As nectar-tap_saprotroph (Põlme et al. 2020); Decay sugar-rich substrates (Põlme et al. 2020) | Põlme S, et al. 2020. Fungal Diversity 105:1-16 ((https://doi.org/10.1007/s13225-020-00466-2)) |
| Otu38405 | 1 | k__Fungi(100);p__Ascomycota(100);c__Saccharomycetes(100);o__Saccharomycetales(100);f__Metschnikowiaceae(100);g__Metschnikowia(100);s__Metschnikowia_pulcherrima(100); | Metschnikowia | 13 | Saprotroph | \|Undefined Saprotroph\| | Yeast | NULL | Probable | As nectar-tap_saprotroph (Põlme et al. 2020); Decay sugar-rich substrates (Põlme et al. 2020) | Põlme S, et al. 2020. Fungal Diversity 105:1-16 ((https://doi.org/10.1007/s13225-020-00466-2)) |
| Otu38406 | 1 | k__Fungi(100);p__Ascomycota(100);c__Saccharomycetes(100);o__Saccharomycetales(100);f__Metschnikowiaceae(100);g__Metschnikowia(100);s__Metschnikowia_pulcherrima(100); | Metschnikowia | 13 | Saprotroph | \|Undefined Saprotroph\| | Yeast | NULL | Probable | As nectar-tap_saprotroph (Põlme et al. 2020); Decay sugar-rich substrates (Põlme et al. 2020) | Põlme S, et al. 2020. Fungal Diversity 105:1-16 ((https://doi.org/10.1007/s13225-020-00466-2)) |
| Otu38407 | 1 | k__Fungi(100);p__Ascomycota(100);c__Saccharomycetes(100);o__Saccharomycetales(100);f__Metschnikowiaceae(100);g__Metschnikowia(100);g__Metschnikowia_unclassified(100); | Metschnikowia | 13 | Saprotroph | \|Undefined Saprotroph\| | Yeast | NULL | Probable | As nectar-tap_saprotroph (Põlme et al. 2020); Decay sugar-rich substrates (Põlme et al. 2020) | Põlme S, et al. 2020. Fungal Diversity 105:1-16 ((https://doi.org/10.1007/s13225-020-00466-2)) |
| Otu38408 | 1 | k__Fungi(100);p__Ascomycota(100);c__Saccharomycetes(100);o__Saccharomycetales(100);f__Metschnikowiaceae(100);g__Metschnikowia(100);s__Metschnikowia_pulcherrima(100); | Metschnikowia | 13 | Saprotroph | \|Undefined Saprotroph\| | Yeast | NULL | Probable | As nectar-tap_saprotroph (Põlme et al. 2020); Decay sugar-rich substrates (Põlme et al. 2020) | Põlme S, et al. 2020. Fungal Diversity 105:1-16 ((https://doi.org/10.1007/s13225-020-00466-2)) |
| Otu38409 | 1 | k__Fungi(100);p__Ascomycota(100);c__Saccharomycetes(100);o__Saccharomycetales(100);f__Metschnikowiaceae(100);g__Metschnikowia(100);g__Metschnikowia_unclassified(100); | Metschnikowia | 13 | Saprotroph | \|Undefined Saprotroph\| | Yeast | NULL | Probable | As nectar-tap_saprotroph (Põlme et al. 2020); Decay sugar-rich substrates (Põlme et al. 2020) | Põlme S, et al. 2020. Fungal Diversity 105:1-16 ((https://doi.org/10.1007/s13225-020-00466-2)) |
| Otu38410 | 1 | k__Fungi(100);p__Ascomycota(100);c__Saccharomycetes(100);o__Saccharomycetales(100);f__Metschnikowiaceae(100);g__Metschnikowia(100);s__Metschnikowia_pulcherrima(100); | Metschnikowia | 13 | Saprotroph | \|Undefined Saprotroph\| | Yeast | NULL | Probable | As nectar-tap_saprotroph (Põlme et al. 2020); Decay sugar-rich substrates (Põlme et al. 2020) | Põlme S, et al. 2020. Fungal Diversity 105:1-16 ((https://doi.org/10.1007/s13225-020-00466-2)) |
| Otu38411 | 1 | k__Fungi(100);p__Ascomycota(100);c__Saccharomycetes(100);o__Saccharomycetales(100);f__Metschnikowiaceae(100);g__Metschnikowia(100);s__Metschnikowia_pulcherrima(100); | Metschnikowia | 13 | Saprotroph | \|Undefined Saprotroph\| | Yeast | NULL | Probable | As nectar-tap_saprotroph (Põlme et al. 2020); Decay sugar-rich substrates (Põlme et al. 2020) | Põlme S, et al. 2020. Fungal Diversity 105:1-16 ((https://doi.org/10.1007/s13225-020-00466-2)) |
| Otu38412 | 1 | k__Fungi(100);p__Ascomycota(100);c__Saccharomycetes(100);o__Saccharomycetales(100);f__Metschnikowiaceae(100);g__Metschnikowia(100);s__Metschnikowia_pulcherrima(100); | Metschnikowia | 13 | Saprotroph | \|Undefined Saprotroph\| | Yeast | NULL | Probable | As nectar-tap_saprotroph (Põlme et al. 2020); Decay sugar-rich substrates (Põlme et al. 2020) | Põlme S, et al. 2020. Fungal Diversity 105:1-16 ((https://doi.org/10.1007/s13225-020-00466-2)) |
| Otu38413 | 1 | k__Fungi(100);p__Ascomycota(100);c__Saccharomycetes(100);o__Saccharomycetales(100);f__Metschnikowiaceae(100);g__Metschnikowia(100);s__Metschnikowia_pulcherrima(100); | Metschnikowia | 13 | Saprotroph | \|Undefined Saprotroph\| | Yeast | NULL | Probable | As nectar-tap_saprotroph (Põlme et al. 2020); Decay sugar-rich substrates (Põlme et al. 2020) | Põlme S, et al. 2020. Fungal Diversity 105:1-16 ((https://doi.org/10.1007/s13225-020-00466-2)) |
| Otu38414 | 1 | k__Fungi(100);p__Ascomycota(100);c__Saccharomycetes(100);o__Saccharomycetales(100);f__Metschnikowiaceae(100);g__Metschnikowia(100);g__Metschnikowia_unclassified(100); | Metschnikowia | 13 | Saprotroph | \|Undefined Saprotroph\| | Yeast | NULL | Probable | As nectar-tap_saprotroph (Põlme et al. 2020); Decay sugar-rich substrates (Põlme et al. 2020) | Põlme S, et al. 2020. Fungal Diversity 105:1-16 ((https://doi.org/10.1007/s13225-020-00466-2)) |
| Otu38415 | 1 | k__Fungi(100);p__Ascomycota(100);c__Saccharomycetes(100);o__Saccharomycetales(100);f__Metschnikowiaceae(100);g__Metschnikowia(100);g__Metschnikowia_unclassified(100); | Metschnikowia | 13 | Saprotroph | \|Undefined Saprotroph\| | Yeast | NULL | Probable | As nectar-tap_saprotroph (Põlme et al. 2020); Decay sugar-rich substrates (Põlme et al. 2020) | Põlme S, et al. 2020. Fungal Diversity 105:1-16 ((https://doi.org/10.1007/s13225-020-00466-2)) |
| Otu38416 | 1 | k__Fungi(100);p__Ascomycota(100);c__Saccharomycetes(100);o__Saccharomycetales(100);f__Metschnikowiaceae(100);g__Metschnikowia(100);s__Metschnikowia_pulcherrima(100); | Metschnikowia | 13 | Saprotroph | \|Undefined Saprotroph\| | Yeast | NULL | Probable | As nectar-tap_saprotroph (Põlme et al. 2020); Decay sugar-rich substrates (Põlme et al. 2020) | Põlme S, et al. 2020. Fungal Diversity 105:1-16 ((https://doi.org/10.1007/s13225-020-00466-2)) |
| Otu38417 | 1 | k__Fungi(100);p__Ascomycota(100);c__Saccharomycetes(100);o__Saccharomycetales(100);f__Metschnikowiaceae(100);g__Metschnikowia(100);g__Metschnikowia_unclassified(100); | Metschnikowia | 13 | Saprotroph | \|Undefined Saprotroph\| | Yeast | NULL | Probable | As nectar-tap_saprotroph (Põlme et al. 2020); Decay sugar-rich substrates (Põlme et al. 2020) | Põlme S, et al. 2020. Fungal Diversity 105:1-16 ((https://doi.org/10.1007/s13225-020-00466-2)) |
| Otu38418 | 1 | k__Fungi(100);p__Ascomycota(100);c__Saccharomycetes(100);o__Saccharomycetales(100);f__Metschnikowiaceae(100);g__Metschnikowia(100);s__Metschnikowia_pulcherrima(100); | Metschnikowia | 13 | Saprotroph | \|Undefined Saprotroph\| | Yeast | NULL | Probable | As nectar-tap_saprotroph (Põlme et al. 2020); Decay sugar-rich substrates (Põlme et al. 2020) | Põlme S, et al. 2020. Fungal Diversity 105:1-16 ((https://doi.org/10.1007/s13225-020-00466-2)) |
| Otu38419 | 1 | k__Fungi(100);p__Ascomycota(100);c__Saccharomycetes(100);o__Saccharomycetales(100);f__Metschnikowiaceae(100);g__Metschnikowia(100);g__Metschnikowia_unclassified(100); | Metschnikowia | 13 | Saprotroph | \|Undefined Saprotroph\| | Yeast | NULL | Probable | As nectar-tap_saprotroph (Põlme et al. 2020); Decay sugar-rich substrates (Põlme et al. 2020) | Põlme S, et al. 2020. Fungal Diversity 105:1-16 ((https://doi.org/10.1007/s13225-020-00466-2)) |
| Otu38420 | 1 | k__Fungi(100);p__Ascomycota(100);c__Saccharomycetes(100);o__Saccharomycetales(100);f__Metschnikowiaceae(100);g__Metschnikowia(100);s__Metschnikowia_pulcherrima(100); | Metschnikowia | 13 | Saprotroph | \|Undefined Saprotroph\| | Yeast | NULL | Probable | As nectar-tap_saprotroph (Põlme et al. 2020); Decay sugar-rich substrates (Põlme et al. 2020) | Põlme S, et al. 2020. Fungal Diversity 105:1-16 ((https://doi.org/10.1007/s13225-020-00466-2)) |
| Otu38421 | 1 | k__Fungi(100);p__Ascomycota(100);c__Saccharomycetes(100);o__Saccharomycetales(100);f__Metschnikowiaceae(100);g__Metschnikowia(100);g__Metschnikowia_unclassified(100); | Metschnikowia | 13 | Saprotroph | \|Undefined Saprotroph\| | Yeast | NULL | Probable | As nectar-tap_saprotroph (Põlme et al. 2020); Decay sugar-rich substrates (Põlme et al. 2020) | Põlme S, et al. 2020. Fungal Diversity 105:1-16 ((https://doi.org/10.1007/s13225-020-00466-2)) |
| Otu38422 | 1 | k__Fungi(100);p__Ascomycota(100);c__Saccharomycetes(100);o__Saccharomycetales(100);f__Metschnikowiaceae(100);g__Metschnikowia(100);g__Metschnikowia_unclassified(100); | Metschnikowia | 13 | Saprotroph | \|Undefined Saprotroph\| | Yeast | NULL | Probable | As nectar-tap_saprotroph (Põlme et al. 2020); Decay sugar-rich substrates (Põlme et al. 2020) | Põlme S, et al. 2020. Fungal Diversity 105:1-16 ((https://doi.org/10.1007/s13225-020-00466-2)) |
| Otu38423 | 1 | k__Fungi(100);p__Ascomycota(100);c__Saccharomycetes(100);o__Saccharomycetales(100);f__Metschnikowiaceae(100);g__Metschnikowia(100);s__Metschnikowia_pulcherrima(100); | Metschnikowia | 13 | Saprotroph | \|Undefined Saprotroph\| | Yeast | NULL | Probable | As nectar-tap_saprotroph (Põlme et al. 2020); Decay sugar-rich substrates (Põlme et al. 2020) | Põlme S, et al. 2020. Fungal Diversity 105:1-16 ((https://doi.org/10.1007/s13225-020-00466-2)) |
| Otu38424 | 1 | k__Fungi(100);p__Ascomycota(100);c__Saccharomycetes(100);o__Saccharomycetales(100);f__Metschnikowiaceae(100);g__Metschnikowia(100);g__Metschnikowia_unclassified(100); | Metschnikowia | 13 | Saprotroph | \|Undefined Saprotroph\| | Yeast | NULL | Probable | As nectar-tap_saprotroph (Põlme et al. 2020); Decay sugar-rich substrates (Põlme et al. 2020) | Põlme S, et al. 2020. Fungal Diversity 105:1-16 ((https://doi.org/10.1007/s13225-020-00466-2)) |
| Otu38425 | 1 | k__Fungi(100);p__Ascomycota(100);c__Saccharomycetes(100);o__Saccharomycetales(100);f__Metschnikowiaceae(100);g__Metschnikowia(100);s__Metschnikowia_pulcherrima(100); | Metschnikowia | 13 | Saprotroph | \|Undefined Saprotroph\| | Yeast | NULL | Probable | As nectar-tap_saprotroph (Põlme et al. 2020); Decay sugar-rich substrates (Põlme et al. 2020) | Põlme S, et al. 2020. Fungal Diversity 105:1-16 ((https://doi.org/10.1007/s13225-020-00466-2)) |
| Otu38426 | 1 | k__Fungi(100);p__Ascomycota(100);c__Saccharomycetes(100);o__Saccharomycetales(100);f__Metschnikowiaceae(100);g__Metschnikowia(100);s__Metschnikowia_pulcherrima(100); | Metschnikowia | 13 | Saprotroph | \|Undefined Saprotroph\| | Yeast | NULL | Probable | As nectar-tap_saprotroph (Põlme et al. 2020); Decay sugar-rich substrates (Põlme et al. 2020) | Põlme S, et al. 2020. Fungal Diversity 105:1-16 ((https://doi.org/10.1007/s13225-020-00466-2)) |
| Otu38427 | 1 | k__Fungi(100);p__Ascomycota(100);c__Saccharomycetes(100);o__Saccharomycetales(100);f__Metschnikowiaceae(100);g__Metschnikowia(100);g__Metschnikowia_unclassified(100); | Metschnikowia | 13 | Saprotroph | \|Undefined Saprotroph\| | Yeast | NULL | Probable | As nectar-tap_saprotroph (Põlme et al. 2020); Decay sugar-rich substrates (Põlme et al. 2020) | Põlme S, et al. 2020. Fungal Diversity 105:1-16 ((https://doi.org/10.1007/s13225-020-00466-2)) |
| Otu38429 | 1 | k__Fungi(100);p__Ascomycota(100);c__Saccharomycetes(100);o__Saccharomycetales(100);f__Metschnikowiaceae(100);g__Metschnikowia(100);s__Metschnikowia_pulcherrima(100); | Metschnikowia | 13 | Saprotroph | \|Undefined Saprotroph\| | Yeast | NULL | Probable | As nectar-tap_saprotroph (Põlme et al. 2020); Decay sugar-rich substrates (Põlme et al. 2020) | Põlme S, et al. 2020. Fungal Diversity 105:1-16 ((https://doi.org/10.1007/s13225-020-00466-2)) |
| Otu38430 | 1 | k__Fungi(100);p__Ascomycota(100);c__Saccharomycetes(100);o__Saccharomycetales(100);f__Metschnikowiaceae(100);g__Metschnikowia(100);g__Metschnikowia_unclassified(100); | Metschnikowia | 13 | Saprotroph | \|Undefined Saprotroph\| | Yeast | NULL | Probable | As nectar-tap_saprotroph (Põlme et al. 2020); Decay sugar-rich substrates (Põlme et al. 2020) | Põlme S, et al. 2020. Fungal Diversity 105:1-16 ((https://doi.org/10.1007/s13225-020-00466-2)) |
| Otu38431 | 1 | k__Fungi(100);p__Ascomycota(100);c__Saccharomycetes(100);o__Saccharomycetales(100);f__Metschnikowiaceae(100);g__Metschnikowia(100);s__Metschnikowia_pulcherrima(100); | Metschnikowia | 13 | Saprotroph | \|Undefined Saprotroph\| | Yeast | NULL | Probable | As nectar-tap_saprotroph (Põlme et al. 2020); Decay sugar-rich substrates (Põlme et al. 2020) | Põlme S, et al. 2020. Fungal Diversity 105:1-16 ((https://doi.org/10.1007/s13225-020-00466-2)) |
| Otu38433 | 1 | k__Fungi(100);p__Ascomycota(100);c__Saccharomycetes(100);o__Saccharomycetales(100);f__Metschnikowiaceae(100);g__Metschnikowia(100);s__Metschnikowia_pulcherrima(100); | Metschnikowia | 13 | Saprotroph | \|Undefined Saprotroph\| | Yeast | NULL | Probable | As nectar-tap_saprotroph (Põlme et al. 2020); Decay sugar-rich substrates (Põlme et al. 2020) | Põlme S, et al. 2020. Fungal Diversity 105:1-16 ((https://doi.org/10.1007/s13225-020-00466-2)) |
| Otu38434 | 1 | k__Fungi(100);p__Ascomycota(100);c__Saccharomycetes(100);o__Saccharomycetales(100);f__Metschnikowiaceae(100);g__Metschnikowia(100);s__Metschnikowia_pulcherrima(100); | Metschnikowia | 13 | Saprotroph | \|Undefined Saprotroph\| | Yeast | NULL | Probable | As nectar-tap_saprotroph (Põlme et al. 2020); Decay sugar-rich substrates (Põlme et al. 2020) | Põlme S, et al. 2020. Fungal Diversity 105:1-16 ((https://doi.org/10.1007/s13225-020-00466-2)) |
| Otu38435 | 1 | k__Fungi(100);p__Ascomycota(100);c__Saccharomycetes(100);o__Saccharomycetales(100);f__Metschnikowiaceae(100);g__Metschnikowia(100);s__Metschnikowia_pulcherrima(100); | Metschnikowia | 13 | Saprotroph | \|Undefined Saprotroph\| | Yeast | NULL | Probable | As nectar-tap_saprotroph (Põlme et al. 2020); Decay sugar-rich substrates (Põlme et al. 2020) | Põlme S, et al. 2020. Fungal Diversity 105:1-16 ((https://doi.org/10.1007/s13225-020-00466-2)) |
| Otu38436 | 1 | k__Fungi(100);p__Ascomycota(100);c__Saccharomycetes(100);o__Saccharomycetales(100);f__Metschnikowiaceae(100);g__Metschnikowia(100);s__Metschnikowia_pulcherrima(100); | Metschnikowia | 13 | Saprotroph | \|Undefined Saprotroph\| | Yeast | NULL | Probable | As nectar-tap_saprotroph (Põlme et al. 2020); Decay sugar-rich substrates (Põlme et al. 2020) | Põlme S, et al. 2020. Fungal Diversity 105:1-16 ((https://doi.org/10.1007/s13225-020-00466-2)) |
| Otu38437 | 1 | k__Fungi(100);p__Ascomycota(100);c__Saccharomycetes(100);o__Saccharomycetales(100);f__Metschnikowiaceae(100);g__Metschnikowia(100);s__Metschnikowia_pulcherrima(100); | Metschnikowia | 13 | Saprotroph | \|Undefined Saprotroph\| | Yeast | NULL | Probable | As nectar-tap_saprotroph (Põlme et al. 2020); Decay sugar-rich substrates (Põlme et al. 2020) | Põlme S, et al. 2020. Fungal Diversity 105:1-16 ((https://doi.org/10.1007/s13225-020-00466-2)) |
| Otu38438 | 1 | k__Fungi(100);p__Ascomycota(100);c__Saccharomycetes(100);o__Saccharomycetales(100);f__Metschnikowiaceae(100);g__Metschnikowia(100);s__Metschnikowia_pulcherrima(100); | Metschnikowia | 13 | Saprotroph | \|Undefined Saprotroph\| | Yeast | NULL | Probable | As nectar-tap_saprotroph (Põlme et al. 2020); Decay sugar-rich substrates (Põlme et al. 2020) | Põlme S, et al. 2020. Fungal Diversity 105:1-16 ((https://doi.org/10.1007/s13225-020-00466-2)) |
| Otu38439 | 1 | k__Fungi(100);p__Ascomycota(100);c__Saccharomycetes(100);o__Saccharomycetales(100);f__Metschnikowiaceae(100);g__Metschnikowia(100);s__Metschnikowia_pulcherrima(100); | Metschnikowia | 13 | Saprotroph | \|Undefined Saprotroph\| | Yeast | NULL | Probable | As nectar-tap_saprotroph (Põlme et al. 2020); Decay sugar-rich substrates (Põlme et al. 2020) | Põlme S, et al. 2020. Fungal Diversity 105:1-16 ((https://doi.org/10.1007/s13225-020-00466-2)) |
| Otu38440 | 1 | k__Fungi(100);p__Ascomycota(100);c__Saccharomycetes(100);o__Saccharomycetales(100);f__Metschnikowiaceae(100);g__Metschnikowia(100);s__Metschnikowia_pulcherrima(100); | Metschnikowia | 13 | Saprotroph | \|Undefined Saprotroph\| | Yeast | NULL | Probable | As nectar-tap_saprotroph (Põlme et al. 2020); Decay sugar-rich substrates (Põlme et al. 2020) | Põlme S, et al. 2020. Fungal Diversity 105:1-16 ((https://doi.org/10.1007/s13225-020-00466-2)) |
| Otu38441 | 1 | k__Fungi(100);p__Ascomycota(100);c__Saccharomycetes(100);o__Saccharomycetales(100);f__Metschnikowiaceae(100);g__Metschnikowia(100);g__Metschnikowia_unclassified(100); | Metschnikowia | 13 | Saprotroph | \|Undefined Saprotroph\| | Yeast | NULL | Probable | As nectar-tap_saprotroph (Põlme et al. 2020); Decay sugar-rich substrates (Põlme et al. 2020) | Põlme S, et al. 2020. Fungal Diversity 105:1-16 ((https://doi.org/10.1007/s13225-020-00466-2)) |
| Otu38443 | 1 | k__Fungi(100);p__Ascomycota(100);c__Saccharomycetes(100);o__Saccharomycetales(100);f__Metschnikowiaceae(100);g__Metschnikowia(100);s__Metschnikowia_pulcherrima(100); | Metschnikowia | 13 | Saprotroph | \|Undefined Saprotroph\| | Yeast | NULL | Probable | As nectar-tap_saprotroph (Põlme et al. 2020); Decay sugar-rich substrates (Põlme et al. 2020) | Põlme S, et al. 2020. Fungal Diversity 105:1-16 ((https://doi.org/10.1007/s13225-020-00466-2)) |
| Otu38444 | 1 | k__Fungi(100);p__Ascomycota(100);c__Saccharomycetes(100);o__Saccharomycetales(100);f__Metschnikowiaceae(100);g__Metschnikowia(100);s__Metschnikowia_pulcherrima(100); | Metschnikowia | 13 | Saprotroph | \|Undefined Saprotroph\| | Yeast | NULL | Probable | As nectar-tap_saprotroph (Põlme et al. 2020); Decay sugar-rich substrates (Põlme et al. 2020) | Põlme S, et al. 2020. Fungal Diversity 105:1-16 ((https://doi.org/10.1007/s13225-020-00466-2)) |
| Otu38445 | 1 | k__Fungi(100);p__Ascomycota(100);c__Saccharomycetes(100);o__Saccharomycetales(100);f__Metschnikowiaceae(100);g__Metschnikowia(100);s__Metschnikowia_pulcherrima(100); | Metschnikowia | 13 | Saprotroph | \|Undefined Saprotroph\| | Yeast | NULL | Probable | As nectar-tap_saprotroph (Põlme et al. 2020); Decay sugar-rich substrates (Põlme et al. 2020) | Põlme S, et al. 2020. Fungal Diversity 105:1-16 ((https://doi.org/10.1007/s13225-020-00466-2)) |
| Otu38446 | 1 | k__Fungi(100);p__Ascomycota(100);c__Saccharomycetes(100);o__Saccharomycetales(100);f__Metschnikowiaceae(100);g__Metschnikowia(100);s__Metschnikowia_pulcherrima(100); | Metschnikowia | 13 | Saprotroph | \|Undefined Saprotroph\| | Yeast | NULL | Probable | As nectar-tap_saprotroph (Põlme et al. 2020); Decay sugar-rich substrates (Põlme et al. 2020) | Põlme S, et al. 2020. Fungal Diversity 105:1-16 ((https://doi.org/10.1007/s13225-020-00466-2)) |
| Otu38447 | 1 | k__Fungi(100);p__Ascomycota(100);c__Saccharomycetes(100);o__Saccharomycetales(100);f__Metschnikowiaceae(100);g__Metschnikowia(100);s__Metschnikowia_pulcherrima(100); | Metschnikowia | 13 | Saprotroph | \|Undefined Saprotroph\| | Yeast | NULL | Probable | As nectar-tap_saprotroph (Põlme et al. 2020); Decay sugar-rich substrates (Põlme et al. 2020) | Põlme S, et al. 2020. Fungal Diversity 105:1-16 ((https://doi.org/10.1007/s13225-020-00466-2)) |
| Otu38448 | 1 | k__Fungi(100);p__Ascomycota(100);c__Saccharomycetes(100);o__Saccharomycetales(100);f__Metschnikowiaceae(100);g__Metschnikowia(100);s__Metschnikowia_pulcherrima(100); | Metschnikowia | 13 | Saprotroph | \|Undefined Saprotroph\| | Yeast | NULL | Probable | As nectar-tap_saprotroph (Põlme et al. 2020); Decay sugar-rich substrates (Põlme et al. 2020) | Põlme S, et al. 2020. Fungal Diversity 105:1-16 ((https://doi.org/10.1007/s13225-020-00466-2)) |
| Otu38449 | 1 | k__Fungi(100);p__Ascomycota(100);c__Saccharomycetes(100);o__Saccharomycetales(100);f__Metschnikowiaceae(100);g__Metschnikowia(100);s__Metschnikowia_pulcherrima(100); | Metschnikowia | 13 | Saprotroph | \|Undefined Saprotroph\| | Yeast | NULL | Probable | As nectar-tap_saprotroph (Põlme et al. 2020); Decay sugar-rich substrates (Põlme et al. 2020) | Põlme S, et al. 2020. Fungal Diversity 105:1-16 ((https://doi.org/10.1007/s13225-020-00466-2)) |
| Otu38450 | 1 | k__Fungi(100);p__Ascomycota(100);c__Saccharomycetes(100);o__Saccharomycetales(100);f__Metschnikowiaceae(100);g__Metschnikowia(100);s__Metschnikowia_pulcherrima(100); | Metschnikowia | 13 | Saprotroph | \|Undefined Saprotroph\| | Yeast | NULL | Probable | As nectar-tap_saprotroph (Põlme et al. 2020); Decay sugar-rich substrates (Põlme et al. 2020) | Põlme S, et al. 2020. Fungal Diversity 105:1-16 ((https://doi.org/10.1007/s13225-020-00466-2)) |
| Otu38451 | 1 | k__Fungi(100);p__Ascomycota(100);c__Saccharomycetes(100);o__Saccharomycetales(100);f__Metschnikowiaceae(100);g__Metschnikowia(100);s__Metschnikowia_pulcherrima(100); | Metschnikowia | 13 | Saprotroph | \|Undefined Saprotroph\| | Yeast | NULL | Probable | As nectar-tap_saprotroph (Põlme et al. 2020); Decay sugar-rich substrates (Põlme et al. 2020) | Põlme S, et al. 2020. Fungal Diversity 105:1-16 ((https://doi.org/10.1007/s13225-020-00466-2)) |
| Otu38452 | 1 | k__Fungi(100);p__Ascomycota(100);c__Saccharomycetes(100);o__Saccharomycetales(100);f__Metschnikowiaceae(100);g__Metschnikowia(100);s__Metschnikowia_pulcherrima(100); | Metschnikowia | 13 | Saprotroph | \|Undefined Saprotroph\| | Yeast | NULL | Probable | As nectar-tap_saprotroph (Põlme et al. 2020); Decay sugar-rich substrates (Põlme et al. 2020) | Põlme S, et al. 2020. Fungal Diversity 105:1-16 ((https://doi.org/10.1007/s13225-020-00466-2)) |
| Otu38453 | 1 | k__Fungi(100);p__Ascomycota(100);c__Saccharomycetes(100);o__Saccharomycetales(100);f__Metschnikowiaceae(100);g__Metschnikowia(100);s__Metschnikowia_pulcherrima(100); | Metschnikowia | 13 | Saprotroph | \|Undefined Saprotroph\| | Yeast | NULL | Probable | As nectar-tap_saprotroph (Põlme et al. 2020); Decay sugar-rich substrates (Põlme et al. 2020) | Põlme S, et al. 2020. Fungal Diversity 105:1-16 ((https://doi.org/10.1007/s13225-020-00466-2)) |
| Otu38457 | 1 | k__Fungi(100);p__Ascomycota(100);c__Saccharomycetes(100);o__Saccharomycetales(100);f__Metschnikowiaceae(100);g__Metschnikowia(100);s__Metschnikowia_pulcherrima(100); | Metschnikowia | 13 | Saprotroph | \|Undefined Saprotroph\| | Yeast | NULL | Probable | As nectar-tap_saprotroph (Põlme et al. 2020); Decay sugar-rich substrates (Põlme et al. 2020) | Põlme S, et al. 2020. Fungal Diversity 105:1-16 ((https://doi.org/10.1007/s13225-020-00466-2)) |
| Otu38458 | 1 | k__Fungi(100);p__Ascomycota(100);c__Saccharomycetes(100);o__Saccharomycetales(100);f__Metschnikowiaceae(100);g__Metschnikowia(100);s__Metschnikowia_pulcherrima(100); | Metschnikowia | 13 | Saprotroph | \|Undefined Saprotroph\| | Yeast | NULL | Probable | As nectar-tap_saprotroph (Põlme et al. 2020); Decay sugar-rich substrates (Põlme et al. 2020) | Põlme S, et al. 2020. Fungal Diversity 105:1-16 ((https://doi.org/10.1007/s13225-020-00466-2)) |
| Otu38459 | 1 | k__Fungi(100);p__Ascomycota(100);c__Saccharomycetes(100);o__Saccharomycetales(100);f__Metschnikowiaceae(100);g__Metschnikowia(100);s__Metschnikowia_pulcherrima(100); | Metschnikowia | 13 | Saprotroph | \|Undefined Saprotroph\| | Yeast | NULL | Probable | As nectar-tap_saprotroph (Põlme et al. 2020); Decay sugar-rich substrates (Põlme et al. 2020) | Põlme S, et al. 2020. Fungal Diversity 105:1-16 ((https://doi.org/10.1007/s13225-020-00466-2)) |
| Otu38460 | 1 | k__Fungi(100);p__Ascomycota(100);c__Saccharomycetes(100);o__Saccharomycetales(100);f__Metschnikowiaceae(100);g__Metschnikowia(100);s__Metschnikowia_pulcherrima(100); | Metschnikowia | 13 | Saprotroph | \|Undefined Saprotroph\| | Yeast | NULL | Probable | As nectar-tap_saprotroph (Põlme et al. 2020); Decay sugar-rich substrates (Põlme et al. 2020) | Põlme S, et al. 2020. Fungal Diversity 105:1-16 ((https://doi.org/10.1007/s13225-020-00466-2)) |
| Otu38461 | 1 | k__Fungi(100);p__Ascomycota(100);c__Saccharomycetes(100);o__Saccharomycetales(100);f__Metschnikowiaceae(100);g__Metschnikowia(100);s__Metschnikowia_pulcherrima(100); | Metschnikowia | 13 | Saprotroph | \|Undefined Saprotroph\| | Yeast | NULL | Probable | As nectar-tap_saprotroph (Põlme et al. 2020); Decay sugar-rich substrates (Põlme et al. 2020) | Põlme S, et al. 2020. Fungal Diversity 105:1-16 ((https://doi.org/10.1007/s13225-020-00466-2)) |
| Otu38462 | 1 | k__Fungi(100);p__Ascomycota(100);c__Saccharomycetes(100);o__Saccharomycetales(100);f__Metschnikowiaceae(100);g__Metschnikowia(100);s__Metschnikowia_pulcherrima(100); | Metschnikowia | 13 | Saprotroph | \|Undefined Saprotroph\| | Yeast | NULL | Probable | As nectar-tap_saprotroph (Põlme et al. 2020); Decay sugar-rich substrates (Põlme et al. 2020) | Põlme S, et al. 2020. Fungal Diversity 105:1-16 ((https://doi.org/10.1007/s13225-020-00466-2)) |
| Otu38463 | 1 | k__Fungi(100);p__Ascomycota(100);c__Saccharomycetes(100);o__Saccharomycetales(100);f__Metschnikowiaceae(100);g__Metschnikowia(100);s__Metschnikowia_pulcherrima(100); | Metschnikowia | 13 | Saprotroph | \|Undefined Saprotroph\| | Yeast | NULL | Probable | As nectar-tap_saprotroph (Põlme et al. 2020); Decay sugar-rich substrates (Põlme et al. 2020) | Põlme S, et al. 2020. Fungal Diversity 105:1-16 ((https://doi.org/10.1007/s13225-020-00466-2)) |

| | | | | | | | | | | | |
|---|---|---|---|---|---|---|---|---|---|---|---|
| Otu38464 | 1 | k__Fungi(100);p__Ascomycota(100);c__Saccharomycetes(100);o__Saccharomycetales(100);f__Metschnikowiaceae(100);g__Metschnikowia(100);s__Metschnikowia_pulcherrima(100); | Metschnikowia | 13 | Saprotroph | \|Undefined Saprotroph\| | Yeast | NULL | Probable | As nectar-tap_saprotroph (Põlme et al. 2020); Decay sugar-rich substrates (Põlme et al. 2020) | Põlme S, et al. 2020. Fungal Diversity 105:1-16 ((https://doi.org/10.1007/s13225-020-00466-2)) |
| Otu38465 | 1 | k__Fungi(100);p__Ascomycota(100);c__Saccharomycetes(100);o__Saccharomycetales(100);f__Metschnikowiaceae(100);g__Metschnikowia(100);s__Metschnikowia_pulcherrima(100); | Metschnikowia | 13 | Saprotroph | \|Undefined Saprotroph\| | Yeast | NULL | Probable | As nectar-tap_saprotroph (Põlme et al. 2020); Decay sugar-rich substrates (Põlme et al. 2020) | Põlme S, et al. 2020. Fungal Diversity 105:1-16 ((https://doi.org/10.1007/s13225-020-00466-2)) |
| Otu38466 | 1 | k__Fungi(100);p__Ascomycota(100);c__Saccharomycetes(100);o__Saccharomycetales(100);f__Metschnikowiaceae(100);g__Metschnikowia(100);s__Metschnikowia_pulcherrima(100); | Metschnikowia | 13 | Saprotroph | \|Undefined Saprotroph\| | Yeast | NULL | Probable | As nectar-tap_saprotroph (Põlme et al. 2020); Decay sugar-rich substrates (Põlme et al. 2020) | Põlme S, et al. 2020. Fungal Diversity 105:1-16 ((https://doi.org/10.1007/s13225-020-00466-2)) |
| Otu38467 | 1 | k__Fungi(100);p__Ascomycota(100);c__Saccharomycetes(100);o__Saccharomycetales(100);f__Metschnikowiaceae(100);g__Metschnikowia(100);s__Metschnikowia_pulcherrima(100); | Metschnikowia | 13 | Saprotroph | \|Undefined Saprotroph\| | Yeast | NULL | Probable | As nectar-tap_saprotroph (Põlme et al. 2020); Decay sugar-rich substrates (Põlme et al. 2020) | Põlme S, et al. 2020. Fungal Diversity 105:1-16 ((https://doi.org/10.1007/s13225-020-00466-2)) |
| Otu38469 | 1 | k__Fungi(100);p__Ascomycota(100);c__Saccharomycetes(100);o__Saccharomycetales(100);f__Metschnikowiaceae(100);g__Metschnikowia(100);s__Metschnikowia_pulcherrima(100); | Metschnikowia | 13 | Saprotroph | \|Undefined Saprotroph\| | Yeast | NULL | Probable | As nectar-tap_saprotroph (Põlme et al. 2020); Decay sugar-rich substrates (Põlme et al. 2020) | Põlme S, et al. 2020. Fungal Diversity 105:1-16 ((https://doi.org/10.1007/s13225-020-00466-2)) |
| Otu38470 | 1 | k__Fungi(100);p__Ascomycota(100);c__Saccharomycetes(100);o__Saccharomycetales(100);f__Metschnikowiaceae(100);g__Metschnikowia(100);s__Metschnikowia_pulcherrima(100); | Metschnikowia | 13 | Saprotroph | \|Undefined Saprotroph\| | Yeast | NULL | Probable | As nectar-tap_saprotroph (Põlme et al. 2020); Decay sugar-rich substrates (Põlme et al. 2020) | Põlme S, et al. 2020. Fungal Diversity 105:1-16 ((https://doi.org/10.1007/s13225-020-00466-2)) |
| Otu38471 | 1 | k__Fungi(100);p__Ascomycota(100);c__Saccharomycetes(100);o__Saccharomycetales(100);f__Metschnikowiaceae(100);g__Metschnikowia(100);s__Metschnikowia_pulcherrima(100); | Metschnikowia | 13 | Saprotroph | \|Undefined Saprotroph\| | Yeast | NULL | Probable | As nectar-tap_saprotroph (Põlme et al. 2020); Decay sugar-rich substrates (Põlme et al. 2020) | Põlme S, et al. 2020. Fungal Diversity 105:1-16 ((https://doi.org/10.1007/s13225-020-00466-2)) |
| Otu38472 | 1 | k__Fungi(100);p__Ascomycota(100);c__Saccharomycetes(100);o__Saccharomycetales(100);f__Metschnikowiaceae(100);g__Metschnikowia(100);s__Metschnikowia_pulcherrima(100); | Metschnikowia | 13 | Saprotroph | \|Undefined Saprotroph\| | Yeast | NULL | Probable | As nectar-tap_saprotroph (Põlme et al. 2020); Decay sugar-rich substrates (Põlme et al. 2020) | Põlme S, et al. 2020. Fungal Diversity 105:1-16 ((https://doi.org/10.1007/s13225-020-00466-2)) |
| Otu38473 | 1 | k__Fungi(100);p__Ascomycota(100);c__Saccharomycetes(100);o__Saccharomycetales(100);f__Metschnikowiaceae(100);g__Metschnikowia(100);g__Metschnikowia_unclassified(100); | Metschnikowia | 13 | Saprotroph | \|Undefined Saprotroph\| | Yeast | NULL | Probable | As nectar-tap_saprotroph (Põlme et al. 2020); Decay sugar-rich substrates (Põlme et al. 2020) | Põlme S, et al. 2020. Fungal Diversity 105:1-16 ((https://doi.org/10.1007/s13225-020-00466-2)) |
| Otu38474 | 1 | k__Fungi(100);p__Ascomycota(100);c__Saccharomycetes(100);o__Saccharomycetales(100);f__Metschnikowiaceae(100);g__Metschnikowia(100);s__Metschnikowia_pulcherrima(100); | Metschnikowia | 13 | Saprotroph | \|Undefined Saprotroph\| | Yeast | NULL | Probable | As nectar-tap_saprotroph (Põlme et al. 2020); Decay sugar-rich substrates (Põlme et al. 2020) | Põlme S, et al. 2020. Fungal Diversity 105:1-16 ((https://doi.org/10.1007/s13225-020-00466-2)) |
| Otu38475 | 1 | k__Fungi(100);p__Ascomycota(100);c__Saccharomycetes(100);o__Saccharomycetales(100);f__Metschnikowiaceae(100);g__Metschnikowia(100);s__Metschnikowia_pulcherrima(100); | Metschnikowia | 13 | Saprotroph | \|Undefined Saprotroph\| | Yeast | NULL | Probable | As nectar-tap_saprotroph (Põlme et al. 2020); Decay sugar-rich substrates (Põlme et al. 2020) | Põlme S, et al. 2020. Fungal Diversity 105:1-16 ((https://doi.org/10.1007/s13225-020-00466-2)) |
| Otu38476 | 1 | k__Fungi(100);p__Ascomycota(100);c__Saccharomycetes(100);o__Saccharomycetales(100);f__Metschnikowiaceae(100);g__Metschnikowia(100);s__Metschnikowia_pulcherrima(100); | Metschnikowia | 13 | Saprotroph | \|Undefined Saprotroph\| | Yeast | NULL | Probable | As nectar-tap_saprotroph (Põlme et al. 2020); Decay sugar-rich substrates (Põlme et al. 2020) | Põlme S, et al. 2020. Fungal Diversity 105:1-16 ((https://doi.org/10.1007/s13225-020-00466-2)) |
| Otu38477 | 1 | k__Fungi(100);p__Ascomycota(100);c__Saccharomycetes(100);o__Saccharomycetales(100);f__Metschnikowiaceae(100);g__Metschnikowia(100);s__Metschnikowia_pulcherrima(100); | Metschnikowia | 13 | Saprotroph | \|Undefined Saprotroph\| | Yeast | NULL | Probable | As nectar-tap_saprotroph (Põlme et al. 2020); Decay sugar-rich substrates (Põlme et al. 2020) | Põlme S, et al. 2020. Fungal Diversity 105:1-16 ((https://doi.org/10.1007/s13225-020-00466-2)) |
| Otu38479 | 1 | k__Fungi(100);p__Ascomycota(100);c__Saccharomycetes(100);o__Saccharomycetales(100);f__Metschnikowiaceae(100);g__Metschnikowia(100);g__Metschnikowia_unclassified(100); | Metschnikowia | 13 | Saprotroph | \|Undefined Saprotroph\| | Yeast | NULL | Probable | As nectar-tap_saprotroph (Põlme et al. 2020); Decay sugar-rich substrates (Põlme et al. 2020) | Põlme S, et al. 2020. Fungal Diversity 105:1-16 ((https://doi.org/10.1007/s13225-020-00466-2)) |
| Otu38480 | 1 | k__Fungi(100);p__Ascomycota(100);c__Saccharomycetes(100);o__Saccharomycetales(100);f__Metschnikowiaceae(100);g__Metschnikowia(100);s__Metschnikowia_pulcherrima(100); | Metschnikowia | 13 | Saprotroph | \|Undefined Saprotroph\| | Yeast | NULL | Probable | As nectar-tap_saprotroph (Põlme et al. 2020); Decay sugar-rich substrates (Põlme et al. 2020) | Põlme S, et al. 2020. Fungal Diversity 105:1-16 ((https://doi.org/10.1007/s13225-020-00466-2)) |
| Otu38481 | 1 | k__Fungi(100);p__Ascomycota(100);c__Saccharomycetes(100);o__Saccharomycetales(100);f__Metschnikowiaceae(100);g__Metschnikowia(100);s__Metschnikowia_pulcherrima(100); | Metschnikowia | 13 | Saprotroph | \|Undefined Saprotroph\| | Yeast | NULL | Probable | As nectar-tap_saprotroph (Põlme et al. 2020); Decay sugar-rich substrates (Põlme et al. 2020) | Põlme S, et al. 2020. Fungal Diversity 105:1-16 ((https://doi.org/10.1007/s13225-020-00466-2)) |
| Otu38482 | 1 | k__Fungi(100);p__Ascomycota(100);c__Saccharomycetes(100);o__Saccharomycetales(100);f__Metschnikowiaceae(100);g__Metschnikowia(100);s__Metschnikowia_pulcherrima(100); | Metschnikowia | 13 | Saprotroph | \|Undefined Saprotroph\| | Yeast | NULL | Probable | As nectar-tap_saprotroph (Põlme et al. 2020); Decay sugar-rich substrates (Põlme et al. 2020) | Põlme S, et al. 2020. Fungal Diversity 105:1-16 ((https://doi.org/10.1007/s13225-020-00466-2)) |
| Otu38483 | 1 | k__Fungi(100);p__Ascomycota(100);c__Saccharomycetes(100);o__Saccharomycetales(100);f__Metschnikowiaceae(100);g__Metschnikowia(100);s__Metschnikowia_pulcherrima(100); | Metschnikowia | 13 | Saprotroph | \|Undefined Saprotroph\| | Yeast | NULL | Probable | As nectar-tap_saprotroph (Põlme et al. 2020); Decay sugar-rich substrates (Põlme et al. 2020) | Põlme S, et al. 2020. Fungal Diversity 105:1-16 ((https://doi.org/10.1007/s13225-020-00466-2)) |
| Otu38484 | 1 | k__Fungi(100);p__Ascomycota(100);c__Saccharomycetes(100);o__Saccharomycetales(100);f__Metschnikowiaceae(100);g__Metschnikowia(100);s__Metschnikowia_pulcherrima(100); | Metschnikowia | 13 | Saprotroph | \|Undefined Saprotroph\| | Yeast | NULL | Probable | As nectar-tap_saprotroph (Põlme et al. 2020); Decay sugar-rich substrates (Põlme et al. 2020) | Põlme S, et al. 2020. Fungal Diversity 105:1-16 ((https://doi.org/10.1007/s13225-020-00466-2)) |
| Otu38485 | 1 | k__Fungi(100);p__Ascomycota(100);c__Saccharomycetes(100);o__Saccharomycetales(100);f__Metschnikowiaceae(100);g__Metschnikowia(100);s__Metschnikowia_pulcherrima(100); | Metschnikowia | 13 | Saprotroph | \|Undefined Saprotroph\| | Yeast | NULL | Probable | As nectar-tap_saprotroph (Põlme et al. 2020); Decay sugar-rich substrates (Põlme et al. 2020) | Põlme S, et al. 2020. Fungal Diversity 105:1-16 ((https://doi.org/10.1007/s13225-020-00466-2)) |
| Otu38486 | 1 | k__Fungi(100);p__Ascomycota(100);c__Saccharomycetes(100);o__Saccharomycetales(100);f__Metschnikowiaceae(100);g__Metschnikowia(100);g__Metschnikowia_unclassified(100); | Metschnikowia | 13 | Saprotroph | \|Undefined Saprotroph\| | Yeast | NULL | Probable | As nectar-tap_saprotroph (Põlme et al. 2020); Decay sugar-rich substrates (Põlme et al. 2020) | Põlme S, et al. 2020. Fungal Diversity 105:1-16 ((https://doi.org/10.1007/s13225-020-00466-2)) |
| Otu38487 | 1 | k__Fungi(100);p__Ascomycota(100);c__Saccharomycetes(100);o__Saccharomycetales(100);f__Metschnikowiaceae(100);g__Metschnikowia(100);s__Metschnikowia_pulcherrima(100); | Metschnikowia | 13 | Saprotroph | \|Undefined Saprotroph\| | Yeast | NULL | Probable | As nectar-tap_saprotroph (Põlme et al. 2020); Decay sugar-rich substrates (Põlme et al. 2020) | Põlme S, et al. 2020. Fungal Diversity 105:1-16 ((https://doi.org/10.1007/s13225-020-00466-2)) |
| Otu38488 | 1 | k__Fungi(100);p__Ascomycota(100);c__Saccharomycetes(100);o__Saccharomycetales(100);f__Metschnikowiaceae(100);g__Metschnikowia(100);s__Metschnikowia_pulcherrima(100); | Metschnikowia | 13 | Saprotroph | \|Undefined Saprotroph\| | Yeast | NULL | Probable | As nectar-tap_saprotroph (Põlme et al. 2020); Decay sugar-rich substrates (Põlme et al. 2020) | Põlme S, et al. 2020. Fungal Diversity 105:1-16 ((https://doi.org/10.1007/s13225-020-00466-2)) |
| Otu38489 | 1 | k__Fungi(100);p__Ascomycota(100);c__Saccharomycetes(100);o__Saccharomycetales(100);f__Metschnikowiaceae(100);g__Metschnikowia(100);s__Metschnikowia_pulcherrima(100); | Metschnikowia | 13 | Saprotroph | \|Undefined Saprotroph\| | Yeast | NULL | Probable | As nectar-tap_saprotroph (Põlme et al. 2020); Decay sugar-rich substrates (Põlme et al. 2020) | Põlme S, et al. 2020. Fungal Diversity 105:1-16 ((https://doi.org/10.1007/s13225-020-00466-2)) |
| Otu38490 | 1 | k__Fungi(100);p__Ascomycota(100);c__Saccharomycetes(100);o__Saccharomycetales(100);f__Metschnikowiaceae(100);g__Metschnikowia(100);s__Metschnikowia_pulcherrima(100); | Metschnikowia | 13 | Saprotroph | \|Undefined Saprotroph\| | Yeast | NULL | Probable | As nectar-tap_saprotroph (Põlme et al. 2020); Decay sugar-rich substrates (Põlme et al. 2020) | Põlme S, et al. 2020. Fungal Diversity 105:1-16 ((https://doi.org/10.1007/s13225-020-00466-2)) |
| Otu38491 | 1 | k__Fungi(100);p__Ascomycota(100);c__Saccharomycetes(100);o__Saccharomycetales(100);f__Metschnikowiaceae(100);g__Metschnikowia(100);g__Metschnikowia_unclassified(100); | Metschnikowia | 13 | Saprotroph | \|Undefined Saprotroph\| | Yeast | NULL | Probable | As nectar-tap_saprotroph (Põlme et al. 2020); Decay sugar-rich substrates (Põlme et al. 2020) | Põlme S, et al. 2020. Fungal Diversity 105:1-16 ((https://doi.org/10.1007/s13225-020-00466-2)) |
| Otu38492 | 1 | k__Fungi(100);p__Ascomycota(100);c__Saccharomycetes(100);o__Saccharomycetales(100);f__Metschnikowiaceae(100);g__Metschnikowia(100);s__Metschnikowia_pulcherrima(100); | Metschnikowia | 13 | Saprotroph | \|Undefined Saprotroph\| | Yeast | NULL | Probable | As nectar-tap_saprotroph (Põlme et al. 2020); Decay sugar-rich substrates (Põlme et al. 2020) | Põlme S, et al. 2020. Fungal Diversity 105:1-16 ((https://doi.org/10.1007/s13225-020-00466-2)) |
| Otu38493 | 1 | k__Fungi(100);p__Ascomycota(100);c__Saccharomycetes(100);o__Saccharomycetales(100);f__Metschnikowiaceae(100);g__Metschnikowia(100);s__Metschnikowia_pulcherrima(100); | Metschnikowia | 13 | Saprotroph | \|Undefined Saprotroph\| | Yeast | NULL | Probable | As nectar-tap_saprotroph (Põlme et al. 2020); Decay sugar-rich substrates (Põlme et al. 2020) | Põlme S, et al. 2020. Fungal Diversity 105:1-16 ((https://doi.org/10.1007/s13225-020-00466-2)) |
| Otu38494 | 1 | k__Fungi(100);p__Ascomycota(100);c__Saccharomycetes(100);o__Saccharomycetales(100);f__Metschnikowiaceae(100);g__Metschnikowia(100);s__Metschnikowia_pulcherrima(100); | Metschnikowia | 13 | Saprotroph | \|Undefined Saprotroph\| | Yeast | NULL | Probable | As nectar-tap_saprotroph (Põlme et al. 2020); Decay sugar-rich substrates (Põlme et al. 2020) | Põlme S, et al. 2020. Fungal Diversity 105:1-16 ((https://doi.org/10.1007/s13225-020-00466-2)) |
| Otu38495 | 1 | k__Fungi(100);p__Ascomycota(100);c__Saccharomycetes(100);o__Saccharomycetales(100);f__Metschnikowiaceae(100);g__Metschnikowia(100);s__Metschnikowia_pulcherrima(100); | Metschnikowia | 13 | Saprotroph | \|Undefined Saprotroph\| | Yeast | NULL | Probable | As nectar-tap_saprotroph (Põlme et al. 2020); Decay sugar-rich substrates (Põlme et al. 2020) | Põlme S, et al. 2020. Fungal Diversity 105:1-16 ((https://doi.org/10.1007/s13225-020-00466-2)) |
| Otu38496 | 1 | k__Fungi(100);p__Ascomycota(100);c__Saccharomycetes(100);o__Saccharomycetales(100);f__Metschnikowiaceae(100);g__Metschnikowia(100);s__Metschnikowia_pulcherrima(100); | Metschnikowia | 13 | Saprotroph | \|Undefined Saprotroph\| | Yeast | NULL | Probable | As nectar-tap_saprotroph (Põlme et al. 2020); Decay sugar-rich substrates (Põlme et al. 2020) | Põlme S, et al. 2020. Fungal Diversity 105:1-16 ((https://doi.org/10.1007/s13225-020-00466-2)) |
| Otu38497 | 1 | k__Fungi(100);p__Ascomycota(100);c__Saccharomycetes(100);o__Saccharomycetales(100);f__Metschnikowiaceae(100);g__Metschnikowia(100);s__Metschnikowia_pulcherrima(100); | Metschnikowia | 13 | Saprotroph | \|Undefined Saprotroph\| | Yeast | NULL | Probable | As nectar-tap_saprotroph (Põlme et al. 2020); Decay sugar-rich substrates (Põlme et al. 2020) | Põlme S, et al. 2020. Fungal Diversity 105:1-16 ((https://doi.org/10.1007/s13225-020-00466-2)) |
| Otu38498 | 1 | k__Fungi(100);p__Ascomycota(100);c__Saccharomycetes(100);o__Saccharomycetales(100);f__Metschnikowiaceae(100);g__Metschnikowia(100);s__Metschnikowia_pulcherrima(100); | Metschnikowia | 13 | Saprotroph | \|Undefined Saprotroph\| | Yeast | NULL | Probable | As nectar-tap_saprotroph (Põlme et al. 2020); Decay sugar-rich substrates (Põlme et al. 2020) | Põlme S, et al. 2020. Fungal Diversity 105:1-16 ((https://doi.org/10.1007/s13225-020-00466-2)) |
| Otu38499 | 1 | k__Fungi(100);p__Ascomycota(100);c__Saccharomycetes(100);o__Saccharomycetales(100);f__Metschnikowiaceae(100);g__Metschnikowia(100);s__Metschnikowia_pulcherrima(100); | Metschnikowia | 13 | Saprotroph | \|Undefined Saprotroph\| | Yeast | NULL | Probable | As nectar-tap_saprotroph (Põlme et al. 2020); Decay sugar-rich substrates (Põlme et al. 2020) | Põlme S, et al. 2020. Fungal Diversity 105:1-16 ((https://doi.org/10.1007/s13225-020-00466-2)) |
| Otu38500 | 1 | k__Fungi(100);p__Ascomycota(100);c__Saccharomycetes(100);o__Saccharomycetales(100);f__Metschnikowiaceae(100);g__Metschnikowia(100);s__Metschnikowia_pulcherrima(100); | Metschnikowia | 13 | Saprotroph | \|Undefined Saprotroph\| | Yeast | NULL | Probable | As nectar-tap_saprotroph (Põlme et al. 2020); Decay sugar-rich substrates (Põlme et al. 2020) | Põlme S, et al. 2020. Fungal Diversity 105:1-16 ((https://doi.org/10.1007/s13225-020-00466-2)) |
| Otu38501 | 1 | k__Fungi(100);p__Ascomycota(100);c__Saccharomycetes(100);o__Saccharomycetales(100);f__Metschnikowiaceae(100);g__Metschnikowia(100);s__Metschnikowia_pulcherrima(100); | Metschnikowia | 13 | Saprotroph | \|Undefined Saprotroph\| | Yeast | NULL | Probable | As nectar-tap_saprotroph (Põlme et al. 2020); Decay sugar-rich substrates (Põlme et al. 2020) | Põlme S, et al. 2020. Fungal Diversity 105:1-16 ((https://doi.org/10.1007/s13225-020-00466-2)) |
| Otu38502 | 1 | k__Fungi(100);p__Ascomycota(100);c__Saccharomycetes(100);o__Saccharomycetales(100);f__Metschnikowiaceae(100);g__Metschnikowia(100);s__Metschnikowia_pulcherrima(100); | Metschnikowia | 13 | Saprotroph | \|Undefined Saprotroph\| | Yeast | NULL | Probable | As nectar-tap_saprotroph (Põlme et al. 2020); Decay sugar-rich substrates (Põlme et al. 2020) | Põlme S, et al. 2020. Fungal Diversity 105:1-16 ((https://doi.org/10.1007/s13225-020-00466-2)) |
| Otu38503 | 1 | k__Fungi(100);p__Ascomycota(100);c__Saccharomycetes(100);o__Saccharomycetales(100);f__Metschnikowiaceae(100);g__Metschnikowia(100);s__Metschnikowia_pulcherrima(100); | Metschnikowia | 13 | Saprotroph | \|Undefined Saprotroph\| | Yeast | NULL | Probable | As nectar-tap_saprotroph (Põlme et al. 2020); Decay sugar-rich substrates (Põlme et al. 2020) | Põlme S, et al. 2020. Fungal Diversity 105:1-16 ((https://doi.org/10.1007/s13225-020-00466-2)) |
| Otu38504 | 1 | k__Fungi(100);p__Ascomycota(100);c__Saccharomycetes(100);o__Saccharomycetales(100);f__Metschnikowiaceae(100);g__Metschnikowia(100);g__Metschnikowia_unclassified(100); | Metschnikowia | 13 | Saprotroph | \|Undefined Saprotroph\| | Yeast | NULL | Probable | As nectar-tap_saprotroph (Põlme et al. 2020); Decay sugar-rich substrates (Põlme et al. 2020) | Põlme S, et al. 2020. Fungal Diversity 105:1-16 ((https://doi.org/10.1007/s13225-020-00466-2)) |
| Otu38506 | 1 | k__Fungi(100);p__Ascomycota(100);c__Saccharomycetes(100);o__Saccharomycetales(100);f__Metschnikowiaceae(100);g__Metschnikowia(100);s__Metschnikowia_pulcherrima(100); | Metschnikowia | 13 | Saprotroph | \|Undefined Saprotroph\| | Yeast | NULL | Probable | As nectar-tap_saprotroph (Põlme et al. 2020); Decay sugar-rich substrates (Põlme et al. 2020) | Põlme S, et al. 2020. Fungal Diversity 105:1-16 ((https://doi.org/10.1007/s13225-020-00466-2)) |
| Otu38507 | 1 | k__Fungi(100);p__Ascomycota(100);c__Saccharomycetes(100);o__Saccharomycetales(100);f__Metschnikowiaceae(100);g__Metschnikowia(100);s__Metschnikowia_pulcherrima(100); | Metschnikowia | 13 | Saprotroph | \|Undefined Saprotroph\| | Yeast | NULL | Probable | As nectar-tap_saprotroph (Põlme et al. 2020); Decay sugar-rich substrates (Põlme et al. 2020) | Põlme S, et al. 2020. Fungal Diversity 105:1-16 ((https://doi.org/10.1007/s13225-020-00466-2)) |
| Otu38508 | 1 | k__Fungi(100);p__Ascomycota(100);c__Saccharomycetes(100);o__Saccharomycetales(100);f__Metschnikowiaceae(100);g__Metschnikowia(100);s__Metschnikowia_pulcherrima(100); | Metschnikowia | 13 | Saprotroph | \|Undefined Saprotroph\| | Yeast | NULL | Probable | As nectar-tap_saprotroph (Põlme et al. 2020); Decay sugar-rich substrates (Põlme et al. 2020) | Põlme S, et al. 2020. Fungal Diversity 105:1-16 ((https://doi.org/10.1007/s13225-020-00466-2)) |
| Otu38509 | 1 | k__Fungi(100);p__Ascomycota(100);c__Saccharomycetes(100);o__Saccharomycetales(100);f__Metschnikowiaceae(100);g__Metschnikowia(100);s__Metschnikowia_pulcherrima(100); | Metschnikowia | 13 | Saprotroph | \|Undefined Saprotroph\| | Yeast | NULL | Probable | As nectar-tap_saprotroph (Põlme et al. 2020); Decay sugar-rich substrates (Põlme et al. 2020) | Põlme S, et al. 2020. Fungal Diversity 105:1-16 ((https://doi.org/10.1007/s13225-020-00466-2)) |
| Otu38510 | 1 | k__Fungi(100);p__Ascomycota(100);c__Saccharomycetes(100);o__Saccharomycetales(100);f__Metschnikowiaceae(100);g__Metschnikowia(100);s__Metschnikowia_pulcherrima(100); | Metschnikowia | 13 | Saprotroph | \|Undefined Saprotroph\| | Yeast | NULL | Probable | As nectar-tap_saprotroph (Põlme et al. 2020); Decay sugar-rich substrates (Põlme et al. 2020) | Põlme S, et al. 2020. Fungal Diversity 105:1-16 ((https://doi.org/10.1007/s13225-020-00466-2)) |
| Otu38511 | 1 | k__Fungi(100);p__Ascomycota(100);c__Saccharomycetes(100);o__Saccharomycetales(100);f__Metschnikowiaceae(100);g__Metschnikowia(100);s__Metschnikowia_pulcherrima(100); | Metschnikowia | 13 | Saprotroph | \|Undefined Saprotroph\| | Yeast | NULL | Probable | As nectar-tap_saprotroph (Põlme et al. 2020); Decay sugar-rich substrates (Põlme et al. 2020) | Põlme S, et al. 2020. Fungal Diversity 105:1-16 ((https://doi.org/10.1007/s13225-020-00466-2)) |
| Otu38512 | 1 | k__Fungi(100);p__Ascomycota(100);c__Saccharomycetes(100);o__Saccharomycetales(100);f__Metschnikowiaceae(100);g__Metschnikowia(100);s__Metschnikowia_pulcherrima(100); | Metschnikowia | 13 | Saprotroph | \|Undefined Saprotroph\| | Yeast | NULL | Probable | As nectar-tap_saprotroph (Põlme et al. 2020); Decay sugar-rich substrates (Põlme et al. 2020) | Põlme S, et al. 2020. Fungal Diversity 105:1-16 ((https://doi.org/10.1007/s13225-020-00466-2)) |
| Otu38513 | 1 | k__Fungi(100);p__Ascomycota(100);c__Saccharomycetes(100);o__Saccharomycetales(100);f__Metschnikowiaceae(100);g__Metschnikowia(100);g__Metschnikowia_unclassified(100); | Metschnikowia | 13 | Saprotroph | \|Undefined Saprotroph\| | Yeast | NULL | Probable | As nectar-tap_saprotroph (Põlme et al. 2020); Decay sugar-rich substrates (Põlme et al. 2020) | Põlme S, et al. 2020. Fungal Diversity 105:1-16 ((https://doi.org/10.1007/s13225-020-00466-2)) |
| Otu38514 | 1 | k__Fungi(100);p__Ascomycota(100);c__Saccharomycetes(100);o__Saccharomycetales(100);f__Metschnikowiaceae(100);g__Metschnikowia(100);s__Metschnikowia_pulcherrima(100); | Metschnikowia | 13 | Saprotroph | \|Undefined Saprotroph\| | Yeast | NULL | Probable | As nectar-tap_saprotroph (Põlme et al. 2020); Decay sugar-rich substrates (Põlme et al. 2020) | Põlme S, et al. 2020. Fungal Diversity 105:1-16 ((https://doi.org/10.1007/s13225-020-00466-2)) |
| Otu38516 | 1 | k__Fungi(100);p__Ascomycota(100);c__Saccharomycetes(100);o__Saccharomycetales(100);f__Metschnikowiaceae(100);g__Metschnikowia(100);g__Metschnikowia_unclassified(100); | Metschnikowia | 13 | Saprotroph | \|Undefined Saprotroph\| | Yeast | NULL | Probable | As nectar-tap_saprotroph (Põlme et al. 2020); Decay sugar-rich substrates (Põlme et al. 2020) | Põlme S, et al. 2020. Fungal Diversity 105:1-16 ((https://doi.org/10.1007/s13225-020-00466-2)) |
| Otu38518 | 1 | k__Fungi(100);p__Ascomycota(100);c__Saccharomycetes(100);o__Saccharomycetales(100);f__Metschnikowiaceae(100);g__Metschnikowia(100);s__Metschnikowia_pulcherrima(100); | Metschnikowia | 13 | Saprotroph | \|Undefined Saprotroph\| | Yeast | NULL | Probable | As nectar-tap_saprotroph (Põlme et al. 2020); Decay sugar-rich substrates (Põlme et al. 2020) | Põlme S, et al. 2020. Fungal Diversity 105:1-16 ((https://doi.org/10.1007/s13225-020-00466-2)) |
| Otu38519 | 1 | k__Fungi(100);p__Ascomycota(100);c__Saccharomycetes(100);o__Saccharomycetales(100);f__Metschnikowiaceae(100);g__Metschnikowia(100);s__Metschnikowia_pulcherrima(100); | Metschnikowia | 13 | Saprotroph | \|Undefined Saprotroph\| | Yeast | NULL | Probable | As nectar-tap_saprotroph (Põlme et al. 2020); Decay sugar-rich substrates (Põlme et al. 2020) | Põlme S, et al. 2020. Fungal Diversity 105:1-16 ((https://doi.org/10.1007/s13225-020-00466-2)) |
| Otu38520 | 1 | k__Fungi(100);p__Ascomycota(100);c__Saccharomycetes(100);o__Saccharomycetales(100);f__Metschnikowiaceae(100);g__Metschnikowia(100);g__Metschnikowia_unclassified(100); | Metschnikowia | 13 | Saprotroph | \|Undefined Saprotroph\| | Yeast | NULL | Probable | As nectar-tap_saprotroph (Põlme et al. 2020); Decay sugar-rich substrates (Põlme et al. 2020) | Põlme S, et al. 2020. Fungal Diversity 105:1-16 ((https://doi.org/10.1007/s13225-020-00466-2)) |
| Otu38521 | 1 | k__Fungi(100);p__Ascomycota(100);c__Saccharomycetes(100);o__Saccharomycetales(100);f__Metschnikowiaceae(100);g__Metschnikowia(100);s__Metschnikowia_pulcherrima(100); | Metschnikowia | 13 | Saprotroph | \|Undefined Saprotroph\| | Yeast | NULL | Probable | As nectar-tap_saprotroph (Põlme et al. 2020); Decay sugar-rich substrates (Põlme et al. 2020) | Põlme S, et al. 2020. Fungal Diversity 105:1-16 ((https://doi.org/10.1007/s13225-020-00466-2)) |
| Otu38522 | 1 | k__Fungi(100);p__Ascomycota(100);c__Saccharomycetes(100);o__Saccharomycetales(100);f__Metschnikowiaceae(100);g__Metschnikowia(100);s__Metschnikowia_pulcherrima(100); | Metschnikowia | 13 | Saprotroph | \|Undefined Saprotroph\| | Yeast | NULL | Probable | As nectar-tap_saprotroph (Põlme et al. 2020); Decay sugar-rich substrates (Põlme et al. 2020) | Põlme S, et al. 2020. Fungal Diversity 105:1-16 ((https://doi.org/10.1007/s13225-020-00466-2)) |
| Otu38523 | 1 | k__Fungi(100);p__Ascomycota(100);c__Saccharomycetes(100);o__Saccharomycetales(100);f__Metschnikowiaceae(100);g__Metschnikowia(100);s__Metschnikowia_pulcherrima(100); | Metschnikowia | 13 | Saprotroph | \|Undefined Saprotroph\| | Yeast | NULL | Probable | As nectar-tap_saprotroph (Põlme et al. 2020); Decay sugar-rich substrates (Põlme et al. 2020) | Põlme S, et al. 2020. Fungal Diversity 105:1-16 ((https://doi.org/10.1007/s13225-020-00466-2)) |
| Otu38524 | 1 | k__Fungi(100);p__Ascomycota(100);c__Saccharomycetes(100);o__Saccharomycetales(100);f__Metschnikowiaceae(100);g__Metschnikowia(100);s__Metschnikowia_pulcherrima(100); | Metschnikowia | 13 | Saprotroph | \|Undefined Saprotroph\| | Yeast | NULL | Probable | As nectar-tap_saprotroph (Põlme et al. 2020); Decay sugar-rich substrates (Põlme et al. 2020) | Põlme S, et al. 2020. Fungal Diversity 105:1-16 ((https://doi.org/10.1007/s13225-020-00466-2)) |
| Otu38525 | 1 | k__Fungi(100);p__Ascomycota(100);c__Saccharomycetes(100);o__Saccharomycetales(100);f__Metschnikowiaceae(100);g__Metschnikowia(100);g__Metschnikowia_unclassified(100); | Metschnikowia | 13 | Saprotroph | \|Undefined Saprotroph\| | Yeast | NULL | Probable | As nectar-tap_saprotroph (Põlme et al. 2020); Decay sugar-rich substrates (Põlme et al. 2020) | Põlme S, et al. 2020. Fungal Diversity 105:1-16 ((https://doi.org/10.1007/s13225-020-00466-2)) |
| Otu38526 | 1 | k__Fungi(100);p__Ascomycota(100);c__Saccharomycetes(100);o__Saccharomycetales(100);f__Metschnikowiaceae(100);g__Metschnikowia(100);g__Metschnikowia_unclassified(100); | Metschnikowia | 13 | Saprotroph | \|Undefined Saprotroph\| | Yeast | NULL | Probable | As nectar-tap_saprotroph (Põlme et al. 2020); Decay sugar-rich substrates (Põlme et al. 2020) | Põlme S, et al. 2020. Fungal Diversity 105:1-16 ((https://doi.org/10.1007/s13225-020-00466-2)) |
| Otu38527 | 1 | k__Fungi(100);p__Ascomycota(100);c__Saccharomycetes(100);o__Saccharomycetales(100);f__Metschnikowiaceae(100);g__Metschnikowia(100);g__Metschnikowia_unclassified(100); | Metschnikowia | 13 | Saprotroph | \|Undefined Saprotroph\| | Yeast | NULL | Probable | As nectar-tap_saprotroph (Põlme et al. 2020); Decay sugar-rich substrates (Põlme et al. 2020) | Põlme S, et al. 2020. Fungal Diversity 105:1-16 ((https://doi.org/10.1007/s13225-020-00466-2)) |
| Otu38528 | 1 | k__Fungi(100);p__Ascomycota(100);c__Saccharomycetes(100);o__Saccharomycetales(100);f__Metschnikowiaceae(100);g__Metschnikowia(100);s__Metschnikowia_pulcherrima(100); | Metschnikowia | 13 | Saprotroph | \|Undefined Saprotroph\| | Yeast | NULL | Probable | As nectar-tap_saprotroph (Põlme et al. 2020); Decay sugar-rich substrates (Põlme et al. 2020) | Põlme S, et al. 2020. Fungal Diversity 105:1-16 ((https://doi.org/10.1007/s13225-020-00466-2)) |
| Otu38529 | 1 | k__Fungi(100);p__Ascomycota(100);c__Saccharomycetes(100);o__Saccharomycetales(100);f__Metschnikowiaceae(100);g__Metschnikowia(100);s__Metschnikowia_pulcherrima(100); | Metschnikowia | 13 | Saprotroph | \|Undefined Saprotroph\| | Yeast | NULL | Probable | As nectar-tap_saprotroph (Põlme et al. 2020); Decay sugar-rich substrates (Põlme et al. 2020) | Põlme S, et al. 2020. Fungal Diversity 105:1-16 ((https://doi.org/10.1007/s13225-020-00466-2)) |
| Otu38530 | 1 | k__Fungi(100);p__Ascomycota(100);c__Saccharomycetes(100);o__Saccharomycetales(100);f__Metschnikowiaceae(100);g__Metschnikowia(100);s__Metschnikowia_pulcherrima(100); | Metschnikowia | 13 | Saprotroph | \|Undefined Saprotroph\| | Yeast | NULL | Probable | As nectar-tap_saprotroph (Põlme et al. 2020); Decay sugar-rich substrates (Põlme et al. 2020) | Põlme S, et al. 2020. Fungal Diversity 105:1-16 ((https://doi.org/10.1007/s13225-020-00466-2)) |
| Otu38531 | 1 | k__Fungi(100);p__Ascomycota(100);c__Saccharomycetes(100);o__Saccharomycetales(100);f__Metschnikowiaceae(100);g__Metschnikowia(100);s__Metschnikowia_pulcherrima(100); | Metschnikowia | 13 | Saprotroph | \|Undefined Saprotroph\| | Yeast | NULL | Probable | As nectar-tap_saprotroph (Põlme et al. 2020); Decay sugar-rich substrates (Põlme et al. 2020) | Põlme S, et al. 2020. Fungal Diversity 105:1-16 ((https://doi.org/10.1007/s13225-020-00466-2)) |
| Otu38532 | 1 | k__Fungi(100);p__Ascomycota(100);c__Saccharomycetes(100);o__Saccharomycetales(100);f__Metschnikowiaceae(100);g__Metschnikowia(100);s__Metschnikowia_pulcherrima(100); | Metschnikowia | 13 | Saprotroph | \|Undefined Saprotroph\| | Yeast | NULL | Probable | As nectar-tap_saprotroph (Põlme et al. 2020); Decay sugar-rich substrates (Põlme et al. 2020) | Põlme S, et al. 2020. Fungal Diversity 105:1-16 ((https://doi.org/10.1007/s13225-020-00466-2)) |
| Otu38533 | 1 | k__Fungi(100);p__Ascomycota(100);c__Saccharomycetes(100);o__Saccharomycetales(100);f__Metschnikowiaceae(100);g__Metschnikowia(100);s__Metschnikowia_pulcherrima(100); | Metschnikowia | 13 | Saprotroph | \|Undefined Saprotroph\| | Yeast | NULL | Probable | As nectar-tap_saprotroph (Põlme et al. 2020); Decay sugar-rich substrates (Põlme et al. 2020) | Põlme S, et al. 2020. Fungal Diversity 105:1-16 ((https://doi.org/10.1007/s13225-020-00466-2)) |
| Otu38534 | 1 | k__Fungi(100);p__Ascomycota(100);c__Saccharomycetes(100);o__Saccharomycetales(100);f__Metschnikowiaceae(100);g__Metschnikowia(100);s__Metschnikowia_pulcherrima(100); | Metschnikowia | 13 | Saprotroph | \|Undefined Saprotroph\| | Yeast | NULL | Probable | As nectar-tap_saprotroph (Põlme et al. 2020); Decay sugar-rich substrates (Põlme et al. 2020) | Põlme S, et al. 2020. Fungal Diversity 105:1-16 ((https://doi.org/10.1007/s13225-020-00466-2)) |
| Otu38536 | 1 | k__Fungi(100);p__Ascomycota(100);c__Saccharomycetes(100);o__Saccharomycetales(100);f__Metschnikowiaceae(100);g__Metschnikowia(100);s__Metschnikowia_pulcherrima(100); | Metschnikowia | 13 | Saprotroph | \|Undefined Saprotroph\| | Yeast | NULL | Probable | As nectar-tap_saprotroph (Põlme et al. 2020); Decay sugar-rich substrates (Põlme et al. 2020) | Põlme S, et al. 2020. Fungal Diversity 105:1-16 ((https://doi.org/10.1007/s13225-020-00466-2)) |
| Otu38541 | 1 | k__Fungi(100);p__Ascomycota(100);c__Saccharomycetes(100);o__Saccharomycetales(100);f__Metschnikowiaceae(100);g__Metschnikowia(100);s__Metschnikowia_pulcherrima(100); | Metschnikowia | 13 | Saprotroph | \|Undefined Saprotroph\| | Yeast | NULL | Probable | As nectar-tap_saprotroph (Põlme et al. 2020); Decay sugar-rich substrates (Põlme et al. 2020) | Põlme S, et al. 2020. Fungal Diversity 105:1-16 ((https://doi.org/10.1007/s13225-020-00466-2)) |
| Otu38542 | 1 | k__Fungi(100);p__Ascomycota(100);c__Eurotiomycetes(100);o__Verrucariales(100);f__Verrucariaceae(100);f__Verrucariaceae_unclassified(100);f__Verrucariaceae_unclassified(100); | Verrucariaceae | 9 | Symbiotroph | Lichenized | Thallus | NULL | Highly Probable | Lichenized with green algae, on rocks, soil, etc., more rarely on wood and bark (Cannon & Kirk 2007) | Cannon PF, Kirk PM. 2007. Fungal Families of the World. CAB International, Cambridge (ISBN: 978-0851998275) |
| Otu38543 | 1 | k__Fungi(100);p__Ascomycota(100);c__Dothideomycetes(100);o__Pleosporales(100);f__Pleosporales_family_Incertae_sedis(100);g__Pyrenochaeta(100);s__Pyrenochaeta_keratinophila(100); | Pyrenochaeta | 13 | Pathotroph-Saprotroph | Plant Pathogen-Undefined Saprotroph-\|Wood Saprotroph\| | Microfungus | Soft Rot | Highly Probable | Decay wood (Põlme et al. 2020) | Seehann G, et al. 1975. List of Fungi in Soft-Rot Tissue Tests. Institute fur Holzbiologie und Holzschutz, Bundesforschungsanstalt fur Forst - und Holzwirtshaft, Hamburg - Lohbrugge, Germany; Costa IPMW, et al. 2012. Mycotaxon 119:e26 ((http://mycotaxon.com/resources/checklists/costa_v119_checklist.pdf)); Tedersoo L, et al. 2014. Science 346:e1256688 ((https://doi.org/10.1126/science.1256688)); Põlme S, et al. 2020. Fungal Diversity 105:1-16 ((https://doi.org/10.1007/s13225-020-00466-2)) |
| Otu38545 | 1 | k__Fungi(100);p__Ascomycota(100);c__Sordariomycetes(100);o__Chaetosphaeriales(100);f__Chaetosphaeriaceae(100);g__Menispora(100);s__Menispora_ciliata(100); | Menispora | 13 | Saprotroph-Symbiotroph | Endophyte-Plant Saprotroph-Undefined Saprotroph-\|Wood Saprotroph\| | Microfungus | NULL | Probable | As foliar_endophyte (Põlme et al. 2020); Decay wood, leaf, fruit, seed (Põlme et al. 2020) | Tedersoo L, et al. 2014. Science 346:e1256688 ((https://doi.org/10.1126/science.1256688)); Christian N, et al. 2016. Microbial Ecology 72:621-632 ((https://doi.org/10.1007/s00248-016-0804-y)) (as endophyte); Põlme S, et al. 2020. Fungal Diversity 105:1-16 ((https://doi.org/10.1007/s13225-020-00466-2)) |
| Otu38547 | 1 | k__Fungi(100);p__Ascomycota(100);c__Saccharomycetes(100);o__Saccharomycetales(100);f__Metschnikowiaceae(100);g__Metschnikowia(100);g__Metschnikowia_unclassified(100); | Metschnikowia | 13 | Saprotroph | \|Undefined Saprotroph\| | Yeast | NULL | Probable | As nectar-tap_saprotroph (Põlme et al. 2020); Decay sugar-rich substrates (Põlme et al. 2020) | Põlme S, et al. 2020. Fungal Diversity 105:1-16 ((https://doi.org/10.1007/s13225-020-00466-2)) |
| Otu38548 | 1 | k__Fungi(100);p__Ascomycota(100);c__Saccharomycetes(100);o__Saccharomycetales(100);f__Metschnikowiaceae(100);g__Metschnikowia(100);s__Metschnikowia_pulcherrima(100); | Metschnikowia | 13 | Saprotroph | \|Undefined Saprotroph\| | Yeast | NULL | Probable | As nectar-tap_saprotroph (Põlme et al. 2020); Decay sugar-rich substrates (Põlme et al. 2020) | Põlme S, et al. 2020. Fungal Diversity 105:1-16 ((https://doi.org/10.1007/s13225-020-00466-2)) |
| Otu38549 | 1 | k__Fungi(100);p__Ascomycota(100);c__Saccharomycetes(100);o__Saccharomycetales(100);f__Metschnikowiaceae(100);g__Metschnikowia(100);s__Metschnikowia_pulcherrima(100); | Metschnikowia | 13 | Saprotroph | \|Undefined Saprotroph\| | Yeast | NULL | Probable | As nectar-tap_saprotroph (Põlme et al. 2020); Decay sugar-rich substrates (Põlme et al. 2020) | Põlme S, et al. 2020. Fungal Diversity 105:1-16 ((https://doi.org/10.1007/s13225-020-00466-2)) |
| Otu38550 | 1 | k__Fungi(100);p__Ascomycota(100);c__Saccharomycetes(100);o__Saccharomycetales(100);f__Metschnikowiaceae(100);g__Metschnikowia(100);g__Metschnikowia_unclassified(100); | Metschnikowia | 13 | Saprotroph | \|Undefined Saprotroph\| | Yeast | NULL | Probable | As nectar-tap_saprotroph (Põlme et al. 2020); Decay sugar-rich substrates (Põlme et al. 2020) | Põlme S, et al. 2020. Fungal Diversity 105:1-16 ((https://doi.org/10.1007/s13225-020-00466-2)) |
| Otu38551 | 1 | k__Fungi(100);p__Ascomycota(100);c__Saccharomycetes(100);o__Saccharomycetales(100);f__Metschnikowiaceae(100);g__Metschnikowia(100);s__Metschnikowia_pulcherrima(100); | Metschnikowia | 13 | Saprotroph | \|Undefined Saprotroph\| | Yeast | NULL | Probable | As nectar-tap_saprotroph (Põlme et al. 2020); Decay sugar-rich substrates (Põlme et al. 2020) | Põlme S, et al. 2020. Fungal Diversity 105:1-16 ((https://doi.org/10.1007/s13225-020-00466-2)) |
| Otu38553 | 1 | k__Fungi(100);p__Ascomycota(100);c__Saccharomycetes(100);o__Saccharomycetales(100);f__Metschnikowiaceae(100);g__Metschnikowia(100);s__Metschnikowia_pulcherrima(100); | Metschnikowia | 13 | Saprotroph | \|Undefined Saprotroph\| | Yeast | NULL | Probable | As nectar-tap_saprotroph (Põlme et al. 2020); Decay sugar-rich substrates (Põlme et al. 2020) | Põlme S, et al. 2020. Fungal Diversity 105:1-16 ((https://doi.org/10.1007/s13225-020-00466-2)) |
| Otu38554 | 1 | k__Fungi(100);p__Ascomycota(100);c__Saccharomycetes(100);o__Saccharomycetales(100);f__Metschnikowiaceae(100);g__Metschnikowia(100);s__Metschnikowia_pulcherrima(100); | Metschnikowia | 13 | Saprotroph | \|Undefined Saprotroph\| | Yeast | NULL | Probable | As nectar-tap_saprotroph (Põlme et al. 2020); Decay sugar-rich substrates (Põlme et al. 2020) | Põlme S, et al. 2020. Fungal Diversity 105:1-16 ((https://doi.org/10.1007/s13225-020-00466-2)) |
| Otu38558 | 1 | k__Fungi(100);p__Ascomycota(100);c__Saccharomycetes(100);o__Saccharomycetales(100);f__Metschnikowiaceae(100);g__Metschnikowia(100);s__Metschnikowia_pulcherrima(100); | Metschnikowia | 13 | Saprotroph | \|Undefined Saprotroph\| | Yeast | NULL | Probable | As nectar-tap_saprotroph (Põlme et al. 2020); Decay sugar-rich substrates (Põlme et al. 2020) | Põlme S, et al. 2020. Fungal Diversity 105:1-16 ((https://doi.org/10.1007/s13225-020-00466-2)) |
| Otu38559 | 1 | k__Fungi(100);p__Ascomycota(100);c__Saccharomycetes(100);o__Saccharomycetales(100);f__Metschnikowiaceae(100);g__Metschnikowia(100);s__Metschnikowia_pulcherrima(100); | Metschnikowia | 13 | Saprotroph | \|Undefined Saprotroph\| | Yeast | NULL | Probable | As nectar-tap_saprotroph (Põlme et al. 2020); Decay sugar-rich substrates (Põlme et al. 2020) | Põlme S, et al. 2020. Fungal Diversity 105:1-16 ((https://doi.org/10.1007/s13225-020-00466-2)) |
| Otu38560 | 1 | k__Fungi(100);p__Ascomycota(100);c__Saccharomycetes(100);o__Saccharomycetales(100);f__Metschnikowiaceae(100);g__Metschnikowia(100);s__Metschnikowia_pulcherrima(100); | Metschnikowia | 13 | Saprotroph | \|Undefined Saprotroph\| | Yeast | NULL | Probable | As nectar-tap_saprotroph (Põlme et al. 2020); Decay sugar-rich substrates (Põlme et al. 2020) | Põlme S, et al. 2020. Fungal Diversity 105:1-16 ((https://doi.org/10.1007/s13225-020-00466-2)) |
| Otu38562 | 1 | k__Fungi(100);p__Ascomycota(100);c__Saccharomycetes(100);o__Saccharomycetales(100);f__Metschnikowiaceae(100);g__Metschnikowia(100);s__Metschnikowia_pulcherrima(100); | Metschnikowia | 13 | Saprotroph | \|Undefined Saprotroph\| | Yeast | NULL | Probable | As nectar-tap_saprotroph (Põlme et al. 2020); Decay sugar-rich substrates (Põlme et al. 2020) | Põlme S, et al. 2020. Fungal Diversity 105:1-16 ((https://doi.org/10.1007/s13225-020-00466-2)) |
| Otu38563 | 1 | k__Fungi(100);p__Ascomycota(100);c__Saccharomycetes(100);o__Saccharomycetales(100);f__Metschnikowiaceae(100);g__Metschnikowia(100);s__Metschnikowia_pulcherrima(100); | Metschnikowia | 13 | Saprotroph | \|Undefined Saprotroph\| | Yeast | NULL | Probable | As nectar-tap_saprotroph (Põlme et al. 2020); Decay sugar-rich substrates (Põlme et al. 2020) | Põlme S, et al. 2020. Fungal Diversity 105:1-16 ((https://doi.org/10.1007/s13225-020-00466-2)) |
| Otu38564 | 1 | k__Fungi(100);p__Ascomycota(100);c__Saccharomycetes(100);o__Saccharomycetales(100);f__Metschnikowiaceae(100);g__Metschnikowia(100);s__Metschnikowia_pulcherrima(100); | Metschnikowia | 13 | Saprotroph | \|Undefined Saprotroph\| | Yeast | NULL | Probable | As nectar-tap_saprotroph (Põlme et al. 2020); Decay sugar-rich substrates (Põlme et al. 2020) | Põlme S, et al. 2020. Fungal Diversity 105:1-16 ((https://doi.org/10.1007/s13225-020-00466-2)) |
| Otu38565 | 1 | k__Fungi(100);p__Ascomycota(100);c__Saccharomycetes(100);o__Saccharomycetales(100);f__Metschnikowiaceae(100);g__Metschnikowia(100);s__Metschnikowia_pulcherrima(100); | Metschnikowia | 13 | Saprotroph | \|Undefined Saprotroph\| | Yeast | NULL | Probable | As nectar-tap_saprotroph (Põlme et al. 2020); Decay sugar-rich substrates (Põlme et al. 2020) | Põlme S, et al. 2020. Fungal Diversity 105:1-16 ((https://doi.org/10.1007/s13225-020-00466-2)) |
| Otu38566 | 1 | k__Fungi(100);p__Ascomycota(100);c__Saccharomycetes(100);o__Saccharomycetales(100);f__Metschnikowiaceae(100);g__Metschnikowia(100);s__Metschnikowia_pulcherrima(100); | Metschnikowia | 13 | Saprotroph | \|Undefined Saprotroph\| | Yeast | NULL | Probable | As nectar-tap_saprotroph (Põlme et al. 2020); Decay sugar-rich substrates (Põlme et al. 2020) | Põlme S, et al. 2020. Fungal Diversity 105:1-16 ((https://doi.org/10.1007/s13225-020-00466-2)) |
| Otu38567 | 1 | k__Fungi(100);p__Ascomycota(100);c__Saccharomycetes(100);o__Saccharomycetales(100);f__Metschnikowiaceae(100);g__Metschnikowia(100);s__Metschnikowia_pulcherrima(100); | Metschnikowia | 13 | Saprotroph | \|Undefined Saprotroph\| | Yeast | NULL | Probable | As nectar-tap_saprotroph (Põlme et al. 2020); Decay sugar-rich substrates (Põlme et al. 2020) | Põlme S, et al. 2020. Fungal Diversity 105:1-16 ((https://doi.org/10.1007/s13225-020-00466-2)) |
| Otu38569 | 1 | k__Fungi(100);p__Ascomycota(100);c__Saccharomycetes(100);o__Saccharomycetales(100);f__Metschnikowiaceae(100);g__Metschnikowia(100);s__Metschnikowia_pulcherrima(100); | Metschnikowia | 13 | Saprotroph | \|Undefined Saprotroph\| | Yeast | NULL | Probable | As nectar-tap_saprotroph (Põlme et al. 2020); Decay sugar-rich substrates (Põlme et al. 2020) | Põlme S, et al. 2020. Fungal Diversity 105:1-16 ((https://doi.org/10.1007/s13225-020-00466-2)) |
| Otu38570 | 1 | k__Fungi(100);p__Ascomycota(100);c__Saccharomycetes(100);o__Saccharomycetales(100);f__Metschnikowiaceae(100);g__Metschnikowia(100);s__Metschnikowia_pulcherrima(100); | Metschnikowia | 13 | Saprotroph | \|Undefined Saprotroph\| | Yeast | NULL | Probable | As nectar-tap_saprotroph (Põlme et al. 2020); Decay sugar-rich substrates (Põlme et al. 2020) | Põlme S, et al. 2020. Fungal Diversity 105:1-16 ((https://doi.org/10.1007/s13225-020-00466-2)) |
| Otu38571 | 1 | k__Fungi(100);p__Ascomycota(100);c__Saccharomycetes(100);o__Saccharomycetales(100);f__Metschnikowiaceae(100);g__Metschnikowia(100);s__Metschnikowia_pulcherrima(100); | Metschnikowia | 13 | Saprotroph | \|Undefined Saprotroph\| | Yeast | NULL | Probable | As nectar-tap_saprotroph (Põlme et al. 2020); Decay sugar-rich substrates (Põlme et al. 2020) | Põlme S, et al. 2020. Fungal Diversity 105:1-16 ((https://doi.org/10.1007/s13225-020-00466-2)) |
| Otu38572 | 1 | k__Fungi(100);p__Ascomycota(100);c__Saccharomycetes(100);o__Saccharomycetales(100);f__Metschnikowiaceae(100);g__Metschnikowia(100);s__Metschnikowia_pulcherrima(100); | Metschnikowia | 13 | Saprotroph | \|Undefined Saprotroph\| | Yeast | NULL | Probable | As nectar-tap_saprotroph (Põlme et al. 2020); Decay sugar-rich substrates (Põlme et al. 2020) | Põlme S, et al. 2020. Fungal Diversity 105:1-16 ((https://doi.org/10.1007/s13225-020-00466-2)) |
| Otu38578 | 1 | k__Fungi(100);p__Ascomycota(100);c__Saccharomycetes(100);o__Saccharomycetales(100);f__Metschnikowiaceae(100);g__Metschnikowia(100);s__Metschnikowia_pulcherrima(100); | Metschnikowia | 13 | Saprotroph | \|Undefined Saprotroph\| | Yeast | NULL | Probable | As nectar-tap_saprotroph (Põlme et al. 2020); Decay sugar-rich substrates (Põlme et al. 2020) | Põlme S, et al. 2020. Fungal Diversity 105:1-16 ((https://doi.org/10.1007/s13225-020-00466-2)) |
| Otu38581 | 1 | k__Fungi(100);p__Ascomycota(100);c__Saccharomycetes(100);o__Saccharomycetales(100);f__Metschnikowiaceae(100);g__Metschnikowia(100);s__Metschnikowia_pulcherrima(100); | Metschnikowia | 13 | Saprotroph | \|Undefined Saprotroph\| | Yeast | NULL | Probable | As nectar-tap_saprotroph (Põlme et al. 2020); Decay sugar-rich substrates (Põlme et al. 2020) | Põlme S, et al. 2020. Fungal Diversity 105:1-16 ((https://doi.org/10.1007/s13225-020-00466-2)) |
| Otu38582 | 1 | k__Fungi(100);p__Ascomycota(100);c__Saccharomycetes(100);o__Saccharomycetales(100);f__Metschnikowiaceae(100);g__Metschnikowia(100);g__Metschnikowia_unclassified(100); | Metschnikowia | 13 | Saprotroph | \|Undefined Saprotroph\| | Yeast | NULL | Probable | As nectar-tap_saprotroph (Põlme et al. 2020); Decay sugar-rich substrates (Põlme et al. 2020) | Põlme S, et al. 2020. Fungal Diversity 105:1-16 ((https://doi.org/10.1007/s13225-020-00466-2)) |
| Otu38583 | 1 | k__Fungi(100);p__Ascomycota(100);c__Saccharomycetes(100);o__Saccharomycetales(100);f__Metschnikowiaceae(100);g__Metschnikowia(100);s__Metschnikowia_pulcherrima(100); | Metschnikowia | 13 | Saprotroph | \|Undefined Saprotroph\| | Yeast | NULL | Probable | As nectar-tap_saprotroph (Põlme et al. 2020); Decay sugar-rich substrates (Põlme et al. 2020) | Põlme S, et al. 2020. Fungal Diversity 105:1-16 ((https://doi.org/10.1007/s13225-020-00466-2)) |
| Otu38584 | 1 | k__Fungi(100);p__Ascomycota(100);c__Saccharomycetes(100);o__Saccharomycetales(100);f__Metschnikowiaceae(100);g__Metschnikowia(100);g__Metschnikowia_unclassified(100); | Metschnikowia | 13 | Saprotroph | \|Undefined Saprotroph\| | Yeast | NULL | Probable | As nectar-tap_saprotroph (Põlme et al. 2020); Decay sugar-rich substrates (Põlme et al. 2020) | Põlme S, et al. 2020. Fungal Diversity 105:1-16 ((https://doi.org/10.1007/s13225-020-00466-2)) |
| Otu38585 | 1 | k__Fungi(100);p__Ascomycota(100);c__Saccharomycetes(100);o__Saccharomycetales(100);f__Metschnikowiaceae(100);g__Metschnikowia(100);s__Metschnikowia_pulcherrima(100); | Metschnikowia | 13 | Saprotroph | \|Undefined Saprotroph\| | Yeast | NULL | Probable | As nectar-tap_saprotroph (Põlme et al. 2020); Decay sugar-rich substrates (Põlme et al. 2020) | Põlme S, et al. 2020. Fungal Diversity 105:1-16 ((https://doi.org/10.1007/s13225-020-00466-2)) |
| Otu38587 | 1 | k__Fungi(100);p__Ascomycota(100);c__Saccharomycetes(100);o__Saccharomycetales(100);f__Metschnikowiaceae(100);g__Metschnikowia(100);g__Metschnikowia_unclassified(100); | Metschnikowia | 13 | Saprotroph | \|Undefined Saprotroph\| | Yeast | NULL | Probable | As nectar-tap_saprotroph (Põlme et al. 2020); Decay sugar-rich substrates (Põlme et al. 2020) | Põlme S, et al. 2020. Fungal Diversity 105:1-16 ((https://doi.org/10.1007/s13225-020-00466-2)) |
| Otu38588 | 1 | k__Fungi(100);p__Ascomycota(100);c__Saccharomycetes(100);o__Saccharomycetales(100);f__Metschnikowiaceae(100);g__Metschnikowia(100);s__Metschnikowia_pulcherrima(100); | Metschnikowia | 13 | Saprotroph | \|Undefined Saprotroph\| | Yeast | NULL | Probable | As nectar-tap_saprotroph (Põlme et al. 2020); Decay sugar-rich substrates (Põlme et al. 2020) | Põlme S, et al. 2020. Fungal Diversity 105:1-16 ((https://doi.org/10.1007/s13225-020-00466-2)) |
| Otu38589 | 1 | k__Fungi(100);p__Ascomycota(100);c__Saccharomycetes(100);o__Saccharomycetales(100);f__Metschnikowiaceae(100);g__Metschnikowia(100);s__Metschnikowia_pulcherrima(100); | Metschnikowia | 13 | Saprotroph | \|Undefined Saprotroph\| | Yeast | NULL | Probable | As nectar-tap_saprotroph (Põlme et al. 2020); Decay sugar-rich substrates (Põlme et al. 2020) | Põlme S, et al. 2020. Fungal Diversity 105:1-16 ((https://doi.org/10.1007/s13225-020-00466-2)) |
| Otu38590 | 1 | k__Fungi(100);p__Ascomycota(100);c__Saccharomycetes(100);o__Saccharomycetales(100);f__Metschnikowiaceae(100);g__Metschnikowia(100);g__Metschnikowia_unclassified(100); | Metschnikowia | 13 | Saprotroph | \|Undefined Saprotroph\| | Yeast | NULL | Probable | As nectar-tap_saprotroph (Põlme et al. 2020); Decay sugar-rich substrates (Põlme et al. 2020) | Põlme S, et al. 2020. Fungal Diversity 105:1-16 ((https://doi.org/10.1007/s13225-020-00466-2)) |
| Otu38591 | 1 | k__Fungi(100);p__Ascomycota(100);c__Saccharomycetes(100);o__Saccharomycetales(100);f__Metschnikowiaceae(100);g__Metschnikowia(100);g__Metschnikowia_unclassified(100); | Metschnikowia | 13 | Saprotroph | \|Undefined Saprotroph\| | Yeast | NULL | Probable | As nectar-tap_saprotroph (Põlme et al. 2020); Decay sugar-rich substrates (Põlme et al. 2020) | Põlme S, et al. 2020. Fungal Diversity 105:1-16 ((https://doi.org/10.1007/s13225-020-00466-2)) |
| Otu38592 | 1 | k__Fungi(100);p__Ascomycota(100);c__Saccharomycetes(100);o__Saccharomycetales(100);f__Metschnikowiaceae(100);g__Metschnikowia(100);s__Metschnikowia_pulcherrima(100); | Metschnikowia | 13 | Saprotroph | \|Undefined Saprotroph\| | Yeast | NULL | Probable | As nectar-tap_saprotroph (Põlme et al. 2020); Decay sugar-rich substrates (Põlme et al. 2020) | Põlme S, et al. 2020. Fungal Diversity 105:1-16 ((https://doi.org/10.1007/s13225-020-00466-2)) |
| Otu38593 | 1 | k__Fungi(100);p__Ascomycota(100);c__Saccharomycetes(100);o__Saccharomycetales(100);f__Metschnikowiaceae(100);g__Metschnikowia(100);s__Metschnikowia_pulcherrima(100); | Metschnikowia | 13 | Saprotroph | \|Undefined Saprotroph\| | Yeast | NULL | Probable | As nectar-tap_saprotroph (Põlme et al. 2020); Decay sugar-rich substrates (Põlme et al. 2020) | Põlme S, et al. 2020. Fungal Diversity 105:1-16 ((https://doi.org/10.1007/s13225-020-00466-2)) |
| Otu38594 | 1 | k__Fungi(100);p__Ascomycota(100);c__Dothideomycetes(100);o__Pleosporales(100);f__Tubeufiaceae(100);f__Tubeufiaceae_unclassified(100);f__Tubeufiaceae_unclassified(100); | Tubeufiaceae | 9 | Pathotroph-Saprotroph | Animal Pathogen-Undefined Saprotroph | NULL | NULL | Probable | Biotrophic or saprobic, especially on scale insects or other fungi. The anamorphic forms are fequently aero-aquatic (Cannon & Kirk 2007) | Cannon PF, Kirk PM. 2007. Fungal Families of the World. CAB International, Cambridge (ISBN: 978-0851998275) |
| Otu38598 | 1 | k__Fungi(100);p__Ascomycota(100);c__Dothideomycetes(100);o__Pleosporales(100);f__Pleosporaceae(100);g__Alternaria(100);s__Alternaria_porri(100); | Alternaria | 13 | Pathotroph-Saprotroph-Symbiotroph | Animal Pathogen-Endophyte-\|Plant Pathogen\|-Plant Saprotroph-Wood Saprotroph | Microfungus | Soft Rot | Possible | As foliar_endophyte (Põlme et al. 2020); As litter_saprotroph (Põlme et al. 2020); Decay leaf, fruit, seed (Põlme et al. 2020) | Duncan CG, Eslyn WE. 1966. Mycologia 58:642-664 ((https://doi.org/10.2307/3757045)); Seehann G, et al. 1975. List of Fungi in Soft-Rot Tissue Tests. Institute fur Holzbiologie und Holzschutz, Bundesforschungsanstalt fur Forst - und Holzwirtshaft, Hamburg - Lohbrugge, Germany ((https://www.irg-wp.com/irgdocs/details.php?d6493033-c9ea-40f3-8e3e-5697f32d640d)); Tedersoo L, et al. 2014. Science 346:e1256688 ((https://doi.org/10.1126/science.1256688)); Põlme S, et al. 2020. Fungal Diversity 105:1-16 ((https://doi.org/10.1007/s13225-020-00466-2)) |
| Otu38599 | 1 | k__Fungi(100);p__Ascomycota(100);c__Lecanoromycetes(100);o__Lecanorales(100);o__Lecanorales_unclassified(100);o__Lecanorales_unclassified(100);o__Lecanorales_unclassified(100); | Lecanorales | 7 | Pathotroph-Symbiotroph | Lichen Parasite-Lichenized | Microfungus-Thallus | NULL | Probable | Lichenized with green algae, occasionally lichenicolous, on rocks, mosses and wood (Cannon & Kirk 2007); ...an order of mostly lichen-forming fungi (Kirk et al. 2008) | Cannon PF, Kirk PM. 2007. Fungal Families of the World. CAB International, Cambridge (ISBN: 978-0851998275); Kirk PM et al. 2008. Dictionary of the Fungi. Tenth Edition. CAB International, Wallingford (ISBN: 978-0851998268); Esslinger TL. 2014. North Dakota State University ((http://www.ndsu.edu/pubweb/~esslinge/chcklst/chcklst7.htm)) |
| Otu38600 | 1 | k__Fungi(100);p__Ascomycota(100);c__Saccharomycetes(100);o__Saccharomycetales(100);f__Metschnikowiaceae(100);g__Metschnikowia(100);s__Metschnikowia_pulcherrima(100); | Metschnikowia | 13 | Saprotroph | \|Undefined Saprotroph\| | Yeast | NULL | Probable | As nectar-tap_saprotroph (Põlme et al. 2020); Decay sugar-rich substrates (Põlme et al. 2020) | Põlme S, et al. 2020. Fungal Diversity 105:1-16 ((https://doi.org/10.1007/s13225-020-00466-2)) |
| Otu38601 | 1 | k__Fungi(100);p__Ascomycota(100);c__Saccharomycetes(100);o__Saccharomycetales(100);f__Metschnikowiaceae(100);g__Metschnikowia(100);g__Metschnikowia_unclassified(100); | Metschnikowia | 13 | Saprotroph | \|Undefined Saprotroph\| | Yeast | NULL | Probable | As nectar-tap_saprotroph (Põlme et al. 2020); Decay sugar-rich substrates (Põlme et al. 2020) | Põlme S, et al. 2020. Fungal Diversity 105:1-16 ((https://doi.org/10.1007/s13225-020-00466-2)) |
| Otu38602 | 1 | k__Fungi(100);p__Ascomycota(100);c__Saccharomycetes(100);o__Saccharomycetales(100);f__Metschnikowiaceae(100);g__Metschnikowia(100);g__Metschnikowia_unclassified(100); | Metschnikowia | 13 | Saprotroph | \|Undefined Saprotroph\| | Yeast | NULL | Probable | As nectar-tap_saprotroph (Põlme et al. 2020); Decay sugar-rich substrates (Põlme et al. 2020) | Põlme S, et al. 2020. Fungal Diversity 105:1-16 ((https://doi.org/10.1007/s13225-020-00466-2)) |
| Otu38603 | 1 | k__Fungi(100);p__Ascomycota(100);c__Saccharomycetes(100);o__Saccharomycetales(100);f__Metschnikowiaceae(100);g__Metschnikowia(100);s__Metschnikowia_pulcherrima(100); | Metschnikowia | 13 | Saprotroph | \|Undefined Saprotroph\| | Yeast | NULL | Probable | As nectar-tap_saprotroph (Põlme et al. 2020); Decay sugar-rich substrates (Põlme et al. 2020) | Põlme S, et al. 2020. Fungal Diversity 105:1-16 ((https://doi.org/10.1007/s13225-020-00466-2)) |
| Otu38604 | 1 | k__Fungi(100);p__Ascomycota(100);c__Saccharomycetes(100);o__Saccharomycetales(100);f__Metschnikowiaceae(100);g__Metschnikowia(100);s__Metschnikowia_pulcherrima(100); | Metschnikowia | 13 | Saprotroph | \|Undefined Saprotroph\| | Yeast | NULL | Probable | As nectar-tap_saprotroph (Põlme et al. 2020); Decay sugar-rich substrates (Põlme et al. 2020) | Põlme S, et al. 2020. Fungal Diversity 105:1-16 ((https://doi.org/10.1007/s13225-020-00466-2)) |
| Otu38605 | 1 | k__Fungi(100);p__Ascomycota(100);c__Saccharomycetes(100);o__Saccharomycetales(100);f__Metschnikowiaceae(100);g__Metschnikowia(100);g__Metschnikowia_unclassified(100); | Metschnikowia | 13 | Saprotroph | \|Undefined Saprotroph\| | Yeast | NULL | Probable | As nectar-tap_saprotroph (Põlme et al. 2020); Decay sugar-rich substrates (Põlme et al. 2020) | Põlme S, et al. 2020. Fungal Diversity 105:1-16 ((https://doi.org/10.1007/s13225-020-00466-2)) |
| Otu38606 | 1 | k__Fungi(100);p__Ascomycota(100);c__Saccharomycetes(100);o__Saccharomycetales(100);f__Metschnikowiaceae(100);g__Metschnikowia(100);s__Metschnikowia_pulcherrima(100); | Metschnikowia | 13 | Saprotroph | \|Undefined Saprotroph\| | Yeast | NULL | Probable | As nectar-tap_saprotroph (Põlme et al. 2020); Decay sugar-rich substrates (Põlme et al. 2020) | Põlme S, et al. 2020. Fungal Diversity 105:1-16 ((https://doi.org/10.1007/s13225-020-00466-2)) |
| Otu38607 | 1 | k__Fungi(100);p__Ascomycota(100);c__Saccharomycetes(100);o__Saccharomycetales(100);f__Metschnikowiaceae(100);g__Metschnikowia(100);s__Metschnikowia_pulcherrima(100); | Metschnikowia | 13 | Saprotroph | \|Undefined Saprotroph\| | Yeast | NULL | Probable | As nectar-tap_saprotroph (Põlme et al. 2020); Decay sugar-rich substrates (Põlme et al. 2020) | Põlme S, et al. 2020. Fungal Diversity 105:1-16 ((https://doi.org/10.1007/s13225-020-00466-2)) |
| Otu38608 | 1 | k__Fungi(100);p__Ascomycota(100);c__Saccharomycetes(100);o__Saccharomycetales(100);f__Metschnikowiaceae(100);g__Metschnikowia(100);s__Metschnikowia_pulcherrima(100); | Metschnikowia | 13 | Saprotroph | \|Undefined Saprotroph\| | Yeast | NULL | Probable | As nectar-tap_saprotroph (Põlme et al. 2020); Decay sugar-rich substrates (Põlme et al. 2020) | Põlme S, et al. 2020. Fungal Diversity 105:1-16 ((https://doi.org/10.1007/s13225-020-00466-2)) |
| Otu38609 | 1 | k__Fungi(100);p__Ascomycota(100);c__Saccharomycetes(100);o__Saccharomycetales(100);f__Metschnikowiaceae(100);g__Metschnikowia(100);g__Metschnikowia_unclassified(100); | Metschnikowia | 13 | Saprotroph | \|Undefined Saprotroph\| | Yeast | NULL | Probable | As nectar-tap_saprotroph (Põlme et al. 2020); Decay sugar-rich substrates (Põlme et al. 2020) | Põlme S, et al. 2020. Fungal Diversity 105:1-16 ((https://doi.org/10.1007/s13225-020-00466-2)) |
| Otu38610 | 1 | k__Fungi(100);p__Ascomycota(100);c__Saccharomycetes(100);o__Saccharomycetales(100);f__Metschnikowiaceae(100);g__Metschnikowia(100);s__Metschnikowia_pulcherrima(100); | Metschnikowia | 13 | Saprotroph | \|Undefined Saprotroph\| | Yeast | NULL | Probable | As nectar-tap_saprotroph (Põlme et al. 2020); Decay sugar-rich substrates (Põlme et al. 2020) | Põlme S, et al. 2020. Fungal Diversity 105:1-16 ((https://doi.org/10.1007/s13225-020-00466-2)) |
| Otu38611 | 1 | k__Fungi(100);p__Ascomycota(100);c__Saccharomycetes(100);o__Saccharomycetales(100);f__Metschnikowiaceae(100);g__Metschnikowia(100);g__Metschnikowia_unclassified(100); | Metschnikowia | 13 | Saprotroph | \|Undefined Saprotroph\| | Yeast | NULL | Probable | As nectar-tap_saprotroph (Põlme et al. 2020); Decay sugar-rich substrates (Põlme et al. 2020) | Põlme S, et al. 2020. Fungal Diversity 105:1-16 ((https://doi.org/10.1007/s13225-020-00466-2)) |
| Otu38612 | 1 | k__Fungi(100);p__Ascomycota(100);c__Saccharomycetes(100);o__Saccharomycetales(100);f__Metschnikowiaceae(100);g__Metschnikowia(100);s__Metschnikowia_pulcherrima(100); | Metschnikowia | 13 | Saprotroph | \|Undefined Saprotroph\| | Yeast | NULL | Probable | As nectar-tap_saprotroph (Põlme et al. 2020); Decay sugar-rich substrates (Põlme et al. 2020) | Põlme S, et al. 2020. Fungal Diversity 105:1-16 ((https://doi.org/10.1007/s13225-020-00466-2)) |
| Otu38613 | 1 | k__Fungi(100);p__Ascomycota(100);c__Saccharomycetes(100);o__Saccharomycetales(100);f__Metschnikowiaceae(100);g__Metschnikowia(100);s__Metschnikowia_pulcherrima(100); | Metschnikowia | 13 | Saprotroph | \|Undefined Saprotroph\| | Yeast | NULL | Probable | As nectar-tap_saprotroph (Põlme et al. 2020); Decay sugar-rich substrates (Põlme et al. 2020) | Põlme S, et al. 2020. Fungal Diversity 105:1-16 ((https://doi.org/10.1007/s13225-020-00466-2)) |
| Otu38614 | 1 | k__Fungi(100);p__Ascomycota(100);c__Saccharomycetes(100);o__Saccharomycetales(100);f__Metschnikowiaceae(100);g__Metschnikowia(100);s__Metschnikowia_pulcherrima(100); | Metschnikowia | 13 | Saprotroph | \|Undefined Saprotroph\| | Yeast | NULL | Probable | As nectar-tap_saprotroph (Põlme et al. 2020); Decay sugar-rich substrates (Põlme et al. 2020) | Põlme S, et al. 2020. Fungal Diversity 105:1-16 ((https://doi.org/10.1007/s13225-020-00466-2)) |
| Otu38615 | 1 | k__Fungi(100);p__Ascomycota(100);c__Saccharomycetes(100);o__Saccharomycetales(100);o__Saccharomycetales_unclassified(100);o__Saccharomycetales_unclassified(100);o__Saccharomycetales_unclassified(100); | Saccharomycetales | 7 | Saprotroph | Undefined Saprotroph | Yeast | NULL | Possible | NULL | Sterkenburg E, et al. 2015. New Phytologist 207:1145-1158 ((https://doi.org/10.1111/nph.13426)) |
| Otu38616 | 1 | k__Fungi(100);p__Ascomycota(100);c__Saccharomycetes(100);o__Saccharomycetales(100);f__Metschnikowiaceae(100);g__Metschnikowia(100);s__Metschnikowia_pulcherrima(100); | Metschnikowia | 13 | Saprotroph | \|Undefined Saprotroph\| | Yeast | NULL | Probable | As nectar-tap_saprotroph (Põlme et al. 2020); Decay sugar-rich substrates (Põlme et al. 2020) | Põlme S, et al. 2020. Fungal Diversity 105:1-16 ((https://doi.org/10.1007/s13225-020-00466-2)) |
| Otu38617 | 1 | k__Fungi(100);p__Ascomycota(100);c__Saccharomycetes(100);o__Saccharomycetales(100);f__Metschnikowiaceae(100);g__Metschnikowia(100);s__Metschnikowia_pulcherrima(100); | Metschnikowia | 13 | Saprotroph | \|Undefined Saprotroph\| | Yeast | NULL | Probable | As nectar-tap_saprotroph (Põlme et al. 2020); Decay sugar-rich substrates (Põlme et al. 2020) | Põlme S, et al. 2020. Fungal Diversity 105:1-16 ((https://doi.org/10.1007/s13225-020-00466-2)) |
| Otu38618 | 1 | k__Fungi(100);p__Ascomycota(100);c__Saccharomycetes(100);o__Saccharomycetales(100);f__Metschnikowiaceae(100);g__Metschnikowia(100);s__Metschnikowia_pulcherrima(100); | Metschnikowia | 13 | Saprotroph | \|Undefined Saprotroph\| | Yeast | NULL | Probable | As nectar-tap_saprotroph (Põlme et al. 2020); Decay sugar-rich substrates (Põlme et al. 2020) | Põlme S, et al. 2020. Fungal Diversity 105:1-16 ((https://doi.org/10.1007/s13225-020-00466-2)) |
| Otu38619 | 1 | k__Fungi(100);p__Ascomycota(100);c__Saccharomycetes(100);o__Saccharomycetales(100);f__Metschnikowiaceae(100);g__Metschnikowia(100);s__Metschnikowia_pulcherrima(100); | Metschnikowia | 13 | Saprotroph | \|Undefined Saprotroph\| | Yeast | NULL | Probable | As nectar-tap_saprotroph (Põlme et al. 2020); Decay sugar-rich substrates (Põlme et al. 2020) | Põlme S, et al. 2020. Fungal Diversity 105:1-16 ((https://doi.org/10.1007/s13225-020-00466-2)) |
| Otu38620 | 1 | k__Fungi(100);p__Ascomycota(100);c__Saccharomycetes(100);o__Saccharomycetales(100);f__Metschnikowiaceae(100);g__Metschnikowia(100);s__Metschnikowia_pulcherrima(100); | Metschnikowia | 13 | Saprotroph | \|Undefined Saprotroph\| | Yeast | NULL | Probable | As nectar-tap_saprotroph (Põlme et al. 2020); Decay sugar-rich substrates (Põlme et al. 2020) | Põlme S, et al. 2020. Fungal Diversity 105:1-16 ((https://doi.org/10.1007/s13225-020-00466-2)) |
| Otu38621 | 1 | k__Fungi(100);p__Ascomycota(100);c__Saccharomycetes(100);o__Saccharomycetales(100);f__Metschnikowiaceae(100);g__Metschnikowia(100);g__Metschnikowia_unclassified(100); | Metschnikowia | 13 | Saprotroph | \|Undefined Saprotroph\| | Yeast | NULL | Probable | As nectar-tap_saprotroph (Põlme et al. 2020); Decay sugar-rich substrates (Põlme et al. 2020) | Põlme S, et al. 2020. Fungal Diversity 105:1-16 ((https://doi.org/10.1007/s13225-020-00466-2)) |
| Otu38622 | 1 | k__Fungi(100);p__Ascomycota(100);c__Saccharomycetes(100);o__Saccharomycetales(100);f__Metschnikowiaceae(100);g__Metschnikowia(100);s__Metschnikowia_pulcherrima(100); | Metschnikowia | 13 | Saprotroph | \|Undefined Saprotroph\| | Yeast | NULL | Probable | As nectar-tap_saprotroph (Põlme et al. 2020); Decay sugar-rich substrates (Põlme et al. 2020) | Põlme S, et al. 2020. Fungal Diversity 105:1-16 ((https://doi.org/10.1007/s13225-020-00466-2)) |
| Otu38624 | 1 | k__Fungi(100);p__Ascomycota(100);c__Saccharomycetes(100);o__Saccharomycetales(100);f__Metschnikowiaceae(100);g__Metschnikowia(100);s__Metschnikowia_pulcherrima(100); | Metschnikowia | 13 | Saprotroph | \|Undefined Saprotroph\| | Yeast | NULL | Probable | As nectar-tap_saprotroph (Põlme et al. 2020); Decay sugar-rich substrates (Põlme et al. 2020) | Põlme S, et al. 2020. Fungal Diversity 105:1-16 ((https://doi.org/10.1007/s13225-020-00466-2)) |
| Otu38625 | 1 | k__Fungi(100);p__Ascomycota(100);c__Saccharomycetes(100);o__Saccharomycetales(100);f__Metschnikowiaceae(100);g__Metschnikowia(100);g__Metschnikowia_unclassified(100); | Metschnikowia | 13 | Saprotroph | \|Undefined Saprotroph\| | Yeast | NULL | Probable | As nectar-tap_saprotroph (Põlme et al. 2020); Decay sugar-rich substrates (Põlme et al. 2020) | Põlme S, et al. 2020. Fungal Diversity 105:1-16 ((https://doi.org/10.1007/s13225-020-00466-2)) |
| Otu38626 | 1 | k__Fungi(100);p__Ascomycota(100);c__Saccharomycetes(100);o__Saccharomycetales(100);f__Metschnikowiaceae(100);g__Metschnikowia(100);s__Metschnikowia_pulcherrima(100); | Metschnikowia | 13 | Saprotroph | \|Undefined Saprotroph\| | Yeast | NULL | Probable | As nectar-tap_saprotroph (Põlme et al. 2020); Decay sugar-rich substrates (Põlme et al. 2020) | Põlme S, et al. 2020. Fungal Diversity 105:1-16 ((https://doi.org/10.1007/s13225-020-00466-2)) |
| Otu38627 | 1 | k__Fungi(100);p__Ascomycota(100);c__Saccharomycetes(100);o__Saccharomycetales(100);f__Metschnikowiaceae(100);g__Metschnikowia(100);s__Metschnikowia_pulcherrima(100); | Metschnikowia | 13 | Saprotroph | \|Undefined Saprotroph\| | Yeast | NULL | Probable | As nectar-tap_saprotroph (Põlme et al. 2020); Decay sugar-rich substrates (Põlme et al. 2020) | Põlme S, et al. 2020. Fungal Diversity 105:1-16 ((https://doi.org/10.1007/s13225-020-00466-2)) |
| Otu38628 | 1 | k__Fungi(100);p__Ascomycota(100);c__Saccharomycetes(100);o__Saccharomycetales(100);f__Metschnikowiaceae(100);g__Metschnikowia(100);s__Metschnikowia_pulcherrima(100); | Metschnikowia | 13 | Saprotroph | \|Undefined Saprotroph\| | Yeast | NULL | Probable | As nectar-tap_saprotroph (Põlme et al. 2020); Decay sugar-rich substrates (Põlme et al. 2020) | Põlme S, et al. 2020. Fungal Diversity 105:1-16 ((https://doi.org/10.1007/s13225-020-00466-2)) |
| Otu38629 | 1 | k__Fungi(100);p__Ascomycota(100);c__Saccharomycetes(100);o__Saccharomycetales(100);f__Metschnikowiaceae(100);g__Metschnikowia(100);s__Metschnikowia_pulcherrima(100); | Metschnikowia | 13 | Saprotroph | \|Undefined Saprotroph\| | Yeast | NULL | Probable | As nectar-tap_saprotroph (Põlme et al. 2020); Decay sugar-rich substrates (Põlme et al. 2020) | Põlme S, et al. 2020. Fungal Diversity 105:1-16 ((https://doi.org/10.1007/s13225-020-00466-2)) |
| Otu38630 | 1 | k__Fungi(100);p__Ascomycota(100);c__Saccharomycetes(100);o__Saccharomycetales(100);f__Metschnikowiaceae(100);g__Metschnikowia(100);s__Metschnikowia_pulcherrima(100); | Metschnikowia | 13 | Saprotroph | \|Undefined Saprotroph\| | Yeast | NULL | Probable | As nectar-tap_saprotroph (Põlme et al. 2020); Decay sugar-rich substrates (Põlme et al. 2020) | Põlme S, et al. 2020. Fungal Diversity 105:1-16 ((https://doi.org/10.1007/s13225-020-00466-2)) |
| Otu38631 | 1 | k__Fungi(100);p__Ascomycota(100);c__Sordariomycetes(100);o__Hypocreales(100);f__Nectriaceae(100);g__Gibberella(100);g__Gibberella_unclassified(100); | Gibberella | 13 | Pathotroph-Saprotroph-Symbiotroph | Endophyte-\|Plant Pathogen\|-Plant Saprotroph-Undefined Saprotroph | Microfungus | Soft Rot | Probable | As foliar_endophyte (Põlme et al. 2020); As litter_saprotroph (Põlme et al. 2020); Causing damping-off disease (Põlme et al. 2020); Decay leaf, fruit, seed, roots, soil, animal material (Põlme et al. 2020) | Tedersoo L, et al. 2014. Science 346:e1256688 ((https://doi.org/10.1126/science.1256688)); Põlme S, et al. 2020. Fungal Diversity 105:1-16 ((https://doi.org/10.1007/s13225-020-00466-2)) |
| Otu38632 | 1 | k__Fungi(100);p__Ascomycota(100);c__Saccharomycetes(100);o__Saccharomycetales(100);f__Metschnikowiaceae(100);g__Metschnikowia(100);s__Metschnikowia_pulcherrima(100); | Metschnikowia | 13 | Saprotroph | \|Undefined Saprotroph\| | Yeast | NULL | Probable | As nectar-tap_saprotroph (Põlme et al. 2020); Decay sugar-rich substrates (Põlme et al. 2020) | Põlme S, et al. 2020. Fungal Diversity 105:1-16 ((https://doi.org/10.1007/s13225-020-00466-2)) |
| Otu38634 | 1 | k__Fungi(100);p__Ascomycota(100);c__Saccharomycetes(100);o__Saccharomycetales(100);f__Metschnikowiaceae(100);g__Metschnikowia(100);g__Metschnikowia_unclassified(100); | Metschnikowia | 13 | Saprotroph | \|Undefined Saprotroph\| | Yeast | NULL | Probable | As nectar-tap_saprotroph (Põlme et al. 2020); Decay sugar-rich substrates (Põlme et al. 2020) | Põlme S, et al. 2020. Fungal Diversity 105:1-16 ((https://doi.org/10.1007/s13225-020-00466-2)) |
| Otu38635 | 1 | k__Fungi(100);p__Ascomycota(100);c__Saccharomycetes(100);o__Saccharomycetales(100);f__Metschnikowiaceae(100);g__Metschnikowia(100);g__Metschnikowia_unclassified(100); | Metschnikowia | 13 | Saprotroph | \|Undefined Saprotroph\| | Yeast | NULL | Probable | As nectar-tap_saprotroph (Põlme et al. 2020); Decay sugar-rich substrates (Põlme et al. 2020) | Põlme S, et al. 2020. Fungal Diversity 105:1-16 ((https://doi.org/10.1007/s13225-020-00466-2)) |
| Otu38637 | 1 | k__Fungi(100);p__Ascomycota(100);c__Saccharomycetes(100);o__Saccharomycetales(100);f__Metschnikowiaceae(100);g__Metschnikowia(100);s__Metschnikowia_pulcherrima(100); | Metschnikowia | 13 | Saprotroph | \|Undefined Saprotroph\| | Yeast | NULL | Probable | As nectar-tap_saprotroph (Põlme et al. 2020); Decay sugar-rich substrates (Põlme et al. 2020) | Põlme S, et al. 2020. Fungal Diversity 105:1-16 ((https://doi.org/10.1007/s13225-020-00466-2)) |
| Otu38638 | 1 | k__Fungi(100);p__Ascomycota(100);c__Saccharomycetes(100);o__Saccharomycetales(100);f__Metschnikowiaceae(100);g__Metschnikowia(100);s__Metschnikowia_pulcherrima(100); | Metschnikowia | 13 | Saprotroph | \|Undefined Saprotroph\| | Yeast | NULL | Probable | As nectar-tap_saprotroph (Põlme et al. 2020); Decay sugar-rich substrates (Põlme et al. 2020) | Põlme S, et al. 2020. Fungal Diversity 105:1-16 ((https://doi.org/10.1007/s13225-020-00466-2)) |
| Otu38639 | 1 | k__Fungi(100);p__Ascomycota(100);c__Saccharomycetes(100);o__Saccharomycetales(100);f__Metschnikowiaceae(100);g__Metschnikowia(100);s__Metschnikowia_pulcherrima(100); | Metschnikowia | 13 | Saprotroph | \|Undefined Saprotroph\| | Yeast | NULL | Probable | As nectar-tap_saprotroph (Põlme et al. 2020); Decay sugar-rich substrates (Põlme et al. 2020) | Põlme S, et al. 2020. Fungal Diversity 105:1-16 ((https://doi.org/10.1007/s13225-020-00466-2)) |
| Otu38640 | 1 | k__Fungi(100);p__Ascomycota(100);c__Saccharomycetes(100);o__Saccharomycetales(100);f__Metschnikowiaceae(100);g__Metschnikowia(100);g__Metschnikowia_unclassified(100); | Metschnikowia | 13 | Saprotroph | \|Undefined Saprotroph\| | Yeast | NULL | Probable | As nectar-tap_saprotroph (Põlme et al. 2020); Decay sugar-rich substrates (Põlme et al. 2020) | Põlme S, et al. 2020. Fungal Diversity 105:1-16 ((https://doi.org/10.1007/s13225-020-00466-2)) |
| Otu38641 | 1 | k__Fungi(100);p__Ascomycota(100);c__Saccharomycetes(100);o__Saccharomycetales(100);f__Metschnikowiaceae(100);g__Metschnikowia(100);s__Metschnikowia_pulcherrima(100); | Metschnikowia | 13 | Saprotroph | \|Undefined Saprotroph\| | Yeast | NULL | Probable | As nectar-tap_saprotroph (Põlme et al. 2020); Decay sugar-rich substrates (Põlme et al. 2020) | Põlme S, et al. 2020. Fungal Diversity 105:1-16 ((https://doi.org/10.1007/s13225-020-00466-2)) |
| Otu38642 | 1 | k__Fungi(100);p__Ascomycota(100);c__Saccharomycetes(100);o__Saccharomycetales(100);f__Metschnikowiaceae(100);g__Metschnikowia(100);s__Metschnikowia_pulcherrima(100); | Metschnikowia | 13 | Saprotroph | \|Undefined Saprotroph\| | Yeast | NULL | Probable | As nectar-tap_saprotroph (Põlme et al. 2020); Decay sugar-rich substrates (Põlme et al. 2020) | Põlme S, et al. 2020. Fungal Diversity 105:1-16 ((https://doi.org/10.1007/s13225-020-00466-2)) |
| Otu38643 | 1 | k__Fungi(100);p__Ascomycota(100);c__Saccharomycetes(100);o__Saccharomycetales(100);f__Metschnikowiaceae(100);g__Metschnikowia(100);g__Metschnikowia_unclassified(100); | Metschnikowia | 13 | Saprotroph | \|Undefined Saprotroph\| | Yeast | NULL | Probable | As nectar-tap_saprotroph (Põlme et al. 2020); Decay sugar-rich substrates (Põlme et al. 2020) | Põlme S, et al. 2020. Fungal Diversity 105:1-16 ((https://doi.org/10.1007/s13225-020-00466-2)) |
| Otu38644 | 1 | k__Fungi(100);p__Ascomycota(100);c__Saccharomycetes(100);o__Saccharomycetales(100);f__Metschnikowiaceae(100);g__Metschnikowia(100);s__Metschnikowia_pulcherrima(100); | Metschnikowia | 13 | Saprotroph | \|Undefined Saprotroph\| | Yeast | NULL | Probable | As nectar-tap_saprotroph (Põlme et al. 2020); Decay sugar-rich substrates (Põlme et al. 2020) | Põlme S, et al. 2020. Fungal Diversity 105:1-16 ((https://doi.org/10.1007/s13225-020-00466-2)) |
| Otu38645 | 1 | k__Fungi(100);p__Ascomycota(100);c__Saccharomycetes(100);o__Saccharomycetales(100);f__Metschnikowiaceae(100);g__Metschnikowia(100);s__Metschnikowia_pulcherrima(100); | Metschnikowia | 13 | Saprotroph | \|Undefined Saprotroph\| | Yeast | NULL | Probable | As nectar-tap_saprotroph (Põlme et al. 2020); Decay sugar-rich substrates (Põlme et al. 2020) | Põlme S, et al. 2020. Fungal Diversity 105:1-16 ((https://doi.org/10.1007/s13225-020-00466-2)) |
| Otu38646 | 1 | k__Fungi(100);p__Ascomycota(100);c__Saccharomycetes(100);o__Saccharomycetales(100);f__Metschnikowiaceae(100);g__Metschnikowia(100);s__Metschnikowia_pulcherrima(100); | Metschnikowia | 13 | Saprotroph | \|Undefined Saprotroph\| | Yeast | NULL | Probable | As nectar-tap_saprotroph (Põlme et al. 2020); Decay sugar-rich substrates (Põlme et al. 2020) | Põlme S, et al. 2020. Fungal Diversity 105:1-16 ((https://doi.org/10.1007/s13225-020-00466-2)) |
| Otu38647 | 1 | k__Fungi(100);p__Ascomycota(100);c__Saccharomycetes(100);o__Saccharomycetales(100);f__Metschnikowiaceae(100);g__Metschnikowia(100);g__Metschnikowia_unclassified(100); | Metschnikowia | 13 | Saprotroph | \|Undefined Saprotroph\| | Yeast | NULL | Probable | As nectar-tap_saprotroph (Põlme et al. 2020); Decay sugar-rich substrates (Põlme et al. 2020) | Põlme S, et al. 2020. Fungal Diversity 105:1-16 ((https://doi.org/10.1007/s13225-020-00466-2)) |
| Otu38648 | 1 | k__Fungi(100);p__Ascomycota(100);c__Saccharomycetes(100);o__Saccharomycetales(100);f__Metschnikowiaceae(100);g__Metschnikowia(100);s__Metschnikowia_pulcherrima(100); | Metschnikowia | 13 | Saprotroph | \|Undefined Saprotroph\| | Yeast | NULL | Probable | As nectar-tap_saprotroph (Põlme et al. 2020); Decay sugar-rich substrates (Põlme et al. 2020) | Põlme S, et al. 2020. Fungal Diversity 105:1-16 ((https://doi.org/10.1007/s13225-020-00466-2)) |
| Otu38649 | 1 | k__Fungi(100);p__Ascomycota(100);c__Saccharomycetes(100);o__Saccharomycetales(100);f__Metschnikowiaceae(100);g__Metschnikowia(100);s__Metschnikowia_pulcherrima(100); | Metschnikowia | 13 | Saprotroph | \|Undefined Saprotroph\| | Yeast | NULL | Probable | As nectar-tap_saprotroph (Põlme et al. 2020); Decay sugar-rich substrates (Põlme et al. 2020) | Põlme S, et al. 2020. Fungal Diversity 105:1-16 ((https://doi.org/10.1007/s13225-020-00466-2)) |
| Otu38650 | 1 | k__Fungi(100);p__Ascomycota(100);c__Saccharomycetes(100);o__Saccharomycetales(100);f__Metschnikowiaceae(100);g__Metschnikowia(100);s__Metschnikowia_pulcherrima(100); | Metschnikowia | 13 | Saprotroph | \|Undefined Saprotroph\| | Yeast | NULL | Probable | As nectar-tap_saprotroph (Põlme et al. 2020); Decay sugar-rich substrates (Põlme et al. 2020) | Põlme S, et al. 2020. Fungal Diversity 105:1-16 ((https://doi.org/10.1007/s13225-020-00466-2)) |
| Otu38651 | 1 | k__Fungi(100);p__Ascomycota(100);c__Saccharomycetes(100);o__Saccharomycetales(100);f__Metschnikowiaceae(100);g__Metschnikowia(100);s__Metschnikowia_pulcherrima(100); | Metschnikowia | 13 | Saprotroph | \|Undefined Saprotroph\| | Yeast | NULL | Probable | As nectar-tap_saprotroph (Põlme et al. 2020); Decay sugar-rich substrates (Põlme et al. 2020) | Põlme S, et al. 2020. Fungal Diversity 105:1-16 ((https://doi.org/10.1007/s13225-020-00466-2)) |
| Otu38652 | 1 | k__Fungi(100);p__Ascomycota(100);c__Saccharomycetes(100);o__Saccharomycetales(100);f__Metschnikowiaceae(100);g__Metschnikowia(100);s__Metschnikowia_pulcherrima(100); | Metschnikowia | 13 | Saprotroph | \|Undefined Saprotroph\| | Yeast | NULL | Probable | As nectar-tap_saprotroph (Põlme et al. 2020); Decay sugar-rich substrates (Põlme et al. 2020) | Põlme S, et al. 2020. Fungal Diversity 105:1-16 ((https://doi.org/10.1007/s13225-020-00466-2)) |
| Otu38653 | 1 | k__Fungi(100);p__Ascomycota(100);c__Saccharomycetes(100);o__Saccharomycetales(100);f__Metschnikowiaceae(100);g__Metschnikowia(100);s__Metschnikowia_pulcherrima(100); | Metschnikowia | 13 | Saprotroph | \|Undefined Saprotroph\| | Yeast | NULL | Probable | As nectar-tap_saprotroph (Põlme et al. 2020); Decay sugar-rich substrates (Põlme et al. 2020) | Põlme S, et al. 2020. Fungal Diversity 105:1-16 ((https://doi.org/10.1007/s13225-020-00466-2)) |
| Otu38654 | 1 | k__Fungi(100);p__Ascomycota(100);c__Saccharomycetes(100);o__Saccharomycetales(100);f__Metschnikowiaceae(100);g__Metschnikowia(100);s__Metschnikowia_pulcherrima(100); | Metschnikowia | 13 | Saprotroph | \|Undefined Saprotroph\| | Yeast | NULL | Probable | As nectar-tap_saprotroph (Põlme et al. 2020); Decay sugar-rich substrates (Põlme et al. 2020) | Põlme S, et al. 2020. Fungal Diversity 105:1-16 ((https://doi.org/10.1007/s13225-020-00466-2)) |
| Otu38655 | 1 | k__Fungi(100);p__Ascomycota(100);c__Saccharomycetes(100);o__Saccharomycetales(100);f__Metschnikowiaceae(100);g__Metschnikowia(100);s__Metschnikowia_pulcherrima(100); | Metschnikowia | 13 | Saprotroph | \|Undefined Saprotroph\| | Yeast | NULL | Probable | As nectar-tap_saprotroph (Põlme et al. 2020); Decay sugar-rich substrates (Põlme et al. 2020) | Põlme S, et al. 2020. Fungal Diversity 105:1-16 ((https://doi.org/10.1007/s13225-020-00466-2)) |
| Otu38656 | 1 | k__Fungi(100);p__Ascomycota(100);c__Saccharomycetes(100);o__Saccharomycetales(100);f__Metschnikowiaceae(100);g__Metschnikowia(100);s__Metschnikowia_pulcherrima(100); | Metschnikowia | 13 | Saprotroph | \|Undefined Saprotroph\| | Yeast | NULL | Probable | As nectar-tap_saprotroph (Põlme et al. 2020); Decay sugar-rich substrates (Põlme et al. 2020) | Põlme S, et al. 2020. Fungal Diversity 105:1-16 ((https://doi.org/10.1007/s13225-020-00466-2)) |
| Otu38657 | 1 | k__Fungi(100);p__Ascomycota(100);c__Saccharomycetes(100);o__Saccharomycetales(100);f__Metschnikowiaceae(100);g__Metschnikowia(100);s__Metschnikowia_pulcherrima(100); | Metschnikowia | 13 | Saprotroph | \|Undefined Saprotroph\| | Yeast | NULL | Probable | As nectar-tap_saprotroph (Põlme et al. 2020); Decay sugar-rich substrates (Põlme et al. 2020) | Põlme S, et al. 2020. Fungal Diversity 105:1-16 ((https://doi.org/10.1007/s13225-020-00466-2)) |
| Otu38658 | 1 | k__Fungi(100);p__Ascomycota(100);c__Saccharomycetes(100);o__Saccharomycetales(100);f__Metschnikowiaceae(100);g__Metschnikowia(100);s__Metschnikowia_pulcherrima(100); | Metschnikowia | 13 | Saprotroph | \|Undefined Saprotroph\| | Yeast | NULL | Probable | As nectar-tap_saprotroph (Põlme et al. 2020); Decay sugar-rich substrates (Põlme et al. 2020) | Põlme S, et al. 2020. Fungal Diversity 105:1-16 ((https://doi.org/10.1007/s13225-020-00466-2)) |
| Otu38659 | 1 | k__Fungi(100);p__Ascomycota(100);c__Saccharomycetes(100);o__Saccharomycetales(100);f__Metschnikowiaceae(100);g__Metschnikowia(100);s__Metschnikowia_pulcherrima(100); | Metschnikowia | 13 | Saprotroph | \|Undefined Saprotroph\| | Yeast | NULL | Probable | As nectar-tap_saprotroph (Põlme et al. 2020); Decay sugar-rich substrates (Põlme et al. 2020) | Põlme S, et al. 2020. Fungal Diversity 105:1-16 ((https://doi.org/10.1007/s13225-020-00466-2)) |
| Otu38660 | 1 | k__Fungi(100);p__Ascomycota(100);c__Saccharomycetes(100);o__Saccharomycetales(100);f__Metschnikowiaceae(100);g__Metschnikowia(100);s__Metschnikowia_pulcherrima(100); | Metschnikowia | 13 | Saprotroph | \|Undefined Saprotroph\| | Yeast | NULL | Probable | As nectar-tap_saprotroph (Põlme et al. 2020); Decay sugar-rich substrates (Põlme et al. 2020) | Põlme S, et al. 2020. Fungal Diversity 105:1-16 ((https://doi.org/10.1007/s13225-020-00466-2)) |
| Otu38661 | 1 | k__Fungi(100);p__Ascomycota(100);c__Saccharomycetes(100);o__Saccharomycetales(100);f__Metschnikowiaceae(100);g__Metschnikowia(100);s__Metschnikowia_pulcherrima(100); | Metschnikowia | 13 | Saprotroph | \|Undefined Saprotroph\| | Yeast | NULL | Probable | As nectar-tap_saprotroph (Põlme et al. 2020); Decay sugar-rich substrates (Põlme et al. 2020) | Põlme S, et al. 2020. Fungal Diversity 105:1-16 ((https://doi.org/10.1007/s13225-020-00466-2)) |
| Otu38662 | 1 | k__Fungi(100);p__Ascomycota(100);c__Saccharomycetes(100);o__Saccharomycetales(100);f__Metschnikowiaceae(100);g__Metschnikowia(100);s__Metschnikowia_pulcherrima(100); | Metschnikowia | 13 | Saprotroph | \|Undefined Saprotroph\| | Yeast | NULL | Probable | As nectar-tap_saprotroph (Põlme et al. 2020); Decay sugar-rich substrates (Põlme et al. 2020) | Põlme S, et al. 2020. Fungal Diversity 105:1-16 ((https://doi.org/10.1007/s13225-020-00466-2)) |
| Otu38663 | 1 | k__Fungi(100);p__Ascomycota(100);c__Saccharomycetes(100);o__Saccharomycetales(100);f__Metschnikowiaceae(100);g__Metschnikowia(100);s__Metschnikowia_pulcherrima(100); | Metschnikowia | 13 | Saprotroph | \|Undefined Saprotroph\| | Yeast | NULL | Probable | As nectar-tap_saprotroph (Põlme et al. 2020); Decay sugar-rich substrates (Põlme et al. 2020) | Põlme S, et al. 2020. Fungal Diversity 105:1-16 ((https://doi.org/10.1007/s13225-020-00466-2)) |
| Otu38664 | 1 | k__Fungi(100);p__Ascomycota(100);c__Saccharomycetes(100);o__Saccharomycetales(100);f__Metschnikowiaceae(100);g__Metschnikowia(100);s__Metschnikowia_pulcherrima(100); | Metschnikowia | 13 | Saprotroph | \|Undefined Saprotroph\| | Yeast | NULL | Probable | As nectar-tap_saprotroph (Põlme et al. 2020); Decay sugar-rich substrates (Põlme et al. 2020) | Põlme S, et al. 2020. Fungal Diversity 105:1-16 ((https://doi.org/10.1007/s13225-020-00466-2)) |
| Otu38665 | 1 | k__Fungi(100);p__Ascomycota(100);c__Saccharomycetes(100);o__Saccharomycetales(100);f__Metschnikowiaceae(100);g__Metschnikowia(100);g__Metschnikowia_unclassified(100); | Metschnikowia | 13 | Saprotroph | \|Undefined Saprotroph\| | Yeast | NULL | Probable | As nectar-tap_saprotroph (Põlme et al. 2020); Decay sugar-rich substrates (Põlme et al. 2020) | Põlme S, et al. 2020. Fungal Diversity 105:1-16 ((https://doi.org/10.1007/s13225-020-00466-2)) |
| Otu38666 | 1 | k__Fungi(100);p__Ascomycota(100);c__Saccharomycetes(100);o__Saccharomycetales(100);f__Metschnikowiaceae(100);g__Metschnikowia(100);g__Metschnikowia_unclassified(100); | Metschnikowia | 13 | Saprotroph | \|Undefined Saprotroph\| | Yeast | NULL | Probable | As nectar-tap_saprotroph (Põlme et al. 2020); Decay sugar-rich substrates (Põlme et al. 2020) | Põlme S, et al. 2020. Fungal Diversity 105:1-16 ((https://doi.org/10.1007/s13225-020-00466-2)) |
| Otu38667 | 1 | k__Fungi(100);p__Ascomycota(100);c__Saccharomycetes(100);o__Saccharomycetales(100);f__Metschnikowiaceae(100);g__Metschnikowia(100);s__Metschnikowia_pulcherrima(100); | Metschnikowia | 13 | Saprotroph | \|Undefined Saprotroph\| | Yeast | NULL | Probable | As nectar-tap_saprotroph (Põlme et al. 2020); Decay sugar-rich substrates (Põlme et al. 2020) | Põlme S, et al. 2020. Fungal Diversity 105:1-16 ((https://doi.org/10.1007/s13225-020-00466-2)) |
| Otu38668 | 1 | k__Fungi(100);p__Ascomycota(100);c__Saccharomycetes(100);o__Saccharomycetales(100);f__Metschnikowiaceae(100);g__Metschnikowia(100);s__Metschnikowia_pulcherrima(100); | Metschnikowia | 13 | Saprotroph | \|Undefined Saprotroph\| | Yeast | NULL | Probable | As nectar-tap_saprotroph (Põlme et al. 2020); Decay sugar-rich substrates (Põlme et al. 2020) | Põlme S, et al. 2020. Fungal Diversity 105:1-16 ((https://doi.org/10.1007/s13225-020-00466-2)) |
| Otu38669 | 1 | k__Fungi(100);p__Ascomycota(100);c__Saccharomycetes(100);o__Saccharomycetales(100);f__Metschnikowiaceae(100);g__Metschnikowia(100);s__Metschnikowia_pulcherrima(100); | Metschnikowia | 13 | Saprotroph | \|Undefined Saprotroph\| | Yeast | NULL | Probable | As nectar-tap_saprotroph (Põlme et al. 2020); Decay sugar-rich substrates (Põlme et al. 2020) | Põlme S, et al. 2020. Fungal Diversity 105:1-16 ((https://doi.org/10.1007/s13225-020-00466-2)) |
| Otu38670 | 1 | k__Fungi(100);p__Ascomycota(100);c__Saccharomycetes(100);o__Saccharomycetales(100);f__Metschnikowiaceae(100);g__Metschnikowia(100);s__Metschnikowia_pulcherrima(100); | Metschnikowia | 13 | Saprotroph | \|Undefined Saprotroph\| | Yeast | NULL | Probable | As nectar-tap_saprotroph (Põlme et al. 2020); Decay sugar-rich substrates (Põlme et al. 2020) | Põlme S, et al. 2020. Fungal Diversity 105:1-16 ((https://doi.org/10.1007/s13225-020-00466-2)) |
| Otu38671 | 1 | k__Fungi(100);p__Ascomycota(100);c__Saccharomycetes(100);o__Saccharomycetales(100);f__Metschnikowiaceae(100);g__Metschnikowia(100);s__Metschnikowia_pulcherrima(100); | Metschnikowia | 13 | Saprotroph | \|Undefined Saprotroph\| | Yeast | NULL | Probable | As nectar-tap_saprotroph (Põlme et al. 2020); Decay sugar-rich substrates (Põlme et al. 2020) | Põlme S, et al. 2020. Fungal Diversity 105:1-16 ((https://doi.org/10.1007/s13225-020-00466-2)) |
| Otu38672 | 1 | k__Fungi(100);p__Ascomycota(100);c__Saccharomycetes(100);o__Saccharomycetales(100);f__Metschnikowiaceae(100);g__Metschnikowia(100);s__Metschnikowia_pulcherrima(100); | Metschnikowia | 13 | Saprotroph | \|Undefined Saprotroph\| | Yeast | NULL | Probable | As nectar-tap_saprotroph (Põlme et al. 2020); Decay sugar-rich substrates (Põlme et al. 2020) | Põlme S, et al. 2020. Fungal Diversity 105:1-16 ((https://doi.org/10.1007/s13225-020-00466-2)) |
| Otu38673 | 1 | k__Fungi(100);p__Ascomycota(100);c__Saccharomycetes(100);o__Saccharomycetales(100);f__Metschnikowiaceae(100);g__Metschnikowia(100);s__Metschnikowia_pulcherrima(100); | Metschnikowia | 13 | Saprotroph | \|Undefined Saprotroph\| | Yeast | NULL | Probable | As nectar-tap_saprotroph (Põlme et al. 2020); Decay sugar-rich substrates (Põlme et al. 2020) | Põlme S, et al. 2020. Fungal Diversity 105:1-16 ((https://doi.org/10.1007/s13225-020-00466-2)) |
| Otu38674 | 1 | k__Fungi(100);p__Ascomycota(100);c__Saccharomycetes(100);o__Saccharomycetales(100);f__Metschnikowiaceae(100);g__Metschnikowia(100);s__Metschnikowia_pulcherrima(100); | Metschnikowia | 13 | Saprotroph | \|Undefined Saprotroph\| | Yeast | NULL | Probable | As nectar-tap_saprotroph (Põlme et al. 2020); Decay sugar-rich substrates (Põlme et al. 2020) | Põlme S, et al. 2020. Fungal Diversity 105:1-16 ((https://doi.org/10.1007/s13225-020-00466-2)) |
| Otu38675 | 1 | k__Fungi(100);p__Ascomycota(100);c__Saccharomycetes(100);o__Saccharomycetales(100);f__Metschnikowiaceae(100);g__Metschnikowia(100);s__Metschnikowia_pulcherrima(100); | Metschnikowia | 13 | Saprotroph | \|Undefined Saprotroph\| | Yeast | NULL | Probable | As nectar-tap_saprotroph (Põlme et al. 2020); Decay sugar-rich substrates (Põlme et al. 2020) | Põlme S, et al. 2020. Fungal Diversity 105:1-16 ((https://doi.org/10.1007/s13225-020-00466-2)) |
| Otu38676 | 1 | k__Fungi(100);p__Ascomycota(100);c__Saccharomycetes(100);o__Saccharomycetales(100);f__Metschnikowiaceae(100);g__Metschnikowia(100);s__Metschnikowia_pulcherrima(100); | Metschnikowia | 13 | Saprotroph | \|Undefined Saprotroph\| | Yeast | NULL | Probable | As nectar-tap_saprotroph (Põlme et al. 2020); Decay sugar-rich substrates (Põlme et al. 2020) | Põlme S, et al. 2020. Fungal Diversity 105:1-16 ((https://doi.org/10.1007/s13225-020-00466-2)) |
| Otu38677 | 1 | k__Fungi(100);p__Ascomycota(100);c__Saccharomycetes(100);o__Saccharomycetales(100);f__Metschnikowiaceae(100);g__Metschnikowia(100);s__Metschnikowia_pulcherrima(100); | Metschnikowia | 13 | Saprotroph | \|Undefined Saprotroph\| | Yeast | NULL | Probable | As nectar-tap_saprotroph (Põlme et al. 2020); Decay sugar-rich substrates (Põlme et al. 2020) | Põlme S, et al. 2020. Fungal Diversity 105:1-16 ((https://doi.org/10.1007/s13225-020-00466-2)) |
| Otu38678 | 1 | k__Fungi(100);p__Ascomycota(100);c__Saccharomycetes(100);o__Saccharomycetales(100);f__Metschnikowiaceae(100);g__Metschnikowia(100);s__Metschnikowia_pulcherrima(100); | Metschnikowia | 13 | Saprotroph | \|Undefined Saprotroph\| | Yeast | NULL | Probable | As nectar-tap_saprotroph (Põlme et al. 2020); Decay sugar-rich substrates (Põlme et al. 2020) | Põlme S, et al. 2020. Fungal Diversity 105:1-16 ((https://doi.org/10.1007/s13225-020-00466-2)) |
| Otu38679 | 1 | k__Fungi(100);p__Ascomycota(100);c__Saccharomycetes(100);o__Saccharomycetales(100);f__Metschnikowiaceae(100);g__Metschnikowia(100);s__Metschnikowia_pulcherrima(100); | Metschnikowia | 13 | Saprotroph | \|Undefined Saprotroph\| | Yeast | NULL | Probable | As nectar-tap_saprotroph (Põlme et al. 2020); Decay sugar-rich substrates (Põlme et al. 2020) | Põlme S, et al. 2020. Fungal Diversity 105:1-16 ((https://doi.org/10.1007/s13225-020-00466-2)) |
| Otu38680 | 1 | k__Fungi(100);p__Ascomycota(100);c__Saccharomycetes(100);o__Saccharomycetales(100);f__Metschnikowiaceae(100);g__Metschnikowia(100);g__Metschnikowia_unclassified(100); | Metschnikowia | 13 | Saprotroph | \|Undefined Saprotroph\| | Yeast | NULL | Probable | As nectar-tap_saprotroph (Põlme et al. 2020); Decay sugar-rich substrates (Põlme et al. 2020) | Põlme S, et al. 2020. Fungal Diversity 105:1-16 ((https://doi.org/10.1007/s13225-020-00466-2)) |
| Otu38681 | 1 | k__Fungi(100);p__Ascomycota(100);c__Saccharomycetes(100);o__Saccharomycetales(100);f__Metschnikowiaceae(100);g__Metschnikowia(100);s__Metschnikowia_pulcherrima(100); | Metschnikowia | 13 | Saprotroph | \|Undefined Saprotroph\| | Yeast | NULL | Probable | As nectar-tap_saprotroph (Põlme et al. 2020); Decay sugar-rich substrates (Põlme et al. 2020) | Põlme S, et al. 2020. Fungal Diversity 105:1-16 ((https://doi.org/10.1007/s13225-020-00466-2)) |
| Otu38682 | 1 | k__Fungi(100);p__Ascomycota(100);c__Saccharomycetes(100);o__Saccharomycetales(100);f__Metschnikowiaceae(100);g__Metschnikowia(100);s__Metschnikowia_pulcherrima(100); | Metschnikowia | 13 | Saprotroph | \|Undefined Saprotroph\| | Yeast | NULL | Probable | As nectar-tap_saprotroph (Põlme et al. 2020); Decay sugar-rich substrates (Põlme et al. 2020) | Põlme S, et al. 2020. Fungal Diversity 105:1-16 ((https://doi.org/10.1007/s13225-020-00466-2)) |
| Otu38683 | 1 | k__Fungi(100);p__Ascomycota(100);c__Saccharomycetes(100);o__Saccharomycetales(100);f__Metschnikowiaceae(100);g__Metschnikowia(100);s__Metschnikowia_pulcherrima(100); | Metschnikowia | 13 | Saprotroph | \|Undefined Saprotroph\| | Yeast | NULL | Probable | As nectar-tap_saprotroph (Põlme et al. 2020); Decay sugar-rich substrates (Põlme et al. 2020) | Põlme S, et al. 2020. Fungal Diversity 105:1-16 ((https://doi.org/10.1007/s13225-020-00466-2)) |
| Otu38684 | 1 | k__Fungi(100);p__Ascomycota(100);c__Saccharomycetes(100);o__Saccharomycetales(100);f__Metschnikowiaceae(100);g__Metschnikowia(100);g__Metschnikowia_unclassified(100); | Metschnikowia | 13 | Saprotroph | \|Undefined Saprotroph\| | Yeast | NULL | Probable | As nectar-tap_saprotroph (Põlme et al. 2020); Decay sugar-rich substrates (Põlme et al. 2020) | Põlme S, et al. 2020. Fungal Diversity 105:1-16 ((https://doi.org/10.1007/s13225-020-00466-2)) |
| Otu38685 | 1 | k__Fungi(100);p__Ascomycota(100);c__Saccharomycetes(100);o__Saccharomycetales(100);f__Metschnikowiaceae(100);g__Metschnikowia(100);g__Metschnikowia_unclassified(100); | Metschnikowia | 13 | Saprotroph | \|Undefined Saprotroph\| | Yeast | NULL | Probable | As nectar-tap_saprotroph (Põlme et al. 2020); Decay sugar-rich substrates (Põlme et al. 2020) | Põlme S, et al. 2020. Fungal Diversity 105:1-16 ((https://doi.org/10.1007/s13225-020-00466-2)) |
| Otu38686 | 1 | k__Fungi(100);p__Ascomycota(100);c__Saccharomycetes(100);o__Saccharomycetales(100);f__Metschnikowiaceae(100);g__Metschnikowia(100);s__Metschnikowia_pulcherrima(100); | Metschnikowia | 13 | Saprotroph | \|Undefined Saprotroph\| | Yeast | NULL | Probable | As nectar-tap_saprotroph (Põlme et al. 2020); Decay sugar-rich substrates (Põlme et al. 2020) | Põlme S, et al. 2020. Fungal Diversity 105:1-16 ((https://doi.org/10.1007/s13225-020-00466-2)) |
| Otu38687 | 1 | k__Fungi(100);p__Ascomycota(100);c__Saccharomycetes(100);o__Saccharomycetales(100);f__Metschnikowiaceae(100);g__Metschnikowia(100);s__Metschnikowia_pulcherrima(100); | Metschnikowia | 13 | Saprotroph | \|Undefined Saprotroph\| | Yeast | NULL | Probable | As nectar-tap_saprotroph (Põlme et al. 2020); Decay sugar-rich substrates (Põlme et al. 2020) | Põlme S, et al. 2020. Fungal Diversity 105:1-16 ((https://doi.org/10.1007/s13225-020-00466-2)) |
| Otu38688 | 1 | k__Fungi(100);p__Ascomycota(100);c__Saccharomycetes(100);o__Saccharomycetales(100);f__Metschnikowiaceae(100);g__Metschnikowia(100);s__Metschnikowia_pulcherrima(100); | Metschnikowia | 13 | Saprotroph | \|Undefined Saprotroph\| | Yeast | NULL | Probable | As nectar-tap_saprotroph (Põlme et al. 2020); Decay sugar-rich substrates (Põlme et al. 2020) | Põlme S, et al. 2020. Fungal Diversity 105:1-16 ((https://doi.org/10.1007/s13225-020-00466-2)) |
| Otu38689 | 1 | k__Fungi(100);p__Ascomycota(100);c__Saccharomycetes(100);o__Saccharomycetales(100);f__Metschnikowiaceae(100);g__Metschnikowia(100);s__Metschnikowia_pulcherrima(100); | Metschnikowia | 13 | Saprotroph | \|Undefined Saprotroph\| | Yeast | NULL | Probable | As nectar-tap_saprotroph (Põlme et al. 2020); Decay sugar-rich substrates (Põlme et al. 2020) | Põlme S, et al. 2020. Fungal Diversity 105:1-16 ((https://doi.org/10.1007/s13225-020-00466-2)) |
| Otu38690 | 1 | k__Fungi(100);p__Ascomycota(100);c__Saccharomycetes(100);o__Saccharomycetales(100);f__Metschnikowiaceae(100);g__Metschnikowia(100);s__Metschnikowia_pulcherrima(100); | Metschnikowia | 13 | Saprotroph | \|Undefined Saprotroph\| | Yeast | NULL | Probable | As nectar-tap_saprotroph (Põlme et al. 2020); Decay sugar-rich substrates (Põlme et al. 2020) | Põlme S, et al. 2020. Fungal Diversity 105:1-16 ((https://doi.org/10.1007/s13225-020-00466-2)) |
| Otu38691 | 1 | k__Fungi(100);p__Ascomycota(100);c__Saccharomycetes(100);o__Saccharomycetales(100);f__Metschnikowiaceae(100);g__Metschnikowia(100);s__Metschnikowia_pulcherrima(100); | Metschnikowia | 13 | Saprotroph | \|Undefined Saprotroph\| | Yeast | NULL | Probable | As nectar-tap_saprotroph (Põlme et al. 2020); Decay sugar-rich substrates (Põlme et al. 2020) | Põlme S, et al. 2020. Fungal Diversity 105:1-16 ((https://doi.org/10.1007/s13225-020-00466-2)) |
| Otu38693 | 1 | k__Fungi(100);p__Ascomycota(100);c__Saccharomycetes(100);o__Saccharomycetales(100);f__Metschnikowiaceae(100);g__Metschnikowia(100);s__Metschnikowia_pulcherrima(100); | Metschnikowia | 13 | Saprotroph | \|Undefined Saprotroph\| | Yeast | NULL | Probable | As nectar-tap_saprotroph (Põlme et al. 2020); Decay sugar-rich substrates (Põlme et al. 2020) | Põlme S, et al. 2020. Fungal Diversity 105:1-16 ((https://doi.org/10.1007/s13225-020-00466-2)) |
| Otu38695 | 1 | k__Fungi(100);p__Ascomycota(100);c__Saccharomycetes(100);o__Saccharomycetales(100);f__Metschnikowiaceae(100);g__Metschnikowia(100);g__Metschnikowia_unclassified(100); | Metschnikowia | 13 | Saprotroph | \|Undefined Saprotroph\| | Yeast | NULL | Probable | As nectar-tap_saprotroph (Põlme et al. 2020); Decay sugar-rich substrates (Põlme et al. 2020) | Põlme S, et al. 2020. Fungal Diversity 105:1-16 ((https://doi.org/10.1007/s13225-020-00466-2)) |
| Otu38697 | 1 | k__Fungi(100);p__Ascomycota(100);c__Saccharomycetes(100);o__Saccharomycetales(100);f__Metschnikowiaceae(100);g__Metschnikowia(100);g__Metschnikowia_unclassified(100); | Metschnikowia | 13 | Saprotroph | \|Undefined Saprotroph\| | Yeast | NULL | Probable | As nectar-tap_saprotroph (Põlme et al. 2020); Decay sugar-rich substrates (Põlme et al. 2020) | Põlme S, et al. 2020. Fungal Diversity 105:1-16 ((https://doi.org/10.1007/s13225-020-00466-2)) |
| Otu38699 | 1 | k__Fungi(100);p__Ascomycota(100);c__Saccharomycetes(100);o__Saccharomycetales(100);f__Metschnikowiaceae(100);g__Metschnikowia(100);g__Metschnikowia_unclassified(100); | Metschnikowia | 13 | Saprotroph | \|Undefined Saprotroph\| | Yeast | NULL | Probable | As nectar-tap_saprotroph (Põlme et al. 2020); Decay sugar-rich substrates (Põlme et al. 2020) | Põlme S, et al. 2020. Fungal Diversity 105:1-16 ((https://doi.org/10.1007/s13225-020-00466-2)) |
| Otu38700 | 1 | k__Fungi(100);p__Ascomycota(100);c__Saccharomycetes(100);o__Saccharomycetales(100);f__Metschnikowiaceae(100);g__Metschnikowia(100);g__Metschnikowia_unclassified(100); | Metschnikowia | 13 | Saprotroph | \|Undefined Saprotroph\| | Yeast | NULL | Probable | As nectar-tap_saprotroph (Põlme et al. 2020); Decay sugar-rich substrates (Põlme et al. 2020) | Põlme S, et al. 2020. Fungal Diversity 105:1-16 ((https://doi.org/10.1007/s13225-020-00466-2)) |
| Otu38701 | 1 | k__Fungi(100);p__Ascomycota(100);c__Saccharomycetes(100);o__Saccharomycetales(100);f__Metschnikowiaceae(100);g__Metschnikowia(100);s__Metschnikowia_pulcherrima(100); | Metschnikowia | 13 | Saprotroph | \|Undefined Saprotroph\| | Yeast | NULL | Probable | As nectar-tap_saprotroph (Põlme et al. 2020); Decay sugar-rich substrates (Põlme et al. 2020) | Põlme S, et al. 2020. Fungal Diversity 105:1-16 ((https://doi.org/10.1007/s13225-020-00466-2)) |
| Otu38702 | 1 | k__Fungi(100);p__Basidiomycota(100);c__Tremellomycetes(100);o__Tremellales(100);f__Tremellales_family_Incertae_sedis(100);g__Fibulobasidium(100);s__Fibulobasidium_inconspicuum(100); | Fibulobasidium | 13 | Saprotroph | Undefined Saprotroph-\|Wood Saprotroph\| | Dimorphic-Facultative Yeast | NULL | Highly Probable | Decay wood (Põlme et al. 2020) | Kurtzman CP, et al. (eds.) 2011. The Yeasts, a Taxonomic Study. Fifth Edition. Vols 1-3. Elsevier, San Diego (ISBN: 9780444521491); Põlme S, et al. 2020. Fungal Diversity 105:1-16 ((https://doi.org/10.1007/s13225-020-00466-2)) |
| Otu38704 | 1 | k__Fungi(100);p__Ascomycota(100);c__Saccharomycetes(100);o__Saccharomycetales(100);f__Metschnikowiaceae(100);g__Metschnikowia(100);s__Metschnikowia_pulcherrima(100); | Metschnikowia | 13 | Saprotroph | \|Undefined Saprotroph\| | Yeast | NULL | Probable | As nectar-tap_saprotroph (Põlme et al. 2020); Decay sugar-rich substrates (Põlme et al. 2020) | Põlme S, et al. 2020. Fungal Diversity 105:1-16 ((https://doi.org/10.1007/s13225-020-00466-2)) |
| Otu38705 | 1 | k__Fungi(100);p__Ascomycota(100);c__Saccharomycetes(100);o__Saccharomycetales(100);f__Metschnikowiaceae(100);g__Metschnikowia(100);s__Metschnikowia_pulcherrima(100); | Metschnikowia | 13 | Saprotroph | \|Undefined Saprotroph\| | Yeast | NULL | Probable | As nectar-tap_saprotroph (Põlme et al. 2020); Decay sugar-rich substrates (Põlme et al. 2020) | Põlme S, et al. 2020. Fungal Diversity 105:1-16 ((https://doi.org/10.1007/s13225-020-00466-2)) |
| Otu38707 | 1 | k__Fungi(100);p__Ascomycota(100);c__Saccharomycetes(100);o__Saccharomycetales(100);f__Metschnikowiaceae(100);g__Metschnikowia(100);g__Metschnikowia_unclassified(100); | Metschnikowia | 13 | Saprotroph | \|Undefined Saprotroph\| | Yeast | NULL | Probable | As nectar-tap_saprotroph (Põlme et al. 2020); Decay sugar-rich substrates (Põlme et al. 2020) | Põlme S, et al. 2020. Fungal Diversity 105:1-16 ((https://doi.org/10.1007/s13225-020-00466-2)) |
| Otu38708 | 1 | k__Fungi(100);p__Ascomycota(100);c__Saccharomycetes(100);o__Saccharomycetales(100);f__Metschnikowiaceae(100);g__Metschnikowia(100);g__Metschnikowia_unclassified(100); | Metschnikowia | 13 | Saprotroph | \|Undefined Saprotroph\| | Yeast | NULL | Probable | As nectar-tap_saprotroph (Põlme et al. 2020); Decay sugar-rich substrates (Põlme et al. 2020) | Põlme S, et al. 2020. Fungal Diversity 105:1-16 ((https://doi.org/10.1007/s13225-020-00466-2)) |

| | | | | | | | | | | | | |
|---|---|---|---|---|---|---|---|---|---|---|---|---|
| Otu38709 | 1 | k__Fungi(100);p__Ascomycota(100);c__Saccharomycetes(100);o__Saccharomycetales(100);f__Metschnikowiaceae(100);g__Metschnikowia(100);g__Metschnikowia_unclassified(100); | Metschnikowia | | 13 | Saprotroph | \|Undefined Saprotroph\| | Yeast | NULL | Probable | As nectar-tap_saprotroph (Põlme et al. 2020); Decay sugar-rich substrates (Põlme et al. 2020) | Põlme S, et al. 2020. Fungal Diversity 105:1-16 ((https://doi.org/10.1007/s13225-020-00466-2)) |
| Otu38710 | 1 | k__Fungi(100);p__Ascomycota(100);c__Saccharomycetes(100);o__Saccharomycetales(100);f__Metschnikowiaceae(100);g__Metschnikowia(100);g__Metschnikowia_unclassified(100); | Metschnikowia | | 13 | Saprotroph | \|Undefined Saprotroph\| | Yeast | NULL | Probable | As nectar-tap_saprotroph (Põlme et al. 2020); Decay sugar-rich substrates (Põlme et al. 2020) | Põlme S, et al. 2020. Fungal Diversity 105:1-16 ((https://doi.org/10.1007/s13225-020-00466-2)) |
| Otu38711 | 1 | k__Fungi(100);p__Ascomycota(100);c__Saccharomycetes(100);o__Saccharomycetales(100);f__Metschnikowiaceae(100);g__Metschnikowia(100);g__Metschnikowia_unclassified(100); | Metschnikowia | | 13 | Saprotroph | \|Undefined Saprotroph\| | Yeast | NULL | Probable | As nectar-tap_saprotroph (Põlme et al. 2020); Decay sugar-rich substrates (Põlme et al. 2020) | Põlme S, et al. 2020. Fungal Diversity 105:1-16 ((https://doi.org/10.1007/s13225-020-00466-2)) |
| Otu38712 | 1 | k__Fungi(100);p__Ascomycota(100);c__Saccharomycetes(100);o__Saccharomycetales(100);f__Metschnikowiaceae(100);g__Metschnikowia(100);g__Metschnikowia_unclassified(100); | Metschnikowia | | 13 | Saprotroph | \|Undefined Saprotroph\| | Yeast | NULL | Probable | As nectar-tap_saprotroph (Põlme et al. 2020); Decay sugar-rich substrates (Põlme et al. 2020) | Põlme S, et al. 2020. Fungal Diversity 105:1-16 ((https://doi.org/10.1007/s13225-020-00466-2)) |
| Otu38713 | 1 | k__Fungi(100);p__Ascomycota(100);c__Saccharomycetes(100);o__Saccharomycetales(100);f__Metschnikowiaceae(100);g__Metschnikowia(100);g__Metschnikowia_unclassified(100); | Metschnikowia | | 13 | Saprotroph | \|Undefined Saprotroph\| | Yeast | NULL | Probable | As nectar-tap_saprotroph (Põlme et al. 2020); Decay sugar-rich substrates (Põlme et al. 2020) | Põlme S, et al. 2020. Fungal Diversity 105:1-16 ((https://doi.org/10.1007/s13225-020-00466-2)) |
| Otu38714 | 1 | k__Fungi(100);p__Ascomycota(100);c__Saccharomycetes(100);o__Saccharomycetales(100);f__Metschnikowiaceae(100);g__Metschnikowia(100);g__Metschnikowia_unclassified(100); | Metschnikowia | | 13 | Saprotroph | \|Undefined Saprotroph\| | Yeast | NULL | Probable | As nectar-tap_saprotroph (Põlme et al. 2020); Decay sugar-rich substrates (Põlme et al. 2020) | Põlme S, et al. 2020. Fungal Diversity 105:1-16 ((https://doi.org/10.1007/s13225-020-00466-2)) |
| Otu38715 | 1 | k__Fungi(100);p__Ascomycota(100);c__Saccharomycetes(100);o__Saccharomycetales(100);f__Metschnikowiaceae(100);g__Metschnikowia(100);g__Metschnikowia_unclassified(100); | Metschnikowia | | 13 | Saprotroph | \|Undefined Saprotroph\| | Yeast | NULL | Probable | As nectar-tap_saprotroph (Põlme et al. 2020); Decay sugar-rich substrates (Põlme et al. 2020) | Põlme S, et al. 2020. Fungal Diversity 105:1-16 ((https://doi.org/10.1007/s13225-020-00466-2)) |
| Otu38716 | 1 | k__Fungi(100);p__Ascomycota(100);c__Saccharomycetes(100);o__Saccharomycetales(100);f__Metschnikowiaceae(100);g__Metschnikowia(100);g__Metschnikowia_unclassified(100); | Metschnikowia | | 13 | Saprotroph | \|Undefined Saprotroph\| | Yeast | NULL | Probable | As nectar-tap_saprotroph (Põlme et al. 2020); Decay sugar-rich substrates (Põlme et al. 2020) | Põlme S, et al. 2020. Fungal Diversity 105:1-16 ((https://doi.org/10.1007/s13225-020-00466-2)) |
| Otu38717 | 1 | k__Fungi(100);p__Ascomycota(100);c__Saccharomycetes(100);o__Saccharomycetales(100);f__Trichomonascaceae(100);g__Blastobotrys(100);g__Blastobotrys_unclassified(100); | Blastobotrys | | 13 | Saprotroph | \|Undefined Saprotroph\| | Yeast | Its unusual metal | Highly Probable | As nectar-tap_saprotroph (Põlme et al. 2020); Decay sugar-rich substrates (Põlme et al. 2020); In the course of identifying alternative microorganisms for biotechnological interest, A. adeninivorans has already proved its strengthened competitiveness as a promising cell factory for many more applications. | Kunze G, et al. 2014. Biotechnology for Biofuels 7:66 ((https://doi.org/10.1186/1754-6834-7-66)); Põlme S, et al. 2020. Fungal Diversity 105:1-16 ((https://doi.org/10.1007/s13225-020-00466-2)) |
| Otu38718 | 1 | k__Fungi(100);p__Ascomycota(100);c__Pezizomycetes(100);o__Pezizales(100);f__Ascobolaceae(100);g__Ascobolus(100);s__Ascobolus_sp(100); | Ascobolus | | 13 | Saprotroph | Dung Saprotroph-\|Plant Saprotroph\|-Wood Saprotroph | Microfungus | Soft Rot | Probable | As litter_saprotroph (Põlme et al. 2020); Decay leaf, fruit, seed, dung (Põlme et al. 2020) | Duncan CG, Eslyn WE. 1966. Mycologia 58:642-645 ((https://doi.org/10.2307/3757045)); Bell A. 1983. Dung Fungi: An Illustrated Guide to Coprophilous Fungi in New Zealand. Victoria University Press, Wellington (ISBN: 978-0864730015); Tedersoo L, et al. 2014. Science 346:e1256688 ((https://doi.org/10.1126/science.1256688)); Põlme S, et al. 2020. Fungal Diversity 105:1-16 ((https://doi.org/10.1007/s13225-020-00466-2)) |
| Otu38719 | 1 | k__Fungi(100);p__Ascomycota(100);c__Dothideomycetes(100);o__Pleosporales(100);f__Pleosporales_family_Incertae_sedis(100);g__Phoma(100);g__Phoma_unclassified(100); | Phoma | | 13 | Pathotroph-Saprotroph-Symbiotroph | Dung Saprotroph-Endophyte-Lichen Parasite-\|Plant Pathogen\|-Plant Saprotroph-Undefined Saprotroph-Wood Saprotroph | Microfungus | Soft Rot | Possible | As foliar_endophyte (Põlme et al. 2020); As litter_saprotroph (Põlme et al. 2020); Decay leaf, fruit, seed, roots, soil (Põlme et al. 2020) | Richardson MJ. 1972. Transactions of the British Mycological Society 58:37-48 ((https://doi.org/10.1016/S0007-1536(72)80069-X)); Seehann G, et al. 1975. List of Fungi in Soft-Rot Tissue Tests. Institute fur Holzbiologie und Holzschutz, Bundesforschungsanstalt fur Forst - und Holzwirtshaft, Hamburg - Lohbrugge, Germany; de Gruyter J, et al. 2010. Mycologia 102:1066-1081 ((https://doi.org/10.3852/09-240)); Costa IPMW, et al. 2012. Mycotaxon 119:e26 ((http://mycotaxon.com/resources/checklists/costa_v119_checklist.pdf)); Tedersoo L, et al. 2014. Science 346:e1256688 ((https://doi.org/10.1126/science.1256688)); Lawrey JD, Diederich P. 2016. http://www.lichenicolous.net (accessed 24 May 2017) ((http://www.lichenicolous.net)); Põlme S, et al. 2020. Fungal Diversity 105:1-16 ((https://doi.org/10.1007/s13225-020-00466-2)) |
| Otu38721 | 1 | k__Fungi(100);p__Ascomycota(100);c__Dothideomycetes(100);o__Patellariales(100);f__Patellariaceae(100);g__Rhytidhysteron(100);s__Rhytidhysteron_rufulum(100); | Rhytidhysteron | | 13 | Pathotroph-Saprotroph | Plant Pathogen-Undefined Saprotroph-\|Wood Saprotroph\| | Microfungus | Soft Rot | Probable | Decay: wood (Põlme et al. 2020) | Tedersoo L, et al. 2014. Science 346:e1256688 ((https://doi.org/10.1126/science.1256688)); Põlme S, et al. 2020. Fungal Diversity 105:1-16 ((https://doi.org/10.1007/s13225-020-00466-2)) |
| Otu38722 | 1 | k__Fungi(100);p__Ascomycota(100);c__Dothideomycetes(100);o__Pleosporales(100);f__Didymosphaeriaceae(100);g__Roussoella(100);s__Roussoella_japanensis(100); | Roussoella | | 13 | Saprotroph | Undefined Saprotroph-\|Wood Saprotroph\| | Microfungus | NULL | Probable | Decay wood (Põlme et al. 2020) | Tedersoo L, et al. 2014. Science 346:e1256688 ((https://doi.org/10.1126/science.1256688)); Põlme S, et al. 2020. Fungal Diversity 105:1-16 ((https://doi.org/10.1007/s13225-020-00466-2)) |
| Otu38724 | 1 | k__Fungi(100);p__Ascomycota(100);c__Dothideomycetes(100);o__Pleosporales(100);f__Pleosporales_family_Incertae_sedis(100);g__Phoma(100);g__Phoma_unclassified(100); | Phoma | | 13 | Pathotroph-Saprotroph-Symbiotroph | Dung Saprotroph-Endophyte-Lichen Parasite-\|Plant Pathogen\|-Plant Saprotroph-Undefined Saprotroph-Wood Saprotroph | Microfungus | Soft Rot | Possible | As foliar_endophyte (Põlme et al. 2020); As litter_saprotroph (Põlme et al. 2020); Decay leaf, fruit, seed, roots, soil (Põlme et al. 2020) | Richardson MJ. 1972. Transactions of the British Mycological Society 58:37-48 ((https://doi.org/10.1016/S0007-1536(72)80069-X)); Seehann G, et al. 1975. List of Fungi in Soft-Rot Tissue Tests. Institute fur Holzbiologie und Holzschutz, Bundesforschungsanstalt fur Forst - und Holzwirtshaft, Hamburg - Lohbrugge, Germany; de Gruyter J, et al. 2010. Mycologia 102:1066-1081 ((https://doi.org/10.3852/09-240)); Costa IPMW, et al. 2012. Mycotaxon 119:e26 ((http://mycotaxon.com/resources/checklists/costa_v119_checklist.pdf)); Tedersoo L, et al. 2014. Science 346:e1256688 ((https://doi.org/10.1126/science.1256688)); Lawrey JD, Diederich P. 2016. http://www.lichenicolous.net (accessed 24 May 2017) ((http://www.lichenicolous.net)); Põlme S, et al. 2020. Fungal Diversity 105:1-16 ((https://doi.org/10.1007/s13225-020-00466-2)) |
| Otu38725 | 1 | k__Fungi(100);p__Ascomycota(100);c__Dothideomycetes(100);o__Pleosporales(100);f__Pleosporales_family_Incertae_sedis(100);g__Pyrenochaeta(100);s__Pyrenochaeta_keratinophila(100); | Pyrenochaeta | | 13 | Pathotroph-Saprotroph | Plant Pathogen-Undefined Saprotroph-\|Wood Saprotroph\| | Microfungus | Soft Rot | Highly Probable | Decay wood (Põlme et al. 2020) | Seehann G, et al. 1975. List of Fungi in Soft-Rot Tissue Tests. Institute fur Holzbiologie und Holzschutz, Bundesforschungsanstalt fur Forst - und Holzwirtshaft, Hamburg - Lohbrugge, Germany; Costa IPMW, et al. 2012. Mycotaxon 119:e26 ((http://mycotaxon.com/resources/checklists/costa_v119_checklist.pdf)); Tedersoo L, et al. 2014. Science 346:e1256688 ((https://doi.org/10.1126/science.1256688)); Põlme S, et al. 2020. Fungal Diversity 105:1-16 ((https://doi.org/10.1007/s13225-020-00466-2)) |
| Otu38726 | 1 | k__Fungi(100);p__Ascomycota(100);c__Dothideomycetes(100);o__Pleosporales(100);f__Pleosporales_family_Incertae_sedis(100);g__Phoma(100);g__Phoma_unclassified(100); | Phoma | | 13 | Pathotroph-Saprotroph-Symbiotroph | Dung Saprotroph-Endophyte-Lichen Parasite-\|Plant Pathogen\|-Plant Saprotroph-Undefined Saprotroph-Wood Saprotroph | Microfungus | Soft Rot | Possible | As foliar_endophyte (Põlme et al. 2020); As litter_saprotroph (Põlme et al. 2020); Decay leaf, fruit, seed, roots, soil (Põlme et al. 2020) | Richardson MJ. 1972. Transactions of the British Mycological Society 58:37-48 ((https://doi.org/10.1016/S0007-1536(72)80069-X)); Seehann G, et al. 1975. List of Fungi in Soft-Rot Tissue Tests. Institute fur Holzbiologie und Holzschutz, Bundesforschungsanstalt fur Forst - und Holzwirtshaft, Hamburg - Lohbrugge, Germany; de Gruyter J, et al. 2010. Mycologia 102:1066-1081 ((https://doi.org/10.3852/09-240)); Costa IPMW, et al. 2012. Mycotaxon 119:e26 ((http://mycotaxon.com/resources/checklists/costa_v119_checklist.pdf)); Tedersoo L, et al. 2014. Science 346:e1256688 ((https://doi.org/10.1126/science.1256688)); Lawrey JD, Diederich P. 2016. http://www.lichenicolous.net (accessed 24 May 2017) ((http://www.lichenicolous.net)); Põlme S, et al. 2020. Fungal Diversity 105:1-16 ((https://doi.org/10.1007/s13225-020-00466-2)) |
| Otu38728 | 1 | k__Fungi(100);p__Ascomycota(100);c__Dothideomycetes(100);o__Pleosporales(100);f__Phaeosphaeriaceae(100);g__Stagonospora(100);s__Stagonospora_convolvuli(100); | Stagonospora | | 13 | Pathotroph | \|Plant Pathogen\| | Microfungus | NULL | Probable | NULL | Tedersoo L, et al. 2014. Science 346:e1256688 ((https://doi.org/10.1126/science.1256688)); Põlme S, et al. 2020. Fungal Diversity 105:1-16 ((https://doi.org/10.1007/s13225-020-00466-2)) |
| Otu38730 | 1 | k__Fungi(100);p__Ascomycota(100);c__Saccharomycetes(100);o__Saccharomycetales(100);f__Metschnikowiaceae(100);g__Metschnikowia(100);g__Metschnikowia_unclassified(100); | Metschnikowia | | 13 | Saprotroph | \|Undefined Saprotroph\| | Yeast | NULL | Probable | As nectar-tap_saprotroph (Põlme et al. 2020); Decay sugar-rich substrates (Põlme et al. 2020) | Põlme S, et al. 2020. Fungal Diversity 105:1-16 ((https://doi.org/10.1007/s13225-020-00466-2)) |
| Otu38731 | 1 | k__Fungi(100);p__Ascomycota(100);c__Dothideomycetes(100);o__Pleosporales(100);f__Pleosporales_family_Incertae_sedis(100);g__Phoma(100);g__Phoma_unclassified(100); | Phoma | | 13 | Pathotroph-Saprotroph-Symbiotroph | Dung Saprotroph-Endophyte-Lichen Parasite-\|Plant Pathogen\|-Plant Saprotroph-Undefined Saprotroph-Wood Saprotroph | Microfungus | Soft Rot | Possible | As foliar_endophyte (Põlme et al. 2020); As litter_saprotroph (Põlme et al. 2020); Decay leaf, fruit, seed, roots, soil (Põlme et al. 2020) | Richardson MJ. 1972. Transactions of the British Mycological Society 58:37-48 ((https://doi.org/10.1016/S0007-1536(72)80069-X)); Seehann G, et al. 1975. List of Fungi in Soft-Rot Tissue Tests. Institute fur Holzbiologie und Holzschutz, Bundesforschungsanstalt fur Forst - und Holzwirtshaft, Hamburg - Lohbrugge, Germany; de Gruyter J, et al. 2010. Mycologia 102:1066-1081 ((https://doi.org/10.3852/09-240)); Costa IPMW, et al. 2012. Mycotaxon 119:e26 ((http://mycotaxon.com/resources/checklists/costa_v119_checklist.pdf)); Tedersoo L, et al. 2014. Science 346:e1256688 ((https://doi.org/10.1126/science.1256688)); Lawrey JD, Diederich P. 2016. http://www.lichenicolous.net (accessed 24 May 2017) ((http://www.lichenicolous.net)); Põlme S, et al. 2020. Fungal Diversity 105:1-16 ((https://doi.org/10.1007/s13225-020-00466-2)) |
| Otu38735 | 1 | k__Fungi(100);p__Ascomycota(100);c__Saccharomycetes(100);o__Saccharomycetales(100);f__Metschnikowiaceae(100);g__Metschnikowia(100);s__Metschnikowia_pulcherrima(100); | Metschnikowia | | 13 | Saprotroph | \|Undefined Saprotroph\| | Yeast | NULL | Probable | As nectar-tap_saprotroph (Põlme et al. 2020); Decay sugar-rich substrates (Põlme et al. 2020) | Põlme S, et al. 2020. Fungal Diversity 105:1-16 ((https://doi.org/10.1007/s13225-020-00466-2)) |
| Otu38741 | 1 | k__Fungi(100);p__Ascomycota(100);c__Saccharomycetes(100);o__Saccharomycetales(100);f__Metschnikowiaceae(100);g__Metschnikowia(100);s__Metschnikowia_pulcherrima(100); | Metschnikowia | | 13 | Saprotroph | \|Undefined Saprotroph\| | Yeast | NULL | Probable | As nectar-tap_saprotroph (Põlme et al. 2020); Decay sugar-rich substrates (Põlme et al. 2020) | Põlme S, et al. 2020. Fungal Diversity 105:1-16 ((https://doi.org/10.1007/s13225-020-00466-2)) |
| Otu38743 | 1 | k__Fungi(100);p__Ascomycota(100);c__Saccharomycetes(100);o__Saccharomycetales(100);f__Metschnikowiaceae(100);g__Metschnikowia(100);s__Metschnikowia_pulcherrima(100); | Metschnikowia | | 13 | Saprotroph | \|Undefined Saprotroph\| | Yeast | NULL | Probable | As nectar-tap_saprotroph (Põlme et al. 2020); Decay sugar-rich substrates (Põlme et al. 2020) | Põlme S, et al. 2020. Fungal Diversity 105:1-16 ((https://doi.org/10.1007/s13225-020-00466-2)) |
| Otu38744 | 1 | k__Fungi(100);p__Ascomycota(100);c__Saccharomycetes(100);o__Saccharomycetales(100);f__Metschnikowiaceae(100);g__Metschnikowia(100);s__Metschnikowia_pulcherrima(100); | Metschnikowia | | 13 | Saprotroph | \|Undefined Saprotroph\| | Yeast | NULL | Probable | As nectar-tap_saprotroph (Põlme et al. 2020); Decay sugar-rich substrates (Põlme et al. 2020) | Põlme S, et al. 2020. Fungal Diversity 105:1-16 ((https://doi.org/10.1007/s13225-020-00466-2)) |
| Otu38745 | 1 | k__Fungi(100);p__Ascomycota(100);c__Saccharomycetes(100);o__Saccharomycetales(100);f__Metschnikowiaceae(100);g__Metschnikowia(100);s__Metschnikowia_pulcherrima(100); | Metschnikowia | | 13 | Saprotroph | \|Undefined Saprotroph\| | Yeast | NULL | Probable | As nectar-tap_saprotroph (Põlme et al. 2020); Decay sugar-rich substrates (Põlme et al. 2020) | Põlme S, et al. 2020. Fungal Diversity 105:1-16 ((https://doi.org/10.1007/s13225-020-00466-2)) |
| Otu38746 | 1 | k__Fungi(100);p__Ascomycota(100);c__Saccharomycetes(100);o__Saccharomycetales(100);f__Metschnikowiaceae(100);g__Metschnikowia(100);g__Metschnikowia_unclassified(100); | Metschnikowia | | 13 | Saprotroph | \|Undefined Saprotroph\| | Yeast | NULL | Probable | As nectar-tap_saprotroph (Põlme et al. 2020); Decay sugar-rich substrates (Põlme et al. 2020) | Põlme S, et al. 2020. Fungal Diversity 105:1-16 ((https://doi.org/10.1007/s13225-020-00466-2)) |
| Otu38747 | 1 | k__Fungi(100);p__Ascomycota(100);c__Saccharomycetes(100);o__Saccharomycetales(100);f__Metschnikowiaceae(100);g__Metschnikowia(100);s__Metschnikowia_pulcherrima(100); | Metschnikowia | | 13 | Saprotroph | \|Undefined Saprotroph\| | Yeast | NULL | Probable | As nectar-tap_saprotroph (Põlme et al. 2020); Decay sugar-rich substrates (Põlme et al. 2020) | Põlme S, et al. 2020. Fungal Diversity 105:1-16 ((https://doi.org/10.1007/s13225-020-00466-2)) |
| Otu38748 | 1 | k__Fungi(100);p__Ascomycota(100);c__Saccharomycetes(100);o__Saccharomycetales(100);f__Metschnikowiaceae(100);g__Metschnikowia(100);g__Metschnikowia_unclassified(100); | Metschnikowia | | 13 | Saprotroph | \|Undefined Saprotroph\| | Yeast | NULL | Probable | As nectar-tap_saprotroph (Põlme et al. 2020); Decay sugar-rich substrates (Põlme et al. 2020) | Põlme S, et al. 2020. Fungal Diversity 105:1-16 ((https://doi.org/10.1007/s13225-020-00466-2)) |
| Otu38749 | 1 | k__Fungi(100);p__Ascomycota(100);c__Saccharomycetes(100);o__Saccharomycetales(100);f__Metschnikowiaceae(100);g__Metschnikowia(100);s__Metschnikowia_pulcherrima(100); | Metschnikowia | | 13 | Saprotroph | \|Undefined Saprotroph\| | Yeast | NULL | Probable | As nectar-tap_saprotroph (Põlme et al. 2020); Decay sugar-rich substrates (Põlme et al. 2020) | Põlme S, et al. 2020. Fungal Diversity 105:1-16 ((https://doi.org/10.1007/s13225-020-00466-2)) |
| Otu38750 | 1 | k__Fungi(100);p__Ascomycota(100);c__Saccharomycetes(100);o__Saccharomycetales(100);f__Metschnikowiaceae(100);g__Metschnikowia(100);g__Metschnikowia_unclassified(100); | Metschnikowia | | 13 | Saprotroph | \|Undefined Saprotroph\| | Yeast | NULL | Probable | As nectar-tap_saprotroph (Põlme et al. 2020); Decay sugar-rich substrates (Põlme et al. 2020) | Põlme S, et al. 2020. Fungal Diversity 105:1-16 ((https://doi.org/10.1007/s13225-020-00466-2)) |
| Otu38751 | 1 | k__Fungi(100);p__Ascomycota(100);c__Saccharomycetes(100);o__Saccharomycetales(100);f__Metschnikowiaceae(100);g__Metschnikowia(100);s__Metschnikowia_pulcherrima(100); | Metschnikowia | | 13 | Saprotroph | \|Undefined Saprotroph\| | Yeast | NULL | Probable | As nectar-tap_saprotroph (Põlme et al. 2020); Decay sugar-rich substrates (Põlme et al. 2020) | Põlme S, et al. 2020. Fungal Diversity 105:1-16 ((https://doi.org/10.1007/s13225-020-00466-2)) |
| Otu38752 | 1 | k__Fungi(100);p__Ascomycota(100);c__Saccharomycetes(100);o__Saccharomycetales(100);f__Metschnikowiaceae(100);g__Metschnikowia(100);g__Metschnikowia_unclassified(100); | Metschnikowia | | 13 | Saprotroph | \|Undefined Saprotroph\| | Yeast | NULL | Probable | As nectar-tap_saprotroph (Põlme et al. 2020); Decay sugar-rich substrates (Põlme et al. 2020) | Põlme S, et al. 2020. Fungal Diversity 105:1-16 ((https://doi.org/10.1007/s13225-020-00466-2)) |
| Otu38753 | 1 | k__Fungi(100);p__Ascomycota(100);c__Saccharomycetes(100);o__Saccharomycetales(100);f__Metschnikowiaceae(100);g__Metschnikowia(100);g__Metschnikowia_unclassified(100); | Metschnikowia | | 13 | Saprotroph | \|Undefined Saprotroph\| | Yeast | NULL | Probable | As nectar-tap_saprotroph (Põlme et al. 2020); Decay sugar-rich substrates (Põlme et al. 2020) | Põlme S, et al. 2020. Fungal Diversity 105:1-16 ((https://doi.org/10.1007/s13225-020-00466-2)) |
| Otu38754 | 1 | k__Fungi(100);p__Ascomycota(100);c__Saccharomycetes(100);o__Saccharomycetales(100);f__Metschnikowiaceae(100);g__Metschnikowia(100);g__Metschnikowia_unclassified(100); | Metschnikowia | | 13 | Saprotroph | \|Undefined Saprotroph\| | Yeast | NULL | Probable | As nectar-tap_saprotroph (Põlme et al. 2020); Decay sugar-rich substrates (Põlme et al. 2020) | Põlme S, et al. 2020. Fungal Diversity 105:1-16 ((https://doi.org/10.1007/s13225-020-00466-2)) |
| Otu38755 | 1 | k__Fungi(100);p__Ascomycota(100);c__Saccharomycetes(100);o__Saccharomycetales(100);f__Metschnikowiaceae(100);g__Metschnikowia(100);g__Metschnikowia_unclassified(100); | Metschnikowia | | 13 | Saprotroph | \|Undefined Saprotroph\| | Yeast | NULL | Probable | As nectar-tap_saprotroph (Põlme et al. 2020); Decay sugar-rich substrates (Põlme et al. 2020) | Põlme S, et al. 2020. Fungal Diversity 105:1-16 ((https://doi.org/10.1007/s13225-020-00466-2)) |
| Otu38756 | 1 | k__Fungi(100);p__Ascomycota(100);c__Saccharomycetes(100);o__Saccharomycetales(100);f__Metschnikowiaceae(100);g__Metschnikowia(100);s__Metschnikowia_pulcherrima(100); | Metschnikowia | | 13 | Saprotroph | \|Undefined Saprotroph\| | Yeast | NULL | Probable | As nectar-tap_saprotroph (Põlme et al. 2020); Decay sugar-rich substrates (Põlme et al. 2020) | Põlme S, et al. 2020. Fungal Diversity 105:1-16 ((https://doi.org/10.1007/s13225-020-00466-2)) |
| Otu38757 | 1 | k__Fungi(100);p__Ascomycota(100);c__Saccharomycetes(100);o__Saccharomycetales(100);f__Metschnikowiaceae(100);g__Metschnikowia(100);s__Metschnikowia_pulcherrima(100); | Metschnikowia | | 13 | Saprotroph | \|Undefined Saprotroph\| | Yeast | NULL | Probable | As nectar-tap_saprotroph (Põlme et al. 2020); Decay sugar-rich substrates (Põlme et al. 2020) | Põlme S, et al. 2020. Fungal Diversity 105:1-16 ((https://doi.org/10.1007/s13225-020-00466-2)) |
| Otu38758 | 1 | k__Fungi(100);p__Ascomycota(100);c__Saccharomycetes(100);o__Saccharomycetales(100);f__Metschnikowiaceae(100);g__Metschnikowia(100);s__Metschnikowia_pulcherrima(100); | Metschnikowia | | 13 | Saprotroph | \|Undefined Saprotroph\| | Yeast | NULL | Probable | As nectar-tap_saprotroph (Põlme et al. 2020); Decay sugar-rich substrates (Põlme et al. 2020) | Põlme S, et al. 2020. Fungal Diversity 105:1-16 ((https://doi.org/10.1007/s13225-020-00466-2)) |
| Otu38759 | 1 | k__Fungi(100);p__Ascomycota(100);c__Saccharomycetes(100);o__Saccharomycetales(100);f__Metschnikowiaceae(100);g__Metschnikowia(100);s__Metschnikowia_pulcherrima(100); | Metschnikowia | | 13 | Saprotroph | \|Undefined Saprotroph\| | Yeast | NULL | Probable | As nectar-tap_saprotroph (Põlme et al. 2020); Decay sugar-rich substrates (Põlme et al. 2020) | Põlme S, et al. 2020. Fungal Diversity 105:1-16 ((https://doi.org/10.1007/s13225-020-00466-2)) |
| Otu38760 | 1 | k__Fungi(100);p__Ascomycota(100);c__Saccharomycetes(100);o__Saccharomycetales(100);f__Metschnikowiaceae(100);g__Metschnikowia(100);s__Metschnikowia_pulcherrima(100); | Metschnikowia | | 13 | Saprotroph | \|Undefined Saprotroph\| | Yeast | NULL | Probable | As nectar-tap_saprotroph (Põlme et al. 2020); Decay sugar-rich substrates (Põlme et al. 2020) | Põlme S, et al. 2020. Fungal Diversity 105:1-16 ((https://doi.org/10.1007/s13225-020-00466-2)) |
| Otu38761 | 1 | k__Fungi(100);p__Ascomycota(100);c__Saccharomycetes(100);o__Saccharomycetales(100);f__Metschnikowiaceae(100);g__Metschnikowia(100);g__Metschnikowia_unclassified(100); | Metschnikowia | | 13 | Saprotroph | \|Undefined Saprotroph\| | Yeast | NULL | Probable | As nectar-tap_saprotroph (Põlme et al. 2020); Decay sugar-rich substrates (Põlme et al. 2020) | Põlme S, et al. 2020. Fungal Diversity 105:1-16 ((https://doi.org/10.1007/s13225-020-00466-2)) |
| Otu38762 | 1 | k__Fungi(100);p__Ascomycota(100);c__Saccharomycetes(100);o__Saccharomycetales(100);f__Metschnikowiaceae(100);g__Metschnikowia(100);s__Metschnikowia_pulcherrima(100); | Metschnikowia | | 13 | Saprotroph | \|Undefined Saprotroph\| | Yeast | NULL | Probable | As nectar-tap_saprotroph (Põlme et al. 2020); Decay sugar-rich substrates (Põlme et al. 2020) | Põlme S, et al. 2020. Fungal Diversity 105:1-16 ((https://doi.org/10.1007/s13225-020-00466-2)) |
| Otu38763 | 1 | k__Fungi(100);p__Ascomycota(100);c__Saccharomycetes(100);o__Saccharomycetales(100);f__Metschnikowiaceae(100);g__Metschnikowia(100);s__Metschnikowia_pulcherrima(100); | Metschnikowia | | 13 | Saprotroph | \|Undefined Saprotroph\| | Yeast | NULL | Probable | As nectar-tap_saprotroph (Põlme et al. 2020); Decay sugar-rich substrates (Põlme et al. 2020) | Põlme S, et al. 2020. Fungal Diversity 105:1-16 ((https://doi.org/10.1007/s13225-020-00466-2)) |
| Otu38764 | 1 | k__Fungi(100);p__Ascomycota(100);c__Saccharomycetes(100);o__Saccharomycetales(100);f__Metschnikowiaceae(100);g__Metschnikowia(100);s__Metschnikowia_pulcherrima(100); | Metschnikowia | | 13 | Saprotroph | \|Undefined Saprotroph\| | Yeast | NULL | Probable | As nectar-tap_saprotroph (Põlme et al. 2020); Decay sugar-rich substrates (Põlme et al. 2020) | Põlme S, et al. 2020. Fungal Diversity 105:1-16 ((https://doi.org/10.1007/s13225-020-00466-2)) |
| Otu38765 | 1 | k__Fungi(100);p__Ascomycota(100);c__Saccharomycetes(100);o__Saccharomycetales(100);f__Metschnikowiaceae(100);g__Metschnikowia(100);s__Metschnikowia_pulcherrima(100); | Metschnikowia | | 13 | Saprotroph | \|Undefined Saprotroph\| | Yeast | NULL | Probable | As nectar-tap_saprotroph (Põlme et al. 2020); Decay sugar-rich substrates (Põlme et al. 2020) | Põlme S, et al. 2020. Fungal Diversity 105:1-16 ((https://doi.org/10.1007/s13225-020-00466-2)) |
| Otu38766 | 1 | k__Fungi(100);p__Ascomycota(100);c__Saccharomycetes(100);o__Saccharomycetales(100);f__Metschnikowiaceae(100);g__Metschnikowia(100);s__Metschnikowia_pulcherrima(100); | Metschnikowia | | 13 | Saprotroph | \|Undefined Saprotroph\| | Yeast | NULL | Probable | As nectar-tap_saprotroph (Põlme et al. 2020); Decay sugar-rich substrates (Põlme et al. 2020) | Põlme S, et al. 2020. Fungal Diversity 105:1-16 ((https://doi.org/10.1007/s13225-020-00466-2)) |
| Otu38767 | 1 | k__Fungi(100);p__Ascomycota(100);c__Saccharomycetes(100);o__Saccharomycetales(100);f__Metschnikowiaceae(100);g__Metschnikowia(100);s__Metschnikowia_pulcherrima(100); | Metschnikowia | | 13 | Saprotroph | \|Undefined Saprotroph\| | Yeast | NULL | Probable | As nectar-tap_saprotroph (Põlme et al. 2020); Decay sugar-rich substrates (Põlme et al. 2020) | Põlme S, et al. 2020. Fungal Diversity 105:1-16 ((https://doi.org/10.1007/s13225-020-00466-2)) |
| Otu38768 | 1 | k__Fungi(100);p__Ascomycota(100);c__Saccharomycetes(100);o__Saccharomycetales(100);f__Metschnikowiaceae(100);g__Metschnikowia(100);s__Metschnikowia_pulcherrima(100); | Metschnikowia | | 13 | Saprotroph | \|Undefined Saprotroph\| | Yeast | NULL | Probable | As nectar-tap_saprotroph (Põlme et al. 2020); Decay sugar-rich substrates (Põlme et al. 2020) | Põlme S, et al. 2020. Fungal Diversity 105:1-16 ((https://doi.org/10.1007/s13225-020-00466-2)) |
| Otu38769 | 1 | k__Fungi(100);p__Ascomycota(100);c__Saccharomycetes(100);o__Saccharomycetales(100);f__Metschnikowiaceae(100);g__Metschnikowia(100);g__Metschnikowia_unclassified(100); | Metschnikowia | | 13 | Saprotroph | \|Undefined Saprotroph\| | Yeast | NULL | Probable | As nectar-tap_saprotroph (Põlme et al. 2020); Decay sugar-rich substrates (Põlme et al. 2020) | Põlme S, et al. 2020. Fungal Diversity 105:1-16 ((https://doi.org/10.1007/s13225-020-00466-2)) |
| Otu38770 | 1 | k__Fungi(100);p__Ascomycota(100);c__Saccharomycetes(100);o__Saccharomycetales(100);f__Metschnikowiaceae(100);g__Metschnikowia(100);g__Metschnikowia_unclassified(100); | Metschnikowia | | 13 | Saprotroph | \|Undefined Saprotroph\| | Yeast | NULL | Probable | As nectar-tap_saprotroph (Põlme et al. 2020); Decay sugar-rich substrates (Põlme et al. 2020) | Põlme S, et al. 2020. Fungal Diversity 105:1-16 ((https://doi.org/10.1007/s13225-020-00466-2)) |
| Otu38772 | 1 | k__Fungi(100);p__Ascomycota(100);c__Saccharomycetes(100);o__Saccharomycetales(100);f__Metschnikowiaceae(100);g__Metschnikowia(100);g__Metschnikowia_unclassified(100); | Metschnikowia | | 13 | Saprotroph | \|Undefined Saprotroph\| | Yeast | NULL | Probable | As nectar-tap_saprotroph (Põlme et al. 2020); Decay sugar-rich substrates (Põlme et al. 2020) | Põlme S, et al. 2020. Fungal Diversity 105:1-16 ((https://doi.org/10.1007/s13225-020-00466-2)) |
| Otu38773 | 1 | k__Fungi(100);p__Ascomycota(100);c__Saccharomycetes(100);o__Saccharomycetales(100);f__Metschnikowiaceae(100);g__Metschnikowia(100);g__Metschnikowia_unclassified(100); | Metschnikowia | | 13 | Saprotroph | \|Undefined Saprotroph\| | Yeast | NULL | Probable | As nectar-tap_saprotroph (Põlme et al. 2020); Decay sugar-rich substrates (Põlme et al. 2020) | Põlme S, et al. 2020. Fungal Diversity 105:1-16 ((https://doi.org/10.1007/s13225-020-00466-2)) |
| Otu38774 | 1 | k__Fungi(100);p__Ascomycota(100);c__Saccharomycetes(100);o__Saccharomycetales(100);f__Metschnikowiaceae(100);g__Metschnikowia(100);g__Metschnikowia_unclassified(100); | Metschnikowia | | 13 | Saprotroph | \|Undefined Saprotroph\| | Yeast | NULL | Probable | As nectar-tap_saprotroph (Põlme et al. 2020); Decay sugar-rich substrates (Põlme et al. 2020) | Põlme S, et al. 2020. Fungal Diversity 105:1-16 ((https://doi.org/10.1007/s13225-020-00466-2)) |
| Otu38776 | 1 | k__Fungi(100);p__Ascomycota(100);c__Saccharomycetes(100);o__Saccharomycetales(100);f__Metschnikowiaceae(100);g__Metschnikowia(100);g__Metschnikowia_unclassified(100); | Metschnikowia | | 13 | Saprotroph | \|Undefined Saprotroph\| | Yeast | NULL | Probable | As nectar-tap_saprotroph (Põlme et al. 2020); Decay sugar-rich substrates (Põlme et al. 2020) | Põlme S, et al. 2020. Fungal Diversity 105:1-16 ((https://doi.org/10.1007/s13225-020-00466-2)) |
| Otu38777 | 1 | k__Fungi(100);p__Ascomycota(100);c__Saccharomycetes(100);o__Saccharomycetales(100);f__Metschnikowiaceae(100);g__Metschnikowia(100);g__Metschnikowia_unclassified(100); | Metschnikowia | | 13 | Saprotroph | \|Undefined Saprotroph\| | Yeast | NULL | Probable | As nectar-tap_saprotroph (Põlme et al. 2020); Decay sugar-rich substrates (Põlme et al. 2020) | Põlme S, et al. 2020. Fungal Diversity 105:1-16 ((https://doi.org/10.1007/s13225-020-00466-2)) |
| Otu38778 | 1 | k__Fungi(100);p__Ascomycota(100);c__Saccharomycetes(100);o__Saccharomycetales(100);f__Metschnikowiaceae(100);g__Metschnikowia(100);g__Metschnikowia_unclassified(100); | Metschnikowia | | 13 | Saprotroph | \|Undefined Saprotroph\| | Yeast | NULL | Probable | As nectar-tap_saprotroph (Põlme et al. 2020); Decay sugar-rich substrates (Põlme et al. 2020) | Põlme S, et al. 2020. Fungal Diversity 105:1-16 ((https://doi.org/10.1007/s13225-020-00466-2)) |
| Otu38779 | 1 | k__Fungi(100);p__Ascomycota(100);c__Saccharomycetes(100);o__Saccharomycetales(100);f__Metschnikowiaceae(100);g__Metschnikowia(100);g__Metschnikowia_unclassified(100); | Metschnikowia | | 13 | Saprotroph | \|Undefined Saprotroph\| | Yeast | NULL | Probable | As nectar-tap_saprotroph (Põlme et al. 2020); Decay sugar-rich substrates (Põlme et al. 2020) | Põlme S, et al. 2020. Fungal Diversity 105:1-16 ((https://doi.org/10.1007/s13225-020-00466-2)) |
| Otu38780 | 1 | k__Fungi(100);p__Ascomycota(100);c__Saccharomycetes(100);o__Saccharomycetales(100);f__Metschnikowiaceae(100);g__Metschnikowia(100);s__Metschnikowia_pulcherrima(100); | Metschnikowia | | 13 | Saprotroph | \|Undefined Saprotroph\| | Yeast | NULL | Probable | As nectar-tap_saprotroph (Põlme et al. 2020); Decay sugar-rich substrates (Põlme et al. 2020) | Põlme S, et al. 2020. Fungal Diversity 105:1-16 ((https://doi.org/10.1007/s13225-020-00466-2)) |
| Otu38782 | 1 | k__Fungi(100);p__Ascomycota(100);c__Saccharomycetes(100);o__Saccharomycetales(100);f__Metschnikowiaceae(100);g__Metschnikowia(100);g__Metschnikowia_unclassified(100); | Metschnikowia | | 13 | Saprotroph | \|Undefined Saprotroph\| | Yeast | NULL | Probable | As nectar-tap_saprotroph (Põlme et al. 2020); Decay sugar-rich substrates (Põlme et al. 2020) | Põlme S, et al. 2020. Fungal Diversity 105:1-16 ((https://doi.org/10.1007/s13225-020-00466-2)) |
| Otu38783 | 1 | k__Fungi(100);p__Ascomycota(100);c__Saccharomycetes(100);o__Saccharomycetales(100);f__Metschnikowiaceae(100);g__Metschnikowia(100);s__Metschnikowia_pulcherrima(100); | Metschnikowia | | 13 | Saprotroph | \|Undefined Saprotroph\| | Yeast | NULL | Probable | As nectar-tap_saprotroph (Põlme et al. 2020); Decay sugar-rich substrates (Põlme et al. 2020) | Põlme S, et al. 2020. Fungal Diversity 105:1-16 ((https://doi.org/10.1007/s13225-020-00466-2)) |
| Otu38784 | 1 | k__Fungi(100);p__Ascomycota(100);c__Saccharomycetes(100);o__Saccharomycetales(100);f__Metschnikowiaceae(100);g__Metschnikowia(100);s__Metschnikowia_pulcherrima(100); | Metschnikowia | | 13 | Saprotroph | \|Undefined Saprotroph\| | Yeast | NULL | Probable | As nectar-tap_saprotroph (Põlme et al. 2020); Decay sugar-rich substrates (Põlme et al. 2020) | Põlme S, et al. 2020. Fungal Diversity 105:1-16 ((https://doi.org/10.1007/s13225-020-00466-2)) |
| Otu38785 | 1 | k__Fungi(100);p__Ascomycota(100);c__Saccharomycetes(100);o__Saccharomycetales(100);f__Metschnikowiaceae(100);g__Metschnikowia(100);s__Metschnikowia_pulcherrima(100); | Metschnikowia | | 13 | Saprotroph | \|Undefined Saprotroph\| | Yeast | NULL | Probable | As nectar-tap_saprotroph (Põlme et al. 2020); Decay sugar-rich substrates (Põlme et al. 2020) | Põlme S, et al. 2020. Fungal Diversity 105:1-16 ((https://doi.org/10.1007/s13225-020-00466-2)) |
| Otu38786 | 1 | k__Fungi(100);p__Ascomycota(100);c__Saccharomycetes(100);o__Saccharomycetales(100);f__Metschnikowiaceae(100);g__Metschnikowia(100);s__Metschnikowia_pulcherrima(100); | Metschnikowia | | 13 | Saprotroph | \|Undefined Saprotroph\| | Yeast | NULL | Probable | As nectar-tap_saprotroph (Põlme et al. 2020); Decay sugar-rich substrates (Põlme et al. 2020) | Põlme S, et al. 2020. Fungal Diversity 105:1-16 ((https://doi.org/10.1007/s13225-020-00466-2)) |
| Otu38787 | 1 | k__Fungi(100);p__Ascomycota(100);c__Saccharomycetes(100);o__Saccharomycetales(100);f__Metschnikowiaceae(100);g__Metschnikowia(100);s__Metschnikowia_pulcherrima(100); | Metschnikowia | | 13 | Saprotroph | \|Undefined Saprotroph\| | Yeast | NULL | Probable | As nectar-tap_saprotroph (Põlme et al. 2020); Decay sugar-rich substrates (Põlme et al. 2020) | Põlme S, et al. 2020. Fungal Diversity 105:1-16 ((https://doi.org/10.1007/s13225-020-00466-2)) |
| Otu38788 | 1 | k__Fungi(100);p__Ascomycota(100);c__Saccharomycetes(100);o__Saccharomycetales(100);f__Metschnikowiaceae(100);g__Metschnikowia(100);s__Metschnikowia_pulcherrima(100); | Metschnikowia | | 13 | Saprotroph | \|Undefined Saprotroph\| | Yeast | NULL | Probable | As nectar-tap_saprotroph (Põlme et al. 2020); Decay sugar-rich substrates (Põlme et al. 2020) | Põlme S, et al. 2020. Fungal Diversity 105:1-16 ((https://doi.org/10.1007/s13225-020-00466-2)) |
| Otu38789 | 1 | k__Fungi(100);p__Ascomycota(100);c__Saccharomycetes(100);o__Saccharomycetales(100);f__Metschnikowiaceae(100);g__Metschnikowia(100);g__Metschnikowia_unclassified(100); | Metschnikowia | | 13 | Saprotroph | \|Undefined Saprotroph\| | Yeast | NULL | Probable | As nectar-tap_saprotroph (Põlme et al. 2020); Decay sugar-rich substrates (Põlme et al. 2020) | Põlme S, et al. 2020. Fungal Diversity 105:1-16 ((https://doi.org/10.1007/s13225-020-00466-2)) |
| Otu38790 | 1 | k__Fungi(100);p__Ascomycota(100);c__Saccharomycetes(100);o__Saccharomycetales(100);f__Metschnikowiaceae(100);g__Metschnikowia(100);g__Metschnikowia_unclassified(100); | Metschnikowia | | 13 | Saprotroph | \|Undefined Saprotroph\| | Yeast | NULL | Probable | As nectar-tap_saprotroph (Põlme et al. 2020); Decay sugar-rich substrates (Põlme et al. 2020) | Põlme S, et al. 2020. Fungal Diversity 105:1-16 ((https://doi.org/10.1007/s13225-020-00466-2)) |
| Otu38791 | 1 | k__Fungi(100);p__Ascomycota(100);c__Saccharomycetes(100);o__Saccharomycetales(100);f__Metschnikowiaceae(100);g__Metschnikowia(100);s__Metschnikowia_pulcherrima(100); | Metschnikowia | | 13 | Saprotroph | \|Undefined Saprotroph\| | Yeast | NULL | Probable | As nectar-tap_saprotroph (Põlme et al. 2020); Decay sugar-rich substrates (Põlme et al. 2020) | Põlme S, et al. 2020. Fungal Diversity 105:1-16 ((https://doi.org/10.1007/s13225-020-00466-2)) |
| Otu38792 | 1 | k__Fungi(100);p__Ascomycota(100);c__Saccharomycetes(100);o__Saccharomycetales(100);f__Metschnikowiaceae(100);g__Metschnikowia(100);s__Metschnikowia_pulcherrima(100); | Metschnikowia | | 13 | Saprotroph | \|Undefined Saprotroph\| | Yeast | NULL | Probable | As nectar-tap_saprotroph (Põlme et al. 2020); Decay sugar-rich substrates (Põlme et al. 2020) | Põlme S, et al. 2020. Fungal Diversity 105:1-16 ((https://doi.org/10.1007/s13225-020-00466-2)) |
| Otu38793 | 1 | k__Fungi(100);p__Ascomycota(100);c__Saccharomycetes(100);o__Saccharomycetales(100);f__Metschnikowiaceae(100);g__Metschnikowia(100);s__Metschnikowia_pulcherrima(100); | Metschnikowia | | 13 | Saprotroph | \|Undefined Saprotroph\| | Yeast | NULL | Probable | As nectar-tap_saprotroph (Põlme et al. 2020); Decay sugar-rich substrates (Põlme et al. 2020) | Põlme S, et al. 2020. Fungal Diversity 105:1-16 ((https://doi.org/10.1007/s13225-020-00466-2)) |
| Otu38794 | 1 | k__Fungi(100);p__Ascomycota(100);c__Saccharomycetes(100);o__Saccharomycetales(100);f__Metschnikowiaceae(100);g__Metschnikowia(100);s__Metschnikowia_pulcherrima(100); | Metschnikowia | | 13 | Saprotroph | \|Undefined Saprotroph\| | Yeast | NULL | Probable | As nectar-tap_saprotroph (Põlme et al. 2020); Decay sugar-rich substrates (Põlme et al. 2020) | Põlme S, et al. 2020. Fungal Diversity 105:1-16 ((https://doi.org/10.1007/s13225-020-00466-2)) |
| Otu38795 | 1 | k__Fungi(100);p__Ascomycota(100);c__Saccharomycetes(100);o__Saccharomycetales(100);f__Metschnikowiaceae(100);g__Metschnikowia(100);s__Metschnikowia_pulcherrima(100); | Metschnikowia | | 13 | Saprotroph | \|Undefined Saprotroph\| | Yeast | NULL | Probable | As nectar-tap_saprotroph (Põlme et al. 2020); Decay sugar-rich substrates (Põlme et al. 2020) | Põlme S, et al. 2020. Fungal Diversity 105:1-16 ((https://doi.org/10.1007/s13225-020-00466-2)) |
| Otu38796 | 1 | k__Fungi(100);p__Ascomycota(100);c__Saccharomycetes(100);o__Saccharomycetales(100);f__Metschnikowiaceae(100);g__Metschnikowia(100);s__Metschnikowia_pulcherrima(100); | Metschnikowia | | 13 | Saprotroph | \|Undefined Saprotroph\| | Yeast | NULL | Probable | As nectar-tap_saprotroph (Põlme et al. 2020); Decay sugar-rich substrates (Põlme et al. 2020) | Põlme S, et al. 2020. Fungal Diversity 105:1-16 ((https://doi.org/10.1007/s13225-020-00466-2)) |
| Otu38797 | 1 | k__Fungi(100);p__Ascomycota(100);c__Saccharomycetes(100);o__Saccharomycetales(100);f__Metschnikowiaceae(100);g__Metschnikowia(100);s__Metschnikowia_pulcherrima(100); | Metschnikowia | | 13 | Saprotroph | \|Undefined Saprotroph\| | Yeast | NULL | Probable | As nectar-tap_saprotroph (Põlme et al. 2020); Decay sugar-rich substrates (Põlme et al. 2020) | Põlme S, et al. 2020. Fungal Diversity 105:1-16 ((https://doi.org/10.1007/s13225-020-00466-2)) |
| Otu38798 | 1 | k__Fungi(100);p__Ascomycota(100);c__Saccharomycetes(100);o__Saccharomycetales(100);f__Metschnikowiaceae(100);g__Metschnikowia(100);g__Metschnikowia_unclassified(100); | Metschnikowia | | 13 | Saprotroph | \|Undefined Saprotroph\| | Yeast | NULL | Probable | As nectar-tap_saprotroph (Põlme et al. 2020); Decay sugar-rich substrates (Põlme et al. 2020) | Põlme S, et al. 2020. Fungal Diversity 105:1-16 ((https://doi.org/10.1007/s13225-020-00466-2)) |
| Otu38800 | 1 | k__Fungi(100);p__Ascomycota(100);c__Saccharomycetes(100);o__Saccharomycetales(100);f__Metschnikowiaceae(100);g__Metschnikowia(100);s__Metschnikowia_pulcherrima(100); | Metschnikowia | | 13 | Saprotroph | \|Undefined Saprotroph\| | Yeast | NULL | Probable | As nectar-tap_saprotroph (Põlme et al. 2020); Decay sugar-rich substrates (Põlme et al. 2020) | Põlme S, et al. 2020. Fungal Diversity 105:1-16 ((https://doi.org/10.1007/s13225-020-00466-2)) |
| Otu38801 | 1 | k__Fungi(100);p__Ascomycota(100);c__Saccharomycetes(100);o__Saccharomycetales(100);f__Metschnikowiaceae(100);g__Metschnikowia(100);s__Metschnikowia_pulcherrima(100); | Metschnikowia | | 13 | Saprotroph | \|Undefined Saprotroph\| | Yeast | NULL | Probable | As nectar-tap_saprotroph (Põlme et al. 2020); Decay sugar-rich substrates (Põlme et al. 2020) | Põlme S, et al. 2020. Fungal Diversity 105:1-16 ((https://doi.org/10.1007/s13225-020-00466-2)) |
| Otu38802 | 1 | k__Fungi(100);p__Ascomycota(100);c__Saccharomycetes(100);o__Saccharomycetales(100);f__Metschnikowiaceae(100);g__Metschnikowia(100);g__Metschnikowia_unclassified(100); | Metschnikowia | | 13 | Saprotroph | \|Undefined Saprotroph\| | Yeast | NULL | Probable | As nectar-tap_saprotroph (Põlme et al. 2020); Decay sugar-rich substrates (Põlme et al. 2020) | Põlme S, et al. 2020. Fungal Diversity 105:1-16 ((https://doi.org/10.1007/s13225-020-00466-2)) |
| Otu38803 | 1 | k__Fungi(100);p__Ascomycota(100);c__Saccharomycetes(100);o__Saccharomycetales(100);f__Metschnikowiaceae(100);g__Metschnikowia(100);g__Metschnikowia_unclassified(100); | Metschnikowia | | 13 | Saprotroph | \|Undefined Saprotroph\| | Yeast | NULL | Probable | As nectar-tap_saprotroph (Põlme et al. 2020); Decay sugar-rich substrates (Põlme et al. 2020) | Põlme S, et al. 2020. Fungal Diversity 105:1-16 ((https://doi.org/10.1007/s13225-020-00466-2)) |
| Otu38804 | 1 | k__Fungi(100);p__Ascomycota(100);c__Saccharomycetes(100);o__Saccharomycetales(100);f__Metschnikowiaceae(100);g__Metschnikowia(100);s__Metschnikowia_pulcherrima(100); | Metschnikowia | | 13 | Saprotroph | \|Undefined Saprotroph\| | Yeast | NULL | Probable | As nectar-tap_saprotroph (Põlme et al. 2020); Decay sugar-rich substrates (Põlme et al. 2020) | Põlme S, et al. 2020. Fungal Diversity 105:1-16 ((https://doi.org/10.1007/s13225-020-00466-2)) |
| Otu38806 | 1 | k__Fungi(100);p__Ascomycota(100);c__Saccharomycetes(100);o__Saccharomycetales(100);f__Metschnikowiaceae(100);g__Metschnikowia(100);g__Metschnikowia_unclassified(100); | Metschnikowia | | 13 | Saprotroph | \|Undefined Saprotroph\| | Yeast | NULL | Probable | As nectar-tap_saprotroph (Põlme et al. 2020); Decay sugar-rich substrates (Põlme et al. 2020) | Põlme S, et al. 2020. Fungal Diversity 105:1-16 ((https://doi.org/10.1007/s13225-020-00466-2)) |
| Otu38808 | 1 | k__Fungi(100);p__Ascomycota(100);c__Saccharomycetes(100);o__Saccharomycetales(100);f__Metschnikowiaceae(100);g__Metschnikowia(100);s__Metschnikowia_pulcherrima(100); | Metschnikowia | | 13 | Saprotroph | \|Undefined Saprotroph\| | Yeast | NULL | Probable | As nectar-tap_saprotroph (Põlme et al. 2020); Decay sugar-rich substrates (Põlme et al. 2020) | Põlme S, et al. 2020. Fungal Diversity 105:1-16 ((https://doi.org/10.1007/s13225-020-00466-2)) |
| Otu38809 | 1 | k__Fungi(100);p__Ascomycota(100);c__Saccharomycetes(100);o__Saccharomycetales(100);f__Metschnikowiaceae(100);g__Metschnikowia(100);s__Metschnikowia_pulcherrima(100); | Metschnikowia | | 13 | Saprotroph | \|Undefined Saprotroph\| | Yeast | NULL | Probable | As nectar-tap_saprotroph (Põlme et al. 2020); Decay sugar-rich substrates (Põlme et al. 2020) | Põlme S, et al. 2020. Fungal Diversity 105:1-16 ((https://doi.org/10.1007/s13225-020-00466-2)) |
| Otu38810 | 1 | k__Fungi(100);p__Ascomycota(100);c__Saccharomycetes(100);o__Saccharomycetales(100);f__Metschnikowiaceae(100);g__Metschnikowia(100);s__Metschnikowia_pulcherrima(100); | Metschnikowia | | 13 | Saprotroph | \|Undefined Saprotroph\| | Yeast | NULL | Probable | As nectar-tap_saprotroph (Põlme et al. 2020); Decay sugar-rich substrates (Põlme et al. 2020) | Põlme S, et al. 2020. Fungal Diversity 105:1-16 ((https://doi.org/10.1007/s13225-020-00466-2)) |
| Otu38811 | 1 | k__Fungi(100);p__Ascomycota(100);c__Saccharomycetes(100);o__Saccharomycetales(100);f__Metschnikowiaceae(100);g__Metschnikowia(100);s__Metschnikowia_pulcherrima(100); | Metschnikowia | | 13 | Saprotroph | \|Undefined Saprotroph\| | Yeast | NULL | Probable | As nectar-tap_saprotroph (Põlme et al. 2020); Decay sugar-rich substrates (Põlme et al. 2020) | Põlme S, et al. 2020. Fungal Diversity 105:1-16 ((https://doi.org/10.1007/s13225-020-00466-2)) |
| Otu38812 | 1 | k__Fungi(100);p__Ascomycota(100);c__Saccharomycetes(100);o__Saccharomycetales(100);f__Metschnikowiaceae(100);g__Metschnikowia(100);s__Metschnikowia_pulcherrima(100); | Metschnikowia | | 13 | Saprotroph | \|Undefined Saprotroph\| | Yeast | NULL | Probable | As nectar-tap_saprotroph (Põlme et al. 2020); Decay sugar-rich substrates (Põlme et al. 2020) | Põlme S, et al. 2020. Fungal Diversity 105:1-16 ((https://doi.org/10.1007/s13225-020-00466-2)) |
| Otu38813 | 1 | k__Fungi(100);p__Ascomycota(100);c__Saccharomycetes(100);o__Saccharomycetales(100);f__Metschnikowiaceae(100);g__Metschnikowia(100);s__Metschnikowia_pulcherrima(100); | Metschnikowia | | 13 | Saprotroph | \|Undefined Saprotroph\| | Yeast | NULL | Probable | As nectar-tap_saprotroph (Põlme et al. 2020); Decay sugar-rich substrates (Põlme et al. 2020) | Põlme S, et al. 2020. Fungal Diversity 105:1-16 ((https://doi.org/10.1007/s13225-020-00466-2)) |
| Otu38814 | 1 | k__Fungi(100);p__Ascomycota(100);c__Saccharomycetes(100);o__Saccharomycetales(100);f__Metschnikowiaceae(100);g__Metschnikowia(100);s__Metschnikowia_pulcherrima(100); | Metschnikowia | | 13 | Saprotroph | \|Undefined Saprotroph\| | Yeast | NULL | Probable | As nectar-tap_saprotroph (Põlme et al. 2020); Decay sugar-rich substrates (Põlme et al. 2020) | Põlme S, et al. 2020. Fungal Diversity 105:1-16 ((https://doi.org/10.1007/s13225-020-00466-2)) |
| Otu38815 | 1 | k__Fungi(100);p__Ascomycota(100);c__Saccharomycetes(100);o__Saccharomycetales(100);f__Metschnikowiaceae(100);g__Metschnikowia(100);s__Metschnikowia_pulcherrima(100); | Metschnikowia | | 13 | Saprotroph | \|Undefined Saprotroph\| | Yeast | NULL | Probable | As nectar-tap_saprotroph (Põlme et al. 2020); Decay sugar-rich substrates (Põlme et al. 2020) | Põlme S, et al. 2020. Fungal Diversity 105:1-16 ((https://doi.org/10.1007/s13225-020-00466-2)) |
| Otu38817 | 1 | k__Fungi(100);p__Ascomycota(100);c__Saccharomycetes(100);o__Saccharomycetales(100);f__Metschnikowiaceae(100);g__Metschnikowia(100);s__Metschnikowia_pulcherrima(100); | Metschnikowia | | 13 | Saprotroph | \|Undefined Saprotroph\| | Yeast | NULL | Probable | As nectar-tap_saprotroph (Põlme et al. 2020); Decay sugar-rich substrates (Põlme et al. 2020) | Põlme S, et al. 2020. Fungal Diversity 105:1-16 ((https://doi.org/10.1007/s13225-020-00466-2)) |
| Otu38818 | 1 | k__Fungi(100);p__Ascomycota(100);c__Saccharomycetes(100);o__Saccharomycetales(100);f__Metschnikowiaceae(100);g__Metschnikowia(100);s__Metschnikowia_pulcherrima(100); | Metschnikowia | | 13 | Saprotroph | \|Undefined Saprotroph\| | Yeast | NULL | Probable | As nectar-tap_saprotroph (Põlme et al. 2020); Decay sugar-rich substrates (Põlme et al. 2020) | Põlme S, et al. 2020. Fungal Diversity 105:1-16 ((https://doi.org/10.1007/s13225-020-00466-2)) |
| Otu38819 | 1 | k__Fungi(100);p__Ascomycota(100);c__Saccharomycetes(100);o__Saccharomycetales(100);f__Metschnikowiaceae(100);g__Metschnikowia(100);s__Metschnikowia_pulcherrima(100); | Metschnikowia | | 13 | Saprotroph | \|Undefined Saprotroph\| | Yeast | NULL | Probable | As nectar-tap_saprotroph (Põlme et al. 2020); Decay sugar-rich substrates (Põlme et al. 2020) | Põlme S, et al. 2020. Fungal Diversity 105:1-16 ((https://doi.org/10.1007/s13225-020-00466-2)) |
| Otu38820 | 1 | k__Fungi(100);p__Ascomycota(100);c__Saccharomycetes(100);o__Saccharomycetales(100);f__Metschnikowiaceae(100);g__Metschnikowia(100);s__Metschnikowia_pulcherrima(100); | Metschnikowia | | 13 | Saprotroph | \|Undefined Saprotroph\| | Yeast | NULL | Probable | As nectar-tap_saprotroph (Põlme et al. 2020); Decay sugar-rich substrates (Põlme et al. 2020) | Põlme S, et al. 2020. Fungal Diversity 105:1-16 ((https://doi.org/10.1007/s13225-020-00466-2)) |
| Otu38821 | 1 | k__Fungi(100);p__Ascomycota(100);c__Saccharomycetes(100);o__Saccharomycetales(100);f__Metschnikowiaceae(100);g__Metschnikowia(100);s__Metschnikowia_pulcherrima(100); | Metschnikowia | | 13 | Saprotroph | \|Undefined Saprotroph\| | Yeast | NULL | Probable | As nectar-tap_saprotroph (Põlme et al. 2020); Decay sugar-rich substrates (Põlme et al. 2020) | Põlme S, et al. 2020. Fungal Diversity 105:1-16 ((https://doi.org/10.1007/s13225-020-00466-2)) |
| Otu38822 | 1 | k__Fungi(100);p__Ascomycota(100);c__Saccharomycetes(100);o__Saccharomycetales(100);f__Metschnikowiaceae(100);g__Metschnikowia(100);s__Metschnikowia_pulcherrima(100); | Metschnikowia | | 13 | Saprotroph | \|Undefined Saprotroph\| | Yeast | NULL | Probable | As nectar-tap_saprotroph (Põlme et al. 2020); Decay sugar-rich substrates (Põlme et al. 2020) | Põlme S, et al. 2020. Fungal Diversity 105:1-16 ((https://doi.org/10.1007/s13225-020-00466-2)) |
| Otu38823 | 1 | k__Fungi(100);p__Ascomycota(100);c__Saccharomycetes(100);o__Saccharomycetales(100);f__Metschnikowiaceae(100);g__Metschnikowia(100);s__Metschnikowia_pulcherrima(100); | Metschnikowia | | 13 | Saprotroph | \|Undefined Saprotroph\| | Yeast | NULL | Probable | As nectar-tap_saprotroph (Põlme et al. 2020); Decay sugar-rich substrates (Põlme et al. 2020) | Põlme S, et al. 2020. Fungal Diversity 105:1-16 ((https://doi.org/10.1007/s13225-020-00466-2)) |
| Otu38824 | 1 | k__Fungi(100);p__Ascomycota(100);c__Saccharomycetes(100);o__Saccharomycetales(100);f__Metschnikowiaceae(100);g__Metschnikowia(100);s__Metschnikowia_pulcherrima(100); | Metschnikowia | | 13 | Saprotroph | \|Undefined Saprotroph\| | Yeast | NULL | Probable | As nectar-tap_saprotroph (Põlme et al. 2020); Decay sugar-rich substrates (Põlme et al. 2020) | Põlme S, et al. 2020. Fungal Diversity 105:1-16 ((https://doi.org/10.1007/s13225-020-00466-2)) |
| Otu38825 | 1 | k__Fungi(100);p__Ascomycota(100);c__Saccharomycetes(100);o__Saccharomycetales(100);f__Metschnikowiaceae(100);g__Metschnikowia(100);s__Metschnikowia_pulcherrima(100); | Metschnikowia | | 13 | Saprotroph | \|Undefined Saprotroph\| | Yeast | NULL | Probable | As nectar-tap_saprotroph (Põlme et al. 2020); Decay sugar-rich substrates (Põlme et al. 2020) | Põlme S, et al. 2020. Fungal Diversity 105:1-16 ((https://doi.org/10.1007/s13225-020-00466-2)) |
| Otu38826 | 1 | k__Fungi(100);p__Ascomycota(100);c__Saccharomycetes(100);o__Saccharomycetales(100);f__Metschnikowiaceae(100);g__Metschnikowia(100);s__Metschnikowia_pulcherrima(100); | Metschnikowia | | 13 | Saprotroph | \|Undefined Saprotroph\| | Yeast | NULL | Probable | As nectar-tap_saprotroph (Põlme et al. 2020); Decay sugar-rich substrates (Põlme et al. 2020) | Põlme S, et al. 2020. Fungal Diversity 105:1-16 ((https://doi.org/10.1007/s13225-020-00466-2)) |
| Otu38827 | 1 | k__Fungi(100);p__Ascomycota(100);c__Saccharomycetes(100);o__Saccharomycetales(100);f__Metschnikowiaceae(100);g__Metschnikowia(100);g__Metschnikowia_unclassified(100); | Metschnikowia | | 13 | Saprotroph | \|Undefined Saprotroph\| | Yeast | NULL | Probable | As nectar-tap_saprotroph (Põlme et al. 2020); Decay sugar-rich substrates (Põlme et al. 2020) | Põlme S, et al. 2020. Fungal Diversity 105:1-16 ((https://doi.org/10.1007/s13225-020-00466-2)) |
| Otu38828 | 1 | k__Fungi(100);p__Ascomycota(100);c__Saccharomycetes(100);o__Saccharomycetales(100);f__Metschnikowiaceae(100);g__Metschnikowia(100);g__Metschnikowia_unclassified(100); | Metschnikowia | | 13 | Saprotroph | \|Undefined Saprotroph\| | Yeast | NULL | Probable | As nectar-tap_saprotroph (Põlme et al. 2020); Decay sugar-rich substrates (Põlme et al. 2020) | Põlme S, et al. 2020. Fungal Diversity 105:1-16 ((https://doi.org/10.1007/s13225-020-00466-2)) |
| Otu38829 | 1 | k__Fungi(100);p__Ascomycota(100);c__Saccharomycetes(100);o__Saccharomycetales(100);f__Metschnikowiaceae(100);g__Metschnikowia(100);s__Metschnikowia_pulcherrima(100); | Metschnikowia | | 13 | Saprotroph | \|Undefined Saprotroph\| | Yeast | NULL | Probable | As nectar-tap_saprotroph (Põlme et al. 2020); Decay sugar-rich substrates (Põlme et al. 2020) | Põlme S, et al. 2020. Fungal Diversity 105:1-16 ((https://doi.org/10.1007/s13225-020-00466-2)) |
| Otu38830 | 1 | k__Fungi(100);p__Ascomycota(100);c__Saccharomycetes(100);o__Saccharomycetales(100);f__Metschnikowiaceae(100);g__Metschnikowia(100);g__Metschnikowia_unclassified(100); | Metschnikowia | | 13 | Saprotroph | \|Undefined Saprotroph\| | Yeast | NULL | Probable | As nectar-tap_saprotroph (Põlme et al. 2020); Decay sugar-rich substrates (Põlme et al. 2020) | Põlme S, et al. 2020. Fungal Diversity 105:1-16 ((https://doi.org/10.1007/s13225-020-00466-2)) |
| Otu38831 | 1 | k__Fungi(100);p__Ascomycota(100);c__Saccharomycetes(100);o__Saccharomycetales(100);f__Metschnikowiaceae(100);g__Metschnikowia(100);s__Metschnikowia_pulcherrima(100); | Metschnikowia | | 13 | Saprotroph | \|Undefined Saprotroph\| | Yeast | NULL | Probable | As nectar-tap_saprotroph (Põlme et al. 2020); Decay sugar-rich substrates (Põlme et al. 2020) | Põlme S, et al. 2020. Fungal Diversity 105:1-16 ((https://doi.org/10.1007/s13225-020-00466-2)) |
| Otu38832 | 1 | k__Fungi(100);p__Ascomycota(100);c__Saccharomycetes(100);o__Saccharomycetales(100);f__Metschnikowiaceae(100);g__Metschnikowia(100);s__Metschnikowia_pulcherrima(100); | Metschnikowia | | 13 | Saprotroph | \|Undefined Saprotroph\| | Yeast | NULL | Probable | As nectar-tap_saprotroph (Põlme et al. 2020); Decay sugar-rich substrates (Põlme et al. 2020) | Põlme S, et al. 2020. Fungal Diversity 105:1-16 ((https://doi.org/10.1007/s13225-020-00466-2)) |
| Otu38833 | 1 | k__Fungi(100);p__Ascomycota(100);c__Saccharomycetes(100);o__Saccharomycetales(100);f__Metschnikowiaceae(100);g__Metschnikowia(100);g__Metschnikowia_unclassified(100); | Metschnikowia | | 13 | Saprotroph | \|Undefined Saprotroph\| | Yeast | NULL | Probable | As nectar-tap_saprotroph (Põlme et al. 2020); Decay sugar-rich substrates (Põlme et al. 2020) | Põlme S, et al. 2020. Fungal Diversity 105:1-16 ((https://doi.org/10.1007/s13225-020-00466-2)) |
| Otu38834 | 1 | k__Fungi(100);p__Ascomycota(100);c__Saccharomycetes(100);o__Saccharomycetales(100);f__Metschnikowiaceae(100);g__Metschnikowia(100);s__Metschnikowia_pulcherrima(100); | Metschnikowia | | 13 | Saprotroph | \|Undefined Saprotroph\| | Yeast | NULL | Probable | As nectar-tap_saprotroph (Põlme et al. 2020); Decay sugar-rich substrates (Põlme et al. 2020) | Põlme S, et al. 2020. Fungal Diversity 105:1-16 ((https://doi.org/10.1007/s13225-020-00466-2)) |
| Otu38835 | 1 | k__Fungi(100);p__Ascomycota(100);c__Saccharomycetes(100);o__Saccharomycetales(100);f__Metschnikowiaceae(100);g__Metschnikowia(100);s__Metschnikowia_pulcherrima(100); | Metschnikowia | | 13 | Saprotroph | \|Undefined Saprotroph\| | Yeast | NULL | Probable | As nectar-tap_saprotroph (Põlme et al. 2020); Decay sugar-rich substrates (Põlme et al. 2020) | Põlme S, et al. 2020. Fungal Diversity 105:1-16 ((https://doi.org/10.1007/s13225-020-00466-2)) |
| Otu38836 | 1 | k__Fungi(100);p__Ascomycota(100);c__Saccharomycetes(100);o__Saccharomycetales(100);f__Metschnikowiaceae(100);g__Metschnikowia(100);g__Metschnikowia_unclassified(100); | Metschnikowia | | 13 | Saprotroph | \|Undefined Saprotroph\| | Yeast | NULL | Probable | As nectar-tap_saprotroph (Põlme et al. 2020); Decay sugar-rich substrates (Põlme et al. 2020) | Põlme S, et al. 2020. Fungal Diversity 105:1-16 ((https://doi.org/10.1007/s13225-020-00466-2)) |
| Otu38837 | 1 | k__Fungi(100);p__Ascomycota(100);c__Saccharomycetes(100);o__Saccharomycetales(100);f__Metschnikowiaceae(100);g__Metschnikowia(100);s__Metschnikowia_pulcherrima(100); | Metschnikowia | | 13 | Saprotroph | \|Undefined Saprotroph\| | Yeast | NULL | Probable | As nectar-tap_saprotroph (Põlme et al. 2020); Decay sugar-rich substrates (Põlme et al. 2020) | Põlme S, et al. 2020. Fungal Diversity 105:1-16 ((https://doi.org/10.1007/s13225-020-00466-2)) |
| Otu38839 | 1 | k__Fungi(100);p__Ascomycota(100);c__Saccharomycetes(100);o__Saccharomycetales(100);f__Metschnikowiaceae(100);g__Metschnikowia(100);s__Metschnikowia_pulcherrima(100); | Metschnikowia | | 13 | Saprotroph | \|Undefined Saprotroph\| | Yeast | NULL | Probable | As nectar-tap_saprotroph (Põlme et al. 2020); Decay sugar-rich substrates (Põlme et al. 2020) | Põlme S, et al. 2020. Fungal Diversity 105:1-16 ((https://doi.org/10.1007/s13225-020-00466-2)) |
| Otu38840 | 1 | k__Fungi(100);p__Ascomycota(100);c__Saccharomycetes(100);o__Saccharomycetales(100);f__Metschnikowiaceae(100);g__Metschnikowia(100);s__Metschnikowia_pulcherrima(100); | Metschnikowia | | 13 | Saprotroph | \|Undefined Saprotroph\| | Yeast | NULL | Probable | As nectar-tap_saprotroph (Põlme et al. 2020); Decay sugar-rich substrates (Põlme et al. 2020) | Põlme S, et al. 2020. Fungal Diversity 105:1-16 ((https://doi.org/10.1007/s13225-020-00466-2)) |
| Otu38842 | 1 | k__Fungi(100);p__Ascomycota(100);c__Saccharomycetes(100);o__Saccharomycetales(100);f__Metschnikowiaceae(100);g__Metschnikowia(100);s__Metschnikowia_pulcherrima(100); | Metschnikowia | | 13 | Saprotroph | \|Undefined Saprotroph\| | Yeast | NULL | Probable | As nectar-tap_saprotroph (Põlme et al. 2020); Decay sugar-rich substrates (Põlme et al. 2020) | Põlme S, et al. 2020. Fungal Diversity 105:1-16 ((https://doi.org/10.1007/s13225-020-00466-2)) |
| Otu38843 | 1 | k__Fungi(100);p__Ascomycota(100);c__Saccharomycetes(100);o__Saccharomycetales(100);f__Metschnikowiaceae(100);g__Metschnikowia(100);s__Metschnikowia_pulcherrima(100); | Metschnikowia | | 13 | Saprotroph | \|Undefined Saprotroph\| | Yeast | NULL | Probable | As nectar-tap_saprotroph (Põlme et al. 2020); Decay sugar-rich substrates (Põlme et al. 2020) | Põlme S, et al. 2020. Fungal Diversity 105:1-16 ((https://doi.org/10.1007/s13225-020-00466-2)) |
| Otu38844 | 1 | k__Fungi(100);p__Ascomycota(100);c__Saccharomycetes(100);o__Saccharomycetales(100);f__Metschnikowiaceae(100);g__Metschnikowia(100);s__Metschnikowia_pulcherrima(100); | Metschnikowia | | 13 | Saprotroph | \|Undefined Saprotroph\| | Yeast | NULL | Probable | As nectar-tap_saprotroph (Põlme et al. 2020); Decay sugar-rich substrates (Põlme et al. 2020) | Põlme S, et al. 2020. Fungal Diversity 105:1-16 ((https://doi.org/10.1007/s13225-020-00466-2)) |
| Otu38845 | 1 | k__Fungi(100);p__Ascomycota(100);c__Saccharomycetes(100);o__Saccharomycetales(100);f__Metschnikowiaceae(100);g__Metschnikowia(100);s__Metschnikowia_pulcherrima(100); | Metschnikowia | | 13 | Saprotroph | \|Undefined Saprotroph\| | Yeast | NULL | Probable | As nectar-tap_saprotroph (Põlme et al. 2020); Decay sugar-rich substrates (Põlme et al. 2020) | Põlme S, et al. 2020. Fungal Diversity 105:1-16 ((https://doi.org/10.1007/s13225-020-00466-2)) |
| Otu38846 | 1 | k__Fungi(100);p__Ascomycota(100);c__Saccharomycetes(100);o__Saccharomycetales(100);f__Metschnikowiaceae(100);g__Metschnikowia(100);s__Metschnikowia_viticola(100); | Metschnikowia | | 13 | Saprotroph | \|Undefined Saprotroph\| | Yeast | NULL | Probable | As nectar-tap_saprotroph (Põlme et al. 2020); Decay sugar-rich substrates (Põlme et al. 2020) | Põlme S, et al. 2020. Fungal Diversity 105:1-16 ((https://doi.org/10.1007/s13225-020-00466-2)) |
| Otu38847 | 1 | k__Fungi(100);p__Ascomycota(100);c__Saccharomycetes(100);o__Saccharomycetales(100);f__Metschnikowiaceae(100);g__Metschnikowia(100);s__Metschnikowia_pulcherrima(100); | Metschnikowia | | 13 | Saprotroph | \|Undefined Saprotroph\| | Yeast | NULL | Probable | As nectar-tap_saprotroph (Põlme et al. 2020); Decay sugar-rich substrates (Põlme et al. 2020) | Põlme S, et al. 2020. Fungal Diversity 105:1-16 ((https://doi.org/10.1007/s13225-020-00466-2)) |
| Otu38848 | 1 | k__Fungi(100);p__Ascomycota(100);c__Saccharomycetes(100);o__Saccharomycetales(100);f__Metschnikowiaceae(100);g__Metschnikowia(100);s__Metschnikowia_pulcherrima(100); | Metschnikowia | | 13 | Saprotroph | \|Undefined Saprotroph\| | Yeast | NULL | Probable | As nectar-tap_saprotroph (Põlme et al. 2020); Decay sugar-rich substrates (Põlme et al. 2020) | Põlme S, et al. 2020. Fungal Diversity 105:1-16 ((https://doi.org/10.1007/s13225-020-00466-2)) |
| Otu38849 | 1 | k__Fungi(100);p__Ascomycota(100);c__Saccharomycetes(100);o__Saccharomycetales(100);f__Metschnikowiaceae(100);g__Metschnikowia(100);s__Metschnikowia_pulcherrima(100); | Metschnikowia | | 13 | Saprotroph | \|Undefined Saprotroph\| | Yeast | NULL | Probable | As nectar-tap_saprotroph (Põlme et al. 2020); Decay sugar-rich substrates (Põlme et al. 2020) | Põlme S, et al. 2020. Fungal Diversity 105:1-16 ((https://doi.org/10.1007/s13225-020-00466-2)) |
| Otu38850 | 1 | k__Fungi(100);p__Ascomycota(100);c__Saccharomycetes(100);o__Saccharomycetales(100);f__Metschnikowiaceae(100);g__Metschnikowia(100);s__Metschnikowia_pulcherrima(100); | Metschnikowia | | 13 | Saprotroph | \|Undefined Saprotroph\| | Yeast | NULL | Probable | As nectar-tap_saprotroph (Põlme et al. 2020); Decay sugar-rich substrates (Põlme et al. 2020) | Põlme S, et al. 2020. Fungal Diversity 105:1-16 ((https://doi.org/10.1007/s13225-020-00466-2)) |
| Otu38851 | 1 | k__Fungi(100);p__Ascomycota(100);c__Saccharomycetes(100);o__Saccharomycetales(100);f__Metschnikowiaceae(100);g__Metschnikowia(100);s__Metschnikowia_pulcherrima(100); | Metschnikowia | | 13 | Saprotroph | \|Undefined Saprotroph\| | Yeast | NULL | Probable | As nectar-tap_saprotroph (Põlme et al. 2020); Decay sugar-rich substrates (Põlme et al. 2020) | Põlme S, et al. 2020. Fungal Diversity 105:1-16 ((https://doi.org/10.1007/s13225-020-00466-2)) |
| Otu38852 | 1 | k__Fungi(100);p__Ascomycota(100);c__Saccharomycetes(100);o__Saccharomycetales(100);f__Metschnikowiaceae(100);g__Metschnikowia(100);s__Metschnikowia_pulcherrima(100); | Metschnikowia | | 13 | Saprotroph | \|Undefined Saprotroph\| | Yeast | NULL | Probable | As nectar-tap_saprotroph (Põlme et al. 2020); Decay sugar-rich substrates (Põlme et al. 2020) | Põlme S, et al. 2020. Fungal Diversity 105:1-16 ((https://doi.org/10.1007/s13225-020-00466-2)) |
| Otu38853 | 1 | k__Fungi(100);p__Ascomycota(100);c__Saccharomycetes(100);o__Saccharomycetales(100);f__Metschnikowiaceae(100);g__Metschnikowia(100);g__Metschnikowia_unclassified(100); | Metschnikowia | | 13 | Saprotroph | \|Undefined Saprotroph\| | Yeast | NULL | Probable | As nectar-tap_saprotroph (Põlme et al. 2020); Decay sugar-rich substrates (Põlme et al. 2020) | Põlme S, et al. 2020. Fungal Diversity 105:1-16 ((https://doi.org/10.1007/s13225-020-00466-2)) |
| Otu38854 | 1 | k__Fungi(100);p__Ascomycota(100);c__Saccharomycetes(100);o__Saccharomycetales(100);f__Metschnikowiaceae(100);g__Metschnikowia(100);g__Metschnikowia_unclassified(100); | Metschnikowia | | 13 | Saprotroph | \|Undefined Saprotroph\| | Yeast | NULL | Probable | As nectar-tap_saprotroph (Põlme et al. 2020); Decay sugar-rich substrates (Põlme et al. 2020) | Põlme S, et al. 2020. Fungal Diversity 105:1-16 ((https://doi.org/10.1007/s13225-020-00466-2)) |
| Otu38855 | 1 | k__Fungi(100);p__Ascomycota(100);c__Saccharomycetes(100);o__Saccharomycetales(100);f__Metschnikowiaceae(100);g__Metschnikowia(100);s__Metschnikowia_pulcherrima(100); | Metschnikowia | | 13 | Saprotroph | \|Undefined Saprotroph\| | Yeast | NULL | Probable | As nectar-tap_saprotroph (Põlme et al. 2020); Decay sugar-rich substrates (Põlme et al. 2020) | Põlme S, et al. 2020. Fungal Diversity 105:1-16 ((https://doi.org/10.1007/s13225-020-00466-2)) |
| Otu38856 | 1 | k__Fungi(100);p__Ascomycota(100);c__Saccharomycetes(100);o__Saccharomycetales(100);f__Metschnikowiaceae(100);g__Metschnikowia(100);s__Metschnikowia_pulcherrima(100); | Metschnikowia | | 13 | Saprotroph | \|Undefined Saprotroph\| | Yeast | NULL | Probable | As nectar-tap_saprotroph (Põlme et al. 2020); Decay sugar-rich substrates (Põlme et al. 2020) | Põlme S, et al. 2020. Fungal Diversity 105:1-16 ((https://doi.org/10.1007/s13225-020-00466-2)) |
| Otu38857 | 1 | k__Fungi(100);p__Ascomycota(100);c__Saccharomycetes(100);o__Saccharomycetales(100);f__Metschnikowiaceae(100);g__Metschnikowia(100);s__Metschnikowia_pulcherrima(100); | Metschnikowia | | 13 | Saprotroph | \|Undefined Saprotroph\| | Yeast | NULL | Probable | As nectar-tap_saprotroph (Põlme et al. 2020); Decay sugar-rich substrates (Põlme et al. 2020) | Põlme S, et al. 2020. Fungal Diversity 105:1-16 ((https://doi.org/10.1007/s13225-020-00466-2)) |
| Otu38858 | 1 | k__Fungi(100);p__Ascomycota(100);c__Saccharomycetes(100);o__Saccharomycetales(100);f__Metschnikowiaceae(100);g__Metschnikowia(100);s__Metschnikowia_pulcherrima(100); | Metschnikowia | | 13 | Saprotroph | \|Undefined Saprotroph\| | Yeast | NULL | Probable | As nectar-tap_saprotroph (Põlme et al. 2020); Decay sugar-rich substrates (Põlme et al. 2020) | Põlme S, et al. 2020. Fungal Diversity 105:1-16 ((https://doi.org/10.1007/s13225-020-00466-2)) |
| Otu38859 | 1 | k__Fungi(100);p__Ascomycota(100);c__Saccharomycetes(100);o__Saccharomycetales(100);f__Metschnikowiaceae(100);g__Metschnikowia(100);s__Metschnikowia_pulcherrima(100); | Metschnikowia | | 13 | Saprotroph | \|Undefined Saprotroph\| | Yeast | NULL | Probable | As nectar-tap_saprotroph (Põlme et al. 2020); Decay sugar-rich substrates (Põlme et al. 2020) | Põlme S, et al. 2020. Fungal Diversity 105:1-16 ((https://doi.org/10.1007/s13225-020-00466-2)) |
| Otu38860 | 1 | k__Fungi(100);p__Ascomycota(100);c__Saccharomycetes(100);o__Saccharomycetales(100);f__Metschnikowiaceae(100);g__Metschnikowia(100);s__Metschnikowia_pulcherrima(100); | Metschnikowia | | 13 | Saprotroph | \|Undefined Saprotroph\| | Yeast | NULL | Probable | As nectar-tap_saprotroph (Põlme et al. 2020); Decay sugar-rich substrates (Põlme et al. 2020) | Põlme S, et al. 2020. Fungal Diversity 105:1-16 ((https://doi.org/10.1007/s13225-020-00466-2)) |
| Otu38861 | 1 | k__Fungi(100);p__Ascomycota(100);c__Saccharomycetes(100);o__Saccharomycetales(100);f__Metschnikowiaceae(100);g__Metschnikowia(100);g__Metschnikowia_unclassified(100); | Metschnikowia | | 13 | Saprotroph | \|Undefined Saprotroph\| | Yeast | NULL | Probable | As nectar-tap_saprotroph (Põlme et al. 2020); Decay sugar-rich substrates (Põlme et al. 2020) | Põlme S, et al. 2020. Fungal Diversity 105:1-16 ((https://doi.org/10.1007/s13225-020-00466-2)) |
| Otu38862 | 1 | k__Fungi(100);p__Ascomycota(100);c__Saccharomycetes(100);o__Saccharomycetales(100);f__Metschnikowiaceae(100);g__Metschnikowia(100);s__Metschnikowia_pulcherrima(100); | Metschnikowia | | 13 | Saprotroph | \|Undefined Saprotroph\| | Yeast | NULL | Probable | As nectar-tap_saprotroph (Põlme et al. 2020); Decay sugar-rich substrates (Põlme et al. 2020) | Põlme S, et al. 2020. Fungal Diversity 105:1-16 ((https://doi.org/10.1007/s13225-020-00466-2)) |
| Otu38863 | 1 | k__Fungi(100);p__Ascomycota(100);c__Saccharomycetes(100);o__Saccharomycetales(100);f__Metschnikowiaceae(100);g__Metschnikowia(100);g__Metschnikowia_unclassified(100); | Metschnikowia | | 13 | Saprotroph | \|Undefined Saprotroph\| | Yeast | NULL | Probable | As nectar-tap_saprotroph (Põlme et al. 2020); Decay sugar-rich substrates (Põlme et al. 2020) | Põlme S, et al. 2020. Fungal Diversity 105:1-16 ((https://doi.org/10.1007/s13225-020-00466-2)) |
| Otu38864 | 1 | k__Fungi(100);p__Ascomycota(100);c__Saccharomycetes(100);o__Saccharomycetales(100);f__Metschnikowiaceae(100);g__Metschnikowia(100);s__Metschnikowia_pulcherrima(100); | Metschnikowia | | 13 | Saprotroph | \|Undefined Saprotroph\| | Yeast | NULL | Probable | As nectar-tap_saprotroph (Põlme et al. 2020); Decay sugar-rich substrates (Põlme et al. 2020) | Põlme S, et al. 2020. Fungal Diversity 105:1-16 ((https://doi.org/10.1007/s13225-020-00466-2)) |
| Otu38865 | 1 | k__Fungi(100);p__Ascomycota(100);c__Saccharomycetes(100);o__Saccharomycetales(100);f__Metschnikowiaceae(100);g__Metschnikowia(100);s__Metschnikowia_pulcherrima(100); | Metschnikowia | | 13 | Saprotroph | \|Undefined Saprotroph\| | Yeast | NULL | Probable | As nectar-tap_saprotroph (Põlme et al. 2020); Decay sugar-rich substrates (Põlme et al. 2020) | Põlme S, et al. 2020. Fungal Diversity 105:1-16 ((https://doi.org/10.1007/s13225-020-00466-2)) |
| Otu38866 | 1 | k__Fungi(100);p__Ascomycota(100);c__Saccharomycetes(100);o__Saccharomycetales(100);f__Metschnikowiaceae(100);g__Metschnikowia(100);g__Metschnikowia_unclassified(100); | Metschnikowia | | 13 | Saprotroph | \|Undefined Saprotroph\| | Yeast | NULL | Probable | As nectar-tap_saprotroph (Põlme et al. 2020); Decay sugar-rich substrates (Põlme et al. 2020) | Põlme S, et al. 2020. Fungal Diversity 105:1-16 ((https://doi.org/10.1007/s13225-020-00466-2)) |
| Otu38867 | 1 | k__Fungi(100);p__Ascomycota(100);c__Saccharomycetes(100);o__Saccharomycetales(100);f__Metschnikowiaceae(100);g__Metschnikowia(100);s__Metschnikowia_pulcherrima(100); | Metschnikowia | | 13 | Saprotroph | \|Undefined Saprotroph\| | Yeast | NULL | Probable | As nectar-tap_saprotroph (Põlme et al. 2020); Decay sugar-rich substrates (Põlme et al. 2020) | Põlme S, et al. 2020. Fungal Diversity 105:1-16 ((https://doi.org/10.1007/s13225-020-00466-2)) |
| Otu38868 | 1 | k__Fungi(100);p__Ascomycota(100);c__Saccharomycetes(100);o__Saccharomycetales(100);f__Metschnikowiaceae(100);g__Metschnikowia(100);g__Metschnikowia_unclassified(100); | Metschnikowia | | 13 | Saprotroph | \|Undefined Saprotroph\| | Yeast | NULL | Probable | As nectar-tap_saprotroph (Põlme et al. 2020); Decay sugar-rich substrates (Põlme et al. 2020) | Põlme S, et al. 2020. Fungal Diversity 105:1-16 ((https://doi.org/10.1007/s13225-020-00466-2)) |
| Otu38869 | 1 | k__Fungi(100);p__Ascomycota(100);c__Saccharomycetes(100);o__Saccharomycetales(100);f__Metschnikowiaceae(100);g__Metschnikowia(100);g__Metschnikowia_unclassified(100); | Metschnikowia | | 13 | Saprotroph | \|Undefined Saprotroph\| | Yeast | NULL | Probable | As nectar-tap_saprotroph (Põlme et al. 2020); Decay sugar-rich substrates (Põlme et al. 2020) | Põlme S, et al. 2020. Fungal Diversity 105:1-16 ((https://doi.org/10.1007/s13225-020-00466-2)) |
| Otu38870 | 1 | k__Fungi(100);p__Ascomycota(100);c__Saccharomycetes(100);o__Saccharomycetales(100);f__Metschnikowiaceae(100);g__Metschnikowia(100);s__Metschnikowia_pulcherrima(100); | Metschnikowia | | 13 | Saprotroph | \|Undefined Saprotroph\| | Yeast | NULL | Probable | As nectar-tap_saprotroph (Põlme et al. 2020); Decay sugar-rich substrates (Põlme et al. 2020) | Põlme S, et al. 2020. Fungal Diversity 105:1-16 ((https://doi.org/10.1007/s13225-020-00466-2)) |
| Otu38871 | 1 | k__Fungi(100);p__Ascomycota(100);c__Saccharomycetes(100);o__Saccharomycetales(100);f__Metschnikowiaceae(100);g__Metschnikowia(100);s__Metschnikowia_pulcherrima(100); | Metschnikowia | | 13 | Saprotroph | \|Undefined Saprotroph\| | Yeast | NULL | Probable | As nectar-tap_saprotroph (Põlme et al. 2020); Decay sugar-rich substrates (Põlme et al. 2020) | Põlme S, et al. 2020. Fungal Diversity 105:1-16 ((https://doi.org/10.1007/s13225-020-00466-2)) |
| Otu38872 | 1 | k__Fungi(100);p__Ascomycota(100);c__Saccharomycetes(100);o__Saccharomycetales(100);f__Metschnikowiaceae(100);g__Metschnikowia(100);s__Metschnikowia_pulcherrima(100); | Metschnikowia | | 13 | Saprotroph | \|Undefined Saprotroph\| | Yeast | NULL | Probable | As nectar-tap_saprotroph (Põlme et al. 2020); Decay sugar-rich substrates (Põlme et al. 2020) | Põlme S, et al. 2020. Fungal Diversity 105:1-16 ((https://doi.org/10.1007/s13225-020-00466-2)) |
| Otu38873 | 1 | k__Fungi(100);p__Ascomycota(100);c__Saccharomycetes(100);o__Saccharomycetales(100);f__Metschnikowiaceae(100);g__Metschnikowia(100);s__Metschnikowia_pulcherrima(100); | Metschnikowia | | 13 | Saprotroph | \|Undefined Saprotroph\| | Yeast | NULL | Probable | As nectar-tap_saprotroph (Põlme et al. 2020); Decay sugar-rich substrates (Põlme et al. 2020) | Põlme S, et al. 2020. Fungal Diversity 105:1-16 ((https://doi.org/10.1007/s13225-020-00466-2)) |
| Otu38874 | 1 | k__Fungi(100);p__Ascomycota(100);c__Saccharomycetes(100);o__Saccharomycetales(100);f__Metschnikowiaceae(100);g__Metschnikowia(100);g__Metschnikowia_unclassified(100); | Metschnikowia | | 13 | Saprotroph | \|Undefined Saprotroph\| | Yeast | NULL | Probable | As nectar-tap_saprotroph (Põlme et al. 2020); Decay sugar-rich substrates (Põlme et al. 2020) | Põlme S, et al. 2020. Fungal Diversity 105:1-16 ((https://doi.org/10.1007/s13225-020-00466-2)) |
| Otu38875 | 1 | k__Fungi(100);p__Ascomycota(100);c__Saccharomycetes(100);o__Saccharomycetales(100);f__Metschnikowiaceae(100);g__Metschnikowia(100);s__Metschnikowia_pulcherrima(100); | Metschnikowia | | 13 | Saprotroph | \|Undefined Saprotroph\| | Yeast | NULL | Probable | As nectar-tap_saprotroph (Põlme et al. 2020); Decay sugar-rich substrates (Põlme et al. 2020) | Põlme S, et al. 2020. Fungal Diversity 105:1-16 ((https://doi.org/10.1007/s13225-020-00466-2)) |
| Otu38878 | 1 | k__Fungi(100);p__Ascomycota(100);c__Saccharomycetes(100);o__Saccharomycetales(100);f__Metschnikowiaceae(100);g__Metschnikowia(100);s__Metschnikowia_pulcherrima(100); | Metschnikowia | | 13 | Saprotroph | \|Undefined Saprotroph\| | Yeast | NULL | Probable | As nectar-tap_saprotroph (Põlme et al. 2020); Decay sugar-rich substrates (Põlme et al. 2020) | Põlme S, et al. 2020. Fungal Diversity 105:1-16 ((https://doi.org/10.1007/s13225-020-00466-2)) |
| Otu38879 | 1 | k__Fungi(100);p__Ascomycota(100);c__Saccharomycetes(100);o__Saccharomycetales(100);f__Metschnikowiaceae(100);g__Metschnikowia(100);s__Metschnikowia_pulcherrima(100); | Metschnikowia | | 13 | Saprotroph | \|Undefined Saprotroph\| | Yeast | NULL | Probable | As nectar-tap_saprotroph (Põlme et al. 2020); Decay sugar-rich substrates (Põlme et al. 2020) | Põlme S, et al. 2020. Fungal Diversity 105:1-16 ((https://doi.org/10.1007/s13225-020-00466-2)) |
| Otu38880 | 1 | k__Fungi(100);p__Ascomycota(100);c__Saccharomycetes(100);o__Saccharomycetales(100);f__Metschnikowiaceae(100);g__Metschnikowia(100);s__Metschnikowia_pulcherrima(100); | Metschnikowia | | 13 | Saprotroph | \|Undefined Saprotroph\| | Yeast | NULL | Probable | As nectar-tap_saprotroph (Põlme et al. 2020); Decay sugar-rich substrates (Põlme et al. 2020) | Põlme S, et al. 2020. Fungal Diversity 105:1-16 ((https://doi.org/10.1007/s13225-020-00466-2)) |
| Otu38881 | 1 | k__Fungi(100);p__Ascomycota(100);c__Saccharomycetes(100);o__Saccharomycetales(100);f__Metschnikowiaceae(100);g__Metschnikowia(100);g__Metschnikowia_unclassified(100); | Metschnikowia | | 13 | Saprotroph | \|Undefined Saprotroph\| | Yeast | NULL | Probable | As nectar-tap_saprotroph (Põlme et al. 2020); Decay sugar-rich substrates (Põlme et al. 2020) | Põlme S, et al. 2020. Fungal Diversity 105:1-16 ((https://doi.org/10.1007/s13225-020-00466-2)) |
| Otu38882 | 1 | k__Fungi(100);p__Ascomycota(100);c__Saccharomycetes(100);o__Saccharomycetales(100);f__Metschnikowiaceae(100);g__Metschnikowia(100);s__Metschnikowia_pulcherrima(100); | Metschnikowia | | 13 | Saprotroph | \|Undefined Saprotroph\| | Yeast | NULL | Probable | As nectar-tap_saprotroph (Põlme et al. 2020); Decay sugar-rich substrates (Põlme et al. 2020) | Põlme S, et al. 2020. Fungal Diversity 105:1-16 ((https://doi.org/10.1007/s13225-020-00466-2)) |
| Otu38883 | 1 | k__Fungi(100);p__Ascomycota(100);c__Saccharomycetes(100);o__Saccharomycetales(100);f__Metschnikowiaceae(100);g__Metschnikowia(100);s__Metschnikowia_pulcherrima(100); | Metschnikowia | | 13 | Saprotroph | \|Undefined Saprotroph\| | Yeast | NULL | Probable | As nectar-tap_saprotroph (Põlme et al. 2020); Decay sugar-rich substrates (Põlme et al. 2020) | Põlme S, et al. 2020. Fungal Diversity 105:1-16 ((https://doi.org/10.1007/s13225-020-00466-2)) |
| Otu38884 | 1 | k__Fungi(100);p__Ascomycota(100);c__Saccharomycetes(100);o__Saccharomycetales(100);f__Metschnikowiaceae(100);g__Metschnikowia(100);s__Metschnikowia_pulcherrima(100); | Metschnikowia | | 13 | Saprotroph | \|Undefined Saprotroph\| | Yeast | NULL | Probable | As nectar-tap_saprotroph (Põlme et al. 2020); Decay sugar-rich substrates (Põlme et al. 2020) | Põlme S, et al. 2020. Fungal Diversity 105:1-16 ((https://doi.org/10.1007/s13225-020-00466-2)) |
| Otu38885 | 1 | k__Fungi(100);p__Ascomycota(100);c__Saccharomycetes(100);o__Saccharomycetales(100);f__Metschnikowiaceae(100);g__Metschnikowia(100);s__Metschnikowia_pulcherrima(100); | Metschnikowia | | 13 | Saprotroph | \|Undefined Saprotroph\| | Yeast | NULL | Probable | As nectar-tap_saprotroph (Põlme et al. 2020); Decay sugar-rich substrates (Põlme et al. 2020) | Põlme S, et al. 2020. Fungal Diversity 105:1-16 ((https://doi.org/10.1007/s13225-020-00466-2)) |
| Otu38886 | 1 | k__Fungi(100);p__Ascomycota(100);c__Saccharomycetes(100);o__Saccharomycetales(100);f__Metschnikowiaceae(100);g__Metschnikowia(100);s__Metschnikowia_pulcherrima(100); | Metschnikowia | | 13 | Saprotroph | \|Undefined Saprotroph\| | Yeast | NULL | Probable | As nectar-tap_saprotroph (Põlme et al. 2020); Decay sugar-rich substrates (Põlme et al. 2020) | Põlme S, et al. 2020. Fungal Diversity 105:1-16 ((https://doi.org/10.1007/s13225-020-00466-2)) |
| Otu38887 | 1 | k__Fungi(100);p__Ascomycota(100);c__Saccharomycetes(100);o__Saccharomycetales(100);f__Metschnikowiaceae(100);g__Metschnikowia(100);g__Metschnikowia_unclassified(100); | Metschnikowia | | 13 | Saprotroph | \|Undefined Saprotroph\| | Yeast | NULL | Probable | As nectar-tap_saprotroph (Põlme et al. 2020); Decay sugar-rich substrates (Põlme et al. 2020) | Põlme S, et al. 2020. Fungal Diversity 105:1-16 ((https://doi.org/10.1007/s13225-020-00466-2)) |
| Otu38888 | 1 | k__Fungi(100);p__Ascomycota(100);c__Saccharomycetes(100);o__Saccharomycetales(100);f__Metschnikowiaceae(100);g__Metschnikowia(100);s__Metschnikowia_pulcherrima(100); | Metschnikowia | | 13 | Saprotroph | \|Undefined Saprotroph\| | Yeast | NULL | Probable | As nectar-tap_saprotroph (Põlme et al. 2020); Decay sugar-rich substrates (Põlme et al. 2020) | Põlme S, et al. 2020. Fungal Diversity 105:1-16 ((https://doi.org/10.1007/s13225-020-00466-2)) |
| Otu38889 | 1 | k__Fungi(100);p__Ascomycota(100);c__Saccharomycetes(100);o__Saccharomycetales(100);f__Metschnikowiaceae(100);g__Metschnikowia(100);g__Metschnikowia_unclassified(100); | Metschnikowia | | 13 | Saprotroph | \|Undefined Saprotroph\| | Yeast | NULL | Probable | As nectar-tap_saprotroph (Põlme et al. 2020); Decay sugar-rich substrates (Põlme et al. 2020) | Põlme S, et al. 2020. Fungal Diversity 105:1-16 ((https://doi.org/10.1007/s13225-020-00466-2)) |
| Otu38890 | 1 | k__Fungi(100);p__Ascomycota(100);c__Saccharomycetes(100);o__Saccharomycetales(100);f__Metschnikowiaceae(100);g__Metschnikowia(100);s__Metschnikowia_pulcherrima(100); | Metschnikowia | | 13 | Saprotroph | \|Undefined Saprotroph\| | Yeast | NULL | Probable | As nectar-tap_saprotroph (Põlme et al. 2020); Decay sugar-rich substrates (Põlme et al. 2020) | Põlme S, et al. 2020. Fungal Diversity 105:1-16 ((https://doi.org/10.1007/s13225-020-00466-2)) |
| Otu38891 | 1 | k__Fungi(100);p__Ascomycota(100);c__Saccharomycetes(100);o__Saccharomycetales(100);f__Metschnikowiaceae(100);g__Metschnikowia(100);g__Metschnikowia_unclassified(100); | Metschnikowia | | 13 | Saprotroph | \|Undefined Saprotroph\| | Yeast | NULL | Probable | As nectar-tap_saprotroph (Põlme et al. 2020); Decay sugar-rich substrates (Põlme et al. 2020) | Põlme S, et al. 2020. Fungal Diversity 105:1-16 ((https://doi.org/10.1007/s13225-020-00466-2)) |
| Otu38892 | 1 | k__Fungi(100);p__Ascomycota(100);c__Saccharomycetes(100);o__Saccharomycetales(100);f__Metschnikowiaceae(100);g__Metschnikowia(100);s__Metschnikowia_pulcherrima(100); | Metschnikowia | | 13 | Saprotroph | \|Undefined Saprotroph\| | Yeast | NULL | Probable | As nectar-tap_saprotroph (Põlme et al. 2020); Decay sugar-rich substrates (Põlme et al. 2020) | Põlme S, et al. 2020. Fungal Diversity 105:1-16 ((https://doi.org/10.1007/s13225-020-00466-2)) |
| Otu38893 | 1 | k__Fungi(100);p__Ascomycota(100);c__Saccharomycetes(100);o__Saccharomycetales(100);f__Metschnikowiaceae(100);g__Metschnikowia(100);s__Metschnikowia_pulcherrima(100); | Metschnikowia | | 13 | Saprotroph | \|Undefined Saprotroph\| | Yeast | NULL | Probable | As nectar-tap_saprotroph (Põlme et al. 2020); Decay sugar-rich substrates (Põlme et al. 2020) | Põlme S, et al. 2020. Fungal Diversity 105:1-16 ((https://doi.org/10.1007/s13225-020-00466-2)) |
| Otu38894 | 1 | k__Fungi(100);p__Ascomycota(100);c__Saccharomycetes(100);o__Saccharomycetales(100);f__Metschnikowiaceae(100);g__Metschnikowia(100);g__Metschnikowia_unclassified(100); | Metschnikowia | | 13 | Saprotroph | \|Undefined Saprotroph\| | Yeast | NULL | Probable | As nectar-tap_saprotroph (Põlme et al. 2020); Decay sugar-rich substrates (Põlme et al. 2020) | Põlme S, et al. 2020. Fungal Diversity 105:1-16 ((https://doi.org/10.1007/s13225-020-00466-2)) |
| Otu38895 | 1 | k__Fungi(100);p__Ascomycota(100);c__Saccharomycetes(100);o__Saccharomycetales(100);f__Metschnikowiaceae(100);g__Metschnikowia(100);g__Metschnikowia_unclassified(100); | Metschnikowia | | 13 | Saprotroph | \|Undefined Saprotroph\| | Yeast | NULL | Probable | As nectar-tap_saprotroph (Põlme et al. 2020); Decay sugar-rich substrates (Põlme et al. 2020) | Põlme S, et al. 2020. Fungal Diversity 105:1-16 ((https://doi.org/10.1007/s13225-020-00466-2)) |
| Otu38896 | 1 | k__Fungi(100);p__Ascomycota(100);c__Saccharomycetes(100);o__Saccharomycetales(100);f__Metschnikowiaceae(100);g__Metschnikowia(100);s__Metschnikowia_pulcherrima(100); | Metschnikowia | | 13 | Saprotroph | \|Undefined Saprotroph\| | Yeast | NULL | Probable | As nectar-tap_saprotroph (Põlme et al. 2020); Decay sugar-rich substrates (Põlme et al. 2020) | Põlme S, et al. 2020. Fungal Diversity 105:1-16 ((https://doi.org/10.1007/s13225-020-00466-2)) |
| Otu38898 | 1 | k__Fungi(100);p__Ascomycota(100);c__Saccharomycetes(100);o__Saccharomycetales(100);f__Metschnikowiaceae(100);g__Metschnikowia(100);s__Metschnikowia_pulcherrima(100); | Metschnikowia | | 13 | Saprotroph | \|Undefined Saprotroph\| | Yeast | NULL | Probable | As nectar-tap_saprotroph (Põlme et al. 2020); Decay sugar-rich substrates (Põlme et al. 2020) | Põlme S, et al. 2020. Fungal Diversity 105:1-16 ((https://doi.org/10.1007/s13225-020-00466-2)) |
| Otu38901 | 1 | k__Fungi(100);p__Ascomycota(100);c__Saccharomycetes(100);o__Saccharomycetales(100);f__Metschnikowiaceae(100);g__Metschnikowia(100);s__Metschnikowia_pulcherrima(100); | Metschnikowia | | 13 | Saprotroph | \|Undefined Saprotroph\| | Yeast | NULL | Probable | As nectar-tap_saprotroph (Põlme et al. 2020); Decay sugar-rich substrates (Põlme et al. 2020) | Põlme S, et al. 2020. Fungal Diversity 105:1-16 ((https://doi.org/10.1007/s13225-020-00466-2)) |
| Otu38902 | 1 | k__Fungi(100);p__Ascomycota(100);c__Saccharomycetes(100);o__Saccharomycetales(100);f__Metschnikowiaceae(100);g__Metschnikowia(100);s__Metschnikowia_pulcherrima(100); | Metschnikowia | | 13 | Saprotroph | \|Undefined Saprotroph\| | Yeast | NULL | Probable | As nectar-tap_saprotroph (Põlme et al. 2020); Decay sugar-rich substrates (Põlme et al. 2020) | Põlme S, et al. 2020. Fungal Diversity 105:1-16 ((https://doi.org/10.1007/s13225-020-00466-2)) |
| Otu38905 | 1 | k__Fungi(100);p__Ascomycota(100);c__Saccharomycetes(100);o__Saccharomycetales(100);f__Metschnikowiaceae(100);g__Metschnikowia(100);g__Metschnikowia_unclassified(100); | Metschnikowia | | 13 | Saprotroph | \|Undefined Saprotroph\| | Yeast | NULL | Probable | As nectar-tap_saprotroph (Põlme et al. 2020); Decay sugar-rich substrates (Põlme et al. 2020) | Põlme S, et al. 2020. Fungal Diversity 105:1-16 ((https://doi.org/10.1007/s13225-020-00466-2)) |
| Otu38906 | 1 | k__Fungi(100);p__Ascomycota(100);c__Saccharomycetes(100);o__Saccharomycetales(100);f__Metschnikowiaceae(100);g__Metschnikowia(100);s__Metschnikowia_pulcherrima(100); | Metschnikowia | | 13 | Saprotroph | \|Undefined Saprotroph\| | Yeast | NULL | Probable | As nectar-tap_saprotroph (Põlme et al. 2020); Decay sugar-rich substrates (Põlme et al. 2020) | Põlme S, et al. 2020. Fungal Diversity 105:1-16 ((https://doi.org/10.1007/s13225-020-00466-2)) |
| Otu38907 | 1 | k__Fungi(100);p__Ascomycota(100);c__Saccharomycetes(100);o__Saccharomycetales(100);f__Metschnikowiaceae(100);g__Metschnikowia(100);g__Metschnikowia_unclassified(100); | Metschnikowia | | 13 | Saprotroph | \|Undefined Saprotroph\| | Yeast | NULL | Probable | As nectar-tap_saprotroph (Põlme et al. 2020); Decay sugar-rich substrates (Põlme et al. 2020) | Põlme S, et al. 2020. Fungal Diversity 105:1-16 ((https://doi.org/10.1007/s13225-020-00466-2)) |
| Otu38909 | 1 | k__Fungi(100);p__Ascomycota(100);c__Saccharomycetes(100);o__Saccharomycetales(100);f__Metschnikowiaceae(100);g__Metschnikowia(100);s__Metschnikowia_pulcherrima(100); | Metschnikowia | | 13 | Saprotroph | \|Undefined Saprotroph\| | Yeast | NULL | Probable | As nectar-tap_saprotroph (Põlme et al. 2020); Decay sugar-rich substrates (Põlme et al. 2020) | Põlme S, et al. 2020. Fungal Diversity 105:1-16 ((https://doi.org/10.1007/s13225-020-00466-2)) |
| Otu38910 | 1 | k__Fungi(100);p__Ascomycota(100);c__Saccharomycetes(100);o__Saccharomycetales(100);f__Metschnikowiaceae(100);g__Metschnikowia(100);g__Metschnikowia_unclassified(100); | Metschnikowia | | 13 | Saprotroph | \|Undefined Saprotroph\| | Yeast | NULL | Probable | As nectar-tap_saprotroph (Põlme et al. 2020); Decay sugar-rich substrates (Põlme et al. 2020) | Põlme S, et al. 2020. Fungal Diversity 105:1-16 ((https://doi.org/10.1007/s13225-020-00466-2)) |
| Otu38912 | 1 | k__Fungi(100);p__Ascomycota(100);c__Saccharomycetes(100);o__Saccharomycetales(100);f__Metschnikowiaceae(100);g__Metschnikowia(100);s__Metschnikowia_pulcherrima(100); | Metschnikowia | | 13 | Saprotroph | \|Undefined Saprotroph\| | Yeast | NULL | Probable | As nectar-tap_saprotroph (Põlme et al. 2020); Decay sugar-rich substrates (Põlme et al. 2020) | Põlme S, et al. 2020. Fungal Diversity 105:1-16 ((https://doi.org/10.1007/s13225-020-00466-2)) |
| Otu38913 | 1 | k__Fungi(100);p__Ascomycota(100);c__Saccharomycetes(100);o__Saccharomycetales(100);f__Metschnikowiaceae(100);g__Metschnikowia(100);s__Metschnikowia_pulcherrima(100); | Metschnikowia | | 13 | Saprotroph | \|Undefined Saprotroph\| | Yeast | NULL | Probable | As nectar-tap_saprotroph (Põlme et al. 2020); Decay sugar-rich substrates (Põlme et al. 2020) | Põlme S, et al. 2020. Fungal Diversity 105:1-16 ((https://doi.org/10.1007/s13225-020-00466-2)) |
| Otu38914 | 1 | k__Fungi(100);p__Ascomycota(100);c__Saccharomycetes(100);o__Saccharomycetales(100);f__Metschnikowiaceae(100);g__Metschnikowia(100);s__Metschnikowia_pulcherrima(100); | Metschnikowia | | 13 | Saprotroph | \|Undefined Saprotroph\| | Yeast | NULL | Probable | As nectar-tap_saprotroph (Põlme et al. 2020); Decay sugar-rich substrates (Põlme et al. 2020) | Põlme S, et al. 2020. Fungal Diversity 105:1-16 ((https://doi.org/10.1007/s13225-020-00466-2)) |
| Otu38915 | 1 | k__Fungi(100);p__Ascomycota(100);c__Saccharomycetes(100);o__Saccharomycetales(100);f__Metschnikowiaceae(100);g__Metschnikowia(100);s__Metschnikowia_pulcherrima(100); | Metschnikowia | | 13 | Saprotroph | \|Undefined Saprotroph\| | Yeast | NULL | Probable | As nectar-tap_saprotroph (Põlme et al. 2020); Decay sugar-rich substrates (Põlme et al. 2020) | Põlme S, et al. 2020. Fungal Diversity 105:1-16 ((https://doi.org/10.1007/s13225-020-00466-2)) |
| [illegible] | [illegible] | [illegible] | [illegible] | | [illegible] | [illegible] | [illegible] | [illegible] | [illegible] | [illegible] | [illegible] | [illegible] |

| | | | | | | | | | | | |
|---|---|---|---|---|---|---|---|---|---|---|---|
| Otu38934 | 1 | k__Fungi(100);p__Ascomycota(100);c__Saccharomycetes(100);o__Saccharomycetales(100);f__Metschnikowiaceae(100);g__Metschnikowia(100);s__Metschnikowia_pulcherrima(100); | Metschnikowia | 13 | Saprotroph | \|Undefined Saprotroph\| | Yeast | NULL | Probable | As nectar-tap_saprotroph (Põlme et al. 2020); Decay sugar-rich substrates (Põlme et al. 2020) | Põlme S, et al. 2020. Fungal Diversity 105:1-16 ((https://doi.org/10.1007/s13225-020-00466-2)) |
| Otu38935 | 1 | k__Fungi(100);p__Ascomycota(100);c__Saccharomycetes(100);o__Saccharomycetales(100);f__Metschnikowiaceae(100);g__Metschnikowia(100);s__Metschnikowia_pulcherrima(100); | Metschnikowia | 13 | Saprotroph | \|Undefined Saprotroph\| | Yeast | NULL | Probable | As nectar-tap_saprotroph (Põlme et al. 2020); Decay sugar-rich substrates (Põlme et al. 2020) | Põlme S, et al. 2020. Fungal Diversity 105:1-16 ((https://doi.org/10.1007/s13225-020-00466-2)) |
| Otu38936 | 1 | k__Fungi(100);p__Ascomycota(100);c__Saccharomycetes(100);o__Saccharomycetales(100);f__Metschnikowiaceae(100);g__Metschnikowia(100);s__Metschnikowia_pulcherrima(100); | Metschnikowia | 13 | Saprotroph | \|Undefined Saprotroph\| | Yeast | NULL | Probable | As nectar-tap_saprotroph (Põlme et al. 2020); Decay sugar-rich substrates (Põlme et al. 2020) | Põlme S, et al. 2020. Fungal Diversity 105:1-16 ((https://doi.org/10.1007/s13225-020-00466-2)) |
| Otu38937 | 1 | k__Fungi(100);p__Ascomycota(100);c__Saccharomycetes(100);o__Saccharomycetales(100);f__Metschnikowiaceae(100);g__Metschnikowia(100);g__Metschnikowia_unclassified(100); | Metschnikowia | 13 | Saprotroph | \|Undefined Saprotroph\| | Yeast | NULL | Probable | As nectar-tap_saprotroph (Põlme et al. 2020); Decay sugar-rich substrates (Põlme et al. 2020) | Põlme S, et al. 2020. Fungal Diversity 105:1-16 ((https://doi.org/10.1007/s13225-020-00466-2)) |
| Otu38944 | 1 | k__Fungi(100);p__Ascomycota(100);c__Saccharomycetes(100);o__Saccharomycetales(100);f__Metschnikowiaceae(100);g__Metschnikowia(100);g__Metschnikowia_unclassified(100); | Metschnikowia | 13 | Saprotroph | \|Undefined Saprotroph\| | Yeast | NULL | Probable | As nectar-tap_saprotroph (Põlme et al. 2020); Decay sugar-rich substrates (Põlme et al. 2020) | Põlme S, et al. 2020. Fungal Diversity 105:1-16 ((https://doi.org/10.1007/s13225-020-00466-2)) |
| Otu38945 | 1 | k__Fungi(100);p__Ascomycota(100);c__Saccharomycetes(100);o__Saccharomycetales(100);f__Metschnikowiaceae(100);g__Metschnikowia(100);g__Metschnikowia_unclassified(100); | Metschnikowia | 13 | Saprotroph | \|Undefined Saprotroph\| | Yeast | NULL | Probable | As nectar-tap_saprotroph (Põlme et al. 2020); Decay sugar-rich substrates (Põlme et al. 2020) | Põlme S, et al. 2020. Fungal Diversity 105:1-16 ((https://doi.org/10.1007/s13225-020-00466-2)) |
| Otu38946 | 1 | k__Fungi(100);p__Ascomycota(100);c__Saccharomycetes(100);o__Saccharomycetales(100);f__Metschnikowiaceae(100);g__Metschnikowia(100);s__Metschnikowia_pulcherrima(100); | Metschnikowia | 13 | Saprotroph | \|Undefined Saprotroph\| | Yeast | NULL | Probable | As nectar-tap_saprotroph (Põlme et al. 2020); Decay sugar-rich substrates (Põlme et al. 2020) | Põlme S, et al. 2020. Fungal Diversity 105:1-16 ((https://doi.org/10.1007/s13225-020-00466-2)) |
| Otu38950 | 1 | k__Fungi(100);p__Ascomycota(100);c__Saccharomycetes(100);o__Saccharomycetales(100);f__Metschnikowiaceae(100);g__Metschnikowia(100);s__Metschnikowia_pulcherrima(100); | Metschnikowia | 13 | Saprotroph | \|Undefined Saprotroph\| | Yeast | NULL | Probable | As nectar-tap_saprotroph (Põlme et al. 2020); Decay sugar-rich substrates (Põlme et al. 2020) | Põlme S, et al. 2020. Fungal Diversity 105:1-16 ((https://doi.org/10.1007/s13225-020-00466-2)) |
| Otu38951 | 1 | k__Fungi(100);p__Ascomycota(100);c__Saccharomycetes(100);o__Saccharomycetales(100);f__Metschnikowiaceae(100);g__Metschnikowia(100);g__Metschnikowia_unclassified(100); | Metschnikowia | 13 | Saprotroph | \|Undefined Saprotroph\| | Yeast | NULL | Probable | As nectar-tap_saprotroph (Põlme et al. 2020); Decay sugar-rich substrates (Põlme et al. 2020) | Põlme S, et al. 2020. Fungal Diversity 105:1-16 ((https://doi.org/10.1007/s13225-020-00466-2)) |
| Otu38952 | 1 | k__Fungi(100);p__Ascomycota(100);c__Saccharomycetes(100);o__Saccharomycetales(100);f__Metschnikowiaceae(100);g__Metschnikowia(100);s__Metschnikowia_pulcherrima(100); | Metschnikowia | 13 | Saprotroph | \|Undefined Saprotroph\| | Yeast | NULL | Probable | As nectar-tap_saprotroph (Põlme et al. 2020); Decay sugar-rich substrates (Põlme et al. 2020) | Põlme S, et al. 2020. Fungal Diversity 105:1-16 ((https://doi.org/10.1007/s13225-020-00466-2)) |
| Otu38953 | 1 | k__Fungi(100);p__Ascomycota(100);c__Saccharomycetes(100);o__Saccharomycetales(100);f__Metschnikowiaceae(100);g__Metschnikowia(100);g__Metschnikowia_unclassified(100); | Metschnikowia | 13 | Saprotroph | \|Undefined Saprotroph\| | Yeast | NULL | Probable | As nectar-tap_saprotroph (Põlme et al. 2020); Decay sugar-rich substrates (Põlme et al. 2020) | Põlme S, et al. 2020. Fungal Diversity 105:1-16 ((https://doi.org/10.1007/s13225-020-00466-2)) |
| Otu38954 | 1 | k__Fungi(100);p__Ascomycota(100);c__Saccharomycetes(100);o__Saccharomycetales(100);f__Metschnikowiaceae(100);g__Metschnikowia(100);s__Metschnikowia_pulcherrima(100); | Metschnikowia | 13 | Saprotroph | \|Undefined Saprotroph\| | Yeast | NULL | Probable | As nectar-tap_saprotroph (Põlme et al. 2020); Decay sugar-rich substrates (Põlme et al. 2020) | Põlme S, et al. 2020. Fungal Diversity 105:1-16 ((https://doi.org/10.1007/s13225-020-00466-2)) |
| Otu38955 | 1 | k__Fungi(100);p__Ascomycota(100);c__Saccharomycetes(100);o__Saccharomycetales(100);f__Metschnikowiaceae(100);g__Metschnikowia(100);s__Metschnikowia_pulcherrima(100); | Metschnikowia | 13 | Saprotroph | \|Undefined Saprotroph\| | Yeast | NULL | Probable | As nectar-tap_saprotroph (Põlme et al. 2020); Decay sugar-rich substrates (Põlme et al. 2020) | Põlme S, et al. 2020. Fungal Diversity 105:1-16 ((https://doi.org/10.1007/s13225-020-00466-2)) |
| Otu38956 | 1 | k__Fungi(100);p__Ascomycota(100);c__Saccharomycetes(100);o__Saccharomycetales(100);f__Metschnikowiaceae(100);g__Metschnikowia(100);s__Metschnikowia_pulcherrima(100); | Metschnikowia | 13 | Saprotroph | \|Undefined Saprotroph\| | Yeast | NULL | Probable | As nectar-tap_saprotroph (Põlme et al. 2020); Decay sugar-rich substrates (Põlme et al. 2020) | Põlme S, et al. 2020. Fungal Diversity 105:1-16 ((https://doi.org/10.1007/s13225-020-00466-2)) |
| Otu38957 | 1 | k__Fungi(100);p__Ascomycota(100);c__Saccharomycetes(100);o__Saccharomycetales(100);f__Metschnikowiaceae(100);g__Metschnikowia(100);s__Metschnikowia_pulcherrima(100); | Metschnikowia | 13 | Saprotroph | \|Undefined Saprotroph\| | Yeast | NULL | Probable | As nectar-tap_saprotroph (Põlme et al. 2020); Decay sugar-rich substrates (Põlme et al. 2020) | Põlme S, et al. 2020. Fungal Diversity 105:1-16 ((https://doi.org/10.1007/s13225-020-00466-2)) |
| Otu38958 | 1 | k__Fungi(100);p__Ascomycota(100);c__Saccharomycetes(100);o__Saccharomycetales(100);f__Metschnikowiaceae(100);g__Metschnikowia(100);g__Metschnikowia_unclassified(100); | Metschnikowia | 13 | Saprotroph | \|Undefined Saprotroph\| | Yeast | NULL | Probable | As nectar-tap_saprotroph (Põlme et al. 2020); Decay sugar-rich substrates (Põlme et al. 2020) | Põlme S, et al. 2020. Fungal Diversity 105:1-16 ((https://doi.org/10.1007/s13225-020-00466-2)) |
| Otu38959 | 1 | k__Fungi(100);p__Ascomycota(100);c__Saccharomycetes(100);o__Saccharomycetales(100);f__Metschnikowiaceae(100);g__Metschnikowia(100);s__Metschnikowia_pulcherrima(100); | Metschnikowia | 13 | Saprotroph | \|Undefined Saprotroph\| | Yeast | NULL | Probable | As nectar-tap_saprotroph (Põlme et al. 2020); Decay sugar-rich substrates (Põlme et al. 2020) | Põlme S, et al. 2020. Fungal Diversity 105:1-16 ((https://doi.org/10.1007/s13225-020-00466-2)) |
| Otu38960 | 1 | k__Fungi(100);p__Ascomycota(100);c__Saccharomycetes(100);o__Saccharomycetales(100);f__Metschnikowiaceae(100);g__Metschnikowia(100);s__Metschnikowia_pulcherrima(100); | Metschnikowia | 13 | Saprotroph | \|Undefined Saprotroph\| | Yeast | NULL | Probable | As nectar-tap_saprotroph (Põlme et al. 2020); Decay sugar-rich substrates (Põlme et al. 2020) | Põlme S, et al. 2020. Fungal Diversity 105:1-16 ((https://doi.org/10.1007/s13225-020-00466-2)) |
| Otu38961 | 1 | k__Fungi(100);p__Ascomycota(100);c__Saccharomycetes(100);o__Saccharomycetales(100);f__Metschnikowiaceae(100);g__Metschnikowia(100);s__Metschnikowia_pulcherrima(100); | Metschnikowia | 13 | Saprotroph | \|Undefined Saprotroph\| | Yeast | NULL | Probable | As nectar-tap_saprotroph (Põlme et al. 2020); Decay sugar-rich substrates (Põlme et al. 2020) | Põlme S, et al. 2020. Fungal Diversity 105:1-16 ((https://doi.org/10.1007/s13225-020-00466-2)) |
| Otu38962 | 1 | k__Fungi(100);p__Ascomycota(100);c__Saccharomycetes(100);o__Saccharomycetales(100);f__Metschnikowiaceae(100);g__Metschnikowia(100);g__Metschnikowia_unclassified(100); | Metschnikowia | 13 | Saprotroph | \|Undefined Saprotroph\| | Yeast | NULL | Probable | As nectar-tap_saprotroph (Põlme et al. 2020); Decay sugar-rich substrates (Põlme et al. 2020) | Põlme S, et al. 2020. Fungal Diversity 105:1-16 ((https://doi.org/10.1007/s13225-020-00466-2)) |
| Otu38963 | 1 | k__Fungi(100);p__Ascomycota(100);c__Saccharomycetes(100);o__Saccharomycetales(100);f__Metschnikowiaceae(100);g__Metschnikowia(100);s__Metschnikowia_pulcherrima(100); | Metschnikowia | 13 | Saprotroph | \|Undefined Saprotroph\| | Yeast | NULL | Probable | As nectar-tap_saprotroph (Põlme et al. 2020); Decay sugar-rich substrates (Põlme et al. 2020) | Põlme S, et al. 2020. Fungal Diversity 105:1-16 ((https://doi.org/10.1007/s13225-020-00466-2)) |
| Otu38964 | 1 | k__Fungi(100);p__Ascomycota(100);c__Saccharomycetes(100);o__Saccharomycetales(100);f__Metschnikowiaceae(100);g__Metschnikowia(100);s__Metschnikowia_pulcherrima(100); | Metschnikowia | 13 | Saprotroph | \|Undefined Saprotroph\| | Yeast | NULL | Probable | As nectar-tap_saprotroph (Põlme et al. 2020); Decay sugar-rich substrates (Põlme et al. 2020) | Põlme S, et al. 2020. Fungal Diversity 105:1-16 ((https://doi.org/10.1007/s13225-020-00466-2)) |
| Otu38965 | 1 | k__Fungi(100);p__Ascomycota(100);c__Saccharomycetes(100);o__Saccharomycetales(100);f__Metschnikowiaceae(100);g__Metschnikowia(100);s__Metschnikowia_pulcherrima(100); | Metschnikowia | 13 | Saprotroph | \|Undefined Saprotroph\| | Yeast | NULL | Probable | As nectar-tap_saprotroph (Põlme et al. 2020); Decay sugar-rich substrates (Põlme et al. 2020) | Põlme S, et al. 2020. Fungal Diversity 105:1-16 ((https://doi.org/10.1007/s13225-020-00466-2)) |
| Otu38967 | 1 | k__Fungi(100);p__Ascomycota(100);c__Saccharomycetes(100);o__Saccharomycetales(100);f__Metschnikowiaceae(100);g__Metschnikowia(100);s__Metschnikowia_pulcherrima(100); | Metschnikowia | 13 | Saprotroph | \|Undefined Saprotroph\| | Yeast | NULL | Probable | As nectar-tap_saprotroph (Põlme et al. 2020); Decay sugar-rich substrates (Põlme et al. 2020) | Põlme S, et al. 2020. Fungal Diversity 105:1-16 ((https://doi.org/10.1007/s13225-020-00466-2)) |
| Otu38968 | 1 | k__Fungi(100);p__Ascomycota(100);c__Saccharomycetes(100);o__Saccharomycetales(100);f__Metschnikowiaceae(100);g__Metschnikowia(100);g__Metschnikowia_unclassified(100); | Metschnikowia | 13 | Saprotroph | \|Undefined Saprotroph\| | Yeast | NULL | Probable | As nectar-tap_saprotroph (Põlme et al. 2020); Decay sugar-rich substrates (Põlme et al. 2020) | Põlme S, et al. 2020. Fungal Diversity 105:1-16 ((https://doi.org/10.1007/s13225-020-00466-2)) |
| Otu38969 | 1 | k__Fungi(100);p__Ascomycota(100);c__Saccharomycetes(100);o__Saccharomycetales(100);f__Metschnikowiaceae(100);g__Metschnikowia(100);s__Metschnikowia_pulcherrima(100); | Metschnikowia | 13 | Saprotroph | \|Undefined Saprotroph\| | Yeast | NULL | Probable | As nectar-tap_saprotroph (Põlme et al. 2020); Decay sugar-rich substrates (Põlme et al. 2020) | Põlme S, et al. 2020. Fungal Diversity 105:1-16 ((https://doi.org/10.1007/s13225-020-00466-2)) |
| Otu38970 | 1 | k__Fungi(100);p__Ascomycota(100);c__Saccharomycetes(100);o__Saccharomycetales(100);f__Metschnikowiaceae(100);g__Metschnikowia(100);s__Metschnikowia_pulcherrima(100); | Metschnikowia | 13 | Saprotroph | \|Undefined Saprotroph\| | Yeast | NULL | Probable | As nectar-tap_saprotroph (Põlme et al. 2020); Decay sugar-rich substrates (Põlme et al. 2020) | Põlme S, et al. 2020. Fungal Diversity 105:1-16 ((https://doi.org/10.1007/s13225-020-00466-2)) |
| Otu38972 | 1 | k__Fungi(100);p__Ascomycota(100);c__Saccharomycetes(100);o__Saccharomycetales(100);f__Metschnikowiaceae(100);g__Metschnikowia(100);s__Metschnikowia_pulcherrima(100); | Metschnikowia | 13 | Saprotroph | \|Undefined Saprotroph\| | Yeast | NULL | Probable | As nectar-tap_saprotroph (Põlme et al. 2020); Decay sugar-rich substrates (Põlme et al. 2020) | Põlme S, et al. 2020. Fungal Diversity 105:1-16 ((https://doi.org/10.1007/s13225-020-00466-2)) |
| Otu38973 | 1 | k__Fungi(100);p__Ascomycota(100);c__Saccharomycetes(100);o__Saccharomycetales(100);f__Metschnikowiaceae(100);g__Metschnikowia(100);s__Metschnikowia_pulcherrima(100); | Metschnikowia | 13 | Saprotroph | \|Undefined Saprotroph\| | Yeast | NULL | Probable | As nectar-tap_saprotroph (Põlme et al. 2020); Decay sugar-rich substrates (Põlme et al. 2020) | Põlme S, et al. 2020. Fungal Diversity 105:1-16 ((https://doi.org/10.1007/s13225-020-00466-2)) |
| Otu38974 | 1 | k__Fungi(100);p__Ascomycota(100);c__Saccharomycetes(100);o__Saccharomycetales(100);f__Metschnikowiaceae(100);g__Metschnikowia(100);g__Metschnikowia_unclassified(100); | Metschnikowia | 13 | Saprotroph | \|Undefined Saprotroph\| | Yeast | NULL | Probable | As nectar-tap_saprotroph (Põlme et al. 2020); Decay sugar-rich substrates (Põlme et al. 2020) | Põlme S, et al. 2020. Fungal Diversity 105:1-16 ((https://doi.org/10.1007/s13225-020-00466-2)) |
| Otu38975 | 1 | k__Fungi(100);p__Ascomycota(100);c__Saccharomycetes(100);o__Saccharomycetales(100);f__Metschnikowiaceae(100);g__Metschnikowia(100);s__Metschnikowia_pulcherrima(100); | Metschnikowia | 13 | Saprotroph | \|Undefined Saprotroph\| | Yeast | NULL | Probable | As nectar-tap_saprotroph (Põlme et al. 2020); Decay sugar-rich substrates (Põlme et al. 2020) | Põlme S, et al. 2020. Fungal Diversity 105:1-16 ((https://doi.org/10.1007/s13225-020-00466-2)) |
| Otu38976 | 1 | k__Fungi(100);p__Ascomycota(100);c__Saccharomycetes(100);o__Saccharomycetales(100);f__Metschnikowiaceae(100);g__Metschnikowia(100);s__Metschnikowia_pulcherrima(100); | Metschnikowia | 13 | Saprotroph | \|Undefined Saprotroph\| | Yeast | NULL | Probable | As nectar-tap_saprotroph (Põlme et al. 2020); Decay sugar-rich substrates (Põlme et al. 2020) | Põlme S, et al. 2020. Fungal Diversity 105:1-16 ((https://doi.org/10.1007/s13225-020-00466-2)) |
| Otu38977 | 1 | k__Fungi(100);p__Ascomycota(100);c__Saccharomycetes(100);o__Saccharomycetales(100);f__Metschnikowiaceae(100);g__Metschnikowia(100);g__Metschnikowia_unclassified(100); | Metschnikowia | 13 | Saprotroph | \|Undefined Saprotroph\| | Yeast | NULL | Probable | As nectar-tap_saprotroph (Põlme et al. 2020); Decay sugar-rich substrates (Põlme et al. 2020) | Põlme S, et al. 2020. Fungal Diversity 105:1-16 ((https://doi.org/10.1007/s13225-020-00466-2)) |
| Otu38978 | 1 | k__Fungi(100);p__Ascomycota(100);c__Saccharomycetes(100);o__Saccharomycetales(100);f__Metschnikowiaceae(100);g__Metschnikowia(100);s__Metschnikowia_pulcherrima(100); | Metschnikowia | 13 | Saprotroph | \|Undefined Saprotroph\| | Yeast | NULL | Probable | As nectar-tap_saprotroph (Põlme et al. 2020); Decay sugar-rich substrates (Põlme et al. 2020) | Põlme S, et al. 2020. Fungal Diversity 105:1-16 ((https://doi.org/10.1007/s13225-020-00466-2)) |
| Otu38979 | 1 | k__Fungi(100);p__Ascomycota(100);c__Saccharomycetes(100);o__Saccharomycetales(100);f__Metschnikowiaceae(100);g__Metschnikowia(100);s__Metschnikowia_pulcherrima(100); | Metschnikowia | 13 | Saprotroph | \|Undefined Saprotroph\| | Yeast | NULL | Probable | As nectar-tap_saprotroph (Põlme et al. 2020); Decay sugar-rich substrates (Põlme et al. 2020) | Põlme S, et al. 2020. Fungal Diversity 105:1-16 ((https://doi.org/10.1007/s13225-020-00466-2)) |
| Otu38981 | 1 | k__Fungi(100);p__Ascomycota(100);c__Saccharomycetes(100);o__Saccharomycetales(100);f__Metschnikowiaceae(100);g__Metschnikowia(100);s__Metschnikowia_pulcherrima(100); | Metschnikowia | 13 | Saprotroph | \|Undefined Saprotroph\| | Yeast | NULL | Probable | As nectar-tap_saprotroph (Põlme et al. 2020); Decay sugar-rich substrates (Põlme et al. 2020) | Põlme S, et al. 2020. Fungal Diversity 105:1-16 ((https://doi.org/10.1007/s13225-020-00466-2)) |
| Otu38982 | 1 | k__Fungi(100);p__Ascomycota(100);c__Saccharomycetes(100);o__Saccharomycetales(100);f__Metschnikowiaceae(100);g__Metschnikowia(100);s__Metschnikowia_pulcherrima(100); | Metschnikowia | 13 | Saprotroph | \|Undefined Saprotroph\| | Yeast | NULL | Probable | As nectar-tap_saprotroph (Põlme et al. 2020); Decay sugar-rich substrates (Põlme et al. 2020) | Põlme S, et al. 2020. Fungal Diversity 105:1-16 ((https://doi.org/10.1007/s13225-020-00466-2)) |
| Otu38983 | 1 | k__Fungi(100);p__Ascomycota(100);c__Saccharomycetes(100);o__Saccharomycetales(100);f__Metschnikowiaceae(100);g__Metschnikowia(100);s__Metschnikowia_pulcherrima(100); | Metschnikowia | 13 | Saprotroph | \|Undefined Saprotroph\| | Yeast | NULL | Probable | As nectar-tap_saprotroph (Põlme et al. 2020); Decay sugar-rich substrates (Põlme et al. 2020) | Põlme S, et al. 2020. Fungal Diversity 105:1-16 ((https://doi.org/10.1007/s13225-020-00466-2)) |
| Otu38984 | 1 | k__Fungi(100);p__Ascomycota(100);c__Saccharomycetes(100);o__Saccharomycetales(100);f__Metschnikowiaceae(100);g__Metschnikowia(100);s__Metschnikowia_pulcherrima(100); | Metschnikowia | 13 | Saprotroph | \|Undefined Saprotroph\| | Yeast | NULL | Probable | As nectar-tap_saprotroph (Põlme et al. 2020); Decay sugar-rich substrates (Põlme et al. 2020) | Põlme S, et al. 2020. Fungal Diversity 105:1-16 ((https://doi.org/10.1007/s13225-020-00466-2)) |
| Otu38985 | 1 | k__Fungi(100);p__Ascomycota(100);c__Saccharomycetes(100);o__Saccharomycetales(100);f__Metschnikowiaceae(100);g__Metschnikowia(100);s__Metschnikowia_pulcherrima(100); | Metschnikowia | 13 | Saprotroph | \|Undefined Saprotroph\| | Yeast | NULL | Probable | As nectar-tap_saprotroph (Põlme et al. 2020); Decay sugar-rich substrates (Põlme et al. 2020) | Põlme S, et al. 2020. Fungal Diversity 105:1-16 ((https://doi.org/10.1007/s13225-020-00466-2)) |
| Otu38986 | 1 | k__Fungi(100);p__Ascomycota(100);c__Saccharomycetes(100);o__Saccharomycetales(100);f__Metschnikowiaceae(100);g__Metschnikowia(100);s__Metschnikowia_pulcherrima(100); | Metschnikowia | 13 | Saprotroph | \|Undefined Saprotroph\| | Yeast | NULL | Probable | As nectar-tap_saprotroph (Põlme et al. 2020); Decay sugar-rich substrates (Põlme et al. 2020) | Põlme S, et al. 2020. Fungal Diversity 105:1-16 ((https://doi.org/10.1007/s13225-020-00466-2)) |
| Otu38987 | 1 | k__Fungi(100);p__Ascomycota(100);c__Saccharomycetes(100);o__Saccharomycetales(100);f__Metschnikowiaceae(100);g__Metschnikowia(100);s__Metschnikowia_pulcherrima(100); | Metschnikowia | 13 | Saprotroph | \|Undefined Saprotroph\| | Yeast | NULL | Probable | As nectar-tap_saprotroph (Põlme et al. 2020); Decay sugar-rich substrates (Põlme et al. 2020) | Põlme S, et al. 2020. Fungal Diversity 105:1-16 ((https://doi.org/10.1007/s13225-020-00466-2)) |
| Otu38988 | 1 | k__Fungi(100);p__Ascomycota(100);c__Saccharomycetes(100);o__Saccharomycetales(100);f__Metschnikowiaceae(100);g__Metschnikowia(100);g__Metschnikowia_unclassified(100); | Metschnikowia | 13 | Saprotroph | \|Undefined Saprotroph\| | Yeast | NULL | Probable | As nectar-tap_saprotroph (Põlme et al. 2020); Decay sugar-rich substrates (Põlme et al. 2020) | Põlme S, et al. 2020. Fungal Diversity 105:1-16 ((https://doi.org/10.1007/s13225-020-00466-2)) |
| Otu38989 | 1 | k__Fungi(100);p__Ascomycota(100);c__Saccharomycetes(100);o__Saccharomycetales(100);f__Metschnikowiaceae(100);g__Metschnikowia(100);s__Metschnikowia_pulcherrima(100); | Metschnikowia | 13 | Saprotroph | \|Undefined Saprotroph\| | Yeast | NULL | Probable | As nectar-tap_saprotroph (Põlme et al. 2020); Decay sugar-rich substrates (Põlme et al. 2020) | Põlme S, et al. 2020. Fungal Diversity 105:1-16 ((https://doi.org/10.1007/s13225-020-00466-2)) |
| Otu38990 | 1 | k__Fungi(100);p__Ascomycota(100);c__Dothideomycetes(100);o__Dothideomycetes_order_Incertae_sedis(100);f__Myxotrichaceae(100);g__Oidiodendron(100);s__Oidiodendron_sp_22MS12(100); | Oidiodendron | 13 | Pathotroph-Saprotroph-Symbiotroph | Endophyte-Ericoid Mycorrhizal-\|Undefined Saprotroph\| | Microfungus | NULL | Probable | As root_endophyte (Põlme et al. 2020); As soil_saprotroph (Põlme et al. 2020); Dark Septate Endophyte; Decay soil (Põlme et al. 2020) | Newsham KK. 2011. New Phytologist 190:783-793 ((https://doi.org/10.1111/j.1469-8137.2010.03611.x)) (as dark septate root endophytes); Põlme S, et al. 2020. Fungal Diversity 105:1-16 ((https://doi.org/10.1007/s13225-020-00466-2)) |
| Otu38991 | 1 | k__Fungi(100);p__Ascomycota(100);c__Saccharomycetes(100);o__Saccharomycetales(100);f__Metschnikowiaceae(100);g__Metschnikowia(100);s__Metschnikowia_pulcherrima(100); | Metschnikowia | 13 | Saprotroph | \|Undefined Saprotroph\| | Yeast | NULL | Probable | As nectar-tap_saprotroph (Põlme et al. 2020); Decay sugar-rich substrates (Põlme et al. 2020) | Põlme S, et al. 2020. Fungal Diversity 105:1-16 ((https://doi.org/10.1007/s13225-020-00466-2)) |
| Otu38992 | 1 | k__Fungi(100);p__Ascomycota(100);c__Saccharomycetes(100);o__Saccharomycetales(100);f__Metschnikowiaceae(100);g__Metschnikowia(100);s__Metschnikowia_pulcherrima(100); | Metschnikowia | 13 | Saprotroph | \|Undefined Saprotroph\| | Yeast | NULL | Probable | As nectar-tap_saprotroph (Põlme et al. 2020); Decay sugar-rich substrates (Põlme et al. 2020) | Põlme S, et al. 2020. Fungal Diversity 105:1-16 ((https://doi.org/10.1007/s13225-020-00466-2)) |
| Otu38993 | 1 | k__Fungi(100);p__Ascomycota(100);c__Saccharomycetes(100);o__Saccharomycetales(100);f__Metschnikowiaceae(100);g__Metschnikowia(100);s__Metschnikowia_pulcherrima(100); | Metschnikowia | 13 | Saprotroph | \|Undefined Saprotroph\| | Yeast | NULL | Probable | As nectar-tap_saprotroph (Põlme et al. 2020); Decay sugar-rich substrates (Põlme et al. 2020) | Põlme S, et al. 2020. Fungal Diversity 105:1-16 ((https://doi.org/10.1007/s13225-020-00466-2)) |
| Otu38995 | 1 | k__Fungi(100);p__Ascomycota(100);c__Saccharomycetes(100);o__Saccharomycetales(100);f__Metschnikowiaceae(100);g__Metschnikowia(100);s__Metschnikowia_pulcherrima(100); | Metschnikowia | 13 | Saprotroph | \|Undefined Saprotroph\| | Yeast | NULL | Probable | As nectar-tap_saprotroph (Põlme et al. 2020); Decay sugar-rich substrates (Põlme et al. 2020) | Põlme S, et al. 2020. Fungal Diversity 105:1-16 ((https://doi.org/10.1007/s13225-020-00466-2)) |
| Otu38996 | 1 | k__Fungi(100);p__Ascomycota(100);c__Saccharomycetes(100);o__Saccharomycetales(100);f__Metschnikowiaceae(100);g__Metschnikowia(100);g__Metschnikowia_unclassified(100); | Metschnikowia | 13 | Saprotroph | \|Undefined Saprotroph\| | Yeast | NULL | Probable | As nectar-tap_saprotroph (Põlme et al. 2020); Decay sugar-rich substrates (Põlme et al. 2020) | Põlme S, et al. 2020. Fungal Diversity 105:1-16 ((https://doi.org/10.1007/s13225-020-00466-2)) |
| Otu38997 | 1 | k__Fungi(100);p__Ascomycota(100);c__Saccharomycetes(100);o__Saccharomycetales(100);f__Metschnikowiaceae(100);g__Metschnikowia(100);s__Metschnikowia_pulcherrima(100); | Metschnikowia | 13 | Saprotroph | \|Undefined Saprotroph\| | Yeast | NULL | Probable | As nectar-tap_saprotroph (Põlme et al. 2020); Decay sugar-rich substrates (Põlme et al. 2020) | Põlme S, et al. 2020. Fungal Diversity 105:1-16 ((https://doi.org/10.1007/s13225-020-00466-2)) |
| Otu38998 | 1 | k__Fungi(100);p__Ascomycota(100);c__Saccharomycetes(100);o__Saccharomycetales(100);f__Metschnikowiaceae(100);g__Metschnikowia(100);g__Metschnikowia_unclassified(100); | Metschnikowia | 13 | Saprotroph | \|Undefined Saprotroph\| | Yeast | NULL | Probable | As nectar-tap_saprotroph (Põlme et al. 2020); Decay sugar-rich substrates (Põlme et al. 2020) | Põlme S, et al. 2020. Fungal Diversity 105:1-16 ((https://doi.org/10.1007/s13225-020-00466-2)) |
| Otu38999 | 1 | k__Fungi(100);p__Ascomycota(100);c__Saccharomycetes(100);o__Saccharomycetales(100);f__Metschnikowiaceae(100);g__Metschnikowia(100);s__Metschnikowia_pulcherrima(100); | Metschnikowia | 13 | Saprotroph | \|Undefined Saprotroph\| | Yeast | NULL | Probable | As nectar-tap_saprotroph (Põlme et al. 2020); Decay sugar-rich substrates (Põlme et al. 2020) | Põlme S, et al. 2020. Fungal Diversity 105:1-16 ((https://doi.org/10.1007/s13225-020-00466-2)) |
| Otu39000 | 1 | k__Fungi(100);p__Ascomycota(100);c__Saccharomycetes(100);o__Saccharomycetales(100);f__Metschnikowiaceae(100);g__Metschnikowia(100);s__Metschnikowia_pulcherrima(100); | Metschnikowia | 13 | Saprotroph | \|Undefined Saprotroph\| | Yeast | NULL | Probable | As nectar-tap_saprotroph (Põlme et al. 2020); Decay sugar-rich substrates (Põlme et al. 2020) | Põlme S, et al. 2020. Fungal Diversity 105:1-16 ((https://doi.org/10.1007/s13225-020-00466-2)) |
| Otu39001 | 1 | k__Fungi(100);p__Ascomycota(100);c__Saccharomycetes(100);o__Saccharomycetales(100);f__Metschnikowiaceae(100);g__Metschnikowia(100);s__Metschnikowia_pulcherrima(100); | Metschnikowia | 13 | Saprotroph | \|Undefined Saprotroph\| | Yeast | NULL | Probable | As nectar-tap_saprotroph (Põlme et al. 2020); Decay sugar-rich substrates (Põlme et al. 2020) | Põlme S, et al. 2020. Fungal Diversity 105:1-16 ((https://doi.org/10.1007/s13225-020-00466-2)) |
| Otu39002 | 1 | k__Fungi(100);p__Ascomycota(100);c__Saccharomycetes(100);o__Saccharomycetales(100);f__Metschnikowiaceae(100);g__Metschnikowia(100);g__Metschnikowia_unclassified(100); | Metschnikowia | 13 | Saprotroph | \|Undefined Saprotroph\| | Yeast | NULL | Probable | As nectar-tap_saprotroph (Põlme et al. 2020); Decay sugar-rich substrates (Põlme et al. 2020) | Põlme S, et al. 2020. Fungal Diversity 105:1-16 ((https://doi.org/10.1007/s13225-020-00466-2)) |
| Otu39003 | 1 | k__Fungi(100);p__Ascomycota(100);c__Saccharomycetes(100);o__Saccharomycetales(100);f__Metschnikowiaceae(100);g__Metschnikowia(100);s__Metschnikowia_pulcherrima(100); | Metschnikowia | 13 | Saprotroph | \|Undefined Saprotroph\| | Yeast | NULL | Probable | As nectar-tap_saprotroph (Põlme et al. 2020); Decay sugar-rich substrates (Põlme et al. 2020) | Põlme S, et al. 2020. Fungal Diversity 105:1-16 ((https://doi.org/10.1007/s13225-020-00466-2)) |
| Otu39004 | 1 | k__Fungi(100);p__Ascomycota(100);c__Saccharomycetes(100);o__Saccharomycetales(100);f__Metschnikowiaceae(100);g__Metschnikowia(100);s__Metschnikowia_pulcherrima(100); | Metschnikowia | 13 | Saprotroph | \|Undefined Saprotroph\| | Yeast | NULL | Probable | As nectar-tap_saprotroph (Põlme et al. 2020); Decay sugar-rich substrates (Põlme et al. 2020) | Põlme S, et al. 2020. Fungal Diversity 105:1-16 ((https://doi.org/10.1007/s13225-020-00466-2)) |
| Otu39005 | 1 | k__Fungi(100);p__Ascomycota(100);c__Saccharomycetes(100);o__Saccharomycetales(100);f__Metschnikowiaceae(100);g__Metschnikowia(100);s__Metschnikowia_pulcherrima(100); | Metschnikowia | 13 | Saprotroph | \|Undefined Saprotroph\| | Yeast | NULL | Probable | As nectar-tap_saprotroph (Põlme et al. 2020); Decay sugar-rich substrates (Põlme et al. 2020) | Põlme S, et al. 2020. Fungal Diversity 105:1-16 ((https://doi.org/10.1007/s13225-020-00466-2)) |
| Otu39006 | 1 | k__Fungi(100);p__Ascomycota(100);c__Saccharomycetes(100);o__Saccharomycetales(100);f__Metschnikowiaceae(100);g__Metschnikowia(100);s__Metschnikowia_pulcherrima(100); | Metschnikowia | 13 | Saprotroph | \|Undefined Saprotroph\| | Yeast | NULL | Probable | As nectar-tap_saprotroph (Põlme et al. 2020); Decay sugar-rich substrates (Põlme et al. 2020) | Põlme S, et al. 2020. Fungal Diversity 105:1-16 ((https://doi.org/10.1007/s13225-020-00466-2)) |
| Otu39007 | 1 | k__Fungi(100);p__Ascomycota(100);c__Saccharomycetes(100);o__Saccharomycetales(100);f__Metschnikowiaceae(100);g__Metschnikowia(100);s__Metschnikowia_pulcherrima(100); | Metschnikowia | 13 | Saprotroph | \|Undefined Saprotroph\| | Yeast | NULL | Probable | As nectar-tap_saprotroph (Põlme et al. 2020); Decay sugar-rich substrates (Põlme et al. 2020) | Põlme S, et al. 2020. Fungal Diversity 105:1-16 ((https://doi.org/10.1007/s13225-020-00466-2)) |
| Otu39008 | 1 | k__Fungi(100);p__Ascomycota(100);c__Saccharomycetes(100);o__Saccharomycetales(100);f__Metschnikowiaceae(100);g__Metschnikowia(100);s__Metschnikowia_pulcherrima(100); | Metschnikowia | 13 | Saprotroph | \|Undefined Saprotroph\| | Yeast | NULL | Probable | As nectar-tap_saprotroph (Põlme et al. 2020); Decay sugar-rich substrates (Põlme et al. 2020) | Põlme S, et al. 2020. Fungal Diversity 105:1-16 ((https://doi.org/10.1007/s13225-020-00466-2)) |
| Otu39009 | 1 | k__Fungi(100);p__Ascomycota(100);c__Saccharomycetes(100);o__Saccharomycetales(100);f__Metschnikowiaceae(100);g__Metschnikowia(100);s__Metschnikowia_pulcherrima(100); | Metschnikowia | 13 | Saprotroph | \|Undefined Saprotroph\| | Yeast | NULL | Probable | As nectar-tap_saprotroph (Põlme et al. 2020); Decay sugar-rich substrates (Põlme et al. 2020) | Põlme S, et al. 2020. Fungal Diversity 105:1-16 ((https://doi.org/10.1007/s13225-020-00466-2)) |
| Otu39010 | 1 | k__Fungi(100);p__Ascomycota(100);c__Saccharomycetes(100);o__Saccharomycetales(100);f__Metschnikowiaceae(100);g__Metschnikowia(100);s__Metschnikowia_pulcherrima(100); | Metschnikowia | 13 | Saprotroph | \|Undefined Saprotroph\| | Yeast | NULL | Probable | As nectar-tap_saprotroph (Põlme et al. 2020); Decay sugar-rich substrates (Põlme et al. 2020) | Põlme S, et al. 2020. Fungal Diversity 105:1-16 ((https://doi.org/10.1007/s13225-020-00466-2)) |
| Otu39011 | 1 | k__Fungi(100);p__Ascomycota(100);c__Saccharomycetes(100);o__Saccharomycetales(100);f__Metschnikowiaceae(100);g__Metschnikowia(100);s__Metschnikowia_pulcherrima(100); | Metschnikowia | 13 | Saprotroph | \|Undefined Saprotroph\| | Yeast | NULL | Probable | As nectar-tap_saprotroph (Põlme et al. 2020); Decay sugar-rich substrates (Põlme et al. 2020) | Põlme S, et al. 2020. Fungal Diversity 105:1-16 ((https://doi.org/10.1007/s13225-020-00466-2)) |
| Otu39012 | 1 | k__Fungi(100);p__Ascomycota(100);c__Saccharomycetes(100);o__Saccharomycetales(100);f__Metschnikowiaceae(100);g__Metschnikowia(100);s__Metschnikowia_pulcherrima(100); | Metschnikowia | 13 | Saprotroph | \|Undefined Saprotroph\| | Yeast | NULL | Probable | As nectar-tap_saprotroph (Põlme et al. 2020); Decay sugar-rich substrates (Põlme et al. 2020) | Põlme S, et al. 2020. Fungal Diversity 105:1-16 ((https://doi.org/10.1007/s13225-020-00466-2)) |
| Otu39013 | 1 | k__Fungi(100);p__Ascomycota(100);c__Saccharomycetes(100);o__Saccharomycetales(100);f__Metschnikowiaceae(100);g__Metschnikowia(100);g__Metschnikowia_unclassified(100); | Metschnikowia | 13 | Saprotroph | \|Undefined Saprotroph\| | Yeast | NULL | Probable | As nectar-tap_saprotroph (Põlme et al. 2020); Decay sugar-rich substrates (Põlme et al. 2020) | Põlme S, et al. 2020. Fungal Diversity 105:1-16 ((https://doi.org/10.1007/s13225-020-00466-2)) |
| Otu39014 | 1 | k__Fungi(100);p__Ascomycota(100);c__Saccharomycetes(100);o__Saccharomycetales(100);f__Metschnikowiaceae(100);g__Metschnikowia(100);s__Metschnikowia_pulcherrima(100); | Metschnikowia | 13 | Saprotroph | \|Undefined Saprotroph\| | Yeast | NULL | Probable | As nectar-tap_saprotroph (Põlme et al. 2020); Decay sugar-rich substrates (Põlme et al. 2020) | Põlme S, et al. 2020. Fungal Diversity 105:1-16 ((https://doi.org/10.1007/s13225-020-00466-2)) |
| Otu39015 | 1 | k__Fungi(100);p__Ascomycota(100);c__Saccharomycetes(100);o__Saccharomycetales(100);f__Metschnikowiaceae(100);g__Metschnikowia(100);g__Metschnikowia_unclassified(100); | Metschnikowia | 13 | Saprotroph | \|Undefined Saprotroph\| | Yeast | NULL | Probable | As nectar-tap_saprotroph (Põlme et al. 2020); Decay sugar-rich substrates (Põlme et al. 2020) | Põlme S, et al. 2020. Fungal Diversity 105:1-16 ((https://doi.org/10.1007/s13225-020-00466-2)) |
| Otu39016 | 1 | k__Fungi(100);p__Ascomycota(100);c__Saccharomycetes(100);o__Saccharomycetales(100);f__Metschnikowiaceae(100);g__Metschnikowia(100);s__Metschnikowia_pulcherrima(100); | Metschnikowia | 13 | Saprotroph | \|Undefined Saprotroph\| | Yeast | NULL | Probable | As nectar-tap_saprotroph (Põlme et al. 2020); Decay sugar-rich substrates (Põlme et al. 2020) | Põlme S, et al. 2020. Fungal Diversity 105:1-16 ((https://doi.org/10.1007/s13225-020-00466-2)) |
| Otu39017 | 1 | k__Fungi(100);p__Ascomycota(100);c__Saccharomycetes(100);o__Saccharomycetales(100);f__Metschnikowiaceae(100);g__Metschnikowia(100);s__Metschnikowia_pulcherrima(100); | Metschnikowia | 13 | Saprotroph | \|Undefined Saprotroph\| | Yeast | NULL | Probable | As nectar-tap_saprotroph (Põlme et al. 2020); Decay sugar-rich substrates (Põlme et al. 2020) | Põlme S, et al. 2020. Fungal Diversity 105:1-16 ((https://doi.org/10.1007/s13225-020-00466-2)) |
| Otu39021 | 1 | k__Fungi(100);p__Ascomycota(100);c__Saccharomycetes(100);o__Saccharomycetales(100);f__Metschnikowiaceae(100);g__Metschnikowia(100);s__Metschnikowia_pulcherrima(100); | Metschnikowia | 13 | Saprotroph | \|Undefined Saprotroph\| | Yeast | NULL | Probable | As nectar-tap_saprotroph (Põlme et al. 2020); Decay sugar-rich substrates (Põlme et al. 2020) | Põlme S, et al. 2020. Fungal Diversity 105:1-16 ((https://doi.org/10.1007/s13225-020-00466-2)) |
| Otu39022 | 1 | k__Fungi(100);p__Ascomycota(100);c__Saccharomycetes(100);o__Saccharomycetales(100);f__Metschnikowiaceae(100);g__Metschnikowia(100);g__Metschnikowia_unclassified(100); | Metschnikowia | 13 | Saprotroph | \|Undefined Saprotroph\| | Yeast | NULL | Probable | As nectar-tap_saprotroph (Põlme et al. 2020); Decay sugar-rich substrates (Põlme et al. 2020) | Põlme S, et al. 2020. Fungal Diversity 105:1-16 ((https://doi.org/10.1007/s13225-020-00466-2)) |
| Otu39023 | 1 | k__Fungi(100);p__Ascomycota(100);c__Saccharomycetes(100);o__Saccharomycetales(100);f__Metschnikowiaceae(100);g__Metschnikowia(100);s__Metschnikowia_pulcherrima(100); | Metschnikowia | 13 | Saprotroph | \|Undefined Saprotroph\| | Yeast | NULL | Probable | As nectar-tap_saprotroph (Põlme et al. 2020); Decay sugar-rich substrates (Põlme et al. 2020) | Põlme S, et al. 2020. Fungal Diversity 105:1-16 ((https://doi.org/10.1007/s13225-020-00466-2)) |
| Otu39024 | 1 | k__Fungi(100);p__Ascomycota(100);c__Saccharomycetes(100);o__Saccharomycetales(100);f__Metschnikowiaceae(100);g__Metschnikowia(100);s__Metschnikowia_pulcherrima(100); | Metschnikowia | 13 | Saprotroph | \|Undefined Saprotroph\| | Yeast | NULL | Probable | As nectar-tap_saprotroph (Põlme et al. 2020); Decay sugar-rich substrates (Põlme et al. 2020) | Põlme S, et al. 2020. Fungal Diversity 105:1-16 ((https://doi.org/10.1007/s13225-020-00466-2)) |
| Otu39025 | 1 | k__Fungi(100);p__Ascomycota(100);c__Saccharomycetes(100);o__Saccharomycetales(100);f__Metschnikowiaceae(100);g__Metschnikowia(100);s__Metschnikowia_pulcherrima(100); | Metschnikowia | 13 | Saprotroph | \|Undefined Saprotroph\| | Yeast | NULL | Probable | As nectar-tap_saprotroph (Põlme et al. 2020); Decay sugar-rich substrates (Põlme et al. 2020) | Põlme S, et al. 2020. Fungal Diversity 105:1-16 ((https://doi.org/10.1007/s13225-020-00466-2)) |
| Otu39026 | 1 | k__Fungi(100);p__Ascomycota(100);c__Saccharomycetes(100);o__Saccharomycetales(100);f__Metschnikowiaceae(100);g__Metschnikowia(100);s__Metschnikowia_pulcherrima(100); | Metschnikowia | 13 | Saprotroph | \|Undefined Saprotroph\| | Yeast | NULL | Probable | As nectar-tap_saprotroph (Põlme et al. 2020); Decay sugar-rich substrates (Põlme et al. 2020) | Põlme S, et al. 2020. Fungal Diversity 105:1-16 ((https://doi.org/10.1007/s13225-020-00466-2)) |
| Otu39027 | 1 | k__Fungi(100);p__Ascomycota(100);c__Saccharomycetes(100);o__Saccharomycetales(100);f__Metschnikowiaceae(100);g__Metschnikowia(100);s__Metschnikowia_pulcherrima(100); | Metschnikowia | 13 | Saprotroph | \|Undefined Saprotroph\| | Yeast | NULL | Probable | As nectar-tap_saprotroph (Põlme et al. 2020); Decay sugar-rich substrates (Põlme et al. 2020) | Põlme S, et al. 2020. Fungal Diversity 105:1-16 ((https://doi.org/10.1007/s13225-020-00466-2)) |
| Otu39028 | 1 | k__Fungi(100);p__Ascomycota(100);c__Saccharomycetes(100);o__Saccharomycetales(100);f__Metschnikowiaceae(100);g__Metschnikowia(100);s__Metschnikowia_pulcherrima(100); | Metschnikowia | 13 | Saprotroph | \|Undefined Saprotroph\| | Yeast | NULL | Probable | As nectar-tap_saprotroph (Põlme et al. 2020); Decay sugar-rich substrates (Põlme et al. 2020) | Põlme S, et al. 2020. Fungal Diversity 105:1-16 ((https://doi.org/10.1007/s13225-020-00466-2)) |
| Otu39029 | 1 | k__Fungi(100);p__Ascomycota(100);c__Saccharomycetes(100);o__Saccharomycetales(100);f__Metschnikowiaceae(100);g__Metschnikowia(100);s__Metschnikowia_pulcherrima(100); | Metschnikowia | 13 | Saprotroph | \|Undefined Saprotroph\| | Yeast | NULL | Probable | As nectar-tap_saprotroph (Põlme et al. 2020); Decay sugar-rich substrates (Põlme et al. 2020) | Põlme S, et al. 2020. Fungal Diversity 105:1-16 ((https://doi.org/10.1007/s13225-020-00466-2)) |
| Otu39030 | 1 | k__Fungi(100);p__Ascomycota(100);c__Saccharomycetes(100);o__Saccharomycetales(100);f__Metschnikowiaceae(100);g__Metschnikowia(100);g__Metschnikowia_unclassified(100); | Metschnikowia | 13 | Saprotroph | \|Undefined Saprotroph\| | Yeast | NULL | Probable | As nectar-tap_saprotroph (Põlme et al. 2020); Decay sugar-rich substrates (Põlme et al. 2020) | Põlme S, et al. 2020. Fungal Diversity 105:1-16 ((https://doi.org/10.1007/s13225-020-00466-2)) |
| Otu39031 | 1 | k__Fungi(100);p__Ascomycota(100);c__Saccharomycetes(100);o__Saccharomycetales(100);f__Metschnikowiaceae(100);g__Metschnikowia(100);g__Metschnikowia_unclassified(100); | Metschnikowia | 13 | Saprotroph | \|Undefined Saprotroph\| | Yeast | NULL | Probable | As nectar-tap_saprotroph (Põlme et al. 2020); Decay sugar-rich substrates (Põlme et al. 2020) | Põlme S, et al. 2020. Fungal Diversity 105:1-16 ((https://doi.org/10.1007/s13225-020-00466-2)) |
| Otu39032 | 1 | k__Fungi(100);p__Ascomycota(100);c__Saccharomycetes(100);o__Saccharomycetales(100);f__Metschnikowiaceae(100);g__Metschnikowia(100);s__Metschnikowia_pulcherrima(100); | Metschnikowia | 13 | Saprotroph | \|Undefined Saprotroph\| | Yeast | NULL | Probable | As nectar-tap_saprotroph (Põlme et al. 2020); Decay sugar-rich substrates (Põlme et al. 2020) | Põlme S, et al. 2020. Fungal Diversity 105:1-16 ((https://doi.org/10.1007/s13225-020-00466-2)) |
| Otu39033 | 1 | k__Fungi(100);p__Ascomycota(100);c__Saccharomycetes(100);o__Saccharomycetales(100);f__Metschnikowiaceae(100);g__Metschnikowia(100);s__Metschnikowia_pulcherrima(100); | Metschnikowia | 13 | Saprotroph | \|Undefined Saprotroph\| | Yeast | NULL | Probable | As nectar-tap_saprotroph (Põlme et al. 2020); Decay sugar-rich substrates (Põlme et al. 2020) | Põlme S, et al. 2020. Fungal Diversity 105:1-16 ((https://doi.org/10.1007/s13225-020-00466-2)) |
| Otu39034 | 1 | k__Fungi(100);p__Ascomycota(100);c__Saccharomycetes(100);o__Saccharomycetales(100);f__Metschnikowiaceae(100);g__Metschnikowia(100);g__Metschnikowia_unclassified(100); | Metschnikowia | 13 | Saprotroph | \|Undefined Saprotroph\| | Yeast | NULL | Probable | As nectar-tap_saprotroph (Põlme et al. 2020); Decay sugar-rich substrates (Põlme et al. 2020) | Põlme S, et al. 2020. Fungal Diversity 105:1-16 ((https://doi.org/10.1007/s13225-020-00466-2)) |
| Otu39035 | 1 | k__Fungi(100);p__Ascomycota(100);c__Saccharomycetes(100);o__Saccharomycetales(100);f__Metschnikowiaceae(100);g__Metschnikowia(100);s__Metschnikowia_pulcherrima(100); | Metschnikowia | 13 | Saprotroph | \|Undefined Saprotroph\| | Yeast | NULL | Probable | As nectar-tap_saprotroph (Põlme et al. 2020); Decay sugar-rich substrates (Põlme et al. 2020) | Põlme S, et al. 2020. Fungal Diversity 105:1-16 ((https://doi.org/10.1007/s13225-020-00466-2)) |
| Otu39036 | 1 | k__Fungi(100);p__Ascomycota(100);c__Saccharomycetes(100);o__Saccharomycetales(100);f__Metschnikowiaceae(100);g__Metschnikowia(100);s__Metschnikowia_pulcherrima(100); | Metschnikowia | 13 | Saprotroph | \|Undefined Saprotroph\| | Yeast | NULL | Probable | As nectar-tap_saprotroph (Põlme et al. 2020); Decay sugar-rich substrates (Põlme et al. 2020) | Põlme S, et al. 2020. Fungal Diversity 105:1-16 ((https://doi.org/10.1007/s13225-020-00466-2)) |
| Otu39037 | 1 | k__Fungi(100);p__Ascomycota(100);c__Saccharomycetes(100);o__Saccharomycetales(100);f__Metschnikowiaceae(100);g__Metschnikowia(100);s__Metschnikowia_pulcherrima(100); | Metschnikowia | 13 | Saprotroph | \|Undefined Saprotroph\| | Yeast | NULL | Probable | As nectar-tap_saprotroph (Põlme et al. 2020); Decay sugar-rich substrates (Põlme et al. 2020) | Põlme S, et al. 2020. Fungal Diversity 105:1-16 ((https://doi.org/10.1007/s13225-020-00466-2)) |
| Otu39038 | 1 | k__Fungi(100);p__Ascomycota(100);c__Saccharomycetes(100);o__Saccharomycetales(100);f__Metschnikowiaceae(100);g__Metschnikowia(100);s__Metschnikowia_pulcherrima(100); | Metschnikowia | 13 | Saprotroph | \|Undefined Saprotroph\| | Yeast | NULL | Probable | As nectar-tap_saprotroph (Põlme et al. 2020); Decay sugar-rich substrates (Põlme et al. 2020) | Põlme S, et al. 2020. Fungal Diversity 105:1-16 ((https://doi.org/10.1007/s13225-020-00466-2)) |
| Otu39039 | 1 | k__Fungi(100);p__Ascomycota(100);c__Saccharomycetes(100);o__Saccharomycetales(100);f__Metschnikowiaceae(100);g__Metschnikowia(100);s__Metschnikowia_pulcherrima(100); | Metschnikowia | 13 | Saprotroph | \|Undefined Saprotroph\| | Yeast | NULL | Probable | As nectar-tap_saprotroph (Põlme et al. 2020); Decay sugar-rich substrates (Põlme et al. 2020) | Põlme S, et al. 2020. Fungal Diversity 105:1-16 ((https://doi.org/10.1007/s13225-020-00466-2)) |
| Otu39040 | 1 | k__Fungi(100);p__Ascomycota(100);c__Saccharomycetes(100);o__Saccharomycetales(100);f__Metschnikowiaceae(100);g__Metschnikowia(100);g__Metschnikowia_unclassified(100); | Metschnikowia | 13 | Saprotroph | \|Undefined Saprotroph\| | Yeast | NULL | Probable | As nectar-tap_saprotroph (Põlme et al. 2020); Decay sugar-rich substrates (Põlme et al. 2020) | Põlme S, et al. 2020. Fungal Diversity 105:1-16 ((https://doi.org/10.1007/s13225-020-00466-2)) |
| Otu39041 | 1 | k__Fungi(100);p__Ascomycota(100);c__Saccharomycetes(100);o__Saccharomycetales(100);f__Metschnikowiaceae(100);g__Metschnikowia(100);g__Metschnikowia_unclassified(100); | Metschnikowia | 13 | Saprotroph | \|Undefined Saprotroph\| | Yeast | NULL | Probable | As nectar-tap_saprotroph (Põlme et al. 2020); Decay sugar-rich substrates (Põlme et al. 2020) | Põlme S, et al. 2020. Fungal Diversity 105:1-16 ((https://doi.org/10.1007/s13225-020-00466-2)) |
| Otu39043 | 1 | k__Fungi(100);p__Ascomycota(100);c__Saccharomycetes(100);o__Saccharomycetales(100);f__Metschnikowiaceae(100);g__Metschnikowia(100);s__Metschnikowia_pulcherrima(100); | Metschnikowia | 13 | Saprotroph | \|Undefined Saprotroph\| | Yeast | NULL | Probable | As nectar-tap_saprotroph (Põlme et al. 2020); Decay sugar-rich substrates (Põlme et al. 2020) | Põlme S, et al. 2020. Fungal Diversity 105:1-16 ((https://doi.org/10.1007/s13225-020-00466-2)) |
| Otu39044 | 1 | k__Fungi(100);p__Ascomycota(100);c__Saccharomycetes(100);o__Saccharomycetales(100);f__Metschnikowiaceae(100);g__Metschnikowia(100);s__Metschnikowia_pulcherrima(100); | Metschnikowia | 13 | Saprotroph | \|Undefined Saprotroph\| | Yeast | NULL | Probable | As nectar-tap_saprotroph (Põlme et al. 2020); Decay sugar-rich substrates (Põlme et al. 2020) | Põlme S, et al. 2020. Fungal Diversity 105:1-16 ((https://doi.org/10.1007/s13225-020-00466-2)) |
| Otu39045 | 1 | k__Fungi(100);p__Ascomycota(100);c__Saccharomycetes(100);o__Saccharomycetales(100);f__Metschnikowiaceae(100);g__Metschnikowia(100);s__Metschnikowia_pulcherrima(100); | Metschnikowia | 13 | Saprotroph | \|Undefined Saprotroph\| | Yeast | NULL | Probable | As nectar-tap_saprotroph (Põlme et al. 2020); Decay sugar-rich substrates (Põlme et al. 2020) | Põlme S, et al. 2020. Fungal Diversity 105:1-16 ((https://doi.org/10.1007/s13225-020-00466-2)) |
| Otu39046 | 1 | k__Fungi(100);p__Ascomycota(100);c__Saccharomycetes(100);o__Saccharomycetales(100);f__Metschnikowiaceae(100);g__Metschnikowia(100);g__Metschnikowia_unclassified(100); | Metschnikowia | 13 | Saprotroph | \|Undefined Saprotroph\| | Yeast | NULL | Probable | As nectar-tap_saprotroph (Põlme et al. 2020); Decay sugar-rich substrates (Põlme et al. 2020) | Põlme S, et al. 2020. Fungal Diversity 105:1-16 ((https://doi.org/10.1007/s13225-020-00466-2)) |
| Otu39047 | 1 | k__Fungi(100);p__Ascomycota(100);c__Saccharomycetes(100);o__Saccharomycetales(100);f__Metschnikowiaceae(100);g__Metschnikowia(100);s__Metschnikowia_pulcherrima(100); | Metschnikowia | 13 | Saprotroph | \|Undefined Saprotroph\| | Yeast | NULL | Probable | As nectar-tap_saprotroph (Põlme et al. 2020); Decay sugar-rich substrates (Põlme et al. 2020) | Põlme S, et al. 2020. Fungal Diversity 105:1-16 ((https://doi.org/10.1007/s13225-020-00466-2)) |
| Otu39048 | 1 | k__Fungi(100);p__Ascomycota(100);c__Saccharomycetes(100);o__Saccharomycetales(100);f__Metschnikowiaceae(100);g__Metschnikowia(100);s__Metschnikowia_pulcherrima(100); | Metschnikowia | 13 | Saprotroph | \|Undefined Saprotroph\| | Yeast | NULL | Probable | As nectar-tap_saprotroph (Põlme et al. 2020); Decay sugar-rich substrates (Põlme et al. 2020) | Põlme S, et al. 2020. Fungal Diversity 105:1-16 ((https://doi.org/10.1007/s13225-020-00466-2)) |
| Otu39049 | 1 | k__Fungi(100);p__Ascomycota(100);c__Saccharomycetes(100);o__Saccharomycetales(100);f__Metschnikowiaceae(100);g__Metschnikowia(100);s__Metschnikowia_pulcherrima(100); | Metschnikowia | 13 | Saprotroph | \|Undefined Saprotroph\| | Yeast | NULL | Probable | As nectar-tap_saprotroph (Põlme et al. 2020); Decay sugar-rich substrates (Põlme et al. 2020) | Põlme S, et al. 2020. Fungal Diversity 105:1-16 ((https://doi.org/10.1007/s13225-020-00466-2)) |
| Otu39050 | 1 | k__Fungi(100);p__Ascomycota(100);c__Saccharomycetes(100);o__Saccharomycetales(100);f__Metschnikowiaceae(100);g__Metschnikowia(100);s__Metschnikowia_pulcherrima(100); | Metschnikowia | 13 | Saprotroph | \|Undefined Saprotroph\| | Yeast | NULL | Probable | As nectar-tap_saprotroph (Põlme et al. 2020); Decay sugar-rich substrates (Põlme et al. 2020) | Põlme S, et al. 2020. Fungal Diversity 105:1-16 ((https://doi.org/10.1007/s13225-020-00466-2)) |
| Otu39052 | 1 | k__Fungi(100);p__Ascomycota(100);c__Saccharomycetes(100);o__Saccharomycetales(100);f__Metschnikowiaceae(100);g__Metschnikowia(100);s__Metschnikowia_pulcherrima(100); | Metschnikowia | 13 | Saprotroph | \|Undefined Saprotroph\| | Yeast | NULL | Probable | As nectar-tap_saprotroph (Põlme et al. 2020); Decay sugar-rich substrates (Põlme et al. 2020) | Põlme S, et al. 2020. Fungal Diversity 105:1-16 ((https://doi.org/10.1007/s13225-020-00466-2)) |
| Otu39053 | 1 | k__Fungi(100);p__Ascomycota(100);c__Saccharomycetes(100);o__Saccharomycetales(100);f__Metschnikowiaceae(100);g__Metschnikowia(100);s__Metschnikowia_pulcherrima(100); | Metschnikowia | 13 | Saprotroph | \|Undefined Saprotroph\| | Yeast | NULL | Probable | As nectar-tap_saprotroph (Põlme et al. 2020); Decay sugar-rich substrates (Põlme et al. 2020) | Põlme S, et al. 2020. Fungal Diversity 105:1-16 ((https://doi.org/10.1007/s13225-020-00466-2)) |
| Otu39056 | 1 | k__Fungi(100);p__Ascomycota(100);c__Saccharomycetes(100);o__Saccharomycetales(100);f__Metschnikowiaceae(100);g__Metschnikowia(100);s__Metschnikowia_pulcherrima(100); | Metschnikowia | 13 | Saprotroph | \|Undefined Saprotroph\| | Yeast | NULL | Probable | As nectar-tap_saprotroph (Põlme et al. 2020); Decay sugar-rich substrates (Põlme et al. 2020) | Põlme S, et al. 2020. Fungal Diversity 105:1-16 ((https://doi.org/10.1007/s13225-020-00466-2)) |
| Otu39057 | 1 | k__Fungi(100);p__Ascomycota(100);c__Saccharomycetes(100);o__Saccharomycetales(100);f__Metschnikowiaceae(100);g__Metschnikowia(100);s__Metschnikowia_pulcherrima(100); | Metschnikowia | 13 | Saprotroph | \|Undefined Saprotroph\| | Yeast | NULL | Probable | As nectar-tap_saprotroph (Põlme et al. 2020); Decay sugar-rich substrates (Põlme et al. 2020) | Põlme S, et al. 2020. Fungal Diversity 105:1-16 ((https://doi.org/10.1007/s13225-020-00466-2)) |
| Otu39058 | 1 | k__Fungi(100);p__Ascomycota(100);c__Saccharomycetes(100);o__Saccharomycetales(100);f__Metschnikowiaceae(100);g__Metschnikowia(100);s__Metschnikowia_pulcherrima(100); | Metschnikowia | 13 | Saprotroph | \|Undefined Saprotroph\| | Yeast | NULL | Probable | As nectar-tap_saprotroph (Põlme et al. 2020); Decay sugar-rich substrates (Põlme et al. 2020) | Põlme S, et al. 2020. Fungal Diversity 105:1-16 ((https://doi.org/10.1007/s13225-020-00466-2)) |
| Otu39059 | 1 | k__Fungi(100);p__Ascomycota(100);c__Saccharomycetes(100);o__Saccharomycetales(100);f__Metschnikowiaceae(100);g__Metschnikowia(100);g__Metschnikowia_unclassified(100); | Metschnikowia | 13 | Saprotroph | \|Undefined Saprotroph\| | Yeast | NULL | Probable | As nectar-tap_saprotroph (Põlme et al. 2020); Decay sugar-rich substrates (Põlme et al. 2020) | Põlme S, et al. 2020. Fungal Diversity 105:1-16 ((https://doi.org/10.1007/s13225-020-00466-2)) |
| Otu39060 | 1 | k__Fungi(100);p__Ascomycota(100);c__Saccharomycetes(100);o__Saccharomycetales(100);f__Metschnikowiaceae(100);g__Metschnikowia(100);s__Metschnikowia_pulcherrima(100); | Metschnikowia | 13 | Saprotroph | \|Undefined Saprotroph\| | Yeast | NULL | Probable | As nectar-tap_saprotroph (Põlme et al. 2020); Decay sugar-rich substrates (Põlme et al. 2020) | Põlme S, et al. 2020. Fungal Diversity 105:1-16 ((https://doi.org/10.1007/s13225-020-00466-2)) |
| Otu39061 | 1 | k__Fungi(100);p__Ascomycota(100);c__Saccharomycetes(100);o__Saccharomycetales(100);f__Metschnikowiaceae(100);g__Metschnikowia(100);s__Metschnikowia_pulcherrima(100); | Metschnikowia | 13 | Saprotroph | \|Undefined Saprotroph\| | Yeast | NULL | Probable | As nectar-tap_saprotroph (Põlme et al. 2020); Decay sugar-rich substrates (Põlme et al. 2020) | Põlme S, et al. 2020. Fungal Diversity 105:1-16 ((https://doi.org/10.1007/s13225-020-00466-2)) |
| Otu39062 | 1 | k__Fungi(100);p__Ascomycota(100);c__Saccharomycetes(100);o__Saccharomycetales(100);f__Metschnikowiaceae(100);g__Metschnikowia(100);s__Metschnikowia_pulcherrima(100); | Metschnikowia | 13 | Saprotroph | \|Undefined Saprotroph\| | Yeast | NULL | Probable | As nectar-tap_saprotroph (Põlme et al. 2020); Decay sugar-rich substrates (Põlme et al. 2020) | Põlme S, et al. 2020. Fungal Diversity 105:1-16 ((https://doi.org/10.1007/s13225-020-00466-2)) |
| Otu39063 | 1 | k__Fungi(100);p__Ascomycota(100);c__Saccharomycetes(100);o__Saccharomycetales(100);f__Metschnikowiaceae(100);g__Metschnikowia(100);s__Metschnikowia_pulcherrima(100); | Metschnikowia | 13 | Saprotroph | \|Undefined Saprotroph\| | Yeast | NULL | Probable | As nectar-tap_saprotroph (Põlme et al. 2020); Decay sugar-rich substrates (Põlme et al. 2020) | Põlme S, et al. 2020. Fungal Diversity 105:1-16 ((https://doi.org/10.1007/s13225-020-00466-2)) |
| Otu39064 | 1 | k__Fungi(100);p__Ascomycota(100);c__Saccharomycetes(100);o__Saccharomycetales(100);f__Metschnikowiaceae(100);g__Metschnikowia(100);s__Metschnikowia_pulcherrima(100); | Metschnikowia | 13 | Saprotroph | \|Undefined Saprotroph\| | Yeast | NULL | Probable | As nectar-tap_saprotroph (Põlme et al. 2020); Decay sugar-rich substrates (Põlme et al. 2020) | Põlme S, et al. 2020. Fungal Diversity 105:1-16 ((https://doi.org/10.1007/s13225-020-00466-2)) |
| Otu39067 | 1 | k__Fungi(100);p__Ascomycota(100);c__Saccharomycetes(100);o__Saccharomycetales(100);f__Metschnikowiaceae(100);g__Metschnikowia(100);g__Metschnikowia_unclassified(100); | Metschnikowia | 13 | Saprotroph | \|Undefined Saprotroph\| | Yeast | NULL | Probable | As nectar-tap_saprotroph (Põlme et al. 2020); Decay sugar-rich substrates (Põlme et al. 2020) | Põlme S, et al. 2020. Fungal Diversity 105:1-16 ((https://doi.org/10.1007/s13225-020-00466-2)) |
| Otu39068 | 1 | k__Fungi(100);p__Ascomycota(100);c__Saccharomycetes(100);o__Saccharomycetales(100);f__Metschnikowiaceae(100);g__Metschnikowia(100);s__Metschnikowia_pulcherrima(100); | Metschnikowia | 13 | Saprotroph | \|Undefined Saprotroph\| | Yeast | NULL | Probable | As nectar-tap_saprotroph (Põlme et al. 2020); Decay sugar-rich substrates (Põlme et al. 2020) | Põlme S, et al. 2020. Fungal Diversity 105:1-16 ((https://doi.org/10.1007/s13225-020-00466-2)) |
| Otu39069 | 1 | k__Fungi(100);p__Ascomycota(100);c__Saccharomycetes(100);o__Saccharomycetales(100);f__Metschnikowiaceae(100);g__Metschnikowia(100);s__Metschnikowia_pulcherrima(100); | Metschnikowia | 13 | Saprotroph | \|Undefined Saprotroph\| | Yeast | NULL | Probable | As nectar-tap_saprotroph (Põlme et al. 2020); Decay sugar-rich substrates (Põlme et al. 2020) | Põlme S, et al. 2020. Fungal Diversity 105:1-16 ((https://doi.org/10.1007/s13225-020-00466-2)) |
| Otu39070 | 1 | k__Fungi(100);p__Ascomycota(100);c__Saccharomycetes(100);o__Saccharomycetales(100);f__Metschnikowiaceae(100);g__Metschnikowia(100);s__Metschnikowia_pulcherrima(100); | Metschnikowia | 13 | Saprotroph | \|Undefined Saprotroph\| | Yeast | NULL | Probable | As nectar-tap_saprotroph (Põlme et al. 2020); Decay sugar-rich substrates (Põlme et al. 2020) | Põlme S, et al. 2020. Fungal Diversity 105:1-16 ((https://doi.org/10.1007/s13225-020-00466-2)) |
| Otu39071 | 1 | k__Fungi(100);p__Ascomycota(100);c__Saccharomycetes(100);o__Saccharomycetales(100);f__Metschnikowiaceae(100);g__Metschnikowia(100);s__Metschnikowia_pulcherrima(100); | Metschnikowia | 13 | Saprotroph | \|Undefined Saprotroph\| | Yeast | NULL | Probable | As nectar-tap_saprotroph (Põlme et al. 2020); Decay sugar-rich substrates (Põlme et al. 2020) | Põlme S, et al. 2020. Fungal Diversity 105:1-16 ((https://doi.org/10.1007/s13225-020-00466-2)) |
| Otu39074 | 1 | k__Fungi(100);p__Ascomycota(100);c__Saccharomycetes(100);o__Saccharomycetales(100);f__Metschnikowiaceae(100);g__Metschnikowia(100);g__Metschnikowia_unclassified(100); | Metschnikowia | 13 | Saprotroph | \|Undefined Saprotroph\| | Yeast | NULL | Probable | As nectar-tap_saprotroph (Põlme et al. 2020); Decay sugar-rich substrates (Põlme et al. 2020) | Põlme S, et al. 2020. Fungal Diversity 105:1-16 ((https://doi.org/10.1007/s13225-020-00466-2)) |
| Otu39075 | 1 | k__Fungi(100);p__Ascomycota(100);c__Saccharomycetes(100);o__Saccharomycetales(100);f__Metschnikowiaceae(100);g__Metschnikowia(100);s__Metschnikowia_pulcherrima(100); | Metschnikowia | 13 | Saprotroph | \|Undefined Saprotroph\| | Yeast | NULL | Probable | As nectar-tap_saprotroph (Põlme et al. 2020); Decay sugar-rich substrates (Põlme et al. 2020) | Põlme S, et al. 2020. Fungal Diversity 105:1-16 ((https://doi.org/10.1007/s13225-020-00466-2)) |
| Otu39077 | 1 | k__Fungi(100);p__Ascomycota(100);c__Saccharomycetes(100);o__Saccharomycetales(100);f__Metschnikowiaceae(100);g__Metschnikowia(100);s__Metschnikowia_pulcherrima(100); | Metschnikowia | 13 | Saprotroph | \|Undefined Saprotroph\| | Yeast | NULL | Probable | As nectar-tap_saprotroph (Põlme et al. 2020); Decay sugar-rich substrates (Põlme et al. 2020) | Põlme S, et al. 2020. Fungal Diversity 105:1-16 ((https://doi.org/10.1007/s13225-020-00466-2)) |
| Otu39078 | 1 | k__Fungi(100);p__Ascomycota(100);c__Saccharomycetes(100);o__Saccharomycetales(100);f__Metschnikowiaceae(100);g__Metschnikowia(100);g__Metschnikowia_unclassified(100); | Metschnikowia | 13 | Saprotroph | \|Undefined Saprotroph\| | Yeast | NULL | Probable | As nectar-tap_saprotroph (Põlme et al. 2020); Decay sugar-rich substrates (Põlme et al. 2020) | Põlme S, et al. 2020. Fungal Diversity 105:1-16 ((https://doi.org/10.1007/s13225-020-00466-2)) |
| Otu39079 | 1 | k__Fungi(100);p__Ascomycota(100);c__Saccharomycetes(100);o__Saccharomycetales(100);f__Metschnikowiaceae(100);g__Metschnikowia(100);s__Metschnikowia_pulcherrima(100); | Metschnikowia | 13 | Saprotroph | \|Undefined Saprotroph\| | Yeast | NULL | Probable | As nectar-tap_saprotroph (Põlme et al. 2020); Decay sugar-rich substrates (Põlme et al. 2020) | Põlme S, et al. 2020. Fungal Diversity 105:1-16 ((https://doi.org/10.1007/s13225-020-00466-2)) |
| Otu39080 | 1 | k__Fungi(100);p__Ascomycota(100);c__Saccharomycetes(100);o__Saccharomycetales(100);f__Metschnikowiaceae(100);g__Metschnikowia(100);s__Metschnikowia_pulcherrima(100); | Metschnikowia | 13 | Saprotroph | \|Undefined Saprotroph\| | Yeast | NULL | Probable | As nectar-tap_saprotroph (Põlme et al. 2020); Decay sugar-rich substrates (Põlme et al. 2020) | Põlme S, et al. 2020. Fungal Diversity 105:1-16 ((https://doi.org/10.1007/s13225-020-00466-2)) |
| Otu39081 | 1 | k__Fungi(100);p__Ascomycota(100);c__Saccharomycetes(100);o__Saccharomycetales(100);f__Metschnikowiaceae(100);g__Metschnikowia(100);s__Metschnikowia_pulcherrima(100); | Metschnikowia | 13 | Saprotroph | \|Undefined Saprotroph\| | Yeast | NULL | Probable | As nectar-tap_saprotroph (Põlme et al. 2020); Decay sugar-rich substrates (Põlme et al. 2020) | Põlme S, et al. 2020. Fungal Diversity 105:1-16 ((https://doi.org/10.1007/s13225-020-00466-2)) |
| Otu39082 | 1 | k__Fungi(100);p__Ascomycota(100);c__Saccharomycetes(100);o__Saccharomycetales(100);f__Metschnikowiaceae(100);g__Metschnikowia(100);s__Metschnikowia_pulcherrima(100); | Metschnikowia | 13 | Saprotroph | \|Undefined Saprotroph\| | Yeast | NULL | Probable | As nectar-tap_saprotroph (Põlme et al. 2020); Decay sugar-rich substrates (Põlme et al. 2020) | Põlme S, et al. 2020. Fungal Diversity 105:1-16 ((https://doi.org/10.1007/s13225-020-00466-2)) |
| Otu39083 | 1 | k__Fungi(100);p__Ascomycota(100);c__Saccharomycetes(100);o__Saccharomycetales(100);f__Metschnikowiaceae(100);g__Metschnikowia(100);s__Metschnikowia_pulcherrima(100); | Metschnikowia | 13 | Saprotroph | \|Undefined Saprotroph\| | Yeast | NULL | Probable | As nectar-tap_saprotroph (Põlme et al. 2020); Decay sugar-rich substrates (Põlme et al. 2020) | Põlme S, et al. 2020. Fungal Diversity 105:1-16 ((https://doi.org/10.1007/s13225-020-00466-2)) |
| Otu39084 | 1 | k__Fungi(100);p__Ascomycota(100);c__Saccharomycetes(100);o__Saccharomycetales(100);f__Metschnikowiaceae(100);g__Metschnikowia(100);s__Metschnikowia_pulcherrima(100); | Metschnikowia | 13 | Saprotroph | \|Undefined Saprotroph\| | Yeast | NULL | Probable | As nectar-tap_saprotroph (Põlme et al. 2020); Decay sugar-rich substrates (Põlme et al. 2020) | Põlme S, et al. 2020. Fungal Diversity 105:1-16 ((https://doi.org/10.1007/s13225-020-00466-2)) |
| Otu39085 | 1 | k__Fungi(100);p__Ascomycota(100);c__Saccharomycetes(100);o__Saccharomycetales(100);f__Metschnikowiaceae(100);g__Metschnikowia(100);g__Metschnikowia_unclassified(100); | Metschnikowia | 13 | Saprotroph | \|Undefined Saprotroph\| | Yeast | NULL | Probable | As nectar-tap_saprotroph (Põlme et al. 2020); Decay sugar-rich substrates (Põlme et al. 2020) | Põlme S, et al. 2020. Fungal Diversity 105:1-16 ((https://doi.org/10.1007/s13225-020-00466-2)) |
| Otu39086 | 1 | k__Fungi(100);p__Ascomycota(100);c__Saccharomycetes(100);o__Saccharomycetales(100);f__Metschnikowiaceae(100);g__Metschnikowia(100);s__Metschnikowia_pulcherrima(100); | Metschnikowia | 13 | Saprotroph | \|Undefined Saprotroph\| | Yeast | NULL | Probable | As nectar-tap_saprotroph (Põlme et al. 2020); Decay sugar-rich substrates (Põlme et al. 2020) | Põlme S, et al. 2020. Fungal Diversity 105:1-16 ((https://doi.org/10.1007/s13225-020-00466-2)) |
| Otu39087 | 1 | k__Fungi(100);p__Ascomycota(100);c__Saccharomycetes(100);o__Saccharomycetales(100);f__Metschnikowiaceae(100);g__Metschnikowia(100);s__Metschnikowia_pulcherrima(100); | Metschnikowia | 13 | Saprotroph | \|Undefined Saprotroph\| | Yeast | NULL | Probable | As nectar-tap_saprotroph (Põlme et al. 2020); Decay sugar-rich substrates (Põlme et al. 2020) | Põlme S, et al. 2020. Fungal Diversity 105:1-16 ((https://doi.org/10.1007/s13225-020-00466-2)) |
| Otu39088 | 1 | k__Fungi(100);p__Ascomycota(100);c__Saccharomycetes(100);o__Saccharomycetales(100);f__Metschnikowiaceae(100);g__Metschnikowia(100);s__Metschnikowia_pulcherrima(100); | Metschnikowia | 13 | Saprotroph | \|Undefined Saprotroph\| | Yeast | NULL | Probable | As nectar-tap_saprotroph (Põlme et al. 2020); Decay sugar-rich substrates (Põlme et al. 2020) | Põlme S, et al. 2020. Fungal Diversity 105:1-16 ((https://doi.org/10.1007/s13225-020-00466-2)) |
| Otu39089 | 1 | k__Fungi(100);p__Ascomycota(100);c__Saccharomycetes(100);o__Saccharomycetales(100);f__Metschnikowiaceae(100);g__Metschnikowia(100);s__Metschnikowia_pulcherrima(100); | Metschnikowia | 13 | Saprotroph | \|Undefined Saprotroph\| | Yeast | NULL | Probable | As nectar-tap_saprotroph (Põlme et al. 2020); Decay sugar-rich substrates (Põlme et al. 2020) | Põlme S, et al. 2020. Fungal Diversity 105:1-16 ((https://doi.org/10.1007/s13225-020-00466-2)) |
| Otu39090 | 1 | k__Fungi(100);p__Ascomycota(100);c__Saccharomycetes(100);o__Saccharomycetales(100);f__Metschnikowiaceae(100);g__Metschnikowia(100);s__Metschnikowia_pulcherrima(100); | Metschnikowia | 13 | Saprotroph | \|Undefined Saprotroph\| | Yeast | NULL | Probable | As nectar-tap_saprotroph (Põlme et al. 2020); Decay sugar-rich substrates (Põlme et al. 2020) | Põlme S, et al. 2020. Fungal Diversity 105:1-16 ((https://doi.org/10.1007/s13225-020-00466-2)) |
| Otu39091 | 1 | k__Fungi(100);p__Ascomycota(100);c__Saccharomycetes(100);o__Saccharomycetales(100);f__Metschnikowiaceae(100);g__Metschnikowia(100);s__Metschnikowia_pulcherrima(100); | Metschnikowia | 13 | Saprotroph | \|Undefined Saprotroph\| | Yeast | NULL | Probable | As nectar-tap_saprotroph (Põlme et al. 2020); Decay sugar-rich substrates (Põlme et al. 2020) | Põlme S, et al. 2020. Fungal Diversity 105:1-16 ((https://doi.org/10.1007/s13225-020-00466-2)) |
| Otu39092 | 1 | k__Fungi(100);p__Ascomycota(100);c__Saccharomycetes(100);o__Saccharomycetales(100);f__Metschnikowiaceae(100);g__Metschnikowia(100);s__Metschnikowia_pulcherrima(100); | Metschnikowia | 13 | Saprotroph | \|Undefined Saprotroph\| | Yeast | NULL | Probable | As nectar-tap_saprotroph (Põlme et al. 2020); Decay sugar-rich substrates (Põlme et al. 2020) | Põlme S, et al. 2020. Fungal Diversity 105:1-16 ((https://doi.org/10.1007/s13225-020-00466-2)) |
| Otu39093 | 1 | k__Fungi(100);p__Ascomycota(100);c__Saccharomycetes(100);o__Saccharomycetales(100);f__Metschnikowiaceae(100);g__Metschnikowia(100);g__Metschnikowia_unclassified(100); | Metschnikowia | 13 | Saprotroph | \|Undefined Saprotroph\| | Yeast | NULL | Probable | As nectar-tap_saprotroph (Põlme et al. 2020); Decay sugar-rich substrates (Põlme et al. 2020) | Põlme S, et al. 2020. Fungal Diversity 105:1-16 ((https://doi.org/10.1007/s13225-020-00466-2)) |
| Otu39094 | 1 | k__Fungi(100);p__Ascomycota(100);c__Saccharomycetes(100);o__Saccharomycetales(100);f__Metschnikowiaceae(100);g__Metschnikowia(100);s__Metschnikowia_pulcherrima(100); | Metschnikowia | 13 | Saprotroph | \|Undefined Saprotroph\| | Yeast | NULL | Probable | As nectar-tap_saprotroph (Põlme et al. 2020); Decay sugar-rich substrates (Põlme et al. 2020) | Põlme S, et al. 2020. Fungal Diversity 105:1-16 ((https://doi.org/10.1007/s13225-020-00466-2)) |
| Otu39095 | 1 | k__Fungi(100);p__Ascomycota(100);c__Saccharomycetes(100);o__Saccharomycetales(100);f__Metschnikowiaceae(100);g__Metschnikowia(100);s__Metschnikowia_pulcherrima(100); | Metschnikowia | 13 | Saprotroph | \|Undefined Saprotroph\| | Yeast | NULL | Probable | As nectar-tap_saprotroph (Põlme et al. 2020); Decay sugar-rich substrates (Põlme et al. 2020) | Põlme S, et al. 2020. Fungal Diversity 105:1-16 ((https://doi.org/10.1007/s13225-020-00466-2)) |
| Otu39096 | 1 | k__Fungi(100);p__Ascomycota(100);c__Saccharomycetes(100);o__Saccharomycetales(100);f__Metschnikowiaceae(100);g__Metschnikowia(100);s__Metschnikowia_pulcherrima(100); | Metschnikowia | 13 | Saprotroph | \|Undefined Saprotroph\| | Yeast | NULL | Probable | As nectar-tap_saprotroph (Põlme et al. 2020); Decay sugar-rich substrates (Põlme et al. 2020) | Põlme S, et al. 2020. Fungal Diversity 105:1-16 ((https://doi.org/10.1007/s13225-020-00466-2)) |
| Otu39097 | 1 | k__Fungi(100);p__Ascomycota(100);c__Saccharomycetes(100);o__Saccharomycetales(100);f__Metschnikowiaceae(100);g__Metschnikowia(100);g__Metschnikowia_unclassified(100); | Metschnikowia | 13 | Saprotroph | \|Undefined Saprotroph\| | Yeast | NULL | Probable | As nectar-tap_saprotroph (Põlme et al. 2020); Decay sugar-rich substrates (Põlme et al. 2020) | Põlme S, et al. 2020. Fungal Diversity 105:1-16 ((https://doi.org/10.1007/s13225-020-00466-2)) |
| Otu39098 | 1 | k__Fungi(100);p__Ascomycota(100);c__Saccharomycetes(100);o__Saccharomycetales(100);f__Metschnikowiaceae(100);g__Metschnikowia(100);s__Metschnikowia_pulcherrima(100); | Metschnikowia | 13 | Saprotroph | \|Undefined Saprotroph\| | Yeast | NULL | Probable | As nectar-tap_saprotroph (Põlme et al. 2020); Decay sugar-rich substrates (Põlme et al. 2020) | Põlme S, et al. 2020. Fungal Diversity 105:1-16 ((https://doi.org/10.1007/s13225-020-00466-2)) |
| Otu39099 | 1 | k__Fungi(100);p__Ascomycota(100);c__Saccharomycetes(100);o__Saccharomycetales(100);f__Metschnikowiaceae(100);g__Metschnikowia(100);s__Metschnikowia_pulcherrima(100); | Metschnikowia | 13 | Saprotroph | \|Undefined Saprotroph\| | Yeast | NULL | Probable | As nectar-tap_saprotroph (Põlme et al. 2020); Decay sugar-rich substrates (Põlme et al. 2020) | Põlme S, et al. 2020. Fungal Diversity 105:1-16 ((https://doi.org/10.1007/s13225-020-00466-2)) |
| Otu39100 | 1 | k__Fungi(100);p__Ascomycota(100);c__Saccharomycetes(100);o__Saccharomycetales(100);f__Metschnikowiaceae(100);g__Metschnikowia(100);s__Metschnikowia_pulcherrima(100); | Metschnikowia | 13 | Saprotroph | \|Undefined Saprotroph\| | Yeast | NULL | Probable | As nectar-tap_saprotroph (Põlme et al. 2020); Decay sugar-rich substrates (Põlme et al. 2020) | Põlme S, et al. 2020. Fungal Diversity 105:1-16 ((https://doi.org/10.1007/s13225-020-00466-2)) |
| Otu39101 | 1 | k__Fungi(100);p__Ascomycota(100);c__Saccharomycetes(100);o__Saccharomycetales(100);f__Metschnikowiaceae(100);g__Metschnikowia(100);s__Metschnikowia_pulcherrima(100); | Metschnikowia | 13 | Saprotroph | \|Undefined Saprotroph\| | Yeast | NULL | Probable | As nectar-tap_saprotroph (Põlme et al. 2020); Decay sugar-rich substrates (Põlme et al. 2020) | Põlme S, et al. 2020. Fungal Diversity 105:1-16 ((https://doi.org/10.1007/s13225-020-00466-2)) |
| Otu39102 | 1 | k__Fungi(100);p__Ascomycota(100);c__Saccharomycetes(100);o__Saccharomycetales(100);f__Metschnikowiaceae(100);g__Metschnikowia(100);s__Metschnikowia_pulcherrima(100); | Metschnikowia | 13 | Saprotroph | \|Undefined Saprotroph\| | Yeast | NULL | Probable | As nectar-tap_saprotroph (Põlme et al. 2020); Decay sugar-rich substrates (Põlme et al. 2020) | Põlme S, et al. 2020. Fungal Diversity 105:1-16 ((https://doi.org/10.1007/s13225-020-00466-2)) |
| Otu39103 | 1 | k__Fungi(100);p__Ascomycota(100);c__Saccharomycetes(100);o__Saccharomycetales(100);f__Metschnikowiaceae(100);g__Metschnikowia(100);s__Metschnikowia_pulcherrima(100); | Metschnikowia | 13 | Saprotroph | \|Undefined Saprotroph\| | Yeast | NULL | Probable | As nectar-tap_saprotroph (Põlme et al. 2020); Decay sugar-rich substrates (Põlme et al. 2020) | Põlme S, et al. 2020. Fungal Diversity 105:1-16 ((https://doi.org/10.1007/s13225-020-00466-2)) |
| Otu39104 | 1 | k__Fungi(100);p__Ascomycota(100);c__Saccharomycetes(100);o__Saccharomycetales(100);f__Metschnikowiaceae(100);g__Metschnikowia(100);g__Metschnikowia_unclassified(100); | Metschnikowia | 13 | Saprotroph | \|Undefined Saprotroph\| | Yeast | NULL | Probable | As nectar-tap_saprotroph (Põlme et al. 2020); Decay sugar-rich substrates (Põlme et al. 2020) | Põlme S, et al. 2020. Fungal Diversity 105:1-16 ((https://doi.org/10.1007/s13225-020-00466-2)) |
| Otu39105 | 1 | k__Fungi(100);p__Ascomycota(100);c__Saccharomycetes(100);o__Saccharomycetales(100);f__Metschnikowiaceae(100);g__Metschnikowia(100);s__Metschnikowia_pulcherrima(100); | Metschnikowia | 13 | Saprotroph | \|Undefined Saprotroph\| | Yeast | NULL | Probable | As nectar-tap_saprotroph (Põlme et al. 2020); Decay sugar-rich substrates (Põlme et al. 2020) | Põlme S, et al. 2020. Fungal Diversity 105:1-16 ((https://doi.org/10.1007/s13225-020-00466-2)) |
| Otu39106 | 1 | k__Fungi(100);p__Ascomycota(100);c__Saccharomycetes(100);o__Saccharomycetales(100);f__Metschnikowiaceae(100);g__Metschnikowia(100);s__Metschnikowia_pulcherrima(100); | Metschnikowia | 13 | Saprotroph | \|Undefined Saprotroph\| | Yeast | NULL | Probable | As nectar-tap_saprotroph (Põlme et al. 2020); Decay sugar-rich substrates (Põlme et al. 2020) | Põlme S, et al. 2020. Fungal Diversity 105:1-16 ((https://doi.org/10.1007/s13225-020-00466-2)) |
| Otu39107 | 1 | k__Fungi(100);p__Ascomycota(100);c__Saccharomycetes(100);o__Saccharomycetales(100);f__Metschnikowiaceae(100);g__Metschnikowia(100);s__Metschnikowia_pulcherrima(100); | Metschnikowia | 13 | Saprotroph | \|Undefined Saprotroph\| | Yeast | NULL | Probable | As nectar-tap_saprotroph (Põlme et al. 2020); Decay sugar-rich substrates (Põlme et al. 2020) | Põlme S, et al. 2020. Fungal Diversity 105:1-16 ((https://doi.org/10.1007/s13225-020-00466-2)) |
| Otu39108 | 1 | k__Fungi(100);p__Ascomycota(100);c__Saccharomycetes(100);o__Saccharomycetales(100);f__Metschnikowiaceae(100);g__Metschnikowia(100);s__Metschnikowia_pulcherrima(100); | Metschnikowia | 13 | Saprotroph | \|Undefined Saprotroph\| | Yeast | NULL | Probable | As nectar-tap_saprotroph (Põlme et al. 2020); Decay sugar-rich substrates (Põlme et al. 2020) | Põlme S, et al. 2020. Fungal Diversity 105:1-16 ((https://doi.org/10.1007/s13225-020-00466-2)) |
| Otu39109 | 1 | k__Fungi(100);p__Ascomycota(100);c__Saccharomycetes(100);o__Saccharomycetales(100);f__Metschnikowiaceae(100);g__Metschnikowia(100);s__Metschnikowia_pulcherrima(100); | Metschnikowia | 13 | Saprotroph | \|Undefined Saprotroph\| | Yeast | NULL | Probable | As nectar-tap_saprotroph (Põlme et al. 2020); Decay sugar-rich substrates (Põlme et al. 2020) | Põlme S, et al. 2020. Fungal Diversity 105:1-16 ((https://doi.org/10.1007/s13225-020-00466-2)) |
| Otu39110 | 1 | k__Fungi(100);p__Ascomycota(100);c__Saccharomycetes(100);o__Saccharomycetales(100);f__Metschnikowiaceae(100);g__Metschnikowia(100);s__Metschnikowia_pulcherrima(100); | Metschnikowia | 13 | Saprotroph | \|Undefined Saprotroph\| | Yeast | NULL | Probable | As nectar-tap_saprotroph (Põlme et al. 2020); Decay sugar-rich substrates (Põlme et al. 2020) | Põlme S, et al. 2020. Fungal Diversity 105:1-16 ((https://doi.org/10.1007/s13225-020-00466-2)) |
| Otu39111 | 1 | k__Fungi(100);p__Ascomycota(100);c__Saccharomycetes(100);o__Saccharomycetales(100);f__Metschnikowiaceae(100);g__Metschnikowia(100);s__Metschnikowia_pulcherrima(100); | Metschnikowia | 13 | Saprotroph | \|Undefined Saprotroph\| | Yeast | NULL | Probable | As nectar-tap_saprotroph (Põlme et al. 2020); Decay sugar-rich substrates (Põlme et al. 2020) | Põlme S, et al. 2020. Fungal Diversity 105:1-16 ((https://doi.org/10.1007/s13225-020-00466-2)) |
| Otu39112 | 1 | k__Fungi(100);p__Ascomycota(100);c__Saccharomycetes(100);o__Saccharomycetales(100);f__Metschnikowiaceae(100);g__Metschnikowia(100);g__Metschnikowia_unclassified(100); | Metschnikowia | 13 | Saprotroph | \|Undefined Saprotroph\| | Yeast | NULL | Probable | As nectar-tap_saprotroph (Põlme et al. 2020); Decay sugar-rich substrates (Põlme et al. 2020) | Põlme S, et al. 2020. Fungal Diversity 105:1-16 ((https://doi.org/10.1007/s13225-020-00466-2)) |
| Otu39113 | 1 | k__Fungi(100);p__Ascomycota(100);c__Saccharomycetes(100);o__Saccharomycetales(100);f__Metschnikowiaceae(100);g__Metschnikowia(100);g__Metschnikowia_unclassified(100); | Metschnikowia | 13 | Saprotroph | \|Undefined Saprotroph\| | Yeast | NULL | Probable | As nectar-tap_saprotroph (Põlme et al. 2020); Decay sugar-rich substrates (Põlme et al. 2020) | Põlme S, et al. 2020. Fungal Diversity 105:1-16 ((https://doi.org/10.1007/s13225-020-00466-2)) |
| Otu39114 | 1 | k__Fungi(100);p__Ascomycota(100);c__Saccharomycetes(100);o__Saccharomycetales(100);f__Metschnikowiaceae(100);g__Metschnikowia(100);s__Metschnikowia_pulcherrima(100); | Metschnikowia | 13 | Saprotroph | \|Undefined Saprotroph\| | Yeast | NULL | Probable | As nectar-tap_saprotroph (Põlme et al. 2020); Decay sugar-rich substrates (Põlme et al. 2020) | Põlme S, et al. 2020. Fungal Diversity 105:1-16 ((https://doi.org/10.1007/s13225-020-00466-2)) |
| Otu39115 | 1 | k__Fungi(100);p__Ascomycota(100);c__Saccharomycetes(100);o__Saccharomycetales(100);f__Metschnikowiaceae(100);g__Metschnikowia(100);s__Metschnikowia_pulcherrima(100); | Metschnikowia | 13 | Saprotroph | \|Undefined Saprotroph\| | Yeast | NULL | Probable | As nectar-tap_saprotroph (Põlme et al. 2020); Decay sugar-rich substrates (Põlme et al. 2020) | Põlme S, et al. 2020. Fungal Diversity 105:1-16 ((https://doi.org/10.1007/s13225-020-00466-2)) |
| Otu39116 | 1 | k__Fungi(100);p__Ascomycota(100);c__Saccharomycetes(100);o__Saccharomycetales(100);f__Metschnikowiaceae(100);g__Metschnikowia(100);s__Metschnikowia_pulcherrima(100); | Metschnikowia | 13 | Saprotroph | \|Undefined Saprotroph\| | Yeast | NULL | Probable | As nectar-tap_saprotroph (Põlme et al. 2020); Decay sugar-rich substrates (Põlme et al. 2020) | Põlme S, et al. 2020. Fungal Diversity 105:1-16 ((https://doi.org/10.1007/s13225-020-00466-2)) |
| Otu39117 | 1 | k__Fungi(100);p__Ascomycota(100);c__Saccharomycetes(100);o__Saccharomycetales(100);f__Metschnikowiaceae(100);g__Metschnikowia(100);s__Metschnikowia_pulcherrima(100); | Metschnikowia | 13 | Saprotroph | \|Undefined Saprotroph\| | Yeast | NULL | Probable | As nectar-tap_saprotroph (Põlme et al. 2020); Decay sugar-rich substrates (Põlme et al. 2020) | Põlme S, et al. 2020. Fungal Diversity 105:1-16 ((https://doi.org/10.1007/s13225-020-00466-2)) |
| Otu39118 | 1 | k__Fungi(100);p__Ascomycota(100);c__Saccharomycetes(100);o__Saccharomycetales(100);f__Metschnikowiaceae(100);g__Metschnikowia(100);s__Metschnikowia_pulcherrima(100); | Metschnikowia | 13 | Saprotroph | \|Undefined Saprotroph\| | Yeast | NULL | Probable | As nectar-tap_saprotroph (Põlme et al. 2020); Decay sugar-rich substrates (Põlme et al. 2020) | Põlme S, et al. 2020. Fungal Diversity 105:1-16 ((https://doi.org/10.1007/s13225-020-00466-2)) |
| Otu39119 | 1 | k__Fungi(100);p__Ascomycota(100);c__Saccharomycetes(100);o__Saccharomycetales(100);f__Metschnikowiaceae(100);g__Metschnikowia(100);s__Metschnikowia_pulcherrima(100); | Metschnikowia | 13 | Saprotroph | \|Undefined Saprotroph\| | Yeast | NULL | Probable | As nectar-tap_saprotroph (Põlme et al. 2020); Decay sugar-rich substrates (Põlme et al. 2020) | Põlme S, et al. 2020. Fungal Diversity 105:1-16 ((https://doi.org/10.1007/s13225-020-00466-2)) |
| Otu39120 | 1 | k__Fungi(100);p__Ascomycota(100);c__Saccharomycetes(100);o__Saccharomycetales(100);f__Metschnikowiaceae(100);g__Metschnikowia(100);s__Metschnikowia_pulcherrima(100); | Metschnikowia | 13 | Saprotroph | \|Undefined Saprotroph\| | Yeast | NULL | Probable | As nectar-tap_saprotroph (Põlme et al. 2020); Decay sugar-rich substrates (Põlme et al. 2020) | Põlme S, et al. 2020. Fungal Diversity 105:1-16 ((https://doi.org/10.1007/s13225-020-00466-2)) |
| Otu39121 | 1 | k__Fungi(100);p__Ascomycota(100);c__Saccharomycetes(100);o__Saccharomycetales(100);f__Metschnikowiaceae(100);g__Metschnikowia(100);s__Metschnikowia_pulcherrima(100); | Metschnikowia | 13 | Saprotroph | \|Undefined Saprotroph\| | Yeast | NULL | Probable | As nectar-tap_saprotroph (Põlme et al. 2020); Decay sugar-rich substrates (Põlme et al. 2020) | Põlme S, et al. 2020. Fungal Diversity 105:1-16 ((https://doi.org/10.1007/s13225-020-00466-2)) |
| Otu39122 | 1 | k__Fungi(100);p__Ascomycota(100);c__Saccharomycetes(100);o__Saccharomycetales(100);f__Metschnikowiaceae(100);g__Metschnikowia(100);s__Metschnikowia_pulcherrima(100); | Metschnikowia | 13 | Saprotroph | \|Undefined Saprotroph\| | Yeast | NULL | Probable | As nectar-tap_saprotroph (Põlme et al. 2020); Decay sugar-rich substrates (Põlme et al. 2020) | Põlme S, et al. 2020. Fungal Diversity 105:1-16 ((https://doi.org/10.1007/s13225-020-00466-2)) |
| Otu39123 | 1 | k__Fungi(100);p__Ascomycota(100);c__Saccharomycetes(100);o__Saccharomycetales(100);f__Metschnikowiaceae(100);g__Metschnikowia(100);g__Metschnikowia_unclassified(100); | Metschnikowia | 13 | Saprotroph | \|Undefined Saprotroph\| | Yeast | NULL | Probable | As nectar-tap_saprotroph (Põlme et al. 2020); Decay sugar-rich substrates (Põlme et al. 2020) | Põlme S, et al. 2020. Fungal Diversity 105:1-16 ((https://doi.org/10.1007/s13225-020-00466-2)) |
| Otu39124 | 1 | k__Fungi(100);p__Ascomycota(100);c__Saccharomycetes(100);o__Saccharomycetales(100);f__Metschnikowiaceae(100);g__Metschnikowia(100);s__Metschnikowia_pulcherrima(100); | Metschnikowia | 13 | Saprotroph | \|Undefined Saprotroph\| | Yeast | NULL | Probable | As nectar-tap_saprotroph (Põlme et al. 2020); Decay sugar-rich substrates (Põlme et al. 2020) | Põlme S, et al. 2020. Fungal Diversity 105:1-16 ((https://doi.org/10.1007/s13225-020-00466-2)) |
| Otu39126 | 1 | k__Fungi(100);p__Ascomycota(100);c__Saccharomycetes(100);o__Saccharomycetales(100);f__Metschnikowiaceae(100);g__Metschnikowia(100);g__Metschnikowia_unclassified(100); | Metschnikowia | 13 | Saprotroph | \|Undefined Saprotroph\| | Yeast | NULL | Probable | As nectar-tap_saprotroph (Põlme et al. 2020); Decay sugar-rich substrates (Põlme et al. 2020) | Põlme S, et al. 2020. Fungal Diversity 105:1-16 ((https://doi.org/10.1007/s13225-020-00466-2)) |
| Otu39127 | 1 | k__Fungi(100);p__Ascomycota(100);c__Saccharomycetes(100);o__Saccharomycetales(100);f__Metschnikowiaceae(100);g__Metschnikowia(100);s__Metschnikowia_pulcherrima(100); | Metschnikowia | 13 | Saprotroph | \|Undefined Saprotroph\| | Yeast | NULL | Probable | As nectar-tap_saprotroph (Põlme et al. 2020); Decay sugar-rich substrates (Põlme et al. 2020) | Põlme S, et al. 2020. Fungal Diversity 105:1-16 ((https://doi.org/10.1007/s13225-020-00466-2)) |
| Otu39128 | 1 | k__Fungi(100);p__Ascomycota(100);c__Saccharomycetes(100);o__Saccharomycetales(100);f__Metschnikowiaceae(100);g__Metschnikowia(100);s__Metschnikowia_pulcherrima(100); | Metschnikowia | 13 | Saprotroph | \|Undefined Saprotroph\| | Yeast | NULL | Probable | As nectar-tap_saprotroph (Põlme et al. 2020); Decay sugar-rich substrates (Põlme et al. 2020) | Põlme S, et al. 2020. Fungal Diversity 105:1-16 ((https://doi.org/10.1007/s13225-020-00466-2)) |
| Otu39129 | 1 | k__Fungi(100);p__Ascomycota(100);c__Saccharomycetes(100);o__Saccharomycetales(100);f__Metschnikowiaceae(100);g__Metschnikowia(100);s__Metschnikowia_pulcherrima(100); | Metschnikowia | 13 | Saprotroph | \|Undefined Saprotroph\| | Yeast | NULL | Probable | As nectar-tap_saprotroph (Põlme et al. 2020); Decay sugar-rich substrates (Põlme et al. 2020) | Põlme S, et al. 2020. Fungal Diversity 105:1-16 ((https://doi.org/10.1007/s13225-020-00466-2)) |
| Otu39130 | 1 | k__Fungi(100);p__Ascomycota(100);c__Saccharomycetes(100);o__Saccharomycetales(100);f__Metschnikowiaceae(100);g__Metschnikowia(100);s__Metschnikowia_pulcherrima(100); | Metschnikowia | 13 | Saprotroph | \|Undefined Saprotroph\| | Yeast | NULL | Probable | As nectar-tap_saprotroph (Põlme et al. 2020); Decay sugar-rich substrates (Põlme et al. 2020) | Põlme S, et al. 2020. Fungal Diversity 105:1-16 ((https://doi.org/10.1007/s13225-020-00466-2)) |
| Otu39131 | 1 | k__Fungi(100);p__Ascomycota(100);c__Saccharomycetes(100);o__Saccharomycetales(100);f__Metschnikowiaceae(100);g__Metschnikowia(100);s__Metschnikowia_pulcherrima(100); | Metschnikowia | 13 | Saprotroph | \|Undefined Saprotroph\| | Yeast | NULL | Probable | As nectar-tap_saprotroph (Põlme et al. 2020); Decay sugar-rich substrates (Põlme et al. 2020) | Põlme S, et al. 2020. Fungal Diversity 105:1-16 ((https://doi.org/10.1007/s13225-020-00466-2)) |
| Otu39132 | 1 | k__Fungi(100);p__Ascomycota(100);c__Saccharomycetes(100);o__Saccharomycetales(100);f__Metschnikowiaceae(100);g__Metschnikowia(100);s__Metschnikowia_pulcherrima(100); | Metschnikowia | 13 | Saprotroph | \|Undefined Saprotroph\| | Yeast | NULL | Probable | As nectar-tap_saprotroph (Põlme et al. 2020); Decay sugar-rich substrates (Põlme et al. 2020) | Põlme S, et al. 2020. Fungal Diversity 105:1-16 ((https://doi.org/10.1007/s13225-020-00466-2)) |
| Otu39133 | 1 | k__Fungi(100);p__Ascomycota(100);c__Saccharomycetes(100);o__Saccharomycetales(100);f__Metschnikowiaceae(100);g__Metschnikowia(100);g__Metschnikowia_unclassified(100); | Metschnikowia | 13 | Saprotroph | \|Undefined Saprotroph\| | Yeast | NULL | Probable | As nectar-tap_saprotroph (Põlme et al. 2020); Decay sugar-rich substrates (Põlme et al. 2020) | Põlme S, et al. 2020. Fungal Diversity 105:1-16 ((https://doi.org/10.1007/s13225-020-00466-2)) |
| Otu39134 | 1 | k__Fungi(100);p__Ascomycota(100);c__Saccharomycetes(100);o__Saccharomycetales(100);f__Metschnikowiaceae(100);g__Metschnikowia(100);s__Metschnikowia_pulcherrima(100); | Metschnikowia | 13 | Saprotroph | \|Undefined Saprotroph\| | Yeast | NULL | Probable | As nectar-tap_saprotroph (Põlme et al. 2020); Decay sugar-rich substrates (Põlme et al. 2020) | Põlme S, et al. 2020. Fungal Diversity 105:1-16 ((https://doi.org/10.1007/s13225-020-00466-2)) |
| Otu39135 | 1 | k__Fungi(100);p__Ascomycota(100);c__Saccharomycetes(100);o__Saccharomycetales(100);f__Metschnikowiaceae(100);g__Metschnikowia(100);s__Metschnikowia_pulcherrima(100); | Metschnikowia | 13 | Saprotroph | \|Undefined Saprotroph\| | Yeast | NULL | Probable | As nectar-tap_saprotroph (Põlme et al. 2020); Decay sugar-rich substrates (Põlme et al. 2020) | Põlme S, et al. 2020. Fungal Diversity 105:1-16 ((https://doi.org/10.1007/s13225-020-00466-2)) |
| Otu39136 | 1 | k__Fungi(100);p__Ascomycota(100);c__Saccharomycetes(100);o__Saccharomycetales(100);f__Metschnikowiaceae(100);g__Metschnikowia(100);g__Metschnikowia_unclassified(100); | Metschnikowia | 13 | Saprotroph | \|Undefined Saprotroph\| | Yeast | NULL | Probable | As nectar-tap_saprotroph (Põlme et al. 2020); Decay sugar-rich substrates (Põlme et al. 2020) | Põlme S, et al. 2020. Fungal Diversity 105:1-16 ((https://doi.org/10.1007/s13225-020-00466-2)) |
| Otu39137 | 1 | k__Fungi(100);p__Ascomycota(100);c__Saccharomycetes(100);o__Saccharomycetales(100);f__Metschnikowiaceae(100);g__Metschnikowia(100);s__Metschnikowia_pulcherrima(100); | Metschnikowia | 13 | Saprotroph | \|Undefined Saprotroph\| | Yeast | NULL | Probable | As nectar-tap_saprotroph (Põlme et al. 2020); Decay sugar-rich substrates (Põlme et al. 2020) | Põlme S, et al. 2020. Fungal Diversity 105:1-16 ((https://doi.org/10.1007/s13225-020-00466-2)) |
| Otu39138 | 1 | k__Fungi(100);p__Ascomycota(100);c__Saccharomycetes(100);o__Saccharomycetales(100);f__Metschnikowiaceae(100);g__Metschnikowia(100);s__Metschnikowia_pulcherrima(100); | Metschnikowia | 13 | Saprotroph | \|Undefined Saprotroph\| | Yeast | NULL | Probable | As nectar-tap_saprotroph (Põlme et al. 2020); Decay sugar-rich substrates (Põlme et al. 2020) | Põlme S, et al. 2020. Fungal Diversity 105:1-16 ((https://doi.org/10.1007/s13225-020-00466-2)) |
| Otu39139 | 1 | k__Fungi(100);p__Ascomycota(100);c__Saccharomycetes(100);o__Saccharomycetales(100);f__Metschnikowiaceae(100);g__Metschnikowia(100);s__Metschnikowia_pulcherrima(100); | Metschnikowia | 13 | Saprotroph | \|Undefined Saprotroph\| | Yeast | NULL | Probable | As nectar-tap_saprotroph (Põlme et al. 2020); Decay sugar-rich substrates (Põlme et al. 2020) | Põlme S, et al. 2020. Fungal Diversity 105:1-16 ((https://doi.org/10.1007/s13225-020-00466-2)) |
| Otu39140 | 1 | k__Fungi(100);p__Ascomycota(100);c__Saccharomycetes(100);o__Saccharomycetales(100);f__Metschnikowiaceae(100);g__Metschnikowia(100);s__Metschnikowia_pulcherrima(100); | Metschnikowia | 13 | Saprotroph | \|Undefined Saprotroph\| | Yeast | NULL | Probable | As nectar-tap_saprotroph (Põlme et al. 2020); Decay sugar-rich substrates (Põlme et al. 2020) | Põlme S, et al. 2020. Fungal Diversity 105:1-16 ((https://doi.org/10.1007/s13225-020-00466-2)) |
| Otu39141 | 1 | k__Fungi(100);p__Ascomycota(100);c__Saccharomycetes(100);o__Saccharomycetales(100);f__Metschnikowiaceae(100);g__Metschnikowia(100);s__Metschnikowia_pulcherrima(100); | Metschnikowia | 13 | Saprotroph | \|Undefined Saprotroph\| | Yeast | NULL | Probable | As nectar-tap_saprotroph (Põlme et al. 2020); Decay sugar-rich substrates (Põlme et al. 2020) | Põlme S, et al. 2020. Fungal Diversity 105:1-16 ((https://doi.org/10.1007/s13225-020-00466-2)) |
| Otu39142 | 1 | k__Fungi(100);p__Ascomycota(100);c__Saccharomycetes(100);o__Saccharomycetales(100);f__Metschnikowiaceae(100);g__Metschnikowia(100);s__Metschnikowia_pulcherrima(100); | Metschnikowia | 13 | Saprotroph | \|Undefined Saprotroph\| | Yeast | NULL | Probable | As nectar-tap_saprotroph (Põlme et al. 2020); Decay sugar-rich substrates (Põlme et al. 2020) | Põlme S, et al. 2020. Fungal Diversity 105:1-16 ((https://doi.org/10.1007/s13225-020-00466-2)) |
| Otu39143 | 1 | k__Fungi(100);p__Ascomycota(100);c__Saccharomycetes(100);o__Saccharomycetales(100);f__Metschnikowiaceae(100);g__Metschnikowia(100);s__Metschnikowia_pulcherrima(100); | Metschnikowia | 13 | Saprotroph | \|Undefined Saprotroph\| | Yeast | NULL | Probable | As nectar-tap_saprotroph (Põlme et al. 2020); Decay sugar-rich substrates (Põlme et al. 2020) | Põlme S, et al. 2020. Fungal Diversity 105:1-16 ((https://doi.org/10.1007/s13225-020-00466-2)) |
| Otu39144 | 1 | k__Fungi(100);p__Ascomycota(100);c__Saccharomycetes(100);o__Saccharomycetales(100);f__Metschnikowiaceae(100);g__Metschnikowia(100);s__Metschnikowia_pulcherrima(100); | Metschnikowia | 13 | Saprotroph | \|Undefined Saprotroph\| | Yeast | NULL | Probable | As nectar-tap_saprotroph (Põlme et al. 2020); Decay sugar-rich substrates (Põlme et al. 2020) | Põlme S, et al. 2020. Fungal Diversity 105:1-16 ((https://doi.org/10.1007/s13225-020-00466-2)) |
| Otu39145 | 1 | k__Fungi(100);p__Ascomycota(100);c__Saccharomycetes(100);o__Saccharomycetales(100);f__Metschnikowiaceae(100);g__Metschnikowia(100);s__Metschnikowia_pulcherrima(100); | Metschnikowia | 13 | Saprotroph | \|Undefined Saprotroph\| | Yeast | NULL | Probable | As nectar-tap_saprotroph (Põlme et al. 2020); Decay sugar-rich substrates (Põlme et al. 2020) | Põlme S, et al. 2020. Fungal Diversity 105:1-16 ((https://doi.org/10.1007/s13225-020-00466-2)) |
| Otu39146 | 1 | k__Fungi(100);p__Ascomycota(100);c__Saccharomycetes(100);o__Saccharomycetales(100);f__Metschnikowiaceae(100);g__Metschnikowia(100);s__Metschnikowia_pulcherrima(100); | Metschnikowia | 13 | Saprotroph | \|Undefined Saprotroph\| | Yeast | NULL | Probable | As nectar-tap_saprotroph (Põlme et al. 2020); Decay sugar-rich substrates (Põlme et al. 2020) | Põlme S, et al. 2020. Fungal Diversity 105:1-16 ((https://doi.org/10.1007/s13225-020-00466-2)) |
| Otu39147 | 1 | k__Fungi(100);p__Ascomycota(100);c__Saccharomycetes(100);o__Saccharomycetales(100);f__Metschnikowiaceae(100);g__Metschnikowia(100);g__Metschnikowia_unclassified(100); | Metschnikowia | 13 | Saprotroph | \|Undefined Saprotroph\| | Yeast | NULL | Probable | As nectar-tap_saprotroph (Põlme et al. 2020); Decay sugar-rich substrates (Põlme et al. 2020) | Põlme S, et al. 2020. Fungal Diversity 105:1-16 ((https://doi.org/10.1007/s13225-020-00466-2)) |
| Otu39148 | 1 | k__Fungi(100);p__Ascomycota(100);c__Saccharomycetes(100);o__Saccharomycetales(100);f__Metschnikowiaceae(100);g__Metschnikowia(100);s__Metschnikowia_pulcherrima(100); | Metschnikowia | 13 | Saprotroph | \|Undefined Saprotroph\| | Yeast | NULL | Probable | As nectar-tap_saprotroph (Põlme et al. 2020); Decay sugar-rich substrates (Põlme et al. 2020) | Põlme S, et al. 2020. Fungal Diversity 105:1-16 ((https://doi.org/10.1007/s13225-020-00466-2)) |
| Otu39149 | 1 | k__Fungi(100);p__Ascomycota(100);c__Saccharomycetes(100);o__Saccharomycetales(100);f__Metschnikowiaceae(100);g__Metschnikowia(100);s__Metschnikowia_pulcherrima(100); | Metschnikowia | 13 | Saprotroph | \|Undefined Saprotroph\| | Yeast | NULL | Probable | As nectar-tap_saprotroph (Põlme et al. 2020); Decay sugar-rich substrates (Põlme et al. 2020) | Põlme S, et al. 2020. Fungal Diversity 105:1-16 ((https://doi.org/10.1007/s13225-020-00466-2)) |
| Otu39150 | 1 | k__Fungi(100);p__Ascomycota(100);c__Saccharomycetes(100);o__Saccharomycetales(100);f__Metschnikowiaceae(100);g__Metschnikowia(100);s__Metschnikowia_pulcherrima(100); | Metschnikowia | 13 | Saprotroph | \|Undefined Saprotroph\| | Yeast | NULL | Probable | As nectar-tap_saprotroph (Põlme et al. 2020); Decay sugar-rich substrates (Põlme et al. 2020) | Põlme S, et al. 2020. Fungal Diversity 105:1-16 ((https://doi.org/10.1007/s13225-020-00466-2)) |
| Otu39151 | 1 | k__Fungi(100);p__Ascomycota(100);c__Saccharomycetes(100);o__Saccharomycetales(100);f__Metschnikowiaceae(100);g__Metschnikowia(100);s__Metschnikowia_pulcherrima(100); | Metschnikowia | 13 | Saprotroph | \|Undefined Saprotroph\| | Yeast | NULL | Probable | As nectar-tap_saprotroph (Põlme et al. 2020); Decay sugar-rich substrates (Põlme et al. 2020) | Põlme S, et al. 2020. Fungal Diversity 105:1-16 ((https://doi.org/10.1007/s13225-020-00466-2)) |
| Otu39152 | 1 | k__Fungi(100);p__Ascomycota(100);c__Saccharomycetes(100);o__Saccharomycetales(100);f__Metschnikowiaceae(100);g__Metschnikowia(100);s__Metschnikowia_pulcherrima(100); | Metschnikowia | 13 | Saprotroph | \|Undefined Saprotroph\| | Yeast | NULL | Probable | As nectar-tap_saprotroph (Põlme et al. 2020); Decay sugar-rich substrates (Põlme et al. 2020) | Põlme S, et al. 2020. Fungal Diversity 105:1-16 ((https://doi.org/10.1007/s13225-020-00466-2)) |
| Otu39153 | 1 | k__Fungi(100);p__Ascomycota(100);c__Saccharomycetes(100);o__Saccharomycetales(100);f__Metschnikowiaceae(100);g__Metschnikowia(100);g__Metschnikowia_unclassified(100); | Metschnikowia | 13 | Saprotroph | \|Undefined Saprotroph\| | Yeast | NULL | Probable | As nectar-tap_saprotroph (Põlme et al. 2020); Decay sugar-rich substrates (Põlme et al. 2020) | Põlme S, et al. 2020. Fungal Diversity 105:1-16 ((https://doi.org/10.1007/s13225-020-00466-2)) |
| Otu39154 | 1 | k__Fungi(100);p__Ascomycota(100);c__Saccharomycetes(100);o__Saccharomycetales(100);f__Metschnikowiaceae(100);g__Metschnikowia(100);s__Metschnikowia_pulcherrima(100); | Metschnikowia | 13 | Saprotroph | \|Undefined Saprotroph\| | Yeast | NULL | Probable | As nectar-tap_saprotroph (Põlme et al. 2020); Decay sugar-rich substrates (Põlme et al. 2020) | Põlme S, et al. 2020. Fungal Diversity 105:1-16 ((https://doi.org/10.1007/s13225-020-00466-2)) |
| Otu39155 | 1 | k__Fungi(100);p__Ascomycota(100);c__Saccharomycetes(100);o__Saccharomycetales(100);f__Metschnikowiaceae(100);g__Metschnikowia(100);g__Metschnikowia_unclassified(100); | Metschnikowia | 13 | Saprotroph | \|Undefined Saprotroph\| | Yeast | NULL | Probable | As nectar-tap_saprotroph (Põlme et al. 2020); Decay sugar-rich substrates (Põlme et al. 2020) | Põlme S, et al. 2020. Fungal Diversity 105:1-16 ((https://doi.org/10.1007/s13225-020-00466-2)) |
| Otu39156 | 1 | k__Fungi(100);p__Ascomycota(100);c__Saccharomycetes(100);o__Saccharomycetales(100);f__Metschnikowiaceae(100);g__Metschnikowia(100);s__Metschnikowia_pulcherrima(100); | Metschnikowia | 13 | Saprotroph | \|Undefined Saprotroph\| | Yeast | NULL | Probable | As nectar-tap_saprotroph (Põlme et al. 2020); Decay sugar-rich substrates (Põlme et al. 2020) | Põlme S, et al. 2020. Fungal Diversity 105:1-16 ((https://doi.org/10.1007/s13225-020-00466-2)) |
| Otu39157 | 1 | k__Fungi(100);p__Ascomycota(100);c__Saccharomycetes(100);o__Saccharomycetales(100);f__Metschnikowiaceae(100);g__Metschnikowia(100);s__Metschnikowia_pulcherrima(100); | Metschnikowia | 13 | Saprotroph | \|Undefined Saprotroph\| | Yeast | NULL | Probable | As nectar-tap_saprotroph (Põlme et al. 2020); Decay sugar-rich substrates (Põlme et al. 2020) | Põlme S, et al. 2020. Fungal Diversity 105:1-16 ((https://doi.org/10.1007/s13225-020-00466-2)) |
| Otu39158 | 1 | k__Fungi(100);p__Ascomycota(100);c__Saccharomycetes(100);o__Saccharomycetales(100);f__Metschnikowiaceae(100);g__Metschnikowia(100);g__Metschnikowia_unclassified(100); | Metschnikowia | 13 | Saprotroph | \|Undefined Saprotroph\| | Yeast | NULL | Probable | As nectar-tap_saprotroph (Põlme et al. 2020); Decay sugar-rich substrates (Põlme et al. 2020) | Põlme S, et al. 2020. Fungal Diversity 105:1-16 ((https://doi.org/10.1007/s13225-020-00466-2)) |
| Otu39160 | 1 | k__Fungi(100);p__Ascomycota(100);c__Saccharomycetes(100);o__Saccharomycetales(100);f__Metschnikowiaceae(100);g__Metschnikowia(100);s__Metschnikowia_pulcherrima(100); | Metschnikowia | 13 | Saprotroph | \|Undefined Saprotroph\| | Yeast | NULL | Probable | As nectar-tap_saprotroph (Põlme et al. 2020); Decay sugar-rich substrates (Põlme et al. 2020) | Põlme S, et al. 2020. Fungal Diversity 105:1-16 ((https://doi.org/10.1007/s13225-020-00466-2)) |
| Otu39161 | 1 | k__Fungi(100);p__Ascomycota(100);c__Saccharomycetes(100);o__Saccharomycetales(100);f__Metschnikowiaceae(100);g__Metschnikowia(100);s__Metschnikowia_pulcherrima(100); | Metschnikowia | 13 | Saprotroph | \|Undefined Saprotroph\| | Yeast | NULL | Probable | As nectar-tap_saprotroph (Põlme et al. 2020); Decay sugar-rich substrates (Põlme et al. 2020) | Põlme S, et al. 2020. Fungal Diversity 105:1-16 ((https://doi.org/10.1007/s13225-020-00466-2)) |
| Otu39162 | 1 | k__Fungi(100);p__Ascomycota(100);c__Saccharomycetes(100);o__Saccharomycetales(100);f__Metschnikowiaceae(100);g__Metschnikowia(100);g__Metschnikowia_unclassified(100); | Metschnikowia | 13 | Saprotroph | \|Undefined Saprotroph\| | Yeast | NULL | Probable | As nectar-tap_saprotroph (Põlme et al. 2020); Decay sugar-rich substrates (Põlme et al. 2020) | Põlme S, et al. 2020. Fungal Diversity 105:1-16 ((https://doi.org/10.1007/s13225-020-00466-2)) |
| Otu39163 | 1 | k__Fungi(100);p__Ascomycota(100);c__Saccharomycetes(100);o__Saccharomycetales(100);f__Metschnikowiaceae(100);g__Metschnikowia(100);s__Metschnikowia_pulcherrima(100); | Metschnikowia | 13 | Saprotroph | \|Undefined Saprotroph\| | Yeast | NULL | Probable | As nectar-tap_saprotroph (Põlme et al. 2020); Decay sugar-rich substrates (Põlme et al. 2020) | Põlme S, et al. 2020. Fungal Diversity 105:1-16 ((https://doi.org/10.1007/s13225-020-00466-2)) |
| Otu39166 | 1 | k__Fungi(100);p__Ascomycota(100);c__Saccharomycetes(100);o__Saccharomycetales(100);f__Metschnikowiaceae(100);g__Metschnikowia(100);s__Metschnikowia_pulcherrima(100); | Metschnikowia | 13 | Saprotroph | \|Undefined Saprotroph\| | Yeast | NULL | Probable | As nectar-tap_saprotroph (Põlme et al. 2020); Decay sugar-rich substrates (Põlme et al. 2020) | Põlme S, et al. 2020. Fungal Diversity 105:1-16 ((https://doi.org/10.1007/s13225-020-00466-2)) |
| Otu39167 | 1 | k__Fungi(100);p__Ascomycota(100);c__Saccharomycetes(100);o__Saccharomycetales(100);f__Metschnikowiaceae(100);g__Metschnikowia(100);s__Metschnikowia_pulcherrima(100); | Metschnikowia | 13 | Saprotroph | \|Undefined Saprotroph\| | Yeast | NULL | Probable | As nectar-tap_saprotroph (Põlme et al. 2020); Decay sugar-rich substrates (Põlme et al. 2020) | Põlme S, et al. 2020. Fungal Diversity 105:1-16 ((https://doi.org/10.1007/s13225-020-00466-2)) |
| Otu39168 | 1 | k__Fungi(100);p__Ascomycota(100);c__Saccharomycetes(100);o__Saccharomycetales(100);f__Metschnikowiaceae(100);g__Metschnikowia(100);s__Metschnikowia_pulcherrima(100); | Metschnikowia | 13 | Saprotroph | \|Undefined Saprotroph\| | Yeast | NULL | Probable | As nectar-tap_saprotroph (Põlme et al. 2020); Decay sugar-rich substrates (Põlme et al. 2020) | Põlme S, et al. 2020. Fungal Diversity 105:1-16 ((https://doi.org/10.1007/s13225-020-00466-2)) |
| Otu39169 | 1 | k__Fungi(100);p__Ascomycota(100);c__Saccharomycetes(100);o__Saccharomycetales(100);f__Metschnikowiaceae(100);g__Metschnikowia(100);s__Metschnikowia_pulcherrima(100); | Metschnikowia | 13 | Saprotroph | \|Undefined Saprotroph\| | Yeast | NULL | Probable | As nectar-tap_saprotroph (Põlme et al. 2020); Decay sugar-rich substrates (Põlme et al. 2020) | Põlme S, et al. 2020. Fungal Diversity 105:1-16 ((https://doi.org/10.1007/s13225-020-00466-2)) |
| Otu39170 | 1 | k__Fungi(100);p__Ascomycota(100);c__Saccharomycetes(100);o__Saccharomycetales(100);f__Metschnikowiaceae(100);g__Metschnikowia(100);s__Metschnikowia_pulcherrima(100); | Metschnikowia | 13 | Saprotroph | \|Undefined Saprotroph\| | Yeast | NULL | Probable | As nectar-tap_saprotroph (Põlme et al. 2020); Decay sugar-rich substrates (Põlme et al. 2020) | Põlme S, et al. 2020. Fungal Diversity 105:1-16 ((https://doi.org/10.1007/s13225-020-00466-2)) |
| Otu39171 | 1 | k__Fungi(100);p__Ascomycota(100);c__Saccharomycetes(100);o__Saccharomycetales(100);f__Metschnikowiaceae(100);g__Metschnikowia(100);s__Metschnikowia_pulcherrima(100); | Metschnikowia | 13 | Saprotroph | \|Undefined Saprotroph\| | Yeast | NULL | Probable | As nectar-tap_saprotroph (Põlme et al. 2020); Decay sugar-rich substrates (Põlme et al. 2020) | Põlme S, et al. 2020. Fungal Diversity 105:1-16 ((https://doi.org/10.1007/s13225-020-00466-2)) |
| Otu39172 | 1 | k__Fungi(100);p__Ascomycota(100);c__Saccharomycetes(100);o__Saccharomycetales(100);f__Metschnikowiaceae(100);g__Metschnikowia(100);s__Metschnikowia_pulcherrima(100); | Metschnikowia | 13 | Saprotroph | \|Undefined Saprotroph\| | Yeast | NULL | Probable | As nectar-tap_saprotroph (Põlme et al. 2020); Decay sugar-rich substrates (Põlme et al. 2020) | Põlme S, et al. 2020. Fungal Diversity 105:1-16 ((https://doi.org/10.1007/s13225-020-00466-2)) |
| Otu39174 | 1 | k__Fungi(100);p__Ascomycota(100);c__Saccharomycetes(100);o__Saccharomycetales(100);f__Metschnikowiaceae(100);g__Metschnikowia(100);s__Metschnikowia_pulcherrima(100); | Metschnikowia | 13 | Saprotroph | \|Undefined Saprotroph\| | Yeast | NULL | Probable | As nectar-tap_saprotroph (Põlme et al. 2020); Decay sugar-rich substrates (Põlme et al. 2020) | Põlme S, et al. 2020. Fungal Diversity 105:1-16 ((https://doi.org/10.1007/s13225-020-00466-2)) |
| Otu39175 | 1 | k__Fungi(100);p__Ascomycota(100);c__Saccharomycetes(100);o__Saccharomycetales(100);f__Metschnikowiaceae(100);g__Metschnikowia(100);s__Metschnikowia_pulcherrima(100); | Metschnikowia | 13 | Saprotroph | \|Undefined Saprotroph\| | Yeast | NULL | Probable | As nectar-tap_saprotroph (Põlme et al. 2020); Decay sugar-rich substrates (Põlme et al. 2020) | Põlme S, et al. 2020. Fungal Diversity 105:1-16 ((https://doi.org/10.1007/s13225-020-00466-2)) |
| Otu39176 | 1 | k__Fungi(100);p__Ascomycota(100);c__Saccharomycetes(100);o__Saccharomycetales(100);f__Metschnikowiaceae(100);g__Metschnikowia(100);g__Metschnikowia_unclassified(100); | Metschnikowia | 13 | Saprotroph | \|Undefined Saprotroph\| | Yeast | NULL | Probable | As nectar-tap_saprotroph (Põlme et al. 2020); Decay sugar-rich substrates (Põlme et al. 2020) | Põlme S, et al. 2020. Fungal Diversity 105:1-16 ((https://doi.org/10.1007/s13225-020-00466-2)) |
| Otu39178 | 1 | k__Fungi(100);p__Ascomycota(100);c__Saccharomycetes(100);o__Saccharomycetales(100);f__Metschnikowiaceae(100);g__Metschnikowia(100);s__Metschnikowia_pulcherrima(100); | Metschnikowia | 13 | Saprotroph | \|Undefined Saprotroph\| | Yeast | NULL | Probable | As nectar-tap_saprotroph (Põlme et al. 2020); Decay sugar-rich substrates (Põlme et al. 2020) | Põlme S, et al. 2020. Fungal Diversity 105:1-16 ((https://doi.org/10.1007/s13225-020-00466-2)) |
| Otu39179 | 1 | k__Fungi(100);p__Ascomycota(100);c__Saccharomycetes(100);o__Saccharomycetales(100);f__Metschnikowiaceae(100);g__Metschnikowia(100);s__Metschnikowia_pulcherrima(100); | Metschnikowia | 13 | Saprotroph | \|Undefined Saprotroph\| | Yeast | NULL | Probable | As nectar-tap_saprotroph (Põlme et al. 2020); Decay sugar-rich substrates (Põlme et al. 2020) | Põlme S, et al. 2020. Fungal Diversity 105:1-16 ((https://doi.org/10.1007/s13225-020-00466-2)) |
| Otu39180 | 1 | k__Fungi(100);p__Ascomycota(100);c__Saccharomycetes(100);o__Saccharomycetales(100);f__Metschnikowiaceae(100);g__Metschnikowia(100);s__Metschnikowia_pulcherrima(100); | Metschnikowia | 13 | Saprotroph | \|Undefined Saprotroph\| | Yeast | NULL | Probable | As nectar-tap_saprotroph (Põlme et al. 2020); Decay sugar-rich substrates (Põlme et al. 2020) | Põlme S, et al. 2020. Fungal Diversity 105:1-16 ((https://doi.org/10.1007/s13225-020-00466-2)) |

| | | | | | | | | | | | |
|---|---|---|---|---|---|---|---|---|---|---|---|
| Otu39181 | 1 | k__Fungi(100);p__Ascomycota(100);c__Saccharomycetes(100);o__Saccharomycetales(100);f__Metschnikowiaceae(100);g__Metschnikowia(100);s__Metschnikowia_pulcherrima(100); | Metschnikowia | 13 | Saprotroph | \|Undefined Saprotroph\| | Yeast | NULL | Probable | As nectar-tap_saprotroph (Põlme et al. 2020); Decay sugar-rich substrates (Põlme et al. 2020) | Põlme S, et al. 2020. Fungal Diversity 105:1-16 ((https://doi.org/10.1007/s13225-020-00466-2)) |
| Otu39182 | 1 | k__Fungi(100);p__Ascomycota(100);c__Saccharomycetes(100);o__Saccharomycetales(100);f__Metschnikowiaceae(100);g__Metschnikowia(100);g__Metschnikowia_unclassified(100); | Metschnikowia | 13 | Saprotroph | \|Undefined Saprotroph\| | Yeast | NULL | Probable | As nectar-tap_saprotroph (Põlme et al. 2020); Decay sugar-rich substrates (Põlme et al. 2020) | Põlme S, et al. 2020. Fungal Diversity 105:1-16 ((https://doi.org/10.1007/s13225-020-00466-2)) |
| Otu39183 | 1 | k__Fungi(100);p__Ascomycota(100);c__Saccharomycetes(100);o__Saccharomycetales(100);f__Metschnikowiaceae(100);g__Metschnikowia(100);s__Metschnikowia_pulcherrima(100); | Metschnikowia | 13 | Saprotroph | \|Undefined Saprotroph\| | Yeast | NULL | Probable | As nectar-tap_saprotroph (Põlme et al. 2020); Decay sugar-rich substrates (Põlme et al. 2020) | Põlme S, et al. 2020. Fungal Diversity 105:1-16 ((https://doi.org/10.1007/s13225-020-00466-2)) |
| Otu39184 | 1 | k__Fungi(100);p__Ascomycota(100);c__Saccharomycetes(100);o__Saccharomycetales(100);f__Metschnikowiaceae(100);g__Metschnikowia(100);s__Metschnikowia_pulcherrima(100); | Metschnikowia | 13 | Saprotroph | \|Undefined Saprotroph\| | Yeast | NULL | Probable | As nectar-tap_saprotroph (Põlme et al. 2020); Decay sugar-rich substrates (Põlme et al. 2020) | Põlme S, et al. 2020. Fungal Diversity 105:1-16 ((https://doi.org/10.1007/s13225-020-00466-2)) |
| Otu39185 | 1 | k__Fungi(100);p__Ascomycota(100);c__Saccharomycetes(100);o__Saccharomycetales(100);f__Metschnikowiaceae(100);g__Metschnikowia(100);s__Metschnikowia_pulcherrima(100); | Metschnikowia | 13 | Saprotroph | \|Undefined Saprotroph\| | Yeast | NULL | Probable | As nectar-tap_saprotroph (Põlme et al. 2020); Decay sugar-rich substrates (Põlme et al. 2020) | Põlme S, et al. 2020. Fungal Diversity 105:1-16 ((https://doi.org/10.1007/s13225-020-00466-2)) |
| Otu39186 | 1 | k__Fungi(100);p__Ascomycota(100);c__Saccharomycetes(100);o__Saccharomycetales(100);f__Metschnikowiaceae(100);g__Metschnikowia(100);s__Metschnikowia_pulcherrima(100); | Metschnikowia | 13 | Saprotroph | \|Undefined Saprotroph\| | Yeast | NULL | Probable | As nectar-tap_saprotroph (Põlme et al. 2020); Decay sugar-rich substrates (Põlme et al. 2020) | Põlme S, et al. 2020. Fungal Diversity 105:1-16 ((https://doi.org/10.1007/s13225-020-00466-2)) |
| Otu39187 | 1 | k__Fungi(100);p__Ascomycota(100);c__Saccharomycetes(100);o__Saccharomycetales(100);f__Metschnikowiaceae(100);g__Metschnikowia(100);g__Metschnikowia_unclassified(100); | Metschnikowia | 13 | Saprotroph | \|Undefined Saprotroph\| | Yeast | NULL | Probable | As nectar-tap_saprotroph (Põlme et al. 2020); Decay sugar-rich substrates (Põlme et al. 2020) | Põlme S, et al. 2020. Fungal Diversity 105:1-16 ((https://doi.org/10.1007/s13225-020-00466-2)) |
| Otu39188 | 1 | k__Fungi(100);p__Ascomycota(100);c__Saccharomycetes(100);o__Saccharomycetales(100);f__Metschnikowiaceae(100);g__Metschnikowia(100);s__Metschnikowia_pulcherrima(100); | Metschnikowia | 13 | Saprotroph | \|Undefined Saprotroph\| | Yeast | NULL | Probable | As nectar-tap_saprotroph (Põlme et al. 2020); Decay sugar-rich substrates (Põlme et al. 2020) | Põlme S, et al. 2020. Fungal Diversity 105:1-16 ((https://doi.org/10.1007/s13225-020-00466-2)) |
| Otu39189 | 1 | k__Fungi(100);p__Ascomycota(100);c__Saccharomycetes(100);o__Saccharomycetales(100);f__Metschnikowiaceae(100);g__Metschnikowia(100);s__Metschnikowia_pulcherrima(100); | Metschnikowia | 13 | Saprotroph | \|Undefined Saprotroph\| | Yeast | NULL | Probable | As nectar-tap_saprotroph (Põlme et al. 2020); Decay sugar-rich substrates (Põlme et al. 2020) | Põlme S, et al. 2020. Fungal Diversity 105:1-16 ((https://doi.org/10.1007/s13225-020-00466-2)) |
| Otu39190 | 1 | k__Fungi(100);p__Ascomycota(100);c__Saccharomycetes(100);o__Saccharomycetales(100);f__Metschnikowiaceae(100);g__Metschnikowia(100);g__Metschnikowia_unclassified(100); | Metschnikowia | 13 | Saprotroph | \|Undefined Saprotroph\| | Yeast | NULL | Probable | As nectar-tap_saprotroph (Põlme et al. 2020); Decay sugar-rich substrates (Põlme et al. 2020) | Põlme S, et al. 2020. Fungal Diversity 105:1-16 ((https://doi.org/10.1007/s13225-020-00466-2)) |
| Otu39191 | 1 | k__Fungi(100);p__Ascomycota(100);c__Saccharomycetes(100);o__Saccharomycetales(100);f__Metschnikowiaceae(100);g__Metschnikowia(100);s__Metschnikowia_pulcherrima(100); | Metschnikowia | 13 | Saprotroph | \|Undefined Saprotroph\| | Yeast | NULL | Probable | As nectar-tap_saprotroph (Põlme et al. 2020); Decay sugar-rich substrates (Põlme et al. 2020) | Põlme S, et al. 2020. Fungal Diversity 105:1-16 ((https://doi.org/10.1007/s13225-020-00466-2)) |
| Otu39192 | 1 | k__Fungi(100);p__Ascomycota(100);c__Saccharomycetes(100);o__Saccharomycetales(100);f__Metschnikowiaceae(100);g__Metschnikowia(100);s__Metschnikowia_pulcherrima(100); | Metschnikowia | 13 | Saprotroph | \|Undefined Saprotroph\| | Yeast | NULL | Probable | As nectar-tap_saprotroph (Põlme et al. 2020); Decay sugar-rich substrates (Põlme et al. 2020) | Põlme S, et al. 2020. Fungal Diversity 105:1-16 ((https://doi.org/10.1007/s13225-020-00466-2)) |
| Otu39194 | 1 | k__Fungi(100);p__Ascomycota(100);c__Saccharomycetes(100);o__Saccharomycetales(100);f__Metschnikowiaceae(100);g__Metschnikowia(100);s__Metschnikowia_pulcherrima(100); | Metschnikowia | 13 | Saprotroph | \|Undefined Saprotroph\| | Yeast | NULL | Probable | As nectar-tap_saprotroph (Põlme et al. 2020); Decay sugar-rich substrates (Põlme et al. 2020) | Põlme S, et al. 2020. Fungal Diversity 105:1-16 ((https://doi.org/10.1007/s13225-020-00466-2)) |
| Otu39195 | 1 | k__Fungi(100);p__Ascomycota(100);c__Saccharomycetes(100);o__Saccharomycetales(100);f__Metschnikowiaceae(100);g__Metschnikowia(100);s__Metschnikowia_pulcherrima(100); | Metschnikowia | 13 | Saprotroph | \|Undefined Saprotroph\| | Yeast | NULL | Probable | As nectar-tap_saprotroph (Põlme et al. 2020); Decay sugar-rich substrates (Põlme et al. 2020) | Põlme S, et al. 2020. Fungal Diversity 105:1-16 ((https://doi.org/10.1007/s13225-020-00466-2)) |
| Otu39196 | 1 | k__Fungi(100);p__Ascomycota(100);c__Saccharomycetes(100);o__Saccharomycetales(100);f__Saccharomycetales_family_Incertae_sedis(100);g__Candida(100);g__Candida_unclassified(100); | Candida | 13 | Pathotroph-Saprotroph-Symbiotroph | Animal Pathogen-Endophyte-Endosymbiont-Epiphyte-\|Undefined Saprotroph\| | Dimorphic Yeast-Yeast | NULL | Probable | As nectar-tap_saprotroph (Põlme et al. 2020); Decay sugar-rich substrates (Põlme et al. 2020); This asexual genus is polyphyletic and contain species that function very differently depend on what they are. Caution should be taken when interpreting guilds for Candida species. | Manolakaki D, et al. 2010. Virulence. 1: 367-375 ((https://doi.org/10.4161/viru.1.5.12796)); Mekbib SB, et al. 2011. Tropical Plant Pathology 36:233-240 ((https://doi.org/10.1590/S1982-56762011000400004)); Põlme S, et al. 2020. Fungal Diversity 105:1-16 ((https://doi.org/10.1007/s13225-020-00466-2)) |
| Otu39197 | 1 | k__Fungi(100);p__Ascomycota(100);c__Saccharomycetes(100);o__Saccharomycetales(100);f__Metschnikowiaceae(100);g__Metschnikowia(100);s__Metschnikowia_pulcherrima(100); | Metschnikowia | 13 | Saprotroph | \|Undefined Saprotroph\| | Yeast | NULL | Probable | As nectar-tap_saprotroph (Põlme et al. 2020); Decay sugar-rich substrates (Põlme et al. 2020) | Põlme S, et al. 2020. Fungal Diversity 105:1-16 ((https://doi.org/10.1007/s13225-020-00466-2)) |
| Otu39198 | 1 | k__Fungi(100);p__Ascomycota(100);c__Saccharomycetes(100);o__Saccharomycetales(100);f__Metschnikowiaceae(100);g__Metschnikowia(100);g__Metschnikowia_unclassified(100); | Metschnikowia | 13 | Saprotroph | \|Undefined Saprotroph\| | Yeast | NULL | Probable | As nectar-tap_saprotroph (Põlme et al. 2020); Decay sugar-rich substrates (Põlme et al. 2020) | Põlme S, et al. 2020. Fungal Diversity 105:1-16 ((https://doi.org/10.1007/s13225-020-00466-2)) |
| Otu39199 | 1 | k__Fungi(100);p__Ascomycota(100);c__Saccharomycetes(100);o__Saccharomycetales(100);f__Metschnikowiaceae(100);g__Metschnikowia(100);g__Metschnikowia_unclassified(100); | Metschnikowia | 13 | Saprotroph | \|Undefined Saprotroph\| | Yeast | NULL | Probable | As nectar-tap_saprotroph (Põlme et al. 2020); Decay sugar-rich substrates (Põlme et al. 2020) | Põlme S, et al. 2020. Fungal Diversity 105:1-16 ((https://doi.org/10.1007/s13225-020-00466-2)) |
| Otu39200 | 1 | k__Fungi(100);p__Ascomycota(100);c__Saccharomycetes(100);o__Saccharomycetales(100);f__Metschnikowiaceae(100);g__Metschnikowia(100);g__Metschnikowia_unclassified(100); | Metschnikowia | 13 | Saprotroph | \|Undefined Saprotroph\| | Yeast | NULL | Probable | As nectar-tap_saprotroph (Põlme et al. 2020); Decay sugar-rich substrates (Põlme et al. 2020) | Põlme S, et al. 2020. Fungal Diversity 105:1-16 ((https://doi.org/10.1007/s13225-020-00466-2)) |
| Otu39201 | 1 | k__Fungi(100);p__Ascomycota(100);c__Saccharomycetes(100);o__Saccharomycetales(100);f__Metschnikowiaceae(100);g__Metschnikowia(100);s__Metschnikowia_pulcherrima(100); | Metschnikowia | 13 | Saprotroph | \|Undefined Saprotroph\| | Yeast | NULL | Probable | As nectar-tap_saprotroph (Põlme et al. 2020); Decay sugar-rich substrates (Põlme et al. 2020) | Põlme S, et al. 2020. Fungal Diversity 105:1-16 ((https://doi.org/10.1007/s13225-020-00466-2)) |
| Otu39202 | 1 | k__Fungi(100);p__Ascomycota(100);c__Saccharomycetes(100);o__Saccharomycetales(100);f__Metschnikowiaceae(100);g__Metschnikowia(100);s__Metschnikowia_pulcherrima(100); | Metschnikowia | 13 | Saprotroph | \|Undefined Saprotroph\| | Yeast | NULL | Probable | As nectar-tap_saprotroph (Põlme et al. 2020); Decay sugar-rich substrates (Põlme et al. 2020) | Põlme S, et al. 2020. Fungal Diversity 105:1-16 ((https://doi.org/10.1007/s13225-020-00466-2)) |
| Otu39203 | 1 | k__Fungi(100);p__Ascomycota(100);c__Saccharomycetes(100);o__Saccharomycetales(100);f__Metschnikowiaceae(100);g__Metschnikowia(100);s__Metschnikowia_pulcherrima(100); | Metschnikowia | 13 | Saprotroph | \|Undefined Saprotroph\| | Yeast | NULL | Probable | As nectar-tap_saprotroph (Põlme et al. 2020); Decay sugar-rich substrates (Põlme et al. 2020) | Põlme S, et al. 2020. Fungal Diversity 105:1-16 ((https://doi.org/10.1007/s13225-020-00466-2)) |
| Otu39204 | 1 | k__Fungi(100);p__Ascomycota(100);c__Saccharomycetes(100);o__Saccharomycetales(100);f__Metschnikowiaceae(100);g__Metschnikowia(100);s__Metschnikowia_pulcherrima(100); | Metschnikowia | 13 | Saprotroph | \|Undefined Saprotroph\| | Yeast | NULL | Probable | As nectar-tap_saprotroph (Põlme et al. 2020); Decay sugar-rich substrates (Põlme et al. 2020) | Põlme S, et al. 2020. Fungal Diversity 105:1-16 ((https://doi.org/10.1007/s13225-020-00466-2)) |
| Otu39205 | 1 | k__Fungi(100);p__Ascomycota(100);c__Saccharomycetes(100);o__Saccharomycetales(100);f__Metschnikowiaceae(100);g__Metschnikowia(100);s__Metschnikowia_pulcherrima(100); | Metschnikowia | 13 | Saprotroph | \|Undefined Saprotroph\| | Yeast | NULL | Probable | As nectar-tap_saprotroph (Põlme et al. 2020); Decay sugar-rich substrates (Põlme et al. 2020) | Põlme S, et al. 2020. Fungal Diversity 105:1-16 ((https://doi.org/10.1007/s13225-020-00466-2)) |
| Otu39206 | 1 | k__Fungi(100);p__Ascomycota(100);c__Saccharomycetes(100);o__Saccharomycetales(100);f__Metschnikowiaceae(100);g__Metschnikowia(100);g__Metschnikowia_unclassified(100); | Metschnikowia | 13 | Saprotroph | \|Undefined Saprotroph\| | Yeast | NULL | Probable | As nectar-tap_saprotroph (Põlme et al. 2020); Decay sugar-rich substrates (Põlme et al. 2020) | Põlme S, et al. 2020. Fungal Diversity 105:1-16 ((https://doi.org/10.1007/s13225-020-00466-2)) |
| Otu39207 | 1 | k__Fungi(100);p__Ascomycota(100);c__Saccharomycetes(100);o__Saccharomycetales(100);f__Metschnikowiaceae(100);g__Metschnikowia(100);s__Metschnikowia_pulcherrima(100); | Metschnikowia | 13 | Saprotroph | \|Undefined Saprotroph\| | Yeast | NULL | Probable | As nectar-tap_saprotroph (Põlme et al. 2020); Decay sugar-rich substrates (Põlme et al. 2020) | Põlme S, et al. 2020. Fungal Diversity 105:1-16 ((https://doi.org/10.1007/s13225-020-00466-2)) |
| Otu39208 | 1 | k__Fungi(100);p__Ascomycota(100);c__Saccharomycetes(100);o__Saccharomycetales(100);f__Metschnikowiaceae(100);g__Metschnikowia(100);s__Metschnikowia_pulcherrima(100); | Metschnikowia | 13 | Saprotroph | \|Undefined Saprotroph\| | Yeast | NULL | Probable | As nectar-tap_saprotroph (Põlme et al. 2020); Decay sugar-rich substrates (Põlme et al. 2020) | Põlme S, et al. 2020. Fungal Diversity 105:1-16 ((https://doi.org/10.1007/s13225-020-00466-2)) |
| Otu39209 | 1 | k__Fungi(100);p__Ascomycota(100);c__Saccharomycetes(100);o__Saccharomycetales(100);f__Metschnikowiaceae(100);g__Metschnikowia(100);g__Metschnikowia_unclassified(100); | Metschnikowia | 13 | Saprotroph | \|Undefined Saprotroph\| | Yeast | NULL | Probable | As nectar-tap_saprotroph (Põlme et al. 2020); Decay sugar-rich substrates (Põlme et al. 2020) | Põlme S, et al. 2020. Fungal Diversity 105:1-16 ((https://doi.org/10.1007/s13225-020-00466-2)) |
| Otu39210 | 1 | k__Fungi(100);p__Ascomycota(100);c__Saccharomycetes(100);o__Saccharomycetales(100);f__Metschnikowiaceae(100);g__Metschnikowia(100);s__Metschnikowia_pulcherrima(100); | Metschnikowia | 13 | Saprotroph | \|Undefined Saprotroph\| | Yeast | NULL | Probable | As nectar-tap_saprotroph (Põlme et al. 2020); Decay sugar-rich substrates (Põlme et al. 2020) | Põlme S, et al. 2020. Fungal Diversity 105:1-16 ((https://doi.org/10.1007/s13225-020-00466-2)) |
| Otu39211 | 1 | k__Fungi(100);p__Ascomycota(100);c__Saccharomycetes(100);o__Saccharomycetales(100);f__Metschnikowiaceae(100);g__Metschnikowia(100);s__Metschnikowia_pulcherrima(100); | Metschnikowia | 13 | Saprotroph | \|Undefined Saprotroph\| | Yeast | NULL | Probable | As nectar-tap_saprotroph (Põlme et al. 2020); Decay sugar-rich substrates (Põlme et al. 2020) | Põlme S, et al. 2020. Fungal Diversity 105:1-16 ((https://doi.org/10.1007/s13225-020-00466-2)) |
| Otu39212 | 1 | k__Fungi(100);p__Ascomycota(100);c__Saccharomycetes(100);o__Saccharomycetales(100);f__Metschnikowiaceae(100);g__Metschnikowia(100);g__Metschnikowia_unclassified(100); | Metschnikowia | 13 | Saprotroph | \|Undefined Saprotroph\| | Yeast | NULL | Probable | As nectar-tap_saprotroph (Põlme et al. 2020); Decay sugar-rich substrates (Põlme et al. 2020) | Põlme S, et al. 2020. Fungal Diversity 105:1-16 ((https://doi.org/10.1007/s13225-020-00466-2)) |
| Otu39213 | 1 | k__Fungi(100);p__Ascomycota(100);c__Saccharomycetes(100);o__Saccharomycetales(100);f__Metschnikowiaceae(100);g__Metschnikowia(100);g__Metschnikowia_unclassified(100); | Metschnikowia | 13 | Saprotroph | \|Undefined Saprotroph\| | Yeast | NULL | Probable | As nectar-tap_saprotroph (Põlme et al. 2020); Decay sugar-rich substrates (Põlme et al. 2020) | Põlme S, et al. 2020. Fungal Diversity 105:1-16 ((https://doi.org/10.1007/s13225-020-00466-2)) |
| Otu39214 | 1 | k__Fungi(100);p__Ascomycota(100);c__Saccharomycetes(100);o__Saccharomycetales(100);f__Metschnikowiaceae(100);g__Metschnikowia(100);g__Metschnikowia_unclassified(100); | Metschnikowia | 13 | Saprotroph | \|Undefined Saprotroph\| | Yeast | NULL | Probable | As nectar-tap_saprotroph (Põlme et al. 2020); Decay sugar-rich substrates (Põlme et al. 2020) | Põlme S, et al. 2020. Fungal Diversity 105:1-16 ((https://doi.org/10.1007/s13225-020-00466-2)) |
| Otu39215 | 1 | k__Fungi(100);p__Ascomycota(100);c__Saccharomycetes(100);o__Saccharomycetales(100);f__Metschnikowiaceae(100);g__Metschnikowia(100);s__Metschnikowia_pulcherrima(100); | Metschnikowia | 13 | Saprotroph | \|Undefined Saprotroph\| | Yeast | NULL | Probable | As nectar-tap_saprotroph (Põlme et al. 2020); Decay sugar-rich substrates (Põlme et al. 2020) | Põlme S, et al. 2020. Fungal Diversity 105:1-16 ((https://doi.org/10.1007/s13225-020-00466-2)) |
| Otu39216 | 1 | k__Fungi(100);p__Ascomycota(100);c__Saccharomycetes(100);o__Saccharomycetales(100);f__Metschnikowiaceae(100);g__Metschnikowia(100);s__Metschnikowia_pulcherrima(100); | Metschnikowia | 13 | Saprotroph | \|Undefined Saprotroph\| | Yeast | NULL | Probable | As nectar-tap_saprotroph (Põlme et al. 2020); Decay sugar-rich substrates (Põlme et al. 2020) | Põlme S, et al. 2020. Fungal Diversity 105:1-16 ((https://doi.org/10.1007/s13225-020-00466-2)) |
| Otu39217 | 1 | k__Fungi(100);p__Ascomycota(100);c__Saccharomycetes(100);o__Saccharomycetales(100);f__Metschnikowiaceae(100);g__Metschnikowia(100);s__Metschnikowia_pulcherrima(100); | Metschnikowia | 13 | Saprotroph | \|Undefined Saprotroph\| | Yeast | NULL | Probable | As nectar-tap_saprotroph (Põlme et al. 2020); Decay sugar-rich substrates (Põlme et al. 2020) | Põlme S, et al. 2020. Fungal Diversity 105:1-16 ((https://doi.org/10.1007/s13225-020-00466-2)) |
| Otu39218 | 1 | k__Fungi(100);p__Ascomycota(100);c__Saccharomycetes(100);o__Saccharomycetales(100);f__Metschnikowiaceae(100);g__Metschnikowia(100);g__Metschnikowia_unclassified(100); | Metschnikowia | 13 | Saprotroph | \|Undefined Saprotroph\| | Yeast | NULL | Probable | As nectar-tap_saprotroph (Põlme et al. 2020); Decay sugar-rich substrates (Põlme et al. 2020) | Põlme S, et al. 2020. Fungal Diversity 105:1-16 ((https://doi.org/10.1007/s13225-020-00466-2)) |
| Otu39219 | 1 | k__Fungi(100);p__Ascomycota(100);c__Saccharomycetes(100);o__Saccharomycetales(100);f__Metschnikowiaceae(100);g__Metschnikowia(100);g__Metschnikowia_unclassified(100); | Metschnikowia | 13 | Saprotroph | \|Undefined Saprotroph\| | Yeast | NULL | Probable | As nectar-tap_saprotroph (Põlme et al. 2020); Decay sugar-rich substrates (Põlme et al. 2020) | Põlme S, et al. 2020. Fungal Diversity 105:1-16 ((https://doi.org/10.1007/s13225-020-00466-2)) |
| Otu39220 | 1 | k__Fungi(100);p__Ascomycota(100);c__Saccharomycetes(100);o__Saccharomycetales(100);f__Metschnikowiaceae(100);g__Metschnikowia(100);g__Metschnikowia_unclassified(100); | Metschnikowia | 13 | Saprotroph | \|Undefined Saprotroph\| | Yeast | NULL | Probable | As nectar-tap_saprotroph (Põlme et al. 2020); Decay sugar-rich substrates (Põlme et al. 2020) | Põlme S, et al. 2020. Fungal Diversity 105:1-16 ((https://doi.org/10.1007/s13225-020-00466-2)) |
| Otu39221 | 1 | k__Fungi(100);p__Ascomycota(100);c__Saccharomycetes(100);o__Saccharomycetales(100);f__Metschnikowiaceae(100);g__Metschnikowia(100);g__Metschnikowia_unclassified(100); | Metschnikowia | 13 | Saprotroph | \|Undefined Saprotroph\| | Yeast | NULL | Probable | As nectar-tap_saprotroph (Põlme et al. 2020); Decay sugar-rich substrates (Põlme et al. 2020) | Põlme S, et al. 2020. Fungal Diversity 105:1-16 ((https://doi.org/10.1007/s13225-020-00466-2)) |
| Otu39222 | 1 | k__Fungi(100);p__Ascomycota(100);c__Saccharomycetes(100);o__Saccharomycetales(100);f__Metschnikowiaceae(100);g__Metschnikowia(100);s__Metschnikowia_pulcherrima(100); | Metschnikowia | 13 | Saprotroph | \|Undefined Saprotroph\| | Yeast | NULL | Probable | As nectar-tap_saprotroph (Põlme et al. 2020); Decay sugar-rich substrates (Põlme et al. 2020) | Põlme S, et al. 2020. Fungal Diversity 105:1-16 ((https://doi.org/10.1007/s13225-020-00466-2)) |
| Otu39223 | 1 | k__Fungi(100);p__Ascomycota(100);c__Saccharomycetes(100);o__Saccharomycetales(100);f__Metschnikowiaceae(100);g__Metschnikowia(100);s__Metschnikowia_pulcherrima(100); | Metschnikowia | 13 | Saprotroph | \|Undefined Saprotroph\| | Yeast | NULL | Probable | As nectar-tap_saprotroph (Põlme et al. 2020); Decay sugar-rich substrates (Põlme et al. 2020) | Põlme S, et al. 2020. Fungal Diversity 105:1-16 ((https://doi.org/10.1007/s13225-020-00466-2)) |
| Otu39224 | 1 | k__Fungi(100);p__Ascomycota(100);c__Saccharomycetes(100);o__Saccharomycetales(100);f__Metschnikowiaceae(100);g__Metschnikowia(100);s__Metschnikowia_pulcherrima(100); | Metschnikowia | 13 | Saprotroph | \|Undefined Saprotroph\| | Yeast | NULL | Probable | As nectar-tap_saprotroph (Põlme et al. 2020); Decay sugar-rich substrates (Põlme et al. 2020) | Põlme S, et al. 2020. Fungal Diversity 105:1-16 ((https://doi.org/10.1007/s13225-020-00466-2)) |
| Otu39225 | 1 | k__Fungi(100);p__Ascomycota(100);c__Saccharomycetes(100);o__Saccharomycetales(100);f__Metschnikowiaceae(100);g__Metschnikowia(100);s__Metschnikowia_pulcherrima(100); | Metschnikowia | 13 | Saprotroph | \|Undefined Saprotroph\| | Yeast | NULL | Probable | As nectar-tap_saprotroph (Põlme et al. 2020); Decay sugar-rich substrates (Põlme et al. 2020) | Põlme S, et al. 2020. Fungal Diversity 105:1-16 ((https://doi.org/10.1007/s13225-020-00466-2)) |
| Otu39226 | 1 | k__Fungi(100);p__Ascomycota(100);c__Saccharomycetes(100);o__Saccharomycetales(100);f__Metschnikowiaceae(100);g__Metschnikowia(100);s__Metschnikowia_pulcherrima(100); | Metschnikowia | 13 | Saprotroph | \|Undefined Saprotroph\| | Yeast | NULL | Probable | As nectar-tap_saprotroph (Põlme et al. 2020); Decay sugar-rich substrates (Põlme et al. 2020) | Põlme S, et al. 2020. Fungal Diversity 105:1-16 ((https://doi.org/10.1007/s13225-020-00466-2)) |
| Otu39227 | 1 | k__Fungi(100);p__Ascomycota(100);c__Saccharomycetes(100);o__Saccharomycetales(100);f__Metschnikowiaceae(100);g__Metschnikowia(100);s__Metschnikowia_pulcherrima(100); | Metschnikowia | 13 | Saprotroph | \|Undefined Saprotroph\| | Yeast | NULL | Probable | As nectar-tap_saprotroph (Põlme et al. 2020); Decay sugar-rich substrates (Põlme et al. 2020) | Põlme S, et al. 2020. Fungal Diversity 105:1-16 ((https://doi.org/10.1007/s13225-020-00466-2)) |
| Otu39228 | 1 | k__Fungi(100);p__Ascomycota(100);c__Saccharomycetes(100);o__Saccharomycetales(100);f__Metschnikowiaceae(100);g__Metschnikowia(100);s__Metschnikowia_pulcherrima(100); | Metschnikowia | 13 | Saprotroph | \|Undefined Saprotroph\| | Yeast | NULL | Probable | As nectar-tap_saprotroph (Põlme et al. 2020); Decay sugar-rich substrates (Põlme et al. 2020) | Põlme S, et al. 2020. Fungal Diversity 105:1-16 ((https://doi.org/10.1007/s13225-020-00466-2)) |
| Otu39229 | 1 | k__Fungi(100);p__Ascomycota(100);c__Saccharomycetes(100);o__Saccharomycetales(100);f__Metschnikowiaceae(100);g__Metschnikowia(100);s__Metschnikowia_pulcherrima(100); | Metschnikowia | 13 | Saprotroph | \|Undefined Saprotroph\| | Yeast | NULL | Probable | As nectar-tap_saprotroph (Põlme et al. 2020); Decay sugar-rich substrates (Põlme et al. 2020) | Põlme S, et al. 2020. Fungal Diversity 105:1-16 ((https://doi.org/10.1007/s13225-020-00466-2)) |
| Otu39230 | 1 | k__Fungi(100);p__Ascomycota(100);c__Saccharomycetes(100);o__Saccharomycetales(100);f__Metschnikowiaceae(100);g__Metschnikowia(100);s__Metschnikowia_pulcherrima(100); | Metschnikowia | 13 | Saprotroph | \|Undefined Saprotroph\| | Yeast | NULL | Probable | As nectar-tap_saprotroph (Põlme et al. 2020); Decay sugar-rich substrates (Põlme et al. 2020) | Põlme S, et al. 2020. Fungal Diversity 105:1-16 ((https://doi.org/10.1007/s13225-020-00466-2)) |
| Otu39231 | 1 | k__Fungi(100);p__Ascomycota(100);c__Saccharomycetes(100);o__Saccharomycetales(100);f__Metschnikowiaceae(100);g__Metschnikowia(100);s__Metschnikowia_pulcherrima(100); | Metschnikowia | 13 | Saprotroph | \|Undefined Saprotroph\| | Yeast | NULL | Probable | As nectar-tap_saprotroph (Põlme et al. 2020); Decay sugar-rich substrates (Põlme et al. 2020) | Põlme S, et al. 2020. Fungal Diversity 105:1-16 ((https://doi.org/10.1007/s13225-020-00466-2)) |
| Otu39232 | 1 | k__Fungi(100);p__Ascomycota(100);c__Saccharomycetes(100);o__Saccharomycetales(100);f__Metschnikowiaceae(100);g__Metschnikowia(100);g__Metschnikowia_unclassified(100); | Metschnikowia | 13 | Saprotroph | \|Undefined Saprotroph\| | Yeast | NULL | Probable | As nectar-tap_saprotroph (Põlme et al. 2020); Decay sugar-rich substrates (Põlme et al. 2020) | Põlme S, et al. 2020. Fungal Diversity 105:1-16 ((https://doi.org/10.1007/s13225-020-00466-2)) |
| Otu39233 | 1 | k__Fungi(100);p__Ascomycota(100);c__Saccharomycetes(100);o__Saccharomycetales(100);f__Metschnikowiaceae(100);g__Metschnikowia(100);s__Metschnikowia_pulcherrima(100); | Metschnikowia | 13 | Saprotroph | \|Undefined Saprotroph\| | Yeast | NULL | Probable | As nectar-tap_saprotroph (Põlme et al. 2020); Decay sugar-rich substrates (Põlme et al. 2020) | Põlme S, et al. 2020. Fungal Diversity 105:1-16 ((https://doi.org/10.1007/s13225-020-00466-2)) |
| Otu39234 | 1 | k__Fungi(100);p__Ascomycota(100);c__Saccharomycetes(100);o__Saccharomycetales(100);f__Metschnikowiaceae(100);g__Metschnikowia(100);s__Metschnikowia_pulcherrima(100); | Metschnikowia | 13 | Saprotroph | \|Undefined Saprotroph\| | Yeast | NULL | Probable | As nectar-tap_saprotroph (Põlme et al. 2020); Decay sugar-rich substrates (Põlme et al. 2020) | Põlme S, et al. 2020. Fungal Diversity 105:1-16 ((https://doi.org/10.1007/s13225-020-00466-2)) |
| Otu39235 | 1 | k__Fungi(100);p__Ascomycota(100);c__Saccharomycetes(100);o__Saccharomycetales(100);f__Metschnikowiaceae(100);g__Metschnikowia(100);s__Metschnikowia_pulcherrima(100); | Metschnikowia | 13 | Saprotroph | \|Undefined Saprotroph\| | Yeast | NULL | Probable | As nectar-tap_saprotroph (Põlme et al. 2020); Decay sugar-rich substrates (Põlme et al. 2020) | Põlme S, et al. 2020. Fungal Diversity 105:1-16 ((https://doi.org/10.1007/s13225-020-00466-2)) |
| Otu39236 | 1 | k__Fungi(100);p__Ascomycota(100);c__Saccharomycetes(100);o__Saccharomycetales(100);f__Metschnikowiaceae(100);g__Metschnikowia(100);s__Metschnikowia_pulcherrima(100); | Metschnikowia | 13 | Saprotroph | \|Undefined Saprotroph\| | Yeast | NULL | Probable | As nectar-tap_saprotroph (Põlme et al. 2020); Decay sugar-rich substrates (Põlme et al. 2020) | Põlme S, et al. 2020. Fungal Diversity 105:1-16 ((https://doi.org/10.1007/s13225-020-00466-2)) |
| Otu39237 | 1 | k__Fungi(100);p__Ascomycota(100);c__Saccharomycetes(100);o__Saccharomycetales(100);f__Metschnikowiaceae(100);g__Metschnikowia(100);s__Metschnikowia_pulcherrima(100); | Metschnikowia | 13 | Saprotroph | \|Undefined Saprotroph\| | Yeast | NULL | Probable | As nectar-tap_saprotroph (Põlme et al. 2020); Decay sugar-rich substrates (Põlme et al. 2020) | Põlme S, et al. 2020. Fungal Diversity 105:1-16 ((https://doi.org/10.1007/s13225-020-00466-2)) |
| Otu39238 | 1 | k__Fungi(100);p__Ascomycota(100);c__Saccharomycetes(100);o__Saccharomycetales(100);f__Metschnikowiaceae(100);g__Metschnikowia(100);s__Metschnikowia_pulcherrima(100); | Metschnikowia | 13 | Saprotroph | \|Undefined Saprotroph\| | Yeast | NULL | Probable | As nectar-tap_saprotroph (Põlme et al. 2020); Decay sugar-rich substrates (Põlme et al. 2020) | Põlme S, et al. 2020. Fungal Diversity 105:1-16 ((https://doi.org/10.1007/s13225-020-00466-2)) |
| Otu39239 | 1 | k__Fungi(100);p__Ascomycota(100);c__Saccharomycetes(100);o__Saccharomycetales(100);f__Metschnikowiaceae(100);g__Metschnikowia(100);g__Metschnikowia_unclassified(100); | Metschnikowia | 13 | Saprotroph | \|Undefined Saprotroph\| | Yeast | NULL | Probable | As nectar-tap_saprotroph (Põlme et al. 2020); Decay sugar-rich substrates (Põlme et al. 2020) | Põlme S, et al. 2020. Fungal Diversity 105:1-16 ((https://doi.org/10.1007/s13225-020-00466-2)) |
| Otu39240 | 1 | k__Fungi(100);p__Ascomycota(100);c__Saccharomycetes(100);o__Saccharomycetales(100);f__Metschnikowiaceae(100);g__Metschnikowia(100);s__Metschnikowia_pulcherrima(100); | Metschnikowia | 13 | Saprotroph | \|Undefined Saprotroph\| | Yeast | NULL | Probable | As nectar-tap_saprotroph (Põlme et al. 2020); Decay sugar-rich substrates (Põlme et al. 2020) | Põlme S, et al. 2020. Fungal Diversity 105:1-16 ((https://doi.org/10.1007/s13225-020-00466-2)) |
| Otu39241 | 1 | k__Fungi(100);p__Ascomycota(100);c__Saccharomycetes(100);o__Saccharomycetales(100);f__Metschnikowiaceae(100);g__Metschnikowia(100);g__Metschnikowia_unclassified(100); | Metschnikowia | 13 | Saprotroph | \|Undefined Saprotroph\| | Yeast | NULL | Probable | As nectar-tap_saprotroph (Põlme et al. 2020); Decay sugar-rich substrates (Põlme et al. 2020) | Põlme S, et al. 2020. Fungal Diversity 105:1-16 ((https://doi.org/10.1007/s13225-020-00466-2)) |
| Otu39242 | 1 | k__Fungi(100);p__Ascomycota(100);c__Saccharomycetes(100);o__Saccharomycetales(100);f__Metschnikowiaceae(100);g__Metschnikowia(100);s__Metschnikowia_pulcherrima(100); | Metschnikowia | 13 | Saprotroph | \|Undefined Saprotroph\| | Yeast | NULL | Probable | As nectar-tap_saprotroph (Põlme et al. 2020); Decay sugar-rich substrates (Põlme et al. 2020) | Põlme S, et al. 2020. Fungal Diversity 105:1-16 ((https://doi.org/10.1007/s13225-020-00466-2)) |
| Otu39243 | 1 | k__Fungi(100);p__Ascomycota(100);c__Saccharomycetes(100);o__Saccharomycetales(100);f__Metschnikowiaceae(100);g__Metschnikowia(100);s__Metschnikowia_pulcherrima(100); | Metschnikowia | 13 | Saprotroph | \|Undefined Saprotroph\| | Yeast | NULL | Probable | As nectar-tap_saprotroph (Põlme et al. 2020); Decay sugar-rich substrates (Põlme et al. 2020) | Põlme S, et al. 2020. Fungal Diversity 105:1-16 ((https://doi.org/10.1007/s13225-020-00466-2)) |
| Otu39245 | 1 | k__Fungi(100);p__Ascomycota(100);c__Saccharomycetes(100);o__Saccharomycetales(100);f__Metschnikowiaceae(100);g__Metschnikowia(100);s__Metschnikowia_pulcherrima(100); | Metschnikowia | 13 | Saprotroph | \|Undefined Saprotroph\| | Yeast | NULL | Probable | As nectar-tap_saprotroph (Põlme et al. 2020); Decay sugar-rich substrates (Põlme et al. 2020) | Põlme S, et al. 2020. Fungal Diversity 105:1-16 ((https://doi.org/10.1007/s13225-020-00466-2)) |
| Otu39246 | 1 | k__Fungi(100);p__Ascomycota(100);c__Saccharomycetes(100);o__Saccharomycetales(100);f__Metschnikowiaceae(100);g__Metschnikowia(100);s__Metschnikowia_pulcherrima(100); | Metschnikowia | 13 | Saprotroph | \|Undefined Saprotroph\| | Yeast | NULL | Probable | As nectar-tap_saprotroph (Põlme et al. 2020); Decay sugar-rich substrates (Põlme et al. 2020) | Põlme S, et al. 2020. Fungal Diversity 105:1-16 ((https://doi.org/10.1007/s13225-020-00466-2)) |
| Otu39247 | 1 | k__Fungi(100);p__Ascomycota(100);c__Saccharomycetes(100);o__Saccharomycetales(100);f__Metschnikowiaceae(100);g__Metschnikowia(100);s__Metschnikowia_pulcherrima(100); | Metschnikowia | 13 | Saprotroph | \|Undefined Saprotroph\| | Yeast | NULL | Probable | As nectar-tap_saprotroph (Põlme et al. 2020); Decay sugar-rich substrates (Põlme et al. 2020) | Põlme S, et al. 2020. Fungal Diversity 105:1-16 ((https://doi.org/10.1007/s13225-020-00466-2)) |
| Otu39248 | 1 | k__Fungi(100);p__Ascomycota(100);c__Saccharomycetes(100);o__Saccharomycetales(100);f__Metschnikowiaceae(100);g__Metschnikowia(100);s__Metschnikowia_pulcherrima(100); | Metschnikowia | 13 | Saprotroph | \|Undefined Saprotroph\| | Yeast | NULL | Probable | As nectar-tap_saprotroph (Põlme et al. 2020); Decay sugar-rich substrates (Põlme et al. 2020) | Põlme S, et al. 2020. Fungal Diversity 105:1-16 ((https://doi.org/10.1007/s13225-020-00466-2)) |
| Otu39249 | 1 | k__Fungi(100);p__Ascomycota(100);c__Saccharomycetes(100);o__Saccharomycetales(100);f__Metschnikowiaceae(100);g__Metschnikowia(100);g__Metschnikowia_unclassified(100); | Metschnikowia | 13 | Saprotroph | \|Undefined Saprotroph\| | Yeast | NULL | Probable | As nectar-tap_saprotroph (Põlme et al. 2020); Decay sugar-rich substrates (Põlme et al. 2020) | Põlme S, et al. 2020. Fungal Diversity 105:1-16 ((https://doi.org/10.1007/s13225-020-00466-2)) |
| Otu39250 | 1 | k__Fungi(100);p__Ascomycota(100);c__Saccharomycetes(100);o__Saccharomycetales(100);f__Metschnikowiaceae(100);g__Metschnikowia(100);g__Metschnikowia_unclassified(100); | Metschnikowia | 13 | Saprotroph | \|Undefined Saprotroph\| | Yeast | NULL | Probable | As nectar-tap_saprotroph (Põlme et al. 2020); Decay sugar-rich substrates (Põlme et al. 2020) | Põlme S, et al. 2020. Fungal Diversity 105:1-16 ((https://doi.org/10.1007/s13225-020-00466-2)) |
| Otu39251 | 1 | k__Fungi(100);p__Ascomycota(100);c__Saccharomycetes(100);o__Saccharomycetales(100);f__Metschnikowiaceae(100);g__Metschnikowia(100);s__Metschnikowia_pulcherrima(100); | Metschnikowia | 13 | Saprotroph | \|Undefined Saprotroph\| | Yeast | NULL | Probable | As nectar-tap_saprotroph (Põlme et al. 2020); Decay sugar-rich substrates (Põlme et al. 2020) | Põlme S, et al. 2020. Fungal Diversity 105:1-16 ((https://doi.org/10.1007/s13225-020-00466-2)) |
| Otu39252 | 1 | k__Fungi(100);p__Ascomycota(100);c__Saccharomycetes(100);o__Saccharomycetales(100);f__Metschnikowiaceae(100);g__Metschnikowia(100);s__Metschnikowia_pulcherrima(100); | Metschnikowia | 13 | Saprotroph | \|Undefined Saprotroph\| | Yeast | NULL | Probable | As nectar-tap_saprotroph (Põlme et al. 2020); Decay sugar-rich substrates (Põlme et al. 2020) | Põlme S, et al. 2020. Fungal Diversity 105:1-16 ((https://doi.org/10.1007/s13225-020-00466-2)) |
| Otu39253 | 1 | k__Fungi(100);p__Ascomycota(100);c__Saccharomycetes(100);o__Saccharomycetales(100);f__Metschnikowiaceae(100);g__Metschnikowia(100);s__Metschnikowia_pulcherrima(100); | Metschnikowia | 13 | Saprotroph | \|Undefined Saprotroph\| | Yeast | NULL | Probable | As nectar-tap_saprotroph (Põlme et al. 2020); Decay sugar-rich substrates (Põlme et al. 2020) | Põlme S, et al. 2020. Fungal Diversity 105:1-16 ((https://doi.org/10.1007/s13225-020-00466-2)) |
| Otu39254 | 1 | k__Fungi(100);p__Ascomycota(100);c__Saccharomycetes(100);o__Saccharomycetales(100);f__Metschnikowiaceae(100);g__Metschnikowia(100);s__Metschnikowia_pulcherrima(100); | Metschnikowia | 13 | Saprotroph | \|Undefined Saprotroph\| | Yeast | NULL | Probable | As nectar-tap_saprotroph (Põlme et al. 2020); Decay sugar-rich substrates (Põlme et al. 2020) | Põlme S, et al. 2020. Fungal Diversity 105:1-16 ((https://doi.org/10.1007/s13225-020-00466-2)) |
| Otu39256 | 1 | k__Fungi(100);p__Ascomycota(100);c__Saccharomycetes(100);o__Saccharomycetales(100);f__Metschnikowiaceae(100);g__Metschnikowia(100);s__Metschnikowia_pulcherrima(100); | Metschnikowia | 13 | Saprotroph | \|Undefined Saprotroph\| | Yeast | NULL | Probable | As nectar-tap_saprotroph (Põlme et al. 2020); Decay sugar-rich substrates (Põlme et al. 2020) | Põlme S, et al. 2020. Fungal Diversity 105:1-16 ((https://doi.org/10.1007/s13225-020-00466-2)) |
| Otu39257 | 1 | k__Fungi(100);p__Ascomycota(100);c__Saccharomycetes(100);o__Saccharomycetales(100);f__Metschnikowiaceae(100);g__Metschnikowia(100);s__Metschnikowia_pulcherrima(100); | Metschnikowia | 13 | Saprotroph | \|Undefined Saprotroph\| | Yeast | NULL | Probable | As nectar-tap_saprotroph (Põlme et al. 2020); Decay sugar-rich substrates (Põlme et al. 2020) | Põlme S, et al. 2020. Fungal Diversity 105:1-16 ((https://doi.org/10.1007/s13225-020-00466-2)) |
| Otu39259 | 1 | k__Fungi(100);p__Ascomycota(100);c__Saccharomycetes(100);o__Saccharomycetales(100);f__Metschnikowiaceae(100);g__Metschnikowia(100);s__Metschnikowia_pulcherrima(100); | Metschnikowia | 13 | Saprotroph | \|Undefined Saprotroph\| | Yeast | NULL | Probable | As nectar-tap_saprotroph (Põlme et al. 2020); Decay sugar-rich substrates (Põlme et al. 2020) | Põlme S, et al. 2020. Fungal Diversity 105:1-16 ((https://doi.org/10.1007/s13225-020-00466-2)) |
| Otu39260 | 1 | k__Fungi(100);p__Ascomycota(100);c__Saccharomycetes(100);o__Saccharomycetales(100);f__Metschnikowiaceae(100);g__Metschnikowia(100);g__Metschnikowia_unclassified(100); | Metschnikowia | 13 | Saprotroph | \|Undefined Saprotroph\| | Yeast | NULL | Probable | As nectar-tap_saprotroph (Põlme et al. 2020); Decay sugar-rich substrates (Põlme et al. 2020) | Põlme S, et al. 2020. Fungal Diversity 105:1-16 ((https://doi.org/10.1007/s13225-020-00466-2)) |
| Otu39261 | 1 | k__Fungi(100);p__Ascomycota(100);c__Saccharomycetes(100);o__Saccharomycetales(100);f__Metschnikowiaceae(100);g__Metschnikowia(100);s__Metschnikowia_pulcherrima(100); | Metschnikowia | 13 | Saprotroph | \|Undefined Saprotroph\| | Yeast | NULL | Probable | As nectar-tap_saprotroph (Põlme et al. 2020); Decay sugar-rich substrates (Põlme et al. 2020) | Põlme S, et al. 2020. Fungal Diversity 105:1-16 ((https://doi.org/10.1007/s13225-020-00466-2)) |
| Otu39262 | 1 | k__Fungi(100);p__Ascomycota(100);c__Saccharomycetes(100);o__Saccharomycetales(100);f__Metschnikowiaceae(100);g__Metschnikowia(100);s__Metschnikowia_pulcherrima(100); | Metschnikowia | 13 | Saprotroph | \|Undefined Saprotroph\| | Yeast | NULL | Probable | As nectar-tap_saprotroph (Põlme et al. 2020); Decay sugar-rich substrates (Põlme et al. 2020) | Põlme S, et al. 2020. Fungal Diversity 105:1-16 ((https://doi.org/10.1007/s13225-020-00466-2)) |
| Otu39263 | 1 | k__Fungi(100);p__Ascomycota(100);c__Saccharomycetes(100);o__Saccharomycetales(100);f__Metschnikowiaceae(100);g__Metschnikowia(100);s__Metschnikowia_pulcherrima(100); | Metschnikowia | 13 | Saprotroph | \|Undefined Saprotroph\| | Yeast | NULL | Probable | As nectar-tap_saprotroph (Põlme et al. 2020); Decay sugar-rich substrates (Põlme et al. 2020) | Põlme S, et al. 2020. Fungal Diversity 105:1-16 ((https://doi.org/10.1007/s13225-020-00466-2)) |
| Otu39264 | 1 | k__Fungi(100);p__Ascomycota(100);c__Saccharomycetes(100);o__Saccharomycetales(100);f__Metschnikowiaceae(100);g__Metschnikowia(100);s__Metschnikowia_pulcherrima(100); | Metschnikowia | 13 | Saprotroph | \|Undefined Saprotroph\| | Yeast | NULL | Probable | As nectar-tap_saprotroph (Põlme et al. 2020); Decay sugar-rich substrates (Põlme et al. 2020) | Põlme S, et al. 2020. Fungal Diversity 105:1-16 ((https://doi.org/10.1007/s13225-020-00466-2)) |
| Otu39265 | 1 | k__Fungi(100);p__Ascomycota(100);c__Saccharomycetes(100);o__Saccharomycetales(100);f__Metschnikowiaceae(100);g__Metschnikowia(100);s__Metschnikowia_pulcherrima(100); | Metschnikowia | 13 | Saprotroph | \|Undefined Saprotroph\| | Yeast | NULL | Probable | As nectar-tap_saprotroph (Põlme et al. 2020); Decay sugar-rich substrates (Põlme et al. 2020) | Põlme S, et al. 2020. Fungal Diversity 105:1-16 ((https://doi.org/10.1007/s13225-020-00466-2)) |
| Otu39266 | 1 | k__Fungi(100);p__Ascomycota(100);c__Saccharomycetes(100);o__Saccharomycetales(100);f__Metschnikowiaceae(100);g__Metschnikowia(100);s__Metschnikowia_pulcherrima(100); | Metschnikowia | 13 | Saprotroph | \|Undefined Saprotroph\| | Yeast | NULL | Probable | As nectar-tap_saprotroph (Põlme et al. 2020); Decay sugar-rich substrates (Põlme et al. 2020) | Põlme S, et al. 2020. Fungal Diversity 105:1-16 ((https://doi.org/10.1007/s13225-020-00466-2)) |
| Otu39267 | 1 | k__Fungi(100);p__Ascomycota(100);c__Saccharomycetes(100);o__Saccharomycetales(100);f__Metschnikowiaceae(100);g__Metschnikowia(100);s__Metschnikowia_pulcherrima(100); | Metschnikowia | 13 | Saprotroph | \|Undefined Saprotroph\| | Yeast | NULL | Probable | As nectar-tap_saprotroph (Põlme et al. 2020); Decay sugar-rich substrates (Põlme et al. 2020) | Põlme S, et al. 2020. Fungal Diversity 105:1-16 ((https://doi.org/10.1007/s13225-020-00466-2)) |
| Otu39268 | 1 | k__Fungi(100);p__Ascomycota(100);c__Saccharomycetes(100);o__Saccharomycetales(100);f__Metschnikowiaceae(100);g__Metschnikowia(100);g__Metschnikowia_unclassified(100); | Metschnikowia | 13 | Saprotroph | \|Undefined Saprotroph\| | Yeast | NULL | Probable | As nectar-tap_saprotroph (Põlme et al. 2020); Decay sugar-rich substrates (Põlme et al. 2020) | Põlme S, et al. 2020. Fungal Diversity 105:1-16 ((https://doi.org/10.1007/s13225-020-00466-2)) |
| Otu39269 | 1 | k__Fungi(100);p__Ascomycota(100);c__Saccharomycetes(100);o__Saccharomycetales(100);f__Metschnikowiaceae(100);g__Metschnikowia(100);s__Metschnikowia_pulcherrima(100); | Metschnikowia | 13 | Saprotroph | \|Undefined Saprotroph\| | Yeast | NULL | Probable | As nectar-tap_saprotroph (Põlme et al. 2020); Decay sugar-rich substrates (Põlme et al. 2020) | Põlme S, et al. 2020. Fungal Diversity 105:1-16 ((https://doi.org/10.1007/s13225-020-00466-2)) |
| Otu39270 | 1 | k__Fungi(100);p__Ascomycota(100);c__Saccharomycetes(100);o__Saccharomycetales(100);f__Metschnikowiaceae(100);g__Metschnikowia(100);g__Metschnikowia_unclassified(100); | Metschnikowia | 13 | Saprotroph | \|Undefined Saprotroph\| | Yeast | NULL | Probable | As nectar-tap_saprotroph (Põlme et al. 2020); Decay sugar-rich substrates (Põlme et al. 2020) | Põlme S, et al. 2020. Fungal Diversity 105:1-16 ((https://doi.org/10.1007/s13225-020-00466-2)) |
| Otu39271 | 1 | k__Fungi(100);p__Ascomycota(100);c__Saccharomycetes(100);o__Saccharomycetales(100);f__Metschnikowiaceae(100);g__Metschnikowia(100);g__Metschnikowia_unclassified(100); | Metschnikowia | 13 | Saprotroph | \|Undefined Saprotroph\| | Yeast | NULL | Probable | As nectar-tap_saprotroph (Põlme et al. 2020); Decay sugar-rich substrates (Põlme et al. 2020) | Põlme S, et al. 2020. Fungal Diversity 105:1-16 ((https://doi.org/10.1007/s13225-020-00466-2)) |
| Otu39272 | 1 | k__Fungi(100);p__Ascomycota(100);c__Saccharomycetes(100);o__Saccharomycetales(100);f__Metschnikowiaceae(100);g__Metschnikowia(100);g__Metschnikowia_unclassified(100); | Metschnikowia | 13 | Saprotroph | \|Undefined Saprotroph\| | Yeast | NULL | Probable | As nectar-tap_saprotroph (Põlme et al. 2020); Decay sugar-rich substrates (Põlme et al. 2020) | Põlme S, et al. 2020. Fungal Diversity 105:1-16 ((https://doi.org/10.1007/s13225-020-00466-2)) |
| Otu39273 | 1 | k__Fungi(100);p__Ascomycota(100);c__Saccharomycetes(100);o__Saccharomycetales(100);f__Metschnikowiaceae(100);g__Metschnikowia(100);s__Metschnikowia_pulcherrima(100); | Metschnikowia | 13 | Saprotroph | \|Undefined Saprotroph\| | Yeast | NULL | Probable | As nectar-tap_saprotroph (Põlme et al. 2020); Decay sugar-rich substrates (Põlme et al. 2020) | Põlme S, et al. 2020. Fungal Diversity 105:1-16 ((https://doi.org/10.1007/s13225-020-00466-2)) |
| Otu39274 | 1 | k__Fungi(100);p__Ascomycota(100);c__Saccharomycetes(100);o__Saccharomycetales(100);f__Metschnikowiaceae(100);g__Metschnikowia(100);s__Metschnikowia_pulcherrima(100); | Metschnikowia | 13 | Saprotroph | \|Undefined Saprotroph\| | Yeast | NULL | Probable | As nectar-tap_saprotroph (Põlme et al. 2020); Decay sugar-rich substrates (Põlme et al. 2020) | Põlme S, et al. 2020. Fungal Diversity 105:1-16 ((https://doi.org/10.1007/s13225-020-00466-2)) |
| Otu39275 | 1 | k__Fungi(100);p__Ascomycota(100);c__Saccharomycetes(100);o__Saccharomycetales(100);f__Metschnikowiaceae(100);g__Metschnikowia(100);s__Metschnikowia_pulcherrima(100); | Metschnikowia | 13 | Saprotroph | \|Undefined Saprotroph\| | Yeast | NULL | Probable | As nectar-tap_saprotroph (Põlme et al. 2020); Decay sugar-rich substrates (Põlme et al. 2020) | Põlme S, et al. 2020. Fungal Diversity 105:1-16 ((https://doi.org/10.1007/s13225-020-00466-2)) |
| Otu39276 | 1 | k__Fungi(100);p__Ascomycota(100);c__Saccharomycetes(100);o__Saccharomycetales(100);f__Metschnikowiaceae(100);g__Metschnikowia(100);s__Metschnikowia_pulcherrima(100); | Metschnikowia | 13 | Saprotroph | \|Undefined Saprotroph\| | Yeast | NULL | Probable | As nectar-tap_saprotroph (Põlme et al. 2020); Decay sugar-rich substrates (Põlme et al. 2020) | Põlme S, et al. 2020. Fungal Diversity 105:1-16 ((https://doi.org/10.1007/s13225-020-00466-2)) |
| Otu39277 | 1 | k__Fungi(100);p__Ascomycota(100);c__Saccharomycetes(100);o__Saccharomycetales(100);f__Metschnikowiaceae(100);g__Metschnikowia(100);g__Metschnikowia_unclassified(100); | Metschnikowia | 13 | Saprotroph | \|Undefined Saprotroph\| | Yeast | NULL | Probable | As nectar-tap_saprotroph (Põlme et al. 2020); Decay sugar-rich substrates (Põlme et al. 2020) | Põlme S, et al. 2020. Fungal Diversity 105:1-16 ((https://doi.org/10.1007/s13225-020-00466-2)) |
| Otu39278 | 1 | k__Fungi(100);p__Ascomycota(100);c__Saccharomycetes(100);o__Saccharomycetales(100);f__Metschnikowiaceae(100);g__Metschnikowia(100);s__Metschnikowia_pulcherrima(100); | Metschnikowia | 13 | Saprotroph | \|Undefined Saprotroph\| | Yeast | NULL | Probable | As nectar-tap_saprotroph (Põlme et al. 2020); Decay sugar-rich substrates (Põlme et al. 2020) | Põlme S, et al. 2020. Fungal Diversity 105:1-16 ((https://doi.org/10.1007/s13225-020-00466-2)) |
| Otu39279 | 1 | k__Fungi(100);p__Ascomycota(100);c__Saccharomycetes(100);o__Saccharomycetales(100);f__Metschnikowiaceae(100);g__Metschnikowia(100);s__Metschnikowia_pulcherrima(100); | Metschnikowia | 13 | Saprotroph | \|Undefined Saprotroph\| | Yeast | NULL | Probable | As nectar-tap_saprotroph (Põlme et al. 2020); Decay sugar-rich substrates (Põlme et al. 2020) | Põlme S, et al. 2020. Fungal Diversity 105:1-16 ((https://doi.org/10.1007/s13225-020-00466-2)) |
| Otu39280 | 1 | k__Fungi(100);p__Ascomycota(100);c__Saccharomycetes(100);o__Saccharomycetales(100);f__Metschnikowiaceae(100);g__Metschnikowia(100);g__Metschnikowia_unclassified(100); | Metschnikowia | 13 | Saprotroph | \|Undefined Saprotroph\| | Yeast | NULL | Probable | As nectar-tap_saprotroph (Põlme et al. 2020); Decay sugar-rich substrates (Põlme et al. 2020) | Põlme S, et al. 2020. Fungal Diversity 105:1-16 ((https://doi.org/10.1007/s13225-020-00466-2)) |
| Otu39281 | 1 | k__Fungi(100);p__Ascomycota(100);c__Saccharomycetes(100);o__Saccharomycetales(100);f__Metschnikowiaceae(100);g__Metschnikowia(100);s__Metschnikowia_pulcherrima(100); | Metschnikowia | 13 | Saprotroph | \|Undefined Saprotroph\| | Yeast | NULL | Probable | As nectar-tap_saprotroph (Põlme et al. 2020); Decay sugar-rich substrates (Põlme et al. 2020) | Põlme S, et al. 2020. Fungal Diversity 105:1-16 ((https://doi.org/10.1007/s13225-020-00466-2)) |
| Otu39282 | 1 | k__Fungi(100);p__Ascomycota(100);c__Saccharomycetes(100);o__Saccharomycetales(100);f__Metschnikowiaceae(100);g__Metschnikowia(100);s__Metschnikowia_pulcherrima(100); | Metschnikowia | 13 | Saprotroph | \|Undefined Saprotroph\| | Yeast | NULL | Probable | As nectar-tap_saprotroph (Põlme et al. 2020); Decay sugar-rich substrates (Põlme et al. 2020) | Põlme S, et al. 2020. Fungal Diversity 105:1-16 ((https://doi.org/10.1007/s13225-020-00466-2)) |
| Otu39283 | 1 | k__Fungi(100);p__Ascomycota(100);c__Saccharomycetes(100);o__Saccharomycetales(100);f__Metschnikowiaceae(100);g__Metschnikowia(100);s__Metschnikowia_pulcherrima(100); | Metschnikowia | 13 | Saprotroph | \|Undefined Saprotroph\| | Yeast | NULL | Probable | As nectar-tap_saprotroph (Põlme et al. 2020); Decay sugar-rich substrates (Põlme et al. 2020) | Põlme S, et al. 2020. Fungal Diversity 105:1-16 ((https://doi.org/10.1007/s13225-020-00466-2)) |
| Otu39284 | 1 | k__Fungi(100);p__Ascomycota(100);c__Saccharomycetes(100);o__Saccharomycetales(100);f__Metschnikowiaceae(100);g__Metschnikowia(100);s__Metschnikowia_pulcherrima(100); | Metschnikowia | 13 | Saprotroph | \|Undefined Saprotroph\| | Yeast | NULL | Probable | As nectar-tap_saprotroph (Põlme et al. 2020); Decay sugar-rich substrates (Põlme et al. 2020) | Põlme S, et al. 2020. Fungal Diversity 105:1-16 ((https://doi.org/10.1007/s13225-020-00466-2)) |
| Otu39285 | 1 | k__Fungi(100);p__Ascomycota(100);c__Saccharomycetes(100);o__Saccharomycetales(100);f__Metschnikowiaceae(100);g__Metschnikowia(100);s__Metschnikowia_pulcherrima(100); | Metschnikowia | 13 | Saprotroph | \|Undefined Saprotroph\| | Yeast | NULL | Probable | As nectar-tap_saprotroph (Põlme et al. 2020); Decay sugar-rich substrates (Põlme et al. 2020) | Põlme S, et al. 2020. Fungal Diversity 105:1-16 ((https://doi.org/10.1007/s13225-020-00466-2)) |
| Otu39286 | 1 | k__Fungi(100);p__Ascomycota(100);c__Saccharomycetes(100);o__Saccharomycetales(100);f__Metschnikowiaceae(100);g__Metschnikowia(100);g__Metschnikowia_unclassified(100); | Metschnikowia | 13 | Saprotroph | \|Undefined Saprotroph\| | Yeast | NULL | Probable | As nectar-tap_saprotroph (Põlme et al. 2020); Decay sugar-rich substrates (Põlme et al. 2020) | Põlme S, et al. 2020. Fungal Diversity 105:1-16 ((https://doi.org/10.1007/s13225-020-00466-2)) |
| Otu39287 | 1 | k__Fungi(100);p__Ascomycota(100);c__Saccharomycetes(100);o__Saccharomycetales(100);f__Metschnikowiaceae(100);g__Metschnikowia(100);g__Metschnikowia_unclassified(100); | Metschnikowia | 13 | Saprotroph | \|Undefined Saprotroph\| | Yeast | NULL | Probable | As nectar-tap_saprotroph (Põlme et al. 2020); Decay sugar-rich substrates (Põlme et al. 2020) | Põlme S, et al. 2020. Fungal Diversity 105:1-16 ((https://doi.org/10.1007/s13225-020-00466-2)) |
| Otu39288 | 1 | k__Fungi(100);p__Ascomycota(100);c__Saccharomycetes(100);o__Saccharomycetales(100);f__Metschnikowiaceae(100);g__Metschnikowia(100);s__Metschnikowia_pulcherrima(100); | Metschnikowia | 13 | Saprotroph | \|Undefined Saprotroph\| | Yeast | NULL | Probable | As nectar-tap_saprotroph (Põlme et al. 2020); Decay sugar-rich substrates (Põlme et al. 2020) | Põlme S, et al. 2020. Fungal Diversity 105:1-16 ((https://doi.org/10.1007/s13225-020-00466-2)) |
| Otu39289 | 1 | k__Fungi(100);p__Ascomycota(100);c__Saccharomycetes(100);o__Saccharomycetales(100);f__Metschnikowiaceae(100);g__Metschnikowia(100);g__Metschnikowia_unclassified(100); | Metschnikowia | 13 | Saprotroph | \|Undefined Saprotroph\| | Yeast | NULL | Probable | As nectar-tap_saprotroph (Põlme et al. 2020); Decay sugar-rich substrates (Põlme et al. 2020) | Põlme S, et al. 2020. Fungal Diversity 105:1-16 ((https://doi.org/10.1007/s13225-020-00466-2)) |
| Otu39290 | 1 | k__Fungi(100);p__Ascomycota(100);c__Saccharomycetes(100);o__Saccharomycetales(100);f__Metschnikowiaceae(100);g__Metschnikowia(100);s__Metschnikowia_pulcherrima(100); | Metschnikowia | 13 | Saprotroph | \|Undefined Saprotroph\| | Yeast | NULL | Probable | As nectar-tap_saprotroph (Põlme et al. 2020); Decay sugar-rich substrates (Põlme et al. 2020) | Põlme S, et al. 2020. Fungal Diversity 105:1-16 ((https://doi.org/10.1007/s13225-020-00466-2)) |
| Otu39291 | 1 | k__Fungi(100);p__Ascomycota(100);c__Saccharomycetes(100);o__Saccharomycetales(100);f__Metschnikowiaceae(100);g__Metschnikowia(100);s__Metschnikowia_pulcherrima(100); | Metschnikowia | 13 | Saprotroph | \|Undefined Saprotroph\| | Yeast | NULL | Probable | As nectar-tap_saprotroph (Põlme et al. 2020); Decay sugar-rich substrates (Põlme et al. 2020) | Põlme S, et al. 2020. Fungal Diversity 105:1-16 ((https://doi.org/10.1007/s13225-020-00466-2)) |
| Otu39292 | 1 | k__Fungi(100);p__Ascomycota(100);c__Saccharomycetes(100);o__Saccharomycetales(100);f__Metschnikowiaceae(100);g__Metschnikowia(100);s__Metschnikowia_pulcherrima(100); | Metschnikowia | 13 | Saprotroph | \|Undefined Saprotroph\| | Yeast | NULL | Probable | As nectar-tap_saprotroph (Põlme et al. 2020); Decay sugar-rich substrates (Põlme et al. 2020) | Põlme S, et al. 2020. Fungal Diversity 105:1-16 ((https://doi.org/10.1007/s13225-020-00466-2)) |
| Otu39294 | 1 | k__Fungi(100);p__Ascomycota(100);c__Saccharomycetes(100);o__Saccharomycetales(100);f__Metschnikowiaceae(100);g__Metschnikowia(100);g__Metschnikowia_unclassified(100); | Metschnikowia | 13 | Saprotroph | \|Undefined Saprotroph\| | Yeast | NULL | Probable | As nectar-tap_saprotroph (Põlme et al. 2020); Decay sugar-rich substrates (Põlme et al. 2020) | Põlme S, et al. 2020. Fungal Diversity 105:1-16 ((https://doi.org/10.1007/s13225-020-00466-2)) |
| Otu39295 | 1 | k__Fungi(100);p__Ascomycota(100);c__Saccharomycetes(100);o__Saccharomycetales(100);f__Metschnikowiaceae(100);g__Metschnikowia(100);s__Metschnikowia_pulcherrima(100); | Metschnikowia | 13 | Saprotroph | \|Undefined Saprotroph\| | Yeast | NULL | Probable | As nectar-tap_saprotroph (Põlme et al. 2020); Decay sugar-rich substrates (Põlme et al. 2020) | Põlme S, et al. 2020. Fungal Diversity 105:1-16 ((https://doi.org/10.1007/s13225-020-00466-2)) |
| Otu39296 | 1 | k__Fungi(100);p__Ascomycota(100);c__Saccharomycetes(100);o__Saccharomycetales(100);f__Metschnikowiaceae(100);g__Metschnikowia(100);s__Metschnikowia_pulcherrima(100); | Metschnikowia | 13 | Saprotroph | \|Undefined Saprotroph\| | Yeast | NULL | Probable | As nectar-tap_saprotroph (Põlme et al. 2020); Decay sugar-rich substrates (Põlme et al. 2020) | Põlme S, et al. 2020. Fungal Diversity 105:1-16 ((https://doi.org/10.1007/s13225-020-00466-2)) |
| Otu39297 | 1 | k__Fungi(100);p__Ascomycota(100);c__Saccharomycetes(100);o__Saccharomycetales(100);f__Metschnikowiaceae(100);g__Metschnikowia(100);g__Metschnikowia_unclassified(100); | Metschnikowia | 13 | Saprotroph | \|Undefined Saprotroph\| | Yeast | NULL | Probable | As nectar-tap_saprotroph (Põlme et al. 2020); Decay sugar-rich substrates (Põlme et al. 2020) | Põlme S, et al. 2020. Fungal Diversity 105:1-16 ((https://doi.org/10.1007/s13225-020-00466-2)) |
| Otu39298 | 1 | k__Fungi(100);p__Ascomycota(100);c__Saccharomycetes(100);o__Saccharomycetales(100);f__Metschnikowiaceae(100);g__Metschnikowia(100);g__Metschnikowia_unclassified(100); | Metschnikowia | 13 | Saprotroph | \|Undefined Saprotroph\| | Yeast | NULL | Probable | As nectar-tap_saprotroph (Põlme et al. 2020); Decay sugar-rich substrates (Põlme et al. 2020) | Põlme S, et al. 2020. Fungal Diversity 105:1-16 ((https://doi.org/10.1007/s13225-020-00466-2)) |
| Otu39299 | 1 | k__Fungi(100);p__Ascomycota(100);c__Saccharomycetes(100);o__Saccharomycetales(100);f__Metschnikowiaceae(100);g__Metschnikowia(100);g__Metschnikowia_unclassified(100); | Metschnikowia | 13 | Saprotroph | \|Undefined Saprotroph\| | Yeast | NULL | Probable | As nectar-tap_saprotroph (Põlme et al. 2020); Decay sugar-rich substrates (Põlme et al. 2020) | Põlme S, et al. 2020. Fungal Diversity 105:1-16 ((https://doi.org/10.1007/s13225-020-00466-2)) |
| Otu39300 | 1 | k__Fungi(100);p__Ascomycota(100);c__Saccharomycetes(100);o__Saccharomycetales(100);f__Metschnikowiaceae(100);g__Metschnikowia(100);s__Metschnikowia_pulcherrima(100); | Metschnikowia | 13 | Saprotroph | \|Undefined Saprotroph\| | Yeast | NULL | Probable | As nectar-tap_saprotroph (Põlme et al. 2020); Decay sugar-rich substrates (Põlme et al. 2020) | Põlme S, et al. 2020. Fungal Diversity 105:1-16 ((https://doi.org/10.1007/s13225-020-00466-2)) |
| Otu39301 | 1 | k__Fungi(100);p__Ascomycota(100);c__Saccharomycetes(100);o__Saccharomycetales(100);f__Metschnikowiaceae(100);g__Metschnikowia(100);g__Metschnikowia_unclassified(100); | Metschnikowia | 13 | Saprotroph | \|Undefined Saprotroph\| | Yeast | NULL | Probable | As nectar-tap_saprotroph (Põlme et al. 2020); Decay sugar-rich substrates (Põlme et al. 2020) | Põlme S, et al. 2020. Fungal Diversity 105:1-16 ((https://doi.org/10.1007/s13225-020-00466-2)) |
| Otu39302 | 1 | k__Fungi(100);p__Ascomycota(100);c__Saccharomycetes(100);o__Saccharomycetales(100);f__Metschnikowiaceae(100);g__Metschnikowia(100);g__Metschnikowia_unclassified(100); | Metschnikowia | 13 | Saprotroph | \|Undefined Saprotroph\| | Yeast | NULL | Probable | As nectar-tap_saprotroph (Põlme et al. 2020); Decay sugar-rich substrates (Põlme et al. 2020) | Põlme S, et al. 2020. Fungal Diversity 105:1-16 ((https://doi.org/10.1007/s13225-020-00466-2)) |
| Otu39303 | 1 | k__Fungi(100);p__Ascomycota(100);c__Saccharomycetes(100);o__Saccharomycetales(100);f__Metschnikowiaceae(100);g__Metschnikowia(100);g__Metschnikowia_unclassified(100); | Metschnikowia | 13 | Saprotroph | \|Undefined Saprotroph\| | Yeast | NULL | Probable | As nectar-tap_saprotroph (Põlme et al. 2020); Decay sugar-rich substrates (Põlme et al. 2020) | Põlme S, et al. 2020. Fungal Diversity 105:1-16 ((https://doi.org/10.1007/s13225-020-00466-2)) |
| Otu39304 | 1 | k__Fungi(100);p__Ascomycota(100);c__Saccharomycetes(100);o__Saccharomycetales(100);f__Metschnikowiaceae(100);g__Metschnikowia(100);g__Metschnikowia_unclassified(100); | Metschnikowia | 13 | Saprotroph | \|Undefined Saprotroph\| | Yeast | NULL | Probable | As nectar-tap_saprotroph (Põlme et al. 2020); Decay sugar-rich substrates (Põlme et al. 2020) | Põlme S, et al. 2020. Fungal Diversity 105:1-16 ((https://doi.org/10.1007/s13225-020-00466-2)) |
| Otu39305 | 1 | k__Fungi(100);p__Ascomycota(100);c__Saccharomycetes(100);o__Saccharomycetales(100);f__Metschnikowiaceae(100);g__Metschnikowia(100);g__Metschnikowia_unclassified(100); | Metschnikowia | 13 | Saprotroph | \|Undefined Saprotroph\| | Yeast | NULL | Probable | As nectar-tap_saprotroph (Põlme et al. 2020); Decay sugar-rich substrates (Põlme et al. 2020) | Põlme S, et al. 2020. Fungal Diversity 105:1-16 ((https://doi.org/10.1007/s13225-020-00466-2)) |
| Otu39306 | 1 | k__Fungi(100);p__Ascomycota(100);c__Saccharomycetes(100);o__Saccharomycetales(100);f__Metschnikowiaceae(100);g__Metschnikowia(100);g__Metschnikowia_unclassified(100); | Metschnikowia | 13 | Saprotroph | \|Undefined Saprotroph\| | Yeast | NULL | Probable | As nectar-tap_saprotroph (Põlme et al. 2020); Decay sugar-rich substrates (Põlme et al. 2020) | Põlme S, et al. 2020. Fungal Diversity 105:1-16 ((https://doi.org/10.1007/s13225-020-00466-2)) |
| Otu39307 | 1 | k__Fungi(100);p__Ascomycota(100);c__Saccharomycetes(100);o__Saccharomycetales(100);f__Metschnikowiaceae(100);g__Metschnikowia(100);g__Metschnikowia_unclassified(100); | Metschnikowia | 13 | Saprotroph | \|Undefined Saprotroph\| | Yeast | NULL | Probable | As nectar-tap_saprotroph (Põlme et al. 2020); Decay sugar-rich substrates (Põlme et al. 2020) | Põlme S, et al. 2020. Fungal Diversity 105:1-16 ((https://doi.org/10.1007/s13225-020-00466-2)) |
| Otu39308 | 1 | k__Fungi(100);p__Ascomycota(100);c__Saccharomycetes(100);o__Saccharomycetales(100);f__Metschnikowiaceae(100);g__Metschnikowia(100);s__Metschnikowia_pulcherrima(100); | Metschnikowia | 13 | Saprotroph | \|Undefined Saprotroph\| | Yeast | NULL | Probable | As nectar-tap_saprotroph (Põlme et al. 2020); Decay sugar-rich substrates (Põlme et al. 2020) | Põlme S, et al. 2020. Fungal Diversity 105:1-16 ((https://doi.org/10.1007/s13225-020-00466-2)) |
| Otu39309 | 1 | k__Fungi(100);p__Ascomycota(100);c__Saccharomycetes(100);o__Saccharomycetales(100);f__Metschnikowiaceae(100);g__Metschnikowia(100);g__Metschnikowia_unclassified(100); | Metschnikowia | 13 | Saprotroph | \|Undefined Saprotroph\| | Yeast | NULL | Probable | As nectar-tap_saprotroph (Põlme et al. 2020); Decay sugar-rich substrates (Põlme et al. 2020) | Põlme S, et al. 2020. Fungal Diversity 105:1-16 ((https://doi.org/10.1007/s13225-020-00466-2)) |
| Otu39310 | 1 | k__Fungi(100);p__Ascomycota(100);c__Saccharomycetes(100);o__Saccharomycetales(100);f__Metschnikowiaceae(100);g__Metschnikowia(100);s__Metschnikowia_pulcherrima(100); | Metschnikowia | 13 | Saprotroph | \|Undefined Saprotroph\| | Yeast | NULL | Probable | As nectar-tap_saprotroph (Põlme et al. 2020); Decay sugar-rich substrates (Põlme et al. 2020) | Põlme S, et al. 2020. Fungal Diversity 105:1-16 ((https://doi.org/10.1007/s13225-020-00466-2)) |
| Otu39311 | 1 | k__Fungi(100);p__Ascomycota(100);c__Saccharomycetes(100);o__Saccharomycetales(100);f__Metschnikowiaceae(100);g__Metschnikowia(100);s__Metschnikowia_pulcherrima(100); | Metschnikowia | 13 | Saprotroph | \|Undefined Saprotroph\| | Yeast | NULL | Probable | As nectar-tap_saprotroph (Põlme et al. 2020); Decay sugar-rich substrates (Põlme et al. 2020) | Põlme S, et al. 2020. Fungal Diversity 105:1-16 ((https://doi.org/10.1007/s13225-020-00466-2)) |
| Otu39312 | 1 | k__Fungi(100);p__Ascomycota(100);c__Saccharomycetes(100);o__Saccharomycetales(100);f__Metschnikowiaceae(100);g__Metschnikowia(100);s__Metschnikowia_pulcherrima(100); | Metschnikowia | 13 | Saprotroph | \|Undefined Saprotroph\| | Yeast | NULL | Probable | As nectar-tap_saprotroph (Põlme et al. 2020); Decay sugar-rich substrates (Põlme et al. 2020) | Põlme S, et al. 2020. Fungal Diversity 105:1-16 ((https://doi.org/10.1007/s13225-020-00466-2)) |
| Otu39313 | 1 | k__Fungi(100);p__Ascomycota(100);c__Saccharomycetes(100);o__Saccharomycetales(100);f__Metschnikowiaceae(100);g__Metschnikowia(100);s__Metschnikowia_pulcherrima(100); | Metschnikowia | 13 | Saprotroph | \|Undefined Saprotroph\| | Yeast | NULL | Probable | As nectar-tap_saprotroph (Põlme et al. 2020); Decay sugar-rich substrates (Põlme et al. 2020) | Põlme S, et al. 2020. Fungal Diversity 105:1-16 ((https://doi.org/10.1007/s13225-020-00466-2)) |
| Otu39315 | 1 | k__Fungi(100);p__Ascomycota(100);c__Saccharomycetes(100);o__Saccharomycetales(100);f__Metschnikowiaceae(100);g__Metschnikowia(100);s__Metschnikowia_pulcherrima(100); | Metschnikowia | 13 | Saprotroph | \|Undefined Saprotroph\| | Yeast | NULL | Probable | As nectar-tap_saprotroph (Põlme et al. 2020); Decay sugar-rich substrates (Põlme et al. 2020) | Põlme S, et al. 2020. Fungal Diversity 105:1-16 ((https://doi.org/10.1007/s13225-020-00466-2)) |
| Otu39316 | 1 | k__Fungi(100);p__Ascomycota(100);c__Saccharomycetes(100);o__Saccharomycetales(100);f__Metschnikowiaceae(100);g__Metschnikowia(100);s__Metschnikowia_pulcherrima(100); | Metschnikowia | 13 | Saprotroph | \|Undefined Saprotroph\| | Yeast | NULL | Probable | As nectar-tap_saprotroph (Põlme et al. 2020); Decay sugar-rich substrates (Põlme et al. 2020) | Põlme S, et al. 2020. Fungal Diversity 105:1-16 ((https://doi.org/10.1007/s13225-020-00466-2)) |
| Otu39317 | 1 | k__Fungi(100);p__Ascomycota(100);c__Saccharomycetes(100);o__Saccharomycetales(100);f__Metschnikowiaceae(100);g__Metschnikowia(100);g__Metschnikowia_unclassified(100); | Metschnikowia | 13 | Saprotroph | \|Undefined Saprotroph\| | Yeast | NULL | Probable | As nectar-tap_saprotroph (Põlme et al. 2020); Decay sugar-rich substrates (Põlme et al. 2020) | Põlme S, et al. 2020. Fungal Diversity 105:1-16 ((https://doi.org/10.1007/s13225-020-00466-2)) |
| Otu39318 | 1 | k__Fungi(100);p__Ascomycota(100);c__Saccharomycetes(100);o__Saccharomycetales(100);f__Metschnikowiaceae(100);g__Metschnikowia(100);s__Metschnikowia_pulcherrima(100); | Metschnikowia | 13 | Saprotroph | \|Undefined Saprotroph\| | Yeast | NULL | Probable | As nectar-tap_saprotroph (Põlme et al. 2020); Decay sugar-rich substrates (Põlme et al. 2020) | Põlme S, et al. 2020. Fungal Diversity 105:1-16 ((https://doi.org/10.1007/s13225-020-00466-2)) |
| Otu39319 | 1 | k__Fungi(100);p__Ascomycota(100);c__Saccharomycetes(100);o__Saccharomycetales(100);f__Metschnikowiaceae(100);g__Metschnikowia(100);g__Metschnikowia_unclassified(100); | Metschnikowia | 13 | Saprotroph | \|Undefined Saprotroph\| | Yeast | NULL | Probable | As nectar-tap_saprotroph (Põlme et al. 2020); Decay sugar-rich substrates (Põlme et al. 2020) | Põlme S, et al. 2020. Fungal Diversity 105:1-16 ((https://doi.org/10.1007/s13225-020-00466-2)) |
| Otu39320 | 1 | k__Fungi(100);p__Ascomycota(100);c__Saccharomycetes(100);o__Saccharomycetales(100);f__Metschnikowiaceae(100);g__Metschnikowia(100);s__Metschnikowia_pulcherrima(100); | Metschnikowia | 13 | Saprotroph | \|Undefined Saprotroph\| | Yeast | NULL | Probable | As nectar-tap_saprotroph (Põlme et al. 2020); Decay sugar-rich substrates (Põlme et al. 2020) | Põlme S, et al. 2020. Fungal Diversity 105:1-16 ((https://doi.org/10.1007/s13225-020-00466-2)) |
| Otu39321 | 1 | k__Fungi(100);p__Ascomycota(100);c__Saccharomycetes(100);o__Saccharomycetales(100);f__Metschnikowiaceae(100);g__Metschnikowia(100);s__Metschnikowia_pulcherrima(100); | Metschnikowia | 13 | Saprotroph | \|Undefined Saprotroph\| | Yeast | NULL | Probable | As nectar-tap_saprotroph (Põlme et al. 2020); Decay sugar-rich substrates (Põlme et al. 2020) | Põlme S, et al. 2020. Fungal Diversity 105:1-16 ((https://doi.org/10.1007/s13225-020-00466-2)) |
| Otu39322 | 1 | k__Fungi(100);p__Ascomycota(100);c__Saccharomycetes(100);o__Saccharomycetales(100);f__Metschnikowiaceae(100);g__Metschnikowia(100);g__Metschnikowia_unclassified(100); | Metschnikowia | 13 | Saprotroph | \|Undefined Saprotroph\| | Yeast | NULL | Probable | As nectar-tap_saprotroph (Põlme et al. 2020); Decay sugar-rich substrates (Põlme et al. 2020) | Põlme S, et al. 2020. Fungal Diversity 105:1-16 ((https://doi.org/10.1007/s13225-020-00466-2)) |
| Otu39323 | 1 | k__Fungi(100);p__Ascomycota(100);c__Saccharomycetes(100);o__Saccharomycetales(100);f__Metschnikowiaceae(100);g__Metschnikowia(100);s__Metschnikowia_pulcherrima(100); | Metschnikowia | 13 | Saprotroph | \|Undefined Saprotroph\| | Yeast | NULL | Probable | As nectar-tap_saprotroph (Põlme et al. 2020); Decay sugar-rich substrates (Põlme et al. 2020) | Põlme S, et al. 2020. Fungal Diversity 105:1-16 ((https://doi.org/10.1007/s13225-020-00466-2)) |
| Otu39324 | 1 | k__Fungi(100);p__Ascomycota(100);c__Saccharomycetes(100);o__Saccharomycetales(100);f__Metschnikowiaceae(100);g__Metschnikowia(100);g__Metschnikowia_unclassified(100); | Metschnikowia | 13 | Saprotroph | \|Undefined Saprotroph\| | Yeast | NULL | Probable | As nectar-tap_saprotroph (Põlme et al. 2020); Decay sugar-rich substrates (Põlme et al. 2020) | Põlme S, et al. 2020. Fungal Diversity 105:1-16 ((https://doi.org/10.1007/s13225-020-00466-2)) |
| Otu39325 | 1 | k__Fungi(100);p__Ascomycota(100);c__Saccharomycetes(100);o__Saccharomycetales(100);f__Metschnikowiaceae(100);g__Metschnikowia(100);s__Metschnikowia_pulcherrima(100); | Metschnikowia | 13 | Saprotroph | \|Undefined Saprotroph\| | Yeast | NULL | Probable | As nectar-tap_saprotroph (Põlme et al. 2020); Decay sugar-rich substrates (Põlme et al. 2020) | Põlme S, et al. 2020. Fungal Diversity 105:1-16 ((https://doi.org/10.1007/s13225-020-00466-2)) |
| Otu39326 | 1 | k__Fungi(100);p__Ascomycota(100);c__Saccharomycetes(100);o__Saccharomycetales(100);f__Metschnikowiaceae(100);g__Metschnikowia(100);s__Metschnikowia_pulcherrima(100); | Metschnikowia | 13 | Saprotroph | \|Undefined Saprotroph\| | Yeast | NULL | Probable | As nectar-tap_saprotroph (Põlme et al. 2020); Decay sugar-rich substrates (Põlme et al. 2020) | Põlme S, et al. 2020. Fungal Diversity 105:1-16 ((https://doi.org/10.1007/s13225-020-00466-2)) |
| Otu39327 | 1 | k__Fungi(100);p__Ascomycota(100);c__Saccharomycetes(100);o__Saccharomycetales(100);f__Metschnikowiaceae(100);g__Metschnikowia(100);s__Metschnikowia_pulcherrima(100); | Metschnikowia | 13 | Saprotroph | \|Undefined Saprotroph\| | Yeast | NULL | Probable | As nectar-tap_saprotroph (Põlme et al. 2020); Decay sugar-rich substrates (Põlme et al. 2020) | Põlme S, et al. 2020. Fungal Diversity 105:1-16 ((https://doi.org/10.1007/s13225-020-00466-2)) |
| Otu39328 | 1 | k__Fungi(100);p__Ascomycota(100);c__Saccharomycetes(100);o__Saccharomycetales(100);f__Metschnikowiaceae(100);g__Metschnikowia(100);s__Metschnikowia_pulcherrima(100); | Metschnikowia | 13 | Saprotroph | \|Undefined Saprotroph\| | Yeast | NULL | Probable | As nectar-tap_saprotroph (Põlme et al. 2020); Decay sugar-rich substrates (Põlme et al. 2020) | Põlme S, et al. 2020. Fungal Diversity 105:1-16 ((https://doi.org/10.1007/s13225-020-00466-2)) |
| Otu39330 | 1 | k__Fungi(100);p__Ascomycota(100);c__Saccharomycetes(100);o__Saccharomycetales(100);f__Metschnikowiaceae(100);g__Metschnikowia(100);g__Metschnikowia_unclassified(100); | Metschnikowia | 13 | Saprotroph | \|Undefined Saprotroph\| | Yeast | NULL | Probable | As nectar-tap_saprotroph (Põlme et al. 2020); Decay sugar-rich substrates (Põlme et al. 2020) | Põlme S, et al. 2020. Fungal Diversity 105:1-16 ((https://doi.org/10.1007/s13225-020-00466-2)) |
| Otu39331 | 1 | k__Fungi(100);p__Ascomycota(100);c__Saccharomycetes(100);o__Saccharomycetales(100);f__Metschnikowiaceae(100);g__Metschnikowia(100);s__Metschnikowia_pulcherrima(100); | Metschnikowia | 13 | Saprotroph | \|Undefined Saprotroph\| | Yeast | NULL | Probable | As nectar-tap_saprotroph (Põlme et al. 2020); Decay sugar-rich substrates (Põlme et al. 2020) | Põlme S, et al. 2020. Fungal Diversity 105:1-16 ((https://doi.org/10.1007/s13225-020-00466-2)) |
| Otu39332 | 1 | k__Fungi(100);p__Ascomycota(100);c__Saccharomycetes(100);o__Saccharomycetales(100);f__Metschnikowiaceae(100);g__Metschnikowia(100);s__Metschnikowia_pulcherrima(100); | Metschnikowia | 13 | Saprotroph | \|Undefined Saprotroph\| | Yeast | NULL | Probable | As nectar-tap_saprotroph (Põlme et al. 2020); Decay sugar-rich substrates (Põlme et al. 2020) | Põlme S, et al. 2020. Fungal Diversity 105:1-16 ((https://doi.org/10.1007/s13225-020-00466-2)) |
| Otu39333 | 1 | k__Fungi(100);p__Ascomycota(100);c__Saccharomycetes(100);o__Saccharomycetales(100);f__Metschnikowiaceae(100);g__Metschnikowia(100);g__Metschnikowia_unclassified(100); | Metschnikowia | 13 | Saprotroph | \|Undefined Saprotroph\| | Yeast | NULL | Probable | As nectar-tap_saprotroph (Põlme et al. 2020); Decay sugar-rich substrates (Põlme et al. 2020) | Põlme S, et al. 2020. Fungal Diversity 105:1-16 ((https://doi.org/10.1007/s13225-020-00466-2)) |
| Otu39334 | 1 | k__Fungi(100);p__Ascomycota(100);c__Saccharomycetes(100);o__Saccharomycetales(100);f__Metschnikowiaceae(100);g__Metschnikowia(100);s__Metschnikowia_pulcherrima(100); | Metschnikowia | 13 | Saprotroph | \|Undefined Saprotroph\| | Yeast | NULL | Probable | As nectar-tap_saprotroph (Põlme et al. 2020); Decay sugar-rich substrates (Põlme et al. 2020) | Põlme S, et al. 2020. Fungal Diversity 105:1-16 ((https://doi.org/10.1007/s13225-020-00466-2)) |
| Otu39335 | 1 | k__Fungi(100);p__Ascomycota(100);c__Saccharomycetes(100);o__Saccharomycetales(100);f__Metschnikowiaceae(100);g__Metschnikowia(100);g__Metschnikowia_unclassified(100); | Metschnikowia | 13 | Saprotroph | \|Undefined Saprotroph\| | Yeast | NULL | Probable | As nectar-tap_saprotroph (Põlme et al. 2020); Decay sugar-rich substrates (Põlme et al. 2020) | Põlme S, et al. 2020. Fungal Diversity 105:1-16 ((https://doi.org/10.1007/s13225-020-00466-2)) |
| Otu39336 | 1 | k__Fungi(100);p__Ascomycota(100);c__Saccharomycetes(100);o__Saccharomycetales(100);f__Metschnikowiaceae(100);g__Metschnikowia(100);s__Metschnikowia_pulcherrima(100); | Metschnikowia | 13 | Saprotroph | \|Undefined Saprotroph\| | Yeast | NULL | Probable | As nectar-tap_saprotroph (Põlme et al. 2020); Decay sugar-rich substrates (Põlme et al. 2020) | Põlme S, et al. 2020. Fungal Diversity 105:1-16 ((https://doi.org/10.1007/s13225-020-00466-2)) |
| Otu39337 | 1 | k__Fungi(100);p__Ascomycota(100);c__Saccharomycetes(100);o__Saccharomycetales(100);f__Metschnikowiaceae(100);g__Metschnikowia(100);s__Metschnikowia_pulcherrima(100); | Metschnikowia | 13 | Saprotroph | \|Undefined Saprotroph\| | Yeast | NULL | Probable | As nectar-tap_saprotroph (Põlme et al. 2020); Decay sugar-rich substrates (Põlme et al. 2020) | Põlme S, et al. 2020. Fungal Diversity 105:1-16 ((https://doi.org/10.1007/s13225-020-00466-2)) |
| Otu39338 | 1 | k__Fungi(100);p__Ascomycota(100);c__Saccharomycetes(100);o__Saccharomycetales(100);f__Metschnikowiaceae(100);g__Metschnikowia(100);s__Metschnikowia_pulcherrima(100); | Metschnikowia | 13 | Saprotroph | \|Undefined Saprotroph\| | Yeast | NULL | Probable | As nectar-tap_saprotroph (Põlme et al. 2020); Decay sugar-rich substrates (Põlme et al. 2020) | Põlme S, et al. 2020. Fungal Diversity 105:1-16 ((https://doi.org/10.1007/s13225-020-00466-2)) |
| Otu39339 | 1 | k__Fungi(100);p__Ascomycota(100);c__Saccharomycetes(100);o__Saccharomycetales(100);f__Metschnikowiaceae(100);g__Metschnikowia(100);s__Metschnikowia_pulcherrima(100); | Metschnikowia | 13 | Saprotroph | \|Undefined Saprotroph\| | Yeast | NULL | Probable | As nectar-tap_saprotroph (Põlme et al. 2020); Decay sugar-rich substrates (Põlme et al. 2020) | Põlme S, et al. 2020. Fungal Diversity 105:1-16 ((https://doi.org/10.1007/s13225-020-00466-2)) |
| Otu39340 | 1 | k__Fungi(100);p__Ascomycota(100);c__Saccharomycetes(100);o__Saccharomycetales(100);f__Metschnikowiaceae(100);g__Metschnikowia(100);g__Metschnikowia_unclassified(100); | Metschnikowia | 13 | Saprotroph | \|Undefined Saprotroph\| | Yeast | NULL | Probable | As nectar-tap_saprotroph (Põlme et al. 2020); Decay sugar-rich substrates (Põlme et al. 2020) | Põlme S, et al. 2020. Fungal Diversity 105:1-16 ((https://doi.org/10.1007/s13225-020-00466-2)) |
| Otu39341 | 1 | k__Fungi(100);p__Ascomycota(100);c__Saccharomycetes(100);o__Saccharomycetales(100);f__Metschnikowiaceae(100);g__Metschnikowia(100);g__Metschnikowia_unclassified(100); | Metschnikowia | 13 | Saprotroph | \|Undefined Saprotroph\| | Yeast | NULL | Probable | As nectar-tap_saprotroph (Põlme et al. 2020); Decay sugar-rich substrates (Põlme et al. 2020) | Põlme S, et al. 2020. Fungal Diversity 105:1-16 ((https://doi.org/10.1007/s13225-020-00466-2)) |
| Otu39342 | 1 | k__Fungi(100);p__Ascomycota(100);c__Saccharomycetes(100);o__Saccharomycetales(100);f__Metschnikowiaceae(100);g__Metschnikowia(100);s__Metschnikowia_pulcherrima(100); | Metschnikowia | 13 | Saprotroph | \|Undefined Saprotroph\| | Yeast | NULL | Probable | As nectar-tap_saprotroph (Põlme et al. 2020); Decay sugar-rich substrates (Põlme et al. 2020) | Põlme S, et al. 2020. Fungal Diversity 105:1-16 ((https://doi.org/10.1007/s13225-020-00466-2)) |
| Otu39343 | 1 | k__Fungi(100);p__Ascomycota(100);c__Saccharomycetes(100);o__Saccharomycetales(100);f__Metschnikowiaceae(100);g__Metschnikowia(100);s__Metschnikowia_pulcherrima(100); | Metschnikowia | 13 | Saprotroph | \|Undefined Saprotroph\| | Yeast | NULL | Probable | As nectar-tap_saprotroph (Põlme et al. 2020); Decay sugar-rich substrates (Põlme et al. 2020) | Põlme S, et al. 2020. Fungal Diversity 105:1-16 ((https://doi.org/10.1007/s13225-020-00466-2)) |
| Otu39346 | 1 | k__Fungi(100);p__Ascomycota(100);c__Saccharomycetes(100);o__Saccharomycetales(100);f__Metschnikowiaceae(100);g__Metschnikowia(100);g__Metschnikowia_unclassified(100); | Metschnikowia | 13 | Saprotroph | \|Undefined Saprotroph\| | Yeast | NULL | Probable | As nectar-tap_saprotroph (Põlme et al. 2020); Decay sugar-rich substrates (Põlme et al. 2020) | Põlme S, et al. 2020. Fungal Diversity 105:1-16 ((https://doi.org/10.1007/s13225-020-00466-2)) |
| Otu39347 | 1 | k__Fungi(100);p__Ascomycota(100);c__Saccharomycetes(100);o__Saccharomycetales(100);f__Metschnikowiaceae(100);g__Metschnikowia(100);s__Metschnikowia_pulcherrima(100); | Metschnikowia | 13 | Saprotroph | \|Undefined Saprotroph\| | Yeast | NULL | Probable | As nectar-tap_saprotroph (Põlme et al. 2020); Decay sugar-rich substrates (Põlme et al. 2020) | Põlme S, et al. 2020. Fungal Diversity 105:1-16 ((https://doi.org/10.1007/s13225-020-00466-2)) |
| Otu39348 | 1 | k__Fungi(100);p__Ascomycota(100);c__Saccharomycetes(100);o__Saccharomycetales(100);f__Metschnikowiaceae(100);g__Metschnikowia(100);s__Metschnikowia_pulcherrima(100); | Metschnikowia | 13 | Saprotroph | \|Undefined Saprotroph\| | Yeast | NULL | Probable | As nectar-tap_saprotroph (Põlme et al. 2020); Decay sugar-rich substrates (Põlme et al. 2020) | Põlme S, et al. 2020. Fungal Diversity 105:1-16 ((https://doi.org/10.1007/s13225-020-00466-2)) |
| Otu39349 | 1 | k__Fungi(100);p__Ascomycota(100);c__Saccharomycetes(100);o__Saccharomycetales(100);f__Metschnikowiaceae(100);g__Metschnikowia(100);g__Metschnikowia_unclassified(100); | Metschnikowia | 13 | Saprotroph | \|Undefined Saprotroph\| | Yeast | NULL | Probable | As nectar-tap_saprotroph (Põlme et al. 2020); Decay sugar-rich substrates (Põlme et al. 2020) | Põlme S, et al. 2020. Fungal Diversity 105:1-16 ((https://doi.org/10.1007/s13225-020-00466-2)) |
| Otu39350 | 1 | k__Fungi(100);p__Ascomycota(100);c__Saccharomycetes(100);o__Saccharomycetales(100);f__Metschnikowiaceae(100);g__Metschnikowia(100);s__Metschnikowia_pulcherrima(100); | Metschnikowia | 13 | Saprotroph | \|Undefined Saprotroph\| | Yeast | NULL | Probable | As nectar-tap_saprotroph (Põlme et al. 2020); Decay sugar-rich substrates (Põlme et al. 2020) | Põlme S, et al. 2020. Fungal Diversity 105:1-16 ((https://doi.org/10.1007/s13225-020-00466-2)) |
| Otu39351 | 1 | k__Fungi(100);p__Ascomycota(100);c__Saccharomycetes(100);o__Saccharomycetales(100);f__Metschnikowiaceae(100);g__Metschnikowia(100);g__Metschnikowia_unclassified(100); | Metschnikowia | 13 | Saprotroph | \|Undefined Saprotroph\| | Yeast | NULL | Probable | As nectar-tap_saprotroph (Põlme et al. 2020); Decay sugar-rich substrates (Põlme et al. 2020) | Põlme S, et al. 2020. Fungal Diversity 105:1-16 ((https://doi.org/10.1007/s13225-020-00466-2)) |
| Otu39352 | 1 | k__Fungi(100);p__Ascomycota(100);c__Saccharomycetes(100);o__Saccharomycetales(100);f__Metschnikowiaceae(100);g__Metschnikowia(100);g__Metschnikowia_unclassified(100); | Metschnikowia | 13 | Saprotroph | \|Undefined Saprotroph\| | Yeast | NULL | Probable | As nectar-tap_saprotroph (Põlme et al. 2020); Decay sugar-rich substrates (Põlme et al. 2020) | Põlme S, et al. 2020. Fungal Diversity 105:1-16 ((https://doi.org/10.1007/s13225-020-00466-2)) |
| Otu39353 | 1 | k__Fungi(100);p__Ascomycota(100);c__Saccharomycetes(100);o__Saccharomycetales(100);f__Metschnikowiaceae(100);g__Metschnikowia(100);s__Metschnikowia_pulcherrima(100); | Metschnikowia | 13 | Saprotroph | \|Undefined Saprotroph\| | Yeast | NULL | Probable | As nectar-tap_saprotroph (Põlme et al. 2020); Decay sugar-rich substrates (Põlme et al. 2020) | Põlme S, et al. 2020. Fungal Diversity 105:1-16 ((https://doi.org/10.1007/s13225-020-00466-2)) |
| Otu39354 | 1 | k__Fungi(100);p__Ascomycota(100);c__Saccharomycetes(100);o__Saccharomycetales(100);f__Metschnikowiaceae(100);g__Metschnikowia(100);s__Metschnikowia_pulcherrima(100); | Metschnikowia | 13 | Saprotroph | \|Undefined Saprotroph\| | Yeast | NULL | Probable | As nectar-tap_saprotroph (Põlme et al. 2020); Decay sugar-rich substrates (Põlme et al. 2020) | Põlme S, et al. 2020. Fungal Diversity 105:1-16 ((https://doi.org/10.1007/s13225-020-00466-2)) |
| Otu39355 | 1 | k__Fungi(100);p__Ascomycota(100);c__Saccharomycetes(100);o__Saccharomycetales(100);f__Metschnikowiaceae(100);g__Metschnikowia(100);s__Metschnikowia_pulcherrima(100); | Metschnikowia | 13 | Saprotroph | \|Undefined Saprotroph\| | Yeast | NULL | Probable | As nectar-tap_saprotroph (Põlme et al. 2020); Decay sugar-rich substrates (Põlme et al. 2020) | Põlme S, et al. 2020. Fungal Diversity 105:1-16 ((https://doi.org/10.1007/s13225-020-00466-2)) |
| Otu39356 | 1 | k__Fungi(100);p__Ascomycota(100);c__Saccharomycetes(100);o__Saccharomycetales(100);f__Metschnikowiaceae(100);g__Metschnikowia(100);s__Metschnikowia_pulcherrima(100); | Metschnikowia | 13 | Saprotroph | \|Undefined Saprotroph\| | Yeast | NULL | Probable | As nectar-tap_saprotroph (Põlme et al. 2020); Decay sugar-rich substrates (Põlme et al. 2020) | Põlme S, et al. 2020. Fungal Diversity 105:1-16 ((https://doi.org/10.1007/s13225-020-00466-2)) |
| Otu39357 | 1 | k__Fungi(100);p__Ascomycota(100);c__Saccharomycetes(100);o__Saccharomycetales(100);f__Metschnikowiaceae(100);g__Metschnikowia(100);s__Metschnikowia_pulcherrima(100); | Metschnikowia | 13 | Saprotroph | \|Undefined Saprotroph\| | Yeast | NULL | Probable | As nectar-tap_saprotroph (Põlme et al. 2020); Decay sugar-rich substrates (Põlme et al. 2020) | Põlme S, et al. 2020. Fungal Diversity 105:1-16 ((https://doi.org/10.1007/s13225-020-00466-2)) |
| Otu39358 | 1 | k__Fungi(100);p__Ascomycota(100);c__Saccharomycetes(100);o__Saccharomycetales(100);f__Metschnikowiaceae(100);g__Metschnikowia(100);s__Metschnikowia_pulcherrima(100); | Metschnikowia | 13 | Saprotroph | \|Undefined Saprotroph\| | Yeast | NULL | Probable | As nectar-tap_saprotroph (Põlme et al. 2020); Decay sugar-rich substrates (Põlme et al. 2020) | Põlme S, et al. 2020. Fungal Diversity 105:1-16 ((https://doi.org/10.1007/s13225-020-00466-2)) |
| Otu39359 | 1 | k__Fungi(100);p__Ascomycota(100);c__Saccharomycetes(100);o__Saccharomycetales(100);f__Metschnikowiaceae(100);g__Metschnikowia(100);s__Metschnikowia_pulcherrima(100); | Metschnikowia | 13 | Saprotroph | \|Undefined Saprotroph\| | Yeast | NULL | Probable | As nectar-tap_saprotroph (Põlme et al. 2020); Decay sugar-rich substrates (Põlme et al. 2020) | Põlme S, et al. 2020. Fungal Diversity 105:1-16 ((https://doi.org/10.1007/s13225-020-00466-2)) |
| Otu39360 | 1 | k__Fungi(100);p__Ascomycota(100);c__Saccharomycetes(100);o__Saccharomycetales(100);f__Metschnikowiaceae(100);g__Metschnikowia(100);s__Metschnikowia_pulcherrima(100); | Metschnikowia | 13 | Saprotroph | \|Undefined Saprotroph\| | Yeast | NULL | Probable | As nectar-tap_saprotroph (Põlme et al. 2020); Decay sugar-rich substrates (Põlme et al. 2020) | Põlme S, et al. 2020. Fungal Diversity 105:1-16 ((https://doi.org/10.1007/s13225-020-00466-2)) |
| Otu39361 | 1 | k__Fungi(100);p__Ascomycota(100);c__Saccharomycetes(100);o__Saccharomycetales(100);f__Metschnikowiaceae(100);g__Metschnikowia(100);s__Metschnikowia_pulcherrima(100); | Metschnikowia | 13 | Saprotroph | \|Undefined Saprotroph\| | Yeast | NULL | Probable | As nectar-tap_saprotroph (Põlme et al. 2020); Decay sugar-rich substrates (Põlme et al. 2020) | Põlme S, et al. 2020. Fungal Diversity 105:1-16 ((https://doi.org/10.1007/s13225-020-00466-2)) |
| Otu39362 | 1 | k__Fungi(100);p__Ascomycota(100);c__Saccharomycetes(100);o__Saccharomycetales(100);f__Metschnikowiaceae(100);g__Metschnikowia(100);s__Metschnikowia_pulcherrima(100); | Metschnikowia | 13 | Saprotroph | \|Undefined Saprotroph\| | Yeast | NULL | Probable | As nectar-tap_saprotroph (Põlme et al. 2020); Decay sugar-rich substrates (Põlme et al. 2020) | Põlme S, et al. 2020. Fungal Diversity 105:1-16 ((https://doi.org/10.1007/s13225-020-00466-2)) |
| Otu39363 | 1 | k__Fungi(100);p__Ascomycota(100);c__Saccharomycetes(100);o__Saccharomycetales(100);f__Metschnikowiaceae(100);g__Metschnikowia(100);g__Metschnikowia_unclassified(100); | Metschnikowia | 13 | Saprotroph | \|Undefined Saprotroph\| | Yeast | NULL | Probable | As nectar-tap_saprotroph (Põlme et al. 2020); Decay sugar-rich substrates (Põlme et al. 2020) | Põlme S, et al. 2020. Fungal Diversity 105:1-16 ((https://doi.org/10.1007/s13225-020-00466-2)) |
| Otu39364 | 1 | k__Fungi(100);p__Ascomycota(100);c__Saccharomycetes(100);o__Saccharomycetales(100);f__Metschnikowiaceae(100);g__Metschnikowia(100);s__Metschnikowia_pulcherrima(100); | Metschnikowia | 13 | Saprotroph | \|Undefined Saprotroph\| | Yeast | NULL | Probable | As nectar-tap_saprotroph (Põlme et al. 2020); Decay sugar-rich substrates (Põlme et al. 2020) | Põlme S, et al. 2020. Fungal Diversity 105:1-16 ((https://doi.org/10.1007/s13225-020-00466-2)) |
| Otu39365 | 1 | k__Fungi(100);p__Ascomycota(100);c__Saccharomycetes(100);o__Saccharomycetales(100);f__Metschnikowiaceae(100);g__Metschnikowia(100);s__Metschnikowia_pulcherrima(100); | Metschnikowia | 13 | Saprotroph | \|Undefined Saprotroph\| | Yeast | NULL | Probable | As nectar-tap_saprotroph (Põlme et al. 2020); Decay sugar-rich substrates (Põlme et al. 2020) | Põlme S, et al. 2020. Fungal Diversity 105:1-16 ((https://doi.org/10.1007/s13225-020-00466-2)) |
| Otu39366 | 1 | k__Fungi(100);p__Ascomycota(100);c__Saccharomycetes(100);o__Saccharomycetales(100);f__Metschnikowiaceae(100);g__Metschnikowia(100);s__Metschnikowia_pulcherrima(100); | Metschnikowia | 13 | Saprotroph | \|Undefined Saprotroph\| | Yeast | NULL | Probable | As nectar-tap_saprotroph (Põlme et al. 2020); Decay sugar-rich substrates (Põlme et al. 2020) | Põlme S, et al. 2020. Fungal Diversity 105:1-16 ((https://doi.org/10.1007/s13225-020-00466-2)) |
| Otu39367 | 1 | k__Fungi(100);p__Ascomycota(100);c__Saccharomycetes(100);o__Saccharomycetales(100);f__Metschnikowiaceae(100);g__Metschnikowia(100);s__Metschnikowia_pulcherrima(100); | Metschnikowia | 13 | Saprotroph | \|Undefined Saprotroph\| | Yeast | NULL | Probable | As nectar-tap_saprotroph (Põlme et al. 2020); Decay sugar-rich substrates (Põlme et al. 2020) | Põlme S, et al. 2020. Fungal Diversity 105:1-16 ((https://doi.org/10.1007/s13225-020-00466-2)) |
| Otu39368 | 1 | k__Fungi(100);p__Ascomycota(100);c__Saccharomycetes(100);o__Saccharomycetales(100);f__Metschnikowiaceae(100);g__Metschnikowia(100);s__Metschnikowia_pulcherrima(100); | Metschnikowia | 13 | Saprotroph | \|Undefined Saprotroph\| | Yeast | NULL | Probable | As nectar-tap_saprotroph (Põlme et al. 2020); Decay sugar-rich substrates (Põlme et al. 2020) | Põlme S, et al. 2020. Fungal Diversity 105:1-16 ((https://doi.org/10.1007/s13225-020-00466-2)) |
| Otu39370 | 1 | k__Fungi(100);p__Ascomycota(100);c__Saccharomycetes(100);o__Saccharomycetales(100);f__Metschnikowiaceae(100);g__Metschnikowia(100);s__Metschnikowia_pulcherrima(100); | Metschnikowia | 13 | Saprotroph | \|Undefined Saprotroph\| | Yeast | NULL | Probable | As nectar-tap_saprotroph (Põlme et al. 2020); Decay sugar-rich substrates (Põlme et al. 2020) | Põlme S, et al. 2020. Fungal Diversity 105:1-16 ((https://doi.org/10.1007/s13225-020-00466-2)) |
| Otu39371 | 1 | k__Fungi(100);p__Ascomycota(100);c__Saccharomycetes(100);o__Saccharomycetales(100);f__Metschnikowiaceae(100);g__Metschnikowia(100);s__Metschnikowia_pulcherrima(100); | Metschnikowia | 13 | Saprotroph | \|Undefined Saprotroph\| | Yeast | NULL | Probable | As nectar-tap_saprotroph (Põlme et al. 2020); Decay sugar-rich substrates (Põlme et al. 2020) | Põlme S, et al. 2020. Fungal Diversity 105:1-16 ((https://doi.org/10.1007/s13225-020-00466-2)) |
| Otu39372 | 1 | k__Fungi(100);p__Ascomycota(100);c__Saccharomycetes(100);o__Saccharomycetales(100);f__Metschnikowiaceae(100);g__Metschnikowia(100);s__Metschnikowia_pulcherrima(100); | Metschnikowia | 13 | Saprotroph | \|Undefined Saprotroph\| | Yeast | NULL | Probable | As nectar-tap_saprotroph (Põlme et al. 2020); Decay sugar-rich substrates (Põlme et al. 2020) | Põlme S, et al. 2020. Fungal Diversity 105:1-16 ((https://doi.org/10.1007/s13225-020-00466-2)) |
| Otu39373 | 1 | k__Fungi(100);p__Ascomycota(100);c__Saccharomycetes(100);o__Saccharomycetales(100);f__Metschnikowiaceae(100);g__Metschnikowia(100);s__Metschnikowia_pulcherrima(100); | Metschnikowia | 13 | Saprotroph | \|Undefined Saprotroph\| | Yeast | NULL | Probable | As nectar-tap_saprotroph (Põlme et al. 2020); Decay sugar-rich substrates (Põlme et al. 2020) | Põlme S, et al. 2020. Fungal Diversity 105:1-16 ((https://doi.org/10.1007/s13225-020-00466-2)) |
| Otu39374 | 1 | k__Fungi(100);p__Ascomycota(100);c__Saccharomycetes(100);o__Saccharomycetales(100);f__Metschnikowiaceae(100);g__Metschnikowia(100);s__Metschnikowia_pulcherrima(100); | Metschnikowia | 13 | Saprotroph | \|Undefined Saprotroph\| | Yeast | NULL | Probable | As nectar-tap_saprotroph (Põlme et al. 2020); Decay sugar-rich substrates (Põlme et al. 2020) | Põlme S, et al. 2020. Fungal Diversity 105:1-16 ((https://doi.org/10.1007/s13225-020-00466-2)) |
| Otu39375 | 1 | k__Fungi(100);p__Ascomycota(100);c__Saccharomycetes(100);o__Saccharomycetales(100);f__Metschnikowiaceae(100);g__Metschnikowia(100);s__Metschnikowia_pulcherrima(100); | Metschnikowia | 13 | Saprotroph | \|Undefined Saprotroph\| | Yeast | NULL | Probable | As nectar-tap_saprotroph (Põlme et al. 2020); Decay sugar-rich substrates (Põlme et al. 2020) | Põlme S, et al. 2020. Fungal Diversity 105:1-16 ((https://doi.org/10.1007/s13225-020-00466-2)) |
| Otu39376 | 1 | k__Fungi(100);p__Ascomycota(100);c__Saccharomycetes(100);o__Saccharomycetales(100);f__Metschnikowiaceae(100);g__Metschnikowia(100);s__Metschnikowia_pulcherrima(100); | Metschnikowia | 13 | Saprotroph | \|Undefined Saprotroph\| | Yeast | NULL | Probable | As nectar-tap_saprotroph (Põlme et al. 2020); Decay sugar-rich substrates (Põlme et al. 2020) | Põlme S, et al. 2020. Fungal Diversity 105:1-16 ((https://doi.org/10.1007/s13225-020-00466-2)) |
| Otu39377 | 1 | k__Fungi(100);p__Ascomycota(100);c__Saccharomycetes(100);o__Saccharomycetales(100);f__Metschnikowiaceae(100);g__Metschnikowia(100);g__Metschnikowia_unclassified(100); | Metschnikowia | 13 | Saprotroph | \|Undefined Saprotroph\| | Yeast | NULL | Probable | As nectar-tap_saprotroph (Põlme et al. 2020); Decay sugar-rich substrates (Põlme et al. 2020) | Põlme S, et al. 2020. Fungal Diversity 105:1-16 ((https://doi.org/10.1007/s13225-020-00466-2)) |
| Otu39378 | 1 | k__Fungi(100);p__Ascomycota(100);c__Saccharomycetes(100);o__Saccharomycetales(100);f__Metschnikowiaceae(100);g__Metschnikowia(100);g__Metschnikowia_unclassified(100); | Metschnikowia | 13 | Saprotroph | \|Undefined Saprotroph\| | Yeast | NULL | Probable | As nectar-tap_saprotroph (Põlme et al. 2020); Decay sugar-rich substrates (Põlme et al. 2020) | Põlme S, et al. 2020. Fungal Diversity 105:1-16 ((https://doi.org/10.1007/s13225-020-00466-2)) |
| Otu39379 | 1 | k__Fungi(100);p__Ascomycota(100);c__Saccharomycetes(100);o__Saccharomycetales(100);f__Metschnikowiaceae(100);g__Metschnikowia(100);s__Metschnikowia_pulcherrima(100); | Metschnikowia | 13 | Saprotroph | \|Undefined Saprotroph\| | Yeast | NULL | Probable | As nectar-tap_saprotroph (Põlme et al. 2020); Decay sugar-rich substrates (Põlme et al. 2020) | Põlme S, et al. 2020. Fungal Diversity 105:1-16 ((https://doi.org/10.1007/s13225-020-00466-2)) |
| Otu39380 | 1 | k__Fungi(100);p__Ascomycota(100);c__Saccharomycetes(100);o__Saccharomycetales(100);f__Metschnikowiaceae(100);g__Metschnikowia(100);s__Metschnikowia_pulcherrima(100); | Metschnikowia | 13 | Saprotroph | \|Undefined Saprotroph\| | Yeast | NULL | Probable | As nectar-tap_saprotroph (Põlme et al. 2020); Decay sugar-rich substrates (Põlme et al. 2020) | Põlme S, et al. 2020. Fungal Diversity 105:1-16 ((https://doi.org/10.1007/s13225-020-00466-2)) |
| Otu39381 | 1 | k__Fungi(100);p__Ascomycota(100);c__Saccharomycetes(100);o__Saccharomycetales(100);f__Metschnikowiaceae(100);g__Metschnikowia(100);s__Metschnikowia_pulcherrima(100); | Metschnikowia | 13 | Saprotroph | \|Undefined Saprotroph\| | Yeast | NULL | Probable | As nectar-tap_saprotroph (Põlme et al. 2020); Decay sugar-rich substrates (Põlme et al. 2020) | Põlme S, et al. 2020. Fungal Diversity 105:1-16 ((https://doi.org/10.1007/s13225-020-00466-2)) |
| Otu39382 | 1 | k__Fungi(100);p__Ascomycota(100);c__Saccharomycetes(100);o__Saccharomycetales(100);f__Metschnikowiaceae(100);g__Metschnikowia(100);s__Metschnikowia_pulcherrima(100); | Metschnikowia | 13 | Saprotroph | \|Undefined Saprotroph\| | Yeast | NULL | Probable | As nectar-tap_saprotroph (Põlme et al. 2020); Decay sugar-rich substrates (Põlme et al. 2020) | Põlme S, et al. 2020. Fungal Diversity 105:1-16 ((https://doi.org/10.1007/s13225-020-00466-2)) |
| Otu39383 | 1 | k__Fungi(100);p__Ascomycota(100);c__Saccharomycetes(100);o__Saccharomycetales(100);f__Metschnikowiaceae(100);g__Metschnikowia(100);s__Metschnikowia_pulcherrima(100); | Metschnikowia | 13 | Saprotroph | \|Undefined Saprotroph\| | Yeast | NULL | Probable | As nectar-tap_saprotroph (Põlme et al. 2020); Decay sugar-rich substrates (Põlme et al. 2020) | Põlme S, et al. 2020. Fungal Diversity 105:1-16 ((https://doi.org/10.1007/s13225-020-00466-2)) |
| Otu39384 | 1 | k__Fungi(100);p__Ascomycota(100);c__Saccharomycetes(100);o__Saccharomycetales(100);f__Metschnikowiaceae(100);g__Metschnikowia(100);g__Metschnikowia_unclassified(100); | Metschnikowia | 13 | Saprotroph | \|Undefined Saprotroph\| | Yeast | NULL | Probable | As nectar-tap_saprotroph (Põlme et al. 2020); Decay sugar-rich substrates (Põlme et al. 2020) | Põlme S, et al. 2020. Fungal Diversity 105:1-16 ((https://doi.org/10.1007/s13225-020-00466-2)) |
| Otu39385 | 1 | k__Fungi(100);p__Ascomycota(100);c__Saccharomycetes(100);o__Saccharomycetales(100);f__Metschnikowiaceae(100);g__Metschnikowia(100);s__Metschnikowia_pulcherrima(100); | Metschnikowia | 13 | Saprotroph | \|Undefined Saprotroph\| | Yeast | NULL | Probable | As nectar-tap_saprotroph (Põlme et al. 2020); Decay sugar-rich substrates (Põlme et al. 2020) | Põlme S, et al. 2020. Fungal Diversity 105:1-16 ((https://doi.org/10.1007/s13225-020-00466-2)) |
| Otu39386 | 1 | k__Fungi(100);p__Ascomycota(100);c__Saccharomycetes(100);o__Saccharomycetales(100);f__Metschnikowiaceae(100);g__Metschnikowia(100);g__Metschnikowia_unclassified(100); | Metschnikowia | 13 | Saprotroph | \|Undefined Saprotroph\| | Yeast | NULL | Probable | As nectar-tap_saprotroph (Põlme et al. 2020); Decay sugar-rich substrates (Põlme et al. 2020) | Põlme S, et al. 2020. Fungal Diversity 105:1-16 ((https://doi.org/10.1007/s13225-020-00466-2)) |
| Otu39387 | 1 | k__Fungi(100);p__Ascomycota(100);c__Saccharomycetes(100);o__Saccharomycetales(100);f__Metschnikowiaceae(100);g__Metschnikowia(100);g__Metschnikowia_unclassified(100); | Metschnikowia | 13 | Saprotroph | \|Undefined Saprotroph\| | Yeast | NULL | Probable | As nectar-tap_saprotroph (Põlme et al. 2020); Decay sugar-rich substrates (Põlme et al. 2020) | Põlme S, et al. 2020. Fungal Diversity 105:1-16 ((https://doi.org/10.1007/s13225-020-00466-2)) |
| Otu39388 | 1 | k__Fungi(100);p__Ascomycota(100);c__Saccharomycetes(100);o__Saccharomycetales(100);f__Metschnikowiaceae(100);g__Metschnikowia(100);g__Metschnikowia_unclassified(100); | Metschnikowia | 13 | Saprotroph | \|Undefined Saprotroph\| | Yeast | NULL | Probable | As nectar-tap_saprotroph (Põlme et al. 2020); Decay sugar-rich substrates (Põlme et al. 2020) | Põlme S, et al. 2020. Fungal Diversity 105:1-16 ((https://doi.org/10.1007/s13225-020-00466-2)) |
| Otu39389 | 1 | k__Fungi(100);p__Ascomycota(100);c__Saccharomycetes(100);o__Saccharomycetales(100);f__Metschnikowiaceae(100);g__Metschnikowia(100);s__Metschnikowia_pulcherrima(100); | Metschnikowia | 13 | Saprotroph | \|Undefined Saprotroph\| | Yeast | NULL | Probable | As nectar-tap_saprotroph (Põlme et al. 2020); Decay sugar-rich substrates (Põlme et al. 2020) | Põlme S, et al. 2020. Fungal Diversity 105:1-16 ((https://doi.org/10.1007/s13225-020-00466-2)) |
| Otu39390 | 1 | k__Fungi(100);p__Ascomycota(100);c__Saccharomycetes(100);o__Saccharomycetales(100);f__Metschnikowiaceae(100);g__Metschnikowia(100);s__Metschnikowia_pulcherrima(100); | Metschnikowia | 13 | Saprotroph | \|Undefined Saprotroph\| | Yeast | NULL | Probable | As nectar-tap_saprotroph (Põlme et al. 2020); Decay sugar-rich substrates (Põlme et al. 2020) | Põlme S, et al. 2020. Fungal Diversity 105:1-16 ((https://doi.org/10.1007/s13225-020-00466-2)) |
| Otu39391 | 1 | k__Fungi(100);p__Ascomycota(100);c__Saccharomycetes(100);o__Saccharomycetales(100);f__Metschnikowiaceae(100);g__Metschnikowia(100);s__Metschnikowia_pulcherrima(100); | Metschnikowia | 13 | Saprotroph | \|Undefined Saprotroph\| | Yeast | NULL | Probable | As nectar-tap_saprotroph (Põlme et al. 2020); Decay sugar-rich substrates (Põlme et al. 2020) | Põlme S, et al. 2020. Fungal Diversity 105:1-16 ((https://doi.org/10.1007/s13225-020-00466-2)) |
| Otu39392 | 1 | k__Fungi(100);p__Ascomycota(100);c__Saccharomycetes(100);o__Saccharomycetales(100);f__Metschnikowiaceae(100);g__Metschnikowia(100);s__Metschnikowia_pulcherrima(100); | Metschnikowia | 13 | Saprotroph | \|Undefined Saprotroph\| | Yeast | NULL | Probable | As nectar-tap_saprotroph (Põlme et al. 2020); Decay sugar-rich substrates (Põlme et al. 2020) | Põlme S, et al. 2020. Fungal Diversity 105:1-16 ((https://doi.org/10.1007/s13225-020-00466-2)) |
| Otu39393 | 1 | k__Fungi(100);p__Ascomycota(100);c__Saccharomycetes(100);o__Saccharomycetales(100);f__Metschnikowiaceae(100);g__Metschnikowia(100);s__Metschnikowia_pulcherrima(100); | Metschnikowia | 13 | Saprotroph | \|Undefined Saprotroph\| | Yeast | NULL | Probable | As nectar-tap_saprotroph (Põlme et al. 2020); Decay sugar-rich substrates (Põlme et al. 2020) | Põlme S, et al. 2020. Fungal Diversity 105:1-16 ((https://doi.org/10.1007/s13225-020-00466-2)) |
| Otu39394 | 1 | k__Fungi(100);p__Ascomycota(100);c__Saccharomycetes(100);o__Saccharomycetales(100);f__Metschnikowiaceae(100);g__Metschnikowia(100);s__Metschnikowia_pulcherrima(100); | Metschnikowia | 13 | Saprotroph | \|Undefined Saprotroph\| | Yeast | NULL | Probable | As nectar-tap_saprotroph (Põlme et al. 2020); Decay sugar-rich substrates (Põlme et al. 2020) | Põlme S, et al. 2020. Fungal Diversity 105:1-16 ((https://doi.org/10.1007/s13225-020-00466-2)) |
| Otu39395 | 1 | k__Fungi(100);p__Ascomycota(100);c__Saccharomycetes(100);o__Saccharomycetales(100);f__Metschnikowiaceae(100);g__Metschnikowia(100);g__Metschnikowia_unclassified(100); | Metschnikowia | 13 | Saprotroph | \|Undefined Saprotroph\| | Yeast | NULL | Probable | As nectar-tap_saprotroph (Põlme et al. 2020); Decay sugar-rich substrates (Põlme et al. 2020) | Põlme S, et al. 2020. Fungal Diversity 105:1-16 ((https://doi.org/10.1007/s13225-020-00466-2)) |
| Otu39396 | 1 | k__Fungi(100);p__Ascomycota(100);c__Saccharomycetes(100);o__Saccharomycetales(100);f__Metschnikowiaceae(100);g__Metschnikowia(100);s__Metschnikowia_pulcherrima(100); | Metschnikowia | 13 | Saprotroph | \|Undefined Saprotroph\| | Yeast | NULL | Probable | As nectar-tap_saprotroph (Põlme et al. 2020); Decay sugar-rich substrates (Põlme et al. 2020) | Põlme S, et al. 2020. Fungal Diversity 105:1-16 ((https://doi.org/10.1007/s13225-020-00466-2)) |
| Otu39397 | 1 | k__Fungi(100);p__Ascomycota(100);c__Saccharomycetes(100);o__Saccharomycetales(100);f__Metschnikowiaceae(100);g__Metschnikowia(100);s__Metschnikowia_pulcherrima(100); | Metschnikowia | 13 | Saprotroph | \|Undefined Saprotroph\| | Yeast | NULL | Probable | As nectar-tap_saprotroph (Põlme et al. 2020); Decay sugar-rich substrates (Põlme et al. 2020) | Põlme S, et al. 2020. Fungal Diversity 105:1-16 ((https://doi.org/10.1007/s13225-020-00466-2)) |
| Otu39399 | 1 | k__Fungi(100);p__Ascomycota(100);c__Saccharomycetes(100);o__Saccharomycetales(100);f__Metschnikowiaceae(100);g__Metschnikowia(100);s__Metschnikowia_pulcherrima(100); | Metschnikowia | 13 | Saprotroph | \|Undefined Saprotroph\| | Yeast | NULL | Probable | As nectar-tap_saprotroph (Põlme et al. 2020); Decay sugar-rich substrates (Põlme et al. 2020) | Põlme S, et al. 2020. Fungal Diversity 105:1-16 ((https://doi.org/10.1007/s13225-020-00466-2)) |
| Otu39400 | 1 | k__Fungi(100);p__Ascomycota(100);c__Saccharomycetes(100);o__Saccharomycetales(100);f__Metschnikowiaceae(100);g__Metschnikowia(100);s__Metschnikowia_pulcherrima(100); | Metschnikowia | 13 | Saprotroph | \|Undefined Saprotroph\| | Yeast | NULL | Probable | As nectar-tap_saprotroph (Põlme et al. 2020); Decay sugar-rich substrates (Põlme et al. 2020) | Põlme S, et al. 2020. Fungal Diversity 105:1-16 ((https://doi.org/10.1007/s13225-020-00466-2)) |
| Otu39401 | 1 | k__Fungi(100);p__Ascomycota(100);c__Saccharomycetes(100);o__Saccharomycetales(100);f__Metschnikowiaceae(100);g__Metschnikowia(100);s__Metschnikowia_pulcherrima(100); | Metschnikowia | 13 | Saprotroph | \|Undefined Saprotroph\| | Yeast | NULL | Probable | As nectar-tap_saprotroph (Põlme et al. 2020); Decay sugar-rich substrates (Põlme et al. 2020) | Põlme S, et al. 2020. Fungal Diversity 105:1-16 ((https://doi.org/10.1007/s13225-020-00466-2)) |
| Otu39403 | 1 | k__Fungi(100);p__Ascomycota(100);c__Saccharomycetes(100);o__Saccharomycetales(100);f__Metschnikowiaceae(100);g__Metschnikowia(100);s__Metschnikowia_pulcherrima(100); | Metschnikowia | 13 | Saprotroph | \|Undefined Saprotroph\| | Yeast | NULL | Probable | As nectar-tap_saprotroph (Põlme et al. 2020); Decay sugar-rich substrates (Põlme et al. 2020) | Põlme S, et al. 2020. Fungal Diversity 105:1-16 ((https://doi.org/10.1007/s13225-020-00466-2)) |
| Otu39404 | 1 | k__Fungi(100);p__Ascomycota(100);c__Saccharomycetes(100);o__Saccharomycetales(100);f__Metschnikowiaceae(100);g__Metschnikowia(100);s__Metschnikowia_pulcherrima(100); | Metschnikowia | 13 | Saprotroph | \|Undefined Saprotroph\| | Yeast | NULL | Probable | As nectar-tap_saprotroph (Põlme et al. 2020); Decay sugar-rich substrates (Põlme et al. 2020) | Põlme S, et al. 2020. Fungal Diversity 105:1-16 ((https://doi.org/10.1007/s13225-020-00466-2)) |
| Otu39405 | 1 | k__Fungi(100);p__Ascomycota(100);c__Saccharomycetes(100);o__Saccharomycetales(100);f__Metschnikowiaceae(100);g__Metschnikowia(100);g__Metschnikowia_unclassified(100); | Metschnikowia | 13 | Saprotroph | \|Undefined Saprotroph\| | Yeast | NULL | Probable | As nectar-tap_saprotroph (Põlme et al. 2020); Decay sugar-rich substrates (Põlme et al. 2020) | Põlme S, et al. 2020. Fungal Diversity 105:1-16 ((https://doi.org/10.1007/s13225-020-00466-2)) |
| Otu39406 | 1 | k__Fungi(100);p__Ascomycota(100);c__Saccharomycetes(100);o__Saccharomycetales(100);f__Metschnikowiaceae(100);g__Metschnikowia(100);g__Metschnikowia_unclassified(100); | Metschnikowia | 13 | Saprotroph | \|Undefined Saprotroph\| | Yeast | NULL | Probable | As nectar-tap_saprotroph (Põlme et al. 2020); Decay sugar-rich substrates (Põlme et al. 2020) | Põlme S, et al. 2020. Fungal Diversity 105:1-16 ((https://doi.org/10.1007/s13225-020-00466-2)) |
| Otu39407 | 1 | k__Fungi(100);p__Ascomycota(100);c__Saccharomycetes(100);o__Saccharomycetales(100);f__Metschnikowiaceae(100);g__Metschnikowia(100);s__Metschnikowia_pulcherrima(100); | Metschnikowia | 13 | Saprotroph | \|Undefined Saprotroph\| | Yeast | NULL | Probable | As nectar-tap_saprotroph (Põlme et al. 2020); Decay sugar-rich substrates (Põlme et al. 2020) | Põlme S, et al. 2020. Fungal Diversity 105:1-16 ((https://doi.org/10.1007/s13225-020-00466-2)) |

| | | | | | | | | | | | |
|---|---|---|---|---|---|---|---|---|---|---|---|
| Otu39408 | 1 | k__Fungi(100);p__Ascomycota(100);c__Saccharomycetes(100);o__Saccharomycetales(100);f__Metschnikowiaceae(100);g__Metschnikowia(100);g__Metschnikowia_unclassified(100); | Metschnikowia | 13 | Saprotroph | \|Undefined Saprotroph\| | Yeast | NULL | Probable | As nectar-tap_saprotroph (Põlme et al. 2020); Decay sugar-rich substrates (Põlme et al. 2020) | Põlme S, et al. 2020. Fungal Diversity 105:1-16 ((https://doi.org/10.1007/s13225-020-00466-2)) |
| Otu39409 | 1 | k__Fungi(100);p__Ascomycota(100);c__Saccharomycetes(100);o__Saccharomycetales(100);f__Metschnikowiaceae(100);g__Metschnikowia(100);s__Metschnikowia_pulcherrima(100); | Metschnikowia | 13 | Saprotroph | \|Undefined Saprotroph\| | Yeast | NULL | Probable | As nectar-tap_saprotroph (Põlme et al. 2020); Decay sugar-rich substrates (Põlme et al. 2020) | Põlme S, et al. 2020. Fungal Diversity 105:1-16 ((https://doi.org/10.1007/s13225-020-00466-2)) |
| Otu39410 | 1 | k__Fungi(100);p__Ascomycota(100);c__Saccharomycetes(100);o__Saccharomycetales(100);f__Metschnikowiaceae(100);g__Metschnikowia(100);s__Metschnikowia_pulcherrima(100); | Metschnikowia | 13 | Saprotroph | \|Undefined Saprotroph\| | Yeast | NULL | Probable | As nectar-tap_saprotroph (Põlme et al. 2020); Decay sugar-rich substrates (Põlme et al. 2020) | Põlme S, et al. 2020. Fungal Diversity 105:1-16 ((https://doi.org/10.1007/s13225-020-00466-2)) |
| Otu39412 | 1 | k__Fungi(100);p__Ascomycota(100);c__Saccharomycetes(100);o__Saccharomycetales(100);f__Metschnikowiaceae(100);g__Metschnikowia(100);g__Metschnikowia_unclassified(100); | Metschnikowia | 13 | Saprotroph | \|Undefined Saprotroph\| | Yeast | NULL | Probable | As nectar-tap_saprotroph (Põlme et al. 2020); Decay sugar-rich substrates (Põlme et al. 2020) | Põlme S, et al. 2020. Fungal Diversity 105:1-16 ((https://doi.org/10.1007/s13225-020-00466-2)) |
| Otu39414 | 1 | k__Fungi(100);p__Ascomycota(100);c__Saccharomycetes(100);o__Saccharomycetales(100);f__Metschnikowiaceae(100);g__Metschnikowia(100);s__Metschnikowia_pulcherrima(100); | Metschnikowia | 13 | Saprotroph | \|Undefined Saprotroph\| | Yeast | NULL | Probable | As nectar-tap_saprotroph (Põlme et al. 2020); Decay sugar-rich substrates (Põlme et al. 2020) | Põlme S, et al. 2020. Fungal Diversity 105:1-16 ((https://doi.org/10.1007/s13225-020-00466-2)) |
| Otu39415 | 1 | k__Fungi(100);p__Ascomycota(100);c__Saccharomycetes(100);o__Saccharomycetales(100);f__Metschnikowiaceae(100);g__Metschnikowia(100);s__Metschnikowia_pulcherrima(100); | Metschnikowia | 13 | Saprotroph | \|Undefined Saprotroph\| | Yeast | NULL | Probable | As nectar-tap_saprotroph (Põlme et al. 2020); Decay sugar-rich substrates (Põlme et al. 2020) | Põlme S, et al. 2020. Fungal Diversity 105:1-16 ((https://doi.org/10.1007/s13225-020-00466-2)) |
| Otu39416 | 1 | k__Fungi(100);p__Ascomycota(100);c__Saccharomycetes(100);o__Saccharomycetales(100);f__Metschnikowiaceae(100);g__Metschnikowia(100);s__Metschnikowia_pulcherrima(100); | Metschnikowia | 13 | Saprotroph | \|Undefined Saprotroph\| | Yeast | NULL | Probable | As nectar-tap_saprotroph (Põlme et al. 2020); Decay sugar-rich substrates (Põlme et al. 2020) | Põlme S, et al. 2020. Fungal Diversity 105:1-16 ((https://doi.org/10.1007/s13225-020-00466-2)) |
| Otu39417 | 1 | k__Fungi(100);p__Ascomycota(100);c__Saccharomycetes(100);o__Saccharomycetales(100);f__Metschnikowiaceae(100);g__Metschnikowia(100);s__Metschnikowia_pulcherrima(100); | Metschnikowia | 13 | Saprotroph | \|Undefined Saprotroph\| | Yeast | NULL | Probable | As nectar-tap_saprotroph (Põlme et al. 2020); Decay sugar-rich substrates (Põlme et al. 2020) | Põlme S, et al. 2020. Fungal Diversity 105:1-16 ((https://doi.org/10.1007/s13225-020-00466-2)) |
| Otu39418 | 1 | k__Fungi(100);p__Ascomycota(100);c__Saccharomycetes(100);o__Saccharomycetales(100);f__Metschnikowiaceae(100);g__Metschnikowia(100);s__Metschnikowia_pulcherrima(100); | Metschnikowia | 13 | Saprotroph | \|Undefined Saprotroph\| | Yeast | NULL | Probable | As nectar-tap_saprotroph (Põlme et al. 2020); Decay sugar-rich substrates (Põlme et al. 2020) | Põlme S, et al. 2020. Fungal Diversity 105:1-16 ((https://doi.org/10.1007/s13225-020-00466-2)) |
| Otu39419 | 1 | k__Fungi(100);p__Ascomycota(100);c__Saccharomycetes(100);o__Saccharomycetales(100);f__Metschnikowiaceae(100);g__Metschnikowia(100);s__Metschnikowia_pulcherrima(100); | Metschnikowia | 13 | Saprotroph | \|Undefined Saprotroph\| | Yeast | NULL | Probable | As nectar-tap_saprotroph (Põlme et al. 2020); Decay sugar-rich substrates (Põlme et al. 2020) | Põlme S, et al. 2020. Fungal Diversity 105:1-16 ((https://doi.org/10.1007/s13225-020-00466-2)) |
| Otu39420 | 1 | k__Fungi(100);p__Ascomycota(100);c__Saccharomycetes(100);o__Saccharomycetales(100);f__Metschnikowiaceae(100);g__Metschnikowia(100);g__Metschnikowia_unclassified(100); | Metschnikowia | 13 | Saprotroph | \|Undefined Saprotroph\| | Yeast | NULL | Probable | As nectar-tap_saprotroph (Põlme et al. 2020); Decay sugar-rich substrates (Põlme et al. 2020) | Põlme S, et al. 2020. Fungal Diversity 105:1-16 ((https://doi.org/10.1007/s13225-020-00466-2)) |
| Otu39421 | 1 | k__Fungi(100);p__Ascomycota(100);c__Saccharomycetes(100);o__Saccharomycetales(100);f__Metschnikowiaceae(100);g__Metschnikowia(100);s__Metschnikowia_pulcherrima(100); | Metschnikowia | 13 | Saprotroph | \|Undefined Saprotroph\| | Yeast | NULL | Probable | As nectar-tap_saprotroph (Põlme et al. 2020); Decay sugar-rich substrates (Põlme et al. 2020) | Põlme S, et al. 2020. Fungal Diversity 105:1-16 ((https://doi.org/10.1007/s13225-020-00466-2)) |
| Otu39422 | 1 | k__Fungi(100);p__Ascomycota(100);c__Saccharomycetes(100);o__Saccharomycetales(100);f__Metschnikowiaceae(100);g__Metschnikowia(100);s__Metschnikowia_pulcherrima(100); | Metschnikowia | 13 | Saprotroph | \|Undefined Saprotroph\| | Yeast | NULL | Probable | As nectar-tap_saprotroph (Põlme et al. 2020); Decay sugar-rich substrates (Põlme et al. 2020) | Põlme S, et al. 2020. Fungal Diversity 105:1-16 ((https://doi.org/10.1007/s13225-020-00466-2)) |
| Otu39423 | 1 | k__Fungi(100);p__Ascomycota(100);c__Saccharomycetes(100);o__Saccharomycetales(100);f__Metschnikowiaceae(100);g__Metschnikowia(100);s__Metschnikowia_pulcherrima(100); | Metschnikowia | 13 | Saprotroph | \|Undefined Saprotroph\| | Yeast | NULL | Probable | As nectar-tap_saprotroph (Põlme et al. 2020); Decay sugar-rich substrates (Põlme et al. 2020) | Põlme S, et al. 2020. Fungal Diversity 105:1-16 ((https://doi.org/10.1007/s13225-020-00466-2)) |
| Otu39424 | 1 | k__Fungi(100);p__Ascomycota(100);c__Saccharomycetes(100);o__Saccharomycetales(100);f__Metschnikowiaceae(100);g__Metschnikowia(100);s__Metschnikowia_pulcherrima(100); | Metschnikowia | 13 | Saprotroph | \|Undefined Saprotroph\| | Yeast | NULL | Probable | As nectar-tap_saprotroph (Põlme et al. 2020); Decay sugar-rich substrates (Põlme et al. 2020) | Põlme S, et al. 2020. Fungal Diversity 105:1-16 ((https://doi.org/10.1007/s13225-020-00466-2)) |
| Otu39425 | 1 | k__Fungi(100);p__Ascomycota(100);c__Saccharomycetes(100);o__Saccharomycetales(100);f__Metschnikowiaceae(100);g__Metschnikowia(100);s__Metschnikowia_pulcherrima(100); | Metschnikowia | 13 | Saprotroph | \|Undefined Saprotroph\| | Yeast | NULL | Probable | As nectar-tap_saprotroph (Põlme et al. 2020); Decay sugar-rich substrates (Põlme et al. 2020) | Põlme S, et al. 2020. Fungal Diversity 105:1-16 ((https://doi.org/10.1007/s13225-020-00466-2)) |
| Otu39426 | 1 | k__Fungi(100);p__Ascomycota(100);c__Saccharomycetes(100);o__Saccharomycetales(100);f__Metschnikowiaceae(100);g__Metschnikowia(100);s__Metschnikowia_pulcherrima(100); | Metschnikowia | 13 | Saprotroph | \|Undefined Saprotroph\| | Yeast | NULL | Probable | As nectar-tap_saprotroph (Põlme et al. 2020); Decay sugar-rich substrates (Põlme et al. 2020) | Põlme S, et al. 2020. Fungal Diversity 105:1-16 ((https://doi.org/10.1007/s13225-020-00466-2)) |
| Otu39427 | 1 | k__Fungi(100);p__Ascomycota(100);c__Saccharomycetes(100);o__Saccharomycetales(100);f__Metschnikowiaceae(100);g__Metschnikowia(100);s__Metschnikowia_pulcherrima(100); | Metschnikowia | 13 | Saprotroph | \|Undefined Saprotroph\| | Yeast | NULL | Probable | As nectar-tap_saprotroph (Põlme et al. 2020); Decay sugar-rich substrates (Põlme et al. 2020) | Põlme S, et al. 2020. Fungal Diversity 105:1-16 ((https://doi.org/10.1007/s13225-020-00466-2)) |
| Otu39428 | 1 | k__Fungi(100);p__Ascomycota(100);c__Saccharomycetes(100);o__Saccharomycetales(100);f__Metschnikowiaceae(100);g__Metschnikowia(100);s__Metschnikowia_pulcherrima(100); | Metschnikowia | 13 | Saprotroph | \|Undefined Saprotroph\| | Yeast | NULL | Probable | As nectar-tap_saprotroph (Põlme et al. 2020); Decay sugar-rich substrates (Põlme et al. 2020) | Põlme S, et al. 2020. Fungal Diversity 105:1-16 ((https://doi.org/10.1007/s13225-020-00466-2)) |
| Otu39430 | 1 | k__Fungi(100);p__Ascomycota(100);c__Saccharomycetes(100);o__Saccharomycetales(100);f__Metschnikowiaceae(100);g__Metschnikowia(100);s__Metschnikowia_pulcherrima(100); | Metschnikowia | 13 | Saprotroph | \|Undefined Saprotroph\| | Yeast | NULL | Probable | As nectar-tap_saprotroph (Põlme et al. 2020); Decay sugar-rich substrates (Põlme et al. 2020) | Põlme S, et al. 2020. Fungal Diversity 105:1-16 ((https://doi.org/10.1007/s13225-020-00466-2)) |
| Otu39431 | 1 | k__Fungi(100);p__Ascomycota(100);c__Saccharomycetes(100);o__Saccharomycetales(100);f__Metschnikowiaceae(100);g__Metschnikowia(100);s__Metschnikowia_pulcherrima(100); | Metschnikowia | 13 | Saprotroph | \|Undefined Saprotroph\| | Yeast | NULL | Probable | As nectar-tap_saprotroph (Põlme et al. 2020); Decay sugar-rich substrates (Põlme et al. 2020) | Põlme S, et al. 2020. Fungal Diversity 105:1-16 ((https://doi.org/10.1007/s13225-020-00466-2)) |
| Otu39432 | 1 | k__Fungi(100);p__Ascomycota(100);c__Saccharomycetes(100);o__Saccharomycetales(100);f__Metschnikowiaceae(100);g__Metschnikowia(100);s__Metschnikowia_pulcherrima(100); | Metschnikowia | 13 | Saprotroph | \|Undefined Saprotroph\| | Yeast | NULL | Probable | As nectar-tap_saprotroph (Põlme et al. 2020); Decay sugar-rich substrates (Põlme et al. 2020) | Põlme S, et al. 2020. Fungal Diversity 105:1-16 ((https://doi.org/10.1007/s13225-020-00466-2)) |
| Otu39433 | 1 | k__Fungi(100);p__Ascomycota(100);c__Saccharomycetes(100);o__Saccharomycetales(100);f__Metschnikowiaceae(100);g__Metschnikowia(100);s__Metschnikowia_pulcherrima(100); | Metschnikowia | 13 | Saprotroph | \|Undefined Saprotroph\| | Yeast | NULL | Probable | As nectar-tap_saprotroph (Põlme et al. 2020); Decay sugar-rich substrates (Põlme et al. 2020) | Põlme S, et al. 2020. Fungal Diversity 105:1-16 ((https://doi.org/10.1007/s13225-020-00466-2)) |
| Otu39434 | 1 | k__Fungi(100);p__Ascomycota(100);c__Saccharomycetes(100);o__Saccharomycetales(100);f__Metschnikowiaceae(100);g__Metschnikowia(100);g__Metschnikowia_unclassified(100); | Metschnikowia | 13 | Saprotroph | \|Undefined Saprotroph\| | Yeast | NULL | Probable | As nectar-tap_saprotroph (Põlme et al. 2020); Decay sugar-rich substrates (Põlme et al. 2020) | Põlme S, et al. 2020. Fungal Diversity 105:1-16 ((https://doi.org/10.1007/s13225-020-00466-2)) |
| Otu39435 | 1 | k__Fungi(100);p__Ascomycota(100);c__Saccharomycetes(100);o__Saccharomycetales(100);f__Metschnikowiaceae(100);g__Metschnikowia(100);g__Metschnikowia_unclassified(100); | Metschnikowia | 13 | Saprotroph | \|Undefined Saprotroph\| | Yeast | NULL | Probable | As nectar-tap_saprotroph (Põlme et al. 2020); Decay sugar-rich substrates (Põlme et al. 2020) | Põlme S, et al. 2020. Fungal Diversity 105:1-16 ((https://doi.org/10.1007/s13225-020-00466-2)) |
| Otu39436 | 1 | k__Fungi(100);p__Ascomycota(100);c__Saccharomycetes(100);o__Saccharomycetales(100);f__Metschnikowiaceae(100);g__Metschnikowia(100);s__Metschnikowia_pulcherrima(100); | Metschnikowia | 13 | Saprotroph | \|Undefined Saprotroph\| | Yeast | NULL | Probable | As nectar-tap_saprotroph (Põlme et al. 2020); Decay sugar-rich substrates (Põlme et al. 2020) | Põlme S, et al. 2020. Fungal Diversity 105:1-16 ((https://doi.org/10.1007/s13225-020-00466-2)) |
| Otu39437 | 1 | k__Fungi(100);p__Ascomycota(100);c__Saccharomycetes(100);o__Saccharomycetales(100);f__Metschnikowiaceae(100);g__Metschnikowia(100);s__Metschnikowia_pulcherrima(100); | Metschnikowia | 13 | Saprotroph | \|Undefined Saprotroph\| | Yeast | NULL | Probable | As nectar-tap_saprotroph (Põlme et al. 2020); Decay sugar-rich substrates (Põlme et al. 2020) | Põlme S, et al. 2020. Fungal Diversity 105:1-16 ((https://doi.org/10.1007/s13225-020-00466-2)) |
| Otu39438 | 1 | k__Fungi(100);p__Ascomycota(100);c__Saccharomycetes(100);o__Saccharomycetales(100);f__Metschnikowiaceae(100);g__Metschnikowia(100);g__Metschnikowia_unclassified(100); | Metschnikowia | 13 | Saprotroph | \|Undefined Saprotroph\| | Yeast | NULL | Probable | As nectar-tap_saprotroph (Põlme et al. 2020); Decay sugar-rich substrates (Põlme et al. 2020) | Põlme S, et al. 2020. Fungal Diversity 105:1-16 ((https://doi.org/10.1007/s13225-020-00466-2)) |
| Otu39439 | 1 | k__Fungi(100);p__Ascomycota(100);c__Saccharomycetes(100);o__Saccharomycetales(100);f__Metschnikowiaceae(100);g__Metschnikowia(100);s__Metschnikowia_pulcherrima(100); | Metschnikowia | 13 | Saprotroph | \|Undefined Saprotroph\| | Yeast | NULL | Probable | As nectar-tap_saprotroph (Põlme et al. 2020); Decay sugar-rich substrates (Põlme et al. 2020) | Põlme S, et al. 2020. Fungal Diversity 105:1-16 ((https://doi.org/10.1007/s13225-020-00466-2)) |
| Otu39440 | 1 | k__Fungi(100);p__Ascomycota(100);c__Saccharomycetes(100);o__Saccharomycetales(100);f__Metschnikowiaceae(100);g__Metschnikowia(100);s__Metschnikowia_pulcherrima(100); | Metschnikowia | 13 | Saprotroph | \|Undefined Saprotroph\| | Yeast | NULL | Probable | As nectar-tap_saprotroph (Põlme et al. 2020); Decay sugar-rich substrates (Põlme et al. 2020) | Põlme S, et al. 2020. Fungal Diversity 105:1-16 ((https://doi.org/10.1007/s13225-020-00466-2)) |
| Otu39441 | 1 | k__Fungi(100);p__Ascomycota(100);c__Saccharomycetes(100);o__Saccharomycetales(100);f__Metschnikowiaceae(100);g__Metschnikowia(100);s__Metschnikowia_pulcherrima(100); | Metschnikowia | 13 | Saprotroph | \|Undefined Saprotroph\| | Yeast | NULL | Probable | As nectar-tap_saprotroph (Põlme et al. 2020); Decay sugar-rich substrates (Põlme et al. 2020) | Põlme S, et al. 2020. Fungal Diversity 105:1-16 ((https://doi.org/10.1007/s13225-020-00466-2)) |
| Otu39442 | 1 | k__Fungi(100);p__Ascomycota(100);c__Saccharomycetes(100);o__Saccharomycetales(100);f__Metschnikowiaceae(100);g__Metschnikowia(100);s__Metschnikowia_pulcherrima(100); | Metschnikowia | 13 | Saprotroph | \|Undefined Saprotroph\| | Yeast | NULL | Probable | As nectar-tap_saprotroph (Põlme et al. 2020); Decay sugar-rich substrates (Põlme et al. 2020) | Põlme S, et al. 2020. Fungal Diversity 105:1-16 ((https://doi.org/10.1007/s13225-020-00466-2)) |
| Otu39443 | 1 | k__Fungi(100);p__Ascomycota(100);c__Saccharomycetes(100);o__Saccharomycetales(100);f__Metschnikowiaceae(100);g__Metschnikowia(100);s__Metschnikowia_pulcherrima(100); | Metschnikowia | 13 | Saprotroph | \|Undefined Saprotroph\| | Yeast | NULL | Probable | As nectar-tap_saprotroph (Põlme et al. 2020); Decay sugar-rich substrates (Põlme et al. 2020) | Põlme S, et al. 2020. Fungal Diversity 105:1-16 ((https://doi.org/10.1007/s13225-020-00466-2)) |
| Otu39444 | 1 | k__Fungi(100);p__Ascomycota(100);c__Saccharomycetes(100);o__Saccharomycetales(100);f__Metschnikowiaceae(100);g__Metschnikowia(100);g__Metschnikowia_unclassified(100); | Metschnikowia | 13 | Saprotroph | \|Undefined Saprotroph\| | Yeast | NULL | Probable | As nectar-tap_saprotroph (Põlme et al. 2020); Decay sugar-rich substrates (Põlme et al. 2020) | Põlme S, et al. 2020. Fungal Diversity 105:1-16 ((https://doi.org/10.1007/s13225-020-00466-2)) |
| Otu39445 | 1 | k__Fungi(100);p__Ascomycota(100);c__Saccharomycetes(100);o__Saccharomycetales(100);f__Metschnikowiaceae(100);g__Metschnikowia(100);g__Metschnikowia_unclassified(100); | Metschnikowia | 13 | Saprotroph | \|Undefined Saprotroph\| | Yeast | NULL | Probable | As nectar-tap_saprotroph (Põlme et al. 2020); Decay sugar-rich substrates (Põlme et al. 2020) | Põlme S, et al. 2020. Fungal Diversity 105:1-16 ((https://doi.org/10.1007/s13225-020-00466-2)) |
| Otu39446 | 1 | k__Fungi(100);p__Ascomycota(100);c__Saccharomycetes(100);o__Saccharomycetales(100);f__Metschnikowiaceae(100);g__Metschnikowia(100);s__Metschnikowia_pulcherrima(100); | Metschnikowia | 13 | Saprotroph | \|Undefined Saprotroph\| | Yeast | NULL | Probable | As nectar-tap_saprotroph (Põlme et al. 2020); Decay sugar-rich substrates (Põlme et al. 2020) | Põlme S, et al. 2020. Fungal Diversity 105:1-16 ((https://doi.org/10.1007/s13225-020-00466-2)) |
| Otu39447 | 1 | k__Fungi(100);p__Ascomycota(100);c__Saccharomycetes(100);o__Saccharomycetales(100);f__Metschnikowiaceae(100);g__Metschnikowia(100);s__Metschnikowia_pulcherrima(100); | Metschnikowia | 13 | Saprotroph | \|Undefined Saprotroph\| | Yeast | NULL | Probable | As nectar-tap_saprotroph (Põlme et al. 2020); Decay sugar-rich substrates (Põlme et al. 2020) | Põlme S, et al. 2020. Fungal Diversity 105:1-16 ((https://doi.org/10.1007/s13225-020-00466-2)) |
| Otu39448 | 1 | k__Fungi(100);p__Ascomycota(100);c__Saccharomycetes(100);o__Saccharomycetales(100);f__Metschnikowiaceae(100);g__Metschnikowia(100);s__Metschnikowia_pulcherrima(100); | Metschnikowia | 13 | Saprotroph | \|Undefined Saprotroph\| | Yeast | NULL | Probable | As nectar-tap_saprotroph (Põlme et al. 2020); Decay sugar-rich substrates (Põlme et al. 2020) | Põlme S, et al. 2020. Fungal Diversity 105:1-16 ((https://doi.org/10.1007/s13225-020-00466-2)) |
| Otu39449 | 1 | k__Fungi(100);p__Ascomycota(100);c__Saccharomycetes(100);o__Saccharomycetales(100);f__Metschnikowiaceae(100);g__Metschnikowia(100);s__Metschnikowia_pulcherrima(100); | Metschnikowia | 13 | Saprotroph | \|Undefined Saprotroph\| | Yeast | NULL | Probable | As nectar-tap_saprotroph (Põlme et al. 2020); Decay sugar-rich substrates (Põlme et al. 2020) | Põlme S, et al. 2020. Fungal Diversity 105:1-16 ((https://doi.org/10.1007/s13225-020-00466-2)) |
| Otu39450 | 1 | k__Fungi(100);p__Ascomycota(100);c__Saccharomycetes(100);o__Saccharomycetales(100);f__Metschnikowiaceae(100);g__Metschnikowia(100);g__Metschnikowia_unclassified(100); | Metschnikowia | 13 | Saprotroph | \|Undefined Saprotroph\| | Yeast | NULL | Probable | As nectar-tap_saprotroph (Põlme et al. 2020); Decay sugar-rich substrates (Põlme et al. 2020) | Põlme S, et al. 2020. Fungal Diversity 105:1-16 ((https://doi.org/10.1007/s13225-020-00466-2)) |
| Otu39451 | 1 | k__Fungi(100);p__Ascomycota(100);c__Saccharomycetes(100);o__Saccharomycetales(100);f__Metschnikowiaceae(100);g__Metschnikowia(100);g__Metschnikowia_unclassified(100); | Metschnikowia | 13 | Saprotroph | \|Undefined Saprotroph\| | Yeast | NULL | Probable | As nectar-tap_saprotroph (Põlme et al. 2020); Decay sugar-rich substrates (Põlme et al. 2020) | Põlme S, et al. 2020. Fungal Diversity 105:1-16 ((https://doi.org/10.1007/s13225-020-00466-2)) |
| Otu39452 | 1 | k__Fungi(100);p__Ascomycota(100);c__Saccharomycetes(100);o__Saccharomycetales(100);f__Metschnikowiaceae(100);g__Metschnikowia(100);s__Metschnikowia_pulcherrima(100); | Metschnikowia | 13 | Saprotroph | \|Undefined Saprotroph\| | Yeast | NULL | Probable | As nectar-tap_saprotroph (Põlme et al. 2020); Decay sugar-rich substrates (Põlme et al. 2020) | Põlme S, et al. 2020. Fungal Diversity 105:1-16 ((https://doi.org/10.1007/s13225-020-00466-2)) |
| Otu39453 | 1 | k__Fungi(100);p__Ascomycota(100);c__Saccharomycetes(100);o__Saccharomycetales(100);f__Metschnikowiaceae(100);g__Metschnikowia(100);g__Metschnikowia_unclassified(100); | Metschnikowia | 13 | Saprotroph | \|Undefined Saprotroph\| | Yeast | NULL | Probable | As nectar-tap_saprotroph (Põlme et al. 2020); Decay sugar-rich substrates (Põlme et al. 2020) | Põlme S, et al. 2020. Fungal Diversity 105:1-16 ((https://doi.org/10.1007/s13225-020-00466-2)) |
| Otu39454 | 1 | k__Fungi(100);p__Ascomycota(100);c__Saccharomycetes(100);o__Saccharomycetales(100);f__Metschnikowiaceae(100);g__Metschnikowia(100);g__Metschnikowia_unclassified(100); | Metschnikowia | 13 | Saprotroph | \|Undefined Saprotroph\| | Yeast | NULL | Probable | As nectar-tap_saprotroph (Põlme et al. 2020); Decay sugar-rich substrates (Põlme et al. 2020) | Põlme S, et al. 2020. Fungal Diversity 105:1-16 ((https://doi.org/10.1007/s13225-020-00466-2)) |
| Otu39455 | 1 | k__Fungi(100);p__Ascomycota(100);c__Saccharomycetes(100);o__Saccharomycetales(100);f__Metschnikowiaceae(100);g__Metschnikowia(100);g__Metschnikowia_unclassified(100); | Metschnikowia | 13 | Saprotroph | \|Undefined Saprotroph\| | Yeast | NULL | Probable | As nectar-tap_saprotroph (Põlme et al. 2020); Decay sugar-rich substrates (Põlme et al. 2020) | Põlme S, et al. 2020. Fungal Diversity 105:1-16 ((https://doi.org/10.1007/s13225-020-00466-2)) |
| Otu39456 | 1 | k__Fungi(100);p__Ascomycota(100);c__Saccharomycetes(100);o__Saccharomycetales(100);f__Metschnikowiaceae(100);g__Metschnikowia(100);s__Metschnikowia_pulcherrima(100); | Metschnikowia | 13 | Saprotroph | \|Undefined Saprotroph\| | Yeast | NULL | Probable | As nectar-tap_saprotroph (Põlme et al. 2020); Decay sugar-rich substrates (Põlme et al. 2020) | Põlme S, et al. 2020. Fungal Diversity 105:1-16 ((https://doi.org/10.1007/s13225-020-00466-2)) |
| Otu39457 | 1 | k__Fungi(100);p__Ascomycota(100);c__Saccharomycetes(100);o__Saccharomycetales(100);f__Metschnikowiaceae(100);g__Metschnikowia(100);g__Metschnikowia_unclassified(100); | Metschnikowia | 13 | Saprotroph | \|Undefined Saprotroph\| | Yeast | NULL | Probable | As nectar-tap_saprotroph (Põlme et al. 2020); Decay sugar-rich substrates (Põlme et al. 2020) | Põlme S, et al. 2020. Fungal Diversity 105:1-16 ((https://doi.org/10.1007/s13225-020-00466-2)) |
| Otu39458 | 1 | k__Fungi(100);p__Ascomycota(100);c__Saccharomycetes(100);o__Saccharomycetales(100);f__Metschnikowiaceae(100);g__Metschnikowia(100);s__Metschnikowia_pulcherrima(100); | Metschnikowia | 13 | Saprotroph | \|Undefined Saprotroph\| | Yeast | NULL | Probable | As nectar-tap_saprotroph (Põlme et al. 2020); Decay sugar-rich substrates (Põlme et al. 2020) | Põlme S, et al. 2020. Fungal Diversity 105:1-16 ((https://doi.org/10.1007/s13225-020-00466-2)) |
| Otu39459 | 1 | k__Fungi(100);p__Ascomycota(100);c__Saccharomycetes(100);o__Saccharomycetales(100);f__Metschnikowiaceae(100);g__Metschnikowia(100);g__Metschnikowia_unclassified(100); | Metschnikowia | 13 | Saprotroph | \|Undefined Saprotroph\| | Yeast | NULL | Probable | As nectar-tap_saprotroph (Põlme et al. 2020); Decay sugar-rich substrates (Põlme et al. 2020) | Põlme S, et al. 2020. Fungal Diversity 105:1-16 ((https://doi.org/10.1007/s13225-020-00466-2)) |
| Otu39460 | 1 | k__Fungi(100);p__Ascomycota(100);c__Saccharomycetes(100);o__Saccharomycetales(100);f__Metschnikowiaceae(100);g__Metschnikowia(100);g__Metschnikowia_unclassified(100); | Metschnikowia | 13 | Saprotroph | \|Undefined Saprotroph\| | Yeast | NULL | Probable | As nectar-tap_saprotroph (Põlme et al. 2020); Decay sugar-rich substrates (Põlme et al. 2020) | Põlme S, et al. 2020. Fungal Diversity 105:1-16 ((https://doi.org/10.1007/s13225-020-00466-2)) |
| Otu39461 | 1 | k__Fungi(100);p__Ascomycota(100);c__Saccharomycetes(100);o__Saccharomycetales(100);f__Metschnikowiaceae(100);g__Metschnikowia(100);s__Metschnikowia_pulcherrima(100); | Metschnikowia | 13 | Saprotroph | \|Undefined Saprotroph\| | Yeast | NULL | Probable | As nectar-tap_saprotroph (Põlme et al. 2020); Decay sugar-rich substrates (Põlme et al. 2020) | Põlme S, et al. 2020. Fungal Diversity 105:1-16 ((https://doi.org/10.1007/s13225-020-00466-2)) |
| Otu39462 | 1 | k__Fungi(100);p__Ascomycota(100);c__Saccharomycetes(100);o__Saccharomycetales(100);f__Metschnikowiaceae(100);g__Metschnikowia(100);s__Metschnikowia_pulcherrima(100); | Metschnikowia | 13 | Saprotroph | \|Undefined Saprotroph\| | Yeast | NULL | Probable | As nectar-tap_saprotroph (Põlme et al. 2020); Decay sugar-rich substrates (Põlme et al. 2020) | Põlme S, et al. 2020. Fungal Diversity 105:1-16 ((https://doi.org/10.1007/s13225-020-00466-2)) |
| Otu39463 | 1 | k__Fungi(100);p__Ascomycota(100);c__Saccharomycetes(100);o__Saccharomycetales(100);f__Metschnikowiaceae(100);g__Metschnikowia(100);g__Metschnikowia_unclassified(100); | Metschnikowia | 13 | Saprotroph | \|Undefined Saprotroph\| | Yeast | NULL | Probable | As nectar-tap_saprotroph (Põlme et al. 2020); Decay sugar-rich substrates (Põlme et al. 2020) | Põlme S, et al. 2020. Fungal Diversity 105:1-16 ((https://doi.org/10.1007/s13225-020-00466-2)) |
| Otu39464 | 1 | k__Fungi(100);p__Ascomycota(100);c__Saccharomycetes(100);o__Saccharomycetales(100);f__Metschnikowiaceae(100);g__Metschnikowia(100);s__Metschnikowia_pulcherrima(100); | Metschnikowia | 13 | Saprotroph | \|Undefined Saprotroph\| | Yeast | NULL | Probable | As nectar-tap_saprotroph (Põlme et al. 2020); Decay sugar-rich substrates (Põlme et al. 2020) | Põlme S, et al. 2020. Fungal Diversity 105:1-16 ((https://doi.org/10.1007/s13225-020-00466-2)) |
| Otu39465 | 1 | k__Fungi(100);p__Ascomycota(100);c__Saccharomycetes(100);o__Saccharomycetales(100);f__Metschnikowiaceae(100);g__Metschnikowia(100);s__Metschnikowia_pulcherrima(100); | Metschnikowia | 13 | Saprotroph | \|Undefined Saprotroph\| | Yeast | NULL | Probable | As nectar-tap_saprotroph (Põlme et al. 2020); Decay sugar-rich substrates (Põlme et al. 2020) | Põlme S, et al. 2020. Fungal Diversity 105:1-16 ((https://doi.org/10.1007/s13225-020-00466-2)) |
| Otu39466 | 1 | k__Fungi(100);p__Ascomycota(100);c__Saccharomycetes(100);o__Saccharomycetales(100);f__Metschnikowiaceae(100);g__Metschnikowia(100);s__Metschnikowia_pulcherrima(100); | Metschnikowia | 13 | Saprotroph | \|Undefined Saprotroph\| | Yeast | NULL | Probable | As nectar-tap_saprotroph (Põlme et al. 2020); Decay sugar-rich substrates (Põlme et al. 2020) | Põlme S, et al. 2020. Fungal Diversity 105:1-16 ((https://doi.org/10.1007/s13225-020-00466-2)) |
| Otu39467 | 1 | k__Fungi(100);p__Ascomycota(100);c__Saccharomycetes(100);o__Saccharomycetales(100);f__Metschnikowiaceae(100);g__Metschnikowia(100);g__Metschnikowia_unclassified(100); | Metschnikowia | 13 | Saprotroph | \|Undefined Saprotroph\| | Yeast | NULL | Probable | As nectar-tap_saprotroph (Põlme et al. 2020); Decay sugar-rich substrates (Põlme et al. 2020) | Põlme S, et al. 2020. Fungal Diversity 105:1-16 ((https://doi.org/10.1007/s13225-020-00466-2)) |
| Otu39468 | 1 | k__Fungi(100);p__Ascomycota(100);c__Saccharomycetes(100);o__Saccharomycetales(100);f__Metschnikowiaceae(100);g__Metschnikowia(100);s__Metschnikowia_pulcherrima(100); | Metschnikowia | 13 | Saprotroph | \|Undefined Saprotroph\| | Yeast | NULL | Probable | As nectar-tap_saprotroph (Põlme et al. 2020); Decay sugar-rich substrates (Põlme et al. 2020) | Põlme S, et al. 2020. Fungal Diversity 105:1-16 ((https://doi.org/10.1007/s13225-020-00466-2)) |
| Otu39469 | 1 | k__Fungi(100);p__Ascomycota(100);c__Saccharomycetes(100);o__Saccharomycetales(100);f__Metschnikowiaceae(100);g__Metschnikowia(100);s__Metschnikowia_pulcherrima(100); | Metschnikowia | 13 | Saprotroph | \|Undefined Saprotroph\| | Yeast | NULL | Probable | As nectar-tap_saprotroph (Põlme et al. 2020); Decay sugar-rich substrates (Põlme et al. 2020) | Põlme S, et al. 2020. Fungal Diversity 105:1-16 ((https://doi.org/10.1007/s13225-020-00466-2)) |
| Otu39470 | 1 | k__Fungi(100);p__Ascomycota(100);c__Saccharomycetes(100);o__Saccharomycetales(100);f__Metschnikowiaceae(100);g__Metschnikowia(100);s__Metschnikowia_pulcherrima(100); | Metschnikowia | 13 | Saprotroph | \|Undefined Saprotroph\| | Yeast | NULL | Probable | As nectar-tap_saprotroph (Põlme et al. 2020); Decay sugar-rich substrates (Põlme et al. 2020) | Põlme S, et al. 2020. Fungal Diversity 105:1-16 ((https://doi.org/10.1007/s13225-020-00466-2)) |
| Otu39471 | 1 | k__Fungi(100);p__Ascomycota(100);c__Saccharomycetes(100);o__Saccharomycetales(100);f__Metschnikowiaceae(100);g__Metschnikowia(100);g__Metschnikowia_unclassified(100); | Metschnikowia | 13 | Saprotroph | \|Undefined Saprotroph\| | Yeast | NULL | Probable | As nectar-tap_saprotroph (Põlme et al. 2020); Decay sugar-rich substrates (Põlme et al. 2020) | Põlme S, et al. 2020. Fungal Diversity 105:1-16 ((https://doi.org/10.1007/s13225-020-00466-2)) |
| Otu39472 | 1 | k__Fungi(100);p__Ascomycota(100);c__Saccharomycetes(100);o__Saccharomycetales(100);f__Metschnikowiaceae(100);g__Metschnikowia(100);s__Metschnikowia_pulcherrima(100); | Metschnikowia | 13 | Saprotroph | \|Undefined Saprotroph\| | Yeast | NULL | Probable | As nectar-tap_saprotroph (Põlme et al. 2020); Decay sugar-rich substrates (Põlme et al. 2020) | Põlme S, et al. 2020. Fungal Diversity 105:1-16 ((https://doi.org/10.1007/s13225-020-00466-2)) |
| Otu39473 | 1 | k__Fungi(100);p__Ascomycota(100);c__Saccharomycetes(100);o__Saccharomycetales(100);f__Metschnikowiaceae(100);g__Metschnikowia(100);s__Metschnikowia_pulcherrima(100); | Metschnikowia | 13 | Saprotroph | \|Undefined Saprotroph\| | Yeast | NULL | Probable | As nectar-tap_saprotroph (Põlme et al. 2020); Decay sugar-rich substrates (Põlme et al. 2020) | Põlme S, et al. 2020. Fungal Diversity 105:1-16 ((https://doi.org/10.1007/s13225-020-00466-2)) |
| Otu39474 | 1 | k__Fungi(100);p__Ascomycota(100);c__Saccharomycetes(100);o__Saccharomycetales(100);f__Metschnikowiaceae(100);g__Metschnikowia(100);s__Metschnikowia_pulcherrima(100); | Metschnikowia | 13 | Saprotroph | \|Undefined Saprotroph\| | Yeast | NULL | Probable | As nectar-tap_saprotroph (Põlme et al. 2020); Decay sugar-rich substrates (Põlme et al. 2020) | Põlme S, et al. 2020. Fungal Diversity 105:1-16 ((https://doi.org/10.1007/s13225-020-00466-2)) |
| Otu39475 | 1 | k__Fungi(100);p__Ascomycota(100);c__Saccharomycetes(100);o__Saccharomycetales(100);f__Metschnikowiaceae(100);g__Metschnikowia(100);s__Metschnikowia_pulcherrima(100); | Metschnikowia | 13 | Saprotroph | \|Undefined Saprotroph\| | Yeast | NULL | Probable | As nectar-tap_saprotroph (Põlme et al. 2020); Decay sugar-rich substrates (Põlme et al. 2020) | Põlme S, et al. 2020. Fungal Diversity 105:1-16 ((https://doi.org/10.1007/s13225-020-00466-2)) |
| Otu39476 | 1 | k__Fungi(100);p__Ascomycota(100);c__Saccharomycetes(100);o__Saccharomycetales(100);f__Metschnikowiaceae(100);g__Metschnikowia(100);s__Metschnikowia_pulcherrima(100); | Metschnikowia | 13 | Saprotroph | \|Undefined Saprotroph\| | Yeast | NULL | Probable | As nectar-tap_saprotroph (Põlme et al. 2020); Decay sugar-rich substrates (Põlme et al. 2020) | Põlme S, et al. 2020. Fungal Diversity 105:1-16 ((https://doi.org/10.1007/s13225-020-00466-2)) |
| Otu39477 | 1 | k__Fungi(100);p__Ascomycota(100);c__Saccharomycetes(100);o__Saccharomycetales(100);f__Metschnikowiaceae(100);g__Metschnikowia(100);s__Metschnikowia_pulcherrima(100); | Metschnikowia | 13 | Saprotroph | \|Undefined Saprotroph\| | Yeast | NULL | Probable | As nectar-tap_saprotroph (Põlme et al. 2020); Decay sugar-rich substrates (Põlme et al. 2020) | Põlme S, et al. 2020. Fungal Diversity 105:1-16 ((https://doi.org/10.1007/s13225-020-00466-2)) |
| Otu39478 | 1 | k__Fungi(100);p__Ascomycota(100);c__Saccharomycetes(100);o__Saccharomycetales(100);f__Metschnikowiaceae(100);g__Metschnikowia(100);s__Metschnikowia_pulcherrima(100); | Metschnikowia | 13 | Saprotroph | \|Undefined Saprotroph\| | Yeast | NULL | Probable | As nectar-tap_saprotroph (Põlme et al. 2020); Decay sugar-rich substrates (Põlme et al. 2020) | Põlme S, et al. 2020. Fungal Diversity 105:1-16 ((https://doi.org/10.1007/s13225-020-00466-2)) |
| Otu39479 | 1 | k__Fungi(100);p__Ascomycota(100);c__Saccharomycetes(100);o__Saccharomycetales(100);f__Metschnikowiaceae(100);g__Metschnikowia(100);g__Metschnikowia_unclassified(100); | Metschnikowia | 13 | Saprotroph | \|Undefined Saprotroph\| | Yeast | NULL | Probable | As nectar-tap_saprotroph (Põlme et al. 2020); Decay sugar-rich substrates (Põlme et al. 2020) | Põlme S, et al. 2020. Fungal Diversity 105:1-16 ((https://doi.org/10.1007/s13225-020-00466-2)) |
| Otu39480 | 1 | k__Fungi(100);p__Ascomycota(100);c__Saccharomycetes(100);o__Saccharomycetales(100);f__Metschnikowiaceae(100);g__Metschnikowia(100);g__Metschnikowia_unclassified(100); | Metschnikowia | 13 | Saprotroph | \|Undefined Saprotroph\| | Yeast | NULL | Probable | As nectar-tap_saprotroph (Põlme et al. 2020); Decay sugar-rich substrates (Põlme et al. 2020) | Põlme S, et al. 2020. Fungal Diversity 105:1-16 ((https://doi.org/10.1007/s13225-020-00466-2)) |
| Otu39481 | 1 | k__Fungi(100);p__Ascomycota(100);c__Saccharomycetes(100);o__Saccharomycetales(100);f__Metschnikowiaceae(100);g__Metschnikowia(100);s__Metschnikowia_pulcherrima(100); | Metschnikowia | 13 | Saprotroph | \|Undefined Saprotroph\| | Yeast | NULL | Probable | As nectar-tap_saprotroph (Põlme et al. 2020); Decay sugar-rich substrates (Põlme et al. 2020) | Põlme S, et al. 2020. Fungal Diversity 105:1-16 ((https://doi.org/10.1007/s13225-020-00466-2)) |
| Otu39482 | 1 | k__Fungi(100);p__Ascomycota(100);c__Saccharomycetes(100);o__Saccharomycetales(100);f__Metschnikowiaceae(100);g__Metschnikowia(100);s__Metschnikowia_pulcherrima(100); | Metschnikowia | 13 | Saprotroph | \|Undefined Saprotroph\| | Yeast | NULL | Probable | As nectar-tap_saprotroph (Põlme et al. 2020); Decay sugar-rich substrates (Põlme et al. 2020) | Põlme S, et al. 2020. Fungal Diversity 105:1-16 ((https://doi.org/10.1007/s13225-020-00466-2)) |
| Otu39483 | 1 | k__Fungi(100);p__Ascomycota(100);c__Saccharomycetes(100);o__Saccharomycetales(100);f__Metschnikowiaceae(100);g__Metschnikowia(100);s__Metschnikowia_pulcherrima(100); | Metschnikowia | 13 | Saprotroph | \|Undefined Saprotroph\| | Yeast | NULL | Probable | As nectar-tap_saprotroph (Põlme et al. 2020); Decay sugar-rich substrates (Põlme et al. 2020) | Põlme S, et al. 2020. Fungal Diversity 105:1-16 ((https://doi.org/10.1007/s13225-020-00466-2)) |
| Otu39484 | 1 | k__Fungi(100);p__Ascomycota(100);c__Saccharomycetes(100);o__Saccharomycetales(100);f__Metschnikowiaceae(100);g__Metschnikowia(100);s__Metschnikowia_pulcherrima(100); | Metschnikowia | 13 | Saprotroph | \|Undefined Saprotroph\| | Yeast | NULL | Probable | As nectar-tap_saprotroph (Põlme et al. 2020); Decay sugar-rich substrates (Põlme et al. 2020) | Põlme S, et al. 2020. Fungal Diversity 105:1-16 ((https://doi.org/10.1007/s13225-020-00466-2)) |
| Otu39485 | 1 | k__Fungi(100);p__Ascomycota(100);c__Saccharomycetes(100);o__Saccharomycetales(100);f__Metschnikowiaceae(100);g__Metschnikowia(100);s__Metschnikowia_pulcherrima(100); | Metschnikowia | 13 | Saprotroph | \|Undefined Saprotroph\| | Yeast | NULL | Probable | As nectar-tap_saprotroph (Põlme et al. 2020); Decay sugar-rich substrates (Põlme et al. 2020) | Põlme S, et al. 2020. Fungal Diversity 105:1-16 ((https://doi.org/10.1007/s13225-020-00466-2)) |
| Otu39486 | 1 | k__Fungi(100);p__Ascomycota(100);c__Sordariomycetes(100);o__Hypocreales(100);f__Nectriaceae(100);f__Nectriaceae_unclassified(100);f__Nectriaceae_unclassified(100); | Nectriaceae | 9 | Pathotroph-Saprotroph-Symbiotroph | Animal Pathogen-Endophyte-Fungal Parasite-Lichen Parasite-Plant Pathogen-Wood Saprotroph | Microfungus | NULL | Possible | Fusarium species are amongst the economically significant plant pathogens (Cannon & Kirk 2007) | Cannon PF, Kirk PM. 2007. Fungal Families of the World. CAB International, Cambridge ((ISBN: 978-0851998275)); Hirooka Y, et al. 2012. Studies in Mycology 71:1-210 ((https://doi.org/10.3114/sim0001)); Tedersoo L, et al. 2014. Science 346:e1256688 ((https://doi.org/10.1126/science.1256688)); The American Phytopathological Society Common Names of Plant Diseases ((http://www.apsnet.org/publications/commonnames/Pages/default.aspx)); https://www.cdc.gov ((https://www.cdc.gov)) |
| Otu39487 | 1 | k__Fungi(100);p__Ascomycota(100);c__Saccharomycetes(100);o__Saccharomycetales(100);f__Metschnikowiaceae(100);g__Metschnikowia(100);g__Metschnikowia_unclassified(100); | Metschnikowia | 13 | Saprotroph | \|Undefined Saprotroph\| | Yeast | NULL | Probable | As nectar-tap_saprotroph (Põlme et al. 2020); Decay sugar-rich substrates (Põlme et al. 2020) | Põlme S, et al. 2020. Fungal Diversity 105:1-16 ((https://doi.org/10.1007/s13225-020-00466-2)) |
| Otu39488 | 1 | k__Fungi(100);p__Ascomycota(100);c__Saccharomycetes(100);o__Saccharomycetales(100);f__Metschnikowiaceae(100);g__Metschnikowia(100);s__Metschnikowia_pulcherrima(100); | Metschnikowia | 13 | Saprotroph | \|Undefined Saprotroph\| | Yeast | NULL | Probable | As nectar-tap_saprotroph (Põlme et al. 2020); Decay sugar-rich substrates (Põlme et al. 2020) | Põlme S, et al. 2020. Fungal Diversity 105:1-16 ((https://doi.org/10.1007/s13225-020-00466-2)) |
| Otu39489 | 1 | k__Fungi(100);p__Ascomycota(100);c__Saccharomycetes(100);o__Saccharomycetales(100);f__Metschnikowiaceae(100);g__Metschnikowia(100);g__Metschnikowia_unclassified(100); | Metschnikowia | 13 | Saprotroph | \|Undefined Saprotroph\| | Yeast | NULL | Probable | As nectar-tap_saprotroph (Põlme et al. 2020); Decay sugar-rich substrates (Põlme et al. 2020) | Põlme S, et al. 2020. Fungal Diversity 105:1-16 ((https://doi.org/10.1007/s13225-020-00466-2)) |
| Otu39490 | 1 | k__Fungi(100);p__Ascomycota(100);c__Saccharomycetes(100);o__Saccharomycetales(100);f__Metschnikowiaceae(100);g__Metschnikowia(100);s__Metschnikowia_pulcherrima(100); | Metschnikowia | 13 | Saprotroph | \|Undefined Saprotroph\| | Yeast | NULL | Probable | As nectar-tap_saprotroph (Põlme et al. 2020); Decay sugar-rich substrates (Põlme et al. 2020) | Põlme S, et al. 2020. Fungal Diversity 105:1-16 ((https://doi.org/10.1007/s13225-020-00466-2)) |
| Otu39491 | 1 | k__Fungi(100);p__Ascomycota(100);c__Saccharomycetes(100);o__Saccharomycetales(100);f__Metschnikowiaceae(100);g__Metschnikowia(100);s__Metschnikowia_pulcherrima(100); | Metschnikowia | 13 | Saprotroph | \|Undefined Saprotroph\| | Yeast | NULL | Probable | As nectar-tap_saprotroph (Põlme et al. 2020); Decay sugar-rich substrates (Põlme et al. 2020) | Põlme S, et al. 2020. Fungal Diversity 105:1-16 ((https://doi.org/10.1007/s13225-020-00466-2)) |
| Otu39492 | 1 | k__Fungi(100);p__Ascomycota(100);c__Saccharomycetes(100);o__Saccharomycetales(100);f__Metschnikowiaceae(100);g__Metschnikowia(100);g__Metschnikowia_unclassified(100); | Metschnikowia | 13 | Saprotroph | \|Undefined Saprotroph\| | Yeast | NULL | Probable | As nectar-tap_saprotroph (Põlme et al. 2020); Decay sugar-rich substrates (Põlme et al. 2020) | Põlme S, et al. 2020. Fungal Diversity 105:1-16 ((https://doi.org/10.1007/s13225-020-00466-2)) |
| Otu39493 | 1 | k__Fungi(100);p__Ascomycota(100);c__Saccharomycetes(100);o__Saccharomycetales(100);f__Metschnikowiaceae(100);g__Metschnikowia(100);g__Metschnikowia_unclassified(100); | Metschnikowia | 13 | Saprotroph | \|Undefined Saprotroph\| | Yeast | NULL | Probable | As nectar-tap_saprotroph (Põlme et al. 2020); Decay sugar-rich substrates (Põlme et al. 2020) | Põlme S, et al. 2020. Fungal Diversity 105:1-16 ((https://doi.org/10.1007/s13225-020-00466-2)) |
| Otu39494 | 1 | k__Fungi(100);p__Ascomycota(100);c__Saccharomycetes(100);o__Saccharomycetales(100);f__Metschnikowiaceae(100);g__Metschnikowia(100);s__Metschnikowia_pulcherrima(100); | Metschnikowia | 13 | Saprotroph | \|Undefined Saprotroph\| | Yeast | NULL | Probable | As nectar-tap_saprotroph (Põlme et al. 2020); Decay sugar-rich substrates (Põlme et al. 2020) | Põlme S, et al. 2020. Fungal Diversity 105:1-16 ((https://doi.org/10.1007/s13225-020-00466-2)) |
| Otu39495 | 1 | k__Fungi(100);p__Ascomycota(100);c__Saccharomycetes(100);o__Saccharomycetales(100);f__Metschnikowiaceae(100);g__Metschnikowia(100);g__Metschnikowia_unclassified(100); | Metschnikowia | 13 | Saprotroph | \|Undefined Saprotroph\| | Yeast | NULL | Probable | As nectar-tap_saprotroph (Põlme et al. 2020); Decay sugar-rich substrates (Põlme et al. 2020) | Põlme S, et al. 2020. Fungal Diversity 105:1-16 ((https://doi.org/10.1007/s13225-020-00466-2)) |
| Otu39496 | 1 | k__Fungi(100);p__Ascomycota(100);c__Saccharomycetes(100);o__Saccharomycetales(100);f__Metschnikowiaceae(100);g__Metschnikowia(100);s__Metschnikowia_pulcherrima(100); | Metschnikowia | 13 | Saprotroph | \|Undefined Saprotroph\| | Yeast | NULL | Probable | As nectar-tap_saprotroph (Põlme et al. 2020); Decay sugar-rich substrates (Põlme et al. 2020) | Põlme S, et al. 2020. Fungal Diversity 105:1-16 ((https://doi.org/10.1007/s13225-020-00466-2)) |
| Otu39497 | 1 | k__Fungi(100);p__Ascomycota(100);c__Saccharomycetes(100);o__Saccharomycetales(100);f__Metschnikowiaceae(100);g__Metschnikowia(100);s__Metschnikowia_pulcherrima(100); | Metschnikowia | 13 | Saprotroph | \|Undefined Saprotroph\| | Yeast | NULL | Probable | As nectar-tap_saprotroph (Põlme et al. 2020); Decay sugar-rich substrates (Põlme et al. 2020) | Põlme S, et al. 2020. Fungal Diversity 105:1-16 ((https://doi.org/10.1007/s13225-020-00466-2)) |
| Otu39498 | 1 | k__Fungi(100);p__Ascomycota(100);c__Saccharomycetes(100);o__Saccharomycetales(100);f__Metschnikowiaceae(100);g__Metschnikowia(100);s__Metschnikowia_pulcherrima(100); | Metschnikowia | 13 | Saprotroph | \|Undefined Saprotroph\| | Yeast | NULL | Probable | As nectar-tap_saprotroph (Põlme et al. 2020); Decay sugar-rich substrates (Põlme et al. 2020) | Põlme S, et al. 2020. Fungal Diversity 105:1-16 ((https://doi.org/10.1007/s13225-020-00466-2)) |
| Otu39499 | 1 | k__Fungi(100);p__Ascomycota(100);c__Saccharomycetes(100);o__Saccharomycetales(100);f__Metschnikowiaceae(100);g__Metschnikowia(100);s__Metschnikowia_pulcherrima(100); | Metschnikowia | 13 | Saprotroph | \|Undefined Saprotroph\| | Yeast | NULL | Probable | As nectar-tap_saprotroph (Põlme et al. 2020); Decay sugar-rich substrates (Põlme et al. 2020) | Põlme S, et al. 2020. Fungal Diversity 105:1-16 ((https://doi.org/10.1007/s13225-020-00466-2)) |
| Otu39500 | 1 | k__Fungi(100);p__Ascomycota(100);c__Saccharomycetes(100);o__Saccharomycetales(100);f__Metschnikowiaceae(100);g__Metschnikowia(100);s__Metschnikowia_pulcherrima(100); | Metschnikowia | 13 | Saprotroph | \|Undefined Saprotroph\| | Yeast | NULL | Probable | As nectar-tap_saprotroph (Põlme et al. 2020); Decay sugar-rich substrates (Põlme et al. 2020) | Põlme S, et al. 2020. Fungal Diversity 105:1-16 ((https://doi.org/10.1007/s13225-020-00466-2)) |
| Otu39501 | 1 | k__Fungi(100);p__Ascomycota(100);c__Saccharomycetes(100);o__Saccharomycetales(100);f__Metschnikowiaceae(100);g__Metschnikowia(100);g__Metschnikowia_unclassified(100); | Metschnikowia | 13 | Saprotroph | \|Undefined Saprotroph\| | Yeast | NULL | Probable | As nectar-tap_saprotroph (Põlme et al. 2020); Decay sugar-rich substrates (Põlme et al. 2020) | Põlme S, et al. 2020. Fungal Diversity 105:1-16 ((https://doi.org/10.1007/s13225-020-00466-2)) |
| Otu39502 | 1 | k__Fungi(100);p__Ascomycota(100);c__Saccharomycetes(100);o__Saccharomycetales(100);f__Metschnikowiaceae(100);g__Metschnikowia(100);s__Metschnikowia_pulcherrima(100); | Metschnikowia | 13 | Saprotroph | \|Undefined Saprotroph\| | Yeast | NULL | Probable | As nectar-tap_saprotroph (Põlme et al. 2020); Decay sugar-rich substrates (Põlme et al. 2020) | Põlme S, et al. 2020. Fungal Diversity 105:1-16 ((https://doi.org/10.1007/s13225-020-00466-2)) |
| Otu39503 | 1 | k__Fungi(100);p__Ascomycota(100);c__Saccharomycetes(100);o__Saccharomycetales(100);f__Metschnikowiaceae(100);g__Metschnikowia(100);s__Metschnikowia_pulcherrima(100); | Metschnikowia | 13 | Saprotroph | \|Undefined Saprotroph\| | Yeast | NULL | Probable | As nectar-tap_saprotroph (Põlme et al. 2020); Decay sugar-rich substrates (Põlme et al. 2020) | Põlme S, et al. 2020. Fungal Diversity 105:1-16 ((https://doi.org/10.1007/s13225-020-00466-2)) |
| Otu39504 | 1 | k__Fungi(100);p__Ascomycota(100);c__Saccharomycetes(100);o__Saccharomycetales(100);f__Metschnikowiaceae(100);g__Metschnikowia(100);s__Metschnikowia_pulcherrima(100); | Metschnikowia | 13 | Saprotroph | \|Undefined Saprotroph\| | Yeast | NULL | Probable | As nectar-tap_saprotroph (Põlme et al. 2020); Decay sugar-rich substrates (Põlme et al. 2020) | Põlme S, et al. 2020. Fungal Diversity 105:1-16 ((https://doi.org/10.1007/s13225-020-00466-2)) |
| Otu39505 | 1 | k__Fungi(100);p__Ascomycota(100);c__Saccharomycetes(100);o__Saccharomycetales(100);f__Metschnikowiaceae(100);g__Metschnikowia(100);s__Metschnikowia_pulcherrima(100); | Metschnikowia | 13 | Saprotroph | \|Undefined Saprotroph\| | Yeast | NULL | Probable | As nectar-tap_saprotroph (Põlme et al. 2020); Decay sugar-rich substrates (Põlme et al. 2020) | Põlme S, et al. 2020. Fungal Diversity 105:1-16 ((https://doi.org/10.1007/s13225-020-00466-2)) |
| Otu39506 | 1 | k__Fungi(100);p__Ascomycota(100);c__Saccharomycetes(100);o__Saccharomycetales(100);f__Metschnikowiaceae(100);g__Metschnikowia(100);s__Metschnikowia_pulcherrima(100); | Metschnikowia | 13 | Saprotroph | \|Undefined Saprotroph\| | Yeast | NULL | Probable | As nectar-tap_saprotroph (Põlme et al. 2020); Decay sugar-rich substrates (Põlme et al. 2020) | Põlme S, et al. 2020. Fungal Diversity 105:1-16 ((https://doi.org/10.1007/s13225-020-00466-2)) |
| Otu39507 | 1 | k__Fungi(100);p__Ascomycota(100);c__Saccharomycetes(100);o__Saccharomycetales(100);f__Metschnikowiaceae(100);g__Metschnikowia(100);s__Metschnikowia_pulcherrima(100); | Metschnikowia | 13 | Saprotroph | \|Undefined Saprotroph\| | Yeast | NULL | Probable | As nectar-tap_saprotroph (Põlme et al. 2020); Decay sugar-rich substrates (Põlme et al. 2020) | Põlme S, et al. 2020. Fungal Diversity 105:1-16 ((https://doi.org/10.1007/s13225-020-00466-2)) |
| Otu39508 | 1 | k__Fungi(100);p__Ascomycota(100);c__Saccharomycetes(100);o__Saccharomycetales(100);f__Metschnikowiaceae(100);g__Metschnikowia(100);g__Metschnikowia_unclassified(100); | Metschnikowia | 13 | Saprotroph | \|Undefined Saprotroph\| | Yeast | NULL | Probable | As nectar-tap_saprotroph (Põlme et al. 2020); Decay sugar-rich substrates (Põlme et al. 2020) | Põlme S, et al. 2020. Fungal Diversity 105:1-16 ((https://doi.org/10.1007/s13225-020-00466-2)) |
| Otu39509 | 1 | k__Fungi(100);p__Ascomycota(100);c__Saccharomycetes(100);o__Saccharomycetales(100);f__Metschnikowiaceae(100);g__Metschnikowia(100);g__Metschnikowia_unclassified(100); | Metschnikowia | 13 | Saprotroph | \|Undefined Saprotroph\| | Yeast | NULL | Probable | As nectar-tap_saprotroph (Põlme et al. 2020); Decay sugar-rich substrates (Põlme et al. 2020) | Põlme S, et al. 2020. Fungal Diversity 105:1-16 ((https://doi.org/10.1007/s13225-020-00466-2)) |
| Otu39510 | 1 | k__Fungi(100);p__Ascomycota(100);c__Saccharomycetes(100);o__Saccharomycetales(100);f__Metschnikowiaceae(100);g__Metschnikowia(100);s__Metschnikowia_pulcherrima(100); | Metschnikowia | 13 | Saprotroph | \|Undefined Saprotroph\| | Yeast | NULL | Probable | As nectar-tap_saprotroph (Põlme et al. 2020); Decay sugar-rich substrates (Põlme et al. 2020) | Põlme S, et al. 2020. Fungal Diversity 105:1-16 ((https://doi.org/10.1007/s13225-020-00466-2)) |
| Otu39511 | 1 | k__Fungi(100);p__Ascomycota(100);c__Saccharomycetes(100);o__Saccharomycetales(100);f__Metschnikowiaceae(100);g__Metschnikowia(100);g__Metschnikowia_unclassified(100); | Metschnikowia | 13 | Saprotroph | \|Undefined Saprotroph\| | Yeast | NULL | Probable | As nectar-tap_saprotroph (Põlme et al. 2020); Decay sugar-rich substrates (Põlme et al. 2020) | Põlme S, et al. 2020. Fungal Diversity 105:1-16 ((https://doi.org/10.1007/s13225-020-00466-2)) |
| Otu39512 | 1 | k__Fungi(100);p__Ascomycota(100);c__Saccharomycetes(100);o__Saccharomycetales(100);f__Metschnikowiaceae(100);g__Metschnikowia(100);g__Metschnikowia_unclassified(100); | Metschnikowia | 13 | Saprotroph | \|Undefined Saprotroph\| | Yeast | NULL | Probable | As nectar-tap_saprotroph (Põlme et al. 2020); Decay sugar-rich substrates (Põlme et al. 2020) | Põlme S, et al. 2020. Fungal Diversity 105:1-16 ((https://doi.org/10.1007/s13225-020-00466-2)) |
| Otu39513 | 1 | k__Fungi(100);p__Ascomycota(100);c__Saccharomycetes(100);o__Saccharomycetales(100);f__Metschnikowiaceae(100);g__Metschnikowia(100);s__Metschnikowia_pulcherrima(100); | Metschnikowia | 13 | Saprotroph | \|Undefined Saprotroph\| | Yeast | NULL | Probable | As nectar-tap_saprotroph (Põlme et al. 2020); Decay sugar-rich substrates (Põlme et al. 2020) | Põlme S, et al. 2020. Fungal Diversity 105:1-16 ((https://doi.org/10.1007/s13225-020-00466-2)) |
| Otu39514 | 1 | k__Fungi(100);p__Ascomycota(100);c__Saccharomycetes(100);o__Saccharomycetales(100);f__Metschnikowiaceae(100);g__Metschnikowia(100);s__Metschnikowia_pulcherrima(100); | Metschnikowia | 13 | Saprotroph | \|Undefined Saprotroph\| | Yeast | NULL | Probable | As nectar-tap_saprotroph (Põlme et al. 2020); Decay sugar-rich substrates (Põlme et al. 2020) | Põlme S, et al. 2020. Fungal Diversity 105:1-16 ((https://doi.org/10.1007/s13225-020-00466-2)) |
| Otu39515 | 1 | k__Fungi(100);p__Ascomycota(100);c__Saccharomycetes(100);o__Saccharomycetales(100);f__Metschnikowiaceae(100);g__Metschnikowia(100);s__Metschnikowia_pulcherrima(100); | Metschnikowia | 13 | Saprotroph | \|Undefined Saprotroph\| | Yeast | NULL | Probable | As nectar-tap_saprotroph (Põlme et al. 2020); Decay sugar-rich substrates (Põlme et al. 2020) | Põlme S, et al. 2020. Fungal Diversity 105:1-16 ((https://doi.org/10.1007/s13225-020-00466-2)) |
| Otu39516 | 1 | k__Fungi(100);p__Ascomycota(100);c__Saccharomycetes(100);o__Saccharomycetales(100);f__Metschnikowiaceae(100);g__Metschnikowia(100);s__Metschnikowia_pulcherrima(100); | Metschnikowia | 13 | Saprotroph | \|Undefined Saprotroph\| | Yeast | NULL | Probable | As nectar-tap_saprotroph (Põlme et al. 2020); Decay sugar-rich substrates (Põlme et al. 2020) | Põlme S, et al. 2020. Fungal Diversity 105:1-16 ((https://doi.org/10.1007/s13225-020-00466-2)) |
| Otu39517 | 1 | k__Fungi(100);p__Ascomycota(100);c__Saccharomycetes(100);o__Saccharomycetales(100);f__Metschnikowiaceae(100);g__Metschnikowia(100);s__Metschnikowia_pulcherrima(100); | Metschnikowia | 13 | Saprotroph | \|Undefined Saprotroph\| | Yeast | NULL | Probable | As nectar-tap_saprotroph (Põlme et al. 2020); Decay sugar-rich substrates (Põlme et al. 2020) | Põlme S, et al. 2020. Fungal Diversity 105:1-16 ((https://doi.org/10.1007/s13225-020-00466-2)) |
| Otu39518 | 1 | k__Fungi(100);p__Ascomycota(100);c__Saccharomycetes(100);o__Saccharomycetales(100);f__Metschnikowiaceae(100);g__Metschnikowia(100);s__Metschnikowia_pulcherrima(100); | Metschnikowia | 13 | Saprotroph | \|Undefined Saprotroph\| | Yeast | NULL | Probable | As nectar-tap_saprotroph (Põlme et al. 2020); Decay sugar-rich substrates (Põlme et al. 2020) | Põlme S, et al. 2020. Fungal Diversity 105:1-16 ((https://doi.org/10.1007/s13225-020-00466-2)) |
| Otu39519 | 1 | k__Fungi(100);p__Ascomycota(100);c__Saccharomycetes(100);o__Saccharomycetales(100);f__Metschnikowiaceae(100);g__Metschnikowia(100);s__Metschnikowia_pulcherrima(100); | Metschnikowia | 13 | Saprotroph | \|Undefined Saprotroph\| | Yeast | NULL | Probable | As nectar-tap_saprotroph (Põlme et al. 2020); Decay sugar-rich substrates (Põlme et al. 2020) | Põlme S, et al. 2020. Fungal Diversity 105:1-16 ((https://doi.org/10.1007/s13225-020-00466-2)) |
| Otu39520 | 1 | k__Fungi(100);p__Ascomycota(100);c__Saccharomycetes(100);o__Saccharomycetales(100);f__Metschnikowiaceae(100);g__Metschnikowia(100);s__Metschnikowia_pulcherrima(100); | Metschnikowia | 13 | Saprotroph | \|Undefined Saprotroph\| | Yeast | NULL | Probable | As nectar-tap_saprotroph (Põlme et al. 2020); Decay sugar-rich substrates (Põlme et al. 2020) | Põlme S, et al. 2020. Fungal Diversity 105:1-16 ((https://doi.org/10.1007/s13225-020-00466-2)) |
| Otu39521 | 1 | k__Fungi(100);p__Ascomycota(100);c__Saccharomycetes(100);o__Saccharomycetales(100);f__Metschnikowiaceae(100);g__Metschnikowia(100);g__Metschnikowia_unclassified(100); | Metschnikowia | 13 | Saprotroph | \|Undefined Saprotroph\| | Yeast | NULL | Probable | As nectar-tap_saprotroph (Põlme et al. 2020); Decay sugar-rich substrates (Põlme et al. 2020) | Põlme S, et al. 2020. Fungal Diversity 105:1-16 ((https://doi.org/10.1007/s13225-020-00466-2)) |
| Otu39522 | 1 | k__Fungi(100);p__Ascomycota(100);c__Saccharomycetes(100);o__Saccharomycetales(100);f__Metschnikowiaceae(100);g__Metschnikowia(100);s__Metschnikowia_pulcherrima(100); | Metschnikowia | 13 | Saprotroph | \|Undefined Saprotroph\| | Yeast | NULL | Probable | As nectar-tap_saprotroph (Põlme et al. 2020); Decay sugar-rich substrates (Põlme et al. 2020) | Põlme S, et al. 2020. Fungal Diversity 105:1-16 ((https://doi.org/10.1007/s13225-020-00466-2)) |
| Otu39523 | 1 | k__Fungi(100);p__Ascomycota(100);c__Saccharomycetes(100);o__Saccharomycetales(100);f__Metschnikowiaceae(100);g__Metschnikowia(100);s__Metschnikowia_pulcherrima(100); | Metschnikowia | 13 | Saprotroph | \|Undefined Saprotroph\| | Yeast | NULL | Probable | As nectar-tap_saprotroph (Põlme et al. 2020); Decay sugar-rich substrates (Põlme et al. 2020) | Põlme S, et al. 2020. Fungal Diversity 105:1-16 ((https://doi.org/10.1007/s13225-020-00466-2)) |
| Otu39524 | 1 | k__Fungi(100);p__Ascomycota(100);c__Saccharomycetes(100);o__Saccharomycetales(100);f__Metschnikowiaceae(100);g__Metschnikowia(100);s__Metschnikowia_pulcherrima(100); | Metschnikowia | 13 | Saprotroph | \|Undefined Saprotroph\| | Yeast | NULL | Probable | As nectar-tap_saprotroph (Põlme et al. 2020); Decay sugar-rich substrates (Põlme et al. 2020) | Põlme S, et al. 2020. Fungal Diversity 105:1-16 ((https://doi.org/10.1007/s13225-020-00466-2)) |
| Otu39525 | 1 | k__Fungi(100);p__Ascomycota(100);c__Saccharomycetes(100);o__Saccharomycetales(100);f__Metschnikowiaceae(100);g__Metschnikowia(100);s__Metschnikowia_pulcherrima(100); | Metschnikowia | 13 | Saprotroph | \|Undefined Saprotroph\| | Yeast | NULL | Probable | As nectar-tap_saprotroph (Põlme et al. 2020); Decay sugar-rich substrates (Põlme et al. 2020) | Põlme S, et al. 2020. Fungal Diversity 105:1-16 ((https://doi.org/10.1007/s13225-020-00466-2)) |
| Otu39526 | 1 | k__Fungi(100);p__Ascomycota(100);c__Saccharomycetes(100);o__Saccharomycetales(100);f__Metschnikowiaceae(100);g__Metschnikowia(100);s__Metschnikowia_pulcherrima(100); | Metschnikowia | 13 | Saprotroph | \|Undefined Saprotroph\| | Yeast | NULL | Probable | As nectar-tap_saprotroph (Põlme et al. 2020); Decay sugar-rich substrates (Põlme et al. 2020) | Põlme S, et al. 2020. Fungal Diversity 105:1-16 ((https://doi.org/10.1007/s13225-020-00466-2)) |
| Otu39527 | 1 | k__Fungi(100);p__Ascomycota(100);c__Saccharomycetes(100);o__Saccharomycetales(100);f__Metschnikowiaceae(100);g__Metschnikowia(100);s__Metschnikowia_pulcherrima(100); | Metschnikowia | 13 | Saprotroph | \|Undefined Saprotroph\| | Yeast | NULL | Probable | As nectar-tap_saprotroph (Põlme et al. 2020); Decay sugar-rich substrates (Põlme et al. 2020) | Põlme S, et al. 2020. Fungal Diversity 105:1-16 ((https://doi.org/10.1007/s13225-020-00466-2)) |
| Otu39528 | 1 | k__Fungi(100);p__Ascomycota(100);c__Saccharomycetes(100);o__Saccharomycetales(100);f__Metschnikowiaceae(100);g__Metschnikowia(100);s__Metschnikowia_pulcherrima(100); | Metschnikowia | 13 | Saprotroph | \|Undefined Saprotroph\| | Yeast | NULL | Probable | As nectar-tap_saprotroph (Põlme et al. 2020); Decay sugar-rich substrates (Põlme et al. 2020) | Põlme S, et al. 2020. Fungal Diversity 105:1-16 ((https://doi.org/10.1007/s13225-020-00466-2)) |
| Otu39529 | 1 | k__Fungi(100);p__Ascomycota(100);c__Saccharomycetes(100);o__Saccharomycetales(100);f__Metschnikowiaceae(100);g__Metschnikowia(100);s__Metschnikowia_pulcherrima(100); | Metschnikowia | 13 | Saprotroph | \|Undefined Saprotroph\| | Yeast | NULL | Probable | As nectar-tap_saprotroph (Põlme et al. 2020); Decay sugar-rich substrates (Põlme et al. 2020) | Põlme S, et al. 2020. Fungal Diversity 105:1-16 ((https://doi.org/10.1007/s13225-020-00466-2)) |
| Otu39530 | 1 | k__Fungi(100);p__Ascomycota(100);c__Saccharomycetes(100);o__Saccharomycetales(100);f__Metschnikowiaceae(100);g__Metschnikowia(100);s__Metschnikowia_pulcherrima(100); | Metschnikowia | 13 | Saprotroph | \|Undefined Saprotroph\| | Yeast | NULL | Probable | As nectar-tap_saprotroph (Põlme et al. 2020); Decay sugar-rich substrates (Põlme et al. 2020) | Põlme S, et al. 2020. Fungal Diversity 105:1-16 ((https://doi.org/10.1007/s13225-020-00466-2)) |
| Otu39531 | 1 | k__Fungi(100);p__Ascomycota(100);c__Saccharomycetes(100);o__Saccharomycetales(100);f__Metschnikowiaceae(100);g__Metschnikowia(100);s__Metschnikowia_pulcherrima(100); | Metschnikowia | 13 | Saprotroph | \|Undefined Saprotroph\| | Yeast | NULL | Probable | As nectar-tap_saprotroph (Põlme et al. 2020); Decay sugar-rich substrates (Põlme et al. 2020) | Põlme S, et al. 2020. Fungal Diversity 105:1-16 ((https://doi.org/10.1007/s13225-020-00466-2)) |
| Otu39532 | 1 | k__Fungi(100);p__Ascomycota(100);c__Saccharomycetes(100);o__Saccharomycetales(100);f__Metschnikowiaceae(100);g__Metschnikowia(100);s__Metschnikowia_pulcherrima(100); | Metschnikowia | 13 | Saprotroph | \|Undefined Saprotroph\| | Yeast | NULL | Probable | As nectar-tap_saprotroph (Põlme et al. 2020); Decay sugar-rich substrates (Põlme et al. 2020) | Põlme S, et al. 2020. Fungal Diversity 105:1-16 ((https://doi.org/10.1007/s13225-020-00466-2)) |
| Otu39533 | 1 | k__Fungi(100);p__Ascomycota(100);c__Saccharomycetes(100);o__Saccharomycetales(100);f__Metschnikowiaceae(100);g__Metschnikowia(100);s__Metschnikowia_pulcherrima(100); | Metschnikowia | 13 | Saprotroph | \|Undefined Saprotroph\| | Yeast | NULL | Probable | As nectar-tap_saprotroph (Põlme et al. 2020); Decay sugar-rich substrates (Põlme et al. 2020) | Põlme S, et al. 2020. Fungal Diversity 105:1-16 ((https://doi.org/10.1007/s13225-020-00466-2)) |
| Otu39534 | 1 | k__Fungi(100);p__Ascomycota(100);c__Saccharomycetes(100);o__Saccharomycetales(100);f__Metschnikowiaceae(100);g__Metschnikowia(100);g__Metschnikowia_unclassified(100); | Metschnikowia | 13 | Saprotroph | \|Undefined Saprotroph\| | Yeast | NULL | Probable | As nectar-tap_saprotroph (Põlme et al. 2020); Decay sugar-rich substrates (Põlme et al. 2020) | Põlme S, et al. 2020. Fungal Diversity 105:1-16 ((https://doi.org/10.1007/s13225-020-00466-2)) |
| Otu39535 | 1 | k__Fungi(100);p__Ascomycota(100);c__Saccharomycetes(100);o__Saccharomycetales(100);f__Metschnikowiaceae(100);g__Metschnikowia(100);s__Metschnikowia_pulcherrima(100); | Metschnikowia | 13 | Saprotroph | \|Undefined Saprotroph\| | Yeast | NULL | Probable | As nectar-tap_saprotroph (Põlme et al. 2020); Decay sugar-rich substrates (Põlme et al. 2020) | Põlme S, et al. 2020. Fungal Diversity 105:1-16 ((https://doi.org/10.1007/s13225-020-00466-2)) |
| Otu39536 | 1 | k__Fungi(100);p__Ascomycota(100);c__Saccharomycetes(100);o__Saccharomycetales(100);f__Metschnikowiaceae(100);g__Metschnikowia(100);s__Metschnikowia_pulcherrima(100); | Metschnikowia | 13 | Saprotroph | \|Undefined Saprotroph\| | Yeast | NULL | Probable | As nectar-tap_saprotroph (Põlme et al. 2020); Decay sugar-rich substrates (Põlme et al. 2020) | Põlme S, et al. 2020. Fungal Diversity 105:1-16 ((https://doi.org/10.1007/s13225-020-00466-2)) |
| Otu39537 | 1 | k__Fungi(100);p__Ascomycota(100);c__Saccharomycetes(100);o__Saccharomycetales(100);f__Metschnikowiaceae(100);g__Metschnikowia(100);s__Metschnikowia_pulcherrima(100); | Metschnikowia | 13 | Saprotroph | \|Undefined Saprotroph\| | Yeast | NULL | Probable | As nectar-tap_saprotroph (Põlme et al. 2020); Decay sugar-rich substrates (Põlme et al. 2020) | Põlme S, et al. 2020. Fungal Diversity 105:1-16 ((https://doi.org/10.1007/s13225-020-00466-2)) |
| Otu39538 | 1 | k__Fungi(100);p__Ascomycota(100);c__Saccharomycetes(100);o__Saccharomycetales(100);f__Metschnikowiaceae(100);g__Metschnikowia(100);s__Metschnikowia_pulcherrima(100); | Metschnikowia | 13 | Saprotroph | \|Undefined Saprotroph\| | Yeast | NULL | Probable | As nectar-tap_saprotroph (Põlme et al. 2020); Decay sugar-rich substrates (Põlme et al. 2020) | Põlme S, et al. 2020. Fungal Diversity 105:1-16 ((https://doi.org/10.1007/s13225-020-00466-2)) |
| Otu39539 | 1 | k__Fungi(100);p__Ascomycota(100);c__Saccharomycetes(100);o__Saccharomycetales(100);f__Metschnikowiaceae(100);g__Metschnikowia(100);g__Metschnikowia_unclassified(100); | Metschnikowia | 13 | Saprotroph | \|Undefined Saprotroph\| | Yeast | NULL | Probable | As nectar-tap_saprotroph (Põlme et al. 2020); Decay sugar-rich substrates (Põlme et al. 2020) | Põlme S, et al. 2020. Fungal Diversity 105:1-16 ((https://doi.org/10.1007/s13225-020-00466-2)) |
| Otu39540 | 1 | k__Fungi(100);p__Ascomycota(100);c__Saccharomycetes(100);o__Saccharomycetales(100);f__Metschnikowiaceae(100);g__Metschnikowia(100);g__Metschnikowia_unclassified(100); | Metschnikowia | 13 | Saprotroph | \|Undefined Saprotroph\| | Yeast | NULL | Probable | As nectar-tap_saprotroph (Põlme et al. 2020); Decay sugar-rich substrates (Põlme et al. 2020) | Põlme S, et al. 2020. Fungal Diversity 105:1-16 ((https://doi.org/10.1007/s13225-020-00466-2)) |
| Otu39541 | 1 | k__Fungi(100);p__Ascomycota(100);c__Saccharomycetes(100);o__Saccharomycetales(100);f__Metschnikowiaceae(100);g__Metschnikowia(100);s__Metschnikowia_pulcherrima(100); | Metschnikowia | 13 | Saprotroph | \|Undefined Saprotroph\| | Yeast | NULL | Probable | As nectar-tap_saprotroph (Põlme et al. 2020); Decay sugar-rich substrates (Põlme et al. 2020) | Põlme S, et al. 2020. Fungal Diversity 105:1-16 ((https://doi.org/10.1007/s13225-020-00466-2)) |
| Otu39542 | 1 | k__Fungi(100);p__Ascomycota(100);c__Saccharomycetes(100);o__Saccharomycetales(100);f__Metschnikowiaceae(100);g__Metschnikowia(100);s__Metschnikowia_pulcherrima(100); | Metschnikowia | 13 | Saprotroph | \|Undefined Saprotroph\| | Yeast | NULL | Probable | As nectar-tap_saprotroph (Põlme et al. 2020); Decay sugar-rich substrates (Põlme et al. 2020) | Põlme S, et al. 2020. Fungal Diversity 105:1-16 ((https://doi.org/10.1007/s13225-020-00466-2)) |
| Otu39543 | 1 | k__Fungi(100);p__Ascomycota(100);c__Saccharomycetes(100);o__Saccharomycetales(100);f__Metschnikowiaceae(100);g__Metschnikowia(100);s__Metschnikowia_pulcherrima(100); | Metschnikowia | 13 | Saprotroph | \|Undefined Saprotroph\| | Yeast | NULL | Probable | As nectar-tap_saprotroph (Põlme et al. 2020); Decay sugar-rich substrates (Põlme et al. 2020) | Põlme S, et al. 2020. Fungal Diversity 105:1-16 ((https://doi.org/10.1007/s13225-020-00466-2)) |
| Otu39544 | 1 | k__Fungi(100);p__Ascomycota(100);c__Saccharomycetes(100);o__Saccharomycetales(100);f__Metschnikowiaceae(100);g__Metschnikowia(100);s__Metschnikowia_pulcherrima(100); | Metschnikowia | 13 | Saprotroph | \|Undefined Saprotroph\| | Yeast | NULL | Probable | As nectar-tap_saprotroph (Põlme et al. 2020); Decay sugar-rich substrates (Põlme et al. 2020) | Põlme S, et al. 2020. Fungal Diversity 105:1-16 ((https://doi.org/10.1007/s13225-020-00466-2)) |
| Otu39545 | 1 | k__Fungi(100);p__Ascomycota(100);c__Saccharomycetes(100);o__Saccharomycetales(100);f__Metschnikowiaceae(100);g__Metschnikowia(100);g__Metschnikowia_unclassified(100); | Metschnikowia | 13 | Saprotroph | \|Undefined Saprotroph\| | Yeast | NULL | Probable | As nectar-tap_saprotroph (Põlme et al. 2020); Decay sugar-rich substrates (Põlme et al. 2020) | Põlme S, et al. 2020. Fungal Diversity 105:1-16 ((https://doi.org/10.1007/s13225-020-00466-2)) |
| Otu39546 | 1 | k__Fungi(100);p__Ascomycota(100);c__Saccharomycetes(100);o__Saccharomycetales(100);f__Metschnikowiaceae(100);g__Metschnikowia(100);s__Metschnikowia_pulcherrima(100); | Metschnikowia | 13 | Saprotroph | \|Undefined Saprotroph\| | Yeast | NULL | Probable | As nectar-tap_saprotroph (Põlme et al. 2020); Decay sugar-rich substrates (Põlme et al. 2020) | Põlme S, et al. 2020. Fungal Diversity 105:1-16 ((https://doi.org/10.1007/s13225-020-00466-2)) |
| Otu39547 | 1 | k__Fungi(100);p__Ascomycota(100);c__Saccharomycetes(100);o__Saccharomycetales(100);f__Metschnikowiaceae(100);g__Metschnikowia(100);s__Metschnikowia_pulcherrima(100); | Metschnikowia | 13 | Saprotroph | \|Undefined Saprotroph\| | Yeast | NULL | Probable | As nectar-tap_saprotroph (Põlme et al. 2020); Decay sugar-rich substrates (Põlme et al. 2020) | Põlme S, et al. 2020. Fungal Diversity 105:1-16 ((https://doi.org/10.1007/s13225-020-00466-2)) |
| Otu39548 | 1 | k__Fungi(100);p__Ascomycota(100);c__Saccharomycetes(100);o__Saccharomycetales(100);f__Metschnikowiaceae(100);g__Metschnikowia(100);s__Metschnikowia_pulcherrima(100); | Metschnikowia | 13 | Saprotroph | \|Undefined Saprotroph\| | Yeast | NULL | Probable | As nectar-tap_saprotroph (Põlme et al. 2020); Decay sugar-rich substrates (Põlme et al. 2020) | Põlme S, et al. 2020. Fungal Diversity 105:1-16 ((https://doi.org/10.1007/s13225-020-00466-2)) |
| Otu39549 | 1 | k__Fungi(100);p__Ascomycota(100);c__Saccharomycetes(100);o__Saccharomycetales(100);f__Metschnikowiaceae(100);g__Metschnikowia(100);g__Metschnikowia_unclassified(100); | Metschnikowia | 13 | Saprotroph | \|Undefined Saprotroph\| | Yeast | NULL | Probable | As nectar-tap_saprotroph (Põlme et al. 2020); Decay sugar-rich substrates (Põlme et al. 2020) | Põlme S, et al. 2020. Fungal Diversity 105:1-16 ((https://doi.org/10.1007/s13225-020-00466-2)) |
| Otu39550 | 1 | k__Fungi(100);p__Ascomycota(100);c__Saccharomycetes(100);o__Saccharomycetales(100);f__Metschnikowiaceae(100);g__Metschnikowia(100);g__Metschnikowia_unclassified(100); | Metschnikowia | 13 | Saprotroph | \|Undefined Saprotroph\| | Yeast | NULL | Probable | As nectar-tap_saprotroph (Põlme et al. 2020); Decay sugar-rich substrates (Põlme et al. 2020) | Põlme S, et al. 2020. Fungal Diversity 105:1-16 ((https://doi.org/10.1007/s13225-020-00466-2)) |
| Otu39551 | 1 | k__Fungi(100);p__Ascomycota(100);c__Saccharomycetes(100);o__Saccharomycetales(100);f__Metschnikowiaceae(100);g__Metschnikowia(100);s__Metschnikowia_pulcherrima(100); | Metschnikowia | 13 | Saprotroph | \|Undefined Saprotroph\| | Yeast | NULL | Probable | As nectar-tap_saprotroph (Põlme et al. 2020); Decay sugar-rich substrates (Põlme et al. 2020) | Põlme S, et al. 2020. Fungal Diversity 105:1-16 ((https://doi.org/10.1007/s13225-020-00466-2)) |
| Otu39552 | 1 | k__Fungi(100);p__Ascomycota(100);c__Saccharomycetes(100);o__Saccharomycetales(100);f__Metschnikowiaceae(100);g__Metschnikowia(100);s__Metschnikowia_pulcherrima(100); | Metschnikowia | 13 | Saprotroph | \|Undefined Saprotroph\| | Yeast | NULL | Probable | As nectar-tap_saprotroph (Põlme et al. 2020); Decay sugar-rich substrates (Põlme et al. 2020) | Põlme S, et al. 2020. Fungal Diversity 105:1-16 ((https://doi.org/10.1007/s13225-020-00466-2)) |
| Otu39553 | 1 | k__Fungi(100);p__Ascomycota(100);c__Saccharomycetes(100);o__Saccharomycetales(100);f__Metschnikowiaceae(100);g__Metschnikowia(100);s__Metschnikowia_pulcherrima(100); | Metschnikowia | 13 | Saprotroph | \|Undefined Saprotroph\| | Yeast | NULL | Probable | As nectar-tap_saprotroph (Põlme et al. 2020); Decay sugar-rich substrates (Põlme et al. 2020) | Põlme S, et al. 2020. Fungal Diversity 105:1-16 ((https://doi.org/10.1007/s13225-020-00466-2)) |
| Otu39554 | 1 | k__Fungi(100);p__Ascomycota(100);c__Saccharomycetes(100);o__Saccharomycetales(100);f__Metschnikowiaceae(100);g__Metschnikowia(100);s__Metschnikowia_pulcherrima(100); | Metschnikowia | 13 | Saprotroph | \|Undefined Saprotroph\| | Yeast | NULL | Probable | As nectar-tap_saprotroph (Põlme et al. 2020); Decay sugar-rich substrates (Põlme et al. 2020) | Põlme S, et al. 2020. Fungal Diversity 105:1-16 ((https://doi.org/10.1007/s13225-020-00466-2)) |
| Otu39555 | 1 | k__Fungi(100);p__Ascomycota(100);c__Saccharomycetes(100);o__Saccharomycetales(100);f__Metschnikowiaceae(100);g__Metschnikowia(100);s__Metschnikowia_pulcherrima(100); | Metschnikowia | 13 | Saprotroph | \|Undefined Saprotroph\| | Yeast | NULL | Probable | As nectar-tap_saprotroph (Põlme et al. 2020); Decay sugar-rich substrates (Põlme et al. 2020) | Põlme S, et al. 2020. Fungal Diversity 105:1-16 ((https://doi.org/10.1007/s13225-020-00466-2)) |
| Otu39556 | 1 | k__Fungi(100);p__Ascomycota(100);c__Saccharomycetes(100);o__Saccharomycetales(100);f__Metschnikowiaceae(100);g__Metschnikowia(100);g__Metschnikowia_unclassified(100); | Metschnikowia | 13 | Saprotroph | \|Undefined Saprotroph\| | Yeast | NULL | Probable | As nectar-tap_saprotroph (Põlme et al. 2020); Decay sugar-rich substrates (Põlme et al. 2020) | Põlme S, et al. 2020. Fungal Diversity 105:1-16 ((https://doi.org/10.1007/s13225-020-00466-2)) |
| Otu39557 | 1 | k__Fungi(100);p__Ascomycota(100);c__Saccharomycetes(100);o__Saccharomycetales(100);f__Metschnikowiaceae(100);g__Metschnikowia(100);s__Metschnikowia_pulcherrima(100); | Metschnikowia | 13 | Saprotroph | \|Undefined Saprotroph\| | Yeast | NULL | Probable | As nectar-tap_saprotroph (Põlme et al. 2020); Decay sugar-rich substrates (Põlme et al. 2020) | Põlme S, et al. 2020. Fungal Diversity 105:1-16 ((https://doi.org/10.1007/s13225-020-00466-2)) |
| Otu39558 | 1 | k__Fungi(100);p__Ascomycota(100);c__Saccharomycetes(100);o__Saccharomycetales(100);f__Metschnikowiaceae(100);g__Metschnikowia(100);s__Metschnikowia_pulcherrima(100); | Metschnikowia | 13 | Saprotroph | \|Undefined Saprotroph\| | Yeast | NULL | Probable | As nectar-tap_saprotroph (Põlme et al. 2020); Decay sugar-rich substrates (Põlme et al. 2020) | Põlme S, et al. 2020. Fungal Diversity 105:1-16 ((https://doi.org/10.1007/s13225-020-00466-2)) |
| Otu39559 | 1 | k__Fungi(100);p__Ascomycota(100);c__Saccharomycetes(100);o__Saccharomycetales(100);f__Metschnikowiaceae(100);g__Metschnikowia(100);s__Metschnikowia_pulcherrima(100); | Metschnikowia | 13 | Saprotroph | \|Undefined Saprotroph\| | Yeast | NULL | Probable | As nectar-tap_saprotroph (Põlme et al. 2020); Decay sugar-rich substrates (Põlme et al. 2020) | Põlme S, et al. 2020. Fungal Diversity 105:1-16 ((https://doi.org/10.1007/s13225-020-00466-2)) |
| Otu39560 | 1 | k__Fungi(100);p__Ascomycota(100);c__Saccharomycetes(100);o__Saccharomycetales(100);f__Metschnikowiaceae(100);g__Metschnikowia(100);s__Metschnikowia_pulcherrima(100); | Metschnikowia | 13 | Saprotroph | \|Undefined Saprotroph\| | Yeast | NULL | Probable | As nectar-tap_saprotroph (Põlme et al. 2020); Decay sugar-rich substrates (Põlme et al. 2020) | Põlme S, et al. 2020. Fungal Diversity 105:1-16 ((https://doi.org/10.1007/s13225-020-00466-2)) |
| Otu39561 | 1 | k__Fungi(100);p__Ascomycota(100);c__Saccharomycetes(100);o__Saccharomycetales(100);f__Metschnikowiaceae(100);g__Metschnikowia(100);s__Metschnikowia_pulcherrima(100); | Metschnikowia | 13 | Saprotroph | \|Undefined Saprotroph\| | Yeast | NULL | Probable | As nectar-tap_saprotroph (Põlme et al. 2020); Decay sugar-rich substrates (Põlme et al. 2020) | Põlme S, et al. 2020. Fungal Diversity 105:1-16 ((https://doi.org/10.1007/s13225-020-00466-2)) |
| Otu39562 | 1 | k__Fungi(100);p__Ascomycota(100);c__Saccharomycetes(100);o__Saccharomycetales(100);f__Metschnikowiaceae(100);g__Metschnikowia(100);s__Metschnikowia_pulcherrima(100); | Metschnikowia | 13 | Saprotroph | \|Undefined Saprotroph\| | Yeast | NULL | Probable | As nectar-tap_saprotroph (Põlme et al. 2020); Decay sugar-rich substrates (Põlme et al. 2020) | Põlme S, et al. 2020. Fungal Diversity 105:1-16 ((https://doi.org/10.1007/s13225-020-00466-2)) |
| Otu39563 | 1 | k__Fungi(100);p__Ascomycota(100);c__Saccharomycetes(100);o__Saccharomycetales(100);f__Metschnikowiaceae(100);g__Metschnikowia(100);s__Metschnikowia_pulcherrima(100); | Metschnikowia | 13 | Saprotroph | \|Undefined Saprotroph\| | Yeast | NULL | Probable | As nectar-tap_saprotroph (Põlme et al. 2020); Decay sugar-rich substrates (Põlme et al. 2020) | Põlme S, et al. 2020. Fungal Diversity 105:1-16 ((https://doi.org/10.1007/s13225-020-00466-2)) |
| Otu39564 | 1 | k__Fungi(100);p__Ascomycota(100);c__Saccharomycetes(100);o__Saccharomycetales(100);f__Metschnikowiaceae(100);g__Metschnikowia(100);s__Metschnikowia_pulcherrima(100); | Metschnikowia | 13 | Saprotroph | \|Undefined Saprotroph\| | Yeast | NULL | Probable | As nectar-tap_saprotroph (Põlme et al. 2020); Decay sugar-rich substrates (Põlme et al. 2020) | Põlme S, et al. 2020. Fungal Diversity 105:1-16 ((https://doi.org/10.1007/s13225-020-00466-2)) |
| Otu39565 | 1 | k__Fungi(100);p__Ascomycota(100);c__Saccharomycetes(100);o__Saccharomycetales(100);f__Metschnikowiaceae(100);g__Metschnikowia(100);s__Metschnikowia_pulcherrima(100); | Metschnikowia | 13 | Saprotroph | \|Undefined Saprotroph\| | Yeast | NULL | Probable | As nectar-tap_saprotroph (Põlme et al. 2020); Decay sugar-rich substrates (Põlme et al. 2020) | Põlme S, et al. 2020. Fungal Diversity 105:1-16 ((https://doi.org/10.1007/s13225-020-00466-2)) |
| Otu39566 | 1 | k__Fungi(100);p__Ascomycota(100);c__Saccharomycetes(100);o__Saccharomycetales(100);f__Metschnikowiaceae(100);g__Metschnikowia(100);s__Metschnikowia_pulcherrima(100); | Metschnikowia | 13 | Saprotroph | \|Undefined Saprotroph\| | Yeast | NULL | Probable | As nectar-tap_saprotroph (Põlme et al. 2020); Decay sugar-rich substrates (Põlme et al. 2020) | Põlme S, et al. 2020. Fungal Diversity 105:1-16 ((https://doi.org/10.1007/s13225-020-00466-2)) |
| Otu39567 | 1 | k__Fungi(100);p__Ascomycota(100);c__Saccharomycetes(100);o__Saccharomycetales(100);f__Metschnikowiaceae(100);g__Metschnikowia(100);g__Metschnikowia_unclassified(100); | Metschnikowia | 13 | Saprotroph | \|Undefined Saprotroph\| | Yeast | NULL | Probable | As nectar-tap_saprotroph (Põlme et al. 2020); Decay sugar-rich substrates (Põlme et al. 2020) | Põlme S, et al. 2020. Fungal Diversity 105:1-16 ((https://doi.org/10.1007/s13225-020-00466-2)) |
| Otu39568 | 1 | k__Fungi(100);p__Ascomycota(100);c__Saccharomycetes(100);o__Saccharomycetales(100);f__Metschnikowiaceae(100);g__Metschnikowia(100);s__Metschnikowia_pulcherrima(100); | Metschnikowia | 13 | Saprotroph | \|Undefined Saprotroph\| | Yeast | NULL | Probable | As nectar-tap_saprotroph (Põlme et al. 2020); Decay sugar-rich substrates (Põlme et al. 2020) | Põlme S, et al. 2020. Fungal Diversity 105:1-16 ((https://doi.org/10.1007/s13225-020-00466-2)) |
| Otu39569 | 1 | k__Fungi(100);p__Ascomycota(100);c__Saccharomycetes(100);o__Saccharomycetales(100);f__Metschnikowiaceae(100);g__Metschnikowia(100);s__Metschnikowia_pulcherrima(100); | Metschnikowia | 13 | Saprotroph | \|Undefined Saprotroph\| | Yeast | NULL | Probable | As nectar-tap_saprotroph (Põlme et al. 2020); Decay sugar-rich substrates (Põlme et al. 2020) | Põlme S, et al. 2020. Fungal Diversity 105:1-16 ((https://doi.org/10.1007/s13225-020-00466-2)) |
| Otu39570 | 1 | k__Fungi(100);p__Ascomycota(100);c__Saccharomycetes(100);o__Saccharomycetales(100);f__Metschnikowiaceae(100);g__Metschnikowia(100);g__Metschnikowia_unclassified(100); | Metschnikowia | 13 | Saprotroph | \|Undefined Saprotroph\| | Yeast | NULL | Probable | As nectar-tap_saprotroph (Põlme et al. 2020); Decay sugar-rich substrates (Põlme et al. 2020) | Põlme S, et al. 2020. Fungal Diversity 105:1-16 ((https://doi.org/10.1007/s13225-020-00466-2)) |
| Otu39571 | 1 | k__Fungi(100);p__Ascomycota(100);c__Saccharomycetes(100);o__Saccharomycetales(100);f__Metschnikowiaceae(100);g__Metschnikowia(100);g__Metschnikowia_unclassified(100); | Metschnikowia | 13 | Saprotroph | \|Undefined Saprotroph\| | Yeast | NULL | Probable | As nectar-tap_saprotroph (Põlme et al. 2020); Decay sugar-rich substrates (Põlme et al. 2020) | Põlme S, et al. 2020. Fungal Diversity 105:1-16 ((https://doi.org/10.1007/s13225-020-00466-2)) |
| Otu39572 | 1 | k__Fungi(100);p__Ascomycota(100);c__Saccharomycetes(100);o__Saccharomycetales(100);f__Metschnikowiaceae(100);g__Metschnikowia(100);g__Metschnikowia_unclassified(100); | Metschnikowia | 13 | Saprotroph | \|Undefined Saprotroph\| | Yeast | NULL | Probable | As nectar-tap_saprotroph (Põlme et al. 2020); Decay sugar-rich substrates (Põlme et al. 2020) | Põlme S, et al. 2020. Fungal Diversity 105:1-16 ((https://doi.org/10.1007/s13225-020-00466-2)) |
| Otu39574 | 1 | k__Fungi(100);p__Ascomycota(100);c__Saccharomycetes(100);o__Saccharomycetales(100);f__Metschnikowiaceae(100);g__Metschnikowia(100);s__Metschnikowia_pulcherrima(100); | Metschnikowia | 13 | Saprotroph | \|Undefined Saprotroph\| | Yeast | NULL | Probable | As nectar-tap_saprotroph (Põlme et al. 2020); Decay sugar-rich substrates (Põlme et al. 2020) | Põlme S, et al. 2020. Fungal Diversity 105:1-16 ((https://doi.org/10.1007/s13225-020-00466-2)) |
| Otu39575 | 1 | k__Fungi(100);p__Ascomycota(100);c__Saccharomycetes(100);o__Saccharomycetales(100);f__Metschnikowiaceae(100);g__Metschnikowia(100);s__Metschnikowia_pulcherrima(100); | Metschnikowia | 13 | Saprotroph | \|Undefined Saprotroph\| | Yeast | NULL | Probable | As nectar-tap_saprotroph (Põlme et al. 2020); Decay sugar-rich substrates (Põlme et al. 2020) | Põlme S, et al. 2020. Fungal Diversity 105:1-16 ((https://doi.org/10.1007/s13225-020-00466-2)) |
| Otu39576 | 1 | k__Fungi(100);p__Ascomycota(100);c__Saccharomycetes(100);o__Saccharomycetales(100);f__Metschnikowiaceae(100);g__Metschnikowia(100);s__Metschnikowia_pulcherrima(100); | Metschnikowia | 13 | Saprotroph | \|Undefined Saprotroph\| | Yeast | NULL | Probable | As nectar-tap_saprotroph (Põlme et al. 2020); Decay sugar-rich substrates (Põlme et al. 2020) | Põlme S, et al. 2020. Fungal Diversity 105:1-16 ((https://doi.org/10.1007/s13225-020-00466-2)) |
| Otu39577 | 1 | k__Fungi(100);p__Ascomycota(100);c__Saccharomycetes(100);o__Saccharomycetales(100);f__Metschnikowiaceae(100);g__Metschnikowia(100);s__Metschnikowia_pulcherrima(100); | Metschnikowia | 13 | Saprotroph | \|Undefined Saprotroph\| | Yeast | NULL | Probable | As nectar-tap_saprotroph (Põlme et al. 2020); Decay sugar-rich substrates (Põlme et al. 2020) | Põlme S, et al. 2020. Fungal Diversity 105:1-16 ((https://doi.org/10.1007/s13225-020-00466-2)) |
| Otu39578 | 1 | k__Fungi(100);p__Ascomycota(100);c__Saccharomycetes(100);o__Saccharomycetales(100);f__Metschnikowiaceae(100);g__Metschnikowia(100);g__Metschnikowia_unclassified(100); | Metschnikowia | 13 | Saprotroph | \|Undefined Saprotroph\| | Yeast | NULL | Probable | As nectar-tap_saprotroph (Põlme et al. 2020); Decay sugar-rich substrates (Põlme et al. 2020) | Põlme S, et al. 2020. Fungal Diversity 105:1-16 ((https://doi.org/10.1007/s13225-020-00466-2)) |
| Otu39579 | 1 | k__Fungi(100);p__Ascomycota(100);c__Saccharomycetes(100);o__Saccharomycetales(100);f__Metschnikowiaceae(100);g__Metschnikowia(100);g__Metschnikowia_unclassified(100); | Metschnikowia | 13 | Saprotroph | \|Undefined Saprotroph\| | Yeast | NULL | Probable | As nectar-tap_saprotroph (Põlme et al. 2020); Decay sugar-rich substrates (Põlme et al. 2020) | Põlme S, et al. 2020. Fungal Diversity 105:1-16 ((https://doi.org/10.1007/s13225-020-00466-2)) |
| Otu39580 | 1 | k__Fungi(100);p__Ascomycota(100);c__Saccharomycetes(100);o__Saccharomycetales(100);f__Metschnikowiaceae(100);g__Metschnikowia(100);s__Metschnikowia_pulcherrima(100); | Metschnikowia | 13 | Saprotroph | \|Undefined Saprotroph\| | Yeast | NULL | Probable | As nectar-tap_saprotroph (Põlme et al. 2020); Decay sugar-rich substrates (Põlme et al. 2020) | Põlme S, et al. 2020. Fungal Diversity 105:1-16 ((https://doi.org/10.1007/s13225-020-00466-2)) |
| Otu39581 | 1 | k__Fungi(100);p__Ascomycota(100);c__Saccharomycetes(100);o__Saccharomycetales(100);f__Metschnikowiaceae(100);g__Metschnikowia(100);s__Metschnikowia_pulcherrima(100); | Metschnikowia | 13 | Saprotroph | \|Undefined Saprotroph\| | Yeast | NULL | Probable | As nectar-tap_saprotroph (Põlme et al. 2020); Decay sugar-rich substrates (Põlme et al. 2020) | Põlme S, et al. 2020. Fungal Diversity 105:1-16 ((https://doi.org/10.1007/s13225-020-00466-2)) |
| Otu39582 | 1 | k__Fungi(100);p__Ascomycota(100);c__Saccharomycetes(100);o__Saccharomycetales(100);f__Metschnikowiaceae(100);g__Metschnikowia(100);g__Metschnikowia_unclassified(100); | Metschnikowia | 13 | Saprotroph | \|Undefined Saprotroph\| | Yeast | NULL | Probable | As nectar-tap_saprotroph (Põlme et al. 2020); Decay sugar-rich substrates (Põlme et al. 2020) | Põlme S, et al. 2020. Fungal Diversity 105:1-16 ((https://doi.org/10.1007/s13225-020-00466-2)) |
| Otu39583 | 1 | k__Fungi(100);p__Ascomycota(100);c__Saccharomycetes(100);o__Saccharomycetales(100);f__Metschnikowiaceae(100);g__Metschnikowia(100);s__Metschnikowia_pulcherrima(100); | Metschnikowia | 13 | Saprotroph | \|Undefined Saprotroph\| | Yeast | NULL | Probable | As nectar-tap_saprotroph (Põlme et al. 2020); Decay sugar-rich substrates (Põlme et al. 2020) | Põlme S, et al. 2020. Fungal Diversity 105:1-16 ((https://doi.org/10.1007/s13225-020-00466-2)) |
| Otu39584 | 1 | k__Fungi(100);p__Ascomycota(100);c__Saccharomycetes(100);o__Saccharomycetales(100);f__Metschnikowiaceae(100);g__Metschnikowia(100);s__Metschnikowia_pulcherrima(100); | Metschnikowia | 13 | Saprotroph | \|Undefined Saprotroph\| | Yeast | NULL | Probable | As nectar-tap_saprotroph (Põlme et al. 2020); Decay sugar-rich substrates (Põlme et al. 2020) | Põlme S, et al. 2020. Fungal Diversity 105:1-16 ((https://doi.org/10.1007/s13225-020-00466-2)) |
| Otu39585 | 1 | k__Fungi(100);p__Ascomycota(100);c__Saccharomycetes(100);o__Saccharomycetales(100);f__Metschnikowiaceae(100);g__Metschnikowia(100);s__Metschnikowia_pulcherrima(100); | Metschnikowia | 13 | Saprotroph | \|Undefined Saprotroph\| | Yeast | NULL | Probable | As nectar-tap_saprotroph (Põlme et al. 2020); Decay sugar-rich substrates (Põlme et al. 2020) | Põlme S, et al. 2020. Fungal Diversity 105:1-16 ((https://doi.org/10.1007/s13225-020-00466-2)) |
| Otu39586 | 1 | k__Fungi(100);p__Ascomycota(100);c__Saccharomycetes(100);o__Saccharomycetales(100);f__Metschnikowiaceae(100);g__Metschnikowia(100);s__Metschnikowia_pulcherrima(100); | Metschnikowia | 13 | Saprotroph | \|Undefined Saprotroph\| | Yeast | NULL | Probable | As nectar-tap_saprotroph (Põlme et al. 2020); Decay sugar-rich substrates (Põlme et al. 2020) | Põlme S, et al. 2020. Fungal Diversity 105:1-16 ((https://doi.org/10.1007/s13225-020-00466-2)) |
| Otu39587 | 1 | k__Fungi(100);p__Ascomycota(100);c__Saccharomycetes(100);o__Saccharomycetales(100);f__Metschnikowiaceae(100);g__Metschnikowia(100);g__Metschnikowia_unclassified(100); | Metschnikowia | 13 | Saprotroph | \|Undefined Saprotroph\| | Yeast | NULL | Probable | As nectar-tap_saprotroph (Põlme et al. 2020); Decay sugar-rich substrates (Põlme et al. 2020) | Põlme S, et al. 2020. Fungal Diversity 105:1-16 ((https://doi.org/10.1007/s13225-020-00466-2)) |
| Otu39588 | 1 | k__Fungi(100);p__Ascomycota(100);c__Saccharomycetes(100);o__Saccharomycetales(100);f__Metschnikowiaceae(100);g__Metschnikowia(100);g__Metschnikowia_unclassified(100); | Metschnikowia | 13 | Saprotroph | \|Undefined Saprotroph\| | Yeast | NULL | Probable | As nectar-tap_saprotroph (Põlme et al. 2020); Decay sugar-rich substrates (Põlme et al. 2020) | Põlme S, et al. 2020. Fungal Diversity 105:1-16 ((https://doi.org/10.1007/s13225-020-00466-2)) |
| Otu39589 | 1 | k__Fungi(100);p__Ascomycota(100);c__Saccharomycetes(100);o__Saccharomycetales(100);f__Metschnikowiaceae(100);g__Metschnikowia(100);s__Metschnikowia_pulcherrima(100); | Metschnikowia | 13 | Saprotroph | \|Undefined Saprotroph\| | Yeast | NULL | Probable | As nectar-tap_saprotroph (Põlme et al. 2020); Decay sugar-rich substrates (Põlme et al. 2020) | Põlme S, et al. 2020. Fungal Diversity 105:1-16 ((https://doi.org/10.1007/s13225-020-00466-2)) |
| Otu39590 | 1 | k__Fungi(100);p__Ascomycota(100);c__Saccharomycetes(100);o__Saccharomycetales(100);f__Metschnikowiaceae(100);g__Metschnikowia(100);s__Metschnikowia_pulcherrima(100); | Metschnikowia | 13 | Saprotroph | \|Undefined Saprotroph\| | Yeast | NULL | Probable | As nectar-tap_saprotroph (Põlme et al. 2020); Decay sugar-rich substrates (Põlme et al. 2020) | Põlme S, et al. 2020. Fungal Diversity 105:1-16 ((https://doi.org/10.1007/s13225-020-00466-2)) |
| Otu39591 | 1 | k__Fungi(100);p__Ascomycota(100);c__Saccharomycetes(100);o__Saccharomycetales(100);f__Metschnikowiaceae(100);g__Metschnikowia(100);g__Metschnikowia_unclassified(100); | Metschnikowia | 13 | Saprotroph | \|Undefined Saprotroph\| | Yeast | NULL | Probable | As nectar-tap_saprotroph (Põlme et al. 2020); Decay sugar-rich substrates (Põlme et al. 2020) | Põlme S, et al. 2020. Fungal Diversity 105:1-16 ((https://doi.org/10.1007/s13225-020-00466-2)) |
| Otu39592 | 1 | k__Fungi(100);p__Ascomycota(100);c__Saccharomycetes(100);o__Saccharomycetales(100);f__Metschnikowiaceae(100);g__Metschnikowia(100);g__Metschnikowia_unclassified(100); | Metschnikowia | 13 | Saprotroph | \|Undefined Saprotroph\| | Yeast | NULL | Probable | As nectar-tap_saprotroph (Põlme et al. 2020); Decay sugar-rich substrates (Põlme et al. 2020) | Põlme S, et al. 2020. Fungal Diversity 105:1-16 ((https://doi.org/10.1007/s13225-020-00466-2)) |
| Otu39593 | 1 | k__Fungi(100);p__Ascomycota(100);c__Saccharomycetes(100);o__Saccharomycetales(100);f__Metschnikowiaceae(100);g__Metschnikowia(100);s__Metschnikowia_pulcherrima(100); | Metschnikowia | 13 | Saprotroph | \|Undefined Saprotroph\| | Yeast | NULL | Probable | As nectar-tap_saprotroph (Põlme et al. 2020); Decay sugar-rich substrates (Põlme et al. 2020) | Põlme S, et al. 2020. Fungal Diversity 105:1-16 ((https://doi.org/10.1007/s13225-020-00466-2)) |
| Otu39594 | 1 | k__Fungi(100);p__Ascomycota(100);c__Saccharomycetes(100);o__Saccharomycetales(100);f__Metschnikowiaceae(100);g__Metschnikowia(100);s__Metschnikowia_pulcherrima(100); | Metschnikowia | 13 | Saprotroph | \|Undefined Saprotroph\| | Yeast | NULL | Probable | As nectar-tap_saprotroph (Põlme et al. 2020); Decay sugar-rich substrates (Põlme et al. 2020) | Põlme S, et al. 2020. Fungal Diversity 105:1-16 ((https://doi.org/10.1007/s13225-020-00466-2)) |
| Otu39595 | 1 | k__Fungi(100);p__Ascomycota(100);c__Saccharomycetes(100);o__Saccharomycetales(100);f__Metschnikowiaceae(100);g__Metschnikowia(100);g__Metschnikowia_unclassified(100); | Metschnikowia | 13 | Saprotroph | \|Undefined Saprotroph\| | Yeast | NULL | Probable | As nectar-tap_saprotroph (Põlme et al. 2020); Decay sugar-rich substrates (Põlme et al. 2020) | Põlme S, et al. 2020. Fungal Diversity 105:1-16 ((https://doi.org/10.1007/s13225-020-00466-2)) |
| Otu39596 | 1 | k__Fungi(100);p__Ascomycota(100);c__Saccharomycetes(100);o__Saccharomycetales(100);f__Metschnikowiaceae(100);g__Metschnikowia(100);s__Metschnikowia_pulcherrima(100); | Metschnikowia | 13 | Saprotroph | \|Undefined Saprotroph\| | Yeast | NULL | Probable | As nectar-tap_saprotroph (Põlme et al. 2020); Decay sugar-rich substrates (Põlme et al. 2020) | Põlme S, et al. 2020. Fungal Diversity 105:1-16 ((https://doi.org/10.1007/s13225-020-00466-2)) |
| Otu39597 | 1 | k__Fungi(100);p__Ascomycota(100);c__Saccharomycetes(100);o__Saccharomycetales(100);f__Metschnikowiaceae(100);g__Metschnikowia(100);s__Metschnikowia_pulcherrima(100); | Metschnikowia | 13 | Saprotroph | \|Undefined Saprotroph\| | Yeast | NULL | Probable | As nectar-tap_saprotroph (Põlme et al. 2020); Decay sugar-rich substrates (Põlme et al. 2020) | Põlme S, et al. 2020. Fungal Diversity 105:1-16 ((https://doi.org/10.1007/s13225-020-00466-2)) |
| Otu39598 | 1 | k__Fungi(100);p__Ascomycota(100);c__Saccharomycetes(100);o__Saccharomycetales(100);f__Metschnikowiaceae(100);g__Metschnikowia(100);s__Metschnikowia_pulcherrima(100); | Metschnikowia | 13 | Saprotroph | \|Undefined Saprotroph\| | Yeast | NULL | Probable | As nectar-tap_saprotroph (Põlme et al. 2020); Decay sugar-rich substrates (Põlme et al. 2020) | Põlme S, et al. 2020. Fungal Diversity 105:1-16 ((https://doi.org/10.1007/s13225-020-00466-2)) |
| Otu39599 | 1 | k__Fungi(100);p__Ascomycota(100);c__Saccharomycetes(100);o__Saccharomycetales(100);f__Metschnikowiaceae(100);g__Metschnikowia(100);s__Metschnikowia_pulcherrima(100); | Metschnikowia | 13 | Saprotroph | \|Undefined Saprotroph\| | Yeast | NULL | Probable | As nectar-tap_saprotroph (Põlme et al. 2020); Decay sugar-rich substrates (Põlme et al. 2020) | Põlme S, et al. 2020. Fungal Diversity 105:1-16 ((https://doi.org/10.1007/s13225-020-00466-2)) |
| Otu39600 | 1 | k__Fungi(100);p__Ascomycota(100);c__Saccharomycetes(100);o__Saccharomycetales(100);f__Metschnikowiaceae(100);g__Metschnikowia(100);s__Metschnikowia_pulcherrima(100); | Metschnikowia | 13 | Saprotroph | \|Undefined Saprotroph\| | Yeast | NULL | Probable | As nectar-tap_saprotroph (Põlme et al. 2020); Decay sugar-rich substrates (Põlme et al. 2020) | Põlme S, et al. 2020. Fungal Diversity 105:1-16 ((https://doi.org/10.1007/s13225-020-00466-2)) |
| Otu39601 | 1 | k__Fungi(100);p__Ascomycota(100);c__Saccharomycetes(100);o__Saccharomycetales(100);f__Metschnikowiaceae(100);g__Metschnikowia(100);s__Metschnikowia_pulcherrima(100); | Metschnikowia | 13 | Saprotroph | \|Undefined Saprotroph\| | Yeast | NULL | Probable | As nectar-tap_saprotroph (Põlme et al. 2020); Decay sugar-rich substrates (Põlme et al. 2020) | Põlme S, et al. 2020. Fungal Diversity 105:1-16 ((https://doi.org/10.1007/s13225-020-00466-2)) |
| Otu39602 | 1 | k__Fungi(100);p__Ascomycota(100);c__Saccharomycetes(100);o__Saccharomycetales(100);f__Metschnikowiaceae(100);g__Metschnikowia(100);s__Metschnikowia_pulcherrima(100); | Metschnikowia | 13 | Saprotroph | \|Undefined Saprotroph\| | Yeast | NULL | Probable | As nectar-tap_saprotroph (Põlme et al. 2020); Decay sugar-rich substrates (Põlme et al. 2020) | Põlme S, et al. 2020. Fungal Diversity 105:1-16 ((https://doi.org/10.1007/s13225-020-00466-2)) |
| Otu39603 | 1 | k__Fungi(100);p__Ascomycota(100);c__Saccharomycetes(100);o__Saccharomycetales(100);f__Metschnikowiaceae(100);g__Metschnikowia(100);s__Metschnikowia_pulcherrima(100); | Metschnikowia | 13 | Saprotroph | \|Undefined Saprotroph\| | Yeast | NULL | Probable | As nectar-tap_saprotroph (Põlme et al. 2020); Decay sugar-rich substrates (Põlme et al. 2020) | Põlme S, et al. 2020. Fungal Diversity 105:1-16 ((https://doi.org/10.1007/s13225-020-00466-2)) |
| Otu39604 | 1 | k__Fungi(100);p__Ascomycota(100);c__Saccharomycetes(100);o__Saccharomycetales(100);f__Metschnikowiaceae(100);g__Metschnikowia(100);g__Metschnikowia_unclassified(100); | Metschnikowia | 13 | Saprotroph | \|Undefined Saprotroph\| | Yeast | NULL | Probable | As nectar-tap_saprotroph (Põlme et al. 2020); Decay sugar-rich substrates (Põlme et al. 2020) | Põlme S, et al. 2020. Fungal Diversity 105:1-16 ((https://doi.org/10.1007/s13225-020-00466-2)) |
| Otu39605 | 1 | k__Fungi(100);p__Ascomycota(100);c__Saccharomycetes(100);o__Saccharomycetales(100);f__Metschnikowiaceae(100);g__Metschnikowia(100);s__Metschnikowia_pulcherrima(100); | Metschnikowia | 13 | Saprotroph | \|Undefined Saprotroph\| | Yeast | NULL | Probable | As nectar-tap_saprotroph (Põlme et al. 2020); Decay sugar-rich substrates (Põlme et al. 2020) | Põlme S, et al. 2020. Fungal Diversity 105:1-16 ((https://doi.org/10.1007/s13225-020-00466-2)) |
| Otu39606 | 1 | k__Fungi(100);p__Ascomycota(100);c__Saccharomycetes(100);o__Saccharomycetales(100);f__Metschnikowiaceae(100);g__Metschnikowia(100);s__Metschnikowia_pulcherrima(100); | Metschnikowia | 13 | Saprotroph | \|Undefined Saprotroph\| | Yeast | NULL | Probable | As nectar-tap_saprotroph (Põlme et al. 2020); Decay sugar-rich substrates (Põlme et al. 2020) | Põlme S, et al. 2020. Fungal Diversity 105:1-16 ((https://doi.org/10.1007/s13225-020-00466-2)) |
| Otu39607 | 1 | k__Fungi(100);p__Ascomycota(100);c__Saccharomycetes(100);o__Saccharomycetales(100);f__Metschnikowiaceae(100);g__Metschnikowia(100);s__Metschnikowia_pulcherrima(100); | Metschnikowia | 13 | Saprotroph | \|Undefined Saprotroph\| | Yeast | NULL | Probable | As nectar-tap_saprotroph (Põlme et al. 2020); Decay sugar-rich substrates (Põlme et al. 2020) | Põlme S, et al. 2020. Fungal Diversity 105:1-16 ((https://doi.org/10.1007/s13225-020-00466-2)) |
| Otu39608 | 1 | k__Fungi(100);p__Ascomycota(100);c__Saccharomycetes(100);o__Saccharomycetales(100);f__Metschnikowiaceae(100);g__Metschnikowia(100);s__Metschnikowia_pulcherrima(100); | Metschnikowia | 13 | Saprotroph | \|Undefined Saprotroph\| | Yeast | NULL | Probable | As nectar-tap_saprotroph (Põlme et al. 2020); Decay sugar-rich substrates (Põlme et al. 2020) | Põlme S, et al. 2020. Fungal Diversity 105:1-16 ((https://doi.org/10.1007/s13225-020-00466-2)) |
| Otu39609 | 1 | k__Fungi(100);p__Ascomycota(100);c__Saccharomycetes(100);o__Saccharomycetales(100);f__Metschnikowiaceae(100);g__Metschnikowia(100);s__Metschnikowia_pulcherrima(100); | Metschnikowia | 13 | Saprotroph | \|Undefined Saprotroph\| | Yeast | NULL | Probable | As nectar-tap_saprotroph (Põlme et al. 2020); Decay sugar-rich substrates (Põlme et al. 2020) | Põlme S, et al. 2020. Fungal Diversity 105:1-16 ((https://doi.org/10.1007/s13225-020-00466-2)) |
| Otu39610 | 1 | k__Fungi(100);p__Ascomycota(100);c__Saccharomycetes(100);o__Saccharomycetales(100);f__Metschnikowiaceae(100);g__Metschnikowia(100);g__Metschnikowia_unclassified(100); | Metschnikowia | 13 | Saprotroph | \|Undefined Saprotroph\| | Yeast | NULL | Probable | As nectar-tap_saprotroph (Põlme et al. 2020); Decay sugar-rich substrates (Põlme et al. 2020) | Põlme S, et al. 2020. Fungal Diversity 105:1-16 ((https://doi.org/10.1007/s13225-020-00466-2)) |
| Otu39611 | 1 | k__Fungi(100);p__Ascomycota(100);c__Saccharomycetes(100);o__Saccharomycetales(100);f__Metschnikowiaceae(100);g__Metschnikowia(100);g__Metschnikowia_unclassified(100); | Metschnikowia | 13 | Saprotroph | \|Undefined Saprotroph\| | Yeast | NULL | Probable | As nectar-tap_saprotroph (Põlme et al. 2020); Decay sugar-rich substrates (Põlme et al. 2020) | Põlme S, et al. 2020. Fungal Diversity 105:1-16 ((https://doi.org/10.1007/s13225-020-00466-2)) |
| Otu39612 | 1 | k__Fungi(100);p__Ascomycota(100);c__Saccharomycetes(100);o__Saccharomycetales(100);f__Metschnikowiaceae(100);g__Metschnikowia(100);s__Metschnikowia_pulcherrima(100); | Metschnikowia | 13 | Saprotroph | \|Undefined Saprotroph\| | Yeast | NULL | Probable | As nectar-tap_saprotroph (Põlme et al. 2020); Decay sugar-rich substrates (Põlme et al. 2020) | Põlme S, et al. 2020. Fungal Diversity 105:1-16 ((https://doi.org/10.1007/s13225-020-00466-2)) |
| Otu39613 | 1 | k__Fungi(100);p__Ascomycota(100);c__Saccharomycetes(100);o__Saccharomycetales(100);f__Metschnikowiaceae(100);g__Metschnikowia(100);s__Metschnikowia_pulcherrima(100); | Metschnikowia | 13 | Saprotroph | \|Undefined Saprotroph\| | Yeast | NULL | Probable | As nectar-tap_saprotroph (Põlme et al. 2020); Decay sugar-rich substrates (Põlme et al. 2020) | Põlme S, et al. 2020. Fungal Diversity 105:1-16 ((https://doi.org/10.1007/s13225-020-00466-2)) |
| Otu39614 | 1 | k__Fungi(100);p__Ascomycota(100);c__Saccharomycetes(100);o__Saccharomycetales(100);f__Metschnikowiaceae(100);g__Metschnikowia(100);g__Metschnikowia_unclassified(100); | Metschnikowia | 13 | Saprotroph | \|Undefined Saprotroph\| | Yeast | NULL | Probable | As nectar-tap_saprotroph (Põlme et al. 2020); Decay sugar-rich substrates (Põlme et al. 2020) | Põlme S, et al. 2020. Fungal Diversity 105:1-16 ((https://doi.org/10.1007/s13225-020-00466-2)) |
| Otu39615 | 1 | k__Fungi(100);p__Ascomycota(100);c__Saccharomycetes(100);o__Saccharomycetales(100);f__Metschnikowiaceae(100);g__Metschnikowia(100);s__Metschnikowia_pulcherrima(100); | Metschnikowia | 13 | Saprotroph | \|Undefined Saprotroph\| | Yeast | NULL | Probable | As nectar-tap_saprotroph (Põlme et al. 2020); Decay sugar-rich substrates (Põlme et al. 2020) | Põlme S, et al. 2020. Fungal Diversity 105:1-16 ((https://doi.org/10.1007/s13225-020-00466-2)) |
| Otu39616 | 1 | k__Fungi(100);p__Ascomycota(100);c__Saccharomycetes(100);o__Saccharomycetales(100);f__Metschnikowiaceae(100);g__Metschnikowia(100);g__Metschnikowia_unclassified(100); | Metschnikowia | 13 | Saprotroph | \|Undefined Saprotroph\| | Yeast | NULL | Probable | As nectar-tap_saprotroph (Põlme et al. 2020); Decay sugar-rich substrates (Põlme et al. 2020) | Põlme S, et al. 2020. Fungal Diversity 105:1-16 ((https://doi.org/10.1007/s13225-020-00466-2)) |
| Otu39617 | 1 | k__Fungi(100);p__Ascomycota(100);c__Saccharomycetes(100);o__Saccharomycetales(100);f__Metschnikowiaceae(100);g__Metschnikowia(100);g__Metschnikowia_unclassified(100); | Metschnikowia | 13 | Saprotroph | \|Undefined Saprotroph\| | Yeast | NULL | Probable | As nectar-tap_saprotroph (Põlme et al. 2020); Decay sugar-rich substrates (Põlme et al. 2020) | Põlme S, et al. 2020. Fungal Diversity 105:1-16 ((https://doi.org/10.1007/s13225-020-00466-2)) |
| Otu39618 | 1 | k__Fungi(100);p__Ascomycota(100);c__Saccharomycetes(100);o__Saccharomycetales(100);f__Metschnikowiaceae(100);g__Metschnikowia(100);s__Metschnikowia_pulcherrima(100); | Metschnikowia | 13 | Saprotroph | \|Undefined Saprotroph\| | Yeast | NULL | Probable | As nectar-tap_saprotroph (Põlme et al. 2020); Decay sugar-rich substrates (Põlme et al. 2020) | Põlme S, et al. 2020. Fungal Diversity 105:1-16 ((https://doi.org/10.1007/s13225-020-00466-2)) |
| Otu39619 | 1 | k__Fungi(100);p__Ascomycota(100);c__Saccharomycetes(100);o__Saccharomycetales(100);f__Metschnikowiaceae(100);g__Metschnikowia(100);g__Metschnikowia_unclassified(100); | Metschnikowia | 13 | Saprotroph | \|Undefined Saprotroph\| | Yeast | NULL | Probable | As nectar-tap_saprotroph (Põlme et al. 2020); Decay sugar-rich substrates (Põlme et al. 2020) | Põlme S, et al. 2020. Fungal Diversity 105:1-16 ((https://doi.org/10.1007/s13225-020-00466-2)) |
| Otu39620 | 1 | k__Fungi(100);p__Ascomycota(100);c__Saccharomycetes(100);o__Saccharomycetales(100);f__Metschnikowiaceae(100);g__Metschnikowia(100);g__Metschnikowia_unclassified(100); | Metschnikowia | 13 | Saprotroph | \|Undefined Saprotroph\| | Yeast | NULL | Probable | As nectar-tap_saprotroph (Põlme et al. 2020); Decay sugar-rich substrates (Põlme et al. 2020) | Põlme S, et al. 2020. Fungal Diversity 105:1-16 ((https://doi.org/10.1007/s13225-020-00466-2)) |
| Otu39621 | 1 | k__Fungi(100);p__Ascomycota(100);c__Saccharomycetes(100);o__Saccharomycetales(100);f__Metschnikowiaceae(100);g__Metschnikowia(100);s__Metschnikowia_pulcherrima(100); | Metschnikowia | 13 | Saprotroph | \|Undefined Saprotroph\| | Yeast | NULL | Probable | As nectar-tap_saprotroph (Põlme et al. 2020); Decay sugar-rich substrates (Põlme et al. 2020) | Põlme S, et al. 2020. Fungal Diversity 105:1-16 ((https://doi.org/10.1007/s13225-020-00466-2)) |
| Otu39622 | 1 | k__Fungi(100);p__Ascomycota(100);c__Saccharomycetes(100);o__Saccharomycetales(100);f__Metschnikowiaceae(100);g__Metschnikowia(100);s__Metschnikowia_pulcherrima(100); | Metschnikowia | 13 | Saprotroph | \|Undefined Saprotroph\| | Yeast | NULL | Probable | As nectar-tap_saprotroph (Põlme et al. 2020); Decay sugar-rich substrates (Põlme et al. 2020) | Põlme S, et al. 2020. Fungal Diversity 105:1-16 ((https://doi.org/10.1007/s13225-020-00466-2)) |
| Otu39623 | 1 | k__Fungi(100);p__Ascomycota(100);c__Saccharomycetes(100);o__Saccharomycetales(100);f__Metschnikowiaceae(100);g__Metschnikowia(100);g__Metschnikowia_unclassified(100); | Metschnikowia | 13 | Saprotroph | \|Undefined Saprotroph\| | Yeast | NULL | Probable | As nectar-tap_saprotroph (Põlme et al. 2020); Decay sugar-rich substrates (Põlme et al. 2020) | Põlme S, et al. 2020. Fungal Diversity 105:1-16 ((https://doi.org/10.1007/s13225-020-00466-2)) |
| Otu39624 | 1 | k__Fungi(100);p__Ascomycota(100);c__Saccharomycetes(100);o__Saccharomycetales(100);f__Metschnikowiaceae(100);g__Metschnikowia(100);s__Metschnikowia_pulcherrima(100); | Metschnikowia | 13 | Saprotroph | \|Undefined Saprotroph\| | Yeast | NULL | Probable | As nectar-tap_saprotroph (Põlme et al. 2020); Decay sugar-rich substrates (Põlme et al. 2020) | Põlme S, et al. 2020. Fungal Diversity 105:1-16 ((https://doi.org/10.1007/s13225-020-00466-2)) |
| Otu39627 | 1 | k__Fungi(100);p__Ascomycota(100);c__Saccharomycetes(100);o__Saccharomycetales(100);f__Metschnikowiaceae(100);g__Metschnikowia(100);g__Metschnikowia_unclassified(100); | Metschnikowia | 13 | Saprotroph | \|Undefined Saprotroph\| | Yeast | NULL | Probable | As nectar-tap_saprotroph (Põlme et al. 2020); Decay sugar-rich substrates (Põlme et al. 2020) | Põlme S, et al. 2020. Fungal Diversity 105:1-16 ((https://doi.org/10.1007/s13225-020-00466-2)) |

| | | | | | | | | | | | |
|---|---|---|---|---|---|---|---|---|---|---|---|
| Otu39630 | 1 | k__Fungi(100);p__Ascomycota(100);c__Saccharomycetes(100);o__Saccharomycetales(100);f__Metschnikowiaceae(100);g__Metschnikowia(100);g__Metschnikowia_unclassified(100); | Metschnikowia | 13 | Saprotroph | \|Undefined Saprotroph\| | Yeast | NULL | Probable | As nectar-tap_saprotroph (Põlme et al. 2020); Decay sugar-rich substrates (Põlme et al. 2020) | Põlme S, et al. 2020. Fungal Diversity 105:1-16 ((https://doi.org/10.1007/s13225-020-00466-2)) |
| Otu39631 | 1 | k__Fungi(100);p__Ascomycota(100);c__Saccharomycetes(100);o__Saccharomycetales(100);f__Metschnikowiaceae(100);g__Metschnikowia(100);s__Metschnikowia_pulcherrima(100); | Metschnikowia | 13 | Saprotroph | \|Undefined Saprotroph\| | Yeast | NULL | Probable | As nectar-tap_saprotroph (Põlme et al. 2020); Decay sugar-rich substrates (Põlme et al. 2020) | Põlme S, et al. 2020. Fungal Diversity 105:1-16 ((https://doi.org/10.1007/s13225-020-00466-2)) |
| Otu39632 | 1 | k__Fungi(100);p__Ascomycota(100);c__Saccharomycetes(100);o__Saccharomycetales(100);f__Metschnikowiaceae(100);g__Metschnikowia(100);s__Metschnikowia_pulcherrima(100); | Metschnikowia | 13 | Saprotroph | \|Undefined Saprotroph\| | Yeast | NULL | Probable | As nectar-tap_saprotroph (Põlme et al. 2020); Decay sugar-rich substrates (Põlme et al. 2020) | Põlme S, et al. 2020. Fungal Diversity 105:1-16 ((https://doi.org/10.1007/s13225-020-00466-2)) |
| Otu39633 | 1 | k__Fungi(100);p__Ascomycota(100);c__Saccharomycetes(100);o__Saccharomycetales(100);f__Metschnikowiaceae(100);g__Metschnikowia(100);s__Metschnikowia_pulcherrima(100); | Metschnikowia | 13 | Saprotroph | \|Undefined Saprotroph\| | Yeast | NULL | Probable | As nectar-tap_saprotroph (Põlme et al. 2020); Decay sugar-rich substrates (Põlme et al. 2020) | Põlme S, et al. 2020. Fungal Diversity 105:1-16 ((https://doi.org/10.1007/s13225-020-00466-2)) |
| Otu39634 | 1 | k__Fungi(100);p__Ascomycota(100);c__Saccharomycetes(100);o__Saccharomycetales(100);f__Metschnikowiaceae(100);g__Metschnikowia(100);s__Metschnikowia_pulcherrima(100); | Metschnikowia | 13 | Saprotroph | \|Undefined Saprotroph\| | Yeast | NULL | Probable | As nectar-tap_saprotroph (Põlme et al. 2020); Decay sugar-rich substrates (Põlme et al. 2020) | Põlme S, et al. 2020. Fungal Diversity 105:1-16 ((https://doi.org/10.1007/s13225-020-00466-2)) |
| Otu39635 | 1 | k__Fungi(100);p__Ascomycota(100);c__Saccharomycetes(100);o__Saccharomycetales(100);f__Metschnikowiaceae(100);g__Metschnikowia(100);s__Metschnikowia_pulcherrima(100); | Metschnikowia | 13 | Saprotroph | \|Undefined Saprotroph\| | Yeast | NULL | Probable | As nectar-tap_saprotroph (Põlme et al. 2020); Decay sugar-rich substrates (Põlme et al. 2020) | Põlme S, et al. 2020. Fungal Diversity 105:1-16 ((https://doi.org/10.1007/s13225-020-00466-2)) |
| Otu39636 | 1 | k__Fungi(100);p__Ascomycota(100);c__Saccharomycetes(100);o__Saccharomycetales(100);f__Metschnikowiaceae(100);g__Metschnikowia(100);s__Metschnikowia_pulcherrima(100); | Metschnikowia | 13 | Saprotroph | \|Undefined Saprotroph\| | Yeast | NULL | Probable | As nectar-tap_saprotroph (Põlme et al. 2020); Decay sugar-rich substrates (Põlme et al. 2020) | Põlme S, et al. 2020. Fungal Diversity 105:1-16 ((https://doi.org/10.1007/s13225-020-00466-2)) |
| Otu39637 | 1 | k__Fungi(100);p__Ascomycota(100);c__Saccharomycetes(100);o__Saccharomycetales(100);f__Metschnikowiaceae(100);g__Metschnikowia(100);s__Metschnikowia_pulcherrima(100); | Metschnikowia | 13 | Saprotroph | \|Undefined Saprotroph\| | Yeast | NULL | Probable | As nectar-tap_saprotroph (Põlme et al. 2020); Decay sugar-rich substrates (Põlme et al. 2020) | Põlme S, et al. 2020. Fungal Diversity 105:1-16 ((https://doi.org/10.1007/s13225-020-00466-2)) |
| Otu39638 | 1 | k__Fungi(100);p__Ascomycota(100);c__Saccharomycetes(100);o__Saccharomycetales(100);f__Metschnikowiaceae(100);g__Metschnikowia(100);s__Metschnikowia_pulcherrima(100); | Metschnikowia | 13 | Saprotroph | \|Undefined Saprotroph\| | Yeast | NULL | Probable | As nectar-tap_saprotroph (Põlme et al. 2020); Decay sugar-rich substrates (Põlme et al. 2020) | Põlme S, et al. 2020. Fungal Diversity 105:1-16 ((https://doi.org/10.1007/s13225-020-00466-2)) |
| Otu39639 | 1 | k__Fungi(100);p__Ascomycota(100);c__Saccharomycetes(100);o__Saccharomycetales(100);f__Metschnikowiaceae(100);g__Metschnikowia(100);s__Metschnikowia_pulcherrima(100); | Metschnikowia | 13 | Saprotroph | \|Undefined Saprotroph\| | Yeast | NULL | Probable | As nectar-tap_saprotroph (Põlme et al. 2020); Decay sugar-rich substrates (Põlme et al. 2020) | Põlme S, et al. 2020. Fungal Diversity 105:1-16 ((https://doi.org/10.1007/s13225-020-00466-2)) |
| Otu39641 | 1 | k__Fungi(100);p__Ascomycota(100);c__Saccharomycetes(100);o__Saccharomycetales(100);f__Metschnikowiaceae(100);g__Metschnikowia(100);s__Metschnikowia_pulcherrima(100); | Metschnikowia | 13 | Saprotroph | \|Undefined Saprotroph\| | Yeast | NULL | Probable | As nectar-tap_saprotroph (Põlme et al. 2020); Decay sugar-rich substrates (Põlme et al. 2020) | Põlme S, et al. 2020. Fungal Diversity 105:1-16 ((https://doi.org/10.1007/s13225-020-00466-2)) |
| Otu39643 | 1 | k__Fungi(100);p__Ascomycota(100);c__Saccharomycetes(100);o__Saccharomycetales(100);f__Metschnikowiaceae(100);g__Metschnikowia(100);g__Metschnikowia_unclassified(100); | Metschnikowia | 13 | Saprotroph | \|Undefined Saprotroph\| | Yeast | NULL | Probable | As nectar-tap_saprotroph (Põlme et al. 2020); Decay sugar-rich substrates (Põlme et al. 2020) | Põlme S, et al. 2020. Fungal Diversity 105:1-16 ((https://doi.org/10.1007/s13225-020-00466-2)) |
| Otu39644 | 1 | k__Fungi(100);p__Ascomycota(100);c__Saccharomycetes(100);o__Saccharomycetales(100);f__Metschnikowiaceae(100);g__Metschnikowia(100);s__Metschnikowia_pulcherrima(100); | Metschnikowia | 13 | Saprotroph | \|Undefined Saprotroph\| | Yeast | NULL | Probable | As nectar-tap_saprotroph (Põlme et al. 2020); Decay sugar-rich substrates (Põlme et al. 2020) | Põlme S, et al. 2020. Fungal Diversity 105:1-16 ((https://doi.org/10.1007/s13225-020-00466-2)) |
| Otu39645 | 1 | k__Fungi(100);p__Ascomycota(100);c__Saccharomycetes(100);o__Saccharomycetales(100);f__Metschnikowiaceae(100);g__Metschnikowia(100);s__Metschnikowia_pulcherrima(100); | Metschnikowia | 13 | Saprotroph | \|Undefined Saprotroph\| | Yeast | NULL | Probable | As nectar-tap_saprotroph (Põlme et al. 2020); Decay sugar-rich substrates (Põlme et al. 2020) | Põlme S, et al. 2020. Fungal Diversity 105:1-16 ((https://doi.org/10.1007/s13225-020-00466-2)) |
| Otu39646 | 1 | k__Fungi(100);p__Ascomycota(100);c__Saccharomycetes(100);o__Saccharomycetales(100);f__Metschnikowiaceae(100);g__Metschnikowia(100);s__Metschnikowia_pulcherrima(100); | Metschnikowia | 13 | Saprotroph | \|Undefined Saprotroph\| | Yeast | NULL | Probable | As nectar-tap_saprotroph (Põlme et al. 2020); Decay sugar-rich substrates (Põlme et al. 2020) | Põlme S, et al. 2020. Fungal Diversity 105:1-16 ((https://doi.org/10.1007/s13225-020-00466-2)) |
| Otu39647 | 1 | k__Fungi(100);p__Ascomycota(100);c__Saccharomycetes(100);o__Saccharomycetales(100);f__Metschnikowiaceae(100);g__Metschnikowia(100);s__Metschnikowia_pulcherrima(100); | Metschnikowia | 13 | Saprotroph | \|Undefined Saprotroph\| | Yeast | NULL | Probable | As nectar-tap_saprotroph (Põlme et al. 2020); Decay sugar-rich substrates (Põlme et al. 2020) | Põlme S, et al. 2020. Fungal Diversity 105:1-16 ((https://doi.org/10.1007/s13225-020-00466-2)) |
| Otu39648 | 1 | k__Fungi(100);p__Ascomycota(100);c__Saccharomycetes(100);o__Saccharomycetales(100);f__Metschnikowiaceae(100);g__Metschnikowia(100);s__Metschnikowia_pulcherrima(100); | Metschnikowia | 13 | Saprotroph | \|Undefined Saprotroph\| | Yeast | NULL | Probable | As nectar-tap_saprotroph (Põlme et al. 2020); Decay sugar-rich substrates (Põlme et al. 2020) | Põlme S, et al. 2020. Fungal Diversity 105:1-16 ((https://doi.org/10.1007/s13225-020-00466-2)) |
| Otu39649 | 1 | k__Fungi(100);p__Ascomycota(100);c__Saccharomycetes(100);o__Saccharomycetales(100);f__Metschnikowiaceae(100);g__Metschnikowia(100);g__Metschnikowia_unclassified(100); | Metschnikowia | 13 | Saprotroph | \|Undefined Saprotroph\| | Yeast | NULL | Probable | As nectar-tap_saprotroph (Põlme et al. 2020); Decay sugar-rich substrates (Põlme et al. 2020) | Põlme S, et al. 2020. Fungal Diversity 105:1-16 ((https://doi.org/10.1007/s13225-020-00466-2)) |
| Otu39650 | 1 | k__Fungi(100);p__Ascomycota(100);c__Saccharomycetes(100);o__Saccharomycetales(100);f__Metschnikowiaceae(100);g__Metschnikowia(100);s__Metschnikowia_pulcherrima(100); | Metschnikowia | 13 | Saprotroph | \|Undefined Saprotroph\| | Yeast | NULL | Probable | As nectar-tap_saprotroph (Põlme et al. 2020); Decay sugar-rich substrates (Põlme et al. 2020) | Põlme S, et al. 2020. Fungal Diversity 105:1-16 ((https://doi.org/10.1007/s13225-020-00466-2)) |
| Otu39651 | 1 | k__Fungi(100);p__Ascomycota(100);c__Saccharomycetes(100);o__Saccharomycetales(100);f__Metschnikowiaceae(100);g__Metschnikowia(100);s__Metschnikowia_pulcherrima(100); | Metschnikowia | 13 | Saprotroph | \|Undefined Saprotroph\| | Yeast | NULL | Probable | As nectar-tap_saprotroph (Põlme et al. 2020); Decay sugar-rich substrates (Põlme et al. 2020) | Põlme S, et al. 2020. Fungal Diversity 105:1-16 ((https://doi.org/10.1007/s13225-020-00466-2)) |
| Otu39652 | 1 | k__Fungi(100);p__Ascomycota(100);c__Saccharomycetes(100);o__Saccharomycetales(100);f__Metschnikowiaceae(100);g__Metschnikowia(100);s__Metschnikowia_pulcherrima(100); | Metschnikowia | 13 | Saprotroph | \|Undefined Saprotroph\| | Yeast | NULL | Probable | As nectar-tap_saprotroph (Põlme et al. 2020); Decay sugar-rich substrates (Põlme et al. 2020) | Põlme S, et al. 2020. Fungal Diversity 105:1-16 ((https://doi.org/10.1007/s13225-020-00466-2)) |
| Otu39653 | 1 | k__Fungi(100);p__Ascomycota(100);c__Saccharomycetes(100);o__Saccharomycetales(100);f__Metschnikowiaceae(100);g__Metschnikowia(100);s__Metschnikowia_pulcherrima(100); | Metschnikowia | 13 | Saprotroph | \|Undefined Saprotroph\| | Yeast | NULL | Probable | As nectar-tap_saprotroph (Põlme et al. 2020); Decay sugar-rich substrates (Põlme et al. 2020) | Põlme S, et al. 2020. Fungal Diversity 105:1-16 ((https://doi.org/10.1007/s13225-020-00466-2)) |
| Otu39654 | 1 | k__Fungi(100);p__Ascomycota(100);c__Saccharomycetes(100);o__Saccharomycetales(100);f__Metschnikowiaceae(100);g__Metschnikowia(100);s__Metschnikowia_pulcherrima(100); | Metschnikowia | 13 | Saprotroph | \|Undefined Saprotroph\| | Yeast | NULL | Probable | As nectar-tap_saprotroph (Põlme et al. 2020); Decay sugar-rich substrates (Põlme et al. 2020) | Põlme S, et al. 2020. Fungal Diversity 105:1-16 ((https://doi.org/10.1007/s13225-020-00466-2)) |
| Otu39655 | 1 | k__Fungi(100);p__Ascomycota(100);c__Saccharomycetes(100);o__Saccharomycetales(100);f__Metschnikowiaceae(100);g__Metschnikowia(100);g__Metschnikowia_unclassified(100); | Metschnikowia | 13 | Saprotroph | \|Undefined Saprotroph\| | Yeast | NULL | Probable | As nectar-tap_saprotroph (Põlme et al. 2020); Decay sugar-rich substrates (Põlme et al. 2020) | Põlme S, et al. 2020. Fungal Diversity 105:1-16 ((https://doi.org/10.1007/s13225-020-00466-2)) |
| Otu39656 | 1 | k__Fungi(100);p__Ascomycota(100);c__Saccharomycetes(100);o__Saccharomycetales(100);f__Metschnikowiaceae(100);g__Metschnikowia(100);g__Metschnikowia_unclassified(100); | Metschnikowia | 13 | Saprotroph | \|Undefined Saprotroph\| | Yeast | NULL | Probable | As nectar-tap_saprotroph (Põlme et al. 2020); Decay sugar-rich substrates (Põlme et al. 2020) | Põlme S, et al. 2020. Fungal Diversity 105:1-16 ((https://doi.org/10.1007/s13225-020-00466-2)) |
| Otu39657 | 1 | k__Fungi(100);p__Ascomycota(100);c__Saccharomycetes(100);o__Saccharomycetales(100);f__Metschnikowiaceae(100);g__Metschnikowia(100);g__Metschnikowia_unclassified(100); | Metschnikowia | 13 | Saprotroph | \|Undefined Saprotroph\| | Yeast | NULL | Probable | As nectar-tap_saprotroph (Põlme et al. 2020); Decay sugar-rich substrates (Põlme et al. 2020) | Põlme S, et al. 2020. Fungal Diversity 105:1-16 ((https://doi.org/10.1007/s13225-020-00466-2)) |
| Otu39658 | 1 | k__Fungi(100);p__Ascomycota(100);c__Saccharomycetes(100);o__Saccharomycetales(100);f__Metschnikowiaceae(100);g__Metschnikowia(100);s__Metschnikowia_pulcherrima(100); | Metschnikowia | 13 | Saprotroph | \|Undefined Saprotroph\| | Yeast | NULL | Probable | As nectar-tap_saprotroph (Põlme et al. 2020); Decay sugar-rich substrates (Põlme et al. 2020) | Põlme S, et al. 2020. Fungal Diversity 105:1-16 ((https://doi.org/10.1007/s13225-020-00466-2)) |
| Otu39659 | 1 | k__Fungi(100);p__Ascomycota(100);c__Saccharomycetes(100);o__Saccharomycetales(100);f__Metschnikowiaceae(100);g__Metschnikowia(100);g__Metschnikowia_unclassified(100); | Metschnikowia | 13 | Saprotroph | \|Undefined Saprotroph\| | Yeast | NULL | Probable | As nectar-tap_saprotroph (Põlme et al. 2020); Decay sugar-rich substrates (Põlme et al. 2020) | Põlme S, et al. 2020. Fungal Diversity 105:1-16 ((https://doi.org/10.1007/s13225-020-00466-2)) |
| Otu39660 | 1 | k__Fungi(100);p__Ascomycota(100);c__Saccharomycetes(100);o__Saccharomycetales(100);f__Metschnikowiaceae(100);g__Metschnikowia(100);s__Metschnikowia_pulcherrima(100); | Metschnikowia | 13 | Saprotroph | \|Undefined Saprotroph\| | Yeast | NULL | Probable | As nectar-tap_saprotroph (Põlme et al. 2020); Decay sugar-rich substrates (Põlme et al. 2020) | Põlme S, et al. 2020. Fungal Diversity 105:1-16 ((https://doi.org/10.1007/s13225-020-00466-2)) |
| Otu39661 | 1 | k__Fungi(100);p__Ascomycota(100);c__Saccharomycetes(100);o__Saccharomycetales(100);f__Metschnikowiaceae(100);g__Metschnikowia(100);s__Metschnikowia_pulcherrima(100); | Metschnikowia | 13 | Saprotroph | \|Undefined Saprotroph\| | Yeast | NULL | Probable | As nectar-tap_saprotroph (Põlme et al. 2020); Decay sugar-rich substrates (Põlme et al. 2020) | Põlme S, et al. 2020. Fungal Diversity 105:1-16 ((https://doi.org/10.1007/s13225-020-00466-2)) |
| Otu39662 | 1 | k__Fungi(100);p__Ascomycota(100);c__Saccharomycetes(100);o__Saccharomycetales(100);f__Metschnikowiaceae(100);g__Metschnikowia(100);s__Metschnikowia_pulcherrima(100); | Metschnikowia | 13 | Saprotroph | \|Undefined Saprotroph\| | Yeast | NULL | Probable | As nectar-tap_saprotroph (Põlme et al. 2020); Decay sugar-rich substrates (Põlme et al. 2020) | Põlme S, et al. 2020. Fungal Diversity 105:1-16 ((https://doi.org/10.1007/s13225-020-00466-2)) |
| Otu39663 | 1 | k__Fungi(100);p__Ascomycota(100);c__Saccharomycetes(100);o__Saccharomycetales(100);f__Metschnikowiaceae(100);g__Metschnikowia(100);s__Metschnikowia_pulcherrima(100); | Metschnikowia | 13 | Saprotroph | \|Undefined Saprotroph\| | Yeast | NULL | Probable | As nectar-tap_saprotroph (Põlme et al. 2020); Decay sugar-rich substrates (Põlme et al. 2020) | Põlme S, et al. 2020. Fungal Diversity 105:1-16 ((https://doi.org/10.1007/s13225-020-00466-2)) |
| Otu39664 | 1 | k__Fungi(100);p__Ascomycota(100);c__Saccharomycetes(100);o__Saccharomycetales(100);f__Metschnikowiaceae(100);g__Metschnikowia(100);s__Metschnikowia_pulcherrima(100); | Metschnikowia | 13 | Saprotroph | \|Undefined Saprotroph\| | Yeast | NULL | Probable | As nectar-tap_saprotroph (Põlme et al. 2020); Decay sugar-rich substrates (Põlme et al. 2020) | Põlme S, et al. 2020. Fungal Diversity 105:1-16 ((https://doi.org/10.1007/s13225-020-00466-2)) |
| Otu39665 | 1 | k__Fungi(100);p__Ascomycota(100);c__Saccharomycetes(100);o__Saccharomycetales(100);f__Metschnikowiaceae(100);g__Metschnikowia(100);s__Metschnikowia_pulcherrima(100); | Metschnikowia | 13 | Saprotroph | \|Undefined Saprotroph\| | Yeast | NULL | Probable | As nectar-tap_saprotroph (Põlme et al. 2020); Decay sugar-rich substrates (Põlme et al. 2020) | Põlme S, et al. 2020. Fungal Diversity 105:1-16 ((https://doi.org/10.1007/s13225-020-00466-2)) |
| Otu39666 | 1 | k__Fungi(100);p__Ascomycota(100);c__Saccharomycetes(100);o__Saccharomycetales(100);f__Metschnikowiaceae(100);g__Metschnikowia(100);s__Metschnikowia_pulcherrima(100); | Metschnikowia | 13 | Saprotroph | \|Undefined Saprotroph\| | Yeast | NULL | Probable | As nectar-tap_saprotroph (Põlme et al. 2020); Decay sugar-rich substrates (Põlme et al. 2020) | Põlme S, et al. 2020. Fungal Diversity 105:1-16 ((https://doi.org/10.1007/s13225-020-00466-2)) |
| Otu39667 | 1 | k__Fungi(100);p__Ascomycota(100);c__Saccharomycetes(100);o__Saccharomycetales(100);f__Metschnikowiaceae(100);g__Metschnikowia(100);s__Metschnikowia_pulcherrima(100); | Metschnikowia | 13 | Saprotroph | \|Undefined Saprotroph\| | Yeast | NULL | Probable | As nectar-tap_saprotroph (Põlme et al. 2020); Decay sugar-rich substrates (Põlme et al. 2020) | Põlme S, et al. 2020. Fungal Diversity 105:1-16 ((https://doi.org/10.1007/s13225-020-00466-2)) |
| Otu39668 | 1 | k__Fungi(100);p__Ascomycota(100);c__Saccharomycetes(100);o__Saccharomycetales(100);f__Metschnikowiaceae(100);g__Metschnikowia(100);g__Metschnikowia_unclassified(100); | Metschnikowia | 13 | Saprotroph | \|Undefined Saprotroph\| | Yeast | NULL | Probable | As nectar-tap_saprotroph (Põlme et al. 2020); Decay sugar-rich substrates (Põlme et al. 2020) | Põlme S, et al. 2020. Fungal Diversity 105:1-16 ((https://doi.org/10.1007/s13225-020-00466-2)) |
| Otu39669 | 1 | k__Fungi(100);p__Ascomycota(100);c__Saccharomycetes(100);o__Saccharomycetales(100);f__Metschnikowiaceae(100);g__Metschnikowia(100);s__Metschnikowia_pulcherrima(100); | Metschnikowia | 13 | Saprotroph | \|Undefined Saprotroph\| | Yeast | NULL | Probable | As nectar-tap_saprotroph (Põlme et al. 2020); Decay sugar-rich substrates (Põlme et al. 2020) | Põlme S, et al. 2020. Fungal Diversity 105:1-16 ((https://doi.org/10.1007/s13225-020-00466-2)) |
| Otu39670 | 1 | k__Fungi(100);p__Ascomycota(100);c__Saccharomycetes(100);o__Saccharomycetales(100);f__Metschnikowiaceae(100);g__Metschnikowia(100);g__Metschnikowia_unclassified(100); | Metschnikowia | 13 | Saprotroph | \|Undefined Saprotroph\| | Yeast | NULL | Probable | As nectar-tap_saprotroph (Põlme et al. 2020); Decay sugar-rich substrates (Põlme et al. 2020) | Põlme S, et al. 2020. Fungal Diversity 105:1-16 ((https://doi.org/10.1007/s13225-020-00466-2)) |
| Otu39671 | 1 | k__Fungi(100);p__Ascomycota(100);c__Saccharomycetes(100);o__Saccharomycetales(100);f__Metschnikowiaceae(100);g__Metschnikowia(100);s__Metschnikowia_pulcherrima(100); | Metschnikowia | 13 | Saprotroph | \|Undefined Saprotroph\| | Yeast | NULL | Probable | As nectar-tap_saprotroph (Põlme et al. 2020); Decay sugar-rich substrates (Põlme et al. 2020) | Põlme S, et al. 2020. Fungal Diversity 105:1-16 ((https://doi.org/10.1007/s13225-020-00466-2)) |
| Otu39672 | 1 | k__Fungi(100);p__Ascomycota(100);c__Saccharomycetes(100);o__Saccharomycetales(100);f__Metschnikowiaceae(100);g__Metschnikowia(100);s__Metschnikowia_pulcherrima(100); | Metschnikowia | 13 | Saprotroph | \|Undefined Saprotroph\| | Yeast | NULL | Probable | As nectar-tap_saprotroph (Põlme et al. 2020); Decay sugar-rich substrates (Põlme et al. 2020) | Põlme S, et al. 2020. Fungal Diversity 105:1-16 ((https://doi.org/10.1007/s13225-020-00466-2)) |
| Otu39673 | 1 | k__Fungi(100);p__Ascomycota(100);c__Saccharomycetes(100);o__Saccharomycetales(100);f__Metschnikowiaceae(100);g__Metschnikowia(100);s__Metschnikowia_pulcherrima(100); | Metschnikowia | 13 | Saprotroph | \|Undefined Saprotroph\| | Yeast | NULL | Probable | As nectar-tap_saprotroph (Põlme et al. 2020); Decay sugar-rich substrates (Põlme et al. 2020) | Põlme S, et al. 2020. Fungal Diversity 105:1-16 ((https://doi.org/10.1007/s13225-020-00466-2)) |
| Otu39674 | 1 | k__Fungi(100);p__Ascomycota(100);c__Saccharomycetes(100);o__Saccharomycetales(100);f__Metschnikowiaceae(100);g__Metschnikowia(100);s__Metschnikowia_pulcherrima(100); | Metschnikowia | 13 | Saprotroph | \|Undefined Saprotroph\| | Yeast | NULL | Probable | As nectar-tap_saprotroph (Põlme et al. 2020); Decay sugar-rich substrates (Põlme et al. 2020) | Põlme S, et al. 2020. Fungal Diversity 105:1-16 ((https://doi.org/10.1007/s13225-020-00466-2)) |
| Otu39676 | 1 | k__Fungi(100);p__Ascomycota(100);c__Saccharomycetes(100);o__Saccharomycetales(100);f__Metschnikowiaceae(100);g__Metschnikowia(100);g__Metschnikowia_unclassified(100); | Metschnikowia | 13 | Saprotroph | \|Undefined Saprotroph\| | Yeast | NULL | Probable | As nectar-tap_saprotroph (Põlme et al. 2020); Decay sugar-rich substrates (Põlme et al. 2020) | Põlme S, et al. 2020. Fungal Diversity 105:1-16 ((https://doi.org/10.1007/s13225-020-00466-2)) |
| Otu39677 | 1 | k__Fungi(100);p__Ascomycota(100);c__Saccharomycetes(100);o__Saccharomycetales(100);f__Metschnikowiaceae(100);g__Metschnikowia(100);g__Metschnikowia_unclassified(100); | Metschnikowia | 13 | Saprotroph | \|Undefined Saprotroph\| | Yeast | NULL | Probable | As nectar-tap_saprotroph (Põlme et al. 2020); Decay sugar-rich substrates (Põlme et al. 2020) | Põlme S, et al. 2020. Fungal Diversity 105:1-16 ((https://doi.org/10.1007/s13225-020-00466-2)) |
| Otu39678 | 1 | k__Fungi(100);p__Ascomycota(100);c__Saccharomycetes(100);o__Saccharomycetales(100);f__Metschnikowiaceae(100);g__Metschnikowia(100);s__Metschnikowia_pulcherrima(100); | Metschnikowia | 13 | Saprotroph | \|Undefined Saprotroph\| | Yeast | NULL | Probable | As nectar-tap_saprotroph (Põlme et al. 2020); Decay sugar-rich substrates (Põlme et al. 2020) | Põlme S, et al. 2020. Fungal Diversity 105:1-16 ((https://doi.org/10.1007/s13225-020-00466-2)) |
| Otu39679 | 1 | k__Fungi(100);p__Ascomycota(100);c__Saccharomycetes(100);o__Saccharomycetales(100);f__Metschnikowiaceae(100);g__Metschnikowia(100);s__Metschnikowia_pulcherrima(100); | Metschnikowia | 13 | Saprotroph | \|Undefined Saprotroph\| | Yeast | NULL | Probable | As nectar-tap_saprotroph (Põlme et al. 2020); Decay sugar-rich substrates (Põlme et al. 2020) | Põlme S, et al. 2020. Fungal Diversity 105:1-16 ((https://doi.org/10.1007/s13225-020-00466-2)) |
| Otu39680 | 1 | k__Fungi(100);p__Ascomycota(100);c__Saccharomycetes(100);o__Saccharomycetales(100);f__Metschnikowiaceae(100);g__Metschnikowia(100);s__Metschnikowia_pulcherrima(100); | Metschnikowia | 13 | Saprotroph | \|Undefined Saprotroph\| | Yeast | NULL | Probable | As nectar-tap_saprotroph (Põlme et al. 2020); Decay sugar-rich substrates (Põlme et al. 2020) | Põlme S, et al. 2020. Fungal Diversity 105:1-16 ((https://doi.org/10.1007/s13225-020-00466-2)) |
| Otu39681 | 1 | k__Fungi(100);p__Ascomycota(100);c__Saccharomycetes(100);o__Saccharomycetales(100);f__Metschnikowiaceae(100);g__Metschnikowia(100);g__Metschnikowia_unclassified(100); | Metschnikowia | 13 | Saprotroph | \|Undefined Saprotroph\| | Yeast | NULL | Probable | As nectar-tap_saprotroph (Põlme et al. 2020); Decay sugar-rich substrates (Põlme et al. 2020) | Põlme S, et al. 2020. Fungal Diversity 105:1-16 ((https://doi.org/10.1007/s13225-020-00466-2)) |
| Otu39682 | 1 | k__Fungi(100);p__Ascomycota(100);c__Saccharomycetes(100);o__Saccharomycetales(100);f__Metschnikowiaceae(100);g__Metschnikowia(100);g__Metschnikowia_unclassified(100); | Metschnikowia | 13 | Saprotroph | \|Undefined Saprotroph\| | Yeast | NULL | Probable | As nectar-tap_saprotroph (Põlme et al. 2020); Decay sugar-rich substrates (Põlme et al. 2020) | Põlme S, et al. 2020. Fungal Diversity 105:1-16 ((https://doi.org/10.1007/s13225-020-00466-2)) |
| Otu39683 | 1 | k__Fungi(100);p__Ascomycota(100);c__Saccharomycetes(100);o__Saccharomycetales(100);o__Saccharomycetales_unclassified(100);o__Saccharomycetales_unclassified(100);o__Saccharomycetales_unclassified(100); | Saccharomycetales | 7 | Saprotroph | Undefined Saprotroph | Yeast | NULL | Possible | NULL | Sterkenburg E, et al. 2015. New Phytologist 207:1145-1158 ((https://doi.org/10.1111/nph.13426)) |
| Otu39684 | 1 | k__Fungi(100);p__Ascomycota(100);c__Saccharomycetes(100);o__Saccharomycetales(100);f__Metschnikowiaceae(100);g__Metschnikowia(100);s__Metschnikowia_pulcherrima(100); | Metschnikowia | 13 | Saprotroph | \|Undefined Saprotroph\| | Yeast | NULL | Probable | As nectar-tap_saprotroph (Põlme et al. 2020); Decay sugar-rich substrates (Põlme et al. 2020) | Põlme S, et al. 2020. Fungal Diversity 105:1-16 ((https://doi.org/10.1007/s13225-020-00466-2)) |
| Otu39685 | 1 | k__Fungi(100);p__Ascomycota(100);c__Saccharomycetes(100);o__Saccharomycetales(100);f__Metschnikowiaceae(100);g__Metschnikowia(100);s__Metschnikowia_pulcherrima(100); | Metschnikowia | 13 | Saprotroph | \|Undefined Saprotroph\| | Yeast | NULL | Probable | As nectar-tap_saprotroph (Põlme et al. 2020); Decay sugar-rich substrates (Põlme et al. 2020) | Põlme S, et al. 2020. Fungal Diversity 105:1-16 ((https://doi.org/10.1007/s13225-020-00466-2)) |
| Otu39686 | 1 | k__Fungi(100);p__Ascomycota(100);c__Saccharomycetes(100);o__Saccharomycetales(100);f__Metschnikowiaceae(100);g__Metschnikowia(100);s__Metschnikowia_pulcherrima(100); | Metschnikowia | 13 | Saprotroph | \|Undefined Saprotroph\| | Yeast | NULL | Probable | As nectar-tap_saprotroph (Põlme et al. 2020); Decay sugar-rich substrates (Põlme et al. 2020) | Põlme S, et al. 2020. Fungal Diversity 105:1-16 ((https://doi.org/10.1007/s13225-020-00466-2)) |
| Otu39687 | 1 | k__Fungi(100);p__Ascomycota(100);c__Saccharomycetes(100);o__Saccharomycetales(100);f__Metschnikowiaceae(100);g__Metschnikowia(100);s__Metschnikowia_pulcherrima(100); | Metschnikowia | 13 | Saprotroph | \|Undefined Saprotroph\| | Yeast | NULL | Probable | As nectar-tap_saprotroph (Põlme et al. 2020); Decay sugar-rich substrates (Põlme et al. 2020) | Põlme S, et al. 2020. Fungal Diversity 105:1-16 ((https://doi.org/10.1007/s13225-020-00466-2)) |
| Otu39688 | 1 | k__Fungi(100);p__Ascomycota(100);c__Saccharomycetes(100);o__Saccharomycetales(100);f__Metschnikowiaceae(100);g__Metschnikowia(100);s__Metschnikowia_pulcherrima(100); | Metschnikowia | 13 | Saprotroph | \|Undefined Saprotroph\| | Yeast | NULL | Probable | As nectar-tap_saprotroph (Põlme et al. 2020); Decay sugar-rich substrates (Põlme et al. 2020) | Põlme S, et al. 2020. Fungal Diversity 105:1-16 ((https://doi.org/10.1007/s13225-020-00466-2)) |
| Otu39689 | 1 | k__Fungi(100);p__Ascomycota(100);c__Saccharomycetes(100);o__Saccharomycetales(100);f__Metschnikowiaceae(100);g__Metschnikowia(100);s__Metschnikowia_pulcherrima(100); | Metschnikowia | 13 | Saprotroph | \|Undefined Saprotroph\| | Yeast | NULL | Probable | As nectar-tap_saprotroph (Põlme et al. 2020); Decay sugar-rich substrates (Põlme et al. 2020) | Põlme S, et al. 2020. Fungal Diversity 105:1-16 ((https://doi.org/10.1007/s13225-020-00466-2)) |
| Otu39690 | 1 | k__Fungi(100);p__Ascomycota(100);c__Saccharomycetes(100);o__Saccharomycetales(100);f__Metschnikowiaceae(100);g__Metschnikowia(100);s__Metschnikowia_pulcherrima(100); | Metschnikowia | 13 | Saprotroph | \|Undefined Saprotroph\| | Yeast | NULL | Probable | As nectar-tap_saprotroph (Põlme et al. 2020); Decay sugar-rich substrates (Põlme et al. 2020) | Põlme S, et al. 2020. Fungal Diversity 105:1-16 ((https://doi.org/10.1007/s13225-020-00466-2)) |
| Otu39691 | 1 | k__Fungi(100);p__Ascomycota(100);c__Saccharomycetes(100);o__Saccharomycetales(100);f__Metschnikowiaceae(100);g__Metschnikowia(100);s__Metschnikowia_pulcherrima(100); | Metschnikowia | 13 | Saprotroph | \|Undefined Saprotroph\| | Yeast | NULL | Probable | As nectar-tap_saprotroph (Põlme et al. 2020); Decay sugar-rich substrates (Põlme et al. 2020) | Põlme S, et al. 2020. Fungal Diversity 105:1-16 ((https://doi.org/10.1007/s13225-020-00466-2)) |
| Otu39692 | 1 | k__Fungi(100);p__Ascomycota(100);c__Saccharomycetes(100);o__Saccharomycetales(100);f__Metschnikowiaceae(100);g__Metschnikowia(100);g__Metschnikowia_unclassified(100); | Metschnikowia | 13 | Saprotroph | \|Undefined Saprotroph\| | Yeast | NULL | Probable | As nectar-tap_saprotroph (Põlme et al. 2020); Decay sugar-rich substrates (Põlme et al. 2020) | Põlme S, et al. 2020. Fungal Diversity 105:1-16 ((https://doi.org/10.1007/s13225-020-00466-2)) |
| Otu39693 | 1 | k__Fungi(100);p__Ascomycota(100);c__Saccharomycetes(100);o__Saccharomycetales(100);f__Metschnikowiaceae(100);g__Metschnikowia(100);s__Metschnikowia_pulcherrima(100); | Metschnikowia | 13 | Saprotroph | \|Undefined Saprotroph\| | Yeast | NULL | Probable | As nectar-tap_saprotroph (Põlme et al. 2020); Decay sugar-rich substrates (Põlme et al. 2020) | Põlme S, et al. 2020. Fungal Diversity 105:1-16 ((https://doi.org/10.1007/s13225-020-00466-2)) |
| Otu39694 | 1 | k__Fungi(100);p__Ascomycota(100);c__Saccharomycetes(100);o__Saccharomycetales(100);f__Metschnikowiaceae(100);g__Metschnikowia(100);s__Metschnikowia_pulcherrima(100); | Metschnikowia | 13 | Saprotroph | \|Undefined Saprotroph\| | Yeast | NULL | Probable | As nectar-tap_saprotroph (Põlme et al. 2020); Decay sugar-rich substrates (Põlme et al. 2020) | Põlme S, et al. 2020. Fungal Diversity 105:1-16 ((https://doi.org/10.1007/s13225-020-00466-2)) |
| Otu39695 | 1 | k__Fungi(100);p__Ascomycota(100);c__Saccharomycetes(100);o__Saccharomycetales(100);f__Metschnikowiaceae(100);g__Metschnikowia(100);s__Metschnikowia_pulcherrima(100); | Metschnikowia | 13 | Saprotroph | \|Undefined Saprotroph\| | Yeast | NULL | Probable | As nectar-tap_saprotroph (Põlme et al. 2020); Decay sugar-rich substrates (Põlme et al. 2020) | Põlme S, et al. 2020. Fungal Diversity 105:1-16 ((https://doi.org/10.1007/s13225-020-00466-2)) |
| Otu39696 | 1 | k__Fungi(100);p__Ascomycota(100);c__Saccharomycetes(100);o__Saccharomycetales(100);f__Metschnikowiaceae(100);g__Metschnikowia(100);s__Metschnikowia_pulcherrima(100); | Metschnikowia | 13 | Saprotroph | \|Undefined Saprotroph\| | Yeast | NULL | Probable | As nectar-tap_saprotroph (Põlme et al. 2020); Decay sugar-rich substrates (Põlme et al. 2020) | Põlme S, et al. 2020. Fungal Diversity 105:1-16 ((https://doi.org/10.1007/s13225-020-00466-2)) |
| Otu39697 | 1 | k__Fungi(100);p__Ascomycota(100);c__Saccharomycetes(100);o__Saccharomycetales(100);f__Metschnikowiaceae(100);g__Metschnikowia(100);g__Metschnikowia_unclassified(100); | Metschnikowia | 13 | Saprotroph | \|Undefined Saprotroph\| | Yeast | NULL | Probable | As nectar-tap_saprotroph (Põlme et al. 2020); Decay sugar-rich substrates (Põlme et al. 2020) | Põlme S, et al. 2020. Fungal Diversity 105:1-16 ((https://doi.org/10.1007/s13225-020-00466-2)) |
| Otu39698 | 1 | k__Fungi(100);p__Ascomycota(100);c__Saccharomycetes(100);o__Saccharomycetales(100);f__Metschnikowiaceae(100);g__Metschnikowia(100);g__Metschnikowia_unclassified(100); | Metschnikowia | 13 | Saprotroph | \|Undefined Saprotroph\| | Yeast | NULL | Probable | As nectar-tap_saprotroph (Põlme et al. 2020); Decay sugar-rich substrates (Põlme et al. 2020) | Põlme S, et al. 2020. Fungal Diversity 105:1-16 ((https://doi.org/10.1007/s13225-020-00466-2)) |
| Otu39699 | 1 | k__Fungi(100);p__Ascomycota(100);c__Saccharomycetes(100);o__Saccharomycetales(100);f__Metschnikowiaceae(100);g__Metschnikowia(100);s__Metschnikowia_pulcherrima(100); | Metschnikowia | 13 | Saprotroph | \|Undefined Saprotroph\| | Yeast | NULL | Probable | As nectar-tap_saprotroph (Põlme et al. 2020); Decay sugar-rich substrates (Põlme et al. 2020) | Põlme S, et al. 2020. Fungal Diversity 105:1-16 ((https://doi.org/10.1007/s13225-020-00466-2)) |
| Otu39700 | 1 | k__Fungi(100);p__Ascomycota(100);c__Saccharomycetes(100);o__Saccharomycetales(100);f__Metschnikowiaceae(100);g__Metschnikowia(100);s__Metschnikowia_pulcherrima(100); | Metschnikowia | 13 | Saprotroph | \|Undefined Saprotroph\| | Yeast | NULL | Probable | As nectar-tap_saprotroph (Põlme et al. 2020); Decay sugar-rich substrates (Põlme et al. 2020) | Põlme S, et al. 2020. Fungal Diversity 105:1-16 ((https://doi.org/10.1007/s13225-020-00466-2)) |
| Otu39701 | 1 | k__Fungi(100);p__Ascomycota(100);c__Saccharomycetes(100);o__Saccharomycetales(100);f__Metschnikowiaceae(100);g__Metschnikowia(100);s__Metschnikowia_pulcherrima(100); | Metschnikowia | 13 | Saprotroph | \|Undefined Saprotroph\| | Yeast | NULL | Probable | As nectar-tap_saprotroph (Põlme et al. 2020); Decay sugar-rich substrates (Põlme et al. 2020) | Põlme S, et al. 2020. Fungal Diversity 105:1-16 ((https://doi.org/10.1007/s13225-020-00466-2)) |
| Otu39702 | 1 | k__Fungi(100);p__Ascomycota(100);c__Saccharomycetes(100);o__Saccharomycetales(100);f__Metschnikowiaceae(100);g__Metschnikowia(100);s__Metschnikowia_pulcherrima(100); | Metschnikowia | 13 | Saprotroph | \|Undefined Saprotroph\| | Yeast | NULL | Probable | As nectar-tap_saprotroph (Põlme et al. 2020); Decay sugar-rich substrates (Põlme et al. 2020) | Põlme S, et al. 2020. Fungal Diversity 105:1-16 ((https://doi.org/10.1007/s13225-020-00466-2)) |
| Otu39703 | 1 | k__Fungi(100);p__Ascomycota(100);c__Saccharomycetes(100);o__Saccharomycetales(100);f__Metschnikowiaceae(100);g__Metschnikowia(100);g__Metschnikowia_unclassified(100); | Metschnikowia | 13 | Saprotroph | \|Undefined Saprotroph\| | Yeast | NULL | Probable | As nectar-tap_saprotroph (Põlme et al. 2020); Decay sugar-rich substrates (Põlme et al. 2020) | Põlme S, et al. 2020. Fungal Diversity 105:1-16 ((https://doi.org/10.1007/s13225-020-00466-2)) |
| Otu39704 | 1 | k__Fungi(100);p__Ascomycota(100);c__Saccharomycetes(100);o__Saccharomycetales(100);f__Metschnikowiaceae(100);g__Metschnikowia(100);g__Metschnikowia_unclassified(100); | Metschnikowia | 13 | Saprotroph | \|Undefined Saprotroph\| | Yeast | NULL | Probable | As nectar-tap_saprotroph (Põlme et al. 2020); Decay sugar-rich substrates (Põlme et al. 2020) | Põlme S, et al. 2020. Fungal Diversity 105:1-16 ((https://doi.org/10.1007/s13225-020-00466-2)) |
| Otu39705 | 1 | k__Fungi(100);p__Ascomycota(100);c__Saccharomycetes(100);o__Saccharomycetales(100);f__Metschnikowiaceae(100);g__Metschnikowia(100);s__Metschnikowia_pulcherrima(100); | Metschnikowia | 13 | Saprotroph | \|Undefined Saprotroph\| | Yeast | NULL | Probable | As nectar-tap_saprotroph (Põlme et al. 2020); Decay sugar-rich substrates (Põlme et al. 2020) | Põlme S, et al. 2020. Fungal Diversity 105:1-16 ((https://doi.org/10.1007/s13225-020-00466-2)) |
| Otu39706 | 1 | k__Fungi(100);p__Ascomycota(100);c__Saccharomycetes(100);o__Saccharomycetales(100);f__Metschnikowiaceae(100);g__Metschnikowia(100);g__Metschnikowia_unclassified(100); | Metschnikowia | 13 | Saprotroph | \|Undefined Saprotroph\| | Yeast | NULL | Probable | As nectar-tap_saprotroph (Põlme et al. 2020); Decay sugar-rich substrates (Põlme et al. 2020) | Põlme S, et al. 2020. Fungal Diversity 105:1-16 ((https://doi.org/10.1007/s13225-020-00466-2)) |
| Otu39707 | 1 | k__Fungi(100);p__Ascomycota(100);c__Saccharomycetes(100);o__Saccharomycetales(100);f__Metschnikowiaceae(100);g__Metschnikowia(100);g__Metschnikowia_unclassified(100); | Metschnikowia | 13 | Saprotroph | \|Undefined Saprotroph\| | Yeast | NULL | Probable | As nectar-tap_saprotroph (Põlme et al. 2020); Decay sugar-rich substrates (Põlme et al. 2020) | Põlme S, et al. 2020. Fungal Diversity 105:1-16 ((https://doi.org/10.1007/s13225-020-00466-2)) |
| Otu39708 | 1 | k__Fungi(100);p__Ascomycota(100);c__Saccharomycetes(100);o__Saccharomycetales(100);f__Metschnikowiaceae(100);g__Metschnikowia(100);s__Metschnikowia_pulcherrima(100); | Metschnikowia | 13 | Saprotroph | \|Undefined Saprotroph\| | Yeast | NULL | Probable | As nectar-tap_saprotroph (Põlme et al. 2020); Decay sugar-rich substrates (Põlme et al. 2020) | Põlme S, et al. 2020. Fungal Diversity 105:1-16 ((https://doi.org/10.1007/s13225-020-00466-2)) |
| Otu39709 | 1 | k__Fungi(100);p__Ascomycota(100);c__Saccharomycetes(100);o__Saccharomycetales(100);f__Metschnikowiaceae(100);g__Metschnikowia(100);s__Metschnikowia_pulcherrima(100); | Metschnikowia | 13 | Saprotroph | \|Undefined Saprotroph\| | Yeast | NULL | Probable | As nectar-tap_saprotroph (Põlme et al. 2020); Decay sugar-rich substrates (Põlme et al. 2020) | Põlme S, et al. 2020. Fungal Diversity 105:1-16 ((https://doi.org/10.1007/s13225-020-00466-2)) |
| Otu39710 | 1 | k__Fungi(100);p__Ascomycota(100);c__Saccharomycetes(100);o__Saccharomycetales(100);f__Metschnikowiaceae(100);g__Metschnikowia(100);s__Metschnikowia_pulcherrima(100); | Metschnikowia | 13 | Saprotroph | \|Undefined Saprotroph\| | Yeast | NULL | Probable | As nectar-tap_saprotroph (Põlme et al. 2020); Decay sugar-rich substrates (Põlme et al. 2020) | Põlme S, et al. 2020. Fungal Diversity 105:1-16 ((https://doi.org/10.1007/s13225-020-00466-2)) |
| Otu39711 | 1 | k__Fungi(100);p__Ascomycota(100);c__Saccharomycetes(100);o__Saccharomycetales(100);f__Metschnikowiaceae(100);g__Metschnikowia(100);s__Metschnikowia_pulcherrima(100); | Metschnikowia | 13 | Saprotroph | \|Undefined Saprotroph\| | Yeast | NULL | Probable | As nectar-tap_saprotroph (Põlme et al. 2020); Decay sugar-rich substrates (Põlme et al. 2020) | Põlme S, et al. 2020. Fungal Diversity 105:1-16 ((https://doi.org/10.1007/s13225-020-00466-2)) |
| Otu39712 | 1 | k__Fungi(100);p__Ascomycota(100);c__Saccharomycetes(100);o__Saccharomycetales(100);f__Metschnikowiaceae(100);g__Metschnikowia(100);s__Metschnikowia_pulcherrima(100); | Metschnikowia | 13 | Saprotroph | \|Undefined Saprotroph\| | Yeast | NULL | Probable | As nectar-tap_saprotroph (Põlme et al. 2020); Decay sugar-rich substrates (Põlme et al. 2020) | Põlme S, et al. 2020. Fungal Diversity 105:1-16 ((https://doi.org/10.1007/s13225-020-00466-2)) |
| Otu39713 | 1 | k__Fungi(100);p__Ascomycota(100);c__Saccharomycetes(100);o__Saccharomycetales(100);f__Metschnikowiaceae(100);g__Metschnikowia(100);g__Metschnikowia_unclassified(100); | Metschnikowia | 13 | Saprotroph | \|Undefined Saprotroph\| | Yeast | NULL | Probable | As nectar-tap_saprotroph (Põlme et al. 2020); Decay sugar-rich substrates (Põlme et al. 2020) | Põlme S, et al. 2020. Fungal Diversity 105:1-16 ((https://doi.org/10.1007/s13225-020-00466-2)) |
| Otu39714 | 1 | k__Fungi(100);p__Ascomycota(100);c__Saccharomycetes(100);o__Saccharomycetales(100);f__Metschnikowiaceae(100);g__Metschnikowia(100);s__Metschnikowia_pulcherrima(100); | Metschnikowia | 13 | Saprotroph | \|Undefined Saprotroph\| | Yeast | NULL | Probable | As nectar-tap_saprotroph (Põlme et al. 2020); Decay sugar-rich substrates (Põlme et al. 2020) | Põlme S, et al. 2020. Fungal Diversity 105:1-16 ((https://doi.org/10.1007/s13225-020-00466-2)) |
| Otu39715 | 1 | k__Fungi(100);p__Ascomycota(100);c__Saccharomycetes(100);o__Saccharomycetales(100);f__Metschnikowiaceae(100);g__Metschnikowia(100);s__Metschnikowia_pulcherrima(100); | Metschnikowia | 13 | Saprotroph | \|Undefined Saprotroph\| | Yeast | NULL | Probable | As nectar-tap_saprotroph (Põlme et al. 2020); Decay sugar-rich substrates (Põlme et al. 2020) | Põlme S, et al. 2020. Fungal Diversity 105:1-16 ((https://doi.org/10.1007/s13225-020-00466-2)) |
| Otu39716 | 1 | k__Fungi(100);p__Ascomycota(100);c__Saccharomycetes(100);o__Saccharomycetales(100);f__Metschnikowiaceae(100);g__Metschnikowia(100);s__Metschnikowia_pulcherrima(100); | Metschnikowia | 13 | Saprotroph | \|Undefined Saprotroph\| | Yeast | NULL | Probable | As nectar-tap_saprotroph (Põlme et al. 2020); Decay sugar-rich substrates (Põlme et al. 2020) | Põlme S, et al. 2020. Fungal Diversity 105:1-16 ((https://doi.org/10.1007/s13225-020-00466-2)) |
| Otu39717 | 1 | k__Fungi(100);p__Ascomycota(100);c__Saccharomycetes(100);o__Saccharomycetales(100);f__Metschnikowiaceae(100);g__Metschnikowia(100);s__Metschnikowia_pulcherrima(100); | Metschnikowia | 13 | Saprotroph | \|Undefined Saprotroph\| | Yeast | NULL | Probable | As nectar-tap_saprotroph (Põlme et al. 2020); Decay sugar-rich substrates (Põlme et al. 2020) | Põlme S, et al. 2020. Fungal Diversity 105:1-16 ((https://doi.org/10.1007/s13225-020-00466-2)) |
| Otu39718 | 1 | k__Fungi(100);p__Ascomycota(100);c__Saccharomycetes(100);o__Saccharomycetales(100);f__Metschnikowiaceae(100);g__Metschnikowia(100);s__Metschnikowia_pulcherrima(100); | Metschnikowia | 13 | Saprotroph | \|Undefined Saprotroph\| | Yeast | NULL | Probable | As nectar-tap_saprotroph (Põlme et al. 2020); Decay sugar-rich substrates (Põlme et al. 2020) | Põlme S, et al. 2020. Fungal Diversity 105:1-16 ((https://doi.org/10.1007/s13225-020-00466-2)) |
| Otu39719 | 1 | k__Fungi(100);p__Ascomycota(100);c__Saccharomycetes(100);o__Saccharomycetales(100);f__Metschnikowiaceae(100);g__Metschnikowia(100);s__Metschnikowia_pulcherrima(100); | Metschnikowia | 13 | Saprotroph | \|Undefined Saprotroph\| | Yeast | NULL | Probable | As nectar-tap_saprotroph (Põlme et al. 2020); Decay sugar-rich substrates (Põlme et al. 2020) | Põlme S, et al. 2020. Fungal Diversity 105:1-16 ((https://doi.org/10.1007/s13225-020-00466-2)) |
| Otu39720 | 1 | k__Fungi(100);p__Ascomycota(100);c__Saccharomycetes(100);o__Saccharomycetales(100);f__Metschnikowiaceae(100);g__Metschnikowia(100);s__Metschnikowia_pulcherrima(100); | Metschnikowia | 13 | Saprotroph | \|Undefined Saprotroph\| | Yeast | NULL | Probable | As nectar-tap_saprotroph (Põlme et al. 2020); Decay sugar-rich substrates (Põlme et al. 2020) | Põlme S, et al. 2020. Fungal Diversity 105:1-16 ((https://doi.org/10.1007/s13225-020-00466-2)) |
| Otu39721 | 1 | k__Fungi(100);p__Ascomycota(100);c__Saccharomycetes(100);o__Saccharomycetales(100);f__Metschnikowiaceae(100);g__Metschnikowia(100);s__Metschnikowia_pulcherrima(100); | Metschnikowia | 13 | Saprotroph | \|Undefined Saprotroph\| | Yeast | NULL | Probable | As nectar-tap_saprotroph (Põlme et al. 2020); Decay sugar-rich substrates (Põlme et al. 2020) | Põlme S, et al. 2020. Fungal Diversity 105:1-16 ((https://doi.org/10.1007/s13225-020-00466-2)) |
| Otu39722 | 1 | k__Fungi(100);p__Ascomycota(100);c__Saccharomycetes(100);o__Saccharomycetales(100);f__Metschnikowiaceae(100);g__Metschnikowia(100);s__Metschnikowia_pulcherrima(100); | Metschnikowia | 13 | Saprotroph | \|Undefined Saprotroph\| | Yeast | NULL | Probable | As nectar-tap_saprotroph (Põlme et al. 2020); Decay sugar-rich substrates (Põlme et al. 2020) | Põlme S, et al. 2020. Fungal Diversity 105:1-16 ((https://doi.org/10.1007/s13225-020-00466-2)) |
| Otu39724 | 1 | k__Fungi(100);p__Ascomycota(100);c__Saccharomycetes(100);o__Saccharomycetales(100);f__Metschnikowiaceae(100);g__Metschnikowia(100);s__Metschnikowia_pulcherrima(100); | Metschnikowia | 13 | Saprotroph | \|Undefined Saprotroph\| | Yeast | NULL | Probable | As nectar-tap_saprotroph (Põlme et al. 2020); Decay sugar-rich substrates (Põlme et al. 2020) | Põlme S, et al. 2020. Fungal Diversity 105:1-16 ((https://doi.org/10.1007/s13225-020-00466-2)) |
| Otu39725 | 1 | k__Fungi(100);p__Ascomycota(100);c__Saccharomycetes(100);o__Saccharomycetales(100);f__Metschnikowiaceae(100);g__Metschnikowia(100);s__Metschnikowia_pulcherrima(100); | Metschnikowia | 13 | Saprotroph | \|Undefined Saprotroph\| | Yeast | NULL | Probable | As nectar-tap_saprotroph (Põlme et al. 2020); Decay sugar-rich substrates (Põlme et al. 2020) | Põlme S, et al. 2020. Fungal Diversity 105:1-16 ((https://doi.org/10.1007/s13225-020-00466-2)) |
| Otu39726 | 1 | k__Fungi(100);p__Ascomycota(100);c__Saccharomycetes(100);o__Saccharomycetales(100);f__Metschnikowiaceae(100);g__Metschnikowia(100);s__Metschnikowia_pulcherrima(100); | Metschnikowia | 13 | Saprotroph | \|Undefined Saprotroph\| | Yeast | NULL | Probable | As nectar-tap_saprotroph (Põlme et al. 2020); Decay sugar-rich substrates (Põlme et al. 2020) | Põlme S, et al. 2020. Fungal Diversity 105:1-16 ((https://doi.org/10.1007/s13225-020-00466-2)) |
| Otu39727 | 1 | k__Fungi(100);p__Ascomycota(100);c__Saccharomycetes(100);o__Saccharomycetales(100);f__Metschnikowiaceae(100);g__Metschnikowia(100);g__Metschnikowia_unclassified(100); | Metschnikowia | 13 | Saprotroph | \|Undefined Saprotroph\| | Yeast | NULL | Probable | As nectar-tap_saprotroph (Põlme et al. 2020); Decay sugar-rich substrates (Põlme et al. 2020) | Põlme S, et al. 2020. Fungal Diversity 105:1-16 ((https://doi.org/10.1007/s13225-020-00466-2)) |
| Otu39728 | 1 | k__Fungi(100);p__Ascomycota(100);c__Saccharomycetes(100);o__Saccharomycetales(100);f__Metschnikowiaceae(100);g__Metschnikowia(100);s__Metschnikowia_pulcherrima(100); | Metschnikowia | 13 | Saprotroph | \|Undefined Saprotroph\| | Yeast | NULL | Probable | As nectar-tap_saprotroph (Põlme et al. 2020); Decay sugar-rich substrates (Põlme et al. 2020) | Põlme S, et al. 2020. Fungal Diversity 105:1-16 ((https://doi.org/10.1007/s13225-020-00466-2)) |
| Otu39729 | 1 | k__Fungi(100);p__Ascomycota(100);c__Saccharomycetes(100);o__Saccharomycetales(100);f__Metschnikowiaceae(100);g__Metschnikowia(100);s__Metschnikowia_pulcherrima(100); | Metschnikowia | 13 | Saprotroph | \|Undefined Saprotroph\| | Yeast | NULL | Probable | As nectar-tap_saprotroph (Põlme et al. 2020); Decay sugar-rich substrates (Põlme et al. 2020) | Põlme S, et al. 2020. Fungal Diversity 105:1-16 ((https://doi.org/10.1007/s13225-020-00466-2)) |
| Otu39730 | 1 | k__Fungi(100);p__Ascomycota(100);c__Saccharomycetes(100);o__Saccharomycetales(100);f__Metschnikowiaceae(100);g__Metschnikowia(100);g__Metschnikowia_unclassified(100); | Metschnikowia | 13 | Saprotroph | \|Undefined Saprotroph\| | Yeast | NULL | Probable | As nectar-tap_saprotroph (Põlme et al. 2020); Decay sugar-rich substrates (Põlme et al. 2020) | Põlme S, et al. 2020. Fungal Diversity 105:1-16 ((https://doi.org/10.1007/s13225-020-00466-2)) |
| Otu39731 | 1 | k__Fungi(100);p__Ascomycota(100);c__Saccharomycetes(100);o__Saccharomycetales(100);f__Metschnikowiaceae(100);g__Metschnikowia(100);s__Metschnikowia_pulcherrima(100); | Metschnikowia | 13 | Saprotroph | \|Undefined Saprotroph\| | Yeast | NULL | Probable | As nectar-tap_saprotroph (Põlme et al. 2020); Decay sugar-rich substrates (Põlme et al. 2020) | Põlme S, et al. 2020. Fungal Diversity 105:1-16 ((https://doi.org/10.1007/s13225-020-00466-2)) |
| Otu39732 | 1 | k__Fungi(100);p__Ascomycota(100);c__Saccharomycetes(100);o__Saccharomycetales(100);f__Metschnikowiaceae(100);g__Metschnikowia(100);s__Metschnikowia_pulcherrima(100); | Metschnikowia | 13 | Saprotroph | \|Undefined Saprotroph\| | Yeast | NULL | Probable | As nectar-tap_saprotroph (Põlme et al. 2020); Decay sugar-rich substrates (Põlme et al. 2020) | Põlme S, et al. 2020. Fungal Diversity 105:1-16 ((https://doi.org/10.1007/s13225-020-00466-2)) |
| Otu39733 | 1 | k__Fungi(100);p__Ascomycota(100);c__Saccharomycetes(100);o__Saccharomycetales(100);f__Metschnikowiaceae(100);g__Metschnikowia(100);g__Metschnikowia_unclassified(100); | Metschnikowia | 13 | Saprotroph | \|Undefined Saprotroph\| | Yeast | NULL | Probable | As nectar-tap_saprotroph (Põlme et al. 2020); Decay sugar-rich substrates (Põlme et al. 2020) | Põlme S, et al. 2020. Fungal Diversity 105:1-16 ((https://doi.org/10.1007/s13225-020-00466-2)) |
| Otu39734 | 1 | k__Fungi(100);p__Ascomycota(100);c__Saccharomycetes(100);o__Saccharomycetales(100);f__Metschnikowiaceae(100);g__Metschnikowia(100);s__Metschnikowia_pulcherrima(100); | Metschnikowia | 13 | Saprotroph | \|Undefined Saprotroph\| | Yeast | NULL | Probable | As nectar-tap_saprotroph (Põlme et al. 2020); Decay sugar-rich substrates (Põlme et al. 2020) | Põlme S, et al. 2020. Fungal Diversity 105:1-16 ((https://doi.org/10.1007/s13225-020-00466-2)) |
| Otu39735 | 1 | k__Fungi(100);p__Ascomycota(100);c__Saccharomycetes(100);o__Saccharomycetales(100);f__Metschnikowiaceae(100);g__Metschnikowia(100);s__Metschnikowia_pulcherrima(100); | Metschnikowia | 13 | Saprotroph | \|Undefined Saprotroph\| | Yeast | NULL | Probable | As nectar-tap_saprotroph (Põlme et al. 2020); Decay sugar-rich substrates (Põlme et al. 2020) | Põlme S, et al. 2020. Fungal Diversity 105:1-16 ((https://doi.org/10.1007/s13225-020-00466-2)) |
| Otu39736 | 1 | k__Fungi(100);p__Ascomycota(100);c__Saccharomycetes(100);o__Saccharomycetales(100);f__Metschnikowiaceae(100);g__Metschnikowia(100);g__Metschnikowia_unclassified(100); | Metschnikowia | 13 | Saprotroph | \|Undefined Saprotroph\| | Yeast | NULL | Probable | As nectar-tap_saprotroph (Põlme et al. 2020); Decay sugar-rich substrates (Põlme et al. 2020) | Põlme S, et al. 2020. Fungal Diversity 105:1-16 ((https://doi.org/10.1007/s13225-020-00466-2)) |
| Otu39737 | 1 | k__Fungi(100);p__Ascomycota(100);c__Saccharomycetes(100);o__Saccharomycetales(100);f__Metschnikowiaceae(100);g__Metschnikowia(100);s__Metschnikowia_pulcherrima(100); | Metschnikowia | 13 | Saprotroph | \|Undefined Saprotroph\| | Yeast | NULL | Probable | As nectar-tap_saprotroph (Põlme et al. 2020); Decay sugar-rich substrates (Põlme et al. 2020) | Põlme S, et al. 2020. Fungal Diversity 105:1-16 ((https://doi.org/10.1007/s13225-020-00466-2)) |
| Otu39738 | 1 | k__Fungi(100);p__Ascomycota(100);c__Saccharomycetes(100);o__Saccharomycetales(100);f__Metschnikowiaceae(100);g__Metschnikowia(100);s__Metschnikowia_pulcherrima(100); | Metschnikowia | 13 | Saprotroph | \|Undefined Saprotroph\| | Yeast | NULL | Probable | As nectar-tap_saprotroph (Põlme et al. 2020); Decay sugar-rich substrates (Põlme et al. 2020) | Põlme S, et al. 2020. Fungal Diversity 105:1-16 ((https://doi.org/10.1007/s13225-020-00466-2)) |
| Otu39739 | 1 | k__Fungi(100);p__Ascomycota(100);c__Saccharomycetes(100);o__Saccharomycetales(100);f__Metschnikowiaceae(100);g__Metschnikowia(100);s__Metschnikowia_pulcherrima(100); | Metschnikowia | 13 | Saprotroph | \|Undefined Saprotroph\| | Yeast | NULL | Probable | As nectar-tap_saprotroph (Põlme et al. 2020); Decay sugar-rich substrates (Põlme et al. 2020) | Põlme S, et al. 2020. Fungal Diversity 105:1-16 ((https://doi.org/10.1007/s13225-020-00466-2)) |
| Otu39740 | 1 | k__Fungi(100);p__Ascomycota(100);c__Saccharomycetes(100);o__Saccharomycetales(100);f__Metschnikowiaceae(100);g__Metschnikowia(100);s__Metschnikowia_pulcherrima(100); | Metschnikowia | 13 | Saprotroph | \|Undefined Saprotroph\| | Yeast | NULL | Probable | As nectar-tap_saprotroph (Põlme et al. 2020); Decay sugar-rich substrates (Põlme et al. 2020) | Põlme S, et al. 2020. Fungal Diversity 105:1-16 ((https://doi.org/10.1007/s13225-020-00466-2)) |
| Otu39741 | 1 | k__Fungi(100);p__Ascomycota(100);c__Saccharomycetes(100);o__Saccharomycetales(100);f__Metschnikowiaceae(100);g__Metschnikowia(100);s__Metschnikowia_pulcherrima(100); | Metschnikowia | 13 | Saprotroph | \|Undefined Saprotroph\| | Yeast | NULL | Probable | As nectar-tap_saprotroph (Põlme et al. 2020); Decay sugar-rich substrates (Põlme et al. 2020) | Põlme S, et al. 2020. Fungal Diversity 105:1-16 ((https://doi.org/10.1007/s13225-020-00466-2)) |
| Otu39742 | 1 | k__Fungi(100);p__Ascomycota(100);c__Sordariomycetes(100);o__Hypocreales(100);f__Nectriaceae(100);g__Gibberella(100);s__Gibberella_intricans(100); | Gibberella | 13 | Pathotroph-Saprotroph-Symbiotroph | Endophyte-\|Plant Pathogen\|-Plant Saprotroph-Undefined Saprotroph | Microfungus | Soft Rot | Probable | As foliar_endophyte (Põlme et al. 2020); As litter_saprotroph (Põlme et al. 2020); Causing damping-off disease (Põlme et al. 2020); Decay leaf, fruit, seed, roots, soil, animal material (Põlme et al. 2020) | Tedersoo L, et al. 2014. Science 346:e1256688 ((https://doi.org/10.1126/science.1256688)); Põlme S, et al. 2020. Fungal Diversity 105:1-16 ((https://doi.org/10.1007/s13225-020-00466-2)) |
| Otu39744 | 1 | k__Fungi(100);p__Ascomycota(100);c__Saccharomycetes(100);o__Saccharomycetales(100);f__Metschnikowiaceae(100);g__Metschnikowia(100);g__Metschnikowia_unclassified(100); | Metschnikowia | 13 | Saprotroph | \|Undefined Saprotroph\| | Yeast | NULL | Probable | As nectar-tap_saprotroph (Põlme et al. 2020); Decay sugar-rich substrates (Põlme et al. 2020) | Põlme S, et al. 2020. Fungal Diversity 105:1-16 ((https://doi.org/10.1007/s13225-020-00466-2)) |
| Otu39745 | 1 | k__Fungi(100);p__Ascomycota(100);c__Saccharomycetes(100);o__Saccharomycetales(100);f__Metschnikowiaceae(100);g__Metschnikowia(100);s__Metschnikowia_pulcherrima(100); | Metschnikowia | 13 | Saprotroph | \|Undefined Saprotroph\| | Yeast | NULL | Probable | As nectar-tap_saprotroph (Põlme et al. 2020); Decay sugar-rich substrates (Põlme et al. 2020) | Põlme S, et al. 2020. Fungal Diversity 105:1-16 ((https://doi.org/10.1007/s13225-020-00466-2)) |
| Otu39746 | 1 | k__Fungi(100);p__Ascomycota(100);c__Saccharomycetes(100);o__Saccharomycetales(100);f__Metschnikowiaceae(100);g__Metschnikowia(100);s__Metschnikowia_pulcherrima(100); | Metschnikowia | 13 | Saprotroph | \|Undefined Saprotroph\| | Yeast | NULL | Probable | As nectar-tap_saprotroph (Põlme et al. 2020); Decay sugar-rich substrates (Põlme et al. 2020) | Põlme S, et al. 2020. Fungal Diversity 105:1-16 ((https://doi.org/10.1007/s13225-020-00466-2)) |
| Otu39747 | 1 | k__Fungi(100);p__Ascomycota(100);c__Saccharomycetes(100);o__Saccharomycetales(100);f__Metschnikowiaceae(100);g__Metschnikowia(100);s__Metschnikowia_pulcherrima(100); | Metschnikowia | 13 | Saprotroph | \|Undefined Saprotroph\| | Yeast | NULL | Probable | As nectar-tap_saprotroph (Põlme et al. 2020); Decay sugar-rich substrates (Põlme et al. 2020) | Põlme S, et al. 2020. Fungal Diversity 105:1-16 ((https://doi.org/10.1007/s13225-020-00466-2)) |
| Otu39748 | 1 | k__Fungi(100);p__Ascomycota(100);c__Saccharomycetes(100);o__Saccharomycetales(100);f__Metschnikowiaceae(100);g__Metschnikowia(100);g__Metschnikowia_unclassified(100); | Metschnikowia | 13 | Saprotroph | \|Undefined Saprotroph\| | Yeast | NULL | Probable | As nectar-tap_saprotroph (Põlme et al. 2020); Decay sugar-rich substrates (Põlme et al. 2020) | Põlme S, et al. 2020. Fungal Diversity 105:1-16 ((https://doi.org/10.1007/s13225-020-00466-2)) |
| Otu39749 | 1 | k__Fungi(100);p__Ascomycota(100);c__Saccharomycetes(100);o__Saccharomycetales(100);f__Metschnikowiaceae(100);g__Metschnikowia(100);s__Metschnikowia_pulcherrima(100); | Metschnikowia | 13 | Saprotroph | \|Undefined Saprotroph\| | Yeast | NULL | Probable | As nectar-tap_saprotroph (Põlme et al. 2020); Decay sugar-rich substrates (Põlme et al. 2020) | Põlme S, et al. 2020. Fungal Diversity 105:1-16 ((https://doi.org/10.1007/s13225-020-00466-2)) |
| Otu39750 | 1 | k__Fungi(100);p__Ascomycota(100);c__Saccharomycetes(100);o__Saccharomycetales(100);f__Metschnikowiaceae(100);g__Metschnikowia(100);s__Metschnikowia_pulcherrima(100); | Metschnikowia | 13 | Saprotroph | \|Undefined Saprotroph\| | Yeast | NULL | Probable | As nectar-tap_saprotroph (Põlme et al. 2020); Decay sugar-rich substrates (Põlme et al. 2020) | Põlme S, et al. 2020. Fungal Diversity 105:1-16 ((https://doi.org/10.1007/s13225-020-00466-2)) |
| Otu39751 | 1 | k__Fungi(100);p__Ascomycota(100);c__Saccharomycetes(100);o__Saccharomycetales(100);f__Metschnikowiaceae(100);g__Metschnikowia(100);s__Metschnikowia_pulcherrima(100); | Metschnikowia | 13 | Saprotroph | \|Undefined Saprotroph\| | Yeast | NULL | Probable | As nectar-tap_saprotroph (Põlme et al. 2020); Decay sugar-rich substrates (Põlme et al. 2020) | Põlme S, et al. 2020. Fungal Diversity 105:1-16 ((https://doi.org/10.1007/s13225-020-00466-2)) |
| Otu39752 | 1 | k__Fungi(100);p__Ascomycota(100);c__Saccharomycetes(100);o__Saccharomycetales(100);f__Metschnikowiaceae(100);g__Metschnikowia(100);s__Metschnikowia_pulcherrima(100); | Metschnikowia | 13 | Saprotroph | \|Undefined Saprotroph\| | Yeast | NULL | Probable | As nectar-tap_saprotroph (Põlme et al. 2020); Decay sugar-rich substrates (Põlme et al. 2020) | Põlme S, et al. 2020. Fungal Diversity 105:1-16 ((https://doi.org/10.1007/s13225-020-00466-2)) |
| Otu39753 | 1 | k__Fungi(100);p__Ascomycota(100);c__Saccharomycetes(100);o__Saccharomycetales(100);f__Metschnikowiaceae(100);g__Metschnikowia(100);s__Metschnikowia_pulcherrima(100); | Metschnikowia | 13 | Saprotroph | \|Undefined Saprotroph\| | Yeast | NULL | Probable | As nectar-tap_saprotroph (Põlme et al. 2020); Decay sugar-rich substrates (Põlme et al. 2020) | Põlme S, et al. 2020. Fungal Diversity 105:1-16 ((https://doi.org/10.1007/s13225-020-00466-2)) |
| Otu39754 | 1 | k__Fungi(100);p__Ascomycota(100);c__Saccharomycetes(100);o__Saccharomycetales(100);f__Metschnikowiaceae(100);g__Metschnikowia(100);s__Metschnikowia_pulcherrima(100); | Metschnikowia | 13 | Saprotroph | \|Undefined Saprotroph\| | Yeast | NULL | Probable | As nectar-tap_saprotroph (Põlme et al. 2020); Decay sugar-rich substrates (Põlme et al. 2020) | Põlme S, et al. 2020. Fungal Diversity 105:1-16 ((https://doi.org/10.1007/s13225-020-00466-2)) |
| Otu39755 | 1 | k__Fungi(100);p__Ascomycota(100);c__Saccharomycetes(100);o__Saccharomycetales(100);f__Metschnikowiaceae(100);g__Metschnikowia(100);g__Metschnikowia_unclassified(100); | Metschnikowia | 13 | Saprotroph | \|Undefined Saprotroph\| | Yeast | NULL | Probable | As nectar-tap_saprotroph (Põlme et al. 2020); Decay sugar-rich substrates (Põlme et al. 2020) | Põlme S, et al. 2020. Fungal Diversity 105:1-16 ((https://doi.org/10.1007/s13225-020-00466-2)) |
| Otu39756 | 1 | k__Fungi(100);p__Ascomycota(100);c__Saccharomycetes(100);o__Saccharomycetales(100);f__Metschnikowiaceae(100);g__Metschnikowia(100);s__Metschnikowia_pulcherrima(100); | Metschnikowia | 13 | Saprotroph | \|Undefined Saprotroph\| | Yeast | NULL | Probable | As nectar-tap_saprotroph (Põlme et al. 2020); Decay sugar-rich substrates (Põlme et al. 2020) | Põlme S, et al. 2020. Fungal Diversity 105:1-16 ((https://doi.org/10.1007/s13225-020-00466-2)) |
| Otu39757 | 1 | k__Fungi(100);p__Ascomycota(100);c__Saccharomycetes(100);o__Saccharomycetales(100);f__Metschnikowiaceae(100);g__Metschnikowia(100);s__Metschnikowia_pulcherrima(100); | Metschnikowia | 13 | Saprotroph | \|Undefined Saprotroph\| | Yeast | NULL | Probable | As nectar-tap_saprotroph (Põlme et al. 2020); Decay sugar-rich substrates (Põlme et al. 2020) | Põlme S, et al. 2020. Fungal Diversity 105:1-16 ((https://doi.org/10.1007/s13225-020-00466-2)) |
| Otu39758 | 1 | k__Fungi(100);p__Ascomycota(100);c__Saccharomycetes(100);o__Saccharomycetales(100);f__Metschnikowiaceae(100);g__Metschnikowia(100);s__Metschnikowia_pulcherrima(100); | Metschnikowia | 13 | Saprotroph | \|Undefined Saprotroph\| | Yeast | NULL | Probable | As nectar-tap_saprotroph (Põlme et al. 2020); Decay sugar-rich substrates (Põlme et al. 2020) | Põlme S, et al. 2020. Fungal Diversity 105:1-16 ((https://doi.org/10.1007/s13225-020-00466-2)) |
| Otu39759 | 1 | k__Fungi(100);p__Ascomycota(100);c__Saccharomycetes(100);o__Saccharomycetales(100);f__Metschnikowiaceae(100);g__Metschnikowia(100);s__Metschnikowia_pulcherrima(100); | Metschnikowia | 13 | Saprotroph | \|Undefined Saprotroph\| | Yeast | NULL | Probable | As nectar-tap_saprotroph (Põlme et al. 2020); Decay sugar-rich substrates (Põlme et al. 2020) | Põlme S, et al. 2020. Fungal Diversity 105:1-16 ((https://doi.org/10.1007/s13225-020-00466-2)) |
| Otu39760 | 1 | k__Fungi(100);p__Ascomycota(100);c__Saccharomycetes(100);o__Saccharomycetales(100);f__Metschnikowiaceae(100);g__Metschnikowia(100);s__Metschnikowia_pulcherrima(100); | Metschnikowia | 13 | Saprotroph | \|Undefined Saprotroph\| | Yeast | NULL | Probable | As nectar-tap_saprotroph (Põlme et al. 2020); Decay sugar-rich substrates (Põlme et al. 2020) | Põlme S, et al. 2020. Fungal Diversity 105:1-16 ((https://doi.org/10.1007/s13225-020-00466-2)) |
| Otu39761 | 1 | k__Fungi(100);p__Ascomycota(100);c__Saccharomycetes(100);o__Saccharomycetales(100);f__Metschnikowiaceae(100);g__Metschnikowia(100);s__Metschnikowia_pulcherrima(100); | Metschnikowia | 13 | Saprotroph | \|Undefined Saprotroph\| | Yeast | NULL | Probable | As nectar-tap_saprotroph (Põlme et al. 2020); Decay sugar-rich substrates (Põlme et al. 2020) | Põlme S, et al. 2020. Fungal Diversity 105:1-16 ((https://doi.org/10.1007/s13225-020-00466-2)) |
| Otu39762 | 1 | k__Fungi(100);p__Ascomycota(100);c__Saccharomycetes(100);o__Saccharomycetales(100);f__Metschnikowiaceae(100);g__Metschnikowia(100);s__Metschnikowia_pulcherrima(100); | Metschnikowia | 13 | Saprotroph | \|Undefined Saprotroph\| | Yeast | NULL | Probable | As nectar-tap_saprotroph (Põlme et al. 2020); Decay sugar-rich substrates (Põlme et al. 2020) | Põlme S, et al. 2020. Fungal Diversity 105:1-16 ((https://doi.org/10.1007/s13225-020-00466-2)) |
| Otu39763 | 1 | k__Fungi(100);p__Ascomycota(100);c__Saccharomycetes(100);o__Saccharomycetales(100);f__Metschnikowiaceae(100);g__Metschnikowia(100);s__Metschnikowia_pulcherrima(100); | Metschnikowia | 13 | Saprotroph | \|Undefined Saprotroph\| | Yeast | NULL | Probable | As nectar-tap_saprotroph (Põlme et al. 2020); Decay sugar-rich substrates (Põlme et al. 2020) | Põlme S, et al. 2020. Fungal Diversity 105:1-16 ((https://doi.org/10.1007/s13225-020-00466-2)) |
| Otu39764 | 1 | k__Fungi(100);p__Ascomycota(100);c__Saccharomycetes(100);o__Saccharomycetales(100);f__Metschnikowiaceae(100);g__Metschnikowia(100);s__Metschnikowia_pulcherrima(100); | Metschnikowia | 13 | Saprotroph | \|Undefined Saprotroph\| | Yeast | NULL | Probable | As nectar-tap_saprotroph (Põlme et al. 2020); Decay sugar-rich substrates (Põlme et al. 2020) | Põlme S, et al. 2020. Fungal Diversity 105:1-16 ((https://doi.org/10.1007/s13225-020-00466-2)) |
| Otu39765 | 1 | k__Fungi(100);p__Ascomycota(100);c__Saccharomycetes(100);o__Saccharomycetales(100);f__Metschnikowiaceae(100);g__Metschnikowia(100);s__Metschnikowia_pulcherrima(100); | Metschnikowia | 13 | Saprotroph | \|Undefined Saprotroph\| | Yeast | NULL | Probable | As nectar-tap_saprotroph (Põlme et al. 2020); Decay sugar-rich substrates (Põlme et al. 2020) | Põlme S, et al. 2020. Fungal Diversity 105:1-16 ((https://doi.org/10.1007/s13225-020-00466-2)) |
| Otu39766 | 1 | k__Fungi(100);p__Ascomycota(100);c__Saccharomycetes(100);o__Saccharomycetales(100);f__Metschnikowiaceae(100);g__Metschnikowia(100);s__Metschnikowia_pulcherrima(100); | Metschnikowia | 13 | Saprotroph | \|Undefined Saprotroph\| | Yeast | NULL | Probable | As nectar-tap_saprotroph (Põlme et al. 2020); Decay sugar-rich substrates (Põlme et al. 2020) | Põlme S, et al. 2020. Fungal Diversity 105:1-16 ((https://doi.org/10.1007/s13225-020-00466-2)) |
| Otu39767 | 1 | k__Fungi(100);p__Ascomycota(100);c__Saccharomycetes(100);o__Saccharomycetales(100);f__Metschnikowiaceae(100);g__Metschnikowia(100);s__Metschnikowia_pulcherrima(100); | Metschnikowia | 13 | Saprotroph | \|Undefined Saprotroph\| | Yeast | NULL | Probable | As nectar-tap_saprotroph (Põlme et al. 2020); Decay sugar-rich substrates (Põlme et al. 2020) | Põlme S, et al. 2020. Fungal Diversity 105:1-16 ((https://doi.org/10.1007/s13225-020-00466-2)) |
| Otu39768 | 1 | k__Fungi(100);p__Ascomycota(100);c__Saccharomycetes(100);o__Saccharomycetales(100);f__Metschnikowiaceae(100);g__Metschnikowia(100);s__Metschnikowia_pulcherrima(100); | Metschnikowia | 13 | Saprotroph | \|Undefined Saprotroph\| | Yeast | NULL | Probable | As nectar-tap_saprotroph (Põlme et al. 2020); Decay sugar-rich substrates (Põlme et al. 2020) | Põlme S, et al. 2020. Fungal Diversity 105:1-16 ((https://doi.org/10.1007/s13225-020-00466-2)) |
| Otu39769 | 1 | k__Fungi(100);p__Ascomycota(100);c__Saccharomycetes(100);o__Saccharomycetales(100);f__Metschnikowiaceae(100);g__Metschnikowia(100);s__Metschnikowia_pulcherrima(100); | Metschnikowia | 13 | Saprotroph | \|Undefined Saprotroph\| | Yeast | NULL | Probable | As nectar-tap_saprotroph (Põlme et al. 2020); Decay sugar-rich substrates (Põlme et al. 2020) | Põlme S, et al. 2020. Fungal Diversity 105:1-16 ((https://doi.org/10.1007/s13225-020-00466-2)) |
| Otu39770 | 1 | k__Fungi(100);p__Ascomycota(100);c__Saccharomycetes(100);o__Saccharomycetales(100);f__Metschnikowiaceae(100);g__Metschnikowia(100);s__Metschnikowia_pulcherrima(100); | Metschnikowia | 13 | Saprotroph | \|Undefined Saprotroph\| | Yeast | NULL | Probable | As nectar-tap_saprotroph (Põlme et al. 2020); Decay sugar-rich substrates (Põlme et al. 2020) | Põlme S, et al. 2020. Fungal Diversity 105:1-16 ((https://doi.org/10.1007/s13225-020-00466-2)) |
| Otu39771 | 1 | k__Fungi(100);p__Ascomycota(100);c__Saccharomycetes(100);o__Saccharomycetales(100);f__Metschnikowiaceae(100);g__Metschnikowia(100);s__Metschnikowia_pulcherrima(100); | Metschnikowia | 13 | Saprotroph | \|Undefined Saprotroph\| | Yeast | NULL | Probable | As nectar-tap_saprotroph (Põlme et al. 2020); Decay sugar-rich substrates (Põlme et al. 2020) | Põlme S, et al. 2020. Fungal Diversity 105:1-16 ((https://doi.org/10.1007/s13225-020-00466-2)) |
| Otu39772 | 1 | k__Fungi(100);p__Ascomycota(100);c__Saccharomycetes(100);o__Saccharomycetales(100);f__Metschnikowiaceae(100);g__Metschnikowia(100);s__Metschnikowia_pulcherrima(100); | Metschnikowia | 13 | Saprotroph | \|Undefined Saprotroph\| | Yeast | NULL | Probable | As nectar-tap_saprotroph (Põlme et al. 2020); Decay sugar-rich substrates (Põlme et al. 2020) | Põlme S, et al. 2020. Fungal Diversity 105:1-16 ((https://doi.org/10.1007/s13225-020-00466-2)) |
| Otu39773 | 1 | k__Fungi(100);p__Ascomycota(100);c__Saccharomycetes(100);o__Saccharomycetales(100);f__Metschnikowiaceae(100);g__Metschnikowia(100);g__Metschnikowia_unclassified(100); | Metschnikowia | 13 | Saprotroph | \|Undefined Saprotroph\| | Yeast | NULL | Probable | As nectar-tap_saprotroph (Põlme et al. 2020); Decay sugar-rich substrates (Põlme et al. 2020) | Põlme S, et al. 2020. Fungal Diversity 105:1-16 ((https://doi.org/10.1007/s13225-020-00466-2)) |
| Otu39774 | 1 | k__Fungi(100);p__Ascomycota(100);c__Saccharomycetes(100);o__Saccharomycetales(100);f__Metschnikowiaceae(100);g__Metschnikowia(100);s__Metschnikowia_pulcherrima(100); | Metschnikowia | 13 | Saprotroph | \|Undefined Saprotroph\| | Yeast | NULL | Probable | As nectar-tap_saprotroph (Põlme et al. 2020); Decay sugar-rich substrates (Põlme et al. 2020) | Põlme S, et al. 2020. Fungal Diversity 105:1-16 ((https://doi.org/10.1007/s13225-020-00466-2)) |
| Otu39775 | 1 | k__Fungi(100);p__Ascomycota(100);c__Saccharomycetes(100);o__Saccharomycetales(100);f__Metschnikowiaceae(100);g__Metschnikowia(100);s__Metschnikowia_pulcherrima(100); | Metschnikowia | 13 | Saprotroph | \|Undefined Saprotroph\| | Yeast | NULL | Probable | As nectar-tap_saprotroph (Põlme et al. 2020); Decay sugar-rich substrates (Põlme et al. 2020) | Põlme S, et al. 2020. Fungal Diversity 105:1-16 ((https://doi.org/10.1007/s13225-020-00466-2)) |
| Otu39776 | 1 | k__Fungi(100);p__Ascomycota(100);c__Saccharomycetes(100);o__Saccharomycetales(100);f__Metschnikowiaceae(100);g__Metschnikowia(100);s__Metschnikowia_pulcherrima(100); | Metschnikowia | 13 | Saprotroph | \|Undefined Saprotroph\| | Yeast | NULL | Probable | As nectar-tap_saprotroph (Põlme et al. 2020); Decay sugar-rich substrates (Põlme et al. 2020) | Põlme S, et al. 2020. Fungal Diversity 105:1-16 ((https://doi.org/10.1007/s13225-020-00466-2)) |
| Otu39777 | 1 | k__Fungi(100);p__Ascomycota(100);c__Saccharomycetes(100);o__Saccharomycetales(100);f__Metschnikowiaceae(100);g__Metschnikowia(100);g__Metschnikowia_unclassified(100); | Metschnikowia | 13 | Saprotroph | \|Undefined Saprotroph\| | Yeast | NULL | Probable | As nectar-tap_saprotroph (Põlme et al. 2020); Decay sugar-rich substrates (Põlme et al. 2020) | Põlme S, et al. 2020. Fungal Diversity 105:1-16 ((https://doi.org/10.1007/s13225-020-00466-2)) |
| Otu39778 | 1 | k__Fungi(100);p__Ascomycota(100);c__Saccharomycetes(100);o__Saccharomycetales(100);f__Metschnikowiaceae(100);g__Metschnikowia(100);s__Metschnikowia_pulcherrima(100); | Metschnikowia | 13 | Saprotroph | \|Undefined Saprotroph\| | Yeast | NULL | Probable | As nectar-tap_saprotroph (Põlme et al. 2020); Decay sugar-rich substrates (Põlme et al. 2020) | Põlme S, et al. 2020. Fungal Diversity 105:1-16 ((https://doi.org/10.1007/s13225-020-00466-2)) |
| Otu39779 | 1 | k__Fungi(100);p__Ascomycota(100);c__Saccharomycetes(100);o__Saccharomycetales(100);f__Metschnikowiaceae(100);g__Metschnikowia(100);s__Metschnikowia_pulcherrima(100); | Metschnikowia | 13 | Saprotroph | \|Undefined Saprotroph\| | Yeast | NULL | Probable | As nectar-tap_saprotroph (Põlme et al. 2020); Decay sugar-rich substrates (Põlme et al. 2020) | Põlme S, et al. 2020. Fungal Diversity 105:1-16 ((https://doi.org/10.1007/s13225-020-00466-2)) |
| Otu39780 | 1 | k__Fungi(100);p__Ascomycota(100);c__Saccharomycetes(100);o__Saccharomycetales(100);f__Metschnikowiaceae(100);g__Metschnikowia(100);g__Metschnikowia_unclassified(100); | Metschnikowia | 13 | Saprotroph | \|Undefined Saprotroph\| | Yeast | NULL | Probable | As nectar-tap_saprotroph (Põlme et al. 2020); Decay sugar-rich substrates (Põlme et al. 2020) | Põlme S, et al. 2020. Fungal Diversity 105:1-16 ((https://doi.org/10.1007/s13225-020-00466-2)) |
| Otu39781 | 1 | k__Fungi(100);p__Ascomycota(100);c__Saccharomycetes(100);o__Saccharomycetales(100);f__Metschnikowiaceae(100);g__Metschnikowia(100);g__Metschnikowia_unclassified(100); | Metschnikowia | 13 | Saprotroph | \|Undefined Saprotroph\| | Yeast | NULL | Probable | As nectar-tap_saprotroph (Põlme et al. 2020); Decay sugar-rich substrates (Põlme et al. 2020) | Põlme S, et al. 2020. Fungal Diversity 105:1-16 ((https://doi.org/10.1007/s13225-020-00466-2)) |
| Otu39782 | 1 | k__Fungi(100);p__Ascomycota(100);c__Saccharomycetes(100);o__Saccharomycetales(100);f__Metschnikowiaceae(100);g__Metschnikowia(100);s__Metschnikowia_pulcherrima(100); | Metschnikowia | 13 | Saprotroph | \|Undefined Saprotroph\| | Yeast | NULL | Probable | As nectar-tap_saprotroph (Põlme et al. 2020); Decay sugar-rich substrates (Põlme et al. 2020) | Põlme S, et al. 2020. Fungal Diversity 105:1-16 ((https://doi.org/10.1007/s13225-020-00466-2)) |
| Otu39783 | 1 | k__Fungi(100);p__Ascomycota(100);c__Saccharomycetes(100);o__Saccharomycetales(100);f__Metschnikowiaceae(100);g__Metschnikowia(100);s__Metschnikowia_pulcherrima(100); | Metschnikowia | 13 | Saprotroph | \|Undefined Saprotroph\| | Yeast | NULL | Probable | As nectar-tap_saprotroph (Põlme et al. 2020); Decay sugar-rich substrates (Põlme et al. 2020) | Põlme S, et al. 2020. Fungal Diversity 105:1-16 ((https://doi.org/10.1007/s13225-020-00466-2)) |
| Otu39784 | 1 | k__Fungi(100);p__Ascomycota(100);c__Saccharomycetes(100);o__Saccharomycetales(100);f__Metschnikowiaceae(100);g__Metschnikowia(100);g__Metschnikowia_unclassified(100); | Metschnikowia | 13 | Saprotroph | \|Undefined Saprotroph\| | Yeast | NULL | Probable | As nectar-tap_saprotroph (Põlme et al. 2020); Decay sugar-rich substrates (Põlme et al. 2020) | Põlme S, et al. 2020. Fungal Diversity 105:1-16 ((https://doi.org/10.1007/s13225-020-00466-2)) |
| Otu39785 | 1 | k__Fungi(100);p__Ascomycota(100);c__Saccharomycetes(100);o__Saccharomycetales(100);f__Metschnikowiaceae(100);g__Metschnikowia(100);s__Metschnikowia_pulcherrima(100); | Metschnikowia | 13 | Saprotroph | \|Undefined Saprotroph\| | Yeast | NULL | Probable | As nectar-tap_saprotroph (Põlme et al. 2020); Decay sugar-rich substrates (Põlme et al. 2020) | Põlme S, et al. 2020. Fungal Diversity 105:1-16 ((https://doi.org/10.1007/s13225-020-00466-2)) |
| Otu39786 | 1 | k__Fungi(100);p__Ascomycota(100);c__Saccharomycetes(100);o__Saccharomycetales(100);f__Metschnikowiaceae(100);g__Metschnikowia(100);s__Metschnikowia_viticola(100); | Metschnikowia | 13 | Saprotroph | \|Undefined Saprotroph\| | Yeast | NULL | Probable | As nectar-tap_saprotroph (Põlme et al. 2020); Decay sugar-rich substrates (Põlme et al. 2020) | Põlme S, et al. 2020. Fungal Diversity 105:1-16 ((https://doi.org/10.1007/s13225-020-00466-2)) |
| Otu39787 | 1 | k__Fungi(100);p__Ascomycota(100);c__Saccharomycetes(100);o__Saccharomycetales(100);f__Metschnikowiaceae(100);g__Metschnikowia(100);g__Metschnikowia_unclassified(100); | Metschnikowia | 13 | Saprotroph | \|Undefined Saprotroph\| | Yeast | NULL | Probable | As nectar-tap_saprotroph (Põlme et al. 2020); Decay sugar-rich substrates (Põlme et al. 2020) | Põlme S, et al. 2020. Fungal Diversity 105:1-16 ((https://doi.org/10.1007/s13225-020-00466-2)) |
| Otu39788 | 1 | k__Fungi(100);p__Ascomycota(100);c__Saccharomycetes(100);o__Saccharomycetales(100);f__Metschnikowiaceae(100);g__Metschnikowia(100);g__Metschnikowia_unclassified(100); | Metschnikowia | 13 | Saprotroph | \|Undefined Saprotroph\| | Yeast | NULL | Probable | As nectar-tap_saprotroph (Põlme et al. 2020); Decay sugar-rich substrates (Põlme et al. 2020) | Põlme S, et al. 2020. Fungal Diversity 105:1-16 ((https://doi.org/10.1007/s13225-020-00466-2)) |
| Otu39789 | 1 | k__Fungi(100);p__Ascomycota(100);c__Saccharomycetes(100);o__Saccharomycetales(100);f__Metschnikowiaceae(100);g__Metschnikowia(100);s__Metschnikowia_pulcherrima(100); | Metschnikowia | 13 | Saprotroph | \|Undefined Saprotroph\| | Yeast | NULL | Probable | As nectar-tap_saprotroph (Põlme et al. 2020); Decay sugar-rich substrates (Põlme et al. 2020) | Põlme S, et al. 2020. Fungal Diversity 105:1-16 ((https://doi.org/10.1007/s13225-020-00466-2)) |
| Otu39790 | 1 | k__Fungi(100);p__Ascomycota(100);c__Saccharomycetes(100);o__Saccharomycetales(100);f__Metschnikowiaceae(100);g__Metschnikowia(100);s__Metschnikowia_pulcherrima(100); | Metschnikowia | 13 | Saprotroph | \|Undefined Saprotroph\| | Yeast | NULL | Probable | As nectar-tap_saprotroph (Põlme et al. 2020); Decay sugar-rich substrates (Põlme et al. 2020) | Põlme S, et al. 2020. Fungal Diversity 105:1-16 ((https://doi.org/10.1007/s13225-020-00466-2)) |
| Otu39791 | 1 | k__Fungi(100);p__Ascomycota(100);c__Saccharomycetes(100);o__Saccharomycetales(100);f__Metschnikowiaceae(100);g__Metschnikowia(100);s__Metschnikowia_pulcherrima(100); | Metschnikowia | 13 | Saprotroph | \|Undefined Saprotroph\| | Yeast | NULL | Probable | As nectar-tap_saprotroph (Põlme et al. 2020); Decay sugar-rich substrates (Põlme et al. 2020) | Põlme S, et al. 2020. Fungal Diversity 105:1-16 ((https://doi.org/10.1007/s13225-020-00466-2)) |
| Otu39792 | 1 | k__Fungi(100);p__Ascomycota(100);c__Saccharomycetes(100);o__Saccharomycetales(100);f__Metschnikowiaceae(100);g__Metschnikowia(100);g__Metschnikowia_unclassified(100); | Metschnikowia | 13 | Saprotroph | \|Undefined Saprotroph\| | Yeast | NULL | Probable | As nectar-tap_saprotroph (Põlme et al. 2020); Decay sugar-rich substrates (Põlme et al. 2020) | Põlme S, et al. 2020. Fungal Diversity 105:1-16 ((https://doi.org/10.1007/s13225-020-00466-2)) |
| Otu39793 | 1 | k__Fungi(100);p__Ascomycota(100);c__Saccharomycetes(100);o__Saccharomycetales(100);f__Metschnikowiaceae(100);g__Metschnikowia(100);s__Metschnikowia_pulcherrima(100); | Metschnikowia | 13 | Saprotroph | \|Undefined Saprotroph\| | Yeast | NULL | Probable | As nectar-tap_saprotroph (Põlme et al. 2020); Decay sugar-rich substrates (Põlme et al. 2020) | Põlme S, et al. 2020. Fungal Diversity 105:1-16 ((https://doi.org/10.1007/s13225-020-00466-2)) |
| Otu39794 | 1 | k__Fungi(100);p__Ascomycota(100);c__Saccharomycetes(100);o__Saccharomycetales(100);f__Metschnikowiaceae(100);g__Metschnikowia(100);s__Metschnikowia_pulcherrima(100); | Metschnikowia | 13 | Saprotroph | \|Undefined Saprotroph\| | Yeast | NULL | Probable | As nectar-tap_saprotroph (Põlme et al. 2020); Decay sugar-rich substrates (Põlme et al. 2020) | Põlme S, et al. 2020. Fungal Diversity 105:1-16 ((https://doi.org/10.1007/s13225-020-00466-2)) |
| Otu39796 | 1 | k__Fungi(100);p__Ascomycota(100);c__Saccharomycetes(100);o__Saccharomycetales(100);f__Metschnikowiaceae(100);g__Metschnikowia(100);s__Metschnikowia_pulcherrima(100); | Metschnikowia | 13 | Saprotroph | \|Undefined Saprotroph\| | Yeast | NULL | Probable | As nectar-tap_saprotroph (Põlme et al. 2020); Decay sugar-rich substrates (Põlme et al. 2020) | Põlme S, et al. 2020. Fungal Diversity 105:1-16 ((https://doi.org/10.1007/s13225-020-00466-2)) |
| Otu39797 | 1 | k__Fungi(100);p__Ascomycota(100);c__Saccharomycetes(100);o__Saccharomycetales(100);f__Metschnikowiaceae(100);g__Metschnikowia(100);s__Metschnikowia_pulcherrima(100); | Metschnikowia | 13 | Saprotroph | \|Undefined Saprotroph\| | Yeast | NULL | Probable | As nectar-tap_saprotroph (Põlme et al. 2020); Decay sugar-rich substrates (Põlme et al. 2020) | Põlme S, et al. 2020. Fungal Diversity 105:1-16 ((https://doi.org/10.1007/s13225-020-00466-2)) |
| Otu39798 | 1 | k__Fungi(100);p__Ascomycota(100);c__Saccharomycetes(100);o__Saccharomycetales(100);f__Metschnikowiaceae(100);g__Metschnikowia(100);s__Metschnikowia_pulcherrima(100); | Metschnikowia | 13 | Saprotroph | \|Undefined Saprotroph\| | Yeast | NULL | Probable | As nectar-tap_saprotroph (Põlme et al. 2020); Decay sugar-rich substrates (Põlme et al. 2020) | Põlme S, et al. 2020. Fungal Diversity 105:1-16 ((https://doi.org/10.1007/s13225-020-00466-2)) |
| Otu39799 | 1 | k__Fungi(100);p__Ascomycota(100);c__Saccharomycetes(100);o__Saccharomycetales(100);f__Metschnikowiaceae(100);g__Metschnikowia(100);g__Metschnikowia_unclassified(100); | Metschnikowia | 13 | Saprotroph | \|Undefined Saprotroph\| | Yeast | NULL | Probable | As nectar-tap_saprotroph (Põlme et al. 2020); Decay sugar-rich substrates (Põlme et al. 2020) | Põlme S, et al. 2020. Fungal Diversity 105:1-16 ((https://doi.org/10.1007/s13225-020-00466-2)) |
| Otu39800 | 1 | k__Fungi(100);p__Ascomycota(100);c__Saccharomycetes(100);o__Saccharomycetales(100);f__Metschnikowiaceae(100);g__Metschnikowia(100);g__Metschnikowia_unclassified(100); | Metschnikowia | 13 | Saprotroph | \|Undefined Saprotroph\| | Yeast | NULL | Probable | As nectar-tap_saprotroph (Põlme et al. 2020); Decay sugar-rich substrates (Põlme et al. 2020) | Põlme S, et al. 2020. Fungal Diversity 105:1-16 ((https://doi.org/10.1007/s13225-020-00466-2)) |
| Otu39801 | 1 | k__Fungi(100);p__Ascomycota(100);c__Saccharomycetes(100);o__Saccharomycetales(100);f__Metschnikowiaceae(100);g__Metschnikowia(100);s__Metschnikowia_pulcherrima(100); | Metschnikowia | 13 | Saprotroph | \|Undefined Saprotroph\| | Yeast | NULL | Probable | As nectar-tap_saprotroph (Põlme et al. 2020); Decay sugar-rich substrates (Põlme et al. 2020) | Põlme S, et al. 2020. Fungal Diversity 105:1-16 ((https://doi.org/10.1007/s13225-020-00466-2)) |
| Otu39802 | 1 | k__Fungi(100);p__Ascomycota(100);c__Saccharomycetes(100);o__Saccharomycetales(100);f__Metschnikowiaceae(100);g__Metschnikowia(100);s__Metschnikowia_pulcherrima(100); | Metschnikowia | 13 | Saprotroph | \|Undefined Saprotroph\| | Yeast | NULL | Probable | As nectar-tap_saprotroph (Põlme et al. 2020); Decay sugar-rich substrates (Põlme et al. 2020) | Põlme S, et al. 2020. Fungal Diversity 105:1-16 ((https://doi.org/10.1007/s13225-020-00466-2)) |
| Otu39803 | 1 | k__Fungi(100);p__Ascomycota(100);c__Saccharomycetes(100);o__Saccharomycetales(100);f__Metschnikowiaceae(100);g__Metschnikowia(100);s__Metschnikowia_pulcherrima(100); | Metschnikowia | 13 | Saprotroph | \|Undefined Saprotroph\| | Yeast | NULL | Probable | As nectar-tap_saprotroph (Põlme et al. 2020); Decay sugar-rich substrates (Põlme et al. 2020) | Põlme S, et al. 2020. Fungal Diversity 105:1-16 ((https://doi.org/10.1007/s13225-020-00466-2)) |
| Otu39804 | 1 | k__Fungi(100);p__Ascomycota(100);c__Saccharomycetes(100);o__Saccharomycetales(100);f__Metschnikowiaceae(100);g__Metschnikowia(100);s__Metschnikowia_pulcherrima(100); | Metschnikowia | 13 | Saprotroph | \|Undefined Saprotroph\| | Yeast | NULL | Probable | As nectar-tap_saprotroph (Põlme et al. 2020); Decay sugar-rich substrates (Põlme et al. 2020) | Põlme S, et al. 2020. Fungal Diversity 105:1-16 ((https://doi.org/10.1007/s13225-020-00466-2)) |
| Otu39805 | 1 | k__Fungi(100);p__Ascomycota(100);c__Saccharomycetes(100);o__Saccharomycetales(100);f__Metschnikowiaceae(100);g__Metschnikowia(100);s__Metschnikowia_pulcherrima(100); | Metschnikowia | 13 | Saprotroph | \|Undefined Saprotroph\| | Yeast | NULL | Probable | As nectar-tap_saprotroph (Põlme et al. 2020); Decay sugar-rich substrates (Põlme et al. 2020) | Põlme S, et al. 2020. Fungal Diversity 105:1-16 ((https://doi.org/10.1007/s13225-020-00466-2)) |
| Otu39806 | 1 | k__Fungi(100);p__Ascomycota(100);c__Saccharomycetes(100);o__Saccharomycetales(100);f__Metschnikowiaceae(100);g__Metschnikowia(100);g__Metschnikowia_unclassified(100); | Metschnikowia | 13 | Saprotroph | \|Undefined Saprotroph\| | Yeast | NULL | Probable | As nectar-tap_saprotroph (Põlme et al. 2020); Decay sugar-rich substrates (Põlme et al. 2020) | Põlme S, et al. 2020. Fungal Diversity 105:1-16 ((https://doi.org/10.1007/s13225-020-00466-2)) |
| Otu39807 | 1 | k__Fungi(100);p__Ascomycota(100);c__Saccharomycetes(100);o__Saccharomycetales(100);f__Metschnikowiaceae(100);g__Metschnikowia(100);g__Metschnikowia_unclassified(100); | Metschnikowia | 13 | Saprotroph | \|Undefined Saprotroph\| | Yeast | NULL | Probable | As nectar-tap_saprotroph (Põlme et al. 2020); Decay sugar-rich substrates (Põlme et al. 2020) | Põlme S, et al. 2020. Fungal Diversity 105:1-16 ((https://doi.org/10.1007/s13225-020-00466-2)) |
| Otu39808 | 1 | k__Fungi(100);p__Ascomycota(100);c__Saccharomycetes(100);o__Saccharomycetales(100);f__Metschnikowiaceae(100);g__Metschnikowia(100);s__Metschnikowia_pulcherrima(100); | Metschnikowia | 13 | Saprotroph | \|Undefined Saprotroph\| | Yeast | NULL | Probable | As nectar-tap_saprotroph (Põlme et al. 2020); Decay sugar-rich substrates (Põlme et al. 2020) | Põlme S, et al. 2020. Fungal Diversity 105:1-16 ((https://doi.org/10.1007/s13225-020-00466-2)) |
| Otu39809 | 1 | k__Fungi(100);p__Ascomycota(100);c__Leotiomycetes(100);o__Helotiales(100);f__Helotiales_family_Incertae_sedis(100);g__Spirosphaera(100);s__Spirosphaera_cupreorufescens(100); | Spirosphaera | 13 | Saprotroph | \|Plant Saprotroph\|-Undefined Saprotroph | Microfungus | NULL | Probable | As litter_saprotroph (Põlme et al. 2020); Decay leaf, fruit, seed (Põlme et al. 2020) | Tedersoo L, et al. 2014. Science 346:e1256688 ((https://doi.org/10.1126/science.1256688)); Põlme S, et al. 2020. Fungal Diversity 105:1-16 ((https://doi.org/10.1007/s13225-020-00466-2)) |
| Otu39810 | 1 | k__Fungi(100);p__Ascomycota(100);c__Saccharomycetes(100);o__Saccharomycetales(100);f__Metschnikowiaceae(100);g__Metschnikowia(100);g__Metschnikowia_unclassified(100); | Metschnikowia | 13 | Saprotroph | \|Undefined Saprotroph\| | Yeast | NULL | Probable | As nectar-tap_saprotroph (Põlme et al. 2020); Decay sugar-rich substrates (Põlme et al. 2020) | Põlme S, et al. 2020. Fungal Diversity 105:1-16 ((https://doi.org/10.1007/s13225-020-00466-2)) |
| Otu39811 | 1 | k__Fungi(100);p__Ascomycota(100);c__Saccharomycetes(100);o__Saccharomycetales(100);f__Metschnikowiaceae(100);g__Metschnikowia(100);g__Metschnikowia_unclassified(100); | Metschnikowia | 13 | Saprotroph | \|Undefined Saprotroph\| | Yeast | NULL | Probable | As nectar-tap_saprotroph (Põlme et al. 2020); Decay sugar-rich substrates (Põlme et al. 2020) | Põlme S, et al. 2020. Fungal Diversity 105:1-16 ((https://doi.org/10.1007/s13225-020-00466-2)) |
| Otu39812 | 1 | k__Fungi(100);p__Ascomycota(100);c__Saccharomycetes(100);o__Saccharomycetales(100);f__Metschnikowiaceae(100);g__Metschnikowia(100);s__Metschnikowia_pulcherrima(100); | Metschnikowia | 13 | Saprotroph | \|Undefined Saprotroph\| | Yeast | NULL | Probable | As nectar-tap_saprotroph (Põlme et al. 2020); Decay sugar-rich substrates (Põlme et al. 2020) | Põlme S, et al. 2020. Fungal Diversity 105:1-16 ((https://doi.org/10.1007/s13225-020-00466-2)) |
| Otu39813 | 1 | k__Fungi(100);p__Ascomycota(100);c__Saccharomycetes(100);o__Saccharomycetales(100);f__Metschnikowiaceae(100);g__Metschnikowia(100);s__Metschnikowia_pulcherrima(100); | Metschnikowia | 13 | Saprotroph | \|Undefined Saprotroph\| | Yeast | NULL | Probable | As nectar-tap_saprotroph (Põlme et al. 2020); Decay sugar-rich substrates (Põlme et al. 2020) | Põlme S, et al. 2020. Fungal Diversity 105:1-16 ((https://doi.org/10.1007/s13225-020-00466-2)) |
| Otu39814 | 1 | k__Fungi(100);p__Ascomycota(100);c__Saccharomycetes(100);o__Saccharomycetales(100);f__Metschnikowiaceae(100);g__Metschnikowia(100);s__Metschnikowia_pulcherrima(100); | Metschnikowia | 13 | Saprotroph | \|Undefined Saprotroph\| | Yeast | NULL | Probable | As nectar-tap_saprotroph (Põlme et al. 2020); Decay sugar-rich substrates (Põlme et al. 2020) | Põlme S, et al. 2020. Fungal Diversity 105:1-16 ((https://doi.org/10.1007/s13225-020-00466-2)) |
| Otu39815 | 1 | k__Fungi(100);p__Ascomycota(100);c__Saccharomycetes(100);o__Saccharomycetales(100);f__Metschnikowiaceae(100);g__Metschnikowia(100);s__Metschnikowia_pulcherrima(100); | Metschnikowia | 13 | Saprotroph | \|Undefined Saprotroph\| | Yeast | NULL | Probable | As nectar-tap_saprotroph (Põlme et al. 2020); Decay sugar-rich substrates (Põlme et al. 2020) | Põlme S, et al. 2020. Fungal Diversity 105:1-16 ((https://doi.org/10.1007/s13225-020-00466-2)) |
| Otu39816 | 1 | k__Fungi(100);p__Ascomycota(100);c__Saccharomycetes(100);o__Saccharomycetales(100);f__Metschnikowiaceae(100);g__Metschnikowia(100);g__Metschnikowia_unclassified(100); | Metschnikowia | 13 | Saprotroph | \|Undefined Saprotroph\| | Yeast | NULL | Probable | As nectar-tap_saprotroph (Põlme et al. 2020); Decay sugar-rich substrates (Põlme et al. 2020) | Põlme S, et al. 2020. Fungal Diversity 105:1-16 ((https://doi.org/10.1007/s13225-020-00466-2)) |
| Otu39817 | 1 | k__Fungi(100);p__Ascomycota(100);c__Saccharomycetes(100);o__Saccharomycetales(100);f__Metschnikowiaceae(100);g__Metschnikowia(100);g__Metschnikowia_unclassified(100); | Metschnikowia | 13 | Saprotroph | \|Undefined Saprotroph\| | Yeast | NULL | Probable | As nectar-tap_saprotroph (Põlme et al. 2020); Decay sugar-rich substrates (Põlme et al. 2020) | Põlme S, et al. 2020. Fungal Diversity 105:1-16 ((https://doi.org/10.1007/s13225-020-00466-2)) |
| Otu39818 | 1 | k__Fungi(100);p__Ascomycota(100);c__Saccharomycetes(100);o__Saccharomycetales(100);f__Metschnikowiaceae(100);g__Metschnikowia(100);g__Metschnikowia_unclassified(100); | Metschnikowia | 13 | Saprotroph | \|Undefined Saprotroph\| | Yeast | NULL | Probable | As nectar-tap_saprotroph (Põlme et al. 2020); Decay sugar-rich substrates (Põlme et al. 2020) | Põlme S, et al. 2020. Fungal Diversity 105:1-16 ((https://doi.org/10.1007/s13225-020-00466-2)) |
| Otu39819 | 1 | k__Fungi(100);p__Ascomycota(100);c__Saccharomycetes(100);o__Saccharomycetales(100);f__Metschnikowiaceae(100);g__Metschnikowia(100);s__Metschnikowia_pulcherrima(100); | Metschnikowia | 13 | Saprotroph | \|Undefined Saprotroph\| | Yeast | NULL | Probable | As nectar-tap_saprotroph (Põlme et al. 2020); Decay sugar-rich substrates (Põlme et al. 2020) | Põlme S, et al. 2020. Fungal Diversity 105:1-16 ((https://doi.org/10.1007/s13225-020-00466-2)) |
| Otu39820 | 1 | k__Fungi(100);p__Ascomycota(100);c__Saccharomycetes(100);o__Saccharomycetales(100);f__Metschnikowiaceae(100);g__Metschnikowia(100);g__Metschnikowia_unclassified(100); | Metschnikowia | 13 | Saprotroph | \|Undefined Saprotroph\| | Yeast | NULL | Probable | As nectar-tap_saprotroph (Põlme et al. 2020); Decay sugar-rich substrates (Põlme et al. 2020) | Põlme S, et al. 2020. Fungal Diversity 105:1-16 ((https://doi.org/10.1007/s13225-020-00466-2)) |
| Otu39821 | 1 | k__Fungi(100);p__Ascomycota(100);c__Saccharomycetes(100);o__Saccharomycetales(100);f__Metschnikowiaceae(100);g__Metschnikowia(100);g__Metschnikowia_unclassified(100); | Metschnikowia | 13 | Saprotroph | \|Undefined Saprotroph\| | Yeast | NULL | Probable | As nectar-tap_saprotroph (Põlme et al. 2020); Decay sugar-rich substrates (Põlme et al. 2020) | Põlme S, et al. 2020. Fungal Diversity 105:1-16 ((https://doi.org/10.1007/s13225-020-00466-2)) |
| Otu39822 | 1 | k__Fungi(100);p__Ascomycota(100);c__Saccharomycetes(100);o__Saccharomycetales(100);f__Metschnikowiaceae(100);g__Metschnikowia(100);g__Metschnikowia_unclassified(100); | Metschnikowia | 13 | Saprotroph | \|Undefined Saprotroph\| | Yeast | NULL | Probable | As nectar-tap_saprotroph (Põlme et al. 2020); Decay sugar-rich substrates (Põlme et al. 2020) | Põlme S, et al. 2020. Fungal Diversity 105:1-16 ((https://doi.org/10.1007/s13225-020-00466-2)) |
| Otu39823 | 1 | k__Fungi(100);p__Ascomycota(100);c__Saccharomycetes(100);o__Saccharomycetales(100);f__Metschnikowiaceae(100);g__Metschnikowia(100);g__Metschnikowia_unclassified(100); | Metschnikowia | 13 | Saprotroph | \|Undefined Saprotroph\| | Yeast | NULL | Probable | As nectar-tap_saprotroph (Põlme et al. 2020); Decay sugar-rich substrates (Põlme et al. 2020) | Põlme S, et al. 2020. Fungal Diversity 105:1-16 ((https://doi.org/10.1007/s13225-020-00466-2)) |
| Otu39824 | 1 | k__Fungi(100);p__Ascomycota(100);c__Saccharomycetes(100);o__Saccharomycetales(100);f__Metschnikowiaceae(100);g__Metschnikowia(100);s__Metschnikowia_pulcherrima(100); | Metschnikowia | 13 | Saprotroph | \|Undefined Saprotroph\| | Yeast | NULL | Probable | As nectar-tap_saprotroph (Põlme et al. 2020); Decay sugar-rich substrates (Põlme et al. 2020) | Põlme S, et al. 2020. Fungal Diversity 105:1-16 ((https://doi.org/10.1007/s13225-020-00466-2)) |
| Otu39825 | 1 | k__Fungi(100);p__Ascomycota(100);c__Saccharomycetes(100);o__Saccharomycetales(100);f__Metschnikowiaceae(100);g__Metschnikowia(100);s__Metschnikowia_pulcherrima(100); | Metschnikowia | 13 | Saprotroph | \|Undefined Saprotroph\| | Yeast | NULL | Probable | As nectar-tap_saprotroph (Põlme et al. 2020); Decay sugar-rich substrates (Põlme et al. 2020) | Põlme S, et al. 2020. Fungal Diversity 105:1-16 ((https://doi.org/10.1007/s13225-020-00466-2)) |
| Otu39826 | 1 | k__Fungi(100);p__Ascomycota(100);c__Saccharomycetes(100);o__Saccharomycetales(100);f__Metschnikowiaceae(100);g__Metschnikowia(100);s__Metschnikowia_pulcherrima(100); | Metschnikowia | 13 | Saprotroph | \|Undefined Saprotroph\| | Yeast | NULL | Probable | As nectar-tap_saprotroph (Põlme et al. 2020); Decay sugar-rich substrates (Põlme et al. 2020) | Põlme S, et al. 2020. Fungal Diversity 105:1-16 ((https://doi.org/10.1007/s13225-020-00466-2)) |
| Otu39827 | 1 | k__Fungi(100);p__Ascomycota(100);c__Saccharomycetes(100);o__Saccharomycetales(100);f__Metschnikowiaceae(100);g__Metschnikowia(100);s__Metschnikowia_pulcherrima(100); | Metschnikowia | 13 | Saprotroph | \|Undefined Saprotroph\| | Yeast | NULL | Probable | As nectar-tap_saprotroph (Põlme et al. 2020); Decay sugar-rich substrates (Põlme et al. 2020) | Põlme S, et al. 2020. Fungal Diversity 105:1-16 ((https://doi.org/10.1007/s13225-020-00466-2)) |
| Otu39828 | 1 | k__Fungi(100);p__Ascomycota(100);c__Saccharomycetes(100);o__Saccharomycetales(100);f__Metschnikowiaceae(100);g__Metschnikowia(100);s__Metschnikowia_pulcherrima(100); | Metschnikowia | 13 | Saprotroph | \|Undefined Saprotroph\| | Yeast | NULL | Probable | As nectar-tap_saprotroph (Põlme et al. 2020); Decay sugar-rich substrates (Põlme et al. 2020) | Põlme S, et al. 2020. Fungal Diversity 105:1-16 ((https://doi.org/10.1007/s13225-020-00466-2)) |
| Otu39829 | 1 | k__Fungi(100);p__Ascomycota(100);c__Saccharomycetes(100);o__Saccharomycetales(100);f__Metschnikowiaceae(100);g__Metschnikowia(100);s__Metschnikowia_pulcherrima(100); | Metschnikowia | 13 | Saprotroph | \|Undefined Saprotroph\| | Yeast | NULL | Probable | As nectar-tap_saprotroph (Põlme et al. 2020); Decay sugar-rich substrates (Põlme et al. 2020) | Põlme S, et al. 2020. Fungal Diversity 105:1-16 ((https://doi.org/10.1007/s13225-020-00466-2)) |
| Otu39830 | 1 | k__Fungi(100);p__Ascomycota(100);c__Saccharomycetes(100);o__Saccharomycetales(100);f__Metschnikowiaceae(100);g__Metschnikowia(100);g__Metschnikowia_unclassified(100); | Metschnikowia | 13 | Saprotroph | \|Undefined Saprotroph\| | Yeast | NULL | Probable | As nectar-tap_saprotroph (Põlme et al. 2020); Decay sugar-rich substrates (Põlme et al. 2020) | Põlme S, et al. 2020. Fungal Diversity 105:1-16 ((https://doi.org/10.1007/s13225-020-00466-2)) |
| Otu39831 | 1 | k__Fungi(100);p__Ascomycota(100);c__Saccharomycetes(100);o__Saccharomycetales(100);f__Metschnikowiaceae(100);g__Metschnikowia(100);g__Metschnikowia_unclassified(100); | Metschnikowia | 13 | Saprotroph | \|Undefined Saprotroph\| | Yeast | NULL | Probable | As nectar-tap_saprotroph (Põlme et al. 2020); Decay sugar-rich substrates (Põlme et al. 2020) | Põlme S, et al. 2020. Fungal Diversity 105:1-16 ((https://doi.org/10.1007/s13225-020-00466-2)) |
| Otu39832 | 1 | k__Fungi(100);p__Ascomycota(100);c__Saccharomycetes(100);o__Saccharomycetales(100);f__Metschnikowiaceae(100);g__Metschnikowia(100);s__Metschnikowia_pulcherrima(100); | Metschnikowia | 13 | Saprotroph | \|Undefined Saprotroph\| | Yeast | NULL | Probable | As nectar-tap_saprotroph (Põlme et al. 2020); Decay sugar-rich substrates (Põlme et al. 2020) | Põlme S, et al. 2020. Fungal Diversity 105:1-16 ((https://doi.org/10.1007/s13225-020-00466-2)) |
| Otu39833 | 1 | k__Fungi(100);p__Ascomycota(100);c__Saccharomycetes(100);o__Saccharomycetales(100);f__Metschnikowiaceae(100);g__Metschnikowia(100);s__Metschnikowia_pulcherrima(100); | Metschnikowia | 13 | Saprotroph | \|Undefined Saprotroph\| | Yeast | NULL | Probable | As nectar-tap_saprotroph (Põlme et al. 2020); Decay sugar-rich substrates (Põlme et al. 2020) | Põlme S, et al. 2020. Fungal Diversity 105:1-16 ((https://doi.org/10.1007/s13225-020-00466-2)) |
| Otu39834 | 1 | k__Fungi(100);p__Ascomycota(100);c__Saccharomycetes(100);o__Saccharomycetales(100);f__Metschnikowiaceae(100);g__Metschnikowia(100);s__Metschnikowia_pulcherrima(100); | Metschnikowia | 13 | Saprotroph | \|Undefined Saprotroph\| | Yeast | NULL | Probable | As nectar-tap_saprotroph (Põlme et al. 2020); Decay sugar-rich substrates (Põlme et al. 2020) | Põlme S, et al. 2020. Fungal Diversity 105:1-16 ((https://doi.org/10.1007/s13225-020-00466-2)) |
| Otu39835 | 1 | k__Fungi(100);p__Ascomycota(100);c__Saccharomycetes(100);o__Saccharomycetales(100);f__Metschnikowiaceae(100);g__Metschnikowia(100);g__Metschnikowia_unclassified(100); | Metschnikowia | 13 | Saprotroph | \|Undefined Saprotroph\| | Yeast | NULL | Probable | As nectar-tap_saprotroph (Põlme et al. 2020); Decay sugar-rich substrates (Põlme et al. 2020) | Põlme S, et al. 2020. Fungal Diversity 105:1-16 ((https://doi.org/10.1007/s13225-020-00466-2)) |
| Otu39836 | 1 | k__Fungi(100);p__Ascomycota(100);c__Saccharomycetes(100);o__Saccharomycetales(100);f__Metschnikowiaceae(100);g__Metschnikowia(100);s__Metschnikowia_pulcherrima(100); | Metschnikowia | 13 | Saprotroph | \|Undefined Saprotroph\| | Yeast | NULL | Probable | As nectar-tap_saprotroph (Põlme et al. 2020); Decay sugar-rich substrates (Põlme et al. 2020) | Põlme S, et al. 2020. Fungal Diversity 105:1-16 ((https://doi.org/10.1007/s13225-020-00466-2)) |
| Otu39837 | 1 | k__Fungi(100);p__Ascomycota(100);c__Saccharomycetes(100);o__Saccharomycetales(100);f__Metschnikowiaceae(100);g__Metschnikowia(100);s__Metschnikowia_pulcherrima(100); | Metschnikowia | 13 | Saprotroph | \|Undefined Saprotroph\| | Yeast | NULL | Probable | As nectar-tap_saprotroph (Põlme et al. 2020); Decay sugar-rich substrates (Põlme et al. 2020) | Põlme S, et al. 2020. Fungal Diversity 105:1-16 ((https://doi.org/10.1007/s13225-020-00466-2)) |
| Otu39838 | 1 | k__Fungi(100);p__Ascomycota(100);c__Saccharomycetes(100);o__Saccharomycetales(100);f__Metschnikowiaceae(100);g__Metschnikowia(100);s__Metschnikowia_pulcherrima(100); | Metschnikowia | 13 | Saprotroph | \|Undefined Saprotroph\| | Yeast | NULL | Probable | As nectar-tap_saprotroph (Põlme et al. 2020); Decay sugar-rich substrates (Põlme et al. 2020) | Põlme S, et al. 2020. Fungal Diversity 105:1-16 ((https://doi.org/10.1007/s13225-020-00466-2)) |
| Otu39839 | 1 | k__Fungi(100);p__Ascomycota(100);c__Saccharomycetes(100);o__Saccharomycetales(100);f__Metschnikowiaceae(100);g__Metschnikowia(100);s__Metschnikowia_pulcherrima(100); | Metschnikowia | 13 | Saprotroph | \|Undefined Saprotroph\| | Yeast | NULL | Probable | As nectar-tap_saprotroph (Põlme et al. 2020); Decay sugar-rich substrates (Põlme et al. 2020) | Põlme S, et al. 2020. Fungal Diversity 105:1-16 ((https://doi.org/10.1007/s13225-020-00466-2)) |
| Otu39840 | 1 | k__Fungi(100);p__Ascomycota(100);c__Saccharomycetes(100);o__Saccharomycetales(100);f__Metschnikowiaceae(100);g__Metschnikowia(100);s__Metschnikowia_pulcherrima(100); | Metschnikowia | 13 | Saprotroph | \|Undefined Saprotroph\| | Yeast | NULL | Probable | As nectar-tap_saprotroph (Põlme et al. 2020); Decay sugar-rich substrates (Põlme et al. 2020) | Põlme S, et al. 2020. Fungal Diversity 105:1-16 ((https://doi.org/10.1007/s13225-020-00466-2)) |
| Otu39841 | 1 | k__Fungi(100);p__Ascomycota(100);c__Saccharomycetes(100);o__Saccharomycetales(100);f__Metschnikowiaceae(100);g__Metschnikowia(100);s__Metschnikowia_pulcherrima(100); | Metschnikowia | 13 | Saprotroph | \|Undefined Saprotroph\| | Yeast | NULL | Probable | As nectar-tap_saprotroph (Põlme et al. 2020); Decay sugar-rich substrates (Põlme et al. 2020) | Põlme S, et al. 2020. Fungal Diversity 105:1-16 ((https://doi.org/10.1007/s13225-020-00466-2)) |
| Otu39842 | 1 | k__Fungi(100);p__Ascomycota(100);c__Saccharomycetes(100);o__Saccharomycetales(100);f__Metschnikowiaceae(100);g__Metschnikowia(100);g__Metschnikowia_unclassified(100); | Metschnikowia | 13 | Saprotroph | \|Undefined Saprotroph\| | Yeast | NULL | Probable | As nectar-tap_saprotroph (Põlme et al. 2020); Decay sugar-rich substrates (Põlme et al. 2020) | Põlme S, et al. 2020. Fungal Diversity 105:1-16 ((https://doi.org/10.1007/s13225-020-00466-2)) |
| Otu39843 | 1 | k__Fungi(100);p__Ascomycota(100);c__Saccharomycetes(100);o__Saccharomycetales(100);f__Metschnikowiaceae(100);g__Metschnikowia(100);s__Metschnikowia_pulcherrima(100); | Metschnikowia | 13 | Saprotroph | \|Undefined Saprotroph\| | Yeast | NULL | Probable | As nectar-tap_saprotroph (Põlme et al. 2020); Decay sugar-rich substrates (Põlme et al. 2020) | Põlme S, et al. 2020. Fungal Diversity 105:1-16 ((https://doi.org/10.1007/s13225-020-00466-2)) |
| Otu39844 | 1 | k__Fungi(100);p__Ascomycota(100);c__Saccharomycetes(100);o__Saccharomycetales(100);f__Metschnikowiaceae(100);g__Metschnikowia(100);s__Metschnikowia_pulcherrima(100); | Metschnikowia | 13 | Saprotroph | \|Undefined Saprotroph\| | Yeast | NULL | Probable | As nectar-tap_saprotroph (Põlme et al. 2020); Decay sugar-rich substrates (Põlme et al. 2020) | Põlme S, et al. 2020. Fungal Diversity 105:1-16 ((https://doi.org/10.1007/s13225-020-00466-2)) |
| Otu39845 | 1 | k__Fungi(100);p__Ascomycota(100);c__Saccharomycetes(100);o__Saccharomycetales(100);f__Metschnikowiaceae(100);g__Metschnikowia(100);s__Metschnikowia_pulcherrima(100); | Metschnikowia | 13 | Saprotroph | \|Undefined Saprotroph\| | Yeast | NULL | Probable | As nectar-tap_saprotroph (Põlme et al. 2020); Decay sugar-rich substrates (Põlme et al. 2020) | Põlme S, et al. 2020. Fungal Diversity 105:1-16 ((https://doi.org/10.1007/s13225-020-00466-2)) |
| Otu39846 | 1 | k__Fungi(100);p__Ascomycota(100);c__Saccharomycetes(100);o__Saccharomycetales(100);f__Metschnikowiaceae(100);g__Metschnikowia(100);s__Metschnikowia_pulcherrima(100); | Metschnikowia | 13 | Saprotroph | \|Undefined Saprotroph\| | Yeast | NULL | Probable | As nectar-tap_saprotroph (Põlme et al. 2020); Decay sugar-rich substrates (Põlme et al. 2020) | Põlme S, et al. 2020. Fungal Diversity 105:1-16 ((https://doi.org/10.1007/s13225-020-00466-2)) |
| Otu39847 | 1 | k__Fungi(100);p__Ascomycota(100);c__Saccharomycetes(100);o__Saccharomycetales(100);f__Metschnikowiaceae(100);g__Metschnikowia(100);s__Metschnikowia_pulcherrima(100); | Metschnikowia | 13 | Saprotroph | \|Undefined Saprotroph\| | Yeast | NULL | Probable | As nectar-tap_saprotroph (Põlme et al. 2020); Decay sugar-rich substrates (Põlme et al. 2020) | Põlme S, et al. 2020. Fungal Diversity 105:1-16 ((https://doi.org/10.1007/s13225-020-00466-2)) |
| Otu39848 | 1 | k__Fungi(100);p__Ascomycota(100);c__Saccharomycetes(100);o__Saccharomycetales(100);f__Metschnikowiaceae(100);g__Metschnikowia(100);s__Metschnikowia_pulcherrima(100); | Metschnikowia | 13 | Saprotroph | \|Undefined Saprotroph\| | Yeast | NULL | Probable | As nectar-tap_saprotroph (Põlme et al. 2020); Decay sugar-rich substrates (Põlme et al. 2020) | Põlme S, et al. 2020. Fungal Diversity 105:1-16 ((https://doi.org/10.1007/s13225-020-00466-2)) |
| Otu39849 | 1 | k__Fungi(100);p__Ascomycota(100);c__Saccharomycetes(100);o__Saccharomycetales(100);f__Metschnikowiaceae(100);g__Metschnikowia(100);s__Metschnikowia_pulcherrima(100); | Metschnikowia | 13 | Saprotroph | \|Undefined Saprotroph\| | Yeast | NULL | Probable | As nectar-tap_saprotroph (Põlme et al. 2020); Decay sugar-rich substrates (Põlme et al. 2020) | Põlme S, et al. 2020. Fungal Diversity 105:1-16 ((https://doi.org/10.1007/s13225-020-00466-2)) |
| Otu39850 | 1 | k__Fungi(100);p__Ascomycota(100);c__Saccharomycetes(100);o__Saccharomycetales(100);f__Metschnikowiaceae(100);g__Metschnikowia(100);s__Metschnikowia_pulcherrima(100); | Metschnikowia | 13 | Saprotroph | \|Undefined Saprotroph\| | Yeast | NULL | Probable | As nectar-tap_saprotroph (Põlme et al. 2020); Decay sugar-rich substrates (Põlme et al. 2020) | Põlme S, et al. 2020. Fungal Diversity 105:1-16 ((https://doi.org/10.1007/s13225-020-00466-2)) |

| | | | | | | | | | | | |
|---|---|---|---|---|---|---|---|---|---|---|---|
| Otu39851 | 1 | k__Fungi(100);p__Ascomycota(100);c__Saccharomycetes(100);o__Saccharomycetales(100);f__Metschnikowiaceae(100);g__Metschnikowia(100);s__Metschnikowia_pulcherrima(100); | Metschnikowia | 13 | Saprotroph | \|Undefined Saprotroph\| | Yeast | NULL | Probable | As nectar-tap_saprotroph (Põlme et al. 2020); Decay sugar-rich substrates (Põlme et al. 2020) | Põlme S, et al. 2020. Fungal Diversity 105:1-16 ((https://doi.org/10.1007/s13225-020-00466-2)) |
| Otu39852 | 1 | k__Fungi(100);p__Ascomycota(100);c__Saccharomycetes(100);o__Saccharomycetales(100);f__Metschnikowiaceae(100);g__Metschnikowia(100);s__Metschnikowia_pulcherrima(100); | Metschnikowia | 13 | Saprotroph | \|Undefined Saprotroph\| | Yeast | NULL | Probable | As nectar-tap_saprotroph (Põlme et al. 2020); Decay sugar-rich substrates (Põlme et al. 2020) | Põlme S, et al. 2020. Fungal Diversity 105:1-16 ((https://doi.org/10.1007/s13225-020-00466-2)) |
| Otu39853 | 1 | k__Fungi(100);p__Ascomycota(100);c__Saccharomycetes(100);o__Saccharomycetales(100);f__Metschnikowiaceae(100);g__Metschnikowia(100);s__Metschnikowia_pulcherrima(100); | Metschnikowia | 13 | Saprotroph | \|Undefined Saprotroph\| | Yeast | NULL | Probable | As nectar-tap_saprotroph (Põlme et al. 2020); Decay sugar-rich substrates (Põlme et al. 2020) | Põlme S, et al. 2020. Fungal Diversity 105:1-16 ((https://doi.org/10.1007/s13225-020-00466-2)) |
| Otu39854 | 1 | k__Fungi(100);p__Ascomycota(100);c__Saccharomycetes(100);o__Saccharomycetales(100);f__Metschnikowiaceae(100);g__Metschnikowia(100);s__Metschnikowia_pulcherrima(100); | Metschnikowia | 13 | Saprotroph | \|Undefined Saprotroph\| | Yeast | NULL | Probable | As nectar-tap_saprotroph (Põlme et al. 2020); Decay sugar-rich substrates (Põlme et al. 2020) | Põlme S, et al. 2020. Fungal Diversity 105:1-16 ((https://doi.org/10.1007/s13225-020-00466-2)) |
| Otu39855 | 1 | k__Fungi(100);p__Ascomycota(100);c__Saccharomycetes(100);o__Saccharomycetales(100);f__Metschnikowiaceae(100);g__Metschnikowia(100);s__Metschnikowia_pulcherrima(100); | Metschnikowia | 13 | Saprotroph | \|Undefined Saprotroph\| | Yeast | NULL | Probable | As nectar-tap_saprotroph (Põlme et al. 2020); Decay sugar-rich substrates (Põlme et al. 2020) | Põlme S, et al. 2020. Fungal Diversity 105:1-16 ((https://doi.org/10.1007/s13225-020-00466-2)) |
| Otu39856 | 1 | k__Fungi(100);p__Ascomycota(100);c__Saccharomycetes(100);o__Saccharomycetales(100);f__Metschnikowiaceae(100);g__Metschnikowia(100);g__Metschnikowia_unclassified(100); | Metschnikowia | 13 | Saprotroph | \|Undefined Saprotroph\| | Yeast | NULL | Probable | As nectar-tap_saprotroph (Põlme et al. 2020); Decay sugar-rich substrates (Põlme et al. 2020) | Põlme S, et al. 2020. Fungal Diversity 105:1-16 ((https://doi.org/10.1007/s13225-020-00466-2)) |
| Otu39857 | 1 | k__Fungi(100);p__Ascomycota(100);c__Saccharomycetes(100);o__Saccharomycetales(100);f__Metschnikowiaceae(100);g__Metschnikowia(100);s__Metschnikowia_pulcherrima(100); | Metschnikowia | 13 | Saprotroph | \|Undefined Saprotroph\| | Yeast | NULL | Probable | As nectar-tap_saprotroph (Põlme et al. 2020); Decay sugar-rich substrates (Põlme et al. 2020) | Põlme S, et al. 2020. Fungal Diversity 105:1-16 ((https://doi.org/10.1007/s13225-020-00466-2)) |
| Otu39858 | 1 | k__Fungi(100);p__Ascomycota(100);c__Saccharomycetes(100);o__Saccharomycetales(100);f__Metschnikowiaceae(100);g__Metschnikowia(100);s__Metschnikowia_pulcherrima(100); | Metschnikowia | 13 | Saprotroph | \|Undefined Saprotroph\| | Yeast | NULL | Probable | As nectar-tap_saprotroph (Põlme et al. 2020); Decay sugar-rich substrates (Põlme et al. 2020) | Põlme S, et al. 2020. Fungal Diversity 105:1-16 ((https://doi.org/10.1007/s13225-020-00466-2)) |
| Otu39859 | 1 | k__Fungi(100);p__Ascomycota(100);c__Saccharomycetes(100);o__Saccharomycetales(100);f__Metschnikowiaceae(100);g__Metschnikowia(100);s__Metschnikowia_pulcherrima(100); | Metschnikowia | 13 | Saprotroph | \|Undefined Saprotroph\| | Yeast | NULL | Probable | As nectar-tap_saprotroph (Põlme et al. 2020); Decay sugar-rich substrates (Põlme et al. 2020) | Põlme S, et al. 2020. Fungal Diversity 105:1-16 ((https://doi.org/10.1007/s13225-020-00466-2)) |
| Otu39860 | 1 | k__Fungi(100);p__Ascomycota(100);c__Saccharomycetes(100);o__Saccharomycetales(100);f__Metschnikowiaceae(100);g__Metschnikowia(100);g__Metschnikowia_unclassified(100); | Metschnikowia | 13 | Saprotroph | \|Undefined Saprotroph\| | Yeast | NULL | Probable | As nectar-tap_saprotroph (Põlme et al. 2020); Decay sugar-rich substrates (Põlme et al. 2020) | Põlme S, et al. 2020. Fungal Diversity 105:1-16 ((https://doi.org/10.1007/s13225-020-00466-2)) |
| Otu39861 | 1 | k__Fungi(100);p__Ascomycota(100);c__Saccharomycetes(100);o__Saccharomycetales(100);f__Metschnikowiaceae(100);g__Metschnikowia(100);s__Metschnikowia_pulcherrima(100); | Metschnikowia | 13 | Saprotroph | \|Undefined Saprotroph\| | Yeast | NULL | Probable | As nectar-tap_saprotroph (Põlme et al. 2020); Decay sugar-rich substrates (Põlme et al. 2020) | Põlme S, et al. 2020. Fungal Diversity 105:1-16 ((https://doi.org/10.1007/s13225-020-00466-2)) |
| Otu39862 | 1 | k__Fungi(100);p__Ascomycota(100);c__Saccharomycetes(100);o__Saccharomycetales(100);f__Metschnikowiaceae(100);g__Metschnikowia(100);s__Metschnikowia_pulcherrima(100); | Metschnikowia | 13 | Saprotroph | \|Undefined Saprotroph\| | Yeast | NULL | Probable | As nectar-tap_saprotroph (Põlme et al. 2020); Decay sugar-rich substrates (Põlme et al. 2020) | Põlme S, et al. 2020. Fungal Diversity 105:1-16 ((https://doi.org/10.1007/s13225-020-00466-2)) |
| Otu39863 | 1 | k__Fungi(100);p__Ascomycota(100);c__Saccharomycetes(100);o__Saccharomycetales(100);f__Metschnikowiaceae(100);g__Metschnikowia(100);s__Metschnikowia_pulcherrima(100); | Metschnikowia | 13 | Saprotroph | \|Undefined Saprotroph\| | Yeast | NULL | Probable | As nectar-tap_saprotroph (Põlme et al. 2020); Decay sugar-rich substrates (Põlme et al. 2020) | Põlme S, et al. 2020. Fungal Diversity 105:1-16 ((https://doi.org/10.1007/s13225-020-00466-2)) |
| Otu39864 | 1 | k__Fungi(100);p__Ascomycota(100);c__Saccharomycetes(100);o__Saccharomycetales(100);f__Metschnikowiaceae(100);g__Metschnikowia(100);s__Metschnikowia_pulcherrima(100); | Metschnikowia | 13 | Saprotroph | \|Undefined Saprotroph\| | Yeast | NULL | Probable | As nectar-tap_saprotroph (Põlme et al. 2020); Decay sugar-rich substrates (Põlme et al. 2020) | Põlme S, et al. 2020. Fungal Diversity 105:1-16 ((https://doi.org/10.1007/s13225-020-00466-2)) |
| Otu39865 | 1 | k__Fungi(100);p__Ascomycota(100);c__Saccharomycetes(100);o__Saccharomycetales(100);f__Metschnikowiaceae(100);g__Metschnikowia(100);g__Metschnikowia_unclassified(100); | Metschnikowia | 13 | Saprotroph | \|Undefined Saprotroph\| | Yeast | NULL | Probable | As nectar-tap_saprotroph (Põlme et al. 2020); Decay sugar-rich substrates (Põlme et al. 2020) | Põlme S, et al. 2020. Fungal Diversity 105:1-16 ((https://doi.org/10.1007/s13225-020-00466-2)) |
| Otu39866 | 1 | k__Fungi(100);p__Ascomycota(100);c__Saccharomycetes(100);o__Saccharomycetales(100);f__Metschnikowiaceae(100);g__Metschnikowia(100);s__Metschnikowia_pulcherrima(100); | Metschnikowia | 13 | Saprotroph | \|Undefined Saprotroph\| | Yeast | NULL | Probable | As nectar-tap_saprotroph (Põlme et al. 2020); Decay sugar-rich substrates (Põlme et al. 2020) | Põlme S, et al. 2020. Fungal Diversity 105:1-16 ((https://doi.org/10.1007/s13225-020-00466-2)) |
| Otu39867 | 1 | k__Fungi(100);p__Ascomycota(100);c__Saccharomycetes(100);o__Saccharomycetales(100);f__Metschnikowiaceae(100);g__Metschnikowia(100);s__Metschnikowia_pulcherrima(100); | Metschnikowia | 13 | Saprotroph | \|Undefined Saprotroph\| | Yeast | NULL | Probable | As nectar-tap_saprotroph (Põlme et al. 2020); Decay sugar-rich substrates (Põlme et al. 2020) | Põlme S, et al. 2020. Fungal Diversity 105:1-16 ((https://doi.org/10.1007/s13225-020-00466-2)) |
| Otu39868 | 1 | k__Fungi(100);p__Ascomycota(100);c__Saccharomycetes(100);o__Saccharomycetales(100);f__Metschnikowiaceae(100);g__Metschnikowia(100);s__Metschnikowia_pulcherrima(100); | Metschnikowia | 13 | Saprotroph | \|Undefined Saprotroph\| | Yeast | NULL | Probable | As nectar-tap_saprotroph (Põlme et al. 2020); Decay sugar-rich substrates (Põlme et al. 2020) | Põlme S, et al. 2020. Fungal Diversity 105:1-16 ((https://doi.org/10.1007/s13225-020-00466-2)) |
| Otu39869 | 1 | k__Fungi(100);p__Ascomycota(100);c__Saccharomycetes(100);o__Saccharomycetales(100);f__Metschnikowiaceae(100);g__Metschnikowia(100);s__Metschnikowia_pulcherrima(100); | Metschnikowia | 13 | Saprotroph | \|Undefined Saprotroph\| | Yeast | NULL | Probable | As nectar-tap_saprotroph (Põlme et al. 2020); Decay sugar-rich substrates (Põlme et al. 2020) | Põlme S, et al. 2020. Fungal Diversity 105:1-16 ((https://doi.org/10.1007/s13225-020-00466-2)) |
| Otu39870 | 1 | k__Fungi(100);p__Ascomycota(100);c__Saccharomycetes(100);o__Saccharomycetales(100);f__Metschnikowiaceae(100);g__Metschnikowia(100);g__Metschnikowia_unclassified(100); | Metschnikowia | 13 | Saprotroph | \|Undefined Saprotroph\| | Yeast | NULL | Probable | As nectar-tap_saprotroph (Põlme et al. 2020); Decay sugar-rich substrates (Põlme et al. 2020) | Põlme S, et al. 2020. Fungal Diversity 105:1-16 ((https://doi.org/10.1007/s13225-020-00466-2)) |
| Otu39871 | 1 | k__Fungi(100);p__Ascomycota(100);c__Saccharomycetes(100);o__Saccharomycetales(100);f__Metschnikowiaceae(100);g__Metschnikowia(100);s__Metschnikowia_pulcherrima(100); | Metschnikowia | 13 | Saprotroph | \|Undefined Saprotroph\| | Yeast | NULL | Probable | As nectar-tap_saprotroph (Põlme et al. 2020); Decay sugar-rich substrates (Põlme et al. 2020) | Põlme S, et al. 2020. Fungal Diversity 105:1-16 ((https://doi.org/10.1007/s13225-020-00466-2)) |
| Otu39872 | 1 | k__Fungi(100);p__Ascomycota(100);c__Saccharomycetes(100);o__Saccharomycetales(100);f__Metschnikowiaceae(100);g__Metschnikowia(100);s__Metschnikowia_pulcherrima(100); | Metschnikowia | 13 | Saprotroph | \|Undefined Saprotroph\| | Yeast | NULL | Probable | As nectar-tap_saprotroph (Põlme et al. 2020); Decay sugar-rich substrates (Põlme et al. 2020) | Põlme S, et al. 2020. Fungal Diversity 105:1-16 ((https://doi.org/10.1007/s13225-020-00466-2)) |
| Otu39873 | 1 | k__Fungi(100);p__Ascomycota(100);c__Saccharomycetes(100);o__Saccharomycetales(100);f__Metschnikowiaceae(100);g__Metschnikowia(100);g__Metschnikowia_unclassified(100); | Metschnikowia | 13 | Saprotroph | \|Undefined Saprotroph\| | Yeast | NULL | Probable | As nectar-tap_saprotroph (Põlme et al. 2020); Decay sugar-rich substrates (Põlme et al. 2020) | Põlme S, et al. 2020. Fungal Diversity 105:1-16 ((https://doi.org/10.1007/s13225-020-00466-2)) |
| Otu39874 | 1 | k__Fungi(100);p__Ascomycota(100);c__Saccharomycetes(100);o__Saccharomycetales(100);f__Metschnikowiaceae(100);g__Metschnikowia(100);g__Metschnikowia_unclassified(100); | Metschnikowia | 13 | Saprotroph | \|Undefined Saprotroph\| | Yeast | NULL | Probable | As nectar-tap_saprotroph (Põlme et al. 2020); Decay sugar-rich substrates (Põlme et al. 2020) | Põlme S, et al. 2020. Fungal Diversity 105:1-16 ((https://doi.org/10.1007/s13225-020-00466-2)) |
| Otu39875 | 1 | k__Fungi(100);p__Ascomycota(100);c__Saccharomycetes(100);o__Saccharomycetales(100);f__Metschnikowiaceae(100);g__Metschnikowia(100);s__Metschnikowia_pulcherrima(100); | Metschnikowia | 13 | Saprotroph | \|Undefined Saprotroph\| | Yeast | NULL | Probable | As nectar-tap_saprotroph (Põlme et al. 2020); Decay sugar-rich substrates (Põlme et al. 2020) | Põlme S, et al. 2020. Fungal Diversity 105:1-16 ((https://doi.org/10.1007/s13225-020-00466-2)) |
| Otu39876 | 1 | k__Fungi(100);p__Ascomycota(100);c__Saccharomycetes(100);o__Saccharomycetales(100);f__Metschnikowiaceae(100);g__Metschnikowia(100);g__Metschnikowia_unclassified(100); | Metschnikowia | 13 | Saprotroph | \|Undefined Saprotroph\| | Yeast | NULL | Probable | As nectar-tap_saprotroph (Põlme et al. 2020); Decay sugar-rich substrates (Põlme et al. 2020) | Põlme S, et al. 2020. Fungal Diversity 105:1-16 ((https://doi.org/10.1007/s13225-020-00466-2)) |
| Otu39877 | 1 | k__Fungi(100);p__Ascomycota(100);c__Saccharomycetes(100);o__Saccharomycetales(100);f__Metschnikowiaceae(100);g__Metschnikowia(100);g__Metschnikowia_unclassified(100); | Metschnikowia | 13 | Saprotroph | \|Undefined Saprotroph\| | Yeast | NULL | Probable | As nectar-tap_saprotroph (Põlme et al. 2020); Decay sugar-rich substrates (Põlme et al. 2020) | Põlme S, et al. 2020. Fungal Diversity 105:1-16 ((https://doi.org/10.1007/s13225-020-00466-2)) |
| Otu39878 | 1 | k__Fungi(100);p__Ascomycota(100);c__Saccharomycetes(100);o__Saccharomycetales(100);f__Metschnikowiaceae(100);g__Metschnikowia(100);g__Metschnikowia_unclassified(100); | Metschnikowia | 13 | Saprotroph | \|Undefined Saprotroph\| | Yeast | NULL | Probable | As nectar-tap_saprotroph (Põlme et al. 2020); Decay sugar-rich substrates (Põlme et al. 2020) | Põlme S, et al. 2020. Fungal Diversity 105:1-16 ((https://doi.org/10.1007/s13225-020-00466-2)) |
| Otu39879 | 1 | k__Fungi(100);p__Ascomycota(100);c__Saccharomycetes(100);o__Saccharomycetales(100);f__Metschnikowiaceae(100);g__Metschnikowia(100);s__Metschnikowia_pulcherrima(100); | Metschnikowia | 13 | Saprotroph | \|Undefined Saprotroph\| | Yeast | NULL | Probable | As nectar-tap_saprotroph (Põlme et al. 2020); Decay sugar-rich substrates (Põlme et al. 2020) | Põlme S, et al. 2020. Fungal Diversity 105:1-16 ((https://doi.org/10.1007/s13225-020-00466-2)) |
| Otu39880 | 1 | k__Fungi(100);p__Ascomycota(100);c__Saccharomycetes(100);o__Saccharomycetales(100);f__Metschnikowiaceae(100);g__Metschnikowia(100);s__Metschnikowia_pulcherrima(100); | Metschnikowia | 13 | Saprotroph | \|Undefined Saprotroph\| | Yeast | NULL | Probable | As nectar-tap_saprotroph (Põlme et al. 2020); Decay sugar-rich substrates (Põlme et al. 2020) | Põlme S, et al. 2020. Fungal Diversity 105:1-16 ((https://doi.org/10.1007/s13225-020-00466-2)) |
| Otu39881 | 1 | k__Fungi(100);p__Ascomycota(100);c__Saccharomycetes(100);o__Saccharomycetales(100);f__Metschnikowiaceae(100);g__Metschnikowia(100);s__Metschnikowia_pulcherrima(100); | Metschnikowia | 13 | Saprotroph | \|Undefined Saprotroph\| | Yeast | NULL | Probable | As nectar-tap_saprotroph (Põlme et al. 2020); Decay sugar-rich substrates (Põlme et al. 2020) | Põlme S, et al. 2020. Fungal Diversity 105:1-16 ((https://doi.org/10.1007/s13225-020-00466-2)) |
| Otu39882 | 1 | k__Fungi(100);p__Ascomycota(100);c__Saccharomycetes(100);o__Saccharomycetales(100);f__Metschnikowiaceae(100);g__Metschnikowia(100);s__Metschnikowia_pulcherrima(100); | Metschnikowia | 13 | Saprotroph | \|Undefined Saprotroph\| | Yeast | NULL | Probable | As nectar-tap_saprotroph (Põlme et al. 2020); Decay sugar-rich substrates (Põlme et al. 2020) | Põlme S, et al. 2020. Fungal Diversity 105:1-16 ((https://doi.org/10.1007/s13225-020-00466-2)) |
| Otu39883 | 1 | k__Fungi(100);p__Ascomycota(100);c__Saccharomycetes(100);o__Saccharomycetales(100);f__Metschnikowiaceae(100);g__Metschnikowia(100);s__Metschnikowia_pulcherrima(100); | Metschnikowia | 13 | Saprotroph | \|Undefined Saprotroph\| | Yeast | NULL | Probable | As nectar-tap_saprotroph (Põlme et al. 2020); Decay sugar-rich substrates (Põlme et al. 2020) | Põlme S, et al. 2020. Fungal Diversity 105:1-16 ((https://doi.org/10.1007/s13225-020-00466-2)) |
| Otu39884 | 1 | k__Fungi(100);p__Ascomycota(100);c__Saccharomycetes(100);o__Saccharomycetales(100);f__Metschnikowiaceae(100);g__Metschnikowia(100);s__Metschnikowia_pulcherrima(100); | Metschnikowia | 13 | Saprotroph | \|Undefined Saprotroph\| | Yeast | NULL | Probable | As nectar-tap_saprotroph (Põlme et al. 2020); Decay sugar-rich substrates (Põlme et al. 2020) | Põlme S, et al. 2020. Fungal Diversity 105:1-16 ((https://doi.org/10.1007/s13225-020-00466-2)) |
| Otu39885 | 1 | k__Fungi(100);p__Ascomycota(100);c__Saccharomycetes(100);o__Saccharomycetales(100);f__Metschnikowiaceae(100);g__Metschnikowia(100);s__Metschnikowia_pulcherrima(100); | Metschnikowia | 13 | Saprotroph | \|Undefined Saprotroph\| | Yeast | NULL | Probable | As nectar-tap_saprotroph (Põlme et al. 2020); Decay sugar-rich substrates (Põlme et al. 2020) | Põlme S, et al. 2020. Fungal Diversity 105:1-16 ((https://doi.org/10.1007/s13225-020-00466-2)) |
| Otu39886 | 1 | k__Fungi(100);p__Ascomycota(100);c__Saccharomycetes(100);o__Saccharomycetales(100);f__Metschnikowiaceae(100);g__Metschnikowia(100);s__Metschnikowia_pulcherrima(100); | Metschnikowia | 13 | Saprotroph | \|Undefined Saprotroph\| | Yeast | NULL | Probable | As nectar-tap_saprotroph (Põlme et al. 2020); Decay sugar-rich substrates (Põlme et al. 2020) | Põlme S, et al. 2020. Fungal Diversity 105:1-16 ((https://doi.org/10.1007/s13225-020-00466-2)) |
| Otu39887 | 1 | k__Fungi(100);p__Ascomycota(100);c__Saccharomycetes(100);o__Saccharomycetales(100);f__Metschnikowiaceae(100);g__Metschnikowia(100);s__Metschnikowia_pulcherrima(100); | Metschnikowia | 13 | Saprotroph | \|Undefined Saprotroph\| | Yeast | NULL | Probable | As nectar-tap_saprotroph (Põlme et al. 2020); Decay sugar-rich substrates (Põlme et al. 2020) | Põlme S, et al. 2020. Fungal Diversity 105:1-16 ((https://doi.org/10.1007/s13225-020-00466-2)) |
| Otu39888 | 1 | k__Fungi(100);p__Ascomycota(100);c__Saccharomycetes(100);o__Saccharomycetales(100);f__Metschnikowiaceae(100);g__Metschnikowia(100);s__Metschnikowia_pulcherrima(100); | Metschnikowia | 13 | Saprotroph | \|Undefined Saprotroph\| | Yeast | NULL | Probable | As nectar-tap_saprotroph (Põlme et al. 2020); Decay sugar-rich substrates (Põlme et al. 2020) | Põlme S, et al. 2020. Fungal Diversity 105:1-16 ((https://doi.org/10.1007/s13225-020-00466-2)) |
| Otu39889 | 1 | k__Fungi(100);p__Ascomycota(100);c__Saccharomycetes(100);o__Saccharomycetales(100);f__Metschnikowiaceae(100);g__Metschnikowia(100);s__Metschnikowia_pulcherrima(100); | Metschnikowia | 13 | Saprotroph | \|Undefined Saprotroph\| | Yeast | NULL | Probable | As nectar-tap_saprotroph (Põlme et al. 2020); Decay sugar-rich substrates (Põlme et al. 2020) | Põlme S, et al. 2020. Fungal Diversity 105:1-16 ((https://doi.org/10.1007/s13225-020-00466-2)) |
| Otu39890 | 1 | k__Fungi(100);p__Ascomycota(100);c__Saccharomycetes(100);o__Saccharomycetales(100);f__Metschnikowiaceae(100);g__Metschnikowia(100);s__Metschnikowia_pulcherrima(100); | Metschnikowia | 13 | Saprotroph | \|Undefined Saprotroph\| | Yeast | NULL | Probable | As nectar-tap_saprotroph (Põlme et al. 2020); Decay sugar-rich substrates (Põlme et al. 2020) | Põlme S, et al. 2020. Fungal Diversity 105:1-16 ((https://doi.org/10.1007/s13225-020-00466-2)) |
| Otu39891 | 1 | k__Fungi(100);p__Ascomycota(100);c__Saccharomycetes(100);o__Saccharomycetales(100);f__Metschnikowiaceae(100);g__Metschnikowia(100);s__Metschnikowia_pulcherrima(100); | Metschnikowia | 13 | Saprotroph | \|Undefined Saprotroph\| | Yeast | NULL | Probable | As nectar-tap_saprotroph (Põlme et al. 2020); Decay sugar-rich substrates (Põlme et al. 2020) | Põlme S, et al. 2020. Fungal Diversity 105:1-16 ((https://doi.org/10.1007/s13225-020-00466-2)) |
| Otu39892 | 1 | k__Fungi(100);p__Ascomycota(100);c__Saccharomycetes(100);o__Saccharomycetales(100);f__Metschnikowiaceae(100);g__Metschnikowia(100);s__Metschnikowia_pulcherrima(100); | Metschnikowia | 13 | Saprotroph | \|Undefined Saprotroph\| | Yeast | NULL | Probable | As nectar-tap_saprotroph (Põlme et al. 2020); Decay sugar-rich substrates (Põlme et al. 2020) | Põlme S, et al. 2020. Fungal Diversity 105:1-16 ((https://doi.org/10.1007/s13225-020-00466-2)) |
| Otu39893 | 1 | k__Fungi(100);p__Ascomycota(100);c__Saccharomycetes(100);o__Saccharomycetales(100);f__Metschnikowiaceae(100);g__Metschnikowia(100);g__Metschnikowia_unclassified(100); | Metschnikowia | 13 | Saprotroph | \|Undefined Saprotroph\| | Yeast | NULL | Probable | As nectar-tap_saprotroph (Põlme et al. 2020); Decay sugar-rich substrates (Põlme et al. 2020) | Põlme S, et al. 2020. Fungal Diversity 105:1-16 ((https://doi.org/10.1007/s13225-020-00466-2)) |
| Otu39894 | 1 | k__Fungi(100);p__Ascomycota(100);c__Saccharomycetes(100);o__Saccharomycetales(100);f__Metschnikowiaceae(100);g__Metschnikowia(100);g__Metschnikowia_unclassified(100); | Metschnikowia | 13 | Saprotroph | \|Undefined Saprotroph\| | Yeast | NULL | Probable | As nectar-tap_saprotroph (Põlme et al. 2020); Decay sugar-rich substrates (Põlme et al. 2020) | Põlme S, et al. 2020. Fungal Diversity 105:1-16 ((https://doi.org/10.1007/s13225-020-00466-2)) |
| Otu39895 | 1 | k__Fungi(100);p__Ascomycota(100);c__Saccharomycetes(100);o__Saccharomycetales(100);f__Metschnikowiaceae(100);g__Metschnikowia(100);g__Metschnikowia_unclassified(100); | Metschnikowia | 13 | Saprotroph | \|Undefined Saprotroph\| | Yeast | NULL | Probable | As nectar-tap_saprotroph (Põlme et al. 2020); Decay sugar-rich substrates (Põlme et al. 2020) | Põlme S, et al. 2020. Fungal Diversity 105:1-16 ((https://doi.org/10.1007/s13225-020-00466-2)) |
| Otu39896 | 1 | k__Fungi(100);p__Ascomycota(100);c__Saccharomycetes(100);o__Saccharomycetales(100);f__Metschnikowiaceae(100);g__Metschnikowia(100);s__Metschnikowia_pulcherrima(100); | Metschnikowia | 13 | Saprotroph | \|Undefined Saprotroph\| | Yeast | NULL | Probable | As nectar-tap_saprotroph (Põlme et al. 2020); Decay sugar-rich substrates (Põlme et al. 2020) | Põlme S, et al. 2020. Fungal Diversity 105:1-16 ((https://doi.org/10.1007/s13225-020-00466-2)) |
| Otu39897 | 1 | k__Fungi(100);p__Ascomycota(100);c__Saccharomycetes(100);o__Saccharomycetales(100);f__Metschnikowiaceae(100);g__Metschnikowia(100);s__Metschnikowia_pulcherrima(100); | Metschnikowia | 13 | Saprotroph | \|Undefined Saprotroph\| | Yeast | NULL | Probable | As nectar-tap_saprotroph (Põlme et al. 2020); Decay sugar-rich substrates (Põlme et al. 2020) | Põlme S, et al. 2020. Fungal Diversity 105:1-16 ((https://doi.org/10.1007/s13225-020-00466-2)) |
| Otu39898 | 1 | k__Fungi(100);p__Ascomycota(100);c__Saccharomycetes(100);o__Saccharomycetales(100);f__Metschnikowiaceae(100);g__Metschnikowia(100);g__Metschnikowia_unclassified(100); | Metschnikowia | 13 | Saprotroph | \|Undefined Saprotroph\| | Yeast | NULL | Probable | As nectar-tap_saprotroph (Põlme et al. 2020); Decay sugar-rich substrates (Põlme et al. 2020) | Põlme S, et al. 2020. Fungal Diversity 105:1-16 ((https://doi.org/10.1007/s13225-020-00466-2)) |
| Otu39899 | 1 | k__Fungi(100);p__Ascomycota(100);c__Saccharomycetes(100);o__Saccharomycetales(100);f__Metschnikowiaceae(100);g__Metschnikowia(100);s__Metschnikowia_pulcherrima(100); | Metschnikowia | 13 | Saprotroph | \|Undefined Saprotroph\| | Yeast | NULL | Probable | As nectar-tap_saprotroph (Põlme et al. 2020); Decay sugar-rich substrates (Põlme et al. 2020) | Põlme S, et al. 2020. Fungal Diversity 105:1-16 ((https://doi.org/10.1007/s13225-020-00466-2)) |
| Otu39900 | 1 | k__Fungi(100);p__Ascomycota(100);c__Saccharomycetes(100);o__Saccharomycetales(100);f__Metschnikowiaceae(100);g__Metschnikowia(100);s__Metschnikowia_pulcherrima(100); | Metschnikowia | 13 | Saprotroph | \|Undefined Saprotroph\| | Yeast | NULL | Probable | As nectar-tap_saprotroph (Põlme et al. 2020); Decay sugar-rich substrates (Põlme et al. 2020) | Põlme S, et al. 2020. Fungal Diversity 105:1-16 ((https://doi.org/10.1007/s13225-020-00466-2)) |
| Otu39901 | 1 | k__Fungi(100);p__Ascomycota(100);c__Saccharomycetes(100);o__Saccharomycetales(100);f__Metschnikowiaceae(100);g__Metschnikowia(100);g__Metschnikowia_unclassified(100); | Metschnikowia | 13 | Saprotroph | \|Undefined Saprotroph\| | Yeast | NULL | Probable | As nectar-tap_saprotroph (Põlme et al. 2020); Decay sugar-rich substrates (Põlme et al. 2020) | Põlme S, et al. 2020. Fungal Diversity 105:1-16 ((https://doi.org/10.1007/s13225-020-00466-2)) |
| Otu39902 | 1 | k__Fungi(100);p__Ascomycota(100);c__Saccharomycetes(100);o__Saccharomycetales(100);f__Metschnikowiaceae(100);g__Metschnikowia(100);s__Metschnikowia_pulcherrima(100); | Metschnikowia | 13 | Saprotroph | \|Undefined Saprotroph\| | Yeast | NULL | Probable | As nectar-tap_saprotroph (Põlme et al. 2020); Decay sugar-rich substrates (Põlme et al. 2020) | Põlme S, et al. 2020. Fungal Diversity 105:1-16 ((https://doi.org/10.1007/s13225-020-00466-2)) |
| Otu39903 | 1 | k__Fungi(100);p__Ascomycota(100);c__Saccharomycetes(100);o__Saccharomycetales(100);f__Metschnikowiaceae(100);g__Metschnikowia(100);g__Metschnikowia_unclassified(100); | Metschnikowia | 13 | Saprotroph | \|Undefined Saprotroph\| | Yeast | NULL | Probable | As nectar-tap_saprotroph (Põlme et al. 2020); Decay sugar-rich substrates (Põlme et al. 2020) | Põlme S, et al. 2020. Fungal Diversity 105:1-16 ((https://doi.org/10.1007/s13225-020-00466-2)) |
| Otu39904 | 1 | k__Fungi(100);p__Ascomycota(100);c__Saccharomycetes(100);o__Saccharomycetales(100);f__Metschnikowiaceae(100);g__Metschnikowia(100);s__Metschnikowia_pulcherrima(100); | Metschnikowia | 13 | Saprotroph | \|Undefined Saprotroph\| | Yeast | NULL | Probable | As nectar-tap_saprotroph (Põlme et al. 2020); Decay sugar-rich substrates (Põlme et al. 2020) | Põlme S, et al. 2020. Fungal Diversity 105:1-16 ((https://doi.org/10.1007/s13225-020-00466-2)) |
| Otu39905 | 1 | k__Fungi(100);p__Ascomycota(100);c__Saccharomycetes(100);o__Saccharomycetales(100);f__Metschnikowiaceae(100);g__Metschnikowia(100);s__Metschnikowia_pulcherrima(100); | Metschnikowia | 13 | Saprotroph | \|Undefined Saprotroph\| | Yeast | NULL | Probable | As nectar-tap_saprotroph (Põlme et al. 2020); Decay sugar-rich substrates (Põlme et al. 2020) | Põlme S, et al. 2020. Fungal Diversity 105:1-16 ((https://doi.org/10.1007/s13225-020-00466-2)) |
| Otu39906 | 1 | k__Fungi(100);p__Ascomycota(100);c__Saccharomycetes(100);o__Saccharomycetales(100);f__Metschnikowiaceae(100);g__Metschnikowia(100);g__Metschnikowia_unclassified(100); | Metschnikowia | 13 | Saprotroph | \|Undefined Saprotroph\| | Yeast | NULL | Probable | As nectar-tap_saprotroph (Põlme et al. 2020); Decay sugar-rich substrates (Põlme et al. 2020) | Põlme S, et al. 2020. Fungal Diversity 105:1-16 ((https://doi.org/10.1007/s13225-020-00466-2)) |
| Otu39907 | 1 | k__Fungi(100);p__Ascomycota(100);c__Saccharomycetes(100);o__Saccharomycetales(100);f__Metschnikowiaceae(100);g__Metschnikowia(100);s__Metschnikowia_pulcherrima(100); | Metschnikowia | 13 | Saprotroph | \|Undefined Saprotroph\| | Yeast | NULL | Probable | As nectar-tap_saprotroph (Põlme et al. 2020); Decay sugar-rich substrates (Põlme et al. 2020) | Põlme S, et al. 2020. Fungal Diversity 105:1-16 ((https://doi.org/10.1007/s13225-020-00466-2)) |
| Otu39908 | 1 | k__Fungi(100);p__Ascomycota(100);c__Saccharomycetes(100);o__Saccharomycetales(100);f__Metschnikowiaceae(100);g__Metschnikowia(100);s__Metschnikowia_pulcherrima(100); | Metschnikowia | 13 | Saprotroph | \|Undefined Saprotroph\| | Yeast | NULL | Probable | As nectar-tap_saprotroph (Põlme et al. 2020); Decay sugar-rich substrates (Põlme et al. 2020) | Põlme S, et al. 2020. Fungal Diversity 105:1-16 ((https://doi.org/10.1007/s13225-020-00466-2)) |
| Otu39909 | 1 | k__Fungi(100);p__Ascomycota(100);c__Saccharomycetes(100);o__Saccharomycetales(100);f__Metschnikowiaceae(100);g__Metschnikowia(100);s__Metschnikowia_pulcherrima(100); | Metschnikowia | 13 | Saprotroph | \|Undefined Saprotroph\| | Yeast | NULL | Probable | As nectar-tap_saprotroph (Põlme et al. 2020); Decay sugar-rich substrates (Põlme et al. 2020) | Põlme S, et al. 2020. Fungal Diversity 105:1-16 ((https://doi.org/10.1007/s13225-020-00466-2)) |
| Otu39910 | 1 | k__Fungi(100);p__Ascomycota(100);c__Saccharomycetes(100);o__Saccharomycetales(100);f__Metschnikowiaceae(100);g__Metschnikowia(100);s__Metschnikowia_pulcherrima(100); | Metschnikowia | 13 | Saprotroph | \|Undefined Saprotroph\| | Yeast | NULL | Probable | As nectar-tap_saprotroph (Põlme et al. 2020); Decay sugar-rich substrates (Põlme et al. 2020) | Põlme S, et al. 2020. Fungal Diversity 105:1-16 ((https://doi.org/10.1007/s13225-020-00466-2)) |
| Otu39911 | 1 | k__Fungi(100);p__Ascomycota(100);c__Saccharomycetes(100);o__Saccharomycetales(100);f__Metschnikowiaceae(100);g__Metschnikowia(100);s__Metschnikowia_pulcherrima(100); | Metschnikowia | 13 | Saprotroph | \|Undefined Saprotroph\| | Yeast | NULL | Probable | As nectar-tap_saprotroph (Põlme et al. 2020); Decay sugar-rich substrates (Põlme et al. 2020) | Põlme S, et al. 2020. Fungal Diversity 105:1-16 ((https://doi.org/10.1007/s13225-020-00466-2)) |
| Otu39912 | 1 | k__Fungi(100);p__Ascomycota(100);c__Saccharomycetes(100);o__Saccharomycetales(100);f__Metschnikowiaceae(100);g__Metschnikowia(100);s__Metschnikowia_pulcherrima(100); | Metschnikowia | 13 | Saprotroph | \|Undefined Saprotroph\| | Yeast | NULL | Probable | As nectar-tap_saprotroph (Põlme et al. 2020); Decay sugar-rich substrates (Põlme et al. 2020) | Põlme S, et al. 2020. Fungal Diversity 105:1-16 ((https://doi.org/10.1007/s13225-020-00466-2)) |
| Otu39913 | 1 | k__Fungi(100);p__Ascomycota(100);c__Saccharomycetes(100);o__Saccharomycetales(100);f__Metschnikowiaceae(100);g__Metschnikowia(100);g__Metschnikowia_unclassified(100); | Metschnikowia | 13 | Saprotroph | \|Undefined Saprotroph\| | Yeast | NULL | Probable | As nectar-tap_saprotroph (Põlme et al. 2020); Decay sugar-rich substrates (Põlme et al. 2020) | Põlme S, et al. 2020. Fungal Diversity 105:1-16 ((https://doi.org/10.1007/s13225-020-00466-2)) |
| Otu39914 | 1 | k__Fungi(100);p__Ascomycota(100);c__Saccharomycetes(100);o__Saccharomycetales(100);f__Metschnikowiaceae(100);g__Metschnikowia(100);s__Metschnikowia_pulcherrima(100); | Metschnikowia | 13 | Saprotroph | \|Undefined Saprotroph\| | Yeast | NULL | Probable | As nectar-tap_saprotroph (Põlme et al. 2020); Decay sugar-rich substrates (Põlme et al. 2020) | Põlme S, et al. 2020. Fungal Diversity 105:1-16 ((https://doi.org/10.1007/s13225-020-00466-2)) |
| Otu39915 | 1 | k__Fungi(100);p__Ascomycota(100);c__Saccharomycetes(100);o__Saccharomycetales(100);f__Metschnikowiaceae(100);g__Metschnikowia(100);g__Metschnikowia_unclassified(100); | Metschnikowia | 13 | Saprotroph | \|Undefined Saprotroph\| | Yeast | NULL | Probable | As nectar-tap_saprotroph (Põlme et al. 2020); Decay sugar-rich substrates (Põlme et al. 2020) | Põlme S, et al. 2020. Fungal Diversity 105:1-16 ((https://doi.org/10.1007/s13225-020-00466-2)) |
| Otu39916 | 1 | k__Fungi(100);p__Ascomycota(100);c__Saccharomycetes(100);o__Saccharomycetales(100);f__Metschnikowiaceae(100);g__Metschnikowia(100);s__Metschnikowia_pulcherrima(100); | Metschnikowia | 13 | Saprotroph | \|Undefined Saprotroph\| | Yeast | NULL | Probable | As nectar-tap_saprotroph (Põlme et al. 2020); Decay sugar-rich substrates (Põlme et al. 2020) | Põlme S, et al. 2020. Fungal Diversity 105:1-16 ((https://doi.org/10.1007/s13225-020-00466-2)) |
| Otu39917 | 1 | k__Fungi(100);p__Ascomycota(100);c__Saccharomycetes(100);o__Saccharomycetales(100);f__Metschnikowiaceae(100);g__Metschnikowia(100);s__Metschnikowia_pulcherrima(100); | Metschnikowia | 13 | Saprotroph | \|Undefined Saprotroph\| | Yeast | NULL | Probable | As nectar-tap_saprotroph (Põlme et al. 2020); Decay sugar-rich substrates (Põlme et al. 2020) | Põlme S, et al. 2020. Fungal Diversity 105:1-16 ((https://doi.org/10.1007/s13225-020-00466-2)) |
| Otu39918 | 1 | k__Fungi(100);p__Ascomycota(100);c__Saccharomycetes(100);o__Saccharomycetales(100);f__Metschnikowiaceae(100);g__Metschnikowia(100);g__Metschnikowia_unclassified(100); | Metschnikowia | 13 | Saprotroph | \|Undefined Saprotroph\| | Yeast | NULL | Probable | As nectar-tap_saprotroph (Põlme et al. 2020); Decay sugar-rich substrates (Põlme et al. 2020) | Põlme S, et al. 2020. Fungal Diversity 105:1-16 ((https://doi.org/10.1007/s13225-020-00466-2)) |
| Otu39919 | 1 | k__Fungi(100);p__Ascomycota(100);c__Saccharomycetes(100);o__Saccharomycetales(100);f__Metschnikowiaceae(100);g__Metschnikowia(100);s__Metschnikowia_pulcherrima(100); | Metschnikowia | 13 | Saprotroph | \|Undefined Saprotroph\| | Yeast | NULL | Probable | As nectar-tap_saprotroph (Põlme et al. 2020); Decay sugar-rich substrates (Põlme et al. 2020) | Põlme S, et al. 2020. Fungal Diversity 105:1-16 ((https://doi.org/10.1007/s13225-020-00466-2)) |
| Otu39920 | 1 | k__Fungi(100);p__Ascomycota(100);c__Saccharomycetes(100);o__Saccharomycetales(100);f__Metschnikowiaceae(100);g__Metschnikowia(100);s__Metschnikowia_pulcherrima(100); | Metschnikowia | 13 | Saprotroph | \|Undefined Saprotroph\| | Yeast | NULL | Probable | As nectar-tap_saprotroph (Põlme et al. 2020); Decay sugar-rich substrates (Põlme et al. 2020) | Põlme S, et al. 2020. Fungal Diversity 105:1-16 ((https://doi.org/10.1007/s13225-020-00466-2)) |
| Otu39921 | 1 | k__Fungi(100);p__Ascomycota(100);c__Saccharomycetes(100);o__Saccharomycetales(100);f__Metschnikowiaceae(100);g__Metschnikowia(100);s__Metschnikowia_pulcherrima(100); | Metschnikowia | 13 | Saprotroph | \|Undefined Saprotroph\| | Yeast | NULL | Probable | As nectar-tap_saprotroph (Põlme et al. 2020); Decay sugar-rich substrates (Põlme et al. 2020) | Põlme S, et al. 2020. Fungal Diversity 105:1-16 ((https://doi.org/10.1007/s13225-020-00466-2)) |
| Otu39922 | 1 | k__Fungi(100);p__Ascomycota(100);c__Saccharomycetes(100);o__Saccharomycetales(100);f__Metschnikowiaceae(100);g__Metschnikowia(100);s__Metschnikowia_pulcherrima(100); | Metschnikowia | 13 | Saprotroph | \|Undefined Saprotroph\| | Yeast | NULL | Probable | As nectar-tap_saprotroph (Põlme et al. 2020); Decay sugar-rich substrates (Põlme et al. 2020) | Põlme S, et al. 2020. Fungal Diversity 105:1-16 ((https://doi.org/10.1007/s13225-020-00466-2)) |
| Otu39923 | 1 | k__Fungi(100);p__Ascomycota(100);c__Saccharomycetes(100);o__Saccharomycetales(100);f__Metschnikowiaceae(100);g__Metschnikowia(100);g__Metschnikowia_unclassified(100); | Metschnikowia | 13 | Saprotroph | \|Undefined Saprotroph\| | Yeast | NULL | Probable | As nectar-tap_saprotroph (Põlme et al. 2020); Decay sugar-rich substrates (Põlme et al. 2020) | Põlme S, et al. 2020. Fungal Diversity 105:1-16 ((https://doi.org/10.1007/s13225-020-00466-2)) |
| Otu39924 | 1 | k__Fungi(100);p__Ascomycota(100);c__Saccharomycetes(100);o__Saccharomycetales(100);f__Metschnikowiaceae(100);g__Metschnikowia(100);s__Metschnikowia_pulcherrima(100); | Metschnikowia | 13 | Saprotroph | \|Undefined Saprotroph\| | Yeast | NULL | Probable | As nectar-tap_saprotroph (Põlme et al. 2020); Decay sugar-rich substrates (Põlme et al. 2020) | Põlme S, et al. 2020. Fungal Diversity 105:1-16 ((https://doi.org/10.1007/s13225-020-00466-2)) |
| Otu39925 | 1 | k__Fungi(100);p__Ascomycota(100);c__Saccharomycetes(100);o__Saccharomycetales(100);f__Metschnikowiaceae(100);g__Metschnikowia(100);s__Metschnikowia_pulcherrima(100); | Metschnikowia | 13 | Saprotroph | \|Undefined Saprotroph\| | Yeast | NULL | Probable | As nectar-tap_saprotroph (Põlme et al. 2020); Decay sugar-rich substrates (Põlme et al. 2020) | Põlme S, et al. 2020. Fungal Diversity 105:1-16 ((https://doi.org/10.1007/s13225-020-00466-2)) |
| Otu39926 | 1 | k__Fungi(100);p__Ascomycota(100);c__Saccharomycetes(100);o__Saccharomycetales(100);f__Metschnikowiaceae(100);g__Metschnikowia(100);g__Metschnikowia_unclassified(100); | Metschnikowia | 13 | Saprotroph | \|Undefined Saprotroph\| | Yeast | NULL | Probable | As nectar-tap_saprotroph (Põlme et al. 2020); Decay sugar-rich substrates (Põlme et al. 2020) | Põlme S, et al. 2020. Fungal Diversity 105:1-16 ((https://doi.org/10.1007/s13225-020-00466-2)) |
| Otu39927 | 1 | k__Fungi(100);p__Ascomycota(100);c__Saccharomycetes(100);o__Saccharomycetales(100);f__Metschnikowiaceae(100);g__Metschnikowia(100);s__Metschnikowia_pulcherrima(100); | Metschnikowia | 13 | Saprotroph | \|Undefined Saprotroph\| | Yeast | NULL | Probable | As nectar-tap_saprotroph (Põlme et al. 2020); Decay sugar-rich substrates (Põlme et al. 2020) | Põlme S, et al. 2020. Fungal Diversity 105:1-16 ((https://doi.org/10.1007/s13225-020-00466-2)) |
| Otu39928 | 1 | k__Fungi(100);p__Ascomycota(100);c__Saccharomycetes(100);o__Saccharomycetales(100);f__Metschnikowiaceae(100);g__Metschnikowia(100);s__Metschnikowia_pulcherrima(100); | Metschnikowia | 13 | Saprotroph | \|Undefined Saprotroph\| | Yeast | NULL | Probable | As nectar-tap_saprotroph (Põlme et al. 2020); Decay sugar-rich substrates (Põlme et al. 2020) | Põlme S, et al. 2020. Fungal Diversity 105:1-16 ((https://doi.org/10.1007/s13225-020-00466-2)) |
| Otu39929 | 1 | k__Fungi(100);p__Ascomycota(100);c__Saccharomycetes(100);o__Saccharomycetales(100);f__Metschnikowiaceae(100);g__Metschnikowia(100);s__Metschnikowia_pulcherrima(100); | Metschnikowia | 13 | Saprotroph | \|Undefined Saprotroph\| | Yeast | NULL | Probable | As nectar-tap_saprotroph (Põlme et al. 2020); Decay sugar-rich substrates (Põlme et al. 2020) | Põlme S, et al. 2020. Fungal Diversity 105:1-16 ((https://doi.org/10.1007/s13225-020-00466-2)) |
| Otu39930 | 1 | k__Fungi(100);p__Ascomycota(100);c__Saccharomycetes(100);o__Saccharomycetales(100);f__Metschnikowiaceae(100);g__Metschnikowia(100);g__Metschnikowia_unclassified(100); | Metschnikowia | 13 | Saprotroph | \|Undefined Saprotroph\| | Yeast | NULL | Probable | As nectar-tap_saprotroph (Põlme et al. 2020); Decay sugar-rich substrates (Põlme et al. 2020) | Põlme S, et al. 2020. Fungal Diversity 105:1-16 ((https://doi.org/10.1007/s13225-020-00466-2)) |
| Otu39931 | 1 | k__Fungi(100);p__Ascomycota(100);c__Saccharomycetes(100);o__Saccharomycetales(100);f__Metschnikowiaceae(100);g__Metschnikowia(100);s__Metschnikowia_pulcherrima(100); | Metschnikowia | 13 | Saprotroph | \|Undefined Saprotroph\| | Yeast | NULL | Probable | As nectar-tap_saprotroph (Põlme et al. 2020); Decay sugar-rich substrates (Põlme et al. 2020) | Põlme S, et al. 2020. Fungal Diversity 105:1-16 ((https://doi.org/10.1007/s13225-020-00466-2)) |
| Otu39932 | 1 | k__Fungi(100);p__Ascomycota(100);c__Saccharomycetes(100);o__Saccharomycetales(100);f__Metschnikowiaceae(100);g__Metschnikowia(100);g__Metschnikowia_unclassified(100); | Metschnikowia | 13 | Saprotroph | \|Undefined Saprotroph\| | Yeast | NULL | Probable | As nectar-tap_saprotroph (Põlme et al. 2020); Decay sugar-rich substrates (Põlme et al. 2020) | Põlme S, et al. 2020. Fungal Diversity 105:1-16 ((https://doi.org/10.1007/s13225-020-00466-2)) |
| Otu39933 | 1 | k__Fungi(100);p__Ascomycota(100);c__Saccharomycetes(100);o__Saccharomycetales(100);f__Metschnikowiaceae(100);g__Metschnikowia(100);s__Metschnikowia_pulcherrima(100); | Metschnikowia | 13 | Saprotroph | \|Undefined Saprotroph\| | Yeast | NULL | Probable | As nectar-tap_saprotroph (Põlme et al. 2020); Decay sugar-rich substrates (Põlme et al. 2020) | Põlme S, et al. 2020. Fungal Diversity 105:1-16 ((https://doi.org/10.1007/s13225-020-00466-2)) |
| Otu39935 | 1 | k__Fungi(100);p__Ascomycota(100);c__Saccharomycetes(100);o__Saccharomycetales(100);f__Metschnikowiaceae(100);g__Metschnikowia(100);s__Metschnikowia_pulcherrima(100); | Metschnikowia | 13 | Saprotroph | \|Undefined Saprotroph\| | Yeast | NULL | Probable | As nectar-tap_saprotroph (Põlme et al. 2020); Decay sugar-rich substrates (Põlme et al. 2020) | Põlme S, et al. 2020. Fungal Diversity 105:1-16 ((https://doi.org/10.1007/s13225-020-00466-2)) |
| Otu39936 | 1 | k__Fungi(100);p__Ascomycota(100);c__Saccharomycetes(100);o__Saccharomycetales(100);f__Metschnikowiaceae(100);g__Metschnikowia(100);g__Metschnikowia_unclassified(100); | Metschnikowia | 13 | Saprotroph | \|Undefined Saprotroph\| | Yeast | NULL | Probable | As nectar-tap_saprotroph (Põlme et al. 2020); Decay sugar-rich substrates (Põlme et al. 2020) | Põlme S, et al. 2020. Fungal Diversity 105:1-16 ((https://doi.org/10.1007/s13225-020-00466-2)) |
| Otu39937 | 1 | k__Fungi(100);p__Ascomycota(100);c__Saccharomycetes(100);o__Saccharomycetales(100);f__Metschnikowiaceae(100);g__Metschnikowia(100);g__Metschnikowia_unclassified(100); | Metschnikowia | 13 | Saprotroph | \|Undefined Saprotroph\| | Yeast | NULL | Probable | As nectar-tap_saprotroph (Põlme et al. 2020); Decay sugar-rich substrates (Põlme et al. 2020) | Põlme S, et al. 2020. Fungal Diversity 105:1-16 ((https://doi.org/10.1007/s13225-020-00466-2)) |
| Otu39938 | 1 | k__Fungi(100);p__Ascomycota(100);c__Saccharomycetes(100);o__Saccharomycetales(100);f__Metschnikowiaceae(100);g__Metschnikowia(100);s__Metschnikowia_pulcherrima(100); | Metschnikowia | 13 | Saprotroph | \|Undefined Saprotroph\| | Yeast | NULL | Probable | As nectar-tap_saprotroph (Põlme et al. 2020); Decay sugar-rich substrates (Põlme et al. 2020) | Põlme S, et al. 2020. Fungal Diversity 105:1-16 ((https://doi.org/10.1007/s13225-020-00466-2)) |
| Otu39939 | 1 | k__Fungi(100);p__Ascomycota(100);c__Saccharomycetes(100);o__Saccharomycetales(100);f__Metschnikowiaceae(100);g__Metschnikowia(100);s__Metschnikowia_pulcherrima(100); | Metschnikowia | 13 | Saprotroph | \|Undefined Saprotroph\| | Yeast | NULL | Probable | As nectar-tap_saprotroph (Põlme et al. 2020); Decay sugar-rich substrates (Põlme et al. 2020) | Põlme S, et al. 2020. Fungal Diversity 105:1-16 ((https://doi.org/10.1007/s13225-020-00466-2)) |
| Otu39940 | 1 | k__Fungi(100);p__Ascomycota(100);c__Saccharomycetes(100);o__Saccharomycetales(100);f__Metschnikowiaceae(100);g__Metschnikowia(100);s__Metschnikowia_pulcherrima(100); | Metschnikowia | 13 | Saprotroph | \|Undefined Saprotroph\| | Yeast | NULL | Probable | As nectar-tap_saprotroph (Põlme et al. 2020); Decay sugar-rich substrates (Põlme et al. 2020) | Põlme S, et al. 2020. Fungal Diversity 105:1-16 ((https://doi.org/10.1007/s13225-020-00466-2)) |
| Otu39941 | 1 | k__Fungi(100);p__Ascomycota(100);c__Saccharomycetes(100);o__Saccharomycetales(100);f__Metschnikowiaceae(100);g__Metschnikowia(100);s__Metschnikowia_pulcherrima(100); | Metschnikowia | 13 | Saprotroph | \|Undefined Saprotroph\| | Yeast | NULL | Probable | As nectar-tap_saprotroph (Põlme et al. 2020); Decay sugar-rich substrates (Põlme et al. 2020) | Põlme S, et al. 2020. Fungal Diversity 105:1-16 ((https://doi.org/10.1007/s13225-020-00466-2)) |
| Otu39942 | 1 | k__Fungi(100);p__Ascomycota(100);c__Saccharomycetes(100);o__Saccharomycetales(100);f__Metschnikowiaceae(100);g__Metschnikowia(100);g__Metschnikowia_unclassified(100); | Metschnikowia | 13 | Saprotroph | \|Undefined Saprotroph\| | Yeast | NULL | Probable | As nectar-tap_saprotroph (Põlme et al. 2020); Decay sugar-rich substrates (Põlme et al. 2020) | Põlme S, et al. 2020. Fungal Diversity 105:1-16 ((https://doi.org/10.1007/s13225-020-00466-2)) |
| Otu39943 | 1 | k__Fungi(100);p__Ascomycota(100);c__Saccharomycetes(100);o__Saccharomycetales(100);f__Metschnikowiaceae(100);g__Metschnikowia(100);s__Metschnikowia_pulcherrima(100); | Metschnikowia | 13 | Saprotroph | \|Undefined Saprotroph\| | Yeast | NULL | Probable | As nectar-tap_saprotroph (Põlme et al. 2020); Decay sugar-rich substrates (Põlme et al. 2020) | Põlme S, et al. 2020. Fungal Diversity 105:1-16 ((https://doi.org/10.1007/s13225-020-00466-2)) |
| Otu39945 | 1 | k__Fungi(100);p__Ascomycota(100);c__Saccharomycetes(100);o__Saccharomycetales(100);f__Metschnikowiaceae(100);g__Metschnikowia(100);s__Metschnikowia_pulcherrima(100); | Metschnikowia | 13 | Saprotroph | \|Undefined Saprotroph\| | Yeast | NULL | Probable | As nectar-tap_saprotroph (Põlme et al. 2020); Decay sugar-rich substrates (Põlme et al. 2020) | Põlme S, et al. 2020. Fungal Diversity 105:1-16 ((https://doi.org/10.1007/s13225-020-00466-2)) |
| Otu39946 | 1 | k__Fungi(100);p__Ascomycota(100);c__Saccharomycetes(100);o__Saccharomycetales(100);f__Metschnikowiaceae(100);g__Metschnikowia(100);s__Metschnikowia_pulcherrima(100); | Metschnikowia | 13 | Saprotroph | \|Undefined Saprotroph\| | Yeast | NULL | Probable | As nectar-tap_saprotroph (Põlme et al. 2020); Decay sugar-rich substrates (Põlme et al. 2020) | Põlme S, et al. 2020. Fungal Diversity 105:1-16 ((https://doi.org/10.1007/s13225-020-00466-2)) |
| Otu39947 | 1 | k__Fungi(100);p__Ascomycota(100);c__Saccharomycetes(100);o__Saccharomycetales(100);f__Metschnikowiaceae(100);g__Metschnikowia(100);g__Metschnikowia_unclassified(100); | Metschnikowia | 13 | Saprotroph | \|Undefined Saprotroph\| | Yeast | NULL | Probable | As nectar-tap_saprotroph (Põlme et al. 2020); Decay sugar-rich substrates (Põlme et al. 2020) | Põlme S, et al. 2020. Fungal Diversity 105:1-16 ((https://doi.org/10.1007/s13225-020-00466-2)) |
| Otu39948 | 1 | k__Fungi(100);p__Ascomycota(100);c__Saccharomycetes(100);o__Saccharomycetales(100);f__Metschnikowiaceae(100);g__Metschnikowia(100);g__Metschnikowia_unclassified(100); | Metschnikowia | 13 | Saprotroph | \|Undefined Saprotroph\| | Yeast | NULL | Probable | As nectar-tap_saprotroph (Põlme et al. 2020); Decay sugar-rich substrates (Põlme et al. 2020) | Põlme S, et al. 2020. Fungal Diversity 105:1-16 ((https://doi.org/10.1007/s13225-020-00466-2)) |
| Otu39949 | 1 | k__Fungi(100);p__Ascomycota(100);c__Saccharomycetes(100);o__Saccharomycetales(100);f__Metschnikowiaceae(100);g__Metschnikowia(100);s__Metschnikowia_pulcherrima(100); | Metschnikowia | 13 | Saprotroph | \|Undefined Saprotroph\| | Yeast | NULL | Probable | As nectar-tap_saprotroph (Põlme et al. 2020); Decay sugar-rich substrates (Põlme et al. 2020) | Põlme S, et al. 2020. Fungal Diversity 105:1-16 ((https://doi.org/10.1007/s13225-020-00466-2)) |
| Otu39950 | 1 | k__Fungi(100);p__Ascomycota(100);c__Saccharomycetes(100);o__Saccharomycetales(100);f__Metschnikowiaceae(100);g__Metschnikowia(100);g__Metschnikowia_unclassified(100); | Metschnikowia | 13 | Saprotroph | \|Undefined Saprotroph\| | Yeast | NULL | Probable | As nectar-tap_saprotroph (Põlme et al. 2020); Decay sugar-rich substrates (Põlme et al. 2020) | Põlme S, et al. 2020. Fungal Diversity 105:1-16 ((https://doi.org/10.1007/s13225-020-00466-2)) |
| Otu39951 | 1 | k__Fungi(100);p__Ascomycota(100);c__Saccharomycetes(100);o__Saccharomycetales(100);f__Metschnikowiaceae(100);g__Metschnikowia(100);s__Metschnikowia_pulcherrima(100); | Metschnikowia | 13 | Saprotroph | \|Undefined Saprotroph\| | Yeast | NULL | Probable | As nectar-tap_saprotroph (Põlme et al. 2020); Decay sugar-rich substrates (Põlme et al. 2020) | Põlme S, et al. 2020. Fungal Diversity 105:1-16 ((https://doi.org/10.1007/s13225-020-00466-2)) |
| Otu39952 | 1 | k__Fungi(100);p__Ascomycota(100);c__Saccharomycetes(100);o__Saccharomycetales(100);f__Metschnikowiaceae(100);g__Metschnikowia(100);s__Metschnikowia_pulcherrima(100); | Metschnikowia | 13 | Saprotroph | \|Undefined Saprotroph\| | Yeast | NULL | Probable | As nectar-tap_saprotroph (Põlme et al. 2020); Decay sugar-rich substrates (Põlme et al. 2020) | Põlme S, et al. 2020. Fungal Diversity 105:1-16 ((https://doi.org/10.1007/s13225-020-00466-2)) |
| Otu39953 | 1 | k__Fungi(100);p__Ascomycota(100);c__Saccharomycetes(100);o__Saccharomycetales(100);f__Metschnikowiaceae(100);g__Metschnikowia(100);g__Metschnikowia_unclassified(100); | Metschnikowia | 13 | Saprotroph | \|Undefined Saprotroph\| | Yeast | NULL | Probable | As nectar-tap_saprotroph (Põlme et al. 2020); Decay sugar-rich substrates (Põlme et al. 2020) | Põlme S, et al. 2020. Fungal Diversity 105:1-16 ((https://doi.org/10.1007/s13225-020-00466-2)) |
| Otu39954 | 1 | k__Fungi(100);p__Ascomycota(100);c__Saccharomycetes(100);o__Saccharomycetales(100);f__Metschnikowiaceae(100);g__Metschnikowia(100);s__Metschnikowia_pulcherrima(100); | Metschnikowia | 13 | Saprotroph | \|Undefined Saprotroph\| | Yeast | NULL | Probable | As nectar-tap_saprotroph (Põlme et al. 2020); Decay sugar-rich substrates (Põlme et al. 2020) | Põlme S, et al. 2020. Fungal Diversity 105:1-16 ((https://doi.org/10.1007/s13225-020-00466-2)) |
| Otu39955 | 1 | k__Fungi(100);p__Ascomycota(100);c__Saccharomycetes(100);o__Saccharomycetales(100);f__Metschnikowiaceae(100);g__Metschnikowia(100);s__Metschnikowia_pulcherrima(100); | Metschnikowia | 13 | Saprotroph | \|Undefined Saprotroph\| | Yeast | NULL | Probable | As nectar-tap_saprotroph (Põlme et al. 2020); Decay sugar-rich substrates (Põlme et al. 2020) | Põlme S, et al. 2020. Fungal Diversity 105:1-16 ((https://doi.org/10.1007/s13225-020-00466-2)) |
| Otu39956 | 1 | k__Fungi(100);p__Ascomycota(100);c__Saccharomycetes(100);o__Saccharomycetales(100);f__Metschnikowiaceae(100);g__Metschnikowia(100);g__Metschnikowia_unclassified(100); | Metschnikowia | 13 | Saprotroph | \|Undefined Saprotroph\| | Yeast | NULL | Probable | As nectar-tap_saprotroph (Põlme et al. 2020); Decay sugar-rich substrates (Põlme et al. 2020) | Põlme S, et al. 2020. Fungal Diversity 105:1-16 ((https://doi.org/10.1007/s13225-020-00466-2)) |
| Otu39957 | 1 | k__Fungi(100);p__Ascomycota(100);c__Saccharomycetes(100);o__Saccharomycetales(100);f__Metschnikowiaceae(100);g__Metschnikowia(100);s__Metschnikowia_pulcherrima(100); | Metschnikowia | 13 | Saprotroph | \|Undefined Saprotroph\| | Yeast | NULL | Probable | As nectar-tap_saprotroph (Põlme et al. 2020); Decay sugar-rich substrates (Põlme et al. 2020) | Põlme S, et al. 2020. Fungal Diversity 105:1-16 ((https://doi.org/10.1007/s13225-020-00466-2)) |
| Otu39958 | 1 | k__Fungi(100);p__Ascomycota(100);c__Saccharomycetes(100);o__Saccharomycetales(100);f__Metschnikowiaceae(100);g__Metschnikowia(100);s__Metschnikowia_pulcherrima(100); | Metschnikowia | 13 | Saprotroph | \|Undefined Saprotroph\| | Yeast | NULL | Probable | As nectar-tap_saprotroph (Põlme et al. 2020); Decay sugar-rich substrates (Põlme et al. 2020) | Põlme S, et al. 2020. Fungal Diversity 105:1-16 ((https://doi.org/10.1007/s13225-020-00466-2)) |
| Otu39959 | 1 | k__Fungi(100);p__Ascomycota(100);c__Saccharomycetes(100);o__Saccharomycetales(100);f__Metschnikowiaceae(100);g__Metschnikowia(100);g__Metschnikowia_unclassified(100); | Metschnikowia | 13 | Saprotroph | \|Undefined Saprotroph\| | Yeast | NULL | Probable | As nectar-tap_saprotroph (Põlme et al. 2020); Decay sugar-rich substrates (Põlme et al. 2020) | Põlme S, et al. 2020. Fungal Diversity 105:1-16 ((https://doi.org/10.1007/s13225-020-00466-2)) |
| Otu39960 | 1 | k__Fungi(100);p__Ascomycota(100);c__Saccharomycetes(100);o__Saccharomycetales(100);f__Metschnikowiaceae(100);g__Metschnikowia(100);s__Metschnikowia_pulcherrima(100); | Metschnikowia | 13 | Saprotroph | \|Undefined Saprotroph\| | Yeast | NULL | Probable | As nectar-tap_saprotroph (Põlme et al. 2020); Decay sugar-rich substrates (Põlme et al. 2020) | Põlme S, et al. 2020. Fungal Diversity 105:1-16 ((https://doi.org/10.1007/s13225-020-00466-2)) |
| Otu39961 | 1 | k__Fungi(100);p__Ascomycota(100);c__Saccharomycetes(100);o__Saccharomycetales(100);f__Metschnikowiaceae(100);g__Metschnikowia(100);g__Metschnikowia_unclassified(100); | Metschnikowia | 13 | Saprotroph | \|Undefined Saprotroph\| | Yeast | NULL | Probable | As nectar-tap_saprotroph (Põlme et al. 2020); Decay sugar-rich substrates (Põlme et al. 2020) | Põlme S, et al. 2020. Fungal Diversity 105:1-16 ((https://doi.org/10.1007/s13225-020-00466-2)) |
| Otu39962 | 1 | k__Fungi(100);p__Ascomycota(100);c__Saccharomycetes(100);o__Saccharomycetales(100);f__Metschnikowiaceae(100);g__Metschnikowia(100);s__Metschnikowia_pulcherrima(100); | Metschnikowia | 13 | Saprotroph | \|Undefined Saprotroph\| | Yeast | NULL | Probable | As nectar-tap_saprotroph (Põlme et al. 2020); Decay sugar-rich substrates (Põlme et al. 2020) | Põlme S, et al. 2020. Fungal Diversity 105:1-16 ((https://doi.org/10.1007/s13225-020-00466-2)) |
| Otu39963 | 1 | k__Fungi(100);p__Ascomycota(100);c__Saccharomycetes(100);o__Saccharomycetales(100);f__Metschnikowiaceae(100);g__Metschnikowia(100);s__Metschnikowia_pulcherrima(100); | Metschnikowia | 13 | Saprotroph | \|Undefined Saprotroph\| | Yeast | NULL | Probable | As nectar-tap_saprotroph (Põlme et al. 2020); Decay sugar-rich substrates (Põlme et al. 2020) | Põlme S, et al. 2020. Fungal Diversity 105:1-16 ((https://doi.org/10.1007/s13225-020-00466-2)) |
| Otu39964 | 1 | k__Fungi(100);p__Ascomycota(100);c__Saccharomycetes(100);o__Saccharomycetales(100);f__Metschnikowiaceae(100);g__Metschnikowia(100);s__Metschnikowia_pulcherrima(100); | Metschnikowia | 13 | Saprotroph | \|Undefined Saprotroph\| | Yeast | NULL | Probable | As nectar-tap_saprotroph (Põlme et al. 2020); Decay sugar-rich substrates (Põlme et al. 2020) | Põlme S, et al. 2020. Fungal Diversity 105:1-16 ((https://doi.org/10.1007/s13225-020-00466-2)) |
| Otu39965 | 1 | k__Fungi(100);p__Ascomycota(100);c__Saccharomycetes(100);o__Saccharomycetales(100);f__Metschnikowiaceae(100);g__Metschnikowia(100);s__Metschnikowia_viticola(100); | Metschnikowia | 13 | Saprotroph | \|Undefined Saprotroph\| | Yeast | NULL | Probable | As nectar-tap_saprotroph (Põlme et al. 2020); Decay sugar-rich substrates (Põlme et al. 2020) | Põlme S, et al. 2020. Fungal Diversity 105:1-16 ((https://doi.org/10.1007/s13225-020-00466-2)) |
| Otu39966 | 1 | k__Fungi(100);p__Ascomycota(100);c__Saccharomycetes(100);o__Saccharomycetales(100);f__Metschnikowiaceae(100);g__Metschnikowia(100);s__Metschnikowia_pulcherrima(100); | Metschnikowia | 13 | Saprotroph | \|Undefined Saprotroph\| | Yeast | NULL | Probable | As nectar-tap_saprotroph (Põlme et al. 2020); Decay sugar-rich substrates (Põlme et al. 2020) | Põlme S, et al. 2020. Fungal Diversity 105:1-16 ((https://doi.org/10.1007/s13225-020-00466-2)) |
| Otu39967 | 1 | k__Fungi(100);p__Ascomycota(100);c__Saccharomycetes(100);o__Saccharomycetales(100);f__Metschnikowiaceae(100);g__Metschnikowia(100);s__Metschnikowia_pulcherrima(100); | Metschnikowia | 13 | Saprotroph | \|Undefined Saprotroph\| | Yeast | NULL | Probable | As nectar-tap_saprotroph (Põlme et al. 2020); Decay sugar-rich substrates (Põlme et al. 2020) | Põlme S, et al. 2020. Fungal Diversity 105:1-16 ((https://doi.org/10.1007/s13225-020-00466-2)) |
| Otu39968 | 1 | k__Fungi(100);p__Ascomycota(100);c__Saccharomycetes(100);o__Saccharomycetales(100);f__Metschnikowiaceae(100);g__Metschnikowia(100);s__Metschnikowia_pulcherrima(100); | Metschnikowia | 13 | Saprotroph | \|Undefined Saprotroph\| | Yeast | NULL | Probable | As nectar-tap_saprotroph (Põlme et al. 2020); Decay sugar-rich substrates (Põlme et al. 2020) | Põlme S, et al. 2020. Fungal Diversity 105:1-16 ((https://doi.org/10.1007/s13225-020-00466-2)) |
| Otu39969 | 1 | k__Fungi(100);p__Ascomycota(100);c__Saccharomycetes(100);o__Saccharomycetales(100);f__Metschnikowiaceae(100);g__Metschnikowia(100);s__Metschnikowia_pulcherrima(100); | Metschnikowia | 13 | Saprotroph | \|Undefined Saprotroph\| | Yeast | NULL | Probable | As nectar-tap_saprotroph (Põlme et al. 2020); Decay sugar-rich substrates (Põlme et al. 2020) | Põlme S, et al. 2020. Fungal Diversity 105:1-16 ((https://doi.org/10.1007/s13225-020-00466-2)) |
| Otu39970 | 1 | k__Fungi(100);p__Ascomycota(100);c__Saccharomycetes(100);o__Saccharomycetales(100);f__Metschnikowiaceae(100);g__Metschnikowia(100);s__Metschnikowia_pulcherrima(100); | Metschnikowia | 13 | Saprotroph | \|Undefined Saprotroph\| | Yeast | NULL | Probable | As nectar-tap_saprotroph (Põlme et al. 2020); Decay sugar-rich substrates (Põlme et al. 2020) | Põlme S, et al. 2020. Fungal Diversity 105:1-16 ((https://doi.org/10.1007/s13225-020-00466-2)) |
| Otu39971 | 1 | k__Fungi(100);p__Ascomycota(100);c__Saccharomycetes(100);o__Saccharomycetales(100);f__Metschnikowiaceae(100);g__Metschnikowia(100);s__Metschnikowia_pulcherrima(100); | Metschnikowia | 13 | Saprotroph | \|Undefined Saprotroph\| | Yeast | NULL | Probable | As nectar-tap_saprotroph (Põlme et al. 2020); Decay sugar-rich substrates (Põlme et al. 2020) | Põlme S, et al. 2020. Fungal Diversity 105:1-16 ((https://doi.org/10.1007/s13225-020-00466-2)) |
| Otu39972 | 1 | k__Fungi(100);p__Ascomycota(100);c__Saccharomycetes(100);o__Saccharomycetales(100);f__Metschnikowiaceae(100);g__Metschnikowia(100);g__Metschnikowia_unclassified(100); | Metschnikowia | 13 | Saprotroph | \|Undefined Saprotroph\| | Yeast | NULL | Probable | As nectar-tap_saprotroph (Põlme et al. 2020); Decay sugar-rich substrates (Põlme et al. 2020) | Põlme S, et al. 2020. Fungal Diversity 105:1-16 ((https://doi.org/10.1007/s13225-020-00466-2)) |
| Otu39973 | 1 | k__Fungi(100);p__Ascomycota(100);c__Saccharomycetes(100);o__Saccharomycetales(100);f__Metschnikowiaceae(100);g__Metschnikowia(100);s__Metschnikowia_pulcherrima(100); | Metschnikowia | 13 | Saprotroph | \|Undefined Saprotroph\| | Yeast | NULL | Probable | As nectar-tap_saprotroph (Põlme et al. 2020); Decay sugar-rich substrates (Põlme et al. 2020) | Põlme S, et al. 2020. Fungal Diversity 105:1-16 ((https://doi.org/10.1007/s13225-020-00466-2)) |
| Otu39974 | 1 | k__Fungi(100);p__Ascomycota(100);c__Saccharomycetes(100);o__Saccharomycetales(100);f__Metschnikowiaceae(100);g__Metschnikowia(100);s__Metschnikowia_pulcherrima(100); | Metschnikowia | 13 | Saprotroph | \|Undefined Saprotroph\| | Yeast | NULL | Probable | As nectar-tap_saprotroph (Põlme et al. 2020); Decay sugar-rich substrates (Põlme et al. 2020) | Põlme S, et al. 2020. Fungal Diversity 105:1-16 ((https://doi.org/10.1007/s13225-020-00466-2)) |
| Otu39975 | 1 | k__Fungi(100);p__Ascomycota(100);c__Saccharomycetes(100);o__Saccharomycetales(100);f__Metschnikowiaceae(100);g__Metschnikowia(100);s__Metschnikowia_pulcherrima(100); | Metschnikowia | 13 | Saprotroph | \|Undefined Saprotroph\| | Yeast | NULL | Probable | As nectar-tap_saprotroph (Põlme et al. 2020); Decay sugar-rich substrates (Põlme et al. 2020) | Põlme S, et al. 2020. Fungal Diversity 105:1-16 ((https://doi.org/10.1007/s13225-020-00466-2)) |
| Otu39976 | 1 | k__Fungi(100);p__Ascomycota(100);c__Saccharomycetes(100);o__Saccharomycetales(100);f__Metschnikowiaceae(100);g__Metschnikowia(100);s__Metschnikowia_pulcherrima(100); | Metschnikowia | 13 | Saprotroph | \|Undefined Saprotroph\| | Yeast | NULL | Probable | As nectar-tap_saprotroph (Põlme et al. 2020); Decay sugar-rich substrates (Põlme et al. 2020) | Põlme S, et al. 2020. Fungal Diversity 105:1-16 ((https://doi.org/10.1007/s13225-020-00466-2)) |
| Otu39977 | 1 | k__Fungi(100);p__Ascomycota(100);c__Saccharomycetes(100);o__Saccharomycetales(100);f__Metschnikowiaceae(100);g__Metschnikowia(100);s__Metschnikowia_pulcherrima(100); | Metschnikowia | 13 | Saprotroph | \|Undefined Saprotroph\| | Yeast | NULL | Probable | As nectar-tap_saprotroph (Põlme et al. 2020); Decay sugar-rich substrates (Põlme et al. 2020) | Põlme S, et al. 2020. Fungal Diversity 105:1-16 ((https://doi.org/10.1007/s13225-020-00466-2)) |
| Otu39978 | 1 | k__Fungi(100);p__Ascomycota(100);c__Saccharomycetes(100);o__Saccharomycetales(100);f__Metschnikowiaceae(100);g__Metschnikowia(100);g__Metschnikowia_unclassified(100); | Metschnikowia | 13 | Saprotroph | \|Undefined Saprotroph\| | Yeast | NULL | Probable | As nectar-tap_saprotroph (Põlme et al. 2020); Decay sugar-rich substrates (Põlme et al. 2020) | Põlme S, et al. 2020. Fungal Diversity 105:1-16 ((https://doi.org/10.1007/s13225-020-00466-2)) |
| Otu39979 | 1 | k__Fungi(100);p__Ascomycota(100);c__Saccharomycetes(100);o__Saccharomycetales(100);f__Metschnikowiaceae(100);g__Metschnikowia(100);s__Metschnikowia_pulcherrima(100); | Metschnikowia | 13 | Saprotroph | \|Undefined Saprotroph\| | Yeast | NULL | Probable | As nectar-tap_saprotroph (Põlme et al. 2020); Decay sugar-rich substrates (Põlme et al. 2020) | Põlme S, et al. 2020. Fungal Diversity 105:1-16 ((https://doi.org/10.1007/s13225-020-00466-2)) |
| Otu39980 | 1 | k__Fungi(100);p__Ascomycota(100);c__Saccharomycetes(100);o__Saccharomycetales(100);f__Metschnikowiaceae(100);g__Metschnikowia(100);s__Metschnikowia_pulcherrima(100); | Metschnikowia | 13 | Saprotroph | \|Undefined Saprotroph\| | Yeast | NULL | Probable | As nectar-tap_saprotroph (Põlme et al. 2020); Decay sugar-rich substrates (Põlme et al. 2020) | Põlme S, et al. 2020. Fungal Diversity 105:1-16 ((https://doi.org/10.1007/s13225-020-00466-2)) |
| Otu39981 | 1 | k__Fungi(100);p__Ascomycota(100);c__Saccharomycetes(100);o__Saccharomycetales(100);f__Metschnikowiaceae(100);g__Metschnikowia(100);s__Metschnikowia_pulcherrima(100); | Metschnikowia | 13 | Saprotroph | \|Undefined Saprotroph\| | Yeast | NULL | Probable | As nectar-tap_saprotroph (Põlme et al. 2020); Decay sugar-rich substrates (Põlme et al. 2020) | Põlme S, et al. 2020. Fungal Diversity 105:1-16 ((https://doi.org/10.1007/s13225-020-00466-2)) |
| Otu39982 | 1 | k__Fungi(100);p__Ascomycota(100);c__Saccharomycetes(100);o__Saccharomycetales(100);f__Metschnikowiaceae(100);g__Metschnikowia(100);g__Metschnikowia_unclassified(100); | Metschnikowia | 13 | Saprotroph | \|Undefined Saprotroph\| | Yeast | NULL | Probable | As nectar-tap_saprotroph (Põlme et al. 2020); Decay sugar-rich substrates (Põlme et al. 2020) | Põlme S, et al. 2020. Fungal Diversity 105:1-16 ((https://doi.org/10.1007/s13225-020-00466-2)) |
| Otu39983 | 1 | k__Fungi(100);p__Ascomycota(100);c__Saccharomycetes(100);o__Saccharomycetales(100);f__Metschnikowiaceae(100);g__Metschnikowia(100);s__Metschnikowia_pulcherrima(100); | Metschnikowia | 13 | Saprotroph | \|Undefined Saprotroph\| | Yeast | NULL | Probable | As nectar-tap_saprotroph (Põlme et al. 2020); Decay sugar-rich substrates (Põlme et al. 2020) | Põlme S, et al. 2020. Fungal Diversity 105:1-16 ((https://doi.org/10.1007/s13225-020-00466-2)) |
| Otu39984 | 1 | k__Fungi(100);p__Ascomycota(100);c__Saccharomycetes(100);o__Saccharomycetales(100);f__Metschnikowiaceae(100);g__Metschnikowia(100);s__Metschnikowia_pulcherrima(100); | Metschnikowia | 13 | Saprotroph | \|Undefined Saprotroph\| | Yeast | NULL | Probable | As nectar-tap_saprotroph (Põlme et al. 2020); Decay sugar-rich substrates (Põlme et al. 2020) | Põlme S, et al. 2020. Fungal Diversity 105:1-16 ((https://doi.org/10.1007/s13225-020-00466-2)) |
| Otu39985 | 1 | k__Fungi(100);p__Ascomycota(100);c__Saccharomycetes(100);o__Saccharomycetales(100);f__Metschnikowiaceae(100);g__Metschnikowia(100);s__Metschnikowia_pulcherrima(100); | Metschnikowia | 13 | Saprotroph | \|Undefined Saprotroph\| | Yeast | NULL | Probable | As nectar-tap_saprotroph (Põlme et al. 2020); Decay sugar-rich substrates (Põlme et al. 2020) | Põlme S, et al. 2020. Fungal Diversity 105:1-16 ((https://doi.org/10.1007/s13225-020-00466-2)) |
| Otu39986 | 1 | k__Fungi(100);p__Ascomycota(100);c__Saccharomycetes(100);o__Saccharomycetales(100);f__Metschnikowiaceae(100);g__Metschnikowia(100);s__Metschnikowia_pulcherrima(100); | Metschnikowia | 13 | Saprotroph | \|Undefined Saprotroph\| | Yeast | NULL | Probable | As nectar-tap_saprotroph (Põlme et al. 2020); Decay sugar-rich substrates (Põlme et al. 2020) | Põlme S, et al. 2020. Fungal Diversity 105:1-16 ((https://doi.org/10.1007/s13225-020-00466-2)) |
| Otu39987 | 1 | k__Fungi(100);p__Ascomycota(100);c__Saccharomycetes(100);o__Saccharomycetales(100);f__Metschnikowiaceae(100);g__Metschnikowia(100);s__Metschnikowia_pulcherrima(100); | Metschnikowia | 13 | Saprotroph | \|Undefined Saprotroph\| | Yeast | NULL | Probable | As nectar-tap_saprotroph (Põlme et al. 2020); Decay sugar-rich substrates (Põlme et al. 2020) | Põlme S, et al. 2020. Fungal Diversity 105:1-16 ((https://doi.org/10.1007/s13225-020-00466-2)) |
| Otu39988 | 1 | k__Fungi(100);p__Ascomycota(100);c__Saccharomycetes(100);o__Saccharomycetales(100);f__Metschnikowiaceae(100);g__Metschnikowia(100);s__Metschnikowia_pulcherrima(100); | Metschnikowia | 13 | Saprotroph | \|Undefined Saprotroph\| | Yeast | NULL | Probable | As nectar-tap_saprotroph (Põlme et al. 2020); Decay sugar-rich substrates (Põlme et al. 2020) | Põlme S, et al. 2020. Fungal Diversity 105:1-16 ((https://doi.org/10.1007/s13225-020-00466-2)) |
| Otu39990 | 1 | k__Fungi(100);p__Ascomycota(100);c__Saccharomycetes(100);o__Saccharomycetales(100);f__Metschnikowiaceae(100);g__Metschnikowia(100);g__Metschnikowia_unclassified(100); | Metschnikowia | 13 | Saprotroph | \|Undefined Saprotroph\| | Yeast | NULL | Probable | As nectar-tap_saprotroph (Põlme et al. 2020); Decay sugar-rich substrates (Põlme et al. 2020) | Põlme S, et al. 2020. Fungal Diversity 105:1-16 ((https://doi.org/10.1007/s13225-020-00466-2)) |
| Otu39991 | 1 | k__Fungi(100);p__Ascomycota(100);c__Saccharomycetes(100);o__Saccharomycetales(100);f__Metschnikowiaceae(100);g__Metschnikowia(100);g__Metschnikowia_unclassified(100); | Metschnikowia | 13 | Saprotroph | \|Undefined Saprotroph\| | Yeast | NULL | Probable | As nectar-tap_saprotroph (Põlme et al. 2020); Decay sugar-rich substrates (Põlme et al. 2020) | Põlme S, et al. 2020. Fungal Diversity 105:1-16 ((https://doi.org/10.1007/s13225-020-00466-2)) |
| Otu39992 | 1 | k__Fungi(100);p__Ascomycota(100);c__Saccharomycetes(100);o__Saccharomycetales(100);f__Metschnikowiaceae(100);g__Metschnikowia(100);g__Metschnikowia_unclassified(100); | Metschnikowia | 13 | Saprotroph | \|Undefined Saprotroph\| | Yeast | NULL | Probable | As nectar-tap_saprotroph (Põlme et al. 2020); Decay sugar-rich substrates (Põlme et al. 2020) | Põlme S, et al. 2020. Fungal Diversity 105:1-16 ((https://doi.org/10.1007/s13225-020-00466-2)) |
| Otu39993 | 1 | k__Fungi(100);p__Ascomycota(100);c__Saccharomycetes(100);o__Saccharomycetales(100);f__Metschnikowiaceae(100);g__Metschnikowia(100);g__Metschnikowia_unclassified(100); | Metschnikowia | 13 | Saprotroph | \|Undefined Saprotroph\| | Yeast | NULL | Probable | As nectar-tap_saprotroph (Põlme et al. 2020); Decay sugar-rich substrates (Põlme et al. 2020) | Põlme S, et al. 2020. Fungal Diversity 105:1-16 ((https://doi.org/10.1007/s13225-020-00466-2)) |
| Otu39994 | 1 | k__Fungi(100);p__Ascomycota(100);c__Saccharomycetes(100);o__Saccharomycetales(100);f__Metschnikowiaceae(100);g__Metschnikowia(100);s__Metschnikowia_pulcherrima(100); | Metschnikowia | 13 | Saprotroph | \|Undefined Saprotroph\| | Yeast | NULL | Probable | As nectar-tap_saprotroph (Põlme et al. 2020); Decay sugar-rich substrates (Põlme et al. 2020) | Põlme S, et al. 2020. Fungal Diversity 105:1-16 ((https://doi.org/10.1007/s13225-020-00466-2)) |
| Otu39995 | 1 | k__Fungi(100);p__Ascomycota(100);c__Saccharomycetes(100);o__Saccharomycetales(100);f__Metschnikowiaceae(100);g__Metschnikowia(100);s__Metschnikowia_pulcherrima(100); | Metschnikowia | 13 | Saprotroph | \|Undefined Saprotroph\| | Yeast | NULL | Probable | As nectar-tap_saprotroph (Põlme et al. 2020); Decay sugar-rich substrates (Põlme et al. 2020) | Põlme S, et al. 2020. Fungal Diversity 105:1-16 ((https://doi.org/10.1007/s13225-020-00466-2)) |
| Otu39996 | 1 | k__Fungi(100);p__Ascomycota(100);c__Saccharomycetes(100);o__Saccharomycetales(100);f__Metschnikowiaceae(100);g__Metschnikowia(100);g__Metschnikowia_unclassified(100); | Metschnikowia | 13 | Saprotroph | \|Undefined Saprotroph\| | Yeast | NULL | Probable | As nectar-tap_saprotroph (Põlme et al. 2020); Decay sugar-rich substrates (Põlme et al. 2020) | Põlme S, et al. 2020. Fungal Diversity 105:1-16 ((https://doi.org/10.1007/s13225-020-00466-2)) |
| Otu39997 | 1 | k__Fungi(100);p__Ascomycota(100);c__Saccharomycetes(100);o__Saccharomycetales(100);f__Metschnikowiaceae(100);g__Metschnikowia(100);s__Metschnikowia_pulcherrima(100); | Metschnikowia | 13 | Saprotroph | \|Undefined Saprotroph\| | Yeast | NULL | Probable | As nectar-tap_saprotroph (Põlme et al. 2020); Decay sugar-rich substrates (Põlme et al. 2020) | Põlme S, et al. 2020. Fungal Diversity 105:1-16 ((https://doi.org/10.1007/s13225-020-00466-2)) |
| Otu39998 | 1 | k__Fungi(100);p__Ascomycota(100);c__Saccharomycetes(100);o__Saccharomycetales(100);f__Metschnikowiaceae(100);g__Metschnikowia(100);s__Metschnikowia_pulcherrima(100); | Metschnikowia | 13 | Saprotroph | \|Undefined Saprotroph\| | Yeast | NULL | Probable | As nectar-tap_saprotroph (Põlme et al. 2020); Decay sugar-rich substrates (Põlme et al. 2020) | Põlme S, et al. 2020. Fungal Diversity 105:1-16 ((https://doi.org/10.1007/s13225-020-00466-2)) |
| Otu39999 | 1 | k__Fungi(100);p__Ascomycota(100);c__Saccharomycetes(100);o__Saccharomycetales(100);f__Metschnikowiaceae(100);g__Metschnikowia(100);s__Metschnikowia_pulcherrima(100); | Metschnikowia | 13 | Saprotroph | \|Undefined Saprotroph\| | Yeast | NULL | Probable | As nectar-tap_saprotroph (Põlme et al. 2020); Decay sugar-rich substrates (Põlme et al. 2020) | Põlme S, et al. 2020. Fungal Diversity 105:1-16 ((https://doi.org/10.1007/s13225-020-00466-2)) |
| Otu40000 | 1 | k__Fungi(100);p__Ascomycota(100);c__Saccharomycetes(100);o__Saccharomycetales(100);f__Metschnikowiaceae(100);g__Metschnikowia(100);s__Metschnikowia_pulcherrima(100); | Metschnikowia | 13 | Saprotroph | \|Undefined Saprotroph\| | Yeast | NULL | Probable | As nectar-tap_saprotroph (Põlme et al. 2020); Decay sugar-rich substrates (Põlme et al. 2020) | Põlme S, et al. 2020. Fungal Diversity 105:1-16 ((https://doi.org/10.1007/s13225-020-00466-2)) |
| Otu40001 | 1 | k__Fungi(100);p__Ascomycota(100);c__Saccharomycetes(100);o__Saccharomycetales(100);f__Metschnikowiaceae(100);g__Metschnikowia(100);g__Metschnikowia_unclassified(100); | Metschnikowia | 13 | Saprotroph | \|Undefined Saprotroph\| | Yeast | NULL | Probable | As nectar-tap_saprotroph (Põlme et al. 2020); Decay sugar-rich substrates (Põlme et al. 2020) | Põlme S, et al. 2020. Fungal Diversity 105:1-16 ((https://doi.org/10.1007/s13225-020-00466-2)) |
| Otu40002 | 1 | k__Fungi(100);p__Ascomycota(100);c__Saccharomycetes(100);o__Saccharomycetales(100);f__Metschnikowiaceae(100);g__Metschnikowia(100);s__Metschnikowia_pulcherrima(100); | Metschnikowia | 13 | Saprotroph | \|Undefined Saprotroph\| | Yeast | NULL | Probable | As nectar-tap_saprotroph (Põlme et al. 2020); Decay sugar-rich substrates (Põlme et al. 2020) | Põlme S, et al. 2020. Fungal Diversity 105:1-16 ((https://doi.org/10.1007/s13225-020-00466-2)) |
| Otu40003 | 1 | k__Fungi(100);p__Ascomycota(100);c__Saccharomycetes(100);o__Saccharomycetales(100);f__Metschnikowiaceae(100);g__Metschnikowia(100);g__Metschnikowia_unclassified(100); | Metschnikowia | 13 | Saprotroph | \|Undefined Saprotroph\| | Yeast | NULL | Probable | As nectar-tap_saprotroph (Põlme et al. 2020); Decay sugar-rich substrates (Põlme et al. 2020) | Põlme S, et al. 2020. Fungal Diversity 105:1-16 ((https://doi.org/10.1007/s13225-020-00466-2)) |
| Otu40004 | 1 | k__Fungi(100);p__Ascomycota(100);c__Saccharomycetes(100);o__Saccharomycetales(100);f__Metschnikowiaceae(100);g__Metschnikowia(100);s__Metschnikowia_pulcherrima(100); | Metschnikowia | 13 | Saprotroph | \|Undefined Saprotroph\| | Yeast | NULL | Probable | As nectar-tap_saprotroph (Põlme et al. 2020); Decay sugar-rich substrates (Põlme et al. 2020) | Põlme S, et al. 2020. Fungal Diversity 105:1-16 ((https://doi.org/10.1007/s13225-020-00466-2)) |
| Otu40005 | 1 | k__Fungi(100);p__Ascomycota(100);c__Saccharomycetes(100);o__Saccharomycetales(100);f__Metschnikowiaceae(100);g__Metschnikowia(100);s__Metschnikowia_pulcherrima(100); | Metschnikowia | 13 | Saprotroph | \|Undefined Saprotroph\| | Yeast | NULL | Probable | As nectar-tap_saprotroph (Põlme et al. 2020); Decay sugar-rich substrates (Põlme et al. 2020) | Põlme S, et al. 2020. Fungal Diversity 105:1-16 ((https://doi.org/10.1007/s13225-020-00466-2)) |
| Otu40006 | 1 | k__Fungi(100);p__Ascomycota(100);c__Saccharomycetes(100);o__Saccharomycetales(100);f__Metschnikowiaceae(100);g__Metschnikowia(100);s__Metschnikowia_pulcherrima(100); | Metschnikowia | 13 | Saprotroph | \|Undefined Saprotroph\| | Yeast | NULL | Probable | As nectar-tap_saprotroph (Põlme et al. 2020); Decay sugar-rich substrates (Põlme et al. 2020) | Põlme S, et al. 2020. Fungal Diversity 105:1-16 ((https://doi.org/10.1007/s13225-020-00466-2)) |
| Otu40007 | 1 | k__Fungi(100);p__Ascomycota(100);c__Saccharomycetes(100);o__Saccharomycetales(100);f__Metschnikowiaceae(100);g__Metschnikowia(100);g__Metschnikowia_unclassified(100); | Metschnikowia | 13 | Saprotroph | \|Undefined Saprotroph\| | Yeast | NULL | Probable | As nectar-tap_saprotroph (Põlme et al. 2020); Decay sugar-rich substrates (Põlme et al. 2020) | Põlme S, et al. 2020. Fungal Diversity 105:1-16 ((https://doi.org/10.1007/s13225-020-00466-2)) |
| Otu40008 | 1 | k__Fungi(100);p__Ascomycota(100);c__Saccharomycetes(100);o__Saccharomycetales(100);f__Metschnikowiaceae(100);g__Metschnikowia(100);s__Metschnikowia_pulcherrima(100); | Metschnikowia | 13 | Saprotroph | \|Undefined Saprotroph\| | Yeast | NULL | Probable | As nectar-tap_saprotroph (Põlme et al. 2020); Decay sugar-rich substrates (Põlme et al. 2020) | Põlme S, et al. 2020. Fungal Diversity 105:1-16 ((https://doi.org/10.1007/s13225-020-00466-2)) |
| Otu40009 | 1 | k__Fungi(100);p__Ascomycota(100);c__Saccharomycetes(100);o__Saccharomycetales(100);f__Metschnikowiaceae(100);g__Metschnikowia(100);s__Metschnikowia_pulcherrima(100); | Metschnikowia | 13 | Saprotroph | \|Undefined Saprotroph\| | Yeast | NULL | Probable | As nectar-tap_saprotroph (Põlme et al. 2020); Decay sugar-rich substrates (Põlme et al. 2020) | Põlme S, et al. 2020. Fungal Diversity 105:1-16 ((https://doi.org/10.1007/s13225-020-00466-2)) |
| Otu40010 | 1 | k__Fungi(100);p__Ascomycota(100);c__Saccharomycetes(100);o__Saccharomycetales(100);f__Metschnikowiaceae(100);g__Metschnikowia(100);s__Metschnikowia_pulcherrima(100); | Metschnikowia | 13 | Saprotroph | \|Undefined Saprotroph\| | Yeast | NULL | Probable | As nectar-tap_saprotroph (Põlme et al. 2020); Decay sugar-rich substrates (Põlme et al. 2020) | Põlme S, et al. 2020. Fungal Diversity 105:1-16 ((https://doi.org/10.1007/s13225-020-00466-2)) |
| Otu40011 | 1 | k__Fungi(100);p__Ascomycota(100);c__Saccharomycetes(100);o__Saccharomycetales(100);f__Metschnikowiaceae(100);g__Metschnikowia(100);g__Metschnikowia_unclassified(100); | Metschnikowia | 13 | Saprotroph | \|Undefined Saprotroph\| | Yeast | NULL | Probable | As nectar-tap_saprotroph (Põlme et al. 2020); Decay sugar-rich substrates (Põlme et al. 2020) | Põlme S, et al. 2020. Fungal Diversity 105:1-16 ((https://doi.org/10.1007/s13225-020-00466-2)) |
| Otu40012 | 1 | k__Fungi(100);p__Ascomycota(100);c__Saccharomycetes(100);o__Saccharomycetales(100);f__Metschnikowiaceae(100);g__Metschnikowia(100);g__Metschnikowia_unclassified(100); | Metschnikowia | 13 | Saprotroph | \|Undefined Saprotroph\| | Yeast | NULL | Probable | As nectar-tap_saprotroph (Põlme et al. 2020); Decay sugar-rich substrates (Põlme et al. 2020) | Põlme S, et al. 2020. Fungal Diversity 105:1-16 ((https://doi.org/10.1007/s13225-020-00466-2)) |
| Otu40013 | 1 | k__Fungi(100);p__Ascomycota(100);c__Saccharomycetes(100);o__Saccharomycetales(100);f__Metschnikowiaceae(100);g__Metschnikowia(100);s__Metschnikowia_pulcherrima(100); | Metschnikowia | 13 | Saprotroph | \|Undefined Saprotroph\| | Yeast | NULL | Probable | As nectar-tap_saprotroph (Põlme et al. 2020); Decay sugar-rich substrates (Põlme et al. 2020) | Põlme S, et al. 2020. Fungal Diversity 105:1-16 ((https://doi.org/10.1007/s13225-020-00466-2)) |
| Otu40014 | 1 | k__Fungi(100);p__Ascomycota(100);c__Saccharomycetes(100);o__Saccharomycetales(100);f__Metschnikowiaceae(100);g__Metschnikowia(100);s__Metschnikowia_pulcherrima(100); | Metschnikowia | 13 | Saprotroph | \|Undefined Saprotroph\| | Yeast | NULL | Probable | As nectar-tap_saprotroph (Põlme et al. 2020); Decay sugar-rich substrates (Põlme et al. 2020) | Põlme S, et al. 2020. Fungal Diversity 105:1-16 ((https://doi.org/10.1007/s13225-020-00466-2)) |
| Otu40015 | 1 | k__Fungi(100);p__Ascomycota(100);c__Saccharomycetes(100);o__Saccharomycetales(100);f__Metschnikowiaceae(100);g__Metschnikowia(100);s__Metschnikowia_pulcherrima(100); | Metschnikowia | 13 | Saprotroph | \|Undefined Saprotroph\| | Yeast | NULL | Probable | As nectar-tap_saprotroph (Põlme et al. 2020); Decay sugar-rich substrates (Põlme et al. 2020) | Põlme S, et al. 2020. Fungal Diversity 105:1-16 ((https://doi.org/10.1007/s13225-020-00466-2)) |
| Otu40016 | 1 | k__Fungi(100);p__Ascomycota(100);c__Saccharomycetes(100);o__Saccharomycetales(100);f__Metschnikowiaceae(100);g__Metschnikowia(100);s__Metschnikowia_pulcherrima(100); | Metschnikowia | 13 | Saprotroph | \|Undefined Saprotroph\| | Yeast | NULL | Probable | As nectar-tap_saprotroph (Põlme et al. 2020); Decay sugar-rich substrates (Põlme et al. 2020) | Põlme S, et al. 2020. Fungal Diversity 105:1-16 ((https://doi.org/10.1007/s13225-020-00466-2)) |
| Otu40017 | 1 | k__Fungi(100);p__Ascomycota(100);c__Saccharomycetes(100);o__Saccharomycetales(100);f__Metschnikowiaceae(100);g__Metschnikowia(100);g__Metschnikowia_unclassified(100); | Metschnikowia | 13 | Saprotroph | \|Undefined Saprotroph\| | Yeast | NULL | Probable | As nectar-tap_saprotroph (Põlme et al. 2020); Decay sugar-rich substrates (Põlme et al. 2020) | Põlme S, et al. 2020. Fungal Diversity 105:1-16 ((https://doi.org/10.1007/s13225-020-00466-2)) |
| Otu40018 | 1 | k__Fungi(100);p__Ascomycota(100);c__Saccharomycetes(100);o__Saccharomycetales(100);f__Metschnikowiaceae(100);g__Metschnikowia(100);g__Metschnikowia_unclassified(100); | Metschnikowia | 13 | Saprotroph | \|Undefined Saprotroph\| | Yeast | NULL | Probable | As nectar-tap_saprotroph (Põlme et al. 2020); Decay sugar-rich substrates (Põlme et al. 2020) | Põlme S, et al. 2020. Fungal Diversity 105:1-16 ((https://doi.org/10.1007/s13225-020-00466-2)) |
| Otu40019 | 1 | k__Fungi(100);p__Ascomycota(100);c__Saccharomycetes(100);o__Saccharomycetales(100);f__Metschnikowiaceae(100);g__Metschnikowia(100);s__Metschnikowia_pulcherrima(100); | Metschnikowia | 13 | Saprotroph | \|Undefined Saprotroph\| | Yeast | NULL | Probable | As nectar-tap_saprotroph (Põlme et al. 2020); Decay sugar-rich substrates (Põlme et al. 2020) | Põlme S, et al. 2020. Fungal Diversity 105:1-16 ((https://doi.org/10.1007/s13225-020-00466-2)) |
| Otu40020 | 1 | k__Fungi(100);p__Ascomycota(100);c__Saccharomycetes(100);o__Saccharomycetales(100);f__Metschnikowiaceae(100);g__Metschnikowia(100);g__Metschnikowia_unclassified(100); | Metschnikowia | 13 | Saprotroph | \|Undefined Saprotroph\| | Yeast | NULL | Probable | As nectar-tap_saprotroph (Põlme et al. 2020); Decay sugar-rich substrates (Põlme et al. 2020) | Põlme S, et al. 2020. Fungal Diversity 105:1-16 ((https://doi.org/10.1007/s13225-020-00466-2)) |
| Otu40021 | 1 | k__Fungi(100);p__Ascomycota(100);c__Saccharomycetes(100);o__Saccharomycetales(100);f__Metschnikowiaceae(100);g__Metschnikowia(100);g__Metschnikowia_unclassified(100); | Metschnikowia | 13 | Saprotroph | \|Undefined Saprotroph\| | Yeast | NULL | Probable | As nectar-tap_saprotroph (Põlme et al. 2020); Decay sugar-rich substrates (Põlme et al. 2020) | Põlme S, et al. 2020. Fungal Diversity 105:1-16 ((https://doi.org/10.1007/s13225-020-00466-2)) |
| Otu40022 | 1 | k__Fungi(100);p__Ascomycota(100);c__Saccharomycetes(100);o__Saccharomycetales(100);f__Metschnikowiaceae(100);g__Metschnikowia(100);s__Metschnikowia_pulcherrima(100); | Metschnikowia | 13 | Saprotroph | \|Undefined Saprotroph\| | Yeast | NULL | Probable | As nectar-tap_saprotroph (Põlme et al. 2020); Decay sugar-rich substrates (Põlme et al. 2020) | Põlme S, et al. 2020. Fungal Diversity 105:1-16 ((https://doi.org/10.1007/s13225-020-00466-2)) |
| Otu40023 | 1 | k__Fungi(100);p__Ascomycota(100);c__Saccharomycetes(100);o__Saccharomycetales(100);f__Metschnikowiaceae(100);g__Metschnikowia(100);g__Metschnikowia_unclassified(100); | Metschnikowia | 13 | Saprotroph | \|Undefined Saprotroph\| | Yeast | NULL | Probable | As nectar-tap_saprotroph (Põlme et al. 2020); Decay sugar-rich substrates (Põlme et al. 2020) | Põlme S, et al. 2020. Fungal Diversity 105:1-16 ((https://doi.org/10.1007/s13225-020-00466-2)) |
| Otu40024 | 1 | k__Fungi(100);p__Ascomycota(100);c__Saccharomycetes(100);o__Saccharomycetales(100);f__Metschnikowiaceae(100);g__Metschnikowia(100);s__Metschnikowia_pulcherrima(100); | Metschnikowia | 13 | Saprotroph | \|Undefined Saprotroph\| | Yeast | NULL | Probable | As nectar-tap_saprotroph (Põlme et al. 2020); Decay sugar-rich substrates (Põlme et al. 2020) | Põlme S, et al. 2020. Fungal Diversity 105:1-16 ((https://doi.org/10.1007/s13225-020-00466-2)) |
| Otu40025 | 1 | k__Fungi(100);p__Ascomycota(100);c__Saccharomycetes(100);o__Saccharomycetales(100);f__Metschnikowiaceae(100);g__Metschnikowia(100);s__Metschnikowia_pulcherrima(100); | Metschnikowia | 13 | Saprotroph | \|Undefined Saprotroph\| | Yeast | NULL | Probable | As nectar-tap_saprotroph (Põlme et al. 2020); Decay sugar-rich substrates (Põlme et al. 2020) | Põlme S, et al. 2020. Fungal Diversity 105:1-16 ((https://doi.org/10.1007/s13225-020-00466-2)) |
| Otu40026 | 1 | k__Fungi(100);p__Ascomycota(100);c__Saccharomycetes(100);o__Saccharomycetales(100);f__Metschnikowiaceae(100);g__Metschnikowia(100);s__Metschnikowia_pulcherrima(100); | Metschnikowia | 13 | Saprotroph | \|Undefined Saprotroph\| | Yeast | NULL | Probable | As nectar-tap_saprotroph (Põlme et al. 2020); Decay sugar-rich substrates (Põlme et al. 2020) | Põlme S, et al. 2020. Fungal Diversity 105:1-16 ((https://doi.org/10.1007/s13225-020-00466-2)) |
| Otu40027 | 1 | k__Fungi(100);p__Ascomycota(100);c__Saccharomycetes(100);o__Saccharomycetales(100);f__Metschnikowiaceae(100);g__Metschnikowia(100);g__Metschnikowia_unclassified(100); | Metschnikowia | 13 | Saprotroph | \|Undefined Saprotroph\| | Yeast | NULL | Probable | As nectar-tap_saprotroph (Põlme et al. 2020); Decay sugar-rich substrates (Põlme et al. 2020) | Põlme S, et al. 2020. Fungal Diversity 105:1-16 ((https://doi.org/10.1007/s13225-020-00466-2)) |
| Otu40028 | 1 | k__Fungi(100);p__Ascomycota(100);c__Saccharomycetes(100);o__Saccharomycetales(100);f__Metschnikowiaceae(100);g__Metschnikowia(100);s__Metschnikowia_pulcherrima(100); | Metschnikowia | 13 | Saprotroph | \|Undefined Saprotroph\| | Yeast | NULL | Probable | As nectar-tap_saprotroph (Põlme et al. 2020); Decay sugar-rich substrates (Põlme et al. 2020) | Põlme S, et al. 2020. Fungal Diversity 105:1-16 ((https://doi.org/10.1007/s13225-020-00466-2)) |
| Otu40029 | 1 | k__Fungi(100);p__Ascomycota(100);c__Saccharomycetes(100);o__Saccharomycetales(100);f__Metschnikowiaceae(100);g__Metschnikowia(100);g__Metschnikowia_unclassified(100); | Metschnikowia | 13 | Saprotroph | \|Undefined Saprotroph\| | Yeast | NULL | Probable | As nectar-tap_saprotroph (Põlme et al. 2020); Decay sugar-rich substrates (Põlme et al. 2020) | Põlme S, et al. 2020. Fungal Diversity 105:1-16 ((https://doi.org/10.1007/s13225-020-00466-2)) |
| Otu40030 | 1 | k__Fungi(100);p__Ascomycota(100);c__Saccharomycetes(100);o__Saccharomycetales(100);f__Metschnikowiaceae(100);g__Metschnikowia(100);s__Metschnikowia_pulcherrima(100); | Metschnikowia | 13 | Saprotroph | \|Undefined Saprotroph\| | Yeast | NULL | Probable | As nectar-tap_saprotroph (Põlme et al. 2020); Decay sugar-rich substrates (Põlme et al. 2020) | Põlme S, et al. 2020. Fungal Diversity 105:1-16 ((https://doi.org/10.1007/s13225-020-00466-2)) |
| Otu40031 | 1 | k__Fungi(100);p__Ascomycota(100);c__Saccharomycetes(100);o__Saccharomycetales(100);f__Metschnikowiaceae(100);g__Metschnikowia(100);s__Metschnikowia_pulcherrima(100); | Metschnikowia | 13 | Saprotroph | \|Undefined Saprotroph\| | Yeast | NULL | Probable | As nectar-tap_saprotroph (Põlme et al. 2020); Decay sugar-rich substrates (Põlme et al. 2020) | Põlme S, et al. 2020. Fungal Diversity 105:1-16 ((https://doi.org/10.1007/s13225-020-00466-2)) |
| Otu40032 | 1 | k__Fungi(100);p__Ascomycota(100);c__Saccharomycetes(100);o__Saccharomycetales(100);f__Metschnikowiaceae(100);g__Metschnikowia(100);s__Metschnikowia_pulcherrima(100); | Metschnikowia | 13 | Saprotroph | \|Undefined Saprotroph\| | Yeast | NULL | Probable | As nectar-tap_saprotroph (Põlme et al. 2020); Decay sugar-rich substrates (Põlme et al. 2020) | Põlme S, et al. 2020. Fungal Diversity 105:1-16 ((https://doi.org/10.1007/s13225-020-00466-2)) |
| Otu40033 | 1 | k__Fungi(100);p__Ascomycota(100);c__Saccharomycetes(100);o__Saccharomycetales(100);f__Metschnikowiaceae(100);g__Metschnikowia(100);s__Metschnikowia_pulcherrima(100); | Metschnikowia | 13 | Saprotroph | \|Undefined Saprotroph\| | Yeast | NULL | Probable | As nectar-tap_saprotroph (Põlme et al. 2020); Decay sugar-rich substrates (Põlme et al. 2020) | Põlme S, et al. 2020. Fungal Diversity 105:1-16 ((https://doi.org/10.1007/s13225-020-00466-2)) |
| Otu40034 | 1 | k__Fungi(100);p__Ascomycota(100);c__Saccharomycetes(100);o__Saccharomycetales(100);f__Metschnikowiaceae(100);g__Metschnikowia(100);s__Metschnikowia_pulcherrima(100); | Metschnikowia | 13 | Saprotroph | \|Undefined Saprotroph\| | Yeast | NULL | Probable | As nectar-tap_saprotroph (Põlme et al. 2020); Decay sugar-rich substrates (Põlme et al. 2020) | Põlme S, et al. 2020. Fungal Diversity 105:1-16 ((https://doi.org/10.1007/s13225-020-00466-2)) |
| Otu40035 | 1 | k__Fungi(100);p__Ascomycota(100);c__Saccharomycetes(100);o__Saccharomycetales(100);f__Metschnikowiaceae(100);g__Metschnikowia(100);s__Metschnikowia_pulcherrima(100); | Metschnikowia | 13 | Saprotroph | \|Undefined Saprotroph\| | Yeast | NULL | Probable | As nectar-tap_saprotroph (Põlme et al. 2020); Decay sugar-rich substrates (Põlme et al. 2020) | Põlme S, et al. 2020. Fungal Diversity 105:1-16 ((https://doi.org/10.1007/s13225-020-00466-2)) |
| Otu40036 | 1 | k__Fungi(100);p__Ascomycota(100);c__Saccharomycetes(100);o__Saccharomycetales(100);f__Metschnikowiaceae(100);g__Metschnikowia(100);g__Metschnikowia_unclassified(100); | Metschnikowia | 13 | Saprotroph | \|Undefined Saprotroph\| | Yeast | NULL | Probable | As nectar-tap_saprotroph (Põlme et al. 2020); Decay sugar-rich substrates (Põlme et al. 2020) | Põlme S, et al. 2020. Fungal Diversity 105:1-16 ((https://doi.org/10.1007/s13225-020-00466-2)) |
| Otu40037 | 1 | k__Fungi(100);p__Ascomycota(100);c__Saccharomycetes(100);o__Saccharomycetales(100);f__Metschnikowiaceae(100);g__Metschnikowia(100);g__Metschnikowia_unclassified(100); | Metschnikowia | 13 | Saprotroph | \|Undefined Saprotroph\| | Yeast | NULL | Probable | As nectar-tap_saprotroph (Põlme et al. 2020); Decay sugar-rich substrates (Põlme et al. 2020) | Põlme S, et al. 2020. Fungal Diversity 105:1-16 ((https://doi.org/10.1007/s13225-020-00466-2)) |
| Otu40038 | 1 | k__Fungi(100);p__Ascomycota(100);c__Saccharomycetes(100);o__Saccharomycetales(100);f__Metschnikowiaceae(100);g__Metschnikowia(100);g__Metschnikowia_unclassified(100); | Metschnikowia | 13 | Saprotroph | \|Undefined Saprotroph\| | Yeast | NULL | Probable | As nectar-tap_saprotroph (Põlme et al. 2020); Decay sugar-rich substrates (Põlme et al. 2020) | Põlme S, et al. 2020. Fungal Diversity 105:1-16 ((https://doi.org/10.1007/s13225-020-00466-2)) |
| Otu40039 | 1 | k__Fungi(100);p__Ascomycota(100);c__Saccharomycetes(100);o__Saccharomycetales(100);f__Metschnikowiaceae(100);g__Metschnikowia(100);s__Metschnikowia_pulcherrima(100); | Metschnikowia | 13 | Saprotroph | \|Undefined Saprotroph\| | Yeast | NULL | Probable | As nectar-tap_saprotroph (Põlme et al. 2020); Decay sugar-rich substrates (Põlme et al. 2020) | Põlme S, et al. 2020. Fungal Diversity 105:1-16 ((https://doi.org/10.1007/s13225-020-00466-2)) |
| Otu40040 | 1 | k__Fungi(100);p__Ascomycota(100);c__Saccharomycetes(100);o__Saccharomycetales(100);f__Metschnikowiaceae(100);g__Metschnikowia(100);g__Metschnikowia_unclassified(100); | Metschnikowia | 13 | Saprotroph | \|Undefined Saprotroph\| | Yeast | NULL | Probable | As nectar-tap_saprotroph (Põlme et al. 2020); Decay sugar-rich substrates (Põlme et al. 2020) | Põlme S, et al. 2020. Fungal Diversity 105:1-16 ((https://doi.org/10.1007/s13225-020-00466-2)) |
| Otu40041 | 1 | k__Fungi(100);p__Ascomycota(100);c__Saccharomycetes(100);o__Saccharomycetales(100);f__Metschnikowiaceae(100);g__Metschnikowia(100);s__Metschnikowia_pulcherrima(100); | Metschnikowia | 13 | Saprotroph | \|Undefined Saprotroph\| | Yeast | NULL | Probable | As nectar-tap_saprotroph (Põlme et al. 2020); Decay sugar-rich substrates (Põlme et al. 2020) | Põlme S, et al. 2020. Fungal Diversity 105:1-16 ((https://doi.org/10.1007/s13225-020-00466-2)) |
| Otu40042 | 1 | k__Fungi(100);p__Ascomycota(100);c__Saccharomycetes(100);o__Saccharomycetales(100);f__Metschnikowiaceae(100);g__Metschnikowia(100);s__Metschnikowia_pulcherrima(100); | Metschnikowia | 13 | Saprotroph | \|Undefined Saprotroph\| | Yeast | NULL | Probable | As nectar-tap_saprotroph (Põlme et al. 2020); Decay sugar-rich substrates (Põlme et al. 2020) | Põlme S, et al. 2020. Fungal Diversity 105:1-16 ((https://doi.org/10.1007/s13225-020-00466-2)) |
| Otu40043 | 1 | k__Fungi(100);p__Ascomycota(100);c__Saccharomycetes(100);o__Saccharomycetales(100);f__Metschnikowiaceae(100);g__Metschnikowia(100);s__Metschnikowia_pulcherrima(100); | Metschnikowia | 13 | Saprotroph | \|Undefined Saprotroph\| | Yeast | NULL | Probable | As nectar-tap_saprotroph (Põlme et al. 2020); Decay sugar-rich substrates (Põlme et al. 2020) | Põlme S, et al. 2020. Fungal Diversity 105:1-16 ((https://doi.org/10.1007/s13225-020-00466-2)) |
| Otu40044 | 1 | k__Fungi(100);p__Ascomycota(100);c__Saccharomycetes(100);o__Saccharomycetales(100);f__Metschnikowiaceae(100);g__Metschnikowia(100);s__Metschnikowia_pulcherrima(100); | Metschnikowia | 13 | Saprotroph | \|Undefined Saprotroph\| | Yeast | NULL | Probable | As nectar-tap_saprotroph (Põlme et al. 2020); Decay sugar-rich substrates (Põlme et al. 2020) | Põlme S, et al. 2020. Fungal Diversity 105:1-16 ((https://doi.org/10.1007/s13225-020-00466-2)) |
| Otu40045 | 1 | k__Fungi(100);p__Ascomycota(100);c__Saccharomycetes(100);o__Saccharomycetales(100);f__Metschnikowiaceae(100);g__Metschnikowia(100);g__Metschnikowia_unclassified(100); | Metschnikowia | 13 | Saprotroph | \|Undefined Saprotroph\| | Yeast | NULL | Probable | As nectar-tap_saprotroph (Põlme et al. 2020); Decay sugar-rich substrates (Põlme et al. 2020) | Põlme S, et al. 2020. Fungal Diversity 105:1-16 ((https://doi.org/10.1007/s13225-020-00466-2)) |
| Otu40046 | 1 | k__Fungi(100);p__Ascomycota(100);c__Saccharomycetes(100);o__Saccharomycetales(100);f__Metschnikowiaceae(100);g__Metschnikowia(100);g__Metschnikowia_unclassified(100); | Metschnikowia | 13 | Saprotroph | \|Undefined Saprotroph\| | Yeast | NULL | Probable | As nectar-tap_saprotroph (Põlme et al. 2020); Decay sugar-rich substrates (Põlme et al. 2020) | Põlme S, et al. 2020. Fungal Diversity 105:1-16 ((https://doi.org/10.1007/s13225-020-00466-2)) |
| Otu40047 | 1 | k__Fungi(100);p__Ascomycota(100);c__Saccharomycetes(100);o__Saccharomycetales(100);f__Metschnikowiaceae(100);g__Metschnikowia(100);s__Metschnikowia_pulcherrima(100); | Metschnikowia | 13 | Saprotroph | \|Undefined Saprotroph\| | Yeast | NULL | Probable | As nectar-tap_saprotroph (Põlme et al. 2020); Decay sugar-rich substrates (Põlme et al. 2020) | Põlme S, et al. 2020. Fungal Diversity 105:1-16 ((https://doi.org/10.1007/s13225-020-00466-2)) |
| Otu40048 | 1 | k__Fungi(100);p__Ascomycota(100);c__Saccharomycetes(100);o__Saccharomycetales(100);f__Metschnikowiaceae(100);g__Metschnikowia(100);s__Metschnikowia_pulcherrima(100); | Metschnikowia | 13 | Saprotroph | \|Undefined Saprotroph\| | Yeast | NULL | Probable | As nectar-tap_saprotroph (Põlme et al. 2020); Decay sugar-rich substrates (Põlme et al. 2020) | Põlme S, et al. 2020. Fungal Diversity 105:1-16 ((https://doi.org/10.1007/s13225-020-00466-2)) |
| Otu40049 | 1 | k__Fungi(100);p__Ascomycota(100);c__Saccharomycetes(100);o__Saccharomycetales(100);f__Metschnikowiaceae(100);g__Metschnikowia(100);s__Metschnikowia_pulcherrima(100); | Metschnikowia | 13 | Saprotroph | \|Undefined Saprotroph\| | Yeast | NULL | Probable | As nectar-tap_saprotroph (Põlme et al. 2020); Decay sugar-rich substrates (Põlme et al. 2020) | Põlme S, et al. 2020. Fungal Diversity 105:1-16 ((https://doi.org/10.1007/s13225-020-00466-2)) |
| Otu40050 | 1 | k__Fungi(100);p__Ascomycota(100);c__Saccharomycetes(100);o__Saccharomycetales(100);f__Metschnikowiaceae(100);g__Metschnikowia(100);s__Metschnikowia_pulcherrima(100); | Metschnikowia | 13 | Saprotroph | \|Undefined Saprotroph\| | Yeast | NULL | Probable | As nectar-tap_saprotroph (Põlme et al. 2020); Decay sugar-rich substrates (Põlme et al. 2020) | Põlme S, et al. 2020. Fungal Diversity 105:1-16 ((https://doi.org/10.1007/s13225-020-00466-2)) |
| Otu40051 | 1 | k__Fungi(100);p__Ascomycota(100);c__Saccharomycetes(100);o__Saccharomycetales(100);f__Metschnikowiaceae(100);g__Metschnikowia(100);s__Metschnikowia_pulcherrima(100); | Metschnikowia | 13 | Saprotroph | \|Undefined Saprotroph\| | Yeast | NULL | Probable | As nectar-tap_saprotroph (Põlme et al. 2020); Decay sugar-rich substrates (Põlme et al. 2020) | Põlme S, et al. 2020. Fungal Diversity 105:1-16 ((https://doi.org/10.1007/s13225-020-00466-2)) |
| Otu40052 | 1 | k__Fungi(100);p__Ascomycota(100);c__Saccharomycetes(100);o__Saccharomycetales(100);f__Metschnikowiaceae(100);g__Metschnikowia(100);g__Metschnikowia_unclassified(100); | Metschnikowia | 13 | Saprotroph | \|Undefined Saprotroph\| | Yeast | NULL | Probable | As nectar-tap_saprotroph (Põlme et al. 2020); Decay sugar-rich substrates (Põlme et al. 2020) | Põlme S, et al. 2020. Fungal Diversity 105:1-16 ((https://doi.org/10.1007/s13225-020-00466-2)) |
| Otu40053 | 1 | k__Fungi(100);p__Ascomycota(100);c__Saccharomycetes(100);o__Saccharomycetales(100);f__Metschnikowiaceae(100);g__Metschnikowia(100);s__Metschnikowia_pulcherrima(100); | Metschnikowia | 13 | Saprotroph | \|Undefined Saprotroph\| | Yeast | NULL | Probable | As nectar-tap_saprotroph (Põlme et al. 2020); Decay sugar-rich substrates (Põlme et al. 2020) | Põlme S, et al. 2020. Fungal Diversity 105:1-16 ((https://doi.org/10.1007/s13225-020-00466-2)) |
| Otu40054 | 1 | k__Fungi(100);p__Ascomycota(100);c__Saccharomycetes(100);o__Saccharomycetales(100);f__Metschnikowiaceae(100);g__Metschnikowia(100);s__Metschnikowia_pulcherrima(100); | Metschnikowia | 13 | Saprotroph | \|Undefined Saprotroph\| | Yeast | NULL | Probable | As nectar-tap_saprotroph (Põlme et al. 2020); Decay sugar-rich substrates (Põlme et al. 2020) | Põlme S, et al. 2020. Fungal Diversity 105:1-16 ((https://doi.org/10.1007/s13225-020-00466-2)) |
| Otu40055 | 1 | k__Fungi(100);p__Ascomycota(100);c__Saccharomycetes(100);o__Saccharomycetales(100);f__Metschnikowiaceae(100);g__Metschnikowia(100);g__Metschnikowia_unclassified(100); | Metschnikowia | 13 | Saprotroph | \|Undefined Saprotroph\| | Yeast | NULL | Probable | As nectar-tap_saprotroph (Põlme et al. 2020); Decay sugar-rich substrates (Põlme et al. 2020) | Põlme S, et al. 2020. Fungal Diversity 105:1-16 ((https://doi.org/10.1007/s13225-020-00466-2)) |
| Otu40056 | 1 | k__Fungi(100);p__Ascomycota(100);c__Saccharomycetes(100);o__Saccharomycetales(100);f__Metschnikowiaceae(100);g__Metschnikowia(100);s__Metschnikowia_pulcherrima(100); | Metschnikowia | 13 | Saprotroph | \|Undefined Saprotroph\| | Yeast | NULL | Probable | As nectar-tap_saprotroph (Põlme et al. 2020); Decay sugar-rich substrates (Põlme et al. 2020) | Põlme S, et al. 2020. Fungal Diversity 105:1-16 ((https://doi.org/10.1007/s13225-020-00466-2)) |
| Otu40057 | 1 | k__Fungi(100);p__Ascomycota(100);c__Saccharomycetes(100);o__Saccharomycetales(100);f__Metschnikowiaceae(100);g__Metschnikowia(100);s__Metschnikowia_pulcherrima(100); | Metschnikowia | 13 | Saprotroph | \|Undefined Saprotroph\| | Yeast | NULL | Probable | As nectar-tap_saprotroph (Põlme et al. 2020); Decay sugar-rich substrates (Põlme et al. 2020) | Põlme S, et al. 2020. Fungal Diversity 105:1-16 ((https://doi.org/10.1007/s13225-020-00466-2)) |
| Otu40058 | 1 | k__Fungi(100);p__Ascomycota(100);c__Saccharomycetes(100);o__Saccharomycetales(100);f__Metschnikowiaceae(100);g__Metschnikowia(100);s__Metschnikowia_pulcherrima(100); | Metschnikowia | 13 | Saprotroph | \|Undefined Saprotroph\| | Yeast | NULL | Probable | As nectar-tap_saprotroph (Põlme et al. 2020); Decay sugar-rich substrates (Põlme et al. 2020) | Põlme S, et al. 2020. Fungal Diversity 105:1-16 ((https://doi.org/10.1007/s13225-020-00466-2)) |
| Otu40059 | 1 | k__Fungi(100);p__Ascomycota(100);c__Saccharomycetes(100);o__Saccharomycetales(100);f__Metschnikowiaceae(100);g__Metschnikowia(100);s__Metschnikowia_pulcherrima(100); | Metschnikowia | 13 | Saprotroph | \|Undefined Saprotroph\| | Yeast | NULL | Probable | As nectar-tap_saprotroph (Põlme et al. 2020); Decay sugar-rich substrates (Põlme et al. 2020) | Põlme S, et al. 2020. Fungal Diversity 105:1-16 ((https://doi.org/10.1007/s13225-020-00466-2)) |
| Otu40060 | 1 | k__Fungi(100);p__Ascomycota(100);c__Saccharomycetes(100);o__Saccharomycetales(100);f__Metschnikowiaceae(100);g__Metschnikowia(100);g__Metschnikowia_unclassified(100); | Metschnikowia | 13 | Saprotroph | \|Undefined Saprotroph\| | Yeast | NULL | Probable | As nectar-tap_saprotroph (Põlme et al. 2020); Decay sugar-rich substrates (Põlme et al. 2020) | Põlme S, et al. 2020. Fungal Diversity 105:1-16 ((https://doi.org/10.1007/s13225-020-00466-2)) |
| Otu40061 | 1 | k__Fungi(100);p__Ascomycota(100);c__Saccharomycetes(100);o__Saccharomycetales(100);f__Metschnikowiaceae(100);g__Metschnikowia(100);g__Metschnikowia_unclassified(100); | Metschnikowia | 13 | Saprotroph | \|Undefined Saprotroph\| | Yeast | NULL | Probable | As nectar-tap_saprotroph (Põlme et al. 2020); Decay sugar-rich substrates (Põlme et al. 2020) | Põlme S, et al. 2020. Fungal Diversity 105:1-16 ((https://doi.org/10.1007/s13225-020-00466-2)) |
| Otu40062 | 1 | k__Fungi(100);p__Ascomycota(100);c__Saccharomycetes(100);o__Saccharomycetales(100);f__Metschnikowiaceae(100);g__Metschnikowia(100);s__Metschnikowia_pulcherrima(100); | Metschnikowia | 13 | Saprotroph | \|Undefined Saprotroph\| | Yeast | NULL | Probable | As nectar-tap_saprotroph (Põlme et al. 2020); Decay sugar-rich substrates (Põlme et al. 2020) | Põlme S, et al. 2020. Fungal Diversity 105:1-16 ((https://doi.org/10.1007/s13225-020-00466-2)) |
| Otu40063 | 1 | k__Fungi(100);p__Ascomycota(100);c__Saccharomycetes(100);o__Saccharomycetales(100);f__Metschnikowiaceae(100);g__Metschnikowia(100);s__Metschnikowia_pulcherrima(100); | Metschnikowia | 13 | Saprotroph | \|Undefined Saprotroph\| | Yeast | NULL | Probable | As nectar-tap_saprotroph (Põlme et al. 2020); Decay sugar-rich substrates (Põlme et al. 2020) | Põlme S, et al. 2020. Fungal Diversity 105:1-16 ((https://doi.org/10.1007/s13225-020-00466-2)) |
| Otu40064 | 1 | k__Fungi(100);p__Ascomycota(100);c__Saccharomycetes(100);o__Saccharomycetales(100);f__Metschnikowiaceae(100);g__Metschnikowia(100);s__Metschnikowia_pulcherrima(100); | Metschnikowia | 13 | Saprotroph | \|Undefined Saprotroph\| | Yeast | NULL | Probable | As nectar-tap_saprotroph (Põlme et al. 2020); Decay sugar-rich substrates (Põlme et al. 2020) | Põlme S, et al. 2020. Fungal Diversity 105:1-16 ((https://doi.org/10.1007/s13225-020-00466-2)) |
| Otu40065 | 1 | k__Fungi(100);p__Ascomycota(100);c__Saccharomycetes(100);o__Saccharomycetales(100);f__Metschnikowiaceae(100);g__Metschnikowia(100);s__Metschnikowia_pulcherrima(100); | Metschnikowia | 13 | Saprotroph | \|Undefined Saprotroph\| | Yeast | NULL | Probable | As nectar-tap_saprotroph (Põlme et al. 2020); Decay sugar-rich substrates (Põlme et al. 2020) | Põlme S, et al. 2020. Fungal Diversity 105:1-16 ((https://doi.org/10.1007/s13225-020-00466-2)) |
| Otu40066 | 1 | k__Fungi(100);p__Ascomycota(100);c__Saccharomycetes(100);o__Saccharomycetales(100);f__Metschnikowiaceae(100);g__Metschnikowia(100);s__Metschnikowia_pulcherrima(100); | Metschnikowia | 13 | Saprotroph | \|Undefined Saprotroph\| | Yeast | NULL | Probable | As nectar-tap_saprotroph (Põlme et al. 2020); Decay sugar-rich substrates (Põlme et al. 2020) | Põlme S, et al. 2020. Fungal Diversity 105:1-16 ((https://doi.org/10.1007/s13225-020-00466-2)) |
| Otu40067 | 1 | k__Fungi(100);p__Ascomycota(100);c__Saccharomycetes(100);o__Saccharomycetales(100);f__Metschnikowiaceae(100);g__Metschnikowia(100);s__Metschnikowia_pulcherrima(100); | Metschnikowia | 13 | Saprotroph | \|Undefined Saprotroph\| | Yeast | NULL | Probable | As nectar-tap_saprotroph (Põlme et al. 2020); Decay sugar-rich substrates (Põlme et al. 2020) | Põlme S, et al. 2020. Fungal Diversity 105:1-16 ((https://doi.org/10.1007/s13225-020-00466-2)) |
| Otu40068 | 1 | k__Fungi(100);p__Ascomycota(100);c__Saccharomycetes(100);o__Saccharomycetales(100);f__Metschnikowiaceae(100);g__Metschnikowia(100);s__Metschnikowia_pulcherrima(100); | Metschnikowia | 13 | Saprotroph | \|Undefined Saprotroph\| | Yeast | NULL | Probable | As nectar-tap_saprotroph (Põlme et al. 2020); Decay sugar-rich substrates (Põlme et al. 2020) | Põlme S, et al. 2020. Fungal Diversity 105:1-16 ((https://doi.org/10.1007/s13225-020-00466-2)) |
| Otu40069 | 1 | k__Fungi(100);p__Ascomycota(100);c__Saccharomycetes(100);o__Saccharomycetales(100);f__Metschnikowiaceae(100);g__Metschnikowia(100);s__Metschnikowia_pulcherrima(100); | Metschnikowia | 13 | Saprotroph | \|Undefined Saprotroph\| | Yeast | NULL | Probable | As nectar-tap_saprotroph (Põlme et al. 2020); Decay sugar-rich substrates (Põlme et al. 2020) | Põlme S, et al. 2020. Fungal Diversity 105:1-16 ((https://doi.org/10.1007/s13225-020-00466-2)) |

| | | | | | | | | | | | |
|---|---|---|---|---|---|---|---|---|---|---|---|
| Otu40070 | 1 | k__Fungi(100);p__Ascomycota(100);c__Saccharomycetes(100);o__Saccharomycetales(100);f__Metschnikowiaceae(100);g__Metschnikowia(100);s__Metschnikowia_pulcherrima(100); | Metschnikowia | 13 | Saprotroph | \|Undefined Saprotroph\| | Yeast | NULL | Probable | As nectar-tap_saprotroph (Põlme et al. 2020); Decay sugar-rich substrates (Põlme et al. 2020) | Põlme S, et al. 2020. Fungal Diversity 105:1-16 ((https://doi.org/10.1007/s13225-020-00466-2)) |
| Otu40071 | 1 | k__Fungi(100);p__Ascomycota(100);c__Saccharomycetes(100);o__Saccharomycetales(100);f__Metschnikowiaceae(100);g__Metschnikowia(100);g__Metschnikowia_unclassified(100); | Metschnikowia | 13 | Saprotroph | \|Undefined Saprotroph\| | Yeast | NULL | Probable | As nectar-tap_saprotroph (Põlme et al. 2020); Decay sugar-rich substrates (Põlme et al. 2020) | Põlme S, et al. 2020. Fungal Diversity 105:1-16 ((https://doi.org/10.1007/s13225-020-00466-2)) |
| Otu40072 | 1 | k__Fungi(100);p__Ascomycota(100);c__Saccharomycetes(100);o__Saccharomycetales(100);f__Metschnikowiaceae(100);g__Metschnikowia(100);g__Metschnikowia_unclassified(100); | Metschnikowia | 13 | Saprotroph | \|Undefined Saprotroph\| | Yeast | NULL | Probable | As nectar-tap_saprotroph (Põlme et al. 2020); Decay sugar-rich substrates (Põlme et al. 2020) | Põlme S, et al. 2020. Fungal Diversity 105:1-16 ((https://doi.org/10.1007/s13225-020-00466-2)) |
| Otu40073 | 1 | k__Fungi(100);p__Ascomycota(100);c__Saccharomycetes(100);o__Saccharomycetales(100);f__Metschnikowiaceae(100);g__Metschnikowia(100);g__Metschnikowia_unclassified(100); | Metschnikowia | 13 | Saprotroph | \|Undefined Saprotroph\| | Yeast | NULL | Probable | As nectar-tap_saprotroph (Põlme et al. 2020); Decay sugar-rich substrates (Põlme et al. 2020) | Põlme S, et al. 2020. Fungal Diversity 105:1-16 ((https://doi.org/10.1007/s13225-020-00466-2)) |
| Otu40074 | 1 | k__Fungi(100);p__Ascomycota(100);c__Saccharomycetes(100);o__Saccharomycetales(100);f__Metschnikowiaceae(100);g__Metschnikowia(100);g__Metschnikowia_unclassified(100); | Metschnikowia | 13 | Saprotroph | \|Undefined Saprotroph\| | Yeast | NULL | Probable | As nectar-tap_saprotroph (Põlme et al. 2020); Decay sugar-rich substrates (Põlme et al. 2020) | Põlme S, et al. 2020. Fungal Diversity 105:1-16 ((https://doi.org/10.1007/s13225-020-00466-2)) |
| Otu40075 | 1 | k__Fungi(100);p__Ascomycota(100);c__Saccharomycetes(100);o__Saccharomycetales(100);f__Metschnikowiaceae(100);g__Metschnikowia(100);s__Metschnikowia_pulcherrima(100); | Metschnikowia | 13 | Saprotroph | \|Undefined Saprotroph\| | Yeast | NULL | Probable | As nectar-tap_saprotroph (Põlme et al. 2020); Decay sugar-rich substrates (Põlme et al. 2020) | Põlme S, et al. 2020. Fungal Diversity 105:1-16 ((https://doi.org/10.1007/s13225-020-00466-2)) |
| Otu40076 | 1 | k__Fungi(100);p__Ascomycota(100);c__Saccharomycetes(100);o__Saccharomycetales(100);f__Metschnikowiaceae(100);g__Metschnikowia(100);s__Metschnikowia_pulcherrima(100); | Metschnikowia | 13 | Saprotroph | \|Undefined Saprotroph\| | Yeast | NULL | Probable | As nectar-tap_saprotroph (Põlme et al. 2020); Decay sugar-rich substrates (Põlme et al. 2020) | Põlme S, et al. 2020. Fungal Diversity 105:1-16 ((https://doi.org/10.1007/s13225-020-00466-2)) |
| Otu40077 | 1 | k__Fungi(100);p__Ascomycota(100);c__Saccharomycetes(100);o__Saccharomycetales(100);f__Metschnikowiaceae(100);g__Metschnikowia(100);s__Metschnikowia_pulcherrima(100); | Metschnikowia | 13 | Saprotroph | \|Undefined Saprotroph\| | Yeast | NULL | Probable | As nectar-tap_saprotroph (Põlme et al. 2020); Decay sugar-rich substrates (Põlme et al. 2020) | Põlme S, et al. 2020. Fungal Diversity 105:1-16 ((https://doi.org/10.1007/s13225-020-00466-2)) |
| Otu40078 | 1 | k__Fungi(100);p__Ascomycota(100);c__Saccharomycetes(100);o__Saccharomycetales(100);f__Metschnikowiaceae(100);g__Metschnikowia(100);s__Metschnikowia_pulcherrima(100); | Metschnikowia | 13 | Saprotroph | \|Undefined Saprotroph\| | Yeast | NULL | Probable | As nectar-tap_saprotroph (Põlme et al. 2020); Decay sugar-rich substrates (Põlme et al. 2020) | Põlme S, et al. 2020. Fungal Diversity 105:1-16 ((https://doi.org/10.1007/s13225-020-00466-2)) |
| Otu40079 | 1 | k__Fungi(100);p__Ascomycota(100);c__Saccharomycetes(100);o__Saccharomycetales(100);f__Metschnikowiaceae(100);g__Metschnikowia(100);s__Metschnikowia_pulcherrima(100); | Metschnikowia | 13 | Saprotroph | \|Undefined Saprotroph\| | Yeast | NULL | Probable | As nectar-tap_saprotroph (Põlme et al. 2020); Decay sugar-rich substrates (Põlme et al. 2020) | Põlme S, et al. 2020. Fungal Diversity 105:1-16 ((https://doi.org/10.1007/s13225-020-00466-2)) |
| Otu40080 | 1 | k__Fungi(100);p__Ascomycota(100);c__Saccharomycetes(100);o__Saccharomycetales(100);f__Metschnikowiaceae(100);g__Metschnikowia(100);s__Metschnikowia_pulcherrima(100); | Metschnikowia | 13 | Saprotroph | \|Undefined Saprotroph\| | Yeast | NULL | Probable | As nectar-tap_saprotroph (Põlme et al. 2020); Decay sugar-rich substrates (Põlme et al. 2020) | Põlme S, et al. 2020. Fungal Diversity 105:1-16 ((https://doi.org/10.1007/s13225-020-00466-2)) |
| Otu40081 | 1 | k__Fungi(100);p__Ascomycota(100);c__Saccharomycetes(100);o__Saccharomycetales(100);f__Metschnikowiaceae(100);g__Metschnikowia(100);s__Metschnikowia_pulcherrima(100); | Metschnikowia | 13 | Saprotroph | \|Undefined Saprotroph\| | Yeast | NULL | Probable | As nectar-tap_saprotroph (Põlme et al. 2020); Decay sugar-rich substrates (Põlme et al. 2020) | Põlme S, et al. 2020. Fungal Diversity 105:1-16 ((https://doi.org/10.1007/s13225-020-00466-2)) |
| Otu40082 | 1 | k__Fungi(100);p__Ascomycota(100);c__Saccharomycetes(100);o__Saccharomycetales(100);f__Metschnikowiaceae(100);g__Metschnikowia(100);g__Metschnikowia_unclassified(100); | Metschnikowia | 13 | Saprotroph | \|Undefined Saprotroph\| | Yeast | NULL | Probable | As nectar-tap_saprotroph (Põlme et al. 2020); Decay sugar-rich substrates (Põlme et al. 2020) | Põlme S, et al. 2020. Fungal Diversity 105:1-16 ((https://doi.org/10.1007/s13225-020-00466-2)) |
| Otu40083 | 1 | k__Fungi(100);p__Ascomycota(100);c__Saccharomycetes(100);o__Saccharomycetales(100);f__Metschnikowiaceae(100);g__Metschnikowia(100);s__Metschnikowia_pulcherrima(100); | Metschnikowia | 13 | Saprotroph | \|Undefined Saprotroph\| | Yeast | NULL | Probable | As nectar-tap_saprotroph (Põlme et al. 2020); Decay sugar-rich substrates (Põlme et al. 2020) | Põlme S, et al. 2020. Fungal Diversity 105:1-16 ((https://doi.org/10.1007/s13225-020-00466-2)) |
| Otu40084 | 1 | k__Fungi(100);p__Ascomycota(100);c__Saccharomycetes(100);o__Saccharomycetales(100);f__Metschnikowiaceae(100);g__Metschnikowia(100);g__Metschnikowia_unclassified(100); | Metschnikowia | 13 | Saprotroph | \|Undefined Saprotroph\| | Yeast | NULL | Probable | As nectar-tap_saprotroph (Põlme et al. 2020); Decay sugar-rich substrates (Põlme et al. 2020) | Põlme S, et al. 2020. Fungal Diversity 105:1-16 ((https://doi.org/10.1007/s13225-020-00466-2)) |
| Otu40085 | 1 | k__Fungi(100);p__Ascomycota(100);c__Saccharomycetes(100);o__Saccharomycetales(100);f__Metschnikowiaceae(100);g__Metschnikowia(100);g__Metschnikowia_unclassified(100); | Metschnikowia | 13 | Saprotroph | \|Undefined Saprotroph\| | Yeast | NULL | Probable | As nectar-tap_saprotroph (Põlme et al. 2020); Decay sugar-rich substrates (Põlme et al. 2020) | Põlme S, et al. 2020. Fungal Diversity 105:1-16 ((https://doi.org/10.1007/s13225-020-00466-2)) |
| Otu40086 | 1 | k__Fungi(100);p__Ascomycota(100);c__Saccharomycetes(100);o__Saccharomycetales(100);f__Metschnikowiaceae(100);g__Metschnikowia(100);s__Metschnikowia_pulcherrima(100); | Metschnikowia | 13 | Saprotroph | \|Undefined Saprotroph\| | Yeast | NULL | Probable | As nectar-tap_saprotroph (Põlme et al. 2020); Decay sugar-rich substrates (Põlme et al. 2020) | Põlme S, et al. 2020. Fungal Diversity 105:1-16 ((https://doi.org/10.1007/s13225-020-00466-2)) |
| Otu40088 | 1 | k__Fungi(100);p__Ascomycota(100);c__Saccharomycetes(100);o__Saccharomycetales(100);f__Metschnikowiaceae(100);g__Metschnikowia(100);s__Metschnikowia_pulcherrima(100); | Metschnikowia | 13 | Saprotroph | \|Undefined Saprotroph\| | Yeast | NULL | Probable | As nectar-tap_saprotroph (Põlme et al. 2020); Decay sugar-rich substrates (Põlme et al. 2020) | Põlme S, et al. 2020. Fungal Diversity 105:1-16 ((https://doi.org/10.1007/s13225-020-00466-2)) |
| Otu40089 | 1 | k__Fungi(100);p__Ascomycota(100);c__Saccharomycetes(100);o__Saccharomycetales(100);f__Metschnikowiaceae(100);g__Metschnikowia(100);s__Metschnikowia_pulcherrima(100); | Metschnikowia | 13 | Saprotroph | \|Undefined Saprotroph\| | Yeast | NULL | Probable | As nectar-tap_saprotroph (Põlme et al. 2020); Decay sugar-rich substrates (Põlme et al. 2020) | Põlme S, et al. 2020. Fungal Diversity 105:1-16 ((https://doi.org/10.1007/s13225-020-00466-2)) |
| Otu40090 | 1 | k__Fungi(100);p__Ascomycota(100);c__Saccharomycetes(100);o__Saccharomycetales(100);f__Metschnikowiaceae(100);g__Metschnikowia(100);s__Metschnikowia_pulcherrima(100); | Metschnikowia | 13 | Saprotroph | \|Undefined Saprotroph\| | Yeast | NULL | Probable | As nectar-tap_saprotroph (Põlme et al. 2020); Decay sugar-rich substrates (Põlme et al. 2020) | Põlme S, et al. 2020. Fungal Diversity 105:1-16 ((https://doi.org/10.1007/s13225-020-00466-2)) |
| Otu40091 | 1 | k__Fungi(100);p__Ascomycota(100);c__Saccharomycetes(100);o__Saccharomycetales(100);f__Metschnikowiaceae(100);g__Metschnikowia(100);s__Metschnikowia_pulcherrima(100); | Metschnikowia | 13 | Saprotroph | \|Undefined Saprotroph\| | Yeast | NULL | Probable | As nectar-tap_saprotroph (Põlme et al. 2020); Decay sugar-rich substrates (Põlme et al. 2020) | Põlme S, et al. 2020. Fungal Diversity 105:1-16 ((https://doi.org/10.1007/s13225-020-00466-2)) |
| Otu40092 | 1 | k__Fungi(100);p__Ascomycota(100);c__Saccharomycetes(100);o__Saccharomycetales(100);f__Metschnikowiaceae(100);g__Metschnikowia(100);s__Metschnikowia_pulcherrima(100); | Metschnikowia | 13 | Saprotroph | \|Undefined Saprotroph\| | Yeast | NULL | Probable | As nectar-tap_saprotroph (Põlme et al. 2020); Decay sugar-rich substrates (Põlme et al. 2020) | Põlme S, et al. 2020. Fungal Diversity 105:1-16 ((https://doi.org/10.1007/s13225-020-00466-2)) |
| Otu40093 | 1 | k__Fungi(100);p__Ascomycota(100);c__Saccharomycetes(100);o__Saccharomycetales(100);f__Metschnikowiaceae(100);g__Metschnikowia(100);s__Metschnikowia_pulcherrima(100); | Metschnikowia | 13 | Saprotroph | \|Undefined Saprotroph\| | Yeast | NULL | Probable | As nectar-tap_saprotroph (Põlme et al. 2020); Decay sugar-rich substrates (Põlme et al. 2020) | Põlme S, et al. 2020. Fungal Diversity 105:1-16 ((https://doi.org/10.1007/s13225-020-00466-2)) |
| Otu40094 | 1 | k__Fungi(100);p__Ascomycota(100);c__Saccharomycetes(100);o__Saccharomycetales(100);f__Metschnikowiaceae(100);g__Metschnikowia(100);s__Metschnikowia_pulcherrima(100); | Metschnikowia | 13 | Saprotroph | \|Undefined Saprotroph\| | Yeast | NULL | Probable | As nectar-tap_saprotroph (Põlme et al. 2020); Decay sugar-rich substrates (Põlme et al. 2020) | Põlme S, et al. 2020. Fungal Diversity 105:1-16 ((https://doi.org/10.1007/s13225-020-00466-2)) |
| Otu40095 | 1 | k__Fungi(100);p__Ascomycota(100);c__Saccharomycetes(100);o__Saccharomycetales(100);f__Metschnikowiaceae(100);g__Metschnikowia(100);g__Metschnikowia_unclassified(100); | Metschnikowia | 13 | Saprotroph | \|Undefined Saprotroph\| | Yeast | NULL | Probable | As nectar-tap_saprotroph (Põlme et al. 2020); Decay sugar-rich substrates (Põlme et al. 2020) | Põlme S, et al. 2020. Fungal Diversity 105:1-16 ((https://doi.org/10.1007/s13225-020-00466-2)) |
| Otu40096 | 1 | k__Fungi(100);p__Ascomycota(100);c__Saccharomycetes(100);o__Saccharomycetales(100);f__Metschnikowiaceae(100);g__Metschnikowia(100);s__Metschnikowia_pulcherrima(100); | Metschnikowia | 13 | Saprotroph | \|Undefined Saprotroph\| | Yeast | NULL | Probable | As nectar-tap_saprotroph (Põlme et al. 2020); Decay sugar-rich substrates (Põlme et al. 2020) | Põlme S, et al. 2020. Fungal Diversity 105:1-16 ((https://doi.org/10.1007/s13225-020-00466-2)) |
| Otu40097 | 1 | k__Fungi(100);p__Ascomycota(100);c__Saccharomycetes(100);o__Saccharomycetales(100);f__Metschnikowiaceae(100);g__Metschnikowia(100);s__Metschnikowia_pulcherrima(100); | Metschnikowia | 13 | Saprotroph | \|Undefined Saprotroph\| | Yeast | NULL | Probable | As nectar-tap_saprotroph (Põlme et al. 2020); Decay sugar-rich substrates (Põlme et al. 2020) | Põlme S, et al. 2020. Fungal Diversity 105:1-16 ((https://doi.org/10.1007/s13225-020-00466-2)) |
| Otu40098 | 1 | k__Fungi(100);p__Ascomycota(100);c__Saccharomycetes(100);o__Saccharomycetales(100);f__Metschnikowiaceae(100);g__Metschnikowia(100);s__Metschnikowia_pulcherrima(100); | Metschnikowia | 13 | Saprotroph | \|Undefined Saprotroph\| | Yeast | NULL | Probable | As nectar-tap_saprotroph (Põlme et al. 2020); Decay sugar-rich substrates (Põlme et al. 2020) | Põlme S, et al. 2020. Fungal Diversity 105:1-16 ((https://doi.org/10.1007/s13225-020-00466-2)) |
| Otu40099 | 1 | k__Fungi(100);p__Ascomycota(100);c__Saccharomycetes(100);o__Saccharomycetales(100);f__Metschnikowiaceae(100);g__Metschnikowia(100);s__Metschnikowia_pulcherrima(100); | Metschnikowia | 13 | Saprotroph | \|Undefined Saprotroph\| | Yeast | NULL | Probable | As nectar-tap_saprotroph (Põlme et al. 2020); Decay sugar-rich substrates (Põlme et al. 2020) | Põlme S, et al. 2020. Fungal Diversity 105:1-16 ((https://doi.org/10.1007/s13225-020-00466-2)) |
| Otu40100 | 1 | k__Fungi(100);p__Ascomycota(100);c__Saccharomycetes(100);o__Saccharomycetales(100);f__Metschnikowiaceae(100);g__Metschnikowia(100);s__Metschnikowia_pulcherrima(100); | Metschnikowia | 13 | Saprotroph | \|Undefined Saprotroph\| | Yeast | NULL | Probable | As nectar-tap_saprotroph (Põlme et al. 2020); Decay sugar-rich substrates (Põlme et al. 2020) | Põlme S, et al. 2020. Fungal Diversity 105:1-16 ((https://doi.org/10.1007/s13225-020-00466-2)) |
| Otu40101 | 1 | k__Fungi(100);p__Ascomycota(100);c__Saccharomycetes(100);o__Saccharomycetales(100);f__Metschnikowiaceae(100);g__Metschnikowia(100);s__Metschnikowia_pulcherrima(100); | Metschnikowia | 13 | Saprotroph | \|Undefined Saprotroph\| | Yeast | NULL | Probable | As nectar-tap_saprotroph (Põlme et al. 2020); Decay sugar-rich substrates (Põlme et al. 2020) | Põlme S, et al. 2020. Fungal Diversity 105:1-16 ((https://doi.org/10.1007/s13225-020-00466-2)) |
| Otu40102 | 1 | k__Fungi(100);p__Ascomycota(100);c__Saccharomycetes(100);o__Saccharomycetales(100);f__Metschnikowiaceae(100);g__Metschnikowia(100);s__Metschnikowia_pulcherrima(100); | Metschnikowia | 13 | Saprotroph | \|Undefined Saprotroph\| | Yeast | NULL | Probable | As nectar-tap_saprotroph (Põlme et al. 2020); Decay sugar-rich substrates (Põlme et al. 2020) | Põlme S, et al. 2020. Fungal Diversity 105:1-16 ((https://doi.org/10.1007/s13225-020-00466-2)) |
| Otu40103 | 1 | k__Fungi(100);p__Ascomycota(100);c__Saccharomycetes(100);o__Saccharomycetales(100);f__Metschnikowiaceae(100);g__Metschnikowia(100);s__Metschnikowia_pulcherrima(100); | Metschnikowia | 13 | Saprotroph | \|Undefined Saprotroph\| | Yeast | NULL | Probable | As nectar-tap_saprotroph (Põlme et al. 2020); Decay sugar-rich substrates (Põlme et al. 2020) | Põlme S, et al. 2020. Fungal Diversity 105:1-16 ((https://doi.org/10.1007/s13225-020-00466-2)) |
| Otu40104 | 1 | k__Fungi(100);p__Ascomycota(100);c__Saccharomycetes(100);o__Saccharomycetales(100);f__Metschnikowiaceae(100);g__Metschnikowia(100);s__Metschnikowia_pulcherrima(100); | Metschnikowia | 13 | Saprotroph | \|Undefined Saprotroph\| | Yeast | NULL | Probable | As nectar-tap_saprotroph (Põlme et al. 2020); Decay sugar-rich substrates (Põlme et al. 2020) | Põlme S, et al. 2020. Fungal Diversity 105:1-16 ((https://doi.org/10.1007/s13225-020-00466-2)) |
| Otu40105 | 1 | k__Fungi(100);p__Ascomycota(100);c__Saccharomycetes(100);o__Saccharomycetales(100);f__Metschnikowiaceae(100);g__Metschnikowia(100);g__Metschnikowia_unclassified(100); | Metschnikowia | 13 | Saprotroph | \|Undefined Saprotroph\| | Yeast | NULL | Probable | As nectar-tap_saprotroph (Põlme et al. 2020); Decay sugar-rich substrates (Põlme et al. 2020) | Põlme S, et al. 2020. Fungal Diversity 105:1-16 ((https://doi.org/10.1007/s13225-020-00466-2)) |
| Otu40106 | 1 | k__Fungi(100);p__Ascomycota(100);c__Saccharomycetes(100);o__Saccharomycetales(100);f__Metschnikowiaceae(100);g__Metschnikowia(100);s__Metschnikowia_pulcherrima(100); | Metschnikowia | 13 | Saprotroph | \|Undefined Saprotroph\| | Yeast | NULL | Probable | As nectar-tap_saprotroph (Põlme et al. 2020); Decay sugar-rich substrates (Põlme et al. 2020) | Põlme S, et al. 2020. Fungal Diversity 105:1-16 ((https://doi.org/10.1007/s13225-020-00466-2)) |
| Otu40107 | 1 | k__Fungi(100);p__Ascomycota(100);c__Saccharomycetes(100);o__Saccharomycetales(100);f__Metschnikowiaceae(100);g__Metschnikowia(100);s__Metschnikowia_pulcherrima(100); | Metschnikowia | 13 | Saprotroph | \|Undefined Saprotroph\| | Yeast | NULL | Probable | As nectar-tap_saprotroph (Põlme et al. 2020); Decay sugar-rich substrates (Põlme et al. 2020) | Põlme S, et al. 2020. Fungal Diversity 105:1-16 ((https://doi.org/10.1007/s13225-020-00466-2)) |
| Otu40108 | 1 | k__Fungi(100);p__Ascomycota(100);c__Saccharomycetes(100);o__Saccharomycetales(100);f__Metschnikowiaceae(100);g__Metschnikowia(100);s__Metschnikowia_pulcherrima(100); | Metschnikowia | 13 | Saprotroph | \|Undefined Saprotroph\| | Yeast | NULL | Probable | As nectar-tap_saprotroph (Põlme et al. 2020); Decay sugar-rich substrates (Põlme et al. 2020) | Põlme S, et al. 2020. Fungal Diversity 105:1-16 ((https://doi.org/10.1007/s13225-020-00466-2)) |
| Otu40109 | 1 | k__Fungi(100);p__Ascomycota(100);c__Saccharomycetes(100);o__Saccharomycetales(100);f__Metschnikowiaceae(100);g__Metschnikowia(100);s__Metschnikowia_pulcherrima(100); | Metschnikowia | 13 | Saprotroph | \|Undefined Saprotroph\| | Yeast | NULL | Probable | As nectar-tap_saprotroph (Põlme et al. 2020); Decay sugar-rich substrates (Põlme et al. 2020) | Põlme S, et al. 2020. Fungal Diversity 105:1-16 ((https://doi.org/10.1007/s13225-020-00466-2)) |
| Otu40110 | 1 | k__Fungi(100);p__Ascomycota(100);c__Saccharomycetes(100);o__Saccharomycetales(100);f__Metschnikowiaceae(100);g__Metschnikowia(100);s__Metschnikowia_pulcherrima(100); | Metschnikowia | 13 | Saprotroph | \|Undefined Saprotroph\| | Yeast | NULL | Probable | As nectar-tap_saprotroph (Põlme et al. 2020); Decay sugar-rich substrates (Põlme et al. 2020) | Põlme S, et al. 2020. Fungal Diversity 105:1-16 ((https://doi.org/10.1007/s13225-020-00466-2)) |
| Otu40111 | 1 | k__Fungi(100);p__Ascomycota(100);c__Saccharomycetes(100);o__Saccharomycetales(100);f__Metschnikowiaceae(100);g__Metschnikowia(100);g__Metschnikowia_unclassified(100); | Metschnikowia | 13 | Saprotroph | \|Undefined Saprotroph\| | Yeast | NULL | Probable | As nectar-tap_saprotroph (Põlme et al. 2020); Decay sugar-rich substrates (Põlme et al. 2020) | Põlme S, et al. 2020. Fungal Diversity 105:1-16 ((https://doi.org/10.1007/s13225-020-00466-2)) |
| Otu40112 | 1 | k__Fungi(100);p__Ascomycota(100);c__Saccharomycetes(100);o__Saccharomycetales(100);f__Metschnikowiaceae(100);g__Metschnikowia(100);s__Metschnikowia_pulcherrima(100); | Metschnikowia | 13 | Saprotroph | \|Undefined Saprotroph\| | Yeast | NULL | Probable | As nectar-tap_saprotroph (Põlme et al. 2020); Decay sugar-rich substrates (Põlme et al. 2020) | Põlme S, et al. 2020. Fungal Diversity 105:1-16 ((https://doi.org/10.1007/s13225-020-00466-2)) |
| Otu40113 | 1 | k__Fungi(100);p__Ascomycota(100);c__Saccharomycetes(100);o__Saccharomycetales(100);f__Metschnikowiaceae(100);g__Metschnikowia(100);s__Metschnikowia_pulcherrima(100); | Metschnikowia | 13 | Saprotroph | \|Undefined Saprotroph\| | Yeast | NULL | Probable | As nectar-tap_saprotroph (Põlme et al. 2020); Decay sugar-rich substrates (Põlme et al. 2020) | Põlme S, et al. 2020. Fungal Diversity 105:1-16 ((https://doi.org/10.1007/s13225-020-00466-2)) |
| Otu40114 | 1 | k__Fungi(100);p__Ascomycota(100);c__Saccharomycetes(100);o__Saccharomycetales(100);f__Metschnikowiaceae(100);g__Metschnikowia(100);s__Metschnikowia_pulcherrima(100); | Metschnikowia | 13 | Saprotroph | \|Undefined Saprotroph\| | Yeast | NULL | Probable | As nectar-tap_saprotroph (Põlme et al. 2020); Decay sugar-rich substrates (Põlme et al. 2020) | Põlme S, et al. 2020. Fungal Diversity 105:1-16 ((https://doi.org/10.1007/s13225-020-00466-2)) |
| Otu40116 | 1 | k__Fungi(100);p__Ascomycota(100);c__Saccharomycetes(100);o__Saccharomycetales(100);f__Metschnikowiaceae(100);g__Metschnikowia(100);s__Metschnikowia_pulcherrima(100); | Metschnikowia | 13 | Saprotroph | \|Undefined Saprotroph\| | Yeast | NULL | Probable | As nectar-tap_saprotroph (Põlme et al. 2020); Decay sugar-rich substrates (Põlme et al. 2020) | Põlme S, et al. 2020. Fungal Diversity 105:1-16 ((https://doi.org/10.1007/s13225-020-00466-2)) |
| Otu40118 | 1 | k__Fungi(100);p__Ascomycota(100);c__Saccharomycetes(100);o__Saccharomycetales(100);f__Metschnikowiaceae(100);g__Metschnikowia(100);s__Metschnikowia_pulcherrima(100); | Metschnikowia | 13 | Saprotroph | \|Undefined Saprotroph\| | Yeast | NULL | Probable | As nectar-tap_saprotroph (Põlme et al. 2020); Decay sugar-rich substrates (Põlme et al. 2020) | Põlme S, et al. 2020. Fungal Diversity 105:1-16 ((https://doi.org/10.1007/s13225-020-00466-2)) |
| Otu40119 | 1 | k__Fungi(100);p__Ascomycota(100);c__Saccharomycetes(100);o__Saccharomycetales(100);f__Metschnikowiaceae(100);g__Metschnikowia(100);s__Metschnikowia_pulcherrima(100); | Metschnikowia | 13 | Saprotroph | \|Undefined Saprotroph\| | Yeast | NULL | Probable | As nectar-tap_saprotroph (Põlme et al. 2020); Decay sugar-rich substrates (Põlme et al. 2020) | Põlme S, et al. 2020. Fungal Diversity 105:1-16 ((https://doi.org/10.1007/s13225-020-00466-2)) |
| Otu40120 | 1 | k__Fungi(100);p__Ascomycota(100);c__Saccharomycetes(100);o__Saccharomycetales(100);f__Metschnikowiaceae(100);g__Metschnikowia(100);g__Metschnikowia_unclassified(100); | Metschnikowia | 13 | Saprotroph | \|Undefined Saprotroph\| | Yeast | NULL | Probable | As nectar-tap_saprotroph (Põlme et al. 2020); Decay sugar-rich substrates (Põlme et al. 2020) | Põlme S, et al. 2020. Fungal Diversity 105:1-16 ((https://doi.org/10.1007/s13225-020-00466-2)) |
| Otu40121 | 1 | k__Fungi(100);p__Ascomycota(100);c__Saccharomycetes(100);o__Saccharomycetales(100);f__Metschnikowiaceae(100);g__Metschnikowia(100);s__Metschnikowia_pulcherrima(100); | Metschnikowia | 13 | Saprotroph | \|Undefined Saprotroph\| | Yeast | NULL | Probable | As nectar-tap_saprotroph (Põlme et al. 2020); Decay sugar-rich substrates (Põlme et al. 2020) | Põlme S, et al. 2020. Fungal Diversity 105:1-16 ((https://doi.org/10.1007/s13225-020-00466-2)) |
| Otu40122 | 1 | k__Fungi(100);p__Ascomycota(100);c__Saccharomycetes(100);o__Saccharomycetales(100);f__Metschnikowiaceae(100);g__Metschnikowia(100);s__Metschnikowia_pulcherrima(100); | Metschnikowia | 13 | Saprotroph | \|Undefined Saprotroph\| | Yeast | NULL | Probable | As nectar-tap_saprotroph (Põlme et al. 2020); Decay sugar-rich substrates (Põlme et al. 2020) | Põlme S, et al. 2020. Fungal Diversity 105:1-16 ((https://doi.org/10.1007/s13225-020-00466-2)) |
| Otu40123 | 1 | k__Fungi(100);p__Ascomycota(100);c__Saccharomycetes(100);o__Saccharomycetales(100);f__Metschnikowiaceae(100);g__Metschnikowia(100);s__Metschnikowia_pulcherrima(100); | Metschnikowia | 13 | Saprotroph | \|Undefined Saprotroph\| | Yeast | NULL | Probable | As nectar-tap_saprotroph (Põlme et al. 2020); Decay sugar-rich substrates (Põlme et al. 2020) | Põlme S, et al. 2020. Fungal Diversity 105:1-16 ((https://doi.org/10.1007/s13225-020-00466-2)) |
| Otu40124 | 1 | k__Fungi(100);p__Ascomycota(100);c__Saccharomycetes(100);o__Saccharomycetales(100);f__Metschnikowiaceae(100);g__Metschnikowia(100);g__Metschnikowia_unclassified(100); | Metschnikowia | 13 | Saprotroph | \|Undefined Saprotroph\| | Yeast | NULL | Probable | As nectar-tap_saprotroph (Põlme et al. 2020); Decay sugar-rich substrates (Põlme et al. 2020) | Põlme S, et al. 2020. Fungal Diversity 105:1-16 ((https://doi.org/10.1007/s13225-020-00466-2)) |
| Otu40125 | 1 | k__Fungi(100);p__Ascomycota(100);c__Saccharomycetes(100);o__Saccharomycetales(100);f__Metschnikowiaceae(100);g__Metschnikowia(100);g__Metschnikowia_unclassified(100); | Metschnikowia | 13 | Saprotroph | \|Undefined Saprotroph\| | Yeast | NULL | Probable | As nectar-tap_saprotroph (Põlme et al. 2020); Decay sugar-rich substrates (Põlme et al. 2020) | Põlme S, et al. 2020. Fungal Diversity 105:1-16 ((https://doi.org/10.1007/s13225-020-00466-2)) |
| Otu40126 | 1 | k__Fungi(100);p__Ascomycota(100);c__Saccharomycetes(100);o__Saccharomycetales(100);f__Metschnikowiaceae(100);g__Metschnikowia(100);s__Metschnikowia_pulcherrima(100); | Metschnikowia | 13 | Saprotroph | \|Undefined Saprotroph\| | Yeast | NULL | Probable | As nectar-tap_saprotroph (Põlme et al. 2020); Decay sugar-rich substrates (Põlme et al. 2020) | Põlme S, et al. 2020. Fungal Diversity 105:1-16 ((https://doi.org/10.1007/s13225-020-00466-2)) |
| Otu40127 | 1 | k__Fungi(100);p__Ascomycota(100);c__Saccharomycetes(100);o__Saccharomycetales(100);f__Metschnikowiaceae(100);g__Metschnikowia(100);s__Metschnikowia_pulcherrima(100); | Metschnikowia | 13 | Saprotroph | \|Undefined Saprotroph\| | Yeast | NULL | Probable | As nectar-tap_saprotroph (Põlme et al. 2020); Decay sugar-rich substrates (Põlme et al. 2020) | Põlme S, et al. 2020. Fungal Diversity 105:1-16 ((https://doi.org/10.1007/s13225-020-00466-2)) |
| Otu40129 | 1 | k__Fungi(100);p__Ascomycota(100);c__Saccharomycetes(100);o__Saccharomycetales(100);f__Metschnikowiaceae(100);g__Metschnikowia(100);g__Metschnikowia_unclassified(100); | Metschnikowia | 13 | Saprotroph | \|Undefined Saprotroph\| | Yeast | NULL | Probable | As nectar-tap_saprotroph (Põlme et al. 2020); Decay sugar-rich substrates (Põlme et al. 2020) | Põlme S, et al. 2020. Fungal Diversity 105:1-16 ((https://doi.org/10.1007/s13225-020-00466-2)) |
| Otu40130 | 1 | k__Fungi(100);p__Ascomycota(100);c__Saccharomycetes(100);o__Saccharomycetales(100);f__Metschnikowiaceae(100);g__Metschnikowia(100);s__Metschnikowia_pulcherrima(100); | Metschnikowia | 13 | Saprotroph | \|Undefined Saprotroph\| | Yeast | NULL | Probable | As nectar-tap_saprotroph (Põlme et al. 2020); Decay sugar-rich substrates (Põlme et al. 2020) | Põlme S, et al. 2020. Fungal Diversity 105:1-16 ((https://doi.org/10.1007/s13225-020-00466-2)) |
| Otu40132 | 1 | k__Fungi(100);p__Ascomycota(100);c__Saccharomycetes(100);o__Saccharomycetales(100);f__Metschnikowiaceae(100);g__Metschnikowia(100);s__Metschnikowia_pulcherrima(100); | Metschnikowia | 13 | Saprotroph | \|Undefined Saprotroph\| | Yeast | NULL | Probable | As nectar-tap_saprotroph (Põlme et al. 2020); Decay sugar-rich substrates (Põlme et al. 2020) | Põlme S, et al. 2020. Fungal Diversity 105:1-16 ((https://doi.org/10.1007/s13225-020-00466-2)) |
| Otu40133 | 1 | k__Fungi(100);p__Ascomycota(100);c__Saccharomycetes(100);o__Saccharomycetales(100);f__Metschnikowiaceae(100);g__Metschnikowia(100);g__Metschnikowia_unclassified(100); | Metschnikowia | 13 | Saprotroph | \|Undefined Saprotroph\| | Yeast | NULL | Probable | As nectar-tap_saprotroph (Põlme et al. 2020); Decay sugar-rich substrates (Põlme et al. 2020) | Põlme S, et al. 2020. Fungal Diversity 105:1-16 ((https://doi.org/10.1007/s13225-020-00466-2)) |
| Otu40134 | 1 | k__Fungi(100);p__Ascomycota(100);c__Saccharomycetes(100);o__Saccharomycetales(100);f__Metschnikowiaceae(100);g__Metschnikowia(100);g__Metschnikowia_unclassified(100); | Metschnikowia | 13 | Saprotroph | \|Undefined Saprotroph\| | Yeast | NULL | Probable | As nectar-tap_saprotroph (Põlme et al. 2020); Decay sugar-rich substrates (Põlme et al. 2020) | Põlme S, et al. 2020. Fungal Diversity 105:1-16 ((https://doi.org/10.1007/s13225-020-00466-2)) |
| Otu40135 | 1 | k__Fungi(100);p__Ascomycota(100);c__Saccharomycetes(100);o__Saccharomycetales(100);f__Metschnikowiaceae(100);g__Metschnikowia(100);s__Metschnikowia_pulcherrima(100); | Metschnikowia | 13 | Saprotroph | \|Undefined Saprotroph\| | Yeast | NULL | Probable | As nectar-tap_saprotroph (Põlme et al. 2020); Decay sugar-rich substrates (Põlme et al. 2020) | Põlme S, et al. 2020. Fungal Diversity 105:1-16 ((https://doi.org/10.1007/s13225-020-00466-2)) |
| Otu40136 | 1 | k__Fungi(100);p__Ascomycota(100);c__Saccharomycetes(100);o__Saccharomycetales(100);f__Metschnikowiaceae(100);g__Metschnikowia(100);s__Metschnikowia_pulcherrima(100); | Metschnikowia | 13 | Saprotroph | \|Undefined Saprotroph\| | Yeast | NULL | Probable | As nectar-tap_saprotroph (Põlme et al. 2020); Decay sugar-rich substrates (Põlme et al. 2020) | Põlme S, et al. 2020. Fungal Diversity 105:1-16 ((https://doi.org/10.1007/s13225-020-00466-2)) |
| Otu40137 | 1 | k__Fungi(100);p__Ascomycota(100);c__Saccharomycetes(100);o__Saccharomycetales(100);f__Metschnikowiaceae(100);g__Metschnikowia(100);s__Metschnikowia_pulcherrima(100); | Metschnikowia | 13 | Saprotroph | \|Undefined Saprotroph\| | Yeast | NULL | Probable | As nectar-tap_saprotroph (Põlme et al. 2020); Decay sugar-rich substrates (Põlme et al. 2020) | Põlme S, et al. 2020. Fungal Diversity 105:1-16 ((https://doi.org/10.1007/s13225-020-00466-2)) |
| Otu40138 | 1 | k__Fungi(100);p__Ascomycota(100);c__Saccharomycetes(100);o__Saccharomycetales(100);f__Metschnikowiaceae(100);g__Metschnikowia(100);s__Metschnikowia_pulcherrima(100); | Metschnikowia | 13 | Saprotroph | \|Undefined Saprotroph\| | Yeast | NULL | Probable | As nectar-tap_saprotroph (Põlme et al. 2020); Decay sugar-rich substrates (Põlme et al. 2020) | Põlme S, et al. 2020. Fungal Diversity 105:1-16 ((https://doi.org/10.1007/s13225-020-00466-2)) |
| Otu40139 | 1 | k__Fungi(100);p__Ascomycota(100);c__Saccharomycetes(100);o__Saccharomycetales(100);f__Metschnikowiaceae(100);g__Metschnikowia(100);g__Metschnikowia_unclassified(100); | Metschnikowia | 13 | Saprotroph | \|Undefined Saprotroph\| | Yeast | NULL | Probable | As nectar-tap_saprotroph (Põlme et al. 2020); Decay sugar-rich substrates (Põlme et al. 2020) | Põlme S, et al. 2020. Fungal Diversity 105:1-16 ((https://doi.org/10.1007/s13225-020-00466-2)) |
| Otu40140 | 1 | k__Fungi(100);p__Ascomycota(100);c__Saccharomycetes(100);o__Saccharomycetales(100);f__Metschnikowiaceae(100);g__Metschnikowia(100);g__Metschnikowia_unclassified(100); | Metschnikowia | 13 | Saprotroph | \|Undefined Saprotroph\| | Yeast | NULL | Probable | As nectar-tap_saprotroph (Põlme et al. 2020); Decay sugar-rich substrates (Põlme et al. 2020) | Põlme S, et al. 2020. Fungal Diversity 105:1-16 ((https://doi.org/10.1007/s13225-020-00466-2)) |
| Otu40141 | 1 | k__Fungi(100);p__Ascomycota(100);c__Saccharomycetes(100);o__Saccharomycetales(100);f__Metschnikowiaceae(100);g__Metschnikowia(100);s__Metschnikowia_pulcherrima(100); | Metschnikowia | 13 | Saprotroph | \|Undefined Saprotroph\| | Yeast | NULL | Probable | As nectar-tap_saprotroph (Põlme et al. 2020); Decay sugar-rich substrates (Põlme et al. 2020) | Põlme S, et al. 2020. Fungal Diversity 105:1-16 ((https://doi.org/10.1007/s13225-020-00466-2)) |
| Otu40142 | 1 | k__Fungi(100);p__Ascomycota(100);c__Saccharomycetes(100);o__Saccharomycetales(100);f__Metschnikowiaceae(100);g__Metschnikowia(100);s__Metschnikowia_pulcherrima(100); | Metschnikowia | 13 | Saprotroph | \|Undefined Saprotroph\| | Yeast | NULL | Probable | As nectar-tap_saprotroph (Põlme et al. 2020); Decay sugar-rich substrates (Põlme et al. 2020) | Põlme S, et al. 2020. Fungal Diversity 105:1-16 ((https://doi.org/10.1007/s13225-020-00466-2)) |
| Otu40143 | 1 | k__Fungi(100);p__Ascomycota(100);c__Saccharomycetes(100);o__Saccharomycetales(100);f__Metschnikowiaceae(100);g__Metschnikowia(100);g__Metschnikowia_unclassified(100); | Metschnikowia | 13 | Saprotroph | \|Undefined Saprotroph\| | Yeast | NULL | Probable | As nectar-tap_saprotroph (Põlme et al. 2020); Decay sugar-rich substrates (Põlme et al. 2020) | Põlme S, et al. 2020. Fungal Diversity 105:1-16 ((https://doi.org/10.1007/s13225-020-00466-2)) |
| Otu40144 | 1 | k__Fungi(100);p__Ascomycota(100);c__Saccharomycetes(100);o__Saccharomycetales(100);f__Metschnikowiaceae(100);g__Metschnikowia(100);g__Metschnikowia_unclassified(100); | Metschnikowia | 13 | Saprotroph | \|Undefined Saprotroph\| | Yeast | NULL | Probable | As nectar-tap_saprotroph (Põlme et al. 2020); Decay sugar-rich substrates (Põlme et al. 2020) | Põlme S, et al. 2020. Fungal Diversity 105:1-16 ((https://doi.org/10.1007/s13225-020-00466-2)) |
| Otu40145 | 1 | k__Fungi(100);p__Ascomycota(100);c__Saccharomycetes(100);o__Saccharomycetales(100);f__Metschnikowiaceae(100);g__Metschnikowia(100);g__Metschnikowia_unclassified(100); | Metschnikowia | 13 | Saprotroph | \|Undefined Saprotroph\| | Yeast | NULL | Probable | As nectar-tap_saprotroph (Põlme et al. 2020); Decay sugar-rich substrates (Põlme et al. 2020) | Põlme S, et al. 2020. Fungal Diversity 105:1-16 ((https://doi.org/10.1007/s13225-020-00466-2)) |
| Otu40146 | 1 | k__Fungi(100);p__Ascomycota(100);c__Saccharomycetes(100);o__Saccharomycetales(100);f__Metschnikowiaceae(100);g__Metschnikowia(100);g__Metschnikowia_unclassified(100); | Metschnikowia | 13 | Saprotroph | \|Undefined Saprotroph\| | Yeast | NULL | Probable | As nectar-tap_saprotroph (Põlme et al. 2020); Decay sugar-rich substrates (Põlme et al. 2020) | Põlme S, et al. 2020. Fungal Diversity 105:1-16 ((https://doi.org/10.1007/s13225-020-00466-2)) |
| Otu40147 | 1 | k__Fungi(100);p__Ascomycota(100);c__Saccharomycetes(100);o__Saccharomycetales(100);f__Metschnikowiaceae(100);g__Metschnikowia(100);g__Metschnikowia_unclassified(100); | Metschnikowia | 13 | Saprotroph | \|Undefined Saprotroph\| | Yeast | NULL | Probable | As nectar-tap_saprotroph (Põlme et al. 2020); Decay sugar-rich substrates (Põlme et al. 2020) | Põlme S, et al. 2020. Fungal Diversity 105:1-16 ((https://doi.org/10.1007/s13225-020-00466-2)) |
| Otu40148 | 1 | k__Fungi(100);p__Ascomycota(100);c__Saccharomycetes(100);o__Saccharomycetales(100);f__Metschnikowiaceae(100);g__Metschnikowia(100);g__Metschnikowia_unclassified(100); | Metschnikowia | 13 | Saprotroph | \|Undefined Saprotroph\| | Yeast | NULL | Probable | As nectar-tap_saprotroph (Põlme et al. 2020); Decay sugar-rich substrates (Põlme et al. 2020) | Põlme S, et al. 2020. Fungal Diversity 105:1-16 ((https://doi.org/10.1007/s13225-020-00466-2)) |
| Otu40149 | 1 | k__Fungi(100);p__Ascomycota(100);c__Saccharomycetes(100);o__Saccharomycetales(100);f__Metschnikowiaceae(100);g__Metschnikowia(100);s__Metschnikowia_pulcherrima(100); | Metschnikowia | 13 | Saprotroph | \|Undefined Saprotroph\| | Yeast | NULL | Probable | As nectar-tap_saprotroph (Põlme et al. 2020); Decay sugar-rich substrates (Põlme et al. 2020) | Põlme S, et al. 2020. Fungal Diversity 105:1-16 ((https://doi.org/10.1007/s13225-020-00466-2)) |
| Otu40150 | 1 | k__Fungi(100);p__Ascomycota(100);c__Saccharomycetes(100);o__Saccharomycetales(100);f__Metschnikowiaceae(100);g__Metschnikowia(100);s__Metschnikowia_pulcherrima(100); | Metschnikowia | 13 | Saprotroph | \|Undefined Saprotroph\| | Yeast | NULL | Probable | As nectar-tap_saprotroph (Põlme et al. 2020); Decay sugar-rich substrates (Põlme et al. 2020) | Põlme S, et al. 2020. Fungal Diversity 105:1-16 ((https://doi.org/10.1007/s13225-020-00466-2)) |
| Otu40151 | 1 | k__Fungi(100);p__Ascomycota(100);c__Saccharomycetes(100);o__Saccharomycetales(100);f__Metschnikowiaceae(100);g__Metschnikowia(100);s__Metschnikowia_pulcherrima(100); | Metschnikowia | 13 | Saprotroph | \|Undefined Saprotroph\| | Yeast | NULL | Probable | As nectar-tap_saprotroph (Põlme et al. 2020); Decay sugar-rich substrates (Põlme et al. 2020) | Põlme S, et al. 2020. Fungal Diversity 105:1-16 ((https://doi.org/10.1007/s13225-020-00466-2)) |
| Otu40152 | 1 | k__Fungi(100);p__Ascomycota(100);c__Saccharomycetes(100);o__Saccharomycetales(100);f__Metschnikowiaceae(100);g__Metschnikowia(100);s__Metschnikowia_pulcherrima(100); | Metschnikowia | 13 | Saprotroph | \|Undefined Saprotroph\| | Yeast | NULL | Probable | As nectar-tap_saprotroph (Põlme et al. 2020); Decay sugar-rich substrates (Põlme et al. 2020) | Põlme S, et al. 2020. Fungal Diversity 105:1-16 ((https://doi.org/10.1007/s13225-020-00466-2)) |
| Otu40153 | 1 | k__Fungi(100);p__Ascomycota(100);c__Saccharomycetes(100);o__Saccharomycetales(100);f__Metschnikowiaceae(100);g__Metschnikowia(100);s__Metschnikowia_pulcherrima(100); | Metschnikowia | 13 | Saprotroph | \|Undefined Saprotroph\| | Yeast | NULL | Probable | As nectar-tap_saprotroph (Põlme et al. 2020); Decay sugar-rich substrates (Põlme et al. 2020) | Põlme S, et al. 2020. Fungal Diversity 105:1-16 ((https://doi.org/10.1007/s13225-020-00466-2)) |
| Otu40154 | 1 | k__Fungi(100);p__Ascomycota(100);c__Saccharomycetes(100);o__Saccharomycetales(100);f__Metschnikowiaceae(100);g__Metschnikowia(100);s__Metschnikowia_pulcherrima(100); | Metschnikowia | 13 | Saprotroph | \|Undefined Saprotroph\| | Yeast | NULL | Probable | As nectar-tap_saprotroph (Põlme et al. 2020); Decay sugar-rich substrates (Põlme et al. 2020) | Põlme S, et al. 2020. Fungal Diversity 105:1-16 ((https://doi.org/10.1007/s13225-020-00466-2)) |
| Otu40155 | 1 | k__Fungi(100);p__Ascomycota(100);c__Saccharomycetes(100);o__Saccharomycetales(100);f__Metschnikowiaceae(100);g__Metschnikowia(100);s__Metschnikowia_pulcherrima(100); | Metschnikowia | 13 | Saprotroph | \|Undefined Saprotroph\| | Yeast | NULL | Probable | As nectar-tap_saprotroph (Põlme et al. 2020); Decay sugar-rich substrates (Põlme et al. 2020) | Põlme S, et al. 2020. Fungal Diversity 105:1-16 ((https://doi.org/10.1007/s13225-020-00466-2)) |
| Otu40156 | 1 | k__Fungi(100);p__Ascomycota(100);c__Saccharomycetes(100);o__Saccharomycetales(100);f__Metschnikowiaceae(100);g__Metschnikowia(100);s__Metschnikowia_pulcherrima(100); | Metschnikowia | 13 | Saprotroph | \|Undefined Saprotroph\| | Yeast | NULL | Probable | As nectar-tap_saprotroph (Põlme et al. 2020); Decay sugar-rich substrates (Põlme et al. 2020) | Põlme S, et al. 2020. Fungal Diversity 105:1-16 ((https://doi.org/10.1007/s13225-020-00466-2)) |
| Otu40157 | 1 | k__Fungi(100);p__Ascomycota(100);c__Saccharomycetes(100);o__Saccharomycetales(100);f__Metschnikowiaceae(100);g__Metschnikowia(100);s__Metschnikowia_pulcherrima(100); | Metschnikowia | 13 | Saprotroph | \|Undefined Saprotroph\| | Yeast | NULL | Probable | As nectar-tap_saprotroph (Põlme et al. 2020); Decay sugar-rich substrates (Põlme et al. 2020) | Põlme S, et al. 2020. Fungal Diversity 105:1-16 ((https://doi.org/10.1007/s13225-020-00466-2)) |
| Otu40158 | 1 | k__Fungi(100);p__Ascomycota(100);c__Saccharomycetes(100);o__Saccharomycetales(100);f__Metschnikowiaceae(100);g__Metschnikowia(100);s__Metschnikowia_pulcherrima(100); | Metschnikowia | 13 | Saprotroph | \|Undefined Saprotroph\| | Yeast | NULL | Probable | As nectar-tap_saprotroph (Põlme et al. 2020); Decay sugar-rich substrates (Põlme et al. 2020) | Põlme S, et al. 2020. Fungal Diversity 105:1-16 ((https://doi.org/10.1007/s13225-020-00466-2)) |
| Otu40159 | 1 | k__Fungi(100);p__Ascomycota(100);c__Saccharomycetes(100);o__Saccharomycetales(100);f__Metschnikowiaceae(100);g__Metschnikowia(100);s__Metschnikowia_pulcherrima(100); | Metschnikowia | 13 | Saprotroph | \|Undefined Saprotroph\| | Yeast | NULL | Probable | As nectar-tap_saprotroph (Põlme et al. 2020); Decay sugar-rich substrates (Põlme et al. 2020) | Põlme S, et al. 2020. Fungal Diversity 105:1-16 ((https://doi.org/10.1007/s13225-020-00466-2)) |
| Otu40160 | 1 | k__Fungi(100);p__Ascomycota(100);c__Saccharomycetes(100);o__Saccharomycetales(100);f__Metschnikowiaceae(100);g__Metschnikowia(100);g__Metschnikowia_unclassified(100); | Metschnikowia | 13 | Saprotroph | \|Undefined Saprotroph\| | Yeast | NULL | Probable | As nectar-tap_saprotroph (Põlme et al. 2020); Decay sugar-rich substrates (Põlme et al. 2020) | Põlme S, et al. 2020. Fungal Diversity 105:1-16 ((https://doi.org/10.1007/s13225-020-00466-2)) |
| Otu40161 | 1 | k__Fungi(100);p__Ascomycota(100);c__Saccharomycetes(100);o__Saccharomycetales(100);f__Metschnikowiaceae(100);g__Metschnikowia(100);g__Metschnikowia_unclassified(100); | Metschnikowia | 13 | Saprotroph | \|Undefined Saprotroph\| | Yeast | NULL | Probable | As nectar-tap_saprotroph (Põlme et al. 2020); Decay sugar-rich substrates (Põlme et al. 2020) | Põlme S, et al. 2020. Fungal Diversity 105:1-16 ((https://doi.org/10.1007/s13225-020-00466-2)) |
| Otu40162 | 1 | k__Fungi(100);p__Ascomycota(100);c__Saccharomycetes(100);o__Saccharomycetales(100);f__Metschnikowiaceae(100);g__Metschnikowia(100);g__Metschnikowia_unclassified(100); | Metschnikowia | 13 | Saprotroph | \|Undefined Saprotroph\| | Yeast | NULL | Probable | As nectar-tap_saprotroph (Põlme et al. 2020); Decay sugar-rich substrates (Põlme et al. 2020) | Põlme S, et al. 2020. Fungal Diversity 105:1-16 ((https://doi.org/10.1007/s13225-020-00466-2)) |
| Otu40163 | 1 | k__Fungi(100);p__Ascomycota(100);c__Saccharomycetes(100);o__Saccharomycetales(100);f__Metschnikowiaceae(100);g__Metschnikowia(100);g__Metschnikowia_unclassified(100); | Metschnikowia | 13 | Saprotroph | \|Undefined Saprotroph\| | Yeast | NULL | Probable | As nectar-tap_saprotroph (Põlme et al. 2020); Decay sugar-rich substrates (Põlme et al. 2020) | Põlme S, et al. 2020. Fungal Diversity 105:1-16 ((https://doi.org/10.1007/s13225-020-00466-2)) |
| Otu40164 | 1 | k__Fungi(100);p__Ascomycota(100);c__Saccharomycetes(100);o__Saccharomycetales(100);f__Metschnikowiaceae(100);g__Metschnikowia(100);s__Metschnikowia_pulcherrima(100); | Metschnikowia | 13 | Saprotroph | \|Undefined Saprotroph\| | Yeast | NULL | Probable | As nectar-tap_saprotroph (Põlme et al. 2020); Decay sugar-rich substrates (Põlme et al. 2020) | Põlme S, et al. 2020. Fungal Diversity 105:1-16 ((https://doi.org/10.1007/s13225-020-00466-2)) |
| Otu40165 | 1 | k__Fungi(100);p__Ascomycota(100);c__Saccharomycetes(100);o__Saccharomycetales(100);f__Metschnikowiaceae(100);g__Metschnikowia(100);g__Metschnikowia_unclassified(100); | Metschnikowia | 13 | Saprotroph | \|Undefined Saprotroph\| | Yeast | NULL | Probable | As nectar-tap_saprotroph (Põlme et al. 2020); Decay sugar-rich substrates (Põlme et al. 2020) | Põlme S, et al. 2020. Fungal Diversity 105:1-16 ((https://doi.org/10.1007/s13225-020-00466-2)) |
| Otu40166 | 1 | k__Fungi(100);p__Ascomycota(100);c__Saccharomycetes(100);o__Saccharomycetales(100);f__Metschnikowiaceae(100);g__Metschnikowia(100);s__Metschnikowia_pulcherrima(100); | Metschnikowia | 13 | Saprotroph | \|Undefined Saprotroph\| | Yeast | NULL | Probable | As nectar-tap_saprotroph (Põlme et al. 2020); Decay sugar-rich substrates (Põlme et al. 2020) | Põlme S, et al. 2020. Fungal Diversity 105:1-16 ((https://doi.org/10.1007/s13225-020-00466-2)) |
| Otu40167 | 1 | k__Fungi(100);p__Ascomycota(100);c__Saccharomycetes(100);o__Saccharomycetales(100);f__Metschnikowiaceae(100);g__Metschnikowia(100);s__Metschnikowia_pulcherrima(100); | Metschnikowia | 13 | Saprotroph | \|Undefined Saprotroph\| | Yeast | NULL | Probable | As nectar-tap_saprotroph (Põlme et al. 2020); Decay sugar-rich substrates (Põlme et al. 2020) | Põlme S, et al. 2020. Fungal Diversity 105:1-16 ((https://doi.org/10.1007/s13225-020-00466-2)) |
| Otu40168 | 1 | k__Fungi(100);p__Ascomycota(100);c__Saccharomycetes(100);o__Saccharomycetales(100);f__Metschnikowiaceae(100);g__Metschnikowia(100);g__Metschnikowia_unclassified(100); | Metschnikowia | 13 | Saprotroph | \|Undefined Saprotroph\| | Yeast | NULL | Probable | As nectar-tap_saprotroph (Põlme et al. 2020); Decay sugar-rich substrates (Põlme et al. 2020) | Põlme S, et al. 2020. Fungal Diversity 105:1-16 ((https://doi.org/10.1007/s13225-020-00466-2)) |
| Otu40169 | 1 | k__Fungi(100);p__Ascomycota(100);c__Saccharomycetes(100);o__Saccharomycetales(100);f__Metschnikowiaceae(100);g__Metschnikowia(100);g__Metschnikowia_unclassified(100); | Metschnikowia | 13 | Saprotroph | \|Undefined Saprotroph\| | Yeast | NULL | Probable | As nectar-tap_saprotroph (Põlme et al. 2020); Decay sugar-rich substrates (Põlme et al. 2020) | Põlme S, et al. 2020. Fungal Diversity 105:1-16 ((https://doi.org/10.1007/s13225-020-00466-2)) |
| Otu40170 | 1 | k__Fungi(100);p__Ascomycota(100);c__Saccharomycetes(100);o__Saccharomycetales(100);f__Metschnikowiaceae(100);g__Metschnikowia(100);s__Metschnikowia_pulcherrima(100); | Metschnikowia | 13 | Saprotroph | \|Undefined Saprotroph\| | Yeast | NULL | Probable | As nectar-tap_saprotroph (Põlme et al. 2020); Decay sugar-rich substrates (Põlme et al. 2020) | Põlme S, et al. 2020. Fungal Diversity 105:1-16 ((https://doi.org/10.1007/s13225-020-00466-2)) |
| Otu40171 | 1 | k__Fungi(100);p__Ascomycota(100);c__Saccharomycetes(100);o__Saccharomycetales(100);f__Metschnikowiaceae(100);g__Metschnikowia(100);s__Metschnikowia_pulcherrima(100); | Metschnikowia | 13 | Saprotroph | \|Undefined Saprotroph\| | Yeast | NULL | Probable | As nectar-tap_saprotroph (Põlme et al. 2020); Decay sugar-rich substrates (Põlme et al. 2020) | Põlme S, et al. 2020. Fungal Diversity 105:1-16 ((https://doi.org/10.1007/s13225-020-00466-2)) |
| Otu40172 | 1 | k__Fungi(100);p__Ascomycota(100);c__Saccharomycetes(100);o__Saccharomycetales(100);f__Metschnikowiaceae(100);g__Metschnikowia(100);s__Metschnikowia_pulcherrima(100); | Metschnikowia | 13 | Saprotroph | \|Undefined Saprotroph\| | Yeast | NULL | Probable | As nectar-tap_saprotroph (Põlme et al. 2020); Decay sugar-rich substrates (Põlme et al. 2020) | Põlme S, et al. 2020. Fungal Diversity 105:1-16 ((https://doi.org/10.1007/s13225-020-00466-2)) |
| Otu40173 | 1 | k__Fungi(100);p__Ascomycota(100);c__Saccharomycetes(100);o__Saccharomycetales(100);f__Metschnikowiaceae(100);g__Metschnikowia(100);s__Metschnikowia_pulcherrima(100); | Metschnikowia | 13 | Saprotroph | \|Undefined Saprotroph\| | Yeast | NULL | Probable | As nectar-tap_saprotroph (Põlme et al. 2020); Decay sugar-rich substrates (Põlme et al. 2020) | Põlme S, et al. 2020. Fungal Diversity 105:1-16 ((https://doi.org/10.1007/s13225-020-00466-2)) |
| Otu40175 | 1 | k__Fungi(100);p__Ascomycota(100);c__Saccharomycetes(100);o__Saccharomycetales(100);f__Metschnikowiaceae(100);g__Metschnikowia(100);s__Metschnikowia_pulcherrima(100); | Metschnikowia | 13 | Saprotroph | \|Undefined Saprotroph\| | Yeast | NULL | Probable | As nectar-tap_saprotroph (Põlme et al. 2020); Decay sugar-rich substrates (Põlme et al. 2020) | Põlme S, et al. 2020. Fungal Diversity 105:1-16 ((https://doi.org/10.1007/s13225-020-00466-2)) |
| Otu40176 | 1 | k__Fungi(100);p__Ascomycota(100);c__Saccharomycetes(100);o__Saccharomycetales(100);f__Metschnikowiaceae(100);g__Metschnikowia(100);s__Metschnikowia_pulcherrima(100); | Metschnikowia | 13 | Saprotroph | \|Undefined Saprotroph\| | Yeast | NULL | Probable | As nectar-tap_saprotroph (Põlme et al. 2020); Decay sugar-rich substrates (Põlme et al. 2020) | Põlme S, et al. 2020. Fungal Diversity 105:1-16 ((https://doi.org/10.1007/s13225-020-00466-2)) |
| Otu40177 | 1 | k__Fungi(100);p__Ascomycota(100);c__Saccharomycetes(100);o__Saccharomycetales(100);f__Metschnikowiaceae(100);g__Metschnikowia(100);s__Metschnikowia_pulcherrima(100); | Metschnikowia | 13 | Saprotroph | \|Undefined Saprotroph\| | Yeast | NULL | Probable | As nectar-tap_saprotroph (Põlme et al. 2020); Decay sugar-rich substrates (Põlme et al. 2020) | Põlme S, et al. 2020. Fungal Diversity 105:1-16 ((https://doi.org/10.1007/s13225-020-00466-2)) |
| Otu40178 | 1 | k__Fungi(100);p__Ascomycota(100);c__Saccharomycetes(100);o__Saccharomycetales(100);f__Metschnikowiaceae(100);g__Metschnikowia(100);g__Metschnikowia_unclassified(100); | Metschnikowia | 13 | Saprotroph | \|Undefined Saprotroph\| | Yeast | NULL | Probable | As nectar-tap_saprotroph (Põlme et al. 2020); Decay sugar-rich substrates (Põlme et al. 2020) | Põlme S, et al. 2020. Fungal Diversity 105:1-16 ((https://doi.org/10.1007/s13225-020-00466-2)) |
| Otu40179 | 1 | k__Fungi(100);p__Ascomycota(100);c__Saccharomycetes(100);o__Saccharomycetales(100);f__Metschnikowiaceae(100);g__Metschnikowia(100);s__Metschnikowia_pulcherrima(100); | Metschnikowia | 13 | Saprotroph | \|Undefined Saprotroph\| | Yeast | NULL | Probable | As nectar-tap_saprotroph (Põlme et al. 2020); Decay sugar-rich substrates (Põlme et al. 2020) | Põlme S, et al. 2020. Fungal Diversity 105:1-16 ((https://doi.org/10.1007/s13225-020-00466-2)) |
| Otu40180 | 1 | k__Fungi(100);p__Ascomycota(100);c__Saccharomycetes(100);o__Saccharomycetales(100);f__Metschnikowiaceae(100);g__Metschnikowia(100);g__Metschnikowia_unclassified(100); | Metschnikowia | 13 | Saprotroph | \|Undefined Saprotroph\| | Yeast | NULL | Probable | As nectar-tap_saprotroph (Põlme et al. 2020); Decay sugar-rich substrates (Põlme et al. 2020) | Põlme S, et al. 2020. Fungal Diversity 105:1-16 ((https://doi.org/10.1007/s13225-020-00466-2)) |
| Otu40181 | 1 | k__Fungi(100);p__Ascomycota(100);c__Saccharomycetes(100);o__Saccharomycetales(100);f__Metschnikowiaceae(100);g__Metschnikowia(100);g__Metschnikowia_unclassified(100); | Metschnikowia | 13 | Saprotroph | \|Undefined Saprotroph\| | Yeast | NULL | Probable | As nectar-tap_saprotroph (Põlme et al. 2020); Decay sugar-rich substrates (Põlme et al. 2020) | Põlme S, et al. 2020. Fungal Diversity 105:1-16 ((https://doi.org/10.1007/s13225-020-00466-2)) |
| Otu40182 | 1 | k__Fungi(100);p__Ascomycota(100);c__Saccharomycetes(100);o__Saccharomycetales(100);f__Metschnikowiaceae(100);g__Metschnikowia(100);g__Metschnikowia_unclassified(100); | Metschnikowia | 13 | Saprotroph | \|Undefined Saprotroph\| | Yeast | NULL | Probable | As nectar-tap_saprotroph (Põlme et al. 2020); Decay sugar-rich substrates (Põlme et al. 2020) | Põlme S, et al. 2020. Fungal Diversity 105:1-16 ((https://doi.org/10.1007/s13225-020-00466-2)) |
| Otu40183 | 1 | k__Fungi(100);p__Ascomycota(100);c__Saccharomycetes(100);o__Saccharomycetales(100);f__Metschnikowiaceae(100);g__Metschnikowia(100);s__Metschnikowia_pulcherrima(100); | Metschnikowia | 13 | Saprotroph | \|Undefined Saprotroph\| | Yeast | NULL | Probable | As nectar-tap_saprotroph (Põlme et al. 2020); Decay sugar-rich substrates (Põlme et al. 2020) | Põlme S, et al. 2020. Fungal Diversity 105:1-16 ((https://doi.org/10.1007/s13225-020-00466-2)) |
| Otu40184 | 1 | k__Fungi(100);p__Ascomycota(100);c__Saccharomycetes(100);o__Saccharomycetales(100);f__Metschnikowiaceae(100);g__Metschnikowia(100);g__Metschnikowia_unclassified(100); | Metschnikowia | 13 | Saprotroph | \|Undefined Saprotroph\| | Yeast | NULL | Probable | As nectar-tap_saprotroph (Põlme et al. 2020); Decay sugar-rich substrates (Põlme et al. 2020) | Põlme S, et al. 2020. Fungal Diversity 105:1-16 ((https://doi.org/10.1007/s13225-020-00466-2)) |
| Otu40185 | 1 | k__Fungi(100);p__Ascomycota(100);c__Saccharomycetes(100);o__Saccharomycetales(100);f__Metschnikowiaceae(100);g__Metschnikowia(100);s__Metschnikowia_pulcherrima(100); | Metschnikowia | 13 | Saprotroph | \|Undefined Saprotroph\| | Yeast | NULL | Probable | As nectar-tap_saprotroph (Põlme et al. 2020); Decay sugar-rich substrates (Põlme et al. 2020) | Põlme S, et al. 2020. Fungal Diversity 105:1-16 ((https://doi.org/10.1007/s13225-020-00466-2)) |
| Otu40186 | 1 | k__Fungi(100);p__Ascomycota(100);c__Saccharomycetes(100);o__Saccharomycetales(100);f__Metschnikowiaceae(100);g__Metschnikowia(100);s__Metschnikowia_pulcherrima(100); | Metschnikowia | 13 | Saprotroph | \|Undefined Saprotroph\| | Yeast | NULL | Probable | As nectar-tap_saprotroph (Põlme et al. 2020); Decay sugar-rich substrates (Põlme et al. 2020) | Põlme S, et al. 2020. Fungal Diversity 105:1-16 ((https://doi.org/10.1007/s13225-020-00466-2)) |
| Otu40187 | 1 | k__Fungi(100);p__Ascomycota(100);c__Saccharomycetes(100);o__Saccharomycetales(100);f__Metschnikowiaceae(100);g__Metschnikowia(100);s__Metschnikowia_pulcherrima(100); | Metschnikowia | 13 | Saprotroph | \|Undefined Saprotroph\| | Yeast | NULL | Probable | As nectar-tap_saprotroph (Põlme et al. 2020); Decay sugar-rich substrates (Põlme et al. 2020) | Põlme S, et al. 2020. Fungal Diversity 105:1-16 ((https://doi.org/10.1007/s13225-020-00466-2)) |
| Otu40189 | 1 | k__Fungi(100);p__Ascomycota(100);c__Saccharomycetes(100);o__Saccharomycetales(100);f__Metschnikowiaceae(100);g__Metschnikowia(100);g__Metschnikowia_unclassified(100); | Metschnikowia | 13 | Saprotroph | \|Undefined Saprotroph\| | Yeast | NULL | Probable | As nectar-tap_saprotroph (Põlme et al. 2020); Decay sugar-rich substrates (Põlme et al. 2020) | Põlme S, et al. 2020. Fungal Diversity 105:1-16 ((https://doi.org/10.1007/s13225-020-00466-2)) |
| Otu40190 | 1 | k__Fungi(100);p__Ascomycota(100);c__Saccharomycetes(100);o__Saccharomycetales(100);f__Metschnikowiaceae(100);g__Metschnikowia(100);s__Metschnikowia_pulcherrima(100); | Metschnikowia | 13 | Saprotroph | \|Undefined Saprotroph\| | Yeast | NULL | Probable | As nectar-tap_saprotroph (Põlme et al. 2020); Decay sugar-rich substrates (Põlme et al. 2020) | Põlme S, et al. 2020. Fungal Diversity 105:1-16 ((https://doi.org/10.1007/s13225-020-00466-2)) |
| Otu40191 | 1 | k__Fungi(100);p__Ascomycota(100);c__Saccharomycetes(100);o__Saccharomycetales(100);f__Metschnikowiaceae(100);g__Metschnikowia(100);s__Metschnikowia_pulcherrima(100); | Metschnikowia | 13 | Saprotroph | \|Undefined Saprotroph\| | Yeast | NULL | Probable | As nectar-tap_saprotroph (Põlme et al. 2020); Decay sugar-rich substrates (Põlme et al. 2020) | Põlme S, et al. 2020. Fungal Diversity 105:1-16 ((https://doi.org/10.1007/s13225-020-00466-2)) |
| Otu40192 | 1 | k__Fungi(100);p__Ascomycota(100);c__Saccharomycetes(100);o__Saccharomycetales(100);f__Metschnikowiaceae(100);g__Metschnikowia(100);s__Metschnikowia_pulcherrima(100); | Metschnikowia | 13 | Saprotroph | \|Undefined Saprotroph\| | Yeast | NULL | Probable | As nectar-tap_saprotroph (Põlme et al. 2020); Decay sugar-rich substrates (Põlme et al. 2020) | Põlme S, et al. 2020. Fungal Diversity 105:1-16 ((https://doi.org/10.1007/s13225-020-00466-2)) |
| Otu40193 | 1 | k__Fungi(100);p__Ascomycota(100);c__Saccharomycetes(100);o__Saccharomycetales(100);f__Metschnikowiaceae(100);g__Metschnikowia(100);s__Metschnikowia_pulcherrima(100); | Metschnikowia | 13 | Saprotroph | \|Undefined Saprotroph\| | Yeast | NULL | Probable | As nectar-tap_saprotroph (Põlme et al. 2020); Decay sugar-rich substrates (Põlme et al. 2020) | Põlme S, et al. 2020. Fungal Diversity 105:1-16 ((https://doi.org/10.1007/s13225-020-00466-2)) |
| Otu40194 | 1 | k__Fungi(100);p__Ascomycota(100);c__Saccharomycetes(100);o__Saccharomycetales(100);f__Metschnikowiaceae(100);g__Metschnikowia(100);s__Metschnikowia_pulcherrima(100); | Metschnikowia | 13 | Saprotroph | \|Undefined Saprotroph\| | Yeast | NULL | Probable | As nectar-tap_saprotroph (Põlme et al. 2020); Decay sugar-rich substrates (Põlme et al. 2020) | Põlme S, et al. 2020. Fungal Diversity 105:1-16 ((https://doi.org/10.1007/s13225-020-00466-2)) |
| Otu40195 | 1 | k__Fungi(100);p__Ascomycota(100);c__Saccharomycetes(100);o__Saccharomycetales(100);f__Metschnikowiaceae(100);g__Metschnikowia(100);g__Metschnikowia_unclassified(100); | Metschnikowia | 13 | Saprotroph | \|Undefined Saprotroph\| | Yeast | NULL | Probable | As nectar-tap_saprotroph (Põlme et al. 2020); Decay sugar-rich substrates (Põlme et al. 2020) | Põlme S, et al. 2020. Fungal Diversity 105:1-16 ((https://doi.org/10.1007/s13225-020-00466-2)) |
| Otu40196 | 1 | k__Fungi(100);p__Ascomycota(100);c__Saccharomycetes(100);o__Saccharomycetales(100);f__Metschnikowiaceae(100);g__Metschnikowia(100);s__Metschnikowia_pulcherrima(100); | Metschnikowia | 13 | Saprotroph | \|Undefined Saprotroph\| | Yeast | NULL | Probable | As nectar-tap_saprotroph (Põlme et al. 2020); Decay sugar-rich substrates (Põlme et al. 2020) | Põlme S, et al. 2020. Fungal Diversity 105:1-16 ((https://doi.org/10.1007/s13225-020-00466-2)) |
| Otu40197 | 1 | k__Fungi(100);p__Ascomycota(100);c__Saccharomycetes(100);o__Saccharomycetales(100);f__Metschnikowiaceae(100);g__Metschnikowia(100);s__Metschnikowia_pulcherrima(100); | Metschnikowia | 13 | Saprotroph | \|Undefined Saprotroph\| | Yeast | NULL | Probable | As nectar-tap_saprotroph (Põlme et al. 2020); Decay sugar-rich substrates (Põlme et al. 2020) | Põlme S, et al. 2020. Fungal Diversity 105:1-16 ((https://doi.org/10.1007/s13225-020-00466-2)) |
| Otu40200 | 1 | k__Fungi(100);p__Ascomycota(100);c__Saccharomycetes(100);o__Saccharomycetales(100);f__Metschnikowiaceae(100);g__Metschnikowia(100);s__Metschnikowia_pulcherrima(100); | Metschnikowia | 13 | Saprotroph | \|Undefined Saprotroph\| | Yeast | NULL | Probable | As nectar-tap_saprotroph (Põlme et al. 2020); Decay sugar-rich substrates (Põlme et al. 2020) | Põlme S, et al. 2020. Fungal Diversity 105:1-16 ((https://doi.org/10.1007/s13225-020-00466-2)) |
| Otu40201 | 1 | k__Fungi(100);p__Ascomycota(100);c__Saccharomycetes(100);o__Saccharomycetales(100);f__Metschnikowiaceae(100);g__Metschnikowia(100);s__Metschnikowia_pulcherrima(100); | Metschnikowia | 13 | Saprotroph | \|Undefined Saprotroph\| | Yeast | NULL | Probable | As nectar-tap_saprotroph (Põlme et al. 2020); Decay sugar-rich substrates (Põlme et al. 2020) | Põlme S, et al. 2020. Fungal Diversity 105:1-16 ((https://doi.org/10.1007/s13225-020-00466-2)) |
| Otu40202 | 1 | k__Fungi(100);p__Ascomycota(100);c__Saccharomycetes(100);o__Saccharomycetales(100);f__Metschnikowiaceae(100);g__Metschnikowia(100);s__Metschnikowia_pulcherrima(100); | Metschnikowia | 13 | Saprotroph | \|Undefined Saprotroph\| | Yeast | NULL | Probable | As nectar-tap_saprotroph (Põlme et al. 2020); Decay sugar-rich substrates (Põlme et al. 2020) | Põlme S, et al. 2020. Fungal Diversity 105:1-16 ((https://doi.org/10.1007/s13225-020-00466-2)) |
| Otu40203 | 1 | k__Fungi(100);p__Ascomycota(100);c__Saccharomycetes(100);o__Saccharomycetales(100);f__Metschnikowiaceae(100);g__Metschnikowia(100);s__Metschnikowia_pulcherrima(100); | Metschnikowia | 13 | Saprotroph | \|Undefined Saprotroph\| | Yeast | NULL | Probable | As nectar-tap_saprotroph (Põlme et al. 2020); Decay sugar-rich substrates (Põlme et al. 2020) | Põlme S, et al. 2020. Fungal Diversity 105:1-16 ((https://doi.org/10.1007/s13225-020-00466-2)) |
| Otu40204 | 1 | k__Fungi(100);p__Ascomycota(100);c__Saccharomycetes(100);o__Saccharomycetales(100);f__Metschnikowiaceae(100);g__Metschnikowia(100);s__Metschnikowia_pulcherrima(100); | Metschnikowia | 13 | Saprotroph | \|Undefined Saprotroph\| | Yeast | NULL | Probable | As nectar-tap_saprotroph (Põlme et al. 2020); Decay sugar-rich substrates (Põlme et al. 2020) | Põlme S, et al. 2020. Fungal Diversity 105:1-16 ((https://doi.org/10.1007/s13225-020-00466-2)) |
| Otu40205 | 1 | k__Fungi(100);p__Ascomycota(100);c__Saccharomycetes(100);o__Saccharomycetales(100);f__Metschnikowiaceae(100);g__Metschnikowia(100);s__Metschnikowia_pulcherrima(100); | Metschnikowia | 13 | Saprotroph | \|Undefined Saprotroph\| | Yeast | NULL | Probable | As nectar-tap_saprotroph (Põlme et al. 2020); Decay sugar-rich substrates (Põlme et al. 2020) | Põlme S, et al. 2020. Fungal Diversity 105:1-16 ((https://doi.org/10.1007/s13225-020-00466-2)) |
| Otu40206 | 1 | k__Fungi(100);p__Ascomycota(100);c__Saccharomycetes(100);o__Saccharomycetales(100);f__Metschnikowiaceae(100);g__Metschnikowia(100);s__Metschnikowia_pulcherrima(100); | Metschnikowia | 13 | Saprotroph | \|Undefined Saprotroph\| | Yeast | NULL | Probable | As nectar-tap_saprotroph (Põlme et al. 2020); Decay sugar-rich substrates (Põlme et al. 2020) | Põlme S, et al. 2020. Fungal Diversity 105:1-16 ((https://doi.org/10.1007/s13225-020-00466-2)) |
| Otu40207 | 1 | k__Fungi(100);p__Ascomycota(100);c__Saccharomycetes(100);o__Saccharomycetales(100);f__Metschnikowiaceae(100);g__Metschnikowia(100);s__Metschnikowia_pulcherrima(100); | Metschnikowia | 13 | Saprotroph | \|Undefined Saprotroph\| | Yeast | NULL | Probable | As nectar-tap_saprotroph (Põlme et al. 2020); Decay sugar-rich substrates (Põlme et al. 2020) | Põlme S, et al. 2020. Fungal Diversity 105:1-16 ((https://doi.org/10.1007/s13225-020-00466-2)) |
| Otu40209 | 1 | k__Fungi(100);p__Ascomycota(100);c__Saccharomycetes(100);o__Saccharomycetales(100);f__Metschnikowiaceae(100);g__Metschnikowia(100);s__Metschnikowia_pulcherrima(100); | Metschnikowia | 13 | Saprotroph | \|Undefined Saprotroph\| | Yeast | NULL | Probable | As nectar-tap_saprotroph (Põlme et al. 2020); Decay sugar-rich substrates (Põlme et al. 2020) | Põlme S, et al. 2020. Fungal Diversity 105:1-16 ((https://doi.org/10.1007/s13225-020-00466-2)) |
| Otu40210 | 1 | k__Fungi(100);p__Ascomycota(100);c__Saccharomycetes(100);o__Saccharomycetales(100);f__Metschnikowiaceae(100);g__Metschnikowia(100);s__Metschnikowia_pulcherrima(100); | Metschnikowia | 13 | Saprotroph | \|Undefined Saprotroph\| | Yeast | NULL | Probable | As nectar-tap_saprotroph (Põlme et al. 2020); Decay sugar-rich substrates (Põlme et al. 2020) | Põlme S, et al. 2020. Fungal Diversity 105:1-16 ((https://doi.org/10.1007/s13225-020-00466-2)) |
| Otu40211 | 1 | k__Fungi(100);p__Ascomycota(100);c__Saccharomycetes(100);o__Saccharomycetales(100);f__Metschnikowiaceae(100);g__Metschnikowia(100);s__Metschnikowia_pulcherrima(100); | Metschnikowia | 13 | Saprotroph | \|Undefined Saprotroph\| | Yeast | NULL | Probable | As nectar-tap_saprotroph (Põlme et al. 2020); Decay sugar-rich substrates (Põlme et al. 2020) | Põlme S, et al. 2020. Fungal Diversity 105:1-16 ((https://doi.org/10.1007/s13225-020-00466-2)) |
| Otu40212 | 1 | k__Fungi(100);p__Ascomycota(100);c__Saccharomycetes(100);o__Saccharomycetales(100);f__Metschnikowiaceae(100);g__Metschnikowia(100);s__Metschnikowia_pulcherrima(100); | Metschnikowia | 13 | Saprotroph | \|Undefined Saprotroph\| | Yeast | NULL | Probable | As nectar-tap_saprotroph (Põlme et al. 2020); Decay sugar-rich substrates (Põlme et al. 2020) | Põlme S, et al. 2020. Fungal Diversity 105:1-16 ((https://doi.org/10.1007/s13225-020-00466-2)) |
| Otu40215 | 1 | k__Fungi(100);p__Ascomycota(100);c__Saccharomycetes(100);o__Saccharomycetales(100);f__Metschnikowiaceae(100);g__Metschnikowia(100);s__Metschnikowia_pulcherrima(100); | Metschnikowia | 13 | Saprotroph | \|Undefined Saprotroph\| | Yeast | NULL | Probable | As nectar-tap_saprotroph (Põlme et al. 2020); Decay sugar-rich substrates (Põlme et al. 2020) | Põlme S, et al. 2020. Fungal Diversity 105:1-16 ((https://doi.org/10.1007/s13225-020-00466-2)) |
| Otu40216 | 1 | k__Fungi(100);p__Ascomycota(100);c__Saccharomycetes(100);o__Saccharomycetales(100);f__Metschnikowiaceae(100);g__Metschnikowia(100);s__Metschnikowia_pulcherrima(100); | Metschnikowia | 13 | Saprotroph | \|Undefined Saprotroph\| | Yeast | NULL | Probable | As nectar-tap_saprotroph (Põlme et al. 2020); Decay sugar-rich substrates (Põlme et al. 2020) | Põlme S, et al. 2020. Fungal Diversity 105:1-16 ((https://doi.org/10.1007/s13225-020-00466-2)) |
| Otu40217 | 1 | k__Fungi(100);p__Ascomycota(100);c__Saccharomycetes(100);o__Saccharomycetales(100);f__Metschnikowiaceae(100);g__Metschnikowia(100);s__Metschnikowia_pulcherrima(100); | Metschnikowia | 13 | Saprotroph | \|Undefined Saprotroph\| | Yeast | NULL | Probable | As nectar-tap_saprotroph (Põlme et al. 2020); Decay sugar-rich substrates (Põlme et al. 2020) | Põlme S, et al. 2020. Fungal Diversity 105:1-16 ((https://doi.org/10.1007/s13225-020-00466-2)) |
| Otu40218 | 1 | k__Fungi(100);p__Ascomycota(100);c__Saccharomycetes(100);o__Saccharomycetales(100);f__Metschnikowiaceae(100);g__Metschnikowia(100);s__Metschnikowia_pulcherrima(100); | Metschnikowia | 13 | Saprotroph | \|Undefined Saprotroph\| | Yeast | NULL | Probable | As nectar-tap_saprotroph (Põlme et al. 2020); Decay sugar-rich substrates (Põlme et al. 2020) | Põlme S, et al. 2020. Fungal Diversity 105:1-16 ((https://doi.org/10.1007/s13225-020-00466-2)) |
| Otu40219 | 1 | k__Fungi(100);p__Ascomycota(100);c__Saccharomycetes(100);o__Saccharomycetales(100);f__Metschnikowiaceae(100);g__Metschnikowia(100);g__Metschnikowia_unclassified(100); | Metschnikowia | 13 | Saprotroph | \|Undefined Saprotroph\| | Yeast | NULL | Probable | As nectar-tap_saprotroph (Põlme et al. 2020); Decay sugar-rich substrates (Põlme et al. 2020) | Põlme S, et al. 2020. Fungal Diversity 105:1-16 ((https://doi.org/10.1007/s13225-020-00466-2)) |
| Otu40220 | 1 | k__Fungi(100);p__Ascomycota(100);c__Saccharomycetes(100);o__Saccharomycetales(100);f__Metschnikowiaceae(100);g__Metschnikowia(100);s__Metschnikowia_pulcherrima(100); | Metschnikowia | 13 | Saprotroph | \|Undefined Saprotroph\| | Yeast | NULL | Probable | As nectar-tap_saprotroph (Põlme et al. 2020); Decay sugar-rich substrates (Põlme et al. 2020) | Põlme S, et al. 2020. Fungal Diversity 105:1-16 ((https://doi.org/10.1007/s13225-020-00466-2)) |
| Otu40221 | 1 | k__Fungi(100);p__Ascomycota(100);c__Saccharomycetes(100);o__Saccharomycetales(100);f__Metschnikowiaceae(100);g__Metschnikowia(100);s__Metschnikowia_pulcherrima(100); | Metschnikowia | 13 | Saprotroph | \|Undefined Saprotroph\| | Yeast | NULL | Probable | As nectar-tap_saprotroph (Põlme et al. 2020); Decay sugar-rich substrates (Põlme et al. 2020) | Põlme S, et al. 2020. Fungal Diversity 105:1-16 ((https://doi.org/10.1007/s13225-020-00466-2)) |
| Otu40222 | 1 | k__Fungi(100);p__Ascomycota(100);c__Saccharomycetes(100);o__Saccharomycetales(100);f__Metschnikowiaceae(100);g__Metschnikowia(100);g__Metschnikowia_unclassified(100); | Metschnikowia | 13 | Saprotroph | \|Undefined Saprotroph\| | Yeast | NULL | Probable | As nectar-tap_saprotroph (Põlme et al. 2020); Decay sugar-rich substrates (Põlme et al. 2020) | Põlme S, et al. 2020. Fungal Diversity 105:1-16 ((https://doi.org/10.1007/s13225-020-00466-2)) |
| Otu40223 | 1 | k__Fungi(100);p__Ascomycota(100);c__Saccharomycetes(100);o__Saccharomycetales(100);f__Metschnikowiaceae(100);g__Metschnikowia(100);s__Metschnikowia_pulcherrima(100); | Metschnikowia | 13 | Saprotroph | \|Undefined Saprotroph\| | Yeast | NULL | Probable | As nectar-tap_saprotroph (Põlme et al. 2020); Decay sugar-rich substrates (Põlme et al. 2020) | Põlme S, et al. 2020. Fungal Diversity 105:1-16 ((https://doi.org/10.1007/s13225-020-00466-2)) |
| Otu40224 | 1 | k__Fungi(100);p__Ascomycota(100);c__Saccharomycetes(100);o__Saccharomycetales(100);f__Metschnikowiaceae(100);g__Metschnikowia(100);s__Metschnikowia_pulcherrima(100); | Metschnikowia | 13 | Saprotroph | \|Undefined Saprotroph\| | Yeast | NULL | Probable | As nectar-tap_saprotroph (Põlme et al. 2020); Decay sugar-rich substrates (Põlme et al. 2020) | Põlme S, et al. 2020. Fungal Diversity 105:1-16 ((https://doi.org/10.1007/s13225-020-00466-2)) |
| Otu40225 | 1 | k__Fungi(100);p__Ascomycota(100);c__Saccharomycetes(100);o__Saccharomycetales(100);f__Metschnikowiaceae(100);g__Metschnikowia(100);s__Metschnikowia_pulcherrima(100); | Metschnikowia | 13 | Saprotroph | \|Undefined Saprotroph\| | Yeast | NULL | Probable | As nectar-tap_saprotroph (Põlme et al. 2020); Decay sugar-rich substrates (Põlme et al. 2020) | Põlme S, et al. 2020. Fungal Diversity 105:1-16 ((https://doi.org/10.1007/s13225-020-00466-2)) |
| Otu40226 | 1 | k__Fungi(100);p__Ascomycota(100);c__Saccharomycetes(100);o__Saccharomycetales(100);f__Metschnikowiaceae(100);g__Metschnikowia(100);g__Metschnikowia_unclassified(100); | Metschnikowia | 13 | Saprotroph | \|Undefined Saprotroph\| | Yeast | NULL | Probable | As nectar-tap_saprotroph (Põlme et al. 2020); Decay sugar-rich substrates (Põlme et al. 2020) | Põlme S, et al. 2020. Fungal Diversity 105:1-16 ((https://doi.org/10.1007/s13225-020-00466-2)) |
| Otu40227 | 1 | k__Fungi(100);p__Ascomycota(100);c__Saccharomycetes(100);o__Saccharomycetales(100);f__Metschnikowiaceae(100);g__Metschnikowia(100);s__Metschnikowia_pulcherrima(100); | Metschnikowia | 13 | Saprotroph | \|Undefined Saprotroph\| | Yeast | NULL | Probable | As nectar-tap_saprotroph (Põlme et al. 2020); Decay sugar-rich substrates (Põlme et al. 2020) | Põlme S, et al. 2020. Fungal Diversity 105:1-16 ((https://doi.org/10.1007/s13225-020-00466-2)) |
| Otu40228 | 1 | k__Fungi(100);p__Ascomycota(100);c__Saccharomycetes(100);o__Saccharomycetales(100);f__Metschnikowiaceae(100);g__Metschnikowia(100);s__Metschnikowia_pulcherrima(100); | Metschnikowia | 13 | Saprotroph | \|Undefined Saprotroph\| | Yeast | NULL | Probable | As nectar-tap_saprotroph (Põlme et al. 2020); Decay sugar-rich substrates (Põlme et al. 2020) | Põlme S, et al. 2020. Fungal Diversity 105:1-16 ((https://doi.org/10.1007/s13225-020-00466-2)) |
| Otu40229 | 1 | k__Fungi(100);p__Ascomycota(100);c__Saccharomycetes(100);o__Saccharomycetales(100);f__Metschnikowiaceae(100);g__Metschnikowia(100);g__Metschnikowia_unclassified(100); | Metschnikowia | 13 | Saprotroph | \|Undefined Saprotroph\| | Yeast | NULL | Probable | As nectar-tap_saprotroph (Põlme et al. 2020); Decay sugar-rich substrates (Põlme et al. 2020) | Põlme S, et al. 2020. Fungal Diversity 105:1-16 ((https://doi.org/10.1007/s13225-020-00466-2)) |
| Otu40230 | 1 | k__Fungi(100);p__Ascomycota(100);c__Saccharomycetes(100);o__Saccharomycetales(100);f__Metschnikowiaceae(100);g__Metschnikowia(100);s__Metschnikowia_pulcherrima(100); | Metschnikowia | 13 | Saprotroph | \|Undefined Saprotroph\| | Yeast | NULL | Probable | As nectar-tap_saprotroph (Põlme et al. 2020); Decay sugar-rich substrates (Põlme et al. 2020) | Põlme S, et al. 2020. Fungal Diversity 105:1-16 ((https://doi.org/10.1007/s13225-020-00466-2)) |
| Otu40231 | 1 | k__Fungi(100);p__Ascomycota(100);c__Saccharomycetes(100);o__Saccharomycetales(100);f__Metschnikowiaceae(100);g__Metschnikowia(100);s__Metschnikowia_pulcherrima(100); | Metschnikowia | 13 | Saprotroph | \|Undefined Saprotroph\| | Yeast | NULL | Probable | As nectar-tap_saprotroph (Põlme et al. 2020); Decay sugar-rich substrates (Põlme et al. 2020) | Põlme S, et al. 2020. Fungal Diversity 105:1-16 ((https://doi.org/10.1007/s13225-020-00466-2)) |
| Otu40232 | 1 | k__Fungi(100);p__Ascomycota(100);c__Saccharomycetes(100);o__Saccharomycetales(100);f__Metschnikowiaceae(100);g__Metschnikowia(100);s__Metschnikowia_pulcherrima(100); | Metschnikowia | 13 | Saprotroph | \|Undefined Saprotroph\| | Yeast | NULL | Probable | As nectar-tap_saprotroph (Põlme et al. 2020); Decay sugar-rich substrates (Põlme et al. 2020) | Põlme S, et al. 2020. Fungal Diversity 105:1-16 ((https://doi.org/10.1007/s13225-020-00466-2)) |
| Otu40234 | 1 | k__Fungi(100);p__Ascomycota(100);c__Saccharomycetes(100);o__Saccharomycetales(100);f__Metschnikowiaceae(100);g__Metschnikowia(100);s__Metschnikowia_pulcherrima(100); | Metschnikowia | 13 | Saprotroph | \|Undefined Saprotroph\| | Yeast | NULL | Probable | As nectar-tap_saprotroph (Põlme et al. 2020); Decay sugar-rich substrates (Põlme et al. 2020) | Põlme S, et al. 2020. Fungal Diversity 105:1-16 ((https://doi.org/10.1007/s13225-020-00466-2)) |
| Otu40235 | 1 | k__Fungi(100);p__Ascomycota(100);c__Saccharomycetes(100);o__Saccharomycetales(100);f__Metschnikowiaceae(100);g__Metschnikowia(100);s__Metschnikowia_pulcherrima(100); | Metschnikowia | 13 | Saprotroph | \|Undefined Saprotroph\| | Yeast | NULL | Probable | As nectar-tap_saprotroph (Põlme et al. 2020); Decay sugar-rich substrates (Põlme et al. 2020) | Põlme S, et al. 2020. Fungal Diversity 105:1-16 ((https://doi.org/10.1007/s13225-020-00466-2)) |
| Otu40236 | 1 | k__Fungi(100);p__Ascomycota(100);c__Saccharomycetes(100);o__Saccharomycetales(100);f__Metschnikowiaceae(100);g__Metschnikowia(100);g__Metschnikowia_unclassified(100); | Metschnikowia | 13 | Saprotroph | \|Undefined Saprotroph\| | Yeast | NULL | Probable | As nectar-tap_saprotroph (Põlme et al. 2020); Decay sugar-rich substrates (Põlme et al. 2020) | Põlme S, et al. 2020. Fungal Diversity 105:1-16 ((https://doi.org/10.1007/s13225-020-00466-2)) |
| Otu40237 | 1 | k__Fungi(100);p__Ascomycota(100);c__Saccharomycetes(100);o__Saccharomycetales(100);f__Metschnikowiaceae(100);g__Metschnikowia(100);s__Metschnikowia_pulcherrima(100); | Metschnikowia | 13 | Saprotroph | \|Undefined Saprotroph\| | Yeast | NULL | Probable | As nectar-tap_saprotroph (Põlme et al. 2020); Decay sugar-rich substrates (Põlme et al. 2020) | Põlme S, et al. 2020. Fungal Diversity 105:1-16 ((https://doi.org/10.1007/s13225-020-00466-2)) |
| Otu40239 | 1 | k__Fungi(100);p__Ascomycota(100);c__Saccharomycetes(100);o__Saccharomycetales(100);f__Metschnikowiaceae(100);g__Metschnikowia(100);s__Metschnikowia_pulcherrima(100); | Metschnikowia | 13 | Saprotroph | \|Undefined Saprotroph\| | Yeast | NULL | Probable | As nectar-tap_saprotroph (Põlme et al. 2020); Decay sugar-rich substrates (Põlme et al. 2020) | Põlme S, et al. 2020. Fungal Diversity 105:1-16 ((https://doi.org/10.1007/s13225-020-00466-2)) |
| Otu40241 | 1 | k__Fungi(100);p__Ascomycota(100);c__Saccharomycetes(100);o__Saccharomycetales(100);f__Metschnikowiaceae(100);g__Metschnikowia(100);s__Metschnikowia_pulcherrima(100); | Metschnikowia | 13 | Saprotroph | \|Undefined Saprotroph\| | Yeast | NULL | Probable | As nectar-tap_saprotroph (Põlme et al. 2020); Decay sugar-rich substrates (Põlme et al. 2020) | Põlme S, et al. 2020. Fungal Diversity 105:1-16 ((https://doi.org/10.1007/s13225-020-00466-2)) |
| Otu40242 | 1 | k__Fungi(100);p__Ascomycota(100);c__Saccharomycetes(100);o__Saccharomycetales(100);f__Metschnikowiaceae(100);g__Metschnikowia(100);g__Metschnikowia_unclassified(100); | Metschnikowia | 13 | Saprotroph | \|Undefined Saprotroph\| | Yeast | NULL | Probable | As nectar-tap_saprotroph (Põlme et al. 2020); Decay sugar-rich substrates (Põlme et al. 2020) | Põlme S, et al. 2020. Fungal Diversity 105:1-16 ((https://doi.org/10.1007/s13225-020-00466-2)) |
| Otu40243 | 1 | k__Fungi(100);p__Ascomycota(100);c__Leotiomycetes(100);o__Helotiales(100);f__Hyaloscyphaceae(100);g__Lachnum(100);g__Lachnum_unclassified(100); | Lachnum | 13 | Saprotroph-Symbiotroph | Endophyte-\|Plant Saprotroph\|-Undefined Saprotroph-Wood Saprotroph | Microfungus | NULL | Probable | As litter_saprotroph (Põlme et al. 2020); As root_endophyte (Põlme et al. 2020); Decay wood, leaf, fruit, seed (Põlme et al. 2020) | Tedersoo L, et al. 2014. Science 346:e1256688 ((https://doi.org/10.1126/science.1256688)); Põlme S, et al. 2020. Fungal Diversity 105:1-16 ((https://doi.org/10.1007/s13225-020-00466-2)) |
| Otu40244 | 1 | k__Fungi(100);p__Ascomycota(100);c__Saccharomycetes(100);o__Saccharomycetales(100);f__Metschnikowiaceae(100);g__Metschnikowia(100);g__Metschnikowia_unclassified(100); | Metschnikowia | 13 | Saprotroph | \|Undefined Saprotroph\| | Yeast | NULL | Probable | As nectar-tap_saprotroph (Põlme et al. 2020); Decay sugar-rich substrates (Põlme et al. 2020) | Põlme S, et al. 2020. Fungal Diversity 105:1-16 ((https://doi.org/10.1007/s13225-020-00466-2)) |
| Otu40246 | 1 | k__Fungi(100);p__Ascomycota(100);c__Saccharomycetes(100);o__Saccharomycetales(100);f__Metschnikowiaceae(100);g__Metschnikowia(100);s__Metschnikowia_pulcherrima(100); | Metschnikowia | 13 | Saprotroph | \|Undefined Saprotroph\| | Yeast | NULL | Probable | As nectar-tap_saprotroph (Põlme et al. 2020); Decay sugar-rich substrates (Põlme et al. 2020) | Põlme S, et al. 2020. Fungal Diversity 105:1-16 ((https://doi.org/10.1007/s13225-020-00466-2)) |
| Otu40247 | 1 | k__Fungi(100);p__Ascomycota(100);c__Saccharomycetes(100);o__Saccharomycetales(100);f__Metschnikowiaceae(100);g__Metschnikowia(100);s__Metschnikowia_pulcherrima(100); | Metschnikowia | 13 | Saprotroph | \|Undefined Saprotroph\| | Yeast | NULL | Probable | As nectar-tap_saprotroph (Põlme et al. 2020); Decay sugar-rich substrates (Põlme et al. 2020) | Põlme S, et al. 2020. Fungal Diversity 105:1-16 ((https://doi.org/10.1007/s13225-020-00466-2)) |
| Otu40248 | 1 | k__Fungi(100);p__Ascomycota(100);c__Saccharomycetes(100);o__Saccharomycetales(100);f__Metschnikowiaceae(100);g__Metschnikowia(100);s__Metschnikowia_pulcherrima(100); | Metschnikowia | 13 | Saprotroph | \|Undefined Saprotroph\| | Yeast | NULL | Probable | As nectar-tap_saprotroph (Põlme et al. 2020); Decay sugar-rich substrates (Põlme et al. 2020) | Põlme S, et al. 2020. Fungal Diversity 105:1-16 ((https://doi.org/10.1007/s13225-020-00466-2)) |
| Otu40249 | 1 | k__Fungi(100);p__Ascomycota(100);c__Saccharomycetes(100);o__Saccharomycetales(100);f__Metschnikowiaceae(100);g__Metschnikowia(100);s__Metschnikowia_pulcherrima(100); | Metschnikowia | 13 | Saprotroph | \|Undefined Saprotroph\| | Yeast | NULL | Probable | As nectar-tap_saprotroph (Põlme et al. 2020); Decay sugar-rich substrates (Põlme et al. 2020) | Põlme S, et al. 2020. Fungal Diversity 105:1-16 ((https://doi.org/10.1007/s13225-020-00466-2)) |
| Otu40250 | 1 | k__Fungi(100);p__Ascomycota(100);c__Saccharomycetes(100);o__Saccharomycetales(100);f__Metschnikowiaceae(100);g__Metschnikowia(100);s__Metschnikowia_pulcherrima(100); | Metschnikowia | 13 | Saprotroph | \|Undefined Saprotroph\| | Yeast | NULL | Probable | As nectar-tap_saprotroph (Põlme et al. 2020); Decay sugar-rich substrates (Põlme et al. 2020) | Põlme S, et al. 2020. Fungal Diversity 105:1-16 ((https://doi.org/10.1007/s13225-020-00466-2)) |
| Otu40251 | 1 | k__Fungi(100);p__Ascomycota(100);c__Saccharomycetes(100);o__Saccharomycetales(100);f__Metschnikowiaceae(100);g__Metschnikowia(100);s__Metschnikowia_pulcherrima(100); | Metschnikowia | 13 | Saprotroph | \|Undefined Saprotroph\| | Yeast | NULL | Probable | As nectar-tap_saprotroph (Põlme et al. 2020); Decay sugar-rich substrates (Põlme et al. 2020) | Põlme S, et al. 2020. Fungal Diversity 105:1-16 ((https://doi.org/10.1007/s13225-020-00466-2)) |
| Otu40252 | 1 | k__Fungi(100);p__Ascomycota(100);c__Saccharomycetes(100);o__Saccharomycetales(100);f__Metschnikowiaceae(100);g__Metschnikowia(100);s__Metschnikowia_pulcherrima(100); | Metschnikowia | 13 | Saprotroph | \|Undefined Saprotroph\| | Yeast | NULL | Probable | As nectar-tap_saprotroph (Põlme et al. 2020); Decay sugar-rich substrates (Põlme et al. 2020) | Põlme S, et al. 2020. Fungal Diversity 105:1-16 ((https://doi.org/10.1007/s13225-020-00466-2)) |
| Otu40253 | 1 | k__Fungi(100);p__Ascomycota(100);c__Saccharomycetes(100);o__Saccharomycetales(100);f__Metschnikowiaceae(100);g__Metschnikowia(100);s__Metschnikowia_pulcherrima(100); | Metschnikowia | 13 | Saprotroph | \|Undefined Saprotroph\| | Yeast | NULL | Probable | As nectar-tap_saprotroph (Põlme et al. 2020); Decay sugar-rich substrates (Põlme et al. 2020) | Põlme S, et al. 2020. Fungal Diversity 105:1-16 ((https://doi.org/10.1007/s13225-020-00466-2)) |
| Otu40254 | 1 | k__Fungi(100);p__Ascomycota(100);c__Saccharomycetes(100);o__Saccharomycetales(100);f__Metschnikowiaceae(100);g__Metschnikowia(100);s__Metschnikowia_pulcherrima(100); | Metschnikowia | 13 | Saprotroph | \|Undefined Saprotroph\| | Yeast | NULL | Probable | As nectar-tap_saprotroph (Põlme et al. 2020); Decay sugar-rich substrates (Põlme et al. 2020) | Põlme S, et al. 2020. Fungal Diversity 105:1-16 ((https://doi.org/10.1007/s13225-020-00466-2)) |
| Otu40255 | 1 | k__Fungi(100);p__Ascomycota(100);c__Saccharomycetes(100);o__Saccharomycetales(100);f__Metschnikowiaceae(100);g__Metschnikowia(100);s__Metschnikowia_pulcherrima(100); | Metschnikowia | 13 | Saprotroph | \|Undefined Saprotroph\| | Yeast | NULL | Probable | As nectar-tap_saprotroph (Põlme et al. 2020); Decay sugar-rich substrates (Põlme et al. 2020) | Põlme S, et al. 2020. Fungal Diversity 105:1-16 ((https://doi.org/10.1007/s13225-020-00466-2)) |
| Otu40256 | 1 | k__Fungi(100);p__Ascomycota(100);c__Saccharomycetes(100);o__Saccharomycetales(100);f__Metschnikowiaceae(100);g__Metschnikowia(100);g__Metschnikowia_unclassified(100); | Metschnikowia | 13 | Saprotroph | \|Undefined Saprotroph\| | Yeast | NULL | Probable | As nectar-tap_saprotroph (Põlme et al. 2020); Decay sugar-rich substrates (Põlme et al. 2020) | Põlme S, et al. 2020. Fungal Diversity 105:1-16 ((https://doi.org/10.1007/s13225-020-00466-2)) |
| Otu40257 | 1 | k__Fungi(100);p__Ascomycota(100);c__Saccharomycetes(100);o__Saccharomycetales(100);f__Metschnikowiaceae(100);g__Metschnikowia(100);s__Metschnikowia_pulcherrima(100); | Metschnikowia | 13 | Saprotroph | \|Undefined Saprotroph\| | Yeast | NULL | Probable | As nectar-tap_saprotroph (Põlme et al. 2020); Decay sugar-rich substrates (Põlme et al. 2020) | Põlme S, et al. 2020. Fungal Diversity 105:1-16 ((https://doi.org/10.1007/s13225-020-00466-2)) |
| Otu40258 | 1 | k__Fungi(100);p__Ascomycota(100);c__Saccharomycetes(100);o__Saccharomycetales(100);f__Metschnikowiaceae(100);g__Metschnikowia(100);g__Metschnikowia_unclassified(100); | Metschnikowia | 13 | Saprotroph | \|Undefined Saprotroph\| | Yeast | NULL | Probable | As nectar-tap_saprotroph (Põlme et al. 2020); Decay sugar-rich substrates (Põlme et al. 2020) | Põlme S, et al. 2020. Fungal Diversity 105:1-16 ((https://doi.org/10.1007/s13225-020-00466-2)) |
| Otu40259 | 1 | k__Fungi(100);p__Ascomycota(100);c__Saccharomycetes(100);o__Saccharomycetales(100);f__Metschnikowiaceae(100);g__Metschnikowia(100);s__Metschnikowia_pulcherrima(100); | Metschnikowia | 13 | Saprotroph | \|Undefined Saprotroph\| | Yeast | NULL | Probable | As nectar-tap_saprotroph (Põlme et al. 2020); Decay sugar-rich substrates (Põlme et al. 2020) | Põlme S, et al. 2020. Fungal Diversity 105:1-16 ((https://doi.org/10.1007/s13225-020-00466-2)) |
| Otu40260 | 1 | k__Fungi(100);p__Ascomycota(100);c__Saccharomycetes(100);o__Saccharomycetales(100);f__Metschnikowiaceae(100);g__Metschnikowia(100);g__Metschnikowia_unclassified(100); | Metschnikowia | 13 | Saprotroph | \|Undefined Saprotroph\| | Yeast | NULL | Probable | As nectar-tap_saprotroph (Põlme et al. 2020); Decay sugar-rich substrates (Põlme et al. 2020) | Põlme S, et al. 2020. Fungal Diversity 105:1-16 ((https://doi.org/10.1007/s13225-020-00466-2)) |
| Otu40261 | 1 | k__Fungi(100);p__Ascomycota(100);c__Saccharomycetes(100);o__Saccharomycetales(100);f__Metschnikowiaceae(100);g__Metschnikowia(100);g__Metschnikowia_unclassified(100); | Metschnikowia | 13 | Saprotroph | \|Undefined Saprotroph\| | Yeast | NULL | Probable | As nectar-tap_saprotroph (Põlme et al. 2020); Decay sugar-rich substrates (Põlme et al. 2020) | Põlme S, et al. 2020. Fungal Diversity 105:1-16 ((https://doi.org/10.1007/s13225-020-00466-2)) |
| Otu40262 | 1 | k__Fungi(100);p__Ascomycota(100);c__Saccharomycetes(100);o__Saccharomycetales(100);f__Metschnikowiaceae(100);g__Metschnikowia(100);s__Metschnikowia_pulcherrima(100); | Metschnikowia | 13 | Saprotroph | \|Undefined Saprotroph\| | Yeast | NULL | Probable | As nectar-tap_saprotroph (Põlme et al. 2020); Decay sugar-rich substrates (Põlme et al. 2020) | Põlme S, et al. 2020. Fungal Diversity 105:1-16 ((https://doi.org/10.1007/s13225-020-00466-2)) |
| Otu40263 | 1 | k__Fungi(100);p__Ascomycota(100);c__Saccharomycetes(100);o__Saccharomycetales(100);f__Metschnikowiaceae(100);g__Metschnikowia(100);g__Metschnikowia_unclassified(100); | Metschnikowia | 13 | Saprotroph | \|Undefined Saprotroph\| | Yeast | NULL | Probable | As nectar-tap_saprotroph (Põlme et al. 2020); Decay sugar-rich substrates (Põlme et al. 2020) | Põlme S, et al. 2020. Fungal Diversity 105:1-16 ((https://doi.org/10.1007/s13225-020-00466-2)) |
| Otu40264 | 1 | k__Fungi(100);p__Ascomycota(100);c__Saccharomycetes(100);o__Saccharomycetales(100);f__Metschnikowiaceae(100);g__Metschnikowia(100);s__Metschnikowia_pulcherrima(100); | Metschnikowia | 13 | Saprotroph | \|Undefined Saprotroph\| | Yeast | NULL | Probable | As nectar-tap_saprotroph (Põlme et al. 2020); Decay sugar-rich substrates (Põlme et al. 2020) | Põlme S, et al. 2020. Fungal Diversity 105:1-16 ((https://doi.org/10.1007/s13225-020-00466-2)) |
| Otu40265 | 1 | k__Fungi(100);p__Ascomycota(100);c__Saccharomycetes(100);o__Saccharomycetales(100);f__Metschnikowiaceae(100);g__Metschnikowia(100);s__Metschnikowia_pulcherrima(100); | Metschnikowia | 13 | Saprotroph | \|Undefined Saprotroph\| | Yeast | NULL | Probable | As nectar-tap_saprotroph (Põlme et al. 2020); Decay sugar-rich substrates (Põlme et al. 2020) | Põlme S, et al. 2020. Fungal Diversity 105:1-16 ((https://doi.org/10.1007/s13225-020-00466-2)) |
| Otu40266 | 1 | k__Fungi(100);p__Ascomycota(100);c__Saccharomycetes(100);o__Saccharomycetales(100);f__Metschnikowiaceae(100);g__Metschnikowia(100);s__Metschnikowia_pulcherrima(100); | Metschnikowia | 13 | Saprotroph | \|Undefined Saprotroph\| | Yeast | NULL | Probable | As nectar-tap_saprotroph (Põlme et al. 2020); Decay sugar-rich substrates (Põlme et al. 2020) | Põlme S, et al. 2020. Fungal Diversity 105:1-16 ((https://doi.org/10.1007/s13225-020-00466-2)) |
| Otu40267 | 1 | k__Fungi(100);p__Ascomycota(100);c__Saccharomycetes(100);o__Saccharomycetales(100);f__Metschnikowiaceae(100);g__Metschnikowia(100);s__Metschnikowia_pulcherrima(100); | Metschnikowia | 13 | Saprotroph | \|Undefined Saprotroph\| | Yeast | NULL | Probable | As nectar-tap_saprotroph (Põlme et al. 2020); Decay sugar-rich substrates (Põlme et al. 2020) | Põlme S, et al. 2020. Fungal Diversity 105:1-16 ((https://doi.org/10.1007/s13225-020-00466-2)) |
| Otu40268 | 1 | k__Fungi(100);p__Ascomycota(100);c__Saccharomycetes(100);o__Saccharomycetales(100);f__Metschnikowiaceae(100);g__Metschnikowia(100);s__Metschnikowia_pulcherrima(100); | Metschnikowia | 13 | Saprotroph | \|Undefined Saprotroph\| | Yeast | NULL | Probable | As nectar-tap_saprotroph (Põlme et al. 2020); Decay sugar-rich substrates (Põlme et al. 2020) | Põlme S, et al. 2020. Fungal Diversity 105:1-16 ((https://doi.org/10.1007/s13225-020-00466-2)) |
| Otu40269 | 1 | k__Fungi(100);p__Ascomycota(100);c__Saccharomycetes(100);o__Saccharomycetales(100);f__Metschnikowiaceae(100);g__Metschnikowia(100);g__Metschnikowia_unclassified(100); | Metschnikowia | 13 | Saprotroph | \|Undefined Saprotroph\| | Yeast | NULL | Probable | As nectar-tap_saprotroph (Põlme et al. 2020); Decay sugar-rich substrates (Põlme et al. 2020) | Põlme S, et al. 2020. Fungal Diversity 105:1-16 ((https://doi.org/10.1007/s13225-020-00466-2)) |
| Otu40270 | 1 | k__Fungi(100);p__Ascomycota(100);c__Saccharomycetes(100);o__Saccharomycetales(100);f__Metschnikowiaceae(100);g__Metschnikowia(100);s__Metschnikowia_pulcherrima(100); | Metschnikowia | 13 | Saprotroph | \|Undefined Saprotroph\| | Yeast | NULL | Probable | As nectar-tap_saprotroph (Põlme et al. 2020); Decay sugar-rich substrates (Põlme et al. 2020) | Põlme S, et al. 2020. Fungal Diversity 105:1-16 ((https://doi.org/10.1007/s13225-020-00466-2)) |
| Otu40271 | 1 | k__Fungi(100);p__Ascomycota(100);c__Saccharomycetes(100);o__Saccharomycetales(100);f__Metschnikowiaceae(100);g__Metschnikowia(100);s__Metschnikowia_pulcherrima(100); | Metschnikowia | 13 | Saprotroph | \|Undefined Saprotroph\| | Yeast | NULL | Probable | As nectar-tap_saprotroph (Põlme et al. 2020); Decay sugar-rich substrates (Põlme et al. 2020) | Põlme S, et al. 2020. Fungal Diversity 105:1-16 ((https://doi.org/10.1007/s13225-020-00466-2)) |
| Otu40272 | 1 | k__Fungi(100);p__Ascomycota(100);c__Saccharomycetes(100);o__Saccharomycetales(100);f__Metschnikowiaceae(100);g__Metschnikowia(100);g__Metschnikowia_unclassified(100); | Metschnikowia | 13 | Saprotroph | \|Undefined Saprotroph\| | Yeast | NULL | Probable | As nectar-tap_saprotroph (Põlme et al. 2020); Decay sugar-rich substrates (Põlme et al. 2020) | Põlme S, et al. 2020. Fungal Diversity 105:1-16 ((https://doi.org/10.1007/s13225-020-00466-2)) |
| Otu40273 | 1 | k__Fungi(100);p__Ascomycota(100);c__Saccharomycetes(100);o__Saccharomycetales(100);f__Metschnikowiaceae(100);g__Metschnikowia(100);s__Metschnikowia_pulcherrima(100); | Metschnikowia | 13 | Saprotroph | \|Undefined Saprotroph\| | Yeast | NULL | Probable | As nectar-tap_saprotroph (Põlme et al. 2020); Decay sugar-rich substrates (Põlme et al. 2020) | Põlme S, et al. 2020. Fungal Diversity 105:1-16 ((https://doi.org/10.1007/s13225-020-00466-2)) |
| Otu40274 | 1 | k__Fungi(100);p__Ascomycota(100);c__Saccharomycetes(100);o__Saccharomycetales(100);f__Metschnikowiaceae(100);g__Metschnikowia(100);g__Metschnikowia_unclassified(100); | Metschnikowia | 13 | Saprotroph | \|Undefined Saprotroph\| | Yeast | NULL | Probable | As nectar-tap_saprotroph (Põlme et al. 2020); Decay sugar-rich substrates (Põlme et al. 2020) | Põlme S, et al. 2020. Fungal Diversity 105:1-16 ((https://doi.org/10.1007/s13225-020-00466-2)) |
| Otu40275 | 1 | k__Fungi(100);p__Ascomycota(100);c__Saccharomycetes(100);o__Saccharomycetales(100);f__Metschnikowiaceae(100);g__Metschnikowia(100);s__Metschnikowia_pulcherrima(100); | Metschnikowia | 13 | Saprotroph | \|Undefined Saprotroph\| | Yeast | NULL | Probable | As nectar-tap_saprotroph (Põlme et al. 2020); Decay sugar-rich substrates (Põlme et al. 2020) | Põlme S, et al. 2020. Fungal Diversity 105:1-16 ((https://doi.org/10.1007/s13225-020-00466-2)) |
| Otu40276 | 1 | k__Fungi(100);p__Ascomycota(100);c__Saccharomycetes(100);o__Saccharomycetales(100);f__Metschnikowiaceae(100);g__Metschnikowia(100);s__Metschnikowia_pulcherrima(100); | Metschnikowia | 13 | Saprotroph | \|Undefined Saprotroph\| | Yeast | NULL | Probable | As nectar-tap_saprotroph (Põlme et al. 2020); Decay sugar-rich substrates (Põlme et al. 2020) | Põlme S, et al. 2020. Fungal Diversity 105:1-16 ((https://doi.org/10.1007/s13225-020-00466-2)) |
| Otu40277 | 1 | k__Fungi(100);p__Ascomycota(100);c__Saccharomycetes(100);o__Saccharomycetales(100);f__Metschnikowiaceae(100);g__Metschnikowia(100);s__Metschnikowia_pulcherrima(100); | Metschnikowia | 13 | Saprotroph | \|Undefined Saprotroph\| | Yeast | NULL | Probable | As nectar-tap_saprotroph (Põlme et al. 2020); Decay sugar-rich substrates (Põlme et al. 2020) | Põlme S, et al. 2020. Fungal Diversity 105:1-16 ((https://doi.org/10.1007/s13225-020-00466-2)) |
| Otu40278 | 1 | k__Fungi(100);p__Ascomycota(100);c__Saccharomycetes(100);o__Saccharomycetales(100);f__Metschnikowiaceae(100);g__Metschnikowia(100);g__Metschnikowia_unclassified(100); | Metschnikowia | 13 | Saprotroph | \|Undefined Saprotroph\| | Yeast | NULL | Probable | As nectar-tap_saprotroph (Põlme et al. 2020); Decay sugar-rich substrates (Põlme et al. 2020) | Põlme S, et al. 2020. Fungal Diversity 105:1-16 ((https://doi.org/10.1007/s13225-020-00466-2)) |
| Otu40279 | 1 | k__Fungi(100);p__Ascomycota(100);c__Saccharomycetes(100);o__Saccharomycetales(100);f__Metschnikowiaceae(100);g__Metschnikowia(100);s__Metschnikowia_pulcherrima(100); | Metschnikowia | 13 | Saprotroph | \|Undefined Saprotroph\| | Yeast | NULL | Probable | As nectar-tap_saprotroph (Põlme et al. 2020); Decay sugar-rich substrates (Põlme et al. 2020) | Põlme S, et al. 2020. Fungal Diversity 105:1-16 ((https://doi.org/10.1007/s13225-020-00466-2)) |
| Otu40280 | 1 | k__Fungi(100);p__Ascomycota(100);c__Saccharomycetes(100);o__Saccharomycetales(100);f__Metschnikowiaceae(100);g__Metschnikowia(100);s__Metschnikowia_pulcherrima(100); | Metschnikowia | 13 | Saprotroph | \|Undefined Saprotroph\| | Yeast | NULL | Probable | As nectar-tap_saprotroph (Põlme et al. 2020); Decay sugar-rich substrates (Põlme et al. 2020) | Põlme S, et al. 2020. Fungal Diversity 105:1-16 ((https://doi.org/10.1007/s13225-020-00466-2)) |
| Otu40281 | 1 | k__Fungi(100);p__Ascomycota(100);c__Saccharomycetes(100);o__Saccharomycetales(100);f__Metschnikowiaceae(100);g__Metschnikowia(100);s__Metschnikowia_pulcherrima(100); | Metschnikowia | 13 | Saprotroph | \|Undefined Saprotroph\| | Yeast | NULL | Probable | As nectar-tap_saprotroph (Põlme et al. 2020); Decay sugar-rich substrates (Põlme et al. 2020) | Põlme S, et al. 2020. Fungal Diversity 105:1-16 ((https://doi.org/10.1007/s13225-020-00466-2)) |
| Otu40282 | 1 | k__Fungi(100);p__Ascomycota(100);c__Saccharomycetes(100);o__Saccharomycetales(100);f__Metschnikowiaceae(100);g__Metschnikowia(100);s__Metschnikowia_pulcherrima(100); | Metschnikowia | 13 | Saprotroph | \|Undefined Saprotroph\| | Yeast | NULL | Probable | As nectar-tap_saprotroph (Põlme et al. 2020); Decay sugar-rich substrates (Põlme et al. 2020) | Põlme S, et al. 2020. Fungal Diversity 105:1-16 ((https://doi.org/10.1007/s13225-020-00466-2)) |
| Otu40284 | 1 | k__Fungi(100);p__Ascomycota(100);c__Saccharomycetes(100);o__Saccharomycetales(100);f__Metschnikowiaceae(100);g__Metschnikowia(100);g__Metschnikowia_unclassified(100); | Metschnikowia | 13 | Saprotroph | \|Undefined Saprotroph\| | Yeast | NULL | Probable | As nectar-tap_saprotroph (Põlme et al. 2020); Decay sugar-rich substrates (Põlme et al. 2020) | Põlme S, et al. 2020. Fungal Diversity 105:1-16 ((https://doi.org/10.1007/s13225-020-00466-2)) |
| Otu40285 | 1 | k__Fungi(100);p__Ascomycota(100);c__Saccharomycetes(100);o__Saccharomycetales(100);f__Metschnikowiaceae(100);g__Metschnikowia(100);s__Metschnikowia_pulcherrima(100); | Metschnikowia | 13 | Saprotroph | \|Undefined Saprotroph\| | Yeast | NULL | Probable | As nectar-tap_saprotroph (Põlme et al. 2020); Decay sugar-rich substrates (Põlme et al. 2020) | Põlme S, et al. 2020. Fungal Diversity 105:1-16 ((https://doi.org/10.1007/s13225-020-00466-2)) |
| Otu40286 | 1 | k__Fungi(100);p__Ascomycota(100);c__Saccharomycetes(100);o__Saccharomycetales(100);f__Metschnikowiaceae(100);g__Metschnikowia(100);g__Metschnikowia_unclassified(100); | Metschnikowia | 13 | Saprotroph | \|Undefined Saprotroph\| | Yeast | NULL | Probable | As nectar-tap_saprotroph (Põlme et al. 2020); Decay sugar-rich substrates (Põlme et al. 2020) | Põlme S, et al. 2020. Fungal Diversity 105:1-16 ((https://doi.org/10.1007/s13225-020-00466-2)) |
| Otu40287 | 1 | k__Fungi(100);p__Ascomycota(100);c__Saccharomycetes(100);o__Saccharomycetales(100);f__Metschnikowiaceae(100);g__Metschnikowia(100);s__Metschnikowia_pulcherrima(100); | Metschnikowia | 13 | Saprotroph | \|Undefined Saprotroph\| | Yeast | NULL | Probable | As nectar-tap_saprotroph (Põlme et al. 2020); Decay sugar-rich substrates (Põlme et al. 2020) | Põlme S, et al. 2020. Fungal Diversity 105:1-16 ((https://doi.org/10.1007/s13225-020-00466-2)) |
| Otu40288 | 1 | k__Fungi(100);p__Ascomycota(100);c__Saccharomycetes(100);o__Saccharomycetales(100);f__Metschnikowiaceae(100);g__Metschnikowia(100);s__Metschnikowia_pulcherrima(100); | Metschnikowia | 13 | Saprotroph | \|Undefined Saprotroph\| | Yeast | NULL | Probable | As nectar-tap_saprotroph (Põlme et al. 2020); Decay sugar-rich substrates (Põlme et al. 2020) | Põlme S, et al. 2020. Fungal Diversity 105:1-16 ((https://doi.org/10.1007/s13225-020-00466-2)) |
| Otu40289 | 1 | k__Fungi(100);p__Ascomycota(100);c__Saccharomycetes(100);o__Saccharomycetales(100);f__Metschnikowiaceae(100);g__Metschnikowia(100);s__Metschnikowia_pulcherrima(100); | Metschnikowia | 13 | Saprotroph | \|Undefined Saprotroph\| | Yeast | NULL | Probable | As nectar-tap_saprotroph (Põlme et al. 2020); Decay sugar-rich substrates (Põlme et al. 2020) | Põlme S, et al. 2020. Fungal Diversity 105:1-16 ((https://doi.org/10.1007/s13225-020-00466-2)) |
| Otu40290 | 1 | k__Fungi(100);p__Ascomycota(100);c__Saccharomycetes(100);o__Saccharomycetales(100);f__Metschnikowiaceae(100);g__Metschnikowia(100);s__Metschnikowia_pulcherrima(100); | Metschnikowia | 13 | Saprotroph | \|Undefined Saprotroph\| | Yeast | NULL | Probable | As nectar-tap_saprotroph (Põlme et al. 2020); Decay sugar-rich substrates (Põlme et al. 2020) | Põlme S, et al. 2020. Fungal Diversity 105:1-16 ((https://doi.org/10.1007/s13225-020-00466-2)) |
| Otu40291 | 1 | k__Fungi(100);p__Ascomycota(100);c__Saccharomycetes(100);o__Saccharomycetales(100);f__Metschnikowiaceae(100);g__Metschnikowia(100);s__Metschnikowia_pulcherrima(100); | Metschnikowia | 13 | Saprotroph | \|Undefined Saprotroph\| | Yeast | NULL | Probable | As nectar-tap_saprotroph (Põlme et al. 2020); Decay sugar-rich substrates (Põlme et al. 2020) | Põlme S, et al. 2020. Fungal Diversity 105:1-16 ((https://doi.org/10.1007/s13225-020-00466-2)) |
| Otu40292 | 1 | k__Fungi(100);p__Ascomycota(100);c__Saccharomycetes(100);o__Saccharomycetales(100);f__Metschnikowiaceae(100);g__Metschnikowia(100);s__Metschnikowia_pulcherrima(100); | Metschnikowia | 13 | Saprotroph | \|Undefined Saprotroph\| | Yeast | NULL | Probable | As nectar-tap_saprotroph (Põlme et al. 2020); Decay sugar-rich substrates (Põlme et al. 2020) | Põlme S, et al. 2020. Fungal Diversity 105:1-16 ((https://doi.org/10.1007/s13225-020-00466-2)) |
| Otu40293 | 1 | k__Fungi(100);p__Ascomycota(100);c__Saccharomycetes(100);o__Saccharomycetales(100);f__Metschnikowiaceae(100);g__Metschnikowia(100);s__Metschnikowia_pulcherrima(100); | Metschnikowia | 13 | Saprotroph | \|Undefined Saprotroph\| | Yeast | NULL | Probable | As nectar-tap_saprotroph (Põlme et al. 2020); Decay sugar-rich substrates (Põlme et al. 2020) | Põlme S, et al. 2020. Fungal Diversity 105:1-16 ((https://doi.org/10.1007/s13225-020-00466-2)) |
| Otu40294 | 1 | k__Fungi(100);p__Ascomycota(100);c__Saccharomycetes(100);o__Saccharomycetales(100);f__Metschnikowiaceae(100);g__Metschnikowia(100);g__Metschnikowia_unclassified(100); | Metschnikowia | 13 | Saprotroph | \|Undefined Saprotroph\| | Yeast | NULL | Probable | As nectar-tap_saprotroph (Põlme et al. 2020); Decay sugar-rich substrates (Põlme et al. 2020) | Põlme S, et al. 2020. Fungal Diversity 105:1-16 ((https://doi.org/10.1007/s13225-020-00466-2)) |
| Otu40295 | 1 | k__Fungi(100);p__Ascomycota(100);c__Saccharomycetes(100);o__Saccharomycetales(100);f__Metschnikowiaceae(100);g__Metschnikowia(100);s__Metschnikowia_pulcherrima(100); | Metschnikowia | 13 | Saprotroph | \|Undefined Saprotroph\| | Yeast | NULL | Probable | As nectar-tap_saprotroph (Põlme et al. 2020); Decay sugar-rich substrates (Põlme et al. 2020) | Põlme S, et al. 2020. Fungal Diversity 105:1-16 ((https://doi.org/10.1007/s13225-020-00466-2)) |
| Otu40296 | 1 | k__Fungi(100);p__Ascomycota(100);c__Saccharomycetes(100);o__Saccharomycetales(100);f__Metschnikowiaceae(100);g__Metschnikowia(100);s__Metschnikowia_pulcherrima(100); | Metschnikowia | 13 | Saprotroph | \|Undefined Saprotroph\| | Yeast | NULL | Probable | As nectar-tap_saprotroph (Põlme et al. 2020); Decay sugar-rich substrates (Põlme et al. 2020) | Põlme S, et al. 2020. Fungal Diversity 105:1-16 ((https://doi.org/10.1007/s13225-020-00466-2)) |
| Otu40297 | 1 | k__Fungi(100);p__Ascomycota(100);c__Saccharomycetes(100);o__Saccharomycetales(100);f__Metschnikowiaceae(100);g__Metschnikowia(100);s__Metschnikowia_pulcherrima(100); | Metschnikowia | 13 | Saprotroph | \|Undefined Saprotroph\| | Yeast | NULL | Probable | As nectar-tap_saprotroph (Põlme et al. 2020); Decay sugar-rich substrates (Põlme et al. 2020) | Põlme S, et al. 2020. Fungal Diversity 105:1-16 ((https://doi.org/10.1007/s13225-020-00466-2)) |
| Otu40298 | 1 | k__Fungi(100);p__Ascomycota(100);c__Saccharomycetes(100);o__Saccharomycetales(100);f__Metschnikowiaceae(100);g__Metschnikowia(100);g__Metschnikowia_unclassified(100); | Metschnikowia | 13 | Saprotroph | \|Undefined Saprotroph\| | Yeast | NULL | Probable | As nectar-tap_saprotroph (Põlme et al. 2020); Decay sugar-rich substrates (Põlme et al. 2020) | Põlme S, et al. 2020. Fungal Diversity 105:1-16 ((https://doi.org/10.1007/s13225-020-00466-2)) |
| Otu40299 | 1 | k__Fungi(100);p__Ascomycota(100);c__Saccharomycetes(100);o__Saccharomycetales(100);f__Metschnikowiaceae(100);g__Metschnikowia(100);s__Metschnikowia_pulcherrima(100); | Metschnikowia | 13 | Saprotroph | \|Undefined Saprotroph\| | Yeast | NULL | Probable | As nectar-tap_saprotroph (Põlme et al. 2020); Decay sugar-rich substrates (Põlme et al. 2020) | Põlme S, et al. 2020. Fungal Diversity 105:1-16 ((https://doi.org/10.1007/s13225-020-00466-2)) |
| Otu40300 | 1 | k__Fungi(100);p__Ascomycota(100);c__Saccharomycetes(100);o__Saccharomycetales(100);f__Metschnikowiaceae(100);g__Metschnikowia(100);s__Metschnikowia_pulcherrima(100); | Metschnikowia | 13 | Saprotroph | \|Undefined Saprotroph\| | Yeast | NULL | Probable | As nectar-tap_saprotroph (Põlme et al. 2020); Decay sugar-rich substrates (Põlme et al. 2020) | Põlme S, et al. 2020. Fungal Diversity 105:1-16 ((https://doi.org/10.1007/s13225-020-00466-2)) |
| Otu40301 | 1 | k__Fungi(100);p__Ascomycota(100);c__Saccharomycetes(100);o__Saccharomycetales(100);f__Metschnikowiaceae(100);g__Metschnikowia(100);s__Metschnikowia_pulcherrima(100); | Metschnikowia | 13 | Saprotroph | \|Undefined Saprotroph\| | Yeast | NULL | Probable | As nectar-tap_saprotroph (Põlme et al. 2020); Decay sugar-rich substrates (Põlme et al. 2020) | Põlme S, et al. 2020. Fungal Diversity 105:1-16 ((https://doi.org/10.1007/s13225-020-00466-2)) |
| Otu40302 | 1 | k__Fungi(100);p__Ascomycota(100);c__Saccharomycetes(100);o__Saccharomycetales(100);f__Metschnikowiaceae(100);g__Metschnikowia(100);s__Metschnikowia_pulcherrima(100); | Metschnikowia | 13 | Saprotroph | \|Undefined Saprotroph\| | Yeast | NULL | Probable | As nectar-tap_saprotroph (Põlme et al. 2020); Decay sugar-rich substrates (Põlme et al. 2020) | Põlme S, et al. 2020. Fungal Diversity 105:1-16 ((https://doi.org/10.1007/s13225-020-00466-2)) |

| | | | | | | | | | | | |
|---|---|---|---|---|---|---|---|---|---|---|---|
| Otu40304 | 1 | k__Fungi(100);p__Ascomycota(100);c__Saccharomycetes(100);o__Saccharomycetales(100);f__Metschnikowiaceae(100);g__Metschnikowia(100);g__Metschnikowia_unclassified(100); | Metschnikowia | 13 | Saprotroph | \|Undefined Saprotroph\| | Yeast | NULL | Probable | As nectar-tap_saprotroph (Põlme et al. 2020); Decay sugar-rich substrates (Põlme et al. 2020) | Põlme S, et al. 2020. Fungal Diversity 105:1-16 ((https://doi.org/10.1007/s13225-020-00466-2)) |
| Otu40305 | 1 | k__Fungi(100);p__Ascomycota(100);c__Saccharomycetes(100);o__Saccharomycetales(100);f__Metschnikowiaceae(100);g__Metschnikowia(100);g__Metschnikowia_unclassified(100); | Metschnikowia | 13 | Saprotroph | \|Undefined Saprotroph\| | Yeast | NULL | Probable | As nectar-tap_saprotroph (Põlme et al. 2020); Decay sugar-rich substrates (Põlme et al. 2020) | Põlme S, et al. 2020. Fungal Diversity 105:1-16 ((https://doi.org/10.1007/s13225-020-00466-2)) |
| Otu40306 | 1 | k__Fungi(100);p__Ascomycota(100);c__Saccharomycetes(100);o__Saccharomycetales(100);f__Metschnikowiaceae(100);g__Metschnikowia(100);s__Metschnikowia_pulcherrima(100); | Metschnikowia | 13 | Saprotroph | \|Undefined Saprotroph\| | Yeast | NULL | Probable | As nectar-tap_saprotroph (Põlme et al. 2020); Decay sugar-rich substrates (Põlme et al. 2020) | Põlme S, et al. 2020. Fungal Diversity 105:1-16 ((https://doi.org/10.1007/s13225-020-00466-2)) |
| Otu40307 | 1 | k__Fungi(100);p__Ascomycota(100);c__Saccharomycetes(100);o__Saccharomycetales(100);f__Metschnikowiaceae(100);g__Metschnikowia(100);s__Metschnikowia_pulcherrima(100); | Metschnikowia | 13 | Saprotroph | \|Undefined Saprotroph\| | Yeast | NULL | Probable | As nectar-tap_saprotroph (Põlme et al. 2020); Decay sugar-rich substrates (Põlme et al. 2020) | Põlme S, et al. 2020. Fungal Diversity 105:1-16 ((https://doi.org/10.1007/s13225-020-00466-2)) |
| Otu40308 | 1 | k__Fungi(100);p__Ascomycota(100);c__Saccharomycetes(100);o__Saccharomycetales(100);f__Metschnikowiaceae(100);g__Metschnikowia(100);s__Metschnikowia_pulcherrima(100); | Metschnikowia | 13 | Saprotroph | \|Undefined Saprotroph\| | Yeast | NULL | Probable | As nectar-tap_saprotroph (Põlme et al. 2020); Decay sugar-rich substrates (Põlme et al. 2020) | Põlme S, et al. 2020. Fungal Diversity 105:1-16 ((https://doi.org/10.1007/s13225-020-00466-2)) |
| Otu40309 | 1 | k__Fungi(100);p__Ascomycota(100);c__Saccharomycetes(100);o__Saccharomycetales(100);f__Metschnikowiaceae(100);g__Metschnikowia(100);g__Metschnikowia_unclassified(100); | Metschnikowia | 13 | Saprotroph | \|Undefined Saprotroph\| | Yeast | NULL | Probable | As nectar-tap_saprotroph (Põlme et al. 2020); Decay sugar-rich substrates (Põlme et al. 2020) | Põlme S, et al. 2020. Fungal Diversity 105:1-16 ((https://doi.org/10.1007/s13225-020-00466-2)) |
| Otu40310 | 1 | k__Fungi(100);p__Ascomycota(100);c__Saccharomycetes(100);o__Saccharomycetales(100);f__Metschnikowiaceae(100);g__Metschnikowia(100);g__Metschnikowia_unclassified(100); | Metschnikowia | 13 | Saprotroph | \|Undefined Saprotroph\| | Yeast | NULL | Probable | As nectar-tap_saprotroph (Põlme et al. 2020); Decay sugar-rich substrates (Põlme et al. 2020) | Põlme S, et al. 2020. Fungal Diversity 105:1-16 ((https://doi.org/10.1007/s13225-020-00466-2)) |
| Otu40311 | 1 | k__Fungi(100);p__Ascomycota(100);c__Saccharomycetes(100);o__Saccharomycetales(100);f__Metschnikowiaceae(100);g__Metschnikowia(100);s__Metschnikowia_pulcherrima(100); | Metschnikowia | 13 | Saprotroph | \|Undefined Saprotroph\| | Yeast | NULL | Probable | As nectar-tap_saprotroph (Põlme et al. 2020); Decay sugar-rich substrates (Põlme et al. 2020) | Põlme S, et al. 2020. Fungal Diversity 105:1-16 ((https://doi.org/10.1007/s13225-020-00466-2)) |
| Otu40312 | 1 | k__Fungi(100);p__Ascomycota(100);c__Saccharomycetes(100);o__Saccharomycetales(100);f__Metschnikowiaceae(100);g__Metschnikowia(100);s__Metschnikowia_pulcherrima(100); | Metschnikowia | 13 | Saprotroph | \|Undefined Saprotroph\| | Yeast | NULL | Probable | As nectar-tap_saprotroph (Põlme et al. 2020); Decay sugar-rich substrates (Põlme et al. 2020) | Põlme S, et al. 2020. Fungal Diversity 105:1-16 ((https://doi.org/10.1007/s13225-020-00466-2)) |
| Otu40315 | 1 | k__Fungi(100);p__Ascomycota(100);c__Saccharomycetes(100);o__Saccharomycetales(100);f__Metschnikowiaceae(100);g__Metschnikowia(100);s__Metschnikowia_pulcherrima(100); | Metschnikowia | 13 | Saprotroph | \|Undefined Saprotroph\| | Yeast | NULL | Probable | As nectar-tap_saprotroph (Põlme et al. 2020); Decay sugar-rich substrates (Põlme et al. 2020) | Põlme S, et al. 2020. Fungal Diversity 105:1-16 ((https://doi.org/10.1007/s13225-020-00466-2)) |
| Otu40317 | 1 | k__Fungi(100);p__Ascomycota(100);c__Saccharomycetes(100);o__Saccharomycetales(100);f__Metschnikowiaceae(100);g__Metschnikowia(100);s__Metschnikowia_pulcherrima(100); | Metschnikowia | 13 | Saprotroph | \|Undefined Saprotroph\| | Yeast | NULL | Probable | As nectar-tap_saprotroph (Põlme et al. 2020); Decay sugar-rich substrates (Põlme et al. 2020) | Põlme S, et al. 2020. Fungal Diversity 105:1-16 ((https://doi.org/10.1007/s13225-020-00466-2)) |
| Otu40319 | 1 | k__Fungi(100);p__Ascomycota(100);c__Saccharomycetes(100);o__Saccharomycetales(100);f__Metschnikowiaceae(100);g__Metschnikowia(100);g__Metschnikowia_unclassified(100); | Metschnikowia | 13 | Saprotroph | \|Undefined Saprotroph\| | Yeast | NULL | Probable | As nectar-tap_saprotroph (Põlme et al. 2020); Decay sugar-rich substrates (Põlme et al. 2020) | Põlme S, et al. 2020. Fungal Diversity 105:1-16 ((https://doi.org/10.1007/s13225-020-00466-2)) |
| Otu40320 | 1 | k__Fungi(100);p__Ascomycota(100);c__Saccharomycetes(100);o__Saccharomycetales(100);f__Metschnikowiaceae(100);g__Metschnikowia(100);s__Metschnikowia_pulcherrima(100); | Metschnikowia | 13 | Saprotroph | \|Undefined Saprotroph\| | Yeast | NULL | Probable | As nectar-tap_saprotroph (Põlme et al. 2020); Decay sugar-rich substrates (Põlme et al. 2020) | Põlme S, et al. 2020. Fungal Diversity 105:1-16 ((https://doi.org/10.1007/s13225-020-00466-2)) |
| Otu40321 | 1 | k__Fungi(100);p__Ascomycota(100);c__Saccharomycetes(100);o__Saccharomycetales(100);f__Metschnikowiaceae(100);g__Metschnikowia(100);g__Metschnikowia_unclassified(100); | Metschnikowia | 13 | Saprotroph | \|Undefined Saprotroph\| | Yeast | NULL | Probable | As nectar-tap_saprotroph (Põlme et al. 2020); Decay sugar-rich substrates (Põlme et al. 2020) | Põlme S, et al. 2020. Fungal Diversity 105:1-16 ((https://doi.org/10.1007/s13225-020-00466-2)) |
| Otu40322 | 1 | k__Fungi(100);p__Ascomycota(100);c__Saccharomycetes(100);o__Saccharomycetales(100);f__Metschnikowiaceae(100);g__Metschnikowia(100);s__Metschnikowia_pulcherrima(100); | Metschnikowia | 13 | Saprotroph | \|Undefined Saprotroph\| | Yeast | NULL | Probable | As nectar-tap_saprotroph (Põlme et al. 2020); Decay sugar-rich substrates (Põlme et al. 2020) | Põlme S, et al. 2020. Fungal Diversity 105:1-16 ((https://doi.org/10.1007/s13225-020-00466-2)) |
| Otu40324 | 1 | k__Fungi(100);p__Ascomycota(100);c__Saccharomycetes(100);o__Saccharomycetales(100);f__Metschnikowiaceae(100);g__Metschnikowia(100);s__Metschnikowia_pulcherrima(100); | Metschnikowia | 13 | Saprotroph | \|Undefined Saprotroph\| | Yeast | NULL | Probable | As nectar-tap_saprotroph (Põlme et al. 2020); Decay sugar-rich substrates (Põlme et al. 2020) | Põlme S, et al. 2020. Fungal Diversity 105:1-16 ((https://doi.org/10.1007/s13225-020-00466-2)) |
| Otu40325 | 1 | k__Fungi(100);p__Ascomycota(100);c__Saccharomycetes(100);o__Saccharomycetales(100);f__Metschnikowiaceae(100);g__Metschnikowia(100);s__Metschnikowia_pulcherrima(100); | Metschnikowia | 13 | Saprotroph | \|Undefined Saprotroph\| | Yeast | NULL | Probable | As nectar-tap_saprotroph (Põlme et al. 2020); Decay sugar-rich substrates (Põlme et al. 2020) | Põlme S, et al. 2020. Fungal Diversity 105:1-16 ((https://doi.org/10.1007/s13225-020-00466-2)) |
| Otu40326 | 1 | k__Fungi(100);p__Ascomycota(100);c__Saccharomycetes(100);o__Saccharomycetales(100);f__Metschnikowiaceae(100);g__Metschnikowia(100);s__Metschnikowia_pulcherrima(100); | Metschnikowia | 13 | Saprotroph | \|Undefined Saprotroph\| | Yeast | NULL | Probable | As nectar-tap_saprotroph (Põlme et al. 2020); Decay sugar-rich substrates (Põlme et al. 2020) | Põlme S, et al. 2020. Fungal Diversity 105:1-16 ((https://doi.org/10.1007/s13225-020-00466-2)) |
| Otu40327 | 1 | k__Fungi(100);p__Ascomycota(100);c__Saccharomycetes(100);o__Saccharomycetales(100);f__Metschnikowiaceae(100);g__Metschnikowia(100);g__Metschnikowia_unclassified(100); | Metschnikowia | 13 | Saprotroph | \|Undefined Saprotroph\| | Yeast | NULL | Probable | As nectar-tap_saprotroph (Põlme et al. 2020); Decay sugar-rich substrates (Põlme et al. 2020) | Põlme S, et al. 2020. Fungal Diversity 105:1-16 ((https://doi.org/10.1007/s13225-020-00466-2)) |
| Otu40328 | 1 | k__Fungi(100);p__Ascomycota(100);c__Saccharomycetes(100);o__Saccharomycetales(100);f__Metschnikowiaceae(100);g__Metschnikowia(100);s__Metschnikowia_pulcherrima(100); | Metschnikowia | 13 | Saprotroph | \|Undefined Saprotroph\| | Yeast | NULL | Probable | As nectar-tap_saprotroph (Põlme et al. 2020); Decay sugar-rich substrates (Põlme et al. 2020) | Põlme S, et al. 2020. Fungal Diversity 105:1-16 ((https://doi.org/10.1007/s13225-020-00466-2)) |
| Otu40329 | 1 | k__Fungi(100);p__Ascomycota(100);c__Saccharomycetes(100);o__Saccharomycetales(100);f__Metschnikowiaceae(100);g__Metschnikowia(100);s__Metschnikowia_pulcherrima(100); | Metschnikowia | 13 | Saprotroph | \|Undefined Saprotroph\| | Yeast | NULL | Probable | As nectar-tap_saprotroph (Põlme et al. 2020); Decay sugar-rich substrates (Põlme et al. 2020) | Põlme S, et al. 2020. Fungal Diversity 105:1-16 ((https://doi.org/10.1007/s13225-020-00466-2)) |
| Otu40330 | 1 | k__Fungi(100);p__Ascomycota(100);c__Saccharomycetes(100);o__Saccharomycetales(100);f__Metschnikowiaceae(100);g__Metschnikowia(100);g__Metschnikowia_unclassified(100); | Metschnikowia | 13 | Saprotroph | \|Undefined Saprotroph\| | Yeast | NULL | Probable | As nectar-tap_saprotroph (Põlme et al. 2020); Decay sugar-rich substrates (Põlme et al. 2020) | Põlme S, et al. 2020. Fungal Diversity 105:1-16 ((https://doi.org/10.1007/s13225-020-00466-2)) |
| Otu40331 | 1 | k__Fungi(100);p__Ascomycota(100);c__Saccharomycetes(100);o__Saccharomycetales(100);f__Metschnikowiaceae(100);g__Metschnikowia(100);s__Metschnikowia_pulcherrima(100); | Metschnikowia | 13 | Saprotroph | \|Undefined Saprotroph\| | Yeast | NULL | Probable | As nectar-tap_saprotroph (Põlme et al. 2020); Decay sugar-rich substrates (Põlme et al. 2020) | Põlme S, et al. 2020. Fungal Diversity 105:1-16 ((https://doi.org/10.1007/s13225-020-00466-2)) |
| Otu40332 | 1 | k__Fungi(100);p__Ascomycota(100);c__Saccharomycetes(100);o__Saccharomycetales(100);f__Metschnikowiaceae(100);g__Metschnikowia(100);g__Metschnikowia_unclassified(100); | Metschnikowia | 13 | Saprotroph | \|Undefined Saprotroph\| | Yeast | NULL | Probable | As nectar-tap_saprotroph (Põlme et al. 2020); Decay sugar-rich substrates (Põlme et al. 2020) | Põlme S, et al. 2020. Fungal Diversity 105:1-16 ((https://doi.org/10.1007/s13225-020-00466-2)) |
| Otu40333 | 1 | k__Fungi(100);p__Ascomycota(100);c__Saccharomycetes(100);o__Saccharomycetales(100);f__Metschnikowiaceae(100);g__Metschnikowia(100);s__Metschnikowia_pulcherrima(100); | Metschnikowia | 13 | Saprotroph | \|Undefined Saprotroph\| | Yeast | NULL | Probable | As nectar-tap_saprotroph (Põlme et al. 2020); Decay sugar-rich substrates (Põlme et al. 2020) | Põlme S, et al. 2020. Fungal Diversity 105:1-16 ((https://doi.org/10.1007/s13225-020-00466-2)) |
| Otu40334 | 1 | k__Fungi(100);p__Ascomycota(100);c__Saccharomycetes(100);o__Saccharomycetales(100);f__Metschnikowiaceae(100);g__Metschnikowia(100);s__Metschnikowia_pulcherrima(100); | Metschnikowia | 13 | Saprotroph | \|Undefined Saprotroph\| | Yeast | NULL | Probable | As nectar-tap_saprotroph (Põlme et al. 2020); Decay sugar-rich substrates (Põlme et al. 2020) | Põlme S, et al. 2020. Fungal Diversity 105:1-16 ((https://doi.org/10.1007/s13225-020-00466-2)) |
| Otu40335 | 1 | k__Fungi(100);p__Ascomycota(100);c__Saccharomycetes(100);o__Saccharomycetales(100);f__Metschnikowiaceae(100);g__Metschnikowia(100);s__Metschnikowia_pulcherrima(100); | Metschnikowia | 13 | Saprotroph | \|Undefined Saprotroph\| | Yeast | NULL | Probable | As nectar-tap_saprotroph (Põlme et al. 2020); Decay sugar-rich substrates (Põlme et al. 2020) | Põlme S, et al. 2020. Fungal Diversity 105:1-16 ((https://doi.org/10.1007/s13225-020-00466-2)) |
| Otu40336 | 1 | k__Fungi(100);p__Ascomycota(100);c__Saccharomycetes(100);o__Saccharomycetales(100);f__Metschnikowiaceae(100);g__Metschnikowia(100);s__Metschnikowia_pulcherrima(100); | Metschnikowia | 13 | Saprotroph | \|Undefined Saprotroph\| | Yeast | NULL | Probable | As nectar-tap_saprotroph (Põlme et al. 2020); Decay sugar-rich substrates (Põlme et al. 2020) | Põlme S, et al. 2020. Fungal Diversity 105:1-16 ((https://doi.org/10.1007/s13225-020-00466-2)) |
| Otu40337 | 1 | k__Fungi(100);p__Ascomycota(100);c__Saccharomycetes(100);o__Saccharomycetales(100);f__Metschnikowiaceae(100);g__Metschnikowia(100);s__Metschnikowia_pulcherrima(100); | Metschnikowia | 13 | Saprotroph | \|Undefined Saprotroph\| | Yeast | NULL | Probable | As nectar-tap_saprotroph (Põlme et al. 2020); Decay sugar-rich substrates (Põlme et al. 2020) | Põlme S, et al. 2020. Fungal Diversity 105:1-16 ((https://doi.org/10.1007/s13225-020-00466-2)) |
| Otu40338 | 1 | k__Fungi(100);p__Ascomycota(100);c__Saccharomycetes(100);o__Saccharomycetales(100);f__Metschnikowiaceae(100);g__Metschnikowia(100);s__Metschnikowia_pulcherrima(100); | Metschnikowia | 13 | Saprotroph | \|Undefined Saprotroph\| | Yeast | NULL | Probable | As nectar-tap_saprotroph (Põlme et al. 2020); Decay sugar-rich substrates (Põlme et al. 2020) | Põlme S, et al. 2020. Fungal Diversity 105:1-16 ((https://doi.org/10.1007/s13225-020-00466-2)) |
| Otu40339 | 1 | k__Fungi(100);p__Ascomycota(100);c__Saccharomycetes(100);o__Saccharomycetales(100);f__Metschnikowiaceae(100);g__Metschnikowia(100);s__Metschnikowia_pulcherrima(100); | Metschnikowia | 13 | Saprotroph | \|Undefined Saprotroph\| | Yeast | NULL | Probable | As nectar-tap_saprotroph (Põlme et al. 2020); Decay sugar-rich substrates (Põlme et al. 2020) | Põlme S, et al. 2020. Fungal Diversity 105:1-16 ((https://doi.org/10.1007/s13225-020-00466-2)) |
| Otu40340 | 1 | k__Fungi(100);p__Ascomycota(100);c__Saccharomycetes(100);o__Saccharomycetales(100);f__Metschnikowiaceae(100);g__Metschnikowia(100);s__Metschnikowia_pulcherrima(100); | Metschnikowia | 13 | Saprotroph | \|Undefined Saprotroph\| | Yeast | NULL | Probable | As nectar-tap_saprotroph (Põlme et al. 2020); Decay sugar-rich substrates (Põlme et al. 2020) | Põlme S, et al. 2020. Fungal Diversity 105:1-16 ((https://doi.org/10.1007/s13225-020-00466-2)) |
| Otu40341 | 1 | k__Fungi(100);p__Ascomycota(100);c__Saccharomycetes(100);o__Saccharomycetales(100);f__Metschnikowiaceae(100);g__Metschnikowia(100);s__Metschnikowia_pulcherrima(100); | Metschnikowia | 13 | Saprotroph | \|Undefined Saprotroph\| | Yeast | NULL | Probable | As nectar-tap_saprotroph (Põlme et al. 2020); Decay sugar-rich substrates (Põlme et al. 2020) | Põlme S, et al. 2020. Fungal Diversity 105:1-16 ((https://doi.org/10.1007/s13225-020-00466-2)) |
| Otu40342 | 1 | k__Fungi(100);p__Ascomycota(100);c__Saccharomycetes(100);o__Saccharomycetales(100);f__Metschnikowiaceae(100);g__Metschnikowia(100);s__Metschnikowia_pulcherrima(100); | Metschnikowia | 13 | Saprotroph | \|Undefined Saprotroph\| | Yeast | NULL | Probable | As nectar-tap_saprotroph (Põlme et al. 2020); Decay sugar-rich substrates (Põlme et al. 2020) | Põlme S, et al. 2020. Fungal Diversity 105:1-16 ((https://doi.org/10.1007/s13225-020-00466-2)) |
| Otu40343 | 1 | k__Fungi(100);p__Ascomycota(100);c__Saccharomycetes(100);o__Saccharomycetales(100);f__Metschnikowiaceae(100);g__Metschnikowia(100);g__Metschnikowia_unclassified(100); | Metschnikowia | 13 | Saprotroph | \|Undefined Saprotroph\| | Yeast | NULL | Probable | As nectar-tap_saprotroph (Põlme et al. 2020); Decay sugar-rich substrates (Põlme et al. 2020) | Põlme S, et al. 2020. Fungal Diversity 105:1-16 ((https://doi.org/10.1007/s13225-020-00466-2)) |
| Otu40344 | 1 | k__Fungi(100);p__Ascomycota(100);c__Saccharomycetes(100);o__Saccharomycetales(100);f__Metschnikowiaceae(100);g__Metschnikowia(100);s__Metschnikowia_pulcherrima(100); | Metschnikowia | 13 | Saprotroph | \|Undefined Saprotroph\| | Yeast | NULL | Probable | As nectar-tap_saprotroph (Põlme et al. 2020); Decay sugar-rich substrates (Põlme et al. 2020) | Põlme S, et al. 2020. Fungal Diversity 105:1-16 ((https://doi.org/10.1007/s13225-020-00466-2)) |
| Otu40345 | 1 | k__Fungi(100);p__Ascomycota(100);c__Saccharomycetes(100);o__Saccharomycetales(100);f__Metschnikowiaceae(100);g__Metschnikowia(100);s__Metschnikowia_pulcherrima(100); | Metschnikowia | 13 | Saprotroph | \|Undefined Saprotroph\| | Yeast | NULL | Probable | As nectar-tap_saprotroph (Põlme et al. 2020); Decay sugar-rich substrates (Põlme et al. 2020) | Põlme S, et al. 2020. Fungal Diversity 105:1-16 ((https://doi.org/10.1007/s13225-020-00466-2)) |
| Otu40346 | 1 | k__Fungi(100);p__Ascomycota(100);c__Saccharomycetes(100);o__Saccharomycetales(100);f__Metschnikowiaceae(100);g__Metschnikowia(100);s__Metschnikowia_pulcherrima(100); | Metschnikowia | 13 | Saprotroph | \|Undefined Saprotroph\| | Yeast | NULL | Probable | As nectar-tap_saprotroph (Põlme et al. 2020); Decay sugar-rich substrates (Põlme et al. 2020) | Põlme S, et al. 2020. Fungal Diversity 105:1-16 ((https://doi.org/10.1007/s13225-020-00466-2)) |
| Otu40347 | 1 | k__Fungi(100);p__Ascomycota(100);c__Saccharomycetes(100);o__Saccharomycetales(100);f__Metschnikowiaceae(100);g__Metschnikowia(100);s__Metschnikowia_pulcherrima(100); | Metschnikowia | 13 | Saprotroph | \|Undefined Saprotroph\| | Yeast | NULL | Probable | As nectar-tap_saprotroph (Põlme et al. 2020); Decay sugar-rich substrates (Põlme et al. 2020) | Põlme S, et al. 2020. Fungal Diversity 105:1-16 ((https://doi.org/10.1007/s13225-020-00466-2)) |
| Otu40348 | 1 | k__Fungi(100);p__Ascomycota(100);c__Saccharomycetes(100);o__Saccharomycetales(100);f__Metschnikowiaceae(100);g__Metschnikowia(100);s__Metschnikowia_pulcherrima(100); | Metschnikowia | 13 | Saprotroph | \|Undefined Saprotroph\| | Yeast | NULL | Probable | As nectar-tap_saprotroph (Põlme et al. 2020); Decay sugar-rich substrates (Põlme et al. 2020) | Põlme S, et al. 2020. Fungal Diversity 105:1-16 ((https://doi.org/10.1007/s13225-020-00466-2)) |
| Otu40349 | 1 | k__Fungi(100);p__Ascomycota(100);c__Saccharomycetes(100);o__Saccharomycetales(100);f__Metschnikowiaceae(100);g__Metschnikowia(100);s__Metschnikowia_pulcherrima(100); | Metschnikowia | 13 | Saprotroph | \|Undefined Saprotroph\| | Yeast | NULL | Probable | As nectar-tap_saprotroph (Põlme et al. 2020); Decay sugar-rich substrates (Põlme et al. 2020) | Põlme S, et al. 2020. Fungal Diversity 105:1-16 ((https://doi.org/10.1007/s13225-020-00466-2)) |
| Otu40350 | 1 | k__Fungi(100);p__Ascomycota(100);c__Saccharomycetes(100);o__Saccharomycetales(100);f__Metschnikowiaceae(100);g__Metschnikowia(100);s__Metschnikowia_pulcherrima(100); | Metschnikowia | 13 | Saprotroph | \|Undefined Saprotroph\| | Yeast | NULL | Probable | As nectar-tap_saprotroph (Põlme et al. 2020); Decay sugar-rich substrates (Põlme et al. 2020) | Põlme S, et al. 2020. Fungal Diversity 105:1-16 ((https://doi.org/10.1007/s13225-020-00466-2)) |
| Otu40351 | 1 | k__Fungi(100);p__Ascomycota(100);c__Saccharomycetes(100);o__Saccharomycetales(100);f__Metschnikowiaceae(100);g__Metschnikowia(100);s__Metschnikowia_pulcherrima(100); | Metschnikowia | 13 | Saprotroph | \|Undefined Saprotroph\| | Yeast | NULL | Probable | As nectar-tap_saprotroph (Põlme et al. 2020); Decay sugar-rich substrates (Põlme et al. 2020) | Põlme S, et al. 2020. Fungal Diversity 105:1-16 ((https://doi.org/10.1007/s13225-020-00466-2)) |
| Otu40352 | 1 | k__Fungi(100);p__Ascomycota(100);c__Saccharomycetes(100);o__Saccharomycetales(100);f__Metschnikowiaceae(100);g__Metschnikowia(100);g__Metschnikowia_unclassified(100); | Metschnikowia | 13 | Saprotroph | \|Undefined Saprotroph\| | Yeast | NULL | Probable | As nectar-tap_saprotroph (Põlme et al. 2020); Decay sugar-rich substrates (Põlme et al. 2020) | Põlme S, et al. 2020. Fungal Diversity 105:1-16 ((https://doi.org/10.1007/s13225-020-00466-2)) |
| Otu40353 | 1 | k__Fungi(100);p__Ascomycota(100);c__Saccharomycetes(100);o__Saccharomycetales(100);f__Metschnikowiaceae(100);g__Metschnikowia(100);s__Metschnikowia_pulcherrima(100); | Metschnikowia | 13 | Saprotroph | \|Undefined Saprotroph\| | Yeast | NULL | Probable | As nectar-tap_saprotroph (Põlme et al. 2020); Decay sugar-rich substrates (Põlme et al. 2020) | Põlme S, et al. 2020. Fungal Diversity 105:1-16 ((https://doi.org/10.1007/s13225-020-00466-2)) |
| Otu40354 | 1 | k__Fungi(100);p__Ascomycota(100);c__Saccharomycetes(100);o__Saccharomycetales(100);f__Metschnikowiaceae(100);g__Metschnikowia(100);s__Metschnikowia_pulcherrima(100); | Metschnikowia | 13 | Saprotroph | \|Undefined Saprotroph\| | Yeast | NULL | Probable | As nectar-tap_saprotroph (Põlme et al. 2020); Decay sugar-rich substrates (Põlme et al. 2020) | Põlme S, et al. 2020. Fungal Diversity 105:1-16 ((https://doi.org/10.1007/s13225-020-00466-2)) |
| Otu40356 | 1 | k__Fungi(100);p__Ascomycota(100);c__Saccharomycetes(100);o__Saccharomycetales(100);f__Metschnikowiaceae(100);g__Metschnikowia(100);g__Metschnikowia_unclassified(100); | Metschnikowia | 13 | Saprotroph | \|Undefined Saprotroph\| | Yeast | NULL | Probable | As nectar-tap_saprotroph (Põlme et al. 2020); Decay sugar-rich substrates (Põlme et al. 2020) | Põlme S, et al. 2020. Fungal Diversity 105:1-16 ((https://doi.org/10.1007/s13225-020-00466-2)) |
| Otu40357 | 1 | k__Fungi(100);p__Ascomycota(100);c__Saccharomycetes(100);o__Saccharomycetales(100);f__Metschnikowiaceae(100);g__Metschnikowia(100);g__Metschnikowia_unclassified(100); | Metschnikowia | 13 | Saprotroph | \|Undefined Saprotroph\| | Yeast | NULL | Probable | As nectar-tap_saprotroph (Põlme et al. 2020); Decay sugar-rich substrates (Põlme et al. 2020) | Põlme S, et al. 2020. Fungal Diversity 105:1-16 ((https://doi.org/10.1007/s13225-020-00466-2)) |
| Otu40358 | 1 | k__Fungi(100);p__Ascomycota(100);c__Saccharomycetes(100);o__Saccharomycetales(100);f__Metschnikowiaceae(100);g__Metschnikowia(100);g__Metschnikowia_unclassified(100); | Metschnikowia | 13 | Saprotroph | \|Undefined Saprotroph\| | Yeast | NULL | Probable | As nectar-tap_saprotroph (Põlme et al. 2020); Decay sugar-rich substrates (Põlme et al. 2020) | Põlme S, et al. 2020. Fungal Diversity 105:1-16 ((https://doi.org/10.1007/s13225-020-00466-2)) |
| Otu40359 | 1 | k__Fungi(100);p__Ascomycota(100);c__Saccharomycetes(100);o__Saccharomycetales(100);f__Metschnikowiaceae(100);g__Metschnikowia(100);s__Metschnikowia_pulcherrima(100); | Metschnikowia | 13 | Saprotroph | \|Undefined Saprotroph\| | Yeast | NULL | Probable | As nectar-tap_saprotroph (Põlme et al. 2020); Decay sugar-rich substrates (Põlme et al. 2020) | Põlme S, et al. 2020. Fungal Diversity 105:1-16 ((https://doi.org/10.1007/s13225-020-00466-2)) |
| Otu40360 | 1 | k__Fungi(100);p__Ascomycota(100);c__Saccharomycetes(100);o__Saccharomycetales(100);f__Metschnikowiaceae(100);g__Metschnikowia(100);s__Metschnikowia_pulcherrima(100); | Metschnikowia | 13 | Saprotroph | \|Undefined Saprotroph\| | Yeast | NULL | Probable | As nectar-tap_saprotroph (Põlme et al. 2020); Decay sugar-rich substrates (Põlme et al. 2020) | Põlme S, et al. 2020. Fungal Diversity 105:1-16 ((https://doi.org/10.1007/s13225-020-00466-2)) |
| Otu40361 | 1 | k__Fungi(100);p__Ascomycota(100);c__Saccharomycetes(100);o__Saccharomycetales(100);f__Metschnikowiaceae(100);g__Metschnikowia(100);s__Metschnikowia_pulcherrima(100); | Metschnikowia | 13 | Saprotroph | \|Undefined Saprotroph\| | Yeast | NULL | Probable | As nectar-tap_saprotroph (Põlme et al. 2020); Decay sugar-rich substrates (Põlme et al. 2020) | Põlme S, et al. 2020. Fungal Diversity 105:1-16 ((https://doi.org/10.1007/s13225-020-00466-2)) |
| Otu40362 | 1 | k__Fungi(100);p__Ascomycota(100);c__Saccharomycetes(100);o__Saccharomycetales(100);f__Metschnikowiaceae(100);g__Metschnikowia(100);s__Metschnikowia_pulcherrima(100); | Metschnikowia | 13 | Saprotroph | \|Undefined Saprotroph\| | Yeast | NULL | Probable | As nectar-tap_saprotroph (Põlme et al. 2020); Decay sugar-rich substrates (Põlme et al. 2020) | Põlme S, et al. 2020. Fungal Diversity 105:1-16 ((https://doi.org/10.1007/s13225-020-00466-2)) |
| Otu40363 | 1 | k__Fungi(100);p__Ascomycota(100);c__Saccharomycetes(100);o__Saccharomycetales(100);f__Metschnikowiaceae(100);g__Metschnikowia(100);g__Metschnikowia_unclassified(100); | Metschnikowia | 13 | Saprotroph | \|Undefined Saprotroph\| | Yeast | NULL | Probable | As nectar-tap_saprotroph (Põlme et al. 2020); Decay sugar-rich substrates (Põlme et al. 2020) | Põlme S, et al. 2020. Fungal Diversity 105:1-16 ((https://doi.org/10.1007/s13225-020-00466-2)) |
| Otu40364 | 1 | k__Fungi(100);p__Ascomycota(100);c__Saccharomycetes(100);o__Saccharomycetales(100);f__Metschnikowiaceae(100);g__Metschnikowia(100);s__Metschnikowia_pulcherrima(100); | Metschnikowia | 13 | Saprotroph | \|Undefined Saprotroph\| | Yeast | NULL | Probable | As nectar-tap_saprotroph (Põlme et al. 2020); Decay sugar-rich substrates (Põlme et al. 2020) | Põlme S, et al. 2020. Fungal Diversity 105:1-16 ((https://doi.org/10.1007/s13225-020-00466-2)) |
| Otu40365 | 1 | k__Fungi(100);p__Ascomycota(100);c__Saccharomycetes(100);o__Saccharomycetales(100);f__Metschnikowiaceae(100);g__Metschnikowia(100);s__Metschnikowia_pulcherrima(100); | Metschnikowia | 13 | Saprotroph | \|Undefined Saprotroph\| | Yeast | NULL | Probable | As nectar-tap_saprotroph (Põlme et al. 2020); Decay sugar-rich substrates (Põlme et al. 2020) | Põlme S, et al. 2020. Fungal Diversity 105:1-16 ((https://doi.org/10.1007/s13225-020-00466-2)) |
| Otu40366 | 1 | k__Fungi(100);p__Ascomycota(100);c__Saccharomycetes(100);o__Saccharomycetales(100);f__Metschnikowiaceae(100);g__Metschnikowia(100);s__Metschnikowia_pulcherrima(100); | Metschnikowia | 13 | Saprotroph | \|Undefined Saprotroph\| | Yeast | NULL | Probable | As nectar-tap_saprotroph (Põlme et al. 2020); Decay sugar-rich substrates (Põlme et al. 2020) | Põlme S, et al. 2020. Fungal Diversity 105:1-16 ((https://doi.org/10.1007/s13225-020-00466-2)) |
| Otu40367 | 1 | k__Fungi(100);p__Ascomycota(100);c__Saccharomycetes(100);o__Saccharomycetales(100);f__Metschnikowiaceae(100);g__Metschnikowia(100);g__Metschnikowia_unclassified(100); | Metschnikowia | 13 | Saprotroph | \|Undefined Saprotroph\| | Yeast | NULL | Probable | As nectar-tap_saprotroph (Põlme et al. 2020); Decay sugar-rich substrates (Põlme et al. 2020) | Põlme S, et al. 2020. Fungal Diversity 105:1-16 ((https://doi.org/10.1007/s13225-020-00466-2)) |
| Otu40368 | 1 | k__Fungi(100);p__Ascomycota(100);c__Saccharomycetes(100);o__Saccharomycetales(100);f__Metschnikowiaceae(100);g__Metschnikowia(100);g__Metschnikowia_unclassified(100); | Metschnikowia | 13 | Saprotroph | \|Undefined Saprotroph\| | Yeast | NULL | Probable | As nectar-tap_saprotroph (Põlme et al. 2020); Decay sugar-rich substrates (Põlme et al. 2020) | Põlme S, et al. 2020. Fungal Diversity 105:1-16 ((https://doi.org/10.1007/s13225-020-00466-2)) |
| Otu40369 | 1 | k__Fungi(100);p__Ascomycota(100);c__Saccharomycetes(100);o__Saccharomycetales(100);f__Metschnikowiaceae(100);g__Metschnikowia(100);s__Metschnikowia_pulcherrima(100); | Metschnikowia | 13 | Saprotroph | \|Undefined Saprotroph\| | Yeast | NULL | Probable | As nectar-tap_saprotroph (Põlme et al. 2020); Decay sugar-rich substrates (Põlme et al. 2020) | Põlme S, et al. 2020. Fungal Diversity 105:1-16 ((https://doi.org/10.1007/s13225-020-00466-2)) |
| Otu40370 | 1 | k__Fungi(100);p__Ascomycota(100);c__Saccharomycetes(100);o__Saccharomycetales(100);f__Metschnikowiaceae(100);g__Metschnikowia(100);s__Metschnikowia_pulcherrima(100); | Metschnikowia | 13 | Saprotroph | \|Undefined Saprotroph\| | Yeast | NULL | Probable | As nectar-tap_saprotroph (Põlme et al. 2020); Decay sugar-rich substrates (Põlme et al. 2020) | Põlme S, et al. 2020. Fungal Diversity 105:1-16 ((https://doi.org/10.1007/s13225-020-00466-2)) |
| Otu40371 | 1 | k__Fungi(100);p__Ascomycota(100);c__Saccharomycetes(100);o__Saccharomycetales(100);f__Metschnikowiaceae(100);g__Metschnikowia(100);s__Metschnikowia_pulcherrima(100); | Metschnikowia | 13 | Saprotroph | \|Undefined Saprotroph\| | Yeast | NULL | Probable | As nectar-tap_saprotroph (Põlme et al. 2020); Decay sugar-rich substrates (Põlme et al. 2020) | Põlme S, et al. 2020. Fungal Diversity 105:1-16 ((https://doi.org/10.1007/s13225-020-00466-2)) |
| Otu40372 | 1 | k__Fungi(100);p__Ascomycota(100);c__Saccharomycetes(100);o__Saccharomycetales(100);f__Metschnikowiaceae(100);g__Metschnikowia(100);s__Metschnikowia_pulcherrima(100); | Metschnikowia | 13 | Saprotroph | \|Undefined Saprotroph\| | Yeast | NULL | Probable | As nectar-tap_saprotroph (Põlme et al. 2020); Decay sugar-rich substrates (Põlme et al. 2020) | Põlme S, et al. 2020. Fungal Diversity 105:1-16 ((https://doi.org/10.1007/s13225-020-00466-2)) |
| Otu40373 | 1 | k__Fungi(100);p__Ascomycota(100);c__Saccharomycetes(100);o__Saccharomycetales(100);f__Metschnikowiaceae(100);g__Metschnikowia(100);s__Metschnikowia_pulcherrima(100); | Metschnikowia | 13 | Saprotroph | \|Undefined Saprotroph\| | Yeast | NULL | Probable | As nectar-tap_saprotroph (Põlme et al. 2020); Decay sugar-rich substrates (Põlme et al. 2020) | Põlme S, et al. 2020. Fungal Diversity 105:1-16 ((https://doi.org/10.1007/s13225-020-00466-2)) |
| Otu40374 | 1 | k__Fungi(100);p__Ascomycota(100);c__Saccharomycetes(100);o__Saccharomycetales(100);f__Metschnikowiaceae(100);g__Metschnikowia(100);s__Metschnikowia_pulcherrima(100); | Metschnikowia | 13 | Saprotroph | \|Undefined Saprotroph\| | Yeast | NULL | Probable | As nectar-tap_saprotroph (Põlme et al. 2020); Decay sugar-rich substrates (Põlme et al. 2020) | Põlme S, et al. 2020. Fungal Diversity 105:1-16 ((https://doi.org/10.1007/s13225-020-00466-2)) |
| Otu40375 | 1 | k__Fungi(100);p__Ascomycota(100);c__Saccharomycetes(100);o__Saccharomycetales(100);f__Metschnikowiaceae(100);g__Metschnikowia(100);s__Metschnikowia_pulcherrima(100); | Metschnikowia | 13 | Saprotroph | \|Undefined Saprotroph\| | Yeast | NULL | Probable | As nectar-tap_saprotroph (Põlme et al. 2020); Decay sugar-rich substrates (Põlme et al. 2020) | Põlme S, et al. 2020. Fungal Diversity 105:1-16 ((https://doi.org/10.1007/s13225-020-00466-2)) |
| Otu40376 | 1 | k__Fungi(100);p__Ascomycota(100);c__Saccharomycetes(100);o__Saccharomycetales(100);f__Metschnikowiaceae(100);g__Metschnikowia(100);s__Metschnikowia_pulcherrima(100); | Metschnikowia | 13 | Saprotroph | \|Undefined Saprotroph\| | Yeast | NULL | Probable | As nectar-tap_saprotroph (Põlme et al. 2020); Decay sugar-rich substrates (Põlme et al. 2020) | Põlme S, et al. 2020. Fungal Diversity 105:1-16 ((https://doi.org/10.1007/s13225-020-00466-2)) |
| Otu40377 | 1 | k__Fungi(100);p__Ascomycota(100);c__Saccharomycetes(100);o__Saccharomycetales(100);f__Metschnikowiaceae(100);g__Metschnikowia(100);s__Metschnikowia_pulcherrima(100); | Metschnikowia | 13 | Saprotroph | \|Undefined Saprotroph\| | Yeast | NULL | Probable | As nectar-tap_saprotroph (Põlme et al. 2020); Decay sugar-rich substrates (Põlme et al. 2020) | Põlme S, et al. 2020. Fungal Diversity 105:1-16 ((https://doi.org/10.1007/s13225-020-00466-2)) |
| Otu40378 | 1 | k__Fungi(100);p__Ascomycota(100);c__Saccharomycetes(100);o__Saccharomycetales(100);f__Metschnikowiaceae(100);g__Metschnikowia(100);s__Metschnikowia_pulcherrima(100); | Metschnikowia | 13 | Saprotroph | \|Undefined Saprotroph\| | Yeast | NULL | Probable | As nectar-tap_saprotroph (Põlme et al. 2020); Decay sugar-rich substrates (Põlme et al. 2020) | Põlme S, et al. 2020. Fungal Diversity 105:1-16 ((https://doi.org/10.1007/s13225-020-00466-2)) |
| Otu40379 | 1 | k__Fungi(100);p__Ascomycota(100);c__Saccharomycetes(100);o__Saccharomycetales(100);f__Metschnikowiaceae(100);g__Metschnikowia(100);s__Metschnikowia_pulcherrima(100); | Metschnikowia | 13 | Saprotroph | \|Undefined Saprotroph\| | Yeast | NULL | Probable | As nectar-tap_saprotroph (Põlme et al. 2020); Decay sugar-rich substrates (Põlme et al. 2020) | Põlme S, et al. 2020. Fungal Diversity 105:1-16 ((https://doi.org/10.1007/s13225-020-00466-2)) |
| Otu40380 | 1 | k__Fungi(100);p__Ascomycota(100);c__Saccharomycetes(100);o__Saccharomycetales(100);f__Metschnikowiaceae(100);g__Metschnikowia(100);s__Metschnikowia_pulcherrima(100); | Metschnikowia | 13 | Saprotroph | \|Undefined Saprotroph\| | Yeast | NULL | Probable | As nectar-tap_saprotroph (Põlme et al. 2020); Decay sugar-rich substrates (Põlme et al. 2020) | Põlme S, et al. 2020. Fungal Diversity 105:1-16 ((https://doi.org/10.1007/s13225-020-00466-2)) |
| Otu40381 | 1 | k__Fungi(100);p__Ascomycota(100);c__Saccharomycetes(100);o__Saccharomycetales(100);f__Metschnikowiaceae(100);g__Metschnikowia(100);g__Metschnikowia_unclassified(100); | Metschnikowia | 13 | Saprotroph | \|Undefined Saprotroph\| | Yeast | NULL | Probable | As nectar-tap_saprotroph (Põlme et al. 2020); Decay sugar-rich substrates (Põlme et al. 2020) | Põlme S, et al. 2020. Fungal Diversity 105:1-16 ((https://doi.org/10.1007/s13225-020-00466-2)) |
| Otu40382 | 1 | k__Fungi(100);p__Ascomycota(100);c__Saccharomycetes(100);o__Saccharomycetales(100);f__Metschnikowiaceae(100);g__Metschnikowia(100);s__Metschnikowia_pulcherrima(100); | Metschnikowia | 13 | Saprotroph | \|Undefined Saprotroph\| | Yeast | NULL | Probable | As nectar-tap_saprotroph (Põlme et al. 2020); Decay sugar-rich substrates (Põlme et al. 2020) | Põlme S, et al. 2020. Fungal Diversity 105:1-16 ((https://doi.org/10.1007/s13225-020-00466-2)) |
| Otu40383 | 1 | k__Fungi(100);p__Ascomycota(100);c__Saccharomycetes(100);o__Saccharomycetales(100);f__Metschnikowiaceae(100);g__Metschnikowia(100);g__Metschnikowia_unclassified(100); | Metschnikowia | 13 | Saprotroph | \|Undefined Saprotroph\| | Yeast | NULL | Probable | As nectar-tap_saprotroph (Põlme et al. 2020); Decay sugar-rich substrates (Põlme et al. 2020) | Põlme S, et al. 2020. Fungal Diversity 105:1-16 ((https://doi.org/10.1007/s13225-020-00466-2)) |
| Otu40384 | 1 | k__Fungi(100);p__Ascomycota(100);c__Saccharomycetes(100);o__Saccharomycetales(100);f__Metschnikowiaceae(100);g__Metschnikowia(100);g__Metschnikowia_unclassified(100); | Metschnikowia | 13 | Saprotroph | \|Undefined Saprotroph\| | Yeast | NULL | Probable | As nectar-tap_saprotroph (Põlme et al. 2020); Decay sugar-rich substrates (Põlme et al. 2020) | Põlme S, et al. 2020. Fungal Diversity 105:1-16 ((https://doi.org/10.1007/s13225-020-00466-2)) |
| Otu40385 | 1 | k__Fungi(100);p__Ascomycota(100);c__Saccharomycetes(100);o__Saccharomycetales(100);f__Metschnikowiaceae(100);g__Metschnikowia(100);g__Metschnikowia_unclassified(100); | Metschnikowia | 13 | Saprotroph | \|Undefined Saprotroph\| | Yeast | NULL | Probable | As nectar-tap_saprotroph (Põlme et al. 2020); Decay sugar-rich substrates (Põlme et al. 2020) | Põlme S, et al. 2020. Fungal Diversity 105:1-16 ((https://doi.org/10.1007/s13225-020-00466-2)) |
| Otu40386 | 1 | k__Fungi(100);p__Ascomycota(100);c__Saccharomycetes(100);o__Saccharomycetales(100);f__Metschnikowiaceae(100);g__Metschnikowia(100);g__Metschnikowia_unclassified(100); | Metschnikowia | 13 | Saprotroph | \|Undefined Saprotroph\| | Yeast | NULL | Probable | As nectar-tap_saprotroph (Põlme et al. 2020); Decay sugar-rich substrates (Põlme et al. 2020) | Põlme S, et al. 2020. Fungal Diversity 105:1-16 ((https://doi.org/10.1007/s13225-020-00466-2)) |
| Otu40387 | 1 | k__Fungi(100);p__Ascomycota(100);c__Saccharomycetes(100);o__Saccharomycetales(100);f__Metschnikowiaceae(100);g__Metschnikowia(100);g__Metschnikowia_unclassified(100); | Metschnikowia | 13 | Saprotroph | \|Undefined Saprotroph\| | Yeast | NULL | Probable | As nectar-tap_saprotroph (Põlme et al. 2020); Decay sugar-rich substrates (Põlme et al. 2020) | Põlme S, et al. 2020. Fungal Diversity 105:1-16 ((https://doi.org/10.1007/s13225-020-00466-2)) |
| Otu40388 | 1 | k__Fungi(100);p__Ascomycota(100);c__Saccharomycetes(100);o__Saccharomycetales(100);f__Metschnikowiaceae(100);g__Metschnikowia(100);s__Metschnikowia_pulcherrima(100); | Metschnikowia | 13 | Saprotroph | \|Undefined Saprotroph\| | Yeast | NULL | Probable | As nectar-tap_saprotroph (Põlme et al. 2020); Decay sugar-rich substrates (Põlme et al. 2020) | Põlme S, et al. 2020. Fungal Diversity 105:1-16 ((https://doi.org/10.1007/s13225-020-00466-2)) |
| Otu40389 | 1 | k__Fungi(100);p__Ascomycota(100);c__Saccharomycetes(100);o__Saccharomycetales(100);f__Metschnikowiaceae(100);g__Metschnikowia(100);g__Metschnikowia_unclassified(100); | Metschnikowia | 13 | Saprotroph | \|Undefined Saprotroph\| | Yeast | NULL | Probable | As nectar-tap_saprotroph (Põlme et al. 2020); Decay sugar-rich substrates (Põlme et al. 2020) | Põlme S, et al. 2020. Fungal Diversity 105:1-16 ((https://doi.org/10.1007/s13225-020-00466-2)) |
| Otu40390 | 1 | k__Fungi(100);p__Ascomycota(100);c__Saccharomycetes(100);o__Saccharomycetales(100);f__Metschnikowiaceae(100);g__Metschnikowia(100);g__Metschnikowia_unclassified(100); | Metschnikowia | 13 | Saprotroph | \|Undefined Saprotroph\| | Yeast | NULL | Probable | As nectar-tap_saprotroph (Põlme et al. 2020); Decay sugar-rich substrates (Põlme et al. 2020) | Põlme S, et al. 2020. Fungal Diversity 105:1-16 ((https://doi.org/10.1007/s13225-020-00466-2)) |
| Otu40391 | 1 | k__Fungi(100);p__Ascomycota(100);c__Saccharomycetes(100);o__Saccharomycetales(100);f__Metschnikowiaceae(100);g__Metschnikowia(100);s__Metschnikowia_pulcherrima(100); | Metschnikowia | 13 | Saprotroph | \|Undefined Saprotroph\| | Yeast | NULL | Probable | As nectar-tap_saprotroph (Põlme et al. 2020); Decay sugar-rich substrates (Põlme et al. 2020) | Põlme S, et al. 2020. Fungal Diversity 105:1-16 ((https://doi.org/10.1007/s13225-020-00466-2)) |
| Otu40392 | 1 | k__Fungi(100);p__Ascomycota(100);c__Saccharomycetes(100);o__Saccharomycetales(100);f__Metschnikowiaceae(100);g__Metschnikowia(100);s__Metschnikowia_pulcherrima(100); | Metschnikowia | 13 | Saprotroph | \|Undefined Saprotroph\| | Yeast | NULL | Probable | As nectar-tap_saprotroph (Põlme et al. 2020); Decay sugar-rich substrates (Põlme et al. 2020) | Põlme S, et al. 2020. Fungal Diversity 105:1-16 ((https://doi.org/10.1007/s13225-020-00466-2)) |
| Otu40394 | 1 | k__Fungi(100);p__Ascomycota(100);c__Saccharomycetes(100);o__Saccharomycetales(100);f__Metschnikowiaceae(100);g__Metschnikowia(100);s__Metschnikowia_pulcherrima(100); | Metschnikowia | 13 | Saprotroph | \|Undefined Saprotroph\| | Yeast | NULL | Probable | As nectar-tap_saprotroph (Põlme et al. 2020); Decay sugar-rich substrates (Põlme et al. 2020) | Põlme S, et al. 2020. Fungal Diversity 105:1-16 ((https://doi.org/10.1007/s13225-020-00466-2)) |
| Otu40396 | 1 | k__Fungi(100);p__Ascomycota(100);c__Saccharomycetes(100);o__Saccharomycetales(100);f__Metschnikowiaceae(100);g__Metschnikowia(100);g__Metschnikowia_unclassified(100); | Metschnikowia | 13 | Saprotroph | \|Undefined Saprotroph\| | Yeast | NULL | Probable | As nectar-tap_saprotroph (Põlme et al. 2020); Decay sugar-rich substrates (Põlme et al. 2020) | Põlme S, et al. 2020. Fungal Diversity 105:1-16 ((https://doi.org/10.1007/s13225-020-00466-2)) |
| Otu40397 | 1 | k__Fungi(100);p__Ascomycota(100);c__Saccharomycetes(100);o__Saccharomycetales(100);f__Metschnikowiaceae(100);g__Metschnikowia(100);s__Metschnikowia_pulcherrima(100); | Metschnikowia | 13 | Saprotroph | \|Undefined Saprotroph\| | Yeast | NULL | Probable | As nectar-tap_saprotroph (Põlme et al. 2020); Decay sugar-rich substrates (Põlme et al. 2020) | Põlme S, et al. 2020. Fungal Diversity 105:1-16 ((https://doi.org/10.1007/s13225-020-00466-2)) |
| Otu40398 | 1 | k__Fungi(100);p__Ascomycota(100);c__Saccharomycetes(100);o__Saccharomycetales(100);f__Metschnikowiaceae(100);g__Metschnikowia(100);g__Metschnikowia_unclassified(100); | Metschnikowia | 13 | Saprotroph | \|Undefined Saprotroph\| | Yeast | NULL | Probable | As nectar-tap_saprotroph (Põlme et al. 2020); Decay sugar-rich substrates (Põlme et al. 2020) | Põlme S, et al. 2020. Fungal Diversity 105:1-16 ((https://doi.org/10.1007/s13225-020-00466-2)) |
| Otu40399 | 1 | k__Fungi(100);p__Ascomycota(100);c__Saccharomycetes(100);o__Saccharomycetales(100);f__Metschnikowiaceae(100);g__Metschnikowia(100);s__Metschnikowia_pulcherrima(100); | Metschnikowia | 13 | Saprotroph | \|Undefined Saprotroph\| | Yeast | NULL | Probable | As nectar-tap_saprotroph (Põlme et al. 2020); Decay sugar-rich substrates (Põlme et al. 2020) | Põlme S, et al. 2020. Fungal Diversity 105:1-16 ((https://doi.org/10.1007/s13225-020-00466-2)) |
| Otu40400 | 1 | k__Fungi(100);p__Ascomycota(100);c__Saccharomycetes(100);o__Saccharomycetales(100);f__Metschnikowiaceae(100);g__Metschnikowia(100);s__Metschnikowia_pulcherrima(100); | Metschnikowia | 13 | Saprotroph | \|Undefined Saprotroph\| | Yeast | NULL | Probable | As nectar-tap_saprotroph (Põlme et al. 2020); Decay sugar-rich substrates (Põlme et al. 2020) | Põlme S, et al. 2020. Fungal Diversity 105:1-16 ((https://doi.org/10.1007/s13225-020-00466-2)) |
| Otu40401 | 1 | k__Fungi(100);p__Ascomycota(100);c__Saccharomycetes(100);o__Saccharomycetales(100);f__Metschnikowiaceae(100);g__Metschnikowia(100);g__Metschnikowia_unclassified(100); | Metschnikowia | 13 | Saprotroph | \|Undefined Saprotroph\| | Yeast | NULL | Probable | As nectar-tap_saprotroph (Põlme et al. 2020); Decay sugar-rich substrates (Põlme et al. 2020) | Põlme S, et al. 2020. Fungal Diversity 105:1-16 ((https://doi.org/10.1007/s13225-020-00466-2)) |
| Otu40402 | 1 | k__Fungi(100);p__Ascomycota(100);c__Saccharomycetes(100);o__Saccharomycetales(100);f__Metschnikowiaceae(100);g__Metschnikowia(100);s__Metschnikowia_pulcherrima(100); | Metschnikowia | 13 | Saprotroph | \|Undefined Saprotroph\| | Yeast | NULL | Probable | As nectar-tap_saprotroph (Põlme et al. 2020); Decay sugar-rich substrates (Põlme et al. 2020) | Põlme S, et al. 2020. Fungal Diversity 105:1-16 ((https://doi.org/10.1007/s13225-020-00466-2)) |
| Otu40403 | 1 | k__Fungi(100);p__Ascomycota(100);c__Saccharomycetes(100);o__Saccharomycetales(100);f__Metschnikowiaceae(100);g__Metschnikowia(100);s__Metschnikowia_pulcherrima(100); | Metschnikowia | 13 | Saprotroph | \|Undefined Saprotroph\| | Yeast | NULL | Probable | As nectar-tap_saprotroph (Põlme et al. 2020); Decay sugar-rich substrates (Põlme et al. 2020) | Põlme S, et al. 2020. Fungal Diversity 105:1-16 ((https://doi.org/10.1007/s13225-020-00466-2)) |
| Otu40404 | 1 | k__Fungi(100);p__Ascomycota(100);c__Saccharomycetes(100);o__Saccharomycetales(100);f__Metschnikowiaceae(100);g__Metschnikowia(100);s__Metschnikowia_pulcherrima(100); | Metschnikowia | 13 | Saprotroph | \|Undefined Saprotroph\| | Yeast | NULL | Probable | As nectar-tap_saprotroph (Põlme et al. 2020); Decay sugar-rich substrates (Põlme et al. 2020) | Põlme S, et al. 2020. Fungal Diversity 105:1-16 ((https://doi.org/10.1007/s13225-020-00466-2)) |
| Otu40405 | 1 | k__Fungi(100);p__Ascomycota(100);c__Saccharomycetes(100);o__Saccharomycetales(100);f__Metschnikowiaceae(100);g__Metschnikowia(100);s__Metschnikowia_pulcherrima(100); | Metschnikowia | 13 | Saprotroph | \|Undefined Saprotroph\| | Yeast | NULL | Probable | As nectar-tap_saprotroph (Põlme et al. 2020); Decay sugar-rich substrates (Põlme et al. 2020) | Põlme S, et al. 2020. Fungal Diversity 105:1-16 ((https://doi.org/10.1007/s13225-020-00466-2)) |
| Otu40406 | 1 | k__Fungi(100);p__Ascomycota(100);c__Saccharomycetes(100);o__Saccharomycetales(100);f__Metschnikowiaceae(100);g__Metschnikowia(100);s__Metschnikowia_pulcherrima(100); | Metschnikowia | 13 | Saprotroph | \|Undefined Saprotroph\| | Yeast | NULL | Probable | As nectar-tap_saprotroph (Põlme et al. 2020); Decay sugar-rich substrates (Põlme et al. 2020) | Põlme S, et al. 2020. Fungal Diversity 105:1-16 ((https://doi.org/10.1007/s13225-020-00466-2)) |
| Otu40407 | 1 | k__Fungi(100);p__Ascomycota(100);c__Saccharomycetes(100);o__Saccharomycetales(100);f__Metschnikowiaceae(100);g__Metschnikowia(100);s__Metschnikowia_pulcherrima(100); | Metschnikowia | 13 | Saprotroph | \|Undefined Saprotroph\| | Yeast | NULL | Probable | As nectar-tap_saprotroph (Põlme et al. 2020); Decay sugar-rich substrates (Põlme et al. 2020) | Põlme S, et al. 2020. Fungal Diversity 105:1-16 ((https://doi.org/10.1007/s13225-020-00466-2)) |
| Otu40408 | 1 | k__Fungi(100);p__Ascomycota(100);c__Saccharomycetes(100);o__Saccharomycetales(100);f__Metschnikowiaceae(100);g__Metschnikowia(100);s__Metschnikowia_pulcherrima(100); | Metschnikowia | 13 | Saprotroph | \|Undefined Saprotroph\| | Yeast | NULL | Probable | As nectar-tap_saprotroph (Põlme et al. 2020); Decay sugar-rich substrates (Põlme et al. 2020) | Põlme S, et al. 2020. Fungal Diversity 105:1-16 ((https://doi.org/10.1007/s13225-020-00466-2)) |
| Otu40409 | 1 | k__Fungi(100);p__Ascomycota(100);c__Saccharomycetes(100);o__Saccharomycetales(100);f__Metschnikowiaceae(100);g__Metschnikowia(100);s__Metschnikowia_pulcherrima(100); | Metschnikowia | 13 | Saprotroph | \|Undefined Saprotroph\| | Yeast | NULL | Probable | As nectar-tap_saprotroph (Põlme et al. 2020); Decay sugar-rich substrates (Põlme et al. 2020) | Põlme S, et al. 2020. Fungal Diversity 105:1-16 ((https://doi.org/10.1007/s13225-020-00466-2)) |
| Otu40410 | 1 | k__Fungi(100);p__Ascomycota(100);c__Saccharomycetes(100);o__Saccharomycetales(100);f__Metschnikowiaceae(100);g__Metschnikowia(100);s__Metschnikowia_pulcherrima(100); | Metschnikowia | 13 | Saprotroph | \|Undefined Saprotroph\| | Yeast | NULL | Probable | As nectar-tap_saprotroph (Põlme et al. 2020); Decay sugar-rich substrates (Põlme et al. 2020) | Põlme S, et al. 2020. Fungal Diversity 105:1-16 ((https://doi.org/10.1007/s13225-020-00466-2)) |
| Otu40411 | 1 | k__Fungi(100);p__Ascomycota(100);c__Saccharomycetes(100);o__Saccharomycetales(100);f__Metschnikowiaceae(100);g__Metschnikowia(100);g__Metschnikowia_unclassified(100); | Metschnikowia | 13 | Saprotroph | \|Undefined Saprotroph\| | Yeast | NULL | Probable | As nectar-tap_saprotroph (Põlme et al. 2020); Decay sugar-rich substrates (Põlme et al. 2020) | Põlme S, et al. 2020. Fungal Diversity 105:1-16 ((https://doi.org/10.1007/s13225-020-00466-2)) |
| Otu40412 | 1 | k__Fungi(100);p__Ascomycota(100);c__Saccharomycetes(100);o__Saccharomycetales(100);f__Metschnikowiaceae(100);g__Metschnikowia(100);s__Metschnikowia_pulcherrima(100); | Metschnikowia | 13 | Saprotroph | \|Undefined Saprotroph\| | Yeast | NULL | Probable | As nectar-tap_saprotroph (Põlme et al. 2020); Decay sugar-rich substrates (Põlme et al. 2020) | Põlme S, et al. 2020. Fungal Diversity 105:1-16 ((https://doi.org/10.1007/s13225-020-00466-2)) |
| Otu40413 | 1 | k__Fungi(100);p__Ascomycota(100);c__Saccharomycetes(100);o__Saccharomycetales(100);f__Metschnikowiaceae(100);g__Metschnikowia(100);s__Metschnikowia_pulcherrima(100); | Metschnikowia | 13 | Saprotroph | \|Undefined Saprotroph\| | Yeast | NULL | Probable | As nectar-tap_saprotroph (Põlme et al. 2020); Decay sugar-rich substrates (Põlme et al. 2020) | Põlme S, et al. 2020. Fungal Diversity 105:1-16 ((https://doi.org/10.1007/s13225-020-00466-2)) |
| Otu40414 | 1 | k__Fungi(100);p__Ascomycota(100);c__Saccharomycetes(100);o__Saccharomycetales(100);f__Metschnikowiaceae(100);g__Metschnikowia(100);s__Metschnikowia_pulcherrima(100); | Metschnikowia | 13 | Saprotroph | \|Undefined Saprotroph\| | Yeast | NULL | Probable | As nectar-tap_saprotroph (Põlme et al. 2020); Decay sugar-rich substrates (Põlme et al. 2020) | Põlme S, et al. 2020. Fungal Diversity 105:1-16 ((https://doi.org/10.1007/s13225-020-00466-2)) |
| Otu40415 | 1 | k__Fungi(100);p__Ascomycota(100);c__Saccharomycetes(100);o__Saccharomycetales(100);f__Metschnikowiaceae(100);g__Metschnikowia(100);s__Metschnikowia_pulcherrima(100); | Metschnikowia | 13 | Saprotroph | \|Undefined Saprotroph\| | Yeast | NULL | Probable | As nectar-tap_saprotroph (Põlme et al. 2020); Decay sugar-rich substrates (Põlme et al. 2020) | Põlme S, et al. 2020. Fungal Diversity 105:1-16 ((https://doi.org/10.1007/s13225-020-00466-2)) |
| Otu40416 | 1 | k__Fungi(100);p__Ascomycota(100);c__Saccharomycetes(100);o__Saccharomycetales(100);f__Metschnikowiaceae(100);g__Metschnikowia(100);s__Metschnikowia_pulcherrima(100); | Metschnikowia | 13 | Saprotroph | \|Undefined Saprotroph\| | Yeast | NULL | Probable | As nectar-tap_saprotroph (Põlme et al. 2020); Decay sugar-rich substrates (Põlme et al. 2020) | Põlme S, et al. 2020. Fungal Diversity 105:1-16 ((https://doi.org/10.1007/s13225-020-00466-2)) |
| Otu40417 | 1 | k__Fungi(100);p__Ascomycota(100);c__Saccharomycetes(100);o__Saccharomycetales(100);f__Metschnikowiaceae(100);g__Metschnikowia(100);s__Metschnikowia_pulcherrima(100); | Metschnikowia | 13 | Saprotroph | \|Undefined Saprotroph\| | Yeast | NULL | Probable | As nectar-tap_saprotroph (Põlme et al. 2020); Decay sugar-rich substrates (Põlme et al. 2020) | Põlme S, et al. 2020. Fungal Diversity 105:1-16 ((https://doi.org/10.1007/s13225-020-00466-2)) |
| Otu40418 | 1 | k__Fungi(100);p__Ascomycota(100);c__Saccharomycetes(100);o__Saccharomycetales(100);f__Metschnikowiaceae(100);g__Metschnikowia(100);s__Metschnikowia_pulcherrima(100); | Metschnikowia | 13 | Saprotroph | \|Undefined Saprotroph\| | Yeast | NULL | Probable | As nectar-tap_saprotroph (Põlme et al. 2020); Decay sugar-rich substrates (Põlme et al. 2020) | Põlme S, et al. 2020. Fungal Diversity 105:1-16 ((https://doi.org/10.1007/s13225-020-00466-2)) |
| Otu40419 | 1 | k__Fungi(100);p__Ascomycota(100);c__Saccharomycetes(100);o__Saccharomycetales(100);f__Metschnikowiaceae(100);g__Metschnikowia(100);s__Metschnikowia_pulcherrima(100); | Metschnikowia | 13 | Saprotroph | \|Undefined Saprotroph\| | Yeast | NULL | Probable | As nectar-tap_saprotroph (Põlme et al. 2020); Decay sugar-rich substrates (Põlme et al. 2020) | Põlme S, et al. 2020. Fungal Diversity 105:1-16 ((https://doi.org/10.1007/s13225-020-00466-2)) |
| Otu40420 | 1 | k__Fungi(100);p__Ascomycota(100);c__Saccharomycetes(100);o__Saccharomycetales(100);f__Metschnikowiaceae(100);g__Metschnikowia(100);s__Metschnikowia_pulcherrima(100); | Metschnikowia | 13 | Saprotroph | \|Undefined Saprotroph\| | Yeast | NULL | Probable | As nectar-tap_saprotroph (Põlme et al. 2020); Decay sugar-rich substrates (Põlme et al. 2020) | Põlme S, et al. 2020. Fungal Diversity 105:1-16 ((https://doi.org/10.1007/s13225-020-00466-2)) |
| Otu40421 | 1 | k__Fungi(100);p__Ascomycota(100);c__Saccharomycetes(100);o__Saccharomycetales(100);f__Metschnikowiaceae(100);g__Metschnikowia(100);s__Metschnikowia_pulcherrima(100); | Metschnikowia | 13 | Saprotroph | \|Undefined Saprotroph\| | Yeast | NULL | Probable | As nectar-tap_saprotroph (Põlme et al. 2020); Decay sugar-rich substrates (Põlme et al. 2020) | Põlme S, et al. 2020. Fungal Diversity 105:1-16 ((https://doi.org/10.1007/s13225-020-00466-2)) |
| Otu40422 | 1 | k__Fungi(100);p__Ascomycota(100);c__Saccharomycetes(100);o__Saccharomycetales(100);f__Metschnikowiaceae(100);g__Metschnikowia(100);s__Metschnikowia_pulcherrima(100); | Metschnikowia | 13 | Saprotroph | \|Undefined Saprotroph\| | Yeast | NULL | Probable | As nectar-tap_saprotroph (Põlme et al. 2020); Decay sugar-rich substrates (Põlme et al. 2020) | Põlme S, et al. 2020. Fungal Diversity 105:1-16 ((https://doi.org/10.1007/s13225-020-00466-2)) |
| Otu40424 | 1 | k__Fungi(100);p__Ascomycota(100);c__Saccharomycetes(100);o__Saccharomycetales(100);f__Metschnikowiaceae(100);g__Metschnikowia(100);s__Metschnikowia_pulcherrima(100); | Metschnikowia | 13 | Saprotroph | \|Undefined Saprotroph\| | Yeast | NULL | Probable | As nectar-tap_saprotroph (Põlme et al. 2020); Decay sugar-rich substrates (Põlme et al. 2020) | Põlme S, et al. 2020. Fungal Diversity 105:1-16 ((https://doi.org/10.1007/s13225-020-00466-2)) |
| Otu40425 | 1 | k__Fungi(100);p__Ascomycota(100);c__Saccharomycetes(100);o__Saccharomycetales(100);f__Metschnikowiaceae(100);g__Metschnikowia(100);s__Metschnikowia_pulcherrima(100); | Metschnikowia | 13 | Saprotroph | \|Undefined Saprotroph\| | Yeast | NULL | Probable | As nectar-tap_saprotroph (Põlme et al. 2020); Decay sugar-rich substrates (Põlme et al. 2020) | Põlme S, et al. 2020. Fungal Diversity 105:1-16 ((https://doi.org/10.1007/s13225-020-00466-2)) |
| Otu40426 | 1 | k__Fungi(100);p__Ascomycota(100);c__Saccharomycetes(100);o__Saccharomycetales(100);f__Metschnikowiaceae(100);g__Metschnikowia(100);s__Metschnikowia_pulcherrima(100); | Metschnikowia | 13 | Saprotroph | \|Undefined Saprotroph\| | Yeast | NULL | Probable | As nectar-tap_saprotroph (Põlme et al. 2020); Decay sugar-rich substrates (Põlme et al. 2020) | Põlme S, et al. 2020. Fungal Diversity 105:1-16 ((https://doi.org/10.1007/s13225-020-00466-2)) |
| Otu40427 | 1 | k__Fungi(100);p__Ascomycota(100);c__Saccharomycetes(100);o__Saccharomycetales(100);f__Metschnikowiaceae(100);g__Metschnikowia(100);g__Metschnikowia_unclassified(100); | Metschnikowia | 13 | Saprotroph | \|Undefined Saprotroph\| | Yeast | NULL | Probable | As nectar-tap_saprotroph (Põlme et al. 2020); Decay sugar-rich substrates (Põlme et al. 2020) | Põlme S, et al. 2020. Fungal Diversity 105:1-16 ((https://doi.org/10.1007/s13225-020-00466-2)) |
| Otu40428 | 1 | k__Fungi(100);p__Ascomycota(100);c__Saccharomycetes(100);o__Saccharomycetales(100);f__Metschnikowiaceae(100);g__Metschnikowia(100);g__Metschnikowia_unclassified(100); | Metschnikowia | 13 | Saprotroph | \|Undefined Saprotroph\| | Yeast | NULL | Probable | As nectar-tap_saprotroph (Põlme et al. 2020); Decay sugar-rich substrates (Põlme et al. 2020) | Põlme S, et al. 2020. Fungal Diversity 105:1-16 ((https://doi.org/10.1007/s13225-020-00466-2)) |
| Otu40429 | 1 | k__Fungi(100);p__Ascomycota(100);c__Saccharomycetes(100);o__Saccharomycetales(100);f__Metschnikowiaceae(100);g__Metschnikowia(100);s__Metschnikowia_pulcherrima(100); | Metschnikowia | 13 | Saprotroph | \|Undefined Saprotroph\| | Yeast | NULL | Probable | As nectar-tap_saprotroph (Põlme et al. 2020); Decay sugar-rich substrates (Põlme et al. 2020) | Põlme S, et al. 2020. Fungal Diversity 105:1-16 ((https://doi.org/10.1007/s13225-020-00466-2)) |
| Otu40430 | 1 | k__Fungi(100);p__Ascomycota(100);c__Saccharomycetes(100);o__Saccharomycetales(100);f__Metschnikowiaceae(100);g__Metschnikowia(100);s__Metschnikowia_pulcherrima(100); | Metschnikowia | 13 | Saprotroph | \|Undefined Saprotroph\| | Yeast | NULL | Probable | As nectar-tap_saprotroph (Põlme et al. 2020); Decay sugar-rich substrates (Põlme et al. 2020) | Põlme S, et al. 2020. Fungal Diversity 105:1-16 ((https://doi.org/10.1007/s13225-020-00466-2)) |
| Otu40431 | 1 | k__Fungi(100);p__Ascomycota(100);c__Saccharomycetes(100);o__Saccharomycetales(100);f__Metschnikowiaceae(100);g__Metschnikowia(100);g__Metschnikowia_unclassified(100); | Metschnikowia | 13 | Saprotroph | \|Undefined Saprotroph\| | Yeast | NULL | Probable | As nectar-tap_saprotroph (Põlme et al. 2020); Decay sugar-rich substrates (Põlme et al. 2020) | Põlme S, et al. 2020. Fungal Diversity 105:1-16 ((https://doi.org/10.1007/s13225-020-00466-2)) |
| Otu40432 | 1 | k__Fungi(100);p__Ascomycota(100);c__Saccharomycetes(100);o__Saccharomycetales(100);f__Metschnikowiaceae(100);g__Metschnikowia(100);g__Metschnikowia_unclassified(100); | Metschnikowia | 13 | Saprotroph | \|Undefined Saprotroph\| | Yeast | NULL | Probable | As nectar-tap_saprotroph (Põlme et al. 2020); Decay sugar-rich substrates (Põlme et al. 2020) | Põlme S, et al. 2020. Fungal Diversity 105:1-16 ((https://doi.org/10.1007/s13225-020-00466-2)) |
| Otu40433 | 1 | k__Fungi(100);p__Ascomycota(100);c__Saccharomycetes(100);o__Saccharomycetales(100);f__Metschnikowiaceae(100);g__Metschnikowia(100);g__Metschnikowia_unclassified(100); | Metschnikowia | 13 | Saprotroph | \|Undefined Saprotroph\| | Yeast | NULL | Probable | As nectar-tap_saprotroph (Põlme et al. 2020); Decay sugar-rich substrates (Põlme et al. 2020) | Põlme S, et al. 2020. Fungal Diversity 105:1-16 ((https://doi.org/10.1007/s13225-020-00466-2)) |
| Otu40434 | 1 | k__Fungi(100);p__Ascomycota(100);c__Saccharomycetes(100);o__Saccharomycetales(100);f__Metschnikowiaceae(100);g__Metschnikowia(100);s__Metschnikowia_pulcherrima(100); | Metschnikowia | 13 | Saprotroph | \|Undefined Saprotroph\| | Yeast | NULL | Probable | As nectar-tap_saprotroph (Põlme et al. 2020); Decay sugar-rich substrates (Põlme et al. 2020) | Põlme S, et al. 2020. Fungal Diversity 105:1-16 ((https://doi.org/10.1007/s13225-020-00466-2)) |
| Otu40435 | 1 | k__Fungi(100);p__Ascomycota(100);c__Saccharomycetes(100);o__Saccharomycetales(100);f__Metschnikowiaceae(100);g__Metschnikowia(100);s__Metschnikowia_pulcherrima(100); | Metschnikowia | 13 | Saprotroph | \|Undefined Saprotroph\| | Yeast | NULL | Probable | As nectar-tap_saprotroph (Põlme et al. 2020); Decay sugar-rich substrates (Põlme et al. 2020) | Põlme S, et al. 2020. Fungal Diversity 105:1-16 ((https://doi.org/10.1007/s13225-020-00466-2)) |
| Otu40436 | 1 | k__Fungi(100);p__Ascomycota(100);c__Saccharomycetes(100);o__Saccharomycetales(100);f__Metschnikowiaceae(100);g__Metschnikowia(100);s__Metschnikowia_pulcherrima(100); | Metschnikowia | 13 | Saprotroph | \|Undefined Saprotroph\| | Yeast | NULL | Probable | As nectar-tap_saprotroph (Põlme et al. 2020); Decay sugar-rich substrates (Põlme et al. 2020) | Põlme S, et al. 2020. Fungal Diversity 105:1-16 ((https://doi.org/10.1007/s13225-020-00466-2)) |
| Otu40437 | 1 | k__Fungi(100);p__Ascomycota(100);c__Saccharomycetes(100);o__Saccharomycetales(100);f__Metschnikowiaceae(100);g__Metschnikowia(100);g__Metschnikowia_unclassified(100); | Metschnikowia | 13 | Saprotroph | \|Undefined Saprotroph\| | Yeast | NULL | Probable | As nectar-tap_saprotroph (Põlme et al. 2020); Decay sugar-rich substrates (Põlme et al. 2020) | Põlme S, et al. 2020. Fungal Diversity 105:1-16 ((https://doi.org/10.1007/s13225-020-00466-2)) |
| Otu40438 | 1 | k__Fungi(100);p__Ascomycota(100);c__Saccharomycetes(100);o__Saccharomycetales(100);f__Metschnikowiaceae(100);g__Metschnikowia(100);s__Metschnikowia_pulcherrima(100); | Metschnikowia | 13 | Saprotroph | \|Undefined Saprotroph\| | Yeast | NULL | Probable | As nectar-tap_saprotroph (Põlme et al. 2020); Decay sugar-rich substrates (Põlme et al. 2020) | Põlme S, et al. 2020. Fungal Diversity 105:1-16 ((https://doi.org/10.1007/s13225-020-00466-2)) |
| Otu40439 | 1 | k__Fungi(100);p__Ascomycota(100);c__Saccharomycetes(100);o__Saccharomycetales(100);f__Metschnikowiaceae(100);g__Metschnikowia(100);s__Metschnikowia_pulcherrima(100); | Metschnikowia | 13 | Saprotroph | \|Undefined Saprotroph\| | Yeast | NULL | Probable | As nectar-tap_saprotroph (Põlme et al. 2020); Decay sugar-rich substrates (Põlme et al. 2020) | Põlme S, et al. 2020. Fungal Diversity 105:1-16 ((https://doi.org/10.1007/s13225-020-00466-2)) |
| Otu40440 | 1 | k__Fungi(100);p__Ascomycota(100);c__Saccharomycetes(100);o__Saccharomycetales(100);f__Metschnikowiaceae(100);g__Metschnikowia(100);s__Metschnikowia_pulcherrima(100); | Metschnikowia | 13 | Saprotroph | \|Undefined Saprotroph\| | Yeast | NULL | Probable | As nectar-tap_saprotroph (Põlme et al. 2020); Decay sugar-rich substrates (Põlme et al. 2020) | Põlme S, et al. 2020. Fungal Diversity 105:1-16 ((https://doi.org/10.1007/s13225-020-00466-2)) |
| Otu40441 | 1 | k__Fungi(100);p__Ascomycota(100);c__Saccharomycetes(100);o__Saccharomycetales(100);f__Metschnikowiaceae(100);g__Metschnikowia(100);s__Metschnikowia_pulcherrima(100); | Metschnikowia | 13 | Saprotroph | \|Undefined Saprotroph\| | Yeast | NULL | Probable | As nectar-tap_saprotroph (Põlme et al. 2020); Decay sugar-rich substrates (Põlme et al. 2020) | Põlme S, et al. 2020. Fungal Diversity 105:1-16 ((https://doi.org/10.1007/s13225-020-00466-2)) |
| Otu40442 | 1 | k__Fungi(100);p__Ascomycota(100);c__Saccharomycetes(100);o__Saccharomycetales(100);f__Metschnikowiaceae(100);g__Metschnikowia(100);s__Metschnikowia_pulcherrima(100); | Metschnikowia | 13 | Saprotroph | \|Undefined Saprotroph\| | Yeast | NULL | Probable | As nectar-tap_saprotroph (Põlme et al. 2020); Decay sugar-rich substrates (Põlme et al. 2020) | Põlme S, et al. 2020. Fungal Diversity 105:1-16 ((https://doi.org/10.1007/s13225-020-00466-2)) |
| Otu40443 | 1 | k__Fungi(100);p__Ascomycota(100);c__Saccharomycetes(100);o__Saccharomycetales(100);f__Metschnikowiaceae(100);g__Metschnikowia(100);s__Metschnikowia_pulcherrima(100); | Metschnikowia | 13 | Saprotroph | \|Undefined Saprotroph\| | Yeast | NULL | Probable | As nectar-tap_saprotroph (Põlme et al. 2020); Decay sugar-rich substrates (Põlme et al. 2020) | Põlme S, et al. 2020. Fungal Diversity 105:1-16 ((https://doi.org/10.1007/s13225-020-00466-2)) |
| Otu40444 | 1 | k__Fungi(100);p__Ascomycota(100);c__Saccharomycetes(100);o__Saccharomycetales(100);f__Metschnikowiaceae(100);g__Metschnikowia(100);s__Metschnikowia_pulcherrima(100); | Metschnikowia | 13 | Saprotroph | \|Undefined Saprotroph\| | Yeast | NULL | Probable | As nectar-tap_saprotroph (Põlme et al. 2020); Decay sugar-rich substrates (Põlme et al. 2020) | Põlme S, et al. 2020. Fungal Diversity 105:1-16 ((https://doi.org/10.1007/s13225-020-00466-2)) |
| Otu40445 | 1 | k__Fungi(100);p__Ascomycota(100);c__Saccharomycetes(100);o__Saccharomycetales(100);f__Metschnikowiaceae(100);g__Metschnikowia(100);s__Metschnikowia_pulcherrima(100); | Metschnikowia | 13 | Saprotroph | \|Undefined Saprotroph\| | Yeast | NULL | Probable | As nectar-tap_saprotroph (Põlme et al. 2020); Decay sugar-rich substrates (Põlme et al. 2020) | Põlme S, et al. 2020. Fungal Diversity 105:1-16 ((https://doi.org/10.1007/s13225-020-00466-2)) |
| Otu40446 | 1 | k__Fungi(100);p__Ascomycota(100);c__Saccharomycetes(100);o__Saccharomycetales(100);f__Metschnikowiaceae(100);g__Metschnikowia(100);s__Metschnikowia_pulcherrima(100); | Metschnikowia | 13 | Saprotroph | \|Undefined Saprotroph\| | Yeast | NULL | Probable | As nectar-tap_saprotroph (Põlme et al. 2020); Decay sugar-rich substrates (Põlme et al. 2020) | Põlme S, et al. 2020. Fungal Diversity 105:1-16 ((https://doi.org/10.1007/s13225-020-00466-2)) |
| Otu40447 | 1 | k__Fungi(100);p__Ascomycota(100);c__Saccharomycetes(100);o__Saccharomycetales(100);f__Metschnikowiaceae(100);g__Metschnikowia(100);s__Metschnikowia_pulcherrima(100); | Metschnikowia | 13 | Saprotroph | \|Undefined Saprotroph\| | Yeast | NULL | Probable | As nectar-tap_saprotroph (Põlme et al. 2020); Decay sugar-rich substrates (Põlme et al. 2020) | Põlme S, et al. 2020. Fungal Diversity 105:1-16 ((https://doi.org/10.1007/s13225-020-00466-2)) |
| Otu40448 | 1 | k__Fungi(100);p__Ascomycota(100);c__Saccharomycetes(100);o__Saccharomycetales(100);f__Metschnikowiaceae(100);g__Metschnikowia(100);s__Metschnikowia_pulcherrima(100); | Metschnikowia | 13 | Saprotroph | \|Undefined Saprotroph\| | Yeast | NULL | Probable | As nectar-tap_saprotroph (Põlme et al. 2020); Decay sugar-rich substrates (Põlme et al. 2020) | Põlme S, et al. 2020. Fungal Diversity 105:1-16 ((https://doi.org/10.1007/s13225-020-00466-2)) |
| Otu40449 | 1 | k__Fungi(100);p__Ascomycota(100);c__Saccharomycetes(100);o__Saccharomycetales(100);f__Metschnikowiaceae(100);g__Metschnikowia(100);s__Metschnikowia_pulcherrima(100); | Metschnikowia | 13 | Saprotroph | \|Undefined Saprotroph\| | Yeast | NULL | Probable | As nectar-tap_saprotroph (Põlme et al. 2020); Decay sugar-rich substrates (Põlme et al. 2020) | Põlme S, et al. 2020. Fungal Diversity 105:1-16 ((https://doi.org/10.1007/s13225-020-00466-2)) |
| Otu40450 | 1 | k__Fungi(100);p__Ascomycota(100);c__Saccharomycetes(100);o__Saccharomycetales(100);f__Metschnikowiaceae(100);g__Metschnikowia(100);g__Metschnikowia_unclassified(100); | Metschnikowia | 13 | Saprotroph | \|Undefined Saprotroph\| | Yeast | NULL | Probable | As nectar-tap_saprotroph (Põlme et al. 2020); Decay sugar-rich substrates (Põlme et al. 2020) | Põlme S, et al. 2020. Fungal Diversity 105:1-16 ((https://doi.org/10.1007/s13225-020-00466-2)) |
| Otu40451 | 1 | k__Fungi(100);p__Ascomycota(100);c__Saccharomycetes(100);o__Saccharomycetales(100);f__Metschnikowiaceae(100);g__Metschnikowia(100);s__Metschnikowia_pulcherrima(100); | Metschnikowia | 13 | Saprotroph | \|Undefined Saprotroph\| | Yeast | NULL | Probable | As nectar-tap_saprotroph (Põlme et al. 2020); Decay sugar-rich substrates (Põlme et al. 2020) | Põlme S, et al. 2020. Fungal Diversity 105:1-16 ((https://doi.org/10.1007/s13225-020-00466-2)) |
| Otu40452 | 1 | k__Fungi(100);p__Ascomycota(100);c__Saccharomycetes(100);o__Saccharomycetales(100);f__Metschnikowiaceae(100);g__Metschnikowia(100);s__Metschnikowia_pulcherrima(100); | Metschnikowia | 13 | Saprotroph | \|Undefined Saprotroph\| | Yeast | NULL | Probable | As nectar-tap_saprotroph (Põlme et al. 2020); Decay sugar-rich substrates (Põlme et al. 2020) | Põlme S, et al. 2020. Fungal Diversity 105:1-16 ((https://doi.org/10.1007/s13225-020-00466-2)) |
| Otu40453 | 1 | k__Fungi(100);p__Ascomycota(100);c__Saccharomycetes(100);o__Saccharomycetales(100);f__Metschnikowiaceae(100);g__Metschnikowia(100);s__Metschnikowia_pulcherrima(100); | Metschnikowia | 13 | Saprotroph | \|Undefined Saprotroph\| | Yeast | NULL | Probable | As nectar-tap_saprotroph (Põlme et al. 2020); Decay sugar-rich substrates (Põlme et al. 2020) | Põlme S, et al. 2020. Fungal Diversity 105:1-16 ((https://doi.org/10.1007/s13225-020-00466-2)) |
| Otu40454 | 1 | k__Fungi(100);p__Ascomycota(100);c__Saccharomycetes(100);o__Saccharomycetales(100);f__Metschnikowiaceae(100);g__Metschnikowia(100);s__Metschnikowia_pulcherrima(100); | Metschnikowia | 13 | Saprotroph | \|Undefined Saprotroph\| | Yeast | NULL | Probable | As nectar-tap_saprotroph (Põlme et al. 2020); Decay sugar-rich substrates (Põlme et al. 2020) | Põlme S, et al. 2020. Fungal Diversity 105:1-16 ((https://doi.org/10.1007/s13225-020-00466-2)) |
| Otu40455 | 1 | k__Fungi(100);p__Ascomycota(100);c__Saccharomycetes(100);o__Saccharomycetales(100);f__Metschnikowiaceae(100);g__Metschnikowia(100);s__Metschnikowia_pulcherrima(100); | Metschnikowia | 13 | Saprotroph | \|Undefined Saprotroph\| | Yeast | NULL | Probable | As nectar-tap_saprotroph (Põlme et al. 2020); Decay sugar-rich substrates (Põlme et al. 2020) | Põlme S, et al. 2020. Fungal Diversity 105:1-16 ((https://doi.org/10.1007/s13225-020-00466-2)) |
| Otu40456 | 1 | k__Fungi(100);p__Ascomycota(100);c__Saccharomycetes(100);o__Saccharomycetales(100);f__Metschnikowiaceae(100);g__Metschnikowia(100);s__Metschnikowia_pulcherrima(100); | Metschnikowia | 13 | Saprotroph | \|Undefined Saprotroph\| | Yeast | NULL | Probable | As nectar-tap_saprotroph (Põlme et al. 2020); Decay sugar-rich substrates (Põlme et al. 2020) | Põlme S, et al. 2020. Fungal Diversity 105:1-16 ((https://doi.org/10.1007/s13225-020-00466-2)) |
| Otu40457 | 1 | k__Fungi(100);p__Ascomycota(100);c__Saccharomycetes(100);o__Saccharomycetales(100);f__Metschnikowiaceae(100);g__Metschnikowia(100);s__Metschnikowia_pulcherrima(100); | Metschnikowia | 13 | Saprotroph | \|Undefined Saprotroph\| | Yeast | NULL | Probable | As nectar-tap_saprotroph (Põlme et al. 2020); Decay sugar-rich substrates (Põlme et al. 2020) | Põlme S, et al. 2020. Fungal Diversity 105:1-16 ((https://doi.org/10.1007/s13225-020-00466-2)) |
| Otu40458 | 1 | k__Fungi(100);p__Ascomycota(100);c__Saccharomycetes(100);o__Saccharomycetales(100);f__Metschnikowiaceae(100);g__Metschnikowia(100);s__Metschnikowia_pulcherrima(100); | Metschnikowia | 13 | Saprotroph | \|Undefined Saprotroph\| | Yeast | NULL | Probable | As nectar-tap_saprotroph (Põlme et al. 2020); Decay sugar-rich substrates (Põlme et al. 2020) | Põlme S, et al. 2020. Fungal Diversity 105:1-16 ((https://doi.org/10.1007/s13225-020-00466-2)) |
| Otu40459 | 1 | k__Fungi(100);p__Ascomycota(100);c__Saccharomycetes(100);o__Saccharomycetales(100);f__Metschnikowiaceae(100);g__Metschnikowia(100);s__Metschnikowia_pulcherrima(100); | Metschnikowia | 13 | Saprotroph | \|Undefined Saprotroph\| | Yeast | NULL | Probable | As nectar-tap_saprotroph (Põlme et al. 2020); Decay sugar-rich substrates (Põlme et al. 2020) | Põlme S, et al. 2020. Fungal Diversity 105:1-16 ((https://doi.org/10.1007/s13225-020-00466-2)) |
| Otu40460 | 1 | k__Fungi(100);p__Ascomycota(100);c__Saccharomycetes(100);o__Saccharomycetales(100);f__Metschnikowiaceae(100);g__Metschnikowia(100);s__Metschnikowia_pulcherrima(100); | Metschnikowia | 13 | Saprotroph | \|Undefined Saprotroph\| | Yeast | NULL | Probable | As nectar-tap_saprotroph (Põlme et al. 2020); Decay sugar-rich substrates (Põlme et al. 2020) | Põlme S, et al. 2020. Fungal Diversity 105:1-16 ((https://doi.org/10.1007/s13225-020-00466-2)) |
| Otu40461 | 1 | k__Fungi(100);p__Ascomycota(100);c__Saccharomycetes(100);o__Saccharomycetales(100);f__Metschnikowiaceae(100);g__Metschnikowia(100);s__Metschnikowia_pulcherrima(100); | Metschnikowia | 13 | Saprotroph | \|Undefined Saprotroph\| | Yeast | NULL | Probable | As nectar-tap_saprotroph (Põlme et al. 2020); Decay sugar-rich substrates (Põlme et al. 2020) | Põlme S, et al. 2020. Fungal Diversity 105:1-16 ((https://doi.org/10.1007/s13225-020-00466-2)) |
| Otu40462 | 1 | k__Fungi(100);p__Ascomycota(100);c__Saccharomycetes(100);o__Saccharomycetales(100);f__Metschnikowiaceae(100);g__Metschnikowia(100);s__Metschnikowia_pulcherrima(100); | Metschnikowia | 13 | Saprotroph | \|Undefined Saprotroph\| | Yeast | NULL | Probable | As nectar-tap_saprotroph (Põlme et al. 2020); Decay sugar-rich substrates (Põlme et al. 2020) | Põlme S, et al. 2020. Fungal Diversity 105:1-16 ((https://doi.org/10.1007/s13225-020-00466-2)) |
| Otu40463 | 1 | k__Fungi(100);p__Ascomycota(100);c__Saccharomycetes(100);o__Saccharomycetales(100);f__Metschnikowiaceae(100);g__Metschnikowia(100);s__Metschnikowia_pulcherrima(100); | Metschnikowia | 13 | Saprotroph | \|Undefined Saprotroph\| | Yeast | NULL | Probable | As nectar-tap_saprotroph (Põlme et al. 2020); Decay sugar-rich substrates (Põlme et al. 2020) | Põlme S, et al. 2020. Fungal Diversity 105:1-16 ((https://doi.org/10.1007/s13225-020-00466-2)) |
| Otu40464 | 1 | k__Fungi(100);p__Ascomycota(100);c__Saccharomycetes(100);o__Saccharomycetales(100);f__Metschnikowiaceae(100);g__Metschnikowia(100);s__Metschnikowia_pulcherrima(100); | Metschnikowia | 13 | Saprotroph | \|Undefined Saprotroph\| | Yeast | NULL | Probable | As nectar-tap_saprotroph (Põlme et al. 2020); Decay sugar-rich substrates (Põlme et al. 2020) | Põlme S, et al. 2020. Fungal Diversity 105:1-16 ((https://doi.org/10.1007/s13225-020-00466-2)) |
| Otu40465 | 1 | k__Fungi(100);p__Ascomycota(100);c__Saccharomycetes(100);o__Saccharomycetales(100);f__Metschnikowiaceae(100);g__Metschnikowia(100);g__Metschnikowia_unclassified(100); | Metschnikowia | 13 | Saprotroph | \|Undefined Saprotroph\| | Yeast | NULL | Probable | As nectar-tap_saprotroph (Põlme et al. 2020); Decay sugar-rich substrates (Põlme et al. 2020) | Põlme S, et al. 2020. Fungal Diversity 105:1-16 ((https://doi.org/10.1007/s13225-020-00466-2)) |
| Otu40466 | 1 | k__Fungi(100);p__Ascomycota(100);c__Saccharomycetes(100);o__Saccharomycetales(100);f__Metschnikowiaceae(100);g__Metschnikowia(100);g__Metschnikowia_unclassified(100); | Metschnikowia | 13 | Saprotroph | \|Undefined Saprotroph\| | Yeast | NULL | Probable | As nectar-tap_saprotroph (Põlme et al. 2020); Decay sugar-rich substrates (Põlme et al. 2020) | Põlme S, et al. 2020. Fungal Diversity 105:1-16 ((https://doi.org/10.1007/s13225-020-00466-2)) |
| Otu40467 | 1 | k__Fungi(100);p__Ascomycota(100);c__Saccharomycetes(100);o__Saccharomycetales(100);f__Metschnikowiaceae(100);g__Metschnikowia(100);s__Metschnikowia_pulcherrima(100); | Metschnikowia | 13 | Saprotroph | \|Undefined Saprotroph\| | Yeast | NULL | Probable | As nectar-tap_saprotroph (Põlme et al. 2020); Decay sugar-rich substrates (Põlme et al. 2020) | Põlme S, et al. 2020. Fungal Diversity 105:1-16 ((https://doi.org/10.1007/s13225-020-00466-2)) |
| Otu40468 | 1 | k__Fungi(100);p__Ascomycota(100);c__Saccharomycetes(100);o__Saccharomycetales(100);f__Metschnikowiaceae(100);g__Metschnikowia(100);s__Metschnikowia_pulcherrima(100); | Metschnikowia | 13 | Saprotroph | \|Undefined Saprotroph\| | Yeast | NULL | Probable | As nectar-tap_saprotroph (Põlme et al. 2020); Decay sugar-rich substrates (Põlme et al. 2020) | Põlme S, et al. 2020. Fungal Diversity 105:1-16 ((https://doi.org/10.1007/s13225-020-00466-2)) |
| Otu40469 | 1 | k__Fungi(100);p__Ascomycota(100);c__Saccharomycetes(100);o__Saccharomycetales(100);f__Metschnikowiaceae(100);g__Metschnikowia(100);g__Metschnikowia_unclassified(100); | Metschnikowia | 13 | Saprotroph | \|Undefined Saprotroph\| | Yeast | NULL | Probable | As nectar-tap_saprotroph (Põlme et al. 2020); Decay sugar-rich substrates (Põlme et al. 2020) | Põlme S, et al. 2020. Fungal Diversity 105:1-16 ((https://doi.org/10.1007/s13225-020-00466-2)) |
| Otu40470 | 1 | k__Fungi(100);p__Ascomycota(100);c__Saccharomycetes(100);o__Saccharomycetales(100);f__Metschnikowiaceae(100);g__Metschnikowia(100);s__Metschnikowia_pulcherrima(100); | Metschnikowia | 13 | Saprotroph | \|Undefined Saprotroph\| | Yeast | NULL | Probable | As nectar-tap_saprotroph (Põlme et al. 2020); Decay sugar-rich substrates (Põlme et al. 2020) | Põlme S, et al. 2020. Fungal Diversity 105:1-16 ((https://doi.org/10.1007/s13225-020-00466-2)) |
| Otu40471 | 1 | k__Fungi(100);p__Ascomycota(100);c__Saccharomycetes(100);o__Saccharomycetales(100);f__Metschnikowiaceae(100);g__Metschnikowia(100);g__Metschnikowia_unclassified(100); | Metschnikowia | 13 | Saprotroph | \|Undefined Saprotroph\| | Yeast | NULL | Probable | As nectar-tap_saprotroph (Põlme et al. 2020); Decay sugar-rich substrates (Põlme et al. 2020) | Põlme S, et al. 2020. Fungal Diversity 105:1-16 ((https://doi.org/10.1007/s13225-020-00466-2)) |
| Otu40472 | 1 | k__Fungi(100);p__Ascomycota(100);c__Saccharomycetes(100);o__Saccharomycetales(100);f__Metschnikowiaceae(100);g__Metschnikowia(100);g__Metschnikowia_unclassified(100); | Metschnikowia | 13 | Saprotroph | \|Undefined Saprotroph\| | Yeast | NULL | Probable | As nectar-tap_saprotroph (Põlme et al. 2020); Decay sugar-rich substrates (Põlme et al. 2020) | Põlme S, et al. 2020. Fungal Diversity 105:1-16 ((https://doi.org/10.1007/s13225-020-00466-2)) |
| Otu40473 | 1 | k__Fungi(100);p__Ascomycota(100);c__Saccharomycetes(100);o__Saccharomycetales(100);f__Metschnikowiaceae(100);g__Metschnikowia(100);s__Metschnikowia_pulcherrima(100); | Metschnikowia | 13 | Saprotroph | \|Undefined Saprotroph\| | Yeast | NULL | Probable | As nectar-tap_saprotroph (Põlme et al. 2020); Decay sugar-rich substrates (Põlme et al. 2020) | Põlme S, et al. 2020. Fungal Diversity 105:1-16 ((https://doi.org/10.1007/s13225-020-00466-2)) |
| Otu40474 | 1 | k__Fungi(100);p__Ascomycota(100);c__Saccharomycetes(100);o__Saccharomycetales(100);f__Metschnikowiaceae(100);g__Metschnikowia(100);s__Metschnikowia_pulcherrima(100); | Metschnikowia | 13 | Saprotroph | \|Undefined Saprotroph\| | Yeast | NULL | Probable | As nectar-tap_saprotroph (Põlme et al. 2020); Decay sugar-rich substrates (Põlme et al. 2020) | Põlme S, et al. 2020. Fungal Diversity 105:1-16 ((https://doi.org/10.1007/s13225-020-00466-2)) |
| Otu40475 | 1 | k__Fungi(100);p__Ascomycota(100);c__Saccharomycetes(100);o__Saccharomycetales(100);f__Metschnikowiaceae(100);g__Metschnikowia(100);g__Metschnikowia_unclassified(100); | Metschnikowia | 13 | Saprotroph | \|Undefined Saprotroph\| | Yeast | NULL | Probable | As nectar-tap_saprotroph (Põlme et al. 2020); Decay sugar-rich substrates (Põlme et al. 2020) | Põlme S, et al. 2020. Fungal Diversity 105:1-16 ((https://doi.org/10.1007/s13225-020-00466-2)) |
| Otu40476 | 1 | k__Fungi(100);p__Ascomycota(100);c__Saccharomycetes(100);o__Saccharomycetales(100);f__Metschnikowiaceae(100);g__Metschnikowia(100);g__Metschnikowia_unclassified(100); | Metschnikowia | 13 | Saprotroph | \|Undefined Saprotroph\| | Yeast | NULL | Probable | As nectar-tap_saprotroph (Põlme et al. 2020); Decay sugar-rich substrates (Põlme et al. 2020) | Põlme S, et al. 2020. Fungal Diversity 105:1-16 ((https://doi.org/10.1007/s13225-020-00466-2)) |
| Otu40477 | 1 | k__Fungi(100);p__Ascomycota(100);c__Saccharomycetes(100);o__Saccharomycetales(100);f__Metschnikowiaceae(100);g__Metschnikowia(100);g__Metschnikowia_unclassified(100); | Metschnikowia | 13 | Saprotroph | \|Undefined Saprotroph\| | Yeast | NULL | Probable | As nectar-tap_saprotroph (Põlme et al. 2020); Decay sugar-rich substrates (Põlme et al. 2020) | Põlme S, et al. 2020. Fungal Diversity 105:1-16 ((https://doi.org/10.1007/s13225-020-00466-2)) |
| Otu40478 | 1 | k__Fungi(100);p__Ascomycota(100);c__Saccharomycetes(100);o__Saccharomycetales(100);f__Metschnikowiaceae(100);g__Metschnikowia(100);s__Metschnikowia_pulcherrima(100); | Metschnikowia | 13 | Saprotroph | \|Undefined Saprotroph\| | Yeast | NULL | Probable | As nectar-tap_saprotroph (Põlme et al. 2020); Decay sugar-rich substrates (Põlme et al. 2020) | Põlme S, et al. 2020. Fungal Diversity 105:1-16 ((https://doi.org/10.1007/s13225-020-00466-2)) |
| Otu40479 | 1 | k__Fungi(100);p__Ascomycota(100);c__Saccharomycetes(100);o__Saccharomycetales(100);f__Metschnikowiaceae(100);g__Metschnikowia(100);s__Metschnikowia_pulcherrima(100); | Metschnikowia | 13 | Saprotroph | \|Undefined Saprotroph\| | Yeast | NULL | Probable | As nectar-tap_saprotroph (Põlme et al. 2020); Decay sugar-rich substrates (Põlme et al. 2020) | Põlme S, et al. 2020. Fungal Diversity 105:1-16 ((https://doi.org/10.1007/s13225-020-00466-2)) |
| Otu40480 | 1 | k__Fungi(100);p__Ascomycota(100);c__Saccharomycetes(100);o__Saccharomycetales(100);f__Metschnikowiaceae(100);g__Metschnikowia(100);s__Metschnikowia_pulcherrima(100); | Metschnikowia | 13 | Saprotroph | \|Undefined Saprotroph\| | Yeast | NULL | Probable | As nectar-tap_saprotroph (Põlme et al. 2020); Decay sugar-rich substrates (Põlme et al. 2020) | Põlme S, et al. 2020. Fungal Diversity 105:1-16 ((https://doi.org/10.1007/s13225-020-00466-2)) |
| Otu40481 | 1 | k__Fungi(100);p__Ascomycota(100);c__Saccharomycetes(100);o__Saccharomycetales(100);f__Metschnikowiaceae(100);g__Metschnikowia(100);s__Metschnikowia_pulcherrima(100); | Metschnikowia | 13 | Saprotroph | \|Undefined Saprotroph\| | Yeast | NULL | Probable | As nectar-tap_saprotroph (Põlme et al. 2020); Decay sugar-rich substrates (Põlme et al. 2020) | Põlme S, et al. 2020. Fungal Diversity 105:1-16 ((https://doi.org/10.1007/s13225-020-00466-2)) |
| Otu40482 | 1 | k__Fungi(100);p__Ascomycota(100);c__Saccharomycetes(100);o__Saccharomycetales(100);f__Metschnikowiaceae(100);g__Metschnikowia(100);g__Metschnikowia_unclassified(100); | Metschnikowia | 13 | Saprotroph | \|Undefined Saprotroph\| | Yeast | NULL | Probable | As nectar-tap_saprotroph (Põlme et al. 2020); Decay sugar-rich substrates (Põlme et al. 2020) | Põlme S, et al. 2020. Fungal Diversity 105:1-16 ((https://doi.org/10.1007/s13225-020-00466-2)) |
| Otu40483 | 1 | k__Fungi(100);p__Ascomycota(100);c__Saccharomycetes(100);o__Saccharomycetales(100);f__Metschnikowiaceae(100);g__Metschnikowia(100);g__Metschnikowia_unclassified(100); | Metschnikowia | 13 | Saprotroph | \|Undefined Saprotroph\| | Yeast | NULL | Probable | As nectar-tap_saprotroph (Põlme et al. 2020); Decay sugar-rich substrates (Põlme et al. 2020) | Põlme S, et al. 2020. Fungal Diversity 105:1-16 ((https://doi.org/10.1007/s13225-020-00466-2)) |
| Otu40484 | 1 | k__Fungi(100);p__Ascomycota(100);c__Saccharomycetes(100);o__Saccharomycetales(100);f__Metschnikowiaceae(100);g__Metschnikowia(100);s__Metschnikowia_pulcherrima(100); | Metschnikowia | 13 | Saprotroph | \|Undefined Saprotroph\| | Yeast | NULL | Probable | As nectar-tap_saprotroph (Põlme et al. 2020); Decay sugar-rich substrates (Põlme et al. 2020) | Põlme S, et al. 2020. Fungal Diversity 105:1-16 ((https://doi.org/10.1007/s13225-020-00466-2)) |
| Otu40485 | 1 | k__Fungi(100);p__Ascomycota(100);c__Saccharomycetes(100);o__Saccharomycetales(100);f__Metschnikowiaceae(100);g__Metschnikowia(100);s__Metschnikowia_pulcherrima(100); | Metschnikowia | 13 | Saprotroph | \|Undefined Saprotroph\| | Yeast | NULL | Probable | As nectar-tap_saprotroph (Põlme et al. 2020); Decay sugar-rich substrates (Põlme et al. 2020) | Põlme S, et al. 2020. Fungal Diversity 105:1-16 ((https://doi.org/10.1007/s13225-020-00466-2)) |
| Otu40486 | 1 | k__Fungi(100);p__Ascomycota(100);c__Saccharomycetes(100);o__Saccharomycetales(100);f__Metschnikowiaceae(100);g__Metschnikowia(100);g__Metschnikowia_unclassified(100); | Metschnikowia | 13 | Saprotroph | \|Undefined Saprotroph\| | Yeast | NULL | Probable | As nectar-tap_saprotroph (Põlme et al. 2020); Decay sugar-rich substrates (Põlme et al. 2020) | Põlme S, et al. 2020. Fungal Diversity 105:1-16 ((https://doi.org/10.1007/s13225-020-00466-2)) |
| Otu40487 | 1 | k__Fungi(100);p__Ascomycota(100);c__Saccharomycetes(100);o__Saccharomycetales(100);f__Metschnikowiaceae(100);g__Metschnikowia(100);s__Metschnikowia_pulcherrima(100); | Metschnikowia | 13 | Saprotroph | \|Undefined Saprotroph\| | Yeast | NULL | Probable | As nectar-tap_saprotroph (Põlme et al. 2020); Decay sugar-rich substrates (Põlme et al. 2020) | Põlme S, et al. 2020. Fungal Diversity 105:1-16 ((https://doi.org/10.1007/s13225-020-00466-2)) |
| Otu40488 | 1 | k__Fungi(100);p__Ascomycota(100);c__Saccharomycetes(100);o__Saccharomycetales(100);f__Metschnikowiaceae(100);g__Metschnikowia(100);s__Metschnikowia_pulcherrima(100); | Metschnikowia | 13 | Saprotroph | \|Undefined Saprotroph\| | Yeast | NULL | Probable | As nectar-tap_saprotroph (Põlme et al. 2020); Decay sugar-rich substrates (Põlme et al. 2020) | Põlme S, et al. 2020. Fungal Diversity 105:1-16 ((https://doi.org/10.1007/s13225-020-00466-2)) |
| Otu40489 | 1 | k__Fungi(100);p__Ascomycota(100);c__Saccharomycetes(100);o__Saccharomycetales(100);f__Metschnikowiaceae(100);g__Metschnikowia(100);s__Metschnikowia_pulcherrima(100); | Metschnikowia | 13 | Saprotroph | \|Undefined Saprotroph\| | Yeast | NULL | Probable | As nectar-tap_saprotroph (Põlme et al. 2020); Decay sugar-rich substrates (Põlme et al. 2020) | Põlme S, et al. 2020. Fungal Diversity 105:1-16 ((https://doi.org/10.1007/s13225-020-00466-2)) |
| Otu40490 | 1 | k__Fungi(100);p__Ascomycota(100);c__Saccharomycetes(100);o__Saccharomycetales(100);f__Metschnikowiaceae(100);g__Metschnikowia(100);s__Metschnikowia_pulcherrima(100); | Metschnikowia | 13 | Saprotroph | \|Undefined Saprotroph\| | Yeast | NULL | Probable | As nectar-tap_saprotroph (Põlme et al. 2020); Decay sugar-rich substrates (Põlme et al. 2020) | Põlme S, et al. 2020. Fungal Diversity 105:1-16 ((https://doi.org/10.1007/s13225-020-00466-2)) |
| Otu40491 | 1 | k__Fungi(100);p__Ascomycota(100);c__Saccharomycetes(100);o__Saccharomycetales(100);f__Metschnikowiaceae(100);g__Metschnikowia(100);s__Metschnikowia_pulcherrima(100); | Metschnikowia | 13 | Saprotroph | \|Undefined Saprotroph\| | Yeast | NULL | Probable | As nectar-tap_saprotroph (Põlme et al. 2020); Decay sugar-rich substrates (Põlme et al. 2020) | Põlme S, et al. 2020. Fungal Diversity 105:1-16 ((https://doi.org/10.1007/s13225-020-00466-2)) |
| Otu40492 | 1 | k__Fungi(100);p__Ascomycota(100);c__Saccharomycetes(100);o__Saccharomycetales(100);f__Metschnikowiaceae(100);g__Metschnikowia(100);s__Metschnikowia_pulcherrima(100); | Metschnikowia | 13 | Saprotroph | \|Undefined Saprotroph\| | Yeast | NULL | Probable | As nectar-tap_saprotroph (Põlme et al. 2020); Decay sugar-rich substrates (Põlme et al. 2020) | Põlme S, et al. 2020. Fungal Diversity 105:1-16 ((https://doi.org/10.1007/s13225-020-00466-2)) |
| Otu40493 | 1 | k__Fungi(100);p__Ascomycota(100);c__Saccharomycetes(100);o__Saccharomycetales(100);f__Metschnikowiaceae(100);g__Metschnikowia(100);s__Metschnikowia_pulcherrima(100); | Metschnikowia | 13 | Saprotroph | \|Undefined Saprotroph\| | Yeast | NULL | Probable | As nectar-tap_saprotroph (Põlme et al. 2020); Decay sugar-rich substrates (Põlme et al. 2020) | Põlme S, et al. 2020. Fungal Diversity 105:1-16 ((https://doi.org/10.1007/s13225-020-00466-2)) |
| Otu40494 | 1 | k__Fungi(100);p__Ascomycota(100);c__Saccharomycetes(100);o__Saccharomycetales(100);f__Metschnikowiaceae(100);g__Metschnikowia(100);s__Metschnikowia_pulcherrima(100); | Metschnikowia | 13 | Saprotroph | \|Undefined Saprotroph\| | Yeast | NULL | Probable | As nectar-tap_saprotroph (Põlme et al. 2020); Decay sugar-rich substrates (Põlme et al. 2020) | Põlme S, et al. 2020. Fungal Diversity 105:1-16 ((https://doi.org/10.1007/s13225-020-00466-2)) |
| Otu40495 | 1 | k__Fungi(100);p__Ascomycota(100);c__Saccharomycetes(100);o__Saccharomycetales(100);f__Metschnikowiaceae(100);g__Metschnikowia(100);g__Metschnikowia_unclassified(100); | Metschnikowia | 13 | Saprotroph | \|Undefined Saprotroph\| | Yeast | NULL | Probable | As nectar-tap_saprotroph (Põlme et al. 2020); Decay sugar-rich substrates (Põlme et al. 2020) | Põlme S, et al. 2020. Fungal Diversity 105:1-16 ((https://doi.org/10.1007/s13225-020-00466-2)) |
| Otu40496 | 1 | k__Fungi(100);p__Ascomycota(100);c__Saccharomycetes(100);o__Saccharomycetales(100);f__Metschnikowiaceae(100);g__Metschnikowia(100);s__Metschnikowia_pulcherrima(100); | Metschnikowia | 13 | Saprotroph | \|Undefined Saprotroph\| | Yeast | NULL | Probable | As nectar-tap_saprotroph (Põlme et al. 2020); Decay sugar-rich substrates (Põlme et al. 2020) | Põlme S, et al. 2020. Fungal Diversity 105:1-16 ((https://doi.org/10.1007/s13225-020-00466-2)) |
| Otu40497 | 1 | k__Fungi(100);p__Ascomycota(100);c__Saccharomycetes(100);o__Saccharomycetales(100);f__Metschnikowiaceae(100);g__Metschnikowia(100);s__Metschnikowia_pulcherrima(100); | Metschnikowia | 13 | Saprotroph | \|Undefined Saprotroph\| | Yeast | NULL | Probable | As nectar-tap_saprotroph (Põlme et al. 2020); Decay sugar-rich substrates (Põlme et al. 2020) | Põlme S, et al. 2020. Fungal Diversity 105:1-16 ((https://doi.org/10.1007/s13225-020-00466-2)) |
| Otu40499 | 1 | k__Fungi(100);p__Ascomycota(100);c__Saccharomycetes(100);o__Saccharomycetales(100);f__Metschnikowiaceae(100);g__Metschnikowia(100);s__Metschnikowia_pulcherrima(100); | Metschnikowia | 13 | Saprotroph | \|Undefined Saprotroph\| | Yeast | NULL | Probable | As nectar-tap_saprotroph (Põlme et al. 2020); Decay sugar-rich substrates (Põlme et al. 2020) | Põlme S, et al. 2020. Fungal Diversity 105:1-16 ((https://doi.org/10.1007/s13225-020-00466-2)) |
| Otu40500 | 1 | k__Fungi(100);p__Ascomycota(100);c__Saccharomycetes(100);o__Saccharomycetales(100);f__Metschnikowiaceae(100);g__Metschnikowia(100);s__Metschnikowia_pulcherrima(100); | Metschnikowia | 13 | Saprotroph | \|Undefined Saprotroph\| | Yeast | NULL | Probable | As nectar-tap_saprotroph (Põlme et al. 2020); Decay sugar-rich substrates (Põlme et al. 2020) | Põlme S, et al. 2020. Fungal Diversity 105:1-16 ((https://doi.org/10.1007/s13225-020-00466-2)) |
| Otu40501 | 1 | k__Fungi(100);p__Ascomycota(100);c__Saccharomycetes(100);o__Saccharomycetales(100);f__Metschnikowiaceae(100);g__Metschnikowia(100);g__Metschnikowia_unclassified(100); | Metschnikowia | 13 | Saprotroph | \|Undefined Saprotroph\| | Yeast | NULL | Probable | As nectar-tap_saprotroph (Põlme et al. 2020); Decay sugar-rich substrates (Põlme et al. 2020) | Põlme S, et al. 2020. Fungal Diversity 105:1-16 ((https://doi.org/10.1007/s13225-020-00466-2)) |
| Otu40502 | 1 | k__Fungi(100);p__Ascomycota(100);c__Saccharomycetes(100);o__Saccharomycetales(100);f__Metschnikowiaceae(100);g__Metschnikowia(100);g__Metschnikowia_unclassified(100); | Metschnikowia | 13 | Saprotroph | \|Undefined Saprotroph\| | Yeast | NULL | Probable | As nectar-tap_saprotroph (Põlme et al. 2020); Decay sugar-rich substrates (Põlme et al. 2020) | Põlme S, et al. 2020. Fungal Diversity 105:1-16 ((https://doi.org/10.1007/s13225-020-00466-2)) |
| Otu40503 | 1 | k__Fungi(100);p__Ascomycota(100);c__Saccharomycetes(100);o__Saccharomycetales(100);f__Metschnikowiaceae(100);g__Metschnikowia(100);s__Metschnikowia_pulcherrima(100); | Metschnikowia | 13 | Saprotroph | \|Undefined Saprotroph\| | Yeast | NULL | Probable | As nectar-tap_saprotroph (Põlme et al. 2020); Decay sugar-rich substrates (Põlme et al. 2020) | Põlme S, et al. 2020. Fungal Diversity 105:1-16 ((https://doi.org/10.1007/s13225-020-00466-2)) |
| Otu40504 | 1 | k__Fungi(100);p__Ascomycota(100);c__Saccharomycetes(100);o__Saccharomycetales(100);f__Metschnikowiaceae(100);g__Metschnikowia(100);s__Metschnikowia_pulcherrima(100); | Metschnikowia | 13 | Saprotroph | \|Undefined Saprotroph\| | Yeast | NULL | Probable | As nectar-tap_saprotroph (Põlme et al. 2020); Decay sugar-rich substrates (Põlme et al. 2020) | Põlme S, et al. 2020. Fungal Diversity 105:1-16 ((https://doi.org/10.1007/s13225-020-00466-2)) |
| Otu40505 | 1 | k__Fungi(100);p__Ascomycota(100);c__Saccharomycetes(100);o__Saccharomycetales(100);f__Metschnikowiaceae(100);g__Metschnikowia(100);s__Metschnikowia_pulcherrima(100); | Metschnikowia | 13 | Saprotroph | \|Undefined Saprotroph\| | Yeast | NULL | Probable | As nectar-tap_saprotroph (Põlme et al. 2020); Decay sugar-rich substrates (Põlme et al. 2020) | Põlme S, et al. 2020. Fungal Diversity 105:1-16 ((https://doi.org/10.1007/s13225-020-00466-2)) |
| Otu40506 | 1 | k__Fungi(100);p__Ascomycota(100);c__Saccharomycetes(100);o__Saccharomycetales(100);f__Metschnikowiaceae(100);g__Metschnikowia(100);s__Metschnikowia_pulcherrima(100); | Metschnikowia | 13 | Saprotroph | \|Undefined Saprotroph\| | Yeast | NULL | Probable | As nectar-tap_saprotroph (Põlme et al. 2020); Decay sugar-rich substrates (Põlme et al. 2020) | Põlme S, et al. 2020. Fungal Diversity 105:1-16 ((https://doi.org/10.1007/s13225-020-00466-2)) |
| Otu40507 | 1 | k__Fungi(100);p__Ascomycota(100);c__Saccharomycetes(100);o__Saccharomycetales(100);f__Metschnikowiaceae(100);g__Metschnikowia(100);s__Metschnikowia_pulcherrima(100); | Metschnikowia | 13 | Saprotroph | \|Undefined Saprotroph\| | Yeast | NULL | Probable | As nectar-tap_saprotroph (Põlme et al. 2020); Decay sugar-rich substrates (Põlme et al. 2020) | Põlme S, et al. 2020. Fungal Diversity 105:1-16 ((https://doi.org/10.1007/s13225-020-00466-2)) |
| Otu40508 | 1 | k__Fungi(100);p__Ascomycota(100);c__Saccharomycetes(100);o__Saccharomycetales(100);f__Metschnikowiaceae(100);g__Metschnikowia(100);s__Metschnikowia_pulcherrima(100); | Metschnikowia | 13 | Saprotroph | \|Undefined Saprotroph\| | Yeast | NULL | Probable | As nectar-tap_saprotroph (Põlme et al. 2020); Decay sugar-rich substrates (Põlme et al. 2020) | Põlme S, et al. 2020. Fungal Diversity 105:1-16 ((https://doi.org/10.1007/s13225-020-00466-2)) |
| Otu40509 | 1 | k__Fungi(100);p__Ascomycota(100);c__Saccharomycetes(100);o__Saccharomycetales(100);f__Metschnikowiaceae(100);g__Metschnikowia(100);s__Metschnikowia_pulcherrima(100); | Metschnikowia | 13 | Saprotroph | \|Undefined Saprotroph\| | Yeast | NULL | Probable | As nectar-tap_saprotroph (Põlme et al. 2020); Decay sugar-rich substrates (Põlme et al. 2020) | Põlme S, et al. 2020. Fungal Diversity 105:1-16 ((https://doi.org/10.1007/s13225-020-00466-2)) |
| Otu40510 | 1 | k__Fungi(100);p__Ascomycota(100);c__Saccharomycetes(100);o__Saccharomycetales(100);f__Metschnikowiaceae(100);g__Metschnikowia(100);g__Metschnikowia_unclassified(100); | Metschnikowia | 13 | Saprotroph | \|Undefined Saprotroph\| | Yeast | NULL | Probable | As nectar-tap_saprotroph (Põlme et al. 2020); Decay sugar-rich substrates (Põlme et al. 2020) | Põlme S, et al. 2020. Fungal Diversity 105:1-16 ((https://doi.org/10.1007/s13225-020-00466-2)) |
| Otu40511 | 1 | k__Fungi(100);p__Ascomycota(100);c__Saccharomycetes(100);o__Saccharomycetales(100);f__Metschnikowiaceae(100);g__Metschnikowia(100);s__Metschnikowia_pulcherrima(100); | Metschnikowia | 13 | Saprotroph | \|Undefined Saprotroph\| | Yeast | NULL | Probable | As nectar-tap_saprotroph (Põlme et al. 2020); Decay sugar-rich substrates (Põlme et al. 2020) | Põlme S, et al. 2020. Fungal Diversity 105:1-16 ((https://doi.org/10.1007/s13225-020-00466-2)) |
| Otu40512 | 1 | k__Fungi(100);p__Ascomycota(100);c__Saccharomycetes(100);o__Saccharomycetales(100);f__Metschnikowiaceae(100);g__Metschnikowia(100);s__Metschnikowia_pulcherrima(100); | Metschnikowia | 13 | Saprotroph | \|Undefined Saprotroph\| | Yeast | NULL | Probable | As nectar-tap_saprotroph (Põlme et al. 2020); Decay sugar-rich substrates (Põlme et al. 2020) | Põlme S, et al. 2020. Fungal Diversity 105:1-16 ((https://doi.org/10.1007/s13225-020-00466-2)) |
| Otu40513 | 1 | k__Fungi(100);p__Ascomycota(100);c__Saccharomycetes(100);o__Saccharomycetales(100);f__Metschnikowiaceae(100);g__Metschnikowia(100);g__Metschnikowia_unclassified(100); | Metschnikowia | 13 | Saprotroph | \|Undefined Saprotroph\| | Yeast | NULL | Probable | As nectar-tap_saprotroph (Põlme et al. 2020); Decay sugar-rich substrates (Põlme et al. 2020) | Põlme S, et al. 2020. Fungal Diversity 105:1-16 ((https://doi.org/10.1007/s13225-020-00466-2)) |
| Otu40514 | 1 | k__Fungi(100);p__Ascomycota(100);c__Saccharomycetes(100);o__Saccharomycetales(100);f__Metschnikowiaceae(100);g__Metschnikowia(100);s__Metschnikowia_pulcherrima(100); | Metschnikowia | 13 | Saprotroph | \|Undefined Saprotroph\| | Yeast | NULL | Probable | As nectar-tap_saprotroph (Põlme et al. 2020); Decay sugar-rich substrates (Põlme et al. 2020) | Põlme S, et al. 2020. Fungal Diversity 105:1-16 ((https://doi.org/10.1007/s13225-020-00466-2)) |
| Otu40515 | 1 | k__Fungi(100);p__Ascomycota(100);c__Saccharomycetes(100);o__Saccharomycetales(100);f__Metschnikowiaceae(100);g__Metschnikowia(100);s__Metschnikowia_pulcherrima(100); | Metschnikowia | 13 | Saprotroph | \|Undefined Saprotroph\| | Yeast | NULL | Probable | As nectar-tap_saprotroph (Põlme et al. 2020); Decay sugar-rich substrates (Põlme et al. 2020) | Põlme S, et al. 2020. Fungal Diversity 105:1-16 ((https://doi.org/10.1007/s13225-020-00466-2)) |
| Otu40516 | 1 | k__Fungi(100);p__Ascomycota(100);c__Saccharomycetes(100);o__Saccharomycetales(100);f__Metschnikowiaceae(100);g__Metschnikowia(100);s__Metschnikowia_pulcherrima(100); | Metschnikowia | 13 | Saprotroph | \|Undefined Saprotroph\| | Yeast | NULL | Probable | As nectar-tap_saprotroph (Põlme et al. 2020); Decay sugar-rich substrates (Põlme et al. 2020) | Põlme S, et al. 2020. Fungal Diversity 105:1-16 ((https://doi.org/10.1007/s13225-020-00466-2)) |
| Otu40517 | 1 | k__Fungi(100);p__Ascomycota(100);c__Saccharomycetes(100);o__Saccharomycetales(100);f__Metschnikowiaceae(100);g__Metschnikowia(100);g__Metschnikowia_unclassified(100); | Metschnikowia | 13 | Saprotroph | \|Undefined Saprotroph\| | Yeast | NULL | Probable | As nectar-tap_saprotroph (Põlme et al. 2020); Decay sugar-rich substrates (Põlme et al. 2020) | Põlme S, et al. 2020. Fungal Diversity 105:1-16 ((https://doi.org/10.1007/s13225-020-00466-2)) |
| Otu40518 | 1 | k__Fungi(100);p__Ascomycota(100);c__Saccharomycetes(100);o__Saccharomycetales(100);f__Metschnikowiaceae(100);g__Metschnikowia(100);s__Metschnikowia_pulcherrima(100); | Metschnikowia | 13 | Saprotroph | \|Undefined Saprotroph\| | Yeast | NULL | Probable | As nectar-tap_saprotroph (Põlme et al. 2020); Decay sugar-rich substrates (Põlme et al. 2020) | Põlme S, et al. 2020. Fungal Diversity 105:1-16 ((https://doi.org/10.1007/s13225-020-00466-2)) |
| Otu40520 | 1 | k__Fungi(100);p__Ascomycota(100);c__Saccharomycetes(100);o__Saccharomycetales(100);f__Metschnikowiaceae(100);g__Metschnikowia(100);s__Metschnikowia_pulcherrima(100); | Metschnikowia | 13 | Saprotroph | \|Undefined Saprotroph\| | Yeast | NULL | Probable | As nectar-tap_saprotroph (Põlme et al. 2020); Decay sugar-rich substrates (Põlme et al. 2020) | Põlme S, et al. 2020. Fungal Diversity 105:1-16 ((https://doi.org/10.1007/s13225-020-00466-2)) |
| Otu40521 | 1 | k__Fungi(100);p__Ascomycota(100);c__Saccharomycetes(100);o__Saccharomycetales(100);f__Metschnikowiaceae(100);g__Metschnikowia(100);s__Metschnikowia_pulcherrima(100); | Metschnikowia | 13 | Saprotroph | \|Undefined Saprotroph\| | Yeast | NULL | Probable | As nectar-tap_saprotroph (Põlme et al. 2020); Decay sugar-rich substrates (Põlme et al. 2020) | Põlme S, et al. 2020. Fungal Diversity 105:1-16 ((https://doi.org/10.1007/s13225-020-00466-2)) |
| Otu40522 | 1 | k__Fungi(100);p__Ascomycota(100);c__Saccharomycetes(100);o__Saccharomycetales(100);f__Metschnikowiaceae(100);g__Metschnikowia(100);s__Metschnikowia_pulcherrima(100); | Metschnikowia | 13 | Saprotroph | \|Undefined Saprotroph\| | Yeast | NULL | Probable | As nectar-tap_saprotroph (Põlme et al. 2020); Decay sugar-rich substrates (Põlme et al. 2020) | Põlme S, et al. 2020. Fungal Diversity 105:1-16 ((https://doi.org/10.1007/s13225-020-00466-2)) |
| Otu40523 | 1 | k__Fungi(100);p__Ascomycota(100);c__Saccharomycetes(100);o__Saccharomycetales(100);f__Metschnikowiaceae(100);g__Metschnikowia(100);s__Metschnikowia_pulcherrima(100); | Metschnikowia | 13 | Saprotroph | \|Undefined Saprotroph\| | Yeast | NULL | Probable | As nectar-tap_saprotroph (Põlme et al. 2020); Decay sugar-rich substrates (Põlme et al. 2020) | Põlme S, et al. 2020. Fungal Diversity 105:1-16 ((https://doi.org/10.1007/s13225-020-00466-2)) |
| Otu40525 | 1 | k__Fungi(100);p__Ascomycota(100);c__Saccharomycetes(100);o__Saccharomycetales(100);f__Metschnikowiaceae(100);g__Metschnikowia(100);s__Metschnikowia_pulcherrima(100); | Metschnikowia | 13 | Saprotroph | \|Undefined Saprotroph\| | Yeast | NULL | Probable | As nectar-tap_saprotroph (Põlme et al. 2020); Decay sugar-rich substrates (Põlme et al. 2020) | Põlme S, et al. 2020. Fungal Diversity 105:1-16 ((https://doi.org/10.1007/s13225-020-00466-2)) |
| Otu40530 | 1 | k__Fungi(100);p__Ascomycota(100);c__Saccharomycetes(100);o__Saccharomycetales(100);f__Metschnikowiaceae(100);g__Metschnikowia(100);s__Metschnikowia_pulcherrima(100); | Metschnikowia | 13 | Saprotroph | \|Undefined Saprotroph\| | Yeast | NULL | Probable | As nectar-tap_saprotroph (Põlme et al. 2020); Decay sugar-rich substrates (Põlme et al. 2020) | Põlme S, et al. 2020. Fungal Diversity 105:1-16 ((https://doi.org/10.1007/s13225-020-00466-2)) |
| Otu40531 | 1 | k__Fungi(100);p__Ascomycota(100);c__Saccharomycetes(100);o__Saccharomycetales(100);f__Metschnikowiaceae(100);g__Metschnikowia(100);s__Metschnikowia_pulcherrima(100); | Metschnikowia | 13 | Saprotroph | \|Undefined Saprotroph\| | Yeast | NULL | Probable | As nectar-tap_saprotroph (Põlme et al. 2020); Decay sugar-rich substrates (Põlme et al. 2020) | Põlme S, et al. 2020. Fungal Diversity 105:1-16 ((https://doi.org/10.1007/s13225-020-00466-2)) |
| Otu40533 | 1 | k__Fungi(100);p__Ascomycota(100);c__Saccharomycetes(100);o__Saccharomycetales(100);f__Metschnikowiaceae(100);g__Metschnikowia(100);s__Metschnikowia_pulcherrima(100); | Metschnikowia | 13 | Saprotroph | \|Undefined Saprotroph\| | Yeast | NULL | Probable | As nectar-tap_saprotroph (Põlme et al. 2020); Decay sugar-rich substrates (Põlme et al. 2020) | Põlme S, et al. 2020. Fungal Diversity 105:1-16 ((https://doi.org/10.1007/s13225-020-00466-2)) |
| Otu40534 | 1 | k__Fungi(100);p__Ascomycota(100);c__Saccharomycetes(100);o__Saccharomycetales(100);f__Metschnikowiaceae(100);g__Metschnikowia(100);s__Metschnikowia_pulcherrima(100); | Metschnikowia | 13 | Saprotroph | \|Undefined Saprotroph\| | Yeast | NULL | Probable | As nectar-tap_saprotroph (Põlme et al. 2020); Decay sugar-rich substrates (Põlme et al. 2020) | Põlme S, et al. 2020. Fungal Diversity 105:1-16 ((https://doi.org/10.1007/s13225-020-00466-2)) |
| Otu40536 | 1 | k__Fungi(100);p__Ascomycota(100);c__Saccharomycetes(100);o__Saccharomycetales(100);f__Metschnikowiaceae(100);g__Metschnikowia(100);s__Metschnikowia_pulcherrima(100); | Metschnikowia | 13 | Saprotroph | \|Undefined Saprotroph\| | Yeast | NULL | Probable | As nectar-tap_saprotroph (Põlme et al. 2020); Decay sugar-rich substrates (Põlme et al. 2020) | Põlme S, et al. 2020. Fungal Diversity 105:1-16 ((https://doi.org/10.1007/s13225-020-00466-2)) |
| Otu40538 | 1 | k__Fungi(100);p__Ascomycota(100);c__Saccharomycetes(100);o__Saccharomycetales(100);f__Metschnikowiaceae(100);g__Metschnikowia(100);s__Metschnikowia_pulcherrima(100); | Metschnikowia | 13 | Saprotroph | \|Undefined Saprotroph\| | Yeast | NULL | Probable | As nectar-tap_saprotroph (Põlme et al. 2020); Decay sugar-rich substrates (Põlme et al. 2020) | Põlme S, et al. 2020. Fungal Diversity 105:1-16 ((https://doi.org/10.1007/s13225-020-00466-2)) |

| | | | | | | | | | | | |
|---|---|---|---|---|---|---|---|---|---|---|---|
| Otu40539 | 1 | k__Fungi(100);p__Ascomycota(100);c__Saccharomycetes(100);o__Saccharomycetales(100);f__Metschnikowiaceae(100);g__Metschnikowia(100);s__Metschnikowia_pulcherrima(100); | Metschnikowia | 13 | Saprotroph | \|Undefined Saprotroph\| | Yeast | NULL | Probable | As nectar-tap_saprotroph (Põlme et al. 2020); Decay sugar-rich substrates (Põlme et al. 2020) | Põlme S, et al. 2020. Fungal Diversity 105:1-16 ((https://doi.org/10.1007/s13225-020-00466-2)) |
| Otu40540 | 1 | k__Fungi(100);p__Ascomycota(100);c__Saccharomycetes(100);o__Saccharomycetales(100);f__Metschnikowiaceae(100);g__Metschnikowia(100);s__Metschnikowia_pulcherrima(100); | Metschnikowia | 13 | Saprotroph | \|Undefined Saprotroph\| | Yeast | NULL | Probable | As nectar-tap_saprotroph (Põlme et al. 2020); Decay sugar-rich substrates (Põlme et al. 2020) | Põlme S, et al. 2020. Fungal Diversity 105:1-16 ((https://doi.org/10.1007/s13225-020-00466-2)) |
| Otu40543 | 1 | k__Fungi(100);p__Ascomycota(100);c__Saccharomycetes(100);o__Saccharomycetales(100);f__Metschnikowiaceae(100);g__Metschnikowia(100);s__Metschnikowia_pulcherrima(100); | Metschnikowia | 13 | Saprotroph | \|Undefined Saprotroph\| | Yeast | NULL | Probable | As nectar-tap_saprotroph (Põlme et al. 2020); Decay sugar-rich substrates (Põlme et al. 2020) | Põlme S, et al. 2020. Fungal Diversity 105:1-16 ((https://doi.org/10.1007/s13225-020-00466-2)) |
| Otu40544 | 1 | k__Fungi(100);p__Ascomycota(100);c__Saccharomycetes(100);o__Saccharomycetales(100);f__Metschnikowiaceae(100);g__Metschnikowia(100);s__Metschnikowia_pulcherrima(100); | Metschnikowia | 13 | Saprotroph | \|Undefined Saprotroph\| | Yeast | NULL | Probable | As nectar-tap_saprotroph (Põlme et al. 2020); Decay sugar-rich substrates (Põlme et al. 2020) | Põlme S, et al. 2020. Fungal Diversity 105:1-16 ((https://doi.org/10.1007/s13225-020-00466-2)) |
| Otu40545 | 1 | k__Fungi(100);p__Ascomycota(100);c__Saccharomycetes(100);o__Saccharomycetales(100);f__Metschnikowiaceae(100);g__Metschnikowia(100);s__Metschnikowia_pulcherrima(100); | Metschnikowia | 13 | Saprotroph | \|Undefined Saprotroph\| | Yeast | NULL | Probable | As nectar-tap_saprotroph (Põlme et al. 2020); Decay sugar-rich substrates (Põlme et al. 2020) | Põlme S, et al. 2020. Fungal Diversity 105:1-16 ((https://doi.org/10.1007/s13225-020-00466-2)) |
| Otu40546 | 1 | k__Fungi(100);p__Ascomycota(100);c__Saccharomycetes(100);o__Saccharomycetales(100);f__Metschnikowiaceae(100);g__Metschnikowia(100);g__Metschnikowia_unclassified(100); | Metschnikowia | 13 | Saprotroph | \|Undefined Saprotroph\| | Yeast | NULL | Probable | As nectar-tap_saprotroph (Põlme et al. 2020); Decay sugar-rich substrates (Põlme et al. 2020) | Põlme S, et al. 2020. Fungal Diversity 105:1-16 ((https://doi.org/10.1007/s13225-020-00466-2)) |
| Otu40547 | 1 | k__Fungi(100);p__Ascomycota(100);c__Saccharomycetes(100);o__Saccharomycetales(100);f__Metschnikowiaceae(100);g__Metschnikowia(100);s__Metschnikowia_pulcherrima(100); | Metschnikowia | 13 | Saprotroph | \|Undefined Saprotroph\| | Yeast | NULL | Probable | As nectar-tap_saprotroph (Põlme et al. 2020); Decay sugar-rich substrates (Põlme et al. 2020) | Põlme S, et al. 2020. Fungal Diversity 105:1-16 ((https://doi.org/10.1007/s13225-020-00466-2)) |
| Otu40550 | 1 | k__Fungi(100);p__Ascomycota(100);c__Saccharomycetes(100);o__Saccharomycetales(100);f__Metschnikowiaceae(100);g__Metschnikowia(100);s__Metschnikowia_pulcherrima(100); | Metschnikowia | 13 | Saprotroph | \|Undefined Saprotroph\| | Yeast | NULL | Probable | As nectar-tap_saprotroph (Põlme et al. 2020); Decay sugar-rich substrates (Põlme et al. 2020) | Põlme S, et al. 2020. Fungal Diversity 105:1-16 ((https://doi.org/10.1007/s13225-020-00466-2)) |
| Otu40551 | 1 | k__Fungi(100);p__Ascomycota(100);c__Saccharomycetes(100);o__Saccharomycetales(100);f__Metschnikowiaceae(100);g__Metschnikowia(100);g__Metschnikowia_unclassified(100); | Metschnikowia | 13 | Saprotroph | \|Undefined Saprotroph\| | Yeast | NULL | Probable | As nectar-tap_saprotroph (Põlme et al. 2020); Decay sugar-rich substrates (Põlme et al. 2020) | Põlme S, et al. 2020. Fungal Diversity 105:1-16 ((https://doi.org/10.1007/s13225-020-00466-2)) |
| Otu40552 | 1 | k__Fungi(100);p__Ascomycota(100);c__Saccharomycetes(100);o__Saccharomycetales(100);f__Metschnikowiaceae(100);g__Metschnikowia(100);s__Metschnikowia_pulcherrima(100); | Metschnikowia | 13 | Saprotroph | \|Undefined Saprotroph\| | Yeast | NULL | Probable | As nectar-tap_saprotroph (Põlme et al. 2020); Decay sugar-rich substrates (Põlme et al. 2020) | Põlme S, et al. 2020. Fungal Diversity 105:1-16 ((https://doi.org/10.1007/s13225-020-00466-2)) |
| Otu40553 | 1 | k__Fungi(100);p__Ascomycota(100);c__Saccharomycetes(100);o__Saccharomycetales(100);f__Metschnikowiaceae(100);g__Metschnikowia(100);s__Metschnikowia_pulcherrima(100); | Metschnikowia | 13 | Saprotroph | \|Undefined Saprotroph\| | Yeast | NULL | Probable | As nectar-tap_saprotroph (Põlme et al. 2020); Decay sugar-rich substrates (Põlme et al. 2020) | Põlme S, et al. 2020. Fungal Diversity 105:1-16 ((https://doi.org/10.1007/s13225-020-00466-2)) |
| Otu40554 | 1 | k__Fungi(100);p__Ascomycota(100);c__Saccharomycetes(100);o__Saccharomycetales(100);f__Metschnikowiaceae(100);g__Metschnikowia(100);s__Metschnikowia_pulcherrima(100); | Metschnikowia | 13 | Saprotroph | \|Undefined Saprotroph\| | Yeast | NULL | Probable | As nectar-tap_saprotroph (Põlme et al. 2020); Decay sugar-rich substrates (Põlme et al. 2020) | Põlme S, et al. 2020. Fungal Diversity 105:1-16 ((https://doi.org/10.1007/s13225-020-00466-2)) |
| Otu40557 | 1 | k__Fungi(100);p__Ascomycota(100);c__Saccharomycetes(100);o__Saccharomycetales(100);f__Metschnikowiaceae(100);g__Metschnikowia(100);s__Metschnikowia_pulcherrima(100); | Metschnikowia | 13 | Saprotroph | \|Undefined Saprotroph\| | Yeast | NULL | Probable | As nectar-tap_saprotroph (Põlme et al. 2020); Decay sugar-rich substrates (Põlme et al. 2020) | Põlme S, et al. 2020. Fungal Diversity 105:1-16 ((https://doi.org/10.1007/s13225-020-00466-2)) |
| Otu40559 | 1 | k__Fungi(100);p__Ascomycota(100);c__Saccharomycetes(100);o__Saccharomycetales(100);f__Metschnikowiaceae(100);g__Metschnikowia(100);g__Metschnikowia_unclassified(100); | Metschnikowia | 13 | Saprotroph | \|Undefined Saprotroph\| | Yeast | NULL | Probable | As nectar-tap_saprotroph (Põlme et al. 2020); Decay sugar-rich substrates (Põlme et al. 2020) | Põlme S, et al. 2020. Fungal Diversity 105:1-16 ((https://doi.org/10.1007/s13225-020-00466-2)) |
| Otu40560 | 1 | k__Fungi(100);p__Ascomycota(100);c__Saccharomycetes(100);o__Saccharomycetales(100);f__Metschnikowiaceae(100);g__Metschnikowia(100);g__Metschnikowia_unclassified(100); | Metschnikowia | 13 | Saprotroph | \|Undefined Saprotroph\| | Yeast | NULL | Probable | As nectar-tap_saprotroph (Põlme et al. 2020); Decay sugar-rich substrates (Põlme et al. 2020) | Põlme S, et al. 2020. Fungal Diversity 105:1-16 ((https://doi.org/10.1007/s13225-020-00466-2)) |
| Otu40561 | 1 | k__Fungi(100);p__Ascomycota(100);c__Saccharomycetes(100);o__Saccharomycetales(100);f__Metschnikowiaceae(100);g__Metschnikowia(100);s__Metschnikowia_pulcherrima(100); | Metschnikowia | 13 | Saprotroph | \|Undefined Saprotroph\| | Yeast | NULL | Probable | As nectar-tap_saprotroph (Põlme et al. 2020); Decay sugar-rich substrates (Põlme et al. 2020) | Põlme S, et al. 2020. Fungal Diversity 105:1-16 ((https://doi.org/10.1007/s13225-020-00466-2)) |
| Otu40562 | 1 | k__Fungi(100);p__Ascomycota(100);c__Saccharomycetes(100);o__Saccharomycetales(100);f__Metschnikowiaceae(100);g__Metschnikowia(100);s__Metschnikowia_pulcherrima(100); | Metschnikowia | 13 | Saprotroph | \|Undefined Saprotroph\| | Yeast | NULL | Probable | As nectar-tap_saprotroph (Põlme et al. 2020); Decay sugar-rich substrates (Põlme et al. 2020) | Põlme S, et al. 2020. Fungal Diversity 105:1-16 ((https://doi.org/10.1007/s13225-020-00466-2)) |
| Otu40563 | 1 | k__Fungi(100);p__Ascomycota(100);c__Saccharomycetes(100);o__Saccharomycetales(100);f__Metschnikowiaceae(100);g__Metschnikowia(100);s__Metschnikowia_pulcherrima(100); | Metschnikowia | 13 | Saprotroph | \|Undefined Saprotroph\| | Yeast | NULL | Probable | As nectar-tap_saprotroph (Põlme et al. 2020); Decay sugar-rich substrates (Põlme et al. 2020) | Põlme S, et al. 2020. Fungal Diversity 105:1-16 ((https://doi.org/10.1007/s13225-020-00466-2)) |
| Otu40564 | 1 | k__Fungi(100);p__Ascomycota(100);c__Saccharomycetes(100);o__Saccharomycetales(100);f__Metschnikowiaceae(100);g__Metschnikowia(100);g__Metschnikowia_unclassified(100); | Metschnikowia | 13 | Saprotroph | \|Undefined Saprotroph\| | Yeast | NULL | Probable | As nectar-tap_saprotroph (Põlme et al. 2020); Decay sugar-rich substrates (Põlme et al. 2020) | Põlme S, et al. 2020. Fungal Diversity 105:1-16 ((https://doi.org/10.1007/s13225-020-00466-2)) |
| Otu40565 | 1 | k__Fungi(100);p__Ascomycota(100);c__Saccharomycetes(100);o__Saccharomycetales(100);f__Metschnikowiaceae(100);g__Metschnikowia(100);s__Metschnikowia_pulcherrima(100); | Metschnikowia | 13 | Saprotroph | \|Undefined Saprotroph\| | Yeast | NULL | Probable | As nectar-tap_saprotroph (Põlme et al. 2020); Decay sugar-rich substrates (Põlme et al. 2020) | Põlme S, et al. 2020. Fungal Diversity 105:1-16 ((https://doi.org/10.1007/s13225-020-00466-2)) |
| Otu40566 | 1 | k__Fungi(100);p__Ascomycota(100);c__Saccharomycetes(100);o__Saccharomycetales(100);f__Metschnikowiaceae(100);g__Metschnikowia(100);s__Metschnikowia_pulcherrima(100); | Metschnikowia | 13 | Saprotroph | \|Undefined Saprotroph\| | Yeast | NULL | Probable | As nectar-tap_saprotroph (Põlme et al. 2020); Decay sugar-rich substrates (Põlme et al. 2020) | Põlme S, et al. 2020. Fungal Diversity 105:1-16 ((https://doi.org/10.1007/s13225-020-00466-2)) |
| Otu40569 | 1 | k__Fungi(100);p__Ascomycota(100);c__Saccharomycetes(100);o__Saccharomycetales(100);f__Metschnikowiaceae(100);g__Metschnikowia(100);s__Metschnikowia_pulcherrima(100); | Metschnikowia | 13 | Saprotroph | \|Undefined Saprotroph\| | Yeast | NULL | Probable | As nectar-tap_saprotroph (Põlme et al. 2020); Decay sugar-rich substrates (Põlme et al. 2020) | Põlme S, et al. 2020. Fungal Diversity 105:1-16 ((https://doi.org/10.1007/s13225-020-00466-2)) |
| Otu40571 | 1 | k__Fungi(100);p__Ascomycota(100);c__Saccharomycetes(100);o__Saccharomycetales(100);f__Metschnikowiaceae(100);g__Metschnikowia(100);g__Metschnikowia_unclassified(100); | Metschnikowia | 13 | Saprotroph | \|Undefined Saprotroph\| | Yeast | NULL | Probable | As nectar-tap_saprotroph (Põlme et al. 2020); Decay sugar-rich substrates (Põlme et al. 2020) | Põlme S, et al. 2020. Fungal Diversity 105:1-16 ((https://doi.org/10.1007/s13225-020-00466-2)) |
| Otu40572 | 1 | k__Fungi(100);p__Ascomycota(100);c__Saccharomycetes(100);o__Saccharomycetales(100);f__Metschnikowiaceae(100);g__Metschnikowia(100);s__Metschnikowia_pulcherrima(100); | Metschnikowia | 13 | Saprotroph | \|Undefined Saprotroph\| | Yeast | NULL | Probable | As nectar-tap_saprotroph (Põlme et al. 2020); Decay sugar-rich substrates (Põlme et al. 2020) | Põlme S, et al. 2020. Fungal Diversity 105:1-16 ((https://doi.org/10.1007/s13225-020-00466-2)) |
| Otu40573 | 1 | k__Fungi(100);p__Ascomycota(100);c__Saccharomycetes(100);o__Saccharomycetales(100);f__Metschnikowiaceae(100);g__Metschnikowia(100);g__Metschnikowia_unclassified(100); | Metschnikowia | 13 | Saprotroph | \|Undefined Saprotroph\| | Yeast | NULL | Probable | As nectar-tap_saprotroph (Põlme et al. 2020); Decay sugar-rich substrates (Põlme et al. 2020) | Põlme S, et al. 2020. Fungal Diversity 105:1-16 ((https://doi.org/10.1007/s13225-020-00466-2)) |
| Otu40574 | 1 | k__Fungi(100);p__Ascomycota(100);c__Saccharomycetes(100);o__Saccharomycetales(100);f__Metschnikowiaceae(100);g__Metschnikowia(100);s__Metschnikowia_pulcherrima(100); | Metschnikowia | 13 | Saprotroph | \|Undefined Saprotroph\| | Yeast | NULL | Probable | As nectar-tap_saprotroph (Põlme et al. 2020); Decay sugar-rich substrates (Põlme et al. 2020) | Põlme S, et al. 2020. Fungal Diversity 105:1-16 ((https://doi.org/10.1007/s13225-020-00466-2)) |
| Otu40575 | 1 | k__Fungi(100);p__Ascomycota(100);c__Saccharomycetes(100);o__Saccharomycetales(100);f__Metschnikowiaceae(100);g__Metschnikowia(100);s__Metschnikowia_pulcherrima(100); | Metschnikowia | 13 | Saprotroph | \|Undefined Saprotroph\| | Yeast | NULL | Probable | As nectar-tap_saprotroph (Põlme et al. 2020); Decay sugar-rich substrates (Põlme et al. 2020) | Põlme S, et al. 2020. Fungal Diversity 105:1-16 ((https://doi.org/10.1007/s13225-020-00466-2)) |
| Otu40576 | 1 | k__Fungi(100);p__Ascomycota(100);c__Saccharomycetes(100);o__Saccharomycetales(100);f__Metschnikowiaceae(100);g__Metschnikowia(100);s__Metschnikowia_pulcherrima(100); | Metschnikowia | 13 | Saprotroph | \|Undefined Saprotroph\| | Yeast | NULL | Probable | As nectar-tap_saprotroph (Põlme et al. 2020); Decay sugar-rich substrates (Põlme et al. 2020) | Põlme S, et al. 2020. Fungal Diversity 105:1-16 ((https://doi.org/10.1007/s13225-020-00466-2)) |
| Otu40577 | 1 | k__Fungi(100);p__Ascomycota(100);c__Saccharomycetes(100);o__Saccharomycetales(100);f__Metschnikowiaceae(100);g__Metschnikowia(100);s__Metschnikowia_pulcherrima(100); | Metschnikowia | 13 | Saprotroph | \|Undefined Saprotroph\| | Yeast | NULL | Probable | As nectar-tap_saprotroph (Põlme et al. 2020); Decay sugar-rich substrates (Põlme et al. 2020) | Põlme S, et al. 2020. Fungal Diversity 105:1-16 ((https://doi.org/10.1007/s13225-020-00466-2)) |
| Otu40580 | 1 | k__Fungi(100);p__Ascomycota(100);c__Saccharomycetes(100);o__Saccharomycetales(100);f__Metschnikowiaceae(100);g__Metschnikowia(100);g__Metschnikowia_unclassified(100); | Metschnikowia | 13 | Saprotroph | \|Undefined Saprotroph\| | Yeast | NULL | Probable | As nectar-tap_saprotroph (Põlme et al. 2020); Decay sugar-rich substrates (Põlme et al. 2020) | Põlme S, et al. 2020. Fungal Diversity 105:1-16 ((https://doi.org/10.1007/s13225-020-00466-2)) |
| Otu40581 | 1 | k__Fungi(100);p__Ascomycota(100);c__Saccharomycetes(100);o__Saccharomycetales(100);f__Metschnikowiaceae(100);g__Metschnikowia(100);s__Metschnikowia_pulcherrima(100); | Metschnikowia | 13 | Saprotroph | \|Undefined Saprotroph\| | Yeast | NULL | Probable | As nectar-tap_saprotroph (Põlme et al. 2020); Decay sugar-rich substrates (Põlme et al. 2020) | Põlme S, et al. 2020. Fungal Diversity 105:1-16 ((https://doi.org/10.1007/s13225-020-00466-2)) |
| Otu40582 | 1 | k__Fungi(100);p__Ascomycota(100);c__Saccharomycetes(100);o__Saccharomycetales(100);f__Metschnikowiaceae(100);g__Metschnikowia(100);s__Metschnikowia_pulcherrima(100); | Metschnikowia | 13 | Saprotroph | \|Undefined Saprotroph\| | Yeast | NULL | Probable | As nectar-tap_saprotroph (Põlme et al. 2020); Decay sugar-rich substrates (Põlme et al. 2020) | Põlme S, et al. 2020. Fungal Diversity 105:1-16 ((https://doi.org/10.1007/s13225-020-00466-2)) |
| Otu40583 | 1 | k__Fungi(100);p__Ascomycota(100);c__Saccharomycetes(100);o__Saccharomycetales(100);f__Metschnikowiaceae(100);g__Metschnikowia(100);g__Metschnikowia_unclassified(100); | Metschnikowia | 13 | Saprotroph | \|Undefined Saprotroph\| | Yeast | NULL | Probable | As nectar-tap_saprotroph (Põlme et al. 2020); Decay sugar-rich substrates (Põlme et al. 2020) | Põlme S, et al. 2020. Fungal Diversity 105:1-16 ((https://doi.org/10.1007/s13225-020-00466-2)) |
| Otu40584 | 1 | k__Fungi(100);p__Ascomycota(100);c__Saccharomycetes(100);o__Saccharomycetales(100);f__Metschnikowiaceae(100);g__Metschnikowia(100);g__Metschnikowia_unclassified(100); | Metschnikowia | 13 | Saprotroph | \|Undefined Saprotroph\| | Yeast | NULL | Probable | As nectar-tap_saprotroph (Põlme et al. 2020); Decay sugar-rich substrates (Põlme et al. 2020) | Põlme S, et al. 2020. Fungal Diversity 105:1-16 ((https://doi.org/10.1007/s13225-020-00466-2)) |
| Otu40585 | 1 | k__Fungi(100);p__Ascomycota(100);c__Saccharomycetes(100);o__Saccharomycetales(100);f__Metschnikowiaceae(100);g__Metschnikowia(100);s__Metschnikowia_pulcherrima(100); | Metschnikowia | 13 | Saprotroph | \|Undefined Saprotroph\| | Yeast | NULL | Probable | As nectar-tap_saprotroph (Põlme et al. 2020); Decay sugar-rich substrates (Põlme et al. 2020) | Põlme S, et al. 2020. Fungal Diversity 105:1-16 ((https://doi.org/10.1007/s13225-020-00466-2)) |
| Otu40586 | 1 | k__Fungi(100);p__Ascomycota(100);c__Saccharomycetes(100);o__Saccharomycetales(100);f__Metschnikowiaceae(100);g__Metschnikowia(100);s__Metschnikowia_pulcherrima(100); | Metschnikowia | 13 | Saprotroph | \|Undefined Saprotroph\| | Yeast | NULL | Probable | As nectar-tap_saprotroph (Põlme et al. 2020); Decay sugar-rich substrates (Põlme et al. 2020) | Põlme S, et al. 2020. Fungal Diversity 105:1-16 ((https://doi.org/10.1007/s13225-020-00466-2)) |
| Otu40588 | 1 | k__Fungi(100);p__Ascomycota(100);c__Saccharomycetes(100);o__Saccharomycetales(100);f__Metschnikowiaceae(100);g__Metschnikowia(100);s__Metschnikowia_pulcherrima(100); | Metschnikowia | 13 | Saprotroph | \|Undefined Saprotroph\| | Yeast | NULL | Probable | As nectar-tap_saprotroph (Põlme et al. 2020); Decay sugar-rich substrates (Põlme et al. 2020) | Põlme S, et al. 2020. Fungal Diversity 105:1-16 ((https://doi.org/10.1007/s13225-020-00466-2)) |
| Otu40589 | 1 | k__Fungi(100);p__Ascomycota(100);c__Saccharomycetes(100);o__Saccharomycetales(100);f__Metschnikowiaceae(100);g__Metschnikowia(100);g__Metschnikowia_unclassified(100); | Metschnikowia | 13 | Saprotroph | \|Undefined Saprotroph\| | Yeast | NULL | Probable | As nectar-tap_saprotroph (Põlme et al. 2020); Decay sugar-rich substrates (Põlme et al. 2020) | Põlme S, et al. 2020. Fungal Diversity 105:1-16 ((https://doi.org/10.1007/s13225-020-00466-2)) |
| Otu40591 | 1 | k__Fungi(100);p__Ascomycota(100);c__Saccharomycetes(100);o__Saccharomycetales(100);f__Metschnikowiaceae(100);g__Metschnikowia(100);s__Metschnikowia_pulcherrima(100); | Metschnikowia | 13 | Saprotroph | \|Undefined Saprotroph\| | Yeast | NULL | Probable | As nectar-tap_saprotroph (Põlme et al. 2020); Decay sugar-rich substrates (Põlme et al. 2020) | Põlme S, et al. 2020. Fungal Diversity 105:1-16 ((https://doi.org/10.1007/s13225-020-00466-2)) |
| Otu40592 | 1 | k__Fungi(100);p__Ascomycota(100);c__Saccharomycetes(100);o__Saccharomycetales(100);f__Metschnikowiaceae(100);g__Metschnikowia(100);s__Metschnikowia_pulcherrima(100); | Metschnikowia | 13 | Saprotroph | \|Undefined Saprotroph\| | Yeast | NULL | Probable | As nectar-tap_saprotroph (Põlme et al. 2020); Decay sugar-rich substrates (Põlme et al. 2020) | Põlme S, et al. 2020. Fungal Diversity 105:1-16 ((https://doi.org/10.1007/s13225-020-00466-2)) |
| Otu40593 | 1 | k__Fungi(100);p__Ascomycota(100);c__Saccharomycetes(100);o__Saccharomycetales(100);f__Metschnikowiaceae(100);g__Metschnikowia(100);s__Metschnikowia_pulcherrima(100); | Metschnikowia | 13 | Saprotroph | \|Undefined Saprotroph\| | Yeast | NULL | Probable | As nectar-tap_saprotroph (Põlme et al. 2020); Decay sugar-rich substrates (Põlme et al. 2020) | Põlme S, et al. 2020. Fungal Diversity 105:1-16 ((https://doi.org/10.1007/s13225-020-00466-2)) |
| Otu40594 | 1 | k__Fungi(100);p__Ascomycota(100);c__Saccharomycetes(100);o__Saccharomycetales(100);f__Metschnikowiaceae(100);g__Metschnikowia(100);s__Metschnikowia_pulcherrima(100); | Metschnikowia | 13 | Saprotroph | \|Undefined Saprotroph\| | Yeast | NULL | Probable | As nectar-tap_saprotroph (Põlme et al. 2020); Decay sugar-rich substrates (Põlme et al. 2020) | Põlme S, et al. 2020. Fungal Diversity 105:1-16 ((https://doi.org/10.1007/s13225-020-00466-2)) |
| Otu40595 | 1 | k__Fungi(100);p__Ascomycota(100);c__Saccharomycetes(100);o__Saccharomycetales(100);f__Metschnikowiaceae(100);g__Metschnikowia(100);s__Metschnikowia_pulcherrima(100); | Metschnikowia | 13 | Saprotroph | \|Undefined Saprotroph\| | Yeast | NULL | Probable | As nectar-tap_saprotroph (Põlme et al. 2020); Decay sugar-rich substrates (Põlme et al. 2020) | Põlme S, et al. 2020. Fungal Diversity 105:1-16 ((https://doi.org/10.1007/s13225-020-00466-2)) |
| Otu40596 | 1 | k__Fungi(100);p__Ascomycota(100);c__Saccharomycetes(100);o__Saccharomycetales(100);f__Metschnikowiaceae(100);g__Metschnikowia(100);g__Metschnikowia_unclassified(100); | Metschnikowia | 13 | Saprotroph | \|Undefined Saprotroph\| | Yeast | NULL | Probable | As nectar-tap_saprotroph (Põlme et al. 2020); Decay sugar-rich substrates (Põlme et al. 2020) | Põlme S, et al. 2020. Fungal Diversity 105:1-16 ((https://doi.org/10.1007/s13225-020-00466-2)) |
| Otu40597 | 1 | k__Fungi(100);p__Ascomycota(100);c__Saccharomycetes(100);o__Saccharomycetales(100);f__Metschnikowiaceae(100);g__Metschnikowia(100);s__Metschnikowia_pulcherrima(100); | Metschnikowia | 13 | Saprotroph | \|Undefined Saprotroph\| | Yeast | NULL | Probable | As nectar-tap_saprotroph (Põlme et al. 2020); Decay sugar-rich substrates (Põlme et al. 2020) | Põlme S, et al. 2020. Fungal Diversity 105:1-16 ((https://doi.org/10.1007/s13225-020-00466-2)) |
| Otu40598 | 1 | k__Fungi(100);p__Ascomycota(100);c__Saccharomycetes(100);o__Saccharomycetales(100);f__Metschnikowiaceae(100);g__Metschnikowia(100);g__Metschnikowia_unclassified(100); | Metschnikowia | 13 | Saprotroph | \|Undefined Saprotroph\| | Yeast | NULL | Probable | As nectar-tap_saprotroph (Põlme et al. 2020); Decay sugar-rich substrates (Põlme et al. 2020) | Põlme S, et al. 2020. Fungal Diversity 105:1-16 ((https://doi.org/10.1007/s13225-020-00466-2)) |
| Otu40599 | 1 | k__Fungi(100);p__Ascomycota(100);c__Saccharomycetes(100);o__Saccharomycetales(100);f__Metschnikowiaceae(100);g__Metschnikowia(100);g__Metschnikowia_unclassified(100); | Metschnikowia | 13 | Saprotroph | \|Undefined Saprotroph\| | Yeast | NULL | Probable | As nectar-tap_saprotroph (Põlme et al. 2020); Decay sugar-rich substrates (Põlme et al. 2020) | Põlme S, et al. 2020. Fungal Diversity 105:1-16 ((https://doi.org/10.1007/s13225-020-00466-2)) |
| Otu40600 | 1 | k__Fungi(100);p__Ascomycota(100);c__Saccharomycetes(100);o__Saccharomycetales(100);f__Metschnikowiaceae(100);g__Metschnikowia(100);g__Metschnikowia_unclassified(100); | Metschnikowia | 13 | Saprotroph | \|Undefined Saprotroph\| | Yeast | NULL | Probable | As nectar-tap_saprotroph (Põlme et al. 2020); Decay sugar-rich substrates (Põlme et al. 2020) | Põlme S, et al. 2020. Fungal Diversity 105:1-16 ((https://doi.org/10.1007/s13225-020-00466-2)) |
| Otu40601 | 1 | k__Fungi(100);p__Ascomycota(100);c__Saccharomycetes(100);o__Saccharomycetales(100);f__Metschnikowiaceae(100);g__Metschnikowia(100);g__Metschnikowia_unclassified(100); | Metschnikowia | 13 | Saprotroph | \|Undefined Saprotroph\| | Yeast | NULL | Probable | As nectar-tap_saprotroph (Põlme et al. 2020); Decay sugar-rich substrates (Põlme et al. 2020) | Põlme S, et al. 2020. Fungal Diversity 105:1-16 ((https://doi.org/10.1007/s13225-020-00466-2)) |
| Otu40602 | 1 | k__Fungi(100);p__Ascomycota(100);c__Saccharomycetes(100);o__Saccharomycetales(100);f__Metschnikowiaceae(100);g__Metschnikowia(100);g__Metschnikowia_unclassified(100); | Metschnikowia | 13 | Saprotroph | \|Undefined Saprotroph\| | Yeast | NULL | Probable | As nectar-tap_saprotroph (Põlme et al. 2020); Decay sugar-rich substrates (Põlme et al. 2020) | Põlme S, et al. 2020. Fungal Diversity 105:1-16 ((https://doi.org/10.1007/s13225-020-00466-2)) |
| Otu40603 | 1 | k__Fungi(100);p__Ascomycota(100);c__Saccharomycetes(100);o__Saccharomycetales(100);f__Metschnikowiaceae(100);g__Metschnikowia(100);s__Metschnikowia_pulcherrima(100); | Metschnikowia | 13 | Saprotroph | \|Undefined Saprotroph\| | Yeast | NULL | Probable | As nectar-tap_saprotroph (Põlme et al. 2020); Decay sugar-rich substrates (Põlme et al. 2020) | Põlme S, et al. 2020. Fungal Diversity 105:1-16 ((https://doi.org/10.1007/s13225-020-00466-2)) |
| Otu40604 | 1 | k__Fungi(100);p__Ascomycota(100);c__Saccharomycetes(100);o__Saccharomycetales(100);f__Metschnikowiaceae(100);g__Metschnikowia(100);s__Metschnikowia_pulcherrima(100); | Metschnikowia | 13 | Saprotroph | \|Undefined Saprotroph\| | Yeast | NULL | Probable | As nectar-tap_saprotroph (Põlme et al. 2020); Decay sugar-rich substrates (Põlme et al. 2020) | Põlme S, et al. 2020. Fungal Diversity 105:1-16 ((https://doi.org/10.1007/s13225-020-00466-2)) |
| Otu40605 | 1 | k__Fungi(100);p__Ascomycota(100);c__Saccharomycetes(100);o__Saccharomycetales(100);f__Metschnikowiaceae(100);g__Metschnikowia(100);s__Metschnikowia_pulcherrima(100); | Metschnikowia | 13 | Saprotroph | \|Undefined Saprotroph\| | Yeast | NULL | Probable | As nectar-tap_saprotroph (Põlme et al. 2020); Decay sugar-rich substrates (Põlme et al. 2020) | Põlme S, et al. 2020. Fungal Diversity 105:1-16 ((https://doi.org/10.1007/s13225-020-00466-2)) |
| Otu40606 | 1 | k__Fungi(100);p__Ascomycota(100);c__Saccharomycetes(100);o__Saccharomycetales(100);f__Metschnikowiaceae(100);g__Metschnikowia(100);s__Metschnikowia_pulcherrima(100); | Metschnikowia | 13 | Saprotroph | \|Undefined Saprotroph\| | Yeast | NULL | Probable | As nectar-tap_saprotroph (Põlme et al. 2020); Decay sugar-rich substrates (Põlme et al. 2020) | Põlme S, et al. 2020. Fungal Diversity 105:1-16 ((https://doi.org/10.1007/s13225-020-00466-2)) |
| Otu40607 | 1 | k__Fungi(100);p__Ascomycota(100);c__Saccharomycetes(100);o__Saccharomycetales(100);f__Metschnikowiaceae(100);g__Metschnikowia(100);s__Metschnikowia_pulcherrima(100); | Metschnikowia | 13 | Saprotroph | \|Undefined Saprotroph\| | Yeast | NULL | Probable | As nectar-tap_saprotroph (Põlme et al. 2020); Decay sugar-rich substrates (Põlme et al. 2020) | Põlme S, et al. 2020. Fungal Diversity 105:1-16 ((https://doi.org/10.1007/s13225-020-00466-2)) |
| Otu40608 | 1 | k__Fungi(100);p__Ascomycota(100);c__Saccharomycetes(100);o__Saccharomycetales(100);f__Metschnikowiaceae(100);g__Metschnikowia(100);s__Metschnikowia_pulcherrima(100); | Metschnikowia | 13 | Saprotroph | \|Undefined Saprotroph\| | Yeast | NULL | Probable | As nectar-tap_saprotroph (Põlme et al. 2020); Decay sugar-rich substrates (Põlme et al. 2020) | Põlme S, et al. 2020. Fungal Diversity 105:1-16 ((https://doi.org/10.1007/s13225-020-00466-2)) |
| Otu40609 | 1 | k__Fungi(100);p__Ascomycota(100);c__Saccharomycetes(100);o__Saccharomycetales(100);f__Metschnikowiaceae(100);g__Metschnikowia(100);s__Metschnikowia_pulcherrima(100); | Metschnikowia | 13 | Saprotroph | \|Undefined Saprotroph\| | Yeast | NULL | Probable | As nectar-tap_saprotroph (Põlme et al. 2020); Decay sugar-rich substrates (Põlme et al. 2020) | Põlme S, et al. 2020. Fungal Diversity 105:1-16 ((https://doi.org/10.1007/s13225-020-00466-2)) |
| Otu40610 | 1 | k__Fungi(100);p__Ascomycota(100);c__Saccharomycetes(100);o__Saccharomycetales(100);f__Metschnikowiaceae(100);g__Metschnikowia(100);s__Metschnikowia_pulcherrima(100); | Metschnikowia | 13 | Saprotroph | \|Undefined Saprotroph\| | Yeast | NULL | Probable | As nectar-tap_saprotroph (Põlme et al. 2020); Decay sugar-rich substrates (Põlme et al. 2020) | Põlme S, et al. 2020. Fungal Diversity 105:1-16 ((https://doi.org/10.1007/s13225-020-00466-2)) |
| Otu40611 | 1 | k__Fungi(100);p__Ascomycota(100);c__Saccharomycetes(100);o__Saccharomycetales(100);f__Metschnikowiaceae(100);g__Metschnikowia(100);s__Metschnikowia_pulcherrima(100); | Metschnikowia | 13 | Saprotroph | \|Undefined Saprotroph\| | Yeast | NULL | Probable | As nectar-tap_saprotroph (Põlme et al. 2020); Decay sugar-rich substrates (Põlme et al. 2020) | Põlme S, et al. 2020. Fungal Diversity 105:1-16 ((https://doi.org/10.1007/s13225-020-00466-2)) |
| Otu40612 | 1 | k__Fungi(100);p__Ascomycota(100);c__Saccharomycetes(100);o__Saccharomycetales(100);f__Metschnikowiaceae(100);g__Metschnikowia(100);s__Metschnikowia_pulcherrima(100); | Metschnikowia | 13 | Saprotroph | \|Undefined Saprotroph\| | Yeast | NULL | Probable | As nectar-tap_saprotroph (Põlme et al. 2020); Decay sugar-rich substrates (Põlme et al. 2020) | Põlme S, et al. 2020. Fungal Diversity 105:1-16 ((https://doi.org/10.1007/s13225-020-00466-2)) |
| Otu40613 | 1 | k__Fungi(100);p__Ascomycota(100);c__Saccharomycetes(100);o__Saccharomycetales(100);f__Metschnikowiaceae(100);g__Metschnikowia(100);s__Metschnikowia_pulcherrima(100); | Metschnikowia | 13 | Saprotroph | \|Undefined Saprotroph\| | Yeast | NULL | Probable | As nectar-tap_saprotroph (Põlme et al. 2020); Decay sugar-rich substrates (Põlme et al. 2020) | Põlme S, et al. 2020. Fungal Diversity 105:1-16 ((https://doi.org/10.1007/s13225-020-00466-2)) |
| Otu40615 | 1 | k__Fungi(100);p__Ascomycota(100);c__Saccharomycetes(100);o__Saccharomycetales(100);f__Metschnikowiaceae(100);g__Metschnikowia(100);s__Metschnikowia_pulcherrima(100); | Metschnikowia | 13 | Saprotroph | \|Undefined Saprotroph\| | Yeast | NULL | Probable | As nectar-tap_saprotroph (Põlme et al. 2020); Decay sugar-rich substrates (Põlme et al. 2020) | Põlme S, et al. 2020. Fungal Diversity 105:1-16 ((https://doi.org/10.1007/s13225-020-00466-2)) |
| Otu40616 | 1 | k__Fungi(100);p__Ascomycota(100);c__Saccharomycetes(100);o__Saccharomycetales(100);f__Metschnikowiaceae(100);g__Metschnikowia(100);s__Metschnikowia_pulcherrima(100); | Metschnikowia | 13 | Saprotroph | \|Undefined Saprotroph\| | Yeast | NULL | Probable | As nectar-tap_saprotroph (Põlme et al. 2020); Decay sugar-rich substrates (Põlme et al. 2020) | Põlme S, et al. 2020. Fungal Diversity 105:1-16 ((https://doi.org/10.1007/s13225-020-00466-2)) |
| Otu40617 | 1 | k__Fungi(100);p__Ascomycota(100);c__Saccharomycetes(100);o__Saccharomycetales(100);f__Metschnikowiaceae(100);g__Metschnikowia(100);s__Metschnikowia_pulcherrima(100); | Metschnikowia | 13 | Saprotroph | \|Undefined Saprotroph\| | Yeast | NULL | Probable | As nectar-tap_saprotroph (Põlme et al. 2020); Decay sugar-rich substrates (Põlme et al. 2020) | Põlme S, et al. 2020. Fungal Diversity 105:1-16 ((https://doi.org/10.1007/s13225-020-00466-2)) |
| Otu40619 | 1 | k__Fungi(100);p__Ascomycota(100);c__Saccharomycetes(100);o__Saccharomycetales(100);f__Metschnikowiaceae(100);g__Metschnikowia(100);g__Metschnikowia_unclassified(100); | Metschnikowia | 13 | Saprotroph | \|Undefined Saprotroph\| | Yeast | NULL | Probable | As nectar-tap_saprotroph (Põlme et al. 2020); Decay sugar-rich substrates (Põlme et al. 2020) | Põlme S, et al. 2020. Fungal Diversity 105:1-16 ((https://doi.org/10.1007/s13225-020-00466-2)) |
| Otu40620 | 1 | k__Fungi(100);p__Ascomycota(100);c__Saccharomycetes(100);o__Saccharomycetales(100);f__Metschnikowiaceae(100);g__Metschnikowia(100);s__Metschnikowia_pulcherrima(100); | Metschnikowia | 13 | Saprotroph | \|Undefined Saprotroph\| | Yeast | NULL | Probable | As nectar-tap_saprotroph (Põlme et al. 2020); Decay sugar-rich substrates (Põlme et al. 2020) | Põlme S, et al. 2020. Fungal Diversity 105:1-16 ((https://doi.org/10.1007/s13225-020-00466-2)) |
| Otu40621 | 1 | k__Fungi(100);p__Ascomycota(100);c__Saccharomycetes(100);o__Saccharomycetales(100);f__Metschnikowiaceae(100);g__Metschnikowia(100);s__Metschnikowia_pulcherrima(100); | Metschnikowia | 13 | Saprotroph | \|Undefined Saprotroph\| | Yeast | NULL | Probable | As nectar-tap_saprotroph (Põlme et al. 2020); Decay sugar-rich substrates (Põlme et al. 2020) | Põlme S, et al. 2020. Fungal Diversity 105:1-16 ((https://doi.org/10.1007/s13225-020-00466-2)) |
| Otu40622 | 1 | k__Fungi(100);p__Ascomycota(100);c__Saccharomycetes(100);o__Saccharomycetales(100);f__Metschnikowiaceae(100);g__Metschnikowia(100);s__Metschnikowia_pulcherrima(100); | Metschnikowia | 13 | Saprotroph | \|Undefined Saprotroph\| | Yeast | NULL | Probable | As nectar-tap_saprotroph (Põlme et al. 2020); Decay sugar-rich substrates (Põlme et al. 2020) | Põlme S, et al. 2020. Fungal Diversity 105:1-16 ((https://doi.org/10.1007/s13225-020-00466-2)) |
| Otu40623 | 1 | k__Fungi(100);p__Ascomycota(100);c__Saccharomycetes(100);o__Saccharomycetales(100);f__Metschnikowiaceae(100);g__Metschnikowia(100);s__Metschnikowia_pulcherrima(100); | Metschnikowia | 13 | Saprotroph | \|Undefined Saprotroph\| | Yeast | NULL | Probable | As nectar-tap_saprotroph (Põlme et al. 2020); Decay sugar-rich substrates (Põlme et al. 2020) | Põlme S, et al. 2020. Fungal Diversity 105:1-16 ((https://doi.org/10.1007/s13225-020-00466-2)) |
| Otu40624 | 1 | k__Fungi(100);p__Ascomycota(100);c__Saccharomycetes(100);o__Saccharomycetales(100);f__Metschnikowiaceae(100);g__Metschnikowia(100);s__Metschnikowia_pulcherrima(100); | Metschnikowia | 13 | Saprotroph | \|Undefined Saprotroph\| | Yeast | NULL | Probable | As nectar-tap_saprotroph (Põlme et al. 2020); Decay sugar-rich substrates (Põlme et al. 2020) | Põlme S, et al. 2020. Fungal Diversity 105:1-16 ((https://doi.org/10.1007/s13225-020-00466-2)) |
| Otu40625 | 1 | k__Fungi(100);p__Ascomycota(100);c__Saccharomycetes(100);o__Saccharomycetales(100);f__Metschnikowiaceae(100);g__Metschnikowia(100);s__Metschnikowia_pulcherrima(100); | Metschnikowia | 13 | Saprotroph | \|Undefined Saprotroph\| | Yeast | NULL | Probable | As nectar-tap_saprotroph (Põlme et al. 2020); Decay sugar-rich substrates (Põlme et al. 2020) | Põlme S, et al. 2020. Fungal Diversity 105:1-16 ((https://doi.org/10.1007/s13225-020-00466-2)) |
| Otu40626 | 1 | k__Fungi(100);p__Ascomycota(100);c__Saccharomycetes(100);o__Saccharomycetales(100);f__Metschnikowiaceae(100);g__Metschnikowia(100);s__Metschnikowia_pulcherrima(100); | Metschnikowia | 13 | Saprotroph | \|Undefined Saprotroph\| | Yeast | NULL | Probable | As nectar-tap_saprotroph (Põlme et al. 2020); Decay sugar-rich substrates (Põlme et al. 2020) | Põlme S, et al. 2020. Fungal Diversity 105:1-16 ((https://doi.org/10.1007/s13225-020-00466-2)) |
| Otu40627 | 1 | k__Fungi(100);p__Ascomycota(100);c__Saccharomycetes(100);o__Saccharomycetales(100);f__Metschnikowiaceae(100);g__Metschnikowia(100);s__Metschnikowia_pulcherrima(100); | Metschnikowia | 13 | Saprotroph | \|Undefined Saprotroph\| | Yeast | NULL | Probable | As nectar-tap_saprotroph (Põlme et al. 2020); Decay sugar-rich substrates (Põlme et al. 2020) | Põlme S, et al. 2020. Fungal Diversity 105:1-16 ((https://doi.org/10.1007/s13225-020-00466-2)) |
| Otu40628 | 1 | k__Fungi(100);p__Ascomycota(100);c__Saccharomycetes(100);o__Saccharomycetales(100);f__Metschnikowiaceae(100);g__Metschnikowia(100);s__Metschnikowia_pulcherrima(100); | Metschnikowia | 13 | Saprotroph | \|Undefined Saprotroph\| | Yeast | NULL | Probable | As nectar-tap_saprotroph (Põlme et al. 2020); Decay sugar-rich substrates (Põlme et al. 2020) | Põlme S, et al. 2020. Fungal Diversity 105:1-16 ((https://doi.org/10.1007/s13225-020-00466-2)) |
| Otu40629 | 1 | k__Fungi(100);p__Ascomycota(100);c__Saccharomycetes(100);o__Saccharomycetales(100);f__Metschnikowiaceae(100);g__Metschnikowia(100);g__Metschnikowia_unclassified(100); | Metschnikowia | 13 | Saprotroph | \|Undefined Saprotroph\| | Yeast | NULL | Probable | As nectar-tap_saprotroph (Põlme et al. 2020); Decay sugar-rich substrates (Põlme et al. 2020) | Põlme S, et al. 2020. Fungal Diversity 105:1-16 ((https://doi.org/10.1007/s13225-020-00466-2)) |
| Otu40630 | 1 | k__Fungi(100);p__Ascomycota(100);c__Saccharomycetes(100);o__Saccharomycetales(100);f__Metschnikowiaceae(100);g__Metschnikowia(100);s__Metschnikowia_pulcherrima(100); | Metschnikowia | 13 | Saprotroph | \|Undefined Saprotroph\| | Yeast | NULL | Probable | As nectar-tap_saprotroph (Põlme et al. 2020); Decay sugar-rich substrates (Põlme et al. 2020) | Põlme S, et al. 2020. Fungal Diversity 105:1-16 ((https://doi.org/10.1007/s13225-020-00466-2)) |
| Otu40631 | 1 | k__Fungi(100);p__Ascomycota(100);c__Saccharomycetes(100);o__Saccharomycetales(100);f__Metschnikowiaceae(100);g__Metschnikowia(100);s__Metschnikowia_pulcherrima(100); | Metschnikowia | 13 | Saprotroph | \|Undefined Saprotroph\| | Yeast | NULL | Probable | As nectar-tap_saprotroph (Põlme et al. 2020); Decay sugar-rich substrates (Põlme et al. 2020) | Põlme S, et al. 2020. Fungal Diversity 105:1-16 ((https://doi.org/10.1007/s13225-020-00466-2)) |
| Otu40632 | 1 | k__Fungi(100);p__Ascomycota(100);c__Saccharomycetes(100);o__Saccharomycetales(100);f__Metschnikowiaceae(100);g__Metschnikowia(100);g__Metschnikowia_unclassified(100); | Metschnikowia | 13 | Saprotroph | \|Undefined Saprotroph\| | Yeast | NULL | Probable | As nectar-tap_saprotroph (Põlme et al. 2020); Decay sugar-rich substrates (Põlme et al. 2020) | Põlme S, et al. 2020. Fungal Diversity 105:1-16 ((https://doi.org/10.1007/s13225-020-00466-2)) |
| Otu40633 | 1 | k__Fungi(100);p__Ascomycota(100);c__Saccharomycetes(100);o__Saccharomycetales(100);f__Metschnikowiaceae(100);g__Metschnikowia(100);s__Metschnikowia_pulcherrima(100); | Metschnikowia | 13 | Saprotroph | \|Undefined Saprotroph\| | Yeast | NULL | Probable | As nectar-tap_saprotroph (Põlme et al. 2020); Decay sugar-rich substrates (Põlme et al. 2020) | Põlme S, et al. 2020. Fungal Diversity 105:1-16 ((https://doi.org/10.1007/s13225-020-00466-2)) |
| Otu40634 | 1 | k__Fungi(100);p__Ascomycota(100);c__Saccharomycetes(100);o__Saccharomycetales(100);f__Metschnikowiaceae(100);g__Metschnikowia(100);s__Metschnikowia_pulcherrima(100); | Metschnikowia | 13 | Saprotroph | \|Undefined Saprotroph\| | Yeast | NULL | Probable | As nectar-tap_saprotroph (Põlme et al. 2020); Decay sugar-rich substrates (Põlme et al. 2020) | Põlme S, et al. 2020. Fungal Diversity 105:1-16 ((https://doi.org/10.1007/s13225-020-00466-2)) |
| Otu40635 | 1 | k__Fungi(100);p__Ascomycota(100);c__Saccharomycetes(100);o__Saccharomycetales(100);f__Metschnikowiaceae(100);g__Metschnikowia(100);s__Metschnikowia_pulcherrima(100); | Metschnikowia | 13 | Saprotroph | \|Undefined Saprotroph\| | Yeast | NULL | Probable | As nectar-tap_saprotroph (Põlme et al. 2020); Decay sugar-rich substrates (Põlme et al. 2020) | Põlme S, et al. 2020. Fungal Diversity 105:1-16 ((https://doi.org/10.1007/s13225-020-00466-2)) |
| Otu40636 | 1 | k__Fungi(100);p__Ascomycota(100);c__Saccharomycetes(100);o__Saccharomycetales(100);f__Metschnikowiaceae(100);g__Metschnikowia(100);g__Metschnikowia_unclassified(100); | Metschnikowia | 13 | Saprotroph | \|Undefined Saprotroph\| | Yeast | NULL | Probable | As nectar-tap_saprotroph (Põlme et al. 2020); Decay sugar-rich substrates (Põlme et al. 2020) | Põlme S, et al. 2020. Fungal Diversity 105:1-16 ((https://doi.org/10.1007/s13225-020-00466-2)) |
| Otu40637 | 1 | k__Fungi(100);p__Ascomycota(100);c__Saccharomycetes(100);o__Saccharomycetales(100);f__Metschnikowiaceae(100);g__Metschnikowia(100);g__Metschnikowia_unclassified(100); | Metschnikowia | 13 | Saprotroph | \|Undefined Saprotroph\| | Yeast | NULL | Probable | As nectar-tap_saprotroph (Põlme et al. 2020); Decay sugar-rich substrates (Põlme et al. 2020) | Põlme S, et al. 2020. Fungal Diversity 105:1-16 ((https://doi.org/10.1007/s13225-020-00466-2)) |
| Otu40638 | 1 | k__Fungi(100);p__Ascomycota(100);c__Saccharomycetes(100);o__Saccharomycetales(100);f__Metschnikowiaceae(100);g__Metschnikowia(100);g__Metschnikowia_unclassified(100); | Metschnikowia | 13 | Saprotroph | \|Undefined Saprotroph\| | Yeast | NULL | Probable | As nectar-tap_saprotroph (Põlme et al. 2020); Decay sugar-rich substrates (Põlme et al. 2020) | Põlme S, et al. 2020. Fungal Diversity 105:1-16 ((https://doi.org/10.1007/s13225-020-00466-2)) |
| Otu40639 | 1 | k__Fungi(100);p__Ascomycota(100);c__Saccharomycetes(100);o__Saccharomycetales(100);f__Metschnikowiaceae(100);g__Metschnikowia(100);s__Metschnikowia_pulcherrima(100); | Metschnikowia | 13 | Saprotroph | \|Undefined Saprotroph\| | Yeast | NULL | Probable | As nectar-tap_saprotroph (Põlme et al. 2020); Decay sugar-rich substrates (Põlme et al. 2020) | Põlme S, et al. 2020. Fungal Diversity 105:1-16 ((https://doi.org/10.1007/s13225-020-00466-2)) |
| Otu40640 | 1 | k__Fungi(100);p__Ascomycota(100);c__Saccharomycetes(100);o__Saccharomycetales(100);f__Metschnikowiaceae(100);g__Metschnikowia(100);s__Metschnikowia_pulcherrima(100); | Metschnikowia | 13 | Saprotroph | \|Undefined Saprotroph\| | Yeast | NULL | Probable | As nectar-tap_saprotroph (Põlme et al. 2020); Decay sugar-rich substrates (Põlme et al. 2020) | Põlme S, et al. 2020. Fungal Diversity 105:1-16 ((https://doi.org/10.1007/s13225-020-00466-2)) |
| Otu40641 | 1 | k__Fungi(100);p__Ascomycota(100);c__Saccharomycetes(100);o__Saccharomycetales(100);f__Metschnikowiaceae(100);g__Metschnikowia(100);s__Metschnikowia_pulcherrima(100); | Metschnikowia | 13 | Saprotroph | \|Undefined Saprotroph\| | Yeast | NULL | Probable | As nectar-tap_saprotroph (Põlme et al. 2020); Decay sugar-rich substrates (Põlme et al. 2020) | Põlme S, et al. 2020. Fungal Diversity 105:1-16 ((https://doi.org/10.1007/s13225-020-00466-2)) |
| Otu40642 | 1 | k__Fungi(100);p__Ascomycota(100);c__Saccharomycetes(100);o__Saccharomycetales(100);f__Metschnikowiaceae(100);g__Metschnikowia(100);s__Metschnikowia_pulcherrima(100); | Metschnikowia | 13 | Saprotroph | \|Undefined Saprotroph\| | Yeast | NULL | Probable | As nectar-tap_saprotroph (Põlme et al. 2020); Decay sugar-rich substrates (Põlme et al. 2020) | Põlme S, et al. 2020. Fungal Diversity 105:1-16 ((https://doi.org/10.1007/s13225-020-00466-2)) |
| Otu40643 | 1 | k__Fungi(100);p__Ascomycota(100);c__Saccharomycetes(100);o__Saccharomycetales(100);f__Metschnikowiaceae(100);g__Metschnikowia(100);s__Metschnikowia_pulcherrima(100); | Metschnikowia | 13 | Saprotroph | \|Undefined Saprotroph\| | Yeast | NULL | Probable | As nectar-tap_saprotroph (Põlme et al. 2020); Decay sugar-rich substrates (Põlme et al. 2020) | Põlme S, et al. 2020. Fungal Diversity 105:1-16 ((https://doi.org/10.1007/s13225-020-00466-2)) |
| Otu40644 | 1 | k__Fungi(100);p__Ascomycota(100);c__Saccharomycetes(100);o__Saccharomycetales(100);f__Metschnikowiaceae(100);g__Metschnikowia(100);s__Metschnikowia_pulcherrima(100); | Metschnikowia | 13 | Saprotroph | \|Undefined Saprotroph\| | Yeast | NULL | Probable | As nectar-tap_saprotroph (Põlme et al. 2020); Decay sugar-rich substrates (Põlme et al. 2020) | Põlme S, et al. 2020. Fungal Diversity 105:1-16 ((https://doi.org/10.1007/s13225-020-00466-2)) |
| Otu40645 | 1 | k__Fungi(100);p__Ascomycota(100);c__Saccharomycetes(100);o__Saccharomycetales(100);f__Metschnikowiaceae(100);g__Metschnikowia(100);s__Metschnikowia_pulcherrima(100); | Metschnikowia | 13 | Saprotroph | \|Undefined Saprotroph\| | Yeast | NULL | Probable | As nectar-tap_saprotroph (Põlme et al. 2020); Decay sugar-rich substrates (Põlme et al. 2020) | Põlme S, et al. 2020. Fungal Diversity 105:1-16 ((https://doi.org/10.1007/s13225-020-00466-2)) |
| Otu40646 | 1 | k__Fungi(100);p__Ascomycota(100);c__Saccharomycetes(100);o__Saccharomycetales(100);f__Metschnikowiaceae(100);g__Metschnikowia(100);s__Metschnikowia_pulcherrima(100); | Metschnikowia | 13 | Saprotroph | \|Undefined Saprotroph\| | Yeast | NULL | Probable | As nectar-tap_saprotroph (Põlme et al. 2020); Decay sugar-rich substrates (Põlme et al. 2020) | Põlme S, et al. 2020. Fungal Diversity 105:1-16 ((https://doi.org/10.1007/s13225-020-00466-2)) |
| Otu40647 | 1 | k__Fungi(100);p__Ascomycota(100);c__Saccharomycetes(100);o__Saccharomycetales(100);f__Metschnikowiaceae(100);g__Metschnikowia(100);s__Metschnikowia_pulcherrima(100); | Metschnikowia | 13 | Saprotroph | \|Undefined Saprotroph\| | Yeast | NULL | Probable | As nectar-tap_saprotroph (Põlme et al. 2020); Decay sugar-rich substrates (Põlme et al. 2020) | Põlme S, et al. 2020. Fungal Diversity 105:1-16 ((https://doi.org/10.1007/s13225-020-00466-2)) |
| Otu40648 | 1 | k__Fungi(100);p__Ascomycota(100);c__Saccharomycetes(100);o__Saccharomycetales(100);f__Metschnikowiaceae(100);g__Metschnikowia(100);g__Metschnikowia_unclassified(100); | Metschnikowia | 13 | Saprotroph | \|Undefined Saprotroph\| | Yeast | NULL | Probable | As nectar-tap_saprotroph (Põlme et al. 2020); Decay sugar-rich substrates (Põlme et al. 2020) | Põlme S, et al. 2020. Fungal Diversity 105:1-16 ((https://doi.org/10.1007/s13225-020-00466-2)) |
| Otu40650 | 1 | k__Fungi(100);p__Ascomycota(100);c__Saccharomycetes(100);o__Saccharomycetales(100);f__Metschnikowiaceae(100);g__Metschnikowia(100);g__Metschnikowia_unclassified(100); | Metschnikowia | 13 | Saprotroph | \|Undefined Saprotroph\| | Yeast | NULL | Probable | As nectar-tap_saprotroph (Põlme et al. 2020); Decay sugar-rich substrates (Põlme et al. 2020) | Põlme S, et al. 2020. Fungal Diversity 105:1-16 ((https://doi.org/10.1007/s13225-020-00466-2)) |
| Otu40651 | 1 | k__Fungi(100);p__Ascomycota(100);c__Saccharomycetes(100);o__Saccharomycetales(100);f__Metschnikowiaceae(100);g__Metschnikowia(100);g__Metschnikowia_unclassified(100); | Metschnikowia | 13 | Saprotroph | \|Undefined Saprotroph\| | Yeast | NULL | Probable | As nectar-tap_saprotroph (Põlme et al. 2020); Decay sugar-rich substrates (Põlme et al. 2020) | Põlme S, et al. 2020. Fungal Diversity 105:1-16 ((https://doi.org/10.1007/s13225-020-00466-2)) |
| Otu40652 | 1 | k__Fungi(100);p__Ascomycota(100);c__Saccharomycetes(100);o__Saccharomycetales(100);f__Metschnikowiaceae(100);g__Metschnikowia(100);g__Metschnikowia_unclassified(100); | Metschnikowia | 13 | Saprotroph | \|Undefined Saprotroph\| | Yeast | NULL | Probable | As nectar-tap_saprotroph (Põlme et al. 2020); Decay sugar-rich substrates (Põlme et al. 2020) | Põlme S, et al. 2020. Fungal Diversity 105:1-16 ((https://doi.org/10.1007/s13225-020-00466-2)) |
| Otu40653 | 1 | k__Fungi(100);p__Ascomycota(100);c__Saccharomycetes(100);o__Saccharomycetales(100);f__Metschnikowiaceae(100);g__Metschnikowia(100);s__Metschnikowia_pulcherrima(100); | Metschnikowia | 13 | Saprotroph | \|Undefined Saprotroph\| | Yeast | NULL | Probable | As nectar-tap_saprotroph (Põlme et al. 2020); Decay sugar-rich substrates (Põlme et al. 2020) | Põlme S, et al. 2020. Fungal Diversity 105:1-16 ((https://doi.org/10.1007/s13225-020-00466-2)) |
| Otu40654 | 1 | k__Fungi(100);p__Ascomycota(100);c__Saccharomycetes(100);o__Saccharomycetales(100);f__Metschnikowiaceae(100);g__Metschnikowia(100);s__Metschnikowia_pulcherrima(100); | Metschnikowia | 13 | Saprotroph | \|Undefined Saprotroph\| | Yeast | NULL | Probable | As nectar-tap_saprotroph (Põlme et al. 2020); Decay sugar-rich substrates (Põlme et al. 2020) | Põlme S, et al. 2020. Fungal Diversity 105:1-16 ((https://doi.org/10.1007/s13225-020-00466-2)) |
| Otu40655 | 1 | k__Fungi(100);p__Ascomycota(100);c__Saccharomycetes(100);o__Saccharomycetales(100);f__Metschnikowiaceae(100);g__Metschnikowia(100);s__Metschnikowia_pulcherrima(100); | Metschnikowia | 13 | Saprotroph | \|Undefined Saprotroph\| | Yeast | NULL | Probable | As nectar-tap_saprotroph (Põlme et al. 2020); Decay sugar-rich substrates (Põlme et al. 2020) | Põlme S, et al. 2020. Fungal Diversity 105:1-16 ((https://doi.org/10.1007/s13225-020-00466-2)) |
| Otu40656 | 1 | k__Fungi(100);p__Ascomycota(100);c__Saccharomycetes(100);o__Saccharomycetales(100);f__Metschnikowiaceae(100);g__Metschnikowia(100);s__Metschnikowia_pulcherrima(100); | Metschnikowia | 13 | Saprotroph | \|Undefined Saprotroph\| | Yeast | NULL | Probable | As nectar-tap_saprotroph (Põlme et al. 2020); Decay sugar-rich substrates (Põlme et al. 2020) | Põlme S, et al. 2020. Fungal Diversity 105:1-16 ((https://doi.org/10.1007/s13225-020-00466-2)) |
| Otu40657 | 1 | k__Fungi(100);p__Ascomycota(100);c__Saccharomycetes(100);o__Saccharomycetales(100);f__Metschnikowiaceae(100);g__Metschnikowia(100);g__Metschnikowia_unclassified(100); | Metschnikowia | 13 | Saprotroph | \|Undefined Saprotroph\| | Yeast | NULL | Probable | As nectar-tap_saprotroph (Põlme et al. 2020); Decay sugar-rich substrates (Põlme et al. 2020) | Põlme S, et al. 2020. Fungal Diversity 105:1-16 ((https://doi.org/10.1007/s13225-020-00466-2)) |
| Otu40658 | 1 | k__Fungi(100);p__Ascomycota(100);c__Saccharomycetes(100);o__Saccharomycetales(100);f__Metschnikowiaceae(100);g__Metschnikowia(100);s__Metschnikowia_pulcherrima(100); | Metschnikowia | 13 | Saprotroph | \|Undefined Saprotroph\| | Yeast | NULL | Probable | As nectar-tap_saprotroph (Põlme et al. 2020); Decay sugar-rich substrates (Põlme et al. 2020) | Põlme S, et al. 2020. Fungal Diversity 105:1-16 ((https://doi.org/10.1007/s13225-020-00466-2)) |
| Otu40659 | 1 | k__Fungi(100);p__Ascomycota(100);c__Saccharomycetes(100);o__Saccharomycetales(100);f__Metschnikowiaceae(100);g__Metschnikowia(100);g__Metschnikowia_unclassified(100); | Metschnikowia | 13 | Saprotroph | \|Undefined Saprotroph\| | Yeast | NULL | Probable | As nectar-tap_saprotroph (Põlme et al. 2020); Decay sugar-rich substrates (Põlme et al. 2020) | Põlme S, et al. 2020. Fungal Diversity 105:1-16 ((https://doi.org/10.1007/s13225-020-00466-2)) |
| Otu40660 | 1 | k__Fungi(100);p__Ascomycota(100);c__Saccharomycetes(100);o__Saccharomycetales(100);f__Metschnikowiaceae(100);g__Metschnikowia(100);s__Metschnikowia_pulcherrima(100); | Metschnikowia | 13 | Saprotroph | \|Undefined Saprotroph\| | Yeast | NULL | Probable | As nectar-tap_saprotroph (Põlme et al. 2020); Decay sugar-rich substrates (Põlme et al. 2020) | Põlme S, et al. 2020. Fungal Diversity 105:1-16 ((https://doi.org/10.1007/s13225-020-00466-2)) |
| Otu40661 | 1 | k__Fungi(100);p__Ascomycota(100);c__Saccharomycetes(100);o__Saccharomycetales(100);f__Metschnikowiaceae(100);g__Metschnikowia(100);g__Metschnikowia_unclassified(100); | Metschnikowia | 13 | Saprotroph | \|Undefined Saprotroph\| | Yeast | NULL | Probable | As nectar-tap_saprotroph (Põlme et al. 2020); Decay sugar-rich substrates (Põlme et al. 2020) | Põlme S, et al. 2020. Fungal Diversity 105:1-16 ((https://doi.org/10.1007/s13225-020-00466-2)) |
| Otu40662 | 1 | k__Fungi(100);p__Ascomycota(100);c__Saccharomycetes(100);o__Saccharomycetales(100);f__Metschnikowiaceae(100);g__Metschnikowia(100);s__Metschnikowia_pulcherrima(100); | Metschnikowia | 13 | Saprotroph | \|Undefined Saprotroph\| | Yeast | NULL | Probable | As nectar-tap_saprotroph (Põlme et al. 2020); Decay sugar-rich substrates (Põlme et al. 2020) | Põlme S, et al. 2020. Fungal Diversity 105:1-16 ((https://doi.org/10.1007/s13225-020-00466-2)) |
| Otu40663 | 1 | k__Fungi(100);p__Ascomycota(100);c__Saccharomycetes(100);o__Saccharomycetales(100);f__Metschnikowiaceae(100);g__Metschnikowia(100);s__Metschnikowia_pulcherrima(100); | Metschnikowia | 13 | Saprotroph | \|Undefined Saprotroph\| | Yeast | NULL | Probable | As nectar-tap_saprotroph (Põlme et al. 2020); Decay sugar-rich substrates (Põlme et al. 2020) | Põlme S, et al. 2020. Fungal Diversity 105:1-16 ((https://doi.org/10.1007/s13225-020-00466-2)) |
| Otu40664 | 1 | k__Fungi(100);p__Ascomycota(100);c__Saccharomycetes(100);o__Saccharomycetales(100);f__Metschnikowiaceae(100);g__Metschnikowia(100);s__Metschnikowia_pulcherrima(100); | Metschnikowia | 13 | Saprotroph | \|Undefined Saprotroph\| | Yeast | NULL | Probable | As nectar-tap_saprotroph (Põlme et al. 2020); Decay sugar-rich substrates (Põlme et al. 2020) | Põlme S, et al. 2020. Fungal Diversity 105:1-16 ((https://doi.org/10.1007/s13225-020-00466-2)) |
| Otu40665 | 1 | k__Fungi(100);p__Ascomycota(100);c__Saccharomycetes(100);o__Saccharomycetales(100);f__Metschnikowiaceae(100);g__Metschnikowia(100);s__Metschnikowia_pulcherrima(100); | Metschnikowia | 13 | Saprotroph | \|Undefined Saprotroph\| | Yeast | NULL | Probable | As nectar-tap_saprotroph (Põlme et al. 2020); Decay sugar-rich substrates (Põlme et al. 2020) | Põlme S, et al. 2020. Fungal Diversity 105:1-16 ((https://doi.org/10.1007/s13225-020-00466-2)) |
| Otu40666 | 1 | k__Fungi(100);p__Ascomycota(100);c__Saccharomycetes(100);o__Saccharomycetales(100);f__Metschnikowiaceae(100);g__Metschnikowia(100);s__Metschnikowia_pulcherrima(100); | Metschnikowia | 13 | Saprotroph | \|Undefined Saprotroph\| | Yeast | NULL | Probable | As nectar-tap_saprotroph (Põlme et al. 2020); Decay sugar-rich substrates (Põlme et al. 2020) | Põlme S, et al. 2020. Fungal Diversity 105:1-16 ((https://doi.org/10.1007/s13225-020-00466-2)) |
| Otu40667 | 1 | k__Fungi(100);p__Ascomycota(100);c__Saccharomycetes(100);o__Saccharomycetales(100);f__Metschnikowiaceae(100);g__Metschnikowia(100);s__Metschnikowia_pulcherrima(100); | Metschnikowia | 13 | Saprotroph | \|Undefined Saprotroph\| | Yeast | NULL | Probable | As nectar-tap_saprotroph (Põlme et al. 2020); Decay sugar-rich substrates (Põlme et al. 2020) | Põlme S, et al. 2020. Fungal Diversity 105:1-16 ((https://doi.org/10.1007/s13225-020-00466-2)) |
| Otu40668 | 1 | k__Fungi(100);p__Ascomycota(100);c__Saccharomycetes(100);o__Saccharomycetales(100);f__Metschnikowiaceae(100);g__Metschnikowia(100);s__Metschnikowia_pulcherrima(100); | Metschnikowia | 13 | Saprotroph | \|Undefined Saprotroph\| | Yeast | NULL | Probable | As nectar-tap_saprotroph (Põlme et al. 2020); Decay sugar-rich substrates (Põlme et al. 2020) | Põlme S, et al. 2020. Fungal Diversity 105:1-16 ((https://doi.org/10.1007/s13225-020-00466-2)) |
| Otu40669 | 1 | k__Fungi(100);p__Ascomycota(100);c__Saccharomycetes(100);o__Saccharomycetales(100);f__Metschnikowiaceae(100);g__Metschnikowia(100);s__Metschnikowia_pulcherrima(100); | Metschnikowia | 13 | Saprotroph | \|Undefined Saprotroph\| | Yeast | NULL | Probable | As nectar-tap_saprotroph (Põlme et al. 2020); Decay sugar-rich substrates (Põlme et al. 2020) | Põlme S, et al. 2020. Fungal Diversity 105:1-16 ((https://doi.org/10.1007/s13225-020-00466-2)) |
| Otu40670 | 1 | k__Fungi(100);p__Ascomycota(100);c__Saccharomycetes(100);o__Saccharomycetales(100);f__Metschnikowiaceae(100);g__Metschnikowia(100);s__Metschnikowia_pulcherrima(100); | Metschnikowia | 13 | Saprotroph | \|Undefined Saprotroph\| | Yeast | NULL | Probable | As nectar-tap_saprotroph (Põlme et al. 2020); Decay sugar-rich substrates (Põlme et al. 2020) | Põlme S, et al. 2020. Fungal Diversity 105:1-16 ((https://doi.org/10.1007/s13225-020-00466-2)) |
| Otu40671 | 1 | k__Fungi(100);p__Ascomycota(100);c__Saccharomycetes(100);o__Saccharomycetales(100);f__Metschnikowiaceae(100);g__Metschnikowia(100);s__Metschnikowia_pulcherrima(100); | Metschnikowia | 13 | Saprotroph | \|Undefined Saprotroph\| | Yeast | NULL | Probable | As nectar-tap_saprotroph (Põlme et al. 2020); Decay sugar-rich substrates (Põlme et al. 2020) | Põlme S, et al. 2020. Fungal Diversity 105:1-16 ((https://doi.org/10.1007/s13225-020-00466-2)) |
| Otu40672 | 1 | k__Fungi(100);p__Ascomycota(100);c__Saccharomycetes(100);o__Saccharomycetales(100);f__Metschnikowiaceae(100);g__Metschnikowia(100);s__Metschnikowia_pulcherrima(100); | Metschnikowia | 13 | Saprotroph | \|Undefined Saprotroph\| | Yeast | NULL | Probable | As nectar-tap_saprotroph (Põlme et al. 2020); Decay sugar-rich substrates (Põlme et al. 2020) | Põlme S, et al. 2020. Fungal Diversity 105:1-16 ((https://doi.org/10.1007/s13225-020-00466-2)) |
| Otu40673 | 1 | k__Fungi(100);p__Ascomycota(100);c__Saccharomycetes(100);o__Saccharomycetales(100);f__Metschnikowiaceae(100);g__Metschnikowia(100);s__Metschnikowia_pulcherrima(100); | Metschnikowia | 13 | Saprotroph | \|Undefined Saprotroph\| | Yeast | NULL | Probable | As nectar-tap_saprotroph (Põlme et al. 2020); Decay sugar-rich substrates (Põlme et al. 2020) | Põlme S, et al. 2020. Fungal Diversity 105:1-16 ((https://doi.org/10.1007/s13225-020-00466-2)) |
| Otu40674 | 1 | k__Fungi(100);p__Ascomycota(100);c__Saccharomycetes(100);o__Saccharomycetales(100);f__Metschnikowiaceae(100);g__Metschnikowia(100);s__Metschnikowia_pulcherrima(100); | Metschnikowia | 13 | Saprotroph | \|Undefined Saprotroph\| | Yeast | NULL | Probable | As nectar-tap_saprotroph (Põlme et al. 2020); Decay sugar-rich substrates (Põlme et al. 2020) | Põlme S, et al. 2020. Fungal Diversity 105:1-16 ((https://doi.org/10.1007/s13225-020-00466-2)) |
| Otu40675 | 1 | k__Fungi(100);p__Ascomycota(100);c__Saccharomycetes(100);o__Saccharomycetales(100);f__Metschnikowiaceae(100);g__Metschnikowia(100);g__Metschnikowia_unclassified(100); | Metschnikowia | 13 | Saprotroph | \|Undefined Saprotroph\| | Yeast | NULL | Probable | As nectar-tap_saprotroph (Põlme et al. 2020); Decay sugar-rich substrates (Põlme et al. 2020) | Põlme S, et al. 2020. Fungal Diversity 105:1-16 ((https://doi.org/10.1007/s13225-020-00466-2)) |
| Otu40676 | 1 | k__Fungi(100);p__Ascomycota(100);c__Saccharomycetes(100);o__Saccharomycetales(100);f__Metschnikowiaceae(100);g__Metschnikowia(100);s__Metschnikowia_pulcherrima(100); | Metschnikowia | 13 | Saprotroph | \|Undefined Saprotroph\| | Yeast | NULL | Probable | As nectar-tap_saprotroph (Põlme et al. 2020); Decay sugar-rich substrates (Põlme et al. 2020) | Põlme S, et al. 2020. Fungal Diversity 105:1-16 ((https://doi.org/10.1007/s13225-020-00466-2)) |
| Otu40677 | 1 | k__Fungi(100);p__Ascomycota(100);c__Saccharomycetes(100);o__Saccharomycetales(100);f__Metschnikowiaceae(100);g__Metschnikowia(100);s__Metschnikowia_pulcherrima(100); | Metschnikowia | 13 | Saprotroph | \|Undefined Saprotroph\| | Yeast | NULL | Probable | As nectar-tap_saprotroph (Põlme et al. 2020); Decay sugar-rich substrates (Põlme et al. 2020) | Põlme S, et al. 2020. Fungal Diversity 105:1-16 ((https://doi.org/10.1007/s13225-020-00466-2)) |
| Otu40678 | 1 | k__Fungi(100);p__Ascomycota(100);c__Saccharomycetes(100);o__Saccharomycetales(100);f__Metschnikowiaceae(100);g__Metschnikowia(100);s__Metschnikowia_pulcherrima(100); | Metschnikowia | 13 | Saprotroph | \|Undefined Saprotroph\| | Yeast | NULL | Probable | As nectar-tap_saprotroph (Põlme et al. 2020); Decay sugar-rich substrates (Põlme et al. 2020) | Põlme S, et al. 2020. Fungal Diversity 105:1-16 ((https://doi.org/10.1007/s13225-020-00466-2)) |
| Otu40679 | 1 | k__Fungi(100);p__Ascomycota(100);c__Saccharomycetes(100);o__Saccharomycetales(100);f__Metschnikowiaceae(100);g__Metschnikowia(100);s__Metschnikowia_pulcherrima(100); | Metschnikowia | 13 | Saprotroph | \|Undefined Saprotroph\| | Yeast | NULL | Probable | As nectar-tap_saprotroph (Põlme et al. 2020); Decay sugar-rich substrates (Põlme et al. 2020) | Põlme S, et al. 2020. Fungal Diversity 105:1-16 ((https://doi.org/10.1007/s13225-020-00466-2)) |
| Otu40680 | 1 | k__Fungi(100);p__Ascomycota(100);c__Saccharomycetes(100);o__Saccharomycetales(100);f__Metschnikowiaceae(100);g__Metschnikowia(100);g__Metschnikowia_unclassified(100); | Metschnikowia | 13 | Saprotroph | \|Undefined Saprotroph\| | Yeast | NULL | Probable | As nectar-tap_saprotroph (Põlme et al. 2020); Decay sugar-rich substrates (Põlme et al. 2020) | Põlme S, et al. 2020. Fungal Diversity 105:1-16 ((https://doi.org/10.1007/s13225-020-00466-2)) |
| Otu40681 | 1 | k__Fungi(100);p__Ascomycota(100);c__Saccharomycetes(100);o__Saccharomycetales(100);f__Metschnikowiaceae(100);g__Metschnikowia(100);s__Metschnikowia_pulcherrima(100); | Metschnikowia | 13 | Saprotroph | \|Undefined Saprotroph\| | Yeast | NULL | Probable | As nectar-tap_saprotroph (Põlme et al. 2020); Decay sugar-rich substrates (Põlme et al. 2020) | Põlme S, et al. 2020. Fungal Diversity 105:1-16 ((https://doi.org/10.1007/s13225-020-00466-2)) |
| Otu40682 | 1 | k__Fungi(100);p__Ascomycota(100);c__Saccharomycetes(100);o__Saccharomycetales(100);f__Metschnikowiaceae(100);g__Metschnikowia(100);s__Metschnikowia_pulcherrima(100); | Metschnikowia | 13 | Saprotroph | \|Undefined Saprotroph\| | Yeast | NULL | Probable | As nectar-tap_saprotroph (Põlme et al. 2020); Decay sugar-rich substrates (Põlme et al. 2020) | Põlme S, et al. 2020. Fungal Diversity 105:1-16 ((https://doi.org/10.1007/s13225-020-00466-2)) |
| Otu40683 | 1 | k__Fungi(100);p__Ascomycota(100);c__Saccharomycetes(100);o__Saccharomycetales(100);f__Metschnikowiaceae(100);g__Metschnikowia(100);s__Metschnikowia_pulcherrima(100); | Metschnikowia | 13 | Saprotroph | \|Undefined Saprotroph\| | Yeast | NULL | Probable | As nectar-tap_saprotroph (Põlme et al. 2020); Decay sugar-rich substrates (Põlme et al. 2020) | Põlme S, et al. 2020. Fungal Diversity 105:1-16 ((https://doi.org/10.1007/s13225-020-00466-2)) |
| Otu40684 | 1 | k__Fungi(100);p__Ascomycota(100);c__Saccharomycetes(100);o__Saccharomycetales(100);f__Metschnikowiaceae(100);g__Metschnikowia(100);s__Metschnikowia_viticola(100); | Metschnikowia | 13 | Saprotroph | \|Undefined Saprotroph\| | Yeast | NULL | Probable | As nectar-tap_saprotroph (Põlme et al. 2020); Decay sugar-rich substrates (Põlme et al. 2020) | Põlme S, et al. 2020. Fungal Diversity 105:1-16 ((https://doi.org/10.1007/s13225-020-00466-2)) |
| Otu40685 | 1 | k__Fungi(100);p__Ascomycota(100);c__Saccharomycetes(100);o__Saccharomycetales(100);f__Metschnikowiaceae(100);g__Metschnikowia(100);s__Metschnikowia_pulcherrima(100); | Metschnikowia | 13 | Saprotroph | \|Undefined Saprotroph\| | Yeast | NULL | Probable | As nectar-tap_saprotroph (Põlme et al. 2020); Decay sugar-rich substrates (Põlme et al. 2020) | Põlme S, et al. 2020. Fungal Diversity 105:1-16 ((https://doi.org/10.1007/s13225-020-00466-2)) |
| Otu40686 | 1 | k__Fungi(100);p__Ascomycota(100);c__Saccharomycetes(100);o__Saccharomycetales(100);f__Metschnikowiaceae(100);g__Metschnikowia(100);s__Metschnikowia_pulcherrima(100); | Metschnikowia | 13 | Saprotroph | \|Undefined Saprotroph\| | Yeast | NULL | Probable | As nectar-tap_saprotroph (Põlme et al. 2020); Decay sugar-rich substrates (Põlme et al. 2020) | Põlme S, et al. 2020. Fungal Diversity 105:1-16 ((https://doi.org/10.1007/s13225-020-00466-2)) |
| Otu40687 | 1 | k__Fungi(100);p__Ascomycota(100);c__Saccharomycetes(100);o__Saccharomycetales(100);f__Metschnikowiaceae(100);g__Metschnikowia(100);s__Metschnikowia_pulcherrima(100); | Metschnikowia | 13 | Saprotroph | \|Undefined Saprotroph\| | Yeast | NULL | Probable | As nectar-tap_saprotroph (Põlme et al. 2020); Decay sugar-rich substrates (Põlme et al. 2020) | Põlme S, et al. 2020. Fungal Diversity 105:1-16 ((https://doi.org/10.1007/s13225-020-00466-2)) |
| Otu40688 | 1 | k__Fungi(100);p__Ascomycota(100);c__Saccharomycetes(100);o__Saccharomycetales(100);f__Metschnikowiaceae(100);g__Metschnikowia(100);g__Metschnikowia_unclassified(100); | Metschnikowia | 13 | Saprotroph | \|Undefined Saprotroph\| | Yeast | NULL | Probable | As nectar-tap_saprotroph (Põlme et al. 2020); Decay sugar-rich substrates (Põlme et al. 2020) | Põlme S, et al. 2020. Fungal Diversity 105:1-16 ((https://doi.org/10.1007/s13225-020-00466-2)) |
| Otu40689 | 1 | k__Fungi(100);p__Ascomycota(100);c__Saccharomycetes(100);o__Saccharomycetales(100);f__Metschnikowiaceae(100);g__Metschnikowia(100);s__Metschnikowia_pulcherrima(100); | Metschnikowia | 13 | Saprotroph | \|Undefined Saprotroph\| | Yeast | NULL | Probable | As nectar-tap_saprotroph (Põlme et al. 2020); Decay sugar-rich substrates (Põlme et al. 2020) | Põlme S, et al. 2020. Fungal Diversity 105:1-16 ((https://doi.org/10.1007/s13225-020-00466-2)) |
| Otu40690 | 1 | k__Fungi(100);p__Ascomycota(100);c__Saccharomycetes(100);o__Saccharomycetales(100);f__Metschnikowiaceae(100);g__Metschnikowia(100);s__Metschnikowia_pulcherrima(100); | Metschnikowia | 13 | Saprotroph | \|Undefined Saprotroph\| | Yeast | NULL | Probable | As nectar-tap_saprotroph (Põlme et al. 2020); Decay sugar-rich substrates (Põlme et al. 2020) | Põlme S, et al. 2020. Fungal Diversity 105:1-16 ((https://doi.org/10.1007/s13225-020-00466-2)) |
| Otu40691 | 1 | k__Fungi(100);p__Ascomycota(100);c__Saccharomycetes(100);o__Saccharomycetales(100);f__Metschnikowiaceae(100);g__Metschnikowia(100);s__Metschnikowia_pulcherrima(100); | Metschnikowia | 13 | Saprotroph | \|Undefined Saprotroph\| | Yeast | NULL | Probable | As nectar-tap_saprotroph (Põlme et al. 2020); Decay sugar-rich substrates (Põlme et al. 2020) | Põlme S, et al. 2020. Fungal Diversity 105:1-16 ((https://doi.org/10.1007/s13225-020-00466-2)) |
| Otu40692 | 1 | k__Fungi(100);p__Ascomycota(100);c__Saccharomycetes(100);o__Saccharomycetales(100);f__Metschnikowiaceae(100);g__Metschnikowia(100);g__Metschnikowia_unclassified(100); | Metschnikowia | 13 | Saprotroph | \|Undefined Saprotroph\| | Yeast | NULL | Probable | As nectar-tap_saprotroph (Põlme et al. 2020); Decay sugar-rich substrates (Põlme et al. 2020) | Põlme S, et al. 2020. Fungal Diversity 105:1-16 ((https://doi.org/10.1007/s13225-020-00466-2)) |
| Otu40693 | 1 | k__Fungi(100);p__Ascomycota(100);c__Saccharomycetes(100);o__Saccharomycetales(100);f__Metschnikowiaceae(100);g__Metschnikowia(100);s__Metschnikowia_pulcherrima(100); | Metschnikowia | 13 | Saprotroph | \|Undefined Saprotroph\| | Yeast | NULL | Probable | As nectar-tap_saprotroph (Põlme et al. 2020); Decay sugar-rich substrates (Põlme et al. 2020) | Põlme S, et al. 2020. Fungal Diversity 105:1-16 ((https://doi.org/10.1007/s13225-020-00466-2)) |
| Otu40694 | 1 | k__Fungi(100);p__Ascomycota(100);c__Saccharomycetes(100);o__Saccharomycetales(100);f__Metschnikowiaceae(100);g__Metschnikowia(100);g__Metschnikowia_unclassified(100); | Metschnikowia | 13 | Saprotroph | \|Undefined Saprotroph\| | Yeast | NULL | Probable | As nectar-tap_saprotroph (Põlme et al. 2020); Decay sugar-rich substrates (Põlme et al. 2020) | Põlme S, et al. 2020. Fungal Diversity 105:1-16 ((https://doi.org/10.1007/s13225-020-00466-2)) |
| Otu40695 | 1 | k__Fungi(100);p__Ascomycota(100);c__Saccharomycetes(100);o__Saccharomycetales(100);f__Metschnikowiaceae(100);g__Metschnikowia(100);s__Metschnikowia_pulcherrima(100); | Metschnikowia | 13 | Saprotroph | \|Undefined Saprotroph\| | Yeast | NULL | Probable | As nectar-tap_saprotroph (Põlme et al. 2020); Decay sugar-rich substrates (Põlme et al. 2020) | Põlme S, et al. 2020. Fungal Diversity 105:1-16 ((https://doi.org/10.1007/s13225-020-00466-2)) |
| Otu40696 | 1 | k__Fungi(100);p__Ascomycota(100);c__Saccharomycetes(100);o__Saccharomycetales(100);f__Metschnikowiaceae(100);g__Metschnikowia(100);s__Metschnikowia_pulcherrima(100); | Metschnikowia | 13 | Saprotroph | \|Undefined Saprotroph\| | Yeast | NULL | Probable | As nectar-tap_saprotroph (Põlme et al. 2020); Decay sugar-rich substrates (Põlme et al. 2020) | Põlme S, et al. 2020. Fungal Diversity 105:1-16 ((https://doi.org/10.1007/s13225-020-00466-2)) |
| Otu40697 | 1 | k__Fungi(100);p__Ascomycota(100);c__Saccharomycetes(100);o__Saccharomycetales(100);f__Metschnikowiaceae(100);g__Metschnikowia(100);s__Metschnikowia_pulcherrima(100); | Metschnikowia | 13 | Saprotroph | \|Undefined Saprotroph\| | Yeast | NULL | Probable | As nectar-tap_saprotroph (Põlme et al. 2020); Decay sugar-rich substrates (Põlme et al. 2020) | Põlme S, et al. 2020. Fungal Diversity 105:1-16 ((https://doi.org/10.1007/s13225-020-00466-2)) |
| Otu40698 | 1 | k__Fungi(100);p__Ascomycota(100);c__Saccharomycetes(100);o__Saccharomycetales(100);f__Metschnikowiaceae(100);g__Metschnikowia(100);s__Metschnikowia_pulcherrima(100); | Metschnikowia | 13 | Saprotroph | \|Undefined Saprotroph\| | Yeast | NULL | Probable | As nectar-tap_saprotroph (Põlme et al. 2020); Decay sugar-rich substrates (Põlme et al. 2020) | Põlme S, et al. 2020. Fungal Diversity 105:1-16 ((https://doi.org/10.1007/s13225-020-00466-2)) |
| Otu40699 | 1 | k__Fungi(100);p__Ascomycota(100);c__Saccharomycetes(100);o__Saccharomycetales(100);f__Metschnikowiaceae(100);g__Metschnikowia(100);s__Metschnikowia_pulcherrima(100); | Metschnikowia | 13 | Saprotroph | \|Undefined Saprotroph\| | Yeast | NULL | Probable | As nectar-tap_saprotroph (Põlme et al. 2020); Decay sugar-rich substrates (Põlme et al. 2020) | Põlme S, et al. 2020. Fungal Diversity 105:1-16 ((https://doi.org/10.1007/s13225-020-00466-2)) |
| Otu40700 | 1 | k__Fungi(100);p__Ascomycota(100);c__Saccharomycetes(100);o__Saccharomycetales(100);f__Metschnikowiaceae(100);g__Metschnikowia(100);s__Metschnikowia_pulcherrima(100); | Metschnikowia | 13 | Saprotroph | \|Undefined Saprotroph\| | Yeast | NULL | Probable | As nectar-tap_saprotroph (Põlme et al. 2020); Decay sugar-rich substrates (Põlme et al. 2020) | Põlme S, et al. 2020. Fungal Diversity 105:1-16 ((https://doi.org/10.1007/s13225-020-00466-2)) |
| Otu40701 | 1 | k__Fungi(100);p__Ascomycota(100);c__Saccharomycetes(100);o__Saccharomycetales(100);f__Metschnikowiaceae(100);g__Metschnikowia(100);s__Metschnikowia_pulcherrima(100); | Metschnikowia | 13 | Saprotroph | \|Undefined Saprotroph\| | Yeast | NULL | Probable | As nectar-tap_saprotroph (Põlme et al. 2020); Decay sugar-rich substrates (Põlme et al. 2020) | Põlme S, et al. 2020. Fungal Diversity 105:1-16 ((https://doi.org/10.1007/s13225-020-00466-2)) |
| Otu40702 | 1 | k__Fungi(100);p__Ascomycota(100);c__Saccharomycetes(100);o__Saccharomycetales(100);f__Metschnikowiaceae(100);g__Metschnikowia(100);g__Metschnikowia_unclassified(100); | Metschnikowia | 13 | Saprotroph | \|Undefined Saprotroph\| | Yeast | NULL | Probable | As nectar-tap_saprotroph (Põlme et al. 2020); Decay sugar-rich substrates (Põlme et al. 2020) | Põlme S, et al. 2020. Fungal Diversity 105:1-16 ((https://doi.org/10.1007/s13225-020-00466-2)) |
| Otu40703 | 1 | k__Fungi(100);p__Ascomycota(100);c__Saccharomycetes(100);o__Saccharomycetales(100);f__Metschnikowiaceae(100);g__Metschnikowia(100);g__Metschnikowia_unclassified(100); | Metschnikowia | 13 | Saprotroph | \|Undefined Saprotroph\| | Yeast | NULL | Probable | As nectar-tap_saprotroph (Põlme et al. 2020); Decay sugar-rich substrates (Põlme et al. 2020) | Põlme S, et al. 2020. Fungal Diversity 105:1-16 ((https://doi.org/10.1007/s13225-020-00466-2)) |
| Otu40704 | 1 | k__Fungi(100);p__Ascomycota(100);c__Saccharomycetes(100);o__Saccharomycetales(100);f__Metschnikowiaceae(100);g__Metschnikowia(100);s__Metschnikowia_pulcherrima(100); | Metschnikowia | 13 | Saprotroph | \|Undefined Saprotroph\| | Yeast | NULL | Probable | As nectar-tap_saprotroph (Põlme et al. 2020); Decay sugar-rich substrates (Põlme et al. 2020) | Põlme S, et al. 2020. Fungal Diversity 105:1-16 ((https://doi.org/10.1007/s13225-020-00466-2)) |
| Otu40705 | 1 | k__Fungi(100);p__Ascomycota(100);c__Saccharomycetes(100);o__Saccharomycetales(100);f__Metschnikowiaceae(100);g__Metschnikowia(100);s__Metschnikowia_pulcherrima(100); | Metschnikowia | 13 | Saprotroph | \|Undefined Saprotroph\| | Yeast | NULL | Probable | As nectar-tap_saprotroph (Põlme et al. 2020); Decay sugar-rich substrates (Põlme et al. 2020) | Põlme S, et al. 2020. Fungal Diversity 105:1-16 ((https://doi.org/10.1007/s13225-020-00466-2)) |
| Otu40706 | 1 | k__Fungi(100);p__Ascomycota(100);c__Saccharomycetes(100);o__Saccharomycetales(100);f__Metschnikowiaceae(100);g__Metschnikowia(100);g__Metschnikowia_unclassified(100); | Metschnikowia | 13 | Saprotroph | \|Undefined Saprotroph\| | Yeast | NULL | Probable | As nectar-tap_saprotroph (Põlme et al. 2020); Decay sugar-rich substrates (Põlme et al. 2020) | Põlme S, et al. 2020. Fungal Diversity 105:1-16 ((https://doi.org/10.1007/s13225-020-00466-2)) |
| Otu40707 | 1 | k__Fungi(100);p__Ascomycota(100);c__Saccharomycetes(100);o__Saccharomycetales(100);f__Metschnikowiaceae(100);g__Metschnikowia(100);s__Metschnikowia_pulcherrima(100); | Metschnikowia | 13 | Saprotroph | \|Undefined Saprotroph\| | Yeast | NULL | Probable | As nectar-tap_saprotroph (Põlme et al. 2020); Decay sugar-rich substrates (Põlme et al. 2020) | Põlme S, et al. 2020. Fungal Diversity 105:1-16 ((https://doi.org/10.1007/s13225-020-00466-2)) |
| Otu40708 | 1 | k__Fungi(100);p__Ascomycota(100);c__Saccharomycetes(100);o__Saccharomycetales(100);f__Metschnikowiaceae(100);g__Metschnikowia(100);g__Metschnikowia_unclassified(100); | Metschnikowia | 13 | Saprotroph | \|Undefined Saprotroph\| | Yeast | NULL | Probable | As nectar-tap_saprotroph (Põlme et al. 2020); Decay sugar-rich substrates (Põlme et al. 2020) | Põlme S, et al. 2020. Fungal Diversity 105:1-16 ((https://doi.org/10.1007/s13225-020-00466-2)) |
| Otu40709 | 1 | k__Fungi(100);p__Ascomycota(100);c__Saccharomycetes(100);o__Saccharomycetales(100);f__Metschnikowiaceae(100);g__Metschnikowia(100);s__Metschnikowia_pulcherrima(100); | Metschnikowia | 13 | Saprotroph | \|Undefined Saprotroph\| | Yeast | NULL | Probable | As nectar-tap_saprotroph (Põlme et al. 2020); Decay sugar-rich substrates (Põlme et al. 2020) | Põlme S, et al. 2020. Fungal Diversity 105:1-16 ((https://doi.org/10.1007/s13225-020-00466-2)) |
| Otu40710 | 1 | k__Fungi(100);p__Ascomycota(100);c__Saccharomycetes(100);o__Saccharomycetales(100);f__Metschnikowiaceae(100);g__Metschnikowia(100);s__Metschnikowia_pulcherrima(100); | Metschnikowia | 13 | Saprotroph | \|Undefined Saprotroph\| | Yeast | NULL | Probable | As nectar-tap_saprotroph (Põlme et al. 2020); Decay sugar-rich substrates (Põlme et al. 2020) | Põlme S, et al. 2020. Fungal Diversity 105:1-16 ((https://doi.org/10.1007/s13225-020-00466-2)) |
| Otu40711 | 1 | k__Fungi(100);p__Ascomycota(100);c__Saccharomycetes(100);o__Saccharomycetales(100);f__Metschnikowiaceae(100);g__Metschnikowia(100);s__Metschnikowia_pulcherrima(100); | Metschnikowia | 13 | Saprotroph | \|Undefined Saprotroph\| | Yeast | NULL | Probable | As nectar-tap_saprotroph (Põlme et al. 2020); Decay sugar-rich substrates (Põlme et al. 2020) | Põlme S, et al. 2020. Fungal Diversity 105:1-16 ((https://doi.org/10.1007/s13225-020-00466-2)) |
| Otu40712 | 1 | k__Fungi(100);p__Ascomycota(100);c__Saccharomycetes(100);o__Saccharomycetales(100);f__Metschnikowiaceae(100);g__Metschnikowia(100);s__Metschnikowia_pulcherrima(100); | Metschnikowia | 13 | Saprotroph | \|Undefined Saprotroph\| | Yeast | NULL | Probable | As nectar-tap_saprotroph (Põlme et al. 2020); Decay sugar-rich substrates (Põlme et al. 2020) | Põlme S, et al. 2020. Fungal Diversity 105:1-16 ((https://doi.org/10.1007/s13225-020-00466-2)) |
| Otu40713 | 1 | k__Fungi(100);p__Ascomycota(100);c__Saccharomycetes(100);o__Saccharomycetales(100);f__Metschnikowiaceae(100);g__Metschnikowia(100);s__Metschnikowia_pulcherrima(100); | Metschnikowia | 13 | Saprotroph | \|Undefined Saprotroph\| | Yeast | NULL | Probable | As nectar-tap_saprotroph (Põlme et al. 2020); Decay sugar-rich substrates (Põlme et al. 2020) | Põlme S, et al. 2020. Fungal Diversity 105:1-16 ((https://doi.org/10.1007/s13225-020-00466-2)) |
| Otu40714 | 1 | k__Fungi(100);p__Ascomycota(100);c__Saccharomycetes(100);o__Saccharomycetales(100);f__Metschnikowiaceae(100);g__Metschnikowia(100);s__Metschnikowia_pulcherrima(100); | Metschnikowia | 13 | Saprotroph | \|Undefined Saprotroph\| | Yeast | NULL | Probable | As nectar-tap_saprotroph (Põlme et al. 2020); Decay sugar-rich substrates (Põlme et al. 2020) | Põlme S, et al. 2020. Fungal Diversity 105:1-16 ((https://doi.org/10.1007/s13225-020-00466-2)) |
| Otu40715 | 1 | k__Fungi(100);p__Ascomycota(100);c__Saccharomycetes(100);o__Saccharomycetales(100);f__Metschnikowiaceae(100);g__Metschnikowia(100);g__Metschnikowia_unclassified(100); | Metschnikowia | 13 | Saprotroph | \|Undefined Saprotroph\| | Yeast | NULL | Probable | As nectar-tap_saprotroph (Põlme et al. 2020); Decay sugar-rich substrates (Põlme et al. 2020) | Põlme S, et al. 2020. Fungal Diversity 105:1-16 ((https://doi.org/10.1007/s13225-020-00466-2)) |
| Otu40716 | 1 | k__Fungi(100);p__Ascomycota(100);c__Saccharomycetes(100);o__Saccharomycetales(100);f__Metschnikowiaceae(100);g__Metschnikowia(100);s__Metschnikowia_pulcherrima(100); | Metschnikowia | 13 | Saprotroph | \|Undefined Saprotroph\| | Yeast | NULL | Probable | As nectar-tap_saprotroph (Põlme et al. 2020); Decay sugar-rich substrates (Põlme et al. 2020) | Põlme S, et al. 2020. Fungal Diversity 105:1-16 ((https://doi.org/10.1007/s13225-020-00466-2)) |
| Otu40717 | 1 | k__Fungi(100);p__Ascomycota(100);c__Saccharomycetes(100);o__Saccharomycetales(100);f__Metschnikowiaceae(100);g__Metschnikowia(100);s__Metschnikowia_pulcherrima(100); | Metschnikowia | 13 | Saprotroph | \|Undefined Saprotroph\| | Yeast | NULL | Probable | As nectar-tap_saprotroph (Põlme et al. 2020); Decay sugar-rich substrates (Põlme et al. 2020) | Põlme S, et al. 2020. Fungal Diversity 105:1-16 ((https://doi.org/10.1007/s13225-020-00466-2)) |
| Otu40718 | 1 | k__Fungi(100);p__Ascomycota(100);c__Saccharomycetes(100);o__Saccharomycetales(100);f__Metschnikowiaceae(100);g__Metschnikowia(100);s__Metschnikowia_pulcherrima(100); | Metschnikowia | 13 | Saprotroph | \|Undefined Saprotroph\| | Yeast | NULL | Probable | As nectar-tap_saprotroph (Põlme et al. 2020); Decay sugar-rich substrates (Põlme et al. 2020) | Põlme S, et al. 2020. Fungal Diversity 105:1-16 ((https://doi.org/10.1007/s13225-020-00466-2)) |
| Otu40719 | 1 | k__Fungi(100);p__Ascomycota(100);c__Saccharomycetes(100);o__Saccharomycetales(100);f__Metschnikowiaceae(100);g__Metschnikowia(100);s__Metschnikowia_pulcherrima(100); | Metschnikowia | 13 | Saprotroph | \|Undefined Saprotroph\| | Yeast | NULL | Probable | As nectar-tap_saprotroph (Põlme et al. 2020); Decay sugar-rich substrates (Põlme et al. 2020) | Põlme S, et al. 2020. Fungal Diversity 105:1-16 ((https://doi.org/10.1007/s13225-020-00466-2)) |
| Otu40720 | 1 | k__Fungi(100);p__Ascomycota(100);c__Saccharomycetes(100);o__Saccharomycetales(100);f__Metschnikowiaceae(100);g__Metschnikowia(100);s__Metschnikowia_pulcherrima(100); | Metschnikowia | 13 | Saprotroph | \|Undefined Saprotroph\| | Yeast | NULL | Probable | As nectar-tap_saprotroph (Põlme et al. 2020); Decay sugar-rich substrates (Põlme et al. 2020) | Põlme S, et al. 2020. Fungal Diversity 105:1-16 ((https://doi.org/10.1007/s13225-020-00466-2)) |
| Otu40721 | 1 | k__Fungi(100);p__Ascomycota(100);c__Saccharomycetes(100);o__Saccharomycetales(100);f__Metschnikowiaceae(100);g__Metschnikowia(100);s__Metschnikowia_pulcherrima(100); | Metschnikowia | 13 | Saprotroph | \|Undefined Saprotroph\| | Yeast | NULL | Probable | As nectar-tap_saprotroph (Põlme et al. 2020); Decay sugar-rich substrates (Põlme et al. 2020) | Põlme S, et al. 2020. Fungal Diversity 105:1-16 ((https://doi.org/10.1007/s13225-020-00466-2)) |
| Otu40722 | 1 | k__Fungi(100);p__Ascomycota(100);c__Saccharomycetes(100);o__Saccharomycetales(100);f__Metschnikowiaceae(100);g__Metschnikowia(100);s__Metschnikowia_pulcherrima(100); | Metschnikowia | 13 | Saprotroph | \|Undefined Saprotroph\| | Yeast | NULL | Probable | As nectar-tap_saprotroph (Põlme et al. 2020); Decay sugar-rich substrates (Põlme et al. 2020) | Põlme S, et al. 2020. Fungal Diversity 105:1-16 ((https://doi.org/10.1007/s13225-020-00466-2)) |
| Otu40723 | 1 | k__Fungi(100);p__Ascomycota(100);c__Saccharomycetes(100);o__Saccharomycetales(100);f__Metschnikowiaceae(100);g__Metschnikowia(100);s__Metschnikowia_pulcherrima(100); | Metschnikowia | 13 | Saprotroph | \|Undefined Saprotroph\| | Yeast | NULL | Probable | As nectar-tap_saprotroph (Põlme et al. 2020); Decay sugar-rich substrates (Põlme et al. 2020) | Põlme S, et al. 2020. Fungal Diversity 105:1-16 ((https://doi.org/10.1007/s13225-020-00466-2)) |
| Otu40724 | 1 | k__Fungi(100);p__Ascomycota(100);c__Saccharomycetes(100);o__Saccharomycetales(100);f__Metschnikowiaceae(100);g__Metschnikowia(100);s__Metschnikowia_pulcherrima(100); | Metschnikowia | 13 | Saprotroph | \|Undefined Saprotroph\| | Yeast | NULL | Probable | As nectar-tap_saprotroph (Põlme et al. 2020); Decay sugar-rich substrates (Põlme et al. 2020) | Põlme S, et al. 2020. Fungal Diversity 105:1-16 ((https://doi.org/10.1007/s13225-020-00466-2)) |
| Otu40725 | 1 | k__Fungi(100);p__Ascomycota(100);c__Saccharomycetes(100);o__Saccharomycetales(100);f__Metschnikowiaceae(100);g__Metschnikowia(100);s__Metschnikowia_pulcherrima(100); | Metschnikowia | 13 | Saprotroph | \|Undefined Saprotroph\| | Yeast | NULL | Probable | As nectar-tap_saprotroph (Põlme et al. 2020); Decay sugar-rich substrates (Põlme et al. 2020) | Põlme S, et al. 2020. Fungal Diversity 105:1-16 ((https://doi.org/10.1007/s13225-020-00466-2)) |
| Otu40726 | 1 | k__Fungi(100);p__Ascomycota(100);c__Saccharomycetes(100);o__Saccharomycetales(100);f__Metschnikowiaceae(100);g__Metschnikowia(100);s__Metschnikowia_pulcherrima(100); | Metschnikowia | 13 | Saprotroph | \|Undefined Saprotroph\| | Yeast | NULL | Probable | As nectar-tap_saprotroph (Põlme et al. 2020); Decay sugar-rich substrates (Põlme et al. 2020) | Põlme S, et al. 2020. Fungal Diversity 105:1-16 ((https://doi.org/10.1007/s13225-020-00466-2)) |
| Otu40727 | 1 | k__Fungi(100);p__Ascomycota(100);c__Saccharomycetes(100);o__Saccharomycetales(100);f__Metschnikowiaceae(100);g__Metschnikowia(100);s__Metschnikowia_pulcherrima(100); | Metschnikowia | 13 | Saprotroph | \|Undefined Saprotroph\| | Yeast | NULL | Probable | As nectar-tap_saprotroph (Põlme et al. 2020); Decay sugar-rich substrates (Põlme et al. 2020) | Põlme S, et al. 2020. Fungal Diversity 105:1-16 ((https://doi.org/10.1007/s13225-020-00466-2)) |
| Otu40728 | 1 | k__Fungi(100);p__Ascomycota(100);c__Saccharomycetes(100);o__Saccharomycetales(100);f__Metschnikowiaceae(100);g__Metschnikowia(100);s__Metschnikowia_pulcherrima(100); | Metschnikowia | 13 | Saprotroph | \|Undefined Saprotroph\| | Yeast | NULL | Probable | As nectar-tap_saprotroph (Põlme et al. 2020); Decay sugar-rich substrates (Põlme et al. 2020) | Põlme S, et al. 2020. Fungal Diversity 105:1-16 ((https://doi.org/10.1007/s13225-020-00466-2)) |
| Otu40729 | 1 | k__Fungi(100);p__Ascomycota(100);c__Saccharomycetes(100);o__Saccharomycetales(100);f__Metschnikowiaceae(100);g__Metschnikowia(100);s__Metschnikowia_pulcherrima(100); | Metschnikowia | 13 | Saprotroph | \|Undefined Saprotroph\| | Yeast | NULL | Probable | As nectar-tap_saprotroph (Põlme et al. 2020); Decay sugar-rich substrates (Põlme et al. 2020) | Põlme S, et al. 2020. Fungal Diversity 105:1-16 ((https://doi.org/10.1007/s13225-020-00466-2)) |
| Otu40730 | 1 | k__Fungi(100);p__Ascomycota(100);c__Saccharomycetes(100);o__Saccharomycetales(100);f__Metschnikowiaceae(100);g__Metschnikowia(100);g__Metschnikowia_unclassified(100); | Metschnikowia | 13 | Saprotroph | \|Undefined Saprotroph\| | Yeast | NULL | Probable | As nectar-tap_saprotroph (Põlme et al. 2020); Decay sugar-rich substrates (Põlme et al. 2020) | Põlme S, et al. 2020. Fungal Diversity 105:1-16 ((https://doi.org/10.1007/s13225-020-00466-2)) |
| Otu40731 | 1 | k__Fungi(100);p__Ascomycota(100);c__Saccharomycetes(100);o__Saccharomycetales(100);f__Metschnikowiaceae(100);g__Metschnikowia(100);g__Metschnikowia_unclassified(100); | Metschnikowia | 13 | Saprotroph | \|Undefined Saprotroph\| | Yeast | NULL | Probable | As nectar-tap_saprotroph (Põlme et al. 2020); Decay sugar-rich substrates (Põlme et al. 2020) | Põlme S, et al. 2020. Fungal Diversity 105:1-16 ((https://doi.org/10.1007/s13225-020-00466-2)) |
| Otu40732 | 1 | k__Fungi(100);p__Ascomycota(100);c__Saccharomycetes(100);o__Saccharomycetales(100);f__Metschnikowiaceae(100);g__Metschnikowia(100);g__Metschnikowia_unclassified(100); | Metschnikowia | 13 | Saprotroph | \|Undefined Saprotroph\| | Yeast | NULL | Probable | As nectar-tap_saprotroph (Põlme et al. 2020); Decay sugar-rich substrates (Põlme et al. 2020) | Põlme S, et al. 2020. Fungal Diversity 105:1-16 ((https://doi.org/10.1007/s13225-020-00466-2)) |
| Otu40733 | 1 | k__Fungi(100);p__Ascomycota(100);c__Saccharomycetes(100);o__Saccharomycetales(100);f__Metschnikowiaceae(100);g__Metschnikowia(100);s__Metschnikowia_pulcherrima(100); | Metschnikowia | 13 | Saprotroph | \|Undefined Saprotroph\| | Yeast | NULL | Probable | As nectar-tap_saprotroph (Põlme et al. 2020); Decay sugar-rich substrates (Põlme et al. 2020) | Põlme S, et al. 2020. Fungal Diversity 105:1-16 ((https://doi.org/10.1007/s13225-020-00466-2)) |
| Otu40734 | 1 | k__Fungi(100);p__Ascomycota(100);c__Saccharomycetes(100);o__Saccharomycetales(100);f__Metschnikowiaceae(100);g__Metschnikowia(100);s__Metschnikowia_pulcherrima(100); | Metschnikowia | 13 | Saprotroph | \|Undefined Saprotroph\| | Yeast | NULL | Probable | As nectar-tap_saprotroph (Põlme et al. 2020); Decay sugar-rich substrates (Põlme et al. 2020) | Põlme S, et al. 2020. Fungal Diversity 105:1-16 ((https://doi.org/10.1007/s13225-020-00466-2)) |
| Otu40735 | 1 | k__Fungi(100);p__Ascomycota(100);c__Saccharomycetes(100);o__Saccharomycetales(100);f__Metschnikowiaceae(100);g__Metschnikowia(100);g__Metschnikowia_unclassified(100); | Metschnikowia | 13 | Saprotroph | \|Undefined Saprotroph\| | Yeast | NULL | Probable | As nectar-tap_saprotroph (Põlme et al. 2020); Decay sugar-rich substrates (Põlme et al. 2020) | Põlme S, et al. 2020. Fungal Diversity 105:1-16 ((https://doi.org/10.1007/s13225-020-00466-2)) |
| Otu40736 | 1 | k__Fungi(100);p__Ascomycota(100);c__Saccharomycetes(100);o__Saccharomycetales(100);f__Metschnikowiaceae(100);g__Metschnikowia(100);g__Metschnikowia_unclassified(100); | Metschnikowia | 13 | Saprotroph | \|Undefined Saprotroph\| | Yeast | NULL | Probable | As nectar-tap_saprotroph (Põlme et al. 2020); Decay sugar-rich substrates (Põlme et al. 2020) | Põlme S, et al. 2020. Fungal Diversity 105:1-16 ((https://doi.org/10.1007/s13225-020-00466-2)) |
| Otu40737 | 1 | k__Fungi(100);p__Ascomycota(100);c__Saccharomycetes(100);o__Saccharomycetales(100);f__Metschnikowiaceae(100);g__Metschnikowia(100);s__Metschnikowia_pulcherrima(100); | Metschnikowia | 13 | Saprotroph | \|Undefined Saprotroph\| | Yeast | NULL | Probable | As nectar-tap_saprotroph (Põlme et al. 2020); Decay sugar-rich substrates (Põlme et al. 2020) | Põlme S, et al. 2020. Fungal Diversity 105:1-16 ((https://doi.org/10.1007/s13225-020-00466-2)) |
| Otu40738 | 1 | k__Fungi(100);p__Ascomycota(100);c__Saccharomycetes(100);o__Saccharomycetales(100);f__Metschnikowiaceae(100);g__Metschnikowia(100);s__Metschnikowia_pulcherrima(100); | Metschnikowia | 13 | Saprotroph | \|Undefined Saprotroph\| | Yeast | NULL | Probable | As nectar-tap_saprotroph (Põlme et al. 2020); Decay sugar-rich substrates (Põlme et al. 2020) | Põlme S, et al. 2020. Fungal Diversity 105:1-16 ((https://doi.org/10.1007/s13225-020-00466-2)) |
| Otu40739 | 1 | k__Fungi(100);p__Ascomycota(100);c__Saccharomycetes(100);o__Saccharomycetales(100);f__Metschnikowiaceae(100);g__Metschnikowia(100);s__Metschnikowia_pulcherrima(100); | Metschnikowia | 13 | Saprotroph | \|Undefined Saprotroph\| | Yeast | NULL | Probable | As nectar-tap_saprotroph (Põlme et al. 2020); Decay sugar-rich substrates (Põlme et al. 2020) | Põlme S, et al. 2020. Fungal Diversity 105:1-16 ((https://doi.org/10.1007/s13225-020-00466-2)) |
| Otu40740 | 1 | k__Fungi(100);p__Ascomycota(100);c__Saccharomycetes(100);o__Saccharomycetales(100);f__Metschnikowiaceae(100);g__Metschnikowia(100);s__Metschnikowia_pulcherrima(100); | Metschnikowia | 13 | Saprotroph | \|Undefined Saprotroph\| | Yeast | NULL | Probable | As nectar-tap_saprotroph (Põlme et al. 2020); Decay sugar-rich substrates (Põlme et al. 2020) | Põlme S, et al. 2020. Fungal Diversity 105:1-16 ((https://doi.org/10.1007/s13225-020-00466-2)) |
| Otu40741 | 1 | k__Fungi(100);p__Ascomycota(100);c__Saccharomycetes(100);o__Saccharomycetales(100);f__Metschnikowiaceae(100);g__Metschnikowia(100);s__Metschnikowia_pulcherrima(100); | Metschnikowia | 13 | Saprotroph | \|Undefined Saprotroph\| | Yeast | NULL | Probable | As nectar-tap_saprotroph (Põlme et al. 2020); Decay sugar-rich substrates (Põlme et al. 2020) | Põlme S, et al. 2020. Fungal Diversity 105:1-16 ((https://doi.org/10.1007/s13225-020-00466-2)) |
| Otu40742 | 1 | k__Fungi(100);p__Ascomycota(100);c__Saccharomycetes(100);o__Saccharomycetales(100);f__Metschnikowiaceae(100);g__Metschnikowia(100);s__Metschnikowia_pulcherrima(100); | Metschnikowia | 13 | Saprotroph | \|Undefined Saprotroph\| | Yeast | NULL | Probable | As nectar-tap_saprotroph (Põlme et al. 2020); Decay sugar-rich substrates (Põlme et al. 2020) | Põlme S, et al. 2020. Fungal Diversity 105:1-16 ((https://doi.org/10.1007/s13225-020-00466-2)) |
| Otu40743 | 1 | k__Fungi(100);p__Ascomycota(100);c__Saccharomycetes(100);o__Saccharomycetales(100);f__Metschnikowiaceae(100);g__Metschnikowia(100);g__Metschnikowia_unclassified(100); | Metschnikowia | 13 | Saprotroph | \|Undefined Saprotroph\| | Yeast | NULL | Probable | As nectar-tap_saprotroph (Põlme et al. 2020); Decay sugar-rich substrates (Põlme et al. 2020) | Põlme S, et al. 2020. Fungal Diversity 105:1-16 ((https://doi.org/10.1007/s13225-020-00466-2)) |
| Otu40744 | 1 | k__Fungi(100);p__Ascomycota(100);c__Saccharomycetes(100);o__Saccharomycetales(100);f__Metschnikowiaceae(100);g__Metschnikowia(100);g__Metschnikowia_unclassified(100); | Metschnikowia | 13 | Saprotroph | \|Undefined Saprotroph\| | Yeast | NULL | Probable | As nectar-tap_saprotroph (Põlme et al. 2020); Decay sugar-rich substrates (Põlme et al. 2020) | Põlme S, et al. 2020. Fungal Diversity 105:1-16 ((https://doi.org/10.1007/s13225-020-00466-2)) |
| Otu40745 | 1 | k__Fungi(100);p__Ascomycota(100);c__Saccharomycetes(100);o__Saccharomycetales(100);f__Metschnikowiaceae(100);g__Metschnikowia(100);g__Metschnikowia_unclassified(100); | Metschnikowia | 13 | Saprotroph | \|Undefined Saprotroph\| | Yeast | NULL | Probable | As nectar-tap_saprotroph (Põlme et al. 2020); Decay sugar-rich substrates (Põlme et al. 2020) | Põlme S, et al. 2020. Fungal Diversity 105:1-16 ((https://doi.org/10.1007/s13225-020-00466-2)) |
| Otu40746 | 1 | k__Fungi(100);p__Ascomycota(100);c__Saccharomycetes(100);o__Saccharomycetales(100);f__Metschnikowiaceae(100);g__Metschnikowia(100);g__Metschnikowia_unclassified(100); | Metschnikowia | 13 | Saprotroph | \|Undefined Saprotroph\| | Yeast | NULL | Probable | As nectar-tap_saprotroph (Põlme et al. 2020); Decay sugar-rich substrates (Põlme et al. 2020) | Põlme S, et al. 2020. Fungal Diversity 105:1-16 ((https://doi.org/10.1007/s13225-020-00466-2)) |
| Otu40747 | 1 | k__Fungi(100);p__Ascomycota(100);c__Saccharomycetes(100);o__Saccharomycetales(100);f__Metschnikowiaceae(100);g__Metschnikowia(100);g__Metschnikowia_unclassified(100); | Metschnikowia | 13 | Saprotroph | \|Undefined Saprotroph\| | Yeast | NULL | Probable | As nectar-tap_saprotroph (Põlme et al. 2020); Decay sugar-rich substrates (Põlme et al. 2020) | Põlme S, et al. 2020. Fungal Diversity 105:1-16 ((https://doi.org/10.1007/s13225-020-00466-2)) |
| Otu40748 | 1 | k__Fungi(100);p__Ascomycota(100);c__Saccharomycetes(100);o__Saccharomycetales(100);f__Metschnikowiaceae(100);g__Metschnikowia(100);g__Metschnikowia_unclassified(100); | Metschnikowia | 13 | Saprotroph | \|Undefined Saprotroph\| | Yeast | NULL | Probable | As nectar-tap_saprotroph (Põlme et al. 2020); Decay sugar-rich substrates (Põlme et al. 2020) | Põlme S, et al. 2020. Fungal Diversity 105:1-16 ((https://doi.org/10.1007/s13225-020-00466-2)) |
| Otu40749 | 1 | k__Fungi(100);p__Ascomycota(100);c__Saccharomycetes(100);o__Saccharomycetales(100);f__Metschnikowiaceae(100);g__Metschnikowia(100);s__Metschnikowia_pulcherrima(100); | Metschnikowia | 13 | Saprotroph | \|Undefined Saprotroph\| | Yeast | NULL | Probable | As nectar-tap_saprotroph (Põlme et al. 2020); Decay sugar-rich substrates (Põlme et al. 2020) | Põlme S, et al. 2020. Fungal Diversity 105:1-16 ((https://doi.org/10.1007/s13225-020-00466-2)) |
| Otu40750 | 1 | k__Fungi(100);p__Ascomycota(100);c__Saccharomycetes(100);o__Saccharomycetales(100);f__Metschnikowiaceae(100);g__Metschnikowia(100);s__Metschnikowia_pulcherrima(100); | Metschnikowia | 13 | Saprotroph | \|Undefined Saprotroph\| | Yeast | NULL | Probable | As nectar-tap_saprotroph (Põlme et al. 2020); Decay sugar-rich substrates (Põlme et al. 2020) | Põlme S, et al. 2020. Fungal Diversity 105:1-16 ((https://doi.org/10.1007/s13225-020-00466-2)) |
| Otu40751 | 1 | k__Fungi(100);p__Ascomycota(100);c__Saccharomycetes(100);o__Saccharomycetales(100);f__Metschnikowiaceae(100);g__Metschnikowia(100);s__Metschnikowia_pulcherrima(100); | Metschnikowia | 13 | Saprotroph | \|Undefined Saprotroph\| | Yeast | NULL | Probable | As nectar-tap_saprotroph (Põlme et al. 2020); Decay sugar-rich substrates (Põlme et al. 2020) | Põlme S, et al. 2020. Fungal Diversity 105:1-16 ((https://doi.org/10.1007/s13225-020-00466-2)) |
| Otu40752 | 1 | k__Fungi(100);p__Ascomycota(100);c__Saccharomycetes(100);o__Saccharomycetales(100);f__Metschnikowiaceae(100);g__Metschnikowia(100);s__Metschnikowia_pulcherrima(100); | Metschnikowia | 13 | Saprotroph | \|Undefined Saprotroph\| | Yeast | NULL | Probable | As nectar-tap_saprotroph (Põlme et al. 2020); Decay sugar-rich substrates (Põlme et al. 2020) | Põlme S, et al. 2020. Fungal Diversity 105:1-16 ((https://doi.org/10.1007/s13225-020-00466-2)) |
| Otu40753 | 1 | k__Fungi(100);p__Ascomycota(100);c__Saccharomycetes(100);o__Saccharomycetales(100);f__Metschnikowiaceae(100);g__Metschnikowia(100);s__Metschnikowia_pulcherrima(100); | Metschnikowia | 13 | Saprotroph | \|Undefined Saprotroph\| | Yeast | NULL | Probable | As nectar-tap_saprotroph (Põlme et al. 2020); Decay sugar-rich substrates (Põlme et al. 2020) | Põlme S, et al. 2020. Fungal Diversity 105:1-16 ((https://doi.org/10.1007/s13225-020-00466-2)) |
| Otu40754 | 1 | k__Fungi(100);p__Ascomycota(100);c__Saccharomycetes(100);o__Saccharomycetales(100);f__Metschnikowiaceae(100);g__Metschnikowia(100);s__Metschnikowia_pulcherrima(100); | Metschnikowia | 13 | Saprotroph | \|Undefined Saprotroph\| | Yeast | NULL | Probable | As nectar-tap_saprotroph (Põlme et al. 2020); Decay sugar-rich substrates (Põlme et al. 2020) | Põlme S, et al. 2020. Fungal Diversity 105:1-16 ((https://doi.org/10.1007/s13225-020-00466-2)) |
| Otu40755 | 1 | k__Fungi(100);p__Ascomycota(100);c__Saccharomycetes(100);o__Saccharomycetales(100);f__Metschnikowiaceae(100);g__Metschnikowia(100);s__Metschnikowia_pulcherrima(100); | Metschnikowia | 13 | Saprotroph | \|Undefined Saprotroph\| | Yeast | NULL | Probable | As nectar-tap_saprotroph (Põlme et al. 2020); Decay sugar-rich substrates (Põlme et al. 2020) | Põlme S, et al. 2020. Fungal Diversity 105:1-16 ((https://doi.org/10.1007/s13225-020-00466-2)) |
| Otu40756 | 1 | k__Fungi(100);p__Ascomycota(100);c__Saccharomycetes(100);o__Saccharomycetales(100);f__Metschnikowiaceae(100);g__Metschnikowia(100);g__Metschnikowia_unclassified(100); | Metschnikowia | 13 | Saprotroph | \|Undefined Saprotroph\| | Yeast | NULL | Probable | As nectar-tap_saprotroph (Põlme et al. 2020); Decay sugar-rich substrates (Põlme et al. 2020) | Põlme S, et al. 2020. Fungal Diversity 105:1-16 ((https://doi.org/10.1007/s13225-020-00466-2)) |
| Otu40757 | 1 | k__Fungi(100);p__Ascomycota(100);c__Saccharomycetes(100);o__Saccharomycetales(100);f__Metschnikowiaceae(100);g__Metschnikowia(100);s__Metschnikowia_pulcherrima(100); | Metschnikowia | 13 | Saprotroph | \|Undefined Saprotroph\| | Yeast | NULL | Probable | As nectar-tap_saprotroph (Põlme et al. 2020); Decay sugar-rich substrates (Põlme et al. 2020) | Põlme S, et al. 2020. Fungal Diversity 105:1-16 ((https://doi.org/10.1007/s13225-020-00466-2)) |
| Otu40758 | 1 | k__Fungi(100);p__Ascomycota(100);c__Saccharomycetes(100);o__Saccharomycetales(100);f__Metschnikowiaceae(100);g__Metschnikowia(100);g__Metschnikowia_unclassified(100); | Metschnikowia | 13 | Saprotroph | \|Undefined Saprotroph\| | Yeast | NULL | Probable | As nectar-tap_saprotroph (Põlme et al. 2020); Decay sugar-rich substrates (Põlme et al. 2020) | Põlme S, et al. 2020. Fungal Diversity 105:1-16 ((https://doi.org/10.1007/s13225-020-00466-2)) |
| Otu40759 | 1 | k__Fungi(100);p__Ascomycota(100);c__Saccharomycetes(100);o__Saccharomycetales(100);f__Metschnikowiaceae(100);g__Metschnikowia(100);s__Metschnikowia_pulcherrima(100); | Metschnikowia | 13 | Saprotroph | \|Undefined Saprotroph\| | Yeast | NULL | Probable | As nectar-tap_saprotroph (Põlme et al. 2020); Decay sugar-rich substrates (Põlme et al. 2020) | Põlme S, et al. 2020. Fungal Diversity 105:1-16 ((https://doi.org/10.1007/s13225-020-00466-2)) |
| Otu40760 | 1 | k__Fungi(100);p__Ascomycota(100);c__Saccharomycetes(100);o__Saccharomycetales(100);f__Metschnikowiaceae(100);g__Metschnikowia(100);s__Metschnikowia_pulcherrima(100); | Metschnikowia | 13 | Saprotroph | \|Undefined Saprotroph\| | Yeast | NULL | Probable | As nectar-tap_saprotroph (Põlme et al. 2020); Decay sugar-rich substrates (Põlme et al. 2020) | Põlme S, et al. 2020. Fungal Diversity 105:1-16 ((https://doi.org/10.1007/s13225-020-00466-2)) |
| Otu40761 | 1 | k__Fungi(100);p__Ascomycota(100);c__Saccharomycetes(100);o__Saccharomycetales(100);f__Metschnikowiaceae(100);g__Metschnikowia(100);s__Metschnikowia_pulcherrima(100); | Metschnikowia | 13 | Saprotroph | \|Undefined Saprotroph\| | Yeast | NULL | Probable | As nectar-tap_saprotroph (Põlme et al. 2020); Decay sugar-rich substrates (Põlme et al. 2020) | Põlme S, et al. 2020. Fungal Diversity 105:1-16 ((https://doi.org/10.1007/s13225-020-00466-2)) |
| Otu40762 | 1 | k__Fungi(100);p__Ascomycota(100);c__Saccharomycetes(100);o__Saccharomycetales(100);f__Metschnikowiaceae(100);g__Metschnikowia(100);s__Metschnikowia_pulcherrima(100); | Metschnikowia | 13 | Saprotroph | \|Undefined Saprotroph\| | Yeast | NULL | Probable | As nectar-tap_saprotroph (Põlme et al. 2020); Decay sugar-rich substrates (Põlme et al. 2020) | Põlme S, et al. 2020. Fungal Diversity 105:1-16 ((https://doi.org/10.1007/s13225-020-00466-2)) |
| Otu40763 | 1 | k__Fungi(100);p__Ascomycota(100);c__Saccharomycetes(100);o__Saccharomycetales(100);f__Metschnikowiaceae(100);g__Metschnikowia(100);s__Metschnikowia_pulcherrima(100); | Metschnikowia | 13 | Saprotroph | \|Undefined Saprotroph\| | Yeast | NULL | Probable | As nectar-tap_saprotroph (Põlme et al. 2020); Decay sugar-rich substrates (Põlme et al. 2020) | Põlme S, et al. 2020. Fungal Diversity 105:1-16 ((https://doi.org/10.1007/s13225-020-00466-2)) |
| Otu40764 | 1 | k__Fungi(100);p__Ascomycota(100);c__Saccharomycetes(100);o__Saccharomycetales(100);f__Metschnikowiaceae(100);g__Metschnikowia(100);g__Metschnikowia_unclassified(100); | Metschnikowia | 13 | Saprotroph | \|Undefined Saprotroph\| | Yeast | NULL | Probable | As nectar-tap_saprotroph (Põlme et al. 2020); Decay sugar-rich substrates (Põlme et al. 2020) | Põlme S, et al. 2020. Fungal Diversity 105:1-16 ((https://doi.org/10.1007/s13225-020-00466-2)) |
| Otu40765 | 1 | k__Fungi(100);p__Ascomycota(100);c__Saccharomycetes(100);o__Saccharomycetales(100);f__Metschnikowiaceae(100);g__Metschnikowia(100);s__Metschnikowia_pulcherrima(100); | Metschnikowia | 13 | Saprotroph | \|Undefined Saprotroph\| | Yeast | NULL | Probable | As nectar-tap_saprotroph (Põlme et al. 2020); Decay sugar-rich substrates (Põlme et al. 2020) | Põlme S, et al. 2020. Fungal Diversity 105:1-16 ((https://doi.org/10.1007/s13225-020-00466-2)) |
| Otu40766 | 1 | k__Fungi(100);p__Ascomycota(100);c__Saccharomycetes(100);o__Saccharomycetales(100);f__Metschnikowiaceae(100);g__Metschnikowia(100);s__Metschnikowia_pulcherrima(100); | Metschnikowia | 13 | Saprotroph | \|Undefined Saprotroph\| | Yeast | NULL | Probable | As nectar-tap_saprotroph (Põlme et al. 2020); Decay sugar-rich substrates (Põlme et al. 2020) | Põlme S, et al. 2020. Fungal Diversity 105:1-16 ((https://doi.org/10.1007/s13225-020-00466-2)) |
| Otu40767 | 1 | k__Fungi(100);p__Ascomycota(100);c__Saccharomycetes(100);o__Saccharomycetales(100);f__Metschnikowiaceae(100);g__Metschnikowia(100);s__Metschnikowia_pulcherrima(100); | Metschnikowia | 13 | Saprotroph | \|Undefined Saprotroph\| | Yeast | NULL | Probable | As nectar-tap_saprotroph (Põlme et al. 2020); Decay sugar-rich substrates (Põlme et al. 2020) | Põlme S, et al. 2020. Fungal Diversity 105:1-16 ((https://doi.org/10.1007/s13225-020-00466-2)) |
| Otu40768 | 1 | k__Fungi(100);p__Ascomycota(100);c__Saccharomycetes(100);o__Saccharomycetales(100);f__Metschnikowiaceae(100);g__Metschnikowia(100);s__Metschnikowia_pulcherrima(100); | Metschnikowia | 13 | Saprotroph | \|Undefined Saprotroph\| | Yeast | NULL | Probable | As nectar-tap_saprotroph (Põlme et al. 2020); Decay sugar-rich substrates (Põlme et al. 2020) | Põlme S, et al. 2020. Fungal Diversity 105:1-16 ((https://doi.org/10.1007/s13225-020-00466-2)) |
| Otu40769 | 1 | k__Fungi(100);p__Ascomycota(100);c__Saccharomycetes(100);o__Saccharomycetales(100);f__Metschnikowiaceae(100);g__Metschnikowia(100);s__Metschnikowia_pulcherrima(100); | Metschnikowia | 13 | Saprotroph | \|Undefined Saprotroph\| | Yeast | NULL | Probable | As nectar-tap_saprotroph (Põlme et al. 2020); Decay sugar-rich substrates (Põlme et al. 2020) | Põlme S, et al. 2020. Fungal Diversity 105:1-16 ((https://doi.org/10.1007/s13225-020-00466-2)) |
| Otu40770 | 1 | k__Fungi(100);p__Ascomycota(100);c__Saccharomycetes(100);o__Saccharomycetales(100);f__Metschnikowiaceae(100);g__Metschnikowia(100);g__Metschnikowia_unclassified(100); | Metschnikowia | 13 | Saprotroph | \|Undefined Saprotroph\| | Yeast | NULL | Probable | As nectar-tap_saprotroph (Põlme et al. 2020); Decay sugar-rich substrates (Põlme et al. 2020) | Põlme S, et al. 2020. Fungal Diversity 105:1-16 ((https://doi.org/10.1007/s13225-020-00466-2)) |
| Otu40771 | 1 | k__Fungi(100);p__Ascomycota(100);c__Saccharomycetes(100);o__Saccharomycetales(100);f__Metschnikowiaceae(100);g__Metschnikowia(100);s__Metschnikowia_pulcherrima(100); | Metschnikowia | 13 | Saprotroph | \|Undefined Saprotroph\| | Yeast | NULL | Probable | As nectar-tap_saprotroph (Põlme et al. 2020); Decay sugar-rich substrates (Põlme et al. 2020) | Põlme S, et al. 2020. Fungal Diversity 105:1-16 ((https://doi.org/10.1007/s13225-020-00466-2)) |

| | | | | | | | | | | | |
|---|---|---|---|---|---|---|---|---|---|---|---|
| Otu40773 | 1 | k__Fungi(100);p__Ascomycota(100);c__Saccharomycetes(100);o__Saccharomycetales(100);f__Metschnikowiaceae(100);g__Metschnikowia(100);s__Metschnikowia_pulcherrima(100); | Metschnikowia | 13 | Saprotroph | Undefined Saprotroph | Yeast | NULL | Probable | As nectar-tap_saprotroph (Põlme et al. 2020); Decay sugar-rich substrates (Põlme et al. 2020) | Põlme S, et al. 2020. Fungal Diversity 105:1-16 (https://doi.org/10.1007/s13225-020-00466-2) |
| Otu40774 | 1 | k__Fungi(100);p__Ascomycota(100);c__Saccharomycetes(100);o__Saccharomycetales(100);f__Metschnikowiaceae(100);g__Metschnikowia(100);s__Metschnikowia_pulcherrima(100); | Metschnikowia | 13 | Saprotroph | Undefined Saprotroph | Yeast | NULL | Probable | As nectar-tap_saprotroph (Põlme et al. 2020); Decay sugar-rich substrates (Põlme et al. 2020) | Põlme S, et al. 2020. Fungal Diversity 105:1-16 (https://doi.org/10.1007/s13225-020-00466-2) |
| Otu40775 | 1 | k__Fungi(100);p__Ascomycota(100);c__Saccharomycetes(100);o__Saccharomycetales(100);f__Metschnikowiaceae(100);g__Metschnikowia(100);s__Metschnikowia_pulcherrima(100); | Metschnikowia | 13 | Saprotroph | Undefined Saprotroph | Yeast | NULL | Probable | As nectar-tap_saprotroph (Põlme et al. 2020); Decay sugar-rich substrates (Põlme et al. 2020) | Põlme S, et al. 2020. Fungal Diversity 105:1-16 (https://doi.org/10.1007/s13225-020-00466-2) |
| Otu40776 | 1 | k__Fungi(100);p__Ascomycota(100);c__Saccharomycetes(100);o__Saccharomycetales(100);f__Metschnikowiaceae(100);g__Metschnikowia(100);g__Metschnikowia_unclassified(100); | Metschnikowia | 13 | Saprotroph | Undefined Saprotroph | Yeast | NULL | Probable | As nectar-tap_saprotroph (Põlme et al. 2020); Decay sugar-rich substrates (Põlme et al. 2020) | Põlme S, et al. 2020. Fungal Diversity 105:1-16 (https://doi.org/10.1007/s13225-020-00466-2) |
| Otu40777 | 1 | k__Fungi(100);p__Ascomycota(100);c__Saccharomycetes(100);o__Saccharomycetales(100);f__Metschnikowiaceae(100);g__Metschnikowia(100);g__Metschnikowia_unclassified(100); | Metschnikowia | 13 | Saprotroph | Undefined Saprotroph | Yeast | NULL | Probable | As nectar-tap_saprotroph (Põlme et al. 2020); Decay sugar-rich substrates (Põlme et al. 2020) | Põlme S, et al. 2020. Fungal Diversity 105:1-16 (https://doi.org/10.1007/s13225-020-00466-2) |
| Otu40778 | 1 | k__Fungi(100);p__Ascomycota(100);c__Saccharomycetes(100);o__Saccharomycetales(100);f__Metschnikowiaceae(100);g__Metschnikowia(100);s__Metschnikowia_pulcherrima(100); | Metschnikowia | 13 | Saprotroph | Undefined Saprotroph | Yeast | NULL | Probable | As nectar-tap_saprotroph (Põlme et al. 2020); Decay sugar-rich substrates (Põlme et al. 2020) | Põlme S, et al. 2020. Fungal Diversity 105:1-16 (https://doi.org/10.1007/s13225-020-00466-2) |
| Otu40779 | 1 | k__Fungi(100);p__Ascomycota(100);c__Saccharomycetes(100);o__Saccharomycetales(100);f__Metschnikowiaceae(100);g__Metschnikowia(100);s__Metschnikowia_pulcherrima(100); | Metschnikowia | 13 | Saprotroph | Undefined Saprotroph | Yeast | NULL | Probable | As nectar-tap_saprotroph (Põlme et al. 2020); Decay sugar-rich substrates (Põlme et al. 2020) | Põlme S, et al. 2020. Fungal Diversity 105:1-16 (https://doi.org/10.1007/s13225-020-00466-2) |
| Otu40781 | 1 | k__Fungi(100);p__Ascomycota(100);c__Saccharomycetes(100);o__Saccharomycetales(100);f__Metschnikowiaceae(100);g__Metschnikowia(100);g__Metschnikowia_unclassified(100); | Metschnikowia | 13 | Saprotroph | Undefined Saprotroph | Yeast | NULL | Probable | As nectar-tap_saprotroph (Põlme et al. 2020); Decay sugar-rich substrates (Põlme et al. 2020) | Põlme S, et al. 2020. Fungal Diversity 105:1-16 (https://doi.org/10.1007/s13225-020-00466-2) |
| Otu40782 | 1 | k__Fungi(100);p__Ascomycota(100);c__Saccharomycetes(100);o__Saccharomycetales(100);f__Metschnikowiaceae(100);g__Metschnikowia(100);g__Metschnikowia_unclassified(100); | Metschnikowia | 13 | Saprotroph | Undefined Saprotroph | Yeast | NULL | Probable | As nectar-tap_saprotroph (Põlme et al. 2020); Decay sugar-rich substrates (Põlme et al. 2020) | Põlme S, et al. 2020. Fungal Diversity 105:1-16 (https://doi.org/10.1007/s13225-020-00466-2) |
| Otu40783 | 1 | k__Fungi(100);p__Ascomycota(100);c__Saccharomycetes(100);o__Saccharomycetales(100);f__Metschnikowiaceae(100);g__Metschnikowia(100);s__Metschnikowia_pulcherrima(100); | Metschnikowia | 13 | Saprotroph | Undefined Saprotroph | Yeast | NULL | Probable | As nectar-tap_saprotroph (Põlme et al. 2020); Decay sugar-rich substrates (Põlme et al. 2020) | Põlme S, et al. 2020. Fungal Diversity 105:1-16 (https://doi.org/10.1007/s13225-020-00466-2) |
| Otu40784 | 1 | k__Fungi(100);p__Ascomycota(100);c__Saccharomycetes(100);o__Saccharomycetales(100);f__Metschnikowiaceae(100);g__Metschnikowia(100);s__Metschnikowia_pulcherrima(100); | Metschnikowia | 13 | Saprotroph | Undefined Saprotroph | Yeast | NULL | Probable | As nectar-tap_saprotroph (Põlme et al. 2020); Decay sugar-rich substrates (Põlme et al. 2020) | Põlme S, et al. 2020. Fungal Diversity 105:1-16 (https://doi.org/10.1007/s13225-020-00466-2) |
| Otu40785 | 1 | k__Fungi(100);p__Ascomycota(100);c__Saccharomycetes(100);o__Saccharomycetales(100);f__Metschnikowiaceae(100);g__Metschnikowia(100);s__Metschnikowia_pulcherrima(100); | Metschnikowia | 13 | Saprotroph | Undefined Saprotroph | Yeast | NULL | Probable | As nectar-tap_saprotroph (Põlme et al. 2020); Decay sugar-rich substrates (Põlme et al. 2020) | Põlme S, et al. 2020. Fungal Diversity 105:1-16 (https://doi.org/10.1007/s13225-020-00466-2) |
| Otu40786 | 1 | k__Fungi(100);p__Ascomycota(100);c__Saccharomycetes(100);o__Saccharomycetales(100);f__Metschnikowiaceae(100);g__Metschnikowia(100);s__Metschnikowia_pulcherrima(100); | Metschnikowia | 13 | Saprotroph | Undefined Saprotroph | Yeast | NULL | Probable | As nectar-tap_saprotroph (Põlme et al. 2020); Decay sugar-rich substrates (Põlme et al. 2020) | Põlme S, et al. 2020. Fungal Diversity 105:1-16 (https://doi.org/10.1007/s13225-020-00466-2) |
| Otu40788 | 1 | k__Fungi(100);p__Ascomycota(100);c__Saccharomycetes(100);o__Saccharomycetales(100);f__Metschnikowiaceae(100);g__Metschnikowia(100);s__Metschnikowia_pulcherrima(100); | Metschnikowia | 13 | Saprotroph | Undefined Saprotroph | Yeast | NULL | Probable | As nectar-tap_saprotroph (Põlme et al. 2020); Decay sugar-rich substrates (Põlme et al. 2020) | Põlme S, et al. 2020. Fungal Diversity 105:1-16 (https://doi.org/10.1007/s13225-020-00466-2) |
| Otu40789 | 1 | k__Fungi(100);p__Ascomycota(100);c__Saccharomycetes(100);o__Saccharomycetales(100);f__Metschnikowiaceae(100);g__Metschnikowia(100);s__Metschnikowia_pulcherrima(100); | Metschnikowia | 13 | Saprotroph | Undefined Saprotroph | Yeast | NULL | Probable | As nectar-tap_saprotroph (Põlme et al. 2020); Decay sugar-rich substrates (Põlme et al. 2020) | Põlme S, et al. 2020. Fungal Diversity 105:1-16 (https://doi.org/10.1007/s13225-020-00466-2) |
| Otu40790 | 1 | k__Fungi(100);p__Ascomycota(100);c__Saccharomycetes(100);o__Saccharomycetales(100);f__Metschnikowiaceae(100);g__Metschnikowia(100);s__Metschnikowia_pulcherrima(100); | Metschnikowia | 13 | Saprotroph | Undefined Saprotroph | Yeast | NULL | Probable | As nectar-tap_saprotroph (Põlme et al. 2020); Decay sugar-rich substrates (Põlme et al. 2020) | Põlme S, et al. 2020. Fungal Diversity 105:1-16 (https://doi.org/10.1007/s13225-020-00466-2) |
| Otu40792 | 1 | k__Fungi(100);p__Ascomycota(100);c__Saccharomycetes(100);o__Saccharomycetales(100);f__Metschnikowiaceae(100);g__Metschnikowia(100);s__Metschnikowia_pulcherrima(100); | Metschnikowia | 13 | Saprotroph | Undefined Saprotroph | Yeast | NULL | Probable | As nectar-tap_saprotroph (Põlme et al. 2020); Decay sugar-rich substrates (Põlme et al. 2020) | Põlme S, et al. 2020. Fungal Diversity 105:1-16 (https://doi.org/10.1007/s13225-020-00466-2) |
| Otu40793 | 1 | k__Fungi(100);p__Ascomycota(100);c__Saccharomycetes(100);o__Saccharomycetales(100);f__Metschnikowiaceae(100);g__Metschnikowia(100);s__Metschnikowia_pulcherrima(100); | Metschnikowia | 13 | Saprotroph | Undefined Saprotroph | Yeast | NULL | Probable | As nectar-tap_saprotroph (Põlme et al. 2020); Decay sugar-rich substrates (Põlme et al. 2020) | Põlme S, et al. 2020. Fungal Diversity 105:1-16 (https://doi.org/10.1007/s13225-020-00466-2) |
| Otu40794 | 1 | k__Fungi(100);p__Ascomycota(100);c__Saccharomycetes(100);o__Saccharomycetales(100);f__Metschnikowiaceae(100);g__Metschnikowia(100);s__Metschnikowia_pulcherrima(100); | Metschnikowia | 13 | Saprotroph | Undefined Saprotroph | Yeast | NULL | Probable | As nectar-tap_saprotroph (Põlme et al. 2020); Decay sugar-rich substrates (Põlme et al. 2020) | Põlme S, et al. 2020. Fungal Diversity 105:1-16 (https://doi.org/10.1007/s13225-020-00466-2) |
| Otu40796 | 1 | k__Fungi(100);p__Ascomycota(100);c__Saccharomycetes(100);o__Saccharomycetales(100);f__Metschnikowiaceae(100);g__Metschnikowia(100);s__Metschnikowia_pulcherrima(100); | Metschnikowia | 13 | Saprotroph | Undefined Saprotroph | Yeast | NULL | Probable | As nectar-tap_saprotroph (Põlme et al. 2020); Decay sugar-rich substrates (Põlme et al. 2020) | Põlme S, et al. 2020. Fungal Diversity 105:1-16 (https://doi.org/10.1007/s13225-020-00466-2) |
| Otu40797 | 1 | k__Fungi(100);p__Ascomycota(100);c__Saccharomycetes(100);o__Saccharomycetales(100);f__Metschnikowiaceae(100);g__Metschnikowia(100);s__Metschnikowia_pulcherrima(100); | Metschnikowia | 13 | Saprotroph | Undefined Saprotroph | Yeast | NULL | Probable | As nectar-tap_saprotroph (Põlme et al. 2020); Decay sugar-rich substrates (Põlme et al. 2020) | Põlme S, et al. 2020. Fungal Diversity 105:1-16 (https://doi.org/10.1007/s13225-020-00466-2) |
| Otu40798 | 1 | k__Fungi(100);p__Ascomycota(100);c__Saccharomycetes(100);o__Saccharomycetales(100);f__Metschnikowiaceae(100);g__Metschnikowia(100);g__Metschnikowia_unclassified(100); | Metschnikowia | 13 | Saprotroph | Undefined Saprotroph | Yeast | NULL | Probable | As nectar-tap_saprotroph (Põlme et al. 2020); Decay sugar-rich substrates (Põlme et al. 2020) | Põlme S, et al. 2020. Fungal Diversity 105:1-16 (https://doi.org/10.1007/s13225-020-00466-2) |
| Otu40799 | 1 | k__Fungi(100);p__Ascomycota(100);c__Saccharomycetes(100);o__Saccharomycetales(100);f__Metschnikowiaceae(100);g__Metschnikowia(100);g__Metschnikowia_unclassified(100); | Metschnikowia | 13 | Saprotroph | Undefined Saprotroph | Yeast | NULL | Probable | As nectar-tap_saprotroph (Põlme et al. 2020); Decay sugar-rich substrates (Põlme et al. 2020) | Põlme S, et al. 2020. Fungal Diversity 105:1-16 (https://doi.org/10.1007/s13225-020-00466-2) |
| Otu40800 | 1 | k__Fungi(100);p__Ascomycota(100);c__Saccharomycetes(100);o__Saccharomycetales(100);f__Metschnikowiaceae(100);g__Metschnikowia(100);g__Metschnikowia_unclassified(100); | Metschnikowia | 13 | Saprotroph | Undefined Saprotroph | Yeast | NULL | Probable | As nectar-tap_saprotroph (Põlme et al. 2020); Decay sugar-rich substrates (Põlme et al. 2020) | Põlme S, et al. 2020. Fungal Diversity 105:1-16 (https://doi.org/10.1007/s13225-020-00466-2) |
| Otu40801 | 1 | k__Fungi(100);p__Ascomycota(100);c__Saccharomycetes(100);o__Saccharomycetales(100);f__Metschnikowiaceae(100);g__Metschnikowia(100);s__Metschnikowia_pulcherrima(100); | Metschnikowia | 13 | Saprotroph | Undefined Saprotroph | Yeast | NULL | Probable | As nectar-tap_saprotroph (Põlme et al. 2020); Decay sugar-rich substrates (Põlme et al. 2020) | Põlme S, et al. 2020. Fungal Diversity 105:1-16 (https://doi.org/10.1007/s13225-020-00466-2) |
| Otu40802 | 1 | k__Fungi(100);p__Ascomycota(100);c__Saccharomycetes(100);o__Saccharomycetales(100);f__Metschnikowiaceae(100);g__Metschnikowia(100);s__Metschnikowia_pulcherrima(100); | Metschnikowia | 13 | Saprotroph | Undefined Saprotroph | Yeast | NULL | Probable | As nectar-tap_saprotroph (Põlme et al. 2020); Decay sugar-rich substrates (Põlme et al. 2020) | Põlme S, et al. 2020. Fungal Diversity 105:1-16 (https://doi.org/10.1007/s13225-020-00466-2) |
| Otu40803 | 1 | k__Fungi(100);p__Ascomycota(100);c__Saccharomycetes(100);o__Saccharomycetales(100);f__Metschnikowiaceae(100);g__Metschnikowia(100);g__Metschnikowia_unclassified(100); | Metschnikowia | 13 | Saprotroph | Undefined Saprotroph | Yeast | NULL | Probable | As nectar-tap_saprotroph (Põlme et al. 2020); Decay sugar-rich substrates (Põlme et al. 2020) | Põlme S, et al. 2020. Fungal Diversity 105:1-16 (https://doi.org/10.1007/s13225-020-00466-2) |
| Otu40805 | 1 | k__Fungi(100);p__Ascomycota(100);c__Saccharomycetes(100);o__Saccharomycetales(100);f__Metschnikowiaceae(100);g__Metschnikowia(100);s__Metschnikowia_pulcherrima(100); | Metschnikowia | 13 | Saprotroph | Undefined Saprotroph | Yeast | NULL | Probable | As nectar-tap_saprotroph (Põlme et al. 2020); Decay sugar-rich substrates (Põlme et al. 2020) | Põlme S, et al. 2020. Fungal Diversity 105:1-16 (https://doi.org/10.1007/s13225-020-00466-2) |
| Otu40807 | 1 | k__Fungi(100);p__Ascomycota(100);c__Saccharomycetes(100);o__Saccharomycetales(100);f__Metschnikowiaceae(100);g__Metschnikowia(100);s__Metschnikowia_pulcherrima(100); | Metschnikowia | 13 | Saprotroph | Undefined Saprotroph | Yeast | NULL | Probable | As nectar-tap_saprotroph (Põlme et al. 2020); Decay sugar-rich substrates (Põlme et al. 2020) | Põlme S, et al. 2020. Fungal Diversity 105:1-16 (https://doi.org/10.1007/s13225-020-00466-2) |
| Otu40808 | 1 | k__Fungi(100);p__Ascomycota(100);c__Saccharomycetes(100);o__Saccharomycetales(100);f__Metschnikowiaceae(100);g__Metschnikowia(100);s__Metschnikowia_pulcherrima(100); | Metschnikowia | 13 | Saprotroph | Undefined Saprotroph | Yeast | NULL | Probable | As nectar-tap_saprotroph (Põlme et al. 2020); Decay sugar-rich substrates (Põlme et al. 2020) | Põlme S, et al. 2020. Fungal Diversity 105:1-16 (https://doi.org/10.1007/s13225-020-00466-2) |
| Otu40811 | 1 | k__Fungi(100);p__Ascomycota(100);c__Saccharomycetes(100);o__Saccharomycetales(100);f__Metschnikowiaceae(100);g__Metschnikowia(100);g__Metschnikowia_unclassified(100); | Metschnikowia | 13 | Saprotroph | Undefined Saprotroph | Yeast | NULL | Probable | As nectar-tap_saprotroph (Põlme et al. 2020); Decay sugar-rich substrates (Põlme et al. 2020) | Põlme S, et al. 2020. Fungal Diversity 105:1-16 (https://doi.org/10.1007/s13225-020-00466-2) |
| Otu40812 | 1 | k__Fungi(100);p__Ascomycota(100);c__Saccharomycetes(100);o__Saccharomycetales(100);f__Metschnikowiaceae(100);g__Metschnikowia(100);s__Metschnikowia_pulcherrima(100); | Metschnikowia | 13 | Saprotroph | Undefined Saprotroph | Yeast | NULL | Probable | As nectar-tap_saprotroph (Põlme et al. 2020); Decay sugar-rich substrates (Põlme et al. 2020) | Põlme S, et al. 2020. Fungal Diversity 105:1-16 (https://doi.org/10.1007/s13225-020-00466-2) |
| Otu40813 | 1 | k__Fungi(100);p__Ascomycota(100);c__Saccharomycetes(100);o__Saccharomycetales(100);f__Metschnikowiaceae(100);g__Metschnikowia(100);g__Metschnikowia_unclassified(100); | Metschnikowia | 13 | Saprotroph | Undefined Saprotroph | Yeast | NULL | Probable | As nectar-tap_saprotroph (Põlme et al. 2020); Decay sugar-rich substrates (Põlme et al. 2020) | Põlme S, et al. 2020. Fungal Diversity 105:1-16 (https://doi.org/10.1007/s13225-020-00466-2) |
| Otu40814 | 1 | k__Fungi(100);p__Ascomycota(100);c__Saccharomycetes(100);o__Saccharomycetales(100);f__Metschnikowiaceae(100);g__Metschnikowia(100);g__Metschnikowia_unclassified(100); | Metschnikowia | 13 | Saprotroph | Undefined Saprotroph | Yeast | NULL | Probable | As nectar-tap_saprotroph (Põlme et al. 2020); Decay sugar-rich substrates (Põlme et al. 2020) | Põlme S, et al. 2020. Fungal Diversity 105:1-16 (https://doi.org/10.1007/s13225-020-00466-2) |
| Otu40815 | 1 | k__Fungi(100);p__Ascomycota(100);c__Saccharomycetes(100);o__Saccharomycetales(100);f__Metschnikowiaceae(100);g__Metschnikowia(100);s__Metschnikowia_pulcherrima(100); | Metschnikowia | 13 | Saprotroph | Undefined Saprotroph | Yeast | NULL | Probable | As nectar-tap_saprotroph (Põlme et al. 2020); Decay sugar-rich substrates (Põlme et al. 2020) | Põlme S, et al. 2020. Fungal Diversity 105:1-16 (https://doi.org/10.1007/s13225-020-00466-2) |
| Otu40816 | 1 | k__Fungi(100);p__Ascomycota(100);c__Saccharomycetes(100);o__Saccharomycetales(100);f__Metschnikowiaceae(100);g__Metschnikowia(100);s__Metschnikowia_pulcherrima(100); | Metschnikowia | 13 | Saprotroph | Undefined Saprotroph | Yeast | NULL | Probable | As nectar-tap_saprotroph (Põlme et al. 2020); Decay sugar-rich substrates (Põlme et al. 2020) | Põlme S, et al. 2020. Fungal Diversity 105:1-16 (https://doi.org/10.1007/s13225-020-00466-2) |
| Otu40817 | 1 | k__Fungi(100);p__Ascomycota(100);c__Saccharomycetes(100);o__Saccharomycetales(100);f__Metschnikowiaceae(100);g__Metschnikowia(100);g__Metschnikowia_unclassified(100); | Metschnikowia | 13 | Saprotroph | Undefined Saprotroph | Yeast | NULL | Probable | As nectar-tap_saprotroph (Põlme et al. 2020); Decay sugar-rich substrates (Põlme et al. 2020) | Põlme S, et al. 2020. Fungal Diversity 105:1-16 (https://doi.org/10.1007/s13225-020-00466-2) |
| Otu40818 | 1 | k__Fungi(100);p__Ascomycota(100);c__Saccharomycetes(100);o__Saccharomycetales(100);f__Metschnikowiaceae(100);g__Metschnikowia(100);s__Metschnikowia_pulcherrima(100); | Metschnikowia | 13 | Saprotroph | Undefined Saprotroph | Yeast | NULL | Probable | As nectar-tap_saprotroph (Põlme et al. 2020); Decay sugar-rich substrates (Põlme et al. 2020) | Põlme S, et al. 2020. Fungal Diversity 105:1-16 (https://doi.org/10.1007/s13225-020-00466-2) |
| Otu40819 | 1 | k__Fungi(100);p__Ascomycota(100);c__Saccharomycetes(100);o__Saccharomycetales(100);f__Metschnikowiaceae(100);g__Metschnikowia(100);g__Metschnikowia_unclassified(100); | Metschnikowia | 13 | Saprotroph | Undefined Saprotroph | Yeast | NULL | Probable | As nectar-tap_saprotroph (Põlme et al. 2020); Decay sugar-rich substrates (Põlme et al. 2020) | Põlme S, et al. 2020. Fungal Diversity 105:1-16 (https://doi.org/10.1007/s13225-020-00466-2) |
| Otu40820 | 1 | k__Fungi(100);p__Ascomycota(100);c__Saccharomycetes(100);o__Saccharomycetales(100);f__Metschnikowiaceae(100);g__Metschnikowia(100);g__Metschnikowia_unclassified(100); | Metschnikowia | 13 | Saprotroph | Undefined Saprotroph | Yeast | NULL | Probable | As nectar-tap_saprotroph (Põlme et al. 2020); Decay sugar-rich substrates (Põlme et al. 2020) | Põlme S, et al. 2020. Fungal Diversity 105:1-16 (https://doi.org/10.1007/s13225-020-00466-2) |
| Otu40822 | 1 | k__Fungi(100);p__Ascomycota(100);c__Saccharomycetes(100);o__Saccharomycetales(100);f__Metschnikowiaceae(100);g__Metschnikowia(100);g__Metschnikowia_unclassified(100); | Metschnikowia | 13 | Saprotroph | Undefined Saprotroph | Yeast | NULL | Probable | As nectar-tap_saprotroph (Põlme et al. 2020); Decay sugar-rich substrates (Põlme et al. 2020) | Põlme S, et al. 2020. Fungal Diversity 105:1-16 (https://doi.org/10.1007/s13225-020-00466-2) |
| Otu40824 | 1 | k__Fungi(100);p__Ascomycota(100);c__Saccharomycetes(100);o__Saccharomycetales(100);f__Metschnikowiaceae(100);g__Metschnikowia(100);g__Metschnikowia_unclassified(100); | Metschnikowia | 13 | Saprotroph | Undefined Saprotroph | Yeast | NULL | Probable | As nectar-tap_saprotroph (Põlme et al. 2020); Decay sugar-rich substrates (Põlme et al. 2020) | Põlme S, et al. 2020. Fungal Diversity 105:1-16 (https://doi.org/10.1007/s13225-020-00466-2) |
| Otu40826 | 1 | k__Fungi(100);p__Ascomycota(100);c__Saccharomycetes(100);o__Saccharomycetales(100);f__Metschnikowiaceae(100);g__Metschnikowia(100);g__Metschnikowia_unclassified(100); | Metschnikowia | 13 | Saprotroph | Undefined Saprotroph | Yeast | NULL | Probable | As nectar-tap_saprotroph (Põlme et al. 2020); Decay sugar-rich substrates (Põlme et al. 2020) | Põlme S, et al. 2020. Fungal Diversity 105:1-16 (https://doi.org/10.1007/s13225-020-00466-2) |
| Otu40827 | 1 | k__Fungi(100);p__Ascomycota(100);c__Saccharomycetes(100);o__Saccharomycetales(100);f__Metschnikowiaceae(100);g__Metschnikowia(100);g__Metschnikowia_unclassified(100); | Metschnikowia | 13 | Saprotroph | Undefined Saprotroph | Yeast | NULL | Probable | As nectar-tap_saprotroph (Põlme et al. 2020); Decay sugar-rich substrates (Põlme et al. 2020) | Põlme S, et al. 2020. Fungal Diversity 105:1-16 (https://doi.org/10.1007/s13225-020-00466-2) |
| Otu40829 | 1 | k__Fungi(100);p__Ascomycota(100);c__Saccharomycetes(100);o__Saccharomycetales(100);f__Metschnikowiaceae(100);g__Metschnikowia(100);g__Metschnikowia_unclassified(100); | Metschnikowia | 13 | Saprotroph | Undefined Saprotroph | Yeast | NULL | Probable | As nectar-tap_saprotroph (Põlme et al. 2020); Decay sugar-rich substrates (Põlme et al. 2020) | Põlme S, et al. 2020. Fungal Diversity 105:1-16 (https://doi.org/10.1007/s13225-020-00466-2) |
| Otu40830 | 1 | k__Fungi(100);p__Ascomycota(100);c__Saccharomycetes(100);o__Saccharomycetales(100);f__Metschnikowiaceae(100);g__Metschnikowia(100);g__Metschnikowia_unclassified(100); | Metschnikowia | 13 | Saprotroph | Undefined Saprotroph | Yeast | NULL | Probable | As nectar-tap_saprotroph (Põlme et al. 2020); Decay sugar-rich substrates (Põlme et al. 2020) | Põlme S, et al. 2020. Fungal Diversity 105:1-16 (https://doi.org/10.1007/s13225-020-00466-2) |
| Otu40831 | 1 | k__Fungi(100);p__Ascomycota(100);c__Saccharomycetes(100);o__Saccharomycetales(100);f__Metschnikowiaceae(100);g__Metschnikowia(100);g__Metschnikowia_unclassified(100); | Metschnikowia | 13 | Saprotroph | Undefined Saprotroph | Yeast | NULL | Probable | As nectar-tap_saprotroph (Põlme et al. 2020); Decay sugar-rich substrates (Põlme et al. 2020) | Põlme S, et al. 2020. Fungal Diversity 105:1-16 (https://doi.org/10.1007/s13225-020-00466-2) |
| Otu40833 | 1 | k__Fungi(100);p__Ascomycota(100);c__Saccharomycetes(100);o__Saccharomycetales(100);f__Metschnikowiaceae(100);g__Metschnikowia(100);g__Metschnikowia_unclassified(100); | Metschnikowia | 13 | Saprotroph | Undefined Saprotroph | Yeast | NULL | Probable | As nectar-tap_saprotroph (Põlme et al. 2020); Decay sugar-rich substrates (Põlme et al. 2020) | Põlme S, et al. 2020. Fungal Diversity 105:1-16 (https://doi.org/10.1007/s13225-020-00466-2) |
| Otu40834 | 1 | k__Fungi(100);p__Ascomycota(100);c__Saccharomycetes(100);o__Saccharomycetales(100);f__Metschnikowiaceae(100);g__Metschnikowia(100);g__Metschnikowia_unclassified(100); | Metschnikowia | 13 | Saprotroph | Undefined Saprotroph | Yeast | NULL | Probable | As nectar-tap_saprotroph (Põlme et al. 2020); Decay sugar-rich substrates (Põlme et al. 2020) | Põlme S, et al. 2020. Fungal Diversity 105:1-16 (https://doi.org/10.1007/s13225-020-00466-2) |
| Otu40835 | 1 | k__Fungi(100);p__Ascomycota(100);c__Saccharomycetes(100);o__Saccharomycetales(100);f__Metschnikowiaceae(100);g__Metschnikowia(100);g__Metschnikowia_unclassified(100); | Metschnikowia | 13 | Saprotroph | Undefined Saprotroph | Yeast | NULL | Probable | As nectar-tap_saprotroph (Põlme et al. 2020); Decay sugar-rich substrates (Põlme et al. 2020) | Põlme S, et al. 2020. Fungal Diversity 105:1-16 (https://doi.org/10.1007/s13225-020-00466-2) |
| Otu40836 | 1 | k__Fungi(100);p__Ascomycota(100);c__Saccharomycetes(100);o__Saccharomycetales(100);f__Metschnikowiaceae(100);g__Metschnikowia(100);s__Metschnikowia_pulcherrima(100); | Metschnikowia | 13 | Saprotroph | Undefined Saprotroph | Yeast | NULL | Probable | As nectar-tap_saprotroph (Põlme et al. 2020); Decay sugar-rich substrates (Põlme et al. 2020) | Põlme S, et al. 2020. Fungal Diversity 105:1-16 (https://doi.org/10.1007/s13225-020-00466-2) |
| Otu40837 | 1 | k__Fungi(100);p__Ascomycota(100);c__Saccharomycetes(100);o__Saccharomycetales(100);f__Metschnikowiaceae(100);g__Metschnikowia(100);s__Metschnikowia_pulcherrima(100); | Metschnikowia | 13 | Saprotroph | Undefined Saprotroph | Yeast | NULL | Probable | As nectar-tap_saprotroph (Põlme et al. 2020); Decay sugar-rich substrates (Põlme et al. 2020) | Põlme S, et al. 2020. Fungal Diversity 105:1-16 (https://doi.org/10.1007/s13225-020-00466-2) |
| Otu40838 | 1 | k__Fungi(100);p__Ascomycota(100);c__Saccharomycetes(100);o__Saccharomycetales(100);f__Metschnikowiaceae(100);g__Metschnikowia(100);g__Metschnikowia_unclassified(100); | Metschnikowia | 13 | Saprotroph | Undefined Saprotroph | Yeast | NULL | Probable | As nectar-tap_saprotroph (Põlme et al. 2020); Decay sugar-rich substrates (Põlme et al. 2020) | Põlme S, et al. 2020. Fungal Diversity 105:1-16 (https://doi.org/10.1007/s13225-020-00466-2) |
| Otu40839 | 1 | k__Fungi(100);p__Ascomycota(100);c__Saccharomycetes(100);o__Saccharomycetales(100);f__Metschnikowiaceae(100);g__Metschnikowia(100);s__Metschnikowia_pulcherrima(100); | Metschnikowia | 13 | Saprotroph | Undefined Saprotroph | Yeast | NULL | Probable | As nectar-tap_saprotroph (Põlme et al. 2020); Decay sugar-rich substrates (Põlme et al. 2020) | Põlme S, et al. 2020. Fungal Diversity 105:1-16 (https://doi.org/10.1007/s13225-020-00466-2) |
| Otu40840 | 1 | k__Fungi(100);p__Ascomycota(100);c__Saccharomycetes(100);o__Saccharomycetales(100);f__Metschnikowiaceae(100);g__Metschnikowia(100);s__Metschnikowia_pulcherrima(100); | Metschnikowia | 13 | Saprotroph | Undefined Saprotroph | Yeast | NULL | Probable | As nectar-tap_saprotroph (Põlme et al. 2020); Decay sugar-rich substrates (Põlme et al. 2020) | Põlme S, et al. 2020. Fungal Diversity 105:1-16 (https://doi.org/10.1007/s13225-020-00466-2) |
| Otu40841 | 1 | k__Fungi(100);p__Ascomycota(100);c__Saccharomycetes(100);o__Saccharomycetales(100);f__Metschnikowiaceae(100);g__Metschnikowia(100);s__Metschnikowia_pulcherrima(100); | Metschnikowia | 13 | Saprotroph | Undefined Saprotroph | Yeast | NULL | Probable | As nectar-tap_saprotroph (Põlme et al. 2020); Decay sugar-rich substrates (Põlme et al. 2020) | Põlme S, et al. 2020. Fungal Diversity 105:1-16 (https://doi.org/10.1007/s13225-020-00466-2) |
| Otu40842 | 1 | k__Fungi(100);p__Ascomycota(100);c__Saccharomycetes(100);o__Saccharomycetales(100);f__Metschnikowiaceae(100);g__Metschnikowia(100);s__Metschnikowia_pulcherrima(100); | Metschnikowia | 13 | Saprotroph | Undefined Saprotroph | Yeast | NULL | Probable | As nectar-tap_saprotroph (Põlme et al. 2020); Decay sugar-rich substrates (Põlme et al. 2020) | Põlme S, et al. 2020. Fungal Diversity 105:1-16 (https://doi.org/10.1007/s13225-020-00466-2) |
| Otu40843 | 1 | k__Fungi(100);p__Ascomycota(100);c__Saccharomycetes(100);o__Saccharomycetales(100);f__Metschnikowiaceae(100);g__Metschnikowia(100);g__Metschnikowia_unclassified(100); | Metschnikowia | 13 | Saprotroph | Undefined Saprotroph | Yeast | NULL | Probable | As nectar-tap_saprotroph (Põlme et al. 2020); Decay sugar-rich substrates (Põlme et al. 2020) | Põlme S, et al. 2020. Fungal Diversity 105:1-16 (https://doi.org/10.1007/s13225-020-00466-2) |
| Otu40845 | 1 | k__Fungi(100);p__Ascomycota(100);c__Saccharomycetes(100);o__Saccharomycetales(100);f__Metschnikowiaceae(100);g__Metschnikowia(100);s__Metschnikowia_pulcherrima(100); | Metschnikowia | 13 | Saprotroph | Undefined Saprotroph | Yeast | NULL | Probable | As nectar-tap_saprotroph (Põlme et al. 2020); Decay sugar-rich substrates (Põlme et al. 2020) | Põlme S, et al. 2020. Fungal Diversity 105:1-16 (https://doi.org/10.1007/s13225-020-00466-2) |
| Otu40846 | 1 | k__Fungi(100);p__Ascomycota(100);c__Saccharomycetes(100);o__Saccharomycetales(100);f__Metschnikowiaceae(100);g__Metschnikowia(100);s__Metschnikowia_pulcherrima(100); | Metschnikowia | 13 | Saprotroph | Undefined Saprotroph | Yeast | NULL | Probable | As nectar-tap_saprotroph (Põlme et al. 2020); Decay sugar-rich substrates (Põlme et al. 2020) | Põlme S, et al. 2020. Fungal Diversity 105:1-16 (https://doi.org/10.1007/s13225-020-00466-2) |
| Otu40847 | 1 | k__Fungi(100);p__Ascomycota(100);c__Saccharomycetes(100);o__Saccharomycetales(100);f__Metschnikowiaceae(100);g__Metschnikowia(100);s__Metschnikowia_pulcherrima(100); | Metschnikowia | 13 | Saprotroph | Undefined Saprotroph | Yeast | NULL | Probable | As nectar-tap_saprotroph (Põlme et al. 2020); Decay sugar-rich substrates (Põlme et al. 2020) | Põlme S, et al. 2020. Fungal Diversity 105:1-16 (https://doi.org/10.1007/s13225-020-00466-2) |
| Otu40848 | 1 | k__Fungi(100);p__Ascomycota(100);c__Saccharomycetes(100);o__Saccharomycetales(100);f__Metschnikowiaceae(100);g__Metschnikowia(100);s__Metschnikowia_pulcherrima(100); | Metschnikowia | 13 | Saprotroph | Undefined Saprotroph | Yeast | NULL | Probable | As nectar-tap_saprotroph (Põlme et al. 2020); Decay sugar-rich substrates (Põlme et al. 2020) | Põlme S, et al. 2020. Fungal Diversity 105:1-16 (https://doi.org/10.1007/s13225-020-00466-2) |
| Otu40849 | 1 | k__Fungi(100);p__Ascomycota(100);c__Saccharomycetes(100);o__Saccharomycetales(100);f__Metschnikowiaceae(100);g__Metschnikowia(100);g__Metschnikowia_unclassified(100); | Metschnikowia | 13 | Saprotroph | Undefined Saprotroph | Yeast | NULL | Probable | As nectar-tap_saprotroph (Põlme et al. 2020); Decay sugar-rich substrates (Põlme et al. 2020) | Põlme S, et al. 2020. Fungal Diversity 105:1-16 (https://doi.org/10.1007/s13225-020-00466-2) |
| Otu40850 | 1 | k__Fungi(100);p__Ascomycota(100);c__Saccharomycetes(100);o__Saccharomycetales(100);f__Metschnikowiaceae(100);g__Metschnikowia(100);s__Metschnikowia_pulcherrima(100); | Metschnikowia | 13 | Saprotroph | Undefined Saprotroph | Yeast | NULL | Probable | As nectar-tap_saprotroph (Põlme et al. 2020); Decay sugar-rich substrates (Põlme et al. 2020) | Põlme S, et al. 2020. Fungal Diversity 105:1-16 (https://doi.org/10.1007/s13225-020-00466-2) |
| Otu40851 | 1 | k__Fungi(100);p__Ascomycota(100);c__Saccharomycetes(100);o__Saccharomycetales(100);f__Metschnikowiaceae(100);g__Metschnikowia(100);s__Metschnikowia_pulcherrima(100); | Metschnikowia | 13 | Saprotroph | Undefined Saprotroph | Yeast | NULL | Probable | As nectar-tap_saprotroph (Põlme et al. 2020); Decay sugar-rich substrates (Põlme et al. 2020) | Põlme S, et al. 2020. Fungal Diversity 105:1-16 (https://doi.org/10.1007/s13225-020-00466-2) |
| Otu40852 | 1 | k__Fungi(100);p__Ascomycota(100);c__Saccharomycetes(100);o__Saccharomycetales(100);f__Metschnikowiaceae(100);g__Metschnikowia(100);s__Metschnikowia_pulcherrima(100); | Metschnikowia | 13 | Saprotroph | Undefined Saprotroph | Yeast | NULL | Probable | As nectar-tap_saprotroph (Põlme et al. 2020); Decay sugar-rich substrates (Põlme et al. 2020) | Põlme S, et al. 2020. Fungal Diversity 105:1-16 (https://doi.org/10.1007/s13225-020-00466-2) |
| Otu40854 | 1 | k__Fungi(100);p__Ascomycota(100);c__Saccharomycetes(100);o__Saccharomycetales(100);f__Metschnikowiaceae(100);g__Metschnikowia(100);s__Metschnikowia_pulcherrima(100); | Metschnikowia | 13 | Saprotroph | Undefined Saprotroph | Yeast | NULL | Probable | As nectar-tap_saprotroph (Põlme et al. 2020); Decay sugar-rich substrates (Põlme et al. 2020) | Põlme S, et al. 2020. Fungal Diversity 105:1-16 (https://doi.org/10.1007/s13225-020-00466-2) |
| Otu40855 | 1 | k__Fungi(100);p__Ascomycota(100);c__Saccharomycetes(100);o__Saccharomycetales(100);f__Metschnikowiaceae(100);g__Metschnikowia(100);g__Metschnikowia_unclassified(100); | Metschnikowia | 13 | Saprotroph | Undefined Saprotroph | Yeast | NULL | Probable | As nectar-tap_saprotroph (Põlme et al. 2020); Decay sugar-rich substrates (Põlme et al. 2020) | Põlme S, et al. 2020. Fungal Diversity 105:1-16 (https://doi.org/10.1007/s13225-020-00466-2) |
| Otu40856 | 1 | k__Fungi(100);p__Ascomycota(100);c__Saccharomycetes(100);o__Saccharomycetales(100);f__Metschnikowiaceae(100);g__Metschnikowia(100);s__Metschnikowia_pulcherrima(100); | Metschnikowia | 13 | Saprotroph | Undefined Saprotroph | Yeast | NULL | Probable | As nectar-tap_saprotroph (Põlme et al. 2020); Decay sugar-rich substrates (Põlme et al. 2020) | Põlme S, et al. 2020. Fungal Diversity 105:1-16 (https://doi.org/10.1007/s13225-020-00466-2) |
| Otu40857 | 1 | k__Fungi(100);p__Ascomycota(100);c__Saccharomycetes(100);o__Saccharomycetales(100);f__Metschnikowiaceae(100);g__Metschnikowia(100);g__Metschnikowia_unclassified(100); | Metschnikowia | 13 | Saprotroph | Undefined Saprotroph | Yeast | NULL | Probable | As nectar-tap_saprotroph (Põlme et al. 2020); Decay sugar-rich substrates (Põlme et al. 2020) | Põlme S, et al. 2020. Fungal Diversity 105:1-16 (https://doi.org/10.1007/s13225-020-00466-2) |
| Otu40858 | 1 | k__Fungi(100);p__Ascomycota(100);c__Saccharomycetes(100);o__Saccharomycetales(100);f__Metschnikowiaceae(100);g__Metschnikowia(100);g__Metschnikowia_unclassified(100); | Metschnikowia | 13 | Saprotroph | Undefined Saprotroph | Yeast | NULL | Probable | As nectar-tap_saprotroph (Põlme et al. 2020); Decay sugar-rich substrates (Põlme et al. 2020) | Põlme S, et al. 2020. Fungal Diversity 105:1-16 (https://doi.org/10.1007/s13225-020-00466-2) |
| Otu40859 | 1 | k__Fungi(100);p__Ascomycota(100);c__Saccharomycetes(100);o__Saccharomycetales(100);f__Metschnikowiaceae(100);g__Metschnikowia(100);s__Metschnikowia_pulcherrima(100); | Metschnikowia | 13 | Saprotroph | Undefined Saprotroph | Yeast | NULL | Probable | As nectar-tap_saprotroph (Põlme et al. 2020); Decay sugar-rich substrates (Põlme et al. 2020) | Põlme S, et al. 2020. Fungal Diversity 105:1-16 (https://doi.org/10.1007/s13225-020-00466-2) |
| Otu40860 | 1 | k__Fungi(100);p__Ascomycota(100);c__Saccharomycetes(100);o__Saccharomycetales(100);f__Metschnikowiaceae(100);g__Metschnikowia(100);s__Metschnikowia_pulcherrima(100); | Metschnikowia | 13 | Saprotroph | Undefined Saprotroph | Yeast | NULL | Probable | As nectar-tap_saprotroph (Põlme et al. 2020); Decay sugar-rich substrates (Põlme et al. 2020) | Põlme S, et al. 2020. Fungal Diversity 105:1-16 (https://doi.org/10.1007/s13225-020-00466-2) |
| Otu40861 | 1 | k__Fungi(100);p__Ascomycota(100);c__Saccharomycetes(100);o__Saccharomycetales(100);f__Metschnikowiaceae(100);g__Metschnikowia(100);g__Metschnikowia_unclassified(100); | Metschnikowia | 13 | Saprotroph | Undefined Saprotroph | Yeast | NULL | Probable | As nectar-tap_saprotroph (Põlme et al. 2020); Decay sugar-rich substrates (Põlme et al. 2020) | Põlme S, et al. 2020. Fungal Diversity 105:1-16 (https://doi.org/10.1007/s13225-020-00466-2) |
| Otu40862 | 1 | k__Fungi(100);p__Ascomycota(100);c__Saccharomycetes(100);o__Saccharomycetales(100);f__Metschnikowiaceae(100);g__Metschnikowia(100);s__Metschnikowia_pulcherrima(100); | Metschnikowia | 13 | Saprotroph | Undefined Saprotroph | Yeast | NULL | Probable | As nectar-tap_saprotroph (Põlme et al. 2020); Decay sugar-rich substrates (Põlme et al. 2020) | Põlme S, et al. 2020. Fungal Diversity 105:1-16 (https://doi.org/10.1007/s13225-020-00466-2) |
| Otu40863 | 1 | k__Fungi(100);p__Ascomycota(100);c__Saccharomycetes(100);o__Saccharomycetales(100);f__Metschnikowiaceae(100);g__Metschnikowia(100);g__Metschnikowia_unclassified(100); | Metschnikowia | 13 | Saprotroph | Undefined Saprotroph | Yeast | NULL | Probable | As nectar-tap_saprotroph (Põlme et al. 2020); Decay sugar-rich substrates (Põlme et al. 2020) | Põlme S, et al. 2020. Fungal Diversity 105:1-16 (https://doi.org/10.1007/s13225-020-00466-2) |
| Otu40864 | 1 | k__Fungi(100);p__Ascomycota(100);c__Saccharomycetes(100);o__Saccharomycetales(100);f__Metschnikowiaceae(100);g__Metschnikowia(100);g__Metschnikowia_unclassified(100); | Metschnikowia | 13 | Saprotroph | Undefined Saprotroph | Yeast | NULL | Probable | As nectar-tap_saprotroph (Põlme et al. 2020); Decay sugar-rich substrates (Põlme et al. 2020) | Põlme S, et al. 2020. Fungal Diversity 105:1-16 (https://doi.org/10.1007/s13225-020-00466-2) |
| Otu40865 | 1 | k__Fungi(100);p__Ascomycota(100);c__Saccharomycetes(100);o__Saccharomycetales(100);f__Metschnikowiaceae(100);g__Metschnikowia(100);g__Metschnikowia_unclassified(100); | Metschnikowia | 13 | Saprotroph | Undefined Saprotroph | Yeast | NULL | Probable | As nectar-tap_saprotroph (Põlme et al. 2020); Decay sugar-rich substrates (Põlme et al. 2020) | Põlme S, et al. 2020. Fungal Diversity 105:1-16 (https://doi.org/10.1007/s13225-020-00466-2) |
| Otu40866 | 1 | k__Fungi(100);p__Ascomycota(100);c__Saccharomycetes(100);o__Saccharomycetales(100);f__Metschnikowiaceae(100);g__Metschnikowia(100);g__Metschnikowia_unclassified(100); | Metschnikowia | 13 | Saprotroph | Undefined Saprotroph | Yeast | NULL | Probable | As nectar-tap_saprotroph (Põlme et al. 2020); Decay sugar-rich substrates (Põlme et al. 2020) | Põlme S, et al. 2020. Fungal Diversity 105:1-16 (https://doi.org/10.1007/s13225-020-00466-2) |
| Otu40867 | 1 | k__Fungi(100);p__Ascomycota(100);c__Saccharomycetes(100);o__Saccharomycetales(100);f__Metschnikowiaceae(100);g__Metschnikowia(100);g__Metschnikowia_unclassified(100); | Metschnikowia | 13 | Saprotroph | Undefined Saprotroph | Yeast | NULL | Probable | As nectar-tap_saprotroph (Põlme et al. 2020); Decay sugar-rich substrates (Põlme et al. 2020) | Põlme S, et al. 2020. Fungal Diversity 105:1-16 (https://doi.org/10.1007/s13225-020-00466-2) |
| Otu40868 | 1 | k__Fungi(100);p__Ascomycota(100);c__Saccharomycetes(100);o__Saccharomycetales(100);f__Metschnikowiaceae(100);g__Metschnikowia(100);g__Metschnikowia_unclassified(100); | Metschnikowia | 13 | Saprotroph | Undefined Saprotroph | Yeast | NULL | Probable | As nectar-tap_saprotroph (Põlme et al. 2020); Decay sugar-rich substrates (Põlme et al. 2020) | Põlme S, et al. 2020. Fungal Diversity 105:1-16 (https://doi.org/10.1007/s13225-020-00466-2) |
| Otu40869 | 1 | k__Fungi(100);p__Ascomycota(100);c__Saccharomycetes(100);o__Saccharomycetales(100);f__Metschnikowiaceae(100);g__Metschnikowia(100);s__Metschnikowia_pulcherrima(100); | Metschnikowia | 13 | Saprotroph | Undefined Saprotroph | Yeast | NULL | Probable | As nectar-tap_saprotroph (Põlme et al. 2020); Decay sugar-rich substrates (Põlme et al. 2020) | Põlme S, et al. 2020. Fungal Diversity 105:1-16 (https://doi.org/10.1007/s13225-020-00466-2) |
| Otu40870 | 1 | k__Fungi(100);p__Ascomycota(100);c__Saccharomycetes(100);o__Saccharomycetales(100);f__Metschnikowiaceae(100);g__Metschnikowia(100);s__Metschnikowia_pulcherrima(100); | Metschnikowia | 13 | Saprotroph | Undefined Saprotroph | Yeast | NULL | Probable | As nectar-tap_saprotroph (Põlme et al. 2020); Decay sugar-rich substrates (Põlme et al. 2020) | Põlme S, et al. 2020. Fungal Diversity 105:1-16 (https://doi.org/10.1007/s13225-020-00466-2) |
| Otu40871 | 1 | k__Fungi(100);p__Ascomycota(100);c__Saccharomycetes(100);o__Saccharomycetales(100);f__Metschnikowiaceae(100);g__Metschnikowia(100);s__Metschnikowia_pulcherrima(100); | Metschnikowia | 13 | Saprotroph | Undefined Saprotroph | Yeast | NULL | Probable | As nectar-tap_saprotroph (Põlme et al. 2020); Decay sugar-rich substrates (Põlme et al. 2020) | Põlme S, et al. 2020. Fungal Diversity 105:1-16 (https://doi.org/10.1007/s13225-020-00466-2) |
| Otu40872 | 1 | k__Fungi(100);p__Ascomycota(100);c__Saccharomycetes(100);o__Saccharomycetales(100);f__Metschnikowiaceae(100);g__Metschnikowia(100);s__Metschnikowia_pulcherrima(100); | Metschnikowia | 13 | Saprotroph | Undefined Saprotroph | Yeast | NULL | Probable | As nectar-tap_saprotroph (Põlme et al. 2020); Decay sugar-rich substrates (Põlme et al. 2020) | Põlme S, et al. 2020. Fungal Diversity 105:1-16 (https://doi.org/10.1007/s13225-020-00466-2) |
| Otu40873 | 1 | k__Fungi(100);p__Ascomycota(100);c__Saccharomycetes(100);o__Saccharomycetales(100);f__Metschnikowiaceae(100);g__Metschnikowia(100);s__Metschnikowia_pulcherrima(100); | Metschnikowia | 13 | Saprotroph | Undefined Saprotroph | Yeast | NULL | Probable | As nectar-tap_saprotroph (Põlme et al. 2020); Decay sugar-rich substrates (Põlme et al. 2020) | Põlme S, et al. 2020. Fungal Diversity 105:1-16 (https://doi.org/10.1007/s13225-020-00466-2) |
| Otu40874 | 1 | k__Fungi(100);p__Ascomycota(100);c__Saccharomycetes(100);o__Saccharomycetales(100);f__Metschnikowiaceae(100);g__Metschnikowia(100);s__Metschnikowia_pulcherrima(100); | Metschnikowia | 13 | Saprotroph | Undefined Saprotroph | Yeast | NULL | Probable | As nectar-tap_saprotroph (Põlme et al. 2020); Decay sugar-rich substrates (Põlme et al. 2020) | Põlme S, et al. 2020. Fungal Diversity 105:1-16 (https://doi.org/10.1007/s13225-020-00466-2) |
| Otu40875 | 1 | k__Fungi(100);p__Ascomycota(100);c__Saccharomycetes(100);o__Saccharomycetales(100);f__Metschnikowiaceae(100);g__Metschnikowia(100);g__Metschnikowia_unclassified(100); | Metschnikowia | 13 | Saprotroph | Undefined Saprotroph | Yeast | NULL | Probable | As nectar-tap_saprotroph (Põlme et al. 2020); Decay sugar-rich substrates (Põlme et al. 2020) | Põlme S, et al. 2020. Fungal Diversity 105:1-16 (https://doi.org/10.1007/s13225-020-00466-2) |
| Otu40876 | 1 | k__Fungi(100);p__Ascomycota(100);c__Saccharomycetes(100);o__Saccharomycetales(100);f__Metschnikowiaceae(100);g__Metschnikowia(100);g__Metschnikowia_unclassified(100); | Metschnikowia | 13 | Saprotroph | Undefined Saprotroph | Yeast | NULL | Probable | As nectar-tap_saprotroph (Põlme et al. 2020); Decay sugar-rich substrates (Põlme et al. 2020) | Põlme S, et al. 2020. Fungal Diversity 105:1-16 (https://doi.org/10.1007/s13225-020-00466-2) |
| Otu40877 | 1 | k__Fungi(100);p__Ascomycota(100);c__Saccharomycetes(100);o__Saccharomycetales(100);f__Metschnikowiaceae(100);g__Metschnikowia(100);s__Metschnikowia_pulcherrima(100); | Metschnikowia | 13 | Saprotroph | Undefined Saprotroph | Yeast | NULL | Probable | As nectar-tap_saprotroph (Põlme et al. 2020); Decay sugar-rich substrates (Põlme et al. 2020) | Põlme S, et al. 2020. Fungal Diversity 105:1-16 (https://doi.org/10.1007/s13225-020-00466-2) |
| Otu40878 | 1 | k__Fungi(100);p__Ascomycota(100);c__Saccharomycetes(100);o__Saccharomycetales(100);f__Metschnikowiaceae(100);g__Metschnikowia(100);g__Metschnikowia_unclassified(100); | Metschnikowia | 13 | Saprotroph | Undefined Saprotroph | Yeast | NULL | Probable | As nectar-tap_saprotroph (Põlme et al. 2020); Decay sugar-rich substrates (Põlme et al. 2020) | Põlme S, et al. 2020. Fungal Diversity 105:1-16 (https://doi.org/10.1007/s13225-020-00466-2) |
| Otu40879 | 1 | k__Fungi(100);p__Ascomycota(100);c__Saccharomycetes(100);o__Saccharomycetales(100);f__Metschnikowiaceae(100);g__Metschnikowia(100);s__Metschnikowia_pulcherrima(100); | Metschnikowia | 13 | Saprotroph | Undefined Saprotroph | Yeast | NULL | Probable | As nectar-tap_saprotroph (Põlme et al. 2020); Decay sugar-rich substrates (Põlme et al. 2020) | Põlme S, et al. 2020. Fungal Diversity 105:1-16 (https://doi.org/10.1007/s13225-020-00466-2) |
| Otu40880 | 1 | k__Fungi(100);p__Ascomycota(100);c__Saccharomycetes(100);o__Saccharomycetales(100);f__Metschnikowiaceae(100);g__Metschnikowia(100);s__Metschnikowia_pulcherrima(100); | Metschnikowia | 13 | Saprotroph | Undefined Saprotroph | Yeast | NULL | Probable | As nectar-tap_saprotroph (Põlme et al. 2020); Decay sugar-rich substrates (Põlme et al. 2020) | Põlme S, et al. 2020. Fungal Diversity 105:1-16 (https://doi.org/10.1007/s13225-020-00466-2) |
| Otu40881 | 1 | k__Fungi(100);p__Ascomycota(100);c__Saccharomycetes(100);o__Saccharomycetales(100);f__Metschnikowiaceae(100);g__Metschnikowia(100);s__Metschnikowia_pulcherrima(100); | Metschnikowia | 13 | Saprotroph | Undefined Saprotroph | Yeast | NULL | Probable | As nectar-tap_saprotroph (Põlme et al. 2020); Decay sugar-rich substrates (Põlme et al. 2020) | Põlme S, et al. 2020. Fungal Diversity 105:1-16 (https://doi.org/10.1007/s13225-020-00466-2) |
| Otu40882 | 1 | k__Fungi(100);p__Ascomycota(100);c__Saccharomycetes(100);o__Saccharomycetales(100);f__Metschnikowiaceae(100);g__Metschnikowia(100);s__Metschnikowia_pulcherrima(100); | Metschnikowia | 13 | Saprotroph | Undefined Saprotroph | Yeast | NULL | Probable | As nectar-tap_saprotroph (Põlme et al. 2020); Decay sugar-rich substrates (Põlme et al. 2020) | Põlme S, et al. 2020. Fungal Diversity 105:1-16 (https://doi.org/10.1007/s13225-020-00466-2) |
| Otu40883 | 1 | k__Fungi(100);p__Ascomycota(100);c__Saccharomycetes(100);o__Saccharomycetales(100);f__Metschnikowiaceae(100);g__Metschnikowia(100);s__Metschnikowia_pulcherrima(100); | Metschnikowia | 13 | Saprotroph | Undefined Saprotroph | Yeast | NULL | Probable | As nectar-tap_saprotroph (Põlme et al. 2020); Decay sugar-rich substrates (Põlme et al. 2020) | Põlme S, et al. 2020. Fungal Diversity 105:1-16 (https://doi.org/10.1007/s13225-020-00466-2) |
| Otu40884 | 1 | k__Fungi(100);p__Ascomycota(100);c__Saccharomycetes(100);o__Saccharomycetales(100);f__Metschnikowiaceae(100);g__Metschnikowia(100);s__Metschnikowia_pulcherrima(100); | Metschnikowia | 13 | Saprotroph | Undefined Saprotroph | Yeast | NULL | Probable | As nectar-tap_saprotroph (Põlme et al. 2020); Decay sugar-rich substrates (Põlme et al. 2020) | Põlme S, et al. 2020. Fungal Diversity 105:1-16 (https://doi.org/10.1007/s13225-020-00466-2) |
| Otu40885 | 1 | k__Fungi(100);p__Ascomycota(100);c__Saccharomycetes(100);o__Saccharomycetales(100);f__Metschnikowiaceae(100);g__Metschnikowia(100);s__Metschnikowia_pulcherrima(100); | Metschnikowia | 13 | Saprotroph | Undefined Saprotroph | Yeast | NULL | Probable | As nectar-tap_saprotroph (Põlme et al. 2020); Decay sugar-rich substrates (Põlme et al. 2020) | Põlme S, et al. 2020. Fungal Diversity 105:1-16 (https://doi.org/10.1007/s13225-020-00466-2) |
| Otu40886 | 1 | k__Fungi(100);p__Ascomycota(100);c__Saccharomycetes(100);o__Saccharomycetales(100);f__Metschnikowiaceae(100);g__Metschnikowia(100);s__Metschnikowia_pulcherrima(100); | Metschnikowia | 13 | Saprotroph | Undefined Saprotroph | Yeast | NULL | Probable | As nectar-tap_saprotroph (Põlme et al. 2020); Decay sugar-rich substrates (Põlme et al. 2020) | Põlme S, et al. 2020. Fungal Diversity 105:1-16 (https://doi.org/10.1007/s13225-020-00466-2) |
| Otu40887 | 1 | k__Fungi(100);p__Ascomycota(100);c__Saccharomycetes(100);o__Saccharomycetales(100);f__Metschnikowiaceae(100);g__Metschnikowia(100);s__Metschnikowia_pulcherrima(100); | Metschnikowia | 13 | Saprotroph | Undefined Saprotroph | Yeast | NULL | Probable | As nectar-tap_saprotroph (Põlme et al. 2020); Decay sugar-rich substrates (Põlme et al. 2020) | Põlme S, et al. 2020. Fungal Diversity 105:1-16 (https://doi.org/10.1007/s13225-020-00466-2) |
| Otu40888 | 1 | k__Fungi(100);p__Ascomycota(100);c__Saccharomycetes(100);o__Saccharomycetales(100);f__Metschnikowiaceae(100);g__Metschnikowia(100);s__Metschnikowia_pulcherrima(100); | Metschnikowia | 13 | Saprotroph | Undefined Saprotroph | Yeast | NULL | Probable | As nectar-tap_saprotroph (Põlme et al. 2020); Decay sugar-rich substrates (Põlme et al. 2020) | Põlme S, et al. 2020. Fungal Diversity 105:1-16 (https://doi.org/10.1007/s13225-020-00466-2) |
| Otu40889 | 1 | k__Fungi(100);p__Ascomycota(100);c__Saccharomycetes(100);o__Saccharomycetales(100);f__Metschnikowiaceae(100);g__Metschnikowia(100);s__Metschnikowia_pulcherrima(100); | Metschnikowia | 13 | Saprotroph | Undefined Saprotroph | Yeast | NULL | Probable | As nectar-tap_saprotroph (Põlme et al. 2020); Decay sugar-rich substrates (Põlme et al. 2020) | Põlme S, et al. 2020. Fungal Diversity 105:1-16 (https://doi.org/10.1007/s13225-020-00466-2) |
| Otu40891 | 1 | k__Fungi(100);p__Ascomycota(100);c__Saccharomycetes(100);o__Saccharomycetales(100);f__Metschnikowiaceae(100);g__Metschnikowia(100);s__Metschnikowia_pulcherrima(100); | Metschnikowia | 13 | Saprotroph | Undefined Saprotroph | Yeast | NULL | Probable | As nectar-tap_saprotroph (Põlme et al. 2020); Decay sugar-rich substrates (Põlme et al. 2020) | Põlme S, et al. 2020. Fungal Diversity 105:1-16 (https://doi.org/10.1007/s13225-020-00466-2) |
| Otu40892 | 1 | k__Fungi(100);p__Ascomycota(100);c__Saccharomycetes(100);o__Saccharomycetales(100);f__Metschnikowiaceae(100);g__Metschnikowia(100);s__Metschnikowia_pulcherrima(100); | Metschnikowia | 13 | Saprotroph | Undefined Saprotroph | Yeast | NULL | Probable | As nectar-tap_saprotroph (Põlme et al. 2020); Decay sugar-rich substrates (Põlme et al. 2020) | Põlme S, et al. 2020. Fungal Diversity 105:1-16 (https://doi.org/10.1007/s13225-020-00466-2) |
| Otu40893 | 1 | k__Fungi(100);p__Ascomycota(100);c__Saccharomycetes(100);o__Saccharomycetales(100);f__Metschnikowiaceae(100);g__Metschnikowia(100);s__Metschnikowia_pulcherrima(100); | Metschnikowia | 13 | Saprotroph | Undefined Saprotroph | Yeast | NULL | Probable | As nectar-tap_saprotroph (Põlme et al. 2020); Decay sugar-rich substrates (Põlme et al. 2020) | Põlme S, et al. 2020. Fungal Diversity 105:1-16 (https://doi.org/10.1007/s13225-020-00466-2) |
| Otu40894 | 1 | k__Fungi(100);p__Ascomycota(100);c__Saccharomycetes(100);o__Saccharomycetales(100);f__Metschnikowiaceae(100);g__Metschnikowia(100);s__Metschnikowia_pulcherrima(100); | Metschnikowia | 13 | Saprotroph | Undefined Saprotroph | Yeast | NULL | Probable | As nectar-tap_saprotroph (Põlme et al. 2020); Decay sugar-rich substrates (Põlme et al. 2020) | Põlme S, et al. 2020. Fungal Diversity 105:1-16 (https://doi.org/10.1007/s13225-020-00466-2) |
| Otu40895 | 1 | k__Fungi(100);p__Ascomycota(100);c__Saccharomycetes(100);o__Saccharomycetales(100);f__Metschnikowiaceae(100);g__Metschnikowia(100);s__Metschnikowia_pulcherrima(100); | Metschnikowia | 13 | Saprotroph | Undefined Saprotroph | Yeast | NULL | Probable | As nectar-tap_saprotroph (Põlme et al. 2020); Decay sugar-rich substrates (Põlme et al. 2020) | Põlme S, et al. 2020. Fungal Diversity 105:1-16 (https://doi.org/10.1007/s13225-020-00466-2) |
| Otu40896 | 1 | k__Fungi(100);p__Ascomycota(100);c__Saccharomycetes(100);o__Saccharomycetales(100);f__Metschnikowiaceae(100);g__Metschnikowia(100);s__Metschnikowia_pulcherrima(100); | Metschnikowia | 13 | Saprotroph | Undefined Saprotroph | Yeast | NULL | Probable | As nectar-tap_saprotroph (Põlme et al. 2020); Decay sugar-rich substrates (Põlme et al. 2020) | Põlme S, et al. 2020. Fungal Diversity 105:1-16 (https://doi.org/10.1007/s13225-020-00466-2) |
| Otu40897 | 1 | k__Fungi(100);p__Ascomycota(100);c__Saccharomycetes(100);o__Saccharomycetales(100);f__Metschnikowiaceae(100);g__Metschnikowia(100);s__Metschnikowia_pulcherrima(100); | Metschnikowia | 13 | Saprotroph | Undefined Saprotroph | Yeast | NULL | Probable | As nectar-tap_saprotroph (Põlme et al. 2020); Decay sugar-rich substrates (Põlme et al. 2020) | Põlme S, et al. 2020. Fungal Diversity 105:1-16 (https://doi.org/10.1007/s13225-020-00466-2) |
| Otu40898 | 1 | k__Fungi(100);p__Ascomycota(100);c__Saccharomycetes(100);o__Saccharomycetales(100);f__Metschnikowiaceae(100);g__Metschnikowia(100);s__Metschnikowia_pulcherrima(100); | Metschnikowia | 13 | Saprotroph | Undefined Saprotroph | Yeast | NULL | Probable | As nectar-tap_saprotroph (Põlme et al. 2020); Decay sugar-rich substrates (Põlme et al. 2020) | Põlme S, et al. 2020. Fungal Diversity 105:1-16 (https://doi.org/10.1007/s13225-020-00466-2) |
| Otu40899 | 1 | k__Fungi(100);p__Ascomycota(100);c__Saccharomycetes(100);o__Saccharomycetales(100);f__Metschnikowiaceae(100);g__Metschnikowia(100);s__Metschnikowia_pulcherrima(100); | Metschnikowia | 13 | Saprotroph | Undefined Saprotroph | Yeast | NULL | Probable | As nectar-tap_saprotroph (Põlme et al. 2020); Decay sugar-rich substrates (Põlme et al. 2020) | Põlme S, et al. 2020. Fungal Diversity 105:1-16 (https://doi.org/10.1007/s13225-020-00466-2) |
| Otu40901 | 1 | k__Fungi(100);p__Ascomycota(100);c__Saccharomycetes(100);o__Saccharomycetales(100);f__Metschnikowiaceae(100);g__Metschnikowia(100);s__Metschnikowia_pulcherrima(100); | Metschnikowia | 13 | Saprotroph | Undefined Saprotroph | Yeast | NULL | Probable | As nectar-tap_saprotroph (Põlme et al. 2020); Decay sugar-rich substrates (Põlme et al. 2020) | Põlme S, et al. 2020. Fungal Diversity 105:1-16 (https://doi.org/10.1007/s13225-020-00466-2) |
| Otu40902 | 1 | k__Fungi(100);p__Ascomycota(100);c__Saccharomycetes(100);o__Saccharomycetales(100);f__Metschnikowiaceae(100);g__Metschnikowia(100);s__Metschnikowia_pulcherrima(100); | Metschnikowia | 13 | Saprotroph | Undefined Saprotroph | Yeast | NULL | Probable | As nectar-tap_saprotroph (Põlme et al. 2020); Decay sugar-rich substrates (Põlme et al. 2020) | Põlme S, et al. 2020. Fungal Diversity 105:1-16 (https://doi.org/10.1007/s13225-020-00466-2) |
| Otu40903 | 1 | k__Fungi(100);p__Ascomycota(100);c__Saccharomycetes(100);o__Saccharomycetales(100);f__Metschnikowiaceae(100);g__Metschnikowia(100);g__Metschnikowia_unclassified(100); | Metschnikowia | 13 | Saprotroph | Undefined Saprotroph | Yeast | NULL | Probable | As nectar-tap_saprotroph (Põlme et al. 2020); Decay sugar-rich substrates (Põlme et al. 2020) | Põlme S, et al. 2020. Fungal Diversity 105:1-16 (https://doi.org/10.1007/s13225-020-00466-2) |
| Otu40905 | 1 | k__Fungi(100);p__Ascomycota(100);c__Saccharomycetes(100);o__Saccharomycetales(100);f__Metschnikowiaceae(100);g__Metschnikowia(100);s__Metschnikowia_pulcherrima(100); | Metschnikowia | 13 | Saprotroph | Undefined Saprotroph | Yeast | NULL | Probable | As nectar-tap_saprotroph (Põlme et al. 2020); Decay sugar-rich substrates (Põlme et al. 2020) | Põlme S, et al. 2020. Fungal Diversity 105:1-16 (https://doi.org/10.1007/s13225-020-00466-2) |
| Otu40906 | 1 | k__Fungi(100);p__Ascomycota(100);c__Saccharomycetes(100);o__Saccharomycetales(100);f__Metschnikowiaceae(100);g__Metschnikowia(100);g__Metschnikowia_unclassified(100); | Metschnikowia | 13 | Saprotroph | Undefined Saprotroph | Yeast | NULL | Probable | As nectar-tap_saprotroph (Põlme et al. 2020); Decay sugar-rich substrates (Põlme et al. 2020) | Põlme S, et al. 2020. Fungal Diversity 105:1-16 (https://doi.org/10.1007/s13225-020-00466-2) |
| Otu40907 | 1 | k__Fungi(100);p__Ascomycota(100);c__Saccharomycetes(100);o__Saccharomycetales(100);f__Metschnikowiaceae(100);g__Metschnikowia(100);s__Metschnikowia_pulcherrima(100); | Metschnikowia | 13 | Saprotroph | Undefined Saprotroph | Yeast | NULL | Probable | As nectar-tap_saprotroph (Põlme et al. 2020); Decay sugar-rich substrates (Põlme et al. 2020) | Põlme S, et al. 2020. Fungal Diversity 105:1-16 (https://doi.org/10.1007/s13225-020-00466-2) |
| Otu40908 | 1 | k__Fungi(100);p__Ascomycota(100);c__Saccharomycetes(100);o__Saccharomycetales(100);f__Metschnikowiaceae(100);g__Metschnikowia(100);s__Metschnikowia_pulcherrima(100); | Metschnikowia | 13 | Saprotroph | Undefined Saprotroph | Yeast | NULL | Probable | As nectar-tap_saprotroph (Põlme et al. 2020); Decay sugar-rich substrates (Põlme et al. 2020) | Põlme S, et al. 2020. Fungal Diversity 105:1-16 (https://doi.org/10.1007/s13225-020-00466-2) |
| Otu40909 | 1 | k__Fungi(100);p__Ascomycota(100);c__Saccharomycetes(100);o__Saccharomycetales(100);f__Metschnikowiaceae(100);g__Metschnikowia(100);s__Metschnikowia_pulcherrima(100); | Metschnikowia | 13 | Saprotroph | Undefined Saprotroph | Yeast | NULL | Probable | As nectar-tap_saprotroph (Põlme et al. 2020); Decay sugar-rich substrates (Põlme et al. 2020) | Põlme S, et al. 2020. Fungal Diversity 105:1-16 (https://doi.org/10.1007/s13225-020-00466-2) |
| Otu40910 | 1 | k__Fungi(100);p__Ascomycota(100);c__Saccharomycetes(100);o__Saccharomycetales(100);f__Metschnikowiaceae(100);g__Metschnikowia(100);s__Metschnikowia_pulcherrima(100); | Metschnikowia | 13 | Saprotroph | Undefined Saprotroph | Yeast | NULL | Probable | As nectar-tap_saprotroph (Põlme et al. 2020); Decay sugar-rich substrates (Põlme et al. 2020) | Põlme S, et al. 2020. Fungal Diversity 105:1-16 (https://doi.org/10.1007/s13225-020-00466-2) |
| Otu40911 | 1 | k__Fungi(100);p__Ascomycota(100);c__Saccharomycetes(100);o__Saccharomycetales(100);f__Metschnikowiaceae(100);g__Metschnikowia(100);s__Metschnikowia_pulcherrima(100); | Metschnikowia | 13 | Saprotroph | Undefined Saprotroph | Yeast | NULL | Probable | As nectar-tap_saprotroph (Põlme et al. 2020); Decay sugar-rich substrates (Põlme et al. 2020) | Põlme S, et al. 2020. Fungal Diversity 105:1-16 (https://doi.org/10.1007/s13225-020-00466-2) |
| Otu40912 | 1 | k__Fungi(100);p__Ascomycota(100);c__Saccharomycetes(100);o__Saccharomycetales(100);f__Metschnikowiaceae(100);g__Metschnikowia(100);s__Metschnikowia_pulcherrima(100); | Metschnikowia | 13 | Saprotroph | Undefined Saprotroph | Yeast | NULL | Probable | As nectar-tap_saprotroph (Põlme et al. 2020); Decay sugar-rich substrates (Põlme et al. 2020) | Põlme S, et al. 2020. Fungal Diversity 105:1-16 (https://doi.org/10.1007/s13225-020-00466-2) |
| Otu40913 | 1 | k__Fungi(100);p__Ascomycota(100);c__Saccharomycetes(100);o__Saccharomycetales(100);f__Metschnikowiaceae(100);g__Metschnikowia(100);s__Metschnikowia_pulcherrima(100); | Metschnikowia | 13 | Saprotroph | Undefined Saprotroph | Yeast | NULL | Probable | As nectar-tap_saprotroph (Põlme et al. 2020); Decay sugar-rich substrates (Põlme et al. 2020) | Põlme S, et al. 2020. Fungal Diversity 105:1-16 (https://doi.org/10.1007/s13225-020-00466-2) |
| Otu40914 | 1 | k__Fungi(100);p__Ascomycota(100);c__Saccharomycetes(100);o__Saccharomycetales(100);f__Metschnikowiaceae(100);g__Metschnikowia(100);s__Metschnikowia_pulcherrima(100); | Metschnikowia | 13 | Saprotroph | Undefined Saprotroph | Yeast | NULL | Probable | As nectar-tap_saprotroph (Põlme et al. 2020); Decay sugar-rich substrates (Põlme et al. 2020) | Põlme S, et al. 2020. Fungal Diversity 105:1-16 (https://doi.org/10.1007/s13225-020-00466-2) |
| Otu40915 | 1 | k__Fungi(100);p__Ascomycota(100);c__Saccharomycetes(100);o__Saccharomycetales(100);f__Metschnikowiaceae(100);g__Metschnikowia(100);s__Metschnikowia_pulcherrima(100); | Metschnikowia | 13 | Saprotroph | Undefined Saprotroph | Yeast | NULL | Probable | As nectar-tap_saprotroph (Põlme et al. 2020); Decay sugar-rich substrates (Põlme et al. 2020) | Põlme S, et al. 2020. Fungal Diversity 105:1-16 (https://doi.org/10.1007/s13225-020-00466-2) |
| Otu40916 | 1 | k__Fungi(100);p__Ascomycota(100);c__Saccharomycetes(100);o__Saccharomycetales(100);f__Metschnikowiaceae(100);g__Metschnikowia(100);s__Metschnikowia_pulcherrima(100); | Metschnikowia | 13 | Saprotroph | Undefined Saprotroph | Yeast | NULL | Probable | As nectar-tap_saprotroph (Põlme et al. 2020); Decay sugar-rich substrates (Põlme et al. 2020) | Põlme S, et al. 2020. Fungal Diversity 105:1-16 (https://doi.org/10.1007/s13225-020-00466-2) |
| Otu40917 | 1 | k__Fungi(100);p__Ascomycota(100);c__Saccharomycetes(100);o__Saccharomycetales(100);f__Metschnikowiaceae(100);g__Metschnikowia(100);s__Metschnikowia_pulcherrima(100); | Metschnikowia | 13 | Saprotroph | Undefined Saprotroph | Yeast | NULL | Probable | As nectar-tap_saprotroph (Põlme et al. 2020); Decay sugar-rich substrates (Põlme et al. 2020) | Põlme S, et al. 2020. Fungal Diversity 105:1-16 (https://doi.org/10.1007/s13225-020-00466-2) |
| Otu40918 | 1 | k__Fungi(100);p__Ascomycota(100);c__Saccharomycetes(100);o__Saccharomycetales(100);f__Metschnikowiaceae(100);g__Metschnikowia(100);s__Metschnikowia_pulcherrima(100); | Metschnikowia | 13 | Saprotroph | Undefined Saprotroph | Yeast | NULL | Probable | As nectar-tap_saprotroph (Põlme et al. 2020); Decay sugar-rich substrates (Põlme et al. 2020) | Põlme S, et al. 2020. Fungal Diversity 105:1-16 (https://doi.org/10.1007/s13225-020-00466-2) |
| Otu40920 | 1 | k__Fungi(100);p__Ascomycota(100);c__Saccharomycetes(100);o__Saccharomycetales(100);f__Metschnikowiaceae(100);g__Metschnikowia(100);g__Metschnikowia_unclassified(100); | Metschnikowia | 13 | Saprotroph | Undefined Saprotroph | Yeast | NULL | Probable | As nectar-tap_saprotroph (Põlme et al. 2020); Decay sugar-rich substrates (Põlme et al. 2020) | Põlme S, et al. 2020. Fungal Diversity 105:1-16 (https://doi.org/10.1007/s13225-020-00466-2) |
| Otu40921 | 1 | k__Fungi(100);p__Ascomycota(100);c__Saccharomycetes(100);o__Saccharomycetales(100);f__Metschnikowiaceae(100);g__Metschnikowia(100);g__Metschnikowia_unclassified(100); | Metschnikowia | 13 | Saprotroph | Undefined Saprotroph | Yeast | NULL | Probable | As nectar-tap_saprotroph (Põlme et al. 2020); Decay sugar-rich substrates (Põlme et al. 2020) | Põlme S, et al. 2020. Fungal Diversity 105:1-16 (https://doi.org/10.1007/s13225-020-00466-2) |
| Otu40922 | 1 | k__Fungi(100);p__Ascomycota(100);c__Saccharomycetes(100);o__Saccharomycetales(100);f__Metschnikowiaceae(100);g__Metschnikowia(100);s__Metschnikowia_pulcherrima(100); | Metschnikowia | 13 | Saprotroph | Undefined Saprotroph | Yeast | NULL | Probable | As nectar-tap_saprotroph (Põlme et al. 2020); Decay sugar-rich substrates (Põlme et al. 2020) | Põlme S, et al. 2020. Fungal Diversity 105:1-16 (https://doi.org/10.1007/s13225-020-00466-2) |
| Otu40923 | 1 | k__Fungi(100);p__Ascomycota(100);c__Saccharomycetes(100);o__Saccharomycetales(100);f__Metschnikowiaceae(100);g__Metschnikowia(100);s__Metschnikowia_pulcherrima(100); | Metschnikowia | 13 | Saprotroph | Undefined Saprotroph | Yeast | NULL | Probable | As nectar-tap_saprotroph (Põlme et al. 2020); Decay sugar-rich substrates (Põlme et al. 2020) | Põlme S, et al. 2020. Fungal Diversity 105:1-16 (https://doi.org/10.1007/s13225-020-00466-2) |
| Otu40924 | 1 | k__Fungi(100);p__Ascomycota(100);c__Saccharomycetes(100);o__Saccharomycetales(100);f__Metschnikowiaceae(100);g__Metschnikowia(100);s__Metschnikowia_pulcherrima(100); | Metschnikowia | 13 | Saprotroph | Undefined Saprotroph | Yeast | NULL | Probable | As nectar-tap_saprotroph (Põlme et al. 2020); Decay sugar-rich substrates (Põlme et al. 2020) | Põlme S, et al. 2020. Fungal Diversity 105:1-16 (https://doi.org/10.1007/s13225-020-00466-2) |
| Otu40925 | 1 | k__Fungi(100);p__Ascomycota(100);c__Saccharomycetes(100);o__Saccharomycetales(100);f__Metschnikowiaceae(100);g__Metschnikowia(100);s__Metschnikowia_pulcherrima(100); | Metschnikowia | 13 | Saprotroph | Undefined Saprotroph | Yeast | NULL | Probable | As nectar-tap_saprotroph (Põlme et al. 2020); Decay sugar-rich substrates (Põlme et al. 2020) | Põlme S, et al. 2020. Fungal Diversity 105:1-16 (https://doi.org/10.1007/s13225-020-00466-2) |
| Otu40926 | 1 | k__Fungi(100);p__Ascomycota(100);c__Saccharomycetes(100);o__Saccharomycetales(100);f__Metschnikowiaceae(100);g__Metschnikowia(100);s__Metschnikowia_pulcherrima(100); | Metschnikowia | 13 | Saprotroph | Undefined Saprotroph | Yeast | NULL | Probable | As nectar-tap_saprotroph (Põlme et al. 2020); Decay sugar-rich substrates (Põlme et al. 2020) | Põlme S, et al. 2020. Fungal Diversity 105:1-16 (https://doi.org/10.1007/s13225-020-00466-2) |
| Otu40927 | 1 | k__Fungi(100);p__Ascomycota(100);c__Saccharomycetes(100);o__Saccharomycetales(100);f__Metschnikowiaceae(100);g__Metschnikowia(100);g__Metschnikowia_unclassified(100); | Metschnikowia | 13 | Saprotroph | Undefined Saprotroph | Yeast | NULL | Probable | As nectar-tap_saprotroph (Põlme et al. 2020); Decay sugar-rich substrates (Põlme et al. 2020) | Põlme S, et al. 2020. Fungal Diversity 105:1-16 (https://doi.org/10.1007/s13225-020-00466-2) |
| Otu40928 | 1 | k__Fungi(100);p__Ascomycota(100);c__Saccharomycetes(100);o__Saccharomycetales(100);f__Metschnikowiaceae(100);g__Metschnikowia(100);s__Metschnikowia_pulcherrima(100); | Metschnikowia | 13 | Saprotroph | Undefined Saprotroph | Yeast | NULL | Probable | As nectar-tap_saprotroph (Põlme et al. 2020); Decay sugar-rich substrates (Põlme et al. 2020) | Põlme S, et al. 2020. Fungal Diversity 105:1-16 (https://doi.org/10.1007/s13225-020-00466-2) |
| Otu40929 | 1 | k__Fungi(100);p__Ascomycota(100);c__Saccharomycetes(100);o__Saccharomycetales(100);f__Metschnikowiaceae(100);g__Metschnikowia(100);s__Metschnikowia_pulcherrima(100); | Metschnikowia | 13 | Saprotroph | Undefined Saprotroph | Yeast | NULL | Probable | As nectar-tap_saprotroph (Põlme et al. 2020); Decay sugar-rich substrates (Põlme et al. 2020) | Põlme S, et al. 2020. Fungal Diversity 105:1-16 (https://doi.org/10.1007/s13225-020-00466-2) |
| Otu40930 | 1 | k__Fungi(100);p__Ascomycota(100);c__Saccharomycetes(100);o__Saccharomycetales(100);f__Metschnikowiaceae(100);g__Metschnikowia(100);s__Metschnikowia_pulcherrima(100); | Metschnikowia | 13 | Saprotroph | Undefined Saprotroph | Yeast | NULL | Probable | As nectar-tap_saprotroph (Põlme et al. 2020); Decay sugar-rich substrates (Põlme et al. 2020) | Põlme S, et al. 2020. Fungal Diversity 105:1-16 (https://doi.org/10.1007/s13225-020-00466-2) |
| Otu40931 | 1 | k__Fungi(100);p__Ascomycota(100);c__Saccharomycetes(100);o__Saccharomycetales(100);f__Metschnikowiaceae(100);g__Metschnikowia(100);g__Metschnikowia_unclassified(100); | Metschnikowia | 13 | Saprotroph | Undefined Saprotroph | Yeast | NULL | Probable | As nectar-tap_saprotroph (Põlme et al. 2020); Decay sugar-rich substrates (Põlme et al. 2020) | Põlme S, et al. 2020. Fungal Diversity 105:1-16 (https://doi.org/10.1007/s13225-020-00466-2) |
| Otu40932 | 1 | k__Fungi(100);p__Ascomycota(100);c__Saccharomycetes(100);o__Saccharomycetales(100);f__Metschnikowiaceae(100);g__Metschnikowia(100);g__Metschnikowia_unclassified(100); | Metschnikowia | 13 | Saprotroph | Undefined Saprotroph | Yeast | NULL | Probable | As nectar-tap_saprotroph (Põlme et al. 2020); Decay sugar-rich substrates (Põlme et al. 2020) | Põlme S, et al. 2020. Fungal Diversity 105:1-16 (https://doi.org/10.1007/s13225-020-00466-2) |
| Otu40933 | 1 | k__Fungi(100);p__Ascomycota(100);c__Saccharomycetes(100);o__Saccharomycetales(100);f__Metschnikowiaceae(100);g__Metschnikowia(100);s__Metschnikowia_pulcherrima(100); | Metschnikowia | 13 | Saprotroph | Undefined Saprotroph | Yeast | NULL | Probable | As nectar-tap_saprotroph (Põlme et al. 2020); Decay sugar-rich substrates (Põlme et al. 2020) | Põlme S, et al. 2020. Fungal Diversity 105:1-16 (https://doi.org/10.1007/s13225-020-00466-2) |
| Otu40934 | 1 | k__Fungi(100);p__Ascomycota(100);c__Saccharomycetes(100);o__Saccharomycetales(100);f__Metschnikowiaceae(100);g__Metschnikowia(100);g__Metschnikowia_unclassified(100); | Metschnikowia | 13 | Saprotroph | Undefined Saprotroph | Yeast | NULL | Probable | As nectar-tap_saprotroph (Põlme et al. 2020); Decay sugar-rich substrates (Põlme et al. 2020) | Põlme S, et al. 2020. Fungal Diversity 105:1-16 (https://doi.org/10.1007/s13225-020-00466-2) |
| Otu40935 | 1 | k__Fungi(100);p__Ascomycota(100);c__Saccharomycetes(100);o__Saccharomycetales(100);f__Metschnikowiaceae(100);g__Metschnikowia(100);g__Metschnikowia_unclassified(100); | Metschnikowia | 13 | Saprotroph | Undefined Saprotroph | Yeast | NULL | Probable | As nectar-tap_saprotroph (Põlme et al. 2020); Decay sugar-rich substrates (Põlme et al. 2020) | Põlme S, et al. 2020. Fungal Diversity 105:1-16 (https://doi.org/10.1007/s13225-020-00466-2) |
| Otu40936 | 1 | k__Fungi(100);p__Ascomycota(100);c__Saccharomycetes(100);o__Saccharomycetales(100);f__Metschnikowiaceae(100);g__Metschnikowia(100);s__Metschnikowia_pulcherrima(100); | Metschnikowia | 13 | Saprotroph | Undefined Saprotroph | Yeast | NULL | Probable | As nectar-tap_saprotroph (Põlme et al. 2020); Decay sugar-rich substrates (Põlme et al. 2020) | Põlme S, et al. 2020. Fungal Diversity 105:1-16 (https://doi.org/10.1007/s13225-020-00466-2) |
| Otu40937 | 1 | k__Fungi(100);p__Ascomycota(100);c__Saccharomycetes(100);o__Saccharomycetales(100);f__Metschnikowiaceae(100);g__Metschnikowia(100);s__Metschnikowia_pulcherrima(100); | Metschnikowia | 13 | Saprotroph | Undefined Saprotroph | Yeast | NULL | Probable | As nectar-tap_saprotroph (Põlme et al. 2020); Decay sugar-rich substrates (Põlme et al. 2020) | Põlme S, et al. 2020. Fungal Diversity 105:1-16 (https://doi.org/10.1007/s13225-020-00466-2) |
| Otu40938 | 1 | k__Fungi(100);p__Ascomycota(100);c__Saccharomycetes(100);o__Saccharomycetales(100);f__Metschnikowiaceae(100);g__Metschnikowia(100);s__Metschnikowia_pulcherrima(100); | Metschnikowia | 13 | Saprotroph | Undefined Saprotroph | Yeast | NULL | Probable | As nectar-tap_saprotroph (Põlme et al. 2020); Decay sugar-rich substrates (Põlme et al. 2020) | Põlme S, et al. 2020. Fungal Diversity 105:1-16 (https://doi.org/10.1007/s13225-020-00466-2) |
| Otu40939 | 1 | k__Fungi(100);p__Ascomycota(100);c__Saccharomycetes(100);o__Saccharomycetales(100);f__Metschnikowiaceae(100);g__Metschnikowia(100);g__Metschnikowia_unclassified(100); | Metschnikowia | 13 | Saprotroph | Undefined Saprotroph | Yeast | NULL | Probable | As nectar-tap_saprotroph (Põlme et al. 2020); Decay sugar-rich substrates (Põlme et al. 2020) | Põlme S, et al. 2020. Fungal Diversity 105:1-16 (https://doi.org/10.1007/s13225-020-00466-2) |
| Otu40940 | 1 | k__Fungi(100);p__Ascomycota(100);c__Saccharomycetes(100);o__Saccharomycetales(100);f__Metschnikowiaceae(100);g__Metschnikowia(100);s__Metschnikowia_pulcherrima(100); | Metschnikowia | 13 | Saprotroph | Undefined Saprotroph | Yeast | NULL | Probable | As nectar-tap_saprotroph (Põlme et al. 2020); Decay sugar-rich substrates (Põlme et al. 2020) | Põlme S, et al. 2020. Fungal Diversity 105:1-16 (https://doi.org/10.1007/s13225-020-00466-2) |
| Otu40941 | 1 | k__Fungi(100);p__Ascomycota(100);c__Saccharomycetes(100);o__Saccharomycetales(100);f__Metschnikowiaceae(100);g__Metschnikowia(100);s__Metschnikowia_pulcherrima(100); | Metschnikowia | 13 | Saprotroph | Undefined Saprotroph | Yeast | NULL | Probable | As nectar-tap_saprotroph (Põlme et al. 2020); Decay sugar-rich substrates (Põlme et al. 2020) | Põlme S, et al. 2020. Fungal Diversity 105:1-16 (https://doi.org/10.1007/s13225-020-00466-2) |
| Otu40942 | 1 | k__Fungi(100);p__Ascomycota(100);c__Saccharomycetes(100);o__Saccharomycetales(100);f__Metschnikowiaceae(100);g__Metschnikowia(100);s__Metschnikowia_pulcherrima(100); | Metschnikowia | 13 | Saprotroph | Undefined Saprotroph | Yeast | NULL | Probable | As nectar-tap_saprotroph (Põlme et al. 2020); Decay sugar-rich substrates (Põlme et al. 2020) | Põlme S, et al. 2020. Fungal Diversity 105:1-16 (https://doi.org/10.1007/s13225-020-00466-2) |
| Otu40943 | 1 | k__Fungi(100);p__Ascomycota(100);c__Saccharomycetes(100);o__Saccharomycetales(100);f__Metschnikowiaceae(100);g__Metschnikowia(100);s__Metschnikowia_pulcherrima(100); | Metschnikowia | 13 | Saprotroph | Undefined Saprotroph | Yeast | NULL | Probable | As nectar-tap_saprotroph (Põlme et al. 2020); Decay sugar-rich substrates (Põlme et al. 2020) | Põlme S, et al. 2020. Fungal Diversity 105:1-16 (https://doi.org/10.1007/s13225-020-00466-2) |
| Otu40944 | 1 | k__Fungi(100);p__Ascomycota(100);c__Saccharomycetes(100);o__Saccharomycetales(100);f__Metschnikowiaceae(100);g__Metschnikowia(100);s__Metschnikowia_pulcherrima(100); | Metschnikowia | 13 | Saprotroph | Undefined Saprotroph | Yeast | NULL | Probable | As nectar-tap_saprotroph (Põlme et al. 2020); Decay sugar-rich substrates (Põlme et al. 2020) | Põlme S, et al. 2020. Fungal Diversity 105:1-16 (https://doi.org/10.1007/s13225-020-00466-2) |
| Otu40945 | 1 | k__Fungi(100);p__Ascomycota(100);c__Saccharomycetes(100);o__Saccharomycetales(100);f__Metschnikowiaceae(100);g__Metschnikowia(100);g__Metschnikowia_unclassified(100); | Metschnikowia | 13 | Saprotroph | Undefined Saprotroph | Yeast | NULL | Probable | As nectar-tap_saprotroph (Põlme et al. 2020); Decay sugar-rich substrates (Põlme et al. 2020) | Põlme S, et al. 2020. Fungal Diversity 105:1-16 (https://doi.org/10.1007/s13225-020-00466-2) |
| Otu40946 | 1 | k__Fungi(100);p__Ascomycota(100);c__Saccharomycetes(100);o__Saccharomycetales(100);f__Metschnikowiaceae(100);g__Metschnikowia(100);s__Metschnikowia_pulcherrima(100); | Metschnikowia | 13 | Saprotroph | Undefined Saprotroph | Yeast | NULL | Probable | As nectar-tap_saprotroph (Põlme et al. 2020); Decay sugar-rich substrates (Põlme et al. 2020) | Põlme S, et al. 2020. Fungal Diversity 105:1-16 (https://doi.org/10.1007/s13225-020-00466-2) |
| Otu40947 | 1 | k__Fungi(100);p__Ascomycota(100);c__Saccharomycetes(100);o__Saccharomycetales(100);f__Metschnikowiaceae(100);g__Metschnikowia(100);g__Metschnikowia_unclassified(100); | Metschnikowia | 13 | Saprotroph | Undefined Saprotroph | Yeast | NULL | Probable | As nectar-tap_saprotroph (Põlme et al. 2020); Decay sugar-rich substrates (Põlme et al. 2020) | Põlme S, et al. 2020. Fungal Diversity 105:1-16 (https://doi.org/10.1007/s13225-020-00466-2) |
| Otu40948 | 1 | k__Fungi(100);p__Ascomycota(100);c__Saccharomycetes(100);o__Saccharomycetales(100);f__Metschnikowiaceae(100);g__Metschnikowia(100);g__Metschnikowia_unclassified(100); | Metschnikowia | 13 | Saprotroph | Undefined Saprotroph | Yeast | NULL | Probable | As nectar-tap_saprotroph (Põlme et al. 2020); Decay sugar-rich substrates (Põlme et al. 2020) | Põlme S, et al. 2020. Fungal Diversity 105:1-16 (https://doi.org/10.1007/s13225-020-00466-2) |
| Otu40949 | 1 | k__Fungi(100);p__Ascomycota(100);c__Saccharomycetes(100);o__Saccharomycetales(100);f__Metschnikowiaceae(100);g__Metschnikowia(100);s__Metschnikowia_pulcherrima(100); | Metschnikowia | 13 | Saprotroph | Undefined Saprotroph | Yeast | NULL | Probable | As nectar-tap_saprotroph (Põlme et al. 2020); Decay sugar-rich substrates (Põlme et al. 2020) | Põlme S, et al. 2020. Fungal Diversity 105:1-16 (https://doi.org/10.1007/s13225-020-00466-2) |
| Otu40950 | 1 | k__Fungi(100);p__Ascomycota(100);c__Saccharomycetes(100);o__Saccharomycetales(100);f__Metschnikowiaceae(100);g__Metschnikowia(100);s__Metschnikowia_pulcherrima(100); | Metschnikowia | 13 | Saprotroph | Undefined Saprotroph | Yeast | NULL | Probable | As nectar-tap_saprotroph (Põlme et al. 2020); Decay sugar-rich substrates (Põlme et al. 2020) | Põlme S, et al. 2020. Fungal Diversity 105:1-16 (https://doi.org/10.1007/s13225-020-00466-2) |
| Otu40951 | 1 | k__Fungi(100);p__Ascomycota(100);c__Saccharomycetes(100);o__Saccharomycetales(100);f__Metschnikowiaceae(100);g__Metschnikowia(100);s__Metschnikowia_pulcherrima(100); | Metschnikowia | 13 | Saprotroph | Undefined Saprotroph | Yeast | NULL | Probable | As nectar-tap_saprotroph (Põlme et al. 2020); Decay sugar-rich substrates (Põlme et al. 2020) | Põlme S, et al. 2020. Fungal Diversity 105:1-16 (https://doi.org/10.1007/s13225-020-00466-2) |
| Otu40952 | 1 | k__Fungi(100);p__Ascomycota(100);c__Saccharomycetes(100);o__Saccharomycetales(100);f__Metschnikowiaceae(100);g__Metschnikowia(100);s__Metschnikowia_pulcherrima(100); | Metschnikowia | 13 | Saprotroph | Undefined Saprotroph | Yeast | NULL | Probable | As nectar-tap_saprotroph (Põlme et al. 2020); Decay sugar-rich substrates (Põlme et al. 2020) | Põlme S, et al. 2020. Fungal Diversity 105:1-16 (https://doi.org/10.1007/s13225-020-00466-2) |
| Otu40953 | 1 | k__Fungi(100);p__Ascomycota(100);c__Saccharomycetes(100);o__Saccharomycetales(100);f__Metschnikowiaceae(100);g__Metschnikowia(100);s__Metschnikowia_pulcherrima(100); | Metschnikowia | 13 | Saprotroph | Undefined Saprotroph | Yeast | NULL | Probable | As nectar-tap_saprotroph (Põlme et al. 2020); Decay sugar-rich substrates (Põlme et al. 2020) | Põlme S, et al. 2020. Fungal Diversity 105:1-16 (https://doi.org/10.1007/s13225-020-00466-2) |
| Otu40954 | 1 | k__Fungi(100);p__Ascomycota(100);c__Saccharomycetes(100);o__Saccharomycetales(100);f__Metschnikowiaceae(100);g__Metschnikowia(100);s__Metschnikowia_pulcherrima(100); | Metschnikowia | 13 | Saprotroph | Undefined Saprotroph | Yeast | NULL | Probable | As nectar-tap_saprotroph (Põlme et al. 2020); Decay sugar-rich substrates (Põlme et al. 2020) | Põlme S, et al. 2020. Fungal Diversity 105:1-16 (https://doi.org/10.1007/s13225-020-00466-2) |
| Otu40955 | 1 | k__Fungi(100);p__Ascomycota(100);c__Saccharomycetes(100);o__Saccharomycetales(100);f__Metschnikowiaceae(100);g__Metschnikowia(100);s__Metschnikowia_pulcherrima(100); | Metschnikowia | 13 | Saprotroph | Undefined Saprotroph | Yeast | NULL | Probable | As nectar-tap_saprotroph (Põlme et al. 2020); Decay sugar-rich substrates (Põlme et al. 2020) | Põlme S, et al. 2020. Fungal Diversity 105:1-16 (https://doi.org/10.1007/s13225-020-00466-2) |
| Otu40956 | 1 | k__Fungi(100);p__Ascomycota(100);c__Saccharomycetes(100);o__Saccharomycetales(100);f__Metschnikowiaceae(100);g__Metschnikowia(100);s__Metschnikowia_pulcherrima(100); | Metschnikowia | 13 | Saprotroph | Undefined Saprotroph | Yeast | NULL | Probable | As nectar-tap_saprotroph (Põlme et al. 2020); Decay sugar-rich substrates (Põlme et al. 2020) | Põlme S, et al. 2020. Fungal Diversity 105:1-16 (https://doi.org/10.1007/s13225-020-00466-2) |
| Otu40957 | 1 | k__Fungi(100);p__Ascomycota(100);c__Saccharomycetes(100);o__Saccharomycetales(100);f__Metschnikowiaceae(100);g__Metschnikowia(100);s__Metschnikowia_pulcherrima(100); | Metschnikowia | 13 | Saprotroph | Undefined Saprotroph | Yeast | NULL | Probable | As nectar-tap_saprotroph (Põlme et al. 2020); Decay sugar-rich substrates (Põlme et al. 2020) | Põlme S, et al. 2020. Fungal Diversity 105:1-16 (https://doi.org/10.1007/s13225-020-00466-2) |
| Otu40958 | 1 | k__Fungi(100);p__Ascomycota(100);c__Saccharomycetes(100);o__Saccharomycetales(100);f__Metschnikowiaceae(100);g__Metschnikowia(100);g__Metschnikowia_unclassified(100); | Metschnikowia | 13 | Saprotroph | Undefined Saprotroph | Yeast | NULL | Probable | As nectar-tap_saprotroph (Põlme et al. 2020); Decay sugar-rich substrates (Põlme et al. 2020) | Põlme S, et al. 2020. Fungal Diversity 105:1-16 (https://doi.org/10.1007/s13225-020-00466-2) |
| Otu40959 | 1 | k__Fungi(100);p__Ascomycota(100);c__Saccharomycetes(100);o__Saccharomycetales(100);f__Metschnikowiaceae(100);g__Metschnikowia(100);g__Metschnikowia_unclassified(100); | Metschnikowia | 13 | Saprotroph | Undefined Saprotroph | Yeast | NULL | Probable | As nectar-tap_saprotroph (Põlme et al. 2020); Decay sugar-rich substrates (Põlme et al. 2020) | Põlme S, et al. 2020. Fungal Diversity 105:1-16 (https://doi.org/10.1007/s13225-020-00466-2) |
| Otu40960 | 1 | k__Fungi(100);p__Ascomycota(100);c__Saccharomycetes(100);o__Saccharomycetales(100);f__Metschnikowiaceae(100);g__Metschnikowia(100);g__Metschnikowia_unclassified(100); | Metschnikowia | 13 | Saprotroph | Undefined Saprotroph | Yeast | NULL | Probable | As nectar-tap_saprotroph (Põlme et al. 2020); Decay sugar-rich substrates (Põlme et al. 2020) | Põlme S, et al. 2020. Fungal Diversity 105:1-16 (https://doi.org/10.1007/s13225-020-00466-2) |
| Otu40961 | 1 | k__Fungi(100);p__Ascomycota(100);c__Saccharomycetes(100);o__Saccharomycetales(100);f__Metschnikowiaceae(100);g__Metschnikowia(100);g__Metschnikowia_unclassified(100); | Metschnikowia | 13 | Saprotroph | Undefined Saprotroph | Yeast | NULL | Probable | As nectar-tap_saprotroph (Põlme et al. 2020); Decay sugar-rich substrates (Põlme et al. 2020) | Põlme S, et al. 2020. Fungal Diversity 105:1-16 (https://doi.org/10.1007/s13225-020-00466-2) |
| Otu40962 | 1 | k__Fungi(100);p__Ascomycota(100);c__Saccharomycetes(100);o__Saccharomycetales(100);f__Metschnikowiaceae(100);g__Metschnikowia(100);g__Metschnikowia_unclassified(100); | Metschnikowia | 13 | Saprotroph | Undefined Saprotroph | Yeast | NULL | Probable | As nectar-tap_saprotroph (Põlme et al. 2020); Decay sugar-rich substrates (Põlme et al. 2020) | Põlme S, et al. 2020. Fungal Diversity 105:1-16 (https://doi.org/10.1007/s13225-020-00466-2) |
| Otu40963 | 1 | k__Fungi(100);p__Ascomycota(100);c__Saccharomycetes(100);o__Saccharomycetales(100);f__Metschnikowiaceae(100);g__Metschnikowia(100);g__Metschnikowia_unclassified(100); | Metschnikowia | 13 | Saprotroph | Undefined Saprotroph | Yeast | NULL | Probable | As nectar-tap_saprotroph (Põlme et al. 2020); Decay sugar-rich substrates (Põlme et al. 2020) | Põlme S, et al. 2020. Fungal Diversity 105:1-16 (https://doi.org/10.1007/s13225-020-00466-2) |
| Otu40964 | 1 | k__Fungi(100);p__Ascomycota(100);c__Saccharomycetes(100);o__Saccharomycetales(100);f__Metschnikowiaceae(100);g__Metschnikowia(100);g__Metschnikowia_unclassified(100); | Metschnikowia | 13 | Saprotroph | Undefined Saprotroph | Yeast | NULL | Probable | As nectar-tap_saprotroph (Põlme et al. 2020); Decay sugar-rich substrates (Põlme et al. 2020) | Põlme S, et al. 2020. Fungal Diversity 105:1-16 (https://doi.org/10.1007/s13225-020-00466-2) |
| Otu40965 | 1 | k__Fungi(100);p__Ascomycota(100);c__Saccharomycetes(100);o__Saccharomycetales(100);f__Metschnikowiaceae(100);g__Metschnikowia(100);g__Metschnikowia_unclassified(100); | Metschnikowia | 13 | Saprotroph | Undefined Saprotroph | Yeast | NULL | Probable | As nectar-tap_saprotroph (Põlme et al. 2020); Decay sugar-rich substrates (Põlme et al. 2020) | Põlme S, et al. 2020. Fungal Diversity 105:1-16 (https://doi.org/10.1007/s13225-020-00466-2) |
| Otu40966 | 1 | k__Fungi(100);p__Ascomycota(100);c__Saccharomycetes(100);o__Saccharomycetales(100);f__Metschnikowiaceae(100);g__Metschnikowia(100);g__Metschnikowia_unclassified(100); | Metschnikowia | 13 | Saprotroph | Undefined Saprotroph | Yeast | NULL | Probable | As nectar-tap_saprotroph (Põlme et al. 2020); Decay sugar-rich substrates (Põlme et al. 2020) | Põlme S, et al. 2020. Fungal Diversity 105:1-16 (https://doi.org/10.1007/s13225-020-00466-2) |
| Otu40967 | 1 | k__Fungi(100);p__Ascomycota(100);c__Saccharomycetes(100);o__Saccharomycetales(100);f__Metschnikowiaceae(100);g__Metschnikowia(100);s__Metschnikowia_pulcherrima(100); | Metschnikowia | 13 | Saprotroph | Undefined Saprotroph | Yeast | NULL | Probable | As nectar-tap_saprotroph (Põlme et al. 2020); Decay sugar-rich substrates (Põlme et al. 2020) | Põlme S, et al. 2020. Fungal Diversity 105:1-16 (https://doi.org/10.1007/s13225-020-00466-2) |
| Otu40968 | 1 | k__Fungi(100);p__Ascomycota(100);c__Saccharomycetes(100);o__Saccharomycetales(100);f__Metschnikowiaceae(100);g__Metschnikowia(100);g__Metschnikowia_unclassified(100); | Metschnikowia | 13 | Saprotroph | Undefined Saprotroph | Yeast | NULL | Probable | As nectar-tap_saprotroph (Põlme et al. 2020); Decay sugar-rich substrates (Põlme et al. 2020) | Põlme S, et al. 2020. Fungal Diversity 105:1-16 (https://doi.org/10.1007/s13225-020-00466-2) |
| Otu40969 | 1 | k__Fungi(100);p__Ascomycota(100);c__Saccharomycetes(100);o__Saccharomycetales(100);f__Metschnikowiaceae(100);g__Metschnikowia(100);s__Metschnikowia_pulcherrima(100); | Metschnikowia | 13 | Saprotroph | Undefined Saprotroph | Yeast | NULL | Probable | As nectar-tap_saprotroph (Põlme et al. 2020); Decay sugar-rich substrates (Põlme et al. 2020) | Põlme S, et al. 2020. Fungal Diversity 105:1-16 (https://doi.org/10.1007/s13225-020-00466-2) |
| Otu40970 | 1 | k__Fungi(100);p__Ascomycota(100);c__Saccharomycetes(100);o__Saccharomycetales(100);f__Metschnikowiaceae(100);g__Metschnikowia(100);s__Metschnikowia_pulcherrima(100); | Metschnikowia | 13 | Saprotroph | Undefined Saprotroph | Yeast | NULL | Probable | As nectar-tap_saprotroph (Põlme et al. 2020); Decay sugar-rich substrates (Põlme et al. 2020) | Põlme S, et al. 2020. Fungal Diversity 105:1-16 (https://doi.org/10.1007/s13225-020-00466-2) |
| Otu40971 | 1 | k__Fungi(100);p__Ascomycota(100);c__Saccharomycetes(100);o__Saccharomycetales(100);f__Metschnikowiaceae(100);g__Metschnikowia(100);s__Metschnikowia_pulcherrima(100); | Metschnikowia | 13 | Saprotroph | Undefined Saprotroph | Yeast | NULL | Probable | As nectar-tap_saprotroph (Põlme et al. 2020); Decay sugar-rich substrates (Põlme et al. 2020) | Põlme S, et al. 2020. Fungal Diversity 105:1-16 (https://doi.org/10.1007/s13225-020-00466-2) |
| Otu40972 | 1 | k__Fungi(100);p__Ascomycota(100);c__Saccharomycetes(100);o__Saccharomycetales(100);f__Metschnikowiaceae(100);g__Metschnikowia(100);s__Metschnikowia_pulcherrima(100); | Metschnikowia | 13 | Saprotroph | Undefined Saprotroph | Yeast | NULL | Probable | As nectar-tap_saprotroph (Põlme et al. 2020); Decay sugar-rich substrates (Põlme et al. 2020) | Põlme S, et al. 2020. Fungal Diversity 105:1-16 (https://doi.org/10.1007/s13225-020-00466-2) |
| Otu40973 | 1 | k__Fungi(100);p__Ascomycota(100);c__Saccharomycetes(100);o__Saccharomycetales(100);f__Metschnikowiaceae(100);g__Metschnikowia(100);s__Metschnikowia_pulcherrima(100); | Metschnikowia | 13 | Saprotroph | Undefined Saprotroph | Yeast | NULL | Probable | As nectar-tap_saprotroph (Põlme et al. 2020); Decay sugar-rich substrates (Põlme et al. 2020) | Põlme S, et al. 2020. Fungal Diversity 105:1-16 (https://doi.org/10.1007/s13225-020-00466-2) |
| Otu40974 | 1 | k__Fungi(100);p__Ascomycota(100);c__Saccharomycetes(100);o__Saccharomycetales(100);f__Metschnikowiaceae(100);g__Metschnikowia(100);s__Metschnikowia_pulcherrima(100); | Metschnikowia | 13 | Saprotroph | Undefined Saprotroph | Yeast | NULL | Probable | As nectar-tap_saprotroph (Põlme et al. 2020); Decay sugar-rich substrates (Põlme et al. 2020) | Põlme S, et al. 2020. Fungal Diversity 105:1-16 (https://doi.org/10.1007/s13225-020-00466-2) |
| Otu40975 | 1 | k__Fungi(100);p__Ascomycota(100);c__Saccharomycetes(100);o__Saccharomycetales(100);f__Metschnikowiaceae(100);g__Metschnikowia(100);s__Metschnikowia_pulcherrima(100); | Metschnikowia | 13 | Saprotroph | Undefined Saprotroph | Yeast | NULL | Probable | As nectar-tap_saprotroph (Põlme et al. 2020); Decay sugar-rich substrates (Põlme et al. 2020) | Põlme S, et al. 2020. Fungal Diversity 105:1-16 (https://doi.org/10.1007/s13225-020-00466-2) |
| Otu40976 | 1 | k__Fungi(100);p__Ascomycota(100);c__Saccharomycetes(100);o__Saccharomycetales(100);f__Metschnikowiaceae(100);g__Metschnikowia(100);s__Metschnikowia_pulcherrima(100); | Metschnikowia | 13 | Saprotroph | Undefined Saprotroph | Yeast | NULL | Probable | As nectar-tap_saprotroph (Põlme et al. 2020); Decay sugar-rich substrates (Põlme et al. 2020) | Põlme S, et al. 2020. Fungal Diversity 105:1-16 (https://doi.org/10.1007/s13225-020-00466-2) |
| Otu40977 | 1 | k__Fungi(100);p__Ascomycota(100);c__Saccharomycetes(100);o__Saccharomycetales(100);f__Metschnikowiaceae(100);g__Metschnikowia(100);s__Metschnikowia_pulcherrima(100); | Metschnikowia | 13 | Saprotroph | Undefined Saprotroph | Yeast | NULL | Probable | As nectar-tap_saprotroph (Põlme et al. 2020); Decay sugar-rich substrates (Põlme et al. 2020) | Põlme S, et al. 2020. Fungal Diversity 105:1-16 (https://doi.org/10.1007/s13225-020-00466-2) |
| Otu40978 | 1 | k__Fungi(100);p__Ascomycota(100);c__Saccharomycetes(100);o__Saccharomycetales(100);f__Metschnikowiaceae(100);g__Metschnikowia(100);g__Metschnikowia_unclassified(100); | Metschnikowia | 13 | Saprotroph | Undefined Saprotroph | Yeast | NULL | Probable | As nectar-tap_saprotroph (Põlme et al. 2020); Decay sugar-rich substrates (Põlme et al. 2020) | Põlme S, et al. 2020. Fungal Diversity 105:1-16 (https://doi.org/10.1007/s13225-020-00466-2) |
| Otu40979 | 1 | k__Fungi(100);p__Ascomycota(100);c__Saccharomycetes(100);o__Saccharomycetales(100);f__Metschnikowiaceae(100);g__Metschnikowia(100);s__Metschnikowia_pulcherrima(100); | Metschnikowia | 13 | Saprotroph | Undefined Saprotroph | Yeast | NULL | Probable | As nectar-tap_saprotroph (Põlme et al. 2020); Decay sugar-rich substrates (Põlme et al. 2020) | Põlme S, et al. 2020. Fungal Diversity 105:1-16 (https://doi.org/10.1007/s13225-020-00466-2) |
| Otu40980 | 1 | k__Fungi(100);p__Ascomycota(100);c__Saccharomycetes(100);o__Saccharomycetales(100);f__Metschnikowiaceae(100);g__Metschnikowia(100);s__Metschnikowia_pulcherrima(100); | Metschnikowia | 13 | Saprotroph | Undefined Saprotroph | Yeast | NULL | Probable | As nectar-tap_saprotroph (Põlme et al. 2020); Decay sugar-rich substrates (Põlme et al. 2020) | Põlme S, et al. 2020. Fungal Diversity 105:1-16 (https://doi.org/10.1007/s13225-020-00466-2) |
| Otu40981 | 1 | k__Fungi(100);p__Ascomycota(100);c__Saccharomycetes(100);o__Saccharomycetales(100);f__Metschnikowiaceae(100);g__Metschnikowia(100);s__Metschnikowia_pulcherrima(100); | Metschnikowia | 13 | Saprotroph | Undefined Saprotroph | Yeast | NULL | Probable | As nectar-tap_saprotroph (Põlme et al. 2020); Decay sugar-rich substrates (Põlme et al. 2020) | Põlme S, et al. 2020. Fungal Diversity 105:1-16 (https://doi.org/10.1007/s13225-020-00466-2) |
| Otu40982 | 1 | k__Fungi(100);p__Ascomycota(100);c__Saccharomycetes(100);o__Saccharomycetales(100);f__Metschnikowiaceae(100);g__Metschnikowia(100);s__Metschnikowia_pulcherrima(100); | Metschnikowia | 13 | Saprotroph | Undefined Saprotroph | Yeast | NULL | Probable | As nectar-tap_saprotroph (Põlme et al. 2020); Decay sugar-rich substrates (Põlme et al. 2020) | Põlme S, et al. 2020. Fungal Diversity 105:1-16 (https://doi.org/10.1007/s13225-020-00466-2) |
| Otu40983 | 1 | k__Fungi(100);p__Ascomycota(100);c__Saccharomycetes(100);o__Saccharomycetales(100);f__Metschnikowiaceae(100);g__Metschnikowia(100);s__Metschnikowia_pulcherrima(100); | Metschnikowia | 13 | Saprotroph | Undefined Saprotroph | Yeast | NULL | Probable | As nectar-tap_saprotroph (Põlme et al. 2020); Decay sugar-rich substrates (Põlme et al. 2020) | Põlme S, et al. 2020. Fungal Diversity 105:1-16 (https://doi.org/10.1007/s13225-020-00466-2) |
| Otu40984 | 1 | k__Fungi(100);p__Ascomycota(100);c__Saccharomycetes(100);o__Saccharomycetales(100);f__Metschnikowiaceae(100);g__Metschnikowia(100);s__Metschnikowia_pulcherrima(100); | Metschnikowia | 13 | Saprotroph | Undefined Saprotroph | Yeast | NULL | Probable | As nectar-tap_saprotroph (Põlme et al. 2020); Decay sugar-rich substrates (Põlme et al. 2020) | Põlme S, et al. 2020. Fungal Diversity 105:1-16 (https://doi.org/10.1007/s13225-020-00466-2) |
| Otu40985 | 1 | k__Fungi(100);p__Ascomycota(100);c__Saccharomycetes(100);o__Saccharomycetales(100);f__Metschnikowiaceae(100);g__Metschnikowia(100);g__Metschnikowia_unclassified(100); | Metschnikowia | 13 | Saprotroph | Undefined Saprotroph | Yeast | NULL | Probable | As nectar-tap_saprotroph (Põlme et al. 2020); Decay sugar-rich substrates (Põlme et al. 2020) | Põlme S, et al. 2020. Fungal Diversity 105:1-16 (https://doi.org/10.1007/s13225-020-00466-2) |
| Otu40986 | 1 | k__Fungi(100);p__Ascomycota(100);c__Saccharomycetes(100);o__Saccharomycetales(100);f__Metschnikowiaceae(100);g__Metschnikowia(100);s__Metschnikowia_pulcherrima(100); | Metschnikowia | 13 | Saprotroph | Undefined Saprotroph | Yeast | NULL | Probable | As nectar-tap_saprotroph (Põlme et al. 2020); Decay sugar-rich substrates (Põlme et al. 2020) | Põlme S, et al. 2020. Fungal Diversity 105:1-16 (https://doi.org/10.1007/s13225-020-00466-2) |
| Otu40987 | 1 | k__Fungi(100);p__Ascomycota(100);c__Saccharomycetes(100);o__Saccharomycetales(100);f__Metschnikowiaceae(100);g__Metschnikowia(100);s__Metschnikowia_pulcherrima(100); | Metschnikowia | 13 | Saprotroph | Undefined Saprotroph | Yeast | NULL | Probable | As nectar-tap_saprotroph (Põlme et al. 2020); Decay sugar-rich substrates (Põlme et al. 2020) | Põlme S, et al. 2020. Fungal Diversity 105:1-16 (https://doi.org/10.1007/s13225-020-00466-2) |
| Otu40988 | 1 | k__Fungi(100);p__Ascomycota(100);c__Saccharomycetes(100);o__Saccharomycetales(100);f__Metschnikowiaceae(100);g__Metschnikowia(100);s__Metschnikowia_pulcherrima(100); | Metschnikowia | 13 | Saprotroph | Undefined Saprotroph | Yeast | NULL | Probable | As nectar-tap_saprotroph (Põlme et al. 2020); Decay sugar-rich substrates (Põlme et al. 2020) | Põlme S, et al. 2020. Fungal Diversity 105:1-16 (https://doi.org/10.1007/s13225-020-00466-2) |
| Otu40989 | 1 | k__Fungi(100);p__Ascomycota(100);c__Saccharomycetes(100);o__Saccharomycetales(100);f__Metschnikowiaceae(100);g__Metschnikowia(100);s__Metschnikowia_pulcherrima(100); | Metschnikowia | 13 | Saprotroph | Undefined Saprotroph | Yeast | NULL | Probable | As nectar-tap_saprotroph (Põlme et al. 2020); Decay sugar-rich substrates (Põlme et al. 2020) | Põlme S, et al. 2020. Fungal Diversity 105:1-16 (https://doi.org/10.1007/s13225-020-00466-2) |
| Otu40990 | 1 | k__Fungi(100);p__Ascomycota(100);c__Saccharomycetes(100);o__Saccharomycetales(100);f__Metschnikowiaceae(100);g__Metschnikowia(100);g__Metschnikowia_unclassified(100); | Metschnikowia | 13 | Saprotroph | Undefined Saprotroph | Yeast | NULL | Probable | As nectar-tap_saprotroph (Põlme et al. 2020); Decay sugar-rich substrates (Põlme et al. 2020) | Põlme S, et al. 2020. Fungal Diversity 105:1-16 (https://doi.org/10.1007/s13225-020-00466-2) |
| Otu40991 | 1 | k__Fungi(100);p__Ascomycota(100);c__Saccharomycetes(100);o__Saccharomycetales(100);f__Metschnikowiaceae(100);g__Metschnikowia(100);s__Metschnikowia_pulcherrima(100); | Metschnikowia | 13 | Saprotroph | Undefined Saprotroph | Yeast | NULL | Probable | As nectar-tap_saprotroph (Põlme et al. 2020); Decay sugar-rich substrates (Põlme et al. 2020) | Põlme S, et al. 2020. Fungal Diversity 105:1-16 (https://doi.org/10.1007/s13225-020-00466-2) |
| Otu40992 | 1 | k__Fungi(100);p__Ascomycota(100);c__Saccharomycetes(100);o__Saccharomycetales(100);f__Metschnikowiaceae(100);g__Metschnikowia(100);g__Metschnikowia_unclassified(100); | Metschnikowia | 13 | Saprotroph | Undefined Saprotroph | Yeast | NULL | Probable | As nectar-tap_saprotroph (Põlme et al. 2020); Decay sugar-rich substrates (Põlme et al. 2020) | Põlme S, et al. 2020. Fungal Diversity 105:1-16 (https://doi.org/10.1007/s13225-020-00466-2) |
| Otu40993 | 1 | k__Fungi(100);p__Ascomycota(100);c__Saccharomycetes(100);o__Saccharomycetales(100);f__Metschnikowiaceae(100);g__Metschnikowia(100);s__Metschnikowia_pulcherrima(100); | Metschnikowia | 13 | Saprotroph | Undefined Saprotroph | Yeast | NULL | Probable | As nectar-tap_saprotroph (Põlme et al. 2020); Decay sugar-rich substrates (Põlme et al. 2020) | Põlme S, et al. 2020. Fungal Diversity 105:1-16 (https://doi.org/10.1007/s13225-020-00466-2) |
| Otu40994 | 1 | k__Fungi(100);p__Ascomycota(100);c__Saccharomycetes(100);o__Saccharomycetales(100);f__Metschnikowiaceae(100);g__Metschnikowia(100);s__Metschnikowia_pulcherrima(100); | Metschnikowia | 13 | Saprotroph | Undefined Saprotroph | Yeast | NULL | Probable | As nectar-tap_saprotroph (Põlme et al. 2020); Decay sugar-rich substrates (Põlme et al. 2020) | Põlme S, et al. 2020. Fungal Diversity 105:1-16 (https://doi.org/10.1007/s13225-020-00466-2) |
| Otu40995 | 1 | k__Fungi(100);p__Ascomycota(100);c__Saccharomycetes(100);o__Saccharomycetales(100);f__Metschnikowiaceae(100);g__Metschnikowia(100);g__Metschnikowia_unclassified(100); | Metschnikowia | 13 | Saprotroph | Undefined Saprotroph | Yeast | NULL | Probable | As nectar-tap_saprotroph (Põlme et al. 2020); Decay sugar-rich substrates (Põlme et al. 2020) | Põlme S, et al. 2020. Fungal Diversity 105:1-16 (https://doi.org/10.1007/s13225-020-00466-2) |
| Otu40996 | 1 | k__Fungi(100);p__Ascomycota(100);c__Saccharomycetes(100);o__Saccharomycetales(100);f__Metschnikowiaceae(100);g__Metschnikowia(100);s__Metschnikowia_pulcherrima(100); | Metschnikowia | 13 | Saprotroph | Undefined Saprotroph | Yeast | NULL | Probable | As nectar-tap_saprotroph (Põlme et al. 2020); Decay sugar-rich substrates (Põlme et al. 2020) | Põlme S, et al. 2020. Fungal Diversity 105:1-16 (https://doi.org/10.1007/s13225-020-00466-2) |
| Otu40997 | 1 | k__Fungi(100);p__Ascomycota(100);c__Saccharomycetes(100);o__Saccharomycetales(100);f__Metschnikowiaceae(100);g__Metschnikowia(100);s__Metschnikowia_pulcherrima(100); | Metschnikowia | 13 | Saprotroph | Undefined Saprotroph | Yeast | NULL | Probable | As nectar-tap_saprotroph (Põlme et al. 2020); Decay sugar-rich substrates (Põlme et al. 2020) | Põlme S, et al. 2020. Fungal Diversity 105:1-16 (https://doi.org/10.1007/s13225-020-00466-2) |
| Otu40998 | 1 | k__Fungi(100);p__Ascomycota(100);c__Saccharomycetes(100);o__Saccharomycetales(100);f__Metschnikowiaceae(100);g__Metschnikowia(100);g__Metschnikowia_unclassified(100); | Metschnikowia | 13 | Saprotroph | Undefined Saprotroph | Yeast | NULL | Probable | As nectar-tap_saprotroph (Põlme et al. 2020); Decay sugar-rich substrates (Põlme et al. 2020) | Põlme S, et al. 2020. Fungal Diversity 105:1-16 (https://doi.org/10.1007/s13225-020-00466-2) |
| Otu40999 | 1 | k__Fungi(100);p__Ascomycota(100);c__Saccharomycetes(100);o__Saccharomycetales(100);f__Metschnikowiaceae(100);g__Metschnikowia(100);g__Metschnikowia_unclassified(100); | Metschnikowia | 13 | Saprotroph | Undefined Saprotroph | Yeast | NULL | Probable | As nectar-tap_saprotroph (Põlme et al. 2020); Decay sugar-rich substrates (Põlme et al. 2020) | Põlme S, et al. 2020. Fungal Diversity 105:1-16 (https://doi.org/10.1007/s13225-020-00466-2) |
| Otu41000 | 1 | k__Fungi(100);p__Ascomycota(100);c__Saccharomycetes(100);o__Saccharomycetales(100);f__Metschnikowiaceae(100);g__Metschnikowia(100);s__Metschnikowia_pulcherrima(100); | Metschnikowia | 13 | Saprotroph | Undefined Saprotroph | Yeast | NULL | Probable | As nectar-tap_saprotroph (Põlme et al. 2020); Decay sugar-rich substrates (Põlme et al. 2020) | Põlme S, et al. 2020. Fungal Diversity 105:1-16 (https://doi.org/10.1007/s13225-020-00466-2) |
| Otu41001 | 1 | k__Fungi(100);p__Ascomycota(100);c__Saccharomycetes(100);o__Saccharomycetales(100);f__Metschnikowiaceae(100);g__Metschnikowia(100);s__Metschnikowia_pulcherrima(100); | Metschnikowia | 13 | Saprotroph | Undefined Saprotroph | Yeast | NULL | Probable | As nectar-tap_saprotroph (Põlme et al. 2020); Decay sugar-rich substrates (Põlme et al. 2020) | Põlme S, et al. 2020. Fungal Diversity 105:1-16 (https://doi.org/10.1007/s13225-020-00466-2) |
| Otu41002 | 1 | k__Fungi(100);p__Ascomycota(100);c__Saccharomycetes(100);o__Saccharomycetales(100);f__Metschnikowiaceae(100);g__Metschnikowia(100);s__Metschnikowia_pulcherrima(100); | Metschnikowia | 13 | Saprotroph | Undefined Saprotroph | Yeast | NULL | Probable | As nectar-tap_saprotroph (Põlme et al. 2020); Decay sugar-rich substrates (Põlme et al. 2020) | Põlme S, et al. 2020. Fungal Diversity 105:1-16 (https://doi.org/10.1007/s13225-020-00466-2) |
| Otu41003 | 1 | k__Fungi(100);p__Ascomycota(100);c__Saccharomycetes(100);o__Saccharomycetales(100);f__Metschnikowiaceae(100);g__Metschnikowia(100);g__Metschnikowia_unclassified(100); | Metschnikowia | 13 | Saprotroph | Undefined Saprotroph | Yeast | NULL | Probable | As nectar-tap_saprotroph (Põlme et al. 2020); Decay sugar-rich substrates (Põlme et al. 2020) | Põlme S, et al. 2020. Fungal Diversity 105:1-16 (https://doi.org/10.1007/s13225-020-00466-2) |
| Otu41005 | 1 | k__Fungi(100);p__Ascomycota(100);c__Saccharomycetes(100);o__Saccharomycetales(100);f__Metschnikowiaceae(100);g__Metschnikowia(100);g__Metschnikowia_unclassified(100); | Metschnikowia | 13 | Saprotroph | Undefined Saprotroph | Yeast | NULL | Probable | As nectar-tap_saprotroph (Põlme et al. 2020); Decay sugar-rich substrates (Põlme et al. 2020) | Põlme S, et al. 2020. Fungal Diversity 105:1-16 (https://doi.org/10.1007/s13225-020-00466-2) |
| Otu41007 | 1 | k__Fungi(100);p__Ascomycota(100);c__Saccharomycetes(100);o__Saccharomycetales(100);f__Metschnikowiaceae(100);g__Metschnikowia(100);s__Metschnikowia_pulcherrima(100); | Metschnikowia | 13 | Saprotroph | Undefined Saprotroph | Yeast | NULL | Probable | As nectar-tap_saprotroph (Põlme et al. 2020); Decay sugar-rich substrates (Põlme et al. 2020) | Põlme S, et al. 2020. Fungal Diversity 105:1-16 (https://doi.org/10.1007/s13225-020-00466-2) |
| Otu41008 | 1 | k__Fungi(100);p__Ascomycota(100);c__Saccharomycetes(100);o__Saccharomycetales(100);f__Metschnikowiaceae(100);g__Metschnikowia(100);g__Metschnikowia_unclassified(100); | Metschnikowia | 13 | Saprotroph | Undefined Saprotroph | Yeast | NULL | Probable | As nectar-tap_saprotroph (Põlme et al. 2020); Decay sugar-rich substrates (Põlme et al. 2020) | Põlme S, et al. 2020. Fungal Diversity 105:1-16 (https://doi.org/10.1007/s13225-020-00466-2) |
| Otu41009 | 1 | k__Fungi(100);p__Ascomycota(100);c__Saccharomycetes(100);o__Saccharomycetales(100);f__Metschnikowiaceae(100);g__Metschnikowia(100);s__Metschnikowia_pulcherrima(100); | Metschnikowia | 13 | Saprotroph | Undefined Saprotroph | Yeast | NULL | Probable | As nectar-tap_saprotroph (Põlme et al. 2020); Decay sugar-rich substrates (Põlme et al. 2020) | Põlme S, et al. 2020. Fungal Diversity 105:1-16 (https://doi.org/10.1007/s13225-020-00466-2) |

| | | | | | | | | | | | |
|---|---|---|---|---|---|---|---|---|---|---|---|
| Otu41010 | 1 | k__Fungi(100);p__Ascomycota(100);c__Saccharomycetes(100);o__Saccharomycetales(100);f__Metschnikowiaceae(100);g__Metschnikowia(100);s__Metschnikowia_pulcherrima(100); | Metschnikowia | 13 | Saprotroph | \|Undefined Saprotroph\| | Yeast | NULL | Probable | As nectar-tap_saprotroph (Põlme et al. 2020); Decay sugar-rich substrates (Põlme et al. 2020) | Põlme S, et al. 2020. Fungal Diversity 105:1-16 ((https://doi.org/10.1007/s13225-020-00466-2)) |
| Otu41011 | 1 | k__Fungi(100);p__Ascomycota(100);c__Saccharomycetes(100);o__Saccharomycetales(100);f__Metschnikowiaceae(100);g__Metschnikowia(100);s__Metschnikowia_pulcherrima(100); | Metschnikowia | 13 | Saprotroph | \|Undefined Saprotroph\| | Yeast | NULL | Probable | As nectar-tap_saprotroph (Põlme et al. 2020); Decay sugar-rich substrates (Põlme et al. 2020) | Põlme S, et al. 2020. Fungal Diversity 105:1-16 ((https://doi.org/10.1007/s13225-020-00466-2)) |
| Otu41012 | 1 | k__Fungi(100);p__Ascomycota(100);c__Saccharomycetes(100);o__Saccharomycetales(100);f__Metschnikowiaceae(100);g__Metschnikowia(100);s__Metschnikowia_pulcherrima(100); | Metschnikowia | 13 | Saprotroph | \|Undefined Saprotroph\| | Yeast | NULL | Probable | As nectar-tap_saprotroph (Põlme et al. 2020); Decay sugar-rich substrates (Põlme et al. 2020) | Põlme S, et al. 2020. Fungal Diversity 105:1-16 ((https://doi.org/10.1007/s13225-020-00466-2)) |
| Otu41013 | 1 | k__Fungi(100);p__Ascomycota(100);c__Saccharomycetes(100);o__Saccharomycetales(100);f__Metschnikowiaceae(100);g__Metschnikowia(100);s__Metschnikowia_pulcherrima(100); | Metschnikowia | 13 | Saprotroph | \|Undefined Saprotroph\| | Yeast | NULL | Probable | As nectar-tap_saprotroph (Põlme et al. 2020); Decay sugar-rich substrates (Põlme et al. 2020) | Põlme S, et al. 2020. Fungal Diversity 105:1-16 ((https://doi.org/10.1007/s13225-020-00466-2)) |
| Otu41014 | 1 | k__Fungi(100);p__Ascomycota(100);c__Saccharomycetes(100);o__Saccharomycetales(100);f__Metschnikowiaceae(100);g__Metschnikowia(100);s__Metschnikowia_pulcherrima(100); | Metschnikowia | 13 | Saprotroph | \|Undefined Saprotroph\| | Yeast | NULL | Probable | As nectar-tap_saprotroph (Põlme et al. 2020); Decay sugar-rich substrates (Põlme et al. 2020) | Põlme S, et al. 2020. Fungal Diversity 105:1-16 ((https://doi.org/10.1007/s13225-020-00466-2)) |
| Otu41016 | 1 | k__Fungi(100);p__Ascomycota(100);c__Saccharomycetes(100);o__Saccharomycetales(100);f__Metschnikowiaceae(100);g__Metschnikowia(100);g__Metschnikowia_unclassified(100); | Metschnikowia | 13 | Saprotroph | \|Undefined Saprotroph\| | Yeast | NULL | Probable | As nectar-tap_saprotroph (Põlme et al. 2020); Decay sugar-rich substrates (Põlme et al. 2020) | Põlme S, et al. 2020. Fungal Diversity 105:1-16 ((https://doi.org/10.1007/s13225-020-00466-2)) |
| Otu41017 | 1 | k__Fungi(100);p__Ascomycota(100);c__Saccharomycetes(100);o__Saccharomycetales(100);f__Metschnikowiaceae(100);g__Metschnikowia(100);g__Metschnikowia_unclassified(100); | Metschnikowia | 13 | Saprotroph | \|Undefined Saprotroph\| | Yeast | NULL | Probable | As nectar-tap_saprotroph (Põlme et al. 2020); Decay sugar-rich substrates (Põlme et al. 2020) | Põlme S, et al. 2020. Fungal Diversity 105:1-16 ((https://doi.org/10.1007/s13225-020-00466-2)) |
| Otu41018 | 1 | k__Fungi(100);p__Ascomycota(100);c__Saccharomycetes(100);o__Saccharomycetales(100);f__Metschnikowiaceae(100);g__Metschnikowia(100);g__Metschnikowia_unclassified(100); | Metschnikowia | 13 | Saprotroph | \|Undefined Saprotroph\| | Yeast | NULL | Probable | As nectar-tap_saprotroph (Põlme et al. 2020); Decay sugar-rich substrates (Põlme et al. 2020) | Põlme S, et al. 2020. Fungal Diversity 105:1-16 ((https://doi.org/10.1007/s13225-020-00466-2)) |
| Otu41019 | 1 | k__Fungi(100);p__Ascomycota(100);c__Saccharomycetes(100);o__Saccharomycetales(100);f__Metschnikowiaceae(100);g__Metschnikowia(100);s__Metschnikowia_pulcherrima(100); | Metschnikowia | 13 | Saprotroph | \|Undefined Saprotroph\| | Yeast | NULL | Probable | As nectar-tap_saprotroph (Põlme et al. 2020); Decay sugar-rich substrates (Põlme et al. 2020) | Põlme S, et al. 2020. Fungal Diversity 105:1-16 ((https://doi.org/10.1007/s13225-020-00466-2)) |
| Otu41020 | 1 | k__Fungi(100);p__Ascomycota(100);c__Saccharomycetes(100);o__Saccharomycetales(100);f__Metschnikowiaceae(100);g__Metschnikowia(100);s__Metschnikowia_pulcherrima(100); | Metschnikowia | 13 | Saprotroph | \|Undefined Saprotroph\| | Yeast | NULL | Probable | As nectar-tap_saprotroph (Põlme et al. 2020); Decay sugar-rich substrates (Põlme et al. 2020) | Põlme S, et al. 2020. Fungal Diversity 105:1-16 ((https://doi.org/10.1007/s13225-020-00466-2)) |
| Otu41021 | 1 | k__Fungi(100);p__Ascomycota(100);c__Saccharomycetes(100);o__Saccharomycetales(100);f__Metschnikowiaceae(100);g__Metschnikowia(100);s__Metschnikowia_pulcherrima(100); | Metschnikowia | 13 | Saprotroph | \|Undefined Saprotroph\| | Yeast | NULL | Probable | As nectar-tap_saprotroph (Põlme et al. 2020); Decay sugar-rich substrates (Põlme et al. 2020) | Põlme S, et al. 2020. Fungal Diversity 105:1-16 ((https://doi.org/10.1007/s13225-020-00466-2)) |
| Otu41022 | 1 | k__Fungi(100);p__Ascomycota(100);c__Saccharomycetes(100);o__Saccharomycetales(100);f__Metschnikowiaceae(100);g__Metschnikowia(100);s__Metschnikowia_pulcherrima(100); | Metschnikowia | 13 | Saprotroph | \|Undefined Saprotroph\| | Yeast | NULL | Probable | As nectar-tap_saprotroph (Põlme et al. 2020); Decay sugar-rich substrates (Põlme et al. 2020) | Põlme S, et al. 2020. Fungal Diversity 105:1-16 ((https://doi.org/10.1007/s13225-020-00466-2)) |
| Otu41023 | 1 | k__Fungi(100);p__Ascomycota(100);c__Saccharomycetes(100);o__Saccharomycetales(100);f__Metschnikowiaceae(100);g__Metschnikowia(100);s__Metschnikowia_pulcherrima(100); | Metschnikowia | 13 | Saprotroph | \|Undefined Saprotroph\| | Yeast | NULL | Probable | As nectar-tap_saprotroph (Põlme et al. 2020); Decay sugar-rich substrates (Põlme et al. 2020) | Põlme S, et al. 2020. Fungal Diversity 105:1-16 ((https://doi.org/10.1007/s13225-020-00466-2)) |
| Otu41024 | 1 | k__Fungi(100);p__Ascomycota(100);c__Saccharomycetes(100);o__Saccharomycetales(100);f__Metschnikowiaceae(100);g__Metschnikowia(100);g__Metschnikowia_unclassified(100); | Metschnikowia | 13 | Saprotroph | \|Undefined Saprotroph\| | Yeast | NULL | Probable | As nectar-tap_saprotroph (Põlme et al. 2020); Decay sugar-rich substrates (Põlme et al. 2020) | Põlme S, et al. 2020. Fungal Diversity 105:1-16 ((https://doi.org/10.1007/s13225-020-00466-2)) |
| Otu41025 | 1 | k__Fungi(100);p__Ascomycota(100);c__Saccharomycetes(100);o__Saccharomycetales(100);f__Metschnikowiaceae(100);g__Metschnikowia(100);s__Metschnikowia_pulcherrima(100); | Metschnikowia | 13 | Saprotroph | \|Undefined Saprotroph\| | Yeast | NULL | Probable | As nectar-tap_saprotroph (Põlme et al. 2020); Decay sugar-rich substrates (Põlme et al. 2020) | Põlme S, et al. 2020. Fungal Diversity 105:1-16 ((https://doi.org/10.1007/s13225-020-00466-2)) |
| Otu41026 | 1 | k__Fungi(100);p__Ascomycota(100);c__Saccharomycetes(100);o__Saccharomycetales(100);f__Metschnikowiaceae(100);g__Metschnikowia(100);s__Metschnikowia_pulcherrima(100); | Metschnikowia | 13 | Saprotroph | \|Undefined Saprotroph\| | Yeast | NULL | Probable | As nectar-tap_saprotroph (Põlme et al. 2020); Decay sugar-rich substrates (Põlme et al. 2020) | Põlme S, et al. 2020. Fungal Diversity 105:1-16 ((https://doi.org/10.1007/s13225-020-00466-2)) |
| Otu41029 | 1 | k__Fungi(100);p__Ascomycota(100);c__Saccharomycetes(100);o__Saccharomycetales(100);f__Metschnikowiaceae(100);g__Metschnikowia(100);s__Metschnikowia_pulcherrima(100); | Metschnikowia | 13 | Saprotroph | \|Undefined Saprotroph\| | Yeast | NULL | Probable | As nectar-tap_saprotroph (Põlme et al. 2020); Decay sugar-rich substrates (Põlme et al. 2020) | Põlme S, et al. 2020. Fungal Diversity 105:1-16 ((https://doi.org/10.1007/s13225-020-00466-2)) |
| Otu41030 | 1 | k__Fungi(100);p__Ascomycota(100);c__Saccharomycetes(100);o__Saccharomycetales(100);f__Metschnikowiaceae(100);g__Metschnikowia(100);s__Metschnikowia_pulcherrima(100); | Metschnikowia | 13 | Saprotroph | \|Undefined Saprotroph\| | Yeast | NULL | Probable | As nectar-tap_saprotroph (Põlme et al. 2020); Decay sugar-rich substrates (Põlme et al. 2020) | Põlme S, et al. 2020. Fungal Diversity 105:1-16 ((https://doi.org/10.1007/s13225-020-00466-2)) |
| Otu41033 | 1 | k__Fungi(100);p__Ascomycota(100);c__Saccharomycetes(100);o__Saccharomycetales(100);f__Metschnikowiaceae(100);g__Metschnikowia(100);s__Metschnikowia_pulcherrima(100); | Metschnikowia | 13 | Saprotroph | \|Undefined Saprotroph\| | Yeast | NULL | Probable | As nectar-tap_saprotroph (Põlme et al. 2020); Decay sugar-rich substrates (Põlme et al. 2020) | Põlme S, et al. 2020. Fungal Diversity 105:1-16 ((https://doi.org/10.1007/s13225-020-00466-2)) |
| Otu41035 | 1 | k__Fungi(100);p__Ascomycota(100);c__Saccharomycetes(100);o__Saccharomycetales(100);f__Metschnikowiaceae(100);g__Metschnikowia(100);s__Metschnikowia_pulcherrima(100); | Metschnikowia | 13 | Saprotroph | \|Undefined Saprotroph\| | Yeast | NULL | Probable | As nectar-tap_saprotroph (Põlme et al. 2020); Decay sugar-rich substrates (Põlme et al. 2020) | Põlme S, et al. 2020. Fungal Diversity 105:1-16 ((https://doi.org/10.1007/s13225-020-00466-2)) |
| Otu41036 | 1 | k__Fungi(100);p__Ascomycota(100);c__Saccharomycetes(100);o__Saccharomycetales(100);f__Metschnikowiaceae(100);g__Metschnikowia(100);s__Metschnikowia_pulcherrima(100); | Metschnikowia | 13 | Saprotroph | \|Undefined Saprotroph\| | Yeast | NULL | Probable | As nectar-tap_saprotroph (Põlme et al. 2020); Decay sugar-rich substrates (Põlme et al. 2020) | Põlme S, et al. 2020. Fungal Diversity 105:1-16 ((https://doi.org/10.1007/s13225-020-00466-2)) |
| Otu41037 | 1 | k__Fungi(100);p__Ascomycota(100);c__Saccharomycetes(100);o__Saccharomycetales(100);f__Metschnikowiaceae(100);g__Metschnikowia(100);s__Metschnikowia_pulcherrima(100); | Metschnikowia | 13 | Saprotroph | \|Undefined Saprotroph\| | Yeast | NULL | Probable | As nectar-tap_saprotroph (Põlme et al. 2020); Decay sugar-rich substrates (Põlme et al. 2020) | Põlme S, et al. 2020. Fungal Diversity 105:1-16 ((https://doi.org/10.1007/s13225-020-00466-2)) |
| Otu41038 | 1 | k__Fungi(100);p__Ascomycota(100);c__Saccharomycetes(100);o__Saccharomycetales(100);f__Metschnikowiaceae(100);g__Metschnikowia(100);s__Metschnikowia_pulcherrima(100); | Metschnikowia | 13 | Saprotroph | \|Undefined Saprotroph\| | Yeast | NULL | Probable | As nectar-tap_saprotroph (Põlme et al. 2020); Decay sugar-rich substrates (Põlme et al. 2020) | Põlme S, et al. 2020. Fungal Diversity 105:1-16 ((https://doi.org/10.1007/s13225-020-00466-2)) |
| Otu41039 | 1 | k__Fungi(100);p__Ascomycota(100);c__Saccharomycetes(100);o__Saccharomycetales(100);f__Metschnikowiaceae(100);g__Metschnikowia(100);g__Metschnikowia_unclassified(100); | Metschnikowia | 13 | Saprotroph | \|Undefined Saprotroph\| | Yeast | NULL | Probable | As nectar-tap_saprotroph (Põlme et al. 2020); Decay sugar-rich substrates (Põlme et al. 2020) | Põlme S, et al. 2020. Fungal Diversity 105:1-16 ((https://doi.org/10.1007/s13225-020-00466-2)) |
| Otu41040 | 1 | k__Fungi(100);p__Ascomycota(100);c__Saccharomycetes(100);o__Saccharomycetales(100);f__Metschnikowiaceae(100);g__Metschnikowia(100);s__Metschnikowia_pulcherrima(100); | Metschnikowia | 13 | Saprotroph | \|Undefined Saprotroph\| | Yeast | NULL | Probable | As nectar-tap_saprotroph (Põlme et al. 2020); Decay sugar-rich substrates (Põlme et al. 2020) | Põlme S, et al. 2020. Fungal Diversity 105:1-16 ((https://doi.org/10.1007/s13225-020-00466-2)) |
| Otu41042 | 1 | k__Fungi(100);p__Ascomycota(100);c__Saccharomycetes(100);o__Saccharomycetales(100);f__Metschnikowiaceae(100);g__Metschnikowia(100);s__Metschnikowia_pulcherrima(100); | Metschnikowia | 13 | Saprotroph | \|Undefined Saprotroph\| | Yeast | NULL | Probable | As nectar-tap_saprotroph (Põlme et al. 2020); Decay sugar-rich substrates (Põlme et al. 2020) | Põlme S, et al. 2020. Fungal Diversity 105:1-16 ((https://doi.org/10.1007/s13225-020-00466-2)) |
| Otu41043 | 1 | k__Fungi(100);p__Ascomycota(100);c__Saccharomycetes(100);o__Saccharomycetales(100);f__Metschnikowiaceae(100);g__Metschnikowia(100);g__Metschnikowia_unclassified(100); | Metschnikowia | 13 | Saprotroph | \|Undefined Saprotroph\| | Yeast | NULL | Probable | As nectar-tap_saprotroph (Põlme et al. 2020); Decay sugar-rich substrates (Põlme et al. 2020) | Põlme S, et al. 2020. Fungal Diversity 105:1-16 ((https://doi.org/10.1007/s13225-020-00466-2)) |
| Otu41044 | 1 | k__Fungi(100);p__Ascomycota(100);c__Saccharomycetes(100);o__Saccharomycetales(100);f__Metschnikowiaceae(100);g__Metschnikowia(100);g__Metschnikowia_unclassified(100); | Metschnikowia | 13 | Saprotroph | \|Undefined Saprotroph\| | Yeast | NULL | Probable | As nectar-tap_saprotroph (Põlme et al. 2020); Decay sugar-rich substrates (Põlme et al. 2020) | Põlme S, et al. 2020. Fungal Diversity 105:1-16 ((https://doi.org/10.1007/s13225-020-00466-2)) |
| Otu41045 | 1 | k__Fungi(100);p__Ascomycota(100);c__Saccharomycetes(100);o__Saccharomycetales(100);f__Metschnikowiaceae(100);g__Metschnikowia(100);g__Metschnikowia_unclassified(100); | Metschnikowia | 13 | Saprotroph | \|Undefined Saprotroph\| | Yeast | NULL | Probable | As nectar-tap_saprotroph (Põlme et al. 2020); Decay sugar-rich substrates (Põlme et al. 2020) | Põlme S, et al. 2020. Fungal Diversity 105:1-16 ((https://doi.org/10.1007/s13225-020-00466-2)) |
| Otu41046 | 1 | k__Fungi(100);p__Ascomycota(100);c__Saccharomycetes(100);o__Saccharomycetales(100);f__Metschnikowiaceae(100);g__Metschnikowia(100);s__Metschnikowia_pulcherrima(100); | Metschnikowia | 13 | Saprotroph | \|Undefined Saprotroph\| | Yeast | NULL | Probable | As nectar-tap_saprotroph (Põlme et al. 2020); Decay sugar-rich substrates (Põlme et al. 2020) | Põlme S, et al. 2020. Fungal Diversity 105:1-16 ((https://doi.org/10.1007/s13225-020-00466-2)) |
| Otu41047 | 1 | k__Fungi(100);p__Ascomycota(100);c__Saccharomycetes(100);o__Saccharomycetales(100);f__Metschnikowiaceae(100);g__Metschnikowia(100);s__Metschnikowia_pulcherrima(100); | Metschnikowia | 13 | Saprotroph | \|Undefined Saprotroph\| | Yeast | NULL | Probable | As nectar-tap_saprotroph (Põlme et al. 2020); Decay sugar-rich substrates (Põlme et al. 2020) | Põlme S, et al. 2020. Fungal Diversity 105:1-16 ((https://doi.org/10.1007/s13225-020-00466-2)) |
| Otu41048 | 1 | k__Fungi(100);p__Ascomycota(100);c__Saccharomycetes(100);o__Saccharomycetales(100);f__Metschnikowiaceae(100);g__Metschnikowia(100);s__Metschnikowia_pulcherrima(100); | Metschnikowia | 13 | Saprotroph | \|Undefined Saprotroph\| | Yeast | NULL | Probable | As nectar-tap_saprotroph (Põlme et al. 2020); Decay sugar-rich substrates (Põlme et al. 2020) | Põlme S, et al. 2020. Fungal Diversity 105:1-16 ((https://doi.org/10.1007/s13225-020-00466-2)) |
| Otu41049 | 1 | k__Fungi(100);p__Ascomycota(100);c__Saccharomycetes(100);o__Saccharomycetales(100);f__Metschnikowiaceae(100);g__Metschnikowia(100);g__Metschnikowia_unclassified(100); | Metschnikowia | 13 | Saprotroph | \|Undefined Saprotroph\| | Yeast | NULL | Probable | As nectar-tap_saprotroph (Põlme et al. 2020); Decay sugar-rich substrates (Põlme et al. 2020) | Põlme S, et al. 2020. Fungal Diversity 105:1-16 ((https://doi.org/10.1007/s13225-020-00466-2)) |
| Otu41050 | 1 | k__Fungi(100);p__Ascomycota(100);c__Saccharomycetes(100);o__Saccharomycetales(100);f__Metschnikowiaceae(100);g__Metschnikowia(100);s__Metschnikowia_pulcherrima(100); | Metschnikowia | 13 | Saprotroph | \|Undefined Saprotroph\| | Yeast | NULL | Probable | As nectar-tap_saprotroph (Põlme et al. 2020); Decay sugar-rich substrates (Põlme et al. 2020) | Põlme S, et al. 2020. Fungal Diversity 105:1-16 ((https://doi.org/10.1007/s13225-020-00466-2)) |
| Otu41051 | 1 | k__Fungi(100);p__Ascomycota(100);c__Saccharomycetes(100);o__Saccharomycetales(100);f__Metschnikowiaceae(100);g__Metschnikowia(100);s__Metschnikowia_pulcherrima(100); | Metschnikowia | 13 | Saprotroph | \|Undefined Saprotroph\| | Yeast | NULL | Probable | As nectar-tap_saprotroph (Põlme et al. 2020); Decay sugar-rich substrates (Põlme et al. 2020) | Põlme S, et al. 2020. Fungal Diversity 105:1-16 ((https://doi.org/10.1007/s13225-020-00466-2)) |
| Otu41052 | 1 | k__Fungi(100);p__Ascomycota(100);c__Saccharomycetes(100);o__Saccharomycetales(100);f__Metschnikowiaceae(100);g__Metschnikowia(100);s__Metschnikowia_pulcherrima(100); | Metschnikowia | 13 | Saprotroph | \|Undefined Saprotroph\| | Yeast | NULL | Probable | As nectar-tap_saprotroph (Põlme et al. 2020); Decay sugar-rich substrates (Põlme et al. 2020) | Põlme S, et al. 2020. Fungal Diversity 105:1-16 ((https://doi.org/10.1007/s13225-020-00466-2)) |
| Otu41054 | 1 | k__Fungi(100);p__Ascomycota(100);c__Saccharomycetes(100);o__Saccharomycetales(100);f__Metschnikowiaceae(100);g__Metschnikowia(100);g__Metschnikowia_unclassified(100); | Metschnikowia | 13 | Saprotroph | \|Undefined Saprotroph\| | Yeast | NULL | Probable | As nectar-tap_saprotroph (Põlme et al. 2020); Decay sugar-rich substrates (Põlme et al. 2020) | Põlme S, et al. 2020. Fungal Diversity 105:1-16 ((https://doi.org/10.1007/s13225-020-00466-2)) |
| Otu41055 | 1 | k__Fungi(100);p__Ascomycota(100);c__Saccharomycetes(100);o__Saccharomycetales(100);f__Metschnikowiaceae(100);g__Metschnikowia(100);g__Metschnikowia_unclassified(100); | Metschnikowia | 13 | Saprotroph | \|Undefined Saprotroph\| | Yeast | NULL | Probable | As nectar-tap_saprotroph (Põlme et al. 2020); Decay sugar-rich substrates (Põlme et al. 2020) | Põlme S, et al. 2020. Fungal Diversity 105:1-16 ((https://doi.org/10.1007/s13225-020-00466-2)) |
| Otu41056 | 1 | k__Fungi(100);p__Ascomycota(100);c__Saccharomycetes(100);o__Saccharomycetales(100);f__Metschnikowiaceae(100);g__Metschnikowia(100);s__Metschnikowia_pulcherrima(100); | Metschnikowia | 13 | Saprotroph | \|Undefined Saprotroph\| | Yeast | NULL | Probable | As nectar-tap_saprotroph (Põlme et al. 2020); Decay sugar-rich substrates (Põlme et al. 2020) | Põlme S, et al. 2020. Fungal Diversity 105:1-16 ((https://doi.org/10.1007/s13225-020-00466-2)) |
| Otu41057 | 1 | k__Fungi(100);p__Ascomycota(100);c__Saccharomycetes(100);o__Saccharomycetales(100);f__Metschnikowiaceae(100);g__Metschnikowia(100);g__Metschnikowia_unclassified(100); | Metschnikowia | 13 | Saprotroph | \|Undefined Saprotroph\| | Yeast | NULL | Probable | As nectar-tap_saprotroph (Põlme et al. 2020); Decay sugar-rich substrates (Põlme et al. 2020) | Põlme S, et al. 2020. Fungal Diversity 105:1-16 ((https://doi.org/10.1007/s13225-020-00466-2)) |
| Otu41058 | 1 | k__Fungi(100);p__Ascomycota(100);c__Saccharomycetes(100);o__Saccharomycetales(100);f__Metschnikowiaceae(100);g__Metschnikowia(100);s__Metschnikowia_pulcherrima(100); | Metschnikowia | 13 | Saprotroph | \|Undefined Saprotroph\| | Yeast | NULL | Probable | As nectar-tap_saprotroph (Põlme et al. 2020); Decay sugar-rich substrates (Põlme et al. 2020) | Põlme S, et al. 2020. Fungal Diversity 105:1-16 ((https://doi.org/10.1007/s13225-020-00466-2)) |
| Otu41059 | 1 | k__Fungi(100);p__Ascomycota(100);c__Saccharomycetes(100);o__Saccharomycetales(100);f__Metschnikowiaceae(100);g__Metschnikowia(100);s__Metschnikowia_pulcherrima(100); | Metschnikowia | 13 | Saprotroph | \|Undefined Saprotroph\| | Yeast | NULL | Probable | As nectar-tap_saprotroph (Põlme et al. 2020); Decay sugar-rich substrates (Põlme et al. 2020) | Põlme S, et al. 2020. Fungal Diversity 105:1-16 ((https://doi.org/10.1007/s13225-020-00466-2)) |
| Otu41060 | 1 | k__Fungi(100);p__Ascomycota(100);c__Saccharomycetes(100);o__Saccharomycetales(100);f__Metschnikowiaceae(100);g__Metschnikowia(100);g__Metschnikowia_unclassified(100); | Metschnikowia | 13 | Saprotroph | \|Undefined Saprotroph\| | Yeast | NULL | Probable | As nectar-tap_saprotroph (Põlme et al. 2020); Decay sugar-rich substrates (Põlme et al. 2020) | Põlme S, et al. 2020. Fungal Diversity 105:1-16 ((https://doi.org/10.1007/s13225-020-00466-2)) |
| Otu41061 | 1 | k__Fungi(100);p__Ascomycota(100);c__Saccharomycetes(100);o__Saccharomycetales(100);f__Metschnikowiaceae(100);g__Metschnikowia(100);s__Metschnikowia_pulcherrima(100); | Metschnikowia | 13 | Saprotroph | \|Undefined Saprotroph\| | Yeast | NULL | Probable | As nectar-tap_saprotroph (Põlme et al. 2020); Decay sugar-rich substrates (Põlme et al. 2020) | Põlme S, et al. 2020. Fungal Diversity 105:1-16 ((https://doi.org/10.1007/s13225-020-00466-2)) |
| Otu41062 | 1 | k__Fungi(100);p__Ascomycota(100);c__Saccharomycetes(100);o__Saccharomycetales(100);f__Metschnikowiaceae(100);g__Metschnikowia(100);s__Metschnikowia_pulcherrima(100); | Metschnikowia | 13 | Saprotroph | \|Undefined Saprotroph\| | Yeast | NULL | Probable | As nectar-tap_saprotroph (Põlme et al. 2020); Decay sugar-rich substrates (Põlme et al. 2020) | Põlme S, et al. 2020. Fungal Diversity 105:1-16 ((https://doi.org/10.1007/s13225-020-00466-2)) |
| Otu41064 | 1 | k__Fungi(100);p__Ascomycota(100);c__Saccharomycetes(100);o__Saccharomycetales(100);f__Metschnikowiaceae(100);g__Metschnikowia(100);s__Metschnikowia_pulcherrima(100); | Metschnikowia | 13 | Saprotroph | \|Undefined Saprotroph\| | Yeast | NULL | Probable | As nectar-tap_saprotroph (Põlme et al. 2020); Decay sugar-rich substrates (Põlme et al. 2020) | Põlme S, et al. 2020. Fungal Diversity 105:1-16 ((https://doi.org/10.1007/s13225-020-00466-2)) |
| Otu41065 | 1 | k__Fungi(100);p__Ascomycota(100);c__Saccharomycetes(100);o__Saccharomycetales(100);f__Metschnikowiaceae(100);g__Metschnikowia(100);s__Metschnikowia_pulcherrima(100); | Metschnikowia | 13 | Saprotroph | \|Undefined Saprotroph\| | Yeast | NULL | Probable | As nectar-tap_saprotroph (Põlme et al. 2020); Decay sugar-rich substrates (Põlme et al. 2020) | Põlme S, et al. 2020. Fungal Diversity 105:1-16 ((https://doi.org/10.1007/s13225-020-00466-2)) |
| Otu41066 | 1 | k__Fungi(100);p__Ascomycota(100);c__Saccharomycetes(100);o__Saccharomycetales(100);f__Metschnikowiaceae(100);g__Metschnikowia(100);s__Metschnikowia_pulcherrima(100); | Metschnikowia | 13 | Saprotroph | \|Undefined Saprotroph\| | Yeast | NULL | Probable | As nectar-tap_saprotroph (Põlme et al. 2020); Decay sugar-rich substrates (Põlme et al. 2020) | Põlme S, et al. 2020. Fungal Diversity 105:1-16 ((https://doi.org/10.1007/s13225-020-00466-2)) |
| Otu41067 | 1 | k__Fungi(100);p__Ascomycota(100);c__Saccharomycetes(100);o__Saccharomycetales(100);f__Metschnikowiaceae(100);g__Metschnikowia(100);s__Metschnikowia_pulcherrima(100); | Metschnikowia | 13 | Saprotroph | \|Undefined Saprotroph\| | Yeast | NULL | Probable | As nectar-tap_saprotroph (Põlme et al. 2020); Decay sugar-rich substrates (Põlme et al. 2020) | Põlme S, et al. 2020. Fungal Diversity 105:1-16 ((https://doi.org/10.1007/s13225-020-00466-2)) |
| Otu41068 | 1 | k__Fungi(100);p__Ascomycota(100);c__Saccharomycetes(100);o__Saccharomycetales(100);f__Metschnikowiaceae(100);g__Metschnikowia(100);g__Metschnikowia_unclassified(100); | Metschnikowia | 13 | Saprotroph | \|Undefined Saprotroph\| | Yeast | NULL | Probable | As nectar-tap_saprotroph (Põlme et al. 2020); Decay sugar-rich substrates (Põlme et al. 2020) | Põlme S, et al. 2020. Fungal Diversity 105:1-16 ((https://doi.org/10.1007/s13225-020-00466-2)) |
| Otu41069 | 1 | k__Fungi(100);p__Ascomycota(100);c__Saccharomycetes(100);o__Saccharomycetales(100);f__Metschnikowiaceae(100);g__Metschnikowia(100);s__Metschnikowia_pulcherrima(100); | Metschnikowia | 13 | Saprotroph | \|Undefined Saprotroph\| | Yeast | NULL | Probable | As nectar-tap_saprotroph (Põlme et al. 2020); Decay sugar-rich substrates (Põlme et al. 2020) | Põlme S, et al. 2020. Fungal Diversity 105:1-16 ((https://doi.org/10.1007/s13225-020-00466-2)) |
| Otu41070 | 1 | k__Fungi(100);p__Ascomycota(100);c__Saccharomycetes(100);o__Saccharomycetales(100);f__Metschnikowiaceae(100);g__Metschnikowia(100);s__Metschnikowia_pulcherrima(100); | Metschnikowia | 13 | Saprotroph | \|Undefined Saprotroph\| | Yeast | NULL | Probable | As nectar-tap_saprotroph (Põlme et al. 2020); Decay sugar-rich substrates (Põlme et al. 2020) | Põlme S, et al. 2020. Fungal Diversity 105:1-16 ((https://doi.org/10.1007/s13225-020-00466-2)) |
| Otu41071 | 1 | k__Fungi(100);p__Ascomycota(100);c__Saccharomycetes(100);o__Saccharomycetales(100);f__Metschnikowiaceae(100);g__Metschnikowia(100);s__Metschnikowia_pulcherrima(100); | Metschnikowia | 13 | Saprotroph | \|Undefined Saprotroph\| | Yeast | NULL | Probable | As nectar-tap_saprotroph (Põlme et al. 2020); Decay sugar-rich substrates (Põlme et al. 2020) | Põlme S, et al. 2020. Fungal Diversity 105:1-16 ((https://doi.org/10.1007/s13225-020-00466-2)) |
| Otu41072 | 1 | k__Fungi(100);p__Ascomycota(100);c__Saccharomycetes(100);o__Saccharomycetales(100);f__Metschnikowiaceae(100);g__Metschnikowia(100);s__Metschnikowia_pulcherrima(100); | Metschnikowia | 13 | Saprotroph | \|Undefined Saprotroph\| | Yeast | NULL | Probable | As nectar-tap_saprotroph (Põlme et al. 2020); Decay sugar-rich substrates (Põlme et al. 2020) | Põlme S, et al. 2020. Fungal Diversity 105:1-16 ((https://doi.org/10.1007/s13225-020-00466-2)) |
| Otu41073 | 1 | k__Fungi(100);p__Ascomycota(100);c__Saccharomycetes(100);o__Saccharomycetales(100);f__Metschnikowiaceae(100);g__Metschnikowia(100);s__Metschnikowia_pulcherrima(100); | Metschnikowia | 13 | Saprotroph | \|Undefined Saprotroph\| | Yeast | NULL | Probable | As nectar-tap_saprotroph (Põlme et al. 2020); Decay sugar-rich substrates (Põlme et al. 2020) | Põlme S, et al. 2020. Fungal Diversity 105:1-16 ((https://doi.org/10.1007/s13225-020-00466-2)) |
| Otu41074 | 1 | k__Fungi(100);p__Ascomycota(100);c__Saccharomycetes(100);o__Saccharomycetales(100);f__Metschnikowiaceae(100);g__Metschnikowia(100);g__Metschnikowia_unclassified(100); | Metschnikowia | 13 | Saprotroph | \|Undefined Saprotroph\| | Yeast | NULL | Probable | As nectar-tap_saprotroph (Põlme et al. 2020); Decay sugar-rich substrates (Põlme et al. 2020) | Põlme S, et al. 2020. Fungal Diversity 105:1-16 ((https://doi.org/10.1007/s13225-020-00466-2)) |
| Otu41075 | 1 | k__Fungi(100);p__Ascomycota(100);c__Saccharomycetes(100);o__Saccharomycetales(100);f__Metschnikowiaceae(100);g__Metschnikowia(100);s__Metschnikowia_pulcherrima(100); | Metschnikowia | 13 | Saprotroph | \|Undefined Saprotroph\| | Yeast | NULL | Probable | As nectar-tap_saprotroph (Põlme et al. 2020); Decay sugar-rich substrates (Põlme et al. 2020) | Põlme S, et al. 2020. Fungal Diversity 105:1-16 ((https://doi.org/10.1007/s13225-020-00466-2)) |
| Otu41076 | 1 | k__Fungi(100);p__Ascomycota(100);c__Saccharomycetes(100);o__Saccharomycetales(100);f__Metschnikowiaceae(100);g__Metschnikowia(100);s__Metschnikowia_pulcherrima(100); | Metschnikowia | 13 | Saprotroph | \|Undefined Saprotroph\| | Yeast | NULL | Probable | As nectar-tap_saprotroph (Põlme et al. 2020); Decay sugar-rich substrates (Põlme et al. 2020) | Põlme S, et al. 2020. Fungal Diversity 105:1-16 ((https://doi.org/10.1007/s13225-020-00466-2)) |
| Otu41077 | 1 | k__Fungi(100);p__Basidiomycota(100);c__Agaricomycetes(100);o__Polyporales(100);f__Fomitopsidaceae(100);g__Piptoporus(100);s__Piptoporus_betulinus(100); | Piptoporus | 13 | Pathotroph-Saprotroph | \|Plant Pathogen\|-Wood Saprotroph | Polyporoid | Brown Rot | Highly Probable | As brown-rot (Põlme et al. 2020); As wood_pathogen (Põlme et al. 2020); Decay wood (Põlme et al. 2020) | Gilbertson RL, Ryvarden L. 1987-1987. North American Polypores. Fungiflora, Oslo (ISBN: 978-0945345060); Põlme S, et al. 2020. Fungal Diversity 105:1-16 ((https://doi.org/10.1007/s13225-020-00466-2)) |
| Otu41078 | 1 | k__Fungi(100);p__Ascomycota(100);c__Saccharomycetes(100);o__Saccharomycetales(100);f__Metschnikowiaceae(100);g__Metschnikowia(100);s__Metschnikowia_pulcherrima(100); | Metschnikowia | 13 | Saprotroph | \|Undefined Saprotroph\| | Yeast | NULL | Probable | As nectar-tap_saprotroph (Põlme et al. 2020); Decay sugar-rich substrates (Põlme et al. 2020) | Põlme S, et al. 2020. Fungal Diversity 105:1-16 ((https://doi.org/10.1007/s13225-020-00466-2)) |
| Otu41079 | 1 | k__Fungi(100);p__Ascomycota(100);c__Saccharomycetes(100);o__Saccharomycetales(100);f__Metschnikowiaceae(100);g__Metschnikowia(100);s__Metschnikowia_pulcherrima(100); | Metschnikowia | 13 | Saprotroph | \|Undefined Saprotroph\| | Yeast | NULL | Probable | As nectar-tap_saprotroph (Põlme et al. 2020); Decay sugar-rich substrates (Põlme et al. 2020) | Põlme S, et al. 2020. Fungal Diversity 105:1-16 ((https://doi.org/10.1007/s13225-020-00466-2)) |
| Otu41080 | 1 | k__Fungi(100);p__Ascomycota(100);c__Saccharomycetes(100);o__Saccharomycetales(100);f__Metschnikowiaceae(100);g__Metschnikowia(100);s__Metschnikowia_pulcherrima(100); | Metschnikowia | 13 | Saprotroph | \|Undefined Saprotroph\| | Yeast | NULL | Probable | As nectar-tap_saprotroph (Põlme et al. 2020); Decay sugar-rich substrates (Põlme et al. 2020) | Põlme S, et al. 2020. Fungal Diversity 105:1-16 ((https://doi.org/10.1007/s13225-020-00466-2)) |
| Otu41081 | 1 | k__Fungi(100);p__Ascomycota(100);c__Saccharomycetes(100);o__Saccharomycetales(100);f__Metschnikowiaceae(100);g__Metschnikowia(100);s__Metschnikowia_pulcherrima(100); | Metschnikowia | 13 | Saprotroph | \|Undefined Saprotroph\| | Yeast | NULL | Probable | As nectar-tap_saprotroph (Põlme et al. 2020); Decay sugar-rich substrates (Põlme et al. 2020) | Põlme S, et al. 2020. Fungal Diversity 105:1-16 ((https://doi.org/10.1007/s13225-020-00466-2)) |
| Otu41082 | 1 | k__Fungi(100);p__Ascomycota(100);c__Saccharomycetes(100);o__Saccharomycetales(100);f__Metschnikowiaceae(100);g__Metschnikowia(100);s__Metschnikowia_pulcherrima(100); | Metschnikowia | 13 | Saprotroph | \|Undefined Saprotroph\| | Yeast | NULL | Probable | As nectar-tap_saprotroph (Põlme et al. 2020); Decay sugar-rich substrates (Põlme et al. 2020) | Põlme S, et al. 2020. Fungal Diversity 105:1-16 ((https://doi.org/10.1007/s13225-020-00466-2)) |
| Otu41083 | 1 | k__Fungi(100);p__Ascomycota(100);c__Saccharomycetes(100);o__Saccharomycetales(100);f__Metschnikowiaceae(100);g__Metschnikowia(100);s__Metschnikowia_pulcherrima(100); | Metschnikowia | 13 | Saprotroph | \|Undefined Saprotroph\| | Yeast | NULL | Probable | As nectar-tap_saprotroph (Põlme et al. 2020); Decay sugar-rich substrates (Põlme et al. 2020) | Põlme S, et al. 2020. Fungal Diversity 105:1-16 ((https://doi.org/10.1007/s13225-020-00466-2)) |
| Otu41084 | 1 | k__Fungi(100);p__Ascomycota(100);c__Saccharomycetes(100);o__Saccharomycetales(100);f__Metschnikowiaceae(100);g__Metschnikowia(100);s__Metschnikowia_pulcherrima(100); | Metschnikowia | 13 | Saprotroph | \|Undefined Saprotroph\| | Yeast | NULL | Probable | As nectar-tap_saprotroph (Põlme et al. 2020); Decay sugar-rich substrates (Põlme et al. 2020) | Põlme S, et al. 2020. Fungal Diversity 105:1-16 ((https://doi.org/10.1007/s13225-020-00466-2)) |
| Otu41085 | 1 | k__Fungi(100);p__Ascomycota(100);c__Saccharomycetes(100);o__Saccharomycetales(100);f__Metschnikowiaceae(100);g__Metschnikowia(100);s__Metschnikowia_pulcherrima(100); | Metschnikowia | 13 | Saprotroph | \|Undefined Saprotroph\| | Yeast | NULL | Probable | As nectar-tap_saprotroph (Põlme et al. 2020); Decay sugar-rich substrates (Põlme et al. 2020) | Põlme S, et al. 2020. Fungal Diversity 105:1-16 ((https://doi.org/10.1007/s13225-020-00466-2)) |
| Otu41086 | 1 | k__Fungi(100);p__Ascomycota(100);c__Saccharomycetes(100);o__Saccharomycetales(100);f__Metschnikowiaceae(100);g__Metschnikowia(100);s__Metschnikowia_pulcherrima(100); | Metschnikowia | 13 | Saprotroph | \|Undefined Saprotroph\| | Yeast | NULL | Probable | As nectar-tap_saprotroph (Põlme et al. 2020); Decay sugar-rich substrates (Põlme et al. 2020) | Põlme S, et al. 2020. Fungal Diversity 105:1-16 ((https://doi.org/10.1007/s13225-020-00466-2)) |
| Otu41087 | 1 | k__Fungi(100);p__Ascomycota(100);c__Saccharomycetes(100);o__Saccharomycetales(100);f__Metschnikowiaceae(100);g__Metschnikowia(100);s__Metschnikowia_pulcherrima(100); | Metschnikowia | 13 | Saprotroph | \|Undefined Saprotroph\| | Yeast | NULL | Probable | As nectar-tap_saprotroph (Põlme et al. 2020); Decay sugar-rich substrates (Põlme et al. 2020) | Põlme S, et al. 2020. Fungal Diversity 105:1-16 ((https://doi.org/10.1007/s13225-020-00466-2)) |
| Otu41088 | 1 | k__Fungi(100);p__Ascomycota(100);c__Saccharomycetes(100);o__Saccharomycetales(100);f__Metschnikowiaceae(100);g__Metschnikowia(100);s__Metschnikowia_pulcherrima(100); | Metschnikowia | 13 | Saprotroph | \|Undefined Saprotroph\| | Yeast | NULL | Probable | As nectar-tap_saprotroph (Põlme et al. 2020); Decay sugar-rich substrates (Põlme et al. 2020) | Põlme S, et al. 2020. Fungal Diversity 105:1-16 ((https://doi.org/10.1007/s13225-020-00466-2)) |
| Otu41089 | 1 | k__Fungi(100);p__Ascomycota(100);c__Saccharomycetes(100);o__Saccharomycetales(100);f__Metschnikowiaceae(100);g__Metschnikowia(100);s__Metschnikowia_pulcherrima(100); | Metschnikowia | 13 | Saprotroph | \|Undefined Saprotroph\| | Yeast | NULL | Probable | As nectar-tap_saprotroph (Põlme et al. 2020); Decay sugar-rich substrates (Põlme et al. 2020) | Põlme S, et al. 2020. Fungal Diversity 105:1-16 ((https://doi.org/10.1007/s13225-020-00466-2)) |
| Otu41090 | 1 | k__Fungi(100);p__Ascomycota(100);c__Saccharomycetes(100);o__Saccharomycetales(100);f__Metschnikowiaceae(100);g__Metschnikowia(100);s__Metschnikowia_pulcherrima(100); | Metschnikowia | 13 | Saprotroph | \|Undefined Saprotroph\| | Yeast | NULL | Probable | As nectar-tap_saprotroph (Põlme et al. 2020); Decay sugar-rich substrates (Põlme et al. 2020) | Põlme S, et al. 2020. Fungal Diversity 105:1-16 ((https://doi.org/10.1007/s13225-020-00466-2)) |
| Otu41091 | 1 | k__Fungi(100);p__Ascomycota(100);c__Saccharomycetes(100);o__Saccharomycetales(100);f__Metschnikowiaceae(100);g__Metschnikowia(100);s__Metschnikowia_pulcherrima(100); | Metschnikowia | 13 | Saprotroph | \|Undefined Saprotroph\| | Yeast | NULL | Probable | As nectar-tap_saprotroph (Põlme et al. 2020); Decay sugar-rich substrates (Põlme et al. 2020) | Põlme S, et al. 2020. Fungal Diversity 105:1-16 ((https://doi.org/10.1007/s13225-020-00466-2)) |
| Otu41092 | 1 | k__Fungi(100);p__Ascomycota(100);c__Saccharomycetes(100);o__Saccharomycetales(100);f__Metschnikowiaceae(100);g__Metschnikowia(100);g__Metschnikowia_unclassified(100); | Metschnikowia | 13 | Saprotroph | \|Undefined Saprotroph\| | Yeast | NULL | Probable | As nectar-tap_saprotroph (Põlme et al. 2020); Decay sugar-rich substrates (Põlme et al. 2020) | Põlme S, et al. 2020. Fungal Diversity 105:1-16 ((https://doi.org/10.1007/s13225-020-00466-2)) |
| Otu41094 | 1 | k__Fungi(100);p__Ascomycota(100);c__Saccharomycetes(100);o__Saccharomycetales(100);f__Metschnikowiaceae(100);g__Metschnikowia(100);s__Metschnikowia_pulcherrima(100); | Metschnikowia | 13 | Saprotroph | \|Undefined Saprotroph\| | Yeast | NULL | Probable | As nectar-tap_saprotroph (Põlme et al. 2020); Decay sugar-rich substrates (Põlme et al. 2020) | Põlme S, et al. 2020. Fungal Diversity 105:1-16 ((https://doi.org/10.1007/s13225-020-00466-2)) |
| Otu41095 | 1 | k__Fungi(100);p__Ascomycota(100);c__Saccharomycetes(100);o__Saccharomycetales(100);f__Metschnikowiaceae(100);g__Metschnikowia(100);g__Metschnikowia_unclassified(100); | Metschnikowia | 13 | Saprotroph | \|Undefined Saprotroph\| | Yeast | NULL | Probable | As nectar-tap_saprotroph (Põlme et al. 2020); Decay sugar-rich substrates (Põlme et al. 2020) | Põlme S, et al. 2020. Fungal Diversity 105:1-16 ((https://doi.org/10.1007/s13225-020-00466-2)) |
| Otu41096 | 1 | k__Fungi(100);p__Ascomycota(100);c__Saccharomycetes(100);o__Saccharomycetales(100);f__Metschnikowiaceae(100);g__Metschnikowia(100);s__Metschnikowia_pulcherrima(100); | Metschnikowia | 13 | Saprotroph | \|Undefined Saprotroph\| | Yeast | NULL | Probable | As nectar-tap_saprotroph (Põlme et al. 2020); Decay sugar-rich substrates (Põlme et al. 2020) | Põlme S, et al. 2020. Fungal Diversity 105:1-16 ((https://doi.org/10.1007/s13225-020-00466-2)) |
| Otu41097 | 1 | k__Fungi(100);p__Ascomycota(100);c__Saccharomycetes(100);o__Saccharomycetales(100);f__Metschnikowiaceae(100);g__Metschnikowia(100);g__Metschnikowia_unclassified(100); | Metschnikowia | 13 | Saprotroph | \|Undefined Saprotroph\| | Yeast | NULL | Probable | As nectar-tap_saprotroph (Põlme et al. 2020); Decay sugar-rich substrates (Põlme et al. 2020) | Põlme S, et al. 2020. Fungal Diversity 105:1-16 ((https://doi.org/10.1007/s13225-020-00466-2)) |
| Otu41098 | 1 | k__Fungi(100);p__Ascomycota(100);c__Saccharomycetes(100);o__Saccharomycetales(100);f__Metschnikowiaceae(100);g__Metschnikowia(100);g__Metschnikowia_unclassified(100); | Metschnikowia | 13 | Saprotroph | \|Undefined Saprotroph\| | Yeast | NULL | Probable | As nectar-tap_saprotroph (Põlme et al. 2020); Decay sugar-rich substrates (Põlme et al. 2020) | Põlme S, et al. 2020. Fungal Diversity 105:1-16 ((https://doi.org/10.1007/s13225-020-00466-2)) |
| Otu41099 | 1 | k__Fungi(100);p__Ascomycota(100);c__Saccharomycetes(100);o__Saccharomycetales(100);f__Metschnikowiaceae(100);g__Metschnikowia(100);g__Metschnikowia_unclassified(100); | Metschnikowia | 13 | Saprotroph | \|Undefined Saprotroph\| | Yeast | NULL | Probable | As nectar-tap_saprotroph (Põlme et al. 2020); Decay sugar-rich substrates (Põlme et al. 2020) | Põlme S, et al. 2020. Fungal Diversity 105:1-16 ((https://doi.org/10.1007/s13225-020-00466-2)) |
| Otu41100 | 1 | k__Fungi(100);p__Ascomycota(100);c__Saccharomycetes(100);o__Saccharomycetales(100);f__Metschnikowiaceae(100);g__Metschnikowia(100);g__Metschnikowia_unclassified(100); | Metschnikowia | 13 | Saprotroph | \|Undefined Saprotroph\| | Yeast | NULL | Probable | As nectar-tap_saprotroph (Põlme et al. 2020); Decay sugar-rich substrates (Põlme et al. 2020) | Põlme S, et al. 2020. Fungal Diversity 105:1-16 ((https://doi.org/10.1007/s13225-020-00466-2)) |
| Otu41101 | 1 | k__Fungi(100);p__Ascomycota(100);c__Saccharomycetes(100);o__Saccharomycetales(100);f__Metschnikowiaceae(100);g__Metschnikowia(100);g__Metschnikowia_unclassified(100); | Metschnikowia | 13 | Saprotroph | \|Undefined Saprotroph\| | Yeast | NULL | Probable | As nectar-tap_saprotroph (Põlme et al. 2020); Decay sugar-rich substrates (Põlme et al. 2020) | Põlme S, et al. 2020. Fungal Diversity 105:1-16 ((https://doi.org/10.1007/s13225-020-00466-2)) |
| Otu41102 | 1 | k__Fungi(100);p__Ascomycota(100);c__Saccharomycetes(100);o__Saccharomycetales(100);f__Metschnikowiaceae(100);g__Metschnikowia(100);s__Metschnikowia_pulcherrima(100); | Metschnikowia | 13 | Saprotroph | \|Undefined Saprotroph\| | Yeast | NULL | Probable | As nectar-tap_saprotroph (Põlme et al. 2020); Decay sugar-rich substrates (Põlme et al. 2020) | Põlme S, et al. 2020. Fungal Diversity 105:1-16 ((https://doi.org/10.1007/s13225-020-00466-2)) |
| Otu41103 | 1 | k__Fungi(100);p__Ascomycota(100);c__Saccharomycetes(100);o__Saccharomycetales(100);f__Metschnikowiaceae(100);g__Metschnikowia(100);s__Metschnikowia_pulcherrima(100); | Metschnikowia | 13 | Saprotroph | \|Undefined Saprotroph\| | Yeast | NULL | Probable | As nectar-tap_saprotroph (Põlme et al. 2020); Decay sugar-rich substrates (Põlme et al. 2020) | Põlme S, et al. 2020. Fungal Diversity 105:1-16 ((https://doi.org/10.1007/s13225-020-00466-2)) |
| Otu41104 | 1 | k__Fungi(100);p__Ascomycota(100);c__Saccharomycetes(100);o__Saccharomycetales(100);f__Metschnikowiaceae(100);g__Metschnikowia(100);g__Metschnikowia_unclassified(100); | Metschnikowia | 13 | Saprotroph | \|Undefined Saprotroph\| | Yeast | NULL | Probable | As nectar-tap_saprotroph (Põlme et al. 2020); Decay sugar-rich substrates (Põlme et al. 2020) | Põlme S, et al. 2020. Fungal Diversity 105:1-16 ((https://doi.org/10.1007/s13225-020-00466-2)) |
| Otu41105 | 1 | k__Fungi(100);p__Ascomycota(100);c__Saccharomycetes(100);o__Saccharomycetales(100);f__Metschnikowiaceae(100);g__Metschnikowia(100);s__Metschnikowia_pulcherrima(100); | Metschnikowia | 13 | Saprotroph | \|Undefined Saprotroph\| | Yeast | NULL | Probable | As nectar-tap_saprotroph (Põlme et al. 2020); Decay sugar-rich substrates (Põlme et al. 2020) | Põlme S, et al. 2020. Fungal Diversity 105:1-16 ((https://doi.org/10.1007/s13225-020-00466-2)) |
| Otu41106 | 1 | k__Fungi(100);p__Ascomycota(100);c__Saccharomycetes(100);o__Saccharomycetales(100);f__Metschnikowiaceae(100);g__Metschnikowia(100);s__Metschnikowia_pulcherrima(100); | Metschnikowia | 13 | Saprotroph | \|Undefined Saprotroph\| | Yeast | NULL | Probable | As nectar-tap_saprotroph (Põlme et al. 2020); Decay sugar-rich substrates (Põlme et al. 2020) | Põlme S, et al. 2020. Fungal Diversity 105:1-16 ((https://doi.org/10.1007/s13225-020-00466-2)) |
| Otu41107 | 1 | k__Fungi(100);p__Ascomycota(100);c__Saccharomycetes(100);o__Saccharomycetales(100);f__Metschnikowiaceae(100);g__Metschnikowia(100);g__Metschnikowia_unclassified(100); | Metschnikowia | 13 | Saprotroph | \|Undefined Saprotroph\| | Yeast | NULL | Probable | As nectar-tap_saprotroph (Põlme et al. 2020); Decay sugar-rich substrates (Põlme et al. 2020) | Põlme S, et al. 2020. Fungal Diversity 105:1-16 ((https://doi.org/10.1007/s13225-020-00466-2)) |
| Otu41108 | 1 | k__Fungi(100);p__Ascomycota(100);c__Saccharomycetes(100);o__Saccharomycetales(100);f__Metschnikowiaceae(100);g__Metschnikowia(100);s__Metschnikowia_pulcherrima(100); | Metschnikowia | 13 | Saprotroph | \|Undefined Saprotroph\| | Yeast | NULL | Probable | As nectar-tap_saprotroph (Põlme et al. 2020); Decay sugar-rich substrates (Põlme et al. 2020) | Põlme S, et al. 2020. Fungal Diversity 105:1-16 ((https://doi.org/10.1007/s13225-020-00466-2)) |
| Otu41109 | 1 | k__Fungi(100);p__Ascomycota(100);c__Saccharomycetes(100);o__Saccharomycetales(100);f__Metschnikowiaceae(100);g__Metschnikowia(100);s__Metschnikowia_pulcherrima(100); | Metschnikowia | 13 | Saprotroph | \|Undefined Saprotroph\| | Yeast | NULL | Probable | As nectar-tap_saprotroph (Põlme et al. 2020); Decay sugar-rich substrates (Põlme et al. 2020) | Põlme S, et al. 2020. Fungal Diversity 105:1-16 ((https://doi.org/10.1007/s13225-020-00466-2)) |
| Otu41110 | 1 | k__Fungi(100);p__Ascomycota(100);c__Saccharomycetes(100);o__Saccharomycetales(100);f__Metschnikowiaceae(100);g__Metschnikowia(100);g__Metschnikowia_unclassified(100); | Metschnikowia | 13 | Saprotroph | \|Undefined Saprotroph\| | Yeast | NULL | Probable | As nectar-tap_saprotroph (Põlme et al. 2020); Decay sugar-rich substrates (Põlme et al. 2020) | Põlme S, et al. 2020. Fungal Diversity 105:1-16 ((https://doi.org/10.1007/s13225-020-00466-2)) |
| Otu41111 | 1 | k__Fungi(100);p__Ascomycota(100);c__Saccharomycetes(100);o__Saccharomycetales(100);f__Metschnikowiaceae(100);g__Metschnikowia(100);s__Metschnikowia_pulcherrima(100); | Metschnikowia | 13 | Saprotroph | \|Undefined Saprotroph\| | Yeast | NULL | Probable | As nectar-tap_saprotroph (Põlme et al. 2020); Decay sugar-rich substrates (Põlme et al. 2020) | Põlme S, et al. 2020. Fungal Diversity 105:1-16 ((https://doi.org/10.1007/s13225-020-00466-2)) |
| Otu41112 | 1 | k__Fungi(100);p__Ascomycota(100);c__Saccharomycetes(100);o__Saccharomycetales(100);f__Metschnikowiaceae(100);g__Metschnikowia(100);s__Metschnikowia_pulcherrima(100); | Metschnikowia | 13 | Saprotroph | \|Undefined Saprotroph\| | Yeast | NULL | Probable | As nectar-tap_saprotroph (Põlme et al. 2020); Decay sugar-rich substrates (Põlme et al. 2020) | Põlme S, et al. 2020. Fungal Diversity 105:1-16 ((https://doi.org/10.1007/s13225-020-00466-2)) |
| Otu41113 | 1 | k__Fungi(100);p__Ascomycota(100);c__Saccharomycetes(100);o__Saccharomycetales(100);f__Metschnikowiaceae(100);g__Metschnikowia(100);g__Metschnikowia_unclassified(100); | Metschnikowia | 13 | Saprotroph | \|Undefined Saprotroph\| | Yeast | NULL | Probable | As nectar-tap_saprotroph (Põlme et al. 2020); Decay sugar-rich substrates (Põlme et al. 2020) | Põlme S, et al. 2020. Fungal Diversity 105:1-16 ((https://doi.org/10.1007/s13225-020-00466-2)) |
| Otu41114 | 1 | k__Fungi(100);p__Ascomycota(100);c__Saccharomycetes(100);o__Saccharomycetales(100);f__Metschnikowiaceae(100);g__Metschnikowia(100);s__Metschnikowia_pulcherrima(100); | Metschnikowia | 13 | Saprotroph | \|Undefined Saprotroph\| | Yeast | NULL | Probable | As nectar-tap_saprotroph (Põlme et al. 2020); Decay sugar-rich substrates (Põlme et al. 2020) | Põlme S, et al. 2020. Fungal Diversity 105:1-16 ((https://doi.org/10.1007/s13225-020-00466-2)) |
| Otu41115 | 1 | k__Fungi(100);p__Ascomycota(100);c__Saccharomycetes(100);o__Saccharomycetales(100);f__Metschnikowiaceae(100);g__Metschnikowia(100);s__Metschnikowia_pulcherrima(100); | Metschnikowia | 13 | Saprotroph | \|Undefined Saprotroph\| | Yeast | NULL | Probable | As nectar-tap_saprotroph (Põlme et al. 2020); Decay sugar-rich substrates (Põlme et al. 2020) | Põlme S, et al. 2020. Fungal Diversity 105:1-16 ((https://doi.org/10.1007/s13225-020-00466-2)) |
| Otu41116 | 1 | k__Fungi(100);p__Ascomycota(100);c__Saccharomycetes(100);o__Saccharomycetales(100);f__Metschnikowiaceae(100);g__Metschnikowia(100);s__Metschnikowia_pulcherrima(100); | Metschnikowia | 13 | Saprotroph | \|Undefined Saprotroph\| | Yeast | NULL | Probable | As nectar-tap_saprotroph (Põlme et al. 2020); Decay sugar-rich substrates (Põlme et al. 2020) | Põlme S, et al. 2020. Fungal Diversity 105:1-16 ((https://doi.org/10.1007/s13225-020-00466-2)) |
| Otu41117 | 1 | k__Fungi(100);p__Ascomycota(100);c__Saccharomycetes(100);o__Saccharomycetales(100);f__Metschnikowiaceae(100);g__Metschnikowia(100);s__Metschnikowia_pulcherrima(100); | Metschnikowia | 13 | Saprotroph | \|Undefined Saprotroph\| | Yeast | NULL | Probable | As nectar-tap_saprotroph (Põlme et al. 2020); Decay sugar-rich substrates (Põlme et al. 2020) | Põlme S, et al. 2020. Fungal Diversity 105:1-16 ((https://doi.org/10.1007/s13225-020-00466-2)) |
| Otu41118 | 1 | k__Fungi(100);p__Ascomycota(100);c__Saccharomycetes(100);o__Saccharomycetales(100);f__Metschnikowiaceae(100);g__Metschnikowia(100);s__Metschnikowia_pulcherrima(100); | Metschnikowia | 13 | Saprotroph | \|Undefined Saprotroph\| | Yeast | NULL | Probable | As nectar-tap_saprotroph (Põlme et al. 2020); Decay sugar-rich substrates (Põlme et al. 2020) | Põlme S, et al. 2020. Fungal Diversity 105:1-16 ((https://doi.org/10.1007/s13225-020-00466-2)) |
| Otu41119 | 1 | k__Fungi(100);p__Ascomycota(100);c__Saccharomycetes(100);o__Saccharomycetales(100);f__Metschnikowiaceae(100);g__Metschnikowia(100);s__Metschnikowia_pulcherrima(100); | Metschnikowia | 13 | Saprotroph | \|Undefined Saprotroph\| | Yeast | NULL | Probable | As nectar-tap_saprotroph (Põlme et al. 2020); Decay sugar-rich substrates (Põlme et al. 2020) | Põlme S, et al. 2020. Fungal Diversity 105:1-16 ((https://doi.org/10.1007/s13225-020-00466-2)) |
| Otu41120 | 1 | k__Fungi(100);p__Ascomycota(100);c__Saccharomycetes(100);o__Saccharomycetales(100);f__Metschnikowiaceae(100);g__Metschnikowia(100);s__Metschnikowia_pulcherrima(100); | Metschnikowia | 13 | Saprotroph | \|Undefined Saprotroph\| | Yeast | NULL | Probable | As nectar-tap_saprotroph (Põlme et al. 2020); Decay sugar-rich substrates (Põlme et al. 2020) | Põlme S, et al. 2020. Fungal Diversity 105:1-16 ((https://doi.org/10.1007/s13225-020-00466-2)) |
| Otu41121 | 1 | k__Fungi(100);p__Ascomycota(100);c__Saccharomycetes(100);o__Saccharomycetales(100);f__Metschnikowiaceae(100);g__Metschnikowia(100);s__Metschnikowia_pulcherrima(100); | Metschnikowia | 13 | Saprotroph | \|Undefined Saprotroph\| | Yeast | NULL | Probable | As nectar-tap_saprotroph (Põlme et al. 2020); Decay sugar-rich substrates (Põlme et al. 2020) | Põlme S, et al. 2020. Fungal Diversity 105:1-16 ((https://doi.org/10.1007/s13225-020-00466-2)) |
| Otu41122 | 1 | k__Fungi(100);p__Ascomycota(100);c__Saccharomycetes(100);o__Saccharomycetales(100);f__Metschnikowiaceae(100);g__Metschnikowia(100);s__Metschnikowia_pulcherrima(100); | Metschnikowia | 13 | Saprotroph | \|Undefined Saprotroph\| | Yeast | NULL | Probable | As nectar-tap_saprotroph (Põlme et al. 2020); Decay sugar-rich substrates (Põlme et al. 2020) | Põlme S, et al. 2020. Fungal Diversity 105:1-16 ((https://doi.org/10.1007/s13225-020-00466-2)) |
| Otu41123 | 1 | k__Fungi(100);p__Ascomycota(100);c__Saccharomycetes(100);o__Saccharomycetales(100);f__Metschnikowiaceae(100);g__Metschnikowia(100);s__Metschnikowia_pulcherrima(100); | Metschnikowia | 13 | Saprotroph | \|Undefined Saprotroph\| | Yeast | NULL | Probable | As nectar-tap_saprotroph (Põlme et al. 2020); Decay sugar-rich substrates (Põlme et al. 2020) | Põlme S, et al. 2020. Fungal Diversity 105:1-16 ((https://doi.org/10.1007/s13225-020-00466-2)) |
| Otu41124 | 1 | k__Fungi(100);p__Ascomycota(100);c__Saccharomycetes(100);o__Saccharomycetales(100);f__Metschnikowiaceae(100);g__Metschnikowia(100);s__Metschnikowia_pulcherrima(100); | Metschnikowia | 13 | Saprotroph | \|Undefined Saprotroph\| | Yeast | NULL | Probable | As nectar-tap_saprotroph (Põlme et al. 2020); Decay sugar-rich substrates (Põlme et al. 2020) | Põlme S, et al. 2020. Fungal Diversity 105:1-16 ((https://doi.org/10.1007/s13225-020-00466-2)) |
| Otu41125 | 1 | k__Fungi(100);p__Ascomycota(100);c__Saccharomycetes(100);o__Saccharomycetales(100);f__Metschnikowiaceae(100);g__Metschnikowia(100);s__Metschnikowia_pulcherrima(100); | Metschnikowia | 13 | Saprotroph | \|Undefined Saprotroph\| | Yeast | NULL | Probable | As nectar-tap_saprotroph (Põlme et al. 2020); Decay sugar-rich substrates (Põlme et al. 2020) | Põlme S, et al. 2020. Fungal Diversity 105:1-16 ((https://doi.org/10.1007/s13225-020-00466-2)) |
| Otu41126 | 1 | k__Fungi(100);p__Ascomycota(100);c__Saccharomycetes(100);o__Saccharomycetales(100);f__Metschnikowiaceae(100);g__Metschnikowia(100);s__Metschnikowia_pulcherrima(100); | Metschnikowia | 13 | Saprotroph | \|Undefined Saprotroph\| | Yeast | NULL | Probable | As nectar-tap_saprotroph (Põlme et al. 2020); Decay sugar-rich substrates (Põlme et al. 2020) | Põlme S, et al. 2020. Fungal Diversity 105:1-16 ((https://doi.org/10.1007/s13225-020-00466-2)) |
| Otu41127 | 1 | k__Fungi(100);p__Ascomycota(100);c__Saccharomycetes(100);o__Saccharomycetales(100);f__Metschnikowiaceae(100);g__Metschnikowia(100);g__Metschnikowia_unclassified(100); | Metschnikowia | 13 | Saprotroph | \|Undefined Saprotroph\| | Yeast | NULL | Probable | As nectar-tap_saprotroph (Põlme et al. 2020); Decay sugar-rich substrates (Põlme et al. 2020) | Põlme S, et al. 2020. Fungal Diversity 105:1-16 ((https://doi.org/10.1007/s13225-020-00466-2)) |
| Otu41128 | 1 | k__Fungi(100);p__Ascomycota(100);c__Saccharomycetes(100);o__Saccharomycetales(100);f__Metschnikowiaceae(100);g__Metschnikowia(100);s__Metschnikowia_pulcherrima(100); | Metschnikowia | 13 | Saprotroph | \|Undefined Saprotroph\| | Yeast | NULL | Probable | As nectar-tap_saprotroph (Põlme et al. 2020); Decay sugar-rich substrates (Põlme et al. 2020) | Põlme S, et al. 2020. Fungal Diversity 105:1-16 ((https://doi.org/10.1007/s13225-020-00466-2)) |
| Otu41129 | 1 | k__Fungi(100);p__Ascomycota(100);c__Saccharomycetes(100);o__Saccharomycetales(100);f__Metschnikowiaceae(100);g__Metschnikowia(100);s__Metschnikowia_pulcherrima(100); | Metschnikowia | 13 | Saprotroph | \|Undefined Saprotroph\| | Yeast | NULL | Probable | As nectar-tap_saprotroph (Põlme et al. 2020); Decay sugar-rich substrates (Põlme et al. 2020) | Põlme S, et al. 2020. Fungal Diversity 105:1-16 ((https://doi.org/10.1007/s13225-020-00466-2)) |
| Otu41130 | 1 | k__Fungi(100);p__Ascomycota(100);c__Saccharomycetes(100);o__Saccharomycetales(100);f__Metschnikowiaceae(100);g__Metschnikowia(100);s__Metschnikowia_pulcherrima(100); | Metschnikowia | 13 | Saprotroph | \|Undefined Saprotroph\| | Yeast | NULL | Probable | As nectar-tap_saprotroph (Põlme et al. 2020); Decay sugar-rich substrates (Põlme et al. 2020) | Põlme S, et al. 2020. Fungal Diversity 105:1-16 ((https://doi.org/10.1007/s13225-020-00466-2)) |
| Otu41131 | 1 | k__Fungi(100);p__Ascomycota(100);c__Saccharomycetes(100);o__Saccharomycetales(100);f__Metschnikowiaceae(100);g__Metschnikowia(100);s__Metschnikowia_pulcherrima(100); | Metschnikowia | 13 | Saprotroph | \|Undefined Saprotroph\| | Yeast | NULL | Probable | As nectar-tap_saprotroph (Põlme et al. 2020); Decay sugar-rich substrates (Põlme et al. 2020) | Põlme S, et al. 2020. Fungal Diversity 105:1-16 ((https://doi.org/10.1007/s13225-020-00466-2)) |
| Otu41132 | 1 | k__Fungi(100);p__Ascomycota(100);c__Saccharomycetes(100);o__Saccharomycetales(100);f__Metschnikowiaceae(100);g__Metschnikowia(100);g__Metschnikowia_unclassified(100); | Metschnikowia | 13 | Saprotroph | \|Undefined Saprotroph\| | Yeast | NULL | Probable | As nectar-tap_saprotroph (Põlme et al. 2020); Decay sugar-rich substrates (Põlme et al. 2020) | Põlme S, et al. 2020. Fungal Diversity 105:1-16 ((https://doi.org/10.1007/s13225-020-00466-2)) |
| Otu41133 | 1 | k__Fungi(100);p__Ascomycota(100);c__Saccharomycetes(100);o__Saccharomycetales(100);f__Metschnikowiaceae(100);g__Metschnikowia(100);s__Metschnikowia_pulcherrima(100); | Metschnikowia | 13 | Saprotroph | \|Undefined Saprotroph\| | Yeast | NULL | Probable | As nectar-tap_saprotroph (Põlme et al. 2020); Decay sugar-rich substrates (Põlme et al. 2020) | Põlme S, et al. 2020. Fungal Diversity 105:1-16 ((https://doi.org/10.1007/s13225-020-00466-2)) |
| Otu41134 | 1 | k__Fungi(100);p__Ascomycota(100);c__Saccharomycetes(100);o__Saccharomycetales(100);f__Metschnikowiaceae(100);g__Metschnikowia(100);s__Metschnikowia_pulcherrima(100); | Metschnikowia | 13 | Saprotroph | \|Undefined Saprotroph\| | Yeast | NULL | Probable | As nectar-tap_saprotroph (Põlme et al. 2020); Decay sugar-rich substrates (Põlme et al. 2020) | Põlme S, et al. 2020. Fungal Diversity 105:1-16 ((https://doi.org/10.1007/s13225-020-00466-2)) |
| Otu41135 | 1 | k__Fungi(100);p__Ascomycota(100);c__Saccharomycetes(100);o__Saccharomycetales(100);f__Metschnikowiaceae(100);g__Metschnikowia(100);s__Metschnikowia_pulcherrima(100); | Metschnikowia | 13 | Saprotroph | \|Undefined Saprotroph\| | Yeast | NULL | Probable | As nectar-tap_saprotroph (Põlme et al. 2020); Decay sugar-rich substrates (Põlme et al. 2020) | Põlme S, et al. 2020. Fungal Diversity 105:1-16 ((https://doi.org/10.1007/s13225-020-00466-2)) |
| Otu41136 | 1 | k__Fungi(100);p__Ascomycota(100);c__Saccharomycetes(100);o__Saccharomycetales(100);f__Metschnikowiaceae(100);g__Metschnikowia(100);s__Metschnikowia_pulcherrima(100); | Metschnikowia | 13 | Saprotroph | \|Undefined Saprotroph\| | Yeast | NULL | Probable | As nectar-tap_saprotroph (Põlme et al. 2020); Decay sugar-rich substrates (Põlme et al. 2020) | Põlme S, et al. 2020. Fungal Diversity 105:1-16 ((https://doi.org/10.1007/s13225-020-00466-2)) |
| Otu41137 | 1 | k__Fungi(100);p__Ascomycota(100);c__Saccharomycetes(100);o__Saccharomycetales(100);f__Metschnikowiaceae(100);g__Metschnikowia(100);s__Metschnikowia_pulcherrima(100); | Metschnikowia | 13 | Saprotroph | \|Undefined Saprotroph\| | Yeast | NULL | Probable | As nectar-tap_saprotroph (Põlme et al. 2020); Decay sugar-rich substrates (Põlme et al. 2020) | Põlme S, et al. 2020. Fungal Diversity 105:1-16 ((https://doi.org/10.1007/s13225-020-00466-2)) |
| Otu41138 | 1 | k__Fungi(100);p__Ascomycota(100);c__Saccharomycetes(100);o__Saccharomycetales(100);f__Metschnikowiaceae(100);g__Metschnikowia(100);s__Metschnikowia_pulcherrima(100); | Metschnikowia | 13 | Saprotroph | \|Undefined Saprotroph\| | Yeast | NULL | Probable | As nectar-tap_saprotroph (Põlme et al. 2020); Decay sugar-rich substrates (Põlme et al. 2020) | Põlme S, et al. 2020. Fungal Diversity 105:1-16 ((https://doi.org/10.1007/s13225-020-00466-2)) |
| Otu41139 | 1 | k__Fungi(100);p__Ascomycota(100);c__Saccharomycetes(100);o__Saccharomycetales(100);f__Metschnikowiaceae(100);g__Metschnikowia(100);s__Metschnikowia_pulcherrima(100); | Metschnikowia | 13 | Saprotroph | \|Undefined Saprotroph\| | Yeast | NULL | Probable | As nectar-tap_saprotroph (Põlme et al. 2020); Decay sugar-rich substrates (Põlme et al. 2020) | Põlme S, et al. 2020. Fungal Diversity 105:1-16 ((https://doi.org/10.1007/s13225-020-00466-2)) |
| Otu41140 | 1 | k__Fungi(100);p__Ascomycota(100);c__Saccharomycetes(100);o__Saccharomycetales(100);f__Metschnikowiaceae(100);g__Metschnikowia(100);s__Metschnikowia_pulcherrima(100); | Metschnikowia | 13 | Saprotroph | \|Undefined Saprotroph\| | Yeast | NULL | Probable | As nectar-tap_saprotroph (Põlme et al. 2020); Decay sugar-rich substrates (Põlme et al. 2020) | Põlme S, et al. 2020. Fungal Diversity 105:1-16 ((https://doi.org/10.1007/s13225-020-00466-2)) |
| Otu41141 | 1 | k__Fungi(100);p__Ascomycota(100);c__Saccharomycetes(100);o__Saccharomycetales(100);f__Metschnikowiaceae(100);g__Metschnikowia(100);s__Metschnikowia_pulcherrima(100); | Metschnikowia | 13 | Saprotroph | \|Undefined Saprotroph\| | Yeast | NULL | Probable | As nectar-tap_saprotroph (Põlme et al. 2020); Decay sugar-rich substrates (Põlme et al. 2020) | Põlme S, et al. 2020. Fungal Diversity 105:1-16 ((https://doi.org/10.1007/s13225-020-00466-2)) |
| Otu41142 | 1 | k__Fungi(100);p__Ascomycota(100);c__Saccharomycetes(100);o__Saccharomycetales(100);f__Metschnikowiaceae(100);g__Metschnikowia(100);s__Metschnikowia_pulcherrima(100); | Metschnikowia | 13 | Saprotroph | \|Undefined Saprotroph\| | Yeast | NULL | Probable | As nectar-tap_saprotroph (Põlme et al. 2020); Decay sugar-rich substrates (Põlme et al. 2020) | Põlme S, et al. 2020. Fungal Diversity 105:1-16 ((https://doi.org/10.1007/s13225-020-00466-2)) |
| Otu41143 | 1 | k__Fungi(100);p__Ascomycota(100);c__Saccharomycetes(100);o__Saccharomycetales(100);f__Metschnikowiaceae(100);g__Metschnikowia(100);g__Metschnikowia_unclassified(100); | Metschnikowia | 13 | Saprotroph | \|Undefined Saprotroph\| | Yeast | NULL | Probable | As nectar-tap_saprotroph (Põlme et al. 2020); Decay sugar-rich substrates (Põlme et al. 2020) | Põlme S, et al. 2020. Fungal Diversity 105:1-16 ((https://doi.org/10.1007/s13225-020-00466-2)) |
| Otu41144 | 1 | k__Fungi(100);p__Ascomycota(100);c__Saccharomycetes(100);o__Saccharomycetales(100);f__Metschnikowiaceae(100);g__Metschnikowia(100);g__Metschnikowia_unclassified(100); | Metschnikowia | 13 | Saprotroph | \|Undefined Saprotroph\| | Yeast | NULL | Probable | As nectar-tap_saprotroph (Põlme et al. 2020); Decay sugar-rich substrates (Põlme et al. 2020) | Põlme S, et al. 2020. Fungal Diversity 105:1-16 ((https://doi.org/10.1007/s13225-020-00466-2)) |
| Otu41145 | 1 | k__Fungi(100);p__Ascomycota(100);c__Saccharomycetes(100);o__Saccharomycetales(100);f__Metschnikowiaceae(100);g__Metschnikowia(100);s__Metschnikowia_pulcherrima(100); | Metschnikowia | 13 | Saprotroph | \|Undefined Saprotroph\| | Yeast | NULL | Probable | As nectar-tap_saprotroph (Põlme et al. 2020); Decay sugar-rich substrates (Põlme et al. 2020) | Põlme S, et al. 2020. Fungal Diversity 105:1-16 ((https://doi.org/10.1007/s13225-020-00466-2)) |
| Otu41146 | 1 | k__Fungi(100);p__Ascomycota(100);c__Saccharomycetes(100);o__Saccharomycetales(100);f__Metschnikowiaceae(100);g__Metschnikowia(100);s__Metschnikowia_pulcherrima(100); | Metschnikowia | 13 | Saprotroph | \|Undefined Saprotroph\| | Yeast | NULL | Probable | As nectar-tap_saprotroph (Põlme et al. 2020); Decay sugar-rich substrates (Põlme et al. 2020) | Põlme S, et al. 2020. Fungal Diversity 105:1-16 ((https://doi.org/10.1007/s13225-020-00466-2)) |
| Otu41147 | 1 | k__Fungi(100);p__Ascomycota(100);c__Saccharomycetes(100);o__Saccharomycetales(100);f__Metschnikowiaceae(100);g__Metschnikowia(100);s__Metschnikowia_pulcherrima(100); | Metschnikowia | 13 | Saprotroph | \|Undefined Saprotroph\| | Yeast | NULL | Probable | As nectar-tap_saprotroph (Põlme et al. 2020); Decay sugar-rich substrates (Põlme et al. 2020) | Põlme S, et al. 2020. Fungal Diversity 105:1-16 ((https://doi.org/10.1007/s13225-020-00466-2)) |
| Otu41148 | 1 | k__Fungi(100);p__Ascomycota(100);c__Saccharomycetes(100);o__Saccharomycetales(100);f__Metschnikowiaceae(100);g__Metschnikowia(100);g__Metschnikowia_unclassified(100); | Metschnikowia | 13 | Saprotroph | \|Undefined Saprotroph\| | Yeast | NULL | Probable | As nectar-tap_saprotroph (Põlme et al. 2020); Decay sugar-rich substrates (Põlme et al. 2020) | Põlme S, et al. 2020. Fungal Diversity 105:1-16 ((https://doi.org/10.1007/s13225-020-00466-2)) |
| Otu41149 | 1 | k__Fungi(100);p__Ascomycota(100);c__Saccharomycetes(100);o__Saccharomycetales(100);f__Metschnikowiaceae(100);g__Metschnikowia(100);g__Metschnikowia_unclassified(100); | Metschnikowia | 13 | Saprotroph | \|Undefined Saprotroph\| | Yeast | NULL | Probable | As nectar-tap_saprotroph (Põlme et al. 2020); Decay sugar-rich substrates (Põlme et al. 2020) | Põlme S, et al. 2020. Fungal Diversity 105:1-16 ((https://doi.org/10.1007/s13225-020-00466-2)) |
| Otu41151 | 1 | k__Fungi(100);p__Ascomycota(100);c__Saccharomycetes(100);o__Saccharomycetales(100);f__Metschnikowiaceae(100);g__Metschnikowia(100);s__Metschnikowia_pulcherrima(100); | Metschnikowia | 13 | Saprotroph | \|Undefined Saprotroph\| | Yeast | NULL | Probable | As nectar-tap_saprotroph (Põlme et al. 2020); Decay sugar-rich substrates (Põlme et al. 2020) | Põlme S, et al. 2020. Fungal Diversity 105:1-16 ((https://doi.org/10.1007/s13225-020-00466-2)) |
| Otu41152 | 1 | k__Fungi(100);p__Ascomycota(100);c__Saccharomycetes(100);o__Saccharomycetales(100);f__Metschnikowiaceae(100);g__Metschnikowia(100);g__Metschnikowia_unclassified(100); | Metschnikowia | 13 | Saprotroph | \|Undefined Saprotroph\| | Yeast | NULL | Probable | As nectar-tap_saprotroph (Põlme et al. 2020); Decay sugar-rich substrates (Põlme et al. 2020) | Põlme S, et al. 2020. Fungal Diversity 105:1-16 ((https://doi.org/10.1007/s13225-020-00466-2)) |
| Otu41153 | 1 | k__Fungi(100);p__Ascomycota(100);c__Saccharomycetes(100);o__Saccharomycetales(100);f__Metschnikowiaceae(100);g__Metschnikowia(100);s__Metschnikowia_pulcherrima(100); | Metschnikowia | 13 | Saprotroph | \|Undefined Saprotroph\| | Yeast | NULL | Probable | As nectar-tap_saprotroph (Põlme et al. 2020); Decay sugar-rich substrates (Põlme et al. 2020) | Põlme S, et al. 2020. Fungal Diversity 105:1-16 ((https://doi.org/10.1007/s13225-020-00466-2)) |
| Otu41154 | 1 | k__Fungi(100);p__Ascomycota(100);c__Saccharomycetes(100);o__Saccharomycetales(100);f__Metschnikowiaceae(100);g__Metschnikowia(100);s__Metschnikowia_pulcherrima(100); | Metschnikowia | 13 | Saprotroph | \|Undefined Saprotroph\| | Yeast | NULL | Probable | As nectar-tap_saprotroph (Põlme et al. 2020); Decay sugar-rich substrates (Põlme et al. 2020) | Põlme S, et al. 2020. Fungal Diversity 105:1-16 ((https://doi.org/10.1007/s13225-020-00466-2)) |
| Otu41155 | 1 | k__Fungi(100);p__Ascomycota(100);c__Saccharomycetes(100);o__Saccharomycetales(100);f__Metschnikowiaceae(100);g__Metschnikowia(100);s__Metschnikowia_pulcherrima(100); | Metschnikowia | 13 | Saprotroph | \|Undefined Saprotroph\| | Yeast | NULL | Probable | As nectar-tap_saprotroph (Põlme et al. 2020); Decay sugar-rich substrates (Põlme et al. 2020) | Põlme S, et al. 2020. Fungal Diversity 105:1-16 ((https://doi.org/10.1007/s13225-020-00466-2)) |
| Otu41156 | 1 | k__Fungi(100);p__Ascomycota(100);c__Saccharomycetes(100);o__Saccharomycetales(100);f__Metschnikowiaceae(100);g__Metschnikowia(100);s__Metschnikowia_pulcherrima(100); | Metschnikowia | 13 | Saprotroph | \|Undefined Saprotroph\| | Yeast | NULL | Probable | As nectar-tap_saprotroph (Põlme et al. 2020); Decay sugar-rich substrates (Põlme et al. 2020) | Põlme S, et al. 2020. Fungal Diversity 105:1-16 ((https://doi.org/10.1007/s13225-020-00466-2)) |
| Otu41157 | 1 | k__Fungi(100);p__Ascomycota(100);c__Saccharomycetes(100);o__Saccharomycetales(100);f__Metschnikowiaceae(100);g__Metschnikowia(100);s__Metschnikowia_pulcherrima(100); | Metschnikowia | 13 | Saprotroph | \|Undefined Saprotroph\| | Yeast | NULL | Probable | As nectar-tap_saprotroph (Põlme et al. 2020); Decay sugar-rich substrates (Põlme et al. 2020) | Põlme S, et al. 2020. Fungal Diversity 105:1-16 ((https://doi.org/10.1007/s13225-020-00466-2)) |
| Otu41158 | 1 | k__Fungi(100);p__Ascomycota(100);c__Saccharomycetes(100);o__Saccharomycetales(100);f__Metschnikowiaceae(100);g__Metschnikowia(100);s__Metschnikowia_pulcherrima(100); | Metschnikowia | 13 | Saprotroph | \|Undefined Saprotroph\| | Yeast | NULL | Probable | As nectar-tap_saprotroph (Põlme et al. 2020); Decay sugar-rich substrates (Põlme et al. 2020) | Põlme S, et al. 2020. Fungal Diversity 105:1-16 ((https://doi.org/10.1007/s13225-020-00466-2)) |
| Otu41159 | 1 | k__Fungi(100);p__Ascomycota(100);c__Saccharomycetes(100);o__Saccharomycetales(100);f__Metschnikowiaceae(100);g__Metschnikowia(100);s__Metschnikowia_pulcherrima(100); | Metschnikowia | 13 | Saprotroph | \|Undefined Saprotroph\| | Yeast | NULL | Probable | As nectar-tap_saprotroph (Põlme et al. 2020); Decay sugar-rich substrates (Põlme et al. 2020) | Põlme S, et al. 2020. Fungal Diversity 105:1-16 ((https://doi.org/10.1007/s13225-020-00466-2)) |
| Otu41160 | 1 | k__Fungi(100);p__Ascomycota(100);c__Saccharomycetes(100);o__Saccharomycetales(100);f__Metschnikowiaceae(100);g__Metschnikowia(100);s__Metschnikowia_pulcherrima(100); | Metschnikowia | 13 | Saprotroph | \|Undefined Saprotroph\| | Yeast | NULL | Probable | As nectar-tap_saprotroph (Põlme et al. 2020); Decay sugar-rich substrates (Põlme et al. 2020) | Põlme S, et al. 2020. Fungal Diversity 105:1-16 ((https://doi.org/10.1007/s13225-020-00466-2)) |
| Otu41161 | 1 | k__Fungi(100);p__Ascomycota(100);c__Saccharomycetes(100);o__Saccharomycetales(100);f__Metschnikowiaceae(100);g__Metschnikowia(100);s__Metschnikowia_pulcherrima(100); | Metschnikowia | 13 | Saprotroph | \|Undefined Saprotroph\| | Yeast | NULL | Probable | As nectar-tap_saprotroph (Põlme et al. 2020); Decay sugar-rich substrates (Põlme et al. 2020) | Põlme S, et al. 2020. Fungal Diversity 105:1-16 ((https://doi.org/10.1007/s13225-020-00466-2)) |
| Otu41162 | 1 | k__Fungi(100);p__Ascomycota(100);c__Saccharomycetes(100);o__Saccharomycetales(100);f__Metschnikowiaceae(100);g__Metschnikowia(100);s__Metschnikowia_pulcherrima(100); | Metschnikowia | 13 | Saprotroph | \|Undefined Saprotroph\| | Yeast | NULL | Probable | As nectar-tap_saprotroph (Põlme et al. 2020); Decay sugar-rich substrates (Põlme et al. 2020) | Põlme S, et al. 2020. Fungal Diversity 105:1-16 ((https://doi.org/10.1007/s13225-020-00466-2)) |
| Otu41163 | 1 | k__Fungi(100);p__Ascomycota(100);c__Saccharomycetes(100);o__Saccharomycetales(100);f__Metschnikowiaceae(100);g__Metschnikowia(100);s__Metschnikowia_pulcherrima(100); | Metschnikowia | 13 | Saprotroph | \|Undefined Saprotroph\| | Yeast | NULL | Probable | As nectar-tap_saprotroph (Põlme et al. 2020); Decay sugar-rich substrates (Põlme et al. 2020) | Põlme S, et al. 2020. Fungal Diversity 105:1-16 ((https://doi.org/10.1007/s13225-020-00466-2)) |
| Otu41164 | 1 | k__Fungi(100);p__Ascomycota(100);c__Saccharomycetes(100);o__Saccharomycetales(100);f__Metschnikowiaceae(100);g__Metschnikowia(100);s__Metschnikowia_pulcherrima(100); | Metschnikowia | 13 | Saprotroph | \|Undefined Saprotroph\| | Yeast | NULL | Probable | As nectar-tap_saprotroph (Põlme et al. 2020); Decay sugar-rich substrates (Põlme et al. 2020) | Põlme S, et al. 2020. Fungal Diversity 105:1-16 ((https://doi.org/10.1007/s13225-020-00466-2)) |
| Otu41165 | 1 | k__Fungi(100);p__Ascomycota(100);c__Saccharomycetes(100);o__Saccharomycetales(100);f__Metschnikowiaceae(100);g__Metschnikowia(100);s__Metschnikowia_pulcherrima(100); | Metschnikowia | 13 | Saprotroph | \|Undefined Saprotroph\| | Yeast | NULL | Probable | As nectar-tap_saprotroph (Põlme et al. 2020); Decay sugar-rich substrates (Põlme et al. 2020) | Põlme S, et al. 2020. Fungal Diversity 105:1-16 ((https://doi.org/10.1007/s13225-020-00466-2)) |
| Otu41166 | 1 | k__Fungi(100);p__Ascomycota(100);c__Saccharomycetes(100);o__Saccharomycetales(100);f__Metschnikowiaceae(100);g__Metschnikowia(100);g__Metschnikowia_unclassified(100); | Metschnikowia | 13 | Saprotroph | \|Undefined Saprotroph\| | Yeast | NULL | Probable | As nectar-tap_saprotroph (Põlme et al. 2020); Decay sugar-rich substrates (Põlme et al. 2020) | Põlme S, et al. 2020. Fungal Diversity 105:1-16 ((https://doi.org/10.1007/s13225-020-00466-2)) |
| Otu41167 | 1 | k__Fungi(100);p__Ascomycota(100);c__Saccharomycetes(100);o__Saccharomycetales(100);f__Metschnikowiaceae(100);g__Metschnikowia(100);g__Metschnikowia_unclassified(100); | Metschnikowia | 13 | Saprotroph | \|Undefined Saprotroph\| | Yeast | NULL | Probable | As nectar-tap_saprotroph (Põlme et al. 2020); Decay sugar-rich substrates (Põlme et al. 2020) | Põlme S, et al. 2020. Fungal Diversity 105:1-16 ((https://doi.org/10.1007/s13225-020-00466-2)) |
| Otu41168 | 1 | k__Fungi(100);p__Ascomycota(100);c__Saccharomycetes(100);o__Saccharomycetales(100);f__Metschnikowiaceae(100);g__Metschnikowia(100);s__Metschnikowia_pulcherrima(100); | Metschnikowia | 13 | Saprotroph | \|Undefined Saprotroph\| | Yeast | NULL | Probable | As nectar-tap_saprotroph (Põlme et al. 2020); Decay sugar-rich substrates (Põlme et al. 2020) | Põlme S, et al. 2020. Fungal Diversity 105:1-16 ((https://doi.org/10.1007/s13225-020-00466-2)) |
| Otu41169 | 1 | k__Fungi(100);p__Ascomycota(100);c__Saccharomycetes(100);o__Saccharomycetales(100);f__Metschnikowiaceae(100);g__Metschnikowia(100);s__Metschnikowia_pulcherrima(100); | Metschnikowia | 13 | Saprotroph | \|Undefined Saprotroph\| | Yeast | NULL | Probable | As nectar-tap_saprotroph (Põlme et al. 2020); Decay sugar-rich substrates (Põlme et al. 2020) | Põlme S, et al. 2020. Fungal Diversity 105:1-16 ((https://doi.org/10.1007/s13225-020-00466-2)) |
| Otu41170 | 1 | k__Fungi(100);p__Ascomycota(100);c__Saccharomycetes(100);o__Saccharomycetales(100);f__Metschnikowiaceae(100);g__Metschnikowia(100);s__Metschnikowia_pulcherrima(100); | Metschnikowia | 13 | Saprotroph | \|Undefined Saprotroph\| | Yeast | NULL | Probable | As nectar-tap_saprotroph (Põlme et al. 2020); Decay sugar-rich substrates (Põlme et al. 2020) | Põlme S, et al. 2020. Fungal Diversity 105:1-16 ((https://doi.org/10.1007/s13225-020-00466-2)) |
| Otu41171 | 1 | k__Fungi(100);p__Ascomycota(100);c__Saccharomycetes(100);o__Saccharomycetales(100);f__Metschnikowiaceae(100);g__Metschnikowia(100);s__Metschnikowia_pulcherrima(100); | Metschnikowia | 13 | Saprotroph | \|Undefined Saprotroph\| | Yeast | NULL | Probable | As nectar-tap_saprotroph (Põlme et al. 2020); Decay sugar-rich substrates (Põlme et al. 2020) | Põlme S, et al. 2020. Fungal Diversity 105:1-16 ((https://doi.org/10.1007/s13225-020-00466-2)) |
| Otu41172 | 1 | k__Fungi(100);p__Ascomycota(100);c__Saccharomycetes(100);o__Saccharomycetales(100);f__Metschnikowiaceae(100);g__Metschnikowia(100);s__Metschnikowia_pulcherrima(100); | Metschnikowia | 13 | Saprotroph | \|Undefined Saprotroph\| | Yeast | NULL | Probable | As nectar-tap_saprotroph (Põlme et al. 2020); Decay sugar-rich substrates (Põlme et al. 2020) | Põlme S, et al. 2020. Fungal Diversity 105:1-16 ((https://doi.org/10.1007/s13225-020-00466-2)) |
| Otu41173 | 1 | k__Fungi(100);p__Ascomycota(100);c__Saccharomycetes(100);o__Saccharomycetales(100);f__Metschnikowiaceae(100);g__Metschnikowia(100);s__Metschnikowia_pulcherrima(100); | Metschnikowia | 13 | Saprotroph | \|Undefined Saprotroph\| | Yeast | NULL | Probable | As nectar-tap_saprotroph (Põlme et al. 2020); Decay sugar-rich substrates (Põlme et al. 2020) | Põlme S, et al. 2020. Fungal Diversity 105:1-16 ((https://doi.org/10.1007/s13225-020-00466-2)) |
| Otu41174 | 1 | k__Fungi(100);p__Ascomycota(100);c__Saccharomycetes(100);o__Saccharomycetales(100);f__Metschnikowiaceae(100);g__Metschnikowia(100);s__Metschnikowia_pulcherrima(100); | Metschnikowia | 13 | Saprotroph | \|Undefined Saprotroph\| | Yeast | NULL | Probable | As nectar-tap_saprotroph (Põlme et al. 2020); Decay sugar-rich substrates (Põlme et al. 2020) | Põlme S, et al. 2020. Fungal Diversity 105:1-16 ((https://doi.org/10.1007/s13225-020-00466-2)) |
| Otu41175 | 1 | k__Fungi(100);p__Ascomycota(100);c__Saccharomycetes(100);o__Saccharomycetales(100);f__Metschnikowiaceae(100);g__Metschnikowia(100);g__Metschnikowia_unclassified(100); | Metschnikowia | 13 | Saprotroph | \|Undefined Saprotroph\| | Yeast | NULL | Probable | As nectar-tap_saprotroph (Põlme et al. 2020); Decay sugar-rich substrates (Põlme et al. 2020) | Põlme S, et al. 2020. Fungal Diversity 105:1-16 ((https://doi.org/10.1007/s13225-020-00466-2)) |
| Otu41176 | 1 | k__Fungi(100);p__Ascomycota(100);c__Saccharomycetes(100);o__Saccharomycetales(100);f__Metschnikowiaceae(100);g__Metschnikowia(100);s__Metschnikowia_pulcherrima(100); | Metschnikowia | 13 | Saprotroph | \|Undefined Saprotroph\| | Yeast | NULL | Probable | As nectar-tap_saprotroph (Põlme et al. 2020); Decay sugar-rich substrates (Põlme et al. 2020) | Põlme S, et al. 2020. Fungal Diversity 105:1-16 ((https://doi.org/10.1007/s13225-020-00466-2)) |
| Otu41177 | 1 | k__Fungi(100);p__Ascomycota(100);c__Saccharomycetes(100);o__Saccharomycetales(100);f__Metschnikowiaceae(100);g__Metschnikowia(100);s__Metschnikowia_pulcherrima(100); | Metschnikowia | 13 | Saprotroph | \|Undefined Saprotroph\| | Yeast | NULL | Probable | As nectar-tap_saprotroph (Põlme et al. 2020); Decay sugar-rich substrates (Põlme et al. 2020) | Põlme S, et al. 2020. Fungal Diversity 105:1-16 ((https://doi.org/10.1007/s13225-020-00466-2)) |
| Otu41178 | 1 | k__Fungi(100);p__Ascomycota(100);c__Saccharomycetes(100);o__Saccharomycetales(100);f__Metschnikowiaceae(100);g__Metschnikowia(100);s__Metschnikowia_pulcherrima(100); | Metschnikowia | 13 | Saprotroph | \|Undefined Saprotroph\| | Yeast | NULL | Probable | As nectar-tap_saprotroph (Põlme et al. 2020); Decay sugar-rich substrates (Põlme et al. 2020) | Põlme S, et al. 2020. Fungal Diversity 105:1-16 ((https://doi.org/10.1007/s13225-020-00466-2)) |
| Otu41179 | 1 | k__Fungi(100);p__Ascomycota(100);c__Saccharomycetes(100);o__Saccharomycetales(100);f__Metschnikowiaceae(100);g__Metschnikowia(100);s__Metschnikowia_pulcherrima(100); | Metschnikowia | 13 | Saprotroph | \|Undefined Saprotroph\| | Yeast | NULL | Probable | As nectar-tap_saprotroph (Põlme et al. 2020); Decay sugar-rich substrates (Põlme et al. 2020) | Põlme S, et al. 2020. Fungal Diversity 105:1-16 ((https://doi.org/10.1007/s13225-020-00466-2)) |
| Otu41180 | 1 | k__Fungi(100);p__Ascomycota(100);c__Saccharomycetes(100);o__Saccharomycetales(100);f__Metschnikowiaceae(100);g__Metschnikowia(100);s__Metschnikowia_pulcherrima(100); | Metschnikowia | 13 | Saprotroph | \|Undefined Saprotroph\| | Yeast | NULL | Probable | As nectar-tap_saprotroph (Põlme et al. 2020); Decay sugar-rich substrates (Põlme et al. 2020) | Põlme S, et al. 2020. Fungal Diversity 105:1-16 ((https://doi.org/10.1007/s13225-020-00466-2)) |
| Otu41181 | 1 | k__Fungi(100);p__Ascomycota(100);c__Saccharomycetes(100);o__Saccharomycetales(100);f__Metschnikowiaceae(100);g__Metschnikowia(100);s__Metschnikowia_pulcherrima(100); | Metschnikowia | 13 | Saprotroph | \|Undefined Saprotroph\| | Yeast | NULL | Probable | As nectar-tap_saprotroph (Põlme et al. 2020); Decay sugar-rich substrates (Põlme et al. 2020) | Põlme S, et al. 2020. Fungal Diversity 105:1-16 ((https://doi.org/10.1007/s13225-020-00466-2)) |
| Otu41182 | 1 | k__Fungi(100);p__Ascomycota(100);c__Saccharomycetes(100);o__Saccharomycetales(100);f__Metschnikowiaceae(100);g__Metschnikowia(100);s__Metschnikowia_pulcherrima(100); | Metschnikowia | 13 | Saprotroph | \|Undefined Saprotroph\| | Yeast | NULL | Probable | As nectar-tap_saprotroph (Põlme et al. 2020); Decay sugar-rich substrates (Põlme et al. 2020) | Põlme S, et al. 2020. Fungal Diversity 105:1-16 ((https://doi.org/10.1007/s13225-020-00466-2)) |
| Otu41183 | 1 | k__Fungi(100);p__Ascomycota(100);c__Saccharomycetes(100);o__Saccharomycetales(100);f__Metschnikowiaceae(100);g__Metschnikowia(100);s__Metschnikowia_pulcherrima(100); | Metschnikowia | 13 | Saprotroph | \|Undefined Saprotroph\| | Yeast | NULL | Probable | As nectar-tap_saprotroph (Põlme et al. 2020); Decay sugar-rich substrates (Põlme et al. 2020) | Põlme S, et al. 2020. Fungal Diversity 105:1-16 ((https://doi.org/10.1007/s13225-020-00466-2)) |
| Otu41184 | 1 | k__Fungi(100);p__Ascomycota(100);c__Saccharomycetes(100);o__Saccharomycetales(100);f__Metschnikowiaceae(100);g__Metschnikowia(100);s__Metschnikowia_pulcherrima(100); | Metschnikowia | 13 | Saprotroph | \|Undefined Saprotroph\| | Yeast | NULL | Probable | As nectar-tap_saprotroph (Põlme et al. 2020); Decay sugar-rich substrates (Põlme et al. 2020) | Põlme S, et al. 2020. Fungal Diversity 105:1-16 ((https://doi.org/10.1007/s13225-020-00466-2)) |
| Otu41186 | 1 | k__Fungi(100);p__Ascomycota(100);c__Saccharomycetes(100);o__Saccharomycetales(100);f__Metschnikowiaceae(100);g__Metschnikowia(100);g__Metschnikowia_unclassified(100); | Metschnikowia | 13 | Saprotroph | \|Undefined Saprotroph\| | Yeast | NULL | Probable | As nectar-tap_saprotroph (Põlme et al. 2020); Decay sugar-rich substrates (Põlme et al. 2020) | Põlme S, et al. 2020. Fungal Diversity 105:1-16 ((https://doi.org/10.1007/s13225-020-00466-2)) |
| Otu41187 | 1 | k__Fungi(100);p__Ascomycota(100);c__Saccharomycetes(100);o__Saccharomycetales(100);f__Metschnikowiaceae(100);g__Metschnikowia(100);s__Metschnikowia_pulcherrima(100); | Metschnikowia | 13 | Saprotroph | \|Undefined Saprotroph\| | Yeast | NULL | Probable | As nectar-tap_saprotroph (Põlme et al. 2020); Decay sugar-rich substrates (Põlme et al. 2020) | Põlme S, et al. 2020. Fungal Diversity 105:1-16 ((https://doi.org/10.1007/s13225-020-00466-2)) |
| Otu41188 | 1 | k__Fungi(100);p__Ascomycota(100);c__Saccharomycetes(100);o__Saccharomycetales(100);f__Metschnikowiaceae(100);g__Metschnikowia(100);s__Metschnikowia_pulcherrima(100); | Metschnikowia | 13 | Saprotroph | \|Undefined Saprotroph\| | Yeast | NULL | Probable | As nectar-tap_saprotroph (Põlme et al. 2020); Decay sugar-rich substrates (Põlme et al. 2020) | Põlme S, et al. 2020. Fungal Diversity 105:1-16 ((https://doi.org/10.1007/s13225-020-00466-2)) |
| Otu41189 | 1 | k__Fungi(100);p__Ascomycota(100);c__Saccharomycetes(100);o__Saccharomycetales(100);f__Metschnikowiaceae(100);g__Metschnikowia(100);s__Metschnikowia_pulcherrima(100); | Metschnikowia | 13 | Saprotroph | \|Undefined Saprotroph\| | Yeast | NULL | Probable | As nectar-tap_saprotroph (Põlme et al. 2020); Decay sugar-rich substrates (Põlme et al. 2020) | Põlme S, et al. 2020. Fungal Diversity 105:1-16 ((https://doi.org/10.1007/s13225-020-00466-2)) |
| Otu41190 | 1 | k__Fungi(100);p__Ascomycota(100);c__Saccharomycetes(100);o__Saccharomycetales(100);f__Metschnikowiaceae(100);g__Metschnikowia(100);s__Metschnikowia_pulcherrima(100); | Metschnikowia | 13 | Saprotroph | \|Undefined Saprotroph\| | Yeast | NULL | Probable | As nectar-tap_saprotroph (Põlme et al. 2020); Decay sugar-rich substrates (Põlme et al. 2020) | Põlme S, et al. 2020. Fungal Diversity 105:1-16 ((https://doi.org/10.1007/s13225-020-00466-2)) |
| Otu41191 | 1 | k__Fungi(100);p__Ascomycota(100);c__Saccharomycetes(100);o__Saccharomycetales(100);f__Metschnikowiaceae(100);g__Metschnikowia(100);s__Metschnikowia_pulcherrima(100); | Metschnikowia | 13 | Saprotroph | \|Undefined Saprotroph\| | Yeast | NULL | Probable | As nectar-tap_saprotroph (Põlme et al. 2020); Decay sugar-rich substrates (Põlme et al. 2020) | Põlme S, et al. 2020. Fungal Diversity 105:1-16 ((https://doi.org/10.1007/s13225-020-00466-2)) |
| Otu41192 | 1 | k__Fungi(100);p__Ascomycota(100);c__Saccharomycetes(100);o__Saccharomycetales(100);f__Metschnikowiaceae(100);g__Metschnikowia(100);s__Metschnikowia_pulcherrima(100); | Metschnikowia | 13 | Saprotroph | \|Undefined Saprotroph\| | Yeast | NULL | Probable | As nectar-tap_saprotroph (Põlme et al. 2020); Decay sugar-rich substrates (Põlme et al. 2020) | Põlme S, et al. 2020. Fungal Diversity 105:1-16 ((https://doi.org/10.1007/s13225-020-00466-2)) |
| Otu41193 | 1 | k__Fungi(100);p__Ascomycota(100);c__Saccharomycetes(100);o__Saccharomycetales(100);f__Metschnikowiaceae(100);g__Metschnikowia(100);s__Metschnikowia_pulcherrima(100); | Metschnikowia | 13 | Saprotroph | \|Undefined Saprotroph\| | Yeast | NULL | Probable | As nectar-tap_saprotroph (Põlme et al. 2020); Decay sugar-rich substrates (Põlme et al. 2020) | Põlme S, et al. 2020. Fungal Diversity 105:1-16 ((https://doi.org/10.1007/s13225-020-00466-2)) |
| Otu41194 | 1 | k__Fungi(100);p__Ascomycota(100);c__Saccharomycetes(100);o__Saccharomycetales(100);f__Metschnikowiaceae(100);g__Metschnikowia(100);s__Metschnikowia_pulcherrima(100); | Metschnikowia | 13 | Saprotroph | \|Undefined Saprotroph\| | Yeast | NULL | Probable | As nectar-tap_saprotroph (Põlme et al. 2020); Decay sugar-rich substrates (Põlme et al. 2020) | Põlme S, et al. 2020. Fungal Diversity 105:1-16 ((https://doi.org/10.1007/s13225-020-00466-2)) |
| Otu41195 | 1 | k__Fungi(100);p__Ascomycota(100);c__Saccharomycetes(100);o__Saccharomycetales(100);f__Metschnikowiaceae(100);g__Metschnikowia(100);s__Metschnikowia_pulcherrima(100); | Metschnikowia | 13 | Saprotroph | \|Undefined Saprotroph\| | Yeast | NULL | Probable | As nectar-tap_saprotroph (Põlme et al. 2020); Decay sugar-rich substrates (Põlme et al. 2020) | Põlme S, et al. 2020. Fungal Diversity 105:1-16 ((https://doi.org/10.1007/s13225-020-00466-2)) |
| Otu41196 | 1 | k__Fungi(100);p__Ascomycota(100);c__Saccharomycetes(100);o__Saccharomycetales(100);f__Metschnikowiaceae(100);g__Metschnikowia(100);g__Metschnikowia_unclassified(100); | Metschnikowia | 13 | Saprotroph | \|Undefined Saprotroph\| | Yeast | NULL | Probable | As nectar-tap_saprotroph (Põlme et al. 2020); Decay sugar-rich substrates (Põlme et al. 2020) | Põlme S, et al. 2020. Fungal Diversity 105:1-16 ((https://doi.org/10.1007/s13225-020-00466-2)) |
| Otu41197 | 1 | k__Fungi(100);p__Ascomycota(100);c__Saccharomycetes(100);o__Saccharomycetales(100);f__Metschnikowiaceae(100);g__Metschnikowia(100);s__Metschnikowia_pulcherrima(100); | Metschnikowia | 13 | Saprotroph | \|Undefined Saprotroph\| | Yeast | NULL | Probable | As nectar-tap_saprotroph (Põlme et al. 2020); Decay sugar-rich substrates (Põlme et al. 2020) | Põlme S, et al. 2020. Fungal Diversity 105:1-16 ((https://doi.org/10.1007/s13225-020-00466-2)) |
| Otu41198 | 1 | k__Fungi(100);p__Ascomycota(100);c__Saccharomycetes(100);o__Saccharomycetales(100);f__Metschnikowiaceae(100);g__Metschnikowia(100);s__Metschnikowia_pulcherrima(100); | Metschnikowia | 13 | Saprotroph | \|Undefined Saprotroph\| | Yeast | NULL | Probable | As nectar-tap_saprotroph (Põlme et al. 2020); Decay sugar-rich substrates (Põlme et al. 2020) | Põlme S, et al. 2020. Fungal Diversity 105:1-16 ((https://doi.org/10.1007/s13225-020-00466-2)) |
| Otu41199 | 1 | k__Fungi(100);p__Ascomycota(100);c__Saccharomycetes(100);o__Saccharomycetales(100);f__Metschnikowiaceae(100);g__Metschnikowia(100);s__Metschnikowia_pulcherrima(100); | Metschnikowia | 13 | Saprotroph | \|Undefined Saprotroph\| | Yeast | NULL | Probable | As nectar-tap_saprotroph (Põlme et al. 2020); Decay sugar-rich substrates (Põlme et al. 2020) | Põlme S, et al. 2020. Fungal Diversity 105:1-16 ((https://doi.org/10.1007/s13225-020-00466-2)) |
| Otu41201 | 1 | k__Fungi(100);p__Ascomycota(100);c__Saccharomycetes(100);o__Saccharomycetales(100);f__Metschnikowiaceae(100);g__Metschnikowia(100);s__Metschnikowia_pulcherrima(100); | Metschnikowia | 13 | Saprotroph | \|Undefined Saprotroph\| | Yeast | NULL | Probable | As nectar-tap_saprotroph (Põlme et al. 2020); Decay sugar-rich substrates (Põlme et al. 2020) | Põlme S, et al. 2020. Fungal Diversity 105:1-16 ((https://doi.org/10.1007/s13225-020-00466-2)) |
| Otu41202 | 1 | k__Fungi(100);p__Ascomycota(100);c__Saccharomycetes(100);o__Saccharomycetales(100);f__Metschnikowiaceae(100);g__Metschnikowia(100);g__Metschnikowia_unclassified(100); | Metschnikowia | 13 | Saprotroph | \|Undefined Saprotroph\| | Yeast | NULL | Probable | As nectar-tap_saprotroph (Põlme et al. 2020); Decay sugar-rich substrates (Põlme et al. 2020) | Põlme S, et al. 2020. Fungal Diversity 105:1-16 ((https://doi.org/10.1007/s13225-020-00466-2)) |
| Otu41203 | 1 | k__Fungi(100);p__Ascomycota(100);c__Saccharomycetes(100);o__Saccharomycetales(100);f__Metschnikowiaceae(100);g__Metschnikowia(100);s__Metschnikowia_pulcherrima(100); | Metschnikowia | 13 | Saprotroph | \|Undefined Saprotroph\| | Yeast | NULL | Probable | As nectar-tap_saprotroph (Põlme et al. 2020); Decay sugar-rich substrates (Põlme et al. 2020) | Põlme S, et al. 2020. Fungal Diversity 105:1-16 ((https://doi.org/10.1007/s13225-020-00466-2)) |
| Otu41204 | 1 | k__Fungi(100);p__Ascomycota(100);c__Saccharomycetes(100);o__Saccharomycetales(100);f__Metschnikowiaceae(100);g__Metschnikowia(100);s__Metschnikowia_pulcherrima(100); | Metschnikowia | 13 | Saprotroph | \|Undefined Saprotroph\| | Yeast | NULL | Probable | As nectar-tap_saprotroph (Põlme et al. 2020); Decay sugar-rich substrates (Põlme et al. 2020) | Põlme S, et al. 2020. Fungal Diversity 105:1-16 ((https://doi.org/10.1007/s13225-020-00466-2)) |
| Otu41205 | 1 | k__Fungi(100);p__Ascomycota(100);c__Saccharomycetes(100);o__Saccharomycetales(100);f__Metschnikowiaceae(100);g__Metschnikowia(100);s__Metschnikowia_pulcherrima(100); | Metschnikowia | 13 | Saprotroph | \|Undefined Saprotroph\| | Yeast | NULL | Probable | As nectar-tap_saprotroph (Põlme et al. 2020); Decay sugar-rich substrates (Põlme et al. 2020) | Põlme S, et al. 2020. Fungal Diversity 105:1-16 ((https://doi.org/10.1007/s13225-020-00466-2)) |
| Otu41207 | 1 | k__Fungi(100);p__Ascomycota(100);c__Saccharomycetes(100);o__Saccharomycetales(100);f__Metschnikowiaceae(100);g__Metschnikowia(100);s__Metschnikowia_pulcherrima(100); | Metschnikowia | 13 | Saprotroph | \|Undefined Saprotroph\| | Yeast | NULL | Probable | As nectar-tap_saprotroph (Põlme et al. 2020); Decay sugar-rich substrates (Põlme et al. 2020) | Põlme S, et al. 2020. Fungal Diversity 105:1-16 ((https://doi.org/10.1007/s13225-020-00466-2)) |
| Otu41208 | 1 | k__Fungi(100);p__Ascomycota(100);c__Saccharomycetes(100);o__Saccharomycetales(100);f__Metschnikowiaceae(100);g__Metschnikowia(100);s__Metschnikowia_pulcherrima(100); | Metschnikowia | 13 | Saprotroph | \|Undefined Saprotroph\| | Yeast | NULL | Probable | As nectar-tap_saprotroph (Põlme et al. 2020); Decay sugar-rich substrates (Põlme et al. 2020) | Põlme S, et al. 2020. Fungal Diversity 105:1-16 ((https://doi.org/10.1007/s13225-020-00466-2)) |
| Otu41209 | 1 | k__Fungi(100);p__Ascomycota(100);c__Saccharomycetes(100);o__Saccharomycetales(100);f__Metschnikowiaceae(100);g__Metschnikowia(100);s__Metschnikowia_pulcherrima(100); | Metschnikowia | 13 | Saprotroph | \|Undefined Saprotroph\| | Yeast | NULL | Probable | As nectar-tap_saprotroph (Põlme et al. 2020); Decay sugar-rich substrates (Põlme et al. 2020) | Põlme S, et al. 2020. Fungal Diversity 105:1-16 ((https://doi.org/10.1007/s13225-020-00466-2)) |
| Otu41210 | 1 | k__Fungi(100);p__Ascomycota(100);c__Saccharomycetes(100);o__Saccharomycetales(100);f__Metschnikowiaceae(100);g__Metschnikowia(100);s__Metschnikowia_pulcherrima(100); | Metschnikowia | 13 | Saprotroph | \|Undefined Saprotroph\| | Yeast | NULL | Probable | As nectar-tap_saprotroph (Põlme et al. 2020); Decay sugar-rich substrates (Põlme et al. 2020) | Põlme S, et al. 2020. Fungal Diversity 105:1-16 ((https://doi.org/10.1007/s13225-020-00466-2)) |
| Otu41211 | 1 | k__Fungi(100);p__Ascomycota(100);c__Saccharomycetes(100);o__Saccharomycetales(100);f__Metschnikowiaceae(100);g__Metschnikowia(100);s__Metschnikowia_pulcherrima(100); | Metschnikowia | 13 | Saprotroph | \|Undefined Saprotroph\| | Yeast | NULL | Probable | As nectar-tap_saprotroph (Põlme et al. 2020); Decay sugar-rich substrates (Põlme et al. 2020) | Põlme S, et al. 2020. Fungal Diversity 105:1-16 ((https://doi.org/10.1007/s13225-020-00466-2)) |
| Otu41212 | 1 | k__Fungi(100);p__Ascomycota(100);c__Saccharomycetes(100);o__Saccharomycetales(100);f__Metschnikowiaceae(100);g__Metschnikowia(100);g__Metschnikowia_unclassified(100); | Metschnikowia | 13 | Saprotroph | \|Undefined Saprotroph\| | Yeast | NULL | Probable | As nectar-tap_saprotroph (Põlme et al. 2020); Decay sugar-rich substrates (Põlme et al. 2020) | Põlme S, et al. 2020. Fungal Diversity 105:1-16 ((https://doi.org/10.1007/s13225-020-00466-2)) |
| Otu41213 | 1 | k__Fungi(100);p__Ascomycota(100);c__Saccharomycetes(100);o__Saccharomycetales(100);f__Metschnikowiaceae(100);g__Metschnikowia(100);g__Metschnikowia_unclassified(100); | Metschnikowia | 13 | Saprotroph | \|Undefined Saprotroph\| | Yeast | NULL | Probable | As nectar-tap_saprotroph (Põlme et al. 2020); Decay sugar-rich substrates (Põlme et al. 2020) | Põlme S, et al. 2020. Fungal Diversity 105:1-16 ((https://doi.org/10.1007/s13225-020-00466-2)) |
| Otu41215 | 1 | k__Fungi(100);p__Ascomycota(100);c__Saccharomycetes(100);o__Saccharomycetales(100);f__Metschnikowiaceae(100);g__Metschnikowia(100);s__Metschnikowia_pulcherrima(100); | Metschnikowia | 13 | Saprotroph | \|Undefined Saprotroph\| | Yeast | NULL | Probable | As nectar-tap_saprotroph (Põlme et al. 2020); Decay sugar-rich substrates (Põlme et al. 2020) | Põlme S, et al. 2020. Fungal Diversity 105:1-16 ((https://doi.org/10.1007/s13225-020-00466-2)) |
| Otu41216 | 1 | k__Fungi(100);p__Ascomycota(100);c__Saccharomycetes(100);o__Saccharomycetales(100);f__Metschnikowiaceae(100);g__Metschnikowia(100);s__Metschnikowia_pulcherrima(100); | Metschnikowia | 13 | Saprotroph | \|Undefined Saprotroph\| | Yeast | NULL | Probable | As nectar-tap_saprotroph (Põlme et al. 2020); Decay sugar-rich substrates (Põlme et al. 2020) | Põlme S, et al. 2020. Fungal Diversity 105:1-16 ((https://doi.org/10.1007/s13225-020-00466-2)) |
| Otu41218 | 1 | k__Fungi(100);p__Ascomycota(100);c__Saccharomycetes(100);o__Saccharomycetales(100);f__Metschnikowiaceae(100);g__Metschnikowia(100);g__Metschnikowia_unclassified(100); | Metschnikowia | 13 | Saprotroph | \|Undefined Saprotroph\| | Yeast | NULL | Probable | As nectar-tap_saprotroph (Põlme et al. 2020); Decay sugar-rich substrates (Põlme et al. 2020) | Põlme S, et al. 2020. Fungal Diversity 105:1-16 ((https://doi.org/10.1007/s13225-020-00466-2)) |
| Otu41219 | 1 | k__Fungi(100);p__Ascomycota(100);c__Saccharomycetes(100);o__Saccharomycetales(100);f__Metschnikowiaceae(100);g__Metschnikowia(100);s__Metschnikowia_pulcherrima(100); | Metschnikowia | 13 | Saprotroph | \|Undefined Saprotroph\| | Yeast | NULL | Probable | As nectar-tap_saprotroph (Põlme et al. 2020); Decay sugar-rich substrates (Põlme et al. 2020) | Põlme S, et al. 2020. Fungal Diversity 105:1-16 ((https://doi.org/10.1007/s13225-020-00466-2)) |
| Otu41220 | 1 | k__Fungi(100);p__Ascomycota(100);c__Saccharomycetes(100);o__Saccharomycetales(100);f__Metschnikowiaceae(100);g__Metschnikowia(100);s__Metschnikowia_pulcherrima(100); | Metschnikowia | 13 | Saprotroph | \|Undefined Saprotroph\| | Yeast | NULL | Probable | As nectar-tap_saprotroph (Põlme et al. 2020); Decay sugar-rich substrates (Põlme et al. 2020) | Põlme S, et al. 2020. Fungal Diversity 105:1-16 ((https://doi.org/10.1007/s13225-020-00466-2)) |
| Otu41221 | 1 | k__Fungi(100);p__Ascomycota(100);c__Saccharomycetes(100);o__Saccharomycetales(100);f__Metschnikowiaceae(100);g__Metschnikowia(100);g__Metschnikowia_unclassified(100); | Metschnikowia | 13 | Saprotroph | \|Undefined Saprotroph\| | Yeast | NULL | Probable | As nectar-tap_saprotroph (Põlme et al. 2020); Decay sugar-rich substrates (Põlme et al. 2020) | Põlme S, et al. 2020. Fungal Diversity 105:1-16 ((https://doi.org/10.1007/s13225-020-00466-2)) |
| Otu41222 | 1 | k__Fungi(100);p__Ascomycota(100);c__Saccharomycetes(100);o__Saccharomycetales(100);f__Metschnikowiaceae(100);g__Metschnikowia(100);s__Metschnikowia_pulcherrima(100); | Metschnikowia | 13 | Saprotroph | \|Undefined Saprotroph\| | Yeast | NULL | Probable | As nectar-tap_saprotroph (Põlme et al. 2020); Decay sugar-rich substrates (Põlme et al. 2020) | Põlme S, et al. 2020. Fungal Diversity 105:1-16 ((https://doi.org/10.1007/s13225-020-00466-2)) |
| Otu41223 | 1 | k__Fungi(100);p__Ascomycota(100);c__Saccharomycetes(100);o__Saccharomycetales(100);f__Metschnikowiaceae(100);g__Metschnikowia(100);s__Metschnikowia_pulcherrima(100); | Metschnikowia | 13 | Saprotroph | \|Undefined Saprotroph\| | Yeast | NULL | Probable | As nectar-tap_saprotroph (Põlme et al. 2020); Decay sugar-rich substrates (Põlme et al. 2020) | Põlme S, et al. 2020. Fungal Diversity 105:1-16 ((https://doi.org/10.1007/s13225-020-00466-2)) |
| Otu41224 | 1 | k__Fungi(100);p__Ascomycota(100);c__Saccharomycetes(100);o__Saccharomycetales(100);f__Metschnikowiaceae(100);g__Metschnikowia(100);g__Metschnikowia_unclassified(100); | Metschnikowia | 13 | Saprotroph | \|Undefined Saprotroph\| | Yeast | NULL | Probable | As nectar-tap_saprotroph (Põlme et al. 2020); Decay sugar-rich substrates (Põlme et al. 2020) | Põlme S, et al. 2020. Fungal Diversity 105:1-16 ((https://doi.org/10.1007/s13225-020-00466-2)) |
| Otu41225 | 1 | k__Fungi(100);p__Ascomycota(100);c__Saccharomycetes(100);o__Saccharomycetales(100);f__Metschnikowiaceae(100);g__Metschnikowia(100);s__Metschnikowia_pulcherrima(100); | Metschnikowia | 13 | Saprotroph | \|Undefined Saprotroph\| | Yeast | NULL | Probable | As nectar-tap_saprotroph (Põlme et al. 2020); Decay sugar-rich substrates (Põlme et al. 2020) | Põlme S, et al. 2020. Fungal Diversity 105:1-16 ((https://doi.org/10.1007/s13225-020-00466-2)) |
| Otu41226 | 1 | k__Fungi(100);p__Ascomycota(100);c__Saccharomycetes(100);o__Saccharomycetales(100);f__Metschnikowiaceae(100);g__Metschnikowia(100);s__Metschnikowia_pulcherrima(100); | Metschnikowia | 13 | Saprotroph | \|Undefined Saprotroph\| | Yeast | NULL | Probable | As nectar-tap_saprotroph (Põlme et al. 2020); Decay sugar-rich substrates (Põlme et al. 2020) | Põlme S, et al. 2020. Fungal Diversity 105:1-16 ((https://doi.org/10.1007/s13225-020-00466-2)) |
| Otu41227 | 1 | k__Fungi(100);p__Ascomycota(100);c__Saccharomycetes(100);o__Saccharomycetales(100);f__Metschnikowiaceae(100);g__Metschnikowia(100);g__Metschnikowia_unclassified(100); | Metschnikowia | 13 | Saprotroph | \|Undefined Saprotroph\| | Yeast | NULL | Probable | As nectar-tap_saprotroph (Põlme et al. 2020); Decay sugar-rich substrates (Põlme et al. 2020) | Põlme S, et al. 2020. Fungal Diversity 105:1-16 ((https://doi.org/10.1007/s13225-020-00466-2)) |
| Otu41228 | 1 | k__Fungi(100);p__Ascomycota(100);c__Saccharomycetes(100);o__Saccharomycetales(100);f__Metschnikowiaceae(100);g__Metschnikowia(100);s__Metschnikowia_pulcherrima(100); | Metschnikowia | 13 | Saprotroph | \|Undefined Saprotroph\| | Yeast | NULL | Probable | As nectar-tap_saprotroph (Põlme et al. 2020); Decay sugar-rich substrates (Põlme et al. 2020) | Põlme S, et al. 2020. Fungal Diversity 105:1-16 ((https://doi.org/10.1007/s13225-020-00466-2)) |
| Otu41229 | 1 | k__Fungi(100);p__Ascomycota(100);c__Saccharomycetes(100);o__Saccharomycetales(100);f__Metschnikowiaceae(100);g__Metschnikowia(100);s__Metschnikowia_pulcherrima(100); | Metschnikowia | 13 | Saprotroph | \|Undefined Saprotroph\| | Yeast | NULL | Probable | As nectar-tap_saprotroph (Põlme et al. 2020); Decay sugar-rich substrates (Põlme et al. 2020) | Põlme S, et al. 2020. Fungal Diversity 105:1-16 ((https://doi.org/10.1007/s13225-020-00466-2)) |
| Otu41230 | 1 | k__Fungi(100);p__Ascomycota(100);c__Saccharomycetes(100);o__Saccharomycetales(100);f__Metschnikowiaceae(100);g__Metschnikowia(100);s__Metschnikowia_pulcherrima(100); | Metschnikowia | 13 | Saprotroph | \|Undefined Saprotroph\| | Yeast | NULL | Probable | As nectar-tap_saprotroph (Põlme et al. 2020); Decay sugar-rich substrates (Põlme et al. 2020) | Põlme S, et al. 2020. Fungal Diversity 105:1-16 ((https://doi.org/10.1007/s13225-020-00466-2)) |
| Otu41231 | 1 | k__Fungi(100);p__Ascomycota(100);c__Saccharomycetes(100);o__Saccharomycetales(100);f__Metschnikowiaceae(100);g__Metschnikowia(100);s__Metschnikowia_pulcherrima(100); | Metschnikowia | 13 | Saprotroph | \|Undefined Saprotroph\| | Yeast | NULL | Probable | As nectar-tap_saprotroph (Põlme et al. 2020); Decay sugar-rich substrates (Põlme et al. 2020) | Põlme S, et al. 2020. Fungal Diversity 105:1-16 ((https://doi.org/10.1007/s13225-020-00466-2)) |
| Otu41232 | 1 | k__Fungi(100);p__Ascomycota(100);c__Saccharomycetes(100);o__Saccharomycetales(100);f__Metschnikowiaceae(100);g__Metschnikowia(100);g__Metschnikowia_unclassified(100); | Metschnikowia | 13 | Saprotroph | \|Undefined Saprotroph\| | Yeast | NULL | Probable | As nectar-tap_saprotroph (Põlme et al. 2020); Decay sugar-rich substrates (Põlme et al. 2020) | Põlme S, et al. 2020. Fungal Diversity 105:1-16 ((https://doi.org/10.1007/s13225-020-00466-2)) |
| Otu41233 | 1 | k__Fungi(100);p__Ascomycota(100);c__Saccharomycetes(100);o__Saccharomycetales(100);f__Metschnikowiaceae(100);g__Metschnikowia(100);s__Metschnikowia_pulcherrima(100); | Metschnikowia | 13 | Saprotroph | \|Undefined Saprotroph\| | Yeast | NULL | Probable | As nectar-tap_saprotroph (Põlme et al. 2020); Decay sugar-rich substrates (Põlme et al. 2020) | Põlme S, et al. 2020. Fungal Diversity 105:1-16 ((https://doi.org/10.1007/s13225-020-00466-2)) |
| Otu41234 | 1 | k__Fungi(100);p__Ascomycota(100);c__Saccharomycetes(100);o__Saccharomycetales(100);f__Metschnikowiaceae(100);g__Metschnikowia(100);[illegible] | Metschnikowia | 13 | Saprotroph | \|Undefined Saprotroph\| | Yeast | NULL | Probable | As nectar-tap_saprotroph (Põlme et al. 2020); Decay sugar-rich substrates (Põlme et al. 2020) | Põlme S, et al. 2020. Fungal Diversity 105:1-16 ((https://doi.org/10.1007/s13225-020-00466-2)) |
| Otu41235 | 1 | k__Fungi(100);p__Ascomycota(100);c__Saccharomycetes(100);o__Saccharomycetales(100);f__Metschnikowiaceae(100);g__Metschnikowia(100);[illegible] | Metschnikowia | 13 | Saprotroph | \|Undefined Saprotroph\| | Yeast | NULL | Probable | As nectar-tap_saprotroph (Põlme et al. 2020); Decay sugar-rich substrates (Põlme et al. 2020) | Põlme S, et al. 2020. Fungal Diversity 105:1-16 ((https://doi.org/10.1007/s13225-020-00466-2)) |
| Otu41236 | 1 | k__Fungi(100);p__Ascomycota(100);c__Saccharomycetes(100);o__Saccharomycetales(100);f__Metschnikowiaceae(100);g__Metschnikowia(100);[illegible] | Metschnikowia | 13 | Saprotroph | \|Undefined Saprotroph\| | Yeast | NULL | Probable | As nectar-tap_saprotroph (Põlme et al. 2020); Decay sugar-rich substrates (Põlme et al. 2020) | Põlme S, et al. 2020. Fungal Diversity 105:1-16 ((https://doi.org/10.1007/s13225-020-00466-2)) |
| Otu41237 | 1 | k__Fungi(100);p__Ascomycota(100);c__Saccharomycetes(100);o__Saccharomycetales(100);f__Metschnikowiaceae(100);g__Metschnikowia(100);[illegible] | Metschnikowia | 13 | Saprotroph | \|Undefined Saprotroph\| | Yeast | NULL | Probable | As nectar-tap_saprotroph (Põlme et al. 2020); Decay sugar-rich substrates (Põlme et al. 2020) | Põlme S, et al. 2020. Fungal Diversity 105:1-16 ((https://doi.org/10.1007/s13225-020-00466-2)) |
| Otu41238 | 1 | k__Fungi(100);p__Ascomycota(100);c__Saccharomycetes(100);o__Saccharomycetales(100);f__Metschnikowiaceae(100);g__Metschnikowia(100);[illegible] | Metschnikowia | 13 | Saprotroph | \|Undefined Saprotroph\| | Yeast | NULL | Probable | As nectar-tap_saprotroph (Põlme et al. 2020); Decay sugar-rich substrates (Põlme et al. 2020) | Põlme S, et al. 2020. Fungal Diversity 105:1-16 ((https://doi.org/10.1007/s13225-020-00466-2)) |
| Otu41239 | 1 | k__Fungi(100);p__Ascomycota(100);c__Saccharomycetes(100);o__Saccharomycetales(100);f__Metschnikowiaceae(100);g__Metschnikowia(100);[illegible] | Metschnikowia | 13 | Saprotroph | \|Undefined Saprotroph\| | Yeast | NULL | Probable | As nectar-tap_saprotroph (Põlme et al. 2020); Decay sugar-rich substrates (Põlme et al. 2020) | Põlme S, et al. 2020. Fungal Diversity 105:1-16 ((https://doi.org/10.1007/s13225-020-00466-2)) |
| Otu41240 | 1 | k__Fungi(100);p__Ascomycota(100);c__Saccharomycetes(100);o__Saccharomycetales(100);f__Metschnikowiaceae(100);g__Metschnikowia(100);[illegible] | Metschnikowia | 13 | Saprotroph | \|Undefined Saprotroph\| | Yeast | NULL | Probable | As nectar-tap_saprotroph (Põlme et al. 2020); Decay sugar-rich substrates (Põlme et al. 2020) | Põlme S, et al. 2020. Fungal Diversity 105:1-16 ((https://doi.org/10.1007/s13225-020-00466-2)) |
| Otu41241 | 1 | k__Fungi(100);p__Ascomycota(100);c__Saccharomycetes(100);o__Saccharomycetales(100);f__Metschnikowiaceae(100);g__Metschnikowia(100);[illegible] | Metschnikowia | 13 | Saprotroph | \|Undefined Saprotroph\| | Yeast | NULL | Probable | As nectar-tap_saprotroph (Põlme et al. 2020); Decay sugar-rich substrates (Põlme et al. 2020) | Põlme S, et al. 2020. Fungal Diversity 105:1-16 ((https://doi.org/10.1007/s13225-020-00466-2)) |

| | | | | | | | | | | | |
|---|---|---|---|---|---|---|---|---|---|---|---|
| Otu41242 | 1 | k__Fungi(100);p__Ascomycota(100);c__Saccharomycetes(100);o__Saccharomycetales(100);f__Metschnikowiaceae(100);g__Metschnikowia(100);s__Metschnikowia_pulcherrima(100); | Metschnikowia | 13 | Saprotroph | \|Undefined Saprotroph\| | Yeast | NULL | Probable | As nectar-tap_saprotroph (Põlme et al. 2020); Decay sugar-rich substrates (Põlme et al. 2020) | Põlme S, et al. 2020. Fungal Diversity 105:1-16 ((https://doi.org/10.1007/s13225-020-00466-2)) |
| Otu41243 | 1 | k__Fungi(100);p__Ascomycota(100);c__Saccharomycetes(100);o__Saccharomycetales(100);f__Metschnikowiaceae(100);g__Metschnikowia(100);s__Metschnikowia_pulcherrima(100); | Metschnikowia | 13 | Saprotroph | \|Undefined Saprotroph\| | Yeast | NULL | Probable | As nectar-tap_saprotroph (Põlme et al. 2020); Decay sugar-rich substrates (Põlme et al. 2020) | Põlme S, et al. 2020. Fungal Diversity 105:1-16 ((https://doi.org/10.1007/s13225-020-00466-2)) |
| Otu41244 | 1 | k__Fungi(100);p__Ascomycota(100);c__Saccharomycetes(100);o__Saccharomycetales(100);f__Metschnikowiaceae(100);g__Metschnikowia(100);s__Metschnikowia_pulcherrima(100); | Metschnikowia | 13 | Saprotroph | \|Undefined Saprotroph\| | Yeast | NULL | Probable | As nectar-tap_saprotroph (Põlme et al. 2020); Decay sugar-rich substrates (Põlme et al. 2020) | Põlme S, et al. 2020. Fungal Diversity 105:1-16 ((https://doi.org/10.1007/s13225-020-00466-2)) |
| Otu41245 | 1 | k__Fungi(100);p__Ascomycota(100);c__Saccharomycetes(100);o__Saccharomycetales(100);f__Metschnikowiaceae(100);g__Metschnikowia(100);s__Metschnikowia_pulcherrima(100); | Metschnikowia | 13 | Saprotroph | \|Undefined Saprotroph\| | Yeast | NULL | Probable | As nectar-tap_saprotroph (Põlme et al. 2020); Decay sugar-rich substrates (Põlme et al. 2020) | Põlme S, et al. 2020. Fungal Diversity 105:1-16 ((https://doi.org/10.1007/s13225-020-00466-2)) |
| Otu41246 | 1 | k__Fungi(100);p__Ascomycota(100);c__Saccharomycetes(100);o__Saccharomycetales(100);f__Metschnikowiaceae(100);g__Metschnikowia(100);s__Metschnikowia_pulcherrima(100); | Metschnikowia | 13 | Saprotroph | \|Undefined Saprotroph\| | Yeast | NULL | Probable | As nectar-tap_saprotroph (Põlme et al. 2020); Decay sugar-rich substrates (Põlme et al. 2020) | Põlme S, et al. 2020. Fungal Diversity 105:1-16 ((https://doi.org/10.1007/s13225-020-00466-2)) |
| Otu41247 | 1 | k__Fungi(100);p__Ascomycota(100);c__Saccharomycetes(100);o__Saccharomycetales(100);f__Metschnikowiaceae(100);g__Metschnikowia(100);s__Metschnikowia_pulcherrima(100); | Metschnikowia | 13 | Saprotroph | \|Undefined Saprotroph\| | Yeast | NULL | Probable | As nectar-tap_saprotroph (Põlme et al. 2020); Decay sugar-rich substrates (Põlme et al. 2020) | Põlme S, et al. 2020. Fungal Diversity 105:1-16 ((https://doi.org/10.1007/s13225-020-00466-2)) |
| Otu41248 | 1 | k__Fungi(100);p__Ascomycota(100);c__Saccharomycetes(100);o__Saccharomycetales(100);f__Metschnikowiaceae(100);g__Metschnikowia(100);g__Metschnikowia_unclassified(100); | Metschnikowia | 13 | Saprotroph | \|Undefined Saprotroph\| | Yeast | NULL | Probable | As nectar-tap_saprotroph (Põlme et al. 2020); Decay sugar-rich substrates (Põlme et al. 2020) | Põlme S, et al. 2020. Fungal Diversity 105:1-16 ((https://doi.org/10.1007/s13225-020-00466-2)) |
| Otu41249 | 1 | k__Fungi(100);p__Ascomycota(100);c__Saccharomycetes(100);o__Saccharomycetales(100);f__Metschnikowiaceae(100);g__Metschnikowia(100);s__Metschnikowia_pulcherrima(100); | Metschnikowia | 13 | Saprotroph | \|Undefined Saprotroph\| | Yeast | NULL | Probable | As nectar-tap_saprotroph (Põlme et al. 2020); Decay sugar-rich substrates (Põlme et al. 2020) | Põlme S, et al. 2020. Fungal Diversity 105:1-16 ((https://doi.org/10.1007/s13225-020-00466-2)) |
| Otu41250 | 1 | k__Fungi(100);p__Ascomycota(100);c__Saccharomycetes(100);o__Saccharomycetales(100);f__Metschnikowiaceae(100);g__Metschnikowia(100);s__Metschnikowia_pulcherrima(100); | Metschnikowia | 13 | Saprotroph | \|Undefined Saprotroph\| | Yeast | NULL | Probable | As nectar-tap_saprotroph (Põlme et al. 2020); Decay sugar-rich substrates (Põlme et al. 2020) | Põlme S, et al. 2020. Fungal Diversity 105:1-16 ((https://doi.org/10.1007/s13225-020-00466-2)) |
| Otu41251 | 1 | k__Fungi(100);p__Ascomycota(100);c__Saccharomycetes(100);o__Saccharomycetales(100);f__Metschnikowiaceae(100);g__Metschnikowia(100);s__Metschnikowia_pulcherrima(100); | Metschnikowia | 13 | Saprotroph | \|Undefined Saprotroph\| | Yeast | NULL | Probable | As nectar-tap_saprotroph (Põlme et al. 2020); Decay sugar-rich substrates (Põlme et al. 2020) | Põlme S, et al. 2020. Fungal Diversity 105:1-16 ((https://doi.org/10.1007/s13225-020-00466-2)) |
| Otu41252 | 1 | k__Fungi(100);p__Ascomycota(100);c__Saccharomycetes(100);o__Saccharomycetales(100);f__Metschnikowiaceae(100);g__Metschnikowia(100);s__Metschnikowia_pulcherrima(100); | Metschnikowia | 13 | Saprotroph | \|Undefined Saprotroph\| | Yeast | NULL | Probable | As nectar-tap_saprotroph (Põlme et al. 2020); Decay sugar-rich substrates (Põlme et al. 2020) | Põlme S, et al. 2020. Fungal Diversity 105:1-16 ((https://doi.org/10.1007/s13225-020-00466-2)) |
| Otu41253 | 1 | k__Fungi(100);p__Ascomycota(100);c__Saccharomycetes(100);o__Saccharomycetales(100);f__Metschnikowiaceae(100);g__Metschnikowia(100);s__Metschnikowia_pulcherrima(100); | Metschnikowia | 13 | Saprotroph | \|Undefined Saprotroph\| | Yeast | NULL | Probable | As nectar-tap_saprotroph (Põlme et al. 2020); Decay sugar-rich substrates (Põlme et al. 2020) | Põlme S, et al. 2020. Fungal Diversity 105:1-16 ((https://doi.org/10.1007/s13225-020-00466-2)) |
| Otu41254 | 1 | k__Fungi(100);p__Ascomycota(100);c__Saccharomycetes(100);o__Saccharomycetales(100);f__Metschnikowiaceae(100);g__Metschnikowia(100);s__Metschnikowia_pulcherrima(100); | Metschnikowia | 13 | Saprotroph | \|Undefined Saprotroph\| | Yeast | NULL | Probable | As nectar-tap_saprotroph (Põlme et al. 2020); Decay sugar-rich substrates (Põlme et al. 2020) | Põlme S, et al. 2020. Fungal Diversity 105:1-16 ((https://doi.org/10.1007/s13225-020-00466-2)) |
| Otu41255 | 1 | k__Fungi(100);p__Ascomycota(100);c__Saccharomycetes(100);o__Saccharomycetales(100);f__Metschnikowiaceae(100);g__Metschnikowia(100);s__Metschnikowia_pulcherrima(100); | Metschnikowia | 13 | Saprotroph | \|Undefined Saprotroph\| | Yeast | NULL | Probable | As nectar-tap_saprotroph (Põlme et al. 2020); Decay sugar-rich substrates (Põlme et al. 2020) | Põlme S, et al. 2020. Fungal Diversity 105:1-16 ((https://doi.org/10.1007/s13225-020-00466-2)) |
| Otu41256 | 1 | k__Fungi(100);p__Ascomycota(100);c__Saccharomycetes(100);o__Saccharomycetales(100);f__Metschnikowiaceae(100);g__Metschnikowia(100);g__Metschnikowia_unclassified(100); | Metschnikowia | 13 | Saprotroph | \|Undefined Saprotroph\| | Yeast | NULL | Probable | As nectar-tap_saprotroph (Põlme et al. 2020); Decay sugar-rich substrates (Põlme et al. 2020) | Põlme S, et al. 2020. Fungal Diversity 105:1-16 ((https://doi.org/10.1007/s13225-020-00466-2)) |
| Otu41257 | 1 | k__Fungi(100);p__Ascomycota(100);c__Saccharomycetes(100);o__Saccharomycetales(100);f__Metschnikowiaceae(100);g__Metschnikowia(100);s__Metschnikowia_pulcherrima(100); | Metschnikowia | 13 | Saprotroph | \|Undefined Saprotroph\| | Yeast | NULL | Probable | As nectar-tap_saprotroph (Põlme et al. 2020); Decay sugar-rich substrates (Põlme et al. 2020) | Põlme S, et al. 2020. Fungal Diversity 105:1-16 ((https://doi.org/10.1007/s13225-020-00466-2)) |
| Otu41258 | 1 | k__Fungi(100);p__Ascomycota(100);c__Saccharomycetes(100);o__Saccharomycetales(100);f__Metschnikowiaceae(100);g__Metschnikowia(100);s__Metschnikowia_pulcherrima(100); | Metschnikowia | 13 | Saprotroph | \|Undefined Saprotroph\| | Yeast | NULL | Probable | As nectar-tap_saprotroph (Põlme et al. 2020); Decay sugar-rich substrates (Põlme et al. 2020) | Põlme S, et al. 2020. Fungal Diversity 105:1-16 ((https://doi.org/10.1007/s13225-020-00466-2)) |
| Otu41259 | 1 | k__Fungi(100);p__Ascomycota(100);c__Saccharomycetes(100);o__Saccharomycetales(100);f__Metschnikowiaceae(100);g__Metschnikowia(100);s__Metschnikowia_pulcherrima(100); | Metschnikowia | 13 | Saprotroph | \|Undefined Saprotroph\| | Yeast | NULL | Probable | As nectar-tap_saprotroph (Põlme et al. 2020); Decay sugar-rich substrates (Põlme et al. 2020) | Põlme S, et al. 2020. Fungal Diversity 105:1-16 ((https://doi.org/10.1007/s13225-020-00466-2)) |
| Otu41260 | 1 | k__Fungi(100);p__Ascomycota(100);c__Saccharomycetes(100);o__Saccharomycetales(100);f__Metschnikowiaceae(100);g__Metschnikowia(100);s__Metschnikowia_pulcherrima(100); | Metschnikowia | 13 | Saprotroph | \|Undefined Saprotroph\| | Yeast | NULL | Probable | As nectar-tap_saprotroph (Põlme et al. 2020); Decay sugar-rich substrates (Põlme et al. 2020) | Põlme S, et al. 2020. Fungal Diversity 105:1-16 ((https://doi.org/10.1007/s13225-020-00466-2)) |
| Otu41261 | 1 | k__Fungi(100);p__Ascomycota(100);c__Saccharomycetes(100);o__Saccharomycetales(100);f__Metschnikowiaceae(100);g__Metschnikowia(100);s__Metschnikowia_pulcherrima(100); | Metschnikowia | 13 | Saprotroph | \|Undefined Saprotroph\| | Yeast | NULL | Probable | As nectar-tap_saprotroph (Põlme et al. 2020); Decay sugar-rich substrates (Põlme et al. 2020) | Põlme S, et al. 2020. Fungal Diversity 105:1-16 ((https://doi.org/10.1007/s13225-020-00466-2)) |
| Otu41262 | 1 | k__Fungi(100);p__Ascomycota(100);c__Saccharomycetes(100);o__Saccharomycetales(100);f__Metschnikowiaceae(100);g__Metschnikowia(100);s__Metschnikowia_pulcherrima(100); | Metschnikowia | 13 | Saprotroph | \|Undefined Saprotroph\| | Yeast | NULL | Probable | As nectar-tap_saprotroph (Põlme et al. 2020); Decay sugar-rich substrates (Põlme et al. 2020) | Põlme S, et al. 2020. Fungal Diversity 105:1-16 ((https://doi.org/10.1007/s13225-020-00466-2)) |
| Otu41263 | 1 | k__Fungi(100);p__Ascomycota(100);c__Saccharomycetes(100);o__Saccharomycetales(100);f__Metschnikowiaceae(100);g__Metschnikowia(100);g__Metschnikowia_unclassified(100); | Metschnikowia | 13 | Saprotroph | \|Undefined Saprotroph\| | Yeast | NULL | Probable | As nectar-tap_saprotroph (Põlme et al. 2020); Decay sugar-rich substrates (Põlme et al. 2020) | Põlme S, et al. 2020. Fungal Diversity 105:1-16 ((https://doi.org/10.1007/s13225-020-00466-2)) |
| Otu41264 | 1 | k__Fungi(100);p__Ascomycota(100);c__Saccharomycetes(100);o__Saccharomycetales(100);f__Metschnikowiaceae(100);g__Metschnikowia(100);s__Metschnikowia_pulcherrima(100); | Metschnikowia | 13 | Saprotroph | \|Undefined Saprotroph\| | Yeast | NULL | Probable | As nectar-tap_saprotroph (Põlme et al. 2020); Decay sugar-rich substrates (Põlme et al. 2020) | Põlme S, et al. 2020. Fungal Diversity 105:1-16 ((https://doi.org/10.1007/s13225-020-00466-2)) |
| Otu41265 | 1 | k__Fungi(100);p__Ascomycota(100);c__Saccharomycetes(100);o__Saccharomycetales(100);f__Metschnikowiaceae(100);g__Metschnikowia(100);s__Metschnikowia_pulcherrima(100); | Metschnikowia | 13 | Saprotroph | \|Undefined Saprotroph\| | Yeast | NULL | Probable | As nectar-tap_saprotroph (Põlme et al. 2020); Decay sugar-rich substrates (Põlme et al. 2020) | Põlme S, et al. 2020. Fungal Diversity 105:1-16 ((https://doi.org/10.1007/s13225-020-00466-2)) |
| Otu41266 | 1 | k__Fungi(100);p__Ascomycota(100);c__Saccharomycetes(100);o__Saccharomycetales(100);f__Metschnikowiaceae(100);g__Metschnikowia(100);s__Metschnikowia_pulcherrima(100); | Metschnikowia | 13 | Saprotroph | \|Undefined Saprotroph\| | Yeast | NULL | Probable | As nectar-tap_saprotroph (Põlme et al. 2020); Decay sugar-rich substrates (Põlme et al. 2020) | Põlme S, et al. 2020. Fungal Diversity 105:1-16 ((https://doi.org/10.1007/s13225-020-00466-2)) |
| Otu41267 | 1 | k__Fungi(100);p__Ascomycota(100);c__Saccharomycetes(100);o__Saccharomycetales(100);f__Metschnikowiaceae(100);g__Metschnikowia(100);s__Metschnikowia_pulcherrima(100); | Metschnikowia | 13 | Saprotroph | \|Undefined Saprotroph\| | Yeast | NULL | Probable | As nectar-tap_saprotroph (Põlme et al. 2020); Decay sugar-rich substrates (Põlme et al. 2020) | Põlme S, et al. 2020. Fungal Diversity 105:1-16 ((https://doi.org/10.1007/s13225-020-00466-2)) |
| Otu41268 | 1 | k__Fungi(100);p__Ascomycota(100);c__Saccharomycetes(100);o__Saccharomycetales(100);f__Metschnikowiaceae(100);g__Metschnikowia(100);s__Metschnikowia_pulcherrima(100); | Metschnikowia | 13 | Saprotroph | \|Undefined Saprotroph\| | Yeast | NULL | Probable | As nectar-tap_saprotroph (Põlme et al. 2020); Decay sugar-rich substrates (Põlme et al. 2020) | Põlme S, et al. 2020. Fungal Diversity 105:1-16 ((https://doi.org/10.1007/s13225-020-00466-2)) |
| Otu41269 | 1 | k__Fungi(100);p__Ascomycota(100);c__Saccharomycetes(100);o__Saccharomycetales(100);f__Metschnikowiaceae(100);g__Metschnikowia(100);s__Metschnikowia_pulcherrima(100); | Metschnikowia | 13 | Saprotroph | \|Undefined Saprotroph\| | Yeast | NULL | Probable | As nectar-tap_saprotroph (Põlme et al. 2020); Decay sugar-rich substrates (Põlme et al. 2020) | Põlme S, et al. 2020. Fungal Diversity 105:1-16 ((https://doi.org/10.1007/s13225-020-00466-2)) |
| Otu41270 | 1 | k__Fungi(100);p__Ascomycota(100);c__Saccharomycetes(100);o__Saccharomycetales(100);f__Metschnikowiaceae(100);g__Metschnikowia(100);s__Metschnikowia_pulcherrima(100); | Metschnikowia | 13 | Saprotroph | \|Undefined Saprotroph\| | Yeast | NULL | Probable | As nectar-tap_saprotroph (Põlme et al. 2020); Decay sugar-rich substrates (Põlme et al. 2020) | Põlme S, et al. 2020. Fungal Diversity 105:1-16 ((https://doi.org/10.1007/s13225-020-00466-2)) |
| Otu41271 | 1 | k__Fungi(100);p__Ascomycota(100);c__Saccharomycetes(100);o__Saccharomycetales(100);f__Metschnikowiaceae(100);g__Metschnikowia(100);g__Metschnikowia_unclassified(100); | Metschnikowia | 13 | Saprotroph | \|Undefined Saprotroph\| | Yeast | NULL | Probable | As nectar-tap_saprotroph (Põlme et al. 2020); Decay sugar-rich substrates (Põlme et al. 2020) | Põlme S, et al. 2020. Fungal Diversity 105:1-16 ((https://doi.org/10.1007/s13225-020-00466-2)) |
| Otu41272 | 1 | k__Fungi(100);p__Ascomycota(100);c__Saccharomycetes(100);o__Saccharomycetales(100);f__Metschnikowiaceae(100);g__Metschnikowia(100);g__Metschnikowia_unclassified(100); | Metschnikowia | 13 | Saprotroph | \|Undefined Saprotroph\| | Yeast | NULL | Probable | As nectar-tap_saprotroph (Põlme et al. 2020); Decay sugar-rich substrates (Põlme et al. 2020) | Põlme S, et al. 2020. Fungal Diversity 105:1-16 ((https://doi.org/10.1007/s13225-020-00466-2)) |
| Otu41273 | 1 | k__Fungi(100);p__Ascomycota(100);c__Saccharomycetes(100);o__Saccharomycetales(100);f__Metschnikowiaceae(100);g__Metschnikowia(100);s__Metschnikowia_pulcherrima(100); | Metschnikowia | 13 | Saprotroph | \|Undefined Saprotroph\| | Yeast | NULL | Probable | As nectar-tap_saprotroph (Põlme et al. 2020); Decay sugar-rich substrates (Põlme et al. 2020) | Põlme S, et al. 2020. Fungal Diversity 105:1-16 ((https://doi.org/10.1007/s13225-020-00466-2)) |
| Otu41274 | 1 | k__Fungi(100);p__Ascomycota(100);c__Saccharomycetes(100);o__Saccharomycetales(100);f__Metschnikowiaceae(100);g__Metschnikowia(100);s__Metschnikowia_pulcherrima(100); | Metschnikowia | 13 | Saprotroph | \|Undefined Saprotroph\| | Yeast | NULL | Probable | As nectar-tap_saprotroph (Põlme et al. 2020); Decay sugar-rich substrates (Põlme et al. 2020) | Põlme S, et al. 2020. Fungal Diversity 105:1-16 ((https://doi.org/10.1007/s13225-020-00466-2)) |
| Otu41275 | 1 | k__Fungi(100);p__Ascomycota(100);c__Saccharomycetes(100);o__Saccharomycetales(100);f__Metschnikowiaceae(100);g__Metschnikowia(100);s__Metschnikowia_pulcherrima(100); | Metschnikowia | 13 | Saprotroph | \|Undefined Saprotroph\| | Yeast | NULL | Probable | As nectar-tap_saprotroph (Põlme et al. 2020); Decay sugar-rich substrates (Põlme et al. 2020) | Põlme S, et al. 2020. Fungal Diversity 105:1-16 ((https://doi.org/10.1007/s13225-020-00466-2)) |
| Otu41276 | 1 | k__Fungi(100);p__Ascomycota(100);c__Saccharomycetes(100);o__Saccharomycetales(100);f__Metschnikowiaceae(100);g__Metschnikowia(100);g__Metschnikowia_unclassified(100); | Metschnikowia | 13 | Saprotroph | \|Undefined Saprotroph\| | Yeast | NULL | Probable | As nectar-tap_saprotroph (Põlme et al. 2020); Decay sugar-rich substrates (Põlme et al. 2020) | Põlme S, et al. 2020. Fungal Diversity 105:1-16 ((https://doi.org/10.1007/s13225-020-00466-2)) |
| Otu41277 | 1 | k__Fungi(100);p__Ascomycota(100);c__Saccharomycetes(100);o__Saccharomycetales(100);f__Metschnikowiaceae(100);g__Metschnikowia(100);s__Metschnikowia_pulcherrima(100); | Metschnikowia | 13 | Saprotroph | \|Undefined Saprotroph\| | Yeast | NULL | Probable | As nectar-tap_saprotroph (Põlme et al. 2020); Decay sugar-rich substrates (Põlme et al. 2020) | Põlme S, et al. 2020. Fungal Diversity 105:1-16 ((https://doi.org/10.1007/s13225-020-00466-2)) |
| Otu41278 | 1 | k__Fungi(100);p__Ascomycota(100);c__Saccharomycetes(100);o__Saccharomycetales(100);f__Metschnikowiaceae(100);g__Metschnikowia(100);g__Metschnikowia_unclassified(100); | Metschnikowia | 13 | Saprotroph | \|Undefined Saprotroph\| | Yeast | NULL | Probable | As nectar-tap_saprotroph (Põlme et al. 2020); Decay sugar-rich substrates (Põlme et al. 2020) | Põlme S, et al. 2020. Fungal Diversity 105:1-16 ((https://doi.org/10.1007/s13225-020-00466-2)) |
| Otu41279 | 1 | k__Fungi(100);p__Ascomycota(100);c__Saccharomycetes(100);o__Saccharomycetales(100);f__Metschnikowiaceae(100);g__Metschnikowia(100);s__Metschnikowia_pulcherrima(100); | Metschnikowia | 13 | Saprotroph | \|Undefined Saprotroph\| | Yeast | NULL | Probable | As nectar-tap_saprotroph (Põlme et al. 2020); Decay sugar-rich substrates (Põlme et al. 2020) | Põlme S, et al. 2020. Fungal Diversity 105:1-16 ((https://doi.org/10.1007/s13225-020-00466-2)) |
| Otu41280 | 1 | k__Fungi(100);p__Ascomycota(100);c__Saccharomycetes(100);o__Saccharomycetales(100);f__Metschnikowiaceae(100);g__Metschnikowia(100);s__Metschnikowia_pulcherrima(100); | Metschnikowia | 13 | Saprotroph | \|Undefined Saprotroph\| | Yeast | NULL | Probable | As nectar-tap_saprotroph (Põlme et al. 2020); Decay sugar-rich substrates (Põlme et al. 2020) | Põlme S, et al. 2020. Fungal Diversity 105:1-16 ((https://doi.org/10.1007/s13225-020-00466-2)) |
| Otu41281 | 1 | k__Fungi(100);p__Ascomycota(100);c__Saccharomycetes(100);o__Saccharomycetales(100);f__Metschnikowiaceae(100);g__Metschnikowia(100);s__Metschnikowia_pulcherrima(100); | Metschnikowia | 13 | Saprotroph | \|Undefined Saprotroph\| | Yeast | NULL | Probable | As nectar-tap_saprotroph (Põlme et al. 2020); Decay sugar-rich substrates (Põlme et al. 2020) | Põlme S, et al. 2020. Fungal Diversity 105:1-16 ((https://doi.org/10.1007/s13225-020-00466-2)) |
| Otu41282 | 1 | k__Fungi(100);p__Ascomycota(100);c__Saccharomycetes(100);o__Saccharomycetales(100);f__Metschnikowiaceae(100);g__Metschnikowia(100);s__Metschnikowia_pulcherrima(100); | Metschnikowia | 13 | Saprotroph | \|Undefined Saprotroph\| | Yeast | NULL | Probable | As nectar-tap_saprotroph (Põlme et al. 2020); Decay sugar-rich substrates (Põlme et al. 2020) | Põlme S, et al. 2020. Fungal Diversity 105:1-16 ((https://doi.org/10.1007/s13225-020-00466-2)) |
| Otu41283 | 1 | k__Fungi(100);p__Ascomycota(100);c__Saccharomycetes(100);o__Saccharomycetales(100);f__Metschnikowiaceae(100);g__Metschnikowia(100);g__Metschnikowia_unclassified(100); | Metschnikowia | 13 | Saprotroph | \|Undefined Saprotroph\| | Yeast | NULL | Probable | As nectar-tap_saprotroph (Põlme et al. 2020); Decay sugar-rich substrates (Põlme et al. 2020) | Põlme S, et al. 2020. Fungal Diversity 105:1-16 ((https://doi.org/10.1007/s13225-020-00466-2)) |
| Otu41284 | 1 | k__Fungi(100);p__Ascomycota(100);c__Saccharomycetes(100);o__Saccharomycetales(100);f__Metschnikowiaceae(100);g__Metschnikowia(100);g__Metschnikowia_unclassified(100); | Metschnikowia | 13 | Saprotroph | \|Undefined Saprotroph\| | Yeast | NULL | Probable | As nectar-tap_saprotroph (Põlme et al. 2020); Decay sugar-rich substrates (Põlme et al. 2020) | Põlme S, et al. 2020. Fungal Diversity 105:1-16 ((https://doi.org/10.1007/s13225-020-00466-2)) |
| Otu41285 | 1 | k__Fungi(100);p__Ascomycota(100);c__Saccharomycetes(100);o__Saccharomycetales(100);f__Metschnikowiaceae(100);g__Metschnikowia(100);s__Metschnikowia_pulcherrima(100); | Metschnikowia | 13 | Saprotroph | \|Undefined Saprotroph\| | Yeast | NULL | Probable | As nectar-tap_saprotroph (Põlme et al. 2020); Decay sugar-rich substrates (Põlme et al. 2020) | Põlme S, et al. 2020. Fungal Diversity 105:1-16 ((https://doi.org/10.1007/s13225-020-00466-2)) |
| Otu41286 | 1 | k__Fungi(100);p__Ascomycota(100);c__Saccharomycetes(100);o__Saccharomycetales(100);f__Metschnikowiaceae(100);g__Metschnikowia(100);s__Metschnikowia_pulcherrima(100); | Metschnikowia | 13 | Saprotroph | \|Undefined Saprotroph\| | Yeast | NULL | Probable | As nectar-tap_saprotroph (Põlme et al. 2020); Decay sugar-rich substrates (Põlme et al. 2020) | Põlme S, et al. 2020. Fungal Diversity 105:1-16 ((https://doi.org/10.1007/s13225-020-00466-2)) |
| Otu41287 | 1 | k__Fungi(100);p__Ascomycota(100);c__Saccharomycetes(100);o__Saccharomycetales(100);f__Metschnikowiaceae(100);g__Metschnikowia(100);g__Metschnikowia_unclassified(100); | Metschnikowia | 13 | Saprotroph | \|Undefined Saprotroph\| | Yeast | NULL | Probable | As nectar-tap_saprotroph (Põlme et al. 2020); Decay sugar-rich substrates (Põlme et al. 2020) | Põlme S, et al. 2020. Fungal Diversity 105:1-16 ((https://doi.org/10.1007/s13225-020-00466-2)) |
| Otu41288 | 1 | k__Fungi(100);p__Ascomycota(100);c__Saccharomycetes(100);o__Saccharomycetales(100);f__Metschnikowiaceae(100);g__Metschnikowia(100);s__Metschnikowia_pulcherrima(100); | Metschnikowia | 13 | Saprotroph | \|Undefined Saprotroph\| | Yeast | NULL | Probable | As nectar-tap_saprotroph (Põlme et al. 2020); Decay sugar-rich substrates (Põlme et al. 2020) | Põlme S, et al. 2020. Fungal Diversity 105:1-16 ((https://doi.org/10.1007/s13225-020-00466-2)) |
| Otu41289 | 1 | k__Fungi(100);p__Ascomycota(100);c__Saccharomycetes(100);o__Saccharomycetales(100);f__Metschnikowiaceae(100);g__Metschnikowia(100);s__Metschnikowia_pulcherrima(100); | Metschnikowia | 13 | Saprotroph | \|Undefined Saprotroph\| | Yeast | NULL | Probable | As nectar-tap_saprotroph (Põlme et al. 2020); Decay sugar-rich substrates (Põlme et al. 2020) | Põlme S, et al. 2020. Fungal Diversity 105:1-16 ((https://doi.org/10.1007/s13225-020-00466-2)) |
| Otu41290 | 1 | k__Fungi(100);p__Ascomycota(100);c__Saccharomycetes(100);o__Saccharomycetales(100);f__Metschnikowiaceae(100);g__Metschnikowia(100);s__Metschnikowia_pulcherrima(100); | Metschnikowia | 13 | Saprotroph | \|Undefined Saprotroph\| | Yeast | NULL | Probable | As nectar-tap_saprotroph (Põlme et al. 2020); Decay sugar-rich substrates (Põlme et al. 2020) | Põlme S, et al. 2020. Fungal Diversity 105:1-16 ((https://doi.org/10.1007/s13225-020-00466-2)) |
| Otu41291 | 1 | k__Fungi(100);p__Ascomycota(100);c__Saccharomycetes(100);o__Saccharomycetales(100);f__Metschnikowiaceae(100);g__Metschnikowia(100);s__Metschnikowia_pulcherrima(100); | Metschnikowia | 13 | Saprotroph | \|Undefined Saprotroph\| | Yeast | NULL | Probable | As nectar-tap_saprotroph (Põlme et al. 2020); Decay sugar-rich substrates (Põlme et al. 2020) | Põlme S, et al. 2020. Fungal Diversity 105:1-16 ((https://doi.org/10.1007/s13225-020-00466-2)) |
| Otu41292 | 1 | k__Fungi(100);p__Ascomycota(100);c__Saccharomycetes(100);o__Saccharomycetales(100);f__Metschnikowiaceae(100);g__Metschnikowia(100);g__Metschnikowia_unclassified(100); | Metschnikowia | 13 | Saprotroph | \|Undefined Saprotroph\| | Yeast | NULL | Probable | As nectar-tap_saprotroph (Põlme et al. 2020); Decay sugar-rich substrates (Põlme et al. 2020) | Põlme S, et al. 2020. Fungal Diversity 105:1-16 ((https://doi.org/10.1007/s13225-020-00466-2)) |
| Otu41293 | 1 | k__Fungi(100);p__Ascomycota(100);c__Saccharomycetes(100);o__Saccharomycetales(100);f__Metschnikowiaceae(100);g__Metschnikowia(100);s__Metschnikowia_pulcherrima(100); | Metschnikowia | 13 | Saprotroph | \|Undefined Saprotroph\| | Yeast | NULL | Probable | As nectar-tap_saprotroph (Põlme et al. 2020); Decay sugar-rich substrates (Põlme et al. 2020) | Põlme S, et al. 2020. Fungal Diversity 105:1-16 ((https://doi.org/10.1007/s13225-020-00466-2)) |
| Otu41294 | 1 | k__Fungi(100);p__Ascomycota(100);c__Saccharomycetes(100);o__Saccharomycetales(100);f__Metschnikowiaceae(100);g__Metschnikowia(100);g__Metschnikowia_unclassified(100); | Metschnikowia | 13 | Saprotroph | \|Undefined Saprotroph\| | Yeast | NULL | Probable | As nectar-tap_saprotroph (Põlme et al. 2020); Decay sugar-rich substrates (Põlme et al. 2020) | Põlme S, et al. 2020. Fungal Diversity 105:1-16 ((https://doi.org/10.1007/s13225-020-00466-2)) |
| Otu41295 | 1 | k__Fungi(100);p__Ascomycota(100);c__Saccharomycetes(100);o__Saccharomycetales(100);f__Metschnikowiaceae(100);g__Metschnikowia(100);s__Metschnikowia_pulcherrima(100); | Metschnikowia | 13 | Saprotroph | \|Undefined Saprotroph\| | Yeast | NULL | Probable | As nectar-tap_saprotroph (Põlme et al. 2020); Decay sugar-rich substrates (Põlme et al. 2020) | Põlme S, et al. 2020. Fungal Diversity 105:1-16 ((https://doi.org/10.1007/s13225-020-00466-2)) |
| Otu41296 | 1 | k__Fungi(100);p__Ascomycota(100);c__Saccharomycetes(100);o__Saccharomycetales(100);f__Metschnikowiaceae(100);g__Metschnikowia(100);g__Metschnikowia_unclassified(100); | Metschnikowia | 13 | Saprotroph | \|Undefined Saprotroph\| | Yeast | NULL | Probable | As nectar-tap_saprotroph (Põlme et al. 2020); Decay sugar-rich substrates (Põlme et al. 2020) | Põlme S, et al. 2020. Fungal Diversity 105:1-16 ((https://doi.org/10.1007/s13225-020-00466-2)) |
| Otu41297 | 1 | k__Fungi(100);p__Ascomycota(100);c__Saccharomycetes(100);o__Saccharomycetales(100);f__Metschnikowiaceae(100);g__Metschnikowia(100);g__Metschnikowia_unclassified(100); | Metschnikowia | 13 | Saprotroph | \|Undefined Saprotroph\| | Yeast | NULL | Probable | As nectar-tap_saprotroph (Põlme et al. 2020); Decay sugar-rich substrates (Põlme et al. 2020) | Põlme S, et al. 2020. Fungal Diversity 105:1-16 ((https://doi.org/10.1007/s13225-020-00466-2)) |
| Otu41298 | 1 | k__Fungi(100);p__Ascomycota(100);c__Saccharomycetes(100);o__Saccharomycetales(100);f__Metschnikowiaceae(100);g__Metschnikowia(100);s__Metschnikowia_pulcherrima(100); | Metschnikowia | 13 | Saprotroph | \|Undefined Saprotroph\| | Yeast | NULL | Probable | As nectar-tap_saprotroph (Põlme et al. 2020); Decay sugar-rich substrates (Põlme et al. 2020) | Põlme S, et al. 2020. Fungal Diversity 105:1-16 ((https://doi.org/10.1007/s13225-020-00466-2)) |
| Otu41299 | 1 | k__Fungi(100);p__Ascomycota(100);c__Saccharomycetes(100);o__Saccharomycetales(100);f__Metschnikowiaceae(100);g__Metschnikowia(100);s__Metschnikowia_pulcherrima(100); | Metschnikowia | 13 | Saprotroph | \|Undefined Saprotroph\| | Yeast | NULL | Probable | As nectar-tap_saprotroph (Põlme et al. 2020); Decay sugar-rich substrates (Põlme et al. 2020) | Põlme S, et al. 2020. Fungal Diversity 105:1-16 ((https://doi.org/10.1007/s13225-020-00466-2)) |
| Otu41300 | 1 | k__Fungi(100);p__Ascomycota(100);c__Saccharomycetes(100);o__Saccharomycetales(100);f__Metschnikowiaceae(100);g__Metschnikowia(100);g__Metschnikowia_unclassified(100); | Metschnikowia | 13 | Saprotroph | \|Undefined Saprotroph\| | Yeast | NULL | Probable | As nectar-tap_saprotroph (Põlme et al. 2020); Decay sugar-rich substrates (Põlme et al. 2020) | Põlme S, et al. 2020. Fungal Diversity 105:1-16 ((https://doi.org/10.1007/s13225-020-00466-2)) |
| Otu41301 | 1 | k__Fungi(100);p__Ascomycota(100);c__Saccharomycetes(100);o__Saccharomycetales(100);f__Metschnikowiaceae(100);g__Metschnikowia(100);s__Metschnikowia_pulcherrima(100); | Metschnikowia | 13 | Saprotroph | \|Undefined Saprotroph\| | Yeast | NULL | Probable | As nectar-tap_saprotroph (Põlme et al. 2020); Decay sugar-rich substrates (Põlme et al. 2020) | Põlme S, et al. 2020. Fungal Diversity 105:1-16 ((https://doi.org/10.1007/s13225-020-00466-2)) |
| Otu41302 | 1 | k__Fungi(100);p__Ascomycota(100);c__Saccharomycetes(100);o__Saccharomycetales(100);f__Metschnikowiaceae(100);g__Metschnikowia(100);s__Metschnikowia_pulcherrima(100); | Metschnikowia | 13 | Saprotroph | \|Undefined Saprotroph\| | Yeast | NULL | Probable | As nectar-tap_saprotroph (Põlme et al. 2020); Decay sugar-rich substrates (Põlme et al. 2020) | Põlme S, et al. 2020. Fungal Diversity 105:1-16 ((https://doi.org/10.1007/s13225-020-00466-2)) |
| Otu41303 | 1 | k__Fungi(100);p__Ascomycota(100);c__Saccharomycetes(100);o__Saccharomycetales(100);f__Metschnikowiaceae(100);g__Metschnikowia(100);s__Metschnikowia_pulcherrima(100); | Metschnikowia | 13 | Saprotroph | \|Undefined Saprotroph\| | Yeast | NULL | Probable | As nectar-tap_saprotroph (Põlme et al. 2020); Decay sugar-rich substrates (Põlme et al. 2020) | Põlme S, et al. 2020. Fungal Diversity 105:1-16 ((https://doi.org/10.1007/s13225-020-00466-2)) |
| Otu41304 | 1 | k__Fungi(100);p__Ascomycota(100);c__Saccharomycetes(100);o__Saccharomycetales(100);f__Metschnikowiaceae(100);g__Metschnikowia(100);s__Metschnikowia_pulcherrima(100); | Metschnikowia | 13 | Saprotroph | \|Undefined Saprotroph\| | Yeast | NULL | Probable | As nectar-tap_saprotroph (Põlme et al. 2020); Decay sugar-rich substrates (Põlme et al. 2020) | Põlme S, et al. 2020. Fungal Diversity 105:1-16 ((https://doi.org/10.1007/s13225-020-00466-2)) |
| Otu41306 | 1 | k__Fungi(100);p__Ascomycota(100);c__Saccharomycetes(100);o__Saccharomycetales(100);f__Metschnikowiaceae(100);g__Metschnikowia(100);s__Metschnikowia_pulcherrima(100); | Metschnikowia | 13 | Saprotroph | \|Undefined Saprotroph\| | Yeast | NULL | Probable | As nectar-tap_saprotroph (Põlme et al. 2020); Decay sugar-rich substrates (Põlme et al. 2020) | Põlme S, et al. 2020. Fungal Diversity 105:1-16 ((https://doi.org/10.1007/s13225-020-00466-2)) |
| Otu41307 | 1 | k__Fungi(100);p__Ascomycota(100);c__Saccharomycetes(100);o__Saccharomycetales(100);f__Metschnikowiaceae(100);g__Metschnikowia(100);g__Metschnikowia_unclassified(100); | Metschnikowia | 13 | Saprotroph | \|Undefined Saprotroph\| | Yeast | NULL | Probable | As nectar-tap_saprotroph (Põlme et al. 2020); Decay sugar-rich substrates (Põlme et al. 2020) | Põlme S, et al. 2020. Fungal Diversity 105:1-16 ((https://doi.org/10.1007/s13225-020-00466-2)) |
| Otu41308 | 1 | k__Fungi(100);p__Ascomycota(100);c__Saccharomycetes(100);o__Saccharomycetales(100);f__Metschnikowiaceae(100);g__Metschnikowia(100);s__Metschnikowia_pulcherrima(100); | Metschnikowia | 13 | Saprotroph | \|Undefined Saprotroph\| | Yeast | NULL | Probable | As nectar-tap_saprotroph (Põlme et al. 2020); Decay sugar-rich substrates (Põlme et al. 2020) | Põlme S, et al. 2020. Fungal Diversity 105:1-16 ((https://doi.org/10.1007/s13225-020-00466-2)) |
| Otu41309 | 1 | k__Fungi(100);p__Ascomycota(100);c__Saccharomycetes(100);o__Saccharomycetales(100);f__Metschnikowiaceae(100);g__Metschnikowia(100);g__Metschnikowia_unclassified(100); | Metschnikowia | 13 | Saprotroph | \|Undefined Saprotroph\| | Yeast | NULL | Probable | As nectar-tap_saprotroph (Põlme et al. 2020); Decay sugar-rich substrates (Põlme et al. 2020) | Põlme S, et al. 2020. Fungal Diversity 105:1-16 ((https://doi.org/10.1007/s13225-020-00466-2)) |
| Otu41310 | 1 | k__Fungi(100);p__Ascomycota(100);c__Saccharomycetes(100);o__Saccharomycetales(100);f__Metschnikowiaceae(100);g__Metschnikowia(100);g__Metschnikowia_unclassified(100); | Metschnikowia | 13 | Saprotroph | \|Undefined Saprotroph\| | Yeast | NULL | Probable | As nectar-tap_saprotroph (Põlme et al. 2020); Decay sugar-rich substrates (Põlme et al. 2020) | Põlme S, et al. 2020. Fungal Diversity 105:1-16 ((https://doi.org/10.1007/s13225-020-00466-2)) |
| Otu41311 | 1 | k__Fungi(100);p__Ascomycota(100);c__Saccharomycetes(100);o__Saccharomycetales(100);f__Metschnikowiaceae(100);g__Metschnikowia(100);s__Metschnikowia_pulcherrima(100); | Metschnikowia | 13 | Saprotroph | \|Undefined Saprotroph\| | Yeast | NULL | Probable | As nectar-tap_saprotroph (Põlme et al. 2020); Decay sugar-rich substrates (Põlme et al. 2020) | Põlme S, et al. 2020. Fungal Diversity 105:1-16 ((https://doi.org/10.1007/s13225-020-00466-2)) |
| Otu41312 | 1 | k__Fungi(100);p__Ascomycota(100);c__Saccharomycetes(100);o__Saccharomycetales(100);f__Metschnikowiaceae(100);g__Metschnikowia(100);g__Metschnikowia_unclassified(100); | Metschnikowia | 13 | Saprotroph | \|Undefined Saprotroph\| | Yeast | NULL | Probable | As nectar-tap_saprotroph (Põlme et al. 2020); Decay sugar-rich substrates (Põlme et al. 2020) | Põlme S, et al. 2020. Fungal Diversity 105:1-16 ((https://doi.org/10.1007/s13225-020-00466-2)) |
| Otu41314 | 1 | k__Fungi(100);p__Ascomycota(100);c__Saccharomycetes(100);o__Saccharomycetales(100);f__Metschnikowiaceae(100);g__Metschnikowia(100);s__Metschnikowia_pulcherrima(100); | Metschnikowia | 13 | Saprotroph | \|Undefined Saprotroph\| | Yeast | NULL | Probable | As nectar-tap_saprotroph (Põlme et al. 2020); Decay sugar-rich substrates (Põlme et al. 2020) | Põlme S, et al. 2020. Fungal Diversity 105:1-16 ((https://doi.org/10.1007/s13225-020-00466-2)) |
| Otu41315 | 1 | k__Fungi(100);p__Ascomycota(100);c__Saccharomycetes(100);o__Saccharomycetales(100);f__Metschnikowiaceae(100);g__Metschnikowia(100);g__Metschnikowia_unclassified(100); | Metschnikowia | 13 | Saprotroph | \|Undefined Saprotroph\| | Yeast | NULL | Probable | As nectar-tap_saprotroph (Põlme et al. 2020); Decay sugar-rich substrates (Põlme et al. 2020) | Põlme S, et al. 2020. Fungal Diversity 105:1-16 ((https://doi.org/10.1007/s13225-020-00466-2)) |
| Otu41316 | 1 | k__Fungi(100);p__Ascomycota(100);c__Saccharomycetes(100);o__Saccharomycetales(100);f__Metschnikowiaceae(100);g__Metschnikowia(100);g__Metschnikowia_unclassified(100); | Metschnikowia | 13 | Saprotroph | \|Undefined Saprotroph\| | Yeast | NULL | Probable | As nectar-tap_saprotroph (Põlme et al. 2020); Decay sugar-rich substrates (Põlme et al. 2020) | Põlme S, et al. 2020. Fungal Diversity 105:1-16 ((https://doi.org/10.1007/s13225-020-00466-2)) |
| Otu41317 | 1 | k__Fungi(100);p__Ascomycota(100);c__Saccharomycetes(100);o__Saccharomycetales(100);f__Metschnikowiaceae(100);g__Metschnikowia(100);g__Metschnikowia_unclassified(100); | Metschnikowia | 13 | Saprotroph | \|Undefined Saprotroph\| | Yeast | NULL | Probable | As nectar-tap_saprotroph (Põlme et al. 2020); Decay sugar-rich substrates (Põlme et al. 2020) | Põlme S, et al. 2020. Fungal Diversity 105:1-16 ((https://doi.org/10.1007/s13225-020-00466-2)) |
| Otu41318 | 1 | k__Fungi(100);p__Ascomycota(100);c__Saccharomycetes(100);o__Saccharomycetales(100);f__Metschnikowiaceae(100);g__Metschnikowia(100);g__Metschnikowia_unclassified(100); | Metschnikowia | 13 | Saprotroph | \|Undefined Saprotroph\| | Yeast | NULL | Probable | As nectar-tap_saprotroph (Põlme et al. 2020); Decay sugar-rich substrates (Põlme et al. 2020) | Põlme S, et al. 2020. Fungal Diversity 105:1-16 ((https://doi.org/10.1007/s13225-020-00466-2)) |
| Otu41319 | 1 | k__Fungi(100);p__Ascomycota(100);c__Saccharomycetes(100);o__Saccharomycetales(100);f__Metschnikowiaceae(100);g__Metschnikowia(100);g__Metschnikowia_unclassified(100); | Metschnikowia | 13 | Saprotroph | \|Undefined Saprotroph\| | Yeast | NULL | Probable | As nectar-tap_saprotroph (Põlme et al. 2020); Decay sugar-rich substrates (Põlme et al. 2020) | Põlme S, et al. 2020. Fungal Diversity 105:1-16 ((https://doi.org/10.1007/s13225-020-00466-2)) |
| Otu41320 | 1 | k__Fungi(100);p__Ascomycota(100);c__Saccharomycetes(100);o__Saccharomycetales(100);f__Metschnikowiaceae(100);g__Metschnikowia(100);g__Metschnikowia_unclassified(100); | Metschnikowia | 13 | Saprotroph | \|Undefined Saprotroph\| | Yeast | NULL | Probable | As nectar-tap_saprotroph (Põlme et al. 2020); Decay sugar-rich substrates (Põlme et al. 2020) | Põlme S, et al. 2020. Fungal Diversity 105:1-16 ((https://doi.org/10.1007/s13225-020-00466-2)) |
| Otu41321 | 1 | k__Fungi(100);p__Ascomycota(100);c__Saccharomycetes(100);o__Saccharomycetales(100);f__Metschnikowiaceae(100);g__Metschnikowia(100);g__Metschnikowia_unclassified(100); | Metschnikowia | 13 | Saprotroph | \|Undefined Saprotroph\| | Yeast | NULL | Probable | As nectar-tap_saprotroph (Põlme et al. 2020); Decay sugar-rich substrates (Põlme et al. 2020) | Põlme S, et al. 2020. Fungal Diversity 105:1-16 ((https://doi.org/10.1007/s13225-020-00466-2)) |
| Otu41322 | 1 | k__Fungi(100);p__Ascomycota(100);c__Saccharomycetes(100);o__Saccharomycetales(100);f__Metschnikowiaceae(100);g__Metschnikowia(100);g__Metschnikowia_unclassified(100); | Metschnikowia | 13 | Saprotroph | \|Undefined Saprotroph\| | Yeast | NULL | Probable | As nectar-tap_saprotroph (Põlme et al. 2020); Decay sugar-rich substrates (Põlme et al. 2020) | Põlme S, et al. 2020. Fungal Diversity 105:1-16 ((https://doi.org/10.1007/s13225-020-00466-2)) |
| Otu41323 | 1 | k__Fungi(100);p__Ascomycota(100);c__Saccharomycetes(100);o__Saccharomycetales(100);f__Metschnikowiaceae(100);g__Metschnikowia(100);g__Metschnikowia_unclassified(100); | Metschnikowia | 13 | Saprotroph | \|Undefined Saprotroph\| | Yeast | NULL | Probable | As nectar-tap_saprotroph (Põlme et al. 2020); Decay sugar-rich substrates (Põlme et al. 2020) | Põlme S, et al. 2020. Fungal Diversity 105:1-16 ((https://doi.org/10.1007/s13225-020-00466-2)) |
| Otu41324 | 1 | k__Fungi(100);p__Ascomycota(100);c__Saccharomycetes(100);o__Saccharomycetales(100);f__Metschnikowiaceae(100);g__Metschnikowia(100);g__Metschnikowia_unclassified(100); | Metschnikowia | 13 | Saprotroph | \|Undefined Saprotroph\| | Yeast | NULL | Probable | As nectar-tap_saprotroph (Põlme et al. 2020); Decay sugar-rich substrates (Põlme et al. 2020) | Põlme S, et al. 2020. Fungal Diversity 105:1-16 ((https://doi.org/10.1007/s13225-020-00466-2)) |
| Otu41325 | 1 | k__Fungi(100);p__Ascomycota(100);c__Saccharomycetes(100);o__Saccharomycetales(100);f__Metschnikowiaceae(100);g__Metschnikowia(100);g__Metschnikowia_unclassified(100); | Metschnikowia | 13 | Saprotroph | \|Undefined Saprotroph\| | Yeast | NULL | Probable | As nectar-tap_saprotroph (Põlme et al. 2020); Decay sugar-rich substrates (Põlme et al. 2020) | Põlme S, et al. 2020. Fungal Diversity 105:1-16 ((https://doi.org/10.1007/s13225-020-00466-2)) |
| Otu41326 | 1 | k__Fungi(100);p__Ascomycota(100);c__Saccharomycetes(100);o__Saccharomycetales(100);f__Metschnikowiaceae(100);g__Metschnikowia(100);g__Metschnikowia_unclassified(100); | Metschnikowia | 13 | Saprotroph | \|Undefined Saprotroph\| | Yeast | NULL | Probable | As nectar-tap_saprotroph (Põlme et al. 2020); Decay sugar-rich substrates (Põlme et al. 2020) | Põlme S, et al. 2020. Fungal Diversity 105:1-16 ((https://doi.org/10.1007/s13225-020-00466-2)) |
| Otu41327 | 1 | k__Fungi(100);p__Ascomycota(100);c__Saccharomycetes(100);o__Saccharomycetales(100);f__Metschnikowiaceae(100);g__Metschnikowia(100);g__Metschnikowia_unclassified(100); | Metschnikowia | 13 | Saprotroph | \|Undefined Saprotroph\| | Yeast | NULL | Probable | As nectar-tap_saprotroph (Põlme et al. 2020); Decay sugar-rich substrates (Põlme et al. 2020) | Põlme S, et al. 2020. Fungal Diversity 105:1-16 ((https://doi.org/10.1007/s13225-020-00466-2)) |
| Otu41328 | 1 | k__Fungi(100);p__Ascomycota(100);c__Saccharomycetes(100);o__Saccharomycetales(100);f__Metschnikowiaceae(100);g__Metschnikowia(100);g__Metschnikowia_unclassified(100); | Metschnikowia | 13 | Saprotroph | \|Undefined Saprotroph\| | Yeast | NULL | Probable | As nectar-tap_saprotroph (Põlme et al. 2020); Decay sugar-rich substrates (Põlme et al. 2020) | Põlme S, et al. 2020. Fungal Diversity 105:1-16 ((https://doi.org/10.1007/s13225-020-00466-2)) |
| Otu41329 | 1 | k__Fungi(100);p__Ascomycota(100);c__Saccharomycetes(100);o__Saccharomycetales(100);f__Metschnikowiaceae(100);g__Metschnikowia(100);g__Metschnikowia_unclassified(100); | Metschnikowia | 13 | Saprotroph | \|Undefined Saprotroph\| | Yeast | NULL | Probable | As nectar-tap_saprotroph (Põlme et al. 2020); Decay sugar-rich substrates (Põlme et al. 2020) | Põlme S, et al. 2020. Fungal Diversity 105:1-16 ((https://doi.org/10.1007/s13225-020-00466-2)) |
| Otu41330 | 1 | k__Fungi(100);p__Ascomycota(100);c__Saccharomycetes(100);o__Saccharomycetales(100);f__Metschnikowiaceae(100);g__Metschnikowia(100);g__Metschnikowia_unclassified(100); | Metschnikowia | 13 | Saprotroph | \|Undefined Saprotroph\| | Yeast | NULL | Probable | As nectar-tap_saprotroph (Põlme et al. 2020); Decay sugar-rich substrates (Põlme et al. 2020) | Põlme S, et al. 2020. Fungal Diversity 105:1-16 ((https://doi.org/10.1007/s13225-020-00466-2)) |
| Otu41331 | 1 | k__Fungi(100);p__Ascomycota(100);c__Saccharomycetes(100);o__Saccharomycetales(100);f__Metschnikowiaceae(100);g__Metschnikowia(100);g__Metschnikowia_unclassified(100); | Metschnikowia | 13 | Saprotroph | \|Undefined Saprotroph\| | Yeast | NULL | Probable | As nectar-tap_saprotroph (Põlme et al. 2020); Decay sugar-rich substrates (Põlme et al. 2020) | Põlme S, et al. 2020. Fungal Diversity 105:1-16 ((https://doi.org/10.1007/s13225-020-00466-2)) |
| Otu41332 | 1 | k__Fungi(100);p__Ascomycota(100);c__Saccharomycetes(100);o__Saccharomycetales(100);f__Metschnikowiaceae(100);g__Metschnikowia(100);g__Metschnikowia_unclassified(100); | Metschnikowia | 13 | Saprotroph | \|Undefined Saprotroph\| | Yeast | NULL | Probable | As nectar-tap_saprotroph (Põlme et al. 2020); Decay sugar-rich substrates (Põlme et al. 2020) | Põlme S, et al. 2020. Fungal Diversity 105:1-16 ((https://doi.org/10.1007/s13225-020-00466-2)) |
| Otu41333 | 1 | k__Fungi(100);p__Ascomycota(100);c__Saccharomycetes(100);o__Saccharomycetales(100);f__Metschnikowiaceae(100);g__Metschnikowia(100);g__Metschnikowia_unclassified(100); | Metschnikowia | 13 | Saprotroph | \|Undefined Saprotroph\| | Yeast | NULL | Probable | As nectar-tap_saprotroph (Põlme et al. 2020); Decay sugar-rich substrates (Põlme et al. 2020) | Põlme S, et al. 2020. Fungal Diversity 105:1-16 ((https://doi.org/10.1007/s13225-020-00466-2)) |
| Otu41334 | 1 | k__Fungi(100);p__Ascomycota(100);c__Saccharomycetes(100);o__Saccharomycetales(100);f__Metschnikowiaceae(100);g__Metschnikowia(100);s__Metschnikowia_pulcherrima(100); | Metschnikowia | 13 | Saprotroph | \|Undefined Saprotroph\| | Yeast | NULL | Probable | As nectar-tap_saprotroph (Põlme et al. 2020); Decay sugar-rich substrates (Põlme et al. 2020) | Põlme S, et al. 2020. Fungal Diversity 105:1-16 ((https://doi.org/10.1007/s13225-020-00466-2)) |
| Otu41335 | 1 | k__Fungi(100);p__Ascomycota(100);c__Saccharomycetes(100);o__Saccharomycetales(100);f__Metschnikowiaceae(100);g__Metschnikowia(100);s__Metschnikowia_pulcherrima(100); | Metschnikowia | 13 | Saprotroph | \|Undefined Saprotroph\| | Yeast | NULL | Probable | As nectar-tap_saprotroph (Põlme et al. 2020); Decay sugar-rich substrates (Põlme et al. 2020) | Põlme S, et al. 2020. Fungal Diversity 105:1-16 ((https://doi.org/10.1007/s13225-020-00466-2)) |
| Otu41336 | 1 | k__Fungi(100);p__Ascomycota(100);c__Saccharomycetes(100);o__Saccharomycetales(100);f__Metschnikowiaceae(100);g__Metschnikowia(100);g__Metschnikowia_unclassified(100); | Metschnikowia | 13 | Saprotroph | \|Undefined Saprotroph\| | Yeast | NULL | Probable | As nectar-tap_saprotroph (Põlme et al. 2020); Decay sugar-rich substrates (Põlme et al. 2020) | Põlme S, et al. 2020. Fungal Diversity 105:1-16 ((https://doi.org/10.1007/s13225-020-00466-2)) |
| Otu41337 | 1 | k__Fungi(100);p__Ascomycota(100);c__Saccharomycetes(100);o__Saccharomycetales(100);f__Metschnikowiaceae(100);g__Metschnikowia(100);s__Metschnikowia_pulcherrima(100); | Metschnikowia | 13 | Saprotroph | \|Undefined Saprotroph\| | Yeast | NULL | Probable | As nectar-tap_saprotroph (Põlme et al. 2020); Decay sugar-rich substrates (Põlme et al. 2020) | Põlme S, et al. 2020. Fungal Diversity 105:1-16 ((https://doi.org/10.1007/s13225-020-00466-2)) |
| Otu41338 | 1 | k__Fungi(100);p__Ascomycota(100);c__Saccharomycetes(100);o__Saccharomycetales(100);f__Metschnikowiaceae(100);g__Metschnikowia(100);s__Metschnikowia_pulcherrima(100); | Metschnikowia | 13 | Saprotroph | \|Undefined Saprotroph\| | Yeast | NULL | Probable | As nectar-tap_saprotroph (Põlme et al. 2020); Decay sugar-rich substrates (Põlme et al. 2020) | Põlme S, et al. 2020. Fungal Diversity 105:1-16 ((https://doi.org/10.1007/s13225-020-00466-2)) |
| Otu41339 | 1 | k__Fungi(100);p__Ascomycota(100);c__Saccharomycetes(100);o__Saccharomycetales(100);f__Metschnikowiaceae(100);g__Metschnikowia(100);s__Metschnikowia_pulcherrima(100); | Metschnikowia | 13 | Saprotroph | \|Undefined Saprotroph\| | Yeast | NULL | Probable | As nectar-tap_saprotroph (Põlme et al. 2020); Decay sugar-rich substrates (Põlme et al. 2020) | Põlme S, et al. 2020. Fungal Diversity 105:1-16 ((https://doi.org/10.1007/s13225-020-00466-2)) |
| Otu41340 | 1 | k__Fungi(100);p__Ascomycota(100);c__Saccharomycetes(100);o__Saccharomycetales(100);f__Metschnikowiaceae(100);g__Metschnikowia(100);s__Metschnikowia_pulcherrima(100); | Metschnikowia | 13 | Saprotroph | \|Undefined Saprotroph\| | Yeast | NULL | Probable | As nectar-tap_saprotroph (Põlme et al. 2020); Decay sugar-rich substrates (Põlme et al. 2020) | Põlme S, et al. 2020. Fungal Diversity 105:1-16 ((https://doi.org/10.1007/s13225-020-00466-2)) |
| Otu41341 | 1 | k__Fungi(100);p__Ascomycota(100);c__Saccharomycetes(100);o__Saccharomycetales(100);f__Metschnikowiaceae(100);g__Metschnikowia(100);s__Metschnikowia_pulcherrima(100); | Metschnikowia | 13 | Saprotroph | \|Undefined Saprotroph\| | Yeast | NULL | Probable | As nectar-tap_saprotroph (Põlme et al. 2020); Decay sugar-rich substrates (Põlme et al. 2020) | Põlme S, et al. 2020. Fungal Diversity 105:1-16 ((https://doi.org/10.1007/s13225-020-00466-2)) |
| Otu41342 | 1 | k__Fungi(100);p__Ascomycota(100);c__Saccharomycetes(100);o__Saccharomycetales(100);f__Metschnikowiaceae(100);g__Metschnikowia(100);s__Metschnikowia_pulcherrima(100); | Metschnikowia | 13 | Saprotroph | \|Undefined Saprotroph\| | Yeast | NULL | Probable | As nectar-tap_saprotroph (Põlme et al. 2020); Decay sugar-rich substrates (Põlme et al. 2020) | Põlme S, et al. 2020. Fungal Diversity 105:1-16 ((https://doi.org/10.1007/s13225-020-00466-2)) |
| Otu41343 | 1 | k__Fungi(100);p__Ascomycota(100);c__Saccharomycetes(100);o__Saccharomycetales(100);f__Metschnikowiaceae(100);g__Metschnikowia(100);s__Metschnikowia_pulcherrima(100); | Metschnikowia | 13 | Saprotroph | \|Undefined Saprotroph\| | Yeast | NULL | Probable | As nectar-tap_saprotroph (Põlme et al. 2020); Decay sugar-rich substrates (Põlme et al. 2020) | Põlme S, et al. 2020. Fungal Diversity 105:1-16 ((https://doi.org/10.1007/s13225-020-00466-2)) |
| Otu41344 | 1 | k__Fungi(100);p__Ascomycota(100);c__Saccharomycetes(100);o__Saccharomycetales(100);f__Metschnikowiaceae(100);g__Metschnikowia(100);s__Metschnikowia_pulcherrima(100); | Metschnikowia | 13 | Saprotroph | \|Undefined Saprotroph\| | Yeast | NULL | Probable | As nectar-tap_saprotroph (Põlme et al. 2020); Decay sugar-rich substrates (Põlme et al. 2020) | Põlme S, et al. 2020. Fungal Diversity 105:1-16 ((https://doi.org/10.1007/s13225-020-00466-2)) |
| Otu41345 | 1 | k__Fungi(100);p__Ascomycota(100);c__Saccharomycetes(100);o__Saccharomycetales(100);f__Metschnikowiaceae(100);g__Metschnikowia(100);s__Metschnikowia_pulcherrima(100); | Metschnikowia | 13 | Saprotroph | \|Undefined Saprotroph\| | Yeast | NULL | Probable | As nectar-tap_saprotroph (Põlme et al. 2020); Decay sugar-rich substrates (Põlme et al. 2020) | Põlme S, et al. 2020. Fungal Diversity 105:1-16 ((https://doi.org/10.1007/s13225-020-00466-2)) |
| Otu41346 | 1 | k__Fungi(100);p__Ascomycota(100);c__Saccharomycetes(100);o__Saccharomycetales(100);f__Metschnikowiaceae(100);g__Metschnikowia(100);s__Metschnikowia_pulcherrima(100); | Metschnikowia | 13 | Saprotroph | \|Undefined Saprotroph\| | Yeast | NULL | Probable | As nectar-tap_saprotroph (Põlme et al. 2020); Decay sugar-rich substrates (Põlme et al. 2020) | Põlme S, et al. 2020. Fungal Diversity 105:1-16 ((https://doi.org/10.1007/s13225-020-00466-2)) |
| Otu41347 | 1 | k__Fungi(100);p__Ascomycota(100);c__Saccharomycetes(100);o__Saccharomycetales(100);f__Metschnikowiaceae(100);g__Metschnikowia(100);s__Metschnikowia_pulcherrima(100); | Metschnikowia | 13 | Saprotroph | \|Undefined Saprotroph\| | Yeast | NULL | Probable | As nectar-tap_saprotroph (Põlme et al. 2020); Decay sugar-rich substrates (Põlme et al. 2020) | Põlme S, et al. 2020. Fungal Diversity 105:1-16 ((https://doi.org/10.1007/s13225-020-00466-2)) |
| Otu41348 | 1 | k__Fungi(100);p__Ascomycota(100);c__Saccharomycetes(100);o__Saccharomycetales(100);f__Metschnikowiaceae(100);g__Metschnikowia(100);s__Metschnikowia_pulcherrima(100); | Metschnikowia | 13 | Saprotroph | \|Undefined Saprotroph\| | Yeast | NULL | Probable | As nectar-tap_saprotroph (Põlme et al. 2020); Decay sugar-rich substrates (Põlme et al. 2020) | Põlme S, et al. 2020. Fungal Diversity 105:1-16 ((https://doi.org/10.1007/s13225-020-00466-2)) |
| Otu41349 | 1 | k__Fungi(100);p__Ascomycota(100);c__Saccharomycetes(100);o__Saccharomycetales(100);f__Metschnikowiaceae(100);g__Metschnikowia(100);s__Metschnikowia_pulcherrima(100); | Metschnikowia | 13 | Saprotroph | \|Undefined Saprotroph\| | Yeast | NULL | Probable | As nectar-tap_saprotroph (Põlme et al. 2020); Decay sugar-rich substrates (Põlme et al. 2020) | Põlme S, et al. 2020. Fungal Diversity 105:1-16 ((https://doi.org/10.1007/s13225-020-00466-2)) |
| Otu41350 | 1 | k__Fungi(100);p__Ascomycota(100);c__Saccharomycetes(100);o__Saccharomycetales(100);f__Metschnikowiaceae(100);g__Metschnikowia(100);s__Metschnikowia_pulcherrima(100); | Metschnikowia | 13 | Saprotroph | \|Undefined Saprotroph\| | Yeast | NULL | Probable | As nectar-tap_saprotroph (Põlme et al. 2020); Decay sugar-rich substrates (Põlme et al. 2020) | Põlme S, et al. 2020. Fungal Diversity 105:1-16 ((https://doi.org/10.1007/s13225-020-00466-2)) |
| Otu41351 | 1 | k__Fungi(100);p__Ascomycota(100);c__Saccharomycetes(100);o__Saccharomycetales(100);f__Metschnikowiaceae(100);g__Metschnikowia(100);s__Metschnikowia_pulcherrima(100); | Metschnikowia | 13 | Saprotroph | \|Undefined Saprotroph\| | Yeast | NULL | Probable | As nectar-tap_saprotroph (Põlme et al. 2020); Decay sugar-rich substrates (Põlme et al. 2020) | Põlme S, et al. 2020. Fungal Diversity 105:1-16 ((https://doi.org/10.1007/s13225-020-00466-2)) |
| Otu41352 | 1 | k__Fungi(100);p__Ascomycota(100);c__Saccharomycetes(100);o__Saccharomycetales(100);f__Metschnikowiaceae(100);g__Metschnikowia(100);s__Metschnikowia_pulcherrima(100); | Metschnikowia | 13 | Saprotroph | \|Undefined Saprotroph\| | Yeast | NULL | Probable | As nectar-tap_saprotroph (Põlme et al. 2020); Decay sugar-rich substrates (Põlme et al. 2020) | Põlme S, et al. 2020. Fungal Diversity 105:1-16 ((https://doi.org/10.1007/s13225-020-00466-2)) |
| Otu41353 | 1 | k__Fungi(100);p__Ascomycota(100);c__Saccharomycetes(100);o__Saccharomycetales(100);f__Metschnikowiaceae(100);g__Metschnikowia(100);s__Metschnikowia_pulcherrima(100); | Metschnikowia | 13 | Saprotroph | \|Undefined Saprotroph\| | Yeast | NULL | Probable | As nectar-tap_saprotroph (Põlme et al. 2020); Decay sugar-rich substrates (Põlme et al. 2020) | Põlme S, et al. 2020. Fungal Diversity 105:1-16 ((https://doi.org/10.1007/s13225-020-00466-2)) |
| Otu41354 | 1 | k__Fungi(100);p__Ascomycota(100);c__Saccharomycetes(100);o__Saccharomycetales(100);f__Metschnikowiaceae(100);g__Metschnikowia(100);s__Metschnikowia_pulcherrima(100); | Metschnikowia | 13 | Saprotroph | \|Undefined Saprotroph\| | Yeast | NULL | Probable | As nectar-tap_saprotroph (Põlme et al. 2020); Decay sugar-rich substrates (Põlme et al. 2020) | Põlme S, et al. 2020. Fungal Diversity 105:1-16 ((https://doi.org/10.1007/s13225-020-00466-2)) |
| Otu41355 | 1 | k__Fungi(100);p__Ascomycota(100);c__Saccharomycetes(100);o__Saccharomycetales(100);f__Metschnikowiaceae(100);g__Metschnikowia(100);s__Metschnikowia_pulcherrima(100); | Metschnikowia | 13 | Saprotroph | \|Undefined Saprotroph\| | Yeast | NULL | Probable | As nectar-tap_saprotroph (Põlme et al. 2020); Decay sugar-rich substrates (Põlme et al. 2020) | Põlme S, et al. 2020. Fungal Diversity 105:1-16 ((https://doi.org/10.1007/s13225-020-00466-2)) |
| Otu41356 | 1 | k__Fungi(100);p__Ascomycota(100);c__Saccharomycetes(100);o__Saccharomycetales(100);f__Metschnikowiaceae(100);g__Metschnikowia(100);s__Metschnikowia_pulcherrima(100); | Metschnikowia | 13 | Saprotroph | \|Undefined Saprotroph\| | Yeast | NULL | Probable | As nectar-tap_saprotroph (Põlme et al. 2020); Decay sugar-rich substrates (Põlme et al. 2020) | Põlme S, et al. 2020. Fungal Diversity 105:1-16 ((https://doi.org/10.1007/s13225-020-00466-2)) |
| Otu41357 | 1 | k__Fungi(100);p__Ascomycota(100);c__Saccharomycetes(100);o__Saccharomycetales(100);f__Metschnikowiaceae(100);g__Metschnikowia(100);s__Metschnikowia_pulcherrima(100); | Metschnikowia | 13 | Saprotroph | \|Undefined Saprotroph\| | Yeast | NULL | Probable | As nectar-tap_saprotroph (Põlme et al. 2020); Decay sugar-rich substrates (Põlme et al. 2020) | Põlme S, et al. 2020. Fungal Diversity 105:1-16 ((https://doi.org/10.1007/s13225-020-00466-2)) |
| Otu41358 | 1 | k__Fungi(100);p__Ascomycota(100);c__Saccharomycetes(100);o__Saccharomycetales(100);f__Metschnikowiaceae(100);g__Metschnikowia(100);g__Metschnikowia_unclassified(100); | Metschnikowia | 13 | Saprotroph | \|Undefined Saprotroph\| | Yeast | NULL | Probable | As nectar-tap_saprotroph (Põlme et al. 2020); Decay sugar-rich substrates (Põlme et al. 2020) | Põlme S, et al. 2020. Fungal Diversity 105:1-16 ((https://doi.org/10.1007/s13225-020-00466-2)) |
| Otu41359 | 1 | k__Fungi(100);p__Ascomycota(100);c__Saccharomycetes(100);o__Saccharomycetales(100);f__Metschnikowiaceae(100);g__Metschnikowia(100);s__Metschnikowia_pulcherrima(100); | Metschnikowia | 13 | Saprotroph | \|Undefined Saprotroph\| | Yeast | NULL | Probable | As nectar-tap_saprotroph (Põlme et al. 2020); Decay sugar-rich substrates (Põlme et al. 2020) | Põlme S, et al. 2020. Fungal Diversity 105:1-16 ((https://doi.org/10.1007/s13225-020-00466-2)) |
| Otu41360 | 1 | k__Fungi(100);p__Ascomycota(100);c__Saccharomycetes(100);o__Saccharomycetales(100);f__Metschnikowiaceae(100);g__Metschnikowia(100);g__Metschnikowia_unclassified(100); | Metschnikowia | 13 | Saprotroph | \|Undefined Saprotroph\| | Yeast | NULL | Probable | As nectar-tap_saprotroph (Põlme et al. 2020); Decay sugar-rich substrates (Põlme et al. 2020) | Põlme S, et al. 2020. Fungal Diversity 105:1-16 ((https://doi.org/10.1007/s13225-020-00466-2)) |
| Otu41361 | 1 | k__Fungi(100);p__Ascomycota(100);c__Saccharomycetes(100);o__Saccharomycetales(100);f__Metschnikowiaceae(100);g__Metschnikowia(100);s__Metschnikowia_pulcherrima(100); | Metschnikowia | 13 | Saprotroph | \|Undefined Saprotroph\| | Yeast | NULL | Probable | As nectar-tap_saprotroph (Põlme et al. 2020); Decay sugar-rich substrates (Põlme et al. 2020) | Põlme S, et al. 2020. Fungal Diversity 105:1-16 ((https://doi.org/10.1007/s13225-020-00466-2)) |
| Otu41362 | 1 | k__Fungi(100);p__Ascomycota(100);c__Saccharomycetes(100);o__Saccharomycetales(100);f__Metschnikowiaceae(100);g__Metschnikowia(100);s__Metschnikowia_pulcherrima(100); | Metschnikowia | 13 | Saprotroph | \|Undefined Saprotroph\| | Yeast | NULL | Probable | As nectar-tap_saprotroph (Põlme et al. 2020); Decay sugar-rich substrates (Põlme et al. 2020) | Põlme S, et al. 2020. Fungal Diversity 105:1-16 ((https://doi.org/10.1007/s13225-020-00466-2)) |
| Otu41363 | 1 | k__Fungi(100);p__Ascomycota(100);c__Saccharomycetes(100);o__Saccharomycetales(100);f__Metschnikowiaceae(100);g__Metschnikowia(100);s__Metschnikowia_pulcherrima(100); | Metschnikowia | 13 | Saprotroph | \|Undefined Saprotroph\| | Yeast | NULL | Probable | As nectar-tap_saprotroph (Põlme et al. 2020); Decay sugar-rich substrates (Põlme et al. 2020) | Põlme S, et al. 2020. Fungal Diversity 105:1-16 ((https://doi.org/10.1007/s13225-020-00466-2)) |
| Otu41364 | 1 | k__Fungi(100);p__Ascomycota(100);c__Saccharomycetes(100);o__Saccharomycetales(100);f__Metschnikowiaceae(100);g__Metschnikowia(100);g__Metschnikowia_unclassified(100); | Metschnikowia | 13 | Saprotroph | \|Undefined Saprotroph\| | Yeast | NULL | Probable | As nectar-tap_saprotroph (Põlme et al. 2020); Decay sugar-rich substrates (Põlme et al. 2020) | Põlme S, et al. 2020. Fungal Diversity 105:1-16 ((https://doi.org/10.1007/s13225-020-00466-2)) |
| Otu41365 | 1 | k__Fungi(100);p__Ascomycota(100);c__Saccharomycetes(100);o__Saccharomycetales(100);f__Metschnikowiaceae(100);g__Metschnikowia(100);s__Metschnikowia_pulcherrima(100); | Metschnikowia | 13 | Saprotroph | \|Undefined Saprotroph\| | Yeast | NULL | Probable | As nectar-tap_saprotroph (Põlme et al. 2020); Decay sugar-rich substrates (Põlme et al. 2020) | Põlme S, et al. 2020. Fungal Diversity 105:1-16 ((https://doi.org/10.1007/s13225-020-00466-2)) |
| Otu41366 | 1 | k__Fungi(100);p__Ascomycota(100);c__Saccharomycetes(100);o__Saccharomycetales(100);f__Metschnikowiaceae(100);g__Metschnikowia(100);g__Metschnikowia_unclassified(100); | Metschnikowia | 13 | Saprotroph | \|Undefined Saprotroph\| | Yeast | NULL | Probable | As nectar-tap_saprotroph (Põlme et al. 2020); Decay sugar-rich substrates (Põlme et al. 2020) | Põlme S, et al. 2020. Fungal Diversity 105:1-16 ((https://doi.org/10.1007/s13225-020-00466-2)) |
| Otu41367 | 1 | k__Fungi(100);p__Ascomycota(100);c__Saccharomycetes(100);o__Saccharomycetales(100);f__Metschnikowiaceae(100);g__Metschnikowia(100);s__Metschnikowia_pulcherrima(100); | Metschnikowia | 13 | Saprotroph | \|Undefined Saprotroph\| | Yeast | NULL | Probable | As nectar-tap_saprotroph (Põlme et al. 2020); Decay sugar-rich substrates (Põlme et al. 2020) | Põlme S, et al. 2020. Fungal Diversity 105:1-16 ((https://doi.org/10.1007/s13225-020-00466-2)) |
| Otu41368 | 1 | k__Fungi(100);p__Ascomycota(100);c__Saccharomycetes(100);o__Saccharomycetales(100);f__Metschnikowiaceae(100);g__Metschnikowia(100);g__Metschnikowia_unclassified(100); | Metschnikowia | 13 | Saprotroph | \|Undefined Saprotroph\| | Yeast | NULL | Probable | As nectar-tap_saprotroph (Põlme et al. 2020); Decay sugar-rich substrates (Põlme et al. 2020) | Põlme S, et al. 2020. Fungal Diversity 105:1-16 ((https://doi.org/10.1007/s13225-020-00466-2)) |
| Otu41369 | 1 | k__Fungi(100);p__Ascomycota(100);c__Saccharomycetes(100);o__Saccharomycetales(100);f__Metschnikowiaceae(100);g__Metschnikowia(100);s__Metschnikowia_pulcherrima(100); | Metschnikowia | 13 | Saprotroph | \|Undefined Saprotroph\| | Yeast | NULL | Probable | As nectar-tap_saprotroph (Põlme et al. 2020); Decay sugar-rich substrates (Põlme et al. 2020) | Põlme S, et al. 2020. Fungal Diversity 105:1-16 ((https://doi.org/10.1007/s13225-020-00466-2)) |
| Otu41370 | 1 | k__Fungi(100);p__Ascomycota(100);c__Saccharomycetes(100);o__Saccharomycetales(100);f__Metschnikowiaceae(100);g__Metschnikowia(100);s__Metschnikowia_pulcherrima(100); | Metschnikowia | 13 | Saprotroph | \|Undefined Saprotroph\| | Yeast | NULL | Probable | As nectar-tap_saprotroph (Põlme et al. 2020); Decay sugar-rich substrates (Põlme et al. 2020) | Põlme S, et al. 2020. Fungal Diversity 105:1-16 ((https://doi.org/10.1007/s13225-020-00466-2)) |
| Otu41371 | 1 | k__Fungi(100);p__Ascomycota(100);c__Saccharomycetes(100);o__Saccharomycetales(100);f__Metschnikowiaceae(100);g__Metschnikowia(100);s__Metschnikowia_pulcherrima(100); | Metschnikowia | 13 | Saprotroph | \|Undefined Saprotroph\| | Yeast | NULL | Probable | As nectar-tap_saprotroph (Põlme et al. 2020); Decay sugar-rich substrates (Põlme et al. 2020) | Põlme S, et al. 2020. Fungal Diversity 105:1-16 ((https://doi.org/10.1007/s13225-020-00466-2)) |
| Otu41372 | 1 | k__Fungi(100);p__Ascomycota(100);c__Saccharomycetes(100);o__Saccharomycetales(100);f__Metschnikowiaceae(100);g__Metschnikowia(100);g__Metschnikowia_unclassified(100); | Metschnikowia | 13 | Saprotroph | \|Undefined Saprotroph\| | Yeast | NULL | Probable | As nectar-tap_saprotroph (Põlme et al. 2020); Decay sugar-rich substrates (Põlme et al. 2020) | Põlme S, et al. 2020. Fungal Diversity 105:1-16 ((https://doi.org/10.1007/s13225-020-00466-2)) |
| Otu41373 | 1 | k__Fungi(100);p__Ascomycota(100);c__Saccharomycetes(100);o__Saccharomycetales(100);f__Metschnikowiaceae(100);g__Metschnikowia(100);s__Metschnikowia_pulcherrima(100); | Metschnikowia | 13 | Saprotroph | \|Undefined Saprotroph\| | Yeast | NULL | Probable | As nectar-tap_saprotroph (Põlme et al. 2020); Decay sugar-rich substrates (Põlme et al. 2020) | Põlme S, et al. 2020. Fungal Diversity 105:1-16 ((https://doi.org/10.1007/s13225-020-00466-2)) |
| Otu41374 | 1 | k__Fungi(100);p__Ascomycota(100);c__Saccharomycetes(100);o__Saccharomycetales(100);f__Metschnikowiaceae(100);g__Metschnikowia(100);s__Metschnikowia_pulcherrima(100); | Metschnikowia | 13 | Saprotroph | \|Undefined Saprotroph\| | Yeast | NULL | Probable | As nectar-tap_saprotroph (Põlme et al. 2020); Decay sugar-rich substrates (Põlme et al. 2020) | Põlme S, et al. 2020. Fungal Diversity 105:1-16 ((https://doi.org/10.1007/s13225-020-00466-2)) |
| Otu41375 | 1 | k__Fungi(100);p__Ascomycota(100);c__Saccharomycetes(100);o__Saccharomycetales(100);f__Metschnikowiaceae(100);g__Metschnikowia(100);s__Metschnikowia_pulcherrima(100); | Metschnikowia | 13 | Saprotroph | \|Undefined Saprotroph\| | Yeast | NULL | Probable | As nectar-tap_saprotroph (Põlme et al. 2020); Decay sugar-rich substrates (Põlme et al. 2020) | Põlme S, et al. 2020. Fungal Diversity 105:1-16 ((https://doi.org/10.1007/s13225-020-00466-2)) |
| Otu41376 | 1 | k__Fungi(100);p__Ascomycota(100);c__Saccharomycetes(100);o__Saccharomycetales(100);f__Metschnikowiaceae(100);g__Metschnikowia(100);s__Metschnikowia_pulcherrima(100); | Metschnikowia | 13 | Saprotroph | \|Undefined Saprotroph\| | Yeast | NULL | Probable | As nectar-tap_saprotroph (Põlme et al. 2020); Decay sugar-rich substrates (Põlme et al. 2020) | Põlme S, et al. 2020. Fungal Diversity 105:1-16 ((https://doi.org/10.1007/s13225-020-00466-2)) |
| Otu41377 | 1 | k__Fungi(100);p__Ascomycota(100);c__Saccharomycetes(100);o__Saccharomycetales(100);f__Metschnikowiaceae(100);g__Metschnikowia(100);s__Metschnikowia_pulcherrima(100); | Metschnikowia | 13 | Saprotroph | \|Undefined Saprotroph\| | Yeast | NULL | Probable | As nectar-tap_saprotroph (Põlme et al. 2020); Decay sugar-rich substrates (Põlme et al. 2020) | Põlme S, et al. 2020. Fungal Diversity 105:1-16 ((https://doi.org/10.1007/s13225-020-00466-2)) |
| Otu41378 | 1 | k__Fungi(100);p__Ascomycota(100);c__Saccharomycetes(100);o__Saccharomycetales(100);f__Metschnikowiaceae(100);g__Metschnikowia(100);s__Metschnikowia_pulcherrima(100); | Metschnikowia | 13 | Saprotroph | \|Undefined Saprotroph\| | Yeast | NULL | Probable | As nectar-tap_saprotroph (Põlme et al. 2020); Decay sugar-rich substrates (Põlme et al. 2020) | Põlme S, et al. 2020. Fungal Diversity 105:1-16 ((https://doi.org/10.1007/s13225-020-00466-2)) |
| Otu41379 | 1 | k__Fungi(100);p__Ascomycota(100);c__Saccharomycetes(100);o__Saccharomycetales(100);f__Metschnikowiaceae(100);g__Metschnikowia(100);g__Metschnikowia_unclassified(100); | Metschnikowia | 13 | Saprotroph | \|Undefined Saprotroph\| | Yeast | NULL | Probable | As nectar-tap_saprotroph (Põlme et al. 2020); Decay sugar-rich substrates (Põlme et al. 2020) | Põlme S, et al. 2020. Fungal Diversity 105:1-16 ((https://doi.org/10.1007/s13225-020-00466-2)) |
| Otu41380 | 1 | k__Fungi(100);p__Ascomycota(100);c__Saccharomycetes(100);o__Saccharomycetales(100);f__Metschnikowiaceae(100);g__Metschnikowia(100);s__Metschnikowia_pulcherrima(100); | Metschnikowia | 13 | Saprotroph | \|Undefined Saprotroph\| | Yeast | NULL | Probable | As nectar-tap_saprotroph (Põlme et al. 2020); Decay sugar-rich substrates (Põlme et al. 2020) | Põlme S, et al. 2020. Fungal Diversity 105:1-16 ((https://doi.org/10.1007/s13225-020-00466-2)) |
| Otu41381 | 1 | k__Fungi(100);p__Ascomycota(100);c__Saccharomycetes(100);o__Saccharomycetales(100);f__Metschnikowiaceae(100);g__Metschnikowia(100);s__Metschnikowia_pulcherrima(100); | Metschnikowia | 13 | Saprotroph | \|Undefined Saprotroph\| | Yeast | NULL | Probable | As nectar-tap_saprotroph (Põlme et al. 2020); Decay sugar-rich substrates (Põlme et al. 2020) | Põlme S, et al. 2020. Fungal Diversity 105:1-16 ((https://doi.org/10.1007/s13225-020-00466-2)) |
| Otu41382 | 1 | k__Fungi(100);p__Ascomycota(100);c__Saccharomycetes(100);o__Saccharomycetales(100);f__Metschnikowiaceae(100);g__Metschnikowia(100);g__Metschnikowia_unclassified(100); | Metschnikowia | 13 | Saprotroph | \|Undefined Saprotroph\| | Yeast | NULL | Probable | As nectar-tap_saprotroph (Põlme et al. 2020); Decay sugar-rich substrates (Põlme et al. 2020) | Põlme S, et al. 2020. Fungal Diversity 105:1-16 ((https://doi.org/10.1007/s13225-020-00466-2)) |
| Otu41383 | 1 | k__Fungi(100);p__Ascomycota(100);c__Saccharomycetes(100);o__Saccharomycetales(100);f__Metschnikowiaceae(100);g__Metschnikowia(100);g__Metschnikowia_unclassified(100); | Metschnikowia | 13 | Saprotroph | \|Undefined Saprotroph\| | Yeast | NULL | Probable | As nectar-tap_saprotroph (Põlme et al. 2020); Decay sugar-rich substrates (Põlme et al. 2020) | Põlme S, et al. 2020. Fungal Diversity 105:1-16 ((https://doi.org/10.1007/s13225-020-00466-2)) |
| Otu41384 | 1 | k__Fungi(100);p__Ascomycota(100);c__Saccharomycetes(100);o__Saccharomycetales(100);f__Metschnikowiaceae(100);g__Metschnikowia(100);s__Metschnikowia_pulcherrima(100); | Metschnikowia | 13 | Saprotroph | \|Undefined Saprotroph\| | Yeast | NULL | Probable | As nectar-tap_saprotroph (Põlme et al. 2020); Decay sugar-rich substrates (Põlme et al. 2020) | Põlme S, et al. 2020. Fungal Diversity 105:1-16 ((https://doi.org/10.1007/s13225-020-00466-2)) |
| Otu41385 | 1 | k__Fungi(100);p__Ascomycota(100);c__Saccharomycetes(100);o__Saccharomycetales(100);f__Metschnikowiaceae(100);g__Metschnikowia(100);g__Metschnikowia_unclassified(100); | Metschnikowia | 13 | Saprotroph | \|Undefined Saprotroph\| | Yeast | NULL | Probable | As nectar-tap_saprotroph (Põlme et al. 2020); Decay sugar-rich substrates (Põlme et al. 2020) | Põlme S, et al. 2020. Fungal Diversity 105:1-16 ((https://doi.org/10.1007/s13225-020-00466-2)) |
| Otu41387 | 1 | k__Fungi(100);p__Ascomycota(100);c__Saccharomycetes(100);o__Saccharomycetales(100);f__Metschnikowiaceae(100);g__Metschnikowia(100);s__Metschnikowia_pulcherrima(100); | Metschnikowia | 13 | Saprotroph | \|Undefined Saprotroph\| | Yeast | NULL | Probable | As nectar-tap_saprotroph (Põlme et al. 2020); Decay sugar-rich substrates (Põlme et al. 2020) | Põlme S, et al. 2020. Fungal Diversity 105:1-16 ((https://doi.org/10.1007/s13225-020-00466-2)) |
| Otu41388 | 1 | k__Fungi(100);p__Ascomycota(100);c__Saccharomycetes(100);o__Saccharomycetales(100);f__Metschnikowiaceae(100);g__Metschnikowia(100);s__Metschnikowia_pulcherrima(100); | Metschnikowia | 13 | Saprotroph | \|Undefined Saprotroph\| | Yeast | NULL | Probable | As nectar-tap_saprotroph (Põlme et al. 2020); Decay sugar-rich substrates (Põlme et al. 2020) | Põlme S, et al. 2020. Fungal Diversity 105:1-16 ((https://doi.org/10.1007/s13225-020-00466-2)) |
| Otu41389 | 1 | k__Fungi(100);p__Ascomycota(100);c__Saccharomycetes(100);o__Saccharomycetales(100);f__Metschnikowiaceae(100);g__Metschnikowia(100);s__Metschnikowia_pulcherrima(100); | Metschnikowia | 13 | Saprotroph | \|Undefined Saprotroph\| | Yeast | NULL | Probable | As nectar-tap_saprotroph (Põlme et al. 2020); Decay sugar-rich substrates (Põlme et al. 2020) | Põlme S, et al. 2020. Fungal Diversity 105:1-16 ((https://doi.org/10.1007/s13225-020-00466-2)) |
| Otu41390 | 1 | k__Fungi(100);p__Ascomycota(100);c__Saccharomycetes(100);o__Saccharomycetales(100);f__Metschnikowiaceae(100);g__Metschnikowia(100);s__Metschnikowia_pulcherrima(100); | Metschnikowia | 13 | Saprotroph | \|Undefined Saprotroph\| | Yeast | NULL | Probable | As nectar-tap_saprotroph (Põlme et al. 2020); Decay sugar-rich substrates (Põlme et al. 2020) | Põlme S, et al. 2020. Fungal Diversity 105:1-16 ((https://doi.org/10.1007/s13225-020-00466-2)) |
| Otu41391 | 1 | k__Fungi(100);p__Ascomycota(100);c__Saccharomycetes(100);o__Saccharomycetales(100);f__Metschnikowiaceae(100);g__Metschnikowia(100);g__Metschnikowia_unclassified(100); | Metschnikowia | 13 | Saprotroph | \|Undefined Saprotroph\| | Yeast | NULL | Probable | As nectar-tap_saprotroph (Põlme et al. 2020); Decay sugar-rich substrates (Põlme et al. 2020) | Põlme S, et al. 2020. Fungal Diversity 105:1-16 ((https://doi.org/10.1007/s13225-020-00466-2)) |
| Otu41392 | 1 | k__Fungi(100);p__Ascomycota(100);c__Saccharomycetes(100);o__Saccharomycetales(100);f__Metschnikowiaceae(100);g__Metschnikowia(100);s__Metschnikowia_pulcherrima(100); | Metschnikowia | 13 | Saprotroph | \|Undefined Saprotroph\| | Yeast | NULL | Probable | As nectar-tap_saprotroph (Põlme et al. 2020); Decay sugar-rich substrates (Põlme et al. 2020) | Põlme S, et al. 2020. Fungal Diversity 105:1-16 ((https://doi.org/10.1007/s13225-020-00466-2)) |
| Otu41393 | 1 | k__Fungi(100);p__Ascomycota(100);c__Saccharomycetes(100);o__Saccharomycetales(100);f__Metschnikowiaceae(100);g__Metschnikowia(100);g__Metschnikowia_unclassified(100); | Metschnikowia | 13 | Saprotroph | \|Undefined Saprotroph\| | Yeast | NULL | Probable | As nectar-tap_saprotroph (Põlme et al. 2020); Decay sugar-rich substrates (Põlme et al. 2020) | Põlme S, et al. 2020. Fungal Diversity 105:1-16 ((https://doi.org/10.1007/s13225-020-00466-2)) |
| Otu41395 | 1 | k__Fungi(100);p__Ascomycota(100);c__Saccharomycetes(100);o__Saccharomycetales(100);f__Metschnikowiaceae(100);g__Metschnikowia(100);s__Metschnikowia_pulcherrima(100); | Metschnikowia | 13 | Saprotroph | \|Undefined Saprotroph\| | Yeast | NULL | Probable | As nectar-tap_saprotroph (Põlme et al. 2020); Decay sugar-rich substrates (Põlme et al. 2020) | Põlme S, et al. 2020. Fungal Diversity 105:1-16 ((https://doi.org/10.1007/s13225-020-00466-2)) |
| Otu41399 | 1 | k__Fungi(100);p__Ascomycota(100);c__Saccharomycetes(100);o__Saccharomycetales(100);f__Metschnikowiaceae(100);g__Metschnikowia(100);s__Metschnikowia_pulcherrima(100); | Metschnikowia | 13 | Saprotroph | \|Undefined Saprotroph\| | Yeast | NULL | Probable | As nectar-tap_saprotroph (Põlme et al. 2020); Decay sugar-rich substrates (Põlme et al. 2020) | Põlme S, et al. 2020. Fungal Diversity 105:1-16 ((https://doi.org/10.1007/s13225-020-00466-2)) |
| Otu41400 | 1 | k__Fungi(100);p__Ascomycota(100);c__Saccharomycetes(100);o__Saccharomycetales(100);f__Metschnikowiaceae(100);g__Metschnikowia(100);s__Metschnikowia_pulcherrima(100); | Metschnikowia | 13 | Saprotroph | \|Undefined Saprotroph\| | Yeast | NULL | Probable | As nectar-tap_saprotroph (Põlme et al. 2020); Decay sugar-rich substrates (Põlme et al. 2020) | Põlme S, et al. 2020. Fungal Diversity 105:1-16 ((https://doi.org/10.1007/s13225-020-00466-2)) |
| Otu41401 | 1 | k__Fungi(100);p__Ascomycota(100);c__Saccharomycetes(100);o__Saccharomycetales(100);f__Metschnikowiaceae(100);g__Metschnikowia(100);s__Metschnikowia_pulcherrima(100); | Metschnikowia | 13 | Saprotroph | \|Undefined Saprotroph\| | Yeast | NULL | Probable | As nectar-tap_saprotroph (Põlme et al. 2020); Decay sugar-rich substrates (Põlme et al. 2020) | Põlme S, et al. 2020. Fungal Diversity 105:1-16 ((https://doi.org/10.1007/s13225-020-00466-2)) |
| Otu41402 | 1 | k__Fungi(100);p__Ascomycota(100);c__Saccharomycetes(100);o__Saccharomycetales(100);f__Metschnikowiaceae(100);g__Metschnikowia(100);s__Metschnikowia_pulcherrima(100); | Metschnikowia | 13 | Saprotroph | \|Undefined Saprotroph\| | Yeast | NULL | Probable | As nectar-tap_saprotroph (Põlme et al. 2020); Decay sugar-rich substrates (Põlme et al. 2020) | Põlme S, et al. 2020. Fungal Diversity 105:1-16 ((https://doi.org/10.1007/s13225-020-00466-2)) |
| Otu41403 | 1 | k__Fungi(100);p__Ascomycota(100);c__Saccharomycetes(100);o__Saccharomycetales(100);f__Metschnikowiaceae(100);g__Metschnikowia(100);s__Metschnikowia_pulcherrima(100); | Metschnikowia | 13 | Saprotroph | \|Undefined Saprotroph\| | Yeast | NULL | Probable | As nectar-tap_saprotroph (Põlme et al. 2020); Decay sugar-rich substrates (Põlme et al. 2020) | Põlme S, et al. 2020. Fungal Diversity 105:1-16 ((https://doi.org/10.1007/s13225-020-00466-2)) |
| Otu41404 | 1 | k__Fungi(100);p__Ascomycota(100);c__Saccharomycetes(100);o__Saccharomycetales(100);f__Metschnikowiaceae(100);g__Metschnikowia(100);s__Metschnikowia_pulcherrima(100); | Metschnikowia | 13 | Saprotroph | \|Undefined Saprotroph\| | Yeast | NULL | Probable | As nectar-tap_saprotroph (Põlme et al. 2020); Decay sugar-rich substrates (Põlme et al. 2020) | Põlme S, et al. 2020. Fungal Diversity 105:1-16 ((https://doi.org/10.1007/s13225-020-00466-2)) |
| Otu41405 | 1 | k__Fungi(100);p__Ascomycota(100);c__Saccharomycetes(100);o__Saccharomycetales(100);f__Metschnikowiaceae(100);g__Metschnikowia(100);g__Metschnikowia_unclassified(100); | Metschnikowia | 13 | Saprotroph | \|Undefined Saprotroph\| | Yeast | NULL | Probable | As nectar-tap_saprotroph (Põlme et al. 2020); Decay sugar-rich substrates (Põlme et al. 2020) | Põlme S, et al. 2020. Fungal Diversity 105:1-16 ((https://doi.org/10.1007/s13225-020-00466-2)) |
| Otu41406 | 1 | k__Fungi(100);p__Ascomycota(100);c__Saccharomycetes(100);o__Saccharomycetales(100);f__Metschnikowiaceae(100);g__Metschnikowia(100);s__Metschnikowia_pulcherrima(100); | Metschnikowia | 13 | Saprotroph | \|Undefined Saprotroph\| | Yeast | NULL | Probable | As nectar-tap_saprotroph (Põlme et al. 2020); Decay sugar-rich substrates (Põlme et al. 2020) | Põlme S, et al. 2020. Fungal Diversity 105:1-16 ((https://doi.org/10.1007/s13225-020-00466-2)) |
| Otu41407 | 1 | k__Fungi(100);p__Ascomycota(100);c__Saccharomycetes(100);o__Saccharomycetales(100);f__Metschnikowiaceae(100);g__Metschnikowia(100);s__Metschnikowia_pulcherrima(100); | Metschnikowia | 13 | Saprotroph | \|Undefined Saprotroph\| | Yeast | NULL | Probable | As nectar-tap_saprotroph (Põlme et al. 2020); Decay sugar-rich substrates (Põlme et al. 2020) | Põlme S, et al. 2020. Fungal Diversity 105:1-16 ((https://doi.org/10.1007/s13225-020-00466-2)) |
| Otu41408 | 1 | k__Fungi(100);p__Ascomycota(100);c__Saccharomycetes(100);o__Saccharomycetales(100);f__Metschnikowiaceae(100);g__Metschnikowia(100);s__Metschnikowia_pulcherrima(100); | Metschnikowia | 13 | Saprotroph | \|Undefined Saprotroph\| | Yeast | NULL | Probable | As nectar-tap_saprotroph (Põlme et al. 2020); Decay sugar-rich substrates (Põlme et al. 2020) | Põlme S, et al. 2020. Fungal Diversity 105:1-16 ((https://doi.org/10.1007/s13225-020-00466-2)) |
| Otu41409 | 1 | k__Fungi(100);p__Ascomycota(100);c__Saccharomycetes(100);o__Saccharomycetales(100);f__Metschnikowiaceae(100);g__Metschnikowia(100);s__Metschnikowia_pulcherrima(100); | Metschnikowia | 13 | Saprotroph | \|Undefined Saprotroph\| | Yeast | NULL | Probable | As nectar-tap_saprotroph (Põlme et al. 2020); Decay sugar-rich substrates (Põlme et al. 2020) | Põlme S, et al. 2020. Fungal Diversity 105:1-16 ((https://doi.org/10.1007/s13225-020-00466-2)) |
| Otu41410 | 1 | k__Fungi(100);p__Ascomycota(100);c__Saccharomycetes(100);o__Saccharomycetales(100);f__Metschnikowiaceae(100);g__Metschnikowia(100);s__Metschnikowia_pulcherrima(100); | Metschnikowia | 13 | Saprotroph | \|Undefined Saprotroph\| | Yeast | NULL | Probable | As nectar-tap_saprotroph (Põlme et al. 2020); Decay sugar-rich substrates (Põlme et al. 2020) | Põlme S, et al. 2020. Fungal Diversity 105:1-16 ((https://doi.org/10.1007/s13225-020-00466-2)) |
| Otu41411 | 1 | k__Fungi(100);p__Ascomycota(100);c__Saccharomycetes(100);o__Saccharomycetales(100);f__Metschnikowiaceae(100);g__Metschnikowia(100);s__Metschnikowia_pulcherrima(100); | Metschnikowia | 13 | Saprotroph | \|Undefined Saprotroph\| | Yeast | NULL | Probable | As nectar-tap_saprotroph (Põlme et al. 2020); Decay sugar-rich substrates (Põlme et al. 2020) | Põlme S, et al. 2020. Fungal Diversity 105:1-16 ((https://doi.org/10.1007/s13225-020-00466-2)) |
| Otu41412 | 1 | k__Fungi(100);p__Ascomycota(100);c__Saccharomycetes(100);o__Saccharomycetales(100);f__Metschnikowiaceae(100);g__Metschnikowia(100);s__Metschnikowia_pulcherrima(100); | Metschnikowia | 13 | Saprotroph | \|Undefined Saprotroph\| | Yeast | NULL | Probable | As nectar-tap_saprotroph (Põlme et al. 2020); Decay sugar-rich substrates (Põlme et al. 2020) | Põlme S, et al. 2020. Fungal Diversity 105:1-16 ((https://doi.org/10.1007/s13225-020-00466-2)) |
| Otu41413 | 1 | k__Fungi(100);p__Ascomycota(100);c__Saccharomycetes(100);o__Saccharomycetales(100);f__Metschnikowiaceae(100);g__Metschnikowia(100);s__Metschnikowia_pulcherrima(100); | Metschnikowia | 13 | Saprotroph | \|Undefined Saprotroph\| | Yeast | NULL | Probable | As nectar-tap_saprotroph (Põlme et al. 2020); Decay sugar-rich substrates (Põlme et al. 2020) | Põlme S, et al. 2020. Fungal Diversity 105:1-16 ((https://doi.org/10.1007/s13225-020-00466-2)) |
| Otu41414 | 1 | k__Fungi(100);p__Ascomycota(100);c__Saccharomycetes(100);o__Saccharomycetales(100);f__Metschnikowiaceae(100);g__Metschnikowia(100);s__Metschnikowia_pulcherrima(100); | Metschnikowia | 13 | Saprotroph | \|Undefined Saprotroph\| | Yeast | NULL | Probable | As nectar-tap_saprotroph (Põlme et al. 2020); Decay sugar-rich substrates (Põlme et al. 2020) | Põlme S, et al. 2020. Fungal Diversity 105:1-16 ((https://doi.org/10.1007/s13225-020-00466-2)) |
| Otu41415 | 1 | k__Fungi(100);p__Ascomycota(100);c__Saccharomycetes(100);o__Saccharomycetales(100);f__Metschnikowiaceae(100);g__Metschnikowia(100);s__Metschnikowia_pulcherrima(100); | Metschnikowia | 13 | Saprotroph | \|Undefined Saprotroph\| | Yeast | NULL | Probable | As nectar-tap_saprotroph (Põlme et al. 2020); Decay sugar-rich substrates (Põlme et al. 2020) | Põlme S, et al. 2020. Fungal Diversity 105:1-16 ((https://doi.org/10.1007/s13225-020-00466-2)) |
| Otu41416 | 1 | k__Fungi(100);p__Ascomycota(100);c__Saccharomycetes(100);o__Saccharomycetales(100);f__Metschnikowiaceae(100);g__Metschnikowia(100);s__Metschnikowia_pulcherrima(100); | Metschnikowia | 13 | Saprotroph | \|Undefined Saprotroph\| | Yeast | NULL | Probable | As nectar-tap_saprotroph (Põlme et al. 2020); Decay sugar-rich substrates (Põlme et al. 2020) | Põlme S, et al. 2020. Fungal Diversity 105:1-16 ((https://doi.org/10.1007/s13225-020-00466-2)) |
| Otu41417 | 1 | k__Fungi(100);p__Ascomycota(100);c__Saccharomycetes(100);o__Saccharomycetales(100);f__Metschnikowiaceae(100);g__Metschnikowia(100);s__Metschnikowia_pulcherrima(100); | Metschnikowia | 13 | Saprotroph | \|Undefined Saprotroph\| | Yeast | NULL | Probable | As nectar-tap_saprotroph (Põlme et al. 2020); Decay sugar-rich substrates (Põlme et al. 2020) | Põlme S, et al. 2020. Fungal Diversity 105:1-16 ((https://doi.org/10.1007/s13225-020-00466-2)) |
| Otu41418 | 1 | k__Fungi(100);p__Ascomycota(100);c__Saccharomycetes(100);o__Saccharomycetales(100);f__Metschnikowiaceae(100);g__Metschnikowia(100);g__Metschnikowia_unclassified(100); | Metschnikowia | 13 | Saprotroph | \|Undefined Saprotroph\| | Yeast | NULL | Probable | As nectar-tap_saprotroph (Põlme et al. 2020); Decay sugar-rich substrates (Põlme et al. 2020) | Põlme S, et al. 2020. Fungal Diversity 105:1-16 ((https://doi.org/10.1007/s13225-020-00466-2)) |
| Otu41419 | 1 | k__Fungi(100);p__Ascomycota(100);c__Saccharomycetes(100);o__Saccharomycetales(100);f__Metschnikowiaceae(100);g__Metschnikowia(100);s__Metschnikowia_pulcherrima(100); | Metschnikowia | 13 | Saprotroph | \|Undefined Saprotroph\| | Yeast | NULL | Probable | As nectar-tap_saprotroph (Põlme et al. 2020); Decay sugar-rich substrates (Põlme et al. 2020) | Põlme S, et al. 2020. Fungal Diversity 105:1-16 ((https://doi.org/10.1007/s13225-020-00466-2)) |
| Otu41420 | 1 | k__Fungi(100);p__Ascomycota(100);c__Saccharomycetes(100);o__Saccharomycetales(100);f__Metschnikowiaceae(100);g__Metschnikowia(100);s__Metschnikowia_pulcherrima(100); | Metschnikowia | 13 | Saprotroph | \|Undefined Saprotroph\| | Yeast | NULL | Probable | As nectar-tap_saprotroph (Põlme et al. 2020); Decay sugar-rich substrates (Põlme et al. 2020) | Põlme S, et al. 2020. Fungal Diversity 105:1-16 ((https://doi.org/10.1007/s13225-020-00466-2)) |
| Otu41421 | 1 | k__Fungi(100);p__Ascomycota(100);c__Saccharomycetes(100);o__Saccharomycetales(100);f__Metschnikowiaceae(100);g__Metschnikowia(100);s__Metschnikowia_pulcherrima(100); | Metschnikowia | 13 | Saprotroph | \|Undefined Saprotroph\| | Yeast | NULL | Probable | As nectar-tap_saprotroph (Põlme et al. 2020); Decay sugar-rich substrates (Põlme et al. 2020) | Põlme S, et al. 2020. Fungal Diversity 105:1-16 ((https://doi.org/10.1007/s13225-020-00466-2)) |
| Otu41422 | 1 | k__Fungi(100);p__Ascomycota(100);c__Saccharomycetes(100);o__Saccharomycetales(100);f__Metschnikowiaceae(100);g__Metschnikowia(100);s__Metschnikowia_pulcherrima(100); | Metschnikowia | 13 | Saprotroph | \|Undefined Saprotroph\| | Yeast | NULL | Probable | As nectar-tap_saprotroph (Põlme et al. 2020); Decay sugar-rich substrates (Põlme et al. 2020) | Põlme S, et al. 2020. Fungal Diversity 105:1-16 ((https://doi.org/10.1007/s13225-020-00466-2)) |
| Otu41427 | 1 | k__Fungi(100);p__Ascomycota(100);c__Saccharomycetes(100);o__Saccharomycetales(100);f__Metschnikowiaceae(100);g__Metschnikowia(100);s__Metschnikowia_pulcherrima(100); | Metschnikowia | 13 | Saprotroph | \|Undefined Saprotroph\| | Yeast | NULL | Probable | As nectar-tap_saprotroph (Põlme et al. 2020); Decay sugar-rich substrates (Põlme et al. 2020) | Põlme S, et al. 2020. Fungal Diversity 105:1-16 ((https://doi.org/10.1007/s13225-020-00466-2)) |
| Otu41430 | 1 | k__Fungi(100);p__Ascomycota(100);c__Saccharomycetes(100);o__Saccharomycetales(100);f__Metschnikowiaceae(100);g__Metschnikowia(100);g__Metschnikowia_unclassified(100); | Metschnikowia | 13 | Saprotroph | \|Undefined Saprotroph\| | Yeast | NULL | Probable | As nectar-tap_saprotroph (Põlme et al. 2020); Decay sugar-rich substrates (Põlme et al. 2020) | Põlme S, et al. 2020. Fungal Diversity 105:1-16 ((https://doi.org/10.1007/s13225-020-00466-2)) |
| Otu41431 | 1 | k__Fungi(100);p__Ascomycota(100);c__Saccharomycetes(100);o__Saccharomycetales(100);f__Metschnikowiaceae(100);g__Metschnikowia(100);g__Metschnikowia_unclassified(100); | Metschnikowia | 13 | Saprotroph | \|Undefined Saprotroph\| | Yeast | NULL | Probable | As nectar-tap_saprotroph (Põlme et al. 2020); Decay sugar-rich substrates (Põlme et al. 2020) | Põlme S, et al. 2020. Fungal Diversity 105:1-16 ((https://doi.org/10.1007/s13225-020-00466-2)) |
| Otu41432 | 1 | k__Fungi(100);p__Ascomycota(100);c__Saccharomycetes(100);o__Saccharomycetales(100);f__Metschnikowiaceae(100);g__Metschnikowia(100);s__Metschnikowia_pulcherrima(100); | Metschnikowia | 13 | Saprotroph | \|Undefined Saprotroph\| | Yeast | NULL | Probable | As nectar-tap_saprotroph (Põlme et al. 2020); Decay sugar-rich substrates (Põlme et al. 2020) | Põlme S, et al. 2020. Fungal Diversity 105:1-16 ((https://doi.org/10.1007/s13225-020-00466-2)) |
| Otu41433 | 1 | k__Fungi(100);p__Ascomycota(100);c__Saccharomycetes(100);o__Saccharomycetales(100);f__Metschnikowiaceae(100);g__Metschnikowia(100);s__Metschnikowia_pulcherrima(100); | Metschnikowia | 13 | Saprotroph | \|Undefined Saprotroph\| | Yeast | NULL | Probable | As nectar-tap_saprotroph (Põlme et al. 2020); Decay sugar-rich substrates (Põlme et al. 2020) | Põlme S, et al. 2020. Fungal Diversity 105:1-16 ((https://doi.org/10.1007/s13225-020-00466-2)) |
| Otu41434 | 1 | k__Fungi(100);p__Ascomycota(100);c__Saccharomycetes(100);o__Saccharomycetales(100);f__Metschnikowiaceae(100);g__Metschnikowia(100);s__Metschnikowia_pulcherrima(100); | Metschnikowia | 13 | Saprotroph | \|Undefined Saprotroph\| | Yeast | NULL | Probable | As nectar-tap_saprotroph (Põlme et al. 2020); Decay sugar-rich substrates (Põlme et al. 2020) | Põlme S, et al. 2020. Fungal Diversity 105:1-16 ((https://doi.org/10.1007/s13225-020-00466-2)) |
| Otu41435 | 1 | k__Fungi(100);p__Ascomycota(100);c__Saccharomycetes(100);o__Saccharomycetales(100);f__Metschnikowiaceae(100);g__Metschnikowia(100);s__Metschnikowia_pulcherrima(100); | Metschnikowia | 13 | Saprotroph | \|Undefined Saprotroph\| | Yeast | NULL | Probable | As nectar-tap_saprotroph (Põlme et al. 2020); Decay sugar-rich substrates (Põlme et al. 2020) | Põlme S, et al. 2020. Fungal Diversity 105:1-16 ((https://doi.org/10.1007/s13225-020-00466-2)) |
| Otu41438 | 1 | k__Fungi(100);p__Ascomycota(100);c__Saccharomycetes(100);o__Saccharomycetales(100);f__Metschnikowiaceae(100);g__Metschnikowia(100);s__Metschnikowia_pulcherrima(100); | Metschnikowia | 13 | Saprotroph | \|Undefined Saprotroph\| | Yeast | NULL | Probable | As nectar-tap_saprotroph (Põlme et al. 2020); Decay sugar-rich substrates (Põlme et al. 2020) | Põlme S, et al. 2020. Fungal Diversity 105:1-16 ((https://doi.org/10.1007/s13225-020-00466-2)) |
| Otu41439 | 1 | k__Fungi(100);p__Ascomycota(100);c__Saccharomycetes(100);o__Saccharomycetales(100);f__Metschnikowiaceae(100);g__Metschnikowia(100);s__Metschnikowia_pulcherrima(100); | Metschnikowia | 13 | Saprotroph | \|Undefined Saprotroph\| | Yeast | NULL | Probable | As nectar-tap_saprotroph (Põlme et al. 2020); Decay sugar-rich substrates (Põlme et al. 2020) | Põlme S, et al. 2020. Fungal Diversity 105:1-16 ((https://doi.org/10.1007/s13225-020-00466-2)) |
| Otu41441 | 1 | k__Fungi(100);p__Ascomycota(100);c__Saccharomycetes(100);o__Saccharomycetales(100);f__Metschnikowiaceae(100);g__Metschnikowia(100);g__Metschnikowia_unclassified(100); | Metschnikowia | 13 | Saprotroph | \|Undefined Saprotroph\| | Yeast | NULL | Probable | As nectar-tap_saprotroph (Põlme et al. 2020); Decay sugar-rich substrates (Põlme et al. 2020) | Põlme S, et al. 2020. Fungal Diversity 105:1-16 ((https://doi.org/10.1007/s13225-020-00466-2)) |
| Otu41442 | 1 | k__Fungi(100);p__Ascomycota(100);c__Saccharomycetes(100);o__Saccharomycetales(100);f__Metschnikowiaceae(100);g__Metschnikowia(100);s__Metschnikowia_pulcherrima(100); | Metschnikowia | 13 | Saprotroph | \|Undefined Saprotroph\| | Yeast | NULL | Probable | As nectar-tap_saprotroph (Põlme et al. 2020); Decay sugar-rich substrates (Põlme et al. 2020) | Põlme S, et al. 2020. Fungal Diversity 105:1-16 ((https://doi.org/10.1007/s13225-020-00466-2)) |
| Otu41443 | 1 | k__Fungi(100);p__Ascomycota(100);c__Saccharomycetes(100);o__Saccharomycetales(100);f__Metschnikowiaceae(100);g__Metschnikowia(100);s__Metschnikowia_pulcherrima(100); | Metschnikowia | 13 | Saprotroph | \|Undefined Saprotroph\| | Yeast | NULL | Probable | As nectar-tap_saprotroph (Põlme et al. 2020); Decay sugar-rich substrates (Põlme et al. 2020) | Põlme S, et al. 2020. Fungal Diversity 105:1-16 ((https://doi.org/10.1007/s13225-020-00466-2)) |
| Otu41444 | 1 | k__Fungi(100);p__Ascomycota(100);c__Saccharomycetes(100);o__Saccharomycetales(100);f__Metschnikowiaceae(100);g__Metschnikowia(100);s__Metschnikowia_pulcherrima(100); | Metschnikowia | 13 | Saprotroph | \|Undefined Saprotroph\| | Yeast | NULL | Probable | As nectar-tap_saprotroph (Põlme et al. 2020); Decay sugar-rich substrates (Põlme et al. 2020) | Põlme S, et al. 2020. Fungal Diversity 105:1-16 ((https://doi.org/10.1007/s13225-020-00466-2)) |
| Otu41447 | 1 | k__Fungi(100);p__Ascomycota(100);c__Saccharomycetes(100);o__Saccharomycetales(100);f__Metschnikowiaceae(100);g__Metschnikowia(100);s__Metschnikowia_pulcherrima(100); | Metschnikowia | 13 | Saprotroph | \|Undefined Saprotroph\| | Yeast | NULL | Probable | As nectar-tap_saprotroph (Põlme et al. 2020); Decay sugar-rich substrates (Põlme et al. 2020) | Põlme S, et al. 2020. Fungal Diversity 105:1-16 ((https://doi.org/10.1007/s13225-020-00466-2)) |
| Otu41448 | 1 | k__Fungi(100);p__Ascomycota(100);c__Saccharomycetes(100);o__Saccharomycetales(100);f__Metschnikowiaceae(100);g__Metschnikowia(100);s__Metschnikowia_pulcherrima(100); | Metschnikowia | 13 | Saprotroph | \|Undefined Saprotroph\| | Yeast | NULL | Probable | As nectar-tap_saprotroph (Põlme et al. 2020); Decay sugar-rich substrates (Põlme et al. 2020) | Põlme S, et al. 2020. Fungal Diversity 105:1-16 ((https://doi.org/10.1007/s13225-020-00466-2)) |
| Otu41449 | 1 | k__Fungi(100);p__Ascomycota(100);c__Saccharomycetes(100);o__Saccharomycetales(100);f__Metschnikowiaceae(100);g__Metschnikowia(100);g__Metschnikowia_unclassified(100); | Metschnikowia | 13 | Saprotroph | \|Undefined Saprotroph\| | Yeast | NULL | Probable | As nectar-tap_saprotroph (Põlme et al. 2020); Decay sugar-rich substrates (Põlme et al. 2020) | Põlme S, et al. 2020. Fungal Diversity 105:1-16 ((https://doi.org/10.1007/s13225-020-00466-2)) |
| Otu41450 | 1 | k__Fungi(100);p__Ascomycota(100);c__Saccharomycetes(100);o__Saccharomycetales(100);f__Metschnikowiaceae(100);g__Metschnikowia(100);g__Metschnikowia_unclassified(100); | Metschnikowia | 13 | Saprotroph | \|Undefined Saprotroph\| | Yeast | NULL | Probable | As nectar-tap_saprotroph (Põlme et al. 2020); Decay sugar-rich substrates (Põlme et al. 2020) | Põlme S, et al. 2020. Fungal Diversity 105:1-16 ((https://doi.org/10.1007/s13225-020-00466-2)) |
| Otu41451 | 1 | k__Fungi(100);p__Ascomycota(100);c__Saccharomycetes(100);o__Saccharomycetales(100);f__Metschnikowiaceae(100);g__Metschnikowia(100);s__Metschnikowia_pulcherrima(100); | Metschnikowia | 13 | Saprotroph | \|Undefined Saprotroph\| | Yeast | NULL | Probable | As nectar-tap_saprotroph (Põlme et al. 2020); Decay sugar-rich substrates (Põlme et al. 2020) | Põlme S, et al. 2020. Fungal Diversity 105:1-16 ((https://doi.org/10.1007/s13225-020-00466-2)) |
| Otu41452 | 1 | k__Fungi(100);p__Ascomycota(100);c__Saccharomycetes(100);o__Saccharomycetales(100);f__Metschnikowiaceae(100);g__Metschnikowia(100);s__Metschnikowia_pulcherrima(100); | Metschnikowia | 13 | Saprotroph | \|Undefined Saprotroph\| | Yeast | NULL | Probable | As nectar-tap_saprotroph (Põlme et al. 2020); Decay sugar-rich substrates (Põlme et al. 2020) | Põlme S, et al. 2020. Fungal Diversity 105:1-16 ((https://doi.org/10.1007/s13225-020-00466-2)) |
| Otu41456 | 1 | k__Fungi(100);p__Ascomycota(100);c__Saccharomycetes(100);o__Saccharomycetales(100);f__Metschnikowiaceae(100);g__Metschnikowia(100);g__Metschnikowia_unclassified(100); | Metschnikowia | 13 | Saprotroph | \|Undefined Saprotroph\| | Yeast | NULL | Probable | As nectar-tap_saprotroph (Põlme et al. 2020); Decay sugar-rich substrates (Põlme et al. 2020) | Põlme S, et al. 2020. Fungal Diversity 105:1-16 ((https://doi.org/10.1007/s13225-020-00466-2)) |
| Otu41457 | 1 | k__Fungi(100);p__Ascomycota(100);c__Saccharomycetes(100);o__Saccharomycetales(100);f__Metschnikowiaceae(100);g__Metschnikowia(100);s__Metschnikowia_pulcherrima(100); | Metschnikowia | 13 | Saprotroph | \|Undefined Saprotroph\| | Yeast | NULL | Probable | As nectar-tap_saprotroph (Põlme et al. 2020); Decay sugar-rich substrates (Põlme et al. 2020) | Põlme S, et al. 2020. Fungal Diversity 105:1-16 ((https://doi.org/10.1007/s13225-020-00466-2)) |
| Otu41458 | 1 | k__Fungi(100);p__Ascomycota(100);c__Saccharomycetes(100);o__Saccharomycetales(100);f__Metschnikowiaceae(100);g__Metschnikowia(100);s__Metschnikowia_pulcherrima(100); | Metschnikowia | 13 | Saprotroph | \|Undefined Saprotroph\| | Yeast | NULL | Probable | As nectar-tap_saprotroph (Põlme et al. 2020); Decay sugar-rich substrates (Põlme et al. 2020) | Põlme S, et al. 2020. Fungal Diversity 105:1-16 ((https://doi.org/10.1007/s13225-020-00466-2)) |
| Otu41459 | 1 | k__Fungi(100);p__Ascomycota(100);c__Saccharomycetes(100);o__Saccharomycetales(100);f__Metschnikowiaceae(100);g__Metschnikowia(100);s__Metschnikowia_pulcherrima(100); | Metschnikowia | 13 | Saprotroph | \|Undefined Saprotroph\| | Yeast | NULL | Probable | As nectar-tap_saprotroph (Põlme et al. 2020); Decay sugar-rich substrates (Põlme et al. 2020) | Põlme S, et al. 2020. Fungal Diversity 105:1-16 ((https://doi.org/10.1007/s13225-020-00466-2)) |
| Otu41460 | 1 | k__Fungi(100);p__Ascomycota(100);c__Saccharomycetes(100);o__Saccharomycetales(100);f__Metschnikowiaceae(100);g__Metschnikowia(100);g__Metschnikowia_unclassified(100); | Metschnikowia | 13 | Saprotroph | \|Undefined Saprotroph\| | Yeast | NULL | Probable | As nectar-tap_saprotroph (Põlme et al. 2020); Decay sugar-rich substrates (Põlme et al. 2020) | Põlme S, et al. 2020. Fungal Diversity 105:1-16 ((https://doi.org/10.1007/s13225-020-00466-2)) |
| Otu41461 | 1 | k__Fungi(100);p__Ascomycota(100);c__Saccharomycetes(100);o__Saccharomycetales(100);f__Metschnikowiaceae(100);g__Metschnikowia(100);s__Metschnikowia_pulcherrima(100); | Metschnikowia | 13 | Saprotroph | \|Undefined Saprotroph\| | Yeast | NULL | Probable | As nectar-tap_saprotroph (Põlme et al. 2020); Decay sugar-rich substrates (Põlme et al. 2020) | Põlme S, et al. 2020. Fungal Diversity 105:1-16 ((https://doi.org/10.1007/s13225-020-00466-2)) |
| Otu41462 | 1 | k__Fungi(100);p__Ascomycota(100);c__Saccharomycetes(100);o__Saccharomycetales(100);f__Metschnikowiaceae(100);g__Metschnikowia(100);s__Metschnikowia_pulcherrima(100); | Metschnikowia | 13 | Saprotroph | \|Undefined Saprotroph\| | Yeast | NULL | Probable | As nectar-tap_saprotroph (Põlme et al. 2020); Decay sugar-rich substrates (Põlme et al. 2020) | Põlme S, et al. 2020. Fungal Diversity 105:1-16 ((https://doi.org/10.1007/s13225-020-00466-2)) |
| Otu41463 | 1 | k__Fungi(100);p__Ascomycota(100);c__Saccharomycetes(100);o__Saccharomycetales(100);f__Metschnikowiaceae(100);g__Metschnikowia(100);s__Metschnikowia_pulcherrima(100); | Metschnikowia | 13 | Saprotroph | \|Undefined Saprotroph\| | Yeast | NULL | Probable | As nectar-tap_saprotroph (Põlme et al. 2020); Decay sugar-rich substrates (Põlme et al. 2020) | Põlme S, et al. 2020. Fungal Diversity 105:1-16 ((https://doi.org/10.1007/s13225-020-00466-2)) |
| Otu41464 | 1 | k__Fungi(100);p__Ascomycota(100);c__Saccharomycetes(100);o__Saccharomycetales(100);f__Metschnikowiaceae(100);g__Metschnikowia(100);s__Metschnikowia_pulcherrima(100); | Metschnikowia | 13 | Saprotroph | \|Undefined Saprotroph\| | Yeast | NULL | Probable | As nectar-tap_saprotroph (Põlme et al. 2020); Decay sugar-rich substrates (Põlme et al. 2020) | Põlme S, et al. 2020. Fungal Diversity 105:1-16 ((https://doi.org/10.1007/s13225-020-00466-2)) |
| Otu41465 | 1 | k__Fungi(100);p__Ascomycota(100);c__Saccharomycetes(100);o__Saccharomycetales(100);f__Metschnikowiaceae(100);g__Metschnikowia(100);s__Metschnikowia_pulcherrima(100); | Metschnikowia | 13 | Saprotroph | \|Undefined Saprotroph\| | Yeast | NULL | Probable | As nectar-tap_saprotroph (Põlme et al. 2020); Decay sugar-rich substrates (Põlme et al. 2020) | Põlme S, et al. 2020. Fungal Diversity 105:1-16 ((https://doi.org/10.1007/s13225-020-00466-2)) |
| Otu41466 | 1 | k__Fungi(100);p__Ascomycota(100);c__Saccharomycetes(100);o__Saccharomycetales(100);f__Metschnikowiaceae(100);g__Metschnikowia(100);s__Metschnikowia_pulcherrima(100); | Metschnikowia | 13 | Saprotroph | \|Undefined Saprotroph\| | Yeast | NULL | Probable | As nectar-tap_saprotroph (Põlme et al. 2020); Decay sugar-rich substrates (Põlme et al. 2020) | Põlme S, et al. 2020. Fungal Diversity 105:1-16 ((https://doi.org/10.1007/s13225-020-00466-2)) |
| Otu41467 | 1 | k__Fungi(100);p__Ascomycota(100);c__Saccharomycetes(100);o__Saccharomycetales(100);f__Metschnikowiaceae(100);g__Metschnikowia(100);g__Metschnikowia_unclassified(100); | Metschnikowia | 13 | Saprotroph | \|Undefined Saprotroph\| | Yeast | NULL | Probable | As nectar-tap_saprotroph (Põlme et al. 2020); Decay sugar-rich substrates (Põlme et al. 2020) | Põlme S, et al. 2020. Fungal Diversity 105:1-16 ((https://doi.org/10.1007/s13225-020-00466-2)) |
| Otu41468 | 1 | k__Fungi(100);p__Ascomycota(100);c__Saccharomycetes(100);o__Saccharomycetales(100);f__Metschnikowiaceae(100);g__Metschnikowia(100);s__Metschnikowia_pulcherrima(100); | Metschnikowia | 13 | Saprotroph | \|Undefined Saprotroph\| | Yeast | NULL | Probable | As nectar-tap_saprotroph (Põlme et al. 2020); Decay sugar-rich substrates (Põlme et al. 2020) | Põlme S, et al. 2020. Fungal Diversity 105:1-16 ((https://doi.org/10.1007/s13225-020-00466-2)) |
| Otu41469 | 1 | k__Fungi(100);p__Ascomycota(100);c__Saccharomycetes(100);o__Saccharomycetales(100);f__Metschnikowiaceae(100);g__Metschnikowia(100);s__Metschnikowia_pulcherrima(100); | Metschnikowia | 13 | Saprotroph | \|Undefined Saprotroph\| | Yeast | NULL | Probable | As nectar-tap_saprotroph (Põlme et al. 2020); Decay sugar-rich substrates (Põlme et al. 2020) | Põlme S, et al. 2020. Fungal Diversity 105:1-16 ((https://doi.org/10.1007/s13225-020-00466-2)) |
| Otu41471 | 1 | k__Fungi(100);p__Ascomycota(100);c__Saccharomycetes(100);o__Saccharomycetales(100);f__Metschnikowiaceae(100);g__Metschnikowia(100);s__Metschnikowia_pulcherrima(100); | Metschnikowia | 13 | Saprotroph | \|Undefined Saprotroph\| | Yeast | NULL | Probable | As nectar-tap_saprotroph (Põlme et al. 2020); Decay sugar-rich substrates (Põlme et al. 2020) | Põlme S, et al. 2020. Fungal Diversity 105:1-16 ((https://doi.org/10.1007/s13225-020-00466-2)) |
| Otu41472 | 1 | k__Fungi(100);p__Ascomycota(100);c__Saccharomycetes(100);o__Saccharomycetales(100);f__Metschnikowiaceae(100);g__Metschnikowia(100);g__Metschnikowia_unclassified(100); | Metschnikowia | 13 | Saprotroph | \|Undefined Saprotroph\| | Yeast | NULL | Probable | As nectar-tap_saprotroph (Põlme et al. 2020); Decay sugar-rich substrates (Põlme et al. 2020) | Põlme S, et al. 2020. Fungal Diversity 105:1-16 ((https://doi.org/10.1007/s13225-020-00466-2)) |
| Otu41473 | 1 | k__Fungi(100);p__Ascomycota(100);c__Saccharomycetes(100);o__Saccharomycetales(100);f__Metschnikowiaceae(100);g__Metschnikowia(100);g__Metschnikowia_unclassified(100); | Metschnikowia | 13 | Saprotroph | \|Undefined Saprotroph\| | Yeast | NULL | Probable | As nectar-tap_saprotroph (Põlme et al. 2020); Decay sugar-rich substrates (Põlme et al. 2020) | Põlme S, et al. 2020. Fungal Diversity 105:1-16 ((https://doi.org/10.1007/s13225-020-00466-2)) |
| Otu41474 | 1 | k__Fungi(100);p__Ascomycota(100);c__Saccharomycetes(100);o__Saccharomycetales(100);f__Metschnikowiaceae(100);g__Metschnikowia(100);s__Metschnikowia_pulcherrima(100); | Metschnikowia | 13 | Saprotroph | \|Undefined Saprotroph\| | Yeast | NULL | Probable | As nectar-tap_saprotroph (Põlme et al. 2020); Decay sugar-rich substrates (Põlme et al. 2020) | Põlme S, et al. 2020. Fungal Diversity 105:1-16 ((https://doi.org/10.1007/s13225-020-00466-2)) |
| Otu41475 | 1 | k__Fungi(100);p__Ascomycota(100);c__Saccharomycetes(100);o__Saccharomycetales(100);f__Metschnikowiaceae(100);g__Metschnikowia(100);s__Metschnikowia_pulcherrima(100); | Metschnikowia | 13 | Saprotroph | \|Undefined Saprotroph\| | Yeast | NULL | Probable | As nectar-tap_saprotroph (Põlme et al. 2020); Decay sugar-rich substrates (Põlme et al. 2020) | Põlme S, et al. 2020. Fungal Diversity 105:1-16 ((https://doi.org/10.1007/s13225-020-00466-2)) |
| Otu41476 | 1 | k__Fungi(100);p__Ascomycota(100);c__Saccharomycetes(100);o__Saccharomycetales(100);f__Metschnikowiaceae(100);g__Metschnikowia(100);g__Metschnikowia_unclassified(100); | Metschnikowia | 13 | Saprotroph | \|Undefined Saprotroph\| | Yeast | NULL | Probable | As nectar-tap_saprotroph (Põlme et al. 2020); Decay sugar-rich substrates (Põlme et al. 2020) | Põlme S, et al. 2020. Fungal Diversity 105:1-16 ((https://doi.org/10.1007/s13225-020-00466-2)) |
| Otu41477 | 1 | k__Fungi(100);p__Ascomycota(100);c__Saccharomycetes(100);o__Saccharomycetales(100);f__Metschnikowiaceae(100);g__Metschnikowia(100);s__Metschnikowia_pulcherrima(100); | Metschnikowia | 13 | Saprotroph | \|Undefined Saprotroph\| | Yeast | NULL | Probable | As nectar-tap_saprotroph (Põlme et al. 2020); Decay sugar-rich substrates (Põlme et al. 2020) | Põlme S, et al. 2020. Fungal Diversity 105:1-16 ((https://doi.org/10.1007/s13225-020-00466-2)) |
| Otu41478 | 1 | k__Fungi(100);p__Ascomycota(100);c__Saccharomycetes(100);o__Saccharomycetales(100);f__Metschnikowiaceae(100);g__Metschnikowia(100);s__Metschnikowia_pulcherrima(100); | Metschnikowia | 13 | Saprotroph | \|Undefined Saprotroph\| | Yeast | NULL | Probable | As nectar-tap_saprotroph (Põlme et al. 2020); Decay sugar-rich substrates (Põlme et al. 2020) | Põlme S, et al. 2020. Fungal Diversity 105:1-16 ((https://doi.org/10.1007/s13225-020-00466-2)) |
| Otu41479 | 1 | k__Fungi(100);p__Ascomycota(100);c__Saccharomycetes(100);o__Saccharomycetales(100);f__Metschnikowiaceae(100);g__Metschnikowia(100);s__Metschnikowia_pulcherrima(100); | Metschnikowia | 13 | Saprotroph | \|Undefined Saprotroph\| | Yeast | NULL | Probable | As nectar-tap_saprotroph (Põlme et al. 2020); Decay sugar-rich substrates (Põlme et al. 2020) | Põlme S, et al. 2020. Fungal Diversity 105:1-16 ((https://doi.org/10.1007/s13225-020-00466-2)) |

| | | | | | | | | | | | |
|---|---|---|---|---|---|---|---|---|---|---|---|
| Otu41738 | 1 | k__Fungi(100);p__Ascomycota(100);c__Saccharomycetes(100);o__Saccharomycetales(100);f__Metschnikowiaceae(100);g__Metschnikowia(100);s__Metschnikowia_pulcherrima(100); | Metschnikowia | 13 | Saprotroph | \|Undefined Saprotroph\| | Yeast | NULL | Probable | As nectar-tap_saprotroph (Põlme et al. 2020); Decay sugar-rich substrates (Põlme et al. 2020) | Põlme S, et al. 2020. Fungal Diversity 105:1-16 ((https://doi.org/10.1007/s13225-020-00466-2)) |
| Otu41739 | 1 | k__Fungi(100);p__Ascomycota(100);c__Saccharomycetes(100);o__Saccharomycetales(100);f__Metschnikowiaceae(100);g__Metschnikowia(100);s__Metschnikowia_pulcherrima(100); | Metschnikowia | 13 | Saprotroph | \|Undefined Saprotroph\| | Yeast | NULL | Probable | As nectar-tap_saprotroph (Põlme et al. 2020); Decay sugar-rich substrates (Põlme et al. 2020) | Põlme S, et al. 2020. Fungal Diversity 105:1-16 ((https://doi.org/10.1007/s13225-020-00466-2)) |
| Otu41741 | 1 | k__Fungi(100);p__Ascomycota(100);c__Saccharomycetes(100);o__Saccharomycetales(100);f__Metschnikowiaceae(100);g__Metschnikowia(100);s__Metschnikowia_pulcherrima(100); | Metschnikowia | 13 | Saprotroph | \|Undefined Saprotroph\| | Yeast | NULL | Probable | As nectar-tap_saprotroph (Põlme et al. 2020); Decay sugar-rich substrates (Põlme et al. 2020) | Põlme S, et al. 2020. Fungal Diversity 105:1-16 ((https://doi.org/10.1007/s13225-020-00466-2)) |
| Otu41742 | 1 | k__Fungi(100);p__Ascomycota(100);c__Saccharomycetes(100);o__Saccharomycetales(100);f__Metschnikowiaceae(100);g__Metschnikowia(100);s__Metschnikowia_pulcherrima(100); | Metschnikowia | 13 | Saprotroph | \|Undefined Saprotroph\| | Yeast | NULL | Probable | As nectar-tap_saprotroph (Põlme et al. 2020); Decay sugar-rich substrates (Põlme et al. 2020) | Põlme S, et al. 2020. Fungal Diversity 105:1-16 ((https://doi.org/10.1007/s13225-020-00466-2)) |
| Otu41743 | 1 | k__Fungi(100);p__Ascomycota(100);c__Saccharomycetes(100);o__Saccharomycetales(100);f__Metschnikowiaceae(100);g__Metschnikowia(100);g__Metschnikowia_unclassified(100); | Metschnikowia | 13 | Saprotroph | \|Undefined Saprotroph\| | Yeast | NULL | Probable | As nectar-tap_saprotroph (Põlme et al. 2020); Decay sugar-rich substrates (Põlme et al. 2020) | Põlme S, et al. 2020. Fungal Diversity 105:1-16 ((https://doi.org/10.1007/s13225-020-00466-2)) |
| Otu41744 | 1 | k__Fungi(100);p__Ascomycota(100);c__Saccharomycetes(100);o__Saccharomycetales(100);f__Metschnikowiaceae(100);g__Metschnikowia(100);s__Metschnikowia_pulcherrima(100); | Metschnikowia | 13 | Saprotroph | \|Undefined Saprotroph\| | Yeast | NULL | Probable | As nectar-tap_saprotroph (Põlme et al. 2020); Decay sugar-rich substrates (Põlme et al. 2020) | Põlme S, et al. 2020. Fungal Diversity 105:1-16 ((https://doi.org/10.1007/s13225-020-00466-2)) |
| Otu41745 | 1 | k__Fungi(100);p__Ascomycota(100);c__Saccharomycetes(100);o__Saccharomycetales(100);f__Metschnikowiaceae(100);g__Metschnikowia(100);s__Metschnikowia_pulcherrima(100); | Metschnikowia | 13 | Saprotroph | \|Undefined Saprotroph\| | Yeast | NULL | Probable | As nectar-tap_saprotroph (Põlme et al. 2020); Decay sugar-rich substrates (Põlme et al. 2020) | Põlme S, et al. 2020. Fungal Diversity 105:1-16 ((https://doi.org/10.1007/s13225-020-00466-2)) |
| Otu41746 | 1 | k__Fungi(100);p__Ascomycota(100);c__Saccharomycetes(100);o__Saccharomycetales(100);f__Metschnikowiaceae(100);g__Metschnikowia(100);g__Metschnikowia_unclassified(100); | Metschnikowia | 13 | Saprotroph | \|Undefined Saprotroph\| | Yeast | NULL | Probable | As nectar-tap_saprotroph (Põlme et al. 2020); Decay sugar-rich substrates (Põlme et al. 2020) | Põlme S, et al. 2020. Fungal Diversity 105:1-16 ((https://doi.org/10.1007/s13225-020-00466-2)) |
| Otu41747 | 1 | k__Fungi(100);p__Ascomycota(100);c__Saccharomycetes(100);o__Saccharomycetales(100);f__Metschnikowiaceae(100);g__Metschnikowia(100);s__Metschnikowia_pulcherrima(100); | Metschnikowia | 13 | Saprotroph | \|Undefined Saprotroph\| | Yeast | NULL | Probable | As nectar-tap_saprotroph (Põlme et al. 2020); Decay sugar-rich substrates (Põlme et al. 2020) | Põlme S, et al. 2020. Fungal Diversity 105:1-16 ((https://doi.org/10.1007/s13225-020-00466-2)) |
| Otu41748 | 1 | k__Fungi(100);p__Ascomycota(100);c__Saccharomycetes(100);o__Saccharomycetales(100);f__Metschnikowiaceae(100);g__Metschnikowia(100);s__Metschnikowia_pulcherrima(100); | Metschnikowia | 13 | Saprotroph | \|Undefined Saprotroph\| | Yeast | NULL | Probable | As nectar-tap_saprotroph (Põlme et al. 2020); Decay sugar-rich substrates (Põlme et al. 2020) | Põlme S, et al. 2020. Fungal Diversity 105:1-16 ((https://doi.org/10.1007/s13225-020-00466-2)) |
| Otu41749 | 1 | k__Fungi(100);p__Ascomycota(100);c__Saccharomycetes(100);o__Saccharomycetales(100);f__Metschnikowiaceae(100);g__Metschnikowia(100);g__Metschnikowia_unclassified(100); | Metschnikowia | 13 | Saprotroph | \|Undefined Saprotroph\| | Yeast | NULL | Probable | As nectar-tap_saprotroph (Põlme et al. 2020); Decay sugar-rich substrates (Põlme et al. 2020) | Põlme S, et al. 2020. Fungal Diversity 105:1-16 ((https://doi.org/10.1007/s13225-020-00466-2)) |
| Otu41750 | 1 | k__Fungi(100);p__Ascomycota(100);c__Saccharomycetes(100);o__Saccharomycetales(100);f__Metschnikowiaceae(100);g__Metschnikowia(100);s__Metschnikowia_pulcherrima(100); | Metschnikowia | 13 | Saprotroph | \|Undefined Saprotroph\| | Yeast | NULL | Probable | As nectar-tap_saprotroph (Põlme et al. 2020); Decay sugar-rich substrates (Põlme et al. 2020) | Põlme S, et al. 2020. Fungal Diversity 105:1-16 ((https://doi.org/10.1007/s13225-020-00466-2)) |
| Otu41751 | 1 | k__Fungi(100);p__Ascomycota(100);c__Saccharomycetes(100);o__Saccharomycetales(100);f__Metschnikowiaceae(100);g__Metschnikowia(100);s__Metschnikowia_pulcherrima(100); | Metschnikowia | 13 | Saprotroph | \|Undefined Saprotroph\| | Yeast | NULL | Probable | As nectar-tap_saprotroph (Põlme et al. 2020); Decay sugar-rich substrates (Põlme et al. 2020) | Põlme S, et al. 2020. Fungal Diversity 105:1-16 ((https://doi.org/10.1007/s13225-020-00466-2)) |
| Otu41752 | 1 | k__Fungi(100);p__Ascomycota(100);c__Saccharomycetes(100);o__Saccharomycetales(100);f__Metschnikowiaceae(100);g__Metschnikowia(100);s__Metschnikowia_pulcherrima(100); | Metschnikowia | 13 | Saprotroph | \|Undefined Saprotroph\| | Yeast | NULL | Probable | As nectar-tap_saprotroph (Põlme et al. 2020); Decay sugar-rich substrates (Põlme et al. 2020) | Põlme S, et al. 2020. Fungal Diversity 105:1-16 ((https://doi.org/10.1007/s13225-020-00466-2)) |
| Otu41753 | 1 | k__Fungi(100);p__Ascomycota(100);c__Saccharomycetes(100);o__Saccharomycetales(100);f__Metschnikowiaceae(100);g__Metschnikowia(100);s__Metschnikowia_pulcherrima(100); | Metschnikowia | 13 | Saprotroph | \|Undefined Saprotroph\| | Yeast | NULL | Probable | As nectar-tap_saprotroph (Põlme et al. 2020); Decay sugar-rich substrates (Põlme et al. 2020) | Põlme S, et al. 2020. Fungal Diversity 105:1-16 ((https://doi.org/10.1007/s13225-020-00466-2)) |
| Otu41754 | 1 | k__Fungi(100);p__Ascomycota(100);c__Saccharomycetes(100);o__Saccharomycetales(100);f__Metschnikowiaceae(100);g__Metschnikowia(100);s__Metschnikowia_pulcherrima(100); | Metschnikowia | 13 | Saprotroph | \|Undefined Saprotroph\| | Yeast | NULL | Probable | As nectar-tap_saprotroph (Põlme et al. 2020); Decay sugar-rich substrates (Põlme et al. 2020) | Põlme S, et al. 2020. Fungal Diversity 105:1-16 ((https://doi.org/10.1007/s13225-020-00466-2)) |
| Otu41755 | 1 | k__Fungi(100);p__Ascomycota(100);c__Saccharomycetes(100);o__Saccharomycetales(100);f__Metschnikowiaceae(100);g__Metschnikowia(100);g__Metschnikowia_unclassified(100); | Metschnikowia | 13 | Saprotroph | \|Undefined Saprotroph\| | Yeast | NULL | Probable | As nectar-tap_saprotroph (Põlme et al. 2020); Decay sugar-rich substrates (Põlme et al. 2020) | Põlme S, et al. 2020. Fungal Diversity 105:1-16 ((https://doi.org/10.1007/s13225-020-00466-2)) |
| Otu41756 | 1 | k__Fungi(100);p__Ascomycota(100);c__Saccharomycetes(100);o__Saccharomycetales(100);f__Metschnikowiaceae(100);g__Metschnikowia(100);s__Metschnikowia_pulcherrima(100); | Metschnikowia | 13 | Saprotroph | \|Undefined Saprotroph\| | Yeast | NULL | Probable | As nectar-tap_saprotroph (Põlme et al. 2020); Decay sugar-rich substrates (Põlme et al. 2020) | Põlme S, et al. 2020. Fungal Diversity 105:1-16 ((https://doi.org/10.1007/s13225-020-00466-2)) |
| Otu41757 | 1 | k__Fungi(100);p__Ascomycota(100);c__Saccharomycetes(100);o__Saccharomycetales(100);f__Metschnikowiaceae(100);g__Metschnikowia(100);s__Metschnikowia_pulcherrima(100); | Metschnikowia | 13 | Saprotroph | \|Undefined Saprotroph\| | Yeast | NULL | Probable | As nectar-tap_saprotroph (Põlme et al. 2020); Decay sugar-rich substrates (Põlme et al. 2020) | Põlme S, et al. 2020. Fungal Diversity 105:1-16 ((https://doi.org/10.1007/s13225-020-00466-2)) |
| Otu41758 | 1 | k__Fungi(100);p__Ascomycota(100);c__Saccharomycetes(100);o__Saccharomycetales(100);f__Metschnikowiaceae(100);g__Metschnikowia(100);s__Metschnikowia_pulcherrima(100); | Metschnikowia | 13 | Saprotroph | \|Undefined Saprotroph\| | Yeast | NULL | Probable | As nectar-tap_saprotroph (Põlme et al. 2020); Decay sugar-rich substrates (Põlme et al. 2020) | Põlme S, et al. 2020. Fungal Diversity 105:1-16 ((https://doi.org/10.1007/s13225-020-00466-2)) |
| Otu41759 | 1 | k__Fungi(100);p__Ascomycota(100);c__Saccharomycetes(100);o__Saccharomycetales(100);f__Metschnikowiaceae(100);g__Metschnikowia(100);s__Metschnikowia_pulcherrima(100); | Metschnikowia | 13 | Saprotroph | \|Undefined Saprotroph\| | Yeast | NULL | Probable | As nectar-tap_saprotroph (Põlme et al. 2020); Decay sugar-rich substrates (Põlme et al. 2020) | Põlme S, et al. 2020. Fungal Diversity 105:1-16 ((https://doi.org/10.1007/s13225-020-00466-2)) |
| Otu41760 | 1 | k__Fungi(100);p__Ascomycota(100);c__Saccharomycetes(100);o__Saccharomycetales(100);f__Metschnikowiaceae(100);g__Metschnikowia(100);s__Metschnikowia_pulcherrima(100); | Metschnikowia | 13 | Saprotroph | \|Undefined Saprotroph\| | Yeast | NULL | Probable | As nectar-tap_saprotroph (Põlme et al. 2020); Decay sugar-rich substrates (Põlme et al. 2020) | Põlme S, et al. 2020. Fungal Diversity 105:1-16 ((https://doi.org/10.1007/s13225-020-00466-2)) |
| Otu41761 | 1 | k__Fungi(100);p__Ascomycota(100);c__Saccharomycetes(100);o__Saccharomycetales(100);f__Metschnikowiaceae(100);g__Metschnikowia(100);s__Metschnikowia_pulcherrima(100); | Metschnikowia | 13 | Saprotroph | \|Undefined Saprotroph\| | Yeast | NULL | Probable | As nectar-tap_saprotroph (Põlme et al. 2020); Decay sugar-rich substrates (Põlme et al. 2020) | Põlme S, et al. 2020. Fungal Diversity 105:1-16 ((https://doi.org/10.1007/s13225-020-00466-2)) |
| Otu41762 | 1 | k__Fungi(100);p__Ascomycota(100);c__Saccharomycetes(100);o__Saccharomycetales(100);f__Metschnikowiaceae(100);g__Metschnikowia(100);s__Metschnikowia_pulcherrima(100); | Metschnikowia | 13 | Saprotroph | \|Undefined Saprotroph\| | Yeast | NULL | Probable | As nectar-tap_saprotroph (Põlme et al. 2020); Decay sugar-rich substrates (Põlme et al. 2020) | Põlme S, et al. 2020. Fungal Diversity 105:1-16 ((https://doi.org/10.1007/s13225-020-00466-2)) |
| Otu41763 | 1 | k__Fungi(100);p__Ascomycota(100);c__Saccharomycetes(100);o__Saccharomycetales(100);f__Metschnikowiaceae(100);g__Metschnikowia(100);s__Metschnikowia_pulcherrima(100); | Metschnikowia | 13 | Saprotroph | \|Undefined Saprotroph\| | Yeast | NULL | Probable | As nectar-tap_saprotroph (Põlme et al. 2020); Decay sugar-rich substrates (Põlme et al. 2020) | Põlme S, et al. 2020. Fungal Diversity 105:1-16 ((https://doi.org/10.1007/s13225-020-00466-2)) |
| Otu41764 | 1 | k__Fungi(100);p__Ascomycota(100);c__Saccharomycetes(100);o__Saccharomycetales(100);f__Metschnikowiaceae(100);g__Metschnikowia(100);s__Metschnikowia_pulcherrima(100); | Metschnikowia | 13 | Saprotroph | \|Undefined Saprotroph\| | Yeast | NULL | Probable | As nectar-tap_saprotroph (Põlme et al. 2020); Decay sugar-rich substrates (Põlme et al. 2020) | Põlme S, et al. 2020. Fungal Diversity 105:1-16 ((https://doi.org/10.1007/s13225-020-00466-2)) |
| Otu41765 | 1 | k__Fungi(100);p__Ascomycota(100);c__Saccharomycetes(100);o__Saccharomycetales(100);f__Metschnikowiaceae(100);g__Metschnikowia(100);s__Metschnikowia_pulcherrima(100); | Metschnikowia | 13 | Saprotroph | \|Undefined Saprotroph\| | Yeast | NULL | Probable | As nectar-tap_saprotroph (Põlme et al. 2020); Decay sugar-rich substrates (Põlme et al. 2020) | Põlme S, et al. 2020. Fungal Diversity 105:1-16 ((https://doi.org/10.1007/s13225-020-00466-2)) |
| Otu41766 | 1 | k__Fungi(100);p__Ascomycota(100);c__Saccharomycetes(100);o__Saccharomycetales(100);f__Metschnikowiaceae(100);g__Metschnikowia(100);s__Metschnikowia_pulcherrima(100); | Metschnikowia | 13 | Saprotroph | \|Undefined Saprotroph\| | Yeast | NULL | Probable | As nectar-tap_saprotroph (Põlme et al. 2020); Decay sugar-rich substrates (Põlme et al. 2020) | Põlme S, et al. 2020. Fungal Diversity 105:1-16 ((https://doi.org/10.1007/s13225-020-00466-2)) |
| Otu41768 | 1 | k__Fungi(100);p__Ascomycota(100);c__Saccharomycetes(100);o__Saccharomycetales(100);f__Metschnikowiaceae(100);g__Metschnikowia(100);g__Metschnikowia_unclassified(100); | Metschnikowia | 13 | Saprotroph | \|Undefined Saprotroph\| | Yeast | NULL | Probable | As nectar-tap_saprotroph (Põlme et al. 2020); Decay sugar-rich substrates (Põlme et al. 2020) | Põlme S, et al. 2020. Fungal Diversity 105:1-16 ((https://doi.org/10.1007/s13225-020-00466-2)) |
| Otu41769 | 1 | k__Fungi(100);p__Ascomycota(100);c__Saccharomycetes(100);o__Saccharomycetales(100);f__Metschnikowiaceae(100);g__Metschnikowia(100);s__Metschnikowia_pulcherrima(100); | Metschnikowia | 13 | Saprotroph | \|Undefined Saprotroph\| | Yeast | NULL | Probable | As nectar-tap_saprotroph (Põlme et al. 2020); Decay sugar-rich substrates (Põlme et al. 2020) | Põlme S, et al. 2020. Fungal Diversity 105:1-16 ((https://doi.org/10.1007/s13225-020-00466-2)) |
| Otu41770 | 1 | k__Fungi(100);p__Ascomycota(100);c__Saccharomycetes(100);o__Saccharomycetales(100);f__Metschnikowiaceae(100);g__Metschnikowia(100);s__Metschnikowia_pulcherrima(100); | Metschnikowia | 13 | Saprotroph | \|Undefined Saprotroph\| | Yeast | NULL | Probable | As nectar-tap_saprotroph (Põlme et al. 2020); Decay sugar-rich substrates (Põlme et al. 2020) | Põlme S, et al. 2020. Fungal Diversity 105:1-16 ((https://doi.org/10.1007/s13225-020-00466-2)) |
| Otu41772 | 1 | k__Fungi(100);p__Ascomycota(100);c__Saccharomycetes(100);o__Saccharomycetales(100);f__Metschnikowiaceae(100);g__Metschnikowia(100);g__Metschnikowia_unclassified(100); | Metschnikowia | 13 | Saprotroph | \|Undefined Saprotroph\| | Yeast | NULL | Probable | As nectar-tap_saprotroph (Põlme et al. 2020); Decay sugar-rich substrates (Põlme et al. 2020) | Põlme S, et al. 2020. Fungal Diversity 105:1-16 ((https://doi.org/10.1007/s13225-020-00466-2)) |
| Otu41773 | 1 | k__Fungi(100);p__Ascomycota(100);c__Saccharomycetes(100);o__Saccharomycetales(100);f__Metschnikowiaceae(100);g__Metschnikowia(100);s__Metschnikowia_pulcherrima(100); | Metschnikowia | 13 | Saprotroph | \|Undefined Saprotroph\| | Yeast | NULL | Probable | As nectar-tap_saprotroph (Põlme et al. 2020); Decay sugar-rich substrates (Põlme et al. 2020) | Põlme S, et al. 2020. Fungal Diversity 105:1-16 ((https://doi.org/10.1007/s13225-020-00466-2)) |
| Otu41774 | 1 | k__Fungi(100);p__Ascomycota(100);c__Saccharomycetes(100);o__Saccharomycetales(100);f__Metschnikowiaceae(100);g__Metschnikowia(100);s__Metschnikowia_pulcherrima(100); | Metschnikowia | 13 | Saprotroph | \|Undefined Saprotroph\| | Yeast | NULL | Probable | As nectar-tap_saprotroph (Põlme et al. 2020); Decay sugar-rich substrates (Põlme et al. 2020) | Põlme S, et al. 2020. Fungal Diversity 105:1-16 ((https://doi.org/10.1007/s13225-020-00466-2)) |
| Otu41775 | 1 | k__Fungi(100);p__Ascomycota(100);c__Saccharomycetes(100);o__Saccharomycetales(100);f__Metschnikowiaceae(100);g__Metschnikowia(100);s__Metschnikowia_pulcherrima(100); | Metschnikowia | 13 | Saprotroph | \|Undefined Saprotroph\| | Yeast | NULL | Probable | As nectar-tap_saprotroph (Põlme et al. 2020); Decay sugar-rich substrates (Põlme et al. 2020) | Põlme S, et al. 2020. Fungal Diversity 105:1-16 ((https://doi.org/10.1007/s13225-020-00466-2)) |
| Otu41776 | 1 | k__Fungi(100);p__Ascomycota(100);c__Saccharomycetes(100);o__Saccharomycetales(100);f__Metschnikowiaceae(100);g__Metschnikowia(100);s__Metschnikowia_pulcherrima(100); | Metschnikowia | 13 | Saprotroph | \|Undefined Saprotroph\| | Yeast | NULL | Probable | As nectar-tap_saprotroph (Põlme et al. 2020); Decay sugar-rich substrates (Põlme et al. 2020) | Põlme S, et al. 2020. Fungal Diversity 105:1-16 ((https://doi.org/10.1007/s13225-020-00466-2)) |
| Otu41777 | 1 | k__Fungi(100);p__Ascomycota(100);c__Saccharomycetes(100);o__Saccharomycetales(100);f__Metschnikowiaceae(100);g__Metschnikowia(100);s__Metschnikowia_pulcherrima(100); | Metschnikowia | 13 | Saprotroph | \|Undefined Saprotroph\| | Yeast | NULL | Probable | As nectar-tap_saprotroph (Põlme et al. 2020); Decay sugar-rich substrates (Põlme et al. 2020) | Põlme S, et al. 2020. Fungal Diversity 105:1-16 ((https://doi.org/10.1007/s13225-020-00466-2)) |
| Otu41778 | 1 | k__Fungi(100);p__Ascomycota(100);c__Saccharomycetes(100);o__Saccharomycetales(100);f__Metschnikowiaceae(100);g__Metschnikowia(100);s__Metschnikowia_pulcherrima(100); | Metschnikowia | 13 | Saprotroph | \|Undefined Saprotroph\| | Yeast | NULL | Probable | As nectar-tap_saprotroph (Põlme et al. 2020); Decay sugar-rich substrates (Põlme et al. 2020) | Põlme S, et al. 2020. Fungal Diversity 105:1-16 ((https://doi.org/10.1007/s13225-020-00466-2)) |
| Otu41779 | 1 | k__Fungi(100);p__Ascomycota(100);c__Saccharomycetes(100);o__Saccharomycetales(100);f__Metschnikowiaceae(100);g__Metschnikowia(100);s__Metschnikowia_pulcherrima(100); | Metschnikowia | 13 | Saprotroph | \|Undefined Saprotroph\| | Yeast | NULL | Probable | As nectar-tap_saprotroph (Põlme et al. 2020); Decay sugar-rich substrates (Põlme et al. 2020) | Põlme S, et al. 2020. Fungal Diversity 105:1-16 ((https://doi.org/10.1007/s13225-020-00466-2)) |
| Otu41780 | 1 | k__Fungi(100);p__Ascomycota(100);c__Saccharomycetes(100);o__Saccharomycetales(100);f__Metschnikowiaceae(100);g__Metschnikowia(100);s__Metschnikowia_pulcherrima(100); | Metschnikowia | 13 | Saprotroph | \|Undefined Saprotroph\| | Yeast | NULL | Probable | As nectar-tap_saprotroph (Põlme et al. 2020); Decay sugar-rich substrates (Põlme et al. 2020) | Põlme S, et al. 2020. Fungal Diversity 105:1-16 ((https://doi.org/10.1007/s13225-020-00466-2)) |
| Otu41781 | 1 | k__Fungi(100);p__Ascomycota(100);c__Saccharomycetes(100);o__Saccharomycetales(100);f__Metschnikowiaceae(100);g__Metschnikowia(100);s__Metschnikowia_pulcherrima(100); | Metschnikowia | 13 | Saprotroph | \|Undefined Saprotroph\| | Yeast | NULL | Probable | As nectar-tap_saprotroph (Põlme et al. 2020); Decay sugar-rich substrates (Põlme et al. 2020) | Põlme S, et al. 2020. Fungal Diversity 105:1-16 ((https://doi.org/10.1007/s13225-020-00466-2)) |
| Otu41782 | 1 | k__Fungi(100);p__Ascomycota(100);c__Saccharomycetes(100);o__Saccharomycetales(100);f__Metschnikowiaceae(100);g__Metschnikowia(100);g__Metschnikowia_unclassified(100); | Metschnikowia | 13 | Saprotroph | \|Undefined Saprotroph\| | Yeast | NULL | Probable | As nectar-tap_saprotroph (Põlme et al. 2020); Decay sugar-rich substrates (Põlme et al. 2020) | Põlme S, et al. 2020. Fungal Diversity 105:1-16 ((https://doi.org/10.1007/s13225-020-00466-2)) |
| Otu41783 | 1 | k__Fungi(100);p__Ascomycota(100);c__Saccharomycetes(100);o__Saccharomycetales(100);f__Metschnikowiaceae(100);g__Metschnikowia(100);s__Metschnikowia_pulcherrima(100); | Metschnikowia | 13 | Saprotroph | \|Undefined Saprotroph\| | Yeast | NULL | Probable | As nectar-tap_saprotroph (Põlme et al. 2020); Decay sugar-rich substrates (Põlme et al. 2020) | Põlme S, et al. 2020. Fungal Diversity 105:1-16 ((https://doi.org/10.1007/s13225-020-00466-2)) |
| Otu41784 | 1 | k__Fungi(100);p__Ascomycota(100);c__Saccharomycetes(100);o__Saccharomycetales(100);f__Metschnikowiaceae(100);g__Metschnikowia(100);s__Metschnikowia_pulcherrima(100); | Metschnikowia | 13 | Saprotroph | \|Undefined Saprotroph\| | Yeast | NULL | Probable | As nectar-tap_saprotroph (Põlme et al. 2020); Decay sugar-rich substrates (Põlme et al. 2020) | Põlme S, et al. 2020. Fungal Diversity 105:1-16 ((https://doi.org/10.1007/s13225-020-00466-2)) |
| Otu41785 | 1 | k__Fungi(100);p__Ascomycota(100);c__Saccharomycetes(100);o__Saccharomycetales(100);f__Metschnikowiaceae(100);g__Metschnikowia(100);s__Metschnikowia_pulcherrima(100); | Metschnikowia | 13 | Saprotroph | \|Undefined Saprotroph\| | Yeast | NULL | Probable | As nectar-tap_saprotroph (Põlme et al. 2020); Decay sugar-rich substrates (Põlme et al. 2020) | Põlme S, et al. 2020. Fungal Diversity 105:1-16 ((https://doi.org/10.1007/s13225-020-00466-2)) |
| Otu41786 | 1 | k__Fungi(100);p__Ascomycota(100);c__Saccharomycetes(100);o__Saccharomycetales(100);f__Metschnikowiaceae(100);g__Metschnikowia(100);s__Metschnikowia_pulcherrima(100); | Metschnikowia | 13 | Saprotroph | \|Undefined Saprotroph\| | Yeast | NULL | Probable | As nectar-tap_saprotroph (Põlme et al. 2020); Decay sugar-rich substrates (Põlme et al. 2020) | Põlme S, et al. 2020. Fungal Diversity 105:1-16 ((https://doi.org/10.1007/s13225-020-00466-2)) |
| Otu41787 | 1 | k__Fungi(100);p__Ascomycota(100);c__Saccharomycetes(100);o__Saccharomycetales(100);f__Metschnikowiaceae(100);g__Metschnikowia(100);s__Metschnikowia_pulcherrima(100); | Metschnikowia | 13 | Saprotroph | \|Undefined Saprotroph\| | Yeast | NULL | Probable | As nectar-tap_saprotroph (Põlme et al. 2020); Decay sugar-rich substrates (Põlme et al. 2020) | Põlme S, et al. 2020. Fungal Diversity 105:1-16 ((https://doi.org/10.1007/s13225-020-00466-2)) |
| Otu41788 | 1 | k__Fungi(100);p__Ascomycota(100);c__Saccharomycetes(100);o__Saccharomycetales(100);f__Metschnikowiaceae(100);g__Metschnikowia(100);s__Metschnikowia_pulcherrima(100); | Metschnikowia | 13 | Saprotroph | \|Undefined Saprotroph\| | Yeast | NULL | Probable | As nectar-tap_saprotroph (Põlme et al. 2020); Decay sugar-rich substrates (Põlme et al. 2020) | Põlme S, et al. 2020. Fungal Diversity 105:1-16 ((https://doi.org/10.1007/s13225-020-00466-2)) |
| Otu41789 | 1 | k__Fungi(100);p__Ascomycota(100);c__Saccharomycetes(100);o__Saccharomycetales(100);f__Metschnikowiaceae(100);g__Metschnikowia(100);s__Metschnikowia_pulcherrima(100); | Metschnikowia | 13 | Saprotroph | \|Undefined Saprotroph\| | Yeast | NULL | Probable | As nectar-tap_saprotroph (Põlme et al. 2020); Decay sugar-rich substrates (Põlme et al. 2020) | Põlme S, et al. 2020. Fungal Diversity 105:1-16 ((https://doi.org/10.1007/s13225-020-00466-2)) |
| Otu41790 | 1 | k__Fungi(100);p__Ascomycota(100);c__Saccharomycetes(100);o__Saccharomycetales(100);f__Metschnikowiaceae(100);g__Metschnikowia(100);s__Metschnikowia_pulcherrima(100); | Metschnikowia | 13 | Saprotroph | \|Undefined Saprotroph\| | Yeast | NULL | Probable | As nectar-tap_saprotroph (Põlme et al. 2020); Decay sugar-rich substrates (Põlme et al. 2020) | Põlme S, et al. 2020. Fungal Diversity 105:1-16 ((https://doi.org/10.1007/s13225-020-00466-2)) |
| Otu41791 | 1 | k__Fungi(100);p__Ascomycota(100);c__Saccharomycetes(100);o__Saccharomycetales(100);f__Metschnikowiaceae(100);g__Metschnikowia(100);s__Metschnikowia_pulcherrima(100); | Metschnikowia | 13 | Saprotroph | \|Undefined Saprotroph\| | Yeast | NULL | Probable | As nectar-tap_saprotroph (Põlme et al. 2020); Decay sugar-rich substrates (Põlme et al. 2020) | Põlme S, et al. 2020. Fungal Diversity 105:1-16 ((https://doi.org/10.1007/s13225-020-00466-2)) |
| Otu41792 | 1 | k__Fungi(100);p__Ascomycota(100);c__Saccharomycetes(100);o__Saccharomycetales(100);f__Metschnikowiaceae(100);g__Metschnikowia(100);g__Metschnikowia_unclassified(100); | Metschnikowia | 13 | Saprotroph | \|Undefined Saprotroph\| | Yeast | NULL | Probable | As nectar-tap_saprotroph (Põlme et al. 2020); Decay sugar-rich substrates (Põlme et al. 2020) | Põlme S, et al. 2020. Fungal Diversity 105:1-16 ((https://doi.org/10.1007/s13225-020-00466-2)) |
| Otu41793 | 1 | k__Fungi(100);p__Ascomycota(100);c__Saccharomycetes(100);o__Saccharomycetales(100);f__Metschnikowiaceae(100);g__Metschnikowia(100);s__Metschnikowia_pulcherrima(100); | Metschnikowia | 13 | Saprotroph | \|Undefined Saprotroph\| | Yeast | NULL | Probable | As nectar-tap_saprotroph (Põlme et al. 2020); Decay sugar-rich substrates (Põlme et al. 2020) | Põlme S, et al. 2020. Fungal Diversity 105:1-16 ((https://doi.org/10.1007/s13225-020-00466-2)) |
| Otu41794 | 1 | k__Fungi(100);p__Ascomycota(100);c__Saccharomycetes(100);o__Saccharomycetales(100);f__Metschnikowiaceae(100);g__Metschnikowia(100);g__Metschnikowia_unclassified(100); | Metschnikowia | 13 | Saprotroph | \|Undefined Saprotroph\| | Yeast | NULL | Probable | As nectar-tap_saprotroph (Põlme et al. 2020); Decay sugar-rich substrates (Põlme et al. 2020) | Põlme S, et al. 2020. Fungal Diversity 105:1-16 ((https://doi.org/10.1007/s13225-020-00466-2)) |
| Otu41795 | 1 | k__Fungi(100);p__Ascomycota(100);c__Saccharomycetes(100);o__Saccharomycetales(100);f__Metschnikowiaceae(100);g__Metschnikowia(100);s__Metschnikowia_pulcherrima(100); | Metschnikowia | 13 | Saprotroph | \|Undefined Saprotroph\| | Yeast | NULL | Probable | As nectar-tap_saprotroph (Põlme et al. 2020); Decay sugar-rich substrates (Põlme et al. 2020) | Põlme S, et al. 2020. Fungal Diversity 105:1-16 ((https://doi.org/10.1007/s13225-020-00466-2)) |
| Otu41796 | 1 | k__Fungi(100);p__Ascomycota(100);c__Saccharomycetes(100);o__Saccharomycetales(100);f__Metschnikowiaceae(100);g__Metschnikowia(100);s__Metschnikowia_pulcherrima(100); | Metschnikowia | 13 | Saprotroph | \|Undefined Saprotroph\| | Yeast | NULL | Probable | As nectar-tap_saprotroph (Põlme et al. 2020); Decay sugar-rich substrates (Põlme et al. 2020) | Põlme S, et al. 2020. Fungal Diversity 105:1-16 ((https://doi.org/10.1007/s13225-020-00466-2)) |
| Otu41797 | 1 | k__Fungi(100);p__Ascomycota(100);c__Saccharomycetes(100);o__Saccharomycetales(100);f__Metschnikowiaceae(100);g__Metschnikowia(100);s__Metschnikowia_viticola(100); | Metschnikowia | 13 | Saprotroph | \|Undefined Saprotroph\| | Yeast | NULL | Probable | As nectar-tap_saprotroph (Põlme et al. 2020); Decay sugar-rich substrates (Põlme et al. 2020) | Põlme S, et al. 2020. Fungal Diversity 105:1-16 ((https://doi.org/10.1007/s13225-020-00466-2)) |
| Otu41798 | 1 | k__Fungi(100);p__Ascomycota(100);c__Saccharomycetes(100);o__Saccharomycetales(100);f__Metschnikowiaceae(100);g__Metschnikowia(100);g__Metschnikowia_unclassified(100); | Metschnikowia | 13 | Saprotroph | \|Undefined Saprotroph\| | Yeast | NULL | Probable | As nectar-tap_saprotroph (Põlme et al. 2020); Decay sugar-rich substrates (Põlme et al. 2020) | Põlme S, et al. 2020. Fungal Diversity 105:1-16 ((https://doi.org/10.1007/s13225-020-00466-2)) |
| Otu41799 | 1 | k__Fungi(100);p__Ascomycota(100);c__Saccharomycetes(100);o__Saccharomycetales(100);f__Metschnikowiaceae(100);g__Metschnikowia(100);g__Metschnikowia_unclassified(100); | Metschnikowia | 13 | Saprotroph | \|Undefined Saprotroph\| | Yeast | NULL | Probable | As nectar-tap_saprotroph (Põlme et al. 2020); Decay sugar-rich substrates (Põlme et al. 2020) | Põlme S, et al. 2020. Fungal Diversity 105:1-16 ((https://doi.org/10.1007/s13225-020-00466-2)) |
| Otu41800 | 1 | k__Fungi(100);p__Ascomycota(100);c__Saccharomycetes(100);o__Saccharomycetales(100);f__Metschnikowiaceae(100);g__Metschnikowia(100);s__Metschnikowia_pulcherrima(100); | Metschnikowia | 13 | Saprotroph | \|Undefined Saprotroph\| | Yeast | NULL | Probable | As nectar-tap_saprotroph (Põlme et al. 2020); Decay sugar-rich substrates (Põlme et al. 2020) | Põlme S, et al. 2020. Fungal Diversity 105:1-16 ((https://doi.org/10.1007/s13225-020-00466-2)) |
| Otu41801 | 1 | k__Fungi(100);p__Ascomycota(100);c__Saccharomycetes(100);o__Saccharomycetales(100);f__Metschnikowiaceae(100);g__Metschnikowia(100);s__Metschnikowia_pulcherrima(100); | Metschnikowia | 13 | Saprotroph | \|Undefined Saprotroph\| | Yeast | NULL | Probable | As nectar-tap_saprotroph (Põlme et al. 2020); Decay sugar-rich substrates (Põlme et al. 2020) | Põlme S, et al. 2020. Fungal Diversity 105:1-16 ((https://doi.org/10.1007/s13225-020-00466-2)) |
| Otu41802 | 1 | k__Fungi(100);p__Ascomycota(100);c__Saccharomycetes(100);o__Saccharomycetales(100);f__Metschnikowiaceae(100);g__Metschnikowia(100);s__Metschnikowia_pulcherrima(100); | Metschnikowia | 13 | Saprotroph | \|Undefined Saprotroph\| | Yeast | NULL | Probable | As nectar-tap_saprotroph (Põlme et al. 2020); Decay sugar-rich substrates (Põlme et al. 2020) | Põlme S, et al. 2020. Fungal Diversity 105:1-16 ((https://doi.org/10.1007/s13225-020-00466-2)) |
| Otu41803 | 1 | k__Fungi(100);p__Ascomycota(100);c__Saccharomycetes(100);o__Saccharomycetales(100);f__Metschnikowiaceae(100);g__Metschnikowia(100);g__Metschnikowia_unclassified(100); | Metschnikowia | 13 | Saprotroph | \|Undefined Saprotroph\| | Yeast | NULL | Probable | As nectar-tap_saprotroph (Põlme et al. 2020); Decay sugar-rich substrates (Põlme et al. 2020) | Põlme S, et al. 2020. Fungal Diversity 105:1-16 ((https://doi.org/10.1007/s13225-020-00466-2)) |
| Otu41804 | 1 | k__Fungi(100);p__Ascomycota(100);c__Saccharomycetes(100);o__Saccharomycetales(100);f__Metschnikowiaceae(100);g__Metschnikowia(100);s__Metschnikowia_pulcherrima(100); | Metschnikowia | 13 | Saprotroph | \|Undefined Saprotroph\| | Yeast | NULL | Probable | As nectar-tap_saprotroph (Põlme et al. 2020); Decay sugar-rich substrates (Põlme et al. 2020) | Põlme S, et al. 2020. Fungal Diversity 105:1-16 ((https://doi.org/10.1007/s13225-020-00466-2)) |
| Otu41805 | 1 | k__Fungi(100);p__Ascomycota(100);c__Saccharomycetes(100);o__Saccharomycetales(100);f__Metschnikowiaceae(100);g__Metschnikowia(100);s__Metschnikowia_pulcherrima(100); | Metschnikowia | 13 | Saprotroph | \|Undefined Saprotroph\| | Yeast | NULL | Probable | As nectar-tap_saprotroph (Põlme et al. 2020); Decay sugar-rich substrates (Põlme et al. 2020) | Põlme S, et al. 2020. Fungal Diversity 105:1-16 ((https://doi.org/10.1007/s13225-020-00466-2)) |
| Otu41806 | 1 | k__Fungi(100);p__Ascomycota(100);c__Saccharomycetes(100);o__Saccharomycetales(100);f__Metschnikowiaceae(100);g__Metschnikowia(100);s__Metschnikowia_pulcherrima(100); | Metschnikowia | 13 | Saprotroph | \|Undefined Saprotroph\| | Yeast | NULL | Probable | As nectar-tap_saprotroph (Põlme et al. 2020); Decay sugar-rich substrates (Põlme et al. 2020) | Põlme S, et al. 2020. Fungal Diversity 105:1-16 ((https://doi.org/10.1007/s13225-020-00466-2)) |
| Otu41807 | 1 | k__Fungi(100);p__Ascomycota(100);c__Saccharomycetes(100);o__Saccharomycetales(100);f__Metschnikowiaceae(100);g__Metschnikowia(100);g__Metschnikowia_unclassified(100); | Metschnikowia | 13 | Saprotroph | \|Undefined Saprotroph\| | Yeast | NULL | Probable | As nectar-tap_saprotroph (Põlme et al. 2020); Decay sugar-rich substrates (Põlme et al. 2020) | Põlme S, et al. 2020. Fungal Diversity 105:1-16 ((https://doi.org/10.1007/s13225-020-00466-2)) |
| Otu41808 | 1 | k__Fungi(100);p__Ascomycota(100);c__Saccharomycetes(100);o__Saccharomycetales(100);f__Metschnikowiaceae(100);g__Metschnikowia(100);s__Metschnikowia_pulcherrima(100); | Metschnikowia | 13 | Saprotroph | \|Undefined Saprotroph\| | Yeast | NULL | Probable | As nectar-tap_saprotroph (Põlme et al. 2020); Decay sugar-rich substrates (Põlme et al. 2020) | Põlme S, et al. 2020. Fungal Diversity 105:1-16 ((https://doi.org/10.1007/s13225-020-00466-2)) |
| Otu41809 | 1 | k__Fungi(100);p__Ascomycota(100);c__Saccharomycetes(100);o__Saccharomycetales(100);f__Metschnikowiaceae(100);g__Metschnikowia(100);s__Metschnikowia_pulcherrima(100); | Metschnikowia | 13 | Saprotroph | \|Undefined Saprotroph\| | Yeast | NULL | Probable | As nectar-tap_saprotroph (Põlme et al. 2020); Decay sugar-rich substrates (Põlme et al. 2020) | Põlme S, et al. 2020. Fungal Diversity 105:1-16 ((https://doi.org/10.1007/s13225-020-00466-2)) |
| Otu41810 | 1 | k__Fungi(100);p__Ascomycota(100);c__Saccharomycetes(100);o__Saccharomycetales(100);f__Metschnikowiaceae(100);g__Metschnikowia(100);s__Metschnikowia_pulcherrima(100); | Metschnikowia | 13 | Saprotroph | \|Undefined Saprotroph\| | Yeast | NULL | Probable | As nectar-tap_saprotroph (Põlme et al. 2020); Decay sugar-rich substrates (Põlme et al. 2020) | Põlme S, et al. 2020. Fungal Diversity 105:1-16 ((https://doi.org/10.1007/s13225-020-00466-2)) |
| Otu41811 | 1 | k__Fungi(100);p__Ascomycota(100);c__Saccharomycetes(100);o__Saccharomycetales(100);f__Metschnikowiaceae(100);g__Metschnikowia(100);s__Metschnikowia_pulcherrima(100); | Metschnikowia | 13 | Saprotroph | \|Undefined Saprotroph\| | Yeast | NULL | Probable | As nectar-tap_saprotroph (Põlme et al. 2020); Decay sugar-rich substrates (Põlme et al. 2020) | Põlme S, et al. 2020. Fungal Diversity 105:1-16 ((https://doi.org/10.1007/s13225-020-00466-2)) |
| Otu41812 | 1 | k__Fungi(100);p__Ascomycota(100);c__Saccharomycetes(100);o__Saccharomycetales(100);f__Metschnikowiaceae(100);g__Metschnikowia(100);s__Metschnikowia_pulcherrima(100); | Metschnikowia | 13 | Saprotroph | \|Undefined Saprotroph\| | Yeast | NULL | Probable | As nectar-tap_saprotroph (Põlme et al. 2020); Decay sugar-rich substrates (Põlme et al. 2020) | Põlme S, et al. 2020. Fungal Diversity 105:1-16 ((https://doi.org/10.1007/s13225-020-00466-2)) |
| Otu41813 | 1 | k__Fungi(100);p__Ascomycota(100);c__Saccharomycetes(100);o__Saccharomycetales(100);f__Metschnikowiaceae(100);g__Metschnikowia(100);s__Metschnikowia_pulcherrima(100); | Metschnikowia | 13 | Saprotroph | \|Undefined Saprotroph\| | Yeast | NULL | Probable | As nectar-tap_saprotroph (Põlme et al. 2020); Decay sugar-rich substrates (Põlme et al. 2020) | Põlme S, et al. 2020. Fungal Diversity 105:1-16 ((https://doi.org/10.1007/s13225-020-00466-2)) |
| Otu41814 | 1 | k__Fungi(100);p__Ascomycota(100);c__Saccharomycetes(100);o__Saccharomycetales(100);f__Metschnikowiaceae(100);g__Metschnikowia(100);g__Metschnikowia_unclassified(100); | Metschnikowia | 13 | Saprotroph | \|Undefined Saprotroph\| | Yeast | NULL | Probable | As nectar-tap_saprotroph (Põlme et al. 2020); Decay sugar-rich substrates (Põlme et al. 2020) | Põlme S, et al. 2020. Fungal Diversity 105:1-16 ((https://doi.org/10.1007/s13225-020-00466-2)) |
| Otu41815 | 1 | k__Fungi(100);p__Ascomycota(100);c__Saccharomycetes(100);o__Saccharomycetales(100);f__Metschnikowiaceae(100);g__Metschnikowia(100);s__Metschnikowia_pulcherrima(100); | Metschnikowia | 13 | Saprotroph | \|Undefined Saprotroph\| | Yeast | NULL | Probable | As nectar-tap_saprotroph (Põlme et al. 2020); Decay sugar-rich substrates (Põlme et al. 2020) | Põlme S, et al. 2020. Fungal Diversity 105:1-16 ((https://doi.org/10.1007/s13225-020-00466-2)) |
| Otu41816 | 1 | k__Fungi(100);p__Ascomycota(100);c__Saccharomycetes(100);o__Saccharomycetales(100);f__Metschnikowiaceae(100);g__Metschnikowia(100);g__Metschnikowia_unclassified(100); | Metschnikowia | 13 | Saprotroph | \|Undefined Saprotroph\| | Yeast | NULL | Probable | As nectar-tap_saprotroph (Põlme et al. 2020); Decay sugar-rich substrates (Põlme et al. 2020) | Põlme S, et al. 2020. Fungal Diversity 105:1-16 ((https://doi.org/10.1007/s13225-020-00466-2)) |
| Otu41817 | 1 | k__Fungi(100);p__Ascomycota(100);c__Saccharomycetes(100);o__Saccharomycetales(100);f__Metschnikowiaceae(100);g__Metschnikowia(100);g__Metschnikowia_unclassified(100); | Metschnikowia | 13 | Saprotroph | \|Undefined Saprotroph\| | Yeast | NULL | Probable | As nectar-tap_saprotroph (Põlme et al. 2020); Decay sugar-rich substrates (Põlme et al. 2020) | Põlme S, et al. 2020. Fungal Diversity 105:1-16 ((https://doi.org/10.1007/s13225-020-00466-2)) |
| Otu41818 | 1 | k__Fungi(100);p__Ascomycota(100);c__Saccharomycetes(100);o__Saccharomycetales(100);f__Metschnikowiaceae(100);g__Metschnikowia(100);g__Metschnikowia_unclassified(100); | Metschnikowia | 13 | Saprotroph | \|Undefined Saprotroph\| | Yeast | NULL | Probable | As nectar-tap_saprotroph (Põlme et al. 2020); Decay sugar-rich substrates (Põlme et al. 2020) | Põlme S, et al. 2020. Fungal Diversity 105:1-16 ((https://doi.org/10.1007/s13225-020-00466-2)) |
| Otu41819 | 1 | k__Fungi(100);p__Ascomycota(100);c__Saccharomycetes(100);o__Saccharomycetales(100);f__Metschnikowiaceae(100);g__Metschnikowia(100);g__Metschnikowia_unclassified(100); | Metschnikowia | 13 | Saprotroph | \|Undefined Saprotroph\| | Yeast | NULL | Probable | As nectar-tap_saprotroph (Põlme et al. 2020); Decay sugar-rich substrates (Põlme et al. 2020) | Põlme S, et al. 2020. Fungal Diversity 105:1-16 ((https://doi.org/10.1007/s13225-020-00466-2)) |
| Otu41820 | 1 | k__Fungi(100);p__Ascomycota(100);c__Saccharomycetes(100);o__Saccharomycetales(100);f__Metschnikowiaceae(100);g__Metschnikowia(100);g__Metschnikowia_unclassified(100); | Metschnikowia | 13 | Saprotroph | \|Undefined Saprotroph\| | Yeast | NULL | Probable | As nectar-tap_saprotroph (Põlme et al. 2020); Decay sugar-rich substrates (Põlme et al. 2020) | Põlme S, et al. 2020. Fungal Diversity 105:1-16 ((https://doi.org/10.1007/s13225-020-00466-2)) |
| Otu41822 | 1 | k__Fungi(100);p__Ascomycota(100);c__Saccharomycetes(100);o__Saccharomycetales(100);f__Metschnikowiaceae(100);g__Metschnikowia(100);s__Metschnikowia_pulcherrima(100); | Metschnikowia | 13 | Saprotroph | \|Undefined Saprotroph\| | Yeast | NULL | Probable | As nectar-tap_saprotroph (Põlme et al. 2020); Decay sugar-rich substrates (Põlme et al. 2020) | Põlme S, et al. 2020. Fungal Diversity 105:1-16 ((https://doi.org/10.1007/s13225-020-00466-2)) |
| Otu41823 | 1 | k__Fungi(100);p__Ascomycota(100);c__Saccharomycetes(100);o__Saccharomycetales(100);f__Metschnikowiaceae(100);g__Metschnikowia(100);s__Metschnikowia_pulcherrima(100); | Metschnikowia | 13 | Saprotroph | \|Undefined Saprotroph\| | Yeast | NULL | Probable | As nectar-tap_saprotroph (Põlme et al. 2020); Decay sugar-rich substrates (Põlme et al. 2020) | Põlme S, et al. 2020. Fungal Diversity 105:1-16 ((https://doi.org/10.1007/s13225-020-00466-2)) |
| Otu41824 | 1 | k__Fungi(100);p__Ascomycota(100);c__Saccharomycetes(100);o__Saccharomycetales(100);f__Metschnikowiaceae(100);g__Metschnikowia(100);s__Metschnikowia_pulcherrima(100); | Metschnikowia | 13 | Saprotroph | \|Undefined Saprotroph\| | Yeast | NULL | Probable | As nectar-tap_saprotroph (Põlme et al. 2020); Decay sugar-rich substrates (Põlme et al. 2020) | Põlme S, et al. 2020. Fungal Diversity 105:1-16 ((https://doi.org/10.1007/s13225-020-00466-2)) |
| Otu41825 | 1 | k__Fungi(100);p__Ascomycota(100);c__Saccharomycetes(100);o__Saccharomycetales(100);f__Metschnikowiaceae(100);g__Metschnikowia(100);s__Metschnikowia_pulcherrima(100); | Metschnikowia | 13 | Saprotroph | \|Undefined Saprotroph\| | Yeast | NULL | Probable | As nectar-tap_saprotroph (Põlme et al. 2020); Decay sugar-rich substrates (Põlme et al. 2020) | Põlme S, et al. 2020. Fungal Diversity 105:1-16 ((https://doi.org/10.1007/s13225-020-00466-2)) |
| Otu41826 | 1 | k__Fungi(100);p__Ascomycota(100);c__Saccharomycetes(100);o__Saccharomycetales(100);f__Saccharomycetales_family_Incertae_sedis(100);g__Candida(100);s__Candida_intermedia(100); | Candida | 13 | Pathotroph-Saprotroph-Symbiotroph | Animal Pathogen-Endophyte-Endosymbiont-Epiphyte-\|Undefined Saprotroph\| | Dimorphic Yeast-Yeast | NULL | Probable | As nectar-tap_saprotroph (Põlme et al. 2020); Decay sugar-rich substrates (Põlme et al. 2020); This asexual genus is polyphyletic and contain species that function very differently depend on what they are. Caution should be taken when interpreting guilds for Candida species. | Manolakaki D, et al. 2010. Virulence. 1: 367-375 ((https://doi.org/10.4161/viru.1.5.12796)); Mekbib SB, et al. 2011. Tropical Plant Pathology 36:233-240 ((https://doi.org/10.1590/S1982-56762011000400004)); Põlme S, et al. 2020. Fungal Diversity 105:1-16 ((https://doi.org/10.1007/s13225-020-00466-2)) |
| Otu41827 | 1 | k__Fungi(100);p__Ascomycota(100);c__Saccharomycetes(100);o__Saccharomycetales(100);f__Metschnikowiaceae(100);g__Metschnikowia(100);s__Metschnikowia_pulcherrima(100); | Metschnikowia | 13 | Saprotroph | \|Undefined Saprotroph\| | Yeast | NULL | Probable | As nectar-tap_saprotroph (Põlme et al. 2020); Decay sugar-rich substrates (Põlme et al. 2020) | Põlme S, et al. 2020. Fungal Diversity 105:1-16 ((https://doi.org/10.1007/s13225-020-00466-2)) |
| Otu41828 | 1 | k__Fungi(100);p__Ascomycota(100);c__Saccharomycetes(100);o__Saccharomycetales(100);f__Metschnikowiaceae(100);g__Metschnikowia(100);g__Metschnikowia_unclassified(100); | Metschnikowia | 13 | Saprotroph | \|Undefined Saprotroph\| | Yeast | NULL | Probable | As nectar-tap_saprotroph (Põlme et al. 2020); Decay sugar-rich substrates (Põlme et al. 2020) | Põlme S, et al. 2020. Fungal Diversity 105:1-16 ((https://doi.org/10.1007/s13225-020-00466-2)) |
| Otu41829 | 1 | k__Fungi(100);p__Ascomycota(100);c__Saccharomycetes(100);o__Saccharomycetales(100);f__Metschnikowiaceae(100);g__Metschnikowia(100);s__Metschnikowia_pulcherrima(100); | Metschnikowia | 13 | Saprotroph | \|Undefined Saprotroph\| | Yeast | NULL | Probable | As nectar-tap_saprotroph (Põlme et al. 2020); Decay sugar-rich substrates (Põlme et al. 2020) | Põlme S, et al. 2020. Fungal Diversity 105:1-16 ((https://doi.org/10.1007/s13225-020-00466-2)) |
| Otu41831 | 1 | k__Fungi(100);p__Ascomycota(100);c__Saccharomycetes(100);o__Saccharomycetales(100);f__Metschnikowiaceae(100);g__Metschnikowia(100);s__Metschnikowia_pulcherrima(100); | Metschnikowia | 13 | Saprotroph | \|Undefined Saprotroph\| | Yeast | NULL | Probable | As nectar-tap_saprotroph (Põlme et al. 2020); Decay sugar-rich substrates (Põlme et al. 2020) | Põlme S, et al. 2020. Fungal Diversity 105:1-16 ((https://doi.org/10.1007/s13225-020-00466-2)) |
| Otu41832 | 1 | k__Fungi(100);p__Ascomycota(100);c__Saccharomycetes(100);o__Saccharomycetales(100);f__Metschnikowiaceae(100);g__Metschnikowia(100);g__Metschnikowia_unclassified(100); | Metschnikowia | 13 | Saprotroph | \|Undefined Saprotroph\| | Yeast | NULL | Probable | As nectar-tap_saprotroph (Põlme et al. 2020); Decay sugar-rich substrates (Põlme et al. 2020) | Põlme S, et al. 2020. Fungal Diversity 105:1-16 ((https://doi.org/10.1007/s13225-020-00466-2)) |
| Otu41833 | 1 | k__Fungi(100);p__Ascomycota(100);c__Saccharomycetes(100);o__Saccharomycetales(100);f__Metschnikowiaceae(100);g__Metschnikowia(100);g__Metschnikowia_unclassified(100); | Metschnikowia | 13 | Saprotroph | \|Undefined Saprotroph\| | Yeast | NULL | Probable | As nectar-tap_saprotroph (Põlme et al. 2020); Decay sugar-rich substrates (Põlme et al. 2020) | Põlme S, et al. 2020. Fungal Diversity 105:1-16 ((https://doi.org/10.1007/s13225-020-00466-2)) |
| Otu41834 | 1 | k__Fungi(100);p__Ascomycota(100);c__Saccharomycetes(100);o__Saccharomycetales(100);f__Metschnikowiaceae(100);g__Metschnikowia(100);g__Metschnikowia_unclassified(100); | Metschnikowia | 13 | Saprotroph | \|Undefined Saprotroph\| | Yeast | NULL | Probable | As nectar-tap_saprotroph (Põlme et al. 2020); Decay sugar-rich substrates (Põlme et al. 2020) | Põlme S, et al. 2020. Fungal Diversity 105:1-16 ((https://doi.org/10.1007/s13225-020-00466-2)) |
| Otu41835 | 1 | k__Fungi(100);p__Ascomycota(100);c__Saccharomycetes(100);o__Saccharomycetales(100);f__Metschnikowiaceae(100);g__Metschnikowia(100);g__Metschnikowia_unclassified(100); | Metschnikowia | 13 | Saprotroph | \|Undefined Saprotroph\| | Yeast | NULL | Probable | As nectar-tap_saprotroph (Põlme et al. 2020); Decay sugar-rich substrates (Põlme et al. 2020) | Põlme S, et al. 2020. Fungal Diversity 105:1-16 ((https://doi.org/10.1007/s13225-020-00466-2)) |
| Otu41837 | 1 | k__Fungi(100);p__Ascomycota(100);c__Saccharomycetes(100);o__Saccharomycetales(100);f__Metschnikowiaceae(100);g__Metschnikowia(100);g__Metschnikowia_unclassified(100); | Metschnikowia | 13 | Saprotroph | \|Undefined Saprotroph\| | Yeast | NULL | Probable | As nectar-tap_saprotroph (Põlme et al. 2020); Decay sugar-rich substrates (Põlme et al. 2020) | Põlme S, et al. 2020. Fungal Diversity 105:1-16 ((https://doi.org/10.1007/s13225-020-00466-2)) |
| Otu41838 | 1 | k__Fungi(100);p__Ascomycota(100);c__Saccharomycetes(100);o__Saccharomycetales(100);f__Metschnikowiaceae(100);g__Metschnikowia(100);s__Metschnikowia_pulcherrima(100); | Metschnikowia | 13 | Saprotroph | \|Undefined Saprotroph\| | Yeast | NULL | Probable | As nectar-tap_saprotroph (Põlme et al. 2020); Decay sugar-rich substrates (Põlme et al. 2020) | Põlme S, et al. 2020. Fungal Diversity 105:1-16 ((https://doi.org/10.1007/s13225-020-00466-2)) |
| Otu41839 | 1 | k__Fungi(100);p__Ascomycota(100);c__Saccharomycetes(100);o__Saccharomycetales(100);f__Metschnikowiaceae(100);g__Metschnikowia(100);s__Metschnikowia_pulcherrima(100); | Metschnikowia | 13 | Saprotroph | \|Undefined Saprotroph\| | Yeast | NULL | Probable | As nectar-tap_saprotroph (Põlme et al. 2020); Decay sugar-rich substrates (Põlme et al. 2020) | Põlme S, et al. 2020. Fungal Diversity 105:1-16 ((https://doi.org/10.1007/s13225-020-00466-2)) |
| Otu41840 | 1 | k__Fungi(100);p__Ascomycota(100);c__Saccharomycetes(100);o__Saccharomycetales(100);f__Metschnikowiaceae(100);g__Metschnikowia(100);s__Metschnikowia_pulcherrima(100); | Metschnikowia | 13 | Saprotroph | \|Undefined Saprotroph\| | Yeast | NULL | Probable | As nectar-tap_saprotroph (Põlme et al. 2020); Decay sugar-rich substrates (Põlme et al. 2020) | Põlme S, et al. 2020. Fungal Diversity 105:1-16 ((https://doi.org/10.1007/s13225-020-00466-2)) |
| Otu41841 | 1 | k__Fungi(100);p__Ascomycota(100);c__Saccharomycetes(100);o__Saccharomycetales(100);f__Metschnikowiaceae(100);g__Metschnikowia(100);s__Metschnikowia_pulcherrima(100); | Metschnikowia | 13 | Saprotroph | \|Undefined Saprotroph\| | Yeast | NULL | Probable | As nectar-tap_saprotroph (Põlme et al. 2020); Decay sugar-rich substrates (Põlme et al. 2020) | Põlme S, et al. 2020. Fungal Diversity 105:1-16 ((https://doi.org/10.1007/s13225-020-00466-2)) |
| Otu41842 | 1 | k__Fungi(100);p__Ascomycota(100);c__Saccharomycetes(100);o__Saccharomycetales(100);f__Metschnikowiaceae(100);g__Metschnikowia(100);s__Metschnikowia_pulcherrima(100); | Metschnikowia | 13 | Saprotroph | \|Undefined Saprotroph\| | Yeast | NULL | Probable | As nectar-tap_saprotroph (Põlme et al. 2020); Decay sugar-rich substrates (Põlme et al. 2020) | Põlme S, et al. 2020. Fungal Diversity 105:1-16 ((https://doi.org/10.1007/s13225-020-00466-2)) |
| Otu41843 | 1 | k__Fungi(100);p__Ascomycota(100);c__Saccharomycetes(100);o__Saccharomycetales(100);f__Metschnikowiaceae(100);g__Metschnikowia(100);s__Metschnikowia_pulcherrima(100); | Metschnikowia | 13 | Saprotroph | \|Undefined Saprotroph\| | Yeast | NULL | Probable | As nectar-tap_saprotroph (Põlme et al. 2020); Decay sugar-rich substrates (Põlme et al. 2020) | Põlme S, et al. 2020. Fungal Diversity 105:1-16 ((https://doi.org/10.1007/s13225-020-00466-2)) |
| Otu41844 | 1 | k__Fungi(100);p__Ascomycota(100);c__Saccharomycetes(100);o__Saccharomycetales(100);f__Metschnikowiaceae(100);g__Metschnikowia(100);s__Metschnikowia_pulcherrima(100); | Metschnikowia | 13 | Saprotroph | \|Undefined Saprotroph\| | Yeast | NULL | Probable | As nectar-tap_saprotroph (Põlme et al. 2020); Decay sugar-rich substrates (Põlme et al. 2020) | Põlme S, et al. 2020. Fungal Diversity 105:1-16 ((https://doi.org/10.1007/s13225-020-00466-2)) |
| Otu41845 | 1 | k__Fungi(100);p__Ascomycota(100);c__Saccharomycetes(100);o__Saccharomycetales(100);f__Metschnikowiaceae(100);g__Metschnikowia(100);s__Metschnikowia_pulcherrima(100); | Metschnikowia | 13 | Saprotroph | \|Undefined Saprotroph\| | Yeast | NULL | Probable | As nectar-tap_saprotroph (Põlme et al. 2020); Decay sugar-rich substrates (Põlme et al. 2020) | Põlme S, et al. 2020. Fungal Diversity 105:1-16 ((https://doi.org/10.1007/s13225-020-00466-2)) |
| Otu41846 | 1 | k__Fungi(100);p__Ascomycota(100);c__Saccharomycetes(100);o__Saccharomycetales(100);f__Metschnikowiaceae(100);g__Metschnikowia(100);s__Metschnikowia_pulcherrima(100); | Metschnikowia | 13 | Saprotroph | \|Undefined Saprotroph\| | Yeast | NULL | Probable | As nectar-tap_saprotroph (Põlme et al. 2020); Decay sugar-rich substrates (Põlme et al. 2020) | Põlme S, et al. 2020. Fungal Diversity 105:1-16 ((https://doi.org/10.1007/s13225-020-00466-2)) |
| Otu41847 | 1 | k__Fungi(100);p__Ascomycota(100);c__Saccharomycetes(100);o__Saccharomycetales(100);f__Metschnikowiaceae(100);g__Metschnikowia(100);s__Metschnikowia_pulcherrima(100); | Metschnikowia | 13 | Saprotroph | \|Undefined Saprotroph\| | Yeast | NULL | Probable | As nectar-tap_saprotroph (Põlme et al. 2020); Decay sugar-rich substrates (Põlme et al. 2020) | Põlme S, et al. 2020. Fungal Diversity 105:1-16 ((https://doi.org/10.1007/s13225-020-00466-2)) |
| Otu41848 | 1 | k__Fungi(100);p__Ascomycota(100);c__Saccharomycetes(100);o__Saccharomycetales(100);f__Metschnikowiaceae(100);g__Metschnikowia(100);s__Metschnikowia_pulcherrima(100); | Metschnikowia | 13 | Saprotroph | \|Undefined Saprotroph\| | Yeast | NULL | Probable | As nectar-tap_saprotroph (Põlme et al. 2020); Decay sugar-rich substrates (Põlme et al. 2020) | Põlme S, et al. 2020. Fungal Diversity 105:1-16 ((https://doi.org/10.1007/s13225-020-00466-2)) |
| Otu41849 | 1 | k__Fungi(100);p__Ascomycota(100);c__Saccharomycetes(100);o__Saccharomycetales(100);f__Metschnikowiaceae(100);g__Metschnikowia(100);g__Metschnikowia_unclassified(100); | Metschnikowia | 13 | Saprotroph | \|Undefined Saprotroph\| | Yeast | NULL | Probable | As nectar-tap_saprotroph (Põlme et al. 2020); Decay sugar-rich substrates (Põlme et al. 2020) | Põlme S, et al. 2020. Fungal Diversity 105:1-16 ((https://doi.org/10.1007/s13225-020-00466-2)) |
| Otu41850 | 1 | k__Fungi(100);p__Ascomycota(100);c__Saccharomycetes(100);o__Saccharomycetales(100);f__Metschnikowiaceae(100);g__Metschnikowia(100);s__Metschnikowia_pulcherrima(100); | Metschnikowia | 13 | Saprotroph | \|Undefined Saprotroph\| | Yeast | NULL | Probable | As nectar-tap_saprotroph (Põlme et al. 2020); Decay sugar-rich substrates (Põlme et al. 2020) | Põlme S, et al. 2020. Fungal Diversity 105:1-16 ((https://doi.org/10.1007/s13225-020-00466-2)) |
| Otu41851 | 1 | k__Fungi(100);p__Ascomycota(100);c__Saccharomycetes(100);o__Saccharomycetales(100);f__Metschnikowiaceae(100);g__Metschnikowia(100);s__Metschnikowia_pulcherrima(100); | Metschnikowia | 13 | Saprotroph | \|Undefined Saprotroph\| | Yeast | NULL | Probable | As nectar-tap_saprotroph (Põlme et al. 2020); Decay sugar-rich substrates (Põlme et al. 2020) | Põlme S, et al. 2020. Fungal Diversity 105:1-16 ((https://doi.org/10.1007/s13225-020-00466-2)) |
| Otu41852 | 1 | k__Fungi(100);p__Ascomycota(100);c__Saccharomycetes(100);o__Saccharomycetales(100);f__Metschnikowiaceae(100);g__Metschnikowia(100);s__Metschnikowia_pulcherrima(100); | Metschnikowia | 13 | Saprotroph | \|Undefined Saprotroph\| | Yeast | NULL | Probable | As nectar-tap_saprotroph (Põlme et al. 2020); Decay sugar-rich substrates (Põlme et al. 2020) | Põlme S, et al. 2020. Fungal Diversity 105:1-16 ((https://doi.org/10.1007/s13225-020-00466-2)) |
| Otu41853 | 1 | k__Fungi(100);p__Ascomycota(100);c__Saccharomycetes(100);o__Saccharomycetales(100);f__Metschnikowiaceae(100);g__Metschnikowia(100);s__Metschnikowia_sp(100); | Metschnikowia | 13 | Saprotroph | \|Undefined Saprotroph\| | Yeast | NULL | Probable | As nectar-tap_saprotroph (Põlme et al. 2020); Decay sugar-rich substrates (Põlme et al. 2020) | Põlme S, et al. 2020. Fungal Diversity 105:1-16 ((https://doi.org/10.1007/s13225-020-00466-2)) |
| Otu41854 | 1 | k__Fungi(100);p__Ascomycota(100);c__Saccharomycetes(100);o__Saccharomycetales(100);f__Metschnikowiaceae(100);g__Metschnikowia(100);g__Metschnikowia_unclassified(100); | Metschnikowia | 13 | Saprotroph | \|Undefined Saprotroph\| | Yeast | NULL | Probable | As nectar-tap_saprotroph (Põlme et al. 2020); Decay sugar-rich substrates (Põlme et al. 2020) | Põlme S, et al. 2020. Fungal Diversity 105:1-16 ((https://doi.org/10.1007/s13225-020-00466-2)) |
| Otu41855 | 1 | k__Fungi(100);p__Ascomycota(100);c__Saccharomycetes(100);o__Saccharomycetales(100);f__Metschnikowiaceae(100);g__Metschnikowia(100);g__Metschnikowia_unclassified(100); | Metschnikowia | 13 | Saprotroph | \|Undefined Saprotroph\| | Yeast | NULL | Probable | As nectar-tap_saprotroph (Põlme et al. 2020); Decay sugar-rich substrates (Põlme et al. 2020) | Põlme S, et al. 2020. Fungal Diversity 105:1-16 ((https://doi.org/10.1007/s13225-020-00466-2)) |
| Otu41856 | 1 | k__Fungi(100);p__Ascomycota(100);c__Saccharomycetes(100);o__Saccharomycetales(100);f__Metschnikowiaceae(100);g__Metschnikowia(100);g__Metschnikowia_unclassified(100); | Metschnikowia | 13 | Saprotroph | \|Undefined Saprotroph\| | Yeast | NULL | Probable | As nectar-tap_saprotroph (Põlme et al. 2020); Decay sugar-rich substrates (Põlme et al. 2020) | Põlme S, et al. 2020. Fungal Diversity 105:1-16 ((https://doi.org/10.1007/s13225-020-00466-2)) |
| Otu41857 | 1 | k__Fungi(100);p__Ascomycota(100);c__Saccharomycetes(100);o__Saccharomycetales(100);f__Metschnikowiaceae(100);g__Metschnikowia(100);s__Metschnikowia_pulcherrima(100); | Metschnikowia | 13 | Saprotroph | \|Undefined Saprotroph\| | Yeast | NULL | Probable | As nectar-tap_saprotroph (Põlme et al. 2020); Decay sugar-rich substrates (Põlme et al. 2020) | Põlme S, et al. 2020. Fungal Diversity 105:1-16 ((https://doi.org/10.1007/s13225-020-00466-2)) |
| Otu41858 | 1 | k__Fungi(100);p__Ascomycota(100);c__Saccharomycetes(100);o__Saccharomycetales(100);f__Metschnikowiaceae(100);g__Metschnikowia(100);g__Metschnikowia_unclassified(100); | Metschnikowia | 13 | Saprotroph | \|Undefined Saprotroph\| | Yeast | NULL | Probable | As nectar-tap_saprotroph (Põlme et al. 2020); Decay sugar-rich substrates (Põlme et al. 2020) | Põlme S, et al. 2020. Fungal Diversity 105:1-16 ((https://doi.org/10.1007/s13225-020-00466-2)) |
| Otu41859 | 1 | k__Fungi(100);p__Ascomycota(100);c__Saccharomycetes(100);o__Saccharomycetales(100);f__Metschnikowiaceae(100);g__Metschnikowia(100);s__Metschnikowia_pulcherrima(100); | Metschnikowia | 13 | Saprotroph | \|Undefined Saprotroph\| | Yeast | NULL | Probable | As nectar-tap_saprotroph (Põlme et al. 2020); Decay sugar-rich substrates (Põlme et al. 2020) | Põlme S, et al. 2020. Fungal Diversity 105:1-16 ((https://doi.org/10.1007/s13225-020-00466-2)) |
| Otu41860 | 1 | k__Fungi(100);p__Ascomycota(100);c__Saccharomycetes(100);o__Saccharomycetales(100);f__Metschnikowiaceae(100);g__Metschnikowia(100);s__Metschnikowia_pulcherrima(100); | Metschnikowia | 13 | Saprotroph | \|Undefined Saprotroph\| | Yeast | NULL | Probable | As nectar-tap_saprotroph (Põlme et al. 2020); Decay sugar-rich substrates (Põlme et al. 2020) | Põlme S, et al. 2020. Fungal Diversity 105:1-16 ((https://doi.org/10.1007/s13225-020-00466-2)) |
| Otu41861 | 1 | k__Fungi(100);p__Ascomycota(100);c__Saccharomycetes(100);o__Saccharomycetales(100);f__Metschnikowiaceae(100);g__Metschnikowia(100);s__Metschnikowia_pulcherrima(100); | Metschnikowia | 13 | Saprotroph | \|Undefined Saprotroph\| | Yeast | NULL | Probable | As nectar-tap_saprotroph (Põlme et al. 2020); Decay sugar-rich substrates (Põlme et al. 2020) | Põlme S, et al. 2020. Fungal Diversity 105:1-16 ((https://doi.org/10.1007/s13225-020-00466-2)) |
| Otu41862 | 1 | k__Fungi(100);p__Ascomycota(100);c__Saccharomycetes(100);o__Saccharomycetales(100);f__Metschnikowiaceae(100);g__Metschnikowia(100);s__Metschnikowia_pulcherrima(100); | Metschnikowia | 13 | Saprotroph | \|Undefined Saprotroph\| | Yeast | NULL | Probable | As nectar-tap_saprotroph (Põlme et al. 2020); Decay sugar-rich substrates (Põlme et al. 2020) | Põlme S, et al. 2020. Fungal Diversity 105:1-16 ((https://doi.org/10.1007/s13225-020-00466-2)) |
| Otu41863 | 1 | k__Fungi(100);p__Ascomycota(100);c__Saccharomycetes(100);o__Saccharomycetales(100);f__Metschnikowiaceae(100);g__Metschnikowia(100);g__Metschnikowia_unclassified(100); | Metschnikowia | 13 | Saprotroph | \|Undefined Saprotroph\| | Yeast | NULL | Probable | As nectar-tap_saprotroph (Põlme et al. 2020); Decay sugar-rich substrates (Põlme et al. 2020) | Põlme S, et al. 2020. Fungal Diversity 105:1-16 ((https://doi.org/10.1007/s13225-020-00466-2)) |
| Otu41864 | 1 | k__Fungi(100);p__Ascomycota(100);c__Saccharomycetes(100);o__Saccharomycetales(100);f__Metschnikowiaceae(100);g__Metschnikowia(100);g__Metschnikowia_unclassified(100); | Metschnikowia | 13 | Saprotroph | \|Undefined Saprotroph\| | Yeast | NULL | Probable | As nectar-tap_saprotroph (Põlme et al. 2020); Decay sugar-rich substrates (Põlme et al. 2020) | Põlme S, et al. 2020. Fungal Diversity 105:1-16 ((https://doi.org/10.1007/s13225-020-00466-2)) |
| Otu41865 | 1 | k__Fungi(100);p__Ascomycota(100);c__Saccharomycetes(100);o__Saccharomycetales(100);f__Metschnikowiaceae(100);g__Metschnikowia(100);g__Metschnikowia_unclassified(100); | Metschnikowia | 13 | Saprotroph | \|Undefined Saprotroph\| | Yeast | NULL | Probable | As nectar-tap_saprotroph (Põlme et al. 2020); Decay sugar-rich substrates (Põlme et al. 2020) | Põlme S, et al. 2020. Fungal Diversity 105:1-16 ((https://doi.org/10.1007/s13225-020-00466-2)) |
| Otu41866 | 1 | k__Fungi(100);p__Ascomycota(100);c__Saccharomycetes(100);o__Saccharomycetales(100);f__Metschnikowiaceae(100);g__Metschnikowia(100);s__Metschnikowia_pulcherrima(100); | Metschnikowia | 13 | Saprotroph | \|Undefined Saprotroph\| | Yeast | NULL | Probable | As nectar-tap_saprotroph (Põlme et al. 2020); Decay sugar-rich substrates (Põlme et al. 2020) | Põlme S, et al. 2020. Fungal Diversity 105:1-16 ((https://doi.org/10.1007/s13225-020-00466-2)) |
| Otu41867 | 1 | k__Fungi(100);p__Ascomycota(100);c__Saccharomycetes(100);o__Saccharomycetales(100);f__Metschnikowiaceae(100);g__Metschnikowia(100);s__Metschnikowia_pulcherrima(100); | Metschnikowia | 13 | Saprotroph | \|Undefined Saprotroph\| | Yeast | NULL | Probable | As nectar-tap_saprotroph (Põlme et al. 2020); Decay sugar-rich substrates (Põlme et al. 2020) | Põlme S, et al. 2020. Fungal Diversity 105:1-16 ((https://doi.org/10.1007/s13225-020-00466-2)) |
| Otu41868 | 1 | k__Fungi(100);p__Ascomycota(100);c__Saccharomycetes(100);o__Saccharomycetales(100);f__Metschnikowiaceae(100);g__Metschnikowia(100);s__Metschnikowia_pulcherrima(100); | Metschnikowia | 13 | Saprotroph | \|Undefined Saprotroph\| | Yeast | NULL | Probable | As nectar-tap_saprotroph (Põlme et al. 2020); Decay sugar-rich substrates (Põlme et al. 2020) | Põlme S, et al. 2020. Fungal Diversity 105:1-16 ((https://doi.org/10.1007/s13225-020-00466-2)) |
| Otu41869 | 1 | k__Fungi(100);p__Ascomycota(100);c__Saccharomycetes(100);o__Saccharomycetales(100);f__Metschnikowiaceae(100);g__Metschnikowia(100);s__Metschnikowia_pulcherrima(100); | Metschnikowia | 13 | Saprotroph | \|Undefined Saprotroph\| | Yeast | NULL | Probable | As nectar-tap_saprotroph (Põlme et al. 2020); Decay sugar-rich substrates (Põlme et al. 2020) | Põlme S, et al. 2020. Fungal Diversity 105:1-16 ((https://doi.org/10.1007/s13225-020-00466-2)) |
| Otu41870 | 1 | k__Fungi(100);p__Ascomycota(100);c__Saccharomycetes(100);o__Saccharomycetales(100);f__Metschnikowiaceae(100);g__Metschnikowia(100);s__Metschnikowia_pulcherrima(100); | Metschnikowia | 13 | Saprotroph | \|Undefined Saprotroph\| | Yeast | NULL | Probable | As nectar-tap_saprotroph (Põlme et al. 2020); Decay sugar-rich substrates (Põlme et al. 2020) | Põlme S, et al. 2020. Fungal Diversity 105:1-16 ((https://doi.org/10.1007/s13225-020-00466-2)) |
| Otu41871 | 1 | k__Fungi(100);p__Ascomycota(100);c__Saccharomycetes(100);o__Saccharomycetales(100);f__Metschnikowiaceae(100);g__Metschnikowia(100);s__Metschnikowia_pulcherrima(100); | Metschnikowia | 13 | Saprotroph | \|Undefined Saprotroph\| | Yeast | NULL | Probable | As nectar-tap_saprotroph (Põlme et al. 2020); Decay sugar-rich substrates (Põlme et al. 2020) | Põlme S, et al. 2020. Fungal Diversity 105:1-16 ((https://doi.org/10.1007/s13225-020-00466-2)) |
| Otu41873 | 1 | k__Fungi(100);p__Ascomycota(100);c__Saccharomycetes(100);o__Saccharomycetales(100);f__Metschnikowiaceae(100);g__Metschnikowia(100);s__Metschnikowia_pulcherrima(100); | Metschnikowia | 13 | Saprotroph | \|Undefined Saprotroph\| | Yeast | NULL | Probable | As nectar-tap_saprotroph (Põlme et al. 2020); Decay sugar-rich substrates (Põlme et al. 2020) | Põlme S, et al. 2020. Fungal Diversity 105:1-16 ((https://doi.org/10.1007/s13225-020-00466-2)) |
| Otu41874 | 1 | k__Fungi(100);p__Ascomycota(100);c__Saccharomycetes(100);o__Saccharomycetales(100);f__Metschnikowiaceae(100);g__Metschnikowia(100);s__Metschnikowia_pulcherrima(100); | Metschnikowia | 13 | Saprotroph | \|Undefined Saprotroph\| | Yeast | NULL | Probable | As nectar-tap_saprotroph (Põlme et al. 2020); Decay sugar-rich substrates (Põlme et al. 2020) | Põlme S, et al. 2020. Fungal Diversity 105:1-16 ((https://doi.org/10.1007/s13225-020-00466-2)) |
| Otu41875 | 1 | k__Fungi(100);p__Ascomycota(100);c__Saccharomycetes(100);o__Saccharomycetales(100);f__Metschnikowiaceae(100);g__Metschnikowia(100);g__Metschnikowia_unclassified(100); | Metschnikowia | 13 | Saprotroph | \|Undefined Saprotroph\| | Yeast | NULL | Probable | As nectar-tap_saprotroph (Põlme et al. 2020); Decay sugar-rich substrates (Põlme et al. 2020) | Põlme S, et al. 2020. Fungal Diversity 105:1-16 ((https://doi.org/10.1007/s13225-020-00466-2)) |
| Otu41876 | 1 | k__Fungi(100);p__Ascomycota(100);c__Saccharomycetes(100);o__Saccharomycetales(100);f__Metschnikowiaceae(100);g__Metschnikowia(100);s__Metschnikowia_pulcherrima(100); | Metschnikowia | 13 | Saprotroph | \|Undefined Saprotroph\| | Yeast | NULL | Probable | As nectar-tap_saprotroph (Põlme et al. 2020); Decay sugar-rich substrates (Põlme et al. 2020) | Põlme S, et al. 2020. Fungal Diversity 105:1-16 ((https://doi.org/10.1007/s13225-020-00466-2)) |
| Otu41877 | 1 | k__Fungi(100);p__Ascomycota(100);c__Saccharomycetes(100);o__Saccharomycetales(100);f__Metschnikowiaceae(100);g__Metschnikowia(100);s__Metschnikowia_pulcherrima(100); | Metschnikowia | 13 | Saprotroph | \|Undefined Saprotroph\| | Yeast | NULL | Probable | As nectar-tap_saprotroph (Põlme et al. 2020); Decay sugar-rich substrates (Põlme et al. 2020) | Põlme S, et al. 2020. Fungal Diversity 105:1-16 ((https://doi.org/10.1007/s13225-020-00466-2)) |
| Otu41878 | 1 | k__Fungi(100);p__Ascomycota(100);c__Saccharomycetes(100);o__Saccharomycetales(100);f__Metschnikowiaceae(100);g__Metschnikowia(100);s__Metschnikowia_pulcherrima(100); | Metschnikowia | 13 | Saprotroph | \|Undefined Saprotroph\| | Yeast | NULL | Probable | As nectar-tap_saprotroph (Põlme et al. 2020); Decay sugar-rich substrates (Põlme et al. 2020) | Põlme S, et al. 2020. Fungal Diversity 105:1-16 ((https://doi.org/10.1007/s13225-020-00466-2)) |
| Otu41879 | 1 | k__Fungi(100);p__Ascomycota(100);c__Saccharomycetes(100);o__Saccharomycetales(100);f__Metschnikowiaceae(100);g__Metschnikowia(100);s__Metschnikowia_pulcherrima(100); | Metschnikowia | 13 | Saprotroph | \|Undefined Saprotroph\| | Yeast | NULL | Probable | As nectar-tap_saprotroph (Põlme et al. 2020); Decay sugar-rich substrates (Põlme et al. 2020) | Põlme S, et al. 2020. Fungal Diversity 105:1-16 ((https://doi.org/10.1007/s13225-020-00466-2)) |
| Otu41880 | 1 | k__Fungi(100);p__Ascomycota(100);c__Saccharomycetes(100);o__Saccharomycetales(100);f__Metschnikowiaceae(100);g__Metschnikowia(100);s__Metschnikowia_pulcherrima(100); | Metschnikowia | 13 | Saprotroph | \|Undefined Saprotroph\| | Yeast | NULL | Probable | As nectar-tap_saprotroph (Põlme et al. 2020); Decay sugar-rich substrates (Põlme et al. 2020) | Põlme S, et al. 2020. Fungal Diversity 105:1-16 ((https://doi.org/10.1007/s13225-020-00466-2)) |
| Otu41881 | 1 | k__Fungi(100);p__Ascomycota(100);c__Saccharomycetes(100);o__Saccharomycetales(100);f__Metschnikowiaceae(100);g__Metschnikowia(100);s__Metschnikowia_pulcherrima(100); | Metschnikowia | 13 | Saprotroph | \|Undefined Saprotroph\| | Yeast | NULL | Probable | As nectar-tap_saprotroph (Põlme et al. 2020); Decay sugar-rich substrates (Põlme et al. 2020) | Põlme S, et al. 2020. Fungal Diversity 105:1-16 ((https://doi.org/10.1007/s13225-020-00466-2)) |
| Otu41882 | 1 | k__Fungi(100);p__Ascomycota(100);c__Saccharomycetes(100);o__Saccharomycetales(100);f__Metschnikowiaceae(100);g__Metschnikowia(100);s__Metschnikowia_pulcherrima(100); | Metschnikowia | 13 | Saprotroph | \|Undefined Saprotroph\| | Yeast | NULL | Probable | As nectar-tap_saprotroph (Põlme et al. 2020); Decay sugar-rich substrates (Põlme et al. 2020) | Põlme S, et al. 2020. Fungal Diversity 105:1-16 ((https://doi.org/10.1007/s13225-020-00466-2)) |
| Otu41883 | 1 | k__Fungi(100);p__Ascomycota(100);c__Saccharomycetes(100);o__Saccharomycetales(100);f__Metschnikowiaceae(100);g__Metschnikowia(100);s__Metschnikowia_pulcherrima(100); | Metschnikowia | 13 | Saprotroph | \|Undefined Saprotroph\| | Yeast | NULL | Probable | As nectar-tap_saprotroph (Põlme et al. 2020); Decay sugar-rich substrates (Põlme et al. 2020) | Põlme S, et al. 2020. Fungal Diversity 105:1-16 ((https://doi.org/10.1007/s13225-020-00466-2)) |
| Otu41884 | 1 | k__Fungi(100);p__Ascomycota(100);c__Saccharomycetes(100);o__Saccharomycetales(100);f__Metschnikowiaceae(100);g__Metschnikowia(100);s__Metschnikowia_pulcherrima(100); | Metschnikowia | 13 | Saprotroph | \|Undefined Saprotroph\| | Yeast | NULL | Probable | As nectar-tap_saprotroph (Põlme et al. 2020); Decay sugar-rich substrates (Põlme et al. 2020) | Põlme S, et al. 2020. Fungal Diversity 105:1-16 ((https://doi.org/10.1007/s13225-020-00466-2)) |
| Otu41885 | 1 | k__Fungi(100);p__Ascomycota(100);c__Saccharomycetes(100);o__Saccharomycetales(100);f__Metschnikowiaceae(100);g__Metschnikowia(100);g__Metschnikowia_unclassified(100); | Metschnikowia | 13 | Saprotroph | \|Undefined Saprotroph\| | Yeast | NULL | Probable | As nectar-tap_saprotroph (Põlme et al. 2020); Decay sugar-rich substrates (Põlme et al. 2020) | Põlme S, et al. 2020. Fungal Diversity 105:1-16 ((https://doi.org/10.1007/s13225-020-00466-2)) |
| Otu41886 | 1 | k__Fungi(100);p__Ascomycota(100);c__Saccharomycetes(100);o__Saccharomycetales(100);f__Metschnikowiaceae(100);g__Metschnikowia(100);s__Metschnikowia_pulcherrima(100); | Metschnikowia | 13 | Saprotroph | \|Undefined Saprotroph\| | Yeast | NULL | Probable | As nectar-tap_saprotroph (Põlme et al. 2020); Decay sugar-rich substrates (Põlme et al. 2020) | Põlme S, et al. 2020. Fungal Diversity 105:1-16 ((https://doi.org/10.1007/s13225-020-00466-2)) |
| Otu41887 | 1 | k__Fungi(100);p__Ascomycota(100);c__Saccharomycetes(100);o__Saccharomycetales(100);f__Metschnikowiaceae(100);g__Metschnikowia(100);s__Metschnikowia_pulcherrima(100); | Metschnikowia | 13 | Saprotroph | \|Undefined Saprotroph\| | Yeast | NULL | Probable | As nectar-tap_saprotroph (Põlme et al. 2020); Decay sugar-rich substrates (Põlme et al. 2020) | Põlme S, et al. 2020. Fungal Diversity 105:1-16 ((https://doi.org/10.1007/s13225-020-00466-2)) |
| Otu41888 | 1 | k__Fungi(100);p__Ascomycota(100);c__Saccharomycetes(100);o__Saccharomycetales(100);f__Metschnikowiaceae(100);g__Metschnikowia(100);g__Metschnikowia_unclassified(100); | Metschnikowia | 13 | Saprotroph | \|Undefined Saprotroph\| | Yeast | NULL | Probable | As nectar-tap_saprotroph (Põlme et al. 2020); Decay sugar-rich substrates (Põlme et al. 2020) | Põlme S, et al. 2020. Fungal Diversity 105:1-16 ((https://doi.org/10.1007/s13225-020-00466-2)) |
| Otu41889 | 1 | k__Fungi(100);p__Ascomycota(100);c__Saccharomycetes(100);o__Saccharomycetales(100);f__Metschnikowiaceae(100);g__Metschnikowia(100);s__Metschnikowia_pulcherrima(100); | Metschnikowia | 13 | Saprotroph | \|Undefined Saprotroph\| | Yeast | NULL | Probable | As nectar-tap_saprotroph (Põlme et al. 2020); Decay sugar-rich substrates (Põlme et al. 2020) | Põlme S, et al. 2020. Fungal Diversity 105:1-16 ((https://doi.org/10.1007/s13225-020-00466-2)) |
| Otu41890 | 1 | k__Fungi(100);p__Ascomycota(100);c__Saccharomycetes(100);o__Saccharomycetales(100);f__Metschnikowiaceae(100);g__Metschnikowia(100);g__Metschnikowia_unclassified(100); | Metschnikowia | 13 | Saprotroph | \|Undefined Saprotroph\| | Yeast | NULL | Probable | As nectar-tap_saprotroph (Põlme et al. 2020); Decay sugar-rich substrates (Põlme et al. 2020) | Põlme S, et al. 2020. Fungal Diversity 105:1-16 ((https://doi.org/10.1007/s13225-020-00466-2)) |
| Otu41891 | 1 | k__Fungi(100);p__Ascomycota(100);c__Saccharomycetes(100);o__Saccharomycetales(100);f__Metschnikowiaceae(100);g__Metschnikowia(100);s__Metschnikowia_pulcherrima(100); | Metschnikowia | 13 | Saprotroph | \|Undefined Saprotroph\| | Yeast | NULL | Probable | As nectar-tap_saprotroph (Põlme et al. 2020); Decay sugar-rich substrates (Põlme et al. 2020) | Põlme S, et al. 2020. Fungal Diversity 105:1-16 ((https://doi.org/10.1007/s13225-020-00466-2)) |
| Otu41892 | 1 | k__Fungi(100);p__Ascomycota(100);c__Saccharomycetes(100);o__Saccharomycetales(100);f__Metschnikowiaceae(100);g__Metschnikowia(100);g__Metschnikowia_unclassified(100); | Metschnikowia | 13 | Saprotroph | \|Undefined Saprotroph\| | Yeast | NULL | Probable | As nectar-tap_saprotroph (Põlme et al. 2020); Decay sugar-rich substrates (Põlme et al. 2020) | Põlme S, et al. 2020. Fungal Diversity 105:1-16 ((https://doi.org/10.1007/s13225-020-00466-2)) |
| Otu41893 | 1 | k__Fungi(100);p__Ascomycota(100);c__Saccharomycetes(100);o__Saccharomycetales(100);f__Metschnikowiaceae(100);g__Metschnikowia(100);g__Metschnikowia_unclassified(100); | Metschnikowia | 13 | Saprotroph | \|Undefined Saprotroph\| | Yeast | NULL | Probable | As nectar-tap_saprotroph (Põlme et al. 2020); Decay sugar-rich substrates (Põlme et al. 2020) | Põlme S, et al. 2020. Fungal Diversity 105:1-16 ((https://doi.org/10.1007/s13225-020-00466-2)) |
| Otu41894 | 1 | k__Fungi(100);p__Ascomycota(100);c__Saccharomycetes(100);o__Saccharomycetales(100);f__Metschnikowiaceae(100);g__Metschnikowia(100);s__Metschnikowia_pulcherrima(100); | Metschnikowia | 13 | Saprotroph | \|Undefined Saprotroph\| | Yeast | NULL | Probable | As nectar-tap_saprotroph (Põlme et al. 2020); Decay sugar-rich substrates (Põlme et al. 2020) | Põlme S, et al. 2020. Fungal Diversity 105:1-16 ((https://doi.org/10.1007/s13225-020-00466-2)) |
| Otu41895 | 1 | k__Fungi(100);p__Ascomycota(100);c__Saccharomycetes(100);o__Saccharomycetales(100);f__Metschnikowiaceae(100);g__Metschnikowia(100);g__Metschnikowia_unclassified(100); | Metschnikowia | 13 | Saprotroph | \|Undefined Saprotroph\| | Yeast | NULL | Probable | As nectar-tap_saprotroph (Põlme et al. 2020); Decay sugar-rich substrates (Põlme et al. 2020) | Põlme S, et al. 2020. Fungal Diversity 105:1-16 ((https://doi.org/10.1007/s13225-020-00466-2)) |
| Otu41896 | 1 | k__Fungi(100);p__Ascomycota(100);c__Saccharomycetes(100);o__Saccharomycetales(100);f__Metschnikowiaceae(100);g__Metschnikowia(100);s__Metschnikowia_pulcherrima(100); | Metschnikowia | 13 | Saprotroph | \|Undefined Saprotroph\| | Yeast | NULL | Probable | As nectar-tap_saprotroph (Põlme et al. 2020); Decay sugar-rich substrates (Põlme et al. 2020) | Põlme S, et al. 2020. Fungal Diversity 105:1-16 ((https://doi.org/10.1007/s13225-020-00466-2)) |
| Otu41897 | 1 | k__Fungi(100);p__Ascomycota(100);c__Saccharomycetes(100);o__Saccharomycetales(100);f__Metschnikowiaceae(100);g__Metschnikowia(100);s__Metschnikowia_pulcherrima(100); | Metschnikowia | 13 | Saprotroph | \|Undefined Saprotroph\| | Yeast | NULL | Probable | As nectar-tap_saprotroph (Põlme et al. 2020); Decay sugar-rich substrates (Põlme et al. 2020) | Põlme S, et al. 2020. Fungal Diversity 105:1-16 ((https://doi.org/10.1007/s13225-020-00466-2)) |
| Otu41898 | 1 | k__Fungi(100);p__Ascomycota(100);c__Saccharomycetes(100);o__Saccharomycetales(100);f__Metschnikowiaceae(100);g__Metschnikowia(100);g__Metschnikowia_unclassified(100); | Metschnikowia | 13 | Saprotroph | \|Undefined Saprotroph\| | Yeast | NULL | Probable | As nectar-tap_saprotroph (Põlme et al. 2020); Decay sugar-rich substrates (Põlme et al. 2020) | Põlme S, et al. 2020. Fungal Diversity 105:1-16 ((https://doi.org/10.1007/s13225-020-00466-2)) |
| Otu41899 | 1 | k__Fungi(100);p__Ascomycota(100);c__Saccharomycetes(100);o__Saccharomycetales(100);f__Metschnikowiaceae(100);g__Metschnikowia(100);s__Metschnikowia_pulcherrima(100); | Metschnikowia | 13 | Saprotroph | \|Undefined Saprotroph\| | Yeast | NULL | Probable | As nectar-tap_saprotroph (Põlme et al. 2020); Decay sugar-rich substrates (Põlme et al. 2020) | Põlme S, et al. 2020. Fungal Diversity 105:1-16 ((https://doi.org/10.1007/s13225-020-00466-2)) |
| Otu41900 | 1 | k__Fungi(100);p__Ascomycota(100);c__Saccharomycetes(100);o__Saccharomycetales(100);f__Metschnikowiaceae(100);g__Metschnikowia(100);s__Metschnikowia_pulcherrima(100); | Metschnikowia | 13 | Saprotroph | \|Undefined Saprotroph\| | Yeast | NULL | Probable | As nectar-tap_saprotroph (Põlme et al. 2020); Decay sugar-rich substrates (Põlme et al. 2020) | Põlme S, et al. 2020. Fungal Diversity 105:1-16 ((https://doi.org/10.1007/s13225-020-00466-2)) |
| Otu41901 | 1 | k__Fungi(100);p__Ascomycota(100);c__Saccharomycetes(100);o__Saccharomycetales(100);f__Metschnikowiaceae(100);g__Metschnikowia(100);s__Metschnikowia_pulcherrima(100); | Metschnikowia | 13 | Saprotroph | \|Undefined Saprotroph\| | Yeast | NULL | Probable | As nectar-tap_saprotroph (Põlme et al. 2020); Decay sugar-rich substrates (Põlme et al. 2020) | Põlme S, et al. 2020. Fungal Diversity 105:1-16 ((https://doi.org/10.1007/s13225-020-00466-2)) |
| Otu41902 | 1 | k__Fungi(100);p__Ascomycota(100);c__Saccharomycetes(100);o__Saccharomycetales(100);f__Metschnikowiaceae(100);g__Metschnikowia(100);g__Metschnikowia_unclassified(100); | Metschnikowia | 13 | Saprotroph | \|Undefined Saprotroph\| | Yeast | NULL | Probable | As nectar-tap_saprotroph (Põlme et al. 2020); Decay sugar-rich substrates (Põlme et al. 2020) | Põlme S, et al. 2020. Fungal Diversity 105:1-16 ((https://doi.org/10.1007/s13225-020-00466-2)) |
| Otu41903 | 1 | k__Fungi(100);p__Ascomycota(100);c__Saccharomycetes(100);o__Saccharomycetales(100);f__Metschnikowiaceae(100);g__Metschnikowia(100);g__Metschnikowia_unclassified(100); | Metschnikowia | 13 | Saprotroph | \|Undefined Saprotroph\| | Yeast | NULL | Probable | As nectar-tap_saprotroph (Põlme et al. 2020); Decay sugar-rich substrates (Põlme et al. 2020) | Põlme S, et al. 2020. Fungal Diversity 105:1-16 ((https://doi.org/10.1007/s13225-020-00466-2)) |
| Otu41904 | 1 | k__Fungi(100);p__Ascomycota(100);c__Saccharomycetes(100);o__Saccharomycetales(100);f__Metschnikowiaceae(100);g__Metschnikowia(100);s__Metschnikowia_pulcherrima(100); | Metschnikowia | 13 | Saprotroph | \|Undefined Saprotroph\| | Yeast | NULL | Probable | As nectar-tap_saprotroph (Põlme et al. 2020); Decay sugar-rich substrates (Põlme et al. 2020) | Põlme S, et al. 2020. Fungal Diversity 105:1-16 ((https://doi.org/10.1007/s13225-020-00466-2)) |
| Otu41905 | 1 | k__Fungi(100);p__Ascomycota(100);c__Saccharomycetes(100);o__Saccharomycetales(100);f__Metschnikowiaceae(100);g__Metschnikowia(100);s__Metschnikowia_pulcherrima(100); | Metschnikowia | 13 | Saprotroph | \|Undefined Saprotroph\| | Yeast | NULL | Probable | As nectar-tap_saprotroph (Põlme et al. 2020); Decay sugar-rich substrates (Põlme et al. 2020) | Põlme S, et al. 2020. Fungal Diversity 105:1-16 ((https://doi.org/10.1007/s13225-020-00466-2)) |
| Otu41906 | 1 | k__Fungi(100);p__Ascomycota(100);c__Saccharomycetes(100);o__Saccharomycetales(100);f__Metschnikowiaceae(100);g__Metschnikowia(100);s__Metschnikowia_pulcherrima(100); | Metschnikowia | 13 | Saprotroph | \|Undefined Saprotroph\| | Yeast | NULL | Probable | As nectar-tap_saprotroph (Põlme et al. 2020); Decay sugar-rich substrates (Põlme et al. 2020) | Põlme S, et al. 2020. Fungal Diversity 105:1-16 ((https://doi.org/10.1007/s13225-020-00466-2)) |
| Otu41907 | 1 | k__Fungi(100);p__Ascomycota(100);c__Saccharomycetes(100);o__Saccharomycetales(100);f__Metschnikowiaceae(100);g__Metschnikowia(100);s__Metschnikowia_pulcherrima(100); | Metschnikowia | 13 | Saprotroph | \|Undefined Saprotroph\| | Yeast | NULL | Probable | As nectar-tap_saprotroph (Põlme et al. 2020); Decay sugar-rich substrates (Põlme et al. 2020) | Põlme S, et al. 2020. Fungal Diversity 105:1-16 ((https://doi.org/10.1007/s13225-020-00466-2)) |
| Otu41908 | 1 | k__Fungi(100);p__Ascomycota(100);c__Dothideomycetes(100);o__Dothideomycetes_order_Incertae_sedis(100);f__Myxotrichaceae(100);g__Oidiodendron(100);s__Oidiodendron_sp(100); | Oidiodendron | 13 | Pathotroph-Saprotroph-Symbiotroph | Endophyte-Ericoid Mycorrhizal-\|Undefined Saprotroph\| | Microfungus | NULL | Probable | As root_endophyte (Põlme et al. 2020); As soil_saprotroph (Põlme et al. 2020); Dark Septate Endophyte; Decay soil (Põlme et al. 2020) | Newsham KK. 2011. New Phytologist 190:783-793 ((https://doi.org/10.1111/j.1469-8137.2010.03611.x)) (as dark septate root endophytes); Põlme S, et al. 2020. Fungal Diversity 105:1-16 ((https://doi.org/10.1007/s13225-020-00466-2)) |
| Otu41911 | 1 | k__Fungi(100);p__Ascomycota(100);c__Saccharomycetes(100);o__Saccharomycetales(100);f__Metschnikowiaceae(100);g__Metschnikowia(100);s__Metschnikowia_pulcherrima(100); | Metschnikowia | 13 | Saprotroph | \|Undefined Saprotroph\| | Yeast | NULL | Probable | As nectar-tap_saprotroph (Põlme et al. 2020); Decay sugar-rich substrates (Põlme et al. 2020) | Põlme S, et al. 2020. Fungal Diversity 105:1-16 ((https://doi.org/10.1007/s13225-020-00466-2)) |
| Otu41912 | 1 | k__Fungi(100);p__Ascomycota(100);c__Saccharomycetes(100);o__Saccharomycetales(100);f__Metschnikowiaceae(100);g__Metschnikowia(100);s__Metschnikowia_pulcherrima(100); | Metschnikowia | 13 | Saprotroph | \|Undefined Saprotroph\| | Yeast | NULL | Probable | As nectar-tap_saprotroph (Põlme et al. 2020); Decay sugar-rich substrates (Põlme et al. 2020) | Põlme S, et al. 2020. Fungal Diversity 105:1-16 ((https://doi.org/10.1007/s13225-020-00466-2)) |
| Otu41913 | 1 | k__Fungi(100);p__Ascomycota(100);c__Saccharomycetes(100);o__Saccharomycetales(100);f__Metschnikowiaceae(100);g__Metschnikowia(100);s__Metschnikowia_pulcherrima(100); | Metschnikowia | 13 | Saprotroph | \|Undefined Saprotroph\| | Yeast | NULL | Probable | As nectar-tap_saprotroph (Põlme et al. 2020); Decay sugar-rich substrates (Põlme et al. 2020) | Põlme S, et al. 2020. Fungal Diversity 105:1-16 ((https://doi.org/10.1007/s13225-020-00466-2)) |
| Otu41914 | 1 | k__Fungi(100);p__Ascomycota(100);c__Saccharomycetes(100);o__Saccharomycetales(100);f__Metschnikowiaceae(100);g__Metschnikowia(100);s__Metschnikowia_pulcherrima(100); | Metschnikowia | 13 | Saprotroph | \|Undefined Saprotroph\| | Yeast | NULL | Probable | As nectar-tap_saprotroph (Põlme et al. 2020); Decay sugar-rich substrates (Põlme et al. 2020) | Põlme S, et al. 2020. Fungal Diversity 105:1-16 ((https://doi.org/10.1007/s13225-020-00466-2)) |
| Otu41915 | 1 | k__Fungi(100);p__Ascomycota(100);c__Saccharomycetes(100);o__Saccharomycetales(100);f__Metschnikowiaceae(100);g__Metschnikowia(100);s__Metschnikowia_pulcherrima(100); | Metschnikowia | 13 | Saprotroph | \|Undefined Saprotroph\| | Yeast | NULL | Probable | As nectar-tap_saprotroph (Põlme et al. 2020); Decay sugar-rich substrates (Põlme et al. 2020) | Põlme S, et al. 2020. Fungal Diversity 105:1-16 ((https://doi.org/10.1007/s13225-020-00466-2)) |
| Otu41916 | 1 | k__Fungi(100);p__Ascomycota(100);c__Saccharomycetes(100);o__Saccharomycetales(100);f__Metschnikowiaceae(100);g__Metschnikowia(100);g__Metschnikowia_unclassified(100); | Metschnikowia | 13 | Saprotroph | \|Undefined Saprotroph\| | Yeast | NULL | Probable | As nectar-tap_saprotroph (Põlme et al. 2020); Decay sugar-rich substrates (Põlme et al. 2020) | Põlme S, et al. 2020. Fungal Diversity 105:1-16 ((https://doi.org/10.1007/s13225-020-00466-2)) |
| Otu41917 | 1 | k__Fungi(100);p__Ascomycota(100);c__Saccharomycetes(100);o__Saccharomycetales(100);f__Metschnikowiaceae(100);g__Metschnikowia(100);s__Metschnikowia_pulcherrima(100); | Metschnikowia | 13 | Saprotroph | \|Undefined Saprotroph\| | Yeast | NULL | Probable | As nectar-tap_saprotroph (Põlme et al. 2020); Decay sugar-rich substrates (Põlme et al. 2020) | Põlme S, et al. 2020. Fungal Diversity 105:1-16 ((https://doi.org/10.1007/s13225-020-00466-2)) |
| Otu41918 | 1 | k__Fungi(100);p__Ascomycota(100);c__Saccharomycetes(100);o__Saccharomycetales(100);f__Metschnikowiaceae(100);g__Metschnikowia(100);s__Metschnikowia_pulcherrima(100); | Metschnikowia | 13 | Saprotroph | \|Undefined Saprotroph\| | Yeast | NULL | Probable | As nectar-tap_saprotroph (Põlme et al. 2020); Decay sugar-rich substrates (Põlme et al. 2020) | Põlme S, et al. 2020. Fungal Diversity 105:1-16 ((https://doi.org/10.1007/s13225-020-00466-2)) |
| Otu41919 | 1 | k__Fungi(100);p__Ascomycota(100);c__Saccharomycetes(100);o__Saccharomycetales(100);f__Metschnikowiaceae(100);g__Metschnikowia(100);g__Metschnikowia_unclassified(100); | Metschnikowia | 13 | Saprotroph | \|Undefined Saprotroph\| | Yeast | NULL | Probable | As nectar-tap_saprotroph (Põlme et al. 2020); Decay sugar-rich substrates (Põlme et al. 2020) | Põlme S, et al. 2020. Fungal Diversity 105:1-16 ((https://doi.org/10.1007/s13225-020-00466-2)) |
| Otu41921 | 1 | k__Fungi(100);p__Ascomycota(100);c__Saccharomycetes(100);o__Saccharomycetales(100);f__Metschnikowiaceae(100);g__Metschnikowia(100);s__Metschnikowia_pulcherrima(100); | Metschnikowia | 13 | Saprotroph | \|Undefined Saprotroph\| | Yeast | NULL | Probable | As nectar-tap_saprotroph (Põlme et al. 2020); Decay sugar-rich substrates (Põlme et al. 2020) | Põlme S, et al. 2020. Fungal Diversity 105:1-16 ((https://doi.org/10.1007/s13225-020-00466-2)) |
| Otu41922 | 1 | k__Fungi(100);p__Ascomycota(100);c__Saccharomycetes(100);o__Saccharomycetales(100);f__Metschnikowiaceae(100);g__Metschnikowia(100);s__Metschnikowia_pulcherrima(100); | Metschnikowia | 13 | Saprotroph | \|Undefined Saprotroph\| | Yeast | NULL | Probable | As nectar-tap_saprotroph (Põlme et al. 2020); Decay sugar-rich substrates (Põlme et al. 2020) | Põlme S, et al. 2020. Fungal Diversity 105:1-16 ((https://doi.org/10.1007/s13225-020-00466-2)) |
| Otu41923 | 1 | k__Fungi(100);p__Ascomycota(100);c__Saccharomycetes(100);o__Saccharomycetales(100);f__Metschnikowiaceae(100);g__Metschnikowia(100);s__Metschnikowia_pulcherrima(100); | Metschnikowia | 13 | Saprotroph | \|Undefined Saprotroph\| | Yeast | NULL | Probable | As nectar-tap_saprotroph (Põlme et al. 2020); Decay sugar-rich substrates (Põlme et al. 2020) | Põlme S, et al. 2020. Fungal Diversity 105:1-16 ((https://doi.org/10.1007/s13225-020-00466-2)) |
| Otu41924 | 1 | k__Fungi(100);p__Ascomycota(100);c__Saccharomycetes(100);o__Saccharomycetales(100);f__Metschnikowiaceae(100);g__Metschnikowia(100);s__Metschnikowia_pulcherrima(100); | Metschnikowia | 13 | Saprotroph | \|Undefined Saprotroph\| | Yeast | NULL | Probable | As nectar-tap_saprotroph (Põlme et al. 2020); Decay sugar-rich substrates (Põlme et al. 2020) | Põlme S, et al. 2020. Fungal Diversity 105:1-16 ((https://doi.org/10.1007/s13225-020-00466-2)) |
| Otu41925 | 1 | k__Fungi(100);p__Ascomycota(100);c__Saccharomycetes(100);o__Saccharomycetales(100);f__Metschnikowiaceae(100);g__Metschnikowia(100);s__Metschnikowia_pulcherrima(100); | Metschnikowia | 13 | Saprotroph | \|Undefined Saprotroph\| | Yeast | NULL | Probable | As nectar-tap_saprotroph (Põlme et al. 2020); Decay sugar-rich substrates (Põlme et al. 2020) | Põlme S, et al. 2020. Fungal Diversity 105:1-16 ((https://doi.org/10.1007/s13225-020-00466-2)) |
| Otu41926 | 1 | k__Fungi(100);p__Ascomycota(100);c__Saccharomycetes(100);o__Saccharomycetales(100);f__Metschnikowiaceae(100);g__Metschnikowia(100);g__Metschnikowia_unclassified(100); | Metschnikowia | 13 | Saprotroph | \|Undefined Saprotroph\| | Yeast | NULL | Probable | As nectar-tap_saprotroph (Põlme et al. 2020); Decay sugar-rich substrates (Põlme et al. 2020) | Põlme S, et al. 2020. Fungal Diversity 105:1-16 ((https://doi.org/10.1007/s13225-020-00466-2)) |
| Otu41927 | 1 | k__Fungi(100);p__Ascomycota(100);c__Saccharomycetes(100);o__Saccharomycetales(100);f__Metschnikowiaceae(100);g__Metschnikowia(100);s__Metschnikowia_pulcherrima(100); | Metschnikowia | 13 | Saprotroph | \|Undefined Saprotroph\| | Yeast | NULL | Probable | As nectar-tap_saprotroph (Põlme et al. 2020); Decay sugar-rich substrates (Põlme et al. 2020) | Põlme S, et al. 2020. Fungal Diversity 105:1-16 ((https://doi.org/10.1007/s13225-020-00466-2)) |
| Otu41928 | 1 | k__Fungi(100);p__Ascomycota(100);c__Saccharomycetes(100);o__Saccharomycetales(100);f__Metschnikowiaceae(100);g__Metschnikowia(100);g__Metschnikowia_unclassified(100); | Metschnikowia | 13 | Saprotroph | \|Undefined Saprotroph\| | Yeast | NULL | Probable | As nectar-tap_saprotroph (Põlme et al. 2020); Decay sugar-rich substrates (Põlme et al. 2020) | Põlme S, et al. 2020. Fungal Diversity 105:1-16 ((https://doi.org/10.1007/s13225-020-00466-2)) |
| Otu41929 | 1 | k__Fungi(100);p__Ascomycota(100);c__Saccharomycetes(100);o__Saccharomycetales(100);f__Metschnikowiaceae(100);g__Metschnikowia(100);g__Metschnikowia_unclassified(100); | Metschnikowia | 13 | Saprotroph | \|Undefined Saprotroph\| | Yeast | NULL | Probable | As nectar-tap_saprotroph (Põlme et al. 2020); Decay sugar-rich substrates (Põlme et al. 2020) | Põlme S, et al. 2020. Fungal Diversity 105:1-16 ((https://doi.org/10.1007/s13225-020-00466-2)) |
| Otu41930 | 1 | k__Fungi(100);p__Ascomycota(100);c__Saccharomycetes(100);o__Saccharomycetales(100);f__Metschnikowiaceae(100);g__Metschnikowia(100);s__Metschnikowia_pulcherrima(100); | Metschnikowia | 13 | Saprotroph | \|Undefined Saprotroph\| | Yeast | NULL | Probable | As nectar-tap_saprotroph (Põlme et al. 2020); Decay sugar-rich substrates (Põlme et al. 2020) | Põlme S, et al. 2020. Fungal Diversity 105:1-16 ((https://doi.org/10.1007/s13225-020-00466-2)) |
| Otu41931 | 1 | k__Fungi(100);p__Ascomycota(100);c__Saccharomycetes(100);o__Saccharomycetales(100);f__Metschnikowiaceae(100);g__Metschnikowia(100);s__Metschnikowia_pulcherrima(100); | Metschnikowia | 13 | Saprotroph | \|Undefined Saprotroph\| | Yeast | NULL | Probable | As nectar-tap_saprotroph (Põlme et al. 2020); Decay sugar-rich substrates (Põlme et al. 2020) | Põlme S, et al. 2020. Fungal Diversity 105:1-16 ((https://doi.org/10.1007/s13225-020-00466-2)) |
| Otu41933 | 1 | k__Fungi(100);p__Ascomycota(100);c__Saccharomycetes(100);o__Saccharomycetales(100);f__Metschnikowiaceae(100);g__Metschnikowia(100);s__Metschnikowia_pulcherrima(100); | Metschnikowia | 13 | Saprotroph | \|Undefined Saprotroph\| | Yeast | NULL | Probable | As nectar-tap_saprotroph (Põlme et al. 2020); Decay sugar-rich substrates (Põlme et al. 2020) | Põlme S, et al. 2020. Fungal Diversity 105:1-16 ((https://doi.org/10.1007/s13225-020-00466-2)) |
| Otu41934 | 1 | k__Fungi(100);p__Ascomycota(100);c__Saccharomycetes(100);o__Saccharomycetales(100);f__Metschnikowiaceae(100);g__Metschnikowia(100);g__Metschnikowia_unclassified(100); | Metschnikowia | 13 | Saprotroph | \|Undefined Saprotroph\| | Yeast | NULL | Probable | As nectar-tap_saprotroph (Põlme et al. 2020); Decay sugar-rich substrates (Põlme et al. 2020) | Põlme S, et al. 2020. Fungal Diversity 105:1-16 ((https://doi.org/10.1007/s13225-020-00466-2)) |
| Otu41935 | 1 | k__Fungi(100);p__Ascomycota(100);c__Saccharomycetes(100);o__Saccharomycetales(100);f__Metschnikowiaceae(100);g__Metschnikowia(100);s__Metschnikowia_pulcherrima(100); | Metschnikowia | 13 | Saprotroph | \|Undefined Saprotroph\| | Yeast | NULL | Probable | As nectar-tap_saprotroph (Põlme et al. 2020); Decay sugar-rich substrates (Põlme et al. 2020) | Põlme S, et al. 2020. Fungal Diversity 105:1-16 ((https://doi.org/10.1007/s13225-020-00466-2)) |
| Otu41936 | 1 | k__Fungi(100);p__Ascomycota(100);c__Saccharomycetes(100);o__Saccharomycetales(100);f__Metschnikowiaceae(100);g__Metschnikowia(100);g__Metschnikowia_unclassified(100); | Metschnikowia | 13 | Saprotroph | \|Undefined Saprotroph\| | Yeast | NULL | Probable | As nectar-tap_saprotroph (Põlme et al. 2020); Decay sugar-rich substrates (Põlme et al. 2020) | Põlme S, et al. 2020. Fungal Diversity 105:1-16 ((https://doi.org/10.1007/s13225-020-00466-2)) |
| Otu41937 | 1 | k__Fungi(100);p__Ascomycota(100);c__Saccharomycetes(100);o__Saccharomycetales(100);f__Metschnikowiaceae(100);g__Metschnikowia(100);s__Metschnikowia_pulcherrima(100); | Metschnikowia | 13 | Saprotroph | \|Undefined Saprotroph\| | Yeast | NULL | Probable | As nectar-tap_saprotroph (Põlme et al. 2020); Decay sugar-rich substrates (Põlme et al. 2020) | Põlme S, et al. 2020. Fungal Diversity 105:1-16 ((https://doi.org/10.1007/s13225-020-00466-2)) |
| Otu41938 | 1 | k__Fungi(100);p__Ascomycota(100);c__Saccharomycetes(100);o__Saccharomycetales(100);f__Metschnikowiaceae(100);g__Metschnikowia(100);s__Metschnikowia_pulcherrima(100); | Metschnikowia | 13 | Saprotroph | \|Undefined Saprotroph\| | Yeast | NULL | Probable | As nectar-tap_saprotroph (Põlme et al. 2020); Decay sugar-rich substrates (Põlme et al. 2020) | Põlme S, et al. 2020. Fungal Diversity 105:1-16 ((https://doi.org/10.1007/s13225-020-00466-2)) |
| Otu41939 | 1 | k__Fungi(100);p__Ascomycota(100);c__Saccharomycetes(100);o__Saccharomycetales(100);f__Metschnikowiaceae(100);g__Metschnikowia(100);g__Metschnikowia_unclassified(100); | Metschnikowia | 13 | Saprotroph | \|Undefined Saprotroph\| | Yeast | NULL | Probable | As nectar-tap_saprotroph (Põlme et al. 2020); Decay sugar-rich substrates (Põlme et al. 2020) | Põlme S, et al. 2020. Fungal Diversity 105:1-16 ((https://doi.org/10.1007/s13225-020-00466-2)) |
| Otu41940 | 1 | k__Fungi(100);p__Ascomycota(100);c__Saccharomycetes(100);o__Saccharomycetales(100);f__Metschnikowiaceae(100);g__Metschnikowia(100);g__Metschnikowia_unclassified(100); | Metschnikowia | 13 | Saprotroph | \|Undefined Saprotroph\| | Yeast | NULL | Probable | As nectar-tap_saprotroph (Põlme et al. 2020); Decay sugar-rich substrates (Põlme et al. 2020) | Põlme S, et al. 2020. Fungal Diversity 105:1-16 ((https://doi.org/10.1007/s13225-020-00466-2)) |
| Otu41941 | 1 | k__Fungi(100);p__Ascomycota(100);c__Saccharomycetes(100);o__Saccharomycetales(100);f__Metschnikowiaceae(100);g__Metschnikowia(100);s__Metschnikowia_pulcherrima(100); | Metschnikowia | 13 | Saprotroph | \|Undefined Saprotroph\| | Yeast | NULL | Probable | As nectar-tap_saprotroph (Põlme et al. 2020); Decay sugar-rich substrates (Põlme et al. 2020) | Põlme S, et al. 2020. Fungal Diversity 105:1-16 ((https://doi.org/10.1007/s13225-020-00466-2)) |
| Otu41942 | 1 | k__Fungi(100);p__Ascomycota(100);c__Saccharomycetes(100);o__Saccharomycetales(100);f__Metschnikowiaceae(100);g__Metschnikowia(100);s__Metschnikowia_pulcherrima(100); | Metschnikowia | 13 | Saprotroph | \|Undefined Saprotroph\| | Yeast | NULL | Probable | As nectar-tap_saprotroph (Põlme et al. 2020); Decay sugar-rich substrates (Põlme et al. 2020) | Põlme S, et al. 2020. Fungal Diversity 105:1-16 ((https://doi.org/10.1007/s13225-020-00466-2)) |
| Otu41943 | 1 | k__Fungi(100);p__Ascomycota(100);c__Saccharomycetes(100);o__Saccharomycetales(100);f__Metschnikowiaceae(100);g__Metschnikowia(100);s__Metschnikowia_pulcherrima(100); | Metschnikowia | 13 | Saprotroph | \|Undefined Saprotroph\| | Yeast | NULL | Probable | As nectar-tap_saprotroph (Põlme et al. 2020); Decay sugar-rich substrates (Põlme et al. 2020) | Põlme S, et al. 2020. Fungal Diversity 105:1-16 ((https://doi.org/10.1007/s13225-020-00466-2)) |
| Otu41944 | 1 | k__Fungi(100);p__Ascomycota(100);c__Saccharomycetes(100);o__Saccharomycetales(100);f__Metschnikowiaceae(100);g__Metschnikowia(100);s__Metschnikowia_pulcherrima(100); | Metschnikowia | 13 | Saprotroph | \|Undefined Saprotroph\| | Yeast | NULL | Probable | As nectar-tap_saprotroph (Põlme et al. 2020); Decay sugar-rich substrates (Põlme et al. 2020) | Põlme S, et al. 2020. Fungal Diversity 105:1-16 ((https://doi.org/10.1007/s13225-020-00466-2)) |
| Otu41945 | 1 | k__Fungi(100);p__Ascomycota(100);c__Saccharomycetes(100);o__Saccharomycetales(100);f__Metschnikowiaceae(100);g__Metschnikowia(100);s__Metschnikowia_pulcherrima(100); | Metschnikowia | 13 | Saprotroph | \|Undefined Saprotroph\| | Yeast | NULL | Probable | As nectar-tap_saprotroph (Põlme et al. 2020); Decay sugar-rich substrates (Põlme et al. 2020) | Põlme S, et al. 2020. Fungal Diversity 105:1-16 ((https://doi.org/10.1007/s13225-020-00466-2)) |
| Otu41947 | 1 | k__Fungi(100);p__Ascomycota(100);c__Saccharomycetes(100);o__Saccharomycetales(100);f__Metschnikowiaceae(100);g__Metschnikowia(100);s__Metschnikowia_pulcherrima(100); | Metschnikowia | 13 | Saprotroph | \|Undefined Saprotroph\| | Yeast | NULL | Probable | As nectar-tap_saprotroph (Põlme et al. 2020); Decay sugar-rich substrates (Põlme et al. 2020) | Põlme S, et al. 2020. Fungal Diversity 105:1-16 ((https://doi.org/10.1007/s13225-020-00466-2)) |
| Otu41948 | 1 | k__Fungi(100);p__Ascomycota(100);c__Saccharomycetes(100);o__Saccharomycetales(100);f__Metschnikowiaceae(100);g__Metschnikowia(100);s__Metschnikowia_pulcherrima(100); | Metschnikowia | 13 | Saprotroph | \|Undefined Saprotroph\| | Yeast | NULL | Probable | As nectar-tap_saprotroph (Põlme et al. 2020); Decay sugar-rich substrates (Põlme et al. 2020) | Põlme S, et al. 2020. Fungal Diversity 105:1-16 ((https://doi.org/10.1007/s13225-020-00466-2)) |
| Otu41949 | 1 | k__Fungi(100);p__Ascomycota(100);c__Saccharomycetes(100);o__Saccharomycetales(100);f__Metschnikowiaceae(100);g__Metschnikowia(100);s__Metschnikowia_pulcherrima(100); | Metschnikowia | 13 | Saprotroph | \|Undefined Saprotroph\| | Yeast | NULL | Probable | As nectar-tap_saprotroph (Põlme et al. 2020); Decay sugar-rich substrates (Põlme et al. 2020) | Põlme S, et al. 2020. Fungal Diversity 105:1-16 ((https://doi.org/10.1007/s13225-020-00466-2)) |
| Otu41952 | 1 | k__Fungi(100);p__Ascomycota(100);c__Saccharomycetes(100);o__Saccharomycetales(100);f__Metschnikowiaceae(100);g__Metschnikowia(100);s__Metschnikowia_pulcherrima(100); | Metschnikowia | 13 | Saprotroph | \|Undefined Saprotroph\| | Yeast | NULL | Probable | As nectar-tap_saprotroph (Põlme et al. 2020); Decay sugar-rich substrates (Põlme et al. 2020) | Põlme S, et al. 2020. Fungal Diversity 105:1-16 ((https://doi.org/10.1007/s13225-020-00466-2)) |
| Otu41953 | 1 | k__Fungi(100);p__Ascomycota(100);c__Saccharomycetes(100);o__Saccharomycetales(100);f__Metschnikowiaceae(100);g__Metschnikowia(100);s__Metschnikowia_pulcherrima(100); | Metschnikowia | 13 | Saprotroph | \|Undefined Saprotroph\| | Yeast | NULL | Probable | As nectar-tap_saprotroph (Põlme et al. 2020); Decay sugar-rich substrates (Põlme et al. 2020) | Põlme S, et al. 2020. Fungal Diversity 105:1-16 ((https://doi.org/10.1007/s13225-020-00466-2)) |
| Otu41954 | 1 | k__Fungi(100);p__Ascomycota(100);c__Saccharomycetes(100);o__Saccharomycetales(100);f__Metschnikowiaceae(100);g__Metschnikowia(100);s__Metschnikowia_pulcherrima(100); | Metschnikowia | 13 | Saprotroph | \|Undefined Saprotroph\| | Yeast | NULL | Probable | As nectar-tap_saprotroph (Põlme et al. 2020); Decay sugar-rich substrates (Põlme et al. 2020) | Põlme S, et al. 2020. Fungal Diversity 105:1-16 ((https://doi.org/10.1007/s13225-020-00466-2)) |
| Otu41957 | 1 | k__Fungi(100);p__Ascomycota(100);c__Saccharomycetes(100);o__Saccharomycetales(100);f__Metschnikowiaceae(100);g__Metschnikowia(100);g__Metschnikowia_unclassified(100); | Metschnikowia | 13 | Saprotroph | \|Undefined Saprotroph\| | Yeast | NULL | Probable | As nectar-tap_saprotroph (Põlme et al. 2020); Decay sugar-rich substrates (Põlme et al. 2020) | Põlme S, et al. 2020. Fungal Diversity 105:1-16 ((https://doi.org/10.1007/s13225-020-00466-2)) |
| Otu41959 | 1 | k__Fungi(100);p__Ascomycota(100);c__Saccharomycetes(100);o__Saccharomycetales(100);f__Saccharomycetales_family_Incertae_sedis(100);g__Candida(100);s__Candida_catenulata(100); | Candida | 13 | Pathotroph-Saprotroph-Symbiotroph | Animal Pathogen-Endophyte-Endosymbiont-Epiphyte-\|Undefined Saprotroph\| | Dimorphic Yeast-Yeast | NULL | Probable | As nectar-tap_saprotroph (Põlme et al. 2020); Decay sugar-rich substrates (Põlme et al. 2020); This asexual genus is polyphyletic and contain species that function very differently depend on what they are. Caution should be taken when interpreting guilds for Candida species. | Manolakaki D, et al. 2010. Virulence. 1: 367-375 ((https://doi.org/10.4161/viru.1.5.12796)); Mekbib SB, et al. 2011. Tropical Plant Pathology 36:233-240 ((https://doi.org/10.1590/S1982-56762011000400004)); Põlme S, et al. 2020. Fungal Diversity 105:1-16 ((https://doi.org/10.1007/s13225-020-00466-2)) |
| Otu41961 | 1 | k__Fungi(100);p__Ascomycota(100);c__Saccharomycetes(100);o__Saccharomycetales(100);f__Metschnikowiaceae(100);g__Metschnikowia(100);s__Metschnikowia_pulcherrima(100); | Metschnikowia | 13 | Saprotroph | \|Undefined Saprotroph\| | Yeast | NULL | Probable | As nectar-tap_saprotroph (Põlme et al. 2020); Decay sugar-rich substrates (Põlme et al. 2020) | Põlme S, et al. 2020. Fungal Diversity 105:1-16 ((https://doi.org/10.1007/s13225-020-00466-2)) |
| Otu41962 | 1 | k__Fungi(100);p__Ascomycota(100);c__Saccharomycetes(100);o__Saccharomycetales(100);f__Metschnikowiaceae(100);g__Metschnikowia(100);s__Metschnikowia_pulcherrima(100); | Metschnikowia | 13 | Saprotroph | \|Undefined Saprotroph\| | Yeast | NULL | Probable | As nectar-tap_saprotroph (Põlme et al. 2020); Decay sugar-rich substrates (Põlme et al. 2020) | Põlme S, et al. 2020. Fungal Diversity 105:1-16 ((https://doi.org/10.1007/s13225-020-00466-2)) |
| Otu41963 | 1 | k__Fungi(100);p__Ascomycota(100);c__Saccharomycetes(100);o__Saccharomycetales(100);f__Metschnikowiaceae(100);g__Metschnikowia(100);s__Metschnikowia_pulcherrima(100); | Metschnikowia | 13 | Saprotroph | \|Undefined Saprotroph\| | Yeast | NULL | Probable | As nectar-tap_saprotroph (Põlme et al. 2020); Decay sugar-rich substrates (Põlme et al. 2020) | Põlme S, et al. 2020. Fungal Diversity 105:1-16 ((https://doi.org/10.1007/s13225-020-00466-2)) |
| Otu41964 | 1 | k__Fungi(100);p__Ascomycota(100);c__Saccharomycetes(100);o__Saccharomycetales(100);f__Metschnikowiaceae(100);g__Metschnikowia(100);s__Metschnikowia_pulcherrima(100); | Metschnikowia | 13 | Saprotroph | \|Undefined Saprotroph\| | Yeast | NULL | Probable | As nectar-tap_saprotroph (Põlme et al. 2020); Decay sugar-rich substrates (Põlme et al. 2020) | Põlme S, et al. 2020. Fungal Diversity 105:1-16 ((https://doi.org/10.1007/s13225-020-00466-2)) |
| Otu41965 | 1 | k__Fungi(100);p__Ascomycota(100);c__Saccharomycetes(100);o__Saccharomycetales(100);f__Metschnikowiaceae(100);g__Metschnikowia(100);s__Metschnikowia_pulcherrima(100); | Metschnikowia | 13 | Saprotroph | \|Undefined Saprotroph\| | Yeast | NULL | Probable | As nectar-tap_saprotroph (Põlme et al. 2020); Decay sugar-rich substrates (Põlme et al. 2020) | Põlme S, et al. 2020. Fungal Diversity 105:1-16 ((https://doi.org/10.1007/s13225-020-00466-2)) |
| Otu41966 | 1 | k__Fungi(100);p__Ascomycota(100);c__Saccharomycetes(100);o__Saccharomycetales(100);f__Metschnikowiaceae(100);g__Metschnikowia(100);s__Metschnikowia_pulcherrima(100); | Metschnikowia | 13 | Saprotroph | \|Undefined Saprotroph\| | Yeast | NULL | Probable | As nectar-tap_saprotroph (Põlme et al. 2020); Decay sugar-rich substrates (Põlme et al. 2020) | Põlme S, et al. 2020. Fungal Diversity 105:1-16 ((https://doi.org/10.1007/s13225-020-00466-2)) |
| Otu41967 | 1 | k__Fungi(100);p__Ascomycota(100);c__Saccharomycetes(100);o__Saccharomycetales(100);f__Metschnikowiaceae(100);g__Metschnikowia(100);s__Metschnikowia_pulcherrima(100); | Metschnikowia | 13 | Saprotroph | \|Undefined Saprotroph\| | Yeast | NULL | Probable | As nectar-tap_saprotroph (Põlme et al. 2020); Decay sugar-rich substrates (Põlme et al. 2020) | Põlme S, et al. 2020. Fungal Diversity 105:1-16 ((https://doi.org/10.1007/s13225-020-00466-2)) |
| Otu41968 | 1 | k__Fungi(100);p__Ascomycota(100);c__Saccharomycetes(100);o__Saccharomycetales(100);f__Metschnikowiaceae(100);g__Metschnikowia(100);s__Metschnikowia_pulcherrima(100); | Metschnikowia | 13 | Saprotroph | \|Undefined Saprotroph\| | Yeast | NULL | Probable | As nectar-tap_saprotroph (Põlme et al. 2020); Decay sugar-rich substrates (Põlme et al. 2020) | Põlme S, et al. 2020. Fungal Diversity 105:1-16 ((https://doi.org/10.1007/s13225-020-00466-2)) |

| | | | | | | | | | | | |
|---|---|---|---|---|---|---|---|---|---|---|---|
| Otu41969 | 1 | k__Fungi(100);p__Ascomycota(100);c__Saccharomycetes(100);o__Saccharomycetales(100);f__Metschnikowiaceae(100);g__Metschnikowia(100);s__Metschnikowia_pulcherrima(100); | Metschnikowia | 13 | Saprotroph | \|Undefined Saprotroph\| | Yeast | NULL | Probable | As nectar-tap_saprotroph (Põlme et al. 2020); Decay sugar-rich substrates (Põlme et al. 2020) | Põlme S, et al. 2020. Fungal Diversity 105:1-16 ((https://doi.org/10.1007/s13225-020-00466-2)) |
| Otu41970 | 1 | k__Fungi(100);p__Ascomycota(100);c__Saccharomycetes(100);o__Saccharomycetales(100);f__Metschnikowiaceae(100);g__Metschnikowia(100);s__Metschnikowia_pulcherrima(100); | Metschnikowia | 13 | Saprotroph | \|Undefined Saprotroph\| | Yeast | NULL | Probable | As nectar-tap_saprotroph (Põlme et al. 2020); Decay sugar-rich substrates (Põlme et al. 2020) | Põlme S, et al. 2020. Fungal Diversity 105:1-16 ((https://doi.org/10.1007/s13225-020-00466-2)) |
| Otu41971 | 1 | k__Fungi(100);p__Ascomycota(100);c__Saccharomycetes(100);o__Saccharomycetales(100);f__Metschnikowiaceae(100);g__Metschnikowia(100);s__Metschnikowia_pulcherrima(100); | Metschnikowia | 13 | Saprotroph | \|Undefined Saprotroph\| | Yeast | NULL | Probable | As nectar-tap_saprotroph (Põlme et al. 2020); Decay sugar-rich substrates (Põlme et al. 2020) | Põlme S, et al. 2020. Fungal Diversity 105:1-16 ((https://doi.org/10.1007/s13225-020-00466-2)) |
| Otu41972 | 1 | k__Fungi(100);p__Ascomycota(100);c__Saccharomycetes(100);o__Saccharomycetales(100);f__Metschnikowiaceae(100);g__Metschnikowia(100);g__Metschnikowia_unclassified(100); | Metschnikowia | 13 | Saprotroph | \|Undefined Saprotroph\| | Yeast | NULL | Probable | As nectar-tap_saprotroph (Põlme et al. 2020); Decay sugar-rich substrates (Põlme et al. 2020) | Põlme S, et al. 2020. Fungal Diversity 105:1-16 ((https://doi.org/10.1007/s13225-020-00466-2)) |
| Otu41973 | 1 | k__Fungi(100);p__Ascomycota(100);c__Saccharomycetes(100);o__Saccharomycetales(100);f__Metschnikowiaceae(100);g__Metschnikowia(100);s__Metschnikowia_pulcherrima(100); | Metschnikowia | 13 | Saprotroph | \|Undefined Saprotroph\| | Yeast | NULL | Probable | As nectar-tap_saprotroph (Põlme et al. 2020); Decay sugar-rich substrates (Põlme et al. 2020) | Põlme S, et al. 2020. Fungal Diversity 105:1-16 ((https://doi.org/10.1007/s13225-020-00466-2)) |
| Otu41974 | 1 | k__Fungi(100);p__Ascomycota(100);c__Saccharomycetes(100);o__Saccharomycetales(100);f__Metschnikowiaceae(100);g__Metschnikowia(100);s__Metschnikowia_pulcherrima(100); | Metschnikowia | 13 | Saprotroph | \|Undefined Saprotroph\| | Yeast | NULL | Probable | As nectar-tap_saprotroph (Põlme et al. 2020); Decay sugar-rich substrates (Põlme et al. 2020) | Põlme S, et al. 2020. Fungal Diversity 105:1-16 ((https://doi.org/10.1007/s13225-020-00466-2)) |
| Otu41975 | 1 | k__Fungi(100);p__Ascomycota(100);c__Saccharomycetes(100);o__Saccharomycetales(100);f__Metschnikowiaceae(100);g__Metschnikowia(100);s__Metschnikowia_pulcherrima(100); | Metschnikowia | 13 | Saprotroph | \|Undefined Saprotroph\| | Yeast | NULL | Probable | As nectar-tap_saprotroph (Põlme et al. 2020); Decay sugar-rich substrates (Põlme et al. 2020) | Põlme S, et al. 2020. Fungal Diversity 105:1-16 ((https://doi.org/10.1007/s13225-020-00466-2)) |
| Otu41976 | 1 | k__Fungi(100);p__Ascomycota(100);c__Saccharomycetes(100);o__Saccharomycetales(100);f__Metschnikowiaceae(100);g__Metschnikowia(100);s__Metschnikowia_pulcherrima(100); | Metschnikowia | 13 | Saprotroph | \|Undefined Saprotroph\| | Yeast | NULL | Probable | As nectar-tap_saprotroph (Põlme et al. 2020); Decay sugar-rich substrates (Põlme et al. 2020) | Põlme S, et al. 2020. Fungal Diversity 105:1-16 ((https://doi.org/10.1007/s13225-020-00466-2)) |
| Otu41977 | 1 | k__Fungi(100);p__Ascomycota(100);c__Saccharomycetes(100);o__Saccharomycetales(100);f__Metschnikowiaceae(100);g__Metschnikowia(100);s__Metschnikowia_pulcherrima(100); | Metschnikowia | 13 | Saprotroph | \|Undefined Saprotroph\| | Yeast | NULL | Probable | As nectar-tap_saprotroph (Põlme et al. 2020); Decay sugar-rich substrates (Põlme et al. 2020) | Põlme S, et al. 2020. Fungal Diversity 105:1-16 ((https://doi.org/10.1007/s13225-020-00466-2)) |
| Otu41978 | 1 | k__Fungi(100);p__Ascomycota(100);c__Saccharomycetes(100);o__Saccharomycetales(100);f__Metschnikowiaceae(100);g__Metschnikowia(100);s__Metschnikowia_pulcherrima(100); | Metschnikowia | 13 | Saprotroph | \|Undefined Saprotroph\| | Yeast | NULL | Probable | As nectar-tap_saprotroph (Põlme et al. 2020); Decay sugar-rich substrates (Põlme et al. 2020) | Põlme S, et al. 2020. Fungal Diversity 105:1-16 ((https://doi.org/10.1007/s13225-020-00466-2)) |
| Otu41979 | 1 | k__Fungi(100);p__Ascomycota(100);c__Saccharomycetes(100);o__Saccharomycetales(100);f__Metschnikowiaceae(100);g__Metschnikowia(100);s__Metschnikowia_pulcherrima(100); | Metschnikowia | 13 | Saprotroph | \|Undefined Saprotroph\| | Yeast | NULL | Probable | As nectar-tap_saprotroph (Põlme et al. 2020); Decay sugar-rich substrates (Põlme et al. 2020) | Põlme S, et al. 2020. Fungal Diversity 105:1-16 ((https://doi.org/10.1007/s13225-020-00466-2)) |
| Otu41980 | 1 | k__Fungi(100);p__Ascomycota(100);c__Saccharomycetes(100);o__Saccharomycetales(100);f__Metschnikowiaceae(100);g__Metschnikowia(100);s__Metschnikowia_pulcherrima(100); | Metschnikowia | 13 | Saprotroph | \|Undefined Saprotroph\| | Yeast | NULL | Probable | As nectar-tap_saprotroph (Põlme et al. 2020); Decay sugar-rich substrates (Põlme et al. 2020) | Põlme S, et al. 2020. Fungal Diversity 105:1-16 ((https://doi.org/10.1007/s13225-020-00466-2)) |
| Otu41981 | 1 | k__Fungi(100);p__Ascomycota(100);c__Saccharomycetes(100);o__Saccharomycetales(100);f__Metschnikowiaceae(100);g__Metschnikowia(100);g__Metschnikowia_unclassified(100); | Metschnikowia | 13 | Saprotroph | \|Undefined Saprotroph\| | Yeast | NULL | Probable | As nectar-tap_saprotroph (Põlme et al. 2020); Decay sugar-rich substrates (Põlme et al. 2020) | Põlme S, et al. 2020. Fungal Diversity 105:1-16 ((https://doi.org/10.1007/s13225-020-00466-2)) |
| Otu41982 | 1 | k__Fungi(100);p__Ascomycota(100);c__Saccharomycetes(100);o__Saccharomycetales(100);f__Metschnikowiaceae(100);g__Metschnikowia(100);g__Metschnikowia_unclassified(100); | Metschnikowia | 13 | Saprotroph | \|Undefined Saprotroph\| | Yeast | NULL | Probable | As nectar-tap_saprotroph (Põlme et al. 2020); Decay sugar-rich substrates (Põlme et al. 2020) | Põlme S, et al. 2020. Fungal Diversity 105:1-16 ((https://doi.org/10.1007/s13225-020-00466-2)) |
| Otu41983 | 1 | k__Fungi(100);p__Ascomycota(100);c__Saccharomycetes(100);o__Saccharomycetales(100);f__Metschnikowiaceae(100);g__Metschnikowia(100);s__Metschnikowia_pulcherrima(100); | Metschnikowia | 13 | Saprotroph | \|Undefined Saprotroph\| | Yeast | NULL | Probable | As nectar-tap_saprotroph (Põlme et al. 2020); Decay sugar-rich substrates (Põlme et al. 2020) | Põlme S, et al. 2020. Fungal Diversity 105:1-16 ((https://doi.org/10.1007/s13225-020-00466-2)) |
| Otu41984 | 1 | k__Fungi(100);p__Ascomycota(100);c__Saccharomycetes(100);o__Saccharomycetales(100);f__Metschnikowiaceae(100);g__Metschnikowia(100);g__Metschnikowia_unclassified(100); | Metschnikowia | 13 | Saprotroph | \|Undefined Saprotroph\| | Yeast | NULL | Probable | As nectar-tap_saprotroph (Põlme et al. 2020); Decay sugar-rich substrates (Põlme et al. 2020) | Põlme S, et al. 2020. Fungal Diversity 105:1-16 ((https://doi.org/10.1007/s13225-020-00466-2)) |
| Otu41985 | 1 | k__Fungi(100);p__Ascomycota(100);c__Saccharomycetes(100);o__Saccharomycetales(100);f__Metschnikowiaceae(100);g__Metschnikowia(100);s__Metschnikowia_pulcherrima(100); | Metschnikowia | 13 | Saprotroph | \|Undefined Saprotroph\| | Yeast | NULL | Probable | As nectar-tap_saprotroph (Põlme et al. 2020); Decay sugar-rich substrates (Põlme et al. 2020) | Põlme S, et al. 2020. Fungal Diversity 105:1-16 ((https://doi.org/10.1007/s13225-020-00466-2)) |
| Otu41988 | 1 | k__Fungi(100);p__Ascomycota(100);c__Saccharomycetes(100);o__Saccharomycetales(100);f__Metschnikowiaceae(100);g__Metschnikowia(100);g__Metschnikowia_unclassified(100); | Metschnikowia | 13 | Saprotroph | \|Undefined Saprotroph\| | Yeast | NULL | Probable | As nectar-tap_saprotroph (Põlme et al. 2020); Decay sugar-rich substrates (Põlme et al. 2020) | Põlme S, et al. 2020. Fungal Diversity 105:1-16 ((https://doi.org/10.1007/s13225-020-00466-2)) |
| Otu41989 | 1 | k__Fungi(100);p__Ascomycota(100);c__Saccharomycetes(100);o__Saccharomycetales(100);f__Metschnikowiaceae(100);g__Metschnikowia(100);g__Metschnikowia_unclassified(100); | Metschnikowia | 13 | Saprotroph | \|Undefined Saprotroph\| | Yeast | NULL | Probable | As nectar-tap_saprotroph (Põlme et al. 2020); Decay sugar-rich substrates (Põlme et al. 2020) | Põlme S, et al. 2020. Fungal Diversity 105:1-16 ((https://doi.org/10.1007/s13225-020-00466-2)) |
| Otu41990 | 1 | k__Fungi(100);p__Ascomycota(100);c__Saccharomycetes(100);o__Saccharomycetales(100);f__Metschnikowiaceae(100);g__Metschnikowia(100);g__Metschnikowia_unclassified(100); | Metschnikowia | 13 | Saprotroph | \|Undefined Saprotroph\| | Yeast | NULL | Probable | As nectar-tap_saprotroph (Põlme et al. 2020); Decay sugar-rich substrates (Põlme et al. 2020) | Põlme S, et al. 2020. Fungal Diversity 105:1-16 ((https://doi.org/10.1007/s13225-020-00466-2)) |
| Otu41991 | 1 | k__Fungi(100);p__Ascomycota(100);c__Saccharomycetes(100);o__Saccharomycetales(100);f__Metschnikowiaceae(100);g__Metschnikowia(100);s__Metschnikowia_pulcherrima(100); | Metschnikowia | 13 | Saprotroph | \|Undefined Saprotroph\| | Yeast | NULL | Probable | As nectar-tap_saprotroph (Põlme et al. 2020); Decay sugar-rich substrates (Põlme et al. 2020) | Põlme S, et al. 2020. Fungal Diversity 105:1-16 ((https://doi.org/10.1007/s13225-020-00466-2)) |
| Otu41992 | 1 | k__Fungi(100);p__Ascomycota(100);c__Saccharomycetes(100);o__Saccharomycetales(100);f__Metschnikowiaceae(100);g__Metschnikowia(100);s__Metschnikowia_pulcherrima(100); | Metschnikowia | 13 | Saprotroph | \|Undefined Saprotroph\| | Yeast | NULL | Probable | As nectar-tap_saprotroph (Põlme et al. 2020); Decay sugar-rich substrates (Põlme et al. 2020) | Põlme S, et al. 2020. Fungal Diversity 105:1-16 ((https://doi.org/10.1007/s13225-020-00466-2)) |
| Otu41993 | 1 | k__Fungi(100);p__Ascomycota(100);c__Saccharomycetes(100);o__Saccharomycetales(100);f__Metschnikowiaceae(100);g__Metschnikowia(100);s__Metschnikowia_pulcherrima(100); | Metschnikowia | 13 | Saprotroph | \|Undefined Saprotroph\| | Yeast | NULL | Probable | As nectar-tap_saprotroph (Põlme et al. 2020); Decay sugar-rich substrates (Põlme et al. 2020) | Põlme S, et al. 2020. Fungal Diversity 105:1-16 ((https://doi.org/10.1007/s13225-020-00466-2)) |
| Otu41994 | 1 | k__Fungi(100);p__Ascomycota(100);c__Saccharomycetes(100);o__Saccharomycetales(100);f__Metschnikowiaceae(100);g__Metschnikowia(100);s__Metschnikowia_pulcherrima(100); | Metschnikowia | 13 | Saprotroph | \|Undefined Saprotroph\| | Yeast | NULL | Probable | As nectar-tap_saprotroph (Põlme et al. 2020); Decay sugar-rich substrates (Põlme et al. 2020) | Põlme S, et al. 2020. Fungal Diversity 105:1-16 ((https://doi.org/10.1007/s13225-020-00466-2)) |
| Otu41995 | 1 | k__Fungi(100);p__Ascomycota(100);c__Saccharomycetes(100);o__Saccharomycetales(100);f__Metschnikowiaceae(100);g__Metschnikowia(100);s__Metschnikowia_pulcherrima(100); | Metschnikowia | 13 | Saprotroph | \|Undefined Saprotroph\| | Yeast | NULL | Probable | As nectar-tap_saprotroph (Põlme et al. 2020); Decay sugar-rich substrates (Põlme et al. 2020) | Põlme S, et al. 2020. Fungal Diversity 105:1-16 ((https://doi.org/10.1007/s13225-020-00466-2)) |
| Otu41996 | 1 | k__Fungi(100);p__Ascomycota(100);c__Saccharomycetes(100);o__Saccharomycetales(100);f__Metschnikowiaceae(100);g__Metschnikowia(100);g__Metschnikowia_unclassified(100); | Metschnikowia | 13 | Saprotroph | \|Undefined Saprotroph\| | Yeast | NULL | Probable | As nectar-tap_saprotroph (Põlme et al. 2020); Decay sugar-rich substrates (Põlme et al. 2020) | Põlme S, et al. 2020. Fungal Diversity 105:1-16 ((https://doi.org/10.1007/s13225-020-00466-2)) |
| Otu41997 | 1 | k__Fungi(100);p__Ascomycota(100);c__Saccharomycetes(100);o__Saccharomycetales(100);f__Metschnikowiaceae(100);g__Metschnikowia(100);s__Metschnikowia_pulcherrima(100); | Metschnikowia | 13 | Saprotroph | \|Undefined Saprotroph\| | Yeast | NULL | Probable | As nectar-tap_saprotroph (Põlme et al. 2020); Decay sugar-rich substrates (Põlme et al. 2020) | Põlme S, et al. 2020. Fungal Diversity 105:1-16 ((https://doi.org/10.1007/s13225-020-00466-2)) |
| Otu41998 | 1 | k__Fungi(100);p__Ascomycota(100);c__Saccharomycetes(100);o__Saccharomycetales(100);f__Metschnikowiaceae(100);g__Metschnikowia(100);g__Metschnikowia_unclassified(100); | Metschnikowia | 13 | Saprotroph | \|Undefined Saprotroph\| | Yeast | NULL | Probable | As nectar-tap_saprotroph (Põlme et al. 2020); Decay sugar-rich substrates (Põlme et al. 2020) | Põlme S, et al. 2020. Fungal Diversity 105:1-16 ((https://doi.org/10.1007/s13225-020-00466-2)) |
| Otu41999 | 1 | k__Fungi(100);p__Ascomycota(100);c__Saccharomycetes(100);o__Saccharomycetales(100);f__Metschnikowiaceae(100);g__Metschnikowia(100);s__Metschnikowia_pulcherrima(100); | Metschnikowia | 13 | Saprotroph | \|Undefined Saprotroph\| | Yeast | NULL | Probable | As nectar-tap_saprotroph (Põlme et al. 2020); Decay sugar-rich substrates (Põlme et al. 2020) | Põlme S, et al. 2020. Fungal Diversity 105:1-16 ((https://doi.org/10.1007/s13225-020-00466-2)) |
| Otu42000 | 1 | k__Fungi(100);p__Ascomycota(100);c__Saccharomycetes(100);o__Saccharomycetales(100);f__Metschnikowiaceae(100);g__Metschnikowia(100);s__Metschnikowia_pulcherrima(100); | Metschnikowia | 13 | Saprotroph | \|Undefined Saprotroph\| | Yeast | NULL | Probable | As nectar-tap_saprotroph (Põlme et al. 2020); Decay sugar-rich substrates (Põlme et al. 2020) | Põlme S, et al. 2020. Fungal Diversity 105:1-16 ((https://doi.org/10.1007/s13225-020-00466-2)) |
| Otu42001 | 1 | k__Fungi(100);p__Ascomycota(100);c__Saccharomycetes(100);o__Saccharomycetales(100);f__Metschnikowiaceae(100);g__Metschnikowia(100);s__Metschnikowia_pulcherrima(100); | Metschnikowia | 13 | Saprotroph | \|Undefined Saprotroph\| | Yeast | NULL | Probable | As nectar-tap_saprotroph (Põlme et al. 2020); Decay sugar-rich substrates (Põlme et al. 2020) | Põlme S, et al. 2020. Fungal Diversity 105:1-16 ((https://doi.org/10.1007/s13225-020-00466-2)) |
| Otu42002 | 1 | k__Fungi(100);p__Ascomycota(100);c__Saccharomycetes(100);o__Saccharomycetales(100);f__Metschnikowiaceae(100);g__Metschnikowia(100);s__Metschnikowia_pulcherrima(100); | Metschnikowia | 13 | Saprotroph | \|Undefined Saprotroph\| | Yeast | NULL | Probable | As nectar-tap_saprotroph (Põlme et al. 2020); Decay sugar-rich substrates (Põlme et al. 2020) | Põlme S, et al. 2020. Fungal Diversity 105:1-16 ((https://doi.org/10.1007/s13225-020-00466-2)) |
| Otu42003 | 1 | k__Fungi(100);p__Ascomycota(100);c__Saccharomycetes(100);o__Saccharomycetales(100);f__Metschnikowiaceae(100);g__Metschnikowia(100);g__Metschnikowia_unclassified(100); | Metschnikowia | 13 | Saprotroph | \|Undefined Saprotroph\| | Yeast | NULL | Probable | As nectar-tap_saprotroph (Põlme et al. 2020); Decay sugar-rich substrates (Põlme et al. 2020) | Põlme S, et al. 2020. Fungal Diversity 105:1-16 ((https://doi.org/10.1007/s13225-020-00466-2)) |
| Otu42004 | 1 | k__Fungi(100);p__Ascomycota(100);c__Saccharomycetes(100);o__Saccharomycetales(100);f__Metschnikowiaceae(100);g__Metschnikowia(100);s__Metschnikowia_pulcherrima(100); | Metschnikowia | 13 | Saprotroph | \|Undefined Saprotroph\| | Yeast | NULL | Probable | As nectar-tap_saprotroph (Põlme et al. 2020); Decay sugar-rich substrates (Põlme et al. 2020) | Põlme S, et al. 2020. Fungal Diversity 105:1-16 ((https://doi.org/10.1007/s13225-020-00466-2)) |
| Otu42005 | 1 | k__Fungi(100);p__Ascomycota(100);c__Saccharomycetes(100);o__Saccharomycetales(100);f__Metschnikowiaceae(100);g__Metschnikowia(100);g__Metschnikowia_unclassified(100); | Metschnikowia | 13 | Saprotroph | \|Undefined Saprotroph\| | Yeast | NULL | Probable | As nectar-tap_saprotroph (Põlme et al. 2020); Decay sugar-rich substrates (Põlme et al. 2020) | Põlme S, et al. 2020. Fungal Diversity 105:1-16 ((https://doi.org/10.1007/s13225-020-00466-2)) |
| Otu42006 | 1 | k__Fungi(100);p__Ascomycota(100);c__Saccharomycetes(100);o__Saccharomycetales(100);f__Metschnikowiaceae(100);g__Metschnikowia(100);g__Metschnikowia_unclassified(100); | Metschnikowia | 13 | Saprotroph | \|Undefined Saprotroph\| | Yeast | NULL | Probable | As nectar-tap_saprotroph (Põlme et al. 2020); Decay sugar-rich substrates (Põlme et al. 2020) | Põlme S, et al. 2020. Fungal Diversity 105:1-16 ((https://doi.org/10.1007/s13225-020-00466-2)) |
| Otu42007 | 1 | k__Fungi(100);p__Ascomycota(100);c__Saccharomycetes(100);o__Saccharomycetales(100);f__Metschnikowiaceae(100);g__Metschnikowia(100);s__Metschnikowia_pulcherrima(100); | Metschnikowia | 13 | Saprotroph | \|Undefined Saprotroph\| | Yeast | NULL | Probable | As nectar-tap_saprotroph (Põlme et al. 2020); Decay sugar-rich substrates (Põlme et al. 2020) | Põlme S, et al. 2020. Fungal Diversity 105:1-16 ((https://doi.org/10.1007/s13225-020-00466-2)) |
| Otu42008 | 1 | k__Fungi(100);p__Ascomycota(100);c__Saccharomycetes(100);o__Saccharomycetales(100);f__Metschnikowiaceae(100);g__Metschnikowia(100);s__Metschnikowia_pulcherrima(100); | Metschnikowia | 13 | Saprotroph | \|Undefined Saprotroph\| | Yeast | NULL | Probable | As nectar-tap_saprotroph (Põlme et al. 2020); Decay sugar-rich substrates (Põlme et al. 2020) | Põlme S, et al. 2020. Fungal Diversity 105:1-16 ((https://doi.org/10.1007/s13225-020-00466-2)) |
| Otu42009 | 1 | k__Fungi(100);p__Ascomycota(100);c__Saccharomycetes(100);o__Saccharomycetales(100);f__Metschnikowiaceae(100);g__Metschnikowia(100);s__Metschnikowia_pulcherrima(100); | Metschnikowia | 13 | Saprotroph | \|Undefined Saprotroph\| | Yeast | NULL | Probable | As nectar-tap_saprotroph (Põlme et al. 2020); Decay sugar-rich substrates (Põlme et al. 2020) | Põlme S, et al. 2020. Fungal Diversity 105:1-16 ((https://doi.org/10.1007/s13225-020-00466-2)) |
| Otu42010 | 1 | k__Fungi(100);p__Ascomycota(100);c__Saccharomycetes(100);o__Saccharomycetales(100);f__Metschnikowiaceae(100);g__Metschnikowia(100);s__Metschnikowia_pulcherrima(100); | Metschnikowia | 13 | Saprotroph | \|Undefined Saprotroph\| | Yeast | NULL | Probable | As nectar-tap_saprotroph (Põlme et al. 2020); Decay sugar-rich substrates (Põlme et al. 2020) | Põlme S, et al. 2020. Fungal Diversity 105:1-16 ((https://doi.org/10.1007/s13225-020-00466-2)) |
| Otu42011 | 1 | k__Fungi(100);p__Ascomycota(100);c__Saccharomycetes(100);o__Saccharomycetales(100);f__Metschnikowiaceae(100);g__Metschnikowia(100);s__Metschnikowia_pulcherrima(100); | Metschnikowia | 13 | Saprotroph | \|Undefined Saprotroph\| | Yeast | NULL | Probable | As nectar-tap_saprotroph (Põlme et al. 2020); Decay sugar-rich substrates (Põlme et al. 2020) | Põlme S, et al. 2020. Fungal Diversity 105:1-16 ((https://doi.org/10.1007/s13225-020-00466-2)) |
| Otu42013 | 1 | k__Fungi(100);p__Ascomycota(100);c__Saccharomycetes(100);o__Saccharomycetales(100);f__Metschnikowiaceae(100);g__Metschnikowia(100);g__Metschnikowia_unclassified(100); | Metschnikowia | 13 | Saprotroph | \|Undefined Saprotroph\| | Yeast | NULL | Probable | As nectar-tap_saprotroph (Põlme et al. 2020); Decay sugar-rich substrates (Põlme et al. 2020) | Põlme S, et al. 2020. Fungal Diversity 105:1-16 ((https://doi.org/10.1007/s13225-020-00466-2)) |
| Otu42014 | 1 | k__Fungi(100);p__Ascomycota(100);c__Saccharomycetes(100);o__Saccharomycetales(100);f__Metschnikowiaceae(100);g__Metschnikowia(100);g__Metschnikowia_unclassified(100); | Metschnikowia | 13 | Saprotroph | \|Undefined Saprotroph\| | Yeast | NULL | Probable | As nectar-tap_saprotroph (Põlme et al. 2020); Decay sugar-rich substrates (Põlme et al. 2020) | Põlme S, et al. 2020. Fungal Diversity 105:1-16 ((https://doi.org/10.1007/s13225-020-00466-2)) |
| Otu42015 | 1 | k__Fungi(100);p__Ascomycota(100);c__Saccharomycetes(100);o__Saccharomycetales(100);f__Metschnikowiaceae(100);g__Metschnikowia(100);g__Metschnikowia_unclassified(100); | Metschnikowia | 13 | Saprotroph | \|Undefined Saprotroph\| | Yeast | NULL | Probable | As nectar-tap_saprotroph (Põlme et al. 2020); Decay sugar-rich substrates (Põlme et al. 2020) | Põlme S, et al. 2020. Fungal Diversity 105:1-16 ((https://doi.org/10.1007/s13225-020-00466-2)) |
| Otu42016 | 1 | k__Fungi(100);p__Ascomycota(100);c__Saccharomycetes(100);o__Saccharomycetales(100);f__Metschnikowiaceae(100);g__Metschnikowia(100);s__Metschnikowia_pulcherrima(100); | Metschnikowia | 13 | Saprotroph | \|Undefined Saprotroph\| | Yeast | NULL | Probable | As nectar-tap_saprotroph (Põlme et al. 2020); Decay sugar-rich substrates (Põlme et al. 2020) | Põlme S, et al. 2020. Fungal Diversity 105:1-16 ((https://doi.org/10.1007/s13225-020-00466-2)) |
| Otu42017 | 1 | k__Fungi(100);p__Ascomycota(100);c__Saccharomycetes(100);o__Saccharomycetales(100);f__Metschnikowiaceae(100);g__Metschnikowia(100);g__Metschnikowia_unclassified(100); | Metschnikowia | 13 | Saprotroph | \|Undefined Saprotroph\| | Yeast | NULL | Probable | As nectar-tap_saprotroph (Põlme et al. 2020); Decay sugar-rich substrates (Põlme et al. 2020) | Põlme S, et al. 2020. Fungal Diversity 105:1-16 ((https://doi.org/10.1007/s13225-020-00466-2)) |
| Otu42018 | 1 | k__Fungi(100);p__Ascomycota(100);c__Saccharomycetes(100);o__Saccharomycetales(100);f__Metschnikowiaceae(100);g__Metschnikowia(100);g__Metschnikowia_unclassified(100); | Metschnikowia | 13 | Saprotroph | \|Undefined Saprotroph\| | Yeast | NULL | Probable | As nectar-tap_saprotroph (Põlme et al. 2020); Decay sugar-rich substrates (Põlme et al. 2020) | Põlme S, et al. 2020. Fungal Diversity 105:1-16 ((https://doi.org/10.1007/s13225-020-00466-2)) |
| Otu42019 | 1 | k__Fungi(100);p__Ascomycota(100);c__Saccharomycetes(100);o__Saccharomycetales(100);f__Metschnikowiaceae(100);g__Metschnikowia(100);s__Metschnikowia_pulcherrima(100); | Metschnikowia | 13 | Saprotroph | \|Undefined Saprotroph\| | Yeast | NULL | Probable | As nectar-tap_saprotroph (Põlme et al. 2020); Decay sugar-rich substrates (Põlme et al. 2020) | Põlme S, et al. 2020. Fungal Diversity 105:1-16 ((https://doi.org/10.1007/s13225-020-00466-2)) |
| Otu42020 | 1 | k__Fungi(100);p__Ascomycota(100);c__Saccharomycetes(100);o__Saccharomycetales(100);f__Metschnikowiaceae(100);g__Metschnikowia(100);g__Metschnikowia_unclassified(100); | Metschnikowia | 13 | Saprotroph | \|Undefined Saprotroph\| | Yeast | NULL | Probable | As nectar-tap_saprotroph (Põlme et al. 2020); Decay sugar-rich substrates (Põlme et al. 2020) | Põlme S, et al. 2020. Fungal Diversity 105:1-16 ((https://doi.org/10.1007/s13225-020-00466-2)) |
| Otu42021 | 1 | k__Fungi(100);p__Ascomycota(100);c__Saccharomycetes(100);o__Saccharomycetales(100);f__Metschnikowiaceae(100);g__Metschnikowia(100);s__Metschnikowia_pulcherrima(100); | Metschnikowia | 13 | Saprotroph | \|Undefined Saprotroph\| | Yeast | NULL | Probable | As nectar-tap_saprotroph (Põlme et al. 2020); Decay sugar-rich substrates (Põlme et al. 2020) | Põlme S, et al. 2020. Fungal Diversity 105:1-16 ((https://doi.org/10.1007/s13225-020-00466-2)) |
| Otu42022 | 1 | k__Fungi(100);p__Ascomycota(100);c__Saccharomycetes(100);o__Saccharomycetales(100);f__Metschnikowiaceae(100);g__Metschnikowia(100);s__Metschnikowia_pulcherrima(100); | Metschnikowia | 13 | Saprotroph | \|Undefined Saprotroph\| | Yeast | NULL | Probable | As nectar-tap_saprotroph (Põlme et al. 2020); Decay sugar-rich substrates (Põlme et al. 2020) | Põlme S, et al. 2020. Fungal Diversity 105:1-16 ((https://doi.org/10.1007/s13225-020-00466-2)) |
| Otu42023 | 1 | k__Fungi(100);p__Ascomycota(100);c__Saccharomycetes(100);o__Saccharomycetales(100);f__Metschnikowiaceae(100);g__Metschnikowia(100);s__Metschnikowia_pulcherrima(100); | Metschnikowia | 13 | Saprotroph | \|Undefined Saprotroph\| | Yeast | NULL | Probable | As nectar-tap_saprotroph (Põlme et al. 2020); Decay sugar-rich substrates (Põlme et al. 2020) | Põlme S, et al. 2020. Fungal Diversity 105:1-16 ((https://doi.org/10.1007/s13225-020-00466-2)) |
| Otu42024 | 1 | k__Fungi(100);p__Ascomycota(100);c__Saccharomycetes(100);o__Saccharomycetales(100);f__Metschnikowiaceae(100);g__Metschnikowia(100);s__Metschnikowia_pulcherrima(100); | Metschnikowia | 13 | Saprotroph | \|Undefined Saprotroph\| | Yeast | NULL | Probable | As nectar-tap_saprotroph (Põlme et al. 2020); Decay sugar-rich substrates (Põlme et al. 2020) | Põlme S, et al. 2020. Fungal Diversity 105:1-16 ((https://doi.org/10.1007/s13225-020-00466-2)) |
| Otu42025 | 1 | k__Fungi(100);p__Ascomycota(100);c__Saccharomycetes(100);o__Saccharomycetales(100);f__Metschnikowiaceae(100);g__Metschnikowia(100);s__Metschnikowia_pulcherrima(100); | Metschnikowia | 13 | Saprotroph | \|Undefined Saprotroph\| | Yeast | NULL | Probable | As nectar-tap_saprotroph (Põlme et al. 2020); Decay sugar-rich substrates (Põlme et al. 2020) | Põlme S, et al. 2020. Fungal Diversity 105:1-16 ((https://doi.org/10.1007/s13225-020-00466-2)) |
| Otu42026 | 1 | k__Fungi(100);p__Ascomycota(100);c__Saccharomycetes(100);o__Saccharomycetales(100);f__Metschnikowiaceae(100);g__Metschnikowia(100);g__Metschnikowia_unclassified(100); | Metschnikowia | 13 | Saprotroph | \|Undefined Saprotroph\| | Yeast | NULL | Probable | As nectar-tap_saprotroph (Põlme et al. 2020); Decay sugar-rich substrates (Põlme et al. 2020) | Põlme S, et al. 2020. Fungal Diversity 105:1-16 ((https://doi.org/10.1007/s13225-020-00466-2)) |
| Otu42027 | 1 | k__Fungi(100);p__Ascomycota(100);c__Saccharomycetes(100);o__Saccharomycetales(100);f__Metschnikowiaceae(100);g__Metschnikowia(100);s__Metschnikowia_pulcherrima(100); | Metschnikowia | 13 | Saprotroph | \|Undefined Saprotroph\| | Yeast | NULL | Probable | As nectar-tap_saprotroph (Põlme et al. 2020); Decay sugar-rich substrates (Põlme et al. 2020) | Põlme S, et al. 2020. Fungal Diversity 105:1-16 ((https://doi.org/10.1007/s13225-020-00466-2)) |
| Otu42028 | 1 | k__Fungi(100);p__Ascomycota(100);c__Saccharomycetes(100);o__Saccharomycetales(100);f__Metschnikowiaceae(100);g__Metschnikowia(100);s__Metschnikowia_pulcherrima(100); | Metschnikowia | 13 | Saprotroph | \|Undefined Saprotroph\| | Yeast | NULL | Probable | As nectar-tap_saprotroph (Põlme et al. 2020); Decay sugar-rich substrates (Põlme et al. 2020) | Põlme S, et al. 2020. Fungal Diversity 105:1-16 ((https://doi.org/10.1007/s13225-020-00466-2)) |
| Otu42029 | 1 | k__Fungi(100);p__Ascomycota(100);c__Saccharomycetes(100);o__Saccharomycetales(100);f__Metschnikowiaceae(100);g__Metschnikowia(100);s__Metschnikowia_pulcherrima(100); | Metschnikowia | 13 | Saprotroph | \|Undefined Saprotroph\| | Yeast | NULL | Probable | As nectar-tap_saprotroph (Põlme et al. 2020); Decay sugar-rich substrates (Põlme et al. 2020) | Põlme S, et al. 2020. Fungal Diversity 105:1-16 ((https://doi.org/10.1007/s13225-020-00466-2)) |
| Otu42030 | 1 | k__Fungi(100);p__Ascomycota(100);c__Saccharomycetes(100);o__Saccharomycetales(100);f__Metschnikowiaceae(100);g__Metschnikowia(100);s__Metschnikowia_pulcherrima(100); | Metschnikowia | 13 | Saprotroph | \|Undefined Saprotroph\| | Yeast | NULL | Probable | As nectar-tap_saprotroph (Põlme et al. 2020); Decay sugar-rich substrates (Põlme et al. 2020) | Põlme S, et al. 2020. Fungal Diversity 105:1-16 ((https://doi.org/10.1007/s13225-020-00466-2)) |
| Otu42031 | 1 | k__Fungi(100);p__Ascomycota(100);c__Saccharomycetes(100);o__Saccharomycetales(100);f__Metschnikowiaceae(100);g__Metschnikowia(100);s__Metschnikowia_pulcherrima(100); | Metschnikowia | 13 | Saprotroph | \|Undefined Saprotroph\| | Yeast | NULL | Probable | As nectar-tap_saprotroph (Põlme et al. 2020); Decay sugar-rich substrates (Põlme et al. 2020) | Põlme S, et al. 2020. Fungal Diversity 105:1-16 ((https://doi.org/10.1007/s13225-020-00466-2)) |
| Otu42032 | 1 | k__Fungi(100);p__Ascomycota(100);c__Saccharomycetes(100);o__Saccharomycetales(100);f__Metschnikowiaceae(100);g__Metschnikowia(100);g__Metschnikowia_unclassified(100); | Metschnikowia | 13 | Saprotroph | \|Undefined Saprotroph\| | Yeast | NULL | Probable | As nectar-tap_saprotroph (Põlme et al. 2020); Decay sugar-rich substrates (Põlme et al. 2020) | Põlme S, et al. 2020. Fungal Diversity 105:1-16 ((https://doi.org/10.1007/s13225-020-00466-2)) |
| Otu42033 | 1 | k__Fungi(100);p__Ascomycota(100);c__Saccharomycetes(100);o__Saccharomycetales(100);f__Metschnikowiaceae(100);g__Metschnikowia(100);s__Metschnikowia_pulcherrima(100); | Metschnikowia | 13 | Saprotroph | \|Undefined Saprotroph\| | Yeast | NULL | Probable | As nectar-tap_saprotroph (Põlme et al. 2020); Decay sugar-rich substrates (Põlme et al. 2020) | Põlme S, et al. 2020. Fungal Diversity 105:1-16 ((https://doi.org/10.1007/s13225-020-00466-2)) |
| Otu42034 | 1 | k__Fungi(100);p__Ascomycota(100);c__Saccharomycetes(100);o__Saccharomycetales(100);f__Metschnikowiaceae(100);g__Metschnikowia(100);s__Metschnikowia_pulcherrima(100); | Metschnikowia | 13 | Saprotroph | \|Undefined Saprotroph\| | Yeast | NULL | Probable | As nectar-tap_saprotroph (Põlme et al. 2020); Decay sugar-rich substrates (Põlme et al. 2020) | Põlme S, et al. 2020. Fungal Diversity 105:1-16 ((https://doi.org/10.1007/s13225-020-00466-2)) |
| Otu42035 | 1 | k__Fungi(100);p__Ascomycota(100);c__Saccharomycetes(100);o__Saccharomycetales(100);f__Metschnikowiaceae(100);g__Metschnikowia(100);s__Metschnikowia_pulcherrima(100); | Metschnikowia | 13 | Saprotroph | \|Undefined Saprotroph\| | Yeast | NULL | Probable | As nectar-tap_saprotroph (Põlme et al. 2020); Decay sugar-rich substrates (Põlme et al. 2020) | Põlme S, et al. 2020. Fungal Diversity 105:1-16 ((https://doi.org/10.1007/s13225-020-00466-2)) |
| Otu42036 | 1 | k__Fungi(100);p__Ascomycota(100);c__Saccharomycetes(100);o__Saccharomycetales(100);f__Metschnikowiaceae(100);g__Metschnikowia(100);g__Metschnikowia_unclassified(100); | Metschnikowia | 13 | Saprotroph | \|Undefined Saprotroph\| | Yeast | NULL | Probable | As nectar-tap_saprotroph (Põlme et al. 2020); Decay sugar-rich substrates (Põlme et al. 2020) | Põlme S, et al. 2020. Fungal Diversity 105:1-16 ((https://doi.org/10.1007/s13225-020-00466-2)) |
| Otu42037 | 1 | k__Fungi(100);p__Ascomycota(100);c__Saccharomycetes(100);o__Saccharomycetales(100);f__Metschnikowiaceae(100);g__Metschnikowia(100);s__Metschnikowia_pulcherrima(100); | Metschnikowia | 13 | Saprotroph | \|Undefined Saprotroph\| | Yeast | NULL | Probable | As nectar-tap_saprotroph (Põlme et al. 2020); Decay sugar-rich substrates (Põlme et al. 2020) | Põlme S, et al. 2020. Fungal Diversity 105:1-16 ((https://doi.org/10.1007/s13225-020-00466-2)) |
| Otu42038 | 1 | k__Fungi(100);p__Ascomycota(100);c__Saccharomycetes(100);o__Saccharomycetales(100);f__Metschnikowiaceae(100);g__Metschnikowia(100);s__Metschnikowia_pulcherrima(100); | Metschnikowia | 13 | Saprotroph | \|Undefined Saprotroph\| | Yeast | NULL | Probable | As nectar-tap_saprotroph (Põlme et al. 2020); Decay sugar-rich substrates (Põlme et al. 2020) | Põlme S, et al. 2020. Fungal Diversity 105:1-16 ((https://doi.org/10.1007/s13225-020-00466-2)) |
| Otu42039 | 1 | k__Fungi(100);p__Ascomycota(100);c__Saccharomycetes(100);o__Saccharomycetales(100);f__Metschnikowiaceae(100);g__Metschnikowia(100);g__Metschnikowia_unclassified(100); | Metschnikowia | 13 | Saprotroph | \|Undefined Saprotroph\| | Yeast | NULL | Probable | As nectar-tap_saprotroph (Põlme et al. 2020); Decay sugar-rich substrates (Põlme et al. 2020) | Põlme S, et al. 2020. Fungal Diversity 105:1-16 ((https://doi.org/10.1007/s13225-020-00466-2)) |
| Otu42041 | 1 | k__Fungi(100);p__Ascomycota(100);c__Saccharomycetes(100);o__Saccharomycetales(100);f__Metschnikowiaceae(100);g__Metschnikowia(100);s__Metschnikowia_pulcherrima(100); | Metschnikowia | 13 | Saprotroph | \|Undefined Saprotroph\| | Yeast | NULL | Probable | As nectar-tap_saprotroph (Põlme et al. 2020); Decay sugar-rich substrates (Põlme et al. 2020) | Põlme S, et al. 2020. Fungal Diversity 105:1-16 ((https://doi.org/10.1007/s13225-020-00466-2)) |
| Otu42042 | 1 | k__Fungi(100);p__Ascomycota(100);c__Saccharomycetes(100);o__Saccharomycetales(100);f__Metschnikowiaceae(100);g__Metschnikowia(100);s__Metschnikowia_pulcherrima(100); | Metschnikowia | 13 | Saprotroph | \|Undefined Saprotroph\| | Yeast | NULL | Probable | As nectar-tap_saprotroph (Põlme et al. 2020); Decay sugar-rich substrates (Põlme et al. 2020) | Põlme S, et al. 2020. Fungal Diversity 105:1-16 ((https://doi.org/10.1007/s13225-020-00466-2)) |
| Otu42043 | 1 | k__Fungi(100);p__Ascomycota(100);c__Saccharomycetes(100);o__Saccharomycetales(100);f__Metschnikowiaceae(100);g__Metschnikowia(100);s__Metschnikowia_pulcherrima(100); | Metschnikowia | 13 | Saprotroph | \|Undefined Saprotroph\| | Yeast | NULL | Probable | As nectar-tap_saprotroph (Põlme et al. 2020); Decay sugar-rich substrates (Põlme et al. 2020) | Põlme S, et al. 2020. Fungal Diversity 105:1-16 ((https://doi.org/10.1007/s13225-020-00466-2)) |
| Otu42044 | 1 | k__Fungi(100);p__Ascomycota(100);c__Saccharomycetes(100);o__Saccharomycetales(100);f__Metschnikowiaceae(100);g__Metschnikowia(100);g__Metschnikowia_unclassified(100); | Metschnikowia | 13 | Saprotroph | \|Undefined Saprotroph\| | Yeast | NULL | Probable | As nectar-tap_saprotroph (Põlme et al. 2020); Decay sugar-rich substrates (Põlme et al. 2020) | Põlme S, et al. 2020. Fungal Diversity 105:1-16 ((https://doi.org/10.1007/s13225-020-00466-2)) |
| Otu42046 | 1 | k__Fungi(100);p__Ascomycota(100);c__Saccharomycetes(100);o__Saccharomycetales(100);f__Metschnikowiaceae(100);g__Metschnikowia(100);s__Metschnikowia_pulcherrima(100); | Metschnikowia | 13 | Saprotroph | \|Undefined Saprotroph\| | Yeast | NULL | Probable | As nectar-tap_saprotroph (Põlme et al. 2020); Decay sugar-rich substrates (Põlme et al. 2020) | Põlme S, et al. 2020. Fungal Diversity 105:1-16 ((https://doi.org/10.1007/s13225-020-00466-2)) |
| Otu42047 | 1 | k__Fungi(100);p__Ascomycota(100);c__Saccharomycetes(100);o__Saccharomycetales(100);f__Metschnikowiaceae(100);g__Metschnikowia(100);g__Metschnikowia_unclassified(100); | Metschnikowia | 13 | Saprotroph | \|Undefined Saprotroph\| | Yeast | NULL | Probable | As nectar-tap_saprotroph (Põlme et al. 2020); Decay sugar-rich substrates (Põlme et al. 2020) | Põlme S, et al. 2020. Fungal Diversity 105:1-16 ((https://doi.org/10.1007/s13225-020-00466-2)) |
| Otu42048 | 1 | k__Fungi(100);p__Ascomycota(100);c__Saccharomycetes(100);o__Saccharomycetales(100);f__Metschnikowiaceae(100);g__Metschnikowia(100);g__Metschnikowia_unclassified(100); | Metschnikowia | 13 | Saprotroph | \|Undefined Saprotroph\| | Yeast | NULL | Probable | As nectar-tap_saprotroph (Põlme et al. 2020); Decay sugar-rich substrates (Põlme et al. 2020) | Põlme S, et al. 2020. Fungal Diversity 105:1-16 ((https://doi.org/10.1007/s13225-020-00466-2)) |
| Otu42049 | 1 | k__Fungi(100);p__Ascomycota(100);c__Saccharomycetes(100);o__Saccharomycetales(100);f__Metschnikowiaceae(100);g__Metschnikowia(100);s__Metschnikowia_pulcherrima(100); | Metschnikowia | 13 | Saprotroph | \|Undefined Saprotroph\| | Yeast | NULL | Probable | As nectar-tap_saprotroph (Põlme et al. 2020); Decay sugar-rich substrates (Põlme et al. 2020) | Põlme S, et al. 2020. Fungal Diversity 105:1-16 ((https://doi.org/10.1007/s13225-020-00466-2)) |
| Otu42050 | 1 | k__Fungi(100);p__Ascomycota(100);c__Saccharomycetes(100);o__Saccharomycetales(100);f__Metschnikowiaceae(100);g__Metschnikowia(100);s__Metschnikowia_pulcherrima(100); | Metschnikowia | 13 | Saprotroph | \|Undefined Saprotroph\| | Yeast | NULL | Probable | As nectar-tap_saprotroph (Põlme et al. 2020); Decay sugar-rich substrates (Põlme et al. 2020) | Põlme S, et al. 2020. Fungal Diversity 105:1-16 ((https://doi.org/10.1007/s13225-020-00466-2)) |
| Otu42051 | 1 | k__Fungi(100);p__Ascomycota(100);c__Saccharomycetes(100);o__Saccharomycetales(100);f__Metschnikowiaceae(100);g__Metschnikowia(100);g__Metschnikowia_unclassified(100); | Metschnikowia | 13 | Saprotroph | \|Undefined Saprotroph\| | Yeast | NULL | Probable | As nectar-tap_saprotroph (Põlme et al. 2020); Decay sugar-rich substrates (Põlme et al. 2020) | Põlme S, et al. 2020. Fungal Diversity 105:1-16 ((https://doi.org/10.1007/s13225-020-00466-2)) |
| Otu42052 | 1 | k__Fungi(100);p__Ascomycota(100);c__Saccharomycetes(100);o__Saccharomycetales(100);f__Metschnikowiaceae(100);g__Metschnikowia(100);s__Metschnikowia_pulcherrima(100); | Metschnikowia | 13 | Saprotroph | \|Undefined Saprotroph\| | Yeast | NULL | Probable | As nectar-tap_saprotroph (Põlme et al. 2020); Decay sugar-rich substrates (Põlme et al. 2020) | Põlme S, et al. 2020. Fungal Diversity 105:1-16 ((https://doi.org/10.1007/s13225-020-00466-2)) |
| Otu42053 | 1 | k__Fungi(100);p__Ascomycota(100);c__Saccharomycetes(100);o__Saccharomycetales(100);f__Metschnikowiaceae(100);g__Metschnikowia(100);s__Metschnikowia_pulcherrima(100); | Metschnikowia | 13 | Saprotroph | \|Undefined Saprotroph\| | Yeast | NULL | Probable | As nectar-tap_saprotroph (Põlme et al. 2020); Decay sugar-rich substrates (Põlme et al. 2020) | Põlme S, et al. 2020. Fungal Diversity 105:1-16 ((https://doi.org/10.1007/s13225-020-00466-2)) |
| Otu42055 | 1 | k__Fungi(100);p__Ascomycota(100);c__Saccharomycetes(100);o__Saccharomycetales(100);f__Metschnikowiaceae(100);g__Metschnikowia(100);s__Metschnikowia_pulcherrima(100); | Metschnikowia | 13 | Saprotroph | \|Undefined Saprotroph\| | Yeast | NULL | Probable | As nectar-tap_saprotroph (Põlme et al. 2020); Decay sugar-rich substrates (Põlme et al. 2020) | Põlme S, et al. 2020. Fungal Diversity 105:1-16 ((https://doi.org/10.1007/s13225-020-00466-2)) |
| Otu42056 | 1 | k__Fungi(100);p__Ascomycota(100);c__Saccharomycetes(100);o__Saccharomycetales(100);f__Metschnikowiaceae(100);g__Metschnikowia(100);s__Metschnikowia_pulcherrima(100); | Metschnikowia | 13 | Saprotroph | \|Undefined Saprotroph\| | Yeast | NULL | Probable | As nectar-tap_saprotroph (Põlme et al. 2020); Decay sugar-rich substrates (Põlme et al. 2020) | Põlme S, et al. 2020. Fungal Diversity 105:1-16 ((https://doi.org/10.1007/s13225-020-00466-2)) |
| Otu42057 | 1 | k__Fungi(100);p__Ascomycota(100);c__Saccharomycetes(100);o__Saccharomycetales(100);f__Metschnikowiaceae(100);g__Metschnikowia(100);s__Metschnikowia_pulcherrima(100); | Metschnikowia | 13 | Saprotroph | \|Undefined Saprotroph\| | Yeast | NULL | Probable | As nectar-tap_saprotroph (Põlme et al. 2020); Decay sugar-rich substrates (Põlme et al. 2020) | Põlme S, et al. 2020. Fungal Diversity 105:1-16 ((https://doi.org/10.1007/s13225-020-00466-2)) |
| Otu42058 | 1 | k__Fungi(100);p__Ascomycota(100);c__Saccharomycetes(100);o__Saccharomycetales(100);f__Metschnikowiaceae(100);g__Metschnikowia(100);s__Metschnikowia_pulcherrima(100); | Metschnikowia | 13 | Saprotroph | \|Undefined Saprotroph\| | Yeast | NULL | Probable | As nectar-tap_saprotroph (Põlme et al. 2020); Decay sugar-rich substrates (Põlme et al. 2020) | Põlme S, et al. 2020. Fungal Diversity 105:1-16 ((https://doi.org/10.1007/s13225-020-00466-2)) |
| Otu42059 | 1 | k__Fungi(100);p__Ascomycota(100);c__Saccharomycetes(100);o__Saccharomycetales(100);f__Metschnikowiaceae(100);g__Metschnikowia(100);s__Metschnikowia_pulcherrima(100); | Metschnikowia | 13 | Saprotroph | \|Undefined Saprotroph\| | Yeast | NULL | Probable | As nectar-tap_saprotroph (Põlme et al. 2020); Decay sugar-rich substrates (Põlme et al. 2020) | Põlme S, et al. 2020. Fungal Diversity 105:1-16 ((https://doi.org/10.1007/s13225-020-00466-2)) |
| Otu42060 | 1 | k__Fungi(100);p__Ascomycota(100);c__Saccharomycetes(100);o__Saccharomycetales(100);f__Metschnikowiaceae(100);g__Metschnikowia(100);s__Metschnikowia_pulcherrima(100); | Metschnikowia | 13 | Saprotroph | \|Undefined Saprotroph\| | Yeast | NULL | Probable | As nectar-tap_saprotroph (Põlme et al. 2020); Decay sugar-rich substrates (Põlme et al. 2020) | Põlme S, et al. 2020. Fungal Diversity 105:1-16 ((https://doi.org/10.1007/s13225-020-00466-2)) |
| Otu42061 | 1 | k__Fungi(100);p__Ascomycota(100);c__Saccharomycetes(100);o__Saccharomycetales(100);f__Metschnikowiaceae(100);g__Metschnikowia(100);s__Metschnikowia_pulcherrima(100); | Metschnikowia | 13 | Saprotroph | \|Undefined Saprotroph\| | Yeast | NULL | Probable | As nectar-tap_saprotroph (Põlme et al. 2020); Decay sugar-rich substrates (Põlme et al. 2020) | Põlme S, et al. 2020. Fungal Diversity 105:1-16 ((https://doi.org/10.1007/s13225-020-00466-2)) |
| Otu42062 | 1 | k__Fungi(100);p__Ascomycota(100);c__Saccharomycetes(100);o__Saccharomycetales(100);f__Metschnikowiaceae(100);g__Metschnikowia(100);s__Metschnikowia_pulcherrima(100); | Metschnikowia | 13 | Saprotroph | \|Undefined Saprotroph\| | Yeast | NULL | Probable | As nectar-tap_saprotroph (Põlme et al. 2020); Decay sugar-rich substrates (Põlme et al. 2020) | Põlme S, et al. 2020. Fungal Diversity 105:1-16 ((https://doi.org/10.1007/s13225-020-00466-2)) |
| Otu42063 | 1 | k__Fungi(100);p__Ascomycota(100);c__Saccharomycetes(100);o__Saccharomycetales(100);f__Metschnikowiaceae(100);g__Metschnikowia(100);s__Metschnikowia_pulcherrima(100); | Metschnikowia | 13 | Saprotroph | \|Undefined Saprotroph\| | Yeast | NULL | Probable | As nectar-tap_saprotroph (Põlme et al. 2020); Decay sugar-rich substrates (Põlme et al. 2020) | Põlme S, et al. 2020. Fungal Diversity 105:1-16 ((https://doi.org/10.1007/s13225-020-00466-2)) |
| Otu42064 | 1 | k__Fungi(100);p__Ascomycota(100);c__Saccharomycetes(100);o__Saccharomycetales(100);f__Metschnikowiaceae(100);g__Metschnikowia(100);s__Metschnikowia_pulcherrima(100); | Metschnikowia | 13 | Saprotroph | \|Undefined Saprotroph\| | Yeast | NULL | Probable | As nectar-tap_saprotroph (Põlme et al. 2020); Decay sugar-rich substrates (Põlme et al. 2020) | Põlme S, et al. 2020. Fungal Diversity 105:1-16 ((https://doi.org/10.1007/s13225-020-00466-2)) |
| Otu42065 | 1 | k__Fungi(100);p__Ascomycota(100);c__Saccharomycetes(100);o__Saccharomycetales(100);f__Metschnikowiaceae(100);g__Metschnikowia(100);s__Metschnikowia_pulcherrima(100); | Metschnikowia | 13 | Saprotroph | \|Undefined Saprotroph\| | Yeast | NULL | Probable | As nectar-tap_saprotroph (Põlme et al. 2020); Decay sugar-rich substrates (Põlme et al. 2020) | Põlme S, et al. 2020. Fungal Diversity 105:1-16 ((https://doi.org/10.1007/s13225-020-00466-2)) |
| Otu42066 | 1 | k__Fungi(100);p__Ascomycota(100);c__Saccharomycetes(100);o__Saccharomycetales(100);f__Metschnikowiaceae(100);g__Metschnikowia(100);s__Metschnikowia_pulcherrima(100); | Metschnikowia | 13 | Saprotroph | \|Undefined Saprotroph\| | Yeast | NULL | Probable | As nectar-tap_saprotroph (Põlme et al. 2020); Decay sugar-rich substrates (Põlme et al. 2020) | Põlme S, et al. 2020. Fungal Diversity 105:1-16 ((https://doi.org/10.1007/s13225-020-00466-2)) |
| Otu42067 | 1 | k__Fungi(100);p__Ascomycota(100);c__Saccharomycetes(100);o__Saccharomycetales(100);f__Metschnikowiaceae(100);g__Metschnikowia(100);s__Metschnikowia_pulcherrima(100); | Metschnikowia | 13 | Saprotroph | \|Undefined Saprotroph\| | Yeast | NULL | Probable | As nectar-tap_saprotroph (Põlme et al. 2020); Decay sugar-rich substrates (Põlme et al. 2020) | Põlme S, et al. 2020. Fungal Diversity 105:1-16 ((https://doi.org/10.1007/s13225-020-00466-2)) |
| Otu42068 | 1 | k__Fungi(100);p__Ascomycota(100);c__Saccharomycetes(100);o__Saccharomycetales(100);f__Metschnikowiaceae(100);g__Metschnikowia(100);s__Metschnikowia_pulcherrima(100); | Metschnikowia | 13 | Saprotroph | \|Undefined Saprotroph\| | Yeast | NULL | Probable | As nectar-tap_saprotroph (Põlme et al. 2020); Decay sugar-rich substrates (Põlme et al. 2020) | Põlme S, et al. 2020. Fungal Diversity 105:1-16 ((https://doi.org/10.1007/s13225-020-00466-2)) |
| Otu42069 | 1 | k__Fungi(100);p__Ascomycota(100);c__Saccharomycetes(100);o__Saccharomycetales(100);f__Metschnikowiaceae(100);g__Metschnikowia(100);g__Metschnikowia_unclassified(100); | Metschnikowia | 13 | Saprotroph | \|Undefined Saprotroph\| | Yeast | NULL | Probable | As nectar-tap_saprotroph (Põlme et al. 2020); Decay sugar-rich substrates (Põlme et al. 2020) | Põlme S, et al. 2020. Fungal Diversity 105:1-16 ((https://doi.org/10.1007/s13225-020-00466-2)) |
| Otu42070 | 1 | k__Fungi(100);p__Ascomycota(100);c__Saccharomycetes(100);o__Saccharomycetales(100);f__Metschnikowiaceae(100);g__Metschnikowia(100);s__Metschnikowia_pulcherrima(100); | Metschnikowia | 13 | Saprotroph | \|Undefined Saprotroph\| | Yeast | NULL | Probable | As nectar-tap_saprotroph (Põlme et al. 2020); Decay sugar-rich substrates (Põlme et al. 2020) | Põlme S, et al. 2020. Fungal Diversity 105:1-16 ((https://doi.org/10.1007/s13225-020-00466-2)) |
| Otu42071 | 1 | k__Fungi(100);p__Ascomycota(100);c__Saccharomycetes(100);o__Saccharomycetales(100);f__Metschnikowiaceae(100);g__Metschnikowia(100);s__Metschnikowia_pulcherrima(100); | Metschnikowia | 13 | Saprotroph | \|Undefined Saprotroph\| | Yeast | NULL | Probable | As nectar-tap_saprotroph (Põlme et al. 2020); Decay sugar-rich substrates (Põlme et al. 2020) | Põlme S, et al. 2020. Fungal Diversity 105:1-16 ((https://doi.org/10.1007/s13225-020-00466-2)) |
| Otu42072 | 1 | k__Fungi(100);p__Ascomycota(100);c__Saccharomycetes(100);o__Saccharomycetales(100);f__Metschnikowiaceae(100);g__Metschnikowia(100);s__Metschnikowia_pulcherrima(100); | Metschnikowia | 13 | Saprotroph | \|Undefined Saprotroph\| | Yeast | NULL | Probable | As nectar-tap_saprotroph (Põlme et al. 2020); Decay sugar-rich substrates (Põlme et al. 2020) | Põlme S, et al. 2020. Fungal Diversity 105:1-16 ((https://doi.org/10.1007/s13225-020-00466-2)) |
| Otu42073 | 1 | k__Fungi(100);p__Ascomycota(100);c__Saccharomycetes(100);o__Saccharomycetales(100);f__Metschnikowiaceae(100);g__Metschnikowia(100);s__Metschnikowia_pulcherrima(100); | Metschnikowia | 13 | Saprotroph | \|Undefined Saprotroph\| | Yeast | NULL | Probable | As nectar-tap_saprotroph (Põlme et al. 2020); Decay sugar-rich substrates (Põlme et al. 2020) | Põlme S, et al. 2020. Fungal Diversity 105:1-16 ((https://doi.org/10.1007/s13225-020-00466-2)) |
| Otu42074 | 1 | k__Fungi(100);p__Ascomycota(100);c__Saccharomycetes(100);o__Saccharomycetales(100);f__Metschnikowiaceae(100);g__Metschnikowia(100);s__Metschnikowia_pulcherrima(100); | Metschnikowia | 13 | Saprotroph | \|Undefined Saprotroph\| | Yeast | NULL | Probable | As nectar-tap_saprotroph (Põlme et al. 2020); Decay sugar-rich substrates (Põlme et al. 2020) | Põlme S, et al. 2020. Fungal Diversity 105:1-16 ((https://doi.org/10.1007/s13225-020-00466-2)) |
| Otu42075 | 1 | k__Fungi(100);p__Ascomycota(100);c__Saccharomycetes(100);o__Saccharomycetales(100);f__Metschnikowiaceae(100);g__Metschnikowia(100);g__Metschnikowia_unclassified(100); | Metschnikowia | 13 | Saprotroph | \|Undefined Saprotroph\| | Yeast | NULL | Probable | As nectar-tap_saprotroph (Põlme et al. 2020); Decay sugar-rich substrates (Põlme et al. 2020) | Põlme S, et al. 2020. Fungal Diversity 105:1-16 ((https://doi.org/10.1007/s13225-020-00466-2)) |
| Otu42076 | 1 | k__Fungi(100);p__Ascomycota(100);c__Saccharomycetes(100);o__Saccharomycetales(100);f__Metschnikowiaceae(100);g__Metschnikowia(100);g__Metschnikowia_unclassified(100); | Metschnikowia | 13 | Saprotroph | \|Undefined Saprotroph\| | Yeast | NULL | Probable | As nectar-tap_saprotroph (Põlme et al. 2020); Decay sugar-rich substrates (Põlme et al. 2020) | Põlme S, et al. 2020. Fungal Diversity 105:1-16 ((https://doi.org/10.1007/s13225-020-00466-2)) |
| Otu42077 | 1 | k__Fungi(100);p__Ascomycota(100);c__Saccharomycetes(100);o__Saccharomycetales(100);f__Metschnikowiaceae(100);g__Metschnikowia(100);s__Metschnikowia_pulcherrima(100); | Metschnikowia | 13 | Saprotroph | \|Undefined Saprotroph\| | Yeast | NULL | Probable | As nectar-tap_saprotroph (Põlme et al. 2020); Decay sugar-rich substrates (Põlme et al. 2020) | Põlme S, et al. 2020. Fungal Diversity 105:1-16 ((https://doi.org/10.1007/s13225-020-00466-2)) |
| Otu42078 | 1 | k__Fungi(100);p__Ascomycota(100);c__Saccharomycetes(100);o__Saccharomycetales(100);f__Metschnikowiaceae(100);g__Metschnikowia(100);s__Metschnikowia_pulcherrima(100); | Metschnikowia | 13 | Saprotroph | \|Undefined Saprotroph\| | Yeast | NULL | Probable | As nectar-tap_saprotroph (Põlme et al. 2020); Decay sugar-rich substrates (Põlme et al. 2020) | Põlme S, et al. 2020. Fungal Diversity 105:1-16 ((https://doi.org/10.1007/s13225-020-00466-2)) |
| Otu42079 | 1 | k__Fungi(100);p__Ascomycota(100);c__Saccharomycetes(100);o__Saccharomycetales(100);f__Metschnikowiaceae(100);g__Metschnikowia(100);s__Metschnikowia_pulcherrima(100); | Metschnikowia | 13 | Saprotroph | \|Undefined Saprotroph\| | Yeast | NULL | Probable | As nectar-tap_saprotroph (Põlme et al. 2020); Decay sugar-rich substrates (Põlme et al. 2020) | Põlme S, et al. 2020. Fungal Diversity 105:1-16 ((https://doi.org/10.1007/s13225-020-00466-2)) |
| Otu42080 | 1 | k__Fungi(100);p__Ascomycota(100);c__Saccharomycetes(100);o__Saccharomycetales(100);f__Metschnikowiaceae(100);g__Metschnikowia(100);s__Metschnikowia_pulcherrima(100); | Metschnikowia | 13 | Saprotroph | \|Undefined Saprotroph\| | Yeast | NULL | Probable | As nectar-tap_saprotroph (Põlme et al. 2020); Decay sugar-rich substrates (Põlme et al. 2020) | Põlme S, et al. 2020. Fungal Diversity 105:1-16 ((https://doi.org/10.1007/s13225-020-00466-2)) |
| Otu42081 | 1 | k__Fungi(100);p__Ascomycota(100);c__Saccharomycetes(100);o__Saccharomycetales(100);f__Metschnikowiaceae(100);g__Metschnikowia(100);s__Metschnikowia_pulcherrima(100); | Metschnikowia | 13 | Saprotroph | \|Undefined Saprotroph\| | Yeast | NULL | Probable | As nectar-tap_saprotroph (Põlme et al. 2020); Decay sugar-rich substrates (Põlme et al. 2020) | Põlme S, et al. 2020. Fungal Diversity 105:1-16 ((https://doi.org/10.1007/s13225-020-00466-2)) |
| Otu42082 | 1 | k__Fungi(100);p__Ascomycota(100);c__Saccharomycetes(100);o__Saccharomycetales(100);f__Metschnikowiaceae(100);g__Metschnikowia(100);s__Metschnikowia_pulcherrima(100); | Metschnikowia | 13 | Saprotroph | \|Undefined Saprotroph\| | Yeast | NULL | Probable | As nectar-tap_saprotroph (Põlme et al. 2020); Decay sugar-rich substrates (Põlme et al. 2020) | Põlme S, et al. 2020. Fungal Diversity 105:1-16 ((https://doi.org/10.1007/s13225-020-00466-2)) |
| Otu42083 | 1 | k__Fungi(100);p__Ascomycota(100);c__Saccharomycetes(100);o__Saccharomycetales(100);f__Metschnikowiaceae(100);g__Metschnikowia(100);s__Metschnikowia_pulcherrima(100); | Metschnikowia | 13 | Saprotroph | \|Undefined Saprotroph\| | Yeast | NULL | Probable | As nectar-tap_saprotroph (Põlme et al. 2020); Decay sugar-rich substrates (Põlme et al. 2020) | Põlme S, et al. 2020. Fungal Diversity 105:1-16 ((https://doi.org/10.1007/s13225-020-00466-2)) |
| Otu42084 | 1 | k__Fungi(100);p__Ascomycota(100);c__Saccharomycetes(100);o__Saccharomycetales(100);f__Metschnikowiaceae(100);g__Metschnikowia(100);g__Metschnikowia_unclassified(100); | Metschnikowia | 13 | Saprotroph | \|Undefined Saprotroph\| | Yeast | NULL | Probable | As nectar-tap_saprotroph (Põlme et al. 2020); Decay sugar-rich substrates (Põlme et al. 2020) | Põlme S, et al. 2020. Fungal Diversity 105:1-16 ((https://doi.org/10.1007/s13225-020-00466-2)) |
| Otu42085 | 1 | k__Fungi(100);p__Ascomycota(100);c__Saccharomycetes(100);o__Saccharomycetales(100);f__Metschnikowiaceae(100);g__Metschnikowia(100);s__Metschnikowia_pulcherrima(100); | Metschnikowia | 13 | Saprotroph | \|Undefined Saprotroph\| | Yeast | NULL | Probable | As nectar-tap_saprotroph (Põlme et al. 2020); Decay sugar-rich substrates (Põlme et al. 2020) | Põlme S, et al. 2020. Fungal Diversity 105:1-16 ((https://doi.org/10.1007/s13225-020-00466-2)) |
| Otu42086 | 1 | k__Fungi(100);p__Ascomycota(100);c__Saccharomycetes(100);o__Saccharomycetales(100);f__Metschnikowiaceae(100);g__Metschnikowia(100);g__Metschnikowia_unclassified(100); | Metschnikowia | 13 | Saprotroph | \|Undefined Saprotroph\| | Yeast | NULL | Probable | As nectar-tap_saprotroph (Põlme et al. 2020); Decay sugar-rich substrates (Põlme et al. 2020) | Põlme S, et al. 2020. Fungal Diversity 105:1-16 ((https://doi.org/10.1007/s13225-020-00466-2)) |
| Otu42087 | 1 | k__Fungi(100);p__Ascomycota(100);c__Saccharomycetes(100);o__Saccharomycetales(100);f__Metschnikowiaceae(100);g__Metschnikowia(100);s__Metschnikowia_pulcherrima(100); | Metschnikowia | 13 | Saprotroph | \|Undefined Saprotroph\| | Yeast | NULL | Probable | As nectar-tap_saprotroph (Põlme et al. 2020); Decay sugar-rich substrates (Põlme et al. 2020) | Põlme S, et al. 2020. Fungal Diversity 105:1-16 ((https://doi.org/10.1007/s13225-020-00466-2)) |
| Otu42088 | 1 | k__Fungi(100);p__Ascomycota(100);c__Saccharomycetes(100);o__Saccharomycetales(100);f__Metschnikowiaceae(100);g__Metschnikowia(100);s__Metschnikowia_pulcherrima(100); | Metschnikowia | 13 | Saprotroph | \|Undefined Saprotroph\| | Yeast | NULL | Probable | As nectar-tap_saprotroph (Põlme et al. 2020); Decay sugar-rich substrates (Põlme et al. 2020) | Põlme S, et al. 2020. Fungal Diversity 105:1-16 ((https://doi.org/10.1007/s13225-020-00466-2)) |
| Otu42089 | 1 | k__Fungi(100);p__Ascomycota(100);c__Saccharomycetes(100);o__Saccharomycetales(100);f__Metschnikowiaceae(100);g__Metschnikowia(100);s__Metschnikowia_pulcherrima(100); | Metschnikowia | 13 | Saprotroph | \|Undefined Saprotroph\| | Yeast | NULL | Probable | As nectar-tap_saprotroph (Põlme et al. 2020); Decay sugar-rich substrates (Põlme et al. 2020) | Põlme S, et al. 2020. Fungal Diversity 105:1-16 ((https://doi.org/10.1007/s13225-020-00466-2)) |
| Otu42090 | 1 | k__Fungi(100);p__Ascomycota(100);c__Saccharomycetes(100);o__Saccharomycetales(100);f__Metschnikowiaceae(100);g__Metschnikowia(100);s__Metschnikowia_pulcherrima(100); | Metschnikowia | 13 | Saprotroph | \|Undefined Saprotroph\| | Yeast | NULL | Probable | As nectar-tap_saprotroph (Põlme et al. 2020); Decay sugar-rich substrates (Põlme et al. 2020) | Põlme S, et al. 2020. Fungal Diversity 105:1-16 ((https://doi.org/10.1007/s13225-020-00466-2)) |
| Otu42091 | 1 | k__Fungi(100);p__Ascomycota(100);c__Saccharomycetes(100);o__Saccharomycetales(100);f__Metschnikowiaceae(100);g__Metschnikowia(100);g__Metschnikowia_unclassified(100); | Metschnikowia | 13 | Saprotroph | \|Undefined Saprotroph\| | Yeast | NULL | Probable | As nectar-tap_saprotroph (Põlme et al. 2020); Decay sugar-rich substrates (Põlme et al. 2020) | Põlme S, et al. 2020. Fungal Diversity 105:1-16 ((https://doi.org/10.1007/s13225-020-00466-2)) |
| Otu42092 | 1 | k__Fungi(100);p__Ascomycota(100);c__Saccharomycetes(100);o__Saccharomycetales(100);f__Metschnikowiaceae(100);g__Metschnikowia(100);s__Metschnikowia_pulcherrima(100); | Metschnikowia | 13 | Saprotroph | \|Undefined Saprotroph\| | Yeast | NULL | Probable | As nectar-tap_saprotroph (Põlme et al. 2020); Decay sugar-rich substrates (Põlme et al. 2020) | Põlme S, et al. 2020. Fungal Diversity 105:1-16 ((https://doi.org/10.1007/s13225-020-00466-2)) |
| Otu42093 | 1 | k__Fungi(100);p__Ascomycota(100);c__Saccharomycetes(100);o__Saccharomycetales(100);f__Metschnikowiaceae(100);g__Metschnikowia(100);s__Metschnikowia_pulcherrima(100); | Metschnikowia | 13 | Saprotroph | \|Undefined Saprotroph\| | Yeast | NULL | Probable | As nectar-tap_saprotroph (Põlme et al. 2020); Decay sugar-rich substrates (Põlme et al. 2020) | Põlme S, et al. 2020. Fungal Diversity 105:1-16 ((https://doi.org/10.1007/s13225-020-00466-2)) |
| Otu42094 | 1 | k__Fungi(100);p__Ascomycota(100);c__Saccharomycetes(100);o__Saccharomycetales(100);f__Metschnikowiaceae(100);g__Metschnikowia(100);s__Metschnikowia_pulcherrima(100); | Metschnikowia | 13 | Saprotroph | \|Undefined Saprotroph\| | Yeast | NULL | Probable | As nectar-tap_saprotroph (Põlme et al. 2020); Decay sugar-rich substrates (Põlme et al. 2020) | Põlme S, et al. 2020. Fungal Diversity 105:1-16 ((https://doi.org/10.1007/s13225-020-00466-2)) |
| Otu42095 | 1 | k__Fungi(100);p__Ascomycota(100);c__Saccharomycetes(100);o__Saccharomycetales(100);f__Metschnikowiaceae(100);g__Metschnikowia(100);s__Metschnikowia_pulcherrima(100); | Metschnikowia | 13 | Saprotroph | \|Undefined Saprotroph\| | Yeast | NULL | Probable | As nectar-tap_saprotroph (Põlme et al. 2020); Decay sugar-rich substrates (Põlme et al. 2020) | Põlme S, et al. 2020. Fungal Diversity 105:1-16 ((https://doi.org/10.1007/s13225-020-00466-2)) |
| Otu42096 | 1 | k__Fungi(100);p__Ascomycota(100);c__Saccharomycetes(100);o__Saccharomycetales(100);f__Metschnikowiaceae(100);g__Metschnikowia(100);g__Metschnikowia_unclassified(100); | Metschnikowia | 13 | Saprotroph | \|Undefined Saprotroph\| | Yeast | NULL | Probable | As nectar-tap_saprotroph (Põlme et al. 2020); Decay sugar-rich substrates (Põlme et al. 2020) | Põlme S, et al. 2020. Fungal Diversity 105:1-16 ((https://doi.org/10.1007/s13225-020-00466-2)) |
| Otu42097 | 1 | k__Fungi(100);p__Ascomycota(100);c__Saccharomycetes(100);o__Saccharomycetales(100);f__Metschnikowiaceae(100);g__Metschnikowia(100);s__Metschnikowia_pulcherrima(100); | Metschnikowia | 13 | Saprotroph | \|Undefined Saprotroph\| | Yeast | NULL | Probable | As nectar-tap_saprotroph (Põlme et al. 2020); Decay sugar-rich substrates (Põlme et al. 2020) | Põlme S, et al. 2020. Fungal Diversity 105:1-16 ((https://doi.org/10.1007/s13225-020-00466-2)) |
| Otu42098 | 1 | k__Fungi(100);p__Ascomycota(100);c__Saccharomycetes(100);o__Saccharomycetales(100);f__Metschnikowiaceae(100);g__Metschnikowia(100);s__Metschnikowia_pulcherrima(100); | Metschnikowia | 13 | Saprotroph | \|Undefined Saprotroph\| | Yeast | NULL | Probable | As nectar-tap_saprotroph (Põlme et al. 2020); Decay sugar-rich substrates (Põlme et al. 2020) | Põlme S, et al. 2020. Fungal Diversity 105:1-16 ((https://doi.org/10.1007/s13225-020-00466-2)) |
| Otu42099 | 1 | k__Fungi(100);p__Ascomycota(100);c__Saccharomycetes(100);o__Saccharomycetales(100);f__Metschnikowiaceae(100);g__Metschnikowia(100);g__Metschnikowia_unclassified(100); | Metschnikowia | 13 | Saprotroph | \|Undefined Saprotroph\| | Yeast | NULL | Probable | As nectar-tap_saprotroph (Põlme et al. 2020); Decay sugar-rich substrates (Põlme et al. 2020) | Põlme S, et al. 2020. Fungal Diversity 105:1-16 ((https://doi.org/10.1007/s13225-020-00466-2)) |
| Otu42100 | 1 | k__Fungi(100);p__Ascomycota(100);c__Saccharomycetes(100);o__Saccharomycetales(100);f__Metschnikowiaceae(100);g__Metschnikowia(100);s__Metschnikowia_pulcherrima(100); | Metschnikowia | 13 | Saprotroph | \|Undefined Saprotroph\| | Yeast | NULL | Probable | As nectar-tap_saprotroph (Põlme et al. 2020); Decay sugar-rich substrates (Põlme et al. 2020) | Põlme S, et al. 2020. Fungal Diversity 105:1-16 ((https://doi.org/10.1007/s13225-020-00466-2)) |
| Otu42101 | 1 | k__Fungi(100);p__Ascomycota(100);c__Saccharomycetes(100);o__Saccharomycetales(100);f__Metschnikowiaceae(100);g__Metschnikowia(100);g__Metschnikowia_unclassified(100); | Metschnikowia | 13 | Saprotroph | \|Undefined Saprotroph\| | Yeast | NULL | Probable | As nectar-tap_saprotroph (Põlme et al. 2020); Decay sugar-rich substrates (Põlme et al. 2020) | Põlme S, et al. 2020. Fungal Diversity 105:1-16 ((https://doi.org/10.1007/s13225-020-00466-2)) |
| Otu42102 | 1 | k__Fungi(100);p__Ascomycota(100);c__Saccharomycetes(100);o__Saccharomycetales(100);f__Metschnikowiaceae(100);g__Metschnikowia(100);s__Metschnikowia_pulcherrima(100); | Metschnikowia | 13 | Saprotroph | \|Undefined Saprotroph\| | Yeast | NULL | Probable | As nectar-tap_saprotroph (Põlme et al. 2020); Decay sugar-rich substrates (Põlme et al. 2020) | Põlme S, et al. 2020. Fungal Diversity 105:1-16 ((https://doi.org/10.1007/s13225-020-00466-2)) |
| Otu42103 | 1 | k__Fungi(100);p__Ascomycota(100);c__Saccharomycetes(100);o__Saccharomycetales(100);f__Metschnikowiaceae(100);g__Metschnikowia(100);s__Metschnikowia_pulcherrima(100); | Metschnikowia | 13 | Saprotroph | \|Undefined Saprotroph\| | Yeast | NULL | Probable | As nectar-tap_saprotroph (Põlme et al. 2020); Decay sugar-rich substrates (Põlme et al. 2020) | Põlme S, et al. 2020. Fungal Diversity 105:1-16 ((https://doi.org/10.1007/s13225-020-00466-2)) |
| Otu42104 | 1 | k__Fungi(100);p__Ascomycota(100);c__Saccharomycetes(100);o__Saccharomycetales(100);f__Metschnikowiaceae(100);g__Metschnikowia(100);g__Metschnikowia_unclassified(100); | Metschnikowia | 13 | Saprotroph | \|Undefined Saprotroph\| | Yeast | NULL | Probable | As nectar-tap_saprotroph (Põlme et al. 2020); Decay sugar-rich substrates (Põlme et al. 2020) | Põlme S, et al. 2020. Fungal Diversity 105:1-16 ((https://doi.org/10.1007/s13225-020-00466-2)) |
| Otu42105 | 1 | k__Fungi(100);p__Ascomycota(100);c__Saccharomycetes(100);o__Saccharomycetales(100);f__Metschnikowiaceae(100);g__Metschnikowia(100);g__Metschnikowia_unclassified(100); | Metschnikowia | 13 | Saprotroph | \|Undefined Saprotroph\| | Yeast | NULL | Probable | As nectar-tap_saprotroph (Põlme et al. 2020); Decay sugar-rich substrates (Põlme et al. 2020) | Põlme S, et al. 2020. Fungal Diversity 105:1-16 ((https://doi.org/10.1007/s13225-020-00466-2)) |
| Otu42106 | 1 | k__Fungi(100);p__Ascomycota(100);c__Saccharomycetes(100);o__Saccharomycetales(100);f__Metschnikowiaceae(100);g__Metschnikowia(100);s__Metschnikowia_pulcherrima(100); | Metschnikowia | 13 | Saprotroph | \|Undefined Saprotroph\| | Yeast | NULL | Probable | As nectar-tap_saprotroph (Põlme et al. 2020); Decay sugar-rich substrates (Põlme et al. 2020) | Põlme S, et al. 2020. Fungal Diversity 105:1-16 ((https://doi.org/10.1007/s13225-020-00466-2)) |
| Otu42107 | 1 | k__Fungi(100);p__Ascomycota(100);c__Saccharomycetes(100);o__Saccharomycetales(100);f__Metschnikowiaceae(100);g__Metschnikowia(100);s__Metschnikowia_pulcherrima(100); | Metschnikowia | 13 | Saprotroph | \|Undefined Saprotroph\| | Yeast | NULL | Probable | As nectar-tap_saprotroph (Põlme et al. 2020); Decay sugar-rich substrates (Põlme et al. 2020) | Põlme S, et al. 2020. Fungal Diversity 105:1-16 ((https://doi.org/10.1007/s13225-020-00466-2)) |
| Otu42108 | 1 | k__Fungi(100);p__Ascomycota(100);c__Saccharomycetes(100);o__Saccharomycetales(100);f__Metschnikowiaceae(100);g__Metschnikowia(100);s__Metschnikowia_pulcherrima(100); | Metschnikowia | 13 | Saprotroph | \|Undefined Saprotroph\| | Yeast | NULL | Probable | As nectar-tap_saprotroph (Põlme et al. 2020); Decay sugar-rich substrates (Põlme et al. 2020) | Põlme S, et al. 2020. Fungal Diversity 105:1-16 ((https://doi.org/10.1007/s13225-020-00466-2)) |
| Otu42109 | 1 | k__Fungi(100);p__Ascomycota(100);c__Saccharomycetes(100);o__Saccharomycetales(100);f__Metschnikowiaceae(100);g__Metschnikowia(100);s__Metschnikowia_pulcherrima(100); | Metschnikowia | 13 | Saprotroph | \|Undefined Saprotroph\| | Yeast | NULL | Probable | As nectar-tap_saprotroph (Põlme et al. 2020); Decay sugar-rich substrates (Põlme et al. 2020) | Põlme S, et al. 2020. Fungal Diversity 105:1-16 ((https://doi.org/10.1007/s13225-020-00466-2)) |
| Otu42110 | 1 | k__Fungi(100);p__Ascomycota(100);c__Saccharomycetes(100);o__Saccharomycetales(100);f__Metschnikowiaceae(100);g__Metschnikowia(100);s__Metschnikowia_pulcherrima(100); | Metschnikowia | 13 | Saprotroph | \|Undefined Saprotroph\| | Yeast | NULL | Probable | As nectar-tap_saprotroph (Põlme et al. 2020); Decay sugar-rich substrates (Põlme et al. 2020) | Põlme S, et al. 2020. Fungal Diversity 105:1-16 ((https://doi.org/10.1007/s13225-020-00466-2)) |
| Otu42111 | 1 | k__Fungi(100);p__Ascomycota(100);c__Saccharomycetes(100);o__Saccharomycetales(100);f__Metschnikowiaceae(100);g__Metschnikowia(100);s__Metschnikowia_pulcherrima(100); | Metschnikowia | 13 | Saprotroph | \|Undefined Saprotroph\| | Yeast | NULL | Probable | As nectar-tap_saprotroph (Põlme et al. 2020); Decay sugar-rich substrates (Põlme et al. 2020) | Põlme S, et al. 2020. Fungal Diversity 105:1-16 ((https://doi.org/10.1007/s13225-020-00466-2)) |
| Otu42112 | 1 | k__Fungi(100);p__Ascomycota(100);c__Saccharomycetes(100);o__Saccharomycetales(100);f__Metschnikowiaceae(100);g__Metschnikowia(100);s__Metschnikowia_pulcherrima(100); | Metschnikowia | 13 | Saprotroph | \|Undefined Saprotroph\| | Yeast | NULL | Probable | As nectar-tap_saprotroph (Põlme et al. 2020); Decay sugar-rich substrates (Põlme et al. 2020) | Põlme S, et al. 2020. Fungal Diversity 105:1-16 ((https://doi.org/10.1007/s13225-020-00466-2)) |
| Otu42113 | 1 | k__Fungi(100);p__Ascomycota(100);c__Saccharomycetes(100);o__Saccharomycetales(100);f__Metschnikowiaceae(100);g__Metschnikowia(100);s__Metschnikowia_pulcherrima(100); | Metschnikowia | 13 | Saprotroph | \|Undefined Saprotroph\| | Yeast | NULL | Probable | As nectar-tap_saprotroph (Põlme et al. 2020); Decay sugar-rich substrates (Põlme et al. 2020) | Põlme S, et al. 2020. Fungal Diversity 105:1-16 ((https://doi.org/10.1007/s13225-020-00466-2)) |
| Otu42114 | 1 | k__Fungi(100);p__Ascomycota(100);c__Saccharomycetes(100);o__Saccharomycetales(100);f__Metschnikowiaceae(100);g__Metschnikowia(100);s__Metschnikowia_pulcherrima(100); | Metschnikowia | 13 | Saprotroph | \|Undefined Saprotroph\| | Yeast | NULL | Probable | As nectar-tap_saprotroph (Põlme et al. 2020); Decay sugar-rich substrates (Põlme et al. 2020) | Põlme S, et al. 2020. Fungal Diversity 105:1-16 ((https://doi.org/10.1007/s13225-020-00466-2)) |
| Otu42115 | 1 | k__Fungi(100);p__Ascomycota(100);c__Saccharomycetes(100);o__Saccharomycetales(100);f__Metschnikowiaceae(100);g__Metschnikowia(100);s__Metschnikowia_pulcherrima(100); | Metschnikowia | 13 | Saprotroph | \|Undefined Saprotroph\| | Yeast | NULL | Probable | As nectar-tap_saprotroph (Põlme et al. 2020); Decay sugar-rich substrates (Põlme et al. 2020) | Põlme S, et al. 2020. Fungal Diversity 105:1-16 ((https://doi.org/10.1007/s13225-020-00466-2)) |
| Otu42116 | 1 | k__Fungi(100);p__Ascomycota(100);c__Saccharomycetes(100);o__Saccharomycetales(100);f__Metschnikowiaceae(100);g__Metschnikowia(100);s__Metschnikowia_pulcherrima(100); | Metschnikowia | 13 | Saprotroph | \|Undefined Saprotroph\| | Yeast | NULL | Probable | As nectar-tap_saprotroph (Põlme et al. 2020); Decay sugar-rich substrates (Põlme et al. 2020) | Põlme S, et al. 2020. Fungal Diversity 105:1-16 ((https://doi.org/10.1007/s13225-020-00466-2)) |
| Otu42117 | 1 | k__Fungi(100);p__Ascomycota(100);c__Saccharomycetes(100);o__Saccharomycetales(100);f__Metschnikowiaceae(100);g__Metschnikowia(100);s__Metschnikowia_pulcherrima(100); | Metschnikowia | 13 | Saprotroph | \|Undefined Saprotroph\| | Yeast | NULL | Probable | As nectar-tap_saprotroph (Põlme et al. 2020); Decay sugar-rich substrates (Põlme et al. 2020) | Põlme S, et al. 2020. Fungal Diversity 105:1-16 ((https://doi.org/10.1007/s13225-020-00466-2)) |
| Otu42118 | 1 | k__Fungi(100);p__Ascomycota(100);c__Saccharomycetes(100);o__Saccharomycetales(100);f__Metschnikowiaceae(100);g__Metschnikowia(100);g__Metschnikowia_unclassified(100); | Metschnikowia | 13 | Saprotroph | \|Undefined Saprotroph\| | Yeast | NULL | Probable | As nectar-tap_saprotroph (Põlme et al. 2020); Decay sugar-rich substrates (Põlme et al. 2020) | Põlme S, et al. 2020. Fungal Diversity 105:1-16 ((https://doi.org/10.1007/s13225-020-00466-2)) |
| Otu42119 | 1 | k__Fungi(100);p__Ascomycota(100);c__Saccharomycetes(100);o__Saccharomycetales(100);f__Metschnikowiaceae(100);g__Metschnikowia(100);s__Metschnikowia_pulcherrima(100); | Metschnikowia | 13 | Saprotroph | \|Undefined Saprotroph\| | Yeast | NULL | Probable | As nectar-tap_saprotroph (Põlme et al. 2020); Decay sugar-rich substrates (Põlme et al. 2020) | Põlme S, et al. 2020. Fungal Diversity 105:1-16 ((https://doi.org/10.1007/s13225-020-00466-2)) |
| Otu42120 | 1 | k__Fungi(100);p__Ascomycota(100);c__Saccharomycetes(100);o__Saccharomycetales(100);f__Metschnikowiaceae(100);g__Metschnikowia(100);s__Metschnikowia_pulcherrima(100); | Metschnikowia | 13 | Saprotroph | \|Undefined Saprotroph\| | Yeast | NULL | Probable | As nectar-tap_saprotroph (Põlme et al. 2020); Decay sugar-rich substrates (Põlme et al. 2020) | Põlme S, et al. 2020. Fungal Diversity 105:1-16 ((https://doi.org/10.1007/s13225-020-00466-2)) |
| Otu42121 | 1 | k__Fungi(100);p__Ascomycota(100);c__Saccharomycetes(100);o__Saccharomycetales(100);f__Metschnikowiaceae(100);g__Metschnikowia(100);s__Metschnikowia_pulcherrima(100); | Metschnikowia | 13 | Saprotroph | \|Undefined Saprotroph\| | Yeast | NULL | Probable | As nectar-tap_saprotroph (Põlme et al. 2020); Decay sugar-rich substrates (Põlme et al. 2020) | Põlme S, et al. 2020. Fungal Diversity 105:1-16 ((https://doi.org/10.1007/s13225-020-00466-2)) |
| Otu42123 | 1 | k__Fungi(100);p__Ascomycota(100);c__Saccharomycetes(100);o__Saccharomycetales(100);f__Metschnikowiaceae(100);g__Metschnikowia(100);s__Metschnikowia_pulcherrima(100); | Metschnikowia | 13 | Saprotroph | \|Undefined Saprotroph\| | Yeast | NULL | Probable | As nectar-tap_saprotroph (Põlme et al. 2020); Decay sugar-rich substrates (Põlme et al. 2020) | Põlme S, et al. 2020. Fungal Diversity 105:1-16 ((https://doi.org/10.1007/s13225-020-00466-2)) |
| Otu42125 | 1 | k__Fungi(100);p__Ascomycota(100);c__Saccharomycetes(100);o__Saccharomycetales(100);f__Metschnikowiaceae(100);g__Metschnikowia(100);s__Metschnikowia_pulcherrima(100); | Metschnikowia | 13 | Saprotroph | \|Undefined Saprotroph\| | Yeast | NULL | Probable | As nectar-tap_saprotroph (Põlme et al. 2020); Decay sugar-rich substrates (Põlme et al. 2020) | Põlme S, et al. 2020. Fungal Diversity 105:1-16 ((https://doi.org/10.1007/s13225-020-00466-2)) |
| Otu42126 | 1 | k__Fungi(100);p__Ascomycota(100);c__Saccharomycetes(100);o__Saccharomycetales(100);f__Metschnikowiaceae(100);g__Metschnikowia(100);s__Metschnikowia_pulcherrima(100); | Metschnikowia | 13 | Saprotroph | \|Undefined Saprotroph\| | Yeast | NULL | Probable | As nectar-tap_saprotroph (Põlme et al. 2020); Decay sugar-rich substrates (Põlme et al. 2020) | Põlme S, et al. 2020. Fungal Diversity 105:1-16 ((https://doi.org/10.1007/s13225-020-00466-2)) |
| Otu42127 | 1 | k__Fungi(100);p__Ascomycota(100);c__Saccharomycetes(100);o__Saccharomycetales(100);f__Metschnikowiaceae(100);g__Metschnikowia(100);s__Metschnikowia_pulcherrima(100); | Metschnikowia | 13 | Saprotroph | \|Undefined Saprotroph\| | Yeast | NULL | Probable | As nectar-tap_saprotroph (Põlme et al. 2020); Decay sugar-rich substrates (Põlme et al. 2020) | Põlme S, et al. 2020. Fungal Diversity 105:1-16 ((https://doi.org/10.1007/s13225-020-00466-2)) |
| Otu42130 | 1 | k__Fungi(100);p__Ascomycota(100);c__Saccharomycetes(100);o__Saccharomycetales(100);f__Metschnikowiaceae(100);g__Metschnikowia(100);s__Metschnikowia_pulcherrima(100); | Metschnikowia | 13 | Saprotroph | \|Undefined Saprotroph\| | Yeast | NULL | Probable | As nectar-tap_saprotroph (Põlme et al. 2020); Decay sugar-rich substrates (Põlme et al. 2020) | Põlme S, et al. 2020. Fungal Diversity 105:1-16 ((https://doi.org/10.1007/s13225-020-00466-2)) |
| Otu42131 | 1 | k__Fungi(100);p__Ascomycota(100);c__Saccharomycetes(100);o__Saccharomycetales(100);f__Metschnikowiaceae(100);g__Metschnikowia(100);g__Metschnikowia_unclassified(100); | Metschnikowia | 13 | Saprotroph | \|Undefined Saprotroph\| | Yeast | NULL | Probable | As nectar-tap_saprotroph (Põlme et al. 2020); Decay sugar-rich substrates (Põlme et al. 2020) | Põlme S, et al. 2020. Fungal Diversity 105:1-16 ((https://doi.org/10.1007/s13225-020-00466-2)) |
| Otu42132 | 1 | k__Fungi(100);p__Ascomycota(100);c__Saccharomycetes(100);o__Saccharomycetales(100);f__Metschnikowiaceae(100);g__Metschnikowia(100);g__Metschnikowia_unclassified(100); | Metschnikowia | 13 | Saprotroph | \|Undefined Saprotroph\| | Yeast | NULL | Probable | As nectar-tap_saprotroph (Põlme et al. 2020); Decay sugar-rich substrates (Põlme et al. 2020) | Põlme S, et al. 2020. Fungal Diversity 105:1-16 ((https://doi.org/10.1007/s13225-020-00466-2)) |
| Otu42133 | 1 | k__Fungi(100);p__Ascomycota(100);c__Saccharomycetes(100);o__Saccharomycetales(100);f__Metschnikowiaceae(100);g__Metschnikowia(100);g__Metschnikowia_unclassified(100); | Metschnikowia | 13 | Saprotroph | \|Undefined Saprotroph\| | Yeast | NULL | Probable | As nectar-tap_saprotroph (Põlme et al. 2020); Decay sugar-rich substrates (Põlme et al. 2020) | Põlme S, et al. 2020. Fungal Diversity 105:1-16 ((https://doi.org/10.1007/s13225-020-00466-2)) |
| Otu42134 | 1 | k__Fungi(100);p__Ascomycota(100);c__Saccharomycetes(100);o__Saccharomycetales(100);f__Metschnikowiaceae(100);g__Metschnikowia(100);s__Metschnikowia_pulcherrima(100); | Metschnikowia | 13 | Saprotroph | \|Undefined Saprotroph\| | Yeast | NULL | Probable | As nectar-tap_saprotroph (Põlme et al. 2020); Decay sugar-rich substrates (Põlme et al. 2020) | Põlme S, et al. 2020. Fungal Diversity 105:1-16 ((https://doi.org/10.1007/s13225-020-00466-2)) |
| Otu42135 | 1 | k__Fungi(100);p__Ascomycota(100);c__Saccharomycetes(100);o__Saccharomycetales(100);f__Metschnikowiaceae(100);g__Metschnikowia(100);s__Metschnikowia_pulcherrima(100); | Metschnikowia | 13 | Saprotroph | \|Undefined Saprotroph\| | Yeast | NULL | Probable | As nectar-tap_saprotroph (Põlme et al. 2020); Decay sugar-rich substrates (Põlme et al. 2020) | Põlme S, et al. 2020. Fungal Diversity 105:1-16 ((https://doi.org/10.1007/s13225-020-00466-2)) |
| Otu42136 | 1 | k__Fungi(100);p__Ascomycota(100);c__Saccharomycetes(100);o__Saccharomycetales(100);f__Metschnikowiaceae(100);g__Metschnikowia(100);s__Metschnikowia_pulcherrima(100); | Metschnikowia | 13 | Saprotroph | \|Undefined Saprotroph\| | Yeast | NULL | Probable | As nectar-tap_saprotroph (Põlme et al. 2020); Decay sugar-rich substrates (Põlme et al. 2020) | Põlme S, et al. 2020. Fungal Diversity 105:1-16 ((https://doi.org/10.1007/s13225-020-00466-2)) |
| Otu42139 | 1 | k__Fungi(100);p__Ascomycota(100);c__Saccharomycetes(100);o__Saccharomycetales(100);f__Metschnikowiaceae(100);g__Metschnikowia(100);g__Metschnikowia_unclassified(100); | Metschnikowia | 13 | Saprotroph | \|Undefined Saprotroph\| | Yeast | NULL | Probable | As nectar-tap_saprotroph (Põlme et al. 2020); Decay sugar-rich substrates (Põlme et al. 2020) | Põlme S, et al. 2020. Fungal Diversity 105:1-16 ((https://doi.org/10.1007/s13225-020-00466-2)) |
| Otu42141 | 1 | k__Fungi(100);p__Ascomycota(100);c__Saccharomycetes(100);o__Saccharomycetales(100);f__Metschnikowiaceae(100);g__Metschnikowia(100);s__Metschnikowia_pulcherrima(100); | Metschnikowia | 13 | Saprotroph | \|Undefined Saprotroph\| | Yeast | NULL | Probable | As nectar-tap_saprotroph (Põlme et al. 2020); Decay sugar-rich substrates (Põlme et al. 2020) | Põlme S, et al. 2020. Fungal Diversity 105:1-16 ((https://doi.org/10.1007/s13225-020-00466-2)) |
| Otu42142 | 1 | k__Fungi(100);p__Ascomycota(100);c__Saccharomycetes(100);o__Saccharomycetales(100);f__Metschnikowiaceae(100);g__Metschnikowia(100);s__Metschnikowia_pulcherrima(100); | Metschnikowia | 13 | Saprotroph | \|Undefined Saprotroph\| | Yeast | NULL | Probable | As nectar-tap_saprotroph (Põlme et al. 2020); Decay sugar-rich substrates (Põlme et al. 2020) | Põlme S, et al. 2020. Fungal Diversity 105:1-16 ((https://doi.org/10.1007/s13225-020-00466-2)) |
| Otu42143 | 1 | k__Fungi(100);p__Ascomycota(100);c__Saccharomycetes(100);o__Saccharomycetales(100);f__Metschnikowiaceae(100);g__Metschnikowia(100);s__Metschnikowia_pulcherrima(100); | Metschnikowia | 13 | Saprotroph | \|Undefined Saprotroph\| | Yeast | NULL | Probable | As nectar-tap_saprotroph (Põlme et al. 2020); Decay sugar-rich substrates (Põlme et al. 2020) | Põlme S, et al. 2020. Fungal Diversity 105:1-16 ((https://doi.org/10.1007/s13225-020-00466-2)) |
| Otu42144 | 1 | k__Fungi(100);p__Ascomycota(100);c__Saccharomycetes(100);o__Saccharomycetales(100);f__Metschnikowiaceae(100);g__Metschnikowia(100);s__Metschnikowia_pulcherrima(100); | Metschnikowia | 13 | Saprotroph | \|Undefined Saprotroph\| | Yeast | NULL | Probable | As nectar-tap_saprotroph (Põlme et al. 2020); Decay sugar-rich substrates (Põlme et al. 2020) | Põlme S, et al. 2020. Fungal Diversity 105:1-16 ((https://doi.org/10.1007/s13225-020-00466-2)) |
| Otu42145 | 1 | k__Fungi(100);p__Ascomycota(100);c__Saccharomycetes(100);o__Saccharomycetales(100);f__Metschnikowiaceae(100);g__Metschnikowia(100);g__Metschnikowia_unclassified(100); | Metschnikowia | 13 | Saprotroph | \|Undefined Saprotroph\| | Yeast | NULL | Probable | As nectar-tap_saprotroph (Põlme et al. 2020); Decay sugar-rich substrates (Põlme et al. 2020) | Põlme S, et al. 2020. Fungal Diversity 105:1-16 ((https://doi.org/10.1007/s13225-020-00466-2)) |
| Otu42146 | 1 | k__Fungi(100);p__Ascomycota(100);c__Saccharomycetes(100);o__Saccharomycetales(100);f__Metschnikowiaceae(100);g__Metschnikowia(100);s__Metschnikowia_pulcherrima(100); | Metschnikowia | 13 | Saprotroph | \|Undefined Saprotroph\| | Yeast | NULL | Probable | As nectar-tap_saprotroph (Põlme et al. 2020); Decay sugar-rich substrates (Põlme et al. 2020) | Põlme S, et al. 2020. Fungal Diversity 105:1-16 ((https://doi.org/10.1007/s13225-020-00466-2)) |
| Otu42147 | 1 | k__Fungi(100);p__Ascomycota(100);c__Saccharomycetes(100);o__Saccharomycetales(100);f__Metschnikowiaceae(100);g__Metschnikowia(100);g__Metschnikowia_unclassified(100); | Metschnikowia | 13 | Saprotroph | \|Undefined Saprotroph\| | Yeast | NULL | Probable | As nectar-tap_saprotroph (Põlme et al. 2020); Decay sugar-rich substrates (Põlme et al. 2020) | Põlme S, et al. 2020. Fungal Diversity 105:1-16 ((https://doi.org/10.1007/s13225-020-00466-2)) |
| Otu42148 | 1 | k__Fungi(100);p__Ascomycota(100);c__Saccharomycetes(100);o__Saccharomycetales(100);f__Metschnikowiaceae(100);g__Metschnikowia(100);s__Metschnikowia_pulcherrima(100); | Metschnikowia | 13 | Saprotroph | \|Undefined Saprotroph\| | Yeast | NULL | Probable | As nectar-tap_saprotroph (Põlme et al. 2020); Decay sugar-rich substrates (Põlme et al. 2020) | Põlme S, et al. 2020. Fungal Diversity 105:1-16 ((https://doi.org/10.1007/s13225-020-00466-2)) |
| Otu42153 | 1 | k__Fungi(100);p__Ascomycota(100);c__Saccharomycetes(100);o__Saccharomycetales(100);f__Metschnikowiaceae(100);g__Metschnikowia(100);s__Metschnikowia_pulcherrima(100); | Metschnikowia | 13 | Saprotroph | \|Undefined Saprotroph\| | Yeast | NULL | Probable | As nectar-tap_saprotroph (Põlme et al. 2020); Decay sugar-rich substrates (Põlme et al. 2020) | Põlme S, et al. 2020. Fungal Diversity 105:1-16 ((https://doi.org/10.1007/s13225-020-00466-2)) |
| Otu42164 | 1 | k__Fungi(100);p__Ascomycota(100);c__Saccharomycetes(100);o__Saccharomycetales(100);f__Metschnikowiaceae(100);g__Metschnikowia(100);g__Metschnikowia_unclassified(100); | Metschnikowia | 13 | Saprotroph | \|Undefined Saprotroph\| | Yeast | NULL | Probable | As nectar-tap_saprotroph (Põlme et al. 2020); Decay sugar-rich substrates (Põlme et al. 2020) | Põlme S, et al. 2020. Fungal Diversity 105:1-16 ((https://doi.org/10.1007/s13225-020-00466-2)) |
| Otu42165 | 1 | k__Fungi(100);p__Ascomycota(100);c__Saccharomycetes(100);o__Saccharomycetales(100);f__Metschnikowiaceae(100);g__Metschnikowia(100);s__Metschnikowia_pulcherrima(100); | Metschnikowia | 13 | Saprotroph | \|Undefined Saprotroph\| | Yeast | NULL | Probable | As nectar-tap_saprotroph (Põlme et al. 2020); Decay sugar-rich substrates (Põlme et al. 2020) | Põlme S, et al. 2020. Fungal Diversity 105:1-16 ((https://doi.org/10.1007/s13225-020-00466-2)) |
| Otu42167 | 1 | k__Fungi(100);p__Ascomycota(100);c__Saccharomycetes(100);o__Saccharomycetales(100);f__Metschnikowiaceae(100);g__Metschnikowia(100);g__Metschnikowia_unclassified(100); | Metschnikowia | 13 | Saprotroph | \|Undefined Saprotroph\| | Yeast | NULL | Probable | As nectar-tap_saprotroph (Põlme et al. 2020); Decay sugar-rich substrates (Põlme et al. 2020) | Põlme S, et al. 2020. Fungal Diversity 105:1-16 ((https://doi.org/10.1007/s13225-020-00466-2)) |
| Otu42169 | 1 | k__Fungi(100);p__Ascomycota(100);c__Saccharomycetes(100);o__Saccharomycetales(100);f__Metschnikowiaceae(100);g__Metschnikowia(100);s__Metschnikowia_pulcherrima(100); | Metschnikowia | 13 | Saprotroph | \|Undefined Saprotroph\| | Yeast | NULL | Probable | As nectar-tap_saprotroph (Põlme et al. 2020); Decay sugar-rich substrates (Põlme et al. 2020) | Põlme S, et al. 2020. Fungal Diversity 105:1-16 ((https://doi.org/10.1007/s13225-020-00466-2)) |
| Otu42170 | 1 | k__Fungi(100);p__Ascomycota(100);c__Saccharomycetes(100);o__Saccharomycetales(100);f__Metschnikowiaceae(100);g__Metschnikowia(100);g__Metschnikowia_unclassified(100); | Metschnikowia | 13 | Saprotroph | \|Undefined Saprotroph\| | Yeast | NULL | Probable | As nectar-tap_saprotroph (Põlme et al. 2020); Decay sugar-rich substrates (Põlme et al. 2020) | Põlme S, et al. 2020. Fungal Diversity 105:1-16 ((https://doi.org/10.1007/s13225-020-00466-2)) |
| Otu42172 | 1 | k__Fungi(100);p__Ascomycota(100);c__Saccharomycetes(100);o__Saccharomycetales(100);f__Metschnikowiaceae(100);g__Metschnikowia(100);s__Metschnikowia_pulcherrima(100); | Metschnikowia | 13 | Saprotroph | \|Undefined Saprotroph\| | Yeast | NULL | Probable | As nectar-tap_saprotroph (Põlme et al. 2020); Decay sugar-rich substrates (Põlme et al. 2020) | Põlme S, et al. 2020. Fungal Diversity 105:1-16 ((https://doi.org/10.1007/s13225-020-00466-2)) |
| Otu42173 | 1 | k__Fungi(100);p__Ascomycota(100);c__Saccharomycetes(100);o__Saccharomycetales(100);f__Metschnikowiaceae(100);g__Metschnikowia(100);g__Metschnikowia_unclassified(100); | Metschnikowia | 13 | Saprotroph | \|Undefined Saprotroph\| | Yeast | NULL | Probable | As nectar-tap_saprotroph (Põlme et al. 2020); Decay sugar-rich substrates (Põlme et al. 2020) | Põlme S, et al. 2020. Fungal Diversity 105:1-16 ((https://doi.org/10.1007/s13225-020-00466-2)) |
| Otu42174 | 1 | k__Fungi(100);p__Ascomycota(100);c__Saccharomycetes(100);o__Saccharomycetales(100);f__Metschnikowiaceae(100);g__Metschnikowia(100);s__Metschnikowia_pulcherrima(100); | Metschnikowia | 13 | Saprotroph | \|Undefined Saprotroph\| | Yeast | NULL | Probable | As nectar-tap_saprotroph (Põlme et al. 2020); Decay sugar-rich substrates (Põlme et al. 2020) | Põlme S, et al. 2020. Fungal Diversity 105:1-16 ((https://doi.org/10.1007/s13225-020-00466-2)) |
| Otu42175 | 1 | k__Fungi(100);p__Ascomycota(100);c__Saccharomycetes(100);o__Saccharomycetales(100);f__Metschnikowiaceae(100);g__Metschnikowia(100);g__Metschnikowia_unclassified(100); | Metschnikowia | 13 | Saprotroph | \|Undefined Saprotroph\| | Yeast | NULL | Probable | As nectar-tap_saprotroph (Põlme et al. 2020); Decay sugar-rich substrates (Põlme et al. 2020) | Põlme S, et al. 2020. Fungal Diversity 105:1-16 ((https://doi.org/10.1007/s13225-020-00466-2)) |
| Otu42176 | 1 | k__Fungi(100);p__Ascomycota(100);c__Saccharomycetes(100);o__Saccharomycetales(100);f__Metschnikowiaceae(100);g__Metschnikowia(100);s__Metschnikowia_pulcherrima(100); | Metschnikowia | 13 | Saprotroph | \|Undefined Saprotroph\| | Yeast | NULL | Probable | As nectar-tap_saprotroph (Põlme et al. 2020); Decay sugar-rich substrates (Põlme et al. 2020) | Põlme S, et al. 2020. Fungal Diversity 105:1-16 ((https://doi.org/10.1007/s13225-020-00466-2)) |
| Otu42177 | 1 | k__Fungi(100);p__Ascomycota(100);c__Saccharomycetes(100);o__Saccharomycetales(100);f__Metschnikowiaceae(100);g__Metschnikowia(100);s__Metschnikowia_pulcherrima(100); | Metschnikowia | 13 | Saprotroph | \|Undefined Saprotroph\| | Yeast | NULL | Probable | As nectar-tap_saprotroph (Põlme et al. 2020); Decay sugar-rich substrates (Põlme et al. 2020) | Põlme S, et al. 2020. Fungal Diversity 105:1-16 ((https://doi.org/10.1007/s13225-020-00466-2)) |
| Otu42178 | 1 | k__Fungi(100);p__Ascomycota(100);c__Saccharomycetes(100);o__Saccharomycetales(100);f__Metschnikowiaceae(100);g__Metschnikowia(100);s__Metschnikowia_pulcherrima(100); | Metschnikowia | 13 | Saprotroph | \|Undefined Saprotroph\| | Yeast | NULL | Probable | As nectar-tap_saprotroph (Põlme et al. 2020); Decay sugar-rich substrates (Põlme et al. 2020) | Põlme S, et al. 2020. Fungal Diversity 105:1-16 ((https://doi.org/10.1007/s13225-020-00466-2)) |
| Otu42179 | 1 | k__Fungi(100);p__Ascomycota(100);c__Saccharomycetes(100);o__Saccharomycetales(100);f__Metschnikowiaceae(100);g__Metschnikowia(100);s__Metschnikowia_pulcherrima(100); | Metschnikowia | 13 | Saprotroph | \|Undefined Saprotroph\| | Yeast | NULL | Probable | As nectar-tap_saprotroph (Põlme et al. 2020); Decay sugar-rich substrates (Põlme et al. 2020) | Põlme S, et al. 2020. Fungal Diversity 105:1-16 ((https://doi.org/10.1007/s13225-020-00466-2)) |
| Otu42180 | 1 | k__Fungi(100);p__Ascomycota(100);c__Saccharomycetes(100);o__Saccharomycetales(100);f__Metschnikowiaceae(100);g__Metschnikowia(100);g__Metschnikowia_unclassified(100); | Metschnikowia | 13 | Saprotroph | \|Undefined Saprotroph\| | Yeast | NULL | Probable | As nectar-tap_saprotroph (Põlme et al. 2020); Decay sugar-rich substrates (Põlme et al. 2020) | Põlme S, et al. 2020. Fungal Diversity 105:1-16 ((https://doi.org/10.1007/s13225-020-00466-2)) |
| Otu42181 | 1 | k__Fungi(100);p__Ascomycota(100);c__Saccharomycetes(100);o__Saccharomycetales(100);f__Metschnikowiaceae(100);g__Metschnikowia(100);s__Metschnikowia_pulcherrima(100); | Metschnikowia | 13 | Saprotroph | \|Undefined Saprotroph\| | Yeast | NULL | Probable | As nectar-tap_saprotroph (Põlme et al. 2020); Decay sugar-rich substrates (Põlme et al. 2020) | Põlme S, et al. 2020. Fungal Diversity 105:1-16 ((https://doi.org/10.1007/s13225-020-00466-2)) |
| Otu42182 | 1 | k__Fungi(100);p__Ascomycota(100);c__Saccharomycetes(100);o__Saccharomycetales(100);f__Metschnikowiaceae(100);g__Metschnikowia(100);s__Metschnikowia_pulcherrima(100); | Metschnikowia | 13 | Saprotroph | \|Undefined Saprotroph\| | Yeast | NULL | Probable | As nectar-tap_saprotroph (Põlme et al. 2020); Decay sugar-rich substrates (Põlme et al. 2020) | Põlme S, et al. 2020. Fungal Diversity 105:1-16 ((https://doi.org/10.1007/s13225-020-00466-2)) |
| Otu42184 | 1 | k__Fungi(100);p__Ascomycota(100);c__Saccharomycetes(100);o__Saccharomycetales(100);f__Metschnikowiaceae(100);g__Metschnikowia(100);s__Metschnikowia_pulcherrima(100); | Metschnikowia | 13 | Saprotroph | \|Undefined Saprotroph\| | Yeast | NULL | Probable | As nectar-tap_saprotroph (Põlme et al. 2020); Decay sugar-rich substrates (Põlme et al. 2020) | Põlme S, et al. 2020. Fungal Diversity 105:1-16 ((https://doi.org/10.1007/s13225-020-00466-2)) |
| Otu42185 | 1 | k__Fungi(100);p__Ascomycota(100);c__Saccharomycetes(100);o__Saccharomycetales(100);f__Metschnikowiaceae(100);g__Metschnikowia(100);g__Metschnikowia_unclassified(100); | Metschnikowia | 13 | Saprotroph | \|Undefined Saprotroph\| | Yeast | NULL | Probable | As nectar-tap_saprotroph (Põlme et al. 2020); Decay sugar-rich substrates (Põlme et al. 2020) | Põlme S, et al. 2020. Fungal Diversity 105:1-16 ((https://doi.org/10.1007/s13225-020-00466-2)) |
| Otu42186 | 1 | k__Fungi(100);p__Ascomycota(100);c__Saccharomycetes(100);o__Saccharomycetales(100);f__Metschnikowiaceae(100);g__Metschnikowia(100);g__Metschnikowia_unclassified(100); | Metschnikowia | 13 | Saprotroph | \|Undefined Saprotroph\| | Yeast | NULL | Probable | As nectar-tap_saprotroph (Põlme et al. 2020); Decay sugar-rich substrates (Põlme et al. 2020) | Põlme S, et al. 2020. Fungal Diversity 105:1-16 ((https://doi.org/10.1007/s13225-020-00466-2)) |
| Otu42187 | 1 | k__Fungi(100);p__Ascomycota(100);c__Saccharomycetes(100);o__Saccharomycetales(100);f__Metschnikowiaceae(100);g__Metschnikowia(100);g__Metschnikowia_unclassified(100); | Metschnikowia | 13 | Saprotroph | \|Undefined Saprotroph\| | Yeast | NULL | Probable | As nectar-tap_saprotroph (Põlme et al. 2020); Decay sugar-rich substrates (Põlme et al. 2020) | Põlme S, et al. 2020. Fungal Diversity 105:1-16 ((https://doi.org/10.1007/s13225-020-00466-2)) |
| Otu42188 | 1 | k__Fungi(100);p__Ascomycota(100);c__Saccharomycetes(100);o__Saccharomycetales(100);f__Metschnikowiaceae(100);g__Metschnikowia(100);g__Metschnikowia_unclassified(100); | Metschnikowia | 13 | Saprotroph | \|Undefined Saprotroph\| | Yeast | NULL | Probable | As nectar-tap_saprotroph (Põlme et al. 2020); Decay sugar-rich substrates (Põlme et al. 2020) | Põlme S, et al. 2020. Fungal Diversity 105:1-16 ((https://doi.org/10.1007/s13225-020-00466-2)) |
| Otu42189 | 1 | k__Fungi(100);p__Ascomycota(100);c__Saccharomycetes(100);o__Saccharomycetales(100);f__Metschnikowiaceae(100);g__Metschnikowia(100);s__Metschnikowia_pulcherrima(100); | Metschnikowia | 13 | Saprotroph | \|Undefined Saprotroph\| | Yeast | NULL | Probable | As nectar-tap_saprotroph (Põlme et al. 2020); Decay sugar-rich substrates (Põlme et al. 2020) | Põlme S, et al. 2020. Fungal Diversity 105:1-16 ((https://doi.org/10.1007/s13225-020-00466-2)) |
| Otu42190 | 1 | k__Fungi(100);p__Ascomycota(100);c__Saccharomycetes(100);o__Saccharomycetales(100);f__Metschnikowiaceae(100);g__Metschnikowia(100);s__Metschnikowia_pulcherrima(100); | Metschnikowia | 13 | Saprotroph | \|Undefined Saprotroph\| | Yeast | NULL | Probable | As nectar-tap_saprotroph (Põlme et al. 2020); Decay sugar-rich substrates (Põlme et al. 2020) | Põlme S, et al. 2020. Fungal Diversity 105:1-16 ((https://doi.org/10.1007/s13225-020-00466-2)) |
| Otu42191 | 1 | k__Fungi(100);p__Ascomycota(100);c__Saccharomycetes(100);o__Saccharomycetales(100);f__Metschnikowiaceae(100);g__Metschnikowia(100);s__Metschnikowia_pulcherrima(100); | Metschnikowia | 13 | Saprotroph | \|Undefined Saprotroph\| | Yeast | NULL | Probable | As nectar-tap_saprotroph (Põlme et al. 2020); Decay sugar-rich substrates (Põlme et al. 2020) | Põlme S, et al. 2020. Fungal Diversity 105:1-16 ((https://doi.org/10.1007/s13225-020-00466-2)) |
| Otu42192 | 1 | k__Fungi(100);p__Ascomycota(100);c__Saccharomycetes(100);o__Saccharomycetales(100);f__Metschnikowiaceae(100);g__Metschnikowia(100);s__Metschnikowia_pulcherrima(100); | Metschnikowia | 13 | Saprotroph | \|Undefined Saprotroph\| | Yeast | NULL | Probable | As nectar-tap_saprotroph (Põlme et al. 2020); Decay sugar-rich substrates (Põlme et al. 2020) | Põlme S, et al. 2020. Fungal Diversity 105:1-16 ((https://doi.org/10.1007/s13225-020-00466-2)) |
| Otu42194 | 1 | k__Fungi(100);p__Ascomycota(100);c__Saccharomycetes(100);o__Saccharomycetales(100);f__Metschnikowiaceae(100);g__Metschnikowia(100);s__Metschnikowia_pulcherrima(100); | Metschnikowia | 13 | Saprotroph | \|Undefined Saprotroph\| | Yeast | NULL | Probable | As nectar-tap_saprotroph (Põlme et al. 2020); Decay sugar-rich substrates (Põlme et al. 2020) | Põlme S, et al. 2020. Fungal Diversity 105:1-16 ((https://doi.org/10.1007/s13225-020-00466-2)) |
| Otu42195 | 1 | k__Fungi(100);p__Ascomycota(100);c__Saccharomycetes(100);o__Saccharomycetales(100);f__Saccharomycetales_family_Incertae_sedis(100);g__Candida(100);g__Candida_unclassified(100); | Candida | 13 | Pathotroph-Saprotroph-Symbiotroph | Animal Pathogen-Endophyte-Endosymbiont-Epiphyte-\|Undefined Saprotroph\| | Dimorphic Yeast-Yeast | NULL | Probable | As nectar-tap_saprotroph (Põlme et al. 2020); Decay sugar-rich substrates (Põlme et al. 2020); This asexual genus is polyphyletic and contain species that function very differently depend on what they are. Caution should be taken when interpreting guilds for Candida species. | Manolakaki D, et al. 2010. Virulence. 1: 367-375 ((https://doi.org/10.4161/viru.1.5.12796)); Mekbib SB, et al. 2011. Tropical Plant Pathology 36:233-240 ((https://doi.org/10.1590/S1982-56762011000400004)); Põlme S, et al. 2020. Fungal Diversity 105:1-16 ((https://doi.org/10.1007/s13225-020-00466-2)) |
| Otu42196 | 1 | k__Fungi(100);p__Ascomycota(100);c__Saccharomycetes(100);o__Saccharomycetales(100);f__Metschnikowiaceae(100);g__Metschnikowia(100);g__Metschnikowia_unclassified(100); | Metschnikowia | 13 | Saprotroph | \|Undefined Saprotroph\| | Yeast | NULL | Probable | As nectar-tap_saprotroph (Põlme et al. 2020); Decay sugar-rich substrates (Põlme et al. 2020) | Põlme S, et al. 2020. Fungal Diversity 105:1-16 ((https://doi.org/10.1007/s13225-020-00466-2)) |
| Otu42197 | 1 | k__Fungi(100);p__Ascomycota(100);c__Saccharomycetes(100);o__Saccharomycetales(100);f__Metschnikowiaceae(100);g__Metschnikowia(100);s__Metschnikowia_pulcherrima(100); | Metschnikowia | 13 | Saprotroph | \|Undefined Saprotroph\| | Yeast | NULL | Probable | As nectar-tap_saprotroph (Põlme et al. 2020); Decay sugar-rich substrates (Põlme et al. 2020) | Põlme S, et al. 2020. Fungal Diversity 105:1-16 ((https://doi.org/10.1007/s13225-020-00466-2)) |
| Otu42198 | 1 | k__Fungi(100);p__Ascomycota(100);c__Saccharomycetes(100);o__Saccharomycetales(100);f__Metschnikowiaceae(100);g__Metschnikowia(100);s__Metschnikowia_pulcherrima(100); | Metschnikowia | 13 | Saprotroph | \|Undefined Saprotroph\| | Yeast | NULL | Probable | As nectar-tap_saprotroph (Põlme et al. 2020); Decay sugar-rich substrates (Põlme et al. 2020) | Põlme S, et al. 2020. Fungal Diversity 105:1-16 ((https://doi.org/10.1007/s13225-020-00466-2)) |
| Otu42199 | 1 | k__Fungi(100);p__Ascomycota(100);c__Saccharomycetes(100);o__Saccharomycetales(100);f__Metschnikowiaceae(100);g__Metschnikowia(100);s__Metschnikowia_pulcherrima(100); | Metschnikowia | 13 | Saprotroph | \|Undefined Saprotroph\| | Yeast | NULL | Probable | As nectar-tap_saprotroph (Põlme et al. 2020); Decay sugar-rich substrates (Põlme et al. 2020) | Põlme S, et al. 2020. Fungal Diversity 105:1-16 ((https://doi.org/10.1007/s13225-020-00466-2)) |
| Otu42200 | 1 | k__Fungi(100);p__Ascomycota(100);c__Saccharomycetes(100);o__Saccharomycetales(100);f__Metschnikowiaceae(100);g__Metschnikowia(100);s__Metschnikowia_pulcherrima(100); | Metschnikowia | 13 | Saprotroph | \|Undefined Saprotroph\| | Yeast | NULL | Probable | As nectar-tap_saprotroph (Põlme et al. 2020); Decay sugar-rich substrates (Põlme et al. 2020) | Põlme S, et al. 2020. Fungal Diversity 105:1-16 ((https://doi.org/10.1007/s13225-020-00466-2)) |
| Otu42201 | 1 | k__Fungi(100);p__Ascomycota(100);c__Saccharomycetes(100);o__Saccharomycetales(100);f__Metschnikowiaceae(100);g__Metschnikowia(100);s__Metschnikowia_pulcherrima(100); | Metschnikowia | 13 | Saprotroph | \|Undefined Saprotroph\| | Yeast | NULL | Probable | As nectar-tap_saprotroph (Põlme et al. 2020); Decay sugar-rich substrates (Põlme et al. 2020) | Põlme S, et al. 2020. Fungal Diversity 105:1-16 ((https://doi.org/10.1007/s13225-020-00466-2)) |
| Otu42202 | 1 | k__Fungi(100);p__Ascomycota(100);c__Saccharomycetes(100);o__Saccharomycetales(100);f__Metschnikowiaceae(100);g__Metschnikowia(100);s__Metschnikowia_pulcherrima(100); | Metschnikowia | 13 | Saprotroph | \|Undefined Saprotroph\| | Yeast | NULL | Probable | As nectar-tap_saprotroph (Põlme et al. 2020); Decay sugar-rich substrates (Põlme et al. 2020) | Põlme S, et al. 2020. Fungal Diversity 105:1-16 ((https://doi.org/10.1007/s13225-020-00466-2)) |
| Otu42203 | 1 | k__Fungi(100);p__Ascomycota(100);c__Saccharomycetes(100);o__Saccharomycetales(100);f__Metschnikowiaceae(100);g__Metschnikowia(100);g__Metschnikowia_unclassified(100); | Metschnikowia | 13 | Saprotroph | \|Undefined Saprotroph\| | Yeast | NULL | Probable | As nectar-tap_saprotroph (Põlme et al. 2020); Decay sugar-rich substrates (Põlme et al. 2020) | Põlme S, et al. 2020. Fungal Diversity 105:1-16 ((https://doi.org/10.1007/s13225-020-00466-2)) |
| Otu42204 | 1 | k__Fungi(100);p__Ascomycota(100);c__Saccharomycetes(100);o__Saccharomycetales(100);f__Metschnikowiaceae(100);g__Metschnikowia(100);s__Metschnikowia_pulcherrima(100); | Metschnikowia | 13 | Saprotroph | \|Undefined Saprotroph\| | Yeast | NULL | Probable | As nectar-tap_saprotroph (Põlme et al. 2020); Decay sugar-rich substrates (Põlme et al. 2020) | Põlme S, et al. 2020. Fungal Diversity 105:1-16 ((https://doi.org/10.1007/s13225-020-00466-2)) |
| Otu42205 | 1 | k__Fungi(100);p__Ascomycota(100);c__Saccharomycetes(100);o__Saccharomycetales(100);f__Metschnikowiaceae(100);g__Metschnikowia(100);g__Metschnikowia_unclassified(100); | Metschnikowia | 13 | Saprotroph | \|Undefined Saprotroph\| | Yeast | NULL | Probable | As nectar-tap_saprotroph (Põlme et al. 2020); Decay sugar-rich substrates (Põlme et al. 2020) | Põlme S, et al. 2020. Fungal Diversity 105:1-16 ((https://doi.org/10.1007/s13225-020-00466-2)) |
| Otu42206 | 1 | k__Fungi(100);p__Ascomycota(100);c__Saccharomycetes(100);o__Saccharomycetales(100);f__Metschnikowiaceae(100);g__Metschnikowia(100);s__Metschnikowia_pulcherrima(100); | Metschnikowia | 13 | Saprotroph | \|Undefined Saprotroph\| | Yeast | NULL | Probable | As nectar-tap_saprotroph (Põlme et al. 2020); Decay sugar-rich substrates (Põlme et al. 2020) | Põlme S, et al. 2020. Fungal Diversity 105:1-16 ((https://doi.org/10.1007/s13225-020-00466-2)) |
| Otu42207 | 1 | k__Fungi(100);p__Ascomycota(100);c__Saccharomycetes(100);o__Saccharomycetales(100);f__Metschnikowiaceae(100);g__Metschnikowia(100);g__Metschnikowia_unclassified(100); | Metschnikowia | 13 | Saprotroph | \|Undefined Saprotroph\| | Yeast | NULL | Probable | As nectar-tap_saprotroph (Põlme et al. 2020); Decay sugar-rich substrates (Põlme et al. 2020) | Põlme S, et al. 2020. Fungal Diversity 105:1-16 ((https://doi.org/10.1007/s13225-020-00466-2)) |
| Otu42208 | 1 | k__Fungi(100);p__Ascomycota(100);c__Saccharomycetes(100);o__Saccharomycetales(100);f__Metschnikowiaceae(100);g__Metschnikowia(100);s__Metschnikowia_pulcherrima(100); | Metschnikowia | 13 | Saprotroph | \|Undefined Saprotroph\| | Yeast | NULL | Probable | As nectar-tap_saprotroph (Põlme et al. 2020); Decay sugar-rich substrates (Põlme et al. 2020) | Põlme S, et al. 2020. Fungal Diversity 105:1-16 ((https://doi.org/10.1007/s13225-020-00466-2)) |
| Otu42209 | 1 | k__Fungi(100);p__Ascomycota(100);c__Saccharomycetes(100);o__Saccharomycetales(100);f__Metschnikowiaceae(100);g__Metschnikowia(100);s__Metschnikowia_pulcherrima(100); | Metschnikowia | 13 | Saprotroph | \|Undefined Saprotroph\| | Yeast | NULL | Probable | As nectar-tap_saprotroph (Põlme et al. 2020); Decay sugar-rich substrates (Põlme et al. 2020) | Põlme S, et al. 2020. Fungal Diversity 105:1-16 ((https://doi.org/10.1007/s13225-020-00466-2)) |
| Otu42210 | 1 | k__Fungi(100);p__Ascomycota(100);c__Saccharomycetes(100);o__Saccharomycetales(100);f__Metschnikowiaceae(100);g__Metschnikowia(100);s__Metschnikowia_pulcherrima(100); | Metschnikowia | 13 | Saprotroph | \|Undefined Saprotroph\| | Yeast | NULL | Probable | As nectar-tap_saprotroph (Põlme et al. 2020); Decay sugar-rich substrates (Põlme et al. 2020) | Põlme S, et al. 2020. Fungal Diversity 105:1-16 ((https://doi.org/10.1007/s13225-020-00466-2)) |
| Otu42211 | 1 | k__Fungi(100);p__Ascomycota(100);c__Saccharomycetes(100);o__Saccharomycetales(100);f__Metschnikowiaceae(100);g__Metschnikowia(100);s__Metschnikowia_pulcherrima(100); | Metschnikowia | 13 | Saprotroph | \|Undefined Saprotroph\| | Yeast | NULL | Probable | As nectar-tap_saprotroph (Põlme et al. 2020); Decay sugar-rich substrates (Põlme et al. 2020) | Põlme S, et al. 2020. Fungal Diversity 105:1-16 ((https://doi.org/10.1007/s13225-020-00466-2)) |
| Otu42212 | 1 | k__Fungi(100);p__Ascomycota(100);c__Saccharomycetes(100);o__Saccharomycetales(100);f__Metschnikowiaceae(100);g__Metschnikowia(100);s__Metschnikowia_pulcherrima(100); | Metschnikowia | 13 | Saprotroph | \|Undefined Saprotroph\| | Yeast | NULL | Probable | As nectar-tap_saprotroph (Põlme et al. 2020); Decay sugar-rich substrates (Põlme et al. 2020) | Põlme S, et al. 2020. Fungal Diversity 105:1-16 ((https://doi.org/10.1007/s13225-020-00466-2)) |
| Otu42213 | 1 | k__Fungi(100);p__Ascomycota(100);c__Saccharomycetes(100);o__Saccharomycetales(100);f__Metschnikowiaceae(100);g__Metschnikowia(100);g__Metschnikowia_unclassified(100); | Metschnikowia | 13 | Saprotroph | \|Undefined Saprotroph\| | Yeast | NULL | Probable | As nectar-tap_saprotroph (Põlme et al. 2020); Decay sugar-rich substrates (Põlme et al. 2020) | Põlme S, et al. 2020. Fungal Diversity 105:1-16 ((https://doi.org/10.1007/s13225-020-00466-2)) |
| Otu42214 | 1 | k__Fungi(100);p__Ascomycota(100);c__Saccharomycetes(100);o__Saccharomycetales(100);f__Metschnikowiaceae(100);g__Metschnikowia(100);g__Metschnikowia_unclassified(100); | Metschnikowia | 13 | Saprotroph | \|Undefined Saprotroph\| | Yeast | NULL | Probable | As nectar-tap_saprotroph (Põlme et al. 2020); Decay sugar-rich substrates (Põlme et al. 2020) | Põlme S, et al. 2020. Fungal Diversity 105:1-16 ((https://doi.org/10.1007/s13225-020-00466-2)) |
| Otu42215 | 1 | k__Fungi(100);p__Ascomycota(100);c__Saccharomycetes(100);o__Saccharomycetales(100);f__Metschnikowiaceae(100);g__Metschnikowia(100);g__Metschnikowia_unclassified(100); | Metschnikowia | 13 | Saprotroph | \|Undefined Saprotroph\| | Yeast | NULL | Probable | As nectar-tap_saprotroph (Põlme et al. 2020); Decay sugar-rich substrates (Põlme et al. 2020) | Põlme S, et al. 2020. Fungal Diversity 105:1-16 ((https://doi.org/10.1007/s13225-020-00466-2)) |

<table>
<tr><td>Otu42216</td><td>1</td><td>k__Fungi(100);p__Ascomycota(100);c__Saccharomycetes(100);o__Saccharomycetales(100);f__Metschnikowiaceae(100);g__Metschnikowia(100);s__Metschnikowia_pulcherrima(100);</td><td>Metschnikowia</td><td>13</td><td>Saprotroph</td><td>|Undefined Saprotroph|</td><td>Yeast</td><td>NULL</td><td>Probable</td><td>As nectar-tap_saprotroph (Põlme et al. 2020); Decay sugar-rich substrates (Põlme et al. 2020)</td><td>Põlme S, et al. 2020. Fungal Diversity 105:1-16 ((https://doi.org/10.1007/s13225-020-00466-2))</td></tr>
<tr><td>Otu42217</td><td>1</td><td>k__Fungi(100);p__Ascomycota(100);c__Saccharomycetes(100);o__Saccharomycetales(100);f__Metschnikowiaceae(100);g__Metschnikowia(100);s__Metschnikowia_pulcherrima(100);</td><td>Metschnikowia</td><td>13</td><td>Saprotroph</td><td>|Undefined Saprotroph|</td><td>Yeast</td><td>NULL</td><td>Probable</td><td>As nectar-tap_saprotroph (Põlme et al. 2020); Decay sugar-rich substrates (Põlme et al. 2020)</td><td>Põlme S, et al. 2020. Fungal Diversity 105:1-16 ((https://doi.org/10.1007/s13225-020-00466-2))</td></tr>
<tr><td>Otu42218</td><td>1</td><td>k__Fungi(100);p__Ascomycota(100);c__Saccharomycetes(100);o__Saccharomycetales(100);f__Metschnikowiaceae(100);g__Metschnikowia(100);s__Metschnikowia_pulcherrima(100);</td><td>Metschnikowia</td><td>13</td><td>Saprotroph</td><td>|Undefined Saprotroph|</td><td>Yeast</td><td>NULL</td><td>Probable</td><td>As nectar-tap_saprotroph (Põlme et al. 2020); Decay sugar-rich substrates (Põlme et al. 2020)</td><td>Põlme S, et al. 2020. Fungal Diversity 105:1-16 ((https://doi.org/10.1007/s13225-020-00466-2))</td></tr>
<tr><td>Otu42220</td><td>1</td><td>k__Fungi(100);p__Ascomycota(100);c__Saccharomycetes(100);o__Saccharomycetales(100);f__Metschnikowiaceae(100);g__Metschnikowia(100);s__Metschnikowia_pulcherrima(100);</td><td>Metschnikowia</td><td>13</td><td>Saprotroph</td><td>|Undefined Saprotroph|</td><td>Yeast</td><td>NULL</td><td>Probable</td><td>As nectar-tap_saprotroph (Põlme et al. 2020); Decay sugar-rich substrates (Põlme et al. 2020)</td><td>Põlme S, et al. 2020. Fungal Diversity 105:1-16 ((https://doi.org/10.1007/s13225-020-00466-2))</td></tr>
<tr><td>Otu42225</td><td>1</td><td>k__Fungi(100);p__Ascomycota(100);c__Saccharomycetes(100);o__Saccharomycetales(100);f__Metschnikowiaceae(100);g__Metschnikowia(100);g__Metschnikowia_unclassified(100);</td><td>Metschnikowia</td><td>13</td><td>Saprotroph</td><td>|Undefined Saprotroph|</td><td>Yeast</td><td>NULL</td><td>Probable</td><td>As nectar-tap_saprotroph (Põlme et al. 2020); Decay sugar-rich substrates (Põlme et al. 2020)</td><td>Põlme S, et al. 2020. Fungal Diversity 105:1-16 ((https://doi.org/10.1007/s13225-020-00466-2))</td></tr>
<tr><td>Otu42226</td><td>1</td><td>k__Fungi(100);p__Ascomycota(100);c__Saccharomycetes(100);o__Saccharomycetales(100);f__Metschnikowiaceae(100);g__Metschnikowia(100);g__Metschnikowia_unclassified(100);</td><td>Metschnikowia</td><td>13</td><td>Saprotroph</td><td>|Undefined Saprotroph|</td><td>Yeast</td><td>NULL</td><td>Probable</td><td>As nectar-tap_saprotroph (Põlme et al. 2020); Decay sugar-rich substrates (Põlme et al. 2020)</td><td>Põlme S, et al. 2020. Fungal Diversity 105:1-16 ((https://doi.org/10.1007/s13225-020-00466-2))</td></tr>
<tr><td>Otu42227</td><td>1</td><td>k__Fungi(100);p__Ascomycota(100);c__Saccharomycetes(100);o__Saccharomycetales(100);f__Metschnikowiaceae(100);g__Metschnikowia(100);s__Metschnikowia_pulcherrima(100);</td><td>Metschnikowia</td><td>13</td><td>Saprotroph</td><td>|Undefined Saprotroph|</td><td>Yeast</td><td>NULL</td><td>Probable</td><td>As nectar-tap_saprotroph (Põlme et al. 2020); Decay sugar-rich substrates (Põlme et al. 2020)</td><td>Põlme S, et al. 2020. Fungal Diversity 105:1-16 ((https://doi.org/10.1007/s13225-020-00466-2))</td></tr>
<tr><td>Otu42228</td><td>1</td><td>k__Fungi(100);p__Ascomycota(100);c__Saccharomycetes(100);o__Saccharomycetales(100);f__Metschnikowiaceae(100);g__Metschnikowia(100);s__Metschnikowia_pulcherrima(100);</td><td>Metschnikowia</td><td>13</td><td>Saprotroph</td><td>|Undefined Saprotroph|</td><td>Yeast</td><td>NULL</td><td>Probable</td><td>As nectar-tap_saprotroph (Põlme et al. 2020); Decay sugar-rich substrates (Põlme et al. 2020)</td><td>Põlme S, et al. 2020. Fungal Diversity 105:1-16 ((https://doi.org/10.1007/s13225-020-00466-2))</td></tr>
<tr><td>Otu42229</td><td>1</td><td>k__Fungi(100);p__Ascomycota(100);c__Saccharomycetes(100);o__Saccharomycetales(100);f__Metschnikowiaceae(100);g__Metschnikowia(100);s__Metschnikowia_pulcherrima(100);</td><td>Metschnikowia</td><td>13</td><td>Saprotroph</td><td>|Undefined Saprotroph|</td><td>Yeast</td><td>NULL</td><td>Probable</td><td>As nectar-tap_saprotroph (Põlme et al. 2020); Decay sugar-rich substrates (Põlme et al. 2020)</td><td>Põlme S, et al. 2020. Fungal Diversity 105:1-16 ((https://doi.org/10.1007/s13225-020-00466-2))</td></tr>
<tr><td>Otu42231</td><td>1</td><td>k__Fungi(100);p__Ascomycota(100);c__Saccharomycetes(100);o__Saccharomycetales(100);f__Metschnikowiaceae(100);g__Metschnikowia(100);s__Metschnikowia_pulcherrima(100);</td><td>Metschnikowia</td><td>13</td><td>Saprotroph</td><td>|Undefined Saprotroph|</td><td>Yeast</td><td>NULL</td><td>Probable</td><td>As nectar-tap_saprotroph (Põlme et al. 2020); Decay sugar-rich substrates (Põlme et al. 2020)</td><td>Põlme S, et al. 2020. Fungal Diversity 105:1-16 ((https://doi.org/10.1007/s13225-020-00466-2))</td></tr>
<tr><td>Otu42232</td><td>1</td><td>k__Fungi(100);p__Ascomycota(100);c__Saccharomycetes(100);o__Saccharomycetales(100);f__Metschnikowiaceae(100);g__Metschnikowia(100);s__Metschnikowia_pulcherrima(100);</td><td>Metschnikowia</td><td>13</td><td>Saprotroph</td><td>|Undefined Saprotroph|</td><td>Yeast</td><td>NULL</td><td>Probable</td><td>As nectar-tap_saprotroph (Põlme et al. 2020); Decay sugar-rich substrates (Põlme et al. 2020)</td><td>Põlme S, et al. 2020. Fungal Diversity 105:1-16 ((https://doi.org/10.1007/s13225-020-00466-2))</td></tr>
<tr><td>Otu42235</td><td>1</td><td>k__Fungi(100);p__Ascomycota(100);c__Saccharomycetes(100);o__Saccharomycetales(100);f__Metschnikowiaceae(100);g__Metschnikowia(100);s__Metschnikowia_pulcherrima(100);</td><td>Metschnikowia</td><td>13</td><td>Saprotroph</td><td>|Undefined Saprotroph|</td><td>Yeast</td><td>NULL</td><td>Probable</td><td>As nectar-tap_saprotroph (Põlme et al. 2020); Decay sugar-rich substrates (Põlme et al. 2020)</td><td>Põlme S, et al. 2020. Fungal Diversity 105:1-16 ((https://doi.org/10.1007/s13225-020-00466-2))</td></tr>
<tr><td>Otu42236</td><td>1</td><td>k__Fungi(100);p__Ascomycota(100);c__Saccharomycetes(100);o__Saccharomycetales(100);f__Metschnikowiaceae(100);g__Metschnikowia(100);s__Metschnikowia_pulcherrima(100);</td><td>Metschnikowia</td><td>13</td><td>Saprotroph</td><td>|Undefined Saprotroph|</td><td>Yeast</td><td>NULL</td><td>Probable</td><td>As nectar-tap_saprotroph (Põlme et al. 2020); Decay sugar-rich substrates (Põlme et al. 2020)</td><td>Põlme S, et al. 2020. Fungal Diversity 105:1-16 ((https://doi.org/10.1007/s13225-020-00466-2))</td></tr>
<tr><td>Otu42240</td><td>1</td><td>k__Fungi(100);p__Ascomycota(100);c__Saccharomycetes(100);o__Saccharomycetales(100);f__Metschnikowiaceae(100);g__Metschnikowia(100);s__Metschnikowia_pulcherrima(100);</td><td>Metschnikowia</td><td>13</td><td>Saprotroph</td><td>|Undefined Saprotroph|</td><td>Yeast</td><td>NULL</td><td>Probable</td><td>As nectar-tap_saprotroph (Põlme et al. 2020); Decay sugar-rich substrates (Põlme et al. 2020)</td><td>Põlme S, et al. 2020. Fungal Diversity 105:1-16 ((https://doi.org/10.1007/s13225-020-00466-2))</td></tr>
<tr><td>Otu42241</td><td>1</td><td>k__Fungi(100);p__Ascomycota(100);c__Saccharomycetes(100);o__Saccharomycetales(100);f__Metschnikowiaceae(100);g__Metschnikowia(100);s__Metschnikowia_pulcherrima(100);</td><td>Metschnikowia</td><td>13</td><td>Saprotroph</td><td>|Undefined Saprotroph|</td><td>Yeast</td><td>NULL</td><td>Probable</td><td>As nectar-tap_saprotroph (Põlme et al. 2020); Decay sugar-rich substrates (Põlme et al. 2020)</td><td>Põlme S, et al. 2020. Fungal Diversity 105:1-16 ((https://doi.org/10.1007/s13225-020-00466-2))</td></tr>
<tr><td>Otu42242</td><td>1</td><td>k__Fungi(100);p__Ascomycota(100);c__Saccharomycetes(100);o__Saccharomycetales(100);f__Metschnikowiaceae(100);g__Metschnikowia(100);g__Metschnikowia_unclassified(100);</td><td>Metschnikowia</td><td>13</td><td>Saprotroph</td><td>|Undefined Saprotroph|</td><td>Yeast</td><td>NULL</td><td>Probable</td><td>As nectar-tap_saprotroph (Põlme et al. 2020); Decay sugar-rich substrates (Põlme et al. 2020)</td><td>Põlme S, et al. 2020. Fungal Diversity 105:1-16 ((https://doi.org/10.1007/s13225-020-00466-2))</td></tr>
<tr><td>Otu42243</td><td>1</td><td>k__Fungi(100);p__Ascomycota(100);c__Saccharomycetes(100);o__Saccharomycetales(100);f__Metschnikowiaceae(100);g__Metschnikowia(100);g__Metschnikowia_unclassified(100);</td><td>Metschnikowia</td><td>13</td><td>Saprotroph</td><td>|Undefined Saprotroph|</td><td>Yeast</td><td>NULL</td><td>Probable</td><td>As nectar-tap_saprotroph (Põlme et al. 2020); Decay sugar-rich substrates (Põlme et al. 2020)</td><td>Põlme S, et al. 2020. Fungal Diversity 105:1-16 ((https://doi.org/10.1007/s13225-020-00466-2))</td></tr>
<tr><td>Otu42245</td><td>1</td><td>k__Fungi(100);p__Ascomycota(100);c__Saccharomycetes(100);o__Saccharomycetales(100);f__Metschnikowiaceae(100);g__Metschnikowia(100);g__Metschnikowia_unclassified(100);</td><td>Metschnikowia</td><td>13</td><td>Saprotroph</td><td>|Undefined Saprotroph|</td><td>Yeast</td><td>NULL</td><td>Probable</td><td>As nectar-tap_saprotroph (Põlme et al. 2020); Decay sugar-rich substrates (Põlme et al. 2020)</td><td>Põlme S, et al. 2020. Fungal Diversity 105:1-16 ((https://doi.org/10.1007/s13225-020-00466-2))</td></tr>
<tr><td>Otu42246</td><td>1</td><td>k__Fungi(100);p__Ascomycota(100);c__Saccharomycetes(100);o__Saccharomycetales(100);f__Metschnikowiaceae(100);g__Metschnikowia(100);s__Metschnikowia_pulcherrima(100);</td><td>Metschnikowia</td><td>13</td><td>Saprotroph</td><td>|Undefined Saprotroph|</td><td>Yeast</td><td>NULL</td><td>Probable</td><td>As nectar-tap_saprotroph (Põlme et al. 2020); Decay sugar-rich substrates (Põlme et al. 2020)</td><td>Põlme S, et al. 2020. Fungal Diversity 105:1-16 ((https://doi.org/10.1007/s13225-020-00466-2))</td></tr>
<tr><td>Otu42247</td><td>1</td><td>k__Fungi(100);p__Ascomycota(100);c__Saccharomycetes(100);o__Saccharomycetales(100);f__Metschnikowiaceae(100);g__Metschnikowia(100);s__Metschnikowia_pulcherrima(100);</td><td>Metschnikowia</td><td>13</td><td>Saprotroph</td><td>|Undefined Saprotroph|</td><td>Yeast</td><td>NULL</td><td>Probable</td><td>As nectar-tap_saprotroph (Põlme et al. 2020); Decay sugar-rich substrates (Põlme et al. 2020)</td><td>Põlme S, et al. 2020. Fungal Diversity 105:1-16 ((https://doi.org/10.1007/s13225-020-00466-2))</td></tr>
<tr><td>Otu42248</td><td>1</td><td>k__Fungi(100);p__Ascomycota(100);c__Saccharomycetes(100);o__Saccharomycetales(100);f__Metschnikowiaceae(100);g__Metschnikowia(100);s__Metschnikowia_pulcherrima(100);</td><td>Metschnikowia</td><td>13</td><td>Saprotroph</td><td>|Undefined Saprotroph|</td><td>Yeast</td><td>NULL</td><td>Probable</td><td>As nectar-tap_saprotroph (Põlme et al. 2020); Decay sugar-rich substrates (Põlme et al. 2020)</td><td>Põlme S, et al. 2020. Fungal Diversity 105:1-16 ((https://doi.org/10.1007/s13225-020-00466-2))</td></tr>
<tr><td>Otu42249</td><td>1</td><td>k__Fungi(100);p__Ascomycota(100);c__Saccharomycetes(100);o__Saccharomycetales(100);f__Metschnikowiaceae(100);g__Metschnikowia(100);g__Metschnikowia_unclassified(100);</td><td>Metschnikowia</td><td>13</td><td>Saprotroph</td><td>|Undefined Saprotroph|</td><td>Yeast</td><td>NULL</td><td>Probable</td><td>As nectar-tap_saprotroph (Põlme et al. 2020); Decay sugar-rich substrates (Põlme et al. 2020)</td><td>Põlme S, et al. 2020. Fungal Diversity 105:1-16 ((https://doi.org/10.1007/s13225-020-00466-2))</td></tr>
<tr><td>Otu42250</td><td>1</td><td>k__Fungi(100);p__Ascomycota(100);c__Saccharomycetes(100);o__Saccharomycetales(100);f__Metschnikowiaceae(100);g__Metschnikowia(100);s__Metschnikowia_pulcherrima(100);</td><td>Metschnikowia</td><td>13</td><td>Saprotroph</td><td>|Undefined Saprotroph|</td><td>Yeast</td><td>NULL</td><td>Probable</td><td>As nectar-tap_saprotroph (Põlme et al. 2020); Decay sugar-rich substrates (Põlme et al. 2020)</td><td>Põlme S, et al. 2020. Fungal Diversity 105:1-16 ((https://doi.org/10.1007/s13225-020-00466-2))</td></tr>
<tr><td>Otu42251</td><td>1</td><td>k__Fungi(100);p__Ascomycota(100);c__Saccharomycetes(100);o__Saccharomycetales(100);f__Metschnikowiaceae(100);g__Metschnikowia(100);s__Metschnikowia_pulcherrima(100);</td><td>Metschnikowia</td><td>13</td><td>Saprotroph</td><td>|Undefined Saprotroph|</td><td>Yeast</td><td>NULL</td><td>Probable</td><td>As nectar-tap_saprotroph (Põlme et al. 2020); Decay sugar-rich substrates (Põlme et al. 2020)</td><td>Põlme S, et al. 2020. Fungal Diversity 105:1-16 ((https://doi.org/10.1007/s13225-020-00466-2))</td></tr>
<tr><td>Otu42257</td><td>1</td><td>k__Fungi(100);p__Ascomycota(100);c__Saccharomycetes(100);o__Saccharomycetales(100);f__Metschnikowiaceae(100);g__Metschnikowia(100);s__Metschnikowia_pulcherrima(100);</td><td>Metschnikowia</td><td>13</td><td>Saprotroph</td><td>|Undefined Saprotroph|</td><td>Yeast</td><td>NULL</td><td>Probable</td><td>As nectar-tap_saprotroph (Põlme et al. 2020); Decay sugar-rich substrates (Põlme et al. 2020)</td><td>Põlme S, et al. 2020. Fungal Diversity 105:1-16 ((https://doi.org/10.1007/s13225-020-00466-2))</td></tr>
<tr><td>Otu42263</td><td>1</td><td>k__Fungi(100);p__Ascomycota(100);c__Saccharomycetes(100);o__Saccharomycetales(100);f__Metschnikowiaceae(100);g__Metschnikowia(100);s__Metschnikowia_pulcherrima(100);</td><td>Metschnikowia</td><td>13</td><td>Saprotroph</td><td>|Undefined Saprotroph|</td><td>Yeast</td><td>NULL</td><td>Probable</td><td>As nectar-tap_saprotroph (Põlme et al. 2020); Decay sugar-rich substrates (Põlme et al. 2020)</td><td>Põlme S, et al. 2020. Fungal Diversity 105:1-16 ((https://doi.org/10.1007/s13225-020-00466-2))</td></tr>
<tr><td>Otu42267</td><td>1</td><td>k__Fungi(100);p__Ascomycota(100);c__Saccharomycetes(100);o__Saccharomycetales(100);f__Metschnikowiaceae(100);g__Metschnikowia(100);s__Metschnikowia_pulcherrima(100);</td><td>Metschnikowia</td><td>13</td><td>Saprotroph</td><td>|Undefined Saprotroph|</td><td>Yeast</td><td>NULL</td><td>Probable</td><td>As nectar-tap_saprotroph (Põlme et al. 2020); Decay sugar-rich substrates (Põlme et al. 2020)</td><td>Põlme S, et al. 2020. Fungal Diversity 105:1-16 ((https://doi.org/10.1007/s13225-020-00466-2))</td></tr>
<tr><td>Otu42269</td><td>1</td><td>k__Fungi(100);p__Ascomycota(100);c__Saccharomycetes(100);o__Saccharomycetales(100);f__Metschnikowiaceae(100);g__Metschnikowia(100);s__Metschnikowia_pulcherrima(100);</td><td>Metschnikowia</td><td>13</td><td>Saprotroph</td><td>|Undefined Saprotroph|</td><td>Yeast</td><td>NULL</td><td>Probable</td><td>As nectar-tap_saprotroph (Põlme et al. 2020); Decay sugar-rich substrates (Põlme et al. 2020)</td><td>Põlme S, et al. 2020. Fungal Diversity 105:1-16 ((https://doi.org/10.1007/s13225-020-00466-2))</td></tr>
<tr><td>Otu42270</td><td>1</td><td>k__Fungi(100);p__Ascomycota(100);c__Saccharomycetes(100);o__Saccharomycetales(100);f__Metschnikowiaceae(100);g__Metschnikowia(100);s__Metschnikowia_pulcherrima(100);</td><td>Metschnikowia</td><td>13</td><td>Saprotroph</td><td>|Undefined Saprotroph|</td><td>Yeast</td><td>NULL</td><td>Probable</td><td>As nectar-tap_saprotroph (Põlme et al. 2020); Decay sugar-rich substrates (Põlme et al. 2020)</td><td>Põlme S, et al. 2020. Fungal Diversity 105:1-16 ((https://doi.org/10.1007/s13225-020-00466-2))</td></tr>
<tr><td>Otu42271</td><td>1</td><td>k__Fungi(100);p__Ascomycota(100);c__Saccharomycetes(100);o__Saccharomycetales(100);f__Metschnikowiaceae(100);g__Metschnikowia(100);s__Metschnikowia_pulcherrima(100);</td><td>Metschnikowia</td><td>13</td><td>Saprotroph</td><td>|Undefined Saprotroph|</td><td>Yeast</td><td>NULL</td><td>Probable</td><td>As nectar-tap_saprotroph (Põlme et al. 2020); Decay sugar-rich substrates (Põlme et al. 2020)</td><td>Põlme S, et al. 2020. Fungal Diversity 105:1-16 ((https://doi.org/10.1007/s13225-020-00466-2))</td></tr>
<tr><td>Otu42272</td><td>1</td><td>k__Fungi(100);p__Ascomycota(100);c__Saccharomycetes(100);o__Saccharomycetales(100);f__Metschnikowiaceae(100);g__Metschnikowia(100);s__Metschnikowia_pulcherrima(100);</td><td>Metschnikowia</td><td>13</td><td>Saprotroph</td><td>|Undefined Saprotroph|</td><td>Yeast</td><td>NULL</td><td>Probable</td><td>As nectar-tap_saprotroph (Põlme et al. 2020); Decay sugar-rich substrates (Põlme et al. 2020)</td><td>Põlme S, et al. 2020. Fungal Diversity 105:1-16 ((https://doi.org/10.1007/s13225-020-00466-2))</td></tr>
<tr><td>Otu42273</td><td>1</td><td>k__Fungi(100);p__Ascomycota(100);c__Saccharomycetes(100);o__Saccharomycetales(100);f__Metschnikowiaceae(100);g__Metschnikowia(100);s__Metschnikowia_pulcherrima(100);</td><td>Metschnikowia</td><td>13</td><td>Saprotroph</td><td>|Undefined Saprotroph|</td><td>Yeast</td><td>NULL</td><td>Probable</td><td>As nectar-tap_saprotroph (Põlme et al. 2020); Decay sugar-rich substrates (Põlme et al. 2020)</td><td>Põlme S, et al. 2020. Fungal Diversity 105:1-16 ((https://doi.org/10.1007/s13225-020-00466-2))</td></tr>
<tr><td>Otu42274</td><td>1</td><td>k__Fungi(100);p__Ascomycota(100);c__Saccharomycetes(100);o__Saccharomycetales(100);f__Metschnikowiaceae(100);g__Metschnikowia(100);s__Metschnikowia_pulcherrima(100);</td><td>Metschnikowia</td><td>13</td><td>Saprotroph</td><td>|Undefined Saprotroph|</td><td>Yeast</td><td>NULL</td><td>Probable</td><td>As nectar-tap_saprotroph (Põlme et al. 2020); Decay sugar-rich substrates (Põlme et al. 2020)</td><td>Põlme S, et al. 2020. Fungal Diversity 105:1-16 ((https://doi.org/10.1007/s13225-020-00466-2))</td></tr>
<tr><td>Otu42275</td><td>1</td><td>k__Fungi(100);p__Ascomycota(100);c__Saccharomycetes(100);o__Saccharomycetales(100);f__Metschnikowiaceae(100);g__Metschnikowia(100);g__Metschnikowia_unclassified(100);</td><td>Metschnikowia</td><td>13</td><td>Saprotroph</td><td>|Undefined Saprotroph|</td><td>Yeast</td><td>NULL</td><td>Probable</td><td>As nectar-tap_saprotroph (Põlme et al. 2020); Decay sugar-rich substrates (Põlme et al. 2020)</td><td>Põlme S, et al. 2020. Fungal Diversity 105:1-16 ((https://doi.org/10.1007/s13225-020-00466-2))</td></tr>
<tr><td>Otu42276</td><td>1</td><td>k__Fungi(100);p__Ascomycota(100);c__Saccharomycetes(100);o__Saccharomycetales(100);f__Metschnikowiaceae(100);g__Metschnikowia(100);s__Metschnikowia_pulcherrima(100);</td><td>Metschnikowia</td><td>13</td><td>Saprotroph</td><td>|Undefined Saprotroph|</td><td>Yeast</td><td>NULL</td><td>Probable</td><td>As nectar-tap_saprotroph (Põlme et al. 2020); Decay sugar-rich substrates (Põlme et al. 2020)</td><td>Põlme S, et al. 2020. Fungal Diversity 105:1-16 ((https://doi.org/10.1007/s13225-020-00466-2))</td></tr>
<tr><td>Otu42277</td><td>1</td><td>k__Fungi(100);p__Ascomycota(100);c__Saccharomycetes(100);o__Saccharomycetales(100);f__Metschnikowiaceae(100);g__Metschnikowia(100);s__Metschnikowia_pulcherrima(100);</td><td>Metschnikowia</td><td>13</td><td>Saprotroph</td><td>|Undefined Saprotroph|</td><td>Yeast</td><td>NULL</td><td>Probable</td><td>As nectar-tap_saprotroph (Põlme et al. 2020); Decay sugar-rich substrates (Põlme et al. 2020)</td><td>Põlme S, et al. 2020. Fungal Diversity 105:1-16 ((https://doi.org/10.1007/s13225-020-00466-2))</td></tr>
<tr><td>Otu42278</td><td>1</td><td>k__Fungi(100);p__Ascomycota(100);c__Saccharomycetes(100);o__Saccharomycetales(100);f__Metschnikowiaceae(100);g__Metschnikowia(100);s__Metschnikowia_pulcherrima(100);</td><td>Metschnikowia</td><td>13</td><td>Saprotroph</td><td>|Undefined Saprotroph|</td><td>Yeast</td><td>NULL</td><td>Probable</td><td>As nectar-tap_saprotroph (Põlme et al. 2020); Decay sugar-rich substrates (Põlme et al. 2020)</td><td>Põlme S, et al. 2020. Fungal Diversity 105:1-16 ((https://doi.org/10.1007/s13225-020-00466-2))</td></tr>
<tr><td>Otu42279</td><td>1</td><td>k__Fungi(100);p__Ascomycota(100);c__Saccharomycetes(100);o__Saccharomycetales(100);f__Metschnikowiaceae(100);g__Metschnikowia(100);s__Metschnikowia_pulcherrima(100);</td><td>Metschnikowia</td><td>13</td><td>Saprotroph</td><td>|Undefined Saprotroph|</td><td>Yeast</td><td>NULL</td><td>Probable</td><td>As nectar-tap_saprotroph (Põlme et al. 2020); Decay sugar-rich substrates (Põlme et al. 2020)</td><td>Põlme S, et al. 2020. Fungal Diversity 105:1-16 ((https://doi.org/10.1007/s13225-020-00466-2))</td></tr>
<tr><td>Otu42280</td><td>1</td><td>k__Fungi(100);p__Ascomycota(100);c__Saccharomycetes(100);o__Saccharomycetales(100);f__Metschnikowiaceae(100);g__Metschnikowia(100);s__Metschnikowia_pulcherrima(100);</td><td>Metschnikowia</td><td>13</td><td>Saprotroph</td><td>|Undefined Saprotroph|</td><td>Yeast</td><td>NULL</td><td>Probable</td><td>As nectar-tap_saprotroph (Põlme et al. 2020); Decay sugar-rich substrates (Põlme et al. 2020)</td><td>Põlme S, et al. 2020. Fungal Diversity 105:1-16 ((https://doi.org/10.1007/s13225-020-00466-2))</td></tr>
<tr><td>Otu42281</td><td>1</td><td>k__Fungi(100);p__Ascomycota(100);c__Saccharomycetes(100);o__Saccharomycetales(100);f__Metschnikowiaceae(100);g__Metschnikowia(100);s__Metschnikowia_pulcherrima(100);</td><td>Metschnikowia</td><td>13</td><td>Saprotroph</td><td>|Undefined Saprotroph|</td><td>Yeast</td><td>NULL</td><td>Probable</td><td>As nectar-tap_saprotroph (Põlme et al. 2020); Decay sugar-rich substrates (Põlme et al. 2020)</td><td>Põlme S, et al. 2020. Fungal Diversity 105:1-16 ((https://doi.org/10.1007/s13225-020-00466-2))</td></tr>
<tr><td>Otu42282</td><td>1</td><td>k__Fungi(100);p__Ascomycota(100);c__Saccharomycetes(100);o__Saccharomycetales(100);f__Metschnikowiaceae(100);g__Metschnikowia(100);g__Metschnikowia_unclassified(100);</td><td>Metschnikowia</td><td>13</td><td>Saprotroph</td><td>|Undefined Saprotroph|</td><td>Yeast</td><td>NULL</td><td>Probable</td><td>As nectar-tap_saprotroph (Põlme et al. 2020); Decay sugar-rich substrates (Põlme et al. 2020)</td><td>Põlme S, et al. 2020. Fungal Diversity 105:1-16 ((https://doi.org/10.1007/s13225-020-00466-2))</td></tr>
<tr><td>Otu42283</td><td>1</td><td>k__Fungi(100);p__Ascomycota(100);c__Saccharomycetes(100);o__Saccharomycetales(100);f__Metschnikowiaceae(100);g__Metschnikowia(100);s__Metschnikowia_pulcherrima(100);</td><td>Metschnikowia</td><td>13</td><td>Saprotroph</td><td>|Undefined Saprotroph|</td><td>Yeast</td><td>NULL</td><td>Probable</td><td>As nectar-tap_saprotroph (Põlme et al. 2020); Decay sugar-rich substrates (Põlme et al. 2020)</td><td>Põlme S, et al. 2020. Fungal Diversity 105:1-16 ((https://doi.org/10.1007/s13225-020-00466-2))</td></tr>
<tr><td>Otu42284</td><td>1</td><td>k__Fungi(100);p__Ascomycota(100);c__Saccharomycetes(100);o__Saccharomycetales(100);f__Metschnikowiaceae(100);g__Metschnikowia(100);s__Metschnikowia_pulcherrima(100);</td><td>Metschnikowia</td><td>13</td><td>Saprotroph</td><td>|Undefined Saprotroph|</td><td>Yeast</td><td>NULL</td><td>Probable</td><td>As nectar-tap_saprotroph (Põlme et al. 2020); Decay sugar-rich substrates (Põlme et al. 2020)</td><td>Põlme S, et al. 2020. Fungal Diversity 105:1-16 ((https://doi.org/10.1007/s13225-020-00466-2))</td></tr>
<tr><td>Otu42285</td><td>1</td><td>k__Fungi(100);p__Ascomycota(100);c__Saccharomycetes(100);o__Saccharomycetales(100);f__Metschnikowiaceae(100);g__Metschnikowia(100);s__Metschnikowia_pulcherrima(100);</td><td>Metschnikowia</td><td>13</td><td>Saprotroph</td><td>|Undefined Saprotroph|</td><td>Yeast</td><td>NULL</td><td>Probable</td><td>As nectar-tap_saprotroph (Põlme et al. 2020); Decay sugar-rich substrates (Põlme et al. 2020)</td><td>Põlme S, et al. 2020. Fungal Diversity 105:1-16 ((https://doi.org/10.1007/s13225-020-00466-2))</td></tr>
<tr><td>Otu42286</td><td>1</td><td>k__Fungi(100);p__Ascomycota(100);c__Saccharomycetes(100);o__Saccharomycetales(100);f__Metschnikowiaceae(100);g__Metschnikowia(100);s__Metschnikowia_pulcherrima(100);</td><td>Metschnikowia</td><td>13</td><td>Saprotroph</td><td>|Undefined Saprotroph|</td><td>Yeast</td><td>NULL</td><td>Probable</td><td>As nectar-tap_saprotroph (Põlme et al. 2020); Decay sugar-rich substrates (Põlme et al. 2020)</td><td>Põlme S, et al. 2020. Fungal Diversity 105:1-16 ((https://doi.org/10.1007/s13225-020-00466-2))</td></tr>
<tr><td>Otu42287</td><td>1</td><td>k__Fungi(100);p__Ascomycota(100);c__Saccharomycetes(100);o__Saccharomycetales(100);f__Metschnikowiaceae(100);g__Metschnikowia(100);s__Metschnikowia_pulcherrima(100);</td><td>Metschnikowia</td><td>13</td><td>Saprotroph</td><td>|Undefined Saprotroph|</td><td>Yeast</td><td>NULL</td><td>Probable</td><td>As nectar-tap_saprotroph (Põlme et al. 2020); Decay sugar-rich substrates (Põlme et al. 2020)</td><td>Põlme S, et al. 2020. Fungal Diversity 105:1-16 ((https://doi.org/10.1007/s13225-020-00466-2))</td></tr>
<tr><td>Otu42288</td><td>1</td><td>k__Fungi(100);p__Ascomycota(100);c__Saccharomycetes(100);o__Saccharomycetales(100);f__Metschnikowiaceae(100);g__Metschnikowia(100);s__Metschnikowia_pulcherrima(100);</td><td>Metschnikowia</td><td>13</td><td>Saprotroph</td><td>|Undefined Saprotroph|</td><td>Yeast</td><td>NULL</td><td>Probable</td><td>As nectar-tap_saprotroph (Põlme et al. 2020); Decay sugar-rich substrates (Põlme et al. 2020)</td><td>Põlme S, et al. 2020. Fungal Diversity 105:1-16 ((https://doi.org/10.1007/s13225-020-00466-2))</td></tr>
<tr><td>Otu42289</td><td>1</td><td>k__Fungi(100);p__Ascomycota(100);c__Saccharomycetes(100);o__Saccharomycetales(100);f__Metschnikowiaceae(100);g__Metschnikowia(100);g__Metschnikowia_unclassified(100);</td><td>Metschnikowia</td><td>13</td><td>Saprotroph</td><td>|Undefined Saprotroph|</td><td>Yeast</td><td>NULL</td><td>Probable</td><td>As nectar-tap_saprotroph (Põlme et al. 2020); Decay sugar-rich substrates (Põlme et al. 2020)</td><td>Põlme S, et al. 2020. Fungal Diversity 105:1-16 ((https://doi.org/10.1007/s13225-020-00466-2))</td></tr>
<tr><td>Otu42290</td><td>1</td><td>k__Fungi(100);p__Ascomycota(100);c__Saccharomycetes(100);o__Saccharomycetales(100);f__Metschnikowiaceae(100);g__Metschnikowia(100);s__Metschnikowia_pulcherrima(100);</td><td>Metschnikowia</td><td>13</td><td>Saprotroph</td><td>|Undefined Saprotroph|</td><td>Yeast</td><td>NULL</td><td>Probable</td><td>As nectar-tap_saprotroph (Põlme et al. 2020); Decay sugar-rich substrates (Põlme et al. 2020)</td><td>Põlme S, et al. 2020. Fungal Diversity 105:1-16 ((https://doi.org/10.1007/s13225-020-00466-2))</td></tr>
<tr><td>Otu42291</td><td>1</td><td>k__Fungi(100);p__Ascomycota(100);c__Saccharomycetes(100);o__Saccharomycetales(100);f__Metschnikowiaceae(100);g__Metschnikowia(100);s__Metschnikowia_pulcherrima(100);</td><td>Metschnikowia</td><td>13</td><td>Saprotroph</td><td>|Undefined Saprotroph|</td><td>Yeast</td><td>NULL</td><td>Probable</td><td>As nectar-tap_saprotroph (Põlme et al. 2020); Decay sugar-rich substrates (Põlme et al. 2020)</td><td>Põlme S, et al. 2020. Fungal Diversity 105:1-16 ((https://doi.org/10.1007/s13225-020-00466-2))</td></tr>
<tr><td>Otu42292</td><td>1</td><td>k__Fungi(100);p__Ascomycota(100);c__Saccharomycetes(100);o__Saccharomycetales(100);f__Metschnikowiaceae(100);g__Metschnikowia(100);s__Metschnikowia_pulcherrima(100);</td><td>Metschnikowia</td><td>13</td><td>Saprotroph</td><td>|Undefined Saprotroph|</td><td>Yeast</td><td>NULL</td><td>Probable</td><td>As nectar-tap_saprotroph (Põlme et al. 2020); Decay sugar-rich substrates (Põlme et al. 2020)</td><td>Põlme S, et al. 2020. Fungal Diversity 105:1-16 ((https://doi.org/10.1007/s13225-020-00466-2))</td></tr>
<tr><td>Otu42293</td><td>1</td><td>k__Fungi(100);p__Ascomycota(100);c__Saccharomycetes(100);o__Saccharomycetales(100);f__Metschnikowiaceae(100);g__Metschnikowia(100);s__Metschnikowia_pulcherrima(100);</td><td>Metschnikowia</td><td>13</td><td>Saprotroph</td><td>|Undefined Saprotroph|</td><td>Yeast</td><td>NULL</td><td>Probable</td><td>As nectar-tap_saprotroph (Põlme et al. 2020); Decay sugar-rich substrates (Põlme et al. 2020)</td><td>Põlme S, et al. 2020. Fungal Diversity 105:1-16 ((https://doi.org/10.1007/s13225-020-00466-2))</td></tr>
<tr><td>Otu42294</td><td>1</td><td>k__Fungi(100);p__Ascomycota(100);c__Saccharomycetes(100);o__Saccharomycetales(100);f__Metschnikowiaceae(100);g__Metschnikowia(100);s__Metschnikowia_pulcherrima(100);</td><td>Metschnikowia</td><td>13</td><td>Saprotroph</td><td>|Undefined Saprotroph|</td><td>Yeast</td><td>NULL</td><td>Probable</td><td>As nectar-tap_saprotroph (Põlme et al. 2020); Decay sugar-rich substrates (Põlme et al. 2020)</td><td>Põlme S, et al. 2020. Fungal Diversity 105:1-16 ((https://doi.org/10.1007/s13225-020-00466-2))</td></tr>
<tr><td>Otu42295</td><td>1</td><td>k__Fungi(100);p__Ascomycota(100);c__Saccharomycetes(100);o__Saccharomycetales(100);f__Metschnikowiaceae(100);g__Metschnikowia(100);g__Metschnikowia_unclassified(100);</td><td>Metschnikowia</td><td>13</td><td>Saprotroph</td><td>|Undefined Saprotroph|</td><td>Yeast</td><td>NULL</td><td>Probable</td><td>As nectar-tap_saprotroph (Põlme et al. 2020); Decay sugar-rich substrates (Põlme et al. 2020)</td><td>Põlme S, et al. 2020. Fungal Diversity 105:1-16 ((https://doi.org/10.1007/s13225-020-00466-2))</td></tr>
<tr><td>Otu42296</td><td>1</td><td>k__Fungi(100);p__Ascomycota(100);c__Saccharomycetes(100);o__Saccharomycetales(100);f__Metschnikowiaceae(100);g__Metschnikowia(100);s__Metschnikowia_pulcherrima(100);</td><td>Metschnikowia</td><td>13</td><td>Saprotroph</td><td>|Undefined Saprotroph|</td><td>Yeast</td><td>NULL</td><td>Probable</td><td>As nectar-tap_saprotroph (Põlme et al. 2020); Decay sugar-rich substrates (Põlme et al. 2020)</td><td>Põlme S, et al. 2020. Fungal Diversity 105:1-16 ((https://doi.org/10.1007/s13225-020-00466-2))</td></tr>
<tr><td>Otu42297</td><td>1</td><td>k__Fungi(100);p__Ascomycota(100);c__Saccharomycetes(100);o__Saccharomycetales(100);f__Metschnikowiaceae(100);g__Metschnikowia(100);s__Metschnikowia_pulcherrima(100);</td><td>Metschnikowia</td><td>13</td><td>Saprotroph</td><td>|Undefined Saprotroph|</td><td>Yeast</td><td>NULL</td><td>Probable</td><td>As nectar-tap_saprotroph (Põlme et al. 2020); Decay sugar-rich substrates (Põlme et al. 2020)</td><td>Põlme S, et al. 2020. Fungal Diversity 105:1-16 ((https://doi.org/10.1007/s13225-020-00466-2))</td></tr>
<tr><td>Otu42298</td><td>1</td><td>k__Fungi(100);p__Ascomycota(100);c__Saccharomycetes(100);o__Saccharomycetales(100);f__Metschnikowiaceae(100);g__Metschnikowia(100);g__Metschnikowia_unclassified(100);</td><td>Metschnikowia</td><td>13</td><td>Saprotroph</td><td>|Undefined Saprotroph|</td><td>Yeast</td><td>NULL</td><td>Probable</td><td>As nectar-tap_saprotroph (Põlme et al. 2020); Decay sugar-rich substrates (Põlme et al. 2020)</td><td>Põlme S, et al. 2020. Fungal Diversity 105:1-16 ((https://doi.org/10.1007/s13225-020-00466-2))</td></tr>
<tr><td>Otu42299</td><td>1</td><td>k__Fungi(100);p__Ascomycota(100);c__Saccharomycetes(100);o__Saccharomycetales(100);f__Metschnikowiaceae(100);g__Metschnikowia(100);g__Metschnikowia_unclassified(100);</td><td>Metschnikowia</td><td>13</td><td>Saprotroph</td><td>|Undefined Saprotroph|</td><td>Yeast</td><td>NULL</td><td>Probable</td><td>As nectar-tap_saprotroph (Põlme et al. 2020); Decay sugar-rich substrates (Põlme et al. 2020)</td><td>Põlme S, et al. 2020. Fungal Diversity 105:1-16 ((https://doi.org/10.1007/s13225-020-00466-2))</td></tr>
<tr><td>Otu42300</td><td>1</td><td>k__Fungi(100);p__Ascomycota(100);c__Saccharomycetes(100);o__Saccharomycetales(100);f__Metschnikowiaceae(100);g__Metschnikowia(100);s__Metschnikowia_pulcherrima(100);</td><td>Metschnikowia</td><td>13</td><td>Saprotroph</td><td>|Undefined Saprotroph|</td><td>Yeast</td><td>NULL</td><td>Probable</td><td>As nectar-tap_saprotroph (Põlme et al. 2020); Decay sugar-rich substrates (Põlme et al. 2020)</td><td>Põlme S, et al. 2020. Fungal Diversity 105:1-16 ((https://doi.org/10.1007/s13225-020-00466-2))</td></tr>
<tr><td>Otu42301</td><td>1</td><td>k__Fungi(100);p__Ascomycota(100);c__Saccharomycetes(100);o__Saccharomycetales(100);f__Metschnikowiaceae(100);g__Metschnikowia(100);s__Metschnikowia_pulcherrima(100);</td><td>Metschnikowia</td><td>13</td><td>Saprotroph</td><td>|Undefined Saprotroph|</td><td>Yeast</td><td>NULL</td><td>Probable</td><td>As nectar-tap_saprotroph (Põlme et al. 2020); Decay sugar-rich substrates (Põlme et al. 2020)</td><td>Põlme S, et al. 2020. Fungal Diversity 105:1-16 ((https://doi.org/10.1007/s13225-020-00466-2))</td></tr>
<tr><td>Otu42302</td><td>1</td><td>k__Fungi(100);p__Ascomycota(100);c__Saccharomycetes(100);o__Saccharomycetales(100);f__Metschnikowiaceae(100);g__Metschnikowia(100);s__Metschnikowia_pulcherrima(100);</td><td>Metschnikowia</td><td>13</td><td>Saprotroph</td><td>|Undefined Saprotroph|</td><td>Yeast</td><td>NULL</td><td>Probable</td><td>As nectar-tap_saprotroph (Põlme et al. 2020); Decay sugar-rich substrates (Põlme et al. 2020)</td><td>Põlme S, et al. 2020. Fungal Diversity 105:1-16 ((https://doi.org/10.1007/s13225-020-00466-2))</td></tr>
<tr><td>Otu42303</td><td>1</td><td>k__Fungi(100);p__Ascomycota(100);c__Saccharomycetes(100);o__Saccharomycetales(100);f__Metschnikowiaceae(100);g__Metschnikowia(100);s__Metschnikowia_pulcherrima(100);</td><td>Metschnikowia</td><td>13</td><td>Saprotroph</td><td>|Undefined Saprotroph|</td><td>Yeast</td><td>NULL</td><td>Probable</td><td>As nectar-tap_saprotroph (Põlme et al. 2020); Decay sugar-rich substrates (Põlme et al. 2020)</td><td>Põlme S, et al. 2020. Fungal Diversity 105:1-16 ((https://doi.org/10.1007/s13225-020-00466-2))</td></tr>
<tr><td>Otu42304</td><td>1</td><td>k__Fungi(100);p__Ascomycota(100);c__Saccharomycetes(100);o__Saccharomycetales(100);f__Metschnikowiaceae(100);g__Metschnikowia(100);s__Metschnikowia_pulcherrima(100);</td><td>Metschnikowia</td><td>13</td><td>Saprotroph</td><td>|Undefined Saprotroph|</td><td>Yeast</td><td>NULL</td><td>Probable</td><td>As nectar-tap_saprotroph (Põlme et al. 2020); Decay sugar-rich substrates (Põlme et al. 2020)</td><td>Põlme S, et al. 2020. Fungal Diversity 105:1-16 ((https://doi.org/10.1007/s13225-020-00466-2))</td></tr>
<tr><td>Otu42305</td><td>1</td><td>k__Fungi(100);p__Ascomycota(100);c__Saccharomycetes(100);o__Saccharomycetales(100);f__Metschnikowiaceae(100);g__Metschnikowia(100);s__Metschnikowia_pulcherrima(100);</td><td>Metschnikowia</td><td>13</td><td>Saprotroph</td><td>|Undefined Saprotroph|</td><td>Yeast</td><td>NULL</td><td>Probable</td><td>As nectar-tap_saprotroph (Põlme et al. 2020); Decay sugar-rich substrates (Põlme et al. 2020)</td><td>Põlme S, et al. 2020. Fungal Diversity 105:1-16 ((https://doi.org/10.1007/s13225-020-00466-2))</td></tr>
<tr><td>Otu42306</td><td>1</td><td>k__Fungi(100);p__Ascomycota(100);c__Saccharomycetes(100);o__Saccharomycetales(100);f__Metschnikowiaceae(100);g__Metschnikowia(100);s__Metschnikowia_pulcherrima(100);</td><td>Metschnikowia</td><td>13</td><td>Saprotroph</td><td>|Undefined Saprotroph|</td><td>Yeast</td><td>NULL</td><td>Probable</td><td>As nectar-tap_saprotroph (Põlme et al. 2020); Decay sugar-rich substrates (Põlme et al. 2020)</td><td>Põlme S, et al. 2020. Fungal Diversity 105:1-16 ((https://doi.org/10.1007/s13225-020-00466-2))</td></tr>
<tr><td>Otu42307</td><td>1</td><td>k__Fungi(100);p__Ascomycota(100);c__Saccharomycetes(100);o__Saccharomycetales(100);f__Metschnikowiaceae(100);g__Metschnikowia(100);s__Metschnikowia_pulcherrima(100);</td><td>Metschnikowia</td><td>13</td><td>Saprotroph</td><td>|Undefined Saprotroph|</td><td>Yeast</td><td>NULL</td><td>Probable</td><td>As nectar-tap_saprotroph (Põlme et al. 2020); Decay sugar-rich substrates (Põlme et al. 2020)</td><td>Põlme S, et al. 2020. Fungal Diversity 105:1-16 ((https://doi.org/10.1007/s13225-020-00466-2))</td></tr>
<tr><td>Otu42308</td><td>1</td><td>k__Fungi(100);p__Ascomycota(100);c__Saccharomycetes(100);o__Saccharomycetales(100);f__Metschnikowiaceae(100);g__Metschnikowia(100);s__Metschnikowia_pulcherrima(100);</td><td>Metschnikowia</td><td>13</td><td>Saprotroph</td><td>|Undefined Saprotroph|</td><td>Yeast</td><td>NULL</td><td>Probable</td><td>As nectar-tap_saprotroph (Põlme et al. 2020); Decay sugar-rich substrates (Põlme et al. 2020)</td><td>Põlme S, et al. 2020. Fungal Diversity 105:1-16 ((https://doi.org/10.1007/s13225-020-00466-2))</td></tr>
<tr><td>Otu42309</td><td>1</td><td>k__Fungi(100);p__Ascomycota(100);c__Saccharomycetes(100);o__Saccharomycetales(100);f__Metschnikowiaceae(100);g__Metschnikowia(100);s__Metschnikowia_pulcherrima(100);</td><td>Metschnikowia</td><td>13</td><td>Saprotroph</td><td>|Undefined Saprotroph|</td><td>Yeast</td><td>NULL</td><td>Probable</td><td>As nectar-tap_saprotroph (Põlme et al. 2020); Decay sugar-rich substrates (Põlme et al. 2020)</td><td>Põlme S, et al. 2020. Fungal Diversity 105:1-16 ((https://doi.org/10.1007/s13225-020-00466-2))</td></tr>
<tr><td>Otu42310</td><td>1</td><td>k__Fungi(100);p__Ascomycota(100);c__Saccharomycetes(100);o__Saccharomycetales(100);f__Metschnikowiaceae(100);g__Metschnikowia(100);s__Metschnikowia_pulcherrima(100);</td><td>Metschnikowia</td><td>13</td><td>Saprotroph</td><td>|Undefined Saprotroph|</td><td>Yeast</td><td>NULL</td><td>Probable</td><td>As nectar-tap_saprotroph (Põlme et al. 2020); Decay sugar-rich substrates (Põlme et al. 2020)</td><td>Põlme S, et al. 2020. Fungal Diversity 105:1-16 ((https://doi.org/10.1007/s13225-020-00466-2))</td></tr>
<tr><td>Otu42311</td><td>1</td><td>k__Fungi(100);p__Ascomycota(100);c__Saccharomycetes(100);o__Saccharomycetales(100);f__Metschnikowiaceae(100);g__Metschnikowia(100);s__Metschnikowia_pulcherrima(100);</td><td>Metschnikowia</td><td>13</td><td>Saprotroph</td><td>|Undefined Saprotroph|</td><td>Yeast</td><td>NULL</td><td>Probable</td><td>As nectar-tap_saprotroph (Põlme et al. 2020); Decay sugar-rich substrates (Põlme et al. 2020)</td><td>Põlme S, et al. 2020. Fungal Diversity 105:1-16 ((https://doi.org/10.1007/s13225-020-00466-2))</td></tr>
<tr><td>Otu42312</td><td>1</td><td>k__Fungi(100);p__Ascomycota(100);c__Saccharomycetes(100);o__Saccharomycetales(100);f__Metschnikowiaceae(100);g__Metschnikowia(100);s__Metschnikowia_pulcherrima(100);</td><td>Metschnikowia</td><td>13</td><td>Saprotroph</td><td>|Undefined Saprotroph|</td><td>Yeast</td><td>NULL</td><td>Probable</td><td>As nectar-tap_saprotroph (Põlme et al. 2020); Decay sugar-rich substrates (Põlme et al. 2020)</td><td>Põlme S, et al. 2020. Fungal Diversity 105:1-16 ((https://doi.org/10.1007/s13225-020-00466-2))</td></tr>
<tr><td>Otu42313</td><td>1</td><td>k__Fungi(100);p__Ascomycota(100);c__Saccharomycetes(100);o__Saccharomycetales(100);f__Metschnikowiaceae(100);g__Metschnikowia(100);g__Metschnikowia_unclassified(100);</td><td>Metschnikowia</td><td>13</td><td>Saprotroph</td><td>|Undefined Saprotroph|</td><td>Yeast</td><td>NULL</td><td>Probable</td><td>As nectar-tap_saprotroph (Põlme et al. 2020); Decay sugar-rich substrates (Põlme et al. 2020)</td><td>Põlme S, et al. 2020. Fungal Diversity 105:1-16 ((https://doi.org/10.1007/s13225-020-00466-2))</td></tr>
<tr><td>Otu42314</td><td>1</td><td>k__Fungi(100);p__Ascomycota(100);c__Saccharomycetes(100);o__Saccharomycetales(100);f__Metschnikowiaceae(100);g__Metschnikowia(100);s__Metschnikowia_pulcherrima(100);</td><td>Metschnikowia</td><td>13</td><td>Saprotroph</td><td>|Undefined Saprotroph|</td><td>Yeast</td><td>NULL</td><td>Probable</td><td>As nectar-tap_saprotroph (Põlme et al. 2020); Decay sugar-rich substrates (Põlme et al. 2020)</td><td>Põlme S, et al. 2020. Fungal Diversity 105:1-16 ((https://doi.org/10.1007/s13225-020-00466-2))</td></tr>
<tr><td>Otu42315</td><td>1</td><td>k__Fungi(100);p__Ascomycota(100);c__Saccharomycetes(100);o__Saccharomycetales(100);f__Metschnikowiaceae(100);g__Metschnikowia(100);s__Metschnikowia_pulcherrima(100);</td><td>Metschnikowia</td><td>13</td><td>Saprotroph</td><td>|Undefined Saprotroph|</td><td>Yeast</td><td>NULL</td><td>Probable</td><td>As nectar-tap_saprotroph (Põlme et al. 2020); Decay sugar-rich substrates (Põlme et al. 2020)</td><td>Põlme S, et al. 2020. Fungal Diversity 105:1-16 ((https://doi.org/10.1007/s13225-020-00466-2))</td></tr>
<tr><td>Otu42316</td><td>1</td><td>k__Fungi(100);p__Ascomycota(100);c__Saccharomycetes(100);o__Saccharomycetales(100);f__Metschnikowiaceae(100);g__Metschnikowia(100);s__Metschnikowia_pulcherrima(100);</td><td>Metschnikowia</td><td>13</td><td>Saprotroph</td><td>|Undefined Saprotroph|</td><td>Yeast</td><td>NULL</td><td>Probable</td><td>As nectar-tap_saprotroph (Põlme et al. 2020); Decay sugar-rich substrates (Põlme et al. 2020)</td><td>Põlme S, et al. 2020. Fungal Diversity 105:1-16 ((https://doi.org/10.1007/s13225-020-00466-2))</td></tr>
<tr><td>Otu42317</td><td>1</td><td>k__Fungi(100);p__Ascomycota(100);c__Saccharomycetes(100);o__Saccharomycetales(100);f__Metschnikowiaceae(100);g__Metschnikowia(100);g__Metschnikowia_unclassified(100);</td><td>Metschnikowia</td><td>13</td><td>Saprotroph</td><td>|Undefined Saprotroph|</td><td>Yeast</td><td>NULL</td><td>Probable</td><td>As nectar-tap_saprotroph (Põlme et al. 2020); Decay sugar-rich substrates (Põlme et al. 2020)</td><td>Põlme S, et al. 2020. Fungal Diversity 105:1-16 ((https://doi.org/10.1007/s13225-020-00466-2))</td></tr>
<tr><td>Otu42318</td><td>1</td><td>k__Fungi(100);p__Ascomycota(100);c__Saccharomycetes(100);o__Saccharomycetales(100);f__Metschnikowiaceae(100);g__Metschnikowia(100);g__Metschnikowia_unclassified(100);</td><td>Metschnikowia</td><td>13</td><td>Saprotroph</td><td>|Undefined Saprotroph|</td><td>Yeast</td><td>NULL</td><td>Probable</td><td>As nectar-tap_saprotroph (Põlme et al. 2020); Decay sugar-rich substrates (Põlme et al. 2020)</td><td>Põlme S, et al. 2020. Fungal Diversity 105:1-16 ((https://doi.org/10.1007/s13225-020-00466-2))</td></tr>
<tr><td>Otu42319</td><td>1</td><td>k__Fungi(100);p__Ascomycota(100);c__Saccharomycetes(100);o__Saccharomycetales(100);f__Metschnikowiaceae(100);g__Metschnikowia(100);s__Metschnikowia_pulcherrima(100);</td><td>Metschnikowia</td><td>13</td><td>Saprotroph</td><td>|Undefined Saprotroph|</td><td>Yeast</td><td>NULL</td><td>Probable</td><td>As nectar-tap_saprotroph (Põlme et al. 2020); Decay sugar-rich substrates (Põlme et al. 2020)</td><td>Põlme S, et al. 2020. Fungal Diversity 105:1-16 ((https://doi.org/10.1007/s13225-020-00466-2))</td></tr>
<tr><td>Otu42320</td><td>1</td><td>k__Fungi(100);p__Ascomycota(100);c__Saccharomycetes(100);o__Saccharomycetales(100);f__Metschnikowiaceae(100);g__Metschnikowia(100);s__Metschnikowia_pulcherrima(100);</td><td>Metschnikowia</td><td>13</td><td>Saprotroph</td><td>|Undefined Saprotroph|</td><td>Yeast</td><td>NULL</td><td>Probable</td><td>As nectar-tap_saprotroph (Põlme et al. 2020); Decay sugar-rich substrates (Põlme et al. 2020)</td><td>Põlme S, et al. 2020. Fungal Diversity 105:1-16 ((https://doi.org/10.1007/s13225-020-00466-2))</td></tr>
<tr><td>Otu42321</td><td>1</td><td>k__Fungi(100);p__Ascomycota(100);c__Saccharomycetes(100);o__Saccharomycetales(100);f__Metschnikowiaceae(100);g__Metschnikowia(100);s__Metschnikowia_pulcherrima(100);</td><td>Metschnikowia</td><td>13</td><td>Saprotroph</td><td>|Undefined Saprotroph|</td><td>Yeast</td><td>NULL</td><td>Probable</td><td>As nectar-tap_saprotroph (Põlme et al. 2020); Decay sugar-rich substrates (Põlme et al. 2020)</td><td>Põlme S, et al. 2020. Fungal Diversity 105:1-16 ((https://doi.org/10.1007/s13225-020-00466-2))</td></tr>
<tr><td>Otu42323</td><td>1</td><td>k__Fungi(100);p__Ascomycota(100);c__Saccharomycetes(100);o__Saccharomycetales(100);f__Metschnikowiaceae(100);g__Metschnikowia(100);s__Metschnikowia_pulcherrima(100);</td><td>Metschnikowia</td><td>13</td><td>Saprotroph</td><td>|Undefined Saprotroph|</td><td>Yeast</td><td>NULL</td><td>Probable</td><td>As nectar-tap_saprotroph (Põlme et al. 2020); Decay sugar-rich substrates (Põlme et al. 2020)</td><td>Põlme S, et al. 2020. Fungal Diversity 105:1-16 ((https://doi.org/10.1007/s13225-020-00466-2))</td></tr>
<tr><td>Otu42324</td><td>1</td><td>k__Fungi(100);p__Ascomycota(100);c__Saccharomycetes(100);o__Saccharomycetales(100);f__Metschnikowiaceae(100);g__Metschnikowia(100);s__Metschnikowia_pulcherrima(100);</td><td>Metschnikowia</td><td>13</td><td>Saprotroph</td><td>|Undefined Saprotroph|</td><td>Yeast</td><td>NULL</td><td>Probable</td><td>As nectar-tap_saprotroph (Põlme et al. 2020); Decay sugar-rich substrates (Põlme et al. 2020)</td><td>Põlme S, et al. 2020. Fungal Diversity 105:1-16 ((https://doi.org/10.1007/s13225-020-00466-2))</td></tr>
<tr><td>Otu42325</td><td>1</td><td>k__Fungi(100);p__Ascomycota(100);c__Saccharomycetes(100);o__Saccharomycetales(100);f__Metschnikowiaceae(100);g__Metschnikowia(100);s__Metschnikowia_pulcherrima(100);</td><td>Metschnikowia</td><td>13</td><td>Saprotroph</td><td>|Undefined Saprotroph|</td><td>Yeast</td><td>NULL</td><td>Probable</td><td>As nectar-tap_saprotroph (Põlme et al. 2020); Decay sugar-rich substrates (Põlme et al. 2020)</td><td>Põlme S, et al. 2020. Fungal Diversity 105:1-16 ((https://doi.org/10.1007/s13225-020-00466-2))</td></tr>
<tr><td>Otu42326</td><td>1</td><td>k__Fungi(100);p__Ascomycota(100);c__Saccharomycetes(100);o__Saccharomycetales(100);f__Metschnikowiaceae(100);g__Metschnikowia(100);s__Metschnikowia_pulcherrima(100);</td><td>Metschnikowia</td><td>13</td><td>Saprotroph</td><td>|Undefined Saprotroph|</td><td>Yeast</td><td>NULL</td><td>Probable</td><td>As nectar-tap_saprotroph (Põlme et al. 2020); Decay sugar-rich substrates (Põlme et al. 2020)</td><td>Põlme S, et al. 2020. Fungal Diversity 105:1-16 ((https://doi.org/10.1007/s13225-020-00466-2))</td></tr>
<tr><td>Otu42327</td><td>1</td><td>k__Fungi(100);p__Ascomycota(100);c__Saccharomycetes(100);o__Saccharomycetales(100);f__Metschnikowiaceae(100);g__Metschnikowia(100);g__Metschnikowia_unclassified(100);</td><td>Metschnikowia</td><td>13</td><td>Saprotroph</td><td>|Undefined Saprotroph|</td><td>Yeast</td><td>NULL</td><td>Probable</td><td>As nectar-tap_saprotroph (Põlme et al. 2020); Decay sugar-rich substrates (Põlme et al. 2020)</td><td>Põlme S, et al. 2020. Fungal Diversity 105:1-16 ((https://doi.org/10.1007/s13225-020-00466-2))</td></tr>
<tr><td>Otu42328</td><td>1</td><td>k__Fungi(100);p__Ascomycota(100);c__Saccharomycetes(100);o__Saccharomycetales(100);f__Metschnikowiaceae(100);g__Metschnikowia(100);s__Metschnikowia_pulcherrima(100);</td><td>Metschnikowia</td><td>13</td><td>Saprotroph</td><td>|Undefined Saprotroph|</td><td>Yeast</td><td>NULL</td><td>Probable</td><td>As nectar-tap_saprotroph (Põlme et al. 2020); Decay sugar-rich substrates (Põlme et al. 2020)</td><td>Põlme S, et al. 2020. Fungal Diversity 105:1-16 ((https://doi.org/10.1007/s13225-020-00466-2))</td></tr>
<tr><td>Otu42329</td><td>1</td><td>k__Fungi(100);p__Ascomycota(100);c__Saccharomycetes(100);o__Saccharomycetales(100);f__Metschnikowiaceae(100);g__Metschnikowia(100);s__Metschnikowia_pulcherrima(100);</td><td>Metschnikowia</td><td>13</td><td>Saprotroph</td><td>|Undefined Saprotroph|</td><td>Yeast</td><td>NULL</td><td>Probable</td><td>As nectar-tap_saprotroph (Põlme et al. 2020); Decay sugar-rich substrates (Põlme et al. 2020)</td><td>Põlme S, et al. 2020. Fungal Diversity 105:1-16 ((https://doi.org/10.1007/s13225-020-00466-2))</td></tr>
<tr><td>Otu42330</td><td>1</td><td>k__Fungi(100);p__Ascomycota(100);c__Saccharomycetes(100);o__Saccharomycetales(100);f__Metschnikowiaceae(100);g__Metschnikowia(100);s__Metschnikowia_pulcherrima(100);</td><td>Metschnikowia</td><td>13</td><td>Saprotroph</td><td>|Undefined Saprotroph|</td><td>Yeast</td><td>NULL</td><td>Probable</td><td>As nectar-tap_saprotroph (Põlme et al. 2020); Decay sugar-rich substrates (Põlme et al. 2020)</td><td>Põlme S, et al. 2020. Fungal Diversity 105:1-16 ((https://doi.org/10.1007/s13225-020-00466-2))</td></tr>
<tr><td>Otu42333</td><td>1</td><td>k__Fungi(100);p__Ascomycota(100);c__Saccharomycetes(100);o__Saccharomycetales(100);f__Metschnikowiaceae(100);g__Metschnikowia(100);s__Metschnikowia_pulcherrima(100);</td><td>Metschnikowia</td><td>13</td><td>Saprotroph</td><td>|Undefined Saprotroph|</td><td>Yeast</td><td>NULL</td><td>Probable</td><td>As nectar-tap_saprotroph (Põlme et al. 2020); Decay sugar-rich substrates (Põlme et al. 2020)</td><td>Põlme S, et al. 2020. Fungal Diversity 105:1-16 ((https://doi.org/10.1007/s13225-020-00466-2))</td></tr>
<tr><td>Otu42335</td><td>1</td><td>k__Fungi(100);p__Ascomycota(100);c__Saccharomycetes(100);o__Saccharomycetales(100);f__Metschnikowiaceae(100);g__Metschnikowia(100);s__Metschnikowia_pulcherrima(100);</td><td>Metschnikowia</td><td>13</td><td>Saprotroph</td><td>|Undefined Saprotroph|</td><td>Yeast</td><td>NULL</td><td>Probable</td><td>As nectar-tap_saprotroph (Põlme et al. 2020); Decay sugar-rich substrates (Põlme et al. 2020)</td><td>Põlme S, et al. 2020. Fungal Diversity 105:1-16 ((https://doi.org/10.1007/s13225-020-00466-2))</td></tr>
<tr><td>Otu42339</td><td>1</td><td>k__Fungi(100);p__Ascomycota(100);c__Saccharomycetes(100);o__Saccharomycetales(100);f__Metschnikowiaceae(100);g__Metschnikowia(100);g__Metschnikowia_unclassified(100);</td><td>Metschnikowia</td><td>13</td><td>Saprotroph</td><td>|Undefined Saprotroph|</td><td>Yeast</td><td>NULL</td><td>Probable</td><td>As nectar-tap_saprotroph (Põlme et al. 2020); Decay sugar-rich substrates (Põlme et al. 2020)</td><td>Põlme S, et al. 2020. Fungal Diversity 105:1-16 ((https://doi.org/10.1007/s13225-020-00466-2))</td></tr>
<tr><td>Otu42341</td><td>1</td><td>k__Fungi(100);p__Ascomycota(100);c__Saccharomycetes(100);o__Saccharomycetales(100);f__Metschnikowiaceae(100);g__Metschnikowia(100);s__Metschnikowia_pulcherrima(100);</td><td>Metschnikowia</td><td>13</td><td>Saprotroph</td><td>|Undefined Saprotroph|</td><td>Yeast</td><td>NULL</td><td>Probable</td><td>As nectar-tap_saprotroph (Põlme et al. 2020); Decay sugar-rich substrates (Põlme et al. 2020)</td><td>Põlme S, et al. 2020. Fungal Diversity 105:1-16 ((https://doi.org/10.1007/s13225-020-00466-2))</td></tr>
<tr><td>Otu42342</td><td>1</td><td>k__Fungi(100);p__Ascomycota(100);c__Saccharomycetes(100);o__Saccharomycetales(100);f__Metschnikowiaceae(100);g__Metschnikowia(100);g__Metschnikowia_unclassified(100);</td><td>Metschnikowia</td><td>13</td><td>Saprotroph</td><td>|Undefined Saprotroph|</td><td>Yeast</td><td>NULL</td><td>Probable</td><td>As nectar-tap_saprotroph (Põlme et al. 2020); Decay sugar-rich substrates (Põlme et al. 2020)</td><td>Põlme S, et al. 2020. Fungal Diversity 105:1-16 ((https://doi.org/10.1007/s13225-020-00466-2))</td></tr>
<tr><td>Otu42343</td><td>1</td><td>k__Fungi(100);p__Ascomycota(100);c__Saccharomycetes(100);o__Saccharomycetales(100);f__Metschnikowiaceae(100);g__Metschnikowia(100);g__Metschnikowia_unclassified(100);</td><td>Metschnikowia</td><td>13</td><td>Saprotroph</td><td>|Undefined Saprotroph|</td><td>Yeast</td><td>NULL</td><td>Probable</td><td>As nectar-tap_saprotroph (Põlme et al. 2020); Decay sugar-rich substrates (Põlme et al. 2020)</td><td>Põlme S, et al. 2020. Fungal Diversity 105:1-16 ((https://doi.org/10.1007/s13225-020-00466-2))</td></tr>
<tr><td>Otu42344</td><td>1</td><td>k__Fungi(100);p__Ascomycota(100);c__Saccharomycetes(100);o__Saccharomycetales(100);f__Metschnikowiaceae(100);g__Metschnikowia(100);g__Metschnikowia_unclassified(100);</td><td>Metschnikowia</td><td>13</td><td>Saprotroph</td><td>|Undefined Saprotroph|</td><td>Yeast</td><td>NULL</td><td>Probable</td><td>As nectar-tap_saprotroph (Põlme et al. 2020); Decay sugar-rich substrates (Põlme et al. 2020)</td><td>Põlme S, et al. 2020. Fungal Diversity 105:1-16 ((https://doi.org/10.1007/s13225-020-00466-2))</td></tr>
<tr><td>Otu42345</td><td>1</td><td>k__Fungi(100);p__Ascomycota(100);c__Saccharomycetes(100);o__Saccharomycetales(100);f__Metschnikowiaceae(100);g__Metschnikowia(100);s__Metschnikowia_pulcherrima(100);</td><td>Metschnikowia</td><td>13</td><td>Saprotroph</td><td>|Undefined Saprotroph|</td><td>Yeast</td><td>NULL</td><td>Probable</td><td>As nectar-tap_saprotroph (Põlme et al. 2020); Decay sugar-rich substrates (Põlme et al. 2020)</td><td>Põlme S, et al. 2020. Fungal Diversity 105:1-16 ((https://doi.org/10.1007/s13225-020-00466-2))</td></tr>
<tr><td>Otu42346</td><td>1</td><td>k__Fungi(100);p__Ascomycota(100);c__Saccharomycetes(100);o__Saccharomycetales(100);f__Metschnikowiaceae(100);g__Metschnikowia(100);s__Metschnikowia_pulcherrima(100);</td><td>Metschnikowia</td><td>13</td><td>Saprotroph</td><td>|Undefined Saprotroph|</td><td>Yeast</td><td>NULL</td><td>Probable</td><td>As nectar-tap_saprotroph (Põlme et al. 2020); Decay sugar-rich substrates (Põlme et al. 2020)</td><td>Põlme S, et al. 2020. Fungal Diversity 105:1-16 ((https://doi.org/10.1007/s13225-020-00466-2))</td></tr>
<tr><td>Otu42347</td><td>1</td><td>k__Fungi(100);p__Ascomycota(100);c__Saccharomycetes(100);o__Saccharomycetales(100);f__Metschnikowiaceae(100);g__Metschnikowia(100);s__Metschnikowia_pulcherrima(100);</td><td>Metschnikowia</td><td>13</td><td>Saprotroph</td><td>|Undefined Saprotroph|</td><td>Yeast</td><td>NULL</td><td>Probable</td><td>As nectar-tap_saprotroph (Põlme et al. 2020); Decay sugar-rich substrates (Põlme et al. 2020)</td><td>Põlme S, et al. 2020. Fungal Diversity 105:1-16 ((https://doi.org/10.1007/s13225-020-00466-2))</td></tr>
<tr><td>Otu42348</td><td>1</td><td>k__Fungi(100);p__Ascomycota(100);c__Saccharomycetes(100);o__Saccharomycetales(100);f__Metschnikowiaceae(100);g__Metschnikowia(100);g__Metschnikowia_unclassified(100);</td><td>Metschnikowia</td><td>13</td><td>Saprotroph</td><td>|Undefined Saprotroph|</td><td>Yeast</td><td>NULL</td><td>Probable</td><td>As nectar-tap_saprotroph (Põlme et al. 2020); Decay sugar-rich substrates (Põlme et al. 2020)</td><td>Põlme S, et al. 2020. Fungal Diversity 105:1-16 ((https://doi.org/10.1007/s13225-020-00466-2))</td></tr>
<tr><td>Otu42349</td><td>1</td><td>k__Fungi(100);p__Ascomycota(100);c__Saccharomycetes(100);o__Saccharomycetales(100);f__Metschnikowiaceae(100);g__Metschnikowia(100);g__Metschnikowia_unclassified(100);</td><td>Metschnikowia</td><td>13</td><td>Saprotroph</td><td>|Undefined Saprotroph|</td><td>Yeast</td><td>NULL</td><td>Probable</td><td>As nectar-tap_saprotroph (Põlme et al. 2020); Decay sugar-rich substrates (Põlme et al. 2020)</td><td>Põlme S, et al. 2020. Fungal Diversity 105:1-16 ((https://doi.org/10.1007/s13225-020-00466-2))</td></tr>
<tr><td>Otu42350</td><td>1</td><td>k__Fungi(100);p__Ascomycota(100);c__Saccharomycetes(100);o__Saccharomycetales(100);f__Metschnikowiaceae(100);g__Metschnikowia(100);s__Metschnikowia_pulcherrima(100);</td><td>Metschnikowia</td><td>13</td><td>Saprotroph</td><td>|Undefined Saprotroph|</td><td>Yeast</td><td>NULL</td><td>Probable</td><td>As nectar-tap_saprotroph (Põlme et al. 2020); Decay sugar-rich substrates (Põlme et al. 2020)</td><td>Põlme S, et al. 2020. Fungal Diversity 105:1-16 ((https://doi.org/10.1007/s13225-020-00466-2))</td></tr>
<tr><td>Otu42351</td><td>1</td><td>k__Fungi(100);p__Ascomycota(100);c__Saccharomycetes(100);o__Saccharomycetales(100);f__Metschnikowiaceae(100);g__Metschnikowia(100);s__Metschnikowia_pulcherrima(100);</td><td>Metschnikowia</td><td>13</td><td>Saprotroph</td><td>|Undefined Saprotroph|</td><td>Yeast</td><td>NULL</td><td>Probable</td><td>As nectar-tap_saprotroph (Põlme et al. 2020); Decay sugar-rich substrates (Põlme et al. 2020)</td><td>Põlme S, et al. 2020. Fungal Diversity 105:1-16 ((https://doi.org/10.1007/s13225-020-00466-2))</td></tr>
<tr><td>Otu42352</td><td>1</td><td>k__Fungi(100);p__Ascomycota(100);c__Saccharomycetes(100);o__Saccharomycetales(100);f__Metschnikowiaceae(100);g__Metschnikowia(100);s__Metschnikowia_pulcherrima(100);</td><td>Metschnikowia</td><td>13</td><td>Saprotroph</td><td>|Undefined Saprotroph|</td><td>Yeast</td><td>NULL</td><td>Probable</td><td>As nectar-tap_saprotroph (Põlme et al. 2020); Decay sugar-rich substrates (Põlme et al. 2020)</td><td>Põlme S, et al. 2020. Fungal Diversity 105:1-16 ((https://doi.org/10.1007/s13225-020-00466-2))</td></tr>
<tr><td>Otu42353</td><td>1</td><td>k__Fungi(100);p__Ascomycota(100);c__Saccharomycetes(100);o__Saccharomycetales(100);f__Metschnikowiaceae(100);g__Metschnikowia(100);g__Metschnikowia_unclassified(100);</td><td>Metschnikowia</td><td>13</td><td>Saprotroph</td><td>|Undefined Saprotroph|</td><td>Yeast</td><td>NULL</td><td>Probable</td><td>As nectar-tap_saprotroph (Põlme et al. 2020); Decay sugar-rich substrates (Põlme et al. 2020)</td><td>Põlme S, et al. 2020. Fungal Diversity 105:1-16 ((https://doi.org/10.1007/s13225-020-00466-2))</td></tr>
<tr><td>Otu42355</td><td>1</td><td>k__Fungi(100);p__Ascomycota(100);c__Saccharomycetes(100);o__Saccharomycetales(100);f__Metschnikowiaceae(100);g__Metschnikowia(100);g__Metschnikowia_unclassified(100);</td><td>Metschnikowia</td><td>13</td><td>Saprotroph</td><td>|Undefined Saprotroph|</td><td>Yeast</td><td>NULL</td><td>Probable</td><td>As nectar-tap_saprotroph (Põlme et al. 2020); Decay sugar-rich substrates (Põlme et al. 2020)</td><td>Põlme S, et al. 2020. Fungal Diversity 105:1-16 ((https://doi.org/10.1007/s13225-020-00466-2))</td></tr>
<tr><td>Otu42356</td><td>1</td><td>k__Fungi(100);p__Ascomycota(100);c__Saccharomycetes(100);o__Saccharomycetales(100);f__Metschnikowiaceae(100);g__Metschnikowia(100);s__Metschnikowia_pulcherrima(100);</td><td>Metschnikowia</td><td>13</td><td>Saprotroph</td><td>|Undefined Saprotroph|</td><td>Yeast</td><td>NULL</td><td>Probable</td><td>As nectar-tap_saprotroph (Põlme et al. 2020); Decay sugar-rich substrates (Põlme et al. 2020)</td><td>Põlme S, et al. 2020. Fungal Diversity 105:1-16 ((https://doi.org/10.1007/s13225-020-00466-2))</td></tr>
<tr><td>Otu42357</td><td>1</td><td>k__Fungi(100);p__Ascomycota(100);c__Saccharomycetes(100);o__Saccharomycetales(100);f__Metschnikowiaceae(100);g__Metschnikowia(100);s__Metschnikowia_pulcherrima(100);</td><td>Metschnikowia</td><td>13</td><td>Saprotroph</td><td>|Undefined Saprotroph|</td><td>Yeast</td><td>NULL</td><td>Probable</td><td>As nectar-tap_saprotroph (Põlme et al. 2020); Decay sugar-rich substrates (Põlme et al. 2020)</td><td>Põlme S, et al. 2020. Fungal Diversity 105:1-16 ((https://doi.org/10.1007/s13225-020-00466-2))</td></tr>
<tr><td>Otu42358</td><td>1</td><td>k__Fungi(100);p__Ascomycota(100);c__Saccharomycetes(100);o__Saccharomycetales(100);f__Metschnikowiaceae(100);g__Metschnikowia(100);s__Metschnikowia_pulcherrima(100);</td><td>Metschnikowia</td><td>13</td><td>Saprotroph</td><td>|Undefined Saprotroph|</td><td>Yeast</td><td>NULL</td><td>Probable</td><td>As nectar-tap_saprotroph (Põlme et al. 2020); Decay sugar-rich substrates (Põlme et al. 2020)</td><td>Põlme S, et al. 2020. Fungal Diversity 105:1-16 ((https://doi.org/10.1007/s13225-020-00466-2))</td></tr>
<tr><td>Otu42360</td><td>1</td><td>k__Fungi(100);p__Ascomycota(100);c__Saccharomycetes(100);o__Saccharomycetales(100);f__Metschnikowiaceae(100);g__Metschnikowia(100);g__Metschnikowia_unclassified(100);</td><td>Metschnikowia</td><td>13</td><td>Saprotroph</td><td>|Undefined Saprotroph|</td><td>Yeast</td><td>NULL</td><td>Probable</td><td>As nectar-tap_saprotroph (Põlme et al. 2020); Decay sugar-rich substrates (Põlme et al. 2020)</td><td>Põlme S, et al. 2020. Fungal Diversity 105:1-16 ((https://doi.org/10.1007/s13225-020-00466-2))</td></tr>
<tr><td>Otu42361</td><td>1</td><td>k__Fungi(100);p__Ascomycota(100);c__Saccharomycetes(100);o__Saccharomycetales(100);f__Metschnikowiaceae(100);g__Metschnikowia(100);s__Metschnikowia_pulcherrima(100);</td><td>Metschnikowia</td><td>13</td><td>Saprotroph</td><td>|Undefined Saprotroph|</td><td>Yeast</td><td>NULL</td><td>Probable</td><td>As nectar-tap_saprotroph (Põlme et al. 2020); Decay sugar-rich substrates (Põlme et al. 2020)</td><td>Põlme S, et al. 2020. Fungal Diversity 105:1-16 ((https://doi.org/10.1007/s13225-020-00466-2))</td></tr>
<tr><td>Otu42362</td><td>1</td><td>k__Fungi(100);p__Ascomycota(100);c__Saccharomycetes(100);o__Saccharomycetales(100);f__Metschnikowiaceae(100);g__Metschnikowia(100);s__Metschnikowia_pulcherrima(100);</td><td>Metschnikowia</td><td>13</td><td>Saprotroph</td><td>|Undefined Saprotroph|</td><td>Yeast</td><td>NULL</td><td>Probable</td><td>As nectar-tap_saprotroph (Põlme et al. 2020); Decay sugar-rich substrates (Põlme et al. 2020)</td><td>Põlme S, et al. 2020. Fungal Diversity 105:1-16 ((https://doi.org/10.1007/s13225-020-00466-2))</td></tr>
<tr><td>Otu42363</td><td>1</td><td>k__Fungi(100);p__Ascomycota(100);c__Saccharomycetes(100);o__Saccharomycetales(100);f__Metschnikowiaceae(100);g__Metschnikowia(100);s__Metschnikowia_pulcherrima(100);</td><td>Metschnikowia</td><td>13</td><td>Saprotroph</td><td>|Undefined Saprotroph|</td><td>Yeast</td><td>NULL</td><td>Probable</td><td>As nectar-tap_saprotroph (Põlme et al. 2020); Decay sugar-rich substrates (Põlme et al. 2020)</td><td>Põlme S, et al. 2020. Fungal Diversity 105:1-16 ((https://doi.org/10.1007/s13225-020-00466-2))</td></tr>
<tr><td>Otu42365</td><td>1</td><td>k__Fungi(100);p__Ascomycota(100);c__Saccharomycetes(100);o__Saccharomycetales(100);f__Metschnikowiaceae(100);g__Metschnikowia(100);s__Metschnikowia_pulcherrima(100);</td><td>Metschnikowia</td><td>13</td><td>Saprotroph</td><td>|Undefined Saprotroph|</td><td>Yeast</td><td>NULL</td><td>Probable</td><td>As nectar-tap_saprotroph (Põlme et al. 2020); Decay sugar-rich substrates (Põlme et al. 2020)</td><td>Põlme S, et al. 2020. Fungal Diversity 105:1-16 ((https://doi.org/10.1007/s13225-020-00466-2))</td></tr>
<tr><td>Otu42366</td><td>1</td><td>k__Fungi(100);p__Ascomycota(100);c__Saccharomycetes(100);o__Saccharomycetales(100);f__Metschnikowiaceae(100);g__Metschnikowia(100);s__Metschnikowia_pulcherrima(100);</td><td>Metschnikowia</td><td>13</td><td>Saprotroph</td><td>|Undefined Saprotroph|</td><td>Yeast</td><td>NULL</td><td>Probable</td><td>As nectar-tap_saprotroph (Põlme et al. 2020); Decay sugar-rich substrates (Põlme et al. 2020)</td><td>Põlme S, et al. 2020. Fungal Diversity 105:1-16 ((https://doi.org/10.1007/s13225-020-00466-2))</td></tr>
<tr><td>Otu42367</td><td>1</td><td>k__Fungi(100);p__Ascomycota(100);c__Saccharomycetes(100);o__Saccharomycetales(100);f__Metschnikowiaceae(100);g__Metschnikowia(100);s__Metschnikowia_pulcherrima(100);</td><td>Metschnikowia</td><td>13</td><td>Saprotroph</td><td>|Undefined Saprotroph|</td><td>Yeast</td><td>NULL</td><td>Probable</td><td>As nectar-tap_saprotroph (Põlme et al. 2020); Decay sugar-rich substrates (Põlme et al. 2020)</td><td>Põlme S, et al. 2020. Fungal Diversity 105:1-16 ((https://doi.org/10.1007/s13225-020-00466-2))</td></tr>
<tr><td>Otu42368</td><td>1</td><td>k__Fungi(100);p__Ascomycota(100);c__Saccharomycetes(100);o__Saccharomycetales(100);f__Metschnikowiaceae(100);g__Metschnikowia(100);g__Metschnikowia_unclassified(100);</td><td>Metschnikowia</td><td>13</td><td>Saprotroph</td><td>|Undefined Saprotroph|</td><td>Yeast</td><td>NULL</td><td>Probable</td><td>As nectar-tap_saprotroph (Põlme et al. 2020); Decay sugar-rich substrates (Põlme et al. 2020)</td><td>Põlme S, et al. 2020. Fungal Diversity 105:1-16 ((https://doi.org/10.1007/s13225-020-00466-2))</td></tr>
<tr><td>Otu42369</td><td>1</td><td>k__Fungi(100);p__Ascomycota(100);c__Saccharomycetes(100);o__Saccharomycetales(100);f__Metschnikowiaceae(100);g__Metschnikowia(100);s__Metschnikowia_pulcherrima(100);</td><td>Metschnikowia</td><td>13</td><td>Saprotroph</td><td>|Undefined Saprotroph|</td><td>Yeast</td><td>NULL</td><td>Probable</td><td>As nectar-tap_saprotroph (Põlme et al. 2020); Decay sugar-rich substrates (Põlme et al. 2020)</td><td>Põlme S, et al. 2020. Fungal Diversity 105:1-16 ((https://doi.org/10.1007/s13225-020-00466-2))</td></tr>
<tr><td>Otu42370</td><td>1</td><td>k__Fungi(100);p__Ascomycota(100);c__Saccharomycetes(100);o__Saccharomycetales(100);f__Metschnikowiaceae(100);g__Metschnikowia(100);s__Metschnikowia_pulcherrima(100);</td><td>Metschnikowia</td><td>13</td><td>Saprotroph</td><td>|Undefined Saprotroph|</td><td>Yeast</td><td>NULL</td><td>Probable</td><td>As nectar-tap_saprotroph (Põlme et al. 2020); Decay sugar-rich substrates (Põlme et al. 2020)</td><td>Põlme S, et al. 2020. Fungal Diversity 105:1-16 ((https://doi.org/10.1007/s13225-020-00466-2))</td></tr>
<tr><td>Otu42371</td><td>1</td><td>k__Fungi(100);p__Ascomycota(100);c__Saccharomycetes(100);o__Saccharomycetales(100);f__Metschnikowiaceae(100);g__Metschnikowia(100);s__Metschnikowia_pulcherrima(100);</td><td>Metschnikowia</td><td>13</td><td>Saprotroph</td><td>|Undefined Saprotroph|</td><td>Yeast</td><td>NULL</td><td>Probable</td><td>As nectar-tap_saprotroph (Põlme et al. 2020); Decay sugar-rich substrates (Põlme et al. 2020)</td><td>Põlme S, et al. 2020. Fungal Diversity 105:1-16 ((https://doi.org/10.1007/s13225-020-00466-2))</td></tr>
<tr><td>Otu42372</td><td>1</td><td>k__Fungi(100);p__Ascomycota(100);c__Saccharomycetes(100);o__Saccharomycetales(100);f__Metschnikowiaceae(100);g__Metschnikowia(100);s__Metschnikowia_pulcherrima(100);</td><td>Metschnikowia</td><td>13</td><td>Saprotroph</td><td>|Undefined Saprotroph|</td><td>Yeast</td><td>NULL</td><td>Probable</td><td>As nectar-tap_saprotroph (Põlme et al. 2020); Decay sugar-rich substrates (Põlme et al. 2020)</td><td>Põlme S, et al. 2020. Fungal Diversity 105:1-16 ((https://doi.org/10.1007/s13225-020-00466-2))</td></tr>
<tr><td>Otu42373</td><td>1</td><td>k__Fungi(100);p__Ascomycota(100);c__Saccharomycetes(100);o__Saccharomycetales(100);f__Metschnikowiaceae(100);g__Metschnikowia(100);s__Metschnikowia_pulcherrima(100);</td><td>Metschnikowia</td><td>13</td><td>Saprotroph</td><td>|Undefined Saprotroph|</td><td>Yeast</td><td>NULL</td><td>Probable</td><td>As nectar-tap_saprotroph (Põlme et al. 2020); Decay sugar-rich substrates (Põlme et al. 2020)</td><td>Põlme S, et al. 2020. Fungal Diversity 105:1-16 ((https://doi.org/10.1007/s13225-020-00466-2))</td></tr>
<tr><td>Otu42374</td><td>1</td><td>k__Fungi(100);p__Ascomycota(100);c__Saccharomycetes(100);o__Saccharomycetales(100);f__Metschnikowiaceae(100);g__Metschnikowia(100);s__Metschnikowia_pulcherrima(100);</td><td>Metschnikowia</td><td>13</td><td>Saprotroph</td><td>|Undefined Saprotroph|</td><td>Yeast</td><td>NULL</td><td>Probable</td><td>As nectar-tap_saprotroph (Põlme et al. 2020); Decay sugar-rich substrates (Põlme et al. 2020)</td><td>Põlme S, et al. 2020. Fungal Diversity 105:1-16 ((https://doi.org/10.1007/s13225-020-00466-2))</td></tr>
<tr><td>Otu42375</td><td>1</td><td>k__Fungi(100);p__Ascomycota(100);c__Saccharomycetes(100);o__Saccharomycetales(100);f__Metschnikowiaceae(100);g__Metschnikowia(100);s__Metschnikowia_pulcherrima(100);</td><td>Metschnikowia</td><td>13</td><td>Saprotroph</td><td>|Undefined Saprotroph|</td><td>Yeast</td><td>NULL</td><td>Probable</td><td>As nectar-tap_saprotroph (Põlme et al. 2020); Decay sugar-rich substrates (Põlme et al. 2020)</td><td>Põlme S, et al. 2020. Fungal Diversity 105:1-16 ((https://doi.org/10.1007/s13225-020-00466-2))</td></tr>
<tr><td>Otu42377</td><td>1</td><td>k__Fungi(100);p__Ascomycota(100);c__Saccharomycetes(100);o__Saccharomycetales(100);f__Metschnikowiaceae(100);g__Metschnikowia(100);g__Metschnikowia_unclassified(100);</td><td>Metschnikowia</td><td>13</td><td>Saprotroph</td><td>|Undefined Saprotroph|</td><td>Yeast</td><td>NULL</td><td>Probable</td><td>As nectar-tap_saprotroph (Põlme et al. 2020); Decay sugar-rich substrates (Põlme et al. 2020)</td><td>Põlme S, et al. 2020. Fungal Diversity 105:1-16 ((https://doi.org/10.1007/s13225-020-00466-2))</td></tr>
<tr><td>Otu42378</td><td>1</td><td>k__Fungi(100);p__Ascomycota(100);c__Saccharomycetes(100);o__Saccharomycetales(100);f__Metschnikowiaceae(100);g__Metschnikowia(100);g__Metschnikowia_unclassified(100);</td><td>Metschnikowia</td><td>13</td><td>Saprotroph</td><td>|Undefined Saprotroph|</td><td>Yeast</td><td>NULL</td><td>Probable</td><td>As nectar-tap_saprotroph (Põlme et al. 2020); Decay sugar-rich substrates (Põlme et al. 2020)</td><td>Põlme S, et al. 2020. Fungal Diversity 105:1-16 ((https://doi.org/10.1007/s13225-020-00466-2))</td></tr>
<tr><td>Otu42383</td><td>1</td><td>k__Fungi(100);p__Ascomycota(100);c__Saccharomycetes(100);o__Saccharomycetales(100);f__Metschnikowiaceae(100);g__Metschnikowia(100);s__Metschnikowia_pulcherrima(100);</td><td>Metschnikowia</td><td>13</td><td>Saprotroph</td><td>|Undefined Saprotroph|</td><td>Yeast</td><td>NULL</td><td>Probable</td><td>As nectar-tap_saprotroph (Põlme et al. 2020); Decay sugar-rich substrates (Põlme et al. 2020)</td><td>Põlme S, et al. 2020. Fungal Diversity 105:1-16 ((https://doi.org/10.1007/s13225-020-00466-2))</td></tr>
<tr><td>Otu42384</td><td>1</td><td>k__Fungi(100);p__Ascomycota(100);c__Saccharomycetes(100);o__Saccharomycetales(100);f__Metschnikowiaceae(100);g__Metschnikowia(100);s__Metschnikowia_pulcherrima(100);</td><td>Metschnikowia</td><td>13</td><td>Saprotroph</td><td>|Undefined Saprotroph|</td><td>Yeast</td><td>NULL</td><td>Probable</td><td>As nectar-tap_saprotroph (Põlme et al. 2020); Decay sugar-rich substrates (Põlme et al. 2020)</td><td>Põlme S, et al. 2020. Fungal Diversity 105:1-16 ((https://doi.org/10.1007/s13225-020-00466-2))</td></tr>
<tr><td>Otu42385</td><td>1</td><td>k__Fungi(100);p__Ascomycota(100);c__Saccharomycetes(100);o__Saccharomycetales(100);f__Metschnikowiaceae(100);g__Metschnikowia(100);g__Metschnikowia_unclassified(100);</td><td>Metschnikowia</td><td>13</td><td>Saprotroph</td><td>|Undefined Saprotroph|</td><td>Yeast</td><td>NULL</td><td>Probable</td><td>As nectar-tap_saprotroph (Põlme et al. 2020); Decay sugar-rich substrates (Põlme et al. 2020)</td><td>Põlme S, et al. 2020. Fungal Diversity 105:1-16 ((https://doi.org/10.1007/s13225-020-00466-2))</td></tr>
<tr><td>Otu42389</td><td>1</td><td>k__Fungi(100);p__Ascomycota(100);c__Saccharomycetes(100);o__Saccharomycetales(100);f__Metschnikowiaceae(100);g__Metschnikowia(100);s__Metschnikowia_pulcherrima(100);</td><td>Metschnikowia</td><td>13</td><td>Saprotroph</td><td>|Undefined Saprotroph|</td><td>Yeast</td><td>NULL</td><td>Probable</td><td>As nectar-tap_saprotroph (Põlme et al. 2020); Decay sugar-rich substrates (Põlme et al. 2020)</td><td>Põlme S, et al. 2020. Fungal Diversity 105:1-16 ((https://doi.org/10.1007/s13225-020-00466-2))</td></tr>
<tr><td>Otu42390</td><td>1</td><td>k__Fungi(100);p__Ascomycota(100);c__Saccharomycetes(100);o__Saccharomycetales(100);f__Metschnikowiaceae(100);g__Metschnikowia(100);s__Metschnikowia_pulcherrima(100);</td><td>Metschnikowia</td><td>13</td><td>Saprotroph</td><td>|Undefined Saprotroph|</td><td>Yeast</td><td>NULL</td><td>Probable</td><td>As nectar-tap_saprotroph (Põlme et al. 2020); Decay sugar-rich substrates (Põlme et al. 2020)</td><td>Põlme S, et al. 2020. Fungal Diversity 105:1-16 ((https://doi.org/10.1007/s13225-020-00466-2))</td></tr>
<tr><td>Otu42391</td><td>1</td><td>k__Fungi(100);p__Ascomycota(100);c__Saccharomycetes(100);o__Saccharomycetales(100);f__Metschnikowiaceae(100);g__Metschnikowia(100);s__Metschnikowia_pulcherrima(100);</td><td>Metschnikowia</td><td>13</td><td>Saprotroph</td><td>|Undefined Saprotroph|</td><td>Yeast</td><td>NULL</td><td>Probable</td><td>As nectar-tap_saprotroph (Põlme et al. 2020); Decay sugar-rich substrates (Põlme et al. 2020)</td><td>Põlme S, et al. 2020. Fungal Diversity 105:1-16 ((https://doi.org/10.1007/s13225-020-00466-2))</td></tr>
<tr><td>Otu42392</td><td>1</td><td>k__Fungi(100);p__Ascomycota(100);c__Saccharomycetes(100);o__Saccharomycetales(100);f__Metschnikowiaceae(100);g__Metschnikowia(100);s__Metschnikowia_pulcherrima(100);</td><td>Metschnikowia</td><td>13</td><td>Saprotroph</td><td>|Undefined Saprotroph|</td><td>Yeast</td><td>NULL</td><td>Probable</td><td>As nectar-tap_saprotroph (Põlme et al. 2020); Decay sugar-rich substrates (Põlme et al. 2020)</td><td>Põlme S, et al. 2020. Fungal Diversity 105:1-16 ((https://doi.org/10.1007/s13225-020-00466-2))</td></tr>
<tr><td>Otu42393</td><td>1</td><td>k__Fungi(100);p__Ascomycota(100);c__Saccharomycetes(100);o__Saccharomycetales(100);f__Metschnikowiaceae(100);g__Metschnikowia(100);s__Metschnikowia_pulcherrima(100);</td><td>Metschnikowia</td><td>13</td><td>Saprotroph</td><td>|Undefined Saprotroph|</td><td>Yeast</td><td>NULL</td><td>Probable</td><td>As nectar-tap_saprotroph (Põlme et al. 2020); Decay sugar-rich substrates (Põlme et al. 2020)</td><td>Põlme S, et al. 2020. Fungal Diversity 105:1-16 ((https://doi.org/10.1007/s13225-020-00466-2))</td></tr>
<tr><td>Otu42394</td><td>1</td><td>k__Fungi(100);p__Ascomycota(100);c__Saccharomycetes(100);o__Saccharomycetales(100);f__Metschnikowiaceae(100);g__Metschnikowia(100);g__Metschnikowia_unclassified(100);</td><td>Metschnikowia</td><td>13</td><td>Saprotroph</td><td>|Undefined Saprotroph|</td><td>Yeast</td><td>NULL</td><td>Probable</td><td>As nectar-tap_saprotroph (Põlme et al. 2020); Decay sugar-rich substrates (Põlme et al. 2020)</td><td>Põlme S, et al. 2020. Fungal Diversity 105:1-16 ((https://doi.org/10.1007/s13225-020-00466-2))</td></tr>
<tr><td>Otu42395</td><td>1</td><td>k__Fungi(100);p__Ascomycota(100);c__Saccharomycetes(100);o__Saccharomycetales(100);f__Metschnikowiaceae(100);g__Metschnikowia(100);g__Metschnikowia_unclassified(100);</td><td>Metschnikowia</td><td>13</td><td>Saprotroph</td><td>|Undefined Saprotroph|</td><td>Yeast</td><td>NULL</td><td>Probable</td><td>As nectar-tap_saprotroph (Põlme et al. 2020); Decay sugar-rich substrates (Põlme et al. 2020)</td><td>Põlme S, et al. 2020. Fungal Diversity 105:1-16 ((https://doi.org/10.1007/s13225-020-00466-2))</td></tr>
<tr><td>Otu42396</td><td>1</td><td>k__Fungi(100);p__Ascomycota(100);c__Saccharomycetes(100);o__Saccharomycetales(100);f__Metschnikowiaceae(100);g__Metschnikowia(100);s__Metschnikowia_pulcherrima(100);</td><td>Metschnikowia</td><td>13</td><td>Saprotroph</td><td>|Undefined Saprotroph|</td><td>Yeast</td><td>NULL</td><td>Probable</td><td>As nectar-tap_saprotroph (Põlme et al. 2020); Decay sugar-rich substrates (Põlme et al. 2020)</td><td>Põlme S, et al. 2020. Fungal Diversity 105:1-16 ((https://doi.org/10.1007/s13225-020-00466-2))</td></tr>
<tr><td>Otu42397</td><td>1</td><td>k__Fungi(100);p__Ascomycota(100);c__Saccharomycetes(100);o__Saccharomycetales(100);f__Metschnikowiaceae(100);g__Metschnikowia(100);s__Metschnikowia_pulcherrima(100);</td><td>Metschnikowia</td><td>13</td><td>Saprotroph</td><td>|Undefined Saprotroph|</td><td>Yeast</td><td>NULL</td><td>Probable</td><td>As nectar-tap_saprotroph (Põlme et al. 2020); Decay sugar-rich substrates (Põlme et al. 2020)</td><td>Põlme S, et al. 2020. Fungal Diversity 105:1-16 ((https://doi.org/10.1007/s13225-020-00466-2))</td></tr>
<tr><td>Otu42398</td><td>1</td><td>k__Fungi(100);p__Ascomycota(100);c__Saccharomycetes(100);o__Saccharomycetales(100);f__Metschnikowiaceae(100);g__Metschnikowia(100);s__Metschnikowia_pulcherrima(100);</td><td>Metschnikowia</td><td>13</td><td>Saprotroph</td><td>|Undefined Saprotroph|</td><td>Yeast</td><td>NULL</td><td>Probable</td><td>As nectar-tap_saprotroph (Põlme et al. 2020); Decay sugar-rich substrates (Põlme et al. 2020)</td><td>Põlme S, et al. 2020. Fungal Diversity 105:1-16 ((https://doi.org/10.1007/s13225-020-00466-2))</td></tr>
<tr><td>Otu42399</td><td>1</td><td>k__Fungi(100);p__Ascomycota(100);c__Saccharomycetes(100);o__Saccharomycetales(100);f__Metschnikowiaceae(100);g__Metschnikowia(100);s__Metschnikowia_pulcherrima(100);</td><td>Metschnikowia</td><td>13</td><td>Saprotroph</td><td>|Undefined Saprotroph|</td><td>Yeast</td><td>NULL</td><td>Probable</td><td>As nectar-tap_saprotroph (Põlme et al. 2020); Decay sugar-rich substrates (Põlme et al. 2020)</td><td>Põlme S, et al. 2020. Fungal Diversity 105:1-16 ((https://doi.org/10.1007/s13225-020-00466-2))</td></tr>
<tr><td>Otu42400</td><td>1</td><td>k__Fungi(100);p__Ascomycota(100);c__Saccharomycetes(100);o__Saccharomycetales(100);f__Metschnikowiaceae(100);g__Metschnikowia(100);g__Metschnikowia_unclassified(100);</td><td>Metschnikowia</td><td>13</td><td>Saprotroph</td><td>|Undefined Saprotroph|</td><td>Yeast</td><td>NULL</td><td>Probable</td><td>As nectar-tap_saprotroph (Põlme et al. 2020); Decay sugar-rich substrates (Põlme et al. 2020)</td><td>Põlme S, et al. 2020. Fungal Diversity 105:1-16 ((https://doi.org/10.1007/s13225-020-00466-2))</td></tr>
<tr><td>Otu42401</td><td>1</td><td>k__Fungi(100);p__Ascomycota(100);c__Saccharomycetes(100);o__Saccharomycetales(100);f__Metschnikowiaceae(100);g__Metschnikowia(100);s__Metschnikowia_pulcherrima(100);</td><td>Metschnikowia</td><td>13</td><td>Saprotroph</td><td>|Undefined Saprotroph|</td><td>Yeast</td><td>NULL</td><td>Probable</td><td>As nectar-tap_saprotroph (Põlme et al. 2020); Decay sugar-rich substrates (Põlme et al. 2020)</td><td>Põlme S, et al. 2020. Fungal Diversity 105:1-16 ((https://doi.org/10.1007/s13225-020-00466-2))</td></tr>
<tr><td>Otu42402</td><td>1</td><td>k__Fungi(100);p__Ascomycota(100);c__Saccharomycetes(100);o__Saccharomycetales(100);f__Metschnikowiaceae(100);g__Metschnikowia(100);s__Metschnikowia_pulcherrima(100);</td><td>Metschnikowia</td><td>13</td><td>Saprotroph</td><td>|Undefined Saprotroph|</td><td>Yeast</td><td>NULL</td><td>Probable</td><td>As nectar-tap_saprotroph (Põlme et al. 2020); Decay sugar-rich substrates (Põlme et al. 2020)</td><td>Põlme S, et al. 2020. Fungal Diversity 105:1-16 ((https://doi.org/10.1007/s13225-020-00466-2))</td></tr>
<tr><td>Otu42403</td><td>1</td><td>k__Fungi(100);p__Ascomycota(100);c__Saccharomycetes(100);o__Saccharomycetales(100);f__Metschnikowiaceae(100);g__Metschnikowia(100);g__Metschnikowia_unclassified(100);</td><td>Metschnikowia</td><td>13</td><td>Saprotroph</td><td>|Undefined Saprotroph|</td><td>Yeast</td><td>NULL</td><td>Probable</td><td>As nectar-tap_saprotroph (Põlme et al. 2020); Decay sugar-rich substrates (Põlme et al. 2020)</td><td>Põlme S, et al. 2020. Fungal Diversity 105:1-16 ((https://doi.org/10.1007/s13225-020-00466-2))</td></tr>
<tr><td>Otu42404</td><td>1</td><td>k__Fungi(100);p__Ascomycota(100);c__Saccharomycetes(100);o__Saccharomycetales(100);f__Metschnikowiaceae(100);g__Metschnikowia(100);s__Metschnikowia_pulcherrima(100);</td><td>Metschnikowia</td><td>13</td><td>Saprotroph</td><td>|Undefined Saprotroph|</td><td>Yeast</td><td>NULL</td><td>Probable</td><td>As nectar-tap_saprotroph (Põlme et al. 2020); Decay sugar-rich substrates (Põlme et al. 2020)</td><td>Põlme S, et al. 2020. Fungal Diversity 105:1-16 ((https://doi.org/10.1007/s13225-020-00466-2))</td></tr>
<tr><td>Otu42406</td><td>1</td><td>k__Fungi(100);p__Ascomycota(100);c__Saccharomycetes(100);o__Saccharomycetales(100);f__Metschnikowiaceae(100);g__Metschnikowia(100);g__Metschnikowia_unclassified(100);</td><td>Metschnikowia</td><td>13</td><td>Saprotroph</td><td>|Undefined Saprotroph|</td><td>Yeast</td><td>NULL</td><td>Probable</td><td>As nectar-tap_saprotroph (Põlme et al. 2020); Decay sugar-rich substrates (Põlme et al. 2020)</td><td>Põlme S, et al. 2020. Fungal Diversity 105:1-16 ((https://doi.org/10.1007/s13225-020-00466-2))</td></tr>
<tr><td>Otu42407</td><td>1</td><td>k__Fungi(100);p__Ascomycota(100);c__Saccharomycetes(100);o__Saccharomycetales(100);f__Metschnikowiaceae(100);g__Metschnikowia(100);s__Metschnikowia_pulcherrima(100);</td><td>Metschnikowia</td><td>13</td><td>Saprotroph</td><td>|Undefined Saprotroph|</td><td>Yeast</td><td>NULL</td><td>Probable</td><td>As nectar-tap_saprotroph (Põlme et al. 2020); Decay sugar-rich substrates (Põlme et al. 2020)</td><td>Põlme S, et al. 2020. Fungal Diversity 105:1-16 ((https://doi.org/10.1007/s13225-020-00466-2))</td></tr>
<tr><td>Otu42410</td><td>1</td><td>k__Fungi(100);p__Ascomycota(100);c__Saccharomycetes(100);o__Saccharomycetales(100);f__Metschnikowiaceae(100);g__Metschnikowia(100);g__Metschnikowia_unclassified(100);</td><td>Metschnikowia</td><td>13</td><td>Saprotroph</td><td>|Undefined Saprotroph|</td><td>Yeast</td><td>NULL</td><td>Probable</td><td>As nectar-tap_saprotroph (Põlme et al. 2020); Decay sugar-rich substrates (Põlme et al. 2020)</td><td>Põlme S, et al. 2020. Fungal Diversity 105:1-16 ((https://doi.org/10.1007/s13225-020-00466-2))</td></tr>
<tr><td>Otu42411</td><td>1</td><td>k__Fungi(100);p__Ascomycota(100);c__Saccharomycetes(100);o__Saccharomycetales(100);f__Metschnikowiaceae(100);g__Metschnikowia(100);s__Metschnikowia_pulcherrima(100);</td><td>Metschnikowia</td><td>13</td><td>Saprotroph</td><td>|Undefined Saprotroph|</td><td>Yeast</td><td>NULL</td><td>Probable</td><td>As nectar-tap_saprotroph (Põlme et al. 2020); Decay sugar-rich substrates (Põlme et al. 2020)</td><td>Põlme S, et al. 2020. Fungal Diversity 105:1-16 ((https://doi.org/10.1007/s13225-020-00466-2))</td></tr>
<tr><td>Otu42412</td><td>1</td><td>k__Fungi(100);p__Ascomycota(100);c__Saccharomycetes(100);o__Saccharomycetales(100);f__Metschnikowiaceae(100);g__Metschnikowia(100);g__Metschnikowia_unclassified(100);</td><td>Metschnikowia</td><td>13</td><td>Saprotroph</td><td>|Undefined Saprotroph|</td><td>Yeast</td><td>NULL</td><td>Probable</td><td>As nectar-tap_saprotroph (Põlme et al. 2020); Decay sugar-rich substrates (Põlme et al. 2020)</td><td>Põlme S, et al. 2020. Fungal Diversity 105:1-16 ((https://doi.org/10.1007/s13225-020-00466-2))</td></tr>
<tr><td>Otu42413</td><td>1</td><td>k__Fungi(100);p__Ascomycota(100);c__Saccharomycetes(100);o__Saccharomycetales(100);f__Metschnikowiaceae(100);g__Metschnikowia(100);s__Metschnikowia_pulcherrima(100);</td><td>Metschnikowia</td><td>13</td><td>Saprotroph</td><td>|Undefined Saprotroph|</td><td>Yeast</td><td>NULL</td><td>Probable</td><td>As nectar-tap_saprotroph (Põlme et al. 2020); Decay sugar-rich substrates (Põlme et al. 2020)</td><td>Põlme S, et al. 2020. Fungal Diversity 105:1-16 ((https://doi.org/10.1007/s13225-020-00466-2))</td></tr>
<tr><td>Otu42414</td><td>1</td><td>k__Fungi(100);p__Ascomycota(100);c__Saccharomycetes(100);o__Saccharomycetales(100);f__Metschnikowiaceae(100);g__Metschnikowia(100);g__Metschnikowia_unclassified(100);</td><td>Metschnikowia</td><td>13</td><td>Saprotroph</td><td>|Undefined Saprotroph|</td><td>Yeast</td><td>NULL</td><td>Probable</td><td>As nectar-tap_saprotroph (Põlme et al. 2020); Decay sugar-rich substrates (Põlme et al. 2020)</td><td>Põlme S, et al. 2020. Fungal Diversity 105:1-16 ((https://doi.org/10.1007/s13225-020-00466-2))</td></tr>
<tr><td>Otu42415</td><td>1</td><td>k__Fungi(100);p__Ascomycota(100);c__Saccharomycetes(100);o__Saccharomycetales(100);f__Metschnikowiaceae(100);g__Metschnikowia(100);s__Metschnikowia_pulcherrima(100);</td><td>Metschnikowia</td><td>13</td><td>Saprotroph</td><td>|Undefined Saprotroph|</td><td>Yeast</td><td>NULL</td><td>Probable</td><td>As nectar-tap_saprotroph (Põlme et al. 2020); Decay sugar-rich substrates (Põlme et al. 2020)</td><td>Põlme S, et al. 2020. Fungal Diversity 105:1-16 ((https://doi.org/10.1007/s13225-020-00466-2))</td></tr>
<tr><td>Otu42418</td><td>1</td><td>k__Fungi(100);p__Ascomycota(100);c__Saccharomycetes(100);o__Saccharomycetales(100);f__Metschnikowiaceae(100);g__Metschnikowia(100);s__Metschnikowia_pulcherrima(100);</td><td>Metschnikowia</td><td>13</td><td>Saprotroph</td><td>|Undefined Saprotroph|</td><td>Yeast</td><td>NULL</td><td>Probable</td><td>As nectar-tap_saprotroph (Põlme et al. 2020); Decay sugar-rich substrates (Põlme et al. 2020)</td><td>Põlme S, et al. 2020. Fungal Diversity 105:1-16 ((https://doi.org/10.1007/s13225-020-00466-2))</td></tr>
<tr><td>Otu42425</td><td>1</td><td>k__Fungi(100);p__Ascomycota(100);c__Saccharomycetes(100);o__Saccharomycetales(100);f__Metschnikowiaceae(100);g__Metschnikowia(100);g__Metschnikowia_unclassified(100);</td><td>Metschnikowia</td><td>13</td><td>Saprotroph</td><td>|Undefined Saprotroph|</td><td>Yeast</td><td>NULL</td><td>Probable</td><td>As nectar-tap_saprotroph (Põlme et al. 2020); Decay sugar-rich substrates (Põlme et al. 2020)</td><td>Põlme S, et al. 2020. Fungal Diversity 105:1-16 ((https://doi.org/10.1007/s13225-020-00466-2))</td></tr>
<tr><td>Otu42426</td><td>1</td><td>k__Fungi(100);p__Ascomycota(100);c__Saccharomycetes(100);o__Saccharomycetales(100);f__Metschnikowiaceae(100);g__Metschnikowia(100);s__Metschnikowia_pulcherrima(100);</td><td>Metschnikowia</td><td>13</td><td>Saprotroph</td><td>|Undefined Saprotroph|</td><td>Yeast</td><td>NULL</td><td>Probable</td><td>As nectar-tap_saprotroph (Põlme et al. 2020); Decay sugar-rich substrates (Põlme et al. 2020)</td><td>Põlme S, et al. 2020. Fungal Diversity 105:1-16 ((https://doi.org/10.1007/s13225-020-00466-2))</td></tr>
<tr><td>Otu42427</td><td>1</td><td>k__Fungi(100);p__Ascomycota(100);c__Saccharomycetes(100);o__Saccharomycetales(100);f__Metschnikowiaceae(100);g__Metschnikowia(100);s__Metschnikowia_pulcherrima(100);</td><td>Metschnikowia</td><td>13</td><td>Saprotroph</td><td>|Undefined Saprotroph|</td><td>Yeast</td><td>NULL</td><td>Probable</td><td>As nectar-tap_saprotroph (Põlme et al. 2020); Decay sugar-rich substrates (Põlme et al. 2020)</td><td>Põlme S, et al. 2020. Fungal Diversity 105:1-16 ((https://doi.org/10.1007/s13225-020-00466-2))</td></tr>
<tr><td>Otu42428</td><td>1</td><td>k__Fungi(100);p__Ascomycota(100);c__Saccharomycetes(100);o__Saccharomycetales(100);f__Metschnikowiaceae(100);g__Metschnikowia(100);g__Metschnikowia_unclassified(100);</td><td>Metschnikowia</td><td>13</td><td>Saprotroph</td><td>|Undefined Saprotroph|</td><td>Yeast</td><td>NULL</td><td>Probable</td><td>As nectar-tap_saprotroph (Põlme et al. 2020); Decay sugar-rich substrates (Põlme et al. 2020)</td><td>Põlme S, et al. 2020. Fungal Diversity 105:1-16 ((https://doi.org/10.1007/s13225-020-00466-2))</td></tr>
<tr><td>Otu42429</td><td>1</td><td>k__Fungi(100);p__Ascomycota(100);c__Saccharomycetes(100);o__Saccharomycetales(100);f__Metschnikowiaceae(100);g__Metschnikowia(100);g__Metschnikowia_unclassified(100);</td><td>Metschnikowia</td><td>13</td><td>Saprotroph</td><td>|Undefined Saprotroph|</td><td>Yeast</td><td>NULL</td><td>Probable</td><td>As nectar-tap_saprotroph (Põlme et al. 2020); Decay sugar-rich substrates (Põlme et al. 2020)</td><td>Põlme S, et al. 2020. Fungal Diversity 105:1-16 ((https://doi.org/10.1007/s13225-020-00466-2))</td></tr>
<tr><td>Otu42430</td><td>1</td><td>k__Fungi(100);p__Ascomycota(100);c__Saccharomycetes(100);o__Saccharomycetales(100);f__Metschnikowiaceae(100);g__Metschnikowia(100);s__Metschnikowia_pulcherrima(100);</td><td>Metschnikowia</td><td>13</td><td>Saprotroph</td><td>|Undefined Saprotroph|</td><td>Yeast</td><td>NULL</td><td>Probable</td><td>As nectar-tap_saprotroph (Põlme et al. 2020); Decay sugar-rich substrates (Põlme et al. 2020)</td><td>Põlme S, et al. 2020. Fungal Diversity 105:1-16 ((https://doi.org/10.1007/s13225-020-00466-2))</td></tr>
<tr><td>Otu42431</td><td>1</td><td>k__Fungi(100);p__Ascomycota(100);c__Saccharomycetes(100);o__Saccharomycetales(100);f__Metschnikowiaceae(100);g__Metschnikowia(100);s__Metschnikowia_pulcherrima(100);</td><td>Metschnikowia</td><td>13</td><td>Saprotroph</td><td>|Undefined Saprotroph|</td><td>Yeast</td><td>NULL</td><td>Probable</td><td>As nectar-tap_saprotroph (Põlme et al. 2020); Decay sugar-rich substrates (Põlme et al. 2020)</td><td>Põlme S, et al. 2020. Fungal Diversity 105:1-16 ((https://doi.org/10.1007/s13225-020-00466-2))</td></tr>
<tr><td>Otu42432</td><td>1</td><td>k__Fungi(100);p__Ascomycota(100);c__Saccharomycetes(100);o__Saccharomycetales(100);f__Metschnikowiaceae(100);g__Metschnikowia(100);g__Metschnikowia_unclassified(100);</td><td>Metschnikowia</td><td>13</td><td>Saprotroph</td><td>|Undefined Saprotroph|</td><td>Yeast</td><td>NULL</td><td>Probable</td><td>As nectar-tap_saprotroph (Põlme et al. 2020); Decay sugar-rich substrates (Põlme et al. 2020)</td><td>Põlme S, et al. 2020. Fungal Diversity 105:1-16 ((https://doi.org/10.1007/s13225-020-00466-2))</td></tr>
<tr><td>Otu42433</td><td>1</td><td>k__Fungi(100);p__Ascomycota(100);c__Saccharomycetes(100);o__Saccharomycetales(100);f__Metschnikowiaceae(100);g__Metschnikowia(100);g__Metschnikowia_unclassified(100);</td><td>Metschnikowia</td><td>13</td><td>Saprotroph</td><td>|Undefined Saprotroph|</td><td>Yeast</td><td>NULL</td><td>Probable</td><td>As nectar-tap_saprotroph (Põlme et al. 2020); Decay sugar-rich substrates (Põlme et al. 2020)</td><td>Põlme S, et al. 2020. Fungal Diversity 105:1-16 ((https://doi.org/10.1007/s13225-020-00466-2))</td></tr>
<tr><td>Otu42434</td><td>1</td><td>k__Fungi(100);p__Ascomycota(100);c__Saccharomycetes(100);o__Saccharomycetales(100);f__Metschnikowiaceae(100);g__Metschnikowia(100);s__Metschnikowia_pulcherrima(100);</td><td>Metschnikowia</td><td>13</td><td>Saprotroph</td><td>|Undefined Saprotroph|</td><td>Yeast</td><td>NULL</td><td>Probable</td><td>As nectar-tap_saprotroph (Põlme et al. 2020); Decay sugar-rich substrates (Põlme et al. 2020)</td><td>Põlme S, et al. 2020. Fungal Diversity 105:1-16 ((https://doi.org/10.1007/s13225-020-00466-2))</td></tr>
<tr><td>Otu42436</td><td>1</td><td>k__Fungi(100);p__Ascomycota(100);c__Saccharomycetes(100);o__Saccharomycetales(100);f__Metschnikowiaceae(100);g__Metschnikowia(100);s__Metschnikowia_pulcherrima(100);</td><td>Metschnikowia</td><td>13</td><td>Saprotroph</td><td>|Undefined Saprotroph|</td><td>Yeast</td><td>NULL</td><td>Probable</td><td>As nectar-tap_saprotroph (Põlme et al. 2020); Decay sugar-rich substrates (Põlme et al. 2020)</td><td>Põlme S, et al. 2020. Fungal Diversity 105:1-16 ((https://doi.org/10.1007/s13225-020-00466-2))</td></tr>
<tr><td>Otu42439</td><td>1</td><td>k__Fungi(100);p__Ascomycota(100);c__Saccharomycetes(100);o__Saccharomycetales(100);f__Metschnikowiaceae(100);g__Metschnikowia(100);g__Metschnikowia_unclassified(100);</td><td>Metschnikowia</td><td>13</td><td>Saprotroph</td><td>|Undefined Saprotroph|</td><td>Yeast</td><td>NULL</td><td>Probable</td><td>As nectar-tap_saprotroph (Põlme et al. 2020); Decay sugar-rich substrates (Põlme et al. 2020)</td><td>Põlme S, et al. 2020. Fungal Diversity 105:1-16 ((https://doi.org/10.1007/s13225-020-00466-2))</td></tr>
<tr><td>Otu42440</td><td>1</td><td>k__Fungi(100);p__Ascomycota(100);c__Saccharomycetes(100);o__Saccharomycetales(100);f__Metschnikowiaceae(100);g__Metschnikowia(100);g__Metschnikowia_unclassified(100);</td><td>Metschnikowia</td><td>13</td><td>Saprotroph</td><td>|Undefined Saprotroph|</td><td>Yeast</td><td>NULL</td><td>Probable</td><td>As nectar-tap_saprotroph (Põlme et al. 2020); Decay sugar-rich substrates (Põlme et al. 2020)</td><td>Põlme S, et al. 2020. Fungal Diversity 105:1-16 ((https://doi.org/10.1007/s13225-020-00466-2))</td></tr>
<tr><td>Otu42441</td><td>1</td><td>k__Fungi(100);p__Ascomycota(100);c__Saccharomycetes(100);o__Saccharomycetales(100);f__Metschnikowiaceae(100);g__Metschnikowia(100);g__Metschnikowia_unclassified(100);</td><td>Metschnikowia</td><td>13</td><td>Saprotroph</td><td>|Undefined Saprotroph|</td><td>Yeast</td><td>NULL</td><td>Probable</td><td>As nectar-tap_saprotroph (Põlme et al. 2020); Decay sugar-rich substrates (Põlme et al. 2020)</td><td>Põlme S, et al. 2020. Fungal Diversity 105:1-16 ((https://doi.org/10.1007/s13225-020-00466-2))</td></tr>
<tr><td>Otu42442</td><td>1</td><td>k__Fungi(100);p__Ascomycota(100);c__Saccharomycetes(100);o__Saccharomycetales(100);f__Metschnikowiaceae(100);g__Metschnikowia(100);g__Metschnikowia_unclassified(100);</td><td>Metschnikowia</td><td>13</td><td>Saprotroph</td><td>|Undefined Saprotroph|</td><td>Yeast</td><td>NULL</td><td>Probable</td><td>As nectar-tap_saprotroph (Põlme et al. 2020); Decay sugar-rich substrates (Põlme et al. 2020)</td><td>Põlme S, et al. 2020. Fungal Diversity 105:1-16 ((https://doi.org/10.1007/s13225-020-00466-2))</td></tr>
<tr><td>Otu42443</td><td>1</td><td>k__Fungi(100);p__Ascomycota(100);c__Saccharomycetes(100);o__Saccharomycetales(100);f__Metschnikowiaceae(100);g__Metschnikowia(100);s__Metschnikowia_pulcherrima(100);</td><td>Metschnikowia</td><td>13</td><td>Saprotroph</td><td>|Undefined Saprotroph|</td><td>Yeast</td><td>NULL</td><td>Probable</td><td>As nectar-tap_saprotroph (Põlme et al. 2020); Decay sugar-rich substrates (Põlme et al. 2020)</td><td>Põlme S, et al. 2020. Fungal Diversity 105:1-16 ((https://doi.org/10.1007/s13225-020-00466-2))</td></tr>
<tr><td>Otu42444</td><td>1</td><td>k__Fungi(100);p__Ascomycota(100);c__Saccharomycetes(100);o__Saccharomycetales(100);f__Metschnikowiaceae(100);g__Metschnikowia(100);s__Metschnikowia_pulcherrima(100);</td><td>Metschnikowia</td><td>13</td><td>Saprotroph</td><td>|Undefined Saprotroph|</td><td>Yeast</td><td>NULL</td><td>Probable</td><td>As nectar-tap_saprotroph (Põlme et al. 2020); Decay sugar-rich substrates (Põlme et al. 2020)</td><td>Põlme S, et al. 2020. Fungal Diversity 105:1-16 ((https://doi.org/10.1007/s13225-020-00466-2))</td></tr>
<tr><td>Otu42445</td><td>1</td><td>k__Fungi(100);p__Ascomycota(100);c__Saccharomycetes(100);o__Saccharomycetales(100);f__Metschnikowiaceae(100);g__Metschnikowia(100);g__Metschnikowia_unclassified(100);</td><td>Metschnikowia</td><td>13</td><td>Saprotroph</td><td>|Undefined Saprotroph|</td><td>Yeast</td><td>NULL</td><td>Probable</td><td>As nectar-tap_saprotroph (Põlme et al. 2020); Decay sugar-rich substrates (Põlme et al. 2020)</td><td>Põlme S, et al. 2020. Fungal Diversity 105:1-16 ((https://doi.org/10.1007/s13225-020-00466-2))</td></tr>
<tr><td>Otu42446</td><td>1</td><td>k__Fungi(100);p__Ascomycota(100);c__Saccharomycetes(100);o__Saccharomycetales(100);f__Metschnikowiaceae(100);g__Metschnikowia(100);g__Metschnikowia_unclassified(100);</td><td>Metschnikowia</td><td>13</td><td>Saprotroph</td><td>|Undefined Saprotroph|</td><td>Yeast</td><td>NULL</td><td>Probable</td><td>As nectar-tap_saprotroph (Põlme et al. 2020); Decay sugar-rich substrates (Põlme et al. 2020)</td><td>Põlme S, et al. 2020. Fungal Diversity 105:1-16 ((https://doi.org/10.1007/s13225-020-00466-2))</td></tr>
<tr><td>Otu42447</td><td>1</td><td>k__Fungi(100);p__Ascomycota(100);c__Saccharomycetes(100);o__Saccharomycetales(100);f__Metschnikowiaceae(100);g__Metschnikowia(100);s__Metschnikowia_pulcherrima(100);</td><td>Metschnikowia</td><td>13</td><td>Saprotroph</td><td>|Undefined Saprotroph|</td><td>Yeast</td><td>NULL</td><td>Probable</td><td>As nectar-tap_saprotroph (Põlme et al. 2020); Decay sugar-rich substrates (Põlme et al. 2020)</td><td>Põlme S, et al. 2020. Fungal Diversity 105:1-16 ((https://doi.org/10.1007/s13225-020-00466-2))</td></tr>
<tr><td>Otu42448</td><td>1</td><td>k__Fungi(100);p__Ascomycota(100);c__Saccharomycetes(100);o__Saccharomycetales(100);f__Metschnikowiaceae(100);g__Metschnikowia(100);s__Metschnikowia_pulcherrima(100);</td><td>Metschnikowia</td><td>13</td><td>Saprotroph</td><td>|Undefined Saprotroph|</td><td>Yeast</td><td>NULL</td><td>Probable</td><td>As nectar-tap_saprotroph (Põlme et al. 2020); Decay sugar-rich substrates (Põlme et al. 2020)</td><td>Põlme S, et al. 2020. Fungal Diversity 105:1-16 ((https://doi.org/10.1007/s13225-020-00466-2))</td></tr>
<tr><td>Otu42449</td><td>1</td><td>k__Fungi(100);p__Ascomycota(100);c__Saccharomycetes(100);o__Saccharomycetales(100);f__Metschnikowiaceae(100);g__Metschnikowia(100);s__Metschnikowia_pulcherrima(100);</td><td>Metschnikowia</td><td>13</td><td>Saprotroph</td><td>|Undefined Saprotroph|</td><td>Yeast</td><td>NULL</td><td>Probable</td><td>As nectar-tap_saprotroph (Põlme et al. 2020); Decay sugar-rich substrates (Põlme et al. 2020)</td><td>Põlme S, et al. 2020. Fungal Diversity 105:1-16 ((https://doi.org/10.1007/s13225-020-00466-2))</td></tr>
<tr><td>Otu42450</td><td>1</td><td>k__Fungi(100);p__Ascomycota(100);c__Saccharomycetes(100);o__Saccharomycetales(100);f__Metschnikowiaceae(100);g__Metschnikowia(100);s__Metschnikowia_pulcherrima(100);</td><td>Metschnikowia</td><td>13</td><td>Saprotroph</td><td>|Undefined Saprotroph|</td><td>Yeast</td><td>NULL</td><td>Probable</td><td>As nectar-tap_saprotroph (Põlme et al. 2020); Decay sugar-rich substrates (Põlme et al. 2020)</td><td>Põlme S, et al. 2020. Fungal Diversity 105:1-16 ((https://doi.org/10.1007/s13225-020-00466-2))</td></tr>
<tr><td>Otu42451</td><td>1</td><td>k__Fungi(100);p__Ascomycota(100);c__Saccharomycetes(100);o__Saccharomycetales(100);f__Metschnikowiaceae(100);g__Metschnikowia(100);s__Metschnikowia_pulcherrima(100);</td><td>Metschnikowia</td><td>13</td><td>Saprotroph</td><td>|Undefined Saprotroph|</td><td>Yeast</td><td>NULL</td><td>Probable</td><td>As nectar-tap_saprotroph (Põlme et al. 2020); Decay sugar-rich substrates (Põlme et al. 2020)</td><td>Põlme S, et al. 2020. Fungal Diversity 105:1-16 ((https://doi.org/10.1007/s13225-020-00466-2))</td></tr>
<tr><td>Otu42452</td><td>1</td><td>k__Fungi(100);p__Ascomycota(100);c__Saccharomycetes(100);o__Saccharomycetales(100);f__Metschnikowiaceae(100);g__Metschnikowia(100);s__Metschnikowia_pulcherrima(100);</td><td>Metschnikowia</td><td>13</td><td>Saprotroph</td><td>|Undefined Saprotroph|</td><td>Yeast</td><td>NULL</td><td>Probable</td><td>As nectar-tap_saprotroph (Põlme et al. 2020); Decay sugar-rich substrates (Põlme et al. 2020)</td><td>Põlme S, et al. 2020. Fungal Diversity 105:1-16 ((https://doi.org/10.1007/s13225-020-00466-2))</td></tr>
<tr><td>Otu42453</td><td>1</td><td>k__Fungi(100);p__Ascomycota(100);c__Saccharomycetes(100);o__Saccharomycetales(100);f__Metschnikowiaceae(100);g__Metschnikowia(100);g__Metschnikowia_unclassified(100);</td><td>Metschnikowia</td><td>13</td><td>Saprotroph</td><td>|Undefined Saprotroph|</td><td>Yeast</td><td>NULL</td><td>Probable</td><td>As nectar-tap_saprotroph (Põlme et al. 2020); Decay sugar-rich substrates (Põlme et al. 2020)</td><td>Põlme S, et al. 2020. Fungal Diversity 105:1-16 ((https://doi.org/10.1007/s13225-020-00466-2))</td></tr>
<tr><td>Otu42454</td><td>1</td><td>k__Fungi(100);p__Ascomycota(100);c__Saccharomycetes(100);o__Saccharomycetales(100);f__Metschnikowiaceae(100);g__Metschnikowia(100);g__Metschnikowia_unclassified(100);</td><td>Metschnikowia</td><td>13</td><td>Saprotroph</td><td>|Undefined Saprotroph|</td><td>Yeast</td><td>NULL</td><td>Probable</td><td>As nectar-tap_saprotroph (Põlme et al. 2020); Decay sugar-rich substrates (Põlme et al. 2020)</td><td>Põlme S, et al. 2020. Fungal Diversity 105:1-16 ((https://doi.org/10.1007/s13225-020-00466-2))</td></tr>
<tr><td>Otu42455</td><td>1</td><td>k__Fungi(100);p__Ascomycota(100);c__Saccharomycetes(100);o__Saccharomycetales(100);f__Metschnikowiaceae(100);g__Metschnikowia(100);s__Metschnikowia_pulcherrima(100);</td><td>Metschnikowia</td><td>13</td><td>Saprotroph</td><td>|Undefined Saprotroph|</td><td>Yeast</td><td>NULL</td><td>Probable</td><td>As nectar-tap_saprotroph (Põlme et al. 2020); Decay sugar-rich substrates (Põlme et al. 2020)</td><td>Põlme S, et al. 2020. Fungal Diversity 105:1-16 ((https://doi.org/10.1007/s13225-020-00466-2))</td></tr>
<tr><td>Otu42456</td><td>1</td><td>k__Fungi(100);p__Ascomycota(100);c__Saccharomycetes(100);o__Saccharomycetales(100);f__Metschnikowiaceae(100);g__Metschnikowia(100);g__Metschnikowia_unclassified(100);</td><td>Metschnikowia</td><td>13</td><td>Saprotroph</td><td>|Undefined Saprotroph|</td><td>Yeast</td><td>NULL</td><td>Probable</td><td>As nectar-tap_saprotroph (Põlme et al. 2020); Decay sugar-rich substrates (Põlme et al. 2020)</td><td>Põlme S, et al. 2020. Fungal Diversity 105:1-16 ((https://doi.org/10.1007/s13225-020-00466-2))</td></tr>
<tr><td>Otu42457</td><td>1</td><td>k__Fungi(100);p__Ascomycota(100);c__Saccharomycetes(100);o__Saccharomycetales(100);f__Metschnikowiaceae(100);g__Metschnikowia(100);g__Metschnikowia_unclassified(100);</td><td>Metschnikowia</td><td>13</td><td>Saprotroph</td><td>|Undefined Saprotroph|</td><td>Yeast</td><td>NULL</td><td>Probable</td><td>As nectar-tap_saprotroph (Põlme et al. 2020); Decay sugar-rich substrates (Põlme et al. 2020)</td><td>Põlme S, et al. 2020. Fungal Diversity 105:1-16 ((https://doi.org/10.1007/s13225-020-00466-2))</td></tr>
<tr><td>Otu42458</td><td>1</td><td>k__Fungi(100);p__Ascomycota(100);c__Saccharomycetes(100);o__Saccharomycetales(100);f__Metschnikowiaceae(100);g__Metschnikowia(100);g__Metschnikowia_unclassified(100);</td><td>Metschnikowia</td><td>13</td><td>Saprotroph</td><td>|Undefined Saprotroph|</td><td>Yeast</td><td>NULL</td><td>Probable</td><td>As nectar-tap_saprotroph (Põlme et al. 2020); Decay sugar-rich substrates (Põlme et al. 2020)</td><td>Põlme S, et al. 2020. Fungal Diversity 105:1-16 ((https://doi.org/10.1007/s13225-020-00466-2))</td></tr>
<tr><td>Otu42459</td><td>1</td><td>k__Fungi(100);p__Ascomycota(100);c__Saccharomycetes(100);o__Saccharomycetales(100);f__Metschnikowiaceae(100);g__Metschnikowia(100);s__Metschnikowia_pulcherrima(100);</td><td>Metschnikowia</td><td>13</td><td>Saprotroph</td><td>|Undefined Saprotroph|</td><td>Yeast</td><td>NULL</td><td>Probable</td><td>As nectar-tap_saprotroph (Põlme et al. 2020); Decay sugar-rich substrates (Põlme et al. 2020)</td><td>Põlme S, et al. 2020. Fungal Diversity 105:1-16 ((https://doi.org/10.1007/s13225-020-00466-2))</td></tr>
<tr><td>Otu42460</td><td>1</td><td>k__Fungi(100);p__Ascomycota(100);c__Saccharomycetes(100);o__Saccharomycetales(100);f__Metschnikowiaceae(100);g__Metschnikowia(100);s__Metschnikowia_pulcherrima(100);</td><td>Metschnikowia</td><td>13</td><td>Saprotroph</td><td>|Undefined Saprotroph|</td><td>Yeast</td><td>NULL</td><td>Probable</td><td>As nectar-tap_saprotroph (Põlme et al. 2020); Decay sugar-rich substrates (Põlme et al. 2020)</td><td>Põlme S, et al. 2020. Fungal Diversity 105:1-16 ((https://doi.org/10.1007/s13225-020-00466-2))</td></tr>
<tr><td>Otu42461</td><td>1</td><td>k__Fungi(100);p__Ascomycota(100);c__Saccharomycetes(100);o__Saccharomycetales(100);f__Metschnikowiaceae(100);g__Metschnikowia(100);s__Metschnikowia_pulcherrima(100);</td><td>Metschnikowia</td><td>13</td><td>Saprotroph</td><td>|Undefined Saprotroph|</td><td>Yeast</td><td>NULL</td><td>Probable</td><td>As nectar-tap_saprotroph (Põlme et al. 2020); Decay sugar-rich substrates (Põlme et al. 2020)</td><td>Põlme S, et al. 2020. Fungal Diversity 105:1-16 ((https://doi.org/10.1007/s13225-020-00466-2))</td></tr>
<tr><td>Otu42462</td><td>1</td><td>k__Fungi(100);p__Ascomycota(100);c__Saccharomycetes(100);o__Saccharomycetales(100);f__Metschnikowiaceae(100);g__Metschnikowia(100);g__Metschnikowia_unclassified(100);</td><td>Metschnikowia</td><td>13</td><td>Saprotroph</td><td>|Undefined Saprotroph|</td><td>Yeast</td><td>NULL</td><td>Probable</td><td>As nectar-tap_saprotroph (Põlme et al. 2020); Decay sugar-rich substrates (Põlme et al. 2020)</td><td>Põlme S, et al. 2020. Fungal Diversity 105:1-16 ((https://doi.org/10.1007/s13225-020-00466-2))</td></tr>
<tr><td>Otu42463</td><td>1</td><td>k__Fungi(100);p__Ascomycota(100);c__Saccharomycetes(100);o__Saccharomycetales(100);f__Metschnikowiaceae(100);g__Metschnikowia(100);s__Metschnikowia_pulcherrima(100);</td><td>Metschnikowia</td><td>13</td><td>Saprotroph</td><td>|Undefined Saprotroph|</td><td>Yeast</td><td>NULL</td><td>Probable</td><td>As nectar-tap_saprotroph (Põlme et al. 2020); Decay sugar-rich substrates (Põlme et al. 2020)</td><td>Põlme S, et al. 2020. Fungal Diversity 105:1-16 ((https://doi.org/10.1007/s13225-020-00466-2))</td></tr>
<tr><td>Otu42464</td><td>1</td><td>k__Fungi(100);p__Ascomycota(100);c__Saccharomycetes(100);o__Saccharomycetales(100);f__Metschnikowiaceae(100);g__Metschnikowia(100);s__Metschnikowia_pulcherrima(100);</td><td>Metschnikowia</td><td>13</td><td>Saprotroph</td><td>|Undefined Saprotroph|</td><td>Yeast</td><td>NULL</td><td>Probable</td><td>As nectar-tap_saprotroph (Põlme et al. 2020); Decay sugar-rich substrates (Põlme et al. 2020)</td><td>Põlme S, et al. 2020. Fungal Diversity 105:1-16 ((https://doi.org/10.1007/s13225-020-00466-2))</td></tr>
<tr><td>Otu42465</td><td>1</td><td>k__Fungi(100);p__Ascomycota(100);c__Saccharomycetes(100);o__Saccharomycetales(100);f__Metschnikowiaceae(100);g__Metschnikowia(100);s__Metschnikowia_pulcherrima(100);</td><td>Metschnikowia</td><td>13</td><td>Saprotroph</td><td>|Undefined Saprotroph|</td><td>Yeast</td><td>NULL</td><td>Probable</td><td>As nectar-tap_saprotroph (Põlme et al. 2020); Decay sugar-rich substrates (Põlme et al. 2020)</td><td>Põlme S, et al. 2020. Fungal Diversity 105:1-16 ((https://doi.org/10.1007/s13225-020-00466-2))</td></tr>
<tr><td>Otu42466</td><td>1</td><td>k__Fungi(100);p__Ascomycota(100);c__Saccharomycetes(100);o__Saccharomycetales(100);f__Metschnikowiaceae(100);g__Metschnikowia(100);s__Metschnikowia_pulcherrima(100);</td><td>Metschnikowia</td><td>13</td><td>Saprotroph</td><td>|Undefined Saprotroph|</td><td>Yeast</td><td>NULL</td><td>Probable</td><td>As nectar-tap_saprotroph (Põlme et al. 2020); Decay sugar-rich substrates (Põlme et al. 2020)</td><td>Põlme S, et al. 2020. Fungal Diversity 105:1-16 ((https://doi.org/10.1007/s13225-020-00466-2))</td></tr>
<tr><td>Otu42467</td><td>1</td><td>k__Fungi(100);p__Ascomycota(100);c__Saccharomycetes(100);o__Saccharomycetales(100);f__Metschnikowiaceae(100);g__Metschnikowia(100);s__Metschnikowia_pulcherrima(100);</td><td>Metschnikowia</td><td>13</td><td>Saprotroph</td><td>|Undefined Saprotroph|</td><td>Yeast</td><td>NULL</td><td>Probable</td><td>As nectar-tap_saprotroph (Põlme et al. 2020); Decay sugar-rich substrates (Põlme et al. 2020)</td><td>Põlme S, et al. 2020. Fungal Diversity 105:1-16 ((https://doi.org/10.1007/s13225-020-00466-2))</td></tr>
<tr><td>Otu42468</td><td>1</td><td>k__Fungi(100);p__Ascomycota(100);c__Saccharomycetes(100);o__Saccharomycetales(100);f__Metschnikowiaceae(100);g__Metschnikowia(100);s__Metschnikowia_pulcherrima(100);</td><td>Metschnikowia</td><td>13</td><td>Saprotroph</td><td>|Undefined Saprotroph|</td><td>Yeast</td><td>NULL</td><td>Probable</td><td>As nectar-tap_saprotroph (Põlme et al. 2020); Decay sugar-rich substrates (Põlme et al. 2020)</td><td>Põlme S, et al. 2020. Fungal Diversity 105:1-16 ((https://doi.org/10.1007/s13225-020-00466-2))</td></tr>
<tr><td>Otu42469</td><td>1</td><td>k__Fungi(100);p__Ascomycota(100);c__Saccharomycetes(100);o__Saccharomycetales(100);f__Metschnikowiaceae(100);g__Metschnikowia(100);s__Metschnikowia_pulcherrima(100);</td><td>Metschnikowia</td><td>13</td><td>Saprotroph</td><td>|Undefined Saprotroph|</td><td>Yeast</td><td>NULL</td><td>Probable</td><td>As nectar-tap_saprotroph (Põlme et al. 2020); Decay sugar-rich substrates (Põlme et al. 2020)</td><td>Põlme S, et al. 2020. Fungal Diversity 105:1-16 ((https://doi.org/10.1007/s13225-020-00466-2))</td></tr>
<tr><td>Otu42470</td><td>1</td><td>k__Fungi(100);p__Ascomycota(100);c__Saccharomycetes(100);o__Saccharomycetales(100);f__Metschnikowiaceae(100);g__Metschnikowia(100);g__Metschnikowia_unclassified(100);</td><td>Metschnikowia</td><td>13</td><td>Saprotroph</td><td>|Undefined Saprotroph|</td><td>Yeast</td><td>NULL</td><td>Probable</td><td>As nectar-tap_saprotroph (Põlme et al. 2020); Decay sugar-rich substrates (Põlme et al. 2020)</td><td>Põlme S, et al. 2020. Fungal Diversity 105:1-16 ((https://doi.org/10.1007/s13225-020-00466-2))</td></tr>
<tr><td>Otu42471</td><td>1</td><td>k__Fungi(100);p__Ascomycota(100);c__Saccharomycetes(100);o__Saccharomycetales(100);f__Metschnikowiaceae(100);g__Metschnikowia(100);s__Metschnikowia_pulcherrima(100);</td><td>Metschnikowia</td><td>13</td><td>Saprotroph</td><td>|Undefined Saprotroph|</td><td>Yeast</td><td>NULL</td><td>Probable</td><td>As nectar-tap_saprotroph (Põlme et al. 2020); Decay sugar-rich substrates (Põlme et al. 2020)</td><td>Põlme S, et al. 2020. Fungal Diversity 105:1-16 ((https://doi.org/10.1007/s13225-020-00466-2))</td></tr>
<tr><td>Otu42472</td><td>1</td><td>k__Fungi(100);p__Ascomycota(100);c__Saccharomycetes(100);o__Saccharomycetales(100);f__Metschnikowiaceae(100);g__Metschnikowia(100);g__Metschnikowia_unclassified(100);</td><td>Metschnikowia</td><td>13</td><td>Saprotroph</td><td>|Undefined Saprotroph|</td><td>Yeast</td><td>NULL</td><td>Probable</td><td>As nectar-tap_saprotroph (Põlme et al. 2020); Decay sugar-rich substrates (Põlme et al. 2020)</td><td>Põlme S, et al. 2020. Fungal Diversity 105:1-16 ((https://doi.org/10.1007/s13225-020-00466-2))</td></tr>
<tr><td>Otu42473</td><td>1</td><td>k__Fungi(100);p__Ascomycota(100);c__Saccharomycetes(100);o__Saccharomycetales(100);f__Metschnikowiaceae(100);g__Metschnikowia(100);s__Metschnikowia_pulcherrima(100);</td><td>Metschnikowia</td><td>13</td><td>Saprotroph</td><td>|Undefined Saprotroph|</td><td>Yeast</td><td>NULL</td><td>Probable</td><td>As nectar-tap_saprotroph (Põlme et al. 2020); Decay sugar-rich substrates (Põlme et al. 2020)</td><td>Põlme S, et al. 2020. Fungal Diversity 105:1-16 ((https://doi.org/10.1007/s13225-020-00466-2))</td></tr>
<tr><td>Otu42474</td><td>1</td><td>k__Fungi(100);p__Ascomycota(100);c__Saccharomycetes(100);o__Saccharomycetales(100);f__Metschnikowiaceae(100);g__Metschnikowia(100);s__Metschnikowia_pulcherrima(100);</td><td>Metschnikowia</td><td>13</td><td>Saprotroph</td><td>|Undefined Saprotroph|</td><td>Yeast</td><td>NULL</td><td>Probable</td><td>As nectar-tap_saprotroph (Põlme et al. 2020); Decay sugar-rich substrates (Põlme et al. 2020)</td><td>Põlme S, et al. 2020. Fungal Diversity 105:1-16 ((https://doi.org/10.1007/s13225-020-00466-2))</td></tr>
<tr><td>Otu42475</td><td>1</td><td>k__Fungi(100);p__Ascomycota(100);c__Saccharomycetes(100);o__Saccharomycetales(100);f__Metschnikowiaceae(100);g__Metschnikowia(100);s__Metschnikowia_pulcherrima(100);</td><td>Metschnikowia</td><td>13</td><td>Saprotroph</td><td>|Undefined Saprotroph|</td><td>Yeast</td><td>NULL</td><td>Probable</td><td>As nectar-tap_saprotroph (Põlme et al. 2020); Decay sugar-rich substrates (Põlme et al. 2020)</td><td>Põlme S, et al. 2020. Fungal Diversity 105:1-16 ((https://doi.org/10.1007/s13225-020-00466-2))</td></tr>
<tr><td>Otu42476</td><td>1</td><td>k__Fungi(100);p__Ascomycota(100);c__Saccharomycetes(100);o__Saccharomycetales(100);f__Metschnikowiaceae(100);g__Metschnikowia(100);s__Metschnikowia_pulcherrima(100);</td><td>Metschnikowia</td><td>13</td><td>Saprotroph</td><td>|Undefined Saprotroph|</td><td>Yeast</td><td>NULL</td><td>Probable</td><td>As nectar-tap_saprotroph (Põlme et al. 2020); Decay sugar-rich substrates (Põlme et al. 2020)</td><td>Põlme S, et al. 2020. Fungal Diversity 105:1-16 ((https://doi.org/10.1007/s13225-020-00466-2))</td></tr>
<tr><td>Otu42477</td><td>1</td><td>k__Fungi(100);p__Ascomycota(100);c__Saccharomycetes(100);o__Saccharomycetales(100);f__Metschnikowiaceae(100);g__Metschnikowia(100);s__Metschnikowia_pulcherrima(100);</td><td>Metschnikowia</td><td>13</td><td>Saprotroph</td><td>|Undefined Saprotroph|</td><td>Yeast</td><td>NULL</td><td>Probable</td><td>As nectar-tap_saprotroph (Põlme et al. 2020); Decay sugar-rich substrates (Põlme et al. 2020)</td><td>Põlme S, et al. 2020. Fungal Diversity 105:1-16 ((https://doi.org/10.1007/s13225-020-00466-2))</td></tr>
<tr><td>Otu42478</td><td>1</td><td>k__Fungi(100);p__Ascomycota(100);c__Saccharomycetes(100);o__Saccharomycetales(100);f__Metschnikowiaceae(100);g__Metschnikowia(100);s__Metschnikowia_pulcherrima(100);</td><td>Metschnikowia</td><td>13</td><td>Saprotroph</td><td>|Undefined Saprotroph|</td><td>Yeast</td><td>NULL</td><td>Probable</td><td>As nectar-tap_saprotroph (Põlme et al. 2020); Decay sugar-rich substrates (Põlme et al. 2020)</td><td>Põlme S, et al. 2020. Fungal Diversity 105:1-16 ((https://doi.org/10.1007/s13225-020-00466-2))</td></tr>
<tr><td>Otu42482</td><td>1</td><td>k__Fungi(100);p__Ascomycota(100);c__Saccharomycetes(100);o__Saccharomycetales(100);f__Metschnikowiaceae(100);g__Metschnikowia(100);s__Metschnikowia_pulcherrima(100);</td><td>Metschnikowia</td><td>13</td><td>Saprotroph</td><td>|Undefined Saprotroph|</td><td>Yeast</td><td>NULL</td><td>Probable</td><td>As nectar-tap_saprotroph (Põlme et al. 2020); Decay sugar-rich substrates (Põlme et al. 2020)</td><td>Põlme S, et al. 2020. Fungal Diversity 105:1-16 ((https://doi.org/10.1007/s13225-020-00466-2))</td></tr>
<tr><td>Otu42483</td><td>1</td><td>k__Fungi(100);p__Ascomycota(100);c__Saccharomycetes(100);o__Saccharomycetales(100);f__Metschnikowiaceae(100);g__Metschnikowia(100);s__Metschnikowia_pulcherrima(100);</td><td>Metschnikowia</td><td>13</td><td>Saprotroph</td><td>|Undefined Saprotroph|</td><td>Yeast</td><td>NULL</td><td>Probable</td><td>As nectar-tap_saprotroph (Põlme et al. 2020); Decay sugar-rich substrates (Põlme et al. 2020)</td><td>Põlme S, et al. 2020. Fungal Diversity 105:1-16 ((https://doi.org/10.1007/s13225-020-00466-2))</td></tr>
<tr><td>Otu42487</td><td>1</td><td>k__Fungi(100);p__Ascomycota(100);c__Saccharomycetes(100);o__Saccharomycetales(100);f__Metschnikowiaceae(100);g__Metschnikowia(100);s__Metschnikowia_pulcherrima(100);</td><td>Metschnikowia</td><td>13</td><td>Saprotroph</td><td>|Undefined Saprotroph|</td><td>Yeast</td><td>NULL</td><td>Probable</td><td>As nectar-tap_saprotroph (Põlme et al. 2020); Decay sugar-rich substrates (Põlme et al. 2020)</td><td>Põlme S, et al. 2020. Fungal Diversity 105:1-16 ((https://doi.org/10.1007/s13225-020-00466-2))</td></tr>
<tr><td>Otu42488</td><td>1</td><td>k__Fungi(100);p__Ascomycota(100);c__Saccharomycetes(100);o__Saccharomycetales(100);f__Metschnikowiaceae(100);g__Metschnikowia(100);s__Metschnikowia_pulcherrima(100);</td><td>Metschnikowia</td><td>13</td><td>Saprotroph</td><td>|Undefined Saprotroph|</td><td>Yeast</td><td>NULL</td><td>Probable</td><td>As nectar-tap_saprotroph (Põlme et al. 2020); Decay sugar-rich substrates (Põlme et al. 2020)</td><td>Põlme S, et al. 2020. Fungal Diversity 105:1-16 ((https://doi.org/10.1007/s13225-020-00466-2))</td></tr>
<tr><td>Otu42489</td><td>1</td><td>k__Fungi(100);p__Ascomycota(100);c__Saccharomycetes(100);o__Saccharomycetales(100);f__Metschnikowiaceae(100);g__Metschnikowia(100);s__Metschnikowia_pulcherrima(100);</td><td>Metschnikowia</td><td>13</td><td>Saprotroph</td><td>|Undefined Saprotroph|</td><td>Yeast</td><td>NULL</td><td>Probable</td><td>As nectar-tap_saprotroph (Põlme et al. 2020); Decay sugar-rich substrates (Põlme et al. 2020)</td><td>Põlme S, et al. 2020. Fungal Diversity 105:1-16 ((https://doi.org/10.1007/s13225-020-00466-2))</td></tr>
<tr><td>Otu42490</td><td>1</td><td>k__Fungi(100);p__Ascomycota(100);c__Saccharomycetes(100);o__Saccharomycetales(100);f__Metschnikowiaceae(100);g__Metschnikowia(100);g__Metschnikowia_unclassified(100);</td><td>Metschnikowia</td><td>13</td><td>Saprotroph</td><td>|Undefined Saprotroph|</td><td>Yeast</td><td>NULL</td><td>Probable</td><td>As nectar-tap_saprotroph (Põlme et al. 2020); Decay sugar-rich substrates (Põlme et al. 2020)</td><td>Põlme S, et al. 2020. Fungal Diversity 105:1-16 ((https://doi.org/10.1007/s13225-020-00466-2))</td></tr>
<tr><td>Otu42492</td><td>1</td><td>k__Fungi(100);p__Ascomycota(100);c__Saccharomycetes(100);o__Saccharomycetales(100);f__Metschnikowiaceae(100);g__Metschnikowia(100);s__Metschnikowia_pulcherrima(100);</td><td>Metschnikowia</td><td>13</td><td>Saprotroph</td><td>|Undefined Saprotroph|</td><td>Yeast</td><td>NULL</td><td>Probable</td><td>As nectar-tap_saprotroph (Põlme et al. 2020); Decay sugar-rich substrates (Põlme et al. 2020)</td><td>Põlme S, et al. 2020. Fungal Diversity 105:1-16 ((https://doi.org/10.1007/s13225-020-00466-2))</td></tr>
<tr><td>Otu42493</td><td>1</td><td>k__Fungi(100);p__Ascomycota(100);c__Saccharomycetes(100);o__Saccharomycetales(100);f__Metschnikowiaceae(100);g__Metschnikowia(100);g__Metschnikowia_unclassified(100);</td><td>Metschnikowia</td><td>13</td><td>Saprotroph</td><td>|Undefined Saprotroph|</td><td>Yeast</td><td>NULL</td><td>Probable</td><td>As nectar-tap_saprotroph (Põlme et al. 2020); Decay sugar-rich substrates (Põlme et al. 2020)</td><td>Põlme S, et al. 2020. Fungal Diversity 105:1-16 ((https://doi.org/10.1007/s13225-020-00466-2))</td></tr>
<tr><td>Otu42494</td><td>1</td><td>k__Fungi(100);p__Ascomycota(100);c__Saccharomycetes(100);o__Saccharomycetales(100);f__Metschnikowiaceae(100);g__Metschnikowia(100);s__Metschnikowia_pulcherrima(100);</td><td>Metschnikowia</td><td>13</td><td>Saprotroph</td><td>|Undefined Saprotroph|</td><td>Yeast</td><td>NULL</td><td>Probable</td><td>As nectar-tap_saprotroph (Põlme et al. 2020); Decay sugar-rich substrates (Põlme et al. 2020)</td><td>Põlme S, et al. 2020. Fungal Diversity 105:1-16 ((https://doi.org/10.1007/s13225-020-00466-2))</td></tr>
<tr><td>Otu42495</td><td>1</td><td>k__Fungi(100);p__Ascomycota(100);c__Saccharomycetes(100);o__Saccharomycetales(100);f__Metschnikowiaceae(100);g__Metschnikowia(100);s__Metschnikowia_pulcherrima(100);</td><td>Metschnikowia</td><td>13</td><td>Saprotroph</td><td>|Undefined Saprotroph|</td><td>Yeast</td><td>NULL</td><td>Probable</td><td>As nectar-tap_saprotroph (Põlme et al. 2020); Decay sugar-rich substrates (Põlme et al. 2020)</td><td>Põlme S, et al. 2020. Fungal Diversity 105:1-16 ((https://doi.org/10.1007/s13225-020-00466-2))</td></tr>
<tr><td>Otu42496</td><td>1</td><td>k__Fungi(100);p__Ascomycota(100);c__Saccharomycetes(100);o__Saccharomycetales(100);f__Metschnikowiaceae(100);g__Metschnikowia(100);s__Metschnikowia_pulcherrima(100);</td><td>Metschnikowia</td><td>13</td><td>Saprotroph</td><td>|Undefined Saprotroph|</td><td>Yeast</td><td>NULL</td><td>Probable</td><td>As nectar-tap_saprotroph (Põlme et al. 2020); Decay sugar-rich substrates (Põlme et al. 2020)</td><td>Põlme S, et al. 2020. Fungal Diversity 105:1-16 ((https://doi.org/10.1007/s13225-020-00466-2))</td></tr>
<tr><td>Otu42497</td><td>1</td><td>k__Fungi(100);p__Ascomycota(100);c__Saccharomycetes(100);o__Saccharomycetales(100);f__Metschnikowiaceae(100);g__Metschnikowia(100);s__Metschnikowia_pulcherrima(100);</td><td>Metschnikowia</td><td>13</td><td>Saprotroph</td><td>|Undefined Saprotroph|</td><td>Yeast</td><td>NULL</td><td>Probable</td><td>As nectar-tap_saprotroph (Põlme et al. 2020); Decay sugar-rich substrates (Põlme et al. 2020)</td><td>Põlme S, et al. 2020. Fungal Diversity 105:1-16 ((https://doi.org/10.1007/s13225-020-00466-2))</td></tr>
</table>

| | | | | | | | | | | | |
|---|---|---|---|---|---|---|---|---|---|---|---|
| Otu42498 | 1 | k__Fungi(100);p__Ascomycota(100);c__Saccharomycetes(100);o__Saccharomycetales(100);f__Metschnikowiaceae(100);g__Metschnikowia(100);s__Metschnikowia_pulcherrima(100); | Metschnikowia | 13 | Saprotroph | \|Undefined Saprotroph\| | Yeast | NULL | Probable | As nectar-tap_saprotroph (Põlme et al. 2020); Decay sugar-rich substrates (Põlme et al. 2020) | Põlme S, et al. 2020. Fungal Diversity 105:1-16 ((https://doi.org/10.1007/s13225-020-00466-2)) |
| Otu42499 | 1 | k__Fungi(100);p__Ascomycota(100);c__Saccharomycetes(100);o__Saccharomycetales(100);f__Metschnikowiaceae(100);g__Metschnikowia(100);s__Metschnikowia_pulcherrima(100); | Metschnikowia | 13 | Saprotroph | \|Undefined Saprotroph\| | Yeast | NULL | Probable | As nectar-tap_saprotroph (Põlme et al. 2020); Decay sugar-rich substrates (Põlme et al. 2020) | Põlme S, et al. 2020. Fungal Diversity 105:1-16 ((https://doi.org/10.1007/s13225-020-00466-2)) |
| Otu42500 | 1 | k__Fungi(100);p__Ascomycota(100);c__Saccharomycetes(100);o__Saccharomycetales(100);f__Metschnikowiaceae(100);g__Metschnikowia(100);g__Metschnikowia_unclassified(100); | Metschnikowia | 13 | Saprotroph | \|Undefined Saprotroph\| | Yeast | NULL | Probable | As nectar-tap_saprotroph (Põlme et al. 2020); Decay sugar-rich substrates (Põlme et al. 2020) | Põlme S, et al. 2020. Fungal Diversity 105:1-16 ((https://doi.org/10.1007/s13225-020-00466-2)) |
| Otu42501 | 1 | k__Fungi(100);p__Ascomycota(100);c__Saccharomycetes(100);o__Saccharomycetales(100);f__Metschnikowiaceae(100);g__Metschnikowia(100);s__Metschnikowia_pulcherrima(100); | Metschnikowia | 13 | Saprotroph | \|Undefined Saprotroph\| | Yeast | NULL | Probable | As nectar-tap_saprotroph (Põlme et al. 2020); Decay sugar-rich substrates (Põlme et al. 2020) | Põlme S, et al. 2020. Fungal Diversity 105:1-16 ((https://doi.org/10.1007/s13225-020-00466-2)) |
| Otu42502 | 1 | k__Fungi(100);p__Ascomycota(100);c__Saccharomycetes(100);o__Saccharomycetales(100);f__Metschnikowiaceae(100);g__Metschnikowia(100);s__Metschnikowia_pulcherrima(100); | Metschnikowia | 13 | Saprotroph | \|Undefined Saprotroph\| | Yeast | NULL | Probable | As nectar-tap_saprotroph (Põlme et al. 2020); Decay sugar-rich substrates (Põlme et al. 2020) | Põlme S, et al. 2020. Fungal Diversity 105:1-16 ((https://doi.org/10.1007/s13225-020-00466-2)) |
| Otu42503 | 1 | k__Fungi(100);p__Ascomycota(100);c__Saccharomycetes(100);o__Saccharomycetales(100);f__Metschnikowiaceae(100);g__Metschnikowia(100);g__Metschnikowia_unclassified(100); | Metschnikowia | 13 | Saprotroph | \|Undefined Saprotroph\| | Yeast | NULL | Probable | As nectar-tap_saprotroph (Põlme et al. 2020); Decay sugar-rich substrates (Põlme et al. 2020) | Põlme S, et al. 2020. Fungal Diversity 105:1-16 ((https://doi.org/10.1007/s13225-020-00466-2)) |
| Otu42504 | 1 | k__Fungi(100);p__Ascomycota(100);c__Saccharomycetes(100);o__Saccharomycetales(100);f__Metschnikowiaceae(100);g__Metschnikowia(100);s__Metschnikowia_pulcherrima(100); | Metschnikowia | 13 | Saprotroph | \|Undefined Saprotroph\| | Yeast | NULL | Probable | As nectar-tap_saprotroph (Põlme et al. 2020); Decay sugar-rich substrates (Põlme et al. 2020) | Põlme S, et al. 2020. Fungal Diversity 105:1-16 ((https://doi.org/10.1007/s13225-020-00466-2)) |
| Otu42505 | 1 | k__Fungi(100);p__Ascomycota(100);c__Saccharomycetes(100);o__Saccharomycetales(100);f__Metschnikowiaceae(100);g__Metschnikowia(100);g__Metschnikowia_unclassified(100); | Metschnikowia | 13 | Saprotroph | \|Undefined Saprotroph\| | Yeast | NULL | Probable | As nectar-tap_saprotroph (Põlme et al. 2020); Decay sugar-rich substrates (Põlme et al. 2020) | Põlme S, et al. 2020. Fungal Diversity 105:1-16 ((https://doi.org/10.1007/s13225-020-00466-2)) |
| Otu42506 | 1 | k__Fungi(100);p__Ascomycota(100);c__Saccharomycetes(100);o__Saccharomycetales(100);f__Metschnikowiaceae(100);g__Metschnikowia(100);s__Metschnikowia_pulcherrima(100); | Metschnikowia | 13 | Saprotroph | \|Undefined Saprotroph\| | Yeast | NULL | Probable | As nectar-tap_saprotroph (Põlme et al. 2020); Decay sugar-rich substrates (Põlme et al. 2020) | Põlme S, et al. 2020. Fungal Diversity 105:1-16 ((https://doi.org/10.1007/s13225-020-00466-2)) |
| Otu42507 | 1 | k__Fungi(100);p__Ascomycota(100);c__Saccharomycetes(100);o__Saccharomycetales(100);f__Metschnikowiaceae(100);g__Metschnikowia(100);s__Metschnikowia_pulcherrima(100); | Metschnikowia | 13 | Saprotroph | \|Undefined Saprotroph\| | Yeast | NULL | Probable | As nectar-tap_saprotroph (Põlme et al. 2020); Decay sugar-rich substrates (Põlme et al. 2020) | Põlme S, et al. 2020. Fungal Diversity 105:1-16 ((https://doi.org/10.1007/s13225-020-00466-2)) |
| Otu42508 | 1 | k__Fungi(100);p__Ascomycota(100);c__Saccharomycetes(100);o__Saccharomycetales(100);f__Metschnikowiaceae(100);g__Metschnikowia(100);s__Metschnikowia_pulcherrima(100); | Metschnikowia | 13 | Saprotroph | \|Undefined Saprotroph\| | Yeast | NULL | Probable | As nectar-tap_saprotroph (Põlme et al. 2020); Decay sugar-rich substrates (Põlme et al. 2020) | Põlme S, et al. 2020. Fungal Diversity 105:1-16 ((https://doi.org/10.1007/s13225-020-00466-2)) |
| Otu42510 | 1 | k__Fungi(100);p__Ascomycota(100);c__Saccharomycetes(100);o__Saccharomycetales(100);f__Metschnikowiaceae(100);g__Metschnikowia(100);g__Metschnikowia_unclassified(100); | Metschnikowia | 13 | Saprotroph | \|Undefined Saprotroph\| | Yeast | NULL | Probable | As nectar-tap_saprotroph (Põlme et al. 2020); Decay sugar-rich substrates (Põlme et al. 2020) | Põlme S, et al. 2020. Fungal Diversity 105:1-16 ((https://doi.org/10.1007/s13225-020-00466-2)) |
| Otu42511 | 1 | k__Fungi(100);p__Ascomycota(100);c__Saccharomycetes(100);o__Saccharomycetales(100);f__Metschnikowiaceae(100);g__Metschnikowia(100);g__Metschnikowia_unclassified(100); | Metschnikowia | 13 | Saprotroph | \|Undefined Saprotroph\| | Yeast | NULL | Probable | As nectar-tap_saprotroph (Põlme et al. 2020); Decay sugar-rich substrates (Põlme et al. 2020) | Põlme S, et al. 2020. Fungal Diversity 105:1-16 ((https://doi.org/10.1007/s13225-020-00466-2)) |
| Otu42512 | 1 | k__Fungi(100);p__Ascomycota(100);c__Saccharomycetes(100);o__Saccharomycetales(100);f__Metschnikowiaceae(100);g__Metschnikowia(100);s__Metschnikowia_pulcherrima(100); | Metschnikowia | 13 | Saprotroph | \|Undefined Saprotroph\| | Yeast | NULL | Probable | As nectar-tap_saprotroph (Põlme et al. 2020); Decay sugar-rich substrates (Põlme et al. 2020) | Põlme S, et al. 2020. Fungal Diversity 105:1-16 ((https://doi.org/10.1007/s13225-020-00466-2)) |
| Otu42513 | 1 | k__Fungi(100);p__Ascomycota(100);c__Saccharomycetes(100);o__Saccharomycetales(100);f__Metschnikowiaceae(100);g__Metschnikowia(100);g__Metschnikowia_unclassified(100); | Metschnikowia | 13 | Saprotroph | \|Undefined Saprotroph\| | Yeast | NULL | Probable | As nectar-tap_saprotroph (Põlme et al. 2020); Decay sugar-rich substrates (Põlme et al. 2020) | Põlme S, et al. 2020. Fungal Diversity 105:1-16 ((https://doi.org/10.1007/s13225-020-00466-2)) |
| Otu42514 | 1 | k__Fungi(100);p__Ascomycota(100);c__Saccharomycetes(100);o__Saccharomycetales(100);f__Metschnikowiaceae(100);g__Metschnikowia(100);s__Metschnikowia_pulcherrima(100); | Metschnikowia | 13 | Saprotroph | \|Undefined Saprotroph\| | Yeast | NULL | Probable | As nectar-tap_saprotroph (Põlme et al. 2020); Decay sugar-rich substrates (Põlme et al. 2020) | Põlme S, et al. 2020. Fungal Diversity 105:1-16 ((https://doi.org/10.1007/s13225-020-00466-2)) |
| Otu42515 | 1 | k__Fungi(100);p__Ascomycota(100);c__Saccharomycetes(100);o__Saccharomycetales(100);f__Metschnikowiaceae(100);g__Metschnikowia(100);s__Metschnikowia_pulcherrima(100); | Metschnikowia | 13 | Saprotroph | \|Undefined Saprotroph\| | Yeast | NULL | Probable | As nectar-tap_saprotroph (Põlme et al. 2020); Decay sugar-rich substrates (Põlme et al. 2020) | Põlme S, et al. 2020. Fungal Diversity 105:1-16 ((https://doi.org/10.1007/s13225-020-00466-2)) |
| Otu42516 | 1 | k__Fungi(100);p__Ascomycota(100);c__Saccharomycetes(100);o__Saccharomycetales(100);f__Metschnikowiaceae(100);g__Metschnikowia(100);s__Metschnikowia_pulcherrima(100); | Metschnikowia | 13 | Saprotroph | \|Undefined Saprotroph\| | Yeast | NULL | Probable | As nectar-tap_saprotroph (Põlme et al. 2020); Decay sugar-rich substrates (Põlme et al. 2020) | Põlme S, et al. 2020. Fungal Diversity 105:1-16 ((https://doi.org/10.1007/s13225-020-00466-2)) |
| Otu42517 | 1 | k__Fungi(100);p__Ascomycota(100);c__Saccharomycetes(100);o__Saccharomycetales(100);f__Metschnikowiaceae(100);g__Metschnikowia(100);s__Metschnikowia_pulcherrima(100); | Metschnikowia | 13 | Saprotroph | \|Undefined Saprotroph\| | Yeast | NULL | Probable | As nectar-tap_saprotroph (Põlme et al. 2020); Decay sugar-rich substrates (Põlme et al. 2020) | Põlme S, et al. 2020. Fungal Diversity 105:1-16 ((https://doi.org/10.1007/s13225-020-00466-2)) |
| Otu42518 | 1 | k__Fungi(100);p__Ascomycota(100);c__Saccharomycetes(100);o__Saccharomycetales(100);f__Metschnikowiaceae(100);g__Metschnikowia(100);s__Metschnikowia_pulcherrima(100); | Metschnikowia | 13 | Saprotroph | \|Undefined Saprotroph\| | Yeast | NULL | Probable | As nectar-tap_saprotroph (Põlme et al. 2020); Decay sugar-rich substrates (Põlme et al. 2020) | Põlme S, et al. 2020. Fungal Diversity 105:1-16 ((https://doi.org/10.1007/s13225-020-00466-2)) |
| Otu42519 | 1 | k__Fungi(100);p__Ascomycota(100);c__Saccharomycetes(100);o__Saccharomycetales(100);f__Metschnikowiaceae(100);g__Metschnikowia(100);s__Metschnikowia_pulcherrima(100); | Metschnikowia | 13 | Saprotroph | \|Undefined Saprotroph\| | Yeast | NULL | Probable | As nectar-tap_saprotroph (Põlme et al. 2020); Decay sugar-rich substrates (Põlme et al. 2020) | Põlme S, et al. 2020. Fungal Diversity 105:1-16 ((https://doi.org/10.1007/s13225-020-00466-2)) |
| Otu42520 | 1 | k__Fungi(100);p__Ascomycota(100);c__Saccharomycetes(100);o__Saccharomycetales(100);f__Metschnikowiaceae(100);g__Metschnikowia(100);s__Metschnikowia_pulcherrima(100); | Metschnikowia | 13 | Saprotroph | \|Undefined Saprotroph\| | Yeast | NULL | Probable | As nectar-tap_saprotroph (Põlme et al. 2020); Decay sugar-rich substrates (Põlme et al. 2020) | Põlme S, et al. 2020. Fungal Diversity 105:1-16 ((https://doi.org/10.1007/s13225-020-00466-2)) |
| Otu42521 | 1 | k__Fungi(100);p__Ascomycota(100);c__Saccharomycetes(100);o__Saccharomycetales(100);f__Metschnikowiaceae(100);g__Metschnikowia(100);s__Metschnikowia_pulcherrima(100); | Metschnikowia | 13 | Saprotroph | \|Undefined Saprotroph\| | Yeast | NULL | Probable | As nectar-tap_saprotroph (Põlme et al. 2020); Decay sugar-rich substrates (Põlme et al. 2020) | Põlme S, et al. 2020. Fungal Diversity 105:1-16 ((https://doi.org/10.1007/s13225-020-00466-2)) |
| Otu42524 | 1 | k__Fungi(100);p__Ascomycota(100);c__Saccharomycetes(100);o__Saccharomycetales(100);f__Metschnikowiaceae(100);g__Metschnikowia(100);s__Metschnikowia_pulcherrima(100); | Metschnikowia | 13 | Saprotroph | \|Undefined Saprotroph\| | Yeast | NULL | Probable | As nectar-tap_saprotroph (Põlme et al. 2020); Decay sugar-rich substrates (Põlme et al. 2020) | Põlme S, et al. 2020. Fungal Diversity 105:1-16 ((https://doi.org/10.1007/s13225-020-00466-2)) |
| Otu42525 | 1 | k__Fungi(100);p__Ascomycota(100);c__Saccharomycetes(100);o__Saccharomycetales(100);f__Metschnikowiaceae(100);g__Metschnikowia(100);s__Metschnikowia_pulcherrima(100); | Metschnikowia | 13 | Saprotroph | \|Undefined Saprotroph\| | Yeast | NULL | Probable | As nectar-tap_saprotroph (Põlme et al. 2020); Decay sugar-rich substrates (Põlme et al. 2020) | Põlme S, et al. 2020. Fungal Diversity 105:1-16 ((https://doi.org/10.1007/s13225-020-00466-2)) |
| Otu42526 | 1 | k__Fungi(100);p__Ascomycota(100);c__Saccharomycetes(100);o__Saccharomycetales(100);f__Metschnikowiaceae(100);g__Metschnikowia(100);s__Metschnikowia_pulcherrima(100); | Metschnikowia | 13 | Saprotroph | \|Undefined Saprotroph\| | Yeast | NULL | Probable | As nectar-tap_saprotroph (Põlme et al. 2020); Decay sugar-rich substrates (Põlme et al. 2020) | Põlme S, et al. 2020. Fungal Diversity 105:1-16 ((https://doi.org/10.1007/s13225-020-00466-2)) |
| Otu42529 | 1 | k__Fungi(100);p__Ascomycota(100);c__Saccharomycetes(100);o__Saccharomycetales(100);f__Metschnikowiaceae(100);g__Metschnikowia(100);g__Metschnikowia_unclassified(100); | Metschnikowia | 13 | Saprotroph | \|Undefined Saprotroph\| | Yeast | NULL | Probable | As nectar-tap_saprotroph (Põlme et al. 2020); Decay sugar-rich substrates (Põlme et al. 2020) | Põlme S, et al. 2020. Fungal Diversity 105:1-16 ((https://doi.org/10.1007/s13225-020-00466-2)) |
| Otu42530 | 1 | k__Fungi(100);p__Ascomycota(100);c__Saccharomycetes(100);o__Saccharomycetales(100);f__Metschnikowiaceae(100);g__Metschnikowia(100);s__Metschnikowia_pulcherrima(100); | Metschnikowia | 13 | Saprotroph | \|Undefined Saprotroph\| | Yeast | NULL | Probable | As nectar-tap_saprotroph (Põlme et al. 2020); Decay sugar-rich substrates (Põlme et al. 2020) | Põlme S, et al. 2020. Fungal Diversity 105:1-16 ((https://doi.org/10.1007/s13225-020-00466-2)) |
| Otu42531 | 1 | k__Fungi(100);p__Ascomycota(100);c__Saccharomycetes(100);o__Saccharomycetales(100);f__Metschnikowiaceae(100);g__Metschnikowia(100);s__Metschnikowia_pulcherrima(100); | Metschnikowia | 13 | Saprotroph | \|Undefined Saprotroph\| | Yeast | NULL | Probable | As nectar-tap_saprotroph (Põlme et al. 2020); Decay sugar-rich substrates (Põlme et al. 2020) | Põlme S, et al. 2020. Fungal Diversity 105:1-16 ((https://doi.org/10.1007/s13225-020-00466-2)) |
| Otu42533 | 1 | k__Fungi(100);p__Ascomycota(100);c__Saccharomycetes(100);o__Saccharomycetales(100);f__Metschnikowiaceae(100);g__Metschnikowia(100);s__Metschnikowia_pulcherrima(100); | Metschnikowia | 13 | Saprotroph | \|Undefined Saprotroph\| | Yeast | NULL | Probable | As nectar-tap_saprotroph (Põlme et al. 2020); Decay sugar-rich substrates (Põlme et al. 2020) | Põlme S, et al. 2020. Fungal Diversity 105:1-16 ((https://doi.org/10.1007/s13225-020-00466-2)) |
| Otu42534 | 1 | k__Fungi(100);p__Ascomycota(100);c__Saccharomycetes(100);o__Saccharomycetales(100);f__Metschnikowiaceae(100);g__Metschnikowia(100);s__Metschnikowia_pulcherrima(100); | Metschnikowia | 13 | Saprotroph | \|Undefined Saprotroph\| | Yeast | NULL | Probable | As nectar-tap_saprotroph (Põlme et al. 2020); Decay sugar-rich substrates (Põlme et al. 2020) | Põlme S, et al. 2020. Fungal Diversity 105:1-16 ((https://doi.org/10.1007/s13225-020-00466-2)) |
| Otu42535 | 1 | k__Fungi(100);p__Ascomycota(100);c__Saccharomycetes(100);o__Saccharomycetales(100);f__Metschnikowiaceae(100);g__Metschnikowia(100);s__Metschnikowia_pulcherrima(100); | Metschnikowia | 13 | Saprotroph | \|Undefined Saprotroph\| | Yeast | NULL | Probable | As nectar-tap_saprotroph (Põlme et al. 2020); Decay sugar-rich substrates (Põlme et al. 2020) | Põlme S, et al. 2020. Fungal Diversity 105:1-16 ((https://doi.org/10.1007/s13225-020-00466-2)) |
| Otu42536 | 1 | k__Fungi(100);p__Ascomycota(100);c__Saccharomycetes(100);o__Saccharomycetales(100);f__Metschnikowiaceae(100);g__Metschnikowia(100);g__Metschnikowia_unclassified(100); | Metschnikowia | 13 | Saprotroph | \|Undefined Saprotroph\| | Yeast | NULL | Probable | As nectar-tap_saprotroph (Põlme et al. 2020); Decay sugar-rich substrates (Põlme et al. 2020) | Põlme S, et al. 2020. Fungal Diversity 105:1-16 ((https://doi.org/10.1007/s13225-020-00466-2)) |
| Otu42537 | 1 | k__Fungi(100);p__Ascomycota(100);c__Saccharomycetes(100);o__Saccharomycetales(100);f__Metschnikowiaceae(100);g__Metschnikowia(100);s__Metschnikowia_pulcherrima(100); | Metschnikowia | 13 | Saprotroph | \|Undefined Saprotroph\| | Yeast | NULL | Probable | As nectar-tap_saprotroph (Põlme et al. 2020); Decay sugar-rich substrates (Põlme et al. 2020) | Põlme S, et al. 2020. Fungal Diversity 105:1-16 ((https://doi.org/10.1007/s13225-020-00466-2)) |
| Otu42538 | 1 | k__Fungi(100);p__Ascomycota(100);c__Saccharomycetes(100);o__Saccharomycetales(100);f__Metschnikowiaceae(100);g__Metschnikowia(100);g__Metschnikowia_unclassified(100); | Metschnikowia | 13 | Saprotroph | \|Undefined Saprotroph\| | Yeast | NULL | Probable | As nectar-tap_saprotroph (Põlme et al. 2020); Decay sugar-rich substrates (Põlme et al. 2020) | Põlme S, et al. 2020. Fungal Diversity 105:1-16 ((https://doi.org/10.1007/s13225-020-00466-2)) |
| Otu42539 | 1 | k__Fungi(100);p__Ascomycota(100);c__Saccharomycetes(100);o__Saccharomycetales(100);f__Metschnikowiaceae(100);g__Metschnikowia(100);s__Metschnikowia_pulcherrima(100); | Metschnikowia | 13 | Saprotroph | \|Undefined Saprotroph\| | Yeast | NULL | Probable | As nectar-tap_saprotroph (Põlme et al. 2020); Decay sugar-rich substrates (Põlme et al. 2020) | Põlme S, et al. 2020. Fungal Diversity 105:1-16 ((https://doi.org/10.1007/s13225-020-00466-2)) |
| Otu42540 | 1 | k__Fungi(100);p__Ascomycota(100);c__Saccharomycetes(100);o__Saccharomycetales(100);f__Metschnikowiaceae(100);g__Metschnikowia(100);g__Metschnikowia_unclassified(100); | Metschnikowia | 13 | Saprotroph | \|Undefined Saprotroph\| | Yeast | NULL | Probable | As nectar-tap_saprotroph (Põlme et al. 2020); Decay sugar-rich substrates (Põlme et al. 2020) | Põlme S, et al. 2020. Fungal Diversity 105:1-16 ((https://doi.org/10.1007/s13225-020-00466-2)) |
| Otu42541 | 1 | k__Fungi(100);p__Ascomycota(100);c__Saccharomycetes(100);o__Saccharomycetales(100);f__Metschnikowiaceae(100);g__Metschnikowia(100);s__Metschnikowia_pulcherrima(100); | Metschnikowia | 13 | Saprotroph | \|Undefined Saprotroph\| | Yeast | NULL | Probable | As nectar-tap_saprotroph (Põlme et al. 2020); Decay sugar-rich substrates (Põlme et al. 2020) | Põlme S, et al. 2020. Fungal Diversity 105:1-16 ((https://doi.org/10.1007/s13225-020-00466-2)) |
| Otu42542 | 1 | k__Fungi(100);p__Ascomycota(100);c__Saccharomycetes(100);o__Saccharomycetales(100);f__Metschnikowiaceae(100);g__Metschnikowia(100);s__Metschnikowia_pulcherrima(100); | Metschnikowia | 13 | Saprotroph | \|Undefined Saprotroph\| | Yeast | NULL | Probable | As nectar-tap_saprotroph (Põlme et al. 2020); Decay sugar-rich substrates (Põlme et al. 2020) | Põlme S, et al. 2020. Fungal Diversity 105:1-16 ((https://doi.org/10.1007/s13225-020-00466-2)) |
| Otu42543 | 1 | k__Fungi(100);p__Ascomycota(100);c__Saccharomycetes(100);o__Saccharomycetales(100);f__Metschnikowiaceae(100);g__Metschnikowia(100);s__Metschnikowia_pulcherrima(100); | Metschnikowia | 13 | Saprotroph | \|Undefined Saprotroph\| | Yeast | NULL | Probable | As nectar-tap_saprotroph (Põlme et al. 2020); Decay sugar-rich substrates (Põlme et al. 2020) | Põlme S, et al. 2020. Fungal Diversity 105:1-16 ((https://doi.org/10.1007/s13225-020-00466-2)) |
| Otu42544 | 1 | k__Fungi(100);p__Ascomycota(100);c__Saccharomycetes(100);o__Saccharomycetales(100);f__Metschnikowiaceae(100);g__Metschnikowia(100);s__Metschnikowia_pulcherrima(100); | Metschnikowia | 13 | Saprotroph | \|Undefined Saprotroph\| | Yeast | NULL | Probable | As nectar-tap_saprotroph (Põlme et al. 2020); Decay sugar-rich substrates (Põlme et al. 2020) | Põlme S, et al. 2020. Fungal Diversity 105:1-16 ((https://doi.org/10.1007/s13225-020-00466-2)) |
| Otu42545 | 1 | k__Fungi(100);p__Ascomycota(100);c__Saccharomycetes(100);o__Saccharomycetales(100);f__Metschnikowiaceae(100);g__Metschnikowia(100);g__Metschnikowia_unclassified(100); | Metschnikowia | 13 | Saprotroph | \|Undefined Saprotroph\| | Yeast | NULL | Probable | As nectar-tap_saprotroph (Põlme et al. 2020); Decay sugar-rich substrates (Põlme et al. 2020) | Põlme S, et al. 2020. Fungal Diversity 105:1-16 ((https://doi.org/10.1007/s13225-020-00466-2)) |
| Otu42546 | 1 | k__Fungi(100);p__Ascomycota(100);c__Saccharomycetes(100);o__Saccharomycetales(100);f__Metschnikowiaceae(100);g__Metschnikowia(100);g__Metschnikowia_unclassified(100); | Metschnikowia | 13 | Saprotroph | \|Undefined Saprotroph\| | Yeast | NULL | Probable | As nectar-tap_saprotroph (Põlme et al. 2020); Decay sugar-rich substrates (Põlme et al. 2020) | Põlme S, et al. 2020. Fungal Diversity 105:1-16 ((https://doi.org/10.1007/s13225-020-00466-2)) |
| Otu42547 | 1 | k__Fungi(100);p__Ascomycota(100);c__Saccharomycetes(100);o__Saccharomycetales(100);f__Metschnikowiaceae(100);g__Metschnikowia(100);s__Metschnikowia_pulcherrima(100); | Metschnikowia | 13 | Saprotroph | \|Undefined Saprotroph\| | Yeast | NULL | Probable | As nectar-tap_saprotroph (Põlme et al. 2020); Decay sugar-rich substrates (Põlme et al. 2020) | Põlme S, et al. 2020. Fungal Diversity 105:1-16 ((https://doi.org/10.1007/s13225-020-00466-2)) |
| Otu42548 | 1 | k__Fungi(100);p__Ascomycota(100);c__Saccharomycetes(100);o__Saccharomycetales(100);f__Metschnikowiaceae(100);g__Metschnikowia(100);g__Metschnikowia_unclassified(100); | Metschnikowia | 13 | Saprotroph | \|Undefined Saprotroph\| | Yeast | NULL | Probable | As nectar-tap_saprotroph (Põlme et al. 2020); Decay sugar-rich substrates (Põlme et al. 2020) | Põlme S, et al. 2020. Fungal Diversity 105:1-16 ((https://doi.org/10.1007/s13225-020-00466-2)) |
| Otu42549 | 1 | k__Fungi(100);p__Ascomycota(100);c__Saccharomycetes(100);o__Saccharomycetales(100);f__Metschnikowiaceae(100);g__Metschnikowia(100);g__Metschnikowia_unclassified(100); | Metschnikowia | 13 | Saprotroph | \|Undefined Saprotroph\| | Yeast | NULL | Probable | As nectar-tap_saprotroph (Põlme et al. 2020); Decay sugar-rich substrates (Põlme et al. 2020) | Põlme S, et al. 2020. Fungal Diversity 105:1-16 ((https://doi.org/10.1007/s13225-020-00466-2)) |
| Otu42550 | 1 | k__Fungi(100);p__Ascomycota(100);c__Saccharomycetes(100);o__Saccharomycetales(100);f__Metschnikowiaceae(100);g__Metschnikowia(100);s__Metschnikowia_pulcherrima(100); | Metschnikowia | 13 | Saprotroph | \|Undefined Saprotroph\| | Yeast | NULL | Probable | As nectar-tap_saprotroph (Põlme et al. 2020); Decay sugar-rich substrates (Põlme et al. 2020) | Põlme S, et al. 2020. Fungal Diversity 105:1-16 ((https://doi.org/10.1007/s13225-020-00466-2)) |
| Otu42551 | 1 | k__Fungi(100);p__Ascomycota(100);c__Saccharomycetes(100);o__Saccharomycetales(100);f__Metschnikowiaceae(100);g__Metschnikowia(100);s__Metschnikowia_pulcherrima(100); | Metschnikowia | 13 | Saprotroph | \|Undefined Saprotroph\| | Yeast | NULL | Probable | As nectar-tap_saprotroph (Põlme et al. 2020); Decay sugar-rich substrates (Põlme et al. 2020) | Põlme S, et al. 2020. Fungal Diversity 105:1-16 ((https://doi.org/10.1007/s13225-020-00466-2)) |
| Otu42552 | 1 | k__Fungi(100);p__Ascomycota(100);c__Saccharomycetes(100);o__Saccharomycetales(100);f__Metschnikowiaceae(100);g__Metschnikowia(100);s__Metschnikowia_pulcherrima(100); | Metschnikowia | 13 | Saprotroph | \|Undefined Saprotroph\| | Yeast | NULL | Probable | As nectar-tap_saprotroph (Põlme et al. 2020); Decay sugar-rich substrates (Põlme et al. 2020) | Põlme S, et al. 2020. Fungal Diversity 105:1-16 ((https://doi.org/10.1007/s13225-020-00466-2)) |
| Otu42553 | 1 | k__Fungi(100);p__Ascomycota(100);c__Saccharomycetes(100);o__Saccharomycetales(100);f__Metschnikowiaceae(100);g__Metschnikowia(100);s__Metschnikowia_pulcherrima(100); | Metschnikowia | 13 | Saprotroph | \|Undefined Saprotroph\| | Yeast | NULL | Probable | As nectar-tap_saprotroph (Põlme et al. 2020); Decay sugar-rich substrates (Põlme et al. 2020) | Põlme S, et al. 2020. Fungal Diversity 105:1-16 ((https://doi.org/10.1007/s13225-020-00466-2)) |
| Otu42554 | 1 | k__Fungi(100);p__Ascomycota(100);c__Saccharomycetes(100);o__Saccharomycetales(100);f__Metschnikowiaceae(100);g__Metschnikowia(100);s__Metschnikowia_pulcherrima(100); | Metschnikowia | 13 | Saprotroph | \|Undefined Saprotroph\| | Yeast | NULL | Probable | As nectar-tap_saprotroph (Põlme et al. 2020); Decay sugar-rich substrates (Põlme et al. 2020) | Põlme S, et al. 2020. Fungal Diversity 105:1-16 ((https://doi.org/10.1007/s13225-020-00466-2)) |
| Otu42555 | 1 | k__Fungi(100);p__Ascomycota(100);c__Saccharomycetes(100);o__Saccharomycetales(100);f__Metschnikowiaceae(100);g__Metschnikowia(100);g__Metschnikowia_unclassified(100); | Metschnikowia | 13 | Saprotroph | \|Undefined Saprotroph\| | Yeast | NULL | Probable | As nectar-tap_saprotroph (Põlme et al. 2020); Decay sugar-rich substrates (Põlme et al. 2020) | Põlme S, et al. 2020. Fungal Diversity 105:1-16 ((https://doi.org/10.1007/s13225-020-00466-2)) |
| Otu42556 | 1 | k__Fungi(100);p__Ascomycota(100);c__Saccharomycetes(100);o__Saccharomycetales(100);f__Metschnikowiaceae(100);g__Metschnikowia(100);s__Metschnikowia_pulcherrima(100); | Metschnikowia | 13 | Saprotroph | \|Undefined Saprotroph\| | Yeast | NULL | Probable | As nectar-tap_saprotroph (Põlme et al. 2020); Decay sugar-rich substrates (Põlme et al. 2020) | Põlme S, et al. 2020. Fungal Diversity 105:1-16 ((https://doi.org/10.1007/s13225-020-00466-2)) |
| Otu42557 | 1 | k__Fungi(100);p__Ascomycota(100);c__Saccharomycetes(100);o__Saccharomycetales(100);f__Metschnikowiaceae(100);g__Metschnikowia(100);g__Metschnikowia_unclassified(100); | Metschnikowia | 13 | Saprotroph | \|Undefined Saprotroph\| | Yeast | NULL | Probable | As nectar-tap_saprotroph (Põlme et al. 2020); Decay sugar-rich substrates (Põlme et al. 2020) | Põlme S, et al. 2020. Fungal Diversity 105:1-16 ((https://doi.org/10.1007/s13225-020-00466-2)) |
| Otu42558 | 1 | k__Fungi(100);p__Ascomycota(100);c__Saccharomycetes(100);o__Saccharomycetales(100);f__Metschnikowiaceae(100);g__Metschnikowia(100);s__Metschnikowia_pulcherrima(100); | Metschnikowia | 13 | Saprotroph | \|Undefined Saprotroph\| | Yeast | NULL | Probable | As nectar-tap_saprotroph (Põlme et al. 2020); Decay sugar-rich substrates (Põlme et al. 2020) | Põlme S, et al. 2020. Fungal Diversity 105:1-16 ((https://doi.org/10.1007/s13225-020-00466-2)) |
| Otu42559 | 1 | k__Fungi(100);p__Ascomycota(100);c__Saccharomycetes(100);o__Saccharomycetales(100);f__Metschnikowiaceae(100);g__Metschnikowia(100);g__Metschnikowia_unclassified(100); | Metschnikowia | 13 | Saprotroph | \|Undefined Saprotroph\| | Yeast | NULL | Probable | As nectar-tap_saprotroph (Põlme et al. 2020); Decay sugar-rich substrates (Põlme et al. 2020) | Põlme S, et al. 2020. Fungal Diversity 105:1-16 ((https://doi.org/10.1007/s13225-020-00466-2)) |
| Otu42560 | 1 | k__Fungi(100);p__Ascomycota(100);c__Saccharomycetes(100);o__Saccharomycetales(100);f__Metschnikowiaceae(100);g__Metschnikowia(100);g__Metschnikowia_unclassified(100); | Metschnikowia | 13 | Saprotroph | \|Undefined Saprotroph\| | Yeast | NULL | Probable | As nectar-tap_saprotroph (Põlme et al. 2020); Decay sugar-rich substrates (Põlme et al. 2020) | Põlme S, et al. 2020. Fungal Diversity 105:1-16 ((https://doi.org/10.1007/s13225-020-00466-2)) |
| Otu42561 | 1 | k__Fungi(100);p__Ascomycota(100);c__Saccharomycetes(100);o__Saccharomycetales(100);f__Metschnikowiaceae(100);g__Metschnikowia(100);s__Metschnikowia_pulcherrima(100); | Metschnikowia | 13 | Saprotroph | \|Undefined Saprotroph\| | Yeast | NULL | Probable | As nectar-tap_saprotroph (Põlme et al. 2020); Decay sugar-rich substrates (Põlme et al. 2020) | Põlme S, et al. 2020. Fungal Diversity 105:1-16 ((https://doi.org/10.1007/s13225-020-00466-2)) |
| Otu42562 | 1 | k__Fungi(100);p__Ascomycota(100);c__Saccharomycetes(100);o__Saccharomycetales(100);f__Metschnikowiaceae(100);g__Metschnikowia(100);g__Metschnikowia_unclassified(100); | Metschnikowia | 13 | Saprotroph | \|Undefined Saprotroph\| | Yeast | NULL | Probable | As nectar-tap_saprotroph (Põlme et al. 2020); Decay sugar-rich substrates (Põlme et al. 2020) | Põlme S, et al. 2020. Fungal Diversity 105:1-16 ((https://doi.org/10.1007/s13225-020-00466-2)) |
| Otu42563 | 1 | k__Fungi(100);p__Ascomycota(100);c__Saccharomycetes(100);o__Saccharomycetales(100);f__Metschnikowiaceae(100);g__Metschnikowia(100);s__Metschnikowia_pulcherrima(100); | Metschnikowia | 13 | Saprotroph | \|Undefined Saprotroph\| | Yeast | NULL | Probable | As nectar-tap_saprotroph (Põlme et al. 2020); Decay sugar-rich substrates (Põlme et al. 2020) | Põlme S, et al. 2020. Fungal Diversity 105:1-16 ((https://doi.org/10.1007/s13225-020-00466-2)) |
| Otu42564 | 1 | k__Fungi(100);p__Ascomycota(100);c__Saccharomycetes(100);o__Saccharomycetales(100);f__Metschnikowiaceae(100);g__Metschnikowia(100);s__Metschnikowia_pulcherrima(100); | Metschnikowia | 13 | Saprotroph | \|Undefined Saprotroph\| | Yeast | NULL | Probable | As nectar-tap_saprotroph (Põlme et al. 2020); Decay sugar-rich substrates (Põlme et al. 2020) | Põlme S, et al. 2020. Fungal Diversity 105:1-16 ((https://doi.org/10.1007/s13225-020-00466-2)) |
| Otu42565 | 1 | k__Fungi(100);p__Ascomycota(100);c__Saccharomycetes(100);o__Saccharomycetales(100);f__Metschnikowiaceae(100);g__Metschnikowia(100);s__Metschnikowia_pulcherrima(100); | Metschnikowia | 13 | Saprotroph | \|Undefined Saprotroph\| | Yeast | NULL | Probable | As nectar-tap_saprotroph (Põlme et al. 2020); Decay sugar-rich substrates (Põlme et al. 2020) | Põlme S, et al. 2020. Fungal Diversity 105:1-16 ((https://doi.org/10.1007/s13225-020-00466-2)) |
| Otu42566 | 1 | k__Fungi(100);p__Ascomycota(100);c__Saccharomycetes(100);o__Saccharomycetales(100);f__Metschnikowiaceae(100);g__Metschnikowia(100);s__Metschnikowia_pulcherrima(100); | Metschnikowia | 13 | Saprotroph | \|Undefined Saprotroph\| | Yeast | NULL | Probable | As nectar-tap_saprotroph (Põlme et al. 2020); Decay sugar-rich substrates (Põlme et al. 2020) | Põlme S, et al. 2020. Fungal Diversity 105:1-16 ((https://doi.org/10.1007/s13225-020-00466-2)) |
| Otu42567 | 1 | k__Fungi(100);p__Ascomycota(100);c__Saccharomycetes(100);o__Saccharomycetales(100);f__Metschnikowiaceae(100);g__Metschnikowia(100);s__Metschnikowia_pulcherrima(100); | Metschnikowia | 13 | Saprotroph | \|Undefined Saprotroph\| | Yeast | NULL | Probable | As nectar-tap_saprotroph (Põlme et al. 2020); Decay sugar-rich substrates (Põlme et al. 2020) | Põlme S, et al. 2020. Fungal Diversity 105:1-16 ((https://doi.org/10.1007/s13225-020-00466-2)) |
| Otu42568 | 1 | k__Fungi(100);p__Ascomycota(100);c__Saccharomycetes(100);o__Saccharomycetales(100);f__Metschnikowiaceae(100);g__Metschnikowia(100);s__Metschnikowia_pulcherrima(100); | Metschnikowia | 13 | Saprotroph | \|Undefined Saprotroph\| | Yeast | NULL | Probable | As nectar-tap_saprotroph (Põlme et al. 2020); Decay sugar-rich substrates (Põlme et al. 2020) | Põlme S, et al. 2020. Fungal Diversity 105:1-16 ((https://doi.org/10.1007/s13225-020-00466-2)) |
| Otu42569 | 1 | k__Fungi(100);p__Ascomycota(100);c__Saccharomycetes(100);o__Saccharomycetales(100);f__Metschnikowiaceae(100);g__Metschnikowia(100);g__Metschnikowia_unclassified(100); | Metschnikowia | 13 | Saprotroph | \|Undefined Saprotroph\| | Yeast | NULL | Probable | As nectar-tap_saprotroph (Põlme et al. 2020); Decay sugar-rich substrates (Põlme et al. 2020) | Põlme S, et al. 2020. Fungal Diversity 105:1-16 ((https://doi.org/10.1007/s13225-020-00466-2)) |
| Otu42570 | 1 | k__Fungi(100);p__Ascomycota(100);c__Saccharomycetes(100);o__Saccharomycetales(100);f__Metschnikowiaceae(100);g__Metschnikowia(100);g__Metschnikowia_unclassified(100); | Metschnikowia | 13 | Saprotroph | \|Undefined Saprotroph\| | Yeast | NULL | Probable | As nectar-tap_saprotroph (Põlme et al. 2020); Decay sugar-rich substrates (Põlme et al. 2020) | Põlme S, et al. 2020. Fungal Diversity 105:1-16 ((https://doi.org/10.1007/s13225-020-00466-2)) |
| Otu42571 | 1 | k__Fungi(100);p__Ascomycota(100);c__Saccharomycetes(100);o__Saccharomycetales(100);f__Metschnikowiaceae(100);g__Metschnikowia(100);s__Metschnikowia_pulcherrima(100); | Metschnikowia | 13 | Saprotroph | \|Undefined Saprotroph\| | Yeast | NULL | Probable | As nectar-tap_saprotroph (Põlme et al. 2020); Decay sugar-rich substrates (Põlme et al. 2020) | Põlme S, et al. 2020. Fungal Diversity 105:1-16 ((https://doi.org/10.1007/s13225-020-00466-2)) |
| Otu42572 | 1 | k__Fungi(100);p__Ascomycota(100);c__Saccharomycetes(100);o__Saccharomycetales(100);f__Metschnikowiaceae(100);g__Metschnikowia(100);g__Metschnikowia_unclassified(100); | Metschnikowia | 13 | Saprotroph | \|Undefined Saprotroph\| | Yeast | NULL | Probable | As nectar-tap_saprotroph (Põlme et al. 2020); Decay sugar-rich substrates (Põlme et al. 2020) | Põlme S, et al. 2020. Fungal Diversity 105:1-16 ((https://doi.org/10.1007/s13225-020-00466-2)) |
| Otu42573 | 1 | k__Fungi(100);p__Ascomycota(100);c__Saccharomycetes(100);o__Saccharomycetales(100);f__Metschnikowiaceae(100);g__Metschnikowia(100);g__Metschnikowia_unclassified(100); | Metschnikowia | 13 | Saprotroph | \|Undefined Saprotroph\| | Yeast | NULL | Probable | As nectar-tap_saprotroph (Põlme et al. 2020); Decay sugar-rich substrates (Põlme et al. 2020) | Põlme S, et al. 2020. Fungal Diversity 105:1-16 ((https://doi.org/10.1007/s13225-020-00466-2)) |
| Otu42574 | 1 | k__Fungi(100);p__Ascomycota(100);c__Saccharomycetes(100);o__Saccharomycetales(100);f__Metschnikowiaceae(100);g__Metschnikowia(100);g__Metschnikowia_unclassified(100); | Metschnikowia | 13 | Saprotroph | \|Undefined Saprotroph\| | Yeast | NULL | Probable | As nectar-tap_saprotroph (Põlme et al. 2020); Decay sugar-rich substrates (Põlme et al. 2020) | Põlme S, et al. 2020. Fungal Diversity 105:1-16 ((https://doi.org/10.1007/s13225-020-00466-2)) |
| Otu42575 | 1 | k__Fungi(100);p__Ascomycota(100);c__Saccharomycetes(100);o__Saccharomycetales(100);f__Metschnikowiaceae(100);g__Metschnikowia(100);s__Metschnikowia_pulcherrima(100); | Metschnikowia | 13 | Saprotroph | \|Undefined Saprotroph\| | Yeast | NULL | Probable | As nectar-tap_saprotroph (Põlme et al. 2020); Decay sugar-rich substrates (Põlme et al. 2020) | Põlme S, et al. 2020. Fungal Diversity 105:1-16 ((https://doi.org/10.1007/s13225-020-00466-2)) |
| Otu42576 | 1 | k__Fungi(100);p__Ascomycota(100);c__Saccharomycetes(100);o__Saccharomycetales(100);f__Metschnikowiaceae(100);g__Metschnikowia(100);s__Metschnikowia_pulcherrima(100); | Metschnikowia | 13 | Saprotroph | \|Undefined Saprotroph\| | Yeast | NULL | Probable | As nectar-tap_saprotroph (Põlme et al. 2020); Decay sugar-rich substrates (Põlme et al. 2020) | Põlme S, et al. 2020. Fungal Diversity 105:1-16 ((https://doi.org/10.1007/s13225-020-00466-2)) |
| Otu42577 | 1 | k__Fungi(100);p__Ascomycota(100);c__Saccharomycetes(100);o__Saccharomycetales(100);f__Metschnikowiaceae(100);g__Metschnikowia(100);s__Metschnikowia_pulcherrima(100); | Metschnikowia | 13 | Saprotroph | \|Undefined Saprotroph\| | Yeast | NULL | Probable | As nectar-tap_saprotroph (Põlme et al. 2020); Decay sugar-rich substrates (Põlme et al. 2020) | Põlme S, et al. 2020. Fungal Diversity 105:1-16 ((https://doi.org/10.1007/s13225-020-00466-2)) |
| Otu42578 | 1 | k__Fungi(100);p__Ascomycota(100);c__Saccharomycetes(100);o__Saccharomycetales(100);f__Metschnikowiaceae(100);g__Metschnikowia(100);s__Metschnikowia_pulcherrima(100); | Metschnikowia | 13 | Saprotroph | \|Undefined Saprotroph\| | Yeast | NULL | Probable | As nectar-tap_saprotroph (Põlme et al. 2020); Decay sugar-rich substrates (Põlme et al. 2020) | Põlme S, et al. 2020. Fungal Diversity 105:1-16 ((https://doi.org/10.1007/s13225-020-00466-2)) |
| Otu42579 | 1 | k__Fungi(100);p__Ascomycota(100);c__Saccharomycetes(100);o__Saccharomycetales(100);f__Metschnikowiaceae(100);g__Metschnikowia(100);s__Metschnikowia_pulcherrima(100); | Metschnikowia | 13 | Saprotroph | \|Undefined Saprotroph\| | Yeast | NULL | Probable | As nectar-tap_saprotroph (Põlme et al. 2020); Decay sugar-rich substrates (Põlme et al. 2020) | Põlme S, et al. 2020. Fungal Diversity 105:1-16 ((https://doi.org/10.1007/s13225-020-00466-2)) |
| Otu42580 | 1 | k__Fungi(100);p__Ascomycota(100);c__Saccharomycetes(100);o__Saccharomycetales(100);f__Metschnikowiaceae(100);g__Metschnikowia(100);s__Metschnikowia_pulcherrima(100); | Metschnikowia | 13 | Saprotroph | \|Undefined Saprotroph\| | Yeast | NULL | Probable | As nectar-tap_saprotroph (Põlme et al. 2020); Decay sugar-rich substrates (Põlme et al. 2020) | Põlme S, et al. 2020. Fungal Diversity 105:1-16 ((https://doi.org/10.1007/s13225-020-00466-2)) |
| Otu42581 | 1 | k__Fungi(100);p__Ascomycota(100);c__Saccharomycetes(100);o__Saccharomycetales(100);f__Metschnikowiaceae(100);g__Metschnikowia(100);s__Metschnikowia_pulcherrima(100); | Metschnikowia | 13 | Saprotroph | \|Undefined Saprotroph\| | Yeast | NULL | Probable | As nectar-tap_saprotroph (Põlme et al. 2020); Decay sugar-rich substrates (Põlme et al. 2020) | Põlme S, et al. 2020. Fungal Diversity 105:1-16 ((https://doi.org/10.1007/s13225-020-00466-2)) |
| Otu42582 | 1 | k__Fungi(100);p__Ascomycota(100);c__Saccharomycetes(100);o__Saccharomycetales(100);f__Metschnikowiaceae(100);g__Metschnikowia(100);s__Metschnikowia_pulcherrima(100); | Metschnikowia | 13 | Saprotroph | \|Undefined Saprotroph\| | Yeast | NULL | Probable | As nectar-tap_saprotroph (Põlme et al. 2020); Decay sugar-rich substrates (Põlme et al. 2020) | Põlme S, et al. 2020. Fungal Diversity 105:1-16 ((https://doi.org/10.1007/s13225-020-00466-2)) |
| Otu42583 | 1 | k__Fungi(100);p__Ascomycota(100);c__Saccharomycetes(100);o__Saccharomycetales(100);f__Metschnikowiaceae(100);g__Metschnikowia(100);s__Metschnikowia_pulcherrima(100); | Metschnikowia | 13 | Saprotroph | \|Undefined Saprotroph\| | Yeast | NULL | Probable | As nectar-tap_saprotroph (Põlme et al. 2020); Decay sugar-rich substrates (Põlme et al. 2020) | Põlme S, et al. 2020. Fungal Diversity 105:1-16 ((https://doi.org/10.1007/s13225-020-00466-2)) |
| Otu42584 | 1 | k__Fungi(100);p__Ascomycota(100);c__Saccharomycetes(100);o__Saccharomycetales(100);f__Metschnikowiaceae(100);g__Metschnikowia(100);s__Metschnikowia_pulcherrima(100); | Metschnikowia | 13 | Saprotroph | \|Undefined Saprotroph\| | Yeast | NULL | Probable | As nectar-tap_saprotroph (Põlme et al. 2020); Decay sugar-rich substrates (Põlme et al. 2020) | Põlme S, et al. 2020. Fungal Diversity 105:1-16 ((https://doi.org/10.1007/s13225-020-00466-2)) |
| Otu42585 | 1 | k__Fungi(100);p__Ascomycota(100);c__Saccharomycetes(100);o__Saccharomycetales(100);f__Metschnikowiaceae(100);g__Metschnikowia(100);s__Metschnikowia_pulcherrima(100); | Metschnikowia | 13 | Saprotroph | \|Undefined Saprotroph\| | Yeast | NULL | Probable | As nectar-tap_saprotroph (Põlme et al. 2020); Decay sugar-rich substrates (Põlme et al. 2020) | Põlme S, et al. 2020. Fungal Diversity 105:1-16 ((https://doi.org/10.1007/s13225-020-00466-2)) |
| Otu42586 | 1 | k__Fungi(100);p__Ascomycota(100);c__Saccharomycetes(100);o__Saccharomycetales(100);f__Metschnikowiaceae(100);g__Metschnikowia(100);s__Metschnikowia_pulcherrima(100); | Metschnikowia | 13 | Saprotroph | \|Undefined Saprotroph\| | Yeast | NULL | Probable | As nectar-tap_saprotroph (Põlme et al. 2020); Decay sugar-rich substrates (Põlme et al. 2020) | Põlme S, et al. 2020. Fungal Diversity 105:1-16 ((https://doi.org/10.1007/s13225-020-00466-2)) |
| Otu42587 | 1 | k__Fungi(100);p__Ascomycota(100);c__Saccharomycetes(100);o__Saccharomycetales(100);f__Metschnikowiaceae(100);g__Metschnikowia(100);s__Metschnikowia_pulcherrima(100); | Metschnikowia | 13 | Saprotroph | \|Undefined Saprotroph\| | Yeast | NULL | Probable | As nectar-tap_saprotroph (Põlme et al. 2020); Decay sugar-rich substrates (Põlme et al. 2020) | Põlme S, et al. 2020. Fungal Diversity 105:1-16 ((https://doi.org/10.1007/s13225-020-00466-2)) |
| Otu42588 | 1 | k__Fungi(100);p__Ascomycota(100);c__Saccharomycetes(100);o__Saccharomycetales(100);f__Metschnikowiaceae(100);g__Metschnikowia(100);g__Metschnikowia_unclassified(100); | Metschnikowia | 13 | Saprotroph | \|Undefined Saprotroph\| | Yeast | NULL | Probable | As nectar-tap_saprotroph (Põlme et al. 2020); Decay sugar-rich substrates (Põlme et al. 2020) | Põlme S, et al. 2020. Fungal Diversity 105:1-16 ((https://doi.org/10.1007/s13225-020-00466-2)) |
| Otu42589 | 1 | k__Fungi(100);p__Ascomycota(100);c__Saccharomycetes(100);o__Saccharomycetales(100);f__Metschnikowiaceae(100);g__Metschnikowia(100);s__Metschnikowia_pulcherrima(100); | Metschnikowia | 13 | Saprotroph | \|Undefined Saprotroph\| | Yeast | NULL | Probable | As nectar-tap_saprotroph (Põlme et al. 2020); Decay sugar-rich substrates (Põlme et al. 2020) | Põlme S, et al. 2020. Fungal Diversity 105:1-16 ((https://doi.org/10.1007/s13225-020-00466-2)) |
| Otu42590 | 1 | k__Fungi(100);p__Ascomycota(100);c__Saccharomycetes(100);o__Saccharomycetales(100);f__Metschnikowiaceae(100);g__Metschnikowia(100);g__Metschnikowia_unclassified(100); | Metschnikowia | 13 | Saprotroph | \|Undefined Saprotroph\| | Yeast | NULL | Probable | As nectar-tap_saprotroph (Põlme et al. 2020); Decay sugar-rich substrates (Põlme et al. 2020) | Põlme S, et al. 2020. Fungal Diversity 105:1-16 ((https://doi.org/10.1007/s13225-020-00466-2)) |
| Otu42591 | 1 | k__Fungi(100);p__Ascomycota(100);c__Saccharomycetes(100);o__Saccharomycetales(100);f__Metschnikowiaceae(100);g__Metschnikowia(100);s__Metschnikowia_pulcherrima(100); | Metschnikowia | 13 | Saprotroph | \|Undefined Saprotroph\| | Yeast | NULL | Probable | As nectar-tap_saprotroph (Põlme et al. 2020); Decay sugar-rich substrates (Põlme et al. 2020) | Põlme S, et al. 2020. Fungal Diversity 105:1-16 ((https://doi.org/10.1007/s13225-020-00466-2)) |
| Otu42592 | 1 | k__Fungi(100);p__Ascomycota(100);c__Saccharomycetes(100);o__Saccharomycetales(100);f__Metschnikowiaceae(100);g__Metschnikowia(100);s__Metschnikowia_pulcherrima(100); | Metschnikowia | 13 | Saprotroph | \|Undefined Saprotroph\| | Yeast | NULL | Probable | As nectar-tap_saprotroph (Põlme et al. 2020); Decay sugar-rich substrates (Põlme et al. 2020) | Põlme S, et al. 2020. Fungal Diversity 105:1-16 ((https://doi.org/10.1007/s13225-020-00466-2)) |
| Otu42593 | 1 | k__Fungi(100);p__Ascomycota(100);c__Saccharomycetes(100);o__Saccharomycetales(100);f__Metschnikowiaceae(100);g__Metschnikowia(100);s__Metschnikowia_pulcherrima(100); | Metschnikowia | 13 | Saprotroph | \|Undefined Saprotroph\| | Yeast | NULL | Probable | As nectar-tap_saprotroph (Põlme et al. 2020); Decay sugar-rich substrates (Põlme et al. 2020) | Põlme S, et al. 2020. Fungal Diversity 105:1-16 ((https://doi.org/10.1007/s13225-020-00466-2)) |
| Otu42594 | 1 | k__Fungi(100);p__Ascomycota(100);c__Saccharomycetes(100);o__Saccharomycetales(100);f__Metschnikowiaceae(100);g__Metschnikowia(100);s__Metschnikowia_pulcherrima(100); | Metschnikowia | 13 | Saprotroph | \|Undefined Saprotroph\| | Yeast | NULL | Probable | As nectar-tap_saprotroph (Põlme et al. 2020); Decay sugar-rich substrates (Põlme et al. 2020) | Põlme S, et al. 2020. Fungal Diversity 105:1-16 ((https://doi.org/10.1007/s13225-020-00466-2)) |
| Otu42595 | 1 | k__Fungi(100);p__Ascomycota(100);c__Saccharomycetes(100);o__Saccharomycetales(100);f__Metschnikowiaceae(100);g__Metschnikowia(100);s__Metschnikowia_pulcherrima(100); | Metschnikowia | 13 | Saprotroph | \|Undefined Saprotroph\| | Yeast | NULL | Probable | As nectar-tap_saprotroph (Põlme et al. 2020); Decay sugar-rich substrates (Põlme et al. 2020) | Põlme S, et al. 2020. Fungal Diversity 105:1-16 ((https://doi.org/10.1007/s13225-020-00466-2)) |
| Otu42596 | 1 | k__Fungi(100);p__Ascomycota(100);c__Saccharomycetes(100);o__Saccharomycetales(100);f__Metschnikowiaceae(100);g__Metschnikowia(100);g__Metschnikowia_unclassified(100); | Metschnikowia | 13 | Saprotroph | \|Undefined Saprotroph\| | Yeast | NULL | Probable | As nectar-tap_saprotroph (Põlme et al. 2020); Decay sugar-rich substrates (Põlme et al. 2020) | Põlme S, et al. 2020. Fungal Diversity 105:1-16 ((https://doi.org/10.1007/s13225-020-00466-2)) |
| Otu42597 | 1 | k__Fungi(100);p__Ascomycota(100);c__Saccharomycetes(100);o__Saccharomycetales(100);f__Metschnikowiaceae(100);g__Metschnikowia(100);s__Metschnikowia_pulcherrima(100); | Metschnikowia | 13 | Saprotroph | \|Undefined Saprotroph\| | Yeast | NULL | Probable | As nectar-tap_saprotroph (Põlme et al. 2020); Decay sugar-rich substrates (Põlme et al. 2020) | Põlme S, et al. 2020. Fungal Diversity 105:1-16 ((https://doi.org/10.1007/s13225-020-00466-2)) |
| Otu42598 | 1 | k__Fungi(100);p__Ascomycota(100);c__Saccharomycetes(100);o__Saccharomycetales(100);f__Metschnikowiaceae(100);g__Metschnikowia(100);s__Metschnikowia_pulcherrima(100); | Metschnikowia | 13 | Saprotroph | \|Undefined Saprotroph\| | Yeast | NULL | Probable | As nectar-tap_saprotroph (Põlme et al. 2020); Decay sugar-rich substrates (Põlme et al. 2020) | Põlme S, et al. 2020. Fungal Diversity 105:1-16 ((https://doi.org/10.1007/s13225-020-00466-2)) |
| Otu42599 | 1 | k__Fungi(100);p__Ascomycota(100);c__Saccharomycetes(100);o__Saccharomycetales(100);f__Metschnikowiaceae(100);g__Metschnikowia(100);g__Metschnikowia_unclassified(100); | Metschnikowia | 13 | Saprotroph | \|Undefined Saprotroph\| | Yeast | NULL | Probable | As nectar-tap_saprotroph (Põlme et al. 2020); Decay sugar-rich substrates (Põlme et al. 2020) | Põlme S, et al. 2020. Fungal Diversity 105:1-16 ((https://doi.org/10.1007/s13225-020-00466-2)) |
| Otu42600 | 1 | k__Fungi(100);p__Ascomycota(100);c__Saccharomycetes(100);o__Saccharomycetales(100);f__Metschnikowiaceae(100);g__Metschnikowia(100);s__Metschnikowia_pulcherrima(100); | Metschnikowia | 13 | Saprotroph | \|Undefined Saprotroph\| | Yeast | NULL | Probable | As nectar-tap_saprotroph (Põlme et al. 2020); Decay sugar-rich substrates (Põlme et al. 2020) | Põlme S, et al. 2020. Fungal Diversity 105:1-16 ((https://doi.org/10.1007/s13225-020-00466-2)) |
| Otu42601 | 1 | k__Fungi(100);p__Ascomycota(100);c__Saccharomycetes(100);o__Saccharomycetales(100);f__Metschnikowiaceae(100);g__Metschnikowia(100);s__Metschnikowia_pulcherrima(100); | Metschnikowia | 13 | Saprotroph | \|Undefined Saprotroph\| | Yeast | NULL | Probable | As nectar-tap_saprotroph (Põlme et al. 2020); Decay sugar-rich substrates (Põlme et al. 2020) | Põlme S, et al. 2020. Fungal Diversity 105:1-16 ((https://doi.org/10.1007/s13225-020-00466-2)) |
| Otu42603 | 1 | k__Fungi(100);p__Ascomycota(100);c__Saccharomycetes(100);o__Saccharomycetales(100);f__Metschnikowiaceae(100);g__Metschnikowia(100);g__Metschnikowia_unclassified(100); | Metschnikowia | 13 | Saprotroph | \|Undefined Saprotroph\| | Yeast | NULL | Probable | As nectar-tap_saprotroph (Põlme et al. 2020); Decay sugar-rich substrates (Põlme et al. 2020) | Põlme S, et al. 2020. Fungal Diversity 105:1-16 ((https://doi.org/10.1007/s13225-020-00466-2)) |
| Otu42604 | 1 | k__Fungi(100);p__Ascomycota(100);c__Saccharomycetes(100);o__Saccharomycetales(100);f__Metschnikowiaceae(100);g__Metschnikowia(100);s__Metschnikowia_pulcherrima(100); | Metschnikowia | 13 | Saprotroph | \|Undefined Saprotroph\| | Yeast | NULL | Probable | As nectar-tap_saprotroph (Põlme et al. 2020); Decay sugar-rich substrates (Põlme et al. 2020) | Põlme S, et al. 2020. Fungal Diversity 105:1-16 ((https://doi.org/10.1007/s13225-020-00466-2)) |
| Otu42605 | 1 | k__Fungi(100);p__Ascomycota(100);c__Saccharomycetes(100);o__Saccharomycetales(100);f__Metschnikowiaceae(100);g__Metschnikowia(100);s__Metschnikowia_pulcherrima(100); | Metschnikowia | 13 | Saprotroph | \|Undefined Saprotroph\| | Yeast | NULL | Probable | As nectar-tap_saprotroph (Põlme et al. 2020); Decay sugar-rich substrates (Põlme et al. 2020) | Põlme S, et al. 2020. Fungal Diversity 105:1-16 ((https://doi.org/10.1007/s13225-020-00466-2)) |
| Otu42606 | 1 | k__Fungi(100);p__Ascomycota(100);c__Saccharomycetes(100);o__Saccharomycetales(100);f__Metschnikowiaceae(100);g__Metschnikowia(100);s__Metschnikowia_pulcherrima(100); | Metschnikowia | 13 | Saprotroph | \|Undefined Saprotroph\| | Yeast | NULL | Probable | As nectar-tap_saprotroph (Põlme et al. 2020); Decay sugar-rich substrates (Põlme et al. 2020) | Põlme S, et al. 2020. Fungal Diversity 105:1-16 ((https://doi.org/10.1007/s13225-020-00466-2)) |
| Otu42607 | 1 | k__Fungi(100);p__Ascomycota(100);c__Saccharomycetes(100);o__Saccharomycetales(100);f__Metschnikowiaceae(100);g__Metschnikowia(100);s__Metschnikowia_pulcherrima(100); | Metschnikowia | 13 | Saprotroph | \|Undefined Saprotroph\| | Yeast | NULL | Probable | As nectar-tap_saprotroph (Põlme et al. 2020); Decay sugar-rich substrates (Põlme et al. 2020) | Põlme S, et al. 2020. Fungal Diversity 105:1-16 ((https://doi.org/10.1007/s13225-020-00466-2)) |
| Otu42608 | 1 | k__Fungi(100);p__Ascomycota(100);c__Saccharomycetes(100);o__Saccharomycetales(100);f__Metschnikowiaceae(100);g__Metschnikowia(100);s__Metschnikowia_pulcherrima(100); | Metschnikowia | 13 | Saprotroph | \|Undefined Saprotroph\| | Yeast | NULL | Probable | As nectar-tap_saprotroph (Põlme et al. 2020); Decay sugar-rich substrates (Põlme et al. 2020) | Põlme S, et al. 2020. Fungal Diversity 105:1-16 ((https://doi.org/10.1007/s13225-020-00466-2)) |
| Otu42609 | 1 | k__Fungi(100);p__Ascomycota(100);c__Saccharomycetes(100);o__Saccharomycetales(100);f__Metschnikowiaceae(100);g__Metschnikowia(100);g__Metschnikowia_unclassified(100); | Metschnikowia | 13 | Saprotroph | \|Undefined Saprotroph\| | Yeast | NULL | Probable | As nectar-tap_saprotroph (Põlme et al. 2020); Decay sugar-rich substrates (Põlme et al. 2020) | Põlme S, et al. 2020. Fungal Diversity 105:1-16 ((https://doi.org/10.1007/s13225-020-00466-2)) |
| Otu42610 | 1 | k__Fungi(100);p__Ascomycota(100);c__Saccharomycetes(100);o__Saccharomycetales(100);f__Metschnikowiaceae(100);g__Metschnikowia(100);s__Metschnikowia_pulcherrima(100); | Metschnikowia | 13 | Saprotroph | \|Undefined Saprotroph\| | Yeast | NULL | Probable | As nectar-tap_saprotroph (Põlme et al. 2020); Decay sugar-rich substrates (Põlme et al. 2020) | Põlme S, et al. 2020. Fungal Diversity 105:1-16 ((https://doi.org/10.1007/s13225-020-00466-2)) |
| Otu42611 | 1 | k__Fungi(100);p__Ascomycota(100);c__Saccharomycetes(100);o__Saccharomycetales(100);f__Metschnikowiaceae(100);g__Metschnikowia(100);g__Metschnikowia_unclassified(100); | Metschnikowia | 13 | Saprotroph | \|Undefined Saprotroph\| | Yeast | NULL | Probable | As nectar-tap_saprotroph (Põlme et al. 2020); Decay sugar-rich substrates (Põlme et al. 2020) | Põlme S, et al. 2020. Fungal Diversity 105:1-16 ((https://doi.org/10.1007/s13225-020-00466-2)) |
| Otu42612 | 1 | k__Fungi(100);p__Ascomycota(100);c__Saccharomycetes(100);o__Saccharomycetales(100);f__Metschnikowiaceae(100);g__Metschnikowia(100);g__Metschnikowia_unclassified(100); | Metschnikowia | 13 | Saprotroph | \|Undefined Saprotroph\| | Yeast | NULL | Probable | As nectar-tap_saprotroph (Põlme et al. 2020); Decay sugar-rich substrates (Põlme et al. 2020) | Põlme S, et al. 2020. Fungal Diversity 105:1-16 ((https://doi.org/10.1007/s13225-020-00466-2)) |
| Otu42613 | 1 | k__Fungi(100);p__Ascomycota(100);c__Saccharomycetes(100);o__Saccharomycetales(100);f__Metschnikowiaceae(100);g__Metschnikowia(100);s__Metschnikowia_pulcherrima(100); | Metschnikowia | 13 | Saprotroph | \|Undefined Saprotroph\| | Yeast | NULL | Probable | As nectar-tap_saprotroph (Põlme et al. 2020); Decay sugar-rich substrates (Põlme et al. 2020) | Põlme S, et al. 2020. Fungal Diversity 105:1-16 ((https://doi.org/10.1007/s13225-020-00466-2)) |
| Otu42614 | 1 | k__Fungi(100);p__Ascomycota(100);c__Saccharomycetes(100);o__Saccharomycetales(100);f__Metschnikowiaceae(100);g__Metschnikowia(100);g__Metschnikowia_unclassified(100); | Metschnikowia | 13 | Saprotroph | \|Undefined Saprotroph\| | Yeast | NULL | Probable | As nectar-tap_saprotroph (Põlme et al. 2020); Decay sugar-rich substrates (Põlme et al. 2020) | Põlme S, et al. 2020. Fungal Diversity 105:1-16 ((https://doi.org/10.1007/s13225-020-00466-2)) |
| Otu42615 | 1 | k__Fungi(100);p__Ascomycota(100);c__Saccharomycetes(100);o__Saccharomycetales(100);f__Metschnikowiaceae(100);g__Metschnikowia(100);s__Metschnikowia_pulcherrima(100); | Metschnikowia | 13 | Saprotroph | \|Undefined Saprotroph\| | Yeast | NULL | Probable | As nectar-tap_saprotroph (Põlme et al. 2020); Decay sugar-rich substrates (Põlme et al. 2020) | Põlme S, et al. 2020. Fungal Diversity 105:1-16 ((https://doi.org/10.1007/s13225-020-00466-2)) |
| Otu42616 | 1 | k__Fungi(100);p__Ascomycota(100);c__Saccharomycetes(100);o__Saccharomycetales(100);f__Metschnikowiaceae(100);g__Metschnikowia(100);s__Metschnikowia_pulcherrima(100); | Metschnikowia | 13 | Saprotroph | \|Undefined Saprotroph\| | Yeast | NULL | Probable | As nectar-tap_saprotroph (Põlme et al. 2020); Decay sugar-rich substrates (Põlme et al. 2020) | Põlme S, et al. 2020. Fungal Diversity 105:1-16 ((https://doi.org/10.1007/s13225-020-00466-2)) |
| Otu42617 | 1 | k__Fungi(100);p__Ascomycota(100);c__Saccharomycetes(100);o__Saccharomycetales(100);f__Metschnikowiaceae(100);g__Metschnikowia(100);s__Metschnikowia_pulcherrima(100); | Metschnikowia | 13 | Saprotroph | \|Undefined Saprotroph\| | Yeast | NULL | Probable | As nectar-tap_saprotroph (Põlme et al. 2020); Decay sugar-rich substrates (Põlme et al. 2020) | Põlme S, et al. 2020. Fungal Diversity 105:1-16 ((https://doi.org/10.1007/s13225-020-00466-2)) |
| Otu42618 | 1 | k__Fungi(100);p__Ascomycota(100);c__Saccharomycetes(100);o__Saccharomycetales(100);f__Metschnikowiaceae(100);g__Metschnikowia(100);s__Metschnikowia_pulcherrima(100); | Metschnikowia | 13 | Saprotroph | \|Undefined Saprotroph\| | Yeast | NULL | Probable | As nectar-tap_saprotroph (Põlme et al. 2020); Decay sugar-rich substrates (Põlme et al. 2020) | Põlme S, et al. 2020. Fungal Diversity 105:1-16 ((https://doi.org/10.1007/s13225-020-00466-2)) |
| Otu42619 | 1 | k__Fungi(100);p__Ascomycota(100);c__Saccharomycetes(100);o__Saccharomycetales(100);f__Metschnikowiaceae(100);g__Metschnikowia(100);s__Metschnikowia_pulcherrima(100); | Metschnikowia | 13 | Saprotroph | \|Undefined Saprotroph\| | Yeast | NULL | Probable | As nectar-tap_saprotroph (Põlme et al. 2020); Decay sugar-rich substrates (Põlme et al. 2020) | Põlme S, et al. 2020. Fungal Diversity 105:1-16 ((https://doi.org/10.1007/s13225-020-00466-2)) |
| Otu42620 | 1 | k__Fungi(100);p__Ascomycota(100);c__Saccharomycetes(100);o__Saccharomycetales(100);f__Metschnikowiaceae(100);g__Metschnikowia(100);s__Metschnikowia_pulcherrima(100); | Metschnikowia | 13 | Saprotroph | \|Undefined Saprotroph\| | Yeast | NULL | Probable | As nectar-tap_saprotroph (Põlme et al. 2020); Decay sugar-rich substrates (Põlme et al. 2020) | Põlme S, et al. 2020. Fungal Diversity 105:1-16 ((https://doi.org/10.1007/s13225-020-00466-2)) |
| Otu42621 | 1 | k__Fungi(100);p__Ascomycota(100);c__Saccharomycetes(100);o__Saccharomycetales(100);f__Metschnikowiaceae(100);g__Metschnikowia(100);s__Metschnikowia_pulcherrima(100); | Metschnikowia | 13 | Saprotroph | \|Undefined Saprotroph\| | Yeast | NULL | Probable | As nectar-tap_saprotroph (Põlme et al. 2020); Decay sugar-rich substrates (Põlme et al. 2020) | Põlme S, et al. 2020. Fungal Diversity 105:1-16 ((https://doi.org/10.1007/s13225-020-00466-2)) |
| Otu42622 | 1 | k__Fungi(100);p__Ascomycota(100);c__Saccharomycetes(100);o__Saccharomycetales(100);f__Metschnikowiaceae(100);g__Metschnikowia(100);s__Metschnikowia_pulcherrima(100); | Metschnikowia | 13 | Saprotroph | \|Undefined Saprotroph\| | Yeast | NULL | Probable | As nectar-tap_saprotroph (Põlme et al. 2020); Decay sugar-rich substrates (Põlme et al. 2020) | Põlme S, et al. 2020. Fungal Diversity 105:1-16 ((https://doi.org/10.1007/s13225-020-00466-2)) |
| Otu42623 | 1 | k__Fungi(100);p__Ascomycota(100);c__Saccharomycetes(100);o__Saccharomycetales(100);f__Metschnikowiaceae(100);g__Metschnikowia(100);g__Metschnikowia_unclassified(100); | Metschnikowia | 13 | Saprotroph | \|Undefined Saprotroph\| | Yeast | NULL | Probable | As nectar-tap_saprotroph (Põlme et al. 2020); Decay sugar-rich substrates (Põlme et al. 2020) | Põlme S, et al. 2020. Fungal Diversity 105:1-16 ((https://doi.org/10.1007/s13225-020-00466-2)) |
| Otu42624 | 1 | k__Fungi(100);p__Ascomycota(100);c__Saccharomycetes(100);o__Saccharomycetales(100);f__Metschnikowiaceae(100);g__Metschnikowia(100);s__Metschnikowia_pulcherrima(100); | Metschnikowia | 13 | Saprotroph | \|Undefined Saprotroph\| | Yeast | NULL | Probable | As nectar-tap_saprotroph (Põlme et al. 2020); Decay sugar-rich substrates (Põlme et al. 2020) | Põlme S, et al. 2020. Fungal Diversity 105:1-16 ((https://doi.org/10.1007/s13225-020-00466-2)) |
| Otu42625 | 1 | k__Fungi(100);p__Ascomycota(100);c__Saccharomycetes(100);o__Saccharomycetales(100);f__Metschnikowiaceae(100);g__Metschnikowia(100);s__Metschnikowia_pulcherrima(100); | Metschnikowia | 13 | Saprotroph | \|Undefined Saprotroph\| | Yeast | NULL | Probable | As nectar-tap_saprotroph (Põlme et al. 2020); Decay sugar-rich substrates (Põlme et al. 2020) | Põlme S, et al. 2020. Fungal Diversity 105:1-16 ((https://doi.org/10.1007/s13225-020-00466-2)) |
| Otu42626 | 1 | k__Fungi(100);p__Ascomycota(100);c__Saccharomycetes(100);o__Saccharomycetales(100);f__Metschnikowiaceae(100);g__Metschnikowia(100);s__Metschnikowia_pulcherrima(100); | Metschnikowia | 13 | Saprotroph | \|Undefined Saprotroph\| | Yeast | NULL | Probable | As nectar-tap_saprotroph (Põlme et al. 2020); Decay sugar-rich substrates (Põlme et al. 2020) | Põlme S, et al. 2020. Fungal Diversity 105:1-16 ((https://doi.org/10.1007/s13225-020-00466-2)) |
| Otu42627 | 1 | k__Fungi(100);p__Ascomycota(100);c__Saccharomycetes(100);o__Saccharomycetales(100);f__Metschnikowiaceae(100);g__Metschnikowia(100);s__Metschnikowia_pulcherrima(100); | Metschnikowia | 13 | Saprotroph | \|Undefined Saprotroph\| | Yeast | NULL | Probable | As nectar-tap_saprotroph (Põlme et al. 2020); Decay sugar-rich substrates (Põlme et al. 2020) | Põlme S, et al. 2020. Fungal Diversity 105:1-16 ((https://doi.org/10.1007/s13225-020-00466-2)) |
| Otu42628 | 1 | k__Fungi(100);p__Ascomycota(100);c__Saccharomycetes(100);o__Saccharomycetales(100);f__Metschnikowiaceae(100);g__Metschnikowia(100);s__Metschnikowia_pulcherrima(100); | Metschnikowia | 13 | Saprotroph | \|Undefined Saprotroph\| | Yeast | NULL | Probable | As nectar-tap_saprotroph (Põlme et al. 2020); Decay sugar-rich substrates (Põlme et al. 2020) | Põlme S, et al. 2020. Fungal Diversity 105:1-16 ((https://doi.org/10.1007/s13225-020-00466-2)) |
| Otu42629 | 1 | k__Fungi(100);p__Ascomycota(100);c__Saccharomycetes(100);o__Saccharomycetales(100);f__Metschnikowiaceae(100);g__Metschnikowia(100);s__Metschnikowia_pulcherrima(100); | Metschnikowia | 13 | Saprotroph | \|Undefined Saprotroph\| | Yeast | NULL | Probable | As nectar-tap_saprotroph (Põlme et al. 2020); Decay sugar-rich substrates (Põlme et al. 2020) | Põlme S, et al. 2020. Fungal Diversity 105:1-16 ((https://doi.org/10.1007/s13225-020-00466-2)) |
| Otu42630 | 1 | k__Fungi(100);p__Ascomycota(100);c__Saccharomycetes(100);o__Saccharomycetales(100);f__Metschnikowiaceae(100);g__Metschnikowia(100);s__Metschnikowia_pulcherrima(100); | Metschnikowia | 13 | Saprotroph | \|Undefined Saprotroph\| | Yeast | NULL | Probable | As nectar-tap_saprotroph (Põlme et al. 2020); Decay sugar-rich substrates (Põlme et al. 2020) | Põlme S, et al. 2020. Fungal Diversity 105:1-16 ((https://doi.org/10.1007/s13225-020-00466-2)) |
| Otu42631 | 1 | k__Fungi(100);p__Ascomycota(100);c__Saccharomycetes(100);o__Saccharomycetales(100);f__Metschnikowiaceae(100);g__Metschnikowia(100);s__Metschnikowia_pulcherrima(100); | Metschnikowia | 13 | Saprotroph | \|Undefined Saprotroph\| | Yeast | NULL | Probable | As nectar-tap_saprotroph (Põlme et al. 2020); Decay sugar-rich substrates (Põlme et al. 2020) | Põlme S, et al. 2020. Fungal Diversity 105:1-16 ((https://doi.org/10.1007/s13225-020-00466-2)) |
| Otu42632 | 1 | k__Fungi(100);p__Ascomycota(100);c__Saccharomycetes(100);o__Saccharomycetales(100);f__Metschnikowiaceae(100);g__Metschnikowia(100);s__Metschnikowia_pulcherrima(100); | Metschnikowia | 13 | Saprotroph | \|Undefined Saprotroph\| | Yeast | NULL | Probable | As nectar-tap_saprotroph (Põlme et al. 2020); Decay sugar-rich substrates (Põlme et al. 2020) | Põlme S, et al. 2020. Fungal Diversity 105:1-16 ((https://doi.org/10.1007/s13225-020-00466-2)) |
| Otu42633 | 1 | k__Fungi(100);p__Ascomycota(100);c__Saccharomycetes(100);o__Saccharomycetales(100);f__Metschnikowiaceae(100);g__Metschnikowia(100);s__Metschnikowia_pulcherrima(100); | Metschnikowia | 13 | Saprotroph | \|Undefined Saprotroph\| | Yeast | NULL | Probable | As nectar-tap_saprotroph (Põlme et al. 2020); Decay sugar-rich substrates (Põlme et al. 2020) | Põlme S, et al. 2020. Fungal Diversity 105:1-16 ((https://doi.org/10.1007/s13225-020-00466-2)) |
| Otu42634 | 1 | k__Fungi(100);p__Ascomycota(100);c__Saccharomycetes(100);o__Saccharomycetales(100);f__Metschnikowiaceae(100);g__Metschnikowia(100);s__Metschnikowia_pulcherrima(100); | Metschnikowia | 13 | Saprotroph | \|Undefined Saprotroph\| | Yeast | NULL | Probable | As nectar-tap_saprotroph (Põlme et al. 2020); Decay sugar-rich substrates (Põlme et al. 2020) | Põlme S, et al. 2020. Fungal Diversity 105:1-16 ((https://doi.org/10.1007/s13225-020-00466-2)) |
| Otu42635 | 1 | k__Fungi(100);p__Ascomycota(100);c__Saccharomycetes(100);o__Saccharomycetales(100);f__Metschnikowiaceae(100);g__Metschnikowia(100);s__Metschnikowia_pulcherrima(100); | Metschnikowia | 13 | Saprotroph | \|Undefined Saprotroph\| | Yeast | NULL | Probable | As nectar-tap_saprotroph (Põlme et al. 2020); Decay sugar-rich substrates (Põlme et al. 2020) | Põlme S, et al. 2020. Fungal Diversity 105:1-16 ((https://doi.org/10.1007/s13225-020-00466-2)) |
| Otu42636 | 1 | k__Fungi(100);p__Ascomycota(100);c__Saccharomycetes(100);o__Saccharomycetales(100);f__Metschnikowiaceae(100);g__Metschnikowia(100);g__Metschnikowia_unclassified(100); | Metschnikowia | 13 | Saprotroph | \|Undefined Saprotroph\| | Yeast | NULL | Probable | As nectar-tap_saprotroph (Põlme et al. 2020); Decay sugar-rich substrates (Põlme et al. 2020) | Põlme S, et al. 2020. Fungal Diversity 105:1-16 ((https://doi.org/10.1007/s13225-020-00466-2)) |
| Otu42637 | 1 | k__Fungi(100);p__Ascomycota(100);c__Saccharomycetes(100);o__Saccharomycetales(100);f__Metschnikowiaceae(100);g__Metschnikowia(100);g__Metschnikowia_unclassified(100); | Metschnikowia | 13 | Saprotroph | \|Undefined Saprotroph\| | Yeast | NULL | Probable | As nectar-tap_saprotroph (Põlme et al. 2020); Decay sugar-rich substrates (Põlme et al. 2020) | Põlme S, et al. 2020. Fungal Diversity 105:1-16 ((https://doi.org/10.1007/s13225-020-00466-2)) |
| Otu42638 | 1 | k__Fungi(100);p__Ascomycota(100);c__Saccharomycetes(100);o__Saccharomycetales(100);f__Metschnikowiaceae(100);g__Metschnikowia(100);s__Metschnikowia_pulcherrima(100); | Metschnikowia | 13 | Saprotroph | \|Undefined Saprotroph\| | Yeast | NULL | Probable | As nectar-tap_saprotroph (Põlme et al. 2020); Decay sugar-rich substrates (Põlme et al. 2020) | Põlme S, et al. 2020. Fungal Diversity 105:1-16 ((https://doi.org/10.1007/s13225-020-00466-2)) |
| Otu42639 | 1 | k__Fungi(100);p__Ascomycota(100);c__Saccharomycetes(100);o__Saccharomycetales(100);f__Metschnikowiaceae(100);g__Metschnikowia(100);s__Metschnikowia_pulcherrima(100); | Metschnikowia | 13 | Saprotroph | \|Undefined Saprotroph\| | Yeast | NULL | Probable | As nectar-tap_saprotroph (Põlme et al. 2020); Decay sugar-rich substrates (Põlme et al. 2020) | Põlme S, et al. 2020. Fungal Diversity 105:1-16 ((https://doi.org/10.1007/s13225-020-00466-2)) |
| Otu42640 | 1 | k__Fungi(100);p__Ascomycota(100);c__Saccharomycetes(100);o__Saccharomycetales(100);f__Metschnikowiaceae(100);g__Metschnikowia(100);s__Metschnikowia_pulcherrima(100); | Metschnikowia | 13 | Saprotroph | \|Undefined Saprotroph\| | Yeast | NULL | Probable | As nectar-tap_saprotroph (Põlme et al. 2020); Decay sugar-rich substrates (Põlme et al. 2020) | Põlme S, et al. 2020. Fungal Diversity 105:1-16 ((https://doi.org/10.1007/s13225-020-00466-2)) |
| Otu42641 | 1 | k__Fungi(100);p__Ascomycota(100);c__Saccharomycetes(100);o__Saccharomycetales(100);f__Metschnikowiaceae(100);g__Metschnikowia(100);s__Metschnikowia_pulcherrima(100); | Metschnikowia | 13 | Saprotroph | \|Undefined Saprotroph\| | Yeast | NULL | Probable | As nectar-tap_saprotroph (Põlme et al. 2020); Decay sugar-rich substrates (Põlme et al. 2020) | Põlme S, et al. 2020. Fungal Diversity 105:1-16 ((https://doi.org/10.1007/s13225-020-00466-2)) |
| Otu42642 | 1 | k__Fungi(100);p__Ascomycota(100);c__Saccharomycetes(100);o__Saccharomycetales(100);f__Metschnikowiaceae(100);g__Metschnikowia(100);s__Metschnikowia_pulcherrima(100); | Metschnikowia | 13 | Saprotroph | \|Undefined Saprotroph\| | Yeast | NULL | Probable | As nectar-tap_saprotroph (Põlme et al. 2020); Decay sugar-rich substrates (Põlme et al. 2020) | Põlme S, et al. 2020. Fungal Diversity 105:1-16 ((https://doi.org/10.1007/s13225-020-00466-2)) |
| Otu42643 | 1 | k__Fungi(100);p__Ascomycota(100);c__Saccharomycetes(100);o__Saccharomycetales(100);f__Metschnikowiaceae(100);g__Metschnikowia(100);s__Metschnikowia_pulcherrima(100); | Metschnikowia | 13 | Saprotroph | \|Undefined Saprotroph\| | Yeast | NULL | Probable | As nectar-tap_saprotroph (Põlme et al. 2020); Decay sugar-rich substrates (Põlme et al. 2020) | Põlme S, et al. 2020. Fungal Diversity 105:1-16 ((https://doi.org/10.1007/s13225-020-00466-2)) |
| Otu42644 | 1 | k__Fungi(100);p__Ascomycota(100);c__Saccharomycetes(100);o__Saccharomycetales(100);f__Metschnikowiaceae(100);g__Metschnikowia(100);s__Metschnikowia_pulcherrima(100); | Metschnikowia | 13 | Saprotroph | \|Undefined Saprotroph\| | Yeast | NULL | Probable | As nectar-tap_saprotroph (Põlme et al. 2020); Decay sugar-rich substrates (Põlme et al. 2020) | Põlme S, et al. 2020. Fungal Diversity 105:1-16 ((https://doi.org/10.1007/s13225-020-00466-2)) |
| Otu42645 | 1 | k__Fungi(100);p__Ascomycota(100);c__Saccharomycetes(100);o__Saccharomycetales(100);f__Metschnikowiaceae(100);g__Metschnikowia(100);s__Metschnikowia_pulcherrima(100); | Metschnikowia | 13 | Saprotroph | \|Undefined Saprotroph\| | Yeast | NULL | Probable | As nectar-tap_saprotroph (Põlme et al. 2020); Decay sugar-rich substrates (Põlme et al. 2020) | Põlme S, et al. 2020. Fungal Diversity 105:1-16 ((https://doi.org/10.1007/s13225-020-00466-2)) |
| Otu42646 | 1 | k__Fungi(100);p__Ascomycota(100);c__Saccharomycetes(100);o__Saccharomycetales(100);f__Metschnikowiaceae(100);g__Metschnikowia(100);g__Metschnikowia_unclassified(100); | Metschnikowia | 13 | Saprotroph | \|Undefined Saprotroph\| | Yeast | NULL | Probable | As nectar-tap_saprotroph (Põlme et al. 2020); Decay sugar-rich substrates (Põlme et al. 2020) | Põlme S, et al. 2020. Fungal Diversity 105:1-16 ((https://doi.org/10.1007/s13225-020-00466-2)) |
| Otu42647 | 1 | k__Fungi(100);p__Ascomycota(100);c__Saccharomycetes(100);o__Saccharomycetales(100);f__Metschnikowiaceae(100);g__Metschnikowia(100);g__Metschnikowia_unclassified(100); | Metschnikowia | 13 | Saprotroph | \|Undefined Saprotroph\| | Yeast | NULL | Probable | As nectar-tap_saprotroph (Põlme et al. 2020); Decay sugar-rich substrates (Põlme et al. 2020) | Põlme S, et al. 2020. Fungal Diversity 105:1-16 ((https://doi.org/10.1007/s13225-020-00466-2)) |
| Otu42649 | 1 | k__Fungi(100);p__Basidiomycota(100);c__Tremellomycetes(100);o__Tremellales(100);f__Tremellales_family_Incertae_sedis(100);g__Hannaella(100);s__Hannaella_luteola(100); | Hannaella | 13 | Pathotroph-Saprotroph | \|Animal Parasite\|-Undefined Saprotroph | Yeast | NULL | Probable | As animal_decomposer (Põlme et al. 2020); Decay animal material (Põlme et al. 2020) | Põlme S, et al. 2020. Fungal Diversity 105:1-16 ((https://doi.org/10.1007/s13225-020-00466-2)) |
| Otu42650 | 1 | k__Fungi(100);p__Ascomycota(100);c__Saccharomycetes(100);o__Saccharomycetales(100);f__Metschnikowiaceae(100);g__Metschnikowia(100);s__Metschnikowia_pulcherrima(100); | Metschnikowia | 13 | Saprotroph | \|Undefined Saprotroph\| | Yeast | NULL | Probable | As nectar-tap_saprotroph (Põlme et al. 2020); Decay sugar-rich substrates (Põlme et al. 2020) | Põlme S, et al. 2020. Fungal Diversity 105:1-16 ((https://doi.org/10.1007/s13225-020-00466-2)) |
| Otu42651 | 1 | k__Fungi(100);p__Ascomycota(100);c__Saccharomycetes(100);o__Saccharomycetales(100);f__Metschnikowiaceae(100);g__Metschnikowia(100);s__Metschnikowia_pulcherrima(100); | Metschnikowia | 13 | Saprotroph | \|Undefined Saprotroph\| | Yeast | NULL | Probable | As nectar-tap_saprotroph (Põlme et al. 2020); Decay sugar-rich substrates (Põlme et al. 2020) | Põlme S, et al. 2020. Fungal Diversity 105:1-16 ((https://doi.org/10.1007/s13225-020-00466-2)) |
| Otu42652 | 1 | k__Fungi(100);p__Ascomycota(100);c__Saccharomycetes(100);o__Saccharomycetales(100);f__Metschnikowiaceae(100);g__Metschnikowia(100);g__Metschnikowia_unclassified(100); | Metschnikowia | 13 | Saprotroph | \|Undefined Saprotroph\| | Yeast | NULL | Probable | As nectar-tap_saprotroph (Põlme et al. 2020); Decay sugar-rich substrates (Põlme et al. 2020) | Põlme S, et al. 2020. Fungal Diversity 105:1-16 ((https://doi.org/10.1007/s13225-020-00466-2)) |
| Otu42653 | 1 | k__Fungi(100);p__Ascomycota(100);c__Saccharomycetes(100);o__Saccharomycetales(100);f__Metschnikowiaceae(100);g__Metschnikowia(100);s__Metschnikowia_pulcherrima(100); | Metschnikowia | 13 | Saprotroph | \|Undefined Saprotroph\| | Yeast | NULL | Probable | As nectar-tap_saprotroph (Põlme et al. 2020); Decay sugar-rich substrates (Põlme et al. 2020) | Põlme S, et al. 2020. Fungal Diversity 105:1-16 ((https://doi.org/10.1007/s13225-020-00466-2)) |
| Otu42654 | 1 | k__Fungi(100);p__Ascomycota(100);c__Saccharomycetes(100);o__Saccharomycetales(100);f__Metschnikowiaceae(100);g__Metschnikowia(100);g__Metschnikowia_unclassified(100); | Metschnikowia | 13 | Saprotroph | \|Undefined Saprotroph\| | Yeast | NULL | Probable | As nectar-tap_saprotroph (Põlme et al. 2020); Decay sugar-rich substrates (Põlme et al. 2020) | Põlme S, et al. 2020. Fungal Diversity 105:1-16 ((https://doi.org/10.1007/s13225-020-00466-2)) |
| Otu42655 | 1 | k__Fungi(100);p__Ascomycota(100);c__Saccharomycetes(100);o__Saccharomycetales(100);f__Metschnikowiaceae(100);g__Metschnikowia(100);s__Metschnikowia_pulcherrima(100); | Metschnikowia | 13 | Saprotroph | \|Undefined Saprotroph\| | Yeast | NULL | Probable | As nectar-tap_saprotroph (Põlme et al. 2020); Decay sugar-rich substrates (Põlme et al. 2020) | Põlme S, et al. 2020. Fungal Diversity 105:1-16 ((https://doi.org/10.1007/s13225-020-00466-2)) |
| Otu42656 | 1 | k__Fungi(100);p__Ascomycota(100);c__Saccharomycetes(100);o__Saccharomycetales(100);f__Metschnikowiaceae(100);g__Metschnikowia(100);s__Metschnikowia_pulcherrima(100); | Metschnikowia | 13 | Saprotroph | \|Undefined Saprotroph\| | Yeast | NULL | Probable | As nectar-tap_saprotroph (Põlme et al. 2020); Decay sugar-rich substrates (Põlme et al. 2020) | Põlme S, et al. 2020. Fungal Diversity 105:1-16 ((https://doi.org/10.1007/s13225-020-00466-2)) |
| Otu42657 | 1 | k__Fungi(100);p__Ascomycota(100);c__Saccharomycetes(100);o__Saccharomycetales(100);f__Metschnikowiaceae(100);g__Metschnikowia(100);s__Metschnikowia_pulcherrima(100); | Metschnikowia | 13 | Saprotroph | \|Undefined Saprotroph\| | Yeast | NULL | Probable | As nectar-tap_saprotroph (Põlme et al. 2020); Decay sugar-rich substrates (Põlme et al. 2020) | Põlme S, et al. 2020. Fungal Diversity 105:1-16 ((https://doi.org/10.1007/s13225-020-00466-2)) |
| Otu42658 | 1 | k__Fungi(100);p__Ascomycota(100);c__Saccharomycetes(100);o__Saccharomycetales(100);f__Metschnikowiaceae(100);g__Metschnikowia(100);s__Metschnikowia_pulcherrima(100); | Metschnikowia | 13 | Saprotroph | \|Undefined Saprotroph\| | Yeast | NULL | Probable | As nectar-tap_saprotroph (Põlme et al. 2020); Decay sugar-rich substrates (Põlme et al. 2020) | Põlme S, et al. 2020. Fungal Diversity 105:1-16 ((https://doi.org/10.1007/s13225-020-00466-2)) |
| Otu42659 | 1 | k__Fungi(100);p__Ascomycota(100);c__Saccharomycetes(100);o__Saccharomycetales(100);f__Metschnikowiaceae(100);g__Metschnikowia(100);s__Metschnikowia_pulcherrima(100); | Metschnikowia | 13 | Saprotroph | \|Undefined Saprotroph\| | Yeast | NULL | Probable | As nectar-tap_saprotroph (Põlme et al. 2020); Decay sugar-rich substrates (Põlme et al. 2020) | Põlme S, et al. 2020. Fungal Diversity 105:1-16 ((https://doi.org/10.1007/s13225-020-00466-2)) |
| Otu42660 | 1 | k__Fungi(100);p__Ascomycota(100);c__Saccharomycetes(100);o__Saccharomycetales(100);f__Metschnikowiaceae(100);g__Metschnikowia(100);g__Metschnikowia_unclassified(100); | Metschnikowia | 13 | Saprotroph | \|Undefined Saprotroph\| | Yeast | NULL | Probable | As nectar-tap_saprotroph (Põlme et al. 2020); Decay sugar-rich substrates (Põlme et al. 2020) | Põlme S, et al. 2020. Fungal Diversity 105:1-16 ((https://doi.org/10.1007/s13225-020-00466-2)) |
| Otu42661 | 1 | k__Fungi(100);p__Ascomycota(100);c__Saccharomycetes(100);o__Saccharomycetales(100);f__Metschnikowiaceae(100);g__Metschnikowia(100);s__Metschnikowia_pulcherrima(100); | Metschnikowia | 13 | Saprotroph | \|Undefined Saprotroph\| | Yeast | NULL | Probable | As nectar-tap_saprotroph (Põlme et al. 2020); Decay sugar-rich substrates (Põlme et al. 2020) | Põlme S, et al. 2020. Fungal Diversity 105:1-16 ((https://doi.org/10.1007/s13225-020-00466-2)) |
| Otu42662 | 1 | k__Fungi(100);p__Ascomycota(100);c__Saccharomycetes(100);o__Saccharomycetales(100);f__Metschnikowiaceae(100);g__Metschnikowia(100);s__Metschnikowia_pulcherrima(100); | Metschnikowia | 13 | Saprotroph | \|Undefined Saprotroph\| | Yeast | NULL | Probable | As nectar-tap_saprotroph (Põlme et al. 2020); Decay sugar-rich substrates (Põlme et al. 2020) | Põlme S, et al. 2020. Fungal Diversity 105:1-16 ((https://doi.org/10.1007/s13225-020-00466-2)) |
| Otu42663 | 1 | k__Fungi(100);p__Ascomycota(100);c__Saccharomycetes(100);o__Saccharomycetales(100);f__Metschnikowiaceae(100);g__Metschnikowia(100);s__Metschnikowia_pulcherrima(100); | Metschnikowia | 13 | Saprotroph | \|Undefined Saprotroph\| | Yeast | NULL | Probable | As nectar-tap_saprotroph (Põlme et al. 2020); Decay sugar-rich substrates (Põlme et al. 2020) | Põlme S, et al. 2020. Fungal Diversity 105:1-16 ((https://doi.org/10.1007/s13225-020-00466-2)) |
| Otu42664 | 1 | k__Fungi(100);p__Ascomycota(100);c__Saccharomycetes(100);o__Saccharomycetales(100);f__Metschnikowiaceae(100);g__Metschnikowia(100);s__Metschnikowia_pulcherrima(100); | Metschnikowia | 13 | Saprotroph | \|Undefined Saprotroph\| | Yeast | NULL | Probable | As nectar-tap_saprotroph (Põlme et al. 2020); Decay sugar-rich substrates (Põlme et al. 2020) | Põlme S, et al. 2020. Fungal Diversity 105:1-16 ((https://doi.org/10.1007/s13225-020-00466-2)) |
| Otu42665 | 1 | k__Fungi(100);p__Ascomycota(100);c__Saccharomycetes(100);o__Saccharomycetales(100);f__Metschnikowiaceae(100);g__Metschnikowia(100);s__Metschnikowia_pulcherrima(100); | Metschnikowia | 13 | Saprotroph | \|Undefined Saprotroph\| | Yeast | NULL | Probable | As nectar-tap_saprotroph (Põlme et al. 2020); Decay sugar-rich substrates (Põlme et al. 2020) | Põlme S, et al. 2020. Fungal Diversity 105:1-16 ((https://doi.org/10.1007/s13225-020-00466-2)) |
| Otu42666 | 1 | k__Fungi(100);p__Ascomycota(100);c__Saccharomycetes(100);o__Saccharomycetales(100);f__Metschnikowiaceae(100);g__Metschnikowia(100);s__Metschnikowia_pulcherrima(100); | Metschnikowia | 13 | Saprotroph | \|Undefined Saprotroph\| | Yeast | NULL | Probable | As nectar-tap_saprotroph (Põlme et al. 2020); Decay sugar-rich substrates (Põlme et al. 2020) | Põlme S, et al. 2020. Fungal Diversity 105:1-16 ((https://doi.org/10.1007/s13225-020-00466-2)) |
| Otu42667 | 1 | k__Fungi(100);p__Ascomycota(100);c__Saccharomycetes(100);o__Saccharomycetales(100);f__Metschnikowiaceae(100);g__Metschnikowia(100);s__Metschnikowia_pulcherrima(100); | Metschnikowia | 13 | Saprotroph | \|Undefined Saprotroph\| | Yeast | NULL | Probable | As nectar-tap_saprotroph (Põlme et al. 2020); Decay sugar-rich substrates (Põlme et al. 2020) | Põlme S, et al. 2020. Fungal Diversity 105:1-16 ((https://doi.org/10.1007/s13225-020-00466-2)) |
| Otu42668 | 1 | k__Fungi(100);p__Ascomycota(100);c__Saccharomycetes(100);o__Saccharomycetales(100);f__Metschnikowiaceae(100);g__Metschnikowia(100);s__Metschnikowia_pulcherrima(100); | Metschnikowia | 13 | Saprotroph | \|Undefined Saprotroph\| | Yeast | NULL | Probable | As nectar-tap_saprotroph (Põlme et al. 2020); Decay sugar-rich substrates (Põlme et al. 2020) | Põlme S, et al. 2020. Fungal Diversity 105:1-16 ((https://doi.org/10.1007/s13225-020-00466-2)) |
| Otu42669 | 1 | k__Fungi(100);p__Ascomycota(100);c__Saccharomycetes(100);o__Saccharomycetales(100);f__Metschnikowiaceae(100);g__Metschnikowia(100);s__Metschnikowia_pulcherrima(100); | Metschnikowia | 13 | Saprotroph | \|Undefined Saprotroph\| | Yeast | NULL | Probable | As nectar-tap_saprotroph (Põlme et al. 2020); Decay sugar-rich substrates (Põlme et al. 2020) | Põlme S, et al. 2020. Fungal Diversity 105:1-16 ((https://doi.org/10.1007/s13225-020-00466-2)) |
| Otu42670 | 1 | k__Fungi(100);p__Ascomycota(100);c__Saccharomycetes(100);o__Saccharomycetales(100);f__Metschnikowiaceae(100);g__Metschnikowia(100);s__Metschnikowia_pulcherrima(100); | Metschnikowia | 13 | Saprotroph | \|Undefined Saprotroph\| | Yeast | NULL | Probable | As nectar-tap_saprotroph (Põlme et al. 2020); Decay sugar-rich substrates (Põlme et al. 2020) | Põlme S, et al. 2020. Fungal Diversity 105:1-16 ((https://doi.org/10.1007/s13225-020-00466-2)) |
| Otu42671 | 1 | k__Fungi(100);p__Ascomycota(100);c__Saccharomycetes(100);o__Saccharomycetales(100);f__Metschnikowiaceae(100);g__Metschnikowia(100);s__Metschnikowia_pulcherrima(100); | Metschnikowia | 13 | Saprotroph | \|Undefined Saprotroph\| | Yeast | NULL | Probable | As nectar-tap_saprotroph (Põlme et al. 2020); Decay sugar-rich substrates (Põlme et al. 2020) | Põlme S, et al. 2020. Fungal Diversity 105:1-16 ((https://doi.org/10.1007/s13225-020-00466-2)) |
| Otu42672 | 1 | k__Fungi(100);p__Ascomycota(100);c__Saccharomycetes(100);o__Saccharomycetales(100);f__Metschnikowiaceae(100);g__Metschnikowia(100);s__Metschnikowia_pulcherrima(100); | Metschnikowia | 13 | Saprotroph | \|Undefined Saprotroph\| | Yeast | NULL | Probable | As nectar-tap_saprotroph (Põlme et al. 2020); Decay sugar-rich substrates (Põlme et al. 2020) | Põlme S, et al. 2020. Fungal Diversity 105:1-16 ((https://doi.org/10.1007/s13225-020-00466-2)) |
| Otu42673 | 1 | k__Fungi(100);p__Ascomycota(100);c__Saccharomycetes(100);o__Saccharomycetales(100);f__Metschnikowiaceae(100);g__Metschnikowia(100);s__Metschnikowia_pulcherrima(100); | Metschnikowia | 13 | Saprotroph | \|Undefined Saprotroph\| | Yeast | NULL | Probable | As nectar-tap_saprotroph (Põlme et al. 2020); Decay sugar-rich substrates (Põlme et al. 2020) | Põlme S, et al. 2020. Fungal Diversity 105:1-16 ((https://doi.org/10.1007/s13225-020-00466-2)) |
| Otu42674 | 1 | k__Fungi(100);p__Ascomycota(100);c__Saccharomycetes(100);o__Saccharomycetales(100);f__Metschnikowiaceae(100);g__Metschnikowia(100);s__Metschnikowia_pulcherrima(100); | Metschnikowia | 13 | Saprotroph | \|Undefined Saprotroph\| | Yeast | NULL | Probable | As nectar-tap_saprotroph (Põlme et al. 2020); Decay sugar-rich substrates (Põlme et al. 2020) | Põlme S, et al. 2020. Fungal Diversity 105:1-16 ((https://doi.org/10.1007/s13225-020-00466-2)) |
| Otu42675 | 1 | k__Fungi(100);p__Ascomycota(100);c__Saccharomycetes(100);o__Saccharomycetales(100);f__Metschnikowiaceae(100);g__Metschnikowia(100);s__Metschnikowia_pulcherrima(100); | Metschnikowia | 13 | Saprotroph | \|Undefined Saprotroph\| | Yeast | NULL | Probable | As nectar-tap_saprotroph (Põlme et al. 2020); Decay sugar-rich substrates (Põlme et al. 2020) | Põlme S, et al. 2020. Fungal Diversity 105:1-16 ((https://doi.org/10.1007/s13225-020-00466-2)) |
| Otu42676 | 1 | k__Fungi(100);p__Ascomycota(100);c__Saccharomycetes(100);o__Saccharomycetales(100);f__Metschnikowiaceae(100);g__Metschnikowia(100);s__Metschnikowia_pulcherrima(100); | Metschnikowia | 13 | Saprotroph | \|Undefined Saprotroph\| | Yeast | NULL | Probable | As nectar-tap_saprotroph (Põlme et al. 2020); Decay sugar-rich substrates (Põlme et al. 2020) | Põlme S, et al. 2020. Fungal Diversity 105:1-16 ((https://doi.org/10.1007/s13225-020-00466-2)) |
| Otu42677 | 1 | k__Fungi(100);p__Ascomycota(100);c__Saccharomycetes(100);o__Saccharomycetales(100);f__Metschnikowiaceae(100);g__Metschnikowia(100);s__Metschnikowia_pulcherrima(100); | Metschnikowia | 13 | Saprotroph | \|Undefined Saprotroph\| | Yeast | NULL | Probable | As nectar-tap_saprotroph (Põlme et al. 2020); Decay sugar-rich substrates (Põlme et al. 2020) | Põlme S, et al. 2020. Fungal Diversity 105:1-16 ((https://doi.org/10.1007/s13225-020-00466-2)) |
| Otu42678 | 1 | k__Fungi(100);p__Ascomycota(100);c__Saccharomycetes(100);o__Saccharomycetales(100);f__Metschnikowiaceae(100);g__Metschnikowia(100);s__Metschnikowia_pulcherrima(100); | Metschnikowia | 13 | Saprotroph | \|Undefined Saprotroph\| | Yeast | NULL | Probable | As nectar-tap_saprotroph (Põlme et al. 2020); Decay sugar-rich substrates (Põlme et al. 2020) | Põlme S, et al. 2020. Fungal Diversity 105:1-16 ((https://doi.org/10.1007/s13225-020-00466-2)) |
| Otu42679 | 1 | k__Fungi(100);p__Ascomycota(100);c__Saccharomycetes(100);o__Saccharomycetales(100);f__Metschnikowiaceae(100);g__Metschnikowia(100);s__Metschnikowia_pulcherrima(100); | Metschnikowia | 13 | Saprotroph | \|Undefined Saprotroph\| | Yeast | NULL | Probable | As nectar-tap_saprotroph (Põlme et al. 2020); Decay sugar-rich substrates (Põlme et al. 2020) | Põlme S, et al. 2020. Fungal Diversity 105:1-16 ((https://doi.org/10.1007/s13225-020-00466-2)) |
| Otu42680 | 1 | k__Fungi(100);p__Ascomycota(100);c__Saccharomycetes(100);o__Saccharomycetales(100);f__Metschnikowiaceae(100);g__Metschnikowia(100);g__Metschnikowia_unclassified(100); | Metschnikowia | 13 | Saprotroph | \|Undefined Saprotroph\| | Yeast | NULL | Probable | As nectar-tap_saprotroph (Põlme et al. 2020); Decay sugar-rich substrates (Põlme et al. 2020) | Põlme S, et al. 2020. Fungal Diversity 105:1-16 ((https://doi.org/10.1007/s13225-020-00466-2)) |
| Otu42681 | 1 | k__Fungi(100);p__Ascomycota(100);c__Saccharomycetes(100);o__Saccharomycetales(100);f__Metschnikowiaceae(100);g__Metschnikowia(100);g__Metschnikowia_unclassified(100); | Metschnikowia | 13 | Saprotroph | \|Undefined Saprotroph\| | Yeast | NULL | Probable | As nectar-tap_saprotroph (Põlme et al. 2020); Decay sugar-rich substrates (Põlme et al. 2020) | Põlme S, et al. 2020. Fungal Diversity 105:1-16 ((https://doi.org/10.1007/s13225-020-00466-2)) |
| Otu42682 | 1 | k__Fungi(100);p__Ascomycota(100);c__Saccharomycetes(100);o__Saccharomycetales(100);f__Metschnikowiaceae(100);g__Metschnikowia(100);s__Metschnikowia_pulcherrima(100); | Metschnikowia | 13 | Saprotroph | \|Undefined Saprotroph\| | Yeast | NULL | Probable | As nectar-tap_saprotroph (Põlme et al. 2020); Decay sugar-rich substrates (Põlme et al. 2020) | Põlme S, et al. 2020. Fungal Diversity 105:1-16 ((https://doi.org/10.1007/s13225-020-00466-2)) |
| Otu42683 | 1 | k__Fungi(100);p__Ascomycota(100);c__Saccharomycetes(100);o__Saccharomycetales(100);f__Metschnikowiaceae(100);g__Metschnikowia(100);s__Metschnikowia_pulcherrima(100); | Metschnikowia | 13 | Saprotroph | \|Undefined Saprotroph\| | Yeast | NULL | Probable | As nectar-tap_saprotroph (Põlme et al. 2020); Decay sugar-rich substrates (Põlme et al. 2020) | Põlme S, et al. 2020. Fungal Diversity 105:1-16 ((https://doi.org/10.1007/s13225-020-00466-2)) |
| Otu42684 | 1 | k__Fungi(100);p__Ascomycota(100);c__Saccharomycetes(100);o__Saccharomycetales(100);f__Metschnikowiaceae(100);g__Metschnikowia(100);s__Metschnikowia_pulcherrima(100); | Metschnikowia | 13 | Saprotroph | \|Undefined Saprotroph\| | Yeast | NULL | Probable | As nectar-tap_saprotroph (Põlme et al. 2020); Decay sugar-rich substrates (Põlme et al. 2020) | Põlme S, et al. 2020. Fungal Diversity 105:1-16 ((https://doi.org/10.1007/s13225-020-00466-2)) |
| Otu42685 | 1 | k__Fungi(100);p__Ascomycota(100);c__Saccharomycetes(100);o__Saccharomycetales(100);f__Metschnikowiaceae(100);g__Metschnikowia(100);g__Metschnikowia_unclassified(100); | Metschnikowia | 13 | Saprotroph | \|Undefined Saprotroph\| | Yeast | NULL | Probable | As nectar-tap_saprotroph (Põlme et al. 2020); Decay sugar-rich substrates (Põlme et al. 2020) | Põlme S, et al. 2020. Fungal Diversity 105:1-16 ((https://doi.org/10.1007/s13225-020-00466-2)) |
| Otu42686 | 1 | k__Fungi(100);p__Ascomycota(100);c__Saccharomycetes(100);o__Saccharomycetales(100);f__Metschnikowiaceae(100);g__Metschnikowia(100);s__Metschnikowia_pulcherrima(100); | Metschnikowia | 13 | Saprotroph | \|Undefined Saprotroph\| | Yeast | NULL | Probable | As nectar-tap_saprotroph (Põlme et al. 2020); Decay sugar-rich substrates (Põlme et al. 2020) | Põlme S, et al. 2020. Fungal Diversity 105:1-16 ((https://doi.org/10.1007/s13225-020-00466-2)) |
| Otu42687 | 1 | k__Fungi(100);p__Ascomycota(100);c__Saccharomycetes(100);o__Saccharomycetales(100);f__Metschnikowiaceae(100);g__Metschnikowia(100);s__Metschnikowia_pulcherrima(100); | Metschnikowia | 13 | Saprotroph | \|Undefined Saprotroph\| | Yeast | NULL | Probable | As nectar-tap_saprotroph (Põlme et al. 2020); Decay sugar-rich substrates (Põlme et al. 2020) | Põlme S, et al. 2020. Fungal Diversity 105:1-16 ((https://doi.org/10.1007/s13225-020-00466-2)) |
| Otu42688 | 1 | k__Fungi(100);p__Ascomycota(100);c__Saccharomycetes(100);o__Saccharomycetales(100);f__Metschnikowiaceae(100);g__Metschnikowia(100);s__Metschnikowia_pulcherrima(100); | Metschnikowia | 13 | Saprotroph | \|Undefined Saprotroph\| | Yeast | NULL | Probable | As nectar-tap_saprotroph (Põlme et al. 2020); Decay sugar-rich substrates (Põlme et al. 2020) | Põlme S, et al. 2020. Fungal Diversity 105:1-16 ((https://doi.org/10.1007/s13225-020-00466-2)) |
| Otu42689 | 1 | k__Fungi(100);p__Ascomycota(100);c__Saccharomycetes(100);o__Saccharomycetales(100);f__Metschnikowiaceae(100);g__Metschnikowia(100);g__Metschnikowia_unclassified(100); | Metschnikowia | 13 | Saprotroph | \|Undefined Saprotroph\| | Yeast | NULL | Probable | As nectar-tap_saprotroph (Põlme et al. 2020); Decay sugar-rich substrates (Põlme et al. 2020) | Põlme S, et al. 2020. Fungal Diversity 105:1-16 ((https://doi.org/10.1007/s13225-020-00466-2)) |
| Otu42690 | 1 | k__Fungi(100);p__Ascomycota(100);c__Saccharomycetes(100);o__Saccharomycetales(100);f__Metschnikowiaceae(100);g__Metschnikowia(100);s__Metschnikowia_pulcherrima(100); | Metschnikowia | 13 | Saprotroph | \|Undefined Saprotroph\| | Yeast | NULL | Probable | As nectar-tap_saprotroph (Põlme et al. 2020); Decay sugar-rich substrates (Põlme et al. 2020) | Põlme S, et al. 2020. Fungal Diversity 105:1-16 ((https://doi.org/10.1007/s13225-020-00466-2)) |
| Otu42691 | 1 | k__Fungi(100);p__Ascomycota(100);c__Saccharomycetes(100);o__Saccharomycetales(100);f__Metschnikowiaceae(100);g__Metschnikowia(100);s__Metschnikowia_pulcherrima(100); | Metschnikowia | 13 | Saprotroph | \|Undefined Saprotroph\| | Yeast | NULL | Probable | As nectar-tap_saprotroph (Põlme et al. 2020); Decay sugar-rich substrates (Põlme et al. 2020) | Põlme S, et al. 2020. Fungal Diversity 105:1-16 ((https://doi.org/10.1007/s13225-020-00466-2)) |
| Otu42692 | 1 | k__Fungi(100);p__Ascomycota(100);c__Saccharomycetes(100);o__Saccharomycetales(100);f__Metschnikowiaceae(100);g__Metschnikowia(100);g__Metschnikowia_unclassified(100); | Metschnikowia | 13 | Saprotroph | \|Undefined Saprotroph\| | Yeast | NULL | Probable | As nectar-tap_saprotroph (Põlme et al. 2020); Decay sugar-rich substrates (Põlme et al. 2020) | Põlme S, et al. 2020. Fungal Diversity 105:1-16 ((https://doi.org/10.1007/s13225-020-00466-2)) |
| Otu42693 | 1 | k__Fungi(100);p__Ascomycota(100);c__Saccharomycetes(100);o__Saccharomycetales(100);f__Metschnikowiaceae(100);g__Metschnikowia(100);s__Metschnikowia_pulcherrima(100); | Metschnikowia | 13 | Saprotroph | \|Undefined Saprotroph\| | Yeast | NULL | Probable | As nectar-tap_saprotroph (Põlme et al. 2020); Decay sugar-rich substrates (Põlme et al. 2020) | Põlme S, et al. 2020. Fungal Diversity 105:1-16 ((https://doi.org/10.1007/s13225-020-00466-2)) |
| Otu42694 | 1 | k__Fungi(100);p__Ascomycota(100);c__Saccharomycetes(100);o__Saccharomycetales(100);f__Metschnikowiaceae(100);g__Metschnikowia(100);s__Metschnikowia_pulcherrima(100); | Metschnikowia | 13 | Saprotroph | \|Undefined Saprotroph\| | Yeast | NULL | Probable | As nectar-tap_saprotroph (Põlme et al. 2020); Decay sugar-rich substrates (Põlme et al. 2020) | Põlme S, et al. 2020. Fungal Diversity 105:1-16 ((https://doi.org/10.1007/s13225-020-00466-2)) |
| Otu42695 | 1 | k__Fungi(100);p__Ascomycota(100);c__Saccharomycetes(100);o__Saccharomycetales(100);f__Metschnikowiaceae(100);g__Metschnikowia(100);s__Metschnikowia_pulcherrima(100); | Metschnikowia | 13 | Saprotroph | \|Undefined Saprotroph\| | Yeast | NULL | Probable | As nectar-tap_saprotroph (Põlme et al. 2020); Decay sugar-rich substrates (Põlme et al. 2020) | Põlme S, et al. 2020. Fungal Diversity 105:1-16 ((https://doi.org/10.1007/s13225-020-00466-2)) |
| Otu42696 | 1 | k__Fungi(100);p__Ascomycota(100);c__Saccharomycetes(100);o__Saccharomycetales(100);f__Metschnikowiaceae(100);g__Metschnikowia(100);s__Metschnikowia_pulcherrima(100); | Metschnikowia | 13 | Saprotroph | \|Undefined Saprotroph\| | Yeast | NULL | Probable | As nectar-tap_saprotroph (Põlme et al. 2020); Decay sugar-rich substrates (Põlme et al. 2020) | Põlme S, et al. 2020. Fungal Diversity 105:1-16 ((https://doi.org/10.1007/s13225-020-00466-2)) |
| Otu42697 | 1 | k__Fungi(100);p__Ascomycota(100);c__Saccharomycetes(100);o__Saccharomycetales(100);f__Metschnikowiaceae(100);g__Metschnikowia(100);s__Metschnikowia_pulcherrima(100); | Metschnikowia | 13 | Saprotroph | \|Undefined Saprotroph\| | Yeast | NULL | Probable | As nectar-tap_saprotroph (Põlme et al. 2020); Decay sugar-rich substrates (Põlme et al. 2020) | Põlme S, et al. 2020. Fungal Diversity 105:1-16 ((https://doi.org/10.1007/s13225-020-00466-2)) |
| Otu42698 | 1 | k__Fungi(100);p__Ascomycota(100);c__Saccharomycetes(100);o__Saccharomycetales(100);f__Metschnikowiaceae(100);g__Metschnikowia(100);g__Metschnikowia_unclassified(100); | Metschnikowia | 13 | Saprotroph | \|Undefined Saprotroph\| | Yeast | NULL | Probable | As nectar-tap_saprotroph (Põlme et al. 2020); Decay sugar-rich substrates (Põlme et al. 2020) | Põlme S, et al. 2020. Fungal Diversity 105:1-16 ((https://doi.org/10.1007/s13225-020-00466-2)) |
| Otu42699 | 1 | k__Fungi(100);p__Ascomycota(100);c__Saccharomycetes(100);o__Saccharomycetales(100);f__Metschnikowiaceae(100);g__Metschnikowia(100);s__Metschnikowia_pulcherrima(100); | Metschnikowia | 13 | Saprotroph | \|Undefined Saprotroph\| | Yeast | NULL | Probable | As nectar-tap_saprotroph (Põlme et al. 2020); Decay sugar-rich substrates (Põlme et al. 2020) | Põlme S, et al. 2020. Fungal Diversity 105:1-16 ((https://doi.org/10.1007/s13225-020-00466-2)) |
| Otu42700 | 1 | k__Fungi(100);p__Ascomycota(100);c__Saccharomycetes(100);o__Saccharomycetales(100);f__Metschnikowiaceae(100);g__Metschnikowia(100);s__Metschnikowia_pulcherrima(100); | Metschnikowia | 13 | Saprotroph | \|Undefined Saprotroph\| | Yeast | NULL | Probable | As nectar-tap_saprotroph (Põlme et al. 2020); Decay sugar-rich substrates (Põlme et al. 2020) | Põlme S, et al. 2020. Fungal Diversity 105:1-16 ((https://doi.org/10.1007/s13225-020-00466-2)) |
| Otu42701 | 1 | k__Fungi(100);p__Ascomycota(100);c__Saccharomycetes(100);o__Saccharomycetales(100);f__Metschnikowiaceae(100);g__Metschnikowia(100);g__Metschnikowia_unclassified(100); | Metschnikowia | 13 | Saprotroph | \|Undefined Saprotroph\| | Yeast | NULL | Probable | As nectar-tap_saprotroph (Põlme et al. 2020); Decay sugar-rich substrates (Põlme et al. 2020) | Põlme S, et al. 2020. Fungal Diversity 105:1-16 ((https://doi.org/10.1007/s13225-020-00466-2)) |
| Otu42702 | 1 | k__Fungi(100);p__Ascomycota(100);c__Saccharomycetes(100);o__Saccharomycetales(100);f__Metschnikowiaceae(100);g__Metschnikowia(100);g__Metschnikowia_unclassified(100); | Metschnikowia | 13 | Saprotroph | \|Undefined Saprotroph\| | Yeast | NULL | Probable | As nectar-tap_saprotroph (Põlme et al. 2020); Decay sugar-rich substrates (Põlme et al. 2020) | Põlme S, et al. 2020. Fungal Diversity 105:1-16 ((https://doi.org/10.1007/s13225-020-00466-2)) |
| Otu42703 | 1 | k__Fungi(100);p__Ascomycota(100);c__Saccharomycetes(100);o__Saccharomycetales(100);f__Metschnikowiaceae(100);g__Metschnikowia(100);s__Metschnikowia_pulcherrima(100); | Metschnikowia | 13 | Saprotroph | \|Undefined Saprotroph\| | Yeast | NULL | Probable | As nectar-tap_saprotroph (Põlme et al. 2020); Decay sugar-rich substrates (Põlme et al. 2020) | Põlme S, et al. 2020. Fungal Diversity 105:1-16 ((https://doi.org/10.1007/s13225-020-00466-2)) |
| Otu42704 | 1 | k__Fungi(100);p__Ascomycota(100);c__Saccharomycetes(100);o__Saccharomycetales(100);f__Metschnikowiaceae(100);g__Metschnikowia(100);s__Metschnikowia_pulcherrima(100); | Metschnikowia | 13 | Saprotroph | \|Undefined Saprotroph\| | Yeast | NULL | Probable | As nectar-tap_saprotroph (Põlme et al. 2020); Decay sugar-rich substrates (Põlme et al. 2020) | Põlme S, et al. 2020. Fungal Diversity 105:1-16 ((https://doi.org/10.1007/s13225-020-00466-2)) |
| Otu42705 | 1 | k__Fungi(100);p__Ascomycota(100);c__Saccharomycetes(100);o__Saccharomycetales(100);f__Metschnikowiaceae(100);g__Metschnikowia(100);s__Metschnikowia_pulcherrima(100); | Metschnikowia | 13 | Saprotroph | \|Undefined Saprotroph\| | Yeast | NULL | Probable | As nectar-tap_saprotroph (Põlme et al. 2020); Decay sugar-rich substrates (Põlme et al. 2020) | Põlme S, et al. 2020. Fungal Diversity 105:1-16 ((https://doi.org/10.1007/s13225-020-00466-2)) |
| Otu42706 | 1 | k__Fungi(100);p__Ascomycota(100);c__Saccharomycetes(100);o__Saccharomycetales(100);f__Metschnikowiaceae(100);g__Metschnikowia(100);s__Metschnikowia_pulcherrima(100); | Metschnikowia | 13 | Saprotroph | \|Undefined Saprotroph\| | Yeast | NULL | Probable | As nectar-tap_saprotroph (Põlme et al. 2020); Decay sugar-rich substrates (Põlme et al. 2020) | Põlme S, et al. 2020. Fungal Diversity 105:1-16 ((https://doi.org/10.1007/s13225-020-00466-2)) |
| Otu42707 | 1 | k__Fungi(100);p__Ascomycota(100);c__Saccharomycetes(100);o__Saccharomycetales(100);f__Metschnikowiaceae(100);g__Metschnikowia(100);s__Metschnikowia_pulcherrima(100); | Metschnikowia | 13 | Saprotroph | \|Undefined Saprotroph\| | Yeast | NULL | Probable | As nectar-tap_saprotroph (Põlme et al. 2020); Decay sugar-rich substrates (Põlme et al. 2020) | Põlme S, et al. 2020. Fungal Diversity 105:1-16 ((https://doi.org/10.1007/s13225-020-00466-2)) |
| Otu42708 | 1 | k__Fungi(100);p__Ascomycota(100);c__Saccharomycetes(100);o__Saccharomycetales(100);f__Metschnikowiaceae(100);g__Metschnikowia(100);s__Metschnikowia_pulcherrima(100); | Metschnikowia | 13 | Saprotroph | \|Undefined Saprotroph\| | Yeast | NULL | Probable | As nectar-tap_saprotroph (Põlme et al. 2020); Decay sugar-rich substrates (Põlme et al. 2020) | Põlme S, et al. 2020. Fungal Diversity 105:1-16 ((https://doi.org/10.1007/s13225-020-00466-2)) |
| Otu42709 | 1 | k__Fungi(100);p__Ascomycota(100);c__Saccharomycetes(100);o__Saccharomycetales(100);f__Metschnikowiaceae(100);g__Metschnikowia(100);s__Metschnikowia_pulcherrima(100); | Metschnikowia | 13 | Saprotroph | \|Undefined Saprotroph\| | Yeast | NULL | Probable | As nectar-tap_saprotroph (Põlme et al. 2020); Decay sugar-rich substrates (Põlme et al. 2020) | Põlme S, et al. 2020. Fungal Diversity 105:1-16 ((https://doi.org/10.1007/s13225-020-00466-2)) |
| Otu42710 | 1 | k__Fungi(100);p__Ascomycota(100);c__Saccharomycetes(100);o__Saccharomycetales(100);f__Metschnikowiaceae(100);g__Metschnikowia(100);s__Metschnikowia_pulcherrima(100); | Metschnikowia | 13 | Saprotroph | \|Undefined Saprotroph\| | Yeast | NULL | Probable | As nectar-tap_saprotroph (Põlme et al. 2020); Decay sugar-rich substrates (Põlme et al. 2020) | Põlme S, et al. 2020. Fungal Diversity 105:1-16 ((https://doi.org/10.1007/s13225-020-00466-2)) |
| Otu42711 | 1 | k__Fungi(100);p__Ascomycota(100);c__Saccharomycetes(100);o__Saccharomycetales(100);f__Metschnikowiaceae(100);g__Metschnikowia(100);s__Metschnikowia_pulcherrima(100); | Metschnikowia | 13 | Saprotroph | \|Undefined Saprotroph\| | Yeast | NULL | Probable | As nectar-tap_saprotroph (Põlme et al. 2020); Decay sugar-rich substrates (Põlme et al. 2020) | Põlme S, et al. 2020. Fungal Diversity 105:1-16 ((https://doi.org/10.1007/s13225-020-00466-2)) |
| Otu42712 | 1 | k__Fungi(100);p__Ascomycota(100);c__Saccharomycetes(100);o__Saccharomycetales(100);f__Metschnikowiaceae(100);g__Metschnikowia(100);s__Metschnikowia_pulcherrima(100); | Metschnikowia | 13 | Saprotroph | \|Undefined Saprotroph\| | Yeast | NULL | Probable | As nectar-tap_saprotroph (Põlme et al. 2020); Decay sugar-rich substrates (Põlme et al. 2020) | Põlme S, et al. 2020. Fungal Diversity 105:1-16 ((https://doi.org/10.1007/s13225-020-00466-2)) |
| Otu42713 | 1 | k__Fungi(100);p__Ascomycota(100);c__Saccharomycetes(100);o__Saccharomycetales(100);f__Metschnikowiaceae(100);g__Metschnikowia(100);s__Metschnikowia_pulcherrima(100); | Metschnikowia | 13 | Saprotroph | \|Undefined Saprotroph\| | Yeast | NULL | Probable | As nectar-tap_saprotroph (Põlme et al. 2020); Decay sugar-rich substrates (Põlme et al. 2020) | Põlme S, et al. 2020. Fungal Diversity 105:1-16 ((https://doi.org/10.1007/s13225-020-00466-2)) |
| Otu42714 | 1 | k__Fungi(100);p__Ascomycota(100);c__Saccharomycetes(100);o__Saccharomycetales(100);f__Metschnikowiaceae(100);g__Metschnikowia(100);s__Metschnikowia_pulcherrima(100); | Metschnikowia | 13 | Saprotroph | \|Undefined Saprotroph\| | Yeast | NULL | Probable | As nectar-tap_saprotroph (Põlme et al. 2020); Decay sugar-rich substrates (Põlme et al. 2020) | Põlme S, et al. 2020. Fungal Diversity 105:1-16 ((https://doi.org/10.1007/s13225-020-00466-2)) |
| Otu42715 | 1 | k__Fungi(100);p__Ascomycota(100);c__Saccharomycetes(100);o__Saccharomycetales(100);f__Metschnikowiaceae(100);g__Metschnikowia(100);s__Metschnikowia_pulcherrima(100); | Metschnikowia | 13 | Saprotroph | \|Undefined Saprotroph\| | Yeast | NULL | Probable | As nectar-tap_saprotroph (Põlme et al. 2020); Decay sugar-rich substrates (Põlme et al. 2020) | Põlme S, et al. 2020. Fungal Diversity 105:1-16 ((https://doi.org/10.1007/s13225-020-00466-2)) |
| Otu42716 | 1 | k__Fungi(100);p__Ascomycota(100);c__Saccharomycetes(100);o__Saccharomycetales(100);f__Metschnikowiaceae(100);g__Metschnikowia(100);s__Metschnikowia_pulcherrima(100); | Metschnikowia | 13 | Saprotroph | \|Undefined Saprotroph\| | Yeast | NULL | Probable | As nectar-tap_saprotroph (Põlme et al. 2020); Decay sugar-rich substrates (Põlme et al. 2020) | Põlme S, et al. 2020. Fungal Diversity 105:1-16 ((https://doi.org/10.1007/s13225-020-00466-2)) |
| Otu42717 | 1 | k__Fungi(100);p__Ascomycota(100);c__Saccharomycetes(100);o__Saccharomycetales(100);f__Metschnikowiaceae(100);g__Metschnikowia(100);s__Metschnikowia_pulcherrima(100); | Metschnikowia | 13 | Saprotroph | \|Undefined Saprotroph\| | Yeast | NULL | Probable | As nectar-tap_saprotroph (Põlme et al. 2020); Decay sugar-rich substrates (Põlme et al. 2020) | Põlme S, et al. 2020. Fungal Diversity 105:1-16 ((https://doi.org/10.1007/s13225-020-00466-2)) |
| Otu42718 | 1 | k__Fungi(100);p__Ascomycota(100);c__Saccharomycetes(100);o__Saccharomycetales(100);f__Metschnikowiaceae(100);g__Metschnikowia(100);g__Metschnikowia_unclassified(100); | Metschnikowia | 13 | Saprotroph | \|Undefined Saprotroph\| | Yeast | NULL | Probable | As nectar-tap_saprotroph (Põlme et al. 2020); Decay sugar-rich substrates (Põlme et al. 2020) | Põlme S, et al. 2020. Fungal Diversity 105:1-16 ((https://doi.org/10.1007/s13225-020-00466-2)) |
| Otu42719 | 1 | k__Fungi(100);p__Ascomycota(100);c__Saccharomycetes(100);o__Saccharomycetales(100);f__Metschnikowiaceae(100);g__Metschnikowia(100);s__Metschnikowia_pulcherrima(100); | Metschnikowia | 13 | Saprotroph | \|Undefined Saprotroph\| | Yeast | NULL | Probable | As nectar-tap_saprotroph (Põlme et al. 2020); Decay sugar-rich substrates (Põlme et al. 2020) | Põlme S, et al. 2020. Fungal Diversity 105:1-16 ((https://doi.org/10.1007/s13225-020-00466-2)) |
| Otu42720 | 1 | k__Fungi(100);p__Ascomycota(100);c__Saccharomycetes(100);o__Saccharomycetales(100);f__Metschnikowiaceae(100);g__Metschnikowia(100);g__Metschnikowia_unclassified(100); | Metschnikowia | 13 | Saprotroph | \|Undefined Saprotroph\| | Yeast | NULL | Probable | As nectar-tap_saprotroph (Põlme et al. 2020); Decay sugar-rich substrates (Põlme et al. 2020) | Põlme S, et al. 2020. Fungal Diversity 105:1-16 ((https://doi.org/10.1007/s13225-020-00466-2)) |
| Otu42721 | 1 | k__Fungi(100);p__Ascomycota(100);c__Saccharomycetes(100);o__Saccharomycetales(100);f__Metschnikowiaceae(100);g__Metschnikowia(100);g__Metschnikowia_unclassified(100); | Metschnikowia | 13 | Saprotroph | \|Undefined Saprotroph\| | Yeast | NULL | Probable | As nectar-tap_saprotroph (Põlme et al. 2020); Decay sugar-rich substrates (Põlme et al. 2020) | Põlme S, et al. 2020. Fungal Diversity 105:1-16 ((https://doi.org/10.1007/s13225-020-00466-2)) |

| | | | | | | | | | | | |
|---|---|---|---|---|---|---|---|---|---|---|---|
| Otu42722 | 1 | k__Fungi(100);p__Ascomycota(100);c__Saccharomycetes(100);o__Saccharomycetales(100);f__Metschnikowiaceae(100);g__Metschnikowia(100);s__Metschnikowia_pulcherrima(100); | Metschnikowia | 13 | Saprotroph | \|Undefined Saprotroph\| | Yeast | NULL | Probable | As nectar-tap_saprotroph (Põlme et al. 2020); Decay sugar-rich substrates (Põlme et al. 2020) | Põlme S, et al. 2020. Fungal Diversity 105:1-16 ((https://doi.org/10.1007/s13225-020-00466-2)) |
| Otu42723 | 1 | k__Fungi(100);p__Ascomycota(100);c__Saccharomycetes(100);o__Saccharomycetales(100);f__Metschnikowiaceae(100);g__Metschnikowia(100);s__Metschnikowia_pulcherrima(100); | Metschnikowia | 13 | Saprotroph | \|Undefined Saprotroph\| | Yeast | NULL | Probable | As nectar-tap_saprotroph (Põlme et al. 2020); Decay sugar-rich substrates (Põlme et al. 2020) | Põlme S, et al. 2020. Fungal Diversity 105:1-16 ((https://doi.org/10.1007/s13225-020-00466-2)) |
| Otu42724 | 1 | k__Fungi(100);p__Ascomycota(100);c__Saccharomycetes(100);o__Saccharomycetales(100);f__Metschnikowiaceae(100);g__Metschnikowia(100);s__Metschnikowia_pulcherrima(100); | Metschnikowia | 13 | Saprotroph | \|Undefined Saprotroph\| | Yeast | NULL | Probable | As nectar-tap_saprotroph (Põlme et al. 2020); Decay sugar-rich substrates (Põlme et al. 2020) | Põlme S, et al. 2020. Fungal Diversity 105:1-16 ((https://doi.org/10.1007/s13225-020-00466-2)) |
| Otu42725 | 1 | k__Fungi(100);p__Ascomycota(100);c__Saccharomycetes(100);o__Saccharomycetales(100);f__Metschnikowiaceae(100);g__Metschnikowia(100);s__Metschnikowia_pulcherrima(100); | Metschnikowia | 13 | Saprotroph | \|Undefined Saprotroph\| | Yeast | NULL | Probable | As nectar-tap_saprotroph (Põlme et al. 2020); Decay sugar-rich substrates (Põlme et al. 2020) | Põlme S, et al. 2020. Fungal Diversity 105:1-16 ((https://doi.org/10.1007/s13225-020-00466-2)) |
| Otu42726 | 1 | k__Fungi(100);p__Ascomycota(100);c__Saccharomycetes(100);o__Saccharomycetales(100);f__Metschnikowiaceae(100);g__Metschnikowia(100);s__Metschnikowia_pulcherrima(100); | Metschnikowia | 13 | Saprotroph | \|Undefined Saprotroph\| | Yeast | NULL | Probable | As nectar-tap_saprotroph (Põlme et al. 2020); Decay sugar-rich substrates (Põlme et al. 2020) | Põlme S, et al. 2020. Fungal Diversity 105:1-16 ((https://doi.org/10.1007/s13225-020-00466-2)) |
| Otu42727 | 1 | k__Fungi(100);p__Ascomycota(100);c__Saccharomycetes(100);o__Saccharomycetales(100);f__Metschnikowiaceae(100);g__Metschnikowia(100);s__Metschnikowia_pulcherrima(100); | Metschnikowia | 13 | Saprotroph | \|Undefined Saprotroph\| | Yeast | NULL | Probable | As nectar-tap_saprotroph (Põlme et al. 2020); Decay sugar-rich substrates (Põlme et al. 2020) | Põlme S, et al. 2020. Fungal Diversity 105:1-16 ((https://doi.org/10.1007/s13225-020-00466-2)) |
| Otu42728 | 1 | k__Fungi(100);p__Ascomycota(100);c__Saccharomycetes(100);o__Saccharomycetales(100);f__Metschnikowiaceae(100);g__Metschnikowia(100);s__Metschnikowia_pulcherrima(100); | Metschnikowia | 13 | Saprotroph | \|Undefined Saprotroph\| | Yeast | NULL | Probable | As nectar-tap_saprotroph (Põlme et al. 2020); Decay sugar-rich substrates (Põlme et al. 2020) | Põlme S, et al. 2020. Fungal Diversity 105:1-16 ((https://doi.org/10.1007/s13225-020-00466-2)) |
| Otu42729 | 1 | k__Fungi(100);p__Ascomycota(100);c__Saccharomycetes(100);o__Saccharomycetales(100);f__Metschnikowiaceae(100);g__Metschnikowia(100);g__Metschnikowia_unclassified(100); | Metschnikowia | 13 | Saprotroph | \|Undefined Saprotroph\| | Yeast | NULL | Probable | As nectar-tap_saprotroph (Põlme et al. 2020); Decay sugar-rich substrates (Põlme et al. 2020) | Põlme S, et al. 2020. Fungal Diversity 105:1-16 ((https://doi.org/10.1007/s13225-020-00466-2)) |
| Otu42730 | 1 | k__Fungi(100);p__Ascomycota(100);c__Saccharomycetes(100);o__Saccharomycetales(100);f__Metschnikowiaceae(100);g__Metschnikowia(100);s__Metschnikowia_pulcherrima(100); | Metschnikowia | 13 | Saprotroph | \|Undefined Saprotroph\| | Yeast | NULL | Probable | As nectar-tap_saprotroph (Põlme et al. 2020); Decay sugar-rich substrates (Põlme et al. 2020) | Põlme S, et al. 2020. Fungal Diversity 105:1-16 ((https://doi.org/10.1007/s13225-020-00466-2)) |
| Otu42731 | 1 | k__Fungi(100);p__Ascomycota(100);c__Saccharomycetes(100);o__Saccharomycetales(100);f__Metschnikowiaceae(100);g__Metschnikowia(100);s__Metschnikowia_pulcherrima(100); | Metschnikowia | 13 | Saprotroph | \|Undefined Saprotroph\| | Yeast | NULL | Probable | As nectar-tap_saprotroph (Põlme et al. 2020); Decay sugar-rich substrates (Põlme et al. 2020) | Põlme S, et al. 2020. Fungal Diversity 105:1-16 ((https://doi.org/10.1007/s13225-020-00466-2)) |
| Otu42732 | 1 | k__Fungi(100);p__Ascomycota(100);c__Saccharomycetes(100);o__Saccharomycetales(100);f__Metschnikowiaceae(100);g__Metschnikowia(100);g__Metschnikowia_unclassified(100); | Metschnikowia | 13 | Saprotroph | \|Undefined Saprotroph\| | Yeast | NULL | Probable | As nectar-tap_saprotroph (Põlme et al. 2020); Decay sugar-rich substrates (Põlme et al. 2020) | Põlme S, et al. 2020. Fungal Diversity 105:1-16 ((https://doi.org/10.1007/s13225-020-00466-2)) |
| Otu42733 | 1 | k__Fungi(100);p__Ascomycota(100);c__Saccharomycetes(100);o__Saccharomycetales(100);f__Metschnikowiaceae(100);g__Metschnikowia(100);g__Metschnikowia_unclassified(100); | Metschnikowia | 13 | Saprotroph | \|Undefined Saprotroph\| | Yeast | NULL | Probable | As nectar-tap_saprotroph (Põlme et al. 2020); Decay sugar-rich substrates (Põlme et al. 2020) | Põlme S, et al. 2020. Fungal Diversity 105:1-16 ((https://doi.org/10.1007/s13225-020-00466-2)) |
| Otu42734 | 1 | k__Fungi(100);p__Ascomycota(100);c__Saccharomycetes(100);o__Saccharomycetales(100);f__Metschnikowiaceae(100);g__Metschnikowia(100);g__Metschnikowia_unclassified(100); | Metschnikowia | 13 | Saprotroph | \|Undefined Saprotroph\| | Yeast | NULL | Probable | As nectar-tap_saprotroph (Põlme et al. 2020); Decay sugar-rich substrates (Põlme et al. 2020) | Põlme S, et al. 2020. Fungal Diversity 105:1-16 ((https://doi.org/10.1007/s13225-020-00466-2)) |
| Otu42735 | 1 | k__Fungi(100);p__Ascomycota(100);c__Saccharomycetes(100);o__Saccharomycetales(100);f__Metschnikowiaceae(100);g__Metschnikowia(100);s__Metschnikowia_pulcherrima(100); | Metschnikowia | 13 | Saprotroph | \|Undefined Saprotroph\| | Yeast | NULL | Probable | As nectar-tap_saprotroph (Põlme et al. 2020); Decay sugar-rich substrates (Põlme et al. 2020) | Põlme S, et al. 2020. Fungal Diversity 105:1-16 ((https://doi.org/10.1007/s13225-020-00466-2)) |
| Otu42736 | 1 | k__Fungi(100);p__Ascomycota(100);c__Saccharomycetes(100);o__Saccharomycetales(100);f__Metschnikowiaceae(100);g__Metschnikowia(100);s__Metschnikowia_pulcherrima(100); | Metschnikowia | 13 | Saprotroph | \|Undefined Saprotroph\| | Yeast | NULL | Probable | As nectar-tap_saprotroph (Põlme et al. 2020); Decay sugar-rich substrates (Põlme et al. 2020) | Põlme S, et al. 2020. Fungal Diversity 105:1-16 ((https://doi.org/10.1007/s13225-020-00466-2)) |
| Otu42737 | 1 | k__Fungi(100);p__Ascomycota(100);c__Saccharomycetes(100);o__Saccharomycetales(100);f__Metschnikowiaceae(100);g__Metschnikowia(100);s__Metschnikowia_pulcherrima(100); | Metschnikowia | 13 | Saprotroph | \|Undefined Saprotroph\| | Yeast | NULL | Probable | As nectar-tap_saprotroph (Põlme et al. 2020); Decay sugar-rich substrates (Põlme et al. 2020) | Põlme S, et al. 2020. Fungal Diversity 105:1-16 ((https://doi.org/10.1007/s13225-020-00466-2)) |
| Otu42738 | 1 | k__Fungi(100);p__Ascomycota(100);c__Saccharomycetes(100);o__Saccharomycetales(100);f__Metschnikowiaceae(100);g__Metschnikowia(100);g__Metschnikowia_unclassified(100); | Metschnikowia | 13 | Saprotroph | \|Undefined Saprotroph\| | Yeast | NULL | Probable | As nectar-tap_saprotroph (Põlme et al. 2020); Decay sugar-rich substrates (Põlme et al. 2020) | Põlme S, et al. 2020. Fungal Diversity 105:1-16 ((https://doi.org/10.1007/s13225-020-00466-2)) |
| Otu42739 | 1 | k__Fungi(100);p__Ascomycota(100);c__Saccharomycetes(100);o__Saccharomycetales(100);f__Metschnikowiaceae(100);g__Metschnikowia(100);s__Metschnikowia_pulcherrima(100); | Metschnikowia | 13 | Saprotroph | \|Undefined Saprotroph\| | Yeast | NULL | Probable | As nectar-tap_saprotroph (Põlme et al. 2020); Decay sugar-rich substrates (Põlme et al. 2020) | Põlme S, et al. 2020. Fungal Diversity 105:1-16 ((https://doi.org/10.1007/s13225-020-00466-2)) |
| Otu42740 | 1 | k__Fungi(100);p__Ascomycota(100);c__Saccharomycetes(100);o__Saccharomycetales(100);f__Metschnikowiaceae(100);g__Metschnikowia(100);s__Metschnikowia_pulcherrima(100); | Metschnikowia | 13 | Saprotroph | \|Undefined Saprotroph\| | Yeast | NULL | Probable | As nectar-tap_saprotroph (Põlme et al. 2020); Decay sugar-rich substrates (Põlme et al. 2020) | Põlme S, et al. 2020. Fungal Diversity 105:1-16 ((https://doi.org/10.1007/s13225-020-00466-2)) |
| Otu42741 | 1 | k__Fungi(100);p__Ascomycota(100);c__Saccharomycetes(100);o__Saccharomycetales(100);f__Metschnikowiaceae(100);g__Metschnikowia(100);g__Metschnikowia_unclassified(100); | Metschnikowia | 13 | Saprotroph | \|Undefined Saprotroph\| | Yeast | NULL | Probable | As nectar-tap_saprotroph (Põlme et al. 2020); Decay sugar-rich substrates (Põlme et al. 2020) | Põlme S, et al. 2020. Fungal Diversity 105:1-16 ((https://doi.org/10.1007/s13225-020-00466-2)) |
| Otu42742 | 1 | k__Fungi(100);p__Ascomycota(100);c__Saccharomycetes(100);o__Saccharomycetales(100);f__Metschnikowiaceae(100);g__Metschnikowia(100);s__Metschnikowia_pulcherrima(100); | Metschnikowia | 13 | Saprotroph | \|Undefined Saprotroph\| | Yeast | NULL | Probable | As nectar-tap_saprotroph (Põlme et al. 2020); Decay sugar-rich substrates (Põlme et al. 2020) | Põlme S, et al. 2020. Fungal Diversity 105:1-16 ((https://doi.org/10.1007/s13225-020-00466-2)) |
| Otu42743 | 1 | k__Fungi(100);p__Ascomycota(100);c__Saccharomycetes(100);o__Saccharomycetales(100);f__Metschnikowiaceae(100);g__Metschnikowia(100);s__Metschnikowia_pulcherrima(100); | Metschnikowia | 13 | Saprotroph | \|Undefined Saprotroph\| | Yeast | NULL | Probable | As nectar-tap_saprotroph (Põlme et al. 2020); Decay sugar-rich substrates (Põlme et al. 2020) | Põlme S, et al. 2020. Fungal Diversity 105:1-16 ((https://doi.org/10.1007/s13225-020-00466-2)) |
| Otu42746 | 1 | k__Fungi(100);p__Ascomycota(100);c__Saccharomycetes(100);o__Saccharomycetales(100);f__Metschnikowiaceae(100);g__Metschnikowia(100);s__Metschnikowia_pulcherrima(100); | Metschnikowia | 13 | Saprotroph | \|Undefined Saprotroph\| | Yeast | NULL | Probable | As nectar-tap_saprotroph (Põlme et al. 2020); Decay sugar-rich substrates (Põlme et al. 2020) | Põlme S, et al. 2020. Fungal Diversity 105:1-16 ((https://doi.org/10.1007/s13225-020-00466-2)) |
| Otu42747 | 1 | k__Fungi(100);p__Ascomycota(100);c__Saccharomycetes(100);o__Saccharomycetales(100);f__Metschnikowiaceae(100);g__Metschnikowia(100);s__Metschnikowia_pulcherrima(100); | Metschnikowia | 13 | Saprotroph | \|Undefined Saprotroph\| | Yeast | NULL | Probable | As nectar-tap_saprotroph (Põlme et al. 2020); Decay sugar-rich substrates (Põlme et al. 2020) | Põlme S, et al. 2020. Fungal Diversity 105:1-16 ((https://doi.org/10.1007/s13225-020-00466-2)) |
| Otu42749 | 1 | k__Fungi(100);p__Ascomycota(100);c__Saccharomycetes(100);o__Saccharomycetales(100);f__Metschnikowiaceae(100);g__Metschnikowia(100);s__Metschnikowia_pulcherrima(100); | Metschnikowia | 13 | Saprotroph | \|Undefined Saprotroph\| | Yeast | NULL | Probable | As nectar-tap_saprotroph (Põlme et al. 2020); Decay sugar-rich substrates (Põlme et al. 2020) | Põlme S, et al. 2020. Fungal Diversity 105:1-16 ((https://doi.org/10.1007/s13225-020-00466-2)) |
| Otu42752 | 1 | k__Fungi(100);p__Ascomycota(100);c__Saccharomycetes(100);o__Saccharomycetales(100);f__Metschnikowiaceae(100);g__Metschnikowia(100);s__Metschnikowia_pulcherrima(100); | Metschnikowia | 13 | Saprotroph | \|Undefined Saprotroph\| | Yeast | NULL | Probable | As nectar-tap_saprotroph (Põlme et al. 2020); Decay sugar-rich substrates (Põlme et al. 2020) | Põlme S, et al. 2020. Fungal Diversity 105:1-16 ((https://doi.org/10.1007/s13225-020-00466-2)) |
| Otu42754 | 1 | k__Fungi(100);p__Ascomycota(100);c__Saccharomycetes(100);o__Saccharomycetales(100);f__Metschnikowiaceae(100);g__Metschnikowia(100);s__Metschnikowia_pulcherrima(100); | Metschnikowia | 13 | Saprotroph | \|Undefined Saprotroph\| | Yeast | NULL | Probable | As nectar-tap_saprotroph (Põlme et al. 2020); Decay sugar-rich substrates (Põlme et al. 2020) | Põlme S, et al. 2020. Fungal Diversity 105:1-16 ((https://doi.org/10.1007/s13225-020-00466-2)) |
| Otu42755 | 1 | k__Fungi(100);p__Ascomycota(100);c__Saccharomycetes(100);o__Saccharomycetales(100);f__Metschnikowiaceae(100);g__Metschnikowia(100);g__Metschnikowia_unclassified(100); | Metschnikowia | 13 | Saprotroph | \|Undefined Saprotroph\| | Yeast | NULL | Probable | As nectar-tap_saprotroph (Põlme et al. 2020); Decay sugar-rich substrates (Põlme et al. 2020) | Põlme S, et al. 2020. Fungal Diversity 105:1-16 ((https://doi.org/10.1007/s13225-020-00466-2)) |
| Otu42756 | 1 | k__Fungi(100);p__Ascomycota(100);c__Saccharomycetes(100);o__Saccharomycetales(100);f__Metschnikowiaceae(100);g__Metschnikowia(100);g__Metschnikowia_unclassified(100); | Metschnikowia | 13 | Saprotroph | \|Undefined Saprotroph\| | Yeast | NULL | Probable | As nectar-tap_saprotroph (Põlme et al. 2020); Decay sugar-rich substrates (Põlme et al. 2020) | Põlme S, et al. 2020. Fungal Diversity 105:1-16 ((https://doi.org/10.1007/s13225-020-00466-2)) |
| Otu42757 | 1 | k__Fungi(100);p__Ascomycota(100);c__Saccharomycetes(100);o__Saccharomycetales(100);f__Metschnikowiaceae(100);g__Metschnikowia(100);s__Metschnikowia_pulcherrima(100); | Metschnikowia | 13 | Saprotroph | \|Undefined Saprotroph\| | Yeast | NULL | Probable | As nectar-tap_saprotroph (Põlme et al. 2020); Decay sugar-rich substrates (Põlme et al. 2020) | Põlme S, et al. 2020. Fungal Diversity 105:1-16 ((https://doi.org/10.1007/s13225-020-00466-2)) |
| Otu42758 | 1 | k__Fungi(100);p__Ascomycota(100);c__Saccharomycetes(100);o__Saccharomycetales(100);f__Metschnikowiaceae(100);g__Metschnikowia(100);s__Metschnikowia_pulcherrima(100); | Metschnikowia | 13 | Saprotroph | \|Undefined Saprotroph\| | Yeast | NULL | Probable | As nectar-tap_saprotroph (Põlme et al. 2020); Decay sugar-rich substrates (Põlme et al. 2020) | Põlme S, et al. 2020. Fungal Diversity 105:1-16 ((https://doi.org/10.1007/s13225-020-00466-2)) |
| Otu42759 | 1 | k__Fungi(100);p__Ascomycota(100);c__Saccharomycetes(100);o__Saccharomycetales(100);f__Metschnikowiaceae(100);g__Metschnikowia(100);s__Metschnikowia_pulcherrima(100); | Metschnikowia | 13 | Saprotroph | \|Undefined Saprotroph\| | Yeast | NULL | Probable | As nectar-tap_saprotroph (Põlme et al. 2020); Decay sugar-rich substrates (Põlme et al. 2020) | Põlme S, et al. 2020. Fungal Diversity 105:1-16 ((https://doi.org/10.1007/s13225-020-00466-2)) |
| Otu42760 | 1 | k__Fungi(100);p__Ascomycota(100);c__Saccharomycetes(100);o__Saccharomycetales(100);f__Metschnikowiaceae(100);g__Metschnikowia(100);s__Metschnikowia_pulcherrima(100); | Metschnikowia | 13 | Saprotroph | \|Undefined Saprotroph\| | Yeast | NULL | Probable | As nectar-tap_saprotroph (Põlme et al. 2020); Decay sugar-rich substrates (Põlme et al. 2020) | Põlme S, et al. 2020. Fungal Diversity 105:1-16 ((https://doi.org/10.1007/s13225-020-00466-2)) |
| Otu42761 | 1 | k__Fungi(100);p__Ascomycota(100);c__Saccharomycetes(100);o__Saccharomycetales(100);f__Metschnikowiaceae(100);g__Metschnikowia(100);s__Metschnikowia_pulcherrima(100); | Metschnikowia | 13 | Saprotroph | \|Undefined Saprotroph\| | Yeast | NULL | Probable | As nectar-tap_saprotroph (Põlme et al. 2020); Decay sugar-rich substrates (Põlme et al. 2020) | Põlme S, et al. 2020. Fungal Diversity 105:1-16 ((https://doi.org/10.1007/s13225-020-00466-2)) |
| Otu42762 | 1 | k__Fungi(100);p__Ascomycota(100);c__Saccharomycetes(100);o__Saccharomycetales(100);f__Metschnikowiaceae(100);g__Metschnikowia(100);s__Metschnikowia_pulcherrima(100); | Metschnikowia | 13 | Saprotroph | \|Undefined Saprotroph\| | Yeast | NULL | Probable | As nectar-tap_saprotroph (Põlme et al. 2020); Decay sugar-rich substrates (Põlme et al. 2020) | Põlme S, et al. 2020. Fungal Diversity 105:1-16 ((https://doi.org/10.1007/s13225-020-00466-2)) |
| Otu42763 | 1 | k__Fungi(100);p__Ascomycota(100);c__Saccharomycetes(100);o__Saccharomycetales(100);f__Metschnikowiaceae(100);g__Metschnikowia(100);g__Metschnikowia_unclassified(100); | Metschnikowia | 13 | Saprotroph | \|Undefined Saprotroph\| | Yeast | NULL | Probable | As nectar-tap_saprotroph (Põlme et al. 2020); Decay sugar-rich substrates (Põlme et al. 2020) | Põlme S, et al. 2020. Fungal Diversity 105:1-16 ((https://doi.org/10.1007/s13225-020-00466-2)) |
| Otu42765 | 1 | k__Fungi(100);p__Ascomycota(100);c__Saccharomycetes(100);o__Saccharomycetales(100);f__Metschnikowiaceae(100);g__Metschnikowia(100);g__Metschnikowia_unclassified(100); | Metschnikowia | 13 | Saprotroph | \|Undefined Saprotroph\| | Yeast | NULL | Probable | As nectar-tap_saprotroph (Põlme et al. 2020); Decay sugar-rich substrates (Põlme et al. 2020) | Põlme S, et al. 2020. Fungal Diversity 105:1-16 ((https://doi.org/10.1007/s13225-020-00466-2)) |
| Otu42766 | 1 | k__Fungi(100);p__Ascomycota(100);c__Saccharomycetes(100);o__Saccharomycetales(100);f__Saccharomycetales_family_Incertae_sedis(100);g__Candida(100);g__Candida_unclassified(100); | Candida | 13 | Pathotroph-Saprotroph-Symbiotroph | Animal Pathogen-Endophyte-Endosymbiont-Epiphyte-\|Undefined Saprotroph\| | Dimorphic Yeast-Yeast | NULL | Probable | As nectar-tap_saprotroph (Põlme et al. 2020); Decay sugar-rich substrates (Põlme et al. 2020); This asexual genus is polyphyletic and contain species that function very differently depend on what they are. Caution should be taken when interpreting guilds for Candida species. | Manolakaki D, et al. 2010. Virulence. 1: 367-375 ((https://doi.org/10.4161/viru.1.5.12796)); Mekbib SB, et al. 2011. Tropical Plant Pathology 36:233-240 ((https://doi.org/10.1590/S1982-56762011000400004)); Põlme S, et al. 2020. Fungal Diversity 105:1-16 ((https://doi.org/10.1007/s13225-020-00466-2)) |
| Otu42767 | 1 | k__Fungi(100);p__Ascomycota(100);c__Saccharomycetes(100);o__Saccharomycetales(100);f__Metschnikowiaceae(100);g__Metschnikowia(100);s__Metschnikowia_pulcherrima(100); | Metschnikowia | 13 | Saprotroph | \|Undefined Saprotroph\| | Yeast | NULL | Probable | As nectar-tap_saprotroph (Põlme et al. 2020); Decay sugar-rich substrates (Põlme et al. 2020) | Põlme S, et al. 2020. Fungal Diversity 105:1-16 ((https://doi.org/10.1007/s13225-020-00466-2)) |
| Otu42768 | 1 | k__Fungi(100);p__Ascomycota(100);c__Saccharomycetes(100);o__Saccharomycetales(100);f__Metschnikowiaceae(100);g__Metschnikowia(100);s__Metschnikowia_pulcherrima(100); | Metschnikowia | 13 | Saprotroph | \|Undefined Saprotroph\| | Yeast | NULL | Probable | As nectar-tap_saprotroph (Põlme et al. 2020); Decay sugar-rich substrates (Põlme et al. 2020) | Põlme S, et al. 2020. Fungal Diversity 105:1-16 ((https://doi.org/10.1007/s13225-020-00466-2)) |
| Otu42769 | 1 | k__Fungi(100);p__Ascomycota(100);c__Saccharomycetes(100);o__Saccharomycetales(100);f__Metschnikowiaceae(100);g__Metschnikowia(100);s__Metschnikowia_pulcherrima(100); | Metschnikowia | 13 | Saprotroph | \|Undefined Saprotroph\| | Yeast | NULL | Probable | As nectar-tap_saprotroph (Põlme et al. 2020); Decay sugar-rich substrates (Põlme et al. 2020) | Põlme S, et al. 2020. Fungal Diversity 105:1-16 ((https://doi.org/10.1007/s13225-020-00466-2)) |
| Otu42770 | 1 | k__Fungi(100);p__Ascomycota(100);c__Saccharomycetes(100);o__Saccharomycetales(100);f__Metschnikowiaceae(100);g__Metschnikowia(100);s__Metschnikowia_pulcherrima(100); | Metschnikowia | 13 | Saprotroph | \|Undefined Saprotroph\| | Yeast | NULL | Probable | As nectar-tap_saprotroph (Põlme et al. 2020); Decay sugar-rich substrates (Põlme et al. 2020) | Põlme S, et al. 2020. Fungal Diversity 105:1-16 ((https://doi.org/10.1007/s13225-020-00466-2)) |
| Otu42771 | 1 | k__Fungi(100);p__Ascomycota(100);c__Saccharomycetes(100);o__Saccharomycetales(100);f__Metschnikowiaceae(100);g__Metschnikowia(100);g__Metschnikowia_unclassified(100); | Metschnikowia | 13 | Saprotroph | \|Undefined Saprotroph\| | Yeast | NULL | Probable | As nectar-tap_saprotroph (Põlme et al. 2020); Decay sugar-rich substrates (Põlme et al. 2020) | Põlme S, et al. 2020. Fungal Diversity 105:1-16 ((https://doi.org/10.1007/s13225-020-00466-2)) |
| Otu42772 | 1 | k__Fungi(100);p__Ascomycota(100);c__Saccharomycetes(100);o__Saccharomycetales(100);f__Metschnikowiaceae(100);g__Metschnikowia(100);s__Metschnikowia_pulcherrima(100); | Metschnikowia | 13 | Saprotroph | \|Undefined Saprotroph\| | Yeast | NULL | Probable | As nectar-tap_saprotroph (Põlme et al. 2020); Decay sugar-rich substrates (Põlme et al. 2020) | Põlme S, et al. 2020. Fungal Diversity 105:1-16 ((https://doi.org/10.1007/s13225-020-00466-2)) |
| Otu42774 | 1 | k__Fungi(100);p__Ascomycota(100);c__Saccharomycetes(100);o__Saccharomycetales(100);f__Metschnikowiaceae(100);g__Metschnikowia(100);s__Metschnikowia_pulcherrima(100); | Metschnikowia | 13 | Saprotroph | \|Undefined Saprotroph\| | Yeast | NULL | Probable | As nectar-tap_saprotroph (Põlme et al. 2020); Decay sugar-rich substrates (Põlme et al. 2020) | Põlme S, et al. 2020. Fungal Diversity 105:1-16 ((https://doi.org/10.1007/s13225-020-00466-2)) |
| Otu42775 | 1 | k__Fungi(100);p__Ascomycota(100);c__Saccharomycetes(100);o__Saccharomycetales(100);f__Metschnikowiaceae(100);g__Metschnikowia(100);g__Metschnikowia_unclassified(100); | Metschnikowia | 13 | Saprotroph | \|Undefined Saprotroph\| | Yeast | NULL | Probable | As nectar-tap_saprotroph (Põlme et al. 2020); Decay sugar-rich substrates (Põlme et al. 2020) | Põlme S, et al. 2020. Fungal Diversity 105:1-16 ((https://doi.org/10.1007/s13225-020-00466-2)) |
| Otu42776 | 1 | k__Fungi(100);p__Ascomycota(100);c__Saccharomycetes(100);o__Saccharomycetales(100);f__Metschnikowiaceae(100);g__Metschnikowia(100);s__Metschnikowia_pulcherrima(100); | Metschnikowia | 13 | Saprotroph | \|Undefined Saprotroph\| | Yeast | NULL | Probable | As nectar-tap_saprotroph (Põlme et al. 2020); Decay sugar-rich substrates (Põlme et al. 2020) | Põlme S, et al. 2020. Fungal Diversity 105:1-16 ((https://doi.org/10.1007/s13225-020-00466-2)) |
| Otu42777 | 1 | k__Fungi(100);p__Ascomycota(100);c__Saccharomycetes(100);o__Saccharomycetales(100);f__Metschnikowiaceae(100);g__Metschnikowia(100);s__Metschnikowia_pulcherrima(100); | Metschnikowia | 13 | Saprotroph | \|Undefined Saprotroph\| | Yeast | NULL | Probable | As nectar-tap_saprotroph (Põlme et al. 2020); Decay sugar-rich substrates (Põlme et al. 2020) | Põlme S, et al. 2020. Fungal Diversity 105:1-16 ((https://doi.org/10.1007/s13225-020-00466-2)) |
| Otu42778 | 1 | k__Fungi(100);p__Ascomycota(100);c__Saccharomycetes(100);o__Saccharomycetales(100);f__Metschnikowiaceae(100);g__Metschnikowia(100);s__Metschnikowia_pulcherrima(100); | Metschnikowia | 13 | Saprotroph | \|Undefined Saprotroph\| | Yeast | NULL | Probable | As nectar-tap_saprotroph (Põlme et al. 2020); Decay sugar-rich substrates (Põlme et al. 2020) | Põlme S, et al. 2020. Fungal Diversity 105:1-16 ((https://doi.org/10.1007/s13225-020-00466-2)) |
| Otu42779 | 1 | k__Fungi(100);p__Ascomycota(100);c__Saccharomycetes(100);o__Saccharomycetales(100);f__Metschnikowiaceae(100);g__Metschnikowia(100);g__Metschnikowia_unclassified(100); | Metschnikowia | 13 | Saprotroph | \|Undefined Saprotroph\| | Yeast | NULL | Probable | As nectar-tap_saprotroph (Põlme et al. 2020); Decay sugar-rich substrates (Põlme et al. 2020) | Põlme S, et al. 2020. Fungal Diversity 105:1-16 ((https://doi.org/10.1007/s13225-020-00466-2)) |
| Otu42780 | 1 | k__Fungi(100);p__Ascomycota(100);c__Saccharomycetes(100);o__Saccharomycetales(100);f__Metschnikowiaceae(100);g__Metschnikowia(100);g__Metschnikowia_unclassified(100); | Metschnikowia | 13 | Saprotroph | \|Undefined Saprotroph\| | Yeast | NULL | Probable | As nectar-tap_saprotroph (Põlme et al. 2020); Decay sugar-rich substrates (Põlme et al. 2020) | Põlme S, et al. 2020. Fungal Diversity 105:1-16 ((https://doi.org/10.1007/s13225-020-00466-2)) |
| Otu42781 | 1 | k__Fungi(100);p__Ascomycota(100);c__Saccharomycetes(100);o__Saccharomycetales(100);f__Metschnikowiaceae(100);g__Metschnikowia(100);g__Metschnikowia_unclassified(100); | Metschnikowia | 13 | Saprotroph | \|Undefined Saprotroph\| | Yeast | NULL | Probable | As nectar-tap_saprotroph (Põlme et al. 2020); Decay sugar-rich substrates (Põlme et al. 2020) | Põlme S, et al. 2020. Fungal Diversity 105:1-16 ((https://doi.org/10.1007/s13225-020-00466-2)) |
| Otu42782 | 1 | k__Fungi(100);p__Ascomycota(100);c__Saccharomycetes(100);o__Saccharomycetales(100);f__Metschnikowiaceae(100);g__Metschnikowia(100);s__Metschnikowia_pulcherrima(100); | Metschnikowia | 13 | Saprotroph | \|Undefined Saprotroph\| | Yeast | NULL | Probable | As nectar-tap_saprotroph (Põlme et al. 2020); Decay sugar-rich substrates (Põlme et al. 2020) | Põlme S, et al. 2020. Fungal Diversity 105:1-16 ((https://doi.org/10.1007/s13225-020-00466-2)) |
| Otu42783 | 1 | k__Fungi(100);p__Ascomycota(100);c__Saccharomycetes(100);o__Saccharomycetales(100);f__Metschnikowiaceae(100);g__Metschnikowia(100);s__Metschnikowia_pulcherrima(100); | Metschnikowia | 13 | Saprotroph | \|Undefined Saprotroph\| | Yeast | NULL | Probable | As nectar-tap_saprotroph (Põlme et al. 2020); Decay sugar-rich substrates (Põlme et al. 2020) | Põlme S, et al. 2020. Fungal Diversity 105:1-16 ((https://doi.org/10.1007/s13225-020-00466-2)) |
| Otu42784 | 1 | k__Fungi(100);p__Ascomycota(100);c__Saccharomycetes(100);o__Saccharomycetales(100);f__Metschnikowiaceae(100);g__Metschnikowia(100);s__Metschnikowia_pulcherrima(100); | Metschnikowia | 13 | Saprotroph | \|Undefined Saprotroph\| | Yeast | NULL | Probable | As nectar-tap_saprotroph (Põlme et al. 2020); Decay sugar-rich substrates (Põlme et al. 2020) | Põlme S, et al. 2020. Fungal Diversity 105:1-16 ((https://doi.org/10.1007/s13225-020-00466-2)) |
| Otu42785 | 1 | k__Fungi(100);p__Ascomycota(100);c__Saccharomycetes(100);o__Saccharomycetales(100);f__Metschnikowiaceae(100);g__Metschnikowia(100);s__Metschnikowia_pulcherrima(100); | Metschnikowia | 13 | Saprotroph | \|Undefined Saprotroph\| | Yeast | NULL | Probable | As nectar-tap_saprotroph (Põlme et al. 2020); Decay sugar-rich substrates (Põlme et al. 2020) | Põlme S, et al. 2020. Fungal Diversity 105:1-16 ((https://doi.org/10.1007/s13225-020-00466-2)) |
| Otu42786 | 1 | k__Fungi(100);p__Ascomycota(100);c__Saccharomycetes(100);o__Saccharomycetales(100);f__Metschnikowiaceae(100);g__Metschnikowia(100);s__Metschnikowia_pulcherrima(100); | Metschnikowia | 13 | Saprotroph | \|Undefined Saprotroph\| | Yeast | NULL | Probable | As nectar-tap_saprotroph (Põlme et al. 2020); Decay sugar-rich substrates (Põlme et al. 2020) | Põlme S, et al. 2020. Fungal Diversity 105:1-16 ((https://doi.org/10.1007/s13225-020-00466-2)) |
| Otu42787 | 1 | k__Fungi(100);p__Ascomycota(100);c__Saccharomycetes(100);o__Saccharomycetales(100);f__Metschnikowiaceae(100);g__Metschnikowia(100);g__Metschnikowia_unclassified(100); | Metschnikowia | 13 | Saprotroph | \|Undefined Saprotroph\| | Yeast | NULL | Probable | As nectar-tap_saprotroph (Põlme et al. 2020); Decay sugar-rich substrates (Põlme et al. 2020) | Põlme S, et al. 2020. Fungal Diversity 105:1-16 ((https://doi.org/10.1007/s13225-020-00466-2)) |
| Otu42788 | 1 | k__Fungi(100);p__Ascomycota(100);c__Saccharomycetes(100);o__Saccharomycetales(100);f__Metschnikowiaceae(100);g__Metschnikowia(100);s__Metschnikowia_pulcherrima(100); | Metschnikowia | 13 | Saprotroph | \|Undefined Saprotroph\| | Yeast | NULL | Probable | As nectar-tap_saprotroph (Põlme et al. 2020); Decay sugar-rich substrates (Põlme et al. 2020) | Põlme S, et al. 2020. Fungal Diversity 105:1-16 ((https://doi.org/10.1007/s13225-020-00466-2)) |
| Otu42789 | 1 | k__Fungi(100);p__Ascomycota(100);c__Saccharomycetes(100);o__Saccharomycetales(100);f__Metschnikowiaceae(100);g__Metschnikowia(100);s__Metschnikowia_pulcherrima(100); | Metschnikowia | 13 | Saprotroph | \|Undefined Saprotroph\| | Yeast | NULL | Probable | As nectar-tap_saprotroph (Põlme et al. 2020); Decay sugar-rich substrates (Põlme et al. 2020) | Põlme S, et al. 2020. Fungal Diversity 105:1-16 ((https://doi.org/10.1007/s13225-020-00466-2)) |
| Otu42790 | 1 | k__Fungi(100);p__Ascomycota(100);c__Saccharomycetes(100);o__Saccharomycetales(100);f__Metschnikowiaceae(100);g__Metschnikowia(100);g__Metschnikowia_unclassified(100); | Metschnikowia | 13 | Saprotroph | \|Undefined Saprotroph\| | Yeast | NULL | Probable | As nectar-tap_saprotroph (Põlme et al. 2020); Decay sugar-rich substrates (Põlme et al. 2020) | Põlme S, et al. 2020. Fungal Diversity 105:1-16 ((https://doi.org/10.1007/s13225-020-00466-2)) |
| Otu42791 | 1 | k__Fungi(100);p__Ascomycota(100);c__Saccharomycetes(100);o__Saccharomycetales(100);f__Metschnikowiaceae(100);g__Metschnikowia(100);s__Metschnikowia_pulcherrima(100); | Metschnikowia | 13 | Saprotroph | \|Undefined Saprotroph\| | Yeast | NULL | Probable | As nectar-tap_saprotroph (Põlme et al. 2020); Decay sugar-rich substrates (Põlme et al. 2020) | Põlme S, et al. 2020. Fungal Diversity 105:1-16 ((https://doi.org/10.1007/s13225-020-00466-2)) |
| Otu42792 | 1 | k__Fungi(100);p__Ascomycota(100);c__Saccharomycetes(100);o__Saccharomycetales(100);f__Metschnikowiaceae(100);g__Metschnikowia(100);s__Metschnikowia_pulcherrima(100); | Metschnikowia | 13 | Saprotroph | \|Undefined Saprotroph\| | Yeast | NULL | Probable | As nectar-tap_saprotroph (Põlme et al. 2020); Decay sugar-rich substrates (Põlme et al. 2020) | Põlme S, et al. 2020. Fungal Diversity 105:1-16 ((https://doi.org/10.1007/s13225-020-00466-2)) |
| Otu42793 | 1 | k__Fungi(100);p__Ascomycota(100);c__Saccharomycetes(100);o__Saccharomycetales(100);f__Metschnikowiaceae(100);g__Metschnikowia(100);s__Metschnikowia_pulcherrima(100); | Metschnikowia | 13 | Saprotroph | \|Undefined Saprotroph\| | Yeast | NULL | Probable | As nectar-tap_saprotroph (Põlme et al. 2020); Decay sugar-rich substrates (Põlme et al. 2020) | Põlme S, et al. 2020. Fungal Diversity 105:1-16 ((https://doi.org/10.1007/s13225-020-00466-2)) |
| Otu42794 | 1 | k__Fungi(100);p__Ascomycota(100);c__Saccharomycetes(100);o__Saccharomycetales(100);f__Metschnikowiaceae(100);g__Metschnikowia(100);s__Metschnikowia_pulcherrima(100); | Metschnikowia | 13 | Saprotroph | \|Undefined Saprotroph\| | Yeast | NULL | Probable | As nectar-tap_saprotroph (Põlme et al. 2020); Decay sugar-rich substrates (Põlme et al. 2020) | Põlme S, et al. 2020. Fungal Diversity 105:1-16 ((https://doi.org/10.1007/s13225-020-00466-2)) |
| Otu42795 | 1 | k__Fungi(100);p__Ascomycota(100);c__Saccharomycetes(100);o__Saccharomycetales(100);f__Metschnikowiaceae(100);g__Metschnikowia(100);s__Metschnikowia_pulcherrima(100); | Metschnikowia | 13 | Saprotroph | \|Undefined Saprotroph\| | Yeast | NULL | Probable | As nectar-tap_saprotroph (Põlme et al. 2020); Decay sugar-rich substrates (Põlme et al. 2020) | Põlme S, et al. 2020. Fungal Diversity 105:1-16 ((https://doi.org/10.1007/s13225-020-00466-2)) |
| Otu42796 | 1 | k__Fungi(100);p__Ascomycota(100);c__Saccharomycetes(100);o__Saccharomycetales(100);f__Metschnikowiaceae(100);g__Metschnikowia(100);g__Metschnikowia_unclassified(100); | Metschnikowia | 13 | Saprotroph | \|Undefined Saprotroph\| | Yeast | NULL | Probable | As nectar-tap_saprotroph (Põlme et al. 2020); Decay sugar-rich substrates (Põlme et al. 2020) | Põlme S, et al. 2020. Fungal Diversity 105:1-16 ((https://doi.org/10.1007/s13225-020-00466-2)) |
| Otu42797 | 1 | k__Fungi(100);p__Ascomycota(100);c__Saccharomycetes(100);o__Saccharomycetales(100);f__Metschnikowiaceae(100);g__Metschnikowia(100);s__Metschnikowia_pulcherrima(100); | Metschnikowia | 13 | Saprotroph | \|Undefined Saprotroph\| | Yeast | NULL | Probable | As nectar-tap_saprotroph (Põlme et al. 2020); Decay sugar-rich substrates (Põlme et al. 2020) | Põlme S, et al. 2020. Fungal Diversity 105:1-16 ((https://doi.org/10.1007/s13225-020-00466-2)) |
| Otu42798 | 1 | k__Fungi(100);p__Ascomycota(100);c__Saccharomycetes(100);o__Saccharomycetales(100);f__Metschnikowiaceae(100);g__Metschnikowia(100);s__Metschnikowia_pulcherrima(100); | Metschnikowia | 13 | Saprotroph | \|Undefined Saprotroph\| | Yeast | NULL | Probable | As nectar-tap_saprotroph (Põlme et al. 2020); Decay sugar-rich substrates (Põlme et al. 2020) | Põlme S, et al. 2020. Fungal Diversity 105:1-16 ((https://doi.org/10.1007/s13225-020-00466-2)) |
| Otu42799 | 1 | k__Fungi(100);p__Ascomycota(100);c__Saccharomycetes(100);o__Saccharomycetales(100);f__Metschnikowiaceae(100);g__Metschnikowia(100);g__Metschnikowia_unclassified(100); | Metschnikowia | 13 | Saprotroph | \|Undefined Saprotroph\| | Yeast | NULL | Probable | As nectar-tap_saprotroph (Põlme et al. 2020); Decay sugar-rich substrates (Põlme et al. 2020) | Põlme S, et al. 2020. Fungal Diversity 105:1-16 ((https://doi.org/10.1007/s13225-020-00466-2)) |
| Otu42800 | 1 | k__Fungi(100);p__Ascomycota(100);c__Saccharomycetes(100);o__Saccharomycetales(100);f__Metschnikowiaceae(100);g__Metschnikowia(100);g__Metschnikowia_unclassified(100); | Metschnikowia | 13 | Saprotroph | \|Undefined Saprotroph\| | Yeast | NULL | Probable | As nectar-tap_saprotroph (Põlme et al. 2020); Decay sugar-rich substrates (Põlme et al. 2020) | Põlme S, et al. 2020. Fungal Diversity 105:1-16 ((https://doi.org/10.1007/s13225-020-00466-2)) |
| Otu42801 | 1 | k__Fungi(100);p__Ascomycota(100);c__Saccharomycetes(100);o__Saccharomycetales(100);f__Metschnikowiaceae(100);g__Metschnikowia(100);s__Metschnikowia_pulcherrima(100); | Metschnikowia | 13 | Saprotroph | \|Undefined Saprotroph\| | Yeast | NULL | Probable | As nectar-tap_saprotroph (Põlme et al. 2020); Decay sugar-rich substrates (Põlme et al. 2020) | Põlme S, et al. 2020. Fungal Diversity 105:1-16 ((https://doi.org/10.1007/s13225-020-00466-2)) |
| Otu42802 | 1 | k__Fungi(100);p__Ascomycota(100);c__Saccharomycetes(100);o__Saccharomycetales(100);f__Metschnikowiaceae(100);g__Metschnikowia(100);s__Metschnikowia_pulcherrima(100); | Metschnikowia | 13 | Saprotroph | \|Undefined Saprotroph\| | Yeast | NULL | Probable | As nectar-tap_saprotroph (Põlme et al. 2020); Decay sugar-rich substrates (Põlme et al. 2020) | Põlme S, et al. 2020. Fungal Diversity 105:1-16 ((https://doi.org/10.1007/s13225-020-00466-2)) |
| Otu42803 | 1 | k__Fungi(100);p__Ascomycota(100);c__Saccharomycetes(100);o__Saccharomycetales(100);f__Metschnikowiaceae(100);g__Metschnikowia(100);g__Metschnikowia_unclassified(100); | Metschnikowia | 13 | Saprotroph | \|Undefined Saprotroph\| | Yeast | NULL | Probable | As nectar-tap_saprotroph (Põlme et al. 2020); Decay sugar-rich substrates (Põlme et al. 2020) | Põlme S, et al. 2020. Fungal Diversity 105:1-16 ((https://doi.org/10.1007/s13225-020-00466-2)) |
| Otu42805 | 1 | k__Fungi(100);p__Ascomycota(100);c__Saccharomycetes(100);o__Saccharomycetales(100);f__Metschnikowiaceae(100);g__Metschnikowia(100);s__Metschnikowia_pulcherrima(100); | Metschnikowia | 13 | Saprotroph | \|Undefined Saprotroph\| | Yeast | NULL | Probable | As nectar-tap_saprotroph (Põlme et al. 2020); Decay sugar-rich substrates (Põlme et al. 2020) | Põlme S, et al. 2020. Fungal Diversity 105:1-16 ((https://doi.org/10.1007/s13225-020-00466-2)) |
| Otu42806 | 1 | k__Fungi(100);p__Ascomycota(100);c__Saccharomycetes(100);o__Saccharomycetales(100);f__Metschnikowiaceae(100);g__Metschnikowia(100);g__Metschnikowia_unclassified(100); | Metschnikowia | 13 | Saprotroph | \|Undefined Saprotroph\| | Yeast | NULL | Probable | As nectar-tap_saprotroph (Põlme et al. 2020); Decay sugar-rich substrates (Põlme et al. 2020) | Põlme S, et al. 2020. Fungal Diversity 105:1-16 ((https://doi.org/10.1007/s13225-020-00466-2)) |
| Otu42807 | 1 | k__Fungi(100);p__Ascomycota(100);c__Saccharomycetes(100);o__Saccharomycetales(100);f__Metschnikowiaceae(100);g__Metschnikowia(100);g__Metschnikowia_unclassified(100); | Metschnikowia | 13 | Saprotroph | \|Undefined Saprotroph\| | Yeast | NULL | Probable | As nectar-tap_saprotroph (Põlme et al. 2020); Decay sugar-rich substrates (Põlme et al. 2020) | Põlme S, et al. 2020. Fungal Diversity 105:1-16 ((https://doi.org/10.1007/s13225-020-00466-2)) |
| Otu42808 | 1 | k__Fungi(100);p__Ascomycota(100);c__Saccharomycetes(100);o__Saccharomycetales(100);f__Metschnikowiaceae(100);g__Metschnikowia(100);s__Metschnikowia_pulcherrima(100); | Metschnikowia | 13 | Saprotroph | \|Undefined Saprotroph\| | Yeast | NULL | Probable | As nectar-tap_saprotroph (Põlme et al. 2020); Decay sugar-rich substrates (Põlme et al. 2020) | Põlme S, et al. 2020. Fungal Diversity 105:1-16 ((https://doi.org/10.1007/s13225-020-00466-2)) |
| Otu42809 | 1 | k__Fungi(100);p__Ascomycota(100);c__Saccharomycetes(100);o__Saccharomycetales(100);f__Metschnikowiaceae(100);g__Metschnikowia(100);g__Metschnikowia_unclassified(100); | Metschnikowia | 13 | Saprotroph | \|Undefined Saprotroph\| | Yeast | NULL | Probable | As nectar-tap_saprotroph (Põlme et al. 2020); Decay sugar-rich substrates (Põlme et al. 2020) | Põlme S, et al. 2020. Fungal Diversity 105:1-16 ((https://doi.org/10.1007/s13225-020-00466-2)) |
| Otu42810 | 1 | k__Fungi(100);p__Ascomycota(100);c__Saccharomycetes(100);o__Saccharomycetales(100);f__Metschnikowiaceae(100);g__Metschnikowia(100);s__Metschnikowia_pulcherrima(100); | Metschnikowia | 13 | Saprotroph | \|Undefined Saprotroph\| | Yeast | NULL | Probable | As nectar-tap_saprotroph (Põlme et al. 2020); Decay sugar-rich substrates (Põlme et al. 2020) | Põlme S, et al. 2020. Fungal Diversity 105:1-16 ((https://doi.org/10.1007/s13225-020-00466-2)) |
| Otu42811 | 1 | k__Fungi(100);p__Ascomycota(100);c__Saccharomycetes(100);o__Saccharomycetales(100);f__Metschnikowiaceae(100);g__Metschnikowia(100);s__Metschnikowia_pulcherrima(100); | Metschnikowia | 13 | Saprotroph | \|Undefined Saprotroph\| | Yeast | NULL | Probable | As nectar-tap_saprotroph (Põlme et al. 2020); Decay sugar-rich substrates (Põlme et al. 2020) | Põlme S, et al. 2020. Fungal Diversity 105:1-16 ((https://doi.org/10.1007/s13225-020-00466-2)) |
| Otu42812 | 1 | k__Fungi(100);p__Ascomycota(100);c__Saccharomycetes(100);o__Saccharomycetales(100);f__Metschnikowiaceae(100);g__Metschnikowia(100);s__Metschnikowia_pulcherrima(100); | Metschnikowia | 13 | Saprotroph | \|Undefined Saprotroph\| | Yeast | NULL | Probable | As nectar-tap_saprotroph (Põlme et al. 2020); Decay sugar-rich substrates (Põlme et al. 2020) | Põlme S, et al. 2020. Fungal Diversity 105:1-16 ((https://doi.org/10.1007/s13225-020-00466-2)) |
| Otu42813 | 1 | k__Fungi(100);p__Ascomycota(100);c__Saccharomycetes(100);o__Saccharomycetales(100);f__Metschnikowiaceae(100);g__Metschnikowia(100);g__Metschnikowia_unclassified(100); | Metschnikowia | 13 | Saprotroph | \|Undefined Saprotroph\| | Yeast | NULL | Probable | As nectar-tap_saprotroph (Põlme et al. 2020); Decay sugar-rich substrates (Põlme et al. 2020) | Põlme S, et al. 2020. Fungal Diversity 105:1-16 ((https://doi.org/10.1007/s13225-020-00466-2)) |
| Otu42814 | 1 | k__Fungi(100);p__Ascomycota(100);c__Saccharomycetes(100);o__Saccharomycetales(100);f__Metschnikowiaceae(100);g__Metschnikowia(100);s__Metschnikowia_pulcherrima(100); | Metschnikowia | 13 | Saprotroph | \|Undefined Saprotroph\| | Yeast | NULL | Probable | As nectar-tap_saprotroph (Põlme et al. 2020); Decay sugar-rich substrates (Põlme et al. 2020) | Põlme S, et al. 2020. Fungal Diversity 105:1-16 ((https://doi.org/10.1007/s13225-020-00466-2)) |
| Otu42815 | 1 | k__Fungi(100);p__Ascomycota(100);c__Saccharomycetes(100);o__Saccharomycetales(100);f__Metschnikowiaceae(100);g__Metschnikowia(100);s__Metschnikowia_pulcherrima(100); | Metschnikowia | 13 | Saprotroph | \|Undefined Saprotroph\| | Yeast | NULL | Probable | As nectar-tap_saprotroph (Põlme et al. 2020); Decay sugar-rich substrates (Põlme et al. 2020) | Põlme S, et al. 2020. Fungal Diversity 105:1-16 ((https://doi.org/10.1007/s13225-020-00466-2)) |
| Otu42816 | 1 | k__Fungi(100);p__Ascomycota(100);c__Saccharomycetes(100);o__Saccharomycetales(100);f__Metschnikowiaceae(100);g__Metschnikowia(100);s__Metschnikowia_pulcherrima(100); | Metschnikowia | 13 | Saprotroph | \|Undefined Saprotroph\| | Yeast | NULL | Probable | As nectar-tap_saprotroph (Põlme et al. 2020); Decay sugar-rich substrates (Põlme et al. 2020) | Põlme S, et al. 2020. Fungal Diversity 105:1-16 ((https://doi.org/10.1007/s13225-020-00466-2)) |
| Otu42817 | 1 | k__Fungi(100);p__Ascomycota(100);c__Saccharomycetes(100);o__Saccharomycetales(100);f__Metschnikowiaceae(100);g__Metschnikowia(100);g__Metschnikowia_unclassified(100); | Metschnikowia | 13 | Saprotroph | \|Undefined Saprotroph\| | Yeast | NULL | Probable | As nectar-tap_saprotroph (Põlme et al. 2020); Decay sugar-rich substrates (Põlme et al. 2020) | Põlme S, et al. 2020. Fungal Diversity 105:1-16 ((https://doi.org/10.1007/s13225-020-00466-2)) |
| Otu42818 | 1 | k__Fungi(100);p__Ascomycota(100);c__Saccharomycetes(100);o__Saccharomycetales(100);f__Metschnikowiaceae(100);g__Metschnikowia(100);g__Metschnikowia_unclassified(100); | Metschnikowia | 13 | Saprotroph | \|Undefined Saprotroph\| | Yeast | NULL | Probable | As nectar-tap_saprotroph (Põlme et al. 2020); Decay sugar-rich substrates (Põlme et al. 2020) | Põlme S, et al. 2020. Fungal Diversity 105:1-16 ((https://doi.org/10.1007/s13225-020-00466-2)) |
| Otu42819 | 1 | k__Fungi(100);p__Ascomycota(100);c__Saccharomycetes(100);o__Saccharomycetales(100);f__Metschnikowiaceae(100);g__Metschnikowia(100);s__Metschnikowia_pulcherrima(100); | Metschnikowia | 13 | Saprotroph | \|Undefined Saprotroph\| | Yeast | NULL | Probable | As nectar-tap_saprotroph (Põlme et al. 2020); Decay sugar-rich substrates (Põlme et al. 2020) | Põlme S, et al. 2020. Fungal Diversity 105:1-16 ((https://doi.org/10.1007/s13225-020-00466-2)) |
| Otu42820 | 1 | k__Fungi(100);p__Ascomycota(100);c__Saccharomycetes(100);o__Saccharomycetales(100);f__Metschnikowiaceae(100);g__Metschnikowia(100);s__Metschnikowia_pulcherrima(100); | Metschnikowia | 13 | Saprotroph | \|Undefined Saprotroph\| | Yeast | NULL | Probable | As nectar-tap_saprotroph (Põlme et al. 2020); Decay sugar-rich substrates (Põlme et al. 2020) | Põlme S, et al. 2020. Fungal Diversity 105:1-16 ((https://doi.org/10.1007/s13225-020-00466-2)) |
| Otu42821 | 1 | k__Fungi(100);p__Ascomycota(100);c__Saccharomycetes(100);o__Saccharomycetales(100);f__Metschnikowiaceae(100);g__Metschnikowia(100);s__Metschnikowia_pulcherrima(100); | Metschnikowia | 13 | Saprotroph | \|Undefined Saprotroph\| | Yeast | NULL | Probable | As nectar-tap_saprotroph (Põlme et al. 2020); Decay sugar-rich substrates (Põlme et al. 2020) | Põlme S, et al. 2020. Fungal Diversity 105:1-16 ((https://doi.org/10.1007/s13225-020-00466-2)) |
| Otu42822 | 1 | k__Fungi(100);p__Ascomycota(100);c__Saccharomycetes(100);o__Saccharomycetales(100);f__Metschnikowiaceae(100);g__Metschnikowia(100);g__Metschnikowia_unclassified(100); | Metschnikowia | 13 | Saprotroph | \|Undefined Saprotroph\| | Yeast | NULL | Probable | As nectar-tap_saprotroph (Põlme et al. 2020); Decay sugar-rich substrates (Põlme et al. 2020) | Põlme S, et al. 2020. Fungal Diversity 105:1-16 ((https://doi.org/10.1007/s13225-020-00466-2)) |
| Otu42823 | 1 | k__Fungi(100);p__Ascomycota(100);c__Saccharomycetes(100);o__Saccharomycetales(100);f__Metschnikowiaceae(100);g__Metschnikowia(100);g__Metschnikowia_unclassified(100); | Metschnikowia | 13 | Saprotroph | \|Undefined Saprotroph\| | Yeast | NULL | Probable | As nectar-tap_saprotroph (Põlme et al. 2020); Decay sugar-rich substrates (Põlme et al. 2020) | Põlme S, et al. 2020. Fungal Diversity 105:1-16 ((https://doi.org/10.1007/s13225-020-00466-2)) |
| Otu42824 | 1 | k__Fungi(100);p__Ascomycota(100);c__Saccharomycetes(100);o__Saccharomycetales(100);f__Metschnikowiaceae(100);g__Metschnikowia(100);g__Metschnikowia_unclassified(100); | Metschnikowia | 13 | Saprotroph | \|Undefined Saprotroph\| | Yeast | NULL | Probable | As nectar-tap_saprotroph (Põlme et al. 2020); Decay sugar-rich substrates (Põlme et al. 2020) | Põlme S, et al. 2020. Fungal Diversity 105:1-16 ((https://doi.org/10.1007/s13225-020-00466-2)) |
| Otu42825 | 1 | k__Fungi(100);p__Ascomycota(100);c__Saccharomycetes(100);o__Saccharomycetales(100);f__Metschnikowiaceae(100);g__Metschnikowia(100);g__Metschnikowia_unclassified(100); | Metschnikowia | 13 | Saprotroph | \|Undefined Saprotroph\| | Yeast | NULL | Probable | As nectar-tap_saprotroph (Põlme et al. 2020); Decay sugar-rich substrates (Põlme et al. 2020) | Põlme S, et al. 2020. Fungal Diversity 105:1-16 ((https://doi.org/10.1007/s13225-020-00466-2)) |
| Otu42826 | 1 | k__Fungi(100);p__Ascomycota(100);c__Saccharomycetes(100);o__Saccharomycetales(100);f__Metschnikowiaceae(100);g__Metschnikowia(100);s__Metschnikowia_pulcherrima(100); | Metschnikowia | 13 | Saprotroph | \|Undefined Saprotroph\| | Yeast | NULL | Probable | As nectar-tap_saprotroph (Põlme et al. 2020); Decay sugar-rich substrates (Põlme et al. 2020) | Põlme S, et al. 2020. Fungal Diversity 105:1-16 ((https://doi.org/10.1007/s13225-020-00466-2)) |
| Otu42827 | 1 | k__Fungi(100);p__Ascomycota(100);c__Saccharomycetes(100);o__Saccharomycetales(100);f__Metschnikowiaceae(100);g__Metschnikowia(100);s__Metschnikowia_pulcherrima(100); | Metschnikowia | 13 | Saprotroph | \|Undefined Saprotroph\| | Yeast | NULL | Probable | As nectar-tap_saprotroph (Põlme et al. 2020); Decay sugar-rich substrates (Põlme et al. 2020) | Põlme S, et al. 2020. Fungal Diversity 105:1-16 ((https://doi.org/10.1007/s13225-020-00466-2)) |
| Otu42829 | 1 | k__Fungi(100);p__Ascomycota(100);c__Saccharomycetes(100);o__Saccharomycetales(100);f__Metschnikowiaceae(100);g__Metschnikowia(100);g__Metschnikowia_unclassified(100); | Metschnikowia | 13 | Saprotroph | \|Undefined Saprotroph\| | Yeast | NULL | Probable | As nectar-tap_saprotroph (Põlme et al. 2020); Decay sugar-rich substrates (Põlme et al. 2020) | Põlme S, et al. 2020. Fungal Diversity 105:1-16 ((https://doi.org/10.1007/s13225-020-00466-2)) |
| Otu42830 | 1 | k__Fungi(100);p__Ascomycota(100);c__Saccharomycetes(100);o__Saccharomycetales(100);f__Metschnikowiaceae(100);g__Metschnikowia(100);s__Metschnikowia_pulcherrima(100); | Metschnikowia | 13 | Saprotroph | \|Undefined Saprotroph\| | Yeast | NULL | Probable | As nectar-tap_saprotroph (Põlme et al. 2020); Decay sugar-rich substrates (Põlme et al. 2020) | Põlme S, et al. 2020. Fungal Diversity 105:1-16 ((https://doi.org/10.1007/s13225-020-00466-2)) |
| Otu42831 | 1 | k__Fungi(100);p__Ascomycota(100);c__Saccharomycetes(100);o__Saccharomycetales(100);f__Metschnikowiaceae(100);g__Metschnikowia(100);s__Metschnikowia_pulcherrima(100); | Metschnikowia | 13 | Saprotroph | \|Undefined Saprotroph\| | Yeast | NULL | Probable | As nectar-tap_saprotroph (Põlme et al. 2020); Decay sugar-rich substrates (Põlme et al. 2020) | Põlme S, et al. 2020. Fungal Diversity 105:1-16 ((https://doi.org/10.1007/s13225-020-00466-2)) |
| Otu42832 | 1 | k__Fungi(100);p__Ascomycota(100);c__Saccharomycetes(100);o__Saccharomycetales(100);f__Metschnikowiaceae(100);g__Metschnikowia(100);s__Metschnikowia_pulcherrima(100); | Metschnikowia | 13 | Saprotroph | \|Undefined Saprotroph\| | Yeast | NULL | Probable | As nectar-tap_saprotroph (Põlme et al. 2020); Decay sugar-rich substrates (Põlme et al. 2020) | Põlme S, et al. 2020. Fungal Diversity 105:1-16 ((https://doi.org/10.1007/s13225-020-00466-2)) |
| Otu42833 | 1 | k__Fungi(100);p__Ascomycota(100);c__Saccharomycetes(100);o__Saccharomycetales(100);f__Metschnikowiaceae(100);g__Metschnikowia(100);s__Metschnikowia_pulcherrima(100); | Metschnikowia | 13 | Saprotroph | \|Undefined Saprotroph\| | Yeast | NULL | Probable | As nectar-tap_saprotroph (Põlme et al. 2020); Decay sugar-rich substrates (Põlme et al. 2020) | Põlme S, et al. 2020. Fungal Diversity 105:1-16 ((https://doi.org/10.1007/s13225-020-00466-2)) |
| Otu42834 | 1 | k__Fungi(100);p__Ascomycota(100);c__Saccharomycetes(100);o__Saccharomycetales(100);f__Metschnikowiaceae(100);g__Metschnikowia(100);g__Metschnikowia_unclassified(100); | Metschnikowia | 13 | Saprotroph | \|Undefined Saprotroph\| | Yeast | NULL | Probable | As nectar-tap_saprotroph (Põlme et al. 2020); Decay sugar-rich substrates (Põlme et al. 2020) | Põlme S, et al. 2020. Fungal Diversity 105:1-16 ((https://doi.org/10.1007/s13225-020-00466-2)) |
| Otu42835 | 1 | k__Fungi(100);p__Ascomycota(100);c__Saccharomycetes(100);o__Saccharomycetales(100);f__Metschnikowiaceae(100);g__Metschnikowia(100);g__Metschnikowia_unclassified(100); | Metschnikowia | 13 | Saprotroph | \|Undefined Saprotroph\| | Yeast | NULL | Probable | As nectar-tap_saprotroph (Põlme et al. 2020); Decay sugar-rich substrates (Põlme et al. 2020) | Põlme S, et al. 2020. Fungal Diversity 105:1-16 ((https://doi.org/10.1007/s13225-020-00466-2)) |
| Otu42836 | 1 | k__Fungi(100);p__Ascomycota(100);c__Saccharomycetes(100);o__Saccharomycetales(100);f__Metschnikowiaceae(100);g__Metschnikowia(100);s__Metschnikowia_pulcherrima(100); | Metschnikowia | 13 | Saprotroph | \|Undefined Saprotroph\| | Yeast | NULL | Probable | As nectar-tap_saprotroph (Põlme et al. 2020); Decay sugar-rich substrates (Põlme et al. 2020) | Põlme S, et al. 2020. Fungal Diversity 105:1-16 ((https://doi.org/10.1007/s13225-020-00466-2)) |
| Otu42837 | 1 | k__Fungi(100);p__Ascomycota(100);c__Saccharomycetes(100);o__Saccharomycetales(100);f__Metschnikowiaceae(100);g__Metschnikowia(100);s__Metschnikowia_pulcherrima(100); | Metschnikowia | 13 | Saprotroph | \|Undefined Saprotroph\| | Yeast | NULL | Probable | As nectar-tap_saprotroph (Põlme et al. 2020); Decay sugar-rich substrates (Põlme et al. 2020) | Põlme S, et al. 2020. Fungal Diversity 105:1-16 ((https://doi.org/10.1007/s13225-020-00466-2)) |
| Otu42838 | 1 | k__Fungi(100);p__Ascomycota(100);c__Saccharomycetes(100);o__Saccharomycetales(100);f__Metschnikowiaceae(100);g__Metschnikowia(100);g__Metschnikowia_unclassified(100); | Metschnikowia | 13 | Saprotroph | \|Undefined Saprotroph\| | Yeast | NULL | Probable | As nectar-tap_saprotroph (Põlme et al. 2020); Decay sugar-rich substrates (Põlme et al. 2020) | Põlme S, et al. 2020. Fungal Diversity 105:1-16 ((https://doi.org/10.1007/s13225-020-00466-2)) |
| Otu42839 | 1 | k__Fungi(100);p__Ascomycota(100);c__Saccharomycetes(100);o__Saccharomycetales(100);f__Metschnikowiaceae(100);g__Metschnikowia(100);s__Metschnikowia_pulcherrima(100); | Metschnikowia | 13 | Saprotroph | \|Undefined Saprotroph\| | Yeast | NULL | Probable | As nectar-tap_saprotroph (Põlme et al. 2020); Decay sugar-rich substrates (Põlme et al. 2020) | Põlme S, et al. 2020. Fungal Diversity 105:1-16 ((https://doi.org/10.1007/s13225-020-00466-2)) |
| Otu42840 | 1 | k__Fungi(100);p__Ascomycota(100);c__Saccharomycetes(100);o__Saccharomycetales(100);f__Metschnikowiaceae(100);g__Metschnikowia(100);s__Metschnikowia_pulcherrima(100); | Metschnikowia | 13 | Saprotroph | \|Undefined Saprotroph\| | Yeast | NULL | Probable | As nectar-tap_saprotroph (Põlme et al. 2020); Decay sugar-rich substrates (Põlme et al. 2020) | Põlme S, et al. 2020. Fungal Diversity 105:1-16 ((https://doi.org/10.1007/s13225-020-00466-2)) |
| Otu42841 | 1 | k__Fungi(100);p__Ascomycota(100);c__Saccharomycetes(100);o__Saccharomycetales(100);f__Metschnikowiaceae(100);g__Metschnikowia(100);g__Metschnikowia_unclassified(100); | Metschnikowia | 13 | Saprotroph | \|Undefined Saprotroph\| | Yeast | NULL | Probable | As nectar-tap_saprotroph (Põlme et al. 2020); Decay sugar-rich substrates (Põlme et al. 2020) | Põlme S, et al. 2020. Fungal Diversity 105:1-16 ((https://doi.org/10.1007/s13225-020-00466-2)) |
| Otu42843 | 1 | k__Fungi(100);p__Ascomycota(100);c__Saccharomycetes(100);o__Saccharomycetales(100);f__Metschnikowiaceae(100);g__Metschnikowia(100);g__Metschnikowia_unclassified(100); | Metschnikowia | 13 | Saprotroph | \|Undefined Saprotroph\| | Yeast | NULL | Probable | As nectar-tap_saprotroph (Põlme et al. 2020); Decay sugar-rich substrates (Põlme et al. 2020) | Põlme S, et al. 2020. Fungal Diversity 105:1-16 ((https://doi.org/10.1007/s13225-020-00466-2)) |
| Otu42844 | 1 | k__Fungi(100);p__Ascomycota(100);c__Saccharomycetes(100);o__Saccharomycetales(100);f__Metschnikowiaceae(100);g__Metschnikowia(100);s__Metschnikowia_pulcherrima(100); | Metschnikowia | 13 | Saprotroph | \|Undefined Saprotroph\| | Yeast | NULL | Probable | As nectar-tap_saprotroph (Põlme et al. 2020); Decay sugar-rich substrates (Põlme et al. 2020) | Põlme S, et al. 2020. Fungal Diversity 105:1-16 ((https://doi.org/10.1007/s13225-020-00466-2)) |
| Otu42845 | 1 | k__Fungi(100);p__Ascomycota(100);c__Saccharomycetes(100);o__Saccharomycetales(100);f__Metschnikowiaceae(100);g__Metschnikowia(100);s__Metschnikowia_pulcherrima(100); | Metschnikowia | 13 | Saprotroph | \|Undefined Saprotroph\| | Yeast | NULL | Probable | As nectar-tap_saprotroph (Põlme et al. 2020); Decay sugar-rich substrates (Põlme et al. 2020) | Põlme S, et al. 2020. Fungal Diversity 105:1-16 ((https://doi.org/10.1007/s13225-020-00466-2)) |
| Otu42847 | 1 | k__Fungi(100);p__Ascomycota(100);c__Saccharomycetes(100);o__Saccharomycetales(100);f__Metschnikowiaceae(100);g__Metschnikowia(100);s__Metschnikowia_pulcherrima(100); | Metschnikowia | 13 | Saprotroph | \|Undefined Saprotroph\| | Yeast | NULL | Probable | As nectar-tap_saprotroph (Põlme et al. 2020); Decay sugar-rich substrates (Põlme et al. 2020) | Põlme S, et al. 2020. Fungal Diversity 105:1-16 ((https://doi.org/10.1007/s13225-020-00466-2)) |
| Otu42848 | 1 | k__Fungi(100);p__Ascomycota(100);c__Saccharomycetes(100);o__Saccharomycetales(100);f__Metschnikowiaceae(100);g__Metschnikowia(100);g__Metschnikowia_unclassified(100); | Metschnikowia | 13 | Saprotroph | \|Undefined Saprotroph\| | Yeast | NULL | Probable | As nectar-tap_saprotroph (Põlme et al. 2020); Decay sugar-rich substrates (Põlme et al. 2020) | Põlme S, et al. 2020. Fungal Diversity 105:1-16 ((https://doi.org/10.1007/s13225-020-00466-2)) |
| Otu42849 | 1 | k__Fungi(100);p__Ascomycota(100);c__Saccharomycetes(100);o__Saccharomycetales(100);f__Metschnikowiaceae(100);g__Metschnikowia(100);s__Metschnikowia_pulcherrima(100); | Metschnikowia | 13 | Saprotroph | \|Undefined Saprotroph\| | Yeast | NULL | Probable | As nectar-tap_saprotroph (Põlme et al. 2020); Decay sugar-rich substrates (Põlme et al. 2020) | Põlme S, et al. 2020. Fungal Diversity 105:1-16 ((https://doi.org/10.1007/s13225-020-00466-2)) |
| Otu42850 | 1 | k__Fungi(100);p__Ascomycota(100);c__Saccharomycetes(100);o__Saccharomycetales(100);f__Metschnikowiaceae(100);g__Metschnikowia(100);g__Metschnikowia_unclassified(100); | Metschnikowia | 13 | Saprotroph | \|Undefined Saprotroph\| | Yeast | NULL | Probable | As nectar-tap_saprotroph (Põlme et al. 2020); Decay sugar-rich substrates (Põlme et al. 2020) | Põlme S, et al. 2020. Fungal Diversity 105:1-16 ((https://doi.org/10.1007/s13225-020-00466-2)) |
| Otu42851 | 1 | k__Fungi(100);p__Ascomycota(100);c__Saccharomycetes(100);o__Saccharomycetales(100);f__Metschnikowiaceae(100);g__Metschnikowia(100);s__Metschnikowia_pulcherrima(100); | Metschnikowia | 13 | Saprotroph | \|Undefined Saprotroph\| | Yeast | NULL | Probable | As nectar-tap_saprotroph (Põlme et al. 2020); Decay sugar-rich substrates (Põlme et al. 2020) | Põlme S, et al. 2020. Fungal Diversity 105:1-16 ((https://doi.org/10.1007/s13225-020-00466-2)) |
| Otu42852 | 1 | k__Fungi(100);p__Ascomycota(100);c__Saccharomycetes(100);o__Saccharomycetales(100);f__Metschnikowiaceae(100);g__Metschnikowia(100);s__Metschnikowia_pulcherrima(100); | Metschnikowia | 13 | Saprotroph | \|Undefined Saprotroph\| | Yeast | NULL | Probable | As nectar-tap_saprotroph (Põlme et al. 2020); Decay sugar-rich substrates (Põlme et al. 2020) | Põlme S, et al. 2020. Fungal Diversity 105:1-16 ((https://doi.org/10.1007/s13225-020-00466-2)) |
| Otu42853 | 1 | k__Fungi(100);p__Ascomycota(100);c__Saccharomycetes(100);o__Saccharomycetales(100);f__Metschnikowiaceae(100);g__Metschnikowia(100);g__Metschnikowia_unclassified(100); | Metschnikowia | 13 | Saprotroph | \|Undefined Saprotroph\| | Yeast | NULL | Probable | As nectar-tap_saprotroph (Põlme et al. 2020); Decay sugar-rich substrates (Põlme et al. 2020) | Põlme S, et al. 2020. Fungal Diversity 105:1-16 ((https://doi.org/10.1007/s13225-020-00466-2)) |
| Otu42854 | 1 | k__Fungi(100);p__Ascomycota(100);c__Saccharomycetes(100);o__Saccharomycetales(100);f__Metschnikowiaceae(100);g__Metschnikowia(100);s__Metschnikowia_pulcherrima(100); | Metschnikowia | 13 | Saprotroph | \|Undefined Saprotroph\| | Yeast | NULL | Probable | As nectar-tap_saprotroph (Põlme et al. 2020); Decay sugar-rich substrates (Põlme et al. 2020) | Põlme S, et al. 2020. Fungal Diversity 105:1-16 ((https://doi.org/10.1007/s13225-020-00466-2)) |
| Otu42855 | 1 | k__Fungi(100);p__Ascomycota(100);c__Saccharomycetes(100);o__Saccharomycetales(100);f__Metschnikowiaceae(100);g__Metschnikowia(100);s__Metschnikowia_pulcherrima(100); | Metschnikowia | 13 | Saprotroph | \|Undefined Saprotroph\| | Yeast | NULL | Probable | As nectar-tap_saprotroph (Põlme et al. 2020); Decay sugar-rich substrates (Põlme et al. 2020) | Põlme S, et al. 2020. Fungal Diversity 105:1-16 ((https://doi.org/10.1007/s13225-020-00466-2)) |
| Otu42856 | 1 | k__Fungi(100);p__Ascomycota(100);c__Saccharomycetes(100);o__Saccharomycetales(100);f__Metschnikowiaceae(100);g__Metschnikowia(100);g__Metschnikowia_unclassified(100); | Metschnikowia | 13 | Saprotroph | \|Undefined Saprotroph\| | Yeast | NULL | Probable | As nectar-tap_saprotroph (Põlme et al. 2020); Decay sugar-rich substrates (Põlme et al. 2020) | Põlme S, et al. 2020. Fungal Diversity 105:1-16 ((https://doi.org/10.1007/s13225-020-00466-2)) |
| Otu42857 | 1 | k__Fungi(100);p__Ascomycota(100);c__Saccharomycetes(100);o__Saccharomycetales(100);f__Metschnikowiaceae(100);g__Metschnikowia(100);s__Metschnikowia_pulcherrima(100); | Metschnikowia | 13 | Saprotroph | \|Undefined Saprotroph\| | Yeast | NULL | Probable | As nectar-tap_saprotroph (Põlme et al. 2020); Decay sugar-rich substrates (Põlme et al. 2020) | Põlme S, et al. 2020. Fungal Diversity 105:1-16 ((https://doi.org/10.1007/s13225-020-00466-2)) |
| Otu42858 | 1 | k__Fungi(100);p__Ascomycota(100);c__Saccharomycetes(100);o__Saccharomycetales(100);f__Metschnikowiaceae(100);g__Metschnikowia(100);s__Metschnikowia_pulcherrima(100); | Metschnikowia | 13 | Saprotroph | \|Undefined Saprotroph\| | Yeast | NULL | Probable | As nectar-tap_saprotroph (Põlme et al. 2020); Decay sugar-rich substrates (Põlme et al. 2020) | Põlme S, et al. 2020. Fungal Diversity 105:1-16 ((https://doi.org/10.1007/s13225-020-00466-2)) |
| Otu42859 | 1 | k__Fungi(100);p__Ascomycota(100);c__Saccharomycetes(100);o__Saccharomycetales(100);f__Metschnikowiaceae(100);g__Metschnikowia(100);s__Metschnikowia_pulcherrima(100); | Metschnikowia | 13 | Saprotroph | \|Undefined Saprotroph\| | Yeast | NULL | Probable | As nectar-tap_saprotroph (Põlme et al. 2020); Decay sugar-rich substrates (Põlme et al. 2020) | Põlme S, et al. 2020. Fungal Diversity 105:1-16 ((https://doi.org/10.1007/s13225-020-00466-2)) |
| Otu42860 | 1 | k__Fungi(100);p__Ascomycota(100);c__Saccharomycetes(100);o__Saccharomycetales(100);f__Metschnikowiaceae(100);g__Metschnikowia(100);s__Metschnikowia_pulcherrima(100); | Metschnikowia | 13 | Saprotroph | \|Undefined Saprotroph\| | Yeast | NULL | Probable | As nectar-tap_saprotroph (Põlme et al. 2020); Decay sugar-rich substrates (Põlme et al. 2020) | Põlme S, et al. 2020. Fungal Diversity 105:1-16 ((https://doi.org/10.1007/s13225-020-00466-2)) |
| Otu42861 | 1 | k__Fungi(100);p__Ascomycota(100);c__Saccharomycetes(100);o__Saccharomycetales(100);f__Metschnikowiaceae(100);g__Metschnikowia(100);s__Metschnikowia_pulcherrima(100); | Metschnikowia | 13 | Saprotroph | \|Undefined Saprotroph\| | Yeast | NULL | Probable | As nectar-tap_saprotroph (Põlme et al. 2020); Decay sugar-rich substrates (Põlme et al. 2020) | Põlme S, et al. 2020. Fungal Diversity 105:1-16 ((https://doi.org/10.1007/s13225-020-00466-2)) |
| Otu42862 | 1 | k__Fungi(100);p__Ascomycota(100);c__Saccharomycetes(100);o__Saccharomycetales(100);f__Metschnikowiaceae(100);g__Metschnikowia(100);s__Metschnikowia_pulcherrima(100); | Metschnikowia | 13 | Saprotroph | \|Undefined Saprotroph\| | Yeast | NULL | Probable | As nectar-tap_saprotroph (Põlme et al. 2020); Decay sugar-rich substrates (Põlme et al. 2020) | Põlme S, et al. 2020. Fungal Diversity 105:1-16 ((https://doi.org/10.1007/s13225-020-00466-2)) |
| Otu42864 | 1 | k__Fungi(100);p__Ascomycota(100);c__Saccharomycetes(100);o__Saccharomycetales(100);f__Metschnikowiaceae(100);g__Metschnikowia(100);g__Metschnikowia_unclassified(100); | Metschnikowia | 13 | Saprotroph | \|Undefined Saprotroph\| | Yeast | NULL | Probable | As nectar-tap_saprotroph (Põlme et al. 2020); Decay sugar-rich substrates (Põlme et al. 2020) | Põlme S, et al. 2020. Fungal Diversity 105:1-16 ((https://doi.org/10.1007/s13225-020-00466-2)) |
| Otu42865 | 1 | k__Fungi(100);p__Ascomycota(100);c__Saccharomycetes(100);o__Saccharomycetales(100);f__Metschnikowiaceae(100);g__Metschnikowia(100);s__Metschnikowia_pulcherrima(100); | Metschnikowia | 13 | Saprotroph | \|Undefined Saprotroph\| | Yeast | NULL | Probable | As nectar-tap_saprotroph (Põlme et al. 2020); Decay sugar-rich substrates (Põlme et al. 2020) | Põlme S, et al. 2020. Fungal Diversity 105:1-16 ((https://doi.org/10.1007/s13225-020-00466-2)) |
| Otu42866 | 1 | k__Fungi(100);p__Ascomycota(100);c__Saccharomycetes(100);o__Saccharomycetales(100);f__Metschnikowiaceae(100);g__Metschnikowia(100);s__Metschnikowia_pulcherrima(100); | Metschnikowia | 13 | Saprotroph | \|Undefined Saprotroph\| | Yeast | NULL | Probable | As nectar-tap_saprotroph (Põlme et al. 2020); Decay sugar-rich substrates (Põlme et al. 2020) | Põlme S, et al. 2020. Fungal Diversity 105:1-16 ((https://doi.org/10.1007/s13225-020-00466-2)) |
| Otu42867 | 1 | k__Fungi(100);p__Ascomycota(100);c__Saccharomycetes(100);o__Saccharomycetales(100);f__Metschnikowiaceae(100);g__Metschnikowia(100);s__Metschnikowia_pulcherrima(100); | Metschnikowia | 13 | Saprotroph | \|Undefined Saprotroph\| | Yeast | NULL | Probable | As nectar-tap_saprotroph (Põlme et al. 2020); Decay sugar-rich substrates (Põlme et al. 2020) | Põlme S, et al. 2020. Fungal Diversity 105:1-16 ((https://doi.org/10.1007/s13225-020-00466-2)) |
| Otu42868 | 1 | k__Fungi(100);p__Ascomycota(100);c__Saccharomycetes(100);o__Saccharomycetales(100);f__Metschnikowiaceae(100);g__Metschnikowia(100);g__Metschnikowia_unclassified(100); | Metschnikowia | 13 | Saprotroph | \|Undefined Saprotroph\| | Yeast | NULL | Probable | As nectar-tap_saprotroph (Põlme et al. 2020); Decay sugar-rich substrates (Põlme et al. 2020) | Põlme S, et al. 2020. Fungal Diversity 105:1-16 ((https://doi.org/10.1007/s13225-020-00466-2)) |
| Otu42869 | 1 | k__Fungi(100);p__Ascomycota(100);c__Saccharomycetes(100);o__Saccharomycetales(100);f__Metschnikowiaceae(100);g__Metschnikowia(100);g__Metschnikowia_unclassified(100); | Metschnikowia | 13 | Saprotroph | \|Undefined Saprotroph\| | Yeast | NULL | Probable | As nectar-tap_saprotroph (Põlme et al. 2020); Decay sugar-rich substrates (Põlme et al. 2020) | Põlme S, et al. 2020. Fungal Diversity 105:1-16 ((https://doi.org/10.1007/s13225-020-00466-2)) |
| Otu42870 | 1 | k__Fungi(100);p__Ascomycota(100);c__Saccharomycetes(100);o__Saccharomycetales(100);f__Metschnikowiaceae(100);g__Metschnikowia(100);g__Metschnikowia_unclassified(100); | Metschnikowia | 13 | Saprotroph | \|Undefined Saprotroph\| | Yeast | NULL | Probable | As nectar-tap_saprotroph (Põlme et al. 2020); Decay sugar-rich substrates (Põlme et al. 2020) | Põlme S, et al. 2020. Fungal Diversity 105:1-16 ((https://doi.org/10.1007/s13225-020-00466-2)) |
| Otu42871 | 1 | k__Fungi(100);p__Ascomycota(100);c__Saccharomycetes(100);o__Saccharomycetales(100);f__Metschnikowiaceae(100);g__Metschnikowia(100);s__Metschnikowia_pulcherrima(100); | Metschnikowia | 13 | Saprotroph | \|Undefined Saprotroph\| | Yeast | NULL | Probable | As nectar-tap_saprotroph (Põlme et al. 2020); Decay sugar-rich substrates (Põlme et al. 2020) | Põlme S, et al. 2020. Fungal Diversity 105:1-16 ((https://doi.org/10.1007/s13225-020-00466-2)) |
| Otu42872 | 1 | k__Fungi(100);p__Ascomycota(100);c__Saccharomycetes(100);o__Saccharomycetales(100);f__Metschnikowiaceae(100);g__Metschnikowia(100);g__Metschnikowia_unclassified(100); | Metschnikowia | 13 | Saprotroph | \|Undefined Saprotroph\| | Yeast | NULL | Probable | As nectar-tap_saprotroph (Põlme et al. 2020); Decay sugar-rich substrates (Põlme et al. 2020) | Põlme S, et al. 2020. Fungal Diversity 105:1-16 ((https://doi.org/10.1007/s13225-020-00466-2)) |
| Otu42873 | 1 | k__Fungi(100);p__Ascomycota(100);c__Saccharomycetes(100);o__Saccharomycetales(100);f__Metschnikowiaceae(100);g__Metschnikowia(100);g__Metschnikowia_unclassified(100); | Metschnikowia | 13 | Saprotroph | \|Undefined Saprotroph\| | Yeast | NULL | Probable | As nectar-tap_saprotroph (Põlme et al. 2020); Decay sugar-rich substrates (Põlme et al. 2020) | Põlme S, et al. 2020. Fungal Diversity 105:1-16 ((https://doi.org/10.1007/s13225-020-00466-2)) |
| Otu42875 | 1 | k__Fungi(100);p__Ascomycota(100);c__Saccharomycetes(100);o__Saccharomycetales(100);f__Metschnikowiaceae(100);g__Metschnikowia(100);s__Metschnikowia_pulcherrima(100); | Metschnikowia | 13 | Saprotroph | \|Undefined Saprotroph\| | Yeast | NULL | Probable | As nectar-tap_saprotroph (Põlme et al. 2020); Decay sugar-rich substrates (Põlme et al. 2020) | Põlme S, et al. 2020. Fungal Diversity 105:1-16 ((https://doi.org/10.1007/s13225-020-00466-2)) |
| Otu42876 | 1 | k__Fungi(100);p__Ascomycota(100);c__Saccharomycetes(100);o__Saccharomycetales(100);f__Metschnikowiaceae(100);g__Metschnikowia(100);s__Metschnikowia_pulcherrima(100); | Metschnikowia | 13 | Saprotroph | \|Undefined Saprotroph\| | Yeast | NULL | Probable | As nectar-tap_saprotroph (Põlme et al. 2020); Decay sugar-rich substrates (Põlme et al. 2020) | Põlme S, et al. 2020. Fungal Diversity 105:1-16 ((https://doi.org/10.1007/s13225-020-00466-2)) |
| Otu42877 | 1 | k__Fungi(100);p__Ascomycota(100);c__Saccharomycetes(100);o__Saccharomycetales(100);f__Metschnikowiaceae(100);g__Metschnikowia(100);s__Metschnikowia_pulcherrima(100); | Metschnikowia | 13 | Saprotroph | \|Undefined Saprotroph\| | Yeast | NULL | Probable | As nectar-tap_saprotroph (Põlme et al. 2020); Decay sugar-rich substrates (Põlme et al. 2020) | Põlme S, et al. 2020. Fungal Diversity 105:1-16 ((https://doi.org/10.1007/s13225-020-00466-2)) |
| Otu42878 | 1 | k__Fungi(100);p__Ascomycota(100);c__Saccharomycetes(100);o__Saccharomycetales(100);f__Metschnikowiaceae(100);g__Metschnikowia(100);s__Metschnikowia_pulcherrima(100); | Metschnikowia | 13 | Saprotroph | \|Undefined Saprotroph\| | Yeast | NULL | Probable | As nectar-tap_saprotroph (Põlme et al. 2020); Decay sugar-rich substrates (Põlme et al. 2020) | Põlme S, et al. 2020. Fungal Diversity 105:1-16 ((https://doi.org/10.1007/s13225-020-00466-2)) |
| Otu42880 | 1 | k__Fungi(100);p__Ascomycota(100);c__Saccharomycetes(100);o__Saccharomycetales(100);f__Metschnikowiaceae(100);g__Metschnikowia(100);s__Metschnikowia_pulcherrima(100); | Metschnikowia | 13 | Saprotroph | \|Undefined Saprotroph\| | Yeast | NULL | Probable | As nectar-tap_saprotroph (Põlme et al. 2020); Decay sugar-rich substrates (Põlme et al. 2020) | Põlme S, et al. 2020. Fungal Diversity 105:1-16 ((https://doi.org/10.1007/s13225-020-00466-2)) |
| Otu42884 | 1 | k__Fungi(100);p__Ascomycota(100);c__Saccharomycetes(100);o__Saccharomycetales(100);f__Metschnikowiaceae(100);g__Metschnikowia(100);s__Metschnikowia_pulcherrima(100); | Metschnikowia | 13 | Saprotroph | \|Undefined Saprotroph\| | Yeast | NULL | Probable | As nectar-tap_saprotroph (Põlme et al. 2020); Decay sugar-rich substrates (Põlme et al. 2020) | Põlme S, et al. 2020. Fungal Diversity 105:1-16 ((https://doi.org/10.1007/s13225-020-00466-2)) |
| Otu42886 | 1 | k__Fungi(100);p__Ascomycota(100);c__Saccharomycetes(100);o__Saccharomycetales(100);f__Metschnikowiaceae(100);g__Metschnikowia(100);s__Metschnikowia_pulcherrima(100); | Metschnikowia | 13 | Saprotroph | \|Undefined Saprotroph\| | Yeast | NULL | Probable | As nectar-tap_saprotroph (Põlme et al. 2020); Decay sugar-rich substrates (Põlme et al. 2020) | Põlme S, et al. 2020. Fungal Diversity 105:1-16 ((https://doi.org/10.1007/s13225-020-00466-2)) |
| Otu42887 | 1 | k__Fungi(100);p__Ascomycota(100);c__Saccharomycetes(100);o__Saccharomycetales(100);f__Metschnikowiaceae(100);g__Metschnikowia(100);s__Metschnikowia_pulcherrima(100); | Metschnikowia | 13 | Saprotroph | \|Undefined Saprotroph\| | Yeast | NULL | Probable | As nectar-tap_saprotroph (Põlme et al. 2020); Decay sugar-rich substrates (Põlme et al. 2020) | Põlme S, et al. 2020. Fungal Diversity 105:1-16 ((https://doi.org/10.1007/s13225-020-00466-2)) |
| Otu42888 | 1 | k__Fungi(100);p__Ascomycota(100);c__Saccharomycetes(100);o__Saccharomycetales(100);f__Metschnikowiaceae(100);g__Metschnikowia(100);s__Metschnikowia_pulcherrima(100); | Metschnikowia | 13 | Saprotroph | \|Undefined Saprotroph\| | Yeast | NULL | Probable | As nectar-tap_saprotroph (Põlme et al. 2020); Decay sugar-rich substrates (Põlme et al. 2020) | Põlme S, et al. 2020. Fungal Diversity 105:1-16 ((https://doi.org/10.1007/s13225-020-00466-2)) |
| Otu42890 | 1 | k__Fungi(100);p__Ascomycota(100);c__Saccharomycetes(100);o__Saccharomycetales(100);f__Metschnikowiaceae(100);g__Metschnikowia(100);s__Metschnikowia_pulcherrima(100); | Metschnikowia | 13 | Saprotroph | \|Undefined Saprotroph\| | Yeast | NULL | Probable | As nectar-tap_saprotroph (Põlme et al. 2020); Decay sugar-rich substrates (Põlme et al. 2020) | Põlme S, et al. 2020. Fungal Diversity 105:1-16 ((https://doi.org/10.1007/s13225-020-00466-2)) |
| Otu42891 | 1 | k__Fungi(100);p__Ascomycota(100);c__Saccharomycetes(100);o__Saccharomycetales(100);f__Metschnikowiaceae(100);g__Metschnikowia(100);g__Metschnikowia_unclassified(100); | Metschnikowia | 13 | Saprotroph | \|Undefined Saprotroph\| | Yeast | NULL | Probable | As nectar-tap_saprotroph (Põlme et al. 2020); Decay sugar-rich substrates (Põlme et al. 2020) | Põlme S, et al. 2020. Fungal Diversity 105:1-16 ((https://doi.org/10.1007/s13225-020-00466-2)) |
| Otu42892 | 1 | k__Fungi(100);p__Ascomycota(100);c__Saccharomycetes(100);o__Saccharomycetales(100);f__Metschnikowiaceae(100);g__Metschnikowia(100);s__Metschnikowia_pulcherrima(100); | Metschnikowia | 13 | Saprotroph | \|Undefined Saprotroph\| | Yeast | NULL | Probable | As nectar-tap_saprotroph (Põlme et al. 2020); Decay sugar-rich substrates (Põlme et al. 2020) | Põlme S, et al. 2020. Fungal Diversity 105:1-16 ((https://doi.org/10.1007/s13225-020-00466-2)) |
| Otu42893 | 1 | k__Fungi(100);p__Ascomycota(100);c__Saccharomycetes(100);o__Saccharomycetales(100);f__Metschnikowiaceae(100);g__Metschnikowia(100);s__Metschnikowia_pulcherrima(100); | Metschnikowia | 13 | Saprotroph | \|Undefined Saprotroph\| | Yeast | NULL | Probable | As nectar-tap_saprotroph (Põlme et al. 2020); Decay sugar-rich substrates (Põlme et al. 2020) | Põlme S, et al. 2020. Fungal Diversity 105:1-16 ((https://doi.org/10.1007/s13225-020-00466-2)) |
| Otu42894 | 1 | k__Fungi(100);p__Ascomycota(100);c__Saccharomycetes(100);o__Saccharomycetales(100);f__Metschnikowiaceae(100);g__Metschnikowia(100);g__Metschnikowia_unclassified(100); | Metschnikowia | 13 | Saprotroph | \|Undefined Saprotroph\| | Yeast | NULL | Probable | As nectar-tap_saprotroph (Põlme et al. 2020); Decay sugar-rich substrates (Põlme et al. 2020) | Põlme S, et al. 2020. Fungal Diversity 105:1-16 ((https://doi.org/10.1007/s13225-020-00466-2)) |
| Otu42897 | 1 | k__Fungi(100);p__Ascomycota(100);c__Saccharomycetes(100);o__Saccharomycetales(100);f__Metschnikowiaceae(100);g__Metschnikowia(100);s__Metschnikowia_pulcherrima(100); | Metschnikowia | 13 | Saprotroph | \|Undefined Saprotroph\| | Yeast | NULL | Probable | As nectar-tap_saprotroph (Põlme et al. 2020); Decay sugar-rich substrates (Põlme et al. 2020) | Põlme S, et al. 2020. Fungal Diversity 105:1-16 ((https://doi.org/10.1007/s13225-020-00466-2)) |
| Otu42900 | 1 | k__Fungi(100);p__Ascomycota(100);c__Saccharomycetes(100);o__Saccharomycetales(100);f__Metschnikowiaceae(100);g__Metschnikowia(100);g__Metschnikowia_unclassified(100); | Metschnikowia | 13 | Saprotroph | \|Undefined Saprotroph\| | Yeast | NULL | Probable | As nectar-tap_saprotroph (Põlme et al. 2020); Decay sugar-rich substrates (Põlme et al. 2020) | Põlme S, et al. 2020. Fungal Diversity 105:1-16 ((https://doi.org/10.1007/s13225-020-00466-2)) |
| Otu42901 | 1 | k__Fungi(100);p__Ascomycota(100);c__Saccharomycetes(100);o__Saccharomycetales(100);f__Metschnikowiaceae(100);g__Metschnikowia(100);g__Metschnikowia_unclassified(100); | Metschnikowia | 13 | Saprotroph | \|Undefined Saprotroph\| | Yeast | NULL | Probable | As nectar-tap_saprotroph (Põlme et al. 2020); Decay sugar-rich substrates (Põlme et al. 2020) | Põlme S, et al. 2020. Fungal Diversity 105:1-16 ((https://doi.org/10.1007/s13225-020-00466-2)) |
| Otu42902 | 1 | k__Fungi(100);p__Ascomycota(100);c__Saccharomycetes(100);o__Saccharomycetales(100);f__Metschnikowiaceae(100);g__Metschnikowia(100);g__Metschnikowia_unclassified(100); | Metschnikowia | 13 | Saprotroph | \|Undefined Saprotroph\| | Yeast | NULL | Probable | As nectar-tap_saprotroph (Põlme et al. 2020); Decay sugar-rich substrates (Põlme et al. 2020) | Põlme S, et al. 2020. Fungal Diversity 105:1-16 ((https://doi.org/10.1007/s13225-020-00466-2)) |
| Otu42903 | 1 | k__Fungi(100);p__Ascomycota(100);c__Saccharomycetes(100);o__Saccharomycetales(100);f__Metschnikowiaceae(100);g__Metschnikowia(100);s__Metschnikowia_pulcherrima(100); | Metschnikowia | 13 | Saprotroph | \|Undefined Saprotroph\| | Yeast | NULL | Probable | As nectar-tap_saprotroph (Põlme et al. 2020); Decay sugar-rich substrates (Põlme et al. 2020) | Põlme S, et al. 2020. Fungal Diversity 105:1-16 ((https://doi.org/10.1007/s13225-020-00466-2)) |
| Otu42904 | 1 | k__Fungi(100);p__Ascomycota(100);c__Saccharomycetes(100);o__Saccharomycetales(100);f__Metschnikowiaceae(100);g__Metschnikowia(100);s__Metschnikowia_pulcherrima(100); | Metschnikowia | 13 | Saprotroph | \|Undefined Saprotroph\| | Yeast | NULL | Probable | As nectar-tap_saprotroph (Põlme et al. 2020); Decay sugar-rich substrates (Põlme et al. 2020) | Põlme S, et al. 2020. Fungal Diversity 105:1-16 ((https://doi.org/10.1007/s13225-020-00466-2)) |
| Otu42905 | 1 | k__Fungi(100);p__Ascomycota(100);c__Saccharomycetes(100);o__Saccharomycetales(100);f__Metschnikowiaceae(100);g__Metschnikowia(100);g__Metschnikowia_unclassified(100); | Metschnikowia | 13 | Saprotroph | \|Undefined Saprotroph\| | Yeast | NULL | Probable | As nectar-tap_saprotroph (Põlme et al. 2020); Decay sugar-rich substrates (Põlme et al. 2020) | Põlme S, et al. 2020. Fungal Diversity 105:1-16 ((https://doi.org/10.1007/s13225-020-00466-2)) |
| Otu42907 | 1 | k__Fungi(100);p__Ascomycota(100);c__Saccharomycetes(100);o__Saccharomycetales(100);f__Metschnikowiaceae(100);g__Metschnikowia(100);g__Metschnikowia_unclassified(100); | Metschnikowia | 13 | Saprotroph | \|Undefined Saprotroph\| | Yeast | NULL | Probable | As nectar-tap_saprotroph (Põlme et al. 2020); Decay sugar-rich substrates (Põlme et al. 2020) | Põlme S, et al. 2020. Fungal Diversity 105:1-16 ((https://doi.org/10.1007/s13225-020-00466-2)) |
| Otu42908 | 1 | k__Fungi(100);p__Ascomycota(100);c__Saccharomycetes(100);o__Saccharomycetales(100);f__Metschnikowiaceae(100);g__Metschnikowia(100);s__Metschnikowia_pulcherrima(100); | Metschnikowia | 13 | Saprotroph | \|Undefined Saprotroph\| | Yeast | NULL | Probable | As nectar-tap_saprotroph (Põlme et al. 2020); Decay sugar-rich substrates (Põlme et al. 2020) | Põlme S, et al. 2020. Fungal Diversity 105:1-16 ((https://doi.org/10.1007/s13225-020-00466-2)) |
| Otu42909 | 1 | k__Fungi(100);p__Ascomycota(100);c__Saccharomycetes(100);o__Saccharomycetales(100);f__Metschnikowiaceae(100);g__Metschnikowia(100);s__Metschnikowia_pulcherrima(100); | Metschnikowia | 13 | Saprotroph | \|Undefined Saprotroph\| | Yeast | NULL | Probable | As nectar-tap_saprotroph (Põlme et al. 2020); Decay sugar-rich substrates (Põlme et al. 2020) | Põlme S, et al. 2020. Fungal Diversity 105:1-16 ((https://doi.org/10.1007/s13225-020-00466-2)) |
| Otu42911 | 1 | k__Fungi(100);p__Ascomycota(100);c__Saccharomycetes(100);o__Saccharomycetales(100);f__Metschnikowiaceae(100);g__Metschnikowia(100);s__Metschnikowia_pulcherrima(100); | Metschnikowia | 13 | Saprotroph | \|Undefined Saprotroph\| | Yeast | NULL | Probable | As nectar-tap_saprotroph (Põlme et al. 2020); Decay sugar-rich substrates (Põlme et al. 2020) | Põlme S, et al. 2020. Fungal Diversity 105:1-16 ((https://doi.org/10.1007/s13225-020-00466-2)) |
| Otu42913 | 1 | k__Fungi(100);p__Ascomycota(100);c__Saccharomycetes(100);o__Saccharomycetales(100);f__Metschnikowiaceae(100);g__Metschnikowia(100);s__Metschnikowia_pulcherrima(100); | Metschnikowia | 13 | Saprotroph | \|Undefined Saprotroph\| | Yeast | NULL | Probable | As nectar-tap_saprotroph (Põlme et al. 2020); Decay sugar-rich substrates (Põlme et al. 2020) | Põlme S, et al. 2020. Fungal Diversity 105:1-16 ((https://doi.org/10.1007/s13225-020-00466-2)) |
| Otu42914 | 1 | k__Fungi(100);p__Ascomycota(100);c__Saccharomycetes(100);o__Saccharomycetales(100);f__Metschnikowiaceae(100);g__Metschnikowia(100);s__Metschnikowia_pulcherrima(100); | Metschnikowia | 13 | Saprotroph | \|Undefined Saprotroph\| | Yeast | NULL | Probable | As nectar-tap_saprotroph (Põlme et al. 2020); Decay sugar-rich substrates (Põlme et al. 2020) | Põlme S, et al. 2020. Fungal Diversity 105:1-16 ((https://doi.org/10.1007/s13225-020-00466-2)) |
| Otu42915 | 1 | k__Fungi(100);p__Ascomycota(100);c__Saccharomycetes(100);o__Saccharomycetales(100);f__Metschnikowiaceae(100);g__Metschnikowia(100);s__Metschnikowia_pulcherrima(100); | Metschnikowia | 13 | Saprotroph | \|Undefined Saprotroph\| | Yeast | NULL | Probable | As nectar-tap_saprotroph (Põlme et al. 2020); Decay sugar-rich substrates (Põlme et al. 2020) | Põlme S, et al. 2020. Fungal Diversity 105:1-16 ((https://doi.org/10.1007/s13225-020-00466-2)) |
| Otu42916 | 1 | k__Fungi(100);p__Ascomycota(100);c__Saccharomycetes(100);o__Saccharomycetales(100);f__Metschnikowiaceae(100);g__Metschnikowia(100);g__Metschnikowia_unclassified(100); | Metschnikowia | 13 | Saprotroph | \|Undefined Saprotroph\| | Yeast | NULL | Probable | As nectar-tap_saprotroph (Põlme et al. 2020); Decay sugar-rich substrates (Põlme et al. 2020) | Põlme S, et al. 2020. Fungal Diversity 105:1-16 ((https://doi.org/10.1007/s13225-020-00466-2)) |
| Otu42917 | 1 | k__Fungi(100);p__Ascomycota(100);c__Saccharomycetes(100);o__Saccharomycetales(100);f__Metschnikowiaceae(100);g__Metschnikowia(100);s__Metschnikowia_pulcherrima(100); | Metschnikowia | 13 | Saprotroph | \|Undefined Saprotroph\| | Yeast | NULL | Probable | As nectar-tap_saprotroph (Põlme et al. 2020); Decay sugar-rich substrates (Põlme et al. 2020) | Põlme S, et al. 2020. Fungal Diversity 105:1-16 ((https://doi.org/10.1007/s13225-020-00466-2)) |
| Otu42919 | 1 | k__Fungi(100);p__Ascomycota(100);c__Saccharomycetes(100);o__Saccharomycetales(100);f__Metschnikowiaceae(100);g__Metschnikowia(100);g__Metschnikowia_unclassified(100); | Metschnikowia | 13 | Saprotroph | \|Undefined Saprotroph\| | Yeast | NULL | Probable | As nectar-tap_saprotroph (Põlme et al. 2020); Decay sugar-rich substrates (Põlme et al. 2020) | Põlme S, et al. 2020. Fungal Diversity 105:1-16 ((https://doi.org/10.1007/s13225-020-00466-2)) |
| Otu42920 | 1 | k__Fungi(100);p__Ascomycota(100);c__Saccharomycetes(100);o__Saccharomycetales(100);f__Metschnikowiaceae(100);g__Metschnikowia(100);s__Metschnikowia_pulcherrima(100); | Metschnikowia | 13 | Saprotroph | \|Undefined Saprotroph\| | Yeast | NULL | Probable | As nectar-tap_saprotroph (Põlme et al. 2020); Decay sugar-rich substrates (Põlme et al. 2020) | Põlme S, et al. 2020. Fungal Diversity 105:1-16 ((https://doi.org/10.1007/s13225-020-00466-2)) |
| Otu42921 | 1 | k__Fungi(100);p__Ascomycota(100);c__Saccharomycetes(100);o__Saccharomycetales(100);f__Metschnikowiaceae(100);g__Metschnikowia(100);s__Metschnikowia_pulcherrima(100); | Metschnikowia | 13 | Saprotroph | \|Undefined Saprotroph\| | Yeast | NULL | Probable | As nectar-tap_saprotroph (Põlme et al. 2020); Decay sugar-rich substrates (Põlme et al. 2020) | Põlme S, et al. 2020. Fungal Diversity 105:1-16 ((https://doi.org/10.1007/s13225-020-00466-2)) |
| Otu42922 | 1 | k__Fungi(100);p__Ascomycota(100);c__Saccharomycetes(100);o__Saccharomycetales(100);f__Metschnikowiaceae(100);g__Metschnikowia(100);s__Metschnikowia_pulcherrima(100); | Metschnikowia | 13 | Saprotroph | \|Undefined Saprotroph\| | Yeast | NULL | Probable | As nectar-tap_saprotroph (Põlme et al. 2020); Decay sugar-rich substrates (Põlme et al. 2020) | Põlme S, et al. 2020. Fungal Diversity 105:1-16 ((https://doi.org/10.1007/s13225-020-00466-2)) |
| Otu42923 | 1 | k__Fungi(100);p__Ascomycota(100);c__Saccharomycetes(100);o__Saccharomycetales(100);f__Metschnikowiaceae(100);g__Metschnikowia(100);s__Metschnikowia_pulcherrima(100); | Metschnikowia | 13 | Saprotroph | \|Undefined Saprotroph\| | Yeast | NULL | Probable | As nectar-tap_saprotroph (Põlme et al. 2020); Decay sugar-rich substrates (Põlme et al. 2020) | Põlme S, et al. 2020. Fungal Diversity 105:1-16 ((https://doi.org/10.1007/s13225-020-00466-2)) |
| Otu42924 | 1 | k__Fungi(100);p__Ascomycota(100);c__Saccharomycetes(100);o__Saccharomycetales(100);f__Metschnikowiaceae(100);g__Metschnikowia(100);s__Metschnikowia_pulcherrima(100); | Metschnikowia | 13 | Saprotroph | \|Undefined Saprotroph\| | Yeast | NULL | Probable | As nectar-tap_saprotroph (Põlme et al. 2020); Decay sugar-rich substrates (Põlme et al. 2020) | Põlme S, et al. 2020. Fungal Diversity 105:1-16 ((https://doi.org/10.1007/s13225-020-00466-2)) |
| Otu42925 | 1 | k__Fungi(100);p__Ascomycota(100);c__Saccharomycetes(100);o__Saccharomycetales(100);f__Metschnikowiaceae(100);g__Metschnikowia(100);s__Metschnikowia_pulcherrima(100); | Metschnikowia | 13 | Saprotroph | \|Undefined Saprotroph\| | Yeast | NULL | Probable | As nectar-tap_saprotroph (Põlme et al. 2020); Decay sugar-rich substrates (Põlme et al. 2020) | Põlme S, et al. 2020. Fungal Diversity 105:1-16 ((https://doi.org/10.1007/s13225-020-00466-2)) |
| Otu42926 | 1 | k__Fungi(100);p__Ascomycota(100);c__Saccharomycetes(100);o__Saccharomycetales(100);f__Metschnikowiaceae(100);g__Metschnikowia(100);s__Metschnikowia_pulcherrima(100); | Metschnikowia | 13 | Saprotroph | \|Undefined Saprotroph\| | Yeast | NULL | Probable | As nectar-tap_saprotroph (Põlme et al. 2020); Decay sugar-rich substrates (Põlme et al. 2020) | Põlme S, et al. 2020. Fungal Diversity 105:1-16 ((https://doi.org/10.1007/s13225-020-00466-2)) |
| Otu42927 | 1 | k__Fungi(100);p__Ascomycota(100);c__Saccharomycetes(100);o__Saccharomycetales(100);f__Metschnikowiaceae(100);g__Metschnikowia(100);s__Metschnikowia_pulcherrima(100); | Metschnikowia | 13 | Saprotroph | \|Undefined Saprotroph\| | Yeast | NULL | Probable | As nectar-tap_saprotroph (Põlme et al. 2020); Decay sugar-rich substrates (Põlme et al. 2020) | Põlme S, et al. 2020. Fungal Diversity 105:1-16 ((https://doi.org/10.1007/s13225-020-00466-2)) |
| Otu42928 | 1 | k__Fungi(100);p__Ascomycota(100);c__Saccharomycetes(100);o__Saccharomycetales(100);f__Metschnikowiaceae(100);g__Metschnikowia(100);g__Metschnikowia_unclassified(100); | Metschnikowia | 13 | Saprotroph | \|Undefined Saprotroph\| | Yeast | NULL | Probable | As nectar-tap_saprotroph (Põlme et al. 2020); Decay sugar-rich substrates (Põlme et al. 2020) | Põlme S, et al. 2020. Fungal Diversity 105:1-16 ((https://doi.org/10.1007/s13225-020-00466-2)) |
| Otu42929 | 1 | k__Fungi(100);p__Ascomycota(100);c__Saccharomycetes(100);o__Saccharomycetales(100);f__Metschnikowiaceae(100);g__Metschnikowia(100);g__Metschnikowia_unclassified(100); | Metschnikowia | 13 | Saprotroph | \|Undefined Saprotroph\| | Yeast | NULL | Probable | As nectar-tap_saprotroph (Põlme et al. 2020); Decay sugar-rich substrates (Põlme et al. 2020) | Põlme S, et al. 2020. Fungal Diversity 105:1-16 ((https://doi.org/10.1007/s13225-020-00466-2)) |
| Otu42930 | 1 | k__Fungi(100);p__Ascomycota(100);c__Saccharomycetes(100);o__Saccharomycetales(100);f__Metschnikowiaceae(100);g__Metschnikowia(100);s__Metschnikowia_pulcherrima(100); | Metschnikowia | 13 | Saprotroph | \|Undefined Saprotroph\| | Yeast | NULL | Probable | As nectar-tap_saprotroph (Põlme et al. 2020); Decay sugar-rich substrates (Põlme et al. 2020) | Põlme S, et al. 2020. Fungal Diversity 105:1-16 ((https://doi.org/10.1007/s13225-020-00466-2)) |
| Otu42931 | 1 | k__Fungi(100);p__Ascomycota(100);c__Saccharomycetes(100);o__Saccharomycetales(100);f__Metschnikowiaceae(100);g__Metschnikowia(100);s__Metschnikowia_pulcherrima(100); | Metschnikowia | 13 | Saprotroph | \|Undefined Saprotroph\| | Yeast | NULL | Probable | As nectar-tap_saprotroph (Põlme et al. 2020); Decay sugar-rich substrates (Põlme et al. 2020) | Põlme S, et al. 2020. Fungal Diversity 105:1-16 ((https://doi.org/10.1007/s13225-020-00466-2)) |
| Otu42932 | 1 | k__Fungi(100);p__Ascomycota(100);c__Saccharomycetes(100);o__Saccharomycetales(100);f__Metschnikowiaceae(100);g__Metschnikowia(100);g__Metschnikowia_unclassified(100); | Metschnikowia | 13 | Saprotroph | \|Undefined Saprotroph\| | Yeast | NULL | Probable | As nectar-tap_saprotroph (Põlme et al. 2020); Decay sugar-rich substrates (Põlme et al. 2020) | Põlme S, et al. 2020. Fungal Diversity 105:1-16 ((https://doi.org/10.1007/s13225-020-00466-2)) |
| Otu42933 | 1 | k__Fungi(100);p__Ascomycota(100);c__Saccharomycetes(100);o__Saccharomycetales(100);f__Metschnikowiaceae(100);g__Metschnikowia(100);g__Metschnikowia_unclassified(100); | Metschnikowia | 13 | Saprotroph | \|Undefined Saprotroph\| | Yeast | NULL | Probable | As nectar-tap_saprotroph (Põlme et al. 2020); Decay sugar-rich substrates (Põlme et al. 2020) | Põlme S, et al. 2020. Fungal Diversity 105:1-16 ((https://doi.org/10.1007/s13225-020-00466-2)) |
| Otu42934 | 1 | k__Fungi(100);p__Ascomycota(100);c__Saccharomycetes(100);o__Saccharomycetales(100);f__Metschnikowiaceae(100);g__Metschnikowia(100);g__Metschnikowia_unclassified(100); | Metschnikowia | 13 | Saprotroph | \|Undefined Saprotroph\| | Yeast | NULL | Probable | As nectar-tap_saprotroph (Põlme et al. 2020); Decay sugar-rich substrates (Põlme et al. 2020) | Põlme S, et al. 2020. Fungal Diversity 105:1-16 ((https://doi.org/10.1007/s13225-020-00466-2)) |
| Otu42935 | 1 | k__Fungi(100);p__Ascomycota(100);c__Saccharomycetes(100);o__Saccharomycetales(100);f__Metschnikowiaceae(100);g__Metschnikowia(100);g__Metschnikowia_unclassified(100); | Metschnikowia | 13 | Saprotroph | \|Undefined Saprotroph\| | Yeast | NULL | Probable | As nectar-tap_saprotroph (Põlme et al. 2020); Decay sugar-rich substrates (Põlme et al. 2020) | Põlme S, et al. 2020. Fungal Diversity 105:1-16 ((https://doi.org/10.1007/s13225-020-00466-2)) |
| Otu42936 | 1 | k__Fungi(100);p__Ascomycota(100);c__Saccharomycetes(100);o__Saccharomycetales(100);f__Metschnikowiaceae(100);g__Metschnikowia(100);s__Metschnikowia_pulcherrima(100); | Metschnikowia | 13 | Saprotroph | \|Undefined Saprotroph\| | Yeast | NULL | Probable | As nectar-tap_saprotroph (Põlme et al. 2020); Decay sugar-rich substrates (Põlme et al. 2020) | Põlme S, et al. 2020. Fungal Diversity 105:1-16 ((https://doi.org/10.1007/s13225-020-00466-2)) |
| Otu42937 | 1 | k__Fungi(100);p__Ascomycota(100);c__Saccharomycetes(100);o__Saccharomycetales(100);f__Metschnikowiaceae(100);g__Metschnikowia(100);s__Metschnikowia_pulcherrima(100); | Metschnikowia | 13 | Saprotroph | \|Undefined Saprotroph\| | Yeast | NULL | Probable | As nectar-tap_saprotroph (Põlme et al. 2020); Decay sugar-rich substrates (Põlme et al. 2020) | Põlme S, et al. 2020. Fungal Diversity 105:1-16 ((https://doi.org/10.1007/s13225-020-00466-2)) |
| Otu42938 | 1 | k__Fungi(100);p__Ascomycota(100);c__Saccharomycetes(100);o__Saccharomycetales(100);f__Metschnikowiaceae(100);g__Metschnikowia(100);s__Metschnikowia_pulcherrima(100); | Metschnikowia | 13 | Saprotroph | \|Undefined Saprotroph\| | Yeast | NULL | Probable | As nectar-tap_saprotroph (Põlme et al. 2020); Decay sugar-rich substrates (Põlme et al. 2020) | Põlme S, et al. 2020. Fungal Diversity 105:1-16 ((https://doi.org/10.1007/s13225-020-00466-2)) |
| Otu42939 | 1 | k__Fungi(100);p__Ascomycota(100);c__Saccharomycetes(100);o__Saccharomycetales(100);f__Metschnikowiaceae(100);g__Metschnikowia(100);s__Metschnikowia_pulcherrima(100); | Metschnikowia | 13 | Saprotroph | \|Undefined Saprotroph\| | Yeast | NULL | Probable | As nectar-tap_saprotroph (Põlme et al. 2020); Decay sugar-rich substrates (Põlme et al. 2020) | Põlme S, et al. 2020. Fungal Diversity 105:1-16 ((https://doi.org/10.1007/s13225-020-00466-2)) |
| Otu42940 | 1 | k__Fungi(100);p__Ascomycota(100);c__Saccharomycetes(100);o__Saccharomycetales(100);f__Metschnikowiaceae(100);g__Metschnikowia(100);s__Metschnikowia_pulcherrima(100); | Metschnikowia | 13 | Saprotroph | \|Undefined Saprotroph\| | Yeast | NULL | Probable | As nectar-tap_saprotroph (Põlme et al. 2020); Decay sugar-rich substrates (Põlme et al. 2020) | Põlme S, et al. 2020. Fungal Diversity 105:1-16 ((https://doi.org/10.1007/s13225-020-00466-2)) |
| Otu42941 | 1 | k__Fungi(100);p__Ascomycota(100);c__Saccharomycetes(100);o__Saccharomycetales(100);f__Metschnikowiaceae(100);g__Metschnikowia(100);s__Metschnikowia_pulcherrima(100); | Metschnikowia | 13 | Saprotroph | \|Undefined Saprotroph\| | Yeast | NULL | Probable | As nectar-tap_saprotroph (Põlme et al. 2020); Decay sugar-rich substrates (Põlme et al. 2020) | Põlme S, et al. 2020. Fungal Diversity 105:1-16 ((https://doi.org/10.1007/s13225-020-00466-2)) |
| Otu42944 | 1 | k__Fungi(100);p__Ascomycota(100);c__Saccharomycetes(100);o__Saccharomycetales(100);f__Metschnikowiaceae(100);g__Metschnikowia(100);s__Metschnikowia_pulcherrima(100); | Metschnikowia | 13 | Saprotroph | \|Undefined Saprotroph\| | Yeast | NULL | Probable | As nectar-tap_saprotroph (Põlme et al. 2020); Decay sugar-rich substrates (Põlme et al. 2020) | Põlme S, et al. 2020. Fungal Diversity 105:1-16 ((https://doi.org/10.1007/s13225-020-00466-2)) |
| Otu42945 | 1 | k__Fungi(100);p__Ascomycota(100);c__Saccharomycetes(100);o__Saccharomycetales(100);f__Metschnikowiaceae(100);g__Metschnikowia(100);s__Metschnikowia_pulcherrima(100); | Metschnikowia | 13 | Saprotroph | \|Undefined Saprotroph\| | Yeast | NULL | Probable | As nectar-tap_saprotroph (Põlme et al. 2020); Decay sugar-rich substrates (Põlme et al. 2020) | Põlme S, et al. 2020. Fungal Diversity 105:1-16 ((https://doi.org/10.1007/s13225-020-00466-2)) |
| Otu42946 | 1 | k__Fungi(100);p__Ascomycota(100);c__Saccharomycetes(100);o__Saccharomycetales(100);f__Metschnikowiaceae(100);g__Metschnikowia(100);s__Metschnikowia_pulcherrima(100); | Metschnikowia | 13 | Saprotroph | \|Undefined Saprotroph\| | Yeast | NULL | Probable | As nectar-tap_saprotroph (Põlme et al. 2020); Decay sugar-rich substrates (Põlme et al. 2020) | Põlme S, et al. 2020. Fungal Diversity 105:1-16 ((https://doi.org/10.1007/s13225-020-00466-2)) |
| Otu42947 | 1 | k__Fungi(100);p__Ascomycota(100);c__Saccharomycetes(100);o__Saccharomycetales(100);f__Metschnikowiaceae(100);g__Metschnikowia(100);g__Metschnikowia_unclassified(100); | Metschnikowia | 13 | Saprotroph | \|Undefined Saprotroph\| | Yeast | NULL | Probable | As nectar-tap_saprotroph (Põlme et al. 2020); Decay sugar-rich substrates (Põlme et al. 2020) | Põlme S, et al. 2020. Fungal Diversity 105:1-16 ((https://doi.org/10.1007/s13225-020-00466-2)) |
| Otu42948 | 1 | k__Fungi(100);p__Ascomycota(100);c__Saccharomycetes(100);o__Saccharomycetales(100);f__Metschnikowiaceae(100);g__Metschnikowia(100);s__Metschnikowia_pulcherrima(100); | Metschnikowia | 13 | Saprotroph | \|Undefined Saprotroph\| | Yeast | NULL | Probable | As nectar-tap_saprotroph (Põlme et al. 2020); Decay sugar-rich substrates (Põlme et al. 2020) | Põlme S, et al. 2020. Fungal Diversity 105:1-16 ((https://doi.org/10.1007/s13225-020-00466-2)) |
| Otu42949 | 1 | k__Fungi(100);p__Ascomycota(100);c__Saccharomycetes(100);o__Saccharomycetales(100);f__Metschnikowiaceae(100);g__Metschnikowia(100);s__Metschnikowia_pulcherrima(100); | Metschnikowia | 13 | Saprotroph | \|Undefined Saprotroph\| | Yeast | NULL | Probable | As nectar-tap_saprotroph (Põlme et al. 2020); Decay sugar-rich substrates (Põlme et al. 2020) | Põlme S, et al. 2020. Fungal Diversity 105:1-16 ((https://doi.org/10.1007/s13225-020-00466-2)) |
| Otu42950 | 1 | k__Fungi(100);p__Ascomycota(100);c__Saccharomycetes(100);o__Saccharomycetales(100);f__Metschnikowiaceae(100);g__Metschnikowia(100);g__Metschnikowia_unclassified(100); | Metschnikowia | 13 | Saprotroph | \|Undefined Saprotroph\| | Yeast | NULL | Probable | As nectar-tap_saprotroph (Põlme et al. 2020); Decay sugar-rich substrates (Põlme et al. 2020) | Põlme S, et al. 2020. Fungal Diversity 105:1-16 ((https://doi.org/10.1007/s13225-020-00466-2)) |
| Otu42951 | 1 | k__Fungi(100);p__Ascomycota(100);c__Saccharomycetes(100);o__Saccharomycetales(100);f__Metschnikowiaceae(100);g__Metschnikowia(100);s__Metschnikowia_pulcherrima(100); | Metschnikowia | 13 | Saprotroph | \|Undefined Saprotroph\| | Yeast | NULL | Probable | As nectar-tap_saprotroph (Põlme et al. 2020); Decay sugar-rich substrates (Põlme et al. 2020) | Põlme S, et al. 2020. Fungal Diversity 105:1-16 ((https://doi.org/10.1007/s13225-020-00466-2)) |
| Otu42952 | 1 | k__Fungi(100);p__Ascomycota(100);c__Saccharomycetes(100);o__Saccharomycetales(100);f__Metschnikowiaceae(100);g__Metschnikowia(100);s__Metschnikowia_pulcherrima(100); | Metschnikowia | 13 | Saprotroph | \|Undefined Saprotroph\| | Yeast | NULL | Probable | As nectar-tap_saprotroph (Põlme et al. 2020); Decay sugar-rich substrates (Põlme et al. 2020) | Põlme S, et al. 2020. Fungal Diversity 105:1-16 ((https://doi.org/10.1007/s13225-020-00466-2)) |
| Otu42953 | 1 | k__Fungi(100);p__Ascomycota(100);c__Saccharomycetes(100);o__Saccharomycetales(100);f__Metschnikowiaceae(100);g__Metschnikowia(100);s__Metschnikowia_pulcherrima(100); | Metschnikowia | 13 | Saprotroph | \|Undefined Saprotroph\| | Yeast | NULL | Probable | As nectar-tap_saprotroph (Põlme et al. 2020); Decay sugar-rich substrates (Põlme et al. 2020) | Põlme S, et al. 2020. Fungal Diversity 105:1-16 ((https://doi.org/10.1007/s13225-020-00466-2)) |
| Otu42954 | 1 | k__Fungi(100);p__Ascomycota(100);c__Saccharomycetes(100);o__Saccharomycetales(100);f__Metschnikowiaceae(100);g__Metschnikowia(100);s__Metschnikowia_pulcherrima(100); | Metschnikowia | 13 | Saprotroph | \|Undefined Saprotroph\| | Yeast | NULL | Probable | As nectar-tap_saprotroph (Põlme et al. 2020); Decay sugar-rich substrates (Põlme et al. 2020) | Põlme S, et al. 2020. Fungal Diversity 105:1-16 ((https://doi.org/10.1007/s13225-020-00466-2)) |
| Otu42956 | 1 | k__Fungi(100);p__Ascomycota(100);c__Saccharomycetes(100);o__Saccharomycetales(100);f__Metschnikowiaceae(100);g__Metschnikowia(100);g__Metschnikowia_unclassified(100); | Metschnikowia | 13 | Saprotroph | \|Undefined Saprotroph\| | Yeast | NULL | Probable | As nectar-tap_saprotroph (Põlme et al. 2020); Decay sugar-rich substrates (Põlme et al. 2020) | Põlme S, et al. 2020. Fungal Diversity 105:1-16 ((https://doi.org/10.1007/s13225-020-00466-2)) |
| Otu42957 | 1 | k__Fungi(100);p__Ascomycota(100);c__Saccharomycetes(100);o__Saccharomycetales(100);f__Metschnikowiaceae(100);g__Metschnikowia(100);s__Metschnikowia_pulcherrima(100); | Metschnikowia | 13 | Saprotroph | \|Undefined Saprotroph\| | Yeast | NULL | Probable | As nectar-tap_saprotroph (Põlme et al. 2020); Decay sugar-rich substrates (Põlme et al. 2020) | Põlme S, et al. 2020. Fungal Diversity 105:1-16 ((https://doi.org/10.1007/s13225-020-00466-2)) |
| Otu42958 | 1 | k__Fungi(100);p__Ascomycota(100);c__Saccharomycetes(100);o__Saccharomycetales(100);f__Metschnikowiaceae(100);g__Metschnikowia(100);g__Metschnikowia_unclassified(100); | Metschnikowia | 13 | Saprotroph | \|Undefined Saprotroph\| | Yeast | NULL | Probable | As nectar-tap_saprotroph (Põlme et al. 2020); Decay sugar-rich substrates (Põlme et al. 2020) | Põlme S, et al. 2020. Fungal Diversity 105:1-16 ((https://doi.org/10.1007/s13225-020-00466-2)) |
| Otu42960 | 1 | k__Fungi(100);p__Ascomycota(100);c__Saccharomycetes(100);o__Saccharomycetales(100);f__Metschnikowiaceae(100);g__Metschnikowia(100);g__Metschnikowia_unclassified(100); | Metschnikowia | 13 | Saprotroph | \|Undefined Saprotroph\| | Yeast | NULL | Probable | As nectar-tap_saprotroph (Põlme et al. 2020); Decay sugar-rich substrates (Põlme et al. 2020) | Põlme S, et al. 2020. Fungal Diversity 105:1-16 ((https://doi.org/10.1007/s13225-020-00466-2)) |
| Otu42961 | 1 | k__Fungi(100);p__Ascomycota(100);c__Saccharomycetes(100);o__Saccharomycetales(100);f__Metschnikowiaceae(100);g__Metschnikowia(100);s__Metschnikowia_pulcherrima(100); | Metschnikowia | 13 | Saprotroph | \|Undefined Saprotroph\| | Yeast | NULL | Probable | As nectar-tap_saprotroph (Põlme et al. 2020); Decay sugar-rich substrates (Põlme et al. 2020) | Põlme S, et al. 2020. Fungal Diversity 105:1-16 ((https://doi.org/10.1007/s13225-020-00466-2)) |
| Otu42962 | 1 | k__Fungi(100);p__Ascomycota(100);c__Saccharomycetes(100);o__Saccharomycetales(100);f__Metschnikowiaceae(100);g__Metschnikowia(100);s__Metschnikowia_pulcherrima(100); | Metschnikowia | 13 | Saprotroph | \|Undefined Saprotroph\| | Yeast | NULL | Probable | As nectar-tap_saprotroph (Põlme et al. 2020); Decay sugar-rich substrates (Põlme et al. 2020) | Põlme S, et al. 2020. Fungal Diversity 105:1-16 ((https://doi.org/10.1007/s13225-020-00466-2)) |
| Otu42963 | 1 | k__Fungi(100);p__Ascomycota(100);c__Saccharomycetes(100);o__Saccharomycetales(100);f__Metschnikowiaceae(100);g__Metschnikowia(100);g__Metschnikowia_unclassified(100); | Metschnikowia | 13 | Saprotroph | \|Undefined Saprotroph\| | Yeast | NULL | Probable | As nectar-tap_saprotroph (Põlme et al. 2020); Decay sugar-rich substrates (Põlme et al. 2020) | Põlme S, et al. 2020. Fungal Diversity 105:1-16 ((https://doi.org/10.1007/s13225-020-00466-2)) |
| Otu42964 | 1 | k__Fungi(100);p__Ascomycota(100);c__Saccharomycetes(100);o__Saccharomycetales(100);f__Metschnikowiaceae(100);g__Metschnikowia(100);g__Metschnikowia_unclassified(100); | Metschnikowia | 13 | Saprotroph | \|Undefined Saprotroph\| | Yeast | NULL | Probable | As nectar-tap_saprotroph (Põlme et al. 2020); Decay sugar-rich substrates (Põlme et al. 2020) | Põlme S, et al. 2020. Fungal Diversity 105:1-16 ((https://doi.org/10.1007/s13225-020-00466-2)) |
| Otu42965 | 1 | k__Fungi(100);p__Ascomycota(100);c__Saccharomycetes(100);o__Saccharomycetales(100);f__Metschnikowiaceae(100);g__Metschnikowia(100);s__Metschnikowia_pulcherrima(100); | Metschnikowia | 13 | Saprotroph | \|Undefined Saprotroph\| | Yeast | NULL | Probable | As nectar-tap_saprotroph (Põlme et al. 2020); Decay sugar-rich substrates (Põlme et al. 2020) | Põlme S, et al. 2020. Fungal Diversity 105:1-16 ((https://doi.org/10.1007/s13225-020-00466-2)) |
| Otu42966 | 1 | k__Fungi(100);p__Ascomycota(100);c__Saccharomycetes(100);o__Saccharomycetales(100);f__Metschnikowiaceae(100);g__Metschnikowia(100);g__Metschnikowia_unclassified(100); | Metschnikowia | 13 | Saprotroph | \|Undefined Saprotroph\| | Yeast | NULL | Probable | As nectar-tap_saprotroph (Põlme et al. 2020); Decay sugar-rich substrates (Põlme et al. 2020) | Põlme S, et al. 2020. Fungal Diversity 105:1-16 ((https://doi.org/10.1007/s13225-020-00466-2)) |
| Otu42967 | 1 | k__Fungi(100);p__Ascomycota(100);c__Saccharomycetes(100);o__Saccharomycetales(100);f__Metschnikowiaceae(100);g__Metschnikowia(100);g__Metschnikowia_unclassified(100); | Metschnikowia | 13 | Saprotroph | \|Undefined Saprotroph\| | Yeast | NULL | Probable | As nectar-tap_saprotroph (Põlme et al. 2020); Decay sugar-rich substrates (Põlme et al. 2020) | Põlme S, et al. 2020. Fungal Diversity 105:1-16 ((https://doi.org/10.1007/s13225-020-00466-2)) |
| Otu42968 | 1 | k__Fungi(100);p__Ascomycota(100);c__Saccharomycetes(100);o__Saccharomycetales(100);f__Metschnikowiaceae(100);g__Metschnikowia(100);s__Metschnikowia_pulcherrima(100); | Metschnikowia | 13 | Saprotroph | \|Undefined Saprotroph\| | Yeast | NULL | Probable | As nectar-tap_saprotroph (Põlme et al. 2020); Decay sugar-rich substrates (Põlme et al. 2020) | Põlme S, et al. 2020. Fungal Diversity 105:1-16 ((https://doi.org/10.1007/s13225-020-00466-2)) |

| | | | | | | | | | | | |
|---|---|---|---|---|---|---|---|---|---|---|---|
| Otu42969 | 1 | k__Fungi(100);p__Ascomycota(100);c__Saccharomycetes(100);o__Saccharomycetales(100);f__Metschnikowiaceae(100);g__Metschnikowia(100);s__Metschnikowia_pulcherrima(100); | Metschnikowia | 13 | Saprotroph | \|Undefined Saprotroph\| | Yeast | NULL | Probable | As nectar-tap_saprotroph (Põlme et al. 2020); Decay sugar-rich substrates (Põlme et al. 2020) | Põlme S, et al. 2020. Fungal Diversity 105:1-16 ((https://doi.org/10.1007/s13225-020-00466-2)) |
| Otu42970 | 1 | k__Fungi(100);p__Ascomycota(100);c__Saccharomycetes(100);o__Saccharomycetales(100);f__Metschnikowiaceae(100);g__Metschnikowia(100);g__Metschnikowia_unclassified(100); | Metschnikowia | 13 | Saprotroph | \|Undefined Saprotroph\| | Yeast | NULL | Probable | As nectar-tap_saprotroph (Põlme et al. 2020); Decay sugar-rich substrates (Põlme et al. 2020) | Põlme S, et al. 2020. Fungal Diversity 105:1-16 ((https://doi.org/10.1007/s13225-020-00466-2)) |
| Otu42971 | 1 | k__Fungi(100);p__Ascomycota(100);c__Saccharomycetes(100);o__Saccharomycetales(100);f__Metschnikowiaceae(100);g__Metschnikowia(100);s__Metschnikowia_pulcherrima(100); | Metschnikowia | 13 | Saprotroph | \|Undefined Saprotroph\| | Yeast | NULL | Probable | As nectar-tap_saprotroph (Põlme et al. 2020); Decay sugar-rich substrates (Põlme et al. 2020) | Põlme S, et al. 2020. Fungal Diversity 105:1-16 ((https://doi.org/10.1007/s13225-020-00466-2)) |
| Otu42972 | 1 | k__Fungi(100);p__Ascomycota(100);c__Saccharomycetes(100);o__Saccharomycetales(100);f__Metschnikowiaceae(100);g__Metschnikowia(100);g__Metschnikowia_unclassified(100); | Metschnikowia | 13 | Saprotroph | \|Undefined Saprotroph\| | Yeast | NULL | Probable | As nectar-tap_saprotroph (Põlme et al. 2020); Decay sugar-rich substrates (Põlme et al. 2020) | Põlme S, et al. 2020. Fungal Diversity 105:1-16 ((https://doi.org/10.1007/s13225-020-00466-2)) |
| Otu42973 | 1 | k__Fungi(100);p__Ascomycota(100);c__Saccharomycetes(100);o__Saccharomycetales(100);f__Metschnikowiaceae(100);g__Metschnikowia(100);s__Metschnikowia_pulcherrima(100); | Metschnikowia | 13 | Saprotroph | \|Undefined Saprotroph\| | Yeast | NULL | Probable | As nectar-tap_saprotroph (Põlme et al. 2020); Decay sugar-rich substrates (Põlme et al. 2020) | Põlme S, et al. 2020. Fungal Diversity 105:1-16 ((https://doi.org/10.1007/s13225-020-00466-2)) |
| Otu42974 | 1 | k__Fungi(100);p__Ascomycota(100);c__Saccharomycetes(100);o__Saccharomycetales(100);f__Metschnikowiaceae(100);g__Metschnikowia(100);g__Metschnikowia_unclassified(100); | Metschnikowia | 13 | Saprotroph | \|Undefined Saprotroph\| | Yeast | NULL | Probable | As nectar-tap_saprotroph (Põlme et al. 2020); Decay sugar-rich substrates (Põlme et al. 2020) | Põlme S, et al. 2020. Fungal Diversity 105:1-16 ((https://doi.org/10.1007/s13225-020-00466-2)) |
| Otu42975 | 1 | k__Fungi(100);p__Ascomycota(100);c__Saccharomycetes(100);o__Saccharomycetales(100);f__Metschnikowiaceae(100);g__Metschnikowia(100);s__Metschnikowia_pulcherrima(100); | Metschnikowia | 13 | Saprotroph | \|Undefined Saprotroph\| | Yeast | NULL | Probable | As nectar-tap_saprotroph (Põlme et al. 2020); Decay sugar-rich substrates (Põlme et al. 2020) | Põlme S, et al. 2020. Fungal Diversity 105:1-16 ((https://doi.org/10.1007/s13225-020-00466-2)) |
| Otu42976 | 1 | k__Fungi(100);p__Ascomycota(100);c__Saccharomycetes(100);o__Saccharomycetales(100);f__Metschnikowiaceae(100);g__Metschnikowia(100);s__Metschnikowia_viticola(100); | Metschnikowia | 13 | Saprotroph | \|Undefined Saprotroph\| | Yeast | NULL | Probable | As nectar-tap_saprotroph (Põlme et al. 2020); Decay sugar-rich substrates (Põlme et al. 2020) | Põlme S, et al. 2020. Fungal Diversity 105:1-16 ((https://doi.org/10.1007/s13225-020-00466-2)) |
| Otu42977 | 1 | k__Fungi(100);p__Ascomycota(100);c__Saccharomycetes(100);o__Saccharomycetales(100);f__Metschnikowiaceae(100);g__Metschnikowia(100);s__Metschnikowia_pulcherrima(100); | Metschnikowia | 13 | Saprotroph | \|Undefined Saprotroph\| | Yeast | NULL | Probable | As nectar-tap_saprotroph (Põlme et al. 2020); Decay sugar-rich substrates (Põlme et al. 2020) | Põlme S, et al. 2020. Fungal Diversity 105:1-16 ((https://doi.org/10.1007/s13225-020-00466-2)) |
| Otu42978 | 1 | k__Fungi(100);p__Ascomycota(100);c__Saccharomycetes(100);o__Saccharomycetales(100);f__Metschnikowiaceae(100);g__Metschnikowia(100);s__Metschnikowia_pulcherrima(100); | Metschnikowia | 13 | Saprotroph | \|Undefined Saprotroph\| | Yeast | NULL | Probable | As nectar-tap_saprotroph (Põlme et al. 2020); Decay sugar-rich substrates (Põlme et al. 2020) | Põlme S, et al. 2020. Fungal Diversity 105:1-16 ((https://doi.org/10.1007/s13225-020-00466-2)) |
| Otu42980 | 1 | k__Fungi(100);p__Ascomycota(100);c__Saccharomycetes(100);o__Saccharomycetales(100);f__Metschnikowiaceae(100);g__Metschnikowia(100);g__Metschnikowia_unclassified(100); | Metschnikowia | 13 | Saprotroph | \|Undefined Saprotroph\| | Yeast | NULL | Probable | As nectar-tap_saprotroph (Põlme et al. 2020); Decay sugar-rich substrates (Põlme et al. 2020) | Põlme S, et al. 2020. Fungal Diversity 105:1-16 ((https://doi.org/10.1007/s13225-020-00466-2)) |
| Otu42981 | 1 | k__Fungi(100);p__Ascomycota(100);c__Saccharomycetes(100);o__Saccharomycetales(100);f__Metschnikowiaceae(100);g__Metschnikowia(100);g__Metschnikowia_unclassified(100); | Metschnikowia | 13 | Saprotroph | \|Undefined Saprotroph\| | Yeast | NULL | Probable | As nectar-tap_saprotroph (Põlme et al. 2020); Decay sugar-rich substrates (Põlme et al. 2020) | Põlme S, et al. 2020. Fungal Diversity 105:1-16 ((https://doi.org/10.1007/s13225-020-00466-2)) |
| Otu42982 | 1 | k__Fungi(100);p__Ascomycota(100);c__Saccharomycetes(100);o__Saccharomycetales(100);f__Metschnikowiaceae(100);g__Metschnikowia(100);s__Metschnikowia_pulcherrima(100); | Metschnikowia | 13 | Saprotroph | \|Undefined Saprotroph\| | Yeast | NULL | Probable | As nectar-tap_saprotroph (Põlme et al. 2020); Decay sugar-rich substrates (Põlme et al. 2020) | Põlme S, et al. 2020. Fungal Diversity 105:1-16 ((https://doi.org/10.1007/s13225-020-00466-2)) |
| Otu42983 | 1 | k__Fungi(100);p__Ascomycota(100);c__Saccharomycetes(100);o__Saccharomycetales(100);f__Metschnikowiaceae(100);g__Metschnikowia(100);g__Metschnikowia_unclassified(100); | Metschnikowia | 13 | Saprotroph | \|Undefined Saprotroph\| | Yeast | NULL | Probable | As nectar-tap_saprotroph (Põlme et al. 2020); Decay sugar-rich substrates (Põlme et al. 2020) | Põlme S, et al. 2020. Fungal Diversity 105:1-16 ((https://doi.org/10.1007/s13225-020-00466-2)) |
| Otu42984 | 1 | k__Fungi(100);p__Ascomycota(100);c__Saccharomycetes(100);o__Saccharomycetales(100);f__Metschnikowiaceae(100);g__Metschnikowia(100);s__Metschnikowia_pulcherrima(100); | Metschnikowia | 13 | Saprotroph | \|Undefined Saprotroph\| | Yeast | NULL | Probable | As nectar-tap_saprotroph (Põlme et al. 2020); Decay sugar-rich substrates (Põlme et al. 2020) | Põlme S, et al. 2020. Fungal Diversity 105:1-16 ((https://doi.org/10.1007/s13225-020-00466-2)) |
| Otu42985 | 1 | k__Fungi(100);p__Ascomycota(100);c__Saccharomycetes(100);o__Saccharomycetales(100);f__Metschnikowiaceae(100);g__Metschnikowia(100);s__Metschnikowia_pulcherrima(100); | Metschnikowia | 13 | Saprotroph | \|Undefined Saprotroph\| | Yeast | NULL | Probable | As nectar-tap_saprotroph (Põlme et al. 2020); Decay sugar-rich substrates (Põlme et al. 2020) | Põlme S, et al. 2020. Fungal Diversity 105:1-16 ((https://doi.org/10.1007/s13225-020-00466-2)) |
| Otu42986 | 1 | k__Fungi(100);p__Ascomycota(100);c__Saccharomycetes(100);o__Saccharomycetales(100);f__Metschnikowiaceae(100);g__Metschnikowia(100);s__Metschnikowia_pulcherrima(100); | Metschnikowia | 13 | Saprotroph | \|Undefined Saprotroph\| | Yeast | NULL | Probable | As nectar-tap_saprotroph (Põlme et al. 2020); Decay sugar-rich substrates (Põlme et al. 2020) | Põlme S, et al. 2020. Fungal Diversity 105:1-16 ((https://doi.org/10.1007/s13225-020-00466-2)) |
| Otu42988 | 1 | k__Fungi(100);p__Ascomycota(100);c__Saccharomycetes(100);o__Saccharomycetales(100);f__Metschnikowiaceae(100);g__Metschnikowia(100);s__Metschnikowia_pulcherrima(100); | Metschnikowia | 13 | Saprotroph | \|Undefined Saprotroph\| | Yeast | NULL | Probable | As nectar-tap_saprotroph (Põlme et al. 2020); Decay sugar-rich substrates (Põlme et al. 2020) | Põlme S, et al. 2020. Fungal Diversity 105:1-16 ((https://doi.org/10.1007/s13225-020-00466-2)) |
| Otu42989 | 1 | k__Fungi(100);p__Ascomycota(100);c__Saccharomycetes(100);o__Saccharomycetales(100);f__Metschnikowiaceae(100);g__Metschnikowia(100);g__Metschnikowia_unclassified(100); | Metschnikowia | 13 | Saprotroph | \|Undefined Saprotroph\| | Yeast | NULL | Probable | As nectar-tap_saprotroph (Põlme et al. 2020); Decay sugar-rich substrates (Põlme et al. 2020) | Põlme S, et al. 2020. Fungal Diversity 105:1-16 ((https://doi.org/10.1007/s13225-020-00466-2)) |
| Otu42990 | 1 | k__Fungi(100);p__Ascomycota(100);c__Saccharomycetes(100);o__Saccharomycetales(100);f__Metschnikowiaceae(100);g__Metschnikowia(100);g__Metschnikowia_unclassified(100); | Metschnikowia | 13 | Saprotroph | \|Undefined Saprotroph\| | Yeast | NULL | Probable | As nectar-tap_saprotroph (Põlme et al. 2020); Decay sugar-rich substrates (Põlme et al. 2020) | Põlme S, et al. 2020. Fungal Diversity 105:1-16 ((https://doi.org/10.1007/s13225-020-00466-2)) |
| Otu42991 | 1 | k__Fungi(100);p__Ascomycota(100);c__Saccharomycetes(100);o__Saccharomycetales(100);f__Metschnikowiaceae(100);g__Metschnikowia(100);g__Metschnikowia_unclassified(100); | Metschnikowia | 13 | Saprotroph | \|Undefined Saprotroph\| | Yeast | NULL | Probable | As nectar-tap_saprotroph (Põlme et al. 2020); Decay sugar-rich substrates (Põlme et al. 2020) | Põlme S, et al. 2020. Fungal Diversity 105:1-16 ((https://doi.org/10.1007/s13225-020-00466-2)) |
| Otu42992 | 1 | k__Fungi(100);p__Ascomycota(100);c__Saccharomycetes(100);o__Saccharomycetales(100);f__Metschnikowiaceae(100);g__Metschnikowia(100);g__Metschnikowia_unclassified(100); | Metschnikowia | 13 | Saprotroph | \|Undefined Saprotroph\| | Yeast | NULL | Probable | As nectar-tap_saprotroph (Põlme et al. 2020); Decay sugar-rich substrates (Põlme et al. 2020) | Põlme S, et al. 2020. Fungal Diversity 105:1-16 ((https://doi.org/10.1007/s13225-020-00466-2)) |
| Otu42993 | 1 | k__Fungi(100);p__Ascomycota(100);c__Saccharomycetes(100);o__Saccharomycetales(100);f__Metschnikowiaceae(100);g__Metschnikowia(100);g__Metschnikowia_unclassified(100); | Metschnikowia | 13 | Saprotroph | \|Undefined Saprotroph\| | Yeast | NULL | Probable | As nectar-tap_saprotroph (Põlme et al. 2020); Decay sugar-rich substrates (Põlme et al. 2020) | Põlme S, et al. 2020. Fungal Diversity 105:1-16 ((https://doi.org/10.1007/s13225-020-00466-2)) |
| Otu42994 | 1 | k__Fungi(100);p__Ascomycota(100);c__Saccharomycetes(100);o__Saccharomycetales(100);f__Metschnikowiaceae(100);g__Metschnikowia(100);s__Metschnikowia_pulcherrima(100); | Metschnikowia | 13 | Saprotroph | \|Undefined Saprotroph\| | Yeast | NULL | Probable | As nectar-tap_saprotroph (Põlme et al. 2020); Decay sugar-rich substrates (Põlme et al. 2020) | Põlme S, et al. 2020. Fungal Diversity 105:1-16 ((https://doi.org/10.1007/s13225-020-00466-2)) |
| Otu42995 | 1 | k__Fungi(100);p__Ascomycota(100);c__Saccharomycetes(100);o__Saccharomycetales(100);f__Metschnikowiaceae(100);g__Metschnikowia(100);s__Metschnikowia_pulcherrima(100); | Metschnikowia | 13 | Saprotroph | \|Undefined Saprotroph\| | Yeast | NULL | Probable | As nectar-tap_saprotroph (Põlme et al. 2020); Decay sugar-rich substrates (Põlme et al. 2020) | Põlme S, et al. 2020. Fungal Diversity 105:1-16 ((https://doi.org/10.1007/s13225-020-00466-2)) |
| Otu42996 | 1 | k__Fungi(100);p__Ascomycota(100);c__Saccharomycetes(100);o__Saccharomycetales(100);f__Metschnikowiaceae(100);g__Metschnikowia(100);g__Metschnikowia_unclassified(100); | Metschnikowia | 13 | Saprotroph | \|Undefined Saprotroph\| | Yeast | NULL | Probable | As nectar-tap_saprotroph (Põlme et al. 2020); Decay sugar-rich substrates (Põlme et al. 2020) | Põlme S, et al. 2020. Fungal Diversity 105:1-16 ((https://doi.org/10.1007/s13225-020-00466-2)) |
| Otu42997 | 1 | k__Fungi(100);p__Ascomycota(100);c__Saccharomycetes(100);o__Saccharomycetales(100);f__Metschnikowiaceae(100);g__Metschnikowia(100);s__Metschnikowia_pulcherrima(100); | Metschnikowia | 13 | Saprotroph | \|Undefined Saprotroph\| | Yeast | NULL | Probable | As nectar-tap_saprotroph (Põlme et al. 2020); Decay sugar-rich substrates (Põlme et al. 2020) | Põlme S, et al. 2020. Fungal Diversity 105:1-16 ((https://doi.org/10.1007/s13225-020-00466-2)) |
| Otu42998 | 1 | k__Fungi(100);p__Ascomycota(100);c__Saccharomycetes(100);o__Saccharomycetales(100);f__Metschnikowiaceae(100);g__Metschnikowia(100);g__Metschnikowia_unclassified(100); | Metschnikowia | 13 | Saprotroph | \|Undefined Saprotroph\| | Yeast | NULL | Probable | As nectar-tap_saprotroph (Põlme et al. 2020); Decay sugar-rich substrates (Põlme et al. 2020) | Põlme S, et al. 2020. Fungal Diversity 105:1-16 ((https://doi.org/10.1007/s13225-020-00466-2)) |
| Otu42999 | 1 | k__Fungi(100);p__Ascomycota(100);c__Saccharomycetes(100);o__Saccharomycetales(100);f__Metschnikowiaceae(100);g__Metschnikowia(100);s__Metschnikowia_pulcherrima(100); | Metschnikowia | 13 | Saprotroph | \|Undefined Saprotroph\| | Yeast | NULL | Probable | As nectar-tap_saprotroph (Põlme et al. 2020); Decay sugar-rich substrates (Põlme et al. 2020) | Põlme S, et al. 2020. Fungal Diversity 105:1-16 ((https://doi.org/10.1007/s13225-020-00466-2)) |
| Otu43000 | 1 | k__Fungi(100);p__Ascomycota(100);c__Saccharomycetes(100);o__Saccharomycetales(100);f__Metschnikowiaceae(100);g__Metschnikowia(100);s__Metschnikowia_pulcherrima(100); | Metschnikowia | 13 | Saprotroph | \|Undefined Saprotroph\| | Yeast | NULL | Probable | As nectar-tap_saprotroph (Põlme et al. 2020); Decay sugar-rich substrates (Põlme et al. 2020) | Põlme S, et al. 2020. Fungal Diversity 105:1-16 ((https://doi.org/10.1007/s13225-020-00466-2)) |
| Otu43001 | 1 | k__Fungi(100);p__Ascomycota(100);c__Saccharomycetes(100);o__Saccharomycetales(100);f__Metschnikowiaceae(100);g__Metschnikowia(100);s__Metschnikowia_pulcherrima(100); | Metschnikowia | 13 | Saprotroph | \|Undefined Saprotroph\| | Yeast | NULL | Probable | As nectar-tap_saprotroph (Põlme et al. 2020); Decay sugar-rich substrates (Põlme et al. 2020) | Põlme S, et al. 2020. Fungal Diversity 105:1-16 ((https://doi.org/10.1007/s13225-020-00466-2)) |
| Otu43002 | 1 | k__Fungi(100);p__Ascomycota(100);c__Saccharomycetes(100);o__Saccharomycetales(100);f__Metschnikowiaceae(100);g__Metschnikowia(100);s__Metschnikowia_pulcherrima(100); | Metschnikowia | 13 | Saprotroph | \|Undefined Saprotroph\| | Yeast | NULL | Probable | As nectar-tap_saprotroph (Põlme et al. 2020); Decay sugar-rich substrates (Põlme et al. 2020) | Põlme S, et al. 2020. Fungal Diversity 105:1-16 ((https://doi.org/10.1007/s13225-020-00466-2)) |
| Otu43003 | 1 | k__Fungi(100);p__Ascomycota(100);c__Saccharomycetes(100);o__Saccharomycetales(100);f__Metschnikowiaceae(100);g__Metschnikowia(100);s__Metschnikowia_pulcherrima(100); | Metschnikowia | 13 | Saprotroph | \|Undefined Saprotroph\| | Yeast | NULL | Probable | As nectar-tap_saprotroph (Põlme et al. 2020); Decay sugar-rich substrates (Põlme et al. 2020) | Põlme S, et al. 2020. Fungal Diversity 105:1-16 ((https://doi.org/10.1007/s13225-020-00466-2)) |
| Otu43004 | 1 | k__Fungi(100);p__Ascomycota(100);c__Saccharomycetes(100);o__Saccharomycetales(100);f__Metschnikowiaceae(100);g__Metschnikowia(100);s__Metschnikowia_pulcherrima(100); | Metschnikowia | 13 | Saprotroph | \|Undefined Saprotroph\| | Yeast | NULL | Probable | As nectar-tap_saprotroph (Põlme et al. 2020); Decay sugar-rich substrates (Põlme et al. 2020) | Põlme S, et al. 2020. Fungal Diversity 105:1-16 ((https://doi.org/10.1007/s13225-020-00466-2)) |
| Otu43005 | 1 | k__Fungi(100);p__Ascomycota(100);c__Saccharomycetes(100);o__Saccharomycetales(100);f__Metschnikowiaceae(100);g__Metschnikowia(100);g__Metschnikowia_unclassified(100); | Metschnikowia | 13 | Saprotroph | \|Undefined Saprotroph\| | Yeast | NULL | Probable | As nectar-tap_saprotroph (Põlme et al. 2020); Decay sugar-rich substrates (Põlme et al. 2020) | Põlme S, et al. 2020. Fungal Diversity 105:1-16 ((https://doi.org/10.1007/s13225-020-00466-2)) |
| Otu43006 | 1 | k__Fungi(100);p__Ascomycota(100);c__Saccharomycetes(100);o__Saccharomycetales(100);f__Metschnikowiaceae(100);g__Metschnikowia(100);s__Metschnikowia_pulcherrima(100); | Metschnikowia | 13 | Saprotroph | \|Undefined Saprotroph\| | Yeast | NULL | Probable | As nectar-tap_saprotroph (Põlme et al. 2020); Decay sugar-rich substrates (Põlme et al. 2020) | Põlme S, et al. 2020. Fungal Diversity 105:1-16 ((https://doi.org/10.1007/s13225-020-00466-2)) |
| Otu43007 | 1 | k__Fungi(100);p__Ascomycota(100);c__Saccharomycetes(100);o__Saccharomycetales(100);f__Metschnikowiaceae(100);g__Metschnikowia(100);s__Metschnikowia_pulcherrima(100); | Metschnikowia | 13 | Saprotroph | \|Undefined Saprotroph\| | Yeast | NULL | Probable | As nectar-tap_saprotroph (Põlme et al. 2020); Decay sugar-rich substrates (Põlme et al. 2020) | Põlme S, et al. 2020. Fungal Diversity 105:1-16 ((https://doi.org/10.1007/s13225-020-00466-2)) |
| Otu43008 | 1 | k__Fungi(100);p__Ascomycota(100);c__Saccharomycetes(100);o__Saccharomycetales(100);f__Metschnikowiaceae(100);g__Metschnikowia(100);g__Metschnikowia_unclassified(100); | Metschnikowia | 13 | Saprotroph | \|Undefined Saprotroph\| | Yeast | NULL | Probable | As nectar-tap_saprotroph (Põlme et al. 2020); Decay sugar-rich substrates (Põlme et al. 2020) | Põlme S, et al. 2020. Fungal Diversity 105:1-16 ((https://doi.org/10.1007/s13225-020-00466-2)) |
| Otu43009 | 1 | k__Fungi(100);p__Ascomycota(100);c__Saccharomycetes(100);o__Saccharomycetales(100);f__Metschnikowiaceae(100);g__Metschnikowia(100);g__Metschnikowia_unclassified(100); | Metschnikowia | 13 | Saprotroph | \|Undefined Saprotroph\| | Yeast | NULL | Probable | As nectar-tap_saprotroph (Põlme et al. 2020); Decay sugar-rich substrates (Põlme et al. 2020) | Põlme S, et al. 2020. Fungal Diversity 105:1-16 ((https://doi.org/10.1007/s13225-020-00466-2)) |
| Otu43010 | 1 | k__Fungi(100);p__Ascomycota(100);c__Saccharomycetes(100);o__Saccharomycetales(100);f__Metschnikowiaceae(100);g__Metschnikowia(100);g__Metschnikowia_unclassified(100); | Metschnikowia | 13 | Saprotroph | \|Undefined Saprotroph\| | Yeast | NULL | Probable | As nectar-tap_saprotroph (Põlme et al. 2020); Decay sugar-rich substrates (Põlme et al. 2020) | Põlme S, et al. 2020. Fungal Diversity 105:1-16 ((https://doi.org/10.1007/s13225-020-00466-2)) |
| Otu43011 | 1 | k__Fungi(100);p__Ascomycota(100);c__Saccharomycetes(100);o__Saccharomycetales(100);f__Metschnikowiaceae(100);g__Metschnikowia(100);s__Metschnikowia_pulcherrima(100); | Metschnikowia | 13 | Saprotroph | \|Undefined Saprotroph\| | Yeast | NULL | Probable | As nectar-tap_saprotroph (Põlme et al. 2020); Decay sugar-rich substrates (Põlme et al. 2020) | Põlme S, et al. 2020. Fungal Diversity 105:1-16 ((https://doi.org/10.1007/s13225-020-00466-2)) |
| Otu43012 | 1 | k__Fungi(100);p__Ascomycota(100);c__Saccharomycetes(100);o__Saccharomycetales(100);f__Metschnikowiaceae(100);g__Metschnikowia(100);s__Metschnikowia_pulcherrima(100); | Metschnikowia | 13 | Saprotroph | \|Undefined Saprotroph\| | Yeast | NULL | Probable | As nectar-tap_saprotroph (Põlme et al. 2020); Decay sugar-rich substrates (Põlme et al. 2020) | Põlme S, et al. 2020. Fungal Diversity 105:1-16 ((https://doi.org/10.1007/s13225-020-00466-2)) |
| Otu43013 | 1 | k__Fungi(100);p__Ascomycota(100);c__Saccharomycetes(100);o__Saccharomycetales(100);f__Metschnikowiaceae(100);g__Metschnikowia(100);g__Metschnikowia_unclassified(100); | Metschnikowia | 13 | Saprotroph | \|Undefined Saprotroph\| | Yeast | NULL | Probable | As nectar-tap_saprotroph (Põlme et al. 2020); Decay sugar-rich substrates (Põlme et al. 2020) | Põlme S, et al. 2020. Fungal Diversity 105:1-16 ((https://doi.org/10.1007/s13225-020-00466-2)) |
| Otu43014 | 1 | k__Fungi(100);p__Ascomycota(100);c__Saccharomycetes(100);o__Saccharomycetales(100);f__Metschnikowiaceae(100);g__Metschnikowia(100);s__Metschnikowia_pulcherrima(100); | Metschnikowia | 13 | Saprotroph | \|Undefined Saprotroph\| | Yeast | NULL | Probable | As nectar-tap_saprotroph (Põlme et al. 2020); Decay sugar-rich substrates (Põlme et al. 2020) | Põlme S, et al. 2020. Fungal Diversity 105:1-16 ((https://doi.org/10.1007/s13225-020-00466-2)) |
| Otu43015 | 1 | k__Fungi(100);p__Ascomycota(100);c__Saccharomycetes(100);o__Saccharomycetales(100);f__Metschnikowiaceae(100);g__Metschnikowia(100);s__Metschnikowia_pulcherrima(100); | Metschnikowia | 13 | Saprotroph | \|Undefined Saprotroph\| | Yeast | NULL | Probable | As nectar-tap_saprotroph (Põlme et al. 2020); Decay sugar-rich substrates (Põlme et al. 2020) | Põlme S, et al. 2020. Fungal Diversity 105:1-16 ((https://doi.org/10.1007/s13225-020-00466-2)) |
| Otu43016 | 1 | k__Fungi(100);p__Ascomycota(100);c__Saccharomycetes(100);o__Saccharomycetales(100);f__Metschnikowiaceae(100);g__Metschnikowia(100);s__Metschnikowia_pulcherrima(100); | Metschnikowia | 13 | Saprotroph | \|Undefined Saprotroph\| | Yeast | NULL | Probable | As nectar-tap_saprotroph (Põlme et al. 2020); Decay sugar-rich substrates (Põlme et al. 2020) | Põlme S, et al. 2020. Fungal Diversity 105:1-16 ((https://doi.org/10.1007/s13225-020-00466-2)) |
| Otu43017 | 1 | k__Fungi(100);p__Ascomycota(100);c__Saccharomycetes(100);o__Saccharomycetales(100);f__Metschnikowiaceae(100);g__Metschnikowia(100);s__Metschnikowia_pulcherrima(100); | Metschnikowia | 13 | Saprotroph | \|Undefined Saprotroph\| | Yeast | NULL | Probable | As nectar-tap_saprotroph (Põlme et al. 2020); Decay sugar-rich substrates (Põlme et al. 2020) | Põlme S, et al. 2020. Fungal Diversity 105:1-16 ((https://doi.org/10.1007/s13225-020-00466-2)) |
| Otu43018 | 1 | k__Fungi(100);p__Ascomycota(100);c__Saccharomycetes(100);o__Saccharomycetales(100);f__Metschnikowiaceae(100);g__Metschnikowia(100);s__Metschnikowia_pulcherrima(100); | Metschnikowia | 13 | Saprotroph | \|Undefined Saprotroph\| | Yeast | NULL | Probable | As nectar-tap_saprotroph (Põlme et al. 2020); Decay sugar-rich substrates (Põlme et al. 2020) | Põlme S, et al. 2020. Fungal Diversity 105:1-16 ((https://doi.org/10.1007/s13225-020-00466-2)) |
| Otu43019 | 1 | k__Fungi(100);p__Ascomycota(100);c__Saccharomycetes(100);o__Saccharomycetales(100);f__Metschnikowiaceae(100);g__Metschnikowia(100);s__Metschnikowia_pulcherrima(100); | Metschnikowia | 13 | Saprotroph | \|Undefined Saprotroph\| | Yeast | NULL | Probable | As nectar-tap_saprotroph (Põlme et al. 2020); Decay sugar-rich substrates (Põlme et al. 2020) | Põlme S, et al. 2020. Fungal Diversity 105:1-16 ((https://doi.org/10.1007/s13225-020-00466-2)) |
| Otu43020 | 1 | k__Fungi(100);p__Ascomycota(100);c__Saccharomycetes(100);o__Saccharomycetales(100);f__Metschnikowiaceae(100);g__Metschnikowia(100);g__Metschnikowia_unclassified(100); | Metschnikowia | 13 | Saprotroph | \|Undefined Saprotroph\| | Yeast | NULL | Probable | As nectar-tap_saprotroph (Põlme et al. 2020); Decay sugar-rich substrates (Põlme et al. 2020) | Põlme S, et al. 2020. Fungal Diversity 105:1-16 ((https://doi.org/10.1007/s13225-020-00466-2)) |
| Otu43021 | 1 | k__Fungi(100);p__Ascomycota(100);c__Saccharomycetes(100);o__Saccharomycetales(100);f__Metschnikowiaceae(100);g__Metschnikowia(100);s__Metschnikowia_pulcherrima(100); | Metschnikowia | 13 | Saprotroph | \|Undefined Saprotroph\| | Yeast | NULL | Probable | As nectar-tap_saprotroph (Põlme et al. 2020); Decay sugar-rich substrates (Põlme et al. 2020) | Põlme S, et al. 2020. Fungal Diversity 105:1-16 ((https://doi.org/10.1007/s13225-020-00466-2)) |
| Otu43022 | 1 | k__Fungi(100);p__Ascomycota(100);c__Saccharomycetes(100);o__Saccharomycetales(100);f__Metschnikowiaceae(100);g__Metschnikowia(100);s__Metschnikowia_pulcherrima(100); | Metschnikowia | 13 | Saprotroph | \|Undefined Saprotroph\| | Yeast | NULL | Probable | As nectar-tap_saprotroph (Põlme et al. 2020); Decay sugar-rich substrates (Põlme et al. 2020) | Põlme S, et al. 2020. Fungal Diversity 105:1-16 ((https://doi.org/10.1007/s13225-020-00466-2)) |
| Otu43023 | 1 | k__Fungi(100);p__Ascomycota(100);c__Saccharomycetes(100);o__Saccharomycetales(100);f__Metschnikowiaceae(100);g__Metschnikowia(100);s__Metschnikowia_pulcherrima(100); | Metschnikowia | 13 | Saprotroph | \|Undefined Saprotroph\| | Yeast | NULL | Probable | As nectar-tap_saprotroph (Põlme et al. 2020); Decay sugar-rich substrates (Põlme et al. 2020) | Põlme S, et al. 2020. Fungal Diversity 105:1-16 ((https://doi.org/10.1007/s13225-020-00466-2)) |
| Otu43024 | 1 | k__Fungi(100);p__Ascomycota(100);c__Saccharomycetes(100);o__Saccharomycetales(100);f__Metschnikowiaceae(100);g__Metschnikowia(100);s__Metschnikowia_pulcherrima(100); | Metschnikowia | 13 | Saprotroph | \|Undefined Saprotroph\| | Yeast | NULL | Probable | As nectar-tap_saprotroph (Põlme et al. 2020); Decay sugar-rich substrates (Põlme et al. 2020) | Põlme S, et al. 2020. Fungal Diversity 105:1-16 ((https://doi.org/10.1007/s13225-020-00466-2)) |
| Otu43025 | 1 | k__Fungi(100);p__Ascomycota(100);c__Saccharomycetes(100);o__Saccharomycetales(100);f__Metschnikowiaceae(100);g__Metschnikowia(100);g__Metschnikowia_unclassified(100); | Metschnikowia | 13 | Saprotroph | \|Undefined Saprotroph\| | Yeast | NULL | Probable | As nectar-tap_saprotroph (Põlme et al. 2020); Decay sugar-rich substrates (Põlme et al. 2020) | Põlme S, et al. 2020. Fungal Diversity 105:1-16 ((https://doi.org/10.1007/s13225-020-00466-2)) |
| Otu43026 | 1 | k__Fungi(100);p__Ascomycota(100);c__Saccharomycetes(100);o__Saccharomycetales(100);f__Metschnikowiaceae(100);g__Metschnikowia(100);s__Metschnikowia_pulcherrima(100); | Metschnikowia | 13 | Saprotroph | \|Undefined Saprotroph\| | Yeast | NULL | Probable | As nectar-tap_saprotroph (Põlme et al. 2020); Decay sugar-rich substrates (Põlme et al. 2020) | Põlme S, et al. 2020. Fungal Diversity 105:1-16 ((https://doi.org/10.1007/s13225-020-00466-2)) |
| Otu43027 | 1 | k__Fungi(100);p__Ascomycota(100);c__Saccharomycetes(100);o__Saccharomycetales(100);f__Metschnikowiaceae(100);g__Metschnikowia(100);s__Metschnikowia_pulcherrima(100); | Metschnikowia | 13 | Saprotroph | \|Undefined Saprotroph\| | Yeast | NULL | Probable | As nectar-tap_saprotroph (Põlme et al. 2020); Decay sugar-rich substrates (Põlme et al. 2020) | Põlme S, et al. 2020. Fungal Diversity 105:1-16 ((https://doi.org/10.1007/s13225-020-00466-2)) |
| Otu43028 | 1 | k__Fungi(100);p__Ascomycota(100);c__Saccharomycetes(100);o__Saccharomycetales(100);f__Metschnikowiaceae(100);g__Metschnikowia(100);g__Metschnikowia_unclassified(100); | Metschnikowia | 13 | Saprotroph | \|Undefined Saprotroph\| | Yeast | NULL | Probable | As nectar-tap_saprotroph (Põlme et al. 2020); Decay sugar-rich substrates (Põlme et al. 2020) | Põlme S, et al. 2020. Fungal Diversity 105:1-16 ((https://doi.org/10.1007/s13225-020-00466-2)) |
| Otu43029 | 1 | k__Fungi(100);p__Ascomycota(100);c__Saccharomycetes(100);o__Saccharomycetales(100);f__Metschnikowiaceae(100);g__Metschnikowia(100);s__Metschnikowia_pulcherrima(100); | Metschnikowia | 13 | Saprotroph | \|Undefined Saprotroph\| | Yeast | NULL | Probable | As nectar-tap_saprotroph (Põlme et al. 2020); Decay sugar-rich substrates (Põlme et al. 2020) | Põlme S, et al. 2020. Fungal Diversity 105:1-16 ((https://doi.org/10.1007/s13225-020-00466-2)) |
| Otu43030 | 1 | k__Fungi(100);p__Ascomycota(100);c__Saccharomycetes(100);o__Saccharomycetales(100);f__Metschnikowiaceae(100);g__Metschnikowia(100);s__Metschnikowia_pulcherrima(100); | Metschnikowia | 13 | Saprotroph | \|Undefined Saprotroph\| | Yeast | NULL | Probable | As nectar-tap_saprotroph (Põlme et al. 2020); Decay sugar-rich substrates (Põlme et al. 2020) | Põlme S, et al. 2020. Fungal Diversity 105:1-16 ((https://doi.org/10.1007/s13225-020-00466-2)) |
| Otu43031 | 1 | k__Fungi(100);p__Ascomycota(100);c__Saccharomycetes(100);o__Saccharomycetales(100);f__Metschnikowiaceae(100);g__Metschnikowia(100);s__Metschnikowia_pulcherrima(100); | Metschnikowia | 13 | Saprotroph | \|Undefined Saprotroph\| | Yeast | NULL | Probable | As nectar-tap_saprotroph (Põlme et al. 2020); Decay sugar-rich substrates (Põlme et al. 2020) | Põlme S, et al. 2020. Fungal Diversity 105:1-16 ((https://doi.org/10.1007/s13225-020-00466-2)) |
| Otu43032 | 1 | k__Fungi(100);p__Ascomycota(100);c__Saccharomycetes(100);o__Saccharomycetales(100);f__Metschnikowiaceae(100);g__Metschnikowia(100);s__Metschnikowia_pulcherrima(100); | Metschnikowia | 13 | Saprotroph | \|Undefined Saprotroph\| | Yeast | NULL | Probable | As nectar-tap_saprotroph (Põlme et al. 2020); Decay sugar-rich substrates (Põlme et al. 2020) | Põlme S, et al. 2020. Fungal Diversity 105:1-16 ((https://doi.org/10.1007/s13225-020-00466-2)) |
| Otu43033 | 1 | k__Fungi(100);p__Ascomycota(100);c__Saccharomycetes(100);o__Saccharomycetales(100);f__Metschnikowiaceae(100);g__Metschnikowia(100);s__Metschnikowia_pulcherrima(100); | Metschnikowia | 13 | Saprotroph | \|Undefined Saprotroph\| | Yeast | NULL | Probable | As nectar-tap_saprotroph (Põlme et al. 2020); Decay sugar-rich substrates (Põlme et al. 2020) | Põlme S, et al. 2020. Fungal Diversity 105:1-16 ((https://doi.org/10.1007/s13225-020-00466-2)) |
| Otu43034 | 1 | k__Fungi(100);p__Ascomycota(100);c__Saccharomycetes(100);o__Saccharomycetales(100);f__Metschnikowiaceae(100);g__Metschnikowia(100);s__Metschnikowia_pulcherrima(100); | Metschnikowia | 13 | Saprotroph | \|Undefined Saprotroph\| | Yeast | NULL | Probable | As nectar-tap_saprotroph (Põlme et al. 2020); Decay sugar-rich substrates (Põlme et al. 2020) | Põlme S, et al. 2020. Fungal Diversity 105:1-16 ((https://doi.org/10.1007/s13225-020-00466-2)) |
| Otu43035 | 1 | k__Fungi(100);p__Ascomycota(100);c__Saccharomycetes(100);o__Saccharomycetales(100);f__Metschnikowiaceae(100);g__Metschnikowia(100);g__Metschnikowia_unclassified(100); | Metschnikowia | 13 | Saprotroph | \|Undefined Saprotroph\| | Yeast | NULL | Probable | As nectar-tap_saprotroph (Põlme et al. 2020); Decay sugar-rich substrates (Põlme et al. 2020) | Põlme S, et al. 2020. Fungal Diversity 105:1-16 ((https://doi.org/10.1007/s13225-020-00466-2)) |
| Otu43036 | 1 | k__Fungi(100);p__Ascomycota(100);c__Saccharomycetes(100);o__Saccharomycetales(100);f__Metschnikowiaceae(100);g__Metschnikowia(100);s__Metschnikowia_pulcherrima(100); | Metschnikowia | 13 | Saprotroph | \|Undefined Saprotroph\| | Yeast | NULL | Probable | As nectar-tap_saprotroph (Põlme et al. 2020); Decay sugar-rich substrates (Põlme et al. 2020) | Põlme S, et al. 2020. Fungal Diversity 105:1-16 ((https://doi.org/10.1007/s13225-020-00466-2)) |
| Otu43037 | 1 | k__Fungi(100);p__Ascomycota(100);c__Saccharomycetes(100);o__Saccharomycetales(100);f__Metschnikowiaceae(100);g__Metschnikowia(100);s__Metschnikowia_pulcherrima(100); | Metschnikowia | 13 | Saprotroph | \|Undefined Saprotroph\| | Yeast | NULL | Probable | As nectar-tap_saprotroph (Põlme et al. 2020); Decay sugar-rich substrates (Põlme et al. 2020) | Põlme S, et al. 2020. Fungal Diversity 105:1-16 ((https://doi.org/10.1007/s13225-020-00466-2)) |
| Otu43038 | 1 | k__Fungi(100);p__Ascomycota(100);c__Saccharomycetes(100);o__Saccharomycetales(100);f__Metschnikowiaceae(100);g__Metschnikowia(100);s__Metschnikowia_pulcherrima(100); | Metschnikowia | 13 | Saprotroph | \|Undefined Saprotroph\| | Yeast | NULL | Probable | As nectar-tap_saprotroph (Põlme et al. 2020); Decay sugar-rich substrates (Põlme et al. 2020) | Põlme S, et al. 2020. Fungal Diversity 105:1-16 ((https://doi.org/10.1007/s13225-020-00466-2)) |
| Otu43039 | 1 | k__Fungi(100);p__Ascomycota(100);c__Saccharomycetes(100);o__Saccharomycetales(100);f__Metschnikowiaceae(100);g__Metschnikowia(100);s__Metschnikowia_pulcherrima(100); | Metschnikowia | 13 | Saprotroph | \|Undefined Saprotroph\| | Yeast | NULL | Probable | As nectar-tap_saprotroph (Põlme et al. 2020); Decay sugar-rich substrates (Põlme et al. 2020) | Põlme S, et al. 2020. Fungal Diversity 105:1-16 ((https://doi.org/10.1007/s13225-020-00466-2)) |
| Otu43040 | 1 | k__Fungi(100);p__Ascomycota(100);c__Saccharomycetes(100);o__Saccharomycetales(100);f__Metschnikowiaceae(100);g__Metschnikowia(100);s__Metschnikowia_pulcherrima(100); | Metschnikowia | 13 | Saprotroph | \|Undefined Saprotroph\| | Yeast | NULL | Probable | As nectar-tap_saprotroph (Põlme et al. 2020); Decay sugar-rich substrates (Põlme et al. 2020) | Põlme S, et al. 2020. Fungal Diversity 105:1-16 ((https://doi.org/10.1007/s13225-020-00466-2)) |
| Otu43041 | 1 | k__Fungi(100);p__Ascomycota(100);c__Saccharomycetes(100);o__Saccharomycetales(100);f__Metschnikowiaceae(100);g__Metschnikowia(100);s__Metschnikowia_pulcherrima(100); | Metschnikowia | 13 | Saprotroph | \|Undefined Saprotroph\| | Yeast | NULL | Probable | As nectar-tap_saprotroph (Põlme et al. 2020); Decay sugar-rich substrates (Põlme et al. 2020) | Põlme S, et al. 2020. Fungal Diversity 105:1-16 ((https://doi.org/10.1007/s13225-020-00466-2)) |
| Otu43042 | 1 | k__Fungi(100);p__Ascomycota(100);c__Saccharomycetes(100);o__Saccharomycetales(100);f__Metschnikowiaceae(100);g__Metschnikowia(100);s__Metschnikowia_pulcherrima(100); | Metschnikowia | 13 | Saprotroph | \|Undefined Saprotroph\| | Yeast | NULL | Probable | As nectar-tap_saprotroph (Põlme et al. 2020); Decay sugar-rich substrates (Põlme et al. 2020) | Põlme S, et al. 2020. Fungal Diversity 105:1-16 ((https://doi.org/10.1007/s13225-020-00466-2)) |
| Otu43043 | 1 | k__Fungi(100);p__Ascomycota(100);c__Saccharomycetes(100);o__Saccharomycetales(100);f__Metschnikowiaceae(100);g__Metschnikowia(100);s__Metschnikowia_pulcherrima(100); | Metschnikowia | 13 | Saprotroph | \|Undefined Saprotroph\| | Yeast | NULL | Probable | As nectar-tap_saprotroph (Põlme et al. 2020); Decay sugar-rich substrates (Põlme et al. 2020) | Põlme S, et al. 2020. Fungal Diversity 105:1-16 ((https://doi.org/10.1007/s13225-020-00466-2)) |
| Otu43044 | 1 | k__Fungi(100);p__Ascomycota(100);c__Saccharomycetes(100);o__Saccharomycetales(100);f__Metschnikowiaceae(100);g__Metschnikowia(100);s__Metschnikowia_pulcherrima(100); | Metschnikowia | 13 | Saprotroph | \|Undefined Saprotroph\| | Yeast | NULL | Probable | As nectar-tap_saprotroph (Põlme et al. 2020); Decay sugar-rich substrates (Põlme et al. 2020) | Põlme S, et al. 2020. Fungal Diversity 105:1-16 ((https://doi.org/10.1007/s13225-020-00466-2)) |
| Otu43045 | 1 | k__Fungi(100);p__Ascomycota(100);c__Saccharomycetes(100);o__Saccharomycetales(100);f__Metschnikowiaceae(100);g__Metschnikowia(100);s__Metschnikowia_pulcherrima(100); | Metschnikowia | 13 | Saprotroph | \|Undefined Saprotroph\| | Yeast | NULL | Probable | As nectar-tap_saprotroph (Põlme et al. 2020); Decay sugar-rich substrates (Põlme et al. 2020) | Põlme S, et al. 2020. Fungal Diversity 105:1-16 ((https://doi.org/10.1007/s13225-020-00466-2)) |
| Otu43046 | 1 | k__Fungi(100);p__Ascomycota(100);c__Saccharomycetes(100);o__Saccharomycetales(100);f__Metschnikowiaceae(100);g__Metschnikowia(100);s__Metschnikowia_pulcherrima(100); | Metschnikowia | 13 | Saprotroph | \|Undefined Saprotroph\| | Yeast | NULL | Probable | As nectar-tap_saprotroph (Põlme et al. 2020); Decay sugar-rich substrates (Põlme et al. 2020) | Põlme S, et al. 2020. Fungal Diversity 105:1-16 ((https://doi.org/10.1007/s13225-020-00466-2)) |
| Otu43047 | 1 | k__Fungi(100);p__Ascomycota(100);c__Saccharomycetes(100);o__Saccharomycetales(100);f__Metschnikowiaceae(100);g__Metschnikowia(100);s__Metschnikowia_pulcherrima(100); | Metschnikowia | 13 | Saprotroph | \|Undefined Saprotroph\| | Yeast | NULL | Probable | As nectar-tap_saprotroph (Põlme et al. 2020); Decay sugar-rich substrates (Põlme et al. 2020) | Põlme S, et al. 2020. Fungal Diversity 105:1-16 ((https://doi.org/10.1007/s13225-020-00466-2)) |
| Otu43048 | 1 | k__Fungi(100);p__Ascomycota(100);c__Saccharomycetes(100);o__Saccharomycetales(100);f__Metschnikowiaceae(100);g__Metschnikowia(100);s__Metschnikowia_pulcherrima(100); | Metschnikowia | 13 | Saprotroph | \|Undefined Saprotroph\| | Yeast | NULL | Probable | As nectar-tap_saprotroph (Põlme et al. 2020); Decay sugar-rich substrates (Põlme et al. 2020) | Põlme S, et al. 2020. Fungal Diversity 105:1-16 ((https://doi.org/10.1007/s13225-020-00466-2)) |
| Otu43049 | 1 | k__Fungi(100);p__Ascomycota(100);c__Saccharomycetes(100);o__Saccharomycetales(100);f__Metschnikowiaceae(100);g__Metschnikowia(100);s__Metschnikowia_pulcherrima(100); | Metschnikowia | 13 | Saprotroph | \|Undefined Saprotroph\| | Yeast | NULL | Probable | As nectar-tap_saprotroph (Põlme et al. 2020); Decay sugar-rich substrates (Põlme et al. 2020) | Põlme S, et al. 2020. Fungal Diversity 105:1-16 ((https://doi.org/10.1007/s13225-020-00466-2)) |
| Otu43050 | 1 | k__Fungi(100);p__Ascomycota(100);c__Saccharomycetes(100);o__Saccharomycetales(100);f__Metschnikowiaceae(100);g__Metschnikowia(100);g__Metschnikowia_unclassified(100); | Metschnikowia | 13 | Saprotroph | \|Undefined Saprotroph\| | Yeast | NULL | Probable | As nectar-tap_saprotroph (Põlme et al. 2020); Decay sugar-rich substrates (Põlme et al. 2020) | Põlme S, et al. 2020. Fungal Diversity 105:1-16 ((https://doi.org/10.1007/s13225-020-00466-2)) |
| Otu43051 | 1 | k__Fungi(100);p__Ascomycota(100);c__Saccharomycetes(100);o__Saccharomycetales(100);f__Metschnikowiaceae(100);g__Metschnikowia(100);g__Metschnikowia_unclassified(100); | Metschnikowia | 13 | Saprotroph | \|Undefined Saprotroph\| | Yeast | NULL | Probable | As nectar-tap_saprotroph (Põlme et al. 2020); Decay sugar-rich substrates (Põlme et al. 2020) | Põlme S, et al. 2020. Fungal Diversity 105:1-16 ((https://doi.org/10.1007/s13225-020-00466-2)) |
| Otu43052 | 1 | k__Fungi(100);p__Ascomycota(100);c__Saccharomycetes(100);o__Saccharomycetales(100);f__Metschnikowiaceae(100);g__Metschnikowia(100);g__Metschnikowia_unclassified(100); | Metschnikowia | 13 | Saprotroph | \|Undefined Saprotroph\| | Yeast | NULL | Probable | As nectar-tap_saprotroph (Põlme et al. 2020); Decay sugar-rich substrates (Põlme et al. 2020) | Põlme S, et al. 2020. Fungal Diversity 105:1-16 ((https://doi.org/10.1007/s13225-020-00466-2)) |
| Otu43053 | 1 | k__Fungi(100);p__Ascomycota(100);c__Saccharomycetes(100);o__Saccharomycetales(100);f__Metschnikowiaceae(100);g__Metschnikowia(100);g__Metschnikowia_unclassified(100); | Metschnikowia | 13 | Saprotroph | \|Undefined Saprotroph\| | Yeast | NULL | Probable | As nectar-tap_saprotroph (Põlme et al. 2020); Decay sugar-rich substrates (Põlme et al. 2020) | Põlme S, et al. 2020. Fungal Diversity 105:1-16 ((https://doi.org/10.1007/s13225-020-00466-2)) |
| Otu43054 | 1 | k__Fungi(100);p__Ascomycota(100);c__Saccharomycetes(100);o__Saccharomycetales(100);f__Metschnikowiaceae(100);g__Metschnikowia(100);g__Metschnikowia_unclassified(100); | Metschnikowia | 13 | Saprotroph | \|Undefined Saprotroph\| | Yeast | NULL | Probable | As nectar-tap_saprotroph (Põlme et al. 2020); Decay sugar-rich substrates (Põlme et al. 2020) | Põlme S, et al. 2020. Fungal Diversity 105:1-16 ((https://doi.org/10.1007/s13225-020-00466-2)) |
| Otu43055 | 1 | k__Fungi(100);p__Ascomycota(100);c__Saccharomycetes(100);o__Saccharomycetales(100);f__Metschnikowiaceae(100);g__Metschnikowia(100);g__Metschnikowia_unclassified(100); | Metschnikowia | 13 | Saprotroph | \|Undefined Saprotroph\| | Yeast | NULL | Probable | As nectar-tap_saprotroph (Põlme et al. 2020); Decay sugar-rich substrates (Põlme et al. 2020) | Põlme S, et al. 2020. Fungal Diversity 105:1-16 ((https://doi.org/10.1007/s13225-020-00466-2)) |
| Otu43056 | 1 | k__Fungi(100);p__Ascomycota(100);c__Saccharomycetes(100);o__Saccharomycetales(100);f__Metschnikowiaceae(100);g__Metschnikowia(100);s__Metschnikowia_pulcherrima(100); | Metschnikowia | 13 | Saprotroph | \|Undefined Saprotroph\| | Yeast | NULL | Probable | As nectar-tap_saprotroph (Põlme et al. 2020); Decay sugar-rich substrates (Põlme et al. 2020) | Põlme S, et al. 2020. Fungal Diversity 105:1-16 ((https://doi.org/10.1007/s13225-020-00466-2)) |
| Otu43057 | 1 | k__Fungi(100);p__Ascomycota(100);c__Saccharomycetes(100);o__Saccharomycetales(100);f__Metschnikowiaceae(100);g__Metschnikowia(100);g__Metschnikowia_unclassified(100); | Metschnikowia | 13 | Saprotroph | \|Undefined Saprotroph\| | Yeast | NULL | Probable | As nectar-tap_saprotroph (Põlme et al. 2020); Decay sugar-rich substrates (Põlme et al. 2020) | Põlme S, et al. 2020. Fungal Diversity 105:1-16 ((https://doi.org/10.1007/s13225-020-00466-2)) |
| Otu43058 | 1 | k__Fungi(100);p__Ascomycota(100);c__Saccharomycetes(100);o__Saccharomycetales(100);f__Metschnikowiaceae(100);g__Metschnikowia(100);s__Metschnikowia_pulcherrima(100); | Metschnikowia | 13 | Saprotroph | \|Undefined Saprotroph\| | Yeast | NULL | Probable | As nectar-tap_saprotroph (Põlme et al. 2020); Decay sugar-rich substrates (Põlme et al. 2020) | Põlme S, et al. 2020. Fungal Diversity 105:1-16 ((https://doi.org/10.1007/s13225-020-00466-2)) |
| Otu43059 | 1 | k__Fungi(100);p__Ascomycota(100);c__Saccharomycetes(100);o__Saccharomycetales(100);f__Metschnikowiaceae(100);g__Metschnikowia(100);s__Metschnikowia_pulcherrima(100); | Metschnikowia | 13 | Saprotroph | \|Undefined Saprotroph\| | Yeast | NULL | Probable | As nectar-tap_saprotroph (Põlme et al. 2020); Decay sugar-rich substrates (Põlme et al. 2020) | Põlme S, et al. 2020. Fungal Diversity 105:1-16 ((https://doi.org/10.1007/s13225-020-00466-2)) |
| Otu43060 | 1 | k__Fungi(100);p__Ascomycota(100);c__Saccharomycetes(100);o__Saccharomycetales(100);f__Metschnikowiaceae(100);g__Metschnikowia(100);s__Metschnikowia_pulcherrima(100); | Metschnikowia | 13 | Saprotroph | \|Undefined Saprotroph\| | Yeast | NULL | Probable | As nectar-tap_saprotroph (Põlme et al. 2020); Decay sugar-rich substrates (Põlme et al. 2020) | Põlme S, et al. 2020. Fungal Diversity 105:1-16 ((https://doi.org/10.1007/s13225-020-00466-2)) |
| Otu43062 | 1 | k__Fungi(100);p__Ascomycota(100);c__Saccharomycetes(100);o__Saccharomycetales(100);f__Metschnikowiaceae(100);g__Metschnikowia(100);s__Metschnikowia_pulcherrima(100); | Metschnikowia | 13 | Saprotroph | \|Undefined Saprotroph\| | Yeast | NULL | Probable | As nectar-tap_saprotroph (Põlme et al. 2020); Decay sugar-rich substrates (Põlme et al. 2020) | Põlme S, et al. 2020. Fungal Diversity 105:1-16 ((https://doi.org/10.1007/s13225-020-00466-2)) |
| Otu43064 | 1 | k__Fungi(100);p__Ascomycota(100);c__Saccharomycetes(100);o__Saccharomycetales(100);f__Metschnikowiaceae(100);g__Metschnikowia(100);s__Metschnikowia_pulcherrima(100); | Metschnikowia | 13 | Saprotroph | \|Undefined Saprotroph\| | Yeast | NULL | Probable | As nectar-tap_saprotroph (Põlme et al. 2020); Decay sugar-rich substrates (Põlme et al. 2020) | Põlme S, et al. 2020. Fungal Diversity 105:1-16 ((https://doi.org/10.1007/s13225-020-00466-2)) |
| Otu43065 | 1 | k__Fungi(100);p__Ascomycota(100);c__Saccharomycetes(100);o__Saccharomycetales(100);f__Metschnikowiaceae(100);g__Metschnikowia(100);s__Metschnikowia_pulcherrima(100); | Metschnikowia | 13 | Saprotroph | \|Undefined Saprotroph\| | Yeast | NULL | Probable | As nectar-tap_saprotroph (Põlme et al. 2020); Decay sugar-rich substrates (Põlme et al. 2020) | Põlme S, et al. 2020. Fungal Diversity 105:1-16 ((https://doi.org/10.1007/s13225-020-00466-2)) |
| Otu43066 | 1 | k__Fungi(100);p__Ascomycota(100);c__Saccharomycetes(100);o__Saccharomycetales(100);f__Metschnikowiaceae(100);g__Metschnikowia(100);s__Metschnikowia_pulcherrima(100); | Metschnikowia | 13 | Saprotroph | \|Undefined Saprotroph\| | Yeast | NULL | Probable | As nectar-tap_saprotroph (Põlme et al. 2020); Decay sugar-rich substrates (Põlme et al. 2020) | Põlme S, et al. 2020. Fungal Diversity 105:1-16 ((https://doi.org/10.1007/s13225-020-00466-2)) |
| Otu43067 | 1 | k__Fungi(100);p__Ascomycota(100);c__Saccharomycetes(100);o__Saccharomycetales(100);f__Metschnikowiaceae(100);g__Metschnikowia(100);s__Metschnikowia_pulcherrima(100); | Metschnikowia | 13 | Saprotroph | \|Undefined Saprotroph\| | Yeast | NULL | Probable | As nectar-tap_saprotroph (Põlme et al. 2020); Decay sugar-rich substrates (Põlme et al. 2020) | Põlme S, et al. 2020. Fungal Diversity 105:1-16 ((https://doi.org/10.1007/s13225-020-00466-2)) |
| Otu43068 | 1 | k__Fungi(100);p__Ascomycota(100);c__Saccharomycetes(100);o__Saccharomycetales(100);f__Metschnikowiaceae(100);g__Metschnikowia(100);s__Metschnikowia_pulcherrima(100); | Metschnikowia | 13 | Saprotroph | \|Undefined Saprotroph\| | Yeast | NULL | Probable | As nectar-tap_saprotroph (Põlme et al. 2020); Decay sugar-rich substrates (Põlme et al. 2020) | Põlme S, et al. 2020. Fungal Diversity 105:1-16 ((https://doi.org/10.1007/s13225-020-00466-2)) |
| Otu43069 | 1 | k__Fungi(100);p__Ascomycota(100);c__Saccharomycetes(100);o__Saccharomycetales(100);f__Metschnikowiaceae(100);g__Metschnikowia(100);g__Metschnikowia_unclassified(100); | Metschnikowia | 13 | Saprotroph | \|Undefined Saprotroph\| | Yeast | NULL | Probable | As nectar-tap_saprotroph (Põlme et al. 2020); Decay sugar-rich substrates (Põlme et al. 2020) | Põlme S, et al. 2020. Fungal Diversity 105:1-16 ((https://doi.org/10.1007/s13225-020-00466-2)) |
| Otu43070 | 1 | k__Fungi(100);p__Ascomycota(100);c__Saccharomycetes(100);o__Saccharomycetales(100);f__Metschnikowiaceae(100);g__Metschnikowia(100);g__Metschnikowia_unclassified(100); | Metschnikowia | 13 | Saprotroph | \|Undefined Saprotroph\| | Yeast | NULL | Probable | As nectar-tap_saprotroph (Põlme et al. 2020); Decay sugar-rich substrates (Põlme et al. 2020) | Põlme S, et al. 2020. Fungal Diversity 105:1-16 ((https://doi.org/10.1007/s13225-020-00466-2)) |
| Otu43071 | 1 | k__Fungi(100);p__Ascomycota(100);c__Saccharomycetes(100);o__Saccharomycetales(100);f__Metschnikowiaceae(100);g__Metschnikowia(100);s__Metschnikowia_pulcherrima(100); | Metschnikowia | 13 | Saprotroph | \|Undefined Saprotroph\| | Yeast | NULL | Probable | As nectar-tap_saprotroph (Põlme et al. 2020); Decay sugar-rich substrates (Põlme et al. 2020) | Põlme S, et al. 2020. Fungal Diversity 105:1-16 ((https://doi.org/10.1007/s13225-020-00466-2)) |
| Otu43072 | 1 | k__Fungi(100);p__Ascomycota(100);c__Saccharomycetes(100);o__Saccharomycetales(100);f__Metschnikowiaceae(100);g__Metschnikowia(100);s__Metschnikowia_pulcherrima(100); | Metschnikowia | 13 | Saprotroph | \|Undefined Saprotroph\| | Yeast | NULL | Probable | As nectar-tap_saprotroph (Põlme et al. 2020); Decay sugar-rich substrates (Põlme et al. 2020) | Põlme S, et al. 2020. Fungal Diversity 105:1-16 ((https://doi.org/10.1007/s13225-020-00466-2)) |
| Otu43073 | 1 | k__Fungi(100);p__Ascomycota(100);c__Saccharomycetes(100);o__Saccharomycetales(100);f__Metschnikowiaceae(100);g__Metschnikowia(100);s__Metschnikowia_pulcherrima(100); | Metschnikowia | 13 | Saprotroph | \|Undefined Saprotroph\| | Yeast | NULL | Probable | As nectar-tap_saprotroph (Põlme et al. 2020); Decay sugar-rich substrates (Põlme et al. 2020) | Põlme S, et al. 2020. Fungal Diversity 105:1-16 ((https://doi.org/10.1007/s13225-020-00466-2)) |
| Otu43074 | 1 | k__Fungi(100);p__Ascomycota(100);c__Saccharomycetes(100);o__Saccharomycetales(100);f__Metschnikowiaceae(100);g__Metschnikowia(100);s__Metschnikowia_pulcherrima(100); | Metschnikowia | 13 | Saprotroph | \|Undefined Saprotroph\| | Yeast | NULL | Probable | As nectar-tap_saprotroph (Põlme et al. 2020); Decay sugar-rich substrates (Põlme et al. 2020) | Põlme S, et al. 2020. Fungal Diversity 105:1-16 ((https://doi.org/10.1007/s13225-020-00466-2)) |
| Otu43075 | 1 | k__Fungi(100);p__Ascomycota(100);c__Saccharomycetes(100);o__Saccharomycetales(100);f__Metschnikowiaceae(100);g__Metschnikowia(100);s__Metschnikowia_pulcherrima(100); | Metschnikowia | 13 | Saprotroph | \|Undefined Saprotroph\| | Yeast | NULL | Probable | As nectar-tap_saprotroph (Põlme et al. 2020); Decay sugar-rich substrates (Põlme et al. 2020) | Põlme S, et al. 2020. Fungal Diversity 105:1-16 ((https://doi.org/10.1007/s13225-020-00466-2)) |
| Otu43076 | 1 | k__Fungi(100);p__Ascomycota(100);c__Saccharomycetes(100);o__Saccharomycetales(100);f__Metschnikowiaceae(100);g__Metschnikowia(100);s__Metschnikowia_pulcherrima(100); | Metschnikowia | 13 | Saprotroph | \|Undefined Saprotroph\| | Yeast | NULL | Probable | As nectar-tap_saprotroph (Põlme et al. 2020); Decay sugar-rich substrates (Põlme et al. 2020) | Põlme S, et al. 2020. Fungal Diversity 105:1-16 ((https://doi.org/10.1007/s13225-020-00466-2)) |
| Otu43077 | 1 | k__Fungi(100);p__Ascomycota(100);c__Saccharomycetes(100);o__Saccharomycetales(100);f__Metschnikowiaceae(100);g__Metschnikowia(100);g__Metschnikowia_unclassified(100); | Metschnikowia | 13 | Saprotroph | \|Undefined Saprotroph\| | Yeast | NULL | Probable | As nectar-tap_saprotroph (Põlme et al. 2020); Decay sugar-rich substrates (Põlme et al. 2020) | Põlme S, et al. 2020. Fungal Diversity 105:1-16 ((https://doi.org/10.1007/s13225-020-00466-2)) |
| Otu43078 | 1 | k__Fungi(100);p__Ascomycota(100);c__Saccharomycetes(100);o__Saccharomycetales(100);f__Metschnikowiaceae(100);g__Metschnikowia(100);s__Metschnikowia_pulcherrima(100); | Metschnikowia | 13 | Saprotroph | \|Undefined Saprotroph\| | Yeast | NULL | Probable | As nectar-tap_saprotroph (Põlme et al. 2020); Decay sugar-rich substrates (Põlme et al. 2020) | Põlme S, et al. 2020. Fungal Diversity 105:1-16 ((https://doi.org/10.1007/s13225-020-00466-2)) |
| Otu43079 | 1 | k__Fungi(100);p__Ascomycota(100);c__Saccharomycetes(100);o__Saccharomycetales(100);f__Metschnikowiaceae(100);g__Metschnikowia(100);s__Metschnikowia_pulcherrima(100); | Metschnikowia | 13 | Saprotroph | \|Undefined Saprotroph\| | Yeast | NULL | Probable | As nectar-tap_saprotroph (Põlme et al. 2020); Decay sugar-rich substrates (Põlme et al. 2020) | Põlme S, et al. 2020. Fungal Diversity 105:1-16 ((https://doi.org/10.1007/s13225-020-00466-2)) |
| Otu43080 | 1 | k__Fungi(100);p__Ascomycota(100);c__Saccharomycetes(100);o__Saccharomycetales(100);f__Metschnikowiaceae(100);g__Metschnikowia(100);g__Metschnikowia_unclassified(100); | Metschnikowia | 13 | Saprotroph | \|Undefined Saprotroph\| | Yeast | NULL | Probable | As nectar-tap_saprotroph (Põlme et al. 2020); Decay sugar-rich substrates (Põlme et al. 2020) | Põlme S, et al. 2020. Fungal Diversity 105:1-16 ((https://doi.org/10.1007/s13225-020-00466-2)) |
| Otu43081 | 1 | k__Fungi(100);p__Ascomycota(100);c__Saccharomycetes(100);o__Saccharomycetales(100);f__Metschnikowiaceae(100);g__Metschnikowia(100);s__Metschnikowia_pulcherrima(100); | Metschnikowia | 13 | Saprotroph | \|Undefined Saprotroph\| | Yeast | NULL | Probable | As nectar-tap_saprotroph (Põlme et al. 2020); Decay sugar-rich substrates (Põlme et al. 2020) | Põlme S, et al. 2020. Fungal Diversity 105:1-16 ((https://doi.org/10.1007/s13225-020-00466-2)) |
| Otu43083 | 1 | k__Fungi(100);p__Ascomycota(100);c__Saccharomycetes(100);o__Saccharomycetales(100);f__Metschnikowiaceae(100);g__Metschnikowia(100);s__Metschnikowia_pulcherrima(100); | Metschnikowia | 13 | Saprotroph | \|Undefined Saprotroph\| | Yeast | NULL | Probable | As nectar-tap_saprotroph (Põlme et al. 2020); Decay sugar-rich substrates (Põlme et al. 2020) | Põlme S, et al. 2020. Fungal Diversity 105:1-16 ((https://doi.org/10.1007/s13225-020-00466-2)) |
| Otu43084 | 1 | k__Fungi(100);p__Ascomycota(100);c__Saccharomycetes(100);o__Saccharomycetales(100);f__Metschnikowiaceae(100);g__Metschnikowia(100);s__Metschnikowia_pulcherrima(100); | Metschnikowia | 13 | Saprotroph | \|Undefined Saprotroph\| | Yeast | NULL | Probable | As nectar-tap_saprotroph (Põlme et al. 2020); Decay sugar-rich substrates (Põlme et al. 2020) | Põlme S, et al. 2020. Fungal Diversity 105:1-16 ((https://doi.org/10.1007/s13225-020-00466-2)) |
| Otu43086 | 1 | k__Fungi(100);p__Ascomycota(100);c__Saccharomycetes(100);o__Saccharomycetales(100);f__Metschnikowiaceae(100);g__Metschnikowia(100);s__Metschnikowia_pulcherrima(100); | Metschnikowia | 13 | Saprotroph | \|Undefined Saprotroph\| | Yeast | NULL | Probable | As nectar-tap_saprotroph (Põlme et al. 2020); Decay sugar-rich substrates (Põlme et al. 2020) | Põlme S, et al. 2020. Fungal Diversity 105:1-16 ((https://doi.org/10.1007/s13225-020-00466-2)) |
| Otu43087 | 1 | k__Fungi(100);p__Ascomycota(100);c__Saccharomycetes(100);o__Saccharomycetales(100);f__Metschnikowiaceae(100);g__Metschnikowia(100);s__Metschnikowia_pulcherrima(100); | Metschnikowia | 13 | Saprotroph | \|Undefined Saprotroph\| | Yeast | NULL | Probable | As nectar-tap_saprotroph (Põlme et al. 2020); Decay sugar-rich substrates (Põlme et al. 2020) | Põlme S, et al. 2020. Fungal Diversity 105:1-16 ((https://doi.org/10.1007/s13225-020-00466-2)) |
| Otu43088 | 1 | k__Fungi(100);p__Ascomycota(100);c__Saccharomycetes(100);o__Saccharomycetales(100);f__Metschnikowiaceae(100);g__Metschnikowia(100);s__Metschnikowia_pulcherrima(100); | Metschnikowia | 13 | Saprotroph | \|Undefined Saprotroph\| | Yeast | NULL | Probable | As nectar-tap_saprotroph (Põlme et al. 2020); Decay sugar-rich substrates (Põlme et al. 2020) | Põlme S, et al. 2020. Fungal Diversity 105:1-16 ((https://doi.org/10.1007/s13225-020-00466-2)) |
| Otu43089 | 1 | k__Fungi(100);p__Ascomycota(100);c__Saccharomycetes(100);o__Saccharomycetales(100);f__Metschnikowiaceae(100);g__Metschnikowia(100);s__Metschnikowia_pulcherrima(100); | Metschnikowia | 13 | Saprotroph | \|Undefined Saprotroph\| | Yeast | NULL | Probable | As nectar-tap_saprotroph (Põlme et al. 2020); Decay sugar-rich substrates (Põlme et al. 2020) | Põlme S, et al. 2020. Fungal Diversity 105:1-16 ((https://doi.org/10.1007/s13225-020-00466-2)) |
| Otu43090 | 1 | k__Fungi(100);p__Ascomycota(100);c__Sordariomycetes(100);o__Hypocreales(100);f__Nectriaceae(100);g__Fusarium(100);g__Fusarium_unclassified(100); | Fusarium | 13 | Pathotroph-Saprotroph-Symbiotroph | Animal Pathogen-Endophyte-Lichen Parasite-\|Plant Pathogen\|-Plant Saprotroph-Undefined Saprotroph-Wood Saprotroph | Microfungus | Soft Rot | Possible | As foliar_endophyte (Põlme et al. 2020); As litter_saprotroph (Põlme et al. 2020); Causing damping-off disease (Põlme et al. 2020); Decay leaf, fruit, seed, roots, soil, animal material (Põlme et al. 2020); Likely opportunistic human pathogen (Irinyi et al. 2015) | Duncan CG, Eslyn WE. 1966. Mycologia 58:642-664 ((https://doi.org/10.2307/3757045)); Seehann G, et al. 1975. List of Fungi in Soft-Rot Tissue Tests. Institute fur Holzbiologie und Holzschutz, Bundesforschungsanstalt fur Forst - und Holzwirtshaft, Hamburg - Lohbrugge, Germany; Costa IPMW, et al. 2012. Mycotaxon 119:e26 ((http://mycotaxon.com/resources/checklists/costa_v119_checklist.pdf)); Tedersoo L, et al. 2014. Science 346:e1256688 ((https://doi.org/10.1126/science.1256688)); Irinyi L, et al. 2015. Medical Mycology 53:313-337 ((https://doi.org/10.1093/mmy/myv008)); Schroers, et al. 2016. Mycologia 108:806-819 ((https://doi.org/10.3852/15-255)); Põlme S, et al. 2020. Fungal Diversity 105:1-16 ((https://doi.org/10.1007/s13225-020-00466-2)); The American Phytopathological Society Common Names of Plant Diseases ((http://www.apsnet.org/publications/commonnames/Pages/default.aspx)); Lawrey JD, Diederich P. 2016. http://www.lichenicolous.net (accessed 24 May 2017) ((http://www.lichenicolous.net)) |
| Otu43091 | 1 | k__Fungi(100);p__Ascomycota(100);c__Saccharomycetes(100);o__Saccharomycetales(100);f__Metschnikowiaceae(100);g__Metschnikowia(100);s__Metschnikowia_pulcherrima(100); | Metschnikowia | 13 | Saprotroph | \|Undefined Saprotroph\| | Yeast | NULL | Probable | As nectar-tap_saprotroph (Põlme et al. 2020); Decay sugar-rich substrates (Põlme et al. 2020) | Põlme S, et al. 2020. Fungal Diversity 105:1-16 ((https://doi.org/10.1007/s13225-020-00466-2)) |
| Otu43092 | 1 | k__Fungi(100);p__Ascomycota(100);c__Saccharomycetes(100);o__Saccharomycetales(100);f__Metschnikowiaceae(100);g__Metschnikowia(100);g__Metschnikowia_unclassified(100); | Metschnikowia | 13 | Saprotroph | \|Undefined Saprotroph\| | Yeast | NULL | Probable | As nectar-tap_saprotroph (Põlme et al. 2020); Decay sugar-rich substrates (Põlme et al. 2020) | Põlme S, et al. 2020. Fungal Diversity 105:1-16 ((https://doi.org/10.1007/s13225-020-00466-2)) |
| Otu43093 | 1 | k__Fungi(100);p__Ascomycota(100);c__Saccharomycetes(100);o__Saccharomycetales(100);f__Metschnikowiaceae(100);g__Metschnikowia(100);s__Metschnikowia_pulcherrima(100); | Metschnikowia | 13 | Saprotroph | \|Undefined Saprotroph\| | Yeast | NULL | Probable | As nectar-tap_saprotroph (Põlme et al. 2020); Decay sugar-rich substrates (Põlme et al. 2020) | Põlme S, et al. 2020. Fungal Diversity 105:1-16 ((https://doi.org/10.1007/s13225-020-00466-2)) |
| Otu43094 | 1 | k__Fungi(100);p__Ascomycota(100);c__Saccharomycetes(100);o__Saccharomycetales(100);f__Metschnikowiaceae(100);g__Metschnikowia(100);s__Metschnikowia_pulcherrima(100); | Metschnikowia | 13 | Saprotroph | \|Undefined Saprotroph\| | Yeast | NULL | Probable | As nectar-tap_saprotroph (Põlme et al. 2020); Decay sugar-rich substrates (Põlme et al. 2020) | Põlme S, et al. 2020. Fungal Diversity 105:1-16 ((https://doi.org/10.1007/s13225-020-00466-2)) |
| Otu43095 | 1 | k__Fungi(100);p__Ascomycota(100);c__Saccharomycetes(100);o__Saccharomycetales(100);f__Metschnikowiaceae(100);g__Metschnikowia(100);s__Metschnikowia_pulcherrima(100); | Metschnikowia | 13 | Saprotroph | \|Undefined Saprotroph\| | Yeast | NULL | Probable | As nectar-tap_saprotroph (Põlme et al. 2020); Decay sugar-rich substrates (Põlme et al. 2020) | Põlme S, et al. 2020. Fungal Diversity 105:1-16 ((https://doi.org/10.1007/s13225-020-00466-2)) |
| Otu43096 | 1 | k__Fungi(100);p__Ascomycota(100);c__Saccharomycetes(100);o__Saccharomycetales(100);f__Metschnikowiaceae(100);g__Metschnikowia(100);s__Metschnikowia_pulcherrima(100); | Metschnikowia | 13 | Saprotroph | \|Undefined Saprotroph\| | Yeast | NULL | Probable | As nectar-tap_saprotroph (Põlme et al. 2020); Decay sugar-rich substrates (Põlme et al. 2020) | Põlme S, et al. 2020. Fungal Diversity 105:1-16 ((https://doi.org/10.1007/s13225-020-00466-2)) |
| Otu43097 | 1 | k__Fungi(100);p__Ascomycota(100);c__Saccharomycetes(100);o__Saccharomycetales(100);f__Metschnikowiaceae(100);g__Metschnikowia(100);s__Metschnikowia_pulcherrima(100); | Metschnikowia | 13 | Saprotroph | \|Undefined Saprotroph\| | Yeast | NULL | Probable | As nectar-tap_saprotroph (Põlme et al. 2020); Decay sugar-rich substrates (Põlme et al. 2020) | Põlme S, et al. 2020. Fungal Diversity 105:1-16 ((https://doi.org/10.1007/s13225-020-00466-2)) |
| Otu43099 | 1 | k__Fungi(100);p__Ascomycota(100);c__Saccharomycetes(100);o__Saccharomycetales(100);f__Metschnikowiaceae(100);g__Metschnikowia(100);s__Metschnikowia_pulcherrima(100); | Metschnikowia | 13 | Saprotroph | \|Undefined Saprotroph\| | Yeast | NULL | Probable | As nectar-tap_saprotroph (Põlme et al. 2020); Decay sugar-rich substrates (Põlme et al. 2020) | Põlme S, et al. 2020. Fungal Diversity 105:1-16 ((https://doi.org/10.1007/s13225-020-00466-2)) |
| Otu43100 | 1 | k__Fungi(100);p__Ascomycota(100);c__Saccharomycetes(100);o__Saccharomycetales(100);f__Metschnikowiaceae(100);g__Metschnikowia(100);g__Metschnikowia_unclassified(100); | Metschnikowia | 13 | Saprotroph | \|Undefined Saprotroph\| | Yeast | NULL | Probable | As nectar-tap_saprotroph (Põlme et al. 2020); Decay sugar-rich substrates (Põlme et al. 2020) | Põlme S, et al. 2020. Fungal Diversity 105:1-16 ((https://doi.org/10.1007/s13225-020-00466-2)) |
| Otu43101 | 1 | k__Fungi(100);p__Ascomycota(100);c__Saccharomycetes(100);o__Saccharomycetales(100);f__Metschnikowiaceae(100);g__Metschnikowia(100);s__Metschnikowia_pulcherrima(100); | Metschnikowia | 13 | Saprotroph | \|Undefined Saprotroph\| | Yeast | NULL | Probable | As nectar-tap_saprotroph (Põlme et al. 2020); Decay sugar-rich substrates (Põlme et al. 2020) | Põlme S, et al. 2020. Fungal Diversity 105:1-16 ((https://doi.org/10.1007/s13225-020-00466-2)) |
| Otu43102 | 1 | k__Fungi(100);p__Ascomycota(100);c__Saccharomycetes(100);o__Saccharomycetales(100);f__Metschnikowiaceae(100);g__Metschnikowia(100);g__Metschnikowia_unclassified(100); | Metschnikowia | 13 | Saprotroph | \|Undefined Saprotroph\| | Yeast | NULL | Probable | As nectar-tap_saprotroph (Põlme et al. 2020); Decay sugar-rich substrates (Põlme et al. 2020) | Põlme S, et al. 2020. Fungal Diversity 105:1-16 ((https://doi.org/10.1007/s13225-020-00466-2)) |
| Otu43103 | 1 | k__Fungi(100);p__Ascomycota(100);c__Saccharomycetes(100);o__Saccharomycetales(100);f__Metschnikowiaceae(100);g__Metschnikowia(100);s__Metschnikowia_pulcherrima(100); | Metschnikowia | 13 | Saprotroph | \|Undefined Saprotroph\| | Yeast | NULL | Probable | As nectar-tap_saprotroph (Põlme et al. 2020); Decay sugar-rich substrates (Põlme et al. 2020) | Põlme S, et al. 2020. Fungal Diversity 105:1-16 ((https://doi.org/10.1007/s13225-020-00466-2)) |
| Otu43104 | 1 | k__Fungi(100);p__Ascomycota(100);c__Saccharomycetes(100);o__Saccharomycetales(100);f__Metschnikowiaceae(100);g__Metschnikowia(100);s__Metschnikowia_pulcherrima(100); | Metschnikowia | 13 | Saprotroph | \|Undefined Saprotroph\| | Yeast | NULL | Probable | As nectar-tap_saprotroph (Põlme et al. 2020); Decay sugar-rich substrates (Põlme et al. 2020) | Põlme S, et al. 2020. Fungal Diversity 105:1-16 ((https://doi.org/10.1007/s13225-020-00466-2)) |
| Otu43105 | 1 | k__Fungi(100);p__Ascomycota(100);c__Saccharomycetes(100);o__Saccharomycetales(100);f__Metschnikowiaceae(100);g__Metschnikowia(100);s__Metschnikowia_pulcherrima(100); | Metschnikowia | 13 | Saprotroph | \|Undefined Saprotroph\| | Yeast | NULL | Probable | As nectar-tap_saprotroph (Põlme et al. 2020); Decay sugar-rich substrates (Põlme et al. 2020) | Põlme S, et al. 2020. Fungal Diversity 105:1-16 ((https://doi.org/10.1007/s13225-020-00466-2)) |
| Otu43106 | 1 | k__Fungi(100);p__Ascomycota(100);c__Saccharomycetes(100);o__Saccharomycetales(100);f__Metschnikowiaceae(100);g__Metschnikowia(100);s__Metschnikowia_pulcherrima(100); | Metschnikowia | 13 | Saprotroph | \|Undefined Saprotroph\| | Yeast | NULL | Probable | As nectar-tap_saprotroph (Põlme et al. 2020); Decay sugar-rich substrates (Põlme et al. 2020) | Põlme S, et al. 2020. Fungal Diversity 105:1-16 ((https://doi.org/10.1007/s13225-020-00466-2)) |
| Otu43107 | 1 | k__Fungi(100);p__Ascomycota(100);c__Saccharomycetes(100);o__Saccharomycetales(100);f__Metschnikowiaceae(100);g__Metschnikowia(100);g__Metschnikowia_unclassified(100); | Metschnikowia | 13 | Saprotroph | \|Undefined Saprotroph\| | Yeast | NULL | Probable | As nectar-tap_saprotroph (Põlme et al. 2020); Decay sugar-rich substrates (Põlme et al. 2020) | Põlme S, et al. 2020. Fungal Diversity 105:1-16 ((https://doi.org/10.1007/s13225-020-00466-2)) |
| Otu43108 | 1 | k__Fungi(100);p__Ascomycota(100);c__Saccharomycetes(100);o__Saccharomycetales(100);f__Metschnikowiaceae(100);g__Metschnikowia(100);s__Metschnikowia_pulcherrima(100); | Metschnikowia | 13 | Saprotroph | \|Undefined Saprotroph\| | Yeast | NULL | Probable | As nectar-tap_saprotroph (Põlme et al. 2020); Decay sugar-rich substrates (Põlme et al. 2020) | Põlme S, et al. 2020. Fungal Diversity 105:1-16 ((https://doi.org/10.1007/s13225-020-00466-2)) |
| Otu43109 | 1 | k__Fungi(100);p__Ascomycota(100);c__Saccharomycetes(100);o__Saccharomycetales(100);f__Metschnikowiaceae(100);g__Metschnikowia(100);s__Metschnikowia_pulcherrima(100); | Metschnikowia | 13 | Saprotroph | \|Undefined Saprotroph\| | Yeast | NULL | Probable | As nectar-tap_saprotroph (Põlme et al. 2020); Decay sugar-rich substrates (Põlme et al. 2020) | Põlme S, et al. 2020. Fungal Diversity 105:1-16 ((https://doi.org/10.1007/s13225-020-00466-2)) |
| Otu43110 | 1 | k__Fungi(100);p__Ascomycota(100);c__Saccharomycetes(100);o__Saccharomycetales(100);f__Metschnikowiaceae(100);g__Metschnikowia(100);s__Metschnikowia_pulcherrima(100); | Metschnikowia | 13 | Saprotroph | \|Undefined Saprotroph\| | Yeast | NULL | Probable | As nectar-tap_saprotroph (Põlme et al. 2020); Decay sugar-rich substrates (Põlme et al. 2020) | Põlme S, et al. 2020. Fungal Diversity 105:1-16 ((https://doi.org/10.1007/s13225-020-00466-2)) |
| Otu43111 | 1 | k__Fungi(100);p__Ascomycota(100);c__Saccharomycetes(100);o__Saccharomycetales(100);f__Metschnikowiaceae(100);g__Metschnikowia(100);s__Metschnikowia_pulcherrima(100); | Metschnikowia | 13 | Saprotroph | \|Undefined Saprotroph\| | Yeast | NULL | Probable | As nectar-tap_saprotroph (Põlme et al. 2020); Decay sugar-rich substrates (Põlme et al. 2020) | Põlme S, et al. 2020. Fungal Diversity 105:1-16 ((https://doi.org/10.1007/s13225-020-00466-2)) |
| Otu43112 | 1 | k__Fungi(100);p__Ascomycota(100);c__Saccharomycetes(100);o__Saccharomycetales(100);f__Metschnikowiaceae(100);g__Metschnikowia(100);s__Metschnikowia_pulcherrima(100); | Metschnikowia | 13 | Saprotroph | \|Undefined Saprotroph\| | Yeast | NULL | Probable | As nectar-tap_saprotroph (Põlme et al. 2020); Decay sugar-rich substrates (Põlme et al. 2020) | Põlme S, et al. 2020. Fungal Diversity 105:1-16 ((https://doi.org/10.1007/s13225-020-00466-2)) |
| Otu43113 | 1 | k__Fungi(100);p__Ascomycota(100);c__Saccharomycetes(100);o__Saccharomycetales(100);f__Metschnikowiaceae(100);g__Metschnikowia(100);g__Metschnikowia_unclassified(100); | Metschnikowia | 13 | Saprotroph | \|Undefined Saprotroph\| | Yeast | NULL | Probable | As nectar-tap_saprotroph (Põlme et al. 2020); Decay sugar-rich substrates (Põlme et al. 2020) | Põlme S, et al. 2020. Fungal Diversity 105:1-16 ((https://doi.org/10.1007/s13225-020-00466-2)) |
| Otu43114 | 1 | k__Fungi(100);p__Ascomycota(100);c__Saccharomycetes(100);o__Saccharomycetales(100);f__Metschnikowiaceae(100);g__Metschnikowia(100);s__Metschnikowia_pulcherrima(100); | Metschnikowia | 13 | Saprotroph | \|Undefined Saprotroph\| | Yeast | NULL | Probable | As nectar-tap_saprotroph (Põlme et al. 2020); Decay sugar-rich substrates (Põlme et al. 2020) | Põlme S, et al. 2020. Fungal Diversity 105:1-16 ((https://doi.org/10.1007/s13225-020-00466-2)) |
| Otu43115 | 1 | k__Fungi(100);p__Ascomycota(100);c__Saccharomycetes(100);o__Saccharomycetales(100);f__Metschnikowiaceae(100);g__Metschnikowia(100);s__Metschnikowia_pulcherrima(100); | Metschnikowia | 13 | Saprotroph | \|Undefined Saprotroph\| | Yeast | NULL | Probable | As nectar-tap_saprotroph (Põlme et al. 2020); Decay sugar-rich substrates (Põlme et al. 2020) | Põlme S, et al. 2020. Fungal Diversity 105:1-16 ((https://doi.org/10.1007/s13225-020-00466-2)) |
| Otu43116 | 1 | k__Fungi(100);p__Ascomycota(100);c__Saccharomycetes(100);o__Saccharomycetales(100);f__Metschnikowiaceae(100);g__Metschnikowia(100);s__Metschnikowia_pulcherrima(100); | Metschnikowia | 13 | Saprotroph | \|Undefined Saprotroph\| | Yeast | NULL | Probable | As nectar-tap_saprotroph (Põlme et al. 2020); Decay sugar-rich substrates (Põlme et al. 2020) | Põlme S, et al. 2020. Fungal Diversity 105:1-16 ((https://doi.org/10.1007/s13225-020-00466-2)) |
| Otu43117 | 1 | k__Fungi(100);p__Ascomycota(100);c__Saccharomycetes(100);o__Saccharomycetales(100);f__Metschnikowiaceae(100);g__Metschnikowia(100);s__Metschnikowia_pulcherrima(100); | Metschnikowia | 13 | Saprotroph | \|Undefined Saprotroph\| | Yeast | NULL | Probable | As nectar-tap_saprotroph (Põlme et al. 2020); Decay sugar-rich substrates (Põlme et al. 2020) | Põlme S, et al. 2020. Fungal Diversity 105:1-16 ((https://doi.org/10.1007/s13225-020-00466-2)) |
| Otu43118 | 1 | k__Fungi(100);p__Ascomycota(100);c__Saccharomycetes(100);o__Saccharomycetales(100);f__Metschnikowiaceae(100);g__Metschnikowia(100);g__Metschnikowia_unclassified(100); | Metschnikowia | 13 | Saprotroph | \|Undefined Saprotroph\| | Yeast | NULL | Probable | As nectar-tap_saprotroph (Põlme et al. 2020); Decay sugar-rich substrates (Põlme et al. 2020) | Põlme S, et al. 2020. Fungal Diversity 105:1-16 ((https://doi.org/10.1007/s13225-020-00466-2)) |
| Otu43119 | 1 | k__Fungi(100);p__Ascomycota(100);c__Saccharomycetes(100);o__Saccharomycetales(100);f__Metschnikowiaceae(100);g__Metschnikowia(100);g__Metschnikowia_unclassified(100); | Metschnikowia | 13 | Saprotroph | \|Undefined Saprotroph\| | Yeast | NULL | Probable | As nectar-tap_saprotroph (Põlme et al. 2020); Decay sugar-rich substrates (Põlme et al. 2020) | Põlme S, et al. 2020. Fungal Diversity 105:1-16 ((https://doi.org/10.1007/s13225-020-00466-2)) |
| Otu43120 | 1 | k__Fungi(100);p__Ascomycota(100);c__Saccharomycetes(100);o__Saccharomycetales(100);f__Metschnikowiaceae(100);g__Metschnikowia(100);s__Metschnikowia_pulcherrima(100); | Metschnikowia | 13 | Saprotroph | \|Undefined Saprotroph\| | Yeast | NULL | Probable | As nectar-tap_saprotroph (Põlme et al. 2020); Decay sugar-rich substrates (Põlme et al. 2020) | Põlme S, et al. 2020. Fungal Diversity 105:1-16 ((https://doi.org/10.1007/s13225-020-00466-2)) |
| Otu43121 | 1 | k__Fungi(100);p__Ascomycota(100);c__Saccharomycetes(100);o__Saccharomycetales(100);f__Metschnikowiaceae(100);g__Metschnikowia(100);s__Metschnikowia_pulcherrima(100); | Metschnikowia | 13 | Saprotroph | \|Undefined Saprotroph\| | Yeast | NULL | Probable | As nectar-tap_saprotroph (Põlme et al. 2020); Decay sugar-rich substrates (Põlme et al. 2020) | Põlme S, et al. 2020. Fungal Diversity 105:1-16 ((https://doi.org/10.1007/s13225-020-00466-2)) |
| Otu43122 | 1 | k__Fungi(100);p__Ascomycota(100);c__Saccharomycetes(100);o__Saccharomycetales(100);f__Metschnikowiaceae(100);g__Metschnikowia(100);s__Metschnikowia_pulcherrima(100); | Metschnikowia | 13 | Saprotroph | \|Undefined Saprotroph\| | Yeast | NULL | Probable | As nectar-tap_saprotroph (Põlme et al. 2020); Decay sugar-rich substrates (Põlme et al. 2020) | Põlme S, et al. 2020. Fungal Diversity 105:1-16 ((https://doi.org/10.1007/s13225-020-00466-2)) |
| Otu43123 | 1 | k__Fungi(100);p__Ascomycota(100);c__Saccharomycetes(100);o__Saccharomycetales(100);f__Metschnikowiaceae(100);g__Metschnikowia(100);s__Metschnikowia_pulcherrima(100); | Metschnikowia | 13 | Saprotroph | \|Undefined Saprotroph\| | Yeast | NULL | Probable | As nectar-tap_saprotroph (Põlme et al. 2020); Decay sugar-rich substrates (Põlme et al. 2020) | Põlme S, et al. 2020. Fungal Diversity 105:1-16 ((https://doi.org/10.1007/s13225-020-00466-2)) |
| Otu43124 | 1 | k__Fungi(100);p__Ascomycota(100);c__Saccharomycetes(100);o__Saccharomycetales(100);f__Metschnikowiaceae(100);g__Metschnikowia(100);s__Metschnikowia_pulcherrima(100); | Metschnikowia | 13 | Saprotroph | \|Undefined Saprotroph\| | Yeast | NULL | Probable | As nectar-tap_saprotroph (Põlme et al. 2020); Decay sugar-rich substrates (Põlme et al. 2020) | Põlme S, et al. 2020. Fungal Diversity 105:1-16 ((https://doi.org/10.1007/s13225-020-00466-2)) |
| Otu43125 | 1 | k__Fungi(100);p__Ascomycota(100);c__Saccharomycetes(100);o__Saccharomycetales(100);f__Metschnikowiaceae(100);g__Metschnikowia(100);s__Metschnikowia_pulcherrima(100); | Metschnikowia | 13 | Saprotroph | \|Undefined Saprotroph\| | Yeast | NULL | Probable | As nectar-tap_saprotroph (Põlme et al. 2020); Decay sugar-rich substrates (Põlme et al. 2020) | Põlme S, et al. 2020. Fungal Diversity 105:1-16 ((https://doi.org/10.1007/s13225-020-00466-2)) |
| Otu43126 | 1 | k__Fungi(100);p__Ascomycota(100);c__Saccharomycetes(100);o__Saccharomycetales(100);f__Metschnikowiaceae(100);g__Metschnikowia(100);s__Metschnikowia_pulcherrima(100); | Metschnikowia | 13 | Saprotroph | \|Undefined Saprotroph\| | Yeast | NULL | Probable | As nectar-tap_saprotroph (Põlme et al. 2020); Decay sugar-rich substrates (Põlme et al. 2020) | Põlme S, et al. 2020. Fungal Diversity 105:1-16 ((https://doi.org/10.1007/s13225-020-00466-2)) |
| Otu43127 | 1 | k__Fungi(100);p__Ascomycota(100);c__Saccharomycetes(100);o__Saccharomycetales(100);f__Metschnikowiaceae(100);g__Metschnikowia(100);s__Metschnikowia_pulcherrima(100); | Metschnikowia | 13 | Saprotroph | \|Undefined Saprotroph\| | Yeast | NULL | Probable | As nectar-tap_saprotroph (Põlme et al. 2020); Decay sugar-rich substrates (Põlme et al. 2020) | Põlme S, et al. 2020. Fungal Diversity 105:1-16 ((https://doi.org/10.1007/s13225-020-00466-2)) |
| Otu43128 | 1 | k__Fungi(100);p__Ascomycota(100);c__Saccharomycetes(100);o__Saccharomycetales(100);f__Metschnikowiaceae(100);g__Metschnikowia(100);s__Metschnikowia_pulcherrima(100); | Metschnikowia | 13 | Saprotroph | \|Undefined Saprotroph\| | Yeast | NULL | Probable | As nectar-tap_saprotroph (Põlme et al. 2020); Decay sugar-rich substrates (Põlme et al. 2020) | Põlme S, et al. 2020. Fungal Diversity 105:1-16 ((https://doi.org/10.1007/s13225-020-00466-2)) |
| Otu43129 | 1 | k__Fungi(100);p__Ascomycota(100);c__Saccharomycetes(100);o__Saccharomycetales(100);f__Metschnikowiaceae(100);g__Metschnikowia(100);s__Metschnikowia_pulcherrima(100); | Metschnikowia | 13 | Saprotroph | \|Undefined Saprotroph\| | Yeast | NULL | Probable | As nectar-tap_saprotroph (Põlme et al. 2020); Decay sugar-rich substrates (Põlme et al. 2020) | Põlme S, et al. 2020. Fungal Diversity 105:1-16 ((https://doi.org/10.1007/s13225-020-00466-2)) |
| Otu43130 | 1 | k__Fungi(100);p__Ascomycota(100);c__Saccharomycetes(100);o__Saccharomycetales(100);f__Metschnikowiaceae(100);g__Metschnikowia(100);s__Metschnikowia_pulcherrima(100); | Metschnikowia | 13 | Saprotroph | \|Undefined Saprotroph\| | Yeast | NULL | Probable | As nectar-tap_saprotroph (Põlme et al. 2020); Decay sugar-rich substrates (Põlme et al. 2020) | Põlme S, et al. 2020. Fungal Diversity 105:1-16 ((https://doi.org/10.1007/s13225-020-00466-2)) |
| Otu43131 | 1 | k__Fungi(100);p__Ascomycota(100);c__Saccharomycetes(100);o__Saccharomycetales(100);f__Metschnikowiaceae(100);g__Metschnikowia(100);s__Metschnikowia_pulcherrima(100); | Metschnikowia | 13 | Saprotroph | \|Undefined Saprotroph\| | Yeast | NULL | Probable | As nectar-tap_saprotroph (Põlme et al. 2020); Decay sugar-rich substrates (Põlme et al. 2020) | Põlme S, et al. 2020. Fungal Diversity 105:1-16 ((https://doi.org/10.1007/s13225-020-00466-2)) |
| Otu43132 | 1 | k__Fungi(100);p__Ascomycota(100);c__Saccharomycetes(100);o__Saccharomycetales(100);f__Metschnikowiaceae(100);g__Metschnikowia(100);g__Metschnikowia_unclassified(100); | Metschnikowia | 13 | Saprotroph | \|Undefined Saprotroph\| | Yeast | NULL | Probable | As nectar-tap_saprotroph (Põlme et al. 2020); Decay sugar-rich substrates (Põlme et al. 2020) | Põlme S, et al. 2020. Fungal Diversity 105:1-16 ((https://doi.org/10.1007/s13225-020-00466-2)) |
| Otu43133 | 1 | k__Fungi(100);p__Ascomycota(100);c__Saccharomycetes(100);o__Saccharomycetales(100);f__Metschnikowiaceae(100);g__Metschnikowia(100);s__Metschnikowia_pulcherrima(100); | Metschnikowia | 13 | Saprotroph | \|Undefined Saprotroph\| | Yeast | NULL | Probable | As nectar-tap_saprotroph (Põlme et al. 2020); Decay sugar-rich substrates (Põlme et al. 2020) | Põlme S, et al. 2020. Fungal Diversity 105:1-16 ((https://doi.org/10.1007/s13225-020-00466-2)) |
| Otu43134 | 1 | k__Fungi(100);p__Ascomycota(100);c__Saccharomycetes(100);o__Saccharomycetales(100);f__Metschnikowiaceae(100);g__Metschnikowia(100);s__Metschnikowia_pulcherrima(100); | Metschnikowia | 13 | Saprotroph | \|Undefined Saprotroph\| | Yeast | NULL | Probable | As nectar-tap_saprotroph (Põlme et al. 2020); Decay sugar-rich substrates (Põlme et al. 2020) | Põlme S, et al. 2020. Fungal Diversity 105:1-16 ((https://doi.org/10.1007/s13225-020-00466-2)) |
| Otu43135 | 1 | k__Fungi(100);p__Ascomycota(100);c__Saccharomycetes(100);o__Saccharomycetales(100);f__Metschnikowiaceae(100);g__Metschnikowia(100);s__Metschnikowia_pulcherrima(100); | Metschnikowia | 13 | Saprotroph | \|Undefined Saprotroph\| | Yeast | NULL | Probable | As nectar-tap_saprotroph (Põlme et al. 2020); Decay sugar-rich substrates (Põlme et al. 2020) | Põlme S, et al. 2020. Fungal Diversity 105:1-16 ((https://doi.org/10.1007/s13225-020-00466-2)) |
| Otu43136 | 1 | k__Fungi(100);p__Ascomycota(100);c__Saccharomycetes(100);o__Saccharomycetales(100);f__Metschnikowiaceae(100);g__Metschnikowia(100);s__Metschnikowia_pulcherrima(100); | Metschnikowia | 13 | Saprotroph | \|Undefined Saprotroph\| | Yeast | NULL | Probable | As nectar-tap_saprotroph (Põlme et al. 2020); Decay sugar-rich substrates (Põlme et al. 2020) | Põlme S, et al. 2020. Fungal Diversity 105:1-16 ((https://doi.org/10.1007/s13225-020-00466-2)) |
| Otu43137 | 1 | k__Fungi(100);p__Ascomycota(100);c__Saccharomycetes(100);o__Saccharomycetales(100);f__Metschnikowiaceae(100);g__Metschnikowia(100);s__Metschnikowia_pulcherrima(100); | Metschnikowia | 13 | Saprotroph | \|Undefined Saprotroph\| | Yeast | NULL | Probable | As nectar-tap_saprotroph (Põlme et al. 2020); Decay sugar-rich substrates (Põlme et al. 2020) | Põlme S, et al. 2020. Fungal Diversity 105:1-16 ((https://doi.org/10.1007/s13225-020-00466-2)) |
| Otu43138 | 1 | k__Fungi(100);p__Ascomycota(100);c__Saccharomycetes(100);o__Saccharomycetales(100);f__Metschnikowiaceae(100);g__Metschnikowia(100);s__Metschnikowia_pulcherrima(100); | Metschnikowia | 13 | Saprotroph | \|Undefined Saprotroph\| | Yeast | NULL | Probable | As nectar-tap_saprotroph (Põlme et al. 2020); Decay sugar-rich substrates (Põlme et al. 2020) | Põlme S, et al. 2020. Fungal Diversity 105:1-16 ((https://doi.org/10.1007/s13225-020-00466-2)) |
| Otu43139 | 1 | k__Fungi(100);p__Ascomycota(100);c__Saccharomycetes(100);o__Saccharomycetales(100);f__Metschnikowiaceae(100);g__Metschnikowia(100);s__Metschnikowia_pulcherrima(100); | Metschnikowia | 13 | Saprotroph | \|Undefined Saprotroph\| | Yeast | NULL | Probable | As nectar-tap_saprotroph (Põlme et al. 2020); Decay sugar-rich substrates (Põlme et al. 2020) | Põlme S, et al. 2020. Fungal Diversity 105:1-16 ((https://doi.org/10.1007/s13225-020-00466-2)) |
| Otu43140 | 1 | k__Fungi(100);p__Ascomycota(100);c__Saccharomycetes(100);o__Saccharomycetales(100);f__Metschnikowiaceae(100);g__Metschnikowia(100);s__Metschnikowia_pulcherrima(100); | Metschnikowia | 13 | Saprotroph | \|Undefined Saprotroph\| | Yeast | NULL | Probable | As nectar-tap_saprotroph (Põlme et al. 2020); Decay sugar-rich substrates (Põlme et al. 2020) | Põlme S, et al. 2020. Fungal Diversity 105:1-16 ((https://doi.org/10.1007/s13225-020-00466-2)) |
| Otu43141 | 1 | k__Fungi(100);p__Ascomycota(100);c__Saccharomycetes(100);o__Saccharomycetales(100);f__Metschnikowiaceae(100);g__Metschnikowia(100);s__Metschnikowia_pulcherrima(100); | Metschnikowia | 13 | Saprotroph | \|Undefined Saprotroph\| | Yeast | NULL | Probable | As nectar-tap_saprotroph (Põlme et al. 2020); Decay sugar-rich substrates (Põlme et al. 2020) | Põlme S, et al. 2020. Fungal Diversity 105:1-16 ((https://doi.org/10.1007/s13225-020-00466-2)) |
| Otu43142 | 1 | k__Fungi(100);p__Ascomycota(100);c__Saccharomycetes(100);o__Saccharomycetales(100);f__Metschnikowiaceae(100);g__Metschnikowia(100);g__Metschnikowia_unclassified(100); | Metschnikowia | 13 | Saprotroph | \|Undefined Saprotroph\| | Yeast | NULL | Probable | As nectar-tap_saprotroph (Põlme et al. 2020); Decay sugar-rich substrates (Põlme et al. 2020) | Põlme S, et al. 2020. Fungal Diversity 105:1-16 ((https://doi.org/10.1007/s13225-020-00466-2)) |
| Otu43143 | 1 | k__Fungi(100);p__Ascomycota(100);c__Saccharomycetes(100);o__Saccharomycetales(100);f__Metschnikowiaceae(100);g__Metschnikowia(100);g__Metschnikowia_unclassified(100); | Metschnikowia | 13 | Saprotroph | \|Undefined Saprotroph\| | Yeast | NULL | Probable | As nectar-tap_saprotroph (Põlme et al. 2020); Decay sugar-rich substrates (Põlme et al. 2020) | Põlme S, et al. 2020. Fungal Diversity 105:1-16 ((https://doi.org/10.1007/s13225-020-00466-2)) |
| Otu43144 | 1 | k__Fungi(100);p__Ascomycota(100);c__Saccharomycetes(100);o__Saccharomycetales(100);f__Metschnikowiaceae(100);g__Metschnikowia(100);s__Metschnikowia_pulcherrima(100); | Metschnikowia | 13 | Saprotroph | \|Undefined Saprotroph\| | Yeast | NULL | Probable | As nectar-tap_saprotroph (Põlme et al. 2020); Decay sugar-rich substrates (Põlme et al. 2020) | Põlme S, et al. 2020. Fungal Diversity 105:1-16 ((https://doi.org/10.1007/s13225-020-00466-2)) |
| Otu43145 | 1 | k__Fungi(100);p__Ascomycota(100);c__Saccharomycetes(100);o__Saccharomycetales(100);f__Metschnikowiaceae(100);g__Metschnikowia(100);g__Metschnikowia_unclassified(100); | Metschnikowia | 13 | Saprotroph | \|Undefined Saprotroph\| | Yeast | NULL | Probable | As nectar-tap_saprotroph (Põlme et al. 2020); Decay sugar-rich substrates (Põlme et al. 2020) | Põlme S, et al. 2020. Fungal Diversity 105:1-16 ((https://doi.org/10.1007/s13225-020-00466-2)) |
| Otu43146 | 1 | k__Fungi(100);p__Ascomycota(100);c__Saccharomycetes(100);o__Saccharomycetales(100);f__Metschnikowiaceae(100);g__Metschnikowia(100);s__Metschnikowia_pulcherrima(100); | Metschnikowia | 13 | Saprotroph | \|Undefined Saprotroph\| | Yeast | NULL | Probable | As nectar-tap_saprotroph (Põlme et al. 2020); Decay sugar-rich substrates (Põlme et al. 2020) | Põlme S, et al. 2020. Fungal Diversity 105:1-16 ((https://doi.org/10.1007/s13225-020-00466-2)) |
| Otu43147 | 1 | k__Fungi(100);p__Ascomycota(100);c__Saccharomycetes(100);o__Saccharomycetales(100);f__Metschnikowiaceae(100);g__Metschnikowia(100);s__Metschnikowia_pulcherrima(100); | Metschnikowia | 13 | Saprotroph | \|Undefined Saprotroph\| | Yeast | NULL | Probable | As nectar-tap_saprotroph (Põlme et al. 2020); Decay sugar-rich substrates (Põlme et al. 2020) | Põlme S, et al. 2020. Fungal Diversity 105:1-16 ((https://doi.org/10.1007/s13225-020-00466-2)) |
| Otu43148 | 1 | k__Fungi(100);p__Ascomycota(100);c__Saccharomycetes(100);o__Saccharomycetales(100);f__Metschnikowiaceae(100);g__Metschnikowia(100);s__Metschnikowia_pulcherrima(100); | Metschnikowia | 13 | Saprotroph | \|Undefined Saprotroph\| | Yeast | NULL | Probable | As nectar-tap_saprotroph (Põlme et al. 2020); Decay sugar-rich substrates (Põlme et al. 2020) | Põlme S, et al. 2020. Fungal Diversity 105:1-16 ((https://doi.org/10.1007/s13225-020-00466-2)) |
| Otu43149 | 1 | k__Fungi(100);p__Ascomycota(100);c__Saccharomycetes(100);o__Saccharomycetales(100);f__Metschnikowiaceae(100);g__Metschnikowia(100);g__Metschnikowia_unclassified(100); | Metschnikowia | 13 | Saprotroph | \|Undefined Saprotroph\| | Yeast | NULL | Probable | As nectar-tap_saprotroph (Põlme et al. 2020); Decay sugar-rich substrates (Põlme et al. 2020) | Põlme S, et al. 2020. Fungal Diversity 105:1-16 ((https://doi.org/10.1007/s13225-020-00466-2)) |
| Otu43150 | 1 | k__Fungi(100);p__Ascomycota(100);c__Saccharomycetes(100);o__Saccharomycetales(100);f__Metschnikowiaceae(100);g__Metschnikowia(100);g__Metschnikowia_unclassified(100); | Metschnikowia | 13 | Saprotroph | \|Undefined Saprotroph\| | Yeast | NULL | Probable | As nectar-tap_saprotroph (Põlme et al. 2020); Decay sugar-rich substrates (Põlme et al. 2020) | Põlme S, et al. 2020. Fungal Diversity 105:1-16 ((https://doi.org/10.1007/s13225-020-00466-2)) |
| Otu43154 | 1 | k__Fungi(100);p__Ascomycota(100);c__Saccharomycetes(100);o__Saccharomycetales(100);f__Metschnikowiaceae(100);g__Metschnikowia(100);s__Metschnikowia_pulcherrima(100); | Metschnikowia | 13 | Saprotroph | \|Undefined Saprotroph\| | Yeast | NULL | Probable | As nectar-tap_saprotroph (Põlme et al. 2020); Decay sugar-rich substrates (Põlme et al. 2020) | Põlme S, et al. 2020. Fungal Diversity 105:1-16 ((https://doi.org/10.1007/s13225-020-00466-2)) |
| Otu43157 | 1 | k__Fungi(100);p__Ascomycota(100);c__Saccharomycetes(100);o__Saccharomycetales(100);f__Metschnikowiaceae(100);g__Metschnikowia(100);g__Metschnikowia_unclassified(100); | Metschnikowia | 13 | Saprotroph | \|Undefined Saprotroph\| | Yeast | NULL | Probable | As nectar-tap_saprotroph (Põlme et al. 2020); Decay sugar-rich substrates (Põlme et al. 2020) | Põlme S, et al. 2020. Fungal Diversity 105:1-16 ((https://doi.org/10.1007/s13225-020-00466-2)) |
| Otu43158 | 1 | k__Fungi(100);p__Ascomycota(100);c__Saccharomycetes(100);o__Saccharomycetales(100);f__Metschnikowiaceae(100);g__Metschnikowia(100);s__Metschnikowia_pulcherrima(100); | Metschnikowia | 13 | Saprotroph | \|Undefined Saprotroph\| | Yeast | NULL | Probable | As nectar-tap_saprotroph (Põlme et al. 2020); Decay sugar-rich substrates (Põlme et al. 2020) | Põlme S, et al. 2020. Fungal Diversity 105:1-16 ((https://doi.org/10.1007/s13225-020-00466-2)) |
| Otu43159 | 1 | k__Fungi(100);p__Ascomycota(100);c__Saccharomycetes(100);o__Saccharomycetales(100);f__Metschnikowiaceae(100);g__Metschnikowia(100);s__Metschnikowia_pulcherrima(100); | Metschnikowia | 13 | Saprotroph | \|Undefined Saprotroph\| | Yeast | NULL | Probable | As nectar-tap_saprotroph (Põlme et al. 2020); Decay sugar-rich substrates (Põlme et al. 2020) | Põlme S, et al. 2020. Fungal Diversity 105:1-16 ((https://doi.org/10.1007/s13225-020-00466-2)) |
| Otu43160 | 1 | k__Fungi(100);p__Ascomycota(100);c__Saccharomycetes(100);o__Saccharomycetales(100);f__Metschnikowiaceae(100);g__Metschnikowia(100);g__Metschnikowia_unclassified(100); | Metschnikowia | 13 | Saprotroph | \|Undefined Saprotroph\| | Yeast | NULL | Probable | As nectar-tap_saprotroph (Põlme et al. 2020); Decay sugar-rich substrates (Põlme et al. 2020) | Põlme S, et al. 2020. Fungal Diversity 105:1-16 ((https://doi.org/10.1007/s13225-020-00466-2)) |
| Otu43161 | 1 | k__Fungi(100);p__Ascomycota(100);c__Saccharomycetes(100);o__Saccharomycetales(100);f__Metschnikowiaceae(100);g__Metschnikowia(100);s__Metschnikowia_pulcherrima(100); | Metschnikowia | 13 | Saprotroph | \|Undefined Saprotroph\| | Yeast | NULL | Probable | As nectar-tap_saprotroph (Põlme et al. 2020); Decay sugar-rich substrates (Põlme et al. 2020) | Põlme S, et al. 2020. Fungal Diversity 105:1-16 ((https://doi.org/10.1007/s13225-020-00466-2)) |
| Otu43166 | 1 | k__Fungi(100);p__Ascomycota(100);c__Saccharomycetes(100);o__Saccharomycetales(100);f__Metschnikowiaceae(100);g__Metschnikowia(100);s__Metschnikowia_pulcherrima(100); | Metschnikowia | 13 | Saprotroph | \|Undefined Saprotroph\| | Yeast | NULL | Probable | As nectar-tap_saprotroph (Põlme et al. 2020); Decay sugar-rich substrates (Põlme et al. 2020) | Põlme S, et al. 2020. Fungal Diversity 105:1-16 ((https://doi.org/10.1007/s13225-020-00466-2)) |
| Otu43168 | 1 | k__Fungi(100);p__Ascomycota(100);c__Saccharomycetes(100);o__Saccharomycetales(100);f__Metschnikowiaceae(100);g__Metschnikowia(100);g__Metschnikowia_unclassified(100); | Metschnikowia | 13 | Saprotroph | \|Undefined Saprotroph\| | Yeast | NULL | Probable | As nectar-tap_saprotroph (Põlme et al. 2020); Decay sugar-rich substrates (Põlme et al. 2020) | Põlme S, et al. 2020. Fungal Diversity 105:1-16 ((https://doi.org/10.1007/s13225-020-00466-2)) |
| Otu43170 | 1 | k__Fungi(100);p__Ascomycota(100);c__Sordariomycetes(100);o__Sordariales(100);f__Cephalothecaceae(100);f__Cephalothecaceae_unclassified(100);f__Cephalothecaceae_unclassified(100); | Cephalothecaceae | 9 | Saprotroph | Fungal Parasite-Wood Saprotroph | Microfungus | NULL | Probable | Typically saprobic, fruiting on very rotten wood or other fungi (Cannon & Kirk 2007) | Cannon PF, Kirk PM. 2007. Fungal Families of the World. CAB International, Cambridge (ISBN: 978-0851998275) |
| Otu43171 | 1 | k__Fungi(100);p__Ascomycota(100);c__Saccharomycetes(100);o__Saccharomycetales(100);f__Metschnikowiaceae(100);g__Metschnikowia(100);s__Metschnikowia_pulcherrima(100); | Metschnikowia | 13 | Saprotroph | \|Undefined Saprotroph\| | Yeast | NULL | Probable | As nectar-tap_saprotroph (Põlme et al. 2020); Decay sugar-rich substrates (Põlme et al. 2020) | Põlme S, et al. 2020. Fungal Diversity 105:1-16 ((https://doi.org/10.1007/s13225-020-00466-2)) |
| Otu43172 | 1 | k__Fungi(100);p__Ascomycota(100);c__Saccharomycetes(100);o__Saccharomycetales(100);f__Metschnikowiaceae(100);g__Metschnikowia(100);s__Metschnikowia_pulcherrima(100); | Metschnikowia | 13 | Saprotroph | \|Undefined Saprotroph\| | Yeast | NULL | Probable | As nectar-tap_saprotroph (Põlme et al. 2020); Decay sugar-rich substrates (Põlme et al. 2020) | Põlme S, et al. 2020. Fungal Diversity 105:1-16 ((https://doi.org/10.1007/s13225-020-00466-2)) |
| Otu43174 | 1 | k__Fungi(100);p__Ascomycota(100);c__Saccharomycetes(100);o__Saccharomycetales(100);f__Metschnikowiaceae(100);g__Metschnikowia(100);s__Metschnikowia_pulcherrima(100); | Metschnikowia | 13 | Saprotroph | \|Undefined Saprotroph\| | Yeast | NULL | Probable | As nectar-tap_saprotroph (Põlme et al. 2020); Decay sugar-rich substrates (Põlme et al. 2020) | Põlme S, et al. 2020. Fungal Diversity 105:1-16 ((https://doi.org/10.1007/s13225-020-00466-2)) |
| Otu43175 | 1 | k__Fungi(100);p__Ascomycota(100);c__Saccharomycetes(100);o__Saccharomycetales(100);f__Metschnikowiaceae(100);g__Metschnikowia(100);s__Metschnikowia_pulcherrima(100); | Metschnikowia | 13 | Saprotroph | \|Undefined Saprotroph\| | Yeast | NULL | Probable | As nectar-tap_saprotroph (Põlme et al. 2020); Decay sugar-rich substrates (Põlme et al. 2020) | Põlme S, et al. 2020. Fungal Diversity 105:1-16 ((https://doi.org/10.1007/s13225-020-00466-2)) |
| Otu43176 | 1 | k__Fungi(100);p__Ascomycota(100);c__Saccharomycetes(100);o__Saccharomycetales(100);f__Metschnikowiaceae(100);g__Metschnikowia(100);s__Metschnikowia_pulcherrima(100); | Metschnikowia | 13 | Saprotroph | \|Undefined Saprotroph\| | Yeast | NULL | Probable | As nectar-tap_saprotroph (Põlme et al. 2020); Decay sugar-rich substrates (Põlme et al. 2020) | Põlme S, et al. 2020. Fungal Diversity 105:1-16 ((https://doi.org/10.1007/s13225-020-00466-2)) |
| Otu43178 | 1 | k__Fungi(100);p__Ascomycota(100);c__Saccharomycetes(100);o__Saccharomycetales(100);f__Metschnikowiaceae(100);g__Metschnikowia(100);s__Metschnikowia_pulcherrima(100); | Metschnikowia | 13 | Saprotroph | \|Undefined Saprotroph\| | Yeast | NULL | Probable | As nectar-tap_saprotroph (Põlme et al. 2020); Decay sugar-rich substrates (Põlme et al. 2020) | Põlme S, et al. 2020. Fungal Diversity 105:1-16 ((https://doi.org/10.1007/s13225-020-00466-2)) |
| Otu43179 | 1 | k__Fungi(100);p__Ascomycota(100);c__Saccharomycetes(100);o__Saccharomycetales(100);f__Metschnikowiaceae(100);g__Metschnikowia(100);g__Metschnikowia_unclassified(100); | Metschnikowia | 13 | Saprotroph | \|Undefined Saprotroph\| | Yeast | NULL | Probable | As nectar-tap_saprotroph (Põlme et al. 2020); Decay sugar-rich substrates (Põlme et al. 2020) | Põlme S, et al. 2020. Fungal Diversity 105:1-16 ((https://doi.org/10.1007/s13225-020-00466-2)) |
| Otu43180 | 1 | k__Fungi(100);p__Ascomycota(100);c__Saccharomycetes(100);o__Saccharomycetales(100);f__Metschnikowiaceae(100);g__Metschnikowia(100);s__Metschnikowia_pulcherrima(100); | Metschnikowia | 13 | Saprotroph | \|Undefined Saprotroph\| | Yeast | NULL | Probable | As nectar-tap_saprotroph (Põlme et al. 2020); Decay sugar-rich substrates (Põlme et al. 2020) | Põlme S, et al. 2020. Fungal Diversity 105:1-16 ((https://doi.org/10.1007/s13225-020-00466-2)) |
| Otu43181 | 1 | k__Fungi(100);p__Ascomycota(100);c__Saccharomycetes(100);o__Saccharomycetales(100);f__Metschnikowiaceae(100);g__Metschnikowia(100);s__Metschnikowia_pulcherrima(100); | Metschnikowia | 13 | Saprotroph | \|Undefined Saprotroph\| | Yeast | NULL | Probable | As nectar-tap_saprotroph (Põlme et al. 2020); Decay sugar-rich substrates (Põlme et al. 2020) | Põlme S, et al. 2020. Fungal Diversity 105:1-16 ((https://doi.org/10.1007/s13225-020-00466-2)) |
| Otu43183 | 1 | k__Fungi(100);p__Ascomycota(100);c__Saccharomycetes(100);o__Saccharomycetales(100);f__Metschnikowiaceae(100);g__Metschnikowia(100);s__Metschnikowia_pulcherrima(100); | Metschnikowia | 13 | Saprotroph | \|Undefined Saprotroph\| | Yeast | NULL | Probable | As nectar-tap_saprotroph (Põlme et al. 2020); Decay sugar-rich substrates (Põlme et al. 2020) | Põlme S, et al. 2020. Fungal Diversity 105:1-16 ((https://doi.org/10.1007/s13225-020-00466-2)) |
| Otu43186 | 1 | k__Fungi(100);p__Ascomycota(100);c__Saccharomycetes(100);o__Saccharomycetales(100);f__Metschnikowiaceae(100);g__Metschnikowia(100);s__Metschnikowia_pulcherrima(100); | Metschnikowia | 13 | Saprotroph | \|Undefined Saprotroph\| | Yeast | NULL | Probable | As nectar-tap_saprotroph (Põlme et al. 2020); Decay sugar-rich substrates (Põlme et al. 2020) | Põlme S, et al. 2020. Fungal Diversity 105:1-16 ((https://doi.org/10.1007/s13225-020-00466-2)) |
| Otu43187 | 1 | k__Fungi(100);p__Ascomycota(100);c__Saccharomycetes(100);o__Saccharomycetales(100);f__Metschnikowiaceae(100);g__Metschnikowia(100);s__Metschnikowia_pulcherrima(100); | Metschnikowia | 13 | Saprotroph | \|Undefined Saprotroph\| | Yeast | NULL | Probable | As nectar-tap_saprotroph (Põlme et al. 2020); Decay sugar-rich substrates (Põlme et al. 2020) | Põlme S, et al. 2020. Fungal Diversity 105:1-16 ((https://doi.org/10.1007/s13225-020-00466-2)) |
| Otu43188 | 1 | k__Fungi(100);p__Ascomycota(100);c__Saccharomycetes(100);o__Saccharomycetales(100);f__Metschnikowiaceae(100);g__Metschnikowia(100);s__Metschnikowia_pulcherrima(100); | Metschnikowia | 13 | Saprotroph | \|Undefined Saprotroph\| | Yeast | NULL | Probable | As nectar-tap_saprotroph (Põlme et al. 2020); Decay sugar-rich substrates (Põlme et al. 2020) | Põlme S, et al. 2020. Fungal Diversity 105:1-16 ((https://doi.org/10.1007/s13225-020-00466-2)) |
| Otu43190 | 1 | k__Fungi(100);p__Ascomycota(100);c__Saccharomycetes(100);o__Saccharomycetales(100);f__Metschnikowiaceae(100);g__Metschnikowia(100);s__Metschnikowia_pulcherrima(100); | Metschnikowia | 13 | Saprotroph | \|Undefined Saprotroph\| | Yeast | NULL | Probable | As nectar-tap_saprotroph (Põlme et al. 2020); Decay sugar-rich substrates (Põlme et al. 2020) | Põlme S, et al. 2020. Fungal Diversity 105:1-16 ((https://doi.org/10.1007/s13225-020-00466-2)) |
| Otu43191 | 1 | k__Fungi(100);p__Ascomycota(100);c__Saccharomycetes(100);o__Saccharomycetales(100);f__Metschnikowiaceae(100);g__Metschnikowia(100);s__Metschnikowia_pulcherrima(100); | Metschnikowia | 13 | Saprotroph | \|Undefined Saprotroph\| | Yeast | NULL | Probable | As nectar-tap_saprotroph (Põlme et al. 2020); Decay sugar-rich substrates (Põlme et al. 2020) | Põlme S, et al. 2020. Fungal Diversity 105:1-16 ((https://doi.org/10.1007/s13225-020-00466-2)) |
| Otu43192 | 1 | k__Fungi(100);p__Ascomycota(100);c__Leotiomycetes(100);o__Helotiales(100);f__Vibrisseaceae(100);g__Acephala(100);s__Acephala_sp_1(100); | Acephala | 13 | Saprotroph-Symbiotroph | Ectomycorrhizal-Endophyte-\|Undefined Saprotroph\| | Microfungus | NULL | Possible | As root_endophyte_dark_septate (Põlme et al. 2020); As soil_saprotroph (Põlme et al. 2020); Decay soil (Põlme et al. 2020) | Tedersoo L, et al. 2010. Mycorrhiza 20:217-263 ((https://doi.org/10.1007/s00572-009-0274-x)) (as ECM/not ECM); Põlme S, et al. 2020. Fungal Diversity 105:1-16 ((https://doi.org/10.1007/s13225-020-00466-2)) |
| Otu43193 | 1 | k__Fungi(100);p__Ascomycota(100);c__Saccharomycetes(100);o__Saccharomycetales(100);f__Metschnikowiaceae(100);g__Metschnikowia(100);s__Metschnikowia_pulcherrima(100); | Metschnikowia | 13 | Saprotroph | \|Undefined Saprotroph\| | Yeast | NULL | Probable | As nectar-tap_saprotroph (Põlme et al. 2020); Decay sugar-rich substrates (Põlme et al. 2020) | Põlme S, et al. 2020. Fungal Diversity 105:1-16 ((https://doi.org/10.1007/s13225-020-00466-2)) |
| Otu43194 | 1 | k__Fungi(100);p__Ascomycota(100);c__Saccharomycetes(100);o__Saccharomycetales(100);f__Metschnikowiaceae(100);g__Metschnikowia(100);s__Metschnikowia_pulcherrima(100); | Metschnikowia | 13 | Saprotroph | \|Undefined Saprotroph\| | Yeast | NULL | Probable | As nectar-tap_saprotroph (Põlme et al. 2020); Decay sugar-rich substrates (Põlme et al. 2020) | Põlme S, et al. 2020. Fungal Diversity 105:1-16 ((https://doi.org/10.1007/s13225-020-00466-2)) |
| Otu43195 | 1 | k__Fungi(100);p__Ascomycota(100);c__Saccharomycetes(100);o__Saccharomycetales(100);f__Metschnikowiaceae(100);g__Metschnikowia(100);s__Metschnikowia_pulcherrima(100); | Metschnikowia | 13 | Saprotroph | \|Undefined Saprotroph\| | Yeast | NULL | Probable | As nectar-tap_saprotroph (Põlme et al. 2020); Decay sugar-rich substrates (Põlme et al. 2020) | Põlme S, et al. 2020. Fungal Diversity 105:1-16 ((https://doi.org/10.1007/s13225-020-00466-2)) |
| Otu43197 | 1 | k__Fungi(100);p__Ascomycota(100);c__Saccharomycetes(100);o__Saccharomycetales(100);f__Metschnikowiaceae(100);g__Metschnikowia(100);s__Metschnikowia_pulcherrima(100); | Metschnikowia | 13 | Saprotroph | \|Undefined Saprotroph\| | Yeast | NULL | Probable | As nectar-tap_saprotroph (Põlme et al. 2020); Decay sugar-rich substrates (Põlme et al. 2020) | Põlme S, et al. 2020. Fungal Diversity 105:1-16 ((https://doi.org/10.1007/s13225-020-00466-2)) |
| Otu43198 | 1 | k__Fungi(100);p__Ascomycota(100);c__Saccharomycetes(100);o__Saccharomycetales(100);f__Metschnikowiaceae(100);g__Metschnikowia(100);s__Metschnikowia_pulcherrima(100); | Metschnikowia | 13 | Saprotroph | \|Undefined Saprotroph\| | Yeast | NULL | Probable | As nectar-tap_saprotroph (Põlme et al. 2020); Decay sugar-rich substrates (Põlme et al. 2020) | Põlme S, et al. 2020. Fungal Diversity 105:1-16 ((https://doi.org/10.1007/s13225-020-00466-2)) |
| Otu43203 | 1 | k__Fungi(100);p__Ascomycota(100);c__Saccharomycetes(100);o__Saccharomycetales(100);f__Metschnikowiaceae(100);g__Metschnikowia(100);g__Metschnikowia_unclassified(100); | Metschnikowia | 13 | Saprotroph | \|Undefined Saprotroph\| | Yeast | NULL | Probable | As nectar-tap_saprotroph (Põlme et al. 2020); Decay sugar-rich substrates (Põlme et al. 2020) | Põlme S, et al. 2020. Fungal Diversity 105:1-16 ((https://doi.org/10.1007/s13225-020-00466-2)) |
| Otu43204 | 1 | k__Fungi(100);p__Ascomycota(100);c__Saccharomycetes(100);o__Saccharomycetales(100);f__Metschnikowiaceae(100);g__Metschnikowia(100);s__Metschnikowia_pulcherrima(100); | Metschnikowia | 13 | Saprotroph | \|Undefined Saprotroph\| | Yeast | NULL | Probable | As nectar-tap_saprotroph (Põlme et al. 2020); Decay sugar-rich substrates (Põlme et al. 2020) | Põlme S, et al. 2020. Fungal Diversity 105:1-16 ((https://doi.org/10.1007/s13225-020-00466-2)) |
| Otu43205 | 1 | k__Fungi(100);p__Ascomycota(100);c__Saccharomycetes(100);o__Saccharomycetales(100);f__Metschnikowiaceae(100);g__Metschnikowia(100);s__Metschnikowia_pulcherrima(100); | Metschnikowia | 13 | Saprotroph | \|Undefined Saprotroph\| | Yeast | NULL | Probable | As nectar-tap_saprotroph (Põlme et al. 2020); Decay sugar-rich substrates (Põlme et al. 2020) | Põlme S, et al. 2020. Fungal Diversity 105:1-16 ((https://doi.org/10.1007/s13225-020-00466-2)) |
| Otu43206 | 1 | k__Fungi(100);p__Ascomycota(100);c__Saccharomycetes(100);o__Saccharomycetales(100);f__Metschnikowiaceae(100);g__Metschnikowia(100);g__Metschnikowia_unclassified(100); | Metschnikowia | 13 | Saprotroph | \|Undefined Saprotroph\| | Yeast | NULL | Probable | As nectar-tap_saprotroph (Põlme et al. 2020); Decay sugar-rich substrates (Põlme et al. 2020) | Põlme S, et al. 2020. Fungal Diversity 105:1-16 ((https://doi.org/10.1007/s13225-020-00466-2)) |
| Otu43207 | 1 | k__Fungi(100);p__Ascomycota(100);c__Saccharomycetes(100);o__Saccharomycetales(100);f__Metschnikowiaceae(100);g__Metschnikowia(100);s__Metschnikowia_pulcherrima(100); | Metschnikowia | 13 | Saprotroph | \|Undefined Saprotroph\| | Yeast | NULL | Probable | As nectar-tap_saprotroph (Põlme et al. 2020); Decay sugar-rich substrates (Põlme et al. 2020) | Põlme S, et al. 2020. Fungal Diversity 105:1-16 ((https://doi.org/10.1007/s13225-020-00466-2)) |
| Otu43208 | 1 | k__Fungi(100);p__Ascomycota(100);c__Saccharomycetes(100);o__Saccharomycetales(100);f__Metschnikowiaceae(100);g__Metschnikowia(100);g__Metschnikowia_unclassified(100); | Metschnikowia | 13 | Saprotroph | \|Undefined Saprotroph\| | Yeast | NULL | Probable | As nectar-tap_saprotroph (Põlme et al. 2020); Decay sugar-rich substrates (Põlme et al. 2020) | Põlme S, et al. 2020. Fungal Diversity 105:1-16 ((https://doi.org/10.1007/s13225-020-00466-2)) |

| | | | | | | | | | | | |
|---|---|---|---|---|---|---|---|---|---|---|---|
| Otu43209 | 1 | k__Fungi(100);p__Ascomycota(100);c__Saccharomycetes(100);o__Saccharomycetales(100);f__Metschnikowiaceae(100);g__Metschnikowia(100);s__Metschnikowia_pulcherrima(100); | Metschnikowia | 13 | Saprotroph | \|Undefined Saprotroph\| | Yeast | NULL | Probable | As nectar-tap_saprotroph (Põlme et al. 2020); Decay sugar-rich substrates (Põlme et al. 2020) | Põlme S, et al. 2020. Fungal Diversity 105:1-16 ((https://doi.org/10.1007/s13225-020-00466-2)) |
| Otu43210 | 1 | k__Fungi(100);p__Ascomycota(100);c__Saccharomycetes(100);o__Saccharomycetales(100);f__Metschnikowiaceae(100);g__Metschnikowia(100);s__Metschnikowia_pulcherrima(100); | Metschnikowia | 13 | Saprotroph | \|Undefined Saprotroph\| | Yeast | NULL | Probable | As nectar-tap_saprotroph (Põlme et al. 2020); Decay sugar-rich substrates (Põlme et al. 2020) | Põlme S, et al. 2020. Fungal Diversity 105:1-16 ((https://doi.org/10.1007/s13225-020-00466-2)) |
| Otu43212 | 1 | k__Fungi(100);p__Ascomycota(100);c__Saccharomycetes(100);o__Saccharomycetales(100);f__Metschnikowiaceae(100);g__Metschnikowia(100);g__Metschnikowia_unclassified(100); | Metschnikowia | 13 | Saprotroph | \|Undefined Saprotroph\| | Yeast | NULL | Probable | As nectar-tap_saprotroph (Põlme et al. 2020); Decay sugar-rich substrates (Põlme et al. 2020) | Põlme S, et al. 2020. Fungal Diversity 105:1-16 ((https://doi.org/10.1007/s13225-020-00466-2)) |
| Otu43213 | 1 | k__Fungi(100);p__Ascomycota(100);c__Saccharomycetes(100);o__Saccharomycetales(100);f__Metschnikowiaceae(100);g__Metschnikowia(100);g__Metschnikowia_unclassified(100); | Metschnikowia | 13 | Saprotroph | \|Undefined Saprotroph\| | Yeast | NULL | Probable | As nectar-tap_saprotroph (Põlme et al. 2020); Decay sugar-rich substrates (Põlme et al. 2020) | Põlme S, et al. 2020. Fungal Diversity 105:1-16 ((https://doi.org/10.1007/s13225-020-00466-2)) |
| Otu43214 | 1 | k__Fungi(100);p__Ascomycota(100);c__Saccharomycetes(100);o__Saccharomycetales(100);f__Metschnikowiaceae(100);g__Metschnikowia(100);g__Metschnikowia_unclassified(100); | Metschnikowia | 13 | Saprotroph | \|Undefined Saprotroph\| | Yeast | NULL | Probable | As nectar-tap_saprotroph (Põlme et al. 2020); Decay sugar-rich substrates (Põlme et al. 2020) | Põlme S, et al. 2020. Fungal Diversity 105:1-16 ((https://doi.org/10.1007/s13225-020-00466-2)) |
| Otu43215 | 1 | k__Fungi(100);p__Ascomycota(100);c__Saccharomycetes(100);o__Saccharomycetales(100);f__Metschnikowiaceae(100);g__Metschnikowia(100);g__Metschnikowia_unclassified(100); | Metschnikowia | 13 | Saprotroph | \|Undefined Saprotroph\| | Yeast | NULL | Probable | As nectar-tap_saprotroph (Põlme et al. 2020); Decay sugar-rich substrates (Põlme et al. 2020) | Põlme S, et al. 2020. Fungal Diversity 105:1-16 ((https://doi.org/10.1007/s13225-020-00466-2)) |
| Otu43217 | 1 | k__Fungi(100);p__Ascomycota(100);c__Saccharomycetes(100);o__Saccharomycetales(100);f__Metschnikowiaceae(100);g__Metschnikowia(100);g__Metschnikowia_unclassified(100); | Metschnikowia | 13 | Saprotroph | \|Undefined Saprotroph\| | Yeast | NULL | Probable | As nectar-tap_saprotroph (Põlme et al. 2020); Decay sugar-rich substrates (Põlme et al. 2020) | Põlme S, et al. 2020. Fungal Diversity 105:1-16 ((https://doi.org/10.1007/s13225-020-00466-2)) |
| Otu43219 | 1 | k__Fungi(100);p__Ascomycota(100);c__Saccharomycetes(100);o__Saccharomycetales(100);f__Metschnikowiaceae(100);g__Metschnikowia(100);s__Metschnikowia_pulcherrima(100); | Metschnikowia | 13 | Saprotroph | \|Undefined Saprotroph\| | Yeast | NULL | Probable | As nectar-tap_saprotroph (Põlme et al. 2020); Decay sugar-rich substrates (Põlme et al. 2020) | Põlme S, et al. 2020. Fungal Diversity 105:1-16 ((https://doi.org/10.1007/s13225-020-00466-2)) |
| Otu43220 | 1 | k__Fungi(100);p__Ascomycota(100);c__Saccharomycetes(100);o__Saccharomycetales(100);f__Metschnikowiaceae(100);g__Metschnikowia(100);s__Metschnikowia_pulcherrima(100); | Metschnikowia | 13 | Saprotroph | \|Undefined Saprotroph\| | Yeast | NULL | Probable | As nectar-tap_saprotroph (Põlme et al. 2020); Decay sugar-rich substrates (Põlme et al. 2020) | Põlme S, et al. 2020. Fungal Diversity 105:1-16 ((https://doi.org/10.1007/s13225-020-00466-2)) |
| Otu43221 | 1 | k__Fungi(100);p__Ascomycota(100);c__Saccharomycetes(100);o__Saccharomycetales(100);f__Metschnikowiaceae(100);g__Metschnikowia(100);g__Metschnikowia_unclassified(100); | Metschnikowia | 13 | Saprotroph | \|Undefined Saprotroph\| | Yeast | NULL | Probable | As nectar-tap_saprotroph (Põlme et al. 2020); Decay sugar-rich substrates (Põlme et al. 2020) | Põlme S, et al. 2020. Fungal Diversity 105:1-16 ((https://doi.org/10.1007/s13225-020-00466-2)) |
| Otu43222 | 1 | k__Fungi(100);p__Ascomycota(100);c__Saccharomycetes(100);o__Saccharomycetales(100);f__Metschnikowiaceae(100);g__Metschnikowia(100);s__Metschnikowia_pulcherrima(100); | Metschnikowia | 13 | Saprotroph | \|Undefined Saprotroph\| | Yeast | NULL | Probable | As nectar-tap_saprotroph (Põlme et al. 2020); Decay sugar-rich substrates (Põlme et al. 2020) | Põlme S, et al. 2020. Fungal Diversity 105:1-16 ((https://doi.org/10.1007/s13225-020-00466-2)) |
| Otu43223 | 1 | k__Fungi(100);p__Ascomycota(100);c__Saccharomycetes(100);o__Saccharomycetales(100);f__Metschnikowiaceae(100);g__Metschnikowia(100);s__Metschnikowia_pulcherrima(100); | Metschnikowia | 13 | Saprotroph | \|Undefined Saprotroph\| | Yeast | NULL | Probable | As nectar-tap_saprotroph (Põlme et al. 2020); Decay sugar-rich substrates (Põlme et al. 2020) | Põlme S, et al. 2020. Fungal Diversity 105:1-16 ((https://doi.org/10.1007/s13225-020-00466-2)) |
| Otu43224 | 1 | k__Fungi(100);p__Ascomycota(100);c__Saccharomycetes(100);o__Saccharomycetales(100);f__Metschnikowiaceae(100);g__Metschnikowia(100);s__Metschnikowia_pulcherrima(100); | Metschnikowia | 13 | Saprotroph | \|Undefined Saprotroph\| | Yeast | NULL | Probable | As nectar-tap_saprotroph (Põlme et al. 2020); Decay sugar-rich substrates (Põlme et al. 2020) | Põlme S, et al. 2020. Fungal Diversity 105:1-16 ((https://doi.org/10.1007/s13225-020-00466-2)) |
| Otu43225 | 1 | k__Fungi(100);p__Ascomycota(100);c__Saccharomycetes(100);o__Saccharomycetales(100);f__Metschnikowiaceae(100);g__Metschnikowia(100);s__Metschnikowia_pulcherrima(100); | Metschnikowia | 13 | Saprotroph | \|Undefined Saprotroph\| | Yeast | NULL | Probable | As nectar-tap_saprotroph (Põlme et al. 2020); Decay sugar-rich substrates (Põlme et al. 2020) | Põlme S, et al. 2020. Fungal Diversity 105:1-16 ((https://doi.org/10.1007/s13225-020-00466-2)) |
| Otu43226 | 1 | k__Fungi(100);p__Ascomycota(100);c__Saccharomycetes(100);o__Saccharomycetales(100);f__Metschnikowiaceae(100);g__Metschnikowia(100);g__Metschnikowia_unclassified(100); | Metschnikowia | 13 | Saprotroph | \|Undefined Saprotroph\| | Yeast | NULL | Probable | As nectar-tap_saprotroph (Põlme et al. 2020); Decay sugar-rich substrates (Põlme et al. 2020) | Põlme S, et al. 2020. Fungal Diversity 105:1-16 ((https://doi.org/10.1007/s13225-020-00466-2)) |
| Otu43227 | 1 | k__Fungi(100);p__Ascomycota(100);c__Saccharomycetes(100);o__Saccharomycetales(100);f__Metschnikowiaceae(100);g__Metschnikowia(100);s__Metschnikowia_pulcherrima(100); | Metschnikowia | 13 | Saprotroph | \|Undefined Saprotroph\| | Yeast | NULL | Probable | As nectar-tap_saprotroph (Põlme et al. 2020); Decay sugar-rich substrates (Põlme et al. 2020) | Põlme S, et al. 2020. Fungal Diversity 105:1-16 ((https://doi.org/10.1007/s13225-020-00466-2)) |
| Otu43228 | 1 | k__Fungi(100);p__Ascomycota(100);c__Saccharomycetes(100);o__Saccharomycetales(100);f__Metschnikowiaceae(100);g__Metschnikowia(100);g__Metschnikowia_unclassified(100); | Metschnikowia | 13 | Saprotroph | \|Undefined Saprotroph\| | Yeast | NULL | Probable | As nectar-tap_saprotroph (Põlme et al. 2020); Decay sugar-rich substrates (Põlme et al. 2020) | Põlme S, et al. 2020. Fungal Diversity 105:1-16 ((https://doi.org/10.1007/s13225-020-00466-2)) |
| Otu43229 | 1 | k__Fungi(100);p__Ascomycota(100);c__Saccharomycetes(100);o__Saccharomycetales(100);f__Metschnikowiaceae(100);g__Metschnikowia(100);s__Metschnikowia_pulcherrima(100); | Metschnikowia | 13 | Saprotroph | \|Undefined Saprotroph\| | Yeast | NULL | Probable | As nectar-tap_saprotroph (Põlme et al. 2020); Decay sugar-rich substrates (Põlme et al. 2020) | Põlme S, et al. 2020. Fungal Diversity 105:1-16 ((https://doi.org/10.1007/s13225-020-00466-2)) |
| Otu43230 | 1 | k__Fungi(100);p__Ascomycota(100);c__Saccharomycetes(100);o__Saccharomycetales(100);f__Metschnikowiaceae(100);g__Metschnikowia(100);s__Metschnikowia_pulcherrima(100); | Metschnikowia | 13 | Saprotroph | \|Undefined Saprotroph\| | Yeast | NULL | Probable | As nectar-tap_saprotroph (Põlme et al. 2020); Decay sugar-rich substrates (Põlme et al. 2020) | Põlme S, et al. 2020. Fungal Diversity 105:1-16 ((https://doi.org/10.1007/s13225-020-00466-2)) |
| Otu43231 | 1 | k__Fungi(100);p__Ascomycota(100);c__Saccharomycetes(100);o__Saccharomycetales(100);f__Metschnikowiaceae(100);g__Metschnikowia(100);g__Metschnikowia_unclassified(100); | Metschnikowia | 13 | Saprotroph | \|Undefined Saprotroph\| | Yeast | NULL | Probable | As nectar-tap_saprotroph (Põlme et al. 2020); Decay sugar-rich substrates (Põlme et al. 2020) | Põlme S, et al. 2020. Fungal Diversity 105:1-16 ((https://doi.org/10.1007/s13225-020-00466-2)) |
| Otu43233 | 1 | k__Fungi(100);p__Ascomycota(100);c__Saccharomycetes(100);o__Saccharomycetales(100);f__Metschnikowiaceae(100);g__Metschnikowia(100);s__Metschnikowia_pulcherrima(100); | Metschnikowia | 13 | Saprotroph | \|Undefined Saprotroph\| | Yeast | NULL | Probable | As nectar-tap_saprotroph (Põlme et al. 2020); Decay sugar-rich substrates (Põlme et al. 2020) | Põlme S, et al. 2020. Fungal Diversity 105:1-16 ((https://doi.org/10.1007/s13225-020-00466-2)) |
| Otu43234 | 1 | k__Fungi(100);p__Ascomycota(100);c__Saccharomycetes(100);o__Saccharomycetales(100);f__Metschnikowiaceae(100);g__Metschnikowia(100);g__Metschnikowia_unclassified(100); | Metschnikowia | 13 | Saprotroph | \|Undefined Saprotroph\| | Yeast | NULL | Probable | As nectar-tap_saprotroph (Põlme et al. 2020); Decay sugar-rich substrates (Põlme et al. 2020) | Põlme S, et al. 2020. Fungal Diversity 105:1-16 ((https://doi.org/10.1007/s13225-020-00466-2)) |
| Otu43235 | 1 | k__Fungi(100);p__Ascomycota(100);c__Saccharomycetes(100);o__Saccharomycetales(100);f__Metschnikowiaceae(100);g__Metschnikowia(100);s__Metschnikowia_pulcherrima(100); | Metschnikowia | 13 | Saprotroph | \|Undefined Saprotroph\| | Yeast | NULL | Probable | As nectar-tap_saprotroph (Põlme et al. 2020); Decay sugar-rich substrates (Põlme et al. 2020) | Põlme S, et al. 2020. Fungal Diversity 105:1-16 ((https://doi.org/10.1007/s13225-020-00466-2)) |
| Otu43236 | 1 | k__Fungi(100);p__Ascomycota(100);c__Saccharomycetes(100);o__Saccharomycetales(100);f__Metschnikowiaceae(100);g__Metschnikowia(100);s__Metschnikowia_pulcherrima(100); | Metschnikowia | 13 | Saprotroph | \|Undefined Saprotroph\| | Yeast | NULL | Probable | As nectar-tap_saprotroph (Põlme et al. 2020); Decay sugar-rich substrates (Põlme et al. 2020) | Põlme S, et al. 2020. Fungal Diversity 105:1-16 ((https://doi.org/10.1007/s13225-020-00466-2)) |
| Otu43237 | 1 | k__Fungi(100);p__Ascomycota(100);c__Saccharomycetes(100);o__Saccharomycetales(100);f__Metschnikowiaceae(100);g__Metschnikowia(100);s__Metschnikowia_pulcherrima(100); | Metschnikowia | 13 | Saprotroph | \|Undefined Saprotroph\| | Yeast | NULL | Probable | As nectar-tap_saprotroph (Põlme et al. 2020); Decay sugar-rich substrates (Põlme et al. 2020) | Põlme S, et al. 2020. Fungal Diversity 105:1-16 ((https://doi.org/10.1007/s13225-020-00466-2)) |
| Otu43238 | 1 | k__Fungi(100);p__Ascomycota(100);c__Saccharomycetes(100);o__Saccharomycetales(100);f__Metschnikowiaceae(100);g__Metschnikowia(100);s__Metschnikowia_pulcherrima(100); | Metschnikowia | 13 | Saprotroph | \|Undefined Saprotroph\| | Yeast | NULL | Probable | As nectar-tap_saprotroph (Põlme et al. 2020); Decay sugar-rich substrates (Põlme et al. 2020) | Põlme S, et al. 2020. Fungal Diversity 105:1-16 ((https://doi.org/10.1007/s13225-020-00466-2)) |
| Otu43239 | 1 | k__Fungi(100);p__Ascomycota(100);c__Saccharomycetes(100);o__Saccharomycetales(100);f__Metschnikowiaceae(100);g__Metschnikowia(100);s__Metschnikowia_pulcherrima(100); | Metschnikowia | 13 | Saprotroph | \|Undefined Saprotroph\| | Yeast | NULL | Probable | As nectar-tap_saprotroph (Põlme et al. 2020); Decay sugar-rich substrates (Põlme et al. 2020) | Põlme S, et al. 2020. Fungal Diversity 105:1-16 ((https://doi.org/10.1007/s13225-020-00466-2)) |
| Otu43240 | 1 | k__Fungi(100);p__Ascomycota(100);c__Saccharomycetes(100);o__Saccharomycetales(100);f__Metschnikowiaceae(100);g__Metschnikowia(100);s__Metschnikowia_pulcherrima(100); | Metschnikowia | 13 | Saprotroph | \|Undefined Saprotroph\| | Yeast | NULL | Probable | As nectar-tap_saprotroph (Põlme et al. 2020); Decay sugar-rich substrates (Põlme et al. 2020) | Põlme S, et al. 2020. Fungal Diversity 105:1-16 ((https://doi.org/10.1007/s13225-020-00466-2)) |
| Otu43242 | 1 | k__Fungi(100);p__Ascomycota(100);c__Saccharomycetes(100);o__Saccharomycetales(100);f__Metschnikowiaceae(100);g__Metschnikowia(100);s__Metschnikowia_pulcherrima(100); | Metschnikowia | 13 | Saprotroph | \|Undefined Saprotroph\| | Yeast | NULL | Probable | As nectar-tap_saprotroph (Põlme et al. 2020); Decay sugar-rich substrates (Põlme et al. 2020) | Põlme S, et al. 2020. Fungal Diversity 105:1-16 ((https://doi.org/10.1007/s13225-020-00466-2)) |
| Otu43243 | 1 | k__Fungi(100);p__Ascomycota(100);c__Saccharomycetes(100);o__Saccharomycetales(100);f__Metschnikowiaceae(100);g__Metschnikowia(100);s__Metschnikowia_pulcherrima(100); | Metschnikowia | 13 | Saprotroph | \|Undefined Saprotroph\| | Yeast | NULL | Probable | As nectar-tap_saprotroph (Põlme et al. 2020); Decay sugar-rich substrates (Põlme et al. 2020) | Põlme S, et al. 2020. Fungal Diversity 105:1-16 ((https://doi.org/10.1007/s13225-020-00466-2)) |
| Otu43244 | 1 | k__Fungi(100);p__Ascomycota(100);c__Saccharomycetes(100);o__Saccharomycetales(100);f__Metschnikowiaceae(100);g__Metschnikowia(100);s__Metschnikowia_pulcherrima(100); | Metschnikowia | 13 | Saprotroph | \|Undefined Saprotroph\| | Yeast | NULL | Probable | As nectar-tap_saprotroph (Põlme et al. 2020); Decay sugar-rich substrates (Põlme et al. 2020) | Põlme S, et al. 2020. Fungal Diversity 105:1-16 ((https://doi.org/10.1007/s13225-020-00466-2)) |
| Otu43245 | 1 | k__Fungi(100);p__Ascomycota(100);c__Saccharomycetes(100);o__Saccharomycetales(100);f__Metschnikowiaceae(100);g__Metschnikowia(100);s__Metschnikowia_pulcherrima(100); | Metschnikowia | 13 | Saprotroph | \|Undefined Saprotroph\| | Yeast | NULL | Probable | As nectar-tap_saprotroph (Põlme et al. 2020); Decay sugar-rich substrates (Põlme et al. 2020) | Põlme S, et al. 2020. Fungal Diversity 105:1-16 ((https://doi.org/10.1007/s13225-020-00466-2)) |
| Otu43246 | 1 | k__Fungi(100);p__Ascomycota(100);c__Saccharomycetes(100);o__Saccharomycetales(100);f__Metschnikowiaceae(100);g__Metschnikowia(100);s__Metschnikowia_pulcherrima(100); | Metschnikowia | 13 | Saprotroph | \|Undefined Saprotroph\| | Yeast | NULL | Probable | As nectar-tap_saprotroph (Põlme et al. 2020); Decay sugar-rich substrates (Põlme et al. 2020) | Põlme S, et al. 2020. Fungal Diversity 105:1-16 ((https://doi.org/10.1007/s13225-020-00466-2)) |
| Otu43247 | 1 | k__Fungi(100);p__Ascomycota(100);c__Saccharomycetes(100);o__Saccharomycetales(100);f__Metschnikowiaceae(100);g__Metschnikowia(100);s__Metschnikowia_pulcherrima(100); | Metschnikowia | 13 | Saprotroph | \|Undefined Saprotroph\| | Yeast | NULL | Probable | As nectar-tap_saprotroph (Põlme et al. 2020); Decay sugar-rich substrates (Põlme et al. 2020) | Põlme S, et al. 2020. Fungal Diversity 105:1-16 ((https://doi.org/10.1007/s13225-020-00466-2)) |
| Otu43252 | 1 | k__Fungi(100);p__Ascomycota(100);c__Saccharomycetes(100);o__Saccharomycetales(100);f__Metschnikowiaceae(100);g__Metschnikowia(100);s__Metschnikowia_pulcherrima(100); | Metschnikowia | 13 | Saprotroph | \|Undefined Saprotroph\| | Yeast | NULL | Probable | As nectar-tap_saprotroph (Põlme et al. 2020); Decay sugar-rich substrates (Põlme et al. 2020) | Põlme S, et al. 2020. Fungal Diversity 105:1-16 ((https://doi.org/10.1007/s13225-020-00466-2)) |
| Otu43253 | 1 | k__Fungi(100);p__Ascomycota(100);c__Saccharomycetes(100);o__Saccharomycetales(100);f__Metschnikowiaceae(100);g__Metschnikowia(100);s__Metschnikowia_pulcherrima(100); | Metschnikowia | 13 | Saprotroph | \|Undefined Saprotroph\| | Yeast | NULL | Probable | As nectar-tap_saprotroph (Põlme et al. 2020); Decay sugar-rich substrates (Põlme et al. 2020) | Põlme S, et al. 2020. Fungal Diversity 105:1-16 ((https://doi.org/10.1007/s13225-020-00466-2)) |
| Otu43254 | 1 | k__Fungi(100);p__Ascomycota(100);c__Saccharomycetes(100);o__Saccharomycetales(100);f__Saccharomycetales_family_Incertae_sedis(100);g__Candida(100);s__Candida_sp_AS23486(100); | Candida | 13 | Pathotroph-Saprotroph-Symbiotroph | Animal Pathogen-Endophyte-Endosymbiont-Epiphyte-\|Undefined Saprotroph\| | Dimorphic Yeast-Yeast | NULL | Probable | As nectar-tap_saprotroph (Põlme et al. 2020); Decay sugar-rich substrates (Põlme et al. 2020); This asexual genus is polyphyletic and contain species that function very differently depend on what they are. Caution should be taken when interpreting guilds for Candida species. | Manolakaki D, et al. 2010. Virulence. 1: 367-375 ((https://doi.org/10.4161/viru.1.5.12796)); Mekbib SB, et al. 2011. Tropical Plant Pathology 36:233-240 ((https://doi.org/10.1590/S1982-56762011000400004)); Põlme S, et al. 2020. Fungal Diversity 105:1-16 ((https://doi.org/10.1007/s13225-020-00466-2)) |
| Otu43255 | 1 | k__Fungi(100);p__Ascomycota(100);c__Saccharomycetes(100);o__Saccharomycetales(100);f__Metschnikowiaceae(100);g__Metschnikowia(100);s__Metschnikowia_pulcherrima(100); | Metschnikowia | 13 | Saprotroph | \|Undefined Saprotroph\| | Yeast | NULL | Probable | As nectar-tap_saprotroph (Põlme et al. 2020); Decay sugar-rich substrates (Põlme et al. 2020) | Põlme S, et al. 2020. Fungal Diversity 105:1-16 ((https://doi.org/10.1007/s13225-020-00466-2)) |
| Otu43256 | 1 | k__Fungi(100);p__Ascomycota(100);c__Saccharomycetes(100);o__Saccharomycetales(100);f__Metschnikowiaceae(100);g__Metschnikowia(100);s__Metschnikowia_pulcherrima(100); | Metschnikowia | 13 | Saprotroph | \|Undefined Saprotroph\| | Yeast | NULL | Probable | As nectar-tap_saprotroph (Põlme et al. 2020); Decay sugar-rich substrates (Põlme et al. 2020) | Põlme S, et al. 2020. Fungal Diversity 105:1-16 ((https://doi.org/10.1007/s13225-020-00466-2)) |
| Otu43257 | 1 | k__Fungi(100);p__Ascomycota(100);c__Saccharomycetes(100);o__Saccharomycetales(100);f__Metschnikowiaceae(100);g__Metschnikowia(100);s__Metschnikowia_pulcherrima(100); | Metschnikowia | 13 | Saprotroph | \|Undefined Saprotroph\| | Yeast | NULL | Probable | As nectar-tap_saprotroph (Põlme et al. 2020); Decay sugar-rich substrates (Põlme et al. 2020) | Põlme S, et al. 2020. Fungal Diversity 105:1-16 ((https://doi.org/10.1007/s13225-020-00466-2)) |
| Otu43258 | 1 | k__Fungi(100);p__Ascomycota(100);c__Saccharomycetes(100);o__Saccharomycetales(100);f__Metschnikowiaceae(100);g__Metschnikowia(100);g__Metschnikowia_unclassified(100); | Metschnikowia | 13 | Saprotroph | \|Undefined Saprotroph\| | Yeast | NULL | Probable | As nectar-tap_saprotroph (Põlme et al. 2020); Decay sugar-rich substrates (Põlme et al. 2020) | Põlme S, et al. 2020. Fungal Diversity 105:1-16 ((https://doi.org/10.1007/s13225-020-00466-2)) |
| Otu43259 | 1 | k__Fungi(100);p__Ascomycota(100);c__Saccharomycetes(100);o__Saccharomycetales(100);f__Metschnikowiaceae(100);g__Metschnikowia(100);s__Metschnikowia_pulcherrima(100); | Metschnikowia | 13 | Saprotroph | \|Undefined Saprotroph\| | Yeast | NULL | Probable | As nectar-tap_saprotroph (Põlme et al. 2020); Decay sugar-rich substrates (Põlme et al. 2020) | Põlme S, et al. 2020. Fungal Diversity 105:1-16 ((https://doi.org/10.1007/s13225-020-00466-2)) |
| Otu43260 | 1 | k__Fungi(100);p__Ascomycota(100);c__Saccharomycetes(100);o__Saccharomycetales(100);f__Metschnikowiaceae(100);g__Metschnikowia(100);s__Metschnikowia_pulcherrima(100); | Metschnikowia | 13 | Saprotroph | \|Undefined Saprotroph\| | Yeast | NULL | Probable | As nectar-tap_saprotroph (Põlme et al. 2020); Decay sugar-rich substrates (Põlme et al. 2020) | Põlme S, et al. 2020. Fungal Diversity 105:1-16 ((https://doi.org/10.1007/s13225-020-00466-2)) |
| Otu43262 | 1 | k__Fungi(100);p__Ascomycota(100);c__Saccharomycetes(100);o__Saccharomycetales(100);f__Metschnikowiaceae(100);g__Metschnikowia(100);s__Metschnikowia_pulcherrima(100); | Metschnikowia | 13 | Saprotroph | \|Undefined Saprotroph\| | Yeast | NULL | Probable | As nectar-tap_saprotroph (Põlme et al. 2020); Decay sugar-rich substrates (Põlme et al. 2020) | Põlme S, et al. 2020. Fungal Diversity 105:1-16 ((https://doi.org/10.1007/s13225-020-00466-2)) |
| Otu43263 | 1 | k__Fungi(100);p__Ascomycota(100);c__Saccharomycetes(100);o__Saccharomycetales(100);f__Metschnikowiaceae(100);g__Metschnikowia(100);s__Metschnikowia_pulcherrima(100); | Metschnikowia | 13 | Saprotroph | \|Undefined Saprotroph\| | Yeast | NULL | Probable | As nectar-tap_saprotroph (Põlme et al. 2020); Decay sugar-rich substrates (Põlme et al. 2020) | Põlme S, et al. 2020. Fungal Diversity 105:1-16 ((https://doi.org/10.1007/s13225-020-00466-2)) |
| Otu43264 | 1 | k__Fungi(100);p__Ascomycota(100);c__Saccharomycetes(100);o__Saccharomycetales(100);f__Metschnikowiaceae(100);g__Metschnikowia(100);s__Metschnikowia_pulcherrima(100); | Metschnikowia | 13 | Saprotroph | \|Undefined Saprotroph\| | Yeast | NULL | Probable | As nectar-tap_saprotroph (Põlme et al. 2020); Decay sugar-rich substrates (Põlme et al. 2020) | Põlme S, et al. 2020. Fungal Diversity 105:1-16 ((https://doi.org/10.1007/s13225-020-00466-2)) |
| Otu43265 | 1 | k__Fungi(100);p__Ascomycota(100);c__Saccharomycetes(100);o__Saccharomycetales(100);f__Metschnikowiaceae(100);g__Metschnikowia(100);g__Metschnikowia_unclassified(100); | Metschnikowia | 13 | Saprotroph | \|Undefined Saprotroph\| | Yeast | NULL | Probable | As nectar-tap_saprotroph (Põlme et al. 2020); Decay sugar-rich substrates (Põlme et al. 2020) | Põlme S, et al. 2020. Fungal Diversity 105:1-16 ((https://doi.org/10.1007/s13225-020-00466-2)) |
| Otu43266 | 1 | k__Fungi(100);p__Ascomycota(100);c__Saccharomycetes(100);o__Saccharomycetales(100);f__Metschnikowiaceae(100);g__Metschnikowia(100);s__Metschnikowia_pulcherrima(100); | Metschnikowia | 13 | Saprotroph | \|Undefined Saprotroph\| | Yeast | NULL | Probable | As nectar-tap_saprotroph (Põlme et al. 2020); Decay sugar-rich substrates (Põlme et al. 2020) | Põlme S, et al. 2020. Fungal Diversity 105:1-16 ((https://doi.org/10.1007/s13225-020-00466-2)) |
| Otu43267 | 1 | k__Fungi(100);p__Ascomycota(100);c__Saccharomycetes(100);o__Saccharomycetales(100);f__Metschnikowiaceae(100);g__Metschnikowia(100);s__Metschnikowia_pulcherrima(100); | Metschnikowia | 13 | Saprotroph | \|Undefined Saprotroph\| | Yeast | NULL | Probable | As nectar-tap_saprotroph (Põlme et al. 2020); Decay sugar-rich substrates (Põlme et al. 2020) | Põlme S, et al. 2020. Fungal Diversity 105:1-16 ((https://doi.org/10.1007/s13225-020-00466-2)) |
| Otu43268 | 1 | k__Fungi(100);p__Ascomycota(100);c__Saccharomycetes(100);o__Saccharomycetales(100);f__Metschnikowiaceae(100);g__Metschnikowia(100);s__Metschnikowia_pulcherrima(100); | Metschnikowia | 13 | Saprotroph | \|Undefined Saprotroph\| | Yeast | NULL | Probable | As nectar-tap_saprotroph (Põlme et al. 2020); Decay sugar-rich substrates (Põlme et al. 2020) | Põlme S, et al. 2020. Fungal Diversity 105:1-16 ((https://doi.org/10.1007/s13225-020-00466-2)) |
| Otu43269 | 1 | k__Fungi(100);p__Ascomycota(100);c__Saccharomycetes(100);o__Saccharomycetales(100);f__Metschnikowiaceae(100);g__Metschnikowia(100);s__Metschnikowia_pulcherrima(100); | Metschnikowia | 13 | Saprotroph | \|Undefined Saprotroph\| | Yeast | NULL | Probable | As nectar-tap_saprotroph (Põlme et al. 2020); Decay sugar-rich substrates (Põlme et al. 2020) | Põlme S, et al. 2020. Fungal Diversity 105:1-16 ((https://doi.org/10.1007/s13225-020-00466-2)) |
| Otu43270 | 1 | k__Fungi(100);p__Ascomycota(100);c__Saccharomycetes(100);o__Saccharomycetales(100);f__Metschnikowiaceae(100);g__Metschnikowia(100);s__Metschnikowia_pulcherrima(100); | Metschnikowia | 13 | Saprotroph | \|Undefined Saprotroph\| | Yeast | NULL | Probable | As nectar-tap_saprotroph (Põlme et al. 2020); Decay sugar-rich substrates (Põlme et al. 2020) | Põlme S, et al. 2020. Fungal Diversity 105:1-16 ((https://doi.org/10.1007/s13225-020-00466-2)) |
| Otu43271 | 1 | k__Fungi(100);p__Ascomycota(100);c__Saccharomycetes(100);o__Saccharomycetales(100);f__Metschnikowiaceae(100);g__Metschnikowia(100);g__Metschnikowia_unclassified(100); | Metschnikowia | 13 | Saprotroph | \|Undefined Saprotroph\| | Yeast | NULL | Probable | As nectar-tap_saprotroph (Põlme et al. 2020); Decay sugar-rich substrates (Põlme et al. 2020) | Põlme S, et al. 2020. Fungal Diversity 105:1-16 ((https://doi.org/10.1007/s13225-020-00466-2)) |
| Otu43272 | 1 | k__Fungi(100);p__Ascomycota(100);c__Saccharomycetes(100);o__Saccharomycetales(100);f__Metschnikowiaceae(100);g__Metschnikowia(100);s__Metschnikowia_pulcherrima(100); | Metschnikowia | 13 | Saprotroph | \|Undefined Saprotroph\| | Yeast | NULL | Probable | As nectar-tap_saprotroph (Põlme et al. 2020); Decay sugar-rich substrates (Põlme et al. 2020) | Põlme S, et al. 2020. Fungal Diversity 105:1-16 ((https://doi.org/10.1007/s13225-020-00466-2)) |
| Otu43275 | 1 | k__Fungi(100);p__Ascomycota(100);c__Saccharomycetes(100);o__Saccharomycetales(100);f__Metschnikowiaceae(100);g__Metschnikowia(100);s__Metschnikowia_pulcherrima(100); | Metschnikowia | 13 | Saprotroph | \|Undefined Saprotroph\| | Yeast | NULL | Probable | As nectar-tap_saprotroph (Põlme et al. 2020); Decay sugar-rich substrates (Põlme et al. 2020) | Põlme S, et al. 2020. Fungal Diversity 105:1-16 ((https://doi.org/10.1007/s13225-020-00466-2)) |
| Otu43276 | 1 | k__Fungi(100);p__Ascomycota(100);c__Saccharomycetes(100);o__Saccharomycetales(100);f__Metschnikowiaceae(100);g__Metschnikowia(100);s__Metschnikowia_pulcherrima(100); | Metschnikowia | 13 | Saprotroph | \|Undefined Saprotroph\| | Yeast | NULL | Probable | As nectar-tap_saprotroph (Põlme et al. 2020); Decay sugar-rich substrates (Põlme et al. 2020) | Põlme S, et al. 2020. Fungal Diversity 105:1-16 ((https://doi.org/10.1007/s13225-020-00466-2)) |
| Otu43277 | 1 | k__Fungi(100);p__Ascomycota(100);c__Saccharomycetes(100);o__Saccharomycetales(100);f__Metschnikowiaceae(100);g__Metschnikowia(100);s__Metschnikowia_pulcherrima(100); | Metschnikowia | 13 | Saprotroph | \|Undefined Saprotroph\| | Yeast | NULL | Probable | As nectar-tap_saprotroph (Põlme et al. 2020); Decay sugar-rich substrates (Põlme et al. 2020) | Põlme S, et al. 2020. Fungal Diversity 105:1-16 ((https://doi.org/10.1007/s13225-020-00466-2)) |
| Otu43278 | 1 | k__Fungi(100);p__Ascomycota(100);c__Saccharomycetes(100);o__Saccharomycetales(100);f__Metschnikowiaceae(100);g__Metschnikowia(100);s__Metschnikowia_pulcherrima(100); | Metschnikowia | 13 | Saprotroph | \|Undefined Saprotroph\| | Yeast | NULL | Probable | As nectar-tap_saprotroph (Põlme et al. 2020); Decay sugar-rich substrates (Põlme et al. 2020) | Põlme S, et al. 2020. Fungal Diversity 105:1-16 ((https://doi.org/10.1007/s13225-020-00466-2)) |
| Otu43279 | 1 | k__Fungi(100);p__Ascomycota(100);c__Saccharomycetes(100);o__Saccharomycetales(100);f__Metschnikowiaceae(100);g__Metschnikowia(100);g__Metschnikowia_unclassified(100); | Metschnikowia | 13 | Saprotroph | \|Undefined Saprotroph\| | Yeast | NULL | Probable | As nectar-tap_saprotroph (Põlme et al. 2020); Decay sugar-rich substrates (Põlme et al. 2020) | Põlme S, et al. 2020. Fungal Diversity 105:1-16 ((https://doi.org/10.1007/s13225-020-00466-2)) |
| Otu43280 | 1 | k__Fungi(100);p__Ascomycota(100);c__Saccharomycetes(100);o__Saccharomycetales(100);f__Metschnikowiaceae(100);g__Metschnikowia(100);s__Metschnikowia_pulcherrima(100); | Metschnikowia | 13 | Saprotroph | \|Undefined Saprotroph\| | Yeast | NULL | Probable | As nectar-tap_saprotroph (Põlme et al. 2020); Decay sugar-rich substrates (Põlme et al. 2020) | Põlme S, et al. 2020. Fungal Diversity 105:1-16 ((https://doi.org/10.1007/s13225-020-00466-2)) |
| Otu43281 | 1 | k__Fungi(100);p__Ascomycota(100);c__Saccharomycetes(100);o__Saccharomycetales(100);f__Metschnikowiaceae(100);g__Metschnikowia(100);s__Metschnikowia_pulcherrima(100); | Metschnikowia | 13 | Saprotroph | \|Undefined Saprotroph\| | Yeast | NULL | Probable | As nectar-tap_saprotroph (Põlme et al. 2020); Decay sugar-rich substrates (Põlme et al. 2020) | Põlme S, et al. 2020. Fungal Diversity 105:1-16 ((https://doi.org/10.1007/s13225-020-00466-2)) |
| Otu43282 | 1 | k__Fungi(100);p__Ascomycota(100);c__Saccharomycetes(100);o__Saccharomycetales(100);f__Metschnikowiaceae(100);g__Metschnikowia(100);s__Metschnikowia_pulcherrima(100); | Metschnikowia | 13 | Saprotroph | \|Undefined Saprotroph\| | Yeast | NULL | Probable | As nectar-tap_saprotroph (Põlme et al. 2020); Decay sugar-rich substrates (Põlme et al. 2020) | Põlme S, et al. 2020. Fungal Diversity 105:1-16 ((https://doi.org/10.1007/s13225-020-00466-2)) |
| Otu43283 | 1 | k__Fungi(100);p__Ascomycota(100);c__Saccharomycetes(100);o__Saccharomycetales(100);f__Metschnikowiaceae(100);g__Metschnikowia(100);g__Metschnikowia_unclassified(100); | Metschnikowia | 13 | Saprotroph | \|Undefined Saprotroph\| | Yeast | NULL | Probable | As nectar-tap_saprotroph (Põlme et al. 2020); Decay sugar-rich substrates (Põlme et al. 2020) | Põlme S, et al. 2020. Fungal Diversity 105:1-16 ((https://doi.org/10.1007/s13225-020-00466-2)) |
| Otu43284 | 1 | k__Fungi(100);p__Ascomycota(100);c__Saccharomycetes(100);o__Saccharomycetales(100);f__Metschnikowiaceae(100);g__Metschnikowia(100);s__Metschnikowia_pulcherrima(100); | Metschnikowia | 13 | Saprotroph | \|Undefined Saprotroph\| | Yeast | NULL | Probable | As nectar-tap_saprotroph (Põlme et al. 2020); Decay sugar-rich substrates (Põlme et al. 2020) | Põlme S, et al. 2020. Fungal Diversity 105:1-16 ((https://doi.org/10.1007/s13225-020-00466-2)) |
| Otu43285 | 1 | k__Fungi(100);p__Ascomycota(100);c__Saccharomycetes(100);o__Saccharomycetales(100);f__Metschnikowiaceae(100);g__Metschnikowia(100);s__Metschnikowia_pulcherrima(100); | Metschnikowia | 13 | Saprotroph | \|Undefined Saprotroph\| | Yeast | NULL | Probable | As nectar-tap_saprotroph (Põlme et al. 2020); Decay sugar-rich substrates (Põlme et al. 2020) | Põlme S, et al. 2020. Fungal Diversity 105:1-16 ((https://doi.org/10.1007/s13225-020-00466-2)) |
| Otu43286 | 1 | k__Fungi(100);p__Ascomycota(100);c__Saccharomycetes(100);o__Saccharomycetales(100);f__Metschnikowiaceae(100);g__Metschnikowia(100);g__Metschnikowia_unclassified(100); | Metschnikowia | 13 | Saprotroph | \|Undefined Saprotroph\| | Yeast | NULL | Probable | As nectar-tap_saprotroph (Põlme et al. 2020); Decay sugar-rich substrates (Põlme et al. 2020) | Põlme S, et al. 2020. Fungal Diversity 105:1-16 ((https://doi.org/10.1007/s13225-020-00466-2)) |
| Otu43287 | 1 | k__Fungi(100);p__Ascomycota(100);c__Saccharomycetes(100);o__Saccharomycetales(100);f__Metschnikowiaceae(100);g__Metschnikowia(100);s__Metschnikowia_pulcherrima(100); | Metschnikowia | 13 | Saprotroph | \|Undefined Saprotroph\| | Yeast | NULL | Probable | As nectar-tap_saprotroph (Põlme et al. 2020); Decay sugar-rich substrates (Põlme et al. 2020) | Põlme S, et al. 2020. Fungal Diversity 105:1-16 ((https://doi.org/10.1007/s13225-020-00466-2)) |
| Otu43288 | 1 | k__Fungi(100);p__Ascomycota(100);c__Saccharomycetes(100);o__Saccharomycetales(100);f__Metschnikowiaceae(100);g__Metschnikowia(100);s__Metschnikowia_pulcherrima(100); | Metschnikowia | 13 | Saprotroph | \|Undefined Saprotroph\| | Yeast | NULL | Probable | As nectar-tap_saprotroph (Põlme et al. 2020); Decay sugar-rich substrates (Põlme et al. 2020) | Põlme S, et al. 2020. Fungal Diversity 105:1-16 ((https://doi.org/10.1007/s13225-020-00466-2)) |
| Otu43289 | 1 | k__Fungi(100);p__Ascomycota(100);c__Saccharomycetes(100);o__Saccharomycetales(100);f__Metschnikowiaceae(100);g__Metschnikowia(100);s__Metschnikowia_pulcherrima(100); | Metschnikowia | 13 | Saprotroph | \|Undefined Saprotroph\| | Yeast | NULL | Probable | As nectar-tap_saprotroph (Põlme et al. 2020); Decay sugar-rich substrates (Põlme et al. 2020) | Põlme S, et al. 2020. Fungal Diversity 105:1-16 ((https://doi.org/10.1007/s13225-020-00466-2)) |
| Otu43290 | 1 | k__Fungi(100);p__Ascomycota(100);c__Saccharomycetes(100);o__Saccharomycetales(100);f__Metschnikowiaceae(100);g__Metschnikowia(100);s__Metschnikowia_pulcherrima(100); | Metschnikowia | 13 | Saprotroph | \|Undefined Saprotroph\| | Yeast | NULL | Probable | As nectar-tap_saprotroph (Põlme et al. 2020); Decay sugar-rich substrates (Põlme et al. 2020) | Põlme S, et al. 2020. Fungal Diversity 105:1-16 ((https://doi.org/10.1007/s13225-020-00466-2)) |
| Otu43291 | 1 | k__Fungi(100);p__Ascomycota(100);c__Saccharomycetes(100);o__Saccharomycetales(100);f__Metschnikowiaceae(100);g__Metschnikowia(100);s__Metschnikowia_pulcherrima(100); | Metschnikowia | 13 | Saprotroph | \|Undefined Saprotroph\| | Yeast | NULL | Probable | As nectar-tap_saprotroph (Põlme et al. 2020); Decay sugar-rich substrates (Põlme et al. 2020) | Põlme S, et al. 2020. Fungal Diversity 105:1-16 ((https://doi.org/10.1007/s13225-020-00466-2)) |
| Otu43292 | 1 | k__Fungi(100);p__Ascomycota(100);c__Saccharomycetes(100);o__Saccharomycetales(100);f__Metschnikowiaceae(100);g__Metschnikowia(100);s__Metschnikowia_pulcherrima(100); | Metschnikowia | 13 | Saprotroph | \|Undefined Saprotroph\| | Yeast | NULL | Probable | As nectar-tap_saprotroph (Põlme et al. 2020); Decay sugar-rich substrates (Põlme et al. 2020) | Põlme S, et al. 2020. Fungal Diversity 105:1-16 ((https://doi.org/10.1007/s13225-020-00466-2)) |
| Otu43293 | 1 | k__Fungi(100);p__Ascomycota(100);c__Saccharomycetes(100);o__Saccharomycetales(100);f__Metschnikowiaceae(100);g__Metschnikowia(100);s__Metschnikowia_pulcherrima(100); | Metschnikowia | 13 | Saprotroph | \|Undefined Saprotroph\| | Yeast | NULL | Probable | As nectar-tap_saprotroph (Põlme et al. 2020); Decay sugar-rich substrates (Põlme et al. 2020) | Põlme S, et al. 2020. Fungal Diversity 105:1-16 ((https://doi.org/10.1007/s13225-020-00466-2)) |
| Otu43294 | 1 | k__Fungi(100);p__Ascomycota(100);c__Saccharomycetes(100);o__Saccharomycetales(100);f__Metschnikowiaceae(100);g__Metschnikowia(100);s__Metschnikowia_pulcherrima(100); | Metschnikowia | 13 | Saprotroph | \|Undefined Saprotroph\| | Yeast | NULL | Probable | As nectar-tap_saprotroph (Põlme et al. 2020); Decay sugar-rich substrates (Põlme et al. 2020) | Põlme S, et al. 2020. Fungal Diversity 105:1-16 ((https://doi.org/10.1007/s13225-020-00466-2)) |
| Otu43295 | 1 | k__Fungi(100);p__Ascomycota(100);c__Saccharomycetes(100);o__Saccharomycetales(100);f__Metschnikowiaceae(100);g__Metschnikowia(100);s__Metschnikowia_pulcherrima(100); | Metschnikowia | 13 | Saprotroph | \|Undefined Saprotroph\| | Yeast | NULL | Probable | As nectar-tap_saprotroph (Põlme et al. 2020); Decay sugar-rich substrates (Põlme et al. 2020) | Põlme S, et al. 2020. Fungal Diversity 105:1-16 ((https://doi.org/10.1007/s13225-020-00466-2)) |
| Otu43296 | 1 | k__Fungi(100);p__Ascomycota(100);c__Saccharomycetes(100);o__Saccharomycetales(100);f__Metschnikowiaceae(100);g__Metschnikowia(100);s__Metschnikowia_pulcherrima(100); | Metschnikowia | 13 | Saprotroph | \|Undefined Saprotroph\| | Yeast | NULL | Probable | As nectar-tap_saprotroph (Põlme et al. 2020); Decay sugar-rich substrates (Põlme et al. 2020) | Põlme S, et al. 2020. Fungal Diversity 105:1-16 ((https://doi.org/10.1007/s13225-020-00466-2)) |
| Otu43297 | 1 | k__Fungi(100);p__Ascomycota(100);c__Saccharomycetes(100);o__Saccharomycetales(100);f__Metschnikowiaceae(100);g__Metschnikowia(100);s__Metschnikowia_pulcherrima(100); | Metschnikowia | 13 | Saprotroph | \|Undefined Saprotroph\| | Yeast | NULL | Probable | As nectar-tap_saprotroph (Põlme et al. 2020); Decay sugar-rich substrates (Põlme et al. 2020) | Põlme S, et al. 2020. Fungal Diversity 105:1-16 ((https://doi.org/10.1007/s13225-020-00466-2)) |
| Otu43298 | 1 | k__Fungi(100);p__Ascomycota(100);c__Saccharomycetes(100);o__Saccharomycetales(100);f__Metschnikowiaceae(100);g__Metschnikowia(100);s__Metschnikowia_pulcherrima(100); | Metschnikowia | 13 | Saprotroph | \|Undefined Saprotroph\| | Yeast | NULL | Probable | As nectar-tap_saprotroph (Põlme et al. 2020); Decay sugar-rich substrates (Põlme et al. 2020) | Põlme S, et al. 2020. Fungal Diversity 105:1-16 ((https://doi.org/10.1007/s13225-020-00466-2)) |
| Otu43299 | 1 | k__Fungi(100);p__Ascomycota(100);c__Saccharomycetes(100);o__Saccharomycetales(100);f__Metschnikowiaceae(100);g__Metschnikowia(100);g__Metschnikowia_unclassified(100); | Metschnikowia | 13 | Saprotroph | \|Undefined Saprotroph\| | Yeast | NULL | Probable | As nectar-tap_saprotroph (Põlme et al. 2020); Decay sugar-rich substrates (Põlme et al. 2020) | Põlme S, et al. 2020. Fungal Diversity 105:1-16 ((https://doi.org/10.1007/s13225-020-00466-2)) |
| Otu43300 | 1 | k__Fungi(100);p__Ascomycota(100);c__Saccharomycetes(100);o__Saccharomycetales(100);f__Metschnikowiaceae(100);g__Metschnikowia(100);s__Metschnikowia_pulcherrima(100); | Metschnikowia | 13 | Saprotroph | \|Undefined Saprotroph\| | Yeast | NULL | Probable | As nectar-tap_saprotroph (Põlme et al. 2020); Decay sugar-rich substrates (Põlme et al. 2020) | Põlme S, et al. 2020. Fungal Diversity 105:1-16 ((https://doi.org/10.1007/s13225-020-00466-2)) |
| Otu43301 | 1 | k__Fungi(100);p__Ascomycota(100);c__Saccharomycetes(100);o__Saccharomycetales(100);f__Metschnikowiaceae(100);g__Metschnikowia(100);s__Metschnikowia_pulcherrima(100); | Metschnikowia | 13 | Saprotroph | \|Undefined Saprotroph\| | Yeast | NULL | Probable | As nectar-tap_saprotroph (Põlme et al. 2020); Decay sugar-rich substrates (Põlme et al. 2020) | Põlme S, et al. 2020. Fungal Diversity 105:1-16 ((https://doi.org/10.1007/s13225-020-00466-2)) |
| Otu43302 | 1 | k__Fungi(100);p__Ascomycota(100);c__Saccharomycetes(100);o__Saccharomycetales(100);f__Metschnikowiaceae(100);g__Metschnikowia(100);s__Metschnikowia_pulcherrima(100); | Metschnikowia | 13 | Saprotroph | \|Undefined Saprotroph\| | Yeast | NULL | Probable | As nectar-tap_saprotroph (Põlme et al. 2020); Decay sugar-rich substrates (Põlme et al. 2020) | Põlme S, et al. 2020. Fungal Diversity 105:1-16 ((https://doi.org/10.1007/s13225-020-00466-2)) |
| Otu43303 | 1 | k__Fungi(100);p__Ascomycota(100);c__Saccharomycetes(100);o__Saccharomycetales(100);f__Metschnikowiaceae(100);g__Metschnikowia(100);g__Metschnikowia_unclassified(100); | Metschnikowia | 13 | Saprotroph | \|Undefined Saprotroph\| | Yeast | NULL | Probable | As nectar-tap_saprotroph (Põlme et al. 2020); Decay sugar-rich substrates (Põlme et al. 2020) | Põlme S, et al. 2020. Fungal Diversity 105:1-16 ((https://doi.org/10.1007/s13225-020-00466-2)) |
| Otu43304 | 1 | k__Fungi(100);p__Ascomycota(100);c__Saccharomycetes(100);o__Saccharomycetales(100);f__Metschnikowiaceae(100);g__Metschnikowia(100);s__Metschnikowia_pulcherrima(100); | Metschnikowia | 13 | Saprotroph | \|Undefined Saprotroph\| | Yeast | NULL | Probable | As nectar-tap_saprotroph (Põlme et al. 2020); Decay sugar-rich substrates (Põlme et al. 2020) | Põlme S, et al. 2020. Fungal Diversity 105:1-16 ((https://doi.org/10.1007/s13225-020-00466-2)) |
| Otu43305 | 1 | k__Fungi(100);p__Ascomycota(100);c__Saccharomycetes(100);o__Saccharomycetales(100);f__Metschnikowiaceae(100);g__Metschnikowia(100);s__Metschnikowia_pulcherrima(100); | Metschnikowia | 13 | Saprotroph | \|Undefined Saprotroph\| | Yeast | NULL | Probable | As nectar-tap_saprotroph (Põlme et al. 2020); Decay sugar-rich substrates (Põlme et al. 2020) | Põlme S, et al. 2020. Fungal Diversity 105:1-16 ((https://doi.org/10.1007/s13225-020-00466-2)) |
| Otu43306 | 1 | k__Fungi(100);p__Ascomycota(100);c__Saccharomycetes(100);o__Saccharomycetales(100);f__Metschnikowiaceae(100);g__Metschnikowia(100);s__Metschnikowia_pulcherrima(100); | Metschnikowia | 13 | Saprotroph | \|Undefined Saprotroph\| | Yeast | NULL | Probable | As nectar-tap_saprotroph (Põlme et al. 2020); Decay sugar-rich substrates (Põlme et al. 2020) | Põlme S, et al. 2020. Fungal Diversity 105:1-16 ((https://doi.org/10.1007/s13225-020-00466-2)) |
| Otu43307 | 1 | k__Fungi(100);p__Ascomycota(100);c__Saccharomycetes(100);o__Saccharomycetales(100);f__Metschnikowiaceae(100);g__Metschnikowia(100);g__Metschnikowia_unclassified(100); | Metschnikowia | 13 | Saprotroph | \|Undefined Saprotroph\| | Yeast | NULL | Probable | As nectar-tap_saprotroph (Põlme et al. 2020); Decay sugar-rich substrates (Põlme et al. 2020) | Põlme S, et al. 2020. Fungal Diversity 105:1-16 ((https://doi.org/10.1007/s13225-020-00466-2)) |
| Otu43309 | 1 | k__Fungi(100);p__Ascomycota(100);c__Saccharomycetes(100);o__Saccharomycetales(100);f__Metschnikowiaceae(100);g__Metschnikowia(100);s__Metschnikowia_pulcherrima(100); | Metschnikowia | 13 | Saprotroph | \|Undefined Saprotroph\| | Yeast | NULL | Probable | As nectar-tap_saprotroph (Põlme et al. 2020); Decay sugar-rich substrates (Põlme et al. 2020) | Põlme S, et al. 2020. Fungal Diversity 105:1-16 ((https://doi.org/10.1007/s13225-020-00466-2)) |
| Otu43312 | 1 | k__Fungi(100);p__Ascomycota(100);c__Saccharomycetes(100);o__Saccharomycetales(100);f__Metschnikowiaceae(100);g__Metschnikowia(100);s__Metschnikowia_pulcherrima(100); | Metschnikowia | 13 | Saprotroph | \|Undefined Saprotroph\| | Yeast | NULL | Probable | As nectar-tap_saprotroph (Põlme et al. 2020); Decay sugar-rich substrates (Põlme et al. 2020) | Põlme S, et al. 2020. Fungal Diversity 105:1-16 ((https://doi.org/10.1007/s13225-020-00466-2)) |
| Otu43313 | 1 | k__Fungi(100);p__Ascomycota(100);c__Saccharomycetes(100);o__Saccharomycetales(100);f__Metschnikowiaceae(100);g__Metschnikowia(100);s__Metschnikowia_pulcherrima(100); | Metschnikowia | 13 | Saprotroph | \|Undefined Saprotroph\| | Yeast | NULL | Probable | As nectar-tap_saprotroph (Põlme et al. 2020); Decay sugar-rich substrates (Põlme et al. 2020) | Põlme S, et al. 2020. Fungal Diversity 105:1-16 ((https://doi.org/10.1007/s13225-020-00466-2)) |
| Otu43315 | 1 | k__Fungi(100);p__Ascomycota(100);c__Saccharomycetes(100);o__Saccharomycetales(100);f__Metschnikowiaceae(100);g__Metschnikowia(100);s__Metschnikowia_pulcherrima(100); | Metschnikowia | 13 | Saprotroph | \|Undefined Saprotroph\| | Yeast | NULL | Probable | As nectar-tap_saprotroph (Põlme et al. 2020); Decay sugar-rich substrates (Põlme et al. 2020) | Põlme S, et al. 2020. Fungal Diversity 105:1-16 ((https://doi.org/10.1007/s13225-020-00466-2)) |
| Otu43316 | 1 | k__Fungi(100);p__Ascomycota(100);c__Saccharomycetes(100);o__Saccharomycetales(100);f__Metschnikowiaceae(100);g__Metschnikowia(100);g__Metschnikowia_unclassified(100); | Metschnikowia | 13 | Saprotroph | \|Undefined Saprotroph\| | Yeast | NULL | Probable | As nectar-tap_saprotroph (Põlme et al. 2020); Decay sugar-rich substrates (Põlme et al. 2020) | Põlme S, et al. 2020. Fungal Diversity 105:1-16 ((https://doi.org/10.1007/s13225-020-00466-2)) |
| Otu43317 | 1 | k__Fungi(100);p__Ascomycota(100);c__Saccharomycetes(100);o__Saccharomycetales(100);f__Metschnikowiaceae(100);g__Metschnikowia(100);s__Metschnikowia_pulcherrima(100); | Metschnikowia | 13 | Saprotroph | \|Undefined Saprotroph\| | Yeast | NULL | Probable | As nectar-tap_saprotroph (Põlme et al. 2020); Decay sugar-rich substrates (Põlme et al. 2020) | Põlme S, et al. 2020. Fungal Diversity 105:1-16 ((https://doi.org/10.1007/s13225-020-00466-2)) |
| Otu43318 | 1 | k__Fungi(100);p__Ascomycota(100);c__Saccharomycetes(100);o__Saccharomycetales(100);f__Metschnikowiaceae(100);g__Metschnikowia(100);s__Metschnikowia_pulcherrima(100); | Metschnikowia | 13 | Saprotroph | \|Undefined Saprotroph\| | Yeast | NULL | Probable | As nectar-tap_saprotroph (Põlme et al. 2020); Decay sugar-rich substrates (Põlme et al. 2020) | Põlme S, et al. 2020. Fungal Diversity 105:1-16 ((https://doi.org/10.1007/s13225-020-00466-2)) |
| Otu43319 | 1 | k__Fungi(100);p__Ascomycota(100);c__Saccharomycetes(100);o__Saccharomycetales(100);f__Metschnikowiaceae(100);g__Metschnikowia(100);s__Metschnikowia_pulcherrima(100); | Metschnikowia | 13 | Saprotroph | \|Undefined Saprotroph\| | Yeast | NULL | Probable | As nectar-tap_saprotroph (Põlme et al. 2020); Decay sugar-rich substrates (Põlme et al. 2020) | Põlme S, et al. 2020. Fungal Diversity 105:1-16 ((https://doi.org/10.1007/s13225-020-00466-2)) |
| Otu43321 | 1 | k__Fungi(100);p__Ascomycota(100);c__Saccharomycetes(100);o__Saccharomycetales(100);f__Metschnikowiaceae(100);g__Metschnikowia(100);s__Metschnikowia_pulcherrima(100); | Metschnikowia | 13 | Saprotroph | \|Undefined Saprotroph\| | Yeast | NULL | Probable | As nectar-tap_saprotroph (Põlme et al. 2020); Decay sugar-rich substrates (Põlme et al. 2020) | Põlme S, et al. 2020. Fungal Diversity 105:1-16 ((https://doi.org/10.1007/s13225-020-00466-2)) |
| Otu43322 | 1 | k__Fungi(100);p__Ascomycota(100);c__Saccharomycetes(100);o__Saccharomycetales(100);f__Metschnikowiaceae(100);g__Metschnikowia(100);s__Metschnikowia_pulcherrima(100); | Metschnikowia | 13 | Saprotroph | \|Undefined Saprotroph\| | Yeast | NULL | Probable | As nectar-tap_saprotroph (Põlme et al. 2020); Decay sugar-rich substrates (Põlme et al. 2020) | Põlme S, et al. 2020. Fungal Diversity 105:1-16 ((https://doi.org/10.1007/s13225-020-00466-2)) |
| Otu43323 | 1 | k__Fungi(100);p__Ascomycota(100);c__Saccharomycetes(100);o__Saccharomycetales(100);f__Metschnikowiaceae(100);g__Metschnikowia(100);g__Metschnikowia_unclassified(100); | Metschnikowia | 13 | Saprotroph | \|Undefined Saprotroph\| | Yeast | NULL | Probable | As nectar-tap_saprotroph (Põlme et al. 2020); Decay sugar-rich substrates (Põlme et al. 2020) | Põlme S, et al. 2020. Fungal Diversity 105:1-16 ((https://doi.org/10.1007/s13225-020-00466-2)) |
| Otu43324 | 1 | k__Fungi(100);p__Ascomycota(100);c__Saccharomycetes(100);o__Saccharomycetales(100);f__Metschnikowiaceae(100);g__Metschnikowia(100);s__Metschnikowia_pulcherrima(100); | Metschnikowia | 13 | Saprotroph | \|Undefined Saprotroph\| | Yeast | NULL | Probable | As nectar-tap_saprotroph (Põlme et al. 2020); Decay sugar-rich substrates (Põlme et al. 2020) | Põlme S, et al. 2020. Fungal Diversity 105:1-16 ((https://doi.org/10.1007/s13225-020-00466-2)) |
| Otu43325 | 1 | k__Fungi(100);p__Ascomycota(100);c__Saccharomycetes(100);o__Saccharomycetales(100);f__Metschnikowiaceae(100);g__Metschnikowia(100);s__Metschnikowia_pulcherrima(100); | Metschnikowia | 13 | Saprotroph | \|Undefined Saprotroph\| | Yeast | NULL | Probable | As nectar-tap_saprotroph (Põlme et al. 2020); Decay sugar-rich substrates (Põlme et al. 2020) | Põlme S, et al. 2020. Fungal Diversity 105:1-16 ((https://doi.org/10.1007/s13225-020-00466-2)) |
| Otu43326 | 1 | k__Fungi(100);p__Ascomycota(100);c__Saccharomycetes(100);o__Saccharomycetales(100);f__Metschnikowiaceae(100);g__Metschnikowia(100);s__Metschnikowia_pulcherrima(100); | Metschnikowia | 13 | Saprotroph | \|Undefined Saprotroph\| | Yeast | NULL | Probable | As nectar-tap_saprotroph (Põlme et al. 2020); Decay sugar-rich substrates (Põlme et al. 2020) | Põlme S, et al. 2020. Fungal Diversity 105:1-16 ((https://doi.org/10.1007/s13225-020-00466-2)) |
| Otu43327 | 1 | k__Fungi(100);p__Ascomycota(100);c__Saccharomycetes(100);o__Saccharomycetales(100);f__Metschnikowiaceae(100);g__Metschnikowia(100);g__Metschnikowia_unclassified(100); | Metschnikowia | 13 | Saprotroph | \|Undefined Saprotroph\| | Yeast | NULL | Probable | As nectar-tap_saprotroph (Põlme et al. 2020); Decay sugar-rich substrates (Põlme et al. 2020) | Põlme S, et al. 2020. Fungal Diversity 105:1-16 ((https://doi.org/10.1007/s13225-020-00466-2)) |
| Otu43328 | 1 | k__Fungi(100);p__Ascomycota(100);c__Saccharomycetes(100);o__Saccharomycetales(100);f__Metschnikowiaceae(100);g__Metschnikowia(100);g__Metschnikowia_unclassified(100); | Metschnikowia | 13 | Saprotroph | \|Undefined Saprotroph\| | Yeast | NULL | Probable | As nectar-tap_saprotroph (Põlme et al. 2020); Decay sugar-rich substrates (Põlme et al. 2020) | Põlme S, et al. 2020. Fungal Diversity 105:1-16 ((https://doi.org/10.1007/s13225-020-00466-2)) |
| Otu43329 | 1 | k__Fungi(100);p__Ascomycota(100);c__Saccharomycetes(100);o__Saccharomycetales(100);f__Metschnikowiaceae(100);g__Metschnikowia(100);s__Metschnikowia_pulcherrima(100); | Metschnikowia | 13 | Saprotroph | \|Undefined Saprotroph\| | Yeast | NULL | Probable | As nectar-tap_saprotroph (Põlme et al. 2020); Decay sugar-rich substrates (Põlme et al. 2020) | Põlme S, et al. 2020. Fungal Diversity 105:1-16 ((https://doi.org/10.1007/s13225-020-00466-2)) |
| Otu43330 | 1 | k__Fungi(100);p__Ascomycota(100);c__Saccharomycetes(100);o__Saccharomycetales(100);f__Metschnikowiaceae(100);g__Metschnikowia(100);s__Metschnikowia_pulcherrima(100); | Metschnikowia | 13 | Saprotroph | \|Undefined Saprotroph\| | Yeast | NULL | Probable | As nectar-tap_saprotroph (Põlme et al. 2020); Decay sugar-rich substrates (Põlme et al. 2020) | Põlme S, et al. 2020. Fungal Diversity 105:1-16 ((https://doi.org/10.1007/s13225-020-00466-2)) |
| Otu43331 | 1 | k__Fungi(100);p__Ascomycota(100);c__Saccharomycetes(100);o__Saccharomycetales(100);f__Metschnikowiaceae(100);g__Metschnikowia(100);g__Metschnikowia_unclassified(100); | Metschnikowia | 13 | Saprotroph | \|Undefined Saprotroph\| | Yeast | NULL | Probable | As nectar-tap_saprotroph (Põlme et al. 2020); Decay sugar-rich substrates (Põlme et al. 2020) | Põlme S, et al. 2020. Fungal Diversity 105:1-16 ((https://doi.org/10.1007/s13225-020-00466-2)) |
| Otu43332 | 1 | k__Fungi(100);p__Ascomycota(100);c__Saccharomycetes(100);o__Saccharomycetales(100);f__Metschnikowiaceae(100);g__Metschnikowia(100);g__Metschnikowia_unclassified(100); | Metschnikowia | 13 | Saprotroph | \|Undefined Saprotroph\| | Yeast | NULL | Probable | As nectar-tap_saprotroph (Põlme et al. 2020); Decay sugar-rich substrates (Põlme et al. 2020) | Põlme S, et al. 2020. Fungal Diversity 105:1-16 ((https://doi.org/10.1007/s13225-020-00466-2)) |
| Otu43333 | 1 | k__Fungi(100);p__Ascomycota(100);c__Saccharomycetes(100);o__Saccharomycetales(100);f__Metschnikowiaceae(100);g__Metschnikowia(100);s__Metschnikowia_pulcherrima(100); | Metschnikowia | 13 | Saprotroph | \|Undefined Saprotroph\| | Yeast | NULL | Probable | As nectar-tap_saprotroph (Põlme et al. 2020); Decay sugar-rich substrates (Põlme et al. 2020) | Põlme S, et al. 2020. Fungal Diversity 105:1-16 ((https://doi.org/10.1007/s13225-020-00466-2)) |
| Otu43334 | 1 | k__Fungi(100);p__Ascomycota(100);c__Saccharomycetes(100);o__Saccharomycetales(100);f__Metschnikowiaceae(100);g__Metschnikowia(100);s__Metschnikowia_pulcherrima(100); | Metschnikowia | 13 | Saprotroph | \|Undefined Saprotroph\| | Yeast | NULL | Probable | As nectar-tap_saprotroph (Põlme et al. 2020); Decay sugar-rich substrates (Põlme et al. 2020) | Põlme S, et al. 2020. Fungal Diversity 105:1-16 ((https://doi.org/10.1007/s13225-020-00466-2)) |
| Otu43335 | 1 | k__Fungi(100);p__Ascomycota(100);c__Saccharomycetes(100);o__Saccharomycetales(100);f__Metschnikowiaceae(100);g__Metschnikowia(100);s__Metschnikowia_pulcherrima(100); | Metschnikowia | 13 | Saprotroph | \|Undefined Saprotroph\| | Yeast | NULL | Probable | As nectar-tap_saprotroph (Põlme et al. 2020); Decay sugar-rich substrates (Põlme et al. 2020) | Põlme S, et al. 2020. Fungal Diversity 105:1-16 ((https://doi.org/10.1007/s13225-020-00466-2)) |
| Otu43336 | 1 | k__Fungi(100);p__Ascomycota(100);c__Saccharomycetes(100);o__Saccharomycetales(100);f__Metschnikowiaceae(100);g__Metschnikowia(100);s__Metschnikowia_pulcherrima(100); | Metschnikowia | 13 | Saprotroph | \|Undefined Saprotroph\| | Yeast | NULL | Probable | As nectar-tap_saprotroph (Põlme et al. 2020); Decay sugar-rich substrates (Põlme et al. 2020) | Põlme S, et al. 2020. Fungal Diversity 105:1-16 ((https://doi.org/10.1007/s13225-020-00466-2)) |
| Otu43337 | 1 | k__Fungi(100);p__Ascomycota(100);c__Saccharomycetes(100);o__Saccharomycetales(100);f__Metschnikowiaceae(100);g__Metschnikowia(100);s__Metschnikowia_pulcherrima(100); | Metschnikowia | 13 | Saprotroph | \|Undefined Saprotroph\| | Yeast | NULL | Probable | As nectar-tap_saprotroph (Põlme et al. 2020); Decay sugar-rich substrates (Põlme et al. 2020) | Põlme S, et al. 2020. Fungal Diversity 105:1-16 ((https://doi.org/10.1007/s13225-020-00466-2)) |
| Otu43338 | 1 | k__Fungi(100);p__Ascomycota(100);c__Saccharomycetes(100);o__Saccharomycetales(100);f__Metschnikowiaceae(100);g__Metschnikowia(100);s__Metschnikowia_pulcherrima(100); | Metschnikowia | 13 | Saprotroph | \|Undefined Saprotroph\| | Yeast | NULL | Probable | As nectar-tap_saprotroph (Põlme et al. 2020); Decay sugar-rich substrates (Põlme et al. 2020) | Põlme S, et al. 2020. Fungal Diversity 105:1-16 ((https://doi.org/10.1007/s13225-020-00466-2)) |
| Otu43339 | 1 | k__Fungi(100);p__Ascomycota(100);c__Saccharomycetes(100);o__Saccharomycetales(100);f__Metschnikowiaceae(100);g__Metschnikowia(100);s__Metschnikowia_pulcherrima(100); | Metschnikowia | 13 | Saprotroph | \|Undefined Saprotroph\| | Yeast | NULL | Probable | As nectar-tap_saprotroph (Põlme et al. 2020); Decay sugar-rich substrates (Põlme et al. 2020) | Põlme S, et al. 2020. Fungal Diversity 105:1-16 ((https://doi.org/10.1007/s13225-020-00466-2)) |
| Otu43340 | 1 | k__Fungi(100);p__Ascomycota(100);c__Saccharomycetes(100);o__Saccharomycetales(100);f__Metschnikowiaceae(100);g__Metschnikowia(100);s__Metschnikowia_pulcherrima(100); | Metschnikowia | 13 | Saprotroph | \|Undefined Saprotroph\| | Yeast | NULL | Probable | As nectar-tap_saprotroph (Põlme et al. 2020); Decay sugar-rich substrates (Põlme et al. 2020) | Põlme S, et al. 2020. Fungal Diversity 105:1-16 ((https://doi.org/10.1007/s13225-020-00466-2)) |
| Otu43341 | 1 | k__Fungi(100);p__Ascomycota(100);c__Saccharomycetes(100);o__Saccharomycetales(100);f__Metschnikowiaceae(100);g__Metschnikowia(100);s__Metschnikowia_pulcherrima(100); | Metschnikowia | 13 | Saprotroph | \|Undefined Saprotroph\| | Yeast | NULL | Probable | As nectar-tap_saprotroph (Põlme et al. 2020); Decay sugar-rich substrates (Põlme et al. 2020) | Põlme S, et al. 2020. Fungal Diversity 105:1-16 ((https://doi.org/10.1007/s13225-020-00466-2)) |
| Otu43342 | 1 | k__Fungi(100);p__Ascomycota(100);c__Saccharomycetes(100);o__Saccharomycetales(100);f__Metschnikowiaceae(100);g__Metschnikowia(100);s__Metschnikowia_pulcherrima(100); | Metschnikowia | 13 | Saprotroph | \|Undefined Saprotroph\| | Yeast | NULL | Probable | As nectar-tap_saprotroph (Põlme et al. 2020); Decay sugar-rich substrates (Põlme et al. 2020) | Põlme S, et al. 2020. Fungal Diversity 105:1-16 ((https://doi.org/10.1007/s13225-020-00466-2)) |
| Otu43343 | 1 | k__Fungi(100);p__Ascomycota(100);c__Saccharomycetes(100);o__Saccharomycetales(100);f__Metschnikowiaceae(100);g__Metschnikowia(100);g__Metschnikowia_unclassified(100); | Metschnikowia | 13 | Saprotroph | \|Undefined Saprotroph\| | Yeast | NULL | Probable | As nectar-tap_saprotroph (Põlme et al. 2020); Decay sugar-rich substrates (Põlme et al. 2020) | Põlme S, et al. 2020. Fungal Diversity 105:1-16 ((https://doi.org/10.1007/s13225-020-00466-2)) |
| Otu43344 | 1 | k__Fungi(100);p__Ascomycota(100);c__Saccharomycetes(100);o__Saccharomycetales(100);f__Metschnikowiaceae(100);g__Metschnikowia(100);g__Metschnikowia_unclassified(100); | Metschnikowia | 13 | Saprotroph | \|Undefined Saprotroph\| | Yeast | NULL | Probable | As nectar-tap_saprotroph (Põlme et al. 2020); Decay sugar-rich substrates (Põlme et al. 2020) | Põlme S, et al. 2020. Fungal Diversity 105:1-16 ((https://doi.org/10.1007/s13225-020-00466-2)) |
| Otu43345 | 1 | k__Fungi(100);p__Ascomycota(100);c__Saccharomycetes(100);o__Saccharomycetales(100);f__Metschnikowiaceae(100);g__Metschnikowia(100);s__Metschnikowia_pulcherrima(100); | Metschnikowia | 13 | Saprotroph | \|Undefined Saprotroph\| | Yeast | NULL | Probable | As nectar-tap_saprotroph (Põlme et al. 2020); Decay sugar-rich substrates (Põlme et al. 2020) | Põlme S, et al. 2020. Fungal Diversity 105:1-16 ((https://doi.org/10.1007/s13225-020-00466-2)) |
| Otu43346 | 1 | k__Fungi(100);p__Ascomycota(100);c__Saccharomycetes(100);o__Saccharomycetales(100);f__Metschnikowiaceae(100);g__Metschnikowia(100);s__Metschnikowia_pulcherrima(100); | Metschnikowia | 13 | Saprotroph | \|Undefined Saprotroph\| | Yeast | NULL | Probable | As nectar-tap_saprotroph (Põlme et al. 2020); Decay sugar-rich substrates (Põlme et al. 2020) | Põlme S, et al. 2020. Fungal Diversity 105:1-16 ((https://doi.org/10.1007/s13225-020-00466-2)) |
| Otu43347 | 1 | k__Fungi(100);p__Ascomycota(100);c__Saccharomycetes(100);o__Saccharomycetales(100);f__Metschnikowiaceae(100);g__Metschnikowia(100);s__Metschnikowia_pulcherrima(100); | Metschnikowia | 13 | Saprotroph | \|Undefined Saprotroph\| | Yeast | NULL | Probable | As nectar-tap_saprotroph (Põlme et al. 2020); Decay sugar-rich substrates (Põlme et al. 2020) | Põlme S, et al. 2020. Fungal Diversity 105:1-16 ((https://doi.org/10.1007/s13225-020-00466-2)) |
| Otu43348 | 1 | k__Fungi(100);p__Ascomycota(100);c__Saccharomycetes(100);o__Saccharomycetales(100);f__Metschnikowiaceae(100);g__Metschnikowia(100);s__Metschnikowia_pulcherrima(100); | Metschnikowia | 13 | Saprotroph | \|Undefined Saprotroph\| | Yeast | NULL | Probable | As nectar-tap_saprotroph (Põlme et al. 2020); Decay sugar-rich substrates (Põlme et al. 2020) | Põlme S, et al. 2020. Fungal Diversity 105:1-16 ((https://doi.org/10.1007/s13225-020-00466-2)) |
| Otu43349 | 1 | k__Fungi(100);p__Ascomycota(100);c__Saccharomycetes(100);o__Saccharomycetales(100);f__Metschnikowiaceae(100);g__Metschnikowia(100);s__Metschnikowia_pulcherrima(100); | Metschnikowia | 13 | Saprotroph | \|Undefined Saprotroph\| | Yeast | NULL | Probable | As nectar-tap_saprotroph (Põlme et al. 2020); Decay sugar-rich substrates (Põlme et al. 2020) | Põlme S, et al. 2020. Fungal Diversity 105:1-16 ((https://doi.org/10.1007/s13225-020-00466-2)) |
| Otu43350 | 1 | k__Fungi(100);p__Ascomycota(100);c__Saccharomycetes(100);o__Saccharomycetales(100);f__Metschnikowiaceae(100);g__Metschnikowia(100);s__Metschnikowia_pulcherrima(100); | Metschnikowia | 13 | Saprotroph | \|Undefined Saprotroph\| | Yeast | NULL | Probable | As nectar-tap_saprotroph (Põlme et al. 2020); Decay sugar-rich substrates (Põlme et al. 2020) | Põlme S, et al. 2020. Fungal Diversity 105:1-16 ((https://doi.org/10.1007/s13225-020-00466-2)) |
| Otu43351 | 1 | k__Fungi(100);p__Ascomycota(100);c__Saccharomycetes(100);o__Saccharomycetales(100);f__Metschnikowiaceae(100);g__Metschnikowia(100);s__Metschnikowia_pulcherrima(100); | Metschnikowia | 13 | Saprotroph | \|Undefined Saprotroph\| | Yeast | NULL | Probable | As nectar-tap_saprotroph (Põlme et al. 2020); Decay sugar-rich substrates (Põlme et al. 2020) | Põlme S, et al. 2020. Fungal Diversity 105:1-16 ((https://doi.org/10.1007/s13225-020-00466-2)) |
| Otu43352 | 1 | k__Fungi(100);p__Ascomycota(100);c__Saccharomycetes(100);o__Saccharomycetales(100);f__Metschnikowiaceae(100);g__Metschnikowia(100);s__Metschnikowia_pulcherrima(100); | Metschnikowia | 13 | Saprotroph | \|Undefined Saprotroph\| | Yeast | NULL | Probable | As nectar-tap_saprotroph (Põlme et al. 2020); Decay sugar-rich substrates (Põlme et al. 2020) | Põlme S, et al. 2020. Fungal Diversity 105:1-16 ((https://doi.org/10.1007/s13225-020-00466-2)) |
| Otu43353 | 1 | k__Fungi(100);p__Ascomycota(100);c__Saccharomycetes(100);o__Saccharomycetales(100);f__Metschnikowiaceae(100);g__Metschnikowia(100);s__Metschnikowia_pulcherrima(100); | Metschnikowia | 13 | Saprotroph | \|Undefined Saprotroph\| | Yeast | NULL | Probable | As nectar-tap_saprotroph (Põlme et al. 2020); Decay sugar-rich substrates (Põlme et al. 2020) | Põlme S, et al. 2020. Fungal Diversity 105:1-16 ((https://doi.org/10.1007/s13225-020-00466-2)) |
| Otu43354 | 1 | k__Fungi(100);p__Ascomycota(100);c__Saccharomycetes(100);o__Saccharomycetales(100);f__Metschnikowiaceae(100);g__Metschnikowia(100);s__Metschnikowia_pulcherrima(100); | Metschnikowia | 13 | Saprotroph | \|Undefined Saprotroph\| | Yeast | NULL | Probable | As nectar-tap_saprotroph (Põlme et al. 2020); Decay sugar-rich substrates (Põlme et al. 2020) | Põlme S, et al. 2020. Fungal Diversity 105:1-16 ((https://doi.org/10.1007/s13225-020-00466-2)) |
| Otu43355 | 1 | k__Fungi(100);p__Ascomycota(100);c__Saccharomycetes(100);o__Saccharomycetales(100);f__Metschnikowiaceae(100);g__Metschnikowia(100);s__Metschnikowia_pulcherrima(100); | Metschnikowia | 13 | Saprotroph | \|Undefined Saprotroph\| | Yeast | NULL | Probable | As nectar-tap_saprotroph (Põlme et al. 2020); Decay sugar-rich substrates (Põlme et al. 2020) | Põlme S, et al. 2020. Fungal Diversity 105:1-16 ((https://doi.org/10.1007/s13225-020-00466-2)) |
| Otu43357 | 1 | k__Fungi(100);p__Ascomycota(100);c__Saccharomycetes(100);o__Saccharomycetales(100);f__Metschnikowiaceae(100);g__Metschnikowia(100);g__Metschnikowia_unclassified(100); | Metschnikowia | 13 | Saprotroph | \|Undefined Saprotroph\| | Yeast | NULL | Probable | As nectar-tap_saprotroph (Põlme et al. 2020); Decay sugar-rich substrates (Põlme et al. 2020) | Põlme S, et al. 2020. Fungal Diversity 105:1-16 ((https://doi.org/10.1007/s13225-020-00466-2)) |
| Otu43358 | 1 | k__Fungi(100);p__Ascomycota(100);c__Saccharomycetes(100);o__Saccharomycetales(100);f__Metschnikowiaceae(100);g__Metschnikowia(100);s__Metschnikowia_pulcherrima(100); | Metschnikowia | 13 | Saprotroph | \|Undefined Saprotroph\| | Yeast | NULL | Probable | As nectar-tap_saprotroph (Põlme et al. 2020); Decay sugar-rich substrates (Põlme et al. 2020) | Põlme S, et al. 2020. Fungal Diversity 105:1-16 ((https://doi.org/10.1007/s13225-020-00466-2)) |
| Otu43359 | 1 | k__Fungi(100);p__Ascomycota(100);c__Saccharomycetes(100);o__Saccharomycetales(100);f__Metschnikowiaceae(100);g__Metschnikowia(100);s__Metschnikowia_pulcherrima(100); | Metschnikowia | 13 | Saprotroph | \|Undefined Saprotroph\| | Yeast | NULL | Probable | As nectar-tap_saprotroph (Põlme et al. 2020); Decay sugar-rich substrates (Põlme et al. 2020) | Põlme S, et al. 2020. Fungal Diversity 105:1-16 ((https://doi.org/10.1007/s13225-020-00466-2)) |
| Otu43360 | 1 | k__Fungi(100);p__Ascomycota(100);c__Saccharomycetes(100);o__Saccharomycetales(100);f__Metschnikowiaceae(100);g__Metschnikowia(100);s__Metschnikowia_pulcherrima(100); | Metschnikowia | 13 | Saprotroph | \|Undefined Saprotroph\| | Yeast | NULL | Probable | As nectar-tap_saprotroph (Põlme et al. 2020); Decay sugar-rich substrates (Põlme et al. 2020) | Põlme S, et al. 2020. Fungal Diversity 105:1-16 ((https://doi.org/10.1007/s13225-020-00466-2)) |
| Otu43361 | 1 | k__Fungi(100);p__Ascomycota(100);c__Saccharomycetes(100);o__Saccharomycetales(100);f__Metschnikowiaceae(100);g__Metschnikowia(100);s__Metschnikowia_pulcherrima(100); | Metschnikowia | 13 | Saprotroph | \|Undefined Saprotroph\| | Yeast | NULL | Probable | As nectar-tap_saprotroph (Põlme et al. 2020); Decay sugar-rich substrates (Põlme et al. 2020) | Põlme S, et al. 2020. Fungal Diversity 105:1-16 ((https://doi.org/10.1007/s13225-020-00466-2)) |
| Otu43362 | 1 | k__Fungi(100);p__Ascomycota(100);c__Saccharomycetes(100);o__Saccharomycetales(100);f__Metschnikowiaceae(100);g__Metschnikowia(100);s__Metschnikowia_pulcherrima(100); | Metschnikowia | 13 | Saprotroph | \|Undefined Saprotroph\| | Yeast | NULL | Probable | As nectar-tap_saprotroph (Põlme et al. 2020); Decay sugar-rich substrates (Põlme et al. 2020) | Põlme S, et al. 2020. Fungal Diversity 105:1-16 ((https://doi.org/10.1007/s13225-020-00466-2)) |
| Otu43363 | 1 | k__Fungi(100);p__Ascomycota(100);c__Saccharomycetes(100);o__Saccharomycetales(100);f__Metschnikowiaceae(100);g__Metschnikowia(100);s__Metschnikowia_pulcherrima(100); | Metschnikowia | 13 | Saprotroph | \|Undefined Saprotroph\| | Yeast | NULL | Probable | As nectar-tap_saprotroph (Põlme et al. 2020); Decay sugar-rich substrates (Põlme et al. 2020) | Põlme S, et al. 2020. Fungal Diversity 105:1-16 ((https://doi.org/10.1007/s13225-020-00466-2)) |
| Otu43364 | 1 | k__Fungi(100);p__Ascomycota(100);c__Saccharomycetes(100);o__Saccharomycetales(100);f__Metschnikowiaceae(100);g__Metschnikowia(100);g__Metschnikowia_unclassified(100); | Metschnikowia | 13 | Saprotroph | \|Undefined Saprotroph\| | Yeast | NULL | Probable | As nectar-tap_saprotroph (Põlme et al. 2020); Decay sugar-rich substrates (Põlme et al. 2020) | Põlme S, et al. 2020. Fungal Diversity 105:1-16 ((https://doi.org/10.1007/s13225-020-00466-2)) |
| Otu43365 | 1 | k__Fungi(100);p__Ascomycota(100);c__Saccharomycetes(100);o__Saccharomycetales(100);f__Metschnikowiaceae(100);g__Metschnikowia(100);s__Metschnikowia_pulcherrima(100); | Metschnikowia | 13 | Saprotroph | \|Undefined Saprotroph\| | Yeast | NULL | Probable | As nectar-tap_saprotroph (Põlme et al. 2020); Decay sugar-rich substrates (Põlme et al. 2020) | Põlme S, et al. 2020. Fungal Diversity 105:1-16 ((https://doi.org/10.1007/s13225-020-00466-2)) |
| Otu43366 | 1 | k__Fungi(100);p__Ascomycota(100);c__Saccharomycetes(100);o__Saccharomycetales(100);f__Metschnikowiaceae(100);g__Metschnikowia(100);g__Metschnikowia_unclassified(100); | Metschnikowia | 13 | Saprotroph | \|Undefined Saprotroph\| | Yeast | NULL | Probable | As nectar-tap_saprotroph (Põlme et al. 2020); Decay sugar-rich substrates (Põlme et al. 2020) | Põlme S, et al. 2020. Fungal Diversity 105:1-16 ((https://doi.org/10.1007/s13225-020-00466-2)) |
| Otu43367 | 1 | k__Fungi(100);p__Ascomycota(100);c__Saccharomycetes(100);o__Saccharomycetales(100);f__Metschnikowiaceae(100);g__Metschnikowia(100);g__Metschnikowia_unclassified(100); | Metschnikowia | 13 | Saprotroph | \|Undefined Saprotroph\| | Yeast | NULL | Probable | As nectar-tap_saprotroph (Põlme et al. 2020); Decay sugar-rich substrates (Põlme et al. 2020) | Põlme S, et al. 2020. Fungal Diversity 105:1-16 ((https://doi.org/10.1007/s13225-020-00466-2)) |
| Otu43368 | 1 | k__Fungi(100);p__Ascomycota(100);c__Saccharomycetes(100);o__Saccharomycetales(100);f__Metschnikowiaceae(100);g__Metschnikowia(100);s__Metschnikowia_pulcherrima(100); | Metschnikowia | 13 | Saprotroph | \|Undefined Saprotroph\| | Yeast | NULL | Probable | As nectar-tap_saprotroph (Põlme et al. 2020); Decay sugar-rich substrates (Põlme et al. 2020) | Põlme S, et al. 2020. Fungal Diversity 105:1-16 ((https://doi.org/10.1007/s13225-020-00466-2)) |
| Otu43369 | 1 | k__Fungi(100);p__Ascomycota(100);c__Saccharomycetes(100);o__Saccharomycetales(100);f__Metschnikowiaceae(100);g__Metschnikowia(100);s__Metschnikowia_pulcherrima(100); | Metschnikowia | 13 | Saprotroph | \|Undefined Saprotroph\| | Yeast | NULL | Probable | As nectar-tap_saprotroph (Põlme et al. 2020); Decay sugar-rich substrates (Põlme et al. 2020) | Põlme S, et al. 2020. Fungal Diversity 105:1-16 ((https://doi.org/10.1007/s13225-020-00466-2)) |
| Otu43370 | 1 | k__Fungi(100);p__Ascomycota(100);c__Saccharomycetes(100);o__Saccharomycetales(100);f__Metschnikowiaceae(100);g__Metschnikowia(100);s__Metschnikowia_pulcherrima(100); | Metschnikowia | 13 | Saprotroph | \|Undefined Saprotroph\| | Yeast | NULL | Probable | As nectar-tap_saprotroph (Põlme et al. 2020); Decay sugar-rich substrates (Põlme et al. 2020) | Põlme S, et al. 2020. Fungal Diversity 105:1-16 ((https://doi.org/10.1007/s13225-020-00466-2)) |
| Otu43371 | 1 | k__Fungi(100);p__Ascomycota(100);c__Saccharomycetes(100);o__Saccharomycetales(100);f__Metschnikowiaceae(100);g__Metschnikowia(100);s__Metschnikowia_pulcherrima(100); | Metschnikowia | 13 | Saprotroph | \|Undefined Saprotroph\| | Yeast | NULL | Probable | As nectar-tap_saprotroph (Põlme et al. 2020); Decay sugar-rich substrates (Põlme et al. 2020) | Põlme S, et al. 2020. Fungal Diversity 105:1-16 ((https://doi.org/10.1007/s13225-020-00466-2)) |
| Otu43372 | 1 | k__Fungi(100);p__Ascomycota(100);c__Saccharomycetes(100);o__Saccharomycetales(100);f__Metschnikowiaceae(100);g__Metschnikowia(100);g__Metschnikowia_unclassified(100); | Metschnikowia | 13 | Saprotroph | \|Undefined Saprotroph\| | Yeast | NULL | Probable | As nectar-tap_saprotroph (Põlme et al. 2020); Decay sugar-rich substrates (Põlme et al. 2020) | Põlme S, et al. 2020. Fungal Diversity 105:1-16 ((https://doi.org/10.1007/s13225-020-00466-2)) |
| Otu43373 | 1 | k__Fungi(100);p__Ascomycota(100);c__Saccharomycetes(100);o__Saccharomycetales(100);f__Metschnikowiaceae(100);g__Metschnikowia(100);s__Metschnikowia_pulcherrima(100); | Metschnikowia | 13 | Saprotroph | \|Undefined Saprotroph\| | Yeast | NULL | Probable | As nectar-tap_saprotroph (Põlme et al. 2020); Decay sugar-rich substrates (Põlme et al. 2020) | Põlme S, et al. 2020. Fungal Diversity 105:1-16 ((https://doi.org/10.1007/s13225-020-00466-2)) |
| Otu43374 | 1 | k__Fungi(100);p__Ascomycota(100);c__Saccharomycetes(100);o__Saccharomycetales(100);f__Metschnikowiaceae(100);g__Metschnikowia(100);s__Metschnikowia_pulcherrima(100); | Metschnikowia | 13 | Saprotroph | \|Undefined Saprotroph\| | Yeast | NULL | Probable | As nectar-tap_saprotroph (Põlme et al. 2020); Decay sugar-rich substrates (Põlme et al. 2020) | Põlme S, et al. 2020. Fungal Diversity 105:1-16 ((https://doi.org/10.1007/s13225-020-00466-2)) |
| Otu43375 | 1 | k__Fungi(100);p__Ascomycota(100);c__Saccharomycetes(100);o__Saccharomycetales(100);f__Metschnikowiaceae(100);g__Metschnikowia(100);s__Metschnikowia_pulcherrima(100); | Metschnikowia | 13 | Saprotroph | \|Undefined Saprotroph\| | Yeast | NULL | Probable | As nectar-tap_saprotroph (Põlme et al. 2020); Decay sugar-rich substrates (Põlme et al. 2020) | Põlme S, et al. 2020. Fungal Diversity 105:1-16 ((https://doi.org/10.1007/s13225-020-00466-2)) |
| Otu43376 | 1 | k__Fungi(100);p__Ascomycota(100);c__Saccharomycetes(100);o__Saccharomycetales(100);f__Metschnikowiaceae(100);g__Metschnikowia(100);s__Metschnikowia_pulcherrima(100); | Metschnikowia | 13 | Saprotroph | \|Undefined Saprotroph\| | Yeast | NULL | Probable | As nectar-tap_saprotroph (Põlme et al. 2020); Decay sugar-rich substrates (Põlme et al. 2020) | Põlme S, et al. 2020. Fungal Diversity 105:1-16 ((https://doi.org/10.1007/s13225-020-00466-2)) |
| Otu43377 | 1 | k__Fungi(100);p__Ascomycota(100);c__Saccharomycetes(100);o__Saccharomycetales(100);f__Metschnikowiaceae(100);g__Metschnikowia(100);s__Metschnikowia_pulcherrima(100); | Metschnikowia | 13 | Saprotroph | \|Undefined Saprotroph\| | Yeast | NULL | Probable | As nectar-tap_saprotroph (Põlme et al. 2020); Decay sugar-rich substrates (Põlme et al. 2020) | Põlme S, et al. 2020. Fungal Diversity 105:1-16 ((https://doi.org/10.1007/s13225-020-00466-2)) |
| Otu43378 | 1 | k__Fungi(100);p__Ascomycota(100);c__Saccharomycetes(100);o__Saccharomycetales(100);f__Metschnikowiaceae(100);g__Metschnikowia(100);g__Metschnikowia_unclassified(100); | Metschnikowia | 13 | Saprotroph | \|Undefined Saprotroph\| | Yeast | NULL | Probable | As nectar-tap_saprotroph (Põlme et al. 2020); Decay sugar-rich substrates (Põlme et al. 2020) | Põlme S, et al. 2020. Fungal Diversity 105:1-16 ((https://doi.org/10.1007/s13225-020-00466-2)) |
| Otu43379 | 1 | k__Fungi(100);p__Ascomycota(100);c__Saccharomycetes(100);o__Saccharomycetales(100);f__Metschnikowiaceae(100);g__Metschnikowia(100);s__Metschnikowia_pulcherrima(100); | Metschnikowia | 13 | Saprotroph | \|Undefined Saprotroph\| | Yeast | NULL | Probable | As nectar-tap_saprotroph (Põlme et al. 2020); Decay sugar-rich substrates (Põlme et al. 2020) | Põlme S, et al. 2020. Fungal Diversity 105:1-16 ((https://doi.org/10.1007/s13225-020-00466-2)) |
| Otu43380 | 1 | k__Fungi(100);p__Ascomycota(100);c__Saccharomycetes(100);o__Saccharomycetales(100);f__Metschnikowiaceae(100);g__Metschnikowia(100);s__Metschnikowia_pulcherrima(100); | Metschnikowia | 13 | Saprotroph | \|Undefined Saprotroph\| | Yeast | NULL | Probable | As nectar-tap_saprotroph (Põlme et al. 2020); Decay sugar-rich substrates (Põlme et al. 2020) | Põlme S, et al. 2020. Fungal Diversity 105:1-16 ((https://doi.org/10.1007/s13225-020-00466-2)) |
| Otu43381 | 1 | k__Fungi(100);p__Ascomycota(100);c__Saccharomycetes(100);o__Saccharomycetales(100);f__Metschnikowiaceae(100);g__Metschnikowia(100);s__Metschnikowia_pulcherrima(100); | Metschnikowia | 13 | Saprotroph | \|Undefined Saprotroph\| | Yeast | NULL | Probable | As nectar-tap_saprotroph (Põlme et al. 2020); Decay sugar-rich substrates (Põlme et al. 2020) | Põlme S, et al. 2020. Fungal Diversity 105:1-16 ((https://doi.org/10.1007/s13225-020-00466-2)) |
| Otu43382 | 1 | k__Fungi(100);p__Ascomycota(100);c__Saccharomycetes(100);o__Saccharomycetales(100);f__Metschnikowiaceae(100);g__Metschnikowia(100);g__Metschnikowia_unclassified(100); | Metschnikowia | 13 | Saprotroph | \|Undefined Saprotroph\| | Yeast | NULL | Probable | As nectar-tap_saprotroph (Põlme et al. 2020); Decay sugar-rich substrates (Põlme et al. 2020) | Põlme S, et al. 2020. Fungal Diversity 105:1-16 ((https://doi.org/10.1007/s13225-020-00466-2)) |
| Otu43383 | 1 | k__Fungi(100);p__Ascomycota(100);c__Saccharomycetes(100);o__Saccharomycetales(100);f__Metschnikowiaceae(100);g__Metschnikowia(100);s__Metschnikowia_pulcherrima(100); | Metschnikowia | 13 | Saprotroph | \|Undefined Saprotroph\| | Yeast | NULL | Probable | As nectar-tap_saprotroph (Põlme et al. 2020); Decay sugar-rich substrates (Põlme et al. 2020) | Põlme S, et al. 2020. Fungal Diversity 105:1-16 ((https://doi.org/10.1007/s13225-020-00466-2)) |
| Otu43384 | 1 | k__Fungi(100);p__Ascomycota(100);c__Saccharomycetes(100);o__Saccharomycetales(100);f__Metschnikowiaceae(100);g__Metschnikowia(100);g__Metschnikowia_unclassified(100); | Metschnikowia | 13 | Saprotroph | \|Undefined Saprotroph\| | Yeast | NULL | Probable | As nectar-tap_saprotroph (Põlme et al. 2020); Decay sugar-rich substrates (Põlme et al. 2020) | Põlme S, et al. 2020. Fungal Diversity 105:1-16 ((https://doi.org/10.1007/s13225-020-00466-2)) |
| Otu43385 | 1 | k__Fungi(100);p__Ascomycota(100);c__Saccharomycetes(100);o__Saccharomycetales(100);f__Metschnikowiaceae(100);g__Metschnikowia(100);s__Metschnikowia_pulcherrima(100); | Metschnikowia | 13 | Saprotroph | \|Undefined Saprotroph\| | Yeast | NULL | Probable | As nectar-tap_saprotroph (Põlme et al. 2020); Decay sugar-rich substrates (Põlme et al. 2020) | Põlme S, et al. 2020. Fungal Diversity 105:1-16 ((https://doi.org/10.1007/s13225-020-00466-2)) |
| Otu43386 | 1 | k__Fungi(100);p__Ascomycota(100);c__Saccharomycetes(100);o__Saccharomycetales(100);f__Metschnikowiaceae(100);g__Metschnikowia(100);s__Metschnikowia_pulcherrima(100); | Metschnikowia | 13 | Saprotroph | \|Undefined Saprotroph\| | Yeast | NULL | Probable | As nectar-tap_saprotroph (Põlme et al. 2020); Decay sugar-rich substrates (Põlme et al. 2020) | Põlme S, et al. 2020. Fungal Diversity 105:1-16 ((https://doi.org/10.1007/s13225-020-00466-2)) |
| Otu43387 | 1 | k__Fungi(100);p__Ascomycota(100);c__Saccharomycetes(100);o__Saccharomycetales(100);f__Metschnikowiaceae(100);g__Metschnikowia(100);s__Metschnikowia_pulcherrima(100); | Metschnikowia | 13 | Saprotroph | \|Undefined Saprotroph\| | Yeast | NULL | Probable | As nectar-tap_saprotroph (Põlme et al. 2020); Decay sugar-rich substrates (Põlme et al. 2020) | Põlme S, et al. 2020. Fungal Diversity 105:1-16 ((https://doi.org/10.1007/s13225-020-00466-2)) |
| Otu43388 | 1 | k__Fungi(100);p__Ascomycota(100);c__Saccharomycetes(100);o__Saccharomycetales(100);f__Metschnikowiaceae(100);g__Metschnikowia(100);g__Metschnikowia_unclassified(100); | Metschnikowia | 13 | Saprotroph | \|Undefined Saprotroph\| | Yeast | NULL | Probable | As nectar-tap_saprotroph (Põlme et al. 2020); Decay sugar-rich substrates (Põlme et al. 2020) | Põlme S, et al. 2020. Fungal Diversity 105:1-16 ((https://doi.org/10.1007/s13225-020-00466-2)) |
| Otu43389 | 1 | k__Fungi(100);p__Ascomycota(100);c__Saccharomycetes(100);o__Saccharomycetales(100);f__Metschnikowiaceae(100);g__Metschnikowia(100);s__Metschnikowia_pulcherrima(100); | Metschnikowia | 13 | Saprotroph | \|Undefined Saprotroph\| | Yeast | NULL | Probable | As nectar-tap_saprotroph (Põlme et al. 2020); Decay sugar-rich substrates (Põlme et al. 2020) | Põlme S, et al. 2020. Fungal Diversity 105:1-16 ((https://doi.org/10.1007/s13225-020-00466-2)) |
| Otu43390 | 1 | k__Fungi(100);p__Ascomycota(100);c__Saccharomycetes(100);o__Saccharomycetales(100);f__Metschnikowiaceae(100);g__Metschnikowia(100);s__Metschnikowia_pulcherrima(100); | Metschnikowia | 13 | Saprotroph | \|Undefined Saprotroph\| | Yeast | NULL | Probable | As nectar-tap_saprotroph (Põlme et al. 2020); Decay sugar-rich substrates (Põlme et al. 2020) | Põlme S, et al. 2020. Fungal Diversity 105:1-16 ((https://doi.org/10.1007/s13225-020-00466-2)) |
| Otu43391 | 1 | k__Fungi(100);p__Ascomycota(100);c__Saccharomycetes(100);o__Saccharomycetales(100);f__Metschnikowiaceae(100);g__Metschnikowia(100);g__Metschnikowia_unclassified(100); | Metschnikowia | 13 | Saprotroph | \|Undefined Saprotroph\| | Yeast | NULL | Probable | As nectar-tap_saprotroph (Põlme et al. 2020); Decay sugar-rich substrates (Põlme et al. 2020) | Põlme S, et al. 2020. Fungal Diversity 105:1-16 ((https://doi.org/10.1007/s13225-020-00466-2)) |
| Otu43392 | 1 | k__Fungi(100);p__Ascomycota(100);c__Saccharomycetes(100);o__Saccharomycetales(100);f__Metschnikowiaceae(100);g__Metschnikowia(100);s__Metschnikowia_pulcherrima(100); | Metschnikowia | 13 | Saprotroph | \|Undefined Saprotroph\| | Yeast | NULL | Probable | As nectar-tap_saprotroph (Põlme et al. 2020); Decay sugar-rich substrates (Põlme et al. 2020) | Põlme S, et al. 2020. Fungal Diversity 105:1-16 ((https://doi.org/10.1007/s13225-020-00466-2)) |
| Otu43394 | 1 | k__Fungi(100);p__Zygomycota(100);c__Zygomycota_class_Incertae_sedis(100);o__Mortierellales(100);f__Mortierellaceae(100);g__Mortierella(100);g__Mortierella_unclassified(100); | Mortierella | 13 | Saprotroph-Symbiotroph | Endophyte-Plant Saprotroph-\|Undefined Saprotroph\| | Microfungus | NULL | Probable | As chitinolytic (Põlme et al. 2020); As soil_saprotroph (Põlme et al. 2020); Decay roots, soil (Põlme et al. 2020); Saprobic in soil and thus frequently isolated from soil and material contaminated with soil such as hay or rotten silage and spoil tips (Cannon & Kirk 2007) | Cannon PF, Kirk PM. 2007. Fungal Families of the World. CAB International, Cambridge (ISBN: 978-0851998275); Tedersoo L, et al. 2014. Science 346:e1256688 ((https://doi.org/10.1126/science.1256688)); Purahong W, et al. 2016. Molecular Ecology 25:4059-4074 ((https://doi.org/10.1111/mec.13739)); Põlme S, et al. 2020. Fungal Diversity 105:1-16 ((https://doi.org/10.1007/s13225-020-00466-2)) |
| Otu43395 | 1 | k__Fungi(100);p__Ascomycota(100);c__Saccharomycetes(100);o__Saccharomycetales(100);f__Metschnikowiaceae(100);g__Metschnikowia(100);s__Metschnikowia_pulcherrima(100); | Metschnikowia | 13 | Saprotroph | \|Undefined Saprotroph\| | Yeast | NULL | Probable | As nectar-tap_saprotroph (Põlme et al. 2020); Decay sugar-rich substrates (Põlme et al. 2020) | Põlme S, et al. 2020. Fungal Diversity 105:1-16 ((https://doi.org/10.1007/s13225-020-00466-2)) |
| Otu43396 | 1 | k__Fungi(100);p__Ascomycota(100);c__Saccharomycetes(100);o__Saccharomycetales(100);f__Metschnikowiaceae(100);g__Metschnikowia(100);s__Metschnikowia_pulcherrima(100); | Metschnikowia | 13 | Saprotroph | \|Undefined Saprotroph\| | Yeast | NULL | Probable | As nectar-tap_saprotroph (Põlme et al. 2020); Decay sugar-rich substrates (Põlme et al. 2020) | Põlme S, et al. 2020. Fungal Diversity 105:1-16 ((https://doi.org/10.1007/s13225-020-00466-2)) |
| Otu43397 | 1 | k__Fungi(100);p__Ascomycota(100);c__Saccharomycetes(100);o__Saccharomycetales(100);f__Metschnikowiaceae(100);g__Metschnikowia(100);s__Metschnikowia_pulcherrima(100); | Metschnikowia | 13 | Saprotroph | \|Undefined Saprotroph\| | Yeast | NULL | Probable | As nectar-tap_saprotroph (Põlme et al. 2020); Decay sugar-rich substrates (Põlme et al. 2020) | Põlme S, et al. 2020. Fungal Diversity 105:1-16 ((https://doi.org/10.1007/s13225-020-00466-2)) |
| Otu43398 | 1 | k__Fungi(100);p__Ascomycota(100);c__Saccharomycetes(100);o__Saccharomycetales(100);f__Metschnikowiaceae(100);g__Metschnikowia(100);s__Metschnikowia_pulcherrima(100); | Metschnikowia | 13 | Saprotroph | \|Undefined Saprotroph\| | Yeast | NULL | Probable | As nectar-tap_saprotroph (Põlme et al. 2020); Decay sugar-rich substrates (Põlme et al. 2020) | Põlme S, et al. 2020. Fungal Diversity 105:1-16 ((https://doi.org/10.1007/s13225-020-00466-2)) |
| Otu43399 | 1 | k__Fungi(100);p__Ascomycota(100);c__Saccharomycetes(100);o__Saccharomycetales(100);f__Metschnikowiaceae(100);g__Metschnikowia(100);g__Metschnikowia_unclassified(100); | Metschnikowia | 13 | Saprotroph | \|Undefined Saprotroph\| | Yeast | NULL | Probable | As nectar-tap_saprotroph (Põlme et al. 2020); Decay sugar-rich substrates (Põlme et al. 2020) | Põlme S, et al. 2020. Fungal Diversity 105:1-16 ((https://doi.org/10.1007/s13225-020-00466-2)) |
| Otu43400 | 1 | k__Fungi(100);p__Ascomycota(100);c__Saccharomycetes(100);o__Saccharomycetales(100);f__Metschnikowiaceae(100);g__Metschnikowia(100);s__Metschnikowia_pulcherrima(100); | Metschnikowia | 13 | Saprotroph | \|Undefined Saprotroph\| | Yeast | NULL | Probable | As nectar-tap_saprotroph (Põlme et al. 2020); Decay sugar-rich substrates (Põlme et al. 2020) | Põlme S, et al. 2020. Fungal Diversity 105:1-16 ((https://doi.org/10.1007/s13225-020-00466-2)) |
| Otu43401 | 1 | k__Fungi(100);p__Ascomycota(100);c__Saccharomycetes(100);o__Saccharomycetales(100);f__Metschnikowiaceae(100);g__Metschnikowia(100);g__Metschnikowia_unclassified(100); | Metschnikowia | 13 | Saprotroph | \|Undefined Saprotroph\| | Yeast | NULL | Probable | As nectar-tap_saprotroph (Põlme et al. 2020); Decay sugar-rich substrates (Põlme et al. 2020) | Põlme S, et al. 2020. Fungal Diversity 105:1-16 ((https://doi.org/10.1007/s13225-020-00466-2)) |
| Otu43402 | 1 | k__Fungi(100);p__Ascomycota(100);c__Saccharomycetes(100);o__Saccharomycetales(100);f__Metschnikowiaceae(100);g__Metschnikowia(100);s__Metschnikowia_pulcherrima(100); | Metschnikowia | 13 | Saprotroph | \|Undefined Saprotroph\| | Yeast | NULL | Probable | As nectar-tap_saprotroph (Põlme et al. 2020); Decay sugar-rich substrates (Põlme et al. 2020) | Põlme S, et al. 2020. Fungal Diversity 105:1-16 ((https://doi.org/10.1007/s13225-020-00466-2)) |
| Otu43403 | 1 | k__Fungi(100);p__Ascomycota(100);c__Saccharomycetes(100);o__Saccharomycetales(100);f__Metschnikowiaceae(100);g__Metschnikowia(100);g__Metschnikowia_unclassified(100); | Metschnikowia | 13 | Saprotroph | \|Undefined Saprotroph\| | Yeast | NULL | Probable | As nectar-tap_saprotroph (Põlme et al. 2020); Decay sugar-rich substrates (Põlme et al. 2020) | Põlme S, et al. 2020. Fungal Diversity 105:1-16 ((https://doi.org/10.1007/s13225-020-00466-2)) |
| Otu43404 | 1 | k__Fungi(100);p__Ascomycota(100);c__Saccharomycetes(100);o__Saccharomycetales(100);f__Metschnikowiaceae(100);g__Metschnikowia(100);s__Metschnikowia_pulcherrima(100); | Metschnikowia | 13 | Saprotroph | \|Undefined Saprotroph\| | Yeast | NULL | Probable | As nectar-tap_saprotroph (Põlme et al. 2020); Decay sugar-rich substrates (Põlme et al. 2020) | Põlme S, et al. 2020. Fungal Diversity 105:1-16 ((https://doi.org/10.1007/s13225-020-00466-2)) |
| Otu43405 | 1 | k__Fungi(100);p__Ascomycota(100);c__Saccharomycetes(100);o__Saccharomycetales(100);f__Metschnikowiaceae(100);g__Metschnikowia(100);s__Metschnikowia_pulcherrima(100); | Metschnikowia | 13 | Saprotroph | \|Undefined Saprotroph\| | Yeast | NULL | Probable | As nectar-tap_saprotroph (Põlme et al. 2020); Decay sugar-rich substrates (Põlme et al. 2020) | Põlme S, et al. 2020. Fungal Diversity 105:1-16 ((https://doi.org/10.1007/s13225-020-00466-2)) |
| Otu43406 | 1 | k__Fungi(100);p__Ascomycota(100);c__Saccharomycetes(100);o__Saccharomycetales(100);f__Metschnikowiaceae(100);g__Metschnikowia(100);s__Metschnikowia_pulcherrima(100); | Metschnikowia | 13 | Saprotroph | \|Undefined Saprotroph\| | Yeast | NULL | Probable | As nectar-tap_saprotroph (Põlme et al. 2020); Decay sugar-rich substrates (Põlme et al. 2020) | Põlme S, et al. 2020. Fungal Diversity 105:1-16 ((https://doi.org/10.1007/s13225-020-00466-2)) |
| Otu43407 | 1 | k__Fungi(100);p__Ascomycota(100);c__Saccharomycetes(100);o__Saccharomycetales(100);f__Metschnikowiaceae(100);g__Metschnikowia(100);s__Metschnikowia_pulcherrima(100); | Metschnikowia | 13 | Saprotroph | \|Undefined Saprotroph\| | Yeast | NULL | Probable | As nectar-tap_saprotroph (Põlme et al. 2020); Decay sugar-rich substrates (Põlme et al. 2020) | Põlme S, et al. 2020. Fungal Diversity 105:1-16 ((https://doi.org/10.1007/s13225-020-00466-2)) |
| Otu43408 | 1 | k__Fungi(100);p__Ascomycota(100);c__Saccharomycetes(100);o__Saccharomycetales(100);f__Metschnikowiaceae(100);g__Metschnikowia(100);s__Metschnikowia_pulcherrima(100); | Metschnikowia | 13 | Saprotroph | \|Undefined Saprotroph\| | Yeast | NULL | Probable | As nectar-tap_saprotroph (Põlme et al. 2020); Decay sugar-rich substrates (Põlme et al. 2020) | Põlme S, et al. 2020. Fungal Diversity 105:1-16 ((https://doi.org/10.1007/s13225-020-00466-2)) |
| Otu43409 | 1 | k__Fungi(100);p__Ascomycota(100);c__Saccharomycetes(100);o__Saccharomycetales(100);f__Metschnikowiaceae(100);g__Metschnikowia(100);s__Metschnikowia_pulcherrima(100); | Metschnikowia | 13 | Saprotroph | \|Undefined Saprotroph\| | Yeast | NULL | Probable | As nectar-tap_saprotroph (Põlme et al. 2020); Decay sugar-rich substrates (Põlme et al. 2020) | Põlme S, et al. 2020. Fungal Diversity 105:1-16 ((https://doi.org/10.1007/s13225-020-00466-2)) |
| Otu43411 | 1 | k__Fungi(100);p__Ascomycota(100);c__Saccharomycetes(100);o__Saccharomycetales(100);f__Metschnikowiaceae(100);g__Metschnikowia(100);s__Metschnikowia_pulcherrima(100); | Metschnikowia | 13 | Saprotroph | \|Undefined Saprotroph\| | Yeast | NULL | Probable | As nectar-tap_saprotroph (Põlme et al. 2020); Decay sugar-rich substrates (Põlme et al. 2020) | Põlme S, et al. 2020. Fungal Diversity 105:1-16 ((https://doi.org/10.1007/s13225-020-00466-2)) |
| Otu43412 | 1 | k__Fungi(100);p__Ascomycota(100);c__Saccharomycetes(100);o__Saccharomycetales(100);f__Metschnikowiaceae(100);g__Metschnikowia(100);s__Metschnikowia_pulcherrima(100); | Metschnikowia | 13 | Saprotroph | \|Undefined Saprotroph\| | Yeast | NULL | Probable | As nectar-tap_saprotroph (Põlme et al. 2020); Decay sugar-rich substrates (Põlme et al. 2020) | Põlme S, et al. 2020. Fungal Diversity 105:1-16 ((https://doi.org/10.1007/s13225-020-00466-2)) |
| Otu43413 | 1 | k__Fungi(100);p__Ascomycota(100);c__Saccharomycetes(100);o__Saccharomycetales(100);f__Metschnikowiaceae(100);g__Metschnikowia(100);s__Metschnikowia_pulcherrima(100); | Metschnikowia | 13 | Saprotroph | \|Undefined Saprotroph\| | Yeast | NULL | Probable | As nectar-tap_saprotroph (Põlme et al. 2020); Decay sugar-rich substrates (Põlme et al. 2020) | Põlme S, et al. 2020. Fungal Diversity 105:1-16 ((https://doi.org/10.1007/s13225-020-00466-2)) |
| Otu43414 | 1 | k__Fungi(100);p__Ascomycota(100);c__Saccharomycetes(100);o__Saccharomycetales(100);f__Metschnikowiaceae(100);g__Metschnikowia(100);g__Metschnikowia_unclassified(100); | Metschnikowia | 13 | Saprotroph | \|Undefined Saprotroph\| | Yeast | NULL | Probable | As nectar-tap_saprotroph (Põlme et al. 2020); Decay sugar-rich substrates (Põlme et al. 2020) | Põlme S, et al. 2020. Fungal Diversity 105:1-16 ((https://doi.org/10.1007/s13225-020-00466-2)) |
| Otu43415 | 1 | k__Fungi(100);p__Ascomycota(100);c__Saccharomycetes(100);o__Saccharomycetales(100);f__Metschnikowiaceae(100);g__Metschnikowia(100);s__Metschnikowia_pulcherrima(100); | Metschnikowia | 13 | Saprotroph | \|Undefined Saprotroph\| | Yeast | NULL | Probable | As nectar-tap_saprotroph (Põlme et al. 2020); Decay sugar-rich substrates (Põlme et al. 2020) | Põlme S, et al. 2020. Fungal Diversity 105:1-16 ((https://doi.org/10.1007/s13225-020-00466-2)) |
| Otu43416 | 1 | k__Fungi(100);p__Ascomycota(100);c__Saccharomycetes(100);o__Saccharomycetales(100);f__Metschnikowiaceae(100);g__Metschnikowia(100);s__Metschnikowia_pulcherrima(100); | Metschnikowia | 13 | Saprotroph | \|Undefined Saprotroph\| | Yeast | NULL | Probable | As nectar-tap_saprotroph (Põlme et al. 2020); Decay sugar-rich substrates (Põlme et al. 2020) | Põlme S, et al. 2020. Fungal Diversity 105:1-16 ((https://doi.org/10.1007/s13225-020-00466-2)) |
| Otu43417 | 1 | k__Fungi(100);p__Ascomycota(100);c__Saccharomycetes(100);o__Saccharomycetales(100);f__Metschnikowiaceae(100);g__Metschnikowia(100);s__Metschnikowia_pulcherrima(100); | Metschnikowia | 13 | Saprotroph | \|Undefined Saprotroph\| | Yeast | NULL | Probable | As nectar-tap_saprotroph (Põlme et al. 2020); Decay sugar-rich substrates (Põlme et al. 2020) | Põlme S, et al. 2020. Fungal Diversity 105:1-16 ((https://doi.org/10.1007/s13225-020-00466-2)) |
| Otu43418 | 1 | k__Fungi(100);p__Ascomycota(100);c__Saccharomycetes(100);o__Saccharomycetales(100);f__Metschnikowiaceae(100);g__Metschnikowia(100);s__Metschnikowia_pulcherrima(100); | Metschnikowia | 13 | Saprotroph | \|Undefined Saprotroph\| | Yeast | NULL | Probable | As nectar-tap_saprotroph (Põlme et al. 2020); Decay sugar-rich substrates (Põlme et al. 2020) | Põlme S, et al. 2020. Fungal Diversity 105:1-16 ((https://doi.org/10.1007/s13225-020-00466-2)) |
| Otu43419 | 1 | k__Fungi(100);p__Ascomycota(100);c__Saccharomycetes(100);o__Saccharomycetales(100);f__Metschnikowiaceae(100);g__Metschnikowia(100);s__Metschnikowia_pulcherrima(100); | Metschnikowia | 13 | Saprotroph | \|Undefined Saprotroph\| | Yeast | NULL | Probable | As nectar-tap_saprotroph (Põlme et al. 2020); Decay sugar-rich substrates (Põlme et al. 2020) | Põlme S, et al. 2020. Fungal Diversity 105:1-16 ((https://doi.org/10.1007/s13225-020-00466-2)) |
| Otu43420 | 1 | k__Fungi(100);p__Ascomycota(100);c__Saccharomycetes(100);o__Saccharomycetales(100);f__Metschnikowiaceae(100);g__Metschnikowia(100);g__Metschnikowia_unclassified(100); | Metschnikowia | 13 | Saprotroph | \|Undefined Saprotroph\| | Yeast | NULL | Probable | As nectar-tap_saprotroph (Põlme et al. 2020); Decay sugar-rich substrates (Põlme et al. 2020) | Põlme S, et al. 2020. Fungal Diversity 105:1-16 ((https://doi.org/10.1007/s13225-020-00466-2)) |
| Otu43421 | 1 | k__Fungi(100);p__Ascomycota(100);c__Saccharomycetes(100);o__Saccharomycetales(100);f__Metschnikowiaceae(100);g__Metschnikowia(100);g__Metschnikowia_unclassified(100); | Metschnikowia | 13 | Saprotroph | \|Undefined Saprotroph\| | Yeast | NULL | Probable | As nectar-tap_saprotroph (Põlme et al. 2020); Decay sugar-rich substrates (Põlme et al. 2020) | Põlme S, et al. 2020. Fungal Diversity 105:1-16 ((https://doi.org/10.1007/s13225-020-00466-2)) |
| Otu43422 | 1 | k__Fungi(100);p__Ascomycota(100);c__Saccharomycetes(100);o__Saccharomycetales(100);f__Metschnikowiaceae(100);g__Metschnikowia(100);s__Metschnikowia_pulcherrima(100); | Metschnikowia | 13 | Saprotroph | \|Undefined Saprotroph\| | Yeast | NULL | Probable | As nectar-tap_saprotroph (Põlme et al. 2020); Decay sugar-rich substrates (Põlme et al. 2020) | Põlme S, et al. 2020. Fungal Diversity 105:1-16 ((https://doi.org/10.1007/s13225-020-00466-2)) |
| Otu43423 | 1 | k__Fungi(100);p__Ascomycota(100);c__Saccharomycetes(100);o__Saccharomycetales(100);f__Metschnikowiaceae(100);g__Metschnikowia(100);g__Metschnikowia_unclassified(100); | Metschnikowia | 13 | Saprotroph | \|Undefined Saprotroph\| | Yeast | NULL | Probable | As nectar-tap_saprotroph (Põlme et al. 2020); Decay sugar-rich substrates (Põlme et al. 2020) | Põlme S, et al. 2020. Fungal Diversity 105:1-16 ((https://doi.org/10.1007/s13225-020-00466-2)) |
| Otu43424 | 1 | k__Fungi(100);p__Ascomycota(100);c__Saccharomycetes(100);o__Saccharomycetales(100);f__Metschnikowiaceae(100);g__Metschnikowia(100);s__Metschnikowia_pulcherrima(100); | Metschnikowia | 13 | Saprotroph | \|Undefined Saprotroph\| | Yeast | NULL | Probable | As nectar-tap_saprotroph (Põlme et al. 2020); Decay sugar-rich substrates (Põlme et al. 2020) | Põlme S, et al. 2020. Fungal Diversity 105:1-16 ((https://doi.org/10.1007/s13225-020-00466-2)) |
| Otu43425 | 1 | k__Fungi(100);p__Ascomycota(100);c__Saccharomycetes(100);o__Saccharomycetales(100);f__Metschnikowiaceae(100);g__Metschnikowia(100);s__Metschnikowia_pulcherrima(100); | Metschnikowia | 13 | Saprotroph | \|Undefined Saprotroph\| | Yeast | NULL | Probable | As nectar-tap_saprotroph (Põlme et al. 2020); Decay sugar-rich substrates (Põlme et al. 2020) | Põlme S, et al. 2020. Fungal Diversity 105:1-16 ((https://doi.org/10.1007/s13225-020-00466-2)) |
| Otu43426 | 1 | k__Fungi(100);p__Ascomycota(100);c__Saccharomycetes(100);o__Saccharomycetales(100);f__Metschnikowiaceae(100);g__Metschnikowia(100);s__Metschnikowia_pulcherrima(100); | Metschnikowia | 13 | Saprotroph | \|Undefined Saprotroph\| | Yeast | NULL | Probable | As nectar-tap_saprotroph (Põlme et al. 2020); Decay sugar-rich substrates (Põlme et al. 2020) | Põlme S, et al. 2020. Fungal Diversity 105:1-16 ((https://doi.org/10.1007/s13225-020-00466-2)) |
| Otu43427 | 1 | k__Fungi(100);p__Ascomycota(100);c__Saccharomycetes(100);o__Saccharomycetales(100);f__Metschnikowiaceae(100);g__Metschnikowia(100);s__Metschnikowia_pulcherrima(100); | Metschnikowia | 13 | Saprotroph | \|Undefined Saprotroph\| | Yeast | NULL | Probable | As nectar-tap_saprotroph (Põlme et al. 2020); Decay sugar-rich substrates (Põlme et al. 2020) | Põlme S, et al. 2020. Fungal Diversity 105:1-16 ((https://doi.org/10.1007/s13225-020-00466-2)) |
| Otu43428 | 1 | k__Fungi(100);p__Ascomycota(100);c__Saccharomycetes(100);o__Saccharomycetales(100);f__Metschnikowiaceae(100);g__Metschnikowia(100);g__Metschnikowia_unclassified(100); | Metschnikowia | 13 | Saprotroph | \|Undefined Saprotroph\| | Yeast | NULL | Probable | As nectar-tap_saprotroph (Põlme et al. 2020); Decay sugar-rich substrates (Põlme et al. 2020) | Põlme S, et al. 2020. Fungal Diversity 105:1-16 ((https://doi.org/10.1007/s13225-020-00466-2)) |
| Otu43429 | 1 | k__Fungi(100);p__Ascomycota(100);c__Saccharomycetes(100);o__Saccharomycetales(100);f__Metschnikowiaceae(100);g__Metschnikowia(100);s__Metschnikowia_pulcherrima(100); | Metschnikowia | 13 | Saprotroph | \|Undefined Saprotroph\| | Yeast | NULL | Probable | As nectar-tap_saprotroph (Põlme et al. 2020); Decay sugar-rich substrates (Põlme et al. 2020) | Põlme S, et al. 2020. Fungal Diversity 105:1-16 ((https://doi.org/10.1007/s13225-020-00466-2)) |
| Otu43430 | 1 | k__Fungi(100);p__Ascomycota(100);c__Saccharomycetes(100);o__Saccharomycetales(100);f__Metschnikowiaceae(100);g__Metschnikowia(100);g__Metschnikowia_unclassified(100); | Metschnikowia | 13 | Saprotroph | \|Undefined Saprotroph\| | Yeast | NULL | Probable | As nectar-tap_saprotroph (Põlme et al. 2020); Decay sugar-rich substrates (Põlme et al. 2020) | Põlme S, et al. 2020. Fungal Diversity 105:1-16 ((https://doi.org/10.1007/s13225-020-00466-2)) |
| Otu43431 | 1 | k__Fungi(100);p__Ascomycota(100);c__Saccharomycetes(100);o__Saccharomycetales(100);f__Metschnikowiaceae(100);g__Metschnikowia(100);s__Metschnikowia_pulcherrima(100); | Metschnikowia | 13 | Saprotroph | \|Undefined Saprotroph\| | Yeast | NULL | Probable | As nectar-tap_saprotroph (Põlme et al. 2020); Decay sugar-rich substrates (Põlme et al. 2020) | Põlme S, et al. 2020. Fungal Diversity 105:1-16 ((https://doi.org/10.1007/s13225-020-00466-2)) |
| Otu43432 | 1 | k__Fungi(100);p__Ascomycota(100);c__Saccharomycetes(100);o__Saccharomycetales(100);f__Metschnikowiaceae(100);g__Metschnikowia(100);s__Metschnikowia_pulcherrima(100); | Metschnikowia | 13 | Saprotroph | \|Undefined Saprotroph\| | Yeast | NULL | Probable | As nectar-tap_saprotroph (Põlme et al. 2020); Decay sugar-rich substrates (Põlme et al. 2020) | Põlme S, et al. 2020. Fungal Diversity 105:1-16 ((https://doi.org/10.1007/s13225-020-00466-2)) |
| Otu43433 | 1 | k__Fungi(100);p__Ascomycota(100);c__Saccharomycetes(100);o__Saccharomycetales(100);f__Metschnikowiaceae(100);g__Metschnikowia(100);s__Metschnikowia_pulcherrima(100); | Metschnikowia | 13 | Saprotroph | \|Undefined Saprotroph\| | Yeast | NULL | Probable | As nectar-tap_saprotroph (Põlme et al. 2020); Decay sugar-rich substrates (Põlme et al. 2020) | Põlme S, et al. 2020. Fungal Diversity 105:1-16 ((https://doi.org/10.1007/s13225-020-00466-2)) |
| Otu43434 | 1 | k__Fungi(100);p__Ascomycota(100);c__Saccharomycetes(100);o__Saccharomycetales(100);f__Metschnikowiaceae(100);g__Metschnikowia(100);g__Metschnikowia_unclassified(100); | Metschnikowia | 13 | Saprotroph | \|Undefined Saprotroph\| | Yeast | NULL | Probable | As nectar-tap_saprotroph (Põlme et al. 2020); Decay sugar-rich substrates (Põlme et al. 2020) | Põlme S, et al. 2020. Fungal Diversity 105:1-16 ((https://doi.org/10.1007/s13225-020-00466-2)) |
| Otu43435 | 1 | k__Fungi(100);p__Ascomycota(100);c__Saccharomycetes(100);o__Saccharomycetales(100);f__Metschnikowiaceae(100);g__Metschnikowia(100);s__Metschnikowia_pulcherrima(100); | Metschnikowia | 13 | Saprotroph | \|Undefined Saprotroph\| | Yeast | NULL | Probable | As nectar-tap_saprotroph (Põlme et al. 2020); Decay sugar-rich substrates (Põlme et al. 2020) | Põlme S, et al. 2020. Fungal Diversity 105:1-16 ((https://doi.org/10.1007/s13225-020-00466-2)) |
| Otu43436 | 1 | k__Fungi(100);p__Ascomycota(100);c__Saccharomycetes(100);o__Saccharomycetales(100);f__Metschnikowiaceae(100);g__Metschnikowia(100);s__Metschnikowia_pulcherrima(100); | Metschnikowia | 13 | Saprotroph | \|Undefined Saprotroph\| | Yeast | NULL | Probable | As nectar-tap_saprotroph (Põlme et al. 2020); Decay sugar-rich substrates (Põlme et al. 2020) | Põlme S, et al. 2020. Fungal Diversity 105:1-16 ((https://doi.org/10.1007/s13225-020-00466-2)) |
| Otu43437 | 1 | k__Fungi(100);p__Ascomycota(100);c__Saccharomycetes(100);o__Saccharomycetales(100);f__Metschnikowiaceae(100);g__Metschnikowia(100);s__Metschnikowia_pulcherrima(100); | Metschnikowia | 13 | Saprotroph | \|Undefined Saprotroph\| | Yeast | NULL | Probable | As nectar-tap_saprotroph (Põlme et al. 2020); Decay sugar-rich substrates (Põlme et al. 2020) | Põlme S, et al. 2020. Fungal Diversity 105:1-16 ((https://doi.org/10.1007/s13225-020-00466-2)) |
| Otu43438 | 1 | k__Fungi(100);p__Ascomycota(100);c__Saccharomycetes(100);o__Saccharomycetales(100);f__Metschnikowiaceae(100);g__Metschnikowia(100);s__Metschnikowia_pulcherrima(100); | Metschnikowia | 13 | Saprotroph | \|Undefined Saprotroph\| | Yeast | NULL | Probable | As nectar-tap_saprotroph (Põlme et al. 2020); Decay sugar-rich substrates (Põlme et al. 2020) | Põlme S, et al. 2020. Fungal Diversity 105:1-16 ((https://doi.org/10.1007/s13225-020-00466-2)) |
| Otu43439 | 1 | k__Fungi(100);p__Ascomycota(100);c__Saccharomycetes(100);o__Saccharomycetales(100);f__Metschnikowiaceae(100);g__Metschnikowia(100);s__Metschnikowia_pulcherrima(100); | Metschnikowia | 13 | Saprotroph | \|Undefined Saprotroph\| | Yeast | NULL | Probable | As nectar-tap_saprotroph (Põlme et al. 2020); Decay sugar-rich substrates (Põlme et al. 2020) | Põlme S, et al. 2020. Fungal Diversity 105:1-16 ((https://doi.org/10.1007/s13225-020-00466-2)) |
| Otu43440 | 1 | k__Fungi(100);p__Ascomycota(100);c__Saccharomycetes(100);o__Saccharomycetales(100);f__Metschnikowiaceae(100);g__Metschnikowia(100);g__Metschnikowia_unclassified(100); | Metschnikowia | 13 | Saprotroph | \|Undefined Saprotroph\| | Yeast | NULL | Probable | As nectar-tap_saprotroph (Põlme et al. 2020); Decay sugar-rich substrates (Põlme et al. 2020) | Põlme S, et al. 2020. Fungal Diversity 105:1-16 ((https://doi.org/10.1007/s13225-020-00466-2)) |
| Otu43441 | 1 | k__Fungi(100);p__Ascomycota(100);c__Saccharomycetes(100);o__Saccharomycetales(100);f__Metschnikowiaceae(100);g__Metschnikowia(100);s__Metschnikowia_pulcherrima(100); | Metschnikowia | 13 | Saprotroph | \|Undefined Saprotroph\| | Yeast | NULL | Probable | As nectar-tap_saprotroph (Põlme et al. 2020); Decay sugar-rich substrates (Põlme et al. 2020) | Põlme S, et al. 2020. Fungal Diversity 105:1-16 ((https://doi.org/10.1007/s13225-020-00466-2)) |
| Otu43442 | 1 | k__Fungi(100);p__Ascomycota(100);c__Saccharomycetes(100);o__Saccharomycetales(100);f__Metschnikowiaceae(100);g__Metschnikowia(100);s__Metschnikowia_pulcherrima(100); | Metschnikowia | 13 | Saprotroph | \|Undefined Saprotroph\| | Yeast | NULL | Probable | As nectar-tap_saprotroph (Põlme et al. 2020); Decay sugar-rich substrates (Põlme et al. 2020) | Põlme S, et al. 2020. Fungal Diversity 105:1-16 ((https://doi.org/10.1007/s13225-020-00466-2)) |

| | | | | | | | | | | | |
|---|---|---|---|---|---|---|---|---|---|---|---|
| Otu43443 | 1 | k__Fungi(100);p__Ascomycota(100);c__Saccharomycetes(100);o__Saccharomycetales(100);f__Metschnikowiaceae(100);g__Metschnikowia(100);s__Metschnikowia_pulcherrima(100); | Metschnikowia | 13 | Saprotroph | \|Undefined Saprotroph\| | Yeast | NULL | Probable | As nectar-tap_saprotroph (Põlme et al. 2020); Decay sugar-rich substrates (Põlme et al. 2020) | Põlme S, et al. 2020. Fungal Diversity 105:1-16 ((https://doi.org/10.1007/s13225-020-00466-2)) |
| Otu43444 | 1 | k__Fungi(100);p__Ascomycota(100);c__Saccharomycetes(100);o__Saccharomycetales(100);f__Metschnikowiaceae(100);g__Metschnikowia(100);s__Metschnikowia_pulcherrima(100); | Metschnikowia | 13 | Saprotroph | \|Undefined Saprotroph\| | Yeast | NULL | Probable | As nectar-tap_saprotroph (Põlme et al. 2020); Decay sugar-rich substrates (Põlme et al. 2020) | Põlme S, et al. 2020. Fungal Diversity 105:1-16 ((https://doi.org/10.1007/s13225-020-00466-2)) |
| Otu43445 | 1 | k__Fungi(100);p__Ascomycota(100);c__Saccharomycetes(100);o__Saccharomycetales(100);f__Metschnikowiaceae(100);g__Metschnikowia(100);g__Metschnikowia_unclassified(100); | Metschnikowia | 13 | Saprotroph | \|Undefined Saprotroph\| | Yeast | NULL | Probable | As nectar-tap_saprotroph (Põlme et al. 2020); Decay sugar-rich substrates (Põlme et al. 2020) | Põlme S, et al. 2020. Fungal Diversity 105:1-16 ((https://doi.org/10.1007/s13225-020-00466-2)) |
| Otu43446 | 1 | k__Fungi(100);p__Ascomycota(100);c__Saccharomycetes(100);o__Saccharomycetales(100);f__Metschnikowiaceae(100);g__Metschnikowia(100);s__Metschnikowia_pulcherrima(100); | Metschnikowia | 13 | Saprotroph | \|Undefined Saprotroph\| | Yeast | NULL | Probable | As nectar-tap_saprotroph (Põlme et al. 2020); Decay sugar-rich substrates (Põlme et al. 2020) | Põlme S, et al. 2020. Fungal Diversity 105:1-16 ((https://doi.org/10.1007/s13225-020-00466-2)) |
| Otu43447 | 1 | k__Fungi(100);p__Ascomycota(100);c__Saccharomycetes(100);o__Saccharomycetales(100);f__Metschnikowiaceae(100);g__Metschnikowia(100);s__Metschnikowia_pulcherrima(100); | Metschnikowia | 13 | Saprotroph | \|Undefined Saprotroph\| | Yeast | NULL | Probable | As nectar-tap_saprotroph (Põlme et al. 2020); Decay sugar-rich substrates (Põlme et al. 2020) | Põlme S, et al. 2020. Fungal Diversity 105:1-16 ((https://doi.org/10.1007/s13225-020-00466-2)) |
| Otu43448 | 1 | k__Fungi(100);p__Ascomycota(100);c__Saccharomycetes(100);o__Saccharomycetales(100);f__Metschnikowiaceae(100);g__Metschnikowia(100);s__Metschnikowia_pulcherrima(100); | Metschnikowia | 13 | Saprotroph | \|Undefined Saprotroph\| | Yeast | NULL | Probable | As nectar-tap_saprotroph (Põlme et al. 2020); Decay sugar-rich substrates (Põlme et al. 2020) | Põlme S, et al. 2020. Fungal Diversity 105:1-16 ((https://doi.org/10.1007/s13225-020-00466-2)) |
| Otu43449 | 1 | k__Fungi(100);p__Ascomycota(100);c__Saccharomycetes(100);o__Saccharomycetales(100);f__Metschnikowiaceae(100);g__Metschnikowia(100);s__Metschnikowia_pulcherrima(100); | Metschnikowia | 13 | Saprotroph | \|Undefined Saprotroph\| | Yeast | NULL | Probable | As nectar-tap_saprotroph (Põlme et al. 2020); Decay sugar-rich substrates (Põlme et al. 2020) | Põlme S, et al. 2020. Fungal Diversity 105:1-16 ((https://doi.org/10.1007/s13225-020-00466-2)) |
| Otu43450 | 1 | k__Fungi(100);p__Ascomycota(100);c__Saccharomycetes(100);o__Saccharomycetales(100);f__Metschnikowiaceae(100);g__Metschnikowia(100);g__Metschnikowia_unclassified(100); | Metschnikowia | 13 | Saprotroph | \|Undefined Saprotroph\| | Yeast | NULL | Probable | As nectar-tap_saprotroph (Põlme et al. 2020); Decay sugar-rich substrates (Põlme et al. 2020) | Põlme S, et al. 2020. Fungal Diversity 105:1-16 ((https://doi.org/10.1007/s13225-020-00466-2)) |
| Otu43451 | 1 | k__Fungi(100);p__Ascomycota(100);c__Saccharomycetes(100);o__Saccharomycetales(100);f__Metschnikowiaceae(100);g__Metschnikowia(100);s__Metschnikowia_pulcherrima(100); | Metschnikowia | 13 | Saprotroph | \|Undefined Saprotroph\| | Yeast | NULL | Probable | As nectar-tap_saprotroph (Põlme et al. 2020); Decay sugar-rich substrates (Põlme et al. 2020) | Põlme S, et al. 2020. Fungal Diversity 105:1-16 ((https://doi.org/10.1007/s13225-020-00466-2)) |
| Otu43452 | 1 | k__Fungi(100);p__Ascomycota(100);c__Saccharomycetes(100);o__Saccharomycetales(100);f__Metschnikowiaceae(100);g__Metschnikowia(100);g__Metschnikowia_unclassified(100); | Metschnikowia | 13 | Saprotroph | \|Undefined Saprotroph\| | Yeast | NULL | Probable | As nectar-tap_saprotroph (Põlme et al. 2020); Decay sugar-rich substrates (Põlme et al. 2020) | Põlme S, et al. 2020. Fungal Diversity 105:1-16 ((https://doi.org/10.1007/s13225-020-00466-2)) |
| Otu43453 | 1 | k__Fungi(100);p__Ascomycota(100);c__Saccharomycetes(100);o__Saccharomycetales(100);f__Metschnikowiaceae(100);g__Metschnikowia(100);s__Metschnikowia_pulcherrima(100); | Metschnikowia | 13 | Saprotroph | \|Undefined Saprotroph\| | Yeast | NULL | Probable | As nectar-tap_saprotroph (Põlme et al. 2020); Decay sugar-rich substrates (Põlme et al. 2020) | Põlme S, et al. 2020. Fungal Diversity 105:1-16 ((https://doi.org/10.1007/s13225-020-00466-2)) |
| Otu43454 | 1 | k__Fungi(100);p__Ascomycota(100);c__Saccharomycetes(100);o__Saccharomycetales(100);f__Metschnikowiaceae(100);g__Metschnikowia(100);s__Metschnikowia_pulcherrima(100); | Metschnikowia | 13 | Saprotroph | \|Undefined Saprotroph\| | Yeast | NULL | Probable | As nectar-tap_saprotroph (Põlme et al. 2020); Decay sugar-rich substrates (Põlme et al. 2020) | Põlme S, et al. 2020. Fungal Diversity 105:1-16 ((https://doi.org/10.1007/s13225-020-00466-2)) |
| Otu43455 | 1 | k__Fungi(100);p__Ascomycota(100);c__Saccharomycetes(100);o__Saccharomycetales(100);f__Metschnikowiaceae(100);g__Metschnikowia(100);g__Metschnikowia_unclassified(100); | Metschnikowia | 13 | Saprotroph | \|Undefined Saprotroph\| | Yeast | NULL | Probable | As nectar-tap_saprotroph (Põlme et al. 2020); Decay sugar-rich substrates (Põlme et al. 2020) | Põlme S, et al. 2020. Fungal Diversity 105:1-16 ((https://doi.org/10.1007/s13225-020-00466-2)) |
| Otu43456 | 1 | k__Fungi(100);p__Ascomycota(100);c__Saccharomycetes(100);o__Saccharomycetales(100);f__Metschnikowiaceae(100);g__Metschnikowia(100);s__Metschnikowia_pulcherrima(100); | Metschnikowia | 13 | Saprotroph | \|Undefined Saprotroph\| | Yeast | NULL | Probable | As nectar-tap_saprotroph (Põlme et al. 2020); Decay sugar-rich substrates (Põlme et al. 2020) | Põlme S, et al. 2020. Fungal Diversity 105:1-16 ((https://doi.org/10.1007/s13225-020-00466-2)) |
| Otu43457 | 1 | k__Fungi(100);p__Ascomycota(100);c__Saccharomycetes(100);o__Saccharomycetales(100);f__Metschnikowiaceae(100);g__Metschnikowia(100);s__Metschnikowia_pulcherrima(100); | Metschnikowia | 13 | Saprotroph | \|Undefined Saprotroph\| | Yeast | NULL | Probable | As nectar-tap_saprotroph (Põlme et al. 2020); Decay sugar-rich substrates (Põlme et al. 2020) | Põlme S, et al. 2020. Fungal Diversity 105:1-16 ((https://doi.org/10.1007/s13225-020-00466-2)) |
| Otu43458 | 1 | k__Fungi(100);p__Ascomycota(100);c__Saccharomycetes(100);o__Saccharomycetales(100);f__Metschnikowiaceae(100);g__Metschnikowia(100);g__Metschnikowia_unclassified(100); | Metschnikowia | 13 | Saprotroph | \|Undefined Saprotroph\| | Yeast | NULL | Probable | As nectar-tap_saprotroph (Põlme et al. 2020); Decay sugar-rich substrates (Põlme et al. 2020) | Põlme S, et al. 2020. Fungal Diversity 105:1-16 ((https://doi.org/10.1007/s13225-020-00466-2)) |
| Otu43459 | 1 | k__Fungi(100);p__Ascomycota(100);c__Saccharomycetes(100);o__Saccharomycetales(100);f__Metschnikowiaceae(100);g__Metschnikowia(100);s__Metschnikowia_pulcherrima(100); | Metschnikowia | 13 | Saprotroph | \|Undefined Saprotroph\| | Yeast | NULL | Probable | As nectar-tap_saprotroph (Põlme et al. 2020); Decay sugar-rich substrates (Põlme et al. 2020) | Põlme S, et al. 2020. Fungal Diversity 105:1-16 ((https://doi.org/10.1007/s13225-020-00466-2)) |
| Otu43460 | 1 | k__Fungi(100);p__Ascomycota(100);c__Saccharomycetes(100);o__Saccharomycetales(100);f__Metschnikowiaceae(100);g__Metschnikowia(100);s__Metschnikowia_pulcherrima(100); | Metschnikowia | 13 | Saprotroph | \|Undefined Saprotroph\| | Yeast | NULL | Probable | As nectar-tap_saprotroph (Põlme et al. 2020); Decay sugar-rich substrates (Põlme et al. 2020) | Põlme S, et al. 2020. Fungal Diversity 105:1-16 ((https://doi.org/10.1007/s13225-020-00466-2)) |
| Otu43461 | 1 | k__Fungi(100);p__Ascomycota(100);c__Saccharomycetes(100);o__Saccharomycetales(100);f__Metschnikowiaceae(100);g__Metschnikowia(100);g__Metschnikowia_unclassified(100); | Metschnikowia | 13 | Saprotroph | \|Undefined Saprotroph\| | Yeast | NULL | Probable | As nectar-tap_saprotroph (Põlme et al. 2020); Decay sugar-rich substrates (Põlme et al. 2020) | Põlme S, et al. 2020. Fungal Diversity 105:1-16 ((https://doi.org/10.1007/s13225-020-00466-2)) |
| Otu43462 | 1 | k__Fungi(100);p__Ascomycota(100);c__Saccharomycetes(100);o__Saccharomycetales(100);f__Metschnikowiaceae(100);g__Metschnikowia(100);s__Metschnikowia_pulcherrima(100); | Metschnikowia | 13 | Saprotroph | \|Undefined Saprotroph\| | Yeast | NULL | Probable | As nectar-tap_saprotroph (Põlme et al. 2020); Decay sugar-rich substrates (Põlme et al. 2020) | Põlme S, et al. 2020. Fungal Diversity 105:1-16 ((https://doi.org/10.1007/s13225-020-00466-2)) |
| Otu43463 | 1 | k__Fungi(100);p__Ascomycota(100);c__Saccharomycetes(100);o__Saccharomycetales(100);f__Metschnikowiaceae(100);g__Metschnikowia(100);g__Metschnikowia_unclassified(100); | Metschnikowia | 13 | Saprotroph | \|Undefined Saprotroph\| | Yeast | NULL | Probable | As nectar-tap_saprotroph (Põlme et al. 2020); Decay sugar-rich substrates (Põlme et al. 2020) | Põlme S, et al. 2020. Fungal Diversity 105:1-16 ((https://doi.org/10.1007/s13225-020-00466-2)) |
| Otu43464 | 1 | k__Fungi(100);p__Ascomycota(100);c__Saccharomycetes(100);o__Saccharomycetales(100);f__Metschnikowiaceae(100);g__Metschnikowia(100);s__Metschnikowia_pulcherrima(100); | Metschnikowia | 13 | Saprotroph | \|Undefined Saprotroph\| | Yeast | NULL | Probable | As nectar-tap_saprotroph (Põlme et al. 2020); Decay sugar-rich substrates (Põlme et al. 2020) | Põlme S, et al. 2020. Fungal Diversity 105:1-16 ((https://doi.org/10.1007/s13225-020-00466-2)) |
| Otu43465 | 1 | k__Fungi(100);p__Ascomycota(100);c__Saccharomycetes(100);o__Saccharomycetales(100);f__Metschnikowiaceae(100);g__Metschnikowia(100);s__Metschnikowia_pulcherrima(100); | Metschnikowia | 13 | Saprotroph | \|Undefined Saprotroph\| | Yeast | NULL | Probable | As nectar-tap_saprotroph (Põlme et al. 2020); Decay sugar-rich substrates (Põlme et al. 2020) | Põlme S, et al. 2020. Fungal Diversity 105:1-16 ((https://doi.org/10.1007/s13225-020-00466-2)) |
| Otu43466 | 1 | k__Fungi(100);p__Ascomycota(100);c__Saccharomycetes(100);o__Saccharomycetales(100);f__Metschnikowiaceae(100);g__Metschnikowia(100);s__Metschnikowia_pulcherrima(100); | Metschnikowia | 13 | Saprotroph | \|Undefined Saprotroph\| | Yeast | NULL | Probable | As nectar-tap_saprotroph (Põlme et al. 2020); Decay sugar-rich substrates (Põlme et al. 2020) | Põlme S, et al. 2020. Fungal Diversity 105:1-16 ((https://doi.org/10.1007/s13225-020-00466-2)) |
| Otu43467 | 1 | k__Fungi(100);p__Ascomycota(100);c__Saccharomycetes(100);o__Saccharomycetales(100);f__Metschnikowiaceae(100);g__Metschnikowia(100);s__Metschnikowia_pulcherrima(100); | Metschnikowia | 13 | Saprotroph | \|Undefined Saprotroph\| | Yeast | NULL | Probable | As nectar-tap_saprotroph (Põlme et al. 2020); Decay sugar-rich substrates (Põlme et al. 2020) | Põlme S, et al. 2020. Fungal Diversity 105:1-16 ((https://doi.org/10.1007/s13225-020-00466-2)) |
| Otu43468 | 1 | k__Fungi(100);p__Ascomycota(100);c__Saccharomycetes(100);o__Saccharomycetales(100);f__Metschnikowiaceae(100);g__Metschnikowia(100);s__Metschnikowia_pulcherrima(100); | Metschnikowia | 13 | Saprotroph | \|Undefined Saprotroph\| | Yeast | NULL | Probable | As nectar-tap_saprotroph (Põlme et al. 2020); Decay sugar-rich substrates (Põlme et al. 2020) | Põlme S, et al. 2020. Fungal Diversity 105:1-16 ((https://doi.org/10.1007/s13225-020-00466-2)) |
| Otu43469 | 1 | k__Fungi(100);p__Ascomycota(100);c__Saccharomycetes(100);o__Saccharomycetales(100);f__Metschnikowiaceae(100);g__Metschnikowia(100);s__Metschnikowia_pulcherrima(100); | Metschnikowia | 13 | Saprotroph | \|Undefined Saprotroph\| | Yeast | NULL | Probable | As nectar-tap_saprotroph (Põlme et al. 2020); Decay sugar-rich substrates (Põlme et al. 2020) | Põlme S, et al. 2020. Fungal Diversity 105:1-16 ((https://doi.org/10.1007/s13225-020-00466-2)) |
| Otu43470 | 1 | k__Fungi(100);p__Ascomycota(100);c__Saccharomycetes(100);o__Saccharomycetales(100);f__Metschnikowiaceae(100);g__Metschnikowia(100);g__Metschnikowia_unclassified(100); | Metschnikowia | 13 | Saprotroph | \|Undefined Saprotroph\| | Yeast | NULL | Probable | As nectar-tap_saprotroph (Põlme et al. 2020); Decay sugar-rich substrates (Põlme et al. 2020) | Põlme S, et al. 2020. Fungal Diversity 105:1-16 ((https://doi.org/10.1007/s13225-020-00466-2)) |
| Otu43471 | 1 | k__Fungi(100);p__Ascomycota(100);c__Saccharomycetes(100);o__Saccharomycetales(100);f__Metschnikowiaceae(100);g__Metschnikowia(100);s__Metschnikowia_pulcherrima(100); | Metschnikowia | 13 | Saprotroph | \|Undefined Saprotroph\| | Yeast | NULL | Probable | As nectar-tap_saprotroph (Põlme et al. 2020); Decay sugar-rich substrates (Põlme et al. 2020) | Põlme S, et al. 2020. Fungal Diversity 105:1-16 ((https://doi.org/10.1007/s13225-020-00466-2)) |
| Otu43472 | 1 | k__Fungi(100);p__Ascomycota(100);c__Saccharomycetes(100);o__Saccharomycetales(100);f__Metschnikowiaceae(100);g__Metschnikowia(100);s__Metschnikowia_pulcherrima(100); | Metschnikowia | 13 | Saprotroph | \|Undefined Saprotroph\| | Yeast | NULL | Probable | As nectar-tap_saprotroph (Põlme et al. 2020); Decay sugar-rich substrates (Põlme et al. 2020) | Põlme S, et al. 2020. Fungal Diversity 105:1-16 ((https://doi.org/10.1007/s13225-020-00466-2)) |
| Otu43473 | 1 | k__Fungi(100);p__Ascomycota(100);c__Saccharomycetes(100);o__Saccharomycetales(100);f__Metschnikowiaceae(100);g__Metschnikowia(100);g__Metschnikowia_unclassified(100); | Metschnikowia | 13 | Saprotroph | \|Undefined Saprotroph\| | Yeast | NULL | Probable | As nectar-tap_saprotroph (Põlme et al. 2020); Decay sugar-rich substrates (Põlme et al. 2020) | Põlme S, et al. 2020. Fungal Diversity 105:1-16 ((https://doi.org/10.1007/s13225-020-00466-2)) |
| Otu43474 | 1 | k__Fungi(100);p__Ascomycota(100);c__Saccharomycetes(100);o__Saccharomycetales(100);f__Metschnikowiaceae(100);g__Metschnikowia(100);s__Metschnikowia_pulcherrima(100); | Metschnikowia | 13 | Saprotroph | \|Undefined Saprotroph\| | Yeast | NULL | Probable | As nectar-tap_saprotroph (Põlme et al. 2020); Decay sugar-rich substrates (Põlme et al. 2020) | Põlme S, et al. 2020. Fungal Diversity 105:1-16 ((https://doi.org/10.1007/s13225-020-00466-2)) |
| Otu43475 | 1 | k__Fungi(100);p__Ascomycota(100);c__Saccharomycetes(100);o__Saccharomycetales(100);f__Metschnikowiaceae(100);g__Metschnikowia(100);s__Metschnikowia_pulcherrima(100); | Metschnikowia | 13 | Saprotroph | \|Undefined Saprotroph\| | Yeast | NULL | Probable | As nectar-tap_saprotroph (Põlme et al. 2020); Decay sugar-rich substrates (Põlme et al. 2020) | Põlme S, et al. 2020. Fungal Diversity 105:1-16 ((https://doi.org/10.1007/s13225-020-00466-2)) |
| Otu43476 | 1 | k__Fungi(100);p__Ascomycota(100);c__Saccharomycetes(100);o__Saccharomycetales(100);f__Metschnikowiaceae(100);g__Metschnikowia(100);s__Metschnikowia_pulcherrima(100); | Metschnikowia | 13 | Saprotroph | \|Undefined Saprotroph\| | Yeast | NULL | Probable | As nectar-tap_saprotroph (Põlme et al. 2020); Decay sugar-rich substrates (Põlme et al. 2020) | Põlme S, et al. 2020. Fungal Diversity 105:1-16 ((https://doi.org/10.1007/s13225-020-00466-2)) |
| Otu43477 | 1 | k__Fungi(100);p__Ascomycota(100);c__Saccharomycetes(100);o__Saccharomycetales(100);f__Metschnikowiaceae(100);g__Metschnikowia(100);s__Metschnikowia_pulcherrima(100); | Metschnikowia | 13 | Saprotroph | \|Undefined Saprotroph\| | Yeast | NULL | Probable | As nectar-tap_saprotroph (Põlme et al. 2020); Decay sugar-rich substrates (Põlme et al. 2020) | Põlme S, et al. 2020. Fungal Diversity 105:1-16 ((https://doi.org/10.1007/s13225-020-00466-2)) |
| Otu43478 | 1 | k__Fungi(100);p__Ascomycota(100);c__Saccharomycetes(100);o__Saccharomycetales(100);f__Metschnikowiaceae(100);g__Metschnikowia(100);s__Metschnikowia_pulcherrima(100); | Metschnikowia | 13 | Saprotroph | \|Undefined Saprotroph\| | Yeast | NULL | Probable | As nectar-tap_saprotroph (Põlme et al. 2020); Decay sugar-rich substrates (Põlme et al. 2020) | Põlme S, et al. 2020. Fungal Diversity 105:1-16 ((https://doi.org/10.1007/s13225-020-00466-2)) |
| Otu43479 | 1 | k__Fungi(100);p__Ascomycota(100);c__Saccharomycetes(100);o__Saccharomycetales(100);f__Metschnikowiaceae(100);g__Metschnikowia(100);s__Metschnikowia_pulcherrima(100); | Metschnikowia | 13 | Saprotroph | \|Undefined Saprotroph\| | Yeast | NULL | Probable | As nectar-tap_saprotroph (Põlme et al. 2020); Decay sugar-rich substrates (Põlme et al. 2020) | Põlme S, et al. 2020. Fungal Diversity 105:1-16 ((https://doi.org/10.1007/s13225-020-00466-2)) |
| Otu43480 | 1 | k__Fungi(100);p__Ascomycota(100);c__Saccharomycetes(100);o__Saccharomycetales(100);f__Metschnikowiaceae(100);g__Metschnikowia(100);s__Metschnikowia_pulcherrima(100); | Metschnikowia | 13 | Saprotroph | \|Undefined Saprotroph\| | Yeast | NULL | Probable | As nectar-tap_saprotroph (Põlme et al. 2020); Decay sugar-rich substrates (Põlme et al. 2020) | Põlme S, et al. 2020. Fungal Diversity 105:1-16 ((https://doi.org/10.1007/s13225-020-00466-2)) |
| Otu43481 | 1 | k__Fungi(100);p__Ascomycota(100);c__Saccharomycetes(100);o__Saccharomycetales(100);f__Metschnikowiaceae(100);g__Metschnikowia(100);s__Metschnikowia_pulcherrima(100); | Metschnikowia | 13 | Saprotroph | \|Undefined Saprotroph\| | Yeast | NULL | Probable | As nectar-tap_saprotroph (Põlme et al. 2020); Decay sugar-rich substrates (Põlme et al. 2020) | Põlme S, et al. 2020. Fungal Diversity 105:1-16 ((https://doi.org/10.1007/s13225-020-00466-2)) |
| Otu43482 | 1 | k__Fungi(100);p__Ascomycota(100);c__Saccharomycetes(100);o__Saccharomycetales(100);f__Metschnikowiaceae(100);g__Metschnikowia(100);g__Metschnikowia_unclassified(100); | Metschnikowia | 13 | Saprotroph | \|Undefined Saprotroph\| | Yeast | NULL | Probable | As nectar-tap_saprotroph (Põlme et al. 2020); Decay sugar-rich substrates (Põlme et al. 2020) | Põlme S, et al. 2020. Fungal Diversity 105:1-16 ((https://doi.org/10.1007/s13225-020-00466-2)) |
| Otu43483 | 1 | k__Fungi(100);p__Ascomycota(100);c__Saccharomycetes(100);o__Saccharomycetales(100);f__Metschnikowiaceae(100);g__Metschnikowia(100);g__Metschnikowia_unclassified(100); | Metschnikowia | 13 | Saprotroph | \|Undefined Saprotroph\| | Yeast | NULL | Probable | As nectar-tap_saprotroph (Põlme et al. 2020); Decay sugar-rich substrates (Põlme et al. 2020) | Põlme S, et al. 2020. Fungal Diversity 105:1-16 ((https://doi.org/10.1007/s13225-020-00466-2)) |
| Otu43484 | 1 | k__Fungi(100);p__Ascomycota(100);c__Saccharomycetes(100);o__Saccharomycetales(100);f__Metschnikowiaceae(100);g__Metschnikowia(100);s__Metschnikowia_pulcherrima(100); | Metschnikowia | 13 | Saprotroph | \|Undefined Saprotroph\| | Yeast | NULL | Probable | As nectar-tap_saprotroph (Põlme et al. 2020); Decay sugar-rich substrates (Põlme et al. 2020) | Põlme S, et al. 2020. Fungal Diversity 105:1-16 ((https://doi.org/10.1007/s13225-020-00466-2)) |
| Otu43485 | 1 | k__Fungi(100);p__Ascomycota(100);c__Saccharomycetes(100);o__Saccharomycetales(100);f__Metschnikowiaceae(100);g__Metschnikowia(100);s__Metschnikowia_pulcherrima(100); | Metschnikowia | 13 | Saprotroph | \|Undefined Saprotroph\| | Yeast | NULL | Probable | As nectar-tap_saprotroph (Põlme et al. 2020); Decay sugar-rich substrates (Põlme et al. 2020) | Põlme S, et al. 2020. Fungal Diversity 105:1-16 ((https://doi.org/10.1007/s13225-020-00466-2)) |
| Otu43486 | 1 | k__Fungi(100);p__Ascomycota(100);c__Saccharomycetes(100);o__Saccharomycetales(100);f__Metschnikowiaceae(100);g__Metschnikowia(100);s__Metschnikowia_pulcherrima(100); | Metschnikowia | 13 | Saprotroph | \|Undefined Saprotroph\| | Yeast | NULL | Probable | As nectar-tap_saprotroph (Põlme et al. 2020); Decay sugar-rich substrates (Põlme et al. 2020) | Põlme S, et al. 2020. Fungal Diversity 105:1-16 ((https://doi.org/10.1007/s13225-020-00466-2)) |
| Otu43487 | 1 | k__Fungi(100);p__Ascomycota(100);c__Saccharomycetes(100);o__Saccharomycetales(100);f__Metschnikowiaceae(100);g__Metschnikowia(100);s__Metschnikowia_pulcherrima(100); | Metschnikowia | 13 | Saprotroph | \|Undefined Saprotroph\| | Yeast | NULL | Probable | As nectar-tap_saprotroph (Põlme et al. 2020); Decay sugar-rich substrates (Põlme et al. 2020) | Põlme S, et al. 2020. Fungal Diversity 105:1-16 ((https://doi.org/10.1007/s13225-020-00466-2)) |
| Otu43488 | 1 | k__Fungi(100);p__Ascomycota(100);c__Saccharomycetes(100);o__Saccharomycetales(100);f__Metschnikowiaceae(100);g__Metschnikowia(100);s__Metschnikowia_pulcherrima(100); | Metschnikowia | 13 | Saprotroph | \|Undefined Saprotroph\| | Yeast | NULL | Probable | As nectar-tap_saprotroph (Põlme et al. 2020); Decay sugar-rich substrates (Põlme et al. 2020) | Põlme S, et al. 2020. Fungal Diversity 105:1-16 ((https://doi.org/10.1007/s13225-020-00466-2)) |
| Otu43489 | 1 | k__Fungi(100);p__Ascomycota(100);c__Saccharomycetes(100);o__Saccharomycetales(100);f__Metschnikowiaceae(100);g__Metschnikowia(100);s__Metschnikowia_pulcherrima(100); | Metschnikowia | 13 | Saprotroph | \|Undefined Saprotroph\| | Yeast | NULL | Probable | As nectar-tap_saprotroph (Põlme et al. 2020); Decay sugar-rich substrates (Põlme et al. 2020) | Põlme S, et al. 2020. Fungal Diversity 105:1-16 ((https://doi.org/10.1007/s13225-020-00466-2)) |
| Otu43490 | 1 | k__Fungi(100);p__Ascomycota(100);c__Saccharomycetes(100);o__Saccharomycetales(100);f__Metschnikowiaceae(100);g__Metschnikowia(100);g__Metschnikowia_unclassified(100); | Metschnikowia | 13 | Saprotroph | \|Undefined Saprotroph\| | Yeast | NULL | Probable | As nectar-tap_saprotroph (Põlme et al. 2020); Decay sugar-rich substrates (Põlme et al. 2020) | Põlme S, et al. 2020. Fungal Diversity 105:1-16 ((https://doi.org/10.1007/s13225-020-00466-2)) |
| Otu43491 | 1 | k__Fungi(100);p__Ascomycota(100);c__Saccharomycetes(100);o__Saccharomycetales(100);f__Metschnikowiaceae(100);g__Metschnikowia(100);g__Metschnikowia_unclassified(100); | Metschnikowia | 13 | Saprotroph | \|Undefined Saprotroph\| | Yeast | NULL | Probable | As nectar-tap_saprotroph (Põlme et al. 2020); Decay sugar-rich substrates (Põlme et al. 2020) | Põlme S, et al. 2020. Fungal Diversity 105:1-16 ((https://doi.org/10.1007/s13225-020-00466-2)) |
| Otu43492 | 1 | k__Fungi(100);p__Ascomycota(100);c__Saccharomycetes(100);o__Saccharomycetales(100);f__Metschnikowiaceae(100);g__Metschnikowia(100);s__Metschnikowia_pulcherrima(100); | Metschnikowia | 13 | Saprotroph | \|Undefined Saprotroph\| | Yeast | NULL | Probable | As nectar-tap_saprotroph (Põlme et al. 2020); Decay sugar-rich substrates (Põlme et al. 2020) | Põlme S, et al. 2020. Fungal Diversity 105:1-16 ((https://doi.org/10.1007/s13225-020-00466-2)) |
| Otu43493 | 1 | k__Fungi(100);p__Ascomycota(100);c__Saccharomycetes(100);o__Saccharomycetales(100);f__Metschnikowiaceae(100);g__Metschnikowia(100);g__Metschnikowia_unclassified(100); | Metschnikowia | 13 | Saprotroph | \|Undefined Saprotroph\| | Yeast | NULL | Probable | As nectar-tap_saprotroph (Põlme et al. 2020); Decay sugar-rich substrates (Põlme et al. 2020) | Põlme S, et al. 2020. Fungal Diversity 105:1-16 ((https://doi.org/10.1007/s13225-020-00466-2)) |
| Otu43494 | 1 | k__Fungi(100);p__Ascomycota(100);c__Saccharomycetes(100);o__Saccharomycetales(100);f__Metschnikowiaceae(100);g__Metschnikowia(100);s__Metschnikowia_pulcherrima(100); | Metschnikowia | 13 | Saprotroph | \|Undefined Saprotroph\| | Yeast | NULL | Probable | As nectar-tap_saprotroph (Põlme et al. 2020); Decay sugar-rich substrates (Põlme et al. 2020) | Põlme S, et al. 2020. Fungal Diversity 105:1-16 ((https://doi.org/10.1007/s13225-020-00466-2)) |
| Otu43495 | 1 | k__Fungi(100);p__Ascomycota(100);c__Saccharomycetes(100);o__Saccharomycetales(100);f__Metschnikowiaceae(100);g__Metschnikowia(100);s__Metschnikowia_pulcherrima(100); | Metschnikowia | 13 | Saprotroph | \|Undefined Saprotroph\| | Yeast | NULL | Probable | As nectar-tap_saprotroph (Põlme et al. 2020); Decay sugar-rich substrates (Põlme et al. 2020) | Põlme S, et al. 2020. Fungal Diversity 105:1-16 ((https://doi.org/10.1007/s13225-020-00466-2)) |
| Otu43496 | 1 | k__Fungi(100);p__Ascomycota(100);c__Saccharomycetes(100);o__Saccharomycetales(100);f__Metschnikowiaceae(100);g__Metschnikowia(100);s__Metschnikowia_pulcherrima(100); | Metschnikowia | 13 | Saprotroph | \|Undefined Saprotroph\| | Yeast | NULL | Probable | As nectar-tap_saprotroph (Põlme et al. 2020); Decay sugar-rich substrates (Põlme et al. 2020) | Põlme S, et al. 2020. Fungal Diversity 105:1-16 ((https://doi.org/10.1007/s13225-020-00466-2)) |
| Otu43497 | 1 | k__Fungi(100);p__Ascomycota(100);c__Saccharomycetes(100);o__Saccharomycetales(100);f__Metschnikowiaceae(100);g__Metschnikowia(100);s__Metschnikowia_pulcherrima(100); | Metschnikowia | 13 | Saprotroph | \|Undefined Saprotroph\| | Yeast | NULL | Probable | As nectar-tap_saprotroph (Põlme et al. 2020); Decay sugar-rich substrates (Põlme et al. 2020) | Põlme S, et al. 2020. Fungal Diversity 105:1-16 ((https://doi.org/10.1007/s13225-020-00466-2)) |
| Otu43498 | 1 | k__Fungi(100);p__Ascomycota(100);c__Saccharomycetes(100);o__Saccharomycetales(100);f__Metschnikowiaceae(100);g__Metschnikowia(100);s__Metschnikowia_pulcherrima(100); | Metschnikowia | 13 | Saprotroph | \|Undefined Saprotroph\| | Yeast | NULL | Probable | As nectar-tap_saprotroph (Põlme et al. 2020); Decay sugar-rich substrates (Põlme et al. 2020) | Põlme S, et al. 2020. Fungal Diversity 105:1-16 ((https://doi.org/10.1007/s13225-020-00466-2)) |
| Otu43499 | 1 | k__Fungi(100);p__Ascomycota(100);c__Saccharomycetes(100);o__Saccharomycetales(100);f__Metschnikowiaceae(100);g__Metschnikowia(100);s__Metschnikowia_pulcherrima(100); | Metschnikowia | 13 | Saprotroph | \|Undefined Saprotroph\| | Yeast | NULL | Probable | As nectar-tap_saprotroph (Põlme et al. 2020); Decay sugar-rich substrates (Põlme et al. 2020) | Põlme S, et al. 2020. Fungal Diversity 105:1-16 ((https://doi.org/10.1007/s13225-020-00466-2)) |
| Otu43500 | 1 | k__Fungi(100);p__Ascomycota(100);c__Saccharomycetes(100);o__Saccharomycetales(100);f__Metschnikowiaceae(100);g__Metschnikowia(100);s__Metschnikowia_pulcherrima(100); | Metschnikowia | 13 | Saprotroph | \|Undefined Saprotroph\| | Yeast | NULL | Probable | As nectar-tap_saprotroph (Põlme et al. 2020); Decay sugar-rich substrates (Põlme et al. 2020) | Põlme S, et al. 2020. Fungal Diversity 105:1-16 ((https://doi.org/10.1007/s13225-020-00466-2)) |
| Otu43501 | 1 | k__Fungi(100);p__Ascomycota(100);c__Saccharomycetes(100);o__Saccharomycetales(100);f__Metschnikowiaceae(100);g__Metschnikowia(100);g__Metschnikowia_unclassified(100); | Metschnikowia | 13 | Saprotroph | \|Undefined Saprotroph\| | Yeast | NULL | Probable | As nectar-tap_saprotroph (Põlme et al. 2020); Decay sugar-rich substrates (Põlme et al. 2020) | Põlme S, et al. 2020. Fungal Diversity 105:1-16 ((https://doi.org/10.1007/s13225-020-00466-2)) |
| Otu43502 | 1 | k__Fungi(100);p__Ascomycota(100);c__Saccharomycetes(100);o__Saccharomycetales(100);f__Metschnikowiaceae(100);g__Metschnikowia(100);s__Metschnikowia_pulcherrima(100); | Metschnikowia | 13 | Saprotroph | \|Undefined Saprotroph\| | Yeast | NULL | Probable | As nectar-tap_saprotroph (Põlme et al. 2020); Decay sugar-rich substrates (Põlme et al. 2020) | Põlme S, et al. 2020. Fungal Diversity 105:1-16 ((https://doi.org/10.1007/s13225-020-00466-2)) |
| Otu43503 | 1 | k__Fungi(100);p__Ascomycota(100);c__Saccharomycetes(100);o__Saccharomycetales(100);f__Metschnikowiaceae(100);g__Metschnikowia(100);s__Metschnikowia_pulcherrima(100); | Metschnikowia | 13 | Saprotroph | \|Undefined Saprotroph\| | Yeast | NULL | Probable | As nectar-tap_saprotroph (Põlme et al. 2020); Decay sugar-rich substrates (Põlme et al. 2020) | Põlme S, et al. 2020. Fungal Diversity 105:1-16 ((https://doi.org/10.1007/s13225-020-00466-2)) |
| Otu43504 | 1 | k__Fungi(100);p__Ascomycota(100);c__Saccharomycetes(100);o__Saccharomycetales(100);f__Metschnikowiaceae(100);g__Metschnikowia(100);g__Metschnikowia_unclassified(100); | Metschnikowia | 13 | Saprotroph | \|Undefined Saprotroph\| | Yeast | NULL | Probable | As nectar-tap_saprotroph (Põlme et al. 2020); Decay sugar-rich substrates (Põlme et al. 2020) | Põlme S, et al. 2020. Fungal Diversity 105:1-16 ((https://doi.org/10.1007/s13225-020-00466-2)) |
| Otu43506 | 1 | k__Fungi(100);p__Ascomycota(100);c__Saccharomycetes(100);o__Saccharomycetales(100);f__Metschnikowiaceae(100);g__Metschnikowia(100);s__Metschnikowia_pulcherrima(100); | Metschnikowia | 13 | Saprotroph | \|Undefined Saprotroph\| | Yeast | NULL | Probable | As nectar-tap_saprotroph (Põlme et al. 2020); Decay sugar-rich substrates (Põlme et al. 2020) | Põlme S, et al. 2020. Fungal Diversity 105:1-16 ((https://doi.org/10.1007/s13225-020-00466-2)) |
| Otu43507 | 1 | k__Fungi(100);p__Ascomycota(100);c__Saccharomycetes(100);o__Saccharomycetales(100);f__Metschnikowiaceae(100);g__Metschnikowia(100);s__Metschnikowia_pulcherrima(100); | Metschnikowia | 13 | Saprotroph | \|Undefined Saprotroph\| | Yeast | NULL | Probable | As nectar-tap_saprotroph (Põlme et al. 2020); Decay sugar-rich substrates (Põlme et al. 2020) | Põlme S, et al. 2020. Fungal Diversity 105:1-16 ((https://doi.org/10.1007/s13225-020-00466-2)) |
| Otu43508 | 1 | k__Fungi(100);p__Ascomycota(100);c__Saccharomycetes(100);o__Saccharomycetales(100);f__Metschnikowiaceae(100);g__Metschnikowia(100);s__Metschnikowia_pulcherrima(100); | Metschnikowia | 13 | Saprotroph | \|Undefined Saprotroph\| | Yeast | NULL | Probable | As nectar-tap_saprotroph (Põlme et al. 2020); Decay sugar-rich substrates (Põlme et al. 2020) | Põlme S, et al. 2020. Fungal Diversity 105:1-16 ((https://doi.org/10.1007/s13225-020-00466-2)) |
| Otu43509 | 1 | k__Fungi(100);p__Ascomycota(100);c__Saccharomycetes(100);o__Saccharomycetales(100);f__Metschnikowiaceae(100);g__Metschnikowia(100);s__Metschnikowia_pulcherrima(100); | Metschnikowia | 13 | Saprotroph | \|Undefined Saprotroph\| | Yeast | NULL | Probable | As nectar-tap_saprotroph (Põlme et al. 2020); Decay sugar-rich substrates (Põlme et al. 2020) | Põlme S, et al. 2020. Fungal Diversity 105:1-16 ((https://doi.org/10.1007/s13225-020-00466-2)) |
| Otu43510 | 1 | k__Fungi(100);p__Ascomycota(100);c__Saccharomycetes(100);o__Saccharomycetales(100);f__Metschnikowiaceae(100);g__Metschnikowia(100);g__Metschnikowia_unclassified(100); | Metschnikowia | 13 | Saprotroph | \|Undefined Saprotroph\| | Yeast | NULL | Probable | As nectar-tap_saprotroph (Põlme et al. 2020); Decay sugar-rich substrates (Põlme et al. 2020) | Põlme S, et al. 2020. Fungal Diversity 105:1-16 ((https://doi.org/10.1007/s13225-020-00466-2)) |
| Otu43511 | 1 | k__Fungi(100);p__Ascomycota(100);c__Saccharomycetes(100);o__Saccharomycetales(100);f__Metschnikowiaceae(100);g__Metschnikowia(100);s__Metschnikowia_pulcherrima(100); | Metschnikowia | 13 | Saprotroph | \|Undefined Saprotroph\| | Yeast | NULL | Probable | As nectar-tap_saprotroph (Põlme et al. 2020); Decay sugar-rich substrates (Põlme et al. 2020) | Põlme S, et al. 2020. Fungal Diversity 105:1-16 ((https://doi.org/10.1007/s13225-020-00466-2)) |
| Otu43512 | 1 | k__Fungi(100);p__Ascomycota(100);c__Saccharomycetes(100);o__Saccharomycetales(100);f__Metschnikowiaceae(100);g__Metschnikowia(100);s__Metschnikowia_pulcherrima(100); | Metschnikowia | 13 | Saprotroph | \|Undefined Saprotroph\| | Yeast | NULL | Probable | As nectar-tap_saprotroph (Põlme et al. 2020); Decay sugar-rich substrates (Põlme et al. 2020) | Põlme S, et al. 2020. Fungal Diversity 105:1-16 ((https://doi.org/10.1007/s13225-020-00466-2)) |
| Otu43513 | 1 | k__Fungi(100);p__Ascomycota(100);c__Saccharomycetes(100);o__Saccharomycetales(100);f__Metschnikowiaceae(100);g__Metschnikowia(100);s__Metschnikowia_pulcherrima(100); | Metschnikowia | 13 | Saprotroph | \|Undefined Saprotroph\| | Yeast | NULL | Probable | As nectar-tap_saprotroph (Põlme et al. 2020); Decay sugar-rich substrates (Põlme et al. 2020) | Põlme S, et al. 2020. Fungal Diversity 105:1-16 ((https://doi.org/10.1007/s13225-020-00466-2)) |
| Otu43514 | 1 | k__Fungi(100);p__Ascomycota(100);c__Saccharomycetes(100);o__Saccharomycetales(100);f__Metschnikowiaceae(100);g__Metschnikowia(100);s__Metschnikowia_pulcherrima(100); | Metschnikowia | 13 | Saprotroph | \|Undefined Saprotroph\| | Yeast | NULL | Probable | As nectar-tap_saprotroph (Põlme et al. 2020); Decay sugar-rich substrates (Põlme et al. 2020) | Põlme S, et al. 2020. Fungal Diversity 105:1-16 ((https://doi.org/10.1007/s13225-020-00466-2)) |
| Otu43515 | 1 | k__Fungi(100);p__Ascomycota(100);c__Saccharomycetes(100);o__Saccharomycetales(100);f__Metschnikowiaceae(100);g__Metschnikowia(100);s__Metschnikowia_pulcherrima(100); | Metschnikowia | 13 | Saprotroph | \|Undefined Saprotroph\| | Yeast | NULL | Probable | As nectar-tap_saprotroph (Põlme et al. 2020); Decay sugar-rich substrates (Põlme et al. 2020) | Põlme S, et al. 2020. Fungal Diversity 105:1-16 ((https://doi.org/10.1007/s13225-020-00466-2)) |
| Otu43516 | 1 | k__Fungi(100);p__Ascomycota(100);c__Saccharomycetes(100);o__Saccharomycetales(100);f__Metschnikowiaceae(100);g__Metschnikowia(100);s__Metschnikowia_pulcherrima(100); | Metschnikowia | 13 | Saprotroph | \|Undefined Saprotroph\| | Yeast | NULL | Probable | As nectar-tap_saprotroph (Põlme et al. 2020); Decay sugar-rich substrates (Põlme et al. 2020) | Põlme S, et al. 2020. Fungal Diversity 105:1-16 ((https://doi.org/10.1007/s13225-020-00466-2)) |
| Otu43517 | 1 | k__Fungi(100);p__Ascomycota(100);c__Saccharomycetes(100);o__Saccharomycetales(100);f__Metschnikowiaceae(100);g__Metschnikowia(100);s__Metschnikowia_pulcherrima(100); | Metschnikowia | 13 | Saprotroph | \|Undefined Saprotroph\| | Yeast | NULL | Probable | As nectar-tap_saprotroph (Põlme et al. 2020); Decay sugar-rich substrates (Põlme et al. 2020) | Põlme S, et al. 2020. Fungal Diversity 105:1-16 ((https://doi.org/10.1007/s13225-020-00466-2)) |
| Otu43518 | 1 | k__Fungi(100);p__Ascomycota(100);c__Saccharomycetes(100);o__Saccharomycetales(100);f__Metschnikowiaceae(100);g__Metschnikowia(100);g__Metschnikowia_unclassified(100); | Metschnikowia | 13 | Saprotroph | \|Undefined Saprotroph\| | Yeast | NULL | Probable | As nectar-tap_saprotroph (Põlme et al. 2020); Decay sugar-rich substrates (Põlme et al. 2020) | Põlme S, et al. 2020. Fungal Diversity 105:1-16 ((https://doi.org/10.1007/s13225-020-00466-2)) |
| Otu43519 | 1 | k__Fungi(100);p__Ascomycota(100);c__Saccharomycetes(100);o__Saccharomycetales(100);f__Metschnikowiaceae(100);g__Metschnikowia(100);g__Metschnikowia_unclassified(100); | Metschnikowia | 13 | Saprotroph | \|Undefined Saprotroph\| | Yeast | NULL | Probable | As nectar-tap_saprotroph (Põlme et al. 2020); Decay sugar-rich substrates (Põlme et al. 2020) | Põlme S, et al. 2020. Fungal Diversity 105:1-16 ((https://doi.org/10.1007/s13225-020-00466-2)) |
| Otu43520 | 1 | k__Fungi(100);p__Ascomycota(100);c__Saccharomycetes(100);o__Saccharomycetales(100);f__Metschnikowiaceae(100);g__Metschnikowia(100);g__Metschnikowia_unclassified(100); | Metschnikowia | 13 | Saprotroph | \|Undefined Saprotroph\| | Yeast | NULL | Probable | As nectar-tap_saprotroph (Põlme et al. 2020); Decay sugar-rich substrates (Põlme et al. 2020) | Põlme S, et al. 2020. Fungal Diversity 105:1-16 ((https://doi.org/10.1007/s13225-020-00466-2)) |
| Otu43521 | 1 | k__Fungi(100);p__Ascomycota(100);c__Saccharomycetes(100);o__Saccharomycetales(100);f__Metschnikowiaceae(100);g__Metschnikowia(100);g__Metschnikowia_unclassified(100); | Metschnikowia | 13 | Saprotroph | \|Undefined Saprotroph\| | Yeast | NULL | Probable | As nectar-tap_saprotroph (Põlme et al. 2020); Decay sugar-rich substrates (Põlme et al. 2020) | Põlme S, et al. 2020. Fungal Diversity 105:1-16 ((https://doi.org/10.1007/s13225-020-00466-2)) |
| Otu43522 | 1 | k__Fungi(100);p__Ascomycota(100);c__Saccharomycetes(100);o__Saccharomycetales(100);f__Metschnikowiaceae(100);g__Metschnikowia(100);g__Metschnikowia_unclassified(100); | Metschnikowia | 13 | Saprotroph | \|Undefined Saprotroph\| | Yeast | NULL | Probable | As nectar-tap_saprotroph (Põlme et al. 2020); Decay sugar-rich substrates (Põlme et al. 2020) | Põlme S, et al. 2020. Fungal Diversity 105:1-16 ((https://doi.org/10.1007/s13225-020-00466-2)) |
| Otu43523 | 1 | k__Fungi(100);p__Ascomycota(100);c__Saccharomycetes(100);o__Saccharomycetales(100);f__Metschnikowiaceae(100);g__Metschnikowia(100);g__Metschnikowia_unclassified(100); | Metschnikowia | 13 | Saprotroph | \|Undefined Saprotroph\| | Yeast | NULL | Probable | As nectar-tap_saprotroph (Põlme et al. 2020); Decay sugar-rich substrates (Põlme et al. 2020) | Põlme S, et al. 2020. Fungal Diversity 105:1-16 ((https://doi.org/10.1007/s13225-020-00466-2)) |
| Otu43524 | 1 | k__Fungi(100);p__Ascomycota(100);c__Saccharomycetes(100);o__Saccharomycetales(100);f__Metschnikowiaceae(100);g__Metschnikowia(100);s__Metschnikowia_pulcherrima(100); | Metschnikowia | 13 | Saprotroph | \|Undefined Saprotroph\| | Yeast | NULL | Probable | As nectar-tap_saprotroph (Põlme et al. 2020); Decay sugar-rich substrates (Põlme et al. 2020) | Põlme S, et al. 2020. Fungal Diversity 105:1-16 ((https://doi.org/10.1007/s13225-020-00466-2)) |
| Otu43526 | 1 | k__Fungi(100);p__Ascomycota(100);c__Saccharomycetes(100);o__Saccharomycetales(100);f__Metschnikowiaceae(100);g__Metschnikowia(100);s__Metschnikowia_pulcherrima(100); | Metschnikowia | 13 | Saprotroph | \|Undefined Saprotroph\| | Yeast | NULL | Probable | As nectar-tap_saprotroph (Põlme et al. 2020); Decay sugar-rich substrates (Põlme et al. 2020) | Põlme S, et al. 2020. Fungal Diversity 105:1-16 ((https://doi.org/10.1007/s13225-020-00466-2)) |
| Otu43527 | 1 | k__Fungi(100);p__Ascomycota(100);c__Saccharomycetes(100);o__Saccharomycetales(100);f__Metschnikowiaceae(100);g__Metschnikowia(100);s__Metschnikowia_pulcherrima(100); | Metschnikowia | 13 | Saprotroph | \|Undefined Saprotroph\| | Yeast | NULL | Probable | As nectar-tap_saprotroph (Põlme et al. 2020); Decay sugar-rich substrates (Põlme et al. 2020) | Põlme S, et al. 2020. Fungal Diversity 105:1-16 ((https://doi.org/10.1007/s13225-020-00466-2)) |
| Otu43528 | 1 | k__Fungi(100);p__Ascomycota(100);c__Saccharomycetes(100);o__Saccharomycetales(100);f__Metschnikowiaceae(100);g__Metschnikowia(100);s__Metschnikowia_pulcherrima(100); | Metschnikowia | 13 | Saprotroph | \|Undefined Saprotroph\| | Yeast | NULL | Probable | As nectar-tap_saprotroph (Põlme et al. 2020); Decay sugar-rich substrates (Põlme et al. 2020) | Põlme S, et al. 2020. Fungal Diversity 105:1-16 ((https://doi.org/10.1007/s13225-020-00466-2)) |
| Otu43529 | 1 | k__Fungi(100);p__Ascomycota(100);c__Saccharomycetes(100);o__Saccharomycetales(100);f__Metschnikowiaceae(100);g__Metschnikowia(100);s__Metschnikowia_pulcherrima(100); | Metschnikowia | 13 | Saprotroph | \|Undefined Saprotroph\| | Yeast | NULL | Probable | As nectar-tap_saprotroph (Põlme et al. 2020); Decay sugar-rich substrates (Põlme et al. 2020) | Põlme S, et al. 2020. Fungal Diversity 105:1-16 ((https://doi.org/10.1007/s13225-020-00466-2)) |
| Otu43530 | 1 | k__Fungi(100);p__Ascomycota(100);c__Saccharomycetes(100);o__Saccharomycetales(100);f__Metschnikowiaceae(100);g__Metschnikowia(100);s__Metschnikowia_pulcherrima(100); | Metschnikowia | 13 | Saprotroph | \|Undefined Saprotroph\| | Yeast | NULL | Probable | As nectar-tap_saprotroph (Põlme et al. 2020); Decay sugar-rich substrates (Põlme et al. 2020) | Põlme S, et al. 2020. Fungal Diversity 105:1-16 ((https://doi.org/10.1007/s13225-020-00466-2)) |
| Otu43531 | 1 | k__Fungi(100);p__Ascomycota(100);c__Saccharomycetes(100);o__Saccharomycetales(100);f__Metschnikowiaceae(100);g__Metschnikowia(100);g__Metschnikowia_unclassified(100); | Metschnikowia | 13 | Saprotroph | \|Undefined Saprotroph\| | Yeast | NULL | Probable | As nectar-tap_saprotroph (Põlme et al. 2020); Decay sugar-rich substrates (Põlme et al. 2020) | Põlme S, et al. 2020. Fungal Diversity 105:1-16 ((https://doi.org/10.1007/s13225-020-00466-2)) |
| Otu43532 | 1 | k__Fungi(100);p__Ascomycota(100);c__Saccharomycetes(100);o__Saccharomycetales(100);f__Metschnikowiaceae(100);g__Metschnikowia(100);s__Metschnikowia_pulcherrima(100); | Metschnikowia | 13 | Saprotroph | \|Undefined Saprotroph\| | Yeast | NULL | Probable | As nectar-tap_saprotroph (Põlme et al. 2020); Decay sugar-rich substrates (Põlme et al. 2020) | Põlme S, et al. 2020. Fungal Diversity 105:1-16 ((https://doi.org/10.1007/s13225-020-00466-2)) |
| Otu43533 | 1 | k__Fungi(100);p__Ascomycota(100);c__Saccharomycetes(100);o__Saccharomycetales(100);f__Metschnikowiaceae(100);g__Metschnikowia(100);s__Metschnikowia_pulcherrima(100); | Metschnikowia | 13 | Saprotroph | \|Undefined Saprotroph\| | Yeast | NULL | Probable | As nectar-tap_saprotroph (Põlme et al. 2020); Decay sugar-rich substrates (Põlme et al. 2020) | Põlme S, et al. 2020. Fungal Diversity 105:1-16 ((https://doi.org/10.1007/s13225-020-00466-2)) |
| Otu43534 | 1 | k__Fungi(100);p__Ascomycota(100);c__Saccharomycetes(100);o__Saccharomycetales(100);f__Metschnikowiaceae(100);g__Metschnikowia(100);s__Metschnikowia_pulcherrima(100); | Metschnikowia | 13 | Saprotroph | \|Undefined Saprotroph\| | Yeast | NULL | Probable | As nectar-tap_saprotroph (Põlme et al. 2020); Decay sugar-rich substrates (Põlme et al. 2020) | Põlme S, et al. 2020. Fungal Diversity 105:1-16 ((https://doi.org/10.1007/s13225-020-00466-2)) |
| Otu43535 | 1 | k__Fungi(100);p__Ascomycota(100);c__Saccharomycetes(100);o__Saccharomycetales(100);f__Metschnikowiaceae(100);g__Metschnikowia(100);s__Metschnikowia_pulcherrima(100); | Metschnikowia | 13 | Saprotroph | \|Undefined Saprotroph\| | Yeast | NULL | Probable | As nectar-tap_saprotroph (Põlme et al. 2020); Decay sugar-rich substrates (Põlme et al. 2020) | Põlme S, et al. 2020. Fungal Diversity 105:1-16 ((https://doi.org/10.1007/s13225-020-00466-2)) |
| Otu43536 | 1 | k__Fungi(100);p__Ascomycota(100);c__Saccharomycetes(100);o__Saccharomycetales(100);f__Metschnikowiaceae(100);g__Metschnikowia(100);g__Metschnikowia_unclassified(100); | Metschnikowia | 13 | Saprotroph | \|Undefined Saprotroph\| | Yeast | NULL | Probable | As nectar-tap_saprotroph (Põlme et al. 2020); Decay sugar-rich substrates (Põlme et al. 2020) | Põlme S, et al. 2020. Fungal Diversity 105:1-16 ((https://doi.org/10.1007/s13225-020-00466-2)) |
| Otu43537 | 1 | k__Fungi(100);p__Ascomycota(100);c__Saccharomycetes(100);o__Saccharomycetales(100);f__Metschnikowiaceae(100);g__Metschnikowia(100);s__Metschnikowia_pulcherrima(100); | Metschnikowia | 13 | Saprotroph | \|Undefined Saprotroph\| | Yeast | NULL | Probable | As nectar-tap_saprotroph (Põlme et al. 2020); Decay sugar-rich substrates (Põlme et al. 2020) | Põlme S, et al. 2020. Fungal Diversity 105:1-16 ((https://doi.org/10.1007/s13225-020-00466-2)) |
| Otu43538 | 1 | k__Fungi(100);p__Ascomycota(100);c__Saccharomycetes(100);o__Saccharomycetales(100);f__Metschnikowiaceae(100);g__Metschnikowia(100);s__Metschnikowia_pulcherrima(100); | Metschnikowia | 13 | Saprotroph | \|Undefined Saprotroph\| | Yeast | NULL | Probable | As nectar-tap_saprotroph (Põlme et al. 2020); Decay sugar-rich substrates (Põlme et al. 2020) | Põlme S, et al. 2020. Fungal Diversity 105:1-16 ((https://doi.org/10.1007/s13225-020-00466-2)) |
| Otu43539 | 1 | k__Fungi(100);p__Ascomycota(100);c__Saccharomycetes(100);o__Saccharomycetales(100);f__Metschnikowiaceae(100);g__Metschnikowia(100);s__Metschnikowia_pulcherrima(100); | Metschnikowia | 13 | Saprotroph | \|Undefined Saprotroph\| | Yeast | NULL | Probable | As nectar-tap_saprotroph (Põlme et al. 2020); Decay sugar-rich substrates (Põlme et al. 2020) | Põlme S, et al. 2020. Fungal Diversity 105:1-16 ((https://doi.org/10.1007/s13225-020-00466-2)) |
| Otu43540 | 1 | k__Fungi(100);p__Ascomycota(100);c__Saccharomycetes(100);o__Saccharomycetales(100);f__Metschnikowiaceae(100);g__Metschnikowia(100);s__Metschnikowia_pulcherrima(100); | Metschnikowia | 13 | Saprotroph | \|Undefined Saprotroph\| | Yeast | NULL | Probable | As nectar-tap_saprotroph (Põlme et al. 2020); Decay sugar-rich substrates (Põlme et al. 2020) | Põlme S, et al. 2020. Fungal Diversity 105:1-16 ((https://doi.org/10.1007/s13225-020-00466-2)) |
| Otu43541 | 1 | k__Fungi(100);p__Ascomycota(100);c__Saccharomycetes(100);o__Saccharomycetales(100);f__Metschnikowiaceae(100);g__Metschnikowia(100);g__Metschnikowia_unclassified(100); | Metschnikowia | 13 | Saprotroph | \|Undefined Saprotroph\| | Yeast | NULL | Probable | As nectar-tap_saprotroph (Põlme et al. 2020); Decay sugar-rich substrates (Põlme et al. 2020) | Põlme S, et al. 2020. Fungal Diversity 105:1-16 ((https://doi.org/10.1007/s13225-020-00466-2)) |
| Otu43542 | 1 | k__Fungi(100);p__Ascomycota(100);c__Saccharomycetes(100);o__Saccharomycetales(100);f__Metschnikowiaceae(100);g__Metschnikowia(100);g__Metschnikowia_unclassified(100); | Metschnikowia | 13 | Saprotroph | \|Undefined Saprotroph\| | Yeast | NULL | Probable | As nectar-tap_saprotroph (Põlme et al. 2020); Decay sugar-rich substrates (Põlme et al. 2020) | Põlme S, et al. 2020. Fungal Diversity 105:1-16 ((https://doi.org/10.1007/s13225-020-00466-2)) |
| Otu43543 | 1 | k__Fungi(100);p__Ascomycota(100);c__Saccharomycetes(100);o__Saccharomycetales(100);f__Metschnikowiaceae(100);g__Metschnikowia(100);s__Metschnikowia_pulcherrima(100); | Metschnikowia | 13 | Saprotroph | \|Undefined Saprotroph\| | Yeast | NULL | Probable | As nectar-tap_saprotroph (Põlme et al. 2020); Decay sugar-rich substrates (Põlme et al. 2020) | Põlme S, et al. 2020. Fungal Diversity 105:1-16 ((https://doi.org/10.1007/s13225-020-00466-2)) |
| Otu43544 | 1 | k__Fungi(100);p__Ascomycota(100);c__Saccharomycetes(100);o__Saccharomycetales(100);f__Metschnikowiaceae(100);g__Metschnikowia(100);g__Metschnikowia_unclassified(100); | Metschnikowia | 13 | Saprotroph | \|Undefined Saprotroph\| | Yeast | NULL | Probable | As nectar-tap_saprotroph (Põlme et al. 2020); Decay sugar-rich substrates (Põlme et al. 2020) | Põlme S, et al. 2020. Fungal Diversity 105:1-16 ((https://doi.org/10.1007/s13225-020-00466-2)) |
| Otu43545 | 1 | k__Fungi(100);p__Ascomycota(100);c__Saccharomycetes(100);o__Saccharomycetales(100);f__Metschnikowiaceae(100);g__Metschnikowia(100);s__Metschnikowia_pulcherrima(100); | Metschnikowia | 13 | Saprotroph | \|Undefined Saprotroph\| | Yeast | NULL | Probable | As nectar-tap_saprotroph (Põlme et al. 2020); Decay sugar-rich substrates (Põlme et al. 2020) | Põlme S, et al. 2020. Fungal Diversity 105:1-16 ((https://doi.org/10.1007/s13225-020-00466-2)) |
| Otu43546 | 1 | k__Fungi(100);p__Ascomycota(100);c__Saccharomycetes(100);o__Saccharomycetales(100);f__Metschnikowiaceae(100);g__Metschnikowia(100);s__Metschnikowia_pulcherrima(100); | Metschnikowia | 13 | Saprotroph | \|Undefined Saprotroph\| | Yeast | NULL | Probable | As nectar-tap_saprotroph (Põlme et al. 2020); Decay sugar-rich substrates (Põlme et al. 2020) | Põlme S, et al. 2020. Fungal Diversity 105:1-16 ((https://doi.org/10.1007/s13225-020-00466-2)) |
| Otu43547 | 1 | k__Fungi(100);p__Ascomycota(100);c__Saccharomycetes(100);o__Saccharomycetales(100);f__Metschnikowiaceae(100);g__Metschnikowia(100);s__Metschnikowia_pulcherrima(100); | Metschnikowia | 13 | Saprotroph | \|Undefined Saprotroph\| | Yeast | NULL | Probable | As nectar-tap_saprotroph (Põlme et al. 2020); Decay sugar-rich substrates (Põlme et al. 2020) | Põlme S, et al. 2020. Fungal Diversity 105:1-16 ((https://doi.org/10.1007/s13225-020-00466-2)) |
| Otu43548 | 1 | k__Fungi(100);p__Ascomycota(100);c__Saccharomycetes(100);o__Saccharomycetales(100);f__Metschnikowiaceae(100);g__Metschnikowia(100);g__Metschnikowia_unclassified(100); | Metschnikowia | 13 | Saprotroph | \|Undefined Saprotroph\| | Yeast | NULL | Probable | As nectar-tap_saprotroph (Põlme et al. 2020); Decay sugar-rich substrates (Põlme et al. 2020) | Põlme S, et al. 2020. Fungal Diversity 105:1-16 ((https://doi.org/10.1007/s13225-020-00466-2)) |
| Otu43549 | 1 | k__Fungi(100);p__Ascomycota(100);c__Saccharomycetes(100);o__Saccharomycetales(100);f__Metschnikowiaceae(100);g__Metschnikowia(100);g__Metschnikowia_unclassified(100); | Metschnikowia | 13 | Saprotroph | \|Undefined Saprotroph\| | Yeast | NULL | Probable | As nectar-tap_saprotroph (Põlme et al. 2020); Decay sugar-rich substrates (Põlme et al. 2020) | Põlme S, et al. 2020. Fungal Diversity 105:1-16 ((https://doi.org/10.1007/s13225-020-00466-2)) |
| Otu43550 | 1 | k__Fungi(100);p__Ascomycota(100);c__Saccharomycetes(100);o__Saccharomycetales(100);f__Metschnikowiaceae(100);g__Metschnikowia(100);g__Metschnikowia_unclassified(100); | Metschnikowia | 13 | Saprotroph | \|Undefined Saprotroph\| | Yeast | NULL | Probable | As nectar-tap_saprotroph (Põlme et al. 2020); Decay sugar-rich substrates (Põlme et al. 2020) | Põlme S, et al. 2020. Fungal Diversity 105:1-16 ((https://doi.org/10.1007/s13225-020-00466-2)) |
| Otu43552 | 1 | k__Fungi(100);p__Ascomycota(100);c__Saccharomycetes(100);o__Saccharomycetales(100);f__Metschnikowiaceae(100);g__Metschnikowia(100);s__Metschnikowia_pulcherrima(100); | Metschnikowia | 13 | Saprotroph | \|Undefined Saprotroph\| | Yeast | NULL | Probable | As nectar-tap_saprotroph (Põlme et al. 2020); Decay sugar-rich substrates (Põlme et al. 2020) | Põlme S, et al. 2020. Fungal Diversity 105:1-16 ((https://doi.org/10.1007/s13225-020-00466-2)) |
| Otu43553 | 1 | k__Fungi(100);p__Ascomycota(100);c__Saccharomycetes(100);o__Saccharomycetales(100);f__Metschnikowiaceae(100);g__Metschnikowia(100);s__Metschnikowia_pulcherrima(100); | Metschnikowia | 13 | Saprotroph | \|Undefined Saprotroph\| | Yeast | NULL | Probable | As nectar-tap_saprotroph (Põlme et al. 2020); Decay sugar-rich substrates (Põlme et al. 2020) | Põlme S, et al. 2020. Fungal Diversity 105:1-16 ((https://doi.org/10.1007/s13225-020-00466-2)) |
| Otu43554 | 1 | k__Fungi(100);p__Ascomycota(100);c__Saccharomycetes(100);o__Saccharomycetales(100);f__Metschnikowiaceae(100);g__Metschnikowia(100);s__Metschnikowia_pulcherrima(100); | Metschnikowia | 13 | Saprotroph | \|Undefined Saprotroph\| | Yeast | NULL | Probable | As nectar-tap_saprotroph (Põlme et al. 2020); Decay sugar-rich substrates (Põlme et al. 2020) | Põlme S, et al. 2020. Fungal Diversity 105:1-16 ((https://doi.org/10.1007/s13225-020-00466-2)) |
| Otu43555 | 1 | k__Fungi(100);p__Ascomycota(100);c__Saccharomycetes(100);o__Saccharomycetales(100);f__Metschnikowiaceae(100);g__Metschnikowia(100);s__Metschnikowia_pulcherrima(100); | Metschnikowia | 13 | Saprotroph | \|Undefined Saprotroph\| | Yeast | NULL | Probable | As nectar-tap_saprotroph (Põlme et al. 2020); Decay sugar-rich substrates (Põlme et al. 2020) | Põlme S, et al. 2020. Fungal Diversity 105:1-16 ((https://doi.org/10.1007/s13225-020-00466-2)) |
| Otu43556 | 1 | k__Fungi(100);p__Ascomycota(100);c__Saccharomycetes(100);o__Saccharomycetales(100);f__Metschnikowiaceae(100);g__Metschnikowia(100);s__Metschnikowia_pulcherrima(100); | Metschnikowia | 13 | Saprotroph | \|Undefined Saprotroph\| | Yeast | NULL | Probable | As nectar-tap_saprotroph (Põlme et al. 2020); Decay sugar-rich substrates (Põlme et al. 2020) | Põlme S, et al. 2020. Fungal Diversity 105:1-16 ((https://doi.org/10.1007/s13225-020-00466-2)) |
| Otu43557 | 1 | k__Fungi(100);p__Ascomycota(100);c__Saccharomycetes(100);o__Saccharomycetales(100);f__Metschnikowiaceae(100);g__Metschnikowia(100);s__Metschnikowia_pulcherrima(100); | Metschnikowia | 13 | Saprotroph | \|Undefined Saprotroph\| | Yeast | NULL | Probable | As nectar-tap_saprotroph (Põlme et al. 2020); Decay sugar-rich substrates (Põlme et al. 2020) | Põlme S, et al. 2020. Fungal Diversity 105:1-16 ((https://doi.org/10.1007/s13225-020-00466-2)) |
| Otu43559 | 1 | k__Fungi(100);p__Ascomycota(100);c__Saccharomycetes(100);o__Saccharomycetales(100);f__Metschnikowiaceae(100);g__Metschnikowia(100);s__Metschnikowia_pulcherrima(100); | Metschnikowia | 13 | Saprotroph | \|Undefined Saprotroph\| | Yeast | NULL | Probable | As nectar-tap_saprotroph (Põlme et al. 2020); Decay sugar-rich substrates (Põlme et al. 2020) | Põlme S, et al. 2020. Fungal Diversity 105:1-16 ((https://doi.org/10.1007/s13225-020-00466-2)) |
| Otu43560 | 1 | k__Fungi(100);p__Ascomycota(100);c__Saccharomycetes(100);o__Saccharomycetales(100);f__Metschnikowiaceae(100);g__Metschnikowia(100);s__Metschnikowia_pulcherrima(100); | Metschnikowia | 13 | Saprotroph | \|Undefined Saprotroph\| | Yeast | NULL | Probable | As nectar-tap_saprotroph (Põlme et al. 2020); Decay sugar-rich substrates (Põlme et al. 2020) | Põlme S, et al. 2020. Fungal Diversity 105:1-16 ((https://doi.org/10.1007/s13225-020-00466-2)) |
| Otu43561 | 1 | k__Fungi(100);p__Ascomycota(100);c__Saccharomycetes(100);o__Saccharomycetales(100);f__Metschnikowiaceae(100);g__Metschnikowia(100);s__Metschnikowia_pulcherrima(100); | Metschnikowia | 13 | Saprotroph | \|Undefined Saprotroph\| | Yeast | NULL | Probable | As nectar-tap_saprotroph (Põlme et al. 2020); Decay sugar-rich substrates (Põlme et al. 2020) | Põlme S, et al. 2020. Fungal Diversity 105:1-16 ((https://doi.org/10.1007/s13225-020-00466-2)) |
| Otu43562 | 1 | k__Fungi(100);p__Ascomycota(100);c__Saccharomycetes(100);o__Saccharomycetales(100);f__Metschnikowiaceae(100);g__Metschnikowia(100);s__Metschnikowia_pulcherrima(100); | Metschnikowia | 13 | Saprotroph | \|Undefined Saprotroph\| | Yeast | NULL | Probable | As nectar-tap_saprotroph (Põlme et al. 2020); Decay sugar-rich substrates (Põlme et al. 2020) | Põlme S, et al. 2020. Fungal Diversity 105:1-16 ((https://doi.org/10.1007/s13225-020-00466-2)) |
| Otu43563 | 1 | k__Fungi(100);p__Ascomycota(100);c__Saccharomycetes(100);o__Saccharomycetales(100);f__Metschnikowiaceae(100);g__Metschnikowia(100);s__Metschnikowia_pulcherrima(100); | Metschnikowia | 13 | Saprotroph | \|Undefined Saprotroph\| | Yeast | NULL | Probable | As nectar-tap_saprotroph (Põlme et al. 2020); Decay sugar-rich substrates (Põlme et al. 2020) | Põlme S, et al. 2020. Fungal Diversity 105:1-16 ((https://doi.org/10.1007/s13225-020-00466-2)) |
| Otu43564 | 1 | k__Fungi(100);p__Ascomycota(100);c__Saccharomycetes(100);o__Saccharomycetales(100);f__Metschnikowiaceae(100);g__Metschnikowia(100);s__Metschnikowia_pulcherrima(100); | Metschnikowia | 13 | Saprotroph | \|Undefined Saprotroph\| | Yeast | NULL | Probable | As nectar-tap_saprotroph (Põlme et al. 2020); Decay sugar-rich substrates (Põlme et al. 2020) | Põlme S, et al. 2020. Fungal Diversity 105:1-16 ((https://doi.org/10.1007/s13225-020-00466-2)) |
| Otu43565 | 1 | k__Fungi(100);p__Ascomycota(100);c__Saccharomycetes(100);o__Saccharomycetales(100);f__Metschnikowiaceae(100);g__Metschnikowia(100);s__Metschnikowia_pulcherrima(100); | Metschnikowia | 13 | Saprotroph | \|Undefined Saprotroph\| | Yeast | NULL | Probable | As nectar-tap_saprotroph (Põlme et al. 2020); Decay sugar-rich substrates (Põlme et al. 2020) | Põlme S, et al. 2020. Fungal Diversity 105:1-16 ((https://doi.org/10.1007/s13225-020-00466-2)) |
| Otu43566 | 1 | k__Fungi(100);p__Ascomycota(100);c__Saccharomycetes(100);o__Saccharomycetales(100);f__Metschnikowiaceae(100);g__Metschnikowia(100);s__Metschnikowia_pulcherrima(100); | Metschnikowia | 13 | Saprotroph | \|Undefined Saprotroph\| | Yeast | NULL | Probable | As nectar-tap_saprotroph (Põlme et al. 2020); Decay sugar-rich substrates (Põlme et al. 2020) | Põlme S, et al. 2020. Fungal Diversity 105:1-16 ((https://doi.org/10.1007/s13225-020-00466-2)) |
| Otu43568 | 1 | k__Fungi(100);p__Ascomycota(100);c__Saccharomycetes(100);o__Saccharomycetales(100);f__Metschnikowiaceae(100);g__Metschnikowia(100);s__Metschnikowia_pulcherrima(100); | Metschnikowia | 13 | Saprotroph | \|Undefined Saprotroph\| | Yeast | NULL | Probable | As nectar-tap_saprotroph (Põlme et al. 2020); Decay sugar-rich substrates (Põlme et al. 2020) | Põlme S, et al. 2020. Fungal Diversity 105:1-16 ((https://doi.org/10.1007/s13225-020-00466-2)) |
| Otu43569 | 1 | k__Fungi(100);p__Ascomycota(100);c__Saccharomycetes(100);o__Saccharomycetales(100);f__Metschnikowiaceae(100);g__Metschnikowia(100);g__Metschnikowia_unclassified(100); | Metschnikowia | 13 | Saprotroph | \|Undefined Saprotroph\| | Yeast | NULL | Probable | As nectar-tap_saprotroph (Põlme et al. 2020); Decay sugar-rich substrates (Põlme et al. 2020) | Põlme S, et al. 2020. Fungal Diversity 105:1-16 ((https://doi.org/10.1007/s13225-020-00466-2)) |
| Otu43570 | 1 | k__Fungi(100);p__Ascomycota(100);c__Saccharomycetes(100);o__Saccharomycetales(100);f__Metschnikowiaceae(100);g__Metschnikowia(100);g__Metschnikowia_unclassified(100); | Metschnikowia | 13 | Saprotroph | \|Undefined Saprotroph\| | Yeast | NULL | Probable | As nectar-tap_saprotroph (Põlme et al. 2020); Decay sugar-rich substrates (Põlme et al. 2020) | Põlme S, et al. 2020. Fungal Diversity 105:1-16 ((https://doi.org/10.1007/s13225-020-00466-2)) |
| Otu43571 | 1 | k__Fungi(100);p__Ascomycota(100);c__Saccharomycetes(100);o__Saccharomycetales(100);f__Metschnikowiaceae(100);g__Metschnikowia(100);s__Metschnikowia_pulcherrima(100); | Metschnikowia | 13 | Saprotroph | \|Undefined Saprotroph\| | Yeast | NULL | Probable | As nectar-tap_saprotroph (Põlme et al. 2020); Decay sugar-rich substrates (Põlme et al. 2020) | Põlme S, et al. 2020. Fungal Diversity 105:1-16 ((https://doi.org/10.1007/s13225-020-00466-2)) |
| Otu43572 | 1 | k__Fungi(100);p__Ascomycota(100);c__Saccharomycetes(100);o__Saccharomycetales(100);f__Metschnikowiaceae(100);g__Metschnikowia(100);s__Metschnikowia_pulcherrima(100); | Metschnikowia | 13 | Saprotroph | \|Undefined Saprotroph\| | Yeast | NULL | Probable | As nectar-tap_saprotroph (Põlme et al. 2020); Decay sugar-rich substrates (Põlme et al. 2020) | Põlme S, et al. 2020. Fungal Diversity 105:1-16 ((https://doi.org/10.1007/s13225-020-00466-2)) |
| Otu43573 | 1 | k__Fungi(100);p__Ascomycota(100);c__Saccharomycetes(100);o__Saccharomycetales(100);f__Metschnikowiaceae(100);g__Metschnikowia(100);g__Metschnikowia_unclassified(100); | Metschnikowia | 13 | Saprotroph | \|Undefined Saprotroph\| | Yeast | NULL | Probable | As nectar-tap_saprotroph (Põlme et al. 2020); Decay sugar-rich substrates (Põlme et al. 2020) | Põlme S, et al. 2020. Fungal Diversity 105:1-16 ((https://doi.org/10.1007/s13225-020-00466-2)) |
| Otu43574 | 1 | k__Fungi(100);p__Ascomycota(100);c__Saccharomycetes(100);o__Saccharomycetales(100);f__Metschnikowiaceae(100);g__Metschnikowia(100);g__Metschnikowia_unclassified(100); | Metschnikowia | 13 | Saprotroph | \|Undefined Saprotroph\| | Yeast | NULL | Probable | As nectar-tap_saprotroph (Põlme et al. 2020); Decay sugar-rich substrates (Põlme et al. 2020) | Põlme S, et al. 2020. Fungal Diversity 105:1-16 ((https://doi.org/10.1007/s13225-020-00466-2)) |
| Otu43575 | 1 | k__Fungi(100);p__Ascomycota(100);c__Saccharomycetes(100);o__Saccharomycetales(100);f__Metschnikowiaceae(100);g__Metschnikowia(100);s__Metschnikowia_pulcherrima(100); | Metschnikowia | 13 | Saprotroph | \|Undefined Saprotroph\| | Yeast | NULL | Probable | As nectar-tap_saprotroph (Põlme et al. 2020); Decay sugar-rich substrates (Põlme et al. 2020) | Põlme S, et al. 2020. Fungal Diversity 105:1-16 ((https://doi.org/10.1007/s13225-020-00466-2)) |
| Otu43576 | 1 | k__Fungi(100);p__Ascomycota(100);c__Saccharomycetes(100);o__Saccharomycetales(100);f__Metschnikowiaceae(100);g__Metschnikowia(100);s__Metschnikowia_pulcherrima(100); | Metschnikowia | 13 | Saprotroph | \|Undefined Saprotroph\| | Yeast | NULL | Probable | As nectar-tap_saprotroph (Põlme et al. 2020); Decay sugar-rich substrates (Põlme et al. 2020) | Põlme S, et al. 2020. Fungal Diversity 105:1-16 ((https://doi.org/10.1007/s13225-020-00466-2)) |
| Otu43577 | 1 | k__Fungi(100);p__Ascomycota(100);c__Saccharomycetes(100);o__Saccharomycetales(100);f__Metschnikowiaceae(100);g__Metschnikowia(100);s__Metschnikowia_pulcherrima(100); | Metschnikowia | 13 | Saprotroph | \|Undefined Saprotroph\| | Yeast | NULL | Probable | As nectar-tap_saprotroph (Põlme et al. 2020); Decay sugar-rich substrates (Põlme et al. 2020) | Põlme S, et al. 2020. Fungal Diversity 105:1-16 ((https://doi.org/10.1007/s13225-020-00466-2)) |
| Otu43578 | 1 | k__Fungi(100);p__Ascomycota(100);c__Saccharomycetes(100);o__Saccharomycetales(100);f__Metschnikowiaceae(100);g__Metschnikowia(100);g__Metschnikowia_unclassified(100); | Metschnikowia | 13 | Saprotroph | \|Undefined Saprotroph\| | Yeast | NULL | Probable | As nectar-tap_saprotroph (Põlme et al. 2020); Decay sugar-rich substrates (Põlme et al. 2020) | Põlme S, et al. 2020. Fungal Diversity 105:1-16 ((https://doi.org/10.1007/s13225-020-00466-2)) |
| Otu43579 | 1 | k__Fungi(100);p__Ascomycota(100);c__Saccharomycetes(100);o__Saccharomycetales(100);f__Metschnikowiaceae(100);g__Metschnikowia(100);g__Metschnikowia_unclassified(100); | Metschnikowia | 13 | Saprotroph | \|Undefined Saprotroph\| | Yeast | NULL | Probable | As nectar-tap_saprotroph (Põlme et al. 2020); Decay sugar-rich substrates (Põlme et al. 2020) | Põlme S, et al. 2020. Fungal Diversity 105:1-16 ((https://doi.org/10.1007/s13225-020-00466-2)) |
| Otu43580 | 1 | k__Fungi(100);p__Ascomycota(100);c__Saccharomycetes(100);o__Saccharomycetales(100);f__Metschnikowiaceae(100);g__Metschnikowia(100);s__Metschnikowia_pulcherrima(100); | Metschnikowia | 13 | Saprotroph | \|Undefined Saprotroph\| | Yeast | NULL | Probable | As nectar-tap_saprotroph (Põlme et al. 2020); Decay sugar-rich substrates (Põlme et al. 2020) | Põlme S, et al. 2020. Fungal Diversity 105:1-16 ((https://doi.org/10.1007/s13225-020-00466-2)) |
| Otu43581 | 1 | k__Fungi(100);p__Ascomycota(100);c__Saccharomycetes(100);o__Saccharomycetales(100);f__Metschnikowiaceae(100);g__Metschnikowia(100);g__Metschnikowia_unclassified(100); | Metschnikowia | 13 | Saprotroph | \|Undefined Saprotroph\| | Yeast | NULL | Probable | As nectar-tap_saprotroph (Põlme et al. 2020); Decay sugar-rich substrates (Põlme et al. 2020) | Põlme S, et al. 2020. Fungal Diversity 105:1-16 ((https://doi.org/10.1007/s13225-020-00466-2)) |
| Otu43582 | 1 | k__Fungi(100);p__Ascomycota(100);c__Saccharomycetes(100);o__Saccharomycetales(100);f__Metschnikowiaceae(100);g__Metschnikowia(100);s__Metschnikowia_pulcherrima(100); | Metschnikowia | 13 | Saprotroph | \|Undefined Saprotroph\| | Yeast | NULL | Probable | As nectar-tap_saprotroph (Põlme et al. 2020); Decay sugar-rich substrates (Põlme et al. 2020) | Põlme S, et al. 2020. Fungal Diversity 105:1-16 ((https://doi.org/10.1007/s13225-020-00466-2)) |
| Otu43583 | 1 | k__Fungi(100);p__Ascomycota(100);c__Saccharomycetes(100);o__Saccharomycetales(100);f__Metschnikowiaceae(100);g__Metschnikowia(100);s__Metschnikowia_pulcherrima(100); | Metschnikowia | 13 | Saprotroph | \|Undefined Saprotroph\| | Yeast | NULL | Probable | As nectar-tap_saprotroph (Põlme et al. 2020); Decay sugar-rich substrates (Põlme et al. 2020) | Põlme S, et al. 2020. Fungal Diversity 105:1-16 ((https://doi.org/10.1007/s13225-020-00466-2)) |
| Otu43584 | 1 | k__Fungi(100);p__Ascomycota(100);c__Saccharomycetes(100);o__Saccharomycetales(100);f__Metschnikowiaceae(100);g__Metschnikowia(100);s__Metschnikowia_pulcherrima(100); | Metschnikowia | 13 | Saprotroph | \|Undefined Saprotroph\| | Yeast | NULL | Probable | As nectar-tap_saprotroph (Põlme et al. 2020); Decay sugar-rich substrates (Põlme et al. 2020) | Põlme S, et al. 2020. Fungal Diversity 105:1-16 ((https://doi.org/10.1007/s13225-020-00466-2)) |
| Otu43586 | 1 | k__Fungi(100);p__Ascomycota(100);c__Saccharomycetes(100);o__Saccharomycetales(100);f__Metschnikowiaceae(100);g__Metschnikowia(100);s__Metschnikowia_pulcherrima(100); | Metschnikowia | 13 | Saprotroph | \|Undefined Saprotroph\| | Yeast | NULL | Probable | As nectar-tap_saprotroph (Põlme et al. 2020); Decay sugar-rich substrates (Põlme et al. 2020) | Põlme S, et al. 2020. Fungal Diversity 105:1-16 ((https://doi.org/10.1007/s13225-020-00466-2)) |
| Otu43587 | 1 | k__Fungi(100);p__Ascomycota(100);c__Saccharomycetes(100);o__Saccharomycetales(100);f__Metschnikowiaceae(100);g__Metschnikowia(100);g__Metschnikowia_unclassified(100); | Metschnikowia | 13 | Saprotroph | \|Undefined Saprotroph\| | Yeast | NULL | Probable | As nectar-tap_saprotroph (Põlme et al. 2020); Decay sugar-rich substrates (Põlme et al. 2020) | Põlme S, et al. 2020. Fungal Diversity 105:1-16 ((https://doi.org/10.1007/s13225-020-00466-2)) |
| Otu43588 | 1 | k__Fungi(100);p__Ascomycota(100);c__Saccharomycetes(100);o__Saccharomycetales(100);f__Metschnikowiaceae(100);g__Metschnikowia(100);s__Metschnikowia_pulcherrima(100); | Metschnikowia | 13 | Saprotroph | \|Undefined Saprotroph\| | Yeast | NULL | Probable | As nectar-tap_saprotroph (Põlme et al. 2020); Decay sugar-rich substrates (Põlme et al. 2020) | Põlme S, et al. 2020. Fungal Diversity 105:1-16 ((https://doi.org/10.1007/s13225-020-00466-2)) |
| Otu43589 | 1 | k__Fungi(100);p__Ascomycota(100);c__Saccharomycetes(100);o__Saccharomycetales(100);f__Metschnikowiaceae(100);g__Metschnikowia(100);s__Metschnikowia_pulcherrima(100); | Metschnikowia | 13 | Saprotroph | \|Undefined Saprotroph\| | Yeast | NULL | Probable | As nectar-tap_saprotroph (Põlme et al. 2020); Decay sugar-rich substrates (Põlme et al. 2020) | Põlme S, et al. 2020. Fungal Diversity 105:1-16 ((https://doi.org/10.1007/s13225-020-00466-2)) |
| Otu43590 | 1 | k__Fungi(100);p__Ascomycota(100);c__Saccharomycetes(100);o__Saccharomycetales(100);f__Metschnikowiaceae(100);g__Metschnikowia(100);s__Metschnikowia_pulcherrima(100); | Metschnikowia | 13 | Saprotroph | \|Undefined Saprotroph\| | Yeast | NULL | Probable | As nectar-tap_saprotroph (Põlme et al. 2020); Decay sugar-rich substrates (Põlme et al. 2020) | Põlme S, et al. 2020. Fungal Diversity 105:1-16 ((https://doi.org/10.1007/s13225-020-00466-2)) |
| Otu43591 | 1 | k__Fungi(100);p__Ascomycota(100);c__Saccharomycetes(100);o__Saccharomycetales(100);f__Metschnikowiaceae(100);g__Metschnikowia(100);g__Metschnikowia_unclassified(100); | Metschnikowia | 13 | Saprotroph | \|Undefined Saprotroph\| | Yeast | NULL | Probable | As nectar-tap_saprotroph (Põlme et al. 2020); Decay sugar-rich substrates (Põlme et al. 2020) | Põlme S, et al. 2020. Fungal Diversity 105:1-16 ((https://doi.org/10.1007/s13225-020-00466-2)) |
| Otu43592 | 1 | k__Fungi(100);p__Ascomycota(100);c__Saccharomycetes(100);o__Saccharomycetales(100);f__Metschnikowiaceae(100);g__Metschnikowia(100);s__Metschnikowia_pulcherrima(100); | Metschnikowia | 13 | Saprotroph | \|Undefined Saprotroph\| | Yeast | NULL | Probable | As nectar-tap_saprotroph (Põlme et al. 2020); Decay sugar-rich substrates (Põlme et al. 2020) | Põlme S, et al. 2020. Fungal Diversity 105:1-16 ((https://doi.org/10.1007/s13225-020-00466-2)) |
| Otu43593 | 1 | k__Fungi(100);p__Ascomycota(100);c__Saccharomycetes(100);o__Saccharomycetales(100);f__Metschnikowiaceae(100);g__Metschnikowia(100);g__Metschnikowia_unclassified(100); | Metschnikowia | 13 | Saprotroph | \|Undefined Saprotroph\| | Yeast | NULL | Probable | As nectar-tap_saprotroph (Põlme et al. 2020); Decay sugar-rich substrates (Põlme et al. 2020) | Põlme S, et al. 2020. Fungal Diversity 105:1-16 ((https://doi.org/10.1007/s13225-020-00466-2)) |
| Otu43594 | 1 | k__Fungi(100);p__Ascomycota(100);c__Saccharomycetes(100);o__Saccharomycetales(100);f__Metschnikowiaceae(100);g__Metschnikowia(100);s__Metschnikowia_pulcherrima(100); | Metschnikowia | 13 | Saprotroph | \|Undefined Saprotroph\| | Yeast | NULL | Probable | As nectar-tap_saprotroph (Põlme et al. 2020); Decay sugar-rich substrates (Põlme et al. 2020) | Põlme S, et al. 2020. Fungal Diversity 105:1-16 ((https://doi.org/10.1007/s13225-020-00466-2)) |
| Otu43595 | 1 | k__Fungi(100);p__Ascomycota(100);c__Saccharomycetes(100);o__Saccharomycetales(100);f__Metschnikowiaceae(100);g__Metschnikowia(100);s__Metschnikowia_pulcherrima(100); | Metschnikowia | 13 | Saprotroph | \|Undefined Saprotroph\| | Yeast | NULL | Probable | As nectar-tap_saprotroph (Põlme et al. 2020); Decay sugar-rich substrates (Põlme et al. 2020) | Põlme S, et al. 2020. Fungal Diversity 105:1-16 ((https://doi.org/10.1007/s13225-020-00466-2)) |
| Otu43596 | 1 | k__Fungi(100);p__Ascomycota(100);c__Saccharomycetes(100);o__Saccharomycetales(100);f__Metschnikowiaceae(100);g__Metschnikowia(100);s__Metschnikowia_pulcherrima(100); | Metschnikowia | 13 | Saprotroph | \|Undefined Saprotroph\| | Yeast | NULL | Probable | As nectar-tap_saprotroph (Põlme et al. 2020); Decay sugar-rich substrates (Põlme et al. 2020) | Põlme S, et al. 2020. Fungal Diversity 105:1-16 ((https://doi.org/10.1007/s13225-020-00466-2)) |
| Otu43597 | 1 | k__Fungi(100);p__Ascomycota(100);c__Saccharomycetes(100);o__Saccharomycetales(100);f__Metschnikowiaceae(100);g__Metschnikowia(100);s__Metschnikowia_pulcherrima(100); | Metschnikowia | 13 | Saprotroph | \|Undefined Saprotroph\| | Yeast | NULL | Probable | As nectar-tap_saprotroph (Põlme et al. 2020); Decay sugar-rich substrates (Põlme et al. 2020) | Põlme S, et al. 2020. Fungal Diversity 105:1-16 ((https://doi.org/10.1007/s13225-020-00466-2)) |
| Otu43598 | 1 | k__Fungi(100);p__Ascomycota(100);c__Saccharomycetes(100);o__Saccharomycetales(100);f__Metschnikowiaceae(100);g__Metschnikowia(100);g__Metschnikowia_unclassified(100); | Metschnikowia | 13 | Saprotroph | \|Undefined Saprotroph\| | Yeast | NULL | Probable | As nectar-tap_saprotroph (Põlme et al. 2020); Decay sugar-rich substrates (Põlme et al. 2020) | Põlme S, et al. 2020. Fungal Diversity 105:1-16 ((https://doi.org/10.1007/s13225-020-00466-2)) |
| Otu43599 | 1 | k__Fungi(100);p__Ascomycota(100);c__Saccharomycetes(100);o__Saccharomycetales(100);f__Metschnikowiaceae(100);g__Metschnikowia(100);g__Metschnikowia_unclassified(100); | Metschnikowia | 13 | Saprotroph | \|Undefined Saprotroph\| | Yeast | NULL | Probable | As nectar-tap_saprotroph (Põlme et al. 2020); Decay sugar-rich substrates (Põlme et al. 2020) | Põlme S, et al. 2020. Fungal Diversity 105:1-16 ((https://doi.org/10.1007/s13225-020-00466-2)) |
| Otu43600 | 1 | k__Fungi(100);p__Ascomycota(100);c__Saccharomycetes(100);o__Saccharomycetales(100);f__Metschnikowiaceae(100);g__Metschnikowia(100);s__Metschnikowia_pulcherrima(100); | Metschnikowia | 13 | Saprotroph | \|Undefined Saprotroph\| | Yeast | NULL | Probable | As nectar-tap_saprotroph (Põlme et al. 2020); Decay sugar-rich substrates (Põlme et al. 2020) | Põlme S, et al. 2020. Fungal Diversity 105:1-16 ((https://doi.org/10.1007/s13225-020-00466-2)) |
| Otu43601 | 1 | k__Fungi(100);p__Ascomycota(100);c__Saccharomycetes(100);o__Saccharomycetales(100);f__Metschnikowiaceae(100);g__Metschnikowia(100);s__Metschnikowia_pulcherrima(100); | Metschnikowia | 13 | Saprotroph | \|Undefined Saprotroph\| | Yeast | NULL | Probable | As nectar-tap_saprotroph (Põlme et al. 2020); Decay sugar-rich substrates (Põlme et al. 2020) | Põlme S, et al. 2020. Fungal Diversity 105:1-16 ((https://doi.org/10.1007/s13225-020-00466-2)) |
| Otu43602 | 1 | k__Fungi(100);p__Ascomycota(100);c__Saccharomycetes(100);o__Saccharomycetales(100);f__Metschnikowiaceae(100);g__Metschnikowia(100);s__Metschnikowia_pulcherrima(100); | Metschnikowia | 13 | Saprotroph | \|Undefined Saprotroph\| | Yeast | NULL | Probable | As nectar-tap_saprotroph (Põlme et al. 2020); Decay sugar-rich substrates (Põlme et al. 2020) | Põlme S, et al. 2020. Fungal Diversity 105:1-16 ((https://doi.org/10.1007/s13225-020-00466-2)) |
| Otu43603 | 1 | k__Fungi(100);p__Ascomycota(100);c__Saccharomycetes(100);o__Saccharomycetales(100);f__Metschnikowiaceae(100);g__Metschnikowia(100);s__Metschnikowia_pulcherrima(100); | Metschnikowia | 13 | Saprotroph | \|Undefined Saprotroph\| | Yeast | NULL | Probable | As nectar-tap_saprotroph (Põlme et al. 2020); Decay sugar-rich substrates (Põlme et al. 2020) | Põlme S, et al. 2020. Fungal Diversity 105:1-16 ((https://doi.org/10.1007/s13225-020-00466-2)) |
| Otu43604 | 1 | k__Fungi(100);p__Ascomycota(100);c__Saccharomycetes(100);o__Saccharomycetales(100);f__Metschnikowiaceae(100);g__Metschnikowia(100);g__Metschnikowia_unclassified(100); | Metschnikowia | 13 | Saprotroph | \|Undefined Saprotroph\| | Yeast | NULL | Probable | As nectar-tap_saprotroph (Põlme et al. 2020); Decay sugar-rich substrates (Põlme et al. 2020) | Põlme S, et al. 2020. Fungal Diversity 105:1-16 ((https://doi.org/10.1007/s13225-020-00466-2)) |
| Otu43605 | 1 | k__Fungi(100);p__Ascomycota(100);c__Saccharomycetes(100);o__Saccharomycetales(100);f__Metschnikowiaceae(100);g__Metschnikowia(100);s__Metschnikowia_pulcherrima(100); | Metschnikowia | 13 | Saprotroph | \|Undefined Saprotroph\| | Yeast | NULL | Probable | As nectar-tap_saprotroph (Põlme et al. 2020); Decay sugar-rich substrates (Põlme et al. 2020) | Põlme S, et al. 2020. Fungal Diversity 105:1-16 ((https://doi.org/10.1007/s13225-020-00466-2)) |
| Otu43606 | 1 | k__Fungi(100);p__Ascomycota(100);c__Saccharomycetes(100);o__Saccharomycetales(100);f__Metschnikowiaceae(100);g__Metschnikowia(100);s__Metschnikowia_pulcherrima(100); | Metschnikowia | 13 | Saprotroph | \|Undefined Saprotroph\| | Yeast | NULL | Probable | As nectar-tap_saprotroph (Põlme et al. 2020); Decay sugar-rich substrates (Põlme et al. 2020) | Põlme S, et al. 2020. Fungal Diversity 105:1-16 ((https://doi.org/10.1007/s13225-020-00466-2)) |
| Otu43607 | 1 | k__Fungi(100);p__Ascomycota(100);c__Saccharomycetes(100);o__Saccharomycetales(100);f__Metschnikowiaceae(100);g__Metschnikowia(100);s__Metschnikowia_pulcherrima(100); | Metschnikowia | 13 | Saprotroph | \|Undefined Saprotroph\| | Yeast | NULL | Probable | As nectar-tap_saprotroph (Põlme et al. 2020); Decay sugar-rich substrates (Põlme et al. 2020) | Põlme S, et al. 2020. Fungal Diversity 105:1-16 ((https://doi.org/10.1007/s13225-020-00466-2)) |
| Otu43608 | 1 | k__Fungi(100);p__Ascomycota(100);c__Saccharomycetes(100);o__Saccharomycetales(100);f__Metschnikowiaceae(100);g__Metschnikowia(100);s__Metschnikowia_pulcherrima(100); | Metschnikowia | 13 | Saprotroph | \|Undefined Saprotroph\| | Yeast | NULL | Probable | As nectar-tap_saprotroph (Põlme et al. 2020); Decay sugar-rich substrates (Põlme et al. 2020) | Põlme S, et al. 2020. Fungal Diversity 105:1-16 ((https://doi.org/10.1007/s13225-020-00466-2)) |
| Otu43609 | 1 | k__Fungi(100);p__Ascomycota(100);c__Saccharomycetes(100);o__Saccharomycetales(100);f__Metschnikowiaceae(100);g__Metschnikowia(100);s__Metschnikowia_pulcherrima(100); | Metschnikowia | 13 | Saprotroph | \|Undefined Saprotroph\| | Yeast | NULL | Probable | As nectar-tap_saprotroph (Põlme et al. 2020); Decay sugar-rich substrates (Põlme et al. 2020) | Põlme S, et al. 2020. Fungal Diversity 105:1-16 ((https://doi.org/10.1007/s13225-020-00466-2)) |
| Otu43610 | 1 | k__Fungi(100);p__Ascomycota(100);c__Saccharomycetes(100);o__Saccharomycetales(100);f__Metschnikowiaceae(100);g__Metschnikowia(100);s__Metschnikowia_pulcherrima(100); | Metschnikowia | 13 | Saprotroph | \|Undefined Saprotroph\| | Yeast | NULL | Probable | As nectar-tap_saprotroph (Põlme et al. 2020); Decay sugar-rich substrates (Põlme et al. 2020) | Põlme S, et al. 2020. Fungal Diversity 105:1-16 ((https://doi.org/10.1007/s13225-020-00466-2)) |
| Otu43611 | 1 | k__Fungi(100);p__Ascomycota(100);c__Saccharomycetes(100);o__Saccharomycetales(100);f__Metschnikowiaceae(100);g__Metschnikowia(100);g__Metschnikowia_unclassified(100); | Metschnikowia | 13 | Saprotroph | \|Undefined Saprotroph\| | Yeast | NULL | Probable | As nectar-tap_saprotroph (Põlme et al. 2020); Decay sugar-rich substrates (Põlme et al. 2020) | Põlme S, et al. 2020. Fungal Diversity 105:1-16 ((https://doi.org/10.1007/s13225-020-00466-2)) |
| Otu43612 | 1 | k__Fungi(100);p__Ascomycota(100);c__Saccharomycetes(100);o__Saccharomycetales(100);f__Metschnikowiaceae(100);g__Metschnikowia(100);s__Metschnikowia_pulcherrima(100); | Metschnikowia | 13 | Saprotroph | \|Undefined Saprotroph\| | Yeast | NULL | Probable | As nectar-tap_saprotroph (Põlme et al. 2020); Decay sugar-rich substrates (Põlme et al. 2020) | Põlme S, et al. 2020. Fungal Diversity 105:1-16 ((https://doi.org/10.1007/s13225-020-00466-2)) |
| Otu43614 | 1 | k__Fungi(100);p__Ascomycota(100);c__Saccharomycetes(100);o__Saccharomycetales(100);f__Metschnikowiaceae(100);g__Metschnikowia(100);s__Metschnikowia_pulcherrima(100); | Metschnikowia | 13 | Saprotroph | \|Undefined Saprotroph\| | Yeast | NULL | Probable | As nectar-tap_saprotroph (Põlme et al. 2020); Decay sugar-rich substrates (Põlme et al. 2020) | Põlme S, et al. 2020. Fungal Diversity 105:1-16 ((https://doi.org/10.1007/s13225-020-00466-2)) |
| Otu43615 | 1 | k__Fungi(100);p__Ascomycota(100);c__Saccharomycetes(100);o__Saccharomycetales(100);f__Metschnikowiaceae(100);g__Metschnikowia(100);s__Metschnikowia_pulcherrima(100); | Metschnikowia | 13 | Saprotroph | \|Undefined Saprotroph\| | Yeast | NULL | Probable | As nectar-tap_saprotroph (Põlme et al. 2020); Decay sugar-rich substrates (Põlme et al. 2020) | Põlme S, et al. 2020. Fungal Diversity 105:1-16 ((https://doi.org/10.1007/s13225-020-00466-2)) |
| Otu43616 | 1 | k__Fungi(100);p__Ascomycota(100);c__Saccharomycetes(100);o__Saccharomycetales(100);f__Metschnikowiaceae(100);g__Metschnikowia(100);g__Metschnikowia_unclassified(100); | Metschnikowia | 13 | Saprotroph | \|Undefined Saprotroph\| | Yeast | NULL | Probable | As nectar-tap_saprotroph (Põlme et al. 2020); Decay sugar-rich substrates (Põlme et al. 2020) | Põlme S, et al. 2020. Fungal Diversity 105:1-16 ((https://doi.org/10.1007/s13225-020-00466-2)) |
| Otu43617 | 1 | k__Fungi(100);p__Ascomycota(100);c__Saccharomycetes(100);o__Saccharomycetales(100);f__Metschnikowiaceae(100);g__Metschnikowia(100);s__Metschnikowia_pulcherrima(100); | Metschnikowia | 13 | Saprotroph | \|Undefined Saprotroph\| | Yeast | NULL | Probable | As nectar-tap_saprotroph (Põlme et al. 2020); Decay sugar-rich substrates (Põlme et al. 2020) | Põlme S, et al. 2020. Fungal Diversity 105:1-16 ((https://doi.org/10.1007/s13225-020-00466-2)) |
| Otu43618 | 1 | k__Fungi(100);p__Ascomycota(100);c__Saccharomycetes(100);o__Saccharomycetales(100);f__Metschnikowiaceae(100);g__Metschnikowia(100);s__Metschnikowia_pulcherrima(100); | Metschnikowia | 13 | Saprotroph | \|Undefined Saprotroph\| | Yeast | NULL | Probable | As nectar-tap_saprotroph (Põlme et al. 2020); Decay sugar-rich substrates (Põlme et al. 2020) | Põlme S, et al. 2020. Fungal Diversity 105:1-16 ((https://doi.org/10.1007/s13225-020-00466-2)) |
| Otu43619 | 1 | k__Fungi(100);p__Ascomycota(100);c__Saccharomycetes(100);o__Saccharomycetales(100);f__Metschnikowiaceae(100);g__Metschnikowia(100);s__Metschnikowia_pulcherrima(100); | Metschnikowia | 13 | Saprotroph | \|Undefined Saprotroph\| | Yeast | NULL | Probable | As nectar-tap_saprotroph (Põlme et al. 2020); Decay sugar-rich substrates (Põlme et al. 2020) | Põlme S, et al. 2020. Fungal Diversity 105:1-16 ((https://doi.org/10.1007/s13225-020-00466-2)) |
| Otu43620 | 1 | k__Fungi(100);p__Ascomycota(100);c__Saccharomycetes(100);o__Saccharomycetales(100);f__Metschnikowiaceae(100);g__Metschnikowia(100);s__Metschnikowia_pulcherrima(100); | Metschnikowia | 13 | Saprotroph | \|Undefined Saprotroph\| | Yeast | NULL | Probable | As nectar-tap_saprotroph (Põlme et al. 2020); Decay sugar-rich substrates (Põlme et al. 2020) | Põlme S, et al. 2020. Fungal Diversity 105:1-16 ((https://doi.org/10.1007/s13225-020-00466-2)) |
| Otu43622 | 1 | k__Fungi(100);p__Ascomycota(100);c__Saccharomycetes(100);o__Saccharomycetales(100);f__Metschnikowiaceae(100);g__Metschnikowia(100);g__Metschnikowia_unclassified(100); | Metschnikowia | 13 | Saprotroph | \|Undefined Saprotroph\| | Yeast | NULL | Probable | As nectar-tap_saprotroph (Põlme et al. 2020); Decay sugar-rich substrates (Põlme et al. 2020) | Põlme S, et al. 2020. Fungal Diversity 105:1-16 ((https://doi.org/10.1007/s13225-020-00466-2)) |
| Otu43624 | 1 | k__Fungi(100);p__Ascomycota(100);c__Saccharomycetes(100);o__Saccharomycetales(100);f__Metschnikowiaceae(100);g__Metschnikowia(100);s__Metschnikowia_pulcherrima(100); | Metschnikowia | 13 | Saprotroph | \|Undefined Saprotroph\| | Yeast | NULL | Probable | As nectar-tap_saprotroph (Põlme et al. 2020); Decay sugar-rich substrates (Põlme et al. 2020) | Põlme S, et al. 2020. Fungal Diversity 105:1-16 ((https://doi.org/10.1007/s13225-020-00466-2)) |
| Otu43625 | 1 | k__Fungi(100);p__Ascomycota(100);c__Saccharomycetes(100);o__Saccharomycetales(100);f__Metschnikowiaceae(100);g__Metschnikowia(100);s__Metschnikowia_pulcherrima(100); | Metschnikowia | 13 | Saprotroph | \|Undefined Saprotroph\| | Yeast | NULL | Probable | As nectar-tap_saprotroph (Põlme et al. 2020); Decay sugar-rich substrates (Põlme et al. 2020) | Põlme S, et al. 2020. Fungal Diversity 105:1-16 ((https://doi.org/10.1007/s13225-020-00466-2)) |
| Otu43626 | 1 | k__Fungi(100);p__Ascomycota(100);c__Saccharomycetes(100);o__Saccharomycetales(100);f__Metschnikowiaceae(100);g__Metschnikowia(100);s__Metschnikowia_pulcherrima(100); | Metschnikowia | 13 | Saprotroph | \|Undefined Saprotroph\| | Yeast | NULL | Probable | As nectar-tap_saprotroph (Põlme et al. 2020); Decay sugar-rich substrates (Põlme et al. 2020) | Põlme S, et al. 2020. Fungal Diversity 105:1-16 ((https://doi.org/10.1007/s13225-020-00466-2)) |
| Otu43627 | 1 | k__Fungi(100);p__Ascomycota(100);c__Saccharomycetes(100);o__Saccharomycetales(100);f__Metschnikowiaceae(100);g__Metschnikowia(100);s__Metschnikowia_pulcherrima(100); | Metschnikowia | 13 | Saprotroph | \|Undefined Saprotroph\| | Yeast | NULL | Probable | As nectar-tap_saprotroph (Põlme et al. 2020); Decay sugar-rich substrates (Põlme et al. 2020) | Põlme S, et al. 2020. Fungal Diversity 105:1-16 ((https://doi.org/10.1007/s13225-020-00466-2)) |
| Otu43628 | 1 | k__Fungi(100);p__Ascomycota(100);c__Saccharomycetes(100);o__Saccharomycetales(100);f__Metschnikowiaceae(100);g__Metschnikowia(100);s__Metschnikowia_pulcherrima(100); | Metschnikowia | 13 | Saprotroph | \|Undefined Saprotroph\| | Yeast | NULL | Probable | As nectar-tap_saprotroph (Põlme et al. 2020); Decay sugar-rich substrates (Põlme et al. 2020) | Põlme S, et al. 2020. Fungal Diversity 105:1-16 ((https://doi.org/10.1007/s13225-020-00466-2)) |
| Otu43629 | 1 | k__Fungi(100);p__Ascomycota(100);c__Saccharomycetes(100);o__Saccharomycetales(100);f__Metschnikowiaceae(100);g__Metschnikowia(100);g__Metschnikowia_unclassified(100); | Metschnikowia | 13 | Saprotroph | \|Undefined Saprotroph\| | Yeast | NULL | Probable | As nectar-tap_saprotroph (Põlme et al. 2020); Decay sugar-rich substrates (Põlme et al. 2020) | Põlme S, et al. 2020. Fungal Diversity 105:1-16 ((https://doi.org/10.1007/s13225-020-00466-2)) |
| Otu43630 | 1 | k__Fungi(100);p__Ascomycota(100);c__Saccharomycetes(100);o__Saccharomycetales(100);f__Metschnikowiaceae(100);g__Metschnikowia(100);g__Metschnikowia_unclassified(100); | Metschnikowia | 13 | Saprotroph | \|Undefined Saprotroph\| | Yeast | NULL | Probable | As nectar-tap_saprotroph (Põlme et al. 2020); Decay sugar-rich substrates (Põlme et al. 2020) | Põlme S, et al. 2020. Fungal Diversity 105:1-16 ((https://doi.org/10.1007/s13225-020-00466-2)) |
| Otu43631 | 1 | k__Fungi(100);p__Ascomycota(100);c__Saccharomycetes(100);o__Saccharomycetales(100);f__Metschnikowiaceae(100);g__Metschnikowia(100);s__Metschnikowia_pulcherrima(100); | Metschnikowia | 13 | Saprotroph | \|Undefined Saprotroph\| | Yeast | NULL | Probable | As nectar-tap_saprotroph (Põlme et al. 2020); Decay sugar-rich substrates (Põlme et al. 2020) | Põlme S, et al. 2020. Fungal Diversity 105:1-16 ((https://doi.org/10.1007/s13225-020-00466-2)) |
| Otu43632 | 1 | k__Fungi(100);p__Ascomycota(100);c__Saccharomycetes(100);o__Saccharomycetales(100);f__Metschnikowiaceae(100);g__Metschnikowia(100);s__Metschnikowia_pulcherrima(100); | Metschnikowia | 13 | Saprotroph | \|Undefined Saprotroph\| | Yeast | NULL | Probable | As nectar-tap_saprotroph (Põlme et al. 2020); Decay sugar-rich substrates (Põlme et al. 2020) | Põlme S, et al. 2020. Fungal Diversity 105:1-16 ((https://doi.org/10.1007/s13225-020-00466-2)) |
| Otu43633 | 1 | k__Fungi(100);p__Ascomycota(100);c__Saccharomycetes(100);o__Saccharomycetales(100);f__Metschnikowiaceae(100);g__Metschnikowia(100);s__Metschnikowia_pulcherrima(100); | Metschnikowia | 13 | Saprotroph | \|Undefined Saprotroph\| | Yeast | NULL | Probable | As nectar-tap_saprotroph (Põlme et al. 2020); Decay sugar-rich substrates (Põlme et al. 2020) | Põlme S, et al. 2020. Fungal Diversity 105:1-16 ((https://doi.org/10.1007/s13225-020-00466-2)) |
| Otu43634 | 1 | k__Fungi(100);p__Ascomycota(100);c__Saccharomycetes(100);o__Saccharomycetales(100);f__Metschnikowiaceae(100);g__Metschnikowia(100);s__Metschnikowia_pulcherrima(100); | Metschnikowia | 13 | Saprotroph | \|Undefined Saprotroph\| | Yeast | NULL | Probable | As nectar-tap_saprotroph (Põlme et al. 2020); Decay sugar-rich substrates (Põlme et al. 2020) | Põlme S, et al. 2020. Fungal Diversity 105:1-16 ((https://doi.org/10.1007/s13225-020-00466-2)) |
| Otu43635 | 1 | k__Fungi(100);p__Ascomycota(100);c__Saccharomycetes(100);o__Saccharomycetales(100);f__Metschnikowiaceae(100);g__Metschnikowia(100);s__Metschnikowia_pulcherrima(100); | Metschnikowia | 13 | Saprotroph | \|Undefined Saprotroph\| | Yeast | NULL | Probable | As nectar-tap_saprotroph (Põlme et al. 2020); Decay sugar-rich substrates (Põlme et al. 2020) | Põlme S, et al. 2020. Fungal Diversity 105:1-16 ((https://doi.org/10.1007/s13225-020-00466-2)) |
| Otu43636 | 1 | k__Fungi(100);p__Ascomycota(100);c__Saccharomycetes(100);o__Saccharomycetales(100);f__Metschnikowiaceae(100);g__Metschnikowia(100);s__Metschnikowia_pulcherrima(100); | Metschnikowia | 13 | Saprotroph | \|Undefined Saprotroph\| | Yeast | NULL | Probable | As nectar-tap_saprotroph (Põlme et al. 2020); Decay sugar-rich substrates (Põlme et al. 2020) | Põlme S, et al. 2020. Fungal Diversity 105:1-16 ((https://doi.org/10.1007/s13225-020-00466-2)) |
| Otu43637 | 1 | k__Fungi(100);p__Ascomycota(100);c__Saccharomycetes(100);o__Saccharomycetales(100);f__Metschnikowiaceae(100);g__Metschnikowia(100);s__Metschnikowia_pulcherrima(100); | Metschnikowia | 13 | Saprotroph | \|Undefined Saprotroph\| | Yeast | NULL | Probable | As nectar-tap_saprotroph (Põlme et al. 2020); Decay sugar-rich substrates (Põlme et al. 2020) | Põlme S, et al. 2020. Fungal Diversity 105:1-16 ((https://doi.org/10.1007/s13225-020-00466-2)) |
| Otu43638 | 1 | k__Fungi(100);p__Ascomycota(100);c__Saccharomycetes(100);o__Saccharomycetales(100);f__Metschnikowiaceae(100);g__Metschnikowia(100);s__Metschnikowia_pulcherrima(100); | Metschnikowia | 13 | Saprotroph | \|Undefined Saprotroph\| | Yeast | NULL | Probable | As nectar-tap_saprotroph (Põlme et al. 2020); Decay sugar-rich substrates (Põlme et al. 2020) | Põlme S, et al. 2020. Fungal Diversity 105:1-16 ((https://doi.org/10.1007/s13225-020-00466-2)) |
| Otu43639 | 1 | k__Fungi(100);p__Ascomycota(100);c__Saccharomycetes(100);o__Saccharomycetales(100);f__Metschnikowiaceae(100);g__Metschnikowia(100);s__Metschnikowia_pulcherrima(100); | Metschnikowia | 13 | Saprotroph | \|Undefined Saprotroph\| | Yeast | NULL | Probable | As nectar-tap_saprotroph (Põlme et al. 2020); Decay sugar-rich substrates (Põlme et al. 2020) | Põlme S, et al. 2020. Fungal Diversity 105:1-16 ((https://doi.org/10.1007/s13225-020-00466-2)) |
| Otu43640 | 1 | k__Fungi(100);p__Ascomycota(100);c__Saccharomycetes(100);o__Saccharomycetales(100);f__Metschnikowiaceae(100);g__Metschnikowia(100);g__Metschnikowia_unclassified(100); | Metschnikowia | 13 | Saprotroph | \|Undefined Saprotroph\| | Yeast | NULL | Probable | As nectar-tap_saprotroph (Põlme et al. 2020); Decay sugar-rich substrates (Põlme et al. 2020) | Põlme S, et al. 2020. Fungal Diversity 105:1-16 ((https://doi.org/10.1007/s13225-020-00466-2)) |
| Otu43641 | 1 | k__Fungi(100);p__Ascomycota(100);c__Saccharomycetes(100);o__Saccharomycetales(100);f__Metschnikowiaceae(100);g__Metschnikowia(100);s__Metschnikowia_pulcherrima(100); | Metschnikowia | 13 | Saprotroph | \|Undefined Saprotroph\| | Yeast | NULL | Probable | As nectar-tap_saprotroph (Põlme et al. 2020); Decay sugar-rich substrates (Põlme et al. 2020) | Põlme S, et al. 2020. Fungal Diversity 105:1-16 ((https://doi.org/10.1007/s13225-020-00466-2)) |
| Otu43642 | 1 | k__Fungi(100);p__Ascomycota(100);c__Saccharomycetes(100);o__Saccharomycetales(100);f__Metschnikowiaceae(100);g__Metschnikowia(100);s__Metschnikowia_pulcherrima(100); | Metschnikowia | 13 | Saprotroph | \|Undefined Saprotroph\| | Yeast | NULL | Probable | As nectar-tap_saprotroph (Põlme et al. 2020); Decay sugar-rich substrates (Põlme et al. 2020) | Põlme S, et al. 2020. Fungal Diversity 105:1-16 ((https://doi.org/10.1007/s13225-020-00466-2)) |
| Otu43644 | 1 | k__Fungi(100);p__Ascomycota(100);c__Saccharomycetes(100);o__Saccharomycetales(100);f__Metschnikowiaceae(100);g__Metschnikowia(100);s__Metschnikowia_pulcherrima(100); | Metschnikowia | 13 | Saprotroph | \|Undefined Saprotroph\| | Yeast | NULL | Probable | As nectar-tap_saprotroph (Põlme et al. 2020); Decay sugar-rich substrates (Põlme et al. 2020) | Põlme S, et al. 2020. Fungal Diversity 105:1-16 ((https://doi.org/10.1007/s13225-020-00466-2)) |
| Otu43645 | 1 | k__Fungi(100);p__Ascomycota(100);c__Saccharomycetes(100);o__Saccharomycetales(100);f__Metschnikowiaceae(100);g__Metschnikowia(100);g__Metschnikowia_unclassified(100); | Metschnikowia | 13 | Saprotroph | \|Undefined Saprotroph\| | Yeast | NULL | Probable | As nectar-tap_saprotroph (Põlme et al. 2020); Decay sugar-rich substrates (Põlme et al. 2020) | Põlme S, et al. 2020. Fungal Diversity 105:1-16 ((https://doi.org/10.1007/s13225-020-00466-2)) |
| Otu43646 | 1 | k__Fungi(100);p__Ascomycota(100);c__Saccharomycetes(100);o__Saccharomycetales(100);f__Metschnikowiaceae(100);g__Metschnikowia(100);s__Metschnikowia_pulcherrima(100); | Metschnikowia | 13 | Saprotroph | \|Undefined Saprotroph\| | Yeast | NULL | Probable | As nectar-tap_saprotroph (Põlme et al. 2020); Decay sugar-rich substrates (Põlme et al. 2020) | Põlme S, et al. 2020. Fungal Diversity 105:1-16 ((https://doi.org/10.1007/s13225-020-00466-2)) |
| Otu43647 | 1 | k__Fungi(100);p__Ascomycota(100);c__Saccharomycetes(100);o__Saccharomycetales(100);f__Metschnikowiaceae(100);g__Metschnikowia(100);s__Metschnikowia_pulcherrima(100); | Metschnikowia | 13 | Saprotroph | \|Undefined Saprotroph\| | Yeast | NULL | Probable | As nectar-tap_saprotroph (Põlme et al. 2020); Decay sugar-rich substrates (Põlme et al. 2020) | Põlme S, et al. 2020. Fungal Diversity 105:1-16 ((https://doi.org/10.1007/s13225-020-00466-2)) |
| Otu43648 | 1 | k__Fungi(100);p__Ascomycota(100);c__Saccharomycetes(100);o__Saccharomycetales(100);f__Metschnikowiaceae(100);g__Metschnikowia(100);g__Metschnikowia_unclassified(100); | Metschnikowia | 13 | Saprotroph | \|Undefined Saprotroph\| | Yeast | NULL | Probable | As nectar-tap_saprotroph (Põlme et al. 2020); Decay sugar-rich substrates (Põlme et al. 2020) | Põlme S, et al. 2020. Fungal Diversity 105:1-16 ((https://doi.org/10.1007/s13225-020-00466-2)) |
| Otu43649 | 1 | k__Fungi(100);p__Ascomycota(100);c__Saccharomycetes(100);o__Saccharomycetales(100);f__Metschnikowiaceae(100);g__Metschnikowia(100);s__Metschnikowia_pulcherrima(100); | Metschnikowia | 13 | Saprotroph | \|Undefined Saprotroph\| | Yeast | NULL | Probable | As nectar-tap_saprotroph (Põlme et al. 2020); Decay sugar-rich substrates (Põlme et al. 2020) | Põlme S, et al. 2020. Fungal Diversity 105:1-16 ((https://doi.org/10.1007/s13225-020-00466-2)) |
| Otu43650 | 1 | k__Fungi(100);p__Ascomycota(100);c__Saccharomycetes(100);o__Saccharomycetales(100);f__Metschnikowiaceae(100);g__Metschnikowia(100);s__Metschnikowia_pulcherrima(100); | Metschnikowia | 13 | Saprotroph | \|Undefined Saprotroph\| | Yeast | NULL | Probable | As nectar-tap_saprotroph (Põlme et al. 2020); Decay sugar-rich substrates (Põlme et al. 2020) | Põlme S, et al. 2020. Fungal Diversity 105:1-16 ((https://doi.org/10.1007/s13225-020-00466-2)) |
| Otu43651 | 1 | k__Fungi(100);p__Ascomycota(100);c__Saccharomycetes(100);o__Saccharomycetales(100);f__Metschnikowiaceae(100);g__Metschnikowia(100);s__Metschnikowia_pulcherrima(100); | Metschnikowia | 13 | Saprotroph | \|Undefined Saprotroph\| | Yeast | NULL | Probable | As nectar-tap_saprotroph (Põlme et al. 2020); Decay sugar-rich substrates (Põlme et al. 2020) | Põlme S, et al. 2020. Fungal Diversity 105:1-16 ((https://doi.org/10.1007/s13225-020-00466-2)) |
| Otu43652 | 1 | k__Fungi(100);p__Ascomycota(100);c__Saccharomycetes(100);o__Saccharomycetales(100);f__Metschnikowiaceae(100);g__Metschnikowia(100);g__Metschnikowia_unclassified(100); | Metschnikowia | 13 | Saprotroph | \|Undefined Saprotroph\| | Yeast | NULL | Probable | As nectar-tap_saprotroph (Põlme et al. 2020); Decay sugar-rich substrates (Põlme et al. 2020) | Põlme S, et al. 2020. Fungal Diversity 105:1-16 ((https://doi.org/10.1007/s13225-020-00466-2)) |
| Otu43654 | 1 | k__Fungi(100);p__Ascomycota(100);c__Saccharomycetes(100);o__Saccharomycetales(100);f__Metschnikowiaceae(100);g__Metschnikowia(100);s__Metschnikowia_pulcherrima(100); | Metschnikowia | 13 | Saprotroph | \|Undefined Saprotroph\| | Yeast | NULL | Probable | As nectar-tap_saprotroph (Põlme et al. 2020); Decay sugar-rich substrates (Põlme et al. 2020) | Põlme S, et al. 2020. Fungal Diversity 105:1-16 ((https://doi.org/10.1007/s13225-020-00466-2)) |
| Otu43655 | 1 | k__Fungi(100);p__Ascomycota(100);c__Saccharomycetes(100);o__Saccharomycetales(100);f__Metschnikowiaceae(100);g__Metschnikowia(100);g__Metschnikowia_unclassified(100); | Metschnikowia | 13 | Saprotroph | \|Undefined Saprotroph\| | Yeast | NULL | Probable | As nectar-tap_saprotroph (Põlme et al. 2020); Decay sugar-rich substrates (Põlme et al. 2020) | Põlme S, et al. 2020. Fungal Diversity 105:1-16 ((https://doi.org/10.1007/s13225-020-00466-2)) |
| Otu43656 | 1 | k__Fungi(100);p__Ascomycota(100);c__Saccharomycetes(100);o__Saccharomycetales(100);f__Metschnikowiaceae(100);g__Metschnikowia(100);s__Metschnikowia_pulcherrima(100); | Metschnikowia | 13 | Saprotroph | \|Undefined Saprotroph\| | Yeast | NULL | Probable | As nectar-tap_saprotroph (Põlme et al. 2020); Decay sugar-rich substrates (Põlme et al. 2020) | Põlme S, et al. 2020. Fungal Diversity 105:1-16 ((https://doi.org/10.1007/s13225-020-00466-2)) |
| Otu43657 | 1 | k__Fungi(100);p__Ascomycota(100);c__Saccharomycetes(100);o__Saccharomycetales(100);f__Metschnikowiaceae(100);g__Metschnikowia(100);s__Metschnikowia_pulcherrima(100); | Metschnikowia | 13 | Saprotroph | \|Undefined Saprotroph\| | Yeast | NULL | Probable | As nectar-tap_saprotroph (Põlme et al. 2020); Decay sugar-rich substrates (Põlme et al. 2020) | Põlme S, et al. 2020. Fungal Diversity 105:1-16 ((https://doi.org/10.1007/s13225-020-00466-2)) |
| Otu43660 | 1 | k__Fungi(100);p__Ascomycota(100);c__Saccharomycetes(100);o__Saccharomycetales(100);f__Metschnikowiaceae(100);g__Metschnikowia(100);s__Metschnikowia_pulcherrima(100); | Metschnikowia | 13 | Saprotroph | \|Undefined Saprotroph\| | Yeast | NULL | Probable | As nectar-tap_saprotroph (Põlme et al. 2020); Decay sugar-rich substrates (Põlme et al. 2020) | Põlme S, et al. 2020. Fungal Diversity 105:1-16 ((https://doi.org/10.1007/s13225-020-00466-2)) |
| Otu43661 | 1 | k__Fungi(100);p__Ascomycota(100);c__Saccharomycetes(100);o__Saccharomycetales(100);f__Metschnikowiaceae(100);g__Metschnikowia(100);g__Metschnikowia_unclassified(100); | Metschnikowia | 13 | Saprotroph | \|Undefined Saprotroph\| | Yeast | NULL | Probable | As nectar-tap_saprotroph (Põlme et al. 2020); Decay sugar-rich substrates (Põlme et al. 2020) | Põlme S, et al. 2020. Fungal Diversity 105:1-16 ((https://doi.org/10.1007/s13225-020-00466-2)) |
| Otu43662 | 1 | k__Fungi(100);p__Ascomycota(100);c__Saccharomycetes(100);o__Saccharomycetales(100);f__Metschnikowiaceae(100);g__Metschnikowia(100);s__Metschnikowia_pulcherrima(100); | Metschnikowia | 13 | Saprotroph | \|Undefined Saprotroph\| | Yeast | NULL | Probable | As nectar-tap_saprotroph (Põlme et al. 2020); Decay sugar-rich substrates (Põlme et al. 2020) | Põlme S, et al. 2020. Fungal Diversity 105:1-16 ((https://doi.org/10.1007/s13225-020-00466-2)) |
| Otu43663 | 1 | k__Fungi(100);p__Ascomycota(100);c__Saccharomycetes(100);o__Saccharomycetales(100);f__Metschnikowiaceae(100);g__Metschnikowia(100);g__Metschnikowia_unclassified(100); | Metschnikowia | 13 | Saprotroph | \|Undefined Saprotroph\| | Yeast | NULL | Probable | As nectar-tap_saprotroph (Põlme et al. 2020); Decay sugar-rich substrates (Põlme et al. 2020) | Põlme S, et al. 2020. Fungal Diversity 105:1-16 ((https://doi.org/10.1007/s13225-020-00466-2)) |
| Otu43664 | 1 | k__Fungi(100);p__Ascomycota(100);c__Saccharomycetes(100);o__Saccharomycetales(100);f__Metschnikowiaceae(100);g__Metschnikowia(100);s__Metschnikowia_pulcherrima(100); | Metschnikowia | 13 | Saprotroph | \|Undefined Saprotroph\| | Yeast | NULL | Probable | As nectar-tap_saprotroph (Põlme et al. 2020); Decay sugar-rich substrates (Põlme et al. 2020) | Põlme S, et al. 2020. Fungal Diversity 105:1-16 ((https://doi.org/10.1007/s13225-020-00466-2)) |
| Otu43665 | 1 | k__Fungi(100);p__Ascomycota(100);c__Saccharomycetes(100);o__Saccharomycetales(100);f__Metschnikowiaceae(100);g__Metschnikowia(100);s__Metschnikowia_pulcherrima(100); | Metschnikowia | 13 | Saprotroph | \|Undefined Saprotroph\| | Yeast | NULL | Probable | As nectar-tap_saprotroph (Põlme et al. 2020); Decay sugar-rich substrates (Põlme et al. 2020) | Põlme S, et al. 2020. Fungal Diversity 105:1-16 ((https://doi.org/10.1007/s13225-020-00466-2)) |
| Otu43666 | 1 | k__Fungi(100);p__Ascomycota(100);c__Saccharomycetes(100);o__Saccharomycetales(100);f__Metschnikowiaceae(100);g__Metschnikowia(100);s__Metschnikowia_pulcherrima(100); | Metschnikowia | 13 | Saprotroph | \|Undefined Saprotroph\| | Yeast | NULL | Probable | As nectar-tap_saprotroph (Põlme et al. 2020); Decay sugar-rich substrates (Põlme et al. 2020) | Põlme S, et al. 2020. Fungal Diversity 105:1-16 ((https://doi.org/10.1007/s13225-020-00466-2)) |
| Otu43667 | 1 | k__Fungi(100);p__Ascomycota(100);c__Saccharomycetes(100);o__Saccharomycetales(100);f__Metschnikowiaceae(100);g__Metschnikowia(100);s__Metschnikowia_pulcherrima(100); | Metschnikowia | 13 | Saprotroph | \|Undefined Saprotroph\| | Yeast | NULL | Probable | As nectar-tap_saprotroph (Põlme et al. 2020); Decay sugar-rich substrates (Põlme et al. 2020) | Põlme S, et al. 2020. Fungal Diversity 105:1-16 ((https://doi.org/10.1007/s13225-020-00466-2)) |
| Otu43668 | 1 | k__Fungi(100);p__Ascomycota(100);c__Saccharomycetes(100);o__Saccharomycetales(100);f__Metschnikowiaceae(100);g__Metschnikowia(100);s__Metschnikowia_pulcherrima(100); | Metschnikowia | 13 | Saprotroph | \|Undefined Saprotroph\| | Yeast | NULL | Probable | As nectar-tap_saprotroph (Põlme et al. 2020); Decay sugar-rich substrates (Põlme et al. 2020) | Põlme S, et al. 2020. Fungal Diversity 105:1-16 ((https://doi.org/10.1007/s13225-020-00466-2)) |
| Otu43669 | 1 | k__Fungi(100);p__Ascomycota(100);c__Saccharomycetes(100);o__Saccharomycetales(100);f__Metschnikowiaceae(100);g__Metschnikowia(100);s__Metschnikowia_pulcherrima(100); | Metschnikowia | 13 | Saprotroph | \|Undefined Saprotroph\| | Yeast | NULL | Probable | As nectar-tap_saprotroph (Põlme et al. 2020); Decay sugar-rich substrates (Põlme et al. 2020) | Põlme S, et al. 2020. Fungal Diversity 105:1-16 ((https://doi.org/10.1007/s13225-020-00466-2)) |
| Otu43670 | 1 | k__Fungi(100);p__Ascomycota(100);c__Saccharomycetes(100);o__Saccharomycetales(100);f__Metschnikowiaceae(100);g__Metschnikowia(100);s__Metschnikowia_pulcherrima(100); | Metschnikowia | 13 | Saprotroph | \|Undefined Saprotroph\| | Yeast | NULL | Probable | As nectar-tap_saprotroph (Põlme et al. 2020); Decay sugar-rich substrates (Põlme et al. 2020) | Põlme S, et al. 2020. Fungal Diversity 105:1-16 ((https://doi.org/10.1007/s13225-020-00466-2)) |
| Otu43671 | 1 | k__Fungi(100);p__Ascomycota(100);c__Saccharomycetes(100);o__Saccharomycetales(100);f__Metschnikowiaceae(100);g__Metschnikowia(100);s__Metschnikowia_pulcherrima(100); | Metschnikowia | 13 | Saprotroph | \|Undefined Saprotroph\| | Yeast | NULL | Probable | As nectar-tap_saprotroph (Põlme et al. 2020); Decay sugar-rich substrates (Põlme et al. 2020) | Põlme S, et al. 2020. Fungal Diversity 105:1-16 ((https://doi.org/10.1007/s13225-020-00466-2)) |

| | | | | | | | | | | | |
|---|---|---|---|---|---|---|---|---|---|---|---|
| Otu43672 | 1 | k__Fungi(100);p__Ascomycota(100);c__Saccharomycetes(100);o__Saccharomycetales(100);f__Metschnikowiaceae(100);g__Metschnikowia(100);g__Metschnikowia_unclassified(100); | Metschnikowia | 13 | Saprotroph | \|Undefined Saprotroph\| | Yeast | NULL | Probable | As nectar-tap_saprotroph (Põlme et al. 2020); Decay sugar-rich substrates (Põlme et al. 2020) | Põlme S, et al. 2020. Fungal Diversity 105:1-16 ((https://doi.org/10.1007/s13225-020-00466-2)) |
| Otu43674 | 1 | k__Fungi(100);p__Ascomycota(100);c__Saccharomycetes(100);o__Saccharomycetales(100);f__Metschnikowiaceae(100);g__Metschnikowia(100);g__Metschnikowia_unclassified(100); | Metschnikowia | 13 | Saprotroph | \|Undefined Saprotroph\| | Yeast | NULL | Probable | As nectar-tap_saprotroph (Põlme et al. 2020); Decay sugar-rich substrates (Põlme et al. 2020) | Põlme S, et al. 2020. Fungal Diversity 105:1-16 ((https://doi.org/10.1007/s13225-020-00466-2)) |
| Otu43675 | 1 | k__Fungi(100);p__Ascomycota(100);c__Saccharomycetes(100);o__Saccharomycetales(100);f__Metschnikowiaceae(100);g__Metschnikowia(100);s__Metschnikowia_pulcherrima(100); | Metschnikowia | 13 | Saprotroph | \|Undefined Saprotroph\| | Yeast | NULL | Probable | As nectar-tap_saprotroph (Põlme et al. 2020); Decay sugar-rich substrates (Põlme et al. 2020) | Põlme S, et al. 2020. Fungal Diversity 105:1-16 ((https://doi.org/10.1007/s13225-020-00466-2)) |
| Otu43676 | 1 | k__Fungi(100);p__Ascomycota(100);c__Saccharomycetes(100);o__Saccharomycetales(100);f__Metschnikowiaceae(100);g__Metschnikowia(100);s__Metschnikowia_pulcherrima(100); | Metschnikowia | 13 | Saprotroph | \|Undefined Saprotroph\| | Yeast | NULL | Probable | As nectar-tap_saprotroph (Põlme et al. 2020); Decay sugar-rich substrates (Põlme et al. 2020) | Põlme S, et al. 2020. Fungal Diversity 105:1-16 ((https://doi.org/10.1007/s13225-020-00466-2)) |
| Otu43677 | 1 | k__Fungi(100);p__Ascomycota(100);c__Saccharomycetes(100);o__Saccharomycetales(100);f__Metschnikowiaceae(100);g__Metschnikowia(100);s__Metschnikowia_pulcherrima(100); | Metschnikowia | 13 | Saprotroph | \|Undefined Saprotroph\| | Yeast | NULL | Probable | As nectar-tap_saprotroph (Põlme et al. 2020); Decay sugar-rich substrates (Põlme et al. 2020) | Põlme S, et al. 2020. Fungal Diversity 105:1-16 ((https://doi.org/10.1007/s13225-020-00466-2)) |
| Otu43678 | 1 | k__Fungi(100);p__Ascomycota(100);c__Saccharomycetes(100);o__Saccharomycetales(100);f__Metschnikowiaceae(100);g__Metschnikowia(100);s__Metschnikowia_pulcherrima(100); | Metschnikowia | 13 | Saprotroph | \|Undefined Saprotroph\| | Yeast | NULL | Probable | As nectar-tap_saprotroph (Põlme et al. 2020); Decay sugar-rich substrates (Põlme et al. 2020) | Põlme S, et al. 2020. Fungal Diversity 105:1-16 ((https://doi.org/10.1007/s13225-020-00466-2)) |
| Otu43679 | 1 | k__Fungi(100);p__Ascomycota(100);c__Saccharomycetes(100);o__Saccharomycetales(100);f__Metschnikowiaceae(100);g__Metschnikowia(100);s__Metschnikowia_pulcherrima(100); | Metschnikowia | 13 | Saprotroph | \|Undefined Saprotroph\| | Yeast | NULL | Probable | As nectar-tap_saprotroph (Põlme et al. 2020); Decay sugar-rich substrates (Põlme et al. 2020) | Põlme S, et al. 2020. Fungal Diversity 105:1-16 ((https://doi.org/10.1007/s13225-020-00466-2)) |
| Otu43680 | 1 | k__Fungi(100);p__Ascomycota(100);c__Saccharomycetes(100);o__Saccharomycetales(100);f__Metschnikowiaceae(100);g__Metschnikowia(100);g__Metschnikowia_unclassified(100); | Metschnikowia | 13 | Saprotroph | \|Undefined Saprotroph\| | Yeast | NULL | Probable | As nectar-tap_saprotroph (Põlme et al. 2020); Decay sugar-rich substrates (Põlme et al. 2020) | Põlme S, et al. 2020. Fungal Diversity 105:1-16 ((https://doi.org/10.1007/s13225-020-00466-2)) |
| Otu43681 | 1 | k__Fungi(100);p__Ascomycota(100);c__Saccharomycetes(100);o__Saccharomycetales(100);f__Metschnikowiaceae(100);g__Metschnikowia(100);s__Metschnikowia_pulcherrima(100); | Metschnikowia | 13 | Saprotroph | \|Undefined Saprotroph\| | Yeast | NULL | Probable | As nectar-tap_saprotroph (Põlme et al. 2020); Decay sugar-rich substrates (Põlme et al. 2020) | Põlme S, et al. 2020. Fungal Diversity 105:1-16 ((https://doi.org/10.1007/s13225-020-00466-2)) |
| Otu43682 | 1 | k__Fungi(100);p__Ascomycota(100);c__Sordariomycetes(100);o__Sordariales(100);f__Chaetomiaceae(100);g__Humicola(100);s__Humicola_grisea_var._grisea(100); | Humicola | 13 | Pathotroph-Saprotroph | Plant Pathogen-Undefined Saprotroph-\|Wood Saprotroph\| | Microfungus | Soft Rot | Probable | Decay wood (Põlme et al. 2020) | Seehann G, et al. 1975. List of Fungi in Soft-Rot Tissue Tests. Institute fur Holzbiologie und Holzschutz, Bundesforschungsanstalt fur Forst - und Holzwirtshaft, Hamburg - Lohbrugge, Germany; Costa IPMW, et al. 2012. Mycotaxon 119:e26 ((http://mycotaxon.com/resources/checklists/costa_v119_checklist.pdf)); Tedersoo L, et al. 2014. Science 346:e1256688 ((https://doi.org/10.1126/science.1256688)); Põlme S, et al. 2020. Fungal Diversity 105:1-16 ((https://doi.org/10.1007/s13225-020-00466-2)) |
| Otu43683 | 1 | k__Fungi(100);p__Ascomycota(100);c__Saccharomycetes(100);o__Saccharomycetales(100);f__Metschnikowiaceae(100);g__Metschnikowia(100);s__Metschnikowia_pulcherrima(100); | Metschnikowia | 13 | Saprotroph | \|Undefined Saprotroph\| | Yeast | NULL | Probable | As nectar-tap_saprotroph (Põlme et al. 2020); Decay sugar-rich substrates (Põlme et al. 2020) | Põlme S, et al. 2020. Fungal Diversity 105:1-16 ((https://doi.org/10.1007/s13225-020-00466-2)) |
| Otu43684 | 1 | k__Fungi(100);p__Ascomycota(100);c__Saccharomycetes(100);o__Saccharomycetales(100);f__Metschnikowiaceae(100);g__Metschnikowia(100);g__Metschnikowia_unclassified(100); | Metschnikowia | 13 | Saprotroph | \|Undefined Saprotroph\| | Yeast | NULL | Probable | As nectar-tap_saprotroph (Põlme et al. 2020); Decay sugar-rich substrates (Põlme et al. 2020) | Põlme S, et al. 2020. Fungal Diversity 105:1-16 ((https://doi.org/10.1007/s13225-020-00466-2)) |
| Otu43685 | 1 | k__Fungi(100);p__Ascomycota(100);c__Saccharomycetes(100);o__Saccharomycetales(100);f__Metschnikowiaceae(100);g__Metschnikowia(100);g__Metschnikowia_unclassified(100); | Metschnikowia | 13 | Saprotroph | \|Undefined Saprotroph\| | Yeast | NULL | Probable | As nectar-tap_saprotroph (Põlme et al. 2020); Decay sugar-rich substrates (Põlme et al. 2020) | Põlme S, et al. 2020. Fungal Diversity 105:1-16 ((https://doi.org/10.1007/s13225-020-00466-2)) |
| Otu43686 | 1 | k__Fungi(100);p__Ascomycota(100);c__Saccharomycetes(100);o__Saccharomycetales(100);f__Metschnikowiaceae(100);g__Metschnikowia(100);g__Metschnikowia_unclassified(100); | Metschnikowia | 13 | Saprotroph | \|Undefined Saprotroph\| | Yeast | NULL | Probable | As nectar-tap_saprotroph (Põlme et al. 2020); Decay sugar-rich substrates (Põlme et al. 2020) | Põlme S, et al. 2020. Fungal Diversity 105:1-16 ((https://doi.org/10.1007/s13225-020-00466-2)) |
| Otu43687 | 1 | k__Fungi(100);p__Ascomycota(100);c__Saccharomycetes(100);o__Saccharomycetales(100);f__Metschnikowiaceae(100);g__Metschnikowia(100);g__Metschnikowia_unclassified(100); | Metschnikowia | 13 | Saprotroph | \|Undefined Saprotroph\| | Yeast | NULL | Probable | As nectar-tap_saprotroph (Põlme et al. 2020); Decay sugar-rich substrates (Põlme et al. 2020) | Põlme S, et al. 2020. Fungal Diversity 105:1-16 ((https://doi.org/10.1007/s13225-020-00466-2)) |
| Otu43688 | 1 | k__Fungi(100);p__Ascomycota(100);c__Saccharomycetes(100);o__Saccharomycetales(100);f__Metschnikowiaceae(100);g__Metschnikowia(100);g__Metschnikowia_unclassified(100); | Metschnikowia | 13 | Saprotroph | \|Undefined Saprotroph\| | Yeast | NULL | Probable | As nectar-tap_saprotroph (Põlme et al. 2020); Decay sugar-rich substrates (Põlme et al. 2020) | Põlme S, et al. 2020. Fungal Diversity 105:1-16 ((https://doi.org/10.1007/s13225-020-00466-2)) |
| Otu43689 | 1 | k__Fungi(100);p__Ascomycota(100);c__Saccharomycetes(100);o__Saccharomycetales(100);f__Metschnikowiaceae(100);g__Metschnikowia(100);s__Metschnikowia_pulcherrima(100); | Metschnikowia | 13 | Saprotroph | \|Undefined Saprotroph\| | Yeast | NULL | Probable | As nectar-tap_saprotroph (Põlme et al. 2020); Decay sugar-rich substrates (Põlme et al. 2020) | Põlme S, et al. 2020. Fungal Diversity 105:1-16 ((https://doi.org/10.1007/s13225-020-00466-2)) |
| Otu43690 | 1 | k__Fungi(100);p__Ascomycota(100);c__Saccharomycetes(100);o__Saccharomycetales(100);f__Metschnikowiaceae(100);g__Metschnikowia(100);s__Metschnikowia_pulcherrima(100); | Metschnikowia | 13 | Saprotroph | \|Undefined Saprotroph\| | Yeast | NULL | Probable | As nectar-tap_saprotroph (Põlme et al. 2020); Decay sugar-rich substrates (Põlme et al. 2020) | Põlme S, et al. 2020. Fungal Diversity 105:1-16 ((https://doi.org/10.1007/s13225-020-00466-2)) |
| Otu43691 | 1 | k__Fungi(100);p__Ascomycota(100);c__Saccharomycetes(100);o__Saccharomycetales(100);f__Metschnikowiaceae(100);g__Metschnikowia(100);s__Metschnikowia_pulcherrima(100); | Metschnikowia | 13 | Saprotroph | \|Undefined Saprotroph\| | Yeast | NULL | Probable | As nectar-tap_saprotroph (Põlme et al. 2020); Decay sugar-rich substrates (Põlme et al. 2020) | Põlme S, et al. 2020. Fungal Diversity 105:1-16 ((https://doi.org/10.1007/s13225-020-00466-2)) |
| Otu43692 | 1 | k__Fungi(100);p__Ascomycota(100);c__Saccharomycetes(100);o__Saccharomycetales(100);f__Metschnikowiaceae(100);g__Metschnikowia(100);s__Metschnikowia_pulcherrima(100); | Metschnikowia | 13 | Saprotroph | \|Undefined Saprotroph\| | Yeast | NULL | Probable | As nectar-tap_saprotroph (Põlme et al. 2020); Decay sugar-rich substrates (Põlme et al. 2020) | Põlme S, et al. 2020. Fungal Diversity 105:1-16 ((https://doi.org/10.1007/s13225-020-00466-2)) |
| Otu43693 | 1 | k__Fungi(100);p__Ascomycota(100);c__Saccharomycetes(100);o__Saccharomycetales(100);f__Metschnikowiaceae(100);g__Metschnikowia(100);s__Metschnikowia_pulcherrima(100); | Metschnikowia | 13 | Saprotroph | \|Undefined Saprotroph\| | Yeast | NULL | Probable | As nectar-tap_saprotroph (Põlme et al. 2020); Decay sugar-rich substrates (Põlme et al. 2020) | Põlme S, et al. 2020. Fungal Diversity 105:1-16 ((https://doi.org/10.1007/s13225-020-00466-2)) |
| Otu43694 | 1 | k__Fungi(100);p__Ascomycota(100);c__Saccharomycetes(100);o__Saccharomycetales(100);f__Metschnikowiaceae(100);g__Metschnikowia(100);s__Metschnikowia_pulcherrima(100); | Metschnikowia | 13 | Saprotroph | \|Undefined Saprotroph\| | Yeast | NULL | Probable | As nectar-tap_saprotroph (Põlme et al. 2020); Decay sugar-rich substrates (Põlme et al. 2020) | Põlme S, et al. 2020. Fungal Diversity 105:1-16 ((https://doi.org/10.1007/s13225-020-00466-2)) |
| Otu43695 | 1 | k__Fungi(100);p__Ascomycota(100);c__Saccharomycetes(100);o__Saccharomycetales(100);f__Metschnikowiaceae(100);g__Metschnikowia(100);s__Metschnikowia_pulcherrima(100); | Metschnikowia | 13 | Saprotroph | \|Undefined Saprotroph\| | Yeast | NULL | Probable | As nectar-tap_saprotroph (Põlme et al. 2020); Decay sugar-rich substrates (Põlme et al. 2020) | Põlme S, et al. 2020. Fungal Diversity 105:1-16 ((https://doi.org/10.1007/s13225-020-00466-2)) |
| Otu43696 | 1 | k__Fungi(100);p__Ascomycota(100);c__Saccharomycetes(100);o__Saccharomycetales(100);f__Metschnikowiaceae(100);g__Metschnikowia(100);s__Metschnikowia_pulcherrima(100); | Metschnikowia | 13 | Saprotroph | \|Undefined Saprotroph\| | Yeast | NULL | Probable | As nectar-tap_saprotroph (Põlme et al. 2020); Decay sugar-rich substrates (Põlme et al. 2020) | Põlme S, et al. 2020. Fungal Diversity 105:1-16 ((https://doi.org/10.1007/s13225-020-00466-2)) |
| Otu43697 | 1 | k__Fungi(100);p__Ascomycota(100);c__Saccharomycetes(100);o__Saccharomycetales(100);f__Metschnikowiaceae(100);g__Metschnikowia(100);s__Metschnikowia_pulcherrima(100); | Metschnikowia | 13 | Saprotroph | \|Undefined Saprotroph\| | Yeast | NULL | Probable | As nectar-tap_saprotroph (Põlme et al. 2020); Decay sugar-rich substrates (Põlme et al. 2020) | Põlme S, et al. 2020. Fungal Diversity 105:1-16 ((https://doi.org/10.1007/s13225-020-00466-2)) |
| Otu43698 | 1 | k__Fungi(100);p__Ascomycota(100);c__Saccharomycetes(100);o__Saccharomycetales(100);f__Metschnikowiaceae(100);g__Metschnikowia(100);s__Metschnikowia_pulcherrima(100); | Metschnikowia | 13 | Saprotroph | \|Undefined Saprotroph\| | Yeast | NULL | Probable | As nectar-tap_saprotroph (Põlme et al. 2020); Decay sugar-rich substrates (Põlme et al. 2020) | Põlme S, et al. 2020. Fungal Diversity 105:1-16 ((https://doi.org/10.1007/s13225-020-00466-2)) |
| Otu43699 | 1 | k__Fungi(100);p__Ascomycota(100);c__Saccharomycetes(100);o__Saccharomycetales(100);f__Metschnikowiaceae(100);g__Metschnikowia(100);s__Metschnikowia_pulcherrima(100); | Metschnikowia | 13 | Saprotroph | \|Undefined Saprotroph\| | Yeast | NULL | Probable | As nectar-tap_saprotroph (Põlme et al. 2020); Decay sugar-rich substrates (Põlme et al. 2020) | Põlme S, et al. 2020. Fungal Diversity 105:1-16 ((https://doi.org/10.1007/s13225-020-00466-2)) |
| Otu43700 | 1 | k__Fungi(100);p__Ascomycota(100);c__Saccharomycetes(100);o__Saccharomycetales(100);f__Metschnikowiaceae(100);g__Metschnikowia(100);s__Metschnikowia_pulcherrima(100); | Metschnikowia | 13 | Saprotroph | \|Undefined Saprotroph\| | Yeast | NULL | Probable | As nectar-tap_saprotroph (Põlme et al. 2020); Decay sugar-rich substrates (Põlme et al. 2020) | Põlme S, et al. 2020. Fungal Diversity 105:1-16 ((https://doi.org/10.1007/s13225-020-00466-2)) |
| Otu43701 | 1 | k__Fungi(100);p__Ascomycota(100);c__Saccharomycetes(100);o__Saccharomycetales(100);f__Metschnikowiaceae(100);g__Metschnikowia(100);s__Metschnikowia_pulcherrima(100); | Metschnikowia | 13 | Saprotroph | \|Undefined Saprotroph\| | Yeast | NULL | Probable | As nectar-tap_saprotroph (Põlme et al. 2020); Decay sugar-rich substrates (Põlme et al. 2020) | Põlme S, et al. 2020. Fungal Diversity 105:1-16 ((https://doi.org/10.1007/s13225-020-00466-2)) |
| Otu43702 | 1 | k__Fungi(100);p__Ascomycota(100);c__Saccharomycetes(100);o__Saccharomycetales(100);f__Metschnikowiaceae(100);g__Metschnikowia(100);g__Metschnikowia_unclassified(100); | Metschnikowia | 13 | Saprotroph | \|Undefined Saprotroph\| | Yeast | NULL | Probable | As nectar-tap_saprotroph (Põlme et al. 2020); Decay sugar-rich substrates (Põlme et al. 2020) | Põlme S, et al. 2020. Fungal Diversity 105:1-16 ((https://doi.org/10.1007/s13225-020-00466-2)) |
| Otu43703 | 1 | k__Fungi(100);p__Ascomycota(100);c__Saccharomycetes(100);o__Saccharomycetales(100);f__Metschnikowiaceae(100);g__Metschnikowia(100);s__Metschnikowia_pulcherrima(100); | Metschnikowia | 13 | Saprotroph | \|Undefined Saprotroph\| | Yeast | NULL | Probable | As nectar-tap_saprotroph (Põlme et al. 2020); Decay sugar-rich substrates (Põlme et al. 2020) | Põlme S, et al. 2020. Fungal Diversity 105:1-16 ((https://doi.org/10.1007/s13225-020-00466-2)) |
| Otu43704 | 1 | k__Fungi(100);p__Ascomycota(100);c__Saccharomycetes(100);o__Saccharomycetales(100);f__Metschnikowiaceae(100);g__Metschnikowia(100);s__Metschnikowia_pulcherrima(100); | Metschnikowia | 13 | Saprotroph | \|Undefined Saprotroph\| | Yeast | NULL | Probable | As nectar-tap_saprotroph (Põlme et al. 2020); Decay sugar-rich substrates (Põlme et al. 2020) | Põlme S, et al. 2020. Fungal Diversity 105:1-16 ((https://doi.org/10.1007/s13225-020-00466-2)) |
| Otu43705 | 1 | k__Fungi(100);p__Ascomycota(100);c__Saccharomycetes(100);o__Saccharomycetales(100);f__Metschnikowiaceae(100);g__Metschnikowia(100);g__Metschnikowia_unclassified(100); | Metschnikowia | 13 | Saprotroph | \|Undefined Saprotroph\| | Yeast | NULL | Probable | As nectar-tap_saprotroph (Põlme et al. 2020); Decay sugar-rich substrates (Põlme et al. 2020) | Põlme S, et al. 2020. Fungal Diversity 105:1-16 ((https://doi.org/10.1007/s13225-020-00466-2)) |
| Otu43706 | 1 | k__Fungi(100);p__Ascomycota(100);c__Saccharomycetes(100);o__Saccharomycetales(100);f__Metschnikowiaceae(100);g__Metschnikowia(100);s__Metschnikowia_pulcherrima(100); | Metschnikowia | 13 | Saprotroph | \|Undefined Saprotroph\| | Yeast | NULL | Probable | As nectar-tap_saprotroph (Põlme et al. 2020); Decay sugar-rich substrates (Põlme et al. 2020) | Põlme S, et al. 2020. Fungal Diversity 105:1-16 ((https://doi.org/10.1007/s13225-020-00466-2)) |
| Otu43707 | 1 | k__Fungi(100);p__Ascomycota(100);c__Saccharomycetes(100);o__Saccharomycetales(100);f__Metschnikowiaceae(100);g__Metschnikowia(100);s__Metschnikowia_pulcherrima(100); | Metschnikowia | 13 | Saprotroph | \|Undefined Saprotroph\| | Yeast | NULL | Probable | As nectar-tap_saprotroph (Põlme et al. 2020); Decay sugar-rich substrates (Põlme et al. 2020) | Põlme S, et al. 2020. Fungal Diversity 105:1-16 ((https://doi.org/10.1007/s13225-020-00466-2)) |
| Otu43708 | 1 | k__Fungi(100);p__Ascomycota(100);c__Saccharomycetes(100);o__Saccharomycetales(100);f__Metschnikowiaceae(100);g__Metschnikowia(100);s__Metschnikowia_pulcherrima(100); | Metschnikowia | 13 | Saprotroph | \|Undefined Saprotroph\| | Yeast | NULL | Probable | As nectar-tap_saprotroph (Põlme et al. 2020); Decay sugar-rich substrates (Põlme et al. 2020) | Põlme S, et al. 2020. Fungal Diversity 105:1-16 ((https://doi.org/10.1007/s13225-020-00466-2)) |
| Otu43709 | 1 | k__Fungi(100);p__Ascomycota(100);c__Saccharomycetes(100);o__Saccharomycetales(100);f__Metschnikowiaceae(100);g__Metschnikowia(100);s__Metschnikowia_pulcherrima(100); | Metschnikowia | 13 | Saprotroph | \|Undefined Saprotroph\| | Yeast | NULL | Probable | As nectar-tap_saprotroph (Põlme et al. 2020); Decay sugar-rich substrates (Põlme et al. 2020) | Põlme S, et al. 2020. Fungal Diversity 105:1-16 ((https://doi.org/10.1007/s13225-020-00466-2)) |
| Otu43710 | 1 | k__Fungi(100);p__Ascomycota(100);c__Saccharomycetes(100);o__Saccharomycetales(100);f__Metschnikowiaceae(100);g__Metschnikowia(100);s__Metschnikowia_pulcherrima(100); | Metschnikowia | 13 | Saprotroph | \|Undefined Saprotroph\| | Yeast | NULL | Probable | As nectar-tap_saprotroph (Põlme et al. 2020); Decay sugar-rich substrates (Põlme et al. 2020) | Põlme S, et al. 2020. Fungal Diversity 105:1-16 ((https://doi.org/10.1007/s13225-020-00466-2)) |
| Otu43711 | 1 | k__Fungi(100);p__Ascomycota(100);c__Saccharomycetes(100);o__Saccharomycetales(100);f__Metschnikowiaceae(100);g__Metschnikowia(100);s__Metschnikowia_pulcherrima(100); | Metschnikowia | 13 | Saprotroph | \|Undefined Saprotroph\| | Yeast | NULL | Probable | As nectar-tap_saprotroph (Põlme et al. 2020); Decay sugar-rich substrates (Põlme et al. 2020) | Põlme S, et al. 2020. Fungal Diversity 105:1-16 ((https://doi.org/10.1007/s13225-020-00466-2)) |
| Otu43712 | 1 | k__Fungi(100);p__Ascomycota(100);c__Saccharomycetes(100);o__Saccharomycetales(100);f__Metschnikowiaceae(100);g__Metschnikowia(100);s__Metschnikowia_pulcherrima(100); | Metschnikowia | 13 | Saprotroph | \|Undefined Saprotroph\| | Yeast | NULL | Probable | As nectar-tap_saprotroph (Põlme et al. 2020); Decay sugar-rich substrates (Põlme et al. 2020) | Põlme S, et al. 2020. Fungal Diversity 105:1-16 ((https://doi.org/10.1007/s13225-020-00466-2)) |
| Otu43713 | 1 | k__Fungi(100);p__Ascomycota(100);c__Saccharomycetes(100);o__Saccharomycetales(100);f__Metschnikowiaceae(100);g__Metschnikowia(100);s__Metschnikowia_pulcherrima(100); | Metschnikowia | 13 | Saprotroph | \|Undefined Saprotroph\| | Yeast | NULL | Probable | As nectar-tap_saprotroph (Põlme et al. 2020); Decay sugar-rich substrates (Põlme et al. 2020) | Põlme S, et al. 2020. Fungal Diversity 105:1-16 ((https://doi.org/10.1007/s13225-020-00466-2)) |
| Otu43714 | 1 | k__Fungi(100);p__Ascomycota(100);c__Saccharomycetes(100);o__Saccharomycetales(100);f__Metschnikowiaceae(100);g__Metschnikowia(100);s__Metschnikowia_pulcherrima(100); | Metschnikowia | 13 | Saprotroph | \|Undefined Saprotroph\| | Yeast | NULL | Probable | As nectar-tap_saprotroph (Põlme et al. 2020); Decay sugar-rich substrates (Põlme et al. 2020) | Põlme S, et al. 2020. Fungal Diversity 105:1-16 ((https://doi.org/10.1007/s13225-020-00466-2)) |
| Otu43715 | 1 | k__Fungi(100);p__Ascomycota(100);c__Saccharomycetes(100);o__Saccharomycetales(100);f__Metschnikowiaceae(100);g__Metschnikowia(100);s__Metschnikowia_pulcherrima(100); | Metschnikowia | 13 | Saprotroph | \|Undefined Saprotroph\| | Yeast | NULL | Probable | As nectar-tap_saprotroph (Põlme et al. 2020); Decay sugar-rich substrates (Põlme et al. 2020) | Põlme S, et al. 2020. Fungal Diversity 105:1-16 ((https://doi.org/10.1007/s13225-020-00466-2)) |
| Otu43716 | 1 | k__Fungi(100);p__Ascomycota(100);c__Saccharomycetes(100);o__Saccharomycetales(100);f__Metschnikowiaceae(100);g__Metschnikowia(100);s__Metschnikowia_pulcherrima(100); | Metschnikowia | 13 | Saprotroph | \|Undefined Saprotroph\| | Yeast | NULL | Probable | As nectar-tap_saprotroph (Põlme et al. 2020); Decay sugar-rich substrates (Põlme et al. 2020) | Põlme S, et al. 2020. Fungal Diversity 105:1-16 ((https://doi.org/10.1007/s13225-020-00466-2)) |
| Otu43717 | 1 | k__Fungi(100);p__Ascomycota(100);c__Saccharomycetes(100);o__Saccharomycetales(100);f__Metschnikowiaceae(100);g__Metschnikowia(100);s__Metschnikowia_pulcherrima(100); | Metschnikowia | 13 | Saprotroph | \|Undefined Saprotroph\| | Yeast | NULL | Probable | As nectar-tap_saprotroph (Põlme et al. 2020); Decay sugar-rich substrates (Põlme et al. 2020) | Põlme S, et al. 2020. Fungal Diversity 105:1-16 ((https://doi.org/10.1007/s13225-020-00466-2)) |
| Otu43718 | 1 | k__Fungi(100);p__Ascomycota(100);c__Saccharomycetes(100);o__Saccharomycetales(100);f__Metschnikowiaceae(100);g__Metschnikowia(100);g__Metschnikowia_unclassified(100); | Metschnikowia | 13 | Saprotroph | \|Undefined Saprotroph\| | Yeast | NULL | Probable | As nectar-tap_saprotroph (Põlme et al. 2020); Decay sugar-rich substrates (Põlme et al. 2020) | Põlme S, et al. 2020. Fungal Diversity 105:1-16 ((https://doi.org/10.1007/s13225-020-00466-2)) |
| Otu43719 | 1 | k__Fungi(100);p__Ascomycota(100);c__Saccharomycetes(100);o__Saccharomycetales(100);f__Metschnikowiaceae(100);g__Metschnikowia(100);g__Metschnikowia_unclassified(100); | Metschnikowia | 13 | Saprotroph | \|Undefined Saprotroph\| | Yeast | NULL | Probable | As nectar-tap_saprotroph (Põlme et al. 2020); Decay sugar-rich substrates (Põlme et al. 2020) | Põlme S, et al. 2020. Fungal Diversity 105:1-16 ((https://doi.org/10.1007/s13225-020-00466-2)) |
| Otu43720 | 1 | k__Fungi(100);p__Ascomycota(100);c__Saccharomycetes(100);o__Saccharomycetales(100);f__Metschnikowiaceae(100);g__Metschnikowia(100);s__Metschnikowia_pulcherrima(100); | Metschnikowia | 13 | Saprotroph | \|Undefined Saprotroph\| | Yeast | NULL | Probable | As nectar-tap_saprotroph (Põlme et al. 2020); Decay sugar-rich substrates (Põlme et al. 2020) | Põlme S, et al. 2020. Fungal Diversity 105:1-16 ((https://doi.org/10.1007/s13225-020-00466-2)) |
| Otu43721 | 1 | k__Fungi(100);p__Ascomycota(100);c__Saccharomycetes(100);o__Saccharomycetales(100);f__Metschnikowiaceae(100);g__Metschnikowia(100);s__Metschnikowia_pulcherrima(100); | Metschnikowia | 13 | Saprotroph | \|Undefined Saprotroph\| | Yeast | NULL | Probable | As nectar-tap_saprotroph (Põlme et al. 2020); Decay sugar-rich substrates (Põlme et al. 2020) | Põlme S, et al. 2020. Fungal Diversity 105:1-16 ((https://doi.org/10.1007/s13225-020-00466-2)) |
| Otu43722 | 1 | k__Fungi(100);p__Ascomycota(100);c__Saccharomycetes(100);o__Saccharomycetales(100);f__Metschnikowiaceae(100);g__Metschnikowia(100);g__Metschnikowia_unclassified(100); | Metschnikowia | 13 | Saprotroph | \|Undefined Saprotroph\| | Yeast | NULL | Probable | As nectar-tap_saprotroph (Põlme et al. 2020); Decay sugar-rich substrates (Põlme et al. 2020) | Põlme S, et al. 2020. Fungal Diversity 105:1-16 ((https://doi.org/10.1007/s13225-020-00466-2)) |
| Otu43723 | 1 | k__Fungi(100);p__Ascomycota(100);c__Saccharomycetes(100);o__Saccharomycetales(100);f__Metschnikowiaceae(100);g__Metschnikowia(100);s__Metschnikowia_pulcherrima(100); | Metschnikowia | 13 | Saprotroph | \|Undefined Saprotroph\| | Yeast | NULL | Probable | As nectar-tap_saprotroph (Põlme et al. 2020); Decay sugar-rich substrates (Põlme et al. 2020) | Põlme S, et al. 2020. Fungal Diversity 105:1-16 ((https://doi.org/10.1007/s13225-020-00466-2)) |
| Otu43724 | 1 | k__Fungi(100);p__Ascomycota(100);c__Saccharomycetes(100);o__Saccharomycetales(100);f__Metschnikowiaceae(100);g__Metschnikowia(100);s__Metschnikowia_pulcherrima(100); | Metschnikowia | 13 | Saprotroph | \|Undefined Saprotroph\| | Yeast | NULL | Probable | As nectar-tap_saprotroph (Põlme et al. 2020); Decay sugar-rich substrates (Põlme et al. 2020) | Põlme S, et al. 2020. Fungal Diversity 105:1-16 ((https://doi.org/10.1007/s13225-020-00466-2)) |
| Otu43725 | 1 | k__Fungi(100);p__Ascomycota(100);c__Saccharomycetes(100);o__Saccharomycetales(100);f__Metschnikowiaceae(100);g__Metschnikowia(100);s__Metschnikowia_pulcherrima(100); | Metschnikowia | 13 | Saprotroph | \|Undefined Saprotroph\| | Yeast | NULL | Probable | As nectar-tap_saprotroph (Põlme et al. 2020); Decay sugar-rich substrates (Põlme et al. 2020) | Põlme S, et al. 2020. Fungal Diversity 105:1-16 ((https://doi.org/10.1007/s13225-020-00466-2)) |
| Otu43726 | 1 | k__Fungi(100);p__Ascomycota(100);c__Saccharomycetes(100);o__Saccharomycetales(100);f__Metschnikowiaceae(100);g__Metschnikowia(100);s__Metschnikowia_pulcherrima(100); | Metschnikowia | 13 | Saprotroph | \|Undefined Saprotroph\| | Yeast | NULL | Probable | As nectar-tap_saprotroph (Põlme et al. 2020); Decay sugar-rich substrates (Põlme et al. 2020) | Põlme S, et al. 2020. Fungal Diversity 105:1-16 ((https://doi.org/10.1007/s13225-020-00466-2)) |
| Otu43727 | 1 | k__Fungi(100);p__Ascomycota(100);c__Saccharomycetes(100);o__Saccharomycetales(100);f__Metschnikowiaceae(100);g__Metschnikowia(100);s__Metschnikowia_pulcherrima(100); | Metschnikowia | 13 | Saprotroph | \|Undefined Saprotroph\| | Yeast | NULL | Probable | As nectar-tap_saprotroph (Põlme et al. 2020); Decay sugar-rich substrates (Põlme et al. 2020) | Põlme S, et al. 2020. Fungal Diversity 105:1-16 ((https://doi.org/10.1007/s13225-020-00466-2)) |
| Otu43728 | 1 | k__Fungi(100);p__Ascomycota(100);c__Saccharomycetes(100);o__Saccharomycetales(100);f__Metschnikowiaceae(100);g__Metschnikowia(100);s__Metschnikowia_pulcherrima(100); | Metschnikowia | 13 | Saprotroph | \|Undefined Saprotroph\| | Yeast | NULL | Probable | As nectar-tap_saprotroph (Põlme et al. 2020); Decay sugar-rich substrates (Põlme et al. 2020) | Põlme S, et al. 2020. Fungal Diversity 105:1-16 ((https://doi.org/10.1007/s13225-020-00466-2)) |
| Otu43729 | 1 | k__Fungi(100);p__Ascomycota(100);c__Saccharomycetes(100);o__Saccharomycetales(100);f__Metschnikowiaceae(100);g__Metschnikowia(100);s__Metschnikowia_pulcherrima(100); | Metschnikowia | 13 | Saprotroph | \|Undefined Saprotroph\| | Yeast | NULL | Probable | As nectar-tap_saprotroph (Põlme et al. 2020); Decay sugar-rich substrates (Põlme et al. 2020) | Põlme S, et al. 2020. Fungal Diversity 105:1-16 ((https://doi.org/10.1007/s13225-020-00466-2)) |
| Otu43730 | 1 | k__Fungi(100);p__Ascomycota(100);c__Saccharomycetes(100);o__Saccharomycetales(100);f__Metschnikowiaceae(100);g__Metschnikowia(100);s__Metschnikowia_pulcherrima(100); | Metschnikowia | 13 | Saprotroph | \|Undefined Saprotroph\| | Yeast | NULL | Probable | As nectar-tap_saprotroph (Põlme et al. 2020); Decay sugar-rich substrates (Põlme et al. 2020) | Põlme S, et al. 2020. Fungal Diversity 105:1-16 ((https://doi.org/10.1007/s13225-020-00466-2)) |
| Otu43731 | 1 | k__Fungi(100);p__Ascomycota(100);c__Saccharomycetes(100);o__Saccharomycetales(100);f__Metschnikowiaceae(100);g__Metschnikowia(100);s__Metschnikowia_pulcherrima(100); | Metschnikowia | 13 | Saprotroph | \|Undefined Saprotroph\| | Yeast | NULL | Probable | As nectar-tap_saprotroph (Põlme et al. 2020); Decay sugar-rich substrates (Põlme et al. 2020) | Põlme S, et al. 2020. Fungal Diversity 105:1-16 ((https://doi.org/10.1007/s13225-020-00466-2)) |
| Otu43732 | 1 | k__Fungi(100);p__Ascomycota(100);c__Saccharomycetes(100);o__Saccharomycetales(100);f__Metschnikowiaceae(100);g__Metschnikowia(100);g__Metschnikowia_unclassified(100); | Metschnikowia | 13 | Saprotroph | \|Undefined Saprotroph\| | Yeast | NULL | Probable | As nectar-tap_saprotroph (Põlme et al. 2020); Decay sugar-rich substrates (Põlme et al. 2020) | Põlme S, et al. 2020. Fungal Diversity 105:1-16 ((https://doi.org/10.1007/s13225-020-00466-2)) |
| Otu43733 | 1 | k__Fungi(100);p__Ascomycota(100);c__Saccharomycetes(100);o__Saccharomycetales(100);f__Metschnikowiaceae(100);g__Metschnikowia(100);s__Metschnikowia_pulcherrima(100); | Metschnikowia | 13 | Saprotroph | \|Undefined Saprotroph\| | Yeast | NULL | Probable | As nectar-tap_saprotroph (Põlme et al. 2020); Decay sugar-rich substrates (Põlme et al. 2020) | Põlme S, et al. 2020. Fungal Diversity 105:1-16 ((https://doi.org/10.1007/s13225-020-00466-2)) |
| Otu43734 | 1 | k__Fungi(100);p__Ascomycota(100);c__Saccharomycetes(100);o__Saccharomycetales(100);f__Metschnikowiaceae(100);g__Metschnikowia(100);s__Metschnikowia_pulcherrima(100); | Metschnikowia | 13 | Saprotroph | \|Undefined Saprotroph\| | Yeast | NULL | Probable | As nectar-tap_saprotroph (Põlme et al. 2020); Decay sugar-rich substrates (Põlme et al. 2020) | Põlme S, et al. 2020. Fungal Diversity 105:1-16 ((https://doi.org/10.1007/s13225-020-00466-2)) |
| Otu43735 | 1 | k__Fungi(100);p__Ascomycota(100);c__Saccharomycetes(100);o__Saccharomycetales(100);f__Metschnikowiaceae(100);g__Metschnikowia(100);s__Metschnikowia_pulcherrima(100); | Metschnikowia | 13 | Saprotroph | \|Undefined Saprotroph\| | Yeast | NULL | Probable | As nectar-tap_saprotroph (Põlme et al. 2020); Decay sugar-rich substrates (Põlme et al. 2020) | Põlme S, et al. 2020. Fungal Diversity 105:1-16 ((https://doi.org/10.1007/s13225-020-00466-2)) |
| Otu43736 | 1 | k__Fungi(100);p__Ascomycota(100);c__Saccharomycetes(100);o__Saccharomycetales(100);f__Metschnikowiaceae(100);g__Metschnikowia(100);s__Metschnikowia_pulcherrima(100); | Metschnikowia | 13 | Saprotroph | \|Undefined Saprotroph\| | Yeast | NULL | Probable | As nectar-tap_saprotroph (Põlme et al. 2020); Decay sugar-rich substrates (Põlme et al. 2020) | Põlme S, et al. 2020. Fungal Diversity 105:1-16 ((https://doi.org/10.1007/s13225-020-00466-2)) |
| Otu43737 | 1 | k__Fungi(100);p__Ascomycota(100);c__Saccharomycetes(100);o__Saccharomycetales(100);f__Metschnikowiaceae(100);g__Metschnikowia(100);s__Metschnikowia_pulcherrima(100); | Metschnikowia | 13 | Saprotroph | \|Undefined Saprotroph\| | Yeast | NULL | Probable | As nectar-tap_saprotroph (Põlme et al. 2020); Decay sugar-rich substrates (Põlme et al. 2020) | Põlme S, et al. 2020. Fungal Diversity 105:1-16 ((https://doi.org/10.1007/s13225-020-00466-2)) |
| Otu43739 | 1 | k__Fungi(100);p__Ascomycota(100);c__Saccharomycetes(100);o__Saccharomycetales(100);f__Metschnikowiaceae(100);g__Metschnikowia(100);s__Metschnikowia_pulcherrima(100); | Metschnikowia | 13 | Saprotroph | \|Undefined Saprotroph\| | Yeast | NULL | Probable | As nectar-tap_saprotroph (Põlme et al. 2020); Decay sugar-rich substrates (Põlme et al. 2020) | Põlme S, et al. 2020. Fungal Diversity 105:1-16 ((https://doi.org/10.1007/s13225-020-00466-2)) |
| Otu43740 | 1 | k__Fungi(100);p__Ascomycota(100);c__Saccharomycetes(100);o__Saccharomycetales(100);f__Metschnikowiaceae(100);g__Metschnikowia(100);g__Metschnikowia_unclassified(100); | Metschnikowia | 13 | Saprotroph | \|Undefined Saprotroph\| | Yeast | NULL | Probable | As nectar-tap_saprotroph (Põlme et al. 2020); Decay sugar-rich substrates (Põlme et al. 2020) | Põlme S, et al. 2020. Fungal Diversity 105:1-16 ((https://doi.org/10.1007/s13225-020-00466-2)) |
| Otu43741 | 1 | k__Fungi(100);p__Zygomycota(100);c__Zygomycota_class_Incertae_sedis(100);o__Mortierellales(100);f__Mortierellaceae(100);g__Mortierella(100);s__Mortierella_umbellata(100); | Mortierella | 13 | Saprotroph-Symbiotroph | Endophyte-Plant Saprotroph-\|Undefined Saprotroph\| | Microfungus | NULL | Probable | As chitinolytic (Põlme et al. 2020); As soil_saprotroph (Põlme et al. 2020); Decay roots, soil (Põlme et al. 2020); Saprobic in soil and thus frequently isolated from soil and material contaminated with soil such as hay or rotten silage and spoil tips (Cannon & Kirk 2007) | Cannon PF, Kirk PM. 2007. Fungal Families of the World. CAB International, Cambridge (ISBN: 978-0851998275); Tedersoo L, et al. 2014. Science 346:e1256688 ((https://doi.org/10.1126/science.1256688)); Purahong W, et al. 2016. Molecular Ecology 25:4059-4074 ((https://doi.org/10.1111/mec.13739)); Põlme S, et al. 2020. Fungal Diversity 105:1-16 ((https://doi.org/10.1007/s13225-020-00466-2)) |
| Otu43743 | 1 | k__Fungi(100);p__Ascomycota(100);c__Saccharomycetes(100);o__Saccharomycetales(100);f__Metschnikowiaceae(100);g__Metschnikowia(100);s__Metschnikowia_pulcherrima(100); | Metschnikowia | 13 | Saprotroph | \|Undefined Saprotroph\| | Yeast | NULL | Probable | As nectar-tap_saprotroph (Põlme et al. 2020); Decay sugar-rich substrates (Põlme et al. 2020) | Põlme S, et al. 2020. Fungal Diversity 105:1-16 ((https://doi.org/10.1007/s13225-020-00466-2)) |
| Otu43744 | 1 | k__Fungi(100);p__Ascomycota(100);c__Saccharomycetes(100);o__Saccharomycetales(100);f__Metschnikowiaceae(100);g__Metschnikowia(100);g__Metschnikowia_unclassified(100); | Metschnikowia | 13 | Saprotroph | \|Undefined Saprotroph\| | Yeast | NULL | Probable | As nectar-tap_saprotroph (Põlme et al. 2020); Decay sugar-rich substrates (Põlme et al. 2020) | Põlme S, et al. 2020. Fungal Diversity 105:1-16 ((https://doi.org/10.1007/s13225-020-00466-2)) |
| Otu43745 | 1 | k__Fungi(100);p__Ascomycota(100);c__Saccharomycetes(100);o__Saccharomycetales(100);f__Metschnikowiaceae(100);g__Metschnikowia(100);s__Metschnikowia_pulcherrima(100); | Metschnikowia | 13 | Saprotroph | \|Undefined Saprotroph\| | Yeast | NULL | Probable | As nectar-tap_saprotroph (Põlme et al. 2020); Decay sugar-rich substrates (Põlme et al. 2020) | Põlme S, et al. 2020. Fungal Diversity 105:1-16 ((https://doi.org/10.1007/s13225-020-00466-2)) |
| Otu43746 | 1 | k__Fungi(100);p__Ascomycota(100);c__Saccharomycetes(100);o__Saccharomycetales(100);f__Metschnikowiaceae(100);g__Metschnikowia(100);s__Metschnikowia_pulcherrima(100); | Metschnikowia | 13 | Saprotroph | \|Undefined Saprotroph\| | Yeast | NULL | Probable | As nectar-tap_saprotroph (Põlme et al. 2020); Decay sugar-rich substrates (Põlme et al. 2020) | Põlme S, et al. 2020. Fungal Diversity 105:1-16 ((https://doi.org/10.1007/s13225-020-00466-2)) |
| Otu43747 | 1 | k__Fungi(100);p__Ascomycota(100);c__Saccharomycetes(100);o__Saccharomycetales(100);f__Metschnikowiaceae(100);g__Metschnikowia(100);s__Metschnikowia_pulcherrima(100); | Metschnikowia | 13 | Saprotroph | \|Undefined Saprotroph\| | Yeast | NULL | Probable | As nectar-tap_saprotroph (Põlme et al. 2020); Decay sugar-rich substrates (Põlme et al. 2020) | Põlme S, et al. 2020. Fungal Diversity 105:1-16 ((https://doi.org/10.1007/s13225-020-00466-2)) |
| Otu43748 | 1 | k__Fungi(100);p__Ascomycota(100);c__Saccharomycetes(100);o__Saccharomycetales(100);f__Metschnikowiaceae(100);g__Metschnikowia(100);s__Metschnikowia_pulcherrima(100); | Metschnikowia | 13 | Saprotroph | \|Undefined Saprotroph\| | Yeast | NULL | Probable | As nectar-tap_saprotroph (Põlme et al. 2020); Decay sugar-rich substrates (Põlme et al. 2020) | Põlme S, et al. 2020. Fungal Diversity 105:1-16 ((https://doi.org/10.1007/s13225-020-00466-2)) |
| Otu43749 | 1 | k__Fungi(100);p__Ascomycota(100);c__Saccharomycetes(100);o__Saccharomycetales(100);f__Metschnikowiaceae(100);g__Metschnikowia(100);s__Metschnikowia_pulcherrima(100); | Metschnikowia | 13 | Saprotroph | \|Undefined Saprotroph\| | Yeast | NULL | Probable | As nectar-tap_saprotroph (Põlme et al. 2020); Decay sugar-rich substrates (Põlme et al. 2020) | Põlme S, et al. 2020. Fungal Diversity 105:1-16 ((https://doi.org/10.1007/s13225-020-00466-2)) |
| Otu43750 | 1 | k__Fungi(100);p__Ascomycota(100);c__Saccharomycetes(100);o__Saccharomycetales(100);f__Metschnikowiaceae(100);g__Metschnikowia(100);g__Metschnikowia_unclassified(100); | Metschnikowia | 13 | Saprotroph | \|Undefined Saprotroph\| | Yeast | NULL | Probable | As nectar-tap_saprotroph (Põlme et al. 2020); Decay sugar-rich substrates (Põlme et al. 2020) | Põlme S, et al. 2020. Fungal Diversity 105:1-16 ((https://doi.org/10.1007/s13225-020-00466-2)) |
| Otu43751 | 1 | k__Fungi(100);p__Ascomycota(100);c__Saccharomycetes(100);o__Saccharomycetales(100);f__Metschnikowiaceae(100);g__Metschnikowia(100);g__Metschnikowia_unclassified(100); | Metschnikowia | 13 | Saprotroph | \|Undefined Saprotroph\| | Yeast | NULL | Probable | As nectar-tap_saprotroph (Põlme et al. 2020); Decay sugar-rich substrates (Põlme et al. 2020) | Põlme S, et al. 2020. Fungal Diversity 105:1-16 ((https://doi.org/10.1007/s13225-020-00466-2)) |
| Otu43752 | 1 | k__Fungi(100);p__Ascomycota(100);c__Saccharomycetes(100);o__Saccharomycetales(100);f__Metschnikowiaceae(100);g__Metschnikowia(100);s__Metschnikowia_pulcherrima(100); | Metschnikowia | 13 | Saprotroph | \|Undefined Saprotroph\| | Yeast | NULL | Probable | As nectar-tap_saprotroph (Põlme et al. 2020); Decay sugar-rich substrates (Põlme et al. 2020) | Põlme S, et al. 2020. Fungal Diversity 105:1-16 ((https://doi.org/10.1007/s13225-020-00466-2)) |
| Otu43753 | 1 | k__Fungi(100);p__Ascomycota(100);c__Saccharomycetes(100);o__Saccharomycetales(100);f__Metschnikowiaceae(100);g__Metschnikowia(100);g__Metschnikowia_unclassified(100); | Metschnikowia | 13 | Saprotroph | \|Undefined Saprotroph\| | Yeast | NULL | Probable | As nectar-tap_saprotroph (Põlme et al. 2020); Decay sugar-rich substrates (Põlme et al. 2020) | Põlme S, et al. 2020. Fungal Diversity 105:1-16 ((https://doi.org/10.1007/s13225-020-00466-2)) |
| Otu43754 | 1 | k__Fungi(100);p__Ascomycota(100);c__Saccharomycetes(100);o__Saccharomycetales(100);f__Metschnikowiaceae(100);g__Metschnikowia(100);s__Metschnikowia_pulcherrima(100); | Metschnikowia | 13 | Saprotroph | \|Undefined Saprotroph\| | Yeast | NULL | Probable | As nectar-tap_saprotroph (Põlme et al. 2020); Decay sugar-rich substrates (Põlme et al. 2020) | Põlme S, et al. 2020. Fungal Diversity 105:1-16 ((https://doi.org/10.1007/s13225-020-00466-2)) |
| Otu43755 | 1 | k__Fungi(100);p__Ascomycota(100);c__Saccharomycetes(100);o__Saccharomycetales(100);f__Metschnikowiaceae(100);g__Metschnikowia(100);s__Metschnikowia_pulcherrima(100); | Metschnikowia | 13 | Saprotroph | \|Undefined Saprotroph\| | Yeast | NULL | Probable | As nectar-tap_saprotroph (Põlme et al. 2020); Decay sugar-rich substrates (Põlme et al. 2020) | Põlme S, et al. 2020. Fungal Diversity 105:1-16 ((https://doi.org/10.1007/s13225-020-00466-2)) |
| Otu43756 | 1 | k__Fungi(100);p__Ascomycota(100);c__Saccharomycetes(100);o__Saccharomycetales(100);f__Metschnikowiaceae(100);g__Metschnikowia(100);s__Metschnikowia_pulcherrima(100); | Metschnikowia | 13 | Saprotroph | \|Undefined Saprotroph\| | Yeast | NULL | Probable | As nectar-tap_saprotroph (Põlme et al. 2020); Decay sugar-rich substrates (Põlme et al. 2020) | Põlme S, et al. 2020. Fungal Diversity 105:1-16 ((https://doi.org/10.1007/s13225-020-00466-2)) |
| Otu43757 | 1 | k__Fungi(100);p__Ascomycota(100);c__Saccharomycetes(100);o__Saccharomycetales(100);f__Metschnikowiaceae(100);g__Metschnikowia(100);s__Metschnikowia_pulcherrima(100); | Metschnikowia | 13 | Saprotroph | \|Undefined Saprotroph\| | Yeast | NULL | Probable | As nectar-tap_saprotroph (Põlme et al. 2020); Decay sugar-rich substrates (Põlme et al. 2020) | Põlme S, et al. 2020. Fungal Diversity 105:1-16 ((https://doi.org/10.1007/s13225-020-00466-2)) |
| Otu43758 | 1 | k__Fungi(100);p__Ascomycota(100);c__Saccharomycetes(100);o__Saccharomycetales(100);f__Metschnikowiaceae(100);g__Metschnikowia(100);g__Metschnikowia_unclassified(100); | Metschnikowia | 13 | Saprotroph | \|Undefined Saprotroph\| | Yeast | NULL | Probable | As nectar-tap_saprotroph (Põlme et al. 2020); Decay sugar-rich substrates (Põlme et al. 2020) | Põlme S, et al. 2020. Fungal Diversity 105:1-16 ((https://doi.org/10.1007/s13225-020-00466-2)) |
| Otu43759 | 1 | k__Fungi(100);p__Ascomycota(100);c__Saccharomycetes(100);o__Saccharomycetales(100);f__Metschnikowiaceae(100);g__Metschnikowia(100);s__Metschnikowia_pulcherrima(100); | Metschnikowia | 13 | Saprotroph | \|Undefined Saprotroph\| | Yeast | NULL | Probable | As nectar-tap_saprotroph (Põlme et al. 2020); Decay sugar-rich substrates (Põlme et al. 2020) | Põlme S, et al. 2020. Fungal Diversity 105:1-16 ((https://doi.org/10.1007/s13225-020-00466-2)) |
| Otu43760 | 1 | k__Fungi(100);p__Ascomycota(100);c__Saccharomycetes(100);o__Saccharomycetales(100);f__Metschnikowiaceae(100);g__Metschnikowia(100);s__Metschnikowia_pulcherrima(100); | Metschnikowia | 13 | Saprotroph | \|Undefined Saprotroph\| | Yeast | NULL | Probable | As nectar-tap_saprotroph (Põlme et al. 2020); Decay sugar-rich substrates (Põlme et al. 2020) | Põlme S, et al. 2020. Fungal Diversity 105:1-16 ((https://doi.org/10.1007/s13225-020-00466-2)) |
| Otu43761 | 1 | k__Fungi(100);p__Ascomycota(100);c__Saccharomycetes(100);o__Saccharomycetales(100);f__Metschnikowiaceae(100);g__Metschnikowia(100);s__Metschnikowia_pulcherrima(100); | Metschnikowia | 13 | Saprotroph | \|Undefined Saprotroph\| | Yeast | NULL | Probable | As nectar-tap_saprotroph (Põlme et al. 2020); Decay sugar-rich substrates (Põlme et al. 2020) | Põlme S, et al. 2020. Fungal Diversity 105:1-16 ((https://doi.org/10.1007/s13225-020-00466-2)) |
| Otu43762 | 1 | k__Fungi(100);p__Ascomycota(100);c__Saccharomycetes(100);o__Saccharomycetales(100);f__Metschnikowiaceae(100);g__Metschnikowia(100);s__Metschnikowia_pulcherrima(100); | Metschnikowia | 13 | Saprotroph | \|Undefined Saprotroph\| | Yeast | NULL | Probable | As nectar-tap_saprotroph (Põlme et al. 2020); Decay sugar-rich substrates (Põlme et al. 2020) | Põlme S, et al. 2020. Fungal Diversity 105:1-16 ((https://doi.org/10.1007/s13225-020-00466-2)) |
| Otu43763 | 1 | k__Fungi(100);p__Ascomycota(100);c__Saccharomycetes(100);o__Saccharomycetales(100);f__Metschnikowiaceae(100);g__Metschnikowia(100);g__Metschnikowia_unclassified(100); | Metschnikowia | 13 | Saprotroph | \|Undefined Saprotroph\| | Yeast | NULL | Probable | As nectar-tap_saprotroph (Põlme et al. 2020); Decay sugar-rich substrates (Põlme et al. 2020) | Põlme S, et al. 2020. Fungal Diversity 105:1-16 ((https://doi.org/10.1007/s13225-020-00466-2)) |
| Otu43764 | 1 | k__Fungi(100);p__Ascomycota(100);c__Saccharomycetes(100);o__Saccharomycetales(100);f__Metschnikowiaceae(100);g__Metschnikowia(100);s__Metschnikowia_pulcherrima(100); | Metschnikowia | 13 | Saprotroph | \|Undefined Saprotroph\| | Yeast | NULL | Probable | As nectar-tap_saprotroph (Põlme et al. 2020); Decay sugar-rich substrates (Põlme et al. 2020) | Põlme S, et al. 2020. Fungal Diversity 105:1-16 ((https://doi.org/10.1007/s13225-020-00466-2)) |
| Otu43765 | 1 | k__Fungi(100);p__Ascomycota(100);c__Saccharomycetes(100);o__Saccharomycetales(100);f__Metschnikowiaceae(100);g__Metschnikowia(100);g__Metschnikowia_unclassified(100); | Metschnikowia | 13 | Saprotroph | \|Undefined Saprotroph\| | Yeast | NULL | Probable | As nectar-tap_saprotroph (Põlme et al. 2020); Decay sugar-rich substrates (Põlme et al. 2020) | Põlme S, et al. 2020. Fungal Diversity 105:1-16 ((https://doi.org/10.1007/s13225-020-00466-2)) |
| Otu43766 | 1 | k__Fungi(100);p__Ascomycota(100);c__Saccharomycetes(100);o__Saccharomycetales(100);f__Metschnikowiaceae(100);g__Metschnikowia(100);s__Metschnikowia_pulcherrima(100); | Metschnikowia | 13 | Saprotroph | \|Undefined Saprotroph\| | Yeast | NULL | Probable | As nectar-tap_saprotroph (Põlme et al. 2020); Decay sugar-rich substrates (Põlme et al. 2020) | Põlme S, et al. 2020. Fungal Diversity 105:1-16 ((https://doi.org/10.1007/s13225-020-00466-2)) |
| Otu43767 | 1 | k__Fungi(100);p__Ascomycota(100);c__Saccharomycetes(100);o__Saccharomycetales(100);f__Metschnikowiaceae(100);g__Metschnikowia(100);s__Metschnikowia_pulcherrima(100); | Metschnikowia | 13 | Saprotroph | \|Undefined Saprotroph\| | Yeast | NULL | Probable | As nectar-tap_saprotroph (Põlme et al. 2020); Decay sugar-rich substrates (Põlme et al. 2020) | Põlme S, et al. 2020. Fungal Diversity 105:1-16 ((https://doi.org/10.1007/s13225-020-00466-2)) |
| Otu43768 | 1 | k__Fungi(100);p__Ascomycota(100);c__Saccharomycetes(100);o__Saccharomycetales(100);f__Metschnikowiaceae(100);g__Metschnikowia(100);s__Metschnikowia_pulcherrima(100); | Metschnikowia | 13 | Saprotroph | \|Undefined Saprotroph\| | Yeast | NULL | Probable | As nectar-tap_saprotroph (Põlme et al. 2020); Decay sugar-rich substrates (Põlme et al. 2020) | Põlme S, et al. 2020. Fungal Diversity 105:1-16 ((https://doi.org/10.1007/s13225-020-00466-2)) |
| Otu43769 | 1 | k__Fungi(100);p__Ascomycota(100);c__Saccharomycetes(100);o__Saccharomycetales(100);f__Metschnikowiaceae(100);g__Metschnikowia(100);g__Metschnikowia_unclassified(100); | Metschnikowia | 13 | Saprotroph | \|Undefined Saprotroph\| | Yeast | NULL | Probable | As nectar-tap_saprotroph (Põlme et al. 2020); Decay sugar-rich substrates (Põlme et al. 2020) | Põlme S, et al. 2020. Fungal Diversity 105:1-16 ((https://doi.org/10.1007/s13225-020-00466-2)) |
| Otu43770 | 1 | k__Fungi(100);p__Ascomycota(100);c__Saccharomycetes(100);o__Saccharomycetales(100);f__Metschnikowiaceae(100);g__Metschnikowia(100);s__Metschnikowia_pulcherrima(100); | Metschnikowia | 13 | Saprotroph | \|Undefined Saprotroph\| | Yeast | NULL | Probable | As nectar-tap_saprotroph (Põlme et al. 2020); Decay sugar-rich substrates (Põlme et al. 2020) | Põlme S, et al. 2020. Fungal Diversity 105:1-16 ((https://doi.org/10.1007/s13225-020-00466-2)) |
| Otu43771 | 1 | k__Fungi(100);p__Ascomycota(100);c__Saccharomycetes(100);o__Saccharomycetales(100);f__Metschnikowiaceae(100);g__Metschnikowia(100);g__Metschnikowia_unclassified(100); | Metschnikowia | 13 | Saprotroph | \|Undefined Saprotroph\| | Yeast | NULL | Probable | As nectar-tap_saprotroph (Põlme et al. 2020); Decay sugar-rich substrates (Põlme et al. 2020) | Põlme S, et al. 2020. Fungal Diversity 105:1-16 ((https://doi.org/10.1007/s13225-020-00466-2)) |
| Otu43772 | 1 | k__Fungi(100);p__Ascomycota(100);c__Saccharomycetes(100);o__Saccharomycetales(100);f__Metschnikowiaceae(100);g__Metschnikowia(100);g__Metschnikowia_unclassified(100); | Metschnikowia | 13 | Saprotroph | \|Undefined Saprotroph\| | Yeast | NULL | Probable | As nectar-tap_saprotroph (Põlme et al. 2020); Decay sugar-rich substrates (Põlme et al. 2020) | Põlme S, et al. 2020. Fungal Diversity 105:1-16 ((https://doi.org/10.1007/s13225-020-00466-2)) |
| Otu43773 | 1 | k__Fungi(100);p__Ascomycota(100);c__Saccharomycetes(100);o__Saccharomycetales(100);f__Metschnikowiaceae(100);g__Metschnikowia(100);g__Metschnikowia_unclassified(100); | Metschnikowia | 13 | Saprotroph | \|Undefined Saprotroph\| | Yeast | NULL | Probable | As nectar-tap_saprotroph (Põlme et al. 2020); Decay sugar-rich substrates (Põlme et al. 2020) | Põlme S, et al. 2020. Fungal Diversity 105:1-16 ((https://doi.org/10.1007/s13225-020-00466-2)) |
| Otu43774 | 1 | k__Fungi(100);p__Ascomycota(100);c__Saccharomycetes(100);o__Saccharomycetales(100);f__Metschnikowiaceae(100);g__Metschnikowia(100);s__Metschnikowia_pulcherrima(100); | Metschnikowia | 13 | Saprotroph | \|Undefined Saprotroph\| | Yeast | NULL | Probable | As nectar-tap_saprotroph (Põlme et al. 2020); Decay sugar-rich substrates (Põlme et al. 2020) | Põlme S, et al. 2020. Fungal Diversity 105:1-16 ((https://doi.org/10.1007/s13225-020-00466-2)) |
| Otu43775 | 1 | k__Fungi(100);p__Ascomycota(100);c__Saccharomycetes(100);o__Saccharomycetales(100);f__Metschnikowiaceae(100);g__Metschnikowia(100);g__Metschnikowia_unclassified(100); | Metschnikowia | 13 | Saprotroph | \|Undefined Saprotroph\| | Yeast | NULL | Probable | As nectar-tap_saprotroph (Põlme et al. 2020); Decay sugar-rich substrates (Põlme et al. 2020) | Põlme S, et al. 2020. Fungal Diversity 105:1-16 ((https://doi.org/10.1007/s13225-020-00466-2)) |
| Otu43776 | 1 | k__Fungi(100);p__Ascomycota(100);c__Saccharomycetes(100);o__Saccharomycetales(100);f__Metschnikowiaceae(100);g__Metschnikowia(100);s__Metschnikowia_pulcherrima(100); | Metschnikowia | 13 | Saprotroph | \|Undefined Saprotroph\| | Yeast | NULL | Probable | As nectar-tap_saprotroph (Põlme et al. 2020); Decay sugar-rich substrates (Põlme et al. 2020) | Põlme S, et al. 2020. Fungal Diversity 105:1-16 ((https://doi.org/10.1007/s13225-020-00466-2)) |
| Otu43777 | 1 | k__Fungi(100);p__Ascomycota(100);c__Saccharomycetes(100);o__Saccharomycetales(100);f__Metschnikowiaceae(100);g__Metschnikowia(100);g__Metschnikowia_unclassified(100); | Metschnikowia | 13 | Saprotroph | \|Undefined Saprotroph\| | Yeast | NULL | Probable | As nectar-tap_saprotroph (Põlme et al. 2020); Decay sugar-rich substrates (Põlme et al. 2020) | Põlme S, et al. 2020. Fungal Diversity 105:1-16 ((https://doi.org/10.1007/s13225-020-00466-2)) |
| Otu43778 | 1 | k__Fungi(100);p__Ascomycota(100);c__Saccharomycetes(100);o__Saccharomycetales(100);f__Metschnikowiaceae(100);g__Metschnikowia(100);s__Metschnikowia_pulcherrima(100); | Metschnikowia | 13 | Saprotroph | \|Undefined Saprotroph\| | Yeast | NULL | Probable | As nectar-tap_saprotroph (Põlme et al. 2020); Decay sugar-rich substrates (Põlme et al. 2020) | Põlme S, et al. 2020. Fungal Diversity 105:1-16 ((https://doi.org/10.1007/s13225-020-00466-2)) |
| Otu43780 | 1 | k__Fungi(100);p__Ascomycota(100);c__Saccharomycetes(100);o__Saccharomycetales(100);f__Metschnikowiaceae(100);g__Metschnikowia(100);g__Metschnikowia_unclassified(100); | Metschnikowia | 13 | Saprotroph | \|Undefined Saprotroph\| | Yeast | NULL | Probable | As nectar-tap_saprotroph (Põlme et al. 2020); Decay sugar-rich substrates (Põlme et al. 2020) | Põlme S, et al. 2020. Fungal Diversity 105:1-16 ((https://doi.org/10.1007/s13225-020-00466-2)) |
| Otu43781 | 1 | k__Fungi(100);p__Ascomycota(100);c__Saccharomycetes(100);o__Saccharomycetales(100);f__Metschnikowiaceae(100);g__Metschnikowia(100);s__Metschnikowia_pulcherrima(100); | Metschnikowia | 13 | Saprotroph | \|Undefined Saprotroph\| | Yeast | NULL | Probable | As nectar-tap_saprotroph (Põlme et al. 2020); Decay sugar-rich substrates (Põlme et al. 2020) | Põlme S, et al. 2020. Fungal Diversity 105:1-16 ((https://doi.org/10.1007/s13225-020-00466-2)) |
| Otu43782 | 1 | k__Fungi(100);p__Ascomycota(100);c__Saccharomycetes(100);o__Saccharomycetales(100);f__Metschnikowiaceae(100);g__Metschnikowia(100);s__Metschnikowia_pulcherrima(100); | Metschnikowia | 13 | Saprotroph | \|Undefined Saprotroph\| | Yeast | NULL | Probable | As nectar-tap_saprotroph (Põlme et al. 2020); Decay sugar-rich substrates (Põlme et al. 2020) | Põlme S, et al. 2020. Fungal Diversity 105:1-16 ((https://doi.org/10.1007/s13225-020-00466-2)) |
| Otu43783 | 1 | k__Fungi(100);p__Ascomycota(100);c__Saccharomycetes(100);o__Saccharomycetales(100);f__Metschnikowiaceae(100);g__Metschnikowia(100);s__Metschnikowia_pulcherrima(100); | Metschnikowia | 13 | Saprotroph | \|Undefined Saprotroph\| | Yeast | NULL | Probable | As nectar-tap_saprotroph (Põlme et al. 2020); Decay sugar-rich substrates (Põlme et al. 2020) | Põlme S, et al. 2020. Fungal Diversity 105:1-16 ((https://doi.org/10.1007/s13225-020-00466-2)) |
| Otu43784 | 1 | k__Fungi(100);p__Ascomycota(100);c__Saccharomycetes(100);o__Saccharomycetales(100);f__Metschnikowiaceae(100);g__Metschnikowia(100);g__Metschnikowia_unclassified(100); | Metschnikowia | 13 | Saprotroph | \|Undefined Saprotroph\| | Yeast | NULL | Probable | As nectar-tap_saprotroph (Põlme et al. 2020); Decay sugar-rich substrates (Põlme et al. 2020) | Põlme S, et al. 2020. Fungal Diversity 105:1-16 ((https://doi.org/10.1007/s13225-020-00466-2)) |
| Otu43785 | 1 | k__Fungi(100);p__Ascomycota(100);c__Saccharomycetes(100);o__Saccharomycetales(100);f__Metschnikowiaceae(100);g__Metschnikowia(100);s__Metschnikowia_pulcherrima(100); | Metschnikowia | 13 | Saprotroph | \|Undefined Saprotroph\| | Yeast | NULL | Probable | As nectar-tap_saprotroph (Põlme et al. 2020); Decay sugar-rich substrates (Põlme et al. 2020) | Põlme S, et al. 2020. Fungal Diversity 105:1-16 ((https://doi.org/10.1007/s13225-020-00466-2)) |
| Otu43786 | 1 | k__Fungi(100);p__Ascomycota(100);c__Saccharomycetes(100);o__Saccharomycetales(100);f__Metschnikowiaceae(100);g__Metschnikowia(100);s__Metschnikowia_pulcherrima(100); | Metschnikowia | 13 | Saprotroph | \|Undefined Saprotroph\| | Yeast | NULL | Probable | As nectar-tap_saprotroph (Põlme et al. 2020); Decay sugar-rich substrates (Põlme et al. 2020) | Põlme S, et al. 2020. Fungal Diversity 105:1-16 ((https://doi.org/10.1007/s13225-020-00466-2)) |
| Otu43787 | 1 | k__Fungi(100);p__Ascomycota(100);c__Saccharomycetes(100);o__Saccharomycetales(100);f__Metschnikowiaceae(100);g__Metschnikowia(100);s__Metschnikowia_pulcherrima(100); | Metschnikowia | 13 | Saprotroph | \|Undefined Saprotroph\| | Yeast | NULL | Probable | As nectar-tap_saprotroph (Põlme et al. 2020); Decay sugar-rich substrates (Põlme et al. 2020) | Põlme S, et al. 2020. Fungal Diversity 105:1-16 ((https://doi.org/10.1007/s13225-020-00466-2)) |
| Otu43788 | 1 | k__Fungi(100);p__Ascomycota(100);c__Saccharomycetes(100);o__Saccharomycetales(100);f__Metschnikowiaceae(100);g__Metschnikowia(100);s__Metschnikowia_pulcherrima(100); | Metschnikowia | 13 | Saprotroph | \|Undefined Saprotroph\| | Yeast | NULL | Probable | As nectar-tap_saprotroph (Põlme et al. 2020); Decay sugar-rich substrates (Põlme et al. 2020) | Põlme S, et al. 2020. Fungal Diversity 105:1-16 ((https://doi.org/10.1007/s13225-020-00466-2)) |
| Otu43789 | 1 | k__Fungi(100);p__Ascomycota(100);c__Saccharomycetes(100);o__Saccharomycetales(100);f__Metschnikowiaceae(100);g__Metschnikowia(100);g__Metschnikowia_unclassified(100); | Metschnikowia | 13 | Saprotroph | \|Undefined Saprotroph\| | Yeast | NULL | Probable | As nectar-tap_saprotroph (Põlme et al. 2020); Decay sugar-rich substrates (Põlme et al. 2020) | Põlme S, et al. 2020. Fungal Diversity 105:1-16 ((https://doi.org/10.1007/s13225-020-00466-2)) |
| Otu43790 | 1 | k__Fungi(100);p__Ascomycota(100);c__Saccharomycetes(100);o__Saccharomycetales(100);f__Metschnikowiaceae(100);g__Metschnikowia(100);s__Metschnikowia_pulcherrima(100); | Metschnikowia | 13 | Saprotroph | \|Undefined Saprotroph\| | Yeast | NULL | Probable | As nectar-tap_saprotroph (Põlme et al. 2020); Decay sugar-rich substrates (Põlme et al. 2020) | Põlme S, et al. 2020. Fungal Diversity 105:1-16 ((https://doi.org/10.1007/s13225-020-00466-2)) |
| Otu43791 | 1 | k__Fungi(100);p__Ascomycota(100);c__Saccharomycetes(100);o__Saccharomycetales(100);f__Metschnikowiaceae(100);g__Metschnikowia(100);g__Metschnikowia_unclassified(100); | Metschnikowia | 13 | Saprotroph | \|Undefined Saprotroph\| | Yeast | NULL | Probable | As nectar-tap_saprotroph (Põlme et al. 2020); Decay sugar-rich substrates (Põlme et al. 2020) | Põlme S, et al. 2020. Fungal Diversity 105:1-16 ((https://doi.org/10.1007/s13225-020-00466-2)) |
| Otu43792 | 1 | k__Fungi(100);p__Ascomycota(100);c__Saccharomycetes(100);o__Saccharomycetales(100);f__Metschnikowiaceae(100);g__Metschnikowia(100);s__Metschnikowia_pulcherrima(100); | Metschnikowia | 13 | Saprotroph | \|Undefined Saprotroph\| | Yeast | NULL | Probable | As nectar-tap_saprotroph (Põlme et al. 2020); Decay sugar-rich substrates (Põlme et al. 2020) | Põlme S, et al. 2020. Fungal Diversity 105:1-16 ((https://doi.org/10.1007/s13225-020-00466-2)) |
| Otu43793 | 1 | k__Fungi(100);p__Ascomycota(100);c__Saccharomycetes(100);o__Saccharomycetales(100);f__Metschnikowiaceae(100);g__Metschnikowia(100);g__Metschnikowia_unclassified(100); | Metschnikowia | 13 | Saprotroph | \|Undefined Saprotroph\| | Yeast | NULL | Probable | As nectar-tap_saprotroph (Põlme et al. 2020); Decay sugar-rich substrates (Põlme et al. 2020) | Põlme S, et al. 2020. Fungal Diversity 105:1-16 ((https://doi.org/10.1007/s13225-020-00466-2)) |
| Otu43794 | 1 | k__Fungi(100);p__Ascomycota(100);c__Saccharomycetes(100);o__Saccharomycetales(100);f__Metschnikowiaceae(100);g__Metschnikowia(100);s__Metschnikowia_pulcherrima(100); | Metschnikowia | 13 | Saprotroph | \|Undefined Saprotroph\| | Yeast | NULL | Probable | As nectar-tap_saprotroph (Põlme et al. 2020); Decay sugar-rich substrates (Põlme et al. 2020) | Põlme S, et al. 2020. Fungal Diversity 105:1-16 ((https://doi.org/10.1007/s13225-020-00466-2)) |
| Otu43795 | 1 | k__Fungi(100);p__Ascomycota(100);c__Saccharomycetes(100);o__Saccharomycetales(100);f__Metschnikowiaceae(100);g__Metschnikowia(100);s__Metschnikowia_pulcherrima(100); | Metschnikowia | 13 | Saprotroph | \|Undefined Saprotroph\| | Yeast | NULL | Probable | As nectar-tap_saprotroph (Põlme et al. 2020); Decay sugar-rich substrates (Põlme et al. 2020) | Põlme S, et al. 2020. Fungal Diversity 105:1-16 ((https://doi.org/10.1007/s13225-020-00466-2)) |
| Otu43796 | 1 | k__Fungi(100);p__Ascomycota(100);c__Saccharomycetes(100);o__Saccharomycetales(100);f__Metschnikowiaceae(100);g__Metschnikowia(100);g__Metschnikowia_unclassified(100); | Metschnikowia | 13 | Saprotroph | \|Undefined Saprotroph\| | Yeast | NULL | Probable | As nectar-tap_saprotroph (Põlme et al. 2020); Decay sugar-rich substrates (Põlme et al. 2020) | Põlme S, et al. 2020. Fungal Diversity 105:1-16 ((https://doi.org/10.1007/s13225-020-00466-2)) |
| Otu43797 | 1 | k__Fungi(100);p__Ascomycota(100);c__Saccharomycetes(100);o__Saccharomycetales(100);f__Metschnikowiaceae(100);g__Metschnikowia(100);s__Metschnikowia_pulcherrima(100); | Metschnikowia | 13 | Saprotroph | \|Undefined Saprotroph\| | Yeast | NULL | Probable | As nectar-tap_saprotroph (Põlme et al. 2020); Decay sugar-rich substrates (Põlme et al. 2020) | Põlme S, et al. 2020. Fungal Diversity 105:1-16 ((https://doi.org/10.1007/s13225-020-00466-2)) |
| Otu43798 | 1 | k__Fungi(100);p__Ascomycota(100);c__Saccharomycetes(100);o__Saccharomycetales(100);f__Metschnikowiaceae(100);g__Metschnikowia(100);s__Metschnikowia_pulcherrima(100); | Metschnikowia | 13 | Saprotroph | \|Undefined Saprotroph\| | Yeast | NULL | Probable | As nectar-tap_saprotroph (Põlme et al. 2020); Decay sugar-rich substrates (Põlme et al. 2020) | Põlme S, et al. 2020. Fungal Diversity 105:1-16 ((https://doi.org/10.1007/s13225-020-00466-2)) |
| Otu43799 | 1 | k__Fungi(100);p__Ascomycota(100);c__Saccharomycetes(100);o__Saccharomycetales(100);f__Metschnikowiaceae(100);g__Metschnikowia(100);g__Metschnikowia_unclassified(100); | Metschnikowia | 13 | Saprotroph | \|Undefined Saprotroph\| | Yeast | NULL | Probable | As nectar-tap_saprotroph (Põlme et al. 2020); Decay sugar-rich substrates (Põlme et al. 2020) | Põlme S, et al. 2020. Fungal Diversity 105:1-16 ((https://doi.org/10.1007/s13225-020-00466-2)) |
| Otu43800 | 1 | k__Fungi(100);p__Ascomycota(100);c__Saccharomycetes(100);o__Saccharomycetales(100);f__Metschnikowiaceae(100);g__Metschnikowia(100);g__Metschnikowia_unclassified(100); | Metschnikowia | 13 | Saprotroph | \|Undefined Saprotroph\| | Yeast | NULL | Probable | As nectar-tap_saprotroph (Põlme et al. 2020); Decay sugar-rich substrates (Põlme et al. 2020) | Põlme S, et al. 2020. Fungal Diversity 105:1-16 ((https://doi.org/10.1007/s13225-020-00466-2)) |
| Otu43801 | 1 | k__Fungi(100);p__Ascomycota(100);c__Saccharomycetes(100);o__Saccharomycetales(100);f__Metschnikowiaceae(100);g__Metschnikowia(100);s__Metschnikowia_pulcherrima(100); | Metschnikowia | 13 | Saprotroph | \|Undefined Saprotroph\| | Yeast | NULL | Probable | As nectar-tap_saprotroph (Põlme et al. 2020); Decay sugar-rich substrates (Põlme et al. 2020) | Põlme S, et al. 2020. Fungal Diversity 105:1-16 ((https://doi.org/10.1007/s13225-020-00466-2)) |
| Otu43802 | 1 | k__Fungi(100);p__Ascomycota(100);c__Saccharomycetes(100);o__Saccharomycetales(100);f__Metschnikowiaceae(100);g__Metschnikowia(100);s__Metschnikowia_pulcherrima(100); | Metschnikowia | 13 | Saprotroph | \|Undefined Saprotroph\| | Yeast | NULL | Probable | As nectar-tap_saprotroph (Põlme et al. 2020); Decay sugar-rich substrates (Põlme et al. 2020) | Põlme S, et al. 2020. Fungal Diversity 105:1-16 ((https://doi.org/10.1007/s13225-020-00466-2)) |
| Otu43803 | 1 | k__Fungi(100);p__Ascomycota(100);c__Saccharomycetes(100);o__Saccharomycetales(100);f__Metschnikowiaceae(100);g__Metschnikowia(100);s__Metschnikowia_viticola(100); | Metschnikowia | 13 | Saprotroph | \|Undefined Saprotroph\| | Yeast | NULL | Probable | As nectar-tap_saprotroph (Põlme et al. 2020); Decay sugar-rich substrates (Põlme et al. 2020) | Põlme S, et al. 2020. Fungal Diversity 105:1-16 ((https://doi.org/10.1007/s13225-020-00466-2)) |
| Otu43804 | 1 | k__Fungi(100);p__Ascomycota(100);c__Saccharomycetes(100);o__Saccharomycetales(100);f__Metschnikowiaceae(100);g__Metschnikowia(100);s__Metschnikowia_pulcherrima(100); | Metschnikowia | 13 | Saprotroph | \|Undefined Saprotroph\| | Yeast | NULL | Probable | As nectar-tap_saprotroph (Põlme et al. 2020); Decay sugar-rich substrates (Põlme et al. 2020) | Põlme S, et al. 2020. Fungal Diversity 105:1-16 ((https://doi.org/10.1007/s13225-020-00466-2)) |
| Otu43805 | 1 | k__Fungi(100);p__Ascomycota(100);c__Saccharomycetes(100);o__Saccharomycetales(100);f__Metschnikowiaceae(100);g__Metschnikowia(100);s__Metschnikowia_pulcherrima(100); | Metschnikowia | 13 | Saprotroph | \|Undefined Saprotroph\| | Yeast | NULL | Probable | As nectar-tap_saprotroph (Põlme et al. 2020); Decay sugar-rich substrates (Põlme et al. 2020) | Põlme S, et al. 2020. Fungal Diversity 105:1-16 ((https://doi.org/10.1007/s13225-020-00466-2)) |
| Otu43806 | 1 | k__Fungi(100);p__Ascomycota(100);c__Saccharomycetes(100);o__Saccharomycetales(100);f__Metschnikowiaceae(100);g__Metschnikowia(100);s__Metschnikowia_pulcherrima(100); | Metschnikowia | 13 | Saprotroph | \|Undefined Saprotroph\| | Yeast | NULL | Probable | As nectar-tap_saprotroph (Põlme et al. 2020); Decay sugar-rich substrates (Põlme et al. 2020) | Põlme S, et al. 2020. Fungal Diversity 105:1-16 ((https://doi.org/10.1007/s13225-020-00466-2)) |
| Otu43807 | 1 | k__Fungi(100);p__Ascomycota(100);c__Saccharomycetes(100);o__Saccharomycetales(100);f__Metschnikowiaceae(100);g__Metschnikowia(100);s__Metschnikowia_pulcherrima(100); | Metschnikowia | 13 | Saprotroph | \|Undefined Saprotroph\| | Yeast | NULL | Probable | As nectar-tap_saprotroph (Põlme et al. 2020); Decay sugar-rich substrates (Põlme et al. 2020) | Põlme S, et al. 2020. Fungal Diversity 105:1-16 ((https://doi.org/10.1007/s13225-020-00466-2)) |
| Otu43808 | 1 | k__Fungi(100);p__Ascomycota(100);c__Saccharomycetes(100);o__Saccharomycetales(100);f__Metschnikowiaceae(100);g__Metschnikowia(100);s__Metschnikowia_pulcherrima(100); | Metschnikowia | 13 | Saprotroph | \|Undefined Saprotroph\| | Yeast | NULL | Probable | As nectar-tap_saprotroph (Põlme et al. 2020); Decay sugar-rich substrates (Põlme et al. 2020) | Põlme S, et al. 2020. Fungal Diversity 105:1-16 ((https://doi.org/10.1007/s13225-020-00466-2)) |
| Otu43809 | 1 | k__Fungi(100);p__Ascomycota(100);c__Saccharomycetes(100);o__Saccharomycetales(100);f__Metschnikowiaceae(100);g__Metschnikowia(100);s__Metschnikowia_pulcherrima(100); | Metschnikowia | 13 | Saprotroph | \|Undefined Saprotroph\| | Yeast | NULL | Probable | As nectar-tap_saprotroph (Põlme et al. 2020); Decay sugar-rich substrates (Põlme et al. 2020) | Põlme S, et al. 2020. Fungal Diversity 105:1-16 ((https://doi.org/10.1007/s13225-020-00466-2)) |
| Otu43810 | 1 | k__Fungi(100);p__Ascomycota(100);c__Saccharomycetes(100);o__Saccharomycetales(100);f__Metschnikowiaceae(100);g__Metschnikowia(100);s__Metschnikowia_pulcherrima(100); | Metschnikowia | 13 | Saprotroph | \|Undefined Saprotroph\| | Yeast | NULL | Probable | As nectar-tap_saprotroph (Põlme et al. 2020); Decay sugar-rich substrates (Põlme et al. 2020) | Põlme S, et al. 2020. Fungal Diversity 105:1-16 ((https://doi.org/10.1007/s13225-020-00466-2)) |
| Otu43811 | 1 | k__Fungi(100);p__Ascomycota(100);c__Saccharomycetes(100);o__Saccharomycetales(100);f__Metschnikowiaceae(100);g__Metschnikowia(100);s__Metschnikowia_pulcherrima(100); | Metschnikowia | 13 | Saprotroph | \|Undefined Saprotroph\| | Yeast | NULL | Probable | As nectar-tap_saprotroph (Põlme et al. 2020); Decay sugar-rich substrates (Põlme et al. 2020) | Põlme S, et al. 2020. Fungal Diversity 105:1-16 ((https://doi.org/10.1007/s13225-020-00466-2)) |
| Otu43812 | 1 | k__Fungi(100);p__Ascomycota(100);c__Saccharomycetes(100);o__Saccharomycetales(100);f__Metschnikowiaceae(100);g__Metschnikowia(100);s__Metschnikowia_pulcherrima(100); | Metschnikowia | 13 | Saprotroph | \|Undefined Saprotroph\| | Yeast | NULL | Probable | As nectar-tap_saprotroph (Põlme et al. 2020); Decay sugar-rich substrates (Põlme et al. 2020) | Põlme S, et al. 2020. Fungal Diversity 105:1-16 ((https://doi.org/10.1007/s13225-020-00466-2)) |
| Otu43813 | 1 | k__Fungi(100);p__Ascomycota(100);c__Saccharomycetes(100);o__Saccharomycetales(100);f__Metschnikowiaceae(100);g__Metschnikowia(100);s__Metschnikowia_pulcherrima(100); | Metschnikowia | 13 | Saprotroph | \|Undefined Saprotroph\| | Yeast | NULL | Probable | As nectar-tap_saprotroph (Põlme et al. 2020); Decay sugar-rich substrates (Põlme et al. 2020) | Põlme S, et al. 2020. Fungal Diversity 105:1-16 ((https://doi.org/10.1007/s13225-020-00466-2)) |
| Otu43814 | 1 | k__Fungi(100);p__Ascomycota(100);c__Saccharomycetes(100);o__Saccharomycetales(100);f__Metschnikowiaceae(100);g__Metschnikowia(100);g__Metschnikowia_unclassified(100); | Metschnikowia | 13 | Saprotroph | \|Undefined Saprotroph\| | Yeast | NULL | Probable | As nectar-tap_saprotroph (Põlme et al. 2020); Decay sugar-rich substrates (Põlme et al. 2020) | Põlme S, et al. 2020. Fungal Diversity 105:1-16 ((https://doi.org/10.1007/s13225-020-00466-2)) |
| Otu43815 | 1 | k__Fungi(100);p__Ascomycota(100);c__Saccharomycetes(100);o__Saccharomycetales(100);f__Metschnikowiaceae(100);g__Metschnikowia(100);g__Metschnikowia_unclassified(100); | Metschnikowia | 13 | Saprotroph | \|Undefined Saprotroph\| | Yeast | NULL | Probable | As nectar-tap_saprotroph (Põlme et al. 2020); Decay sugar-rich substrates (Põlme et al. 2020) | Põlme S, et al. 2020. Fungal Diversity 105:1-16 ((https://doi.org/10.1007/s13225-020-00466-2)) |
| Otu43816 | 1 | k__Fungi(100);p__Ascomycota(100);c__Saccharomycetes(100);o__Saccharomycetales(100);f__Metschnikowiaceae(100);g__Metschnikowia(100);g__Metschnikowia_unclassified(100); | Metschnikowia | 13 | Saprotroph | \|Undefined Saprotroph\| | Yeast | NULL | Probable | As nectar-tap_saprotroph (Põlme et al. 2020); Decay sugar-rich substrates (Põlme et al. 2020) | Põlme S, et al. 2020. Fungal Diversity 105:1-16 ((https://doi.org/10.1007/s13225-020-00466-2)) |
| Otu43817 | 1 | k__Fungi(100);p__Ascomycota(100);c__Saccharomycetes(100);o__Saccharomycetales(100);f__Metschnikowiaceae(100);g__Metschnikowia(100);s__Metschnikowia_pulcherrima(100); | Metschnikowia | 13 | Saprotroph | \|Undefined Saprotroph\| | Yeast | NULL | Probable | As nectar-tap_saprotroph (Põlme et al. 2020); Decay sugar-rich substrates (Põlme et al. 2020) | Põlme S, et al. 2020. Fungal Diversity 105:1-16 ((https://doi.org/10.1007/s13225-020-00466-2)) |
| Otu43818 | 1 | k__Fungi(100);p__Ascomycota(100);c__Saccharomycetes(100);o__Saccharomycetales(100);f__Metschnikowiaceae(100);g__Metschnikowia(100);s__Metschnikowia_pulcherrima(100); | Metschnikowia | 13 | Saprotroph | \|Undefined Saprotroph\| | Yeast | NULL | Probable | As nectar-tap_saprotroph (Põlme et al. 2020); Decay sugar-rich substrates (Põlme et al. 2020) | Põlme S, et al. 2020. Fungal Diversity 105:1-16 ((https://doi.org/10.1007/s13225-020-00466-2)) |
| Otu43819 | 1 | k__Fungi(100);p__Ascomycota(100);c__Saccharomycetes(100);o__Saccharomycetales(100);f__Metschnikowiaceae(100);g__Metschnikowia(100);s__Metschnikowia_pulcherrima(100); | Metschnikowia | 13 | Saprotroph | \|Undefined Saprotroph\| | Yeast | NULL | Probable | As nectar-tap_saprotroph (Põlme et al. 2020); Decay sugar-rich substrates (Põlme et al. 2020) | Põlme S, et al. 2020. Fungal Diversity 105:1-16 ((https://doi.org/10.1007/s13225-020-00466-2)) |
| Otu43820 | 1 | k__Fungi(100);p__Ascomycota(100);c__Saccharomycetes(100);o__Saccharomycetales(100);f__Metschnikowiaceae(100);g__Metschnikowia(100);s__Metschnikowia_pulcherrima(100); | Metschnikowia | 13 | Saprotroph | \|Undefined Saprotroph\| | Yeast | NULL | Probable | As nectar-tap_saprotroph (Põlme et al. 2020); Decay sugar-rich substrates (Põlme et al. 2020) | Põlme S, et al. 2020. Fungal Diversity 105:1-16 ((https://doi.org/10.1007/s13225-020-00466-2)) |
| Otu43821 | 1 | k__Fungi(100);p__Ascomycota(100);c__Saccharomycetes(100);o__Saccharomycetales(100);f__Metschnikowiaceae(100);g__Metschnikowia(100);s__Metschnikowia_pulcherrima(100); | Metschnikowia | 13 | Saprotroph | \|Undefined Saprotroph\| | Yeast | NULL | Probable | As nectar-tap_saprotroph (Põlme et al. 2020); Decay sugar-rich substrates (Põlme et al. 2020) | Põlme S, et al. 2020. Fungal Diversity 105:1-16 ((https://doi.org/10.1007/s13225-020-00466-2)) |
| Otu43822 | 1 | k__Fungi(100);p__Ascomycota(100);c__Saccharomycetes(100);o__Saccharomycetales(100);f__Metschnikowiaceae(100);g__Metschnikowia(100);s__Metschnikowia_pulcherrima(100); | Metschnikowia | 13 | Saprotroph | \|Undefined Saprotroph\| | Yeast | NULL | Probable | As nectar-tap_saprotroph (Põlme et al. 2020); Decay sugar-rich substrates (Põlme et al. 2020) | Põlme S, et al. 2020. Fungal Diversity 105:1-16 ((https://doi.org/10.1007/s13225-020-00466-2)) |
| Otu43823 | 1 | k__Fungi(100);p__Ascomycota(100);c__Saccharomycetes(100);o__Saccharomycetales(100);f__Metschnikowiaceae(100);g__Metschnikowia(100);s__Metschnikowia_pulcherrima(100); | Metschnikowia | 13 | Saprotroph | \|Undefined Saprotroph\| | Yeast | NULL | Probable | As nectar-tap_saprotroph (Põlme et al. 2020); Decay sugar-rich substrates (Põlme et al. 2020) | Põlme S, et al. 2020. Fungal Diversity 105:1-16 ((https://doi.org/10.1007/s13225-020-00466-2)) |
| Otu43824 | 1 | k__Fungi(100);p__Ascomycota(100);c__Saccharomycetes(100);o__Saccharomycetales(100);f__Metschnikowiaceae(100);g__Metschnikowia(100);s__Metschnikowia_pulcherrima(100); | Metschnikowia | 13 | Saprotroph | \|Undefined Saprotroph\| | Yeast | NULL | Probable | As nectar-tap_saprotroph (Põlme et al. 2020); Decay sugar-rich substrates (Põlme et al. 2020) | Põlme S, et al. 2020. Fungal Diversity 105:1-16 ((https://doi.org/10.1007/s13225-020-00466-2)) |
| Otu43825 | 1 | k__Fungi(100);p__Ascomycota(100);c__Saccharomycetes(100);o__Saccharomycetales(100);f__Metschnikowiaceae(100);g__Metschnikowia(100);s__Metschnikowia_pulcherrima(100); | Metschnikowia | 13 | Saprotroph | \|Undefined Saprotroph\| | Yeast | NULL | Probable | As nectar-tap_saprotroph (Põlme et al. 2020); Decay sugar-rich substrates (Põlme et al. 2020) | Põlme S, et al. 2020. Fungal Diversity 105:1-16 ((https://doi.org/10.1007/s13225-020-00466-2)) |
| Otu43826 | 1 | k__Fungi(100);p__Ascomycota(100);c__Saccharomycetes(100);o__Saccharomycetales(100);f__Metschnikowiaceae(100);g__Metschnikowia(100);g__Metschnikowia_unclassified(100); | Metschnikowia | 13 | Saprotroph | \|Undefined Saprotroph\| | Yeast | NULL | Probable | As nectar-tap_saprotroph (Põlme et al. 2020); Decay sugar-rich substrates (Põlme et al. 2020) | Põlme S, et al. 2020. Fungal Diversity 105:1-16 ((https://doi.org/10.1007/s13225-020-00466-2)) |
| Otu43827 | 1 | k__Fungi(100);p__Ascomycota(100);c__Saccharomycetes(100);o__Saccharomycetales(100);f__Metschnikowiaceae(100);g__Metschnikowia(100);s__Metschnikowia_pulcherrima(100); | Metschnikowia | 13 | Saprotroph | \|Undefined Saprotroph\| | Yeast | NULL | Probable | As nectar-tap_saprotroph (Põlme et al. 2020); Decay sugar-rich substrates (Põlme et al. 2020) | Põlme S, et al. 2020. Fungal Diversity 105:1-16 ((https://doi.org/10.1007/s13225-020-00466-2)) |
| Otu43828 | 1 | k__Fungi(100);p__Ascomycota(100);c__Saccharomycetes(100);o__Saccharomycetales(100);f__Metschnikowiaceae(100);g__Metschnikowia(100);s__Metschnikowia_pulcherrima(100); | Metschnikowia | 13 | Saprotroph | \|Undefined Saprotroph\| | Yeast | NULL | Probable | As nectar-tap_saprotroph (Põlme et al. 2020); Decay sugar-rich substrates (Põlme et al. 2020) | Põlme S, et al. 2020. Fungal Diversity 105:1-16 ((https://doi.org/10.1007/s13225-020-00466-2)) |
| Otu43829 | 1 | k__Fungi(100);p__Ascomycota(100);c__Saccharomycetes(100);o__Saccharomycetales(100);f__Metschnikowiaceae(100);g__Metschnikowia(100);s__Metschnikowia_pulcherrima(100); | Metschnikowia | 13 | Saprotroph | \|Undefined Saprotroph\| | Yeast | NULL | Probable | As nectar-tap_saprotroph (Põlme et al. 2020); Decay sugar-rich substrates (Põlme et al. 2020) | Põlme S, et al. 2020. Fungal Diversity 105:1-16 ((https://doi.org/10.1007/s13225-020-00466-2)) |
| Otu43830 | 1 | k__Fungi(100);p__Ascomycota(100);c__Saccharomycetes(100);o__Saccharomycetales(100);f__Metschnikowiaceae(100);g__Metschnikowia(100);s__Metschnikowia_pulcherrima(100); | Metschnikowia | 13 | Saprotroph | \|Undefined Saprotroph\| | Yeast | NULL | Probable | As nectar-tap_saprotroph (Põlme et al. 2020); Decay sugar-rich substrates (Põlme et al. 2020) | Põlme S, et al. 2020. Fungal Diversity 105:1-16 ((https://doi.org/10.1007/s13225-020-00466-2)) |
| Otu43831 | 1 | k__Fungi(100);p__Ascomycota(100);c__Saccharomycetes(100);o__Saccharomycetales(100);f__Metschnikowiaceae(100);g__Metschnikowia(100);g__Metschnikowia_unclassified(100); | Metschnikowia | 13 | Saprotroph | \|Undefined Saprotroph\| | Yeast | NULL | Probable | As nectar-tap_saprotroph (Põlme et al. 2020); Decay sugar-rich substrates (Põlme et al. 2020) | Põlme S, et al. 2020. Fungal Diversity 105:1-16 ((https://doi.org/10.1007/s13225-020-00466-2)) |
| Otu43832 | 1 | k__Fungi(100);p__Ascomycota(100);c__Saccharomycetes(100);o__Saccharomycetales(100);f__Metschnikowiaceae(100);g__Metschnikowia(100);s__Metschnikowia_pulcherrima(100); | Metschnikowia | 13 | Saprotroph | \|Undefined Saprotroph\| | Yeast | NULL | Probable | As nectar-tap_saprotroph (Põlme et al. 2020); Decay sugar-rich substrates (Põlme et al. 2020) | Põlme S, et al. 2020. Fungal Diversity 105:1-16 ((https://doi.org/10.1007/s13225-020-00466-2)) |
| Otu43833 | 1 | k__Fungi(100);p__Ascomycota(100);c__Saccharomycetes(100);o__Saccharomycetales(100);f__Metschnikowiaceae(100);g__Metschnikowia(100);s__Metschnikowia_pulcherrima(100); | Metschnikowia | 13 | Saprotroph | \|Undefined Saprotroph\| | Yeast | NULL | Probable | As nectar-tap_saprotroph (Põlme et al. 2020); Decay sugar-rich substrates (Põlme et al. 2020) | Põlme S, et al. 2020. Fungal Diversity 105:1-16 ((https://doi.org/10.1007/s13225-020-00466-2)) |
| Otu43834 | 1 | k__Fungi(100);p__Ascomycota(100);c__Saccharomycetes(100);o__Saccharomycetales(100);f__Metschnikowiaceae(100);g__Metschnikowia(100);s__Metschnikowia_pulcherrima(100); | Metschnikowia | 13 | Saprotroph | \|Undefined Saprotroph\| | Yeast | NULL | Probable | As nectar-tap_saprotroph (Põlme et al. 2020); Decay sugar-rich substrates (Põlme et al. 2020) | Põlme S, et al. 2020. Fungal Diversity 105:1-16 ((https://doi.org/10.1007/s13225-020-00466-2)) |
| Otu43835 | 1 | k__Fungi(100);p__Ascomycota(100);c__Saccharomycetes(100);o__Saccharomycetales(100);f__Metschnikowiaceae(100);g__Metschnikowia(100);s__Metschnikowia_pulcherrima(100); | Metschnikowia | 13 | Saprotroph | \|Undefined Saprotroph\| | Yeast | NULL | Probable | As nectar-tap_saprotroph (Põlme et al. 2020); Decay sugar-rich substrates (Põlme et al. 2020) | Põlme S, et al. 2020. Fungal Diversity 105:1-16 ((https://doi.org/10.1007/s13225-020-00466-2)) |
| Otu43836 | 1 | k__Fungi(100);p__Ascomycota(100);c__Saccharomycetes(100);o__Saccharomycetales(100);f__Metschnikowiaceae(100);g__Metschnikowia(100);s__Metschnikowia_pulcherrima(100); | Metschnikowia | 13 | Saprotroph | \|Undefined Saprotroph\| | Yeast | NULL | Probable | As nectar-tap_saprotroph (Põlme et al. 2020); Decay sugar-rich substrates (Põlme et al. 2020) | Põlme S, et al. 2020. Fungal Diversity 105:1-16 ((https://doi.org/10.1007/s13225-020-00466-2)) |
| Otu43837 | 1 | k__Fungi(100);p__Ascomycota(100);c__Saccharomycetes(100);o__Saccharomycetales(100);f__Metschnikowiaceae(100);g__Metschnikowia(100);g__Metschnikowia_unclassified(100); | Metschnikowia | 13 | Saprotroph | \|Undefined Saprotroph\| | Yeast | NULL | Probable | As nectar-tap_saprotroph (Põlme et al. 2020); Decay sugar-rich substrates (Põlme et al. 2020) | Põlme S, et al. 2020. Fungal Diversity 105:1-16 ((https://doi.org/10.1007/s13225-020-00466-2)) |
| Otu43838 | 1 | k__Fungi(100);p__Ascomycota(100);c__Saccharomycetes(100);o__Saccharomycetales(100);f__Metschnikowiaceae(100);g__Metschnikowia(100);g__Metschnikowia_unclassified(100); | Metschnikowia | 13 | Saprotroph | \|Undefined Saprotroph\| | Yeast | NULL | Probable | As nectar-tap_saprotroph (Põlme et al. 2020); Decay sugar-rich substrates (Põlme et al. 2020) | Põlme S, et al. 2020. Fungal Diversity 105:1-16 ((https://doi.org/10.1007/s13225-020-00466-2)) |
| Otu43839 | 1 | k__Fungi(100);p__Ascomycota(100);c__Saccharomycetes(100);o__Saccharomycetales(100);f__Metschnikowiaceae(100);g__Metschnikowia(100);s__Metschnikowia_pulcherrima(100); | Metschnikowia | 13 | Saprotroph | \|Undefined Saprotroph\| | Yeast | NULL | Probable | As nectar-tap_saprotroph (Põlme et al. 2020); Decay sugar-rich substrates (Põlme et al. 2020) | Põlme S, et al. 2020. Fungal Diversity 105:1-16 ((https://doi.org/10.1007/s13225-020-00466-2)) |
| Otu43840 | 1 | k__Fungi(100);p__Ascomycota(100);c__Saccharomycetes(100);o__Saccharomycetales(100);f__Metschnikowiaceae(100);g__Metschnikowia(100);s__Metschnikowia_pulcherrima(100); | Metschnikowia | 13 | Saprotroph | \|Undefined Saprotroph\| | Yeast | NULL | Probable | As nectar-tap_saprotroph (Põlme et al. 2020); Decay sugar-rich substrates (Põlme et al. 2020) | Põlme S, et al. 2020. Fungal Diversity 105:1-16 ((https://doi.org/10.1007/s13225-020-00466-2)) |
| Otu43841 | 1 | k__Fungi(100);p__Ascomycota(100);c__Saccharomycetes(100);o__Saccharomycetales(100);f__Metschnikowiaceae(100);g__Metschnikowia(100);s__Metschnikowia_pulcherrima(100); | Metschnikowia | 13 | Saprotroph | \|Undefined Saprotroph\| | Yeast | NULL | Probable | As nectar-tap_saprotroph (Põlme et al. 2020); Decay sugar-rich substrates (Põlme et al. 2020) | Põlme S, et al. 2020. Fungal Diversity 105:1-16 ((https://doi.org/10.1007/s13225-020-00466-2)) |
| Otu43842 | 1 | k__Fungi(100);p__Ascomycota(100);c__Saccharomycetes(100);o__Saccharomycetales(100);f__Metschnikowiaceae(100);g__Metschnikowia(100);s__Metschnikowia_pulcherrima(100); | Metschnikowia | 13 | Saprotroph | \|Undefined Saprotroph\| | Yeast | NULL | Probable | As nectar-tap_saprotroph (Põlme et al. 2020); Decay sugar-rich substrates (Põlme et al. 2020) | Põlme S, et al. 2020. Fungal Diversity 105:1-16 ((https://doi.org/10.1007/s13225-020-00466-2)) |
| Otu43843 | 1 | k__Fungi(100);p__Ascomycota(100);c__Saccharomycetes(100);o__Saccharomycetales(100);f__Metschnikowiaceae(100);g__Metschnikowia(100);s__Metschnikowia_pulcherrima(100); | Metschnikowia | 13 | Saprotroph | \|Undefined Saprotroph\| | Yeast | NULL | Probable | As nectar-tap_saprotroph (Põlme et al. 2020); Decay sugar-rich substrates (Põlme et al. 2020) | Põlme S, et al. 2020. Fungal Diversity 105:1-16 ((https://doi.org/10.1007/s13225-020-00466-2)) |
| Otu43844 | 1 | k__Fungi(100);p__Ascomycota(100);c__Saccharomycetes(100);o__Saccharomycetales(100);f__Metschnikowiaceae(100);g__Metschnikowia(100);g__Metschnikowia_unclassified(100); | Metschnikowia | 13 | Saprotroph | \|Undefined Saprotroph\| | Yeast | NULL | Probable | As nectar-tap_saprotroph (Põlme et al. 2020); Decay sugar-rich substrates (Põlme et al. 2020) | Põlme S, et al. 2020. Fungal Diversity 105:1-16 ((https://doi.org/10.1007/s13225-020-00466-2)) |
| Otu43846 | 1 | k__Fungi(100);p__Ascomycota(100);c__Saccharomycetes(100);o__Saccharomycetales(100);f__Metschnikowiaceae(100);g__Metschnikowia(100);g__Metschnikowia_unclassified(100); | Metschnikowia | 13 | Saprotroph | \|Undefined Saprotroph\| | Yeast | NULL | Probable | As nectar-tap_saprotroph (Põlme et al. 2020); Decay sugar-rich substrates (Põlme et al. 2020) | Põlme S, et al. 2020. Fungal Diversity 105:1-16 ((https://doi.org/10.1007/s13225-020-00466-2)) |
| Otu43847 | 1 | k__Fungi(100);p__Ascomycota(100);c__Saccharomycetes(100);o__Saccharomycetales(100);f__Metschnikowiaceae(100);g__Metschnikowia(100);s__Metschnikowia_pulcherrima(100); | Metschnikowia | 13 | Saprotroph | \|Undefined Saprotroph\| | Yeast | NULL | Probable | As nectar-tap_saprotroph (Põlme et al. 2020); Decay sugar-rich substrates (Põlme et al. 2020) | Põlme S, et al. 2020. Fungal Diversity 105:1-16 ((https://doi.org/10.1007/s13225-020-00466-2)) |
| Otu43849 | 1 | k__Fungi(100);p__Ascomycota(100);c__Saccharomycetes(100);o__Saccharomycetales(100);f__Metschnikowiaceae(100);g__Metschnikowia(100);s__Metschnikowia_pulcherrima(100); | Metschnikowia | 13 | Saprotroph | \|Undefined Saprotroph\| | Yeast | NULL | Probable | As nectar-tap_saprotroph (Põlme et al. 2020); Decay sugar-rich substrates (Põlme et al. 2020) | Põlme S, et al. 2020. Fungal Diversity 105:1-16 ((https://doi.org/10.1007/s13225-020-00466-2)) |
| Otu43850 | 1 | k__Fungi(100);p__Ascomycota(100);c__Saccharomycetes(100);o__Saccharomycetales(100);f__Metschnikowiaceae(100);g__Metschnikowia(100);s__Metschnikowia_pulcherrima(100); | Metschnikowia | 13 | Saprotroph | \|Undefined Saprotroph\| | Yeast | NULL | Probable | As nectar-tap_saprotroph (Põlme et al. 2020); Decay sugar-rich substrates (Põlme et al. 2020) | Põlme S, et al. 2020. Fungal Diversity 105:1-16 ((https://doi.org/10.1007/s13225-020-00466-2)) |
| Otu43851 | 1 | k__Fungi(100);p__Ascomycota(100);c__Saccharomycetes(100);o__Saccharomycetales(100);f__Metschnikowiaceae(100);g__Metschnikowia(100);s__Metschnikowia_pulcherrima(100); | Metschnikowia | 13 | Saprotroph | \|Undefined Saprotroph\| | Yeast | NULL | Probable | As nectar-tap_saprotroph (Põlme et al. 2020); Decay sugar-rich substrates (Põlme et al. 2020) | Põlme S, et al. 2020. Fungal Diversity 105:1-16 ((https://doi.org/10.1007/s13225-020-00466-2)) |
| Otu43852 | 1 | k__Fungi(100);p__Ascomycota(100);c__Saccharomycetes(100);o__Saccharomycetales(100);f__Metschnikowiaceae(100);g__Metschnikowia(100);s__Metschnikowia_pulcherrima(100); | Metschnikowia | 13 | Saprotroph | \|Undefined Saprotroph\| | Yeast | NULL | Probable | As nectar-tap_saprotroph (Põlme et al. 2020); Decay sugar-rich substrates (Põlme et al. 2020) | Põlme S, et al. 2020. Fungal Diversity 105:1-16 ((https://doi.org/10.1007/s13225-020-00466-2)) |
| Otu43853 | 1 | k__Fungi(100);p__Ascomycota(100);c__Saccharomycetes(100);o__Saccharomycetales(100);f__Metschnikowiaceae(100);g__Metschnikowia(100);s__Metschnikowia_pulcherrima(100); | Metschnikowia | 13 | Saprotroph | \|Undefined Saprotroph\| | Yeast | NULL | Probable | As nectar-tap_saprotroph (Põlme et al. 2020); Decay sugar-rich substrates (Põlme et al. 2020) | Põlme S, et al. 2020. Fungal Diversity 105:1-16 ((https://doi.org/10.1007/s13225-020-00466-2)) |
| Otu43854 | 1 | k__Fungi(100);p__Ascomycota(100);c__Saccharomycetes(100);o__Saccharomycetales(100);f__Metschnikowiaceae(100);g__Metschnikowia(100);s__Metschnikowia_pulcherrima(100); | Metschnikowia | 13 | Saprotroph | \|Undefined Saprotroph\| | Yeast | NULL | Probable | As nectar-tap_saprotroph (Põlme et al. 2020); Decay sugar-rich substrates (Põlme et al. 2020) | Põlme S, et al. 2020. Fungal Diversity 105:1-16 ((https://doi.org/10.1007/s13225-020-00466-2)) |
| Otu43855 | 1 | k__Fungi(100);p__Ascomycota(100);c__Saccharomycetes(100);o__Saccharomycetales(100);f__Metschnikowiaceae(100);g__Metschnikowia(100);g__Metschnikowia_unclassified(100); | Metschnikowia | 13 | Saprotroph | \|Undefined Saprotroph\| | Yeast | NULL | Probable | As nectar-tap_saprotroph (Põlme et al. 2020); Decay sugar-rich substrates (Põlme et al. 2020) | Põlme S, et al. 2020. Fungal Diversity 105:1-16 ((https://doi.org/10.1007/s13225-020-00466-2)) |
| Otu43856 | 1 | k__Fungi(100);p__Ascomycota(100);c__Saccharomycetes(100);o__Saccharomycetales(100);f__Metschnikowiaceae(100);g__Metschnikowia(100);s__Metschnikowia_viticola(100); | Metschnikowia | 13 | Saprotroph | \|Undefined Saprotroph\| | Yeast | NULL | Probable | As nectar-tap_saprotroph (Põlme et al. 2020); Decay sugar-rich substrates (Põlme et al. 2020) | Põlme S, et al. 2020. Fungal Diversity 105:1-16 ((https://doi.org/10.1007/s13225-020-00466-2)) |
| Otu43857 | 1 | k__Fungi(100);p__Ascomycota(100);c__Saccharomycetes(100);o__Saccharomycetales(100);f__Metschnikowiaceae(100);g__Metschnikowia(100);s__Metschnikowia_pulcherrima(100); | Metschnikowia | 13 | Saprotroph | \|Undefined Saprotroph\| | Yeast | NULL | Probable | As nectar-tap_saprotroph (Põlme et al. 2020); Decay sugar-rich substrates (Põlme et al. 2020) | Põlme S, et al. 2020. Fungal Diversity 105:1-16 ((https://doi.org/10.1007/s13225-020-00466-2)) |
| Otu43858 | 1 | k__Fungi(100);p__Ascomycota(100);c__Saccharomycetes(100);o__Saccharomycetales(100);f__Metschnikowiaceae(100);g__Metschnikowia(100);s__Metschnikowia_pulcherrima(100); | Metschnikowia | 13 | Saprotroph | \|Undefined Saprotroph\| | Yeast | NULL | Probable | As nectar-tap_saprotroph (Põlme et al. 2020); Decay sugar-rich substrates (Põlme et al. 2020) | Põlme S, et al. 2020. Fungal Diversity 105:1-16 ((https://doi.org/10.1007/s13225-020-00466-2)) |
| Otu43859 | 1 | k__Fungi(100);p__Ascomycota(100);c__Saccharomycetes(100);o__Saccharomycetales(100);f__Metschnikowiaceae(100);g__Metschnikowia(100);s__Metschnikowia_pulcherrima(100); | Metschnikowia | 13 | Saprotroph | \|Undefined Saprotroph\| | Yeast | NULL | Probable | As nectar-tap_saprotroph (Põlme et al. 2020); Decay sugar-rich substrates (Põlme et al. 2020) | Põlme S, et al. 2020. Fungal Diversity 105:1-16 ((https://doi.org/10.1007/s13225-020-00466-2)) |
| Otu43860 | 1 | k__Fungi(100);p__Ascomycota(100);c__Saccharomycetes(100);o__Saccharomycetales(100);f__Metschnikowiaceae(100);g__Metschnikowia(100);s__Metschnikowia_pulcherrima(100); | Metschnikowia | 13 | Saprotroph | \|Undefined Saprotroph\| | Yeast | NULL | Probable | As nectar-tap_saprotroph (Põlme et al. 2020); Decay sugar-rich substrates (Põlme et al. 2020) | Põlme S, et al. 2020. Fungal Diversity 105:1-16 ((https://doi.org/10.1007/s13225-020-00466-2)) |
| Otu43861 | 1 | k__Fungi(100);p__Ascomycota(100);c__Saccharomycetes(100);o__Saccharomycetales(100);f__Metschnikowiaceae(100);g__Metschnikowia(100);s__Metschnikowia_pulcherrima(100); | Metschnikowia | 13 | Saprotroph | \|Undefined Saprotroph\| | Yeast | NULL | Probable | As nectar-tap_saprotroph (Põlme et al. 2020); Decay sugar-rich substrates (Põlme et al. 2020) | Põlme S, et al. 2020. Fungal Diversity 105:1-16 ((https://doi.org/10.1007/s13225-020-00466-2)) |
| Otu43862 | 1 | k__Fungi(100);p__Ascomycota(100);c__Saccharomycetes(100);o__Saccharomycetales(100);f__Metschnikowiaceae(100);g__Metschnikowia(100);s__Metschnikowia_pulcherrima(100); | Metschnikowia | 13 | Saprotroph | \|Undefined Saprotroph\| | Yeast | NULL | Probable | As nectar-tap_saprotroph (Põlme et al. 2020); Decay sugar-rich substrates (Põlme et al. 2020) | Põlme S, et al. 2020. Fungal Diversity 105:1-16 ((https://doi.org/10.1007/s13225-020-00466-2)) |
| Otu43863 | 1 | k__Fungi(100);p__Ascomycota(100);c__Saccharomycetes(100);o__Saccharomycetales(100);f__Metschnikowiaceae(100);g__Metschnikowia(100);s__Metschnikowia_pulcherrima(100); | Metschnikowia | 13 | Saprotroph | \|Undefined Saprotroph\| | Yeast | NULL | Probable | As nectar-tap_saprotroph (Põlme et al. 2020); Decay sugar-rich substrates (Põlme et al. 2020) | Põlme S, et al. 2020. Fungal Diversity 105:1-16 ((https://doi.org/10.1007/s13225-020-00466-2)) |
| Otu43864 | 1 | k__Fungi(100);p__Ascomycota(100);c__Saccharomycetes(100);o__Saccharomycetales(100);f__Metschnikowiaceae(100);g__Metschnikowia(100);g__Metschnikowia_unclassified(100); | Metschnikowia | 13 | Saprotroph | \|Undefined Saprotroph\| | Yeast | NULL | Probable | As nectar-tap_saprotroph (Põlme et al. 2020); Decay sugar-rich substrates (Põlme et al. 2020) | Põlme S, et al. 2020. Fungal Diversity 105:1-16 ((https://doi.org/10.1007/s13225-020-00466-2)) |
| Otu43865 | 1 | k__Fungi(100);p__Ascomycota(100);c__Saccharomycetes(100);o__Saccharomycetales(100);f__Metschnikowiaceae(100);g__Metschnikowia(100);s__Metschnikowia_pulcherrima(100); | Metschnikowia | 13 | Saprotroph | \|Undefined Saprotroph\| | Yeast | NULL | Probable | As nectar-tap_saprotroph (Põlme et al. 2020); Decay sugar-rich substrates (Põlme et al. 2020) | Põlme S, et al. 2020. Fungal Diversity 105:1-16 ((https://doi.org/10.1007/s13225-020-00466-2)) |
| Otu43866 | 1 | k__Fungi(100);p__Ascomycota(100);c__Saccharomycetes(100);o__Saccharomycetales(100);f__Metschnikowiaceae(100);g__Metschnikowia(100);g__Metschnikowia_unclassified(100); | Metschnikowia | 13 | Saprotroph | \|Undefined Saprotroph\| | Yeast | NULL | Probable | As nectar-tap_saprotroph (Põlme et al. 2020); Decay sugar-rich substrates (Põlme et al. 2020) | Põlme S, et al. 2020. Fungal Diversity 105:1-16 ((https://doi.org/10.1007/s13225-020-00466-2)) |
| Otu43867 | 1 | k__Fungi(100);p__Ascomycota(100);c__Saccharomycetes(100);o__Saccharomycetales(100);f__Metschnikowiaceae(100);g__Metschnikowia(100);s__Metschnikowia_pulcherrima(100); | Metschnikowia | 13 | Saprotroph | \|Undefined Saprotroph\| | Yeast | NULL | Probable | As nectar-tap_saprotroph (Põlme et al. 2020); Decay sugar-rich substrates (Põlme et al. 2020) | Põlme S, et al. 2020. Fungal Diversity 105:1-16 ((https://doi.org/10.1007/s13225-020-00466-2)) |
| Otu43868 | 1 | k__Fungi(100);p__Ascomycota(100);c__Saccharomycetes(100);o__Saccharomycetales(100);f__Metschnikowiaceae(100);g__Metschnikowia(100);s__Metschnikowia_pulcherrima(100); | Metschnikowia | 13 | Saprotroph | \|Undefined Saprotroph\| | Yeast | NULL | Probable | As nectar-tap_saprotroph (Põlme et al. 2020); Decay sugar-rich substrates (Põlme et al. 2020) | Põlme S, et al. 2020. Fungal Diversity 105:1-16 ((https://doi.org/10.1007/s13225-020-00466-2)) |
| Otu43869 | 1 | k__Fungi(100);p__Ascomycota(100);c__Saccharomycetes(100);o__Saccharomycetales(100);f__Metschnikowiaceae(100);g__Metschnikowia(100);s__Metschnikowia_pulcherrima(100); | Metschnikowia | 13 | Saprotroph | \|Undefined Saprotroph\| | Yeast | NULL | Probable | As nectar-tap_saprotroph (Põlme et al. 2020); Decay sugar-rich substrates (Põlme et al. 2020) | Põlme S, et al. 2020. Fungal Diversity 105:1-16 ((https://doi.org/10.1007/s13225-020-00466-2)) |
| Otu43870 | 1 | k__Fungi(100);p__Ascomycota(100);c__Saccharomycetes(100);o__Saccharomycetales(100);f__Metschnikowiaceae(100);g__Metschnikowia(100);s__Metschnikowia_pulcherrima(100); | Metschnikowia | 13 | Saprotroph | \|Undefined Saprotroph\| | Yeast | NULL | Probable | As nectar-tap_saprotroph (Põlme et al. 2020); Decay sugar-rich substrates (Põlme et al. 2020) | Põlme S, et al. 2020. Fungal Diversity 105:1-16 ((https://doi.org/10.1007/s13225-020-00466-2)) |
| Otu43871 | 1 | k__Fungi(100);p__Ascomycota(100);c__Saccharomycetes(100);o__Saccharomycetales(100);f__Metschnikowiaceae(100);g__Metschnikowia(100);s__Metschnikowia_pulcherrima(100); | Metschnikowia | 13 | Saprotroph | \|Undefined Saprotroph\| | Yeast | NULL | Probable | As nectar-tap_saprotroph (Põlme et al. 2020); Decay sugar-rich substrates (Põlme et al. 2020) | Põlme S, et al. 2020. Fungal Diversity 105:1-16 ((https://doi.org/10.1007/s13225-020-00466-2)) |
| Otu43872 | 1 | k__Fungi(100);p__Ascomycota(100);c__Saccharomycetes(100);o__Saccharomycetales(100);f__Metschnikowiaceae(100);g__Metschnikowia(100);s__Metschnikowia_pulcherrima(100); | Metschnikowia | 13 | Saprotroph | \|Undefined Saprotroph\| | Yeast | NULL | Probable | As nectar-tap_saprotroph (Põlme et al. 2020); Decay sugar-rich substrates (Põlme et al. 2020) | Põlme S, et al. 2020. Fungal Diversity 105:1-16 ((https://doi.org/10.1007/s13225-020-00466-2)) |
| Otu43873 | 1 | k__Fungi(100);p__Ascomycota(100);c__Saccharomycetes(100);o__Saccharomycetales(100);f__Metschnikowiaceae(100);g__Metschnikowia(100);s__Metschnikowia_pulcherrima(100); | Metschnikowia | 13 | Saprotroph | \|Undefined Saprotroph\| | Yeast | NULL | Probable | As nectar-tap_saprotroph (Põlme et al. 2020); Decay sugar-rich substrates (Põlme et al. 2020) | Põlme S, et al. 2020. Fungal Diversity 105:1-16 ((https://doi.org/10.1007/s13225-020-00466-2)) |
| Otu43874 | 1 | k__Fungi(100);p__Ascomycota(100);c__Saccharomycetes(100);o__Saccharomycetales(100);f__Metschnikowiaceae(100);g__Metschnikowia(100);s__Metschnikowia_pulcherrima(100); | Metschnikowia | 13 | Saprotroph | \|Undefined Saprotroph\| | Yeast | NULL | Probable | As nectar-tap_saprotroph (Põlme et al. 2020); Decay sugar-rich substrates (Põlme et al. 2020) | Põlme S, et al. 2020. Fungal Diversity 105:1-16 ((https://doi.org/10.1007/s13225-020-00466-2)) |
| Otu43875 | 1 | k__Fungi(100);p__Ascomycota(100);c__Saccharomycetes(100);o__Saccharomycetales(100);f__Metschnikowiaceae(100);g__Metschnikowia(100);s__Metschnikowia_pulcherrima(100); | Metschnikowia | 13 | Saprotroph | \|Undefined Saprotroph\| | Yeast | NULL | Probable | As nectar-tap_saprotroph (Põlme et al. 2020); Decay sugar-rich substrates (Põlme et al. 2020) | Põlme S, et al. 2020. Fungal Diversity 105:1-16 ((https://doi.org/10.1007/s13225-020-00466-2)) |
| Otu43876 | 1 | k__Fungi(100);p__Ascomycota(100);c__Saccharomycetes(100);o__Saccharomycetales(100);f__Metschnikowiaceae(100);g__Metschnikowia(100);s__Metschnikowia_pulcherrima(100); | Metschnikowia | 13 | Saprotroph | \|Undefined Saprotroph\| | Yeast | NULL | Probable | As nectar-tap_saprotroph (Põlme et al. 2020); Decay sugar-rich substrates (Põlme et al. 2020) | Põlme S, et al. 2020. Fungal Diversity 105:1-16 ((https://doi.org/10.1007/s13225-020-00466-2)) |
| Otu43877 | 1 | k__Fungi(100);p__Ascomycota(100);c__Saccharomycetes(100);o__Saccharomycetales(100);f__Metschnikowiaceae(100);g__Metschnikowia(100);g__Metschnikowia_unclassified(100); | Metschnikowia | 13 | Saprotroph | \|Undefined Saprotroph\| | Yeast | NULL | Probable | As nectar-tap_saprotroph (Põlme et al. 2020); Decay sugar-rich substrates (Põlme et al. 2020) | Põlme S, et al. 2020. Fungal Diversity 105:1-16 ((https://doi.org/10.1007/s13225-020-00466-2)) |
| Otu43878 | 1 | k__Fungi(100);p__Ascomycota(100);c__Saccharomycetes(100);o__Saccharomycetales(100);f__Metschnikowiaceae(100);g__Metschnikowia(100);s__Metschnikowia_pulcherrima(100); | Metschnikowia | 13 | Saprotroph | \|Undefined Saprotroph\| | Yeast | NULL | Probable | As nectar-tap_saprotroph (Põlme et al. 2020); Decay sugar-rich substrates (Põlme et al. 2020) | Põlme S, et al. 2020. Fungal Diversity 105:1-16 ((https://doi.org/10.1007/s13225-020-00466-2)) |
| Otu43879 | 1 | k__Fungi(100);p__Ascomycota(100);c__Saccharomycetes(100);o__Saccharomycetales(100);f__Metschnikowiaceae(100);g__Metschnikowia(100);g__Metschnikowia_unclassified(100); | Metschnikowia | 13 | Saprotroph | \|Undefined Saprotroph\| | Yeast | NULL | Probable | As nectar-tap_saprotroph (Põlme et al. 2020); Decay sugar-rich substrates (Põlme et al. 2020) | Põlme S, et al. 2020. Fungal Diversity 105:1-16 ((https://doi.org/10.1007/s13225-020-00466-2)) |
| Otu43880 | 1 | k__Fungi(100);p__Ascomycota(100);c__Saccharomycetes(100);o__Saccharomycetales(100);f__Metschnikowiaceae(100);g__Metschnikowia(100);g__Metschnikowia_unclassified(100); | Metschnikowia | 13 | Saprotroph | \|Undefined Saprotroph\| | Yeast | NULL | Probable | As nectar-tap_saprotroph (Põlme et al. 2020); Decay sugar-rich substrates (Põlme et al. 2020) | Põlme S, et al. 2020. Fungal Diversity 105:1-16 ((https://doi.org/10.1007/s13225-020-00466-2)) |
| Otu43881 | 1 | k__Fungi(100);p__Ascomycota(100);c__Saccharomycetes(100);o__Saccharomycetales(100);f__Metschnikowiaceae(100);g__Metschnikowia(100);g__Metschnikowia_unclassified(100); | Metschnikowia | 13 | Saprotroph | \|Undefined Saprotroph\| | Yeast | NULL | Probable | As nectar-tap_saprotroph (Põlme et al. 2020); Decay sugar-rich substrates (Põlme et al. 2020) | Põlme S, et al. 2020. Fungal Diversity 105:1-16 ((https://doi.org/10.1007/s13225-020-00466-2)) |
| Otu43882 | 1 | k__Fungi(100);p__Ascomycota(100);c__Saccharomycetes(100);o__Saccharomycetales(100);f__Metschnikowiaceae(100);g__Metschnikowia(100);g__Metschnikowia_unclassified(100); | Metschnikowia | 13 | Saprotroph | \|Undefined Saprotroph\| | Yeast | NULL | Probable | As nectar-tap_saprotroph (Põlme et al. 2020); Decay sugar-rich substrates (Põlme et al. 2020) | Põlme S, et al. 2020. Fungal Diversity 105:1-16 ((https://doi.org/10.1007/s13225-020-00466-2)) |
| Otu43883 | 1 | k__Fungi(100);p__Ascomycota(100);c__Saccharomycetes(100);o__Saccharomycetales(100);f__Metschnikowiaceae(100);g__Metschnikowia(100);g__Metschnikowia_unclassified(100); | Metschnikowia | 13 | Saprotroph | \|Undefined Saprotroph\| | Yeast | NULL | Probable | As nectar-tap_saprotroph (Põlme et al. 2020); Decay sugar-rich substrates (Põlme et al. 2020) | Põlme S, et al. 2020. Fungal Diversity 105:1-16 ((https://doi.org/10.1007/s13225-020-00466-2)) |
| Otu43884 | 1 | k__Fungi(100);p__Ascomycota(100);c__Saccharomycetes(100);o__Saccharomycetales(100);f__Metschnikowiaceae(100);g__Metschnikowia(100);g__Metschnikowia_unclassified(100); | Metschnikowia | 13 | Saprotroph | \|Undefined Saprotroph\| | Yeast | NULL | Probable | As nectar-tap_saprotroph (Põlme et al. 2020); Decay sugar-rich substrates (Põlme et al. 2020) | Põlme S, et al. 2020. Fungal Diversity 105:1-16 ((https://doi.org/10.1007/s13225-020-00466-2)) |
| Otu43885 | 1 | k__Fungi(100);p__Ascomycota(100);c__Saccharomycetes(100);o__Saccharomycetales(100);f__Metschnikowiaceae(100);g__Metschnikowia(100);g__Metschnikowia_unclassified(100); | Metschnikowia | 13 | Saprotroph | \|Undefined Saprotroph\| | Yeast | NULL | Probable | As nectar-tap_saprotroph (Põlme et al. 2020); Decay sugar-rich substrates (Põlme et al. 2020) | Põlme S, et al. 2020. Fungal Diversity 105:1-16 ((https://doi.org/10.1007/s13225-020-00466-2)) |
| Otu43886 | 1 | k__Fungi(100);p__Ascomycota(100);c__Saccharomycetes(100);o__Saccharomycetales(100);f__Metschnikowiaceae(100);g__Metschnikowia(100);g__Metschnikowia_unclassified(100); | Metschnikowia | 13 | Saprotroph | \|Undefined Saprotroph\| | Yeast | NULL | Probable | As nectar-tap_saprotroph (Põlme et al. 2020); Decay sugar-rich substrates (Põlme et al. 2020) | Põlme S, et al. 2020. Fungal Diversity 105:1-16 ((https://doi.org/10.1007/s13225-020-00466-2)) |
| Otu43887 | 1 | k__Fungi(100);p__Ascomycota(100);c__Saccharomycetes(100);o__Saccharomycetales(100);f__Metschnikowiaceae(100);g__Metschnikowia(100);g__Metschnikowia_unclassified(100); | Metschnikowia | 13 | Saprotroph | \|Undefined Saprotroph\| | Yeast | NULL | Probable | As nectar-tap_saprotroph (Põlme et al. 2020); Decay sugar-rich substrates (Põlme et al. 2020) | Põlme S, et al. 2020. Fungal Diversity 105:1-16 ((https://doi.org/10.1007/s13225-020-00466-2)) |
| Otu43888 | 1 | k__Fungi(100);p__Ascomycota(100);c__Saccharomycetes(100);o__Saccharomycetales(100);f__Metschnikowiaceae(100);g__Metschnikowia(100);g__Metschnikowia_unclassified(100); | Metschnikowia | 13 | Saprotroph | \|Undefined Saprotroph\| | Yeast | NULL | Probable | As nectar-tap_saprotroph (Põlme et al. 2020); Decay sugar-rich substrates (Põlme et al. 2020) | Põlme S, et al. 2020. Fungal Diversity 105:1-16 ((https://doi.org/10.1007/s13225-020-00466-2)) |
| Otu43889 | 1 | k__Fungi(100);p__Ascomycota(100);c__Saccharomycetes(100);o__Saccharomycetales(100);f__Metschnikowiaceae(100);g__Metschnikowia(100);g__Metschnikowia_unclassified(100); | Metschnikowia | 13 | Saprotroph | \|Undefined Saprotroph\| | Yeast | NULL | Probable | As nectar-tap_saprotroph (Põlme et al. 2020); Decay sugar-rich substrates (Põlme et al. 2020) | Põlme S, et al. 2020. Fungal Diversity 105:1-16 ((https://doi.org/10.1007/s13225-020-00466-2)) |
| Otu43890 | 1 | k__Fungi(100);p__Ascomycota(100);c__Saccharomycetes(100);o__Saccharomycetales(100);f__Metschnikowiaceae(100);g__Metschnikowia(100);s__Metschnikowia_pulcherrima(100); | Metschnikowia | 13 | Saprotroph | \|Undefined Saprotroph\| | Yeast | NULL | Probable | As nectar-tap_saprotroph (Põlme et al. 2020); Decay sugar-rich substrates (Põlme et al. 2020) | Põlme S, et al. 2020. Fungal Diversity 105:1-16 ((https://doi.org/10.1007/s13225-020-00466-2)) |

| | | | | | | | | | | | |
|---|---|---|---|---|---|---|---|---|---|---|---|
| Otu43891 | 1 | k__Fungi(100);p__Ascomycota(100);c__Saccharomycetes(100);o__Saccharomycetales(100);f__Metschnikowiaceae(100);g__Metschnikowia(100);g__Metschnikowia_unclassified(100); | Metschnikowia | 13 | Saprotroph | \|Undefined Saprotroph\| | Yeast | NULL | Probable | As nectar-tap_saprotroph (Põlme et al. 2020); Decay sugar-rich substrates (Põlme et al. 2020) | Põlme S, et al. 2020. Fungal Diversity 105:1-16 ((https://doi.org/10.1007/s13225-020-00466-2)) |
| Otu43892 | 1 | k__Fungi(100);p__Ascomycota(100);c__Saccharomycetes(100);o__Saccharomycetales(100);f__Metschnikowiaceae(100);g__Metschnikowia(100);s__Metschnikowia_pulcherrima(100); | Metschnikowia | 13 | Saprotroph | \|Undefined Saprotroph\| | Yeast | NULL | Probable | As nectar-tap_saprotroph (Põlme et al. 2020); Decay sugar-rich substrates (Põlme et al. 2020) | Põlme S, et al. 2020. Fungal Diversity 105:1-16 ((https://doi.org/10.1007/s13225-020-00466-2)) |
| Otu43893 | 1 | k__Fungi(100);p__Ascomycota(100);c__Saccharomycetes(100);o__Saccharomycetales(100);f__Metschnikowiaceae(100);g__Metschnikowia(100);g__Metschnikowia_unclassified(100); | Metschnikowia | 13 | Saprotroph | \|Undefined Saprotroph\| | Yeast | NULL | Probable | As nectar-tap_saprotroph (Põlme et al. 2020); Decay sugar-rich substrates (Põlme et al. 2020) | Põlme S, et al. 2020. Fungal Diversity 105:1-16 ((https://doi.org/10.1007/s13225-020-00466-2)) |
| Otu43896 | 1 | k__Fungi(100);p__Ascomycota(100);c__Saccharomycetes(100);o__Saccharomycetales(100);f__Metschnikowiaceae(100);g__Metschnikowia(100);s__Metschnikowia_pulcherrima(100); | Metschnikowia | 13 | Saprotroph | \|Undefined Saprotroph\| | Yeast | NULL | Probable | As nectar-tap_saprotroph (Põlme et al. 2020); Decay sugar-rich substrates (Põlme et al. 2020) | Põlme S, et al. 2020. Fungal Diversity 105:1-16 ((https://doi.org/10.1007/s13225-020-00466-2)) |
| Otu43897 | 1 | k__Fungi(100);p__Ascomycota(100);c__Saccharomycetes(100);o__Saccharomycetales(100);f__Metschnikowiaceae(100);g__Metschnikowia(100);s__Metschnikowia_pulcherrima(100); | Metschnikowia | 13 | Saprotroph | \|Undefined Saprotroph\| | Yeast | NULL | Probable | As nectar-tap_saprotroph (Põlme et al. 2020); Decay sugar-rich substrates (Põlme et al. 2020) | Põlme S, et al. 2020. Fungal Diversity 105:1-16 ((https://doi.org/10.1007/s13225-020-00466-2)) |
| Otu43898 | 1 | k__Fungi(100);p__Ascomycota(100);c__Saccharomycetes(100);o__Saccharomycetales(100);f__Metschnikowiaceae(100);g__Metschnikowia(100);g__Metschnikowia_unclassified(100); | Metschnikowia | 13 | Saprotroph | \|Undefined Saprotroph\| | Yeast | NULL | Probable | As nectar-tap_saprotroph (Põlme et al. 2020); Decay sugar-rich substrates (Põlme et al. 2020) | Põlme S, et al. 2020. Fungal Diversity 105:1-16 ((https://doi.org/10.1007/s13225-020-00466-2)) |
| Otu43899 | 1 | k__Fungi(100);p__Ascomycota(100);c__Saccharomycetes(100);o__Saccharomycetales(100);f__Metschnikowiaceae(100);g__Metschnikowia(100);g__Metschnikowia_unclassified(100); | Metschnikowia | 13 | Saprotroph | \|Undefined Saprotroph\| | Yeast | NULL | Probable | As nectar-tap_saprotroph (Põlme et al. 2020); Decay sugar-rich substrates (Põlme et al. 2020) | Põlme S, et al. 2020. Fungal Diversity 105:1-16 ((https://doi.org/10.1007/s13225-020-00466-2)) |
| Otu43901 | 1 | k__Fungi(100);p__Ascomycota(100);c__Saccharomycetes(100);o__Saccharomycetales(100);f__Metschnikowiaceae(100);g__Metschnikowia(100);s__Metschnikowia_pulcherrima(100); | Metschnikowia | 13 | Saprotroph | \|Undefined Saprotroph\| | Yeast | NULL | Probable | As nectar-tap_saprotroph (Põlme et al. 2020); Decay sugar-rich substrates (Põlme et al. 2020) | Põlme S, et al. 2020. Fungal Diversity 105:1-16 ((https://doi.org/10.1007/s13225-020-00466-2)) |
| Otu43902 | 1 | k__Fungi(100);p__Ascomycota(100);c__Saccharomycetes(100);o__Saccharomycetales(100);f__Metschnikowiaceae(100);g__Metschnikowia(100);s__Metschnikowia_pulcherrima(100); | Metschnikowia | 13 | Saprotroph | \|Undefined Saprotroph\| | Yeast | NULL | Probable | As nectar-tap_saprotroph (Põlme et al. 2020); Decay sugar-rich substrates (Põlme et al. 2020) | Põlme S, et al. 2020. Fungal Diversity 105:1-16 ((https://doi.org/10.1007/s13225-020-00466-2)) |
| Otu43903 | 1 | k__Fungi(100);p__Ascomycota(100);c__Saccharomycetes(100);o__Saccharomycetales(100);f__Metschnikowiaceae(100);g__Metschnikowia(100);s__Metschnikowia_pulcherrima(100); | Metschnikowia | 13 | Saprotroph | \|Undefined Saprotroph\| | Yeast | NULL | Probable | As nectar-tap_saprotroph (Põlme et al. 2020); Decay sugar-rich substrates (Põlme et al. 2020) | Põlme S, et al. 2020. Fungal Diversity 105:1-16 ((https://doi.org/10.1007/s13225-020-00466-2)) |
| Otu43904 | 1 | k__Fungi(100);p__Ascomycota(100);c__Saccharomycetes(100);o__Saccharomycetales(100);f__Metschnikowiaceae(100);g__Metschnikowia(100);s__Metschnikowia_pulcherrima(100); | Metschnikowia | 13 | Saprotroph | \|Undefined Saprotroph\| | Yeast | NULL | Probable | As nectar-tap_saprotroph (Põlme et al. 2020); Decay sugar-rich substrates (Põlme et al. 2020) | Põlme S, et al. 2020. Fungal Diversity 105:1-16 ((https://doi.org/10.1007/s13225-020-00466-2)) |
| Otu43905 | 1 | k__Fungi(100);p__Ascomycota(100);c__Saccharomycetes(100);o__Saccharomycetales(100);f__Metschnikowiaceae(100);g__Metschnikowia(100);s__Metschnikowia_pulcherrima(100); | Metschnikowia | 13 | Saprotroph | \|Undefined Saprotroph\| | Yeast | NULL | Probable | As nectar-tap_saprotroph (Põlme et al. 2020); Decay sugar-rich substrates (Põlme et al. 2020) | Põlme S, et al. 2020. Fungal Diversity 105:1-16 ((https://doi.org/10.1007/s13225-020-00466-2)) |
| Otu43906 | 1 | k__Fungi(100);p__Ascomycota(100);c__Saccharomycetes(100);o__Saccharomycetales(100);f__Metschnikowiaceae(100);g__Metschnikowia(100);s__Metschnikowia_pulcherrima(100); | Metschnikowia | 13 | Saprotroph | \|Undefined Saprotroph\| | Yeast | NULL | Probable | As nectar-tap_saprotroph (Põlme et al. 2020); Decay sugar-rich substrates (Põlme et al. 2020) | Põlme S, et al. 2020. Fungal Diversity 105:1-16 ((https://doi.org/10.1007/s13225-020-00466-2)) |
| Otu43907 | 1 | k__Fungi(100);p__Ascomycota(100);c__Saccharomycetes(100);o__Saccharomycetales(100);f__Metschnikowiaceae(100);g__Metschnikowia(100);s__Metschnikowia_pulcherrima(100); | Metschnikowia | 13 | Saprotroph | \|Undefined Saprotroph\| | Yeast | NULL | Probable | As nectar-tap_saprotroph (Põlme et al. 2020); Decay sugar-rich substrates (Põlme et al. 2020) | Põlme S, et al. 2020. Fungal Diversity 105:1-16 ((https://doi.org/10.1007/s13225-020-00466-2)) |
| Otu43908 | 1 | k__Fungi(100);p__Ascomycota(100);c__Saccharomycetes(100);o__Saccharomycetales(100);f__Metschnikowiaceae(100);g__Metschnikowia(100);s__Metschnikowia_pulcherrima(100); | Metschnikowia | 13 | Saprotroph | \|Undefined Saprotroph\| | Yeast | NULL | Probable | As nectar-tap_saprotroph (Põlme et al. 2020); Decay sugar-rich substrates (Põlme et al. 2020) | Põlme S, et al. 2020. Fungal Diversity 105:1-16 ((https://doi.org/10.1007/s13225-020-00466-2)) |
| Otu43909 | 1 | k__Fungi(100);p__Ascomycota(100);c__Saccharomycetes(100);o__Saccharomycetales(100);f__Metschnikowiaceae(100);g__Metschnikowia(100);s__Metschnikowia_pulcherrima(100); | Metschnikowia | 13 | Saprotroph | \|Undefined Saprotroph\| | Yeast | NULL | Probable | As nectar-tap_saprotroph (Põlme et al. 2020); Decay sugar-rich substrates (Põlme et al. 2020) | Põlme S, et al. 2020. Fungal Diversity 105:1-16 ((https://doi.org/10.1007/s13225-020-00466-2)) |
| Otu43910 | 1 | k__Fungi(100);p__Ascomycota(100);c__Saccharomycetes(100);o__Saccharomycetales(100);f__Metschnikowiaceae(100);g__Metschnikowia(100);s__Metschnikowia_pulcherrima(100); | Metschnikowia | 13 | Saprotroph | \|Undefined Saprotroph\| | Yeast | NULL | Probable | As nectar-tap_saprotroph (Põlme et al. 2020); Decay sugar-rich substrates (Põlme et al. 2020) | Põlme S, et al. 2020. Fungal Diversity 105:1-16 ((https://doi.org/10.1007/s13225-020-00466-2)) |
| Otu43911 | 1 | k__Fungi(100);p__Ascomycota(100);c__Saccharomycetes(100);o__Saccharomycetales(100);f__Metschnikowiaceae(100);g__Metschnikowia(100);s__Metschnikowia_pulcherrima(100); | Metschnikowia | 13 | Saprotroph | \|Undefined Saprotroph\| | Yeast | NULL | Probable | As nectar-tap_saprotroph (Põlme et al. 2020); Decay sugar-rich substrates (Põlme et al. 2020) | Põlme S, et al. 2020. Fungal Diversity 105:1-16 ((https://doi.org/10.1007/s13225-020-00466-2)) |
| Otu43912 | 1 | k__Fungi(100);p__Ascomycota(100);c__Saccharomycetes(100);o__Saccharomycetales(100);f__Metschnikowiaceae(100);g__Metschnikowia(100);s__Metschnikowia_pulcherrima(100); | Metschnikowia | 13 | Saprotroph | \|Undefined Saprotroph\| | Yeast | NULL | Probable | As nectar-tap_saprotroph (Põlme et al. 2020); Decay sugar-rich substrates (Põlme et al. 2020) | Põlme S, et al. 2020. Fungal Diversity 105:1-16 ((https://doi.org/10.1007/s13225-020-00466-2)) |
| Otu43913 | 1 | k__Fungi(100);p__Ascomycota(100);c__Saccharomycetes(100);o__Saccharomycetales(100);f__Metschnikowiaceae(100);g__Metschnikowia(100);s__Metschnikowia_pulcherrima(100); | Metschnikowia | 13 | Saprotroph | \|Undefined Saprotroph\| | Yeast | NULL | Probable | As nectar-tap_saprotroph (Põlme et al. 2020); Decay sugar-rich substrates (Põlme et al. 2020) | Põlme S, et al. 2020. Fungal Diversity 105:1-16 ((https://doi.org/10.1007/s13225-020-00466-2)) |
| Otu43914 | 1 | k__Fungi(100);p__Ascomycota(100);c__Saccharomycetes(100);o__Saccharomycetales(100);f__Metschnikowiaceae(100);g__Metschnikowia(100);g__Metschnikowia_unclassified(100); | Metschnikowia | 13 | Saprotroph | \|Undefined Saprotroph\| | Yeast | NULL | Probable | As nectar-tap_saprotroph (Põlme et al. 2020); Decay sugar-rich substrates (Põlme et al. 2020) | Põlme S, et al. 2020. Fungal Diversity 105:1-16 ((https://doi.org/10.1007/s13225-020-00466-2)) |
| Otu43916 | 1 | k__Fungi(100);p__Ascomycota(100);c__Saccharomycetes(100);o__Saccharomycetales(100);f__Metschnikowiaceae(100);g__Metschnikowia(100);s__Metschnikowia_pulcherrima(100); | Metschnikowia | 13 | Saprotroph | \|Undefined Saprotroph\| | Yeast | NULL | Probable | As nectar-tap_saprotroph (Põlme et al. 2020); Decay sugar-rich substrates (Põlme et al. 2020) | Põlme S, et al. 2020. Fungal Diversity 105:1-16 ((https://doi.org/10.1007/s13225-020-00466-2)) |
| Otu43917 | 1 | k__Fungi(100);p__Ascomycota(100);c__Saccharomycetes(100);o__Saccharomycetales(100);f__Metschnikowiaceae(100);g__Metschnikowia(100);s__Metschnikowia_pulcherrima(100); | Metschnikowia | 13 | Saprotroph | \|Undefined Saprotroph\| | Yeast | NULL | Probable | As nectar-tap_saprotroph (Põlme et al. 2020); Decay sugar-rich substrates (Põlme et al. 2020) | Põlme S, et al. 2020. Fungal Diversity 105:1-16 ((https://doi.org/10.1007/s13225-020-00466-2)) |
| Otu43920 | 1 | k__Fungi(100);p__Ascomycota(100);c__Saccharomycetes(100);o__Saccharomycetales(100);f__Metschnikowiaceae(100);g__Metschnikowia(100);s__Metschnikowia_pulcherrima(100); | Metschnikowia | 13 | Saprotroph | \|Undefined Saprotroph\| | Yeast | NULL | Probable | As nectar-tap_saprotroph (Põlme et al. 2020); Decay sugar-rich substrates (Põlme et al. 2020) | Põlme S, et al. 2020. Fungal Diversity 105:1-16 ((https://doi.org/10.1007/s13225-020-00466-2)) |
| Otu43921 | 1 | k__Fungi(100);p__Ascomycota(100);c__Saccharomycetes(100);o__Saccharomycetales(100);f__Metschnikowiaceae(100);g__Metschnikowia(100);g__Metschnikowia_unclassified(100); | Metschnikowia | 13 | Saprotroph | \|Undefined Saprotroph\| | Yeast | NULL | Probable | As nectar-tap_saprotroph (Põlme et al. 2020); Decay sugar-rich substrates (Põlme et al. 2020) | Põlme S, et al. 2020. Fungal Diversity 105:1-16 ((https://doi.org/10.1007/s13225-020-00466-2)) |
| Otu43922 | 1 | k__Fungi(100);p__Ascomycota(100);c__Saccharomycetes(100);o__Saccharomycetales(100);f__Metschnikowiaceae(100);g__Metschnikowia(100);s__Metschnikowia_pulcherrima(100); | Metschnikowia | 13 | Saprotroph | \|Undefined Saprotroph\| | Yeast | NULL | Probable | As nectar-tap_saprotroph (Põlme et al. 2020); Decay sugar-rich substrates (Põlme et al. 2020) | Põlme S, et al. 2020. Fungal Diversity 105:1-16 ((https://doi.org/10.1007/s13225-020-00466-2)) |
| Otu43923 | 1 | k__Fungi(100);p__Ascomycota(100);c__Saccharomycetes(100);o__Saccharomycetales(100);f__Metschnikowiaceae(100);g__Metschnikowia(100);s__Metschnikowia_pulcherrima(100); | Metschnikowia | 13 | Saprotroph | \|Undefined Saprotroph\| | Yeast | NULL | Probable | As nectar-tap_saprotroph (Põlme et al. 2020); Decay sugar-rich substrates (Põlme et al. 2020) | Põlme S, et al. 2020. Fungal Diversity 105:1-16 ((https://doi.org/10.1007/s13225-020-00466-2)) |
| Otu43924 | 1 | k__Fungi(100);p__Ascomycota(100);c__Saccharomycetes(100);o__Saccharomycetales(100);f__Metschnikowiaceae(100);g__Metschnikowia(100);g__Metschnikowia_unclassified(100); | Metschnikowia | 13 | Saprotroph | \|Undefined Saprotroph\| | Yeast | NULL | Probable | As nectar-tap_saprotroph (Põlme et al. 2020); Decay sugar-rich substrates (Põlme et al. 2020) | Põlme S, et al. 2020. Fungal Diversity 105:1-16 ((https://doi.org/10.1007/s13225-020-00466-2)) |
| Otu43925 | 1 | k__Fungi(100);p__Ascomycota(100);c__Saccharomycetes(100);o__Saccharomycetales(100);f__Metschnikowiaceae(100);g__Metschnikowia(100);s__Metschnikowia_pulcherrima(100); | Metschnikowia | 13 | Saprotroph | \|Undefined Saprotroph\| | Yeast | NULL | Probable | As nectar-tap_saprotroph (Põlme et al. 2020); Decay sugar-rich substrates (Põlme et al. 2020) | Põlme S, et al. 2020. Fungal Diversity 105:1-16 ((https://doi.org/10.1007/s13225-020-00466-2)) |
| Otu43926 | 1 | k__Fungi(100);p__Ascomycota(100);c__Saccharomycetes(100);o__Saccharomycetales(100);f__Metschnikowiaceae(100);g__Metschnikowia(100);s__Metschnikowia_pulcherrima(100); | Metschnikowia | 13 | Saprotroph | \|Undefined Saprotroph\| | Yeast | NULL | Probable | As nectar-tap_saprotroph (Põlme et al. 2020); Decay sugar-rich substrates (Põlme et al. 2020) | Põlme S, et al. 2020. Fungal Diversity 105:1-16 ((https://doi.org/10.1007/s13225-020-00466-2)) |
| Otu43927 | 1 | k__Fungi(100);p__Ascomycota(100);c__Saccharomycetes(100);o__Saccharomycetales(100);f__Metschnikowiaceae(100);g__Metschnikowia(100);s__Metschnikowia_pulcherrima(100); | Metschnikowia | 13 | Saprotroph | \|Undefined Saprotroph\| | Yeast | NULL | Probable | As nectar-tap_saprotroph (Põlme et al. 2020); Decay sugar-rich substrates (Põlme et al. 2020) | Põlme S, et al. 2020. Fungal Diversity 105:1-16 ((https://doi.org/10.1007/s13225-020-00466-2)) |
| Otu43928 | 1 | k__Fungi(100);p__Ascomycota(100);c__Sordariomycetes(100);o__Sordariales(100);f__Chaetomiaceae(100);f__Chaetomiaceae_unclassified(100);f__Chaetomiaceae_unclassified(100); | Chaetomiaceae | 9 | Pathotroph-Saprotroph-Symbiotroph | Animal Pathogen-Dung Saprotroph-Endophyte-Epiphyte-Plant Saprotroph-Wood Saprotroph | Microfungus | Soft Rot | Possible | Saprobic, acting as key decay organisms of plant materials and plant-derived commodities (Cannon & Kirk 2007); Epiphyte detection method-culture (Massimo et al. 2015); Endophyte detection method-culture (Busby et al. 2016, David et al. 2016); Likely opportunistic human pathogen (Irinyi et al. 2015) | Seehann G, et al. 1975. List of Fungi in Soft-Rot Tissue Tests. Institute fur Holzbiologie und Holzschutz, Bundesforschungsanstalt fur Forst - und Holzwirtshaft, Hamburg - Lohbrugge, Germany; Cannon PF, Kirk PM. 2007. Fungal Families of the World. CAB International, Cambridge ((https://isbnsearch.org/isbn/9780851998275)); Bills GF, et al. 2013. Current Opinion in Microbiology 16:549-565 ((https://doi.org/10.1016/j.mib.2013.08.001)); Tedersoo L, et al. 2014. Science 346:e1256688 ((https://doi.org/10.1126/science.1256688)); Irinyi L, et al. 2015. Medical Mycology 53:313-337 ((https://doi.org/10.1093/mmy/myv008)); Massimo NC, et al. 2015. Microbial Ecology 70:61-76 ((https://doi.org/10.1007/s00248-014-0563-6)); Busby PE, et al. 2016. New Phytologist 209:1681-1692 ((https://doi.org/10.1111/nph.13742)); David AS, et al. 2016. Microbial Ecology 71:912-926 ((https://doi.org/10.1007/s00248-015-0712-6)) |
| Otu43929 | 1 | k__Fungi(100);p__Ascomycota(100);c__Saccharomycetes(100);o__Saccharomycetales(100);f__Metschnikowiaceae(100);g__Metschnikowia(100);s__Metschnikowia_pulcherrima(100); | Metschnikowia | 13 | Saprotroph | \|Undefined Saprotroph\| | Yeast | NULL | Probable | As nectar-tap_saprotroph (Põlme et al. 2020); Decay sugar-rich substrates (Põlme et al. 2020) | Põlme S, et al. 2020. Fungal Diversity 105:1-16 ((https://doi.org/10.1007/s13225-020-00466-2)) |
| Otu43930 | 1 | k__Fungi(100);p__Ascomycota(100);c__Saccharomycetes(100);o__Saccharomycetales(100);f__Metschnikowiaceae(100);g__Metschnikowia(100);s__Metschnikowia_pulcherrima(100); | Metschnikowia | 13 | Saprotroph | \|Undefined Saprotroph\| | Yeast | NULL | Probable | As nectar-tap_saprotroph (Põlme et al. 2020); Decay sugar-rich substrates (Põlme et al. 2020) | Põlme S, et al. 2020. Fungal Diversity 105:1-16 ((https://doi.org/10.1007/s13225-020-00466-2)) |
| Otu43931 | 1 | k__Fungi(100);p__Ascomycota(100);c__Saccharomycetes(100);o__Saccharomycetales(100);f__Metschnikowiaceae(100);g__Metschnikowia(100);s__Metschnikowia_pulcherrima(100); | Metschnikowia | 13 | Saprotroph | \|Undefined Saprotroph\| | Yeast | NULL | Probable | As nectar-tap_saprotroph (Põlme et al. 2020); Decay sugar-rich substrates (Põlme et al. 2020) | Põlme S, et al. 2020. Fungal Diversity 105:1-16 ((https://doi.org/10.1007/s13225-020-00466-2)) |
| Otu43932 | 1 | k__Fungi(100);p__Ascomycota(100);c__Saccharomycetes(100);o__Saccharomycetales(100);f__Metschnikowiaceae(100);g__Metschnikowia(100);s__Metschnikowia_pulcherrima(100); | Metschnikowia | 13 | Saprotroph | \|Undefined Saprotroph\| | Yeast | NULL | Probable | As nectar-tap_saprotroph (Põlme et al. 2020); Decay sugar-rich substrates (Põlme et al. 2020) | Põlme S, et al. 2020. Fungal Diversity 105:1-16 ((https://doi.org/10.1007/s13225-020-00466-2)) |
| Otu43933 | 1 | k__Fungi(100);p__Ascomycota(100);c__Saccharomycetes(100);o__Saccharomycetales(100);f__Metschnikowiaceae(100);g__Metschnikowia(100);g__Metschnikowia_unclassified(100); | Metschnikowia | 13 | Saprotroph | \|Undefined Saprotroph\| | Yeast | NULL | Probable | As nectar-tap_saprotroph (Põlme et al. 2020); Decay sugar-rich substrates (Põlme et al. 2020) | Põlme S, et al. 2020. Fungal Diversity 105:1-16 ((https://doi.org/10.1007/s13225-020-00466-2)) |
| Otu43934 | 1 | k__Fungi(100);p__Ascomycota(100);c__Saccharomycetes(100);o__Saccharomycetales(100);f__Metschnikowiaceae(100);g__Metschnikowia(100);s__Metschnikowia_pulcherrima(100); | Metschnikowia | 13 | Saprotroph | \|Undefined Saprotroph\| | Yeast | NULL | Probable | As nectar-tap_saprotroph (Põlme et al. 2020); Decay sugar-rich substrates (Põlme et al. 2020) | Põlme S, et al. 2020. Fungal Diversity 105:1-16 ((https://doi.org/10.1007/s13225-020-00466-2)) |
| Otu43935 | 1 | k__Fungi(100);p__Ascomycota(100);c__Saccharomycetes(100);o__Saccharomycetales(100);f__Metschnikowiaceae(100);g__Metschnikowia(100);s__Metschnikowia_pulcherrima(100); | Metschnikowia | 13 | Saprotroph | \|Undefined Saprotroph\| | Yeast | NULL | Probable | As nectar-tap_saprotroph (Põlme et al. 2020); Decay sugar-rich substrates (Põlme et al. 2020) | Põlme S, et al. 2020. Fungal Diversity 105:1-16 ((https://doi.org/10.1007/s13225-020-00466-2)) |
| Otu43936 | 1 | k__Fungi(100);p__Ascomycota(100);c__Saccharomycetes(100);o__Saccharomycetales(100);f__Metschnikowiaceae(100);g__Metschnikowia(100);s__Metschnikowia_pulcherrima(100); | Metschnikowia | 13 | Saprotroph | \|Undefined Saprotroph\| | Yeast | NULL | Probable | As nectar-tap_saprotroph (Põlme et al. 2020); Decay sugar-rich substrates (Põlme et al. 2020) | Põlme S, et al. 2020. Fungal Diversity 105:1-16 ((https://doi.org/10.1007/s13225-020-00466-2)) |
| Otu43937 | 1 | k__Fungi(100);p__Ascomycota(100);c__Saccharomycetes(100);o__Saccharomycetales(100);f__Metschnikowiaceae(100);g__Metschnikowia(100);s__Metschnikowia_pulcherrima(100); | Metschnikowia | 13 | Saprotroph | \|Undefined Saprotroph\| | Yeast | NULL | Probable | As nectar-tap_saprotroph (Põlme et al. 2020); Decay sugar-rich substrates (Põlme et al. 2020) | Põlme S, et al. 2020. Fungal Diversity 105:1-16 ((https://doi.org/10.1007/s13225-020-00466-2)) |
| Otu43938 | 1 | k__Fungi(100);p__Ascomycota(100);c__Saccharomycetes(100);o__Saccharomycetales(100);f__Metschnikowiaceae(100);g__Metschnikowia(100);s__Metschnikowia_pulcherrima(100); | Metschnikowia | 13 | Saprotroph | \|Undefined Saprotroph\| | Yeast | NULL | Probable | As nectar-tap_saprotroph (Põlme et al. 2020); Decay sugar-rich substrates (Põlme et al. 2020) | Põlme S, et al. 2020. Fungal Diversity 105:1-16 ((https://doi.org/10.1007/s13225-020-00466-2)) |
| Otu43939 | 1 | k__Fungi(100);p__Ascomycota(100);c__Saccharomycetes(100);o__Saccharomycetales(100);f__Metschnikowiaceae(100);g__Metschnikowia(100);s__Metschnikowia_pulcherrima(100); | Metschnikowia | 13 | Saprotroph | \|Undefined Saprotroph\| | Yeast | NULL | Probable | As nectar-tap_saprotroph (Põlme et al. 2020); Decay sugar-rich substrates (Põlme et al. 2020) | Põlme S, et al. 2020. Fungal Diversity 105:1-16 ((https://doi.org/10.1007/s13225-020-00466-2)) |
| Otu43940 | 1 | k__Fungi(100);p__Ascomycota(100);c__Saccharomycetes(100);o__Saccharomycetales(100);f__Metschnikowiaceae(100);g__Metschnikowia(100);s__Metschnikowia_pulcherrima(100); | Metschnikowia | 13 | Saprotroph | \|Undefined Saprotroph\| | Yeast | NULL | Probable | As nectar-tap_saprotroph (Põlme et al. 2020); Decay sugar-rich substrates (Põlme et al. 2020) | Põlme S, et al. 2020. Fungal Diversity 105:1-16 ((https://doi.org/10.1007/s13225-020-00466-2)) |
| Otu43941 | 1 | k__Fungi(100);p__Ascomycota(100);c__Saccharomycetes(100);o__Saccharomycetales(100);f__Metschnikowiaceae(100);g__Metschnikowia(100);s__Metschnikowia_pulcherrima(100); | Metschnikowia | 13 | Saprotroph | \|Undefined Saprotroph\| | Yeast | NULL | Probable | As nectar-tap_saprotroph (Põlme et al. 2020); Decay sugar-rich substrates (Põlme et al. 2020) | Põlme S, et al. 2020. Fungal Diversity 105:1-16 ((https://doi.org/10.1007/s13225-020-00466-2)) |
| Otu43942 | 1 | k__Fungi(100);p__Ascomycota(100);c__Saccharomycetes(100);o__Saccharomycetales(100);f__Metschnikowiaceae(100);g__Metschnikowia(100);s__Metschnikowia_pulcherrima(100); | Metschnikowia | 13 | Saprotroph | \|Undefined Saprotroph\| | Yeast | NULL | Probable | As nectar-tap_saprotroph (Põlme et al. 2020); Decay sugar-rich substrates (Põlme et al. 2020) | Põlme S, et al. 2020. Fungal Diversity 105:1-16 ((https://doi.org/10.1007/s13225-020-00466-2)) |
| Otu43943 | 1 | k__Fungi(100);p__Ascomycota(100);c__Saccharomycetes(100);o__Saccharomycetales(100);f__Metschnikowiaceae(100);g__Metschnikowia(100);g__Metschnikowia_unclassified(100); | Metschnikowia | 13 | Saprotroph | \|Undefined Saprotroph\| | Yeast | NULL | Probable | As nectar-tap_saprotroph (Põlme et al. 2020); Decay sugar-rich substrates (Põlme et al. 2020) | Põlme S, et al. 2020. Fungal Diversity 105:1-16 ((https://doi.org/10.1007/s13225-020-00466-2)) |
| Otu43944 | 1 | k__Fungi(100);p__Ascomycota(100);c__Saccharomycetes(100);o__Saccharomycetales(100);f__Metschnikowiaceae(100);g__Metschnikowia(100);s__Metschnikowia_pulcherrima(100); | Metschnikowia | 13 | Saprotroph | \|Undefined Saprotroph\| | Yeast | NULL | Probable | As nectar-tap_saprotroph (Põlme et al. 2020); Decay sugar-rich substrates (Põlme et al. 2020) | Põlme S, et al. 2020. Fungal Diversity 105:1-16 ((https://doi.org/10.1007/s13225-020-00466-2)) |
| Otu43945 | 1 | k__Fungi(100);p__Ascomycota(100);c__Saccharomycetes(100);o__Saccharomycetales(100);f__Metschnikowiaceae(100);g__Metschnikowia(100);g__Metschnikowia_unclassified(100); | Metschnikowia | 13 | Saprotroph | \|Undefined Saprotroph\| | Yeast | NULL | Probable | As nectar-tap_saprotroph (Põlme et al. 2020); Decay sugar-rich substrates (Põlme et al. 2020) | Põlme S, et al. 2020. Fungal Diversity 105:1-16 ((https://doi.org/10.1007/s13225-020-00466-2)) |
| Otu43946 | 1 | k__Fungi(100);p__Ascomycota(100);c__Saccharomycetes(100);o__Saccharomycetales(100);f__Metschnikowiaceae(100);g__Metschnikowia(100);s__Metschnikowia_pulcherrima(100); | Metschnikowia | 13 | Saprotroph | \|Undefined Saprotroph\| | Yeast | NULL | Probable | As nectar-tap_saprotroph (Põlme et al. 2020); Decay sugar-rich substrates (Põlme et al. 2020) | Põlme S, et al. 2020. Fungal Diversity 105:1-16 ((https://doi.org/10.1007/s13225-020-00466-2)) |
| Otu43947 | 1 | k__Fungi(100);p__Ascomycota(100);c__Saccharomycetes(100);o__Saccharomycetales(100);f__Metschnikowiaceae(100);g__Metschnikowia(100);s__Metschnikowia_viticola(100); | Metschnikowia | 13 | Saprotroph | \|Undefined Saprotroph\| | Yeast | NULL | Probable | As nectar-tap_saprotroph (Põlme et al. 2020); Decay sugar-rich substrates (Põlme et al. 2020) | Põlme S, et al. 2020. Fungal Diversity 105:1-16 ((https://doi.org/10.1007/s13225-020-00466-2)) |
| Otu43948 | 1 | k__Fungi(100);p__Ascomycota(100);c__Saccharomycetes(100);o__Saccharomycetales(100);f__Metschnikowiaceae(100);g__Metschnikowia(100);g__Metschnikowia_unclassified(100); | Metschnikowia | 13 | Saprotroph | \|Undefined Saprotroph\| | Yeast | NULL | Probable | As nectar-tap_saprotroph (Põlme et al. 2020); Decay sugar-rich substrates (Põlme et al. 2020) | Põlme S, et al. 2020. Fungal Diversity 105:1-16 ((https://doi.org/10.1007/s13225-020-00466-2)) |
| Otu43949 | 1 | k__Fungi(100);p__Ascomycota(100);c__Saccharomycetes(100);o__Saccharomycetales(100);f__Metschnikowiaceae(100);g__Metschnikowia(100);s__Metschnikowia_pulcherrima(100); | Metschnikowia | 13 | Saprotroph | \|Undefined Saprotroph\| | Yeast | NULL | Probable | As nectar-tap_saprotroph (Põlme et al. 2020); Decay sugar-rich substrates (Põlme et al. 2020) | Põlme S, et al. 2020. Fungal Diversity 105:1-16 ((https://doi.org/10.1007/s13225-020-00466-2)) |
| Otu43950 | 1 | k__Fungi(100);p__Ascomycota(100);c__Saccharomycetes(100);o__Saccharomycetales(100);f__Metschnikowiaceae(100);g__Metschnikowia(100);s__Metschnikowia_pulcherrima(100); | Metschnikowia | 13 | Saprotroph | \|Undefined Saprotroph\| | Yeast | NULL | Probable | As nectar-tap_saprotroph (Põlme et al. 2020); Decay sugar-rich substrates (Põlme et al. 2020) | Põlme S, et al. 2020. Fungal Diversity 105:1-16 ((https://doi.org/10.1007/s13225-020-00466-2)) |
| Otu43951 | 1 | k__Fungi(100);p__Ascomycota(100);c__Saccharomycetes(100);o__Saccharomycetales(100);f__Metschnikowiaceae(100);g__Metschnikowia(100);s__Metschnikowia_pulcherrima(100); | Metschnikowia | 13 | Saprotroph | \|Undefined Saprotroph\| | Yeast | NULL | Probable | As nectar-tap_saprotroph (Põlme et al. 2020); Decay sugar-rich substrates (Põlme et al. 2020) | Põlme S, et al. 2020. Fungal Diversity 105:1-16 ((https://doi.org/10.1007/s13225-020-00466-2)) |
| Otu43952 | 1 | k__Fungi(100);p__Ascomycota(100);c__Saccharomycetes(100);o__Saccharomycetales(100);f__Metschnikowiaceae(100);g__Metschnikowia(100);g__Metschnikowia_unclassified(100); | Metschnikowia | 13 | Saprotroph | \|Undefined Saprotroph\| | Yeast | NULL | Probable | As nectar-tap_saprotroph (Põlme et al. 2020); Decay sugar-rich substrates (Põlme et al. 2020) | Põlme S, et al. 2020. Fungal Diversity 105:1-16 ((https://doi.org/10.1007/s13225-020-00466-2)) |
| Otu43953 | 1 | k__Fungi(100);p__Ascomycota(100);c__Saccharomycetes(100);o__Saccharomycetales(100);f__Metschnikowiaceae(100);g__Metschnikowia(100);s__Metschnikowia_pulcherrima(100); | Metschnikowia | 13 | Saprotroph | \|Undefined Saprotroph\| | Yeast | NULL | Probable | As nectar-tap_saprotroph (Põlme et al. 2020); Decay sugar-rich substrates (Põlme et al. 2020) | Põlme S, et al. 2020. Fungal Diversity 105:1-16 ((https://doi.org/10.1007/s13225-020-00466-2)) |
| Otu43954 | 1 | k__Fungi(100);p__Ascomycota(100);c__Saccharomycetes(100);o__Saccharomycetales(100);f__Metschnikowiaceae(100);g__Metschnikowia(100);s__Metschnikowia_pulcherrima(100); | Metschnikowia | 13 | Saprotroph | \|Undefined Saprotroph\| | Yeast | NULL | Probable | As nectar-tap_saprotroph (Põlme et al. 2020); Decay sugar-rich substrates (Põlme et al. 2020) | Põlme S, et al. 2020. Fungal Diversity 105:1-16 ((https://doi.org/10.1007/s13225-020-00466-2)) |
| Otu43955 | 1 | k__Fungi(100);p__Ascomycota(100);c__Saccharomycetes(100);o__Saccharomycetales(100);f__Metschnikowiaceae(100);g__Metschnikowia(100);s__Metschnikowia_pulcherrima(100); | Metschnikowia | 13 | Saprotroph | \|Undefined Saprotroph\| | Yeast | NULL | Probable | As nectar-tap_saprotroph (Põlme et al. 2020); Decay sugar-rich substrates (Põlme et al. 2020) | Põlme S, et al. 2020. Fungal Diversity 105:1-16 ((https://doi.org/10.1007/s13225-020-00466-2)) |
| Otu43957 | 1 | k__Fungi(100);p__Ascomycota(100);c__Saccharomycetes(100);o__Saccharomycetales(100);f__Metschnikowiaceae(100);g__Metschnikowia(100);g__Metschnikowia_unclassified(100); | Metschnikowia | 13 | Saprotroph | \|Undefined Saprotroph\| | Yeast | NULL | Probable | As nectar-tap_saprotroph (Põlme et al. 2020); Decay sugar-rich substrates (Põlme et al. 2020) | Põlme S, et al. 2020. Fungal Diversity 105:1-16 ((https://doi.org/10.1007/s13225-020-00466-2)) |
| Otu43958 | 1 | k__Fungi(100);p__Ascomycota(100);c__Saccharomycetes(100);o__Saccharomycetales(100);f__Metschnikowiaceae(100);g__Metschnikowia(100);s__Metschnikowia_viticola(100); | Metschnikowia | 13 | Saprotroph | \|Undefined Saprotroph\| | Yeast | NULL | Probable | As nectar-tap_saprotroph (Põlme et al. 2020); Decay sugar-rich substrates (Põlme et al. 2020) | Põlme S, et al. 2020. Fungal Diversity 105:1-16 ((https://doi.org/10.1007/s13225-020-00466-2)) |
| Otu43959 | 1 | k__Fungi(100);p__Ascomycota(100);c__Saccharomycetes(100);o__Saccharomycetales(100);f__Metschnikowiaceae(100);g__Metschnikowia(100);s__Metschnikowia_pulcherrima(100); | Metschnikowia | 13 | Saprotroph | \|Undefined Saprotroph\| | Yeast | NULL | Probable | As nectar-tap_saprotroph (Põlme et al. 2020); Decay sugar-rich substrates (Põlme et al. 2020) | Põlme S, et al. 2020. Fungal Diversity 105:1-16 ((https://doi.org/10.1007/s13225-020-00466-2)) |
| Otu43960 | 1 | k__Fungi(100);p__Ascomycota(100);c__Saccharomycetes(100);o__Saccharomycetales(100);f__Metschnikowiaceae(100);g__Metschnikowia(100);s__Metschnikowia_pulcherrima(100); | Metschnikowia | 13 | Saprotroph | \|Undefined Saprotroph\| | Yeast | NULL | Probable | As nectar-tap_saprotroph (Põlme et al. 2020); Decay sugar-rich substrates (Põlme et al. 2020) | Põlme S, et al. 2020. Fungal Diversity 105:1-16 ((https://doi.org/10.1007/s13225-020-00466-2)) |
| Otu43961 | 1 | k__Fungi(100);p__Ascomycota(100);c__Saccharomycetes(100);o__Saccharomycetales(100);f__Metschnikowiaceae(100);g__Metschnikowia(100);s__Metschnikowia_pulcherrima(100); | Metschnikowia | 13 | Saprotroph | \|Undefined Saprotroph\| | Yeast | NULL | Probable | As nectar-tap_saprotroph (Põlme et al. 2020); Decay sugar-rich substrates (Põlme et al. 2020) | Põlme S, et al. 2020. Fungal Diversity 105:1-16 ((https://doi.org/10.1007/s13225-020-00466-2)) |
| Otu43962 | 1 | k__Fungi(100);p__Ascomycota(100);c__Saccharomycetes(100);o__Saccharomycetales(100);f__Metschnikowiaceae(100);g__Metschnikowia(100);g__Metschnikowia_unclassified(100); | Metschnikowia | 13 | Saprotroph | \|Undefined Saprotroph\| | Yeast | NULL | Probable | As nectar-tap_saprotroph (Põlme et al. 2020); Decay sugar-rich substrates (Põlme et al. 2020) | Põlme S, et al. 2020. Fungal Diversity 105:1-16 ((https://doi.org/10.1007/s13225-020-00466-2)) |
| Otu43963 | 1 | k__Fungi(100);p__Ascomycota(100);c__Saccharomycetes(100);o__Saccharomycetales(100);f__Metschnikowiaceae(100);g__Metschnikowia(100);s__Metschnikowia_pulcherrima(100); | Metschnikowia | 13 | Saprotroph | \|Undefined Saprotroph\| | Yeast | NULL | Probable | As nectar-tap_saprotroph (Põlme et al. 2020); Decay sugar-rich substrates (Põlme et al. 2020) | Põlme S, et al. 2020. Fungal Diversity 105:1-16 ((https://doi.org/10.1007/s13225-020-00466-2)) |
| Otu43964 | 1 | k__Fungi(100);p__Ascomycota(100);c__Saccharomycetes(100);o__Saccharomycetales(100);f__Metschnikowiaceae(100);g__Metschnikowia(100);s__Metschnikowia_pulcherrima(100); | Metschnikowia | 13 | Saprotroph | \|Undefined Saprotroph\| | Yeast | NULL | Probable | As nectar-tap_saprotroph (Põlme et al. 2020); Decay sugar-rich substrates (Põlme et al. 2020) | Põlme S, et al. 2020. Fungal Diversity 105:1-16 ((https://doi.org/10.1007/s13225-020-00466-2)) |
| Otu43965 | 1 | k__Fungi(100);p__Ascomycota(100);c__Saccharomycetes(100);o__Saccharomycetales(100);f__Metschnikowiaceae(100);g__Metschnikowia(100);s__Metschnikowia_pulcherrima(100); | Metschnikowia | 13 | Saprotroph | \|Undefined Saprotroph\| | Yeast | NULL | Probable | As nectar-tap_saprotroph (Põlme et al. 2020); Decay sugar-rich substrates (Põlme et al. 2020) | Põlme S, et al. 2020. Fungal Diversity 105:1-16 ((https://doi.org/10.1007/s13225-020-00466-2)) |
| Otu43966 | 1 | k__Fungi(100);p__Ascomycota(100);c__Saccharomycetes(100);o__Saccharomycetales(100);f__Metschnikowiaceae(100);g__Metschnikowia(100);s__Metschnikowia_pulcherrima(100); | Metschnikowia | 13 | Saprotroph | \|Undefined Saprotroph\| | Yeast | NULL | Probable | As nectar-tap_saprotroph (Põlme et al. 2020); Decay sugar-rich substrates (Põlme et al. 2020) | Põlme S, et al. 2020. Fungal Diversity 105:1-16 ((https://doi.org/10.1007/s13225-020-00466-2)) |
| Otu43967 | 1 | k__Fungi(100);p__Ascomycota(100);c__Saccharomycetes(100);o__Saccharomycetales(100);f__Metschnikowiaceae(100);g__Metschnikowia(100);s__Metschnikowia_pulcherrima(100); | Metschnikowia | 13 | Saprotroph | \|Undefined Saprotroph\| | Yeast | NULL | Probable | As nectar-tap_saprotroph (Põlme et al. 2020); Decay sugar-rich substrates (Põlme et al. 2020) | Põlme S, et al. 2020. Fungal Diversity 105:1-16 ((https://doi.org/10.1007/s13225-020-00466-2)) |
| Otu43968 | 1 | k__Fungi(100);p__Ascomycota(100);c__Saccharomycetes(100);o__Saccharomycetales(100);f__Metschnikowiaceae(100);g__Metschnikowia(100);s__Metschnikowia_pulcherrima(100); | Metschnikowia | 13 | Saprotroph | \|Undefined Saprotroph\| | Yeast | NULL | Probable | As nectar-tap_saprotroph (Põlme et al. 2020); Decay sugar-rich substrates (Põlme et al. 2020) | Põlme S, et al. 2020. Fungal Diversity 105:1-16 ((https://doi.org/10.1007/s13225-020-00466-2)) |
| Otu43969 | 1 | k__Fungi(100);p__Ascomycota(100);c__Saccharomycetes(100);o__Saccharomycetales(100);f__Metschnikowiaceae(100);g__Metschnikowia(100);g__Metschnikowia_unclassified(100); | Metschnikowia | 13 | Saprotroph | \|Undefined Saprotroph\| | Yeast | NULL | Probable | As nectar-tap_saprotroph (Põlme et al. 2020); Decay sugar-rich substrates (Põlme et al. 2020) | Põlme S, et al. 2020. Fungal Diversity 105:1-16 ((https://doi.org/10.1007/s13225-020-00466-2)) |
| Otu43970 | 1 | k__Fungi(100);p__Ascomycota(100);c__Saccharomycetes(100);o__Saccharomycetales(100);f__Metschnikowiaceae(100);g__Metschnikowia(100);s__Metschnikowia_pulcherrima(100); | Metschnikowia | 13 | Saprotroph | \|Undefined Saprotroph\| | Yeast | NULL | Probable | As nectar-tap_saprotroph (Põlme et al. 2020); Decay sugar-rich substrates (Põlme et al. 2020) | Põlme S, et al. 2020. Fungal Diversity 105:1-16 ((https://doi.org/10.1007/s13225-020-00466-2)) |
| Otu43971 | 1 | k__Fungi(100);p__Ascomycota(100);c__Saccharomycetes(100);o__Saccharomycetales(100);f__Metschnikowiaceae(100);g__Metschnikowia(100);s__Metschnikowia_pulcherrima(100); | Metschnikowia | 13 | Saprotroph | \|Undefined Saprotroph\| | Yeast | NULL | Probable | As nectar-tap_saprotroph (Põlme et al. 2020); Decay sugar-rich substrates (Põlme et al. 2020) | Põlme S, et al. 2020. Fungal Diversity 105:1-16 ((https://doi.org/10.1007/s13225-020-00466-2)) |
| Otu43973 | 1 | k__Fungi(100);p__Ascomycota(100);c__Saccharomycetes(100);o__Saccharomycetales(100);f__Metschnikowiaceae(100);g__Metschnikowia(100);s__Metschnikowia_pulcherrima(100); | Metschnikowia | 13 | Saprotroph | \|Undefined Saprotroph\| | Yeast | NULL | Probable | As nectar-tap_saprotroph (Põlme et al. 2020); Decay sugar-rich substrates (Põlme et al. 2020) | Põlme S, et al. 2020. Fungal Diversity 105:1-16 ((https://doi.org/10.1007/s13225-020-00466-2)) |
| Otu43974 | 1 | k__Fungi(100);p__Ascomycota(100);c__Saccharomycetes(100);o__Saccharomycetales(100);f__Metschnikowiaceae(100);g__Metschnikowia(100);s__Metschnikowia_pulcherrima(100); | Metschnikowia | 13 | Saprotroph | \|Undefined Saprotroph\| | Yeast | NULL | Probable | As nectar-tap_saprotroph (Põlme et al. 2020); Decay sugar-rich substrates (Põlme et al. 2020) | Põlme S, et al. 2020. Fungal Diversity 105:1-16 ((https://doi.org/10.1007/s13225-020-00466-2)) |
| Otu43975 | 1 | k__Fungi(100);p__Ascomycota(100);c__Saccharomycetes(100);o__Saccharomycetales(100);f__Metschnikowiaceae(100);g__Metschnikowia(100);s__Metschnikowia_pulcherrima(100); | Metschnikowia | 13 | Saprotroph | \|Undefined Saprotroph\| | Yeast | NULL | Probable | As nectar-tap_saprotroph (Põlme et al. 2020); Decay sugar-rich substrates (Põlme et al. 2020) | Põlme S, et al. 2020. Fungal Diversity 105:1-16 ((https://doi.org/10.1007/s13225-020-00466-2)) |
| Otu43976 | 1 | k__Fungi(100);p__Ascomycota(100);c__Saccharomycetes(100);o__Saccharomycetales(100);f__Metschnikowiaceae(100);g__Metschnikowia(100);g__Metschnikowia_unclassified(100); | Metschnikowia | 13 | Saprotroph | \|Undefined Saprotroph\| | Yeast | NULL | Probable | As nectar-tap_saprotroph (Põlme et al. 2020); Decay sugar-rich substrates (Põlme et al. 2020) | Põlme S, et al. 2020. Fungal Diversity 105:1-16 ((https://doi.org/10.1007/s13225-020-00466-2)) |
| Otu43977 | 1 | k__Fungi(100);p__Ascomycota(100);c__Saccharomycetes(100);o__Saccharomycetales(100);f__Metschnikowiaceae(100);g__Metschnikowia(100);s__Metschnikowia_pulcherrima(100); | Metschnikowia | 13 | Saprotroph | \|Undefined Saprotroph\| | Yeast | NULL | Probable | As nectar-tap_saprotroph (Põlme et al. 2020); Decay sugar-rich substrates (Põlme et al. 2020) | Põlme S, et al. 2020. Fungal Diversity 105:1-16 ((https://doi.org/10.1007/s13225-020-00466-2)) |
| Otu43978 | 1 | k__Fungi(100);p__Ascomycota(100);c__Saccharomycetes(100);o__Saccharomycetales(100);f__Metschnikowiaceae(100);g__Metschnikowia(100);s__Metschnikowia_pulcherrima(100); | Metschnikowia | 13 | Saprotroph | \|Undefined Saprotroph\| | Yeast | NULL | Probable | As nectar-tap_saprotroph (Põlme et al. 2020); Decay sugar-rich substrates (Põlme et al. 2020) | Põlme S, et al. 2020. Fungal Diversity 105:1-16 ((https://doi.org/10.1007/s13225-020-00466-2)) |
| Otu43979 | 1 | k__Fungi(100);p__Ascomycota(100);c__Saccharomycetes(100);o__Saccharomycetales(100);f__Metschnikowiaceae(100);g__Metschnikowia(100);s__Metschnikowia_pulcherrima(100); | Metschnikowia | 13 | Saprotroph | \|Undefined Saprotroph\| | Yeast | NULL | Probable | As nectar-tap_saprotroph (Põlme et al. 2020); Decay sugar-rich substrates (Põlme et al. 2020) | Põlme S, et al. 2020. Fungal Diversity 105:1-16 ((https://doi.org/10.1007/s13225-020-00466-2)) |
| Otu43980 | 1 | k__Fungi(100);p__Ascomycota(100);c__Saccharomycetes(100);o__Saccharomycetales(100);f__Metschnikowiaceae(100);g__Metschnikowia(100);s__Metschnikowia_pulcherrima(100); | Metschnikowia | 13 | Saprotroph | \|Undefined Saprotroph\| | Yeast | NULL | Probable | As nectar-tap_saprotroph (Põlme et al. 2020); Decay sugar-rich substrates (Põlme et al. 2020) | Põlme S, et al. 2020. Fungal Diversity 105:1-16 ((https://doi.org/10.1007/s13225-020-00466-2)) |
| Otu43981 | 1 | k__Fungi(100);p__Ascomycota(100);c__Saccharomycetes(100);o__Saccharomycetales(100);f__Metschnikowiaceae(100);g__Metschnikowia(100);s__Metschnikowia_pulcherrima(100); | Metschnikowia | 13 | Saprotroph | \|Undefined Saprotroph\| | Yeast | NULL | Probable | As nectar-tap_saprotroph (Põlme et al. 2020); Decay sugar-rich substrates (Põlme et al. 2020) | Põlme S, et al. 2020. Fungal Diversity 105:1-16 ((https://doi.org/10.1007/s13225-020-00466-2)) |
| Otu43982 | 1 | k__Fungi(100);p__Ascomycota(100);c__Saccharomycetes(100);o__Saccharomycetales(100);f__Metschnikowiaceae(100);g__Metschnikowia(100);s__Metschnikowia_pulcherrima(100); | Metschnikowia | 13 | Saprotroph | \|Undefined Saprotroph\| | Yeast | NULL | Probable | As nectar-tap_saprotroph (Põlme et al. 2020); Decay sugar-rich substrates (Põlme et al. 2020) | Põlme S, et al. 2020. Fungal Diversity 105:1-16 ((https://doi.org/10.1007/s13225-020-00466-2)) |
| Otu43983 | 1 | k__Fungi(100);p__Ascomycota(100);c__Saccharomycetes(100);o__Saccharomycetales(100);f__Metschnikowiaceae(100);g__Metschnikowia(100);s__Metschnikowia_pulcherrima(100); | Metschnikowia | 13 | Saprotroph | \|Undefined Saprotroph\| | Yeast | NULL | Probable | As nectar-tap_saprotroph (Põlme et al. 2020); Decay sugar-rich substrates (Põlme et al. 2020) | Põlme S, et al. 2020. Fungal Diversity 105:1-16 ((https://doi.org/10.1007/s13225-020-00466-2)) |
| Otu43984 | 1 | k__Fungi(100);p__Ascomycota(100);c__Saccharomycetes(100);o__Saccharomycetales(100);f__Metschnikowiaceae(100);g__Metschnikowia(100);s__Metschnikowia_pulcherrima(100); | Metschnikowia | 13 | Saprotroph | \|Undefined Saprotroph\| | Yeast | NULL | Probable | As nectar-tap_saprotroph (Põlme et al. 2020); Decay sugar-rich substrates (Põlme et al. 2020) | Põlme S, et al. 2020. Fungal Diversity 105:1-16 ((https://doi.org/10.1007/s13225-020-00466-2)) |
| Otu43985 | 1 | k__Fungi(100);p__Ascomycota(100);c__Saccharomycetes(100);o__Saccharomycetales(100);f__Metschnikowiaceae(100);g__Metschnikowia(100);g__Metschnikowia_unclassified(100); | Metschnikowia | 13 | Saprotroph | \|Undefined Saprotroph\| | Yeast | NULL | Probable | As nectar-tap_saprotroph (Põlme et al. 2020); Decay sugar-rich substrates (Põlme et al. 2020) | Põlme S, et al. 2020. Fungal Diversity 105:1-16 ((https://doi.org/10.1007/s13225-020-00466-2)) |
| Otu43986 | 1 | k__Fungi(100);p__Ascomycota(100);c__Saccharomycetes(100);o__Saccharomycetales(100);f__Metschnikowiaceae(100);g__Metschnikowia(100);s__Metschnikowia_pulcherrima(100); | Metschnikowia | 13 | Saprotroph | \|Undefined Saprotroph\| | Yeast | NULL | Probable | As nectar-tap_saprotroph (Põlme et al. 2020); Decay sugar-rich substrates (Põlme et al. 2020) | Põlme S, et al. 2020. Fungal Diversity 105:1-16 ((https://doi.org/10.1007/s13225-020-00466-2)) |
| Otu43987 | 1 | k__Fungi(100);p__Ascomycota(100);c__Saccharomycetes(100);o__Saccharomycetales(100);f__Metschnikowiaceae(100);g__Metschnikowia(100);s__Metschnikowia_pulcherrima(100); | Metschnikowia | 13 | Saprotroph | \|Undefined Saprotroph\| | Yeast | NULL | Probable | As nectar-tap_saprotroph (Põlme et al. 2020); Decay sugar-rich substrates (Põlme et al. 2020) | Põlme S, et al. 2020. Fungal Diversity 105:1-16 ((https://doi.org/10.1007/s13225-020-00466-2)) |
| Otu43988 | 1 | k__Fungi(100);p__Ascomycota(100);c__Saccharomycetes(100);o__Saccharomycetales(100);f__Metschnikowiaceae(100);g__Metschnikowia(100);s__Metschnikowia_pulcherrima(100); | Metschnikowia | 13 | Saprotroph | \|Undefined Saprotroph\| | Yeast | NULL | Probable | As nectar-tap_saprotroph (Põlme et al. 2020); Decay sugar-rich substrates (Põlme et al. 2020) | Põlme S, et al. 2020. Fungal Diversity 105:1-16 ((https://doi.org/10.1007/s13225-020-00466-2)) |
| Otu43989 | 1 | k__Fungi(100);p__Ascomycota(100);c__Saccharomycetes(100);o__Saccharomycetales(100);f__Metschnikowiaceae(100);g__Metschnikowia(100);s__Metschnikowia_pulcherrima(100); | Metschnikowia | 13 | Saprotroph | \|Undefined Saprotroph\| | Yeast | NULL | Probable | As nectar-tap_saprotroph (Põlme et al. 2020); Decay sugar-rich substrates (Põlme et al. 2020) | Põlme S, et al. 2020. Fungal Diversity 105:1-16 ((https://doi.org/10.1007/s13225-020-00466-2)) |
| Otu43990 | 1 | k__Fungi(100);p__Ascomycota(100);c__Saccharomycetes(100);o__Saccharomycetales(100);f__Metschnikowiaceae(100);g__Metschnikowia(100);s__Metschnikowia_pulcherrima(100); | Metschnikowia | 13 | Saprotroph | \|Undefined Saprotroph\| | Yeast | NULL | Probable | As nectar-tap_saprotroph (Põlme et al. 2020); Decay sugar-rich substrates (Põlme et al. 2020) | Põlme S, et al. 2020. Fungal Diversity 105:1-16 ((https://doi.org/10.1007/s13225-020-00466-2)) |
| Otu43991 | 1 | k__Fungi(100);p__Ascomycota(100);c__Saccharomycetes(100);o__Saccharomycetales(100);f__Metschnikowiaceae(100);g__Metschnikowia(100);s__Metschnikowia_pulcherrima(100); | Metschnikowia | 13 | Saprotroph | \|Undefined Saprotroph\| | Yeast | NULL | Probable | As nectar-tap_saprotroph (Põlme et al. 2020); Decay sugar-rich substrates (Põlme et al. 2020) | Põlme S, et al. 2020. Fungal Diversity 105:1-16 ((https://doi.org/10.1007/s13225-020-00466-2)) |
| Otu43992 | 1 | k__Fungi(100);p__Ascomycota(100);c__Saccharomycetes(100);o__Saccharomycetales(100);f__Metschnikowiaceae(100);g__Metschnikowia(100);s__Metschnikowia_pulcherrima(100); | Metschnikowia | 13 | Saprotroph | \|Undefined Saprotroph\| | Yeast | NULL | Probable | As nectar-tap_saprotroph (Põlme et al. 2020); Decay sugar-rich substrates (Põlme et al. 2020) | Põlme S, et al. 2020. Fungal Diversity 105:1-16 ((https://doi.org/10.1007/s13225-020-00466-2)) |
| Otu43993 | 1 | k__Fungi(100);p__Ascomycota(100);c__Saccharomycetes(100);o__Saccharomycetales(100);f__Metschnikowiaceae(100);g__Metschnikowia(100);g__Metschnikowia_unclassified(100); | Metschnikowia | 13 | Saprotroph | \|Undefined Saprotroph\| | Yeast | NULL | Probable | As nectar-tap_saprotroph (Põlme et al. 2020); Decay sugar-rich substrates (Põlme et al. 2020) | Põlme S, et al. 2020. Fungal Diversity 105:1-16 ((https://doi.org/10.1007/s13225-020-00466-2)) |
| Otu43994 | 1 | k__Fungi(100);p__Ascomycota(100);c__Saccharomycetes(100);o__Saccharomycetales(100);f__Metschnikowiaceae(100);g__Metschnikowia(100);s__Metschnikowia_pulcherrima(100); | Metschnikowia | 13 | Saprotroph | \|Undefined Saprotroph\| | Yeast | NULL | Probable | As nectar-tap_saprotroph (Põlme et al. 2020); Decay sugar-rich substrates (Põlme et al. 2020) | Põlme S, et al. 2020. Fungal Diversity 105:1-16 ((https://doi.org/10.1007/s13225-020-00466-2)) |
| Otu43995 | 1 | k__Fungi(100);p__Ascomycota(100);c__Saccharomycetes(100);o__Saccharomycetales(100);f__Metschnikowiaceae(100);g__Metschnikowia(100);s__Metschnikowia_pulcherrima(100); | Metschnikowia | 13 | Saprotroph | \|Undefined Saprotroph\| | Yeast | NULL | Probable | As nectar-tap_saprotroph (Põlme et al. 2020); Decay sugar-rich substrates (Põlme et al. 2020) | Põlme S, et al. 2020. Fungal Diversity 105:1-16 ((https://doi.org/10.1007/s13225-020-00466-2)) |
| Otu43996 | 1 | k__Fungi(100);p__Ascomycota(100);c__Saccharomycetes(100);o__Saccharomycetales(100);f__Metschnikowiaceae(100);g__Metschnikowia(100);s__Metschnikowia_pulcherrima(100); | Metschnikowia | 13 | Saprotroph | \|Undefined Saprotroph\| | Yeast | NULL | Probable | As nectar-tap_saprotroph (Põlme et al. 2020); Decay sugar-rich substrates (Põlme et al. 2020) | Põlme S, et al. 2020. Fungal Diversity 105:1-16 ((https://doi.org/10.1007/s13225-020-00466-2)) |
| Otu43997 | 1 | k__Fungi(100);p__Ascomycota(100);c__Saccharomycetes(100);o__Saccharomycetales(100);f__Metschnikowiaceae(100);g__Metschnikowia(100);g__Metschnikowia_unclassified(100); | Metschnikowia | 13 | Saprotroph | \|Undefined Saprotroph\| | Yeast | NULL | Probable | As nectar-tap_saprotroph (Põlme et al. 2020); Decay sugar-rich substrates (Põlme et al. 2020) | Põlme S, et al. 2020. Fungal Diversity 105:1-16 ((https://doi.org/10.1007/s13225-020-00466-2)) |
| Otu43998 | 1 | k__Fungi(100);p__Ascomycota(100);c__Saccharomycetes(100);o__Saccharomycetales(100);f__Metschnikowiaceae(100);g__Metschnikowia(100);s__Metschnikowia_pulcherrima(100); | Metschnikowia | 13 | Saprotroph | \|Undefined Saprotroph\| | Yeast | NULL | Probable | As nectar-tap_saprotroph (Põlme et al. 2020); Decay sugar-rich substrates (Põlme et al. 2020) | Põlme S, et al. 2020. Fungal Diversity 105:1-16 ((https://doi.org/10.1007/s13225-020-00466-2)) |
| Otu43999 | 1 | k__Fungi(100);p__Ascomycota(100);c__Saccharomycetes(100);o__Saccharomycetales(100);f__Metschnikowiaceae(100);g__Metschnikowia(100);s__Metschnikowia_pulcherrima(100); | Metschnikowia | 13 | Saprotroph | \|Undefined Saprotroph\| | Yeast | NULL | Probable | As nectar-tap_saprotroph (Põlme et al. 2020); Decay sugar-rich substrates (Põlme et al. 2020) | Põlme S, et al. 2020. Fungal Diversity 105:1-16 ((https://doi.org/10.1007/s13225-020-00466-2)) |
| Otu44000 | 1 | k__Fungi(100);p__Ascomycota(100);c__Saccharomycetes(100);o__Saccharomycetales(100);f__Metschnikowiaceae(100);g__Metschnikowia(100);s__Metschnikowia_pulcherrima(100); | Metschnikowia | 13 | Saprotroph | \|Undefined Saprotroph\| | Yeast | NULL | Probable | As nectar-tap_saprotroph (Põlme et al. 2020); Decay sugar-rich substrates (Põlme et al. 2020) | Põlme S, et al. 2020. Fungal Diversity 105:1-16 ((https://doi.org/10.1007/s13225-020-00466-2)) |
| Otu44001 | 1 | k__Fungi(100);p__Ascomycota(100);c__Saccharomycetes(100);o__Saccharomycetales(100);f__Metschnikowiaceae(100);g__Metschnikowia(100);s__Metschnikowia_pulcherrima(100); | Metschnikowia | 13 | Saprotroph | \|Undefined Saprotroph\| | Yeast | NULL | Probable | As nectar-tap_saprotroph (Põlme et al. 2020); Decay sugar-rich substrates (Põlme et al. 2020) | Põlme S, et al. 2020. Fungal Diversity 105:1-16 ((https://doi.org/10.1007/s13225-020-00466-2)) |
| Otu44003 | 1 | k__Fungi(100);p__Ascomycota(100);c__Saccharomycetes(100);o__Saccharomycetales(100);f__Metschnikowiaceae(100);g__Metschnikowia(100);s__Metschnikowia_pulcherrima(100); | Metschnikowia | 13 | Saprotroph | \|Undefined Saprotroph\| | Yeast | NULL | Probable | As nectar-tap_saprotroph (Põlme et al. 2020); Decay sugar-rich substrates (Põlme et al. 2020) | Põlme S, et al. 2020. Fungal Diversity 105:1-16 ((https://doi.org/10.1007/s13225-020-00466-2)) |
| Otu44004 | 1 | k__Fungi(100);p__Ascomycota(100);c__Saccharomycetes(100);o__Saccharomycetales(100);f__Metschnikowiaceae(100);g__Metschnikowia(100);s__Metschnikowia_pulcherrima(100); | Metschnikowia | 13 | Saprotroph | \|Undefined Saprotroph\| | Yeast | NULL | Probable | As nectar-tap_saprotroph (Põlme et al. 2020); Decay sugar-rich substrates (Põlme et al. 2020) | Põlme S, et al. 2020. Fungal Diversity 105:1-16 ((https://doi.org/10.1007/s13225-020-00466-2)) |
| Otu44005 | 1 | k__Fungi(100);p__Ascomycota(100);c__Saccharomycetes(100);o__Saccharomycetales(100);f__Metschnikowiaceae(100);g__Metschnikowia(100);s__Metschnikowia_pulcherrima(100); | Metschnikowia | 13 | Saprotroph | \|Undefined Saprotroph\| | Yeast | NULL | Probable | As nectar-tap_saprotroph (Põlme et al. 2020); Decay sugar-rich substrates (Põlme et al. 2020) | Põlme S, et al. 2020. Fungal Diversity 105:1-16 ((https://doi.org/10.1007/s13225-020-00466-2)) |
| Otu44006 | 1 | k__Fungi(100);p__Ascomycota(100);c__Saccharomycetes(100);o__Saccharomycetales(100);f__Metschnikowiaceae(100);g__Metschnikowia(100);s__Metschnikowia_pulcherrima(100); | Metschnikowia | 13 | Saprotroph | \|Undefined Saprotroph\| | Yeast | NULL | Probable | As nectar-tap_saprotroph (Põlme et al. 2020); Decay sugar-rich substrates (Põlme et al. 2020) | Põlme S, et al. 2020. Fungal Diversity 105:1-16 ((https://doi.org/10.1007/s13225-020-00466-2)) |
| Otu44007 | 1 | k__Fungi(100);p__Ascomycota(100);c__Saccharomycetes(100);o__Saccharomycetales(100);f__Metschnikowiaceae(100);g__Metschnikowia(100);s__Metschnikowia_pulcherrima(100); | Metschnikowia | 13 | Saprotroph | \|Undefined Saprotroph\| | Yeast | NULL | Probable | As nectar-tap_saprotroph (Põlme et al. 2020); Decay sugar-rich substrates (Põlme et al. 2020) | Põlme S, et al. 2020. Fungal Diversity 105:1-16 ((https://doi.org/10.1007/s13225-020-00466-2)) |
| Otu44010 | 1 | k__Fungi(100);p__Ascomycota(100);c__Saccharomycetes(100);o__Saccharomycetales(100);f__Metschnikowiaceae(100);g__Metschnikowia(100);s__Metschnikowia_pulcherrima(100); | Metschnikowia | 13 | Saprotroph | \|Undefined Saprotroph\| | Yeast | NULL | Probable | As nectar-tap_saprotroph (Põlme et al. 2020); Decay sugar-rich substrates (Põlme et al. 2020) | Põlme S, et al. 2020. Fungal Diversity 105:1-16 ((https://doi.org/10.1007/s13225-020-00466-2)) |
| Otu44011 | 1 | k__Fungi(100);p__Ascomycota(100);c__Saccharomycetes(100);o__Saccharomycetales(100);f__Metschnikowiaceae(100);g__Metschnikowia(100);s__Metschnikowia_pulcherrima(100); | Metschnikowia | 13 | Saprotroph | \|Undefined Saprotroph\| | Yeast | NULL | Probable | As nectar-tap_saprotroph (Põlme et al. 2020); Decay sugar-rich substrates (Põlme et al. 2020) | Põlme S, et al. 2020. Fungal Diversity 105:1-16 ((https://doi.org/10.1007/s13225-020-00466-2)) |
| Otu44012 | 1 | k__Fungi(100);p__Ascomycota(100);c__Saccharomycetes(100);o__Saccharomycetales(100);f__Metschnikowiaceae(100);g__Metschnikowia(100);g__Metschnikowia_unclassified(100); | Metschnikowia | 13 | Saprotroph | \|Undefined Saprotroph\| | Yeast | NULL | Probable | As nectar-tap_saprotroph (Põlme et al. 2020); Decay sugar-rich substrates (Põlme et al. 2020) | Põlme S, et al. 2020. Fungal Diversity 105:1-16 ((https://doi.org/10.1007/s13225-020-00466-2)) |
| Otu44013 | 1 | k__Fungi(100);p__Ascomycota(100);c__Saccharomycetes(100);o__Saccharomycetales(100);f__Metschnikowiaceae(100);g__Metschnikowia(100);s__Metschnikowia_pulcherrima(100); | Metschnikowia | 13 | Saprotroph | \|Undefined Saprotroph\| | Yeast | NULL | Probable | As nectar-tap_saprotroph (Põlme et al. 2020); Decay sugar-rich substrates (Põlme et al. 2020) | Põlme S, et al. 2020. Fungal Diversity 105:1-16 ((https://doi.org/10.1007/s13225-020-00466-2)) |
| Otu44014 | 1 | k__Fungi(100);p__Ascomycota(100);c__Saccharomycetes(100);o__Saccharomycetales(100);f__Metschnikowiaceae(100);g__Metschnikowia(100);s__Metschnikowia_pulcherrima(100); | Metschnikowia | 13 | Saprotroph | \|Undefined Saprotroph\| | Yeast | NULL | Probable | As nectar-tap_saprotroph (Põlme et al. 2020); Decay sugar-rich substrates (Põlme et al. 2020) | Põlme S, et al. 2020. Fungal Diversity 105:1-16 ((https://doi.org/10.1007/s13225-020-00466-2)) |
| Otu44015 | 1 | k__Fungi(100);p__Ascomycota(100);c__Saccharomycetes(100);o__Saccharomycetales(100);f__Metschnikowiaceae(100);g__Metschnikowia(100);s__Metschnikowia_pulcherrima(100); | Metschnikowia | 13 | Saprotroph | \|Undefined Saprotroph\| | Yeast | NULL | Probable | As nectar-tap_saprotroph (Põlme et al. 2020); Decay sugar-rich substrates (Põlme et al. 2020) | Põlme S, et al. 2020. Fungal Diversity 105:1-16 ((https://doi.org/10.1007/s13225-020-00466-2)) |
| Otu44016 | 1 | k__Fungi(100);p__Ascomycota(100);c__Saccharomycetes(100);o__Saccharomycetales(100);f__Metschnikowiaceae(100);g__Metschnikowia(100);s__Metschnikowia_pulcherrima(100); | Metschnikowia | 13 | Saprotroph | \|Undefined Saprotroph\| | Yeast | NULL | Probable | As nectar-tap_saprotroph (Põlme et al. 2020); Decay sugar-rich substrates (Põlme et al. 2020) | Põlme S, et al. 2020. Fungal Diversity 105:1-16 ((https://doi.org/10.1007/s13225-020-00466-2)) |
| Otu44017 | 1 | k__Fungi(100);p__Ascomycota(100);c__Saccharomycetes(100);o__Saccharomycetales(100);f__Metschnikowiaceae(100);g__Metschnikowia(100);s__Metschnikowia_pulcherrima(100); | Metschnikowia | 13 | Saprotroph | \|Undefined Saprotroph\| | Yeast | NULL | Probable | As nectar-tap_saprotroph (Põlme et al. 2020); Decay sugar-rich substrates (Põlme et al. 2020) | Põlme S, et al. 2020. Fungal Diversity 105:1-16 ((https://doi.org/10.1007/s13225-020-00466-2)) |
| Otu44018 | 1 | k__Fungi(100);p__Ascomycota(100);c__Saccharomycetes(100);o__Saccharomycetales(100);f__Metschnikowiaceae(100);g__Metschnikowia(100);s__Metschnikowia_pulcherrima(100); | Metschnikowia | 13 | Saprotroph | \|Undefined Saprotroph\| | Yeast | NULL | Probable | As nectar-tap_saprotroph (Põlme et al. 2020); Decay sugar-rich substrates (Põlme et al. 2020) | Põlme S, et al. 2020. Fungal Diversity 105:1-16 ((https://doi.org/10.1007/s13225-020-00466-2)) |
| Otu44019 | 1 | k__Fungi(100);p__Ascomycota(100);c__Saccharomycetes(100);o__Saccharomycetales(100);f__Metschnikowiaceae(100);g__Metschnikowia(100);s__Metschnikowia_pulcherrima(100); | Metschnikowia | 13 | Saprotroph | \|Undefined Saprotroph\| | Yeast | NULL | Probable | As nectar-tap_saprotroph (Põlme et al. 2020); Decay sugar-rich substrates (Põlme et al. 2020) | Põlme S, et al. 2020. Fungal Diversity 105:1-16 ((https://doi.org/10.1007/s13225-020-00466-2)) |
| Otu44020 | 1 | k__Fungi(100);p__Ascomycota(100);c__Saccharomycetes(100);o__Saccharomycetales(100);f__Metschnikowiaceae(100);g__Metschnikowia(100);s__Metschnikowia_viticola(100); | Metschnikowia | 13 | Saprotroph | \|Undefined Saprotroph\| | Yeast | NULL | Probable | As nectar-tap_saprotroph (Põlme et al. 2020); Decay sugar-rich substrates (Põlme et al. 2020) | Põlme S, et al. 2020. Fungal Diversity 105:1-16 ((https://doi.org/10.1007/s13225-020-00466-2)) |
| Otu44021 | 1 | k__Fungi(100);p__Ascomycota(100);c__Saccharomycetes(100);o__Saccharomycetales(100);f__Metschnikowiaceae(100);g__Metschnikowia(100);s__Metschnikowia_pulcherrima(100); | Metschnikowia | 13 | Saprotroph | \|Undefined Saprotroph\| | Yeast | NULL | Probable | As nectar-tap_saprotroph (Põlme et al. 2020); Decay sugar-rich substrates (Põlme et al. 2020) | Põlme S, et al. 2020. Fungal Diversity 105:1-16 ((https://doi.org/10.1007/s13225-020-00466-2)) |
| Otu44022 | 1 | k__Fungi(100);p__Ascomycota(100);c__Saccharomycetes(100);o__Saccharomycetales(100);f__Metschnikowiaceae(100);g__Metschnikowia(100);g__Metschnikowia_unclassified(100); | Metschnikowia | 13 | Saprotroph | \|Undefined Saprotroph\| | Yeast | NULL | Probable | As nectar-tap_saprotroph (Põlme et al. 2020); Decay sugar-rich substrates (Põlme et al. 2020) | Põlme S, et al. 2020. Fungal Diversity 105:1-16 ((https://doi.org/10.1007/s13225-020-00466-2)) |
| Otu44023 | 1 | k__Fungi(100);p__Ascomycota(100);c__Saccharomycetes(100);o__Saccharomycetales(100);f__Metschnikowiaceae(100);g__Metschnikowia(100);s__Metschnikowia_pulcherrima(100); | Metschnikowia | 13 | Saprotroph | \|Undefined Saprotroph\| | Yeast | NULL | Probable | As nectar-tap_saprotroph (Põlme et al. 2020); Decay sugar-rich substrates (Põlme et al. 2020) | Põlme S, et al. 2020. Fungal Diversity 105:1-16 ((https://doi.org/10.1007/s13225-020-00466-2)) |
| Otu44024 | 1 | k__Fungi(100);p__Ascomycota(100);c__Saccharomycetes(100);o__Saccharomycetales(100);f__Metschnikowiaceae(100);g__Metschnikowia(100);s__Metschnikowia_pulcherrima(100); | Metschnikowia | 13 | Saprotroph | \|Undefined Saprotroph\| | Yeast | NULL | Probable | As nectar-tap_saprotroph (Põlme et al. 2020); Decay sugar-rich substrates (Põlme et al. 2020) | Põlme S, et al. 2020. Fungal Diversity 105:1-16 ((https://doi.org/10.1007/s13225-020-00466-2)) |
| Otu44025 | 1 | k__Fungi(100);p__Ascomycota(100);c__Saccharomycetes(100);o__Saccharomycetales(100);f__Metschnikowiaceae(100);g__Metschnikowia(100);s__Metschnikowia_pulcherrima(100); | Metschnikowia | 13 | Saprotroph | \|Undefined Saprotroph\| | Yeast | NULL | Probable | As nectar-tap_saprotroph (Põlme et al. 2020); Decay sugar-rich substrates (Põlme et al. 2020) | Põlme S, et al. 2020. Fungal Diversity 105:1-16 ((https://doi.org/10.1007/s13225-020-00466-2)) |
| Otu44026 | 1 | k__Fungi(100);p__Ascomycota(100);c__Saccharomycetes(100);o__Saccharomycetales(100);f__Metschnikowiaceae(100);g__Metschnikowia(100);s__Metschnikowia_pulcherrima(100); | Metschnikowia | 13 | Saprotroph | \|Undefined Saprotroph\| | Yeast | NULL | Probable | As nectar-tap_saprotroph (Põlme et al. 2020); Decay sugar-rich substrates (Põlme et al. 2020) | Põlme S, et al. 2020. Fungal Diversity 105:1-16 ((https://doi.org/10.1007/s13225-020-00466-2)) |
| Otu44027 | 1 | k__Fungi(100);p__Ascomycota(100);c__Saccharomycetes(100);o__Saccharomycetales(100);f__Metschnikowiaceae(100);g__Metschnikowia(100);s__Metschnikowia_pulcherrima(100); | Metschnikowia | 13 | Saprotroph | \|Undefined Saprotroph\| | Yeast | NULL | Probable | As nectar-tap_saprotroph (Põlme et al. 2020); Decay sugar-rich substrates (Põlme et al. 2020) | Põlme S, et al. 2020. Fungal Diversity 105:1-16 ((https://doi.org/10.1007/s13225-020-00466-2)) |
| Otu44028 | 1 | k__Fungi(100);p__Ascomycota(100);c__Saccharomycetes(100);o__Saccharomycetales(100);f__Metschnikowiaceae(100);g__Metschnikowia(100);s__Metschnikowia_pulcherrima(100); | Metschnikowia | 13 | Saprotroph | \|Undefined Saprotroph\| | Yeast | NULL | Probable | As nectar-tap_saprotroph (Põlme et al. 2020); Decay sugar-rich substrates (Põlme et al. 2020) | Põlme S, et al. 2020. Fungal Diversity 105:1-16 ((https://doi.org/10.1007/s13225-020-00466-2)) |
| Otu44029 | 1 | k__Fungi(100);p__Ascomycota(100);c__Saccharomycetes(100);o__Saccharomycetales(100);f__Metschnikowiaceae(100);g__Metschnikowia(100);g__Metschnikowia_unclassified(100); | Metschnikowia | 13 | Saprotroph | \|Undefined Saprotroph\| | Yeast | NULL | Probable | As nectar-tap_saprotroph (Põlme et al. 2020); Decay sugar-rich substrates (Põlme et al. 2020) | Põlme S, et al. 2020. Fungal Diversity 105:1-16 ((https://doi.org/10.1007/s13225-020-00466-2)) |
| Otu44030 | 1 | k__Fungi(100);p__Ascomycota(100);c__Saccharomycetes(100);o__Saccharomycetales(100);f__Metschnikowiaceae(100);g__Metschnikowia(100);s__Metschnikowia_pulcherrima(100); | Metschnikowia | 13 | Saprotroph | \|Undefined Saprotroph\| | Yeast | NULL | Probable | As nectar-tap_saprotroph (Põlme et al. 2020); Decay sugar-rich substrates (Põlme et al. 2020) | Põlme S, et al. 2020. Fungal Diversity 105:1-16 ((https://doi.org/10.1007/s13225-020-00466-2)) |
| Otu44032 | 1 | k__Fungi(100);p__Ascomycota(100);c__Saccharomycetes(100);o__Saccharomycetales(100);f__Metschnikowiaceae(100);g__Metschnikowia(100);g__Metschnikowia_unclassified(100); | Metschnikowia | 13 | Saprotroph | \|Undefined Saprotroph\| | Yeast | NULL | Probable | As nectar-tap_saprotroph (Põlme et al. 2020); Decay sugar-rich substrates (Põlme et al. 2020) | Põlme S, et al. 2020. Fungal Diversity 105:1-16 ((https://doi.org/10.1007/s13225-020-00466-2)) |
| Otu44033 | 1 | k__Fungi(100);p__Ascomycota(100);c__Saccharomycetes(100);o__Saccharomycetales(100);f__Metschnikowiaceae(100);g__Metschnikowia(100);s__Metschnikowia_pulcherrima(100); | Metschnikowia | 13 | Saprotroph | \|Undefined Saprotroph\| | Yeast | NULL | Probable | As nectar-tap_saprotroph (Põlme et al. 2020); Decay sugar-rich substrates (Põlme et al. 2020) | Põlme S, et al. 2020. Fungal Diversity 105:1-16 ((https://doi.org/10.1007/s13225-020-00466-2)) |
| Otu44034 | 1 | k__Fungi(100);p__Ascomycota(100);c__Saccharomycetes(100);o__Saccharomycetales(100);f__Metschnikowiaceae(100);g__Metschnikowia(100);s__Metschnikowia_pulcherrima(100); | Metschnikowia | 13 | Saprotroph | \|Undefined Saprotroph\| | Yeast | NULL | Probable | As nectar-tap_saprotroph (Põlme et al. 2020); Decay sugar-rich substrates (Põlme et al. 2020) | Põlme S, et al. 2020. Fungal Diversity 105:1-16 ((https://doi.org/10.1007/s13225-020-00466-2)) |
| Otu44035 | 1 | k__Fungi(100);p__Ascomycota(100);c__Saccharomycetes(100);o__Saccharomycetales(100);f__Metschnikowiaceae(100);g__Metschnikowia(100);g__Metschnikowia_unclassified(100); | Metschnikowia | 13 | Saprotroph | \|Undefined Saprotroph\| | Yeast | NULL | Probable | As nectar-tap_saprotroph (Põlme et al. 2020); Decay sugar-rich substrates (Põlme et al. 2020) | Põlme S, et al. 2020. Fungal Diversity 105:1-16 ((https://doi.org/10.1007/s13225-020-00466-2)) |
| Otu44036 | 1 | k__Fungi(100);p__Ascomycota(100);c__Saccharomycetes(100);o__Saccharomycetales(100);f__Metschnikowiaceae(100);g__Metschnikowia(100);s__Metschnikowia_pulcherrima(100); | Metschnikowia | 13 | Saprotroph | \|Undefined Saprotroph\| | Yeast | NULL | Probable | As nectar-tap_saprotroph (Põlme et al. 2020); Decay sugar-rich substrates (Põlme et al. 2020) | Põlme S, et al. 2020. Fungal Diversity 105:1-16 ((https://doi.org/10.1007/s13225-020-00466-2)) |
| Otu44037 | 1 | k__Fungi(100);p__Ascomycota(100);c__Saccharomycetes(100);o__Saccharomycetales(100);f__Metschnikowiaceae(100);g__Metschnikowia(100);s__Metschnikowia_pulcherrima(100); | Metschnikowia | 13 | Saprotroph | \|Undefined Saprotroph\| | Yeast | NULL | Probable | As nectar-tap_saprotroph (Põlme et al. 2020); Decay sugar-rich substrates (Põlme et al. 2020) | Põlme S, et al. 2020. Fungal Diversity 105:1-16 ((https://doi.org/10.1007/s13225-020-00466-2)) |
| Otu44038 | 1 | k__Fungi(100);p__Ascomycota(100);c__Saccharomycetes(100);o__Saccharomycetales(100);f__Metschnikowiaceae(100);g__Metschnikowia(100);s__Metschnikowia_pulcherrima(100); | Metschnikowia | 13 | Saprotroph | \|Undefined Saprotroph\| | Yeast | NULL | Probable | As nectar-tap_saprotroph (Põlme et al. 2020); Decay sugar-rich substrates (Põlme et al. 2020) | Põlme S, et al. 2020. Fungal Diversity 105:1-16 ((https://doi.org/10.1007/s13225-020-00466-2)) |
| Otu44039 | 1 | k__Fungi(100);p__Ascomycota(100);c__Saccharomycetes(100);o__Saccharomycetales(100);f__Metschnikowiaceae(100);g__Metschnikowia(100);s__Metschnikowia_pulcherrima(100); | Metschnikowia | 13 | Saprotroph | \|Undefined Saprotroph\| | Yeast | NULL | Probable | As nectar-tap_saprotroph (Põlme et al. 2020); Decay sugar-rich substrates (Põlme et al. 2020) | Põlme S, et al. 2020. Fungal Diversity 105:1-16 ((https://doi.org/10.1007/s13225-020-00466-2)) |
| Otu44040 | 1 | k__Fungi(100);p__Ascomycota(100);c__Saccharomycetes(100);o__Saccharomycetales(100);f__Metschnikowiaceae(100);g__Metschnikowia(100);s__Metschnikowia_pulcherrima(100); | Metschnikowia | 13 | Saprotroph | \|Undefined Saprotroph\| | Yeast | NULL | Probable | As nectar-tap_saprotroph (Põlme et al. 2020); Decay sugar-rich substrates (Põlme et al. 2020) | Põlme S, et al. 2020. Fungal Diversity 105:1-16 ((https://doi.org/10.1007/s13225-020-00466-2)) |
| Otu44041 | 1 | k__Fungi(100);p__Ascomycota(100);c__Saccharomycetes(100);o__Saccharomycetales(100);f__Metschnikowiaceae(100);g__Metschnikowia(100);g__Metschnikowia_unclassified(100); | Metschnikowia | 13 | Saprotroph | \|Undefined Saprotroph\| | Yeast | NULL | Probable | As nectar-tap_saprotroph (Põlme et al. 2020); Decay sugar-rich substrates (Põlme et al. 2020) | Põlme S, et al. 2020. Fungal Diversity 105:1-16 ((https://doi.org/10.1007/s13225-020-00466-2)) |
| Otu44042 | 1 | k__Fungi(100);p__Ascomycota(100);c__Saccharomycetes(100);o__Saccharomycetales(100);f__Metschnikowiaceae(100);g__Metschnikowia(100);s__Metschnikowia_pulcherrima(100); | Metschnikowia | 13 | Saprotroph | \|Undefined Saprotroph\| | Yeast | NULL | Probable | As nectar-tap_saprotroph (Põlme et al. 2020); Decay sugar-rich substrates (Põlme et al. 2020) | Põlme S, et al. 2020. Fungal Diversity 105:1-16 ((https://doi.org/10.1007/s13225-020-00466-2)) |
| Otu44043 | 1 | k__Fungi(100);p__Ascomycota(100);c__Saccharomycetes(100);o__Saccharomycetales(100);f__Metschnikowiaceae(100);g__Metschnikowia(100);s__Metschnikowia_pulcherrima(100); | Metschnikowia | 13 | Saprotroph | \|Undefined Saprotroph\| | Yeast | NULL | Probable | As nectar-tap_saprotroph (Põlme et al. 2020); Decay sugar-rich substrates (Põlme et al. 2020) | Põlme S, et al. 2020. Fungal Diversity 105:1-16 ((https://doi.org/10.1007/s13225-020-00466-2)) |
| Otu44044 | 1 | k__Fungi(100);p__Ascomycota(100);c__Saccharomycetes(100);o__Saccharomycetales(100);f__Metschnikowiaceae(100);g__Metschnikowia(100);g__Metschnikowia_unclassified(100); | Metschnikowia | 13 | Saprotroph | \|Undefined Saprotroph\| | Yeast | NULL | Probable | As nectar-tap_saprotroph (Põlme et al. 2020); Decay sugar-rich substrates (Põlme et al. 2020) | Põlme S, et al. 2020. Fungal Diversity 105:1-16 ((https://doi.org/10.1007/s13225-020-00466-2)) |
| Otu44045 | 1 | k__Fungi(100);p__Ascomycota(100);c__Saccharomycetes(100);o__Saccharomycetales(100);f__Metschnikowiaceae(100);g__Metschnikowia(100);s__Metschnikowia_pulcherrima(100); | Metschnikowia | 13 | Saprotroph | \|Undefined Saprotroph\| | Yeast | NULL | Probable | As nectar-tap_saprotroph (Põlme et al. 2020); Decay sugar-rich substrates (Põlme et al. 2020) | Põlme S, et al. 2020. Fungal Diversity 105:1-16 ((https://doi.org/10.1007/s13225-020-00466-2)) |
| Otu44046 | 1 | k__Fungi(100);p__Ascomycota(100);c__Saccharomycetes(100);o__Saccharomycetales(100);f__Metschnikowiaceae(100);g__Metschnikowia(100);g__Metschnikowia_unclassified(100); | Metschnikowia | 13 | Saprotroph | \|Undefined Saprotroph\| | Yeast | NULL | Probable | As nectar-tap_saprotroph (Põlme et al. 2020); Decay sugar-rich substrates (Põlme et al. 2020) | Põlme S, et al. 2020. Fungal Diversity 105:1-16 ((https://doi.org/10.1007/s13225-020-00466-2)) |
| Otu44047 | 1 | k__Fungi(100);p__Ascomycota(100);c__Saccharomycetes(100);o__Saccharomycetales(100);f__Metschnikowiaceae(100);g__Metschnikowia(100);s__Metschnikowia_pulcherrima(100); | Metschnikowia | 13 | Saprotroph | \|Undefined Saprotroph\| | Yeast | NULL | Probable | As nectar-tap_saprotroph (Põlme et al. 2020); Decay sugar-rich substrates (Põlme et al. 2020) | Põlme S, et al. 2020. Fungal Diversity 105:1-16 ((https://doi.org/10.1007/s13225-020-00466-2)) |
| Otu44048 | 1 | k__Fungi(100);p__Ascomycota(100);c__Saccharomycetes(100);o__Saccharomycetales(100);f__Metschnikowiaceae(100);g__Metschnikowia(100);g__Metschnikowia_unclassified(100); | Metschnikowia | 13 | Saprotroph | \|Undefined Saprotroph\| | Yeast | NULL | Probable | As nectar-tap_saprotroph (Põlme et al. 2020); Decay sugar-rich substrates (Põlme et al. 2020) | Põlme S, et al. 2020. Fungal Diversity 105:1-16 ((https://doi.org/10.1007/s13225-020-00466-2)) |
| Otu44049 | 1 | k__Fungi(100);p__Ascomycota(100);c__Saccharomycetes(100);o__Saccharomycetales(100);f__Metschnikowiaceae(100);g__Metschnikowia(100);s__Metschnikowia_pulcherrima(100); | Metschnikowia | 13 | Saprotroph | \|Undefined Saprotroph\| | Yeast | NULL | Probable | As nectar-tap_saprotroph (Põlme et al. 2020); Decay sugar-rich substrates (Põlme et al. 2020) | Põlme S, et al. 2020. Fungal Diversity 105:1-16 ((https://doi.org/10.1007/s13225-020-00466-2)) |
| Otu44050 | 1 | k__Fungi(100);p__Ascomycota(100);c__Saccharomycetes(100);o__Saccharomycetales(100);f__Metschnikowiaceae(100);g__Metschnikowia(100);s__Metschnikowia_pulcherrima(100); | Metschnikowia | 13 | Saprotroph | \|Undefined Saprotroph\| | Yeast | NULL | Probable | As nectar-tap_saprotroph (Põlme et al. 2020); Decay sugar-rich substrates (Põlme et al. 2020) | Põlme S, et al. 2020. Fungal Diversity 105:1-16 ((https://doi.org/10.1007/s13225-020-00466-2)) |
| Otu44051 | 1 | k__Fungi(100);p__Ascomycota(100);c__Saccharomycetes(100);o__Saccharomycetales(100);f__Metschnikowiaceae(100);g__Metschnikowia(100);s__Metschnikowia_pulcherrima(100); | Metschnikowia | 13 | Saprotroph | \|Undefined Saprotroph\| | Yeast | NULL | Probable | As nectar-tap_saprotroph (Põlme et al. 2020); Decay sugar-rich substrates (Põlme et al. 2020) | Põlme S, et al. 2020. Fungal Diversity 105:1-16 ((https://doi.org/10.1007/s13225-020-00466-2)) |
| Otu44052 | 1 | k__Fungi(100);p__Ascomycota(100);c__Saccharomycetes(100);o__Saccharomycetales(100);f__Metschnikowiaceae(100);g__Metschnikowia(100);g__Metschnikowia_unclassified(100); | Metschnikowia | 13 | Saprotroph | \|Undefined Saprotroph\| | Yeast | NULL | Probable | As nectar-tap_saprotroph (Põlme et al. 2020); Decay sugar-rich substrates (Põlme et al. 2020) | Põlme S, et al. 2020. Fungal Diversity 105:1-16 ((https://doi.org/10.1007/s13225-020-00466-2)) |
| Otu44053 | 1 | k__Fungi(100);p__Ascomycota(100);c__Saccharomycetes(100);o__Saccharomycetales(100);f__Metschnikowiaceae(100);g__Metschnikowia(100);s__Metschnikowia_pulcherrima(100); | Metschnikowia | 13 | Saprotroph | \|Undefined Saprotroph\| | Yeast | NULL | Probable | As nectar-tap_saprotroph (Põlme et al. 2020); Decay sugar-rich substrates (Põlme et al. 2020) | Põlme S, et al. 2020. Fungal Diversity 105:1-16 ((https://doi.org/10.1007/s13225-020-00466-2)) |
| Otu44054 | 1 | k__Fungi(100);p__Ascomycota(100);c__Saccharomycetes(100);o__Saccharomycetales(100);f__Metschnikowiaceae(100);g__Metschnikowia(100);s__Metschnikowia_pulcherrima(100); | Metschnikowia | 13 | Saprotroph | \|Undefined Saprotroph\| | Yeast | NULL | Probable | As nectar-tap_saprotroph (Põlme et al. 2020); Decay sugar-rich substrates (Põlme et al. 2020) | Põlme S, et al. 2020. Fungal Diversity 105:1-16 ((https://doi.org/10.1007/s13225-020-00466-2)) |
| Otu44055 | 1 | k__Fungi(100);p__Ascomycota(100);c__Saccharomycetes(100);o__Saccharomycetales(100);f__Metschnikowiaceae(100);g__Metschnikowia(100);s__Metschnikowia_pulcherrima(100); | Metschnikowia | 13 | Saprotroph | \|Undefined Saprotroph\| | Yeast | NULL | Probable | As nectar-tap_saprotroph (Põlme et al. 2020); Decay sugar-rich substrates (Põlme et al. 2020) | Põlme S, et al. 2020. Fungal Diversity 105:1-16 ((https://doi.org/10.1007/s13225-020-00466-2)) |
| Otu44056 | 1 | k__Fungi(100);p__Ascomycota(100);c__Saccharomycetes(100);o__Saccharomycetales(100);f__Metschnikowiaceae(100);g__Metschnikowia(100);g__Metschnikowia_unclassified(100); | Metschnikowia | 13 | Saprotroph | \|Undefined Saprotroph\| | Yeast | NULL | Probable | As nectar-tap_saprotroph (Põlme et al. 2020); Decay sugar-rich substrates (Põlme et al. 2020) | Põlme S, et al. 2020. Fungal Diversity 105:1-16 ((https://doi.org/10.1007/s13225-020-00466-2)) |
| Otu44057 | 1 | k__Fungi(100);p__Ascomycota(100);c__Saccharomycetes(100);o__Saccharomycetales(100);f__Metschnikowiaceae(100);g__Metschnikowia(100);g__Metschnikowia_unclassified(100); | Metschnikowia | 13 | Saprotroph | \|Undefined Saprotroph\| | Yeast | NULL | Probable | As nectar-tap_saprotroph (Põlme et al. 2020); Decay sugar-rich substrates (Põlme et al. 2020) | Põlme S, et al. 2020. Fungal Diversity 105:1-16 ((https://doi.org/10.1007/s13225-020-00466-2)) |
| Otu44058 | 1 | k__Fungi(100);p__Ascomycota(100);c__Saccharomycetes(100);o__Saccharomycetales(100);f__Metschnikowiaceae(100);g__Metschnikowia(100);s__Metschnikowia_pulcherrima(100); | Metschnikowia | 13 | Saprotroph | \|Undefined Saprotroph\| | Yeast | NULL | Probable | As nectar-tap_saprotroph (Põlme et al. 2020); Decay sugar-rich substrates (Põlme et al. 2020) | Põlme S, et al. 2020. Fungal Diversity 105:1-16 ((https://doi.org/10.1007/s13225-020-00466-2)) |
| Otu44059 | 1 | k__Fungi(100);p__Ascomycota(100);c__Saccharomycetes(100);o__Saccharomycetales(100);f__Metschnikowiaceae(100);g__Metschnikowia(100);g__Metschnikowia_unclassified(100); | Metschnikowia | 13 | Saprotroph | \|Undefined Saprotroph\| | Yeast | NULL | Probable | As nectar-tap_saprotroph (Põlme et al. 2020); Decay sugar-rich substrates (Põlme et al. 2020) | Põlme S, et al. 2020. Fungal Diversity 105:1-16 ((https://doi.org/10.1007/s13225-020-00466-2)) |
| Otu44060 | 1 | k__Fungi(100);p__Ascomycota(100);c__Saccharomycetes(100);o__Saccharomycetales(100);f__Metschnikowiaceae(100);g__Metschnikowia(100);s__Metschnikowia_viticola(100); | Metschnikowia | 13 | Saprotroph | \|Undefined Saprotroph\| | Yeast | NULL | Probable | As nectar-tap_saprotroph (Põlme et al. 2020); Decay sugar-rich substrates (Põlme et al. 2020) | Põlme S, et al. 2020. Fungal Diversity 105:1-16 ((https://doi.org/10.1007/s13225-020-00466-2)) |
| Otu44061 | 1 | k__Fungi(100);p__Ascomycota(100);c__Saccharomycetes(100);o__Saccharomycetales(100);f__Metschnikowiaceae(100);g__Metschnikowia(100);s__Metschnikowia_viticola(100); | Metschnikowia | 13 | Saprotroph | \|Undefined Saprotroph\| | Yeast | NULL | Probable | As nectar-tap_saprotroph (Põlme et al. 2020); Decay sugar-rich substrates (Põlme et al. 2020) | Põlme S, et al. 2020. Fungal Diversity 105:1-16 ((https://doi.org/10.1007/s13225-020-00466-2)) |
| Otu44062 | 1 | k__Fungi(100);p__Ascomycota(100);c__Saccharomycetes(100);o__Saccharomycetales(100);f__Metschnikowiaceae(100);g__Metschnikowia(100);g__Metschnikowia_unclassified(100); | Metschnikowia | 13 | Saprotroph | \|Undefined Saprotroph\| | Yeast | NULL | Probable | As nectar-tap_saprotroph (Põlme et al. 2020); Decay sugar-rich substrates (Põlme et al. 2020) | Põlme S, et al. 2020. Fungal Diversity 105:1-16 ((https://doi.org/10.1007/s13225-020-00466-2)) |
| Otu44063 | 1 | k__Fungi(100);p__Ascomycota(100);c__Saccharomycetes(100);o__Saccharomycetales(100);f__Metschnikowiaceae(100);g__Metschnikowia(100);g__Metschnikowia_unclassified(100); | Metschnikowia | 13 | Saprotroph | \|Undefined Saprotroph\| | Yeast | NULL | Probable | As nectar-tap_saprotroph (Põlme et al. 2020); Decay sugar-rich substrates (Põlme et al. 2020) | Põlme S, et al. 2020. Fungal Diversity 105:1-16 ((https://doi.org/10.1007/s13225-020-00466-2)) |
| Otu44064 | 1 | k__Fungi(100);p__Ascomycota(100);c__Saccharomycetes(100);o__Saccharomycetales(100);f__Metschnikowiaceae(100);g__Metschnikowia(100);g__Metschnikowia_unclassified(100); | Metschnikowia | 13 | Saprotroph | \|Undefined Saprotroph\| | Yeast | NULL | Probable | As nectar-tap_saprotroph (Põlme et al. 2020); Decay sugar-rich substrates (Põlme et al. 2020) | Põlme S, et al. 2020. Fungal Diversity 105:1-16 ((https://doi.org/10.1007/s13225-020-00466-2)) |
| Otu44065 | 1 | k__Fungi(100);p__Ascomycota(100);c__Saccharomycetes(100);o__Saccharomycetales(100);f__Metschnikowiaceae(100);g__Metschnikowia(100);s__Metschnikowia_pulcherrima(100); | Metschnikowia | 13 | Saprotroph | \|Undefined Saprotroph\| | Yeast | NULL | Probable | As nectar-tap_saprotroph (Põlme et al. 2020); Decay sugar-rich substrates (Põlme et al. 2020) | Põlme S, et al. 2020. Fungal Diversity 105:1-16 ((https://doi.org/10.1007/s13225-020-00466-2)) |
| Otu44066 | 1 | k__Fungi(100);p__Ascomycota(100);c__Saccharomycetes(100);o__Saccharomycetales(100);f__Metschnikowiaceae(100);g__Metschnikowia(100);s__Metschnikowia_pulcherrima(100); | Metschnikowia | 13 | Saprotroph | \|Undefined Saprotroph\| | Yeast | NULL | Probable | As nectar-tap_saprotroph (Põlme et al. 2020); Decay sugar-rich substrates (Põlme et al. 2020) | Põlme S, et al. 2020. Fungal Diversity 105:1-16 ((https://doi.org/10.1007/s13225-020-00466-2)) |
| Otu44067 | 1 | k__Fungi(100);p__Ascomycota(100);c__Saccharomycetes(100);o__Saccharomycetales(100);f__Metschnikowiaceae(100);g__Metschnikowia(100);s__Metschnikowia_pulcherrima(100); | Metschnikowia | 13 | Saprotroph | \|Undefined Saprotroph\| | Yeast | NULL | Probable | As nectar-tap_saprotroph (Põlme et al. 2020); Decay sugar-rich substrates (Põlme et al. 2020) | Põlme S, et al. 2020. Fungal Diversity 105:1-16 ((https://doi.org/10.1007/s13225-020-00466-2)) |
| Otu44068 | 1 | k__Fungi(100);p__Ascomycota(100);c__Saccharomycetes(100);o__Saccharomycetales(100);f__Metschnikowiaceae(100);g__Metschnikowia(100);s__Metschnikowia_pulcherrima(100); | Metschnikowia | 13 | Saprotroph | \|Undefined Saprotroph\| | Yeast | NULL | Probable | As nectar-tap_saprotroph (Põlme et al. 2020); Decay sugar-rich substrates (Põlme et al. 2020) | Põlme S, et al. 2020. Fungal Diversity 105:1-16 ((https://doi.org/10.1007/s13225-020-00466-2)) |
| Otu44069 | 1 | k__Fungi(100);p__Ascomycota(100);c__Saccharomycetes(100);o__Saccharomycetales(100);f__Metschnikowiaceae(100);g__Metschnikowia(100);g__Metschnikowia_unclassified(100); | Metschnikowia | 13 | Saprotroph | \|Undefined Saprotroph\| | Yeast | NULL | Probable | As nectar-tap_saprotroph (Põlme et al. 2020); Decay sugar-rich substrates (Põlme et al. 2020) | Põlme S, et al. 2020. Fungal Diversity 105:1-16 ((https://doi.org/10.1007/s13225-020-00466-2)) |
| Otu44070 | 1 | k__Fungi(100);p__Ascomycota(100);c__Saccharomycetes(100);o__Saccharomycetales(100);f__Metschnikowiaceae(100);g__Metschnikowia(100);s__Metschnikowia_pulcherrima(100); | Metschnikowia | 13 | Saprotroph | \|Undefined Saprotroph\| | Yeast | NULL | Probable | As nectar-tap_saprotroph (Põlme et al. 2020); Decay sugar-rich substrates (Põlme et al. 2020) | Põlme S, et al. 2020. Fungal Diversity 105:1-16 ((https://doi.org/10.1007/s13225-020-00466-2)) |
| Otu44071 | 1 | k__Fungi(100);p__Ascomycota(100);c__Saccharomycetes(100);o__Saccharomycetales(100);f__Metschnikowiaceae(100);g__Metschnikowia(100);s__Metschnikowia_pulcherrima(100); | Metschnikowia | 13 | Saprotroph | \|Undefined Saprotroph\| | Yeast | NULL | Probable | As nectar-tap_saprotroph (Põlme et al. 2020); Decay sugar-rich substrates (Põlme et al. 2020) | Põlme S, et al. 2020. Fungal Diversity 105:1-16 ((https://doi.org/10.1007/s13225-020-00466-2)) |
| Otu44073 | 1 | k__Fungi(100);p__Ascomycota(100);c__Saccharomycetes(100);o__Saccharomycetales(100);f__Metschnikowiaceae(100);g__Metschnikowia(100);s__Metschnikowia_pulcherrima(100); | Metschnikowia | 13 | Saprotroph | \|Undefined Saprotroph\| | Yeast | NULL | Probable | As nectar-tap_saprotroph (Põlme et al. 2020); Decay sugar-rich substrates (Põlme et al. 2020) | Põlme S, et al. 2020. Fungal Diversity 105:1-16 ((https://doi.org/10.1007/s13225-020-00466-2)) |
| Otu44074 | 1 | k__Fungi(100);p__Ascomycota(100);c__Saccharomycetes(100);o__Saccharomycetales(100);f__Metschnikowiaceae(100);g__Metschnikowia(100);s__Metschnikowia_pulcherrima(100); | Metschnikowia | 13 | Saprotroph | \|Undefined Saprotroph\| | Yeast | NULL | Probable | As nectar-tap_saprotroph (Põlme et al. 2020); Decay sugar-rich substrates (Põlme et al. 2020) | Põlme S, et al. 2020. Fungal Diversity 105:1-16 ((https://doi.org/10.1007/s13225-020-00466-2)) |
| Otu44075 | 1 | k__Fungi(100);p__Ascomycota(100);c__Saccharomycetes(100);o__Saccharomycetales(100);f__Metschnikowiaceae(100);g__Metschnikowia(100);s__Metschnikowia_pulcherrima(100); | Metschnikowia | 13 | Saprotroph | \|Undefined Saprotroph\| | Yeast | NULL | Probable | As nectar-tap_saprotroph (Põlme et al. 2020); Decay sugar-rich substrates (Põlme et al. 2020) | Põlme S, et al. 2020. Fungal Diversity 105:1-16 ((https://doi.org/10.1007/s13225-020-00466-2)) |
| Otu44076 | 1 | k__Fungi(100);p__Ascomycota(100);c__Saccharomycetes(100);o__Saccharomycetales(100);f__Metschnikowiaceae(100);g__Metschnikowia(100);s__Metschnikowia_pulcherrima(100); | Metschnikowia | 13 | Saprotroph | \|Undefined Saprotroph\| | Yeast | NULL | Probable | As nectar-tap_saprotroph (Põlme et al. 2020); Decay sugar-rich substrates (Põlme et al. 2020) | Põlme S, et al. 2020. Fungal Diversity 105:1-16 ((https://doi.org/10.1007/s13225-020-00466-2)) |
| Otu44077 | 1 | k__Fungi(100);p__Ascomycota(100);c__Saccharomycetes(100);o__Saccharomycetales(100);f__Metschnikowiaceae(100);g__Metschnikowia(100);s__Metschnikowia_pulcherrima(100); | Metschnikowia | 13 | Saprotroph | \|Undefined Saprotroph\| | Yeast | NULL | Probable | As nectar-tap_saprotroph (Põlme et al. 2020); Decay sugar-rich substrates (Põlme et al. 2020) | Põlme S, et al. 2020. Fungal Diversity 105:1-16 ((https://doi.org/10.1007/s13225-020-00466-2)) |
| Otu44078 | 1 | k__Fungi(100);p__Ascomycota(100);c__Saccharomycetes(100);o__Saccharomycetales(100);f__Metschnikowiaceae(100);g__Metschnikowia(100);s__Metschnikowia_pulcherrima(100); | Metschnikowia | 13 | Saprotroph | \|Undefined Saprotroph\| | Yeast | NULL | Probable | As nectar-tap_saprotroph (Põlme et al. 2020); Decay sugar-rich substrates (Põlme et al. 2020) | Põlme S, et al. 2020. Fungal Diversity 105:1-16 ((https://doi.org/10.1007/s13225-020-00466-2)) |
| Otu44079 | 1 | k__Fungi(100);p__Ascomycota(100);c__Saccharomycetes(100);o__Saccharomycetales(100);f__Metschnikowiaceae(100);g__Metschnikowia(100);s__Metschnikowia_pulcherrima(100); | Metschnikowia | 13 | Saprotroph | \|Undefined Saprotroph\| | Yeast | NULL | Probable | As nectar-tap_saprotroph (Põlme et al. 2020); Decay sugar-rich substrates (Põlme et al. 2020) | Põlme S, et al. 2020. Fungal Diversity 105:1-16 ((https://doi.org/10.1007/s13225-020-00466-2)) |
| Otu44080 | 1 | k__Fungi(100);p__Ascomycota(100);c__Saccharomycetes(100);o__Saccharomycetales(100);f__Metschnikowiaceae(100);g__Metschnikowia(100);s__Metschnikowia_pulcherrima(100); | Metschnikowia | 13 | Saprotroph | \|Undefined Saprotroph\| | Yeast | NULL | Probable | As nectar-tap_saprotroph (Põlme et al. 2020); Decay sugar-rich substrates (Põlme et al. 2020) | Põlme S, et al. 2020. Fungal Diversity 105:1-16 ((https://doi.org/10.1007/s13225-020-00466-2)) |
| Otu44081 | 1 | k__Fungi(100);p__Ascomycota(100);c__Saccharomycetes(100);o__Saccharomycetales(100);f__Metschnikowiaceae(100);g__Metschnikowia(100);s__Metschnikowia_pulcherrima(100); | Metschnikowia | 13 | Saprotroph | \|Undefined Saprotroph\| | Yeast | NULL | Probable | As nectar-tap_saprotroph (Põlme et al. 2020); Decay sugar-rich substrates (Põlme et al. 2020) | Põlme S, et al. 2020. Fungal Diversity 105:1-16 ((https://doi.org/10.1007/s13225-020-00466-2)) |
| Otu44082 | 1 | k__Fungi(100);p__Ascomycota(100);c__Saccharomycetes(100);o__Saccharomycetales(100);f__Metschnikowiaceae(100);g__Metschnikowia(100);g__Metschnikowia_unclassified(100); | Metschnikowia | 13 | Saprotroph | \|Undefined Saprotroph\| | Yeast | NULL | Probable | As nectar-tap_saprotroph (Põlme et al. 2020); Decay sugar-rich substrates (Põlme et al. 2020) | Põlme S, et al. 2020. Fungal Diversity 105:1-16 ((https://doi.org/10.1007/s13225-020-00466-2)) |
| Otu44083 | 1 | k__Fungi(100);p__Ascomycota(100);c__Saccharomycetes(100);o__Saccharomycetales(100);f__Metschnikowiaceae(100);g__Metschnikowia(100);g__Metschnikowia_unclassified(100); | Metschnikowia | 13 | Saprotroph | \|Undefined Saprotroph\| | Yeast | NULL | Probable | As nectar-tap_saprotroph (Põlme et al. 2020); Decay sugar-rich substrates (Põlme et al. 2020) | Põlme S, et al. 2020. Fungal Diversity 105:1-16 ((https://doi.org/10.1007/s13225-020-00466-2)) |
| Otu44084 | 1 | k__Fungi(100);p__Ascomycota(100);c__Saccharomycetes(100);o__Saccharomycetales(100);f__Metschnikowiaceae(100);g__Metschnikowia(100);s__Metschnikowia_pulcherrima(100); | Metschnikowia | 13 | Saprotroph | \|Undefined Saprotroph\| | Yeast | NULL | Probable | As nectar-tap_saprotroph (Põlme et al. 2020); Decay sugar-rich substrates (Põlme et al. 2020) | Põlme S, et al. 2020. Fungal Diversity 105:1-16 ((https://doi.org/10.1007/s13225-020-00466-2)) |
| Otu44085 | 1 | k__Fungi(100);p__Ascomycota(100);c__Saccharomycetes(100);o__Saccharomycetales(100);f__Metschnikowiaceae(100);g__Metschnikowia(100);s__Metschnikowia_pulcherrima(100); | Metschnikowia | 13 | Saprotroph | \|Undefined Saprotroph\| | Yeast | NULL | Probable | As nectar-tap_saprotroph (Põlme et al. 2020); Decay sugar-rich substrates (Põlme et al. 2020) | Põlme S, et al. 2020. Fungal Diversity 105:1-16 ((https://doi.org/10.1007/s13225-020-00466-2)) |
| Otu44086 | 1 | k__Fungi(100);p__Ascomycota(100);c__Saccharomycetes(100);o__Saccharomycetales(100);f__Metschnikowiaceae(100);g__Metschnikowia(100);s__Metschnikowia_pulcherrima(100); | Metschnikowia | 13 | Saprotroph | \|Undefined Saprotroph\| | Yeast | NULL | Probable | As nectar-tap_saprotroph (Põlme et al. 2020); Decay sugar-rich substrates (Põlme et al. 2020) | Põlme S, et al. 2020. Fungal Diversity 105:1-16 ((https://doi.org/10.1007/s13225-020-00466-2)) |
| Otu44087 | 1 | k__Fungi(100);p__Ascomycota(100);c__Saccharomycetes(100);o__Saccharomycetales(100);f__Metschnikowiaceae(100);g__Metschnikowia(100);s__Metschnikowia_pulcherrima(100); | Metschnikowia | 13 | Saprotroph | \|Undefined Saprotroph\| | Yeast | NULL | Probable | As nectar-tap_saprotroph (Põlme et al. 2020); Decay sugar-rich substrates (Põlme et al. 2020) | Põlme S, et al. 2020. Fungal Diversity 105:1-16 ((https://doi.org/10.1007/s13225-020-00466-2)) |
| Otu44089 | 1 | k__Fungi(100);p__Ascomycota(100);c__Saccharomycetes(100);o__Saccharomycetales(100);f__Metschnikowiaceae(100);g__Metschnikowia(100);g__Metschnikowia_unclassified(100); | Metschnikowia | 13 | Saprotroph | \|Undefined Saprotroph\| | Yeast | NULL | Probable | As nectar-tap_saprotroph (Põlme et al. 2020); Decay sugar-rich substrates (Põlme et al. 2020) | Põlme S, et al. 2020. Fungal Diversity 105:1-16 ((https://doi.org/10.1007/s13225-020-00466-2)) |
| Otu44090 | 1 | k__Fungi(100);p__Ascomycota(100);c__Saccharomycetes(100);o__Saccharomycetales(100);f__Metschnikowiaceae(100);g__Metschnikowia(100);g__Metschnikowia_unclassified(100); | Metschnikowia | 13 | Saprotroph | \|Undefined Saprotroph\| | Yeast | NULL | Probable | As nectar-tap_saprotroph (Põlme et al. 2020); Decay sugar-rich substrates (Põlme et al. 2020) | Põlme S, et al. 2020. Fungal Diversity 105:1-16 ((https://doi.org/10.1007/s13225-020-00466-2)) |
| Otu44091 | 1 | k__Fungi(100);p__Ascomycota(100);c__Saccharomycetes(100);o__Saccharomycetales(100);f__Metschnikowiaceae(100);g__Metschnikowia(100);s__Metschnikowia_pulcherrima(100); | Metschnikowia | 13 | Saprotroph | \|Undefined Saprotroph\| | Yeast | NULL | Probable | As nectar-tap_saprotroph (Põlme et al. 2020); Decay sugar-rich substrates (Põlme et al. 2020) | Põlme S, et al. 2020. Fungal Diversity 105:1-16 ((https://doi.org/10.1007/s13225-020-00466-2)) |
| Otu44092 | 1 | k__Fungi(100);p__Ascomycota(100);c__Saccharomycetes(100);o__Saccharomycetales(100);f__Metschnikowiaceae(100);g__Metschnikowia(100);s__Metschnikowia_pulcherrima(100); | Metschnikowia | 13 | Saprotroph | \|Undefined Saprotroph\| | Yeast | NULL | Probable | As nectar-tap_saprotroph (Põlme et al. 2020); Decay sugar-rich substrates (Põlme et al. 2020) | Põlme S, et al. 2020. Fungal Diversity 105:1-16 ((https://doi.org/10.1007/s13225-020-00466-2)) |
| Otu44093 | 1 | k__Fungi(100);p__Ascomycota(100);c__Saccharomycetes(100);o__Saccharomycetales(100);f__Metschnikowiaceae(100);g__Metschnikowia(100);s__Metschnikowia_pulcherrima(100); | Metschnikowia | 13 | Saprotroph | \|Undefined Saprotroph\| | Yeast | NULL | Probable | As nectar-tap_saprotroph (Põlme et al. 2020); Decay sugar-rich substrates (Põlme et al. 2020) | Põlme S, et al. 2020. Fungal Diversity 105:1-16 ((https://doi.org/10.1007/s13225-020-00466-2)) |
| Otu44094 | 1 | k__Fungi(100);p__Ascomycota(100);c__Saccharomycetes(100);o__Saccharomycetales(100);f__Metschnikowiaceae(100);g__Metschnikowia(100);s__Metschnikowia_pulcherrima(100); | Metschnikowia | 13 | Saprotroph | \|Undefined Saprotroph\| | Yeast | NULL | Probable | As nectar-tap_saprotroph (Põlme et al. 2020); Decay sugar-rich substrates (Põlme et al. 2020) | Põlme S, et al. 2020. Fungal Diversity 105:1-16 ((https://doi.org/10.1007/s13225-020-00466-2)) |
| Otu44095 | 1 | k__Fungi(100);p__Ascomycota(100);c__Saccharomycetes(100);o__Saccharomycetales(100);f__Metschnikowiaceae(100);g__Metschnikowia(100);s__Metschnikowia_pulcherrima(100); | Metschnikowia | 13 | Saprotroph | \|Undefined Saprotroph\| | Yeast | NULL | Probable | As nectar-tap_saprotroph (Põlme et al. 2020); Decay sugar-rich substrates (Põlme et al. 2020) | Põlme S, et al. 2020. Fungal Diversity 105:1-16 ((https://doi.org/10.1007/s13225-020-00466-2)) |
| Otu44096 | 1 | k__Fungi(100);p__Ascomycota(100);c__Saccharomycetes(100);o__Saccharomycetales(100);f__Metschnikowiaceae(100);g__Metschnikowia(100);s__Metschnikowia_pulcherrima(100); | Metschnikowia | 13 | Saprotroph | \|Undefined Saprotroph\| | Yeast | NULL | Probable | As nectar-tap_saprotroph (Põlme et al. 2020); Decay sugar-rich substrates (Põlme et al. 2020) | Põlme S, et al. 2020. Fungal Diversity 105:1-16 ((https://doi.org/10.1007/s13225-020-00466-2)) |
| Otu44097 | 1 | k__Fungi(100);p__Ascomycota(100);c__Saccharomycetes(100);o__Saccharomycetales(100);f__Metschnikowiaceae(100);g__Metschnikowia(100);s__Metschnikowia_pulcherrima(100); | Metschnikowia | 13 | Saprotroph | \|Undefined Saprotroph\| | Yeast | NULL | Probable | As nectar-tap_saprotroph (Põlme et al. 2020); Decay sugar-rich substrates (Põlme et al. 2020) | Põlme S, et al. 2020. Fungal Diversity 105:1-16 ((https://doi.org/10.1007/s13225-020-00466-2)) |
| Otu44098 | 1 | k__Fungi(100);p__Ascomycota(100);c__Saccharomycetes(100);o__Saccharomycetales(100);f__Metschnikowiaceae(100);g__Metschnikowia(100);s__Metschnikowia_pulcherrima(100); | Metschnikowia | 13 | Saprotroph | \|Undefined Saprotroph\| | Yeast | NULL | Probable | As nectar-tap_saprotroph (Põlme et al. 2020); Decay sugar-rich substrates (Põlme et al. 2020) | Põlme S, et al. 2020. Fungal Diversity 105:1-16 ((https://doi.org/10.1007/s13225-020-00466-2)) |
| Otu44099 | 1 | k__Fungi(100);p__Ascomycota(100);c__Saccharomycetes(100);o__Saccharomycetales(100);f__Metschnikowiaceae(100);g__Metschnikowia(100);s__Metschnikowia_pulcherrima(100); | Metschnikowia | 13 | Saprotroph | \|Undefined Saprotroph\| | Yeast | NULL | Probable | As nectar-tap_saprotroph (Põlme et al. 2020); Decay sugar-rich substrates (Põlme et al. 2020) | Põlme S, et al. 2020. Fungal Diversity 105:1-16 ((https://doi.org/10.1007/s13225-020-00466-2)) |
| Otu44100 | 1 | k__Fungi(100);p__Ascomycota(100);c__Saccharomycetes(100);o__Saccharomycetales(100);f__Metschnikowiaceae(100);g__Metschnikowia(100);s__Metschnikowia_pulcherrima(100); | Metschnikowia | 13 | Saprotroph | \|Undefined Saprotroph\| | Yeast | NULL | Probable | As nectar-tap_saprotroph (Põlme et al. 2020); Decay sugar-rich substrates (Põlme et al. 2020) | Põlme S, et al. 2020. Fungal Diversity 105:1-16 ((https://doi.org/10.1007/s13225-020-00466-2)) |
| Otu44101 | 1 | k__Fungi(100);p__Ascomycota(100);c__Saccharomycetes(100);o__Saccharomycetales(100);f__Metschnikowiaceae(100);g__Metschnikowia(100);s__Metschnikowia_pulcherrima(100); | Metschnikowia | 13 | Saprotroph | \|Undefined Saprotroph\| | Yeast | NULL | Probable | As nectar-tap_saprotroph (Põlme et al. 2020); Decay sugar-rich substrates (Põlme et al. 2020) | Põlme S, et al. 2020. Fungal Diversity 105:1-16 ((https://doi.org/10.1007/s13225-020-00466-2)) |
| Otu44102 | 1 | k__Fungi(100);p__Ascomycota(100);c__Saccharomycetes(100);o__Saccharomycetales(100);f__Metschnikowiaceae(100);g__Metschnikowia(100);s__Metschnikowia_pulcherrima(100); | Metschnikowia | 13 | Saprotroph | \|Undefined Saprotroph\| | Yeast | NULL | Probable | As nectar-tap_saprotroph (Põlme et al. 2020); Decay sugar-rich substrates (Põlme et al. 2020) | Põlme S, et al. 2020. Fungal Diversity 105:1-16 ((https://doi.org/10.1007/s13225-020-00466-2)) |
| Otu44103 | 1 | k__Fungi(100);p__Ascomycota(100);c__Saccharomycetes(100);o__Saccharomycetales(100);f__Metschnikowiaceae(100);g__Metschnikowia(100);s__Metschnikowia_pulcherrima(100); | Metschnikowia | 13 | Saprotroph | \|Undefined Saprotroph\| | Yeast | NULL | Probable | As nectar-tap_saprotroph (Põlme et al. 2020); Decay sugar-rich substrates (Põlme et al. 2020) | Põlme S, et al. 2020. Fungal Diversity 105:1-16 ((https://doi.org/10.1007/s13225-020-00466-2)) |
| Otu44104 | 1 | k__Fungi(100);p__Ascomycota(100);c__Saccharomycetes(100);o__Saccharomycetales(100);f__Metschnikowiaceae(100);g__Metschnikowia(100);s__Metschnikowia_pulcherrima(100); | Metschnikowia | 13 | Saprotroph | \|Undefined Saprotroph\| | Yeast | NULL | Probable | As nectar-tap_saprotroph (Põlme et al. 2020); Decay sugar-rich substrates (Põlme et al. 2020) | Põlme S, et al. 2020. Fungal Diversity 105:1-16 ((https://doi.org/10.1007/s13225-020-00466-2)) |
| Otu44105 | 1 | k__Fungi(100);p__Ascomycota(100);c__Saccharomycetes(100);o__Saccharomycetales(100);f__Metschnikowiaceae(100);g__Metschnikowia(100);s__Metschnikowia_pulcherrima(100); | Metschnikowia | 13 | Saprotroph | \|Undefined Saprotroph\| | Yeast | NULL | Probable | As nectar-tap_saprotroph (Põlme et al. 2020); Decay sugar-rich substrates (Põlme et al. 2020) | Põlme S, et al. 2020. Fungal Diversity 105:1-16 ((https://doi.org/10.1007/s13225-020-00466-2)) |
| Otu44106 | 1 | k__Fungi(100);p__Ascomycota(100);c__Saccharomycetes(100);o__Saccharomycetales(100);f__Metschnikowiaceae(100);g__Metschnikowia(100);g__Metschnikowia_unclassified(100); | Metschnikowia | 13 | Saprotroph | \|Undefined Saprotroph\| | Yeast | NULL | Probable | As nectar-tap_saprotroph (Põlme et al. 2020); Decay sugar-rich substrates (Põlme et al. 2020) | Põlme S, et al. 2020. Fungal Diversity 105:1-16 ((https://doi.org/10.1007/s13225-020-00466-2)) |
| Otu44107 | 1 | k__Fungi(100);p__Ascomycota(100);c__Saccharomycetes(100);o__Saccharomycetales(100);f__Metschnikowiaceae(100);g__Metschnikowia(100);s__Metschnikowia_pulcherrima(100); | Metschnikowia | 13 | Saprotroph | \|Undefined Saprotroph\| | Yeast | NULL | Probable | As nectar-tap_saprotroph (Põlme et al. 2020); Decay sugar-rich substrates (Põlme et al. 2020) | Põlme S, et al. 2020. Fungal Diversity 105:1-16 ((https://doi.org/10.1007/s13225-020-00466-2)) |
| Otu44108 | 1 | k__Fungi(100);p__Ascomycota(100);c__Saccharomycetes(100);o__Saccharomycetales(100);f__Metschnikowiaceae(100);g__Metschnikowia(100);g__Metschnikowia_unclassified(100); | Metschnikowia | 13 | Saprotroph | \|Undefined Saprotroph\| | Yeast | NULL | Probable | As nectar-tap_saprotroph (Põlme et al. 2020); Decay sugar-rich substrates (Põlme et al. 2020) | Põlme S, et al. 2020. Fungal Diversity 105:1-16 ((https://doi.org/10.1007/s13225-020-00466-2)) |
| Otu44109 | 1 | k__Fungi(100);p__Ascomycota(100);c__Saccharomycetes(100);o__Saccharomycetales(100);f__Metschnikowiaceae(100);g__Metschnikowia(100);s__Metschnikowia_pulcherrima(100); | Metschnikowia | 13 | Saprotroph | \|Undefined Saprotroph\| | Yeast | NULL | Probable | As nectar-tap_saprotroph (Põlme et al. 2020); Decay sugar-rich substrates (Põlme et al. 2020) | Põlme S, et al. 2020. Fungal Diversity 105:1-16 ((https://doi.org/10.1007/s13225-020-00466-2)) |
| Otu44110 | 1 | k__Fungi(100);p__Ascomycota(100);c__Saccharomycetes(100);o__Saccharomycetales(100);f__Metschnikowiaceae(100);g__Metschnikowia(100);s__Metschnikowia_pulcherrima(100); | Metschnikowia | 13 | Saprotroph | \|Undefined Saprotroph\| | Yeast | NULL | Probable | As nectar-tap_saprotroph (Põlme et al. 2020); Decay sugar-rich substrates (Põlme et al. 2020) | Põlme S, et al. 2020. Fungal Diversity 105:1-16 ((https://doi.org/10.1007/s13225-020-00466-2)) |
| Otu44111 | 1 | k__Fungi(100);p__Ascomycota(100);c__Saccharomycetes(100);o__Saccharomycetales(100);f__Metschnikowiaceae(100);g__Metschnikowia(100);s__Metschnikowia_pulcherrima(100); | Metschnikowia | 13 | Saprotroph | \|Undefined Saprotroph\| | Yeast | NULL | Probable | As nectar-tap_saprotroph (Põlme et al. 2020); Decay sugar-rich substrates (Põlme et al. 2020) | Põlme S, et al. 2020. Fungal Diversity 105:1-16 ((https://doi.org/10.1007/s13225-020-00466-2)) |
| Otu44112 | 1 | k__Fungi(100);p__Ascomycota(100);c__Saccharomycetes(100);o__Saccharomycetales(100);f__Metschnikowiaceae(100);g__Metschnikowia(100);s__Metschnikowia_pulcherrima(100); | Metschnikowia | 13 | Saprotroph | \|Undefined Saprotroph\| | Yeast | NULL | Probable | As nectar-tap_saprotroph (Põlme et al. 2020); Decay sugar-rich substrates (Põlme et al. 2020) | Põlme S, et al. 2020. Fungal Diversity 105:1-16 ((https://doi.org/10.1007/s13225-020-00466-2)) |
| Otu44113 | 1 | k__Fungi(100);p__Ascomycota(100);c__Saccharomycetes(100);o__Saccharomycetales(100);f__Metschnikowiaceae(100);g__Metschnikowia(100);s__Metschnikowia_pulcherrima(100); | Metschnikowia | 13 | Saprotroph | \|Undefined Saprotroph\| | Yeast | NULL | Probable | As nectar-tap_saprotroph (Põlme et al. 2020); Decay sugar-rich substrates (Põlme et al. 2020) | Põlme S, et al. 2020. Fungal Diversity 105:1-16 ((https://doi.org/10.1007/s13225-020-00466-2)) |
| Otu44114 | 1 | k__Fungi(100);p__Ascomycota(100);c__Saccharomycetes(100);o__Saccharomycetales(100);f__Metschnikowiaceae(100);g__Metschnikowia(100);s__Metschnikowia_pulcherrima(100); | Metschnikowia | 13 | Saprotroph | \|Undefined Saprotroph\| | Yeast | NULL | Probable | As nectar-tap_saprotroph (Põlme et al. 2020); Decay sugar-rich substrates (Põlme et al. 2020) | Põlme S, et al. 2020. Fungal Diversity 105:1-16 ((https://doi.org/10.1007/s13225-020-00466-2)) |
| Otu44115 | 1 | k__Fungi(100);p__Ascomycota(100);c__Saccharomycetes(100);o__Saccharomycetales(100);f__Metschnikowiaceae(100);g__Metschnikowia(100);g__Metschnikowia_unclassified(100); | Metschnikowia | 13 | Saprotroph | \|Undefined Saprotroph\| | Yeast | NULL | Probable | As nectar-tap_saprotroph (Põlme et al. 2020); Decay sugar-rich substrates (Põlme et al. 2020) | Põlme S, et al. 2020. Fungal Diversity 105:1-16 ((https://doi.org/10.1007/s13225-020-00466-2)) |
| Otu44116 | 1 | k__Fungi(100);p__Ascomycota(100);c__Saccharomycetes(100);o__Saccharomycetales(100);f__Metschnikowiaceae(100);g__Metschnikowia(100);s__Metschnikowia_pulcherrima(100); | Metschnikowia | 13 | Saprotroph | \|Undefined Saprotroph\| | Yeast | NULL | Probable | As nectar-tap_saprotroph (Põlme et al. 2020); Decay sugar-rich substrates (Põlme et al. 2020) | Põlme S, et al. 2020. Fungal Diversity 105:1-16 ((https://doi.org/10.1007/s13225-020-00466-2)) |

| | | | | | | | | | | | | |
|---|---|---|---|---|---|---|---|---|---|---|---|---|
| Otu44117 | 1 | k__Fungi(100);p__Ascomycota(100);c__Saccharomycetes(100);o__Saccharomycetales(100);f__Metschnikowiaceae(100);g__Metschnikowia(100);s__Metschnikowia_pulcherrima(100); | Metschnikowia | | 13 | Saprotroph | \|Undefined Saprotroph\| | Yeast | NULL | Probable | As nectar-tap_saprotroph (Põlme et al. 2020); Decay sugar-rich substrates (Põlme et al. 2020) | Põlme S, et al. 2020. Fungal Diversity 105:1-16 ((https://doi.org/10.1007/s13225-020-00466-2)) |
| Otu44118 | 1 | k__Fungi(100);p__Ascomycota(100);c__Saccharomycetes(100);o__Saccharomycetales(100);f__Metschnikowiaceae(100);g__Metschnikowia(100);s__Metschnikowia_pulcherrima(100); | Metschnikowia | | 13 | Saprotroph | \|Undefined Saprotroph\| | Yeast | NULL | Probable | As nectar-tap_saprotroph (Põlme et al. 2020); Decay sugar-rich substrates (Põlme et al. 2020) | Põlme S, et al. 2020. Fungal Diversity 105:1-16 ((https://doi.org/10.1007/s13225-020-00466-2)) |
| Otu44119 | 1 | k__Fungi(100);p__Ascomycota(100);c__Saccharomycetes(100);o__Saccharomycetales(100);f__Metschnikowiaceae(100);g__Metschnikowia(100);s__Metschnikowia_pulcherrima(100); | Metschnikowia | | 13 | Saprotroph | \|Undefined Saprotroph\| | Yeast | NULL | Probable | As nectar-tap_saprotroph (Põlme et al. 2020); Decay sugar-rich substrates (Põlme et al. 2020) | Põlme S, et al. 2020. Fungal Diversity 105:1-16 ((https://doi.org/10.1007/s13225-020-00466-2)) |
| Otu44120 | 1 | k__Fungi(100);p__Ascomycota(100);c__Saccharomycetes(100);o__Saccharomycetales(100);f__Metschnikowiaceae(100);g__Metschnikowia(100);s__Metschnikowia_pulcherrima(100); | Metschnikowia | | 13 | Saprotroph | \|Undefined Saprotroph\| | Yeast | NULL | Probable | As nectar-tap_saprotroph (Põlme et al. 2020); Decay sugar-rich substrates (Põlme et al. 2020) | Põlme S, et al. 2020. Fungal Diversity 105:1-16 ((https://doi.org/10.1007/s13225-020-00466-2)) |
| Otu44121 | 1 | k__Fungi(100);p__Ascomycota(100);c__Saccharomycetes(100);o__Saccharomycetales(100);f__Metschnikowiaceae(100);g__Metschnikowia(100);g__Metschnikowia_unclassified(100); | Metschnikowia | | 13 | Saprotroph | \|Undefined Saprotroph\| | Yeast | NULL | Probable | As nectar-tap_saprotroph (Põlme et al. 2020); Decay sugar-rich substrates (Põlme et al. 2020) | Põlme S, et al. 2020. Fungal Diversity 105:1-16 ((https://doi.org/10.1007/s13225-020-00466-2)) |
| Otu44122 | 1 | k__Fungi(100);p__Ascomycota(100);c__Saccharomycetes(100);o__Saccharomycetales(100);f__Metschnikowiaceae(100);g__Metschnikowia(100);s__Metschnikowia_pulcherrima(100); | Metschnikowia | | 13 | Saprotroph | \|Undefined Saprotroph\| | Yeast | NULL | Probable | As nectar-tap_saprotroph (Põlme et al. 2020); Decay sugar-rich substrates (Põlme et al. 2020) | Põlme S, et al. 2020. Fungal Diversity 105:1-16 ((https://doi.org/10.1007/s13225-020-00466-2)) |
| Otu44124 | 1 | k__Fungi(100);p__Ascomycota(100);c__Saccharomycetes(100);o__Saccharomycetales(100);f__Metschnikowiaceae(100);g__Metschnikowia(100);g__Metschnikowia_unclassified(100); | Metschnikowia | | 13 | Saprotroph | \|Undefined Saprotroph\| | Yeast | NULL | Probable | As nectar-tap_saprotroph (Põlme et al. 2020); Decay sugar-rich substrates (Põlme et al. 2020) | Põlme S, et al. 2020. Fungal Diversity 105:1-16 ((https://doi.org/10.1007/s13225-020-00466-2)) |
| Otu44125 | 1 | k__Fungi(100);p__Ascomycota(100);c__Saccharomycetes(100);o__Saccharomycetales(100);f__Metschnikowiaceae(100);g__Metschnikowia(100);s__Metschnikowia_pulcherrima(100); | Metschnikowia | | 13 | Saprotroph | \|Undefined Saprotroph\| | Yeast | NULL | Probable | As nectar-tap_saprotroph (Põlme et al. 2020); Decay sugar-rich substrates (Põlme et al. 2020) | Põlme S, et al. 2020. Fungal Diversity 105:1-16 ((https://doi.org/10.1007/s13225-020-00466-2)) |
| Otu44126 | 1 | k__Fungi(100);p__Ascomycota(100);c__Saccharomycetes(100);o__Saccharomycetales(100);f__Metschnikowiaceae(100);g__Metschnikowia(100);s__Metschnikowia_pulcherrima(100); | Metschnikowia | | 13 | Saprotroph | \|Undefined Saprotroph\| | Yeast | NULL | Probable | As nectar-tap_saprotroph (Põlme et al. 2020); Decay sugar-rich substrates (Põlme et al. 2020) | Põlme S, et al. 2020. Fungal Diversity 105:1-16 ((https://doi.org/10.1007/s13225-020-00466-2)) |
| Otu44127 | 1 | k__Fungi(100);p__Ascomycota(100);c__Saccharomycetes(100);o__Saccharomycetales(100);f__Metschnikowiaceae(100);g__Metschnikowia(100);s__Metschnikowia_pulcherrima(100); | Metschnikowia | | 13 | Saprotroph | \|Undefined Saprotroph\| | Yeast | NULL | Probable | As nectar-tap_saprotroph (Põlme et al. 2020); Decay sugar-rich substrates (Põlme et al. 2020) | Põlme S, et al. 2020. Fungal Diversity 105:1-16 ((https://doi.org/10.1007/s13225-020-00466-2)) |
| Otu44128 | 1 | k__Fungi(100);p__Ascomycota(100);c__Saccharomycetes(100);o__Saccharomycetales(100);f__Metschnikowiaceae(100);g__Metschnikowia(100);g__Metschnikowia_unclassified(100); | Metschnikowia | | 13 | Saprotroph | \|Undefined Saprotroph\| | Yeast | NULL | Probable | As nectar-tap_saprotroph (Põlme et al. 2020); Decay sugar-rich substrates (Põlme et al. 2020) | Põlme S, et al. 2020. Fungal Diversity 105:1-16 ((https://doi.org/10.1007/s13225-020-00466-2)) |
| Otu44129 | 1 | k__Fungi(100);p__Ascomycota(100);c__Saccharomycetes(100);o__Saccharomycetales(100);f__Metschnikowiaceae(100);g__Metschnikowia(100);g__Metschnikowia_unclassified(100); | Metschnikowia | | 13 | Saprotroph | \|Undefined Saprotroph\| | Yeast | NULL | Probable | As nectar-tap_saprotroph (Põlme et al. 2020); Decay sugar-rich substrates (Põlme et al. 2020) | Põlme S, et al. 2020. Fungal Diversity 105:1-16 ((https://doi.org/10.1007/s13225-020-00466-2)) |
| Otu44130 | 1 | k__Fungi(100);p__Ascomycota(100);c__Saccharomycetes(100);o__Saccharomycetales(100);f__Metschnikowiaceae(100);g__Metschnikowia(100);s__Metschnikowia_pulcherrima(100); | Metschnikowia | | 13 | Saprotroph | \|Undefined Saprotroph\| | Yeast | NULL | Probable | As nectar-tap_saprotroph (Põlme et al. 2020); Decay sugar-rich substrates (Põlme et al. 2020) | Põlme S, et al. 2020. Fungal Diversity 105:1-16 ((https://doi.org/10.1007/s13225-020-00466-2)) |
| Otu44132 | 1 | k__Fungi(100);p__Ascomycota(100);c__Saccharomycetes(100);o__Saccharomycetales(100);f__Metschnikowiaceae(100);g__Metschnikowia(100);g__Metschnikowia_unclassified(100); | Metschnikowia | | 13 | Saprotroph | \|Undefined Saprotroph\| | Yeast | NULL | Probable | As nectar-tap_saprotroph (Põlme et al. 2020); Decay sugar-rich substrates (Põlme et al. 2020) | Põlme S, et al. 2020. Fungal Diversity 105:1-16 ((https://doi.org/10.1007/s13225-020-00466-2)) |
| Otu44133 | 1 | k__Fungi(100);p__Ascomycota(100);c__Saccharomycetes(100);o__Saccharomycetales(100);f__Metschnikowiaceae(100);g__Metschnikowia(100);g__Metschnikowia_unclassified(100); | Metschnikowia | | 13 | Saprotroph | \|Undefined Saprotroph\| | Yeast | NULL | Probable | As nectar-tap_saprotroph (Põlme et al. 2020); Decay sugar-rich substrates (Põlme et al. 2020) | Põlme S, et al. 2020. Fungal Diversity 105:1-16 ((https://doi.org/10.1007/s13225-020-00466-2)) |
| Otu44134 | 1 | k__Fungi(100);p__Ascomycota(100);c__Saccharomycetes(100);o__Saccharomycetales(100);f__Metschnikowiaceae(100);g__Metschnikowia(100);s__Metschnikowia_pulcherrima(100); | Metschnikowia | | 13 | Saprotroph | \|Undefined Saprotroph\| | Yeast | NULL | Probable | As nectar-tap_saprotroph (Põlme et al. 2020); Decay sugar-rich substrates (Põlme et al. 2020) | Põlme S, et al. 2020. Fungal Diversity 105:1-16 ((https://doi.org/10.1007/s13225-020-00466-2)) |
| Otu44135 | 1 | k__Fungi(100);p__Ascomycota(100);c__Saccharomycetes(100);o__Saccharomycetales(100);f__Metschnikowiaceae(100);g__Metschnikowia(100);s__Metschnikowia_pulcherrima(100); | Metschnikowia | | 13 | Saprotroph | \|Undefined Saprotroph\| | Yeast | NULL | Probable | As nectar-tap_saprotroph (Põlme et al. 2020); Decay sugar-rich substrates (Põlme et al. 2020) | Põlme S, et al. 2020. Fungal Diversity 105:1-16 ((https://doi.org/10.1007/s13225-020-00466-2)) |
| Otu44136 | 1 | k__Fungi(100);p__Ascomycota(100);c__Saccharomycetes(100);o__Saccharomycetales(100);f__Metschnikowiaceae(100);g__Metschnikowia(100);s__Metschnikowia_pulcherrima(100); | Metschnikowia | | 13 | Saprotroph | \|Undefined Saprotroph\| | Yeast | NULL | Probable | As nectar-tap_saprotroph (Põlme et al. 2020); Decay sugar-rich substrates (Põlme et al. 2020) | Põlme S, et al. 2020. Fungal Diversity 105:1-16 ((https://doi.org/10.1007/s13225-020-00466-2)) |
| Otu44137 | 1 | k__Fungi(100);p__Ascomycota(100);c__Saccharomycetes(100);o__Saccharomycetales(100);f__Metschnikowiaceae(100);g__Metschnikowia(100);s__Metschnikowia_pulcherrima(100); | Metschnikowia | | 13 | Saprotroph | \|Undefined Saprotroph\| | Yeast | NULL | Probable | As nectar-tap_saprotroph (Põlme et al. 2020); Decay sugar-rich substrates (Põlme et al. 2020) | Põlme S, et al. 2020. Fungal Diversity 105:1-16 ((https://doi.org/10.1007/s13225-020-00466-2)) |
| Otu44138 | 1 | k__Fungi(100);p__Ascomycota(100);c__Saccharomycetes(100);o__Saccharomycetales(100);f__Metschnikowiaceae(100);g__Metschnikowia(100);g__Metschnikowia_unclassified(100); | Metschnikowia | | 13 | Saprotroph | \|Undefined Saprotroph\| | Yeast | NULL | Probable | As nectar-tap_saprotroph (Põlme et al. 2020); Decay sugar-rich substrates (Põlme et al. 2020) | Põlme S, et al. 2020. Fungal Diversity 105:1-16 ((https://doi.org/10.1007/s13225-020-00466-2)) |
| Otu44139 | 1 | k__Fungi(100);p__Ascomycota(100);c__Saccharomycetes(100);o__Saccharomycetales(100);f__Metschnikowiaceae(100);g__Metschnikowia(100);s__Metschnikowia_pulcherrima(100); | Metschnikowia | | 13 | Saprotroph | \|Undefined Saprotroph\| | Yeast | NULL | Probable | As nectar-tap_saprotroph (Põlme et al. 2020); Decay sugar-rich substrates (Põlme et al. 2020) | Põlme S, et al. 2020. Fungal Diversity 105:1-16 ((https://doi.org/10.1007/s13225-020-00466-2)) |
| Otu44140 | 1 | k__Fungi(100);p__Ascomycota(100);c__Saccharomycetes(100);o__Saccharomycetales(100);f__Metschnikowiaceae(100);g__Metschnikowia(100);s__Metschnikowia_pulcherrima(100); | Metschnikowia | | 13 | Saprotroph | \|Undefined Saprotroph\| | Yeast | NULL | Probable | As nectar-tap_saprotroph (Põlme et al. 2020); Decay sugar-rich substrates (Põlme et al. 2020) | Põlme S, et al. 2020. Fungal Diversity 105:1-16 ((https://doi.org/10.1007/s13225-020-00466-2)) |
| Otu44141 | 1 | k__Fungi(100);p__Ascomycota(100);c__Saccharomycetes(100);o__Saccharomycetales(100);f__Metschnikowiaceae(100);g__Metschnikowia(100);s__Metschnikowia_pulcherrima(100); | Metschnikowia | | 13 | Saprotroph | \|Undefined Saprotroph\| | Yeast | NULL | Probable | As nectar-tap_saprotroph (Põlme et al. 2020); Decay sugar-rich substrates (Põlme et al. 2020) | Põlme S, et al. 2020. Fungal Diversity 105:1-16 ((https://doi.org/10.1007/s13225-020-00466-2)) |
| Otu44142 | 1 | k__Fungi(100);p__Ascomycota(100);c__Saccharomycetes(100);o__Saccharomycetales(100);f__Metschnikowiaceae(100);g__Metschnikowia(100);g__Metschnikowia_unclassified(100); | Metschnikowia | | 13 | Saprotroph | \|Undefined Saprotroph\| | Yeast | NULL | Probable | As nectar-tap_saprotroph (Põlme et al. 2020); Decay sugar-rich substrates (Põlme et al. 2020) | Põlme S, et al. 2020. Fungal Diversity 105:1-16 ((https://doi.org/10.1007/s13225-020-00466-2)) |
| Otu44143 | 1 | k__Fungi(100);p__Ascomycota(100);c__Saccharomycetes(100);o__Saccharomycetales(100);f__Metschnikowiaceae(100);g__Metschnikowia(100);s__Metschnikowia_pulcherrima(100); | Metschnikowia | | 13 | Saprotroph | \|Undefined Saprotroph\| | Yeast | NULL | Probable | As nectar-tap_saprotroph (Põlme et al. 2020); Decay sugar-rich substrates (Põlme et al. 2020) | Põlme S, et al. 2020. Fungal Diversity 105:1-16 ((https://doi.org/10.1007/s13225-020-00466-2)) |
| Otu44144 | 1 | k__Fungi(100);p__Ascomycota(100);c__Saccharomycetes(100);o__Saccharomycetales(100);f__Metschnikowiaceae(100);g__Metschnikowia(100);s__Metschnikowia_pulcherrima(100); | Metschnikowia | | 13 | Saprotroph | \|Undefined Saprotroph\| | Yeast | NULL | Probable | As nectar-tap_saprotroph (Põlme et al. 2020); Decay sugar-rich substrates (Põlme et al. 2020) | Põlme S, et al. 2020. Fungal Diversity 105:1-16 ((https://doi.org/10.1007/s13225-020-00466-2)) |
| Otu44145 | 1 | k__Fungi(100);p__Ascomycota(100);c__Saccharomycetes(100);o__Saccharomycetales(100);f__Metschnikowiaceae(100);g__Metschnikowia(100);s__Metschnikowia_pulcherrima(100); | Metschnikowia | | 13 | Saprotroph | \|Undefined Saprotroph\| | Yeast | NULL | Probable | As nectar-tap_saprotroph (Põlme et al. 2020); Decay sugar-rich substrates (Põlme et al. 2020) | Põlme S, et al. 2020. Fungal Diversity 105:1-16 ((https://doi.org/10.1007/s13225-020-00466-2)) |
| Otu44146 | 1 | k__Fungi(100);p__Ascomycota(100);c__Saccharomycetes(100);o__Saccharomycetales(100);f__Metschnikowiaceae(100);g__Metschnikowia(100);s__Metschnikowia_pulcherrima(100); | Metschnikowia | | 13 | Saprotroph | \|Undefined Saprotroph\| | Yeast | NULL | Probable | As nectar-tap_saprotroph (Põlme et al. 2020); Decay sugar-rich substrates (Põlme et al. 2020) | Põlme S, et al. 2020. Fungal Diversity 105:1-16 ((https://doi.org/10.1007/s13225-020-00466-2)) |
| Otu44147 | 1 | k__Fungi(100);p__Ascomycota(100);c__Saccharomycetes(100);o__Saccharomycetales(100);f__Metschnikowiaceae(100);g__Metschnikowia(100);s__Metschnikowia_pulcherrima(100); | Metschnikowia | | 13 | Saprotroph | \|Undefined Saprotroph\| | Yeast | NULL | Probable | As nectar-tap_saprotroph (Põlme et al. 2020); Decay sugar-rich substrates (Põlme et al. 2020) | Põlme S, et al. 2020. Fungal Diversity 105:1-16 ((https://doi.org/10.1007/s13225-020-00466-2)) |
| Otu44148 | 1 | k__Fungi(100);p__Ascomycota(100);c__Saccharomycetes(100);o__Saccharomycetales(100);f__Metschnikowiaceae(100);g__Metschnikowia(100);s__Metschnikowia_pulcherrima(100); | Metschnikowia | | 13 | Saprotroph | \|Undefined Saprotroph\| | Yeast | NULL | Probable | As nectar-tap_saprotroph (Põlme et al. 2020); Decay sugar-rich substrates (Põlme et al. 2020) | Põlme S, et al. 2020. Fungal Diversity 105:1-16 ((https://doi.org/10.1007/s13225-020-00466-2)) |
| Otu44149 | 1 | k__Fungi(100);p__Ascomycota(100);c__Saccharomycetes(100);o__Saccharomycetales(100);f__Metschnikowiaceae(100);g__Metschnikowia(100);s__Metschnikowia_pulcherrima(100); | Metschnikowia | | 13 | Saprotroph | \|Undefined Saprotroph\| | Yeast | NULL | Probable | As nectar-tap_saprotroph (Põlme et al. 2020); Decay sugar-rich substrates (Põlme et al. 2020) | Põlme S, et al. 2020. Fungal Diversity 105:1-16 ((https://doi.org/10.1007/s13225-020-00466-2)) |
| Otu44150 | 1 | k__Fungi(100);p__Ascomycota(100);c__Saccharomycetes(100);o__Saccharomycetales(100);f__Metschnikowiaceae(100);g__Metschnikowia(100);g__Metschnikowia_unclassified(100); | Metschnikowia | | 13 | Saprotroph | \|Undefined Saprotroph\| | Yeast | NULL | Probable | As nectar-tap_saprotroph (Põlme et al. 2020); Decay sugar-rich substrates (Põlme et al. 2020) | Põlme S, et al. 2020. Fungal Diversity 105:1-16 ((https://doi.org/10.1007/s13225-020-00466-2)) |
| Otu44151 | 1 | k__Fungi(100);p__Ascomycota(100);c__Saccharomycetes(100);o__Saccharomycetales(100);f__Metschnikowiaceae(100);g__Metschnikowia(100);s__Metschnikowia_pulcherrima(100); | Metschnikowia | | 13 | Saprotroph | \|Undefined Saprotroph\| | Yeast | NULL | Probable | As nectar-tap_saprotroph (Põlme et al. 2020); Decay sugar-rich substrates (Põlme et al. 2020) | Põlme S, et al. 2020. Fungal Diversity 105:1-16 ((https://doi.org/10.1007/s13225-020-00466-2)) |
| Otu44152 | 1 | k__Fungi(100);p__Ascomycota(100);c__Saccharomycetes(100);o__Saccharomycetales(100);f__Metschnikowiaceae(100);g__Metschnikowia(100);g__Metschnikowia_unclassified(100); | Metschnikowia | | 13 | Saprotroph | \|Undefined Saprotroph\| | Yeast | NULL | Probable | As nectar-tap_saprotroph (Põlme et al. 2020); Decay sugar-rich substrates (Põlme et al. 2020) | Põlme S, et al. 2020. Fungal Diversity 105:1-16 ((https://doi.org/10.1007/s13225-020-00466-2)) |
| Otu44153 | 1 | k__Fungi(100);p__Ascomycota(100);c__Saccharomycetes(100);o__Saccharomycetales(100);f__Metschnikowiaceae(100);g__Metschnikowia(100);g__Metschnikowia_unclassified(100); | Metschnikowia | | 13 | Saprotroph | \|Undefined Saprotroph\| | Yeast | NULL | Probable | As nectar-tap_saprotroph (Põlme et al. 2020); Decay sugar-rich substrates (Põlme et al. 2020) | Põlme S, et al. 2020. Fungal Diversity 105:1-16 ((https://doi.org/10.1007/s13225-020-00466-2)) |
| Otu44154 | 1 | k__Fungi(100);p__Ascomycota(100);c__Saccharomycetes(100);o__Saccharomycetales(100);f__Metschnikowiaceae(100);g__Metschnikowia(100);s__Metschnikowia_pulcherrima(100); | Metschnikowia | | 13 | Saprotroph | \|Undefined Saprotroph\| | Yeast | NULL | Probable | As nectar-tap_saprotroph (Põlme et al. 2020); Decay sugar-rich substrates (Põlme et al. 2020) | Põlme S, et al. 2020. Fungal Diversity 105:1-16 ((https://doi.org/10.1007/s13225-020-00466-2)) |
| Otu44155 | 1 | k__Fungi(100);p__Ascomycota(100);c__Saccharomycetes(100);o__Saccharomycetales(100);f__Metschnikowiaceae(100);g__Metschnikowia(100);s__Metschnikowia_pulcherrima(100); | Metschnikowia | | 13 | Saprotroph | \|Undefined Saprotroph\| | Yeast | NULL | Probable | As nectar-tap_saprotroph (Põlme et al. 2020); Decay sugar-rich substrates (Põlme et al. 2020) | Põlme S, et al. 2020. Fungal Diversity 105:1-16 ((https://doi.org/10.1007/s13225-020-00466-2)) |
| Otu44156 | 1 | k__Fungi(100);p__Ascomycota(100);c__Saccharomycetes(100);o__Saccharomycetales(100);f__Metschnikowiaceae(100);g__Metschnikowia(100);s__Metschnikowia_pulcherrima(100); | Metschnikowia | | 13 | Saprotroph | \|Undefined Saprotroph\| | Yeast | NULL | Probable | As nectar-tap_saprotroph (Põlme et al. 2020); Decay sugar-rich substrates (Põlme et al. 2020) | Põlme S, et al. 2020. Fungal Diversity 105:1-16 ((https://doi.org/10.1007/s13225-020-00466-2)) |
| Otu44157 | 1 | k__Fungi(100);p__Ascomycota(100);c__Sordariomycetes(100);o__Hypocreales(100);f__Nectriaceae(100);g__Fusarium(100);g__Fusarium_unclassified(100); | Fusarium | | 13 | Pathotroph-Saprotroph-Symbiotroph | Animal Pathogen-Endophyte-Lichen Parasite-\|Plant Pathogen\|-Plant Saprotroph-Undefined Saprotroph-Wood Saprotroph | Microfungus | Soft Rot | Possible | As foliar_endophyte (Põlme et al. 2020); As litter_saprotroph (Põlme et al. 2020); Causing damping-off disease (Põlme et al. 2020); Decay leaf, fruit, seed, roots, soil, animal material (Põlme et al. 2020); Likely opportunistic human pathogen (Irinyi et al. 2015) | Duncan CG, Eslyn WE. 1966. Mycologia 58:642-664 ((https://doi.org/10.2307/3757045)); Seehann G, et al. 1975. List of Fungi in Soft-Rot Tissue Tests. Institute fur Holzbiologie und Holzschutz, Bundesforschungsanstalt fur Forst - und Holzwirtshaft, Hamburg - Lohbrugge, Germany; Costa IPMW, et al. 2012. Mycotaxon 119:e26 ((http://mycotaxon.com/resources/checklists/costa_v119_checklist.pdf)); Tedersoo L, et al. 2014. Science 346:e1256688 ((https://doi.org/10.1126/science.1256688)); Irinyi L, et al. 2015. Medical Mycology 53:313-337 ((https://doi.org/10.1093/mmy/myv008)); Schroers, et al. 2016. Mycologia 108:806-819 ((https://doi.org/10.3852/15-255)); Põlme S, et al. 2020. Fungal Diversity 105:1-16 ((https://doi.org/10.1007/s13225-020-00466-2)); The American Phytopathological Society Common Names of Plant Diseases ((http://www.apsnet.org/publications/commonnames/Pages/default.aspx)); Lawrey JD, Diederich P. 2016. http://www.lichenicolous.net (accessed 24 May 2017) ((http://www.lichenicolous.net)) |
| Otu44158 | 1 | k__Fungi(100);p__Ascomycota(100);c__Saccharomycetes(100);o__Saccharomycetales(100);f__Metschnikowiaceae(100);g__Metschnikowia(100);s__Metschnikowia_pulcherrima(100); | Metschnikowia | | 13 | Saprotroph | \|Undefined Saprotroph\| | Yeast | NULL | Probable | As nectar-tap_saprotroph (Põlme et al. 2020); Decay sugar-rich substrates (Põlme et al. 2020) | Põlme S, et al. 2020. Fungal Diversity 105:1-16 ((https://doi.org/10.1007/s13225-020-00466-2)) |
| Otu44159 | 1 | k__Fungi(100);p__Ascomycota(100);c__Saccharomycetes(100);o__Saccharomycetales(100);f__Metschnikowiaceae(100);g__Metschnikowia(100);s__Metschnikowia_pulcherrima(100); | Metschnikowia | | 13 | Saprotroph | \|Undefined Saprotroph\| | Yeast | NULL | Probable | As nectar-tap_saprotroph (Põlme et al. 2020); Decay sugar-rich substrates (Põlme et al. 2020) | Põlme S, et al. 2020. Fungal Diversity 105:1-16 ((https://doi.org/10.1007/s13225-020-00466-2)) |
| Otu44160 | 1 | k__Fungi(100);p__Ascomycota(100);c__Saccharomycetes(100);o__Saccharomycetales(100);f__Metschnikowiaceae(100);g__Metschnikowia(100);s__Metschnikowia_pulcherrima(100); | Metschnikowia | | 13 | Saprotroph | \|Undefined Saprotroph\| | Yeast | NULL | Probable | As nectar-tap_saprotroph (Põlme et al. 2020); Decay sugar-rich substrates (Põlme et al. 2020) | Põlme S, et al. 2020. Fungal Diversity 105:1-16 ((https://doi.org/10.1007/s13225-020-00466-2)) |
| Otu44161 | 1 | k__Fungi(100);p__Ascomycota(100);c__Saccharomycetes(100);o__Saccharomycetales(100);f__Metschnikowiaceae(100);g__Metschnikowia(100);s__Metschnikowia_pulcherrima(100); | Metschnikowia | | 13 | Saprotroph | \|Undefined Saprotroph\| | Yeast | NULL | Probable | As nectar-tap_saprotroph (Põlme et al. 2020); Decay sugar-rich substrates (Põlme et al. 2020) | Põlme S, et al. 2020. Fungal Diversity 105:1-16 ((https://doi.org/10.1007/s13225-020-00466-2)) |
| Otu44163 | 1 | k__Fungi(100);p__Ascomycota(100);c__Saccharomycetes(100);o__Saccharomycetales(100);f__Metschnikowiaceae(100);g__Metschnikowia(100);s__Metschnikowia_pulcherrima(100); | Metschnikowia | | 13 | Saprotroph | \|Undefined Saprotroph\| | Yeast | NULL | Probable | As nectar-tap_saprotroph (Põlme et al. 2020); Decay sugar-rich substrates (Põlme et al. 2020) | Põlme S, et al. 2020. Fungal Diversity 105:1-16 ((https://doi.org/10.1007/s13225-020-00466-2)) |
| Otu44164 | 1 | k__Fungi(100);p__Ascomycota(100);c__Saccharomycetes(100);o__Saccharomycetales(100);f__Metschnikowiaceae(100);g__Metschnikowia(100);g__Metschnikowia_unclassified(100); | Metschnikowia | | 13 | Saprotroph | \|Undefined Saprotroph\| | Yeast | NULL | Probable | As nectar-tap_saprotroph (Põlme et al. 2020); Decay sugar-rich substrates (Põlme et al. 2020) | Põlme S, et al. 2020. Fungal Diversity 105:1-16 ((https://doi.org/10.1007/s13225-020-00466-2)) |
| Otu44165 | 1 | k__Fungi(100);p__Ascomycota(100);c__Saccharomycetes(100);o__Saccharomycetales(100);f__Metschnikowiaceae(100);g__Metschnikowia(100);s__Metschnikowia_pulcherrima(100); | Metschnikowia | | 13 | Saprotroph | \|Undefined Saprotroph\| | Yeast | NULL | Probable | As nectar-tap_saprotroph (Põlme et al. 2020); Decay sugar-rich substrates (Põlme et al. 2020) | Põlme S, et al. 2020. Fungal Diversity 105:1-16 ((https://doi.org/10.1007/s13225-020-00466-2)) |
| Otu44166 | 1 | k__Fungi(100);p__Ascomycota(100);c__Saccharomycetes(100);o__Saccharomycetales(100);f__Metschnikowiaceae(100);g__Metschnikowia(100);g__Metschnikowia_unclassified(100); | Metschnikowia | | 13 | Saprotroph | \|Undefined Saprotroph\| | Yeast | NULL | Probable | As nectar-tap_saprotroph (Põlme et al. 2020); Decay sugar-rich substrates (Põlme et al. 2020) | Põlme S, et al. 2020. Fungal Diversity 105:1-16 ((https://doi.org/10.1007/s13225-020-00466-2)) |
| Otu44167 | 1 | k__Fungi(100);p__Ascomycota(100);c__Saccharomycetes(100);o__Saccharomycetales(100);f__Metschnikowiaceae(100);g__Metschnikowia(100);g__Metschnikowia_unclassified(100); | Metschnikowia | | 13 | Saprotroph | \|Undefined Saprotroph\| | Yeast | NULL | Probable | As nectar-tap_saprotroph (Põlme et al. 2020); Decay sugar-rich substrates (Põlme et al. 2020) | Põlme S, et al. 2020. Fungal Diversity 105:1-16 ((https://doi.org/10.1007/s13225-020-00466-2)) |
| Otu44168 | 1 | k__Fungi(100);p__Ascomycota(100);c__Saccharomycetes(100);o__Saccharomycetales(100);f__Metschnikowiaceae(100);g__Metschnikowia(100);g__Metschnikowia_unclassified(100); | Metschnikowia | | 13 | Saprotroph | \|Undefined Saprotroph\| | Yeast | NULL | Probable | As nectar-tap_saprotroph (Põlme et al. 2020); Decay sugar-rich substrates (Põlme et al. 2020) | Põlme S, et al. 2020. Fungal Diversity 105:1-16 ((https://doi.org/10.1007/s13225-020-00466-2)) |
| Otu44169 | 1 | k__Fungi(100);p__Ascomycota(100);c__Saccharomycetes(100);o__Saccharomycetales(100);f__Metschnikowiaceae(100);g__Metschnikowia(100);g__Metschnikowia_unclassified(100); | Metschnikowia | | 13 | Saprotroph | \|Undefined Saprotroph\| | Yeast | NULL | Probable | As nectar-tap_saprotroph (Põlme et al. 2020); Decay sugar-rich substrates (Põlme et al. 2020) | Põlme S, et al. 2020. Fungal Diversity 105:1-16 ((https://doi.org/10.1007/s13225-020-00466-2)) |
| Otu44170 | 1 | k__Fungi(100);p__Ascomycota(100);c__Saccharomycetes(100);o__Saccharomycetales(100);f__Metschnikowiaceae(100);g__Metschnikowia(100);g__Metschnikowia_unclassified(100); | Metschnikowia | | 13 | Saprotroph | \|Undefined Saprotroph\| | Yeast | NULL | Probable | As nectar-tap_saprotroph (Põlme et al. 2020); Decay sugar-rich substrates (Põlme et al. 2020) | Põlme S, et al. 2020. Fungal Diversity 105:1-16 ((https://doi.org/10.1007/s13225-020-00466-2)) |
| Otu44171 | 1 | k__Fungi(100);p__Ascomycota(100);c__Saccharomycetes(100);o__Saccharomycetales(100);f__Metschnikowiaceae(100);g__Metschnikowia(100);s__Metschnikowia_pulcherrima(100); | Metschnikowia | | 13 | Saprotroph | \|Undefined Saprotroph\| | Yeast | NULL | Probable | As nectar-tap_saprotroph (Põlme et al. 2020); Decay sugar-rich substrates (Põlme et al. 2020) | Põlme S, et al. 2020. Fungal Diversity 105:1-16 ((https://doi.org/10.1007/s13225-020-00466-2)) |
| Otu44172 | 1 | k__Fungi(100);p__Ascomycota(100);c__Saccharomycetes(100);o__Saccharomycetales(100);f__Metschnikowiaceae(100);g__Metschnikowia(100);s__Metschnikowia_pulcherrima(100); | Metschnikowia | | 13 | Saprotroph | \|Undefined Saprotroph\| | Yeast | NULL | Probable | As nectar-tap_saprotroph (Põlme et al. 2020); Decay sugar-rich substrates (Põlme et al. 2020) | Põlme S, et al. 2020. Fungal Diversity 105:1-16 ((https://doi.org/10.1007/s13225-020-00466-2)) |
| Otu44173 | 1 | k__Fungi(100);p__Ascomycota(100);c__Saccharomycetes(100);o__Saccharomycetales(100);f__Metschnikowiaceae(100);g__Metschnikowia(100);g__Metschnikowia_unclassified(100); | Metschnikowia | | 13 | Saprotroph | \|Undefined Saprotroph\| | Yeast | NULL | Probable | As nectar-tap_saprotroph (Põlme et al. 2020); Decay sugar-rich substrates (Põlme et al. 2020) | Põlme S, et al. 2020. Fungal Diversity 105:1-16 ((https://doi.org/10.1007/s13225-020-00466-2)) |
| Otu44174 | 1 | k__Fungi(100);p__Ascomycota(100);c__Saccharomycetes(100);o__Saccharomycetales(100);f__Metschnikowiaceae(100);g__Metschnikowia(100);s__Metschnikowia_pulcherrima(100); | Metschnikowia | | 13 | Saprotroph | \|Undefined Saprotroph\| | Yeast | NULL | Probable | As nectar-tap_saprotroph (Põlme et al. 2020); Decay sugar-rich substrates (Põlme et al. 2020) | Põlme S, et al. 2020. Fungal Diversity 105:1-16 ((https://doi.org/10.1007/s13225-020-00466-2)) |
| Otu44175 | 1 | k__Fungi(100);p__Ascomycota(100);c__Saccharomycetes(100);o__Saccharomycetales(100);f__Metschnikowiaceae(100);g__Metschnikowia(100);s__Metschnikowia_pulcherrima(100); | Metschnikowia | | 13 | Saprotroph | \|Undefined Saprotroph\| | Yeast | NULL | Probable | As nectar-tap_saprotroph (Põlme et al. 2020); Decay sugar-rich substrates (Põlme et al. 2020) | Põlme S, et al. 2020. Fungal Diversity 105:1-16 ((https://doi.org/10.1007/s13225-020-00466-2)) |
| Otu44177 | 1 | k__Fungi(100);p__Ascomycota(100);c__Saccharomycetes(100);o__Saccharomycetales(100);f__Metschnikowiaceae(100);g__Metschnikowia(100);s__Metschnikowia_pulcherrima(100); | Metschnikowia | | 13 | Saprotroph | \|Undefined Saprotroph\| | Yeast | NULL | Probable | As nectar-tap_saprotroph (Põlme et al. 2020); Decay sugar-rich substrates (Põlme et al. 2020) | Põlme S, et al. 2020. Fungal Diversity 105:1-16 ((https://doi.org/10.1007/s13225-020-00466-2)) |
| Otu44178 | 1 | k__Fungi(100);p__Ascomycota(100);c__Saccharomycetes(100);o__Saccharomycetales(100);f__Metschnikowiaceae(100);g__Metschnikowia(100);s__Metschnikowia_pulcherrima(100); | Metschnikowia | | 13 | Saprotroph | \|Undefined Saprotroph\| | Yeast | NULL | Probable | As nectar-tap_saprotroph (Põlme et al. 2020); Decay sugar-rich substrates (Põlme et al. 2020) | Põlme S, et al. 2020. Fungal Diversity 105:1-16 ((https://doi.org/10.1007/s13225-020-00466-2)) |
| Otu44179 | 1 | k__Fungi(100);p__Ascomycota(100);c__Saccharomycetes(100);o__Saccharomycetales(100);f__Metschnikowiaceae(100);g__Metschnikowia(100);s__Metschnikowia_pulcherrima(100); | Metschnikowia | | 13 | Saprotroph | \|Undefined Saprotroph\| | Yeast | NULL | Probable | As nectar-tap_saprotroph (Põlme et al. 2020); Decay sugar-rich substrates (Põlme et al. 2020) | Põlme S, et al. 2020. Fungal Diversity 105:1-16 ((https://doi.org/10.1007/s13225-020-00466-2)) |
| Otu44180 | 1 | k__Fungi(100);p__Ascomycota(100);c__Saccharomycetes(100);o__Saccharomycetales(100);f__Metschnikowiaceae(100);g__Metschnikowia(100);s__Metschnikowia_pulcherrima(100); | Metschnikowia | | 13 | Saprotroph | \|Undefined Saprotroph\| | Yeast | NULL | Probable | As nectar-tap_saprotroph (Põlme et al. 2020); Decay sugar-rich substrates (Põlme et al. 2020) | Põlme S, et al. 2020. Fungal Diversity 105:1-16 ((https://doi.org/10.1007/s13225-020-00466-2)) |
| Otu44181 | 1 | k__Fungi(100);p__Ascomycota(100);c__Saccharomycetes(100);o__Saccharomycetales(100);f__Metschnikowiaceae(100);g__Metschnikowia(100);s__Metschnikowia_pulcherrima(100); | Metschnikowia | | 13 | Saprotroph | \|Undefined Saprotroph\| | Yeast | NULL | Probable | As nectar-tap_saprotroph (Põlme et al. 2020); Decay sugar-rich substrates (Põlme et al. 2020) | Põlme S, et al. 2020. Fungal Diversity 105:1-16 ((https://doi.org/10.1007/s13225-020-00466-2)) |
| Otu44182 | 1 | k__Fungi(100);p__Ascomycota(100);c__Saccharomycetes(100);o__Saccharomycetales(100);f__Metschnikowiaceae(100);g__Metschnikowia(100);g__Metschnikowia_unclassified(100); | Metschnikowia | | 13 | Saprotroph | \|Undefined Saprotroph\| | Yeast | NULL | Probable | As nectar-tap_saprotroph (Põlme et al. 2020); Decay sugar-rich substrates (Põlme et al. 2020) | Põlme S, et al. 2020. Fungal Diversity 105:1-16 ((https://doi.org/10.1007/s13225-020-00466-2)) |
| Otu44184 | 1 | k__Fungi(100);p__Ascomycota(100);c__Saccharomycetes(100);o__Saccharomycetales(100);f__Metschnikowiaceae(100);g__Metschnikowia(100);s__Metschnikowia_pulcherrima(100); | Metschnikowia | | 13 | Saprotroph | \|Undefined Saprotroph\| | Yeast | NULL | Probable | As nectar-tap_saprotroph (Põlme et al. 2020); Decay sugar-rich substrates (Põlme et al. 2020) | Põlme S, et al. 2020. Fungal Diversity 105:1-16 ((https://doi.org/10.1007/s13225-020-00466-2)) |
| Otu44185 | 1 | k__Fungi(100);p__Ascomycota(100);c__Saccharomycetes(100);o__Saccharomycetales(100);f__Metschnikowiaceae(100);g__Metschnikowia(100);s__Metschnikowia_pulcherrima(100); | Metschnikowia | | 13 | Saprotroph | \|Undefined Saprotroph\| | Yeast | NULL | Probable | As nectar-tap_saprotroph (Põlme et al. 2020); Decay sugar-rich substrates (Põlme et al. 2020) | Põlme S, et al. 2020. Fungal Diversity 105:1-16 ((https://doi.org/10.1007/s13225-020-00466-2)) |
| Otu44186 | 1 | k__Fungi(100);p__Ascomycota(100);c__Saccharomycetes(100);o__Saccharomycetales(100);f__Metschnikowiaceae(100);g__Metschnikowia(100);s__Metschnikowia_pulcherrima(100); | Metschnikowia | | 13 | Saprotroph | \|Undefined Saprotroph\| | Yeast | NULL | Probable | As nectar-tap_saprotroph (Põlme et al. 2020); Decay sugar-rich substrates (Põlme et al. 2020) | Põlme S, et al. 2020. Fungal Diversity 105:1-16 ((https://doi.org/10.1007/s13225-020-00466-2)) |
| Otu44187 | 1 | k__Fungi(100);p__Ascomycota(100);c__Saccharomycetes(100);o__Saccharomycetales(100);f__Metschnikowiaceae(100);g__Metschnikowia(100);s__Metschnikowia_pulcherrima(100); | Metschnikowia | | 13 | Saprotroph | \|Undefined Saprotroph\| | Yeast | NULL | Probable | As nectar-tap_saprotroph (Põlme et al. 2020); Decay sugar-rich substrates (Põlme et al. 2020) | Põlme S, et al. 2020. Fungal Diversity 105:1-16 ((https://doi.org/10.1007/s13225-020-00466-2)) |
| Otu44188 | 1 | k__Fungi(100);p__Ascomycota(100);c__Saccharomycetes(100);o__Saccharomycetales(100);f__Metschnikowiaceae(100);g__Metschnikowia(100);s__Metschnikowia_pulcherrima(100); | Metschnikowia | | 13 | Saprotroph | \|Undefined Saprotroph\| | Yeast | NULL | Probable | As nectar-tap_saprotroph (Põlme et al. 2020); Decay sugar-rich substrates (Põlme et al. 2020) | Põlme S, et al. 2020. Fungal Diversity 105:1-16 ((https://doi.org/10.1007/s13225-020-00466-2)) |
| Otu44189 | 1 | k__Fungi(100);p__Ascomycota(100);c__Saccharomycetes(100);o__Saccharomycetales(100);f__Metschnikowiaceae(100);g__Metschnikowia(100);s__Metschnikowia_pulcherrima(100); | Metschnikowia | | 13 | Saprotroph | \|Undefined Saprotroph\| | Yeast | NULL | Probable | As nectar-tap_saprotroph (Põlme et al. 2020); Decay sugar-rich substrates (Põlme et al. 2020) | Põlme S, et al. 2020. Fungal Diversity 105:1-16 ((https://doi.org/10.1007/s13225-020-00466-2)) |
| Otu44190 | 1 | k__Fungi(100);p__Ascomycota(100);c__Saccharomycetes(100);o__Saccharomycetales(100);f__Metschnikowiaceae(100);g__Metschnikowia(100);g__Metschnikowia_unclassified(100); | Metschnikowia | | 13 | Saprotroph | \|Undefined Saprotroph\| | Yeast | NULL | Probable | As nectar-tap_saprotroph (Põlme et al. 2020); Decay sugar-rich substrates (Põlme et al. 2020) | Põlme S, et al. 2020. Fungal Diversity 105:1-16 ((https://doi.org/10.1007/s13225-020-00466-2)) |
| Otu44191 | 1 | k__Fungi(100);p__Ascomycota(100);c__Saccharomycetes(100);o__Saccharomycetales(100);f__Metschnikowiaceae(100);g__Metschnikowia(100);s__Metschnikowia_pulcherrima(100); | Metschnikowia | | 13 | Saprotroph | \|Undefined Saprotroph\| | Yeast | NULL | Probable | As nectar-tap_saprotroph (Põlme et al. 2020); Decay sugar-rich substrates (Põlme et al. 2020) | Põlme S, et al. 2020. Fungal Diversity 105:1-16 ((https://doi.org/10.1007/s13225-020-00466-2)) |
| Otu44192 | 1 | k__Fungi(100);p__Ascomycota(100);c__Saccharomycetes(100);o__Saccharomycetales(100);f__Metschnikowiaceae(100);g__Metschnikowia(100);s__Metschnikowia_pulcherrima(100); | Metschnikowia | | 13 | Saprotroph | \|Undefined Saprotroph\| | Yeast | NULL | Probable | As nectar-tap_saprotroph (Põlme et al. 2020); Decay sugar-rich substrates (Põlme et al. 2020) | Põlme S, et al. 2020. Fungal Diversity 105:1-16 ((https://doi.org/10.1007/s13225-020-00466-2)) |
| Otu44193 | 1 | k__Fungi(100);p__Ascomycota(100);c__Saccharomycetes(100);o__Saccharomycetales(100);f__Metschnikowiaceae(100);g__Metschnikowia(100);s__Metschnikowia_pulcherrima(100); | Metschnikowia | | 13 | Saprotroph | \|Undefined Saprotroph\| | Yeast | NULL | Probable | As nectar-tap_saprotroph (Põlme et al. 2020); Decay sugar-rich substrates (Põlme et al. 2020) | Põlme S, et al. 2020. Fungal Diversity 105:1-16 ((https://doi.org/10.1007/s13225-020-00466-2)) |
| Otu44194 | 1 | k__Fungi(100);p__Ascomycota(100);c__Saccharomycetes(100);o__Saccharomycetales(100);f__Metschnikowiaceae(100);g__Metschnikowia(100);s__Metschnikowia_pulcherrima(100); | Metschnikowia | | 13 | Saprotroph | \|Undefined Saprotroph\| | Yeast | NULL | Probable | As nectar-tap_saprotroph (Põlme et al. 2020); Decay sugar-rich substrates (Põlme et al. 2020) | Põlme S, et al. 2020. Fungal Diversity 105:1-16 ((https://doi.org/10.1007/s13225-020-00466-2)) |
| Otu44195 | 1 | k__Fungi(100);p__Ascomycota(100);c__Saccharomycetes(100);o__Saccharomycetales(100);f__Metschnikowiaceae(100);g__Metschnikowia(100);s__Metschnikowia_pulcherrima(100); | Metschnikowia | | 13 | Saprotroph | \|Undefined Saprotroph\| | Yeast | NULL | Probable | As nectar-tap_saprotroph (Põlme et al. 2020); Decay sugar-rich substrates (Põlme et al. 2020) | Põlme S, et al. 2020. Fungal Diversity 105:1-16 ((https://doi.org/10.1007/s13225-020-00466-2)) |
| Otu44196 | 1 | k__Fungi(100);p__Ascomycota(100);c__Saccharomycetes(100);o__Saccharomycetales(100);f__Metschnikowiaceae(100);g__Metschnikowia(100);s__Metschnikowia_pulcherrima(100); | Metschnikowia | | 13 | Saprotroph | \|Undefined Saprotroph\| | Yeast | NULL | Probable | As nectar-tap_saprotroph (Põlme et al. 2020); Decay sugar-rich substrates (Põlme et al. 2020) | Põlme S, et al. 2020. Fungal Diversity 105:1-16 ((https://doi.org/10.1007/s13225-020-00466-2)) |
| Otu44197 | 1 | k__Fungi(100);p__Ascomycota(100);c__Saccharomycetes(100);o__Saccharomycetales(100);f__Metschnikowiaceae(100);g__Metschnikowia(100);g__Metschnikowia_unclassified(100); | Metschnikowia | | 13 | Saprotroph | \|Undefined Saprotroph\| | Yeast | NULL | Probable | As nectar-tap_saprotroph (Põlme et al. 2020); Decay sugar-rich substrates (Põlme et al. 2020) | Põlme S, et al. 2020. Fungal Diversity 105:1-16 ((https://doi.org/10.1007/s13225-020-00466-2)) |
| Otu44198 | 1 | k__Fungi(100);p__Ascomycota(100);c__Saccharomycetes(100);o__Saccharomycetales(100);f__Metschnikowiaceae(100);g__Metschnikowia(100);g__Metschnikowia_unclassified(100); | Metschnikowia | | 13 | Saprotroph | \|Undefined Saprotroph\| | Yeast | NULL | Probable | As nectar-tap_saprotroph (Põlme et al. 2020); Decay sugar-rich substrates (Põlme et al. 2020) | Põlme S, et al. 2020. Fungal Diversity 105:1-16 ((https://doi.org/10.1007/s13225-020-00466-2)) |
| Otu44199 | 1 | k__Fungi(100);p__Ascomycota(100);c__Saccharomycetes(100);o__Saccharomycetales(100);f__Metschnikowiaceae(100);g__Metschnikowia(100);g__Metschnikowia_unclassified(100); | Metschnikowia | | 13 | Saprotroph | \|Undefined Saprotroph\| | Yeast | NULL | Probable | As nectar-tap_saprotroph (Põlme et al. 2020); Decay sugar-rich substrates (Põlme et al. 2020) | Põlme S, et al. 2020. Fungal Diversity 105:1-16 ((https://doi.org/10.1007/s13225-020-00466-2)) |
| Otu44200 | 1 | k__Fungi(100);p__Ascomycota(100);c__Saccharomycetes(100);o__Saccharomycetales(100);f__Metschnikowiaceae(100);g__Metschnikowia(100);s__Metschnikowia_pulcherrima(100); | Metschnikowia | | 13 | Saprotroph | \|Undefined Saprotroph\| | Yeast | NULL | Probable | As nectar-tap_saprotroph (Põlme et al. 2020); Decay sugar-rich substrates (Põlme et al. 2020) | Põlme S, et al. 2020. Fungal Diversity 105:1-16 ((https://doi.org/10.1007/s13225-020-00466-2)) |
| Otu44201 | 1 | k__Fungi(100);p__Ascomycota(100);c__Saccharomycetes(100);o__Saccharomycetales(100);f__Metschnikowiaceae(100);g__Metschnikowia(100);s__Metschnikowia_pulcherrima(100); | Metschnikowia | | 13 | Saprotroph | \|Undefined Saprotroph\| | Yeast | NULL | Probable | As nectar-tap_saprotroph (Põlme et al. 2020); Decay sugar-rich substrates (Põlme et al. 2020) | Põlme S, et al. 2020. Fungal Diversity 105:1-16 ((https://doi.org/10.1007/s13225-020-00466-2)) |
| Otu44202 | 1 | k__Fungi(100);p__Ascomycota(100);c__Sordariomycetes(100);o__Hypocreales(100);f__Nectriaceae(100);g__Fusarium(100);g__Fusarium_unclassified(100); | Fusarium | | 13 | Pathotroph-Saprotroph-Symbiotroph | Animal Pathogen-Endophyte-Lichen Parasite-\|Plant Pathogen\|-Plant Saprotroph-Undefined Saprotroph-Wood Saprotroph | Microfungus | Soft Rot | Possible | As foliar_endophyte (Põlme et al. 2020); As litter_saprotroph (Põlme et al. 2020); Causing damping-off disease (Põlme et al. 2020); Decay leaf, fruit, seed, roots, soil, animal material (Põlme et al. 2020); Likely opportunistic human pathogen (Irinyi et al. 2015) | Duncan CG, Eslyn WE. 1966. Mycologia 58:642-664 ((https://doi.org/10.2307/3757045)); Seehann G, et al. 1975. List of Fungi in Soft-Rot Tissue Tests. Institute fur Holzbiologie und Holzschutz, Bundesforschungsanstalt fur Forst - und Holzwirtshaft, Hamburg - Lohbrugge, Germany; Costa IPMW, et al. 2012. Mycotaxon 119:e26 ((http://mycotaxon.com/resources/checklists/costa_v119_checklist.pdf)); Tedersoo L, et al. 2014. Science 346:e1256688 ((https://doi.org/10.1126/science.1256688)); Irinyi L, et al. 2015. Medical Mycology 53:313-337 ((https://doi.org/10.1093/mmy/myv008)); Schroers, et al. 2016. Mycologia 108:806-819 ((https://doi.org/10.3852/15-255)); Põlme S, et al. 2020. Fungal Diversity 105:1-16 ((https://doi.org/10.1007/s13225-020-00466-2)); The American Phytopathological Society Common Names of Plant Diseases ((http://www.apsnet.org/publications/commonnames/Pages/default.aspx)); Lawrey JD, Diederich P. 2016. http://www.lichenicolous.net (accessed 24 May 2017) ((http://www.lichenicolous.net)) |
| Otu44203 | 1 | k__Fungi(100);p__Ascomycota(100);c__Saccharomycetes(100);o__Saccharomycetales(100);f__Metschnikowiaceae(100);g__Metschnikowia(100);g__Metschnikowia_unclassified(100); | Metschnikowia | | 13 | Saprotroph | \|Undefined Saprotroph\| | Yeast | NULL | Probable | As nectar-tap_saprotroph (Põlme et al. 2020); Decay sugar-rich substrates (Põlme et al. 2020) | Põlme S, et al. 2020. Fungal Diversity 105:1-16 ((https://doi.org/10.1007/s13225-020-00466-2)) |
| Otu44204 | 1 | k__Fungi(100);p__Ascomycota(100);c__Saccharomycetes(100);o__Saccharomycetales(100);f__Metschnikowiaceae(100);g__Metschnikowia(100);s__Metschnikowia_pulcherrima(100); | Metschnikowia | | 13 | Saprotroph | \|Undefined Saprotroph\| | Yeast | NULL | Probable | As nectar-tap_saprotroph (Põlme et al. 2020); Decay sugar-rich substrates (Põlme et al. 2020) | Põlme S, et al. 2020. Fungal Diversity 105:1-16 ((https://doi.org/10.1007/s13225-020-00466-2)) |
| Otu44205 | 1 | k__Fungi(100);p__Ascomycota(100);c__Saccharomycetes(100);o__Saccharomycetales(100);f__Metschnikowiaceae(100);g__Metschnikowia(100);s__Metschnikowia_pulcherrima(100); | Metschnikowia | | 13 | Saprotroph | \|Undefined Saprotroph\| | Yeast | NULL | Probable | As nectar-tap_saprotroph (Põlme et al. 2020); Decay sugar-rich substrates (Põlme et al. 2020) | Põlme S, et al. 2020. Fungal Diversity 105:1-16 ((https://doi.org/10.1007/s13225-020-00466-2)) |
| Otu44206 | 1 | k__Fungi(100);p__Ascomycota(100);c__Saccharomycetes(100);o__Saccharomycetales(100);f__Metschnikowiaceae(100);g__Metschnikowia(100);s__Metschnikowia_pulcherrima(100); | Metschnikowia | | 13 | Saprotroph | \|Undefined Saprotroph\| | Yeast | NULL | Probable | As nectar-tap_saprotroph (Põlme et al. 2020); Decay sugar-rich substrates (Põlme et al. 2020) | Põlme S, et al. 2020. Fungal Diversity 105:1-16 ((https://doi.org/10.1007/s13225-020-00466-2)) |
| Otu44207 | 1 | k__Fungi(100);p__Ascomycota(100);c__Saccharomycetes(100);o__Saccharomycetales(100);f__Metschnikowiaceae(100);g__Metschnikowia(100);g__Metschnikowia_unclassified(100); | Metschnikowia | | 13 | Saprotroph | \|Undefined Saprotroph\| | Yeast | NULL | Probable | As nectar-tap_saprotroph (Põlme et al. 2020); Decay sugar-rich substrates (Põlme et al. 2020) | Põlme S, et al. 2020. Fungal Diversity 105:1-16 ((https://doi.org/10.1007/s13225-020-00466-2)) |
| Otu44208 | 1 | k__Fungi(100);p__Ascomycota(100);c__Saccharomycetes(100);o__Saccharomycetales(100);f__Metschnikowiaceae(100);g__Metschnikowia(100);s__Metschnikowia_pulcherrima(100); | Metschnikowia | | 13 | Saprotroph | \|Undefined Saprotroph\| | Yeast | NULL | Probable | As nectar-tap_saprotroph (Põlme et al. 2020); Decay sugar-rich substrates (Põlme et al. 2020) | Põlme S, et al. 2020. Fungal Diversity 105:1-16 ((https://doi.org/10.1007/s13225-020-00466-2)) |
| Otu44209 | 1 | k__Fungi(100);p__Ascomycota(100);c__Saccharomycetes(100);o__Saccharomycetales(100);f__Metschnikowiaceae(100);g__Metschnikowia(100);s__Metschnikowia_pulcherrima(100); | Metschnikowia | | 13 | Saprotroph | \|Undefined Saprotroph\| | Yeast | NULL | Probable | As nectar-tap_saprotroph (Põlme et al. 2020); Decay sugar-rich substrates (Põlme et al. 2020) | Põlme S, et al. 2020. Fungal Diversity 105:1-16 ((https://doi.org/10.1007/s13225-020-00466-2)) |
| Otu44210 | 1 | k__Fungi(100);p__Ascomycota(100);c__Saccharomycetes(100);o__Saccharomycetales(100);f__Metschnikowiaceae(100);g__Metschnikowia(100);s__Metschnikowia_pulcherrima(100); | Metschnikowia | | 13 | Saprotroph | \|Undefined Saprotroph\| | Yeast | NULL | Probable | As nectar-tap_saprotroph (Põlme et al. 2020); Decay sugar-rich substrates (Põlme et al. 2020) | Põlme S, et al. 2020. Fungal Diversity 105:1-16 ((https://doi.org/10.1007/s13225-020-00466-2)) |
| Otu44211 | 1 | k__Fungi(100);p__Ascomycota(100);c__Saccharomycetes(100);o__Saccharomycetales(100);f__Metschnikowiaceae(100);g__Metschnikowia(100);g__Metschnikowia_unclassified(100); | Metschnikowia | | 13 | Saprotroph | \|Undefined Saprotroph\| | Yeast | NULL | Probable | As nectar-tap_saprotroph (Põlme et al. 2020); Decay sugar-rich substrates (Põlme et al. 2020) | Põlme S, et al. 2020. Fungal Diversity 105:1-16 ((https://doi.org/10.1007/s13225-020-00466-2)) |
| Otu44212 | 1 | k__Fungi(100);p__Ascomycota(100);c__Saccharomycetes(100);o__Saccharomycetales(100);f__Metschnikowiaceae(100);g__Metschnikowia(100);s__Metschnikowia_pulcherrima(100); | Metschnikowia | | 13 | Saprotroph | \|Undefined Saprotroph\| | Yeast | NULL | Probable | As nectar-tap_saprotroph (Põlme et al. 2020); Decay sugar-rich substrates (Põlme et al. 2020) | Põlme S, et al. 2020. Fungal Diversity 105:1-16 ((https://doi.org/10.1007/s13225-020-00466-2)) |
| Otu44213 | 1 | k__Fungi(100);p__Ascomycota(100);c__Saccharomycetes(100);o__Saccharomycetales(100);f__Metschnikowiaceae(100);g__Metschnikowia(100);s__Metschnikowia_pulcherrima(100); | Metschnikowia | | 13 | Saprotroph | \|Undefined Saprotroph\| | Yeast | NULL | Probable | As nectar-tap_saprotroph (Põlme et al. 2020); Decay sugar-rich substrates (Põlme et al. 2020) | Põlme S, et al. 2020. Fungal Diversity 105:1-16 ((https://doi.org/10.1007/s13225-020-00466-2)) |
| Otu44214 | 1 | k__Fungi(100);p__Ascomycota(100);c__Saccharomycetes(100);o__Saccharomycetales(100);f__Metschnikowiaceae(100);g__Metschnikowia(100);s__Metschnikowia_pulcherrima(100); | Metschnikowia | | 13 | Saprotroph | \|Undefined Saprotroph\| | Yeast | NULL | Probable | As nectar-tap_saprotroph (Põlme et al. 2020); Decay sugar-rich substrates (Põlme et al. 2020) | Põlme S, et al. 2020. Fungal Diversity 105:1-16 ((https://doi.org/10.1007/s13225-020-00466-2)) |
| Otu44215 | 1 | k__Fungi(100);p__Ascomycota(100);c__Saccharomycetes(100);o__Saccharomycetales(100);f__Metschnikowiaceae(100);g__Metschnikowia(100);g__Metschnikowia_unclassified(100); | Metschnikowia | | 13 | Saprotroph | \|Undefined Saprotroph\| | Yeast | NULL | Probable | As nectar-tap_saprotroph (Põlme et al. 2020); Decay sugar-rich substrates (Põlme et al. 2020) | Põlme S, et al. 2020. Fungal Diversity 105:1-16 ((https://doi.org/10.1007/s13225-020-00466-2)) |
| Otu44216 | 1 | k__Fungi(100);p__Ascomycota(100);c__Saccharomycetes(100);o__Saccharomycetales(100);f__Metschnikowiaceae(100);g__Metschnikowia(100);s__Metschnikowia_pulcherrima(100); | Metschnikowia | | 13 | Saprotroph | \|Undefined Saprotroph\| | Yeast | NULL | Probable | As nectar-tap_saprotroph (Põlme et al. 2020); Decay sugar-rich substrates (Põlme et al. 2020) | Põlme S, et al. 2020. Fungal Diversity 105:1-16 ((https://doi.org/10.1007/s13225-020-00466-2)) |
| Otu44217 | 1 | k__Fungi(100);p__Ascomycota(100);c__Saccharomycetes(100);o__Saccharomycetales(100);f__Metschnikowiaceae(100);g__Metschnikowia(100);g__Metschnikowia_unclassified(100); | Metschnikowia | | 13 | Saprotroph | \|Undefined Saprotroph\| | Yeast | NULL | Probable | As nectar-tap_saprotroph (Põlme et al. 2020); Decay sugar-rich substrates (Põlme et al. 2020) | Põlme S, et al. 2020. Fungal Diversity 105:1-16 ((https://doi.org/10.1007/s13225-020-00466-2)) |
| Otu44218 | 1 | k__Fungi(100);p__Ascomycota(100);c__Saccharomycetes(100);o__Saccharomycetales(100);f__Metschnikowiaceae(100);g__Metschnikowia(100);s__Metschnikowia_pulcherrima(100); | Metschnikowia | | 13 | Saprotroph | \|Undefined Saprotroph\| | Yeast | NULL | Probable | As nectar-tap_saprotroph (Põlme et al. 2020); Decay sugar-rich substrates (Põlme et al. 2020) | Põlme S, et al. 2020. Fungal Diversity 105:1-16 ((https://doi.org/10.1007/s13225-020-00466-2)) |
| Otu44219 | 1 | k__Fungi(100);p__Ascomycota(100);c__Saccharomycetes(100);o__Saccharomycetales(100);f__Metschnikowiaceae(100);g__Metschnikowia(100);s__Metschnikowia_pulcherrima(100); | Metschnikowia | | 13 | Saprotroph | \|Undefined Saprotroph\| | Yeast | NULL | Probable | As nectar-tap_saprotroph (Põlme et al. 2020); Decay sugar-rich substrates (Põlme et al. 2020) | Põlme S, et al. 2020. Fungal Diversity 105:1-16 ((https://doi.org/10.1007/s13225-020-00466-2)) |
| Otu44220 | 1 | k__Fungi(100);p__Ascomycota(100);c__Saccharomycetes(100);o__Saccharomycetales(100);f__Metschnikowiaceae(100);g__Metschnikowia(100);s__Metschnikowia_pulcherrima(100); | Metschnikowia | | 13 | Saprotroph | \|Undefined Saprotroph\| | Yeast | NULL | Probable | As nectar-tap_saprotroph (Põlme et al. 2020); Decay sugar-rich substrates (Põlme et al. 2020) | Põlme S, et al. 2020. Fungal Diversity 105:1-16 ((https://doi.org/10.1007/s13225-020-00466-2)) |
| Otu44221 | 1 | k__Fungi(100);p__Ascomycota(100);c__Saccharomycetes(100);o__Saccharomycetales(100);f__Metschnikowiaceae(100);g__Metschnikowia(100);s__Metschnikowia_pulcherrima(100); | Metschnikowia | | 13 | Saprotroph | \|Undefined Saprotroph\| | Yeast | NULL | Probable | As nectar-tap_saprotroph (Põlme et al. 2020); Decay sugar-rich substrates (Põlme et al. 2020) | Põlme S, et al. 2020. Fungal Diversity 105:1-16 ((https://doi.org/10.1007/s13225-020-00466-2)) |
| Otu44222 | 1 | k__Fungi(100);p__Ascomycota(100);c__Saccharomycetes(100);o__Saccharomycetales(100);f__Metschnikowiaceae(100);g__Metschnikowia(100);s__Metschnikowia_pulcherrima(100); | Metschnikowia | | 13 | Saprotroph | \|Undefined Saprotroph\| | Yeast | NULL | Probable | As nectar-tap_saprotroph (Põlme et al. 2020); Decay sugar-rich substrates (Põlme et al. 2020) | Põlme S, et al. 2020. Fungal Diversity 105:1-16 ((https://doi.org/10.1007/s13225-020-00466-2)) |
| Otu44223 | 1 | k__Fungi(100);p__Ascomycota(100);c__Saccharomycetes(100);o__Saccharomycetales(100);f__Metschnikowiaceae(100);g__Metschnikowia(100);s__Metschnikowia_pulcherrima(100); | Metschnikowia | | 13 | Saprotroph | \|Undefined Saprotroph\| | Yeast | NULL | Probable | As nectar-tap_saprotroph (Põlme et al. 2020); Decay sugar-rich substrates (Põlme et al. 2020) | Põlme S, et al. 2020. Fungal Diversity 105:1-16 ((https://doi.org/10.1007/s13225-020-00466-2)) |
| Otu44224 | 1 | k__Fungi(100);p__Ascomycota(100);c__Saccharomycetes(100);o__Saccharomycetales(100);f__Metschnikowiaceae(100);g__Metschnikowia(100);s__Metschnikowia_pulcherrima(100); | Metschnikowia | | 13 | Saprotroph | \|Undefined Saprotroph\| | Yeast | NULL | Probable | As nectar-tap_saprotroph (Põlme et al. 2020); Decay sugar-rich substrates (Põlme et al. 2020) | Põlme S, et al. 2020. Fungal Diversity 105:1-16 ((https://doi.org/10.1007/s13225-020-00466-2)) |
| Otu44225 | 1 | k__Fungi(100);p__Ascomycota(100);c__Saccharomycetes(100);o__Saccharomycetales(100);f__Metschnikowiaceae(100);g__Metschnikowia(100);s__Metschnikowia_pulcherrima(100); | Metschnikowia | | 13 | Saprotroph | \|Undefined Saprotroph\| | Yeast | NULL | Probable | As nectar-tap_saprotroph (Põlme et al. 2020); Decay sugar-rich substrates (Põlme et al. 2020) | Põlme S, et al. 2020. Fungal Diversity 105:1-16 ((https://doi.org/10.1007/s13225-020-00466-2)) |
| Otu44226 | 1 | k__Fungi(100);p__Ascomycota(100);c__Saccharomycetes(100);o__Saccharomycetales(100);f__Metschnikowiaceae(100);g__Metschnikowia(100);s__Metschnikowia_pulcherrima(100); | Metschnikowia | | 13 | Saprotroph | \|Undefined Saprotroph\| | Yeast | NULL | Probable | As nectar-tap_saprotroph (Põlme et al. 2020); Decay sugar-rich substrates (Põlme et al. 2020) | Põlme S, et al. 2020. Fungal Diversity 105:1-16 ((https://doi.org/10.1007/s13225-020-00466-2)) |
| Otu44227 | 1 | k__Fungi(100);p__Ascomycota(100);c__Saccharomycetes(100);o__Saccharomycetales(100);f__Metschnikowiaceae(100);g__Metschnikowia(100);g__Metschnikowia_unclassified(100); | Metschnikowia | | 13 | Saprotroph | \|Undefined Saprotroph\| | Yeast | NULL | Probable | As nectar-tap_saprotroph (Põlme et al. 2020); Decay sugar-rich substrates (Põlme et al. 2020) | Põlme S, et al. 2020. Fungal Diversity 105:1-16 ((https://doi.org/10.1007/s13225-020-00466-2)) |
| Otu44228 | 1 | k__Fungi(100);p__Ascomycota(100);c__Saccharomycetes(100);o__Saccharomycetales(100);f__Metschnikowiaceae(100);g__Metschnikowia(100);g__Metschnikowia_unclassified(100); | Metschnikowia | | 13 | Saprotroph | \|Undefined Saprotroph\| | Yeast | NULL | Probable | As nectar-tap_saprotroph (Põlme et al. 2020); Decay sugar-rich substrates (Põlme et al. 2020) | Põlme S, et al. 2020. Fungal Diversity 105:1-16 ((https://doi.org/10.1007/s13225-020-00466-2)) |
| Otu44229 | 1 | k__Fungi(100);p__Ascomycota(100);c__Saccharomycetes(100);o__Saccharomycetales(100);f__Metschnikowiaceae(100);g__Metschnikowia(100);s__Metschnikowia_pulcherrima(100); | Metschnikowia | | 13 | Saprotroph | \|Undefined Saprotroph\| | Yeast | NULL | Probable | As nectar-tap_saprotroph (Põlme et al. 2020); Decay sugar-rich substrates (Põlme et al. 2020) | Põlme S, et al. 2020. Fungal Diversity 105:1-16 ((https://doi.org/10.1007/s13225-020-00466-2)) |
| Otu44230 | 1 | k__Fungi(100);p__Ascomycota(100);c__Saccharomycetes(100);o__Saccharomycetales(100);f__Metschnikowiaceae(100);g__Metschnikowia(100);s__Metschnikowia_pulcherrima(100); | Metschnikowia | | 13 | Saprotroph | \|Undefined Saprotroph\| | Yeast | NULL | Probable | As nectar-tap_saprotroph (Põlme et al. 2020); Decay sugar-rich substrates (Põlme et al. 2020) | Põlme S, et al. 2020. Fungal Diversity 105:1-16 ((https://doi.org/10.1007/s13225-020-00466-2)) |
| Otu44231 | 1 | k__Fungi(100);p__Ascomycota(100);c__Saccharomycetes(100);o__Saccharomycetales(100);f__Metschnikowiaceae(100);g__Metschnikowia(100);s__Metschnikowia_pulcherrima(100); | Metschnikowia | | 13 | Saprotroph | \|Undefined Saprotroph\| | Yeast | NULL | Probable | As nectar-tap_saprotroph (Põlme et al. 2020); Decay sugar-rich substrates (Põlme et al. 2020) | Põlme S, et al. 2020. Fungal Diversity 105:1-16 ((https://doi.org/10.1007/s13225-020-00466-2)) |
| Otu44232 | 1 | k__Fungi(100);p__Ascomycota(100);c__Saccharomycetes(100);o__Saccharomycetales(100);f__Metschnikowiaceae(100);g__Metschnikowia(100);g__Metschnikowia_unclassified(100); | Metschnikowia | | 13 | Saprotroph | \|Undefined Saprotroph\| | Yeast | NULL | Probable | As nectar-tap_saprotroph (Põlme et al. 2020); Decay sugar-rich substrates (Põlme et al. 2020) | Põlme S, et al. 2020. Fungal Diversity 105:1-16 ((https://doi.org/10.1007/s13225-020-00466-2)) |
| Otu44233 | 1 | k__Fungi(100);p__Ascomycota(100);c__Saccharomycetes(100);o__Saccharomycetales(100);f__Metschnikowiaceae(100);g__Metschnikowia(100);g__Metschnikowia_unclassified(100); | Metschnikowia | | 13 | Saprotroph | \|Undefined Saprotroph\| | Yeast | NULL | Probable | As nectar-tap_saprotroph (Põlme et al. 2020); Decay sugar-rich substrates (Põlme et al. 2020) | Põlme S, et al. 2020. Fungal Diversity 105:1-16 ((https://doi.org/10.1007/s13225-020-00466-2)) |
| Otu44234 | 1 | k__Fungi(100);p__Ascomycota(100);c__Saccharomycetes(100);o__Saccharomycetales(100);f__Metschnikowiaceae(100);g__Metschnikowia(100);g__Metschnikowia_unclassified(100); | Metschnikowia | | 13 | Saprotroph | \|Undefined Saprotroph\| | Yeast | NULL | Probable | As nectar-tap_saprotroph (Põlme et al. 2020); Decay sugar-rich substrates (Põlme et al. 2020) | Põlme S, et al. 2020. Fungal Diversity 105:1-16 ((https://doi.org/10.1007/s13225-020-00466-2)) |
| Otu44235 | 1 | k__Fungi(100);p__Ascomycota(100);c__Saccharomycetes(100);o__Saccharomycetales(100);f__Metschnikowiaceae(100);g__Metschnikowia(100);s__Metschnikowia_pulcherrima(100); | Metschnikowia | | 13 | Saprotroph | \|Undefined Saprotroph\| | Yeast | NULL | Probable | As nectar-tap_saprotroph (Põlme et al. 2020); Decay sugar-rich substrates (Põlme et al. 2020) | Põlme S, et al. 2020. Fungal Diversity 105:1-16 ((https://doi.org/10.1007/s13225-020-00466-2)) |
| Otu44236 | 1 | k__Fungi(100);p__Ascomycota(100);c__Saccharomycetes(100);o__Saccharomycetales(100);f__Metschnikowiaceae(100);g__Metschnikowia(100);s__Metschnikowia_pulcherrima(100); | Metschnikowia | | 13 | Saprotroph | \|Undefined Saprotroph\| | Yeast | NULL | Probable | As nectar-tap_saprotroph (Põlme et al. 2020); Decay sugar-rich substrates (Põlme et al. 2020) | Põlme S, et al. 2020. Fungal Diversity 105:1-16 ((https://doi.org/10.1007/s13225-020-00466-2)) |
| Otu44237 | 1 | k__Fungi(100);p__Ascomycota(100);c__Saccharomycetes(100);o__Saccharomycetales(100);f__Metschnikowiaceae(100);g__Metschnikowia(100);g__Metschnikowia_unclassified(100); | Metschnikowia | | 13 | Saprotroph | \|Undefined Saprotroph\| | Yeast | NULL | Probable | As nectar-tap_saprotroph (Põlme et al. 2020); Decay sugar-rich substrates (Põlme et al. 2020) | Põlme S, et al. 2020. Fungal Diversity 105:1-16 ((https://doi.org/10.1007/s13225-020-00466-2)) |
| Otu44238 | 1 | k__Fungi(100);p__Ascomycota(100);c__Saccharomycetes(100);o__Saccharomycetales(100);f__Metschnikowiaceae(100);g__Metschnikowia(100);s__Metschnikowia_pulcherrima(100); | Metschnikowia | | 13 | Saprotroph | \|Undefined Saprotroph\| | Yeast | NULL | Probable | As nectar-tap_saprotroph (Põlme et al. 2020); Decay sugar-rich substrates (Põlme et al. 2020) | Põlme S, et al. 2020. Fungal Diversity 105:1-16 ((https://doi.org/10.1007/s13225-020-00466-2)) |
| Otu44239 | 1 | k__Fungi(100);p__Ascomycota(100);c__Saccharomycetes(100);o__Saccharomycetales(100);f__Metschnikowiaceae(100);g__Metschnikowia(100);s__Metschnikowia_pulcherrima(100); | Metschnikowia | | 13 | Saprotroph | \|Undefined Saprotroph\| | Yeast | NULL | Probable | As nectar-tap_saprotroph (Põlme et al. 2020); Decay sugar-rich substrates (Põlme et al. 2020) | Põlme S, et al. 2020. Fungal Diversity 105:1-16 ((https://doi.org/10.1007/s13225-020-00466-2)) |
| Otu44240 | 1 | k__Fungi(100);p__Ascomycota(100);c__Saccharomycetes(100);o__Saccharomycetales(100);f__Metschnikowiaceae(100);g__Metschnikowia(100);s__Metschnikowia_pulcherrima(100); | Metschnikowia | | 13 | Saprotroph | \|Undefined Saprotroph\| | Yeast | NULL | Probable | As nectar-tap_saprotroph (Põlme et al. 2020); Decay sugar-rich substrates (Põlme et al. 2020) | Põlme S, et al. 2020. Fungal Diversity 105:1-16 ((https://doi.org/10.1007/s13225-020-00466-2)) |
| Otu44241 | 1 | k__Fungi(100);p__Ascomycota(100);c__Saccharomycetes(100);o__Saccharomycetales(100);f__Metschnikowiaceae(100);g__Metschnikowia(100);s__Metschnikowia_pulcherrima(100); | Metschnikowia | | 13 | Saprotroph | \|Undefined Saprotroph\| | Yeast | NULL | Probable | As nectar-tap_saprotroph (Põlme et al. 2020); Decay sugar-rich substrates (Põlme et al. 2020) | Põlme S, et al. 2020. Fungal Diversity 105:1-16 ((https://doi.org/10.1007/s13225-020-00466-2)) |
| Otu44242 | 1 | k__Fungi(100);p__Ascomycota(100);c__Saccharomycetes(100);o__Saccharomycetales(100);f__Metschnikowiaceae(100);g__Metschnikowia(100);g__Metschnikowia_unclassified(100); | Metschnikowia | | 13 | Saprotroph | \|Undefined Saprotroph\| | Yeast | NULL | Probable | As nectar-tap_saprotroph (Põlme et al. 2020); Decay sugar-rich substrates (Põlme et al. 2020) | Põlme S, et al. 2020. Fungal Diversity 105:1-16 ((https://doi.org/10.1007/s13225-020-00466-2)) |
| Otu44243 | 1 | k__Fungi(100);p__Ascomycota(100);c__Saccharomycetes(100);o__Saccharomycetales(100);f__Metschnikowiaceae(100);g__Metschnikowia(100);s__Metschnikowia_pulcherrima(100); | Metschnikowia | | 13 | Saprotroph | \|Undefined Saprotroph\| | Yeast | NULL | Probable | As nectar-tap_saprotroph (Põlme et al. 2020); Decay sugar-rich substrates (Põlme et al. 2020) | Põlme S, et al. 2020. Fungal Diversity 105:1-16 ((https://doi.org/10.1007/s13225-020-00466-2)) |
| Otu44244 | 1 | k__Fungi(100);p__Ascomycota(100);c__Saccharomycetes(100);o__Saccharomycetales(100);f__Metschnikowiaceae(100);g__Metschnikowia(100);s__Metschnikowia_pulcherrima(100); | Metschnikowia | | 13 | Saprotroph | \|Undefined Saprotroph\| | Yeast | NULL | Probable | As nectar-tap_saprotroph (Põlme et al. 2020); Decay sugar-rich substrates (Põlme et al. 2020) | Põlme S, et al. 2020. Fungal Diversity 105:1-16 ((https://doi.org/10.1007/s13225-020-00466-2)) |
| Otu44245 | 1 | k__Fungi(100);p__Ascomycota(100);c__Saccharomycetes(100);o__Saccharomycetales(100);f__Metschnikowiaceae(100);g__Metschnikowia(100);s__Metschnikowia_pulcherrima(100); | Metschnikowia | | 13 | Saprotroph | \|Undefined Saprotroph\| | Yeast | NULL | Probable | As nectar-tap_saprotroph (Põlme et al. 2020); Decay sugar-rich substrates (Põlme et al. 2020) | Põlme S, et al. 2020. Fungal Diversity 105:1-16 ((https://doi.org/10.1007/s13225-020-00466-2)) |
| Otu44246 | 1 | k__Fungi(100);p__Ascomycota(100);c__Saccharomycetes(100);o__Saccharomycetales(100);f__Metschnikowiaceae(100);g__Metschnikowia(100);s__Metschnikowia_pulcherrima(100); | Metschnikowia | | 13 | Saprotroph | \|Undefined Saprotroph\| | Yeast | NULL | Probable | As nectar-tap_saprotroph (Põlme et al. 2020); Decay sugar-rich substrates (Põlme et al. 2020) | Põlme S, et al. 2020. Fungal Diversity 105:1-16 ((https://doi.org/10.1007/s13225-020-00466-2)) |
| Otu44247 | 1 | k__Fungi(100);p__Ascomycota(100);c__Saccharomycetes(100);o__Saccharomycetales(100);f__Metschnikowiaceae(100);g__Metschnikowia(100);s__Metschnikowia_pulcherrima(100); | Metschnikowia | | 13 | Saprotroph | \|Undefined Saprotroph\| | Yeast | NULL | Probable | As nectar-tap_saprotroph (Põlme et al. 2020); Decay sugar-rich substrates (Põlme et al. 2020) | Põlme S, et al. 2020. Fungal Diversity 105:1-16 ((https://doi.org/10.1007/s13225-020-00466-2)) |
| Otu44248 | 1 | k__Fungi(100);p__Ascomycota(100);c__Saccharomycetes(100);o__Saccharomycetales(100);f__Metschnikowiaceae(100);g__Metschnikowia(100);s__Metschnikowia_pulcherrima(100); | Metschnikowia | | 13 | Saprotroph | \|Undefined Saprotroph\| | Yeast | NULL | Probable | As nectar-tap_saprotroph (Põlme et al. 2020); Decay sugar-rich substrates (Põlme et al. 2020) | Põlme S, et al. 2020. Fungal Diversity 105:1-16 ((https://doi.org/10.1007/s13225-020-00466-2)) |
| Otu44249 | 1 | k__Fungi(100);p__Ascomycota(100);c__Saccharomycetes(100);o__Saccharomycetales(100);f__Metschnikowiaceae(100);g__Metschnikowia(100);s__Metschnikowia_pulcherrima(100); | Metschnikowia | | 13 | Saprotroph | \|Undefined Saprotroph\| | Yeast | NULL | Probable | As nectar-tap_saprotroph (Põlme et al. 2020); Decay sugar-rich substrates (Põlme et al. 2020) | Põlme S, et al. 2020. Fungal Diversity 105:1-16 ((https://doi.org/10.1007/s13225-020-00466-2)) |
| Otu44250 | 1 | k__Fungi(100);p__Ascomycota(100);c__Saccharomycetes(100);o__Saccharomycetales(100);f__Metschnikowiaceae(100);g__Metschnikowia(100);s__Metschnikowia_pulcherrima(100); | Metschnikowia | | 13 | Saprotroph | \|Undefined Saprotroph\| | Yeast | NULL | Probable | As nectar-tap_saprotroph (Põlme et al. 2020); Decay sugar-rich substrates (Põlme et al. 2020) | Põlme S, et al. 2020. Fungal Diversity 105:1-16 ((https://doi.org/10.1007/s13225-020-00466-2)) |
| Otu44251 | 1 | k__Fungi(100);p__Ascomycota(100);c__Saccharomycetes(100);o__Saccharomycetales(100);f__Metschnikowiaceae(100);g__Metschnikowia(100);s__Metschnikowia_pulcherrima(100); | Metschnikowia | | 13 | Saprotroph | \|Undefined Saprotroph\| | Yeast | NULL | Probable | As nectar-tap_saprotroph (Põlme et al. 2020); Decay sugar-rich substrates (Põlme et al. 2020) | Põlme S, et al. 2020. Fungal Diversity 105:1-16 ((https://doi.org/10.1007/s13225-020-00466-2)) |
| Otu44252 | 1 | k__Fungi(100);p__Ascomycota(100);c__Saccharomycetes(100);o__Saccharomycetales(100);f__Metschnikowiaceae(100);g__Metschnikowia(100);g__Metschnikowia_unclassified(100); | Metschnikowia | | 13 | Saprotroph | \|Undefined Saprotroph\| | Yeast | NULL | Probable | As nectar-tap_saprotroph (Põlme et al. 2020); Decay sugar-rich substrates (Põlme et al. 2020) | Põlme S, et al. 2020. Fungal Diversity 105:1-16 ((https://doi.org/10.1007/s13225-020-00466-2)) |
| Otu44253 | 1 | k__Fungi(100);p__Ascomycota(100);c__Saccharomycetes(100);o__Saccharomycetales(100);f__Metschnikowiaceae(100);g__Metschnikowia(100);g__Metschnikowia_unclassified(100); | Metschnikowia | | 13 | Saprotroph | \|Undefined Saprotroph\| | Yeast | NULL | Probable | As nectar-tap_saprotroph (Põlme et al. 2020); Decay sugar-rich substrates (Põlme et al. 2020) | Põlme S, et al. 2020. Fungal Diversity 105:1-16 ((https://doi.org/10.1007/s13225-020-00466-2)) |
| Otu44254 | 1 | k__Fungi(100);p__Ascomycota(100);c__Saccharomycetes(100);o__Saccharomycetales(100);f__Metschnikowiaceae(100);g__Metschnikowia(100);s__Metschnikowia_pulcherrima(100); | Metschnikowia | | 13 | Saprotroph | \|Undefined Saprotroph\| | Yeast | NULL | Probable | As nectar-tap_saprotroph (Põlme et al. 2020); Decay sugar-rich substrates (Põlme et al. 2020) | Põlme S, et al. 2020. Fungal Diversity 105:1-16 ((https://doi.org/10.1007/s13225-020-00466-2)) |
| Otu44255 | 1 | k__Fungi(100);p__Ascomycota(100);c__Saccharomycetes(100);o__Saccharomycetales(100);f__Metschnikowiaceae(100);g__Metschnikowia(100);s__Metschnikowia_pulcherrima(100); | Metschnikowia | | 13 | Saprotroph | \|Undefined Saprotroph\| | Yeast | NULL | Probable | As nectar-tap_saprotroph (Põlme et al. 2020); Decay sugar-rich substrates (Põlme et al. 2020) | Põlme S, et al. 2020. Fungal Diversity 105:1-16 ((https://doi.org/10.1007/s13225-020-00466-2)) |
| Otu44256 | 1 | k__Fungi(100);p__Ascomycota(100);c__Saccharomycetes(100);o__Saccharomycetales(100);f__Metschnikowiaceae(100);g__Metschnikowia(100);s__Metschnikowia_pulcherrima(100); | Metschnikowia | | 13 | Saprotroph | \|Undefined Saprotroph\| | Yeast | NULL | Probable | As nectar-tap_saprotroph (Põlme et al. 2020); Decay sugar-rich substrates (Põlme et al. 2020) | Põlme S, et al. 2020. Fungal Diversity 105:1-16 ((https://doi.org/10.1007/s13225-020-00466-2)) |
| Otu44257 | 1 | k__Fungi(100);p__Ascomycota(100);c__Saccharomycetes(100);o__Saccharomycetales(100);f__Metschnikowiaceae(100);g__Metschnikowia(100);s__Metschnikowia_pulcherrima(100); | Metschnikowia | | 13 | Saprotroph | \|Undefined Saprotroph\| | Yeast | NULL | Probable | As nectar-tap_saprotroph (Põlme et al. 2020); Decay sugar-rich substrates (Põlme et al. 2020) | Põlme S, et al. 2020. Fungal Diversity 105:1-16 ((https://doi.org/10.1007/s13225-020-00466-2)) |
| Otu44258 | 1 | k__Fungi(100);p__Ascomycota(100);c__Saccharomycetes(100);o__Saccharomycetales(100);f__Metschnikowiaceae(100);g__Metschnikowia(100);s__Metschnikowia_pulcherrima(100); | Metschnikowia | | 13 | Saprotroph | \|Undefined Saprotroph\| | Yeast | NULL | Probable | As nectar-tap_saprotroph (Põlme et al. 2020); Decay sugar-rich substrates (Põlme et al. 2020) | Põlme S, et al. 2020. Fungal Diversity 105:1-16 ((https://doi.org/10.1007/s13225-020-00466-2)) |
| Otu44259 | 1 | k__Fungi(100);p__Ascomycota(100);c__Saccharomycetes(100);o__Saccharomycetales(100);f__Metschnikowiaceae(100);g__Metschnikowia(100);s__Metschnikowia_pulcherrima(100); | Metschnikowia | | 13 | Saprotroph | \|Undefined Saprotroph\| | Yeast | NULL | Probable | As nectar-tap_saprotroph (Põlme et al. 2020); Decay sugar-rich substrates (Põlme et al. 2020) | Põlme S, et al. 2020. Fungal Diversity 105:1-16 ((https://doi.org/10.1007/s13225-020-00466-2)) |
| Otu44260 | 1 | k__Fungi(100);p__Ascomycota(100);c__Saccharomycetes(100);o__Saccharomycetales(100);f__Metschnikowiaceae(100);g__Metschnikowia(100);s__Metschnikowia_pulcherrima(100); | Metschnikowia | | 13 | Saprotroph | \|Undefined Saprotroph\| | Yeast | NULL | Probable | As nectar-tap_saprotroph (Põlme et al. 2020); Decay sugar-rich substrates (Põlme et al. 2020) | Põlme S, et al. 2020. Fungal Diversity 105:1-16 ((https://doi.org/10.1007/s13225-020-00466-2)) |
| Otu44261 | 1 | k__Fungi(100);p__Ascomycota(100);c__Saccharomycetes(100);o__Saccharomycetales(100);f__Metschnikowiaceae(100);g__Metschnikowia(100);s__Metschnikowia_pulcherrima(100); | Metschnikowia | | 13 | Saprotroph | \|Undefined Saprotroph\| | Yeast | NULL | Probable | As nectar-tap_saprotroph (Põlme et al. 2020); Decay sugar-rich substrates (Põlme et al. 2020) | Põlme S, et al. 2020. Fungal Diversity 105:1-16 ((https://doi.org/10.1007/s13225-020-00466-2)) |
| Otu44262 | 1 | k__Fungi(100);p__Ascomycota(100);c__Saccharomycetes(100);o__Saccharomycetales(100);f__Metschnikowiaceae(100);g__Metschnikowia(100);s__Metschnikowia_pulcherrima(100); | Metschnikowia | | 13 | Saprotroph | \|Undefined Saprotroph\| | Yeast | NULL | Probable | As nectar-tap_saprotroph (Põlme et al. 2020); Decay sugar-rich substrates (Põlme et al. 2020) | Põlme S, et al. 2020. Fungal Diversity 105:1-16 ((https://doi.org/10.1007/s13225-020-00466-2)) |
| Otu44263 | 1 | k__Fungi(100);p__Ascomycota(100);c__Saccharomycetes(100);o__Saccharomycetales(100);f__Metschnikowiaceae(100);g__Metschnikowia(100);g__Metschnikowia_unclassified(100); | Metschnikowia | | 13 | Saprotroph | \|Undefined Saprotroph\| | Yeast | NULL | Probable | As nectar-tap_saprotroph (Põlme et al. 2020); Decay sugar-rich substrates (Põlme et al. 2020) | Põlme S, et al. 2020. Fungal Diversity 105:1-16 ((https://doi.org/10.1007/s13225-020-00466-2)) |
| Otu44264 | 1 | k__Fungi(100);p__Ascomycota(100);c__Saccharomycetes(100);o__Saccharomycetales(100);f__Metschnikowiaceae(100);g__Metschnikowia(100);s__Metschnikowia_pulcherrima(100); | Metschnikowia | | 13 | Saprotroph | \|Undefined Saprotroph\| | Yeast | NULL | Probable | As nectar-tap_saprotroph (Põlme et al. 2020); Decay sugar-rich substrates (Põlme et al. 2020) | Põlme S, et al. 2020. Fungal Diversity 105:1-16 ((https://doi.org/10.1007/s13225-020-00466-2)) |
| Otu44265 | 1 | k__Fungi(100);p__Ascomycota(100);c__Saccharomycetes(100);o__Saccharomycetales(100);f__Metschnikowiaceae(100);g__Metschnikowia(100);s__Metschnikowia_pulcherrima(100); | Metschnikowia | | 13 | Saprotroph | \|Undefined Saprotroph\| | Yeast | NULL | Probable | As nectar-tap_saprotroph (Põlme et al. 2020); Decay sugar-rich substrates (Põlme et al. 2020) | Põlme S, et al. 2020. Fungal Diversity 105:1-16 ((https://doi.org/10.1007/s13225-020-00466-2)) |
| Otu44266 | 1 | k__Fungi(100);p__Ascomycota(100);c__Saccharomycetes(100);o__Saccharomycetales(100);f__Metschnikowiaceae(100);g__Metschnikowia(100);s__Metschnikowia_pulcherrima(100); | Metschnikowia | | 13 | Saprotroph | \|Undefined Saprotroph\| | Yeast | NULL | Probable | As nectar-tap_saprotroph (Põlme et al. 2020); Decay sugar-rich substrates (Põlme et al. 2020) | Põlme S, et al. 2020. Fungal Diversity 105:1-16 ((https://doi.org/10.1007/s13225-020-00466-2)) |
| Otu44267 | 1 | k__Fungi(100);p__Ascomycota(100);c__Saccharomycetes(100);o__Saccharomycetales(100);f__Metschnikowiaceae(100);g__Metschnikowia(100);s__Metschnikowia_pulcherrima(100); | Metschnikowia | | 13 | Saprotroph | \|Undefined Saprotroph\| | Yeast | NULL | Probable | As nectar-tap_saprotroph (Põlme et al. 2020); Decay sugar-rich substrates (Põlme et al. 2020) | Põlme S, et al. 2020. Fungal Diversity 105:1-16 ((https://doi.org/10.1007/s13225-020-00466-2)) |
| Otu44268 | 1 | k__Fungi(100);p__Ascomycota(100);c__Saccharomycetes(100);o__Saccharomycetales(100);f__Metschnikowiaceae(100);g__Metschnikowia(100);s__Metschnikowia_pulcherrima(100); | Metschnikowia | | 13 | Saprotroph | \|Undefined Saprotroph\| | Yeast | NULL | Probable | As nectar-tap_saprotroph (Põlme et al. 2020); Decay sugar-rich substrates (Põlme et al. 2020) | Põlme S, et al. 2020. Fungal Diversity 105:1-16 ((https://doi.org/10.1007/s13225-020-00466-2)) |
| Otu44269 | 1 | k__Fungi(100);p__Ascomycota(100);c__Saccharomycetes(100);o__Saccharomycetales(100);f__Metschnikowiaceae(100);g__Metschnikowia(100);s__Metschnikowia_pulcherrima(100); | Metschnikowia | | 13 | Saprotroph | \|Undefined Saprotroph\| | Yeast | NULL | Probable | As nectar-tap_saprotroph (Põlme et al. 2020); Decay sugar-rich substrates (Põlme et al. 2020) | Põlme S, et al. 2020. Fungal Diversity 105:1-16 ((https://doi.org/10.1007/s13225-020-00466-2)) |
| Otu44270 | 1 | k__Fungi(100);p__Ascomycota(100);c__Saccharomycetes(100);o__Saccharomycetales(100);f__Metschnikowiaceae(100);g__Metschnikowia(100);s__Metschnikowia_pulcherrima(100); | Metschnikowia | | 13 | Saprotroph | \|Undefined Saprotroph\| | Yeast | NULL | Probable | As nectar-tap_saprotroph (Põlme et al. 2020); Decay sugar-rich substrates (Põlme et al. 2020) | Põlme S, et al. 2020. Fungal Diversity 105:1-16 ((https://doi.org/10.1007/s13225-020-00466-2)) |
| Otu44271 | 1 | k__Fungi(100);p__Ascomycota(100);c__Saccharomycetes(100);o__Saccharomycetales(100);f__Metschnikowiaceae(100);g__Metschnikowia(100);g__Metschnikowia_unclassified(100); | Metschnikowia | | 13 | Saprotroph | \|Undefined Saprotroph\| | Yeast | NULL | Probable | As nectar-tap_saprotroph (Põlme et al. 2020); Decay sugar-rich substrates (Põlme et al. 2020) | Põlme S, et al. 2020. Fungal Diversity 105:1-16 ((https://doi.org/10.1007/s13225-020-00466-2)) |
| Otu44272 | 1 | k__Fungi(100);p__Ascomycota(100);c__Saccharomycetes(100);o__Saccharomycetales(100);f__Metschnikowiaceae(100);g__Metschnikowia(100);g__Metschnikowia_unclassified(100); | Metschnikowia | | 13 | Saprotroph | \|Undefined Saprotroph\| | Yeast | NULL | Probable | As nectar-tap_saprotroph (Põlme et al. 2020); Decay sugar-rich substrates (Põlme et al. 2020) | Põlme S, et al. 2020. Fungal Diversity 105:1-16 ((https://doi.org/10.1007/s13225-020-00466-2)) |
| Otu44273 | 1 | k__Fungi(100);p__Ascomycota(100);c__Saccharomycetes(100);o__Saccharomycetales(100);f__Metschnikowiaceae(100);g__Metschnikowia(100);g__Metschnikowia_unclassified(100); | Metschnikowia | | 13 | Saprotroph | \|Undefined Saprotroph\| | Yeast | NULL | Probable | As nectar-tap_saprotroph (Põlme et al. 2020); Decay sugar-rich substrates (Põlme et al. 2020) | Põlme S, et al. 2020. Fungal Diversity 105:1-16 ((https://doi.org/10.1007/s13225-020-00466-2)) |
| Otu44274 | 1 | k__Fungi(100);p__Ascomycota(100);c__Saccharomycetes(100);o__Saccharomycetales(100);f__Metschnikowiaceae(100);g__Metschnikowia(100);g__Metschnikowia_unclassified(100); | Metschnikowia | | 13 | Saprotroph | \|Undefined Saprotroph\| | Yeast | NULL | Probable | As nectar-tap_saprotroph (Põlme et al. 2020); Decay sugar-rich substrates (Põlme et al. 2020) | Põlme S, et al. 2020. Fungal Diversity 105:1-16 ((https://doi.org/10.1007/s13225-020-00466-2)) |
| Otu44275 | 1 | k__Fungi(100);p__Ascomycota(100);c__Saccharomycetes(100);o__Saccharomycetales(100);f__Metschnikowiaceae(100);g__Metschnikowia(100);s__Metschnikowia_pulcherrima(100); | Metschnikowia | | 13 | Saprotroph | \|Undefined Saprotroph\| | Yeast | NULL | Probable | As nectar-tap_saprotroph (Põlme et al. 2020); Decay sugar-rich substrates (Põlme et al. 2020) | Põlme S, et al. 2020. Fungal Diversity 105:1-16 ((https://doi.org/10.1007/s13225-020-00466-2)) |
| Otu44276 | 1 | k__Fungi(100);p__Ascomycota(100);c__Saccharomycetes(100);o__Saccharomycetales(100);f__Metschnikowiaceae(100);g__Metschnikowia(100);g__Metschnikowia_unclassified(100); | Metschnikowia | | 13 | Saprotroph | \|Undefined Saprotroph\| | Yeast | NULL | Probable | As nectar-tap_saprotroph (Põlme et al. 2020); Decay sugar-rich substrates (Põlme et al. 2020) | Põlme S, et al. 2020. Fungal Diversity 105:1-16 ((https://doi.org/10.1007/s13225-020-00466-2)) |
| Otu44277 | 1 | k__Fungi(100);p__Ascomycota(100);c__Saccharomycetes(100);o__Saccharomycetales(100);f__Metschnikowiaceae(100);g__Metschnikowia(100);g__Metschnikowia_unclassified(100); | Metschnikowia | | 13 | Saprotroph | \|Undefined Saprotroph\| | Yeast | NULL | Probable | As nectar-tap_saprotroph (Põlme et al. 2020); Decay sugar-rich substrates (Põlme et al. 2020) | Põlme S, et al. 2020. Fungal Diversity 105:1-16 ((https://doi.org/10.1007/s13225-020-00466-2)) |
| Otu44278 | 1 | k__Fungi(100);p__Ascomycota(100);c__Saccharomycetes(100);o__Saccharomycetales(100);f__Metschnikowiaceae(100);g__Metschnikowia(100);s__Metschnikowia_pulcherrima(100); | Metschnikowia | | 13 | Saprotroph | \|Undefined Saprotroph\| | Yeast | NULL | Probable | As nectar-tap_saprotroph (Põlme et al. 2020); Decay sugar-rich substrates (Põlme et al. 2020) | Põlme S, et al. 2020. Fungal Diversity 105:1-16 ((https://doi.org/10.1007/s13225-020-00466-2)) |
| Otu44279 | 1 | k__Fungi(100);p__Ascomycota(100);c__Saccharomycetes(100);o__Saccharomycetales(100);f__Metschnikowiaceae(100);g__Metschnikowia(100);s__Metschnikowia_pulcherrima(100); | Metschnikowia | | 13 | Saprotroph | \|Undefined Saprotroph\| | Yeast | NULL | Probable | As nectar-tap_saprotroph (Põlme et al. 2020); Decay sugar-rich substrates (Põlme et al. 2020) | Põlme S, et al. 2020. Fungal Diversity 105:1-16 ((https://doi.org/10.1007/s13225-020-00466-2)) |
| Otu44280 | 1 | k__Fungi(100);p__Ascomycota(100);c__Saccharomycetes(100);o__Saccharomycetales(100);f__Metschnikowiaceae(100);g__Metschnikowia(100);s__Metschnikowia_pulcherrima(100); | Metschnikowia | | 13 | Saprotroph | \|Undefined Saprotroph\| | Yeast | NULL | Probable | As nectar-tap_saprotroph (Põlme et al. 2020); Decay sugar-rich substrates (Põlme et al. 2020) | Põlme S, et al. 2020. Fungal Diversity 105:1-16 ((https://doi.org/10.1007/s13225-020-00466-2)) |
| Otu44281 | 1 | k__Fungi(100);p__Ascomycota(100);c__Saccharomycetes(100);o__Saccharomycetales(100);f__Metschnikowiaceae(100);g__Metschnikowia(100);s__Metschnikowia_pulcherrima(100); | Metschnikowia | | 13 | Saprotroph | \|Undefined Saprotroph\| | Yeast | NULL | Probable | As nectar-tap_saprotroph (Põlme et al. 2020); Decay sugar-rich substrates (Põlme et al. 2020) | Põlme S, et al. 2020. Fungal Diversity 105:1-16 ((https://doi.org/10.1007/s13225-020-00466-2)) |
| Otu44282 | 1 | k__Fungi(100);p__Ascomycota(100);c__Saccharomycetes(100);o__Saccharomycetales(100);f__Metschnikowiaceae(100);g__Metschnikowia(100);s__Metschnikowia_pulcherrima(100); | Metschnikowia | | 13 | Saprotroph | \|Undefined Saprotroph\| | Yeast | NULL | Probable | As nectar-tap_saprotroph (Põlme et al. 2020); Decay sugar-rich substrates (Põlme et al. 2020) | Põlme S, et al. 2020. Fungal Diversity 105:1-16 ((https://doi.org/10.1007/s13225-020-00466-2)) |
| Otu44283 | 1 | k__Fungi(100);p__Ascomycota(100);c__Saccharomycetes(100);o__Saccharomycetales(100);f__Metschnikowiaceae(100);g__Metschnikowia(100);s__Metschnikowia_pulcherrima(100); | Metschnikowia | | 13 | Saprotroph | \|Undefined Saprotroph\| | Yeast | NULL | Probable | As nectar-tap_saprotroph (Põlme et al. 2020); Decay sugar-rich substrates (Põlme et al. 2020) | Põlme S, et al. 2020. Fungal Diversity 105:1-16 ((https://doi.org/10.1007/s13225-020-00466-2)) |
| Otu44284 | 1 | k__Fungi(100);p__Ascomycota(100);c__Saccharomycetes(100);o__Saccharomycetales(100);f__Metschnikowiaceae(100);g__Metschnikowia(100);s__Metschnikowia_pulcherrima(100); | Metschnikowia | | 13 | Saprotroph | \|Undefined Saprotroph\| | Yeast | NULL | Probable | As nectar-tap_saprotroph (Põlme et al. 2020); Decay sugar-rich substrates (Põlme et al. 2020) | Põlme S, et al. 2020. Fungal Diversity 105:1-16 ((https://doi.org/10.1007/s13225-020-00466-2)) |
| Otu44285 | 1 | k__Fungi(100);p__Ascomycota(100);c__Saccharomycetes(100);o__Saccharomycetales(100);f__Metschnikowiaceae(100);g__Metschnikowia(100);s__Metschnikowia_pulcherrima(100); | Metschnikowia | | 13 | Saprotroph | \|Undefined Saprotroph\| | Yeast | NULL | Probable | As nectar-tap_saprotroph (Põlme et al. 2020); Decay sugar-rich substrates (Põlme et al. 2020) | Põlme S, et al. 2020. Fungal Diversity 105:1-16 ((https://doi.org/10.1007/s13225-020-00466-2)) |
| Otu44286 | 1 | k__Fungi(100);p__Ascomycota(100);c__Saccharomycetes(100);o__Saccharomycetales(100);f__Metschnikowiaceae(100);g__Metschnikowia(100);s__Metschnikowia_pulcherrima(100); | Metschnikowia | | 13 | Saprotroph | \|Undefined Saprotroph\| | Yeast | NULL | Probable | As nectar-tap_saprotroph (Põlme et al. 2020); Decay sugar-rich substrates (Põlme et al. 2020) | Põlme S, et al. 2020. Fungal Diversity 105:1-16 ((https://doi.org/10.1007/s13225-020-00466-2)) |
| Otu44287 | 1 | k__Fungi(100);p__Ascomycota(100);c__Saccharomycetes(100);o__Saccharomycetales(100);f__Metschnikowiaceae(100);g__Metschnikowia(100);g__Metschnikowia_unclassified(100); | Metschnikowia | | 13 | Saprotroph | \|Undefined Saprotroph\| | Yeast | NULL | Probable | As nectar-tap_saprotroph (Põlme et al. 2020); Decay sugar-rich substrates (Põlme et al. 2020) | Põlme S, et al. 2020. Fungal Diversity 105:1-16 ((https://doi.org/10.1007/s13225-020-00466-2)) |
| Otu44288 | 1 | k__Fungi(100);p__Ascomycota(100);c__Saccharomycetes(100);o__Saccharomycetales(100);f__Metschnikowiaceae(100);g__Metschnikowia(100);s__Metschnikowia_pulcherrima(100); | Metschnikowia | | 13 | Saprotroph | \|Undefined Saprotroph\| | Yeast | NULL | Probable | As nectar-tap_saprotroph (Põlme et al. 2020); Decay sugar-rich substrates (Põlme et al. 2020) | Põlme S, et al. 2020. Fungal Diversity 105:1-16 ((https://doi.org/10.1007/s13225-020-00466-2)) |
| Otu44289 | 1 | k__Fungi(100);p__Ascomycota(100);c__Saccharomycetes(100);o__Saccharomycetales(100);f__Metschnikowiaceae(100);g__Metschnikowia(100);s__Metschnikowia_pulcherrima(100); | Metschnikowia | | 13 | Saprotroph | \|Undefined Saprotroph\| | Yeast | NULL | Probable | As nectar-tap_saprotroph (Põlme et al. 2020); Decay sugar-rich substrates (Põlme et al. 2020) | Põlme S, et al. 2020. Fungal Diversity 105:1-16 ((https://doi.org/10.1007/s13225-020-00466-2)) |
| Otu44290 | 1 | k__Fungi(100);p__Ascomycota(100);c__Saccharomycetes(100);o__Saccharomycetales(100);f__Metschnikowiaceae(100);g__Metschnikowia(100);s__Metschnikowia_pulcherrima(100); | Metschnikowia | | 13 | Saprotroph | \|Undefined Saprotroph\| | Yeast | NULL | Probable | As nectar-tap_saprotroph (Põlme et al. 2020); Decay sugar-rich substrates (Põlme et al. 2020) | Põlme S, et al. 2020. Fungal Diversity 105:1-16 ((https://doi.org/10.1007/s13225-020-00466-2)) |
| Otu44291 | 1 | k__Fungi(100);p__Ascomycota(100);c__Saccharomycetes(100);o__Saccharomycetales(100);f__Metschnikowiaceae(100);g__Metschnikowia(100);g__Metschnikowia_unclassified(100); | Metschnikowia | | 13 | Saprotroph | \|Undefined Saprotroph\| | Yeast | NULL | Probable | As nectar-tap_saprotroph (Põlme et al. 2020); Decay sugar-rich substrates (Põlme et al. 2020) | Põlme S, et al. 2020. Fungal Diversity 105:1-16 ((https://doi.org/10.1007/s13225-020-00466-2)) |
| Otu44292 | 1 | k__Fungi(100);p__Ascomycota(100);c__Saccharomycetes(100);o__Saccharomycetales(100);f__Metschnikowiaceae(100);g__Metschnikowia(100);g__Metschnikowia_unclassified(100); | Metschnikowia | | 13 | Saprotroph | \|Undefined Saprotroph\| | Yeast | NULL | Probable | As nectar-tap_saprotroph (Põlme et al. 2020); Decay sugar-rich substrates (Põlme et al. 2020) | Põlme S, et al. 2020. Fungal Diversity 105:1-16 ((https://doi.org/10.1007/s13225-020-00466-2)) |
| Otu44293 | 1 | k__Fungi(100);p__Ascomycota(100);c__Saccharomycetes(100);o__Saccharomycetales(100);f__Metschnikowiaceae(100);g__Metschnikowia(100);g__Metschnikowia_unclassified(100); | Metschnikowia | | 13 | Saprotroph | \|Undefined Saprotroph\| | Yeast | NULL | Probable | As nectar-tap_saprotroph (Põlme et al. 2020); Decay sugar-rich substrates (Põlme et al. 2020) | Põlme S, et al. 2020. Fungal Diversity 105:1-16 ((https://doi.org/10.1007/s13225-020-00466-2)) |
| Otu44294 | 1 | k__Fungi(100);p__Ascomycota(100);c__Saccharomycetes(100);o__Saccharomycetales(100);f__Metschnikowiaceae(100);g__Metschnikowia(100);s__Metschnikowia_pulcherrima(100); | Metschnikowia | | 13 | Saprotroph | \|Undefined Saprotroph\| | Yeast | NULL | Probable | As nectar-tap_saprotroph (Põlme et al. 2020); Decay sugar-rich substrates (Põlme et al. 2020) | Põlme S, et al. 2020. Fungal Diversity 105:1-16 ((https://doi.org/10.1007/s13225-020-00466-2)) |
| Otu44295 | 1 | k__Fungi(100);p__Ascomycota(100);c__Saccharomycetes(100);o__Saccharomycetales(100);f__Metschnikowiaceae(100);g__Metschnikowia(100);s__Metschnikowia_pulcherrima(100); | Metschnikowia | | 13 | Saprotroph | \|Undefined Saprotroph\| | Yeast | NULL | Probable | As nectar-tap_saprotroph (Põlme et al. 2020); Decay sugar-rich substrates (Põlme et al. 2020) | Põlme S, et al. 2020. Fungal Diversity 105:1-16 ((https://doi.org/10.1007/s13225-020-00466-2)) |
| Otu44296 | 1 | k__Fungi(100);p__Ascomycota(100);c__Saccharomycetes(100);o__Saccharomycetales(100);f__Metschnikowiaceae(100);g__Metschnikowia(100);s__Metschnikowia_pulcherrima(100); | Metschnikowia | | 13 | Saprotroph | \|Undefined Saprotroph\| | Yeast | NULL | Probable | As nectar-tap_saprotroph (Põlme et al. 2020); Decay sugar-rich substrates (Põlme et al. 2020) | Põlme S, et al. 2020. Fungal Diversity 105:1-16 ((https://doi.org/10.1007/s13225-020-00466-2)) |
| Otu44297 | 1 | k__Fungi(100);p__Ascomycota(100);c__Saccharomycetes(100);o__Saccharomycetales(100);f__Metschnikowiaceae(100);g__Metschnikowia(100);g__Metschnikowia_unclassified(100); | Metschnikowia | | 13 | Saprotroph | \|Undefined Saprotroph\| | Yeast | NULL | Probable | As nectar-tap_saprotroph (Põlme et al. 2020); Decay sugar-rich substrates (Põlme et al. 2020) | Põlme S, et al. 2020. Fungal Diversity 105:1-16 ((https://doi.org/10.1007/s13225-020-00466-2)) |
| Otu44298 | 1 | k__Fungi(100);p__Ascomycota(100);c__Saccharomycetes(100);o__Saccharomycetales(100);f__Metschnikowiaceae(100);g__Metschnikowia(100);g__Metschnikowia_unclassified(100); | Metschnikowia | | 13 | Saprotroph | \|Undefined Saprotroph\| | Yeast | NULL | Probable | As nectar-tap_saprotroph (Põlme et al. 2020); Decay sugar-rich substrates (Põlme et al. 2020) | Põlme S, et al. 2020. Fungal Diversity 105:1-16 ((https://doi.org/10.1007/s13225-020-00466-2)) |
| Otu44299 | 1 | k__Fungi(100);p__Ascomycota(100);c__Saccharomycetes(100);o__Saccharomycetales(100);f__Metschnikowiaceae(100);g__Metschnikowia(100);g__Metschnikowia_unclassified(100); | Metschnikowia | | 13 | Saprotroph | \|Undefined Saprotroph\| | Yeast | NULL | Probable | As nectar-tap_saprotroph (Põlme et al. 2020); Decay sugar-rich substrates (Põlme et al. 2020) | Põlme S, et al. 2020. Fungal Diversity 105:1-16 ((https://doi.org/10.1007/s13225-020-00466-2)) |
| Otu44300 | 1 | k__Fungi(100);p__Ascomycota(100);c__Saccharomycetes(100);o__Saccharomycetales(100);f__Metschnikowiaceae(100);g__Metschnikowia(100);s__Metschnikowia_pulcherrima(100); | Metschnikowia | | 13 | Saprotroph | \|Undefined Saprotroph\| | Yeast | NULL | Probable | As nectar-tap_saprotroph (Põlme et al. 2020); Decay sugar-rich substrates (Põlme et al. 2020) | Põlme S, et al. 2020. Fungal Diversity 105:1-16 ((https://doi.org/10.1007/s13225-020-00466-2)) |
| Otu44302 | 1 | k__Fungi(100);p__Ascomycota(100);c__Saccharomycetes(100);o__Saccharomycetales(100);f__Metschnikowiaceae(100);g__Metschnikowia(100);s__Metschnikowia_pulcherrima(100); | Metschnikowia | | 13 | Saprotroph | \|Undefined Saprotroph\| | Yeast | NULL | Probable | As nectar-tap_saprotroph (Põlme et al. 2020); Decay sugar-rich substrates (Põlme et al. 2020) | Põlme S, et al. 2020. Fungal Diversity 105:1-16 ((https://doi.org/10.1007/s13225-020-00466-2)) |
| Otu44303 | 1 | k__Fungi(100);p__Ascomycota(100);c__Saccharomycetes(100);o__Saccharomycetales(100);f__Metschnikowiaceae(100);g__Metschnikowia(100);s__Metschnikowia_pulcherrima(100); | Metschnikowia | | 13 | Saprotroph | \|Undefined Saprotroph\| | Yeast | NULL | Probable | As nectar-tap_saprotroph (Põlme et al. 2020); Decay sugar-rich substrates (Põlme et al. 2020) | Põlme S, et al. 2020. Fungal Diversity 105:1-16 ((https://doi.org/10.1007/s13225-020-00466-2)) |
| Otu44304 | 1 | k__Fungi(100);p__Ascomycota(100);c__Saccharomycetes(100);o__Saccharomycetales(100);f__Metschnikowiaceae(100);g__Metschnikowia(100);s__Metschnikowia_pulcherrima(100); | Metschnikowia | | 13 | Saprotroph | \|Undefined Saprotroph\| | Yeast | NULL | Probable | As nectar-tap_saprotroph (Põlme et al. 2020); Decay sugar-rich substrates (Põlme et al. 2020) | Põlme S, et al. 2020. Fungal Diversity 105:1-16 ((https://doi.org/10.1007/s13225-020-00466-2)) |
| Otu44305 | 1 | k__Fungi(100);p__Ascomycota(100);c__Saccharomycetes(100);o__Saccharomycetales(100);f__Metschnikowiaceae(100);g__Metschnikowia(100);s__Metschnikowia_pulcherrima(100); | Metschnikowia | | 13 | Saprotroph | \|Undefined Saprotroph\| | Yeast | NULL | Probable | As nectar-tap_saprotroph (Põlme et al. 2020); Decay sugar-rich substrates (Põlme et al. 2020) | Põlme S, et al. 2020. Fungal Diversity 105:1-16 ((https://doi.org/10.1007/s13225-020-00466-2)) |
| Otu44306 | 1 | k__Fungi(100);p__Ascomycota(100);c__Saccharomycetes(100);o__Saccharomycetales(100);f__Metschnikowiaceae(100);g__Metschnikowia(100);s__Metschnikowia_pulcherrima(100); | Metschnikowia | | 13 | Saprotroph | \|Undefined Saprotroph\| | Yeast | NULL | Probable | As nectar-tap_saprotroph (Põlme et al. 2020); Decay sugar-rich substrates (Põlme et al. 2020) | Põlme S, et al. 2020. Fungal Diversity 105:1-16 ((https://doi.org/10.1007/s13225-020-00466-2)) |
| Otu44307 | 1 | k__Fungi(100);p__Ascomycota(100);c__Saccharomycetes(100);o__Saccharomycetales(100);f__Metschnikowiaceae(100);g__Metschnikowia(100);s__Metschnikowia_pulcherrima(100); | Metschnikowia | | 13 | Saprotroph | \|Undefined Saprotroph\| | Yeast | NULL | Probable | As nectar-tap_saprotroph (Põlme et al. 2020); Decay sugar-rich substrates (Põlme et al. 2020) | Põlme S, et al. 2020. Fungal Diversity 105:1-16 ((https://doi.org/10.1007/s13225-020-00466-2)) |
| Otu44308 | 1 | k__Fungi(100);p__Ascomycota(100);c__Saccharomycetes(100);o__Saccharomycetales(100);f__Metschnikowiaceae(100);g__Metschnikowia(100);g__Metschnikowia_unclassified(100); | Metschnikowia | | 13 | Saprotroph | \|Undefined Saprotroph\| | Yeast | NULL | Probable | As nectar-tap_saprotroph (Põlme et al. 2020); Decay sugar-rich substrates (Põlme et al. 2020) | Põlme S, et al. 2020. Fungal Diversity 105:1-16 ((https://doi.org/10.1007/s13225-020-00466-2)) |
| Otu44309 | 1 | k__Fungi(100);p__Ascomycota(100);c__Saccharomycetes(100);o__Saccharomycetales(100);f__Metschnikowiaceae(100);g__Metschnikowia(100);s__Metschnikowia_pulcherrima(100); | Metschnikowia | | 13 | Saprotroph | \|Undefined Saprotroph\| | Yeast | NULL | Probable | As nectar-tap_saprotroph (Põlme et al. 2020); Decay sugar-rich substrates (Põlme et al. 2020) | Põlme S, et al. 2020. Fungal Diversity 105:1-16 ((https://doi.org/10.1007/s13225-020-00466-2)) |
| Otu44311 | 1 | k__Fungi(100);p__Ascomycota(100);c__Saccharomycetes(100);o__Saccharomycetales(100);f__Metschnikowiaceae(100);g__Metschnikowia(100);g__Metschnikowia_unclassified(100); | Metschnikowia | | 13 | Saprotroph | \|Undefined Saprotroph\| | Yeast | NULL | Probable | As nectar-tap_saprotroph (Põlme et al. 2020); Decay sugar-rich substrates (Põlme et al. 2020) | Põlme S, et al. 2020. Fungal Diversity 105:1-16 ((https://doi.org/10.1007/s13225-020-00466-2)) |
| Otu44312 | 1 | k__Fungi(100);p__Ascomycota(100);c__Saccharomycetes(100);o__Saccharomycetales(100);f__Metschnikowiaceae(100);g__Metschnikowia(100);s__Metschnikowia_pulcherrima(100); | Metschnikowia | | 13 | Saprotroph | \|Undefined Saprotroph\| | Yeast | NULL | Probable | As nectar-tap_saprotroph (Põlme et al. 2020); Decay sugar-rich substrates (Põlme et al. 2020) | Põlme S, et al. 2020. Fungal Diversity 105:1-16 ((https://doi.org/10.1007/s13225-020-00466-2)) |
| Otu44313 | 1 | k__Fungi(100);p__Ascomycota(100);c__Saccharomycetes(100);o__Saccharomycetales(100);f__Metschnikowiaceae(100);g__Metschnikowia(100);s__Metschnikowia_pulcherrima(100); | Metschnikowia | | 13 | Saprotroph | \|Undefined Saprotroph\| | Yeast | NULL | Probable | As nectar-tap_saprotroph (Põlme et al. 2020); Decay sugar-rich substrates (Põlme et al. 2020) | Põlme S, et al. 2020. Fungal Diversity 105:1-16 ((https://doi.org/10.1007/s13225-020-00466-2)) |
| Otu44314 | 1 | k__Fungi(100);p__Ascomycota(100);c__Saccharomycetes(100);o__Saccharomycetales(100);f__Metschnikowiaceae(100);g__Metschnikowia(100);s__Metschnikowia_pulcherrima(100); | Metschnikowia | | 13 | Saprotroph | \|Undefined Saprotroph\| | Yeast | NULL | Probable | As nectar-tap_saprotroph (Põlme et al. 2020); Decay sugar-rich substrates (Põlme et al. 2020) | Põlme S, et al. 2020. Fungal Diversity 105:1-16 ((https://doi.org/10.1007/s13225-020-00466-2)) |
| Otu44316 | 1 | k__Fungi(100);p__Ascomycota(100);c__Saccharomycetes(100);o__Saccharomycetales(100);f__Metschnikowiaceae(100);g__Metschnikowia(100);s__Metschnikowia_pulcherrima(100); | Metschnikowia | | 13 | Saprotroph | \|Undefined Saprotroph\| | Yeast | NULL | Probable | As nectar-tap_saprotroph (Põlme et al. 2020); Decay sugar-rich substrates (Põlme et al. 2020) | Põlme S, et al. 2020. Fungal Diversity 105:1-16 ((https://doi.org/10.1007/s13225-020-00466-2)) |
| Otu44317 | 1 | k__Fungi(100);p__Ascomycota(100);c__Saccharomycetes(100);o__Saccharomycetales(100);f__Metschnikowiaceae(100);g__Metschnikowia(100);s__Metschnikowia_pulcherrima(100); | Metschnikowia | | 13 | Saprotroph | \|Undefined Saprotroph\| | Yeast | NULL | Probable | As nectar-tap_saprotroph (Põlme et al. 2020); Decay sugar-rich substrates (Põlme et al. 2020) | Põlme S, et al. 2020. Fungal Diversity 105:1-16 ((https://doi.org/10.1007/s13225-020-00466-2)) |
| Otu44318 | 1 | k__Fungi(100);p__Ascomycota(100);c__Saccharomycetes(100);o__Saccharomycetales(100);f__Metschnikowiaceae(100);g__Metschnikowia(100);s__Metschnikowia_pulcherrima(100); | Metschnikowia | | 13 | Saprotroph | \|Undefined Saprotroph\| | Yeast | NULL | Probable | As nectar-tap_saprotroph (Põlme et al. 2020); Decay sugar-rich substrates (Põlme et al. 2020) | Põlme S, et al. 2020. Fungal Diversity 105:1-16 ((https://doi.org/10.1007/s13225-020-00466-2)) |
| Otu44319 | 1 | k__Fungi(100);p__Ascomycota(100);c__Saccharomycetes(100);o__Saccharomycetales(100);f__Metschnikowiaceae(100);g__Metschnikowia(100);s__Metschnikowia_pulcherrima(100); | Metschnikowia | | 13 | Saprotroph | \|Undefined Saprotroph\| | Yeast | NULL | Probable | As nectar-tap_saprotroph (Põlme et al. 2020); Decay sugar-rich substrates (Põlme et al. 2020) | Põlme S, et al. 2020. Fungal Diversity 105:1-16 ((https://doi.org/10.1007/s13225-020-00466-2)) |
| Otu44320 | 1 | k__Fungi(100);p__Ascomycota(100);c__Saccharomycetes(100);o__Saccharomycetales(100);f__Metschnikowiaceae(100);g__Metschnikowia(100);g__Metschnikowia_unclassified(100); | Metschnikowia | | 13 | Saprotroph | \|Undefined Saprotroph\| | Yeast | NULL | Probable | As nectar-tap_saprotroph (Põlme et al. 2020); Decay sugar-rich substrates (Põlme et al. 2020) | Põlme S, et al. 2020. Fungal Diversity 105:1-16 ((https://doi.org/10.1007/s13225-020-00466-2)) |
| Otu44321 | 1 | k__Fungi(100);p__Ascomycota(100);c__Saccharomycetes(100);o__Saccharomycetales(100);f__Metschnikowiaceae(100);g__Metschnikowia(100);g__Metschnikowia_unclassified(100); | Metschnikowia | | 13 | Saprotroph | \|Undefined Saprotroph\| | Yeast | NULL | Probable | As nectar-tap_saprotroph (Põlme et al. 2020); Decay sugar-rich substrates (Põlme et al. 2020) | Põlme S, et al. 2020. Fungal Diversity 105:1-16 ((https://doi.org/10.1007/s13225-020-00466-2)) |
| Otu44322 | 1 | k__Fungi(100);p__Ascomycota(100);c__Saccharomycetes(100);o__Saccharomycetales(100);f__Metschnikowiaceae(100);g__Metschnikowia(100);s__Metschnikowia_pulcherrima(100); | Metschnikowia | | 13 | Saprotroph | \|Undefined Saprotroph\| | Yeast | NULL | Probable | As nectar-tap_saprotroph (Põlme et al. 2020); Decay sugar-rich substrates (Põlme et al. 2020) | Põlme S, et al. 2020. Fungal Diversity 105:1-16 ((https://doi.org/10.1007/s13225-020-00466-2)) |
| Otu44323 | 1 | k__Fungi(100);p__Ascomycota(100);c__Saccharomycetes(100);o__Saccharomycetales(100);f__Metschnikowiaceae(100);g__Metschnikowia(100);s__Metschnikowia_pulcherrima(100); | Metschnikowia | | 13 | Saprotroph | \|Undefined Saprotroph\| | Yeast | NULL | Probable | As nectar-tap_saprotroph (Põlme et al. 2020); Decay sugar-rich substrates (Põlme et al. 2020) | Põlme S, et al. 2020. Fungal Diversity 105:1-16 ((https://doi.org/10.1007/s13225-020-00466-2)) |
| Otu44325 | 1 | k__Fungi(100);p__Ascomycota(100);c__Saccharomycetes(100);o__Saccharomycetales(100);f__Metschnikowiaceae(100);g__Metschnikowia(100);s__Metschnikowia_pulcherrima(100); | Metschnikowia | | 13 | Saprotroph | \|Undefined Saprotroph\| | Yeast | NULL | Probable | As nectar-tap_saprotroph (Põlme et al. 2020); Decay sugar-rich substrates (Põlme et al. 2020) | Põlme S, et al. 2020. Fungal Diversity 105:1-16 ((https://doi.org/10.1007/s13225-020-00466-2)) |
| Otu44326 | 1 | k__Fungi(100);p__Ascomycota(100);c__Saccharomycetes(100);o__Saccharomycetales(100);f__Saccharomycetaceae(100);g__Kazachstania(100);s__Kazachstania_intestinalis(100); | Kazachstania | | 13 | Saprotroph | \|Undefined Saprotroph\| | Yeast | NULL | Probable | As nectar-tap_saprotroph (Põlme et al. 2020); Decay sugar-rich substrates (Põlme et al. 2020) | Põlme S, et al. 2020. Fungal Diversity 105:1-16 ((https://doi.org/10.1007/s13225-020-00466-2)) |
| Otu44328 | 1 | k__Fungi(100);p__Ascomycota(100);c__Saccharomycetes(100);o__Saccharomycetales(100);f__Metschnikowiaceae(100);g__Metschnikowia(100);s__Metschnikowia_pulcherrima(100); | Metschnikowia | | 13 | Saprotroph | \|Undefined Saprotroph\| | Yeast | NULL | Probable | As nectar-tap_saprotroph (Põlme et al. 2020); Decay sugar-rich substrates (Põlme et al. 2020) | Põlme S, et al. 2020. Fungal Diversity 105:1-16 ((https://doi.org/10.1007/s13225-020-00466-2)) |
| Otu44329 | 1 | k__Fungi(100);p__Ascomycota(100);c__Saccharomycetes(100);o__Saccharomycetales(100);f__Metschnikowiaceae(100);g__Metschnikowia(100);s__Metschnikowia_pulcherrima(100); | Metschnikowia | | 13 | Saprotroph | \|Undefined Saprotroph\| | Yeast | NULL | Probable | As nectar-tap_saprotroph (Põlme et al. 2020); Decay sugar-rich substrates (Põlme et al. 2020) | Põlme S, et al. 2020. Fungal Diversity 105:1-16 ((https://doi.org/10.1007/s13225-020-00466-2)) |
| Otu44330 | 1 | k__Fungi(100);p__Ascomycota(100);c__Saccharomycetes(100);o__Saccharomycetales(100);f__Metschnikowiaceae(100);g__Metschnikowia(100);s__Metschnikowia_pulcherrima(100); | Metschnikowia | | 13 | Saprotroph | \|Undefined Saprotroph\| | Yeast | NULL | Probable | As nectar-tap_saprotroph (Põlme et al. 2020); Decay sugar-rich substrates (Põlme et al. 2020) | Põlme S, et al. 2020. Fungal Diversity 105:1-16 ((https://doi.org/10.1007/s13225-020-00466-2)) |
| Otu44331 | 1 | k__Fungi(100);p__Ascomycota(100);c__Saccharomycetes(100);o__Saccharomycetales(100);f__Metschnikowiaceae(100);g__Metschnikowia(100);s__Metschnikowia_pulcherrima(100); | Metschnikowia | | 13 | Saprotroph | \|Undefined Saprotroph\| | Yeast | NULL | Probable | As nectar-tap_saprotroph (Põlme et al. 2020); Decay sugar-rich substrates (Põlme et al. 2020) | Põlme S, et al. 2020. Fungal Diversity 105:1-16 ((https://doi.org/10.1007/s13225-020-00466-2)) |
| Otu44332 | 1 | k__Fungi(100);p__Ascomycota(100);c__Saccharomycetes(100);o__Saccharomycetales(100);f__Metschnikowiaceae(100);g__Metschnikowia(100);s__Metschnikowia_pulcherrima(100); | Metschnikowia | | 13 | Saprotroph | \|Undefined Saprotroph\| | Yeast | NULL | Probable | As nectar-tap_saprotroph (Põlme et al. 2020); Decay sugar-rich substrates (Põlme et al. 2020) | Põlme S, et al. 2020. Fungal Diversity 105:1-16 ((https://doi.org/10.1007/s13225-020-00466-2)) |

| | | | | | | | | | | | |
|---|---|---|---|---|---|---|---|---|---|---|---|
| Otu44333 | 1 | k__Fungi(100);p__Ascomycota(100);c__Saccharomycetes(100);o__Saccharomycetales(100);f__Metschnikowiaceae(100);g__Metschnikowia(100);s__Metschnikowia_pulcherrima(100); | Metschnikowia | 13 | Saprotroph | \|Undefined Saprotroph\| | Yeast | NULL | Probable | As nectar-tap_saprotroph (Põlme et al. 2020); Decay sugar-rich substrates (Põlme et al. 2020) | Põlme S, et al. 2020. Fungal Diversity 105:1-16 ((https://doi.org/10.1007/s13225-020-00466-2)) |
| Otu44334 | 1 | k__Fungi(100);p__Ascomycota(100);c__Saccharomycetes(100);o__Saccharomycetales(100);f__Metschnikowiaceae(100);g__Metschnikowia(100);s__Metschnikowia_pulcherrima(100); | Metschnikowia | 13 | Saprotroph | \|Undefined Saprotroph\| | Yeast | NULL | Probable | As nectar-tap_saprotroph (Põlme et al. 2020); Decay sugar-rich substrates (Põlme et al. 2020) | Põlme S, et al. 2020. Fungal Diversity 105:1-16 ((https://doi.org/10.1007/s13225-020-00466-2)) |
| Otu44335 | 1 | k__Fungi(100);p__Ascomycota(100);c__Saccharomycetes(100);o__Saccharomycetales(100);f__Metschnikowiaceae(100);g__Metschnikowia(100);s__Metschnikowia_pulcherrima(100); | Metschnikowia | 13 | Saprotroph | \|Undefined Saprotroph\| | Yeast | NULL | Probable | As nectar-tap_saprotroph (Põlme et al. 2020); Decay sugar-rich substrates (Põlme et al. 2020) | Põlme S, et al. 2020. Fungal Diversity 105:1-16 ((https://doi.org/10.1007/s13225-020-00466-2)) |
| Otu44336 | 1 | k__Fungi(100);p__Ascomycota(100);c__Saccharomycetes(100);o__Saccharomycetales(100);f__Metschnikowiaceae(100);g__Metschnikowia(100);g__Metschnikowia_unclassified(100); | Metschnikowia | 13 | Saprotroph | \|Undefined Saprotroph\| | Yeast | NULL | Probable | As nectar-tap_saprotroph (Põlme et al. 2020); Decay sugar-rich substrates (Põlme et al. 2020) | Põlme S, et al. 2020. Fungal Diversity 105:1-16 ((https://doi.org/10.1007/s13225-020-00466-2)) |
| Otu44337 | 1 | k__Fungi(100);p__Ascomycota(100);c__Saccharomycetes(100);o__Saccharomycetales(100);f__Metschnikowiaceae(100);g__Metschnikowia(100);s__Metschnikowia_pulcherrima(100); | Metschnikowia | 13 | Saprotroph | \|Undefined Saprotroph\| | Yeast | NULL | Probable | As nectar-tap_saprotroph (Põlme et al. 2020); Decay sugar-rich substrates (Põlme et al. 2020) | Põlme S, et al. 2020. Fungal Diversity 105:1-16 ((https://doi.org/10.1007/s13225-020-00466-2)) |
| Otu44338 | 1 | k__Fungi(100);p__Ascomycota(100);c__Saccharomycetes(100);o__Saccharomycetales(100);f__Metschnikowiaceae(100);g__Metschnikowia(100);s__Metschnikowia_pulcherrima(100); | Metschnikowia | 13 | Saprotroph | \|Undefined Saprotroph\| | Yeast | NULL | Probable | As nectar-tap_saprotroph (Põlme et al. 2020); Decay sugar-rich substrates (Põlme et al. 2020) | Põlme S, et al. 2020. Fungal Diversity 105:1-16 ((https://doi.org/10.1007/s13225-020-00466-2)) |
| Otu44339 | 1 | k__Fungi(100);p__Ascomycota(100);c__Saccharomycetes(100);o__Saccharomycetales(100);f__Metschnikowiaceae(100);g__Metschnikowia(100);g__Metschnikowia_unclassified(100); | Metschnikowia | 13 | Saprotroph | \|Undefined Saprotroph\| | Yeast | NULL | Probable | As nectar-tap_saprotroph (Põlme et al. 2020); Decay sugar-rich substrates (Põlme et al. 2020) | Põlme S, et al. 2020. Fungal Diversity 105:1-16 ((https://doi.org/10.1007/s13225-020-00466-2)) |
| Otu44340 | 1 | k__Fungi(100);p__Ascomycota(100);c__Saccharomycetes(100);o__Saccharomycetales(100);f__Metschnikowiaceae(100);g__Metschnikowia(100);s__Metschnikowia_pulcherrima(100); | Metschnikowia | 13 | Saprotroph | \|Undefined Saprotroph\| | Yeast | NULL | Probable | As nectar-tap_saprotroph (Põlme et al. 2020); Decay sugar-rich substrates (Põlme et al. 2020) | Põlme S, et al. 2020. Fungal Diversity 105:1-16 ((https://doi.org/10.1007/s13225-020-00466-2)) |
| Otu44341 | 1 | k__Fungi(100);p__Ascomycota(100);c__Saccharomycetes(100);o__Saccharomycetales(100);f__Metschnikowiaceae(100);g__Metschnikowia(100);g__Metschnikowia_unclassified(100); | Metschnikowia | 13 | Saprotroph | \|Undefined Saprotroph\| | Yeast | NULL | Probable | As nectar-tap_saprotroph (Põlme et al. 2020); Decay sugar-rich substrates (Põlme et al. 2020) | Põlme S, et al. 2020. Fungal Diversity 105:1-16 ((https://doi.org/10.1007/s13225-020-00466-2)) |
| Otu44342 | 1 | k__Fungi(100);p__Ascomycota(100);c__Saccharomycetes(100);o__Saccharomycetales(100);f__Metschnikowiaceae(100);g__Metschnikowia(100);s__Metschnikowia_pulcherrima(100); | Metschnikowia | 13 | Saprotroph | \|Undefined Saprotroph\| | Yeast | NULL | Probable | As nectar-tap_saprotroph (Põlme et al. 2020); Decay sugar-rich substrates (Põlme et al. 2020) | Põlme S, et al. 2020. Fungal Diversity 105:1-16 ((https://doi.org/10.1007/s13225-020-00466-2)) |
| Otu44343 | 1 | k__Fungi(100);p__Ascomycota(100);c__Saccharomycetes(100);o__Saccharomycetales(100);f__Metschnikowiaceae(100);g__Metschnikowia(100);s__Metschnikowia_pulcherrima(100); | Metschnikowia | 13 | Saprotroph | \|Undefined Saprotroph\| | Yeast | NULL | Probable | As nectar-tap_saprotroph (Põlme et al. 2020); Decay sugar-rich substrates (Põlme et al. 2020) | Põlme S, et al. 2020. Fungal Diversity 105:1-16 ((https://doi.org/10.1007/s13225-020-00466-2)) |
| Otu44344 | 1 | k__Fungi(100);p__Ascomycota(100);c__Saccharomycetes(100);o__Saccharomycetales(100);f__Metschnikowiaceae(100);g__Metschnikowia(100);s__Metschnikowia_pulcherrima(100); | Metschnikowia | 13 | Saprotroph | \|Undefined Saprotroph\| | Yeast | NULL | Probable | As nectar-tap_saprotroph (Põlme et al. 2020); Decay sugar-rich substrates (Põlme et al. 2020) | Põlme S, et al. 2020. Fungal Diversity 105:1-16 ((https://doi.org/10.1007/s13225-020-00466-2)) |
| Otu44345 | 1 | k__Fungi(100);p__Ascomycota(100);c__Saccharomycetes(100);o__Saccharomycetales(100);f__Metschnikowiaceae(100);g__Metschnikowia(100);s__Metschnikowia_pulcherrima(100); | Metschnikowia | 13 | Saprotroph | \|Undefined Saprotroph\| | Yeast | NULL | Probable | As nectar-tap_saprotroph (Põlme et al. 2020); Decay sugar-rich substrates (Põlme et al. 2020) | Põlme S, et al. 2020. Fungal Diversity 105:1-16 ((https://doi.org/10.1007/s13225-020-00466-2)) |
| Otu44346 | 1 | k__Fungi(100);p__Ascomycota(100);c__Saccharomycetes(100);o__Saccharomycetales(100);f__Metschnikowiaceae(100);g__Metschnikowia(100);g__Metschnikowia_unclassified(100); | Metschnikowia | 13 | Saprotroph | \|Undefined Saprotroph\| | Yeast | NULL | Probable | As nectar-tap_saprotroph (Põlme et al. 2020); Decay sugar-rich substrates (Põlme et al. 2020) | Põlme S, et al. 2020. Fungal Diversity 105:1-16 ((https://doi.org/10.1007/s13225-020-00466-2)) |
| Otu44347 | 1 | k__Fungi(100);p__Ascomycota(100);c__Saccharomycetes(100);o__Saccharomycetales(100);f__Metschnikowiaceae(100);g__Metschnikowia(100);s__Metschnikowia_pulcherrima(100); | Metschnikowia | 13 | Saprotroph | \|Undefined Saprotroph\| | Yeast | NULL | Probable | As nectar-tap_saprotroph (Põlme et al. 2020); Decay sugar-rich substrates (Põlme et al. 2020) | Põlme S, et al. 2020. Fungal Diversity 105:1-16 ((https://doi.org/10.1007/s13225-020-00466-2)) |
| Otu44348 | 1 | k__Fungi(100);p__Ascomycota(100);c__Saccharomycetes(100);o__Saccharomycetales(100);f__Metschnikowiaceae(100);g__Metschnikowia(100);s__Metschnikowia_pulcherrima(100); | Metschnikowia | 13 | Saprotroph | \|Undefined Saprotroph\| | Yeast | NULL | Probable | As nectar-tap_saprotroph (Põlme et al. 2020); Decay sugar-rich substrates (Põlme et al. 2020) | Põlme S, et al. 2020. Fungal Diversity 105:1-16 ((https://doi.org/10.1007/s13225-020-00466-2)) |
| Otu44349 | 1 | k__Fungi(100);p__Ascomycota(100);c__Saccharomycetes(100);o__Saccharomycetales(100);f__Metschnikowiaceae(100);g__Metschnikowia(100);s__Metschnikowia_pulcherrima(100); | Metschnikowia | 13 | Saprotroph | \|Undefined Saprotroph\| | Yeast | NULL | Probable | As nectar-tap_saprotroph (Põlme et al. 2020); Decay sugar-rich substrates (Põlme et al. 2020) | Põlme S, et al. 2020. Fungal Diversity 105:1-16 ((https://doi.org/10.1007/s13225-020-00466-2)) |
| Otu44350 | 1 | k__Fungi(100);p__Ascomycota(100);c__Saccharomycetes(100);o__Saccharomycetales(100);f__Metschnikowiaceae(100);g__Metschnikowia(100);s__Metschnikowia_pulcherrima(100); | Metschnikowia | 13 | Saprotroph | \|Undefined Saprotroph\| | Yeast | NULL | Probable | As nectar-tap_saprotroph (Põlme et al. 2020); Decay sugar-rich substrates (Põlme et al. 2020) | Põlme S, et al. 2020. Fungal Diversity 105:1-16 ((https://doi.org/10.1007/s13225-020-00466-2)) |
| Otu44351 | 1 | k__Fungi(100);p__Ascomycota(100);c__Saccharomycetes(100);o__Saccharomycetales(100);f__Metschnikowiaceae(100);g__Metschnikowia(100);s__Metschnikowia_pulcherrima(100); | Metschnikowia | 13 | Saprotroph | \|Undefined Saprotroph\| | Yeast | NULL | Probable | As nectar-tap_saprotroph (Põlme et al. 2020); Decay sugar-rich substrates (Põlme et al. 2020) | Põlme S, et al. 2020. Fungal Diversity 105:1-16 ((https://doi.org/10.1007/s13225-020-00466-2)) |
| Otu44352 | 1 | k__Fungi(100);p__Ascomycota(100);c__Saccharomycetes(100);o__Saccharomycetales(100);f__Metschnikowiaceae(100);g__Metschnikowia(100);s__Metschnikowia_pulcherrima(100); | Metschnikowia | 13 | Saprotroph | \|Undefined Saprotroph\| | Yeast | NULL | Probable | As nectar-tap_saprotroph (Põlme et al. 2020); Decay sugar-rich substrates (Põlme et al. 2020) | Põlme S, et al. 2020. Fungal Diversity 105:1-16 ((https://doi.org/10.1007/s13225-020-00466-2)) |
| Otu44353 | 1 | k__Fungi(100);p__Ascomycota(100);c__Saccharomycetes(100);o__Saccharomycetales(100);f__Metschnikowiaceae(100);g__Metschnikowia(100);s__Metschnikowia_pulcherrima(100); | Metschnikowia | 13 | Saprotroph | \|Undefined Saprotroph\| | Yeast | NULL | Probable | As nectar-tap_saprotroph (Põlme et al. 2020); Decay sugar-rich substrates (Põlme et al. 2020) | Põlme S, et al. 2020. Fungal Diversity 105:1-16 ((https://doi.org/10.1007/s13225-020-00466-2)) |
| Otu44354 | 1 | k__Fungi(100);p__Ascomycota(100);c__Saccharomycetes(100);o__Saccharomycetales(100);f__Metschnikowiaceae(100);g__Metschnikowia(100);s__Metschnikowia_pulcherrima(100); | Metschnikowia | 13 | Saprotroph | \|Undefined Saprotroph\| | Yeast | NULL | Probable | As nectar-tap_saprotroph (Põlme et al. 2020); Decay sugar-rich substrates (Põlme et al. 2020) | Põlme S, et al. 2020. Fungal Diversity 105:1-16 ((https://doi.org/10.1007/s13225-020-00466-2)) |
| Otu44355 | 1 | k__Fungi(100);p__Ascomycota(100);c__Saccharomycetes(100);o__Saccharomycetales(100);f__Metschnikowiaceae(100);g__Metschnikowia(100);s__Metschnikowia_pulcherrima(100); | Metschnikowia | 13 | Saprotroph | \|Undefined Saprotroph\| | Yeast | NULL | Probable | As nectar-tap_saprotroph (Põlme et al. 2020); Decay sugar-rich substrates (Põlme et al. 2020) | Põlme S, et al. 2020. Fungal Diversity 105:1-16 ((https://doi.org/10.1007/s13225-020-00466-2)) |
| Otu44356 | 1 | k__Fungi(100);p__Ascomycota(100);c__Saccharomycetes(100);o__Saccharomycetales(100);f__Metschnikowiaceae(100);g__Metschnikowia(100);s__Metschnikowia_pulcherrima(100); | Metschnikowia | 13 | Saprotroph | \|Undefined Saprotroph\| | Yeast | NULL | Probable | As nectar-tap_saprotroph (Põlme et al. 2020); Decay sugar-rich substrates (Põlme et al. 2020) | Põlme S, et al. 2020. Fungal Diversity 105:1-16 ((https://doi.org/10.1007/s13225-020-00466-2)) |
| Otu44357 | 1 | k__Fungi(100);p__Ascomycota(100);c__Saccharomycetes(100);o__Saccharomycetales(100);f__Metschnikowiaceae(100);g__Metschnikowia(100);g__Metschnikowia_unclassified(100); | Metschnikowia | 13 | Saprotroph | \|Undefined Saprotroph\| | Yeast | NULL | Probable | As nectar-tap_saprotroph (Põlme et al. 2020); Decay sugar-rich substrates (Põlme et al. 2020) | Põlme S, et al. 2020. Fungal Diversity 105:1-16 ((https://doi.org/10.1007/s13225-020-00466-2)) |
| Otu44358 | 1 | k__Fungi(100);p__Ascomycota(100);c__Saccharomycetes(100);o__Saccharomycetales(100);f__Metschnikowiaceae(100);g__Metschnikowia(100);s__Metschnikowia_pulcherrima(100); | Metschnikowia | 13 | Saprotroph | \|Undefined Saprotroph\| | Yeast | NULL | Probable | As nectar-tap_saprotroph (Põlme et al. 2020); Decay sugar-rich substrates (Põlme et al. 2020) | Põlme S, et al. 2020. Fungal Diversity 105:1-16 ((https://doi.org/10.1007/s13225-020-00466-2)) |
| Otu44359 | 1 | k__Fungi(100);p__Ascomycota(100);c__Saccharomycetes(100);o__Saccharomycetales(100);f__Metschnikowiaceae(100);g__Metschnikowia(100);s__Metschnikowia_pulcherrima(100); | Metschnikowia | 13 | Saprotroph | \|Undefined Saprotroph\| | Yeast | NULL | Probable | As nectar-tap_saprotroph (Põlme et al. 2020); Decay sugar-rich substrates (Põlme et al. 2020) | Põlme S, et al. 2020. Fungal Diversity 105:1-16 ((https://doi.org/10.1007/s13225-020-00466-2)) |
| Otu44360 | 1 | k__Fungi(100);p__Ascomycota(100);c__Saccharomycetes(100);o__Saccharomycetales(100);f__Metschnikowiaceae(100);g__Metschnikowia(100);g__Metschnikowia_unclassified(100); | Metschnikowia | 13 | Saprotroph | \|Undefined Saprotroph\| | Yeast | NULL | Probable | As nectar-tap_saprotroph (Põlme et al. 2020); Decay sugar-rich substrates (Põlme et al. 2020) | Põlme S, et al. 2020. Fungal Diversity 105:1-16 ((https://doi.org/10.1007/s13225-020-00466-2)) |
| Otu44361 | 1 | k__Fungi(100);p__Ascomycota(100);c__Saccharomycetes(100);o__Saccharomycetales(100);f__Metschnikowiaceae(100);g__Metschnikowia(100);s__Metschnikowia_pulcherrima(100); | Metschnikowia | 13 | Saprotroph | \|Undefined Saprotroph\| | Yeast | NULL | Probable | As nectar-tap_saprotroph (Põlme et al. 2020); Decay sugar-rich substrates (Põlme et al. 2020) | Põlme S, et al. 2020. Fungal Diversity 105:1-16 ((https://doi.org/10.1007/s13225-020-00466-2)) |
| Otu44362 | 1 | k__Fungi(100);p__Ascomycota(100);c__Saccharomycetes(100);o__Saccharomycetales(100);f__Metschnikowiaceae(100);g__Metschnikowia(100);g__Metschnikowia_unclassified(100); | Metschnikowia | 13 | Saprotroph | \|Undefined Saprotroph\| | Yeast | NULL | Probable | As nectar-tap_saprotroph (Põlme et al. 2020); Decay sugar-rich substrates (Põlme et al. 2020) | Põlme S, et al. 2020. Fungal Diversity 105:1-16 ((https://doi.org/10.1007/s13225-020-00466-2)) |
| Otu44363 | 1 | k__Fungi(100);p__Ascomycota(100);c__Saccharomycetes(100);o__Saccharomycetales(100);f__Metschnikowiaceae(100);g__Metschnikowia(100);s__Metschnikowia_pulcherrima(100); | Metschnikowia | 13 | Saprotroph | \|Undefined Saprotroph\| | Yeast | NULL | Probable | As nectar-tap_saprotroph (Põlme et al. 2020); Decay sugar-rich substrates (Põlme et al. 2020) | Põlme S, et al. 2020. Fungal Diversity 105:1-16 ((https://doi.org/10.1007/s13225-020-00466-2)) |
| Otu44364 | 1 | k__Fungi(100);p__Ascomycota(100);c__Saccharomycetes(100);o__Saccharomycetales(100);f__Metschnikowiaceae(100);g__Metschnikowia(100);s__Metschnikowia_pulcherrima(100); | Metschnikowia | 13 | Saprotroph | \|Undefined Saprotroph\| | Yeast | NULL | Probable | As nectar-tap_saprotroph (Põlme et al. 2020); Decay sugar-rich substrates (Põlme et al. 2020) | Põlme S, et al. 2020. Fungal Diversity 105:1-16 ((https://doi.org/10.1007/s13225-020-00466-2)) |
| Otu44365 | 1 | k__Fungi(100);p__Ascomycota(100);c__Saccharomycetes(100);o__Saccharomycetales(100);f__Metschnikowiaceae(100);g__Metschnikowia(100);s__Metschnikowia_pulcherrima(100); | Metschnikowia | 13 | Saprotroph | \|Undefined Saprotroph\| | Yeast | NULL | Probable | As nectar-tap_saprotroph (Põlme et al. 2020); Decay sugar-rich substrates (Põlme et al. 2020) | Põlme S, et al. 2020. Fungal Diversity 105:1-16 ((https://doi.org/10.1007/s13225-020-00466-2)) |
| Otu44366 | 1 | k__Fungi(100);p__Ascomycota(100);c__Saccharomycetes(100);o__Saccharomycetales(100);f__Metschnikowiaceae(100);g__Metschnikowia(100);g__Metschnikowia_unclassified(100); | Metschnikowia | 13 | Saprotroph | \|Undefined Saprotroph\| | Yeast | NULL | Probable | As nectar-tap_saprotroph (Põlme et al. 2020); Decay sugar-rich substrates (Põlme et al. 2020) | Põlme S, et al. 2020. Fungal Diversity 105:1-16 ((https://doi.org/10.1007/s13225-020-00466-2)) |
| Otu44367 | 1 | k__Fungi(100);p__Ascomycota(100);c__Saccharomycetes(100);o__Saccharomycetales(100);f__Metschnikowiaceae(100);g__Metschnikowia(100);s__Metschnikowia_pulcherrima(100); | Metschnikowia | 13 | Saprotroph | \|Undefined Saprotroph\| | Yeast | NULL | Probable | As nectar-tap_saprotroph (Põlme et al. 2020); Decay sugar-rich substrates (Põlme et al. 2020) | Põlme S, et al. 2020. Fungal Diversity 105:1-16 ((https://doi.org/10.1007/s13225-020-00466-2)) |
| Otu44368 | 1 | k__Fungi(100);p__Ascomycota(100);c__Saccharomycetes(100);o__Saccharomycetales(100);f__Metschnikowiaceae(100);g__Metschnikowia(100);s__Metschnikowia_pulcherrima(100); | Metschnikowia | 13 | Saprotroph | \|Undefined Saprotroph\| | Yeast | NULL | Probable | As nectar-tap_saprotroph (Põlme et al. 2020); Decay sugar-rich substrates (Põlme et al. 2020) | Põlme S, et al. 2020. Fungal Diversity 105:1-16 ((https://doi.org/10.1007/s13225-020-00466-2)) |
| Otu44369 | 1 | k__Fungi(100);p__Ascomycota(100);c__Saccharomycetes(100);o__Saccharomycetales(100);f__Metschnikowiaceae(100);g__Metschnikowia(100);s__Metschnikowia_pulcherrima(100); | Metschnikowia | 13 | Saprotroph | \|Undefined Saprotroph\| | Yeast | NULL | Probable | As nectar-tap_saprotroph (Põlme et al. 2020); Decay sugar-rich substrates (Põlme et al. 2020) | Põlme S, et al. 2020. Fungal Diversity 105:1-16 ((https://doi.org/10.1007/s13225-020-00466-2)) |
| Otu44370 | 1 | k__Fungi(100);p__Ascomycota(100);c__Saccharomycetes(100);o__Saccharomycetales(100);f__Metschnikowiaceae(100);g__Metschnikowia(100);s__Metschnikowia_pulcherrima(100); | Metschnikowia | 13 | Saprotroph | \|Undefined Saprotroph\| | Yeast | NULL | Probable | As nectar-tap_saprotroph (Põlme et al. 2020); Decay sugar-rich substrates (Põlme et al. 2020) | Põlme S, et al. 2020. Fungal Diversity 105:1-16 ((https://doi.org/10.1007/s13225-020-00466-2)) |
| Otu44371 | 1 | k__Fungi(100);p__Ascomycota(100);c__Saccharomycetes(100);o__Saccharomycetales(100);f__Metschnikowiaceae(100);g__Metschnikowia(100);g__Metschnikowia_unclassified(100); | Metschnikowia | 13 | Saprotroph | \|Undefined Saprotroph\| | Yeast | NULL | Probable | As nectar-tap_saprotroph (Põlme et al. 2020); Decay sugar-rich substrates (Põlme et al. 2020) | Põlme S, et al. 2020. Fungal Diversity 105:1-16 ((https://doi.org/10.1007/s13225-020-00466-2)) |
| Otu44372 | 1 | k__Fungi(100);p__Ascomycota(100);c__Saccharomycetes(100);o__Saccharomycetales(100);f__Metschnikowiaceae(100);g__Metschnikowia(100);s__Metschnikowia_pulcherrima(100); | Metschnikowia | 13 | Saprotroph | \|Undefined Saprotroph\| | Yeast | NULL | Probable | As nectar-tap_saprotroph (Põlme et al. 2020); Decay sugar-rich substrates (Põlme et al. 2020) | Põlme S, et al. 2020. Fungal Diversity 105:1-16 ((https://doi.org/10.1007/s13225-020-00466-2)) |
| Otu44373 | 1 | k__Fungi(100);p__Ascomycota(100);c__Saccharomycetes(100);o__Saccharomycetales(100);f__Metschnikowiaceae(100);g__Metschnikowia(100);s__Metschnikowia_pulcherrima(100); | Metschnikowia | 13 | Saprotroph | \|Undefined Saprotroph\| | Yeast | NULL | Probable | As nectar-tap_saprotroph (Põlme et al. 2020); Decay sugar-rich substrates (Põlme et al. 2020) | Põlme S, et al. 2020. Fungal Diversity 105:1-16 ((https://doi.org/10.1007/s13225-020-00466-2)) |
| Otu44374 | 1 | k__Fungi(100);p__Ascomycota(100);c__Saccharomycetes(100);o__Saccharomycetales(100);f__Metschnikowiaceae(100);g__Metschnikowia(100);s__Metschnikowia_pulcherrima(100); | Metschnikowia | 13 | Saprotroph | \|Undefined Saprotroph\| | Yeast | NULL | Probable | As nectar-tap_saprotroph (Põlme et al. 2020); Decay sugar-rich substrates (Põlme et al. 2020) | Põlme S, et al. 2020. Fungal Diversity 105:1-16 ((https://doi.org/10.1007/s13225-020-00466-2)) |
| Otu44375 | 1 | k__Fungi(100);p__Ascomycota(100);c__Saccharomycetes(100);o__Saccharomycetales(100);f__Metschnikowiaceae(100);g__Metschnikowia(100);s__Metschnikowia_pulcherrima(100); | Metschnikowia | 13 | Saprotroph | \|Undefined Saprotroph\| | Yeast | NULL | Probable | As nectar-tap_saprotroph (Põlme et al. 2020); Decay sugar-rich substrates (Põlme et al. 2020) | Põlme S, et al. 2020. Fungal Diversity 105:1-16 ((https://doi.org/10.1007/s13225-020-00466-2)) |
| Otu44376 | 1 | k__Fungi(100);p__Ascomycota(100);c__Saccharomycetes(100);o__Saccharomycetales(100);f__Metschnikowiaceae(100);g__Metschnikowia(100);s__Metschnikowia_pulcherrima(100); | Metschnikowia | 13 | Saprotroph | \|Undefined Saprotroph\| | Yeast | NULL | Probable | As nectar-tap_saprotroph (Põlme et al. 2020); Decay sugar-rich substrates (Põlme et al. 2020) | Põlme S, et al. 2020. Fungal Diversity 105:1-16 ((https://doi.org/10.1007/s13225-020-00466-2)) |
| Otu44377 | 1 | k__Fungi(100);p__Ascomycota(100);c__Saccharomycetes(100);o__Saccharomycetales(100);f__Metschnikowiaceae(100);g__Metschnikowia(100);s__Metschnikowia_pulcherrima(100); | Metschnikowia | 13 | Saprotroph | \|Undefined Saprotroph\| | Yeast | NULL | Probable | As nectar-tap_saprotroph (Põlme et al. 2020); Decay sugar-rich substrates (Põlme et al. 2020) | Põlme S, et al. 2020. Fungal Diversity 105:1-16 ((https://doi.org/10.1007/s13225-020-00466-2)) |
| Otu44378 | 1 | k__Fungi(100);p__Ascomycota(100);c__Saccharomycetes(100);o__Saccharomycetales(100);f__Metschnikowiaceae(100);g__Metschnikowia(100);s__Metschnikowia_pulcherrima(100); | Metschnikowia | 13 | Saprotroph | \|Undefined Saprotroph\| | Yeast | NULL | Probable | As nectar-tap_saprotroph (Põlme et al. 2020); Decay sugar-rich substrates (Põlme et al. 2020) | Põlme S, et al. 2020. Fungal Diversity 105:1-16 ((https://doi.org/10.1007/s13225-020-00466-2)) |
| Otu44379 | 1 | k__Fungi(100);p__Ascomycota(100);c__Saccharomycetes(100);o__Saccharomycetales(100);f__Metschnikowiaceae(100);g__Metschnikowia(100);g__Metschnikowia_unclassified(100); | Metschnikowia | 13 | Saprotroph | \|Undefined Saprotroph\| | Yeast | NULL | Probable | As nectar-tap_saprotroph (Põlme et al. 2020); Decay sugar-rich substrates (Põlme et al. 2020) | Põlme S, et al. 2020. Fungal Diversity 105:1-16 ((https://doi.org/10.1007/s13225-020-00466-2)) |
| Otu44380 | 1 | k__Fungi(100);p__Ascomycota(100);c__Saccharomycetes(100);o__Saccharomycetales(100);f__Metschnikowiaceae(100);g__Metschnikowia(100);g__Metschnikowia_unclassified(100); | Metschnikowia | 13 | Saprotroph | \|Undefined Saprotroph\| | Yeast | NULL | Probable | As nectar-tap_saprotroph (Põlme et al. 2020); Decay sugar-rich substrates (Põlme et al. 2020) | Põlme S, et al. 2020. Fungal Diversity 105:1-16 ((https://doi.org/10.1007/s13225-020-00466-2)) |
| Otu44381 | 1 | k__Fungi(100);p__Ascomycota(100);c__Saccharomycetes(100);o__Saccharomycetales(100);f__Metschnikowiaceae(100);g__Metschnikowia(100);g__Metschnikowia_unclassified(100); | Metschnikowia | 13 | Saprotroph | \|Undefined Saprotroph\| | Yeast | NULL | Probable | As nectar-tap_saprotroph (Põlme et al. 2020); Decay sugar-rich substrates (Põlme et al. 2020) | Põlme S, et al. 2020. Fungal Diversity 105:1-16 ((https://doi.org/10.1007/s13225-020-00466-2)) |
| Otu44382 | 1 | k__Fungi(100);p__Ascomycota(100);c__Saccharomycetes(100);o__Saccharomycetales(100);f__Metschnikowiaceae(100);g__Metschnikowia(100);s__Metschnikowia_pulcherrima(100); | Metschnikowia | 13 | Saprotroph | \|Undefined Saprotroph\| | Yeast | NULL | Probable | As nectar-tap_saprotroph (Põlme et al. 2020); Decay sugar-rich substrates (Põlme et al. 2020) | Põlme S, et al. 2020. Fungal Diversity 105:1-16 ((https://doi.org/10.1007/s13225-020-00466-2)) |
| Otu44383 | 1 | k__Fungi(100);p__Ascomycota(100);c__Saccharomycetes(100);o__Saccharomycetales(100);f__Metschnikowiaceae(100);g__Metschnikowia(100);g__Metschnikowia_unclassified(100); | Metschnikowia | 13 | Saprotroph | \|Undefined Saprotroph\| | Yeast | NULL | Probable | As nectar-tap_saprotroph (Põlme et al. 2020); Decay sugar-rich substrates (Põlme et al. 2020) | Põlme S, et al. 2020. Fungal Diversity 105:1-16 ((https://doi.org/10.1007/s13225-020-00466-2)) |
| Otu44384 | 1 | k__Fungi(100);p__Ascomycota(100);c__Saccharomycetes(100);o__Saccharomycetales(100);f__Metschnikowiaceae(100);g__Metschnikowia(100);s__Metschnikowia_pulcherrima(100); | Metschnikowia | 13 | Saprotroph | \|Undefined Saprotroph\| | Yeast | NULL | Probable | As nectar-tap_saprotroph (Põlme et al. 2020); Decay sugar-rich substrates (Põlme et al. 2020) | Põlme S, et al. 2020. Fungal Diversity 105:1-16 ((https://doi.org/10.1007/s13225-020-00466-2)) |
| Otu44385 | 1 | k__Fungi(100);p__Ascomycota(100);c__Saccharomycetes(100);o__Saccharomycetales(100);f__Metschnikowiaceae(100);g__Metschnikowia(100);s__Metschnikowia_pulcherrima(100); | Metschnikowia | 13 | Saprotroph | \|Undefined Saprotroph\| | Yeast | NULL | Probable | As nectar-tap_saprotroph (Põlme et al. 2020); Decay sugar-rich substrates (Põlme et al. 2020) | Põlme S, et al. 2020. Fungal Diversity 105:1-16 ((https://doi.org/10.1007/s13225-020-00466-2)) |
| Otu44386 | 1 | k__Fungi(100);p__Ascomycota(100);c__Saccharomycetes(100);o__Saccharomycetales(100);f__Metschnikowiaceae(100);g__Metschnikowia(100);s__Metschnikowia_pulcherrima(100); | Metschnikowia | 13 | Saprotroph | \|Undefined Saprotroph\| | Yeast | NULL | Probable | As nectar-tap_saprotroph (Põlme et al. 2020); Decay sugar-rich substrates (Põlme et al. 2020) | Põlme S, et al. 2020. Fungal Diversity 105:1-16 ((https://doi.org/10.1007/s13225-020-00466-2)) |
| Otu44387 | 1 | k__Fungi(100);p__Ascomycota(100);c__Saccharomycetes(100);o__Saccharomycetales(100);f__Metschnikowiaceae(100);g__Metschnikowia(100);s__Metschnikowia_pulcherrima(100); | Metschnikowia | 13 | Saprotroph | \|Undefined Saprotroph\| | Yeast | NULL | Probable | As nectar-tap_saprotroph (Põlme et al. 2020); Decay sugar-rich substrates (Põlme et al. 2020) | Põlme S, et al. 2020. Fungal Diversity 105:1-16 ((https://doi.org/10.1007/s13225-020-00466-2)) |
| Otu44388 | 1 | k__Fungi(100);p__Ascomycota(100);c__Saccharomycetes(100);o__Saccharomycetales(100);f__Metschnikowiaceae(100);g__Metschnikowia(100);s__Metschnikowia_pulcherrima(100); | Metschnikowia | 13 | Saprotroph | \|Undefined Saprotroph\| | Yeast | NULL | Probable | As nectar-tap_saprotroph (Põlme et al. 2020); Decay sugar-rich substrates (Põlme et al. 2020) | Põlme S, et al. 2020. Fungal Diversity 105:1-16 ((https://doi.org/10.1007/s13225-020-00466-2)) |
| Otu44389 | 1 | k__Fungi(100);p__Ascomycota(100);c__Saccharomycetes(100);o__Saccharomycetales(100);f__Metschnikowiaceae(100);g__Metschnikowia(100);s__Metschnikowia_pulcherrima(100); | Metschnikowia | 13 | Saprotroph | \|Undefined Saprotroph\| | Yeast | NULL | Probable | As nectar-tap_saprotroph (Põlme et al. 2020); Decay sugar-rich substrates (Põlme et al. 2020) | Põlme S, et al. 2020. Fungal Diversity 105:1-16 ((https://doi.org/10.1007/s13225-020-00466-2)) |
| Otu44390 | 1 | k__Fungi(100);p__Ascomycota(100);c__Saccharomycetes(100);o__Saccharomycetales(100);f__Metschnikowiaceae(100);g__Metschnikowia(100);s__Metschnikowia_pulcherrima(100); | Metschnikowia | 13 | Saprotroph | \|Undefined Saprotroph\| | Yeast | NULL | Probable | As nectar-tap_saprotroph (Põlme et al. 2020); Decay sugar-rich substrates (Põlme et al. 2020) | Põlme S, et al. 2020. Fungal Diversity 105:1-16 ((https://doi.org/10.1007/s13225-020-00466-2)) |
| Otu44391 | 1 | k__Fungi(100);p__Ascomycota(100);c__Saccharomycetes(100);o__Saccharomycetales(100);f__Metschnikowiaceae(100);g__Metschnikowia(100);g__Metschnikowia_unclassified(100); | Metschnikowia | 13 | Saprotroph | \|Undefined Saprotroph\| | Yeast | NULL | Probable | As nectar-tap_saprotroph (Põlme et al. 2020); Decay sugar-rich substrates (Põlme et al. 2020) | Põlme S, et al. 2020. Fungal Diversity 105:1-16 ((https://doi.org/10.1007/s13225-020-00466-2)) |
| Otu44392 | 1 | k__Fungi(100);p__Ascomycota(100);c__Saccharomycetes(100);o__Saccharomycetales(100);f__Metschnikowiaceae(100);g__Metschnikowia(100);s__Metschnikowia_pulcherrima(100); | Metschnikowia | 13 | Saprotroph | \|Undefined Saprotroph\| | Yeast | NULL | Probable | As nectar-tap_saprotroph (Põlme et al. 2020); Decay sugar-rich substrates (Põlme et al. 2020) | Põlme S, et al. 2020. Fungal Diversity 105:1-16 ((https://doi.org/10.1007/s13225-020-00466-2)) |
| Otu44393 | 1 | k__Fungi(100);p__Ascomycota(100);c__Saccharomycetes(100);o__Saccharomycetales(100);f__Metschnikowiaceae(100);g__Metschnikowia(100);s__Metschnikowia_pulcherrima(100); | Metschnikowia | 13 | Saprotroph | \|Undefined Saprotroph\| | Yeast | NULL | Probable | As nectar-tap_saprotroph (Põlme et al. 2020); Decay sugar-rich substrates (Põlme et al. 2020) | Põlme S, et al. 2020. Fungal Diversity 105:1-16 ((https://doi.org/10.1007/s13225-020-00466-2)) |
| Otu44394 | 1 | k__Fungi(100);p__Ascomycota(100);c__Saccharomycetes(100);o__Saccharomycetales(100);f__Metschnikowiaceae(100);g__Metschnikowia(100);g__Metschnikowia_unclassified(100); | Metschnikowia | 13 | Saprotroph | \|Undefined Saprotroph\| | Yeast | NULL | Probable | As nectar-tap_saprotroph (Põlme et al. 2020); Decay sugar-rich substrates (Põlme et al. 2020) | Põlme S, et al. 2020. Fungal Diversity 105:1-16 ((https://doi.org/10.1007/s13225-020-00466-2)) |
| Otu44395 | 1 | k__Fungi(100);p__Ascomycota(100);c__Saccharomycetes(100);o__Saccharomycetales(100);f__Metschnikowiaceae(100);g__Metschnikowia(100);s__Metschnikowia_pulcherrima(100); | Metschnikowia | 13 | Saprotroph | \|Undefined Saprotroph\| | Yeast | NULL | Probable | As nectar-tap_saprotroph (Põlme et al. 2020); Decay sugar-rich substrates (Põlme et al. 2020) | Põlme S, et al. 2020. Fungal Diversity 105:1-16 ((https://doi.org/10.1007/s13225-020-00466-2)) |
| Otu44396 | 1 | k__Fungi(100);p__Ascomycota(100);c__Saccharomycetes(100);o__Saccharomycetales(100);f__Metschnikowiaceae(100);g__Metschnikowia(100);s__Metschnikowia_pulcherrima(100); | Metschnikowia | 13 | Saprotroph | \|Undefined Saprotroph\| | Yeast | NULL | Probable | As nectar-tap_saprotroph (Põlme et al. 2020); Decay sugar-rich substrates (Põlme et al. 2020) | Põlme S, et al. 2020. Fungal Diversity 105:1-16 ((https://doi.org/10.1007/s13225-020-00466-2)) |
| Otu44397 | 1 | k__Fungi(100);p__Ascomycota(100);c__Saccharomycetes(100);o__Saccharomycetales(100);f__Metschnikowiaceae(100);g__Metschnikowia(100);g__Metschnikowia_unclassified(100); | Metschnikowia | 13 | Saprotroph | \|Undefined Saprotroph\| | Yeast | NULL | Probable | As nectar-tap_saprotroph (Põlme et al. 2020); Decay sugar-rich substrates (Põlme et al. 2020) | Põlme S, et al. 2020. Fungal Diversity 105:1-16 ((https://doi.org/10.1007/s13225-020-00466-2)) |
| Otu44398 | 1 | k__Fungi(100);p__Ascomycota(100);c__Saccharomycetes(100);o__Saccharomycetales(100);f__Metschnikowiaceae(100);g__Metschnikowia(100);s__Metschnikowia_pulcherrima(100); | Metschnikowia | 13 | Saprotroph | \|Undefined Saprotroph\| | Yeast | NULL | Probable | As nectar-tap_saprotroph (Põlme et al. 2020); Decay sugar-rich substrates (Põlme et al. 2020) | Põlme S, et al. 2020. Fungal Diversity 105:1-16 ((https://doi.org/10.1007/s13225-020-00466-2)) |
| Otu44399 | 1 | k__Fungi(100);p__Ascomycota(100);c__Saccharomycetes(100);o__Saccharomycetales(100);f__Metschnikowiaceae(100);g__Metschnikowia(100);s__Metschnikowia_pulcherrima(100); | Metschnikowia | 13 | Saprotroph | \|Undefined Saprotroph\| | Yeast | NULL | Probable | As nectar-tap_saprotroph (Põlme et al. 2020); Decay sugar-rich substrates (Põlme et al. 2020) | Põlme S, et al. 2020. Fungal Diversity 105:1-16 ((https://doi.org/10.1007/s13225-020-00466-2)) |
| Otu44400 | 1 | k__Fungi(100);p__Ascomycota(100);c__Saccharomycetes(100);o__Saccharomycetales(100);f__Metschnikowiaceae(100);g__Metschnikowia(100);g__Metschnikowia_unclassified(100); | Metschnikowia | 13 | Saprotroph | \|Undefined Saprotroph\| | Yeast | NULL | Probable | As nectar-tap_saprotroph (Põlme et al. 2020); Decay sugar-rich substrates (Põlme et al. 2020) | Põlme S, et al. 2020. Fungal Diversity 105:1-16 ((https://doi.org/10.1007/s13225-020-00466-2)) |
| Otu44401 | 1 | k__Fungi(100);p__Ascomycota(100);c__Saccharomycetes(100);o__Saccharomycetales(100);f__Metschnikowiaceae(100);g__Metschnikowia(100);g__Metschnikowia_unclassified(100); | Metschnikowia | 13 | Saprotroph | \|Undefined Saprotroph\| | Yeast | NULL | Probable | As nectar-tap_saprotroph (Põlme et al. 2020); Decay sugar-rich substrates (Põlme et al. 2020) | Põlme S, et al. 2020. Fungal Diversity 105:1-16 ((https://doi.org/10.1007/s13225-020-00466-2)) |
| Otu44402 | 1 | k__Fungi(100);p__Ascomycota(100);c__Saccharomycetes(100);o__Saccharomycetales(100);f__Metschnikowiaceae(100);g__Metschnikowia(100);s__Metschnikowia_pulcherrima(100); | Metschnikowia | 13 | Saprotroph | \|Undefined Saprotroph\| | Yeast | NULL | Probable | As nectar-tap_saprotroph (Põlme et al. 2020); Decay sugar-rich substrates (Põlme et al. 2020) | Põlme S, et al. 2020. Fungal Diversity 105:1-16 ((https://doi.org/10.1007/s13225-020-00466-2)) |
| Otu44403 | 1 | k__Fungi(100);p__Ascomycota(100);c__Saccharomycetes(100);o__Saccharomycetales(100);f__Metschnikowiaceae(100);g__Metschnikowia(100);g__Metschnikowia_unclassified(100); | Metschnikowia | 13 | Saprotroph | \|Undefined Saprotroph\| | Yeast | NULL | Probable | As nectar-tap_saprotroph (Põlme et al. 2020); Decay sugar-rich substrates (Põlme et al. 2020) | Põlme S, et al. 2020. Fungal Diversity 105:1-16 ((https://doi.org/10.1007/s13225-020-00466-2)) |
| Otu44404 | 1 | k__Fungi(100);p__Ascomycota(100);c__Saccharomycetes(100);o__Saccharomycetales(100);f__Metschnikowiaceae(100);g__Metschnikowia(100);g__Metschnikowia_unclassified(100); | Metschnikowia | 13 | Saprotroph | \|Undefined Saprotroph\| | Yeast | NULL | Probable | As nectar-tap_saprotroph (Põlme et al. 2020); Decay sugar-rich substrates (Põlme et al. 2020) | Põlme S, et al. 2020. Fungal Diversity 105:1-16 ((https://doi.org/10.1007/s13225-020-00466-2)) |
| Otu44405 | 1 | k__Fungi(100);p__Ascomycota(100);c__Saccharomycetes(100);o__Saccharomycetales(100);f__Metschnikowiaceae(100);g__Metschnikowia(100);s__Metschnikowia_pulcherrima(100); | Metschnikowia | 13 | Saprotroph | \|Undefined Saprotroph\| | Yeast | NULL | Probable | As nectar-tap_saprotroph (Põlme et al. 2020); Decay sugar-rich substrates (Põlme et al. 2020) | Põlme S, et al. 2020. Fungal Diversity 105:1-16 ((https://doi.org/10.1007/s13225-020-00466-2)) |
| Otu44406 | 1 | k__Fungi(100);p__Ascomycota(100);c__Saccharomycetes(100);o__Saccharomycetales(100);f__Metschnikowiaceae(100);g__Metschnikowia(100);g__Metschnikowia_unclassified(100); | Metschnikowia | 13 | Saprotroph | \|Undefined Saprotroph\| | Yeast | NULL | Probable | As nectar-tap_saprotroph (Põlme et al. 2020); Decay sugar-rich substrates (Põlme et al. 2020) | Põlme S, et al. 2020. Fungal Diversity 105:1-16 ((https://doi.org/10.1007/s13225-020-00466-2)) |
| Otu44407 | 1 | k__Fungi(100);p__Ascomycota(100);c__Saccharomycetes(100);o__Saccharomycetales(100);f__Metschnikowiaceae(100);g__Metschnikowia(100);s__Metschnikowia_pulcherrima(100); | Metschnikowia | 13 | Saprotroph | \|Undefined Saprotroph\| | Yeast | NULL | Probable | As nectar-tap_saprotroph (Põlme et al. 2020); Decay sugar-rich substrates (Põlme et al. 2020) | Põlme S, et al. 2020. Fungal Diversity 105:1-16 ((https://doi.org/10.1007/s13225-020-00466-2)) |
| Otu44408 | 1 | k__Fungi(100);p__Ascomycota(100);c__Saccharomycetes(100);o__Saccharomycetales(100);f__Metschnikowiaceae(100);g__Metschnikowia(100);s__Metschnikowia_pulcherrima(100); | Metschnikowia | 13 | Saprotroph | \|Undefined Saprotroph\| | Yeast | NULL | Probable | As nectar-tap_saprotroph (Põlme et al. 2020); Decay sugar-rich substrates (Põlme et al. 2020) | Põlme S, et al. 2020. Fungal Diversity 105:1-16 ((https://doi.org/10.1007/s13225-020-00466-2)) |
| Otu44409 | 1 | k__Fungi(100);p__Ascomycota(100);c__Saccharomycetes(100);o__Saccharomycetales(100);f__Metschnikowiaceae(100);g__Metschnikowia(100);s__Metschnikowia_pulcherrima(100); | Metschnikowia | 13 | Saprotroph | \|Undefined Saprotroph\| | Yeast | NULL | Probable | As nectar-tap_saprotroph (Põlme et al. 2020); Decay sugar-rich substrates (Põlme et al. 2020) | Põlme S, et al. 2020. Fungal Diversity 105:1-16 ((https://doi.org/10.1007/s13225-020-00466-2)) |
| Otu44410 | 1 | k__Fungi(100);p__Ascomycota(100);c__Saccharomycetes(100);o__Saccharomycetales(100);f__Metschnikowiaceae(100);g__Metschnikowia(100);s__Metschnikowia_pulcherrima(100); | Metschnikowia | 13 | Saprotroph | \|Undefined Saprotroph\| | Yeast | NULL | Probable | As nectar-tap_saprotroph (Põlme et al. 2020); Decay sugar-rich substrates (Põlme et al. 2020) | Põlme S, et al. 2020. Fungal Diversity 105:1-16 ((https://doi.org/10.1007/s13225-020-00466-2)) |
| Otu44411 | 1 | k__Fungi(100);p__Ascomycota(100);c__Saccharomycetes(100);o__Saccharomycetales(100);f__Metschnikowiaceae(100);g__Metschnikowia(100);s__Metschnikowia_pulcherrima(100); | Metschnikowia | 13 | Saprotroph | \|Undefined Saprotroph\| | Yeast | NULL | Probable | As nectar-tap_saprotroph (Põlme et al. 2020); Decay sugar-rich substrates (Põlme et al. 2020) | Põlme S, et al. 2020. Fungal Diversity 105:1-16 ((https://doi.org/10.1007/s13225-020-00466-2)) |
| Otu44412 | 1 | k__Fungi(100);p__Ascomycota(100);c__Saccharomycetes(100);o__Saccharomycetales(100);f__Metschnikowiaceae(100);g__Metschnikowia(100);g__Metschnikowia_unclassified(100); | Metschnikowia | 13 | Saprotroph | \|Undefined Saprotroph\| | Yeast | NULL | Probable | As nectar-tap_saprotroph (Põlme et al. 2020); Decay sugar-rich substrates (Põlme et al. 2020) | Põlme S, et al. 2020. Fungal Diversity 105:1-16 ((https://doi.org/10.1007/s13225-020-00466-2)) |
| Otu44413 | 1 | k__Fungi(100);p__Ascomycota(100);c__Saccharomycetes(100);o__Saccharomycetales(100);f__Metschnikowiaceae(100);g__Metschnikowia(100);s__Metschnikowia_pulcherrima(100); | Metschnikowia | 13 | Saprotroph | \|Undefined Saprotroph\| | Yeast | NULL | Probable | As nectar-tap_saprotroph (Põlme et al. 2020); Decay sugar-rich substrates (Põlme et al. 2020) | Põlme S, et al. 2020. Fungal Diversity 105:1-16 ((https://doi.org/10.1007/s13225-020-00466-2)) |
| Otu44414 | 1 | k__Fungi(100);p__Ascomycota(100);c__Saccharomycetes(100);o__Saccharomycetales(100);f__Metschnikowiaceae(100);g__Metschnikowia(100);s__Metschnikowia_pulcherrima(100); | Metschnikowia | 13 | Saprotroph | \|Undefined Saprotroph\| | Yeast | NULL | Probable | As nectar-tap_saprotroph (Põlme et al. 2020); Decay sugar-rich substrates (Põlme et al. 2020) | Põlme S, et al. 2020. Fungal Diversity 105:1-16 ((https://doi.org/10.1007/s13225-020-00466-2)) |
| Otu44415 | 1 | k__Fungi(100);p__Ascomycota(100);c__Saccharomycetes(100);o__Saccharomycetales(100);f__Metschnikowiaceae(100);g__Metschnikowia(100);s__Metschnikowia_pulcherrima(100); | Metschnikowia | 13 | Saprotroph | \|Undefined Saprotroph\| | Yeast | NULL | Probable | As nectar-tap_saprotroph (Põlme et al. 2020); Decay sugar-rich substrates (Põlme et al. 2020) | Põlme S, et al. 2020. Fungal Diversity 105:1-16 ((https://doi.org/10.1007/s13225-020-00466-2)) |
| Otu44416 | 1 | k__Fungi(100);p__Ascomycota(100);c__Saccharomycetes(100);o__Saccharomycetales(100);f__Metschnikowiaceae(100);g__Metschnikowia(100);s__Metschnikowia_pulcherrima(100); | Metschnikowia | 13 | Saprotroph | \|Undefined Saprotroph\| | Yeast | NULL | Probable | As nectar-tap_saprotroph (Põlme et al. 2020); Decay sugar-rich substrates (Põlme et al. 2020) | Põlme S, et al. 2020. Fungal Diversity 105:1-16 ((https://doi.org/10.1007/s13225-020-00466-2)) |
| Otu44417 | 1 | k__Fungi(100);p__Ascomycota(100);c__Saccharomycetes(100);o__Saccharomycetales(100);f__Metschnikowiaceae(100);g__Metschnikowia(100);s__Metschnikowia_pulcherrima(100); | Metschnikowia | 13 | Saprotroph | \|Undefined Saprotroph\| | Yeast | NULL | Probable | As nectar-tap_saprotroph (Põlme et al. 2020); Decay sugar-rich substrates (Põlme et al. 2020) | Põlme S, et al. 2020. Fungal Diversity 105:1-16 ((https://doi.org/10.1007/s13225-020-00466-2)) |
| Otu44419 | 1 | k__Fungi(100);p__Ascomycota(100);c__Saccharomycetes(100);o__Saccharomycetales(100);f__Metschnikowiaceae(100);g__Metschnikowia(100);s__Metschnikowia_pulcherrima(100); | Metschnikowia | 13 | Saprotroph | \|Undefined Saprotroph\| | Yeast | NULL | Probable | As nectar-tap_saprotroph (Põlme et al. 2020); Decay sugar-rich substrates (Põlme et al. 2020) | Põlme S, et al. 2020. Fungal Diversity 105:1-16 ((https://doi.org/10.1007/s13225-020-00466-2)) |
| Otu44420 | 1 | k__Fungi(100);p__Ascomycota(100);c__Saccharomycetes(100);o__Saccharomycetales(100);f__Metschnikowiaceae(100);g__Metschnikowia(100);s__Metschnikowia_pulcherrima(100); | Metschnikowia | 13 | Saprotroph | \|Undefined Saprotroph\| | Yeast | NULL | Probable | As nectar-tap_saprotroph (Põlme et al. 2020); Decay sugar-rich substrates (Põlme et al. 2020) | Põlme S, et al. 2020. Fungal Diversity 105:1-16 ((https://doi.org/10.1007/s13225-020-00466-2)) |
| Otu44423 | 1 | k__Fungi(100);p__Ascomycota(100);c__Saccharomycetes(100);o__Saccharomycetales(100);f__Metschnikowiaceae(100);g__Metschnikowia(100);g__Metschnikowia_unclassified(100); | Metschnikowia | 13 | Saprotroph | \|Undefined Saprotroph\| | Yeast | NULL | Probable | As nectar-tap_saprotroph (Põlme et al. 2020); Decay sugar-rich substrates (Põlme et al. 2020) | Põlme S, et al. 2020. Fungal Diversity 105:1-16 ((https://doi.org/10.1007/s13225-020-00466-2)) |
| Otu44424 | 1 | k__Fungi(100);p__Ascomycota(100);c__Saccharomycetes(100);o__Saccharomycetales(100);f__Metschnikowiaceae(100);g__Metschnikowia(100);g__Metschnikowia_unclassified(100); | Metschnikowia | 13 | Saprotroph | \|Undefined Saprotroph\| | Yeast | NULL | Probable | As nectar-tap_saprotroph (Põlme et al. 2020); Decay sugar-rich substrates (Põlme et al. 2020) | Põlme S, et al. 2020. Fungal Diversity 105:1-16 ((https://doi.org/10.1007/s13225-020-00466-2)) |
| Otu44425 | 1 | k__Fungi(100);p__Ascomycota(100);c__Saccharomycetes(100);o__Saccharomycetales(100);f__Metschnikowiaceae(100);g__Metschnikowia(100);g__Metschnikowia_unclassified(100); | Metschnikowia | 13 | Saprotroph | \|Undefined Saprotroph\| | Yeast | NULL | Probable | As nectar-tap_saprotroph (Põlme et al. 2020); Decay sugar-rich substrates (Põlme et al. 2020) | Põlme S, et al. 2020. Fungal Diversity 105:1-16 ((https://doi.org/10.1007/s13225-020-00466-2)) |
| Otu44426 | 1 | k__Fungi(100);p__Ascomycota(100);c__Saccharomycetes(100);o__Saccharomycetales(100);f__Metschnikowiaceae(100);g__Metschnikowia(100);s__Metschnikowia_pulcherrima(100); | Metschnikowia | 13 | Saprotroph | \|Undefined Saprotroph\| | Yeast | NULL | Probable | As nectar-tap_saprotroph (Põlme et al. 2020); Decay sugar-rich substrates (Põlme et al. 2020) | Põlme S, et al. 2020. Fungal Diversity 105:1-16 ((https://doi.org/10.1007/s13225-020-00466-2)) |
| Otu44427 | 1 | k__Fungi(100);p__Ascomycota(100);c__Saccharomycetes(100);o__Saccharomycetales(100);f__Metschnikowiaceae(100);g__Metschnikowia(100);s__Metschnikowia_pulcherrima(100); | Metschnikowia | 13 | Saprotroph | \|Undefined Saprotroph\| | Yeast | NULL | Probable | As nectar-tap_saprotroph (Põlme et al. 2020); Decay sugar-rich substrates (Põlme et al. 2020) | Põlme S, et al. 2020. Fungal Diversity 105:1-16 ((https://doi.org/10.1007/s13225-020-00466-2)) |
| Otu44428 | 1 | k__Fungi(100);p__Ascomycota(100);c__Saccharomycetes(100);o__Saccharomycetales(100);f__Metschnikowiaceae(100);g__Metschnikowia(100);g__Metschnikowia_unclassified(100); | Metschnikowia | 13 | Saprotroph | \|Undefined Saprotroph\| | Yeast | NULL | Probable | As nectar-tap_saprotroph (Põlme et al. 2020); Decay sugar-rich substrates (Põlme et al. 2020) | Põlme S, et al. 2020. Fungal Diversity 105:1-16 ((https://doi.org/10.1007/s13225-020-00466-2)) |
| Otu44429 | 1 | k__Fungi(100);p__Ascomycota(100);c__Saccharomycetes(100);o__Saccharomycetales(100);f__Metschnikowiaceae(100);g__Metschnikowia(100);s__Metschnikowia_pulcherrima(100); | Metschnikowia | 13 | Saprotroph | \|Undefined Saprotroph\| | Yeast | NULL | Probable | As nectar-tap_saprotroph (Põlme et al. 2020); Decay sugar-rich substrates (Põlme et al. 2020) | Põlme S, et al. 2020. Fungal Diversity 105:1-16 ((https://doi.org/10.1007/s13225-020-00466-2)) |
| Otu44430 | 1 | k__Fungi(100);p__Ascomycota(100);c__Saccharomycetes(100);o__Saccharomycetales(100);f__Metschnikowiaceae(100);g__Metschnikowia(100);g__Metschnikowia_unclassified(100); | Metschnikowia | 13 | Saprotroph | \|Undefined Saprotroph\| | Yeast | NULL | Probable | As nectar-tap_saprotroph (Põlme et al. 2020); Decay sugar-rich substrates (Põlme et al. 2020) | Põlme S, et al. 2020. Fungal Diversity 105:1-16 ((https://doi.org/10.1007/s13225-020-00466-2)) |
| Otu44431 | 1 | k__Fungi(100);p__Ascomycota(100);c__Saccharomycetes(100);o__Saccharomycetales(100);f__Metschnikowiaceae(100);g__Metschnikowia(100);s__Metschnikowia_pulcherrima(100); | Metschnikowia | 13 | Saprotroph | \|Undefined Saprotroph\| | Yeast | NULL | Probable | As nectar-tap_saprotroph (Põlme et al. 2020); Decay sugar-rich substrates (Põlme et al. 2020) | Põlme S, et al. 2020. Fungal Diversity 105:1-16 ((https://doi.org/10.1007/s13225-020-00466-2)) |
| Otu44434 | 1 | k__Fungi(100);p__Ascomycota(100);c__Saccharomycetes(100);o__Saccharomycetales(100);f__Metschnikowiaceae(100);g__Metschnikowia(100);s__Metschnikowia_pulcherrima(100); | Metschnikowia | 13 | Saprotroph | \|Undefined Saprotroph\| | Yeast | NULL | Probable | As nectar-tap_saprotroph (Põlme et al. 2020); Decay sugar-rich substrates (Põlme et al. 2020) | Põlme S, et al. 2020. Fungal Diversity 105:1-16 ((https://doi.org/10.1007/s13225-020-00466-2)) |
| Otu44435 | 1 | k__Fungi(100);p__Ascomycota(100);c__Saccharomycetes(100);o__Saccharomycetales(100);f__Metschnikowiaceae(100);g__Metschnikowia(100);g__Metschnikowia_unclassified(100); | Metschnikowia | 13 | Saprotroph | \|Undefined Saprotroph\| | Yeast | NULL | Probable | As nectar-tap_saprotroph (Põlme et al. 2020); Decay sugar-rich substrates (Põlme et al. 2020) | Põlme S, et al. 2020. Fungal Diversity 105:1-16 ((https://doi.org/10.1007/s13225-020-00466-2)) |
| Otu44436 | 1 | k__Fungi(100);p__Ascomycota(100);c__Saccharomycetes(100);o__Saccharomycetales(100);f__Metschnikowiaceae(100);g__Metschnikowia(100);s__Metschnikowia_pulcherrima(100); | Metschnikowia | 13 | Saprotroph | \|Undefined Saprotroph\| | Yeast | NULL | Probable | As nectar-tap_saprotroph (Põlme et al. 2020); Decay sugar-rich substrates (Põlme et al. 2020) | Põlme S, et al. 2020. Fungal Diversity 105:1-16 ((https://doi.org/10.1007/s13225-020-00466-2)) |
| Otu44437 | 1 | k__Fungi(100);p__Ascomycota(100);c__Saccharomycetes(100);o__Saccharomycetales(100);f__Metschnikowiaceae(100);g__Metschnikowia(100);s__Metschnikowia_pulcherrima(100); | Metschnikowia | 13 | Saprotroph | \|Undefined Saprotroph\| | Yeast | NULL | Probable | As nectar-tap_saprotroph (Põlme et al. 2020); Decay sugar-rich substrates (Põlme et al. 2020) | Põlme S, et al. 2020. Fungal Diversity 105:1-16 ((https://doi.org/10.1007/s13225-020-00466-2)) |
| Otu44438 | 1 | k__Fungi(100);p__Ascomycota(100);c__Saccharomycetes(100);o__Saccharomycetales(100);f__Metschnikowiaceae(100);g__Metschnikowia(100);s__Metschnikowia_pulcherrima(100); | Metschnikowia | 13 | Saprotroph | \|Undefined Saprotroph\| | Yeast | NULL | Probable | As nectar-tap_saprotroph (Põlme et al. 2020); Decay sugar-rich substrates (Põlme et al. 2020) | Põlme S, et al. 2020. Fungal Diversity 105:1-16 ((https://doi.org/10.1007/s13225-020-00466-2)) |
| Otu44439 | 1 | k__Fungi(100);p__Ascomycota(100);c__Saccharomycetes(100);o__Saccharomycetales(100);f__Metschnikowiaceae(100);g__Metschnikowia(100);s__Metschnikowia_pulcherrima(100); | Metschnikowia | 13 | Saprotroph | \|Undefined Saprotroph\| | Yeast | NULL | Probable | As nectar-tap_saprotroph (Põlme et al. 2020); Decay sugar-rich substrates (Põlme et al. 2020) | Põlme S, et al. 2020. Fungal Diversity 105:1-16 ((https://doi.org/10.1007/s13225-020-00466-2)) |
| Otu44440 | 1 | k__Fungi(100);p__Ascomycota(100);c__Saccharomycetes(100);o__Saccharomycetales(100);f__Metschnikowiaceae(100);g__Metschnikowia(100);g__Metschnikowia_unclassified(100); | Metschnikowia | 13 | Saprotroph | \|Undefined Saprotroph\| | Yeast | NULL | Probable | As nectar-tap_saprotroph (Põlme et al. 2020); Decay sugar-rich substrates (Põlme et al. 2020) | Põlme S, et al. 2020. Fungal Diversity 105:1-16 ((https://doi.org/10.1007/s13225-020-00466-2)) |
| Otu44441 | 1 | k__Fungi(100);p__Ascomycota(100);c__Saccharomycetes(100);o__Saccharomycetales(100);f__Metschnikowiaceae(100);g__Metschnikowia(100);g__Metschnikowia_unclassified(100); | Metschnikowia | 13 | Saprotroph | \|Undefined Saprotroph\| | Yeast | NULL | Probable | As nectar-tap_saprotroph (Põlme et al. 2020); Decay sugar-rich substrates (Põlme et al. 2020) | Põlme S, et al. 2020. Fungal Diversity 105:1-16 ((https://doi.org/10.1007/s13225-020-00466-2)) |
| Otu44442 | 1 | k__Fungi(100);p__Ascomycota(100);c__Saccharomycetes(100);o__Saccharomycetales(100);f__Metschnikowiaceae(100);g__Metschnikowia(100);s__Metschnikowia_pulcherrima(100); | Metschnikowia | 13 | Saprotroph | \|Undefined Saprotroph\| | Yeast | NULL | Probable | As nectar-tap_saprotroph (Põlme et al. 2020); Decay sugar-rich substrates (Põlme et al. 2020) | Põlme S, et al. 2020. Fungal Diversity 105:1-16 ((https://doi.org/10.1007/s13225-020-00466-2)) |
| Otu44443 | 1 | k__Fungi(100);p__Ascomycota(100);c__Saccharomycetes(100);o__Saccharomycetales(100);f__Metschnikowiaceae(100);g__Metschnikowia(100);g__Metschnikowia_unclassified(100); | Metschnikowia | 13 | Saprotroph | \|Undefined Saprotroph\| | Yeast | NULL | Probable | As nectar-tap_saprotroph (Põlme et al. 2020); Decay sugar-rich substrates (Põlme et al. 2020) | Põlme S, et al. 2020. Fungal Diversity 105:1-16 ((https://doi.org/10.1007/s13225-020-00466-2)) |
| Otu44444 | 1 | k__Fungi(100);p__Ascomycota(100);c__Saccharomycetes(100);o__Saccharomycetales(100);f__Metschnikowiaceae(100);g__Metschnikowia(100);s__Metschnikowia_pulcherrima(100); | Metschnikowia | 13 | Saprotroph | \|Undefined Saprotroph\| | Yeast | NULL | Probable | As nectar-tap_saprotroph (Põlme et al. 2020); Decay sugar-rich substrates (Põlme et al. 2020) | Põlme S, et al. 2020. Fungal Diversity 105:1-16 ((https://doi.org/10.1007/s13225-020-00466-2)) |
| Otu44445 | 1 | k__Fungi(100);p__Ascomycota(100);c__Saccharomycetes(100);o__Saccharomycetales(100);f__Metschnikowiaceae(100);g__Metschnikowia(100);s__Metschnikowia_pulcherrima(100); | Metschnikowia | 13 | Saprotroph | \|Undefined Saprotroph\| | Yeast | NULL | Probable | As nectar-tap_saprotroph (Põlme et al. 2020); Decay sugar-rich substrates (Põlme et al. 2020) | Põlme S, et al. 2020. Fungal Diversity 105:1-16 ((https://doi.org/10.1007/s13225-020-00466-2)) |
| Otu44446 | 1 | k__Fungi(100);p__Ascomycota(100);c__Saccharomycetes(100);o__Saccharomycetales(100);f__Metschnikowiaceae(100);g__Metschnikowia(100);s__Metschnikowia_pulcherrima(100); | Metschnikowia | 13 | Saprotroph | \|Undefined Saprotroph\| | Yeast | NULL | Probable | As nectar-tap_saprotroph (Põlme et al. 2020); Decay sugar-rich substrates (Põlme et al. 2020) | Põlme S, et al. 2020. Fungal Diversity 105:1-16 ((https://doi.org/10.1007/s13225-020-00466-2)) |
| Otu44447 | 1 | k__Fungi(100);p__Ascomycota(100);c__Saccharomycetes(100);o__Saccharomycetales(100);f__Metschnikowiaceae(100);g__Metschnikowia(100);s__Metschnikowia_pulcherrima(100); | Metschnikowia | 13 | Saprotroph | \|Undefined Saprotroph\| | Yeast | NULL | Probable | As nectar-tap_saprotroph (Põlme et al. 2020); Decay sugar-rich substrates (Põlme et al. 2020) | Põlme S, et al. 2020. Fungal Diversity 105:1-16 ((https://doi.org/10.1007/s13225-020-00466-2)) |
| Otu44448 | 1 | k__Fungi(100);p__Ascomycota(100);c__Saccharomycetes(100);o__Saccharomycetales(100);f__Metschnikowiaceae(100);g__Metschnikowia(100);s__Metschnikowia_pulcherrima(100); | Metschnikowia | 13 | Saprotroph | \|Undefined Saprotroph\| | Yeast | NULL | Probable | As nectar-tap_saprotroph (Põlme et al. 2020); Decay sugar-rich substrates (Põlme et al. 2020) | Põlme S, et al. 2020. Fungal Diversity 105:1-16 ((https://doi.org/10.1007/s13225-020-00466-2)) |
| Otu44449 | 1 | k__Fungi(100);p__Ascomycota(100);c__Saccharomycetes(100);o__Saccharomycetales(100);f__Metschnikowiaceae(100);g__Metschnikowia(100);s__Metschnikowia_pulcherrima(100); | Metschnikowia | 13 | Saprotroph | \|Undefined Saprotroph\| | Yeast | NULL | Probable | As nectar-tap_saprotroph (Põlme et al. 2020); Decay sugar-rich substrates (Põlme et al. 2020) | Põlme S, et al. 2020. Fungal Diversity 105:1-16 ((https://doi.org/10.1007/s13225-020-00466-2)) |
| Otu44450 | 1 | k__Fungi(100);p__Ascomycota(100);c__Saccharomycetes(100);o__Saccharomycetales(100);f__Metschnikowiaceae(100);g__Metschnikowia(100);s__Metschnikowia_pulcherrima(100); | Metschnikowia | 13 | Saprotroph | \|Undefined Saprotroph\| | Yeast | NULL | Probable | As nectar-tap_saprotroph (Põlme et al. 2020); Decay sugar-rich substrates (Põlme et al. 2020) | Põlme S, et al. 2020. Fungal Diversity 105:1-16 ((https://doi.org/10.1007/s13225-020-00466-2)) |
| Otu44451 | 1 | k__Fungi(100);p__Ascomycota(100);c__Saccharomycetes(100);o__Saccharomycetales(100);f__Metschnikowiaceae(100);g__Metschnikowia(100);s__Metschnikowia_pulcherrima(100); | Metschnikowia | 13 | Saprotroph | \|Undefined Saprotroph\| | Yeast | NULL | Probable | As nectar-tap_saprotroph (Põlme et al. 2020); Decay sugar-rich substrates (Põlme et al. 2020) | Põlme S, et al. 2020. Fungal Diversity 105:1-16 ((https://doi.org/10.1007/s13225-020-00466-2)) |
| Otu44452 | 1 | k__Fungi(100);p__Ascomycota(100);c__Saccharomycetes(100);o__Saccharomycetales(100);f__Metschnikowiaceae(100);g__Metschnikowia(100);s__Metschnikowia_pulcherrima(100); | Metschnikowia | 13 | Saprotroph | \|Undefined Saprotroph\| | Yeast | NULL | Probable | As nectar-tap_saprotroph (Põlme et al. 2020); Decay sugar-rich substrates (Põlme et al. 2020) | Põlme S, et al. 2020. Fungal Diversity 105:1-16 ((https://doi.org/10.1007/s13225-020-00466-2)) |
| Otu44453 | 1 | k__Fungi(100);p__Ascomycota(100);c__Saccharomycetes(100);o__Saccharomycetales(100);f__Metschnikowiaceae(100);g__Metschnikowia(100);s__Metschnikowia_pulcherrima(100); | Metschnikowia | 13 | Saprotroph | \|Undefined Saprotroph\| | Yeast | NULL | Probable | As nectar-tap_saprotroph (Põlme et al. 2020); Decay sugar-rich substrates (Põlme et al. 2020) | Põlme S, et al. 2020. Fungal Diversity 105:1-16 ((https://doi.org/10.1007/s13225-020-00466-2)) |
| Otu44454 | 1 | k__Fungi(100);p__Ascomycota(100);c__Saccharomycetes(100);o__Saccharomycetales(100);f__Metschnikowiaceae(100);g__Metschnikowia(100);g__Metschnikowia_unclassified(100); | Metschnikowia | 13 | Saprotroph | \|Undefined Saprotroph\| | Yeast | NULL | Probable | As nectar-tap_saprotroph (Põlme et al. 2020); Decay sugar-rich substrates (Põlme et al. 2020) | Põlme S, et al. 2020. Fungal Diversity 105:1-16 ((https://doi.org/10.1007/s13225-020-00466-2)) |
| Otu44455 | 1 | k__Fungi(100);p__Ascomycota(100);c__Saccharomycetes(100);o__Saccharomycetales(100);f__Metschnikowiaceae(100);g__Metschnikowia(100);s__Metschnikowia_pulcherrima(100); | Metschnikowia | 13 | Saprotroph | \|Undefined Saprotroph\| | Yeast | NULL | Probable | As nectar-tap_saprotroph (Põlme et al. 2020); Decay sugar-rich substrates (Põlme et al. 2020) | Põlme S, et al. 2020. Fungal Diversity 105:1-16 ((https://doi.org/10.1007/s13225-020-00466-2)) |
| Otu44456 | 1 | k__Fungi(100);p__Ascomycota(100);c__Saccharomycetes(100);o__Saccharomycetales(100);f__Metschnikowiaceae(100);g__Metschnikowia(100);g__Metschnikowia_unclassified(100); | Metschnikowia | 13 | Saprotroph | \|Undefined Saprotroph\| | Yeast | NULL | Probable | As nectar-tap_saprotroph (Põlme et al. 2020); Decay sugar-rich substrates (Põlme et al. 2020) | Põlme S, et al. 2020. Fungal Diversity 105:1-16 ((https://doi.org/10.1007/s13225-020-00466-2)) |
| Otu44457 | 1 | k__Fungi(100);p__Ascomycota(100);c__Saccharomycetes(100);o__Saccharomycetales(100);f__Metschnikowiaceae(100);g__Metschnikowia(100);s__Metschnikowia_pulcherrima(100); | Metschnikowia | 13 | Saprotroph | \|Undefined Saprotroph\| | Yeast | NULL | Probable | As nectar-tap_saprotroph (Põlme et al. 2020); Decay sugar-rich substrates (Põlme et al. 2020) | Põlme S, et al. 2020. Fungal Diversity 105:1-16 ((https://doi.org/10.1007/s13225-020-00466-2)) |
| Otu44458 | 1 | k__Fungi(100);p__Ascomycota(100);c__Saccharomycetes(100);o__Saccharomycetales(100);f__Metschnikowiaceae(100);g__Metschnikowia(100);g__Metschnikowia_unclassified(100); | Metschnikowia | 13 | Saprotroph | \|Undefined Saprotroph\| | Yeast | NULL | Probable | As nectar-tap_saprotroph (Põlme et al. 2020); Decay sugar-rich substrates (Põlme et al. 2020) | Põlme S, et al. 2020. Fungal Diversity 105:1-16 ((https://doi.org/10.1007/s13225-020-00466-2)) |
| Otu44459 | 1 | k__Fungi(100);p__Ascomycota(100);c__Saccharomycetes(100);o__Saccharomycetales(100);f__Metschnikowiaceae(100);g__Metschnikowia(100);g__Metschnikowia_unclassified(100); | Metschnikowia | 13 | Saprotroph | \|Undefined Saprotroph\| | Yeast | NULL | Probable | As nectar-tap_saprotroph (Põlme et al. 2020); Decay sugar-rich substrates (Põlme et al. 2020) | Põlme S, et al. 2020. Fungal Diversity 105:1-16 ((https://doi.org/10.1007/s13225-020-00466-2)) |
| Otu44460 | 1 | k__Fungi(100);p__Ascomycota(100);c__Saccharomycetes(100);o__Saccharomycetales(100);f__Metschnikowiaceae(100);g__Metschnikowia(100);g__Metschnikowia_unclassified(100); | Metschnikowia | 13 | Saprotroph | \|Undefined Saprotroph\| | Yeast | NULL | Probable | As nectar-tap_saprotroph (Põlme et al. 2020); Decay sugar-rich substrates (Põlme et al. 2020) | Põlme S, et al. 2020. Fungal Diversity 105:1-16 ((https://doi.org/10.1007/s13225-020-00466-2)) |
| Otu44461 | 1 | k__Fungi(100);p__Ascomycota(100);c__Saccharomycetes(100);o__Saccharomycetales(100);f__Metschnikowiaceae(100);g__Metschnikowia(100);s__Metschnikowia_pulcherrima(100); | Metschnikowia | 13 | Saprotroph | \|Undefined Saprotroph\| | Yeast | NULL | Probable | As nectar-tap_saprotroph (Põlme et al. 2020); Decay sugar-rich substrates (Põlme et al. 2020) | Põlme S, et al. 2020. Fungal Diversity 105:1-16 ((https://doi.org/10.1007/s13225-020-00466-2)) |
| Otu44462 | 1 | k__Fungi(100);p__Ascomycota(100);c__Saccharomycetes(100);o__Saccharomycetales(100);f__Metschnikowiaceae(100);g__Metschnikowia(100);s__Metschnikowia_pulcherrima(100); | Metschnikowia | 13 | Saprotroph | \|Undefined Saprotroph\| | Yeast | NULL | Probable | As nectar-tap_saprotroph (Põlme et al. 2020); Decay sugar-rich substrates (Põlme et al. 2020) | Põlme S, et al. 2020. Fungal Diversity 105:1-16 ((https://doi.org/10.1007/s13225-020-00466-2)) |
| Otu44463 | 1 | k__Fungi(100);p__Ascomycota(100);c__Saccharomycetes(100);o__Saccharomycetales(100);f__Metschnikowiaceae(100);g__Metschnikowia(100);s__Metschnikowia_pulcherrima(100); | Metschnikowia | 13 | Saprotroph | \|Undefined Saprotroph\| | Yeast | NULL | Probable | As nectar-tap_saprotroph (Põlme et al. 2020); Decay sugar-rich substrates (Põlme et al. 2020) | Põlme S, et al. 2020. Fungal Diversity 105:1-16 ((https://doi.org/10.1007/s13225-020-00466-2)) |
| Otu44464 | 1 | k__Fungi(100);p__Ascomycota(100);c__Saccharomycetes(100);o__Saccharomycetales(100);f__Metschnikowiaceae(100);g__Metschnikowia(100);s__Metschnikowia_pulcherrima(100); | Metschnikowia | 13 | Saprotroph | \|Undefined Saprotroph\| | Yeast | NULL | Probable | As nectar-tap_saprotroph (Põlme et al. 2020); Decay sugar-rich substrates (Põlme et al. 2020) | Põlme S, et al. 2020. Fungal Diversity 105:1-16 ((https://doi.org/10.1007/s13225-020-00466-2)) |
| Otu44465 | 1 | k__Fungi(100);p__Ascomycota(100);c__Saccharomycetes(100);o__Saccharomycetales(100);f__Metschnikowiaceae(100);g__Metschnikowia(100);s__Metschnikowia_pulcherrima(100); | Metschnikowia | 13 | Saprotroph | \|Undefined Saprotroph\| | Yeast | NULL | Probable | As nectar-tap_saprotroph (Põlme et al. 2020); Decay sugar-rich substrates (Põlme et al. 2020) | Põlme S, et al. 2020. Fungal Diversity 105:1-16 ((https://doi.org/10.1007/s13225-020-00466-2)) |
| Otu44467 | 1 | k__Fungi(100);p__Ascomycota(100);c__Saccharomycetes(100);o__Saccharomycetales(100);f__Metschnikowiaceae(100);g__Metschnikowia(100);s__Metschnikowia_pulcherrima(100); | Metschnikowia | 13 | Saprotroph | \|Undefined Saprotroph\| | Yeast | NULL | Probable | As nectar-tap_saprotroph (Põlme et al. 2020); Decay sugar-rich substrates (Põlme et al. 2020) | Põlme S, et al. 2020. Fungal Diversity 105:1-16 ((https://doi.org/10.1007/s13225-020-00466-2)) |
| Otu44468 | 1 | k__Fungi(100);p__Ascomycota(100);c__Saccharomycetes(100);o__Saccharomycetales(100);f__Metschnikowiaceae(100);g__Metschnikowia(100);s__Metschnikowia_pulcherrima(100); | Metschnikowia | 13 | Saprotroph | \|Undefined Saprotroph\| | Yeast | NULL | Probable | As nectar-tap_saprotroph (Põlme et al. 2020); Decay sugar-rich substrates (Põlme et al. 2020) | Põlme S, et al. 2020. Fungal Diversity 105:1-16 ((https://doi.org/10.1007/s13225-020-00466-2)) |
| Otu44469 | 1 | k__Fungi(100);p__Ascomycota(100);c__Saccharomycetes(100);o__Saccharomycetales(100);f__Metschnikowiaceae(100);g__Metschnikowia(100);g__Metschnikowia_unclassified(100); | Metschnikowia | 13 | Saprotroph | \|Undefined Saprotroph\| | Yeast | NULL | Probable | As nectar-tap_saprotroph (Põlme et al. 2020); Decay sugar-rich substrates (Põlme et al. 2020) | Põlme S, et al. 2020. Fungal Diversity 105:1-16 ((https://doi.org/10.1007/s13225-020-00466-2)) |
| Otu44470 | 1 | k__Fungi(100);p__Ascomycota(100);c__Saccharomycetes(100);o__Saccharomycetales(100);f__Metschnikowiaceae(100);g__Metschnikowia(100);s__Metschnikowia_pulcherrima(100); | Metschnikowia | 13 | Saprotroph | \|Undefined Saprotroph\| | Yeast | NULL | Probable | As nectar-tap_saprotroph (Põlme et al. 2020); Decay sugar-rich substrates (Põlme et al. 2020) | Põlme S, et al. 2020. Fungal Diversity 105:1-16 ((https://doi.org/10.1007/s13225-020-00466-2)) |
| Otu44471 | 1 | k__Fungi(100);p__Ascomycota(100);c__Saccharomycetes(100);o__Saccharomycetales(100);f__Metschnikowiaceae(100);g__Metschnikowia(100);s__Metschnikowia_pulcherrima(100); | Metschnikowia | 13 | Saprotroph | \|Undefined Saprotroph\| | Yeast | NULL | Probable | As nectar-tap_saprotroph (Põlme et al. 2020); Decay sugar-rich substrates (Põlme et al. 2020) | Põlme S, et al. 2020. Fungal Diversity 105:1-16 ((https://doi.org/10.1007/s13225-020-00466-2)) |
| Otu44472 | 1 | k__Fungi(100);p__Ascomycota(100);c__Saccharomycetes(100);o__Saccharomycetales(100);f__Metschnikowiaceae(100);g__Metschnikowia(100);s__Metschnikowia_pulcherrima(100); | Metschnikowia | 13 | Saprotroph | \|Undefined Saprotroph\| | Yeast | NULL | Probable | As nectar-tap_saprotroph (Põlme et al. 2020); Decay sugar-rich substrates (Põlme et al. 2020) | Põlme S, et al. 2020. Fungal Diversity 105:1-16 ((https://doi.org/10.1007/s13225-020-00466-2)) |
| Otu44473 | 1 | k__Fungi(100);p__Ascomycota(100);c__Saccharomycetes(100);o__Saccharomycetales(100);f__Metschnikowiaceae(100);g__Metschnikowia(100);g__Metschnikowia_unclassified(100); | Metschnikowia | 13 | Saprotroph | \|Undefined Saprotroph\| | Yeast | NULL | Probable | As nectar-tap_saprotroph (Põlme et al. 2020); Decay sugar-rich substrates (Põlme et al. 2020) | Põlme S, et al. 2020. Fungal Diversity 105:1-16 ((https://doi.org/10.1007/s13225-020-00466-2)) |
| Otu44474 | 1 | k__Fungi(100);p__Ascomycota(100);c__Saccharomycetes(100);o__Saccharomycetales(100);f__Metschnikowiaceae(100);g__Metschnikowia(100);g__Metschnikowia_unclassified(100); | Metschnikowia | 13 | Saprotroph | \|Undefined Saprotroph\| | Yeast | NULL | Probable | As nectar-tap_saprotroph (Põlme et al. 2020); Decay sugar-rich substrates (Põlme et al. 2020) | Põlme S, et al. 2020. Fungal Diversity 105:1-16 ((https://doi.org/10.1007/s13225-020-00466-2)) |
| Otu44475 | 1 | k__Fungi(100);p__Ascomycota(100);c__Saccharomycetes(100);o__Saccharomycetales(100);f__Metschnikowiaceae(100);g__Metschnikowia(100);g__Metschnikowia_unclassified(100); | Metschnikowia | 13 | Saprotroph | \|Undefined Saprotroph\| | Yeast | NULL | Probable | As nectar-tap_saprotroph (Põlme et al. 2020); Decay sugar-rich substrates (Põlme et al. 2020) | Põlme S, et al. 2020. Fungal Diversity 105:1-16 ((https://doi.org/10.1007/s13225-020-00466-2)) |
| Otu44476 | 1 | k__Fungi(100);p__Ascomycota(100);c__Saccharomycetes(100);o__Saccharomycetales(100);f__Metschnikowiaceae(100);g__Metschnikowia(100);g__Metschnikowia_unclassified(100); | Metschnikowia | 13 | Saprotroph | \|Undefined Saprotroph\| | Yeast | NULL | Probable | As nectar-tap_saprotroph (Põlme et al. 2020); Decay sugar-rich substrates (Põlme et al. 2020) | Põlme S, et al. 2020. Fungal Diversity 105:1-16 ((https://doi.org/10.1007/s13225-020-00466-2)) |
| Otu44477 | 1 | k__Fungi(100);p__Ascomycota(100);c__Saccharomycetes(100);o__Saccharomycetales(100);f__Metschnikowiaceae(100);g__Metschnikowia(100);g__Metschnikowia_unclassified(100); | Metschnikowia | 13 | Saprotroph | \|Undefined Saprotroph\| | Yeast | NULL | Probable | As nectar-tap_saprotroph (Põlme et al. 2020); Decay sugar-rich substrates (Põlme et al. 2020) | Põlme S, et al. 2020. Fungal Diversity 105:1-16 ((https://doi.org/10.1007/s13225-020-00466-2)) |
| Otu44478 | 1 | k__Fungi(100);p__Ascomycota(100);c__Saccharomycetes(100);o__Saccharomycetales(100);f__Metschnikowiaceae(100);g__Metschnikowia(100);g__Metschnikowia_unclassified(100); | Metschnikowia | 13 | Saprotroph | \|Undefined Saprotroph\| | Yeast | NULL | Probable | As nectar-tap_saprotroph (Põlme et al. 2020); Decay sugar-rich substrates (Põlme et al. 2020) | Põlme S, et al. 2020. Fungal Diversity 105:1-16 ((https://doi.org/10.1007/s13225-020-00466-2)) |
| Otu44479 | 1 | k__Fungi(100);p__Ascomycota(100);c__Saccharomycetes(100);o__Saccharomycetales(100);f__Metschnikowiaceae(100);g__Metschnikowia(100);g__Metschnikowia_unclassified(100); | Metschnikowia | 13 | Saprotroph | \|Undefined Saprotroph\| | Yeast | NULL | Probable | As nectar-tap_saprotroph (Põlme et al. 2020); Decay sugar-rich substrates (Põlme et al. 2020) | Põlme S, et al. 2020. Fungal Diversity 105:1-16 ((https://doi.org/10.1007/s13225-020-00466-2)) |
| Otu44480 | 1 | k__Fungi(100);p__Ascomycota(100);c__Saccharomycetes(100);o__Saccharomycetales(100);f__Metschnikowiaceae(100);g__Metschnikowia(100);g__Metschnikowia_unclassified(100); | Metschnikowia | 13 | Saprotroph | \|Undefined Saprotroph\| | Yeast | NULL | Probable | As nectar-tap_saprotroph (Põlme et al. 2020); Decay sugar-rich substrates (Põlme et al. 2020) | Põlme S, et al. 2020. Fungal Diversity 105:1-16 ((https://doi.org/10.1007/s13225-020-00466-2)) |
| Otu44481 | 1 | k__Fungi(100);p__Ascomycota(100);c__Saccharomycetes(100);o__Saccharomycetales(100);f__Metschnikowiaceae(100);g__Metschnikowia(100);g__Metschnikowia_unclassified(100); | Metschnikowia | 13 | Saprotroph | \|Undefined Saprotroph\| | Yeast | NULL | Probable | As nectar-tap_saprotroph (Põlme et al. 2020); Decay sugar-rich substrates (Põlme et al. 2020) | Põlme S, et al. 2020. Fungal Diversity 105:1-16 ((https://doi.org/10.1007/s13225-020-00466-2)) |
| Otu44482 | 1 | k__Fungi(100);p__Ascomycota(100);c__Saccharomycetes(100);o__Saccharomycetales(100);f__Metschnikowiaceae(100);g__Metschnikowia(100);g__Metschnikowia_unclassified(100); | Metschnikowia | 13 | Saprotroph | \|Undefined Saprotroph\| | Yeast | NULL | Probable | As nectar-tap_saprotroph (Põlme et al. 2020); Decay sugar-rich substrates (Põlme et al. 2020) | Põlme S, et al. 2020. Fungal Diversity 105:1-16 ((https://doi.org/10.1007/s13225-020-00466-2)) |
| Otu44483 | 1 | k__Fungi(100);p__Ascomycota(100);c__Saccharomycetes(100);o__Saccharomycetales(100);f__Metschnikowiaceae(100);g__Metschnikowia(100);s__Metschnikowia_pulcherrima(100); | Metschnikowia | 13 | Saprotroph | \|Undefined Saprotroph\| | Yeast | NULL | Probable | As nectar-tap_saprotroph (Põlme et al. 2020); Decay sugar-rich substrates (Põlme et al. 2020) | Põlme S, et al. 2020. Fungal Diversity 105:1-16 ((https://doi.org/10.1007/s13225-020-00466-2)) |
| Otu44484 | 1 | k__Fungi(100);p__Ascomycota(100);c__Saccharomycetes(100);o__Saccharomycetales(100);f__Metschnikowiaceae(100);g__Metschnikowia(100);s__Metschnikowia_pulcherrima(100); | Metschnikowia | 13 | Saprotroph | \|Undefined Saprotroph\| | Yeast | NULL | Probable | As nectar-tap_saprotroph (Põlme et al. 2020); Decay sugar-rich substrates (Põlme et al. 2020) | Põlme S, et al. 2020. Fungal Diversity 105:1-16 ((https://doi.org/10.1007/s13225-020-00466-2)) |
| Otu44485 | 1 | k__Fungi(100);p__Ascomycota(100);c__Saccharomycetes(100);o__Saccharomycetales(100);f__Metschnikowiaceae(100);g__Metschnikowia(100);g__Metschnikowia_unclassified(100); | Metschnikowia | 13 | Saprotroph | \|Undefined Saprotroph\| | Yeast | NULL | Probable | As nectar-tap_saprotroph (Põlme et al. 2020); Decay sugar-rich substrates (Põlme et al. 2020) | Põlme S, et al. 2020. Fungal Diversity 105:1-16 ((https://doi.org/10.1007/s13225-020-00466-2)) |
| Otu44486 | 1 | k__Fungi(100);p__Ascomycota(100);c__Saccharomycetes(100);o__Saccharomycetales(100);f__Metschnikowiaceae(100);g__Metschnikowia(100);s__Metschnikowia_pulcherrima(100); | Metschnikowia | 13 | Saprotroph | \|Undefined Saprotroph\| | Yeast | NULL | Probable | As nectar-tap_saprotroph (Põlme et al. 2020); Decay sugar-rich substrates (Põlme et al. 2020) | Põlme S, et al. 2020. Fungal Diversity 105:1-16 ((https://doi.org/10.1007/s13225-020-00466-2)) |
| Otu44487 | 1 | k__Fungi(100);p__Ascomycota(100);c__Saccharomycetes(100);o__Saccharomycetales(100);f__Metschnikowiaceae(100);g__Metschnikowia(100);g__Metschnikowia_unclassified(100); | Metschnikowia | 13 | Saprotroph | \|Undefined Saprotroph\| | Yeast | NULL | Probable | As nectar-tap_saprotroph (Põlme et al. 2020); Decay sugar-rich substrates (Põlme et al. 2020) | Põlme S, et al. 2020. Fungal Diversity 105:1-16 ((https://doi.org/10.1007/s13225-020-00466-2)) |
| Otu44488 | 1 | k__Fungi(100);p__Ascomycota(100);c__Saccharomycetes(100);o__Saccharomycetales(100);f__Metschnikowiaceae(100);g__Metschnikowia(100);s__Metschnikowia_pulcherrima(100); | Metschnikowia | 13 | Saprotroph | \|Undefined Saprotroph\| | Yeast | NULL | Probable | As nectar-tap_saprotroph (Põlme et al. 2020); Decay sugar-rich substrates (Põlme et al. 2020) | Põlme S, et al. 2020. Fungal Diversity 105:1-16 ((https://doi.org/10.1007/s13225-020-00466-2)) |
| Otu44489 | 1 | k__Fungi(100);p__Ascomycota(100);c__Saccharomycetes(100);o__Saccharomycetales(100);f__Metschnikowiaceae(100);g__Metschnikowia(100);s__Metschnikowia_pulcherrima(100); | Metschnikowia | 13 | Saprotroph | \|Undefined Saprotroph\| | Yeast | NULL | Probable | As nectar-tap_saprotroph (Põlme et al. 2020); Decay sugar-rich substrates (Põlme et al. 2020) | Põlme S, et al. 2020. Fungal Diversity 105:1-16 ((https://doi.org/10.1007/s13225-020-00466-2)) |
| Otu44490 | 1 | k__Fungi(100);p__Ascomycota(100);c__Saccharomycetes(100);o__Saccharomycetales(100);f__Metschnikowiaceae(100);g__Metschnikowia(100);g__Metschnikowia_unclassified(100); | Metschnikowia | 13 | Saprotroph | \|Undefined Saprotroph\| | Yeast | NULL | Probable | As nectar-tap_saprotroph (Põlme et al. 2020); Decay sugar-rich substrates (Põlme et al. 2020) | Põlme S, et al. 2020. Fungal Diversity 105:1-16 ((https://doi.org/10.1007/s13225-020-00466-2)) |
| Otu44491 | 1 | k__Fungi(100);p__Ascomycota(100);c__Saccharomycetes(100);o__Saccharomycetales(100);f__Metschnikowiaceae(100);g__Metschnikowia(100);g__Metschnikowia_unclassified(100); | Metschnikowia | 13 | Saprotroph | \|Undefined Saprotroph\| | Yeast | NULL | Probable | As nectar-tap_saprotroph (Põlme et al. 2020); Decay sugar-rich substrates (Põlme et al. 2020) | Põlme S, et al. 2020. Fungal Diversity 105:1-16 ((https://doi.org/10.1007/s13225-020-00466-2)) |
| Otu44492 | 1 | k__Fungi(100);p__Ascomycota(100);c__Saccharomycetes(100);o__Saccharomycetales(100);f__Metschnikowiaceae(100);g__Metschnikowia(100);g__Metschnikowia_unclassified(100); | Metschnikowia | 13 | Saprotroph | \|Undefined Saprotroph\| | Yeast | NULL | Probable | As nectar-tap_saprotroph (Põlme et al. 2020); Decay sugar-rich substrates (Põlme et al. 2020) | Põlme S, et al. 2020. Fungal Diversity 105:1-16 ((https://doi.org/10.1007/s13225-020-00466-2)) |
| Otu44493 | 1 | k__Fungi(100);p__Ascomycota(100);c__Saccharomycetes(100);o__Saccharomycetales(100);f__Metschnikowiaceae(100);g__Metschnikowia(100);s__Metschnikowia_pulcherrima(100); | Metschnikowia | 13 | Saprotroph | \|Undefined Saprotroph\| | Yeast | NULL | Probable | As nectar-tap_saprotroph (Põlme et al. 2020); Decay sugar-rich substrates (Põlme et al. 2020) | Põlme S, et al. 2020. Fungal Diversity 105:1-16 ((https://doi.org/10.1007/s13225-020-00466-2)) |
| Otu44494 | 1 | k__Fungi(100);p__Ascomycota(100);c__Saccharomycetes(100);o__Saccharomycetales(100);f__Metschnikowiaceae(100);g__Metschnikowia(100);s__Metschnikowia_pulcherrima(100); | Metschnikowia | 13 | Saprotroph | \|Undefined Saprotroph\| | Yeast | NULL | Probable | As nectar-tap_saprotroph (Põlme et al. 2020); Decay sugar-rich substrates (Põlme et al. 2020) | Põlme S, et al. 2020. Fungal Diversity 105:1-16 ((https://doi.org/10.1007/s13225-020-00466-2)) |
| Otu44495 | 1 | k__Fungi(100);p__Ascomycota(100);c__Saccharomycetes(100);o__Saccharomycetales(100);f__Metschnikowiaceae(100);g__Metschnikowia(100);g__Metschnikowia_unclassified(100); | Metschnikowia | 13 | Saprotroph | \|Undefined Saprotroph\| | Yeast | NULL | Probable | As nectar-tap_saprotroph (Põlme et al. 2020); Decay sugar-rich substrates (Põlme et al. 2020) | Põlme S, et al. 2020. Fungal Diversity 105:1-16 ((https://doi.org/10.1007/s13225-020-00466-2)) |
| Otu44496 | 1 | k__Fungi(100);p__Ascomycota(100);c__Saccharomycetes(100);o__Saccharomycetales(100);f__Metschnikowiaceae(100);g__Metschnikowia(100);g__Metschnikowia_unclassified(100); | Metschnikowia | 13 | Saprotroph | \|Undefined Saprotroph\| | Yeast | NULL | Probable | As nectar-tap_saprotroph (Põlme et al. 2020); Decay sugar-rich substrates (Põlme et al. 2020) | Põlme S, et al. 2020. Fungal Diversity 105:1-16 ((https://doi.org/10.1007/s13225-020-00466-2)) |
| Otu44497 | 1 | k__Fungi(100);p__Ascomycota(100);c__Saccharomycetes(100);o__Saccharomycetales(100);f__Metschnikowiaceae(100);g__Metschnikowia(100);g__Metschnikowia_unclassified(100); | Metschnikowia | 13 | Saprotroph | \|Undefined Saprotroph\| | Yeast | NULL | Probable | As nectar-tap_saprotroph (Põlme et al. 2020); Decay sugar-rich substrates (Põlme et al. 2020) | Põlme S, et al. 2020. Fungal Diversity 105:1-16 ((https://doi.org/10.1007/s13225-020-00466-2)) |
| Otu44498 | 1 | k__Fungi(100);p__Ascomycota(100);c__Saccharomycetes(100);o__Saccharomycetales(100);f__Metschnikowiaceae(100);g__Metschnikowia(100);s__Metschnikowia_pulcherrima(100); | Metschnikowia | 13 | Saprotroph | \|Undefined Saprotroph\| | Yeast | NULL | Probable | As nectar-tap_saprotroph (Põlme et al. 2020); Decay sugar-rich substrates (Põlme et al. 2020) | Põlme S, et al. 2020. Fungal Diversity 105:1-16 ((https://doi.org/10.1007/s13225-020-00466-2)) |
| Otu44499 | 1 | k__Fungi(100);p__Ascomycota(100);c__Saccharomycetes(100);o__Saccharomycetales(100);f__Metschnikowiaceae(100);g__Metschnikowia(100);s__Metschnikowia_pulcherrima(100); | Metschnikowia | 13 | Saprotroph | \|Undefined Saprotroph\| | Yeast | NULL | Probable | As nectar-tap_saprotroph (Põlme et al. 2020); Decay sugar-rich substrates (Põlme et al. 2020) | Põlme S, et al. 2020. Fungal Diversity 105:1-16 ((https://doi.org/10.1007/s13225-020-00466-2)) |
| Otu44500 | 1 | k__Fungi(100);p__Ascomycota(100);c__Saccharomycetes(100);o__Saccharomycetales(100);f__Metschnikowiaceae(100);g__Metschnikowia(100);s__Metschnikowia_pulcherrima(100); | Metschnikowia | 13 | Saprotroph | \|Undefined Saprotroph\| | Yeast | NULL | Probable | As nectar-tap_saprotroph (Põlme et al. 2020); Decay sugar-rich substrates (Põlme et al. 2020) | Põlme S, et al. 2020. Fungal Diversity 105:1-16 ((https://doi.org/10.1007/s13225-020-00466-2)) |
| Otu44501 | 1 | k__Fungi(100);p__Ascomycota(100);c__Saccharomycetes(100);o__Saccharomycetales(100);f__Metschnikowiaceae(100);g__Metschnikowia(100);g__Metschnikowia_unclassified(100); | Metschnikowia | 13 | Saprotroph | \|Undefined Saprotroph\| | Yeast | NULL | Probable | As nectar-tap_saprotroph (Põlme et al. 2020); Decay sugar-rich substrates (Põlme et al. 2020) | Põlme S, et al. 2020. Fungal Diversity 105:1-16 ((https://doi.org/10.1007/s13225-020-00466-2)) |
| Otu44502 | 1 | k__Fungi(100);p__Ascomycota(100);c__Saccharomycetes(100);o__Saccharomycetales(100);f__Metschnikowiaceae(100);g__Metschnikowia(100);g__Metschnikowia_unclassified(100); | Metschnikowia | 13 | Saprotroph | \|Undefined Saprotroph\| | Yeast | NULL | Probable | As nectar-tap_saprotroph (Põlme et al. 2020); Decay sugar-rich substrates (Põlme et al. 2020) | Põlme S, et al. 2020. Fungal Diversity 105:1-16 ((https://doi.org/10.1007/s13225-020-00466-2)) |
| Otu44503 | 1 | k__Fungi(100);p__Ascomycota(100);c__Saccharomycetes(100);o__Saccharomycetales(100);f__Metschnikowiaceae(100);g__Metschnikowia(100);s__Metschnikowia_pulcherrima(100); | Metschnikowia | 13 | Saprotroph | \|Undefined Saprotroph\| | Yeast | NULL | Probable | As nectar-tap_saprotroph (Põlme et al. 2020); Decay sugar-rich substrates (Põlme et al. 2020) | Põlme S, et al. 2020. Fungal Diversity 105:1-16 ((https://doi.org/10.1007/s13225-020-00466-2)) |
| Otu44504 | 1 | k__Fungi(100);p__Ascomycota(100);c__Saccharomycetes(100);o__Saccharomycetales(100);f__Metschnikowiaceae(100);g__Metschnikowia(100);s__Metschnikowia_pulcherrima(100); | Metschnikowia | 13 | Saprotroph | \|Undefined Saprotroph\| | Yeast | NULL | Probable | As nectar-tap_saprotroph (Põlme et al. 2020); Decay sugar-rich substrates (Põlme et al. 2020) | Põlme S, et al. 2020. Fungal Diversity 105:1-16 ((https://doi.org/10.1007/s13225-020-00466-2)) |
| Otu44505 | 1 | k__Fungi(100);p__Ascomycota(100);c__Saccharomycetes(100);o__Saccharomycetales(100);f__Metschnikowiaceae(100);g__Metschnikowia(100);s__Metschnikowia_pulcherrima(100); | Metschnikowia | 13 | Saprotroph | \|Undefined Saprotroph\| | Yeast | NULL | Probable | As nectar-tap_saprotroph (Põlme et al. 2020); Decay sugar-rich substrates (Põlme et al. 2020) | Põlme S, et al. 2020. Fungal Diversity 105:1-16 ((https://doi.org/10.1007/s13225-020-00466-2)) |
| Otu44506 | 1 | k__Fungi(100);p__Ascomycota(100);c__Saccharomycetes(100);o__Saccharomycetales(100);f__Metschnikowiaceae(100);g__Metschnikowia(100);g__Metschnikowia_unclassified(100); | Metschnikowia | 13 | Saprotroph | \|Undefined Saprotroph\| | Yeast | NULL | Probable | As nectar-tap_saprotroph (Põlme et al. 2020); Decay sugar-rich substrates (Põlme et al. 2020) | Põlme S, et al. 2020. Fungal Diversity 105:1-16 ((https://doi.org/10.1007/s13225-020-00466-2)) |
| Otu44507 | 1 | k__Fungi(100);p__Ascomycota(100);c__Saccharomycetes(100);o__Saccharomycetales(100);f__Metschnikowiaceae(100);g__Metschnikowia(100);s__Metschnikowia_pulcherrima(100); | Metschnikowia | 13 | Saprotroph | \|Undefined Saprotroph\| | Yeast | NULL | Probable | As nectar-tap_saprotroph (Põlme et al. 2020); Decay sugar-rich substrates (Põlme et al. 2020) | Põlme S, et al. 2020. Fungal Diversity 105:1-16 ((https://doi.org/10.1007/s13225-020-00466-2)) |
| Otu44508 | 1 | k__Fungi(100);p__Ascomycota(100);c__Saccharomycetes(100);o__Saccharomycetales(100);f__Metschnikowiaceae(100);g__Metschnikowia(100);s__Metschnikowia_pulcherrima(100); | Metschnikowia | 13 | Saprotroph | \|Undefined Saprotroph\| | Yeast | NULL | Probable | As nectar-tap_saprotroph (Põlme et al. 2020); Decay sugar-rich substrates (Põlme et al. 2020) | Põlme S, et al. 2020. Fungal Diversity 105:1-16 ((https://doi.org/10.1007/s13225-020-00466-2)) |
| Otu44509 | 1 | k__Fungi(100);p__Ascomycota(100);c__Saccharomycetes(100);o__Saccharomycetales(100);f__Metschnikowiaceae(100);g__Metschnikowia(100);s__Metschnikowia_pulcherrima(100); | Metschnikowia | 13 | Saprotroph | \|Undefined Saprotroph\| | Yeast | NULL | Probable | As nectar-tap_saprotroph (Põlme et al. 2020); Decay sugar-rich substrates (Põlme et al. 2020) | Põlme S, et al. 2020. Fungal Diversity 105:1-16 ((https://doi.org/10.1007/s13225-020-00466-2)) |
| Otu44511 | 1 | k__Fungi(100);p__Ascomycota(100);c__Saccharomycetes(100);o__Saccharomycetales(100);f__Metschnikowiaceae(100);g__Metschnikowia(100);s__Metschnikowia_pulcherrima(100); | Metschnikowia | 13 | Saprotroph | \|Undefined Saprotroph\| | Yeast | NULL | Probable | As nectar-tap_saprotroph (Põlme et al. 2020); Decay sugar-rich substrates (Põlme et al. 2020) | Põlme S, et al. 2020. Fungal Diversity 105:1-16 ((https://doi.org/10.1007/s13225-020-00466-2)) |
| Otu44512 | 1 | k__Fungi(100);p__Ascomycota(100);c__Saccharomycetes(100);o__Saccharomycetales(100);f__Metschnikowiaceae(100);g__Metschnikowia(100);s__Metschnikowia_pulcherrima(100); | Metschnikowia | 13 | Saprotroph | \|Undefined Saprotroph\| | Yeast | NULL | Probable | As nectar-tap_saprotroph (Põlme et al. 2020); Decay sugar-rich substrates (Põlme et al. 2020) | Põlme S, et al. 2020. Fungal Diversity 105:1-16 ((https://doi.org/10.1007/s13225-020-00466-2)) |
| Otu44513 | 1 | k__Fungi(100);p__Ascomycota(100);c__Saccharomycetes(100);o__Saccharomycetales(100);f__Metschnikowiaceae(100);g__Metschnikowia(100);s__Metschnikowia_pulcherrima(100); | Metschnikowia | 13 | Saprotroph | \|Undefined Saprotroph\| | Yeast | NULL | Probable | As nectar-tap_saprotroph (Põlme et al. 2020); Decay sugar-rich substrates (Põlme et al. 2020) | Põlme S, et al. 2020. Fungal Diversity 105:1-16 ((https://doi.org/10.1007/s13225-020-00466-2)) |
| Otu44514 | 1 | k__Fungi(100);p__Ascomycota(100);c__Saccharomycetes(100);o__Saccharomycetales(100);f__Metschnikowiaceae(100);g__Metschnikowia(100);g__Metschnikowia_unclassified(100); | Metschnikowia | 13 | Saprotroph | \|Undefined Saprotroph\| | Yeast | NULL | Probable | As nectar-tap_saprotroph (Põlme et al. 2020); Decay sugar-rich substrates (Põlme et al. 2020) | Põlme S, et al. 2020. Fungal Diversity 105:1-16 ((https://doi.org/10.1007/s13225-020-00466-2)) |
| Otu44515 | 1 | k__Fungi(100);p__Ascomycota(100);c__Saccharomycetes(100);o__Saccharomycetales(100);f__Metschnikowiaceae(100);g__Metschnikowia(100);s__Metschnikowia_pulcherrima(100); | Metschnikowia | 13 | Saprotroph | \|Undefined Saprotroph\| | Yeast | NULL | Probable | As nectar-tap_saprotroph (Põlme et al. 2020); Decay sugar-rich substrates (Põlme et al. 2020) | Põlme S, et al. 2020. Fungal Diversity 105:1-16 ((https://doi.org/10.1007/s13225-020-00466-2)) |
| Otu44516 | 1 | k__Fungi(100);p__Ascomycota(100);c__Saccharomycetes(100);o__Saccharomycetales(100);f__Metschnikowiaceae(100);g__Metschnikowia(100);s__Metschnikowia_pulcherrima(100); | Metschnikowia | 13 | Saprotroph | \|Undefined Saprotroph\| | Yeast | NULL | Probable | As nectar-tap_saprotroph (Põlme et al. 2020); Decay sugar-rich substrates (Põlme et al. 2020) | Põlme S, et al. 2020. Fungal Diversity 105:1-16 ((https://doi.org/10.1007/s13225-020-00466-2)) |
| Otu44517 | 1 | k__Fungi(100);p__Ascomycota(100);c__Saccharomycetes(100);o__Saccharomycetales(100);f__Metschnikowiaceae(100);g__Metschnikowia(100);s__Metschnikowia_pulcherrima(100); | Metschnikowia | 13 | Saprotroph | \|Undefined Saprotroph\| | Yeast | NULL | Probable | As nectar-tap_saprotroph (Põlme et al. 2020); Decay sugar-rich substrates (Põlme et al. 2020) | Põlme S, et al. 2020. Fungal Diversity 105:1-16 ((https://doi.org/10.1007/s13225-020-00466-2)) |
| Otu44518 | 1 | k__Fungi(100);p__Ascomycota(100);c__Saccharomycetes(100);o__Saccharomycetales(100);f__Metschnikowiaceae(100);g__Metschnikowia(100);s__Metschnikowia_pulcherrima(100); | Metschnikowia | 13 | Saprotroph | \|Undefined Saprotroph\| | Yeast | NULL | Probable | As nectar-tap_saprotroph (Põlme et al. 2020); Decay sugar-rich substrates (Põlme et al. 2020) | Põlme S, et al. 2020. Fungal Diversity 105:1-16 ((https://doi.org/10.1007/s13225-020-00466-2)) |
| Otu44519 | 1 | k__Fungi(100);p__Ascomycota(100);c__Saccharomycetes(100);o__Saccharomycetales(100);f__Metschnikowiaceae(100);g__Metschnikowia(100);g__Metschnikowia_unclassified(100); | Metschnikowia | 13 | Saprotroph | \|Undefined Saprotroph\| | Yeast | NULL | Probable | As nectar-tap_saprotroph (Põlme et al. 2020); Decay sugar-rich substrates (Põlme et al. 2020) | Põlme S, et al. 2020. Fungal Diversity 105:1-16 ((https://doi.org/10.1007/s13225-020-00466-2)) |
| Otu44520 | 1 | k__Fungi(100);p__Ascomycota(100);c__Saccharomycetes(100);o__Saccharomycetales(100);f__Metschnikowiaceae(100);g__Metschnikowia(100);s__Metschnikowia_pulcherrima(100); | Metschnikowia | 13 | Saprotroph | \|Undefined Saprotroph\| | Yeast | NULL | Probable | As nectar-tap_saprotroph (Põlme et al. 2020); Decay sugar-rich substrates (Põlme et al. 2020) | Põlme S, et al. 2020. Fungal Diversity 105:1-16 ((https://doi.org/10.1007/s13225-020-00466-2)) |
| Otu44521 | 1 | k__Fungi(100);p__Ascomycota(100);c__Saccharomycetes(100);o__Saccharomycetales(100);f__Metschnikowiaceae(100);g__Metschnikowia(100);s__Metschnikowia_pulcherrima(100); | Metschnikowia | 13 | Saprotroph | \|Undefined Saprotroph\| | Yeast | NULL | Probable | As nectar-tap_saprotroph (Põlme et al. 2020); Decay sugar-rich substrates (Põlme et al. 2020) | Põlme S, et al. 2020. Fungal Diversity 105:1-16 ((https://doi.org/10.1007/s13225-020-00466-2)) |
| Otu44522 | 1 | k__Fungi(100);p__Ascomycota(100);c__Saccharomycetes(100);o__Saccharomycetales(100);f__Metschnikowiaceae(100);g__Metschnikowia(100);s__Metschnikowia_pulcherrima(100); | Metschnikowia | 13 | Saprotroph | \|Undefined Saprotroph\| | Yeast | NULL | Probable | As nectar-tap_saprotroph (Põlme et al. 2020); Decay sugar-rich substrates (Põlme et al. 2020) | Põlme S, et al. 2020. Fungal Diversity 105:1-16 ((https://doi.org/10.1007/s13225-020-00466-2)) |
| Otu44523 | 1 | k__Fungi(100);p__Ascomycota(100);c__Saccharomycetes(100);o__Saccharomycetales(100);f__Metschnikowiaceae(100);g__Metschnikowia(100);s__Metschnikowia_pulcherrima(100); | Metschnikowia | 13 | Saprotroph | \|Undefined Saprotroph\| | Yeast | NULL | Probable | As nectar-tap_saprotroph (Põlme et al. 2020); Decay sugar-rich substrates (Põlme et al. 2020) | Põlme S, et al. 2020. Fungal Diversity 105:1-16 ((https://doi.org/10.1007/s13225-020-00466-2)) |
| Otu44524 | 1 | k__Fungi(100);p__Ascomycota(100);c__Saccharomycetes(100);o__Saccharomycetales(100);f__Metschnikowiaceae(100);g__Metschnikowia(100);s__Metschnikowia_pulcherrima(100); | Metschnikowia | 13 | Saprotroph | \|Undefined Saprotroph\| | Yeast | NULL | Probable | As nectar-tap_saprotroph (Põlme et al. 2020); Decay sugar-rich substrates (Põlme et al. 2020) | Põlme S, et al. 2020. Fungal Diversity 105:1-16 ((https://doi.org/10.1007/s13225-020-00466-2)) |
| Otu44525 | 1 | k__Fungi(100);p__Ascomycota(100);c__Saccharomycetes(100);o__Saccharomycetales(100);f__Metschnikowiaceae(100);g__Metschnikowia(100);s__Metschnikowia_pulcherrima(100); | Metschnikowia | 13 | Saprotroph | \|Undefined Saprotroph\| | Yeast | NULL | Probable | As nectar-tap_saprotroph (Põlme et al. 2020); Decay sugar-rich substrates (Põlme et al. 2020) | Põlme S, et al. 2020. Fungal Diversity 105:1-16 ((https://doi.org/10.1007/s13225-020-00466-2)) |
| Otu44526 | 1 | k__Fungi(100);p__Ascomycota(100);c__Saccharomycetes(100);o__Saccharomycetales(100);f__Metschnikowiaceae(100);g__Metschnikowia(100);s__Metschnikowia_viticola(100); | Metschnikowia | 13 | Saprotroph | \|Undefined Saprotroph\| | Yeast | NULL | Probable | As nectar-tap_saprotroph (Põlme et al. 2020); Decay sugar-rich substrates (Põlme et al. 2020) | Põlme S, et al. 2020. Fungal Diversity 105:1-16 ((https://doi.org/10.1007/s13225-020-00466-2)) |
| Otu44527 | 1 | k__Fungi(100);p__Ascomycota(100);c__Saccharomycetes(100);o__Saccharomycetales(100);f__Metschnikowiaceae(100);g__Metschnikowia(100);g__Metschnikowia_unclassified(100); | Metschnikowia | 13 | Saprotroph | \|Undefined Saprotroph\| | Yeast | NULL | Probable | As nectar-tap_saprotroph (Põlme et al. 2020); Decay sugar-rich substrates (Põlme et al. 2020) | Põlme S, et al. 2020. Fungal Diversity 105:1-16 ((https://doi.org/10.1007/s13225-020-00466-2)) |
| Otu44528 | 1 | k__Fungi(100);p__Ascomycota(100);c__Saccharomycetes(100);o__Saccharomycetales(100);f__Metschnikowiaceae(100);g__Metschnikowia(100);s__Metschnikowia_pulcherrima(100); | Metschnikowia | 13 | Saprotroph | \|Undefined Saprotroph\| | Yeast | NULL | Probable | As nectar-tap_saprotroph (Põlme et al. 2020); Decay sugar-rich substrates (Põlme et al. 2020) | Põlme S, et al. 2020. Fungal Diversity 105:1-16 ((https://doi.org/10.1007/s13225-020-00466-2)) |
| Otu44529 | 1 | k__Fungi(100);p__Ascomycota(100);c__Saccharomycetes(100);o__Saccharomycetales(100);f__Metschnikowiaceae(100);g__Metschnikowia(100);s__Metschnikowia_pulcherrima(100); | Metschnikowia | 13 | Saprotroph | \|Undefined Saprotroph\| | Yeast | NULL | Probable | As nectar-tap_saprotroph (Põlme et al. 2020); Decay sugar-rich substrates (Põlme et al. 2020) | Põlme S, et al. 2020. Fungal Diversity 105:1-16 ((https://doi.org/10.1007/s13225-020-00466-2)) |
| Otu44530 | 1 | k__Fungi(100);p__Ascomycota(100);c__Saccharomycetes(100);o__Saccharomycetales(100);f__Metschnikowiaceae(100);g__Metschnikowia(100);s__Metschnikowia_pulcherrima(100); | Metschnikowia | 13 | Saprotroph | \|Undefined Saprotroph\| | Yeast | NULL | Probable | As nectar-tap_saprotroph (Põlme et al. 2020); Decay sugar-rich substrates (Põlme et al. 2020) | Põlme S, et al. 2020. Fungal Diversity 105:1-16 ((https://doi.org/10.1007/s13225-020-00466-2)) |
| Otu44531 | 1 | k__Fungi(100);p__Ascomycota(100);c__Saccharomycetes(100);o__Saccharomycetales(100);f__Metschnikowiaceae(100);g__Metschnikowia(100);g__Metschnikowia_unclassified(100); | Metschnikowia | 13 | Saprotroph | \|Undefined Saprotroph\| | Yeast | NULL | Probable | As nectar-tap_saprotroph (Põlme et al. 2020); Decay sugar-rich substrates (Põlme et al. 2020) | Põlme S, et al. 2020. Fungal Diversity 105:1-16 ((https://doi.org/10.1007/s13225-020-00466-2)) |
| Otu44532 | 1 | k__Fungi(100);p__Ascomycota(100);c__Saccharomycetes(100);o__Saccharomycetales(100);f__Metschnikowiaceae(100);g__Metschnikowia(100);g__Metschnikowia_unclassified(100); | Metschnikowia | 13 | Saprotroph | \|Undefined Saprotroph\| | Yeast | NULL | Probable | As nectar-tap_saprotroph (Põlme et al. 2020); Decay sugar-rich substrates (Põlme et al. 2020) | Põlme S, et al. 2020. Fungal Diversity 105:1-16 ((https://doi.org/10.1007/s13225-020-00466-2)) |
| Otu44533 | 1 | k__Fungi(100);p__Ascomycota(100);c__Saccharomycetes(100);o__Saccharomycetales(100);f__Metschnikowiaceae(100);g__Metschnikowia(100);s__Metschnikowia_pulcherrima(100); | Metschnikowia | 13 | Saprotroph | \|Undefined Saprotroph\| | Yeast | NULL | Probable | As nectar-tap_saprotroph (Põlme et al. 2020); Decay sugar-rich substrates (Põlme et al. 2020) | Põlme S, et al. 2020. Fungal Diversity 105:1-16 ((https://doi.org/10.1007/s13225-020-00466-2)) |
| Otu44534 | 1 | k__Fungi(100);p__Ascomycota(100);c__Saccharomycetes(100);o__Saccharomycetales(100);f__Metschnikowiaceae(100);g__Metschnikowia(100);g__Metschnikowia_unclassified(100); | Metschnikowia | 13 | Saprotroph | \|Undefined Saprotroph\| | Yeast | NULL | Probable | As nectar-tap_saprotroph (Põlme et al. 2020); Decay sugar-rich substrates (Põlme et al. 2020) | Põlme S, et al. 2020. Fungal Diversity 105:1-16 ((https://doi.org/10.1007/s13225-020-00466-2)) |
| Otu44535 | 1 | k__Fungi(100);p__Ascomycota(100);c__Saccharomycetes(100);o__Saccharomycetales(100);f__Metschnikowiaceae(100);g__Metschnikowia(100);s__Metschnikowia_pulcherrima(100); | Metschnikowia | 13 | Saprotroph | \|Undefined Saprotroph\| | Yeast | NULL | Probable | As nectar-tap_saprotroph (Põlme et al. 2020); Decay sugar-rich substrates (Põlme et al. 2020) | Põlme S, et al. 2020. Fungal Diversity 105:1-16 ((https://doi.org/10.1007/s13225-020-00466-2)) |
| Otu44536 | 1 | k__Fungi(100);p__Ascomycota(100);c__Saccharomycetes(100);o__Saccharomycetales(100);f__Metschnikowiaceae(100);g__Metschnikowia(100);s__Metschnikowia_pulcherrima(100); | Metschnikowia | 13 | Saprotroph | \|Undefined Saprotroph\| | Yeast | NULL | Probable | As nectar-tap_saprotroph (Põlme et al. 2020); Decay sugar-rich substrates (Põlme et al. 2020) | Põlme S, et al. 2020. Fungal Diversity 105:1-16 ((https://doi.org/10.1007/s13225-020-00466-2)) |
| Otu44537 | 1 | k__Fungi(100);p__Ascomycota(100);c__Saccharomycetes(100);o__Saccharomycetales(100);f__Metschnikowiaceae(100);g__Metschnikowia(100);s__Metschnikowia_pulcherrima(100); | Metschnikowia | 13 | Saprotroph | \|Undefined Saprotroph\| | Yeast | NULL | Probable | As nectar-tap_saprotroph (Põlme et al. 2020); Decay sugar-rich substrates (Põlme et al. 2020) | Põlme S, et al. 2020. Fungal Diversity 105:1-16 ((https://doi.org/10.1007/s13225-020-00466-2)) |
| Otu44538 | 1 | k__Fungi(100);p__Ascomycota(100);c__Saccharomycetes(100);o__Saccharomycetales(100);f__Metschnikowiaceae(100);g__Metschnikowia(100);s__Metschnikowia_pulcherrima(100); | Metschnikowia | 13 | Saprotroph | \|Undefined Saprotroph\| | Yeast | NULL | Probable | As nectar-tap_saprotroph (Põlme et al. 2020); Decay sugar-rich substrates (Põlme et al. 2020) | Põlme S, et al. 2020. Fungal Diversity 105:1-16 ((https://doi.org/10.1007/s13225-020-00466-2)) |
| Otu44539 | 1 | k__Fungi(100);p__Ascomycota(100);c__Saccharomycetes(100);o__Saccharomycetales(100);f__Metschnikowiaceae(100);g__Metschnikowia(100);s__Metschnikowia_pulcherrima(100); | Metschnikowia | 13 | Saprotroph | \|Undefined Saprotroph\| | Yeast | NULL | Probable | As nectar-tap_saprotroph (Põlme et al. 2020); Decay sugar-rich substrates (Põlme et al. 2020) | Põlme S, et al. 2020. Fungal Diversity 105:1-16 ((https://doi.org/10.1007/s13225-020-00466-2)) |
| Otu44540 | 1 | k__Fungi(100);p__Ascomycota(100);c__Saccharomycetes(100);o__Saccharomycetales(100);f__Metschnikowiaceae(100);g__Metschnikowia(100);s__Metschnikowia_pulcherrima(100); | Metschnikowia | 13 | Saprotroph | \|Undefined Saprotroph\| | Yeast | NULL | Probable | As nectar-tap_saprotroph (Põlme et al. 2020); Decay sugar-rich substrates (Põlme et al. 2020) | Põlme S, et al. 2020. Fungal Diversity 105:1-16 ((https://doi.org/10.1007/s13225-020-00466-2)) |
| Otu44541 | 1 | k__Fungi(100);p__Ascomycota(100);c__Saccharomycetes(100);o__Saccharomycetales(100);f__Metschnikowiaceae(100);g__Metschnikowia(100);s__Metschnikowia_pulcherrima(100); | Metschnikowia | 13 | Saprotroph | \|Undefined Saprotroph\| | Yeast | NULL | Probable | As nectar-tap_saprotroph (Põlme et al. 2020); Decay sugar-rich substrates (Põlme et al. 2020) | Põlme S, et al. 2020. Fungal Diversity 105:1-16 ((https://doi.org/10.1007/s13225-020-00466-2)) |
| Otu44542 | 1 | k__Fungi(100);p__Ascomycota(100);c__Saccharomycetes(100);o__Saccharomycetales(100);f__Metschnikowiaceae(100);g__Metschnikowia(100);s__Metschnikowia_pulcherrima(100); | Metschnikowia | 13 | Saprotroph | \|Undefined Saprotroph\| | Yeast | NULL | Probable | As nectar-tap_saprotroph (Põlme et al. 2020); Decay sugar-rich substrates (Põlme et al. 2020) | Põlme S, et al. 2020. Fungal Diversity 105:1-16 ((https://doi.org/10.1007/s13225-020-00466-2)) |
| Otu44543 | 1 | k__Fungi(100);p__Ascomycota(100);c__Saccharomycetes(100);o__Saccharomycetales(100);f__Metschnikowiaceae(100);g__Metschnikowia(100);s__Metschnikowia_pulcherrima(100); | Metschnikowia | 13 | Saprotroph | \|Undefined Saprotroph\| | Yeast | NULL | Probable | As nectar-tap_saprotroph (Põlme et al. 2020); Decay sugar-rich substrates (Põlme et al. 2020) | Põlme S, et al. 2020. Fungal Diversity 105:1-16 ((https://doi.org/10.1007/s13225-020-00466-2)) |
| Otu44544 | 1 | k__Fungi(100);p__Ascomycota(100);c__Saccharomycetes(100);o__Saccharomycetales(100);f__Metschnikowiaceae(100);g__Metschnikowia(100);s__Metschnikowia_pulcherrima(100); | Metschnikowia | 13 | Saprotroph | \|Undefined Saprotroph\| | Yeast | NULL | Probable | As nectar-tap_saprotroph (Põlme et al. 2020); Decay sugar-rich substrates (Põlme et al. 2020) | Põlme S, et al. 2020. Fungal Diversity 105:1-16 ((https://doi.org/10.1007/s13225-020-00466-2)) |
| Otu44545 | 1 | k__Fungi(100);p__Ascomycota(100);c__Saccharomycetes(100);o__Saccharomycetales(100);f__Metschnikowiaceae(100);g__Metschnikowia(100);s__Metschnikowia_pulcherrima(100); | Metschnikowia | 13 | Saprotroph | \|Undefined Saprotroph\| | Yeast | NULL | Probable | As nectar-tap_saprotroph (Põlme et al. 2020); Decay sugar-rich substrates (Põlme et al. 2020) | Põlme S, et al. 2020. Fungal Diversity 105:1-16 ((https://doi.org/10.1007/s13225-020-00466-2)) |
| Otu44546 | 1 | k__Fungi(100);p__Ascomycota(100);c__Saccharomycetes(100);o__Saccharomycetales(100);f__Metschnikowiaceae(100);g__Metschnikowia(100);g__Metschnikowia_unclassified(100); | Metschnikowia | 13 | Saprotroph | \|Undefined Saprotroph\| | Yeast | NULL | Probable | As nectar-tap_saprotroph (Põlme et al. 2020); Decay sugar-rich substrates (Põlme et al. 2020) | Põlme S, et al. 2020. Fungal Diversity 105:1-16 ((https://doi.org/10.1007/s13225-020-00466-2)) |
| Otu44547 | 1 | k__Fungi(100);p__Ascomycota(100);c__Saccharomycetes(100);o__Saccharomycetales(100);f__Metschnikowiaceae(100);g__Metschnikowia(100);g__Metschnikowia_unclassified(100); | Metschnikowia | 13 | Saprotroph | \|Undefined Saprotroph\| | Yeast | NULL | Probable | As nectar-tap_saprotroph (Põlme et al. 2020); Decay sugar-rich substrates (Põlme et al. 2020) | Põlme S, et al. 2020. Fungal Diversity 105:1-16 ((https://doi.org/10.1007/s13225-020-00466-2)) |
| Otu44548 | 1 | k__Fungi(100);p__Ascomycota(100);c__Saccharomycetes(100);o__Saccharomycetales(100);f__Metschnikowiaceae(100);g__Metschnikowia(100);s__Metschnikowia_pulcherrima(100); | Metschnikowia | 13 | Saprotroph | \|Undefined Saprotroph\| | Yeast | NULL | Probable | As nectar-tap_saprotroph (Põlme et al. 2020); Decay sugar-rich substrates (Põlme et al. 2020) | Põlme S, et al. 2020. Fungal Diversity 105:1-16 ((https://doi.org/10.1007/s13225-020-00466-2)) |
| Otu44549 | 1 | k__Fungi(100);p__Ascomycota(100);c__Saccharomycetes(100);o__Saccharomycetales(100);f__Metschnikowiaceae(100);g__Metschnikowia(100);g__Metschnikowia_unclassified(100); | Metschnikowia | 13 | Saprotroph | \|Undefined Saprotroph\| | Yeast | NULL | Probable | As nectar-tap_saprotroph (Põlme et al. 2020); Decay sugar-rich substrates (Põlme et al. 2020) | Põlme S, et al. 2020. Fungal Diversity 105:1-16 ((https://doi.org/10.1007/s13225-020-00466-2)) |
| Otu44550 | 1 | k__Fungi(100);p__Ascomycota(100);c__Saccharomycetes(100);o__Saccharomycetales(100);f__Metschnikowiaceae(100);g__Metschnikowia(100);s__Metschnikowia_pulcherrima(100); | Metschnikowia | 13 | Saprotroph | \|Undefined Saprotroph\| | Yeast | NULL | Probable | As nectar-tap_saprotroph (Põlme et al. 2020); Decay sugar-rich substrates (Põlme et al. 2020) | Põlme S, et al. 2020. Fungal Diversity 105:1-16 ((https://doi.org/10.1007/s13225-020-00466-2)) |
| Otu44551 | 1 | k__Fungi(100);p__Ascomycota(100);c__Saccharomycetes(100);o__Saccharomycetales(100);f__Metschnikowiaceae(100);g__Metschnikowia(100);s__Metschnikowia_pulcherrima(100); | Metschnikowia | 13 | Saprotroph | \|Undefined Saprotroph\| | Yeast | NULL | Probable | As nectar-tap_saprotroph (Põlme et al. 2020); Decay sugar-rich substrates (Põlme et al. 2020) | Põlme S, et al. 2020. Fungal Diversity 105:1-16 ((https://doi.org/10.1007/s13225-020-00466-2)) |
| Otu44552 | 1 | k__Fungi(100);p__Ascomycota(100);c__Saccharomycetes(100);o__Saccharomycetales(100);f__Metschnikowiaceae(100);g__Metschnikowia(100);s__Metschnikowia_pulcherrima(100); | Metschnikowia | 13 | Saprotroph | \|Undefined Saprotroph\| | Yeast | NULL | Probable | As nectar-tap_saprotroph (Põlme et al. 2020); Decay sugar-rich substrates (Põlme et al. 2020) | Põlme S, et al. 2020. Fungal Diversity 105:1-16 ((https://doi.org/10.1007/s13225-020-00466-2)) |
| Otu44553 | 1 | k__Fungi(100);p__Ascomycota(100);c__Saccharomycetes(100);o__Saccharomycetales(100);f__Metschnikowiaceae(100);g__Metschnikowia(100);g__Metschnikowia_unclassified(100); | Metschnikowia | 13 | Saprotroph | \|Undefined Saprotroph\| | Yeast | NULL | Probable | As nectar-tap_saprotroph (Põlme et al. 2020); Decay sugar-rich substrates (Põlme et al. 2020) | Põlme S, et al. 2020. Fungal Diversity 105:1-16 ((https://doi.org/10.1007/s13225-020-00466-2)) |
| Otu44554 | 1 | k__Fungi(100);p__Ascomycota(100);c__Saccharomycetes(100);o__Saccharomycetales(100);f__Metschnikowiaceae(100);g__Metschnikowia(100);s__Metschnikowia_pulcherrima(100); | Metschnikowia | 13 | Saprotroph | \|Undefined Saprotroph\| | Yeast | NULL | Probable | As nectar-tap_saprotroph (Põlme et al. 2020); Decay sugar-rich substrates (Põlme et al. 2020) | Põlme S, et al. 2020. Fungal Diversity 105:1-16 ((https://doi.org/10.1007/s13225-020-00466-2)) |
| Otu44556 | 1 | k__Fungi(100);p__Ascomycota(100);c__Saccharomycetes(100);o__Saccharomycetales(100);f__Metschnikowiaceae(100);g__Metschnikowia(100);s__Metschnikowia_pulcherrima(100); | Metschnikowia | 13 | Saprotroph | \|Undefined Saprotroph\| | Yeast | NULL | Probable | As nectar-tap_saprotroph (Põlme et al. 2020); Decay sugar-rich substrates (Põlme et al. 2020) | Põlme S, et al. 2020. Fungal Diversity 105:1-16 ((https://doi.org/10.1007/s13225-020-00466-2)) |

| | | | | | | | | | | | |
|---|---|---|---|---|---|---|---|---|---|---|---|
| Otu44557 | 1 | k__Fungi(100);p__Ascomycota(100);c__Saccharomycetes(100);o__Saccharomycetales(100);f__Metschnikowiaceae(100);g__Metschnikowia(100);s__Metschnikowia_pulcherrima(100); | Metschnikowia | 13 | Saprotroph | \|Undefined Saprotroph\| | Yeast | NULL | Probable | As nectar-tap_saprotroph (Põlme et al. 2020); Decay sugar-rich substrates (Põlme et al. 2020) | Põlme S, et al. 2020. Fungal Diversity 105:1-16 ((https://doi.org/10.1007/s13225-020-00466-2)) |
| Otu44558 | 1 | k__Fungi(100);p__Ascomycota(100);c__Saccharomycetes(100);o__Saccharomycetales(100);f__Metschnikowiaceae(100);g__Metschnikowia(100);s__Metschnikowia_pulcherrima(100); | Metschnikowia | 13 | Saprotroph | \|Undefined Saprotroph\| | Yeast | NULL | Probable | As nectar-tap_saprotroph (Põlme et al. 2020); Decay sugar-rich substrates (Põlme et al. 2020) | Põlme S, et al. 2020. Fungal Diversity 105:1-16 ((https://doi.org/10.1007/s13225-020-00466-2)) |
| Otu44559 | 1 | k__Fungi(100);p__Ascomycota(100);c__Saccharomycetes(100);o__Saccharomycetales(100);f__Metschnikowiaceae(100);g__Metschnikowia(100);s__Metschnikowia_pulcherrima(100); | Metschnikowia | 13 | Saprotroph | \|Undefined Saprotroph\| | Yeast | NULL | Probable | As nectar-tap_saprotroph (Põlme et al. 2020); Decay sugar-rich substrates (Põlme et al. 2020) | Põlme S, et al. 2020. Fungal Diversity 105:1-16 ((https://doi.org/10.1007/s13225-020-00466-2)) |
| Otu44560 | 1 | k__Fungi(100);p__Ascomycota(100);c__Saccharomycetes(100);o__Saccharomycetales(100);f__Metschnikowiaceae(100);g__Metschnikowia(100);s__Metschnikowia_pulcherrima(100); | Metschnikowia | 13 | Saprotroph | \|Undefined Saprotroph\| | Yeast | NULL | Probable | As nectar-tap_saprotroph (Põlme et al. 2020); Decay sugar-rich substrates (Põlme et al. 2020) | Põlme S, et al. 2020. Fungal Diversity 105:1-16 ((https://doi.org/10.1007/s13225-020-00466-2)) |
| Otu44561 | 1 | k__Fungi(100);p__Ascomycota(100);c__Saccharomycetes(100);o__Saccharomycetales(100);f__Metschnikowiaceae(100);g__Metschnikowia(100);s__Metschnikowia_pulcherrima(100); | Metschnikowia | 13 | Saprotroph | \|Undefined Saprotroph\| | Yeast | NULL | Probable | As nectar-tap_saprotroph (Põlme et al. 2020); Decay sugar-rich substrates (Põlme et al. 2020) | Põlme S, et al. 2020. Fungal Diversity 105:1-16 ((https://doi.org/10.1007/s13225-020-00466-2)) |
| Otu44562 | 1 | k__Fungi(100);p__Ascomycota(100);c__Saccharomycetes(100);o__Saccharomycetales(100);f__Metschnikowiaceae(100);g__Metschnikowia(100);s__Metschnikowia_pulcherrima(100); | Metschnikowia | 13 | Saprotroph | \|Undefined Saprotroph\| | Yeast | NULL | Probable | As nectar-tap_saprotroph (Põlme et al. 2020); Decay sugar-rich substrates (Põlme et al. 2020) | Põlme S, et al. 2020. Fungal Diversity 105:1-16 ((https://doi.org/10.1007/s13225-020-00466-2)) |
| Otu44564 | 1 | k__Fungi(100);p__Ascomycota(100);c__Saccharomycetes(100);o__Saccharomycetales(100);f__Metschnikowiaceae(100);g__Metschnikowia(100);s__Metschnikowia_pulcherrima(100); | Metschnikowia | 13 | Saprotroph | \|Undefined Saprotroph\| | Yeast | NULL | Probable | As nectar-tap_saprotroph (Põlme et al. 2020); Decay sugar-rich substrates (Põlme et al. 2020) | Põlme S, et al. 2020. Fungal Diversity 105:1-16 ((https://doi.org/10.1007/s13225-020-00466-2)) |
| Otu44565 | 1 | k__Fungi(100);p__Ascomycota(100);c__Saccharomycetes(100);o__Saccharomycetales(100);f__Metschnikowiaceae(100);g__Metschnikowia(100);s__Metschnikowia_pulcherrima(100); | Metschnikowia | 13 | Saprotroph | \|Undefined Saprotroph\| | Yeast | NULL | Probable | As nectar-tap_saprotroph (Põlme et al. 2020); Decay sugar-rich substrates (Põlme et al. 2020) | Põlme S, et al. 2020. Fungal Diversity 105:1-16 ((https://doi.org/10.1007/s13225-020-00466-2)) |
| Otu44569 | 1 | k__Fungi(100);p__Ascomycota(100);c__Saccharomycetes(100);o__Saccharomycetales(100);f__Metschnikowiaceae(100);g__Metschnikowia(100);s__Metschnikowia_pulcherrima(100); | Metschnikowia | 13 | Saprotroph | \|Undefined Saprotroph\| | Yeast | NULL | Probable | As nectar-tap_saprotroph (Põlme et al. 2020); Decay sugar-rich substrates (Põlme et al. 2020) | Põlme S, et al. 2020. Fungal Diversity 105:1-16 ((https://doi.org/10.1007/s13225-020-00466-2)) |
| Otu44570 | 1 | k__Fungi(100);p__Ascomycota(100);c__Saccharomycetes(100);o__Saccharomycetales(100);f__Metschnikowiaceae(100);g__Metschnikowia(100);g__Metschnikowia_unclassified(100); | Metschnikowia | 13 | Saprotroph | \|Undefined Saprotroph\| | Yeast | NULL | Probable | As nectar-tap_saprotroph (Põlme et al. 2020); Decay sugar-rich substrates (Põlme et al. 2020) | Põlme S, et al. 2020. Fungal Diversity 105:1-16 ((https://doi.org/10.1007/s13225-020-00466-2)) |
| Otu44571 | 1 | k__Fungi(100);p__Ascomycota(100);c__Saccharomycetes(100);o__Saccharomycetales(100);f__Metschnikowiaceae(100);g__Metschnikowia(100);g__Metschnikowia_unclassified(100); | Metschnikowia | 13 | Saprotroph | \|Undefined Saprotroph\| | Yeast | NULL | Probable | As nectar-tap_saprotroph (Põlme et al. 2020); Decay sugar-rich substrates (Põlme et al. 2020) | Põlme S, et al. 2020. Fungal Diversity 105:1-16 ((https://doi.org/10.1007/s13225-020-00466-2)) |
| Otu44572 | 1 | k__Fungi(100);p__Ascomycota(100);c__Saccharomycetes(100);o__Saccharomycetales(100);f__Metschnikowiaceae(100);g__Metschnikowia(100);s__Metschnikowia_pulcherrima(100); | Metschnikowia | 13 | Saprotroph | \|Undefined Saprotroph\| | Yeast | NULL | Probable | As nectar-tap_saprotroph (Põlme et al. 2020); Decay sugar-rich substrates (Põlme et al. 2020) | Põlme S, et al. 2020. Fungal Diversity 105:1-16 ((https://doi.org/10.1007/s13225-020-00466-2)) |
| Otu44573 | 1 | k__Fungi(100);p__Ascomycota(100);c__Saccharomycetes(100);o__Saccharomycetales(100);f__Metschnikowiaceae(100);g__Metschnikowia(100);g__Metschnikowia_unclassified(100); | Metschnikowia | 13 | Saprotroph | \|Undefined Saprotroph\| | Yeast | NULL | Probable | As nectar-tap_saprotroph (Põlme et al. 2020); Decay sugar-rich substrates (Põlme et al. 2020) | Põlme S, et al. 2020. Fungal Diversity 105:1-16 ((https://doi.org/10.1007/s13225-020-00466-2)) |
| Otu44574 | 1 | k__Fungi(100);p__Ascomycota(100);c__Saccharomycetes(100);o__Saccharomycetales(100);f__Metschnikowiaceae(100);g__Metschnikowia(100);g__Metschnikowia_unclassified(100); | Metschnikowia | 13 | Saprotroph | \|Undefined Saprotroph\| | Yeast | NULL | Probable | As nectar-tap_saprotroph (Põlme et al. 2020); Decay sugar-rich substrates (Põlme et al. 2020) | Põlme S, et al. 2020. Fungal Diversity 105:1-16 ((https://doi.org/10.1007/s13225-020-00466-2)) |
| Otu44575 | 1 | k__Fungi(100);p__Ascomycota(100);c__Saccharomycetes(100);o__Saccharomycetales(100);f__Metschnikowiaceae(100);g__Metschnikowia(100);g__Metschnikowia_unclassified(100); | Metschnikowia | 13 | Saprotroph | \|Undefined Saprotroph\| | Yeast | NULL | Probable | As nectar-tap_saprotroph (Põlme et al. 2020); Decay sugar-rich substrates (Põlme et al. 2020) | Põlme S, et al. 2020. Fungal Diversity 105:1-16 ((https://doi.org/10.1007/s13225-020-00466-2)) |
| Otu44576 | 1 | k__Fungi(100);p__Ascomycota(100);c__Saccharomycetes(100);o__Saccharomycetales(100);f__Metschnikowiaceae(100);g__Metschnikowia(100);g__Metschnikowia_unclassified(100); | Metschnikowia | 13 | Saprotroph | \|Undefined Saprotroph\| | Yeast | NULL | Probable | As nectar-tap_saprotroph (Põlme et al. 2020); Decay sugar-rich substrates (Põlme et al. 2020) | Põlme S, et al. 2020. Fungal Diversity 105:1-16 ((https://doi.org/10.1007/s13225-020-00466-2)) |
| Otu44577 | 1 | k__Fungi(100);p__Ascomycota(100);c__Saccharomycetes(100);o__Saccharomycetales(100);f__Metschnikowiaceae(100);g__Metschnikowia(100);s__Metschnikowia_pulcherrima(100); | Metschnikowia | 13 | Saprotroph | \|Undefined Saprotroph\| | Yeast | NULL | Probable | As nectar-tap_saprotroph (Põlme et al. 2020); Decay sugar-rich substrates (Põlme et al. 2020) | Põlme S, et al. 2020. Fungal Diversity 105:1-16 ((https://doi.org/10.1007/s13225-020-00466-2)) |
| Otu44578 | 1 | k__Fungi(100);p__Ascomycota(100);c__Saccharomycetes(100);o__Saccharomycetales(100);f__Metschnikowiaceae(100);g__Metschnikowia(100);s__Metschnikowia_pulcherrima(100); | Metschnikowia | 13 | Saprotroph | \|Undefined Saprotroph\| | Yeast | NULL | Probable | As nectar-tap_saprotroph (Põlme et al. 2020); Decay sugar-rich substrates (Põlme et al. 2020) | Põlme S, et al. 2020. Fungal Diversity 105:1-16 ((https://doi.org/10.1007/s13225-020-00466-2)) |
| Otu44579 | 1 | k__Fungi(100);p__Ascomycota(100);c__Saccharomycetes(100);o__Saccharomycetales(100);f__Metschnikowiaceae(100);g__Metschnikowia(100);s__Metschnikowia_pulcherrima(100); | Metschnikowia | 13 | Saprotroph | \|Undefined Saprotroph\| | Yeast | NULL | Probable | As nectar-tap_saprotroph (Põlme et al. 2020); Decay sugar-rich substrates (Põlme et al. 2020) | Põlme S, et al. 2020. Fungal Diversity 105:1-16 ((https://doi.org/10.1007/s13225-020-00466-2)) |
| Otu44580 | 1 | k__Fungi(100);p__Ascomycota(100);c__Saccharomycetes(100);o__Saccharomycetales(100);f__Metschnikowiaceae(100);g__Metschnikowia(100);s__Metschnikowia_pulcherrima(100); | Metschnikowia | 13 | Saprotroph | \|Undefined Saprotroph\| | Yeast | NULL | Probable | As nectar-tap_saprotroph (Põlme et al. 2020); Decay sugar-rich substrates (Põlme et al. 2020) | Põlme S, et al. 2020. Fungal Diversity 105:1-16 ((https://doi.org/10.1007/s13225-020-00466-2)) |
| Otu44581 | 1 | k__Fungi(100);p__Ascomycota(100);c__Saccharomycetes(100);o__Saccharomycetales(100);f__Metschnikowiaceae(100);g__Metschnikowia(100);s__Metschnikowia_pulcherrima(100); | Metschnikowia | 13 | Saprotroph | \|Undefined Saprotroph\| | Yeast | NULL | Probable | As nectar-tap_saprotroph (Põlme et al. 2020); Decay sugar-rich substrates (Põlme et al. 2020) | Põlme S, et al. 2020. Fungal Diversity 105:1-16 ((https://doi.org/10.1007/s13225-020-00466-2)) |
| Otu44582 | 1 | k__Fungi(100);p__Ascomycota(100);c__Saccharomycetes(100);o__Saccharomycetales(100);f__Metschnikowiaceae(100);g__Metschnikowia(100);s__Metschnikowia_pulcherrima(100); | Metschnikowia | 13 | Saprotroph | \|Undefined Saprotroph\| | Yeast | NULL | Probable | As nectar-tap_saprotroph (Põlme et al. 2020); Decay sugar-rich substrates (Põlme et al. 2020) | Põlme S, et al. 2020. Fungal Diversity 105:1-16 ((https://doi.org/10.1007/s13225-020-00466-2)) |
| Otu44583 | 1 | k__Fungi(100);p__Ascomycota(100);c__Saccharomycetes(100);o__Saccharomycetales(100);f__Metschnikowiaceae(100);g__Metschnikowia(100);s__Metschnikowia_pulcherrima(100); | Metschnikowia | 13 | Saprotroph | \|Undefined Saprotroph\| | Yeast | NULL | Probable | As nectar-tap_saprotroph (Põlme et al. 2020); Decay sugar-rich substrates (Põlme et al. 2020) | Põlme S, et al. 2020. Fungal Diversity 105:1-16 ((https://doi.org/10.1007/s13225-020-00466-2)) |
| Otu44584 | 1 | k__Fungi(100);p__Ascomycota(100);c__Saccharomycetes(100);o__Saccharomycetales(100);f__Metschnikowiaceae(100);g__Metschnikowia(100);s__Metschnikowia_pulcherrima(100); | Metschnikowia | 13 | Saprotroph | \|Undefined Saprotroph\| | Yeast | NULL | Probable | As nectar-tap_saprotroph (Põlme et al. 2020); Decay sugar-rich substrates (Põlme et al. 2020) | Põlme S, et al. 2020. Fungal Diversity 105:1-16 ((https://doi.org/10.1007/s13225-020-00466-2)) |
| Otu44585 | 1 | k__Fungi(100);p__Ascomycota(100);c__Saccharomycetes(100);o__Saccharomycetales(100);f__Metschnikowiaceae(100);g__Metschnikowia(100);s__Metschnikowia_pulcherrima(100); | Metschnikowia | 13 | Saprotroph | \|Undefined Saprotroph\| | Yeast | NULL | Probable | As nectar-tap_saprotroph (Põlme et al. 2020); Decay sugar-rich substrates (Põlme et al. 2020) | Põlme S, et al. 2020. Fungal Diversity 105:1-16 ((https://doi.org/10.1007/s13225-020-00466-2)) |
| Otu44586 | 1 | k__Fungi(100);p__Ascomycota(100);c__Saccharomycetes(100);o__Saccharomycetales(100);f__Metschnikowiaceae(100);g__Metschnikowia(100);s__Metschnikowia_pulcherrima(100); | Metschnikowia | 13 | Saprotroph | \|Undefined Saprotroph\| | Yeast | NULL | Probable | As nectar-tap_saprotroph (Põlme et al. 2020); Decay sugar-rich substrates (Põlme et al. 2020) | Põlme S, et al. 2020. Fungal Diversity 105:1-16 ((https://doi.org/10.1007/s13225-020-00466-2)) |
| Otu44587 | 1 | k__Fungi(100);p__Ascomycota(100);c__Saccharomycetes(100);o__Saccharomycetales(100);f__Metschnikowiaceae(100);g__Metschnikowia(100);g__Metschnikowia_unclassified(100); | Metschnikowia | 13 | Saprotroph | \|Undefined Saprotroph\| | Yeast | NULL | Probable | As nectar-tap_saprotroph (Põlme et al. 2020); Decay sugar-rich substrates (Põlme et al. 2020) | Põlme S, et al. 2020. Fungal Diversity 105:1-16 ((https://doi.org/10.1007/s13225-020-00466-2)) |
| Otu44589 | 1 | k__Fungi(100);p__Ascomycota(100);c__Saccharomycetes(100);o__Saccharomycetales(100);f__Metschnikowiaceae(100);g__Metschnikowia(100);s__Metschnikowia_pulcherrima(100); | Metschnikowia | 13 | Saprotroph | \|Undefined Saprotroph\| | Yeast | NULL | Probable | As nectar-tap_saprotroph (Põlme et al. 2020); Decay sugar-rich substrates (Põlme et al. 2020) | Põlme S, et al. 2020. Fungal Diversity 105:1-16 ((https://doi.org/10.1007/s13225-020-00466-2)) |
| Otu44590 | 1 | k__Fungi(100);p__Ascomycota(100);c__Saccharomycetes(100);o__Saccharomycetales(100);f__Metschnikowiaceae(100);g__Metschnikowia(100);s__Metschnikowia_pulcherrima(100); | Metschnikowia | 13 | Saprotroph | \|Undefined Saprotroph\| | Yeast | NULL | Probable | As nectar-tap_saprotroph (Põlme et al. 2020); Decay sugar-rich substrates (Põlme et al. 2020) | Põlme S, et al. 2020. Fungal Diversity 105:1-16 ((https://doi.org/10.1007/s13225-020-00466-2)) |
| Otu44591 | 1 | k__Fungi(100);p__Ascomycota(100);c__Saccharomycetes(100);o__Saccharomycetales(100);f__Metschnikowiaceae(100);g__Metschnikowia(100);s__Metschnikowia_pulcherrima(100); | Metschnikowia | 13 | Saprotroph | \|Undefined Saprotroph\| | Yeast | NULL | Probable | As nectar-tap_saprotroph (Põlme et al. 2020); Decay sugar-rich substrates (Põlme et al. 2020) | Põlme S, et al. 2020. Fungal Diversity 105:1-16 ((https://doi.org/10.1007/s13225-020-00466-2)) |
| Otu44592 | 1 | k__Fungi(100);p__Ascomycota(100);c__Saccharomycetes(100);o__Saccharomycetales(100);f__Metschnikowiaceae(100);g__Metschnikowia(100);s__Metschnikowia_pulcherrima(100); | Metschnikowia | 13 | Saprotroph | \|Undefined Saprotroph\| | Yeast | NULL | Probable | As nectar-tap_saprotroph (Põlme et al. 2020); Decay sugar-rich substrates (Põlme et al. 2020) | Põlme S, et al. 2020. Fungal Diversity 105:1-16 ((https://doi.org/10.1007/s13225-020-00466-2)) |
| Otu44593 | 1 | k__Fungi(100);p__Ascomycota(100);c__Saccharomycetes(100);o__Saccharomycetales(100);f__Metschnikowiaceae(100);g__Metschnikowia(100);s__Metschnikowia_pulcherrima(100); | Metschnikowia | 13 | Saprotroph | \|Undefined Saprotroph\| | Yeast | NULL | Probable | As nectar-tap_saprotroph (Põlme et al. 2020); Decay sugar-rich substrates (Põlme et al. 2020) | Põlme S, et al. 2020. Fungal Diversity 105:1-16 ((https://doi.org/10.1007/s13225-020-00466-2)) |
| Otu44594 | 1 | k__Fungi(100);p__Ascomycota(100);c__Saccharomycetes(100);o__Saccharomycetales(100);f__Metschnikowiaceae(100);g__Metschnikowia(100);s__Metschnikowia_pulcherrima(100); | Metschnikowia | 13 | Saprotroph | \|Undefined Saprotroph\| | Yeast | NULL | Probable | As nectar-tap_saprotroph (Põlme et al. 2020); Decay sugar-rich substrates (Põlme et al. 2020) | Põlme S, et al. 2020. Fungal Diversity 105:1-16 ((https://doi.org/10.1007/s13225-020-00466-2)) |
| Otu44595 | 1 | k__Fungi(100);p__Ascomycota(100);c__Saccharomycetes(100);o__Saccharomycetales(100);f__Metschnikowiaceae(100);g__Metschnikowia(100);g__Metschnikowia_unclassified(100); | Metschnikowia | 13 | Saprotroph | \|Undefined Saprotroph\| | Yeast | NULL | Probable | As nectar-tap_saprotroph (Põlme et al. 2020); Decay sugar-rich substrates (Põlme et al. 2020) | Põlme S, et al. 2020. Fungal Diversity 105:1-16 ((https://doi.org/10.1007/s13225-020-00466-2)) |
| Otu44596 | 1 | k__Fungi(100);p__Ascomycota(100);c__Saccharomycetes(100);o__Saccharomycetales(100);f__Metschnikowiaceae(100);g__Metschnikowia(100);g__Metschnikowia_unclassified(100); | Metschnikowia | 13 | Saprotroph | \|Undefined Saprotroph\| | Yeast | NULL | Probable | As nectar-tap_saprotroph (Põlme et al. 2020); Decay sugar-rich substrates (Põlme et al. 2020) | Põlme S, et al. 2020. Fungal Diversity 105:1-16 ((https://doi.org/10.1007/s13225-020-00466-2)) |
| Otu44597 | 1 | k__Fungi(100);p__Ascomycota(100);c__Saccharomycetes(100);o__Saccharomycetales(100);f__Metschnikowiaceae(100);g__Metschnikowia(100);s__Metschnikowia_pulcherrima(100); | Metschnikowia | 13 | Saprotroph | \|Undefined Saprotroph\| | Yeast | NULL | Probable | As nectar-tap_saprotroph (Põlme et al. 2020); Decay sugar-rich substrates (Põlme et al. 2020) | Põlme S, et al. 2020. Fungal Diversity 105:1-16 ((https://doi.org/10.1007/s13225-020-00466-2)) |
| Otu44598 | 1 | k__Fungi(100);p__Ascomycota(100);c__Saccharomycetes(100);o__Saccharomycetales(100);f__Metschnikowiaceae(100);g__Metschnikowia(100);s__Metschnikowia_pulcherrima(100); | Metschnikowia | 13 | Saprotroph | \|Undefined Saprotroph\| | Yeast | NULL | Probable | As nectar-tap_saprotroph (Põlme et al. 2020); Decay sugar-rich substrates (Põlme et al. 2020) | Põlme S, et al. 2020. Fungal Diversity 105:1-16 ((https://doi.org/10.1007/s13225-020-00466-2)) |
| Otu44599 | 1 | k__Fungi(100);p__Ascomycota(100);c__Saccharomycetes(100);o__Saccharomycetales(100);f__Metschnikowiaceae(100);g__Metschnikowia(100);g__Metschnikowia_unclassified(100); | Metschnikowia | 13 | Saprotroph | \|Undefined Saprotroph\| | Yeast | NULL | Probable | As nectar-tap_saprotroph (Põlme et al. 2020); Decay sugar-rich substrates (Põlme et al. 2020) | Põlme S, et al. 2020. Fungal Diversity 105:1-16 ((https://doi.org/10.1007/s13225-020-00466-2)) |
| Otu44600 | 1 | k__Fungi(100);p__Ascomycota(100);c__Saccharomycetes(100);o__Saccharomycetales(100);f__Metschnikowiaceae(100);g__Metschnikowia(100);g__Metschnikowia_unclassified(100); | Metschnikowia | 13 | Saprotroph | \|Undefined Saprotroph\| | Yeast | NULL | Probable | As nectar-tap_saprotroph (Põlme et al. 2020); Decay sugar-rich substrates (Põlme et al. 2020) | Põlme S, et al. 2020. Fungal Diversity 105:1-16 ((https://doi.org/10.1007/s13225-020-00466-2)) |
| Otu44601 | 1 | k__Fungi(100);p__Ascomycota(100);c__Saccharomycetes(100);o__Saccharomycetales(100);f__Metschnikowiaceae(100);g__Metschnikowia(100);s__Metschnikowia_pulcherrima(100); | Metschnikowia | 13 | Saprotroph | \|Undefined Saprotroph\| | Yeast | NULL | Probable | As nectar-tap_saprotroph (Põlme et al. 2020); Decay sugar-rich substrates (Põlme et al. 2020) | Põlme S, et al. 2020. Fungal Diversity 105:1-16 ((https://doi.org/10.1007/s13225-020-00466-2)) |
| Otu44602 | 1 | k__Fungi(100);p__Ascomycota(100);c__Saccharomycetes(100);o__Saccharomycetales(100);f__Metschnikowiaceae(100);g__Metschnikowia(100);s__Metschnikowia_pulcherrima(100); | Metschnikowia | 13 | Saprotroph | \|Undefined Saprotroph\| | Yeast | NULL | Probable | As nectar-tap_saprotroph (Põlme et al. 2020); Decay sugar-rich substrates (Põlme et al. 2020) | Põlme S, et al. 2020. Fungal Diversity 105:1-16 ((https://doi.org/10.1007/s13225-020-00466-2)) |
| Otu44603 | 1 | k__Fungi(100);p__Ascomycota(100);c__Saccharomycetes(100);o__Saccharomycetales(100);f__Metschnikowiaceae(100);g__Metschnikowia(100);g__Metschnikowia_unclassified(100); | Metschnikowia | 13 | Saprotroph | \|Undefined Saprotroph\| | Yeast | NULL | Probable | As nectar-tap_saprotroph (Põlme et al. 2020); Decay sugar-rich substrates (Põlme et al. 2020) | Põlme S, et al. 2020. Fungal Diversity 105:1-16 ((https://doi.org/10.1007/s13225-020-00466-2)) |
| Otu44604 | 1 | k__Fungi(100);p__Ascomycota(100);c__Saccharomycetes(100);o__Saccharomycetales(100);f__Metschnikowiaceae(100);g__Metschnikowia(100);s__Metschnikowia_pulcherrima(100); | Metschnikowia | 13 | Saprotroph | \|Undefined Saprotroph\| | Yeast | NULL | Probable | As nectar-tap_saprotroph (Põlme et al. 2020); Decay sugar-rich substrates (Põlme et al. 2020) | Põlme S, et al. 2020. Fungal Diversity 105:1-16 ((https://doi.org/10.1007/s13225-020-00466-2)) |
| Otu44605 | 1 | k__Fungi(100);p__Ascomycota(100);c__Saccharomycetes(100);o__Saccharomycetales(100);f__Metschnikowiaceae(100);g__Metschnikowia(100);s__Metschnikowia_pulcherrima(100); | Metschnikowia | 13 | Saprotroph | \|Undefined Saprotroph\| | Yeast | NULL | Probable | As nectar-tap_saprotroph (Põlme et al. 2020); Decay sugar-rich substrates (Põlme et al. 2020) | Põlme S, et al. 2020. Fungal Diversity 105:1-16 ((https://doi.org/10.1007/s13225-020-00466-2)) |
| Otu44606 | 1 | k__Fungi(100);p__Ascomycota(100);c__Saccharomycetes(100);o__Saccharomycetales(100);f__Metschnikowiaceae(100);g__Metschnikowia(100);s__Metschnikowia_pulcherrima(100); | Metschnikowia | 13 | Saprotroph | \|Undefined Saprotroph\| | Yeast | NULL | Probable | As nectar-tap_saprotroph (Põlme et al. 2020); Decay sugar-rich substrates (Põlme et al. 2020) | Põlme S, et al. 2020. Fungal Diversity 105:1-16 ((https://doi.org/10.1007/s13225-020-00466-2)) |
| Otu44607 | 1 | k__Fungi(100);p__Ascomycota(100);c__Saccharomycetes(100);o__Saccharomycetales(100);f__Metschnikowiaceae(100);g__Metschnikowia(100);s__Metschnikowia_pulcherrima(100); | Metschnikowia | 13 | Saprotroph | \|Undefined Saprotroph\| | Yeast | NULL | Probable | As nectar-tap_saprotroph (Põlme et al. 2020); Decay sugar-rich substrates (Põlme et al. 2020) | Põlme S, et al. 2020. Fungal Diversity 105:1-16 ((https://doi.org/10.1007/s13225-020-00466-2)) |
| Otu44608 | 1 | k__Fungi(100);p__Ascomycota(100);c__Saccharomycetes(100);o__Saccharomycetales(100);f__Metschnikowiaceae(100);g__Metschnikowia(100);g__Metschnikowia_unclassified(100); | Metschnikowia | 13 | Saprotroph | \|Undefined Saprotroph\| | Yeast | NULL | Probable | As nectar-tap_saprotroph (Põlme et al. 2020); Decay sugar-rich substrates (Põlme et al. 2020) | Põlme S, et al. 2020. Fungal Diversity 105:1-16 ((https://doi.org/10.1007/s13225-020-00466-2)) |
| Otu44610 | 1 | k__Fungi(100);p__Ascomycota(100);c__Saccharomycetes(100);o__Saccharomycetales(100);f__Metschnikowiaceae(100);g__Metschnikowia(100);g__Metschnikowia_unclassified(100); | Metschnikowia | 13 | Saprotroph | \|Undefined Saprotroph\| | Yeast | NULL | Probable | As nectar-tap_saprotroph (Põlme et al. 2020); Decay sugar-rich substrates (Põlme et al. 2020) | Põlme S, et al. 2020. Fungal Diversity 105:1-16 ((https://doi.org/10.1007/s13225-020-00466-2)) |
| Otu44611 | 1 | k__Fungi(100);p__Ascomycota(100);c__Saccharomycetes(100);o__Saccharomycetales(100);f__Metschnikowiaceae(100);g__Metschnikowia(100);g__Metschnikowia_unclassified(100); | Metschnikowia | 13 | Saprotroph | \|Undefined Saprotroph\| | Yeast | NULL | Probable | As nectar-tap_saprotroph (Põlme et al. 2020); Decay sugar-rich substrates (Põlme et al. 2020) | Põlme S, et al. 2020. Fungal Diversity 105:1-16 ((https://doi.org/10.1007/s13225-020-00466-2)) |
| Otu44612 | 1 | k__Fungi(100);p__Ascomycota(100);c__Saccharomycetes(100);o__Saccharomycetales(100);f__Metschnikowiaceae(100);g__Metschnikowia(100);g__Metschnikowia_unclassified(100); | Metschnikowia | 13 | Saprotroph | \|Undefined Saprotroph\| | Yeast | NULL | Probable | As nectar-tap_saprotroph (Põlme et al. 2020); Decay sugar-rich substrates (Põlme et al. 2020) | Põlme S, et al. 2020. Fungal Diversity 105:1-16 ((https://doi.org/10.1007/s13225-020-00466-2)) |
| Otu44613 | 1 | k__Fungi(100);p__Ascomycota(100);c__Saccharomycetes(100);o__Saccharomycetales(100);f__Metschnikowiaceae(100);g__Metschnikowia(100);s__Metschnikowia_pulcherrima(100); | Metschnikowia | 13 | Saprotroph | \|Undefined Saprotroph\| | Yeast | NULL | Probable | As nectar-tap_saprotroph (Põlme et al. 2020); Decay sugar-rich substrates (Põlme et al. 2020) | Põlme S, et al. 2020. Fungal Diversity 105:1-16 ((https://doi.org/10.1007/s13225-020-00466-2)) |
| Otu44614 | 1 | k__Fungi(100);p__Ascomycota(100);c__Saccharomycetes(100);o__Saccharomycetales(100);f__Metschnikowiaceae(100);g__Metschnikowia(100);s__Metschnikowia_pulcherrima(100); | Metschnikowia | 13 | Saprotroph | \|Undefined Saprotroph\| | Yeast | NULL | Probable | As nectar-tap_saprotroph (Põlme et al. 2020); Decay sugar-rich substrates (Põlme et al. 2020) | Põlme S, et al. 2020. Fungal Diversity 105:1-16 ((https://doi.org/10.1007/s13225-020-00466-2)) |
| Otu44615 | 1 | k__Fungi(100);p__Ascomycota(100);c__Saccharomycetes(100);o__Saccharomycetales(100);f__Metschnikowiaceae(100);g__Metschnikowia(100);s__Metschnikowia_pulcherrima(100); | Metschnikowia | 13 | Saprotroph | \|Undefined Saprotroph\| | Yeast | NULL | Probable | As nectar-tap_saprotroph (Põlme et al. 2020); Decay sugar-rich substrates (Põlme et al. 2020) | Põlme S, et al. 2020. Fungal Diversity 105:1-16 ((https://doi.org/10.1007/s13225-020-00466-2)) |
| Otu44616 | 1 | k__Fungi(100);p__Ascomycota(100);c__Saccharomycetes(100);o__Saccharomycetales(100);f__Metschnikowiaceae(100);g__Metschnikowia(100);s__Metschnikowia_pulcherrima(100); | Metschnikowia | 13 | Saprotroph | \|Undefined Saprotroph\| | Yeast | NULL | Probable | As nectar-tap_saprotroph (Põlme et al. 2020); Decay sugar-rich substrates (Põlme et al. 2020) | Põlme S, et al. 2020. Fungal Diversity 105:1-16 ((https://doi.org/10.1007/s13225-020-00466-2)) |
| Otu44617 | 1 | k__Fungi(100);p__Ascomycota(100);c__Saccharomycetes(100);o__Saccharomycetales(100);f__Metschnikowiaceae(100);g__Metschnikowia(100);s__Metschnikowia_pulcherrima(100); | Metschnikowia | 13 | Saprotroph | \|Undefined Saprotroph\| | Yeast | NULL | Probable | As nectar-tap_saprotroph (Põlme et al. 2020); Decay sugar-rich substrates (Põlme et al. 2020) | Põlme S, et al. 2020. Fungal Diversity 105:1-16 ((https://doi.org/10.1007/s13225-020-00466-2)) |
| Otu44618 | 1 | k__Fungi(100);p__Ascomycota(100);c__Saccharomycetes(100);o__Saccharomycetales(100);f__Metschnikowiaceae(100);g__Metschnikowia(100);s__Metschnikowia_pulcherrima(100); | Metschnikowia | 13 | Saprotroph | \|Undefined Saprotroph\| | Yeast | NULL | Probable | As nectar-tap_saprotroph (Põlme et al. 2020); Decay sugar-rich substrates (Põlme et al. 2020) | Põlme S, et al. 2020. Fungal Diversity 105:1-16 ((https://doi.org/10.1007/s13225-020-00466-2)) |
| Otu44619 | 1 | k__Fungi(100);p__Ascomycota(100);c__Saccharomycetes(100);o__Saccharomycetales(100);f__Metschnikowiaceae(100);g__Metschnikowia(100);s__Metschnikowia_pulcherrima(100); | Metschnikowia | 13 | Saprotroph | \|Undefined Saprotroph\| | Yeast | NULL | Probable | As nectar-tap_saprotroph (Põlme et al. 2020); Decay sugar-rich substrates (Põlme et al. 2020) | Põlme S, et al. 2020. Fungal Diversity 105:1-16 ((https://doi.org/10.1007/s13225-020-00466-2)) |
| Otu44620 | 1 | k__Fungi(100);p__Ascomycota(100);c__Saccharomycetes(100);o__Saccharomycetales(100);f__Metschnikowiaceae(100);g__Metschnikowia(100);g__Metschnikowia_unclassified(100); | Metschnikowia | 13 | Saprotroph | \|Undefined Saprotroph\| | Yeast | NULL | Probable | As nectar-tap_saprotroph (Põlme et al. 2020); Decay sugar-rich substrates (Põlme et al. 2020) | Põlme S, et al. 2020. Fungal Diversity 105:1-16 ((https://doi.org/10.1007/s13225-020-00466-2)) |
| Otu44621 | 1 | k__Fungi(100);p__Ascomycota(100);c__Saccharomycetes(100);o__Saccharomycetales(100);f__Metschnikowiaceae(100);g__Metschnikowia(100);s__Metschnikowia_pulcherrima(100); | Metschnikowia | 13 | Saprotroph | \|Undefined Saprotroph\| | Yeast | NULL | Probable | As nectar-tap_saprotroph (Põlme et al. 2020); Decay sugar-rich substrates (Põlme et al. 2020) | Põlme S, et al. 2020. Fungal Diversity 105:1-16 ((https://doi.org/10.1007/s13225-020-00466-2)) |
| Otu44622 | 1 | k__Fungi(100);p__Ascomycota(100);c__Saccharomycetes(100);o__Saccharomycetales(100);f__Metschnikowiaceae(100);g__Metschnikowia(100);s__Metschnikowia_pulcherrima(100); | Metschnikowia | 13 | Saprotroph | \|Undefined Saprotroph\| | Yeast | NULL | Probable | As nectar-tap_saprotroph (Põlme et al. 2020); Decay sugar-rich substrates (Põlme et al. 2020) | Põlme S, et al. 2020. Fungal Diversity 105:1-16 ((https://doi.org/10.1007/s13225-020-00466-2)) |
| Otu44623 | 1 | k__Fungi(100);p__Ascomycota(100);c__Saccharomycetes(100);o__Saccharomycetales(100);f__Metschnikowiaceae(100);g__Metschnikowia(100);s__Metschnikowia_pulcherrima(100); | Metschnikowia | 13 | Saprotroph | \|Undefined Saprotroph\| | Yeast | NULL | Probable | As nectar-tap_saprotroph (Põlme et al. 2020); Decay sugar-rich substrates (Põlme et al. 2020) | Põlme S, et al. 2020. Fungal Diversity 105:1-16 ((https://doi.org/10.1007/s13225-020-00466-2)) |
| Otu44624 | 1 | k__Fungi(100);p__Ascomycota(100);c__Saccharomycetes(100);o__Saccharomycetales(100);f__Metschnikowiaceae(100);g__Metschnikowia(100);g__Metschnikowia_unclassified(100); | Metschnikowia | 13 | Saprotroph | \|Undefined Saprotroph\| | Yeast | NULL | Probable | As nectar-tap_saprotroph (Põlme et al. 2020); Decay sugar-rich substrates (Põlme et al. 2020) | Põlme S, et al. 2020. Fungal Diversity 105:1-16 ((https://doi.org/10.1007/s13225-020-00466-2)) |
| Otu44625 | 1 | k__Fungi(100);p__Ascomycota(100);c__Saccharomycetes(100);o__Saccharomycetales(100);f__Metschnikowiaceae(100);g__Metschnikowia(100);s__Metschnikowia_pulcherrima(100); | Metschnikowia | 13 | Saprotroph | \|Undefined Saprotroph\| | Yeast | NULL | Probable | As nectar-tap_saprotroph (Põlme et al. 2020); Decay sugar-rich substrates (Põlme et al. 2020) | Põlme S, et al. 2020. Fungal Diversity 105:1-16 ((https://doi.org/10.1007/s13225-020-00466-2)) |
| Otu44626 | 1 | k__Fungi(100);p__Ascomycota(100);c__Saccharomycetes(100);o__Saccharomycetales(100);f__Metschnikowiaceae(100);g__Metschnikowia(100);s__Metschnikowia_pulcherrima(100); | Metschnikowia | 13 | Saprotroph | \|Undefined Saprotroph\| | Yeast | NULL | Probable | As nectar-tap_saprotroph (Põlme et al. 2020); Decay sugar-rich substrates (Põlme et al. 2020) | Põlme S, et al. 2020. Fungal Diversity 105:1-16 ((https://doi.org/10.1007/s13225-020-00466-2)) |
| Otu44627 | 1 | k__Fungi(100);p__Ascomycota(100);c__Saccharomycetes(100);o__Saccharomycetales(100);f__Metschnikowiaceae(100);g__Metschnikowia(100);g__Metschnikowia_unclassified(100); | Metschnikowia | 13 | Saprotroph | \|Undefined Saprotroph\| | Yeast | NULL | Probable | As nectar-tap_saprotroph (Põlme et al. 2020); Decay sugar-rich substrates (Põlme et al. 2020) | Põlme S, et al. 2020. Fungal Diversity 105:1-16 ((https://doi.org/10.1007/s13225-020-00466-2)) |
| Otu44628 | 1 | k__Fungi(100);p__Ascomycota(100);c__Saccharomycetes(100);o__Saccharomycetales(100);f__Metschnikowiaceae(100);g__Metschnikowia(100);s__Metschnikowia_pulcherrima(100); | Metschnikowia | 13 | Saprotroph | \|Undefined Saprotroph\| | Yeast | NULL | Probable | As nectar-tap_saprotroph (Põlme et al. 2020); Decay sugar-rich substrates (Põlme et al. 2020) | Põlme S, et al. 2020. Fungal Diversity 105:1-16 ((https://doi.org/10.1007/s13225-020-00466-2)) |
| Otu44629 | 1 | k__Fungi(100);p__Ascomycota(100);c__Saccharomycetes(100);o__Saccharomycetales(100);f__Metschnikowiaceae(100);g__Metschnikowia(100);g__Metschnikowia_unclassified(100); | Metschnikowia | 13 | Saprotroph | \|Undefined Saprotroph\| | Yeast | NULL | Probable | As nectar-tap_saprotroph (Põlme et al. 2020); Decay sugar-rich substrates (Põlme et al. 2020) | Põlme S, et al. 2020. Fungal Diversity 105:1-16 ((https://doi.org/10.1007/s13225-020-00466-2)) |
| Otu44630 | 1 | k__Fungi(100);p__Ascomycota(100);c__Saccharomycetes(100);o__Saccharomycetales(100);f__Metschnikowiaceae(100);g__Metschnikowia(100);s__Metschnikowia_pulcherrima(100); | Metschnikowia | 13 | Saprotroph | \|Undefined Saprotroph\| | Yeast | NULL | Probable | As nectar-tap_saprotroph (Põlme et al. 2020); Decay sugar-rich substrates (Põlme et al. 2020) | Põlme S, et al. 2020. Fungal Diversity 105:1-16 ((https://doi.org/10.1007/s13225-020-00466-2)) |
| Otu44631 | 1 | k__Fungi(100);p__Ascomycota(100);c__Saccharomycetes(100);o__Saccharomycetales(100);f__Metschnikowiaceae(100);g__Metschnikowia(100);s__Metschnikowia_pulcherrima(100); | Metschnikowia | 13 | Saprotroph | \|Undefined Saprotroph\| | Yeast | NULL | Probable | As nectar-tap_saprotroph (Põlme et al. 2020); Decay sugar-rich substrates (Põlme et al. 2020) | Põlme S, et al. 2020. Fungal Diversity 105:1-16 ((https://doi.org/10.1007/s13225-020-00466-2)) |
| Otu44632 | 1 | k__Fungi(100);p__Ascomycota(100);c__Saccharomycetes(100);o__Saccharomycetales(100);f__Metschnikowiaceae(100);g__Metschnikowia(100);s__Metschnikowia_pulcherrima(100); | Metschnikowia | 13 | Saprotroph | \|Undefined Saprotroph\| | Yeast | NULL | Probable | As nectar-tap_saprotroph (Põlme et al. 2020); Decay sugar-rich substrates (Põlme et al. 2020) | Põlme S, et al. 2020. Fungal Diversity 105:1-16 ((https://doi.org/10.1007/s13225-020-00466-2)) |
| Otu44633 | 1 | k__Fungi(100);p__Ascomycota(100);c__Saccharomycetes(100);o__Saccharomycetales(100);f__Metschnikowiaceae(100);g__Metschnikowia(100);g__Metschnikowia_unclassified(100); | Metschnikowia | 13 | Saprotroph | \|Undefined Saprotroph\| | Yeast | NULL | Probable | As nectar-tap_saprotroph (Põlme et al. 2020); Decay sugar-rich substrates (Põlme et al. 2020) | Põlme S, et al. 2020. Fungal Diversity 105:1-16 ((https://doi.org/10.1007/s13225-020-00466-2)) |
| Otu44634 | 1 | k__Fungi(100);p__Ascomycota(100);c__Saccharomycetes(100);o__Saccharomycetales(100);f__Metschnikowiaceae(100);g__Metschnikowia(100);s__Metschnikowia_pulcherrima(100); | Metschnikowia | 13 | Saprotroph | \|Undefined Saprotroph\| | Yeast | NULL | Probable | As nectar-tap_saprotroph (Põlme et al. 2020); Decay sugar-rich substrates (Põlme et al. 2020) | Põlme S, et al. 2020. Fungal Diversity 105:1-16 ((https://doi.org/10.1007/s13225-020-00466-2)) |
| Otu44635 | 1 | k__Fungi(100);p__Ascomycota(100);c__Saccharomycetes(100);o__Saccharomycetales(100);f__Metschnikowiaceae(100);g__Metschnikowia(100);s__Metschnikowia_pulcherrima(100); | Metschnikowia | 13 | Saprotroph | \|Undefined Saprotroph\| | Yeast | NULL | Probable | As nectar-tap_saprotroph (Põlme et al. 2020); Decay sugar-rich substrates (Põlme et al. 2020) | Põlme S, et al. 2020. Fungal Diversity 105:1-16 ((https://doi.org/10.1007/s13225-020-00466-2)) |
| Otu44636 | 1 | k__Fungi(100);p__Ascomycota(100);c__Saccharomycetes(100);o__Saccharomycetales(100);f__Metschnikowiaceae(100);g__Metschnikowia(100);s__Metschnikowia_pulcherrima(100); | Metschnikowia | 13 | Saprotroph | \|Undefined Saprotroph\| | Yeast | NULL | Probable | As nectar-tap_saprotroph (Põlme et al. 2020); Decay sugar-rich substrates (Põlme et al. 2020) | Põlme S, et al. 2020. Fungal Diversity 105:1-16 ((https://doi.org/10.1007/s13225-020-00466-2)) |
| Otu44637 | 1 | k__Fungi(100);p__Ascomycota(100);c__Saccharomycetes(100);o__Saccharomycetales(100);f__Metschnikowiaceae(100);g__Metschnikowia(100);g__Metschnikowia_unclassified(100); | Metschnikowia | 13 | Saprotroph | \|Undefined Saprotroph\| | Yeast | NULL | Probable | As nectar-tap_saprotroph (Põlme et al. 2020); Decay sugar-rich substrates (Põlme et al. 2020) | Põlme S, et al. 2020. Fungal Diversity 105:1-16 ((https://doi.org/10.1007/s13225-020-00466-2)) |
| Otu44638 | 1 | k__Fungi(100);p__Ascomycota(100);c__Saccharomycetes(100);o__Saccharomycetales(100);f__Metschnikowiaceae(100);g__Metschnikowia(100);s__Metschnikowia_pulcherrima(100); | Metschnikowia | 13 | Saprotroph | \|Undefined Saprotroph\| | Yeast | NULL | Probable | As nectar-tap_saprotroph (Põlme et al. 2020); Decay sugar-rich substrates (Põlme et al. 2020) | Põlme S, et al. 2020. Fungal Diversity 105:1-16 ((https://doi.org/10.1007/s13225-020-00466-2)) |
| Otu44639 | 1 | k__Fungi(100);p__Ascomycota(100);c__Saccharomycetes(100);o__Saccharomycetales(100);f__Metschnikowiaceae(100);g__Metschnikowia(100);s__Metschnikowia_pulcherrima(100); | Metschnikowia | 13 | Saprotroph | \|Undefined Saprotroph\| | Yeast | NULL | Probable | As nectar-tap_saprotroph (Põlme et al. 2020); Decay sugar-rich substrates (Põlme et al. 2020) | Põlme S, et al. 2020. Fungal Diversity 105:1-16 ((https://doi.org/10.1007/s13225-020-00466-2)) |
| Otu44640 | 1 | k__Fungi(100);p__Ascomycota(100);c__Saccharomycetes(100);o__Saccharomycetales(100);f__Metschnikowiaceae(100);g__Metschnikowia(100);s__Metschnikowia_pulcherrima(100); | Metschnikowia | 13 | Saprotroph | \|Undefined Saprotroph\| | Yeast | NULL | Probable | As nectar-tap_saprotroph (Põlme et al. 2020); Decay sugar-rich substrates (Põlme et al. 2020) | Põlme S, et al. 2020. Fungal Diversity 105:1-16 ((https://doi.org/10.1007/s13225-020-00466-2)) |
| Otu44641 | 1 | k__Fungi(100);p__Ascomycota(100);c__Saccharomycetes(100);o__Saccharomycetales(100);f__Metschnikowiaceae(100);g__Metschnikowia(100);s__Metschnikowia_pulcherrima(100); | Metschnikowia | 13 | Saprotroph | \|Undefined Saprotroph\| | Yeast | NULL | Probable | As nectar-tap_saprotroph (Põlme et al. 2020); Decay sugar-rich substrates (Põlme et al. 2020) | Põlme S, et al. 2020. Fungal Diversity 105:1-16 ((https://doi.org/10.1007/s13225-020-00466-2)) |
| Otu44642 | 1 | k__Fungi(100);p__Ascomycota(100);c__Saccharomycetes(100);o__Saccharomycetales(100);f__Metschnikowiaceae(100);g__Metschnikowia(100);s__Metschnikowia_pulcherrima(100); | Metschnikowia | 13 | Saprotroph | \|Undefined Saprotroph\| | Yeast | NULL | Probable | As nectar-tap_saprotroph (Põlme et al. 2020); Decay sugar-rich substrates (Põlme et al. 2020) | Põlme S, et al. 2020. Fungal Diversity 105:1-16 ((https://doi.org/10.1007/s13225-020-00466-2)) |
| Otu44643 | 1 | k__Fungi(100);p__Ascomycota(100);c__Saccharomycetes(100);o__Saccharomycetales(100);f__Metschnikowiaceae(100);g__Metschnikowia(100);s__Metschnikowia_pulcherrima(100); | Metschnikowia | 13 | Saprotroph | \|Undefined Saprotroph\| | Yeast | NULL | Probable | As nectar-tap_saprotroph (Põlme et al. 2020); Decay sugar-rich substrates (Põlme et al. 2020) | Põlme S, et al. 2020. Fungal Diversity 105:1-16 ((https://doi.org/10.1007/s13225-020-00466-2)) |
| Otu44644 | 1 | k__Fungi(100);p__Ascomycota(100);c__Saccharomycetes(100);o__Saccharomycetales(100);f__Metschnikowiaceae(100);g__Metschnikowia(100);s__Metschnikowia_pulcherrima(100); | Metschnikowia | 13 | Saprotroph | \|Undefined Saprotroph\| | Yeast | NULL | Probable | As nectar-tap_saprotroph (Põlme et al. 2020); Decay sugar-rich substrates (Põlme et al. 2020) | Põlme S, et al. 2020. Fungal Diversity 105:1-16 ((https://doi.org/10.1007/s13225-020-00466-2)) |
| Otu44645 | 1 | k__Fungi(100);p__Ascomycota(100);c__Saccharomycetes(100);o__Saccharomycetales(100);f__Metschnikowiaceae(100);g__Metschnikowia(100);s__Metschnikowia_pulcherrima(100); | Metschnikowia | 13 | Saprotroph | \|Undefined Saprotroph\| | Yeast | NULL | Probable | As nectar-tap_saprotroph (Põlme et al. 2020); Decay sugar-rich substrates (Põlme et al. 2020) | Põlme S, et al. 2020. Fungal Diversity 105:1-16 ((https://doi.org/10.1007/s13225-020-00466-2)) |
| Otu44646 | 1 | k__Fungi(100);p__Ascomycota(100);c__Saccharomycetes(100);o__Saccharomycetales(100);f__Metschnikowiaceae(100);g__Metschnikowia(100);s__Metschnikowia_pulcherrima(100); | Metschnikowia | 13 | Saprotroph | \|Undefined Saprotroph\| | Yeast | NULL | Probable | As nectar-tap_saprotroph (Põlme et al. 2020); Decay sugar-rich substrates (Põlme et al. 2020) | Põlme S, et al. 2020. Fungal Diversity 105:1-16 ((https://doi.org/10.1007/s13225-020-00466-2)) |
| Otu44647 | 1 | k__Fungi(100);p__Ascomycota(100);c__Saccharomycetes(100);o__Saccharomycetales(100);f__Metschnikowiaceae(100);g__Metschnikowia(100);s__Metschnikowia_pulcherrima(100); | Metschnikowia | 13 | Saprotroph | \|Undefined Saprotroph\| | Yeast | NULL | Probable | As nectar-tap_saprotroph (Põlme et al. 2020); Decay sugar-rich substrates (Põlme et al. 2020) | Põlme S, et al. 2020. Fungal Diversity 105:1-16 ((https://doi.org/10.1007/s13225-020-00466-2)) |
| Otu44648 | 1 | k__Fungi(100);p__Ascomycota(100);c__Saccharomycetes(100);o__Saccharomycetales(100);f__Metschnikowiaceae(100);g__Metschnikowia(100);s__Metschnikowia_pulcherrima(100); | Metschnikowia | 13 | Saprotroph | \|Undefined Saprotroph\| | Yeast | NULL | Probable | As nectar-tap_saprotroph (Põlme et al. 2020); Decay sugar-rich substrates (Põlme et al. 2020) | Põlme S, et al. 2020. Fungal Diversity 105:1-16 ((https://doi.org/10.1007/s13225-020-00466-2)) |
| Otu44649 | 1 | k__Fungi(100);p__Ascomycota(100);c__Saccharomycetes(100);o__Saccharomycetales(100);f__Metschnikowiaceae(100);g__Metschnikowia(100);s__Metschnikowia_pulcherrima(100); | Metschnikowia | 13 | Saprotroph | \|Undefined Saprotroph\| | Yeast | NULL | Probable | As nectar-tap_saprotroph (Põlme et al. 2020); Decay sugar-rich substrates (Põlme et al. 2020) | Põlme S, et al. 2020. Fungal Diversity 105:1-16 ((https://doi.org/10.1007/s13225-020-00466-2)) |
| Otu44650 | 1 | k__Fungi(100);p__Ascomycota(100);c__Saccharomycetes(100);o__Saccharomycetales(100);f__Metschnikowiaceae(100);g__Metschnikowia(100);s__Metschnikowia_pulcherrima(100); | Metschnikowia | 13 | Saprotroph | \|Undefined Saprotroph\| | Yeast | NULL | Probable | As nectar-tap_saprotroph (Põlme et al. 2020); Decay sugar-rich substrates (Põlme et al. 2020) | Põlme S, et al. 2020. Fungal Diversity 105:1-16 ((https://doi.org/10.1007/s13225-020-00466-2)) |
| Otu44651 | 1 | k__Fungi(100);p__Ascomycota(100);c__Saccharomycetes(100);o__Saccharomycetales(100);f__Metschnikowiaceae(100);g__Metschnikowia(100);g__Metschnikowia_unclassified(100); | Metschnikowia | 13 | Saprotroph | \|Undefined Saprotroph\| | Yeast | NULL | Probable | As nectar-tap_saprotroph (Põlme et al. 2020); Decay sugar-rich substrates (Põlme et al. 2020) | Põlme S, et al. 2020. Fungal Diversity 105:1-16 ((https://doi.org/10.1007/s13225-020-00466-2)) |
| Otu44652 | 1 | k__Fungi(100);p__Ascomycota(100);c__Saccharomycetes(100);o__Saccharomycetales(100);f__Metschnikowiaceae(100);g__Metschnikowia(100);g__Metschnikowia_unclassified(100); | Metschnikowia | 13 | Saprotroph | \|Undefined Saprotroph\| | Yeast | NULL | Probable | As nectar-tap_saprotroph (Põlme et al. 2020); Decay sugar-rich substrates (Põlme et al. 2020) | Põlme S, et al. 2020. Fungal Diversity 105:1-16 ((https://doi.org/10.1007/s13225-020-00466-2)) |
| Otu44653 | 1 | k__Fungi(100);p__Ascomycota(100);c__Saccharomycetes(100);o__Saccharomycetales(100);f__Metschnikowiaceae(100);g__Metschnikowia(100);s__Metschnikowia_pulcherrima(100); | Metschnikowia | 13 | Saprotroph | \|Undefined Saprotroph\| | Yeast | NULL | Probable | As nectar-tap_saprotroph (Põlme et al. 2020); Decay sugar-rich substrates (Põlme et al. 2020) | Põlme S, et al. 2020. Fungal Diversity 105:1-16 ((https://doi.org/10.1007/s13225-020-00466-2)) |
| Otu44654 | 1 | k__Fungi(100);p__Ascomycota(100);c__Saccharomycetes(100);o__Saccharomycetales(100);f__Metschnikowiaceae(100);g__Metschnikowia(100);s__Metschnikowia_pulcherrima(100); | Metschnikowia | 13 | Saprotroph | \|Undefined Saprotroph\| | Yeast | NULL | Probable | As nectar-tap_saprotroph (Põlme et al. 2020); Decay sugar-rich substrates (Põlme et al. 2020) | Põlme S, et al. 2020. Fungal Diversity 105:1-16 ((https://doi.org/10.1007/s13225-020-00466-2)) |
| Otu44655 | 1 | k__Fungi(100);p__Ascomycota(100);c__Saccharomycetes(100);o__Saccharomycetales(100);f__Metschnikowiaceae(100);g__Metschnikowia(100);s__Metschnikowia_pulcherrima(100); | Metschnikowia | 13 | Saprotroph | \|Undefined Saprotroph\| | Yeast | NULL | Probable | As nectar-tap_saprotroph (Põlme et al. 2020); Decay sugar-rich substrates (Põlme et al. 2020) | Põlme S, et al. 2020. Fungal Diversity 105:1-16 ((https://doi.org/10.1007/s13225-020-00466-2)) |
| Otu44656 | 1 | k__Fungi(100);p__Ascomycota(100);c__Saccharomycetes(100);o__Saccharomycetales(100);f__Metschnikowiaceae(100);g__Metschnikowia(100);s__Metschnikowia_pulcherrima(100); | Metschnikowia | 13 | Saprotroph | \|Undefined Saprotroph\| | Yeast | NULL | Probable | As nectar-tap_saprotroph (Põlme et al. 2020); Decay sugar-rich substrates (Põlme et al. 2020) | Põlme S, et al. 2020. Fungal Diversity 105:1-16 ((https://doi.org/10.1007/s13225-020-00466-2)) |
| Otu44657 | 1 | k__Fungi(100);p__Ascomycota(100);c__Saccharomycetes(100);o__Saccharomycetales(100);f__Metschnikowiaceae(100);g__Metschnikowia(100);s__Metschnikowia_pulcherrima(100); | Metschnikowia | 13 | Saprotroph | \|Undefined Saprotroph\| | Yeast | NULL | Probable | As nectar-tap_saprotroph (Põlme et al. 2020); Decay sugar-rich substrates (Põlme et al. 2020) | Põlme S, et al. 2020. Fungal Diversity 105:1-16 ((https://doi.org/10.1007/s13225-020-00466-2)) |
| Otu44658 | 1 | k__Fungi(100);p__Ascomycota(100);c__Saccharomycetes(100);o__Saccharomycetales(100);f__Metschnikowiaceae(100);g__Metschnikowia(100);g__Metschnikowia_unclassified(100); | Metschnikowia | 13 | Saprotroph | \|Undefined Saprotroph\| | Yeast | NULL | Probable | As nectar-tap_saprotroph (Põlme et al. 2020); Decay sugar-rich substrates (Põlme et al. 2020) | Põlme S, et al. 2020. Fungal Diversity 105:1-16 ((https://doi.org/10.1007/s13225-020-00466-2)) |
| Otu44659 | 1 | k__Fungi(100);p__Ascomycota(100);c__Saccharomycetes(100);o__Saccharomycetales(100);f__Metschnikowiaceae(100);g__Metschnikowia(100);s__Metschnikowia_pulcherrima(100); | Metschnikowia | 13 | Saprotroph | \|Undefined Saprotroph\| | Yeast | NULL | Probable | As nectar-tap_saprotroph (Põlme et al. 2020); Decay sugar-rich substrates (Põlme et al. 2020) | Põlme S, et al. 2020. Fungal Diversity 105:1-16 ((https://doi.org/10.1007/s13225-020-00466-2)) |
| Otu44660 | 1 | k__Fungi(100);p__Ascomycota(100);c__Sordariomycetes(100);o__Hypocreales(100);f__Nectriaceae(100);f__Nectriaceae_unclassified(100);f__Nectriaceae_unclassified(100); | Nectriaceae | 9 | Pathotroph-Saprotroph-Symbiotroph | Animal Pathogen-Endophyte-Fungal Parasite-Lichen Parasite-Plant Pathogen-Wood Saprotroph | Microfungus | NULL | Possible | Fusarium species are amongst the economically significant plant pathogens (Cannon & Kirk 2007) | Cannon PF, Kirk PM. 2007. Fungal Families of the World. CAB International, Cambridge (ISBN: 978-0851998275); Hirooka Y, et al. 2012. Studies in Mycology 71:1-210 ((https://doi.org/10.3114/sim0001)); Tedersoo L, et al. 2014. Science 346:e1256688 ((https://doi.org/10.1126/science.1256688)); The American Phytopathological Society Common Names of Plant Diseases ((http://www.apsnet.org/publications/commonnames/Pages/default.aspx)); https://www.cdc.gov ((https://www.cdc.gov)) |
| Otu44661 | 1 | k__Fungi(100);p__Ascomycota(100);c__Saccharomycetes(100);o__Saccharomycetales(100);f__Metschnikowiaceae(100);g__Metschnikowia(100);s__Metschnikowia_pulcherrima(100); | Metschnikowia | 13 | Saprotroph | \|Undefined Saprotroph\| | Yeast | NULL | Probable | As nectar-tap_saprotroph (Põlme et al. 2020); Decay sugar-rich substrates (Põlme et al. 2020) | Põlme S, et al. 2020. Fungal Diversity 105:1-16 ((https://doi.org/10.1007/s13225-020-00466-2)) |
| Otu44662 | 1 | k__Fungi(100);p__Ascomycota(100);c__Saccharomycetes(100);o__Saccharomycetales(100);f__Metschnikowiaceae(100);g__Metschnikowia(100);s__Metschnikowia_pulcherrima(100); | Metschnikowia | 13 | Saprotroph | \|Undefined Saprotroph\| | Yeast | NULL | Probable | As nectar-tap_saprotroph (Põlme et al. 2020); Decay sugar-rich substrates (Põlme et al. 2020) | Põlme S, et al. 2020. Fungal Diversity 105:1-16 ((https://doi.org/10.1007/s13225-020-00466-2)) |
| Otu44663 | 1 | k__Fungi(100);p__Ascomycota(100);c__Saccharomycetes(100);o__Saccharomycetales(100);f__Metschnikowiaceae(100);g__Metschnikowia(100);s__Metschnikowia_pulcherrima(100); | Metschnikowia | 13 | Saprotroph | \|Undefined Saprotroph\| | Yeast | NULL | Probable | As nectar-tap_saprotroph (Põlme et al. 2020); Decay sugar-rich substrates (Põlme et al. 2020) | Põlme S, et al. 2020. Fungal Diversity 105:1-16 ((https://doi.org/10.1007/s13225-020-00466-2)) |
| Otu44664 | 1 | k__Fungi(100);p__Ascomycota(100);c__Saccharomycetes(100);o__Saccharomycetales(100);f__Metschnikowiaceae(100);g__Metschnikowia(100);s__Metschnikowia_pulcherrima(100); | Metschnikowia | 13 | Saprotroph | \|Undefined Saprotroph\| | Yeast | NULL | Probable | As nectar-tap_saprotroph (Põlme et al. 2020); Decay sugar-rich substrates (Põlme et al. 2020) | Põlme S, et al. 2020. Fungal Diversity 105:1-16 ((https://doi.org/10.1007/s13225-020-00466-2)) |
| Otu44665 | 1 | k__Fungi(100);p__Ascomycota(100);c__Saccharomycetes(100);o__Saccharomycetales(100);f__Metschnikowiaceae(100);g__Metschnikowia(100);g__Metschnikowia_unclassified(100); | Metschnikowia | 13 | Saprotroph | \|Undefined Saprotroph\| | Yeast | NULL | Probable | As nectar-tap_saprotroph (Põlme et al. 2020); Decay sugar-rich substrates (Põlme et al. 2020) | Põlme S, et al. 2020. Fungal Diversity 105:1-16 ((https://doi.org/10.1007/s13225-020-00466-2)) |
| Otu44666 | 1 | k__Fungi(100);p__Ascomycota(100);c__Saccharomycetes(100);o__Saccharomycetales(100);f__Metschnikowiaceae(100);g__Metschnikowia(100);g__Metschnikowia_unclassified(100); | Metschnikowia | 13 | Saprotroph | \|Undefined Saprotroph\| | Yeast | NULL | Probable | As nectar-tap_saprotroph (Põlme et al. 2020); Decay sugar-rich substrates (Põlme et al. 2020) | Põlme S, et al. 2020. Fungal Diversity 105:1-16 ((https://doi.org/10.1007/s13225-020-00466-2)) |
| Otu44667 | 1 | k__Fungi(100);p__Ascomycota(100);c__Saccharomycetes(100);o__Saccharomycetales(100);f__Metschnikowiaceae(100);g__Metschnikowia(100);s__Metschnikowia_pulcherrima(100); | Metschnikowia | 13 | Saprotroph | \|Undefined Saprotroph\| | Yeast | NULL | Probable | As nectar-tap_saprotroph (Põlme et al. 2020); Decay sugar-rich substrates (Põlme et al. 2020) | Põlme S, et al. 2020. Fungal Diversity 105:1-16 ((https://doi.org/10.1007/s13225-020-00466-2)) |
| Otu44668 | 1 | k__Fungi(100);p__Ascomycota(100);c__Saccharomycetes(100);o__Saccharomycetales(100);f__Metschnikowiaceae(100);g__Metschnikowia(100);s__Metschnikowia_pulcherrima(100); | Metschnikowia | 13 | Saprotroph | \|Undefined Saprotroph\| | Yeast | NULL | Probable | As nectar-tap_saprotroph (Põlme et al. 2020); Decay sugar-rich substrates (Põlme et al. 2020) | Põlme S, et al. 2020. Fungal Diversity 105:1-16 ((https://doi.org/10.1007/s13225-020-00466-2)) |
| Otu44669 | 1 | k__Fungi(100);p__Ascomycota(100);c__Saccharomycetes(100);o__Saccharomycetales(100);f__Metschnikowiaceae(100);g__Metschnikowia(100);g__Metschnikowia_unclassified(100); | Metschnikowia | 13 | Saprotroph | \|Undefined Saprotroph\| | Yeast | NULL | Probable | As nectar-tap_saprotroph (Põlme et al. 2020); Decay sugar-rich substrates (Põlme et al. 2020) | Põlme S, et al. 2020. Fungal Diversity 105:1-16 ((https://doi.org/10.1007/s13225-020-00466-2)) |
| Otu44670 | 1 | k__Fungi(100);p__Ascomycota(100);c__Saccharomycetes(100);o__Saccharomycetales(100);f__Metschnikowiaceae(100);g__Metschnikowia(100);s__Metschnikowia_pulcherrima(100); | Metschnikowia | 13 | Saprotroph | \|Undefined Saprotroph\| | Yeast | NULL | Probable | As nectar-tap_saprotroph (Põlme et al. 2020); Decay sugar-rich substrates (Põlme et al. 2020) | Põlme S, et al. 2020. Fungal Diversity 105:1-16 ((https://doi.org/10.1007/s13225-020-00466-2)) |
| Otu44671 | 1 | k__Fungi(100);p__Ascomycota(100);c__Saccharomycetes(100);o__Saccharomycetales(100);f__Metschnikowiaceae(100);g__Metschnikowia(100);s__Metschnikowia_pulcherrima(100); | Metschnikowia | 13 | Saprotroph | \|Undefined Saprotroph\| | Yeast | NULL | Probable | As nectar-tap_saprotroph (Põlme et al. 2020); Decay sugar-rich substrates (Põlme et al. 2020) | Põlme S, et al. 2020. Fungal Diversity 105:1-16 ((https://doi.org/10.1007/s13225-020-00466-2)) |
| Otu44672 | 1 | k__Fungi(100);p__Ascomycota(100);c__Saccharomycetes(100);o__Saccharomycetales(100);f__Metschnikowiaceae(100);g__Metschnikowia(100);s__Metschnikowia_pulcherrima(100); | Metschnikowia | 13 | Saprotroph | \|Undefined Saprotroph\| | Yeast | NULL | Probable | As nectar-tap_saprotroph (Põlme et al. 2020); Decay sugar-rich substrates (Põlme et al. 2020) | Põlme S, et al. 2020. Fungal Diversity 105:1-16 ((https://doi.org/10.1007/s13225-020-00466-2)) |
| Otu44673 | 1 | k__Fungi(100);p__Ascomycota(100);c__Saccharomycetes(100);o__Saccharomycetales(100);f__Metschnikowiaceae(100);g__Metschnikowia(100);s__Metschnikowia_pulcherrima(100); | Metschnikowia | 13 | Saprotroph | \|Undefined Saprotroph\| | Yeast | NULL | Probable | As nectar-tap_saprotroph (Põlme et al. 2020); Decay sugar-rich substrates (Põlme et al. 2020) | Põlme S, et al. 2020. Fungal Diversity 105:1-16 ((https://doi.org/10.1007/s13225-020-00466-2)) |
| Otu44674 | 1 | k__Fungi(100);p__Ascomycota(100);c__Saccharomycetes(100);o__Saccharomycetales(100);f__Metschnikowiaceae(100);g__Metschnikowia(100);g__Metschnikowia_unclassified(100); | Metschnikowia | 13 | Saprotroph | \|Undefined Saprotroph\| | Yeast | NULL | Probable | As nectar-tap_saprotroph (Põlme et al. 2020); Decay sugar-rich substrates (Põlme et al. 2020) | Põlme S, et al. 2020. Fungal Diversity 105:1-16 ((https://doi.org/10.1007/s13225-020-00466-2)) |
| Otu44676 | 1 | k__Fungi(100);p__Ascomycota(100);c__Saccharomycetes(100);o__Saccharomycetales(100);f__Metschnikowiaceae(100);g__Metschnikowia(100);s__Metschnikowia_pulcherrima(100); | Metschnikowia | 13 | Saprotroph | \|Undefined Saprotroph\| | Yeast | NULL | Probable | As nectar-tap_saprotroph (Põlme et al. 2020); Decay sugar-rich substrates (Põlme et al. 2020) | Põlme S, et al. 2020. Fungal Diversity 105:1-16 ((https://doi.org/10.1007/s13225-020-00466-2)) |
| Otu44677 | 1 | k__Fungi(100);p__Ascomycota(100);c__Saccharomycetes(100);o__Saccharomycetales(100);f__Metschnikowiaceae(100);g__Metschnikowia(100);s__Metschnikowia_pulcherrima(100); | Metschnikowia | 13 | Saprotroph | \|Undefined Saprotroph\| | Yeast | NULL | Probable | As nectar-tap_saprotroph (Põlme et al. 2020); Decay sugar-rich substrates (Põlme et al. 2020) | Põlme S, et al. 2020. Fungal Diversity 105:1-16 ((https://doi.org/10.1007/s13225-020-00466-2)) |
| Otu44678 | 1 | k__Fungi(100);p__Ascomycota(100);c__Saccharomycetes(100);o__Saccharomycetales(100);f__Metschnikowiaceae(100);g__Metschnikowia(100);s__Metschnikowia_pulcherrima(100); | Metschnikowia | 13 | Saprotroph | \|Undefined Saprotroph\| | Yeast | NULL | Probable | As nectar-tap_saprotroph (Põlme et al. 2020); Decay sugar-rich substrates (Põlme et al. 2020) | Põlme S, et al. 2020. Fungal Diversity 105:1-16 ((https://doi.org/10.1007/s13225-020-00466-2)) |
| Otu44679 | 1 | k__Fungi(100);p__Ascomycota(100);c__Saccharomycetes(100);o__Saccharomycetales(100);f__Metschnikowiaceae(100);g__Metschnikowia(100);s__Metschnikowia_pulcherrima(100); | Metschnikowia | 13 | Saprotroph | \|Undefined Saprotroph\| | Yeast | NULL | Probable | As nectar-tap_saprotroph (Põlme et al. 2020); Decay sugar-rich substrates (Põlme et al. 2020) | Põlme S, et al. 2020. Fungal Diversity 105:1-16 ((https://doi.org/10.1007/s13225-020-00466-2)) |
| Otu44680 | 1 | k__Fungi(100);p__Ascomycota(100);c__Saccharomycetes(100);o__Saccharomycetales(100);f__Metschnikowiaceae(100);g__Metschnikowia(100);s__Metschnikowia_pulcherrima(100); | Metschnikowia | 13 | Saprotroph | \|Undefined Saprotroph\| | Yeast | NULL | Probable | As nectar-tap_saprotroph (Põlme et al. 2020); Decay sugar-rich substrates (Põlme et al. 2020) | Põlme S, et al. 2020. Fungal Diversity 105:1-16 ((https://doi.org/10.1007/s13225-020-00466-2)) |
| Otu44681 | 1 | k__Fungi(100);p__Ascomycota(100);c__Saccharomycetes(100);o__Saccharomycetales(100);f__Metschnikowiaceae(100);g__Metschnikowia(100);s__Metschnikowia_pulcherrima(100); | Metschnikowia | 13 | Saprotroph | \|Undefined Saprotroph\| | Yeast | NULL | Probable | As nectar-tap_saprotroph (Põlme et al. 2020); Decay sugar-rich substrates (Põlme et al. 2020) | Põlme S, et al. 2020. Fungal Diversity 105:1-16 ((https://doi.org/10.1007/s13225-020-00466-2)) |
| Otu44682 | 1 | k__Fungi(100);p__Ascomycota(100);c__Saccharomycetes(100);o__Saccharomycetales(100);f__Metschnikowiaceae(100);g__Metschnikowia(100);g__Metschnikowia_unclassified(100); | Metschnikowia | 13 | Saprotroph | \|Undefined Saprotroph\| | Yeast | NULL | Probable | As nectar-tap_saprotroph (Põlme et al. 2020); Decay sugar-rich substrates (Põlme et al. 2020) | Põlme S, et al. 2020. Fungal Diversity 105:1-16 ((https://doi.org/10.1007/s13225-020-00466-2)) |
| Otu44683 | 1 | k__Fungi(100);p__Ascomycota(100);c__Saccharomycetes(100);o__Saccharomycetales(100);f__Metschnikowiaceae(100);g__Metschnikowia(100);g__Metschnikowia_unclassified(100); | Metschnikowia | 13 | Saprotroph | \|Undefined Saprotroph\| | Yeast | NULL | Probable | As nectar-tap_saprotroph (Põlme et al. 2020); Decay sugar-rich substrates (Põlme et al. 2020) | Põlme S, et al. 2020. Fungal Diversity 105:1-16 ((https://doi.org/10.1007/s13225-020-00466-2)) |
| Otu44684 | 1 | k__Fungi(100);p__Ascomycota(100);c__Saccharomycetes(100);o__Saccharomycetales(100);f__Metschnikowiaceae(100);g__Metschnikowia(100);s__Metschnikowia_pulcherrima(100); | Metschnikowia | 13 | Saprotroph | \|Undefined Saprotroph\| | Yeast | NULL | Probable | As nectar-tap_saprotroph (Põlme et al. 2020); Decay sugar-rich substrates (Põlme et al. 2020) | Põlme S, et al. 2020. Fungal Diversity 105:1-16 ((https://doi.org/10.1007/s13225-020-00466-2)) |
| Otu44685 | 1 | k__Fungi(100);p__Ascomycota(100);c__Saccharomycetes(100);o__Saccharomycetales(100);f__Metschnikowiaceae(100);g__Metschnikowia(100);s__Metschnikowia_pulcherrima(100); | Metschnikowia | 13 | Saprotroph | \|Undefined Saprotroph\| | Yeast | NULL | Probable | As nectar-tap_saprotroph (Põlme et al. 2020); Decay sugar-rich substrates (Põlme et al. 2020) | Põlme S, et al. 2020. Fungal Diversity 105:1-16 ((https://doi.org/10.1007/s13225-020-00466-2)) |
| Otu44686 | 1 | k__Fungi(100);p__Ascomycota(100);c__Saccharomycetes(100);o__Saccharomycetales(100);f__Metschnikowiaceae(100);g__Metschnikowia(100);g__Metschnikowia_unclassified(100); | Metschnikowia | 13 | Saprotroph | \|Undefined Saprotroph\| | Yeast | NULL | Probable | As nectar-tap_saprotroph (Põlme et al. 2020); Decay sugar-rich substrates (Põlme et al. 2020) | Põlme S, et al. 2020. Fungal Diversity 105:1-16 ((https://doi.org/10.1007/s13225-020-00466-2)) |
| Otu44687 | 1 | k__Fungi(100);p__Ascomycota(100);c__Saccharomycetes(100);o__Saccharomycetales(100);f__Metschnikowiaceae(100);g__Metschnikowia(100);s__Metschnikowia_pulcherrima(100); | Metschnikowia | 13 | Saprotroph | \|Undefined Saprotroph\| | Yeast | NULL | Probable | As nectar-tap_saprotroph (Põlme et al. 2020); Decay sugar-rich substrates (Põlme et al. 2020) | Põlme S, et al. 2020. Fungal Diversity 105:1-16 ((https://doi.org/10.1007/s13225-020-00466-2)) |
| Otu44688 | 1 | k__Fungi(100);p__Ascomycota(100);c__Saccharomycetes(100);o__Saccharomycetales(100);f__Metschnikowiaceae(100);g__Metschnikowia(100);s__Metschnikowia_pulcherrima(100); | Metschnikowia | 13 | Saprotroph | \|Undefined Saprotroph\| | Yeast | NULL | Probable | As nectar-tap_saprotroph (Põlme et al. 2020); Decay sugar-rich substrates (Põlme et al. 2020) | Põlme S, et al. 2020. Fungal Diversity 105:1-16 ((https://doi.org/10.1007/s13225-020-00466-2)) |
| Otu44689 | 1 | k__Fungi(100);p__Ascomycota(100);c__Saccharomycetes(100);o__Saccharomycetales(100);f__Metschnikowiaceae(100);g__Metschnikowia(100);s__Metschnikowia_pulcherrima(100); | Metschnikowia | 13 | Saprotroph | \|Undefined Saprotroph\| | Yeast | NULL | Probable | As nectar-tap_saprotroph (Põlme et al. 2020); Decay sugar-rich substrates (Põlme et al. 2020) | Põlme S, et al. 2020. Fungal Diversity 105:1-16 ((https://doi.org/10.1007/s13225-020-00466-2)) |
| Otu44690 | 1 | k__Fungi(100);p__Ascomycota(100);c__Saccharomycetes(100);o__Saccharomycetales(100);f__Metschnikowiaceae(100);g__Metschnikowia(100);s__Metschnikowia_pulcherrima(100); | Metschnikowia | 13 | Saprotroph | \|Undefined Saprotroph\| | Yeast | NULL | Probable | As nectar-tap_saprotroph (Põlme et al. 2020); Decay sugar-rich substrates (Põlme et al. 2020) | Põlme S, et al. 2020. Fungal Diversity 105:1-16 ((https://doi.org/10.1007/s13225-020-00466-2)) |
| Otu44691 | 1 | k__Fungi(100);p__Ascomycota(100);c__Saccharomycetes(100);o__Saccharomycetales(100);f__Metschnikowiaceae(100);g__Metschnikowia(100);s__Metschnikowia_pulcherrima(100); | Metschnikowia | 13 | Saprotroph | \|Undefined Saprotroph\| | Yeast | NULL | Probable | As nectar-tap_saprotroph (Põlme et al. 2020); Decay sugar-rich substrates (Põlme et al. 2020) | Põlme S, et al. 2020. Fungal Diversity 105:1-16 ((https://doi.org/10.1007/s13225-020-00466-2)) |
| Otu44692 | 1 | k__Fungi(100);p__Ascomycota(100);c__Saccharomycetes(100);o__Saccharomycetales(100);f__Metschnikowiaceae(100);g__Metschnikowia(100);g__Metschnikowia_unclassified(100); | Metschnikowia | 13 | Saprotroph | \|Undefined Saprotroph\| | Yeast | NULL | Probable | As nectar-tap_saprotroph (Põlme et al. 2020); Decay sugar-rich substrates (Põlme et al. 2020) | Põlme S, et al. 2020. Fungal Diversity 105:1-16 ((https://doi.org/10.1007/s13225-020-00466-2)) |
| Otu44693 | 1 | k__Fungi(100);p__Ascomycota(100);c__Saccharomycetes(100);o__Saccharomycetales(100);f__Metschnikowiaceae(100);g__Metschnikowia(100);s__Metschnikowia_pulcherrima(100); | Metschnikowia | 13 | Saprotroph | \|Undefined Saprotroph\| | Yeast | NULL | Probable | As nectar-tap_saprotroph (Põlme et al. 2020); Decay sugar-rich substrates (Põlme et al. 2020) | Põlme S, et al. 2020. Fungal Diversity 105:1-16 ((https://doi.org/10.1007/s13225-020-00466-2)) |
| Otu44694 | 1 | k__Fungi(100);p__Ascomycota(100);c__Saccharomycetes(100);o__Saccharomycetales(100);f__Metschnikowiaceae(100);g__Metschnikowia(100);s__Metschnikowia_pulcherrima(100); | Metschnikowia | 13 | Saprotroph | \|Undefined Saprotroph\| | Yeast | NULL | Probable | As nectar-tap_saprotroph (Põlme et al. 2020); Decay sugar-rich substrates (Põlme et al. 2020) | Põlme S, et al. 2020. Fungal Diversity 105:1-16 ((https://doi.org/10.1007/s13225-020-00466-2)) |
| Otu44695 | 1 | k__Fungi(100);p__Ascomycota(100);c__Saccharomycetes(100);o__Saccharomycetales(100);f__Metschnikowiaceae(100);g__Metschnikowia(100);s__Metschnikowia_pulcherrima(100); | Metschnikowia | 13 | Saprotroph | \|Undefined Saprotroph\| | Yeast | NULL | Probable | As nectar-tap_saprotroph (Põlme et al. 2020); Decay sugar-rich substrates (Põlme et al. 2020) | Põlme S, et al. 2020. Fungal Diversity 105:1-16 ((https://doi.org/10.1007/s13225-020-00466-2)) |
| Otu44696 | 1 | k__Fungi(100);p__Ascomycota(100);c__Saccharomycetes(100);o__Saccharomycetales(100);f__Metschnikowiaceae(100);g__Metschnikowia(100);s__Metschnikowia_pulcherrima(100); | Metschnikowia | 13 | Saprotroph | \|Undefined Saprotroph\| | Yeast | NULL | Probable | As nectar-tap_saprotroph (Põlme et al. 2020); Decay sugar-rich substrates (Põlme et al. 2020) | Põlme S, et al. 2020. Fungal Diversity 105:1-16 ((https://doi.org/10.1007/s13225-020-00466-2)) |
| Otu44697 | 1 | k__Fungi(100);p__Ascomycota(100);c__Saccharomycetes(100);o__Saccharomycetales(100);f__Metschnikowiaceae(100);g__Metschnikowia(100);g__Metschnikowia_unclassified(100); | Metschnikowia | 13 | Saprotroph | \|Undefined Saprotroph\| | Yeast | NULL | Probable | As nectar-tap_saprotroph (Põlme et al. 2020); Decay sugar-rich substrates (Põlme et al. 2020) | Põlme S, et al. 2020. Fungal Diversity 105:1-16 ((https://doi.org/10.1007/s13225-020-00466-2)) |
| Otu44698 | 1 | k__Fungi(100);p__Ascomycota(100);c__Saccharomycetes(100);o__Saccharomycetales(100);f__Metschnikowiaceae(100);g__Metschnikowia(100);s__Metschnikowia_pulcherrima(100); | Metschnikowia | 13 | Saprotroph | \|Undefined Saprotroph\| | Yeast | NULL | Probable | As nectar-tap_saprotroph (Põlme et al. 2020); Decay sugar-rich substrates (Põlme et al. 2020) | Põlme S, et al. 2020. Fungal Diversity 105:1-16 ((https://doi.org/10.1007/s13225-020-00466-2)) |
| Otu44699 | 1 | k__Fungi(100);p__Ascomycota(100);c__Saccharomycetes(100);o__Saccharomycetales(100);f__Metschnikowiaceae(100);g__Metschnikowia(100);s__Metschnikowia_pulcherrima(100); | Metschnikowia | 13 | Saprotroph | \|Undefined Saprotroph\| | Yeast | NULL | Probable | As nectar-tap_saprotroph (Põlme et al. 2020); Decay sugar-rich substrates (Põlme et al. 2020) | Põlme S, et al. 2020. Fungal Diversity 105:1-16 ((https://doi.org/10.1007/s13225-020-00466-2)) |
| Otu44700 | 1 | k__Fungi(100);p__Ascomycota(100);c__Saccharomycetes(100);o__Saccharomycetales(100);f__Metschnikowiaceae(100);g__Metschnikowia(100);s__Metschnikowia_pulcherrima(100); | Metschnikowia | 13 | Saprotroph | \|Undefined Saprotroph\| | Yeast | NULL | Probable | As nectar-tap_saprotroph (Põlme et al. 2020); Decay sugar-rich substrates (Põlme et al. 2020) | Põlme S, et al. 2020. Fungal Diversity 105:1-16 ((https://doi.org/10.1007/s13225-020-00466-2)) |
| Otu44701 | 1 | k__Fungi(100);p__Ascomycota(100);c__Saccharomycetes(100);o__Saccharomycetales(100);f__Metschnikowiaceae(100);g__Metschnikowia(100);s__Metschnikowia_pulcherrima(100); | Metschnikowia | 13 | Saprotroph | \|Undefined Saprotroph\| | Yeast | NULL | Probable | As nectar-tap_saprotroph (Põlme et al. 2020); Decay sugar-rich substrates (Põlme et al. 2020) | Põlme S, et al. 2020. Fungal Diversity 105:1-16 ((https://doi.org/10.1007/s13225-020-00466-2)) |
| Otu44702 | 1 | k__Fungi(100);p__Ascomycota(100);c__Saccharomycetes(100);o__Saccharomycetales(100);f__Metschnikowiaceae(100);g__Metschnikowia(100);s__Metschnikowia_pulcherrima(100); | Metschnikowia | 13 | Saprotroph | \|Undefined Saprotroph\| | Yeast | NULL | Probable | As nectar-tap_saprotroph (Põlme et al. 2020); Decay sugar-rich substrates (Põlme et al. 2020) | Põlme S, et al. 2020. Fungal Diversity 105:1-16 ((https://doi.org/10.1007/s13225-020-00466-2)) |
| Otu44703 | 1 | k__Fungi(100);p__Ascomycota(100);c__Saccharomycetes(100);o__Saccharomycetales(100);f__Metschnikowiaceae(100);g__Metschnikowia(100);s__Metschnikowia_pulcherrima(100); | Metschnikowia | 13 | Saprotroph | \|Undefined Saprotroph\| | Yeast | NULL | Probable | As nectar-tap_saprotroph (Põlme et al. 2020); Decay sugar-rich substrates (Põlme et al. 2020) | Põlme S, et al. 2020. Fungal Diversity 105:1-16 ((https://doi.org/10.1007/s13225-020-00466-2)) |
| Otu44704 | 1 | k__Fungi(100);p__Ascomycota(100);c__Saccharomycetes(100);o__Saccharomycetales(100);f__Metschnikowiaceae(100);g__Metschnikowia(100);s__Metschnikowia_pulcherrima(100); | Metschnikowia | 13 | Saprotroph | \|Undefined Saprotroph\| | Yeast | NULL | Probable | As nectar-tap_saprotroph (Põlme et al. 2020); Decay sugar-rich substrates (Põlme et al. 2020) | Põlme S, et al. 2020. Fungal Diversity 105:1-16 ((https://doi.org/10.1007/s13225-020-00466-2)) |
| Otu44706 | 1 | k__Fungi(100);p__Ascomycota(100);c__Saccharomycetes(100);o__Saccharomycetales(100);f__Metschnikowiaceae(100);g__Metschnikowia(100);s__Metschnikowia_pulcherrima(100); | Metschnikowia | 13 | Saprotroph | \|Undefined Saprotroph\| | Yeast | NULL | Probable | As nectar-tap_saprotroph (Põlme et al. 2020); Decay sugar-rich substrates (Põlme et al. 2020) | Põlme S, et al. 2020. Fungal Diversity 105:1-16 ((https://doi.org/10.1007/s13225-020-00466-2)) |
| Otu44707 | 1 | k__Fungi(100);p__Ascomycota(100);c__Saccharomycetes(100);o__Saccharomycetales(100);f__Metschnikowiaceae(100);g__Metschnikowia(100);s__Metschnikowia_pulcherrima(100); | Metschnikowia | 13 | Saprotroph | \|Undefined Saprotroph\| | Yeast | NULL | Probable | As nectar-tap_saprotroph (Põlme et al. 2020); Decay sugar-rich substrates (Põlme et al. 2020) | Põlme S, et al. 2020. Fungal Diversity 105:1-16 ((https://doi.org/10.1007/s13225-020-00466-2)) |
| Otu44708 | 1 | k__Fungi(100);p__Ascomycota(100);c__Saccharomycetes(100);o__Saccharomycetales(100);f__Metschnikowiaceae(100);g__Metschnikowia(100);s__Metschnikowia_pulcherrima(100); | Metschnikowia | 13 | Saprotroph | \|Undefined Saprotroph\| | Yeast | NULL | Probable | As nectar-tap_saprotroph (Põlme et al. 2020); Decay sugar-rich substrates (Põlme et al. 2020) | Põlme S, et al. 2020. Fungal Diversity 105:1-16 ((https://doi.org/10.1007/s13225-020-00466-2)) |
| Otu44709 | 1 | k__Fungi(100);p__Ascomycota(100);c__Saccharomycetes(100);o__Saccharomycetales(100);f__Metschnikowiaceae(100);g__Metschnikowia(100);s__Metschnikowia_pulcherrima(100); | Metschnikowia | 13 | Saprotroph | \|Undefined Saprotroph\| | Yeast | NULL | Probable | As nectar-tap_saprotroph (Põlme et al. 2020); Decay sugar-rich substrates (Põlme et al. 2020) | Põlme S, et al. 2020. Fungal Diversity 105:1-16 ((https://doi.org/10.1007/s13225-020-00466-2)) |
| Otu44710 | 1 | k__Fungi(100);p__Ascomycota(100);c__Saccharomycetes(100);o__Saccharomycetales(100);f__Metschnikowiaceae(100);g__Metschnikowia(100);s__Metschnikowia_pulcherrima(100); | Metschnikowia | 13 | Saprotroph | \|Undefined Saprotroph\| | Yeast | NULL | Probable | As nectar-tap_saprotroph (Põlme et al. 2020); Decay sugar-rich substrates (Põlme et al. 2020) | Põlme S, et al. 2020. Fungal Diversity 105:1-16 ((https://doi.org/10.1007/s13225-020-00466-2)) |
| Otu44711 | 1 | k__Fungi(100);p__Ascomycota(100);c__Saccharomycetes(100);o__Saccharomycetales(100);f__Metschnikowiaceae(100);g__Metschnikowia(100);s__Metschnikowia_pulcherrima(100); | Metschnikowia | 13 | Saprotroph | \|Undefined Saprotroph\| | Yeast | NULL | Probable | As nectar-tap_saprotroph (Põlme et al. 2020); Decay sugar-rich substrates (Põlme et al. 2020) | Põlme S, et al. 2020. Fungal Diversity 105:1-16 ((https://doi.org/10.1007/s13225-020-00466-2)) |
| Otu44712 | 1 | k__Fungi(100);p__Ascomycota(100);c__Saccharomycetes(100);o__Saccharomycetales(100);f__Metschnikowiaceae(100);g__Metschnikowia(100);s__Metschnikowia_pulcherrima(100); | Metschnikowia | 13 | Saprotroph | \|Undefined Saprotroph\| | Yeast | NULL | Probable | As nectar-tap_saprotroph (Põlme et al. 2020); Decay sugar-rich substrates (Põlme et al. 2020) | Põlme S, et al. 2020. Fungal Diversity 105:1-16 ((https://doi.org/10.1007/s13225-020-00466-2)) |
| Otu44713 | 1 | k__Fungi(100);p__Ascomycota(100);c__Saccharomycetes(100);o__Saccharomycetales(100);f__Metschnikowiaceae(100);g__Metschnikowia(100);s__Metschnikowia_pulcherrima(100); | Metschnikowia | 13 | Saprotroph | \|Undefined Saprotroph\| | Yeast | NULL | Probable | As nectar-tap_saprotroph (Põlme et al. 2020); Decay sugar-rich substrates (Põlme et al. 2020) | Põlme S, et al. 2020. Fungal Diversity 105:1-16 ((https://doi.org/10.1007/s13225-020-00466-2)) |
| Otu44714 | 1 | k__Fungi(100);p__Ascomycota(100);c__Saccharomycetes(100);o__Saccharomycetales(100);f__Metschnikowiaceae(100);g__Metschnikowia(100);s__Metschnikowia_pulcherrima(100); | Metschnikowia | 13 | Saprotroph | \|Undefined Saprotroph\| | Yeast | NULL | Probable | As nectar-tap_saprotroph (Põlme et al. 2020); Decay sugar-rich substrates (Põlme et al. 2020) | Põlme S, et al. 2020. Fungal Diversity 105:1-16 ((https://doi.org/10.1007/s13225-020-00466-2)) |
| Otu44715 | 1 | k__Fungi(100);p__Ascomycota(100);c__Saccharomycetes(100);o__Saccharomycetales(100);f__Metschnikowiaceae(100);g__Metschnikowia(100);s__Metschnikowia_pulcherrima(100); | Metschnikowia | 13 | Saprotroph | \|Undefined Saprotroph\| | Yeast | NULL | Probable | As nectar-tap_saprotroph (Põlme et al. 2020); Decay sugar-rich substrates (Põlme et al. 2020) | Põlme S, et al. 2020. Fungal Diversity 105:1-16 ((https://doi.org/10.1007/s13225-020-00466-2)) |
| Otu44716 | 1 | k__Fungi(100);p__Ascomycota(100);c__Saccharomycetes(100);o__Saccharomycetales(100);f__Metschnikowiaceae(100);g__Metschnikowia(100);s__Metschnikowia_pulcherrima(100); | Metschnikowia | 13 | Saprotroph | \|Undefined Saprotroph\| | Yeast | NULL | Probable | As nectar-tap_saprotroph (Põlme et al. 2020); Decay sugar-rich substrates (Põlme et al. 2020) | Põlme S, et al. 2020. Fungal Diversity 105:1-16 ((https://doi.org/10.1007/s13225-020-00466-2)) |
| Otu44717 | 1 | k__Fungi(100);p__Ascomycota(100);c__Saccharomycetes(100);o__Saccharomycetales(100);f__Metschnikowiaceae(100);g__Metschnikowia(100);g__Metschnikowia_unclassified(100); | Metschnikowia | 13 | Saprotroph | \|Undefined Saprotroph\| | Yeast | NULL | Probable | As nectar-tap_saprotroph (Põlme et al. 2020); Decay sugar-rich substrates (Põlme et al. 2020) | Põlme S, et al. 2020. Fungal Diversity 105:1-16 ((https://doi.org/10.1007/s13225-020-00466-2)) |
| Otu44718 | 1 | k__Fungi(100);p__Ascomycota(100);c__Saccharomycetes(100);o__Saccharomycetales(100);f__Metschnikowiaceae(100);g__Metschnikowia(100);s__Metschnikowia_pulcherrima(100); | Metschnikowia | 13 | Saprotroph | \|Undefined Saprotroph\| | Yeast | NULL | Probable | As nectar-tap_saprotroph (Põlme et al. 2020); Decay sugar-rich substrates (Põlme et al. 2020) | Põlme S, et al. 2020. Fungal Diversity 105:1-16 ((https://doi.org/10.1007/s13225-020-00466-2)) |
| Otu44719 | 1 | k__Fungi(100);p__Ascomycota(100);c__Saccharomycetes(100);o__Saccharomycetales(100);f__Metschnikowiaceae(100);g__Metschnikowia(100);s__Metschnikowia_pulcherrima(100); | Metschnikowia | 13 | Saprotroph | \|Undefined Saprotroph\| | Yeast | NULL | Probable | As nectar-tap_saprotroph (Põlme et al. 2020); Decay sugar-rich substrates (Põlme et al. 2020) | Põlme S, et al. 2020. Fungal Diversity 105:1-16 ((https://doi.org/10.1007/s13225-020-00466-2)) |
| Otu44720 | 1 | k__Fungi(100);p__Ascomycota(100);c__Saccharomycetes(100);o__Saccharomycetales(100);f__Metschnikowiaceae(100);g__Metschnikowia(100);s__Metschnikowia_pulcherrima(100); | Metschnikowia | 13 | Saprotroph | \|Undefined Saprotroph\| | Yeast | NULL | Probable | As nectar-tap_saprotroph (Põlme et al. 2020); Decay sugar-rich substrates (Põlme et al. 2020) | Põlme S, et al. 2020. Fungal Diversity 105:1-16 ((https://doi.org/10.1007/s13225-020-00466-2)) |
| Otu44721 | 1 | k__Fungi(100);p__Ascomycota(100);c__Saccharomycetes(100);o__Saccharomycetales(100);f__Metschnikowiaceae(100);g__Metschnikowia(100);g__Metschnikowia_unclassified(100); | Metschnikowia | 13 | Saprotroph | \|Undefined Saprotroph\| | Yeast | NULL | Probable | As nectar-tap_saprotroph (Põlme et al. 2020); Decay sugar-rich substrates (Põlme et al. 2020) | Põlme S, et al. 2020. Fungal Diversity 105:1-16 ((https://doi.org/10.1007/s13225-020-00466-2)) |
| Otu44722 | 1 | k__Fungi(100);p__Ascomycota(100);c__Saccharomycetes(100);o__Saccharomycetales(100);f__Metschnikowiaceae(100);g__Metschnikowia(100);g__Metschnikowia_unclassified(100); | Metschnikowia | 13 | Saprotroph | \|Undefined Saprotroph\| | Yeast | NULL | Probable | As nectar-tap_saprotroph (Põlme et al. 2020); Decay sugar-rich substrates (Põlme et al. 2020) | Põlme S, et al. 2020. Fungal Diversity 105:1-16 ((https://doi.org/10.1007/s13225-020-00466-2)) |
| Otu44724 | 1 | k__Fungi(100);p__Ascomycota(100);c__Saccharomycetes(100);o__Saccharomycetales(100);f__Metschnikowiaceae(100);g__Metschnikowia(100);s__Metschnikowia_pulcherrima(100); | Metschnikowia | 13 | Saprotroph | \|Undefined Saprotroph\| | Yeast | NULL | Probable | As nectar-tap_saprotroph (Põlme et al. 2020); Decay sugar-rich substrates (Põlme et al. 2020) | Põlme S, et al. 2020. Fungal Diversity 105:1-16 ((https://doi.org/10.1007/s13225-020-00466-2)) |
| Otu44725 | 1 | k__Fungi(100);p__Ascomycota(100);c__Saccharomycetes(100);o__Saccharomycetales(100);f__Metschnikowiaceae(100);g__Metschnikowia(100);s__Metschnikowia_pulcherrima(100); | Metschnikowia | 13 | Saprotroph | \|Undefined Saprotroph\| | Yeast | NULL | Probable | As nectar-tap_saprotroph (Põlme et al. 2020); Decay sugar-rich substrates (Põlme et al. 2020) | Põlme S, et al. 2020. Fungal Diversity 105:1-16 ((https://doi.org/10.1007/s13225-020-00466-2)) |
| Otu44726 | 1 | k__Fungi(100);p__Ascomycota(100);c__Saccharomycetes(100);o__Saccharomycetales(100);f__Metschnikowiaceae(100);g__Metschnikowia(100);s__Metschnikowia_pulcherrima(100); | Metschnikowia | 13 | Saprotroph | \|Undefined Saprotroph\| | Yeast | NULL | Probable | As nectar-tap_saprotroph (Põlme et al. 2020); Decay sugar-rich substrates (Põlme et al. 2020) | Põlme S, et al. 2020. Fungal Diversity 105:1-16 ((https://doi.org/10.1007/s13225-020-00466-2)) |
| Otu44729 | 1 | k__Fungi(100);p__Ascomycota(100);c__Dothideomycetes(100);o__Pleosporales(100);f__Montagnulaceae(100);f__Montagnulaceae_unclassified(100);f__Montagnulaceae_unclassified(100); | Montagnulaceae | 9 | Pathotroph-Saprotroph | Fungal Parasite-Plant Pathogen-Undefined Saprotroph | Microfungus | NULL | Probable | Species in this clade can be saprobic, plant pathogenic, or mycoparasitic (Zhang et al. 2009) | Zhang Y, et al. 2009. Studies in Mycology 64:85-102 ((https://doi.org/10.3114/sim.2009.64.04)) |
| Otu44733 | 1 | k__Fungi(100);p__Ascomycota(100);c__Saccharomycetes(100);o__Saccharomycetales(100);f__Metschnikowiaceae(100);g__Metschnikowia(100);s__Metschnikowia_pulcherrima(100); | Metschnikowia | 13 | Saprotroph | \|Undefined Saprotroph\| | Yeast | NULL | Probable | As nectar-tap_saprotroph (Põlme et al. 2020); Decay sugar-rich substrates (Põlme et al. 2020) | Põlme S, et al. 2020. Fungal Diversity 105:1-16 ((https://doi.org/10.1007/s13225-020-00466-2)) |
| Otu44734 | 1 | k__Fungi(100);p__Ascomycota(100);c__Saccharomycetes(100);o__Saccharomycetales(100);f__Metschnikowiaceae(100);g__Metschnikowia(100);s__Metschnikowia_pulcherrima(100); | Metschnikowia | 13 | Saprotroph | \|Undefined Saprotroph\| | Yeast | NULL | Probable | As nectar-tap_saprotroph (Põlme et al. 2020); Decay sugar-rich substrates (Põlme et al. 2020) | Põlme S, et al. 2020. Fungal Diversity 105:1-16 ((https://doi.org/10.1007/s13225-020-00466-2)) |
| Otu44735 | 1 | k__Fungi(100);p__Ascomycota(100);c__Saccharomycetes(100);o__Saccharomycetales(100);f__Metschnikowiaceae(100);g__Metschnikowia(100);s__Metschnikowia_pulcherrima(100); | Metschnikowia | 13 | Saprotroph | \|Undefined Saprotroph\| | Yeast | NULL | Probable | As nectar-tap_saprotroph (Põlme et al. 2020); Decay sugar-rich substrates (Põlme et al. 2020) | Põlme S, et al. 2020. Fungal Diversity 105:1-16 ((https://doi.org/10.1007/s13225-020-00466-2)) |
| Otu44736 | 1 | k__Fungi(100);p__Ascomycota(100);c__Saccharomycetes(100);o__Saccharomycetales(100);f__Metschnikowiaceae(100);g__Metschnikowia(100);s__Metschnikowia_pulcherrima(100); | Metschnikowia | 13 | Saprotroph | \|Undefined Saprotroph\| | Yeast | NULL | Probable | As nectar-tap_saprotroph (Põlme et al. 2020); Decay sugar-rich substrates (Põlme et al. 2020) | Põlme S, et al. 2020. Fungal Diversity 105:1-16 ((https://doi.org/10.1007/s13225-020-00466-2)) |
| Otu44737 | 1 | k__Fungi(100);p__Ascomycota(100);c__Saccharomycetes(100);o__Saccharomycetales(100);f__Metschnikowiaceae(100);g__Metschnikowia(100);s__Metschnikowia_pulcherrima(100); | Metschnikowia | 13 | Saprotroph | \|Undefined Saprotroph\| | Yeast | NULL | Probable | As nectar-tap_saprotroph (Põlme et al. 2020); Decay sugar-rich substrates (Põlme et al. 2020) | Põlme S, et al. 2020. Fungal Diversity 105:1-16 ((https://doi.org/10.1007/s13225-020-00466-2)) |
| Otu44738 | 1 | k__Fungi(100);p__Ascomycota(100);c__Saccharomycetes(100);o__Saccharomycetales(100);f__Metschnikowiaceae(100);g__Metschnikowia(100);g__Metschnikowia_unclassified(100); | Metschnikowia | 13 | Saprotroph | \|Undefined Saprotroph\| | Yeast | NULL | Probable | As nectar-tap_saprotroph (Põlme et al. 2020); Decay sugar-rich substrates (Põlme et al. 2020) | Põlme S, et al. 2020. Fungal Diversity 105:1-16 ((https://doi.org/10.1007/s13225-020-00466-2)) |
| Otu44739 | 1 | k__Fungi(100);p__Ascomycota(100);c__Saccharomycetes(100);o__Saccharomycetales(100);f__Metschnikowiaceae(100);g__Metschnikowia(100);s__Metschnikowia_pulcherrima(100); | Metschnikowia | 13 | Saprotroph | \|Undefined Saprotroph\| | Yeast | NULL | Probable | As nectar-tap_saprotroph (Põlme et al. 2020); Decay sugar-rich substrates (Põlme et al. 2020) | Põlme S, et al. 2020. Fungal Diversity 105:1-16 ((https://doi.org/10.1007/s13225-020-00466-2)) |
| Otu44740 | 1 | k__Fungi(100);p__Ascomycota(100);c__Saccharomycetes(100);o__Saccharomycetales(100);f__Metschnikowiaceae(100);g__Metschnikowia(100);g__Metschnikowia_unclassified(100); | Metschnikowia | 13 | Saprotroph | \|Undefined Saprotroph\| | Yeast | NULL | Probable | As nectar-tap_saprotroph (Põlme et al. 2020); Decay sugar-rich substrates (Põlme et al. 2020) | Põlme S, et al. 2020. Fungal Diversity 105:1-16 ((https://doi.org/10.1007/s13225-020-00466-2)) |
| Otu44741 | 1 | k__Fungi(100);p__Ascomycota(100);c__Saccharomycetes(100);o__Saccharomycetales(100);f__Metschnikowiaceae(100);g__Metschnikowia(100);s__Metschnikowia_pulcherrima(100); | Metschnikowia | 13 | Saprotroph | \|Undefined Saprotroph\| | Yeast | NULL | Probable | As nectar-tap_saprotroph (Põlme et al. 2020); Decay sugar-rich substrates (Põlme et al. 2020) | Põlme S, et al. 2020. Fungal Diversity 105:1-16 ((https://doi.org/10.1007/s13225-020-00466-2)) |
| Otu44742 | 1 | k__Fungi(100);p__Ascomycota(100);c__Saccharomycetes(100);o__Saccharomycetales(100);f__Metschnikowiaceae(100);g__Metschnikowia(100);s__Metschnikowia_pulcherrima(100); | Metschnikowia | 13 | Saprotroph | \|Undefined Saprotroph\| | Yeast | NULL | Probable | As nectar-tap_saprotroph (Põlme et al. 2020); Decay sugar-rich substrates (Põlme et al. 2020) | Põlme S, et al. 2020. Fungal Diversity 105:1-16 ((https://doi.org/10.1007/s13225-020-00466-2)) |
| Otu44743 | 1 | k__Fungi(100);p__Ascomycota(100);c__Saccharomycetes(100);o__Saccharomycetales(100);f__Metschnikowiaceae(100);g__Metschnikowia(100);s__Metschnikowia_pulcherrima(100); | Metschnikowia | 13 | Saprotroph | \|Undefined Saprotroph\| | Yeast | NULL | Probable | As nectar-tap_saprotroph (Põlme et al. 2020); Decay sugar-rich substrates (Põlme et al. 2020) | Põlme S, et al. 2020. Fungal Diversity 105:1-16 ((https://doi.org/10.1007/s13225-020-00466-2)) |
| Otu44744 | 1 | k__Fungi(100);p__Ascomycota(100);c__Saccharomycetes(100);o__Saccharomycetales(100);f__Metschnikowiaceae(100);g__Metschnikowia(100);g__Metschnikowia_unclassified(100); | Metschnikowia | 13 | Saprotroph | \|Undefined Saprotroph\| | Yeast | NULL | Probable | As nectar-tap_saprotroph (Põlme et al. 2020); Decay sugar-rich substrates (Põlme et al. 2020) | Põlme S, et al. 2020. Fungal Diversity 105:1-16 ((https://doi.org/10.1007/s13225-020-00466-2)) |
| Otu44745 | 1 | k__Fungi(100);p__Ascomycota(100);c__Saccharomycetes(100);o__Saccharomycetales(100);f__Metschnikowiaceae(100);g__Metschnikowia(100);s__Metschnikowia_pulcherrima(100); | Metschnikowia | 13 | Saprotroph | \|Undefined Saprotroph\| | Yeast | NULL | Probable | As nectar-tap_saprotroph (Põlme et al. 2020); Decay sugar-rich substrates (Põlme et al. 2020) | Põlme S, et al. 2020. Fungal Diversity 105:1-16 ((https://doi.org/10.1007/s13225-020-00466-2)) |
| Otu44746 | 1 | k__Fungi(100);p__Ascomycota(100);c__Dothideomycetes(100);o__Capnodiales(100);f__Teratosphaeriaceae(100);g__Catenulostroma(100);s__Catenulostroma_hermanusense(100); | Catenulostroma | 13 | Pathotroph | \|Plant Pathogen\| | Microfungus | NULL | Probable | NULL | Tedersoo L, et al. 2014. Science 346:e1256688 ((https://doi.org/10.1126/science.1256688)); Põlme S, et al. 2020. Fungal Diversity 105:1-16 ((https://doi.org/10.1007/s13225-020-00466-2)) |
| Otu44747 | 1 | k__Fungi(100);p__Ascomycota(100);c__Saccharomycetes(100);o__Saccharomycetales(100);f__Metschnikowiaceae(100);g__Metschnikowia(100);s__Metschnikowia_pulcherrima(100); | Metschnikowia | 13 | Saprotroph | \|Undefined Saprotroph\| | Yeast | NULL | Probable | As nectar-tap_saprotroph (Põlme et al. 2020); Decay sugar-rich substrates (Põlme et al. 2020) | Põlme S, et al. 2020. Fungal Diversity 105:1-16 ((https://doi.org/10.1007/s13225-020-00466-2)) |
| Otu44749 | 1 | k__Fungi(100);p__Ascomycota(100);c__Saccharomycetes(100);o__Saccharomycetales(100);f__Metschnikowiaceae(100);g__Metschnikowia(100);g__Metschnikowia_unclassified(100); | Metschnikowia | 13 | Saprotroph | \|Undefined Saprotroph\| | Yeast | NULL | Probable | As nectar-tap_saprotroph (Põlme et al. 2020); Decay sugar-rich substrates (Põlme et al. 2020) | Põlme S, et al. 2020. Fungal Diversity 105:1-16 ((https://doi.org/10.1007/s13225-020-00466-2)) |
| Otu44750 | 1 | k__Fungi(100);p__Ascomycota(100);c__Saccharomycetes(100);o__Saccharomycetales(100);f__Metschnikowiaceae(100);g__Metschnikowia(100);s__Metschnikowia_pulcherrima(100); | Metschnikowia | 13 | Saprotroph | \|Undefined Saprotroph\| | Yeast | NULL | Probable | As nectar-tap_saprotroph (Põlme et al. 2020); Decay sugar-rich substrates (Põlme et al. 2020) | Põlme S, et al. 2020. Fungal Diversity 105:1-16 ((https://doi.org/10.1007/s13225-020-00466-2)) |
| Otu44751 | 1 | k__Fungi(100);p__Ascomycota(100);c__Saccharomycetes(100);o__Saccharomycetales(100);f__Metschnikowiaceae(100);g__Metschnikowia(100);s__Metschnikowia_pulcherrima(100); | Metschnikowia | 13 | Saprotroph | \|Undefined Saprotroph\| | Yeast | NULL | Probable | As nectar-tap_saprotroph (Põlme et al. 2020); Decay sugar-rich substrates (Põlme et al. 2020) | Põlme S, et al. 2020. Fungal Diversity 105:1-16 ((https://doi.org/10.1007/s13225-020-00466-2)) |
| Otu44752 | 1 | k__Fungi(100);p__Ascomycota(100);c__Saccharomycetes(100);o__Saccharomycetales(100);f__Metschnikowiaceae(100);g__Metschnikowia(100);s__Metschnikowia_pulcherrima(100); | Metschnikowia | 13 | Saprotroph | \|Undefined Saprotroph\| | Yeast | NULL | Probable | As nectar-tap_saprotroph (Põlme et al. 2020); Decay sugar-rich substrates (Põlme et al. 2020) | Põlme S, et al. 2020. Fungal Diversity 105:1-16 ((https://doi.org/10.1007/s13225-020-00466-2)) |
| Otu44753 | 1 | k__Fungi(100);p__Ascomycota(100);c__Saccharomycetes(100);o__Saccharomycetales(100);f__Metschnikowiaceae(100);g__Metschnikowia(100);s__Metschnikowia_pulcherrima(100); | Metschnikowia | 13 | Saprotroph | \|Undefined Saprotroph\| | Yeast | NULL | Probable | As nectar-tap_saprotroph (Põlme et al. 2020); Decay sugar-rich substrates (Põlme et al. 2020) | Põlme S, et al. 2020. Fungal Diversity 105:1-16 ((https://doi.org/10.1007/s13225-020-00466-2)) |
| Otu44754 | 1 | k__Fungi(100);p__Ascomycota(100);c__Saccharomycetes(100);o__Saccharomycetales(100);f__Metschnikowiaceae(100);g__Metschnikowia(100);s__Metschnikowia_pulcherrima(100); | Metschnikowia | 13 | Saprotroph | \|Undefined Saprotroph\| | Yeast | NULL | Probable | As nectar-tap_saprotroph (Põlme et al. 2020); Decay sugar-rich substrates (Põlme et al. 2020) | Põlme S, et al. 2020. Fungal Diversity 105:1-16 ((https://doi.org/10.1007/s13225-020-00466-2)) |
| Otu44756 | 1 | k__Fungi(100);p__Ascomycota(100);c__Saccharomycetes(100);o__Saccharomycetales(100);f__Metschnikowiaceae(100);g__Metschnikowia(100);s__Metschnikowia_pulcherrima(100); | Metschnikowia | 13 | Saprotroph | \|Undefined Saprotroph\| | Yeast | NULL | Probable | As nectar-tap_saprotroph (Põlme et al. 2020); Decay sugar-rich substrates (Põlme et al. 2020) | Põlme S, et al. 2020. Fungal Diversity 105:1-16 ((https://doi.org/10.1007/s13225-020-00466-2)) |
| Otu44757 | 1 | k__Fungi(100);p__Ascomycota(100);c__Saccharomycetes(100);o__Saccharomycetales(100);f__Metschnikowiaceae(100);g__Metschnikowia(100);s__Metschnikowia_pulcherrima(100); | Metschnikowia | 13 | Saprotroph | \|Undefined Saprotroph\| | Yeast | NULL | Probable | As nectar-tap_saprotroph (Põlme et al. 2020); Decay sugar-rich substrates (Põlme et al. 2020) | Põlme S, et al. 2020. Fungal Diversity 105:1-16 ((https://doi.org/10.1007/s13225-020-00466-2)) |
| Otu44758 | 1 | k__Fungi(100);p__Ascomycota(100);c__Saccharomycetes(100);o__Saccharomycetales(100);f__Metschnikowiaceae(100);g__Metschnikowia(100);s__Metschnikowia_pulcherrima(100); | Metschnikowia | 13 | Saprotroph | \|Undefined Saprotroph\| | Yeast | NULL | Probable | As nectar-tap_saprotroph (Põlme et al. 2020); Decay sugar-rich substrates (Põlme et al. 2020) | Põlme S, et al. 2020. Fungal Diversity 105:1-16 ((https://doi.org/10.1007/s13225-020-00466-2)) |
| Otu44759 | 1 | k__Fungi(100);p__Ascomycota(100);c__Saccharomycetes(100);o__Saccharomycetales(100);f__Metschnikowiaceae(100);g__Metschnikowia(100);g__Metschnikowia_unclassified(100); | Metschnikowia | 13 | Saprotroph | \|Undefined Saprotroph\| | Yeast | NULL | Probable | As nectar-tap_saprotroph (Põlme et al. 2020); Decay sugar-rich substrates (Põlme et al. 2020) | Põlme S, et al. 2020. Fungal Diversity 105:1-16 ((https://doi.org/10.1007/s13225-020-00466-2)) |
| Otu44760 | 1 | k__Fungi(100);p__Ascomycota(100);c__Saccharomycetes(100);o__Saccharomycetales(100);f__Metschnikowiaceae(100);g__Metschnikowia(100);g__Metschnikowia_unclassified(100); | Metschnikowia | 13 | Saprotroph | \|Undefined Saprotroph\| | Yeast | NULL | Probable | As nectar-tap_saprotroph (Põlme et al. 2020); Decay sugar-rich substrates (Põlme et al. 2020) | Põlme S, et al. 2020. Fungal Diversity 105:1-16 ((https://doi.org/10.1007/s13225-020-00466-2)) |
| Otu44761 | 1 | k__Fungi(100);p__Ascomycota(100);c__Saccharomycetes(100);o__Saccharomycetales(100);f__Metschnikowiaceae(100);g__Metschnikowia(100);g__Metschnikowia_unclassified(100); | Metschnikowia | 13 | Saprotroph | \|Undefined Saprotroph\| | Yeast | NULL | Probable | As nectar-tap_saprotroph (Põlme et al. 2020); Decay sugar-rich substrates (Põlme et al. 2020) | Põlme S, et al. 2020. Fungal Diversity 105:1-16 ((https://doi.org/10.1007/s13225-020-00466-2)) |
| Otu44763 | 1 | k__Fungi(100);p__Ascomycota(100);c__Saccharomycetes(100);o__Saccharomycetales(100);f__Metschnikowiaceae(100);g__Metschnikowia(100);g__Metschnikowia_unclassified(100); | Metschnikowia | 13 | Saprotroph | \|Undefined Saprotroph\| | Yeast | NULL | Probable | As nectar-tap_saprotroph (Põlme et al. 2020); Decay sugar-rich substrates (Põlme et al. 2020) | Põlme S, et al. 2020. Fungal Diversity 105:1-16 ((https://doi.org/10.1007/s13225-020-00466-2)) |
| Otu44764 | 1 | k__Fungi(100);p__Ascomycota(100);c__Saccharomycetes(100);o__Saccharomycetales(100);f__Metschnikowiaceae(100);g__Metschnikowia(100);g__Metschnikowia_unclassified(100); | Metschnikowia | 13 | Saprotroph | \|Undefined Saprotroph\| | Yeast | NULL | Probable | As nectar-tap_saprotroph (Põlme et al. 2020); Decay sugar-rich substrates (Põlme et al. 2020) | Põlme S, et al. 2020. Fungal Diversity 105:1-16 ((https://doi.org/10.1007/s13225-020-00466-2)) |
| Otu44765 | 1 | k__Fungi(100);p__Ascomycota(100);c__Saccharomycetes(100);o__Saccharomycetales(100);f__Metschnikowiaceae(100);g__Metschnikowia(100);g__Metschnikowia_unclassified(100); | Metschnikowia | 13 | Saprotroph | \|Undefined Saprotroph\| | Yeast | NULL | Probable | As nectar-tap_saprotroph (Põlme et al. 2020); Decay sugar-rich substrates (Põlme et al. 2020) | Põlme S, et al. 2020. Fungal Diversity 105:1-16 ((https://doi.org/10.1007/s13225-020-00466-2)) |
| Otu44766 | 1 | k__Fungi(100);p__Ascomycota(100);c__Saccharomycetes(100);o__Saccharomycetales(100);f__Metschnikowiaceae(100);g__Metschnikowia(100);s__Metschnikowia_pulcherrima(100); | Metschnikowia | 13 | Saprotroph | \|Undefined Saprotroph\| | Yeast | NULL | Probable | As nectar-tap_saprotroph (Põlme et al. 2020); Decay sugar-rich substrates (Põlme et al. 2020) | Põlme S, et al. 2020. Fungal Diversity 105:1-16 ((https://doi.org/10.1007/s13225-020-00466-2)) |
| Otu44767 | 1 | k__Fungi(100);p__Ascomycota(100);c__Saccharomycetes(100);o__Saccharomycetales(100);f__Metschnikowiaceae(100);g__Metschnikowia(100);s__Metschnikowia_pulcherrima(100); | Metschnikowia | 13 | Saprotroph | \|Undefined Saprotroph\| | Yeast | NULL | Probable | As nectar-tap_saprotroph (Põlme et al. 2020); Decay sugar-rich substrates (Põlme et al. 2020) | Põlme S, et al. 2020. Fungal Diversity 105:1-16 ((https://doi.org/10.1007/s13225-020-00466-2)) |
| Otu44768 | 1 | k__Fungi(100);p__Ascomycota(100);c__Saccharomycetes(100);o__Saccharomycetales(100);f__Metschnikowiaceae(100);g__Metschnikowia(100);s__Metschnikowia_pulcherrima(100); | Metschnikowia | 13 | Saprotroph | \|Undefined Saprotroph\| | Yeast | NULL | Probable | As nectar-tap_saprotroph (Põlme et al. 2020); Decay sugar-rich substrates (Põlme et al. 2020) | Põlme S, et al. 2020. Fungal Diversity 105:1-16 ((https://doi.org/10.1007/s13225-020-00466-2)) |
| Otu44769 | 1 | k__Fungi(100);p__Ascomycota(100);c__Saccharomycetes(100);o__Saccharomycetales(100);f__Metschnikowiaceae(100);g__Metschnikowia(100);s__Metschnikowia_pulcherrima(100); | Metschnikowia | 13 | Saprotroph | \|Undefined Saprotroph\| | Yeast | NULL | Probable | As nectar-tap_saprotroph (Põlme et al. 2020); Decay sugar-rich substrates (Põlme et al. 2020) | Põlme S, et al. 2020. Fungal Diversity 105:1-16 ((https://doi.org/10.1007/s13225-020-00466-2)) |
| Otu44770 | 1 | k__Fungi(100);p__Ascomycota(100);c__Saccharomycetes(100);o__Saccharomycetales(100);f__Metschnikowiaceae(100);g__Metschnikowia(100);s__Metschnikowia_pulcherrima(100); | Metschnikowia | 13 | Saprotroph | \|Undefined Saprotroph\| | Yeast | NULL | Probable | As nectar-tap_saprotroph (Põlme et al. 2020); Decay sugar-rich substrates (Põlme et al. 2020) | Põlme S, et al. 2020. Fungal Diversity 105:1-16 ((https://doi.org/10.1007/s13225-020-00466-2)) |
| Otu44771 | 1 | k__Fungi(100);p__Ascomycota(100);c__Saccharomycetes(100);o__Saccharomycetales(100);f__Metschnikowiaceae(100);g__Metschnikowia(100);g__Metschnikowia_unclassified(100); | Metschnikowia | 13 | Saprotroph | \|Undefined Saprotroph\| | Yeast | NULL | Probable | As nectar-tap_saprotroph (Põlme et al. 2020); Decay sugar-rich substrates (Põlme et al. 2020) | Põlme S, et al. 2020. Fungal Diversity 105:1-16 ((https://doi.org/10.1007/s13225-020-00466-2)) |
| Otu44772 | 1 | k__Fungi(100);p__Ascomycota(100);c__Saccharomycetes(100);o__Saccharomycetales(100);f__Metschnikowiaceae(100);g__Metschnikowia(100);g__Metschnikowia_unclassified(100); | Metschnikowia | 13 | Saprotroph | \|Undefined Saprotroph\| | Yeast | NULL | Probable | As nectar-tap_saprotroph (Põlme et al. 2020); Decay sugar-rich substrates (Põlme et al. 2020) | Põlme S, et al. 2020. Fungal Diversity 105:1-16 ((https://doi.org/10.1007/s13225-020-00466-2)) |
| Otu44773 | 1 | k__Fungi(100);p__Ascomycota(100);c__Saccharomycetes(100);o__Saccharomycetales(100);f__Metschnikowiaceae(100);g__Metschnikowia(100);s__Metschnikowia_pulcherrima(100); | Metschnikowia | 13 | Saprotroph | \|Undefined Saprotroph\| | Yeast | NULL | Probable | As nectar-tap_saprotroph (Põlme et al. 2020); Decay sugar-rich substrates (Põlme et al. 2020) | Põlme S, et al. 2020. Fungal Diversity 105:1-16 ((https://doi.org/10.1007/s13225-020-00466-2)) |
| Otu44774 | 1 | k__Fungi(100);p__Ascomycota(100);c__Saccharomycetes(100);o__Saccharomycetales(100);f__Metschnikowiaceae(100);g__Metschnikowia(100);s__Metschnikowia_pulcherrima(100); | Metschnikowia | 13 | Saprotroph | \|Undefined Saprotroph\| | Yeast | NULL | Probable | As nectar-tap_saprotroph (Põlme et al. 2020); Decay sugar-rich substrates (Põlme et al. 2020) | Põlme S, et al. 2020. Fungal Diversity 105:1-16 ((https://doi.org/10.1007/s13225-020-00466-2)) |
| Otu44775 | 1 | k__Fungi(100);p__Ascomycota(100);c__Saccharomycetes(100);o__Saccharomycetales(100);f__Metschnikowiaceae(100);g__Metschnikowia(100);s__Metschnikowia_pulcherrima(100); | Metschnikowia | 13 | Saprotroph | \|Undefined Saprotroph\| | Yeast | NULL | Probable | As nectar-tap_saprotroph (Põlme et al. 2020); Decay sugar-rich substrates (Põlme et al. 2020) | Põlme S, et al. 2020. Fungal Diversity 105:1-16 ((https://doi.org/10.1007/s13225-020-00466-2)) |
| Otu44776 | 1 | k__Fungi(100);p__Ascomycota(100);c__Saccharomycetes(100);o__Saccharomycetales(100);f__Metschnikowiaceae(100);g__Metschnikowia(100);s__Metschnikowia_pulcherrima(100); | Metschnikowia | 13 | Saprotroph | \|Undefined Saprotroph\| | Yeast | NULL | Probable | As nectar-tap_saprotroph (Põlme et al. 2020); Decay sugar-rich substrates (Põlme et al. 2020) | Põlme S, et al. 2020. Fungal Diversity 105:1-16 ((https://doi.org/10.1007/s13225-020-00466-2)) |
| Otu44777 | 1 | k__Fungi(100);p__Ascomycota(100);c__Saccharomycetes(100);o__Saccharomycetales(100);f__Metschnikowiaceae(100);g__Metschnikowia(100);g__Metschnikowia_unclassified(100); | Metschnikowia | 13 | Saprotroph | \|Undefined Saprotroph\| | Yeast | NULL | Probable | As nectar-tap_saprotroph (Põlme et al. 2020); Decay sugar-rich substrates (Põlme et al. 2020) | Põlme S, et al. 2020. Fungal Diversity 105:1-16 ((https://doi.org/10.1007/s13225-020-00466-2)) |
| Otu44778 | 1 | k__Fungi(100);p__Ascomycota(100);c__Saccharomycetes(100);o__Saccharomycetales(100);f__Metschnikowiaceae(100);g__Metschnikowia(100);s__Metschnikowia_pulcherrima(100); | Metschnikowia | 13 | Saprotroph | \|Undefined Saprotroph\| | Yeast | NULL | Probable | As nectar-tap_saprotroph (Põlme et al. 2020); Decay sugar-rich substrates (Põlme et al. 2020) | Põlme S, et al. 2020. Fungal Diversity 105:1-16 ((https://doi.org/10.1007/s13225-020-00466-2)) |
| Otu44779 | 1 | k__Fungi(100);p__Ascomycota(100);c__Saccharomycetes(100);o__Saccharomycetales(100);f__Metschnikowiaceae(100);g__Metschnikowia(100);s__Metschnikowia_pulcherrima(100); | Metschnikowia | 13 | Saprotroph | \|Undefined Saprotroph\| | Yeast | NULL | Probable | As nectar-tap_saprotroph (Põlme et al. 2020); Decay sugar-rich substrates (Põlme et al. 2020) | Põlme S, et al. 2020. Fungal Diversity 105:1-16 ((https://doi.org/10.1007/s13225-020-00466-2)) |
| Otu44780 | 1 | k__Fungi(100);p__Ascomycota(100);c__Saccharomycetes(100);o__Saccharomycetales(100);f__Metschnikowiaceae(100);g__Metschnikowia(100);g__Metschnikowia_unclassified(100); | Metschnikowia | 13 | Saprotroph | \|Undefined Saprotroph\| | Yeast | NULL | Probable | As nectar-tap_saprotroph (Põlme et al. 2020); Decay sugar-rich substrates (Põlme et al. 2020) | Põlme S, et al. 2020. Fungal Diversity 105:1-16 ((https://doi.org/10.1007/s13225-020-00466-2)) |
| Otu44781 | 1 | k__Fungi(100);p__Ascomycota(100);c__Saccharomycetes(100);o__Saccharomycetales(100);f__Metschnikowiaceae(100);g__Metschnikowia(100);g__Metschnikowia_unclassified(100); | Metschnikowia | 13 | Saprotroph | \|Undefined Saprotroph\| | Yeast | NULL | Probable | As nectar-tap_saprotroph (Põlme et al. 2020); Decay sugar-rich substrates (Põlme et al. 2020) | Põlme S, et al. 2020. Fungal Diversity 105:1-16 ((https://doi.org/10.1007/s13225-020-00466-2)) |
| Otu44782 | 1 | k__Fungi(100);p__Ascomycota(100);c__Saccharomycetes(100);o__Saccharomycetales(100);f__Metschnikowiaceae(100);g__Metschnikowia(100);s__Metschnikowia_pulcherrima(100); | Metschnikowia | 13 | Saprotroph | \|Undefined Saprotroph\| | Yeast | NULL | Probable | As nectar-tap_saprotroph (Põlme et al. 2020); Decay sugar-rich substrates (Põlme et al. 2020) | Põlme S, et al. 2020. Fungal Diversity 105:1-16 ((https://doi.org/10.1007/s13225-020-00466-2)) |
| Otu44783 | 1 | k__Fungi(100);p__Ascomycota(100);c__Saccharomycetes(100);o__Saccharomycetales(100);f__Metschnikowiaceae(100);g__Metschnikowia(100);s__Metschnikowia_pulcherrima(100); | Metschnikowia | 13 | Saprotroph | \|Undefined Saprotroph\| | Yeast | NULL | Probable | As nectar-tap_saprotroph (Põlme et al. 2020); Decay sugar-rich substrates (Põlme et al. 2020) | Põlme S, et al. 2020. Fungal Diversity 105:1-16 ((https://doi.org/10.1007/s13225-020-00466-2)) |
| Otu44784 | 1 | k__Fungi(100);p__Ascomycota(100);c__Saccharomycetes(100);o__Saccharomycetales(100);f__Metschnikowiaceae(100);g__Metschnikowia(100);s__Metschnikowia_pulcherrima(100); | Metschnikowia | 13 | Saprotroph | \|Undefined Saprotroph\| | Yeast | NULL | Probable | As nectar-tap_saprotroph (Põlme et al. 2020); Decay sugar-rich substrates (Põlme et al. 2020) | Põlme S, et al. 2020. Fungal Diversity 105:1-16 ((https://doi.org/10.1007/s13225-020-00466-2)) |
| Otu44785 | 1 | k__Fungi(100);p__Ascomycota(100);c__Saccharomycetes(100);o__Saccharomycetales(100);f__Metschnikowiaceae(100);g__Metschnikowia(100);s__Metschnikowia_pulcherrima(100); | Metschnikowia | 13 | Saprotroph | \|Undefined Saprotroph\| | Yeast | NULL | Probable | As nectar-tap_saprotroph (Põlme et al. 2020); Decay sugar-rich substrates (Põlme et al. 2020) | Põlme S, et al. 2020. Fungal Diversity 105:1-16 ((https://doi.org/10.1007/s13225-020-00466-2)) |
| Otu44786 | 1 | k__Fungi(100);p__Ascomycota(100);c__Sordariomycetes(100);o__Hypocreales(100);f__Hypocreales_family_Incertae_sedis(100);g__Geosmithia(100);s__Geosmithia_sp_NRRL_35616(100); | Geosmithia | 13 | Saprotroph | \|Undefined Saprotroph\| | Microfungus | NULL | Probable | Decay leaf, fruit, seed, soil, dung, animal material (Põlme et al. 2020) | Põlme S, et al. 2020. Fungal Diversity 105:1-16 ((https://doi.org/10.1007/s13225-020-00466-2)) |

| | | | | | | | | | | | |
|---|---|---|---|---|---|---|---|---|---|---|---|
| Otu45009 | 1 | k__Fungi(100);p__Ascomycota(100);c__Saccharomycetes(100);o__Saccharomycetales(100);f__Metschnikowiaceae(100);g__Metschnikowia(100);s__Metschnikowia_pulcherrima(100); | Metschnikowia | 13 | Saprotroph | \|Undefined Saprotroph\| | Yeast | NULL | Probable | As nectar-tap_saprotroph (Põlme et al. 2020); Decay sugar-rich substrates (Põlme et al. 2020) | Põlme S, et al. 2020. Fungal Diversity 105:1-16 ((https://doi.org/10.1007/s13225-020-00466-2)) |
| Otu45010 | 1 | k__Fungi(100);p__Ascomycota(100);c__Saccharomycetes(100);o__Saccharomycetales(100);f__Metschnikowiaceae(100);g__Metschnikowia(100);g__Metschnikowia_unclassified(100); | Metschnikowia | 13 | Saprotroph | \|Undefined Saprotroph\| | Yeast | NULL | Probable | As nectar-tap_saprotroph (Põlme et al. 2020); Decay sugar-rich substrates (Põlme et al. 2020) | Põlme S, et al. 2020. Fungal Diversity 105:1-16 ((https://doi.org/10.1007/s13225-020-00466-2)) |
| Otu45011 | 1 | k__Fungi(100);p__Ascomycota(100);c__Saccharomycetes(100);o__Saccharomycetales(100);f__Metschnikowiaceae(100);g__Metschnikowia(100);s__Metschnikowia_pulcherrima(100); | Metschnikowia | 13 | Saprotroph | \|Undefined Saprotroph\| | Yeast | NULL | Probable | As nectar-tap_saprotroph (Põlme et al. 2020); Decay sugar-rich substrates (Põlme et al. 2020) | Põlme S, et al. 2020. Fungal Diversity 105:1-16 ((https://doi.org/10.1007/s13225-020-00466-2)) |
| Otu45012 | 1 | k__Fungi(100);p__Ascomycota(100);c__Saccharomycetes(100);o__Saccharomycetales(100);f__Metschnikowiaceae(100);g__Metschnikowia(100);s__Metschnikowia_pulcherrima(100); | Metschnikowia | 13 | Saprotroph | \|Undefined Saprotroph\| | Yeast | NULL | Probable | As nectar-tap_saprotroph (Põlme et al. 2020); Decay sugar-rich substrates (Põlme et al. 2020) | Põlme S, et al. 2020. Fungal Diversity 105:1-16 ((https://doi.org/10.1007/s13225-020-00466-2)) |
| Otu45013 | 1 | k__Fungi(100);p__Ascomycota(100);c__Saccharomycetes(100);o__Saccharomycetales(100);f__Metschnikowiaceae(100);g__Metschnikowia(100);s__Metschnikowia_pulcherrima(100); | Metschnikowia | 13 | Saprotroph | \|Undefined Saprotroph\| | Yeast | NULL | Probable | As nectar-tap_saprotroph (Põlme et al. 2020); Decay sugar-rich substrates (Põlme et al. 2020) | Põlme S, et al. 2020. Fungal Diversity 105:1-16 ((https://doi.org/10.1007/s13225-020-00466-2)) |
| Otu45014 | 1 | k__Fungi(100);p__Ascomycota(100);c__Saccharomycetes(100);o__Saccharomycetales(100);f__Metschnikowiaceae(100);g__Metschnikowia(100);s__Metschnikowia_pulcherrima(100); | Metschnikowia | 13 | Saprotroph | \|Undefined Saprotroph\| | Yeast | NULL | Probable | As nectar-tap_saprotroph (Põlme et al. 2020); Decay sugar-rich substrates (Põlme et al. 2020) | Põlme S, et al. 2020. Fungal Diversity 105:1-16 ((https://doi.org/10.1007/s13225-020-00466-2)) |
| Otu45015 | 1 | k__Fungi(100);p__Ascomycota(100);c__Saccharomycetes(100);o__Saccharomycetales(100);f__Metschnikowiaceae(100);g__Metschnikowia(100);s__Metschnikowia_pulcherrima(100); | Metschnikowia | 13 | Saprotroph | \|Undefined Saprotroph\| | Yeast | NULL | Probable | As nectar-tap_saprotroph (Põlme et al. 2020); Decay sugar-rich substrates (Põlme et al. 2020) | Põlme S, et al. 2020. Fungal Diversity 105:1-16 ((https://doi.org/10.1007/s13225-020-00466-2)) |
| Otu45016 | 1 | k__Fungi(100);p__Ascomycota(100);c__Saccharomycetes(100);o__Saccharomycetales(100);f__Metschnikowiaceae(100);g__Metschnikowia(100);g__Metschnikowia_unclassified(100); | Metschnikowia | 13 | Saprotroph | \|Undefined Saprotroph\| | Yeast | NULL | Probable | As nectar-tap_saprotroph (Põlme et al. 2020); Decay sugar-rich substrates (Põlme et al. 2020) | Põlme S, et al. 2020. Fungal Diversity 105:1-16 ((https://doi.org/10.1007/s13225-020-00466-2)) |
| Otu45017 | 1 | k__Fungi(100);p__Ascomycota(100);c__Saccharomycetes(100);o__Saccharomycetales(100);f__Metschnikowiaceae(100);g__Metschnikowia(100);g__Metschnikowia_unclassified(100); | Metschnikowia | 13 | Saprotroph | \|Undefined Saprotroph\| | Yeast | NULL | Probable | As nectar-tap_saprotroph (Põlme et al. 2020); Decay sugar-rich substrates (Põlme et al. 2020) | Põlme S, et al. 2020. Fungal Diversity 105:1-16 ((https://doi.org/10.1007/s13225-020-00466-2)) |
| Otu45018 | 1 | k__Fungi(100);p__Ascomycota(100);c__Saccharomycetes(100);o__Saccharomycetales(100);f__Metschnikowiaceae(100);g__Metschnikowia(100);s__Metschnikowia_pulcherrima(100); | Metschnikowia | 13 | Saprotroph | \|Undefined Saprotroph\| | Yeast | NULL | Probable | As nectar-tap_saprotroph (Põlme et al. 2020); Decay sugar-rich substrates (Põlme et al. 2020) | Põlme S, et al. 2020. Fungal Diversity 105:1-16 ((https://doi.org/10.1007/s13225-020-00466-2)) |
| Otu45019 | 1 | k__Fungi(100);p__Ascomycota(100);c__Saccharomycetes(100);o__Saccharomycetales(100);f__Metschnikowiaceae(100);g__Metschnikowia(100);g__Metschnikowia_unclassified(100); | Metschnikowia | 13 | Saprotroph | \|Undefined Saprotroph\| | Yeast | NULL | Probable | As nectar-tap_saprotroph (Põlme et al. 2020); Decay sugar-rich substrates (Põlme et al. 2020) | Põlme S, et al. 2020. Fungal Diversity 105:1-16 ((https://doi.org/10.1007/s13225-020-00466-2)) |
| Otu45020 | 1 | k__Fungi(100);p__Ascomycota(100);c__Saccharomycetes(100);o__Saccharomycetales(100);f__Metschnikowiaceae(100);g__Metschnikowia(100);s__Metschnikowia_viticola(100); | Metschnikowia | 13 | Saprotroph | \|Undefined Saprotroph\| | Yeast | NULL | Probable | As nectar-tap_saprotroph (Põlme et al. 2020); Decay sugar-rich substrates (Põlme et al. 2020) | Põlme S, et al. 2020. Fungal Diversity 105:1-16 ((https://doi.org/10.1007/s13225-020-00466-2)) |
| Otu45021 | 1 | k__Fungi(100);p__Ascomycota(100);c__Saccharomycetes(100);o__Saccharomycetales(100);f__Metschnikowiaceae(100);g__Metschnikowia(100);s__Metschnikowia_pulcherrima(100); | Metschnikowia | 13 | Saprotroph | \|Undefined Saprotroph\| | Yeast | NULL | Probable | As nectar-tap_saprotroph (Põlme et al. 2020); Decay sugar-rich substrates (Põlme et al. 2020) | Põlme S, et al. 2020. Fungal Diversity 105:1-16 ((https://doi.org/10.1007/s13225-020-00466-2)) |
| Otu45022 | 1 | k__Fungi(100);p__Ascomycota(100);c__Saccharomycetes(100);o__Saccharomycetales(100);f__Metschnikowiaceae(100);g__Metschnikowia(100);s__Metschnikowia_pulcherrima(100); | Metschnikowia | 13 | Saprotroph | \|Undefined Saprotroph\| | Yeast | NULL | Probable | As nectar-tap_saprotroph (Põlme et al. 2020); Decay sugar-rich substrates (Põlme et al. 2020) | Põlme S, et al. 2020. Fungal Diversity 105:1-16 ((https://doi.org/10.1007/s13225-020-00466-2)) |
| Otu45023 | 1 | k__Fungi(100);p__Ascomycota(100);c__Saccharomycetes(100);o__Saccharomycetales(100);f__Metschnikowiaceae(100);g__Metschnikowia(100);s__Metschnikowia_pulcherrima(100); | Metschnikowia | 13 | Saprotroph | \|Undefined Saprotroph\| | Yeast | NULL | Probable | As nectar-tap_saprotroph (Põlme et al. 2020); Decay sugar-rich substrates (Põlme et al. 2020) | Põlme S, et al. 2020. Fungal Diversity 105:1-16 ((https://doi.org/10.1007/s13225-020-00466-2)) |
| Otu45024 | 1 | k__Fungi(100);p__Ascomycota(100);c__Saccharomycetes(100);o__Saccharomycetales(100);f__Metschnikowiaceae(100);g__Metschnikowia(100);s__Metschnikowia_pulcherrima(100); | Metschnikowia | 13 | Saprotroph | \|Undefined Saprotroph\| | Yeast | NULL | Probable | As nectar-tap_saprotroph (Põlme et al. 2020); Decay sugar-rich substrates (Põlme et al. 2020) | Põlme S, et al. 2020. Fungal Diversity 105:1-16 ((https://doi.org/10.1007/s13225-020-00466-2)) |
| Otu45025 | 1 | k__Fungi(100);p__Ascomycota(100);c__Saccharomycetes(100);o__Saccharomycetales(100);f__Metschnikowiaceae(100);g__Metschnikowia(100);s__Metschnikowia_pulcherrima(100); | Metschnikowia | 13 | Saprotroph | \|Undefined Saprotroph\| | Yeast | NULL | Probable | As nectar-tap_saprotroph (Põlme et al. 2020); Decay sugar-rich substrates (Põlme et al. 2020) | Põlme S, et al. 2020. Fungal Diversity 105:1-16 ((https://doi.org/10.1007/s13225-020-00466-2)) |
| Otu45026 | 1 | k__Fungi(100);p__Ascomycota(100);c__Saccharomycetes(100);o__Saccharomycetales(100);f__Metschnikowiaceae(100);g__Metschnikowia(100);s__Metschnikowia_pulcherrima(100); | Metschnikowia | 13 | Saprotroph | \|Undefined Saprotroph\| | Yeast | NULL | Probable | As nectar-tap_saprotroph (Põlme et al. 2020); Decay sugar-rich substrates (Põlme et al. 2020) | Põlme S, et al. 2020. Fungal Diversity 105:1-16 ((https://doi.org/10.1007/s13225-020-00466-2)) |
| Otu45027 | 1 | k__Fungi(100);p__Ascomycota(100);c__Saccharomycetes(100);o__Saccharomycetales(100);f__Metschnikowiaceae(100);g__Metschnikowia(100);s__Metschnikowia_pulcherrima(100); | Metschnikowia | 13 | Saprotroph | \|Undefined Saprotroph\| | Yeast | NULL | Probable | As nectar-tap_saprotroph (Põlme et al. 2020); Decay sugar-rich substrates (Põlme et al. 2020) | Põlme S, et al. 2020. Fungal Diversity 105:1-16 ((https://doi.org/10.1007/s13225-020-00466-2)) |
| Otu45028 | 1 | k__Fungi(100);p__Ascomycota(100);c__Saccharomycetes(100);o__Saccharomycetales(100);f__Metschnikowiaceae(100);g__Metschnikowia(100);s__Metschnikowia_pulcherrima(100); | Metschnikowia | 13 | Saprotroph | \|Undefined Saprotroph\| | Yeast | NULL | Probable | As nectar-tap_saprotroph (Põlme et al. 2020); Decay sugar-rich substrates (Põlme et al. 2020) | Põlme S, et al. 2020. Fungal Diversity 105:1-16 ((https://doi.org/10.1007/s13225-020-00466-2)) |
| Otu45029 | 1 | k__Fungi(100);p__Ascomycota(100);c__Saccharomycetes(100);o__Saccharomycetales(100);f__Metschnikowiaceae(100);g__Metschnikowia(100);s__Metschnikowia_viticola(100); | Metschnikowia | 13 | Saprotroph | \|Undefined Saprotroph\| | Yeast | NULL | Probable | As nectar-tap_saprotroph (Põlme et al. 2020); Decay sugar-rich substrates (Põlme et al. 2020) | Põlme S, et al. 2020. Fungal Diversity 105:1-16 ((https://doi.org/10.1007/s13225-020-00466-2)) |
| Otu45030 | 1 | k__Fungi(100);p__Ascomycota(100);c__Saccharomycetes(100);o__Saccharomycetales(100);f__Metschnikowiaceae(100);g__Metschnikowia(100);g__Metschnikowia_unclassified(100); | Metschnikowia | 13 | Saprotroph | \|Undefined Saprotroph\| | Yeast | NULL | Probable | As nectar-tap_saprotroph (Põlme et al. 2020); Decay sugar-rich substrates (Põlme et al. 2020) | Põlme S, et al. 2020. Fungal Diversity 105:1-16 ((https://doi.org/10.1007/s13225-020-00466-2)) |
| Otu45031 | 1 | k__Fungi(100);p__Ascomycota(100);c__Saccharomycetes(100);o__Saccharomycetales(100);f__Metschnikowiaceae(100);g__Metschnikowia(100);s__Metschnikowia_pulcherrima(100); | Metschnikowia | 13 | Saprotroph | \|Undefined Saprotroph\| | Yeast | NULL | Probable | As nectar-tap_saprotroph (Põlme et al. 2020); Decay sugar-rich substrates (Põlme et al. 2020) | Põlme S, et al. 2020. Fungal Diversity 105:1-16 ((https://doi.org/10.1007/s13225-020-00466-2)) |
| Otu45032 | 1 | k__Fungi(100);p__Ascomycota(100);c__Saccharomycetes(100);o__Saccharomycetales(100);f__Metschnikowiaceae(100);g__Metschnikowia(100);g__Metschnikowia_unclassified(100); | Metschnikowia | 13 | Saprotroph | \|Undefined Saprotroph\| | Yeast | NULL | Probable | As nectar-tap_saprotroph (Põlme et al. 2020); Decay sugar-rich substrates (Põlme et al. 2020) | Põlme S, et al. 2020. Fungal Diversity 105:1-16 ((https://doi.org/10.1007/s13225-020-00466-2)) |
| Otu45034 | 1 | k__Fungi(100);p__Ascomycota(100);c__Saccharomycetes(100);o__Saccharomycetales(100);f__Metschnikowiaceae(100);g__Metschnikowia(100);s__Metschnikowia_pulcherrima(100); | Metschnikowia | 13 | Saprotroph | \|Undefined Saprotroph\| | Yeast | NULL | Probable | As nectar-tap_saprotroph (Põlme et al. 2020); Decay sugar-rich substrates (Põlme et al. 2020) | Põlme S, et al. 2020. Fungal Diversity 105:1-16 ((https://doi.org/10.1007/s13225-020-00466-2)) |
| Otu45035 | 1 | k__Fungi(100);p__Ascomycota(100);c__Saccharomycetes(100);o__Saccharomycetales(100);f__Metschnikowiaceae(100);g__Metschnikowia(100);s__Metschnikowia_pulcherrima(100); | Metschnikowia | 13 | Saprotroph | \|Undefined Saprotroph\| | Yeast | NULL | Probable | As nectar-tap_saprotroph (Põlme et al. 2020); Decay sugar-rich substrates (Põlme et al. 2020) | Põlme S, et al. 2020. Fungal Diversity 105:1-16 ((https://doi.org/10.1007/s13225-020-00466-2)) |
| Otu45036 | 1 | k__Fungi(100);p__Ascomycota(100);c__Saccharomycetes(100);o__Saccharomycetales(100);f__Metschnikowiaceae(100);g__Metschnikowia(100);g__Metschnikowia_unclassified(100); | Metschnikowia | 13 | Saprotroph | \|Undefined Saprotroph\| | Yeast | NULL | Probable | As nectar-tap_saprotroph (Põlme et al. 2020); Decay sugar-rich substrates (Põlme et al. 2020) | Põlme S, et al. 2020. Fungal Diversity 105:1-16 ((https://doi.org/10.1007/s13225-020-00466-2)) |
| Otu45037 | 1 | k__Fungi(100);p__Ascomycota(100);c__Saccharomycetes(100);o__Saccharomycetales(100);f__Metschnikowiaceae(100);g__Metschnikowia(100);g__Metschnikowia_unclassified(100); | Metschnikowia | 13 | Saprotroph | \|Undefined Saprotroph\| | Yeast | NULL | Probable | As nectar-tap_saprotroph (Põlme et al. 2020); Decay sugar-rich substrates (Põlme et al. 2020) | Põlme S, et al. 2020. Fungal Diversity 105:1-16 ((https://doi.org/10.1007/s13225-020-00466-2)) |
| Otu45038 | 1 | k__Fungi(100);p__Ascomycota(100);c__Saccharomycetes(100);o__Saccharomycetales(100);f__Metschnikowiaceae(100);g__Metschnikowia(100);s__Metschnikowia_pulcherrima(100); | Metschnikowia | 13 | Saprotroph | \|Undefined Saprotroph\| | Yeast | NULL | Probable | As nectar-tap_saprotroph (Põlme et al. 2020); Decay sugar-rich substrates (Põlme et al. 2020) | Põlme S, et al. 2020. Fungal Diversity 105:1-16 ((https://doi.org/10.1007/s13225-020-00466-2)) |
| Otu45039 | 1 | k__Fungi(100);p__Ascomycota(100);c__Saccharomycetes(100);o__Saccharomycetales(100);f__Metschnikowiaceae(100);g__Metschnikowia(100);g__Metschnikowia_unclassified(100); | Metschnikowia | 13 | Saprotroph | \|Undefined Saprotroph\| | Yeast | NULL | Probable | As nectar-tap_saprotroph (Põlme et al. 2020); Decay sugar-rich substrates (Põlme et al. 2020) | Põlme S, et al. 2020. Fungal Diversity 105:1-16 ((https://doi.org/10.1007/s13225-020-00466-2)) |
| Otu45040 | 1 | k__Fungi(100);p__Ascomycota(100);c__Saccharomycetes(100);o__Saccharomycetales(100);f__Metschnikowiaceae(100);g__Metschnikowia(100);s__Metschnikowia_pulcherrima(100); | Metschnikowia | 13 | Saprotroph | \|Undefined Saprotroph\| | Yeast | NULL | Probable | As nectar-tap_saprotroph (Põlme et al. 2020); Decay sugar-rich substrates (Põlme et al. 2020) | Põlme S, et al. 2020. Fungal Diversity 105:1-16 ((https://doi.org/10.1007/s13225-020-00466-2)) |
| Otu45041 | 1 | k__Fungi(100);p__Ascomycota(100);c__Saccharomycetes(100);o__Saccharomycetales(100);f__Metschnikowiaceae(100);g__Metschnikowia(100);s__Metschnikowia_pulcherrima(100); | Metschnikowia | 13 | Saprotroph | \|Undefined Saprotroph\| | Yeast | NULL | Probable | As nectar-tap_saprotroph (Põlme et al. 2020); Decay sugar-rich substrates (Põlme et al. 2020) | Põlme S, et al. 2020. Fungal Diversity 105:1-16 ((https://doi.org/10.1007/s13225-020-00466-2)) |
| Otu45042 | 1 | k__Fungi(100);p__Ascomycota(100);c__Saccharomycetes(100);o__Saccharomycetales(100);f__Metschnikowiaceae(100);g__Metschnikowia(100);s__Metschnikowia_pulcherrima(100); | Metschnikowia | 13 | Saprotroph | \|Undefined Saprotroph\| | Yeast | NULL | Probable | As nectar-tap_saprotroph (Põlme et al. 2020); Decay sugar-rich substrates (Põlme et al. 2020) | Põlme S, et al. 2020. Fungal Diversity 105:1-16 ((https://doi.org/10.1007/s13225-020-00466-2)) |
| Otu45043 | 1 | k__Fungi(100);p__Ascomycota(100);c__Saccharomycetes(100);o__Saccharomycetales(100);f__Metschnikowiaceae(100);g__Metschnikowia(100);s__Metschnikowia_pulcherrima(100); | Metschnikowia | 13 | Saprotroph | \|Undefined Saprotroph\| | Yeast | NULL | Probable | As nectar-tap_saprotroph (Põlme et al. 2020); Decay sugar-rich substrates (Põlme et al. 2020) | Põlme S, et al. 2020. Fungal Diversity 105:1-16 ((https://doi.org/10.1007/s13225-020-00466-2)) |
| Otu45044 | 1 | k__Fungi(100);p__Ascomycota(100);c__Saccharomycetes(100);o__Saccharomycetales(100);f__Metschnikowiaceae(100);g__Metschnikowia(100);s__Metschnikowia_pulcherrima(100); | Metschnikowia | 13 | Saprotroph | \|Undefined Saprotroph\| | Yeast | NULL | Probable | As nectar-tap_saprotroph (Põlme et al. 2020); Decay sugar-rich substrates (Põlme et al. 2020) | Põlme S, et al. 2020. Fungal Diversity 105:1-16 ((https://doi.org/10.1007/s13225-020-00466-2)) |
| Otu45045 | 1 | k__Fungi(100);p__Ascomycota(100);c__Saccharomycetes(100);o__Saccharomycetales(100);f__Metschnikowiaceae(100);g__Metschnikowia(100);g__Metschnikowia_unclassified(100); | Metschnikowia | 13 | Saprotroph | \|Undefined Saprotroph\| | Yeast | NULL | Probable | As nectar-tap_saprotroph (Põlme et al. 2020); Decay sugar-rich substrates (Põlme et al. 2020) | Põlme S, et al. 2020. Fungal Diversity 105:1-16 ((https://doi.org/10.1007/s13225-020-00466-2)) |
| Otu45046 | 1 | k__Fungi(100);p__Ascomycota(100);c__Saccharomycetes(100);o__Saccharomycetales(100);f__Metschnikowiaceae(100);g__Metschnikowia(100);g__Metschnikowia_unclassified(100); | Metschnikowia | 13 | Saprotroph | \|Undefined Saprotroph\| | Yeast | NULL | Probable | As nectar-tap_saprotroph (Põlme et al. 2020); Decay sugar-rich substrates (Põlme et al. 2020) | Põlme S, et al. 2020. Fungal Diversity 105:1-16 ((https://doi.org/10.1007/s13225-020-00466-2)) |
| Otu45047 | 1 | k__Fungi(100);p__Ascomycota(100);c__Saccharomycetes(100);o__Saccharomycetales(100);f__Metschnikowiaceae(100);g__Metschnikowia(100);g__Metschnikowia_unclassified(100); | Metschnikowia | 13 | Saprotroph | \|Undefined Saprotroph\| | Yeast | NULL | Probable | As nectar-tap_saprotroph (Põlme et al. 2020); Decay sugar-rich substrates (Põlme et al. 2020) | Põlme S, et al. 2020. Fungal Diversity 105:1-16 ((https://doi.org/10.1007/s13225-020-00466-2)) |
| Otu45048 | 1 | k__Fungi(100);p__Ascomycota(100);c__Saccharomycetes(100);o__Saccharomycetales(100);f__Metschnikowiaceae(100);g__Metschnikowia(100);s__Metschnikowia_pulcherrima(100); | Metschnikowia | 13 | Saprotroph | \|Undefined Saprotroph\| | Yeast | NULL | Probable | As nectar-tap_saprotroph (Põlme et al. 2020); Decay sugar-rich substrates (Põlme et al. 2020) | Põlme S, et al. 2020. Fungal Diversity 105:1-16 ((https://doi.org/10.1007/s13225-020-00466-2)) |
| Otu45049 | 1 | k__Fungi(100);p__Ascomycota(100);c__Saccharomycetes(100);o__Saccharomycetales(100);f__Metschnikowiaceae(100);g__Metschnikowia(100);s__Metschnikowia_pulcherrima(100); | Metschnikowia | 13 | Saprotroph | \|Undefined Saprotroph\| | Yeast | NULL | Probable | As nectar-tap_saprotroph (Põlme et al. 2020); Decay sugar-rich substrates (Põlme et al. 2020) | Põlme S, et al. 2020. Fungal Diversity 105:1-16 ((https://doi.org/10.1007/s13225-020-00466-2)) |
| Otu45050 | 1 | k__Fungi(100);p__Ascomycota(100);c__Saccharomycetes(100);o__Saccharomycetales(100);f__Metschnikowiaceae(100);g__Metschnikowia(100);g__Metschnikowia_unclassified(100); | Metschnikowia | 13 | Saprotroph | \|Undefined Saprotroph\| | Yeast | NULL | Probable | As nectar-tap_saprotroph (Põlme et al. 2020); Decay sugar-rich substrates (Põlme et al. 2020) | Põlme S, et al. 2020. Fungal Diversity 105:1-16 ((https://doi.org/10.1007/s13225-020-00466-2)) |
| Otu45051 | 1 | k__Fungi(100);p__Ascomycota(100);c__Saccharomycetes(100);o__Saccharomycetales(100);f__Metschnikowiaceae(100);g__Metschnikowia(100);s__Metschnikowia_pulcherrima(100); | Metschnikowia | 13 | Saprotroph | \|Undefined Saprotroph\| | Yeast | NULL | Probable | As nectar-tap_saprotroph (Põlme et al. 2020); Decay sugar-rich substrates (Põlme et al. 2020) | Põlme S, et al. 2020. Fungal Diversity 105:1-16 ((https://doi.org/10.1007/s13225-020-00466-2)) |
| Otu45052 | 1 | k__Fungi(100);p__Ascomycota(100);c__Saccharomycetes(100);o__Saccharomycetales(100);f__Metschnikowiaceae(100);g__Metschnikowia(100);s__Metschnikowia_pulcherrima(100); | Metschnikowia | 13 | Saprotroph | \|Undefined Saprotroph\| | Yeast | NULL | Probable | As nectar-tap_saprotroph (Põlme et al. 2020); Decay sugar-rich substrates (Põlme et al. 2020) | Põlme S, et al. 2020. Fungal Diversity 105:1-16 ((https://doi.org/10.1007/s13225-020-00466-2)) |
| Otu45053 | 1 | k__Fungi(100);p__Ascomycota(100);c__Saccharomycetes(100);o__Saccharomycetales(100);f__Metschnikowiaceae(100);g__Metschnikowia(100);s__Metschnikowia_pulcherrima(100); | Metschnikowia | 13 | Saprotroph | \|Undefined Saprotroph\| | Yeast | NULL | Probable | As nectar-tap_saprotroph (Põlme et al. 2020); Decay sugar-rich substrates (Põlme et al. 2020) | Põlme S, et al. 2020. Fungal Diversity 105:1-16 ((https://doi.org/10.1007/s13225-020-00466-2)) |
| Otu45054 | 1 | k__Fungi(100);p__Ascomycota(100);c__Saccharomycetes(100);o__Saccharomycetales(100);f__Metschnikowiaceae(100);g__Metschnikowia(100);s__Metschnikowia_pulcherrima(100); | Metschnikowia | 13 | Saprotroph | \|Undefined Saprotroph\| | Yeast | NULL | Probable | As nectar-tap_saprotroph (Põlme et al. 2020); Decay sugar-rich substrates (Põlme et al. 2020) | Põlme S, et al. 2020. Fungal Diversity 105:1-16 ((https://doi.org/10.1007/s13225-020-00466-2)) |
| Otu45055 | 1 | k__Fungi(100);p__Ascomycota(100);c__Saccharomycetes(100);o__Saccharomycetales(100);f__Metschnikowiaceae(100);g__Metschnikowia(100);s__Metschnikowia_pulcherrima(100); | Metschnikowia | 13 | Saprotroph | \|Undefined Saprotroph\| | Yeast | NULL | Probable | As nectar-tap_saprotroph (Põlme et al. 2020); Decay sugar-rich substrates (Põlme et al. 2020) | Põlme S, et al. 2020. Fungal Diversity 105:1-16 ((https://doi.org/10.1007/s13225-020-00466-2)) |
| Otu45056 | 1 | k__Fungi(100);p__Ascomycota(100);c__Saccharomycetes(100);o__Saccharomycetales(100);f__Metschnikowiaceae(100);g__Metschnikowia(100);s__Metschnikowia_pulcherrima(100); | Metschnikowia | 13 | Saprotroph | \|Undefined Saprotroph\| | Yeast | NULL | Probable | As nectar-tap_saprotroph (Põlme et al. 2020); Decay sugar-rich substrates (Põlme et al. 2020) | Põlme S, et al. 2020. Fungal Diversity 105:1-16 ((https://doi.org/10.1007/s13225-020-00466-2)) |
| Otu45057 | 1 | k__Fungi(100);p__Ascomycota(100);c__Saccharomycetes(100);o__Saccharomycetales(100);f__Metschnikowiaceae(100);g__Metschnikowia(100);s__Metschnikowia_pulcherrima(100); | Metschnikowia | 13 | Saprotroph | \|Undefined Saprotroph\| | Yeast | NULL | Probable | As nectar-tap_saprotroph (Põlme et al. 2020); Decay sugar-rich substrates (Põlme et al. 2020) | Põlme S, et al. 2020. Fungal Diversity 105:1-16 ((https://doi.org/10.1007/s13225-020-00466-2)) |
| Otu45058 | 1 | k__Fungi(100);p__Ascomycota(100);c__Saccharomycetes(100);o__Saccharomycetales(100);f__Metschnikowiaceae(100);g__Metschnikowia(100);s__Metschnikowia_pulcherrima(100); | Metschnikowia | 13 | Saprotroph | \|Undefined Saprotroph\| | Yeast | NULL | Probable | As nectar-tap_saprotroph (Põlme et al. 2020); Decay sugar-rich substrates (Põlme et al. 2020) | Põlme S, et al. 2020. Fungal Diversity 105:1-16 ((https://doi.org/10.1007/s13225-020-00466-2)) |
| Otu45059 | 1 | k__Fungi(100);p__Ascomycota(100);c__Saccharomycetes(100);o__Saccharomycetales(100);f__Metschnikowiaceae(100);g__Metschnikowia(100);g__Metschnikowia_unclassified(100); | Metschnikowia | 13 | Saprotroph | \|Undefined Saprotroph\| | Yeast | NULL | Probable | As nectar-tap_saprotroph (Põlme et al. 2020); Decay sugar-rich substrates (Põlme et al. 2020) | Põlme S, et al. 2020. Fungal Diversity 105:1-16 ((https://doi.org/10.1007/s13225-020-00466-2)) |
| Otu45060 | 1 | k__Fungi(100);p__Ascomycota(100);c__Saccharomycetes(100);o__Saccharomycetales(100);f__Metschnikowiaceae(100);g__Metschnikowia(100);s__Metschnikowia_pulcherrima(100); | Metschnikowia | 13 | Saprotroph | \|Undefined Saprotroph\| | Yeast | NULL | Probable | As nectar-tap_saprotroph (Põlme et al. 2020); Decay sugar-rich substrates (Põlme et al. 2020) | Põlme S, et al. 2020. Fungal Diversity 105:1-16 ((https://doi.org/10.1007/s13225-020-00466-2)) |
| Otu45061 | 1 | k__Fungi(100);p__Ascomycota(100);c__Saccharomycetes(100);o__Saccharomycetales(100);f__Metschnikowiaceae(100);g__Metschnikowia(100);s__Metschnikowia_pulcherrima(100); | Metschnikowia | 13 | Saprotroph | \|Undefined Saprotroph\| | Yeast | NULL | Probable | As nectar-tap_saprotroph (Põlme et al. 2020); Decay sugar-rich substrates (Põlme et al. 2020) | Põlme S, et al. 2020. Fungal Diversity 105:1-16 ((https://doi.org/10.1007/s13225-020-00466-2)) |
| Otu45062 | 1 | k__Fungi(100);p__Ascomycota(100);c__Saccharomycetes(100);o__Saccharomycetales(100);f__Metschnikowiaceae(100);g__Metschnikowia(100);g__Metschnikowia_unclassified(100); | Metschnikowia | 13 | Saprotroph | \|Undefined Saprotroph\| | Yeast | NULL | Probable | As nectar-tap_saprotroph (Põlme et al. 2020); Decay sugar-rich substrates (Põlme et al. 2020) | Põlme S, et al. 2020. Fungal Diversity 105:1-16 ((https://doi.org/10.1007/s13225-020-00466-2)) |
| Otu45063 | 1 | k__Fungi(100);p__Ascomycota(100);c__Saccharomycetes(100);o__Saccharomycetales(100);f__Metschnikowiaceae(100);g__Metschnikowia(100);s__Metschnikowia_pulcherrima(100); | Metschnikowia | 13 | Saprotroph | \|Undefined Saprotroph\| | Yeast | NULL | Probable | As nectar-tap_saprotroph (Põlme et al. 2020); Decay sugar-rich substrates (Põlme et al. 2020) | Põlme S, et al. 2020. Fungal Diversity 105:1-16 ((https://doi.org/10.1007/s13225-020-00466-2)) |
| Otu45064 | 1 | k__Fungi(100);p__Ascomycota(100);c__Saccharomycetes(100);o__Saccharomycetales(100);f__Metschnikowiaceae(100);g__Metschnikowia(100);s__Metschnikowia_pulcherrima(100); | Metschnikowia | 13 | Saprotroph | \|Undefined Saprotroph\| | Yeast | NULL | Probable | As nectar-tap_saprotroph (Põlme et al. 2020); Decay sugar-rich substrates (Põlme et al. 2020) | Põlme S, et al. 2020. Fungal Diversity 105:1-16 ((https://doi.org/10.1007/s13225-020-00466-2)) |
| Otu45065 | 1 | k__Fungi(100);p__Ascomycota(100);c__Saccharomycetes(100);o__Saccharomycetales(100);f__Metschnikowiaceae(100);g__Metschnikowia(100);s__Metschnikowia_pulcherrima(100); | Metschnikowia | 13 | Saprotroph | \|Undefined Saprotroph\| | Yeast | NULL | Probable | As nectar-tap_saprotroph (Põlme et al. 2020); Decay sugar-rich substrates (Põlme et al. 2020) | Põlme S, et al. 2020. Fungal Diversity 105:1-16 ((https://doi.org/10.1007/s13225-020-00466-2)) |
| Otu45066 | 1 | k__Fungi(100);p__Ascomycota(100);c__Saccharomycetes(100);o__Saccharomycetales(100);f__Metschnikowiaceae(100);g__Metschnikowia(100);g__Metschnikowia_unclassified(100); | Metschnikowia | 13 | Saprotroph | \|Undefined Saprotroph\| | Yeast | NULL | Probable | As nectar-tap_saprotroph (Põlme et al. 2020); Decay sugar-rich substrates (Põlme et al. 2020) | Põlme S, et al. 2020. Fungal Diversity 105:1-16 ((https://doi.org/10.1007/s13225-020-00466-2)) |
| Otu45067 | 1 | k__Fungi(100);p__Ascomycota(100);c__Saccharomycetes(100);o__Saccharomycetales(100);f__Metschnikowiaceae(100);g__Metschnikowia(100);g__Metschnikowia_unclassified(100); | Metschnikowia | 13 | Saprotroph | \|Undefined Saprotroph\| | Yeast | NULL | Probable | As nectar-tap_saprotroph (Põlme et al. 2020); Decay sugar-rich substrates (Põlme et al. 2020) | Põlme S, et al. 2020. Fungal Diversity 105:1-16 ((https://doi.org/10.1007/s13225-020-00466-2)) |
| Otu45068 | 1 | k__Fungi(100);p__Ascomycota(100);c__Saccharomycetes(100);o__Saccharomycetales(100);f__Metschnikowiaceae(100);g__Metschnikowia(100);s__Metschnikowia_pulcherrima(100); | Metschnikowia | 13 | Saprotroph | \|Undefined Saprotroph\| | Yeast | NULL | Probable | As nectar-tap_saprotroph (Põlme et al. 2020); Decay sugar-rich substrates (Põlme et al. 2020) | Põlme S, et al. 2020. Fungal Diversity 105:1-16 ((https://doi.org/10.1007/s13225-020-00466-2)) |
| Otu45069 | 1 | k__Fungi(100);p__Ascomycota(100);c__Saccharomycetes(100);o__Saccharomycetales(100);f__Metschnikowiaceae(100);g__Metschnikowia(100);s__Metschnikowia_pulcherrima(100); | Metschnikowia | 13 | Saprotroph | \|Undefined Saprotroph\| | Yeast | NULL | Probable | As nectar-tap_saprotroph (Põlme et al. 2020); Decay sugar-rich substrates (Põlme et al. 2020) | Põlme S, et al. 2020. Fungal Diversity 105:1-16 ((https://doi.org/10.1007/s13225-020-00466-2)) |
| Otu45070 | 1 | k__Fungi(100);p__Ascomycota(100);c__Saccharomycetes(100);o__Saccharomycetales(100);f__Metschnikowiaceae(100);g__Metschnikowia(100);s__Metschnikowia_pulcherrima(100); | Metschnikowia | 13 | Saprotroph | \|Undefined Saprotroph\| | Yeast | NULL | Probable | As nectar-tap_saprotroph (Põlme et al. 2020); Decay sugar-rich substrates (Põlme et al. 2020) | Põlme S, et al. 2020. Fungal Diversity 105:1-16 ((https://doi.org/10.1007/s13225-020-00466-2)) |
| Otu45071 | 1 | k__Fungi(100);p__Ascomycota(100);c__Saccharomycetes(100);o__Saccharomycetales(100);f__Metschnikowiaceae(100);g__Metschnikowia(100);g__Metschnikowia_unclassified(100); | Metschnikowia | 13 | Saprotroph | \|Undefined Saprotroph\| | Yeast | NULL | Probable | As nectar-tap_saprotroph (Põlme et al. 2020); Decay sugar-rich substrates (Põlme et al. 2020) | Põlme S, et al. 2020. Fungal Diversity 105:1-16 ((https://doi.org/10.1007/s13225-020-00466-2)) |
| Otu45072 | 1 | k__Fungi(100);p__Ascomycota(100);c__Saccharomycetes(100);o__Saccharomycetales(100);f__Metschnikowiaceae(100);g__Metschnikowia(100);g__Metschnikowia_unclassified(100); | Metschnikowia | 13 | Saprotroph | \|Undefined Saprotroph\| | Yeast | NULL | Probable | As nectar-tap_saprotroph (Põlme et al. 2020); Decay sugar-rich substrates (Põlme et al. 2020) | Põlme S, et al. 2020. Fungal Diversity 105:1-16 ((https://doi.org/10.1007/s13225-020-00466-2)) |
| Otu45073 | 1 | k__Fungi(100);p__Ascomycota(100);c__Saccharomycetes(100);o__Saccharomycetales(100);f__Metschnikowiaceae(100);g__Metschnikowia(100);g__Metschnikowia_unclassified(100); | Metschnikowia | 13 | Saprotroph | \|Undefined Saprotroph\| | Yeast | NULL | Probable | As nectar-tap_saprotroph (Põlme et al. 2020); Decay sugar-rich substrates (Põlme et al. 2020) | Põlme S, et al. 2020. Fungal Diversity 105:1-16 ((https://doi.org/10.1007/s13225-020-00466-2)) |
| Otu45074 | 1 | k__Fungi(100);p__Ascomycota(100);c__Saccharomycetes(100);o__Saccharomycetales(100);f__Metschnikowiaceae(100);g__Metschnikowia(100);s__Metschnikowia_pulcherrima(100); | Metschnikowia | 13 | Saprotroph | \|Undefined Saprotroph\| | Yeast | NULL | Probable | As nectar-tap_saprotroph (Põlme et al. 2020); Decay sugar-rich substrates (Põlme et al. 2020) | Põlme S, et al. 2020. Fungal Diversity 105:1-16 ((https://doi.org/10.1007/s13225-020-00466-2)) |
| Otu45075 | 1 | k__Fungi(100);p__Ascomycota(100);c__Saccharomycetes(100);o__Saccharomycetales(100);f__Metschnikowiaceae(100);g__Metschnikowia(100);s__Metschnikowia_pulcherrima(100); | Metschnikowia | 13 | Saprotroph | \|Undefined Saprotroph\| | Yeast | NULL | Probable | As nectar-tap_saprotroph (Põlme et al. 2020); Decay sugar-rich substrates (Põlme et al. 2020) | Põlme S, et al. 2020. Fungal Diversity 105:1-16 ((https://doi.org/10.1007/s13225-020-00466-2)) |
| Otu45076 | 1 | k__Fungi(100);p__Ascomycota(100);c__Saccharomycetes(100);o__Saccharomycetales(100);f__Metschnikowiaceae(100);g__Metschnikowia(100);s__Metschnikowia_pulcherrima(100); | Metschnikowia | 13 | Saprotroph | \|Undefined Saprotroph\| | Yeast | NULL | Probable | As nectar-tap_saprotroph (Põlme et al. 2020); Decay sugar-rich substrates (Põlme et al. 2020) | Põlme S, et al. 2020. Fungal Diversity 105:1-16 ((https://doi.org/10.1007/s13225-020-00466-2)) |
| Otu45077 | 1 | k__Fungi(100);p__Ascomycota(100);c__Saccharomycetes(100);o__Saccharomycetales(100);f__Metschnikowiaceae(100);g__Metschnikowia(100);s__Metschnikowia_pulcherrima(100); | Metschnikowia | 13 | Saprotroph | \|Undefined Saprotroph\| | Yeast | NULL | Probable | As nectar-tap_saprotroph (Põlme et al. 2020); Decay sugar-rich substrates (Põlme et al. 2020) | Põlme S, et al. 2020. Fungal Diversity 105:1-16 ((https://doi.org/10.1007/s13225-020-00466-2)) |
| Otu45078 | 1 | k__Fungi(100);p__Ascomycota(100);c__Saccharomycetes(100);o__Saccharomycetales(100);f__Metschnikowiaceae(100);g__Metschnikowia(100);s__Metschnikowia_pulcherrima(100); | Metschnikowia | 13 | Saprotroph | \|Undefined Saprotroph\| | Yeast | NULL | Probable | As nectar-tap_saprotroph (Põlme et al. 2020); Decay sugar-rich substrates (Põlme et al. 2020) | Põlme S, et al. 2020. Fungal Diversity 105:1-16 ((https://doi.org/10.1007/s13225-020-00466-2)) |
| Otu45079 | 1 | k__Fungi(100);p__Ascomycota(100);c__Saccharomycetes(100);o__Saccharomycetales(100);f__Metschnikowiaceae(100);g__Metschnikowia(100);g__Metschnikowia_unclassified(100); | Metschnikowia | 13 | Saprotroph | \|Undefined Saprotroph\| | Yeast | NULL | Probable | As nectar-tap_saprotroph (Põlme et al. 2020); Decay sugar-rich substrates (Põlme et al. 2020) | Põlme S, et al. 2020. Fungal Diversity 105:1-16 ((https://doi.org/10.1007/s13225-020-00466-2)) |
| Otu45080 | 1 | k__Fungi(100);p__Ascomycota(100);c__Saccharomycetes(100);o__Saccharomycetales(100);f__Metschnikowiaceae(100);g__Metschnikowia(100);s__Metschnikowia_pulcherrima(100); | Metschnikowia | 13 | Saprotroph | \|Undefined Saprotroph\| | Yeast | NULL | Probable | As nectar-tap_saprotroph (Põlme et al. 2020); Decay sugar-rich substrates (Põlme et al. 2020) | Põlme S, et al. 2020. Fungal Diversity 105:1-16 ((https://doi.org/10.1007/s13225-020-00466-2)) |
| Otu45081 | 1 | k__Fungi(100);p__Ascomycota(100);c__Saccharomycetes(100);o__Saccharomycetales(100);f__Metschnikowiaceae(100);g__Metschnikowia(100);s__Metschnikowia_pulcherrima(100); | Metschnikowia | 13 | Saprotroph | \|Undefined Saprotroph\| | Yeast | NULL | Probable | As nectar-tap_saprotroph (Põlme et al. 2020); Decay sugar-rich substrates (Põlme et al. 2020) | Põlme S, et al. 2020. Fungal Diversity 105:1-16 ((https://doi.org/10.1007/s13225-020-00466-2)) |
| Otu45082 | 1 | k__Fungi(100);p__Ascomycota(100);c__Saccharomycetes(100);o__Saccharomycetales(100);f__Metschnikowiaceae(100);g__Metschnikowia(100);s__Metschnikowia_pulcherrima(100); | Metschnikowia | 13 | Saprotroph | \|Undefined Saprotroph\| | Yeast | NULL | Probable | As nectar-tap_saprotroph (Põlme et al. 2020); Decay sugar-rich substrates (Põlme et al. 2020) | Põlme S, et al. 2020. Fungal Diversity 105:1-16 ((https://doi.org/10.1007/s13225-020-00466-2)) |
| Otu45083 | 1 | k__Fungi(100);p__Ascomycota(100);c__Saccharomycetes(100);o__Saccharomycetales(100);f__Metschnikowiaceae(100);g__Metschnikowia(100);s__Metschnikowia_pulcherrima(100); | Metschnikowia | 13 | Saprotroph | \|Undefined Saprotroph\| | Yeast | NULL | Probable | As nectar-tap_saprotroph (Põlme et al. 2020); Decay sugar-rich substrates (Põlme et al. 2020) | Põlme S, et al. 2020. Fungal Diversity 105:1-16 ((https://doi.org/10.1007/s13225-020-00466-2)) |
| Otu45084 | 1 | k__Fungi(100);p__Ascomycota(100);c__Saccharomycetes(100);o__Saccharomycetales(100);f__Metschnikowiaceae(100);g__Metschnikowia(100);s__Metschnikowia_pulcherrima(100); | Metschnikowia | 13 | Saprotroph | \|Undefined Saprotroph\| | Yeast | NULL | Probable | As nectar-tap_saprotroph (Põlme et al. 2020); Decay sugar-rich substrates (Põlme et al. 2020) | Põlme S, et al. 2020. Fungal Diversity 105:1-16 ((https://doi.org/10.1007/s13225-020-00466-2)) |
| Otu45085 | 1 | k__Fungi(100);p__Ascomycota(100);c__Saccharomycetes(100);o__Saccharomycetales(100);f__Metschnikowiaceae(100);g__Metschnikowia(100);s__Metschnikowia_pulcherrima(100); | Metschnikowia | 13 | Saprotroph | \|Undefined Saprotroph\| | Yeast | NULL | Probable | As nectar-tap_saprotroph (Põlme et al. 2020); Decay sugar-rich substrates (Põlme et al. 2020) | Põlme S, et al. 2020. Fungal Diversity 105:1-16 ((https://doi.org/10.1007/s13225-020-00466-2)) |
| Otu45086 | 1 | k__Fungi(100);p__Ascomycota(100);c__Saccharomycetes(100);o__Saccharomycetales(100);f__Metschnikowiaceae(100);g__Metschnikowia(100);g__Metschnikowia_unclassified(100); | Metschnikowia | 13 | Saprotroph | \|Undefined Saprotroph\| | Yeast | NULL | Probable | As nectar-tap_saprotroph (Põlme et al. 2020); Decay sugar-rich substrates (Põlme et al. 2020) | Põlme S, et al. 2020. Fungal Diversity 105:1-16 ((https://doi.org/10.1007/s13225-020-00466-2)) |
| Otu45087 | 1 | k__Fungi(100);p__Ascomycota(100);c__Saccharomycetes(100);o__Saccharomycetales(100);f__Metschnikowiaceae(100);g__Metschnikowia(100);s__Metschnikowia_pulcherrima(100); | Metschnikowia | 13 | Saprotroph | \|Undefined Saprotroph\| | Yeast | NULL | Probable | As nectar-tap_saprotroph (Põlme et al. 2020); Decay sugar-rich substrates (Põlme et al. 2020) | Põlme S, et al. 2020. Fungal Diversity 105:1-16 ((https://doi.org/10.1007/s13225-020-00466-2)) |
| Otu45088 | 1 | k__Fungi(100);p__Ascomycota(100);c__Saccharomycetes(100);o__Saccharomycetales(100);f__Saccharomycetaceae(100);g__Kazachstania(100);s__Kazachstania_intestinalis(100); | Kazachstania | 13 | Saprotroph | \|Undefined Saprotroph\| | Yeast | NULL | Probable | As nectar-tap_saprotroph (Põlme et al. 2020); Decay sugar-rich substrates (Põlme et al. 2020) | Põlme S, et al. 2020. Fungal Diversity 105:1-16 ((https://doi.org/10.1007/s13225-020-00466-2)) |
| Otu45089 | 1 | k__Fungi(100);p__Ascomycota(100);c__Saccharomycetes(100);o__Saccharomycetales(100);f__Metschnikowiaceae(100);g__Metschnikowia(100);g__Metschnikowia_unclassified(100); | Metschnikowia | 13 | Saprotroph | \|Undefined Saprotroph\| | Yeast | NULL | Probable | As nectar-tap_saprotroph (Põlme et al. 2020); Decay sugar-rich substrates (Põlme et al. 2020) | Põlme S, et al. 2020. Fungal Diversity 105:1-16 ((https://doi.org/10.1007/s13225-020-00466-2)) |
| Otu45090 | 1 | k__Fungi(100);p__Ascomycota(100);c__Saccharomycetes(100);o__Saccharomycetales(100);f__Metschnikowiaceae(100);g__Metschnikowia(100);s__Metschnikowia_pulcherrima(100); | Metschnikowia | 13 | Saprotroph | \|Undefined Saprotroph\| | Yeast | NULL | Probable | As nectar-tap_saprotroph (Põlme et al. 2020); Decay sugar-rich substrates (Põlme et al. 2020) | Põlme S, et al. 2020. Fungal Diversity 105:1-16 ((https://doi.org/10.1007/s13225-020-00466-2)) |
| Otu45091 | 1 | k__Fungi(100);p__Ascomycota(100);c__Saccharomycetes(100);o__Saccharomycetales(100);f__Metschnikowiaceae(100);g__Metschnikowia(100);s__Metschnikowia_pulcherrima(100); | Metschnikowia | 13 | Saprotroph | \|Undefined Saprotroph\| | Yeast | NULL | Probable | As nectar-tap_saprotroph (Põlme et al. 2020); Decay sugar-rich substrates (Põlme et al. 2020) | Põlme S, et al. 2020. Fungal Diversity 105:1-16 ((https://doi.org/10.1007/s13225-020-00466-2)) |
| Otu45092 | 1 | k__Fungi(100);p__Ascomycota(100);c__Saccharomycetes(100);o__Saccharomycetales(100);f__Metschnikowiaceae(100);g__Metschnikowia(100);s__Metschnikowia_pulcherrima(100); | Metschnikowia | 13 | Saprotroph | \|Undefined Saprotroph\| | Yeast | NULL | Probable | As nectar-tap_saprotroph (Põlme et al. 2020); Decay sugar-rich substrates (Põlme et al. 2020) | Põlme S, et al. 2020. Fungal Diversity 105:1-16 ((https://doi.org/10.1007/s13225-020-00466-2)) |
| Otu45093 | 1 | k__Fungi(100);p__Ascomycota(100);c__Saccharomycetes(100);o__Saccharomycetales(100);f__Metschnikowiaceae(100);g__Metschnikowia(100);s__Metschnikowia_pulcherrima(100); | Metschnikowia | 13 | Saprotroph | \|Undefined Saprotroph\| | Yeast | NULL | Probable | As nectar-tap_saprotroph (Põlme et al. 2020); Decay sugar-rich substrates (Põlme et al. 2020) | Põlme S, et al. 2020. Fungal Diversity 105:1-16 ((https://doi.org/10.1007/s13225-020-00466-2)) |
| Otu45094 | 1 | k__Fungi(100);p__Ascomycota(100);c__Saccharomycetes(100);o__Saccharomycetales(100);f__Metschnikowiaceae(100);g__Metschnikowia(100);s__Metschnikowia_pulcherrima(100); | Metschnikowia | 13 | Saprotroph | \|Undefined Saprotroph\| | Yeast | NULL | Probable | As nectar-tap_saprotroph (Põlme et al. 2020); Decay sugar-rich substrates (Põlme et al. 2020) | Põlme S, et al. 2020. Fungal Diversity 105:1-16 ((https://doi.org/10.1007/s13225-020-00466-2)) |
| Otu45095 | 1 | k__Fungi(100);p__Ascomycota(100);c__Saccharomycetes(100);o__Saccharomycetales(100);f__Metschnikowiaceae(100);g__Metschnikowia(100);s__Metschnikowia_pulcherrima(100); | Metschnikowia | 13 | Saprotroph | \|Undefined Saprotroph\| | Yeast | NULL | Probable | As nectar-tap_saprotroph (Põlme et al. 2020); Decay sugar-rich substrates (Põlme et al. 2020) | Põlme S, et al. 2020. Fungal Diversity 105:1-16 ((https://doi.org/10.1007/s13225-020-00466-2)) |
| Otu45096 | 1 | k__Fungi(100);p__Ascomycota(100);c__Saccharomycetes(100);o__Saccharomycetales(100);f__Metschnikowiaceae(100);g__Metschnikowia(100);g__Metschnikowia_unclassified(100); | Metschnikowia | 13 | Saprotroph | \|Undefined Saprotroph\| | Yeast | NULL | Probable | As nectar-tap_saprotroph (Põlme et al. 2020); Decay sugar-rich substrates (Põlme et al. 2020) | Põlme S, et al. 2020. Fungal Diversity 105:1-16 ((https://doi.org/10.1007/s13225-020-00466-2)) |
| Otu45097 | 1 | k__Fungi(100);p__Ascomycota(100);c__Saccharomycetes(100);o__Saccharomycetales(100);f__Metschnikowiaceae(100);g__Metschnikowia(100);s__Metschnikowia_pulcherrima(100); | Metschnikowia | 13 | Saprotroph | \|Undefined Saprotroph\| | Yeast | NULL | Probable | As nectar-tap_saprotroph (Põlme et al. 2020); Decay sugar-rich substrates (Põlme et al. 2020) | Põlme S, et al. 2020. Fungal Diversity 105:1-16 ((https://doi.org/10.1007/s13225-020-00466-2)) |
| Otu45098 | 1 | k__Fungi(100);p__Ascomycota(100);c__Saccharomycetes(100);o__Saccharomycetales(100);f__Metschnikowiaceae(100);g__Metschnikowia(100);s__Metschnikowia_pulcherrima(100); | Metschnikowia | 13 | Saprotroph | \|Undefined Saprotroph\| | Yeast | NULL | Probable | As nectar-tap_saprotroph (Põlme et al. 2020); Decay sugar-rich substrates (Põlme et al. 2020) | Põlme S, et al. 2020. Fungal Diversity 105:1-16 ((https://doi.org/10.1007/s13225-020-00466-2)) |
| Otu45099 | 1 | k__Fungi(100);p__Ascomycota(100);c__Saccharomycetes(100);o__Saccharomycetales(100);f__Metschnikowiaceae(100);g__Metschnikowia(100);g__Metschnikowia_unclassified(100); | Metschnikowia | 13 | Saprotroph | \|Undefined Saprotroph\| | Yeast | NULL | Probable | As nectar-tap_saprotroph (Põlme et al. 2020); Decay sugar-rich substrates (Põlme et al. 2020) | Põlme S, et al. 2020. Fungal Diversity 105:1-16 ((https://doi.org/10.1007/s13225-020-00466-2)) |
| Otu45100 | 1 | k__Fungi(100);p__Ascomycota(100);c__Saccharomycetes(100);o__Saccharomycetales(100);f__Metschnikowiaceae(100);g__Metschnikowia(100);s__Metschnikowia_pulcherrima(100); | Metschnikowia | 13 | Saprotroph | \|Undefined Saprotroph\| | Yeast | NULL | Probable | As nectar-tap_saprotroph (Põlme et al. 2020); Decay sugar-rich substrates (Põlme et al. 2020) | Põlme S, et al. 2020. Fungal Diversity 105:1-16 ((https://doi.org/10.1007/s13225-020-00466-2)) |
| Otu45101 | 1 | k__Fungi(100);p__Ascomycota(100);c__Saccharomycetes(100);o__Saccharomycetales(100);f__Metschnikowiaceae(100);g__Metschnikowia(100);g__Metschnikowia_unclassified(100); | Metschnikowia | 13 | Saprotroph | \|Undefined Saprotroph\| | Yeast | NULL | Probable | As nectar-tap_saprotroph (Põlme et al. 2020); Decay sugar-rich substrates (Põlme et al. 2020) | Põlme S, et al. 2020. Fungal Diversity 105:1-16 ((https://doi.org/10.1007/s13225-020-00466-2)) |
| Otu45102 | 1 | k__Fungi(100);p__Ascomycota(100);c__Saccharomycetes(100);o__Saccharomycetales(100);f__Metschnikowiaceae(100);g__Metschnikowia(100);g__Metschnikowia_unclassified(100); | Metschnikowia | 13 | Saprotroph | \|Undefined Saprotroph\| | Yeast | NULL | Probable | As nectar-tap_saprotroph (Põlme et al. 2020); Decay sugar-rich substrates (Põlme et al. 2020) | Põlme S, et al. 2020. Fungal Diversity 105:1-16 ((https://doi.org/10.1007/s13225-020-00466-2)) |
| Otu45103 | 1 | k__Fungi(100);p__Ascomycota(100);c__Saccharomycetes(100);o__Saccharomycetales(100);f__Metschnikowiaceae(100);g__Metschnikowia(100);g__Metschnikowia_unclassified(100); | Metschnikowia | 13 | Saprotroph | \|Undefined Saprotroph\| | Yeast | NULL | Probable | As nectar-tap_saprotroph (Põlme et al. 2020); Decay sugar-rich substrates (Põlme et al. 2020) | Põlme S, et al. 2020. Fungal Diversity 105:1-16 ((https://doi.org/10.1007/s13225-020-00466-2)) |
| Otu45104 | 1 | k__Fungi(100);p__Ascomycota(100);c__Saccharomycetes(100);o__Saccharomycetales(100);f__Metschnikowiaceae(100);g__Metschnikowia(100);g__Metschnikowia_unclassified(100); | Metschnikowia | 13 | Saprotroph | \|Undefined Saprotroph\| | Yeast | NULL | Probable | As nectar-tap_saprotroph (Põlme et al. 2020); Decay sugar-rich substrates (Põlme et al. 2020) | Põlme S, et al. 2020. Fungal Diversity 105:1-16 ((https://doi.org/10.1007/s13225-020-00466-2)) |
| Otu45105 | 1 | k__Fungi(100);p__Ascomycota(100);c__Saccharomycetes(100);o__Saccharomycetales(100);f__Metschnikowiaceae(100);g__Metschnikowia(100);s__Metschnikowia_pulcherrima(100); | Metschnikowia | 13 | Saprotroph | \|Undefined Saprotroph\| | Yeast | NULL | Probable | As nectar-tap_saprotroph (Põlme et al. 2020); Decay sugar-rich substrates (Põlme et al. 2020) | Põlme S, et al. 2020. Fungal Diversity 105:1-16 ((https://doi.org/10.1007/s13225-020-00466-2)) |
| Otu45106 | 1 | k__Fungi(100);p__Ascomycota(100);c__Saccharomycetes(100);o__Saccharomycetales(100);f__Metschnikowiaceae(100);g__Metschnikowia(100);g__Metschnikowia_unclassified(100); | Metschnikowia | 13 | Saprotroph | \|Undefined Saprotroph\| | Yeast | NULL | Probable | As nectar-tap_saprotroph (Põlme et al. 2020); Decay sugar-rich substrates (Põlme et al. 2020) | Põlme S, et al. 2020. Fungal Diversity 105:1-16 ((https://doi.org/10.1007/s13225-020-00466-2)) |
| Otu45107 | 1 | k__Fungi(100);p__Ascomycota(100);c__Saccharomycetes(100);o__Saccharomycetales(100);f__Metschnikowiaceae(100);g__Metschnikowia(100);s__Metschnikowia_pulcherrima(100); | Metschnikowia | 13 | Saprotroph | \|Undefined Saprotroph\| | Yeast | NULL | Probable | As nectar-tap_saprotroph (Põlme et al. 2020); Decay sugar-rich substrates (Põlme et al. 2020) | Põlme S, et al. 2020. Fungal Diversity 105:1-16 ((https://doi.org/10.1007/s13225-020-00466-2)) |
| Otu45108 | 1 | k__Fungi(100);p__Ascomycota(100);c__Saccharomycetes(100);o__Saccharomycetales(100);f__Metschnikowiaceae(100);g__Metschnikowia(100);s__Metschnikowia_pulcherrima(100); | Metschnikowia | 13 | Saprotroph | \|Undefined Saprotroph\| | Yeast | NULL | Probable | As nectar-tap_saprotroph (Põlme et al. 2020); Decay sugar-rich substrates (Põlme et al. 2020) | Põlme S, et al. 2020. Fungal Diversity 105:1-16 ((https://doi.org/10.1007/s13225-020-00466-2)) |
| Otu45109 | 1 | k__Fungi(100);p__Ascomycota(100);c__Saccharomycetes(100);o__Saccharomycetales(100);f__Metschnikowiaceae(100);g__Metschnikowia(100);g__Metschnikowia_unclassified(100); | Metschnikowia | 13 | Saprotroph | \|Undefined Saprotroph\| | Yeast | NULL | Probable | As nectar-tap_saprotroph (Põlme et al. 2020); Decay sugar-rich substrates (Põlme et al. 2020) | Põlme S, et al. 2020. Fungal Diversity 105:1-16 ((https://doi.org/10.1007/s13225-020-00466-2)) |
| Otu45110 | 1 | k__Fungi(100);p__Ascomycota(100);c__Saccharomycetes(100);o__Saccharomycetales(100);f__Metschnikowiaceae(100);g__Metschnikowia(100);g__Metschnikowia_unclassified(100); | Metschnikowia | 13 | Saprotroph | \|Undefined Saprotroph\| | Yeast | NULL | Probable | As nectar-tap_saprotroph (Põlme et al. 2020); Decay sugar-rich substrates (Põlme et al. 2020) | Põlme S, et al. 2020. Fungal Diversity 105:1-16 ((https://doi.org/10.1007/s13225-020-00466-2)) |
| Otu45111 | 1 | k__Fungi(100);p__Ascomycota(100);c__Saccharomycetes(100);o__Saccharomycetales(100);f__Metschnikowiaceae(100);g__Metschnikowia(100);s__Metschnikowia_pulcherrima(100); | Metschnikowia | 13 | Saprotroph | \|Undefined Saprotroph\| | Yeast | NULL | Probable | As nectar-tap_saprotroph (Põlme et al. 2020); Decay sugar-rich substrates (Põlme et al. 2020) | Põlme S, et al. 2020. Fungal Diversity 105:1-16 ((https://doi.org/10.1007/s13225-020-00466-2)) |
| Otu45112 | 1 | k__Fungi(100);p__Ascomycota(100);c__Saccharomycetes(100);o__Saccharomycetales(100);f__Metschnikowiaceae(100);g__Metschnikowia(100);s__Metschnikowia_pulcherrima(100); | Metschnikowia | 13 | Saprotroph | \|Undefined Saprotroph\| | Yeast | NULL | Probable | As nectar-tap_saprotroph (Põlme et al. 2020); Decay sugar-rich substrates (Põlme et al. 2020) | Põlme S, et al. 2020. Fungal Diversity 105:1-16 ((https://doi.org/10.1007/s13225-020-00466-2)) |
| Otu45113 | 1 | k__Fungi(100);p__Ascomycota(100);c__Saccharomycetes(100);o__Saccharomycetales(100);f__Metschnikowiaceae(100);g__Metschnikowia(100);s__Metschnikowia_pulcherrima(100); | Metschnikowia | 13 | Saprotroph | \|Undefined Saprotroph\| | Yeast | NULL | Probable | As nectar-tap_saprotroph (Põlme et al. 2020); Decay sugar-rich substrates (Põlme et al. 2020) | Põlme S, et al. 2020. Fungal Diversity 105:1-16 ((https://doi.org/10.1007/s13225-020-00466-2)) |
| Otu45114 | 1 | k__Fungi(100);p__Ascomycota(100);c__Saccharomycetes(100);o__Saccharomycetales(100);f__Metschnikowiaceae(100);g__Metschnikowia(100);s__Metschnikowia_pulcherrima(100); | Metschnikowia | 13 | Saprotroph | \|Undefined Saprotroph\| | Yeast | NULL | Probable | As nectar-tap_saprotroph (Põlme et al. 2020); Decay sugar-rich substrates (Põlme et al. 2020) | Põlme S, et al. 2020. Fungal Diversity 105:1-16 ((https://doi.org/10.1007/s13225-020-00466-2)) |
| Otu45115 | 1 | k__Fungi(100);p__Ascomycota(100);c__Saccharomycetes(100);o__Saccharomycetales(100);f__Metschnikowiaceae(100);g__Metschnikowia(100);s__Metschnikowia_pulcherrima(100); | Metschnikowia | 13 | Saprotroph | \|Undefined Saprotroph\| | Yeast | NULL | Probable | As nectar-tap_saprotroph (Põlme et al. 2020); Decay sugar-rich substrates (Põlme et al. 2020) | Põlme S, et al. 2020. Fungal Diversity 105:1-16 ((https://doi.org/10.1007/s13225-020-00466-2)) |
| Otu45116 | 1 | k__Fungi(100);p__Ascomycota(100);c__Saccharomycetes(100);o__Saccharomycetales(100);f__Metschnikowiaceae(100);g__Metschnikowia(100);s__Metschnikowia_pulcherrima(100); | Metschnikowia | 13 | Saprotroph | \|Undefined Saprotroph\| | Yeast | NULL | Probable | As nectar-tap_saprotroph (Põlme et al. 2020); Decay sugar-rich substrates (Põlme et al. 2020) | Põlme S, et al. 2020. Fungal Diversity 105:1-16 ((https://doi.org/10.1007/s13225-020-00466-2)) |
| Otu45117 | 1 | k__Fungi(100);p__Ascomycota(100);c__Saccharomycetes(100);o__Saccharomycetales(100);f__Metschnikowiaceae(100);g__Metschnikowia(100);g__Metschnikowia_unclassified(100); | Metschnikowia | 13 | Saprotroph | \|Undefined Saprotroph\| | Yeast | NULL | Probable | As nectar-tap_saprotroph (Põlme et al. 2020); Decay sugar-rich substrates (Põlme et al. 2020) | Põlme S, et al. 2020. Fungal Diversity 105:1-16 ((https://doi.org/10.1007/s13225-020-00466-2)) |
| Otu45118 | 1 | k__Fungi(100);p__Ascomycota(100);c__Saccharomycetes(100);o__Saccharomycetales(100);f__Metschnikowiaceae(100);g__Metschnikowia(100);s__Metschnikowia_pulcherrima(100); | Metschnikowia | 13 | Saprotroph | \|Undefined Saprotroph\| | Yeast | NULL | Probable | As nectar-tap_saprotroph (Põlme et al. 2020); Decay sugar-rich substrates (Põlme et al. 2020) | Põlme S, et al. 2020. Fungal Diversity 105:1-16 ((https://doi.org/10.1007/s13225-020-00466-2)) |
| Otu45119 | 1 | k__Fungi(100);p__Ascomycota(100);c__Saccharomycetes(100);o__Saccharomycetales(100);f__Metschnikowiaceae(100);g__Metschnikowia(100);s__Metschnikowia_pulcherrima(100); | Metschnikowia | 13 | Saprotroph | \|Undefined Saprotroph\| | Yeast | NULL | Probable | As nectar-tap_saprotroph (Põlme et al. 2020); Decay sugar-rich substrates (Põlme et al. 2020) | Põlme S, et al. 2020. Fungal Diversity 105:1-16 ((https://doi.org/10.1007/s13225-020-00466-2)) |
| Otu45120 | 1 | k__Fungi(100);p__Ascomycota(100);c__Saccharomycetes(100);o__Saccharomycetales(100);f__Metschnikowiaceae(100);g__Metschnikowia(100);s__Metschnikowia_pulcherrima(100); | Metschnikowia | 13 | Saprotroph | \|Undefined Saprotroph\| | Yeast | NULL | Probable | As nectar-tap_saprotroph (Põlme et al. 2020); Decay sugar-rich substrates (Põlme et al. 2020) | Põlme S, et al. 2020. Fungal Diversity 105:1-16 ((https://doi.org/10.1007/s13225-020-00466-2)) |
| Otu45121 | 1 | k__Fungi(100);p__Ascomycota(100);c__Saccharomycetes(100);o__Saccharomycetales(100);f__Metschnikowiaceae(100);g__Metschnikowia(100);s__Metschnikowia_pulcherrima(100); | Metschnikowia | 13 | Saprotroph | \|Undefined Saprotroph\| | Yeast | NULL | Probable | As nectar-tap_saprotroph (Põlme et al. 2020); Decay sugar-rich substrates (Põlme et al. 2020) | Põlme S, et al. 2020. Fungal Diversity 105:1-16 ((https://doi.org/10.1007/s13225-020-00466-2)) |
| Otu45122 | 1 | k__Fungi(100);p__Ascomycota(100);c__Saccharomycetes(100);o__Saccharomycetales(100);f__Metschnikowiaceae(100);g__Metschnikowia(100);s__Metschnikowia_pulcherrima(100); | Metschnikowia | 13 | Saprotroph | \|Undefined Saprotroph\| | Yeast | NULL | Probable | As nectar-tap_saprotroph (Põlme et al. 2020); Decay sugar-rich substrates (Põlme et al. 2020) | Põlme S, et al. 2020. Fungal Diversity 105:1-16 ((https://doi.org/10.1007/s13225-020-00466-2)) |
| Otu45123 | 1 | k__Fungi(100);p__Ascomycota(100);c__Saccharomycetes(100);o__Saccharomycetales(100);f__Metschnikowiaceae(100);g__Metschnikowia(100);g__Metschnikowia_unclassified(100); | Metschnikowia | 13 | Saprotroph | \|Undefined Saprotroph\| | Yeast | NULL | Probable | As nectar-tap_saprotroph (Põlme et al. 2020); Decay sugar-rich substrates (Põlme et al. 2020) | Põlme S, et al. 2020. Fungal Diversity 105:1-16 ((https://doi.org/10.1007/s13225-020-00466-2)) |
| Otu45124 | 1 | k__Fungi(100);p__Ascomycota(100);c__Saccharomycetes(100);o__Saccharomycetales(100);f__Metschnikowiaceae(100);g__Metschnikowia(100);s__Metschnikowia_pulcherrima(100); | Metschnikowia | 13 | Saprotroph | \|Undefined Saprotroph\| | Yeast | NULL | Probable | As nectar-tap_saprotroph (Põlme et al. 2020); Decay sugar-rich substrates (Põlme et al. 2020) | Põlme S, et al. 2020. Fungal Diversity 105:1-16 ((https://doi.org/10.1007/s13225-020-00466-2)) |
| Otu45125 | 1 | k__Fungi(100);p__Ascomycota(100);c__Saccharomycetes(100);o__Saccharomycetales(100);f__Metschnikowiaceae(100);g__Metschnikowia(100);s__Metschnikowia_pulcherrima(100); | Metschnikowia | 13 | Saprotroph | \|Undefined Saprotroph\| | Yeast | NULL | Probable | As nectar-tap_saprotroph (Põlme et al. 2020); Decay sugar-rich substrates (Põlme et al. 2020) | Põlme S, et al. 2020. Fungal Diversity 105:1-16 ((https://doi.org/10.1007/s13225-020-00466-2)) |
| Otu45126 | 1 | k__Fungi(100);p__Ascomycota(100);c__Saccharomycetes(100);o__Saccharomycetales(100);f__Metschnikowiaceae(100);g__Metschnikowia(100);s__Metschnikowia_pulcherrima(100); | Metschnikowia | 13 | Saprotroph | \|Undefined Saprotroph\| | Yeast | NULL | Probable | As nectar-tap_saprotroph (Põlme et al. 2020); Decay sugar-rich substrates (Põlme et al. 2020) | Põlme S, et al. 2020. Fungal Diversity 105:1-16 ((https://doi.org/10.1007/s13225-020-00466-2)) |
| Otu45127 | 1 | k__Fungi(100);p__Ascomycota(100);c__Saccharomycetes(100);o__Saccharomycetales(100);f__Metschnikowiaceae(100);g__Metschnikowia(100);s__Metschnikowia_pulcherrima(100); | Metschnikowia | 13 | Saprotroph | \|Undefined Saprotroph\| | Yeast | NULL | Probable | As nectar-tap_saprotroph (Põlme et al. 2020); Decay sugar-rich substrates (Põlme et al. 2020) | Põlme S, et al. 2020. Fungal Diversity 105:1-16 ((https://doi.org/10.1007/s13225-020-00466-2)) |
| Otu45128 | 1 | k__Fungi(100);p__Ascomycota(100);c__Saccharomycetes(100);o__Saccharomycetales(100);f__Metschnikowiaceae(100);g__Metschnikowia(100);s__Metschnikowia_pulcherrima(100); | Metschnikowia | 13 | Saprotroph | \|Undefined Saprotroph\| | Yeast | NULL | Probable | As nectar-tap_saprotroph (Põlme et al. 2020); Decay sugar-rich substrates (Põlme et al. 2020) | Põlme S, et al. 2020. Fungal Diversity 105:1-16 ((https://doi.org/10.1007/s13225-020-00466-2)) |
| Otu45129 | 1 | k__Fungi(100);p__Ascomycota(100);c__Saccharomycetes(100);o__Saccharomycetales(100);f__Metschnikowiaceae(100);g__Metschnikowia(100);s__Metschnikowia_pulcherrima(100); | Metschnikowia | 13 | Saprotroph | \|Undefined Saprotroph\| | Yeast | NULL | Probable | As nectar-tap_saprotroph (Põlme et al. 2020); Decay sugar-rich substrates (Põlme et al. 2020) | Põlme S, et al. 2020. Fungal Diversity 105:1-16 ((https://doi.org/10.1007/s13225-020-00466-2)) |
| Otu45130 | 1 | k__Fungi(100);p__Ascomycota(100);c__Saccharomycetes(100);o__Saccharomycetales(100);f__Metschnikowiaceae(100);g__Metschnikowia(100);s__Metschnikowia_pulcherrima(100); | Metschnikowia | 13 | Saprotroph | \|Undefined Saprotroph\| | Yeast | NULL | Probable | As nectar-tap_saprotroph (Põlme et al. 2020); Decay sugar-rich substrates (Põlme et al. 2020) | Põlme S, et al. 2020. Fungal Diversity 105:1-16 ((https://doi.org/10.1007/s13225-020-00466-2)) |
| Otu45131 | 1 | k__Fungi(100);p__Ascomycota(100);c__Saccharomycetes(100);o__Saccharomycetales(100);f__Metschnikowiaceae(100);g__Metschnikowia(100);s__Metschnikowia_pulcherrima(100); | Metschnikowia | 13 | Saprotroph | \|Undefined Saprotroph\| | Yeast | NULL | Probable | As nectar-tap_saprotroph (Põlme et al. 2020); Decay sugar-rich substrates (Põlme et al. 2020) | Põlme S, et al. 2020. Fungal Diversity 105:1-16 ((https://doi.org/10.1007/s13225-020-00466-2)) |
| Otu45132 | 1 | k__Fungi(100);p__Ascomycota(100);c__Saccharomycetes(100);o__Saccharomycetales(100);f__Metschnikowiaceae(100);g__Metschnikowia(100);s__Metschnikowia_pulcherrima(100); | Metschnikowia | 13 | Saprotroph | \|Undefined Saprotroph\| | Yeast | NULL | Probable | As nectar-tap_saprotroph (Põlme et al. 2020); Decay sugar-rich substrates (Põlme et al. 2020) | Põlme S, et al. 2020. Fungal Diversity 105:1-16 ((https://doi.org/10.1007/s13225-020-00466-2)) |
| Otu45133 | 1 | k__Fungi(100);p__Ascomycota(100);c__Saccharomycetes(100);o__Saccharomycetales(100);f__Metschnikowiaceae(100);g__Metschnikowia(100);g__Metschnikowia_unclassified(100); | Metschnikowia | 13 | Saprotroph | \|Undefined Saprotroph\| | Yeast | NULL | Probable | As nectar-tap_saprotroph (Põlme et al. 2020); Decay sugar-rich substrates (Põlme et al. 2020) | Põlme S, et al. 2020. Fungal Diversity 105:1-16 ((https://doi.org/10.1007/s13225-020-00466-2)) |
| Otu45134 | 1 | k__Fungi(100);p__Ascomycota(100);c__Saccharomycetes(100);o__Saccharomycetales(100);f__Metschnikowiaceae(100);g__Metschnikowia(100);g__Metschnikowia_unclassified(100); | Metschnikowia | 13 | Saprotroph | \|Undefined Saprotroph\| | Yeast | NULL | Probable | As nectar-tap_saprotroph (Põlme et al. 2020); Decay sugar-rich substrates (Põlme et al. 2020) | Põlme S, et al. 2020. Fungal Diversity 105:1-16 ((https://doi.org/10.1007/s13225-020-00466-2)) |
| Otu45135 | 1 | k__Fungi(100);p__Ascomycota(100);c__Saccharomycetes(100);o__Saccharomycetales(100);f__Metschnikowiaceae(100);g__Metschnikowia(100);s__Metschnikowia_pulcherrima(100); | Metschnikowia | 13 | Saprotroph | \|Undefined Saprotroph\| | Yeast | NULL | Probable | As nectar-tap_saprotroph (Põlme et al. 2020); Decay sugar-rich substrates (Põlme et al. 2020) | Põlme S, et al. 2020. Fungal Diversity 105:1-16 ((https://doi.org/10.1007/s13225-020-00466-2)) |
| Otu45136 | 1 | k__Fungi(100);p__Ascomycota(100);c__Saccharomycetes(100);o__Saccharomycetales(100);f__Metschnikowiaceae(100);g__Metschnikowia(100);s__Metschnikowia_pulcherrima(100); | Metschnikowia | 13 | Saprotroph | \|Undefined Saprotroph\| | Yeast | NULL | Probable | As nectar-tap_saprotroph (Põlme et al. 2020); Decay sugar-rich substrates (Põlme et al. 2020) | Põlme S, et al. 2020. Fungal Diversity 105:1-16 ((https://doi.org/10.1007/s13225-020-00466-2)) |
| Otu45137 | 1 | k__Fungi(100);p__Ascomycota(100);c__Saccharomycetes(100);o__Saccharomycetales(100);f__Metschnikowiaceae(100);g__Metschnikowia(100);s__Metschnikowia_pulcherrima(100); | Metschnikowia | 13 | Saprotroph | \|Undefined Saprotroph\| | Yeast | NULL | Probable | As nectar-tap_saprotroph (Põlme et al. 2020); Decay sugar-rich substrates (Põlme et al. 2020) | Põlme S, et al. 2020. Fungal Diversity 105:1-16 ((https://doi.org/10.1007/s13225-020-00466-2)) |
| Otu45138 | 1 | k__Fungi(100);p__Ascomycota(100);c__Saccharomycetes(100);o__Saccharomycetales(100);f__Metschnikowiaceae(100);g__Metschnikowia(100);g__Metschnikowia_unclassified(100); | Metschnikowia | 13 | Saprotroph | \|Undefined Saprotroph\| | Yeast | NULL | Probable | As nectar-tap_saprotroph (Põlme et al. 2020); Decay sugar-rich substrates (Põlme et al. 2020) | Põlme S, et al. 2020. Fungal Diversity 105:1-16 ((https://doi.org/10.1007/s13225-020-00466-2)) |
| Otu45139 | 1 | k__Fungi(100);p__Ascomycota(100);c__Saccharomycetes(100);o__Saccharomycetales(100);f__Metschnikowiaceae(100);g__Metschnikowia(100);s__Metschnikowia_pulcherrima(100); | Metschnikowia | 13 | Saprotroph | \|Undefined Saprotroph\| | Yeast | NULL | Probable | As nectar-tap_saprotroph (Põlme et al. 2020); Decay sugar-rich substrates (Põlme et al. 2020) | Põlme S, et al. 2020. Fungal Diversity 105:1-16 ((https://doi.org/10.1007/s13225-020-00466-2)) |
| Otu45140 | 1 | k__Fungi(100);p__Ascomycota(100);c__Saccharomycetes(100);o__Saccharomycetales(100);f__Metschnikowiaceae(100);g__Metschnikowia(100);s__Metschnikowia_pulcherrima(100); | Metschnikowia | 13 | Saprotroph | \|Undefined Saprotroph\| | Yeast | NULL | Probable | As nectar-tap_saprotroph (Põlme et al. 2020); Decay sugar-rich substrates (Põlme et al. 2020) | Põlme S, et al. 2020. Fungal Diversity 105:1-16 ((https://doi.org/10.1007/s13225-020-00466-2)) |
| Otu45141 | 1 | k__Fungi(100);p__Ascomycota(100);c__Saccharomycetes(100);o__Saccharomycetales(100);f__Metschnikowiaceae(100);g__Metschnikowia(100);s__Metschnikowia_pulcherrima(100); | Metschnikowia | 13 | Saprotroph | \|Undefined Saprotroph\| | Yeast | NULL | Probable | As nectar-tap_saprotroph (Põlme et al. 2020); Decay sugar-rich substrates (Põlme et al. 2020) | Põlme S, et al. 2020. Fungal Diversity 105:1-16 ((https://doi.org/10.1007/s13225-020-00466-2)) |
| Otu45142 | 1 | k__Fungi(100);p__Ascomycota(100);c__Saccharomycetes(100);o__Saccharomycetales(100);f__Metschnikowiaceae(100);g__Metschnikowia(100);s__Metschnikowia_pulcherrima(100); | Metschnikowia | 13 | Saprotroph | \|Undefined Saprotroph\| | Yeast | NULL | Probable | As nectar-tap_saprotroph (Põlme et al. 2020); Decay sugar-rich substrates (Põlme et al. 2020) | Põlme S, et al. 2020. Fungal Diversity 105:1-16 ((https://doi.org/10.1007/s13225-020-00466-2)) |
| Otu45143 | 1 | k__Fungi(100);p__Ascomycota(100);c__Saccharomycetes(100);o__Saccharomycetales(100);f__Metschnikowiaceae(100);g__Metschnikowia(100);g__Metschnikowia_unclassified(100); | Metschnikowia | 13 | Saprotroph | \|Undefined Saprotroph\| | Yeast | NULL | Probable | As nectar-tap_saprotroph (Põlme et al. 2020); Decay sugar-rich substrates (Põlme et al. 2020) | Põlme S, et al. 2020. Fungal Diversity 105:1-16 ((https://doi.org/10.1007/s13225-020-00466-2)) |
| Otu45144 | 1 | k__Fungi(100);p__Ascomycota(100);c__Saccharomycetes(100);o__Saccharomycetales(100);f__Metschnikowiaceae(100);g__Metschnikowia(100);s__Metschnikowia_pulcherrima(100); | Metschnikowia | 13 | Saprotroph | \|Undefined Saprotroph\| | Yeast | NULL | Probable | As nectar-tap_saprotroph (Põlme et al. 2020); Decay sugar-rich substrates (Põlme et al. 2020) | Põlme S, et al. 2020. Fungal Diversity 105:1-16 ((https://doi.org/10.1007/s13225-020-00466-2)) |
| Otu45145 | 1 | k__Fungi(100);p__Ascomycota(100);c__Saccharomycetes(100);o__Saccharomycetales(100);f__Metschnikowiaceae(100);g__Metschnikowia(100);s__Metschnikowia_pulcherrima(100); | Metschnikowia | 13 | Saprotroph | \|Undefined Saprotroph\| | Yeast | NULL | Probable | As nectar-tap_saprotroph (Põlme et al. 2020); Decay sugar-rich substrates (Põlme et al. 2020) | Põlme S, et al. 2020. Fungal Diversity 105:1-16 ((https://doi.org/10.1007/s13225-020-00466-2)) |
| Otu45146 | 1 | k__Fungi(100);p__Ascomycota(100);c__Saccharomycetes(100);o__Saccharomycetales(100);f__Metschnikowiaceae(100);g__Metschnikowia(100);s__Metschnikowia_pulcherrima(100); | Metschnikowia | 13 | Saprotroph | \|Undefined Saprotroph\| | Yeast | NULL | Probable | As nectar-tap_saprotroph (Põlme et al. 2020); Decay sugar-rich substrates (Põlme et al. 2020) | Põlme S, et al. 2020. Fungal Diversity 105:1-16 ((https://doi.org/10.1007/s13225-020-00466-2)) |
| Otu45147 | 1 | k__Fungi(100);p__Ascomycota(100);c__Saccharomycetes(100);o__Saccharomycetales(100);f__Metschnikowiaceae(100);g__Metschnikowia(100);g__Metschnikowia_unclassified(100); | Metschnikowia | 13 | Saprotroph | \|Undefined Saprotroph\| | Yeast | NULL | Probable | As nectar-tap_saprotroph (Põlme et al. 2020); Decay sugar-rich substrates (Põlme et al. 2020) | Põlme S, et al. 2020. Fungal Diversity 105:1-16 ((https://doi.org/10.1007/s13225-020-00466-2)) |
| Otu45148 | 1 | k__Fungi(100);p__Ascomycota(100);c__Saccharomycetes(100);o__Saccharomycetales(100);f__Metschnikowiaceae(100);g__Metschnikowia(100);s__Metschnikowia_pulcherrima(100); | Metschnikowia | 13 | Saprotroph | \|Undefined Saprotroph\| | Yeast | NULL | Probable | As nectar-tap_saprotroph (Põlme et al. 2020); Decay sugar-rich substrates (Põlme et al. 2020) | Põlme S, et al. 2020. Fungal Diversity 105:1-16 ((https://doi.org/10.1007/s13225-020-00466-2)) |
| Otu45149 | 1 | k__Fungi(100);p__Ascomycota(100);c__Saccharomycetes(100);o__Saccharomycetales(100);f__Metschnikowiaceae(100);g__Metschnikowia(100);s__Metschnikowia_pulcherrima(100); | Metschnikowia | 13 | Saprotroph | \|Undefined Saprotroph\| | Yeast | NULL | Probable | As nectar-tap_saprotroph (Põlme et al. 2020); Decay sugar-rich substrates (Põlme et al. 2020) | Põlme S, et al. 2020. Fungal Diversity 105:1-16 ((https://doi.org/10.1007/s13225-020-00466-2)) |
| Otu45150 | 1 | k__Fungi(100);p__Ascomycota(100);c__Saccharomycetes(100);o__Saccharomycetales(100);f__Metschnikowiaceae(100);g__Metschnikowia(100);s__Metschnikowia_viticola(100); | Metschnikowia | 13 | Saprotroph | \|Undefined Saprotroph\| | Yeast | NULL | Probable | As nectar-tap_saprotroph (Põlme et al. 2020); Decay sugar-rich substrates (Põlme et al. 2020) | Põlme S, et al. 2020. Fungal Diversity 105:1-16 ((https://doi.org/10.1007/s13225-020-00466-2)) |
| Otu45151 | 1 | k__Fungi(100);p__Ascomycota(100);c__Saccharomycetes(100);o__Saccharomycetales(100);f__Metschnikowiaceae(100);g__Metschnikowia(100);s__Metschnikowia_pulcherrima(100); | Metschnikowia | 13 | Saprotroph | \|Undefined Saprotroph\| | Yeast | NULL | Probable | As nectar-tap_saprotroph (Põlme et al. 2020); Decay sugar-rich substrates (Põlme et al. 2020) | Põlme S, et al. 2020. Fungal Diversity 105:1-16 ((https://doi.org/10.1007/s13225-020-00466-2)) |
| Otu45152 | 1 | k__Fungi(100);p__Ascomycota(100);c__Saccharomycetes(100);o__Saccharomycetales(100);f__Metschnikowiaceae(100);g__Metschnikowia(100);s__Metschnikowia_pulcherrima(100); | Metschnikowia | 13 | Saprotroph | \|Undefined Saprotroph\| | Yeast | NULL | Probable | As nectar-tap_saprotroph (Põlme et al. 2020); Decay sugar-rich substrates (Põlme et al. 2020) | Põlme S, et al. 2020. Fungal Diversity 105:1-16 ((https://doi.org/10.1007/s13225-020-00466-2)) |
| Otu45153 | 1 | k__Fungi(100);p__Ascomycota(100);c__Saccharomycetes(100);o__Saccharomycetales(100);f__Metschnikowiaceae(100);g__Metschnikowia(100);s__Metschnikowia_pulcherrima(100); | Metschnikowia | 13 | Saprotroph | \|Undefined Saprotroph\| | Yeast | NULL | Probable | As nectar-tap_saprotroph (Põlme et al. 2020); Decay sugar-rich substrates (Põlme et al. 2020) | Põlme S, et al. 2020. Fungal Diversity 105:1-16 ((https://doi.org/10.1007/s13225-020-00466-2)) |
| Otu45154 | 1 | k__Fungi(100);p__Ascomycota(100);c__Saccharomycetes(100);o__Saccharomycetales(100);f__Metschnikowiaceae(100);g__Metschnikowia(100);s__Metschnikowia_pulcherrima(100); | Metschnikowia | 13 | Saprotroph | \|Undefined Saprotroph\| | Yeast | NULL | Probable | As nectar-tap_saprotroph (Põlme et al. 2020); Decay sugar-rich substrates (Põlme et al. 2020) | Põlme S, et al. 2020. Fungal Diversity 105:1-16 ((https://doi.org/10.1007/s13225-020-00466-2)) |
| Otu45155 | 1 | k__Fungi(100);p__Ascomycota(100);c__Saccharomycetes(100);o__Saccharomycetales(100);f__Metschnikowiaceae(100);g__Metschnikowia(100);s__Metschnikowia_viticola(100); | Metschnikowia | 13 | Saprotroph | \|Undefined Saprotroph\| | Yeast | NULL | Probable | As nectar-tap_saprotroph (Põlme et al. 2020); Decay sugar-rich substrates (Põlme et al. 2020) | Põlme S, et al. 2020. Fungal Diversity 105:1-16 ((https://doi.org/10.1007/s13225-020-00466-2)) |
| Otu45156 | 1 | k__Fungi(100);p__Ascomycota(100);c__Saccharomycetes(100);o__Saccharomycetales(100);f__Metschnikowiaceae(100);g__Metschnikowia(100);s__Metschnikowia_pulcherrima(100); | Metschnikowia | 13 | Saprotroph | \|Undefined Saprotroph\| | Yeast | NULL | Probable | As nectar-tap_saprotroph (Põlme et al. 2020); Decay sugar-rich substrates (Põlme et al. 2020) | Põlme S, et al. 2020. Fungal Diversity 105:1-16 ((https://doi.org/10.1007/s13225-020-00466-2)) |
| Otu45157 | 1 | k__Fungi(100);p__Ascomycota(100);c__Saccharomycetes(100);o__Saccharomycetales(100);f__Metschnikowiaceae(100);g__Metschnikowia(100);s__Metschnikowia_pulcherrima(100); | Metschnikowia | 13 | Saprotroph | \|Undefined Saprotroph\| | Yeast | NULL | Probable | As nectar-tap_saprotroph (Põlme et al. 2020); Decay sugar-rich substrates (Põlme et al. 2020) | Põlme S, et al. 2020. Fungal Diversity 105:1-16 ((https://doi.org/10.1007/s13225-020-00466-2)) |
| Otu45158 | 1 | k__Fungi(100);p__Ascomycota(100);c__Saccharomycetes(100);o__Saccharomycetales(100);f__Metschnikowiaceae(100);g__Metschnikowia(100);s__Metschnikowia_pulcherrima(100); | Metschnikowia | 13 | Saprotroph | \|Undefined Saprotroph\| | Yeast | NULL | Probable | As nectar-tap_saprotroph (Põlme et al. 2020); Decay sugar-rich substrates (Põlme et al. 2020) | Põlme S, et al. 2020. Fungal Diversity 105:1-16 ((https://doi.org/10.1007/s13225-020-00466-2)) |
| Otu45159 | 1 | k__Fungi(100);p__Ascomycota(100);c__Saccharomycetes(100);o__Saccharomycetales(100);f__Metschnikowiaceae(100);g__Metschnikowia(100);s__Metschnikowia_pulcherrima(100); | Metschnikowia | 13 | Saprotroph | \|Undefined Saprotroph\| | Yeast | NULL | Probable | As nectar-tap_saprotroph (Põlme et al. 2020); Decay sugar-rich substrates (Põlme et al. 2020) | Põlme S, et al. 2020. Fungal Diversity 105:1-16 ((https://doi.org/10.1007/s13225-020-00466-2)) |
| Otu45160 | 1 | k__Fungi(100);p__Ascomycota(100);c__Saccharomycetes(100);o__Saccharomycetales(100);f__Metschnikowiaceae(100);g__Metschnikowia(100);s__Metschnikowia_pulcherrima(100); | Metschnikowia | 13 | Saprotroph | \|Undefined Saprotroph\| | Yeast | NULL | Probable | As nectar-tap_saprotroph (Põlme et al. 2020); Decay sugar-rich substrates (Põlme et al. 2020) | Põlme S, et al. 2020. Fungal Diversity 105:1-16 ((https://doi.org/10.1007/s13225-020-00466-2)) |
| Otu45161 | 1 | k__Fungi(100);p__Ascomycota(100);c__Saccharomycetes(100);o__Saccharomycetales(100);f__Metschnikowiaceae(100);g__Metschnikowia(100);s__Metschnikowia_pulcherrima(100); | Metschnikowia | 13 | Saprotroph | \|Undefined Saprotroph\| | Yeast | NULL | Probable | As nectar-tap_saprotroph (Põlme et al. 2020); Decay sugar-rich substrates (Põlme et al. 2020) | Põlme S, et al. 2020. Fungal Diversity 105:1-16 ((https://doi.org/10.1007/s13225-020-00466-2)) |
| Otu45162 | 1 | k__Fungi(100);p__Ascomycota(100);c__Saccharomycetes(100);o__Saccharomycetales(100);f__Metschnikowiaceae(100);g__Metschnikowia(100);g__Metschnikowia_unclassified(100); | Metschnikowia | 13 | Saprotroph | \|Undefined Saprotroph\| | Yeast | NULL | Probable | As nectar-tap_saprotroph (Põlme et al. 2020); Decay sugar-rich substrates (Põlme et al. 2020) | Põlme S, et al. 2020. Fungal Diversity 105:1-16 ((https://doi.org/10.1007/s13225-020-00466-2)) |
| Otu45163 | 1 | k__Fungi(100);p__Ascomycota(100);c__Saccharomycetes(100);o__Saccharomycetales(100);f__Metschnikowiaceae(100);g__Metschnikowia(100);g__Metschnikowia_unclassified(100); | Metschnikowia | 13 | Saprotroph | \|Undefined Saprotroph\| | Yeast | NULL | Probable | As nectar-tap_saprotroph (Põlme et al. 2020); Decay sugar-rich substrates (Põlme et al. 2020) | Põlme S, et al. 2020. Fungal Diversity 105:1-16 ((https://doi.org/10.1007/s13225-020-00466-2)) |
| Otu45164 | 1 | k__Fungi(100);p__Ascomycota(100);c__Saccharomycetes(100);o__Saccharomycetales(100);f__Metschnikowiaceae(100);g__Metschnikowia(100);s__Metschnikowia_pulcherrima(100); | Metschnikowia | 13 | Saprotroph | \|Undefined Saprotroph\| | Yeast | NULL | Probable | As nectar-tap_saprotroph (Põlme et al. 2020); Decay sugar-rich substrates (Põlme et al. 2020) | Põlme S, et al. 2020. Fungal Diversity 105:1-16 ((https://doi.org/10.1007/s13225-020-00466-2)) |
| Otu45165 | 1 | k__Fungi(100);p__Ascomycota(100);c__Saccharomycetes(100);o__Saccharomycetales(100);f__Metschnikowiaceae(100);g__Metschnikowia(100);s__Metschnikowia_pulcherrima(100); | Metschnikowia | 13 | Saprotroph | \|Undefined Saprotroph\| | Yeast | NULL | Probable | As nectar-tap_saprotroph (Põlme et al. 2020); Decay sugar-rich substrates (Põlme et al. 2020) | Põlme S, et al. 2020. Fungal Diversity 105:1-16 ((https://doi.org/10.1007/s13225-020-00466-2)) |
| Otu45166 | 1 | k__Fungi(100);p__Ascomycota(100);c__Saccharomycetes(100);o__Saccharomycetales(100);f__Metschnikowiaceae(100);g__Metschnikowia(100);g__Metschnikowia_unclassified(100); | Metschnikowia | 13 | Saprotroph | \|Undefined Saprotroph\| | Yeast | NULL | Probable | As nectar-tap_saprotroph (Põlme et al. 2020); Decay sugar-rich substrates (Põlme et al. 2020) | Põlme S, et al. 2020. Fungal Diversity 105:1-16 ((https://doi.org/10.1007/s13225-020-00466-2)) |
| Otu45167 | 1 | k__Fungi(100);p__Ascomycota(100);c__Saccharomycetes(100);o__Saccharomycetales(100);f__Metschnikowiaceae(100);g__Metschnikowia(100);s__Metschnikowia_pulcherrima(100); | Metschnikowia | 13 | Saprotroph | \|Undefined Saprotroph\| | Yeast | NULL | Probable | As nectar-tap_saprotroph (Põlme et al. 2020); Decay sugar-rich substrates (Põlme et al. 2020) | Põlme S, et al. 2020. Fungal Diversity 105:1-16 ((https://doi.org/10.1007/s13225-020-00466-2)) |
| Otu45168 | 1 | k__Fungi(100);p__Ascomycota(100);c__Saccharomycetes(100);o__Saccharomycetales(100);f__Metschnikowiaceae(100);g__Metschnikowia(100);g__Metschnikowia_unclassified(100); | Metschnikowia | 13 | Saprotroph | \|Undefined Saprotroph\| | Yeast | NULL | Probable | As nectar-tap_saprotroph (Põlme et al. 2020); Decay sugar-rich substrates (Põlme et al. 2020) | Põlme S, et al. 2020. Fungal Diversity 105:1-16 ((https://doi.org/10.1007/s13225-020-00466-2)) |
| Otu45169 | 1 | k__Fungi(100);p__Ascomycota(100);c__Saccharomycetes(100);o__Saccharomycetales(100);f__Metschnikowiaceae(100);g__Metschnikowia(100);s__Metschnikowia_pulcherrima(100); | Metschnikowia | 13 | Saprotroph | \|Undefined Saprotroph\| | Yeast | NULL | Probable | As nectar-tap_saprotroph (Põlme et al. 2020); Decay sugar-rich substrates (Põlme et al. 2020) | Põlme S, et al. 2020. Fungal Diversity 105:1-16 ((https://doi.org/10.1007/s13225-020-00466-2)) |
| Otu45170 | 1 | k__Fungi(100);p__Ascomycota(100);c__Saccharomycetes(100);o__Saccharomycetales(100);f__Metschnikowiaceae(100);g__Metschnikowia(100);s__Metschnikowia_pulcherrima(100); | Metschnikowia | 13 | Saprotroph | \|Undefined Saprotroph\| | Yeast | NULL | Probable | As nectar-tap_saprotroph (Põlme et al. 2020); Decay sugar-rich substrates (Põlme et al. 2020) | Põlme S, et al. 2020. Fungal Diversity 105:1-16 ((https://doi.org/10.1007/s13225-020-00466-2)) |
| Otu45171 | 1 | k__Fungi(100);p__Ascomycota(100);c__Saccharomycetes(100);o__Saccharomycetales(100);f__Metschnikowiaceae(100);g__Metschnikowia(100);s__Metschnikowia_pulcherrima(100); | Metschnikowia | 13 | Saprotroph | \|Undefined Saprotroph\| | Yeast | NULL | Probable | As nectar-tap_saprotroph (Põlme et al. 2020); Decay sugar-rich substrates (Põlme et al. 2020) | Põlme S, et al. 2020. Fungal Diversity 105:1-16 ((https://doi.org/10.1007/s13225-020-00466-2)) |
| Otu45172 | 1 | k__Fungi(100);p__Ascomycota(100);c__Saccharomycetes(100);o__Saccharomycetales(100);f__Metschnikowiaceae(100);g__Metschnikowia(100);s__Metschnikowia_viticola(100); | Metschnikowia | 13 | Saprotroph | \|Undefined Saprotroph\| | Yeast | NULL | Probable | As nectar-tap_saprotroph (Põlme et al. 2020); Decay sugar-rich substrates (Põlme et al. 2020) | Põlme S, et al. 2020. Fungal Diversity 105:1-16 ((https://doi.org/10.1007/s13225-020-00466-2)) |
| Otu45173 | 1 | k__Fungi(100);p__Ascomycota(100);c__Saccharomycetes(100);o__Saccharomycetales(100);f__Metschnikowiaceae(100);g__Metschnikowia(100);s__Metschnikowia_pulcherrima(100); | Metschnikowia | 13 | Saprotroph | \|Undefined Saprotroph\| | Yeast | NULL | Probable | As nectar-tap_saprotroph (Põlme et al. 2020); Decay sugar-rich substrates (Põlme et al. 2020) | Põlme S, et al. 2020. Fungal Diversity 105:1-16 ((https://doi.org/10.1007/s13225-020-00466-2)) |
| Otu45174 | 1 | k__Fungi(100);p__Ascomycota(100);c__Saccharomycetes(100);o__Saccharomycetales(100);f__Metschnikowiaceae(100);g__Metschnikowia(100);g__Metschnikowia_unclassified(100); | Metschnikowia | 13 | Saprotroph | \|Undefined Saprotroph\| | Yeast | NULL | Probable | As nectar-tap_saprotroph (Põlme et al. 2020); Decay sugar-rich substrates (Põlme et al. 2020) | Põlme S, et al. 2020. Fungal Diversity 105:1-16 ((https://doi.org/10.1007/s13225-020-00466-2)) |
| Otu45175 | 1 | k__Fungi(100);p__Ascomycota(100);c__Saccharomycetes(100);o__Saccharomycetales(100);f__Metschnikowiaceae(100);g__Metschnikowia(100);s__Metschnikowia_pulcherrima(100); | Metschnikowia | 13 | Saprotroph | \|Undefined Saprotroph\| | Yeast | NULL | Probable | As nectar-tap_saprotroph (Põlme et al. 2020); Decay sugar-rich substrates (Põlme et al. 2020) | Põlme S, et al. 2020. Fungal Diversity 105:1-16 ((https://doi.org/10.1007/s13225-020-00466-2)) |
| Otu45176 | 1 | k__Fungi(100);p__Ascomycota(100);c__Saccharomycetes(100);o__Saccharomycetales(100);f__Metschnikowiaceae(100);g__Metschnikowia(100);s__Metschnikowia_pulcherrima(100); | Metschnikowia | 13 | Saprotroph | \|Undefined Saprotroph\| | Yeast | NULL | Probable | As nectar-tap_saprotroph (Põlme et al. 2020); Decay sugar-rich substrates (Põlme et al. 2020) | Põlme S, et al. 2020. Fungal Diversity 105:1-16 ((https://doi.org/10.1007/s13225-020-00466-2)) |
| Otu45177 | 1 | k__Fungi(100);p__Ascomycota(100);c__Saccharomycetes(100);o__Saccharomycetales(100);f__Metschnikowiaceae(100);g__Metschnikowia(100);s__Metschnikowia_pulcherrima(100); | Metschnikowia | 13 | Saprotroph | \|Undefined Saprotroph\| | Yeast | NULL | Probable | As nectar-tap_saprotroph (Põlme et al. 2020); Decay sugar-rich substrates (Põlme et al. 2020) | Põlme S, et al. 2020. Fungal Diversity 105:1-16 ((https://doi.org/10.1007/s13225-020-00466-2)) |
| Otu45178 | 1 | k__Fungi(100);p__Ascomycota(100);c__Saccharomycetes(100);o__Saccharomycetales(100);f__Metschnikowiaceae(100);g__Metschnikowia(100);s__Metschnikowia_pulcherrima(100); | Metschnikowia | 13 | Saprotroph | \|Undefined Saprotroph\| | Yeast | NULL | Probable | As nectar-tap_saprotroph (Põlme et al. 2020); Decay sugar-rich substrates (Põlme et al. 2020) | Põlme S, et al. 2020. Fungal Diversity 105:1-16 ((https://doi.org/10.1007/s13225-020-00466-2)) |
| Otu45179 | 1 | k__Fungi(100);p__Ascomycota(100);c__Saccharomycetes(100);o__Saccharomycetales(100);f__Metschnikowiaceae(100);g__Metschnikowia(100);g__Metschnikowia_unclassified(100); | Metschnikowia | 13 | Saprotroph | \|Undefined Saprotroph\| | Yeast | NULL | Probable | As nectar-tap_saprotroph (Põlme et al. 2020); Decay sugar-rich substrates (Põlme et al. 2020) | Põlme S, et al. 2020. Fungal Diversity 105:1-16 ((https://doi.org/10.1007/s13225-020-00466-2)) |
| Otu45180 | 1 | k__Fungi(100);p__Ascomycota(100);c__Saccharomycetes(100);o__Saccharomycetales(100);f__Metschnikowiaceae(100);g__Metschnikowia(100);s__Metschnikowia_pulcherrima(100); | Metschnikowia | 13 | Saprotroph | \|Undefined Saprotroph\| | Yeast | NULL | Probable | As nectar-tap_saprotroph (Põlme et al. 2020); Decay sugar-rich substrates (Põlme et al. 2020) | Põlme S, et al. 2020. Fungal Diversity 105:1-16 ((https://doi.org/10.1007/s13225-020-00466-2)) |
| Otu45181 | 1 | k__Fungi(100);p__Ascomycota(100);c__Saccharomycetes(100);o__Saccharomycetales(100);f__Metschnikowiaceae(100);g__Metschnikowia(100);s__Metschnikowia_pulcherrima(100); | Metschnikowia | 13 | Saprotroph | \|Undefined Saprotroph\| | Yeast | NULL | Probable | As nectar-tap_saprotroph (Põlme et al. 2020); Decay sugar-rich substrates (Põlme et al. 2020) | Põlme S, et al. 2020. Fungal Diversity 105:1-16 ((https://doi.org/10.1007/s13225-020-00466-2)) |
| Otu45182 | 1 | k__Fungi(100);p__Ascomycota(100);c__Saccharomycetes(100);o__Saccharomycetales(100);f__Metschnikowiaceae(100);g__Metschnikowia(100);s__Metschnikowia_pulcherrima(100); | Metschnikowia | 13 | Saprotroph | \|Undefined Saprotroph\| | Yeast | NULL | Probable | As nectar-tap_saprotroph (Põlme et al. 2020); Decay sugar-rich substrates (Põlme et al. 2020) | Põlme S, et al. 2020. Fungal Diversity 105:1-16 ((https://doi.org/10.1007/s13225-020-00466-2)) |
| Otu45183 | 1 | k__Fungi(100);p__Ascomycota(100);c__Saccharomycetes(100);o__Saccharomycetales(100);f__Metschnikowiaceae(100);g__Metschnikowia(100);s__Metschnikowia_pulcherrima(100); | Metschnikowia | 13 | Saprotroph | \|Undefined Saprotroph\| | Yeast | NULL | Probable | As nectar-tap_saprotroph (Põlme et al. 2020); Decay sugar-rich substrates (Põlme et al. 2020) | Põlme S, et al. 2020. Fungal Diversity 105:1-16 ((https://doi.org/10.1007/s13225-020-00466-2)) |
| Otu45184 | 1 | k__Fungi(100);p__Ascomycota(100);c__Saccharomycetes(100);o__Saccharomycetales(100);f__Metschnikowiaceae(100);g__Metschnikowia(100);s__Metschnikowia_pulcherrima(100); | Metschnikowia | 13 | Saprotroph | \|Undefined Saprotroph\| | Yeast | NULL | Probable | As nectar-tap_saprotroph (Põlme et al. 2020); Decay sugar-rich substrates (Põlme et al. 2020) | Põlme S, et al. 2020. Fungal Diversity 105:1-16 ((https://doi.org/10.1007/s13225-020-00466-2)) |
| Otu45185 | 1 | k__Fungi(100);p__Ascomycota(100);c__Saccharomycetes(100);o__Saccharomycetales(100);f__Metschnikowiaceae(100);g__Metschnikowia(100);g__Metschnikowia_unclassified(100); | Metschnikowia | 13 | Saprotroph | \|Undefined Saprotroph\| | Yeast | NULL | Probable | As nectar-tap_saprotroph (Põlme et al. 2020); Decay sugar-rich substrates (Põlme et al. 2020) | Põlme S, et al. 2020. Fungal Diversity 105:1-16 ((https://doi.org/10.1007/s13225-020-00466-2)) |
| Otu45186 | 1 | k__Fungi(100);p__Ascomycota(100);c__Saccharomycetes(100);o__Saccharomycetales(100);f__Metschnikowiaceae(100);g__Metschnikowia(100);s__Metschnikowia_pulcherrima(100); | Metschnikowia | 13 | Saprotroph | \|Undefined Saprotroph\| | Yeast | NULL | Probable | As nectar-tap_saprotroph (Põlme et al. 2020); Decay sugar-rich substrates (Põlme et al. 2020) | Põlme S, et al. 2020. Fungal Diversity 105:1-16 ((https://doi.org/10.1007/s13225-020-00466-2)) |
| Otu45187 | 1 | k__Fungi(100);p__Ascomycota(100);c__Saccharomycetes(100);o__Saccharomycetales(100);f__Metschnikowiaceae(100);g__Metschnikowia(100);s__Metschnikowia_pulcherrima(100); | Metschnikowia | 13 | Saprotroph | \|Undefined Saprotroph\| | Yeast | NULL | Probable | As nectar-tap_saprotroph (Põlme et al. 2020); Decay sugar-rich substrates (Põlme et al. 2020) | Põlme S, et al. 2020. Fungal Diversity 105:1-16 ((https://doi.org/10.1007/s13225-020-00466-2)) |
| Otu45188 | 1 | k__Fungi(100);p__Ascomycota(100);c__Saccharomycetes(100);o__Saccharomycetales(100);f__Metschnikowiaceae(100);g__Metschnikowia(100);s__Metschnikowia_pulcherrima(100); | Metschnikowia | 13 | Saprotroph | \|Undefined Saprotroph\| | Yeast | NULL | Probable | As nectar-tap_saprotroph (Põlme et al. 2020); Decay sugar-rich substrates (Põlme et al. 2020) | Põlme S, et al. 2020. Fungal Diversity 105:1-16 ((https://doi.org/10.1007/s13225-020-00466-2)) |
| Otu45189 | 1 | k__Fungi(100);p__Ascomycota(100);c__Saccharomycetes(100);o__Saccharomycetales(100);f__Metschnikowiaceae(100);g__Metschnikowia(100);g__Metschnikowia_unclassified(100); | Metschnikowia | 13 | Saprotroph | \|Undefined Saprotroph\| | Yeast | NULL | Probable | As nectar-tap_saprotroph (Põlme et al. 2020); Decay sugar-rich substrates (Põlme et al. 2020) | Põlme S, et al. 2020. Fungal Diversity 105:1-16 ((https://doi.org/10.1007/s13225-020-00466-2)) |
| Otu45190 | 1 | k__Fungi(100);p__Ascomycota(100);c__Saccharomycetes(100);o__Saccharomycetales(100);f__Metschnikowiaceae(100);g__Metschnikowia(100);s__Metschnikowia_pulcherrima(100); | Metschnikowia | 13 | Saprotroph | \|Undefined Saprotroph\| | Yeast | NULL | Probable | As nectar-tap_saprotroph (Põlme et al. 2020); Decay sugar-rich substrates (Põlme et al. 2020) | Põlme S, et al. 2020. Fungal Diversity 105:1-16 ((https://doi.org/10.1007/s13225-020-00466-2)) |
| Otu45191 | 1 | k__Fungi(100);p__Ascomycota(100);c__Saccharomycetes(100);o__Saccharomycetales(100);f__Metschnikowiaceae(100);g__Metschnikowia(100);g__Metschnikowia_unclassified(100); | Metschnikowia | 13 | Saprotroph | \|Undefined Saprotroph\| | Yeast | NULL | Probable | As nectar-tap_saprotroph (Põlme et al. 2020); Decay sugar-rich substrates (Põlme et al. 2020) | Põlme S, et al. 2020. Fungal Diversity 105:1-16 ((https://doi.org/10.1007/s13225-020-00466-2)) |
| Otu45192 | 1 | k__Fungi(100);p__Ascomycota(100);c__Saccharomycetes(100);o__Saccharomycetales(100);f__Metschnikowiaceae(100);g__Metschnikowia(100);g__Metschnikowia_unclassified(100); | Metschnikowia | 13 | Saprotroph | \|Undefined Saprotroph\| | Yeast | NULL | Probable | As nectar-tap_saprotroph (Põlme et al. 2020); Decay sugar-rich substrates (Põlme et al. 2020) | Põlme S, et al. 2020. Fungal Diversity 105:1-16 ((https://doi.org/10.1007/s13225-020-00466-2)) |
| Otu45193 | 1 | k__Fungi(100);p__Ascomycota(100);c__Saccharomycetes(100);o__Saccharomycetales(100);f__Metschnikowiaceae(100);g__Metschnikowia(100);s__Metschnikowia_pulcherrima(100); | Metschnikowia | 13 | Saprotroph | \|Undefined Saprotroph\| | Yeast | NULL | Probable | As nectar-tap_saprotroph (Põlme et al. 2020); Decay sugar-rich substrates (Põlme et al. 2020) | Põlme S, et al. 2020. Fungal Diversity 105:1-16 ((https://doi.org/10.1007/s13225-020-00466-2)) |
| Otu45194 | 1 | k__Fungi(100);p__Ascomycota(100);c__Saccharomycetes(100);o__Saccharomycetales(100);f__Metschnikowiaceae(100);g__Metschnikowia(100);g__Metschnikowia_unclassified(100); | Metschnikowia | 13 | Saprotroph | \|Undefined Saprotroph\| | Yeast | NULL | Probable | As nectar-tap_saprotroph (Põlme et al. 2020); Decay sugar-rich substrates (Põlme et al. 2020) | Põlme S, et al. 2020. Fungal Diversity 105:1-16 ((https://doi.org/10.1007/s13225-020-00466-2)) |
| Otu45195 | 1 | k__Fungi(100);p__Ascomycota(100);c__Saccharomycetes(100);o__Saccharomycetales(100);f__Metschnikowiaceae(100);g__Metschnikowia(100);g__Metschnikowia_unclassified(100); | Metschnikowia | 13 | Saprotroph | \|Undefined Saprotroph\| | Yeast | NULL | Probable | As nectar-tap_saprotroph (Põlme et al. 2020); Decay sugar-rich substrates (Põlme et al. 2020) | Põlme S, et al. 2020. Fungal Diversity 105:1-16 ((https://doi.org/10.1007/s13225-020-00466-2)) |
| Otu45196 | 1 | k__Fungi(100);p__Ascomycota(100);c__Saccharomycetes(100);o__Saccharomycetales(100);f__Metschnikowiaceae(100);g__Metschnikowia(100);g__Metschnikowia_unclassified(100); | Metschnikowia | 13 | Saprotroph | \|Undefined Saprotroph\| | Yeast | NULL | Probable | As nectar-tap_saprotroph (Põlme et al. 2020); Decay sugar-rich substrates (Põlme et al. 2020) | Põlme S, et al. 2020. Fungal Diversity 105:1-16 ((https://doi.org/10.1007/s13225-020-00466-2)) |
| Otu45197 | 1 | k__Fungi(100);p__Ascomycota(100);c__Saccharomycetes(100);o__Saccharomycetales(100);f__Metschnikowiaceae(100);g__Metschnikowia(100);g__Metschnikowia_unclassified(100); | Metschnikowia | 13 | Saprotroph | \|Undefined Saprotroph\| | Yeast | NULL | Probable | As nectar-tap_saprotroph (Põlme et al. 2020); Decay sugar-rich substrates (Põlme et al. 2020) | Põlme S, et al. 2020. Fungal Diversity 105:1-16 ((https://doi.org/10.1007/s13225-020-00466-2)) |
| Otu45198 | 1 | k__Fungi(100);p__Ascomycota(100);c__Saccharomycetes(100);o__Saccharomycetales(100);f__Metschnikowiaceae(100);g__Metschnikowia(100);g__Metschnikowia_unclassified(100); | Metschnikowia | 13 | Saprotroph | \|Undefined Saprotroph\| | Yeast | NULL | Probable | As nectar-tap_saprotroph (Põlme et al. 2020); Decay sugar-rich substrates (Põlme et al. 2020) | Põlme S, et al. 2020. Fungal Diversity 105:1-16 ((https://doi.org/10.1007/s13225-020-00466-2)) |
| Otu45199 | 1 | k__Fungi(100);p__Ascomycota(100);c__Saccharomycetes(100);o__Saccharomycetales(100);f__Metschnikowiaceae(100);g__Metschnikowia(100);g__Metschnikowia_unclassified(100); | Metschnikowia | 13 | Saprotroph | \|Undefined Saprotroph\| | Yeast | NULL | Probable | As nectar-tap_saprotroph (Põlme et al. 2020); Decay sugar-rich substrates (Põlme et al. 2020) | Põlme S, et al. 2020. Fungal Diversity 105:1-16 ((https://doi.org/10.1007/s13225-020-00466-2)) |
| Otu45200 | 1 | k__Fungi(100);p__Ascomycota(100);c__Saccharomycetes(100);o__Saccharomycetales(100);f__Metschnikowiaceae(100);g__Metschnikowia(100);s__Metschnikowia_pulcherrima(100); | Metschnikowia | 13 | Saprotroph | \|Undefined Saprotroph\| | Yeast | NULL | Probable | As nectar-tap_saprotroph (Põlme et al. 2020); Decay sugar-rich substrates (Põlme et al. 2020) | Põlme S, et al. 2020. Fungal Diversity 105:1-16 ((https://doi.org/10.1007/s13225-020-00466-2)) |
| Otu45201 | 1 | k__Fungi(100);p__Ascomycota(100);c__Saccharomycetes(100);o__Saccharomycetales(100);f__Metschnikowiaceae(100);g__Metschnikowia(100);s__Metschnikowia_pulcherrima(100); | Metschnikowia | 13 | Saprotroph | \|Undefined Saprotroph\| | Yeast | NULL | Probable | As nectar-tap_saprotroph (Põlme et al. 2020); Decay sugar-rich substrates (Põlme et al. 2020) | Põlme S, et al. 2020. Fungal Diversity 105:1-16 ((https://doi.org/10.1007/s13225-020-00466-2)) |
| Otu45202 | 1 | k__Fungi(100);p__Ascomycota(100);c__Saccharomycetes(100);o__Saccharomycetales(100);f__Metschnikowiaceae(100);g__Metschnikowia(100);s__Metschnikowia_pulcherrima(100); | Metschnikowia | 13 | Saprotroph | \|Undefined Saprotroph\| | Yeast | NULL | Probable | As nectar-tap_saprotroph (Põlme et al. 2020); Decay sugar-rich substrates (Põlme et al. 2020) | Põlme S, et al. 2020. Fungal Diversity 105:1-16 ((https://doi.org/10.1007/s13225-020-00466-2)) |
| Otu45203 | 1 | k__Fungi(100);p__Ascomycota(100);c__Saccharomycetes(100);o__Saccharomycetales(100);f__Metschnikowiaceae(100);g__Metschnikowia(100);g__Metschnikowia_unclassified(100); | Metschnikowia | 13 | Saprotroph | \|Undefined Saprotroph\| | Yeast | NULL | Probable | As nectar-tap_saprotroph (Põlme et al. 2020); Decay sugar-rich substrates (Põlme et al. 2020) | Põlme S, et al. 2020. Fungal Diversity 105:1-16 ((https://doi.org/10.1007/s13225-020-00466-2)) |
| Otu45204 | 1 | k__Fungi(100);p__Ascomycota(100);c__Saccharomycetes(100);o__Saccharomycetales(100);f__Metschnikowiaceae(100);g__Metschnikowia(100);s__Metschnikowia_pulcherrima(100); | Metschnikowia | 13 | Saprotroph | \|Undefined Saprotroph\| | Yeast | NULL | Probable | As nectar-tap_saprotroph (Põlme et al. 2020); Decay sugar-rich substrates (Põlme et al. 2020) | Põlme S, et al. 2020. Fungal Diversity 105:1-16 ((https://doi.org/10.1007/s13225-020-00466-2)) |
| Otu45205 | 1 | k__Fungi(100);p__Ascomycota(100);c__Saccharomycetes(100);o__Saccharomycetales(100);f__Metschnikowiaceae(100);g__Metschnikowia(100);s__Metschnikowia_viticola(100); | Metschnikowia | 13 | Saprotroph | \|Undefined Saprotroph\| | Yeast | NULL | Probable | As nectar-tap_saprotroph (Põlme et al. 2020); Decay sugar-rich substrates (Põlme et al. 2020) | Põlme S, et al. 2020. Fungal Diversity 105:1-16 ((https://doi.org/10.1007/s13225-020-00466-2)) |
| Otu45206 | 1 | k__Fungi(100);p__Ascomycota(100);c__Saccharomycetes(100);o__Saccharomycetales(100);f__Metschnikowiaceae(100);g__Metschnikowia(100);s__Metschnikowia_pulcherrima(100); | Metschnikowia | 13 | Saprotroph | \|Undefined Saprotroph\| | Yeast | NULL | Probable | As nectar-tap_saprotroph (Põlme et al. 2020); Decay sugar-rich substrates (Põlme et al. 2020) | Põlme S, et al. 2020. Fungal Diversity 105:1-16 ((https://doi.org/10.1007/s13225-020-00466-2)) |
| Otu45207 | 1 | k__Fungi(100);p__Ascomycota(100);c__Saccharomycetes(100);o__Saccharomycetales(100);f__Metschnikowiaceae(100);g__Metschnikowia(100);g__Metschnikowia_unclassified(100); | Metschnikowia | 13 | Saprotroph | \|Undefined Saprotroph\| | Yeast | NULL | Probable | As nectar-tap_saprotroph (Põlme et al. 2020); Decay sugar-rich substrates (Põlme et al. 2020) | Põlme S, et al. 2020. Fungal Diversity 105:1-16 ((https://doi.org/10.1007/s13225-020-00466-2)) |
| Otu45208 | 1 | k__Fungi(100);p__Ascomycota(100);c__Saccharomycetes(100);o__Saccharomycetales(100);f__Metschnikowiaceae(100);g__Metschnikowia(100);s__Metschnikowia_pulcherrima(100); | Metschnikowia | 13 | Saprotroph | \|Undefined Saprotroph\| | Yeast | NULL | Probable | As nectar-tap_saprotroph (Põlme et al. 2020); Decay sugar-rich substrates (Põlme et al. 2020) | Põlme S, et al. 2020. Fungal Diversity 105:1-16 ((https://doi.org/10.1007/s13225-020-00466-2)) |
| Otu45209 | 1 | k__Fungi(100);p__Ascomycota(100);c__Saccharomycetes(100);o__Saccharomycetales(100);f__Metschnikowiaceae(100);g__Metschnikowia(100);s__Metschnikowia_pulcherrima(100); | Metschnikowia | 13 | Saprotroph | \|Undefined Saprotroph\| | Yeast | NULL | Probable | As nectar-tap_saprotroph (Põlme et al. 2020); Decay sugar-rich substrates (Põlme et al. 2020) | Põlme S, et al. 2020. Fungal Diversity 105:1-16 ((https://doi.org/10.1007/s13225-020-00466-2)) |
| Otu45210 | 1 | k__Fungi(100);p__Ascomycota(100);c__Saccharomycetes(100);o__Saccharomycetales(100);f__Metschnikowiaceae(100);g__Metschnikowia(100);s__Metschnikowia_pulcherrima(100); | Metschnikowia | 13 | Saprotroph | \|Undefined Saprotroph\| | Yeast | NULL | Probable | As nectar-tap_saprotroph (Põlme et al. 2020); Decay sugar-rich substrates (Põlme et al. 2020) | Põlme S, et al. 2020. Fungal Diversity 105:1-16 ((https://doi.org/10.1007/s13225-020-00466-2)) |
| Otu45211 | 1 | k__Fungi(100);p__Ascomycota(100);c__Saccharomycetes(100);o__Saccharomycetales(100);f__Metschnikowiaceae(100);g__Metschnikowia(100);s__Metschnikowia_pulcherrima(100); | Metschnikowia | 13 | Saprotroph | \|Undefined Saprotroph\| | Yeast | NULL | Probable | As nectar-tap_saprotroph (Põlme et al. 2020); Decay sugar-rich substrates (Põlme et al. 2020) | Põlme S, et al. 2020. Fungal Diversity 105:1-16 ((https://doi.org/10.1007/s13225-020-00466-2)) |
| Otu45212 | 1 | k__Fungi(100);p__Ascomycota(100);c__Saccharomycetes(100);o__Saccharomycetales(100);f__Metschnikowiaceae(100);g__Metschnikowia(100);s__Metschnikowia_pulcherrima(100); | Metschnikowia | 13 | Saprotroph | \|Undefined Saprotroph\| | Yeast | NULL | Probable | As nectar-tap_saprotroph (Põlme et al. 2020); Decay sugar-rich substrates (Põlme et al. 2020) | Põlme S, et al. 2020. Fungal Diversity 105:1-16 ((https://doi.org/10.1007/s13225-020-00466-2)) |
| Otu45213 | 1 | k__Fungi(100);p__Ascomycota(100);c__Saccharomycetes(100);o__Saccharomycetales(100);f__Metschnikowiaceae(100);g__Metschnikowia(100);s__Metschnikowia_pulcherrima(100); | Metschnikowia | 13 | Saprotroph | \|Undefined Saprotroph\| | Yeast | NULL | Probable | As nectar-tap_saprotroph (Põlme et al. 2020); Decay sugar-rich substrates (Põlme et al. 2020) | Põlme S, et al. 2020. Fungal Diversity 105:1-16 ((https://doi.org/10.1007/s13225-020-00466-2)) |
| Otu45214 | 1 | k__Fungi(100);p__Ascomycota(100);c__Saccharomycetes(100);o__Saccharomycetales(100);f__Metschnikowiaceae(100);g__Metschnikowia(100);g__Metschnikowia_unclassified(100); | Metschnikowia | 13 | Saprotroph | \|Undefined Saprotroph\| | Yeast | NULL | Probable | As nectar-tap_saprotroph (Põlme et al. 2020); Decay sugar-rich substrates (Põlme et al. 2020) | Põlme S, et al. 2020. Fungal Diversity 105:1-16 ((https://doi.org/10.1007/s13225-020-00466-2)) |
| Otu45216 | 1 | k__Fungi(100);p__Ascomycota(100);c__Saccharomycetes(100);o__Saccharomycetales(100);f__Metschnikowiaceae(100);g__Metschnikowia(100);s__Metschnikowia_pulcherrima(100); | Metschnikowia | 13 | Saprotroph | \|Undefined Saprotroph\| | Yeast | NULL | Probable | As nectar-tap_saprotroph (Põlme et al. 2020); Decay sugar-rich substrates (Põlme et al. 2020) | Põlme S, et al. 2020. Fungal Diversity 105:1-16 ((https://doi.org/10.1007/s13225-020-00466-2)) |
| Otu45217 | 1 | k__Fungi(100);p__Ascomycota(100);c__Saccharomycetes(100);o__Saccharomycetales(100);f__Metschnikowiaceae(100);g__Metschnikowia(100);s__Metschnikowia_pulcherrima(100); | Metschnikowia | 13 | Saprotroph | \|Undefined Saprotroph\| | Yeast | NULL | Probable | As nectar-tap_saprotroph (Põlme et al. 2020); Decay sugar-rich substrates (Põlme et al. 2020) | Põlme S, et al. 2020. Fungal Diversity 105:1-16 ((https://doi.org/10.1007/s13225-020-00466-2)) |
| Otu45218 | 1 | k__Fungi(100);p__Ascomycota(100);c__Saccharomycetes(100);o__Saccharomycetales(100);f__Metschnikowiaceae(100);g__Metschnikowia(100);s__Metschnikowia_pulcherrima(100); | Metschnikowia | 13 | Saprotroph | \|Undefined Saprotroph\| | Yeast | NULL | Probable | As nectar-tap_saprotroph (Põlme et al. 2020); Decay sugar-rich substrates (Põlme et al. 2020) | Põlme S, et al. 2020. Fungal Diversity 105:1-16 ((https://doi.org/10.1007/s13225-020-00466-2)) |
| Otu45219 | 1 | k__Fungi(100);p__Ascomycota(100);c__Saccharomycetes(100);o__Saccharomycetales(100);f__Metschnikowiaceae(100);g__Metschnikowia(100);s__Metschnikowia_pulcherrima(100); | Metschnikowia | 13 | Saprotroph | \|Undefined Saprotroph\| | Yeast | NULL | Probable | As nectar-tap_saprotroph (Põlme et al. 2020); Decay sugar-rich substrates (Põlme et al. 2020) | Põlme S, et al. 2020. Fungal Diversity 105:1-16 ((https://doi.org/10.1007/s13225-020-00466-2)) |
| Otu45220 | 1 | k__Fungi(100);p__Ascomycota(100);c__Saccharomycetes(100);o__Saccharomycetales(100);f__Metschnikowiaceae(100);g__Metschnikowia(100);s__Metschnikowia_pulcherrima(100); | Metschnikowia | 13 | Saprotroph | \|Undefined Saprotroph\| | Yeast | NULL | Probable | As nectar-tap_saprotroph (Põlme et al. 2020); Decay sugar-rich substrates (Põlme et al. 2020) | Põlme S, et al. 2020. Fungal Diversity 105:1-16 ((https://doi.org/10.1007/s13225-020-00466-2)) |
| Otu45221 | 1 | k__Fungi(100);p__Ascomycota(100);c__Saccharomycetes(100);o__Saccharomycetales(100);f__Metschnikowiaceae(100);g__Metschnikowia(100);s__Metschnikowia_pulcherrima(100); | Metschnikowia | 13 | Saprotroph | \|Undefined Saprotroph\| | Yeast | NULL | Probable | As nectar-tap_saprotroph (Põlme et al. 2020); Decay sugar-rich substrates (Põlme et al. 2020) | Põlme S, et al. 2020. Fungal Diversity 105:1-16 ((https://doi.org/10.1007/s13225-020-00466-2)) |
| Otu45222 | 1 | k__Fungi(100);p__Ascomycota(100);c__Saccharomycetes(100);o__Saccharomycetales(100);f__Saccharomycetales_family_Incertae_sedis(100);g__Candida(100);s__Candida_dosseyi(100); | Candida | 13 | Pathotroph-Saprotroph-Symbiotroph | Animal Pathogen-Endophyte-Endosymbiont-Epiphyte-\|Undefined Saprotroph\| | Dimorphic Yeast-Yeast | NULL | Probable | As nectar-tap_saprotroph (Põlme et al. 2020); Decay sugar-rich substrates (Põlme et al. 2020). This asexual genus is polyphyletic and contain species that function very differently depend on what they are. Caution should be taken when interpreting guilds for Candida species. | Manolakaki D, et al. 2010. Virulence. 1: 367-375 ((https://doi.org/10.4161/viru.1.5.12796)); Mekbib SB, et al. 2011. Tropical Plant Pathology 36:233-240 ((https://doi.org/10.1590/S1982-56762011000400004)); Põlme S, et al. 2020. Fungal Diversity 105:1-16 ((https://doi.org/10.1007/s13225-020-00466-2)) |
| Otu45223 | 1 | k__Fungi(100);p__Ascomycota(100);c__Saccharomycetes(100);o__Saccharomycetales(100);f__Metschnikowiaceae(100);g__Metschnikowia(100);g__Metschnikowia_unclassified(100); | Metschnikowia | 13 | Saprotroph | \|Undefined Saprotroph\| | Yeast | NULL | Probable | As nectar-tap_saprotroph (Põlme et al. 2020); Decay sugar-rich substrates (Põlme et al. 2020) | Põlme S, et al. 2020. Fungal Diversity 105:1-16 ((https://doi.org/10.1007/s13225-020-00466-2)) |
| Otu45224 | 1 | k__Fungi(100);p__Ascomycota(100);c__Saccharomycetes(100);o__Saccharomycetales(100);f__Metschnikowiaceae(100);g__Metschnikowia(100);s__Metschnikowia_pulcherrima(100); | Metschnikowia | 13 | Saprotroph | \|Undefined Saprotroph\| | Yeast | NULL | Probable | As nectar-tap_saprotroph (Põlme et al. 2020); Decay sugar-rich substrates (Põlme et al. 2020) | Põlme S, et al. 2020. Fungal Diversity 105:1-16 ((https://doi.org/10.1007/s13225-020-00466-2)) |
| Otu45225 | 1 | k__Fungi(100);p__Ascomycota(100);c__Saccharomycetes(100);o__Saccharomycetales(100);f__Metschnikowiaceae(100);g__Metschnikowia(100);s__Metschnikowia_pulcherrima(100); | Metschnikowia | 13 | Saprotroph | \|Undefined Saprotroph\| | Yeast | NULL | Probable | As nectar-tap_saprotroph (Põlme et al. 2020); Decay sugar-rich substrates (Põlme et al. 2020) | Põlme S, et al. 2020. Fungal Diversity 105:1-16 ((https://doi.org/10.1007/s13225-020-00466-2)) |

| | | | | | | | | | | | |
|---|---|---|---|---|---|---|---|---|---|---|---|
| Otu45226 | 1 | k__Fungi(100);p__Ascomycota(100);c__Saccharomycetes(100);o__Saccharomycetales(100);f__Metschnikowiaceae(100);g__Metschnikowia(100);s__Metschnikowia_pulcherrima(100); | Metschnikowia | 13 | Saprotroph | \|Undefined Saprotroph\| | Yeast | NULL | Probable | As nectar-tap_saprotroph (Põlme et al. 2020); Decay sugar-rich substrates (Põlme et al. 2020) | Põlme S, et al. 2020. Fungal Diversity 105:1-16 ((https://doi.org/10.1007/s13225-020-00466-2)) |
| Otu45227 | 1 | k__Fungi(100);p__Ascomycota(100);c__Saccharomycetes(100);o__Saccharomycetales(100);f__Metschnikowiaceae(100);g__Metschnikowia(100);s__Metschnikowia_pulcherrima(100); | Metschnikowia | 13 | Saprotroph | \|Undefined Saprotroph\| | Yeast | NULL | Probable | As nectar-tap_saprotroph (Põlme et al. 2020); Decay sugar-rich substrates (Põlme et al. 2020) | Põlme S, et al. 2020. Fungal Diversity 105:1-16 ((https://doi.org/10.1007/s13225-020-00466-2)) |
| Otu45229 | 1 | k__Fungi(100);p__Ascomycota(100);c__Saccharomycetes(100);o__Saccharomycetales(100);f__Metschnikowiaceae(100);g__Metschnikowia(100);s__Metschnikowia_pulcherrima(100); | Metschnikowia | 13 | Saprotroph | \|Undefined Saprotroph\| | Yeast | NULL | Probable | As nectar-tap_saprotroph (Põlme et al. 2020); Decay sugar-rich substrates (Põlme et al. 2020) | Põlme S, et al. 2020. Fungal Diversity 105:1-16 ((https://doi.org/10.1007/s13225-020-00466-2)) |
| Otu45230 | 1 | k__Fungi(100);p__Ascomycota(100);c__Saccharomycetes(100);o__Saccharomycetales(100);f__Metschnikowiaceae(100);g__Metschnikowia(100);s__Metschnikowia_pulcherrima(100); | Metschnikowia | 13 | Saprotroph | \|Undefined Saprotroph\| | Yeast | NULL | Probable | As nectar-tap_saprotroph (Põlme et al. 2020); Decay sugar-rich substrates (Põlme et al. 2020) | Põlme S, et al. 2020. Fungal Diversity 105:1-16 ((https://doi.org/10.1007/s13225-020-00466-2)) |
| Otu45231 | 1 | k__Fungi(100);p__Ascomycota(100);c__Saccharomycetes(100);o__Saccharomycetales(100);f__Metschnikowiaceae(100);g__Metschnikowia(100);s__Metschnikowia_pulcherrima(100); | Metschnikowia | 13 | Saprotroph | \|Undefined Saprotroph\| | Yeast | NULL | Probable | As nectar-tap_saprotroph (Põlme et al. 2020); Decay sugar-rich substrates (Põlme et al. 2020) | Põlme S, et al. 2020. Fungal Diversity 105:1-16 ((https://doi.org/10.1007/s13225-020-00466-2)) |
| Otu45232 | 1 | k__Fungi(100);p__Ascomycota(100);c__Saccharomycetes(100);o__Saccharomycetales(100);f__Metschnikowiaceae(100);g__Metschnikowia(100);s__Metschnikowia_pulcherrima(100); | Metschnikowia | 13 | Saprotroph | \|Undefined Saprotroph\| | Yeast | NULL | Probable | As nectar-tap_saprotroph (Põlme et al. 2020); Decay sugar-rich substrates (Põlme et al. 2020) | Põlme S, et al. 2020. Fungal Diversity 105:1-16 ((https://doi.org/10.1007/s13225-020-00466-2)) |
| Otu45233 | 1 | k__Fungi(100);p__Ascomycota(100);c__Saccharomycetes(100);o__Saccharomycetales(100);f__Metschnikowiaceae(100);g__Metschnikowia(100);s__Metschnikowia_pulcherrima(100); | Metschnikowia | 13 | Saprotroph | \|Undefined Saprotroph\| | Yeast | NULL | Probable | As nectar-tap_saprotroph (Põlme et al. 2020); Decay sugar-rich substrates (Põlme et al. 2020) | Põlme S, et al. 2020. Fungal Diversity 105:1-16 ((https://doi.org/10.1007/s13225-020-00466-2)) |
| Otu45234 | 1 | k__Fungi(100);p__Ascomycota(100);c__Saccharomycetes(100);o__Saccharomycetales(100);f__Metschnikowiaceae(100);g__Metschnikowia(100);s__Metschnikowia_pulcherrima(100); | Metschnikowia | 13 | Saprotroph | \|Undefined Saprotroph\| | Yeast | NULL | Probable | As nectar-tap_saprotroph (Põlme et al. 2020); Decay sugar-rich substrates (Põlme et al. 2020) | Põlme S, et al. 2020. Fungal Diversity 105:1-16 ((https://doi.org/10.1007/s13225-020-00466-2)) |
| Otu45235 | 1 | k__Fungi(100);p__Ascomycota(100);c__Saccharomycetes(100);o__Saccharomycetales(100);f__Metschnikowiaceae(100);g__Metschnikowia(100);g__Metschnikowia_unclassified(100); | Metschnikowia | 13 | Saprotroph | \|Undefined Saprotroph\| | Yeast | NULL | Probable | As nectar-tap_saprotroph (Põlme et al. 2020); Decay sugar-rich substrates (Põlme et al. 2020) | Põlme S, et al. 2020. Fungal Diversity 105:1-16 ((https://doi.org/10.1007/s13225-020-00466-2)) |
| Otu45236 | 1 | k__Fungi(100);p__Ascomycota(100);c__Saccharomycetes(100);o__Saccharomycetales(100);f__Metschnikowiaceae(100);g__Metschnikowia(100);s__Metschnikowia_pulcherrima(100); | Metschnikowia | 13 | Saprotroph | \|Undefined Saprotroph\| | Yeast | NULL | Probable | As nectar-tap_saprotroph (Põlme et al. 2020); Decay sugar-rich substrates (Põlme et al. 2020) | Põlme S, et al. 2020. Fungal Diversity 105:1-16 ((https://doi.org/10.1007/s13225-020-00466-2)) |
| Otu45237 | 1 | k__Fungi(100);p__Ascomycota(100);c__Saccharomycetes(100);o__Saccharomycetales(100);f__Metschnikowiaceae(100);g__Metschnikowia(100);s__Metschnikowia_pulcherrima(100); | Metschnikowia | 13 | Saprotroph | \|Undefined Saprotroph\| | Yeast | NULL | Probable | As nectar-tap_saprotroph (Põlme et al. 2020); Decay sugar-rich substrates (Põlme et al. 2020) | Põlme S, et al. 2020. Fungal Diversity 105:1-16 ((https://doi.org/10.1007/s13225-020-00466-2)) |
| Otu45238 | 1 | k__Fungi(100);p__Ascomycota(100);c__Saccharomycetes(100);o__Saccharomycetales(100);f__Metschnikowiaceae(100);g__Metschnikowia(100);s__Metschnikowia_pulcherrima(100); | Metschnikowia | 13 | Saprotroph | \|Undefined Saprotroph\| | Yeast | NULL | Probable | As nectar-tap_saprotroph (Põlme et al. 2020); Decay sugar-rich substrates (Põlme et al. 2020) | Põlme S, et al. 2020. Fungal Diversity 105:1-16 ((https://doi.org/10.1007/s13225-020-00466-2)) |
| Otu45239 | 1 | k__Fungi(100);p__Ascomycota(100);c__Saccharomycetes(100);o__Saccharomycetales(100);f__Metschnikowiaceae(100);g__Metschnikowia(100);s__Metschnikowia_pulcherrima(100); | Metschnikowia | 13 | Saprotroph | \|Undefined Saprotroph\| | Yeast | NULL | Probable | As nectar-tap_saprotroph (Põlme et al. 2020); Decay sugar-rich substrates (Põlme et al. 2020) | Põlme S, et al. 2020. Fungal Diversity 105:1-16 ((https://doi.org/10.1007/s13225-020-00466-2)) |
| Otu45240 | 1 | k__Fungi(100);p__Ascomycota(100);c__Saccharomycetes(100);o__Saccharomycetales(100);f__Metschnikowiaceae(100);g__Metschnikowia(100);s__Metschnikowia_pulcherrima(100); | Metschnikowia | 13 | Saprotroph | \|Undefined Saprotroph\| | Yeast | NULL | Probable | As nectar-tap_saprotroph (Põlme et al. 2020); Decay sugar-rich substrates (Põlme et al. 2020) | Põlme S, et al. 2020. Fungal Diversity 105:1-16 ((https://doi.org/10.1007/s13225-020-00466-2)) |
| Otu45241 | 1 | k__Fungi(100);p__Ascomycota(100);c__Saccharomycetes(100);o__Saccharomycetales(100);f__Metschnikowiaceae(100);g__Metschnikowia(100);s__Metschnikowia_pulcherrima(100); | Metschnikowia | 13 | Saprotroph | \|Undefined Saprotroph\| | Yeast | NULL | Probable | As nectar-tap_saprotroph (Põlme et al. 2020); Decay sugar-rich substrates (Põlme et al. 2020) | Põlme S, et al. 2020. Fungal Diversity 105:1-16 ((https://doi.org/10.1007/s13225-020-00466-2)) |
| Otu45242 | 1 | k__Fungi(100);p__Ascomycota(100);c__Saccharomycetes(100);o__Saccharomycetales(100);f__Metschnikowiaceae(100);g__Metschnikowia(100);s__Metschnikowia_pulcherrima(100); | Metschnikowia | 13 | Saprotroph | \|Undefined Saprotroph\| | Yeast | NULL | Probable | As nectar-tap_saprotroph (Põlme et al. 2020); Decay sugar-rich substrates (Põlme et al. 2020) | Põlme S, et al. 2020. Fungal Diversity 105:1-16 ((https://doi.org/10.1007/s13225-020-00466-2)) |
| Otu45243 | 1 | k__Fungi(100);p__Ascomycota(100);c__Saccharomycetes(100);o__Saccharomycetales(100);f__Metschnikowiaceae(100);g__Metschnikowia(100);s__Metschnikowia_pulcherrima(100); | Metschnikowia | 13 | Saprotroph | \|Undefined Saprotroph\| | Yeast | NULL | Probable | As nectar-tap_saprotroph (Põlme et al. 2020); Decay sugar-rich substrates (Põlme et al. 2020) | Põlme S, et al. 2020. Fungal Diversity 105:1-16 ((https://doi.org/10.1007/s13225-020-00466-2)) |
| Otu45244 | 1 | k__Fungi(100);p__Ascomycota(100);c__Saccharomycetes(100);o__Saccharomycetales(100);f__Metschnikowiaceae(100);g__Metschnikowia(100);s__Metschnikowia_pulcherrima(100); | Metschnikowia | 13 | Saprotroph | \|Undefined Saprotroph\| | Yeast | NULL | Probable | As nectar-tap_saprotroph (Põlme et al. 2020); Decay sugar-rich substrates (Põlme et al. 2020) | Põlme S, et al. 2020. Fungal Diversity 105:1-16 ((https://doi.org/10.1007/s13225-020-00466-2)) |
| Otu45245 | 1 | k__Fungi(100);p__Ascomycota(100);c__Saccharomycetes(100);o__Saccharomycetales(100);f__Metschnikowiaceae(100);g__Metschnikowia(100);g__Metschnikowia_unclassified(100); | Metschnikowia | 13 | Saprotroph | \|Undefined Saprotroph\| | Yeast | NULL | Probable | As nectar-tap_saprotroph (Põlme et al. 2020); Decay sugar-rich substrates (Põlme et al. 2020) | Põlme S, et al. 2020. Fungal Diversity 105:1-16 ((https://doi.org/10.1007/s13225-020-00466-2)) |
| Otu45246 | 1 | k__Fungi(100);p__Ascomycota(100);c__Saccharomycetes(100);o__Saccharomycetales(100);f__Metschnikowiaceae(100);g__Metschnikowia(100);s__Metschnikowia_pulcherrima(100); | Metschnikowia | 13 | Saprotroph | \|Undefined Saprotroph\| | Yeast | NULL | Probable | As nectar-tap_saprotroph (Põlme et al. 2020); Decay sugar-rich substrates (Põlme et al. 2020) | Põlme S, et al. 2020. Fungal Diversity 105:1-16 ((https://doi.org/10.1007/s13225-020-00466-2)) |
| Otu45247 | 1 | k__Fungi(100);p__Ascomycota(100);c__Saccharomycetes(100);o__Saccharomycetales(100);f__Metschnikowiaceae(100);g__Metschnikowia(100);g__Metschnikowia_unclassified(100); | Metschnikowia | 13 | Saprotroph | \|Undefined Saprotroph\| | Yeast | NULL | Probable | As nectar-tap_saprotroph (Põlme et al. 2020); Decay sugar-rich substrates (Põlme et al. 2020) | Põlme S, et al. 2020. Fungal Diversity 105:1-16 ((https://doi.org/10.1007/s13225-020-00466-2)) |
| Otu45248 | 1 | k__Fungi(100);p__Ascomycota(100);c__Saccharomycetes(100);o__Saccharomycetales(100);f__Metschnikowiaceae(100);g__Metschnikowia(100);s__Metschnikowia_pulcherrima(100); | Metschnikowia | 13 | Saprotroph | \|Undefined Saprotroph\| | Yeast | NULL | Probable | As nectar-tap_saprotroph (Põlme et al. 2020); Decay sugar-rich substrates (Põlme et al. 2020) | Põlme S, et al. 2020. Fungal Diversity 105:1-16 ((https://doi.org/10.1007/s13225-020-00466-2)) |
| Otu45249 | 1 | k__Fungi(100);p__Ascomycota(100);c__Saccharomycetes(100);o__Saccharomycetales(100);f__Metschnikowiaceae(100);g__Metschnikowia(100);s__Metschnikowia_pulcherrima(100); | Metschnikowia | 13 | Saprotroph | \|Undefined Saprotroph\| | Yeast | NULL | Probable | As nectar-tap_saprotroph (Põlme et al. 2020); Decay sugar-rich substrates (Põlme et al. 2020) | Põlme S, et al. 2020. Fungal Diversity 105:1-16 ((https://doi.org/10.1007/s13225-020-00466-2)) |
| Otu45250 | 1 | k__Fungi(100);p__Ascomycota(100);c__Saccharomycetes(100);o__Saccharomycetales(100);f__Metschnikowiaceae(100);g__Metschnikowia(100);s__Metschnikowia_pulcherrima(100); | Metschnikowia | 13 | Saprotroph | \|Undefined Saprotroph\| | Yeast | NULL | Probable | As nectar-tap_saprotroph (Põlme et al. 2020); Decay sugar-rich substrates (Põlme et al. 2020) | Põlme S, et al. 2020. Fungal Diversity 105:1-16 ((https://doi.org/10.1007/s13225-020-00466-2)) |
| Otu45251 | 1 | k__Fungi(100);p__Ascomycota(100);c__Saccharomycetes(100);o__Saccharomycetales(100);f__Metschnikowiaceae(100);g__Metschnikowia(100);g__Metschnikowia_unclassified(100); | Metschnikowia | 13 | Saprotroph | \|Undefined Saprotroph\| | Yeast | NULL | Probable | As nectar-tap_saprotroph (Põlme et al. 2020); Decay sugar-rich substrates (Põlme et al. 2020) | Põlme S, et al. 2020. Fungal Diversity 105:1-16 ((https://doi.org/10.1007/s13225-020-00466-2)) |
| Otu45252 | 1 | k__Fungi(100);p__Ascomycota(100);c__Saccharomycetes(100);o__Saccharomycetales(100);f__Metschnikowiaceae(100);g__Metschnikowia(100);s__Metschnikowia_pulcherrima(100); | Metschnikowia | 13 | Saprotroph | \|Undefined Saprotroph\| | Yeast | NULL | Probable | As nectar-tap_saprotroph (Põlme et al. 2020); Decay sugar-rich substrates (Põlme et al. 2020) | Põlme S, et al. 2020. Fungal Diversity 105:1-16 ((https://doi.org/10.1007/s13225-020-00466-2)) |
| Otu45253 | 1 | k__Fungi(100);p__Ascomycota(100);c__Saccharomycetes(100);o__Saccharomycetales(100);f__Metschnikowiaceae(100);g__Metschnikowia(100);s__Metschnikowia_pulcherrima(100); | Metschnikowia | 13 | Saprotroph | \|Undefined Saprotroph\| | Yeast | NULL | Probable | As nectar-tap_saprotroph (Põlme et al. 2020); Decay sugar-rich substrates (Põlme et al. 2020) | Põlme S, et al. 2020. Fungal Diversity 105:1-16 ((https://doi.org/10.1007/s13225-020-00466-2)) |
| Otu45254 | 1 | k__Fungi(100);p__Ascomycota(100);c__Saccharomycetes(100);o__Saccharomycetales(100);f__Metschnikowiaceae(100);g__Metschnikowia(100);g__Metschnikowia_unclassified(100); | Metschnikowia | 13 | Saprotroph | \|Undefined Saprotroph\| | Yeast | NULL | Probable | As nectar-tap_saprotroph (Põlme et al. 2020); Decay sugar-rich substrates (Põlme et al. 2020) | Põlme S, et al. 2020. Fungal Diversity 105:1-16 ((https://doi.org/10.1007/s13225-020-00466-2)) |
| Otu45255 | 1 | k__Fungi(100);p__Ascomycota(100);c__Saccharomycetes(100);o__Saccharomycetales(100);f__Metschnikowiaceae(100);g__Metschnikowia(100);s__Metschnikowia_pulcherrima(100); | Metschnikowia | 13 | Saprotroph | \|Undefined Saprotroph\| | Yeast | NULL | Probable | As nectar-tap_saprotroph (Põlme et al. 2020); Decay sugar-rich substrates (Põlme et al. 2020) | Põlme S, et al. 2020. Fungal Diversity 105:1-16 ((https://doi.org/10.1007/s13225-020-00466-2)) |
| Otu45256 | 1 | k__Fungi(100);p__Ascomycota(100);c__Saccharomycetes(100);o__Saccharomycetales(100);f__Metschnikowiaceae(100);g__Metschnikowia(100);s__Metschnikowia_pulcherrima(100); | Metschnikowia | 13 | Saprotroph | \|Undefined Saprotroph\| | Yeast | NULL | Probable | As nectar-tap_saprotroph (Põlme et al. 2020); Decay sugar-rich substrates (Põlme et al. 2020) | Põlme S, et al. 2020. Fungal Diversity 105:1-16 ((https://doi.org/10.1007/s13225-020-00466-2)) |
| Otu45257 | 1 | k__Fungi(100);p__Ascomycota(100);c__Saccharomycetes(100);o__Saccharomycetales(100);f__Metschnikowiaceae(100);g__Metschnikowia(100);s__Metschnikowia_pulcherrima(100); | Metschnikowia | 13 | Saprotroph | \|Undefined Saprotroph\| | Yeast | NULL | Probable | As nectar-tap_saprotroph (Põlme et al. 2020); Decay sugar-rich substrates (Põlme et al. 2020) | Põlme S, et al. 2020. Fungal Diversity 105:1-16 ((https://doi.org/10.1007/s13225-020-00466-2)) |
| Otu45258 | 1 | k__Fungi(100);p__Ascomycota(100);c__Saccharomycetes(100);o__Saccharomycetales(100);f__Metschnikowiaceae(100);g__Metschnikowia(100);s__Metschnikowia_pulcherrima(100); | Metschnikowia | 13 | Saprotroph | \|Undefined Saprotroph\| | Yeast | NULL | Probable | As nectar-tap_saprotroph (Põlme et al. 2020); Decay sugar-rich substrates (Põlme et al. 2020) | Põlme S, et al. 2020. Fungal Diversity 105:1-16 ((https://doi.org/10.1007/s13225-020-00466-2)) |
| Otu45259 | 1 | k__Fungi(100);p__Ascomycota(100);c__Saccharomycetes(100);o__Saccharomycetales(100);f__Metschnikowiaceae(100);g__Metschnikowia(100);s__Metschnikowia_pulcherrima(100); | Metschnikowia | 13 | Saprotroph | \|Undefined Saprotroph\| | Yeast | NULL | Probable | As nectar-tap_saprotroph (Põlme et al. 2020); Decay sugar-rich substrates (Põlme et al. 2020) | Põlme S, et al. 2020. Fungal Diversity 105:1-16 ((https://doi.org/10.1007/s13225-020-00466-2)) |
| Otu45260 | 1 | k__Fungi(100);p__Ascomycota(100);c__Saccharomycetes(100);o__Saccharomycetales(100);f__Metschnikowiaceae(100);g__Metschnikowia(100);s__Metschnikowia_pulcherrima(100); | Metschnikowia | 13 | Saprotroph | \|Undefined Saprotroph\| | Yeast | NULL | Probable | As nectar-tap_saprotroph (Põlme et al. 2020); Decay sugar-rich substrates (Põlme et al. 2020) | Põlme S, et al. 2020. Fungal Diversity 105:1-16 ((https://doi.org/10.1007/s13225-020-00466-2)) |
| Otu45262 | 1 | k__Fungi(100);p__Ascomycota(100);c__Saccharomycetes(100);o__Saccharomycetales(100);f__Metschnikowiaceae(100);g__Metschnikowia(100);g__Metschnikowia_unclassified(100); | Metschnikowia | 13 | Saprotroph | \|Undefined Saprotroph\| | Yeast | NULL | Probable | As nectar-tap_saprotroph (Põlme et al. 2020); Decay sugar-rich substrates (Põlme et al. 2020) | Põlme S, et al. 2020. Fungal Diversity 105:1-16 ((https://doi.org/10.1007/s13225-020-00466-2)) |
| Otu45263 | 1 | k__Fungi(100);p__Ascomycota(100);c__Saccharomycetes(100);o__Saccharomycetales(100);f__Metschnikowiaceae(100);g__Metschnikowia(100);g__Metschnikowia_unclassified(100); | Metschnikowia | 13 | Saprotroph | \|Undefined Saprotroph\| | Yeast | NULL | Probable | As nectar-tap_saprotroph (Põlme et al. 2020); Decay sugar-rich substrates (Põlme et al. 2020) | Põlme S, et al. 2020. Fungal Diversity 105:1-16 ((https://doi.org/10.1007/s13225-020-00466-2)) |
| Otu45264 | 1 | k__Fungi(100);p__Ascomycota(100);c__Saccharomycetes(100);o__Saccharomycetales(100);f__Metschnikowiaceae(100);g__Metschnikowia(100);g__Metschnikowia_unclassified(100); | Metschnikowia | 13 | Saprotroph | \|Undefined Saprotroph\| | Yeast | NULL | Probable | As nectar-tap_saprotroph (Põlme et al. 2020); Decay sugar-rich substrates (Põlme et al. 2020) | Põlme S, et al. 2020. Fungal Diversity 105:1-16 ((https://doi.org/10.1007/s13225-020-00466-2)) |
| Otu45265 | 1 | k__Fungi(100);p__Ascomycota(100);c__Saccharomycetes(100);o__Saccharomycetales(100);f__Metschnikowiaceae(100);g__Metschnikowia(100);s__Metschnikowia_pulcherrima(100); | Metschnikowia | 13 | Saprotroph | \|Undefined Saprotroph\| | Yeast | NULL | Probable | As nectar-tap_saprotroph (Põlme et al. 2020); Decay sugar-rich substrates (Põlme et al. 2020) | Põlme S, et al. 2020. Fungal Diversity 105:1-16 ((https://doi.org/10.1007/s13225-020-00466-2)) |
| Otu45266 | 1 | k__Fungi(100);p__Ascomycota(100);c__Saccharomycetes(100);o__Saccharomycetales(100);f__Metschnikowiaceae(100);g__Metschnikowia(100);s__Metschnikowia_pulcherrima(100); | Metschnikowia | 13 | Saprotroph | \|Undefined Saprotroph\| | Yeast | NULL | Probable | As nectar-tap_saprotroph (Põlme et al. 2020); Decay sugar-rich substrates (Põlme et al. 2020) | Põlme S, et al. 2020. Fungal Diversity 105:1-16 ((https://doi.org/10.1007/s13225-020-00466-2)) |
| Otu45267 | 1 | k__Fungi(100);p__Ascomycota(100);c__Saccharomycetes(100);o__Saccharomycetales(100);f__Metschnikowiaceae(100);g__Metschnikowia(100);s__Metschnikowia_pulcherrima(100); | Metschnikowia | 13 | Saprotroph | \|Undefined Saprotroph\| | Yeast | NULL | Probable | As nectar-tap_saprotroph (Põlme et al. 2020); Decay sugar-rich substrates (Põlme et al. 2020) | Põlme S, et al. 2020. Fungal Diversity 105:1-16 ((https://doi.org/10.1007/s13225-020-00466-2)) |
| Otu45268 | 1 | k__Fungi(100);p__Ascomycota(100);c__Saccharomycetes(100);o__Saccharomycetales(100);f__Metschnikowiaceae(100);g__Metschnikowia(100);g__Metschnikowia_unclassified(100); | Metschnikowia | 13 | Saprotroph | \|Undefined Saprotroph\| | Yeast | NULL | Probable | As nectar-tap_saprotroph (Põlme et al. 2020); Decay sugar-rich substrates (Põlme et al. 2020) | Põlme S, et al. 2020. Fungal Diversity 105:1-16 ((https://doi.org/10.1007/s13225-020-00466-2)) |
| Otu45270 | 1 | k__Fungi(100);p__Ascomycota(100);c__Saccharomycetes(100);o__Saccharomycetales(100);f__Metschnikowiaceae(100);g__Metschnikowia(100);s__Metschnikowia_pulcherrima(100); | Metschnikowia | 13 | Saprotroph | \|Undefined Saprotroph\| | Yeast | NULL | Probable | As nectar-tap_saprotroph (Põlme et al. 2020); Decay sugar-rich substrates (Põlme et al. 2020) | Põlme S, et al. 2020. Fungal Diversity 105:1-16 ((https://doi.org/10.1007/s13225-020-00466-2)) |
| Otu45271 | 1 | k__Fungi(100);p__Ascomycota(100);c__Saccharomycetes(100);o__Saccharomycetales(100);f__Metschnikowiaceae(100);g__Metschnikowia(100);s__Metschnikowia_pulcherrima(100); | Metschnikowia | 13 | Saprotroph | \|Undefined Saprotroph\| | Yeast | NULL | Probable | As nectar-tap_saprotroph (Põlme et al. 2020); Decay sugar-rich substrates (Põlme et al. 2020) | Põlme S, et al. 2020. Fungal Diversity 105:1-16 ((https://doi.org/10.1007/s13225-020-00466-2)) |
| Otu45272 | 1 | k__Fungi(100);p__Ascomycota(100);c__Saccharomycetes(100);o__Saccharomycetales(100);f__Metschnikowiaceae(100);g__Metschnikowia(100);s__Metschnikowia_pulcherrima(100); | Metschnikowia | 13 | Saprotroph | \|Undefined Saprotroph\| | Yeast | NULL | Probable | As nectar-tap_saprotroph (Põlme et al. 2020); Decay sugar-rich substrates (Põlme et al. 2020) | Põlme S, et al. 2020. Fungal Diversity 105:1-16 ((https://doi.org/10.1007/s13225-020-00466-2)) |
| Otu45274 | 1 | k__Fungi(100);p__Ascomycota(100);c__Saccharomycetes(100);o__Saccharomycetales(100);f__Metschnikowiaceae(100);g__Metschnikowia(100);s__Metschnikowia_pulcherrima(100); | Metschnikowia | 13 | Saprotroph | \|Undefined Saprotroph\| | Yeast | NULL | Probable | As nectar-tap_saprotroph (Põlme et al. 2020); Decay sugar-rich substrates (Põlme et al. 2020) | Põlme S, et al. 2020. Fungal Diversity 105:1-16 ((https://doi.org/10.1007/s13225-020-00466-2)) |
| Otu45275 | 1 | k__Fungi(100);p__Ascomycota(100);c__Saccharomycetes(100);o__Saccharomycetales(100);f__Metschnikowiaceae(100);g__Metschnikowia(100);s__Metschnikowia_pulcherrima(100); | Metschnikowia | 13 | Saprotroph | \|Undefined Saprotroph\| | Yeast | NULL | Probable | As nectar-tap_saprotroph (Põlme et al. 2020); Decay sugar-rich substrates (Põlme et al. 2020) | Põlme S, et al. 2020. Fungal Diversity 105:1-16 ((https://doi.org/10.1007/s13225-020-00466-2)) |
| Otu45276 | 1 | k__Fungi(100);p__Ascomycota(100);c__Saccharomycetes(100);o__Saccharomycetales(100);f__Metschnikowiaceae(100);g__Metschnikowia(100);s__Metschnikowia_pulcherrima(100); | Metschnikowia | 13 | Saprotroph | \|Undefined Saprotroph\| | Yeast | NULL | Probable | As nectar-tap_saprotroph (Põlme et al. 2020); Decay sugar-rich substrates (Põlme et al. 2020) | Põlme S, et al. 2020. Fungal Diversity 105:1-16 ((https://doi.org/10.1007/s13225-020-00466-2)) |
| Otu45277 | 1 | k__Fungi(100);p__Ascomycota(100);c__Saccharomycetes(100);o__Saccharomycetales(100);f__Metschnikowiaceae(100);g__Metschnikowia(100);g__Metschnikowia_unclassified(100); | Metschnikowia | 13 | Saprotroph | \|Undefined Saprotroph\| | Yeast | NULL | Probable | As nectar-tap_saprotroph (Põlme et al. 2020); Decay sugar-rich substrates (Põlme et al. 2020) | Põlme S, et al. 2020. Fungal Diversity 105:1-16 ((https://doi.org/10.1007/s13225-020-00466-2)) |
| Otu45278 | 1 | k__Fungi(100);p__Ascomycota(100);c__Saccharomycetes(100);o__Saccharomycetales(100);f__Metschnikowiaceae(100);g__Metschnikowia(100);s__Metschnikowia_pulcherrima(100); | Metschnikowia | 13 | Saprotroph | \|Undefined Saprotroph\| | Yeast | NULL | Probable | As nectar-tap_saprotroph (Põlme et al. 2020); Decay sugar-rich substrates (Põlme et al. 2020) | Põlme S, et al. 2020. Fungal Diversity 105:1-16 ((https://doi.org/10.1007/s13225-020-00466-2)) |
| Otu45279 | 1 | k__Fungi(100);p__Ascomycota(100);c__Saccharomycetes(100);o__Saccharomycetales(100);f__Metschnikowiaceae(100);g__Metschnikowia(100);s__Metschnikowia_pulcherrima(100); | Metschnikowia | 13 | Saprotroph | \|Undefined Saprotroph\| | Yeast | NULL | Probable | As nectar-tap_saprotroph (Põlme et al. 2020); Decay sugar-rich substrates (Põlme et al. 2020) | Põlme S, et al. 2020. Fungal Diversity 105:1-16 ((https://doi.org/10.1007/s13225-020-00466-2)) |
| Otu45280 | 1 | k__Fungi(100);p__Ascomycota(100);c__Saccharomycetes(100);o__Saccharomycetales(100);f__Metschnikowiaceae(100);g__Metschnikowia(100);s__Metschnikowia_pulcherrima(100); | Metschnikowia | 13 | Saprotroph | \|Undefined Saprotroph\| | Yeast | NULL | Probable | As nectar-tap_saprotroph (Põlme et al. 2020); Decay sugar-rich substrates (Põlme et al. 2020) | Põlme S, et al. 2020. Fungal Diversity 105:1-16 ((https://doi.org/10.1007/s13225-020-00466-2)) |
| Otu45281 | 1 | k__Fungi(100);p__Ascomycota(100);c__Saccharomycetes(100);o__Saccharomycetales(100);f__Metschnikowiaceae(100);g__Metschnikowia(100);g__Metschnikowia_unclassified(100); | Metschnikowia | 13 | Saprotroph | \|Undefined Saprotroph\| | Yeast | NULL | Probable | As nectar-tap_saprotroph (Põlme et al. 2020); Decay sugar-rich substrates (Põlme et al. 2020) | Põlme S, et al. 2020. Fungal Diversity 105:1-16 ((https://doi.org/10.1007/s13225-020-00466-2)) |
| Otu45282 | 1 | k__Fungi(100);p__Ascomycota(100);c__Saccharomycetes(100);o__Saccharomycetales(100);f__Metschnikowiaceae(100);g__Metschnikowia(100);s__Metschnikowia_pulcherrima(100); | Metschnikowia | 13 | Saprotroph | \|Undefined Saprotroph\| | Yeast | NULL | Probable | As nectar-tap_saprotroph (Põlme et al. 2020); Decay sugar-rich substrates (Põlme et al. 2020) | Põlme S, et al. 2020. Fungal Diversity 105:1-16 ((https://doi.org/10.1007/s13225-020-00466-2)) |
| Otu45283 | 1 | k__Fungi(100);p__Ascomycota(100);c__Saccharomycetes(100);o__Saccharomycetales(100);f__Metschnikowiaceae(100);g__Metschnikowia(100);s__Metschnikowia_pulcherrima(100); | Metschnikowia | 13 | Saprotroph | \|Undefined Saprotroph\| | Yeast | NULL | Probable | As nectar-tap_saprotroph (Põlme et al. 2020); Decay sugar-rich substrates (Põlme et al. 2020) | Põlme S, et al. 2020. Fungal Diversity 105:1-16 ((https://doi.org/10.1007/s13225-020-00466-2)) |
| Otu45284 | 1 | k__Fungi(100);p__Ascomycota(100);c__Saccharomycetes(100);o__Saccharomycetales(100);f__Metschnikowiaceae(100);g__Metschnikowia(100);g__Metschnikowia_unclassified(100); | Metschnikowia | 13 | Saprotroph | \|Undefined Saprotroph\| | Yeast | NULL | Probable | As nectar-tap_saprotroph (Põlme et al. 2020); Decay sugar-rich substrates (Põlme et al. 2020) | Põlme S, et al. 2020. Fungal Diversity 105:1-16 ((https://doi.org/10.1007/s13225-020-00466-2)) |
| Otu45285 | 1 | k__Fungi(100);p__Ascomycota(100);c__Saccharomycetes(100);o__Saccharomycetales(100);f__Metschnikowiaceae(100);g__Metschnikowia(100);s__Metschnikowia_pulcherrima(100); | Metschnikowia | 13 | Saprotroph | \|Undefined Saprotroph\| | Yeast | NULL | Probable | As nectar-tap_saprotroph (Põlme et al. 2020); Decay sugar-rich substrates (Põlme et al. 2020) | Põlme S, et al. 2020. Fungal Diversity 105:1-16 ((https://doi.org/10.1007/s13225-020-00466-2)) |
| Otu45286 | 1 | k__Fungi(100);p__Ascomycota(100);c__Saccharomycetes(100);o__Saccharomycetales(100);f__Metschnikowiaceae(100);g__Metschnikowia(100);s__Metschnikowia_pulcherrima(100); | Metschnikowia | 13 | Saprotroph | \|Undefined Saprotroph\| | Yeast | NULL | Probable | As nectar-tap_saprotroph (Põlme et al. 2020); Decay sugar-rich substrates (Põlme et al. 2020) | Põlme S, et al. 2020. Fungal Diversity 105:1-16 ((https://doi.org/10.1007/s13225-020-00466-2)) |
| Otu45287 | 1 | k__Fungi(100);p__Ascomycota(100);c__Saccharomycetes(100);o__Saccharomycetales(100);f__Metschnikowiaceae(100);g__Metschnikowia(100);s__Metschnikowia_pulcherrima(100); | Metschnikowia | 13 | Saprotroph | \|Undefined Saprotroph\| | Yeast | NULL | Probable | As nectar-tap_saprotroph (Põlme et al. 2020); Decay sugar-rich substrates (Põlme et al. 2020) | Põlme S, et al. 2020. Fungal Diversity 105:1-16 ((https://doi.org/10.1007/s13225-020-00466-2)) |
| Otu45288 | 1 | k__Fungi(100);p__Ascomycota(100);c__Saccharomycetes(100);o__Saccharomycetales(100);f__Metschnikowiaceae(100);g__Metschnikowia(100);g__Metschnikowia_unclassified(100); | Metschnikowia | 13 | Saprotroph | \|Undefined Saprotroph\| | Yeast | NULL | Probable | As nectar-tap_saprotroph (Põlme et al. 2020); Decay sugar-rich substrates (Põlme et al. 2020) | Põlme S, et al. 2020. Fungal Diversity 105:1-16 ((https://doi.org/10.1007/s13225-020-00466-2)) |
| Otu45289 | 1 | k__Fungi(100);p__Ascomycota(100);c__Saccharomycetes(100);o__Saccharomycetales(100);f__Metschnikowiaceae(100);g__Metschnikowia(100);g__Metschnikowia_unclassified(100); | Metschnikowia | 13 | Saprotroph | \|Undefined Saprotroph\| | Yeast | NULL | Probable | As nectar-tap_saprotroph (Põlme et al. 2020); Decay sugar-rich substrates (Põlme et al. 2020) | Põlme S, et al. 2020. Fungal Diversity 105:1-16 ((https://doi.org/10.1007/s13225-020-00466-2)) |
| Otu45290 | 1 | k__Fungi(100);p__Ascomycota(100);c__Saccharomycetes(100);o__Saccharomycetales(100);f__Metschnikowiaceae(100);g__Metschnikowia(100);s__Metschnikowia_pulcherrima(100); | Metschnikowia | 13 | Saprotroph | \|Undefined Saprotroph\| | Yeast | NULL | Probable | As nectar-tap_saprotroph (Põlme et al. 2020); Decay sugar-rich substrates (Põlme et al. 2020) | Põlme S, et al. 2020. Fungal Diversity 105:1-16 ((https://doi.org/10.1007/s13225-020-00466-2)) |
| Otu45291 | 1 | k__Fungi(100);p__Ascomycota(100);c__Saccharomycetes(100);o__Saccharomycetales(100);f__Metschnikowiaceae(100);g__Metschnikowia(100);s__Metschnikowia_pulcherrima(100); | Metschnikowia | 13 | Saprotroph | \|Undefined Saprotroph\| | Yeast | NULL | Probable | As nectar-tap_saprotroph (Põlme et al. 2020); Decay sugar-rich substrates (Põlme et al. 2020) | Põlme S, et al. 2020. Fungal Diversity 105:1-16 ((https://doi.org/10.1007/s13225-020-00466-2)) |
| Otu45292 | 1 | k__Fungi(100);p__Ascomycota(100);c__Saccharomycetes(100);o__Saccharomycetales(100);f__Metschnikowiaceae(100);g__Metschnikowia(100);s__Metschnikowia_pulcherrima(100); | Metschnikowia | 13 | Saprotroph | \|Undefined Saprotroph\| | Yeast | NULL | Probable | As nectar-tap_saprotroph (Põlme et al. 2020); Decay sugar-rich substrates (Põlme et al. 2020) | Põlme S, et al. 2020. Fungal Diversity 105:1-16 ((https://doi.org/10.1007/s13225-020-00466-2)) |
| Otu45293 | 1 | k__Fungi(100);p__Ascomycota(100);c__Saccharomycetes(100);o__Saccharomycetales(100);f__Metschnikowiaceae(100);g__Metschnikowia(100);s__Metschnikowia_pulcherrima(100); | Metschnikowia | 13 | Saprotroph | \|Undefined Saprotroph\| | Yeast | NULL | Probable | As nectar-tap_saprotroph (Põlme et al. 2020); Decay sugar-rich substrates (Põlme et al. 2020) | Põlme S, et al. 2020. Fungal Diversity 105:1-16 ((https://doi.org/10.1007/s13225-020-00466-2)) |
| Otu45294 | 1 | k__Fungi(100);p__Basidiomycota(100);c__Exobasidiomycetes(100);o__Exobasidiomycetes_order_Incertae_sedis(100);f__Exobasidiomycetes_family_Incertae_sedis(100);g__Acaromyces(100);s__Acaromyces_ingoldii(100); | Acaromyces | 13 | Pathotroph | \|Animal Parasite\|-Animal Pathogen-Plant Pathogen | Dimorphic | NULL | Highly Probable | An anamorphic fungus isolated from cadavers of the citrus rust mite (P. oleivora) infesting grapefruit (C. paradisi) leaves (Boekhout et al. 2003); Decay leaf, fruit, seed, animal material (Põlme et al. 2020); Host - mites (Põlme et al. 2020); Similar to the yeast-like fungi classified in the Ustilaginales, such as Pseudozyma species (Boekhout et al. 2003) | Boekhout T, et al. 2003. International Journal of Systematic and Evolutionary Microbiology 53:1655-1664 ((https://doi.org/10.1099/ijs.0.02434-0)); Põlme S, et al. 2020. Fungal Diversity 105:1-16 ((https://doi.org/10.1007/s13225-020-00466-2)) |
| Otu45295 | 1 | k__Fungi(100);p__Ascomycota(100);c__Dothideomycetes(100);o__Pleosporales(100);f__Pleosporaceae(100);g__Alternaria(100);g__Alternaria_unclassified(100); | Alternaria | 13 | Pathotroph-Saprotroph-Symbiotroph | Animal Pathogen-Endophyte-\|Plant Pathogen\|-Plant Saprotroph-Wood Saprotroph | Microfungus | Soft Rot | Possible | As foliar_endophyte (Põlme et al. 2020); As litter_saprotroph (Põlme et al. 2020); Decay leaf, fruit, seed (Põlme et al. 2020) | Duncan CG, Eslyn WE. 1966. Mycologia 58:642-664 ((https://doi.org/10.2307/3757045)); Seehann G, et al. 1975. List of Fungi in Soft-Rot Tissue Tests. Institute fur Holzbiologie und Holzschutz, Bundesforschungsanstalt fur Forst - und Holzwirtshaft, Hamburg - Lohbrugge, Germany ((https://www.irg-wp.com/irgdocs/details.php?d6493033-c9ea-40f3-8e3e-5697f32d640d)); Tedersoo L, et al. 2014. Science 346:e1256688 ((https://doi.org/10.1126/science.1256688)); Põlme S, et al. 2020. Fungal Diversity 105:1-16 ((https://doi.org/10.1007/s13225-020-00466-2)) |
| Otu45296 | 1 | k__Fungi(100);p__Ascomycota(100);c__Dothideomycetes(100);o__Pleosporales(100);f__Sporormiaceae(100);g__Preussia(100);s__Preussia_sp_GP_2006a(100); | Preussia | 13 | Saprotroph | \|Dung Saprotroph\|-Undefined Saprotroph | Microfungus | NULL | Probable | Decay dung (Põlme et al. 2020) | Tedersoo L, et al. 2014. Science 346:e1256688 ((https://doi.org/10.1126/science.1256688)); Põlme S, et al. 2020. Fungal Diversity 105:1-16 ((https://doi.org/10.1007/s13225-020-00466-2)) |
| Otu45297 | 1 | k__Fungi(100);p__Ascomycota(100);c__Dothideomycetes(100);o__Pleosporales(100);f__Phaeosphaeriaceae(100);f__Phaeosphaeriaceae_unclassified(100);f__Phaeosphaeriaceae_unclassified(100); | Phaeosphaeriaceae | 9 | Pathotroph-Saprotroph | Fungal Parasite-Plant Pathogen-Plant Saprotroph | Microfungus | NULL | Probable | Necrotrophic or saprobic on a wide range of plants, especially monocotyledons; a few species parasitic on other fungi (Cannon & Kirk 2007) | Cannon PF, Kirk PM. 2007. Fungal Families of the World. CAB International, Cambridge (ISBN: 978-0851998275) |
| Otu45299 | 1 | k__Fungi(100);p__Ascomycota(100);c__Dothideomycetes(100);o__Pleosporales(100);f__Phaeosphaeriaceae(100);f__Phaeosphaeriaceae_unclassified(100);f__Phaeosphaeriaceae_unclassified(100); | Phaeosphaeriaceae | 9 | Pathotroph-Saprotroph | Fungal Parasite-Plant Pathogen-Plant Saprotroph | Microfungus | NULL | Probable | Necrotrophic or saprobic on a wide range of plants, especially monocotyledons; a few species parasitic on other fungi (Cannon & Kirk 2007) | Cannon PF, Kirk PM. 2007. Fungal Families of the World. CAB International, Cambridge (ISBN: 978-0851998275) |
| Otu45300 | 1 | k__Fungi(100);p__Ascomycota(100);c__Dothideomycetes(100);o__Pleosporales(100);f__Tubeufiaceae(100);g__unclassified_Tubeufiaceae(100);s__Tubeufiaceae_sp(100); | Tubeufiaceae | 9 | Pathotroph-Saprotroph | Animal Pathogen-Undefined Saprotroph | NULL | NULL | Probable | Biotrophic or saprobic, especially on scale insects or other fungi. The anamorphic forms are fequently aero-aquatic (Cannon & Kirk 2007) | Cannon PF, Kirk PM. 2007. Fungal Families of the World. CAB International, Cambridge (ISBN: 978-0851998275) |
| Otu45301 | 1 | k__Fungi(100);p__Ascomycota(100);c__Saccharomycetes(100);o__Saccharomycetales(100);f__Metschnikowiaceae(100);g__Metschnikowia(100);g__Metschnikowia_unclassified(100); | Metschnikowia | 13 | Saprotroph | \|Undefined Saprotroph\| | Yeast | NULL | Probable | As nectar-tap_saprotroph (Põlme et al. 2020); Decay sugar-rich substrates (Põlme et al. 2020) | Põlme S, et al. 2020. Fungal Diversity 105:1-16 ((https://doi.org/10.1007/s13225-020-00466-2)) |
| Otu45302 | 1 | k__Fungi(100);p__Ascomycota(100);c__Saccharomycetes(100);o__Saccharomycetales(100);f__Metschnikowiaceae(100);g__Metschnikowia(100);g__Metschnikowia_unclassified(100); | Metschnikowia | 13 | Saprotroph | \|Undefined Saprotroph\| | Yeast | NULL | Probable | As nectar-tap_saprotroph (Põlme et al. 2020); Decay sugar-rich substrates (Põlme et al. 2020) | Põlme S, et al. 2020. Fungal Diversity 105:1-16 ((https://doi.org/10.1007/s13225-020-00466-2)) |
| Otu45303 | 1 | k__Fungi(100);p__Ascomycota(100);c__Saccharomycetes(100);o__Saccharomycetales(100);f__Metschnikowiaceae(100);g__Metschnikowia(100);s__Metschnikowia_pulcherrima(100); | Metschnikowia | 13 | Saprotroph | \|Undefined Saprotroph\| | Yeast | NULL | Probable | As nectar-tap_saprotroph (Põlme et al. 2020); Decay sugar-rich substrates (Põlme et al. 2020) | Põlme S, et al. 2020. Fungal Diversity 105:1-16 ((https://doi.org/10.1007/s13225-020-00466-2)) |
| Otu45304 | 1 | k__Fungi(100);p__Ascomycota(100);c__Saccharomycetes(100);o__Saccharomycetales(100);f__Metschnikowiaceae(100);g__Metschnikowia(100);g__Metschnikowia_unclassified(100); | Metschnikowia | 13 | Saprotroph | \|Undefined Saprotroph\| | Yeast | NULL | Probable | As nectar-tap_saprotroph (Põlme et al. 2020); Decay sugar-rich substrates (Põlme et al. 2020) | Põlme S, et al. 2020. Fungal Diversity 105:1-16 ((https://doi.org/10.1007/s13225-020-00466-2)) |
| Otu45305 | 1 | k__Fungi(100);p__Ascomycota(100);c__Saccharomycetes(100);o__Saccharomycetales(100);f__Metschnikowiaceae(100);g__Metschnikowia(100);s__Metschnikowia_pulcherrima(100); | Metschnikowia | 13 | Saprotroph | \|Undefined Saprotroph\| | Yeast | NULL | Probable | As nectar-tap_saprotroph (Põlme et al. 2020); Decay sugar-rich substrates (Põlme et al. 2020) | Põlme S, et al. 2020. Fungal Diversity 105:1-16 ((https://doi.org/10.1007/s13225-020-00466-2)) |
| Otu45306 | 1 | k__Fungi(100);p__Ascomycota(100);c__Saccharomycetes(100);o__Saccharomycetales(100);f__Metschnikowiaceae(100);g__Metschnikowia(100);g__Metschnikowia_unclassified(100); | Metschnikowia | 13 | Saprotroph | \|Undefined Saprotroph\| | Yeast | NULL | Probable | As nectar-tap_saprotroph (Põlme et al. 2020); Decay sugar-rich substrates (Põlme et al. 2020) | Põlme S, et al. 2020. Fungal Diversity 105:1-16 ((https://doi.org/10.1007/s13225-020-00466-2)) |
| Otu45307 | 1 | k__Fungi(100);p__Ascomycota(100);c__Saccharomycetes(100);o__Saccharomycetales(100);f__Metschnikowiaceae(100);g__Metschnikowia(100);g__Metschnikowia_unclassified(100); | Metschnikowia | 13 | Saprotroph | \|Undefined Saprotroph\| | Yeast | NULL | Probable | As nectar-tap_saprotroph (Põlme et al. 2020); Decay sugar-rich substrates (Põlme et al. 2020) | Põlme S, et al. 2020. Fungal Diversity 105:1-16 ((https://doi.org/10.1007/s13225-020-00466-2)) |
| Otu45308 | 1 | k__Fungi(100);p__Ascomycota(100);c__Saccharomycetes(100);o__Saccharomycetales(100);f__Metschnikowiaceae(100);g__Metschnikowia(100);s__Metschnikowia_pulcherrima(100); | Metschnikowia | 13 | Saprotroph | \|Undefined Saprotroph\| | Yeast | NULL | Probable | As nectar-tap_saprotroph (Põlme et al. 2020); Decay sugar-rich substrates (Põlme et al. 2020) | Põlme S, et al. 2020. Fungal Diversity 105:1-16 ((https://doi.org/10.1007/s13225-020-00466-2)) |
| Otu45309 | 1 | k__Fungi(100);p__Ascomycota(100);c__Saccharomycetes(100);o__Saccharomycetales(100);f__Metschnikowiaceae(100);g__Metschnikowia(100);s__Metschnikowia_pulcherrima(100); | Metschnikowia | 13 | Saprotroph | \|Undefined Saprotroph\| | Yeast | NULL | Probable | As nectar-tap_saprotroph (Põlme et al. 2020); Decay sugar-rich substrates (Põlme et al. 2020) | Põlme S, et al. 2020. Fungal Diversity 105:1-16 ((https://doi.org/10.1007/s13225-020-00466-2)) |
| Otu45310 | 1 | k__Fungi(100);p__Ascomycota(100);c__Saccharomycetes(100);o__Saccharomycetales(100);f__Metschnikowiaceae(100);g__Metschnikowia(100);s__Metschnikowia_pulcherrima(100); | Metschnikowia | 13 | Saprotroph | \|Undefined Saprotroph\| | Yeast | NULL | Probable | As nectar-tap_saprotroph (Põlme et al. 2020); Decay sugar-rich substrates (Põlme et al. 2020) | Põlme S, et al. 2020. Fungal Diversity 105:1-16 ((https://doi.org/10.1007/s13225-020-00466-2)) |
| Otu45311 | 1 | k__Fungi(100);p__Ascomycota(100);c__Saccharomycetes(100);o__Saccharomycetales(100);f__Metschnikowiaceae(100);g__Metschnikowia(100);s__Metschnikowia_pulcherrima(100); | Metschnikowia | 13 | Saprotroph | \|Undefined Saprotroph\| | Yeast | NULL | Probable | As nectar-tap_saprotroph (Põlme et al. 2020); Decay sugar-rich substrates (Põlme et al. 2020) | Põlme S, et al. 2020. Fungal Diversity 105:1-16 ((https://doi.org/10.1007/s13225-020-00466-2)) |
| Otu45312 | 1 | k__Fungi(100);p__Ascomycota(100);c__Saccharomycetes(100);o__Saccharomycetales(100);f__Metschnikowiaceae(100);g__Metschnikowia(100);s__Metschnikowia_pulcherrima(100); | Metschnikowia | 13 | Saprotroph | \|Undefined Saprotroph\| | Yeast | NULL | Probable | As nectar-tap_saprotroph (Põlme et al. 2020); Decay sugar-rich substrates (Põlme et al. 2020) | Põlme S, et al. 2020. Fungal Diversity 105:1-16 ((https://doi.org/10.1007/s13225-020-00466-2)) |
| Otu45313 | 1 | k__Fungi(100);p__Ascomycota(100);c__Saccharomycetes(100);o__Saccharomycetales(100);f__Metschnikowiaceae(100);g__Metschnikowia(100);s__Metschnikowia_pulcherrima(100); | Metschnikowia | 13 | Saprotroph | \|Undefined Saprotroph\| | Yeast | NULL | Probable | As nectar-tap_saprotroph (Põlme et al. 2020); Decay sugar-rich substrates (Põlme et al. 2020) | Põlme S, et al. 2020. Fungal Diversity 105:1-16 ((https://doi.org/10.1007/s13225-020-00466-2)) |
| Otu45314 | 1 | k__Fungi(100);p__Ascomycota(100);c__Saccharomycetes(100);o__Saccharomycetales(100);f__Metschnikowiaceae(100);g__Metschnikowia(100);g__Metschnikowia_unclassified(100); | Metschnikowia | 13 | Saprotroph | \|Undefined Saprotroph\| | Yeast | NULL | Probable | As nectar-tap_saprotroph (Põlme et al. 2020); Decay sugar-rich substrates (Põlme et al. 2020) | Põlme S, et al. 2020. Fungal Diversity 105:1-16 ((https://doi.org/10.1007/s13225-020-00466-2)) |
| Otu45315 | 1 | k__Fungi(100);p__Ascomycota(100);c__Saccharomycetes(100);o__Saccharomycetales(100);f__Metschnikowiaceae(100);g__Metschnikowia(100);s__Metschnikowia_pulcherrima(100); | Metschnikowia | 13 | Saprotroph | \|Undefined Saprotroph\| | Yeast | NULL | Probable | As nectar-tap_saprotroph (Põlme et al. 2020); Decay sugar-rich substrates (Põlme et al. 2020) | Põlme S, et al. 2020. Fungal Diversity 105:1-16 ((https://doi.org/10.1007/s13225-020-00466-2)) |
| Otu45316 | 1 | k__Fungi(100);p__Ascomycota(100);c__Saccharomycetes(100);o__Saccharomycetales(100);f__Metschnikowiaceae(100);g__Metschnikowia(100);s__Metschnikowia_pulcherrima(100); | Metschnikowia | 13 | Saprotroph | \|Undefined Saprotroph\| | Yeast | NULL | Probable | As nectar-tap_saprotroph (Põlme et al. 2020); Decay sugar-rich substrates (Põlme et al. 2020) | Põlme S, et al. 2020. Fungal Diversity 105:1-16 ((https://doi.org/10.1007/s13225-020-00466-2)) |
| Otu45317 | 1 | k__Fungi(100);p__Ascomycota(100);c__Saccharomycetes(100);o__Saccharomycetales(100);f__Metschnikowiaceae(100);g__Metschnikowia(100);s__Metschnikowia_pulcherrima(100); | Metschnikowia | 13 | Saprotroph | \|Undefined Saprotroph\| | Yeast | NULL | Probable | As nectar-tap_saprotroph (Põlme et al. 2020); Decay sugar-rich substrates (Põlme et al. 2020) | Põlme S, et al. 2020. Fungal Diversity 105:1-16 ((https://doi.org/10.1007/s13225-020-00466-2)) |
| Otu45318 | 1 | k__Fungi(100);p__Ascomycota(100);c__Saccharomycetes(100);o__Saccharomycetales(100);f__Metschnikowiaceae(100);g__Metschnikowia(100);s__Metschnikowia_pulcherrima(100); | Metschnikowia | 13 | Saprotroph | \|Undefined Saprotroph\| | Yeast | NULL | Probable | As nectar-tap_saprotroph (Põlme et al. 2020); Decay sugar-rich substrates (Põlme et al. 2020) | Põlme S, et al. 2020. Fungal Diversity 105:1-16 ((https://doi.org/10.1007/s13225-020-00466-2)) |
| Otu45319 | 1 | k__Fungi(100);p__Ascomycota(100);c__Saccharomycetes(100);o__Saccharomycetales(100);f__Metschnikowiaceae(100);g__Metschnikowia(100);s__Metschnikowia_pulcherrima(100); | Metschnikowia | 13 | Saprotroph | \|Undefined Saprotroph\| | Yeast | NULL | Probable | As nectar-tap_saprotroph (Põlme et al. 2020); Decay sugar-rich substrates (Põlme et al. 2020) | Põlme S, et al. 2020. Fungal Diversity 105:1-16 ((https://doi.org/10.1007/s13225-020-00466-2)) |
| Otu45320 | 1 | k__Fungi(100);p__Ascomycota(100);c__Saccharomycetes(100);o__Saccharomycetales(100);f__Metschnikowiaceae(100);g__Metschnikowia(100);s__Metschnikowia_pulcherrima(100); | Metschnikowia | 13 | Saprotroph | \|Undefined Saprotroph\| | Yeast | NULL | Probable | As nectar-tap_saprotroph (Põlme et al. 2020); Decay sugar-rich substrates (Põlme et al. 2020) | Põlme S, et al. 2020. Fungal Diversity 105:1-16 ((https://doi.org/10.1007/s13225-020-00466-2)) |
| Otu45321 | 1 | k__Fungi(100);p__Ascomycota(100);c__Saccharomycetes(100);o__Saccharomycetales(100);f__Metschnikowiaceae(100);g__Metschnikowia(100);s__Metschnikowia_pulcherrima(100); | Metschnikowia | 13 | Saprotroph | \|Undefined Saprotroph\| | Yeast | NULL | Probable | As nectar-tap_saprotroph (Põlme et al. 2020); Decay sugar-rich substrates (Põlme et al. 2020) | Põlme S, et al. 2020. Fungal Diversity 105:1-16 ((https://doi.org/10.1007/s13225-020-00466-2)) |
| Otu45322 | 1 | k__Fungi(100);p__Basidiomycota(100);c__Saccharomycetes(100);o__Saccharomycetales(100);f__Metschnikowiaceae(100);g__Metschnikowia(100);s__Metschnikowia_pulcherrima(100); | Metschnikowia | 13 | Saprotroph | \|Undefined Saprotroph\| | Yeast | NULL | Probable | As nectar-tap_saprotroph (Põlme et al. 2020); Decay sugar-rich substrates (Põlme et al. 2020) | Põlme S, et al. 2020. Fungal Diversity 105:1-16 ((https://doi.org/10.1007/s13225-020-00466-2)) |
| Otu45323 | 1 | k__Fungi(100);p__Ascomycota(100);c__Saccharomycetes(100);o__Saccharomycetales(100);f__Metschnikowiaceae(100);g__Metschnikowia(100);g__Metschnikowia_unclassified(100); | Metschnikowia | 13 | Saprotroph | \|Undefined Saprotroph\| | Yeast | NULL | Probable | As nectar-tap_saprotroph (Põlme et al. 2020); Decay sugar-rich substrates (Põlme et al. 2020) | Põlme S, et al. 2020. Fungal Diversity 105:1-16 ((https://doi.org/10.1007/s13225-020-00466-2)) |
| Otu45324 | 1 | k__Fungi(100);p__Ascomycota(100);c__Saccharomycetes(100);o__Saccharomycetales(100);f__Metschnikowiaceae(100);g__Metschnikowia(100);s__Metschnikowia_pulcherrima(100); | Metschnikowia | 13 | Saprotroph | \|Undefined Saprotroph\| | Yeast | NULL | Probable | As nectar-tap_saprotroph (Põlme et al. 2020); Decay sugar-rich substrates (Põlme et al. 2020) | Põlme S, et al. 2020. Fungal Diversity 105:1-16 ((https://doi.org/10.1007/s13225-020-00466-2)) |
| Otu45325 | 1 | k__Fungi(100);p__Ascomycota(100);c__Saccharomycetes(100);o__Saccharomycetales(100);f__Metschnikowiaceae(100);g__Metschnikowia(100);s__Metschnikowia_pulcherrima(100); | Metschnikowia | 13 | Saprotroph | \|Undefined Saprotroph\| | Yeast | NULL | Probable | As nectar-tap_saprotroph (Põlme et al. 2020); Decay sugar-rich substrates (Põlme et al. 2020) | Põlme S, et al. 2020. Fungal Diversity 105:1-16 ((https://doi.org/10.1007/s13225-020-00466-2)) |
| Otu45326 | 1 | k__Fungi(100);p__Ascomycota(100);c__Saccharomycetes(100);o__Saccharomycetales(100);f__Metschnikowiaceae(100);g__Metschnikowia(100);g__Metschnikowia_unclassified(100); | Metschnikowia | 13 | Saprotroph | \|Undefined Saprotroph\| | Yeast | NULL | Probable | As nectar-tap_saprotroph (Põlme et al. 2020); Decay sugar-rich substrates (Põlme et al. 2020) | Põlme S, et al. 2020. Fungal Diversity 105:1-16 ((https://doi.org/10.1007/s13225-020-00466-2)) |
| Otu45327 | 1 | k__Fungi(100);p__Ascomycota(100);c__Saccharomycetes(100);o__Saccharomycetales(100);f__Metschnikowiaceae(100);g__Metschnikowia(100);s__Metschnikowia_pulcherrima(100); | Metschnikowia | 13 | Saprotroph | \|Undefined Saprotroph\| | Yeast | NULL | Probable | As nectar-tap_saprotroph (Põlme et al. 2020); Decay sugar-rich substrates (Põlme et al. 2020) | Põlme S, et al. 2020. Fungal Diversity 105:1-16 ((https://doi.org/10.1007/s13225-020-00466-2)) |
| Otu45328 | 1 | k__Fungi(100);p__Ascomycota(100);c__Saccharomycetes(100);o__Saccharomycetales(100);f__Metschnikowiaceae(100);g__Metschnikowia(100);s__Metschnikowia_pulcherrima(100); | Metschnikowia | 13 | Saprotroph | \|Undefined Saprotroph\| | Yeast | NULL | Probable | As nectar-tap_saprotroph (Põlme et al. 2020); Decay sugar-rich substrates (Põlme et al. 2020) | Põlme S, et al. 2020. Fungal Diversity 105:1-16 ((https://doi.org/10.1007/s13225-020-00466-2)) |
| Otu45330 | 1 | k__Fungi(100);p__Ascomycota(100);c__Saccharomycetes(100);o__Saccharomycetales(100);f__Metschnikowiaceae(100);g__Metschnikowia(100);s__Metschnikowia_pulcherrima(100); | Metschnikowia | 13 | Saprotroph | \|Undefined Saprotroph\| | Yeast | NULL | Probable | As nectar-tap_saprotroph (Põlme et al. 2020); Decay sugar-rich substrates (Põlme et al. 2020) | Põlme S, et al. 2020. Fungal Diversity 105:1-16 ((https://doi.org/10.1007/s13225-020-00466-2)) |
| Otu45331 | 1 | k__Fungi(100);p__Ascomycota(100);c__Saccharomycetes(100);o__Saccharomycetales(100);f__Metschnikowiaceae(100);g__Metschnikowia(100);s__Metschnikowia_pulcherrima(100); | Metschnikowia | 13 | Saprotroph | \|Undefined Saprotroph\| | Yeast | NULL | Probable | As nectar-tap_saprotroph (Põlme et al. 2020); Decay sugar-rich substrates (Põlme et al. 2020) | Põlme S, et al. 2020. Fungal Diversity 105:1-16 ((https://doi.org/10.1007/s13225-020-00466-2)) |
| Otu45332 | 1 | k__Fungi(100);p__Ascomycota(100);c__Saccharomycetes(100);o__Saccharomycetales(100);f__Metschnikowiaceae(100);g__Metschnikowia(100);s__Metschnikowia_pulcherrima(100); | Metschnikowia | 13 | Saprotroph | \|Undefined Saprotroph\| | Yeast | NULL | Probable | As nectar-tap_saprotroph (Põlme et al. 2020); Decay sugar-rich substrates (Põlme et al. 2020) | Põlme S, et al. 2020. Fungal Diversity 105:1-16 ((https://doi.org/10.1007/s13225-020-00466-2)) |
| Otu45333 | 1 | k__Fungi(100);p__Ascomycota(100);c__Saccharomycetes(100);o__Saccharomycetales(100);f__Metschnikowiaceae(100);g__Metschnikowia(100);s__Metschnikowia_pulcherrima(100); | Metschnikowia | 13 | Saprotroph | \|Undefined Saprotroph\| | Yeast | NULL | Probable | As nectar-tap_saprotroph (Põlme et al. 2020); Decay sugar-rich substrates (Põlme et al. 2020) | Põlme S, et al. 2020. Fungal Diversity 105:1-16 ((https://doi.org/10.1007/s13225-020-00466-2)) |
| Otu45334 | 1 | k__Fungi(100);p__Ascomycota(100);c__Saccharomycetes(100);o__Saccharomycetales(100);f__Metschnikowiaceae(100);g__Metschnikowia(100);s__Metschnikowia_pulcherrima(100); | Metschnikowia | 13 | Saprotroph | \|Undefined Saprotroph\| | Yeast | NULL | Probable | As nectar-tap_saprotroph (Põlme et al. 2020); Decay sugar-rich substrates (Põlme et al. 2020) | Põlme S, et al. 2020. Fungal Diversity 105:1-16 ((https://doi.org/10.1007/s13225-020-00466-2)) |
| Otu45335 | 1 | k__Fungi(100);p__Ascomycota(100);c__Saccharomycetes(100);o__Saccharomycetales(100);f__Metschnikowiaceae(100);g__Metschnikowia(100);g__Metschnikowia_unclassified(100); | Metschnikowia | 13 | Saprotroph | \|Undefined Saprotroph\| | Yeast | NULL | Probable | As nectar-tap_saprotroph (Põlme et al. 2020); Decay sugar-rich substrates (Põlme et al. 2020) | Põlme S, et al. 2020. Fungal Diversity 105:1-16 ((https://doi.org/10.1007/s13225-020-00466-2)) |
| Otu45336 | 1 | k__Fungi(100);p__Ascomycota(100);c__Saccharomycetes(100);o__Saccharomycetales(100);f__Metschnikowiaceae(100);g__Metschnikowia(100);s__Metschnikowia_pulcherrima(100); | Metschnikowia | 13 | Saprotroph | \|Undefined Saprotroph\| | Yeast | NULL | Probable | As nectar-tap_saprotroph (Põlme et al. 2020); Decay sugar-rich substrates (Põlme et al. 2020) | Põlme S, et al. 2020. Fungal Diversity 105:1-16 ((https://doi.org/10.1007/s13225-020-00466-2)) |
| Otu45337 | 1 | k__Fungi(100);p__Ascomycota(100);c__Saccharomycetes(100);o__Saccharomycetales(100);f__Metschnikowiaceae(100);g__Metschnikowia(100);s__Metschnikowia_pulcherrima(100); | Metschnikowia | 13 | Saprotroph | \|Undefined Saprotroph\| | Yeast | NULL | Probable | As nectar-tap_saprotroph (Põlme et al. 2020); Decay sugar-rich substrates (Põlme et al. 2020) | Põlme S, et al. 2020. Fungal Diversity 105:1-16 ((https://doi.org/10.1007/s13225-020-00466-2)) |
| Otu45338 | 1 | k__Fungi(100);p__Ascomycota(100);c__Saccharomycetes(100);o__Saccharomycetales(100);f__Metschnikowiaceae(100);g__Metschnikowia(100);s__Metschnikowia_pulcherrima(100); | Metschnikowia | 13 | Saprotroph | \|Undefined Saprotroph\| | Yeast | NULL | Probable | As nectar-tap_saprotroph (Põlme et al. 2020); Decay sugar-rich substrates (Põlme et al. 2020) | Põlme S, et al. 2020. Fungal Diversity 105:1-16 ((https://doi.org/10.1007/s13225-020-00466-2)) |
| Otu45339 | 1 | k__Fungi(100);p__Ascomycota(100);c__Saccharomycetes(100);o__Saccharomycetales(100);f__Metschnikowiaceae(100);g__Metschnikowia(100);s__Metschnikowia_pulcherrima(100); | Metschnikowia | 13 | Saprotroph | \|Undefined Saprotroph\| | Yeast | NULL | Probable | As nectar-tap_saprotroph (Põlme et al. 2020); Decay sugar-rich substrates (Põlme et al. 2020) | Põlme S, et al. 2020. Fungal Diversity 105:1-16 ((https://doi.org/10.1007/s13225-020-00466-2)) |
| Otu45340 | 1 | k__Fungi(100);p__Ascomycota(100);c__Saccharomycetes(100);o__Saccharomycetales(100);f__Metschnikowiaceae(100);g__Metschnikowia(100);s__Metschnikowia_pulcherrima(100); | Metschnikowia | 13 | Saprotroph | \|Undefined Saprotroph\| | Yeast | NULL | Probable | As nectar-tap_saprotroph (Põlme et al. 2020); Decay sugar-rich substrates (Põlme et al. 2020) | Põlme S, et al. 2020. Fungal Diversity 105:1-16 ((https://doi.org/10.1007/s13225-020-00466-2)) |
| Otu45341 | 1 | k__Fungi(100);p__Ascomycota(100);c__Saccharomycetes(100);o__Saccharomycetales(100);f__Metschnikowiaceae(100);g__Metschnikowia(100);s__Metschnikowia_pulcherrima(100); | Metschnikowia | 13 | Saprotroph | \|Undefined Saprotroph\| | Yeast | NULL | Probable | As nectar-tap_saprotroph (Põlme et al. 2020); Decay sugar-rich substrates (Põlme et al. 2020) | Põlme S, et al. 2020. Fungal Diversity 105:1-16 ((https://doi.org/10.1007/s13225-020-00466-2)) |
| Otu45342 | 1 | k__Fungi(100);p__Ascomycota(100);c__Saccharomycetes(100);o__Saccharomycetales(100);f__Metschnikowiaceae(100);g__Metschnikowia(100);g__Metschnikowia_unclassified(100); | Metschnikowia | 13 | Saprotroph | \|Undefined Saprotroph\| | Yeast | NULL | Probable | As nectar-tap_saprotroph (Põlme et al. 2020); Decay sugar-rich substrates (Põlme et al. 2020) | Põlme S, et al. 2020. Fungal Diversity 105:1-16 ((https://doi.org/10.1007/s13225-020-00466-2)) |
| Otu45343 | 1 | k__Fungi(100);p__Ascomycota(100);c__Saccharomycetes(100);o__Saccharomycetales(100);f__Metschnikowiaceae(100);g__Metschnikowia(100);s__Metschnikowia_pulcherrima(100); | Metschnikowia | 13 | Saprotroph | \|Undefined Saprotroph\| | Yeast | NULL | Probable | As nectar-tap_saprotroph (Põlme et al. 2020); Decay sugar-rich substrates (Põlme et al. 2020) | Põlme S, et al. 2020. Fungal Diversity 105:1-16 ((https://doi.org/10.1007/s13225-020-00466-2)) |
| Otu45344 | 1 | k__Fungi(100);p__Ascomycota(100);c__Saccharomycetes(100);o__Saccharomycetales(100);f__Metschnikowiaceae(100);g__Metschnikowia(100);s__Metschnikowia_pulcherrima(100); | Metschnikowia | 13 | Saprotroph | \|Undefined Saprotroph\| | Yeast | NULL | Probable | As nectar-tap_saprotroph (Põlme et al. 2020); Decay sugar-rich substrates (Põlme et al. 2020) | Põlme S, et al. 2020. Fungal Diversity 105:1-16 ((https://doi.org/10.1007/s13225-020-00466-2)) |
| Otu45345 | 1 | k__Fungi(100);p__Ascomycota(100);c__Saccharomycetes(100);o__Saccharomycetales(100);f__Metschnikowiaceae(100);g__Metschnikowia(100);s__Metschnikowia_pulcherrima(100); | Metschnikowia | 13 | Saprotroph | \|Undefined Saprotroph\| | Yeast | NULL | Probable | As nectar-tap_saprotroph (Põlme et al. 2020); Decay sugar-rich substrates (Põlme et al. 2020) | Põlme S, et al. 2020. Fungal Diversity 105:1-16 ((https://doi.org/10.1007/s13225-020-00466-2)) |
| Otu45346 | 1 | k__Fungi(100);p__Ascomycota(100);c__Saccharomycetes(100);o__Saccharomycetales(100);f__Metschnikowiaceae(100);g__Metschnikowia(100);g__Metschnikowia_unclassified(100); | Metschnikowia | 13 | Saprotroph | \|Undefined Saprotroph\| | Yeast | NULL | Probable | As nectar-tap_saprotroph (Põlme et al. 2020); Decay sugar-rich substrates (Põlme et al. 2020) | Põlme S, et al. 2020. Fungal Diversity 105:1-16 ((https://doi.org/10.1007/s13225-020-00466-2)) |
| Otu45347 | 1 | k__Fungi(100);p__Ascomycota(100);c__Saccharomycetes(100);o__Saccharomycetales(100);f__Metschnikowiaceae(100);g__Metschnikowia(100);s__Metschnikowia_pulcherrima(100); | Metschnikowia | 13 | Saprotroph | \|Undefined Saprotroph\| | Yeast | NULL | Probable | As nectar-tap_saprotroph (Põlme et al. 2020); Decay sugar-rich substrates (Põlme et al. 2020) | Põlme S, et al. 2020. Fungal Diversity 105:1-16 ((https://doi.org/10.1007/s13225-020-00466-2)) |
| Otu45348 | 1 | k__Fungi(100);p__Ascomycota(100);c__Saccharomycetes(100);o__Saccharomycetales(100);f__Metschnikowiaceae(100);g__Metschnikowia(100);g__Metschnikowia_unclassified(100); | Metschnikowia | 13 | Saprotroph | \|Undefined Saprotroph\| | Yeast | NULL | Probable | As nectar-tap_saprotroph (Põlme et al. 2020); Decay sugar-rich substrates (Põlme et al. 2020) | Põlme S, et al. 2020. Fungal Diversity 105:1-16 ((https://doi.org/10.1007/s13225-020-00466-2)) |
| Otu45349 | 1 | k__Fungi(100);p__Ascomycota(100);c__Saccharomycetes(100);o__Saccharomycetales(100);f__Metschnikowiaceae(100);g__Metschnikowia(100);s__Metschnikowia_pulcherrima(100); | Metschnikowia | 13 | Saprotroph | \|Undefined Saprotroph\| | Yeast | NULL | Probable | As nectar-tap_saprotroph (Põlme et al. 2020); Decay sugar-rich substrates (Põlme et al. 2020) | Põlme S, et al. 2020. Fungal Diversity 105:1-16 ((https://doi.org/10.1007/s13225-020-00466-2)) |
| Otu45350 | 1 | k__Fungi(100);p__Ascomycota(100);c__Saccharomycetes(100);o__Saccharomycetales(100);f__Metschnikowiaceae(100);g__Metschnikowia(100);s__Metschnikowia_pulcherrima(100); | Metschnikowia | 13 | Saprotroph | \|Undefined Saprotroph\| | Yeast | NULL | Probable | As nectar-tap_saprotroph (Põlme et al. 2020); Decay sugar-rich substrates (Põlme et al. 2020) | Põlme S, et al. 2020. Fungal Diversity 105:1-16 ((https://doi.org/10.1007/s13225-020-00466-2)) |
| Otu45351 | 1 | k__Fungi(100);p__Ascomycota(100);c__Saccharomycetes(100);o__Saccharomycetales(100);f__Metschnikowiaceae(100);g__Metschnikowia(100);s__Metschnikowia_pulcherrima(100); | Metschnikowia | 13 | Saprotroph | \|Undefined Saprotroph\| | Yeast | NULL | Probable | As nectar-tap_saprotroph (Põlme et al. 2020); Decay sugar-rich substrates (Põlme et al. 2020) | Põlme S, et al. 2020. Fungal Diversity 105:1-16 ((https://doi.org/10.1007/s13225-020-00466-2)) |
| Otu45352 | 1 | k__Fungi(100);p__Ascomycota(100);c__Saccharomycetes(100);o__Saccharomycetales(100);f__Metschnikowiaceae(100);g__Metschnikowia(100);s__Metschnikowia_pulcherrima(100); | Metschnikowia | 13 | Saprotroph | \|Undefined Saprotroph\| | Yeast | NULL | Probable | As nectar-tap_saprotroph (Põlme et al. 2020); Decay sugar-rich substrates (Põlme et al. 2020) | Põlme S, et al. 2020. Fungal Diversity 105:1-16 ((https://doi.org/10.1007/s13225-020-00466-2)) |
| Otu45353 | 1 | k__Fungi(100);p__Ascomycota(100);c__Saccharomycetes(100);o__Saccharomycetales(100);f__Metschnikowiaceae(100);g__Metschnikowia(100);s__Metschnikowia_pulcherrima(100); | Metschnikowia | 13 | Saprotroph | \|Undefined Saprotroph\| | Yeast | NULL | Probable | As nectar-tap_saprotroph (Põlme et al. 2020); Decay sugar-rich substrates (Põlme et al. 2020) | Põlme S, et al. 2020. Fungal Diversity 105:1-16 ((https://doi.org/10.1007/s13225-020-00466-2)) |
| Otu45354 | 1 | k__Fungi(100);p__Ascomycota(100);c__Saccharomycetes(100);o__Saccharomycetales(100);f__Metschnikowiaceae(100);g__Metschnikowia(100);s__Metschnikowia_pulcherrima(100); | Metschnikowia | 13 | Saprotroph | \|Undefined Saprotroph\| | Yeast | NULL | Probable | As nectar-tap_saprotroph (Põlme et al. 2020); Decay sugar-rich substrates (Põlme et al. 2020) | Põlme S, et al. 2020. Fungal Diversity 105:1-16 ((https://doi.org/10.1007/s13225-020-00466-2)) |
| Otu45355 | 1 | k__Fungi(100);p__Ascomycota(100);c__Saccharomycetes(100);o__Saccharomycetales(100);f__Metschnikowiaceae(100);g__Metschnikowia(100);s__Metschnikowia_pulcherrima(100); | Metschnikowia | 13 | Saprotroph | \|Undefined Saprotroph\| | Yeast | NULL | Probable | As nectar-tap_saprotroph (Põlme et al. 2020); Decay sugar-rich substrates (Põlme et al. 2020) | Põlme S, et al. 2020. Fungal Diversity 105:1-16 ((https://doi.org/10.1007/s13225-020-00466-2)) |
| Otu45356 | 1 | k__Fungi(100);p__Ascomycota(100);c__Saccharomycetes(100);o__Saccharomycetales(100);f__Metschnikowiaceae(100);g__Metschnikowia(100);s__Metschnikowia_pulcherrima(100); | Metschnikowia | 13 | Saprotroph | \|Undefined Saprotroph\| | Yeast | NULL | Probable | As nectar-tap_saprotroph (Põlme et al. 2020); Decay sugar-rich substrates (Põlme et al. 2020) | Põlme S, et al. 2020. Fungal Diversity 105:1-16 ((https://doi.org/10.1007/s13225-020-00466-2)) |
| Otu45357 | 1 | k__Fungi(100);p__Ascomycota(100);c__Saccharomycetes(100);o__Saccharomycetales(100);f__Metschnikowiaceae(100);g__Metschnikowia(100);g__Metschnikowia_unclassified(100); | Metschnikowia | 13 | Saprotroph | \|Undefined Saprotroph\| | Yeast | NULL | Probable | As nectar-tap_saprotroph (Põlme et al. 2020); Decay sugar-rich substrates (Põlme et al. 2020) | Põlme S, et al. 2020. Fungal Diversity 105:1-16 ((https://doi.org/10.1007/s13225-020-00466-2)) |
| Otu45358 | 1 | k__Fungi(100);p__Ascomycota(100);c__Saccharomycetes(100);o__Saccharomycetales(100);f__Metschnikowiaceae(100);g__Metschnikowia(100);s__Metschnikowia_pulcherrima(100); | Metschnikowia | 13 | Saprotroph | \|Undefined Saprotroph\| | Yeast | NULL | Probable | As nectar-tap_saprotroph (Põlme et al. 2020); Decay sugar-rich substrates (Põlme et al. 2020) | Põlme S, et al. 2020. Fungal Diversity 105:1-16 ((https://doi.org/10.1007/s13225-020-00466-2)) |
| Otu45359 | 1 | k__Fungi(100);p__Ascomycota(100);c__Saccharomycetes(100);o__Saccharomycetales(100);f__Metschnikowiaceae(100);g__Metschnikowia(100);s__Metschnikowia_pulcherrima(100); | Metschnikowia | 13 | Saprotroph | \|Undefined Saprotroph\| | Yeast | NULL | Probable | As nectar-tap_saprotroph (Põlme et al. 2020); Decay sugar-rich substrates (Põlme et al. 2020) | Põlme S, et al. 2020. Fungal Diversity 105:1-16 ((https://doi.org/10.1007/s13225-020-00466-2)) |
| Otu45360 | 1 | k__Fungi(100);p__Ascomycota(100);c__Saccharomycetes(100);o__Saccharomycetales(100);f__Metschnikowiaceae(100);g__Metschnikowia(100);s__Metschnikowia_pulcherrima(100); | Metschnikowia | 13 | Saprotroph | \|Undefined Saprotroph\| | Yeast | NULL | Probable | As nectar-tap_saprotroph (Põlme et al. 2020); Decay sugar-rich substrates (Põlme et al. 2020) | Põlme S, et al. 2020. Fungal Diversity 105:1-16 ((https://doi.org/10.1007/s13225-020-00466-2)) |
| Otu45361 | 1 | k__Fungi(100);p__Ascomycota(100);c__Saccharomycetes(100);o__Saccharomycetales(100);f__Metschnikowiaceae(100);g__Metschnikowia(100);g__Metschnikowia_unclassified(100); | Metschnikowia | 13 | Saprotroph | \|Undefined Saprotroph\| | Yeast | NULL | Probable | As nectar-tap_saprotroph (Põlme et al. 2020); Decay sugar-rich substrates (Põlme et al. 2020) | Põlme S, et al. 2020. Fungal Diversity 105:1-16 ((https://doi.org/10.1007/s13225-020-00466-2)) |
| Otu45362 | 1 | k__Fungi(100);p__Ascomycota(100);c__Saccharomycetes(100);o__Saccharomycetales(100);f__Metschnikowiaceae(100);g__Metschnikowia(100);g__Metschnikowia_unclassified(100); | Metschnikowia | 13 | Saprotroph | \|Undefined Saprotroph\| | Yeast | NULL | Probable | As nectar-tap_saprotroph (Põlme et al. 2020); Decay sugar-rich substrates (Põlme et al. 2020) | Põlme S, et al. 2020. Fungal Diversity 105:1-16 ((https://doi.org/10.1007/s13225-020-00466-2)) |
| Otu45363 | 1 | k__Fungi(100);p__Ascomycota(100);c__Saccharomycetes(100);o__Saccharomycetales(100);f__Metschnikowiaceae(100);g__Metschnikowia(100);s__Metschnikowia_pulcherrima(100); | Metschnikowia | 13 | Saprotroph | \|Undefined Saprotroph\| | Yeast | NULL | Probable | As nectar-tap_saprotroph (Põlme et al. 2020); Decay sugar-rich substrates (Põlme et al. 2020) | Põlme S, et al. 2020. Fungal Diversity 105:1-16 ((https://doi.org/10.1007/s13225-020-00466-2)) |
| Otu45364 | 1 | k__Fungi(100);p__Ascomycota(100);c__Saccharomycetes(100);o__Saccharomycetales(100);f__Metschnikowiaceae(100);g__Metschnikowia(100);s__Metschnikowia_pulcherrima(100); | Metschnikowia | 13 | Saprotroph | \|Undefined Saprotroph\| | Yeast | NULL | Probable | As nectar-tap_saprotroph (Põlme et al. 2020); Decay sugar-rich substrates (Põlme et al. 2020) | Põlme S, et al. 2020. Fungal Diversity 105:1-16 ((https://doi.org/10.1007/s13225-020-00466-2)) |
| Otu45365 | 1 | k__Fungi(100);p__Ascomycota(100);c__Saccharomycetes(100);o__Saccharomycetales(100);f__Metschnikowiaceae(100);g__Metschnikowia(100);g__Metschnikowia_unclassified(100); | Metschnikowia | 13 | Saprotroph | \|Undefined Saprotroph\| | Yeast | NULL | Probable | As nectar-tap_saprotroph (Põlme et al. 2020); Decay sugar-rich substrates (Põlme et al. 2020) | Põlme S, et al. 2020. Fungal Diversity 105:1-16 ((https://doi.org/10.1007/s13225-020-00466-2)) |
| Otu45366 | 1 | k__Fungi(100);p__Ascomycota(100);c__Saccharomycetes(100);o__Saccharomycetales(100);f__Metschnikowiaceae(100);g__Metschnikowia(100);g__Metschnikowia_unclassified(100); | Metschnikowia | 13 | Saprotroph | \|Undefined Saprotroph\| | Yeast | NULL | Probable | As nectar-tap_saprotroph (Põlme et al. 2020); Decay sugar-rich substrates (Põlme et al. 2020) | Põlme S, et al. 2020. Fungal Diversity 105:1-16 ((https://doi.org/10.1007/s13225-020-00466-2)) |
| Otu45367 | 1 | k__Fungi(100);p__Ascomycota(100);c__Saccharomycetes(100);o__Saccharomycetales(100);f__Metschnikowiaceae(100);g__Metschnikowia(100);s__Metschnikowia_pulcherrima(100); | Metschnikowia | 13 | Saprotroph | \|Undefined Saprotroph\| | Yeast | NULL | Probable | As nectar-tap_saprotroph (Põlme et al. 2020); Decay sugar-rich substrates (Põlme et al. 2020) | Põlme S, et al. 2020. Fungal Diversity 105:1-16 ((https://doi.org/10.1007/s13225-020-00466-2)) |
| Otu45368 | 1 | k__Fungi(100);p__Ascomycota(100);c__Saccharomycetes(100);o__Saccharomycetales(100);f__Metschnikowiaceae(100);g__Metschnikowia(100);s__Metschnikowia_pulcherrima(100); | Metschnikowia | 13 | Saprotroph | \|Undefined Saprotroph\| | Yeast | NULL | Probable | As nectar-tap_saprotroph (Põlme et al. 2020); Decay sugar-rich substrates (Põlme et al. 2020) | Põlme S, et al. 2020. Fungal Diversity 105:1-16 ((https://doi.org/10.1007/s13225-020-00466-2)) |
| Otu45369 | 1 | k__Fungi(100);p__Ascomycota(100);c__Saccharomycetes(100);o__Saccharomycetales(100);f__Metschnikowiaceae(100);g__Metschnikowia(100);s__Metschnikowia_pulcherrima(100); | Metschnikowia | 13 | Saprotroph | \|Undefined Saprotroph\| | Yeast | NULL | Probable | As nectar-tap_saprotroph (Põlme et al. 2020); Decay sugar-rich substrates (Põlme et al. 2020) | Põlme S, et al. 2020. Fungal Diversity 105:1-16 ((https://doi.org/10.1007/s13225-020-00466-2)) |
| Otu45370 | 1 | k__Fungi(100);p__Ascomycota(100);c__Saccharomycetes(100);o__Saccharomycetales(100);f__Metschnikowiaceae(100);g__Metschnikowia(100);g__Metschnikowia_unclassified(100); | Metschnikowia | 13 | Saprotroph | \|Undefined Saprotroph\| | Yeast | NULL | Probable | As nectar-tap_saprotroph (Põlme et al. 2020); Decay sugar-rich substrates (Põlme et al. 2020) | Põlme S, et al. 2020. Fungal Diversity 105:1-16 ((https://doi.org/10.1007/s13225-020-00466-2)) |
| Otu45371 | 1 | k__Fungi(100);p__Ascomycota(100);c__Saccharomycetes(100);o__Saccharomycetales(100);f__Metschnikowiaceae(100);g__Metschnikowia(100);s__Metschnikowia_pulcherrima(100); | Metschnikowia | 13 | Saprotroph | \|Undefined Saprotroph\| | Yeast | NULL | Probable | As nectar-tap_saprotroph (Põlme et al. 2020); Decay sugar-rich substrates (Põlme et al. 2020) | Põlme S, et al. 2020. Fungal Diversity 105:1-16 ((https://doi.org/10.1007/s13225-020-00466-2)) |
| Otu45372 | 1 | k__Fungi(100);p__Ascomycota(100);c__Saccharomycetes(100);o__Saccharomycetales(100);f__Metschnikowiaceae(100);g__Metschnikowia(100);s__Metschnikowia_pulcherrima(100); | Metschnikowia | 13 | Saprotroph | \|Undefined Saprotroph\| | Yeast | NULL | Probable | As nectar-tap_saprotroph (Põlme et al. 2020); Decay sugar-rich substrates (Põlme et al. 2020) | Põlme S, et al. 2020. Fungal Diversity 105:1-16 ((https://doi.org/10.1007/s13225-020-00466-2)) |
| Otu45373 | 1 | k__Fungi(100);p__Ascomycota(100);c__Saccharomycetes(100);o__Saccharomycetales(100);f__Metschnikowiaceae(100);g__Metschnikowia(100);g__Metschnikowia_unclassified(100); | Metschnikowia | 13 | Saprotroph | \|Undefined Saprotroph\| | Yeast | NULL | Probable | As nectar-tap_saprotroph (Põlme et al. 2020); Decay sugar-rich substrates (Põlme et al. 2020) | Põlme S, et al. 2020. Fungal Diversity 105:1-16 ((https://doi.org/10.1007/s13225-020-00466-2)) |
| Otu45374 | 1 | k__Fungi(100);p__Ascomycota(100);c__Saccharomycetes(100);o__Saccharomycetales(100);f__Metschnikowiaceae(100);g__Metschnikowia(100);s__Metschnikowia_pulcherrima(100); | Metschnikowia | 13 | Saprotroph | \|Undefined Saprotroph\| | Yeast | NULL | Probable | As nectar-tap_saprotroph (Põlme et al. 2020); Decay sugar-rich substrates (Põlme et al. 2020) | Põlme S, et al. 2020. Fungal Diversity 105:1-16 ((https://doi.org/10.1007/s13225-020-00466-2)) |
| Otu45375 | 1 | k__Fungi(100);p__Ascomycota(100);c__Saccharomycetes(100);o__Saccharomycetales(100);f__Metschnikowiaceae(100);g__Metschnikowia(100);s__Metschnikowia_pulcherrima(100); | Metschnikowia | 13 | Saprotroph | \|Undefined Saprotroph\| | Yeast | NULL | Probable | As nectar-tap_saprotroph (Põlme et al. 2020); Decay sugar-rich substrates (Põlme et al. 2020) | Põlme S, et al. 2020. Fungal Diversity 105:1-16 ((https://doi.org/10.1007/s13225-020-00466-2)) |
| Otu45376 | 1 | k__Fungi(100);p__Ascomycota(100);c__Saccharomycetes(100);o__Saccharomycetales(100);f__Metschnikowiaceae(100);g__Metschnikowia(100);s__Metschnikowia_pulcherrima(100); | Metschnikowia | 13 | Saprotroph | \|Undefined Saprotroph\| | Yeast | NULL | Probable | As nectar-tap_saprotroph (Põlme et al. 2020); Decay sugar-rich substrates (Põlme et al. 2020) | Põlme S, et al. 2020. Fungal Diversity 105:1-16 ((https://doi.org/10.1007/s13225-020-00466-2)) |
| Otu45377 | 1 | k__Fungi(100);p__Ascomycota(100);c__Saccharomycetes(100);o__Saccharomycetales(100);f__Metschnikowiaceae(100);g__Metschnikowia(100);s__Metschnikowia_pulcherrima(100); | Metschnikowia | 13 | Saprotroph | \|Undefined Saprotroph\| | Yeast | NULL | Probable | As nectar-tap_saprotroph (Põlme et al. 2020); Decay sugar-rich substrates (Põlme et al. 2020) | Põlme S, et al. 2020. Fungal Diversity 105:1-16 ((https://doi.org/10.1007/s13225-020-00466-2)) |
| Otu45378 | 1 | k__Fungi(100);p__Ascomycota(100);c__Saccharomycetes(100);o__Saccharomycetales(100);f__Metschnikowiaceae(100);g__Metschnikowia(100);s__Metschnikowia_pulcherrima(100); | Metschnikowia | 13 | Saprotroph | \|Undefined Saprotroph\| | Yeast | NULL | Probable | As nectar-tap_saprotroph (Põlme et al. 2020); Decay sugar-rich substrates (Põlme et al. 2020) | Põlme S, et al. 2020. Fungal Diversity 105:1-16 ((https://doi.org/10.1007/s13225-020-00466-2)) |
| Otu45379 | 1 | k__Fungi(100);p__Ascomycota(100);c__Saccharomycetes(100);o__Saccharomycetales(100);f__Metschnikowiaceae(100);g__Metschnikowia(100);s__Metschnikowia_pulcherrima(100); | Metschnikowia | 13 | Saprotroph | \|Undefined Saprotroph\| | Yeast | NULL | Probable | As nectar-tap_saprotroph (Põlme et al. 2020); Decay sugar-rich substrates (Põlme et al. 2020) | Põlme S, et al. 2020. Fungal Diversity 105:1-16 ((https://doi.org/10.1007/s13225-020-00466-2)) |
| Otu45380 | 1 | k__Fungi(100);p__Ascomycota(100);c__Saccharomycetes(100);o__Saccharomycetales(100);f__Metschnikowiaceae(100);g__Metschnikowia(100);s__Metschnikowia_pulcherrima(100); | Metschnikowia | 13 | Saprotroph | \|Undefined Saprotroph\| | Yeast | NULL | Probable | As nectar-tap_saprotroph (Põlme et al. 2020); Decay sugar-rich substrates (Põlme et al. 2020) | Põlme S, et al. 2020. Fungal Diversity 105:1-16 ((https://doi.org/10.1007/s13225-020-00466-2)) |
| Otu45381 | 1 | k__Fungi(100);p__Ascomycota(100);c__Saccharomycetes(100);o__Saccharomycetales(100);f__Metschnikowiaceae(100);g__Metschnikowia(100);s__Metschnikowia_pulcherrima(100); | Metschnikowia | 13 | Saprotroph | \|Undefined Saprotroph\| | Yeast | NULL | Probable | As nectar-tap_saprotroph (Põlme et al. 2020); Decay sugar-rich substrates (Põlme et al. 2020) | Põlme S, et al. 2020. Fungal Diversity 105:1-16 ((https://doi.org/10.1007/s13225-020-00466-2)) |
| Otu45382 | 1 | k__Fungi(100);p__Ascomycota(100);c__Saccharomycetes(100);o__Saccharomycetales(100);f__Metschnikowiaceae(100);g__Metschnikowia(100);s__Metschnikowia_pulcherrima(100); | Metschnikowia | 13 | Saprotroph | \|Undefined Saprotroph\| | Yeast | NULL | Probable | As nectar-tap_saprotroph (Põlme et al. 2020); Decay sugar-rich substrates (Põlme et al. 2020) | Põlme S, et al. 2020. Fungal Diversity 105:1-16 ((https://doi.org/10.1007/s13225-020-00466-2)) |
| Otu45383 | 1 | k__Fungi(100);p__Ascomycota(100);c__Saccharomycetes(100);o__Saccharomycetales(100);f__Metschnikowiaceae(100);g__Metschnikowia(100);s__Metschnikowia_pulcherrima(100); | Metschnikowia | 13 | Saprotroph | \|Undefined Saprotroph\| | Yeast | NULL | Probable | As nectar-tap_saprotroph (Põlme et al. 2020); Decay sugar-rich substrates (Põlme et al. 2020) | Põlme S, et al. 2020. Fungal Diversity 105:1-16 ((https://doi.org/10.1007/s13225-020-00466-2)) |
| Otu45384 | 1 | k__Fungi(100);p__Ascomycota(100);c__Saccharomycetes(100);o__Saccharomycetales(100);f__Metschnikowiaceae(100);g__Metschnikowia(100);s__Metschnikowia_pulcherrima(100); | Metschnikowia | 13 | Saprotroph | \|Undefined Saprotroph\| | Yeast | NULL | Probable | As nectar-tap_saprotroph (Põlme et al. 2020); Decay sugar-rich substrates (Põlme et al. 2020) | Põlme S, et al. 2020. Fungal Diversity 105:1-16 ((https://doi.org/10.1007/s13225-020-00466-2)) |
| Otu45385 | 1 | k__Fungi(100);p__Ascomycota(100);c__Saccharomycetes(100);o__Saccharomycetales(100);f__Metschnikowiaceae(100);g__Metschnikowia(100);s__Metschnikowia_pulcherrima(100); | Metschnikowia | 13 | Saprotroph | \|Undefined Saprotroph\| | Yeast | NULL | Probable | As nectar-tap_saprotroph (Põlme et al. 2020); Decay sugar-rich substrates (Põlme et al. 2020) | Põlme S, et al. 2020. Fungal Diversity 105:1-16 ((https://doi.org/10.1007/s13225-020-00466-2)) |
| Otu45386 | 1 | k__Fungi(100);p__Ascomycota(100);c__Saccharomycetes(100);o__Saccharomycetales(100);f__Metschnikowiaceae(100);g__Metschnikowia(100);s__Metschnikowia_pulcherrima(100); | Metschnikowia | 13 | Saprotroph | \|Undefined Saprotroph\| | Yeast | NULL | Probable | As nectar-tap_saprotroph (Põlme et al. 2020); Decay sugar-rich substrates (Põlme et al. 2020) | Põlme S, et al. 2020. Fungal Diversity 105:1-16 ((https://doi.org/10.1007/s13225-020-00466-2)) |
| Otu45387 | 1 | k__Fungi(100);p__Ascomycota(100);c__Saccharomycetes(100);o__Saccharomycetales(100);f__Metschnikowiaceae(100);g__Metschnikowia(100);s__Metschnikowia_pulcherrima(100); | Metschnikowia | 13 | Saprotroph | \|Undefined Saprotroph\| | Yeast | NULL | Probable | As nectar-tap_saprotroph (Põlme et al. 2020); Decay sugar-rich substrates (Põlme et al. 2020) | Põlme S, et al. 2020. Fungal Diversity 105:1-16 ((https://doi.org/10.1007/s13225-020-00466-2)) |
| Otu45388 | 1 | k__Fungi(100);p__Ascomycota(100);c__Saccharomycetes(100);o__Saccharomycetales(100);f__Metschnikowiaceae(100);g__Metschnikowia(100);g__Metschnikowia_unclassified(100); | Metschnikowia | 13 | Saprotroph | \|Undefined Saprotroph\| | Yeast | NULL | Probable | As nectar-tap_saprotroph (Põlme et al. 2020); Decay sugar-rich substrates (Põlme et al. 2020) | Põlme S, et al. 2020. Fungal Diversity 105:1-16 ((https://doi.org/10.1007/s13225-020-00466-2)) |
| Otu45389 | 1 | k__Fungi(100);p__Ascomycota(100);c__Saccharomycetes(100);o__Saccharomycetales(100);f__Metschnikowiaceae(100);g__Metschnikowia(100);s__Metschnikowia_pulcherrima(100); | Metschnikowia | 13 | Saprotroph | \|Undefined Saprotroph\| | Yeast | NULL | Probable | As nectar-tap_saprotroph (Põlme et al. 2020); Decay sugar-rich substrates (Põlme et al. 2020) | Põlme S, et al. 2020. Fungal Diversity 105:1-16 ((https://doi.org/10.1007/s13225-020-00466-2)) |
| Otu45390 | 1 | k__Fungi(100);p__Ascomycota(100);c__Saccharomycetes(100);o__Saccharomycetales(100);f__Metschnikowiaceae(100);g__Metschnikowia(100);s__Metschnikowia_pulcherrima(100); | Metschnikowia | 13 | Saprotroph | \|Undefined Saprotroph\| | Yeast | NULL | Probable | As nectar-tap_saprotroph (Põlme et al. 2020); Decay sugar-rich substrates (Põlme et al. 2020) | Põlme S, et al. 2020. Fungal Diversity 105:1-16 ((https://doi.org/10.1007/s13225-020-00466-2)) |
| Otu45391 | 1 | k__Fungi(100);p__Ascomycota(100);c__Saccharomycetes(100);o__Saccharomycetales(100);f__Metschnikowiaceae(100);g__Metschnikowia(100);s__Metschnikowia_pulcherrima(100); | Metschnikowia | 13 | Saprotroph | \|Undefined Saprotroph\| | Yeast | NULL | Probable | As nectar-tap_saprotroph (Põlme et al. 2020); Decay sugar-rich substrates (Põlme et al. 2020) | Põlme S, et al. 2020. Fungal Diversity 105:1-16 ((https://doi.org/10.1007/s13225-020-00466-2)) |
| Otu45392 | 1 | k__Fungi(100);p__Ascomycota(100);c__Saccharomycetes(100);o__Saccharomycetales(100);f__Metschnikowiaceae(100);g__Metschnikowia(100);s__Metschnikowia_pulcherrima(100); | Metschnikowia | 13 | Saprotroph | \|Undefined Saprotroph\| | Yeast | NULL | Probable | As nectar-tap_saprotroph (Põlme et al. 2020); Decay sugar-rich substrates (Põlme et al. 2020) | Põlme S, et al. 2020. Fungal Diversity 105:1-16 ((https://doi.org/10.1007/s13225-020-00466-2)) |
| Otu45393 | 1 | k__Fungi(100);p__Ascomycota(100);c__Saccharomycetes(100);o__Saccharomycetales(100);f__Metschnikowiaceae(100);g__Metschnikowia(100);g__Metschnikowia_unclassified(100); | Metschnikowia | 13 | Saprotroph | \|Undefined Saprotroph\| | Yeast | NULL | Probable | As nectar-tap_saprotroph (Põlme et al. 2020); Decay sugar-rich substrates (Põlme et al. 2020) | Põlme S, et al. 2020. Fungal Diversity 105:1-16 ((https://doi.org/10.1007/s13225-020-00466-2)) |
| Otu45394 | 1 | k__Fungi(100);p__Ascomycota(100);c__Saccharomycetes(100);o__Saccharomycetales(100);f__Metschnikowiaceae(100);g__Metschnikowia(100);s__Metschnikowia_pulcherrima(100); | Metschnikowia | 13 | Saprotroph | \|Undefined Saprotroph\| | Yeast | NULL | Probable | As nectar-tap_saprotroph (Põlme et al. 2020); Decay sugar-rich substrates (Põlme et al. 2020) | Põlme S, et al. 2020. Fungal Diversity 105:1-16 ((https://doi.org/10.1007/s13225-020-00466-2)) |
| Otu45395 | 1 | k__Fungi(100);p__Ascomycota(100);c__Saccharomycetes(100);o__Saccharomycetales(100);f__Metschnikowiaceae(100);g__Metschnikowia(100);s__Metschnikowia_pulcherrima(100); | Metschnikowia | 13 | Saprotroph | \|Undefined Saprotroph\| | Yeast | NULL | Probable | As nectar-tap_saprotroph (Põlme et al. 2020); Decay sugar-rich substrates (Põlme et al. 2020) | Põlme S, et al. 2020. Fungal Diversity 105:1-16 ((https://doi.org/10.1007/s13225-020-00466-2)) |
| Otu45396 | 1 | k__Fungi(100);p__Ascomycota(100);c__Saccharomycetes(100);o__Saccharomycetales(100);f__Metschnikowiaceae(100);g__Metschnikowia(100);g__Metschnikowia_unclassified(100); | Metschnikowia | 13 | Saprotroph | \|Undefined Saprotroph\| | Yeast | NULL | Probable | As nectar-tap_saprotroph (Põlme et al. 2020); Decay sugar-rich substrates (Põlme et al. 2020) | Põlme S, et al. 2020. Fungal Diversity 105:1-16 ((https://doi.org/10.1007/s13225-020-00466-2)) |
| Otu45397 | 1 | k__Fungi(100);p__Ascomycota(100);c__Saccharomycetes(100);o__Saccharomycetales(100);f__Metschnikowiaceae(100);g__Metschnikowia(100);s__Metschnikowia_pulcherrima(100); | Metschnikowia | 13 | Saprotroph | \|Undefined Saprotroph\| | Yeast | NULL | Probable | As nectar-tap_saprotroph (Põlme et al. 2020); Decay sugar-rich substrates (Põlme et al. 2020) | Põlme S, et al. 2020. Fungal Diversity 105:1-16 ((https://doi.org/10.1007/s13225-020-00466-2)) |
| Otu45398 | 1 | k__Fungi(100);p__Ascomycota(100);c__Saccharomycetes(100);o__Saccharomycetales(100);f__Metschnikowiaceae(100);g__Metschnikowia(100);s__Metschnikowia_pulcherrima(100); | Metschnikowia | 13 | Saprotroph | \|Undefined Saprotroph\| | Yeast | NULL | Probable | As nectar-tap_saprotroph (Põlme et al. 2020); Decay sugar-rich substrates (Põlme et al. 2020) | Põlme S, et al. 2020. Fungal Diversity 105:1-16 ((https://doi.org/10.1007/s13225-020-00466-2)) |
| Otu45399 | 1 | k__Fungi(100);p__Ascomycota(100);c__Saccharomycetes(100);o__Saccharomycetales(100);f__Metschnikowiaceae(100);g__Metschnikowia(100);s__Metschnikowia_pulcherrima(100); | Metschnikowia | 13 | Saprotroph | \|Undefined Saprotroph\| | Yeast | NULL | Probable | As nectar-tap_saprotroph (Põlme et al. 2020); Decay sugar-rich substrates (Põlme et al. 2020) | Põlme S, et al. 2020. Fungal Diversity 105:1-16 ((https://doi.org/10.1007/s13225-020-00466-2)) |
| Otu45400 | 1 | k__Fungi(100);p__Ascomycota(100);c__Saccharomycetes(100);o__Saccharomycetales(100);f__Metschnikowiaceae(100);g__Metschnikowia(100);s__Metschnikowia_pulcherrima(100); | Metschnikowia | 13 | Saprotroph | \|Undefined Saprotroph\| | Yeast | NULL | Probable | As nectar-tap_saprotroph (Põlme et al. 2020); Decay sugar-rich substrates (Põlme et al. 2020) | Põlme S, et al. 2020. Fungal Diversity 105:1-16 ((https://doi.org/10.1007/s13225-020-00466-2)) |
| Otu45401 | 1 | k__Fungi(100);p__Ascomycota(100);c__Saccharomycetes(100);o__Saccharomycetales(100);f__Metschnikowiaceae(100);g__Metschnikowia(100);s__Metschnikowia_pulcherrima(100); | Metschnikowia | 13 | Saprotroph | \|Undefined Saprotroph\| | Yeast | NULL | Probable | As nectar-tap_saprotroph (Põlme et al. 2020); Decay sugar-rich substrates (Põlme et al. 2020) | Põlme S, et al. 2020. Fungal Diversity 105:1-16 ((https://doi.org/10.1007/s13225-020-00466-2)) |
| Otu45402 | 1 | k__Fungi(100);p__Ascomycota(100);c__Saccharomycetes(100);o__Saccharomycetales(100);f__Metschnikowiaceae(100);g__Metschnikowia(100);s__Metschnikowia_pulcherrima(100); | Metschnikowia | 13 | Saprotroph | \|Undefined Saprotroph\| | Yeast | NULL | Probable | As nectar-tap_saprotroph (Põlme et al. 2020); Decay sugar-rich substrates (Põlme et al. 2020) | Põlme S, et al. 2020. Fungal Diversity 105:1-16 ((https://doi.org/10.1007/s13225-020-00466-2)) |
| Otu45403 | 1 | k__Fungi(100);p__Ascomycota(100);c__Saccharomycetes(100);o__Saccharomycetales(100);f__Metschnikowiaceae(100);g__Metschnikowia(100);s__Metschnikowia_pulcherrima(100); | Metschnikowia | 13 | Saprotroph | \|Undefined Saprotroph\| | Yeast | NULL | Probable | As nectar-tap_saprotroph (Põlme et al. 2020); Decay sugar-rich substrates (Põlme et al. 2020) | Põlme S, et al. 2020. Fungal Diversity 105:1-16 ((https://doi.org/10.1007/s13225-020-00466-2)) |
| Otu45404 | 1 | k__Fungi(100);p__Ascomycota(100);c__Saccharomycetes(100);o__Saccharomycetales(100);f__Metschnikowiaceae(100);g__Metschnikowia(100);s__Metschnikowia_pulcherrima(100); | Metschnikowia | 13 | Saprotroph | \|Undefined Saprotroph\| | Yeast | NULL | Probable | As nectar-tap_saprotroph (Põlme et al. 2020); Decay sugar-rich substrates (Põlme et al. 2020) | Põlme S, et al. 2020. Fungal Diversity 105:1-16 ((https://doi.org/10.1007/s13225-020-00466-2)) |
| Otu45405 | 1 | k__Fungi(100);p__Ascomycota(100);c__Saccharomycetes(100);o__Saccharomycetales(100);f__Metschnikowiaceae(100);g__Metschnikowia(100);g__Metschnikowia_unclassified(100); | Metschnikowia | 13 | Saprotroph | \|Undefined Saprotroph\| | Yeast | NULL | Probable | As nectar-tap_saprotroph (Põlme et al. 2020); Decay sugar-rich substrates (Põlme et al. 2020) | Põlme S, et al. 2020. Fungal Diversity 105:1-16 ((https://doi.org/10.1007/s13225-020-00466-2)) |
| Otu45406 | 1 | k__Fungi(100);p__Ascomycota(100);c__Saccharomycetes(100);o__Saccharomycetales(100);f__Metschnikowiaceae(100);g__Metschnikowia(100);g__Metschnikowia_unclassified(100); | Metschnikowia | 13 | Saprotroph | \|Undefined Saprotroph\| | Yeast | NULL | Probable | As nectar-tap_saprotroph (Põlme et al. 2020); Decay sugar-rich substrates (Põlme et al. 2020) | Põlme S, et al. 2020. Fungal Diversity 105:1-16 ((https://doi.org/10.1007/s13225-020-00466-2)) |
| Otu45407 | 1 | k__Fungi(100);p__Ascomycota(100);c__Saccharomycetes(100);o__Saccharomycetales(100);f__Metschnikowiaceae(100);g__Metschnikowia(100);s__Metschnikowia_pulcherrima(100); | Metschnikowia | 13 | Saprotroph | \|Undefined Saprotroph\| | Yeast | NULL | Probable | As nectar-tap_saprotroph (Põlme et al. 2020); Decay sugar-rich substrates (Põlme et al. 2020) | Põlme S, et al. 2020. Fungal Diversity 105:1-16 ((https://doi.org/10.1007/s13225-020-00466-2)) |
| Otu45408 | 1 | k__Fungi(100);p__Ascomycota(100);c__Dothideomycetes(100);o__Pleosporales(100);f__Pleosporaceae(100);g__Alternaria(100);g__Alternaria_unclassified(100); | Alternaria | 13 | Pathotroph-Saprotroph-Symbiotroph | Animal Pathogen-Endophyte-\|Plant Pathogen\|-Plant Saprotroph-Wood Saprotroph | Microfungus | Soft Rot | Possible | As foliar_endophyte (Põlme et al. 2020); As litter_saprotroph (Põlme et al. 2020); Decay leaf, fruit, seed (Põlme et al. 2020) | Duncan CG, Eslyn WE. 1966. Mycologia 58:642-664 ((https://doi.org/10.2307/3757045)); Seehann G, et al. 1975. List of Fungi in Soft-Rot Tissue Tests. Institute fur Holzbiologie und Holzschutz, Bundesforschungsanstalt fur Forst - und Holzwirtshaft, Hamburg - Lohbrugge, Germany ((https://www.irg-wp.com/irgdocs/details.php?d6493033-c9ea-40f3-8e3e-5697f32d640d)); Tedersoo L, et al. 2014. Science 346:e1256688 ((https://doi.org/10.1126/science.1256688)); Põlme S, et al. 2020. Fungal Diversity 105:1-16 ((https://doi.org/10.1007/s13225-020-00466-2)) |
| Otu45409 | 1 | k__Fungi(100);p__Ascomycota(100);c__Saccharomycetes(100);o__Saccharomycetales(100);f__Metschnikowiaceae(100);g__Metschnikowia(100);s__Metschnikowia_pulcherrima(100); | Metschnikowia | 13 | Saprotroph | \|Undefined Saprotroph\| | Yeast | NULL | Probable | As nectar-tap_saprotroph (Põlme et al. 2020); Decay sugar-rich substrates (Põlme et al. 2020) | Põlme S, et al. 2020. Fungal Diversity 105:1-16 ((https://doi.org/10.1007/s13225-020-00466-2)) |
| Otu45410 | 1 | k__Fungi(100);p__Ascomycota(100);c__Saccharomycetes(100);o__Saccharomycetales(100);f__Metschnikowiaceae(100);g__Metschnikowia(100);s__Metschnikowia_pulcherrima(100); | Metschnikowia | 13 | Saprotroph | \|Undefined Saprotroph\| | Yeast | NULL | Probable | As nectar-tap_saprotroph (Põlme et al. 2020); Decay sugar-rich substrates (Põlme et al. 2020) | Põlme S, et al. 2020. Fungal Diversity 105:1-16 ((https://doi.org/10.1007/s13225-020-00466-2)) |
| Otu45412 | 1 | k__Fungi(100);p__Ascomycota(100);c__Saccharomycetes(100);o__Saccharomycetales(100);f__Metschnikowiaceae(100);g__Metschnikowia(100);s__Metschnikowia_pulcherrima(100); | Metschnikowia | 13 | Saprotroph | \|Undefined Saprotroph\| | Yeast | NULL | Probable | As nectar-tap_saprotroph (Põlme et al. 2020); Decay sugar-rich substrates (Põlme et al. 2020) | Põlme S, et al. 2020. Fungal Diversity 105:1-16 ((https://doi.org/10.1007/s13225-020-00466-2)) |
| Otu45413 | 1 | k__Fungi(100);p__Ascomycota(100);c__Saccharomycetes(100);o__Saccharomycetales(100);f__Metschnikowiaceae(100);g__Metschnikowia(100);s__Metschnikowia_pulcherrima(100); | Metschnikowia | 13 | Saprotroph | \|Undefined Saprotroph\| | Yeast | NULL | Probable | As nectar-tap_saprotroph (Põlme et al. 2020); Decay sugar-rich substrates (Põlme et al. 2020) | Põlme S, et al. 2020. Fungal Diversity 105:1-16 ((https://doi.org/10.1007/s13225-020-00466-2)) |
| Otu45414 | 1 | k__Fungi(100);p__Ascomycota(100);c__Saccharomycetes(100);o__Saccharomycetales(100);f__Metschnikowiaceae(100);g__Metschnikowia(100);s__Metschnikowia_pulcherrima(100); | Metschnikowia | 13 | Saprotroph | \|Undefined Saprotroph\| | Yeast | NULL | Probable | As nectar-tap_saprotroph (Põlme et al. 2020); Decay sugar-rich substrates (Põlme et al. 2020) | Põlme S, et al. 2020. Fungal Diversity 105:1-16 ((https://doi.org/10.1007/s13225-020-00466-2)) |
| Otu45415 | 1 | k__Fungi(100);p__Ascomycota(100);c__Saccharomycetes(100);o__Saccharomycetales(100);f__Metschnikowiaceae(100);g__Metschnikowia(100);g__Metschnikowia_unclassified(100); | Metschnikowia | 13 | Saprotroph | \|Undefined Saprotroph\| | Yeast | NULL | Probable | As nectar-tap_saprotroph (Põlme et al. 2020); Decay sugar-rich substrates (Põlme et al. 2020) | Põlme S, et al. 2020. Fungal Diversity 105:1-16 ((https://doi.org/10.1007/s13225-020-00466-2)) |
| Otu45416 | 1 | k__Fungi(100);p__Ascomycota(100);c__Saccharomycetes(100);o__Saccharomycetales(100);f__Metschnikowiaceae(100);g__Metschnikowia(100);s__Metschnikowia_pulcherrima(100); | Metschnikowia | 13 | Saprotroph | \|Undefined Saprotroph\| | Yeast | NULL | Probable | As nectar-tap_saprotroph (Põlme et al. 2020); Decay sugar-rich substrates (Põlme et al. 2020) | Põlme S, et al. 2020. Fungal Diversity 105:1-16 ((https://doi.org/10.1007/s13225-020-00466-2)) |
| Otu45417 | 1 | k__Fungi(100);p__Ascomycota(100);c__Saccharomycetes(100);o__Saccharomycetales(100);f__Metschnikowiaceae(100);g__Metschnikowia(100);s__Metschnikowia_pulcherrima(100); | Metschnikowia | 13 | Saprotroph | \|Undefined Saprotroph\| | Yeast | NULL | Probable | As nectar-tap_saprotroph (Põlme et al. 2020); Decay sugar-rich substrates (Põlme et al. 2020) | Põlme S, et al. 2020. Fungal Diversity 105:1-16 ((https://doi.org/10.1007/s13225-020-00466-2)) |
| Otu45418 | 1 | k__Fungi(100);p__Ascomycota(100);c__Saccharomycetes(100);o__Saccharomycetales(100);f__Metschnikowiaceae(100);g__Metschnikowia(100);s__Metschnikowia_pulcherrima(100); | Metschnikowia | 13 | Saprotroph | \|Undefined Saprotroph\| | Yeast | NULL | Probable | As nectar-tap_saprotroph (Põlme et al. 2020); Decay sugar-rich substrates (Põlme et al. 2020) | Põlme S, et al. 2020. Fungal Diversity 105:1-16 ((https://doi.org/10.1007/s13225-020-00466-2)) |
| Otu45419 | 1 | k__Fungi(100);p__Ascomycota(100);c__Saccharomycetes(100);o__Saccharomycetales(100);f__Metschnikowiaceae(100);g__Metschnikowia(100);g__Metschnikowia_unclassified(100); | Metschnikowia | 13 | Saprotroph | \|Undefined Saprotroph\| | Yeast | NULL | Probable | As nectar-tap_saprotroph (Põlme et al. 2020); Decay sugar-rich substrates (Põlme et al. 2020) | Põlme S, et al. 2020. Fungal Diversity 105:1-16 ((https://doi.org/10.1007/s13225-020-00466-2)) |
| Otu45420 | 1 | k__Fungi(100);p__Ascomycota(100);c__Saccharomycetes(100);o__Saccharomycetales(100);f__Metschnikowiaceae(100);g__Metschnikowia(100);s__Metschnikowia_pulcherrima(100); | Metschnikowia | 13 | Saprotroph | \|Undefined Saprotroph\| | Yeast | NULL | Probable | As nectar-tap_saprotroph (Põlme et al. 2020); Decay sugar-rich substrates (Põlme et al. 2020) | Põlme S, et al. 2020. Fungal Diversity 105:1-16 ((https://doi.org/10.1007/s13225-020-00466-2)) |
| Otu45421 | 1 | k__Fungi(100);p__Ascomycota(100);c__Saccharomycetes(100);o__Saccharomycetales(100);f__Metschnikowiaceae(100);g__Metschnikowia(100);g__Metschnikowia_unclassified(100); | Metschnikowia | 13 | Saprotroph | \|Undefined Saprotroph\| | Yeast | NULL | Probable | As nectar-tap_saprotroph (Põlme et al. 2020); Decay sugar-rich substrates (Põlme et al. 2020) | Põlme S, et al. 2020. Fungal Diversity 105:1-16 ((https://doi.org/10.1007/s13225-020-00466-2)) |
| Otu45422 | 1 | k__Fungi(100);p__Ascomycota(100);c__Saccharomycetes(100);o__Saccharomycetales(100);f__Metschnikowiaceae(100);g__Metschnikowia(100);s__Metschnikowia_pulcherrima(100); | Metschnikowia | 13 | Saprotroph | \|Undefined Saprotroph\| | Yeast | NULL | Probable | As nectar-tap_saprotroph (Põlme et al. 2020); Decay sugar-rich substrates (Põlme et al. 2020) | Põlme S, et al. 2020. Fungal Diversity 105:1-16 ((https://doi.org/10.1007/s13225-020-00466-2)) |
| Otu45423 | 1 | k__Fungi(100);p__Ascomycota(100);c__Saccharomycetes(100);o__Saccharomycetales(100);f__Metschnikowiaceae(100);g__Metschnikowia(100);s__Metschnikowia_pulcherrima(100); | Metschnikowia | 13 | Saprotroph | \|Undefined Saprotroph\| | Yeast | NULL | Probable | As nectar-tap_saprotroph (Põlme et al. 2020); Decay sugar-rich substrates (Põlme et al. 2020) | Põlme S, et al. 2020. Fungal Diversity 105:1-16 ((https://doi.org/10.1007/s13225-020-00466-2)) |
| Otu45424 | 1 | k__Fungi(100);p__Ascomycota(100);c__Saccharomycetes(100);o__Saccharomycetales(100);f__Metschnikowiaceae(100);g__Metschnikowia(100);s__Metschnikowia_pulcherrima(100); | Metschnikowia | 13 | Saprotroph | \|Undefined Saprotroph\| | Yeast | NULL | Probable | As nectar-tap_saprotroph (Põlme et al. 2020); Decay sugar-rich substrates (Põlme et al. 2020) | Põlme S, et al. 2020. Fungal Diversity 105:1-16 ((https://doi.org/10.1007/s13225-020-00466-2)) |
| Otu45425 | 1 | k__Fungi(100);p__Ascomycota(100);c__Saccharomycetes(100);o__Saccharomycetales(100);f__Metschnikowiaceae(100);g__Metschnikowia(100);g__Metschnikowia_unclassified(100); | Metschnikowia | 13 | Saprotroph | \|Undefined Saprotroph\| | Yeast | NULL | Probable | As nectar-tap_saprotroph (Põlme et al. 2020); Decay sugar-rich substrates (Põlme et al. 2020) | Põlme S, et al. 2020. Fungal Diversity 105:1-16 ((https://doi.org/10.1007/s13225-020-00466-2)) |
| Otu45426 | 1 | k__Fungi(100);p__Ascomycota(100);c__Saccharomycetes(100);o__Saccharomycetales(100);f__Metschnikowiaceae(100);g__Metschnikowia(100);s__Metschnikowia_pulcherrima(100); | Metschnikowia | 13 | Saprotroph | \|Undefined Saprotroph\| | Yeast | NULL | Probable | As nectar-tap_saprotroph (Põlme et al. 2020); Decay sugar-rich substrates (Põlme et al. 2020) | Põlme S, et al. 2020. Fungal Diversity 105:1-16 ((https://doi.org/10.1007/s13225-020-00466-2)) |
| Otu45427 | 1 | k__Fungi(100);p__Ascomycota(100);c__Saccharomycetes(100);o__Saccharomycetales(100);f__Metschnikowiaceae(100);g__Metschnikowia(100);s__Metschnikowia_pulcherrima(100); | Metschnikowia | 13 | Saprotroph | \|Undefined Saprotroph\| | Yeast | NULL | Probable | As nectar-tap_saprotroph (Põlme et al. 2020); Decay sugar-rich substrates (Põlme et al. 2020) | Põlme S, et al. 2020. Fungal Diversity 105:1-16 ((https://doi.org/10.1007/s13225-020-00466-2)) |
| Otu45428 | 1 | k__Fungi(100);p__Ascomycota(100);c__Saccharomycetes(100);o__Saccharomycetales(100);f__Metschnikowiaceae(100);g__Metschnikowia(100);s__Metschnikowia_pulcherrima(100); | Metschnikowia | 13 | Saprotroph | \|Undefined Saprotroph\| | Yeast | NULL | Probable | As nectar-tap_saprotroph (Põlme et al. 2020); Decay sugar-rich substrates (Põlme et al. 2020) | Põlme S, et al. 2020. Fungal Diversity 105:1-16 ((https://doi.org/10.1007/s13225-020-00466-2)) |
| Otu45429 | 1 | k__Fungi(100);p__Ascomycota(100);c__Saccharomycetes(100);o__Saccharomycetales(100);f__Metschnikowiaceae(100);g__Metschnikowia(100);s__Metschnikowia_pulcherrima(100); | Metschnikowia | 13 | Saprotroph | \|Undefined Saprotroph\| | Yeast | NULL | Probable | As nectar-tap_saprotroph (Põlme et al. 2020); Decay sugar-rich substrates (Põlme et al. 2020) | Põlme S, et al. 2020. Fungal Diversity 105:1-16 ((https://doi.org/10.1007/s13225-020-00466-2)) |
| Otu45430 | 1 | k__Fungi(100);p__Ascomycota(100);c__Saccharomycetes(100);o__Saccharomycetales(100);f__Metschnikowiaceae(100);g__Metschnikowia(100);s__Metschnikowia_pulcherrima(100); | Metschnikowia | 13 | Saprotroph | \|Undefined Saprotroph\| | Yeast | NULL | Probable | As nectar-tap_saprotroph (Põlme et al. 2020); Decay sugar-rich substrates (Põlme et al. 2020) | Põlme S, et al. 2020. Fungal Diversity 105:1-16 ((https://doi.org/10.1007/s13225-020-00466-2)) |
| Otu45432 | 1 | k__Fungi(100);p__Ascomycota(100);c__Saccharomycetes(100);o__Saccharomycetales(100);f__Metschnikowiaceae(100);g__Metschnikowia(100);s__Metschnikowia_pulcherrima(100); | Metschnikowia | 13 | Saprotroph | \|Undefined Saprotroph\| | Yeast | NULL | Probable | As nectar-tap_saprotroph (Põlme et al. 2020); Decay sugar-rich substrates (Põlme et al. 2020) | Põlme S, et al. 2020. Fungal Diversity 105:1-16 ((https://doi.org/10.1007/s13225-020-00466-2)) |
| Otu45433 | 1 | k__Fungi(100);p__Ascomycota(100);c__Saccharomycetes(100);o__Saccharomycetales(100);f__Metschnikowiaceae(100);g__Metschnikowia(100);s__Metschnikowia_pulcherrima(100); | Metschnikowia | 13 | Saprotroph | \|Undefined Saprotroph\| | Yeast | NULL | Probable | As nectar-tap_saprotroph (Põlme et al. 2020); Decay sugar-rich substrates (Põlme et al. 2020) | Põlme S, et al. 2020. Fungal Diversity 105:1-16 ((https://doi.org/10.1007/s13225-020-00466-2)) |
| Otu45434 | 1 | k__Fungi(100);p__Ascomycota(100);c__Saccharomycetes(100);o__Saccharomycetales(100);f__Metschnikowiaceae(100);g__Metschnikowia(100);s__Metschnikowia_pulcherrima(100); | Metschnikowia | 13 | Saprotroph | \|Undefined Saprotroph\| | Yeast | NULL | Probable | As nectar-tap_saprotroph (Põlme et al. 2020); Decay sugar-rich substrates (Põlme et al. 2020) | Põlme S, et al. 2020. Fungal Diversity 105:1-16 ((https://doi.org/10.1007/s13225-020-00466-2)) |
| Otu45435 | 1 | k__Fungi(100);p__Ascomycota(100);c__Saccharomycetes(100);o__Saccharomycetales(100);f__Metschnikowiaceae(100);g__Metschnikowia(100);s__Metschnikowia_pulcherrima(100); | Metschnikowia | 13 | Saprotroph | \|Undefined Saprotroph\| | Yeast | NULL | Probable | As nectar-tap_saprotroph (Põlme et al. 2020); Decay sugar-rich substrates (Põlme et al. 2020) | Põlme S, et al. 2020. Fungal Diversity 105:1-16 ((https://doi.org/10.1007/s13225-020-00466-2)) |
| Otu45436 | 1 | k__Fungi(100);p__Ascomycota(100);c__Saccharomycetes(100);o__Saccharomycetales(100);f__Metschnikowiaceae(100);g__Metschnikowia(100);s__Metschnikowia_pulcherrima(100); | Metschnikowia | 13 | Saprotroph | \|Undefined Saprotroph\| | Yeast | NULL | Probable | As nectar-tap_saprotroph (Põlme et al. 2020); Decay sugar-rich substrates (Põlme et al. 2020) | Põlme S, et al. 2020. Fungal Diversity 105:1-16 ((https://doi.org/10.1007/s13225-020-00466-2)) |
| Otu45437 | 1 | k__Fungi(100);p__Ascomycota(100);c__Saccharomycetes(100);o__Saccharomycetales(100);f__Metschnikowiaceae(100);g__Metschnikowia(100);s__Metschnikowia_pulcherrima(100); | Metschnikowia | 13 | Saprotroph | \|Undefined Saprotroph\| | Yeast | NULL | Probable | As nectar-tap_saprotroph (Põlme et al. 2020); Decay sugar-rich substrates (Põlme et al. 2020) | Põlme S, et al. 2020. Fungal Diversity 105:1-16 ((https://doi.org/10.1007/s13225-020-00466-2)) |
| Otu45438 | 1 | k__Fungi(100);p__Ascomycota(100);c__Saccharomycetes(100);o__Saccharomycetales(100);f__Metschnikowiaceae(100);g__Metschnikowia(100);s__Metschnikowia_pulcherrima(100); | Metschnikowia | 13 | Saprotroph | \|Undefined Saprotroph\| | Yeast | NULL | Probable | As nectar-tap_saprotroph (Põlme et al. 2020); Decay sugar-rich substrates (Põlme et al. 2020) | Põlme S, et al. 2020. Fungal Diversity 105:1-16 ((https://doi.org/10.1007/s13225-020-00466-2)) |
| Otu45439 | 1 | k__Fungi(100);p__Ascomycota(100);c__Saccharomycetes(100);o__Saccharomycetales(100);f__Metschnikowiaceae(100);g__Metschnikowia(100);s__Metschnikowia_pulcherrima(100); | Metschnikowia | 13 | Saprotroph | \|Undefined Saprotroph\| | Yeast | NULL | Probable | As nectar-tap_saprotroph (Põlme et al. 2020); Decay sugar-rich substrates (Põlme et al. 2020) | Põlme S, et al. 2020. Fungal Diversity 105:1-16 ((https://doi.org/10.1007/s13225-020-00466-2)) |
| Otu45440 | 1 | k__Fungi(100);p__Ascomycota(100);c__Saccharomycetes(100);o__Saccharomycetales(100);f__Metschnikowiaceae(100);g__Metschnikowia(100);s__Metschnikowia_pulcherrima(100); | Metschnikowia | 13 | Saprotroph | \|Undefined Saprotroph\| | Yeast | NULL | Probable | As nectar-tap_saprotroph (Põlme et al. 2020); Decay sugar-rich substrates (Põlme et al. 2020) | Põlme S, et al. 2020. Fungal Diversity 105:1-16 ((https://doi.org/10.1007/s13225-020-00466-2)) |
| Otu45441 | 1 | k__Fungi(100);p__Ascomycota(100);c__Saccharomycetes(100);o__Saccharomycetales(100);f__Metschnikowiaceae(100);g__Metschnikowia(100);s__Metschnikowia_pulcherrima(100); | Metschnikowia | 13 | Saprotroph | \|Undefined Saprotroph\| | Yeast | NULL | Probable | As nectar-tap_saprotroph (Põlme et al. 2020); Decay sugar-rich substrates (Põlme et al. 2020) | Põlme S, et al. 2020. Fungal Diversity 105:1-16 ((https://doi.org/10.1007/s13225-020-00466-2)) |
| Otu45442 | 1 | k__Fungi(100);p__Ascomycota(100);c__Saccharomycetes(100);o__Saccharomycetales(100);f__Metschnikowiaceae(100);g__Metschnikowia(100);s__Metschnikowia_pulcherrima(100); | Metschnikowia | 13 | Saprotroph | \|Undefined Saprotroph\| | Yeast | NULL | Probable | As nectar-tap_saprotroph (Põlme et al. 2020); Decay sugar-rich substrates (Põlme et al. 2020) | Põlme S, et al. 2020. Fungal Diversity 105:1-16 ((https://doi.org/10.1007/s13225-020-00466-2)) |

| | | | | | | | | | | | |
|---|---|---|---|---|---|---|---|---|---|---|---|
| Otu45443 | 1 | k__Fungi(100);p__Ascomycota(100);c__Saccharomycetes(100);o__Saccharomycetales(100);f__Metschnikowiaceae(100);g__Metschnikowia(100);s__Metschnikowia_pulcherrima(100); | Metschnikowia | 13 | Saprotroph | \|Undefined Saprotroph\| | Yeast | NULL | Probable | As nectar-tap_saprotroph (Põlme et al. 2020); Decay sugar-rich substrates (Põlme et al. 2020) | Põlme S, et al. 2020. Fungal Diversity 105:1-16 ((https://doi.org/10.1007/s13225-020-00466-2)) |
| Otu45444 | 1 | k__Fungi(100);p__Ascomycota(100);c__Saccharomycetes(100);o__Saccharomycetales(100);f__Metschnikowiaceae(100);g__Metschnikowia(100);g__Metschnikowia_unclassified(100); | Metschnikowia | 13 | Saprotroph | \|Undefined Saprotroph\| | Yeast | NULL | Probable | As nectar-tap_saprotroph (Põlme et al. 2020); Decay sugar-rich substrates (Põlme et al. 2020) | Põlme S, et al. 2020. Fungal Diversity 105:1-16 ((https://doi.org/10.1007/s13225-020-00466-2)) |
| Otu45445 | 1 | k__Fungi(100);p__Ascomycota(100);c__Saccharomycetes(100);o__Saccharomycetales(100);f__Metschnikowiaceae(100);g__Metschnikowia(100);s__Metschnikowia_pulcherrima(100); | Metschnikowia | 13 | Saprotroph | \|Undefined Saprotroph\| | Yeast | NULL | Probable | As nectar-tap_saprotroph (Põlme et al. 2020); Decay sugar-rich substrates (Põlme et al. 2020) | Põlme S, et al. 2020. Fungal Diversity 105:1-16 ((https://doi.org/10.1007/s13225-020-00466-2)) |
| Otu45446 | 1 | k__Fungi(100);p__Ascomycota(100);c__Saccharomycetes(100);o__Saccharomycetales(100);f__Metschnikowiaceae(100);g__Metschnikowia(100);s__Metschnikowia_pulcherrima(100); | Metschnikowia | 13 | Saprotroph | \|Undefined Saprotroph\| | Yeast | NULL | Probable | As nectar-tap_saprotroph (Põlme et al. 2020); Decay sugar-rich substrates (Põlme et al. 2020) | Põlme S, et al. 2020. Fungal Diversity 105:1-16 ((https://doi.org/10.1007/s13225-020-00466-2)) |
| Otu45447 | 1 | k__Fungi(100);p__Ascomycota(100);c__Saccharomycetes(100);o__Saccharomycetales(100);f__Metschnikowiaceae(100);g__Metschnikowia(100);s__Metschnikowia_pulcherrima(100); | Metschnikowia | 13 | Saprotroph | \|Undefined Saprotroph\| | Yeast | NULL | Probable | As nectar-tap_saprotroph (Põlme et al. 2020); Decay sugar-rich substrates (Põlme et al. 2020) | Põlme S, et al. 2020. Fungal Diversity 105:1-16 ((https://doi.org/10.1007/s13225-020-00466-2)) |
| Otu45448 | 1 | k__Fungi(100);p__Ascomycota(100);c__Saccharomycetes(100);o__Saccharomycetales(100);f__Metschnikowiaceae(100);g__Metschnikowia(100);s__Metschnikowia_pulcherrima(100); | Metschnikowia | 13 | Saprotroph | \|Undefined Saprotroph\| | Yeast | NULL | Probable | As nectar-tap_saprotroph (Põlme et al. 2020); Decay sugar-rich substrates (Põlme et al. 2020) | Põlme S, et al. 2020. Fungal Diversity 105:1-16 ((https://doi.org/10.1007/s13225-020-00466-2)) |
| Otu45449 | 1 | k__Fungi(100);p__Ascomycota(100);c__Saccharomycetes(100);o__Saccharomycetales(100);f__Metschnikowiaceae(100);g__Metschnikowia(100);s__Metschnikowia_pulcherrima(100); | Metschnikowia | 13 | Saprotroph | \|Undefined Saprotroph\| | Yeast | NULL | Probable | As nectar-tap_saprotroph (Põlme et al. 2020); Decay sugar-rich substrates (Põlme et al. 2020) | Põlme S, et al. 2020. Fungal Diversity 105:1-16 ((https://doi.org/10.1007/s13225-020-00466-2)) |
| Otu45450 | 1 | k__Fungi(100);p__Ascomycota(100);c__Saccharomycetes(100);o__Saccharomycetales(100);f__Metschnikowiaceae(100);g__Metschnikowia(100);s__Metschnikowia_pulcherrima(100); | Metschnikowia | 13 | Saprotroph | \|Undefined Saprotroph\| | Yeast | NULL | Probable | As nectar-tap_saprotroph (Põlme et al. 2020); Decay sugar-rich substrates (Põlme et al. 2020) | Põlme S, et al. 2020. Fungal Diversity 105:1-16 ((https://doi.org/10.1007/s13225-020-00466-2)) |
| Otu45451 | 1 | k__Fungi(100);p__Ascomycota(100);c__Saccharomycetes(100);o__Saccharomycetales(100);f__Metschnikowiaceae(100);g__Metschnikowia(100);s__Metschnikowia_pulcherrima(100); | Metschnikowia | 13 | Saprotroph | \|Undefined Saprotroph\| | Yeast | NULL | Probable | As nectar-tap_saprotroph (Põlme et al. 2020); Decay sugar-rich substrates (Põlme et al. 2020) | Põlme S, et al. 2020. Fungal Diversity 105:1-16 ((https://doi.org/10.1007/s13225-020-00466-2)) |
| Otu45452 | 1 | k__Fungi(100);p__Ascomycota(100);c__Saccharomycetes(100);o__Saccharomycetales(100);f__Metschnikowiaceae(100);g__Metschnikowia(100);s__Metschnikowia_pulcherrima(100); | Metschnikowia | 13 | Saprotroph | \|Undefined Saprotroph\| | Yeast | NULL | Probable | As nectar-tap_saprotroph (Põlme et al. 2020); Decay sugar-rich substrates (Põlme et al. 2020) | Põlme S, et al. 2020. Fungal Diversity 105:1-16 ((https://doi.org/10.1007/s13225-020-00466-2)) |
| Otu45453 | 1 | k__Fungi(100);p__Ascomycota(100);c__Saccharomycetes(100);o__Saccharomycetales(100);f__Metschnikowiaceae(100);g__Metschnikowia(100);s__Metschnikowia_pulcherrima(100); | Metschnikowia | 13 | Saprotroph | \|Undefined Saprotroph\| | Yeast | NULL | Probable | As nectar-tap_saprotroph (Põlme et al. 2020); Decay sugar-rich substrates (Põlme et al. 2020) | Põlme S, et al. 2020. Fungal Diversity 105:1-16 ((https://doi.org/10.1007/s13225-020-00466-2)) |
| Otu45454 | 1 | k__Fungi(100);p__Ascomycota(100);c__Saccharomycetes(100);o__Saccharomycetales(100);f__Metschnikowiaceae(100);g__Metschnikowia(100);s__Metschnikowia_pulcherrima(100); | Metschnikowia | 13 | Saprotroph | \|Undefined Saprotroph\| | Yeast | NULL | Probable | As nectar-tap_saprotroph (Põlme et al. 2020); Decay sugar-rich substrates (Põlme et al. 2020) | Põlme S, et al. 2020. Fungal Diversity 105:1-16 ((https://doi.org/10.1007/s13225-020-00466-2)) |
| Otu45455 | 1 | k__Fungi(100);p__Ascomycota(100);c__Saccharomycetes(100);o__Saccharomycetales(100);f__Metschnikowiaceae(100);g__Metschnikowia(100);s__Metschnikowia_pulcherrima(100); | Metschnikowia | 13 | Saprotroph | \|Undefined Saprotroph\| | Yeast | NULL | Probable | As nectar-tap_saprotroph (Põlme et al. 2020); Decay sugar-rich substrates (Põlme et al. 2020) | Põlme S, et al. 2020. Fungal Diversity 105:1-16 ((https://doi.org/10.1007/s13225-020-00466-2)) |
| Otu45456 | 1 | k__Fungi(100);p__Ascomycota(100);c__Saccharomycetes(100);o__Saccharomycetales(100);f__Metschnikowiaceae(100);g__Metschnikowia(100);s__Metschnikowia_pulcherrima(100); | Metschnikowia | 13 | Saprotroph | \|Undefined Saprotroph\| | Yeast | NULL | Probable | As nectar-tap_saprotroph (Põlme et al. 2020); Decay sugar-rich substrates (Põlme et al. 2020) | Põlme S, et al. 2020. Fungal Diversity 105:1-16 ((https://doi.org/10.1007/s13225-020-00466-2)) |
| Otu45457 | 1 | k__Fungi(100);p__Ascomycota(100);c__Saccharomycetes(100);o__Saccharomycetales(100);f__Metschnikowiaceae(100);g__Metschnikowia(100);s__Metschnikowia_pulcherrima(100); | Metschnikowia | 13 | Saprotroph | \|Undefined Saprotroph\| | Yeast | NULL | Probable | As nectar-tap_saprotroph (Põlme et al. 2020); Decay sugar-rich substrates (Põlme et al. 2020) | Põlme S, et al. 2020. Fungal Diversity 105:1-16 ((https://doi.org/10.1007/s13225-020-00466-2)) |
| Otu45458 | 1 | k__Fungi(100);p__Ascomycota(100);c__Saccharomycetes(100);o__Saccharomycetales(100);f__Metschnikowiaceae(100);g__Metschnikowia(100);s__Metschnikowia_pulcherrima(100); | Metschnikowia | 13 | Saprotroph | \|Undefined Saprotroph\| | Yeast | NULL | Probable | As nectar-tap_saprotroph (Põlme et al. 2020); Decay sugar-rich substrates (Põlme et al. 2020) | Põlme S, et al. 2020. Fungal Diversity 105:1-16 ((https://doi.org/10.1007/s13225-020-00466-2)) |
| Otu45459 | 1 | k__Fungi(100);p__Ascomycota(100);c__Saccharomycetes(100);o__Saccharomycetales(100);f__Metschnikowiaceae(100);g__Metschnikowia(100);s__Metschnikowia_pulcherrima(100); | Metschnikowia | 13 | Saprotroph | \|Undefined Saprotroph\| | Yeast | NULL | Probable | As nectar-tap_saprotroph (Põlme et al. 2020); Decay sugar-rich substrates (Põlme et al. 2020) | Põlme S, et al. 2020. Fungal Diversity 105:1-16 ((https://doi.org/10.1007/s13225-020-00466-2)) |
| Otu45460 | 1 | k__Fungi(100);p__Ascomycota(100);c__Saccharomycetes(100);o__Saccharomycetales(100);f__Metschnikowiaceae(100);g__Metschnikowia(100);g__Metschnikowia_unclassified(100); | Metschnikowia | 13 | Saprotroph | \|Undefined Saprotroph\| | Yeast | NULL | Probable | As nectar-tap_saprotroph (Põlme et al. 2020); Decay sugar-rich substrates (Põlme et al. 2020) | Põlme S, et al. 2020. Fungal Diversity 105:1-16 ((https://doi.org/10.1007/s13225-020-00466-2)) |
| Otu45461 | 1 | k__Fungi(100);p__Ascomycota(100);c__Saccharomycetes(100);o__Saccharomycetales(100);f__Metschnikowiaceae(100);g__Metschnikowia(100);s__Metschnikowia_pulcherrima(100); | Metschnikowia | 13 | Saprotroph | \|Undefined Saprotroph\| | Yeast | NULL | Probable | As nectar-tap_saprotroph (Põlme et al. 2020); Decay sugar-rich substrates (Põlme et al. 2020) | Põlme S, et al. 2020. Fungal Diversity 105:1-16 ((https://doi.org/10.1007/s13225-020-00466-2)) |
| Otu45462 | 1 | k__Fungi(100);p__Ascomycota(100);c__Saccharomycetes(100);o__Saccharomycetales(100);f__Metschnikowiaceae(100);g__Metschnikowia(100);s__Metschnikowia_pulcherrima(100); | Metschnikowia | 13 | Saprotroph | \|Undefined Saprotroph\| | Yeast | NULL | Probable | As nectar-tap_saprotroph (Põlme et al. 2020); Decay sugar-rich substrates (Põlme et al. 2020) | Põlme S, et al. 2020. Fungal Diversity 105:1-16 ((https://doi.org/10.1007/s13225-020-00466-2)) |
| Otu45463 | 1 | k__Fungi(100);p__Ascomycota(100);c__Saccharomycetes(100);o__Saccharomycetales(100);f__Metschnikowiaceae(100);g__Metschnikowia(100);s__Metschnikowia_pulcherrima(100); | Metschnikowia | 13 | Saprotroph | \|Undefined Saprotroph\| | Yeast | NULL | Probable | As nectar-tap_saprotroph (Põlme et al. 2020); Decay sugar-rich substrates (Põlme et al. 2020) | Põlme S, et al. 2020. Fungal Diversity 105:1-16 ((https://doi.org/10.1007/s13225-020-00466-2)) |
| Otu45464 | 1 | k__Fungi(100);p__Ascomycota(100);c__Saccharomycetes(100);o__Saccharomycetales(100);f__Metschnikowiaceae(100);g__Metschnikowia(100);s__Metschnikowia_pulcherrima(100); | Metschnikowia | 13 | Saprotroph | \|Undefined Saprotroph\| | Yeast | NULL | Probable | As nectar-tap_saprotroph (Põlme et al. 2020); Decay sugar-rich substrates (Põlme et al. 2020) | Põlme S, et al. 2020. Fungal Diversity 105:1-16 ((https://doi.org/10.1007/s13225-020-00466-2)) |
| Otu45465 | 1 | k__Fungi(100);p__Ascomycota(100);c__Saccharomycetes(100);o__Saccharomycetales(100);f__Metschnikowiaceae(100);g__Metschnikowia(100);g__Metschnikowia_unclassified(100); | Metschnikowia | 13 | Saprotroph | \|Undefined Saprotroph\| | Yeast | NULL | Probable | As nectar-tap_saprotroph (Põlme et al. 2020); Decay sugar-rich substrates (Põlme et al. 2020) | Põlme S, et al. 2020. Fungal Diversity 105:1-16 ((https://doi.org/10.1007/s13225-020-00466-2)) |
| Otu45466 | 1 | k__Fungi(100);p__Ascomycota(100);c__Saccharomycetes(100);o__Saccharomycetales(100);f__Metschnikowiaceae(100);g__Metschnikowia(100);s__Metschnikowia_pulcherrima(100); | Metschnikowia | 13 | Saprotroph | \|Undefined Saprotroph\| | Yeast | NULL | Probable | As nectar-tap_saprotroph (Põlme et al. 2020); Decay sugar-rich substrates (Põlme et al. 2020) | Põlme S, et al. 2020. Fungal Diversity 105:1-16 ((https://doi.org/10.1007/s13225-020-00466-2)) |
| Otu45467 | 1 | k__Fungi(100);p__Ascomycota(100);c__Saccharomycetes(100);o__Saccharomycetales(100);f__Metschnikowiaceae(100);g__Metschnikowia(100);s__Metschnikowia_pulcherrima(100); | Metschnikowia | 13 | Saprotroph | \|Undefined Saprotroph\| | Yeast | NULL | Probable | As nectar-tap_saprotroph (Põlme et al. 2020); Decay sugar-rich substrates (Põlme et al. 2020) | Põlme S, et al. 2020. Fungal Diversity 105:1-16 ((https://doi.org/10.1007/s13225-020-00466-2)) |
| Otu45468 | 1 | k__Fungi(100);p__Ascomycota(100);c__Saccharomycetes(100);o__Saccharomycetales(100);f__Metschnikowiaceae(100);g__Metschnikowia(100);s__Metschnikowia_pulcherrima(100); | Metschnikowia | 13 | Saprotroph | \|Undefined Saprotroph\| | Yeast | NULL | Probable | As nectar-tap_saprotroph (Põlme et al. 2020); Decay sugar-rich substrates (Põlme et al. 2020) | Põlme S, et al. 2020. Fungal Diversity 105:1-16 ((https://doi.org/10.1007/s13225-020-00466-2)) |
| Otu45469 | 1 | k__Fungi(100);p__Ascomycota(100);c__Saccharomycetes(100);o__Saccharomycetales(100);f__Metschnikowiaceae(100);g__Metschnikowia(100);s__Metschnikowia_pulcherrima(100); | Metschnikowia | 13 | Saprotroph | \|Undefined Saprotroph\| | Yeast | NULL | Probable | As nectar-tap_saprotroph (Põlme et al. 2020); Decay sugar-rich substrates (Põlme et al. 2020) | Põlme S, et al. 2020. Fungal Diversity 105:1-16 ((https://doi.org/10.1007/s13225-020-00466-2)) |
| Otu45470 | 1 | k__Fungi(100);p__Ascomycota(100);c__Saccharomycetes(100);o__Saccharomycetales(100);f__Metschnikowiaceae(100);g__Metschnikowia(100);g__Metschnikowia_unclassified(100); | Metschnikowia | 13 | Saprotroph | \|Undefined Saprotroph\| | Yeast | NULL | Probable | As nectar-tap_saprotroph (Põlme et al. 2020); Decay sugar-rich substrates (Põlme et al. 2020) | Põlme S, et al. 2020. Fungal Diversity 105:1-16 ((https://doi.org/10.1007/s13225-020-00466-2)) |
| Otu45471 | 1 | k__Fungi(100);p__Ascomycota(100);c__Saccharomycetes(100);o__Saccharomycetales(100);f__Metschnikowiaceae(100);g__Metschnikowia(100);s__Metschnikowia_pulcherrima(100); | Metschnikowia | 13 | Saprotroph | \|Undefined Saprotroph\| | Yeast | NULL | Probable | As nectar-tap_saprotroph (Põlme et al. 2020); Decay sugar-rich substrates (Põlme et al. 2020) | Põlme S, et al. 2020. Fungal Diversity 105:1-16 ((https://doi.org/10.1007/s13225-020-00466-2)) |
| Otu45472 | 1 | k__Fungi(100);p__Ascomycota(100);c__Saccharomycetes(100);o__Saccharomycetales(100);f__Metschnikowiaceae(100);g__Metschnikowia(100);s__Metschnikowia_pulcherrima(100); | Metschnikowia | 13 | Saprotroph | \|Undefined Saprotroph\| | Yeast | NULL | Probable | As nectar-tap_saprotroph (Põlme et al. 2020); Decay sugar-rich substrates (Põlme et al. 2020) | Põlme S, et al. 2020. Fungal Diversity 105:1-16 ((https://doi.org/10.1007/s13225-020-00466-2)) |
| Otu45473 | 1 | k__Fungi(100);p__Ascomycota(100);c__Saccharomycetes(100);o__Saccharomycetales(100);f__Metschnikowiaceae(100);g__Metschnikowia(100);s__Metschnikowia_pulcherrima(100); | Metschnikowia | 13 | Saprotroph | \|Undefined Saprotroph\| | Yeast | NULL | Probable | As nectar-tap_saprotroph (Põlme et al. 2020); Decay sugar-rich substrates (Põlme et al. 2020) | Põlme S, et al. 2020. Fungal Diversity 105:1-16 ((https://doi.org/10.1007/s13225-020-00466-2)) |
| Otu45474 | 1 | k__Fungi(100);p__Ascomycota(100);c__Saccharomycetes(100);o__Saccharomycetales(100);f__Metschnikowiaceae(100);g__Metschnikowia(100);s__Metschnikowia_pulcherrima(100); | Metschnikowia | 13 | Saprotroph | \|Undefined Saprotroph\| | Yeast | NULL | Probable | As nectar-tap_saprotroph (Põlme et al. 2020); Decay sugar-rich substrates (Põlme et al. 2020) | Põlme S, et al. 2020. Fungal Diversity 105:1-16 ((https://doi.org/10.1007/s13225-020-00466-2)) |
| Otu45475 | 1 | k__Fungi(100);p__Ascomycota(100);c__Saccharomycetes(100);o__Saccharomycetales(100);f__Metschnikowiaceae(100);g__Metschnikowia(100);g__Metschnikowia_unclassified(100); | Metschnikowia | 13 | Saprotroph | \|Undefined Saprotroph\| | Yeast | NULL | Probable | As nectar-tap_saprotroph (Põlme et al. 2020); Decay sugar-rich substrates (Põlme et al. 2020) | Põlme S, et al. 2020. Fungal Diversity 105:1-16 ((https://doi.org/10.1007/s13225-020-00466-2)) |
| Otu45476 | 1 | k__Fungi(100);p__Ascomycota(100);c__Saccharomycetes(100);o__Saccharomycetales(100);f__Metschnikowiaceae(100);g__Metschnikowia(100);s__Metschnikowia_pulcherrima(100); | Metschnikowia | 13 | Saprotroph | \|Undefined Saprotroph\| | Yeast | NULL | Probable | As nectar-tap_saprotroph (Põlme et al. 2020); Decay sugar-rich substrates (Põlme et al. 2020) | Põlme S, et al. 2020. Fungal Diversity 105:1-16 ((https://doi.org/10.1007/s13225-020-00466-2)) |
| Otu45477 | 1 | k__Fungi(100);p__Ascomycota(100);c__Saccharomycetes(100);o__Saccharomycetales(100);f__Metschnikowiaceae(100);g__Metschnikowia(100);s__Metschnikowia_pulcherrima(100); | Metschnikowia | 13 | Saprotroph | \|Undefined Saprotroph\| | Yeast | NULL | Probable | As nectar-tap_saprotroph (Põlme et al. 2020); Decay sugar-rich substrates (Põlme et al. 2020) | Põlme S, et al. 2020. Fungal Diversity 105:1-16 ((https://doi.org/10.1007/s13225-020-00466-2)) |
| Otu45478 | 1 | k__Fungi(100);p__Ascomycota(100);c__Saccharomycetes(100);o__Saccharomycetales(100);f__Metschnikowiaceae(100);g__Metschnikowia(100);s__Metschnikowia_pulcherrima(100); | Metschnikowia | 13 | Saprotroph | \|Undefined Saprotroph\| | Yeast | NULL | Probable | As nectar-tap_saprotroph (Põlme et al. 2020); Decay sugar-rich substrates (Põlme et al. 2020) | Põlme S, et al. 2020. Fungal Diversity 105:1-16 ((https://doi.org/10.1007/s13225-020-00466-2)) |
| Otu45479 | 1 | k__Fungi(100);p__Ascomycota(100);c__Saccharomycetes(100);o__Saccharomycetales(100);f__Metschnikowiaceae(100);g__Metschnikowia(100);s__Metschnikowia_pulcherrima(100); | Metschnikowia | 13 | Saprotroph | \|Undefined Saprotroph\| | Yeast | NULL | Probable | As nectar-tap_saprotroph (Põlme et al. 2020); Decay sugar-rich substrates (Põlme et al. 2020) | Põlme S, et al. 2020. Fungal Diversity 105:1-16 ((https://doi.org/10.1007/s13225-020-00466-2)) |
| Otu45480 | 1 | k__Fungi(100);p__Ascomycota(100);c__Saccharomycetes(100);o__Saccharomycetales(100);f__Metschnikowiaceae(100);g__Metschnikowia(100);g__Metschnikowia_unclassified(100); | Metschnikowia | 13 | Saprotroph | \|Undefined Saprotroph\| | Yeast | NULL | Probable | As nectar-tap_saprotroph (Põlme et al. 2020); Decay sugar-rich substrates (Põlme et al. 2020) | Põlme S, et al. 2020. Fungal Diversity 105:1-16 ((https://doi.org/10.1007/s13225-020-00466-2)) |
| Otu45481 | 1 | k__Fungi(100);p__Ascomycota(100);c__Saccharomycetes(100);o__Saccharomycetales(100);f__Metschnikowiaceae(100);g__Metschnikowia(100);g__Metschnikowia_unclassified(100); | Metschnikowia | 13 | Saprotroph | \|Undefined Saprotroph\| | Yeast | NULL | Probable | As nectar-tap_saprotroph (Põlme et al. 2020); Decay sugar-rich substrates (Põlme et al. 2020) | Põlme S, et al. 2020. Fungal Diversity 105:1-16 ((https://doi.org/10.1007/s13225-020-00466-2)) |
| Otu45482 | 1 | k__Fungi(100);p__Ascomycota(100);c__Saccharomycetes(100);o__Saccharomycetales(100);f__Metschnikowiaceae(100);g__Metschnikowia(100);s__Metschnikowia_pulcherrima(100); | Metschnikowia | 13 | Saprotroph | \|Undefined Saprotroph\| | Yeast | NULL | Probable | As nectar-tap_saprotroph (Põlme et al. 2020); Decay sugar-rich substrates (Põlme et al. 2020) | Põlme S, et al. 2020. Fungal Diversity 105:1-16 ((https://doi.org/10.1007/s13225-020-00466-2)) |
| Otu45483 | 1 | k__Fungi(100);p__Ascomycota(100);c__Saccharomycetes(100);o__Saccharomycetales(100);f__Metschnikowiaceae(100);g__Metschnikowia(100);s__Metschnikowia_viticola(100); | Metschnikowia | 13 | Saprotroph | \|Undefined Saprotroph\| | Yeast | NULL | Probable | As nectar-tap_saprotroph (Põlme et al. 2020); Decay sugar-rich substrates (Põlme et al. 2020) | Põlme S, et al. 2020. Fungal Diversity 105:1-16 ((https://doi.org/10.1007/s13225-020-00466-2)) |
| Otu45484 | 1 | k__Fungi(100);p__Ascomycota(100);c__Saccharomycetes(100);o__Saccharomycetales(100);f__Metschnikowiaceae(100);g__Metschnikowia(100);s__Metschnikowia_pulcherrima(100); | Metschnikowia | 13 | Saprotroph | \|Undefined Saprotroph\| | Yeast | NULL | Probable | As nectar-tap_saprotroph (Põlme et al. 2020); Decay sugar-rich substrates (Põlme et al. 2020) | Põlme S, et al. 2020. Fungal Diversity 105:1-16 ((https://doi.org/10.1007/s13225-020-00466-2)) |
| Otu45485 | 1 | k__Fungi(100);p__Ascomycota(100);c__Saccharomycetes(100);o__Saccharomycetales(100);f__Metschnikowiaceae(100);g__Metschnikowia(100);g__Metschnikowia_unclassified(100); | Metschnikowia | 13 | Saprotroph | \|Undefined Saprotroph\| | Yeast | NULL | Probable | As nectar-tap_saprotroph (Põlme et al. 2020); Decay sugar-rich substrates (Põlme et al. 2020) | Põlme S, et al. 2020. Fungal Diversity 105:1-16 ((https://doi.org/10.1007/s13225-020-00466-2)) |
| Otu45486 | 1 | k__Fungi(100);p__Ascomycota(100);c__Saccharomycetes(100);o__Saccharomycetales(100);f__Metschnikowiaceae(100);g__Metschnikowia(100);g__Metschnikowia_unclassified(100); | Metschnikowia | 13 | Saprotroph | \|Undefined Saprotroph\| | Yeast | NULL | Probable | As nectar-tap_saprotroph (Põlme et al. 2020); Decay sugar-rich substrates (Põlme et al. 2020) | Põlme S, et al. 2020. Fungal Diversity 105:1-16 ((https://doi.org/10.1007/s13225-020-00466-2)) |
| Otu45487 | 1 | k__Fungi(100);p__Ascomycota(100);c__Saccharomycetes(100);o__Saccharomycetales(100);f__Metschnikowiaceae(100);g__Metschnikowia(100);g__Metschnikowia_unclassified(100); | Metschnikowia | 13 | Saprotroph | \|Undefined Saprotroph\| | Yeast | NULL | Probable | As nectar-tap_saprotroph (Põlme et al. 2020); Decay sugar-rich substrates (Põlme et al. 2020) | Põlme S, et al. 2020. Fungal Diversity 105:1-16 ((https://doi.org/10.1007/s13225-020-00466-2)) |
| Otu45488 | 1 | k__Fungi(100);p__Ascomycota(100);c__Saccharomycetes(100);o__Saccharomycetales(100);f__Metschnikowiaceae(100);g__Metschnikowia(100);s__Metschnikowia_pulcherrima(100); | Metschnikowia | 13 | Saprotroph | \|Undefined Saprotroph\| | Yeast | NULL | Probable | As nectar-tap_saprotroph (Põlme et al. 2020); Decay sugar-rich substrates (Põlme et al. 2020) | Põlme S, et al. 2020. Fungal Diversity 105:1-16 ((https://doi.org/10.1007/s13225-020-00466-2)) |
| Otu45489 | 1 | k__Fungi(100);p__Ascomycota(100);c__Saccharomycetes(100);o__Saccharomycetales(100);f__Metschnikowiaceae(100);g__Metschnikowia(100);s__Metschnikowia_pulcherrima(100); | Metschnikowia | 13 | Saprotroph | \|Undefined Saprotroph\| | Yeast | NULL | Probable | As nectar-tap_saprotroph (Põlme et al. 2020); Decay sugar-rich substrates (Põlme et al. 2020) | Põlme S, et al. 2020. Fungal Diversity 105:1-16 ((https://doi.org/10.1007/s13225-020-00466-2)) |
| Otu45490 | 1 | k__Fungi(100);p__Ascomycota(100);c__Saccharomycetes(100);o__Saccharomycetales(100);f__Metschnikowiaceae(100);g__Metschnikowia(100);g__Metschnikowia_unclassified(100); | Metschnikowia | 13 | Saprotroph | \|Undefined Saprotroph\| | Yeast | NULL | Probable | As nectar-tap_saprotroph (Põlme et al. 2020); Decay sugar-rich substrates (Põlme et al. 2020) | Põlme S, et al. 2020. Fungal Diversity 105:1-16 ((https://doi.org/10.1007/s13225-020-00466-2)) |
| Otu45492 | 1 | k__Fungi(100);p__Ascomycota(100);c__Saccharomycetes(100);o__Saccharomycetales(100);f__Metschnikowiaceae(100);g__Metschnikowia(100);s__Metschnikowia_pulcherrima(100); | Metschnikowia | 13 | Saprotroph | \|Undefined Saprotroph\| | Yeast | NULL | Probable | As nectar-tap_saprotroph (Põlme et al. 2020); Decay sugar-rich substrates (Põlme et al. 2020) | Põlme S, et al. 2020. Fungal Diversity 105:1-16 ((https://doi.org/10.1007/s13225-020-00466-2)) |
| Otu45493 | 1 | k__Fungi(100);p__Ascomycota(100);c__Saccharomycetes(100);o__Saccharomycetales(100);f__Metschnikowiaceae(100);g__Metschnikowia(100);s__Metschnikowia_viticola(100); | Metschnikowia | 13 | Saprotroph | \|Undefined Saprotroph\| | Yeast | NULL | Probable | As nectar-tap_saprotroph (Põlme et al. 2020); Decay sugar-rich substrates (Põlme et al. 2020) | Põlme S, et al. 2020. Fungal Diversity 105:1-16 ((https://doi.org/10.1007/s13225-020-00466-2)) |
| Otu45494 | 1 | k__Fungi(100);p__Ascomycota(100);c__Saccharomycetes(100);o__Saccharomycetales(100);f__Metschnikowiaceae(100);g__Metschnikowia(100);s__Metschnikowia_viticola(100); | Metschnikowia | 13 | Saprotroph | \|Undefined Saprotroph\| | Yeast | NULL | Probable | As nectar-tap_saprotroph (Põlme et al. 2020); Decay sugar-rich substrates (Põlme et al. 2020) | Põlme S, et al. 2020. Fungal Diversity 105:1-16 ((https://doi.org/10.1007/s13225-020-00466-2)) |
| Otu45495 | 1 | k__Fungi(100);p__Ascomycota(100);c__Saccharomycetes(100);o__Saccharomycetales(100);f__Metschnikowiaceae(100);g__Metschnikowia(100);g__Metschnikowia_unclassified(100); | Metschnikowia | 13 | Saprotroph | \|Undefined Saprotroph\| | Yeast | NULL | Probable | As nectar-tap_saprotroph (Põlme et al. 2020); Decay sugar-rich substrates (Põlme et al. 2020) | Põlme S, et al. 2020. Fungal Diversity 105:1-16 ((https://doi.org/10.1007/s13225-020-00466-2)) |
| Otu45496 | 1 | k__Fungi(100);p__Ascomycota(100);c__Saccharomycetes(100);o__Saccharomycetales(100);f__Metschnikowiaceae(100);g__Metschnikowia(100);s__Metschnikowia_pulcherrima(100); | Metschnikowia | 13 | Saprotroph | \|Undefined Saprotroph\| | Yeast | NULL | Probable | As nectar-tap_saprotroph (Põlme et al. 2020); Decay sugar-rich substrates (Põlme et al. 2020) | Põlme S, et al. 2020. Fungal Diversity 105:1-16 ((https://doi.org/10.1007/s13225-020-00466-2)) |
| Otu45497 | 1 | k__Fungi(100);p__Ascomycota(100);c__Saccharomycetes(100);o__Saccharomycetales(100);f__Metschnikowiaceae(100);g__Metschnikowia(100);s__Metschnikowia_pulcherrima(100); | Metschnikowia | 13 | Saprotroph | \|Undefined Saprotroph\| | Yeast | NULL | Probable | As nectar-tap_saprotroph (Põlme et al. 2020); Decay sugar-rich substrates (Põlme et al. 2020) | Põlme S, et al. 2020. Fungal Diversity 105:1-16 ((https://doi.org/10.1007/s13225-020-00466-2)) |
| Otu45498 | 1 | k__Fungi(100);p__Ascomycota(100);c__Saccharomycetes(100);o__Saccharomycetales(100);f__Metschnikowiaceae(100);g__Metschnikowia(100);g__Metschnikowia_unclassified(100); | Metschnikowia | 13 | Saprotroph | \|Undefined Saprotroph\| | Yeast | NULL | Probable | As nectar-tap_saprotroph (Põlme et al. 2020); Decay sugar-rich substrates (Põlme et al. 2020) | Põlme S, et al. 2020. Fungal Diversity 105:1-16 ((https://doi.org/10.1007/s13225-020-00466-2)) |
| Otu45499 | 1 | k__Fungi(100);p__Ascomycota(100);c__Saccharomycetes(100);o__Saccharomycetales(100);f__Metschnikowiaceae(100);g__Metschnikowia(100);g__Metschnikowia_unclassified(100); | Metschnikowia | 13 | Saprotroph | \|Undefined Saprotroph\| | Yeast | NULL | Probable | As nectar-tap_saprotroph (Põlme et al. 2020); Decay sugar-rich substrates (Põlme et al. 2020) | Põlme S, et al. 2020. Fungal Diversity 105:1-16 ((https://doi.org/10.1007/s13225-020-00466-2)) |
| Otu45500 | 1 | k__Fungi(100);p__Ascomycota(100);c__Saccharomycetes(100);o__Saccharomycetales(100);f__Metschnikowiaceae(100);g__Metschnikowia(100);s__Metschnikowia_pulcherrima(100); | Metschnikowia | 13 | Saprotroph | \|Undefined Saprotroph\| | Yeast | NULL | Probable | As nectar-tap_saprotroph (Põlme et al. 2020); Decay sugar-rich substrates (Põlme et al. 2020) | Põlme S, et al. 2020. Fungal Diversity 105:1-16 ((https://doi.org/10.1007/s13225-020-00466-2)) |
| Otu45501 | 1 | k__Fungi(100);p__Ascomycota(100);c__Saccharomycetes(100);o__Saccharomycetales(100);f__Metschnikowiaceae(100);g__Metschnikowia(100);s__Metschnikowia_pulcherrima(100); | Metschnikowia | 13 | Saprotroph | \|Undefined Saprotroph\| | Yeast | NULL | Probable | As nectar-tap_saprotroph (Põlme et al. 2020); Decay sugar-rich substrates (Põlme et al. 2020) | Põlme S, et al. 2020. Fungal Diversity 105:1-16 ((https://doi.org/10.1007/s13225-020-00466-2)) |
| Otu45502 | 1 | k__Fungi(100);p__Ascomycota(100);c__Saccharomycetes(100);o__Saccharomycetales(100);f__Metschnikowiaceae(100);g__Metschnikowia(100);s__Metschnikowia_pulcherrima(100); | Metschnikowia | 13 | Saprotroph | \|Undefined Saprotroph\| | Yeast | NULL | Probable | As nectar-tap_saprotroph (Põlme et al. 2020); Decay sugar-rich substrates (Põlme et al. 2020) | Põlme S, et al. 2020. Fungal Diversity 105:1-16 ((https://doi.org/10.1007/s13225-020-00466-2)) |
| Otu45503 | 1 | k__Fungi(100);p__Ascomycota(100);c__Saccharomycetes(100);o__Saccharomycetales(100);f__Metschnikowiaceae(100);g__Metschnikowia(100);s__Metschnikowia_pulcherrima(100); | Metschnikowia | 13 | Saprotroph | \|Undefined Saprotroph\| | Yeast | NULL | Probable | As nectar-tap_saprotroph (Põlme et al. 2020); Decay sugar-rich substrates (Põlme et al. 2020) | Põlme S, et al. 2020. Fungal Diversity 105:1-16 ((https://doi.org/10.1007/s13225-020-00466-2)) |
| Otu45504 | 1 | k__Fungi(100);p__Ascomycota(100);c__Saccharomycetes(100);o__Saccharomycetales(100);f__Metschnikowiaceae(100);g__Metschnikowia(100);s__Metschnikowia_pulcherrima(100); | Metschnikowia | 13 | Saprotroph | \|Undefined Saprotroph\| | Yeast | NULL | Probable | As nectar-tap_saprotroph (Põlme et al. 2020); Decay sugar-rich substrates (Põlme et al. 2020) | Põlme S, et al. 2020. Fungal Diversity 105:1-16 ((https://doi.org/10.1007/s13225-020-00466-2)) |
| Otu45505 | 1 | k__Fungi(100);p__Ascomycota(100);c__Saccharomycetes(100);o__Saccharomycetales(100);f__Metschnikowiaceae(100);g__Metschnikowia(100);g__Metschnikowia_unclassified(100); | Metschnikowia | 13 | Saprotroph | \|Undefined Saprotroph\| | Yeast | NULL | Probable | As nectar-tap_saprotroph (Põlme et al. 2020); Decay sugar-rich substrates (Põlme et al. 2020) | Põlme S, et al. 2020. Fungal Diversity 105:1-16 ((https://doi.org/10.1007/s13225-020-00466-2)) |
| Otu45506 | 1 | k__Fungi(100);p__Ascomycota(100);c__Saccharomycetes(100);o__Saccharomycetales(100);f__Metschnikowiaceae(100);g__Metschnikowia(100);s__Metschnikowia_pulcherrima(100); | Metschnikowia | 13 | Saprotroph | \|Undefined Saprotroph\| | Yeast | NULL | Probable | As nectar-tap_saprotroph (Põlme et al. 2020); Decay sugar-rich substrates (Põlme et al. 2020) | Põlme S, et al. 2020. Fungal Diversity 105:1-16 ((https://doi.org/10.1007/s13225-020-00466-2)) |
| Otu45507 | 1 | k__Fungi(100);p__Ascomycota(100);c__Saccharomycetes(100);o__Saccharomycetales(100);f__Metschnikowiaceae(100);g__Metschnikowia(100);s__Metschnikowia_pulcherrima(100); | Metschnikowia | 13 | Saprotroph | \|Undefined Saprotroph\| | Yeast | NULL | Probable | As nectar-tap_saprotroph (Põlme et al. 2020); Decay sugar-rich substrates (Põlme et al. 2020) | Põlme S, et al. 2020. Fungal Diversity 105:1-16 ((https://doi.org/10.1007/s13225-020-00466-2)) |
| Otu45508 | 1 | k__Fungi(100);p__Ascomycota(100);c__Saccharomycetes(100);o__Saccharomycetales(100);f__Metschnikowiaceae(100);g__Metschnikowia(100);s__Metschnikowia_pulcherrima(100); | Metschnikowia | 13 | Saprotroph | \|Undefined Saprotroph\| | Yeast | NULL | Probable | As nectar-tap_saprotroph (Põlme et al. 2020); Decay sugar-rich substrates (Põlme et al. 2020) | Põlme S, et al. 2020. Fungal Diversity 105:1-16 ((https://doi.org/10.1007/s13225-020-00466-2)) |
| Otu45509 | 1 | k__Fungi(100);p__Ascomycota(100);c__Saccharomycetes(100);o__Saccharomycetales(100);f__Metschnikowiaceae(100);g__Metschnikowia(100);s__Metschnikowia_pulcherrima(100); | Metschnikowia | 13 | Saprotroph | \|Undefined Saprotroph\| | Yeast | NULL | Probable | As nectar-tap_saprotroph (Põlme et al. 2020); Decay sugar-rich substrates (Põlme et al. 2020) | Põlme S, et al. 2020. Fungal Diversity 105:1-16 ((https://doi.org/10.1007/s13225-020-00466-2)) |
| Otu45510 | 1 | k__Fungi(100);p__Ascomycota(100);c__Saccharomycetes(100);o__Saccharomycetales(100);f__Metschnikowiaceae(100);g__Metschnikowia(100);s__Metschnikowia_pulcherrima(100); | Metschnikowia | 13 | Saprotroph | \|Undefined Saprotroph\| | Yeast | NULL | Probable | As nectar-tap_saprotroph (Põlme et al. 2020); Decay sugar-rich substrates (Põlme et al. 2020) | Põlme S, et al. 2020. Fungal Diversity 105:1-16 ((https://doi.org/10.1007/s13225-020-00466-2)) |
| Otu45511 | 1 | k__Fungi(100);p__Ascomycota(100);c__Saccharomycetes(100);o__Saccharomycetales(100);f__Metschnikowiaceae(100);g__Metschnikowia(100);s__Metschnikowia_pulcherrima(100); | Metschnikowia | 13 | Saprotroph | \|Undefined Saprotroph\| | Yeast | NULL | Probable | As nectar-tap_saprotroph (Põlme et al. 2020); Decay sugar-rich substrates (Põlme et al. 2020) | Põlme S, et al. 2020. Fungal Diversity 105:1-16 ((https://doi.org/10.1007/s13225-020-00466-2)) |
| Otu45513 | 1 | k__Fungi(100);p__Ascomycota(100);c__Saccharomycetes(100);o__Saccharomycetales(100);f__Metschnikowiaceae(100);g__Metschnikowia(100);s__Metschnikowia_pulcherrima(100); | Metschnikowia | 13 | Saprotroph | \|Undefined Saprotroph\| | Yeast | NULL | Probable | As nectar-tap_saprotroph (Põlme et al. 2020); Decay sugar-rich substrates (Põlme et al. 2020) | Põlme S, et al. 2020. Fungal Diversity 105:1-16 ((https://doi.org/10.1007/s13225-020-00466-2)) |
| Otu45514 | 1 | k__Fungi(100);p__Ascomycota(100);c__Saccharomycetes(100);o__Saccharomycetales(100);f__Metschnikowiaceae(100);g__Metschnikowia(100);g__Metschnikowia_unclassified(100); | Metschnikowia | 13 | Saprotroph | \|Undefined Saprotroph\| | Yeast | NULL | Probable | As nectar-tap_saprotroph (Põlme et al. 2020); Decay sugar-rich substrates (Põlme et al. 2020) | Põlme S, et al. 2020. Fungal Diversity 105:1-16 ((https://doi.org/10.1007/s13225-020-00466-2)) |
| Otu45515 | 1 | k__Fungi(100);p__Ascomycota(100);c__Saccharomycetes(100);o__Saccharomycetales(100);f__Metschnikowiaceae(100);g__Metschnikowia(100);s__Metschnikowia_pulcherrima(100); | Metschnikowia | 13 | Saprotroph | \|Undefined Saprotroph\| | Yeast | NULL | Probable | As nectar-tap_saprotroph (Põlme et al. 2020); Decay sugar-rich substrates (Põlme et al. 2020) | Põlme S, et al. 2020. Fungal Diversity 105:1-16 ((https://doi.org/10.1007/s13225-020-00466-2)) |
| Otu45516 | 1 | k__Fungi(100);p__Ascomycota(100);c__Saccharomycetes(100);o__Saccharomycetales(100);f__Metschnikowiaceae(100);g__Metschnikowia(100);s__Metschnikowia_viticola(100); | Metschnikowia | 13 | Saprotroph | \|Undefined Saprotroph\| | Yeast | NULL | Probable | As nectar-tap_saprotroph (Põlme et al. 2020); Decay sugar-rich substrates (Põlme et al. 2020) | Põlme S, et al. 2020. Fungal Diversity 105:1-16 ((https://doi.org/10.1007/s13225-020-00466-2)) |
| Otu45517 | 1 | k__Fungi(100);p__Ascomycota(100);c__Saccharomycetes(100);o__Saccharomycetales(100);f__Metschnikowiaceae(100);g__Metschnikowia(100);g__Metschnikowia_unclassified(100); | Metschnikowia | 13 | Saprotroph | \|Undefined Saprotroph\| | Yeast | NULL | Probable | As nectar-tap_saprotroph (Põlme et al. 2020); Decay sugar-rich substrates (Põlme et al. 2020) | Põlme S, et al. 2020. Fungal Diversity 105:1-16 ((https://doi.org/10.1007/s13225-020-00466-2)) |
| Otu45518 | 1 | k__Fungi(100);p__Ascomycota(100);c__Saccharomycetes(100);o__Saccharomycetales(100);f__Metschnikowiaceae(100);g__Metschnikowia(100);s__Metschnikowia_pulcherrima(100); | Metschnikowia | 13 | Saprotroph | \|Undefined Saprotroph\| | Yeast | NULL | Probable | As nectar-tap_saprotroph (Põlme et al. 2020); Decay sugar-rich substrates (Põlme et al. 2020) | Põlme S, et al. 2020. Fungal Diversity 105:1-16 ((https://doi.org/10.1007/s13225-020-00466-2)) |
| Otu45519 | 1 | k__Fungi(100);p__Ascomycota(100);c__Saccharomycetes(100);o__Saccharomycetales(100);f__Metschnikowiaceae(100);g__Metschnikowia(100);g__Metschnikowia_unclassified(100); | Metschnikowia | 13 | Saprotroph | \|Undefined Saprotroph\| | Yeast | NULL | Probable | As nectar-tap_saprotroph (Põlme et al. 2020); Decay sugar-rich substrates (Põlme et al. 2020) | Põlme S, et al. 2020. Fungal Diversity 105:1-16 ((https://doi.org/10.1007/s13225-020-00466-2)) |
| Otu45520 | 1 | k__Fungi(100);p__Ascomycota(100);c__Saccharomycetes(100);o__Saccharomycetales(100);f__Metschnikowiaceae(100);g__Metschnikowia(100);s__Metschnikowia_pulcherrima(100); | Metschnikowia | 13 | Saprotroph | \|Undefined Saprotroph\| | Yeast | NULL | Probable | As nectar-tap_saprotroph (Põlme et al. 2020); Decay sugar-rich substrates (Põlme et al. 2020) | Põlme S, et al. 2020. Fungal Diversity 105:1-16 ((https://doi.org/10.1007/s13225-020-00466-2)) |
| Otu45521 | 1 | k__Fungi(100);p__Ascomycota(100);c__Saccharomycetes(100);o__Saccharomycetales(100);f__Metschnikowiaceae(100);g__Metschnikowia(100);s__Metschnikowia_pulcherrima(100); | Metschnikowia | 13 | Saprotroph | \|Undefined Saprotroph\| | Yeast | NULL | Probable | As nectar-tap_saprotroph (Põlme et al. 2020); Decay sugar-rich substrates (Põlme et al. 2020) | Põlme S, et al. 2020. Fungal Diversity 105:1-16 ((https://doi.org/10.1007/s13225-020-00466-2)) |
| Otu45522 | 1 | k__Fungi(100);p__Ascomycota(100);c__Sordariomycetes(100);o__Hypocreales(100);f__Nectriaceae(100);f__Nectriaceae_unclassified(100);f__Nectriaceae_unclassified(100); | Nectriaceae | 9 | Pathotroph-Saprotroph-Symbiotroph | Animal Pathogen-Endophyte-Fungal Parasite-Lichen Parasite-Plant Pathogen-Wood Saprotroph | Microfungus | NULL | Possible | Fusarium species are amongst the economically significant plant pathogens (Cannon & Kirk 2007) | Cannon PF, Kirk PM. 2007. Fungal Families of the World. CAB International, Cambridge (ISBN: 978-0851998275); Hirooka Y, et al. 2012. Studies in Mycology 711-210 ((https://doi.org/10.3114/sim0001)); Tedersoo L, et al. 2014. Science 346:e1256688 ((https://doi.org/10.1126/science.1256688)); The American Phytopathological Society Common Names of Plant Diseases ((http://www.apsnet.org/publications/commonnames/Pages/default.aspx)); https://www.cdc.gov ((https://www.cdc.gov)) |
| Otu45523 | 1 | k__Fungi(100);p__Ascomycota(100);c__Saccharomycetes(100);o__Saccharomycetales(100);f__Metschnikowiaceae(100);g__Metschnikowia(100);s__Metschnikowia_pulcherrima(100); | Metschnikowia | 13 | Saprotroph | \|Undefined Saprotroph\| | Yeast | NULL | Probable | As nectar-tap_saprotroph (Põlme et al. 2020); Decay sugar-rich substrates (Põlme et al. 2020) | Põlme S, et al. 2020. Fungal Diversity 105:1-16 ((https://doi.org/10.1007/s13225-020-00466-2)) |
| Otu45524 | 1 | k__Fungi(100);p__Ascomycota(100);c__Saccharomycetes(100);o__Saccharomycetales(100);f__Metschnikowiaceae(100);g__Metschnikowia(100);s__Metschnikowia_viticola(100); | Metschnikowia | 13 | Saprotroph | \|Undefined Saprotroph\| | Yeast | NULL | Probable | As nectar-tap_saprotroph (Põlme et al. 2020); Decay sugar-rich substrates (Põlme et al. 2020) | Põlme S, et al. 2020. Fungal Diversity 105:1-16 ((https://doi.org/10.1007/s13225-020-00466-2)) |
| Otu45525 | 1 | k__Fungi(100);p__Ascomycota(100);c__Saccharomycetes(100);o__Saccharomycetales(100);f__Metschnikowiaceae(100);g__Metschnikowia(100);s__Metschnikowia_pulcherrima(100); | Metschnikowia | 13 | Saprotroph | \|Undefined Saprotroph\| | Yeast | NULL | Probable | As nectar-tap_saprotroph (Põlme et al. 2020); Decay sugar-rich substrates (Põlme et al. 2020) | Põlme S, et al. 2020. Fungal Diversity 105:1-16 ((https://doi.org/10.1007/s13225-020-00466-2)) |
| Otu45526 | 1 | k__Fungi(100);p__Ascomycota(100);c__Saccharomycetes(100);o__Saccharomycetales(100);f__Metschnikowiaceae(100);g__Metschnikowia(100);s__Metschnikowia_pulcherrima(100); | Metschnikowia | 13 | Saprotroph | \|Undefined Saprotroph\| | Yeast | NULL | Probable | As nectar-tap_saprotroph (Põlme et al. 2020); Decay sugar-rich substrates (Põlme et al. 2020) | Põlme S, et al. 2020. Fungal Diversity 105:1-16 ((https://doi.org/10.1007/s13225-020-00466-2)) |
| Otu45527 | 1 | k__Fungi(100);p__Ascomycota(100);c__Saccharomycetes(100);o__Saccharomycetales(100);f__Metschnikowiaceae(100);g__Metschnikowia(100);s__Metschnikowia_pulcherrima(100); | Metschnikowia | 13 | Saprotroph | \|Undefined Saprotroph\| | Yeast | NULL | Probable | As nectar-tap_saprotroph (Põlme et al. 2020); Decay sugar-rich substrates (Põlme et al. 2020) | Põlme S, et al. 2020. Fungal Diversity 105:1-16 ((https://doi.org/10.1007/s13225-020-00466-2)) |
| Otu45528 | 1 | k__Fungi(100);p__Ascomycota(100);c__Saccharomycetes(100);o__Saccharomycetales(100);f__Metschnikowiaceae(100);g__Metschnikowia(100);s__Metschnikowia_pulcherrima(100); | Metschnikowia | 13 | Saprotroph | \|Undefined Saprotroph\| | Yeast | NULL | Probable | As nectar-tap_saprotroph (Põlme et al. 2020); Decay sugar-rich substrates (Põlme et al. 2020) | Põlme S, et al. 2020. Fungal Diversity 105:1-16 ((https://doi.org/10.1007/s13225-020-00466-2)) |
| Otu45529 | 1 | k__Fungi(100);p__Ascomycota(100);c__Saccharomycetes(100);o__Saccharomycetales(100);f__Metschnikowiaceae(100);g__Metschnikowia(100);s__Metschnikowia_pulcherrima(100); | Metschnikowia | 13 | Saprotroph | \|Undefined Saprotroph\| | Yeast | NULL | Probable | As nectar-tap_saprotroph (Põlme et al. 2020); Decay sugar-rich substrates (Põlme et al. 2020) | Põlme S, et al. 2020. Fungal Diversity 105:1-16 ((https://doi.org/10.1007/s13225-020-00466-2)) |
| Otu45530 | 1 | k__Fungi(100);p__Ascomycota(100);c__Saccharomycetes(100);o__Saccharomycetales(100);f__Metschnikowiaceae(100);g__Metschnikowia(100);s__Metschnikowia_pulcherrima(100); | Metschnikowia | 13 | Saprotroph | \|Undefined Saprotroph\| | Yeast | NULL | Probable | As nectar-tap_saprotroph (Põlme et al. 2020); Decay sugar-rich substrates (Põlme et al. 2020) | Põlme S, et al. 2020. Fungal Diversity 105:1-16 ((https://doi.org/10.1007/s13225-020-00466-2)) |
| Otu45531 | 1 | k__Fungi(100);p__Ascomycota(100);c__Saccharomycetes(100);o__Saccharomycetales(100);f__Metschnikowiaceae(100);g__Metschnikowia(100);s__Metschnikowia_pulcherrima(100); | Metschnikowia | 13 | Saprotroph | \|Undefined Saprotroph\| | Yeast | NULL | Probable | As nectar-tap_saprotroph (Põlme et al. 2020); Decay sugar-rich substrates (Põlme et al. 2020) | Põlme S, et al. 2020. Fungal Diversity 105:1-16 ((https://doi.org/10.1007/s13225-020-00466-2)) |
| Otu45532 | 1 | k__Fungi(100);p__Ascomycota(100);c__Saccharomycetes(100);o__Saccharomycetales(100);f__Metschnikowiaceae(100);g__Metschnikowia(100);s__Metschnikowia_pulcherrima(100); | Metschnikowia | 13 | Saprotroph | \|Undefined Saprotroph\| | Yeast | NULL | Probable | As nectar-tap_saprotroph (Põlme et al. 2020); Decay sugar-rich substrates (Põlme et al. 2020) | Põlme S, et al. 2020. Fungal Diversity 105:1-16 ((https://doi.org/10.1007/s13225-020-00466-2)) |
| Otu45533 | 1 | k__Fungi(100);p__Ascomycota(100);c__Saccharomycetes(100);o__Saccharomycetales(100);f__Metschnikowiaceae(100);g__Metschnikowia(100);g__Metschnikowia_unclassified(100); | Metschnikowia | 13 | Saprotroph | \|Undefined Saprotroph\| | Yeast | NULL | Probable | As nectar-tap_saprotroph (Põlme et al. 2020); Decay sugar-rich substrates (Põlme et al. 2020) | Põlme S, et al. 2020. Fungal Diversity 105:1-16 ((https://doi.org/10.1007/s13225-020-00466-2)) |
| Otu45534 | 1 | k__Fungi(100);p__Ascomycota(100);c__Saccharomycetes(100);o__Saccharomycetales(100);f__Metschnikowiaceae(100);g__Metschnikowia(100);g__Metschnikowia_unclassified(100); | Metschnikowia | 13 | Saprotroph | \|Undefined Saprotroph\| | Yeast | NULL | Probable | As nectar-tap_saprotroph (Põlme et al. 2020); Decay sugar-rich substrates (Põlme et al. 2020) | Põlme S, et al. 2020. Fungal Diversity 105:1-16 ((https://doi.org/10.1007/s13225-020-00466-2)) |
| Otu45535 | 1 | k__Fungi(100);p__Ascomycota(100);c__Saccharomycetes(100);o__Saccharomycetales(100);f__Metschnikowiaceae(100);g__Metschnikowia(100);g__Metschnikowia_unclassified(100); | Metschnikowia | 13 | Saprotroph | \|Undefined Saprotroph\| | Yeast | NULL | Probable | As nectar-tap_saprotroph (Põlme et al. 2020); Decay sugar-rich substrates (Põlme et al. 2020) | Põlme S, et al. 2020. Fungal Diversity 105:1-16 ((https://doi.org/10.1007/s13225-020-00466-2)) |
| Otu45536 | 1 | k__Fungi(100);p__Ascomycota(100);c__Saccharomycetes(100);o__Saccharomycetales(100);f__Metschnikowiaceae(100);g__Metschnikowia(100);g__Metschnikowia_unclassified(100); | Metschnikowia | 13 | Saprotroph | \|Undefined Saprotroph\| | Yeast | NULL | Probable | As nectar-tap_saprotroph (Põlme et al. 2020); Decay sugar-rich substrates (Põlme et al. 2020) | Põlme S, et al. 2020. Fungal Diversity 105:1-16 ((https://doi.org/10.1007/s13225-020-00466-2)) |
| Otu45537 | 1 | k__Fungi(100);p__Ascomycota(100);c__Saccharomycetes(100);o__Saccharomycetales(100);f__Metschnikowiaceae(100);g__Metschnikowia(100);g__Metschnikowia_unclassified(100); | Metschnikowia | 13 | Saprotroph | \|Undefined Saprotroph\| | Yeast | NULL | Probable | As nectar-tap_saprotroph (Põlme et al. 2020); Decay sugar-rich substrates (Põlme et al. 2020) | Põlme S, et al. 2020. Fungal Diversity 105:1-16 ((https://doi.org/10.1007/s13225-020-00466-2)) |
| Otu45538 | 1 | k__Fungi(100);p__Ascomycota(100);c__Saccharomycetes(100);o__Saccharomycetales(100);f__Metschnikowiaceae(100);g__Metschnikowia(100);g__Metschnikowia_unclassified(100); | Metschnikowia | 13 | Saprotroph | \|Undefined Saprotroph\| | Yeast | NULL | Probable | As nectar-tap_saprotroph (Põlme et al. 2020); Decay sugar-rich substrates (Põlme et al. 2020) | Põlme S, et al. 2020. Fungal Diversity 105:1-16 ((https://doi.org/10.1007/s13225-020-00466-2)) |
| Otu45539 | 1 | k__Fungi(100);p__Ascomycota(100);c__Saccharomycetes(100);o__Saccharomycetales(100);f__Metschnikowiaceae(100);g__Metschnikowia(100);g__Metschnikowia_unclassified(100); | Metschnikowia | 13 | Saprotroph | \|Undefined Saprotroph\| | Yeast | NULL | Probable | As nectar-tap_saprotroph (Põlme et al. 2020); Decay sugar-rich substrates (Põlme et al. 2020) | Põlme S, et al. 2020. Fungal Diversity 105:1-16 ((https://doi.org/10.1007/s13225-020-00466-2)) |
| Otu45540 | 1 | k__Fungi(100);p__Ascomycota(100);c__Saccharomycetes(100);o__Saccharomycetales(100);f__Metschnikowiaceae(100);g__Metschnikowia(100);g__Metschnikowia_unclassified(100); | Metschnikowia | 13 | Saprotroph | \|Undefined Saprotroph\| | Yeast | NULL | Probable | As nectar-tap_saprotroph (Põlme et al. 2020); Decay sugar-rich substrates (Põlme et al. 2020) | Põlme S, et al. 2020. Fungal Diversity 105:1-16 ((https://doi.org/10.1007/s13225-020-00466-2)) |
| Otu45541 | 1 | k__Fungi(100);p__Ascomycota(100);c__Saccharomycetes(100);o__Saccharomycetales(100);f__Metschnikowiaceae(100);g__Metschnikowia(100);g__Metschnikowia_unclassified(100); | Metschnikowia | 13 | Saprotroph | \|Undefined Saprotroph\| | Yeast | NULL | Probable | As nectar-tap_saprotroph (Põlme et al. 2020); Decay sugar-rich substrates (Põlme et al. 2020) | Põlme S, et al. 2020. Fungal Diversity 105:1-16 ((https://doi.org/10.1007/s13225-020-00466-2)) |
| Otu45542 | 1 | k__Fungi(100);p__Ascomycota(100);c__Saccharomycetes(100);o__Saccharomycetales(100);f__Metschnikowiaceae(100);g__Metschnikowia(100);g__Metschnikowia_unclassified(100); | Metschnikowia | 13 | Saprotroph | \|Undefined Saprotroph\| | Yeast | NULL | Probable | As nectar-tap_saprotroph (Põlme et al. 2020); Decay sugar-rich substrates (Põlme et al. 2020) | Põlme S, et al. 2020. Fungal Diversity 105:1-16 ((https://doi.org/10.1007/s13225-020-00466-2)) |
| Otu45543 | 1 | k__Fungi(100);p__Ascomycota(100);c__Saccharomycetes(100);o__Saccharomycetales(100);f__Metschnikowiaceae(100);g__Metschnikowia(100);g__Metschnikowia_unclassified(100); | Metschnikowia | 13 | Saprotroph | \|Undefined Saprotroph\| | Yeast | NULL | Probable | As nectar-tap_saprotroph (Põlme et al. 2020); Decay sugar-rich substrates (Põlme et al. 2020) | Põlme S, et al. 2020. Fungal Diversity 105:1-16 ((https://doi.org/10.1007/s13225-020-00466-2)) |
| Otu45544 | 1 | k__Fungi(100);p__Ascomycota(100);c__Saccharomycetes(100);o__Saccharomycetales(100);f__Metschnikowiaceae(100);g__Metschnikowia(100);g__Metschnikowia_unclassified(100); | Metschnikowia | 13 | Saprotroph | \|Undefined Saprotroph\| | Yeast | NULL | Probable | As nectar-tap_saprotroph (Põlme et al. 2020); Decay sugar-rich substrates (Põlme et al. 2020) | Põlme S, et al. 2020. Fungal Diversity 105:1-16 ((https://doi.org/10.1007/s13225-020-00466-2)) |
| Otu45545 | 1 | k__Fungi(100);p__Ascomycota(100);c__Saccharomycetes(100);o__Saccharomycetales(100);f__Metschnikowiaceae(100);g__Metschnikowia(100);g__Metschnikowia_unclassified(100); | Metschnikowia | 13 | Saprotroph | \|Undefined Saprotroph\| | Yeast | NULL | Probable | As nectar-tap_saprotroph (Põlme et al. 2020); Decay sugar-rich substrates (Põlme et al. 2020) | Põlme S, et al. 2020. Fungal Diversity 105:1-16 ((https://doi.org/10.1007/s13225-020-00466-2)) |
| Otu45546 | 1 | k__Fungi(100);p__Ascomycota(100);c__Saccharomycetes(100);o__Saccharomycetales(100);f__Metschnikowiaceae(100);g__Metschnikowia(100);g__Metschnikowia_unclassified(100); | Metschnikowia | 13 | Saprotroph | \|Undefined Saprotroph\| | Yeast | NULL | Probable | As nectar-tap_saprotroph (Põlme et al. 2020); Decay sugar-rich substrates (Põlme et al. 2020) | Põlme S, et al. 2020. Fungal Diversity 105:1-16 ((https://doi.org/10.1007/s13225-020-00466-2)) |
| Otu45547 | 1 | k__Fungi(100);p__Ascomycota(100);c__Saccharomycetes(100);o__Saccharomycetales(100);f__Metschnikowiaceae(100);g__Metschnikowia(100);g__Metschnikowia_unclassified(100); | Metschnikowia | 13 | Saprotroph | \|Undefined Saprotroph\| | Yeast | NULL | Probable | As nectar-tap_saprotroph (Põlme et al. 2020); Decay sugar-rich substrates (Põlme et al. 2020) | Põlme S, et al. 2020. Fungal Diversity 105:1-16 ((https://doi.org/10.1007/s13225-020-00466-2)) |
| Otu45548 | 1 | k__Fungi(100);p__Ascomycota(100);c__Saccharomycetes(100);o__Saccharomycetales(100);f__Metschnikowiaceae(100);g__Metschnikowia(100);g__Metschnikowia_unclassified(100); | Metschnikowia | 13 | Saprotroph | \|Undefined Saprotroph\| | Yeast | NULL | Probable | As nectar-tap_saprotroph (Põlme et al. 2020); Decay sugar-rich substrates (Põlme et al. 2020) | Põlme S, et al. 2020. Fungal Diversity 105:1-16 ((https://doi.org/10.1007/s13225-020-00466-2)) |
| Otu45549 | 1 | k__Fungi(100);p__Ascomycota(100);c__Saccharomycetes(100);o__Saccharomycetales(100);f__Metschnikowiaceae(100);g__Metschnikowia(100);g__Metschnikowia_unclassified(100); | Metschnikowia | 13 | Saprotroph | \|Undefined Saprotroph\| | Yeast | NULL | Probable | As nectar-tap_saprotroph (Põlme et al. 2020); Decay sugar-rich substrates (Põlme et al. 2020) | Põlme S, et al. 2020. Fungal Diversity 105:1-16 ((https://doi.org/10.1007/s13225-020-00466-2)) |
| Otu45550 | 1 | k__Fungi(100);p__Ascomycota(100);c__Saccharomycetes(100);o__Saccharomycetales(100);f__Metschnikowiaceae(100);g__Metschnikowia(100);g__Metschnikowia_unclassified(100); | Metschnikowia | 13 | Saprotroph | \|Undefined Saprotroph\| | Yeast | NULL | Probable | As nectar-tap_saprotroph (Põlme et al. 2020); Decay sugar-rich substrates (Põlme et al. 2020) | Põlme S, et al. 2020. Fungal Diversity 105:1-16 ((https://doi.org/10.1007/s13225-020-00466-2)) |
| Otu45552 | 1 | k__Fungi(100);p__Ascomycota(100);c__Saccharomycetes(100);o__Saccharomycetales(100);f__Metschnikowiaceae(100);g__Metschnikowia(100);g__Metschnikowia_unclassified(100); | Metschnikowia | 13 | Saprotroph | \|Undefined Saprotroph\| | Yeast | NULL | Probable | As nectar-tap_saprotroph (Põlme et al. 2020); Decay sugar-rich substrates (Põlme et al. 2020) | Põlme S, et al. 2020. Fungal Diversity 105:1-16 ((https://doi.org/10.1007/s13225-020-00466-2)) |
| Otu45553 | 1 | k__Fungi(100);p__Ascomycota(100);c__Saccharomycetes(100);o__Saccharomycetales(100);f__Metschnikowiaceae(100);g__Metschnikowia(100);g__Metschnikowia_unclassified(100); | Metschnikowia | 13 | Saprotroph | \|Undefined Saprotroph\| | Yeast | NULL | Probable | As nectar-tap_saprotroph (Põlme et al. 2020); Decay sugar-rich substrates (Põlme et al. 2020) | Põlme S, et al. 2020. Fungal Diversity 105:1-16 ((https://doi.org/10.1007/s13225-020-00466-2)) |
| Otu45554 | 1 | k__Fungi(100);p__Ascomycota(100);c__Saccharomycetes(100);o__Saccharomycetales(100);f__Metschnikowiaceae(100);g__Metschnikowia(100);g__Metschnikowia_unclassified(100); | Metschnikowia | 13 | Saprotroph | \|Undefined Saprotroph\| | Yeast | NULL | Probable | As nectar-tap_saprotroph (Põlme et al. 2020); Decay sugar-rich substrates (Põlme et al. 2020) | Põlme S, et al. 2020. Fungal Diversity 105:1-16 ((https://doi.org/10.1007/s13225-020-00466-2)) |
| Otu45555 | 1 | k__Fungi(100);p__Ascomycota(100);c__Saccharomycetes(100);o__Saccharomycetales(100);f__Metschnikowiaceae(100);g__Metschnikowia(100);g__Metschnikowia_unclassified(100); | Metschnikowia | 13 | Saprotroph | \|Undefined Saprotroph\| | Yeast | NULL | Probable | As nectar-tap_saprotroph (Põlme et al. 2020); Decay sugar-rich substrates (Põlme et al. 2020) | Põlme S, et al. 2020. Fungal Diversity 105:1-16 ((https://doi.org/10.1007/s13225-020-00466-2)) |
| Otu45556 | 1 | k__Fungi(100);p__Ascomycota(100);c__Saccharomycetes(100);o__Saccharomycetales(100);f__Metschnikowiaceae(100);g__Metschnikowia(100);g__Metschnikowia_unclassified(100); | Metschnikowia | 13 | Saprotroph | \|Undefined Saprotroph\| | Yeast | NULL | Probable | As nectar-tap_saprotroph (Põlme et al. 2020); Decay sugar-rich substrates (Põlme et al. 2020) | Põlme S, et al. 2020. Fungal Diversity 105:1-16 ((https://doi.org/10.1007/s13225-020-00466-2)) |
| Otu45557 | 1 | k__Fungi(100);p__Ascomycota(100);c__Saccharomycetes(100);o__Saccharomycetales(100);f__Metschnikowiaceae(100);g__Metschnikowia(100);g__Metschnikowia_unclassified(100); | Metschnikowia | 13 | Saprotroph | \|Undefined Saprotroph\| | Yeast | NULL | Probable | As nectar-tap_saprotroph (Põlme et al. 2020); Decay sugar-rich substrates (Põlme et al. 2020) | Põlme S, et al. 2020. Fungal Diversity 105:1-16 ((https://doi.org/10.1007/s13225-020-00466-2)) |
| Otu45558 | 1 | k__Fungi(100);p__Ascomycota(100);c__Saccharomycetes(100);o__Saccharomycetales(100);f__Metschnikowiaceae(100);g__Metschnikowia(100);g__Metschnikowia_unclassified(100); | Metschnikowia | 13 | Saprotroph | \|Undefined Saprotroph\| | Yeast | NULL | Probable | As nectar-tap_saprotroph (Põlme et al. 2020); Decay sugar-rich substrates (Põlme et al. 2020) | Põlme S, et al. 2020. Fungal Diversity 105:1-16 ((https://doi.org/10.1007/s13225-020-00466-2)) |
| Otu45559 | 1 | k__Fungi(100);p__Ascomycota(100);c__Saccharomycetes(100);o__Saccharomycetales(100);f__Metschnikowiaceae(100);g__Metschnikowia(100);g__Metschnikowia_unclassified(100); | Metschnikowia | 13 | Saprotroph | \|Undefined Saprotroph\| | Yeast | NULL | Probable | As nectar-tap_saprotroph (Põlme et al. 2020); Decay sugar-rich substrates (Põlme et al. 2020) | Põlme S, et al. 2020. Fungal Diversity 105:1-16 ((https://doi.org/10.1007/s13225-020-00466-2)) |
| Otu45560 | 1 | k__Fungi(100);p__Ascomycota(100);c__Saccharomycetes(100);o__Saccharomycetales(100);f__Metschnikowiaceae(100);g__Metschnikowia(100);g__Metschnikowia_unclassified(100); | Metschnikowia | 13 | Saprotroph | \|Undefined Saprotroph\| | Yeast | NULL | Probable | As nectar-tap_saprotroph (Põlme et al. 2020); Decay sugar-rich substrates (Põlme et al. 2020) | Põlme S, et al. 2020. Fungal Diversity 105:1-16 ((https://doi.org/10.1007/s13225-020-00466-2)) |
| Otu45561 | 1 | k__Fungi(100);p__Ascomycota(100);c__Saccharomycetes(100);o__Saccharomycetales(100);f__Metschnikowiaceae(100);g__Metschnikowia(100);g__Metschnikowia_unclassified(100); | Metschnikowia | 13 | Saprotroph | \|Undefined Saprotroph\| | Yeast | NULL | Probable | As nectar-tap_saprotroph (Põlme et al. 2020); Decay sugar-rich substrates (Põlme et al. 2020) | Põlme S, et al. 2020. Fungal Diversity 105:1-16 ((https://doi.org/10.1007/s13225-020-00466-2)) |
| Otu45562 | 1 | k__Fungi(100);p__Ascomycota(100);c__Saccharomycetes(100);o__Saccharomycetales(100);f__Metschnikowiaceae(100);g__Metschnikowia(100);s__Metschnikowia_pulcherrima(100); | Metschnikowia | 13 | Saprotroph | \|Undefined Saprotroph\| | Yeast | NULL | Probable | As nectar-tap_saprotroph (Põlme et al. 2020); Decay sugar-rich substrates (Põlme et al. 2020) | Põlme S, et al. 2020. Fungal Diversity 105:1-16 ((https://doi.org/10.1007/s13225-020-00466-2)) |
| Otu45563 | 1 | k__Fungi(100);p__Ascomycota(100);c__Saccharomycetes(100);o__Saccharomycetales(100);f__Metschnikowiaceae(100);g__Metschnikowia(100);g__Metschnikowia_unclassified(100); | Metschnikowia | 13 | Saprotroph | \|Undefined Saprotroph\| | Yeast | NULL | Probable | As nectar-tap_saprotroph (Põlme et al. 2020); Decay sugar-rich substrates (Põlme et al. 2020) | Põlme S, et al. 2020. Fungal Diversity 105:1-16 ((https://doi.org/10.1007/s13225-020-00466-2)) |
| Otu45565 | 1 | k__Fungi(100);p__Ascomycota(100);c__Saccharomycetes(100);o__Saccharomycetales(100);f__Metschnikowiaceae(100);g__Metschnikowia(100);g__Metschnikowia_unclassified(100); | Metschnikowia | 13 | Saprotroph | \|Undefined Saprotroph\| | Yeast | NULL | Probable | As nectar-tap_saprotroph (Põlme et al. 2020); Decay sugar-rich substrates (Põlme et al. 2020) | Põlme S, et al. 2020. Fungal Diversity 105:1-16 ((https://doi.org/10.1007/s13225-020-00466-2)) |
| Otu45566 | 1 | k__Fungi(100);p__Ascomycota(100);c__Saccharomycetes(100);o__Saccharomycetales(100);f__Metschnikowiaceae(100);g__Metschnikowia(100);g__Metschnikowia_unclassified(100); | Metschnikowia | 13 | Saprotroph | \|Undefined Saprotroph\| | Yeast | NULL | Probable | As nectar-tap_saprotroph (Põlme et al. 2020); Decay sugar-rich substrates (Põlme et al. 2020) | Põlme S, et al. 2020. Fungal Diversity 105:1-16 ((https://doi.org/10.1007/s13225-020-00466-2)) |
| Otu45567 | 1 | k__Fungi(100);p__Ascomycota(100);c__Saccharomycetes(100);o__Saccharomycetales(100);f__Metschnikowiaceae(100);g__Metschnikowia(100);g__Metschnikowia_unclassified(100); | Metschnikowia | 13 | Saprotroph | \|Undefined Saprotroph\| | Yeast | NULL | Probable | As nectar-tap_saprotroph (Põlme et al. 2020); Decay sugar-rich substrates (Põlme et al. 2020) | Põlme S, et al. 2020. Fungal Diversity 105:1-16 ((https://doi.org/10.1007/s13225-020-00466-2)) |
| Otu45568 | 1 | k__Fungi(100);p__Ascomycota(100);c__Saccharomycetes(100);o__Saccharomycetales(100);f__Metschnikowiaceae(100);g__Metschnikowia(100);g__Metschnikowia_unclassified(100); | Metschnikowia | 13 | Saprotroph | \|Undefined Saprotroph\| | Yeast | NULL | Probable | As nectar-tap_saprotroph (Põlme et al. 2020); Decay sugar-rich substrates (Põlme et al. 2020) | Põlme S, et al. 2020. Fungal Diversity 105:1-16 ((https://doi.org/10.1007/s13225-020-00466-2)) |
| Otu45569 | 1 | k__Fungi(100);p__Ascomycota(100);c__Saccharomycetes(100);o__Saccharomycetales(100);f__Metschnikowiaceae(100);g__Metschnikowia(100);g__Metschnikowia_unclassified(100); | Metschnikowia | 13 | Saprotroph | \|Undefined Saprotroph\| | Yeast | NULL | Probable | As nectar-tap_saprotroph (Põlme et al. 2020); Decay sugar-rich substrates (Põlme et al. 2020) | Põlme S, et al. 2020. Fungal Diversity 105:1-16 ((https://doi.org/10.1007/s13225-020-00466-2)) |
| Otu45570 | 1 | k__Fungi(100);p__Ascomycota(100);c__Saccharomycetes(100);o__Saccharomycetales(100);f__Metschnikowiaceae(100);g__Metschnikowia(100);g__Metschnikowia_unclassified(100); | Metschnikowia | 13 | Saprotroph | \|Undefined Saprotroph\| | Yeast | NULL | Probable | As nectar-tap_saprotroph (Põlme et al. 2020); Decay sugar-rich substrates (Põlme et al. 2020) | Põlme S, et al. 2020. Fungal Diversity 105:1-16 ((https://doi.org/10.1007/s13225-020-00466-2)) |
| Otu45571 | 1 | k__Fungi(100);p__Ascomycota(100);c__Saccharomycetes(100);o__Saccharomycetales(100);f__Metschnikowiaceae(100);g__Metschnikowia(100);g__Metschnikowia_unclassified(100); | Metschnikowia | 13 | Saprotroph | \|Undefined Saprotroph\| | Yeast | NULL | Probable | As nectar-tap_saprotroph (Põlme et al. 2020); Decay sugar-rich substrates (Põlme et al. 2020) | Põlme S, et al. 2020. Fungal Diversity 105:1-16 ((https://doi.org/10.1007/s13225-020-00466-2)) |
| Otu45572 | 1 | k__Fungi(100);p__Ascomycota(100);c__Saccharomycetes(100);o__Saccharomycetales(100);f__Metschnikowiaceae(100);g__Metschnikowia(100);g__Metschnikowia_unclassified(100); | Metschnikowia | 13 | Saprotroph | \|Undefined Saprotroph\| | Yeast | NULL | Probable | As nectar-tap_saprotroph (Põlme et al. 2020); Decay sugar-rich substrates (Põlme et al. 2020) | Põlme S, et al. 2020. Fungal Diversity 105:1-16 ((https://doi.org/10.1007/s13225-020-00466-2)) |
| Otu45573 | 1 | k__Fungi(100);p__Ascomycota(100);c__Saccharomycetes(100);o__Saccharomycetales(100);f__Metschnikowiaceae(100);g__Metschnikowia(100);g__Metschnikowia_unclassified(100); | Metschnikowia | 13 | Saprotroph | \|Undefined Saprotroph\| | Yeast | NULL | Probable | As nectar-tap_saprotroph (Põlme et al. 2020); Decay sugar-rich substrates (Põlme et al. 2020) | Põlme S, et al. 2020. Fungal Diversity 105:1-16 ((https://doi.org/10.1007/s13225-020-00466-2)) |
| Otu45574 | 1 | k__Fungi(100);p__Ascomycota(100);c__Saccharomycetes(100);o__Saccharomycetales(100);f__Metschnikowiaceae(100);g__Metschnikowia(100);g__Metschnikowia_unclassified(100); | Metschnikowia | 13 | Saprotroph | \|Undefined Saprotroph\| | Yeast | NULL | Probable | As nectar-tap_saprotroph (Põlme et al. 2020); Decay sugar-rich substrates (Põlme et al. 2020) | Põlme S, et al. 2020. Fungal Diversity 105:1-16 ((https://doi.org/10.1007/s13225-020-00466-2)) |
| Otu45575 | 1 | k__Fungi(100);p__Ascomycota(100);c__Saccharomycetes(100);o__Saccharomycetales(100);f__Metschnikowiaceae(100);g__Metschnikowia(100);g__Metschnikowia_unclassified(100); | Metschnikowia | 13 | Saprotroph | \|Undefined Saprotroph\| | Yeast | NULL | Probable | As nectar-tap_saprotroph (Põlme et al. 2020); Decay sugar-rich substrates (Põlme et al. 2020) | Põlme S, et al. 2020. Fungal Diversity 105:1-16 ((https://doi.org/10.1007/s13225-020-00466-2)) |
| Otu45576 | 1 | k__Fungi(100);p__Ascomycota(100);c__Saccharomycetes(100);o__Saccharomycetales(100);f__Metschnikowiaceae(100);g__Metschnikowia(100);s__Metschnikowia_pulcherrima(100); | Metschnikowia | 13 | Saprotroph | \|Undefined Saprotroph\| | Yeast | NULL | Probable | As nectar-tap_saprotroph (Põlme et al. 2020); Decay sugar-rich substrates (Põlme et al. 2020) | Põlme S, et al. 2020. Fungal Diversity 105:1-16 ((https://doi.org/10.1007/s13225-020-00466-2)) |
| Otu45577 | 1 | k__Fungi(100);p__Ascomycota(100);c__Saccharomycetes(100);o__Saccharomycetales(100);f__Metschnikowiaceae(100);g__Metschnikowia(100);g__Metschnikowia_unclassified(100); | Metschnikowia | 13 | Saprotroph | \|Undefined Saprotroph\| | Yeast | NULL | Probable | As nectar-tap_saprotroph (Põlme et al. 2020); Decay sugar-rich substrates (Põlme et al. 2020) | Põlme S, et al. 2020. Fungal Diversity 105:1-16 ((https://doi.org/10.1007/s13225-020-00466-2)) |
| Otu45578 | 1 | k__Fungi(100);p__Ascomycota(100);c__Saccharomycetes(100);o__Saccharomycetales(100);f__Metschnikowiaceae(100);g__Metschnikowia(100);g__Metschnikowia_unclassified(100); | Metschnikowia | 13 | Saprotroph | \|Undefined Saprotroph\| | Yeast | NULL | Probable | As nectar-tap_saprotroph (Põlme et al. 2020); Decay sugar-rich substrates (Põlme et al. 2020) | Põlme S, et al. 2020. Fungal Diversity 105:1-16 ((https://doi.org/10.1007/s13225-020-00466-2)) |
| Otu45580 | 1 | k__Fungi(100);p__Ascomycota(100);c__Saccharomycetes(100);o__Saccharomycetales(100);f__Metschnikowiaceae(100);g__Metschnikowia(100);g__Metschnikowia_unclassified(100); | Metschnikowia | 13 | Saprotroph | \|Undefined Saprotroph\| | Yeast | NULL | Probable | As nectar-tap_saprotroph (Põlme et al. 2020); Decay sugar-rich substrates (Põlme et al. 2020) | Põlme S, et al. 2020. Fungal Diversity 105:1-16 ((https://doi.org/10.1007/s13225-020-00466-2)) |
| Otu45581 | 1 | k__Fungi(100);p__Ascomycota(100);c__Saccharomycetes(100);o__Saccharomycetales(100);f__Metschnikowiaceae(100);g__Metschnikowia(100);g__Metschnikowia_unclassified(100); | Metschnikowia | 13 | Saprotroph | \|Undefined Saprotroph\| | Yeast | NULL | Probable | As nectar-tap_saprotroph (Põlme et al. 2020); Decay sugar-rich substrates (Põlme et al. 2020) | Põlme S, et al. 2020. Fungal Diversity 105:1-16 ((https://doi.org/10.1007/s13225-020-00466-2)) |
| Otu45582 | 1 | k__Fungi(100);p__Ascomycota(100);c__Saccharomycetes(100);o__Saccharomycetales(100);f__Metschnikowiaceae(100);g__Metschnikowia(100);g__Metschnikowia_unclassified(100); | Metschnikowia | 13 | Saprotroph | \|Undefined Saprotroph\| | Yeast | NULL | Probable | As nectar-tap_saprotroph (Põlme et al. 2020); Decay sugar-rich substrates (Põlme et al. 2020) | Põlme S, et al. 2020. Fungal Diversity 105:1-16 ((https://doi.org/10.1007/s13225-020-00466-2)) |
| Otu45583 | 1 | k__Fungi(100);p__Ascomycota(100);c__Saccharomycetes(100);o__Saccharomycetales(100);f__Metschnikowiaceae(100);g__Metschnikowia(100);g__Metschnikowia_unclassified(100); | Metschnikowia | 13 | Saprotroph | \|Undefined Saprotroph\| | Yeast | NULL | Probable | As nectar-tap_saprotroph (Põlme et al. 2020); Decay sugar-rich substrates (Põlme et al. 2020) | Põlme S, et al. 2020. Fungal Diversity 105:1-16 ((https://doi.org/10.1007/s13225-020-00466-2)) |
| Otu45584 | 1 | k__Fungi(100);p__Ascomycota(100);c__Saccharomycetes(100);o__Saccharomycetales(100);f__Metschnikowiaceae(100);g__Metschnikowia(100);g__Metschnikowia_unclassified(100); | Metschnikowia | 13 | Saprotroph | \|Undefined Saprotroph\| | Yeast | NULL | Probable | As nectar-tap_saprotroph (Põlme et al. 2020); Decay sugar-rich substrates (Põlme et al. 2020) | Põlme S, et al. 2020. Fungal Diversity 105:1-16 ((https://doi.org/10.1007/s13225-020-00466-2)) |
| Otu45585 | 1 | k__Fungi(100);p__Ascomycota(100);c__Saccharomycetes(100);o__Saccharomycetales(100);f__Metschnikowiaceae(100);g__Metschnikowia(100);g__Metschnikowia_unclassified(100); | Metschnikowia | 13 | Saprotroph | \|Undefined Saprotroph\| | Yeast | NULL | Probable | As nectar-tap_saprotroph (Põlme et al. 2020); Decay sugar-rich substrates (Põlme et al. 2020) | Põlme S, et al. 2020. Fungal Diversity 105:1-16 ((https://doi.org/10.1007/s13225-020-00466-2)) |
| Otu45587 | 1 | k__Fungi(100);p__Ascomycota(100);c__Saccharomycetes(100);o__Saccharomycetales(100);f__Metschnikowiaceae(100);g__Metschnikowia(100);g__Metschnikowia_unclassified(100); | Metschnikowia | 13 | Saprotroph | \|Undefined Saprotroph\| | Yeast | NULL | Probable | As nectar-tap_saprotroph (Põlme et al. 2020); Decay sugar-rich substrates (Põlme et al. 2020) | Põlme S, et al. 2020. Fungal Diversity 105:1-16 ((https://doi.org/10.1007/s13225-020-00466-2)) |
| Otu45588 | 1 | k__Fungi(100);p__Ascomycota(100);c__Saccharomycetes(100);o__Saccharomycetales(100);f__Metschnikowiaceae(100);g__Metschnikowia(100);g__Metschnikowia_unclassified(100); | Metschnikowia | 13 | Saprotroph | \|Undefined Saprotroph\| | Yeast | NULL | Probable | As nectar-tap_saprotroph (Põlme et al. 2020); Decay sugar-rich substrates (Põlme et al. 2020) | Põlme S, et al. 2020. Fungal Diversity 105:1-16 ((https://doi.org/10.1007/s13225-020-00466-2)) |
| Otu45589 | 1 | k__Fungi(100);p__Ascomycota(100);c__Saccharomycetes(100);o__Saccharomycetales(100);f__Metschnikowiaceae(100);g__Metschnikowia(100);g__Metschnikowia_unclassified(100); | Metschnikowia | 13 | Saprotroph | \|Undefined Saprotroph\| | Yeast | NULL | Probable | As nectar-tap_saprotroph (Põlme et al. 2020); Decay sugar-rich substrates (Põlme et al. 2020) | Põlme S, et al. 2020. Fungal Diversity 105:1-16 ((https://doi.org/10.1007/s13225-020-00466-2)) |
| Otu45591 | 1 | k__Fungi(100);p__Ascomycota(100);c__Saccharomycetes(100);o__Saccharomycetales(100);f__Metschnikowiaceae(100);g__Metschnikowia(100);g__Metschnikowia_unclassified(100); | Metschnikowia | 13 | Saprotroph | \|Undefined Saprotroph\| | Yeast | NULL | Probable | As nectar-tap_saprotroph (Põlme et al. 2020); Decay sugar-rich substrates (Põlme et al. 2020) | Põlme S, et al. 2020. Fungal Diversity 105:1-16 ((https://doi.org/10.1007/s13225-020-00466-2)) |
| Otu45592 | 1 | k__Fungi(100);p__Ascomycota(100);c__Saccharomycetes(100);o__Saccharomycetales(100);f__Metschnikowiaceae(100);g__Metschnikowia(100);s__Metschnikowia_pulcherrima(100); | Metschnikowia | 13 | Saprotroph | \|Undefined Saprotroph\| | Yeast | NULL | Probable | As nectar-tap_saprotroph (Põlme et al. 2020); Decay sugar-rich substrates (Põlme et al. 2020) | Põlme S, et al. 2020. Fungal Diversity 105:1-16 ((https://doi.org/10.1007/s13225-020-00466-2)) |
| Otu45593 | 1 | k__Fungi(100);p__Ascomycota(100);c__Saccharomycetes(100);o__Saccharomycetales(100);f__Metschnikowiaceae(100);g__Metschnikowia(100);s__Metschnikowia_pulcherrima(100); | Metschnikowia | 13 | Saprotroph | \|Undefined Saprotroph\| | Yeast | NULL | Probable | As nectar-tap_saprotroph (Põlme et al. 2020); Decay sugar-rich substrates (Põlme et al. 2020) | Põlme S, et al. 2020. Fungal Diversity 105:1-16 ((https://doi.org/10.1007/s13225-020-00466-2)) |
| Otu45594 | 1 | k__Fungi(100);p__Ascomycota(100);c__Saccharomycetes(100);o__Saccharomycetales(100);f__Metschnikowiaceae(100);g__Metschnikowia(100);g__Metschnikowia_unclassified(100); | Metschnikowia | 13 | Saprotroph | \|Undefined Saprotroph\| | Yeast | NULL | Probable | As nectar-tap_saprotroph (Põlme et al. 2020); Decay sugar-rich substrates (Põlme et al. 2020) | Põlme S, et al. 2020. Fungal Diversity 105:1-16 ((https://doi.org/10.1007/s13225-020-00466-2)) |
| Otu45595 | 1 | k__Fungi(100);p__Ascomycota(100);c__Saccharomycetes(100);o__Saccharomycetales(100);f__Metschnikowiaceae(100);g__Metschnikowia(100);s__Metschnikowia_pulcherrima(100); | Metschnikowia | 13 | Saprotroph | \|Undefined Saprotroph\| | Yeast | NULL | Probable | As nectar-tap_saprotroph (Põlme et al. 2020); Decay sugar-rich substrates (Põlme et al. 2020) | Põlme S, et al. 2020. Fungal Diversity 105:1-16 ((https://doi.org/10.1007/s13225-020-00466-2)) |
| Otu45596 | 1 | k__Fungi(100);p__Ascomycota(100);c__Saccharomycetes(100);o__Saccharomycetales(100);f__Metschnikowiaceae(100);g__Metschnikowia(100);s__Metschnikowia_pulcherrima(100); | Metschnikowia | 13 | Saprotroph | \|Undefined Saprotroph\| | Yeast | NULL | Probable | As nectar-tap_saprotroph (Põlme et al. 2020); Decay sugar-rich substrates (Põlme et al. 2020) | Põlme S, et al. 2020. Fungal Diversity 105:1-16 ((https://doi.org/10.1007/s13225-020-00466-2)) |
| Otu45597 | 1 | k__Fungi(100);p__Ascomycota(100);c__Saccharomycetes(100);o__Saccharomycetales(100);f__Metschnikowiaceae(100);g__Metschnikowia(100);s__Metschnikowia_pulcherrima(100); | Metschnikowia | 13 | Saprotroph | \|Undefined Saprotroph\| | Yeast | NULL | Probable | As nectar-tap_saprotroph (Põlme et al. 2020); Decay sugar-rich substrates (Põlme et al. 2020) | Põlme S, et al. 2020. Fungal Diversity 105:1-16 ((https://doi.org/10.1007/s13225-020-00466-2)) |
| Otu45598 | 1 | k__Fungi(100);p__Ascomycota(100);c__Saccharomycetes(100);o__Saccharomycetales(100);f__Metschnikowiaceae(100);g__Metschnikowia(100);s__Metschnikowia_pulcherrima(100); | Metschnikowia | 13 | Saprotroph | \|Undefined Saprotroph\| | Yeast | NULL | Probable | As nectar-tap_saprotroph (Põlme et al. 2020); Decay sugar-rich substrates (Põlme et al. 2020) | Põlme S, et al. 2020. Fungal Diversity 105:1-16 ((https://doi.org/10.1007/s13225-020-00466-2)) |
| Otu45599 | 1 | k__Fungi(100);p__Ascomycota(100);c__Saccharomycetes(100);o__Saccharomycetales(100);f__Metschnikowiaceae(100);g__Metschnikowia(100);g__Metschnikowia_unclassified(100); | Metschnikowia | 13 | Saprotroph | \|Undefined Saprotroph\| | Yeast | NULL | Probable | As nectar-tap_saprotroph (Põlme et al. 2020); Decay sugar-rich substrates (Põlme et al. 2020) | Põlme S, et al. 2020. Fungal Diversity 105:1-16 ((https://doi.org/10.1007/s13225-020-00466-2)) |
| Otu45600 | 1 | k__Fungi(100);p__Ascomycota(100);c__Saccharomycetes(100);o__Saccharomycetales(100);f__Metschnikowiaceae(100);g__Metschnikowia(100);g__Metschnikowia_unclassified(100); | Metschnikowia | 13 | Saprotroph | \|Undefined Saprotroph\| | Yeast | NULL | Probable | As nectar-tap_saprotroph (Põlme et al. 2020); Decay sugar-rich substrates (Põlme et al. 2020) | Põlme S, et al. 2020. Fungal Diversity 105:1-16 ((https://doi.org/10.1007/s13225-020-00466-2)) |
| Otu45601 | 1 | k__Fungi(100);p__Ascomycota(100);c__Saccharomycetes(100);o__Saccharomycetales(100);f__Metschnikowiaceae(100);g__Metschnikowia(100);g__Metschnikowia_unclassified(100); | Metschnikowia | 13 | Saprotroph | \|Undefined Saprotroph\| | Yeast | NULL | Probable | As nectar-tap_saprotroph (Põlme et al. 2020); Decay sugar-rich substrates (Põlme et al. 2020) | Põlme S, et al. 2020. Fungal Diversity 105:1-16 ((https://doi.org/10.1007/s13225-020-00466-2)) |
| Otu45602 | 1 | k__Fungi(100);p__Ascomycota(100);c__Saccharomycetes(100);o__Saccharomycetales(100);f__Metschnikowiaceae(100);g__Metschnikowia(100);s__Metschnikowia_pulcherrima(100); | Metschnikowia | 13 | Saprotroph | \|Undefined Saprotroph\| | Yeast | NULL | Probable | As nectar-tap_saprotroph (Põlme et al. 2020); Decay sugar-rich substrates (Põlme et al. 2020) | Põlme S, et al. 2020. Fungal Diversity 105:1-16 ((https://doi.org/10.1007/s13225-020-00466-2)) |
| Otu45603 | 1 | k__Fungi(100);p__Ascomycota(100);c__Saccharomycetes(100);o__Saccharomycetales(100);f__Metschnikowiaceae(100);g__Metschnikowia(100);g__Metschnikowia_unclassified(100); | Metschnikowia | 13 | Saprotroph | \|Undefined Saprotroph\| | Yeast | NULL | Probable | As nectar-tap_saprotroph (Põlme et al. 2020); Decay sugar-rich substrates (Põlme et al. 2020) | Põlme S, et al. 2020. Fungal Diversity 105:1-16 ((https://doi.org/10.1007/s13225-020-00466-2)) |
| Otu45604 | 1 | k__Fungi(100);p__Ascomycota(100);c__Saccharomycetes(100);o__Saccharomycetales(100);f__Metschnikowiaceae(100);g__Metschnikowia(100);s__Metschnikowia_pulcherrima(100); | Metschnikowia | 13 | Saprotroph | \|Undefined Saprotroph\| | Yeast | NULL | Probable | As nectar-tap_saprotroph (Põlme et al. 2020); Decay sugar-rich substrates (Põlme et al. 2020) | Põlme S, et al. 2020. Fungal Diversity 105:1-16 ((https://doi.org/10.1007/s13225-020-00466-2)) |
| Otu45605 | 1 | k__Fungi(100);p__Ascomycota(100);c__Saccharomycetes(100);o__Saccharomycetales(100);f__Metschnikowiaceae(100);g__Metschnikowia(100);s__Metschnikowia_pulcherrima(100); | Metschnikowia | 13 | Saprotroph | \|Undefined Saprotroph\| | Yeast | NULL | Probable | As nectar-tap_saprotroph (Põlme et al. 2020); Decay sugar-rich substrates (Põlme et al. 2020) | Põlme S, et al. 2020. Fungal Diversity 105:1-16 ((https://doi.org/10.1007/s13225-020-00466-2)) |
| Otu45606 | 1 | k__Fungi(100);p__Ascomycota(100);c__Saccharomycetes(100);o__Saccharomycetales(100);f__Metschnikowiaceae(100);g__Metschnikowia(100);s__Metschnikowia_pulcherrima(100); | Metschnikowia | 13 | Saprotroph | \|Undefined Saprotroph\| | Yeast | NULL | Probable | As nectar-tap_saprotroph (Põlme et al. 2020); Decay sugar-rich substrates (Põlme et al. 2020) | Põlme S, et al. 2020. Fungal Diversity 105:1-16 ((https://doi.org/10.1007/s13225-020-00466-2)) |
| Otu45607 | 1 | k__Fungi(100);p__Ascomycota(100);c__Saccharomycetes(100);o__Saccharomycetales(100);f__Metschnikowiaceae(100);g__Metschnikowia(100);s__Metschnikowia_pulcherrima(100); | Metschnikowia | 13 | Saprotroph | \|Undefined Saprotroph\| | Yeast | NULL | Probable | As nectar-tap_saprotroph (Põlme et al. 2020); Decay sugar-rich substrates (Põlme et al. 2020) | Põlme S, et al. 2020. Fungal Diversity 105:1-16 ((https://doi.org/10.1007/s13225-020-00466-2)) |
| Otu45608 | 1 | k__Fungi(100);p__Ascomycota(100);c__Saccharomycetes(100);o__Saccharomycetales(100);f__Metschnikowiaceae(100);g__Metschnikowia(100);s__Metschnikowia_pulcherrima(100); | Metschnikowia | 13 | Saprotroph | \|Undefined Saprotroph\| | Yeast | NULL | Probable | As nectar-tap_saprotroph (Põlme et al. 2020); Decay sugar-rich substrates (Põlme et al. 2020) | Põlme S, et al. 2020. Fungal Diversity 105:1-16 ((https://doi.org/10.1007/s13225-020-00466-2)) |
| Otu45609 | 1 | k__Fungi(100);p__Ascomycota(100);c__Saccharomycetes(100);o__Saccharomycetales(100);f__Metschnikowiaceae(100);g__Metschnikowia(100);s__Metschnikowia_viticola(100); | Metschnikowia | 13 | Saprotroph | \|Undefined Saprotroph\| | Yeast | NULL | Probable | As nectar-tap_saprotroph (Põlme et al. 2020); Decay sugar-rich substrates (Põlme et al. 2020) | Põlme S, et al. 2020. Fungal Diversity 105:1-16 ((https://doi.org/10.1007/s13225-020-00466-2)) |
| Otu45610 | 1 | k__Fungi(100);p__Ascomycota(100);c__Saccharomycetes(100);o__Saccharomycetales(100);f__Metschnikowiaceae(100);g__Metschnikowia(100);s__Metschnikowia_pulcherrima(100); | Metschnikowia | 13 | Saprotroph | \|Undefined Saprotroph\| | Yeast | NULL | Probable | As nectar-tap_saprotroph (Põlme et al. 2020); Decay sugar-rich substrates (Põlme et al. 2020) | Põlme S, et al. 2020. Fungal Diversity 105:1-16 ((https://doi.org/10.1007/s13225-020-00466-2)) |
| Otu45611 | 1 | k__Fungi(100);p__Ascomycota(100);c__Saccharomycetes(100);o__Saccharomycetales(100);f__Metschnikowiaceae(100);g__Metschnikowia(100);g__Metschnikowia_unclassified(100); | Metschnikowia | 13 | Saprotroph | \|Undefined Saprotroph\| | Yeast | NULL | Probable | As nectar-tap_saprotroph (Põlme et al. 2020); Decay sugar-rich substrates (Põlme et al. 2020) | Põlme S, et al. 2020. Fungal Diversity 105:1-16 ((https://doi.org/10.1007/s13225-020-00466-2)) |
| Otu45612 | 1 | k__Fungi(100);p__Ascomycota(100);c__Saccharomycetes(100);o__Saccharomycetales(100);f__Metschnikowiaceae(100);g__Metschnikowia(100);s__Metschnikowia_pulcherrima(100); | Metschnikowia | 13 | Saprotroph | \|Undefined Saprotroph\| | Yeast | NULL | Probable | As nectar-tap_saprotroph (Põlme et al. 2020); Decay sugar-rich substrates (Põlme et al. 2020) | Põlme S, et al. 2020. Fungal Diversity 105:1-16 ((https://doi.org/10.1007/s13225-020-00466-2)) |
| Otu45613 | 1 | k__Fungi(100);p__Ascomycota(100);c__Saccharomycetes(100);o__Saccharomycetales(100);f__Metschnikowiaceae(100);g__Metschnikowia(100);s__Metschnikowia_pulcherrima(100); | Metschnikowia | 13 | Saprotroph | \|Undefined Saprotroph\| | Yeast | NULL | Probable | As nectar-tap_saprotroph (Põlme et al. 2020); Decay sugar-rich substrates (Põlme et al. 2020) | Põlme S, et al. 2020. Fungal Diversity 105:1-16 ((https://doi.org/10.1007/s13225-020-00466-2)) |
| Otu45614 | 1 | k__Fungi(100);p__Ascomycota(100);c__Saccharomycetes(100);o__Saccharomycetales(100);f__Metschnikowiaceae(100);g__Metschnikowia(100);s__Metschnikowia_pulcherrima(100); | Metschnikowia | 13 | Saprotroph | \|Undefined Saprotroph\| | Yeast | NULL | Probable | As nectar-tap_saprotroph (Põlme et al. 2020); Decay sugar-rich substrates (Põlme et al. 2020) | Põlme S, et al. 2020. Fungal Diversity 105:1-16 ((https://doi.org/10.1007/s13225-020-00466-2)) |
| Otu45615 | 1 | k__Fungi(100);p__Ascomycota(100);c__Saccharomycetes(100);o__Saccharomycetales(100);f__Metschnikowiaceae(100);g__Metschnikowia(100);s__Metschnikowia_pulcherrima(100); | Metschnikowia | 13 | Saprotroph | \|Undefined Saprotroph\| | Yeast | NULL | Probable | As nectar-tap_saprotroph (Põlme et al. 2020); Decay sugar-rich substrates (Põlme et al. 2020) | Põlme S, et al. 2020. Fungal Diversity 105:1-16 ((https://doi.org/10.1007/s13225-020-00466-2)) |
| Otu45616 | 1 | k__Fungi(100);p__Ascomycota(100);c__Saccharomycetes(100);o__Saccharomycetales(100);f__Metschnikowiaceae(100);g__Metschnikowia(100);s__Metschnikowia_pulcherrima(100); | Metschnikowia | 13 | Saprotroph | \|Undefined Saprotroph\| | Yeast | NULL | Probable | As nectar-tap_saprotroph (Põlme et al. 2020); Decay sugar-rich substrates (Põlme et al. 2020) | Põlme S, et al. 2020. Fungal Diversity 105:1-16 ((https://doi.org/10.1007/s13225-020-00466-2)) |
| Otu45617 | 1 | k__Fungi(100);p__Ascomycota(100);c__Saccharomycetes(100);o__Saccharomycetales(100);f__Metschnikowiaceae(100);g__Metschnikowia(100);s__Metschnikowia_pulcherrima(100); | Metschnikowia | 13 | Saprotroph | \|Undefined Saprotroph\| | Yeast | NULL | Probable | As nectar-tap_saprotroph (Põlme et al. 2020); Decay sugar-rich substrates (Põlme et al. 2020) | Põlme S, et al. 2020. Fungal Diversity 105:1-16 ((https://doi.org/10.1007/s13225-020-00466-2)) |
| Otu45618 | 1 | k__Fungi(100);p__Ascomycota(100);c__Saccharomycetes(100);o__Saccharomycetales(100);f__Metschnikowiaceae(100);g__Metschnikowia(100);g__Metschnikowia_unclassified(100); | Metschnikowia | 13 | Saprotroph | \|Undefined Saprotroph\| | Yeast | NULL | Probable | As nectar-tap_saprotroph (Põlme et al. 2020); Decay sugar-rich substrates (Põlme et al. 2020) | Põlme S, et al. 2020. Fungal Diversity 105:1-16 ((https://doi.org/10.1007/s13225-020-00466-2)) |
| Otu45619 | 1 | k__Fungi(100);p__Ascomycota(100);c__Saccharomycetes(100);o__Saccharomycetales(100);f__Metschnikowiaceae(100);g__Metschnikowia(100);s__Metschnikowia_pulcherrima(100); | Metschnikowia | 13 | Saprotroph | \|Undefined Saprotroph\| | Yeast | NULL | Probable | As nectar-tap_saprotroph (Põlme et al. 2020); Decay sugar-rich substrates (Põlme et al. 2020) | Põlme S, et al. 2020. Fungal Diversity 105:1-16 ((https://doi.org/10.1007/s13225-020-00466-2)) |
| Otu45620 | 1 | k__Fungi(100);p__Ascomycota(100);c__Saccharomycetes(100);o__Saccharomycetales(100);f__Metschnikowiaceae(100);g__Metschnikowia(100);s__Metschnikowia_pulcherrima(100); | Metschnikowia | 13 | Saprotroph | \|Undefined Saprotroph\| | Yeast | NULL | Probable | As nectar-tap_saprotroph (Põlme et al. 2020); Decay sugar-rich substrates (Põlme et al. 2020) | Põlme S, et al. 2020. Fungal Diversity 105:1-16 ((https://doi.org/10.1007/s13225-020-00466-2)) |
| Otu45621 | 1 | k__Fungi(100);p__Ascomycota(100);c__Saccharomycetes(100);o__Saccharomycetales(100);f__Metschnikowiaceae(100);g__Metschnikowia(100);g__Metschnikowia_unclassified(100); | Metschnikowia | 13 | Saprotroph | \|Undefined Saprotroph\| | Yeast | NULL | Probable | As nectar-tap_saprotroph (Põlme et al. 2020); Decay sugar-rich substrates (Põlme et al. 2020) | Põlme S, et al. 2020. Fungal Diversity 105:1-16 ((https://doi.org/10.1007/s13225-020-00466-2)) |
| Otu45622 | 1 | k__Fungi(100);p__Ascomycota(100);c__Saccharomycetes(100);o__Saccharomycetales(100);f__Metschnikowiaceae(100);g__Metschnikowia(100);g__Metschnikowia_unclassified(100); | Metschnikowia | 13 | Saprotroph | \|Undefined Saprotroph\| | Yeast | NULL | Probable | As nectar-tap_saprotroph (Põlme et al. 2020); Decay sugar-rich substrates (Põlme et al. 2020) | Põlme S, et al. 2020. Fungal Diversity 105:1-16 ((https://doi.org/10.1007/s13225-020-00466-2)) |
| Otu45623 | 1 | k__Fungi(100);p__Ascomycota(100);c__Saccharomycetes(100);o__Saccharomycetales(100);f__Metschnikowiaceae(100);g__Metschnikowia(100);g__Metschnikowia_unclassified(100); | Metschnikowia | 13 | Saprotroph | \|Undefined Saprotroph\| | Yeast | NULL | Probable | As nectar-tap_saprotroph (Põlme et al. 2020); Decay sugar-rich substrates (Põlme et al. 2020) | Põlme S, et al. 2020. Fungal Diversity 105:1-16 ((https://doi.org/10.1007/s13225-020-00466-2)) |
| Otu45624 | 1 | k__Fungi(100);p__Ascomycota(100);c__Saccharomycetes(100);o__Saccharomycetales(100);f__Metschnikowiaceae(100);g__Metschnikowia(100);g__Metschnikowia_unclassified(100); | Metschnikowia | 13 | Saprotroph | \|Undefined Saprotroph\| | Yeast | NULL | Probable | As nectar-tap_saprotroph (Põlme et al. 2020); Decay sugar-rich substrates (Põlme et al. 2020) | Põlme S, et al. 2020. Fungal Diversity 105:1-16 ((https://doi.org/10.1007/s13225-020-00466-2)) |
| Otu45625 | 1 | k__Fungi(100);p__Ascomycota(100);c__Saccharomycetes(100);o__Saccharomycetales(100);f__Metschnikowiaceae(100);g__Metschnikowia(100);g__Metschnikowia_unclassified(100); | Metschnikowia | 13 | Saprotroph | \|Undefined Saprotroph\| | Yeast | NULL | Probable | As nectar-tap_saprotroph (Põlme et al. 2020); Decay sugar-rich substrates (Põlme et al. 2020) | Põlme S, et al. 2020. Fungal Diversity 105:1-16 ((https://doi.org/10.1007/s13225-020-00466-2)) |
| Otu45626 | 1 | k__Fungi(100);p__Ascomycota(100);c__Saccharomycetes(100);o__Saccharomycetales(100);f__Metschnikowiaceae(100);g__Metschnikowia(100);s__Metschnikowia_viticola(100); | Metschnikowia | 13 | Saprotroph | \|Undefined Saprotroph\| | Yeast | NULL | Probable | As nectar-tap_saprotroph (Põlme et al. 2020); Decay sugar-rich substrates (Põlme et al. 2020) | Põlme S, et al. 2020. Fungal Diversity 105:1-16 ((https://doi.org/10.1007/s13225-020-00466-2)) |
| Otu45627 | 1 | k__Fungi(100);p__Ascomycota(100);c__Saccharomycetes(100);o__Saccharomycetales(100);f__Metschnikowiaceae(100);g__Metschnikowia(100);s__Metschnikowia_pulcherrima(100); | Metschnikowia | 13 | Saprotroph | \|Undefined Saprotroph\| | Yeast | NULL | Probable | As nectar-tap_saprotroph (Põlme et al. 2020); Decay sugar-rich substrates (Põlme et al. 2020) | Põlme S, et al. 2020. Fungal Diversity 105:1-16 ((https://doi.org/10.1007/s13225-020-00466-2)) |
| Otu45628 | 1 | k__Fungi(100);p__Ascomycota(100);c__Saccharomycetes(100);o__Saccharomycetales(100);f__Metschnikowiaceae(100);g__Metschnikowia(100);s__Metschnikowia_pulcherrima(100); | Metschnikowia | 13 | Saprotroph | \|Undefined Saprotroph\| | Yeast | NULL | Probable | As nectar-tap_saprotroph (Põlme et al. 2020); Decay sugar-rich substrates (Põlme et al. 2020) | Põlme S, et al. 2020. Fungal Diversity 105:1-16 ((https://doi.org/10.1007/s13225-020-00466-2)) |
| Otu45629 | 1 | k__Fungi(100);p__Ascomycota(100);c__Saccharomycetes(100);o__Saccharomycetales(100);f__Metschnikowiaceae(100);g__Metschnikowia(100);s__Metschnikowia_viticola(100); | Metschnikowia | 13 | Saprotroph | \|Undefined Saprotroph\| | Yeast | NULL | Probable | As nectar-tap_saprotroph (Põlme et al. 2020); Decay sugar-rich substrates (Põlme et al. 2020) | Põlme S, et al. 2020. Fungal Diversity 105:1-16 ((https://doi.org/10.1007/s13225-020-00466-2)) |
| Otu45630 | 1 | k__Fungi(100);p__Ascomycota(100);c__Saccharomycetes(100);o__Saccharomycetales(100);f__Metschnikowiaceae(100);g__Metschnikowia(100);s__Metschnikowia_viticola(100); | Metschnikowia | 13 | Saprotroph | \|Undefined Saprotroph\| | Yeast | NULL | Probable | As nectar-tap_saprotroph (Põlme et al. 2020); Decay sugar-rich substrates (Põlme et al. 2020) | Põlme S, et al. 2020. Fungal Diversity 105:1-16 ((https://doi.org/10.1007/s13225-020-00466-2)) |
| Otu45631 | 1 | k__Fungi(100);p__Ascomycota(100);c__Saccharomycetes(100);o__Saccharomycetales(100);f__Metschnikowiaceae(100);g__Metschnikowia(100);g__Metschnikowia_unclassified(100); | Metschnikowia | 13 | Saprotroph | \|Undefined Saprotroph\| | Yeast | NULL | Probable | As nectar-tap_saprotroph (Põlme et al. 2020); Decay sugar-rich substrates (Põlme et al. 2020) | Põlme S, et al. 2020. Fungal Diversity 105:1-16 ((https://doi.org/10.1007/s13225-020-00466-2)) |
| Otu45632 | 1 | k__Fungi(100);p__Ascomycota(100);c__Saccharomycetes(100);o__Saccharomycetales(100);f__Metschnikowiaceae(100);g__Metschnikowia(100);s__Metschnikowia_pulcherrima(100); | Metschnikowia | 13 | Saprotroph | \|Undefined Saprotroph\| | Yeast | NULL | Probable | As nectar-tap_saprotroph (Põlme et al. 2020); Decay sugar-rich substrates (Põlme et al. 2020) | Põlme S, et al. 2020. Fungal Diversity 105:1-16 ((https://doi.org/10.1007/s13225-020-00466-2)) |
| Otu45633 | 1 | k__Fungi(100);p__Ascomycota(100);c__Saccharomycetes(100);o__Saccharomycetales(100);f__Metschnikowiaceae(100);g__Metschnikowia(100);s__Metschnikowia_pulcherrima(100); | Metschnikowia | 13 | Saprotroph | \|Undefined Saprotroph\| | Yeast | NULL | Probable | As nectar-tap_saprotroph (Põlme et al. 2020); Decay sugar-rich substrates (Põlme et al. 2020) | Põlme S, et al. 2020. Fungal Diversity 105:1-16 ((https://doi.org/10.1007/s13225-020-00466-2)) |
| Otu45634 | 1 | k__Fungi(100);p__Ascomycota(100);c__Saccharomycetes(100);o__Saccharomycetales(100);f__Metschnikowiaceae(100);g__Metschnikowia(100);s__Metschnikowia_pulcherrima(100); | Metschnikowia | 13 | Saprotroph | \|Undefined Saprotroph\| | Yeast | NULL | Probable | As nectar-tap_saprotroph (Põlme et al. 2020); Decay sugar-rich substrates (Põlme et al. 2020) | Põlme S, et al. 2020. Fungal Diversity 105:1-16 ((https://doi.org/10.1007/s13225-020-00466-2)) |
| Otu45635 | 1 | k__Fungi(100);p__Ascomycota(100);c__Saccharomycetes(100);o__Saccharomycetales(100);f__Metschnikowiaceae(100);g__Metschnikowia(100);g__Metschnikowia_unclassified(100); | Metschnikowia | 13 | Saprotroph | \|Undefined Saprotroph\| | Yeast | NULL | Probable | As nectar-tap_saprotroph (Põlme et al. 2020); Decay sugar-rich substrates (Põlme et al. 2020) | Põlme S, et al. 2020. Fungal Diversity 105:1-16 ((https://doi.org/10.1007/s13225-020-00466-2)) |
| Otu45636 | 1 | k__Fungi(100);p__Ascomycota(100);c__Saccharomycetes(100);o__Saccharomycetales(100);f__Metschnikowiaceae(100);g__Metschnikowia(100);s__Metschnikowia_pulcherrima(100); | Metschnikowia | 13 | Saprotroph | \|Undefined Saprotroph\| | Yeast | NULL | Probable | As nectar-tap_saprotroph (Põlme et al. 2020); Decay sugar-rich substrates (Põlme et al. 2020) | Põlme S, et al. 2020. Fungal Diversity 105:1-16 ((https://doi.org/10.1007/s13225-020-00466-2)) |
| Otu45637 | 1 | k__Fungi(100);p__Ascomycota(100);c__Saccharomycetes(100);o__Saccharomycetales(100);f__Metschnikowiaceae(100);g__Metschnikowia(100);s__Metschnikowia_pulcherrima(100); | Metschnikowia | 13 | Saprotroph | \|Undefined Saprotroph\| | Yeast | NULL | Probable | As nectar-tap_saprotroph (Põlme et al. 2020); Decay sugar-rich substrates (Põlme et al. 2020) | Põlme S, et al. 2020. Fungal Diversity 105:1-16 ((https://doi.org/10.1007/s13225-020-00466-2)) |
| Otu45638 | 1 | k__Fungi(100);p__Ascomycota(100);c__Saccharomycetes(100);o__Saccharomycetales(100);f__Metschnikowiaceae(100);g__Metschnikowia(100);s__Metschnikowia_pulcherrima(100); | Metschnikowia | 13 | Saprotroph | \|Undefined Saprotroph\| | Yeast | NULL | Probable | As nectar-tap_saprotroph (Põlme et al. 2020); Decay sugar-rich substrates (Põlme et al. 2020) | Põlme S, et al. 2020. Fungal Diversity 105:1-16 ((https://doi.org/10.1007/s13225-020-00466-2)) |
| Otu45639 | 1 | k__Fungi(100);p__Ascomycota(100);c__Saccharomycetes(100);o__Saccharomycetales(100);f__Metschnikowiaceae(100);g__Metschnikowia(100);s__Metschnikowia_viticola(100); | Metschnikowia | 13 | Saprotroph | \|Undefined Saprotroph\| | Yeast | NULL | Probable | As nectar-tap_saprotroph (Põlme et al. 2020); Decay sugar-rich substrates (Põlme et al. 2020) | Põlme S, et al. 2020. Fungal Diversity 105:1-16 ((https://doi.org/10.1007/s13225-020-00466-2)) |
| Otu45640 | 1 | k__Fungi(100);p__Ascomycota(100);c__Saccharomycetes(100);o__Saccharomycetales(100);f__Metschnikowiaceae(100);g__Metschnikowia(100);g__Metschnikowia_unclassified(100); | Metschnikowia | 13 | Saprotroph | \|Undefined Saprotroph\| | Yeast | NULL | Probable | As nectar-tap_saprotroph (Põlme et al. 2020); Decay sugar-rich substrates (Põlme et al. 2020) | Põlme S, et al. 2020. Fungal Diversity 105:1-16 ((https://doi.org/10.1007/s13225-020-00466-2)) |
| Otu45641 | 1 | k__Fungi(100);p__Ascomycota(100);c__Saccharomycetes(100);o__Saccharomycetales(100);f__Metschnikowiaceae(100);g__Metschnikowia(100);s__Metschnikowia_pulcherrima(100); | Metschnikowia | 13 | Saprotroph | \|Undefined Saprotroph\| | Yeast | NULL | Probable | As nectar-tap_saprotroph (Põlme et al. 2020); Decay sugar-rich substrates (Põlme et al. 2020) | Põlme S, et al. 2020. Fungal Diversity 105:1-16 ((https://doi.org/10.1007/s13225-020-00466-2)) |
| Otu45642 | 1 | k__Fungi(100);p__Ascomycota(100);c__Saccharomycetes(100);o__Saccharomycetales(100);f__Metschnikowiaceae(100);g__Metschnikowia(100);s__Metschnikowia_pulcherrima(100); | Metschnikowia | 13 | Saprotroph | \|Undefined Saprotroph\| | Yeast | NULL | Probable | As nectar-tap_saprotroph (Põlme et al. 2020); Decay sugar-rich substrates (Põlme et al. 2020) | Põlme S, et al. 2020. Fungal Diversity 105:1-16 ((https://doi.org/10.1007/s13225-020-00466-2)) |
| Otu45643 | 1 | k__Fungi(100);p__Ascomycota(100);c__Saccharomycetes(100);o__Saccharomycetales(100);f__Metschnikowiaceae(100);g__Metschnikowia(100);s__Metschnikowia_pulcherrima(100); | Metschnikowia | 13 | Saprotroph | \|Undefined Saprotroph\| | Yeast | NULL | Probable | As nectar-tap_saprotroph (Põlme et al. 2020); Decay sugar-rich substrates (Põlme et al. 2020) | Põlme S, et al. 2020. Fungal Diversity 105:1-16 ((https://doi.org/10.1007/s13225-020-00466-2)) |
| Otu45644 | 1 | k__Fungi(100);p__Ascomycota(100);c__Saccharomycetes(100);o__Saccharomycetales(100);f__Metschnikowiaceae(100);g__Metschnikowia(100);g__Metschnikowia_unclassified(100); | Metschnikowia | 13 | Saprotroph | \|Undefined Saprotroph\| | Yeast | NULL | Probable | As nectar-tap_saprotroph (Põlme et al. 2020); Decay sugar-rich substrates (Põlme et al. 2020) | Põlme S, et al. 2020. Fungal Diversity 105:1-16 ((https://doi.org/10.1007/s13225-020-00466-2)) |
| Otu45645 | 1 | k__Fungi(100);p__Ascomycota(100);c__Saccharomycetes(100);o__Saccharomycetales(100);f__Metschnikowiaceae(100);g__Metschnikowia(100);g__Metschnikowia_unclassified(100); | Metschnikowia | 13 | Saprotroph | \|Undefined Saprotroph\| | Yeast | NULL | Probable | As nectar-tap_saprotroph (Põlme et al. 2020); Decay sugar-rich substrates (Põlme et al. 2020) | Põlme S, et al. 2020. Fungal Diversity 105:1-16 ((https://doi.org/10.1007/s13225-020-00466-2)) |
| Otu45646 | 1 | k__Fungi(100);p__Ascomycota(100);c__Saccharomycetes(100);o__Saccharomycetales(100);f__Metschnikowiaceae(100);g__Metschnikowia(100);s__Metschnikowia_pulcherrima(100); | Metschnikowia | 13 | Saprotroph | \|Undefined Saprotroph\| | Yeast | NULL | Probable | As nectar-tap_saprotroph (Põlme et al. 2020); Decay sugar-rich substrates (Põlme et al. 2020) | Põlme S, et al. 2020. Fungal Diversity 105:1-16 ((https://doi.org/10.1007/s13225-020-00466-2)) |
| Otu45647 | 1 | k__Fungi(100);p__Ascomycota(100);c__Saccharomycetes(100);o__Saccharomycetales(100);f__Metschnikowiaceae(100);g__Metschnikowia(100);s__Metschnikowia_pulcherrima(100); | Metschnikowia | 13 | Saprotroph | \|Undefined Saprotroph\| | Yeast | NULL | Probable | As nectar-tap_saprotroph (Põlme et al. 2020); Decay sugar-rich substrates (Põlme et al. 2020) | Põlme S, et al. 2020. Fungal Diversity 105:1-16 ((https://doi.org/10.1007/s13225-020-00466-2)) |
| Otu45648 | 1 | k__Fungi(100);p__Ascomycota(100);c__Saccharomycetes(100);o__Saccharomycetales(100);f__Metschnikowiaceae(100);g__Metschnikowia(100);s__Metschnikowia_pulcherrima(100); | Metschnikowia | 13 | Saprotroph | \|Undefined Saprotroph\| | Yeast | NULL | Probable | As nectar-tap_saprotroph (Põlme et al. 2020); Decay sugar-rich substrates (Põlme et al. 2020) | Põlme S, et al. 2020. Fungal Diversity 105:1-16 ((https://doi.org/10.1007/s13225-020-00466-2)) |
| Otu45649 | 1 | k__Fungi(100);p__Ascomycota(100);c__Saccharomycetes(100);o__Saccharomycetales(100);f__Metschnikowiaceae(100);g__Metschnikowia(100);s__Metschnikowia_pulcherrima(100); | Metschnikowia | 13 | Saprotroph | \|Undefined Saprotroph\| | Yeast | NULL | Probable | As nectar-tap_saprotroph (Põlme et al. 2020); Decay sugar-rich substrates (Põlme et al. 2020) | Põlme S, et al. 2020. Fungal Diversity 105:1-16 ((https://doi.org/10.1007/s13225-020-00466-2)) |
| Otu45650 | 1 | k__Fungi(100);p__Ascomycota(100);c__Saccharomycetes(100);o__Saccharomycetales(100);f__Metschnikowiaceae(100);g__Metschnikowia(100);s__Metschnikowia_pulcherrima(100); | Metschnikowia | 13 | Saprotroph | \|Undefined Saprotroph\| | Yeast | NULL | Probable | As nectar-tap_saprotroph (Põlme et al. 2020); Decay sugar-rich substrates (Põlme et al. 2020) | Põlme S, et al. 2020. Fungal Diversity 105:1-16 ((https://doi.org/10.1007/s13225-020-00466-2)) |
| Otu45651 | 1 | k__Fungi(100);p__Ascomycota(100);c__Saccharomycetes(100);o__Saccharomycetales(100);f__Metschnikowiaceae(100);g__Metschnikowia(100);s__Metschnikowia_pulcherrima(100); | Metschnikowia | 13 | Saprotroph | \|Undefined Saprotroph\| | Yeast | NULL | Probable | As nectar-tap_saprotroph (Põlme et al. 2020); Decay sugar-rich substrates (Põlme et al. 2020) | Põlme S, et al. 2020. Fungal Diversity 105:1-16 ((https://doi.org/10.1007/s13225-020-00466-2)) |
| Otu45652 | 1 | k__Fungi(100);p__Ascomycota(100);c__Saccharomycetes(100);o__Saccharomycetales(100);f__Metschnikowiaceae(100);g__Metschnikowia(100);s__Metschnikowia_pulcherrima(100); | Metschnikowia | 13 | Saprotroph | \|Undefined Saprotroph\| | Yeast | NULL | Probable | As nectar-tap_saprotroph (Põlme et al. 2020); Decay sugar-rich substrates (Põlme et al. 2020) | Põlme S, et al. 2020. Fungal Diversity 105:1-16 ((https://doi.org/10.1007/s13225-020-00466-2)) |
| Otu45653 | 1 | k__Fungi(100);p__Ascomycota(100);c__Saccharomycetes(100);o__Saccharomycetales(100);f__Metschnikowiaceae(100);g__Metschnikowia(100);s__Metschnikowia_pulcherrima(100); | Metschnikowia | 13 | Saprotroph | \|Undefined Saprotroph\| | Yeast | NULL | Probable | As nectar-tap_saprotroph (Põlme et al. 2020); Decay sugar-rich substrates (Põlme et al. 2020) | Põlme S, et al. 2020. Fungal Diversity 105:1-16 ((https://doi.org/10.1007/s13225-020-00466-2)) |
| Otu45654 | 1 | k__Fungi(100);p__Ascomycota(100);c__Saccharomycetes(100);o__Saccharomycetales(100);f__Metschnikowiaceae(100);g__Metschnikowia(100);s__Metschnikowia_pulcherrima(100); | Metschnikowia | 13 | Saprotroph | \|Undefined Saprotroph\| | Yeast | NULL | Probable | As nectar-tap_saprotroph (Põlme et al. 2020); Decay sugar-rich substrates (Põlme et al. 2020) | Põlme S, et al. 2020. Fungal Diversity 105:1-16 ((https://doi.org/10.1007/s13225-020-00466-2)) |
| Otu45655 | 1 | k__Fungi(100);p__Ascomycota(100);c__Saccharomycetes(100);o__Saccharomycetales(100);f__Metschnikowiaceae(100);g__Metschnikowia(100);s__Metschnikowia_pulcherrima(100); | Metschnikowia | 13 | Saprotroph | \|Undefined Saprotroph\| | Yeast | NULL | Probable | As nectar-tap_saprotroph (Põlme et al. 2020); Decay sugar-rich substrates (Põlme et al. 2020) | Põlme S, et al. 2020. Fungal Diversity 105:1-16 ((https://doi.org/10.1007/s13225-020-00466-2)) |
| Otu45656 | 1 | k__Fungi(100);p__Ascomycota(100);c__Saccharomycetes(100);o__Saccharomycetales(100);f__Metschnikowiaceae(100);g__Metschnikowia(100);s__Metschnikowia_pulcherrima(100); | Metschnikowia | 13 | Saprotroph | \|Undefined Saprotroph\| | Yeast | NULL | Probable | As nectar-tap_saprotroph (Põlme et al. 2020); Decay sugar-rich substrates (Põlme et al. 2020) | Põlme S, et al. 2020. Fungal Diversity 105:1-16 ((https://doi.org/10.1007/s13225-020-00466-2)) |
| Otu45657 | 1 | k__Fungi(100);p__Ascomycota(100);c__Saccharomycetes(100);o__Saccharomycetales(100);f__Metschnikowiaceae(100);g__Metschnikowia(100);g__Metschnikowia_unclassified(100); | Metschnikowia | 13 | Saprotroph | \|Undefined Saprotroph\| | Yeast | NULL | Probable | As nectar-tap_saprotroph (Põlme et al. 2020); Decay sugar-rich substrates (Põlme et al. 2020) | Põlme S, et al. 2020. Fungal Diversity 105:1-16 ((https://doi.org/10.1007/s13225-020-00466-2)) |
| Otu45659 | 1 | k__Fungi(100);p__Ascomycota(100);c__Saccharomycetes(100);o__Saccharomycetales(100);f__Metschnikowiaceae(100);g__Metschnikowia(100);s__Metschnikowia_pulcherrima(100); | Metschnikowia | 13 | Saprotroph | \|Undefined Saprotroph\| | Yeast | NULL | Probable | As nectar-tap_saprotroph (Põlme et al. 2020); Decay sugar-rich substrates (Põlme et al. 2020) | Põlme S, et al. 2020. Fungal Diversity 105:1-16 ((https://doi.org/10.1007/s13225-020-00466-2)) |
| Otu45661 | 1 | k__Fungi(100);p__Ascomycota(100);c__Saccharomycetes(100);o__Saccharomycetales(100);f__Metschnikowiaceae(100);g__Metschnikowia(100);s__Metschnikowia_pulcherrima(100); | Metschnikowia | 13 | Saprotroph | \|Undefined Saprotroph\| | Yeast | NULL | Probable | As nectar-tap_saprotroph (Põlme et al. 2020); Decay sugar-rich substrates (Põlme et al. 2020) | Põlme S, et al. 2020. Fungal Diversity 105:1-16 ((https://doi.org/10.1007/s13225-020-00466-2)) |
| Otu45662 | 1 | k__Fungi(100);p__Ascomycota(100);c__Saccharomycetes(100);o__Saccharomycetales(100);f__Metschnikowiaceae(100);g__Metschnikowia(100);s__Metschnikowia_pulcherrima(100); | Metschnikowia | 13 | Saprotroph | \|Undefined Saprotroph\| | Yeast | NULL | Probable | As nectar-tap_saprotroph (Põlme et al. 2020); Decay sugar-rich substrates (Põlme et al. 2020) | Põlme S, et al. 2020. Fungal Diversity 105:1-16 ((https://doi.org/10.1007/s13225-020-00466-2)) |
| Otu45663 | 1 | k__Fungi(100);p__Ascomycota(100);c__Saccharomycetes(100);o__Saccharomycetales(100);f__Metschnikowiaceae(100);g__Metschnikowia(100);s__Metschnikowia_pulcherrima(100); | Metschnikowia | 13 | Saprotroph | \|Undefined Saprotroph\| | Yeast | NULL | Probable | As nectar-tap_saprotroph (Põlme et al. 2020); Decay sugar-rich substrates (Põlme et al. 2020) | Põlme S, et al. 2020. Fungal Diversity 105:1-16 ((https://doi.org/10.1007/s13225-020-00466-2)) |
| Otu45664 | 1 | k__Fungi(100);p__Ascomycota(100);c__Saccharomycetes(100);o__Saccharomycetales(100);f__Metschnikowiaceae(100);g__Metschnikowia(100);s__Metschnikowia_pulcherrima(100); | Metschnikowia | 13 | Saprotroph | \|Undefined Saprotroph\| | Yeast | NULL | Probable | As nectar-tap_saprotroph (Põlme et al. 2020); Decay sugar-rich substrates (Põlme et al. 2020) | Põlme S, et al. 2020. Fungal Diversity 105:1-16 ((https://doi.org/10.1007/s13225-020-00466-2)) |
| Otu45665 | 1 | k__Fungi(100);p__Ascomycota(100);c__Saccharomycetes(100);o__Saccharomycetales(100);f__Metschnikowiaceae(100);g__Metschnikowia(100);s__Metschnikowia_pulcherrima(100); | Metschnikowia | 13 | Saprotroph | \|Undefined Saprotroph\| | Yeast | NULL | Probable | As nectar-tap_saprotroph (Põlme et al. 2020); Decay sugar-rich substrates (Põlme et al. 2020) | Põlme S, et al. 2020. Fungal Diversity 105:1-16 ((https://doi.org/10.1007/s13225-020-00466-2)) |

| | | | | | | | | | | | |
|---|---|---|---|---|---|---|---|---|---|---|---|
| Otu45666 | 1 | k__Fungi(100);p__Ascomycota(100);c__Saccharomycetes(100);o__Saccharomycetales(100);f__Metschnikowiaceae(100);g__Metschnikowia(100);s__Metschnikowia_pulcherrima(100); | Metschnikowia | 13 | Saprotroph | \|Undefined Saprotroph\| | Yeast | NULL | Probable | As nectar-tap_saprotroph (Põlme et al. 2020); Decay sugar-rich substrates (Põlme et al. 2020) | Põlme S, et al. 2020. Fungal Diversity 105:1-16 ((https://doi.org/10.1007/s13225-020-00466-2)) |
| Otu45667 | 1 | k__Fungi(100);p__Ascomycota(100);c__Saccharomycetes(100);o__Saccharomycetales(100);f__Metschnikowiaceae(100);g__Metschnikowia(100);s__Metschnikowia_pulcherrima(100); | Metschnikowia | 13 | Saprotroph | \|Undefined Saprotroph\| | Yeast | NULL | Probable | As nectar-tap_saprotroph (Põlme et al. 2020); Decay sugar-rich substrates (Põlme et al. 2020) | Põlme S, et al. 2020. Fungal Diversity 105:1-16 ((https://doi.org/10.1007/s13225-020-00466-2)) |
| Otu45668 | 1 | k__Fungi(100);p__Ascomycota(100);c__Saccharomycetes(100);o__Saccharomycetales(100);f__Metschnikowiaceae(100);g__Metschnikowia(100);s__Metschnikowia_pulcherrima(100); | Metschnikowia | 13 | Saprotroph | \|Undefined Saprotroph\| | Yeast | NULL | Probable | As nectar-tap_saprotroph (Põlme et al. 2020); Decay sugar-rich substrates (Põlme et al. 2020) | Põlme S, et al. 2020. Fungal Diversity 105:1-16 ((https://doi.org/10.1007/s13225-020-00466-2)) |
| Otu45669 | 1 | k__Fungi(100);p__Ascomycota(100);c__Saccharomycetes(100);o__Saccharomycetales(100);f__Metschnikowiaceae(100);g__Metschnikowia(100);s__Metschnikowia_pulcherrima(100); | Metschnikowia | 13 | Saprotroph | \|Undefined Saprotroph\| | Yeast | NULL | Probable | As nectar-tap_saprotroph (Põlme et al. 2020); Decay sugar-rich substrates (Põlme et al. 2020) | Põlme S, et al. 2020. Fungal Diversity 105:1-16 ((https://doi.org/10.1007/s13225-020-00466-2)) |
| Otu45670 | 1 | k__Fungi(100);p__Ascomycota(100);c__Saccharomycetes(100);o__Saccharomycetales(100);f__Metschnikowiaceae(100);g__Metschnikowia(100);s__Metschnikowia_pulcherrima(100); | Metschnikowia | 13 | Saprotroph | \|Undefined Saprotroph\| | Yeast | NULL | Probable | As nectar-tap_saprotroph (Põlme et al. 2020); Decay sugar-rich substrates (Põlme et al. 2020) | Põlme S, et al. 2020. Fungal Diversity 105:1-16 ((https://doi.org/10.1007/s13225-020-00466-2)) |
| Otu45671 | 1 | k__Fungi(100);p__Ascomycota(100);c__Saccharomycetes(100);o__Saccharomycetales(100);f__Metschnikowiaceae(100);g__Metschnikowia(100);s__Metschnikowia_pulcherrima(100); | Metschnikowia | 13 | Saprotroph | \|Undefined Saprotroph\| | Yeast | NULL | Probable | As nectar-tap_saprotroph (Põlme et al. 2020); Decay sugar-rich substrates (Põlme et al. 2020) | Põlme S, et al. 2020. Fungal Diversity 105:1-16 ((https://doi.org/10.1007/s13225-020-00466-2)) |
| Otu45672 | 1 | k__Fungi(100);p__Ascomycota(100);c__Saccharomycetes(100);o__Saccharomycetales(100);f__Metschnikowiaceae(100);g__Metschnikowia(100);s__Metschnikowia_pulcherrima(100); | Metschnikowia | 13 | Saprotroph | \|Undefined Saprotroph\| | Yeast | NULL | Probable | As nectar-tap_saprotroph (Põlme et al. 2020); Decay sugar-rich substrates (Põlme et al. 2020) | Põlme S, et al. 2020. Fungal Diversity 105:1-16 ((https://doi.org/10.1007/s13225-020-00466-2)) |
| Otu45673 | 1 | k__Fungi(100);p__Ascomycota(100);c__Saccharomycetes(100);o__Saccharomycetales(100);f__Metschnikowiaceae(100);g__Metschnikowia(100);g__Metschnikowia_unclassified(100); | Metschnikowia | 13 | Saprotroph | \|Undefined Saprotroph\| | Yeast | NULL | Probable | As nectar-tap_saprotroph (Põlme et al. 2020); Decay sugar-rich substrates (Põlme et al. 2020) | Põlme S, et al. 2020. Fungal Diversity 105:1-16 ((https://doi.org/10.1007/s13225-020-00466-2)) |
| Otu45674 | 1 | k__Fungi(100);p__Ascomycota(100);c__Saccharomycetes(100);o__Saccharomycetales(100);f__Metschnikowiaceae(100);g__Metschnikowia(100);s__Metschnikowia_pulcherrima(100); | Metschnikowia | 13 | Saprotroph | \|Undefined Saprotroph\| | Yeast | NULL | Probable | As nectar-tap_saprotroph (Põlme et al. 2020); Decay sugar-rich substrates (Põlme et al. 2020) | Põlme S, et al. 2020. Fungal Diversity 105:1-16 ((https://doi.org/10.1007/s13225-020-00466-2)) |
| Otu45675 | 1 | k__Fungi(100);p__Ascomycota(100);c__Saccharomycetes(100);o__Saccharomycetales(100);f__Metschnikowiaceae(100);g__Metschnikowia(100);s__Metschnikowia_pulcherrima(100); | Metschnikowia | 13 | Saprotroph | \|Undefined Saprotroph\| | Yeast | NULL | Probable | As nectar-tap_saprotroph (Põlme et al. 2020); Decay sugar-rich substrates (Põlme et al. 2020) | Põlme S, et al. 2020. Fungal Diversity 105:1-16 ((https://doi.org/10.1007/s13225-020-00466-2)) |
| Otu45676 | 1 | k__Fungi(100);p__Ascomycota(100);c__Saccharomycetes(100);o__Saccharomycetales(100);f__Metschnikowiaceae(100);g__Metschnikowia(100);g__Metschnikowia_unclassified(100); | Metschnikowia | 13 | Saprotroph | \|Undefined Saprotroph\| | Yeast | NULL | Probable | As nectar-tap_saprotroph (Põlme et al. 2020); Decay sugar-rich substrates (Põlme et al. 2020) | Põlme S, et al. 2020. Fungal Diversity 105:1-16 ((https://doi.org/10.1007/s13225-020-00466-2)) |
| Otu45677 | 1 | k__Fungi(100);p__Ascomycota(100);c__Saccharomycetes(100);o__Saccharomycetales(100);f__Metschnikowiaceae(100);g__Metschnikowia(100);s__Metschnikowia_pulcherrima(100); | Metschnikowia | 13 | Saprotroph | \|Undefined Saprotroph\| | Yeast | NULL | Probable | As nectar-tap_saprotroph (Põlme et al. 2020); Decay sugar-rich substrates (Põlme et al. 2020) | Põlme S, et al. 2020. Fungal Diversity 105:1-16 ((https://doi.org/10.1007/s13225-020-00466-2)) |
| Otu45678 | 1 | k__Fungi(100);p__Ascomycota(100);c__Saccharomycetes(100);o__Saccharomycetales(100);f__Metschnikowiaceae(100);g__Metschnikowia(100);s__Metschnikowia_pulcherrima(100); | Metschnikowia | 13 | Saprotroph | \|Undefined Saprotroph\| | Yeast | NULL | Probable | As nectar-tap_saprotroph (Põlme et al. 2020); Decay sugar-rich substrates (Põlme et al. 2020) | Põlme S, et al. 2020. Fungal Diversity 105:1-16 ((https://doi.org/10.1007/s13225-020-00466-2)) |
| Otu45679 | 1 | k__Fungi(100);p__Ascomycota(100);c__Saccharomycetes(100);o__Saccharomycetales(100);f__Metschnikowiaceae(100);g__Metschnikowia(100);s__Metschnikowia_pulcherrima(100); | Metschnikowia | 13 | Saprotroph | \|Undefined Saprotroph\| | Yeast | NULL | Probable | As nectar-tap_saprotroph (Põlme et al. 2020); Decay sugar-rich substrates (Põlme et al. 2020) | Põlme S, et al. 2020. Fungal Diversity 105:1-16 ((https://doi.org/10.1007/s13225-020-00466-2)) |
| Otu45680 | 1 | k__Fungi(100);p__Ascomycota(100);c__Saccharomycetes(100);o__Saccharomycetales(100);f__Metschnikowiaceae(100);g__Metschnikowia(100);s__Metschnikowia_pulcherrima(100); | Metschnikowia | 13 | Saprotroph | \|Undefined Saprotroph\| | Yeast | NULL | Probable | As nectar-tap_saprotroph (Põlme et al. 2020); Decay sugar-rich substrates (Põlme et al. 2020) | Põlme S, et al. 2020. Fungal Diversity 105:1-16 ((https://doi.org/10.1007/s13225-020-00466-2)) |
| Otu45681 | 1 | k__Fungi(100);p__Ascomycota(100);c__Saccharomycetes(100);o__Saccharomycetales(100);f__Metschnikowiaceae(100);g__Metschnikowia(100);s__Metschnikowia_pulcherrima(100); | Metschnikowia | 13 | Saprotroph | \|Undefined Saprotroph\| | Yeast | NULL | Probable | As nectar-tap_saprotroph (Põlme et al. 2020); Decay sugar-rich substrates (Põlme et al. 2020) | Põlme S, et al. 2020. Fungal Diversity 105:1-16 ((https://doi.org/10.1007/s13225-020-00466-2)) |
| Otu45682 | 1 | k__Fungi(100);p__Ascomycota(100);c__Saccharomycetes(100);o__Saccharomycetales(100);f__Metschnikowiaceae(100);g__Metschnikowia(100);g__Metschnikowia_unclassified(100); | Metschnikowia | 13 | Saprotroph | \|Undefined Saprotroph\| | Yeast | NULL | Probable | As nectar-tap_saprotroph (Põlme et al. 2020); Decay sugar-rich substrates (Põlme et al. 2020) | Põlme S, et al. 2020. Fungal Diversity 105:1-16 ((https://doi.org/10.1007/s13225-020-00466-2)) |
| Otu45683 | 1 | k__Fungi(100);p__Ascomycota(100);c__Saccharomycetes(100);o__Saccharomycetales(100);f__Metschnikowiaceae(100);g__Metschnikowia(100);s__Metschnikowia_pulcherrima(100); | Metschnikowia | 13 | Saprotroph | \|Undefined Saprotroph\| | Yeast | NULL | Probable | As nectar-tap_saprotroph (Põlme et al. 2020); Decay sugar-rich substrates (Põlme et al. 2020) | Põlme S, et al. 2020. Fungal Diversity 105:1-16 ((https://doi.org/10.1007/s13225-020-00466-2)) |
| Otu45684 | 1 | k__Fungi(100);p__Ascomycota(100);c__Saccharomycetes(100);o__Saccharomycetales(100);f__Metschnikowiaceae(100);g__Metschnikowia(100);g__Metschnikowia_unclassified(100); | Metschnikowia | 13 | Saprotroph | \|Undefined Saprotroph\| | Yeast | NULL | Probable | As nectar-tap_saprotroph (Põlme et al. 2020); Decay sugar-rich substrates (Põlme et al. 2020) | Põlme S, et al. 2020. Fungal Diversity 105:1-16 ((https://doi.org/10.1007/s13225-020-00466-2)) |
| Otu45685 | 1 | k__Fungi(100);p__Ascomycota(100);c__Saccharomycetes(100);o__Saccharomycetales(100);f__Metschnikowiaceae(100);g__Metschnikowia(100);s__Metschnikowia_pulcherrima(100); | Metschnikowia | 13 | Saprotroph | \|Undefined Saprotroph\| | Yeast | NULL | Probable | As nectar-tap_saprotroph (Põlme et al. 2020); Decay sugar-rich substrates (Põlme et al. 2020) | Põlme S, et al. 2020. Fungal Diversity 105:1-16 ((https://doi.org/10.1007/s13225-020-00466-2)) |
| Otu45686 | 1 | k__Fungi(100);p__Ascomycota(100);c__Saccharomycetes(100);o__Saccharomycetales(100);f__Metschnikowiaceae(100);g__Metschnikowia(100);s__Metschnikowia_pulcherrima(100); | Metschnikowia | 13 | Saprotroph | \|Undefined Saprotroph\| | Yeast | NULL | Probable | As nectar-tap_saprotroph (Põlme et al. 2020); Decay sugar-rich substrates (Põlme et al. 2020) | Põlme S, et al. 2020. Fungal Diversity 105:1-16 ((https://doi.org/10.1007/s13225-020-00466-2)) |
| Otu45687 | 1 | k__Fungi(100);p__Ascomycota(100);c__Saccharomycetes(100);o__Saccharomycetales(100);f__Metschnikowiaceae(100);g__Metschnikowia(100);g__Metschnikowia_unclassified(100); | Metschnikowia | 13 | Saprotroph | \|Undefined Saprotroph\| | Yeast | NULL | Probable | As nectar-tap_saprotroph (Põlme et al. 2020); Decay sugar-rich substrates (Põlme et al. 2020) | Põlme S, et al. 2020. Fungal Diversity 105:1-16 ((https://doi.org/10.1007/s13225-020-00466-2)) |
| Otu45688 | 1 | k__Fungi(100);p__Ascomycota(100);c__Saccharomycetes(100);o__Saccharomycetales(100);f__Metschnikowiaceae(100);g__Metschnikowia(100);s__Metschnikowia_pulcherrima(100); | Metschnikowia | 13 | Saprotroph | \|Undefined Saprotroph\| | Yeast | NULL | Probable | As nectar-tap_saprotroph (Põlme et al. 2020); Decay sugar-rich substrates (Põlme et al. 2020) | Põlme S, et al. 2020. Fungal Diversity 105:1-16 ((https://doi.org/10.1007/s13225-020-00466-2)) |
| Otu45689 | 1 | k__Fungi(100);p__Ascomycota(100);c__Saccharomycetes(100);o__Saccharomycetales(100);f__Metschnikowiaceae(100);g__Metschnikowia(100);s__Metschnikowia_pulcherrima(100); | Metschnikowia | 13 | Saprotroph | \|Undefined Saprotroph\| | Yeast | NULL | Probable | As nectar-tap_saprotroph (Põlme et al. 2020); Decay sugar-rich substrates (Põlme et al. 2020) | Põlme S, et al. 2020. Fungal Diversity 105:1-16 ((https://doi.org/10.1007/s13225-020-00466-2)) |
| Otu45690 | 1 | k__Fungi(100);p__Ascomycota(100);c__Saccharomycetes(100);o__Saccharomycetales(100);f__Metschnikowiaceae(100);g__Metschnikowia(100);g__Metschnikowia_unclassified(100); | Metschnikowia | 13 | Saprotroph | \|Undefined Saprotroph\| | Yeast | NULL | Probable | As nectar-tap_saprotroph (Põlme et al. 2020); Decay sugar-rich substrates (Põlme et al. 2020) | Põlme S, et al. 2020. Fungal Diversity 105:1-16 ((https://doi.org/10.1007/s13225-020-00466-2)) |
| Otu45691 | 1 | k__Fungi(100);p__Ascomycota(100);c__Saccharomycetes(100);o__Saccharomycetales(100);f__Metschnikowiaceae(100);g__Metschnikowia(100);g__Metschnikowia_unclassified(100); | Metschnikowia | 13 | Saprotroph | \|Undefined Saprotroph\| | Yeast | NULL | Probable | As nectar-tap_saprotroph (Põlme et al. 2020); Decay sugar-rich substrates (Põlme et al. 2020) | Põlme S, et al. 2020. Fungal Diversity 105:1-16 ((https://doi.org/10.1007/s13225-020-00466-2)) |
| Otu45692 | 1 | k__Fungi(100);p__Ascomycota(100);c__Saccharomycetes(100);o__Saccharomycetales(100);f__Metschnikowiaceae(100);g__Metschnikowia(100);g__Metschnikowia_unclassified(100); | Metschnikowia | 13 | Saprotroph | \|Undefined Saprotroph\| | Yeast | NULL | Probable | As nectar-tap_saprotroph (Põlme et al. 2020); Decay sugar-rich substrates (Põlme et al. 2020) | Põlme S, et al. 2020. Fungal Diversity 105:1-16 ((https://doi.org/10.1007/s13225-020-00466-2)) |
| Otu45693 | 1 | k__Fungi(100);p__Ascomycota(100);c__Saccharomycetes(100);o__Saccharomycetales(100);f__Metschnikowiaceae(100);g__Metschnikowia(100);s__Metschnikowia_pulcherrima(100); | Metschnikowia | 13 | Saprotroph | \|Undefined Saprotroph\| | Yeast | NULL | Probable | As nectar-tap_saprotroph (Põlme et al. 2020); Decay sugar-rich substrates (Põlme et al. 2020) | Põlme S, et al. 2020. Fungal Diversity 105:1-16 ((https://doi.org/10.1007/s13225-020-00466-2)) |
| Otu45694 | 1 | k__Fungi(100);p__Ascomycota(100);c__Saccharomycetes(100);o__Saccharomycetales(100);f__Metschnikowiaceae(100);g__Metschnikowia(100);s__Metschnikowia_pulcherrima(100); | Metschnikowia | 13 | Saprotroph | \|Undefined Saprotroph\| | Yeast | NULL | Probable | As nectar-tap_saprotroph (Põlme et al. 2020); Decay sugar-rich substrates (Põlme et al. 2020) | Põlme S, et al. 2020. Fungal Diversity 105:1-16 ((https://doi.org/10.1007/s13225-020-00466-2)) |
| Otu45695 | 1 | k__Fungi(100);p__Ascomycota(100);c__Saccharomycetes(100);o__Saccharomycetales(100);f__Metschnikowiaceae(100);g__Metschnikowia(100);g__Metschnikowia_unclassified(100); | Metschnikowia | 13 | Saprotroph | \|Undefined Saprotroph\| | Yeast | NULL | Probable | As nectar-tap_saprotroph (Põlme et al. 2020); Decay sugar-rich substrates (Põlme et al. 2020) | Põlme S, et al. 2020. Fungal Diversity 105:1-16 ((https://doi.org/10.1007/s13225-020-00466-2)) |
| Otu45696 | 1 | k__Fungi(100);p__Ascomycota(100);c__Saccharomycetes(100);o__Saccharomycetales(100);f__Metschnikowiaceae(100);g__Metschnikowia(100);s__Metschnikowia_pulcherrima(100); | Metschnikowia | 13 | Saprotroph | \|Undefined Saprotroph\| | Yeast | NULL | Probable | As nectar-tap_saprotroph (Põlme et al. 2020); Decay sugar-rich substrates (Põlme et al. 2020) | Põlme S, et al. 2020. Fungal Diversity 105:1-16 ((https://doi.org/10.1007/s13225-020-00466-2)) |
| Otu45697 | 1 | k__Fungi(100);p__Ascomycota(100);c__Saccharomycetes(100);o__Saccharomycetales(100);f__Metschnikowiaceae(100);g__Metschnikowia(100);s__Metschnikowia_pulcherrima(100); | Metschnikowia | 13 | Saprotroph | \|Undefined Saprotroph\| | Yeast | NULL | Probable | As nectar-tap_saprotroph (Põlme et al. 2020); Decay sugar-rich substrates (Põlme et al. 2020) | Põlme S, et al. 2020. Fungal Diversity 105:1-16 ((https://doi.org/10.1007/s13225-020-00466-2)) |
| Otu45698 | 1 | k__Fungi(100);p__Ascomycota(100);c__Saccharomycetes(100);o__Saccharomycetales(100);f__Metschnikowiaceae(100);g__Metschnikowia(100);s__Metschnikowia_pulcherrima(100); | Metschnikowia | 13 | Saprotroph | \|Undefined Saprotroph\| | Yeast | NULL | Probable | As nectar-tap_saprotroph (Põlme et al. 2020); Decay sugar-rich substrates (Põlme et al. 2020) | Põlme S, et al. 2020. Fungal Diversity 105:1-16 ((https://doi.org/10.1007/s13225-020-00466-2)) |
| Otu45699 | 1 | k__Fungi(100);p__Ascomycota(100);c__Saccharomycetes(100);o__Saccharomycetales(100);f__Metschnikowiaceae(100);g__Metschnikowia(100);s__Metschnikowia_pulcherrima(100); | Metschnikowia | 13 | Saprotroph | \|Undefined Saprotroph\| | Yeast | NULL | Probable | As nectar-tap_saprotroph (Põlme et al. 2020); Decay sugar-rich substrates (Põlme et al. 2020) | Põlme S, et al. 2020. Fungal Diversity 105:1-16 ((https://doi.org/10.1007/s13225-020-00466-2)) |
| Otu45700 | 1 | k__Fungi(100);p__Ascomycota(100);c__Saccharomycetes(100);o__Saccharomycetales(100);f__Metschnikowiaceae(100);g__Metschnikowia(100);g__Metschnikowia_unclassified(100); | Metschnikowia | 13 | Saprotroph | \|Undefined Saprotroph\| | Yeast | NULL | Probable | As nectar-tap_saprotroph (Põlme et al. 2020); Decay sugar-rich substrates (Põlme et al. 2020) | Põlme S, et al. 2020. Fungal Diversity 105:1-16 ((https://doi.org/10.1007/s13225-020-00466-2)) |
| Otu45701 | 1 | k__Fungi(100);p__Ascomycota(100);c__Saccharomycetes(100);o__Saccharomycetales(100);f__Metschnikowiaceae(100);g__Metschnikowia(100);s__Metschnikowia_pulcherrima(100); | Metschnikowia | 13 | Saprotroph | \|Undefined Saprotroph\| | Yeast | NULL | Probable | As nectar-tap_saprotroph (Põlme et al. 2020); Decay sugar-rich substrates (Põlme et al. 2020) | Põlme S, et al. 2020. Fungal Diversity 105:1-16 ((https://doi.org/10.1007/s13225-020-00466-2)) |
| Otu45702 | 1 | k__Fungi(100);p__Ascomycota(100);c__Saccharomycetes(100);o__Saccharomycetales(100);f__Metschnikowiaceae(100);g__Metschnikowia(100);s__Metschnikowia_pulcherrima(100); | Metschnikowia | 13 | Saprotroph | \|Undefined Saprotroph\| | Yeast | NULL | Probable | As nectar-tap_saprotroph (Põlme et al. 2020); Decay sugar-rich substrates (Põlme et al. 2020) | Põlme S, et al. 2020. Fungal Diversity 105:1-16 ((https://doi.org/10.1007/s13225-020-00466-2)) |
| Otu45703 | 1 | k__Fungi(100);p__Ascomycota(100);c__Saccharomycetes(100);o__Saccharomycetales(100);f__Metschnikowiaceae(100);g__Metschnikowia(100);g__Metschnikowia_unclassified(100); | Metschnikowia | 13 | Saprotroph | \|Undefined Saprotroph\| | Yeast | NULL | Probable | As nectar-tap_saprotroph (Põlme et al. 2020); Decay sugar-rich substrates (Põlme et al. 2020) | Põlme S, et al. 2020. Fungal Diversity 105:1-16 ((https://doi.org/10.1007/s13225-020-00466-2)) |
| Otu45704 | 1 | k__Fungi(100);p__Ascomycota(100);c__Saccharomycetes(100);o__Saccharomycetales(100);f__Metschnikowiaceae(100);g__Metschnikowia(100);g__Metschnikowia_unclassified(100); | Metschnikowia | 13 | Saprotroph | \|Undefined Saprotroph\| | Yeast | NULL | Probable | As nectar-tap_saprotroph (Põlme et al. 2020); Decay sugar-rich substrates (Põlme et al. 2020) | Põlme S, et al. 2020. Fungal Diversity 105:1-16 ((https://doi.org/10.1007/s13225-020-00466-2)) |
| Otu45705 | 1 | k__Fungi(100);p__Ascomycota(100);c__Saccharomycetes(100);o__Saccharomycetales(100);f__Metschnikowiaceae(100);g__Metschnikowia(100);s__Metschnikowia_pulcherrima(100); | Metschnikowia | 13 | Saprotroph | \|Undefined Saprotroph\| | Yeast | NULL | Probable | As nectar-tap_saprotroph (Põlme et al. 2020); Decay sugar-rich substrates (Põlme et al. 2020) | Põlme S, et al. 2020. Fungal Diversity 105:1-16 ((https://doi.org/10.1007/s13225-020-00466-2)) |
| Otu45706 | 1 | k__Fungi(100);p__Ascomycota(100);c__Saccharomycetes(100);o__Saccharomycetales(100);f__Metschnikowiaceae(100);g__Metschnikowia(100);s__Metschnikowia_pulcherrima(100); | Metschnikowia | 13 | Saprotroph | \|Undefined Saprotroph\| | Yeast | NULL | Probable | As nectar-tap_saprotroph (Põlme et al. 2020); Decay sugar-rich substrates (Põlme et al. 2020) | Põlme S, et al. 2020. Fungal Diversity 105:1-16 ((https://doi.org/10.1007/s13225-020-00466-2)) |
| Otu45707 | 1 | k__Fungi(100);p__Ascomycota(100);c__Saccharomycetes(100);o__Saccharomycetales(100);f__Metschnikowiaceae(100);g__Metschnikowia(100);s__Metschnikowia_pulcherrima(100); | Metschnikowia | 13 | Saprotroph | \|Undefined Saprotroph\| | Yeast | NULL | Probable | As nectar-tap_saprotroph (Põlme et al. 2020); Decay sugar-rich substrates (Põlme et al. 2020) | Põlme S, et al. 2020. Fungal Diversity 105:1-16 ((https://doi.org/10.1007/s13225-020-00466-2)) |
| Otu45708 | 1 | k__Fungi(100);p__Ascomycota(100);c__Saccharomycetes(100);o__Saccharomycetales(100);f__Metschnikowiaceae(100);g__Metschnikowia(100);s__Metschnikowia_pulcherrima(100); | Metschnikowia | 13 | Saprotroph | \|Undefined Saprotroph\| | Yeast | NULL | Probable | As nectar-tap_saprotroph (Põlme et al. 2020); Decay sugar-rich substrates (Põlme et al. 2020) | Põlme S, et al. 2020. Fungal Diversity 105:1-16 ((https://doi.org/10.1007/s13225-020-00466-2)) |
| Otu45709 | 1 | k__Fungi(100);p__Ascomycota(100);c__Saccharomycetes(100);o__Saccharomycetales(100);f__Metschnikowiaceae(100);g__Metschnikowia(100);s__Metschnikowia_pulcherrima(100); | Metschnikowia | 13 | Saprotroph | \|Undefined Saprotroph\| | Yeast | NULL | Probable | As nectar-tap_saprotroph (Põlme et al. 2020); Decay sugar-rich substrates (Põlme et al. 2020) | Põlme S, et al. 2020. Fungal Diversity 105:1-16 ((https://doi.org/10.1007/s13225-020-00466-2)) |
| Otu45710 | 1 | k__Fungi(100);p__Ascomycota(100);c__Saccharomycetes(100);o__Saccharomycetales(100);f__Metschnikowiaceae(100);g__Metschnikowia(100);s__Metschnikowia_pulcherrima(100); | Metschnikowia | 13 | Saprotroph | \|Undefined Saprotroph\| | Yeast | NULL | Probable | As nectar-tap_saprotroph (Põlme et al. 2020); Decay sugar-rich substrates (Põlme et al. 2020) | Põlme S, et al. 2020. Fungal Diversity 105:1-16 ((https://doi.org/10.1007/s13225-020-00466-2)) |
| Otu45711 | 1 | k__Fungi(100);p__Ascomycota(100);c__Saccharomycetes(100);o__Saccharomycetales(100);f__Metschnikowiaceae(100);g__Metschnikowia(100);s__Metschnikowia_pulcherrima(100); | Metschnikowia | 13 | Saprotroph | \|Undefined Saprotroph\| | Yeast | NULL | Probable | As nectar-tap_saprotroph (Põlme et al. 2020); Decay sugar-rich substrates (Põlme et al. 2020) | Põlme S, et al. 2020. Fungal Diversity 105:1-16 ((https://doi.org/10.1007/s13225-020-00466-2)) |
| Otu45712 | 1 | k__Fungi(100);p__Ascomycota(100);c__Saccharomycetes(100);o__Saccharomycetales(100);f__Metschnikowiaceae(100);g__Metschnikowia(100);s__Metschnikowia_pulcherrima(100); | Metschnikowia | 13 | Saprotroph | \|Undefined Saprotroph\| | Yeast | NULL | Probable | As nectar-tap_saprotroph (Põlme et al. 2020); Decay sugar-rich substrates (Põlme et al. 2020) | Põlme S, et al. 2020. Fungal Diversity 105:1-16 ((https://doi.org/10.1007/s13225-020-00466-2)) |
| Otu45713 | 1 | k__Fungi(100);p__Ascomycota(100);c__Saccharomycetes(100);o__Saccharomycetales(100);f__Metschnikowiaceae(100);g__Metschnikowia(100);s__Metschnikowia_pulcherrima(100); | Metschnikowia | 13 | Saprotroph | \|Undefined Saprotroph\| | Yeast | NULL | Probable | As nectar-tap_saprotroph (Põlme et al. 2020); Decay sugar-rich substrates (Põlme et al. 2020) | Põlme S, et al. 2020. Fungal Diversity 105:1-16 ((https://doi.org/10.1007/s13225-020-00466-2)) |
| Otu45714 | 1 | k__Fungi(100);p__Ascomycota(100);c__Saccharomycetes(100);o__Saccharomycetales(100);f__Metschnikowiaceae(100);g__Metschnikowia(100);s__Metschnikowia_viticola(100); | Metschnikowia | 13 | Saprotroph | \|Undefined Saprotroph\| | Yeast | NULL | Probable | As nectar-tap_saprotroph (Põlme et al. 2020); Decay sugar-rich substrates (Põlme et al. 2020) | Põlme S, et al. 2020. Fungal Diversity 105:1-16 ((https://doi.org/10.1007/s13225-020-00466-2)) |
| Otu45716 | 1 | k__Fungi(100);p__Ascomycota(100);c__Saccharomycetes(100);o__Saccharomycetales(100);f__Metschnikowiaceae(100);g__Metschnikowia(100);s__Metschnikowia_viticola(100); | Metschnikowia | 13 | Saprotroph | \|Undefined Saprotroph\| | Yeast | NULL | Probable | As nectar-tap_saprotroph (Põlme et al. 2020); Decay sugar-rich substrates (Põlme et al. 2020) | Põlme S, et al. 2020. Fungal Diversity 105:1-16 ((https://doi.org/10.1007/s13225-020-00466-2)) |
| Otu45717 | 1 | k__Fungi(100);p__Ascomycota(100);c__Saccharomycetes(100);o__Saccharomycetales(100);f__Metschnikowiaceae(100);g__Metschnikowia(100);s__Metschnikowia_viticola(100); | Metschnikowia | 13 | Saprotroph | \|Undefined Saprotroph\| | Yeast | NULL | Probable | As nectar-tap_saprotroph (Põlme et al. 2020); Decay sugar-rich substrates (Põlme et al. 2020) | Põlme S, et al. 2020. Fungal Diversity 105:1-16 ((https://doi.org/10.1007/s13225-020-00466-2)) |
| Otu45718 | 1 | k__Fungi(100);p__Ascomycota(100);c__Saccharomycetes(100);o__Saccharomycetales(100);f__Metschnikowiaceae(100);g__Metschnikowia(100);s__Metschnikowia_pulcherrima(100); | Metschnikowia | 13 | Saprotroph | \|Undefined Saprotroph\| | Yeast | NULL | Probable | As nectar-tap_saprotroph (Põlme et al. 2020); Decay sugar-rich substrates (Põlme et al. 2020) | Põlme S, et al. 2020. Fungal Diversity 105:1-16 ((https://doi.org/10.1007/s13225-020-00466-2)) |
| Otu45719 | 1 | k__Fungi(100);p__Ascomycota(100);c__Saccharomycetes(100);o__Saccharomycetales(100);f__Metschnikowiaceae(100);g__Metschnikowia(100);s__Metschnikowia_pulcherrima(100); | Metschnikowia | 13 | Saprotroph | \|Undefined Saprotroph\| | Yeast | NULL | Probable | As nectar-tap_saprotroph (Põlme et al. 2020); Decay sugar-rich substrates (Põlme et al. 2020) | Põlme S, et al. 2020. Fungal Diversity 105:1-16 ((https://doi.org/10.1007/s13225-020-00466-2)) |
| Otu45720 | 1 | k__Fungi(100);p__Ascomycota(100);c__Saccharomycetes(100);o__Saccharomycetales(100);f__Metschnikowiaceae(100);g__Metschnikowia(100);g__Metschnikowia_unclassified(100); | Metschnikowia | 13 | Saprotroph | \|Undefined Saprotroph\| | Yeast | NULL | Probable | As nectar-tap_saprotroph (Põlme et al. 2020); Decay sugar-rich substrates (Põlme et al. 2020) | Põlme S, et al. 2020. Fungal Diversity 105:1-16 ((https://doi.org/10.1007/s13225-020-00466-2)) |
| Otu45721 | 1 | k__Fungi(100);p__Ascomycota(100);c__Saccharomycetes(100);o__Saccharomycetales(100);f__Metschnikowiaceae(100);g__Metschnikowia(100);s__Metschnikowia_pulcherrima(100); | Metschnikowia | 13 | Saprotroph | \|Undefined Saprotroph\| | Yeast | NULL | Probable | As nectar-tap_saprotroph (Põlme et al. 2020); Decay sugar-rich substrates (Põlme et al. 2020) | Põlme S, et al. 2020. Fungal Diversity 105:1-16 ((https://doi.org/10.1007/s13225-020-00466-2)) |
| Otu45723 | 1 | k__Fungi(100);p__Ascomycota(100);c__Saccharomycetes(100);o__Saccharomycetales(100);f__Metschnikowiaceae(100);g__Metschnikowia(100);s__Metschnikowia_pulcherrima(100); | Metschnikowia | 13 | Saprotroph | \|Undefined Saprotroph\| | Yeast | NULL | Probable | As nectar-tap_saprotroph (Põlme et al. 2020); Decay sugar-rich substrates (Põlme et al. 2020) | Põlme S, et al. 2020. Fungal Diversity 105:1-16 ((https://doi.org/10.1007/s13225-020-00466-2)) |
| Otu45724 | 1 | k__Fungi(100);p__Ascomycota(100);c__Saccharomycetes(100);o__Saccharomycetales(100);f__Metschnikowiaceae(100);g__Metschnikowia(100);s__Metschnikowia_pulcherrima(100); | Metschnikowia | 13 | Saprotroph | \|Undefined Saprotroph\| | Yeast | NULL | Probable | As nectar-tap_saprotroph (Põlme et al. 2020); Decay sugar-rich substrates (Põlme et al. 2020) | Põlme S, et al. 2020. Fungal Diversity 105:1-16 ((https://doi.org/10.1007/s13225-020-00466-2)) |
| Otu45725 | 1 | k__Fungi(100);p__Ascomycota(100);c__Saccharomycetes(100);o__Saccharomycetales(100);f__Metschnikowiaceae(100);g__Metschnikowia(100);s__Metschnikowia_pulcherrima(100); | Metschnikowia | 13 | Saprotroph | \|Undefined Saprotroph\| | Yeast | NULL | Probable | As nectar-tap_saprotroph (Põlme et al. 2020); Decay sugar-rich substrates (Põlme et al. 2020) | Põlme S, et al. 2020. Fungal Diversity 105:1-16 ((https://doi.org/10.1007/s13225-020-00466-2)) |
| Otu45726 | 1 | k__Fungi(100);p__Ascomycota(100);c__Saccharomycetes(100);o__Saccharomycetales(100);f__Metschnikowiaceae(100);g__Metschnikowia(100);s__Metschnikowia_pulcherrima(100); | Metschnikowia | 13 | Saprotroph | \|Undefined Saprotroph\| | Yeast | NULL | Probable | As nectar-tap_saprotroph (Põlme et al. 2020); Decay sugar-rich substrates (Põlme et al. 2020) | Põlme S, et al. 2020. Fungal Diversity 105:1-16 ((https://doi.org/10.1007/s13225-020-00466-2)) |
| Otu45727 | 1 | k__Fungi(100);p__Ascomycota(100);c__Saccharomycetes(100);o__Saccharomycetales(100);f__Metschnikowiaceae(100);g__Metschnikowia(100);s__Metschnikowia_pulcherrima(100); | Metschnikowia | 13 | Saprotroph | \|Undefined Saprotroph\| | Yeast | NULL | Probable | As nectar-tap_saprotroph (Põlme et al. 2020); Decay sugar-rich substrates (Põlme et al. 2020) | Põlme S, et al. 2020. Fungal Diversity 105:1-16 ((https://doi.org/10.1007/s13225-020-00466-2)) |
| Otu45728 | 1 | k__Fungi(100);p__Ascomycota(100);c__Saccharomycetes(100);o__Saccharomycetales(100);f__Metschnikowiaceae(100);g__Metschnikowia(100);s__Metschnikowia_pulcherrima(100); | Metschnikowia | 13 | Saprotroph | \|Undefined Saprotroph\| | Yeast | NULL | Probable | As nectar-tap_saprotroph (Põlme et al. 2020); Decay sugar-rich substrates (Põlme et al. 2020) | Põlme S, et al. 2020. Fungal Diversity 105:1-16 ((https://doi.org/10.1007/s13225-020-00466-2)) |
| Otu45729 | 1 | k__Fungi(100);p__Ascomycota(100);c__Saccharomycetes(100);o__Saccharomycetales(100);f__Metschnikowiaceae(100);g__Metschnikowia(100);g__Metschnikowia_unclassified(100); | Metschnikowia | 13 | Saprotroph | \|Undefined Saprotroph\| | Yeast | NULL | Probable | As nectar-tap_saprotroph (Põlme et al. 2020); Decay sugar-rich substrates (Põlme et al. 2020) | Põlme S, et al. 2020. Fungal Diversity 105:1-16 ((https://doi.org/10.1007/s13225-020-00466-2)) |
| Otu45730 | 1 | k__Fungi(100);p__Ascomycota(100);c__Saccharomycetes(100);o__Saccharomycetales(100);f__Metschnikowiaceae(100);g__Metschnikowia(100);s__Metschnikowia_pulcherrima(100); | Metschnikowia | 13 | Saprotroph | \|Undefined Saprotroph\| | Yeast | NULL | Probable | As nectar-tap_saprotroph (Põlme et al. 2020); Decay sugar-rich substrates (Põlme et al. 2020) | Põlme S, et al. 2020. Fungal Diversity 105:1-16 ((https://doi.org/10.1007/s13225-020-00466-2)) |
| Otu45731 | 1 | k__Fungi(100);p__Ascomycota(100);c__Saccharomycetes(100);o__Saccharomycetales(100);f__Metschnikowiaceae(100);g__Metschnikowia(100);s__Metschnikowia_pulcherrima(100); | Metschnikowia | 13 | Saprotroph | \|Undefined Saprotroph\| | Yeast | NULL | Probable | As nectar-tap_saprotroph (Põlme et al. 2020); Decay sugar-rich substrates (Põlme et al. 2020) | Põlme S, et al. 2020. Fungal Diversity 105:1-16 ((https://doi.org/10.1007/s13225-020-00466-2)) |
| Otu45732 | 1 | k__Fungi(100);p__Ascomycota(100);c__Saccharomycetes(100);o__Saccharomycetales(100);f__Metschnikowiaceae(100);g__Metschnikowia(100);g__Metschnikowia_unclassified(100); | Metschnikowia | 13 | Saprotroph | \|Undefined Saprotroph\| | Yeast | NULL | Probable | As nectar-tap_saprotroph (Põlme et al. 2020); Decay sugar-rich substrates (Põlme et al. 2020) | Põlme S, et al. 2020. Fungal Diversity 105:1-16 ((https://doi.org/10.1007/s13225-020-00466-2)) |
| Otu45733 | 1 | k__Fungi(100);p__Ascomycota(100);c__Saccharomycetes(100);o__Saccharomycetales(100);f__Metschnikowiaceae(100);g__Metschnikowia(100);s__Metschnikowia_pulcherrima(100); | Metschnikowia | 13 | Saprotroph | \|Undefined Saprotroph\| | Yeast | NULL | Probable | As nectar-tap_saprotroph (Põlme et al. 2020); Decay sugar-rich substrates (Põlme et al. 2020) | Põlme S, et al. 2020. Fungal Diversity 105:1-16 ((https://doi.org/10.1007/s13225-020-00466-2)) |
| Otu45734 | 1 | k__Fungi(100);p__Ascomycota(100);c__Saccharomycetes(100);o__Saccharomycetales(100);f__Metschnikowiaceae(100);g__Metschnikowia(100);s__Metschnikowia_pulcherrima(100); | Metschnikowia | 13 | Saprotroph | \|Undefined Saprotroph\| | Yeast | NULL | Probable | As nectar-tap_saprotroph (Põlme et al. 2020); Decay sugar-rich substrates (Põlme et al. 2020) | Põlme S, et al. 2020. Fungal Diversity 105:1-16 ((https://doi.org/10.1007/s13225-020-00466-2)) |
| Otu45735 | 1 | k__Fungi(100);p__Ascomycota(100);c__Saccharomycetes(100);o__Saccharomycetales(100);f__Metschnikowiaceae(100);g__Metschnikowia(100);g__Metschnikowia_unclassified(100); | Metschnikowia | 13 | Saprotroph | \|Undefined Saprotroph\| | Yeast | NULL | Probable | As nectar-tap_saprotroph (Põlme et al. 2020); Decay sugar-rich substrates (Põlme et al. 2020) | Põlme S, et al. 2020. Fungal Diversity 105:1-16 ((https://doi.org/10.1007/s13225-020-00466-2)) |
| Otu45736 | 1 | k__Fungi(100);p__Ascomycota(100);c__Saccharomycetes(100);o__Saccharomycetales(100);f__Metschnikowiaceae(100);g__Metschnikowia(100);s__Metschnikowia_pulcherrima(100); | Metschnikowia | 13 | Saprotroph | \|Undefined Saprotroph\| | Yeast | NULL | Probable | As nectar-tap_saprotroph (Põlme et al. 2020); Decay sugar-rich substrates (Põlme et al. 2020) | Põlme S, et al. 2020. Fungal Diversity 105:1-16 ((https://doi.org/10.1007/s13225-020-00466-2)) |
| Otu45737 | 1 | k__Fungi(100);p__Ascomycota(100);c__Saccharomycetes(100);o__Saccharomycetales(100);f__Metschnikowiaceae(100);g__Metschnikowia(100);s__Metschnikowia_pulcherrima(100); | Metschnikowia | 13 | Saprotroph | \|Undefined Saprotroph\| | Yeast | NULL | Probable | As nectar-tap_saprotroph (Põlme et al. 2020); Decay sugar-rich substrates (Põlme et al. 2020) | Põlme S, et al. 2020. Fungal Diversity 105:1-16 ((https://doi.org/10.1007/s13225-020-00466-2)) |
| Otu45738 | 1 | k__Fungi(100);p__Ascomycota(100);c__Saccharomycetes(100);o__Saccharomycetales(100);f__Metschnikowiaceae(100);g__Metschnikowia(100);s__Metschnikowia_pulcherrima(100); | Metschnikowia | 13 | Saprotroph | \|Undefined Saprotroph\| | Yeast | NULL | Probable | As nectar-tap_saprotroph (Põlme et al. 2020); Decay sugar-rich substrates (Põlme et al. 2020) | Põlme S, et al. 2020. Fungal Diversity 105:1-16 ((https://doi.org/10.1007/s13225-020-00466-2)) |
| Otu45739 | 1 | k__Fungi(100);p__Ascomycota(100);c__Saccharomycetes(100);o__Saccharomycetales(100);f__Metschnikowiaceae(100);g__Metschnikowia(100);s__Metschnikowia_pulcherrima(100); | Metschnikowia | 13 | Saprotroph | \|Undefined Saprotroph\| | Yeast | NULL | Probable | As nectar-tap_saprotroph (Põlme et al. 2020); Decay sugar-rich substrates (Põlme et al. 2020) | Põlme S, et al. 2020. Fungal Diversity 105:1-16 ((https://doi.org/10.1007/s13225-020-00466-2)) |
| Otu45740 | 1 | k__Fungi(100);p__Ascomycota(100);c__Saccharomycetes(100);o__Saccharomycetales(100);f__Metschnikowiaceae(100);g__Metschnikowia(100);s__Metschnikowia_pulcherrima(100); | Metschnikowia | 13 | Saprotroph | \|Undefined Saprotroph\| | Yeast | NULL | Probable | As nectar-tap_saprotroph (Põlme et al. 2020); Decay sugar-rich substrates (Põlme et al. 2020) | Põlme S, et al. 2020. Fungal Diversity 105:1-16 ((https://doi.org/10.1007/s13225-020-00466-2)) |
| Otu45741 | 1 | k__Fungi(100);p__Ascomycota(100);c__Saccharomycetes(100);o__Saccharomycetales(100);f__Metschnikowiaceae(100);g__Metschnikowia(100);s__Metschnikowia_pulcherrima(100); | Metschnikowia | 13 | Saprotroph | \|Undefined Saprotroph\| | Yeast | NULL | Probable | As nectar-tap_saprotroph (Põlme et al. 2020); Decay sugar-rich substrates (Põlme et al. 2020) | Põlme S, et al. 2020. Fungal Diversity 105:1-16 ((https://doi.org/10.1007/s13225-020-00466-2)) |
| Otu45743 | 1 | k__Fungi(100);p__Ascomycota(100);c__Saccharomycetes(100);o__Saccharomycetales(100);f__Metschnikowiaceae(100);g__Metschnikowia(100);s__Metschnikowia_pulcherrima(100); | Metschnikowia | 13 | Saprotroph | \|Undefined Saprotroph\| | Yeast | NULL | Probable | As nectar-tap_saprotroph (Põlme et al. 2020); Decay sugar-rich substrates (Põlme et al. 2020) | Põlme S, et al. 2020. Fungal Diversity 105:1-16 ((https://doi.org/10.1007/s13225-020-00466-2)) |
| Otu45744 | 1 | k__Fungi(100);p__Ascomycota(100);c__Saccharomycetes(100);o__Saccharomycetales(100);f__Metschnikowiaceae(100);g__Metschnikowia(100);g__Metschnikowia_unclassified(100); | Metschnikowia | 13 | Saprotroph | \|Undefined Saprotroph\| | Yeast | NULL | Probable | As nectar-tap_saprotroph (Põlme et al. 2020); Decay sugar-rich substrates (Põlme et al. 2020) | Põlme S, et al. 2020. Fungal Diversity 105:1-16 ((https://doi.org/10.1007/s13225-020-00466-2)) |
| Otu45745 | 1 | k__Fungi(100);p__Ascomycota(100);c__Sordariomycetes(100);o__Hypocreales(100);f__Nectriaceae(100);g__Gibberella(100);s__Gibberella_intricans(100); | Gibberella | 13 | Pathotroph-Saprotroph-Symbiotroph | Endophyte-\|Plant Pathogen\|-Plant Saprotroph-Undefined Saprotroph | Microfungus | Soft Rot | Probable | As foliar_endophyte (Põlme et al. 2020); As litter_saprotroph (Põlme et al. 2020); Causing damping-off disease (Põlme et al. 2020); Decay leaf, fruit, seed, roots, soil, animal material (Põlme et al. 2020) | Tedersoo L, et al. 2014. Science 346:e1256688 ((https://doi.org/10.1126/science.1256688)); Põlme S, et al. 2020. Fungal Diversity 105:1-16 ((https://doi.org/10.1007/s13225-020-00466-2)) |
| Otu45746 | 1 | k__Fungi(100);p__Ascomycota(100);c__Saccharomycetes(100);o__Saccharomycetales(100);f__Metschnikowiaceae(100);g__Metschnikowia(100);s__Metschnikowia_pulcherrima(100); | Metschnikowia | 13 | Saprotroph | \|Undefined Saprotroph\| | Yeast | NULL | Probable | As nectar-tap_saprotroph (Põlme et al. 2020); Decay sugar-rich substrates (Põlme et al. 2020) | Põlme S, et al. 2020. Fungal Diversity 105:1-16 ((https://doi.org/10.1007/s13225-020-00466-2)) |
| Otu45747 | 1 | k__Fungi(100);p__Ascomycota(100);c__Saccharomycetes(100);o__Saccharomycetales(100);f__Metschnikowiaceae(100);g__Metschnikowia(100);s__Metschnikowia_pulcherrima(100); | Metschnikowia | 13 | Saprotroph | \|Undefined Saprotroph\| | Yeast | NULL | Probable | As nectar-tap_saprotroph (Põlme et al. 2020); Decay sugar-rich substrates (Põlme et al. 2020) | Põlme S, et al. 2020. Fungal Diversity 105:1-16 ((https://doi.org/10.1007/s13225-020-00466-2)) |
| Otu45748 | 1 | k__Fungi(100);p__Ascomycota(100);c__Saccharomycetes(100);o__Saccharomycetales(100);f__Metschnikowiaceae(100);g__Metschnikowia(100);s__Metschnikowia_pulcherrima(100); | Metschnikowia | 13 | Saprotroph | \|Undefined Saprotroph\| | Yeast | NULL | Probable | As nectar-tap_saprotroph (Põlme et al. 2020); Decay sugar-rich substrates (Põlme et al. 2020) | Põlme S, et al. 2020. Fungal Diversity 105:1-16 ((https://doi.org/10.1007/s13225-020-00466-2)) |
| Otu45749 | 1 | k__Fungi(100);p__Ascomycota(100);c__Saccharomycetes(100);o__Saccharomycetales(100);f__Metschnikowiaceae(100);g__Metschnikowia(100);s__Metschnikowia_pulcherrima(100); | Metschnikowia | 13 | Saprotroph | \|Undefined Saprotroph\| | Yeast | NULL | Probable | As nectar-tap_saprotroph (Põlme et al. 2020); Decay sugar-rich substrates (Põlme et al. 2020) | Põlme S, et al. 2020. Fungal Diversity 105:1-16 ((https://doi.org/10.1007/s13225-020-00466-2)) |
| Otu45750 | 1 | k__Fungi(100);p__Ascomycota(100);c__Saccharomycetes(100);o__Saccharomycetales(100);f__Metschnikowiaceae(100);g__Metschnikowia(100);s__Metschnikowia_pulcherrima(100); | Metschnikowia | 13 | Saprotroph | \|Undefined Saprotroph\| | Yeast | NULL | Probable | As nectar-tap_saprotroph (Põlme et al. 2020); Decay sugar-rich substrates (Põlme et al. 2020) | Põlme S, et al. 2020. Fungal Diversity 105:1-16 ((https://doi.org/10.1007/s13225-020-00466-2)) |
| Otu45751 | 1 | k__Fungi(100);p__Ascomycota(100);c__Saccharomycetes(100);o__Saccharomycetales(100);f__Metschnikowiaceae(100);g__Metschnikowia(100);s__Metschnikowia_pulcherrima(100); | Metschnikowia | 13 | Saprotroph | \|Undefined Saprotroph\| | Yeast | NULL | Probable | As nectar-tap_saprotroph (Põlme et al. 2020); Decay sugar-rich substrates (Põlme et al. 2020) | Põlme S, et al. 2020. Fungal Diversity 105:1-16 ((https://doi.org/10.1007/s13225-020-00466-2)) |
| Otu45752 | 1 | k__Fungi(100);p__Ascomycota(100);c__Saccharomycetes(100);o__Saccharomycetales(100);f__Metschnikowiaceae(100);g__Metschnikowia(100);s__Metschnikowia_pulcherrima(100); | Metschnikowia | 13 | Saprotroph | \|Undefined Saprotroph\| | Yeast | NULL | Probable | As nectar-tap_saprotroph (Põlme et al. 2020); Decay sugar-rich substrates (Põlme et al. 2020) | Põlme S, et al. 2020. Fungal Diversity 105:1-16 ((https://doi.org/10.1007/s13225-020-00466-2)) |
| Otu45753 | 1 | k__Fungi(100);p__Ascomycota(100);c__Saccharomycetes(100);o__Saccharomycetales(100);f__Metschnikowiaceae(100);g__Metschnikowia(100);s__Metschnikowia_pulcherrima(100); | Metschnikowia | 13 | Saprotroph | \|Undefined Saprotroph\| | Yeast | NULL | Probable | As nectar-tap_saprotroph (Põlme et al. 2020); Decay sugar-rich substrates (Põlme et al. 2020) | Põlme S, et al. 2020. Fungal Diversity 105:1-16 ((https://doi.org/10.1007/s13225-020-00466-2)) |
| Otu45754 | 1 | k__Fungi(100);p__Ascomycota(100);c__Saccharomycetes(100);o__Saccharomycetales(100);f__Metschnikowiaceae(100);g__Metschnikowia(100);s__Metschnikowia_pulcherrima(100); | Metschnikowia | 13 | Saprotroph | \|Undefined Saprotroph\| | Yeast | NULL | Probable | As nectar-tap_saprotroph (Põlme et al. 2020); Decay sugar-rich substrates (Põlme et al. 2020) | Põlme S, et al. 2020. Fungal Diversity 105:1-16 ((https://doi.org/10.1007/s13225-020-00466-2)) |
| Otu45755 | 1 | k__Fungi(100);p__Ascomycota(100);c__Saccharomycetes(100);o__Saccharomycetales(100);f__Metschnikowiaceae(100);g__Metschnikowia(100);s__Metschnikowia_pulcherrima(100); | Metschnikowia | 13 | Saprotroph | \|Undefined Saprotroph\| | Yeast | NULL | Probable | As nectar-tap_saprotroph (Põlme et al. 2020); Decay sugar-rich substrates (Põlme et al. 2020) | Põlme S, et al. 2020. Fungal Diversity 105:1-16 ((https://doi.org/10.1007/s13225-020-00466-2)) |
| Otu45756 | 1 | k__Fungi(100);p__Ascomycota(100);c__Saccharomycetes(100);o__Saccharomycetales(100);f__Metschnikowiaceae(100);g__Metschnikowia(100);s__Metschnikowia_pulcherrima(100); | Metschnikowia | 13 | Saprotroph | \|Undefined Saprotroph\| | Yeast | NULL | Probable | As nectar-tap_saprotroph (Põlme et al. 2020); Decay sugar-rich substrates (Põlme et al. 2020) | Põlme S, et al. 2020. Fungal Diversity 105:1-16 ((https://doi.org/10.1007/s13225-020-00466-2)) |
| Otu45757 | 1 | k__Fungi(100);p__Ascomycota(100);c__Saccharomycetes(100);o__Saccharomycetales(100);f__Metschnikowiaceae(100);g__Metschnikowia(100);s__Metschnikowia_pulcherrima(100); | Metschnikowia | 13 | Saprotroph | \|Undefined Saprotroph\| | Yeast | NULL | Probable | As nectar-tap_saprotroph (Põlme et al. 2020); Decay sugar-rich substrates (Põlme et al. 2020) | Põlme S, et al. 2020. Fungal Diversity 105:1-16 ((https://doi.org/10.1007/s13225-020-00466-2)) |
| Otu45758 | 1 | k__Fungi(100);p__Ascomycota(100);c__Saccharomycetes(100);o__Saccharomycetales(100);f__Metschnikowiaceae(100);g__Metschnikowia(100);s__Metschnikowia_pulcherrima(100); | Metschnikowia | 13 | Saprotroph | \|Undefined Saprotroph\| | Yeast | NULL | Probable | As nectar-tap_saprotroph (Põlme et al. 2020); Decay sugar-rich substrates (Põlme et al. 2020) | Põlme S, et al. 2020. Fungal Diversity 105:1-16 ((https://doi.org/10.1007/s13225-020-00466-2)) |
| Otu45759 | 1 | k__Fungi(100);p__Ascomycota(100);c__Saccharomycetes(100);o__Saccharomycetales(100);f__Metschnikowiaceae(100);g__Metschnikowia(100);g__Metschnikowia_unclassified(100); | Metschnikowia | 13 | Saprotroph | \|Undefined Saprotroph\| | Yeast | NULL | Probable | As nectar-tap_saprotroph (Põlme et al. 2020); Decay sugar-rich substrates (Põlme et al. 2020) | Põlme S, et al. 2020. Fungal Diversity 105:1-16 ((https://doi.org/10.1007/s13225-020-00466-2)) |
| Otu45760 | 1 | k__Fungi(100);p__Ascomycota(100);c__Saccharomycetes(100);o__Saccharomycetales(100);f__Metschnikowiaceae(100);g__Metschnikowia(100);s__Metschnikowia_pulcherrima(100); | Metschnikowia | 13 | Saprotroph | \|Undefined Saprotroph\| | Yeast | NULL | Probable | As nectar-tap_saprotroph (Põlme et al. 2020); Decay sugar-rich substrates (Põlme et al. 2020) | Põlme S, et al. 2020. Fungal Diversity 105:1-16 ((https://doi.org/10.1007/s13225-020-00466-2)) |
| Otu45761 | 1 | k__Fungi(100);p__Ascomycota(100);c__Saccharomycetes(100);o__Saccharomycetales(100);f__Metschnikowiaceae(100);g__Metschnikowia(100);s__Metschnikowia_pulcherrima(100); | Metschnikowia | 13 | Saprotroph | \|Undefined Saprotroph\| | Yeast | NULL | Probable | As nectar-tap_saprotroph (Põlme et al. 2020); Decay sugar-rich substrates (Põlme et al. 2020) | Põlme S, et al. 2020. Fungal Diversity 105:1-16 ((https://doi.org/10.1007/s13225-020-00466-2)) |
| Otu45762 | 1 | k__Fungi(100);p__Ascomycota(100);c__Saccharomycetes(100);o__Saccharomycetales(100);f__Metschnikowiaceae(100);g__Metschnikowia(100);s__Metschnikowia_pulcherrima(100); | Metschnikowia | 13 | Saprotroph | \|Undefined Saprotroph\| | Yeast | NULL | Probable | As nectar-tap_saprotroph (Põlme et al. 2020); Decay sugar-rich substrates (Põlme et al. 2020) | Põlme S, et al. 2020. Fungal Diversity 105:1-16 ((https://doi.org/10.1007/s13225-020-00466-2)) |
| Otu45763 | 1 | k__Fungi(100);p__Ascomycota(100);c__Saccharomycetes(100);o__Saccharomycetales(100);f__Metschnikowiaceae(100);g__Metschnikowia(100);g__Metschnikowia_unclassified(100); | Metschnikowia | 13 | Saprotroph | \|Undefined Saprotroph\| | Yeast | NULL | Probable | As nectar-tap_saprotroph (Põlme et al. 2020); Decay sugar-rich substrates (Põlme et al. 2020) | Põlme S, et al. 2020. Fungal Diversity 105:1-16 ((https://doi.org/10.1007/s13225-020-00466-2)) |
| Otu45764 | 1 | k__Fungi(100);p__Ascomycota(100);c__Saccharomycetes(100);o__Saccharomycetales(100);f__Metschnikowiaceae(100);g__Metschnikowia(100);g__Metschnikowia_unclassified(100); | Metschnikowia | 13 | Saprotroph | \|Undefined Saprotroph\| | Yeast | NULL | Probable | As nectar-tap_saprotroph (Põlme et al. 2020); Decay sugar-rich substrates (Põlme et al. 2020) | Põlme S, et al. 2020. Fungal Diversity 105:1-16 ((https://doi.org/10.1007/s13225-020-00466-2)) |
| Otu45766 | 1 | k__Fungi(100);p__Ascomycota(100);c__Saccharomycetes(100);o__Saccharomycetales(100);f__Metschnikowiaceae(100);g__Metschnikowia(100);s__Metschnikowia_pulcherrima(100); | Metschnikowia | 13 | Saprotroph | \|Undefined Saprotroph\| | Yeast | NULL | Probable | As nectar-tap_saprotroph (Põlme et al. 2020); Decay sugar-rich substrates (Põlme et al. 2020) | Põlme S, et al. 2020. Fungal Diversity 105:1-16 ((https://doi.org/10.1007/s13225-020-00466-2)) |
| Otu45767 | 1 | k__Fungi(100);p__Ascomycota(100);c__Saccharomycetes(100);o__Saccharomycetales(100);f__Metschnikowiaceae(100);g__Metschnikowia(100);s__Metschnikowia_pulcherrima(100); | Metschnikowia | 13 | Saprotroph | \|Undefined Saprotroph\| | Yeast | NULL | Probable | As nectar-tap_saprotroph (Põlme et al. 2020); Decay sugar-rich substrates (Põlme et al. 2020) | Põlme S, et al. 2020. Fungal Diversity 105:1-16 ((https://doi.org/10.1007/s13225-020-00466-2)) |
| Otu45768 | 1 | k__Fungi(100);p__Ascomycota(100);c__Saccharomycetes(100);o__Saccharomycetales(100);f__Metschnikowiaceae(100);g__Metschnikowia(100);s__Metschnikowia_pulcherrima(100); | Metschnikowia | 13 | Saprotroph | \|Undefined Saprotroph\| | Yeast | NULL | Probable | As nectar-tap_saprotroph (Põlme et al. 2020); Decay sugar-rich substrates (Põlme et al. 2020) | Põlme S, et al. 2020. Fungal Diversity 105:1-16 ((https://doi.org/10.1007/s13225-020-00466-2)) |
| Otu45769 | 1 | k__Fungi(100);p__Ascomycota(100);c__Saccharomycetes(100);o__Saccharomycetales(100);f__Metschnikowiaceae(100);g__Metschnikowia(100);s__Metschnikowia_pulcherrima(100); | Metschnikowia | 13 | Saprotroph | \|Undefined Saprotroph\| | Yeast | NULL | Probable | As nectar-tap_saprotroph (Põlme et al. 2020); Decay sugar-rich substrates (Põlme et al. 2020) | Põlme S, et al. 2020. Fungal Diversity 105:1-16 ((https://doi.org/10.1007/s13225-020-00466-2)) |
| Otu45770 | 1 | k__Fungi(100);p__Ascomycota(100);c__Saccharomycetes(100);o__Saccharomycetales(100);f__Metschnikowiaceae(100);g__Metschnikowia(100);s__Metschnikowia_pulcherrima(100); | Metschnikowia | 13 | Saprotroph | \|Undefined Saprotroph\| | Yeast | NULL | Probable | As nectar-tap_saprotroph (Põlme et al. 2020); Decay sugar-rich substrates (Põlme et al. 2020) | Põlme S, et al. 2020. Fungal Diversity 105:1-16 ((https://doi.org/10.1007/s13225-020-00466-2)) |
| Otu45771 | 1 | k__Fungi(100);p__Ascomycota(100);c__Saccharomycetes(100);o__Saccharomycetales(100);f__Metschnikowiaceae(100);g__Metschnikowia(100);s__Metschnikowia_pulcherrima(100); | Metschnikowia | 13 | Saprotroph | \|Undefined Saprotroph\| | Yeast | NULL | Probable | As nectar-tap_saprotroph (Põlme et al. 2020); Decay sugar-rich substrates (Põlme et al. 2020) | Põlme S, et al. 2020. Fungal Diversity 105:1-16 ((https://doi.org/10.1007/s13225-020-00466-2)) |
| Otu45772 | 1 | k__Fungi(100);p__Ascomycota(100);c__Saccharomycetes(100);o__Saccharomycetales(100);f__Metschnikowiaceae(100);g__Metschnikowia(100);s__Metschnikowia_pulcherrima(100); | Metschnikowia | 13 | Saprotroph | \|Undefined Saprotroph\| | Yeast | NULL | Probable | As nectar-tap_saprotroph (Põlme et al. 2020); Decay sugar-rich substrates (Põlme et al. 2020) | Põlme S, et al. 2020. Fungal Diversity 105:1-16 ((https://doi.org/10.1007/s13225-020-00466-2)) |
| Otu45773 | 1 | k__Fungi(100);p__Ascomycota(100);c__Saccharomycetes(100);o__Saccharomycetales(100);f__Metschnikowiaceae(100);g__Metschnikowia(100);s__Metschnikowia_pulcherrima(100); | Metschnikowia | 13 | Saprotroph | \|Undefined Saprotroph\| | Yeast | NULL | Probable | As nectar-tap_saprotroph (Põlme et al. 2020); Decay sugar-rich substrates (Põlme et al. 2020) | Põlme S, et al. 2020. Fungal Diversity 105:1-16 ((https://doi.org/10.1007/s13225-020-00466-2)) |
| Otu45774 | 1 | k__Fungi(100);p__Ascomycota(100);c__Saccharomycetes(100);o__Saccharomycetales(100);f__Metschnikowiaceae(100);g__Metschnikowia(100);s__Metschnikowia_pulcherrima(100); | Metschnikowia | 13 | Saprotroph | \|Undefined Saprotroph\| | Yeast | NULL | Probable | As nectar-tap_saprotroph (Põlme et al. 2020); Decay sugar-rich substrates (Põlme et al. 2020) | Põlme S, et al. 2020. Fungal Diversity 105:1-16 ((https://doi.org/10.1007/s13225-020-00466-2)) |
| Otu45775 | 1 | k__Fungi(100);p__Ascomycota(100);c__Saccharomycetes(100);o__Saccharomycetales(100);f__Metschnikowiaceae(100);g__Metschnikowia(100);g__Metschnikowia_unclassified(100); | Metschnikowia | 13 | Saprotroph | \|Undefined Saprotroph\| | Yeast | NULL | Probable | As nectar-tap_saprotroph (Põlme et al. 2020); Decay sugar-rich substrates (Põlme et al. 2020) | Põlme S, et al. 2020. Fungal Diversity 105:1-16 ((https://doi.org/10.1007/s13225-020-00466-2)) |
| Otu45776 | 1 | k__Fungi(100);p__Ascomycota(100);c__Saccharomycetes(100);o__Saccharomycetales(100);f__Metschnikowiaceae(100);g__Metschnikowia(100);s__Metschnikowia_pulcherrima(100); | Metschnikowia | 13 | Saprotroph | \|Undefined Saprotroph\| | Yeast | NULL | Probable | As nectar-tap_saprotroph (Põlme et al. 2020); Decay sugar-rich substrates (Põlme et al. 2020) | Põlme S, et al. 2020. Fungal Diversity 105:1-16 ((https://doi.org/10.1007/s13225-020-00466-2)) |
| Otu45777 | 1 | k__Fungi(100);p__Ascomycota(100);c__Saccharomycetes(100);o__Saccharomycetales(100);f__Metschnikowiaceae(100);g__Metschnikowia(100);s__Metschnikowia_pulcherrima(100); | Metschnikowia | 13 | Saprotroph | \|Undefined Saprotroph\| | Yeast | NULL | Probable | As nectar-tap_saprotroph (Põlme et al. 2020); Decay sugar-rich substrates (Põlme et al. 2020) | Põlme S, et al. 2020. Fungal Diversity 105:1-16 ((https://doi.org/10.1007/s13225-020-00466-2)) |
| Otu45778 | 1 | k__Fungi(100);p__Ascomycota(100);c__Saccharomycetes(100);o__Saccharomycetales(100);f__Metschnikowiaceae(100);g__Metschnikowia(100);s__Metschnikowia_pulcherrima(100); | Metschnikowia | 13 | Saprotroph | \|Undefined Saprotroph\| | Yeast | NULL | Probable | As nectar-tap_saprotroph (Põlme et al. 2020); Decay sugar-rich substrates (Põlme et al. 2020) | Põlme S, et al. 2020. Fungal Diversity 105:1-16 ((https://doi.org/10.1007/s13225-020-00466-2)) |
| Otu45779 | 1 | k__Fungi(100);p__Ascomycota(100);c__Saccharomycetes(100);o__Saccharomycetales(100);f__Metschnikowiaceae(100);g__Metschnikowia(100);s__Metschnikowia_pulcherrima(100); | Metschnikowia | 13 | Saprotroph | \|Undefined Saprotroph\| | Yeast | NULL | Probable | As nectar-tap_saprotroph (Põlme et al. 2020); Decay sugar-rich substrates (Põlme et al. 2020) | Põlme S, et al. 2020. Fungal Diversity 105:1-16 ((https://doi.org/10.1007/s13225-020-00466-2)) |
| Otu45780 | 1 | k__Fungi(100);p__Ascomycota(100);c__Saccharomycetes(100);o__Saccharomycetales(100);f__Metschnikowiaceae(100);g__Metschnikowia(100);s__Metschnikowia_pulcherrima(100); | Metschnikowia | 13 | Saprotroph | \|Undefined Saprotroph\| | Yeast | NULL | Probable | As nectar-tap_saprotroph (Põlme et al. 2020); Decay sugar-rich substrates (Põlme et al. 2020) | Põlme S, et al. 2020. Fungal Diversity 105:1-16 ((https://doi.org/10.1007/s13225-020-00466-2)) |
| Otu45781 | 1 | k__Fungi(100);p__Ascomycota(100);c__Saccharomycetes(100);o__Saccharomycetales(100);f__Metschnikowiaceae(100);g__Metschnikowia(100);g__Metschnikowia_unclassified(100); | Metschnikowia | 13 | Saprotroph | \|Undefined Saprotroph\| | Yeast | NULL | Probable | As nectar-tap_saprotroph (Põlme et al. 2020); Decay sugar-rich substrates (Põlme et al. 2020) | Põlme S, et al. 2020. Fungal Diversity 105:1-16 ((https://doi.org/10.1007/s13225-020-00466-2)) |
| Otu45782 | 1 | k__Fungi(100);p__Ascomycota(100);c__Saccharomycetes(100);o__Saccharomycetales(100);f__Metschnikowiaceae(100);g__Metschnikowia(100);g__Metschnikowia_unclassified(100); | Metschnikowia | 13 | Saprotroph | \|Undefined Saprotroph\| | Yeast | NULL | Probable | As nectar-tap_saprotroph (Põlme et al. 2020); Decay sugar-rich substrates (Põlme et al. 2020) | Põlme S, et al. 2020. Fungal Diversity 105:1-16 ((https://doi.org/10.1007/s13225-020-00466-2)) |
| Otu45783 | 1 | k__Fungi(100);p__Ascomycota(100);c__Saccharomycetes(100);o__Saccharomycetales(100);f__Metschnikowiaceae(100);g__Metschnikowia(100);s__Metschnikowia_pulcherrima(100); | Metschnikowia | 13 | Saprotroph | \|Undefined Saprotroph\| | Yeast | NULL | Probable | As nectar-tap_saprotroph (Põlme et al. 2020); Decay sugar-rich substrates (Põlme et al. 2020) | Põlme S, et al. 2020. Fungal Diversity 105:1-16 ((https://doi.org/10.1007/s13225-020-00466-2)) |
| Otu45784 | 1 | k__Fungi(100);p__Ascomycota(100);c__Saccharomycetes(100);o__Saccharomycetales(100);f__Metschnikowiaceae(100);g__Metschnikowia(100);s__Metschnikowia_pulcherrima(100); | Metschnikowia | 13 | Saprotroph | \|Undefined Saprotroph\| | Yeast | NULL | Probable | As nectar-tap_saprotroph (Põlme et al. 2020); Decay sugar-rich substrates (Põlme et al. 2020) | Põlme S, et al. 2020. Fungal Diversity 105:1-16 ((https://doi.org/10.1007/s13225-020-00466-2)) |
| Otu45785 | 1 | k__Fungi(100);p__Ascomycota(100);c__Saccharomycetes(100);o__Saccharomycetales(100);f__Metschnikowiaceae(100);g__Metschnikowia(100);s__Metschnikowia_pulcherrima(100); | Metschnikowia | 13 | Saprotroph | \|Undefined Saprotroph\| | Yeast | NULL | Probable | As nectar-tap_saprotroph (Põlme et al. 2020); Decay sugar-rich substrates (Põlme et al. 2020) | Põlme S, et al. 2020. Fungal Diversity 105:1-16 ((https://doi.org/10.1007/s13225-020-00466-2)) |
| Otu45786 | 1 | k__Fungi(100);p__Ascomycota(100);c__Saccharomycetes(100);o__Saccharomycetales(100);f__Metschnikowiaceae(100);g__Metschnikowia(100);s__Metschnikowia_pulcherrima(100); | Metschnikowia | 13 | Saprotroph | \|Undefined Saprotroph\| | Yeast | NULL | Probable | As nectar-tap_saprotroph (Põlme et al. 2020); Decay sugar-rich substrates (Põlme et al. 2020) | Põlme S, et al. 2020. Fungal Diversity 105:1-16 ((https://doi.org/10.1007/s13225-020-00466-2)) |
| Otu45787 | 1 | k__Fungi(100);p__Ascomycota(100);c__Saccharomycetes(100);o__Saccharomycetales(100);f__Metschnikowiaceae(100);g__Metschnikowia(100);s__Metschnikowia_pulcherrima(100); | Metschnikowia | 13 | Saprotroph | \|Undefined Saprotroph\| | Yeast | NULL | Probable | As nectar-tap_saprotroph (Põlme et al. 2020); Decay sugar-rich substrates (Põlme et al. 2020) | Põlme S, et al. 2020. Fungal Diversity 105:1-16 ((https://doi.org/10.1007/s13225-020-00466-2)) |
| Otu45788 | 1 | k__Fungi(100);p__Ascomycota(100);c__Saccharomycetes(100);o__Saccharomycetales(100);f__Metschnikowiaceae(100);g__Metschnikowia(100);s__Metschnikowia_pulcherrima(100); | Metschnikowia | 13 | Saprotroph | \|Undefined Saprotroph\| | Yeast | NULL | Probable | As nectar-tap_saprotroph (Põlme et al. 2020); Decay sugar-rich substrates (Põlme et al. 2020) | Põlme S, et al. 2020. Fungal Diversity 105:1-16 ((https://doi.org/10.1007/s13225-020-00466-2)) |
| Otu45789 | 1 | k__Fungi(100);p__Ascomycota(100);c__Saccharomycetes(100);o__Saccharomycetales(100);f__Metschnikowiaceae(100);g__Metschnikowia(100);s__Metschnikowia_pulcherrima(100); | Metschnikowia | 13 | Saprotroph | \|Undefined Saprotroph\| | Yeast | NULL | Probable | As nectar-tap_saprotroph (Põlme et al. 2020); Decay sugar-rich substrates (Põlme et al. 2020) | Põlme S, et al. 2020. Fungal Diversity 105:1-16 ((https://doi.org/10.1007/s13225-020-00466-2)) |
| Otu45790 | 1 | k__Fungi(100);p__Ascomycota(100);c__Saccharomycetes(100);o__Saccharomycetales(100);f__Metschnikowiaceae(100);g__Metschnikowia(100);s__Metschnikowia_pulcherrima(100); | Metschnikowia | 13 | Saprotroph | \|Undefined Saprotroph\| | Yeast | NULL | Probable | As nectar-tap_saprotroph (Põlme et al. 2020); Decay sugar-rich substrates (Põlme et al. 2020) | Põlme S, et al. 2020. Fungal Diversity 105:1-16 ((https://doi.org/10.1007/s13225-020-00466-2)) |
| Otu45791 | 1 | k__Fungi(100);p__Ascomycota(100);c__Saccharomycetes(100);o__Saccharomycetales(100);f__Metschnikowiaceae(100);g__Metschnikowia(100);s__Metschnikowia_pulcherrima(100); | Metschnikowia | 13 | Saprotroph | \|Undefined Saprotroph\| | Yeast | NULL | Probable | As nectar-tap_saprotroph (Põlme et al. 2020); Decay sugar-rich substrates (Põlme et al. 2020) | Põlme S, et al. 2020. Fungal Diversity 105:1-16 ((https://doi.org/10.1007/s13225-020-00466-2)) |
| Otu45792 | 1 | k__Fungi(100);p__Ascomycota(100);c__Saccharomycetes(100);o__Saccharomycetales(100);f__Metschnikowiaceae(100);g__Metschnikowia(100);s__Metschnikowia_pulcherrima(100); | Metschnikowia | 13 | Saprotroph | \|Undefined Saprotroph\| | Yeast | NULL | Probable | As nectar-tap_saprotroph (Põlme et al. 2020); Decay sugar-rich substrates (Põlme et al. 2020) | Põlme S, et al. 2020. Fungal Diversity 105:1-16 ((https://doi.org/10.1007/s13225-020-00466-2)) |
| Otu45793 | 1 | k__Fungi(100);p__Ascomycota(100);c__Saccharomycetes(100);o__Saccharomycetales(100);f__Metschnikowiaceae(100);g__Metschnikowia(100);s__Metschnikowia_pulcherrima(100); | Metschnikowia | 13 | Saprotroph | \|Undefined Saprotroph\| | Yeast | NULL | Probable | As nectar-tap_saprotroph (Põlme et al. 2020); Decay sugar-rich substrates (Põlme et al. 2020) | Põlme S, et al. 2020. Fungal Diversity 105:1-16 ((https://doi.org/10.1007/s13225-020-00466-2)) |
| Otu45794 | 1 | k__Fungi(100);p__Ascomycota(100);c__Saccharomycetes(100);o__Saccharomycetales(100);f__Metschnikowiaceae(100);g__Metschnikowia(100);s__Metschnikowia_pulcherrima(100); | Metschnikowia | 13 | Saprotroph | \|Undefined Saprotroph\| | Yeast | NULL | Probable | As nectar-tap_saprotroph (Põlme et al. 2020); Decay sugar-rich substrates (Põlme et al. 2020) | Põlme S, et al. 2020. Fungal Diversity 105:1-16 ((https://doi.org/10.1007/s13225-020-00466-2)) |
| Otu45795 | 1 | k__Fungi(100);p__Ascomycota(100);c__Saccharomycetes(100);o__Saccharomycetales(100);f__Metschnikowiaceae(100);g__Metschnikowia(100);s__Metschnikowia_pulcherrima(100); | Metschnikowia | 13 | Saprotroph | \|Undefined Saprotroph\| | Yeast | NULL | Probable | As nectar-tap_saprotroph (Põlme et al. 2020); Decay sugar-rich substrates (Põlme et al. 2020) | Põlme S, et al. 2020. Fungal Diversity 105:1-16 ((https://doi.org/10.1007/s13225-020-00466-2)) |
| Otu45796 | 1 | k__Fungi(100);p__Ascomycota(100);c__Saccharomycetes(100);o__Saccharomycetales(100);f__Metschnikowiaceae(100);g__Metschnikowia(100);s__Metschnikowia_pulcherrima(100); | Metschnikowia | 13 | Saprotroph | \|Undefined Saprotroph\| | Yeast | NULL | Probable | As nectar-tap_saprotroph (Põlme et al. 2020); Decay sugar-rich substrates (Põlme et al. 2020) | Põlme S, et al. 2020. Fungal Diversity 105:1-16 ((https://doi.org/10.1007/s13225-020-00466-2)) |
| Otu45797 | 1 | k__Fungi(100);p__Ascomycota(100);c__Saccharomycetes(100);o__Saccharomycetales(100);f__Metschnikowiaceae(100);g__Metschnikowia(100);s__Metschnikowia_pulcherrima(100); | Metschnikowia | 13 | Saprotroph | \|Undefined Saprotroph\| | Yeast | NULL | Probable | As nectar-tap_saprotroph (Põlme et al. 2020); Decay sugar-rich substrates (Põlme et al. 2020) | Põlme S, et al. 2020. Fungal Diversity 105:1-16 ((https://doi.org/10.1007/s13225-020-00466-2)) |
| Otu45798 | 1 | k__Fungi(100);p__Ascomycota(100);c__Saccharomycetes(100);o__Saccharomycetales(100);f__Metschnikowiaceae(100);g__Metschnikowia(100);s__Metschnikowia_pulcherrima(100); | Metschnikowia | 13 | Saprotroph | \|Undefined Saprotroph\| | Yeast | NULL | Probable | As nectar-tap_saprotroph (Põlme et al. 2020); Decay sugar-rich substrates (Põlme et al. 2020) | Põlme S, et al. 2020. Fungal Diversity 105:1-16 ((https://doi.org/10.1007/s13225-020-00466-2)) |
| Otu45799 | 1 | k__Fungi(100);p__Ascomycota(100);c__Saccharomycetes(100);o__Saccharomycetales(100);f__Metschnikowiaceae(100);g__Metschnikowia(100);s__Metschnikowia_pulcherrima(100); | Metschnikowia | 13 | Saprotroph | \|Undefined Saprotroph\| | Yeast | NULL | Probable | As nectar-tap_saprotroph (Põlme et al. 2020); Decay sugar-rich substrates (Põlme et al. 2020) | Põlme S, et al. 2020. Fungal Diversity 105:1-16 ((https://doi.org/10.1007/s13225-020-00466-2)) |
| Otu45800 | 1 | k__Fungi(100);p__Ascomycota(100);c__Saccharomycetes(100);o__Saccharomycetales(100);f__Metschnikowiaceae(100);g__Metschnikowia(100);s__Metschnikowia_pulcherrima(100); | Metschnikowia | 13 | Saprotroph | \|Undefined Saprotroph\| | Yeast | NULL | Probable | As nectar-tap_saprotroph (Põlme et al. 2020); Decay sugar-rich substrates (Põlme et al. 2020) | Põlme S, et al. 2020. Fungal Diversity 105:1-16 ((https://doi.org/10.1007/s13225-020-00466-2)) |
| Otu45801 | 1 | k__Fungi(100);p__Ascomycota(100);c__Saccharomycetes(100);o__Saccharomycetales(100);f__Metschnikowiaceae(100);g__Metschnikowia(100);s__Metschnikowia_pulcherrima(100); | Metschnikowia | 13 | Saprotroph | \|Undefined Saprotroph\| | Yeast | NULL | Probable | As nectar-tap_saprotroph (Põlme et al. 2020); Decay sugar-rich substrates (Põlme et al. 2020) | Põlme S, et al. 2020. Fungal Diversity 105:1-16 ((https://doi.org/10.1007/s13225-020-00466-2)) |
| Otu45802 | 1 | k__Fungi(100);p__Ascomycota(100);c__Saccharomycetes(100);o__Saccharomycetales(100);f__Metschnikowiaceae(100);g__Metschnikowia(100);s__Metschnikowia_pulcherrima(100); | Metschnikowia | 13 | Saprotroph | \|Undefined Saprotroph\| | Yeast | NULL | Probable | As nectar-tap_saprotroph (Põlme et al. 2020); Decay sugar-rich substrates (Põlme et al. 2020) | Põlme S, et al. 2020. Fungal Diversity 105:1-16 ((https://doi.org/10.1007/s13225-020-00466-2)) |
| Otu45803 | 1 | k__Fungi(100);p__Ascomycota(100);c__Saccharomycetes(100);o__Saccharomycetales(100);f__Metschnikowiaceae(100);g__Metschnikowia(100);s__Metschnikowia_pulcherrima(100); | Metschnikowia | 13 | Saprotroph | \|Undefined Saprotroph\| | Yeast | NULL | Probable | As nectar-tap_saprotroph (Põlme et al. 2020); Decay sugar-rich substrates (Põlme et al. 2020) | Põlme S, et al. 2020. Fungal Diversity 105:1-16 ((https://doi.org/10.1007/s13225-020-00466-2)) |
| Otu45804 | 1 | k__Fungi(100);p__Ascomycota(100);c__Saccharomycetes(100);o__Saccharomycetales(100);f__Metschnikowiaceae(100);g__Metschnikowia(100);s__Metschnikowia_pulcherrima(100); | Metschnikowia | 13 | Saprotroph | \|Undefined Saprotroph\| | Yeast | NULL | Probable | As nectar-tap_saprotroph (Põlme et al. 2020); Decay sugar-rich substrates (Põlme et al. 2020) | Põlme S, et al. 2020. Fungal Diversity 105:1-16 ((https://doi.org/10.1007/s13225-020-00466-2)) |
| Otu45805 | 1 | k__Fungi(100);p__Ascomycota(100);c__Saccharomycetes(100);o__Saccharomycetales(100);f__Metschnikowiaceae(100);g__Metschnikowia(100);s__Metschnikowia_pulcherrima(100); | Metschnikowia | 13 | Saprotroph | \|Undefined Saprotroph\| | Yeast | NULL | Probable | As nectar-tap_saprotroph (Põlme et al. 2020); Decay sugar-rich substrates (Põlme et al. 2020) | Põlme S, et al. 2020. Fungal Diversity 105:1-16 ((https://doi.org/10.1007/s13225-020-00466-2)) |
| Otu45806 | 1 | k__Fungi(100);p__Ascomycota(100);c__Saccharomycetes(100);o__Saccharomycetales(100);f__Metschnikowiaceae(100);g__Metschnikowia(100);s__Metschnikowia_pulcherrima(100); | Metschnikowia | 13 | Saprotroph | \|Undefined Saprotroph\| | Yeast | NULL | Probable | As nectar-tap_saprotroph (Põlme et al. 2020); Decay sugar-rich substrates (Põlme et al. 2020) | Põlme S, et al. 2020. Fungal Diversity 105:1-16 ((https://doi.org/10.1007/s13225-020-00466-2)) |
| Otu45807 | 1 | k__Fungi(100);p__Ascomycota(100);c__Saccharomycetes(100);o__Saccharomycetales(100);f__Metschnikowiaceae(100);g__Metschnikowia(100);s__Metschnikowia_pulcherrima(100); | Metschnikowia | 13 | Saprotroph | \|Undefined Saprotroph\| | Yeast | NULL | Probable | As nectar-tap_saprotroph (Põlme et al. 2020); Decay sugar-rich substrates (Põlme et al. 2020) | Põlme S, et al. 2020. Fungal Diversity 105:1-16 ((https://doi.org/10.1007/s13225-020-00466-2)) |
| Otu45808 | 1 | k__Fungi(100);p__Ascomycota(100);c__Saccharomycetes(100);o__Saccharomycetales(100);f__Metschnikowiaceae(100);g__Metschnikowia(100);s__Metschnikowia_pulcherrima(100); | Metschnikowia | 13 | Saprotroph | \|Undefined Saprotroph\| | Yeast | NULL | Probable | As nectar-tap_saprotroph (Põlme et al. 2020); Decay sugar-rich substrates (Põlme et al. 2020) | Põlme S, et al. 2020. Fungal Diversity 105:1-16 ((https://doi.org/10.1007/s13225-020-00466-2)) |
| Otu45809 | 1 | k__Fungi(100);p__Ascomycota(100);c__Saccharomycetes(100);o__Saccharomycetales(100);f__Metschnikowiaceae(100);g__Metschnikowia(100);s__Metschnikowia_pulcherrima(100); | Metschnikowia | 13 | Saprotroph | \|Undefined Saprotroph\| | Yeast | NULL | Probable | As nectar-tap_saprotroph (Põlme et al. 2020); Decay sugar-rich substrates (Põlme et al. 2020) | Põlme S, et al. 2020. Fungal Diversity 105:1-16 ((https://doi.org/10.1007/s13225-020-00466-2)) |
| Otu45810 | 1 | k__Fungi(100);p__Ascomycota(100);c__Saccharomycetes(100);o__Saccharomycetales(100);f__Metschnikowiaceae(100);g__Metschnikowia(100);s__Metschnikowia_pulcherrima(100); | Metschnikowia | 13 | Saprotroph | \|Undefined Saprotroph\| | Yeast | NULL | Probable | As nectar-tap_saprotroph (Põlme et al. 2020); Decay sugar-rich substrates (Põlme et al. 2020) | Põlme S, et al. 2020. Fungal Diversity 105:1-16 ((https://doi.org/10.1007/s13225-020-00466-2)) |
| Otu45811 | 1 | k__Fungi(100);p__Basidiomycota(100);c__Tremellomycetes(100);o__Tremellales(100);o__Tremellales_unclassified(100);o__Tremellales_unclassified(100);o__Tremellales_unclassified(100); | Tremellales | 7 | Pathotroph-Saprotroph-Symbiotroph | Fungal Parasite-Undefined Saprotroph | Tremelloid-Yeast | NULL | Possible | The order contains both teleomorphic and anamorphic species, most of the latter being yeasts | Stenkenburg E, et al. 2015. New Phytologist 207:1145-1158 ((https://doi.org/10.1111/nph.13426)) |
| Otu45812 | 1 | k__Fungi(100);p__Ascomycota(100);c__Dothideomycetes(100);o__Pleosporales(100);f__Pleosporales_family_Incertae_sedis(100);g__Massariosphaeria(100);s__Massariosphaeria_sp_FAEII23a(100); | Massariosphaeria | 13 | Saprotroph | Undefined Saprotroph-\|Wood Saprotroph\| | Microfungus | NULL | Probable | Decay wood (Põlme et al. 2020) | Tedersoo L, et al. 2014. Science 346:e1256688 ((https://doi.org/10.1126/science.1256688)); Põlme S, et al. 2020. Fungal Diversity 105:1-16 ((https://doi.org/10.1007/s13225-020-00466-2)) |
| Otu45815 | 1 | k__Fungi(100);p__Ascomycota(100);c__Lecanoromycetes(100);o__Lecanorales(100);f__Stereocaulaceae(100);g__Lepraria(100);s__Lepraria_caesiella(100); | Lepraria | 13 | Symbiotroph | \|Lichenized\| | Thallus | NULL | Highly Probable | NULL | Esslinger TL. 2014. North Dakota State University ((http://www.ndsu.edu/pubweb/~esslinge/chcklst/chcklst7.htm)); Põlme S, et al. 2020. Fungal Diversity 105:1-16 ((https://doi.org/10.1007/s13225-020-00466-2)) |
| Otu45817 | 1 | k__Fungi(100);p__Ascomycota(100);c__Saccharomycetes(100);o__Saccharomycetales(100);f__Metschnikowiaceae(100);g__Metschnikowia(100);s__Metschnikowia_pulcherrima(100); | Metschnikowia | 13 | Saprotroph | \|Undefined Saprotroph\| | Yeast | NULL | Probable | As nectar-tap_saprotroph (Põlme et al. 2020); Decay sugar-rich substrates (Põlme et al. 2020) | Põlme S, et al. 2020. Fungal Diversity 105:1-16 ((https://doi.org/10.1007/s13225-020-00466-2)) |
| Otu45818 | 1 | k__Fungi(100);p__Ascomycota(100);c__Saccharomycetes(100);o__Saccharomycetales(100);f__Metschnikowiaceae(100);g__Metschnikowia(100);s__Metschnikowia_pulcherrima(100); | Metschnikowia | 13 | Saprotroph | \|Undefined Saprotroph\| | Yeast | NULL | Probable | As nectar-tap_saprotroph (Põlme et al. 2020); Decay sugar-rich substrates (Põlme et al. 2020) | Põlme S, et al. 2020. Fungal Diversity 105:1-16 ((https://doi.org/10.1007/s13225-020-00466-2)) |
| Otu45819 | 1 | k__Fungi(100);p__Ascomycota(100);c__Saccharomycetes(100);o__Saccharomycetales(100);f__Metschnikowiaceae(100);g__Metschnikowia(100);s__Metschnikowia_pulcherrima(100); | Metschnikowia | 13 | Saprotroph | \|Undefined Saprotroph\| | Yeast | NULL | Probable | As nectar-tap_saprotroph (Põlme et al. 2020); Decay sugar-rich substrates (Põlme et al. 2020) | Põlme S, et al. 2020. Fungal Diversity 105:1-16 ((https://doi.org/10.1007/s13225-020-00466-2)) |
| Otu45820 | 1 | k__Fungi(100);p__Ascomycota(100);c__Saccharomycetes(100);o__Saccharomycetales(100);f__Metschnikowiaceae(100);g__Metschnikowia(100);s__Metschnikowia_pulcherrima(100); | Metschnikowia | 13 | Saprotroph | \|Undefined Saprotroph\| | Yeast | NULL | Probable | As nectar-tap_saprotroph (Põlme et al. 2020); Decay sugar-rich substrates (Põlme et al. 2020) | Põlme S, et al. 2020. Fungal Diversity 105:1-16 ((https://doi.org/10.1007/s13225-020-00466-2)) |
| Otu45821 | 1 | k__Fungi(100);p__Ascomycota(100);c__Saccharomycetes(100);o__Saccharomycetales(100);f__Metschnikowiaceae(100);g__Metschnikowia(100);s__Metschnikowia_pulcherrima(100); | Metschnikowia | 13 | Saprotroph | \|Undefined Saprotroph\| | Yeast | NULL | Probable | As nectar-tap_saprotroph (Põlme et al. 2020); Decay sugar-rich substrates (Põlme et al. 2020) | Põlme S, et al. 2020. Fungal Diversity 105:1-16 ((https://doi.org/10.1007/s13225-020-00466-2)) |
| Otu45822 | 1 | k__Fungi(100);p__Ascomycota(100);c__Saccharomycetes(100);o__Saccharomycetales(100);f__Metschnikowiaceae(100);g__Metschnikowia(100);s__Metschnikowia_pulcherrima(100); | Metschnikowia | 13 | Saprotroph | \|Undefined Saprotroph\| | Yeast | NULL | Probable | As nectar-tap_saprotroph (Põlme et al. 2020); Decay sugar-rich substrates (Põlme et al. 2020) | Põlme S, et al. 2020. Fungal Diversity 105:1-16 ((https://doi.org/10.1007/s13225-020-00466-2)) |
| Otu45824 | 1 | k__Fungi(100);p__Ascomycota(100);c__Saccharomycetes(100);o__Saccharomycetales(100);f__Metschnikowiaceae(100);g__Metschnikowia(100);s__Metschnikowia_pulcherrima(100); | Metschnikowia | 13 | Saprotroph | \|Undefined Saprotroph\| | Yeast | NULL | Probable | As nectar-tap_saprotroph (Põlme et al. 2020); Decay sugar-rich substrates (Põlme et al. 2020) | Põlme S, et al. 2020. Fungal Diversity 105:1-16 ((https://doi.org/10.1007/s13225-020-00466-2)) |
| Otu45825 | 1 | k__Fungi(100);p__Ascomycota(100);c__Saccharomycetes(100);o__Saccharomycetales(100);f__Metschnikowiaceae(100);g__Metschnikowia(100);s__Metschnikowia_pulcherrima(100); | Metschnikowia | 13 | Saprotroph | \|Undefined Saprotroph\| | Yeast | NULL | Probable | As nectar-tap_saprotroph (Põlme et al. 2020); Decay sugar-rich substrates (Põlme et al. 2020) | Põlme S, et al. 2020. Fungal Diversity 105:1-16 ((https://doi.org/10.1007/s13225-020-00466-2)) |
| Otu45827 | 1 | k__Fungi(100);p__Ascomycota(100);c__Saccharomycetes(100);o__Saccharomycetales(100);f__Metschnikowiaceae(100);g__Metschnikowia(100);s__Metschnikowia_pulcherrima(100); | Metschnikowia | 13 | Saprotroph | \|Undefined Saprotroph\| | Yeast | NULL | Probable | As nectar-tap_saprotroph (Põlme et al. 2020); Decay sugar-rich substrates (Põlme et al. 2020) | Põlme S, et al. 2020. Fungal Diversity 105:1-16 ((https://doi.org/10.1007/s13225-020-00466-2)) |
| Otu45829 | 1 | k__Fungi(100);p__Ascomycota(100);c__Saccharomycetes(100);o__Saccharomycetales(100);f__Metschnikowiaceae(100);g__Metschnikowia(100);s__Metschnikowia_pulcherrima(100); | Metschnikowia | 13 | Saprotroph | \|Undefined Saprotroph\| | Yeast | NULL | Probable | As nectar-tap_saprotroph (Põlme et al. 2020); Decay sugar-rich substrates (Põlme et al. 2020) | Põlme S, et al. 2020. Fungal Diversity 105:1-16 ((https://doi.org/10.1007/s13225-020-00466-2)) |
| Otu45830 | 1 | k__Fungi(100);p__Ascomycota(100);c__Saccharomycetes(100);o__Saccharomycetales(100);f__Metschnikowiaceae(100);g__Metschnikowia(100);s__Metschnikowia_pulcherrima(100); | Metschnikowia | 13 | Saprotroph | \|Undefined Saprotroph\| | Yeast | NULL | Probable | As nectar-tap_saprotroph (Põlme et al. 2020); Decay sugar-rich substrates (Põlme et al. 2020) | Põlme S, et al. 2020. Fungal Diversity 105:1-16 ((https://doi.org/10.1007/s13225-020-00466-2)) |
| Otu45831 | 1 | k__Fungi(100);p__Ascomycota(100);c__Saccharomycetes(100);o__Saccharomycetales(100);f__Metschnikowiaceae(100);g__Metschnikowia(100);s__Metschnikowia_pulcherrima(100); | Metschnikowia | 13 | Saprotroph | \|Undefined Saprotroph\| | Yeast | NULL | Probable | As nectar-tap_saprotroph (Põlme et al. 2020); Decay sugar-rich substrates (Põlme et al. 2020) | Põlme S, et al. 2020. Fungal Diversity 105:1-16 ((https://doi.org/10.1007/s13225-020-00466-2)) |
| Otu45832 | 1 | k__Fungi(100);p__Ascomycota(100);c__Saccharomycetes(100);o__Saccharomycetales(100);f__Metschnikowiaceae(100);g__Metschnikowia(100);g__Metschnikowia_unclassified(100); | Metschnikowia | 13 | Saprotroph | \|Undefined Saprotroph\| | Yeast | NULL | Probable | As nectar-tap_saprotroph (Põlme et al. 2020); Decay sugar-rich substrates (Põlme et al. 2020) | Põlme S, et al. 2020. Fungal Diversity 105:1-16 ((https://doi.org/10.1007/s13225-020-00466-2)) |
| Otu45833 | 1 | k__Fungi(100);p__Ascomycota(100);c__Saccharomycetes(100);o__Saccharomycetales(100);f__Metschnikowiaceae(100);g__Metschnikowia(100);s__Metschnikowia_pulcherrima(100); | Metschnikowia | 13 | Saprotroph | \|Undefined Saprotroph\| | Yeast | NULL | Probable | As nectar-tap_saprotroph (Põlme et al. 2020); Decay sugar-rich substrates (Põlme et al. 2020) | Põlme S, et al. 2020. Fungal Diversity 105:1-16 ((https://doi.org/10.1007/s13225-020-00466-2)) |
| Otu45834 | 1 | k__Fungi(100);p__Ascomycota(100);c__Saccharomycetes(100);o__Saccharomycetales(100);f__Metschnikowiaceae(100);g__Metschnikowia(100);s__Metschnikowia_pulcherrima(100); | Metschnikowia | 13 | Saprotroph | \|Undefined Saprotroph\| | Yeast | NULL | Probable | As nectar-tap_saprotroph (Põlme et al. 2020); Decay sugar-rich substrates (Põlme et al. 2020) | Põlme S, et al. 2020. Fungal Diversity 105:1-16 ((https://doi.org/10.1007/s13225-020-00466-2)) |
| Otu45835 | 1 | k__Fungi(100);p__Ascomycota(100);c__Saccharomycetes(100);o__Saccharomycetales(100);f__Metschnikowiaceae(100);g__Metschnikowia(100);s__Metschnikowia_pulcherrima(100); | Metschnikowia | 13 | Saprotroph | \|Undefined Saprotroph\| | Yeast | NULL | Probable | As nectar-tap_saprotroph (Põlme et al. 2020); Decay sugar-rich substrates (Põlme et al. 2020) | Põlme S, et al. 2020. Fungal Diversity 105:1-16 ((https://doi.org/10.1007/s13225-020-00466-2)) |
| Otu45836 | 1 | k__Fungi(100);p__Ascomycota(100);c__Saccharomycetes(100);o__Saccharomycetales(100);f__Metschnikowiaceae(100);g__Metschnikowia(100);s__Metschnikowia_pulcherrima(100); | Metschnikowia | 13 | Saprotroph | \|Undefined Saprotroph\| | Yeast | NULL | Probable | As nectar-tap_saprotroph (Põlme et al. 2020); Decay sugar-rich substrates (Põlme et al. 2020) | Põlme S, et al. 2020. Fungal Diversity 105:1-16 ((https://doi.org/10.1007/s13225-020-00466-2)) |
| Otu45837 | 1 | k__Fungi(100);p__Ascomycota(100);c__Saccharomycetes(100);o__Saccharomycetales(100);f__Metschnikowiaceae(100);g__Metschnikowia(100);s__Metschnikowia_pulcherrima(100); | Metschnikowia | 13 | Saprotroph | \|Undefined Saprotroph\| | Yeast | NULL | Probable | As nectar-tap_saprotroph (Põlme et al. 2020); Decay sugar-rich substrates (Põlme et al. 2020) | Põlme S, et al. 2020. Fungal Diversity 105:1-16 ((https://doi.org/10.1007/s13225-020-00466-2)) |
| Otu45838 | 1 | k__Fungi(100);p__Ascomycota(100);c__Saccharomycetes(100);o__Saccharomycetales(100);f__Metschnikowiaceae(100);g__Metschnikowia(100);s__Metschnikowia_pulcherrima(100); | Metschnikowia | 13 | Saprotroph | \|Undefined Saprotroph\| | Yeast | NULL | Probable | As nectar-tap_saprotroph (Põlme et al. 2020); Decay sugar-rich substrates (Põlme et al. 2020) | Põlme S, et al. 2020. Fungal Diversity 105:1-16 ((https://doi.org/10.1007/s13225-020-00466-2)) |
| Otu45839 | 1 | k__Fungi(100);p__Ascomycota(100);c__Saccharomycetes(100);o__Saccharomycetales(100);f__Metschnikowiaceae(100);g__Metschnikowia(100);s__Metschnikowia_pulcherrima(100); | Metschnikowia | 13 | Saprotroph | \|Undefined Saprotroph\| | Yeast | NULL | Probable | As nectar-tap_saprotroph (Põlme et al. 2020); Decay sugar-rich substrates (Põlme et al. 2020) | Põlme S, et al. 2020. Fungal Diversity 105:1-16 ((https://doi.org/10.1007/s13225-020-00466-2)) |
| Otu45840 | 1 | k__Fungi(100);p__Ascomycota(100);c__Saccharomycetes(100);o__Saccharomycetales(100);f__Metschnikowiaceae(100);g__Metschnikowia(100);s__Metschnikowia_pulcherrima(100); | Metschnikowia | 13 | Saprotroph | \|Undefined Saprotroph\| | Yeast | NULL | Probable | As nectar-tap_saprotroph (Põlme et al. 2020); Decay sugar-rich substrates (Põlme et al. 2020) | Põlme S, et al. 2020. Fungal Diversity 105:1-16 ((https://doi.org/10.1007/s13225-020-00466-2)) |
| Otu45841 | 1 | k__Fungi(100);p__Ascomycota(100);c__Saccharomycetes(100);o__Saccharomycetales(100);f__Metschnikowiaceae(100);g__Metschnikowia(100);s__Metschnikowia_pulcherrima(100); | Metschnikowia | 13 | Saprotroph | \|Undefined Saprotroph\| | Yeast | NULL | Probable | As nectar-tap_saprotroph (Põlme et al. 2020); Decay sugar-rich substrates (Põlme et al. 2020) | Põlme S, et al. 2020. Fungal Diversity 105:1-16 ((https://doi.org/10.1007/s13225-020-00466-2)) |
| Otu45842 | 1 | k__Fungi(100);p__Ascomycota(100);c__Saccharomycetes(100);o__Saccharomycetales(100);f__Metschnikowiaceae(100);g__Metschnikowia(100);s__Metschnikowia_pulcherrima(100); | Metschnikowia | 13 | Saprotroph | \|Undefined Saprotroph\| | Yeast | NULL | Probable | As nectar-tap_saprotroph (Põlme et al. 2020); Decay sugar-rich substrates (Põlme et al. 2020) | Põlme S, et al. 2020. Fungal Diversity 105:1-16 ((https://doi.org/10.1007/s13225-020-00466-2)) |
| Otu45843 | 1 | k__Fungi(100);p__Ascomycota(100);c__Saccharomycetes(100);o__Saccharomycetales(100);f__Metschnikowiaceae(100);g__Metschnikowia(100);s__Metschnikowia_pulcherrima(100); | Metschnikowia | 13 | Saprotroph | \|Undefined Saprotroph\| | Yeast | NULL | Probable | As nectar-tap_saprotroph (Põlme et al. 2020); Decay sugar-rich substrates (Põlme et al. 2020) | Põlme S, et al. 2020. Fungal Diversity 105:1-16 ((https://doi.org/10.1007/s13225-020-00466-2)) |
| Otu45844 | 1 | k__Fungi(100);p__Ascomycota(100);c__Saccharomycetes(100);o__Saccharomycetales(100);f__Metschnikowiaceae(100);g__Metschnikowia(100);s__Metschnikowia_pulcherrima(100); | Metschnikowia | 13 | Saprotroph | \|Undefined Saprotroph\| | Yeast | NULL | Probable | As nectar-tap_saprotroph (Põlme et al. 2020); Decay sugar-rich substrates (Põlme et al. 2020) | Põlme S, et al. 2020. Fungal Diversity 105:1-16 ((https://doi.org/10.1007/s13225-020-00466-2)) |
| Otu45845 | 1 | k__Fungi(100);p__Ascomycota(100);c__Saccharomycetes(100);o__Saccharomycetales(100);f__Metschnikowiaceae(100);g__Metschnikowia(100);g__Metschnikowia_unclassified(100); | Metschnikowia | 13 | Saprotroph | \|Undefined Saprotroph\| | Yeast | NULL | Probable | As nectar-tap_saprotroph (Põlme et al. 2020); Decay sugar-rich substrates (Põlme et al. 2020) | Põlme S, et al. 2020. Fungal Diversity 105:1-16 ((https://doi.org/10.1007/s13225-020-00466-2)) |
| Otu45846 | 1 | k__Fungi(100);p__Ascomycota(100);c__Saccharomycetes(100);o__Saccharomycetales(100);f__Metschnikowiaceae(100);g__Metschnikowia(100);s__Metschnikowia_pulcherrima(100); | Metschnikowia | 13 | Saprotroph | \|Undefined Saprotroph\| | Yeast | NULL | Probable | As nectar-tap_saprotroph (Põlme et al. 2020); Decay sugar-rich substrates (Põlme et al. 2020) | Põlme S, et al. 2020. Fungal Diversity 105:1-16 ((https://doi.org/10.1007/s13225-020-00466-2)) |
| Otu45847 | 1 | k__Fungi(100);p__Ascomycota(100);c__Saccharomycetes(100);o__Saccharomycetales(100);f__Metschnikowiaceae(100);g__Metschnikowia(100);s__Metschnikowia_pulcherrima(100); | Metschnikowia | 13 | Saprotroph | \|Undefined Saprotroph\| | Yeast | NULL | Probable | As nectar-tap_saprotroph (Põlme et al. 2020); Decay sugar-rich substrates (Põlme et al. 2020) | Põlme S, et al. 2020. Fungal Diversity 105:1-16 ((https://doi.org/10.1007/s13225-020-00466-2)) |
| Otu45848 | 1 | k__Fungi(100);p__Ascomycota(100);c__Saccharomycetes(100);o__Saccharomycetales(100);f__Metschnikowiaceae(100);g__Metschnikowia(100);g__Metschnikowia_unclassified(100); | Metschnikowia | 13 | Saprotroph | \|Undefined Saprotroph\| | Yeast | NULL | Probable | As nectar-tap_saprotroph (Põlme et al. 2020); Decay sugar-rich substrates (Põlme et al. 2020) | Põlme S, et al. 2020. Fungal Diversity 105:1-16 ((https://doi.org/10.1007/s13225-020-00466-2)) |
| Otu45849 | 1 | k__Fungi(100);p__Ascomycota(100);c__Saccharomycetes(100);o__Saccharomycetales(100);f__Metschnikowiaceae(100);g__Metschnikowia(100);s__Metschnikowia_pulcherrima(100); | Metschnikowia | 13 | Saprotroph | \|Undefined Saprotroph\| | Yeast | NULL | Probable | As nectar-tap_saprotroph (Põlme et al. 2020); Decay sugar-rich substrates (Põlme et al. 2020) | Põlme S, et al. 2020. Fungal Diversity 105:1-16 ((https://doi.org/10.1007/s13225-020-00466-2)) |
| Otu45850 | 1 | k__Fungi(100);p__Ascomycota(100);c__Saccharomycetes(100);o__Saccharomycetales(100);f__Metschnikowiaceae(100);g__Metschnikowia(100);s__Metschnikowia_pulcherrima(100); | Metschnikowia | 13 | Saprotroph | \|Undefined Saprotroph\| | Yeast | NULL | Probable | As nectar-tap_saprotroph (Põlme et al. 2020); Decay sugar-rich substrates (Põlme et al. 2020) | Põlme S, et al. 2020. Fungal Diversity 105:1-16 ((https://doi.org/10.1007/s13225-020-00466-2)) |
| Otu45851 | 1 | k__Fungi(100);p__Ascomycota(100);c__Saccharomycetes(100);o__Saccharomycetales(100);f__Metschnikowiaceae(100);g__Metschnikowia(100);s__Metschnikowia_pulcherrima(100); | Metschnikowia | 13 | Saprotroph | \|Undefined Saprotroph\| | Yeast | NULL | Probable | As nectar-tap_saprotroph (Põlme et al. 2020); Decay sugar-rich substrates (Põlme et al. 2020) | Põlme S, et al. 2020. Fungal Diversity 105:1-16 ((https://doi.org/10.1007/s13225-020-00466-2)) |
| Otu45852 | 1 | k__Fungi(100);p__Ascomycota(100);c__Saccharomycetes(100);o__Saccharomycetales(100);f__Metschnikowiaceae(100);g__Metschnikowia(100);s__Metschnikowia_pulcherrima(100); | Metschnikowia | 13 | Saprotroph | \|Undefined Saprotroph\| | Yeast | NULL | Probable | As nectar-tap_saprotroph (Põlme et al. 2020); Decay sugar-rich substrates (Põlme et al. 2020) | Põlme S, et al. 2020. Fungal Diversity 105:1-16 ((https://doi.org/10.1007/s13225-020-00466-2)) |
| Otu45853 | 1 | k__Fungi(100);p__Ascomycota(100);c__Saccharomycetes(100);o__Saccharomycetales(100);f__Metschnikowiaceae(100);g__Metschnikowia(100);s__Metschnikowia_pulcherrima(100); | Metschnikowia | 13 | Saprotroph | \|Undefined Saprotroph\| | Yeast | NULL | Probable | As nectar-tap_saprotroph (Põlme et al. 2020); Decay sugar-rich substrates (Põlme et al. 2020) | Põlme S, et al. 2020. Fungal Diversity 105:1-16 ((https://doi.org/10.1007/s13225-020-00466-2)) |
| Otu45854 | 1 | k__Fungi(100);p__Ascomycota(100);c__Saccharomycetes(100);o__Saccharomycetales(100);f__Metschnikowiaceae(100);g__Metschnikowia(100);s__Metschnikowia_pulcherrima(100); | Metschnikowia | 13 | Saprotroph | \|Undefined Saprotroph\| | Yeast | NULL | Probable | As nectar-tap_saprotroph (Põlme et al. 2020); Decay sugar-rich substrates (Põlme et al. 2020) | Põlme S, et al. 2020. Fungal Diversity 105:1-16 ((https://doi.org/10.1007/s13225-020-00466-2)) |
| Otu45855 | 1 | k__Fungi(100);p__Ascomycota(100);c__Saccharomycetes(100);o__Saccharomycetales(100);f__Metschnikowiaceae(100);g__Metschnikowia(100);s__Metschnikowia_pulcherrima(100); | Metschnikowia | 13 | Saprotroph | \|Undefined Saprotroph\| | Yeast | NULL | Probable | As nectar-tap_saprotroph (Põlme et al. 2020); Decay sugar-rich substrates (Põlme et al. 2020) | Põlme S, et al. 2020. Fungal Diversity 105:1-16 ((https://doi.org/10.1007/s13225-020-00466-2)) |
| Otu45856 | 1 | k__Fungi(100);p__Ascomycota(100);c__Saccharomycetes(100);o__Saccharomycetales(100);f__Metschnikowiaceae(100);g__Metschnikowia(100);g__Metschnikowia_unclassified(100); | Metschnikowia | 13 | Saprotroph | \|Undefined Saprotroph\| | Yeast | NULL | Probable | As nectar-tap_saprotroph (Põlme et al. 2020); Decay sugar-rich substrates (Põlme et al. 2020) | Põlme S, et al. 2020. Fungal Diversity 105:1-16 ((https://doi.org/10.1007/s13225-020-00466-2)) |
| Otu45857 | 1 | k__Fungi(100);p__Ascomycota(100);c__Saccharomycetes(100);o__Saccharomycetales(100);f__Metschnikowiaceae(100);g__Metschnikowia(100);s__Metschnikowia_pulcherrima(100); | Metschnikowia | 13 | Saprotroph | \|Undefined Saprotroph\| | Yeast | NULL | Probable | As nectar-tap_saprotroph (Põlme et al. 2020); Decay sugar-rich substrates (Põlme et al. 2020) | Põlme S, et al. 2020. Fungal Diversity 105:1-16 ((https://doi.org/10.1007/s13225-020-00466-2)) |
| Otu45858 | 1 | k__Fungi(100);p__Ascomycota(100);c__Saccharomycetes(100);o__Saccharomycetales(100);f__Metschnikowiaceae(100);g__Metschnikowia(100);s__Metschnikowia_pulcherrima(100); | Metschnikowia | 13 | Saprotroph | \|Undefined Saprotroph\| | Yeast | NULL | Probable | As nectar-tap_saprotroph (Põlme et al. 2020); Decay sugar-rich substrates (Põlme et al. 2020) | Põlme S, et al. 2020. Fungal Diversity 105:1-16 ((https://doi.org/10.1007/s13225-020-00466-2)) |
| Otu45859 | 1 | k__Fungi(100);p__Ascomycota(100);c__Saccharomycetes(100);o__Saccharomycetales(100);f__Metschnikowiaceae(100);g__Metschnikowia(100);s__Metschnikowia_pulcherrima(100); | Metschnikowia | 13 | Saprotroph | \|Undefined Saprotroph\| | Yeast | NULL | Probable | As nectar-tap_saprotroph (Põlme et al. 2020); Decay sugar-rich substrates (Põlme et al. 2020) | Põlme S, et al. 2020. Fungal Diversity 105:1-16 ((https://doi.org/10.1007/s13225-020-00466-2)) |
| Otu45860 | 1 | k__Fungi(100);p__Ascomycota(100);c__Saccharomycetes(100);o__Saccharomycetales(100);f__Metschnikowiaceae(100);g__Metschnikowia(100);g__Metschnikowia_unclassified(100); | Metschnikowia | 13 | Saprotroph | \|Undefined Saprotroph\| | Yeast | NULL | Probable | As nectar-tap_saprotroph (Põlme et al. 2020); Decay sugar-rich substrates (Põlme et al. 2020) | Põlme S, et al. 2020. Fungal Diversity 105:1-16 ((https://doi.org/10.1007/s13225-020-00466-2)) |
| Otu45861 | 1 | k__Fungi(100);p__Ascomycota(100);c__Saccharomycetes(100);o__Saccharomycetales(100);f__Metschnikowiaceae(100);g__Metschnikowia(100);g__Metschnikowia_unclassified(100); | Metschnikowia | 13 | Saprotroph | \|Undefined Saprotroph\| | Yeast | NULL | Probable | As nectar-tap_saprotroph (Põlme et al. 2020); Decay sugar-rich substrates (Põlme et al. 2020) | Põlme S, et al. 2020. Fungal Diversity 105:1-16 ((https://doi.org/10.1007/s13225-020-00466-2)) |
| Otu45862 | 1 | k__Fungi(100);p__Ascomycota(100);c__Saccharomycetes(100);o__Saccharomycetales(100);f__Metschnikowiaceae(100);g__Metschnikowia(100);s__Metschnikowia_pulcherrima(100); | Metschnikowia | 13 | Saprotroph | \|Undefined Saprotroph\| | Yeast | NULL | Probable | As nectar-tap_saprotroph (Põlme et al. 2020); Decay sugar-rich substrates (Põlme et al. 2020) | Põlme S, et al. 2020. Fungal Diversity 105:1-16 ((https://doi.org/10.1007/s13225-020-00466-2)) |
| Otu45863 | 1 | k__Fungi(100);p__Ascomycota(100);c__Saccharomycetes(100);o__Saccharomycetales(100);f__Metschnikowiaceae(100);g__Metschnikowia(100);s__Metschnikowia_pulcherrima(100); | Metschnikowia | 13 | Saprotroph | \|Undefined Saprotroph\| | Yeast | NULL | Probable | As nectar-tap_saprotroph (Põlme et al. 2020); Decay sugar-rich substrates (Põlme et al. 2020) | Põlme S, et al. 2020. Fungal Diversity 105:1-16 ((https://doi.org/10.1007/s13225-020-00466-2)) |
| Otu45864 | 1 | k__Fungi(100);p__Ascomycota(100);c__Saccharomycetes(100);o__Saccharomycetales(100);f__Metschnikowiaceae(100);g__Metschnikowia(100);s__Metschnikowia_pulcherrima(100); | Metschnikowia | 13 | Saprotroph | \|Undefined Saprotroph\| | Yeast | NULL | Probable | As nectar-tap_saprotroph (Põlme et al. 2020); Decay sugar-rich substrates (Põlme et al. 2020) | Põlme S, et al. 2020. Fungal Diversity 105:1-16 ((https://doi.org/10.1007/s13225-020-00466-2)) |
| Otu45865 | 1 | k__Fungi(100);p__Ascomycota(100);c__Saccharomycetes(100);o__Saccharomycetales(100);f__Metschnikowiaceae(100);g__Metschnikowia(100);s__Metschnikowia_pulcherrima(100); | Metschnikowia | 13 | Saprotroph | \|Undefined Saprotroph\| | Yeast | NULL | Probable | As nectar-tap_saprotroph (Põlme et al. 2020); Decay sugar-rich substrates (Põlme et al. 2020) | Põlme S, et al. 2020. Fungal Diversity 105:1-16 ((https://doi.org/10.1007/s13225-020-00466-2)) |
| Otu45866 | 1 | k__Fungi(100);p__Ascomycota(100);c__Saccharomycetes(100);o__Saccharomycetales(100);f__Metschnikowiaceae(100);g__Metschnikowia(100);s__Metschnikowia_pulcherrima(100); | Metschnikowia | 13 | Saprotroph | \|Undefined Saprotroph\| | Yeast | NULL | Probable | As nectar-tap_saprotroph (Põlme et al. 2020); Decay sugar-rich substrates (Põlme et al. 2020) | Põlme S, et al. 2020. Fungal Diversity 105:1-16 ((https://doi.org/10.1007/s13225-020-00466-2)) |
| Otu45867 | 1 | k__Fungi(100);p__Ascomycota(100);c__Saccharomycetes(100);o__Saccharomycetales(100);f__Metschnikowiaceae(100);g__Metschnikowia(100);s__Metschnikowia_pulcherrima(100); | Metschnikowia | 13 | Saprotroph | \|Undefined Saprotroph\| | Yeast | NULL | Probable | As nectar-tap_saprotroph (Põlme et al. 2020); Decay sugar-rich substrates (Põlme et al. 2020) | Põlme S, et al. 2020. Fungal Diversity 105:1-16 ((https://doi.org/10.1007/s13225-020-00466-2)) |
| Otu45868 | 1 | k__Fungi(100);p__Ascomycota(100);c__Saccharomycetes(100);o__Saccharomycetales(100);f__Metschnikowiaceae(100);g__Metschnikowia(100);s__Metschnikowia_pulcherrima(100); | Metschnikowia | 13 | Saprotroph | \|Undefined Saprotroph\| | Yeast | NULL | Probable | As nectar-tap_saprotroph (Põlme et al. 2020); Decay sugar-rich substrates (Põlme et al. 2020) | Põlme S, et al. 2020. Fungal Diversity 105:1-16 ((https://doi.org/10.1007/s13225-020-00466-2)) |
| Otu45869 | 1 | k__Fungi(100);p__Ascomycota(100);c__Saccharomycetes(100);o__Saccharomycetales(100);f__Metschnikowiaceae(100);g__Metschnikowia(100);s__Metschnikowia_pulcherrima(100); | Metschnikowia | 13 | Saprotroph | \|Undefined Saprotroph\| | Yeast | NULL | Probable | As nectar-tap_saprotroph (Põlme et al. 2020); Decay sugar-rich substrates (Põlme et al. 2020) | Põlme S, et al. 2020. Fungal Diversity 105:1-16 ((https://doi.org/10.1007/s13225-020-00466-2)) |
| Otu45870 | 1 | k__Fungi(100);p__Ascomycota(100);c__Saccharomycetes(100);o__Saccharomycetales(100);f__Metschnikowiaceae(100);g__Metschnikowia(100);s__Metschnikowia_pulcherrima(100); | Metschnikowia | 13 | Saprotroph | \|Undefined Saprotroph\| | Yeast | NULL | Probable | As nectar-tap_saprotroph (Põlme et al. 2020); Decay sugar-rich substrates (Põlme et al. 2020) | Põlme S, et al. 2020. Fungal Diversity 105:1-16 ((https://doi.org/10.1007/s13225-020-00466-2)) |
| Otu45871 | 1 | k__Fungi(100);p__Ascomycota(100);c__Saccharomycetes(100);o__Saccharomycetales(100);f__Metschnikowiaceae(100);g__Metschnikowia(100);s__Metschnikowia_pulcherrima(100); | Metschnikowia | 13 | Saprotroph | \|Undefined Saprotroph\| | Yeast | NULL | Probable | As nectar-tap_saprotroph (Põlme et al. 2020); Decay sugar-rich substrates (Põlme et al. 2020) | Põlme S, et al. 2020. Fungal Diversity 105:1-16 ((https://doi.org/10.1007/s13225-020-00466-2)) |
| Otu45872 | 1 | k__Fungi(100);p__Ascomycota(100);c__Saccharomycetes(100);o__Saccharomycetales(100);f__Metschnikowiaceae(100);g__Metschnikowia(100);s__Metschnikowia_pulcherrima(100); | Metschnikowia | 13 | Saprotroph | \|Undefined Saprotroph\| | Yeast | NULL | Probable | As nectar-tap_saprotroph (Põlme et al. 2020); Decay sugar-rich substrates (Põlme et al. 2020) | Põlme S, et al. 2020. Fungal Diversity 105:1-16 ((https://doi.org/10.1007/s13225-020-00466-2)) |
| Otu45873 | 1 | k__Fungi(100);p__Ascomycota(100);c__Saccharomycetes(100);o__Saccharomycetales(100);f__Metschnikowiaceae(100);g__Metschnikowia(100);s__Metschnikowia_pulcherrima(100); | Metschnikowia | 13 | Saprotroph | \|Undefined Saprotroph\| | Yeast | NULL | Probable | As nectar-tap_saprotroph (Põlme et al. 2020); Decay sugar-rich substrates (Põlme et al. 2020) | Põlme S, et al. 2020. Fungal Diversity 105:1-16 ((https://doi.org/10.1007/s13225-020-00466-2)) |
| Otu45874 | 1 | k__Fungi(100);p__Ascomycota(100);c__Saccharomycetes(100);o__Saccharomycetales(100);f__Metschnikowiaceae(100);g__Metschnikowia(100);s__Metschnikowia_pulcherrima(100); | Metschnikowia | 13 | Saprotroph | \|Undefined Saprotroph\| | Yeast | NULL | Probable | As nectar-tap_saprotroph (Põlme et al. 2020); Decay sugar-rich substrates (Põlme et al. 2020) | Põlme S, et al. 2020. Fungal Diversity 105:1-16 ((https://doi.org/10.1007/s13225-020-00466-2)) |
| Otu45875 | 1 | k__Fungi(100);p__Ascomycota(100);c__Saccharomycetes(100);o__Saccharomycetales(100);f__Metschnikowiaceae(100);g__Metschnikowia(100);s__Metschnikowia_pulcherrima(100); | Metschnikowia | 13 | Saprotroph | \|Undefined Saprotroph\| | Yeast | NULL | Probable | As nectar-tap_saprotroph (Põlme et al. 2020); Decay sugar-rich substrates (Põlme et al. 2020) | Põlme S, et al. 2020. Fungal Diversity 105:1-16 ((https://doi.org/10.1007/s13225-020-00466-2)) |
| Otu45877 | 1 | k__Fungi(100);p__Ascomycota(100);c__Saccharomycetes(100);o__Saccharomycetales(100);f__Metschnikowiaceae(100);g__Metschnikowia(100);s__Metschnikowia_pulcherrima(100); | Metschnikowia | 13 | Saprotroph | \|Undefined Saprotroph\| | Yeast | NULL | Probable | As nectar-tap_saprotroph (Põlme et al. 2020); Decay sugar-rich substrates (Põlme et al. 2020) | Põlme S, et al. 2020. Fungal Diversity 105:1-16 ((https://doi.org/10.1007/s13225-020-00466-2)) |
| Otu45878 | 1 | k__Fungi(100);p__Ascomycota(100);c__Saccharomycetes(100);o__Saccharomycetales(100);f__Metschnikowiaceae(100);g__Metschnikowia(100);g__Metschnikowia_unclassified(100); | Metschnikowia | 13 | Saprotroph | \|Undefined Saprotroph\| | Yeast | NULL | Probable | As nectar-tap_saprotroph (Põlme et al. 2020); Decay sugar-rich substrates (Põlme et al. 2020) | Põlme S, et al. 2020. Fungal Diversity 105:1-16 ((https://doi.org/10.1007/s13225-020-00466-2)) |
| Otu45879 | 1 | k__Fungi(100);p__Ascomycota(100);c__Saccharomycetes(100);o__Saccharomycetales(100);f__Metschnikowiaceae(100);g__Metschnikowia(100);s__Metschnikowia_pulcherrima(100); | Metschnikowia | 13 | Saprotroph | \|Undefined Saprotroph\| | Yeast | NULL | Probable | As nectar-tap_saprotroph (Põlme et al. 2020); Decay sugar-rich substrates (Põlme et al. 2020) | Põlme S, et al. 2020. Fungal Diversity 105:1-16 ((https://doi.org/10.1007/s13225-020-00466-2)) |
| Otu45880 | 1 | k__Fungi(100);p__Ascomycota(100);c__Saccharomycetes(100);o__Saccharomycetales(100);f__Metschnikowiaceae(100);g__Metschnikowia(100);s__Metschnikowia_pulcherrima(100); | Metschnikowia | 13 | Saprotroph | \|Undefined Saprotroph\| | Yeast | NULL | Probable | As nectar-tap_saprotroph (Põlme et al. 2020); Decay sugar-rich substrates (Põlme et al. 2020) | Põlme S, et al. 2020. Fungal Diversity 105:1-16 ((https://doi.org/10.1007/s13225-020-00466-2)) |
| Otu45881 | 1 | k__Fungi(100);p__Ascomycota(100);c__Saccharomycetes(100);o__Saccharomycetales(100);f__Metschnikowiaceae(100);g__Metschnikowia(100);s__Metschnikowia_pulcherrima(100); | Metschnikowia | 13 | Saprotroph | \|Undefined Saprotroph\| | Yeast | NULL | Probable | As nectar-tap_saprotroph (Põlme et al. 2020); Decay sugar-rich substrates (Põlme et al. 2020) | Põlme S, et al. 2020. Fungal Diversity 105:1-16 ((https://doi.org/10.1007/s13225-020-00466-2)) |
| Otu45882 | 1 | k__Fungi(100);p__Ascomycota(100);c__Saccharomycetes(100);o__Saccharomycetales(100);f__Metschnikowiaceae(100);g__Metschnikowia(100);s__Metschnikowia_pulcherrima(100); | Metschnikowia | 13 | Saprotroph | \|Undefined Saprotroph\| | Yeast | NULL | Probable | As nectar-tap_saprotroph (Põlme et al. 2020); Decay sugar-rich substrates (Põlme et al. 2020) | Põlme S, et al. 2020. Fungal Diversity 105:1-16 ((https://doi.org/10.1007/s13225-020-00466-2)) |
| Otu45883 | 1 | k__Fungi(100);p__Ascomycota(100);c__Saccharomycetes(100);o__Saccharomycetales(100);f__Metschnikowiaceae(100);g__Metschnikowia(100);s__Metschnikowia_pulcherrima(100); | Metschnikowia | 13 | Saprotroph | \|Undefined Saprotroph\| | Yeast | NULL | Probable | As nectar-tap_saprotroph (Põlme et al. 2020); Decay sugar-rich substrates (Põlme et al. 2020) | Põlme S, et al. 2020. Fungal Diversity 105:1-16 ((https://doi.org/10.1007/s13225-020-00466-2)) |
| Otu45884 | 1 | k__Fungi(100);p__Ascomycota(100);c__Saccharomycetes(100);o__Saccharomycetales(100);f__Metschnikowiaceae(100);g__Metschnikowia(100);s__Metschnikowia_pulcherrima(100); | Metschnikowia | 13 | Saprotroph | \|Undefined Saprotroph\| | Yeast | NULL | Probable | As nectar-tap_saprotroph (Põlme et al. 2020); Decay sugar-rich substrates (Põlme et al. 2020) | Põlme S, et al. 2020. Fungal Diversity 105:1-16 ((https://doi.org/10.1007/s13225-020-00466-2)) |
| Otu45885 | 1 | k__Fungi(100);p__Ascomycota(100);c__Saccharomycetes(100);o__Saccharomycetales(100);f__Metschnikowiaceae(100);g__Metschnikowia(100);g__Metschnikowia_unclassified(100); | Metschnikowia | 13 | Saprotroph | \|Undefined Saprotroph\| | Yeast | NULL | Probable | As nectar-tap_saprotroph (Põlme et al. 2020); Decay sugar-rich substrates (Põlme et al. 2020) | Põlme S, et al. 2020. Fungal Diversity 105:1-16 ((https://doi.org/10.1007/s13225-020-00466-2)) |
| Otu45886 | 1 | k__Fungi(100);p__Ascomycota(100);c__Saccharomycetes(100);o__Saccharomycetales(100);f__Metschnikowiaceae(100);g__Metschnikowia(100);s__Metschnikowia_pulcherrima(100); | Metschnikowia | 13 | Saprotroph | \|Undefined Saprotroph\| | Yeast | NULL | Probable | As nectar-tap_saprotroph (Põlme et al. 2020); Decay sugar-rich substrates (Põlme et al. 2020) | Põlme S, et al. 2020. Fungal Diversity 105:1-16 ((https://doi.org/10.1007/s13225-020-00466-2)) |
| Otu45887 | 1 | k__Fungi(100);p__Ascomycota(100);c__Saccharomycetes(100);o__Saccharomycetales(100);f__Metschnikowiaceae(100);g__Metschnikowia(100);s__Metschnikowia_pulcherrima(100); | Metschnikowia | 13 | Saprotroph | \|Undefined Saprotroph\| | Yeast | NULL | Probable | As nectar-tap_saprotroph (Põlme et al. 2020); Decay sugar-rich substrates (Põlme et al. 2020) | Põlme S, et al. 2020. Fungal Diversity 105:1-16 ((https://doi.org/10.1007/s13225-020-00466-2)) |
| Otu45888 | 1 | k__Fungi(100);p__Ascomycota(100);c__Saccharomycetes(100);o__Saccharomycetales(100);f__Metschnikowiaceae(100);g__Metschnikowia(100);s__Metschnikowia_pulcherrima(100); | Metschnikowia | 13 | Saprotroph | \|Undefined Saprotroph\| | Yeast | NULL | Probable | As nectar-tap_saprotroph (Põlme et al. 2020); Decay sugar-rich substrates (Põlme et al. 2020) | Põlme S, et al. 2020. Fungal Diversity 105:1-16 ((https://doi.org/10.1007/s13225-020-00466-2)) |
| Otu45889 | 1 | k__Fungi(100);p__Ascomycota(100);c__Saccharomycetes(100);o__Saccharomycetales(100);f__Metschnikowiaceae(100);g__Metschnikowia(100);g__Metschnikowia_unclassified(100); | Metschnikowia | 13 | Saprotroph | \|Undefined Saprotroph\| | Yeast | NULL | Probable | As nectar-tap_saprotroph (Põlme et al. 2020); Decay sugar-rich substrates (Põlme et al. 2020) | Põlme S, et al. 2020. Fungal Diversity 105:1-16 ((https://doi.org/10.1007/s13225-020-00466-2)) |
| Otu45890 | 1 | k__Fungi(100);p__Ascomycota(100);c__Saccharomycetes(100);o__Saccharomycetales(100);f__Metschnikowiaceae(100);g__Metschnikowia(100);s__Metschnikowia_pulcherrima(100); | Metschnikowia | 13 | Saprotroph | \|Undefined Saprotroph\| | Yeast | NULL | Probable | As nectar-tap_saprotroph (Põlme et al. 2020); Decay sugar-rich substrates (Põlme et al. 2020) | Põlme S, et al. 2020. Fungal Diversity 105:1-16 ((https://doi.org/10.1007/s13225-020-00466-2)) |

| | | | | | | | | | | | |
|---|---|---|---|---|---|---|---|---|---|---|---|
| Otu45891 | 1 | k__Fungi(100);p__Ascomycota(100);c__Saccharomycetes(100);o__Saccharomycetales(100);f__Metschnikowiaceae(100);g__Metschnikowia(100);s__Metschnikowia_pulcherrima(100); | Metschnikowia | 13 | Saprotroph | \|Undefined Saprotroph\| | Yeast | NULL | Probable | As nectar-tap_saprotroph (Põlme et al. 2020); Decay sugar-rich substrates (Põlme et al. 2020) | Põlme S, et al. 2020. Fungal Diversity 105:1-16 ((https://doi.org/10.1007/s13225-020-00466-2)) |
| Otu45892 | 1 | k__Fungi(100);p__Ascomycota(100);c__Saccharomycetes(100);o__Saccharomycetales(100);f__Metschnikowiaceae(100);g__Metschnikowia(100);s__Metschnikowia_pulcherrima(100); | Metschnikowia | 13 | Saprotroph | \|Undefined Saprotroph\| | Yeast | NULL | Probable | As nectar-tap_saprotroph (Põlme et al. 2020); Decay sugar-rich substrates (Põlme et al. 2020) | Põlme S, et al. 2020. Fungal Diversity 105:1-16 ((https://doi.org/10.1007/s13225-020-00466-2)) |
| Otu45893 | 1 | k__Fungi(100);p__Ascomycota(100);c__Saccharomycetes(100);o__Saccharomycetales(100);f__Metschnikowiaceae(100);g__Metschnikowia(100);s__Metschnikowia_pulcherrima(100); | Metschnikowia | 13 | Saprotroph | \|Undefined Saprotroph\| | Yeast | NULL | Probable | As nectar-tap_saprotroph (Põlme et al. 2020); Decay sugar-rich substrates (Põlme et al. 2020) | Põlme S, et al. 2020. Fungal Diversity 105:1-16 ((https://doi.org/10.1007/s13225-020-00466-2)) |
| Otu45894 | 1 | k__Fungi(100);p__Ascomycota(100);c__Saccharomycetes(100);o__Saccharomycetales(100);f__Metschnikowiaceae(100);g__Metschnikowia(100);s__Metschnikowia_pulcherrima(100); | Metschnikowia | 13 | Saprotroph | \|Undefined Saprotroph\| | Yeast | NULL | Probable | As nectar-tap_saprotroph (Põlme et al. 2020); Decay sugar-rich substrates (Põlme et al. 2020) | Põlme S, et al. 2020. Fungal Diversity 105:1-16 ((https://doi.org/10.1007/s13225-020-00466-2)) |
| Otu45895 | 1 | k__Fungi(100);p__Ascomycota(100);c__Saccharomycetes(100);o__Saccharomycetales(100);f__Metschnikowiaceae(100);g__Metschnikowia(100);s__Metschnikowia_pulcherrima(100); | Metschnikowia | 13 | Saprotroph | \|Undefined Saprotroph\| | Yeast | NULL | Probable | As nectar-tap_saprotroph (Põlme et al. 2020); Decay sugar-rich substrates (Põlme et al. 2020) | Põlme S, et al. 2020. Fungal Diversity 105:1-16 ((https://doi.org/10.1007/s13225-020-00466-2)) |
| Otu45896 | 1 | k__Fungi(100);p__Ascomycota(100);c__Saccharomycetes(100);o__Saccharomycetales(100);f__Metschnikowiaceae(100);g__Metschnikowia(100);s__Metschnikowia_pulcherrima(100); | Metschnikowia | 13 | Saprotroph | \|Undefined Saprotroph\| | Yeast | NULL | Probable | As nectar-tap_saprotroph (Põlme et al. 2020); Decay sugar-rich substrates (Põlme et al. 2020) | Põlme S, et al. 2020. Fungal Diversity 105:1-16 ((https://doi.org/10.1007/s13225-020-00466-2)) |
| Otu45897 | 1 | k__Fungi(100);p__Ascomycota(100);c__Saccharomycetes(100);o__Saccharomycetales(100);f__Metschnikowiaceae(100);g__Metschnikowia(100);s__Metschnikowia_pulcherrima(100); | Metschnikowia | 13 | Saprotroph | \|Undefined Saprotroph\| | Yeast | NULL | Probable | As nectar-tap_saprotroph (Põlme et al. 2020); Decay sugar-rich substrates (Põlme et al. 2020) | Põlme S, et al. 2020. Fungal Diversity 105:1-16 ((https://doi.org/10.1007/s13225-020-00466-2)) |
| Otu45898 | 1 | k__Fungi(100);p__Ascomycota(100);c__Saccharomycetes(100);o__Saccharomycetales(100);f__Metschnikowiaceae(100);g__Metschnikowia(100);g__Metschnikowia_unclassified(100); | Metschnikowia | 13 | Saprotroph | \|Undefined Saprotroph\| | Yeast | NULL | Probable | As nectar-tap_saprotroph (Põlme et al. 2020); Decay sugar-rich substrates (Põlme et al. 2020) | Põlme S, et al. 2020. Fungal Diversity 105:1-16 ((https://doi.org/10.1007/s13225-020-00466-2)) |
| Otu45899 | 1 | k__Fungi(100);p__Ascomycota(100);c__Saccharomycetes(100);o__Saccharomycetales(100);f__Metschnikowiaceae(100);g__Metschnikowia(100);s__Metschnikowia_pulcherrima(100); | Metschnikowia | 13 | Saprotroph | \|Undefined Saprotroph\| | Yeast | NULL | Probable | As nectar-tap_saprotroph (Põlme et al. 2020); Decay sugar-rich substrates (Põlme et al. 2020) | Põlme S, et al. 2020. Fungal Diversity 105:1-16 ((https://doi.org/10.1007/s13225-020-00466-2)) |
| Otu45900 | 1 | k__Fungi(100);p__Ascomycota(100);c__Saccharomycetes(100);o__Saccharomycetales(100);f__Metschnikowiaceae(100);g__Metschnikowia(100);s__Metschnikowia_pulcherrima(100); | Metschnikowia | 13 | Saprotroph | \|Undefined Saprotroph\| | Yeast | NULL | Probable | As nectar-tap_saprotroph (Põlme et al. 2020); Decay sugar-rich substrates (Põlme et al. 2020) | Põlme S, et al. 2020. Fungal Diversity 105:1-16 ((https://doi.org/10.1007/s13225-020-00466-2)) |
| Otu45901 | 1 | k__Fungi(100);p__Ascomycota(100);c__Saccharomycetes(100);o__Saccharomycetales(100);f__Metschnikowiaceae(100);g__Metschnikowia(100);s__Metschnikowia_pulcherrima(100); | Metschnikowia | 13 | Saprotroph | \|Undefined Saprotroph\| | Yeast | NULL | Probable | As nectar-tap_saprotroph (Põlme et al. 2020); Decay sugar-rich substrates (Põlme et al. 2020) | Põlme S, et al. 2020. Fungal Diversity 105:1-16 ((https://doi.org/10.1007/s13225-020-00466-2)) |
| Otu45902 | 1 | k__Fungi(100);p__Ascomycota(100);c__Saccharomycetes(100);o__Saccharomycetales(100);f__Metschnikowiaceae(100);g__Metschnikowia(100);g__Metschnikowia_unclassified(100); | Metschnikowia | 13 | Saprotroph | \|Undefined Saprotroph\| | Yeast | NULL | Probable | As nectar-tap_saprotroph (Põlme et al. 2020); Decay sugar-rich substrates (Põlme et al. 2020) | Põlme S, et al. 2020. Fungal Diversity 105:1-16 ((https://doi.org/10.1007/s13225-020-00466-2)) |
| Otu45903 | 1 | k__Fungi(100);p__Ascomycota(100);c__Saccharomycetes(100);o__Saccharomycetales(100);f__Metschnikowiaceae(100);g__Metschnikowia(100);s__Metschnikowia_pulcherrima(100); | Metschnikowia | 13 | Saprotroph | \|Undefined Saprotroph\| | Yeast | NULL | Probable | As nectar-tap_saprotroph (Põlme et al. 2020); Decay sugar-rich substrates (Põlme et al. 2020) | Põlme S, et al. 2020. Fungal Diversity 105:1-16 ((https://doi.org/10.1007/s13225-020-00466-2)) |
| Otu45904 | 1 | k__Fungi(100);p__Ascomycota(100);c__Saccharomycetes(100);o__Saccharomycetales(100);f__Metschnikowiaceae(100);g__Metschnikowia(100);s__Metschnikowia_pulcherrima(100); | Metschnikowia | 13 | Saprotroph | \|Undefined Saprotroph\| | Yeast | NULL | Probable | As nectar-tap_saprotroph (Põlme et al. 2020); Decay sugar-rich substrates (Põlme et al. 2020) | Põlme S, et al. 2020. Fungal Diversity 105:1-16 ((https://doi.org/10.1007/s13225-020-00466-2)) |
| Otu45905 | 1 | k__Fungi(100);p__Ascomycota(100);c__Saccharomycetes(100);o__Saccharomycetales(100);f__Metschnikowiaceae(100);g__Metschnikowia(100);s__Metschnikowia_pulcherrima(100); | Metschnikowia | 13 | Saprotroph | \|Undefined Saprotroph\| | Yeast | NULL | Probable | As nectar-tap_saprotroph (Põlme et al. 2020); Decay sugar-rich substrates (Põlme et al. 2020) | Põlme S, et al. 2020. Fungal Diversity 105:1-16 ((https://doi.org/10.1007/s13225-020-00466-2)) |
| Otu45906 | 1 | k__Fungi(100);p__Ascomycota(100);c__Saccharomycetes(100);o__Saccharomycetales(100);f__Metschnikowiaceae(100);g__Metschnikowia(100);s__Metschnikowia_pulcherrima(100); | Metschnikowia | 13 | Saprotroph | \|Undefined Saprotroph\| | Yeast | NULL | Probable | As nectar-tap_saprotroph (Põlme et al. 2020); Decay sugar-rich substrates (Põlme et al. 2020) | Põlme S, et al. 2020. Fungal Diversity 105:1-16 ((https://doi.org/10.1007/s13225-020-00466-2)) |
| Otu45907 | 1 | k__Fungi(100);p__Ascomycota(100);c__Saccharomycetes(100);o__Saccharomycetales(100);f__Metschnikowiaceae(100);g__Metschnikowia(100);s__Metschnikowia_pulcherrima(100); | Metschnikowia | 13 | Saprotroph | \|Undefined Saprotroph\| | Yeast | NULL | Probable | As nectar-tap_saprotroph (Põlme et al. 2020); Decay sugar-rich substrates (Põlme et al. 2020) | Põlme S, et al. 2020. Fungal Diversity 105:1-16 ((https://doi.org/10.1007/s13225-020-00466-2)) |
| Otu45908 | 1 | k__Fungi(100);p__Ascomycota(100);c__Saccharomycetes(100);o__Saccharomycetales(100);f__Metschnikowiaceae(100);g__Metschnikowia(100);s__Metschnikowia_pulcherrima(100); | Metschnikowia | 13 | Saprotroph | \|Undefined Saprotroph\| | Yeast | NULL | Probable | As nectar-tap_saprotroph (Põlme et al. 2020); Decay sugar-rich substrates (Põlme et al. 2020) | Põlme S, et al. 2020. Fungal Diversity 105:1-16 ((https://doi.org/10.1007/s13225-020-00466-2)) |
| Otu45909 | 1 | k__Fungi(100);p__Ascomycota(100);c__Saccharomycetes(100);o__Saccharomycetales(100);f__Metschnikowiaceae(100);g__Metschnikowia(100);g__Metschnikowia_unclassified(100); | Metschnikowia | 13 | Saprotroph | \|Undefined Saprotroph\| | Yeast | NULL | Probable | As nectar-tap_saprotroph (Põlme et al. 2020); Decay sugar-rich substrates (Põlme et al. 2020) | Põlme S, et al. 2020. Fungal Diversity 105:1-16 ((https://doi.org/10.1007/s13225-020-00466-2)) |
| Otu45910 | 1 | k__Fungi(100);p__Ascomycota(100);c__Saccharomycetes(100);o__Saccharomycetales(100);f__Metschnikowiaceae(100);g__Metschnikowia(100);g__Metschnikowia_unclassified(100); | Metschnikowia | 13 | Saprotroph | \|Undefined Saprotroph\| | Yeast | NULL | Probable | As nectar-tap_saprotroph (Põlme et al. 2020); Decay sugar-rich substrates (Põlme et al. 2020) | Põlme S, et al. 2020. Fungal Diversity 105:1-16 ((https://doi.org/10.1007/s13225-020-00466-2)) |
| Otu45911 | 1 | k__Fungi(100);p__Ascomycota(100);c__Saccharomycetes(100);o__Saccharomycetales(100);f__Metschnikowiaceae(100);g__Metschnikowia(100);g__Metschnikowia_unclassified(100); | Metschnikowia | 13 | Saprotroph | \|Undefined Saprotroph\| | Yeast | NULL | Probable | As nectar-tap_saprotroph (Põlme et al. 2020); Decay sugar-rich substrates (Põlme et al. 2020) | Põlme S, et al. 2020. Fungal Diversity 105:1-16 ((https://doi.org/10.1007/s13225-020-00466-2)) |
| Otu45912 | 1 | k__Fungi(100);p__Ascomycota(100);c__Saccharomycetes(100);o__Saccharomycetales(100);f__Metschnikowiaceae(100);g__Metschnikowia(100);g__Metschnikowia_unclassified(100); | Metschnikowia | 13 | Saprotroph | \|Undefined Saprotroph\| | Yeast | NULL | Probable | As nectar-tap_saprotroph (Põlme et al. 2020); Decay sugar-rich substrates (Põlme et al. 2020) | Põlme S, et al. 2020. Fungal Diversity 105:1-16 ((https://doi.org/10.1007/s13225-020-00466-2)) |
| Otu45913 | 1 | k__Fungi(100);p__Ascomycota(100);c__Saccharomycetes(100);o__Saccharomycetales(100);f__Metschnikowiaceae(100);g__Metschnikowia(100);s__Metschnikowia_pulcherrima(100); | Metschnikowia | 13 | Saprotroph | \|Undefined Saprotroph\| | Yeast | NULL | Probable | As nectar-tap_saprotroph (Põlme et al. 2020); Decay sugar-rich substrates (Põlme et al. 2020) | Põlme S, et al. 2020. Fungal Diversity 105:1-16 ((https://doi.org/10.1007/s13225-020-00466-2)) |
| Otu45914 | 1 | k__Fungi(100);p__Ascomycota(100);c__Saccharomycetes(100);o__Saccharomycetales(100);f__Metschnikowiaceae(100);g__Metschnikowia(100);s__Metschnikowia_pulcherrima(100); | Metschnikowia | 13 | Saprotroph | \|Undefined Saprotroph\| | Yeast | NULL | Probable | As nectar-tap_saprotroph (Põlme et al. 2020); Decay sugar-rich substrates (Põlme et al. 2020) | Põlme S, et al. 2020. Fungal Diversity 105:1-16 ((https://doi.org/10.1007/s13225-020-00466-2)) |
| Otu45915 | 1 | k__Fungi(100);p__Ascomycota(100);c__Saccharomycetes(100);o__Saccharomycetales(100);f__Metschnikowiaceae(100);g__Metschnikowia(100);g__Metschnikowia_unclassified(100); | Metschnikowia | 13 | Saprotroph | \|Undefined Saprotroph\| | Yeast | NULL | Probable | As nectar-tap_saprotroph (Põlme et al. 2020); Decay sugar-rich substrates (Põlme et al. 2020) | Põlme S, et al. 2020. Fungal Diversity 105:1-16 ((https://doi.org/10.1007/s13225-020-00466-2)) |
| Otu45916 | 1 | k__Fungi(100);p__Ascomycota(100);c__Saccharomycetes(100);o__Saccharomycetales(100);f__Metschnikowiaceae(100);g__Metschnikowia(100);g__Metschnikowia_unclassified(100); | Metschnikowia | 13 | Saprotroph | \|Undefined Saprotroph\| | Yeast | NULL | Probable | As nectar-tap_saprotroph (Põlme et al. 2020); Decay sugar-rich substrates (Põlme et al. 2020) | Põlme S, et al. 2020. Fungal Diversity 105:1-16 ((https://doi.org/10.1007/s13225-020-00466-2)) |
| Otu45917 | 1 | k__Fungi(100);p__Ascomycota(100);c__Saccharomycetes(100);o__Saccharomycetales(100);f__Metschnikowiaceae(100);g__Metschnikowia(100);s__Metschnikowia_pulcherrima(100); | Metschnikowia | 13 | Saprotroph | \|Undefined Saprotroph\| | Yeast | NULL | Probable | As nectar-tap_saprotroph (Põlme et al. 2020); Decay sugar-rich substrates (Põlme et al. 2020) | Põlme S, et al. 2020. Fungal Diversity 105:1-16 ((https://doi.org/10.1007/s13225-020-00466-2)) |
| Otu45918 | 1 | k__Fungi(100);p__Ascomycota(100);c__Saccharomycetes(100);o__Saccharomycetales(100);f__Metschnikowiaceae(100);g__Metschnikowia(100);s__Metschnikowia_pulcherrima(100); | Metschnikowia | 13 | Saprotroph | \|Undefined Saprotroph\| | Yeast | NULL | Probable | As nectar-tap_saprotroph (Põlme et al. 2020); Decay sugar-rich substrates (Põlme et al. 2020) | Põlme S, et al. 2020. Fungal Diversity 105:1-16 ((https://doi.org/10.1007/s13225-020-00466-2)) |
| Otu45919 | 1 | k__Fungi(100);p__Ascomycota(100);c__Saccharomycetes(100);o__Saccharomycetales(100);f__Metschnikowiaceae(100);g__Metschnikowia(100);s__Metschnikowia_pulcherrima(100); | Metschnikowia | 13 | Saprotroph | \|Undefined Saprotroph\| | Yeast | NULL | Probable | As nectar-tap_saprotroph (Põlme et al. 2020); Decay sugar-rich substrates (Põlme et al. 2020) | Põlme S, et al. 2020. Fungal Diversity 105:1-16 ((https://doi.org/10.1007/s13225-020-00466-2)) |
| Otu45920 | 1 | k__Fungi(100);p__Ascomycota(100);c__Saccharomycetes(100);o__Saccharomycetales(100);f__Metschnikowiaceae(100);g__Metschnikowia(100);s__Metschnikowia_pulcherrima(100); | Metschnikowia | 13 | Saprotroph | \|Undefined Saprotroph\| | Yeast | NULL | Probable | As nectar-tap_saprotroph (Põlme et al. 2020); Decay sugar-rich substrates (Põlme et al. 2020) | Põlme S, et al. 2020. Fungal Diversity 105:1-16 ((https://doi.org/10.1007/s13225-020-00466-2)) |
| Otu45921 | 1 | k__Fungi(100);p__Ascomycota(100);c__Saccharomycetes(100);o__Saccharomycetales(100);f__Metschnikowiaceae(100);g__Metschnikowia(100);g__Metschnikowia_unclassified(100); | Metschnikowia | 13 | Saprotroph | \|Undefined Saprotroph\| | Yeast | NULL | Probable | As nectar-tap_saprotroph (Põlme et al. 2020); Decay sugar-rich substrates (Põlme et al. 2020) | Põlme S, et al. 2020. Fungal Diversity 105:1-16 ((https://doi.org/10.1007/s13225-020-00466-2)) |
| Otu45922 | 1 | k__Fungi(100);p__Ascomycota(100);c__Saccharomycetes(100);o__Saccharomycetales(100);f__Metschnikowiaceae(100);g__Metschnikowia(100);s__Metschnikowia_pulcherrima(100); | Metschnikowia | 13 | Saprotroph | \|Undefined Saprotroph\| | Yeast | NULL | Probable | As nectar-tap_saprotroph (Põlme et al. 2020); Decay sugar-rich substrates (Põlme et al. 2020) | Põlme S, et al. 2020. Fungal Diversity 105:1-16 ((https://doi.org/10.1007/s13225-020-00466-2)) |
| Otu45923 | 1 | k__Fungi(100);p__Ascomycota(100);c__Saccharomycetes(100);o__Saccharomycetales(100);f__Metschnikowiaceae(100);g__Metschnikowia(100);s__Metschnikowia_pulcherrima(100); | Metschnikowia | 13 | Saprotroph | \|Undefined Saprotroph\| | Yeast | NULL | Probable | As nectar-tap_saprotroph (Põlme et al. 2020); Decay sugar-rich substrates (Põlme et al. 2020) | Põlme S, et al. 2020. Fungal Diversity 105:1-16 ((https://doi.org/10.1007/s13225-020-00466-2)) |
| Otu45924 | 1 | k__Fungi(100);p__Ascomycota(100);c__Saccharomycetes(100);o__Saccharomycetales(100);f__Metschnikowiaceae(100);g__Metschnikowia(100);s__Metschnikowia_pulcherrima(100); | Metschnikowia | 13 | Saprotroph | \|Undefined Saprotroph\| | Yeast | NULL | Probable | As nectar-tap_saprotroph (Põlme et al. 2020); Decay sugar-rich substrates (Põlme et al. 2020) | Põlme S, et al. 2020. Fungal Diversity 105:1-16 ((https://doi.org/10.1007/s13225-020-00466-2)) |
| Otu45925 | 1 | k__Fungi(100);p__Ascomycota(100);c__Saccharomycetes(100);o__Saccharomycetales(100);f__Metschnikowiaceae(100);g__Metschnikowia(100);s__Metschnikowia_pulcherrima(100); | Metschnikowia | 13 | Saprotroph | \|Undefined Saprotroph\| | Yeast | NULL | Probable | As nectar-tap_saprotroph (Põlme et al. 2020); Decay sugar-rich substrates (Põlme et al. 2020) | Põlme S, et al. 2020. Fungal Diversity 105:1-16 ((https://doi.org/10.1007/s13225-020-00466-2)) |
| Otu45926 | 1 | k__Fungi(100);p__Ascomycota(100);c__Saccharomycetes(100);o__Saccharomycetales(100);f__Metschnikowiaceae(100);g__Metschnikowia(100);g__Metschnikowia_unclassified(100); | Metschnikowia | 13 | Saprotroph | \|Undefined Saprotroph\| | Yeast | NULL | Probable | As nectar-tap_saprotroph (Põlme et al. 2020); Decay sugar-rich substrates (Põlme et al. 2020) | Põlme S, et al. 2020. Fungal Diversity 105:1-16 ((https://doi.org/10.1007/s13225-020-00466-2)) |
| Otu45927 | 1 | k__Fungi(100);p__Ascomycota(100);c__Saccharomycetes(100);o__Saccharomycetales(100);f__Metschnikowiaceae(100);g__Metschnikowia(100);g__Metschnikowia_unclassified(100); | Metschnikowia | 13 | Saprotroph | \|Undefined Saprotroph\| | Yeast | NULL | Probable | As nectar-tap_saprotroph (Põlme et al. 2020); Decay sugar-rich substrates (Põlme et al. 2020) | Põlme S, et al. 2020. Fungal Diversity 105:1-16 ((https://doi.org/10.1007/s13225-020-00466-2)) |
| Otu45928 | 1 | k__Fungi(100);p__Ascomycota(100);c__Saccharomycetes(100);o__Saccharomycetales(100);f__Metschnikowiaceae(100);g__Metschnikowia(100);s__Metschnikowia_pulcherrima(100); | Metschnikowia | 13 | Saprotroph | \|Undefined Saprotroph\| | Yeast | NULL | Probable | As nectar-tap_saprotroph (Põlme et al. 2020); Decay sugar-rich substrates (Põlme et al. 2020) | Põlme S, et al. 2020. Fungal Diversity 105:1-16 ((https://doi.org/10.1007/s13225-020-00466-2)) |
| Otu45930 | 1 | k__Fungi(100);p__Ascomycota(100);c__Saccharomycetes(100);o__Saccharomycetales(100);f__Metschnikowiaceae(100);g__Metschnikowia(100);g__Metschnikowia_unclassified(100); | Metschnikowia | 13 | Saprotroph | \|Undefined Saprotroph\| | Yeast | NULL | Probable | As nectar-tap_saprotroph (Põlme et al. 2020); Decay sugar-rich substrates (Põlme et al. 2020) | Põlme S, et al. 2020. Fungal Diversity 105:1-16 ((https://doi.org/10.1007/s13225-020-00466-2)) |
| Otu45931 | 1 | k__Fungi(100);p__Ascomycota(100);c__Saccharomycetes(100);o__Saccharomycetales(100);f__Metschnikowiaceae(100);g__Metschnikowia(100);s__Metschnikowia_pulcherrima(100); | Metschnikowia | 13 | Saprotroph | \|Undefined Saprotroph\| | Yeast | NULL | Probable | As nectar-tap_saprotroph (Põlme et al. 2020); Decay sugar-rich substrates (Põlme et al. 2020) | Põlme S, et al. 2020. Fungal Diversity 105:1-16 ((https://doi.org/10.1007/s13225-020-00466-2)) |
| Otu45932 | 1 | k__Fungi(100);p__Ascomycota(100);c__Saccharomycetes(100);o__Saccharomycetales(100);f__Metschnikowiaceae(100);g__Metschnikowia(100);s__Metschnikowia_pulcherrima(100); | Metschnikowia | 13 | Saprotroph | \|Undefined Saprotroph\| | Yeast | NULL | Probable | As nectar-tap_saprotroph (Põlme et al. 2020); Decay sugar-rich substrates (Põlme et al. 2020) | Põlme S, et al. 2020. Fungal Diversity 105:1-16 ((https://doi.org/10.1007/s13225-020-00466-2)) |
| Otu45933 | 1 | k__Fungi(100);p__Ascomycota(100);c__Saccharomycetes(100);o__Saccharomycetales(100);f__Metschnikowiaceae(100);g__Metschnikowia(100);s__Metschnikowia_pulcherrima(100); | Metschnikowia | 13 | Saprotroph | \|Undefined Saprotroph\| | Yeast | NULL | Probable | As nectar-tap_saprotroph (Põlme et al. 2020); Decay sugar-rich substrates (Põlme et al. 2020) | Põlme S, et al. 2020. Fungal Diversity 105:1-16 ((https://doi.org/10.1007/s13225-020-00466-2)) |
| Otu45934 | 1 | k__Fungi(100);p__Ascomycota(100);c__Saccharomycetes(100);o__Saccharomycetales(100);f__Metschnikowiaceae(100);g__Metschnikowia(100);s__Metschnikowia_pulcherrima(100); | Metschnikowia | 13 | Saprotroph | \|Undefined Saprotroph\| | Yeast | NULL | Probable | As nectar-tap_saprotroph (Põlme et al. 2020); Decay sugar-rich substrates (Põlme et al. 2020) | Põlme S, et al. 2020. Fungal Diversity 105:1-16 ((https://doi.org/10.1007/s13225-020-00466-2)) |
| Otu45935 | 1 | k__Fungi(100);p__Ascomycota(100);c__Saccharomycetes(100);o__Saccharomycetales(100);f__Metschnikowiaceae(100);g__Metschnikowia(100);s__Metschnikowia_pulcherrima(100); | Metschnikowia | 13 | Saprotroph | \|Undefined Saprotroph\| | Yeast | NULL | Probable | As nectar-tap_saprotroph (Põlme et al. 2020); Decay sugar-rich substrates (Põlme et al. 2020) | Põlme S, et al. 2020. Fungal Diversity 105:1-16 ((https://doi.org/10.1007/s13225-020-00466-2)) |
| Otu45936 | 1 | k__Fungi(100);p__Ascomycota(100);c__Saccharomycetes(100);o__Saccharomycetales(100);f__Metschnikowiaceae(100);g__Metschnikowia(100);s__Metschnikowia_pulcherrima(100); | Metschnikowia | 13 | Saprotroph | \|Undefined Saprotroph\| | Yeast | NULL | Probable | As nectar-tap_saprotroph (Põlme et al. 2020); Decay sugar-rich substrates (Põlme et al. 2020) | Põlme S, et al. 2020. Fungal Diversity 105:1-16 ((https://doi.org/10.1007/s13225-020-00466-2)) |
| Otu45937 | 1 | k__Fungi(100);p__Ascomycota(100);c__Saccharomycetes(100);o__Saccharomycetales(100);f__Metschnikowiaceae(100);g__Metschnikowia(100);s__Metschnikowia_pulcherrima(100); | Metschnikowia | 13 | Saprotroph | \|Undefined Saprotroph\| | Yeast | NULL | Probable | As nectar-tap_saprotroph (Põlme et al. 2020); Decay sugar-rich substrates (Põlme et al. 2020) | Põlme S, et al. 2020. Fungal Diversity 105:1-16 ((https://doi.org/10.1007/s13225-020-00466-2)) |
| Otu45938 | 1 | k__Fungi(100);p__Ascomycota(100);c__Saccharomycetes(100);o__Saccharomycetales(100);f__Metschnikowiaceae(100);g__Metschnikowia(100);s__Metschnikowia_pulcherrima(100); | Metschnikowia | 13 | Saprotroph | \|Undefined Saprotroph\| | Yeast | NULL | Probable | As nectar-tap_saprotroph (Põlme et al. 2020); Decay sugar-rich substrates (Põlme et al. 2020) | Põlme S, et al. 2020. Fungal Diversity 105:1-16 ((https://doi.org/10.1007/s13225-020-00466-2)) |
| Otu45939 | 1 | k__Fungi(100);p__Ascomycota(100);c__Saccharomycetes(100);o__Saccharomycetales(100);f__Metschnikowiaceae(100);g__Metschnikowia(100);s__Metschnikowia_pulcherrima(100); | Metschnikowia | 13 | Saprotroph | \|Undefined Saprotroph\| | Yeast | NULL | Probable | As nectar-tap_saprotroph (Põlme et al. 2020); Decay sugar-rich substrates (Põlme et al. 2020) | Põlme S, et al. 2020. Fungal Diversity 105:1-16 ((https://doi.org/10.1007/s13225-020-00466-2)) |
| Otu45940 | 1 | k__Fungi(100);p__Ascomycota(100);c__Saccharomycetes(100);o__Saccharomycetales(100);f__Metschnikowiaceae(100);g__Metschnikowia(100);g__Metschnikowia_unclassified(100); | Metschnikowia | 13 | Saprotroph | \|Undefined Saprotroph\| | Yeast | NULL | Probable | As nectar-tap_saprotroph (Põlme et al. 2020); Decay sugar-rich substrates (Põlme et al. 2020) | Põlme S, et al. 2020. Fungal Diversity 105:1-16 ((https://doi.org/10.1007/s13225-020-00466-2)) |
| Otu45941 | 1 | k__Fungi(100);p__Ascomycota(100);c__Saccharomycetes(100);o__Saccharomycetales(100);f__Metschnikowiaceae(100);g__Metschnikowia(100);s__Metschnikowia_pulcherrima(100); | Metschnikowia | 13 | Saprotroph | \|Undefined Saprotroph\| | Yeast | NULL | Probable | As nectar-tap_saprotroph (Põlme et al. 2020); Decay sugar-rich substrates (Põlme et al. 2020) | Põlme S, et al. 2020. Fungal Diversity 105:1-16 ((https://doi.org/10.1007/s13225-020-00466-2)) |
| Otu45942 | 1 | k__Fungi(100);p__Ascomycota(100);c__Saccharomycetes(100);o__Saccharomycetales(100);f__Metschnikowiaceae(100);g__Metschnikowia(100);s__Metschnikowia_pulcherrima(100); | Metschnikowia | 13 | Saprotroph | \|Undefined Saprotroph\| | Yeast | NULL | Probable | As nectar-tap_saprotroph (Põlme et al. 2020); Decay sugar-rich substrates (Põlme et al. 2020) | Põlme S, et al. 2020. Fungal Diversity 105:1-16 ((https://doi.org/10.1007/s13225-020-00466-2)) |
| Otu45943 | 1 | k__Fungi(100);p__Ascomycota(100);c__Saccharomycetes(100);o__Saccharomycetales(100);f__Metschnikowiaceae(100);g__Metschnikowia(100);s__Metschnikowia_pulcherrima(100); | Metschnikowia | 13 | Saprotroph | \|Undefined Saprotroph\| | Yeast | NULL | Probable | As nectar-tap_saprotroph (Põlme et al. 2020); Decay sugar-rich substrates (Põlme et al. 2020) | Põlme S, et al. 2020. Fungal Diversity 105:1-16 ((https://doi.org/10.1007/s13225-020-00466-2)) |
| Otu45944 | 1 | k__Fungi(100);p__Ascomycota(100);c__Saccharomycetes(100);o__Saccharomycetales(100);f__Metschnikowiaceae(100);g__Metschnikowia(100);s__Metschnikowia_pulcherrima(100); | Metschnikowia | 13 | Saprotroph | \|Undefined Saprotroph\| | Yeast | NULL | Probable | As nectar-tap_saprotroph (Põlme et al. 2020); Decay sugar-rich substrates (Põlme et al. 2020) | Põlme S, et al. 2020. Fungal Diversity 105:1-16 ((https://doi.org/10.1007/s13225-020-00466-2)) |
| Otu45945 | 1 | k__Fungi(100);p__Ascomycota(100);c__Saccharomycetes(100);o__Saccharomycetales(100);f__Metschnikowiaceae(100);g__Metschnikowia(100);g__Metschnikowia_unclassified(100); | Metschnikowia | 13 | Saprotroph | \|Undefined Saprotroph\| | Yeast | NULL | Probable | As nectar-tap_saprotroph (Põlme et al. 2020); Decay sugar-rich substrates (Põlme et al. 2020) | Põlme S, et al. 2020. Fungal Diversity 105:1-16 ((https://doi.org/10.1007/s13225-020-00466-2)) |
| Otu45946 | 1 | k__Fungi(100);p__Ascomycota(100);c__Saccharomycetes(100);o__Saccharomycetales(100);f__Metschnikowiaceae(100);g__Metschnikowia(100);s__Metschnikowia_pulcherrima(100); | Metschnikowia | 13 | Saprotroph | \|Undefined Saprotroph\| | Yeast | NULL | Probable | As nectar-tap_saprotroph (Põlme et al. 2020); Decay sugar-rich substrates (Põlme et al. 2020) | Põlme S, et al. 2020. Fungal Diversity 105:1-16 ((https://doi.org/10.1007/s13225-020-00466-2)) |
| Otu45947 | 1 | k__Fungi(100);p__Ascomycota(100);c__Saccharomycetes(100);o__Saccharomycetales(100);f__Metschnikowiaceae(100);g__Metschnikowia(100);s__Metschnikowia_pulcherrima(100); | Metschnikowia | 13 | Saprotroph | \|Undefined Saprotroph\| | Yeast | NULL | Probable | As nectar-tap_saprotroph (Põlme et al. 2020); Decay sugar-rich substrates (Põlme et al. 2020) | Põlme S, et al. 2020. Fungal Diversity 105:1-16 ((https://doi.org/10.1007/s13225-020-00466-2)) |
| Otu45948 | 1 | k__Fungi(100);p__Ascomycota(100);c__Saccharomycetes(100);o__Saccharomycetales(100);f__Metschnikowiaceae(100);g__Metschnikowia(100);s__Metschnikowia_pulcherrima(100); | Metschnikowia | 13 | Saprotroph | \|Undefined Saprotroph\| | Yeast | NULL | Probable | As nectar-tap_saprotroph (Põlme et al. 2020); Decay sugar-rich substrates (Põlme et al. 2020) | Põlme S, et al. 2020. Fungal Diversity 105:1-16 ((https://doi.org/10.1007/s13225-020-00466-2)) |
| Otu45949 | 1 | k__Fungi(100);p__Ascomycota(100);c__Saccharomycetes(100);o__Saccharomycetales(100);f__Metschnikowiaceae(100);g__Metschnikowia(100);g__Metschnikowia_unclassified(100); | Metschnikowia | 13 | Saprotroph | \|Undefined Saprotroph\| | Yeast | NULL | Probable | As nectar-tap_saprotroph (Põlme et al. 2020); Decay sugar-rich substrates (Põlme et al. 2020) | Põlme S, et al. 2020. Fungal Diversity 105:1-16 ((https://doi.org/10.1007/s13225-020-00466-2)) |
| Otu45950 | 1 | k__Fungi(100);p__Ascomycota(100);c__Saccharomycetes(100);o__Saccharomycetales(100);f__Metschnikowiaceae(100);g__Metschnikowia(100);s__Metschnikowia_pulcherrima(100); | Metschnikowia | 13 | Saprotroph | \|Undefined Saprotroph\| | Yeast | NULL | Probable | As nectar-tap_saprotroph (Põlme et al. 2020); Decay sugar-rich substrates (Põlme et al. 2020) | Põlme S, et al. 2020. Fungal Diversity 105:1-16 ((https://doi.org/10.1007/s13225-020-00466-2)) |
| Otu45951 | 1 | k__Fungi(100);p__Ascomycota(100);c__Saccharomycetes(100);o__Saccharomycetales(100);f__Metschnikowiaceae(100);g__Metschnikowia(100);s__Metschnikowia_pulcherrima(100); | Metschnikowia | 13 | Saprotroph | \|Undefined Saprotroph\| | Yeast | NULL | Probable | As nectar-tap_saprotroph (Põlme et al. 2020); Decay sugar-rich substrates (Põlme et al. 2020) | Põlme S, et al. 2020. Fungal Diversity 105:1-16 ((https://doi.org/10.1007/s13225-020-00466-2)) |
| Otu45952 | 1 | k__Fungi(100);p__Ascomycota(100);c__Saccharomycetes(100);o__Saccharomycetales(100);f__Metschnikowiaceae(100);g__Metschnikowia(100);g__Metschnikowia_unclassified(100); | Metschnikowia | 13 | Saprotroph | \|Undefined Saprotroph\| | Yeast | NULL | Probable | As nectar-tap_saprotroph (Põlme et al. 2020); Decay sugar-rich substrates (Põlme et al. 2020) | Põlme S, et al. 2020. Fungal Diversity 105:1-16 ((https://doi.org/10.1007/s13225-020-00466-2)) |
| Otu45953 | 1 | k__Fungi(100);p__Ascomycota(100);c__Saccharomycetes(100);o__Saccharomycetales(100);f__Metschnikowiaceae(100);g__Metschnikowia(100);s__Metschnikowia_pulcherrima(100); | Metschnikowia | 13 | Saprotroph | \|Undefined Saprotroph\| | Yeast | NULL | Probable | As nectar-tap_saprotroph (Põlme et al. 2020); Decay sugar-rich substrates (Põlme et al. 2020) | Põlme S, et al. 2020. Fungal Diversity 105:1-16 ((https://doi.org/10.1007/s13225-020-00466-2)) |
| Otu45954 | 1 | k__Fungi(100);p__Ascomycota(100);c__Saccharomycetes(100);o__Saccharomycetales(100);f__Metschnikowiaceae(100);g__Metschnikowia(100);s__Metschnikowia_pulcherrima(100); | Metschnikowia | 13 | Saprotroph | \|Undefined Saprotroph\| | Yeast | NULL | Probable | As nectar-tap_saprotroph (Põlme et al. 2020); Decay sugar-rich substrates (Põlme et al. 2020) | Põlme S, et al. 2020. Fungal Diversity 105:1-16 ((https://doi.org/10.1007/s13225-020-00466-2)) |
| Otu45955 | 1 | k__Fungi(100);p__Ascomycota(100);c__Saccharomycetes(100);o__Saccharomycetales(100);f__Metschnikowiaceae(100);g__Metschnikowia(100);g__Metschnikowia_unclassified(100); | Metschnikowia | 13 | Saprotroph | \|Undefined Saprotroph\| | Yeast | NULL | Probable | As nectar-tap_saprotroph (Põlme et al. 2020); Decay sugar-rich substrates (Põlme et al. 2020) | Põlme S, et al. 2020. Fungal Diversity 105:1-16 ((https://doi.org/10.1007/s13225-020-00466-2)) |
| Otu45956 | 1 | k__Fungi(100);p__Ascomycota(100);c__Saccharomycetes(100);o__Saccharomycetales(100);f__Metschnikowiaceae(100);g__Metschnikowia(100);s__Metschnikowia_pulcherrima(100); | Metschnikowia | 13 | Saprotroph | \|Undefined Saprotroph\| | Yeast | NULL | Probable | As nectar-tap_saprotroph (Põlme et al. 2020); Decay sugar-rich substrates (Põlme et al. 2020) | Põlme S, et al. 2020. Fungal Diversity 105:1-16 ((https://doi.org/10.1007/s13225-020-00466-2)) |
| Otu45957 | 1 | k__Fungi(100);p__Ascomycota(100);c__Saccharomycetes(100);o__Saccharomycetales(100);f__Metschnikowiaceae(100);g__Metschnikowia(100);s__Metschnikowia_pulcherrima(100); | Metschnikowia | 13 | Saprotroph | \|Undefined Saprotroph\| | Yeast | NULL | Probable | As nectar-tap_saprotroph (Põlme et al. 2020); Decay sugar-rich substrates (Põlme et al. 2020) | Põlme S, et al. 2020. Fungal Diversity 105:1-16 ((https://doi.org/10.1007/s13225-020-00466-2)) |
| Otu45958 | 1 | k__Fungi(100);p__Ascomycota(100);c__Saccharomycetes(100);o__Saccharomycetales(100);f__Metschnikowiaceae(100);g__Metschnikowia(100);s__Metschnikowia_pulcherrima(100); | Metschnikowia | 13 | Saprotroph | \|Undefined Saprotroph\| | Yeast | NULL | Probable | As nectar-tap_saprotroph (Põlme et al. 2020); Decay sugar-rich substrates (Põlme et al. 2020) | Põlme S, et al. 2020. Fungal Diversity 105:1-16 ((https://doi.org/10.1007/s13225-020-00466-2)) |
| Otu45959 | 1 | k__Fungi(100);p__Ascomycota(100);c__Saccharomycetes(100);o__Saccharomycetales(100);f__Metschnikowiaceae(100);g__Metschnikowia(100);s__Metschnikowia_pulcherrima(100); | Metschnikowia | 13 | Saprotroph | \|Undefined Saprotroph\| | Yeast | NULL | Probable | As nectar-tap_saprotroph (Põlme et al. 2020); Decay sugar-rich substrates (Põlme et al. 2020) | Põlme S, et al. 2020. Fungal Diversity 105:1-16 ((https://doi.org/10.1007/s13225-020-00466-2)) |
| Otu45960 | 1 | k__Fungi(100);p__Ascomycota(100);c__Saccharomycetes(100);o__Saccharomycetales(100);f__Metschnikowiaceae(100);g__Metschnikowia(100);s__Metschnikowia_pulcherrima(100); | Metschnikowia | 13 | Saprotroph | \|Undefined Saprotroph\| | Yeast | NULL | Probable | As nectar-tap_saprotroph (Põlme et al. 2020); Decay sugar-rich substrates (Põlme et al. 2020) | Põlme S, et al. 2020. Fungal Diversity 105:1-16 ((https://doi.org/10.1007/s13225-020-00466-2)) |
| Otu45961 | 1 | k__Fungi(100);p__Ascomycota(100);c__Saccharomycetes(100);o__Saccharomycetales(100);f__Metschnikowiaceae(100);g__Metschnikowia(100);s__Metschnikowia_pulcherrima(100); | Metschnikowia | 13 | Saprotroph | \|Undefined Saprotroph\| | Yeast | NULL | Probable | As nectar-tap_saprotroph (Põlme et al. 2020); Decay sugar-rich substrates (Põlme et al. 2020) | Põlme S, et al. 2020. Fungal Diversity 105:1-16 ((https://doi.org/10.1007/s13225-020-00466-2)) |
| Otu45962 | 1 | k__Fungi(100);p__Ascomycota(100);c__Saccharomycetes(100);o__Saccharomycetales(100);f__Metschnikowiaceae(100);g__Metschnikowia(100);s__Metschnikowia_pulcherrima(100); | Metschnikowia | 13 | Saprotroph | \|Undefined Saprotroph\| | Yeast | NULL | Probable | As nectar-tap_saprotroph (Põlme et al. 2020); Decay sugar-rich substrates (Põlme et al. 2020) | Põlme S, et al. 2020. Fungal Diversity 105:1-16 ((https://doi.org/10.1007/s13225-020-00466-2)) |
| Otu45963 | 1 | k__Fungi(100);p__Ascomycota(100);c__Saccharomycetes(100);o__Saccharomycetales(100);f__Metschnikowiaceae(100);g__Metschnikowia(100);s__Metschnikowia_pulcherrima(100); | Metschnikowia | 13 | Saprotroph | \|Undefined Saprotroph\| | Yeast | NULL | Probable | As nectar-tap_saprotroph (Põlme et al. 2020); Decay sugar-rich substrates (Põlme et al. 2020) | Põlme S, et al. 2020. Fungal Diversity 105:1-16 ((https://doi.org/10.1007/s13225-020-00466-2)) |
| Otu45964 | 1 | k__Fungi(100);p__Ascomycota(100);c__Saccharomycetes(100);o__Saccharomycetales(100);f__Metschnikowiaceae(100);g__Metschnikowia(100);g__Metschnikowia_unclassified(100); | Metschnikowia | 13 | Saprotroph | \|Undefined Saprotroph\| | Yeast | NULL | Probable | As nectar-tap_saprotroph (Põlme et al. 2020); Decay sugar-rich substrates (Põlme et al. 2020) | Põlme S, et al. 2020. Fungal Diversity 105:1-16 ((https://doi.org/10.1007/s13225-020-00466-2)) |
| Otu45965 | 1 | k__Fungi(100);p__Ascomycota(100);c__Saccharomycetes(100);o__Saccharomycetales(100);f__Metschnikowiaceae(100);g__Metschnikowia(100);s__Metschnikowia_pulcherrima(100); | Metschnikowia | 13 | Saprotroph | \|Undefined Saprotroph\| | Yeast | NULL | Probable | As nectar-tap_saprotroph (Põlme et al. 2020); Decay sugar-rich substrates (Põlme et al. 2020) | Põlme S, et al. 2020. Fungal Diversity 105:1-16 ((https://doi.org/10.1007/s13225-020-00466-2)) |
| Otu45966 | 1 | k__Fungi(100);p__Ascomycota(100);c__Saccharomycetes(100);o__Saccharomycetales(100);f__Metschnikowiaceae(100);g__Metschnikowia(100);s__Metschnikowia_pulcherrima(100); | Metschnikowia | 13 | Saprotroph | \|Undefined Saprotroph\| | Yeast | NULL | Probable | As nectar-tap_saprotroph (Põlme et al. 2020); Decay sugar-rich substrates (Põlme et al. 2020) | Põlme S, et al. 2020. Fungal Diversity 105:1-16 ((https://doi.org/10.1007/s13225-020-00466-2)) |
| Otu45967 | 1 | k__Fungi(100);p__Ascomycota(100);c__Saccharomycetes(100);o__Saccharomycetales(100);f__Metschnikowiaceae(100);g__Metschnikowia(100);s__Metschnikowia_pulcherrima(100); | Metschnikowia | 13 | Saprotroph | \|Undefined Saprotroph\| | Yeast | NULL | Probable | As nectar-tap_saprotroph (Põlme et al. 2020); Decay sugar-rich substrates (Põlme et al. 2020) | Põlme S, et al. 2020. Fungal Diversity 105:1-16 ((https://doi.org/10.1007/s13225-020-00466-2)) |
| Otu45968 | 1 | k__Fungi(100);p__Ascomycota(100);c__Saccharomycetes(100);o__Saccharomycetales(100);f__Metschnikowiaceae(100);g__Metschnikowia(100);s__Metschnikowia_pulcherrima(100); | Metschnikowia | 13 | Saprotroph | \|Undefined Saprotroph\| | Yeast | NULL | Probable | As nectar-tap_saprotroph (Põlme et al. 2020); Decay sugar-rich substrates (Põlme et al. 2020) | Põlme S, et al. 2020. Fungal Diversity 105:1-16 ((https://doi.org/10.1007/s13225-020-00466-2)) |
| Otu45969 | 1 | k__Fungi(100);p__Ascomycota(100);c__Saccharomycetes(100);o__Saccharomycetales(100);f__Metschnikowiaceae(100);g__Metschnikowia(100);s__Metschnikowia_pulcherrima(100); | Metschnikowia | 13 | Saprotroph | \|Undefined Saprotroph\| | Yeast | NULL | Probable | As nectar-tap_saprotroph (Põlme et al. 2020); Decay sugar-rich substrates (Põlme et al. 2020) | Põlme S, et al. 2020. Fungal Diversity 105:1-16 ((https://doi.org/10.1007/s13225-020-00466-2)) |
| Otu45970 | 1 | k__Fungi(100);p__Ascomycota(100);c__Saccharomycetes(100);o__Saccharomycetales(100);f__Metschnikowiaceae(100);g__Metschnikowia(100);s__Metschnikowia_pulcherrima(100); | Metschnikowia | 13 | Saprotroph | \|Undefined Saprotroph\| | Yeast | NULL | Probable | As nectar-tap_saprotroph (Põlme et al. 2020); Decay sugar-rich substrates (Põlme et al. 2020) | Põlme S, et al. 2020. Fungal Diversity 105:1-16 ((https://doi.org/10.1007/s13225-020-00466-2)) |
| Otu45971 | 1 | k__Fungi(100);p__Ascomycota(100);c__Saccharomycetes(100);o__Saccharomycetales(100);f__Metschnikowiaceae(100);g__Metschnikowia(100);g__Metschnikowia_unclassified(100); | Metschnikowia | 13 | Saprotroph | \|Undefined Saprotroph\| | Yeast | NULL | Probable | As nectar-tap_saprotroph (Põlme et al. 2020); Decay sugar-rich substrates (Põlme et al. 2020) | Põlme S, et al. 2020. Fungal Diversity 105:1-16 ((https://doi.org/10.1007/s13225-020-00466-2)) |
| Otu45972 | 1 | k__Fungi(100);p__Ascomycota(100);c__Saccharomycetes(100);o__Saccharomycetales(100);f__Metschnikowiaceae(100);g__Metschnikowia(100);s__Metschnikowia_pulcherrima(100); | Metschnikowia | 13 | Saprotroph | \|Undefined Saprotroph\| | Yeast | NULL | Probable | As nectar-tap_saprotroph (Põlme et al. 2020); Decay sugar-rich substrates (Põlme et al. 2020) | Põlme S, et al. 2020. Fungal Diversity 105:1-16 ((https://doi.org/10.1007/s13225-020-00466-2)) |
| Otu45973 | 1 | k__Fungi(100);p__Ascomycota(100);c__Saccharomycetes(100);o__Saccharomycetales(100);f__Metschnikowiaceae(100);g__Metschnikowia(100);s__Metschnikowia_pulcherrima(100); | Metschnikowia | 13 | Saprotroph | \|Undefined Saprotroph\| | Yeast | NULL | Probable | As nectar-tap_saprotroph (Põlme et al. 2020); Decay sugar-rich substrates (Põlme et al. 2020) | Põlme S, et al. 2020. Fungal Diversity 105:1-16 ((https://doi.org/10.1007/s13225-020-00466-2)) |
| Otu45974 | 1 | k__Fungi(100);p__Ascomycota(100);c__Saccharomycetes(100);o__Saccharomycetales(100);f__Metschnikowiaceae(100);g__Metschnikowia(100);s__Metschnikowia_pulcherrima(100); | Metschnikowia | 13 | Saprotroph | \|Undefined Saprotroph\| | Yeast | NULL | Probable | As nectar-tap_saprotroph (Põlme et al. 2020); Decay sugar-rich substrates (Põlme et al. 2020) | Põlme S, et al. 2020. Fungal Diversity 105:1-16 ((https://doi.org/10.1007/s13225-020-00466-2)) |
| Otu45975 | 1 | k__Fungi(100);p__Ascomycota(100);c__Saccharomycetes(100);o__Saccharomycetales(100);f__Metschnikowiaceae(100);g__Metschnikowia(100);g__Metschnikowia_unclassified(100); | Metschnikowia | 13 | Saprotroph | \|Undefined Saprotroph\| | Yeast | NULL | Probable | As nectar-tap_saprotroph (Põlme et al. 2020); Decay sugar-rich substrates (Põlme et al. 2020) | Põlme S, et al. 2020. Fungal Diversity 105:1-16 ((https://doi.org/10.1007/s13225-020-00466-2)) |
| Otu45976 | 1 | k__Fungi(100);p__Ascomycota(100);c__Saccharomycetes(100);o__Saccharomycetales(100);f__Metschnikowiaceae(100);g__Metschnikowia(100);s__Metschnikowia_pulcherrima(100); | Metschnikowia | 13 | Saprotroph | \|Undefined Saprotroph\| | Yeast | NULL | Probable | As nectar-tap_saprotroph (Põlme et al. 2020); Decay sugar-rich substrates (Põlme et al. 2020) | Põlme S, et al. 2020. Fungal Diversity 105:1-16 ((https://doi.org/10.1007/s13225-020-00466-2)) |
| Otu45977 | 1 | k__Fungi(100);p__Ascomycota(100);c__Saccharomycetes(100);o__Saccharomycetales(100);f__Metschnikowiaceae(100);g__Metschnikowia(100);s__Metschnikowia_pulcherrima(100); | Metschnikowia | 13 | Saprotroph | \|Undefined Saprotroph\| | Yeast | NULL | Probable | As nectar-tap_saprotroph (Põlme et al. 2020); Decay sugar-rich substrates (Põlme et al. 2020) | Põlme S, et al. 2020. Fungal Diversity 105:1-16 ((https://doi.org/10.1007/s13225-020-00466-2)) |
| Otu45978 | 1 | k__Fungi(100);p__Ascomycota(100);c__Saccharomycetes(100);o__Saccharomycetales(100);f__Metschnikowiaceae(100);g__Metschnikowia(100);g__Metschnikowia_unclassified(100); | Metschnikowia | 13 | Saprotroph | \|Undefined Saprotroph\| | Yeast | NULL | Probable | As nectar-tap_saprotroph (Põlme et al. 2020); Decay sugar-rich substrates (Põlme et al. 2020) | Põlme S, et al. 2020. Fungal Diversity 105:1-16 ((https://doi.org/10.1007/s13225-020-00466-2)) |
| Otu45979 | 1 | k__Fungi(100);p__Ascomycota(100);c__Saccharomycetes(100);o__Saccharomycetales(100);f__Metschnikowiaceae(100);g__Metschnikowia(100);s__Metschnikowia_pulcherrima(100); | Metschnikowia | 13 | Saprotroph | \|Undefined Saprotroph\| | Yeast | NULL | Probable | As nectar-tap_saprotroph (Põlme et al. 2020); Decay sugar-rich substrates (Põlme et al. 2020) | Põlme S, et al. 2020. Fungal Diversity 105:1-16 ((https://doi.org/10.1007/s13225-020-00466-2)) |
| Otu45980 | 1 | k__Fungi(100);p__Ascomycota(100);c__Saccharomycetes(100);o__Saccharomycetales(100);f__Metschnikowiaceae(100);g__Metschnikowia(100);s__Metschnikowia_pulcherrima(100); | Metschnikowia | 13 | Saprotroph | \|Undefined Saprotroph\| | Yeast | NULL | Probable | As nectar-tap_saprotroph (Põlme et al. 2020); Decay sugar-rich substrates (Põlme et al. 2020) | Põlme S, et al. 2020. Fungal Diversity 105:1-16 ((https://doi.org/10.1007/s13225-020-00466-2)) |
| Otu45982 | 1 | k__Fungi(100);p__Ascomycota(100);c__Saccharomycetes(100);o__Saccharomycetales(100);f__Metschnikowiaceae(100);g__Metschnikowia(100);s__Metschnikowia_pulcherrima(100); | Metschnikowia | 13 | Saprotroph | \|Undefined Saprotroph\| | Yeast | NULL | Probable | As nectar-tap_saprotroph (Põlme et al. 2020); Decay sugar-rich substrates (Põlme et al. 2020) | Põlme S, et al. 2020. Fungal Diversity 105:1-16 ((https://doi.org/10.1007/s13225-020-00466-2)) |
| Otu45983 | 1 | k__Fungi(100);p__Ascomycota(100);c__Saccharomycetes(100);o__Saccharomycetales(100);f__Metschnikowiaceae(100);g__Metschnikowia(100);s__Metschnikowia_pulcherrima(100); | Metschnikowia | 13 | Saprotroph | \|Undefined Saprotroph\| | Yeast | NULL | Probable | As nectar-tap_saprotroph (Põlme et al. 2020); Decay sugar-rich substrates (Põlme et al. 2020) | Põlme S, et al. 2020. Fungal Diversity 105:1-16 ((https://doi.org/10.1007/s13225-020-00466-2)) |
| Otu45984 | 1 | k__Fungi(100);p__Ascomycota(100);c__Saccharomycetes(100);o__Saccharomycetales(100);f__Metschnikowiaceae(100);g__Metschnikowia(100);g__Metschnikowia_unclassified(100); | Metschnikowia | 13 | Saprotroph | \|Undefined Saprotroph\| | Yeast | NULL | Probable | As nectar-tap_saprotroph (Põlme et al. 2020); Decay sugar-rich substrates (Põlme et al. 2020) | Põlme S, et al. 2020. Fungal Diversity 105:1-16 ((https://doi.org/10.1007/s13225-020-00466-2)) |
| Otu45985 | 1 | k__Fungi(100);p__Ascomycota(100);c__Saccharomycetes(100);o__Saccharomycetales(100);f__Metschnikowiaceae(100);g__Metschnikowia(100);s__Metschnikowia_pulcherrima(100); | Metschnikowia | 13 | Saprotroph | \|Undefined Saprotroph\| | Yeast | NULL | Probable | As nectar-tap_saprotroph (Põlme et al. 2020); Decay sugar-rich substrates (Põlme et al. 2020) | Põlme S, et al. 2020. Fungal Diversity 105:1-16 ((https://doi.org/10.1007/s13225-020-00466-2)) |
| Otu45986 | 1 | k__Fungi(100);p__Ascomycota(100);c__Saccharomycetes(100);o__Saccharomycetales(100);f__Metschnikowiaceae(100);g__Metschnikowia(100);s__Metschnikowia_pulcherrima(100); | Metschnikowia | 13 | Saprotroph | \|Undefined Saprotroph\| | Yeast | NULL | Probable | As nectar-tap_saprotroph (Põlme et al. 2020); Decay sugar-rich substrates (Põlme et al. 2020) | Põlme S, et al. 2020. Fungal Diversity 105:1-16 ((https://doi.org/10.1007/s13225-020-00466-2)) |
| Otu45987 | 1 | k__Fungi(100);p__Ascomycota(100);c__Saccharomycetes(100);o__Saccharomycetales(100);f__Metschnikowiaceae(100);g__Metschnikowia(100);g__Metschnikowia_unclassified(100); | Metschnikowia | 13 | Saprotroph | \|Undefined Saprotroph\| | Yeast | NULL | Probable | As nectar-tap_saprotroph (Põlme et al. 2020); Decay sugar-rich substrates (Põlme et al. 2020) | Põlme S, et al. 2020. Fungal Diversity 105:1-16 ((https://doi.org/10.1007/s13225-020-00466-2)) |
| Otu45988 | 1 | k__Fungi(100);p__Ascomycota(100);c__Saccharomycetes(100);o__Saccharomycetales(100);f__Metschnikowiaceae(100);g__Metschnikowia(100);s__Metschnikowia_pulcherrima(100); | Metschnikowia | 13 | Saprotroph | \|Undefined Saprotroph\| | Yeast | NULL | Probable | As nectar-tap_saprotroph (Põlme et al. 2020); Decay sugar-rich substrates (Põlme et al. 2020) | Põlme S, et al. 2020. Fungal Diversity 105:1-16 ((https://doi.org/10.1007/s13225-020-00466-2)) |
| Otu45989 | 1 | k__Fungi(100);p__Ascomycota(100);c__Saccharomycetes(100);o__Saccharomycetales(100);f__Metschnikowiaceae(100);g__Metschnikowia(100);s__Metschnikowia_pulcherrima(100); | Metschnikowia | 13 | Saprotroph | \|Undefined Saprotroph\| | Yeast | NULL | Probable | As nectar-tap_saprotroph (Põlme et al. 2020); Decay sugar-rich substrates (Põlme et al. 2020) | Põlme S, et al. 2020. Fungal Diversity 105:1-16 ((https://doi.org/10.1007/s13225-020-00466-2)) |
| Otu45990 | 1 | k__Fungi(100);p__Ascomycota(100);c__Saccharomycetes(100);o__Saccharomycetales(100);f__Metschnikowiaceae(100);g__Metschnikowia(100);g__Metschnikowia_unclassified(100); | Metschnikowia | 13 | Saprotroph | \|Undefined Saprotroph\| | Yeast | NULL | Probable | As nectar-tap_saprotroph (Põlme et al. 2020); Decay sugar-rich substrates (Põlme et al. 2020) | Põlme S, et al. 2020. Fungal Diversity 105:1-16 ((https://doi.org/10.1007/s13225-020-00466-2)) |
| Otu45991 | 1 | k__Fungi(100);p__Ascomycota(100);c__Saccharomycetes(100);o__Saccharomycetales(100);f__Metschnikowiaceae(100);g__Metschnikowia(100);s__Metschnikowia_pulcherrima(100); | Metschnikowia | 13 | Saprotroph | \|Undefined Saprotroph\| | Yeast | NULL | Probable | As nectar-tap_saprotroph (Põlme et al. 2020); Decay sugar-rich substrates (Põlme et al. 2020) | Põlme S, et al. 2020. Fungal Diversity 105:1-16 ((https://doi.org/10.1007/s13225-020-00466-2)) |
| Otu45992 | 1 | k__Fungi(100);p__Ascomycota(100);c__Saccharomycetes(100);o__Saccharomycetales(100);f__Metschnikowiaceae(100);g__Metschnikowia(100);s__Metschnikowia_pulcherrima(100); | Metschnikowia | 13 | Saprotroph | \|Undefined Saprotroph\| | Yeast | NULL | Probable | As nectar-tap_saprotroph (Põlme et al. 2020); Decay sugar-rich substrates (Põlme et al. 2020) | Põlme S, et al. 2020. Fungal Diversity 105:1-16 ((https://doi.org/10.1007/s13225-020-00466-2)) |
| Otu45993 | 1 | k__Fungi(100);p__Ascomycota(100);c__Saccharomycetes(100);o__Saccharomycetales(100);f__Metschnikowiaceae(100);g__Metschnikowia(100);s__Metschnikowia_pulcherrima(100); | Metschnikowia | 13 | Saprotroph | \|Undefined Saprotroph\| | Yeast | NULL | Probable | As nectar-tap_saprotroph (Põlme et al. 2020); Decay sugar-rich substrates (Põlme et al. 2020) | Põlme S, et al. 2020. Fungal Diversity 105:1-16 ((https://doi.org/10.1007/s13225-020-00466-2)) |
| Otu45994 | 1 | k__Fungi(100);p__Ascomycota(100);c__Saccharomycetes(100);o__Saccharomycetales(100);f__Metschnikowiaceae(100);g__Metschnikowia(100);s__Metschnikowia_pulcherrima(100); | Metschnikowia | 13 | Saprotroph | \|Undefined Saprotroph\| | Yeast | NULL | Probable | As nectar-tap_saprotroph (Põlme et al. 2020); Decay sugar-rich substrates (Põlme et al. 2020) | Põlme S, et al. 2020. Fungal Diversity 105:1-16 ((https://doi.org/10.1007/s13225-020-00466-2)) |
| Otu45995 | 1 | k__Fungi(100);p__Ascomycota(100);c__Saccharomycetes(100);o__Saccharomycetales(100);f__Metschnikowiaceae(100);g__Metschnikowia(100);s__Metschnikowia_pulcherrima(100); | Metschnikowia | 13 | Saprotroph | \|Undefined Saprotroph\| | Yeast | NULL | Probable | As nectar-tap_saprotroph (Põlme et al. 2020); Decay sugar-rich substrates (Põlme et al. 2020) | Põlme S, et al. 2020. Fungal Diversity 105:1-16 ((https://doi.org/10.1007/s13225-020-00466-2)) |
| Otu45996 | 1 | k__Fungi(100);p__Ascomycota(100);c__Saccharomycetes(100);o__Saccharomycetales(100);f__Metschnikowiaceae(100);g__Metschnikowia(100);s__Metschnikowia_pulcherrima(100); | Metschnikowia | 13 | Saprotroph | \|Undefined Saprotroph\| | Yeast | NULL | Probable | As nectar-tap_saprotroph (Põlme et al. 2020); Decay sugar-rich substrates (Põlme et al. 2020) | Põlme S, et al. 2020. Fungal Diversity 105:1-16 ((https://doi.org/10.1007/s13225-020-00466-2)) |
| Otu45998 | 1 | k__Fungi(100);p__Ascomycota(100);c__Saccharomycetes(100);o__Saccharomycetales(100);f__Metschnikowiaceae(100);g__Metschnikowia(100);s__Metschnikowia_pulcherrima(100); | Metschnikowia | 13 | Saprotroph | \|Undefined Saprotroph\| | Yeast | NULL | Probable | As nectar-tap_saprotroph (Põlme et al. 2020); Decay sugar-rich substrates (Põlme et al. 2020) | Põlme S, et al. 2020. Fungal Diversity 105:1-16 ((https://doi.org/10.1007/s13225-020-00466-2)) |
| Otu45999 | 1 | k__Fungi(100);p__Ascomycota(100);c__Saccharomycetes(100);o__Saccharomycetales(100);f__Metschnikowiaceae(100);g__Metschnikowia(100);s__Metschnikowia_pulcherrima(100); | Metschnikowia | 13 | Saprotroph | \|Undefined Saprotroph\| | Yeast | NULL | Probable | As nectar-tap_saprotroph (Põlme et al. 2020); Decay sugar-rich substrates (Põlme et al. 2020) | Põlme S, et al. 2020. Fungal Diversity 105:1-16 ((https://doi.org/10.1007/s13225-020-00466-2)) |
| Otu46000 | 1 | k__Fungi(100);p__Ascomycota(100);c__Saccharomycetes(100);o__Saccharomycetales(100);f__Metschnikowiaceae(100);g__Metschnikowia(100);g__Metschnikowia_unclassified(100); | Metschnikowia | 13 | Saprotroph | \|Undefined Saprotroph\| | Yeast | NULL | Probable | As nectar-tap_saprotroph (Põlme et al. 2020); Decay sugar-rich substrates (Põlme et al. 2020) | Põlme S, et al. 2020. Fungal Diversity 105:1-16 ((https://doi.org/10.1007/s13225-020-00466-2)) |
| Otu46001 | 1 | k__Fungi(100);p__Ascomycota(100);c__Saccharomycetes(100);o__Saccharomycetales(100);f__Metschnikowiaceae(100);g__Metschnikowia(100);s__Metschnikowia_pulcherrima(100); | Metschnikowia | 13 | Saprotroph | \|Undefined Saprotroph\| | Yeast | NULL | Probable | As nectar-tap_saprotroph (Põlme et al. 2020); Decay sugar-rich substrates (Põlme et al. 2020) | Põlme S, et al. 2020. Fungal Diversity 105:1-16 ((https://doi.org/10.1007/s13225-020-00466-2)) |
| Otu46002 | 1 | k__Fungi(100);p__Ascomycota(100);c__Saccharomycetes(100);o__Saccharomycetales(100);f__Metschnikowiaceae(100);g__Metschnikowia(100);s__Metschnikowia_pulcherrima(100); | Metschnikowia | 13 | Saprotroph | \|Undefined Saprotroph\| | Yeast | NULL | Probable | As nectar-tap_saprotroph (Põlme et al. 2020); Decay sugar-rich substrates (Põlme et al. 2020) | Põlme S, et al. 2020. Fungal Diversity 105:1-16 ((https://doi.org/10.1007/s13225-020-00466-2)) |
| Otu46003 | 1 | k__Fungi(100);p__Ascomycota(100);c__Saccharomycetes(100);o__Saccharomycetales(100);f__Metschnikowiaceae(100);g__Metschnikowia(100);s__Metschnikowia_pulcherrima(100); | Metschnikowia | 13 | Saprotroph | \|Undefined Saprotroph\| | Yeast | NULL | Probable | As nectar-tap_saprotroph (Põlme et al. 2020); Decay sugar-rich substrates (Põlme et al. 2020) | Põlme S, et al. 2020. Fungal Diversity 105:1-16 ((https://doi.org/10.1007/s13225-020-00466-2)) |
| Otu46004 | 1 | k__Fungi(100);p__Ascomycota(100);c__Saccharomycetes(100);o__Saccharomycetales(100);f__Metschnikowiaceae(100);g__Metschnikowia(100);s__Metschnikowia_pulcherrima(100); | Metschnikowia | 13 | Saprotroph | \|Undefined Saprotroph\| | Yeast | NULL | Probable | As nectar-tap_saprotroph (Põlme et al. 2020); Decay sugar-rich substrates (Põlme et al. 2020) | Põlme S, et al. 2020. Fungal Diversity 105:1-16 ((https://doi.org/10.1007/s13225-020-00466-2)) |
| Otu46005 | 1 | k__Fungi(100);p__Ascomycota(100);c__Saccharomycetes(100);o__Saccharomycetales(100);f__Metschnikowiaceae(100);g__Metschnikowia(100);s__Metschnikowia_pulcherrima(100); | Metschnikowia | 13 | Saprotroph | \|Undefined Saprotroph\| | Yeast | NULL | Probable | As nectar-tap_saprotroph (Põlme et al. 2020); Decay sugar-rich substrates (Põlme et al. 2020) | Põlme S, et al. 2020. Fungal Diversity 105:1-16 ((https://doi.org/10.1007/s13225-020-00466-2)) |
| Otu46006 | 1 | k__Fungi(100);p__Ascomycota(100);c__Saccharomycetes(100);o__Saccharomycetales(100);f__Metschnikowiaceae(100);g__Metschnikowia(100);s__Metschnikowia_pulcherrima(100); | Metschnikowia | 13 | Saprotroph | \|Undefined Saprotroph\| | Yeast | NULL | Probable | As nectar-tap_saprotroph (Põlme et al. 2020); Decay sugar-rich substrates (Põlme et al. 2020) | Põlme S, et al. 2020. Fungal Diversity 105:1-16 ((https://doi.org/10.1007/s13225-020-00466-2)) |
| Otu46007 | 1 | k__Fungi(100);p__Ascomycota(100);c__Saccharomycetes(100);o__Saccharomycetales(100);f__Metschnikowiaceae(100);g__Metschnikowia(100);s__Metschnikowia_pulcherrima(100); | Metschnikowia | 13 | Saprotroph | \|Undefined Saprotroph\| | Yeast | NULL | Probable | As nectar-tap_saprotroph (Põlme et al. 2020); Decay sugar-rich substrates (Põlme et al. 2020) | Põlme S, et al. 2020. Fungal Diversity 105:1-16 ((https://doi.org/10.1007/s13225-020-00466-2)) |
| Otu46008 | 1 | k__Fungi(100);p__Ascomycota(100);c__Saccharomycetes(100);o__Saccharomycetales(100);f__Metschnikowiaceae(100);g__Metschnikowia(100);g__Metschnikowia_unclassified(100); | Metschnikowia | 13 | Saprotroph | \|Undefined Saprotroph\| | Yeast | NULL | Probable | As nectar-tap_saprotroph (Põlme et al. 2020); Decay sugar-rich substrates (Põlme et al. 2020) | Põlme S, et al. 2020. Fungal Diversity 105:1-16 ((https://doi.org/10.1007/s13225-020-00466-2)) |
| Otu46009 | 1 | k__Fungi(100);p__Ascomycota(100);c__Saccharomycetes(100);o__Saccharomycetales(100);f__Metschnikowiaceae(100);g__Metschnikowia(100);s__Metschnikowia_pulcherrima(100); | Metschnikowia | 13 | Saprotroph | \|Undefined Saprotroph\| | Yeast | NULL | Probable | As nectar-tap_saprotroph (Põlme et al. 2020); Decay sugar-rich substrates (Põlme et al. 2020) | Põlme S, et al. 2020. Fungal Diversity 105:1-16 ((https://doi.org/10.1007/s13225-020-00466-2)) |
| Otu46010 | 1 | k__Fungi(100);p__Ascomycota(100);c__Saccharomycetes(100);o__Saccharomycetales(100);f__Metschnikowiaceae(100);g__Metschnikowia(100);s__Metschnikowia_pulcherrima(100); | Metschnikowia | 13 | Saprotroph | \|Undefined Saprotroph\| | Yeast | NULL | Probable | As nectar-tap_saprotroph (Põlme et al. 2020); Decay sugar-rich substrates (Põlme et al. 2020) | Põlme S, et al. 2020. Fungal Diversity 105:1-16 ((https://doi.org/10.1007/s13225-020-00466-2)) |
| Otu46011 | 1 | k__Fungi(100);p__Ascomycota(100);c__Saccharomycetes(100);o__Saccharomycetales(100);f__Metschnikowiaceae(100);g__Metschnikowia(100);s__Metschnikowia_pulcherrima(100); | Metschnikowia | 13 | Saprotroph | \|Undefined Saprotroph\| | Yeast | NULL | Probable | As nectar-tap_saprotroph (Põlme et al. 2020); Decay sugar-rich substrates (Põlme et al. 2020) | Põlme S, et al. 2020. Fungal Diversity 105:1-16 ((https://doi.org/10.1007/s13225-020-00466-2)) |
| Otu46012 | 1 | k__Fungi(100);p__Ascomycota(100);c__Saccharomycetes(100);o__Saccharomycetales(100);f__Metschnikowiaceae(100);g__Metschnikowia(100);g__Metschnikowia_unclassified(100); | Metschnikowia | 13 | Saprotroph | \|Undefined Saprotroph\| | Yeast | NULL | Probable | As nectar-tap_saprotroph (Põlme et al. 2020); Decay sugar-rich substrates (Põlme et al. 2020) | Põlme S, et al. 2020. Fungal Diversity 105:1-16 ((https://doi.org/10.1007/s13225-020-00466-2)) |
| Otu46013 | 1 | k__Fungi(100);p__Ascomycota(100);c__Saccharomycetes(100);o__Saccharomycetales(100);f__Metschnikowiaceae(100);g__Metschnikowia(100);s__Metschnikowia_pulcherrima(100); | Metschnikowia | 13 | Saprotroph | \|Undefined Saprotroph\| | Yeast | NULL | Probable | As nectar-tap_saprotroph (Põlme et al. 2020); Decay sugar-rich substrates (Põlme et al. 2020) | Põlme S, et al. 2020. Fungal Diversity 105:1-16 ((https://doi.org/10.1007/s13225-020-00466-2)) |
| Otu46014 | 1 | k__Fungi(100);p__Ascomycota(100);c__Saccharomycetes(100);o__Saccharomycetales(100);f__Metschnikowiaceae(100);g__Metschnikowia(100);s__Metschnikowia_pulcherrima(100); | Metschnikowia | 13 | Saprotroph | \|Undefined Saprotroph\| | Yeast | NULL | Probable | As nectar-tap_saprotroph (Põlme et al. 2020); Decay sugar-rich substrates (Põlme et al. 2020) | Põlme S, et al. 2020. Fungal Diversity 105:1-16 ((https://doi.org/10.1007/s13225-020-00466-2)) |
| Otu46015 | 1 | k__Fungi(100);p__Ascomycota(100);c__Saccharomycetes(100);o__Saccharomycetales(100);f__Metschnikowiaceae(100);g__Metschnikowia(100);s__Metschnikowia_pulcherrima(100); | Metschnikowia | 13 | Saprotroph | \|Undefined Saprotroph\| | Yeast | NULL | Probable | As nectar-tap_saprotroph (Põlme et al. 2020); Decay sugar-rich substrates (Põlme et al. 2020) | Põlme S, et al. 2020. Fungal Diversity 105:1-16 ((https://doi.org/10.1007/s13225-020-00466-2)) |
| Otu46016 | 1 | k__Fungi(100);p__Ascomycota(100);c__Saccharomycetes(100);o__Saccharomycetales(100);f__Metschnikowiaceae(100);g__Metschnikowia(100);s__Metschnikowia_pulcherrima(100); | Metschnikowia | 13 | Saprotroph | \|Undefined Saprotroph\| | Yeast | NULL | Probable | As nectar-tap_saprotroph (Põlme et al. 2020); Decay sugar-rich substrates (Põlme et al. 2020) | Põlme S, et al. 2020. Fungal Diversity 105:1-16 ((https://doi.org/10.1007/s13225-020-00466-2)) |
| Otu46017 | 1 | k__Fungi(100);p__Ascomycota(100);c__Saccharomycetes(100);o__Saccharomycetales(100);f__Metschnikowiaceae(100);g__Metschnikowia(100);g__Metschnikowia_unclassified(100); | Metschnikowia | 13 | Saprotroph | \|Undefined Saprotroph\| | Yeast | NULL | Probable | As nectar-tap_saprotroph (Põlme et al. 2020); Decay sugar-rich substrates (Põlme et al. 2020) | Põlme S, et al. 2020. Fungal Diversity 105:1-16 ((https://doi.org/10.1007/s13225-020-00466-2)) |
| Otu46018 | 1 | k__Fungi(100);p__Ascomycota(100);c__Saccharomycetes(100);o__Saccharomycetales(100);f__Metschnikowiaceae(100);g__Metschnikowia(100);g__Metschnikowia_unclassified(100); | Metschnikowia | 13 | Saprotroph | \|Undefined Saprotroph\| | Yeast | NULL | Probable | As nectar-tap_saprotroph (Põlme et al. 2020); Decay sugar-rich substrates (Põlme et al. 2020) | Põlme S, et al. 2020. Fungal Diversity 105:1-16 ((https://doi.org/10.1007/s13225-020-00466-2)) |
| Otu46019 | 1 | k__Fungi(100);p__Ascomycota(100);c__Saccharomycetes(100);o__Saccharomycetales(100);f__Metschnikowiaceae(100);g__Metschnikowia(100);s__Metschnikowia_pulcherrima(100); | Metschnikowia | 13 | Saprotroph | \|Undefined Saprotroph\| | Yeast | NULL | Probable | As nectar-tap_saprotroph (Põlme et al. 2020); Decay sugar-rich substrates (Põlme et al. 2020) | Põlme S, et al. 2020. Fungal Diversity 105:1-16 ((https://doi.org/10.1007/s13225-020-00466-2)) |
| Otu46021 | 1 | k__Fungi(100);p__Ascomycota(100);c__Saccharomycetes(100);o__Saccharomycetales(100);f__Metschnikowiaceae(100);g__Metschnikowia(100);s__Metschnikowia_pulcherrima(100); | Metschnikowia | 13 | Saprotroph | \|Undefined Saprotroph\| | Yeast | NULL | Probable | As nectar-tap_saprotroph (Põlme et al. 2020); Decay sugar-rich substrates (Põlme et al. 2020) | Põlme S, et al. 2020. Fungal Diversity 105:1-16 ((https://doi.org/10.1007/s13225-020-00466-2)) |
| Otu46022 | 1 | k__Fungi(100);p__Ascomycota(100);c__Saccharomycetes(100);o__Saccharomycetales(100);f__Metschnikowiaceae(100);g__Metschnikowia(100);s__Metschnikowia_pulcherrima(100); | Metschnikowia | 13 | Saprotroph | \|Undefined Saprotroph\| | Yeast | NULL | Probable | As nectar-tap_saprotroph (Põlme et al. 2020); Decay sugar-rich substrates (Põlme et al. 2020) | Põlme S, et al. 2020. Fungal Diversity 105:1-16 ((https://doi.org/10.1007/s13225-020-00466-2)) |
| Otu46023 | 1 | k__Fungi(100);p__Ascomycota(100);c__Saccharomycetes(100);o__Saccharomycetales(100);f__Metschnikowiaceae(100);g__Metschnikowia(100);s__Metschnikowia_pulcherrima(100); | Metschnikowia | 13 | Saprotroph | \|Undefined Saprotroph\| | Yeast | NULL | Probable | As nectar-tap_saprotroph (Põlme et al. 2020); Decay sugar-rich substrates (Põlme et al. 2020) | Põlme S, et al. 2020. Fungal Diversity 105:1-16 ((https://doi.org/10.1007/s13225-020-00466-2)) |
| Otu46024 | 1 | k__Fungi(100);p__Ascomycota(100);c__Saccharomycetes(100);o__Saccharomycetales(100);f__Metschnikowiaceae(100);g__Metschnikowia(100);s__Metschnikowia_pulcherrima(100); | Metschnikowia | 13 | Saprotroph | \|Undefined Saprotroph\| | Yeast | NULL | Probable | As nectar-tap_saprotroph (Põlme et al. 2020); Decay sugar-rich substrates (Põlme et al. 2020) | Põlme S, et al. 2020. Fungal Diversity 105:1-16 ((https://doi.org/10.1007/s13225-020-00466-2)) |
| Otu46025 | 1 | k__Fungi(100);p__Ascomycota(100);c__Saccharomycetes(100);o__Saccharomycetales(100);f__Metschnikowiaceae(100);g__Metschnikowia(100);s__Metschnikowia_pulcherrima(100); | Metschnikowia | 13 | Saprotroph | \|Undefined Saprotroph\| | Yeast | NULL | Probable | As nectar-tap_saprotroph (Põlme et al. 2020); Decay sugar-rich substrates (Põlme et al. 2020) | Põlme S, et al. 2020. Fungal Diversity 105:1-16 ((https://doi.org/10.1007/s13225-020-00466-2)) |
| Otu46026 | 1 | k__Fungi(100);p__Ascomycota(100);c__Saccharomycetes(100);o__Saccharomycetales(100);f__Metschnikowiaceae(100);g__Metschnikowia(100);g__Metschnikowia_unclassified(100); | Metschnikowia | 13 | Saprotroph | \|Undefined Saprotroph\| | Yeast | NULL | Probable | As nectar-tap_saprotroph (Põlme et al. 2020); Decay sugar-rich substrates (Põlme et al. 2020) | Põlme S, et al. 2020. Fungal Diversity 105:1-16 ((https://doi.org/10.1007/s13225-020-00466-2)) |
| Otu46027 | 1 | k__Fungi(100);p__Ascomycota(100);c__Saccharomycetes(100);o__Saccharomycetales(100);f__Metschnikowiaceae(100);g__Metschnikowia(100);s__Metschnikowia_pulcherrima(100); | Metschnikowia | 13 | Saprotroph | \|Undefined Saprotroph\| | Yeast | NULL | Probable | As nectar-tap_saprotroph (Põlme et al. 2020); Decay sugar-rich substrates (Põlme et al. 2020) | Põlme S, et al. 2020. Fungal Diversity 105:1-16 ((https://doi.org/10.1007/s13225-020-00466-2)) |
| Otu46028 | 1 | k__Fungi(100);p__Ascomycota(100);c__Saccharomycetes(100);o__Saccharomycetales(100);f__Metschnikowiaceae(100);g__Metschnikowia(100);s__Metschnikowia_pulcherrima(100); | Metschnikowia | 13 | Saprotroph | \|Undefined Saprotroph\| | Yeast | NULL | Probable | As nectar-tap_saprotroph (Põlme et al. 2020); Decay sugar-rich substrates (Põlme et al. 2020) | Põlme S, et al. 2020. Fungal Diversity 105:1-16 ((https://doi.org/10.1007/s13225-020-00466-2)) |
| Otu46029 | 1 | k__Fungi(100);p__Ascomycota(100);c__Saccharomycetes(100);o__Saccharomycetales(100);f__Metschnikowiaceae(100);g__Metschnikowia(100);s__Metschnikowia_pulcherrima(100); | Metschnikowia | 13 | Saprotroph | \|Undefined Saprotroph\| | Yeast | NULL | Probable | As nectar-tap_saprotroph (Põlme et al. 2020); Decay sugar-rich substrates (Põlme et al. 2020) | Põlme S, et al. 2020. Fungal Diversity 105:1-16 ((https://doi.org/10.1007/s13225-020-00466-2)) |
| Otu46030 | 1 | k__Fungi(100);p__Ascomycota(100);c__Saccharomycetes(100);o__Saccharomycetales(100);f__Metschnikowiaceae(100);g__Metschnikowia(100);s__Metschnikowia_pulcherrima(100); | Metschnikowia | 13 | Saprotroph | \|Undefined Saprotroph\| | Yeast | NULL | Probable | As nectar-tap_saprotroph (Põlme et al. 2020); Decay sugar-rich substrates (Põlme et al. 2020) | Põlme S, et al. 2020. Fungal Diversity 105:1-16 ((https://doi.org/10.1007/s13225-020-00466-2)) |
| Otu46031 | 1 | k__Fungi(100);p__Ascomycota(100);c__Saccharomycetes(100);o__Saccharomycetales(100);f__Metschnikowiaceae(100);g__Metschnikowia(100);s__Metschnikowia_pulcherrima(100); | Metschnikowia | 13 | Saprotroph | \|Undefined Saprotroph\| | Yeast | NULL | Probable | As nectar-tap_saprotroph (Põlme et al. 2020); Decay sugar-rich substrates (Põlme et al. 2020) | Põlme S, et al. 2020. Fungal Diversity 105:1-16 ((https://doi.org/10.1007/s13225-020-00466-2)) |
| Otu46032 | 1 | k__Fungi(100);p__Ascomycota(100);c__Saccharomycetes(100);o__Saccharomycetales(100);f__Metschnikowiaceae(100);g__Metschnikowia(100);g__Metschnikowia_unclassified(100); | Metschnikowia | 13 | Saprotroph | \|Undefined Saprotroph\| | Yeast | NULL | Probable | As nectar-tap_saprotroph (Põlme et al. 2020); Decay sugar-rich substrates (Põlme et al. 2020) | Põlme S, et al. 2020. Fungal Diversity 105:1-16 ((https://doi.org/10.1007/s13225-020-00466-2)) |
| Otu46033 | 1 | k__Fungi(100);p__Ascomycota(100);c__Saccharomycetes(100);o__Saccharomycetales(100);f__Metschnikowiaceae(100);g__Metschnikowia(100);s__Metschnikowia_pulcherrima(100); | Metschnikowia | 13 | Saprotroph | \|Undefined Saprotroph\| | Yeast | NULL | Probable | As nectar-tap_saprotroph (Põlme et al. 2020); Decay sugar-rich substrates (Põlme et al. 2020) | Põlme S, et al. 2020. Fungal Diversity 105:1-16 ((https://doi.org/10.1007/s13225-020-00466-2)) |
| Otu46034 | 1 | k__Fungi(100);p__Ascomycota(100);c__Saccharomycetes(100);o__Saccharomycetales(100);f__Metschnikowiaceae(100);g__Metschnikowia(100);s__Metschnikowia_pulcherrima(100); | Metschnikowia | 13 | Saprotroph | \|Undefined Saprotroph\| | Yeast | NULL | Probable | As nectar-tap_saprotroph (Põlme et al. 2020); Decay sugar-rich substrates (Põlme et al. 2020) | Põlme S, et al. 2020. Fungal Diversity 105:1-16 ((https://doi.org/10.1007/s13225-020-00466-2)) |
| Otu46035 | 1 | k__Fungi(100);p__Ascomycota(100);c__Saccharomycetes(100);o__Saccharomycetales(100);f__Metschnikowiaceae(100);g__Metschnikowia(100);g__Metschnikowia_unclassified(100); | Metschnikowia | 13 | Saprotroph | \|Undefined Saprotroph\| | Yeast | NULL | Probable | As nectar-tap_saprotroph (Põlme et al. 2020); Decay sugar-rich substrates (Põlme et al. 2020) | Põlme S, et al. 2020. Fungal Diversity 105:1-16 ((https://doi.org/10.1007/s13225-020-00466-2)) |
| Otu46036 | 1 | k__Fungi(100);p__Ascomycota(100);c__Saccharomycetes(100);o__Saccharomycetales(100);f__Metschnikowiaceae(100);g__Metschnikowia(100);g__Metschnikowia_unclassified(100); | Metschnikowia | 13 | Saprotroph | \|Undefined Saprotroph\| | Yeast | NULL | Probable | As nectar-tap_saprotroph (Põlme et al. 2020); Decay sugar-rich substrates (Põlme et al. 2020) | Põlme S, et al. 2020. Fungal Diversity 105:1-16 ((https://doi.org/10.1007/s13225-020-00466-2)) |
| Otu46037 | 1 | k__Fungi(100);p__Ascomycota(100);c__Saccharomycetes(100);o__Saccharomycetales(100);f__Metschnikowiaceae(100);g__Metschnikowia(100);g__Metschnikowia_unclassified(100); | Metschnikowia | 13 | Saprotroph | \|Undefined Saprotroph\| | Yeast | NULL | Probable | As nectar-tap_saprotroph (Põlme et al. 2020); Decay sugar-rich substrates (Põlme et al. 2020) | Põlme S, et al. 2020. Fungal Diversity 105:1-16 ((https://doi.org/10.1007/s13225-020-00466-2)) |
| Otu46038 | 1 | k__Fungi(100);p__Ascomycota(100);c__Saccharomycetes(100);o__Saccharomycetales(100);f__Metschnikowiaceae(100);g__Metschnikowia(100);g__Metschnikowia_unclassified(100); | Metschnikowia | 13 | Saprotroph | \|Undefined Saprotroph\| | Yeast | NULL | Probable | As nectar-tap_saprotroph (Põlme et al. 2020); Decay sugar-rich substrates (Põlme et al. 2020) | Põlme S, et al. 2020. Fungal Diversity 105:1-16 ((https://doi.org/10.1007/s13225-020-00466-2)) |
| Otu46039 | 1 | k__Fungi(100);p__Ascomycota(100);c__Saccharomycetes(100);o__Saccharomycetales(100);f__Metschnikowiaceae(100);g__Metschnikowia(100);s__Metschnikowia_pulcherrima(100); | Metschnikowia | 13 | Saprotroph | \|Undefined Saprotroph\| | Yeast | NULL | Probable | As nectar-tap_saprotroph (Põlme et al. 2020); Decay sugar-rich substrates (Põlme et al. 2020) | Põlme S, et al. 2020. Fungal Diversity 105:1-16 ((https://doi.org/10.1007/s13225-020-00466-2)) |
| Otu46040 | 1 | k__Fungi(100);p__Ascomycota(100);c__Saccharomycetes(100);o__Saccharomycetales(100);f__Metschnikowiaceae(100);g__Metschnikowia(100);g__Metschnikowia_unclassified(100); | Metschnikowia | 13 | Saprotroph | \|Undefined Saprotroph\| | Yeast | NULL | Probable | As nectar-tap_saprotroph (Põlme et al. 2020); Decay sugar-rich substrates (Põlme et al. 2020) | Põlme S, et al. 2020. Fungal Diversity 105:1-16 ((https://doi.org/10.1007/s13225-020-00466-2)) |
| Otu46041 | 1 | k__Fungi(100);p__Ascomycota(100);c__Saccharomycetes(100);o__Saccharomycetales(100);f__Metschnikowiaceae(100);g__Metschnikowia(100);s__Metschnikowia_pulcherrima(100); | Metschnikowia | 13 | Saprotroph | \|Undefined Saprotroph\| | Yeast | NULL | Probable | As nectar-tap_saprotroph (Põlme et al. 2020); Decay sugar-rich substrates (Põlme et al. 2020) | Põlme S, et al. 2020. Fungal Diversity 105:1-16 ((https://doi.org/10.1007/s13225-020-00466-2)) |
| Otu46042 | 1 | k__Fungi(100);p__Ascomycota(100);c__Saccharomycetes(100);o__Saccharomycetales(100);f__Metschnikowiaceae(100);g__Metschnikowia(100);s__Metschnikowia_pulcherrima(100); | Metschnikowia | 13 | Saprotroph | \|Undefined Saprotroph\| | Yeast | NULL | Probable | As nectar-tap_saprotroph (Põlme et al. 2020); Decay sugar-rich substrates (Põlme et al. 2020) | Põlme S, et al. 2020. Fungal Diversity 105:1-16 ((https://doi.org/10.1007/s13225-020-00466-2)) |
| Otu46043 | 1 | k__Fungi(100);p__Ascomycota(100);c__Saccharomycetes(100);o__Saccharomycetales(100);f__Metschnikowiaceae(100);g__Metschnikowia(100);s__Metschnikowia_pulcherrima(100); | Metschnikowia | 13 | Saprotroph | \|Undefined Saprotroph\| | Yeast | NULL | Probable | As nectar-tap_saprotroph (Põlme et al. 2020); Decay sugar-rich substrates (Põlme et al. 2020) | Põlme S, et al. 2020. Fungal Diversity 105:1-16 ((https://doi.org/10.1007/s13225-020-00466-2)) |
| Otu46044 | 1 | k__Fungi(100);p__Ascomycota(100);c__Saccharomycetes(100);o__Saccharomycetales(100);f__Metschnikowiaceae(100);g__Metschnikowia(100);s__Metschnikowia_pulcherrima(100); | Metschnikowia | 13 | Saprotroph | \|Undefined Saprotroph\| | Yeast | NULL | Probable | As nectar-tap_saprotroph (Põlme et al. 2020); Decay sugar-rich substrates (Põlme et al. 2020) | Põlme S, et al. 2020. Fungal Diversity 105:1-16 ((https://doi.org/10.1007/s13225-020-00466-2)) |
| Otu46045 | 1 | k__Fungi(100);p__Ascomycota(100);c__Saccharomycetes(100);o__Saccharomycetales(100);f__Metschnikowiaceae(100);g__Metschnikowia(100);s__Metschnikowia_pulcherrima(100); | Metschnikowia | 13 | Saprotroph | \|Undefined Saprotroph\| | Yeast | NULL | Probable | As nectar-tap_saprotroph (Põlme et al. 2020); Decay sugar-rich substrates (Põlme et al. 2020) | Põlme S, et al. 2020. Fungal Diversity 105:1-16 ((https://doi.org/10.1007/s13225-020-00466-2)) |
| Otu46046 | 1 | k__Fungi(100);p__Ascomycota(100);c__Saccharomycetes(100);o__Saccharomycetales(100);f__Metschnikowiaceae(100);g__Metschnikowia(100);g__Metschnikowia_unclassified(100); | Metschnikowia | 13 | Saprotroph | \|Undefined Saprotroph\| | Yeast | NULL | Probable | As nectar-tap_saprotroph (Põlme et al. 2020); Decay sugar-rich substrates (Põlme et al. 2020) | Põlme S, et al. 2020. Fungal Diversity 105:1-16 ((https://doi.org/10.1007/s13225-020-00466-2)) |
| Otu46047 | 1 | k__Fungi(100);p__Ascomycota(100);c__Saccharomycetes(100);o__Saccharomycetales(100);f__Metschnikowiaceae(100);g__Metschnikowia(100);s__Metschnikowia_pulcherrima(100); | Metschnikowia | 13 | Saprotroph | \|Undefined Saprotroph\| | Yeast | NULL | Probable | As nectar-tap_saprotroph (Põlme et al. 2020); Decay sugar-rich substrates (Põlme et al. 2020) | Põlme S, et al. 2020. Fungal Diversity 105:1-16 ((https://doi.org/10.1007/s13225-020-00466-2)) |
| Otu46048 | 1 | k__Fungi(100);p__Ascomycota(100);c__Saccharomycetes(100);o__Saccharomycetales(100);f__Metschnikowiaceae(100);g__Metschnikowia(100);s__Metschnikowia_pulcherrima(100); | Metschnikowia | 13 | Saprotroph | \|Undefined Saprotroph\| | Yeast | NULL | Probable | As nectar-tap_saprotroph (Põlme et al. 2020); Decay sugar-rich substrates (Põlme et al. 2020) | Põlme S, et al. 2020. Fungal Diversity 105:1-16 ((https://doi.org/10.1007/s13225-020-00466-2)) |
| Otu46049 | 1 | k__Fungi(100);p__Ascomycota(100);c__Saccharomycetes(100);o__Saccharomycetales(100);f__Metschnikowiaceae(100);g__Metschnikowia(100);s__Metschnikowia_pulcherrima(100); | Metschnikowia | 13 | Saprotroph | \|Undefined Saprotroph\| | Yeast | NULL | Probable | As nectar-tap_saprotroph (Põlme et al. 2020); Decay sugar-rich substrates (Põlme et al. 2020) | Põlme S, et al. 2020. Fungal Diversity 105:1-16 ((https://doi.org/10.1007/s13225-020-00466-2)) |
| Otu46050 | 1 | k__Fungi(100);p__Ascomycota(100);c__Saccharomycetes(100);o__Saccharomycetales(100);f__Metschnikowiaceae(100);g__Metschnikowia(100);g__Metschnikowia_unclassified(100); | Metschnikowia | 13 | Saprotroph | \|Undefined Saprotroph\| | Yeast | NULL | Probable | As nectar-tap_saprotroph (Põlme et al. 2020); Decay sugar-rich substrates (Põlme et al. 2020) | Põlme S, et al. 2020. Fungal Diversity 105:1-16 ((https://doi.org/10.1007/s13225-020-00466-2)) |
| Otu46051 | 1 | k__Fungi(100);p__Ascomycota(100);c__Saccharomycetes(100);o__Saccharomycetales(100);f__Metschnikowiaceae(100);g__Metschnikowia(100);s__Metschnikowia_pulcherrima(100); | Metschnikowia | 13 | Saprotroph | \|Undefined Saprotroph\| | Yeast | NULL | Probable | As nectar-tap_saprotroph (Põlme et al. 2020); Decay sugar-rich substrates (Põlme et al. 2020) | Põlme S, et al. 2020. Fungal Diversity 105:1-16 ((https://doi.org/10.1007/s13225-020-00466-2)) |
| Otu46052 | 1 | k__Fungi(100);p__Ascomycota(100);c__Saccharomycetes(100);o__Saccharomycetales(100);f__Metschnikowiaceae(100);g__Metschnikowia(100);s__Metschnikowia_pulcherrima(100); | Metschnikowia | 13 | Saprotroph | \|Undefined Saprotroph\| | Yeast | NULL | Probable | As nectar-tap_saprotroph (Põlme et al. 2020); Decay sugar-rich substrates (Põlme et al. 2020) | Põlme S, et al. 2020. Fungal Diversity 105:1-16 ((https://doi.org/10.1007/s13225-020-00466-2)) |
| Otu46053 | 1 | k__Fungi(100);p__Ascomycota(100);c__Saccharomycetes(100);o__Saccharomycetales(100);f__Metschnikowiaceae(100);g__Metschnikowia(100);s__Metschnikowia_pulcherrima(100); | Metschnikowia | 13 | Saprotroph | \|Undefined Saprotroph\| | Yeast | NULL | Probable | As nectar-tap_saprotroph (Põlme et al. 2020); Decay sugar-rich substrates (Põlme et al. 2020) | Põlme S, et al. 2020. Fungal Diversity 105:1-16 ((https://doi.org/10.1007/s13225-020-00466-2)) |
| Otu46054 | 1 | k__Fungi(100);p__Ascomycota(100);c__Saccharomycetes(100);o__Saccharomycetales(100);f__Metschnikowiaceae(100);g__Metschnikowia(100);s__Metschnikowia_pulcherrima(100); | Metschnikowia | 13 | Saprotroph | \|Undefined Saprotroph\| | Yeast | NULL | Probable | As nectar-tap_saprotroph (Põlme et al. 2020); Decay sugar-rich substrates (Põlme et al. 2020) | Põlme S, et al. 2020. Fungal Diversity 105:1-16 ((https://doi.org/10.1007/s13225-020-00466-2)) |
| Otu46055 | 1 | k__Fungi(100);p__Ascomycota(100);c__Saccharomycetes(100);o__Saccharomycetales(100);f__Metschnikowiaceae(100);g__Metschnikowia(100);g__Metschnikowia_unclassified(100); | Metschnikowia | 13 | Saprotroph | \|Undefined Saprotroph\| | Yeast | NULL | Probable | As nectar-tap_saprotroph (Põlme et al. 2020); Decay sugar-rich substrates (Põlme et al. 2020) | Põlme S, et al. 2020. Fungal Diversity 105:1-16 ((https://doi.org/10.1007/s13225-020-00466-2)) |
| Otu46056 | 1 | k__Fungi(100);p__Ascomycota(100);c__Saccharomycetes(100);o__Saccharomycetales(100);f__Metschnikowiaceae(100);g__Metschnikowia(100);s__Metschnikowia_pulcherrima(100); | Metschnikowia | 13 | Saprotroph | \|Undefined Saprotroph\| | Yeast | NULL | Probable | As nectar-tap_saprotroph (Põlme et al. 2020); Decay sugar-rich substrates (Põlme et al. 2020) | Põlme S, et al. 2020. Fungal Diversity 105:1-16 ((https://doi.org/10.1007/s13225-020-00466-2)) |
| Otu46057 | 1 | k__Fungi(100);p__Ascomycota(100);c__Saccharomycetes(100);o__Saccharomycetales(100);f__Metschnikowiaceae(100);g__Metschnikowia(100);s__Metschnikowia_pulcherrima(100); | Metschnikowia | 13 | Saprotroph | \|Undefined Saprotroph\| | Yeast | NULL | Probable | As nectar-tap_saprotroph (Põlme et al. 2020); Decay sugar-rich substrates (Põlme et al. 2020) | Põlme S, et al. 2020. Fungal Diversity 105:1-16 ((https://doi.org/10.1007/s13225-020-00466-2)) |
| Otu46058 | 1 | k__Fungi(100);p__Ascomycota(100);c__Saccharomycetes(100);o__Saccharomycetales(100);f__Metschnikowiaceae(100);g__Metschnikowia(100);s__Metschnikowia_pulcherrima(100); | Metschnikowia | 13 | Saprotroph | \|Undefined Saprotroph\| | Yeast | NULL | Probable | As nectar-tap_saprotroph (Põlme et al. 2020); Decay sugar-rich substrates (Põlme et al. 2020) | Põlme S, et al. 2020. Fungal Diversity 105:1-16 ((https://doi.org/10.1007/s13225-020-00466-2)) |
| Otu46059 | 1 | k__Fungi(100);p__Ascomycota(100);c__Saccharomycetes(100);o__Saccharomycetales(100);f__Metschnikowiaceae(100);g__Metschnikowia(100);g__Metschnikowia_unclassified(100); | Metschnikowia | 13 | Saprotroph | \|Undefined Saprotroph\| | Yeast | NULL | Probable | As nectar-tap_saprotroph (Põlme et al. 2020); Decay sugar-rich substrates (Põlme et al. 2020) | Põlme S, et al. 2020. Fungal Diversity 105:1-16 ((https://doi.org/10.1007/s13225-020-00466-2)) |
| Otu46060 | 1 | k__Fungi(100);p__Ascomycota(100);c__Saccharomycetes(100);o__Saccharomycetales(100);f__Metschnikowiaceae(100);g__Metschnikowia(100);g__Metschnikowia_unclassified(100); | Metschnikowia | 13 | Saprotroph | \|Undefined Saprotroph\| | Yeast | NULL | Probable | As nectar-tap_saprotroph (Põlme et al. 2020); Decay sugar-rich substrates (Põlme et al. 2020) | Põlme S, et al. 2020. Fungal Diversity 105:1-16 ((https://doi.org/10.1007/s13225-020-00466-2)) |
| Otu46061 | 1 | k__Fungi(100);p__Ascomycota(100);c__Saccharomycetes(100);o__Saccharomycetales(100);f__Metschnikowiaceae(100);g__Metschnikowia(100);s__Metschnikowia_pulcherrima(100); | Metschnikowia | 13 | Saprotroph | \|Undefined Saprotroph\| | Yeast | NULL | Probable | As nectar-tap_saprotroph (Põlme et al. 2020); Decay sugar-rich substrates (Põlme et al. 2020) | Põlme S, et al. 2020. Fungal Diversity 105:1-16 ((https://doi.org/10.1007/s13225-020-00466-2)) |
| Otu46062 | 1 | k__Fungi(100);p__Ascomycota(100);c__Saccharomycetes(100);o__Saccharomycetales(100);f__Metschnikowiaceae(100);g__Metschnikowia(100);g__Metschnikowia_unclassified(100); | Metschnikowia | 13 | Saprotroph | \|Undefined Saprotroph\| | Yeast | NULL | Probable | As nectar-tap_saprotroph (Põlme et al. 2020); Decay sugar-rich substrates (Põlme et al. 2020) | Põlme S, et al. 2020. Fungal Diversity 105:1-16 ((https://doi.org/10.1007/s13225-020-00466-2)) |
| Otu46063 | 1 | k__Fungi(100);p__Ascomycota(100);c__Saccharomycetes(100);o__Saccharomycetales(100);f__Metschnikowiaceae(100);g__Metschnikowia(100);s__Metschnikowia_pulcherrima(100); | Metschnikowia | 13 | Saprotroph | \|Undefined Saprotroph\| | Yeast | NULL | Probable | As nectar-tap_saprotroph (Põlme et al. 2020); Decay sugar-rich substrates (Põlme et al. 2020) | Põlme S, et al. 2020. Fungal Diversity 105:1-16 ((https://doi.org/10.1007/s13225-020-00466-2)) |
| Otu46064 | 1 | k__Fungi(100);p__Ascomycota(100);c__Saccharomycetes(100);o__Saccharomycetales(100);f__Metschnikowiaceae(100);g__Metschnikowia(100);g__Metschnikowia_unclassified(100); | Metschnikowia | 13 | Saprotroph | \|Undefined Saprotroph\| | Yeast | NULL | Probable | As nectar-tap_saprotroph (Põlme et al. 2020); Decay sugar-rich substrates (Põlme et al. 2020) | Põlme S, et al. 2020. Fungal Diversity 105:1-16 ((https://doi.org/10.1007/s13225-020-00466-2)) |
| Otu46065 | 1 | k__Fungi(100);p__Ascomycota(100);c__Saccharomycetes(100);o__Saccharomycetales(100);f__Metschnikowiaceae(100);g__Metschnikowia(100);s__Metschnikowia_pulcherrima(100); | Metschnikowia | 13 | Saprotroph | \|Undefined Saprotroph\| | Yeast | NULL | Probable | As nectar-tap_saprotroph (Põlme et al. 2020); Decay sugar-rich substrates (Põlme et al. 2020) | Põlme S, et al. 2020. Fungal Diversity 105:1-16 ((https://doi.org/10.1007/s13225-020-00466-2)) |
| Otu46066 | 1 | k__Fungi(100);p__Ascomycota(100);c__Saccharomycetes(100);o__Saccharomycetales(100);f__Metschnikowiaceae(100);g__Metschnikowia(100);s__Metschnikowia_pulcherrima(100); | Metschnikowia | 13 | Saprotroph | \|Undefined Saprotroph\| | Yeast | NULL | Probable | As nectar-tap_saprotroph (Põlme et al. 2020); Decay sugar-rich substrates (Põlme et al. 2020) | Põlme S, et al. 2020. Fungal Diversity 105:1-16 ((https://doi.org/10.1007/s13225-020-00466-2)) |
| Otu46067 | 1 | k__Fungi(100);p__Ascomycota(100);c__Saccharomycetes(100);o__Saccharomycetales(100);f__Metschnikowiaceae(100);g__Metschnikowia(100);s__Metschnikowia_pulcherrima(100); | Metschnikowia | 13 | Saprotroph | \|Undefined Saprotroph\| | Yeast | NULL | Probable | As nectar-tap_saprotroph (Põlme et al. 2020); Decay sugar-rich substrates (Põlme et al. 2020) | Põlme S, et al. 2020. Fungal Diversity 105:1-16 ((https://doi.org/10.1007/s13225-020-00466-2)) |
| Otu46068 | 1 | k__Fungi(100);p__Ascomycota(100);c__Saccharomycetes(100);o__Saccharomycetales(100);f__Metschnikowiaceae(100);g__Metschnikowia(100);s__Metschnikowia_pulcherrima(100); | Metschnikowia | 13 | Saprotroph | \|Undefined Saprotroph\| | Yeast | NULL | Probable | As nectar-tap_saprotroph (Põlme et al. 2020); Decay sugar-rich substrates (Põlme et al. 2020) | Põlme S, et al. 2020. Fungal Diversity 105:1-16 ((https://doi.org/10.1007/s13225-020-00466-2)) |
| Otu46069 | 1 | k__Fungi(100);p__Ascomycota(100);c__Saccharomycetes(100);o__Saccharomycetales(100);f__Metschnikowiaceae(100);g__Metschnikowia(100);s__Metschnikowia_pulcherrima(100); | Metschnikowia | 13 | Saprotroph | \|Undefined Saprotroph\| | Yeast | NULL | Probable | As nectar-tap_saprotroph (Põlme et al. 2020); Decay sugar-rich substrates (Põlme et al. 2020) | Põlme S, et al. 2020. Fungal Diversity 105:1-16 ((https://doi.org/10.1007/s13225-020-00466-2)) |
| Otu46070 | 1 | k__Fungi(100);p__Ascomycota(100);c__Saccharomycetes(100);o__Saccharomycetales(100);f__Metschnikowiaceae(100);g__Metschnikowia(100);s__Metschnikowia_pulcherrima(100); | Metschnikowia | 13 | Saprotroph | \|Undefined Saprotroph\| | Yeast | NULL | Probable | As nectar-tap_saprotroph (Põlme et al. 2020); Decay sugar-rich substrates (Põlme et al. 2020) | Põlme S, et al. 2020. Fungal Diversity 105:1-16 ((https://doi.org/10.1007/s13225-020-00466-2)) |
| Otu46071 | 1 | k__Fungi(100);p__Ascomycota(100);c__Saccharomycetes(100);o__Saccharomycetales(100);f__Metschnikowiaceae(100);g__Metschnikowia(100);s__Metschnikowia_pulcherrima(100); | Metschnikowia | 13 | Saprotroph | \|Undefined Saprotroph\| | Yeast | NULL | Probable | As nectar-tap_saprotroph (Põlme et al. 2020); Decay sugar-rich substrates (Põlme et al. 2020) | Põlme S, et al. 2020. Fungal Diversity 105:1-16 ((https://doi.org/10.1007/s13225-020-00466-2)) |
| Otu46072 | 1 | k__Fungi(100);p__Ascomycota(100);c__Saccharomycetes(100);o__Saccharomycetales(100);f__Metschnikowiaceae(100);g__Metschnikowia(100);s__Metschnikowia_pulcherrima(100); | Metschnikowia | 13 | Saprotroph | \|Undefined Saprotroph\| | Yeast | NULL | Probable | As nectar-tap_saprotroph (Põlme et al. 2020); Decay sugar-rich substrates (Põlme et al. 2020) | Põlme S, et al. 2020. Fungal Diversity 105:1-16 ((https://doi.org/10.1007/s13225-020-00466-2)) |
| Otu46073 | 1 | k__Fungi(100);p__Ascomycota(100);c__Saccharomycetes(100);o__Saccharomycetales(100);f__Metschnikowiaceae(100);g__Metschnikowia(100);s__Metschnikowia_pulcherrima(100); | Metschnikowia | 13 | Saprotroph | \|Undefined Saprotroph\| | Yeast | NULL | Probable | As nectar-tap_saprotroph (Põlme et al. 2020); Decay sugar-rich substrates (Põlme et al. 2020) | Põlme S, et al. 2020. Fungal Diversity 105:1-16 ((https://doi.org/10.1007/s13225-020-00466-2)) |
| Otu46074 | 1 | k__Fungi(100);p__Ascomycota(100);c__Saccharomycetes(100);o__Saccharomycetales(100);f__Metschnikowiaceae(100);g__Metschnikowia(100);s__Metschnikowia_pulcherrima(100); | Metschnikowia | 13 | Saprotroph | \|Undefined Saprotroph\| | Yeast | NULL | Probable | As nectar-tap_saprotroph (Põlme et al. 2020); Decay sugar-rich substrates (Põlme et al. 2020) | Põlme S, et al. 2020. Fungal Diversity 105:1-16 ((https://doi.org/10.1007/s13225-020-00466-2)) |
| Otu46076 | 1 | k__Fungi(100);p__Ascomycota(100);c__Saccharomycetes(100);o__Saccharomycetales(100);f__Metschnikowiaceae(100);g__Metschnikowia(100);s__Metschnikowia_pulcherrima(100); | Metschnikowia | 13 | Saprotroph | \|Undefined Saprotroph\| | Yeast | NULL | Probable | As nectar-tap_saprotroph (Põlme et al. 2020); Decay sugar-rich substrates (Põlme et al. 2020) | Põlme S, et al. 2020. Fungal Diversity 105:1-16 ((https://doi.org/10.1007/s13225-020-00466-2)) |
| Otu46077 | 1 | k__Fungi(100);p__Ascomycota(100);c__Saccharomycetes(100);o__Saccharomycetales(100);f__Metschnikowiaceae(100);g__Metschnikowia(100);s__Metschnikowia_pulcherrima(100); | Metschnikowia | 13 | Saprotroph | \|Undefined Saprotroph\| | Yeast | NULL | Probable | As nectar-tap_saprotroph (Põlme et al. 2020); Decay sugar-rich substrates (Põlme et al. 2020) | Põlme S, et al. 2020. Fungal Diversity 105:1-16 ((https://doi.org/10.1007/s13225-020-00466-2)) |
| Otu46078 | 1 | k__Fungi(100);p__Ascomycota(100);c__Saccharomycetes(100);o__Saccharomycetales(100);f__Metschnikowiaceae(100);g__Metschnikowia(100);s__Metschnikowia_pulcherrima(100); | Metschnikowia | 13 | Saprotroph | \|Undefined Saprotroph\| | Yeast | NULL | Probable | As nectar-tap_saprotroph (Põlme et al. 2020); Decay sugar-rich substrates (Põlme et al. 2020) | Põlme S, et al. 2020. Fungal Diversity 105:1-16 ((https://doi.org/10.1007/s13225-020-00466-2)) |
| Otu46079 | 1 | k__Fungi(100);p__Ascomycota(100);c__Saccharomycetes(100);o__Saccharomycetales(100);f__Metschnikowiaceae(100);g__Metschnikowia(100);s__Metschnikowia_pulcherrima(100); | Metschnikowia | 13 | Saprotroph | \|Undefined Saprotroph\| | Yeast | NULL | Probable | As nectar-tap_saprotroph (Põlme et al. 2020); Decay sugar-rich substrates (Põlme et al. 2020) | Põlme S, et al. 2020. Fungal Diversity 105:1-16 ((https://doi.org/10.1007/s13225-020-00466-2)) |
| Otu46080 | 1 | k__Fungi(100);p__Ascomycota(100);c__Saccharomycetes(100);o__Saccharomycetales(100);f__Metschnikowiaceae(100);g__Metschnikowia(100);s__Metschnikowia_pulcherrima(100); | Metschnikowia | 13 | Saprotroph | \|Undefined Saprotroph\| | Yeast | NULL | Probable | As nectar-tap_saprotroph (Põlme et al. 2020); Decay sugar-rich substrates (Põlme et al. 2020) | Põlme S, et al. 2020. Fungal Diversity 105:1-16 ((https://doi.org/10.1007/s13225-020-00466-2)) |
| Otu46081 | 1 | k__Fungi(100);p__Ascomycota(100);c__Saccharomycetes(100);o__Saccharomycetales(100);f__Metschnikowiaceae(100);g__Metschnikowia(100);g__Metschnikowia_unclassified(100); | Metschnikowia | 13 | Saprotroph | \|Undefined Saprotroph\| | Yeast | NULL | Probable | As nectar-tap_saprotroph (Põlme et al. 2020); Decay sugar-rich substrates (Põlme et al. 2020) | Põlme S, et al. 2020. Fungal Diversity 105:1-16 ((https://doi.org/10.1007/s13225-020-00466-2)) |
| Otu46082 | 1 | k__Fungi(100);p__Ascomycota(100);c__Saccharomycetes(100);o__Saccharomycetales(100);f__Metschnikowiaceae(100);g__Metschnikowia(100);s__Metschnikowia_pulcherrima(100); | Metschnikowia | 13 | Saprotroph | \|Undefined Saprotroph\| | Yeast | NULL | Probable | As nectar-tap_saprotroph (Põlme et al. 2020); Decay sugar-rich substrates (Põlme et al. 2020) | Põlme S, et al. 2020. Fungal Diversity 105:1-16 ((https://doi.org/10.1007/s13225-020-00466-2)) |
| Otu46083 | 1 | k__Fungi(100);p__Ascomycota(100);c__Saccharomycetes(100);o__Saccharomycetales(100);f__Metschnikowiaceae(100);g__Metschnikowia(100);s__Metschnikowia_pulcherrima(100); | Metschnikowia | 13 | Saprotroph | \|Undefined Saprotroph\| | Yeast | NULL | Probable | As nectar-tap_saprotroph (Põlme et al. 2020); Decay sugar-rich substrates (Põlme et al. 2020) | Põlme S, et al. 2020. Fungal Diversity 105:1-16 ((https://doi.org/10.1007/s13225-020-00466-2)) |
| Otu46084 | 1 | k__Fungi(100);p__Ascomycota(100);c__Saccharomycetes(100);o__Saccharomycetales(100);f__Metschnikowiaceae(100);g__Metschnikowia(100);s__Metschnikowia_pulcherrima(100); | Metschnikowia | 13 | Saprotroph | \|Undefined Saprotroph\| | Yeast | NULL | Probable | As nectar-tap_saprotroph (Põlme et al. 2020); Decay sugar-rich substrates (Põlme et al. 2020) | Põlme S, et al. 2020. Fungal Diversity 105:1-16 ((https://doi.org/10.1007/s13225-020-00466-2)) |
| Otu46085 | 1 | k__Fungi(100);p__Ascomycota(100);c__Saccharomycetes(100);o__Saccharomycetales(100);f__Metschnikowiaceae(100);g__Metschnikowia(100);g__Metschnikowia_unclassified(100); | Metschnikowia | 13 | Saprotroph | \|Undefined Saprotroph\| | Yeast | NULL | Probable | As nectar-tap_saprotroph (Põlme et al. 2020); Decay sugar-rich substrates (Põlme et al. 2020) | Põlme S, et al. 2020. Fungal Diversity 105:1-16 ((https://doi.org/10.1007/s13225-020-00466-2)) |
| Otu46086 | 1 | k__Fungi(100);p__Ascomycota(100);c__Saccharomycetes(100);o__Saccharomycetales(100);f__Metschnikowiaceae(100);g__Metschnikowia(100);s__Metschnikowia_pulcherrima(100); | Metschnikowia | 13 | Saprotroph | \|Undefined Saprotroph\| | Yeast | NULL | Probable | As nectar-tap_saprotroph (Põlme et al. 2020); Decay sugar-rich substrates (Põlme et al. 2020) | Põlme S, et al. 2020. Fungal Diversity 105:1-16 ((https://doi.org/10.1007/s13225-020-00466-2)) |
| Otu46087 | 1 | k__Fungi(100);p__Ascomycota(100);c__Saccharomycetes(100);o__Saccharomycetales(100);f__Metschnikowiaceae(100);g__Metschnikowia(100);s__Metschnikowia_pulcherrima(100); | Metschnikowia | 13 | Saprotroph | \|Undefined Saprotroph\| | Yeast | NULL | Probable | As nectar-tap_saprotroph (Põlme et al. 2020); Decay sugar-rich substrates (Põlme et al. 2020) | Põlme S, et al. 2020. Fungal Diversity 105:1-16 ((https://doi.org/10.1007/s13225-020-00466-2)) |
| Otu46088 | 1 | k__Fungi(100);p__Ascomycota(100);c__Saccharomycetes(100);o__Saccharomycetales(100);f__Metschnikowiaceae(100);g__Metschnikowia(100);g__Metschnikowia_unclassified(100); | Metschnikowia | 13 | Saprotroph | \|Undefined Saprotroph\| | Yeast | NULL | Probable | As nectar-tap_saprotroph (Põlme et al. 2020); Decay sugar-rich substrates (Põlme et al. 2020) | Põlme S, et al. 2020. Fungal Diversity 105:1-16 ((https://doi.org/10.1007/s13225-020-00466-2)) |
| Otu46089 | 1 | k__Fungi(100);p__Ascomycota(100);c__Saccharomycetes(100);o__Saccharomycetales(100);f__Metschnikowiaceae(100);g__Metschnikowia(100);g__Metschnikowia_unclassified(100); | Metschnikowia | 13 | Saprotroph | \|Undefined Saprotroph\| | Yeast | NULL | Probable | As nectar-tap_saprotroph (Põlme et al. 2020); Decay sugar-rich substrates (Põlme et al. 2020) | Põlme S, et al. 2020. Fungal Diversity 105:1-16 ((https://doi.org/10.1007/s13225-020-00466-2)) |
| Otu46090 | 1 | k__Fungi(100);p__Ascomycota(100);c__Saccharomycetes(100);o__Saccharomycetales(100);f__Metschnikowiaceae(100);g__Metschnikowia(100);s__Metschnikowia_pulcherrima(100); | Metschnikowia | 13 | Saprotroph | \|Undefined Saprotroph\| | Yeast | NULL | Probable | As nectar-tap_saprotroph (Põlme et al. 2020); Decay sugar-rich substrates (Põlme et al. 2020) | Põlme S, et al. 2020. Fungal Diversity 105:1-16 ((https://doi.org/10.1007/s13225-020-00466-2)) |
| Otu46091 | 1 | k__Fungi(100);p__Ascomycota(100);c__Saccharomycetes(100);o__Saccharomycetales(100);f__Metschnikowiaceae(100);g__Metschnikowia(100);s__Metschnikowia_pulcherrima(100); | Metschnikowia | 13 | Saprotroph | \|Undefined Saprotroph\| | Yeast | NULL | Probable | As nectar-tap_saprotroph (Põlme et al. 2020); Decay sugar-rich substrates (Põlme et al. 2020) | Põlme S, et al. 2020. Fungal Diversity 105:1-16 ((https://doi.org/10.1007/s13225-020-00466-2)) |
| Otu46092 | 1 | k__Fungi(100);p__Ascomycota(100);c__Saccharomycetes(100);o__Saccharomycetales(100);f__Metschnikowiaceae(100);g__Metschnikowia(100);s__Metschnikowia_pulcherrima(100); | Metschnikowia | 13 | Saprotroph | \|Undefined Saprotroph\| | Yeast | NULL | Probable | As nectar-tap_saprotroph (Põlme et al. 2020); Decay sugar-rich substrates (Põlme et al. 2020) | Põlme S, et al. 2020. Fungal Diversity 105:1-16 ((https://doi.org/10.1007/s13225-020-00466-2)) |
| Otu46093 | 1 | k__Fungi(100);p__Ascomycota(100);c__Saccharomycetes(100);o__Saccharomycetales(100);f__Metschnikowiaceae(100);g__Metschnikowia(100);g__Metschnikowia_unclassified(100); | Metschnikowia | 13 | Saprotroph | \|Undefined Saprotroph\| | Yeast | NULL | Probable | As nectar-tap_saprotroph (Põlme et al. 2020); Decay sugar-rich substrates (Põlme et al. 2020) | Põlme S, et al. 2020. Fungal Diversity 105:1-16 ((https://doi.org/10.1007/s13225-020-00466-2)) |
| Otu46094 | 1 | k__Fungi(100);p__Ascomycota(100);c__Saccharomycetes(100);o__Saccharomycetales(100);f__Metschnikowiaceae(100);g__Metschnikowia(100);s__Metschnikowia_pulcherrima(100); | Metschnikowia | 13 | Saprotroph | \|Undefined Saprotroph\| | Yeast | NULL | Probable | As nectar-tap_saprotroph (Põlme et al. 2020); Decay sugar-rich substrates (Põlme et al. 2020) | Põlme S, et al. 2020. Fungal Diversity 105:1-16 ((https://doi.org/10.1007/s13225-020-00466-2)) |
| Otu46096 | 1 | k__Fungi(100);p__Ascomycota(100);c__Saccharomycetes(100);o__Saccharomycetales(100);f__Metschnikowiaceae(100);g__Metschnikowia(100);s__Metschnikowia_pulcherrima(100); | Metschnikowia | 13 | Saprotroph | \|Undefined Saprotroph\| | Yeast | NULL | Probable | As nectar-tap_saprotroph (Põlme et al. 2020); Decay sugar-rich substrates (Põlme et al. 2020) | Põlme S, et al. 2020. Fungal Diversity 105:1-16 ((https://doi.org/10.1007/s13225-020-00466-2)) |
| Otu46097 | 1 | k__Fungi(100);p__Ascomycota(100);c__Saccharomycetes(100);o__Saccharomycetales(100);f__Metschnikowiaceae(100);g__Metschnikowia(100);s__Metschnikowia_pulcherrima(100); | Metschnikowia | 13 | Saprotroph | \|Undefined Saprotroph\| | Yeast | NULL | Probable | As nectar-tap_saprotroph (Põlme et al. 2020); Decay sugar-rich substrates (Põlme et al. 2020) | Põlme S, et al. 2020. Fungal Diversity 105:1-16 ((https://doi.org/10.1007/s13225-020-00466-2)) |
| Otu46098 | 1 | k__Fungi(100);p__Ascomycota(100);c__Saccharomycetes(100);o__Saccharomycetales(100);f__Metschnikowiaceae(100);g__Metschnikowia(100);s__Metschnikowia_pulcherrima(100); | Metschnikowia | 13 | Saprotroph | \|Undefined Saprotroph\| | Yeast | NULL | Probable | As nectar-tap_saprotroph (Põlme et al. 2020); Decay sugar-rich substrates (Põlme et al. 2020) | Põlme S, et al. 2020. Fungal Diversity 105:1-16 ((https://doi.org/10.1007/s13225-020-00466-2)) |
| Otu46099 | 1 | k__Fungi(100);p__Ascomycota(100);c__Saccharomycetes(100);o__Saccharomycetales(100);f__Metschnikowiaceae(100);g__Metschnikowia(100);s__Metschnikowia_pulcherrima(100); | Metschnikowia | 13 | Saprotroph | \|Undefined Saprotroph\| | Yeast | NULL | Probable | As nectar-tap_saprotroph (Põlme et al. 2020); Decay sugar-rich substrates (Põlme et al. 2020) | Põlme S, et al. 2020. Fungal Diversity 105:1-16 ((https://doi.org/10.1007/s13225-020-00466-2)) |
| Otu46100 | 1 | k__Fungi(100);p__Ascomycota(100);c__Saccharomycetes(100);o__Saccharomycetales(100);f__Metschnikowiaceae(100);g__Metschnikowia(100);g__Metschnikowia_unclassified(100); | Metschnikowia | 13 | Saprotroph | \|Undefined Saprotroph\| | Yeast | NULL | Probable | As nectar-tap_saprotroph (Põlme et al. 2020); Decay sugar-rich substrates (Põlme et al. 2020) | Põlme S, et al. 2020. Fungal Diversity 105:1-16 ((https://doi.org/10.1007/s13225-020-00466-2)) |
| Otu46101 | 1 | k__Fungi(100);p__Ascomycota(100);c__Saccharomycetes(100);o__Saccharomycetales(100);f__Metschnikowiaceae(100);g__Metschnikowia(100);s__Metschnikowia_pulcherrima(100); | Metschnikowia | 13 | Saprotroph | \|Undefined Saprotroph\| | Yeast | NULL | Probable | As nectar-tap_saprotroph (Põlme et al. 2020); Decay sugar-rich substrates (Põlme et al. 2020) | Põlme S, et al. 2020. Fungal Diversity 105:1-16 ((https://doi.org/10.1007/s13225-020-00466-2)) |
| Otu46102 | 1 | k__Fungi(100);p__Ascomycota(100);c__Saccharomycetes(100);o__Saccharomycetales(100);f__Metschnikowiaceae(100);g__Metschnikowia(100);g__Metschnikowia_unclassified(100); | Metschnikowia | 13 | Saprotroph | \|Undefined Saprotroph\| | Yeast | NULL | Probable | As nectar-tap_saprotroph (Põlme et al. 2020); Decay sugar-rich substrates (Põlme et al. 2020) | Põlme S, et al. 2020. Fungal Diversity 105:1-16 ((https://doi.org/10.1007/s13225-020-00466-2)) |
| Otu46103 | 1 | k__Fungi(100);p__Ascomycota(100);c__Saccharomycetes(100);o__Saccharomycetales(100);f__Metschnikowiaceae(100);g__Metschnikowia(100);g__Metschnikowia_unclassified(100); | Metschnikowia | 13 | Saprotroph | \|Undefined Saprotroph\| | Yeast | NULL | Probable | As nectar-tap_saprotroph (Põlme et al. 2020); Decay sugar-rich substrates (Põlme et al. 2020) | Põlme S, et al. 2020. Fungal Diversity 105:1-16 ((https://doi.org/10.1007/s13225-020-00466-2)) |
| Otu46104 | 1 | k__Fungi(100);p__Ascomycota(100);c__Saccharomycetes(100);o__Saccharomycetales(100);f__Metschnikowiaceae(100);g__Metschnikowia(100);s__Metschnikowia_pulcherrima(100); | Metschnikowia | 13 | Saprotroph | \|Undefined Saprotroph\| | Yeast | NULL | Probable | As nectar-tap_saprotroph (Põlme et al. 2020); Decay sugar-rich substrates (Põlme et al. 2020) | Põlme S, et al. 2020. Fungal Diversity 105:1-16 ((https://doi.org/10.1007/s13225-020-00466-2)) |
| Otu46105 | 1 | k__Fungi(100);p__Ascomycota(100);c__Saccharomycetes(100);o__Saccharomycetales(100);f__Metschnikowiaceae(100);g__Metschnikowia(100);s__Metschnikowia_pulcherrima(100); | Metschnikowia | 13 | Saprotroph | \|Undefined Saprotroph\| | Yeast | NULL | Probable | As nectar-tap_saprotroph (Põlme et al. 2020); Decay sugar-rich substrates (Põlme et al. 2020) | Põlme S, et al. 2020. Fungal Diversity 105:1-16 ((https://doi.org/10.1007/s13225-020-00466-2)) |
| Otu46106 | 1 | k__Fungi(100);p__Ascomycota(100);c__Saccharomycetes(100);o__Saccharomycetales(100);f__Metschnikowiaceae(100);g__Metschnikowia(100);s__Metschnikowia_pulcherrima(100); | Metschnikowia | 13 | Saprotroph | \|Undefined Saprotroph\| | Yeast | NULL | Probable | As nectar-tap_saprotroph (Põlme et al. 2020); Decay sugar-rich substrates (Põlme et al. 2020) | Põlme S, et al. 2020. Fungal Diversity 105:1-16 ((https://doi.org/10.1007/s13225-020-00466-2)) |
| Otu46107 | 1 | k__Fungi(100);p__Ascomycota(100);c__Saccharomycetes(100);o__Saccharomycetales(100);f__Metschnikowiaceae(100);g__Metschnikowia(100);s__Metschnikowia_pulcherrima(100); | Metschnikowia | 13 | Saprotroph | \|Undefined Saprotroph\| | Yeast | NULL | Probable | As nectar-tap_saprotroph (Põlme et al. 2020); Decay sugar-rich substrates (Põlme et al. 2020) | Põlme S, et al. 2020. Fungal Diversity 105:1-16 ((https://doi.org/10.1007/s13225-020-00466-2)) |
| Otu46108 | 1 | k__Fungi(100);p__Ascomycota(100);c__Saccharomycetes(100);o__Saccharomycetales(100);f__Metschnikowiaceae(100);g__Metschnikowia(100);s__Metschnikowia_pulcherrima(100); | Metschnikowia | 13 | Saprotroph | \|Undefined Saprotroph\| | Yeast | NULL | Probable | As nectar-tap_saprotroph (Põlme et al. 2020); Decay sugar-rich substrates (Põlme et al. 2020) | Põlme S, et al. 2020. Fungal Diversity 105:1-16 ((https://doi.org/10.1007/s13225-020-00466-2)) |
| Otu46110 | 1 | k__Fungi(100);p__Ascomycota(100);c__Saccharomycetes(100);o__Saccharomycetales(100);f__Metschnikowiaceae(100);g__Metschnikowia(100);s__Metschnikowia_pulcherrima(100); | Metschnikowia | 13 | Saprotroph | \|Undefined Saprotroph\| | Yeast | NULL | Probable | As nectar-tap_saprotroph (Põlme et al. 2020); Decay sugar-rich substrates (Põlme et al. 2020) | Põlme S, et al. 2020. Fungal Diversity 105:1-16 ((https://doi.org/10.1007/s13225-020-00466-2)) |
| Otu46111 | 1 | k__Fungi(100);p__Ascomycota(100);c__Saccharomycetes(100);o__Saccharomycetales(100);f__Metschnikowiaceae(100);g__Metschnikowia(100);s__Metschnikowia_pulcherrima(100); | Metschnikowia | 13 | Saprotroph | \|Undefined Saprotroph\| | Yeast | NULL | Probable | As nectar-tap_saprotroph (Põlme et al. 2020); Decay sugar-rich substrates (Põlme et al. 2020) | Põlme S, et al. 2020. Fungal Diversity 105:1-16 ((https://doi.org/10.1007/s13225-020-00466-2)) |
| Otu46112 | 1 | k__Fungi(100);p__Ascomycota(100);c__Saccharomycetes(100);o__Saccharomycetales(100);f__Metschnikowiaceae(100);g__Metschnikowia(100);g__Metschnikowia_unclassified(100); | Metschnikowia | 13 | Saprotroph | \|Undefined Saprotroph\| | Yeast | NULL | Probable | As nectar-tap_saprotroph (Põlme et al. 2020); Decay sugar-rich substrates (Põlme et al. 2020) | Põlme S, et al. 2020. Fungal Diversity 105:1-16 ((https://doi.org/10.1007/s13225-020-00466-2)) |
| Otu46113 | 1 | k__Fungi(100);p__Ascomycota(100);c__Saccharomycetes(100);o__Saccharomycetales(100);f__Metschnikowiaceae(100);g__Metschnikowia(100);s__Metschnikowia_pulcherrima(100); | Metschnikowia | 13 | Saprotroph | \|Undefined Saprotroph\| | Yeast | NULL | Probable | As nectar-tap_saprotroph (Põlme et al. 2020); Decay sugar-rich substrates (Põlme et al. 2020) | Põlme S, et al. 2020. Fungal Diversity 105:1-16 ((https://doi.org/10.1007/s13225-020-00466-2)) |

| | | | | | | | | | | | |
|---|---|---|---|---|---|---|---|---|---|---|---|
| Otu46115 | 1 | k__Fungi(100);p__Ascomycota(100);c__Saccharomycetes(100);o__Saccharomycetales(100);f__Metschnikowiaceae(100);g__Metschnikowia(100);s__Metschnikowia_pulcherrima(100); | Metschnikowia | 13 | Saprotroph | \|Undefined Saprotroph\| | Yeast | NULL | Probable | As nectar-tap_saprotroph (Põlme et al. 2020); Decay sugar-rich substrates (Põlme et al. 2020) | Põlme S, et al. 2020. Fungal Diversity 105:1-16 ((https://doi.org/10.1007/s13225-020-00466-2)) |
| Otu46116 | 1 | k__Fungi(100);p__Ascomycota(100);c__Saccharomycetes(100);o__Saccharomycetales(100);f__Metschnikowiaceae(100);g__Metschnikowia(100);g__Metschnikowia_unclassified(100); | Metschnikowia | 13 | Saprotroph | \|Undefined Saprotroph\| | Yeast | NULL | Probable | As nectar-tap_saprotroph (Põlme et al. 2020); Decay sugar-rich substrates (Põlme et al. 2020) | Põlme S, et al. 2020. Fungal Diversity 105:1-16 ((https://doi.org/10.1007/s13225-020-00466-2)) |
| Otu46117 | 1 | k__Fungi(100);p__Ascomycota(100);c__Saccharomycetes(100);o__Saccharomycetales(100);f__Metschnikowiaceae(100);g__Metschnikowia(100);g__Metschnikowia_unclassified(100); | Metschnikowia | 13 | Saprotroph | \|Undefined Saprotroph\| | Yeast | NULL | Probable | As nectar-tap_saprotroph (Põlme et al. 2020); Decay sugar-rich substrates (Põlme et al. 2020) | Põlme S, et al. 2020. Fungal Diversity 105:1-16 ((https://doi.org/10.1007/s13225-020-00466-2)) |
| Otu46118 | 1 | k__Fungi(100);p__Ascomycota(100);c__Saccharomycetes(100);o__Saccharomycetales(100);f__Metschnikowiaceae(100);g__Metschnikowia(100);g__Metschnikowia_unclassified(100); | Metschnikowia | 13 | Saprotroph | \|Undefined Saprotroph\| | Yeast | NULL | Probable | As nectar-tap_saprotroph (Põlme et al. 2020); Decay sugar-rich substrates (Põlme et al. 2020) | Põlme S, et al. 2020. Fungal Diversity 105:1-16 ((https://doi.org/10.1007/s13225-020-00466-2)) |
| Otu46119 | 1 | k__Fungi(100);p__Ascomycota(100);c__Saccharomycetes(100);o__Saccharomycetales(100);f__Metschnikowiaceae(100);g__Metschnikowia(100);g__Metschnikowia_unclassified(100); | Metschnikowia | 13 | Saprotroph | \|Undefined Saprotroph\| | Yeast | NULL | Probable | As nectar-tap_saprotroph (Põlme et al. 2020); Decay sugar-rich substrates (Põlme et al. 2020) | Põlme S, et al. 2020. Fungal Diversity 105:1-16 ((https://doi.org/10.1007/s13225-020-00466-2)) |
| Otu46120 | 1 | k__Fungi(100);p__Ascomycota(100);c__Saccharomycetes(100);o__Saccharomycetales(100);f__Metschnikowiaceae(100);g__Metschnikowia(100);g__Metschnikowia_unclassified(100); | Metschnikowia | 13 | Saprotroph | \|Undefined Saprotroph\| | Yeast | NULL | Probable | As nectar-tap_saprotroph (Põlme et al. 2020); Decay sugar-rich substrates (Põlme et al. 2020) | Põlme S, et al. 2020. Fungal Diversity 105:1-16 ((https://doi.org/10.1007/s13225-020-00466-2)) |
| Otu46121 | 1 | k__Fungi(100);p__Ascomycota(100);c__Saccharomycetes(100);o__Saccharomycetales(100);f__Metschnikowiaceae(100);g__Metschnikowia(100);g__Metschnikowia_unclassified(100); | Metschnikowia | 13 | Saprotroph | \|Undefined Saprotroph\| | Yeast | NULL | Probable | As nectar-tap_saprotroph (Põlme et al. 2020); Decay sugar-rich substrates (Põlme et al. 2020) | Põlme S, et al. 2020. Fungal Diversity 105:1-16 ((https://doi.org/10.1007/s13225-020-00466-2)) |
| Otu46122 | 1 | k__Fungi(100);p__Ascomycota(100);c__Saccharomycetes(100);o__Saccharomycetales(100);f__Metschnikowiaceae(100);g__Metschnikowia(100);s__Metschnikowia_pulcherrima(100); | Metschnikowia | 13 | Saprotroph | \|Undefined Saprotroph\| | Yeast | NULL | Probable | As nectar-tap_saprotroph (Põlme et al. 2020); Decay sugar-rich substrates (Põlme et al. 2020) | Põlme S, et al. 2020. Fungal Diversity 105:1-16 ((https://doi.org/10.1007/s13225-020-00466-2)) |
| Otu46123 | 1 | k__Fungi(100);p__Ascomycota(100);c__Saccharomycetes(100);o__Saccharomycetales(100);f__Metschnikowiaceae(100);g__Metschnikowia(100);s__Metschnikowia_pulcherrima(100); | Metschnikowia | 13 | Saprotroph | \|Undefined Saprotroph\| | Yeast | NULL | Probable | As nectar-tap_saprotroph (Põlme et al. 2020); Decay sugar-rich substrates (Põlme et al. 2020) | Põlme S, et al. 2020. Fungal Diversity 105:1-16 ((https://doi.org/10.1007/s13225-020-00466-2)) |
| Otu46124 | 1 | k__Fungi(100);p__Ascomycota(100);c__Saccharomycetes(100);o__Saccharomycetales(100);f__Metschnikowiaceae(100);g__Metschnikowia(100);s__Metschnikowia_pulcherrima(100); | Metschnikowia | 13 | Saprotroph | \|Undefined Saprotroph\| | Yeast | NULL | Probable | As nectar-tap_saprotroph (Põlme et al. 2020); Decay sugar-rich substrates (Põlme et al. 2020) | Põlme S, et al. 2020. Fungal Diversity 105:1-16 ((https://doi.org/10.1007/s13225-020-00466-2)) |
| Otu46125 | 1 | k__Fungi(100);p__Ascomycota(100);c__Saccharomycetes(100);o__Saccharomycetales(100);f__Metschnikowiaceae(100);g__Metschnikowia(100);s__Metschnikowia_pulcherrima(100); | Metschnikowia | 13 | Saprotroph | \|Undefined Saprotroph\| | Yeast | NULL | Probable | As nectar-tap_saprotroph (Põlme et al. 2020); Decay sugar-rich substrates (Põlme et al. 2020) | Põlme S, et al. 2020. Fungal Diversity 105:1-16 ((https://doi.org/10.1007/s13225-020-00466-2)) |
| Otu46127 | 1 | k__Fungi(100);p__Ascomycota(100);c__Saccharomycetes(100);o__Saccharomycetales(100);f__Metschnikowiaceae(100);g__Metschnikowia(100);s__Metschnikowia_pulcherrima(100); | Metschnikowia | 13 | Saprotroph | \|Undefined Saprotroph\| | Yeast | NULL | Probable | As nectar-tap_saprotroph (Põlme et al. 2020); Decay sugar-rich substrates (Põlme et al. 2020) | Põlme S, et al. 2020. Fungal Diversity 105:1-16 ((https://doi.org/10.1007/s13225-020-00466-2)) |
| Otu46128 | 1 | k__Fungi(100);p__Ascomycota(100);c__Saccharomycetes(100);o__Saccharomycetales(100);f__Metschnikowiaceae(100);g__Metschnikowia(100);s__Metschnikowia_pulcherrima(100); | Metschnikowia | 13 | Saprotroph | \|Undefined Saprotroph\| | Yeast | NULL | Probable | As nectar-tap_saprotroph (Põlme et al. 2020); Decay sugar-rich substrates (Põlme et al. 2020) | Põlme S, et al. 2020. Fungal Diversity 105:1-16 ((https://doi.org/10.1007/s13225-020-00466-2)) |
| Otu46129 | 1 | k__Fungi(100);p__Ascomycota(100);c__Saccharomycetes(100);o__Saccharomycetales(100);f__Metschnikowiaceae(100);g__Metschnikowia(100);s__Metschnikowia_pulcherrima(100); | Metschnikowia | 13 | Saprotroph | \|Undefined Saprotroph\| | Yeast | NULL | Probable | As nectar-tap_saprotroph (Põlme et al. 2020); Decay sugar-rich substrates (Põlme et al. 2020) | Põlme S, et al. 2020. Fungal Diversity 105:1-16 ((https://doi.org/10.1007/s13225-020-00466-2)) |
| Otu46130 | 1 | k__Fungi(100);p__Ascomycota(100);c__Saccharomycetes(100);o__Saccharomycetales(100);f__Metschnikowiaceae(100);g__Metschnikowia(100);s__Metschnikowia_pulcherrima(100); | Metschnikowia | 13 | Saprotroph | \|Undefined Saprotroph\| | Yeast | NULL | Probable | As nectar-tap_saprotroph (Põlme et al. 2020); Decay sugar-rich substrates (Põlme et al. 2020) | Põlme S, et al. 2020. Fungal Diversity 105:1-16 ((https://doi.org/10.1007/s13225-020-00466-2)) |
| Otu46131 | 1 | k__Fungi(100);p__Ascomycota(100);c__Saccharomycetes(100);o__Saccharomycetales(100);f__Metschnikowiaceae(100);g__Metschnikowia(100);s__Metschnikowia_pulcherrima(100); | Metschnikowia | 13 | Saprotroph | \|Undefined Saprotroph\| | Yeast | NULL | Probable | As nectar-tap_saprotroph (Põlme et al. 2020); Decay sugar-rich substrates (Põlme et al. 2020) | Põlme S, et al. 2020. Fungal Diversity 105:1-16 ((https://doi.org/10.1007/s13225-020-00466-2)) |
| Otu46132 | 1 | k__Fungi(100);p__Ascomycota(100);c__Saccharomycetes(100);o__Saccharomycetales(100);f__Metschnikowiaceae(100);g__Metschnikowia(100);s__Metschnikowia_pulcherrima(100); | Metschnikowia | 13 | Saprotroph | \|Undefined Saprotroph\| | Yeast | NULL | Probable | As nectar-tap_saprotroph (Põlme et al. 2020); Decay sugar-rich substrates (Põlme et al. 2020) | Põlme S, et al. 2020. Fungal Diversity 105:1-16 ((https://doi.org/10.1007/s13225-020-00466-2)) |
| Otu46133 | 1 | k__Fungi(100);p__Ascomycota(100);c__Saccharomycetes(100);o__Saccharomycetales(100);f__Metschnikowiaceae(100);g__Metschnikowia(100);s__Metschnikowia_pulcherrima(100); | Metschnikowia | 13 | Saprotroph | \|Undefined Saprotroph\| | Yeast | NULL | Probable | As nectar-tap_saprotroph (Põlme et al. 2020); Decay sugar-rich substrates (Põlme et al. 2020) | Põlme S, et al. 2020. Fungal Diversity 105:1-16 ((https://doi.org/10.1007/s13225-020-00466-2)) |
| Otu46134 | 1 | k__Fungi(100);p__Ascomycota(100);c__Saccharomycetes(100);o__Saccharomycetales(100);f__Metschnikowiaceae(100);g__Metschnikowia(100);s__Metschnikowia_pulcherrima(100); | Metschnikowia | 13 | Saprotroph | \|Undefined Saprotroph\| | Yeast | NULL | Probable | As nectar-tap_saprotroph (Põlme et al. 2020); Decay sugar-rich substrates (Põlme et al. 2020) | Põlme S, et al. 2020. Fungal Diversity 105:1-16 ((https://doi.org/10.1007/s13225-020-00466-2)) |
| Otu46135 | 1 | k__Fungi(100);p__Ascomycota(100);c__Saccharomycetes(100);o__Saccharomycetales(100);f__Metschnikowiaceae(100);g__Metschnikowia(100);s__Metschnikowia_pulcherrima(100); | Metschnikowia | 13 | Saprotroph | \|Undefined Saprotroph\| | Yeast | NULL | Probable | As nectar-tap_saprotroph (Põlme et al. 2020); Decay sugar-rich substrates (Põlme et al. 2020) | Põlme S, et al. 2020. Fungal Diversity 105:1-16 ((https://doi.org/10.1007/s13225-020-00466-2)) |
| Otu46136 | 1 | k__Fungi(100);p__Ascomycota(100);c__Saccharomycetes(100);o__Saccharomycetales(100);f__Metschnikowiaceae(100);g__Metschnikowia(100);g__Metschnikowia_unclassified(100); | Metschnikowia | 13 | Saprotroph | \|Undefined Saprotroph\| | Yeast | NULL | Probable | As nectar-tap_saprotroph (Põlme et al. 2020); Decay sugar-rich substrates (Põlme et al. 2020) | Põlme S, et al. 2020. Fungal Diversity 105:1-16 ((https://doi.org/10.1007/s13225-020-00466-2)) |
| Otu46138 | 1 | k__Fungi(100);p__Ascomycota(100);c__Saccharomycetes(100);o__Saccharomycetales(100);f__Metschnikowiaceae(100);g__Metschnikowia(100);s__Metschnikowia_pulcherrima(100); | Metschnikowia | 13 | Saprotroph | \|Undefined Saprotroph\| | Yeast | NULL | Probable | As nectar-tap_saprotroph (Põlme et al. 2020); Decay sugar-rich substrates (Põlme et al. 2020) | Põlme S, et al. 2020. Fungal Diversity 105:1-16 ((https://doi.org/10.1007/s13225-020-00466-2)) |
| Otu46140 | 1 | k__Fungi(100);p__Ascomycota(100);c__Saccharomycetes(100);o__Saccharomycetales(100);f__Metschnikowiaceae(100);g__Metschnikowia(100);g__Metschnikowia_unclassified(100); | Metschnikowia | 13 | Saprotroph | \|Undefined Saprotroph\| | Yeast | NULL | Probable | As nectar-tap_saprotroph (Põlme et al. 2020); Decay sugar-rich substrates (Põlme et al. 2020) | Põlme S, et al. 2020. Fungal Diversity 105:1-16 ((https://doi.org/10.1007/s13225-020-00466-2)) |
| Otu46141 | 1 | k__Fungi(100);p__Ascomycota(100);c__Saccharomycetes(100);o__Saccharomycetales(100);f__Metschnikowiaceae(100);g__Metschnikowia(100);s__Metschnikowia_pulcherrima(100); | Metschnikowia | 13 | Saprotroph | \|Undefined Saprotroph\| | Yeast | NULL | Probable | As nectar-tap_saprotroph (Põlme et al. 2020); Decay sugar-rich substrates (Põlme et al. 2020) | Põlme S, et al. 2020. Fungal Diversity 105:1-16 ((https://doi.org/10.1007/s13225-020-00466-2)) |
| Otu46142 | 1 | k__Fungi(100);p__Ascomycota(100);c__Saccharomycetes(100);o__Saccharomycetales(100);f__Metschnikowiaceae(100);g__Metschnikowia(100);s__Metschnikowia_pulcherrima(100); | Metschnikowia | 13 | Saprotroph | \|Undefined Saprotroph\| | Yeast | NULL | Probable | As nectar-tap_saprotroph (Põlme et al. 2020); Decay sugar-rich substrates (Põlme et al. 2020) | Põlme S, et al. 2020. Fungal Diversity 105:1-16 ((https://doi.org/10.1007/s13225-020-00466-2)) |
| Otu46143 | 1 | k__Fungi(100);p__Ascomycota(100);c__Saccharomycetes(100);o__Saccharomycetales(100);f__Metschnikowiaceae(100);g__Metschnikowia(100);s__Metschnikowia_pulcherrima(100); | Metschnikowia | 13 | Saprotroph | \|Undefined Saprotroph\| | Yeast | NULL | Probable | As nectar-tap_saprotroph (Põlme et al. 2020); Decay sugar-rich substrates (Põlme et al. 2020) | Põlme S, et al. 2020. Fungal Diversity 105:1-16 ((https://doi.org/10.1007/s13225-020-00466-2)) |
| Otu46144 | 1 | k__Fungi(100);p__Ascomycota(100);c__Saccharomycetes(100);o__Saccharomycetales(100);f__Metschnikowiaceae(100);g__Metschnikowia(100);s__Metschnikowia_pulcherrima(100); | Metschnikowia | 13 | Saprotroph | \|Undefined Saprotroph\| | Yeast | NULL | Probable | As nectar-tap_saprotroph (Põlme et al. 2020); Decay sugar-rich substrates (Põlme et al. 2020) | Põlme S, et al. 2020. Fungal Diversity 105:1-16 ((https://doi.org/10.1007/s13225-020-00466-2)) |
| Otu46146 | 1 | k__Fungi(100);p__Ascomycota(100);c__Saccharomycetes(100);o__Saccharomycetales(100);f__Metschnikowiaceae(100);g__Metschnikowia(100);s__Metschnikowia_pulcherrima(100); | Metschnikowia | 13 | Saprotroph | \|Undefined Saprotroph\| | Yeast | NULL | Probable | As nectar-tap_saprotroph (Põlme et al. 2020); Decay sugar-rich substrates (Põlme et al. 2020) | Põlme S, et al. 2020. Fungal Diversity 105:1-16 ((https://doi.org/10.1007/s13225-020-00466-2)) |
| Otu46147 | 1 | k__Fungi(100);p__Ascomycota(100);c__Saccharomycetes(100);o__Saccharomycetales(100);f__Metschnikowiaceae(100);g__Metschnikowia(100);g__Metschnikowia_unclassified(100); | Metschnikowia | 13 | Saprotroph | \|Undefined Saprotroph\| | Yeast | NULL | Probable | As nectar-tap_saprotroph (Põlme et al. 2020); Decay sugar-rich substrates (Põlme et al. 2020) | Põlme S, et al. 2020. Fungal Diversity 105:1-16 ((https://doi.org/10.1007/s13225-020-00466-2)) |
| Otu46148 | 1 | k__Fungi(100);p__Ascomycota(100);c__Saccharomycetes(100);o__Saccharomycetales(100);f__Metschnikowiaceae(100);g__Metschnikowia(100);s__Metschnikowia_pulcherrima(100); | Metschnikowia | 13 | Saprotroph | \|Undefined Saprotroph\| | Yeast | NULL | Probable | As nectar-tap_saprotroph (Põlme et al. 2020); Decay sugar-rich substrates (Põlme et al. 2020) | Põlme S, et al. 2020. Fungal Diversity 105:1-16 ((https://doi.org/10.1007/s13225-020-00466-2)) |
| Otu46149 | 1 | k__Fungi(100);p__Ascomycota(100);c__Saccharomycetes(100);o__Saccharomycetales(100);f__Metschnikowiaceae(100);g__Metschnikowia(100);s__Metschnikowia_pulcherrima(100); | Metschnikowia | 13 | Saprotroph | \|Undefined Saprotroph\| | Yeast | NULL | Probable | As nectar-tap_saprotroph (Põlme et al. 2020); Decay sugar-rich substrates (Põlme et al. 2020) | Põlme S, et al. 2020. Fungal Diversity 105:1-16 ((https://doi.org/10.1007/s13225-020-00466-2)) |
| Otu46150 | 1 | k__Fungi(100);p__Ascomycota(100);c__Saccharomycetes(100);o__Saccharomycetales(100);f__Metschnikowiaceae(100);g__Metschnikowia(100);s__Metschnikowia_pulcherrima(100); | Metschnikowia | 13 | Saprotroph | \|Undefined Saprotroph\| | Yeast | NULL | Probable | As nectar-tap_saprotroph (Põlme et al. 2020); Decay sugar-rich substrates (Põlme et al. 2020) | Põlme S, et al. 2020. Fungal Diversity 105:1-16 ((https://doi.org/10.1007/s13225-020-00466-2)) |
| Otu46151 | 1 | k__Fungi(100);p__Ascomycota(100);c__Saccharomycetes(100);o__Saccharomycetales(100);f__Metschnikowiaceae(100);g__Metschnikowia(100);s__Metschnikowia_pulcherrima(100); | Metschnikowia | 13 | Saprotroph | \|Undefined Saprotroph\| | Yeast | NULL | Probable | As nectar-tap_saprotroph (Põlme et al. 2020); Decay sugar-rich substrates (Põlme et al. 2020) | Põlme S, et al. 2020. Fungal Diversity 105:1-16 ((https://doi.org/10.1007/s13225-020-00466-2)) |
| Otu46152 | 1 | k__Fungi(100);p__Ascomycota(100);c__Saccharomycetes(100);o__Saccharomycetales(100);f__Metschnikowiaceae(100);g__Metschnikowia(100);g__Metschnikowia_unclassified(100); | Metschnikowia | 13 | Saprotroph | \|Undefined Saprotroph\| | Yeast | NULL | Probable | As nectar-tap_saprotroph (Põlme et al. 2020); Decay sugar-rich substrates (Põlme et al. 2020) | Põlme S, et al. 2020. Fungal Diversity 105:1-16 ((https://doi.org/10.1007/s13225-020-00466-2)) |
| Otu46153 | 1 | k__Fungi(100);p__Ascomycota(100);c__Saccharomycetes(100);o__Saccharomycetales(100);f__Metschnikowiaceae(100);g__Metschnikowia(100);g__Metschnikowia_unclassified(100); | Metschnikowia | 13 | Saprotroph | \|Undefined Saprotroph\| | Yeast | NULL | Probable | As nectar-tap_saprotroph (Põlme et al. 2020); Decay sugar-rich substrates (Põlme et al. 2020) | Põlme S, et al. 2020. Fungal Diversity 105:1-16 ((https://doi.org/10.1007/s13225-020-00466-2)) |
| Otu46154 | 1 | k__Fungi(100);p__Ascomycota(100);c__Saccharomycetes(100);o__Saccharomycetales(100);f__Metschnikowiaceae(100);g__Metschnikowia(100);s__Metschnikowia_pulcherrima(100); | Metschnikowia | 13 | Saprotroph | \|Undefined Saprotroph\| | Yeast | NULL | Probable | As nectar-tap_saprotroph (Põlme et al. 2020); Decay sugar-rich substrates (Põlme et al. 2020) | Põlme S, et al. 2020. Fungal Diversity 105:1-16 ((https://doi.org/10.1007/s13225-020-00466-2)) |
| Otu46155 | 1 | k__Fungi(100);p__Ascomycota(100);c__Saccharomycetes(100);o__Saccharomycetales(100);f__Metschnikowiaceae(100);g__Metschnikowia(100);s__Metschnikowia_pulcherrima(100); | Metschnikowia | 13 | Saprotroph | \|Undefined Saprotroph\| | Yeast | NULL | Probable | As nectar-tap_saprotroph (Põlme et al. 2020); Decay sugar-rich substrates (Põlme et al. 2020) | Põlme S, et al. 2020. Fungal Diversity 105:1-16 ((https://doi.org/10.1007/s13225-020-00466-2)) |
| Otu46156 | 1 | k__Fungi(100);p__Ascomycota(100);c__Saccharomycetes(100);o__Saccharomycetales(100);f__Metschnikowiaceae(100);g__Metschnikowia(100);s__Metschnikowia_pulcherrima(100); | Metschnikowia | 13 | Saprotroph | \|Undefined Saprotroph\| | Yeast | NULL | Probable | As nectar-tap_saprotroph (Põlme et al. 2020); Decay sugar-rich substrates (Põlme et al. 2020) | Põlme S, et al. 2020. Fungal Diversity 105:1-16 ((https://doi.org/10.1007/s13225-020-00466-2)) |
| Otu46157 | 1 | k__Fungi(100);p__Ascomycota(100);c__Saccharomycetes(100);o__Saccharomycetales(100);f__Metschnikowiaceae(100);g__Metschnikowia(100);s__Metschnikowia_pulcherrima(100); | Metschnikowia | 13 | Saprotroph | \|Undefined Saprotroph\| | Yeast | NULL | Probable | As nectar-tap_saprotroph (Põlme et al. 2020); Decay sugar-rich substrates (Põlme et al. 2020) | Põlme S, et al. 2020. Fungal Diversity 105:1-16 ((https://doi.org/10.1007/s13225-020-00466-2)) |
| Otu46158 | 1 | k__Fungi(100);p__Ascomycota(100);c__Saccharomycetes(100);o__Saccharomycetales(100);f__Metschnikowiaceae(100);g__Metschnikowia(100);g__Metschnikowia_unclassified(100); | Metschnikowia | 13 | Saprotroph | \|Undefined Saprotroph\| | Yeast | NULL | Probable | As nectar-tap_saprotroph (Põlme et al. 2020); Decay sugar-rich substrates (Põlme et al. 2020) | Põlme S, et al. 2020. Fungal Diversity 105:1-16 ((https://doi.org/10.1007/s13225-020-00466-2)) |
| Otu46159 | 1 | k__Fungi(100);p__Ascomycota(100);c__Saccharomycetes(100);o__Saccharomycetales(100);f__Metschnikowiaceae(100);g__Metschnikowia(100);g__Metschnikowia_unclassified(100); | Metschnikowia | 13 | Saprotroph | \|Undefined Saprotroph\| | Yeast | NULL | Probable | As nectar-tap_saprotroph (Põlme et al. 2020); Decay sugar-rich substrates (Põlme et al. 2020) | Põlme S, et al. 2020. Fungal Diversity 105:1-16 ((https://doi.org/10.1007/s13225-020-00466-2)) |
| Otu46160 | 1 | k__Fungi(100);p__Ascomycota(100);c__Saccharomycetes(100);o__Saccharomycetales(100);f__Metschnikowiaceae(100);g__Metschnikowia(100);g__Metschnikowia_unclassified(100); | Metschnikowia | 13 | Saprotroph | \|Undefined Saprotroph\| | Yeast | NULL | Probable | As nectar-tap_saprotroph (Põlme et al. 2020); Decay sugar-rich substrates (Põlme et al. 2020) | Põlme S, et al. 2020. Fungal Diversity 105:1-16 ((https://doi.org/10.1007/s13225-020-00466-2)) |
| Otu46161 | 1 | k__Fungi(100);p__Ascomycota(100);c__Saccharomycetes(100);o__Saccharomycetales(100);f__Metschnikowiaceae(100);g__Metschnikowia(100);s__Metschnikowia_pulcherrima(100); | Metschnikowia | 13 | Saprotroph | \|Undefined Saprotroph\| | Yeast | NULL | Probable | As nectar-tap_saprotroph (Põlme et al. 2020); Decay sugar-rich substrates (Põlme et al. 2020) | Põlme S, et al. 2020. Fungal Diversity 105:1-16 ((https://doi.org/10.1007/s13225-020-00466-2)) |
| Otu46162 | 1 | k__Fungi(100);p__Ascomycota(100);c__Saccharomycetes(100);o__Saccharomycetales(100);f__Metschnikowiaceae(100);g__Metschnikowia(100);s__Metschnikowia_pulcherrima(100); | Metschnikowia | 13 | Saprotroph | \|Undefined Saprotroph\| | Yeast | NULL | Probable | As nectar-tap_saprotroph (Põlme et al. 2020); Decay sugar-rich substrates (Põlme et al. 2020) | Põlme S, et al. 2020. Fungal Diversity 105:1-16 ((https://doi.org/10.1007/s13225-020-00466-2)) |
| Otu46163 | 1 | k__Fungi(100);p__Ascomycota(100);c__Saccharomycetes(100);o__Saccharomycetales(100);f__Metschnikowiaceae(100);g__Metschnikowia(100);s__Metschnikowia_pulcherrima(100); | Metschnikowia | 13 | Saprotroph | \|Undefined Saprotroph\| | Yeast | NULL | Probable | As nectar-tap_saprotroph (Põlme et al. 2020); Decay sugar-rich substrates (Põlme et al. 2020) | Põlme S, et al. 2020. Fungal Diversity 105:1-16 ((https://doi.org/10.1007/s13225-020-00466-2)) |
| Otu46164 | 1 | k__Fungi(100);p__Ascomycota(100);c__Saccharomycetes(100);o__Saccharomycetales(100);f__Metschnikowiaceae(100);g__Metschnikowia(100);s__Metschnikowia_pulcherrima(100); | Metschnikowia | 13 | Saprotroph | \|Undefined Saprotroph\| | Yeast | NULL | Probable | As nectar-tap_saprotroph (Põlme et al. 2020); Decay sugar-rich substrates (Põlme et al. 2020) | Põlme S, et al. 2020. Fungal Diversity 105:1-16 ((https://doi.org/10.1007/s13225-020-00466-2)) |
| Otu46165 | 1 | k__Fungi(100);p__Ascomycota(100);c__Saccharomycetes(100);o__Saccharomycetales(100);f__Metschnikowiaceae(100);g__Metschnikowia(100);g__Metschnikowia_unclassified(100); | Metschnikowia | 13 | Saprotroph | \|Undefined Saprotroph\| | Yeast | NULL | Probable | As nectar-tap_saprotroph (Põlme et al. 2020); Decay sugar-rich substrates (Põlme et al. 2020) | Põlme S, et al. 2020. Fungal Diversity 105:1-16 ((https://doi.org/10.1007/s13225-020-00466-2)) |
| Otu46166 | 1 | k__Fungi(100);p__Ascomycota(100);c__Saccharomycetes(100);o__Saccharomycetales(100);f__Metschnikowiaceae(100);g__Metschnikowia(100);s__Metschnikowia_pulcherrima(100); | Metschnikowia | 13 | Saprotroph | \|Undefined Saprotroph\| | Yeast | NULL | Probable | As nectar-tap_saprotroph (Põlme et al. 2020); Decay sugar-rich substrates (Põlme et al. 2020) | Põlme S, et al. 2020. Fungal Diversity 105:1-16 ((https://doi.org/10.1007/s13225-020-00466-2)) |
| Otu46167 | 1 | k__Fungi(100);p__Ascomycota(100);c__Saccharomycetes(100);o__Saccharomycetales(100);f__Metschnikowiaceae(100);g__Metschnikowia(100);s__Metschnikowia_pulcherrima(100); | Metschnikowia | 13 | Saprotroph | \|Undefined Saprotroph\| | Yeast | NULL | Probable | As nectar-tap_saprotroph (Põlme et al. 2020); Decay sugar-rich substrates (Põlme et al. 2020) | Põlme S, et al. 2020. Fungal Diversity 105:1-16 ((https://doi.org/10.1007/s13225-020-00466-2)) |
| Otu46168 | 1 | k__Fungi(100);p__Ascomycota(100);c__Saccharomycetes(100);o__Saccharomycetales(100);f__Metschnikowiaceae(100);g__Metschnikowia(100);s__Metschnikowia_pulcherrima(100); | Metschnikowia | 13 | Saprotroph | \|Undefined Saprotroph\| | Yeast | NULL | Probable | As nectar-tap_saprotroph (Põlme et al. 2020); Decay sugar-rich substrates (Põlme et al. 2020) | Põlme S, et al. 2020. Fungal Diversity 105:1-16 ((https://doi.org/10.1007/s13225-020-00466-2)) |
| Otu46169 | 1 | k__Fungi(100);p__Ascomycota(100);c__Saccharomycetes(100);o__Saccharomycetales(100);f__Metschnikowiaceae(100);g__Metschnikowia(100);s__Metschnikowia_pulcherrima(100); | Metschnikowia | 13 | Saprotroph | \|Undefined Saprotroph\| | Yeast | NULL | Probable | As nectar-tap_saprotroph (Põlme et al. 2020); Decay sugar-rich substrates (Põlme et al. 2020) | Põlme S, et al. 2020. Fungal Diversity 105:1-16 ((https://doi.org/10.1007/s13225-020-00466-2)) |
| Otu46170 | 1 | k__Fungi(100);p__Ascomycota(100);c__Saccharomycetes(100);o__Saccharomycetales(100);f__Metschnikowiaceae(100);g__Metschnikowia(100);s__Metschnikowia_pulcherrima(100); | Metschnikowia | 13 | Saprotroph | \|Undefined Saprotroph\| | Yeast | NULL | Probable | As nectar-tap_saprotroph (Põlme et al. 2020); Decay sugar-rich substrates (Põlme et al. 2020) | Põlme S, et al. 2020. Fungal Diversity 105:1-16 ((https://doi.org/10.1007/s13225-020-00466-2)) |
| Otu46171 | 1 | k__Fungi(100);p__Ascomycota(100);c__Saccharomycetes(100);o__Saccharomycetales(100);f__Metschnikowiaceae(100);g__Metschnikowia(100);s__Metschnikowia_pulcherrima(100); | Metschnikowia | 13 | Saprotroph | \|Undefined Saprotroph\| | Yeast | NULL | Probable | As nectar-tap_saprotroph (Põlme et al. 2020); Decay sugar-rich substrates (Põlme et al. 2020) | Põlme S, et al. 2020. Fungal Diversity 105:1-16 ((https://doi.org/10.1007/s13225-020-00466-2)) |
| Otu46172 | 1 | k__Fungi(100);p__Ascomycota(100);c__Saccharomycetes(100);o__Saccharomycetales(100);f__Metschnikowiaceae(100);g__Metschnikowia(100);g__Metschnikowia_unclassified(100); | Metschnikowia | 13 | Saprotroph | \|Undefined Saprotroph\| | Yeast | NULL | Probable | As nectar-tap_saprotroph (Põlme et al. 2020); Decay sugar-rich substrates (Põlme et al. 2020) | Põlme S, et al. 2020. Fungal Diversity 105:1-16 ((https://doi.org/10.1007/s13225-020-00466-2)) |
| Otu46173 | 1 | k__Fungi(100);p__Ascomycota(100);c__Saccharomycetes(100);o__Saccharomycetales(100);f__Metschnikowiaceae(100);g__Metschnikowia(100);g__Metschnikowia_unclassified(100); | Metschnikowia | 13 | Saprotroph | \|Undefined Saprotroph\| | Yeast | NULL | Probable | As nectar-tap_saprotroph (Põlme et al. 2020); Decay sugar-rich substrates (Põlme et al. 2020) | Põlme S, et al. 2020. Fungal Diversity 105:1-16 ((https://doi.org/10.1007/s13225-020-00466-2)) |
| Otu46174 | 1 | k__Fungi(100);p__Ascomycota(100);c__Saccharomycetes(100);o__Saccharomycetales(100);f__Metschnikowiaceae(100);g__Metschnikowia(100);s__Metschnikowia_pulcherrima(100); | Metschnikowia | 13 | Saprotroph | \|Undefined Saprotroph\| | Yeast | NULL | Probable | As nectar-tap_saprotroph (Põlme et al. 2020); Decay sugar-rich substrates (Põlme et al. 2020) | Põlme S, et al. 2020. Fungal Diversity 105:1-16 ((https://doi.org/10.1007/s13225-020-00466-2)) |
| Otu46175 | 1 | k__Fungi(100);p__Ascomycota(100);c__Saccharomycetes(100);o__Saccharomycetales(100);f__Metschnikowiaceae(100);g__Metschnikowia(100);s__Metschnikowia_pulcherrima(100); | Metschnikowia | 13 | Saprotroph | \|Undefined Saprotroph\| | Yeast | NULL | Probable | As nectar-tap_saprotroph (Põlme et al. 2020); Decay sugar-rich substrates (Põlme et al. 2020) | Põlme S, et al. 2020. Fungal Diversity 105:1-16 ((https://doi.org/10.1007/s13225-020-00466-2)) |
| Otu46176 | 1 | k__Fungi(100);p__Ascomycota(100);c__Saccharomycetes(100);o__Saccharomycetales(100);f__Metschnikowiaceae(100);g__Metschnikowia(100);s__Metschnikowia_pulcherrima(100); | Metschnikowia | 13 | Saprotroph | \|Undefined Saprotroph\| | Yeast | NULL | Probable | As nectar-tap_saprotroph (Põlme et al. 2020); Decay sugar-rich substrates (Põlme et al. 2020) | Põlme S, et al. 2020. Fungal Diversity 105:1-16 ((https://doi.org/10.1007/s13225-020-00466-2)) |
| Otu46177 | 1 | k__Fungi(100);p__Ascomycota(100);c__Saccharomycetes(100);o__Saccharomycetales(100);f__Metschnikowiaceae(100);g__Metschnikowia(100);s__Metschnikowia_pulcherrima(100); | Metschnikowia | 13 | Saprotroph | \|Undefined Saprotroph\| | Yeast | NULL | Probable | As nectar-tap_saprotroph (Põlme et al. 2020); Decay sugar-rich substrates (Põlme et al. 2020) | Põlme S, et al. 2020. Fungal Diversity 105:1-16 ((https://doi.org/10.1007/s13225-020-00466-2)) |
| Otu46178 | 1 | k__Fungi(100);p__Ascomycota(100);c__Saccharomycetes(100);o__Saccharomycetales(100);f__Metschnikowiaceae(100);g__Metschnikowia(100);s__Metschnikowia_pulcherrima(100); | Metschnikowia | 13 | Saprotroph | \|Undefined Saprotroph\| | Yeast | NULL | Probable | As nectar-tap_saprotroph (Põlme et al. 2020); Decay sugar-rich substrates (Põlme et al. 2020) | Põlme S, et al. 2020. Fungal Diversity 105:1-16 ((https://doi.org/10.1007/s13225-020-00466-2)) |
| Otu46179 | 1 | k__Fungi(100);p__Ascomycota(100);c__Saccharomycetes(100);o__Saccharomycetales(100);f__Metschnikowiaceae(100);g__Metschnikowia(100);g__Metschnikowia_unclassified(100); | Metschnikowia | 13 | Saprotroph | \|Undefined Saprotroph\| | Yeast | NULL | Probable | As nectar-tap_saprotroph (Põlme et al. 2020); Decay sugar-rich substrates (Põlme et al. 2020) | Põlme S, et al. 2020. Fungal Diversity 105:1-16 ((https://doi.org/10.1007/s13225-020-00466-2)) |
| Otu46180 | 1 | k__Fungi(100);p__Ascomycota(100);c__Saccharomycetes(100);o__Saccharomycetales(100);f__Metschnikowiaceae(100);g__Metschnikowia(100);s__Metschnikowia_pulcherrima(100); | Metschnikowia | 13 | Saprotroph | \|Undefined Saprotroph\| | Yeast | NULL | Probable | As nectar-tap_saprotroph (Põlme et al. 2020); Decay sugar-rich substrates (Põlme et al. 2020) | Põlme S, et al. 2020. Fungal Diversity 105:1-16 ((https://doi.org/10.1007/s13225-020-00466-2)) |
| Otu46181 | 1 | k__Fungi(100);p__Ascomycota(100);c__Saccharomycetes(100);o__Saccharomycetales(100);f__Metschnikowiaceae(100);g__Metschnikowia(100);s__Metschnikowia_pulcherrima(100); | Metschnikowia | 13 | Saprotroph | \|Undefined Saprotroph\| | Yeast | NULL | Probable | As nectar-tap_saprotroph (Põlme et al. 2020); Decay sugar-rich substrates (Põlme et al. 2020) | Põlme S, et al. 2020. Fungal Diversity 105:1-16 ((https://doi.org/10.1007/s13225-020-00466-2)) |
| Otu46182 | 1 | k__Fungi(100);p__Ascomycota(100);c__Saccharomycetes(100);o__Saccharomycetales(100);f__Metschnikowiaceae(100);g__Metschnikowia(100);s__Metschnikowia_pulcherrima(100); | Metschnikowia | 13 | Saprotroph | \|Undefined Saprotroph\| | Yeast | NULL | Probable | As nectar-tap_saprotroph (Põlme et al. 2020); Decay sugar-rich substrates (Põlme et al. 2020) | Põlme S, et al. 2020. Fungal Diversity 105:1-16 ((https://doi.org/10.1007/s13225-020-00466-2)) |
| Otu46183 | 1 | k__Fungi(100);p__Ascomycota(100);c__Saccharomycetes(100);o__Saccharomycetales(100);f__Metschnikowiaceae(100);g__Metschnikowia(100);s__Metschnikowia_pulcherrima(100); | Metschnikowia | 13 | Saprotroph | \|Undefined Saprotroph\| | Yeast | NULL | Probable | As nectar-tap_saprotroph (Põlme et al. 2020); Decay sugar-rich substrates (Põlme et al. 2020) | Põlme S, et al. 2020. Fungal Diversity 105:1-16 ((https://doi.org/10.1007/s13225-020-00466-2)) |
| Otu46184 | 1 | k__Fungi(100);p__Ascomycota(100);c__Saccharomycetes(100);o__Saccharomycetales(100);f__Metschnikowiaceae(100);g__Metschnikowia(100);s__Metschnikowia_pulcherrima(100); | Metschnikowia | 13 | Saprotroph | \|Undefined Saprotroph\| | Yeast | NULL | Probable | As nectar-tap_saprotroph (Põlme et al. 2020); Decay sugar-rich substrates (Põlme et al. 2020) | Põlme S, et al. 2020. Fungal Diversity 105:1-16 ((https://doi.org/10.1007/s13225-020-00466-2)) |
| Otu46185 | 1 | k__Fungi(100);p__Ascomycota(100);c__Saccharomycetes(100);o__Saccharomycetales(100);f__Metschnikowiaceae(100);g__Metschnikowia(100);s__Metschnikowia_pulcherrima(100); | Metschnikowia | 13 | Saprotroph | \|Undefined Saprotroph\| | Yeast | NULL | Probable | As nectar-tap_saprotroph (Põlme et al. 2020); Decay sugar-rich substrates (Põlme et al. 2020) | Põlme S, et al. 2020. Fungal Diversity 105:1-16 ((https://doi.org/10.1007/s13225-020-00466-2)) |
| Otu46186 | 1 | k__Fungi(100);p__Ascomycota(100);c__Saccharomycetes(100);o__Saccharomycetales(100);f__Metschnikowiaceae(100);g__Metschnikowia(100);s__Metschnikowia_pulcherrima(100); | Metschnikowia | 13 | Saprotroph | \|Undefined Saprotroph\| | Yeast | NULL | Probable | As nectar-tap_saprotroph (Põlme et al. 2020); Decay sugar-rich substrates (Põlme et al. 2020) | Põlme S, et al. 2020. Fungal Diversity 105:1-16 ((https://doi.org/10.1007/s13225-020-00466-2)) |
| Otu46187 | 1 | k__Fungi(100);p__Ascomycota(100);c__Saccharomycetes(100);o__Saccharomycetales(100);f__Metschnikowiaceae(100);g__Metschnikowia(100);g__Metschnikowia_unclassified(100); | Metschnikowia | 13 | Saprotroph | \|Undefined Saprotroph\| | Yeast | NULL | Probable | As nectar-tap_saprotroph (Põlme et al. 2020); Decay sugar-rich substrates (Põlme et al. 2020) | Põlme S, et al. 2020. Fungal Diversity 105:1-16 ((https://doi.org/10.1007/s13225-020-00466-2)) |
| Otu46188 | 1 | k__Fungi(100);p__Ascomycota(100);c__Saccharomycetes(100);o__Saccharomycetales(100);f__Metschnikowiaceae(100);g__Metschnikowia(100);g__Metschnikowia_unclassified(100); | Metschnikowia | 13 | Saprotroph | \|Undefined Saprotroph\| | Yeast | NULL | Probable | As nectar-tap_saprotroph (Põlme et al. 2020); Decay sugar-rich substrates (Põlme et al. 2020) | Põlme S, et al. 2020. Fungal Diversity 105:1-16 ((https://doi.org/10.1007/s13225-020-00466-2)) |
| Otu46189 | 1 | k__Fungi(100);p__Ascomycota(100);c__Saccharomycetes(100);o__Saccharomycetales(100);f__Metschnikowiaceae(100);g__Metschnikowia(100);s__Metschnikowia_pulcherrima(100); | Metschnikowia | 13 | Saprotroph | \|Undefined Saprotroph\| | Yeast | NULL | Probable | As nectar-tap_saprotroph (Põlme et al. 2020); Decay sugar-rich substrates (Põlme et al. 2020) | Põlme S, et al. 2020. Fungal Diversity 105:1-16 ((https://doi.org/10.1007/s13225-020-00466-2)) |
| Otu46190 | 1 | k__Fungi(100);p__Ascomycota(100);c__Saccharomycetes(100);o__Saccharomycetales(100);f__Metschnikowiaceae(100);g__Metschnikowia(100);s__Metschnikowia_pulcherrima(100); | Metschnikowia | 13 | Saprotroph | \|Undefined Saprotroph\| | Yeast | NULL | Probable | As nectar-tap_saprotroph (Põlme et al. 2020); Decay sugar-rich substrates (Põlme et al. 2020) | Põlme S, et al. 2020. Fungal Diversity 105:1-16 ((https://doi.org/10.1007/s13225-020-00466-2)) |
| Otu46191 | 1 | k__Fungi(100);p__Ascomycota(100);c__Saccharomycetes(100);o__Saccharomycetales(100);f__Metschnikowiaceae(100);g__Metschnikowia(100);s__Metschnikowia_pulcherrima(100); | Metschnikowia | 13 | Saprotroph | \|Undefined Saprotroph\| | Yeast | NULL | Probable | As nectar-tap_saprotroph (Põlme et al. 2020); Decay sugar-rich substrates (Põlme et al. 2020) | Põlme S, et al. 2020. Fungal Diversity 105:1-16 ((https://doi.org/10.1007/s13225-020-00466-2)) |
| Otu46192 | 1 | k__Fungi(100);p__Ascomycota(100);c__Saccharomycetes(100);o__Saccharomycetales(100);f__Metschnikowiaceae(100);g__Metschnikowia(100);s__Metschnikowia_pulcherrima(100); | Metschnikowia | 13 | Saprotroph | \|Undefined Saprotroph\| | Yeast | NULL | Probable | As nectar-tap_saprotroph (Põlme et al. 2020); Decay sugar-rich substrates (Põlme et al. 2020) | Põlme S, et al. 2020. Fungal Diversity 105:1-16 ((https://doi.org/10.1007/s13225-020-00466-2)) |
| Otu46193 | 1 | k__Fungi(100);p__Ascomycota(100);c__Saccharomycetes(100);o__Saccharomycetales(100);f__Metschnikowiaceae(100);g__Metschnikowia(100);s__Metschnikowia_pulcherrima(100); | Metschnikowia | 13 | Saprotroph | \|Undefined Saprotroph\| | Yeast | NULL | Probable | As nectar-tap_saprotroph (Põlme et al. 2020); Decay sugar-rich substrates (Põlme et al. 2020) | Põlme S, et al. 2020. Fungal Diversity 105:1-16 ((https://doi.org/10.1007/s13225-020-00466-2)) |
| Otu46194 | 1 | k__Fungi(100);p__Ascomycota(100);c__Saccharomycetes(100);o__Saccharomycetales(100);f__Metschnikowiaceae(100);g__Metschnikowia(100);s__Metschnikowia_pulcherrima(100); | Metschnikowia | 13 | Saprotroph | \|Undefined Saprotroph\| | Yeast | NULL | Probable | As nectar-tap_saprotroph (Põlme et al. 2020); Decay sugar-rich substrates (Põlme et al. 2020) | Põlme S, et al. 2020. Fungal Diversity 105:1-16 ((https://doi.org/10.1007/s13225-020-00466-2)) |
| Otu46195 | 1 | k__Fungi(100);p__Ascomycota(100);c__Saccharomycetes(100);o__Saccharomycetales(100);f__Metschnikowiaceae(100);g__Metschnikowia(100);s__Metschnikowia_pulcherrima(100); | Metschnikowia | 13 | Saprotroph | \|Undefined Saprotroph\| | Yeast | NULL | Probable | As nectar-tap_saprotroph (Põlme et al. 2020); Decay sugar-rich substrates (Põlme et al. 2020) | Põlme S, et al. 2020. Fungal Diversity 105:1-16 ((https://doi.org/10.1007/s13225-020-00466-2)) |
| Otu46196 | 1 | k__Fungi(100);p__Ascomycota(100);c__Saccharomycetes(100);o__Saccharomycetales(100);f__Metschnikowiaceae(100);g__Metschnikowia(100);s__Metschnikowia_pulcherrima(100); | Metschnikowia | 13 | Saprotroph | \|Undefined Saprotroph\| | Yeast | NULL | Probable | As nectar-tap_saprotroph (Põlme et al. 2020); Decay sugar-rich substrates (Põlme et al. 2020) | Põlme S, et al. 2020. Fungal Diversity 105:1-16 ((https://doi.org/10.1007/s13225-020-00466-2)) |
| Otu46197 | 1 | k__Fungi(100);p__Ascomycota(100);c__Saccharomycetes(100);o__Saccharomycetales(100);f__Metschnikowiaceae(100);g__Metschnikowia(100);s__Metschnikowia_pulcherrima(100); | Metschnikowia | 13 | Saprotroph | \|Undefined Saprotroph\| | Yeast | NULL | Probable | As nectar-tap_saprotroph (Põlme et al. 2020); Decay sugar-rich substrates (Põlme et al. 2020) | Põlme S, et al. 2020. Fungal Diversity 105:1-16 ((https://doi.org/10.1007/s13225-020-00466-2)) |
| Otu46198 | 1 | k__Fungi(100);p__Ascomycota(100);c__Saccharomycetes(100);o__Saccharomycetales(100);f__Metschnikowiaceae(100);g__Metschnikowia(100);s__Metschnikowia_pulcherrima(100); | Metschnikowia | 13 | Saprotroph | \|Undefined Saprotroph\| | Yeast | NULL | Probable | As nectar-tap_saprotroph (Põlme et al. 2020); Decay sugar-rich substrates (Põlme et al. 2020) | Põlme S, et al. 2020. Fungal Diversity 105:1-16 ((https://doi.org/10.1007/s13225-020-00466-2)) |
| Otu46199 | 1 | k__Fungi(100);p__Ascomycota(100);c__Saccharomycetes(100);o__Saccharomycetales(100);f__Metschnikowiaceae(100);g__Metschnikowia(100);s__Metschnikowia_pulcherrima(100); | Metschnikowia | 13 | Saprotroph | \|Undefined Saprotroph\| | Yeast | NULL | Probable | As nectar-tap_saprotroph (Põlme et al. 2020); Decay sugar-rich substrates (Põlme et al. 2020) | Põlme S, et al. 2020. Fungal Diversity 105:1-16 ((https://doi.org/10.1007/s13225-020-00466-2)) |
| Otu46200 | 1 | k__Fungi(100);p__Ascomycota(100);c__Saccharomycetes(100);o__Saccharomycetales(100);f__Metschnikowiaceae(100);g__Metschnikowia(100);s__Metschnikowia_pulcherrima(100); | Metschnikowia | 13 | Saprotroph | \|Undefined Saprotroph\| | Yeast | NULL | Probable | As nectar-tap_saprotroph (Põlme et al. 2020); Decay sugar-rich substrates (Põlme et al. 2020) | Põlme S, et al. 2020. Fungal Diversity 105:1-16 ((https://doi.org/10.1007/s13225-020-00466-2)) |
| Otu46201 | 1 | k__Fungi(100);p__Ascomycota(100);c__Saccharomycetes(100);o__Saccharomycetales(100);f__Metschnikowiaceae(100);g__Metschnikowia(100);g__Metschnikowia_unclassified(100); | Metschnikowia | 13 | Saprotroph | \|Undefined Saprotroph\| | Yeast | NULL | Probable | As nectar-tap_saprotroph (Põlme et al. 2020); Decay sugar-rich substrates (Põlme et al. 2020) | Põlme S, et al. 2020. Fungal Diversity 105:1-16 ((https://doi.org/10.1007/s13225-020-00466-2)) |
| Otu46202 | 1 | k__Fungi(100);p__Ascomycota(100);c__Saccharomycetes(100);o__Saccharomycetales(100);f__Metschnikowiaceae(100);g__Metschnikowia(100);s__Metschnikowia_pulcherrima(100); | Metschnikowia | 13 | Saprotroph | \|Undefined Saprotroph\| | Yeast | NULL | Probable | As nectar-tap_saprotroph (Põlme et al. 2020); Decay sugar-rich substrates (Põlme et al. 2020) | Põlme S, et al. 2020. Fungal Diversity 105:1-16 ((https://doi.org/10.1007/s13225-020-00466-2)) |
| Otu46203 | 1 | k__Fungi(100);p__Ascomycota(100);c__Saccharomycetes(100);o__Saccharomycetales(100);f__Metschnikowiaceae(100);g__Metschnikowia(100);s__Metschnikowia_pulcherrima(100); | Metschnikowia | 13 | Saprotroph | \|Undefined Saprotroph\| | Yeast | NULL | Probable | As nectar-tap_saprotroph (Põlme et al. 2020); Decay sugar-rich substrates (Põlme et al. 2020) | Põlme S, et al. 2020. Fungal Diversity 105:1-16 ((https://doi.org/10.1007/s13225-020-00466-2)) |
| Otu46204 | 1 | k__Fungi(100);p__Ascomycota(100);c__Saccharomycetes(100);o__Saccharomycetales(100);f__Metschnikowiaceae(100);g__Metschnikowia(100);s__Metschnikowia_pulcherrima(100); | Metschnikowia | 13 | Saprotroph | \|Undefined Saprotroph\| | Yeast | NULL | Probable | As nectar-tap_saprotroph (Põlme et al. 2020); Decay sugar-rich substrates (Põlme et al. 2020) | Põlme S, et al. 2020. Fungal Diversity 105:1-16 ((https://doi.org/10.1007/s13225-020-00466-2)) |
| Otu46205 | 1 | k__Fungi(100);p__Ascomycota(100);c__Saccharomycetes(100);o__Saccharomycetales(100);f__Metschnikowiaceae(100);g__Metschnikowia(100);s__Metschnikowia_pulcherrima(100); | Metschnikowia | 13 | Saprotroph | \|Undefined Saprotroph\| | Yeast | NULL | Probable | As nectar-tap_saprotroph (Põlme et al. 2020); Decay sugar-rich substrates (Põlme et al. 2020) | Põlme S, et al. 2020. Fungal Diversity 105:1-16 ((https://doi.org/10.1007/s13225-020-00466-2)) |
| Otu46206 | 1 | k__Fungi(100);p__Ascomycota(100);c__Saccharomycetes(100);o__Saccharomycetales(100);f__Metschnikowiaceae(100);g__Metschnikowia(100);g__Metschnikowia_unclassified(100); | Metschnikowia | 13 | Saprotroph | \|Undefined Saprotroph\| | Yeast | NULL | Probable | As nectar-tap_saprotroph (Põlme et al. 2020); Decay sugar-rich substrates (Põlme et al. 2020) | Põlme S, et al. 2020. Fungal Diversity 105:1-16 ((https://doi.org/10.1007/s13225-020-00466-2)) |
| Otu46207 | 1 | k__Fungi(100);p__Ascomycota(100);c__Saccharomycetes(100);o__Saccharomycetales(100);f__Metschnikowiaceae(100);g__Metschnikowia(100);g__Metschnikowia_unclassified(100); | Metschnikowia | 13 | Saprotroph | \|Undefined Saprotroph\| | Yeast | NULL | Probable | As nectar-tap_saprotroph (Põlme et al. 2020); Decay sugar-rich substrates (Põlme et al. 2020) | Põlme S, et al. 2020. Fungal Diversity 105:1-16 ((https://doi.org/10.1007/s13225-020-00466-2)) |
| Otu46208 | 1 | k__Fungi(100);p__Ascomycota(100);c__Saccharomycetes(100);o__Saccharomycetales(100);f__Metschnikowiaceae(100);g__Metschnikowia(100);s__Metschnikowia_pulcherrima(100); | Metschnikowia | 13 | Saprotroph | \|Undefined Saprotroph\| | Yeast | NULL | Probable | As nectar-tap_saprotroph (Põlme et al. 2020); Decay sugar-rich substrates (Põlme et al. 2020) | Põlme S, et al. 2020. Fungal Diversity 105:1-16 ((https://doi.org/10.1007/s13225-020-00466-2)) |
| Otu46209 | 1 | k__Fungi(100);p__Ascomycota(100);c__Saccharomycetes(100);o__Saccharomycetales(100);f__Metschnikowiaceae(100);g__Metschnikowia(100);s__Metschnikowia_pulcherrima(100); | Metschnikowia | 13 | Saprotroph | \|Undefined Saprotroph\| | Yeast | NULL | Probable | As nectar-tap_saprotroph (Põlme et al. 2020); Decay sugar-rich substrates (Põlme et al. 2020) | Põlme S, et al. 2020. Fungal Diversity 105:1-16 ((https://doi.org/10.1007/s13225-020-00466-2)) |
| Otu46210 | 1 | k__Fungi(100);p__Ascomycota(100);c__Saccharomycetes(100);o__Saccharomycetales(100);f__Metschnikowiaceae(100);g__Metschnikowia(100);g__Metschnikowia_unclassified(100); | Metschnikowia | 13 | Saprotroph | \|Undefined Saprotroph\| | Yeast | NULL | Probable | As nectar-tap_saprotroph (Põlme et al. 2020); Decay sugar-rich substrates (Põlme et al. 2020) | Põlme S, et al. 2020. Fungal Diversity 105:1-16 ((https://doi.org/10.1007/s13225-020-00466-2)) |
| Otu46211 | 1 | k__Fungi(100);p__Ascomycota(100);c__Saccharomycetes(100);o__Saccharomycetales(100);f__Metschnikowiaceae(100);g__Metschnikowia(100);g__Metschnikowia_unclassified(100); | Metschnikowia | 13 | Saprotroph | \|Undefined Saprotroph\| | Yeast | NULL | Probable | As nectar-tap_saprotroph (Põlme et al. 2020); Decay sugar-rich substrates (Põlme et al. 2020) | Põlme S, et al. 2020. Fungal Diversity 105:1-16 ((https://doi.org/10.1007/s13225-020-00466-2)) |
| Otu46212 | 1 | k__Fungi(100);p__Ascomycota(100);c__Saccharomycetes(100);o__Saccharomycetales(100);f__Metschnikowiaceae(100);g__Metschnikowia(100);s__Metschnikowia_pulcherrima(100); | Metschnikowia | 13 | Saprotroph | \|Undefined Saprotroph\| | Yeast | NULL | Probable | As nectar-tap_saprotroph (Põlme et al. 2020); Decay sugar-rich substrates (Põlme et al. 2020) | Põlme S, et al. 2020. Fungal Diversity 105:1-16 ((https://doi.org/10.1007/s13225-020-00466-2)) |
| Otu46213 | 1 | k__Fungi(100);p__Ascomycota(100);c__Saccharomycetes(100);o__Saccharomycetales(100);f__Metschnikowiaceae(100);g__Metschnikowia(100);g__Metschnikowia_unclassified(100); | Metschnikowia | 13 | Saprotroph | \|Undefined Saprotroph\| | Yeast | NULL | Probable | As nectar-tap_saprotroph (Põlme et al. 2020); Decay sugar-rich substrates (Põlme et al. 2020) | Põlme S, et al. 2020. Fungal Diversity 105:1-16 ((https://doi.org/10.1007/s13225-020-00466-2)) |
| Otu46214 | 1 | k__Fungi(100);p__Ascomycota(100);c__Saccharomycetes(100);o__Saccharomycetales(100);f__Metschnikowiaceae(100);g__Metschnikowia(100);s__Metschnikowia_pulcherrima(100); | Metschnikowia | 13 | Saprotroph | \|Undefined Saprotroph\| | Yeast | NULL | Probable | As nectar-tap_saprotroph (Põlme et al. 2020); Decay sugar-rich substrates (Põlme et al. 2020) | Põlme S, et al. 2020. Fungal Diversity 105:1-16 ((https://doi.org/10.1007/s13225-020-00466-2)) |
| Otu46215 | 1 | k__Fungi(100);p__Ascomycota(100);c__Saccharomycetes(100);o__Saccharomycetales(100);f__Metschnikowiaceae(100);g__Metschnikowia(100);s__Metschnikowia_pulcherrima(100); | Metschnikowia | 13 | Saprotroph | \|Undefined Saprotroph\| | Yeast | NULL | Probable | As nectar-tap_saprotroph (Põlme et al. 2020); Decay sugar-rich substrates (Põlme et al. 2020) | Põlme S, et al. 2020. Fungal Diversity 105:1-16 ((https://doi.org/10.1007/s13225-020-00466-2)) |
| Otu46216 | 1 | k__Fungi(100);p__Ascomycota(100);c__Saccharomycetes(100);o__Saccharomycetales(100);f__Metschnikowiaceae(100);g__Metschnikowia(100);s__Metschnikowia_pulcherrima(100); | Metschnikowia | 13 | Saprotroph | \|Undefined Saprotroph\| | Yeast | NULL | Probable | As nectar-tap_saprotroph (Põlme et al. 2020); Decay sugar-rich substrates (Põlme et al. 2020) | Põlme S, et al. 2020. Fungal Diversity 105:1-16 ((https://doi.org/10.1007/s13225-020-00466-2)) |
| Otu46217 | 1 | k__Fungi(100);p__Ascomycota(100);c__Saccharomycetes(100);o__Saccharomycetales(100);f__Metschnikowiaceae(100);g__Metschnikowia(100);s__Metschnikowia_pulcherrima(100); | Metschnikowia | 13 | Saprotroph | \|Undefined Saprotroph\| | Yeast | NULL | Probable | As nectar-tap_saprotroph (Põlme et al. 2020); Decay sugar-rich substrates (Põlme et al. 2020) | Põlme S, et al. 2020. Fungal Diversity 105:1-16 ((https://doi.org/10.1007/s13225-020-00466-2)) |
| Otu46218 | 1 | k__Fungi(100);p__Ascomycota(100);c__Saccharomycetes(100);o__Saccharomycetales(100);f__Metschnikowiaceae(100);g__Metschnikowia(100);g__Metschnikowia_unclassified(100); | Metschnikowia | 13 | Saprotroph | \|Undefined Saprotroph\| | Yeast | NULL | Probable | As nectar-tap_saprotroph (Põlme et al. 2020); Decay sugar-rich substrates (Põlme et al. 2020) | Põlme S, et al. 2020. Fungal Diversity 105:1-16 ((https://doi.org/10.1007/s13225-020-00466-2)) |
| Otu46219 | 1 | k__Fungi(100);p__Ascomycota(100);c__Saccharomycetes(100);o__Saccharomycetales(100);f__Metschnikowiaceae(100);g__Metschnikowia(100);s__Metschnikowia_pulcherrima(100); | Metschnikowia | 13 | Saprotroph | \|Undefined Saprotroph\| | Yeast | NULL | Probable | As nectar-tap_saprotroph (Põlme et al. 2020); Decay sugar-rich substrates (Põlme et al. 2020) | Põlme S, et al. 2020. Fungal Diversity 105:1-16 ((https://doi.org/10.1007/s13225-020-00466-2)) |
| Otu46220 | 1 | k__Fungi(100);p__Ascomycota(100);c__Saccharomycetes(100);o__Saccharomycetales(100);f__Metschnikowiaceae(100);g__Metschnikowia(100);g__Metschnikowia_unclassified(100); | Metschnikowia | 13 | Saprotroph | \|Undefined Saprotroph\| | Yeast | NULL | Probable | As nectar-tap_saprotroph (Põlme et al. 2020); Decay sugar-rich substrates (Põlme et al. 2020) | Põlme S, et al. 2020. Fungal Diversity 105:1-16 ((https://doi.org/10.1007/s13225-020-00466-2)) |
| Otu46221 | 1 | k__Fungi(100);p__Ascomycota(100);c__Saccharomycetes(100);o__Saccharomycetales(100);f__Metschnikowiaceae(100);g__Metschnikowia(100);s__Metschnikowia_pulcherrima(100); | Metschnikowia | 13 | Saprotroph | \|Undefined Saprotroph\| | Yeast | NULL | Probable | As nectar-tap_saprotroph (Põlme et al. 2020); Decay sugar-rich substrates (Põlme et al. 2020) | Põlme S, et al. 2020. Fungal Diversity 105:1-16 ((https://doi.org/10.1007/s13225-020-00466-2)) |
| Otu46222 | 1 | k__Fungi(100);p__Ascomycota(100);c__Saccharomycetes(100);o__Saccharomycetales(100);f__Metschnikowiaceae(100);g__Metschnikowia(100);s__Metschnikowia_pulcherrima(100); | Metschnikowia | 13 | Saprotroph | \|Undefined Saprotroph\| | Yeast | NULL | Probable | As nectar-tap_saprotroph (Põlme et al. 2020); Decay sugar-rich substrates (Põlme et al. 2020) | Põlme S, et al. 2020. Fungal Diversity 105:1-16 ((https://doi.org/10.1007/s13225-020-00466-2)) |
| Otu46223 | 1 | k__Fungi(100);p__Ascomycota(100);c__Saccharomycetes(100);o__Saccharomycetales(100);f__Metschnikowiaceae(100);g__Metschnikowia(100);g__Metschnikowia_unclassified(100); | Metschnikowia | 13 | Saprotroph | \|Undefined Saprotroph\| | Yeast | NULL | Probable | As nectar-tap_saprotroph (Põlme et al. 2020); Decay sugar-rich substrates (Põlme et al. 2020) | Põlme S, et al. 2020. Fungal Diversity 105:1-16 ((https://doi.org/10.1007/s13225-020-00466-2)) |
| Otu46224 | 1 | k__Fungi(100);p__Ascomycota(100);c__Saccharomycetes(100);o__Saccharomycetales(100);f__Metschnikowiaceae(100);g__Metschnikowia(100);s__Metschnikowia_pulcherrima(100); | Metschnikowia | 13 | Saprotroph | \|Undefined Saprotroph\| | Yeast | NULL | Probable | As nectar-tap_saprotroph (Põlme et al. 2020); Decay sugar-rich substrates (Põlme et al. 2020) | Põlme S, et al. 2020. Fungal Diversity 105:1-16 ((https://doi.org/10.1007/s13225-020-00466-2)) |
| Otu46225 | 1 | k__Fungi(100);p__Ascomycota(100);c__Saccharomycetes(100);o__Saccharomycetales(100);f__Metschnikowiaceae(100);g__Metschnikowia(100);g__Metschnikowia_unclassified(100); | Metschnikowia | 13 | Saprotroph | \|Undefined Saprotroph\| | Yeast | NULL | Probable | As nectar-tap_saprotroph (Põlme et al. 2020); Decay sugar-rich substrates (Põlme et al. 2020) | Põlme S, et al. 2020. Fungal Diversity 105:1-16 ((https://doi.org/10.1007/s13225-020-00466-2)) |
| Otu46226 | 1 | k__Fungi(100);p__Ascomycota(100);c__Saccharomycetes(100);o__Saccharomycetales(100);f__Metschnikowiaceae(100);g__Metschnikowia(100);s__Metschnikowia_pulcherrima(100); | Metschnikowia | 13 | Saprotroph | \|Undefined Saprotroph\| | Yeast | NULL | Probable | As nectar-tap_saprotroph (Põlme et al. 2020); Decay sugar-rich substrates (Põlme et al. 2020) | Põlme S, et al. 2020. Fungal Diversity 105:1-16 ((https://doi.org/10.1007/s13225-020-00466-2)) |
| Otu46227 | 1 | k__Fungi(100);p__Ascomycota(100);c__Saccharomycetes(100);o__Saccharomycetales(100);f__Metschnikowiaceae(100);g__Metschnikowia(100);s__Metschnikowia_pulcherrima(100); | Metschnikowia | 13 | Saprotroph | \|Undefined Saprotroph\| | Yeast | NULL | Probable | As nectar-tap_saprotroph (Põlme et al. 2020); Decay sugar-rich substrates (Põlme et al. 2020) | Põlme S, et al. 2020. Fungal Diversity 105:1-16 ((https://doi.org/10.1007/s13225-020-00466-2)) |
| Otu46228 | 1 | k__Fungi(100);p__Ascomycota(100);c__Saccharomycetes(100);o__Saccharomycetales(100);f__Metschnikowiaceae(100);g__Metschnikowia(100);s__Metschnikowia_pulcherrima(100); | Metschnikowia | 13 | Saprotroph | \|Undefined Saprotroph\| | Yeast | NULL | Probable | As nectar-tap_saprotroph (Põlme et al. 2020); Decay sugar-rich substrates (Põlme et al. 2020) | Põlme S, et al. 2020. Fungal Diversity 105:1-16 ((https://doi.org/10.1007/s13225-020-00466-2)) |
| Otu46229 | 1 | k__Fungi(100);p__Ascomycota(100);c__Saccharomycetes(100);o__Saccharomycetales(100);f__Metschnikowiaceae(100);g__Metschnikowia(100);s__Metschnikowia_pulcherrima(100); | Metschnikowia | 13 | Saprotroph | \|Undefined Saprotroph\| | Yeast | NULL | Probable | As nectar-tap_saprotroph (Põlme et al. 2020); Decay sugar-rich substrates (Põlme et al. 2020) | Põlme S, et al. 2020. Fungal Diversity 105:1-16 ((https://doi.org/10.1007/s13225-020-00466-2)) |
| Otu46230 | 1 | k__Fungi(100);p__Ascomycota(100);c__Saccharomycetes(100);o__Saccharomycetales(100);f__Metschnikowiaceae(100);g__Metschnikowia(100);s__Metschnikowia_pulcherrima(100); | Metschnikowia | 13 | Saprotroph | \|Undefined Saprotroph\| | Yeast | NULL | Probable | As nectar-tap_saprotroph (Põlme et al. 2020); Decay sugar-rich substrates (Põlme et al. 2020) | Põlme S, et al. 2020. Fungal Diversity 105:1-16 ((https://doi.org/10.1007/s13225-020-00466-2)) |
| Otu46231 | 1 | k__Fungi(100);p__Ascomycota(100);c__Saccharomycetes(100);o__Saccharomycetales(100);f__Metschnikowiaceae(100);g__Metschnikowia(100);s__Metschnikowia_pulcherrima(100); | Metschnikowia | 13 | Saprotroph | \|Undefined Saprotroph\| | Yeast | NULL | Probable | As nectar-tap_saprotroph (Põlme et al. 2020); Decay sugar-rich substrates (Põlme et al. 2020) | Põlme S, et al. 2020. Fungal Diversity 105:1-16 ((https://doi.org/10.1007/s13225-020-00466-2)) |
| Otu46232 | 1 | k__Fungi(100);p__Ascomycota(100);c__Saccharomycetes(100);o__Saccharomycetales(100);f__Metschnikowiaceae(100);g__Metschnikowia(100);s__Metschnikowia_pulcherrima(100); | Metschnikowia | 13 | Saprotroph | \|Undefined Saprotroph\| | Yeast | NULL | Probable | As nectar-tap_saprotroph (Põlme et al. 2020); Decay sugar-rich substrates (Põlme et al. 2020) | Põlme S, et al. 2020. Fungal Diversity 105:1-16 ((https://doi.org/10.1007/s13225-020-00466-2)) |
| Otu46233 | 1 | k__Fungi(100);p__Ascomycota(100);c__Saccharomycetes(100);o__Saccharomycetales(100);f__Metschnikowiaceae(100);g__Metschnikowia(100);s__Metschnikowia_pulcherrima(100); | Metschnikowia | 13 | Saprotroph | \|Undefined Saprotroph\| | Yeast | NULL | Probable | As nectar-tap_saprotroph (Põlme et al. 2020); Decay sugar-rich substrates (Põlme et al. 2020) | Põlme S, et al. 2020. Fungal Diversity 105:1-16 ((https://doi.org/10.1007/s13225-020-00466-2)) |
| Otu46234 | 1 | k__Fungi(100);p__Ascomycota(100);c__Saccharomycetes(100);o__Saccharomycetales(100);f__Metschnikowiaceae(100);g__Metschnikowia(100);g__Metschnikowia_unclassified(100); | Metschnikowia | 13 | Saprotroph | \|Undefined Saprotroph\| | Yeast | NULL | Probable | As nectar-tap_saprotroph (Põlme et al. 2020); Decay sugar-rich substrates (Põlme et al. 2020) | Põlme S, et al. 2020. Fungal Diversity 105:1-16 ((https://doi.org/10.1007/s13225-020-00466-2)) |
| Otu46235 | 1 | k__Fungi(100);p__Ascomycota(100);c__Saccharomycetes(100);o__Saccharomycetales(100);f__Metschnikowiaceae(100);g__Metschnikowia(100);g__Metschnikowia_unclassified(100); | Metschnikowia | 13 | Saprotroph | \|Undefined Saprotroph\| | Yeast | NULL | Probable | As nectar-tap_saprotroph (Põlme et al. 2020); Decay sugar-rich substrates (Põlme et al. 2020) | Põlme S, et al. 2020. Fungal Diversity 105:1-16 ((https://doi.org/10.1007/s13225-020-00466-2)) |
| Otu46236 | 1 | k__Fungi(100);p__Ascomycota(100);c__Saccharomycetes(100);o__Saccharomycetales(100);f__Metschnikowiaceae(100);g__Metschnikowia(100);s__Metschnikowia_pulcherrima(100); | Metschnikowia | 13 | Saprotroph | \|Undefined Saprotroph\| | Yeast | NULL | Probable | As nectar-tap_saprotroph (Põlme et al. 2020); Decay sugar-rich substrates (Põlme et al. 2020) | Põlme S, et al. 2020. Fungal Diversity 105:1-16 ((https://doi.org/10.1007/s13225-020-00466-2)) |
| Otu46237 | 1 | k__Fungi(100);p__Ascomycota(100);c__Saccharomycetes(100);o__Saccharomycetales(100);f__Metschnikowiaceae(100);g__Metschnikowia(100);g__Metschnikowia_unclassified(100); | Metschnikowia | 13 | Saprotroph | \|Undefined Saprotroph\| | Yeast | NULL | Probable | As nectar-tap_saprotroph (Põlme et al. 2020); Decay sugar-rich substrates (Põlme et al. 2020) | Põlme S, et al. 2020. Fungal Diversity 105:1-16 ((https://doi.org/10.1007/s13225-020-00466-2)) |
| Otu46238 | 1 | k__Fungi(100);p__Ascomycota(100);c__Saccharomycetes(100);o__Saccharomycetales(100);f__Metschnikowiaceae(100);g__Metschnikowia(100);s__Metschnikowia_pulcherrima(100); | Metschnikowia | 13 | Saprotroph | \|Undefined Saprotroph\| | Yeast | NULL | Probable | As nectar-tap_saprotroph (Põlme et al. 2020); Decay sugar-rich substrates (Põlme et al. 2020) | Põlme S, et al. 2020. Fungal Diversity 105:1-16 ((https://doi.org/10.1007/s13225-020-00466-2)) |
| Otu46239 | 1 | k__Fungi(100);p__Ascomycota(100);c__Saccharomycetes(100);o__Saccharomycetales(100);f__Metschnikowiaceae(100);g__Metschnikowia(100);s__Metschnikowia_pulcherrima(100); | Metschnikowia | 13 | Saprotroph | \|Undefined Saprotroph\| | Yeast | NULL | Probable | As nectar-tap_saprotroph (Põlme et al. 2020); Decay sugar-rich substrates (Põlme et al. 2020) | Põlme S, et al. 2020. Fungal Diversity 105:1-16 ((https://doi.org/10.1007/s13225-020-00466-2)) |
| Otu46241 | 1 | k__Fungi(100);p__Ascomycota(100);c__Saccharomycetes(100);o__Saccharomycetales(100);f__Metschnikowiaceae(100);g__Metschnikowia(100);s__Metschnikowia_pulcherrima(100); | Metschnikowia | 13 | Saprotroph | \|Undefined Saprotroph\| | Yeast | NULL | Probable | As nectar-tap_saprotroph (Põlme et al. 2020); Decay sugar-rich substrates (Põlme et al. 2020) | Põlme S, et al. 2020. Fungal Diversity 105:1-16 ((https://doi.org/10.1007/s13225-020-00466-2)) |
| Otu46242 | 1 | k__Fungi(100);p__Ascomycota(100);c__Saccharomycetes(100);o__Saccharomycetales(100);f__Metschnikowiaceae(100);g__Metschnikowia(100);s__Metschnikowia_pulcherrima(100); | Metschnikowia | 13 | Saprotroph | \|Undefined Saprotroph\| | Yeast | NULL | Probable | As nectar-tap_saprotroph (Põlme et al. 2020); Decay sugar-rich substrates (Põlme et al. 2020) | Põlme S, et al. 2020. Fungal Diversity 105:1-16 ((https://doi.org/10.1007/s13225-020-00466-2)) |
| Otu46243 | 1 | k__Fungi(100);p__Ascomycota(100);c__Saccharomycetes(100);o__Saccharomycetales(100);f__Metschnikowiaceae(100);g__Metschnikowia(100);s__Metschnikowia_pulcherrima(100); | Metschnikowia | 13 | Saprotroph | \|Undefined Saprotroph\| | Yeast | NULL | Probable | As nectar-tap_saprotroph (Põlme et al. 2020); Decay sugar-rich substrates (Põlme et al. 2020) | Põlme S, et al. 2020. Fungal Diversity 105:1-16 ((https://doi.org/10.1007/s13225-020-00466-2)) |
| Otu46244 | 1 | k__Fungi(100);p__Ascomycota(100);c__Saccharomycetes(100);o__Saccharomycetales(100);f__Metschnikowiaceae(100);g__Metschnikowia(100);g__Metschnikowia_unclassified(100); | Metschnikowia | 13 | Saprotroph | \|Undefined Saprotroph\| | Yeast | NULL | Probable | As nectar-tap_saprotroph (Põlme et al. 2020); Decay sugar-rich substrates (Põlme et al. 2020) | Põlme S, et al. 2020. Fungal Diversity 105:1-16 ((https://doi.org/10.1007/s13225-020-00466-2)) |
| Otu46245 | 1 | k__Fungi(100);p__Ascomycota(100);c__Saccharomycetes(100);o__Saccharomycetales(100);f__Metschnikowiaceae(100);g__Metschnikowia(100);s__Metschnikowia_pulcherrima(100); | Metschnikowia | 13 | Saprotroph | \|Undefined Saprotroph\| | Yeast | NULL | Probable | As nectar-tap_saprotroph (Põlme et al. 2020); Decay sugar-rich substrates (Põlme et al. 2020) | Põlme S, et al. 2020. Fungal Diversity 105:1-16 ((https://doi.org/10.1007/s13225-020-00466-2)) |
| Otu46246 | 1 | k__Fungi(100);p__Ascomycota(100);c__Saccharomycetes(100);o__Saccharomycetales(100);f__Metschnikowiaceae(100);g__Metschnikowia(100);g__Metschnikowia_unclassified(100); | Metschnikowia | 13 | Saprotroph | \|Undefined Saprotroph\| | Yeast | NULL | Probable | As nectar-tap_saprotroph (Põlme et al. 2020); Decay sugar-rich substrates (Põlme et al. 2020) | Põlme S, et al. 2020. Fungal Diversity 105:1-16 ((https://doi.org/10.1007/s13225-020-00466-2)) |
| Otu46247 | 1 | k__Fungi(100);p__Ascomycota(100);c__Saccharomycetes(100);o__Saccharomycetales(100);f__Metschnikowiaceae(100);g__Metschnikowia(100);s__Metschnikowia_pulcherrima(100); | Metschnikowia | 13 | Saprotroph | \|Undefined Saprotroph\| | Yeast | NULL | Probable | As nectar-tap_saprotroph (Põlme et al. 2020); Decay sugar-rich substrates (Põlme et al. 2020) | Põlme S, et al. 2020. Fungal Diversity 105:1-16 ((https://doi.org/10.1007/s13225-020-00466-2)) |
| Otu46248 | 1 | k__Fungi(100);p__Ascomycota(100);c__Saccharomycetes(100);o__Saccharomycetales(100);f__Metschnikowiaceae(100);g__Metschnikowia(100);s__Metschnikowia_pulcherrima(100); | Metschnikowia | 13 | Saprotroph | \|Undefined Saprotroph\| | Yeast | NULL | Probable | As nectar-tap_saprotroph (Põlme et al. 2020); Decay sugar-rich substrates (Põlme et al. 2020) | Põlme S, et al. 2020. Fungal Diversity 105:1-16 ((https://doi.org/10.1007/s13225-020-00466-2)) |
| Otu46249 | 1 | k__Fungi(100);p__Ascomycota(100);c__Saccharomycetes(100);o__Saccharomycetales(100);f__Metschnikowiaceae(100);g__Metschnikowia(100);g__Metschnikowia_unclassified(100); | Metschnikowia | 13 | Saprotroph | \|Undefined Saprotroph\| | Yeast | NULL | Probable | As nectar-tap_saprotroph (Põlme et al. 2020); Decay sugar-rich substrates (Põlme et al. 2020) | Põlme S, et al. 2020. Fungal Diversity 105:1-16 ((https://doi.org/10.1007/s13225-020-00466-2)) |
| Otu46250 | 1 | k__Fungi(100);p__Ascomycota(100);c__Saccharomycetes(100);o__Saccharomycetales(100);f__Metschnikowiaceae(100);g__Metschnikowia(100);g__Metschnikowia_unclassified(100); | Metschnikowia | 13 | Saprotroph | \|Undefined Saprotroph\| | Yeast | NULL | Probable | As nectar-tap_saprotroph (Põlme et al. 2020); Decay sugar-rich substrates (Põlme et al. 2020) | Põlme S, et al. 2020. Fungal Diversity 105:1-16 ((https://doi.org/10.1007/s13225-020-00466-2)) |
| Otu46252 | 1 | k__Fungi(100);p__Ascomycota(100);c__Saccharomycetes(100);o__Saccharomycetales(100);f__Metschnikowiaceae(100);g__Metschnikowia(100);s__Metschnikowia_pulcherrima(100); | Metschnikowia | 13 | Saprotroph | \|Undefined Saprotroph\| | Yeast | NULL | Probable | As nectar-tap_saprotroph (Põlme et al. 2020); Decay sugar-rich substrates (Põlme et al. 2020) | Põlme S, et al. 2020. Fungal Diversity 105:1-16 ((https://doi.org/10.1007/s13225-020-00466-2)) |
| Otu46253 | 1 | k__Fungi(100);p__Ascomycota(100);c__Saccharomycetes(100);o__Saccharomycetales(100);f__Metschnikowiaceae(100);g__Metschnikowia(100);s__Metschnikowia_pulcherrima(100); | Metschnikowia | 13 | Saprotroph | \|Undefined Saprotroph\| | Yeast | NULL | Probable | As nectar-tap_saprotroph (Põlme et al. 2020); Decay sugar-rich substrates (Põlme et al. 2020) | Põlme S, et al. 2020. Fungal Diversity 105:1-16 ((https://doi.org/10.1007/s13225-020-00466-2)) |
| Otu46254 | 1 | k__Fungi(100);p__Ascomycota(100);c__Saccharomycetes(100);o__Saccharomycetales(100);f__Metschnikowiaceae(100);g__Metschnikowia(100);s__Metschnikowia_pulcherrima(100); | Metschnikowia | 13 | Saprotroph | \|Undefined Saprotroph\| | Yeast | NULL | Probable | As nectar-tap_saprotroph (Põlme et al. 2020); Decay sugar-rich substrates (Põlme et al. 2020) | Põlme S, et al. 2020. Fungal Diversity 105:1-16 ((https://doi.org/10.1007/s13225-020-00466-2)) |
| Otu46255 | 1 | k__Fungi(100);p__Ascomycota(100);c__Saccharomycetes(100);o__Saccharomycetales(100);f__Metschnikowiaceae(100);g__Metschnikowia(100);s__Metschnikowia_pulcherrima(100); | Metschnikowia | 13 | Saprotroph | \|Undefined Saprotroph\| | Yeast | NULL | Probable | As nectar-tap_saprotroph (Põlme et al. 2020); Decay sugar-rich substrates (Põlme et al. 2020) | Põlme S, et al. 2020. Fungal Diversity 105:1-16 ((https://doi.org/10.1007/s13225-020-00466-2)) |
| Otu46256 | 1 | k__Fungi(100);p__Ascomycota(100);c__Saccharomycetes(100);o__Saccharomycetales(100);f__Metschnikowiaceae(100);g__Metschnikowia(100);s__Metschnikowia_pulcherrima(100); | Metschnikowia | 13 | Saprotroph | \|Undefined Saprotroph\| | Yeast | NULL | Probable | As nectar-tap_saprotroph (Põlme et al. 2020); Decay sugar-rich substrates (Põlme et al. 2020) | Põlme S, et al. 2020. Fungal Diversity 105:1-16 ((https://doi.org/10.1007/s13225-020-00466-2)) |
| Otu46257 | 1 | k__Fungi(100);p__Ascomycota(100);c__Saccharomycetes(100);o__Saccharomycetales(100);f__Metschnikowiaceae(100);g__Metschnikowia(100);s__Metschnikowia_pulcherrima(100); | Metschnikowia | 13 | Saprotroph | \|Undefined Saprotroph\| | Yeast | NULL | Probable | As nectar-tap_saprotroph (Põlme et al. 2020); Decay sugar-rich substrates (Põlme et al. 2020) | Põlme S, et al. 2020. Fungal Diversity 105:1-16 ((https://doi.org/10.1007/s13225-020-00466-2)) |
| Otu46260 | 1 | k__Fungi(100);p__Ascomycota(100);c__Saccharomycetes(100);o__Saccharomycetales(100);f__Metschnikowiaceae(100);g__Metschnikowia(100);s__Metschnikowia_pulcherrima(100); | Metschnikowia | 13 | Saprotroph | \|Undefined Saprotroph\| | Yeast | NULL | Probable | As nectar-tap_saprotroph (Põlme et al. 2020); Decay sugar-rich substrates (Põlme et al. 2020) | Põlme S, et al. 2020. Fungal Diversity 105:1-16 ((https://doi.org/10.1007/s13225-020-00466-2)) |
| Otu46261 | 1 | k__Fungi(100);p__Ascomycota(100);c__Saccharomycetes(100);o__Saccharomycetales(100);f__Metschnikowiaceae(100);g__Metschnikowia(100);s__Metschnikowia_pulcherrima(100); | Metschnikowia | 13 | Saprotroph | \|Undefined Saprotroph\| | Yeast | NULL | Probable | As nectar-tap_saprotroph (Põlme et al. 2020); Decay sugar-rich substrates (Põlme et al. 2020) | Põlme S, et al. 2020. Fungal Diversity 105:1-16 ((https://doi.org/10.1007/s13225-020-00466-2)) |
| Otu46262 | 1 | k__Fungi(100);p__Ascomycota(100);c__Saccharomycetes(100);o__Saccharomycetales(100);f__Metschnikowiaceae(100);g__Metschnikowia(100);s__Metschnikowia_pulcherrima(100); | Metschnikowia | 13 | Saprotroph | \|Undefined Saprotroph\| | Yeast | NULL | Probable | As nectar-tap_saprotroph (Põlme et al. 2020); Decay sugar-rich substrates (Põlme et al. 2020) | Põlme S, et al. 2020. Fungal Diversity 105:1-16 ((https://doi.org/10.1007/s13225-020-00466-2)) |
| Otu46263 | 1 | k__Fungi(100);p__Ascomycota(100);c__Saccharomycetes(100);o__Saccharomycetales(100);f__Metschnikowiaceae(100);g__Metschnikowia(100);s__Metschnikowia_pulcherrima(100); | Metschnikowia | 13 | Saprotroph | \|Undefined Saprotroph\| | Yeast | NULL | Probable | As nectar-tap_saprotroph (Põlme et al. 2020); Decay sugar-rich substrates (Põlme et al. 2020) | Põlme S, et al. 2020. Fungal Diversity 105:1-16 ((https://doi.org/10.1007/s13225-020-00466-2)) |
| Otu46264 | 1 | k__Fungi(100);p__Ascomycota(100);c__Saccharomycetes(100);o__Saccharomycetales(100);f__Metschnikowiaceae(100);g__Metschnikowia(100);s__Metschnikowia_pulcherrima(100); | Metschnikowia | 13 | Saprotroph | \|Undefined Saprotroph\| | Yeast | NULL | Probable | As nectar-tap_saprotroph (Põlme et al. 2020); Decay sugar-rich substrates (Põlme et al. 2020) | Põlme S, et al. 2020. Fungal Diversity 105:1-16 ((https://doi.org/10.1007/s13225-020-00466-2)) |
| Otu46265 | 1 | k__Fungi(100);p__Ascomycota(100);c__Saccharomycetes(100);o__Saccharomycetales(100);f__Metschnikowiaceae(100);g__Metschnikowia(100);g__Metschnikowia_unclassified(100); | Metschnikowia | 13 | Saprotroph | \|Undefined Saprotroph\| | Yeast | NULL | Probable | As nectar-tap_saprotroph (Põlme et al. 2020); Decay sugar-rich substrates (Põlme et al. 2020) | Põlme S, et al. 2020. Fungal Diversity 105:1-16 ((https://doi.org/10.1007/s13225-020-00466-2)) |
| Otu46266 | 1 | k__Fungi(100);p__Ascomycota(100);c__Saccharomycetes(100);o__Saccharomycetales(100);f__Metschnikowiaceae(100);g__Metschnikowia(100);s__Metschnikowia_pulcherrima(100); | Metschnikowia | 13 | Saprotroph | \|Undefined Saprotroph\| | Yeast | NULL | Probable | As nectar-tap_saprotroph (Põlme et al. 2020); Decay sugar-rich substrates (Põlme et al. 2020) | Põlme S, et al. 2020. Fungal Diversity 105:1-16 ((https://doi.org/10.1007/s13225-020-00466-2)) |
| Otu46267 | 1 | k__Fungi(100);p__Ascomycota(100);c__Saccharomycetes(100);o__Saccharomycetales(100);f__Metschnikowiaceae(100);g__Metschnikowia(100);s__Metschnikowia_pulcherrima(100); | Metschnikowia | 13 | Saprotroph | \|Undefined Saprotroph\| | Yeast | NULL | Probable | As nectar-tap_saprotroph (Põlme et al. 2020); Decay sugar-rich substrates (Põlme et al. 2020) | Põlme S, et al. 2020. Fungal Diversity 105:1-16 ((https://doi.org/10.1007/s13225-020-00466-2)) |
| Otu46268 | 1 | k__Fungi(100);p__Ascomycota(100);c__Saccharomycetes(100);o__Saccharomycetales(100);f__Metschnikowiaceae(100);g__Metschnikowia(100);s__Metschnikowia_pulcherrima(100); | Metschnikowia | 13 | Saprotroph | \|Undefined Saprotroph\| | Yeast | NULL | Probable | As nectar-tap_saprotroph (Põlme et al. 2020); Decay sugar-rich substrates (Põlme et al. 2020) | Põlme S, et al. 2020. Fungal Diversity 105:1-16 ((https://doi.org/10.1007/s13225-020-00466-2)) |
| Otu46269 | 1 | k__Fungi(100);p__Ascomycota(100);c__Saccharomycetes(100);o__Saccharomycetales(100);f__Metschnikowiaceae(100);g__Metschnikowia(100);s__Metschnikowia_pulcherrima(100); | Metschnikowia | 13 | Saprotroph | \|Undefined Saprotroph\| | Yeast | NULL | Probable | As nectar-tap_saprotroph (Põlme et al. 2020); Decay sugar-rich substrates (Põlme et al. 2020) | Põlme S, et al. 2020. Fungal Diversity 105:1-16 ((https://doi.org/10.1007/s13225-020-00466-2)) |
| Otu46271 | 1 | k__Fungi(100);p__Ascomycota(100);c__Saccharomycetes(100);o__Saccharomycetales(100);f__Metschnikowiaceae(100);g__Metschnikowia(100);s__Metschnikowia_pulcherrima(100); | Metschnikowia | 13 | Saprotroph | \|Undefined Saprotroph\| | Yeast | NULL | Probable | As nectar-tap_saprotroph (Põlme et al. 2020); Decay sugar-rich substrates (Põlme et al. 2020) | Põlme S, et al. 2020. Fungal Diversity 105:1-16 ((https://doi.org/10.1007/s13225-020-00466-2)) |
| Otu46272 | 1 | k__Fungi(100);p__Ascomycota(100);c__Saccharomycetes(100);o__Saccharomycetales(100);f__Metschnikowiaceae(100);g__Metschnikowia(100);g__Metschnikowia_unclassified(100); | Metschnikowia | 13 | Saprotroph | \|Undefined Saprotroph\| | Yeast | NULL | Probable | As nectar-tap_saprotroph (Põlme et al. 2020); Decay sugar-rich substrates (Põlme et al. 2020) | Põlme S, et al. 2020. Fungal Diversity 105:1-16 ((https://doi.org/10.1007/s13225-020-00466-2)) |
| Otu46273 | 1 | k__Fungi(100);p__Ascomycota(100);c__Saccharomycetes(100);o__Saccharomycetales(100);f__Metschnikowiaceae(100);g__Metschnikowia(100);s__Metschnikowia_pulcherrima(100); | Metschnikowia | 13 | Saprotroph | \|Undefined Saprotroph\| | Yeast | NULL | Probable | As nectar-tap_saprotroph (Põlme et al. 2020); Decay sugar-rich substrates (Põlme et al. 2020) | Põlme S, et al. 2020. Fungal Diversity 105:1-16 ((https://doi.org/10.1007/s13225-020-00466-2)) |
| Otu46274 | 1 | k__Fungi(100);p__Ascomycota(100);c__Saccharomycetes(100);o__Saccharomycetales(100);f__Metschnikowiaceae(100);g__Metschnikowia(100);s__Metschnikowia_pulcherrima(100); | Metschnikowia | 13 | Saprotroph | \|Undefined Saprotroph\| | Yeast | NULL | Probable | As nectar-tap_saprotroph (Põlme et al. 2020); Decay sugar-rich substrates (Põlme et al. 2020) | Põlme S, et al. 2020. Fungal Diversity 105:1-16 ((https://doi.org/10.1007/s13225-020-00466-2)) |
| Otu46275 | 1 | k__Fungi(100);p__Ascomycota(100);c__Saccharomycetes(100);o__Saccharomycetales(100);f__Metschnikowiaceae(100);g__Metschnikowia(100);g__Metschnikowia_unclassified(100); | Metschnikowia | 13 | Saprotroph | \|Undefined Saprotroph\| | Yeast | NULL | Probable | As nectar-tap_saprotroph (Põlme et al. 2020); Decay sugar-rich substrates (Põlme et al. 2020) | Põlme S, et al. 2020. Fungal Diversity 105:1-16 ((https://doi.org/10.1007/s13225-020-00466-2)) |
| Otu46276 | 1 | k__Fungi(100);p__Ascomycota(100);c__Saccharomycetes(100);o__Saccharomycetales(100);f__Metschnikowiaceae(100);g__Metschnikowia(100);g__Metschnikowia_unclassified(100); | Metschnikowia | 13 | Saprotroph | \|Undefined Saprotroph\| | Yeast | NULL | Probable | As nectar-tap_saprotroph (Põlme et al. 2020); Decay sugar-rich substrates (Põlme et al. 2020) | Põlme S, et al. 2020. Fungal Diversity 105:1-16 ((https://doi.org/10.1007/s13225-020-00466-2)) |
| Otu46277 | 1 | k__Fungi(100);p__Ascomycota(100);c__Saccharomycetes(100);o__Saccharomycetales(100);f__Metschnikowiaceae(100);g__Metschnikowia(100);g__Metschnikowia_unclassified(100); | Metschnikowia | 13 | Saprotroph | \|Undefined Saprotroph\| | Yeast | NULL | Probable | As nectar-tap_saprotroph (Põlme et al. 2020); Decay sugar-rich substrates (Põlme et al. 2020) | Põlme S, et al. 2020. Fungal Diversity 105:1-16 ((https://doi.org/10.1007/s13225-020-00466-2)) |
| Otu46278 | 1 | k__Fungi(100);p__Ascomycota(100);c__Saccharomycetes(100);o__Saccharomycetales(100);f__Metschnikowiaceae(100);g__Metschnikowia(100);s__Metschnikowia_pulcherrima(100); | Metschnikowia | 13 | Saprotroph | \|Undefined Saprotroph\| | Yeast | NULL | Probable | As nectar-tap_saprotroph (Põlme et al. 2020); Decay sugar-rich substrates (Põlme et al. 2020) | Põlme S, et al. 2020. Fungal Diversity 105:1-16 ((https://doi.org/10.1007/s13225-020-00466-2)) |
| Otu46279 | 1 | k__Fungi(100);p__Ascomycota(100);c__Saccharomycetes(100);o__Saccharomycetales(100);f__Metschnikowiaceae(100);g__Metschnikowia(100);g__Metschnikowia_unclassified(100); | Metschnikowia | 13 | Saprotroph | \|Undefined Saprotroph\| | Yeast | NULL | Probable | As nectar-tap_saprotroph (Põlme et al. 2020); Decay sugar-rich substrates (Põlme et al. 2020) | Põlme S, et al. 2020. Fungal Diversity 105:1-16 ((https://doi.org/10.1007/s13225-020-00466-2)) |
| Otu46280 | 1 | k__Fungi(100);p__Ascomycota(100);c__Saccharomycetes(100);o__Saccharomycetales(100);f__Metschnikowiaceae(100);g__Metschnikowia(100);s__Metschnikowia_pulcherrima(100); | Metschnikowia | 13 | Saprotroph | \|Undefined Saprotroph\| | Yeast | NULL | Probable | As nectar-tap_saprotroph (Põlme et al. 2020); Decay sugar-rich substrates (Põlme et al. 2020) | Põlme S, et al. 2020. Fungal Diversity 105:1-16 ((https://doi.org/10.1007/s13225-020-00466-2)) |
| Otu46281 | 1 | k__Fungi(100);p__Ascomycota(100);c__Saccharomycetes(100);o__Saccharomycetales(100);f__Metschnikowiaceae(100);g__Metschnikowia(100);s__Metschnikowia_pulcherrima(100); | Metschnikowia | 13 | Saprotroph | \|Undefined Saprotroph\| | Yeast | NULL | Probable | As nectar-tap_saprotroph (Põlme et al. 2020); Decay sugar-rich substrates (Põlme et al. 2020) | Põlme S, et al. 2020. Fungal Diversity 105:1-16 ((https://doi.org/10.1007/s13225-020-00466-2)) |
| Otu46282 | 1 | k__Fungi(100);p__Ascomycota(100);c__Saccharomycetes(100);o__Saccharomycetales(100);f__Metschnikowiaceae(100);g__Metschnikowia(100);s__Metschnikowia_pulcherrima(100); | Metschnikowia | 13 | Saprotroph | \|Undefined Saprotroph\| | Yeast | NULL | Probable | As nectar-tap_saprotroph (Põlme et al. 2020); Decay sugar-rich substrates (Põlme et al. 2020) | Põlme S, et al. 2020. Fungal Diversity 105:1-16 ((https://doi.org/10.1007/s13225-020-00466-2)) |
| Otu46283 | 1 | k__Fungi(100);p__Ascomycota(100);c__Saccharomycetes(100);o__Saccharomycetales(100);f__Metschnikowiaceae(100);g__Metschnikowia(100);s__Metschnikowia_pulcherrima(100); | Metschnikowia | 13 | Saprotroph | \|Undefined Saprotroph\| | Yeast | NULL | Probable | As nectar-tap_saprotroph (Põlme et al. 2020); Decay sugar-rich substrates (Põlme et al. 2020) | Põlme S, et al. 2020. Fungal Diversity 105:1-16 ((https://doi.org/10.1007/s13225-020-00466-2)) |
| Otu46284 | 1 | k__Fungi(100);p__Ascomycota(100);c__Saccharomycetes(100);o__Saccharomycetales(100);f__Metschnikowiaceae(100);g__Metschnikowia(100);s__Metschnikowia_pulcherrima(100); | Metschnikowia | 13 | Saprotroph | \|Undefined Saprotroph\| | Yeast | NULL | Probable | As nectar-tap_saprotroph (Põlme et al. 2020); Decay sugar-rich substrates (Põlme et al. 2020) | Põlme S, et al. 2020. Fungal Diversity 105:1-16 ((https://doi.org/10.1007/s13225-020-00466-2)) |
| Otu46285 | 1 | k__Fungi(100);p__Ascomycota(100);c__Saccharomycetes(100);o__Saccharomycetales(100);f__Metschnikowiaceae(100);g__Metschnikowia(100);s__Metschnikowia_pulcherrima(100); | Metschnikowia | 13 | Saprotroph | \|Undefined Saprotroph\| | Yeast | NULL | Probable | As nectar-tap_saprotroph (Põlme et al. 2020); Decay sugar-rich substrates (Põlme et al. 2020) | Põlme S, et al. 2020. Fungal Diversity 105:1-16 ((https://doi.org/10.1007/s13225-020-00466-2)) |
| Otu46286 | 1 | k__Fungi(100);p__Ascomycota(100);c__Saccharomycetes(100);o__Saccharomycetales(100);f__Metschnikowiaceae(100);g__Metschnikowia(100);s__Metschnikowia_pulcherrima(100); | Metschnikowia | 13 | Saprotroph | \|Undefined Saprotroph\| | Yeast | NULL | Probable | As nectar-tap_saprotroph (Põlme et al. 2020); Decay sugar-rich substrates (Põlme et al. 2020) | Põlme S, et al. 2020. Fungal Diversity 105:1-16 ((https://doi.org/10.1007/s13225-020-00466-2)) |
| Otu46287 | 1 | k__Fungi(100);p__Ascomycota(100);c__Saccharomycetes(100);o__Saccharomycetales(100);f__Metschnikowiaceae(100);g__Metschnikowia(100);s__Metschnikowia_pulcherrima(100); | Metschnikowia | 13 | Saprotroph | \|Undefined Saprotroph\| | Yeast | NULL | Probable | As nectar-tap_saprotroph (Põlme et al. 2020); Decay sugar-rich substrates (Põlme et al. 2020) | Põlme S, et al. 2020. Fungal Diversity 105:1-16 ((https://doi.org/10.1007/s13225-020-00466-2)) |
| Otu46288 | 1 | k__Fungi(100);p__Ascomycota(100);c__Saccharomycetes(100);o__Saccharomycetales(100);f__Metschnikowiaceae(100);g__Metschnikowia(100);s__Metschnikowia_pulcherrima(100); | Metschnikowia | 13 | Saprotroph | \|Undefined Saprotroph\| | Yeast | NULL | Probable | As nectar-tap_saprotroph (Põlme et al. 2020); Decay sugar-rich substrates (Põlme et al. 2020) | Põlme S, et al. 2020. Fungal Diversity 105:1-16 ((https://doi.org/10.1007/s13225-020-00466-2)) |
| Otu46289 | 1 | k__Fungi(100);p__Ascomycota(100);c__Saccharomycetes(100);o__Saccharomycetales(100);f__Metschnikowiaceae(100);g__Metschnikowia(100);s__Metschnikowia_pulcherrima(100); | Metschnikowia | 13 | Saprotroph | \|Undefined Saprotroph\| | Yeast | NULL | Probable | As nectar-tap_saprotroph (Põlme et al. 2020); Decay sugar-rich substrates (Põlme et al. 2020) | Põlme S, et al. 2020. Fungal Diversity 105:1-16 ((https://doi.org/10.1007/s13225-020-00466-2)) |
| Otu46290 | 1 | k__Fungi(100);p__Ascomycota(100);c__Saccharomycetes(100);o__Saccharomycetales(100);f__Metschnikowiaceae(100);g__Metschnikowia(100);s__Metschnikowia_pulcherrima(100); | Metschnikowia | 13 | Saprotroph | \|Undefined Saprotroph\| | Yeast | NULL | Probable | As nectar-tap_saprotroph (Põlme et al. 2020); Decay sugar-rich substrates (Põlme et al. 2020) | Põlme S, et al. 2020. Fungal Diversity 105:1-16 ((https://doi.org/10.1007/s13225-020-00466-2)) |
| Otu46291 | 1 | k__Fungi(100);p__Ascomycota(100);c__Saccharomycetes(100);o__Saccharomycetales(100);f__Metschnikowiaceae(100);g__Metschnikowia(100);s__Metschnikowia_pulcherrima(100); | Metschnikowia | 13 | Saprotroph | \|Undefined Saprotroph\| | Yeast | NULL | Probable | As nectar-tap_saprotroph (Põlme et al. 2020); Decay sugar-rich substrates (Põlme et al. 2020) | Põlme S, et al. 2020. Fungal Diversity 105:1-16 ((https://doi.org/10.1007/s13225-020-00466-2)) |
| Otu46292 | 1 | k__Fungi(100);p__Ascomycota(100);c__Saccharomycetes(100);o__Saccharomycetales(100);f__Metschnikowiaceae(100);g__Metschnikowia(100);g__Metschnikowia_unclassified(100); | Metschnikowia | 13 | Saprotroph | \|Undefined Saprotroph\| | Yeast | NULL | Probable | As nectar-tap_saprotroph (Põlme et al. 2020); Decay sugar-rich substrates (Põlme et al. 2020) | Põlme S, et al. 2020. Fungal Diversity 105:1-16 ((https://doi.org/10.1007/s13225-020-00466-2)) |
| Otu46293 | 1 | k__Fungi(100);p__Ascomycota(100);c__Saccharomycetes(100);o__Saccharomycetales(100);f__Metschnikowiaceae(100);g__Metschnikowia(100);s__Metschnikowia_pulcherrima(100); | Metschnikowia | 13 | Saprotroph | \|Undefined Saprotroph\| | Yeast | NULL | Probable | As nectar-tap_saprotroph (Põlme et al. 2020); Decay sugar-rich substrates (Põlme et al. 2020) | Põlme S, et al. 2020. Fungal Diversity 105:1-16 ((https://doi.org/10.1007/s13225-020-00466-2)) |
| Otu46294 | 1 | k__Fungi(100);p__Ascomycota(100);c__Saccharomycetes(100);o__Saccharomycetales(100);f__Metschnikowiaceae(100);g__Metschnikowia(100);s__Metschnikowia_pulcherrima(100); | Metschnikowia | 13 | Saprotroph | \|Undefined Saprotroph\| | Yeast | NULL | Probable | As nectar-tap_saprotroph (Põlme et al. 2020); Decay sugar-rich substrates (Põlme et al. 2020) | Põlme S, et al. 2020. Fungal Diversity 105:1-16 ((https://doi.org/10.1007/s13225-020-00466-2)) |
| Otu46295 | 1 | k__Fungi(100);p__Ascomycota(100);c__Saccharomycetes(100);o__Saccharomycetales(100);f__Metschnikowiaceae(100);g__Metschnikowia(100);s__Metschnikowia_pulcherrima(100); | Metschnikowia | 13 | Saprotroph | \|Undefined Saprotroph\| | Yeast | NULL | Probable | As nectar-tap_saprotroph (Põlme et al. 2020); Decay sugar-rich substrates (Põlme et al. 2020) | Põlme S, et al. 2020. Fungal Diversity 105:1-16 ((https://doi.org/10.1007/s13225-020-00466-2)) |
| Otu46296 | 1 | k__Fungi(100);p__Ascomycota(100);c__Saccharomycetes(100);o__Saccharomycetales(100);f__Metschnikowiaceae(100);g__Metschnikowia(100);s__Metschnikowia_pulcherrima(100); | Metschnikowia | 13 | Saprotroph | \|Undefined Saprotroph\| | Yeast | NULL | Probable | As nectar-tap_saprotroph (Põlme et al. 2020); Decay sugar-rich substrates (Põlme et al. 2020) | Põlme S, et al. 2020. Fungal Diversity 105:1-16 ((https://doi.org/10.1007/s13225-020-00466-2)) |
| Otu46297 | 1 | k__Fungi(100);p__Ascomycota(100);c__Saccharomycetes(100);o__Saccharomycetales(100);f__Metschnikowiaceae(100);g__Metschnikowia(100);s__Metschnikowia_pulcherrima(100); | Metschnikowia | 13 | Saprotroph | \|Undefined Saprotroph\| | Yeast | NULL | Probable | As nectar-tap_saprotroph (Põlme et al. 2020); Decay sugar-rich substrates (Põlme et al. 2020) | Põlme S, et al. 2020. Fungal Diversity 105:1-16 ((https://doi.org/10.1007/s13225-020-00466-2)) |
| Otu46298 | 1 | k__Fungi(100);p__Ascomycota(100);c__Saccharomycetes(100);o__Saccharomycetales(100);f__Metschnikowiaceae(100);g__Metschnikowia(100);s__Metschnikowia_pulcherrima(100); | Metschnikowia | 13 | Saprotroph | \|Undefined Saprotroph\| | Yeast | NULL | Probable | As nectar-tap_saprotroph (Põlme et al. 2020); Decay sugar-rich substrates (Põlme et al. 2020) | Põlme S, et al. 2020. Fungal Diversity 105:1-16 ((https://doi.org/10.1007/s13225-020-00466-2)) |
| Otu46299 | 1 | k__Fungi(100);p__Ascomycota(100);c__Saccharomycetes(100);o__Saccharomycetales(100);f__Metschnikowiaceae(100);g__Metschnikowia(100);s__Metschnikowia_pulcherrima(100); | Metschnikowia | 13 | Saprotroph | \|Undefined Saprotroph\| | Yeast | NULL | Probable | As nectar-tap_saprotroph (Põlme et al. 2020); Decay sugar-rich substrates (Põlme et al. 2020) | Põlme S, et al. 2020. Fungal Diversity 105:1-16 ((https://doi.org/10.1007/s13225-020-00466-2)) |
| Otu46300 | 1 | k__Fungi(100);p__Ascomycota(100);c__Saccharomycetes(100);o__Saccharomycetales(100);f__Metschnikowiaceae(100);g__Metschnikowia(100);s__Metschnikowia_pulcherrima(100); | Metschnikowia | 13 | Saprotroph | \|Undefined Saprotroph\| | Yeast | NULL | Probable | As nectar-tap_saprotroph (Põlme et al. 2020); Decay sugar-rich substrates (Põlme et al. 2020) | Põlme S, et al. 2020. Fungal Diversity 105:1-16 ((https://doi.org/10.1007/s13225-020-00466-2)) |
| Otu46301 | 1 | k__Fungi(100);p__Ascomycota(100);c__Saccharomycetes(100);o__Saccharomycetales(100);f__Metschnikowiaceae(100);g__Metschnikowia(100);g__Metschnikowia_unclassified(100); | Metschnikowia | 13 | Saprotroph | \|Undefined Saprotroph\| | Yeast | NULL | Probable | As nectar-tap_saprotroph (Põlme et al. 2020); Decay sugar-rich substrates (Põlme et al. 2020) | Põlme S, et al. 2020. Fungal Diversity 105:1-16 ((https://doi.org/10.1007/s13225-020-00466-2)) |
| Otu46302 | 1 | k__Fungi(100);p__Ascomycota(100);c__Saccharomycetes(100);o__Saccharomycetales(100);f__Metschnikowiaceae(100);g__Metschnikowia(100);s__Metschnikowia_pulcherrima(100); | Metschnikowia | 13 | Saprotroph | \|Undefined Saprotroph\| | Yeast | NULL | Probable | As nectar-tap_saprotroph (Põlme et al. 2020); Decay sugar-rich substrates (Põlme et al. 2020) | Põlme S, et al. 2020. Fungal Diversity 105:1-16 ((https://doi.org/10.1007/s13225-020-00466-2)) |
| Otu46303 | 1 | k__Fungi(100);p__Ascomycota(100);c__Saccharomycetes(100);o__Saccharomycetales(100);f__Metschnikowiaceae(100);g__Metschnikowia(100);s__Metschnikowia_pulcherrima(100); | Metschnikowia | 13 | Saprotroph | \|Undefined Saprotroph\| | Yeast | NULL | Probable | As nectar-tap_saprotroph (Põlme et al. 2020); Decay sugar-rich substrates (Põlme et al. 2020) | Põlme S, et al. 2020. Fungal Diversity 105:1-16 ((https://doi.org/10.1007/s13225-020-00466-2)) |
| Otu46304 | 1 | k__Fungi(100);p__Ascomycota(100);c__Saccharomycetes(100);o__Saccharomycetales(100);f__Metschnikowiaceae(100);g__Metschnikowia(100);s__Metschnikowia_pulcherrima(100); | Metschnikowia | 13 | Saprotroph | \|Undefined Saprotroph\| | Yeast | NULL | Probable | As nectar-tap_saprotroph (Põlme et al. 2020); Decay sugar-rich substrates (Põlme et al. 2020) | Põlme S, et al. 2020. Fungal Diversity 105:1-16 ((https://doi.org/10.1007/s13225-020-00466-2)) |
| Otu46305 | 1 | k__Fungi(100);p__Ascomycota(100);c__Saccharomycetes(100);o__Saccharomycetales(100);f__Metschnikowiaceae(100);g__Metschnikowia(100);s__Metschnikowia_pulcherrima(100); | Metschnikowia | 13 | Saprotroph | \|Undefined Saprotroph\| | Yeast | NULL | Probable | As nectar-tap_saprotroph (Põlme et al. 2020); Decay sugar-rich substrates (Põlme et al. 2020) | Põlme S, et al. 2020. Fungal Diversity 105:1-16 ((https://doi.org/10.1007/s13225-020-00466-2)) |
| Otu46306 | 1 | k__Fungi(100);p__Ascomycota(100);c__Saccharomycetes(100);o__Saccharomycetales(100);f__Metschnikowiaceae(100);g__Metschnikowia(100);s__Metschnikowia_pulcherrima(100); | Metschnikowia | 13 | Saprotroph | \|Undefined Saprotroph\| | Yeast | NULL | Probable | As nectar-tap_saprotroph (Põlme et al. 2020); Decay sugar-rich substrates (Põlme et al. 2020) | Põlme S, et al. 2020. Fungal Diversity 105:1-16 ((https://doi.org/10.1007/s13225-020-00466-2)) |
| Otu46307 | 1 | k__Fungi(100);p__Ascomycota(100);c__Saccharomycetes(100);o__Saccharomycetales(100);f__Metschnikowiaceae(100);g__Metschnikowia(100);s__Metschnikowia_pulcherrima(100); | Metschnikowia | 13 | Saprotroph | \|Undefined Saprotroph\| | Yeast | NULL | Probable | As nectar-tap_saprotroph (Põlme et al. 2020); Decay sugar-rich substrates (Põlme et al. 2020) | Põlme S, et al. 2020. Fungal Diversity 105:1-16 ((https://doi.org/10.1007/s13225-020-00466-2)) |
| Otu46308 | 1 | k__Fungi(100);p__Ascomycota(100);c__Saccharomycetes(100);o__Saccharomycetales(100);f__Metschnikowiaceae(100);g__Metschnikowia(100);s__Metschnikowia_pulcherrima(100); | Metschnikowia | 13 | Saprotroph | \|Undefined Saprotroph\| | Yeast | NULL | Probable | As nectar-tap_saprotroph (Põlme et al. 2020); Decay sugar-rich substrates (Põlme et al. 2020) | Põlme S, et al. 2020. Fungal Diversity 105:1-16 ((https://doi.org/10.1007/s13225-020-00466-2)) |
| Otu46309 | 1 | k__Fungi(100);p__Ascomycota(100);c__Saccharomycetes(100);o__Saccharomycetales(100);f__Metschnikowiaceae(100);g__Metschnikowia(100);s__Metschnikowia_pulcherrima(100); | Metschnikowia | 13 | Saprotroph | \|Undefined Saprotroph\| | Yeast | NULL | Probable | As nectar-tap_saprotroph (Põlme et al. 2020); Decay sugar-rich substrates (Põlme et al. 2020) | Põlme S, et al. 2020. Fungal Diversity 105:1-16 ((https://doi.org/10.1007/s13225-020-00466-2)) |
| Otu46310 | 1 | k__Fungi(100);p__Ascomycota(100);c__Saccharomycetes(100);o__Saccharomycetales(100);f__Metschnikowiaceae(100);g__Metschnikowia(100);s__Metschnikowia_pulcherrima(100); | Metschnikowia | 13 | Saprotroph | \|Undefined Saprotroph\| | Yeast | NULL | Probable | As nectar-tap_saprotroph (Põlme et al. 2020); Decay sugar-rich substrates (Põlme et al. 2020) | Põlme S, et al. 2020. Fungal Diversity 105:1-16 ((https://doi.org/10.1007/s13225-020-00466-2)) |
| Otu46311 | 1 | k__Fungi(100);p__Ascomycota(100);c__Saccharomycetes(100);o__Saccharomycetales(100);f__Metschnikowiaceae(100);g__Metschnikowia(100);s__Metschnikowia_pulcherrima(100); | Metschnikowia | 13 | Saprotroph | \|Undefined Saprotroph\| | Yeast | NULL | Probable | As nectar-tap_saprotroph (Põlme et al. 2020); Decay sugar-rich substrates (Põlme et al. 2020) | Põlme S, et al. 2020. Fungal Diversity 105:1-16 ((https://doi.org/10.1007/s13225-020-00466-2)) |
| Otu46312 | 1 | k__Fungi(100);p__Ascomycota(100);c__Saccharomycetes(100);o__Saccharomycetales(100);f__Metschnikowiaceae(100);g__Metschnikowia(100);s__Metschnikowia_pulcherrima(100); | Metschnikowia | 13 | Saprotroph | \|Undefined Saprotroph\| | Yeast | NULL | Probable | As nectar-tap_saprotroph (Põlme et al. 2020); Decay sugar-rich substrates (Põlme et al. 2020) | Põlme S, et al. 2020. Fungal Diversity 105:1-16 ((https://doi.org/10.1007/s13225-020-00466-2)) |
| Otu46313 | 1 | k__Fungi(100);p__Ascomycota(100);c__Saccharomycetes(100);o__Saccharomycetales(100);f__Metschnikowiaceae(100);g__Metschnikowia(100);s__Metschnikowia_pulcherrima(100); | Metschnikowia | 13 | Saprotroph | \|Undefined Saprotroph\| | Yeast | NULL | Probable | As nectar-tap_saprotroph (Põlme et al. 2020); Decay sugar-rich substrates (Põlme et al. 2020) | Põlme S, et al. 2020. Fungal Diversity 105:1-16 ((https://doi.org/10.1007/s13225-020-00466-2)) |
| Otu46314 | 1 | k__Fungi(100);p__Ascomycota(100);c__Saccharomycetes(100);o__Saccharomycetales(100);f__Metschnikowiaceae(100);g__Metschnikowia(100);s__Metschnikowia_pulcherrima(100); | Metschnikowia | 13 | Saprotroph | \|Undefined Saprotroph\| | Yeast | NULL | Probable | As nectar-tap_saprotroph (Põlme et al. 2020); Decay sugar-rich substrates (Põlme et al. 2020) | Põlme S, et al. 2020. Fungal Diversity 105:1-16 ((https://doi.org/10.1007/s13225-020-00466-2)) |
| Otu46315 | 1 | k__Fungi(100);p__Ascomycota(100);c__Saccharomycetes(100);o__Saccharomycetales(100);f__Metschnikowiaceae(100);g__Metschnikowia(100);s__Metschnikowia_pulcherrima(100); | Metschnikowia | 13 | Saprotroph | \|Undefined Saprotroph\| | Yeast | NULL | Probable | As nectar-tap_saprotroph (Põlme et al. 2020); Decay sugar-rich substrates (Põlme et al. 2020) | Põlme S, et al. 2020. Fungal Diversity 105:1-16 ((https://doi.org/10.1007/s13225-020-00466-2)) |
| Otu46316 | 1 | k__Fungi(100);p__Ascomycota(100);c__Saccharomycetes(100);o__Saccharomycetales(100);f__Metschnikowiaceae(100);g__Metschnikowia(100);s__Metschnikowia_pulcherrima(100); | Metschnikowia | 13 | Saprotroph | \|Undefined Saprotroph\| | Yeast | NULL | Probable | As nectar-tap_saprotroph (Põlme et al. 2020); Decay sugar-rich substrates (Põlme et al. 2020) | Põlme S, et al. 2020. Fungal Diversity 105:1-16 ((https://doi.org/10.1007/s13225-020-00466-2)) |
| Otu46317 | 1 | k__Fungi(100);p__Ascomycota(100);c__Saccharomycetes(100);o__Saccharomycetales(100);f__Metschnikowiaceae(100);g__Metschnikowia(100);g__Metschnikowia_unclassified(100); | Metschnikowia | 13 | Saprotroph | \|Undefined Saprotroph\| | Yeast | NULL | Probable | As nectar-tap_saprotroph (Põlme et al. 2020); Decay sugar-rich substrates (Põlme et al. 2020) | Põlme S, et al. 2020. Fungal Diversity 105:1-16 ((https://doi.org/10.1007/s13225-020-00466-2)) |
| Otu46318 | 1 | k__Fungi(100);p__Ascomycota(100);c__Saccharomycetes(100);o__Saccharomycetales(100);f__Metschnikowiaceae(100);g__Metschnikowia(100);g__Metschnikowia_unclassified(100); | Metschnikowia | 13 | Saprotroph | \|Undefined Saprotroph\| | Yeast | NULL | Probable | As nectar-tap_saprotroph (Põlme et al. 2020); Decay sugar-rich substrates (Põlme et al. 2020) | Põlme S, et al. 2020. Fungal Diversity 105:1-16 ((https://doi.org/10.1007/s13225-020-00466-2)) |
| Otu46319 | 1 | k__Fungi(100);p__Ascomycota(100);c__Saccharomycetes(100);o__Saccharomycetales(100);f__Metschnikowiaceae(100);g__Metschnikowia(100);s__Metschnikowia_pulcherrima(100); | Metschnikowia | 13 | Saprotroph | \|Undefined Saprotroph\| | Yeast | NULL | Probable | As nectar-tap_saprotroph (Põlme et al. 2020); Decay sugar-rich substrates (Põlme et al. 2020) | Põlme S, et al. 2020. Fungal Diversity 105:1-16 ((https://doi.org/10.1007/s13225-020-00466-2)) |
| Otu46320 | 1 | k__Fungi(100);p__Ascomycota(100);c__Saccharomycetes(100);o__Saccharomycetales(100);f__Metschnikowiaceae(100);g__Metschnikowia(100);s__Metschnikowia_pulcherrima(100); | Metschnikowia | 13 | Saprotroph | \|Undefined Saprotroph\| | Yeast | NULL | Probable | As nectar-tap_saprotroph (Põlme et al. 2020); Decay sugar-rich substrates (Põlme et al. 2020) | Põlme S, et al. 2020. Fungal Diversity 105:1-16 ((https://doi.org/10.1007/s13225-020-00466-2)) |
| Otu46321 | 1 | k__Fungi(100);p__Ascomycota(100);c__Saccharomycetes(100);o__Saccharomycetales(100);f__Metschnikowiaceae(100);g__Metschnikowia(100);s__Metschnikowia_pulcherrima(100); | Metschnikowia | 13 | Saprotroph | \|Undefined Saprotroph\| | Yeast | NULL | Probable | As nectar-tap_saprotroph (Põlme et al. 2020); Decay sugar-rich substrates (Põlme et al. 2020) | Põlme S, et al. 2020. Fungal Diversity 105:1-16 ((https://doi.org/10.1007/s13225-020-00466-2)) |
| Otu46322 | 1 | k__Fungi(100);p__Ascomycota(100);c__Saccharomycetes(100);o__Saccharomycetales(100);f__Metschnikowiaceae(100);g__Metschnikowia(100);s__Metschnikowia_pulcherrima(100); | Metschnikowia | 13 | Saprotroph | \|Undefined Saprotroph\| | Yeast | NULL | Probable | As nectar-tap_saprotroph (Põlme et al. 2020); Decay sugar-rich substrates (Põlme et al. 2020) | Põlme S, et al. 2020. Fungal Diversity 105:1-16 ((https://doi.org/10.1007/s13225-020-00466-2)) |
| Otu46323 | 1 | k__Fungi(100);p__Ascomycota(100);c__Saccharomycetes(100);o__Saccharomycetales(100);f__Metschnikowiaceae(100);g__Metschnikowia(100);s__Metschnikowia_pulcherrima(100); | Metschnikowia | 13 | Saprotroph | \|Undefined Saprotroph\| | Yeast | NULL | Probable | As nectar-tap_saprotroph (Põlme et al. 2020); Decay sugar-rich substrates (Põlme et al. 2020) | Põlme S, et al. 2020. Fungal Diversity 105:1-16 ((https://doi.org/10.1007/s13225-020-00466-2)) |
| Otu46324 | 1 | k__Fungi(100);p__Ascomycota(100);c__Saccharomycetes(100);o__Saccharomycetales(100);f__Metschnikowiaceae(100);g__Metschnikowia(100);g__Metschnikowia_unclassified(100); | Metschnikowia | 13 | Saprotroph | \|Undefined Saprotroph\| | Yeast | NULL | Probable | As nectar-tap_saprotroph (Põlme et al. 2020); Decay sugar-rich substrates (Põlme et al. 2020) | Põlme S, et al. 2020. Fungal Diversity 105:1-16 ((https://doi.org/10.1007/s13225-020-00466-2)) |
| Otu46325 | 1 | k__Fungi(100);p__Ascomycota(100);c__Saccharomycetes(100);o__Saccharomycetales(100);f__Metschnikowiaceae(100);g__Metschnikowia(100);s__Metschnikowia_pulcherrima(100); | Metschnikowia | 13 | Saprotroph | \|Undefined Saprotroph\| | Yeast | NULL | Probable | As nectar-tap_saprotroph (Põlme et al. 2020); Decay sugar-rich substrates (Põlme et al. 2020) | Põlme S, et al. 2020. Fungal Diversity 105:1-16 ((https://doi.org/10.1007/s13225-020-00466-2)) |
| Otu46326 | 1 | k__Fungi(100);p__Ascomycota(100);c__Saccharomycetes(100);o__Saccharomycetales(100);f__Metschnikowiaceae(100);g__Metschnikowia(100);s__Metschnikowia_pulcherrima(100); | Metschnikowia | 13 | Saprotroph | \|Undefined Saprotroph\| | Yeast | NULL | Probable | As nectar-tap_saprotroph (Põlme et al. 2020); Decay sugar-rich substrates (Põlme et al. 2020) | Põlme S, et al. 2020. Fungal Diversity 105:1-16 ((https://doi.org/10.1007/s13225-020-00466-2)) |
| Otu46327 | 1 | k__Fungi(100);p__Ascomycota(100);c__Saccharomycetes(100);o__Saccharomycetales(100);f__Metschnikowiaceae(100);g__Metschnikowia(100);s__Metschnikowia_pulcherrima(100); | Metschnikowia | 13 | Saprotroph | \|Undefined Saprotroph\| | Yeast | NULL | Probable | As nectar-tap_saprotroph (Põlme et al. 2020); Decay sugar-rich substrates (Põlme et al. 2020) | Põlme S, et al. 2020. Fungal Diversity 105:1-16 ((https://doi.org/10.1007/s13225-020-00466-2)) |
| Otu46328 | 1 | k__Fungi(100);p__Ascomycota(100);c__Saccharomycetes(100);o__Saccharomycetales(100);f__Metschnikowiaceae(100);g__Metschnikowia(100);s__Metschnikowia_pulcherrima(100); | Metschnikowia | 13 | Saprotroph | \|Undefined Saprotroph\| | Yeast | NULL | Probable | As nectar-tap_saprotroph (Põlme et al. 2020); Decay sugar-rich substrates (Põlme et al. 2020) | Põlme S, et al. 2020. Fungal Diversity 105:1-16 ((https://doi.org/10.1007/s13225-020-00466-2)) |
| Otu46329 | 1 | k__Fungi(100);p__Ascomycota(100);c__Saccharomycetes(100);o__Saccharomycetales(100);f__Metschnikowiaceae(100);g__Metschnikowia(100);g__Metschnikowia_unclassified(100); | Metschnikowia | 13 | Saprotroph | \|Undefined Saprotroph\| | Yeast | NULL | Probable | As nectar-tap_saprotroph (Põlme et al. 2020); Decay sugar-rich substrates (Põlme et al. 2020) | Põlme S, et al. 2020. Fungal Diversity 105:1-16 ((https://doi.org/10.1007/s13225-020-00466-2)) |
| Otu46330 | 1 | k__Fungi(100);p__Ascomycota(100);c__Saccharomycetes(100);o__Saccharomycetales(100);f__Metschnikowiaceae(100);g__Metschnikowia(100);s__Metschnikowia_pulcherrima(100); | Metschnikowia | 13 | Saprotroph | \|Undefined Saprotroph\| | Yeast | NULL | Probable | As nectar-tap_saprotroph (Põlme et al. 2020); Decay sugar-rich substrates (Põlme et al. 2020) | Põlme S, et al. 2020. Fungal Diversity 105:1-16 ((https://doi.org/10.1007/s13225-020-00466-2)) |
| Otu46331 | 1 | k__Fungi(100);p__Ascomycota(100);c__Saccharomycetes(100);o__Saccharomycetales(100);f__Metschnikowiaceae(100);g__Metschnikowia(100);s__Metschnikowia_pulcherrima(100); | Metschnikowia | 13 | Saprotroph | \|Undefined Saprotroph\| | Yeast | NULL | Probable | As nectar-tap_saprotroph (Põlme et al. 2020); Decay sugar-rich substrates (Põlme et al. 2020) | Põlme S, et al. 2020. Fungal Diversity 105:1-16 ((https://doi.org/10.1007/s13225-020-00466-2)) |
| Otu46332 | 1 | k__Fungi(100);p__Ascomycota(100);c__Saccharomycetes(100);o__Saccharomycetales(100);f__Metschnikowiaceae(100);g__Metschnikowia(100);s__Metschnikowia_pulcherrima(100); | Metschnikowia | 13 | Saprotroph | \|Undefined Saprotroph\| | Yeast | NULL | Probable | As nectar-tap_saprotroph (Põlme et al. 2020); Decay sugar-rich substrates (Põlme et al. 2020) | Põlme S, et al. 2020. Fungal Diversity 105:1-16 ((https://doi.org/10.1007/s13225-020-00466-2)) |
| Otu46333 | 1 | k__Fungi(100);p__Ascomycota(100);c__Saccharomycetes(100);o__Saccharomycetales(100);f__Metschnikowiaceae(100);g__Metschnikowia(100);s__Metschnikowia_pulcherrima(100); | Metschnikowia | 13 | Saprotroph | \|Undefined Saprotroph\| | Yeast | NULL | Probable | As nectar-tap_saprotroph (Põlme et al. 2020); Decay sugar-rich substrates (Põlme et al. 2020) | Põlme S, et al. 2020. Fungal Diversity 105:1-16 ((https://doi.org/10.1007/s13225-020-00466-2)) |
| Otu46334 | 1 | k__Fungi(100);p__Ascomycota(100);c__Saccharomycetes(100);o__Saccharomycetales(100);f__Metschnikowiaceae(100);g__Metschnikowia(100);s__Metschnikowia_pulcherrima(100); | Metschnikowia | 13 | Saprotroph | \|Undefined Saprotroph\| | Yeast | NULL | Probable | As nectar-tap_saprotroph (Põlme et al. 2020); Decay sugar-rich substrates (Põlme et al. 2020) | Põlme S, et al. 2020. Fungal Diversity 105:1-16 ((https://doi.org/10.1007/s13225-020-00466-2)) |
| Otu46335 | 1 | k__Fungi(100);p__Ascomycota(100);c__Saccharomycetes(100);o__Saccharomycetales(100);f__Metschnikowiaceae(100);g__Metschnikowia(100);s__Metschnikowia_pulcherrima(100); | Metschnikowia | 13 | Saprotroph | \|Undefined Saprotroph\| | Yeast | NULL | Probable | As nectar-tap_saprotroph (Põlme et al. 2020); Decay sugar-rich substrates (Põlme et al. 2020) | Põlme S, et al. 2020. Fungal Diversity 105:1-16 ((https://doi.org/10.1007/s13225-020-00466-2)) |
| Otu46336 | 1 | k__Fungi(100);p__Ascomycota(100);c__Saccharomycetes(100);o__Saccharomycetales(100);f__Metschnikowiaceae(100);g__Metschnikowia(100);s__Metschnikowia_pulcherrima(100); | Metschnikowia | 13 | Saprotroph | \|Undefined Saprotroph\| | Yeast | NULL | Probable | As nectar-tap_saprotroph (Põlme et al. 2020); Decay sugar-rich substrates (Põlme et al. 2020) | Põlme S, et al. 2020. Fungal Diversity 105:1-16 ((https://doi.org/10.1007/s13225-020-00466-2)) |
| Otu46337 | 1 | k__Fungi(100);p__Ascomycota(100);c__Saccharomycetes(100);o__Saccharomycetales(100);f__Metschnikowiaceae(100);g__Metschnikowia(100);s__Metschnikowia_pulcherrima(100); | Metschnikowia | 13 | Saprotroph | \|Undefined Saprotroph\| | Yeast | NULL | Probable | As nectar-tap_saprotroph (Põlme et al. 2020); Decay sugar-rich substrates (Põlme et al. 2020) | Põlme S, et al. 2020. Fungal Diversity 105:1-16 ((https://doi.org/10.1007/s13225-020-00466-2)) |
| Otu46338 | 1 | k__Fungi(100);p__Ascomycota(100);c__Saccharomycetes(100);o__Saccharomycetales(100);f__Metschnikowiaceae(100);g__Metschnikowia(100);g__Metschnikowia_unclassified(100); | Metschnikowia | 13 | Saprotroph | \|Undefined Saprotroph\| | Yeast | NULL | Probable | As nectar-tap_saprotroph (Põlme et al. 2020); Decay sugar-rich substrates (Põlme et al. 2020) | Põlme S, et al. 2020. Fungal Diversity 105:1-16 ((https://doi.org/10.1007/s13225-020-00466-2)) |
| Otu46339 | 1 | k__Fungi(100);p__Ascomycota(100);c__Saccharomycetes(100);o__Saccharomycetales(100);f__Metschnikowiaceae(100);g__Metschnikowia(100);s__Metschnikowia_pulcherrima(100); | Metschnikowia | 13 | Saprotroph | \|Undefined Saprotroph\| | Yeast | NULL | Probable | As nectar-tap_saprotroph (Põlme et al. 2020); Decay sugar-rich substrates (Põlme et al. 2020) | Põlme S, et al. 2020. Fungal Diversity 105:1-16 ((https://doi.org/10.1007/s13225-020-00466-2)) |

| | | | | | | | | | | | |
|---|---|---|---|---|---|---|---|---|---|---|---|
| Otu46340 | 1 | k__Fungi(100);p__Ascomycota(100);c__Saccharomycetes(100);o__Saccharomycetales(100);f__Metschnikowiaceae(100);g__Metschnikowia(100);g__Metschnikowia_unclassified(100); | Metschnikowia | 13 | Saprotroph | \|Undefined Saprotroph\| | Yeast | NULL | Probable | As nectar-tap_saprotroph (Põlme et al. 2020); Decay sugar-rich substrates (Põlme et al. 2020) | Põlme S, et al. 2020. Fungal Diversity 105:1-16 ((https://doi.org/10.1007/s13225-020-00466-2)) |
| Otu46341 | 1 | k__Fungi(100);p__Ascomycota(100);c__Saccharomycetes(100);o__Saccharomycetales(100);f__Metschnikowiaceae(100);g__Metschnikowia(100);g__Metschnikowia_unclassified(100); | Metschnikowia | 13 | Saprotroph | \|Undefined Saprotroph\| | Yeast | NULL | Probable | As nectar-tap_saprotroph (Põlme et al. 2020); Decay sugar-rich substrates (Põlme et al. 2020) | Põlme S, et al. 2020. Fungal Diversity 105:1-16 ((https://doi.org/10.1007/s13225-020-00466-2)) |
| Otu46342 | 1 | k__Fungi(100);p__Ascomycota(100);c__Saccharomycetes(100);o__Saccharomycetales(100);f__Metschnikowiaceae(100);g__Metschnikowia(100);s__Metschnikowia_pulcherrima(100); | Metschnikowia | 13 | Saprotroph | \|Undefined Saprotroph\| | Yeast | NULL | Probable | As nectar-tap_saprotroph (Põlme et al. 2020); Decay sugar-rich substrates (Põlme et al. 2020) | Põlme S, et al. 2020. Fungal Diversity 105:1-16 ((https://doi.org/10.1007/s13225-020-00466-2)) |
| Otu46343 | 1 | k__Fungi(100);p__Ascomycota(100);c__Saccharomycetes(100);o__Saccharomycetales(100);f__Metschnikowiaceae(100);g__Metschnikowia(100);g__Metschnikowia_unclassified(100); | Metschnikowia | 13 | Saprotroph | \|Undefined Saprotroph\| | Yeast | NULL | Probable | As nectar-tap_saprotroph (Põlme et al. 2020); Decay sugar-rich substrates (Põlme et al. 2020) | Põlme S, et al. 2020. Fungal Diversity 105:1-16 ((https://doi.org/10.1007/s13225-020-00466-2)) |
| Otu46344 | 1 | k__Fungi(100);p__Ascomycota(100);c__Saccharomycetes(100);o__Saccharomycetales(100);f__Metschnikowiaceae(100);g__Metschnikowia(100);s__Metschnikowia_pulcherrima(100); | Metschnikowia | 13 | Saprotroph | \|Undefined Saprotroph\| | Yeast | NULL | Probable | As nectar-tap_saprotroph (Põlme et al. 2020); Decay sugar-rich substrates (Põlme et al. 2020) | Põlme S, et al. 2020. Fungal Diversity 105:1-16 ((https://doi.org/10.1007/s13225-020-00466-2)) |
| Otu46345 | 1 | k__Fungi(100);p__Ascomycota(100);c__Saccharomycetes(100);o__Saccharomycetales(100);f__Metschnikowiaceae(100);g__Metschnikowia(100);g__Metschnikowia_unclassified(100); | Metschnikowia | 13 | Saprotroph | \|Undefined Saprotroph\| | Yeast | NULL | Probable | As nectar-tap_saprotroph (Põlme et al. 2020); Decay sugar-rich substrates (Põlme et al. 2020) | Põlme S, et al. 2020. Fungal Diversity 105:1-16 ((https://doi.org/10.1007/s13225-020-00466-2)) |
| Otu46346 | 1 | k__Fungi(100);p__Ascomycota(100);c__Saccharomycetes(100);o__Saccharomycetales(100);f__Metschnikowiaceae(100);g__Metschnikowia(100);s__Metschnikowia_pulcherrima(100); | Metschnikowia | 13 | Saprotroph | \|Undefined Saprotroph\| | Yeast | NULL | Probable | As nectar-tap_saprotroph (Põlme et al. 2020); Decay sugar-rich substrates (Põlme et al. 2020) | Põlme S, et al. 2020. Fungal Diversity 105:1-16 ((https://doi.org/10.1007/s13225-020-00466-2)) |
| Otu46347 | 1 | k__Fungi(100);p__Ascomycota(100);c__Saccharomycetes(100);o__Saccharomycetales(100);f__Metschnikowiaceae(100);g__Metschnikowia(100);s__Metschnikowia_pulcherrima(100); | Metschnikowia | 13 | Saprotroph | \|Undefined Saprotroph\| | Yeast | NULL | Probable | As nectar-tap_saprotroph (Põlme et al. 2020); Decay sugar-rich substrates (Põlme et al. 2020) | Põlme S, et al. 2020. Fungal Diversity 105:1-16 ((https://doi.org/10.1007/s13225-020-00466-2)) |
| Otu46348 | 1 | k__Fungi(100);p__Ascomycota(100);c__Saccharomycetes(100);o__Saccharomycetales(100);f__Metschnikowiaceae(100);g__Metschnikowia(100);s__Metschnikowia_pulcherrima(100); | Metschnikowia | 13 | Saprotroph | \|Undefined Saprotroph\| | Yeast | NULL | Probable | As nectar-tap_saprotroph (Põlme et al. 2020); Decay sugar-rich substrates (Põlme et al. 2020) | Põlme S, et al. 2020. Fungal Diversity 105:1-16 ((https://doi.org/10.1007/s13225-020-00466-2)) |
| Otu46349 | 1 | k__Fungi(100);p__Ascomycota(100);c__Saccharomycetes(100);o__Saccharomycetales(100);f__Metschnikowiaceae(100);g__Metschnikowia(100);g__Metschnikowia_unclassified(100); | Metschnikowia | 13 | Saprotroph | \|Undefined Saprotroph\| | Yeast | NULL | Probable | As nectar-tap_saprotroph (Põlme et al. 2020); Decay sugar-rich substrates (Põlme et al. 2020) | Põlme S, et al. 2020. Fungal Diversity 105:1-16 ((https://doi.org/10.1007/s13225-020-00466-2)) |
| Otu46350 | 1 | k__Fungi(100);p__Ascomycota(100);c__Saccharomycetes(100);o__Saccharomycetales(100);f__Metschnikowiaceae(100);g__Metschnikowia(100);s__Metschnikowia_pulcherrima(100); | Metschnikowia | 13 | Saprotroph | \|Undefined Saprotroph\| | Yeast | NULL | Probable | As nectar-tap_saprotroph (Põlme et al. 2020); Decay sugar-rich substrates (Põlme et al. 2020) | Põlme S, et al. 2020. Fungal Diversity 105:1-16 ((https://doi.org/10.1007/s13225-020-00466-2)) |
| Otu46351 | 1 | k__Fungi(100);p__Ascomycota(100);c__Saccharomycetes(100);o__Saccharomycetales(100);f__Metschnikowiaceae(100);g__Metschnikowia(100);s__Metschnikowia_pulcherrima(100); | Metschnikowia | 13 | Saprotroph | \|Undefined Saprotroph\| | Yeast | NULL | Probable | As nectar-tap_saprotroph (Põlme et al. 2020); Decay sugar-rich substrates (Põlme et al. 2020) | Põlme S, et al. 2020. Fungal Diversity 105:1-16 ((https://doi.org/10.1007/s13225-020-00466-2)) |
| Otu46352 | 1 | k__Fungi(100);p__Ascomycota(100);c__Saccharomycetes(100);o__Saccharomycetales(100);f__Metschnikowiaceae(100);g__Metschnikowia(100);s__Metschnikowia_pulcherrima(100); | Metschnikowia | 13 | Saprotroph | \|Undefined Saprotroph\| | Yeast | NULL | Probable | As nectar-tap_saprotroph (Põlme et al. 2020); Decay sugar-rich substrates (Põlme et al. 2020) | Põlme S, et al. 2020. Fungal Diversity 105:1-16 ((https://doi.org/10.1007/s13225-020-00466-2)) |
| Otu46353 | 1 | k__Fungi(100);p__Ascomycota(100);c__Saccharomycetes(100);o__Saccharomycetales(100);f__Metschnikowiaceae(100);g__Metschnikowia(100);g__Metschnikowia_unclassified(100); | Metschnikowia | 13 | Saprotroph | \|Undefined Saprotroph\| | Yeast | NULL | Probable | As nectar-tap_saprotroph (Põlme et al. 2020); Decay sugar-rich substrates (Põlme et al. 2020) | Põlme S, et al. 2020. Fungal Diversity 105:1-16 ((https://doi.org/10.1007/s13225-020-00466-2)) |
| Otu46354 | 1 | k__Fungi(100);p__Ascomycota(100);c__Saccharomycetes(100);o__Saccharomycetales(100);f__Metschnikowiaceae(100);g__Metschnikowia(100);g__Metschnikowia_unclassified(100); | Metschnikowia | 13 | Saprotroph | \|Undefined Saprotroph\| | Yeast | NULL | Probable | As nectar-tap_saprotroph (Põlme et al. 2020); Decay sugar-rich substrates (Põlme et al. 2020) | Põlme S, et al. 2020. Fungal Diversity 105:1-16 ((https://doi.org/10.1007/s13225-020-00466-2)) |
| Otu46355 | 1 | k__Fungi(100);p__Ascomycota(100);c__Saccharomycetes(100);o__Saccharomycetales(100);f__Metschnikowiaceae(100);g__Metschnikowia(100);s__Metschnikowia_pulcherrima(100); | Metschnikowia | 13 | Saprotroph | \|Undefined Saprotroph\| | Yeast | NULL | Probable | As nectar-tap_saprotroph (Põlme et al. 2020); Decay sugar-rich substrates (Põlme et al. 2020) | Põlme S, et al. 2020. Fungal Diversity 105:1-16 ((https://doi.org/10.1007/s13225-020-00466-2)) |
| Otu46356 | 1 | k__Fungi(100);p__Ascomycota(100);c__Saccharomycetes(100);o__Saccharomycetales(100);f__Metschnikowiaceae(100);g__Metschnikowia(100);s__Metschnikowia_pulcherrima(100); | Metschnikowia | 13 | Saprotroph | \|Undefined Saprotroph\| | Yeast | NULL | Probable | As nectar-tap_saprotroph (Põlme et al. 2020); Decay sugar-rich substrates (Põlme et al. 2020) | Põlme S, et al. 2020. Fungal Diversity 105:1-16 ((https://doi.org/10.1007/s13225-020-00466-2)) |
| Otu46357 | 1 | k__Fungi(100);p__Ascomycota(100);c__Saccharomycetes(100);o__Saccharomycetales(100);f__Metschnikowiaceae(100);g__Metschnikowia(100);s__Metschnikowia_pulcherrima(100); | Metschnikowia | 13 | Saprotroph | \|Undefined Saprotroph\| | Yeast | NULL | Probable | As nectar-tap_saprotroph (Põlme et al. 2020); Decay sugar-rich substrates (Põlme et al. 2020) | Põlme S, et al. 2020. Fungal Diversity 105:1-16 ((https://doi.org/10.1007/s13225-020-00466-2)) |
| Otu46358 | 1 | k__Fungi(100);p__Ascomycota(100);c__Saccharomycetes(100);o__Saccharomycetales(100);f__Metschnikowiaceae(100);g__Metschnikowia(100);s__Metschnikowia_pulcherrima(100); | Metschnikowia | 13 | Saprotroph | \|Undefined Saprotroph\| | Yeast | NULL | Probable | As nectar-tap_saprotroph (Põlme et al. 2020); Decay sugar-rich substrates (Põlme et al. 2020) | Põlme S, et al. 2020. Fungal Diversity 105:1-16 ((https://doi.org/10.1007/s13225-020-00466-2)) |
| Otu46359 | 1 | k__Fungi(100);p__Ascomycota(100);c__Saccharomycetes(100);o__Saccharomycetales(100);f__Metschnikowiaceae(100);g__Metschnikowia(100);s__Metschnikowia_pulcherrima(100); | Metschnikowia | 13 | Saprotroph | \|Undefined Saprotroph\| | Yeast | NULL | Probable | As nectar-tap_saprotroph (Põlme et al. 2020); Decay sugar-rich substrates (Põlme et al. 2020) | Põlme S, et al. 2020. Fungal Diversity 105:1-16 ((https://doi.org/10.1007/s13225-020-00466-2)) |
| Otu46360 | 1 | k__Fungi(100);p__Ascomycota(100);c__Saccharomycetes(100);o__Saccharomycetales(100);f__Metschnikowiaceae(100);g__Metschnikowia(100);s__Metschnikowia_pulcherrima(100); | Metschnikowia | 13 | Saprotroph | \|Undefined Saprotroph\| | Yeast | NULL | Probable | As nectar-tap_saprotroph (Põlme et al. 2020); Decay sugar-rich substrates (Põlme et al. 2020) | Põlme S, et al. 2020. Fungal Diversity 105:1-16 ((https://doi.org/10.1007/s13225-020-00466-2)) |
| Otu46361 | 1 | k__Fungi(100);p__Ascomycota(100);c__Saccharomycetes(100);o__Saccharomycetales(100);f__Metschnikowiaceae(100);g__Metschnikowia(100);g__Metschnikowia_unclassified(100); | Metschnikowia | 13 | Saprotroph | \|Undefined Saprotroph\| | Yeast | NULL | Probable | As nectar-tap_saprotroph (Põlme et al. 2020); Decay sugar-rich substrates (Põlme et al. 2020) | Põlme S, et al. 2020. Fungal Diversity 105:1-16 ((https://doi.org/10.1007/s13225-020-00466-2)) |
| Otu46362 | 1 | k__Fungi(100);p__Ascomycota(100);c__Saccharomycetes(100);o__Saccharomycetales(100);f__Metschnikowiaceae(100);g__Metschnikowia(100);s__Metschnikowia_pulcherrima(100); | Metschnikowia | 13 | Saprotroph | \|Undefined Saprotroph\| | Yeast | NULL | Probable | As nectar-tap_saprotroph (Põlme et al. 2020); Decay sugar-rich substrates (Põlme et al. 2020) | Põlme S, et al. 2020. Fungal Diversity 105:1-16 ((https://doi.org/10.1007/s13225-020-00466-2)) |
| Otu46363 | 1 | k__Fungi(100);p__Ascomycota(100);c__Saccharomycetes(100);o__Saccharomycetales(100);f__Metschnikowiaceae(100);g__Metschnikowia(100);s__Metschnikowia_pulcherrima(100); | Metschnikowia | 13 | Saprotroph | \|Undefined Saprotroph\| | Yeast | NULL | Probable | As nectar-tap_saprotroph (Põlme et al. 2020); Decay sugar-rich substrates (Põlme et al. 2020) | Põlme S, et al. 2020. Fungal Diversity 105:1-16 ((https://doi.org/10.1007/s13225-020-00466-2)) |
| Otu46364 | 1 | k__Fungi(100);p__Ascomycota(100);c__Saccharomycetes(100);o__Saccharomycetales(100);f__Metschnikowiaceae(100);g__Metschnikowia(100);s__Metschnikowia_pulcherrima(100); | Metschnikowia | 13 | Saprotroph | \|Undefined Saprotroph\| | Yeast | NULL | Probable | As nectar-tap_saprotroph (Põlme et al. 2020); Decay sugar-rich substrates (Põlme et al. 2020) | Põlme S, et al. 2020. Fungal Diversity 105:1-16 ((https://doi.org/10.1007/s13225-020-00466-2)) |
| Otu46365 | 1 | k__Fungi(100);p__Ascomycota(100);c__Saccharomycetes(100);o__Saccharomycetales(100);f__Metschnikowiaceae(100);g__Metschnikowia(100);g__Metschnikowia_unclassified(100); | Metschnikowia | 13 | Saprotroph | \|Undefined Saprotroph\| | Yeast | NULL | Probable | As nectar-tap_saprotroph (Põlme et al. 2020); Decay sugar-rich substrates (Põlme et al. 2020) | Põlme S, et al. 2020. Fungal Diversity 105:1-16 ((https://doi.org/10.1007/s13225-020-00466-2)) |
| Otu46366 | 1 | k__Fungi(100);p__Ascomycota(100);c__Saccharomycetes(100);o__Saccharomycetales(100);f__Metschnikowiaceae(100);g__Metschnikowia(100);g__Metschnikowia_unclassified(100); | Metschnikowia | 13 | Saprotroph | \|Undefined Saprotroph\| | Yeast | NULL | Probable | As nectar-tap_saprotroph (Põlme et al. 2020); Decay sugar-rich substrates (Põlme et al. 2020) | Põlme S, et al. 2020. Fungal Diversity 105:1-16 ((https://doi.org/10.1007/s13225-020-00466-2)) |
| Otu46367 | 1 | k__Fungi(100);p__Ascomycota(100);c__Saccharomycetes(100);o__Saccharomycetales(100);f__Metschnikowiaceae(100);g__Metschnikowia(100);s__Metschnikowia_pulcherrima(100); | Metschnikowia | 13 | Saprotroph | \|Undefined Saprotroph\| | Yeast | NULL | Probable | As nectar-tap_saprotroph (Põlme et al. 2020); Decay sugar-rich substrates (Põlme et al. 2020) | Põlme S, et al. 2020. Fungal Diversity 105:1-16 ((https://doi.org/10.1007/s13225-020-00466-2)) |
| Otu46368 | 1 | k__Fungi(100);p__Ascomycota(100);c__Saccharomycetes(100);o__Saccharomycetales(100);f__Metschnikowiaceae(100);g__Metschnikowia(100);s__Metschnikowia_pulcherrima(100); | Metschnikowia | 13 | Saprotroph | \|Undefined Saprotroph\| | Yeast | NULL | Probable | As nectar-tap_saprotroph (Põlme et al. 2020); Decay sugar-rich substrates (Põlme et al. 2020) | Põlme S, et al. 2020. Fungal Diversity 105:1-16 ((https://doi.org/10.1007/s13225-020-00466-2)) |
| Otu46369 | 1 | k__Fungi(100);p__Ascomycota(100);c__Saccharomycetes(100);o__Saccharomycetales(100);f__Metschnikowiaceae(100);g__Metschnikowia(100);s__Metschnikowia_pulcherrima(100); | Metschnikowia | 13 | Saprotroph | \|Undefined Saprotroph\| | Yeast | NULL | Probable | As nectar-tap_saprotroph (Põlme et al. 2020); Decay sugar-rich substrates (Põlme et al. 2020) | Põlme S, et al. 2020. Fungal Diversity 105:1-16 ((https://doi.org/10.1007/s13225-020-00466-2)) |
| Otu46370 | 1 | k__Fungi(100);p__Ascomycota(100);c__Saccharomycetes(100);o__Saccharomycetales(100);f__Metschnikowiaceae(100);g__Metschnikowia(100);s__Metschnikowia_pulcherrima(100); | Metschnikowia | 13 | Saprotroph | \|Undefined Saprotroph\| | Yeast | NULL | Probable | As nectar-tap_saprotroph (Põlme et al. 2020); Decay sugar-rich substrates (Põlme et al. 2020) | Põlme S, et al. 2020. Fungal Diversity 105:1-16 ((https://doi.org/10.1007/s13225-020-00466-2)) |
| Otu46372 | 1 | k__Fungi(100);p__Ascomycota(100);c__Saccharomycetes(100);o__Saccharomycetales(100);f__Metschnikowiaceae(100);g__Metschnikowia(100);s__Metschnikowia_pulcherrima(100); | Metschnikowia | 13 | Saprotroph | \|Undefined Saprotroph\| | Yeast | NULL | Probable | As nectar-tap_saprotroph (Põlme et al. 2020); Decay sugar-rich substrates (Põlme et al. 2020) | Põlme S, et al. 2020. Fungal Diversity 105:1-16 ((https://doi.org/10.1007/s13225-020-00466-2)) |
| Otu46373 | 1 | k__Fungi(100);p__Ascomycota(100);c__Saccharomycetes(100);o__Saccharomycetales(100);f__Metschnikowiaceae(100);g__Metschnikowia(100);s__Metschnikowia_pulcherrima(100); | Metschnikowia | 13 | Saprotroph | \|Undefined Saprotroph\| | Yeast | NULL | Probable | As nectar-tap_saprotroph (Põlme et al. 2020); Decay sugar-rich substrates (Põlme et al. 2020) | Põlme S, et al. 2020. Fungal Diversity 105:1-16 ((https://doi.org/10.1007/s13225-020-00466-2)) |
| Otu46374 | 1 | k__Fungi(100);p__Ascomycota(100);c__Saccharomycetes(100);o__Saccharomycetales(100);f__Metschnikowiaceae(100);g__Metschnikowia(100);s__Metschnikowia_pulcherrima(100); | Metschnikowia | 13 | Saprotroph | \|Undefined Saprotroph\| | Yeast | NULL | Probable | As nectar-tap_saprotroph (Põlme et al. 2020); Decay sugar-rich substrates (Põlme et al. 2020) | Põlme S, et al. 2020. Fungal Diversity 105:1-16 ((https://doi.org/10.1007/s13225-020-00466-2)) |
| Otu46375 | 1 | k__Fungi(100);p__Ascomycota(100);c__Saccharomycetes(100);o__Saccharomycetales(100);f__Metschnikowiaceae(100);g__Metschnikowia(100);s__Metschnikowia_pulcherrima(100); | Metschnikowia | 13 | Saprotroph | \|Undefined Saprotroph\| | Yeast | NULL | Probable | As nectar-tap_saprotroph (Põlme et al. 2020); Decay sugar-rich substrates (Põlme et al. 2020) | Põlme S, et al. 2020. Fungal Diversity 105:1-16 ((https://doi.org/10.1007/s13225-020-00466-2)) |
| Otu46376 | 1 | k__Fungi(100);p__Ascomycota(100);c__Saccharomycetes(100);o__Saccharomycetales(100);f__Metschnikowiaceae(100);g__Metschnikowia(100);s__Metschnikowia_pulcherrima(100); | Metschnikowia | 13 | Saprotroph | \|Undefined Saprotroph\| | Yeast | NULL | Probable | As nectar-tap_saprotroph (Põlme et al. 2020); Decay sugar-rich substrates (Põlme et al. 2020) | Põlme S, et al. 2020. Fungal Diversity 105:1-16 ((https://doi.org/10.1007/s13225-020-00466-2)) |
| Otu46377 | 1 | k__Fungi(100);p__Ascomycota(100);c__Saccharomycetes(100);o__Saccharomycetales(100);f__Metschnikowiaceae(100);g__Metschnikowia(100);s__Metschnikowia_pulcherrima(100); | Metschnikowia | 13 | Saprotroph | \|Undefined Saprotroph\| | Yeast | NULL | Probable | As nectar-tap_saprotroph (Põlme et al. 2020); Decay sugar-rich substrates (Põlme et al. 2020) | Põlme S, et al. 2020. Fungal Diversity 105:1-16 ((https://doi.org/10.1007/s13225-020-00466-2)) |
| Otu46378 | 1 | k__Fungi(100);p__Ascomycota(100);c__Saccharomycetes(100);o__Saccharomycetales(100);f__Metschnikowiaceae(100);g__Metschnikowia(100);s__Metschnikowia_pulcherrima(100); | Metschnikowia | 13 | Saprotroph | \|Undefined Saprotroph\| | Yeast | NULL | Probable | As nectar-tap_saprotroph (Põlme et al. 2020); Decay sugar-rich substrates (Põlme et al. 2020) | Põlme S, et al. 2020. Fungal Diversity 105:1-16 ((https://doi.org/10.1007/s13225-020-00466-2)) |
| Otu46379 | 1 | k__Fungi(100);p__Ascomycota(100);c__Saccharomycetes(100);o__Saccharomycetales(100);f__Metschnikowiaceae(100);g__Metschnikowia(100);s__Metschnikowia_pulcherrima(100); | Metschnikowia | 13 | Saprotroph | \|Undefined Saprotroph\| | Yeast | NULL | Probable | As nectar-tap_saprotroph (Põlme et al. 2020); Decay sugar-rich substrates (Põlme et al. 2020) | Põlme S, et al. 2020. Fungal Diversity 105:1-16 ((https://doi.org/10.1007/s13225-020-00466-2)) |
| Otu46380 | 1 | k__Fungi(100);p__Ascomycota(100);c__Saccharomycetes(100);o__Saccharomycetales(100);f__Metschnikowiaceae(100);g__Metschnikowia(100);s__Metschnikowia_pulcherrima(100); | Metschnikowia | 13 | Saprotroph | \|Undefined Saprotroph\| | Yeast | NULL | Probable | As nectar-tap_saprotroph (Põlme et al. 2020); Decay sugar-rich substrates (Põlme et al. 2020) | Põlme S, et al. 2020. Fungal Diversity 105:1-16 ((https://doi.org/10.1007/s13225-020-00466-2)) |
| Otu46381 | 1 | k__Fungi(100);p__Ascomycota(100);c__Saccharomycetes(100);o__Saccharomycetales(100);f__Metschnikowiaceae(100);g__Metschnikowia(100);g__Metschnikowia_unclassified(100); | Metschnikowia | 13 | Saprotroph | \|Undefined Saprotroph\| | Yeast | NULL | Probable | As nectar-tap_saprotroph (Põlme et al. 2020); Decay sugar-rich substrates (Põlme et al. 2020) | Põlme S, et al. 2020. Fungal Diversity 105:1-16 ((https://doi.org/10.1007/s13225-020-00466-2)) |
| Otu46382 | 1 | k__Fungi(100);p__Ascomycota(100);c__Saccharomycetes(100);o__Saccharomycetales(100);f__Metschnikowiaceae(100);g__Metschnikowia(100);g__Metschnikowia_unclassified(100); | Metschnikowia | 13 | Saprotroph | \|Undefined Saprotroph\| | Yeast | NULL | Probable | As nectar-tap_saprotroph (Põlme et al. 2020); Decay sugar-rich substrates (Põlme et al. 2020) | Põlme S, et al. 2020. Fungal Diversity 105:1-16 ((https://doi.org/10.1007/s13225-020-00466-2)) |
| Otu46383 | 1 | k__Fungi(100);p__Ascomycota(100);c__Saccharomycetes(100);o__Saccharomycetales(100);f__Metschnikowiaceae(100);g__Metschnikowia(100);s__Metschnikowia_pulcherrima(100); | Metschnikowia | 13 | Saprotroph | \|Undefined Saprotroph\| | Yeast | NULL | Probable | As nectar-tap_saprotroph (Põlme et al. 2020); Decay sugar-rich substrates (Põlme et al. 2020) | Põlme S, et al. 2020. Fungal Diversity 105:1-16 ((https://doi.org/10.1007/s13225-020-00466-2)) |
| Otu46384 | 1 | k__Fungi(100);p__Ascomycota(100);c__Saccharomycetes(100);o__Saccharomycetales(100);f__Metschnikowiaceae(100);g__Metschnikowia(100);g__Metschnikowia_unclassified(100); | Metschnikowia | 13 | Saprotroph | \|Undefined Saprotroph\| | Yeast | NULL | Probable | As nectar-tap_saprotroph (Põlme et al. 2020); Decay sugar-rich substrates (Põlme et al. 2020) | Põlme S, et al. 2020. Fungal Diversity 105:1-16 ((https://doi.org/10.1007/s13225-020-00466-2)) |
| Otu46385 | 1 | k__Fungi(100);p__Ascomycota(100);c__Saccharomycetes(100);o__Saccharomycetales(100);f__Metschnikowiaceae(100);g__Metschnikowia(100);g__Metschnikowia_unclassified(100); | Metschnikowia | 13 | Saprotroph | \|Undefined Saprotroph\| | Yeast | NULL | Probable | As nectar-tap_saprotroph (Põlme et al. 2020); Decay sugar-rich substrates (Põlme et al. 2020) | Põlme S, et al. 2020. Fungal Diversity 105:1-16 ((https://doi.org/10.1007/s13225-020-00466-2)) |
| Otu46386 | 1 | k__Fungi(100);p__Ascomycota(100);c__Saccharomycetes(100);o__Saccharomycetales(100);f__Metschnikowiaceae(100);g__Metschnikowia(100);s__Metschnikowia_pulcherrima(100); | Metschnikowia | 13 | Saprotroph | \|Undefined Saprotroph\| | Yeast | NULL | Probable | As nectar-tap_saprotroph (Põlme et al. 2020); Decay sugar-rich substrates (Põlme et al. 2020) | Põlme S, et al. 2020. Fungal Diversity 105:1-16 ((https://doi.org/10.1007/s13225-020-00466-2)) |
| Otu46387 | 1 | k__Fungi(100);p__Ascomycota(100);c__Saccharomycetes(100);o__Saccharomycetales(100);f__Metschnikowiaceae(100);g__Metschnikowia(100);s__Metschnikowia_pulcherrima(100); | Metschnikowia | 13 | Saprotroph | \|Undefined Saprotroph\| | Yeast | NULL | Probable | As nectar-tap_saprotroph (Põlme et al. 2020); Decay sugar-rich substrates (Põlme et al. 2020) | Põlme S, et al. 2020. Fungal Diversity 105:1-16 ((https://doi.org/10.1007/s13225-020-00466-2)) |
| Otu46388 | 1 | k__Fungi(100);p__Ascomycota(100);c__Saccharomycetes(100);o__Saccharomycetales(100);f__Metschnikowiaceae(100);g__Metschnikowia(100);s__Metschnikowia_pulcherrima(100); | Metschnikowia | 13 | Saprotroph | \|Undefined Saprotroph\| | Yeast | NULL | Probable | As nectar-tap_saprotroph (Põlme et al. 2020); Decay sugar-rich substrates (Põlme et al. 2020) | Põlme S, et al. 2020. Fungal Diversity 105:1-16 ((https://doi.org/10.1007/s13225-020-00466-2)) |
| Otu46389 | 1 | k__Fungi(100);p__Ascomycota(100);c__Saccharomycetes(100);o__Saccharomycetales(100);f__Metschnikowiaceae(100);g__Metschnikowia(100);s__Metschnikowia_pulcherrima(100); | Metschnikowia | 13 | Saprotroph | \|Undefined Saprotroph\| | Yeast | NULL | Probable | As nectar-tap_saprotroph (Põlme et al. 2020); Decay sugar-rich substrates (Põlme et al. 2020) | Põlme S, et al. 2020. Fungal Diversity 105:1-16 ((https://doi.org/10.1007/s13225-020-00466-2)) |
| Otu46390 | 1 | k__Fungi(100);p__Ascomycota(100);c__Saccharomycetes(100);o__Saccharomycetales(100);f__Metschnikowiaceae(100);g__Metschnikowia(100);s__Metschnikowia_pulcherrima(100); | Metschnikowia | 13 | Saprotroph | \|Undefined Saprotroph\| | Yeast | NULL | Probable | As nectar-tap_saprotroph (Põlme et al. 2020); Decay sugar-rich substrates (Põlme et al. 2020) | Põlme S, et al. 2020. Fungal Diversity 105:1-16 ((https://doi.org/10.1007/s13225-020-00466-2)) |
| Otu46391 | 1 | k__Fungi(100);p__Ascomycota(100);c__Saccharomycetes(100);o__Saccharomycetales(100);f__Metschnikowiaceae(100);g__Metschnikowia(100);s__Metschnikowia_pulcherrima(100); | Metschnikowia | 13 | Saprotroph | \|Undefined Saprotroph\| | Yeast | NULL | Probable | As nectar-tap_saprotroph (Põlme et al. 2020); Decay sugar-rich substrates (Põlme et al. 2020) | Põlme S, et al. 2020. Fungal Diversity 105:1-16 ((https://doi.org/10.1007/s13225-020-00466-2)) |
| Otu46392 | 1 | k__Fungi(100);p__Ascomycota(100);c__Saccharomycetes(100);o__Saccharomycetales(100);f__Metschnikowiaceae(100);g__Metschnikowia(100);s__Metschnikowia_pulcherrima(100); | Metschnikowia | 13 | Saprotroph | \|Undefined Saprotroph\| | Yeast | NULL | Probable | As nectar-tap_saprotroph (Põlme et al. 2020); Decay sugar-rich substrates (Põlme et al. 2020) | Põlme S, et al. 2020. Fungal Diversity 105:1-16 ((https://doi.org/10.1007/s13225-020-00466-2)) |
| Otu46393 | 1 | k__Fungi(100);p__Ascomycota(100);c__Saccharomycetes(100);o__Saccharomycetales(100);f__Metschnikowiaceae(100);g__Metschnikowia(100);g__Metschnikowia_unclassified(100); | Metschnikowia | 13 | Saprotroph | \|Undefined Saprotroph\| | Yeast | NULL | Probable | As nectar-tap_saprotroph (Põlme et al. 2020); Decay sugar-rich substrates (Põlme et al. 2020) | Põlme S, et al. 2020. Fungal Diversity 105:1-16 ((https://doi.org/10.1007/s13225-020-00466-2)) |
| Otu46395 | 1 | k__Fungi(100);p__Ascomycota(100);c__Saccharomycetes(100);o__Saccharomycetales(100);f__Metschnikowiaceae(100);g__Metschnikowia(100);s__Metschnikowia_pulcherrima(100); | Metschnikowia | 13 | Saprotroph | \|Undefined Saprotroph\| | Yeast | NULL | Probable | As nectar-tap_saprotroph (Põlme et al. 2020); Decay sugar-rich substrates (Põlme et al. 2020) | Põlme S, et al. 2020. Fungal Diversity 105:1-16 ((https://doi.org/10.1007/s13225-020-00466-2)) |
| Otu46396 | 1 | k__Fungi(100);p__Ascomycota(100);c__Saccharomycetes(100);o__Saccharomycetales(100);f__Metschnikowiaceae(100);g__Metschnikowia(100);s__Metschnikowia_pulcherrima(100); | Metschnikowia | 13 | Saprotroph | \|Undefined Saprotroph\| | Yeast | NULL | Probable | As nectar-tap_saprotroph (Põlme et al. 2020); Decay sugar-rich substrates (Põlme et al. 2020) | Põlme S, et al. 2020. Fungal Diversity 105:1-16 ((https://doi.org/10.1007/s13225-020-00466-2)) |
| Otu46397 | 1 | k__Fungi(100);p__Ascomycota(100);c__Saccharomycetes(100);o__Saccharomycetales(100);f__Metschnikowiaceae(100);g__Metschnikowia(100);g__Metschnikowia_unclassified(100); | Metschnikowia | 13 | Saprotroph | \|Undefined Saprotroph\| | Yeast | NULL | Probable | As nectar-tap_saprotroph (Põlme et al. 2020); Decay sugar-rich substrates (Põlme et al. 2020) | Põlme S, et al. 2020. Fungal Diversity 105:1-16 ((https://doi.org/10.1007/s13225-020-00466-2)) |
| Otu46398 | 1 | k__Fungi(100);p__Ascomycota(100);c__Saccharomycetes(100);o__Saccharomycetales(100);f__Metschnikowiaceae(100);g__Metschnikowia(100);g__Metschnikowia_unclassified(100); | Metschnikowia | 13 | Saprotroph | \|Undefined Saprotroph\| | Yeast | NULL | Probable | As nectar-tap_saprotroph (Põlme et al. 2020); Decay sugar-rich substrates (Põlme et al. 2020) | Põlme S, et al. 2020. Fungal Diversity 105:1-16 ((https://doi.org/10.1007/s13225-020-00466-2)) |
| Otu46399 | 1 | k__Fungi(100);p__Ascomycota(100);c__Saccharomycetes(100);o__Saccharomycetales(100);f__Metschnikowiaceae(100);g__Metschnikowia(100);g__Metschnikowia_unclassified(100); | Metschnikowia | 13 | Saprotroph | \|Undefined Saprotroph\| | Yeast | NULL | Probable | As nectar-tap_saprotroph (Põlme et al. 2020); Decay sugar-rich substrates (Põlme et al. 2020) | Põlme S, et al. 2020. Fungal Diversity 105:1-16 ((https://doi.org/10.1007/s13225-020-00466-2)) |
| Otu46400 | 1 | k__Fungi(100);p__Ascomycota(100);c__Saccharomycetes(100);o__Saccharomycetales(100);f__Metschnikowiaceae(100);g__Metschnikowia(100);g__Metschnikowia_unclassified(100); | Metschnikowia | 13 | Saprotroph | \|Undefined Saprotroph\| | Yeast | NULL | Probable | As nectar-tap_saprotroph (Põlme et al. 2020); Decay sugar-rich substrates (Põlme et al. 2020) | Põlme S, et al. 2020. Fungal Diversity 105:1-16 ((https://doi.org/10.1007/s13225-020-00466-2)) |
| Otu46401 | 1 | k__Fungi(100);p__Ascomycota(100);c__Saccharomycetes(100);o__Saccharomycetales(100);f__Metschnikowiaceae(100);g__Metschnikowia(100);g__Metschnikowia_unclassified(100); | Metschnikowia | 13 | Saprotroph | \|Undefined Saprotroph\| | Yeast | NULL | Probable | As nectar-tap_saprotroph (Põlme et al. 2020); Decay sugar-rich substrates (Põlme et al. 2020) | Põlme S, et al. 2020. Fungal Diversity 105:1-16 ((https://doi.org/10.1007/s13225-020-00466-2)) |
| Otu46402 | 1 | k__Fungi(100);p__Ascomycota(100);c__Saccharomycetes(100);o__Saccharomycetales(100);f__Metschnikowiaceae(100);g__Metschnikowia(100);g__Metschnikowia_unclassified(100); | Metschnikowia | 13 | Saprotroph | \|Undefined Saprotroph\| | Yeast | NULL | Probable | As nectar-tap_saprotroph (Põlme et al. 2020); Decay sugar-rich substrates (Põlme et al. 2020) | Põlme S, et al. 2020. Fungal Diversity 105:1-16 ((https://doi.org/10.1007/s13225-020-00466-2)) |
| Otu46403 | 1 | k__Fungi(100);p__Ascomycota(100);c__Saccharomycetes(100);o__Saccharomycetales(100);f__Metschnikowiaceae(100);g__Metschnikowia(100);g__Metschnikowia_unclassified(100); | Metschnikowia | 13 | Saprotroph | \|Undefined Saprotroph\| | Yeast | NULL | Probable | As nectar-tap_saprotroph (Põlme et al. 2020); Decay sugar-rich substrates (Põlme et al. 2020) | Põlme S, et al. 2020. Fungal Diversity 105:1-16 ((https://doi.org/10.1007/s13225-020-00466-2)) |
| Otu46404 | 1 | k__Fungi(100);p__Ascomycota(100);c__Saccharomycetes(100);o__Saccharomycetales(100);f__Metschnikowiaceae(100);g__Metschnikowia(100);g__Metschnikowia_unclassified(100); | Metschnikowia | 13 | Saprotroph | \|Undefined Saprotroph\| | Yeast | NULL | Probable | As nectar-tap_saprotroph (Põlme et al. 2020); Decay sugar-rich substrates (Põlme et al. 2020) | Põlme S, et al. 2020. Fungal Diversity 105:1-16 ((https://doi.org/10.1007/s13225-020-00466-2)) |
| Otu46405 | 1 | k__Fungi(100);p__Ascomycota(100);c__Saccharomycetes(100);o__Saccharomycetales(100);f__Metschnikowiaceae(100);g__Metschnikowia(100);g__Metschnikowia_unclassified(100); | Metschnikowia | 13 | Saprotroph | \|Undefined Saprotroph\| | Yeast | NULL | Probable | As nectar-tap_saprotroph (Põlme et al. 2020); Decay sugar-rich substrates (Põlme et al. 2020) | Põlme S, et al. 2020. Fungal Diversity 105:1-16 ((https://doi.org/10.1007/s13225-020-00466-2)) |
| Otu46406 | 1 | k__Fungi(100);p__Ascomycota(100);c__Saccharomycetes(100);o__Saccharomycetales(100);f__Metschnikowiaceae(100);g__Metschnikowia(100);g__Metschnikowia_unclassified(100); | Metschnikowia | 13 | Saprotroph | \|Undefined Saprotroph\| | Yeast | NULL | Probable | As nectar-tap_saprotroph (Põlme et al. 2020); Decay sugar-rich substrates (Põlme et al. 2020) | Põlme S, et al. 2020. Fungal Diversity 105:1-16 ((https://doi.org/10.1007/s13225-020-00466-2)) |
| Otu46407 | 1 | k__Fungi(100);p__Ascomycota(100);c__Saccharomycetes(100);o__Saccharomycetales(100);f__Metschnikowiaceae(100);g__Metschnikowia(100);g__Metschnikowia_unclassified(100); | Metschnikowia | 13 | Saprotroph | \|Undefined Saprotroph\| | Yeast | NULL | Probable | As nectar-tap_saprotroph (Põlme et al. 2020); Decay sugar-rich substrates (Põlme et al. 2020) | Põlme S, et al. 2020. Fungal Diversity 105:1-16 ((https://doi.org/10.1007/s13225-020-00466-2)) |
| Otu46408 | 1 | k__Fungi(100);p__Ascomycota(100);c__Saccharomycetes(100);o__Saccharomycetales(100);f__Metschnikowiaceae(100);g__Metschnikowia(100);g__Metschnikowia_unclassified(100); | Metschnikowia | 13 | Saprotroph | \|Undefined Saprotroph\| | Yeast | NULL | Probable | As nectar-tap_saprotroph (Põlme et al. 2020); Decay sugar-rich substrates (Põlme et al. 2020) | Põlme S, et al. 2020. Fungal Diversity 105:1-16 ((https://doi.org/10.1007/s13225-020-00466-2)) |
| Otu46409 | 1 | k__Fungi(100);p__Ascomycota(100);c__Saccharomycetes(100);o__Saccharomycetales(100);f__Metschnikowiaceae(100);g__Metschnikowia(100);g__Metschnikowia_unclassified(100); | Metschnikowia | 13 | Saprotroph | \|Undefined Saprotroph\| | Yeast | NULL | Probable | As nectar-tap_saprotroph (Põlme et al. 2020); Decay sugar-rich substrates (Põlme et al. 2020) | Põlme S, et al. 2020. Fungal Diversity 105:1-16 ((https://doi.org/10.1007/s13225-020-00466-2)) |
| Otu46410 | 1 | k__Fungi(100);p__Ascomycota(100);c__Saccharomycetes(100);o__Saccharomycetales(100);f__Metschnikowiaceae(100);g__Metschnikowia(100);g__Metschnikowia_unclassified(100); | Metschnikowia | 13 | Saprotroph | \|Undefined Saprotroph\| | Yeast | NULL | Probable | As nectar-tap_saprotroph (Põlme et al. 2020); Decay sugar-rich substrates (Põlme et al. 2020) | Põlme S, et al. 2020. Fungal Diversity 105:1-16 ((https://doi.org/10.1007/s13225-020-00466-2)) |
| Otu46411 | 1 | k__Fungi(100);p__Ascomycota(100);c__Saccharomycetes(100);o__Saccharomycetales(100);f__Metschnikowiaceae(100);g__Metschnikowia(100);g__Metschnikowia_unclassified(100); | Metschnikowia | 13 | Saprotroph | \|Undefined Saprotroph\| | Yeast | NULL | Probable | As nectar-tap_saprotroph (Põlme et al. 2020); Decay sugar-rich substrates (Põlme et al. 2020) | Põlme S, et al. 2020. Fungal Diversity 105:1-16 ((https://doi.org/10.1007/s13225-020-00466-2)) |
| Otu46412 | 1 | k__Fungi(100);p__Ascomycota(100);c__Saccharomycetes(100);o__Saccharomycetales(100);f__Metschnikowiaceae(100);g__Metschnikowia(100);s__Metschnikowia_pulcherrima(100); | Metschnikowia | 13 | Saprotroph | \|Undefined Saprotroph\| | Yeast | NULL | Probable | As nectar-tap_saprotroph (Põlme et al. 2020); Decay sugar-rich substrates (Põlme et al. 2020) | Põlme S, et al. 2020. Fungal Diversity 105:1-16 ((https://doi.org/10.1007/s13225-020-00466-2)) |
| Otu46413 | 1 | k__Fungi(100);p__Ascomycota(100);c__Saccharomycetes(100);o__Saccharomycetales(100);f__Metschnikowiaceae(100);g__Metschnikowia(100);s__Metschnikowia_pulcherrima(100); | Metschnikowia | 13 | Saprotroph | \|Undefined Saprotroph\| | Yeast | NULL | Probable | As nectar-tap_saprotroph (Põlme et al. 2020); Decay sugar-rich substrates (Põlme et al. 2020) | Põlme S, et al. 2020. Fungal Diversity 105:1-16 ((https://doi.org/10.1007/s13225-020-00466-2)) |
| Otu46415 | 1 | k__Fungi(100);p__Ascomycota(100);c__Saccharomycetes(100);o__Saccharomycetales(100);f__Metschnikowiaceae(100);g__Metschnikowia(100);s__Metschnikowia_pulcherrima(100); | Metschnikowia | 13 | Saprotroph | \|Undefined Saprotroph\| | Yeast | NULL | Probable | As nectar-tap_saprotroph (Põlme et al. 2020); Decay sugar-rich substrates (Põlme et al. 2020) | Põlme S, et al. 2020. Fungal Diversity 105:1-16 ((https://doi.org/10.1007/s13225-020-00466-2)) |
| Otu46416 | 1 | k__Fungi(100);p__Ascomycota(100);c__Saccharomycetes(100);o__Saccharomycetales(100);f__Metschnikowiaceae(100);g__Metschnikowia(100);g__Metschnikowia_unclassified(100); | Metschnikowia | 13 | Saprotroph | \|Undefined Saprotroph\| | Yeast | NULL | Probable | As nectar-tap_saprotroph (Põlme et al. 2020); Decay sugar-rich substrates (Põlme et al. 2020) | Põlme S, et al. 2020. Fungal Diversity 105:1-16 ((https://doi.org/10.1007/s13225-020-00466-2)) |
| Otu46417 | 1 | k__Fungi(100);p__Ascomycota(100);c__Saccharomycetes(100);o__Saccharomycetales(100);f__Metschnikowiaceae(100);g__Metschnikowia(100);s__Metschnikowia_pulcherrima(100); | Metschnikowia | 13 | Saprotroph | \|Undefined Saprotroph\| | Yeast | NULL | Probable | As nectar-tap_saprotroph (Põlme et al. 2020); Decay sugar-rich substrates (Põlme et al. 2020) | Põlme S, et al. 2020. Fungal Diversity 105:1-16 ((https://doi.org/10.1007/s13225-020-00466-2)) |
| Otu46418 | 1 | k__Fungi(100);p__Ascomycota(100);c__Saccharomycetes(100);o__Saccharomycetales(100);f__Metschnikowiaceae(100);g__Metschnikowia(100);g__Metschnikowia_unclassified(100); | Metschnikowia | 13 | Saprotroph | \|Undefined Saprotroph\| | Yeast | NULL | Probable | As nectar-tap_saprotroph (Põlme et al. 2020); Decay sugar-rich substrates (Põlme et al. 2020) | Põlme S, et al. 2020. Fungal Diversity 105:1-16 ((https://doi.org/10.1007/s13225-020-00466-2)) |
| Otu46419 | 1 | k__Fungi(100);p__Ascomycota(100);c__Saccharomycetes(100);o__Saccharomycetales(100);f__Metschnikowiaceae(100);g__Metschnikowia(100);s__Metschnikowia_pulcherrima(100); | Metschnikowia | 13 | Saprotroph | \|Undefined Saprotroph\| | Yeast | NULL | Probable | As nectar-tap_saprotroph (Põlme et al. 2020); Decay sugar-rich substrates (Põlme et al. 2020) | Põlme S, et al. 2020. Fungal Diversity 105:1-16 ((https://doi.org/10.1007/s13225-020-00466-2)) |
| Otu46420 | 1 | k__Fungi(100);p__Ascomycota(100);c__Saccharomycetes(100);o__Saccharomycetales(100);f__Metschnikowiaceae(100);g__Metschnikowia(100);s__Metschnikowia_pulcherrima(100); | Metschnikowia | 13 | Saprotroph | \|Undefined Saprotroph\| | Yeast | NULL | Probable | As nectar-tap_saprotroph (Põlme et al. 2020); Decay sugar-rich substrates (Põlme et al. 2020) | Põlme S, et al. 2020. Fungal Diversity 105:1-16 ((https://doi.org/10.1007/s13225-020-00466-2)) |
| Otu46421 | 1 | k__Fungi(100);p__Ascomycota(100);c__Saccharomycetes(100);o__Saccharomycetales(100);f__Metschnikowiaceae(100);g__Metschnikowia(100);s__Metschnikowia_pulcherrima(100); | Metschnikowia | 13 | Saprotroph | \|Undefined Saprotroph\| | Yeast | NULL | Probable | As nectar-tap_saprotroph (Põlme et al. 2020); Decay sugar-rich substrates (Põlme et al. 2020) | Põlme S, et al. 2020. Fungal Diversity 105:1-16 ((https://doi.org/10.1007/s13225-020-00466-2)) |
| Otu46422 | 1 | k__Fungi(100);p__Ascomycota(100);c__Saccharomycetes(100);o__Saccharomycetales(100);f__Metschnikowiaceae(100);g__Metschnikowia(100);s__Metschnikowia_pulcherrima(100); | Metschnikowia | 13 | Saprotroph | \|Undefined Saprotroph\| | Yeast | NULL | Probable | As nectar-tap_saprotroph (Põlme et al. 2020); Decay sugar-rich substrates (Põlme et al. 2020) | Põlme S, et al. 2020. Fungal Diversity 105:1-16 ((https://doi.org/10.1007/s13225-020-00466-2)) |
| Otu46423 | 1 | k__Fungi(100);p__Ascomycota(100);c__Saccharomycetes(100);o__Saccharomycetales(100);f__Metschnikowiaceae(100);g__Metschnikowia(100);g__Metschnikowia_unclassified(100); | Metschnikowia | 13 | Saprotroph | \|Undefined Saprotroph\| | Yeast | NULL | Probable | As nectar-tap_saprotroph (Põlme et al. 2020); Decay sugar-rich substrates (Põlme et al. 2020) | Põlme S, et al. 2020. Fungal Diversity 105:1-16 ((https://doi.org/10.1007/s13225-020-00466-2)) |
| Otu46424 | 1 | k__Fungi(100);p__Ascomycota(100);c__Saccharomycetes(100);o__Saccharomycetales(100);f__Metschnikowiaceae(100);g__Metschnikowia(100);s__Metschnikowia_pulcherrima(100); | Metschnikowia | 13 | Saprotroph | \|Undefined Saprotroph\| | Yeast | NULL | Probable | As nectar-tap_saprotroph (Põlme et al. 2020); Decay sugar-rich substrates (Põlme et al. 2020) | Põlme S, et al. 2020. Fungal Diversity 105:1-16 ((https://doi.org/10.1007/s13225-020-00466-2)) |
| Otu46425 | 1 | k__Fungi(100);p__Ascomycota(100);c__Saccharomycetes(100);o__Saccharomycetales(100);f__Metschnikowiaceae(100);g__Metschnikowia(100);s__Metschnikowia_pulcherrima(100); | Metschnikowia | 13 | Saprotroph | \|Undefined Saprotroph\| | Yeast | NULL | Probable | As nectar-tap_saprotroph (Põlme et al. 2020); Decay sugar-rich substrates (Põlme et al. 2020) | Põlme S, et al. 2020. Fungal Diversity 105:1-16 ((https://doi.org/10.1007/s13225-020-00466-2)) |
| Otu46426 | 1 | k__Fungi(100);p__Ascomycota(100);c__Saccharomycetes(100);o__Saccharomycetales(100);f__Metschnikowiaceae(100);g__Metschnikowia(100);s__Metschnikowia_pulcherrima(100); | Metschnikowia | 13 | Saprotroph | \|Undefined Saprotroph\| | Yeast | NULL | Probable | As nectar-tap_saprotroph (Põlme et al. 2020); Decay sugar-rich substrates (Põlme et al. 2020) | Põlme S, et al. 2020. Fungal Diversity 105:1-16 ((https://doi.org/10.1007/s13225-020-00466-2)) |
| Otu46427 | 1 | k__Fungi(100);p__Ascomycota(100);c__Saccharomycetes(100);o__Saccharomycetales(100);f__Metschnikowiaceae(100);g__Metschnikowia(100);s__Metschnikowia_pulcherrima(100); | Metschnikowia | 13 | Saprotroph | \|Undefined Saprotroph\| | Yeast | NULL | Probable | As nectar-tap_saprotroph (Põlme et al. 2020); Decay sugar-rich substrates (Põlme et al. 2020) | Põlme S, et al. 2020. Fungal Diversity 105:1-16 ((https://doi.org/10.1007/s13225-020-00466-2)) |
| Otu46428 | 1 | k__Fungi(100);p__Ascomycota(100);c__Saccharomycetes(100);o__Saccharomycetales(100);f__Metschnikowiaceae(100);g__Metschnikowia(100);s__Metschnikowia_pulcherrima(100); | Metschnikowia | 13 | Saprotroph | \|Undefined Saprotroph\| | Yeast | NULL | Probable | As nectar-tap_saprotroph (Põlme et al. 2020); Decay sugar-rich substrates (Põlme et al. 2020) | Põlme S, et al. 2020. Fungal Diversity 105:1-16 ((https://doi.org/10.1007/s13225-020-00466-2)) |
| Otu46429 | 1 | k__Fungi(100);p__Ascomycota(100);c__Saccharomycetes(100);o__Saccharomycetales(100);f__Metschnikowiaceae(100);g__Metschnikowia(100);s__Metschnikowia_pulcherrima(100); | Metschnikowia | 13 | Saprotroph | \|Undefined Saprotroph\| | Yeast | NULL | Probable | As nectar-tap_saprotroph (Põlme et al. 2020); Decay sugar-rich substrates (Põlme et al. 2020) | Põlme S, et al. 2020. Fungal Diversity 105:1-16 ((https://doi.org/10.1007/s13225-020-00466-2)) |
| Otu46430 | 1 | k__Fungi(100);p__Ascomycota(100);c__Saccharomycetes(100);o__Saccharomycetales(100);f__Metschnikowiaceae(100);g__Metschnikowia(100);s__Metschnikowia_pulcherrima(100); | Metschnikowia | 13 | Saprotroph | \|Undefined Saprotroph\| | Yeast | NULL | Probable | As nectar-tap_saprotroph (Põlme et al. 2020); Decay sugar-rich substrates (Põlme et al. 2020) | Põlme S, et al. 2020. Fungal Diversity 105:1-16 ((https://doi.org/10.1007/s13225-020-00466-2)) |
| Otu46431 | 1 | k__Fungi(100);p__Ascomycota(100);c__Saccharomycetes(100);o__Saccharomycetales(100);f__Metschnikowiaceae(100);g__Metschnikowia(100);s__Metschnikowia_pulcherrima(100); | Metschnikowia | 13 | Saprotroph | \|Undefined Saprotroph\| | Yeast | NULL | Probable | As nectar-tap_saprotroph (Põlme et al. 2020); Decay sugar-rich substrates (Põlme et al. 2020) | Põlme S, et al. 2020. Fungal Diversity 105:1-16 ((https://doi.org/10.1007/s13225-020-00466-2)) |
| Otu46432 | 1 | k__Fungi(100);p__Ascomycota(100);c__Saccharomycetes(100);o__Saccharomycetales(100);f__Metschnikowiaceae(100);g__Metschnikowia(100);s__Metschnikowia_pulcherrima(100); | Metschnikowia | 13 | Saprotroph | \|Undefined Saprotroph\| | Yeast | NULL | Probable | As nectar-tap_saprotroph (Põlme et al. 2020); Decay sugar-rich substrates (Põlme et al. 2020) | Põlme S, et al. 2020. Fungal Diversity 105:1-16 ((https://doi.org/10.1007/s13225-020-00466-2)) |
| Otu46433 | 1 | k__Fungi(100);p__Ascomycota(100);c__Saccharomycetes(100);o__Saccharomycetales(100);f__Metschnikowiaceae(100);g__Metschnikowia(100);s__Metschnikowia_pulcherrima(100); | Metschnikowia | 13 | Saprotroph | \|Undefined Saprotroph\| | Yeast | NULL | Probable | As nectar-tap_saprotroph (Põlme et al. 2020); Decay sugar-rich substrates (Põlme et al. 2020) | Põlme S, et al. 2020. Fungal Diversity 105:1-16 ((https://doi.org/10.1007/s13225-020-00466-2)) |
| Otu46434 | 1 | k__Fungi(100);p__Ascomycota(100);c__Saccharomycetes(100);o__Saccharomycetales(100);f__Metschnikowiaceae(100);g__Metschnikowia(100);s__Metschnikowia_pulcherrima(100); | Metschnikowia | 13 | Saprotroph | \|Undefined Saprotroph\| | Yeast | NULL | Probable | As nectar-tap_saprotroph (Põlme et al. 2020); Decay sugar-rich substrates (Põlme et al. 2020) | Põlme S, et al. 2020. Fungal Diversity 105:1-16 ((https://doi.org/10.1007/s13225-020-00466-2)) |
| Otu46435 | 1 | k__Fungi(100);p__Ascomycota(100);c__Saccharomycetes(100);o__Saccharomycetales(100);f__Metschnikowiaceae(100);g__Metschnikowia(100);s__Metschnikowia_pulcherrima(100); | Metschnikowia | 13 | Saprotroph | \|Undefined Saprotroph\| | Yeast | NULL | Probable | As nectar-tap_saprotroph (Põlme et al. 2020); Decay sugar-rich substrates (Põlme et al. 2020) | Põlme S, et al. 2020. Fungal Diversity 105:1-16 ((https://doi.org/10.1007/s13225-020-00466-2)) |
| Otu46436 | 1 | k__Fungi(100);p__Ascomycota(100);c__Saccharomycetes(100);o__Saccharomycetales(100);f__Metschnikowiaceae(100);g__Metschnikowia(100);g__Metschnikowia_unclassified(100); | Metschnikowia | 13 | Saprotroph | \|Undefined Saprotroph\| | Yeast | NULL | Probable | As nectar-tap_saprotroph (Põlme et al. 2020); Decay sugar-rich substrates (Põlme et al. 2020) | Põlme S, et al. 2020. Fungal Diversity 105:1-16 ((https://doi.org/10.1007/s13225-020-00466-2)) |
| Otu46437 | 1 | k__Fungi(100);p__Ascomycota(100);c__Saccharomycetes(100);o__Saccharomycetales(100);f__Metschnikowiaceae(100);g__Metschnikowia(100);s__Metschnikowia_pulcherrima(100); | Metschnikowia | 13 | Saprotroph | \|Undefined Saprotroph\| | Yeast | NULL | Probable | As nectar-tap_saprotroph (Põlme et al. 2020); Decay sugar-rich substrates (Põlme et al. 2020) | Põlme S, et al. 2020. Fungal Diversity 105:1-16 ((https://doi.org/10.1007/s13225-020-00466-2)) |
| Otu46438 | 1 | k__Fungi(100);p__Ascomycota(100);c__Saccharomycetes(100);o__Saccharomycetales(100);f__Metschnikowiaceae(100);g__Metschnikowia(100);s__Metschnikowia_pulcherrima(100); | Metschnikowia | 13 | Saprotroph | \|Undefined Saprotroph\| | Yeast | NULL | Probable | As nectar-tap_saprotroph (Põlme et al. 2020); Decay sugar-rich substrates (Põlme et al. 2020) | Põlme S, et al. 2020. Fungal Diversity 105:1-16 ((https://doi.org/10.1007/s13225-020-00466-2)) |
| Otu46439 | 1 | k__Fungi(100);p__Ascomycota(100);c__Saccharomycetes(100);o__Saccharomycetales(100);f__Metschnikowiaceae(100);g__Metschnikowia(100);s__Metschnikowia_pulcherrima(100); | Metschnikowia | 13 | Saprotroph | \|Undefined Saprotroph\| | Yeast | NULL | Probable | As nectar-tap_saprotroph (Põlme et al. 2020); Decay sugar-rich substrates (Põlme et al. 2020) | Põlme S, et al. 2020. Fungal Diversity 105:1-16 ((https://doi.org/10.1007/s13225-020-00466-2)) |
| Otu46440 | 1 | k__Fungi(100);p__Ascomycota(100);c__Saccharomycetes(100);o__Saccharomycetales(100);f__Metschnikowiaceae(100);g__Metschnikowia(100);s__Metschnikowia_pulcherrima(100); | Metschnikowia | 13 | Saprotroph | \|Undefined Saprotroph\| | Yeast | NULL | Probable | As nectar-tap_saprotroph (Põlme et al. 2020); Decay sugar-rich substrates (Põlme et al. 2020) | Põlme S, et al. 2020. Fungal Diversity 105:1-16 ((https://doi.org/10.1007/s13225-020-00466-2)) |
| Otu46441 | 1 | k__Fungi(100);p__Ascomycota(100);c__Saccharomycetes(100);o__Saccharomycetales(100);f__Metschnikowiaceae(100);g__Metschnikowia(100);s__Metschnikowia_pulcherrima(100); | Metschnikowia | 13 | Saprotroph | \|Undefined Saprotroph\| | Yeast | NULL | Probable | As nectar-tap_saprotroph (Põlme et al. 2020); Decay sugar-rich substrates (Põlme et al. 2020) | Põlme S, et al. 2020. Fungal Diversity 105:1-16 ((https://doi.org/10.1007/s13225-020-00466-2)) |
| Otu46442 | 1 | k__Fungi(100);p__Ascomycota(100);c__Saccharomycetes(100);o__Saccharomycetales(100);f__Metschnikowiaceae(100);g__Metschnikowia(100);s__Metschnikowia_pulcherrima(100); | Metschnikowia | 13 | Saprotroph | \|Undefined Saprotroph\| | Yeast | NULL | Probable | As nectar-tap_saprotroph (Põlme et al. 2020); Decay sugar-rich substrates (Põlme et al. 2020) | Põlme S, et al. 2020. Fungal Diversity 105:1-16 ((https://doi.org/10.1007/s13225-020-00466-2)) |
| Otu46443 | 1 | k__Fungi(100);p__Ascomycota(100);c__Saccharomycetes(100);o__Saccharomycetales(100);f__Metschnikowiaceae(100);g__Metschnikowia(100);s__Metschnikowia_pulcherrima(100); | Metschnikowia | 13 | Saprotroph | \|Undefined Saprotroph\| | Yeast | NULL | Probable | As nectar-tap_saprotroph (Põlme et al. 2020); Decay sugar-rich substrates (Põlme et al. 2020) | Põlme S, et al. 2020. Fungal Diversity 105:1-16 ((https://doi.org/10.1007/s13225-020-00466-2)) |
| Otu46444 | 1 | k__Fungi(100);p__Ascomycota(100);c__Saccharomycetes(100);o__Saccharomycetales(100);f__Metschnikowiaceae(100);g__Metschnikowia(100);s__Metschnikowia_pulcherrima(100); | Metschnikowia | 13 | Saprotroph | \|Undefined Saprotroph\| | Yeast | NULL | Probable | As nectar-tap_saprotroph (Põlme et al. 2020); Decay sugar-rich substrates (Põlme et al. 2020) | Põlme S, et al. 2020. Fungal Diversity 105:1-16 ((https://doi.org/10.1007/s13225-020-00466-2)) |
| Otu46445 | 1 | k__Fungi(100);p__Ascomycota(100);c__Saccharomycetes(100);o__Saccharomycetales(100);f__Metschnikowiaceae(100);g__Metschnikowia(100);s__Metschnikowia_pulcherrima(100); | Metschnikowia | 13 | Saprotroph | \|Undefined Saprotroph\| | Yeast | NULL | Probable | As nectar-tap_saprotroph (Põlme et al. 2020); Decay sugar-rich substrates (Põlme et al. 2020) | Põlme S, et al. 2020. Fungal Diversity 105:1-16 ((https://doi.org/10.1007/s13225-020-00466-2)) |
| Otu46446 | 1 | k__Fungi(100);p__Ascomycota(100);c__Saccharomycetes(100);o__Saccharomycetales(100);f__Metschnikowiaceae(100);g__Metschnikowia(100);s__Metschnikowia_pulcherrima(100); | Metschnikowia | 13 | Saprotroph | \|Undefined Saprotroph\| | Yeast | NULL | Probable | As nectar-tap_saprotroph (Põlme et al. 2020); Decay sugar-rich substrates (Põlme et al. 2020) | Põlme S, et al. 2020. Fungal Diversity 105:1-16 ((https://doi.org/10.1007/s13225-020-00466-2)) |
| Otu46447 | 1 | k__Fungi(100);p__Ascomycota(100);c__Saccharomycetes(100);o__Saccharomycetales(100);f__Metschnikowiaceae(100);g__Metschnikowia(100);g__Metschnikowia_unclassified(100); | Metschnikowia | 13 | Saprotroph | \|Undefined Saprotroph\| | Yeast | NULL | Probable | As nectar-tap_saprotroph (Põlme et al. 2020); Decay sugar-rich substrates (Põlme et al. 2020) | Põlme S, et al. 2020. Fungal Diversity 105:1-16 ((https://doi.org/10.1007/s13225-020-00466-2)) |
| Otu46448 | 1 | k__Fungi(100);p__Ascomycota(100);c__Saccharomycetes(100);o__Saccharomycetales(100);f__Metschnikowiaceae(100);g__Metschnikowia(100);g__Metschnikowia_unclassified(100); | Metschnikowia | 13 | Saprotroph | \|Undefined Saprotroph\| | Yeast | NULL | Probable | As nectar-tap_saprotroph (Põlme et al. 2020); Decay sugar-rich substrates (Põlme et al. 2020) | Põlme S, et al. 2020. Fungal Diversity 105:1-16 ((https://doi.org/10.1007/s13225-020-00466-2)) |
| Otu46449 | 1 | k__Fungi(100);p__Ascomycota(100);c__Saccharomycetes(100);o__Saccharomycetales(100);f__Metschnikowiaceae(100);g__Metschnikowia(100);s__Metschnikowia_pulcherrima(100); | Metschnikowia | 13 | Saprotroph | \|Undefined Saprotroph\| | Yeast | NULL | Probable | As nectar-tap_saprotroph (Põlme et al. 2020); Decay sugar-rich substrates (Põlme et al. 2020) | Põlme S, et al. 2020. Fungal Diversity 105:1-16 ((https://doi.org/10.1007/s13225-020-00466-2)) |
| Otu46450 | 1 | k__Fungi(100);p__Ascomycota(100);c__Saccharomycetes(100);o__Saccharomycetales(100);f__Metschnikowiaceae(100);g__Metschnikowia(100);s__Metschnikowia_pulcherrima(100); | Metschnikowia | 13 | Saprotroph | \|Undefined Saprotroph\| | Yeast | NULL | Probable | As nectar-tap_saprotroph (Põlme et al. 2020); Decay sugar-rich substrates (Põlme et al. 2020) | Põlme S, et al. 2020. Fungal Diversity 105:1-16 ((https://doi.org/10.1007/s13225-020-00466-2)) |
| Otu46451 | 1 | k__Fungi(100);p__Ascomycota(100);c__Saccharomycetes(100);o__Saccharomycetales(100);f__Metschnikowiaceae(100);g__Metschnikowia(100);s__Metschnikowia_pulcherrima(100); | Metschnikowia | 13 | Saprotroph | \|Undefined Saprotroph\| | Yeast | NULL | Probable | As nectar-tap_saprotroph (Põlme et al. 2020); Decay sugar-rich substrates (Põlme et al. 2020) | Põlme S, et al. 2020. Fungal Diversity 105:1-16 ((https://doi.org/10.1007/s13225-020-00466-2)) |
| Otu46452 | 1 | k__Fungi(100);p__Ascomycota(100);c__Saccharomycetes(100);o__Saccharomycetales(100);f__Metschnikowiaceae(100);g__Metschnikowia(100);s__Metschnikowia_pulcherrima(100); | Metschnikowia | 13 | Saprotroph | \|Undefined Saprotroph\| | Yeast | NULL | Probable | As nectar-tap_saprotroph (Põlme et al. 2020); Decay sugar-rich substrates (Põlme et al. 2020) | Põlme S, et al. 2020. Fungal Diversity 105:1-16 ((https://doi.org/10.1007/s13225-020-00466-2)) |
| Otu46453 | 1 | k__Fungi(100);p__Ascomycota(100);c__Saccharomycetes(100);o__Saccharomycetales(100);f__Metschnikowiaceae(100);g__Metschnikowia(100);g__Metschnikowia_unclassified(100); | Metschnikowia | 13 | Saprotroph | \|Undefined Saprotroph\| | Yeast | NULL | Probable | As nectar-tap_saprotroph (Põlme et al. 2020); Decay sugar-rich substrates (Põlme et al. 2020) | Põlme S, et al. 2020. Fungal Diversity 105:1-16 ((https://doi.org/10.1007/s13225-020-00466-2)) |
| Otu46454 | 1 | k__Fungi(100);p__Ascomycota(100);c__Saccharomycetes(100);o__Saccharomycetales(100);f__Metschnikowiaceae(100);g__Metschnikowia(100);s__Metschnikowia_pulcherrima(100); | Metschnikowia | 13 | Saprotroph | \|Undefined Saprotroph\| | Yeast | NULL | Probable | As nectar-tap_saprotroph (Põlme et al. 2020); Decay sugar-rich substrates (Põlme et al. 2020) | Põlme S, et al. 2020. Fungal Diversity 105:1-16 ((https://doi.org/10.1007/s13225-020-00466-2)) |
| Otu46455 | 1 | k__Fungi(100);p__Ascomycota(100);c__Saccharomycetes(100);o__Saccharomycetales(100);f__Metschnikowiaceae(100);g__Metschnikowia(100);s__Metschnikowia_pulcherrima(100); | Metschnikowia | 13 | Saprotroph | \|Undefined Saprotroph\| | Yeast | NULL | Probable | As nectar-tap_saprotroph (Põlme et al. 2020); Decay sugar-rich substrates (Põlme et al. 2020) | Põlme S, et al. 2020. Fungal Diversity 105:1-16 ((https://doi.org/10.1007/s13225-020-00466-2)) |
| Otu46456 | 1 | k__Fungi(100);p__Ascomycota(100);c__Saccharomycetes(100);o__Saccharomycetales(100);f__Metschnikowiaceae(100);g__Metschnikowia(100);s__Metschnikowia_pulcherrima(100); | Metschnikowia | 13 | Saprotroph | \|Undefined Saprotroph\| | Yeast | NULL | Probable | As nectar-tap_saprotroph (Põlme et al. 2020); Decay sugar-rich substrates (Põlme et al. 2020) | Põlme S, et al. 2020. Fungal Diversity 105:1-16 ((https://doi.org/10.1007/s13225-020-00466-2)) |
| Otu46457 | 1 | k__Fungi(100);p__Ascomycota(100);c__Saccharomycetes(100);o__Saccharomycetales(100);f__Metschnikowiaceae(100);g__Metschnikowia(100);s__Metschnikowia_pulcherrima(100); | Metschnikowia | 13 | Saprotroph | \|Undefined Saprotroph\| | Yeast | NULL | Probable | As nectar-tap_saprotroph (Põlme et al. 2020); Decay sugar-rich substrates (Põlme et al. 2020) | Põlme S, et al. 2020. Fungal Diversity 105:1-16 ((https://doi.org/10.1007/s13225-020-00466-2)) |
| Otu46458 | 1 | k__Fungi(100);p__Ascomycota(100);c__Saccharomycetes(100);o__Saccharomycetales(100);f__Metschnikowiaceae(100);g__Metschnikowia(100);s__Metschnikowia_pulcherrima(100); | Metschnikowia | 13 | Saprotroph | \|Undefined Saprotroph\| | Yeast | NULL | Probable | As nectar-tap_saprotroph (Põlme et al. 2020); Decay sugar-rich substrates (Põlme et al. 2020) | Põlme S, et al. 2020. Fungal Diversity 105:1-16 ((https://doi.org/10.1007/s13225-020-00466-2)) |
| Otu46459 | 1 | k__Fungi(100);p__Ascomycota(100);c__Saccharomycetes(100);o__Saccharomycetales(100);f__Metschnikowiaceae(100);g__Metschnikowia(100);s__Metschnikowia_pulcherrima(100); | Metschnikowia | 13 | Saprotroph | \|Undefined Saprotroph\| | Yeast | NULL | Probable | As nectar-tap_saprotroph (Põlme et al. 2020); Decay sugar-rich substrates (Põlme et al. 2020) | Põlme S, et al. 2020. Fungal Diversity 105:1-16 ((https://doi.org/10.1007/s13225-020-00466-2)) |
| Otu46460 | 1 | k__Fungi(100);p__Ascomycota(100);c__Saccharomycetes(100);o__Saccharomycetales(100);f__Metschnikowiaceae(100);g__Metschnikowia(100);s__Metschnikowia_pulcherrima(100); | Metschnikowia | 13 | Saprotroph | \|Undefined Saprotroph\| | Yeast | NULL | Probable | As nectar-tap_saprotroph (Põlme et al. 2020); Decay sugar-rich substrates (Põlme et al. 2020) | Põlme S, et al. 2020. Fungal Diversity 105:1-16 ((https://doi.org/10.1007/s13225-020-00466-2)) |
| Otu46461 | 1 | k__Fungi(100);p__Ascomycota(100);c__Saccharomycetes(100);o__Saccharomycetales(100);f__Metschnikowiaceae(100);g__Metschnikowia(100);s__Metschnikowia_pulcherrima(100); | Metschnikowia | 13 | Saprotroph | \|Undefined Saprotroph\| | Yeast | NULL | Probable | As nectar-tap_saprotroph (Põlme et al. 2020); Decay sugar-rich substrates (Põlme et al. 2020) | Põlme S, et al. 2020. Fungal Diversity 105:1-16 ((https://doi.org/10.1007/s13225-020-00466-2)) |
| Otu46462 | 1 | k__Fungi(100);p__Ascomycota(100);c__Saccharomycetes(100);o__Saccharomycetales(100);f__Metschnikowiaceae(100);g__Metschnikowia(100);s__Metschnikowia_pulcherrima(100); | Metschnikowia | 13 | Saprotroph | \|Undefined Saprotroph\| | Yeast | NULL | Probable | As nectar-tap_saprotroph (Põlme et al. 2020); Decay sugar-rich substrates (Põlme et al. 2020) | Põlme S, et al. 2020. Fungal Diversity 105:1-16 ((https://doi.org/10.1007/s13225-020-00466-2)) |
| Otu46463 | 1 | k__Fungi(100);p__Ascomycota(100);c__Saccharomycetes(100);o__Saccharomycetales(100);f__Metschnikowiaceae(100);g__Metschnikowia(100);g__Metschnikowia_unclassified(100); | Metschnikowia | 13 | Saprotroph | \|Undefined Saprotroph\| | Yeast | NULL | Probable | As nectar-tap_saprotroph (Põlme et al. 2020); Decay sugar-rich substrates (Põlme et al. 2020) | Põlme S, et al. 2020. Fungal Diversity 105:1-16 ((https://doi.org/10.1007/s13225-020-00466-2)) |
| Otu46464 | 1 | k__Fungi(100);p__Ascomycota(100);c__Saccharomycetes(100);o__Saccharomycetales(100);f__Metschnikowiaceae(100);g__Metschnikowia(100);g__Metschnikowia_unclassified(100); | Metschnikowia | 13 | Saprotroph | \|Undefined Saprotroph\| | Yeast | NULL | Probable | As nectar-tap_saprotroph (Põlme et al. 2020); Decay sugar-rich substrates (Põlme et al. 2020) | Põlme S, et al. 2020. Fungal Diversity 105:1-16 ((https://doi.org/10.1007/s13225-020-00466-2)) |
| Otu46465 | 1 | k__Fungi(100);p__Ascomycota(100);c__Saccharomycetes(100);o__Saccharomycetales(100);f__Metschnikowiaceae(100);g__Metschnikowia(100);g__Metschnikowia_unclassified(100); | Metschnikowia | 13 | Saprotroph | \|Undefined Saprotroph\| | Yeast | NULL | Probable | As nectar-tap_saprotroph (Põlme et al. 2020); Decay sugar-rich substrates (Põlme et al. 2020) | Põlme S, et al. 2020. Fungal Diversity 105:1-16 ((https://doi.org/10.1007/s13225-020-00466-2)) |
| Otu46466 | 1 | k__Fungi(100);p__Ascomycota(100);c__Saccharomycetes(100);o__Saccharomycetales(100);f__Metschnikowiaceae(100);g__Metschnikowia(100);s__Metschnikowia_pulcherrima(100); | Metschnikowia | 13 | Saprotroph | \|Undefined Saprotroph\| | Yeast | NULL | Probable | As nectar-tap_saprotroph (Põlme et al. 2020); Decay sugar-rich substrates (Põlme et al. 2020) | Põlme S, et al. 2020. Fungal Diversity 105:1-16 ((https://doi.org/10.1007/s13225-020-00466-2)) |
| Otu46467 | 1 | k__Fungi(100);p__Ascomycota(100);c__Saccharomycetes(100);o__Saccharomycetales(100);f__Metschnikowiaceae(100);g__Metschnikowia(100);s__Metschnikowia_pulcherrima(100); | Metschnikowia | 13 | Saprotroph | \|Undefined Saprotroph\| | Yeast | NULL | Probable | As nectar-tap_saprotroph (Põlme et al. 2020); Decay sugar-rich substrates (Põlme et al. 2020) | Põlme S, et al. 2020. Fungal Diversity 105:1-16 ((https://doi.org/10.1007/s13225-020-00466-2)) |
| Otu46468 | 1 | k__Fungi(100);p__Ascomycota(100);c__Saccharomycetes(100);o__Saccharomycetales(100);f__Metschnikowiaceae(100);g__Metschnikowia(100);s__Metschnikowia_viticola(100); | Metschnikowia | 13 | Saprotroph | \|Undefined Saprotroph\| | Yeast | NULL | Probable | As nectar-tap_saprotroph (Põlme et al. 2020); Decay sugar-rich substrates (Põlme et al. 2020) | Põlme S, et al. 2020. Fungal Diversity 105:1-16 ((https://doi.org/10.1007/s13225-020-00466-2)) |
| Otu46469 | 1 | k__Fungi(100);p__Ascomycota(100);c__Saccharomycetes(100);o__Saccharomycetales(100);f__Metschnikowiaceae(100);g__Metschnikowia(100);g__Metschnikowia_unclassified(100); | Metschnikowia | 13 | Saprotroph | \|Undefined Saprotroph\| | Yeast | NULL | Probable | As nectar-tap_saprotroph (Põlme et al. 2020); Decay sugar-rich substrates (Põlme et al. 2020) | Põlme S, et al. 2020. Fungal Diversity 105:1-16 ((https://doi.org/10.1007/s13225-020-00466-2)) |
| Otu46470 | 1 | k__Fungi(100);p__Ascomycota(100);c__Saccharomycetes(100);o__Saccharomycetales(100);f__Metschnikowiaceae(100);g__Metschnikowia(100);g__Metschnikowia_unclassified(100); | Metschnikowia | 13 | Saprotroph | \|Undefined Saprotroph\| | Yeast | NULL | Probable | As nectar-tap_saprotroph (Põlme et al. 2020); Decay sugar-rich substrates (Põlme et al. 2020) | Põlme S, et al. 2020. Fungal Diversity 105:1-16 ((https://doi.org/10.1007/s13225-020-00466-2)) |
| Otu46471 | 1 | k__Fungi(100);p__Ascomycota(100);c__Saccharomycetes(100);o__Saccharomycetales(100);f__Metschnikowiaceae(100);g__Metschnikowia(100);s__Metschnikowia_pulcherrima(100); | Metschnikowia | 13 | Saprotroph | \|Undefined Saprotroph\| | Yeast | NULL | Probable | As nectar-tap_saprotroph (Põlme et al. 2020); Decay sugar-rich substrates (Põlme et al. 2020) | Põlme S, et al. 2020. Fungal Diversity 105:1-16 ((https://doi.org/10.1007/s13225-020-00466-2)) |
| Otu46472 | 1 | k__Fungi(100);p__Ascomycota(100);c__Saccharomycetes(100);o__Saccharomycetales(100);f__Metschnikowiaceae(100);g__Metschnikowia(100);s__Metschnikowia_pulcherrima(100); | Metschnikowia | 13 | Saprotroph | \|Undefined Saprotroph\| | Yeast | NULL | Probable | As nectar-tap_saprotroph (Põlme et al. 2020); Decay sugar-rich substrates (Põlme et al. 2020) | Põlme S, et al. 2020. Fungal Diversity 105:1-16 ((https://doi.org/10.1007/s13225-020-00466-2)) |
| Otu46473 | 1 | k__Fungi(100);p__Ascomycota(100);c__Saccharomycetes(100);o__Saccharomycetales(100);f__Metschnikowiaceae(100);g__Metschnikowia(100);g__Metschnikowia_unclassified(100); | Metschnikowia | 13 | Saprotroph | \|Undefined Saprotroph\| | Yeast | NULL | Probable | As nectar-tap_saprotroph (Põlme et al. 2020); Decay sugar-rich substrates (Põlme et al. 2020) | Põlme S, et al. 2020. Fungal Diversity 105:1-16 ((https://doi.org/10.1007/s13225-020-00466-2)) |
| Otu46474 | 1 | k__Fungi(100);p__Ascomycota(100);c__Saccharomycetes(100);o__Saccharomycetales(100);f__Metschnikowiaceae(100);g__Metschnikowia(100);s__Metschnikowia_pulcherrima(100); | Metschnikowia | 13 | Saprotroph | \|Undefined Saprotroph\| | Yeast | NULL | Probable | As nectar-tap_saprotroph (Põlme et al. 2020); Decay sugar-rich substrates (Põlme et al. 2020) | Põlme S, et al. 2020. Fungal Diversity 105:1-16 ((https://doi.org/10.1007/s13225-020-00466-2)) |
| Otu46475 | 1 | k__Fungi(100);p__Ascomycota(100);c__Saccharomycetes(100);o__Saccharomycetales(100);f__Metschnikowiaceae(100);g__Metschnikowia(100);s__Metschnikowia_pulcherrima(100); | Metschnikowia | 13 | Saprotroph | \|Undefined Saprotroph\| | Yeast | NULL | Probable | As nectar-tap_saprotroph (Põlme et al. 2020); Decay sugar-rich substrates (Põlme et al. 2020) | Põlme S, et al. 2020. Fungal Diversity 105:1-16 ((https://doi.org/10.1007/s13225-020-00466-2)) |
| Otu46476 | 1 | k__Fungi(100);p__Ascomycota(100);c__Saccharomycetes(100);o__Saccharomycetales(100);f__Metschnikowiaceae(100);g__Metschnikowia(100);s__Metschnikowia_pulcherrima(100); | Metschnikowia | 13 | Saprotroph | \|Undefined Saprotroph\| | Yeast | NULL | Probable | As nectar-tap_saprotroph (Põlme et al. 2020); Decay sugar-rich substrates (Põlme et al. 2020) | Põlme S, et al. 2020. Fungal Diversity 105:1-16 ((https://doi.org/10.1007/s13225-020-00466-2)) |
| Otu46477 | 1 | k__Fungi(100);p__Ascomycota(100);c__Saccharomycetes(100);o__Saccharomycetales(100);f__Metschnikowiaceae(100);g__Metschnikowia(100);s__Metschnikowia_pulcherrima(100); | Metschnikowia | 13 | Saprotroph | \|Undefined Saprotroph\| | Yeast | NULL | Probable | As nectar-tap_saprotroph (Põlme et al. 2020); Decay sugar-rich substrates (Põlme et al. 2020) | Põlme S, et al. 2020. Fungal Diversity 105:1-16 ((https://doi.org/10.1007/s13225-020-00466-2)) |
| Otu46479 | 1 | k__Fungi(100);p__Ascomycota(100);c__Saccharomycetes(100);o__Saccharomycetales(100);f__Metschnikowiaceae(100);g__Metschnikowia(100);g__Metschnikowia_unclassified(100); | Metschnikowia | 13 | Saprotroph | \|Undefined Saprotroph\| | Yeast | NULL | Probable | As nectar-tap_saprotroph (Põlme et al. 2020); Decay sugar-rich substrates (Põlme et al. 2020) | Põlme S, et al. 2020. Fungal Diversity 105:1-16 ((https://doi.org/10.1007/s13225-020-00466-2)) |
| Otu46480 | 1 | k__Fungi(100);p__Ascomycota(100);c__Saccharomycetes(100);o__Saccharomycetales(100);f__Metschnikowiaceae(100);g__Metschnikowia(100);g__Metschnikowia_unclassified(100); | Metschnikowia | 13 | Saprotroph | \|Undefined Saprotroph\| | Yeast | NULL | Probable | As nectar-tap_saprotroph (Põlme et al. 2020); Decay sugar-rich substrates (Põlme et al. 2020) | Põlme S, et al. 2020. Fungal Diversity 105:1-16 ((https://doi.org/10.1007/s13225-020-00466-2)) |
| Otu46481 | 1 | k__Fungi(100);p__Ascomycota(100);c__Saccharomycetes(100);o__Saccharomycetales(100);f__Metschnikowiaceae(100);g__Metschnikowia(100);g__Metschnikowia_unclassified(100); | Metschnikowia | 13 | Saprotroph | \|Undefined Saprotroph\| | Yeast | NULL | Probable | As nectar-tap_saprotroph (Põlme et al. 2020); Decay sugar-rich substrates (Põlme et al. 2020) | Põlme S, et al. 2020. Fungal Diversity 105:1-16 ((https://doi.org/10.1007/s13225-020-00466-2)) |
| Otu46482 | 1 | k__Fungi(100);p__Ascomycota(100);c__Saccharomycetes(100);o__Saccharomycetales(100);f__Metschnikowiaceae(100);g__Metschnikowia(100);s__Metschnikowia_pulcherrima(100); | Metschnikowia | 13 | Saprotroph | \|Undefined Saprotroph\| | Yeast | NULL | Probable | As nectar-tap_saprotroph (Põlme et al. 2020); Decay sugar-rich substrates (Põlme et al. 2020) | Põlme S, et al. 2020. Fungal Diversity 105:1-16 ((https://doi.org/10.1007/s13225-020-00466-2)) |
| Otu46483 | 1 | k__Fungi(100);p__Ascomycota(100);c__Saccharomycetes(100);o__Saccharomycetales(100);f__Metschnikowiaceae(100);g__Metschnikowia(100);s__Metschnikowia_pulcherrima(100); | Metschnikowia | 13 | Saprotroph | \|Undefined Saprotroph\| | Yeast | NULL | Probable | As nectar-tap_saprotroph (Põlme et al. 2020); Decay sugar-rich substrates (Põlme et al. 2020) | Põlme S, et al. 2020. Fungal Diversity 105:1-16 ((https://doi.org/10.1007/s13225-020-00466-2)) |
| Otu46484 | 1 | k__Fungi(100);p__Ascomycota(100);c__Saccharomycetes(100);o__Saccharomycetales(100);f__Metschnikowiaceae(100);g__Metschnikowia(100);s__Metschnikowia_pulcherrima(100); | Metschnikowia | 13 | Saprotroph | \|Undefined Saprotroph\| | Yeast | NULL | Probable | As nectar-tap_saprotroph (Põlme et al. 2020); Decay sugar-rich substrates (Põlme et al. 2020) | Põlme S, et al. 2020. Fungal Diversity 105:1-16 ((https://doi.org/10.1007/s13225-020-00466-2)) |
| Otu46485 | 1 | k__Fungi(100);p__Ascomycota(100);c__Saccharomycetes(100);o__Saccharomycetales(100);f__Metschnikowiaceae(100);g__Metschnikowia(100);s__Metschnikowia_pulcherrima(100); | Metschnikowia | 13 | Saprotroph | \|Undefined Saprotroph\| | Yeast | NULL | Probable | As nectar-tap_saprotroph (Põlme et al. 2020); Decay sugar-rich substrates (Põlme et al. 2020) | Põlme S, et al. 2020. Fungal Diversity 105:1-16 ((https://doi.org/10.1007/s13225-020-00466-2)) |
| Otu46486 | 1 | k__Fungi(100);p__Ascomycota(100);c__Saccharomycetes(100);o__Saccharomycetales(100);f__Metschnikowiaceae(100);g__Metschnikowia(100);g__Metschnikowia_unclassified(100); | Metschnikowia | 13 | Saprotroph | \|Undefined Saprotroph\| | Yeast | NULL | Probable | As nectar-tap_saprotroph (Põlme et al. 2020); Decay sugar-rich substrates (Põlme et al. 2020) | Põlme S, et al. 2020. Fungal Diversity 105:1-16 ((https://doi.org/10.1007/s13225-020-00466-2)) |
| Otu46487 | 1 | k__Fungi(100);p__Ascomycota(100);c__Saccharomycetes(100);o__Saccharomycetales(100);f__Metschnikowiaceae(100);g__Metschnikowia(100);g__Metschnikowia_unclassified(100); | Metschnikowia | 13 | Saprotroph | \|Undefined Saprotroph\| | Yeast | NULL | Probable | As nectar-tap_saprotroph (Põlme et al. 2020); Decay sugar-rich substrates (Põlme et al. 2020) | Põlme S, et al. 2020. Fungal Diversity 105:1-16 ((https://doi.org/10.1007/s13225-020-00466-2)) |
| Otu46488 | 1 | k__Fungi(100);p__Ascomycota(100);c__Saccharomycetes(100);o__Saccharomycetales(100);f__Metschnikowiaceae(100);g__Metschnikowia(100);s__Metschnikowia_pulcherrima(100); | Metschnikowia | 13 | Saprotroph | \|Undefined Saprotroph\| | Yeast | NULL | Probable | As nectar-tap_saprotroph (Põlme et al. 2020); Decay sugar-rich substrates (Põlme et al. 2020) | Põlme S, et al. 2020. Fungal Diversity 105:1-16 ((https://doi.org/10.1007/s13225-020-00466-2)) |
| Otu46489 | 1 | k__Fungi(100);p__Ascomycota(100);c__Saccharomycetes(100);o__Saccharomycetales(100);f__Metschnikowiaceae(100);g__Metschnikowia(100);s__Metschnikowia_pulcherrima(100); | Metschnikowia | 13 | Saprotroph | \|Undefined Saprotroph\| | Yeast | NULL | Probable | As nectar-tap_saprotroph (Põlme et al. 2020); Decay sugar-rich substrates (Põlme et al. 2020) | Põlme S, et al. 2020. Fungal Diversity 105:1-16 ((https://doi.org/10.1007/s13225-020-00466-2)) |
| Otu46490 | 1 | k__Fungi(100);p__Ascomycota(100);c__Saccharomycetes(100);o__Saccharomycetales(100);f__Metschnikowiaceae(100);g__Metschnikowia(100);s__Metschnikowia_pulcherrima(100); | Metschnikowia | 13 | Saprotroph | \|Undefined Saprotroph\| | Yeast | NULL | Probable | As nectar-tap_saprotroph (Põlme et al. 2020); Decay sugar-rich substrates (Põlme et al. 2020) | Põlme S, et al. 2020. Fungal Diversity 105:1-16 ((https://doi.org/10.1007/s13225-020-00466-2)) |
| Otu46491 | 1 | k__Fungi(100);p__Ascomycota(100);c__Saccharomycetes(100);o__Saccharomycetales(100);f__Metschnikowiaceae(100);g__Metschnikowia(100);s__Metschnikowia_pulcherrima(100); | Metschnikowia | 13 | Saprotroph | \|Undefined Saprotroph\| | Yeast | NULL | Probable | As nectar-tap_saprotroph (Põlme et al. 2020); Decay sugar-rich substrates (Põlme et al. 2020) | Põlme S, et al. 2020. Fungal Diversity 105:1-16 ((https://doi.org/10.1007/s13225-020-00466-2)) |
| Otu46492 | 1 | k__Fungi(100);p__Ascomycota(100);c__Saccharomycetes(100);o__Saccharomycetales(100);f__Metschnikowiaceae(100);g__Metschnikowia(100);g__Metschnikowia_unclassified(100); | Metschnikowia | 13 | Saprotroph | \|Undefined Saprotroph\| | Yeast | NULL | Probable | As nectar-tap_saprotroph (Põlme et al. 2020); Decay sugar-rich substrates (Põlme et al. 2020) | Põlme S, et al. 2020. Fungal Diversity 105:1-16 ((https://doi.org/10.1007/s13225-020-00466-2)) |
| Otu46493 | 1 | k__Fungi(100);p__Ascomycota(100);c__Saccharomycetes(100);o__Saccharomycetales(100);f__Metschnikowiaceae(100);g__Metschnikowia(100);s__Metschnikowia_pulcherrima(100); | Metschnikowia | 13 | Saprotroph | \|Undefined Saprotroph\| | Yeast | NULL | Probable | As nectar-tap_saprotroph (Põlme et al. 2020); Decay sugar-rich substrates (Põlme et al. 2020) | Põlme S, et al. 2020. Fungal Diversity 105:1-16 ((https://doi.org/10.1007/s13225-020-00466-2)) |
| Otu46494 | 1 | k__Fungi(100);p__Ascomycota(100);c__Saccharomycetes(100);o__Saccharomycetales(100);f__Metschnikowiaceae(100);g__Metschnikowia(100);s__Metschnikowia_pulcherrima(100); | Metschnikowia | 13 | Saprotroph | \|Undefined Saprotroph\| | Yeast | NULL | Probable | As nectar-tap_saprotroph (Põlme et al. 2020); Decay sugar-rich substrates (Põlme et al. 2020) | Põlme S, et al. 2020. Fungal Diversity 105:1-16 ((https://doi.org/10.1007/s13225-020-00466-2)) |
| Otu46495 | 1 | k__Fungi(100);p__Ascomycota(100);c__Saccharomycetes(100);o__Saccharomycetales(100);f__Metschnikowiaceae(100);g__Metschnikowia(100);g__Metschnikowia_unclassified(100); | Metschnikowia | 13 | Saprotroph | \|Undefined Saprotroph\| | Yeast | NULL | Probable | As nectar-tap_saprotroph (Põlme et al. 2020); Decay sugar-rich substrates (Põlme et al. 2020) | Põlme S, et al. 2020. Fungal Diversity 105:1-16 ((https://doi.org/10.1007/s13225-020-00466-2)) |
| Otu46496 | 1 | k__Fungi(100);p__Ascomycota(100);c__Saccharomycetes(100);o__Saccharomycetales(100);f__Metschnikowiaceae(100);g__Metschnikowia(100);s__Metschnikowia_pulcherrima(100); | Metschnikowia | 13 | Saprotroph | \|Undefined Saprotroph\| | Yeast | NULL | Probable | As nectar-tap_saprotroph (Põlme et al. 2020); Decay sugar-rich substrates (Põlme et al. 2020) | Põlme S, et al. 2020. Fungal Diversity 105:1-16 ((https://doi.org/10.1007/s13225-020-00466-2)) |
| Otu46497 | 1 | k__Fungi(100);p__Ascomycota(100);c__Saccharomycetes(100);o__Saccharomycetales(100);f__Metschnikowiaceae(100);g__Metschnikowia(100);g__Metschnikowia_unclassified(100); | Metschnikowia | 13 | Saprotroph | \|Undefined Saprotroph\| | Yeast | NULL | Probable | As nectar-tap_saprotroph (Põlme et al. 2020); Decay sugar-rich substrates (Põlme et al. 2020) | Põlme S, et al. 2020. Fungal Diversity 105:1-16 ((https://doi.org/10.1007/s13225-020-00466-2)) |
| Otu46498 | 1 | k__Fungi(100);p__Ascomycota(100);c__Saccharomycetes(100);o__Saccharomycetales(100);f__Metschnikowiaceae(100);g__Metschnikowia(100);s__Metschnikowia_pulcherrima(100); | Metschnikowia | 13 | Saprotroph | \|Undefined Saprotroph\| | Yeast | NULL | Probable | As nectar-tap_saprotroph (Põlme et al. 2020); Decay sugar-rich substrates (Põlme et al. 2020) | Põlme S, et al. 2020. Fungal Diversity 105:1-16 ((https://doi.org/10.1007/s13225-020-00466-2)) |
| Otu46499 | 1 | k__Fungi(100);p__Ascomycota(100);c__Saccharomycetes(100);o__Saccharomycetales(100);f__Metschnikowiaceae(100);g__Metschnikowia(100);g__Metschnikowia_unclassified(100); | Metschnikowia | 13 | Saprotroph | \|Undefined Saprotroph\| | Yeast | NULL | Probable | As nectar-tap_saprotroph (Põlme et al. 2020); Decay sugar-rich substrates (Põlme et al. 2020) | Põlme S, et al. 2020. Fungal Diversity 105:1-16 ((https://doi.org/10.1007/s13225-020-00466-2)) |
| Otu46500 | 1 | k__Fungi(100);p__Ascomycota(100);c__Saccharomycetes(100);o__Saccharomycetales(100);f__Metschnikowiaceae(100);g__Metschnikowia(100);s__Metschnikowia_pulcherrima(100); | Metschnikowia | 13 | Saprotroph | \|Undefined Saprotroph\| | Yeast | NULL | Probable | As nectar-tap_saprotroph (Põlme et al. 2020); Decay sugar-rich substrates (Põlme et al. 2020) | Põlme S, et al. 2020. Fungal Diversity 105:1-16 ((https://doi.org/10.1007/s13225-020-00466-2)) |
| Otu46501 | 1 | k__Fungi(100);p__Ascomycota(100);c__Saccharomycetes(100);o__Saccharomycetales(100);f__Metschnikowiaceae(100);g__Metschnikowia(100);s__Metschnikowia_pulcherrima(100); | Metschnikowia | 13 | Saprotroph | \|Undefined Saprotroph\| | Yeast | NULL | Probable | As nectar-tap_saprotroph (Põlme et al. 2020); Decay sugar-rich substrates (Põlme et al. 2020) | Põlme S, et al. 2020. Fungal Diversity 105:1-16 ((https://doi.org/10.1007/s13225-020-00466-2)) |
| Otu46502 | 1 | k__Fungi(100);p__Ascomycota(100);c__Saccharomycetes(100);o__Saccharomycetales(100);f__Metschnikowiaceae(100);g__Metschnikowia(100);s__Metschnikowia_pulcherrima(100); | Metschnikowia | 13 | Saprotroph | \|Undefined Saprotroph\| | Yeast | NULL | Probable | As nectar-tap_saprotroph (Põlme et al. 2020); Decay sugar-rich substrates (Põlme et al. 2020) | Põlme S, et al. 2020. Fungal Diversity 105:1-16 ((https://doi.org/10.1007/s13225-020-00466-2)) |
| Otu46503 | 1 | k__Fungi(100);p__Ascomycota(100);c__Saccharomycetes(100);o__Saccharomycetales(100);f__Metschnikowiaceae(100);g__Metschnikowia(100);s__Metschnikowia_pulcherrima(100); | Metschnikowia | 13 | Saprotroph | \|Undefined Saprotroph\| | Yeast | NULL | Probable | As nectar-tap_saprotroph (Põlme et al. 2020); Decay sugar-rich substrates (Põlme et al. 2020) | Põlme S, et al. 2020. Fungal Diversity 105:1-16 ((https://doi.org/10.1007/s13225-020-00466-2)) |
| Otu46504 | 1 | k__Fungi(100);p__Ascomycota(100);c__Saccharomycetes(100);o__Saccharomycetales(100);f__Metschnikowiaceae(100);g__Metschnikowia(100);s__Metschnikowia_pulcherrima(100); | Metschnikowia | 13 | Saprotroph | \|Undefined Saprotroph\| | Yeast | NULL | Probable | As nectar-tap_saprotroph (Põlme et al. 2020); Decay sugar-rich substrates (Põlme et al. 2020) | Põlme S, et al. 2020. Fungal Diversity 105:1-16 ((https://doi.org/10.1007/s13225-020-00466-2)) |
| Otu46505 | 1 | k__Fungi(100);p__Ascomycota(100);c__Saccharomycetes(100);o__Saccharomycetales(100);f__Metschnikowiaceae(100);g__Metschnikowia(100);g__Metschnikowia_unclassified(100); | Metschnikowia | 13 | Saprotroph | \|Undefined Saprotroph\| | Yeast | NULL | Probable | As nectar-tap_saprotroph (Põlme et al. 2020); Decay sugar-rich substrates (Põlme et al. 2020) | Põlme S, et al. 2020. Fungal Diversity 105:1-16 ((https://doi.org/10.1007/s13225-020-00466-2)) |
| Otu46506 | 1 | k__Fungi(100);p__Ascomycota(100);c__Saccharomycetes(100);o__Saccharomycetales(100);f__Metschnikowiaceae(100);g__Metschnikowia(100);s__Metschnikowia_pulcherrima(100); | Metschnikowia | 13 | Saprotroph | \|Undefined Saprotroph\| | Yeast | NULL | Probable | As nectar-tap_saprotroph (Põlme et al. 2020); Decay sugar-rich substrates (Põlme et al. 2020) | Põlme S, et al. 2020. Fungal Diversity 105:1-16 ((https://doi.org/10.1007/s13225-020-00466-2)) |
| Otu46507 | 1 | k__Fungi(100);p__Ascomycota(100);c__Saccharomycetes(100);o__Saccharomycetales(100);f__Metschnikowiaceae(100);g__Metschnikowia(100);s__Metschnikowia_pulcherrima(100); | Metschnikowia | 13 | Saprotroph | \|Undefined Saprotroph\| | Yeast | NULL | Probable | As nectar-tap_saprotroph (Põlme et al. 2020); Decay sugar-rich substrates (Põlme et al. 2020) | Põlme S, et al. 2020. Fungal Diversity 105:1-16 ((https://doi.org/10.1007/s13225-020-00466-2)) |
| Otu46508 | 1 | k__Fungi(100);p__Ascomycota(100);c__Saccharomycetes(100);o__Saccharomycetales(100);f__Metschnikowiaceae(100);g__Metschnikowia(100);s__Metschnikowia_pulcherrima(100); | Metschnikowia | 13 | Saprotroph | \|Undefined Saprotroph\| | Yeast | NULL | Probable | As nectar-tap_saprotroph (Põlme et al. 2020); Decay sugar-rich substrates (Põlme et al. 2020) | Põlme S, et al. 2020. Fungal Diversity 105:1-16 ((https://doi.org/10.1007/s13225-020-00466-2)) |
| Otu46509 | 1 | k__Fungi(100);p__Ascomycota(100);c__Saccharomycetes(100);o__Saccharomycetales(100);f__Metschnikowiaceae(100);g__Metschnikowia(100);s__Metschnikowia_pulcherrima(100); | Metschnikowia | 13 | Saprotroph | \|Undefined Saprotroph\| | Yeast | NULL | Probable | As nectar-tap_saprotroph (Põlme et al. 2020); Decay sugar-rich substrates (Põlme et al. 2020) | Põlme S, et al. 2020. Fungal Diversity 105:1-16 ((https://doi.org/10.1007/s13225-020-00466-2)) |
| Otu46510 | 1 | k__Fungi(100);p__Basidiomycota(100);c__Tremellomycetes(100);o__Tremellales(100);f__Tremellales_family_Incertae_sedis(100);g__Hannaella(100);s__Hannaella_sp(100); | Hannaella | 13 | Pathotroph-Saprotroph | \|Animal Parasite\|-Undefined Saprotroph | Yeast | NULL | Probable | As animal_decomposer (Põlme et al. 2020); Decay animal material (Põlme et al. 2020) | Põlme S, et al. 2020. Fungal Diversity 105:1-16 ((https://doi.org/10.1007/s13225-020-00466-2)) |
| Otu46511 | 1 | k__Fungi(100);p__Ascomycota(100);c__Saccharomycetes(100);o__Saccharomycetales(100);f__Metschnikowiaceae(100);g__Metschnikowia(100);s__Metschnikowia_pulcherrima(100); | Metschnikowia | 13 | Saprotroph | \|Undefined Saprotroph\| | Yeast | NULL | Probable | As nectar-tap_saprotroph (Põlme et al. 2020); Decay sugar-rich substrates (Põlme et al. 2020) | Põlme S, et al. 2020. Fungal Diversity 105:1-16 ((https://doi.org/10.1007/s13225-020-00466-2)) |
| Otu46512 | 1 | k__Fungi(100);p__Ascomycota(100);c__Saccharomycetes(100);o__Saccharomycetales(100);f__Metschnikowiaceae(100);g__Metschnikowia(100);s__Metschnikowia_pulcherrima(100); | Metschnikowia | 13 | Saprotroph | \|Undefined Saprotroph\| | Yeast | NULL | Probable | As nectar-tap_saprotroph (Põlme et al. 2020); Decay sugar-rich substrates (Põlme et al. 2020) | Põlme S, et al. 2020. Fungal Diversity 105:1-16 ((https://doi.org/10.1007/s13225-020-00466-2)) |
| Otu46513 | 1 | k__Fungi(100);p__Ascomycota(100);c__Saccharomycetes(100);o__Saccharomycetales(100);f__Metschnikowiaceae(100);g__Metschnikowia(100);s__Metschnikowia_pulcherrima(100); | Metschnikowia | 13 | Saprotroph | \|Undefined Saprotroph\| | Yeast | NULL | Probable | As nectar-tap_saprotroph (Põlme et al. 2020); Decay sugar-rich substrates (Põlme et al. 2020) | Põlme S, et al. 2020. Fungal Diversity 105:1-16 ((https://doi.org/10.1007/s13225-020-00466-2)) |
| Otu46514 | 1 | k__Fungi(100);p__Ascomycota(100);c__Saccharomycetes(100);o__Saccharomycetales(100);f__Metschnikowiaceae(100);g__Metschnikowia(100);s__Metschnikowia_pulcherrima(100); | Metschnikowia | 13 | Saprotroph | \|Undefined Saprotroph\| | Yeast | NULL | Probable | As nectar-tap_saprotroph (Põlme et al. 2020); Decay sugar-rich substrates (Põlme et al. 2020) | Põlme S, et al. 2020. Fungal Diversity 105:1-16 ((https://doi.org/10.1007/s13225-020-00466-2)) |
| Otu46515 | 1 | k__Fungi(100);p__Ascomycota(100);c__Saccharomycetes(100);o__Saccharomycetales(100);f__Metschnikowiaceae(100);g__Metschnikowia(100);s__Metschnikowia_viticola(100); | Metschnikowia | 13 | Saprotroph | \|Undefined Saprotroph\| | Yeast | NULL | Probable | As nectar-tap_saprotroph (Põlme et al. 2020); Decay sugar-rich substrates (Põlme et al. 2020) | Põlme S, et al. 2020. Fungal Diversity 105:1-16 ((https://doi.org/10.1007/s13225-020-00466-2)) |
| Otu46516 | 1 | k__Fungi(100);p__Ascomycota(100);c__Saccharomycetes(100);o__Saccharomycetales(100);f__Metschnikowiaceae(100);g__Metschnikowia(100);s__Metschnikowia_pulcherrima(100); | Metschnikowia | 13 | Saprotroph | \|Undefined Saprotroph\| | Yeast | NULL | Probable | As nectar-tap_saprotroph (Põlme et al. 2020); Decay sugar-rich substrates (Põlme et al. 2020) | Põlme S, et al. 2020. Fungal Diversity 105:1-16 ((https://doi.org/10.1007/s13225-020-00466-2)) |
| Otu46517 | 1 | k__Fungi(100);p__Ascomycota(100);c__Saccharomycetes(100);o__Saccharomycetales(100);f__Metschnikowiaceae(100);g__Metschnikowia(100);s__Metschnikowia_pulcherrima(100); | Metschnikowia | 13 | Saprotroph | \|Undefined Saprotroph\| | Yeast | NULL | Probable | As nectar-tap_saprotroph (Põlme et al. 2020); Decay sugar-rich substrates (Põlme et al. 2020) | Põlme S, et al. 2020. Fungal Diversity 105:1-16 ((https://doi.org/10.1007/s13225-020-00466-2)) |
| Otu46518 | 1 | k__Fungi(100);p__Ascomycota(100);c__Saccharomycetes(100);o__Saccharomycetales(100);f__Metschnikowiaceae(100);g__Metschnikowia(100);s__Metschnikowia_pulcherrima(100); | Metschnikowia | 13 | Saprotroph | \|Undefined Saprotroph\| | Yeast | NULL | Probable | As nectar-tap_saprotroph (Põlme et al. 2020); Decay sugar-rich substrates (Põlme et al. 2020) | Põlme S, et al. 2020. Fungal Diversity 105:1-16 ((https://doi.org/10.1007/s13225-020-00466-2)) |
| Otu46519 | 1 | k__Fungi(100);p__Ascomycota(100);c__Saccharomycetes(100);o__Saccharomycetales(100);f__Metschnikowiaceae(100);g__Metschnikowia(100);s__Metschnikowia_pulcherrima(100); | Metschnikowia | 13 | Saprotroph | \|Undefined Saprotroph\| | Yeast | NULL | Probable | As nectar-tap_saprotroph (Põlme et al. 2020); Decay sugar-rich substrates (Põlme et al. 2020) | Põlme S, et al. 2020. Fungal Diversity 105:1-16 ((https://doi.org/10.1007/s13225-020-00466-2)) |
| Otu46520 | 1 | k__Fungi(100);p__Ascomycota(100);c__Saccharomycetes(100);o__Saccharomycetales(100);f__Metschnikowiaceae(100);g__Metschnikowia(100);s__Metschnikowia_pulcherrima(100); | Metschnikowia | 13 | Saprotroph | \|Undefined Saprotroph\| | Yeast | NULL | Probable | As nectar-tap_saprotroph (Põlme et al. 2020); Decay sugar-rich substrates (Põlme et al. 2020) | Põlme S, et al. 2020. Fungal Diversity 105:1-16 ((https://doi.org/10.1007/s13225-020-00466-2)) |
| Otu46521 | 1 | k__Fungi(100);p__Ascomycota(100);c__Saccharomycetes(100);o__Saccharomycetales(100);f__Metschnikowiaceae(100);g__Metschnikowia(100);s__Metschnikowia_pulcherrima(100); | Metschnikowia | 13 | Saprotroph | \|Undefined Saprotroph\| | Yeast | NULL | Probable | As nectar-tap_saprotroph (Põlme et al. 2020); Decay sugar-rich substrates (Põlme et al. 2020) | Põlme S, et al. 2020. Fungal Diversity 105:1-16 ((https://doi.org/10.1007/s13225-020-00466-2)) |
| Otu46522 | 1 | k__Fungi(100);p__Ascomycota(100);c__Saccharomycetes(100);o__Saccharomycetales(100);f__Metschnikowiaceae(100);g__Metschnikowia(100);s__Metschnikowia_pulcherrima(100); | Metschnikowia | 13 | Saprotroph | \|Undefined Saprotroph\| | Yeast | NULL | Probable | As nectar-tap_saprotroph (Põlme et al. 2020); Decay sugar-rich substrates (Põlme et al. 2020) | Põlme S, et al. 2020. Fungal Diversity 105:1-16 ((https://doi.org/10.1007/s13225-020-00466-2)) |
| Otu46523 | 1 | k__Fungi(100);p__Ascomycota(100);c__Saccharomycetes(100);o__Saccharomycetales(100);f__Metschnikowiaceae(100);g__Metschnikowia(100);g__Metschnikowia_unclassified(100); | Metschnikowia | 13 | Saprotroph | \|Undefined Saprotroph\| | Yeast | NULL | Probable | As nectar-tap_saprotroph (Põlme et al. 2020); Decay sugar-rich substrates (Põlme et al. 2020) | Põlme S, et al. 2020. Fungal Diversity 105:1-16 ((https://doi.org/10.1007/s13225-020-00466-2)) |
| Otu46524 | 1 | k__Fungi(100);p__Ascomycota(100);c__Saccharomycetes(100);o__Saccharomycetales(100);f__Metschnikowiaceae(100);g__Metschnikowia(100);s__Metschnikowia_pulcherrima(100); | Metschnikowia | 13 | Saprotroph | \|Undefined Saprotroph\| | Yeast | NULL | Probable | As nectar-tap_saprotroph (Põlme et al. 2020); Decay sugar-rich substrates (Põlme et al. 2020) | Põlme S, et al. 2020. Fungal Diversity 105:1-16 ((https://doi.org/10.1007/s13225-020-00466-2)) |
| Otu46525 | 1 | k__Fungi(100);p__Ascomycota(100);c__Saccharomycetes(100);o__Saccharomycetales(100);f__Metschnikowiaceae(100);g__Metschnikowia(100);g__Metschnikowia_unclassified(100); | Metschnikowia | 13 | Saprotroph | \|Undefined Saprotroph\| | Yeast | NULL | Probable | As nectar-tap_saprotroph (Põlme et al. 2020); Decay sugar-rich substrates (Põlme et al. 2020) | Põlme S, et al. 2020. Fungal Diversity 105:1-16 ((https://doi.org/10.1007/s13225-020-00466-2)) |
| Otu46526 | 1 | k__Fungi(100);p__Ascomycota(100);c__Saccharomycetes(100);o__Saccharomycetales(100);f__Metschnikowiaceae(100);g__Metschnikowia(100);g__Metschnikowia_unclassified(100); | Metschnikowia | 13 | Saprotroph | \|Undefined Saprotroph\| | Yeast | NULL | Probable | As nectar-tap_saprotroph (Põlme et al. 2020); Decay sugar-rich substrates (Põlme et al. 2020) | Põlme S, et al. 2020. Fungal Diversity 105:1-16 ((https://doi.org/10.1007/s13225-020-00466-2)) |
| Otu46527 | 1 | k__Fungi(100);p__Ascomycota(100);c__Saccharomycetes(100);o__Saccharomycetales(100);f__Metschnikowiaceae(100);g__Metschnikowia(100);g__Metschnikowia_unclassified(100); | Metschnikowia | 13 | Saprotroph | \|Undefined Saprotroph\| | Yeast | NULL | Probable | As nectar-tap_saprotroph (Põlme et al. 2020); Decay sugar-rich substrates (Põlme et al. 2020) | Põlme S, et al. 2020. Fungal Diversity 105:1-16 ((https://doi.org/10.1007/s13225-020-00466-2)) |
| Otu46528 | 1 | k__Fungi(100);p__Ascomycota(100);c__Saccharomycetes(100);o__Saccharomycetales(100);f__Metschnikowiaceae(100);g__Metschnikowia(100);g__Metschnikowia_unclassified(100); | Metschnikowia | 13 | Saprotroph | \|Undefined Saprotroph\| | Yeast | NULL | Probable | As nectar-tap_saprotroph (Põlme et al. 2020); Decay sugar-rich substrates (Põlme et al. 2020) | Põlme S, et al. 2020. Fungal Diversity 105:1-16 ((https://doi.org/10.1007/s13225-020-00466-2)) |
| Otu46529 | 1 | k__Fungi(100);p__Ascomycota(100);c__Saccharomycetes(100);o__Saccharomycetales(100);f__Metschnikowiaceae(100);g__Metschnikowia(100);g__Metschnikowia_unclassified(100); | Metschnikowia | 13 | Saprotroph | \|Undefined Saprotroph\| | Yeast | NULL | Probable | As nectar-tap_saprotroph (Põlme et al. 2020); Decay sugar-rich substrates (Põlme et al. 2020) | Põlme S, et al. 2020. Fungal Diversity 105:1-16 ((https://doi.org/10.1007/s13225-020-00466-2)) |
| Otu46530 | 1 | k__Fungi(100);p__Ascomycota(100);c__Saccharomycetes(100);o__Saccharomycetales(100);f__Metschnikowiaceae(100);g__Metschnikowia(100);g__Metschnikowia_unclassified(100); | Metschnikowia | 13 | Saprotroph | \|Undefined Saprotroph\| | Yeast | NULL | Probable | As nectar-tap_saprotroph (Põlme et al. 2020); Decay sugar-rich substrates (Põlme et al. 2020) | Põlme S, et al. 2020. Fungal Diversity 105:1-16 ((https://doi.org/10.1007/s13225-020-00466-2)) |
| Otu46531 | 1 | k__Fungi(100);p__Ascomycota(100);c__Saccharomycetes(100);o__Saccharomycetales(100);f__Metschnikowiaceae(100);g__Metschnikowia(100);g__Metschnikowia_unclassified(100); | Metschnikowia | 13 | Saprotroph | \|Undefined Saprotroph\| | Yeast | NULL | Probable | As nectar-tap_saprotroph (Põlme et al. 2020); Decay sugar-rich substrates (Põlme et al. 2020) | Põlme S, et al. 2020. Fungal Diversity 105:1-16 ((https://doi.org/10.1007/s13225-020-00466-2)) |
| Otu46532 | 1 | k__Fungi(100);p__Ascomycota(100);c__Saccharomycetes(100);o__Saccharomycetales(100);f__Metschnikowiaceae(100);g__Metschnikowia(100);s__Metschnikowia_pulcherrima(100); | Metschnikowia | 13 | Saprotroph | \|Undefined Saprotroph\| | Yeast | NULL | Probable | As nectar-tap_saprotroph (Põlme et al. 2020); Decay sugar-rich substrates (Põlme et al. 2020) | Põlme S, et al. 2020. Fungal Diversity 105:1-16 ((https://doi.org/10.1007/s13225-020-00466-2)) |
| Otu46533 | 1 | k__Fungi(100);p__Ascomycota(100);c__Saccharomycetes(100);o__Saccharomycetales(100);f__Metschnikowiaceae(100);g__Metschnikowia(100);g__Metschnikowia_unclassified(100); | Metschnikowia | 13 | Saprotroph | \|Undefined Saprotroph\| | Yeast | NULL | Probable | As nectar-tap_saprotroph (Põlme et al. 2020); Decay sugar-rich substrates (Põlme et al. 2020) | Põlme S, et al. 2020. Fungal Diversity 105:1-16 ((https://doi.org/10.1007/s13225-020-00466-2)) |
| Otu46534 | 1 | k__Fungi(100);p__Ascomycota(100);c__Saccharomycetes(100);o__Saccharomycetales(100);f__Metschnikowiaceae(100);g__Metschnikowia(100);s__Metschnikowia_pulcherrima(100); | Metschnikowia | 13 | Saprotroph | \|Undefined Saprotroph\| | Yeast | NULL | Probable | As nectar-tap_saprotroph (Põlme et al. 2020); Decay sugar-rich substrates (Põlme et al. 2020) | Põlme S, et al. 2020. Fungal Diversity 105:1-16 ((https://doi.org/10.1007/s13225-020-00466-2)) |
| Otu46535 | 1 | k__Fungi(100);p__Ascomycota(100);c__Saccharomycetes(100);o__Saccharomycetales(100);f__Metschnikowiaceae(100);g__Metschnikowia(100);s__Metschnikowia_pulcherrima(100); | Metschnikowia | 13 | Saprotroph | \|Undefined Saprotroph\| | Yeast | NULL | Probable | As nectar-tap_saprotroph (Põlme et al. 2020); Decay sugar-rich substrates (Põlme et al. 2020) | Põlme S, et al. 2020. Fungal Diversity 105:1-16 ((https://doi.org/10.1007/s13225-020-00466-2)) |
| Otu46536 | 1 | k__Fungi(100);p__Ascomycota(100);c__Saccharomycetes(100);o__Saccharomycetales(100);f__Metschnikowiaceae(100);g__Metschnikowia(100);g__Metschnikowia_unclassified(100); | Metschnikowia | 13 | Saprotroph | \|Undefined Saprotroph\| | Yeast | NULL | Probable | As nectar-tap_saprotroph (Põlme et al. 2020); Decay sugar-rich substrates (Põlme et al. 2020) | Põlme S, et al. 2020. Fungal Diversity 105:1-16 ((https://doi.org/10.1007/s13225-020-00466-2)) |
| Otu46537 | 1 | k__Fungi(100);p__Ascomycota(100);c__Saccharomycetes(100);o__Saccharomycetales(100);f__Metschnikowiaceae(100);g__Metschnikowia(100);g__Metschnikowia_unclassified(100); | Metschnikowia | 13 | Saprotroph | \|Undefined Saprotroph\| | Yeast | NULL | Probable | As nectar-tap_saprotroph (Põlme et al. 2020); Decay sugar-rich substrates (Põlme et al. 2020) | Põlme S, et al. 2020. Fungal Diversity 105:1-16 ((https://doi.org/10.1007/s13225-020-00466-2)) |
| Otu46538 | 1 | k__Fungi(100);p__Ascomycota(100);c__Saccharomycetes(100);o__Saccharomycetales(100);f__Metschnikowiaceae(100);g__Metschnikowia(100);s__Metschnikowia_pulcherrima(100); | Metschnikowia | 13 | Saprotroph | \|Undefined Saprotroph\| | Yeast | NULL | Probable | As nectar-tap_saprotroph (Põlme et al. 2020); Decay sugar-rich substrates (Põlme et al. 2020) | Põlme S, et al. 2020. Fungal Diversity 105:1-16 ((https://doi.org/10.1007/s13225-020-00466-2)) |
| Otu46539 | 1 | k__Fungi(100);p__Ascomycota(100);c__Saccharomycetes(100);o__Saccharomycetales(100);f__Metschnikowiaceae(100);g__Metschnikowia(100);s__Metschnikowia_pulcherrima(100); | Metschnikowia | 13 | Saprotroph | \|Undefined Saprotroph\| | Yeast | NULL | Probable | As nectar-tap_saprotroph (Põlme et al. 2020); Decay sugar-rich substrates (Põlme et al. 2020) | Põlme S, et al. 2020. Fungal Diversity 105:1-16 ((https://doi.org/10.1007/s13225-020-00466-2)) |
| Otu46540 | 1 | k__Fungi(100);p__Ascomycota(100);c__Saccharomycetes(100);o__Saccharomycetales(100);f__Metschnikowiaceae(100);g__Metschnikowia(100);s__Metschnikowia_pulcherrima(100); | Metschnikowia | 13 | Saprotroph | \|Undefined Saprotroph\| | Yeast | NULL | Probable | As nectar-tap_saprotroph (Põlme et al. 2020); Decay sugar-rich substrates (Põlme et al. 2020) | Põlme S, et al. 2020. Fungal Diversity 105:1-16 ((https://doi.org/10.1007/s13225-020-00466-2)) |
| Otu46541 | 1 | k__Fungi(100);p__Ascomycota(100);c__Saccharomycetes(100);o__Saccharomycetales(100);f__Metschnikowiaceae(100);g__Metschnikowia(100);g__Metschnikowia_unclassified(100); | Metschnikowia | 13 | Saprotroph | \|Undefined Saprotroph\| | Yeast | NULL | Probable | As nectar-tap_saprotroph (Põlme et al. 2020); Decay sugar-rich substrates (Põlme et al. 2020) | Põlme S, et al. 2020. Fungal Diversity 105:1-16 ((https://doi.org/10.1007/s13225-020-00466-2)) |
| Otu46542 | 1 | k__Fungi(100);p__Ascomycota(100);c__Saccharomycetes(100);o__Saccharomycetales(100);f__Metschnikowiaceae(100);g__Metschnikowia(100);s__Metschnikowia_pulcherrima(100); | Metschnikowia | 13 | Saprotroph | \|Undefined Saprotroph\| | Yeast | NULL | Probable | As nectar-tap_saprotroph (Põlme et al. 2020); Decay sugar-rich substrates (Põlme et al. 2020) | Põlme S, et al. 2020. Fungal Diversity 105:1-16 ((https://doi.org/10.1007/s13225-020-00466-2)) |
| Otu46543 | 1 | k__Fungi(100);p__Ascomycota(100);c__Saccharomycetes(100);o__Saccharomycetales(100);f__Metschnikowiaceae(100);g__Metschnikowia(100);s__Metschnikowia_pulcherrima(100); | Metschnikowia | 13 | Saprotroph | \|Undefined Saprotroph\| | Yeast | NULL | Probable | As nectar-tap_saprotroph (Põlme et al. 2020); Decay sugar-rich substrates (Põlme et al. 2020) | Põlme S, et al. 2020. Fungal Diversity 105:1-16 ((https://doi.org/10.1007/s13225-020-00466-2)) |
| Otu46544 | 1 | k__Fungi(100);p__Ascomycota(100);c__Saccharomycetes(100);o__Saccharomycetales(100);f__Metschnikowiaceae(100);g__Metschnikowia(100);g__Metschnikowia_unclassified(100); | Metschnikowia | 13 | Saprotroph | \|Undefined Saprotroph\| | Yeast | NULL | Probable | As nectar-tap_saprotroph (Põlme et al. 2020); Decay sugar-rich substrates (Põlme et al. 2020) | Põlme S, et al. 2020. Fungal Diversity 105:1-16 ((https://doi.org/10.1007/s13225-020-00466-2)) |
| Otu46545 | 1 | k__Fungi(100);p__Ascomycota(100);c__Saccharomycetes(100);o__Saccharomycetales(100);f__Metschnikowiaceae(100);g__Metschnikowia(100);s__Metschnikowia_pulcherrima(100); | Metschnikowia | 13 | Saprotroph | \|Undefined Saprotroph\| | Yeast | NULL | Probable | As nectar-tap_saprotroph (Põlme et al. 2020); Decay sugar-rich substrates (Põlme et al. 2020) | Põlme S, et al. 2020. Fungal Diversity 105:1-16 ((https://doi.org/10.1007/s13225-020-00466-2)) |
| Otu46546 | 1 | k__Fungi(100);p__Ascomycota(100);c__Saccharomycetes(100);o__Saccharomycetales(100);f__Metschnikowiaceae(100);g__Metschnikowia(100);s__Metschnikowia_pulcherrima(100); | Metschnikowia | 13 | Saprotroph | \|Undefined Saprotroph\| | Yeast | NULL | Probable | As nectar-tap_saprotroph (Põlme et al. 2020); Decay sugar-rich substrates (Põlme et al. 2020) | Põlme S, et al. 2020. Fungal Diversity 105:1-16 ((https://doi.org/10.1007/s13225-020-00466-2)) |
| Otu46547 | 1 | k__Fungi(100);p__Ascomycota(100);c__Saccharomycetes(100);o__Saccharomycetales(100);f__Metschnikowiaceae(100);g__Metschnikowia(100);s__Metschnikowia_pulcherrima(100); | Metschnikowia | 13 | Saprotroph | \|Undefined Saprotroph\| | Yeast | NULL | Probable | As nectar-tap_saprotroph (Põlme et al. 2020); Decay sugar-rich substrates (Põlme et al. 2020) | Põlme S, et al. 2020. Fungal Diversity 105:1-16 ((https://doi.org/10.1007/s13225-020-00466-2)) |
| Otu46548 | 1 | k__Fungi(100);p__Ascomycota(100);c__Saccharomycetes(100);o__Saccharomycetales(100);f__Metschnikowiaceae(100);g__Metschnikowia(100);s__Metschnikowia_pulcherrima(100); | Metschnikowia | 13 | Saprotroph | \|Undefined Saprotroph\| | Yeast | NULL | Probable | As nectar-tap_saprotroph (Põlme et al. 2020); Decay sugar-rich substrates (Põlme et al. 2020) | Põlme S, et al. 2020. Fungal Diversity 105:1-16 ((https://doi.org/10.1007/s13225-020-00466-2)) |
| Otu46549 | 1 | k__Fungi(100);p__Ascomycota(100);c__Sordariomycetes(100);o__Sordariales(100);f__Cephalothecaceae(100);f__Cephalothecaceae_unclassified(100);f__Cephalothecaceae_unclassified(100); | Cephalothecaceae | 9 | Saprotroph | Fungal Parasite-Wood Saprotroph | Microfungus | NULL | Probable | Typically saprobic, fruiting on very rotten wood or other fungi (Cannon & Kirk 2007) | Cannon PF, Kirk PM. 2007. Fungal Families of the World. CAB International, Cambridge (ISBN: 978-0851998275) |
| Otu46550 | 1 | k__Fungi(100);p__Ascomycota(100);c__Saccharomycetes(100);o__Saccharomycetales(100);f__Metschnikowiaceae(100);g__Metschnikowia(100);s__Metschnikowia_pulcherrima(100); | Metschnikowia | 13 | Saprotroph | \|Undefined Saprotroph\| | Yeast | NULL | Probable | As nectar-tap_saprotroph (Põlme et al. 2020); Decay sugar-rich substrates (Põlme et al. 2020) | Põlme S, et al. 2020. Fungal Diversity 105:1-16 ((https://doi.org/10.1007/s13225-020-00466-2)) |
| Otu46551 | 1 | k__Fungi(100);p__Ascomycota(100);c__Saccharomycetes(100);o__Saccharomycetales(100);f__Metschnikowiaceae(100);g__Metschnikowia(100);s__Metschnikowia_pulcherrima(100); | Metschnikowia | 13 | Saprotroph | \|Undefined Saprotroph\| | Yeast | NULL | Probable | As nectar-tap_saprotroph (Põlme et al. 2020); Decay sugar-rich substrates (Põlme et al. 2020) | Põlme S, et al. 2020. Fungal Diversity 105:1-16 ((https://doi.org/10.1007/s13225-020-00466-2)) |
| Otu46552 | 1 | k__Fungi(100);p__Ascomycota(100);c__Saccharomycetes(100);o__Saccharomycetales(100);f__Metschnikowiaceae(100);g__Metschnikowia(100);s__Metschnikowia_pulcherrima(100); | Metschnikowia | 13 | Saprotroph | \|Undefined Saprotroph\| | Yeast | NULL | Probable | As nectar-tap_saprotroph (Põlme et al. 2020); Decay sugar-rich substrates (Põlme et al. 2020) | Põlme S, et al. 2020. Fungal Diversity 105:1-16 ((https://doi.org/10.1007/s13225-020-00466-2)) |
| Otu46553 | 1 | k__Fungi(100);p__Ascomycota(100);c__Saccharomycetes(100);o__Saccharomycetales(100);f__Metschnikowiaceae(100);g__Metschnikowia(100);g__Metschnikowia_unclassified(100); | Metschnikowia | 13 | Saprotroph | \|Undefined Saprotroph\| | Yeast | NULL | Probable | As nectar-tap_saprotroph (Põlme et al. 2020); Decay sugar-rich substrates (Põlme et al. 2020) | Põlme S, et al. 2020. Fungal Diversity 105:1-16 ((https://doi.org/10.1007/s13225-020-00466-2)) |
| Otu46554 | 1 | k__Fungi(100);p__Ascomycota(100);c__Saccharomycetes(100);o__Saccharomycetales(100);f__Metschnikowiaceae(100);g__Metschnikowia(100);g__Metschnikowia_unclassified(100); | Metschnikowia | 13 | Saprotroph | \|Undefined Saprotroph\| | Yeast | NULL | Probable | As nectar-tap_saprotroph (Põlme et al. 2020); Decay sugar-rich substrates (Põlme et al. 2020) | Põlme S, et al. 2020. Fungal Diversity 105:1-16 ((https://doi.org/10.1007/s13225-020-00466-2)) |
| Otu46555 | 1 | k__Fungi(100);p__Ascomycota(100);c__Saccharomycetes(100);o__Saccharomycetales(100);f__Metschnikowiaceae(100);g__Metschnikowia(100);s__Metschnikowia_pulcherrima(100); | Metschnikowia | 13 | Saprotroph | \|Undefined Saprotroph\| | Yeast | NULL | Probable | As nectar-tap_saprotroph (Põlme et al. 2020); Decay sugar-rich substrates (Põlme et al. 2020) | Põlme S, et al. 2020. Fungal Diversity 105:1-16 ((https://doi.org/10.1007/s13225-020-00466-2)) |
| Otu46556 | 1 | k__Fungi(100);p__Ascomycota(100);c__Saccharomycetes(100);o__Saccharomycetales(100);f__Metschnikowiaceae(100);g__Metschnikowia(100);s__Metschnikowia_pulcherrima(100); | Metschnikowia | 13 | Saprotroph | \|Undefined Saprotroph\| | Yeast | NULL | Probable | As nectar-tap_saprotroph (Põlme et al. 2020); Decay sugar-rich substrates (Põlme et al. 2020) | Põlme S, et al. 2020. Fungal Diversity 105:1-16 ((https://doi.org/10.1007/s13225-020-00466-2)) |
| Otu46557 | 1 | k__Fungi(100);p__Ascomycota(100);c__Saccharomycetes(100);o__Saccharomycetales(100);f__Metschnikowiaceae(100);g__Metschnikowia(100);g__Metschnikowia_unclassified(100); | Metschnikowia | 13 | Saprotroph | \|Undefined Saprotroph\| | Yeast | NULL | Probable | As nectar-tap_saprotroph (Põlme et al. 2020); Decay sugar-rich substrates (Põlme et al. 2020) | Põlme S, et al. 2020. Fungal Diversity 105:1-16 ((https://doi.org/10.1007/s13225-020-00466-2)) |
| Otu46558 | 1 | k__Fungi(100);p__Ascomycota(100);c__Saccharomycetes(100);o__Saccharomycetales(100);f__Metschnikowiaceae(100);g__Metschnikowia(100);s__Metschnikowia_pulcherrima(100); | Metschnikowia | 13 | Saprotroph | \|Undefined Saprotroph\| | Yeast | NULL | Probable | As nectar-tap_saprotroph (Põlme et al. 2020); Decay sugar-rich substrates (Põlme et al. 2020) | Põlme S, et al. 2020. Fungal Diversity 105:1-16 ((https://doi.org/10.1007/s13225-020-00466-2)) |
| Otu46559 | 1 | k__Fungi(100);p__Ascomycota(100);c__Saccharomycetes(100);o__Saccharomycetales(100);f__Metschnikowiaceae(100);g__Metschnikowia(100);s__Metschnikowia_pulcherrima(100); | Metschnikowia | 13 | Saprotroph | \|Undefined Saprotroph\| | Yeast | NULL | Probable | As nectar-tap_saprotroph (Põlme et al. 2020); Decay sugar-rich substrates (Põlme et al. 2020) | Põlme S, et al. 2020. Fungal Diversity 105:1-16 ((https://doi.org/10.1007/s13225-020-00466-2)) |

| | | | | | | | | | | | |
|---|---|---|---|---|---|---|---|---|---|---|---|
| Otu46560 | 1 | k__Fungi(100);p__Ascomycota(100);c__Saccharomycetes(100);o__Saccharomycetales(100);f__Metschnikowiaceae(100);g__Metschnikowia(100);g__Metschnikowia_unclassified(100); | Metschnikowia | 13 | Saprotroph | \|Undefined Saprotroph\| | Yeast | NULL | Probable | As nectar-tap_saprotroph (Põlme et al. 2020); Decay sugar-rich substrates (Põlme et al. 2020) | Põlme S, et al. 2020. Fungal Diversity 105:1-16 ((https://doi.org/10.1007/s13225-020-00466-2)) |
| Otu46561 | 1 | k__Fungi(100);p__Ascomycota(100);c__Saccharomycetes(100);o__Saccharomycetales(100);f__Metschnikowiaceae(100);g__Metschnikowia(100);s__Metschnikowia_pulcherrima(100); | Metschnikowia | 13 | Saprotroph | \|Undefined Saprotroph\| | Yeast | NULL | Probable | As nectar-tap_saprotroph (Põlme et al. 2020); Decay sugar-rich substrates (Põlme et al. 2020) | Põlme S, et al. 2020. Fungal Diversity 105:1-16 ((https://doi.org/10.1007/s13225-020-00466-2)) |
| Otu46562 | 1 | k__Fungi(100);p__Ascomycota(100);c__Saccharomycetes(100);o__Saccharomycetales(100);f__Metschnikowiaceae(100);g__Metschnikowia(100);g__Metschnikowia_unclassified(100); | Metschnikowia | 13 | Saprotroph | \|Undefined Saprotroph\| | Yeast | NULL | Probable | As nectar-tap_saprotroph (Põlme et al. 2020); Decay sugar-rich substrates (Põlme et al. 2020) | Põlme S, et al. 2020. Fungal Diversity 105:1-16 ((https://doi.org/10.1007/s13225-020-00466-2)) |
| Otu46563 | 1 | k__Fungi(100);p__Ascomycota(100);c__Saccharomycetes(100);o__Saccharomycetales(100);f__Metschnikowiaceae(100);g__Metschnikowia(100);s__Metschnikowia_pulcherrima(100); | Metschnikowia | 13 | Saprotroph | \|Undefined Saprotroph\| | Yeast | NULL | Probable | As nectar-tap_saprotroph (Põlme et al. 2020); Decay sugar-rich substrates (Põlme et al. 2020) | Põlme S, et al. 2020. Fungal Diversity 105:1-16 ((https://doi.org/10.1007/s13225-020-00466-2)) |
| Otu46565 | 1 | k__Fungi(100);p__Ascomycota(100);c__Saccharomycetes(100);o__Saccharomycetales(100);f__Metschnikowiaceae(100);g__Metschnikowia(100);g__Metschnikowia_unclassified(100); | Metschnikowia | 13 | Saprotroph | \|Undefined Saprotroph\| | Yeast | NULL | Probable | As nectar-tap_saprotroph (Põlme et al. 2020); Decay sugar-rich substrates (Põlme et al. 2020) | Põlme S, et al. 2020. Fungal Diversity 105:1-16 ((https://doi.org/10.1007/s13225-020-00466-2)) |
| Otu46566 | 1 | k__Fungi(100);p__Ascomycota(100);c__Saccharomycetes(100);o__Saccharomycetales(100);f__Metschnikowiaceae(100);g__Metschnikowia(100);s__Metschnikowia_pulcherrima(100); | Metschnikowia | 13 | Saprotroph | \|Undefined Saprotroph\| | Yeast | NULL | Probable | As nectar-tap_saprotroph (Põlme et al. 2020); Decay sugar-rich substrates (Põlme et al. 2020) | Põlme S, et al. 2020. Fungal Diversity 105:1-16 ((https://doi.org/10.1007/s13225-020-00466-2)) |
| Otu46567 | 1 | k__Fungi(100);p__Ascomycota(100);c__Saccharomycetes(100);o__Saccharomycetales(100);f__Metschnikowiaceae(100);g__Metschnikowia(100);s__Metschnikowia_pulcherrima(100); | Metschnikowia | 13 | Saprotroph | \|Undefined Saprotroph\| | Yeast | NULL | Probable | As nectar-tap_saprotroph (Põlme et al. 2020); Decay sugar-rich substrates (Põlme et al. 2020) | Põlme S, et al. 2020. Fungal Diversity 105:1-16 ((https://doi.org/10.1007/s13225-020-00466-2)) |
| Otu46568 | 1 | k__Fungi(100);p__Ascomycota(100);c__Saccharomycetes(100);o__Saccharomycetales(100);f__Metschnikowiaceae(100);g__Metschnikowia(100);g__Metschnikowia_unclassified(100); | Metschnikowia | 13 | Saprotroph | \|Undefined Saprotroph\| | Yeast | NULL | Probable | As nectar-tap_saprotroph (Põlme et al. 2020); Decay sugar-rich substrates (Põlme et al. 2020) | Põlme S, et al. 2020. Fungal Diversity 105:1-16 ((https://doi.org/10.1007/s13225-020-00466-2)) |
| Otu46569 | 1 | k__Fungi(100);p__Ascomycota(100);c__Saccharomycetes(100);o__Saccharomycetales(100);f__Metschnikowiaceae(100);g__Metschnikowia(100);g__Metschnikowia_unclassified(100); | Metschnikowia | 13 | Saprotroph | \|Undefined Saprotroph\| | Yeast | NULL | Probable | As nectar-tap_saprotroph (Põlme et al. 2020); Decay sugar-rich substrates (Põlme et al. 2020) | Põlme S, et al. 2020. Fungal Diversity 105:1-16 ((https://doi.org/10.1007/s13225-020-00466-2)) |
| Otu46570 | 1 | k__Fungi(100);p__Ascomycota(100);c__Saccharomycetes(100);o__Saccharomycetales(100);f__Metschnikowiaceae(100);g__Metschnikowia(100);g__Metschnikowia_unclassified(100); | Metschnikowia | 13 | Saprotroph | \|Undefined Saprotroph\| | Yeast | NULL | Probable | As nectar-tap_saprotroph (Põlme et al. 2020); Decay sugar-rich substrates (Põlme et al. 2020) | Põlme S, et al. 2020. Fungal Diversity 105:1-16 ((https://doi.org/10.1007/s13225-020-00466-2)) |
| Otu46571 | 1 | k__Fungi(100);p__Ascomycota(100);c__Saccharomycetes(100);o__Saccharomycetales(100);f__Metschnikowiaceae(100);g__Metschnikowia(100);g__Metschnikowia_unclassified(100); | Metschnikowia | 13 | Saprotroph | \|Undefined Saprotroph\| | Yeast | NULL | Probable | As nectar-tap_saprotroph (Põlme et al. 2020); Decay sugar-rich substrates (Põlme et al. 2020) | Põlme S, et al. 2020. Fungal Diversity 105:1-16 ((https://doi.org/10.1007/s13225-020-00466-2)) |
| Otu46572 | 1 | k__Fungi(100);p__Ascomycota(100);c__Saccharomycetes(100);o__Saccharomycetales(100);f__Metschnikowiaceae(100);g__Metschnikowia(100);g__Metschnikowia_unclassified(100); | Metschnikowia | 13 | Saprotroph | \|Undefined Saprotroph\| | Yeast | NULL | Probable | As nectar-tap_saprotroph (Põlme et al. 2020); Decay sugar-rich substrates (Põlme et al. 2020) | Põlme S, et al. 2020. Fungal Diversity 105:1-16 ((https://doi.org/10.1007/s13225-020-00466-2)) |
| Otu46573 | 1 | k__Fungi(100);p__Ascomycota(100);c__Saccharomycetes(100);o__Saccharomycetales(100);f__Metschnikowiaceae(100);g__Metschnikowia(100);g__Metschnikowia_unclassified(100); | Metschnikowia | 13 | Saprotroph | \|Undefined Saprotroph\| | Yeast | NULL | Probable | As nectar-tap_saprotroph (Põlme et al. 2020); Decay sugar-rich substrates (Põlme et al. 2020) | Põlme S, et al. 2020. Fungal Diversity 105:1-16 ((https://doi.org/10.1007/s13225-020-00466-2)) |
| Otu46574 | 1 | k__Fungi(100);p__Ascomycota(100);c__Saccharomycetes(100);o__Saccharomycetales(100);f__Metschnikowiaceae(100);g__Metschnikowia(100);g__Metschnikowia_unclassified(100); | Metschnikowia | 13 | Saprotroph | \|Undefined Saprotroph\| | Yeast | NULL | Probable | As nectar-tap_saprotroph (Põlme et al. 2020); Decay sugar-rich substrates (Põlme et al. 2020) | Põlme S, et al. 2020. Fungal Diversity 105:1-16 ((https://doi.org/10.1007/s13225-020-00466-2)) |
| Otu46575 | 1 | k__Fungi(100);p__Ascomycota(100);c__Saccharomycetes(100);o__Saccharomycetales(100);f__Metschnikowiaceae(100);g__Metschnikowia(100);s__Metschnikowia_pulcherrima(100); | Metschnikowia | 13 | Saprotroph | \|Undefined Saprotroph\| | Yeast | NULL | Probable | As nectar-tap_saprotroph (Põlme et al. 2020); Decay sugar-rich substrates (Põlme et al. 2020) | Põlme S, et al. 2020. Fungal Diversity 105:1-16 ((https://doi.org/10.1007/s13225-020-00466-2)) |
| Otu46576 | 1 | k__Fungi(100);p__Ascomycota(100);c__Saccharomycetes(100);o__Saccharomycetales(100);f__Metschnikowiaceae(100);g__Metschnikowia(100);s__Metschnikowia_pulcherrima(100); | Metschnikowia | 13 | Saprotroph | \|Undefined Saprotroph\| | Yeast | NULL | Probable | As nectar-tap_saprotroph (Põlme et al. 2020); Decay sugar-rich substrates (Põlme et al. 2020) | Põlme S, et al. 2020. Fungal Diversity 105:1-16 ((https://doi.org/10.1007/s13225-020-00466-2)) |
| Otu46577 | 1 | k__Fungi(100);p__Ascomycota(100);c__Saccharomycetes(100);o__Saccharomycetales(100);f__Metschnikowiaceae(100);g__Metschnikowia(100);s__Metschnikowia_pulcherrima(100); | Metschnikowia | 13 | Saprotroph | \|Undefined Saprotroph\| | Yeast | NULL | Probable | As nectar-tap_saprotroph (Põlme et al. 2020); Decay sugar-rich substrates (Põlme et al. 2020) | Põlme S, et al. 2020. Fungal Diversity 105:1-16 ((https://doi.org/10.1007/s13225-020-00466-2)) |
| Otu46578 | 1 | k__Fungi(100);p__Ascomycota(100);c__Saccharomycetes(100);o__Saccharomycetales(100);f__Metschnikowiaceae(100);g__Metschnikowia(100);s__Metschnikowia_pulcherrima(100); | Metschnikowia | 13 | Saprotroph | \|Undefined Saprotroph\| | Yeast | NULL | Probable | As nectar-tap_saprotroph (Põlme et al. 2020); Decay sugar-rich substrates (Põlme et al. 2020) | Põlme S, et al. 2020. Fungal Diversity 105:1-16 ((https://doi.org/10.1007/s13225-020-00466-2)) |
| Otu46579 | 1 | k__Fungi(100);p__Ascomycota(100);c__Saccharomycetes(100);o__Saccharomycetales(100);f__Metschnikowiaceae(100);g__Metschnikowia(100);s__Metschnikowia_pulcherrima(100); | Metschnikowia | 13 | Saprotroph | \|Undefined Saprotroph\| | Yeast | NULL | Probable | As nectar-tap_saprotroph (Põlme et al. 2020); Decay sugar-rich substrates (Põlme et al. 2020) | Põlme S, et al. 2020. Fungal Diversity 105:1-16 ((https://doi.org/10.1007/s13225-020-00466-2)) |
| Otu46580 | 1 | k__Fungi(100);p__Ascomycota(100);c__Saccharomycetes(100);o__Saccharomycetales(100);f__Metschnikowiaceae(100);g__Metschnikowia(100);s__Metschnikowia_pulcherrima(100); | Metschnikowia | 13 | Saprotroph | \|Undefined Saprotroph\| | Yeast | NULL | Probable | As nectar-tap_saprotroph (Põlme et al. 2020); Decay sugar-rich substrates (Põlme et al. 2020) | Põlme S, et al. 2020. Fungal Diversity 105:1-16 ((https://doi.org/10.1007/s13225-020-00466-2)) |
| Otu46581 | 1 | k__Fungi(100);p__Ascomycota(100);c__Saccharomycetes(100);o__Saccharomycetales(100);f__Metschnikowiaceae(100);g__Metschnikowia(100);s__Metschnikowia_pulcherrima(100); | Metschnikowia | 13 | Saprotroph | \|Undefined Saprotroph\| | Yeast | NULL | Probable | As nectar-tap_saprotroph (Põlme et al. 2020); Decay sugar-rich substrates (Põlme et al. 2020) | Põlme S, et al. 2020. Fungal Diversity 105:1-16 ((https://doi.org/10.1007/s13225-020-00466-2)) |
| Otu46582 | 1 | k__Fungi(100);p__Ascomycota(100);c__Saccharomycetes(100);o__Saccharomycetales(100);f__Metschnikowiaceae(100);g__Metschnikowia(100);s__Metschnikowia_pulcherrima(100); | Metschnikowia | 13 | Saprotroph | \|Undefined Saprotroph\| | Yeast | NULL | Probable | As nectar-tap_saprotroph (Põlme et al. 2020); Decay sugar-rich substrates (Põlme et al. 2020) | Põlme S, et al. 2020. Fungal Diversity 105:1-16 ((https://doi.org/10.1007/s13225-020-00466-2)) |
| Otu46583 | 1 | k__Fungi(100);p__Ascomycota(100);c__Saccharomycetes(100);o__Saccharomycetales(100);f__Metschnikowiaceae(100);g__Metschnikowia(100);s__Metschnikowia_pulcherrima(100); | Metschnikowia | 13 | Saprotroph | \|Undefined Saprotroph\| | Yeast | NULL | Probable | As nectar-tap_saprotroph (Põlme et al. 2020); Decay sugar-rich substrates (Põlme et al. 2020) | Põlme S, et al. 2020. Fungal Diversity 105:1-16 ((https://doi.org/10.1007/s13225-020-00466-2)) |
| Otu46584 | 1 | k__Fungi(100);p__Ascomycota(100);c__Saccharomycetes(100);o__Saccharomycetales(100);f__Metschnikowiaceae(100);g__Metschnikowia(100);s__Metschnikowia_pulcherrima(100); | Metschnikowia | 13 | Saprotroph | \|Undefined Saprotroph\| | Yeast | NULL | Probable | As nectar-tap_saprotroph (Põlme et al. 2020); Decay sugar-rich substrates (Põlme et al. 2020) | Põlme S, et al. 2020. Fungal Diversity 105:1-16 ((https://doi.org/10.1007/s13225-020-00466-2)) |
| Otu46585 | 1 | k__Fungi(100);p__Ascomycota(100);c__Saccharomycetes(100);o__Saccharomycetales(100);f__Metschnikowiaceae(100);g__Metschnikowia(100);s__Metschnikowia_pulcherrima(100); | Metschnikowia | 13 | Saprotroph | \|Undefined Saprotroph\| | Yeast | NULL | Probable | As nectar-tap_saprotroph (Põlme et al. 2020); Decay sugar-rich substrates (Põlme et al. 2020) | Põlme S, et al. 2020. Fungal Diversity 105:1-16 ((https://doi.org/10.1007/s13225-020-00466-2)) |
| Otu46586 | 1 | k__Fungi(100);p__Ascomycota(100);c__Saccharomycetes(100);o__Saccharomycetales(100);f__Metschnikowiaceae(100);g__Metschnikowia(100);s__Metschnikowia_pulcherrima(100); | Metschnikowia | 13 | Saprotroph | \|Undefined Saprotroph\| | Yeast | NULL | Probable | As nectar-tap_saprotroph (Põlme et al. 2020); Decay sugar-rich substrates (Põlme et al. 2020) | Põlme S, et al. 2020. Fungal Diversity 105:1-16 ((https://doi.org/10.1007/s13225-020-00466-2)) |
| Otu46587 | 1 | k__Fungi(100);p__Ascomycota(100);c__Saccharomycetes(100);o__Saccharomycetales(100);f__Metschnikowiaceae(100);g__Metschnikowia(100);g__Metschnikowia_unclassified(100); | Metschnikowia | 13 | Saprotroph | \|Undefined Saprotroph\| | Yeast | NULL | Probable | As nectar-tap_saprotroph (Põlme et al. 2020); Decay sugar-rich substrates (Põlme et al. 2020) | Põlme S, et al. 2020. Fungal Diversity 105:1-16 ((https://doi.org/10.1007/s13225-020-00466-2)) |
| Otu46588 | 1 | k__Fungi(100);p__Ascomycota(100);c__Saccharomycetes(100);o__Saccharomycetales(100);f__Metschnikowiaceae(100);g__Metschnikowia(100);s__Metschnikowia_pulcherrima(100); | Metschnikowia | 13 | Saprotroph | \|Undefined Saprotroph\| | Yeast | NULL | Probable | As nectar-tap_saprotroph (Põlme et al. 2020); Decay sugar-rich substrates (Põlme et al. 2020) | Põlme S, et al. 2020. Fungal Diversity 105:1-16 ((https://doi.org/10.1007/s13225-020-00466-2)) |
| Otu46589 | 1 | k__Fungi(100);p__Ascomycota(100);c__Saccharomycetes(100);o__Saccharomycetales(100);f__Metschnikowiaceae(100);g__Metschnikowia(100);s__Metschnikowia_pulcherrima(100); | Metschnikowia | 13 | Saprotroph | \|Undefined Saprotroph\| | Yeast | NULL | Probable | As nectar-tap_saprotroph (Põlme et al. 2020); Decay sugar-rich substrates (Põlme et al. 2020) | Põlme S, et al. 2020. Fungal Diversity 105:1-16 ((https://doi.org/10.1007/s13225-020-00466-2)) |
| Otu46590 | 1 | k__Fungi(100);p__Ascomycota(100);c__Saccharomycetes(100);o__Saccharomycetales(100);f__Metschnikowiaceae(100);g__Metschnikowia(100);g__Metschnikowia_unclassified(100); | Metschnikowia | 13 | Saprotroph | \|Undefined Saprotroph\| | Yeast | NULL | Probable | As nectar-tap_saprotroph (Põlme et al. 2020); Decay sugar-rich substrates (Põlme et al. 2020) | Põlme S, et al. 2020. Fungal Diversity 105:1-16 ((https://doi.org/10.1007/s13225-020-00466-2)) |
| Otu46591 | 1 | k__Fungi(100);p__Ascomycota(100);c__Saccharomycetes(100);o__Saccharomycetales(100);f__Metschnikowiaceae(100);g__Metschnikowia(100);g__Metschnikowia_unclassified(100); | Metschnikowia | 13 | Saprotroph | \|Undefined Saprotroph\| | Yeast | NULL | Probable | As nectar-tap_saprotroph (Põlme et al. 2020); Decay sugar-rich substrates (Põlme et al. 2020) | Põlme S, et al. 2020. Fungal Diversity 105:1-16 ((https://doi.org/10.1007/s13225-020-00466-2)) |
| Otu46592 | 1 | k__Fungi(100);p__Ascomycota(100);c__Saccharomycetes(100);o__Saccharomycetales(100);f__Metschnikowiaceae(100);g__Metschnikowia(100);s__Metschnikowia_pulcherrima(100); | Metschnikowia | 13 | Saprotroph | \|Undefined Saprotroph\| | Yeast | NULL | Probable | As nectar-tap_saprotroph (Põlme et al. 2020); Decay sugar-rich substrates (Põlme et al. 2020) | Põlme S, et al. 2020. Fungal Diversity 105:1-16 ((https://doi.org/10.1007/s13225-020-00466-2)) |
| Otu46593 | 1 | k__Fungi(100);p__Ascomycota(100);c__Saccharomycetes(100);o__Saccharomycetales(100);f__Metschnikowiaceae(100);g__Metschnikowia(100);g__Metschnikowia_unclassified(100); | Metschnikowia | 13 | Saprotroph | \|Undefined Saprotroph\| | Yeast | NULL | Probable | As nectar-tap_saprotroph (Põlme et al. 2020); Decay sugar-rich substrates (Põlme et al. 2020) | Põlme S, et al. 2020. Fungal Diversity 105:1-16 ((https://doi.org/10.1007/s13225-020-00466-2)) |
| Otu46594 | 1 | k__Fungi(100);p__Ascomycota(100);c__Saccharomycetes(100);o__Saccharomycetales(100);f__Metschnikowiaceae(100);g__Metschnikowia(100);s__Metschnikowia_pulcherrima(100); | Metschnikowia | 13 | Saprotroph | \|Undefined Saprotroph\| | Yeast | NULL | Probable | As nectar-tap_saprotroph (Põlme et al. 2020); Decay sugar-rich substrates (Põlme et al. 2020) | Põlme S, et al. 2020. Fungal Diversity 105:1-16 ((https://doi.org/10.1007/s13225-020-00466-2)) |
| Otu46595 | 1 | k__Fungi(100);p__Ascomycota(100);c__Saccharomycetes(100);o__Saccharomycetales(100);f__Metschnikowiaceae(100);g__Metschnikowia(100);g__Metschnikowia_unclassified(100); | Metschnikowia | 13 | Saprotroph | \|Undefined Saprotroph\| | Yeast | NULL | Probable | As nectar-tap_saprotroph (Põlme et al. 2020); Decay sugar-rich substrates (Põlme et al. 2020) | Põlme S, et al. 2020. Fungal Diversity 105:1-16 ((https://doi.org/10.1007/s13225-020-00466-2)) |
| Otu46596 | 1 | k__Fungi(100);p__Ascomycota(100);c__Saccharomycetes(100);o__Saccharomycetales(100);f__Metschnikowiaceae(100);g__Metschnikowia(100);g__Metschnikowia_unclassified(100); | Metschnikowia | 13 | Saprotroph | \|Undefined Saprotroph\| | Yeast | NULL | Probable | As nectar-tap_saprotroph (Põlme et al. 2020); Decay sugar-rich substrates (Põlme et al. 2020) | Põlme S, et al. 2020. Fungal Diversity 105:1-16 ((https://doi.org/10.1007/s13225-020-00466-2)) |
| Otu46597 | 1 | k__Fungi(100);p__Ascomycota(100);c__Saccharomycetes(100);o__Saccharomycetales(100);f__Metschnikowiaceae(100);g__Metschnikowia(100);g__Metschnikowia_unclassified(100); | Metschnikowia | 13 | Saprotroph | \|Undefined Saprotroph\| | Yeast | NULL | Probable | As nectar-tap_saprotroph (Põlme et al. 2020); Decay sugar-rich substrates (Põlme et al. 2020) | Põlme S, et al. 2020. Fungal Diversity 105:1-16 ((https://doi.org/10.1007/s13225-020-00466-2)) |
| Otu46598 | 1 | k__Fungi(100);p__Ascomycota(100);c__Saccharomycetes(100);o__Saccharomycetales(100);f__Metschnikowiaceae(100);g__Metschnikowia(100);s__Metschnikowia_pulcherrima(100); | Metschnikowia | 13 | Saprotroph | \|Undefined Saprotroph\| | Yeast | NULL | Probable | As nectar-tap_saprotroph (Põlme et al. 2020); Decay sugar-rich substrates (Põlme et al. 2020) | Põlme S, et al. 2020. Fungal Diversity 105:1-16 ((https://doi.org/10.1007/s13225-020-00466-2)) |
| Otu46599 | 1 | k__Fungi(100);p__Ascomycota(100);c__Saccharomycetes(100);o__Saccharomycetales(100);f__Metschnikowiaceae(100);g__Metschnikowia(100);s__Metschnikowia_pulcherrima(100); | Metschnikowia | 13 | Saprotroph | \|Undefined Saprotroph\| | Yeast | NULL | Probable | As nectar-tap_saprotroph (Põlme et al. 2020); Decay sugar-rich substrates (Põlme et al. 2020) | Põlme S, et al. 2020. Fungal Diversity 105:1-16 ((https://doi.org/10.1007/s13225-020-00466-2)) |
| Otu46600 | 1 | k__Fungi(100);p__Ascomycota(100);c__Saccharomycetes(100);o__Saccharomycetales(100);f__Metschnikowiaceae(100);g__Metschnikowia(100);s__Metschnikowia_pulcherrima(100); | Metschnikowia | 13 | Saprotroph | \|Undefined Saprotroph\| | Yeast | NULL | Probable | As nectar-tap_saprotroph (Põlme et al. 2020); Decay sugar-rich substrates (Põlme et al. 2020) | Põlme S, et al. 2020. Fungal Diversity 105:1-16 ((https://doi.org/10.1007/s13225-020-00466-2)) |
| Otu46601 | 1 | k__Fungi(100);p__Ascomycota(100);c__Saccharomycetes(100);o__Saccharomycetales(100);f__Metschnikowiaceae(100);g__Metschnikowia(100);s__Metschnikowia_pulcherrima(100); | Metschnikowia | 13 | Saprotroph | \|Undefined Saprotroph\| | Yeast | NULL | Probable | As nectar-tap_saprotroph (Põlme et al. 2020); Decay sugar-rich substrates (Põlme et al. 2020) | Põlme S, et al. 2020. Fungal Diversity 105:1-16 ((https://doi.org/10.1007/s13225-020-00466-2)) |
| Otu46602 | 1 | k__Fungi(100);p__Ascomycota(100);c__Saccharomycetes(100);o__Saccharomycetales(100);f__Metschnikowiaceae(100);g__Metschnikowia(100);s__Metschnikowia_pulcherrima(100); | Metschnikowia | 13 | Saprotroph | \|Undefined Saprotroph\| | Yeast | NULL | Probable | As nectar-tap_saprotroph (Põlme et al. 2020); Decay sugar-rich substrates (Põlme et al. 2020) | Põlme S, et al. 2020. Fungal Diversity 105:1-16 ((https://doi.org/10.1007/s13225-020-00466-2)) |
| Otu46603 | 1 | k__Fungi(100);p__Ascomycota(100);c__Saccharomycetes(100);o__Saccharomycetales(100);f__Metschnikowiaceae(100);g__Metschnikowia(100);g__Metschnikowia_unclassified(100); | Metschnikowia | 13 | Saprotroph | \|Undefined Saprotroph\| | Yeast | NULL | Probable | As nectar-tap_saprotroph (Põlme et al. 2020); Decay sugar-rich substrates (Põlme et al. 2020) | Põlme S, et al. 2020. Fungal Diversity 105:1-16 ((https://doi.org/10.1007/s13225-020-00466-2)) |
| Otu46604 | 1 | k__Fungi(100);p__Ascomycota(100);c__Saccharomycetes(100);o__Saccharomycetales(100);f__Metschnikowiaceae(100);g__Metschnikowia(100);s__Metschnikowia_pulcherrima(100); | Metschnikowia | 13 | Saprotroph | \|Undefined Saprotroph\| | Yeast | NULL | Probable | As nectar-tap_saprotroph (Põlme et al. 2020); Decay sugar-rich substrates (Põlme et al. 2020) | Põlme S, et al. 2020. Fungal Diversity 105:1-16 ((https://doi.org/10.1007/s13225-020-00466-2)) |
| Otu46605 | 1 | k__Fungi(100);p__Ascomycota(100);c__Saccharomycetes(100);o__Saccharomycetales(100);f__Metschnikowiaceae(100);g__Metschnikowia(100);s__Metschnikowia_pulcherrima(100); | Metschnikowia | 13 | Saprotroph | \|Undefined Saprotroph\| | Yeast | NULL | Probable | As nectar-tap_saprotroph (Põlme et al. 2020); Decay sugar-rich substrates (Põlme et al. 2020) | Põlme S, et al. 2020. Fungal Diversity 105:1-16 ((https://doi.org/10.1007/s13225-020-00466-2)) |
| Otu46606 | 1 | k__Fungi(100);p__Ascomycota(100);c__Saccharomycetes(100);o__Saccharomycetales(100);f__Metschnikowiaceae(100);g__Metschnikowia(100);s__Metschnikowia_pulcherrima(100); | Metschnikowia | 13 | Saprotroph | \|Undefined Saprotroph\| | Yeast | NULL | Probable | As nectar-tap_saprotroph (Põlme et al. 2020); Decay sugar-rich substrates (Põlme et al. 2020) | Põlme S, et al. 2020. Fungal Diversity 105:1-16 ((https://doi.org/10.1007/s13225-020-00466-2)) |
| Otu46607 | 1 | k__Fungi(100);p__Ascomycota(100);c__Saccharomycetes(100);o__Saccharomycetales(100);f__Metschnikowiaceae(100);g__Metschnikowia(100);s__Metschnikowia_pulcherrima(100); | Metschnikowia | 13 | Saprotroph | \|Undefined Saprotroph\| | Yeast | NULL | Probable | As nectar-tap_saprotroph (Põlme et al. 2020); Decay sugar-rich substrates (Põlme et al. 2020) | Põlme S, et al. 2020. Fungal Diversity 105:1-16 ((https://doi.org/10.1007/s13225-020-00466-2)) |
| Otu46608 | 1 | k__Fungi(100);p__Ascomycota(100);c__Saccharomycetes(100);o__Saccharomycetales(100);f__Metschnikowiaceae(100);g__Metschnikowia(100);s__Metschnikowia_pulcherrima(100); | Metschnikowia | 13 | Saprotroph | \|Undefined Saprotroph\| | Yeast | NULL | Probable | As nectar-tap_saprotroph (Põlme et al. 2020); Decay sugar-rich substrates (Põlme et al. 2020) | Põlme S, et al. 2020. Fungal Diversity 105:1-16 ((https://doi.org/10.1007/s13225-020-00466-2)) |
| Otu46609 | 1 | k__Fungi(100);p__Ascomycota(100);c__Saccharomycetes(100);o__Saccharomycetales(100);f__Metschnikowiaceae(100);g__Metschnikowia(100);s__Metschnikowia_pulcherrima(100); | Metschnikowia | 13 | Saprotroph | \|Undefined Saprotroph\| | Yeast | NULL | Probable | As nectar-tap_saprotroph (Põlme et al. 2020); Decay sugar-rich substrates (Põlme et al. 2020) | Põlme S, et al. 2020. Fungal Diversity 105:1-16 ((https://doi.org/10.1007/s13225-020-00466-2)) |
| Otu46610 | 1 | k__Fungi(100);p__Ascomycota(100);c__Saccharomycetes(100);o__Saccharomycetales(100);f__Metschnikowiaceae(100);g__Metschnikowia(100);s__Metschnikowia_pulcherrima(100); | Metschnikowia | 13 | Saprotroph | \|Undefined Saprotroph\| | Yeast | NULL | Probable | As nectar-tap_saprotroph (Põlme et al. 2020); Decay sugar-rich substrates (Põlme et al. 2020) | Põlme S, et al. 2020. Fungal Diversity 105:1-16 ((https://doi.org/10.1007/s13225-020-00466-2)) |
| Otu46611 | 1 | k__Fungi(100);p__Ascomycota(100);c__Saccharomycetes(100);o__Saccharomycetales(100);f__Metschnikowiaceae(100);g__Metschnikowia(100);s__Metschnikowia_pulcherrima(100); | Metschnikowia | 13 | Saprotroph | \|Undefined Saprotroph\| | Yeast | NULL | Probable | As nectar-tap_saprotroph (Põlme et al. 2020); Decay sugar-rich substrates (Põlme et al. 2020) | Põlme S, et al. 2020. Fungal Diversity 105:1-16 ((https://doi.org/10.1007/s13225-020-00466-2)) |
| Otu46612 | 1 | k__Fungi(100);p__Ascomycota(100);c__Saccharomycetes(100);o__Saccharomycetales(100);f__Metschnikowiaceae(100);g__Metschnikowia(100);s__Metschnikowia_pulcherrima(100); | Metschnikowia | 13 | Saprotroph | \|Undefined Saprotroph\| | Yeast | NULL | Probable | As nectar-tap_saprotroph (Põlme et al. 2020); Decay sugar-rich substrates (Põlme et al. 2020) | Põlme S, et al. 2020. Fungal Diversity 105:1-16 ((https://doi.org/10.1007/s13225-020-00466-2)) |
| Otu46613 | 1 | k__Fungi(100);p__Ascomycota(100);c__Saccharomycetes(100);o__Saccharomycetales(100);f__Metschnikowiaceae(100);g__Metschnikowia(100);s__Metschnikowia_pulcherrima(100); | Metschnikowia | 13 | Saprotroph | \|Undefined Saprotroph\| | Yeast | NULL | Probable | As nectar-tap_saprotroph (Põlme et al. 2020); Decay sugar-rich substrates (Põlme et al. 2020) | Põlme S, et al. 2020. Fungal Diversity 105:1-16 ((https://doi.org/10.1007/s13225-020-00466-2)) |
| Otu46614 | 1 | k__Fungi(100);p__Ascomycota(100);c__Saccharomycetes(100);o__Saccharomycetales(100);f__Metschnikowiaceae(100);g__Metschnikowia(100);s__Metschnikowia_pulcherrima(100); | Metschnikowia | 13 | Saprotroph | \|Undefined Saprotroph\| | Yeast | NULL | Probable | As nectar-tap_saprotroph (Põlme et al. 2020); Decay sugar-rich substrates (Põlme et al. 2020) | Põlme S, et al. 2020. Fungal Diversity 105:1-16 ((https://doi.org/10.1007/s13225-020-00466-2)) |
| Otu46615 | 1 | k__Fungi(100);p__Ascomycota(100);c__Saccharomycetes(100);o__Saccharomycetales(100);f__Metschnikowiaceae(100);g__Metschnikowia(100);s__Metschnikowia_pulcherrima(100); | Metschnikowia | 13 | Saprotroph | \|Undefined Saprotroph\| | Yeast | NULL | Probable | As nectar-tap_saprotroph (Põlme et al. 2020); Decay sugar-rich substrates (Põlme et al. 2020) | Põlme S, et al. 2020. Fungal Diversity 105:1-16 ((https://doi.org/10.1007/s13225-020-00466-2)) |
| Otu46616 | 1 | k__Fungi(100);p__Ascomycota(100);c__Saccharomycetes(100);o__Saccharomycetales(100);f__Metschnikowiaceae(100);g__Metschnikowia(100);g__Metschnikowia_unclassified(100); | Metschnikowia | 13 | Saprotroph | \|Undefined Saprotroph\| | Yeast | NULL | Probable | As nectar-tap_saprotroph (Põlme et al. 2020); Decay sugar-rich substrates (Põlme et al. 2020) | Põlme S, et al. 2020. Fungal Diversity 105:1-16 ((https://doi.org/10.1007/s13225-020-00466-2)) |
| Otu46617 | 1 | k__Fungi(100);p__Ascomycota(100);c__Saccharomycetes(100);o__Saccharomycetales(100);f__Metschnikowiaceae(100);g__Metschnikowia(100);s__Metschnikowia_pulcherrima(100); | Metschnikowia | 13 | Saprotroph | \|Undefined Saprotroph\| | Yeast | NULL | Probable | As nectar-tap_saprotroph (Põlme et al. 2020); Decay sugar-rich substrates (Põlme et al. 2020) | Põlme S, et al. 2020. Fungal Diversity 105:1-16 ((https://doi.org/10.1007/s13225-020-00466-2)) |
| Otu46618 | 1 | k__Fungi(100);p__Ascomycota(100);c__Saccharomycetes(100);o__Saccharomycetales(100);f__Metschnikowiaceae(100);g__Metschnikowia(100);s__Metschnikowia_pulcherrima(100); | Metschnikowia | 13 | Saprotroph | \|Undefined Saprotroph\| | Yeast | NULL | Probable | As nectar-tap_saprotroph (Põlme et al. 2020); Decay sugar-rich substrates (Põlme et al. 2020) | Põlme S, et al. 2020. Fungal Diversity 105:1-16 ((https://doi.org/10.1007/s13225-020-00466-2)) |
| Otu46619 | 1 | k__Fungi(100);p__Ascomycota(100);c__Saccharomycetes(100);o__Saccharomycetales(100);f__Metschnikowiaceae(100);g__Metschnikowia(100);g__Metschnikowia_unclassified(100); | Metschnikowia | 13 | Saprotroph | \|Undefined Saprotroph\| | Yeast | NULL | Probable | As nectar-tap_saprotroph (Põlme et al. 2020); Decay sugar-rich substrates (Põlme et al. 2020) | Põlme S, et al. 2020. Fungal Diversity 105:1-16 ((https://doi.org/10.1007/s13225-020-00466-2)) |
| Otu46620 | 1 | k__Fungi(100);p__Ascomycota(100);c__Saccharomycetes(100);o__Saccharomycetales(100);f__Metschnikowiaceae(100);g__Metschnikowia(100);s__Metschnikowia_pulcherrima(100); | Metschnikowia | 13 | Saprotroph | \|Undefined Saprotroph\| | Yeast | NULL | Probable | As nectar-tap_saprotroph (Põlme et al. 2020); Decay sugar-rich substrates (Põlme et al. 2020) | Põlme S, et al. 2020. Fungal Diversity 105:1-16 ((https://doi.org/10.1007/s13225-020-00466-2)) |
| Otu46621 | 1 | k__Fungi(100);p__Ascomycota(100);c__Saccharomycetes(100);o__Saccharomycetales(100);f__Metschnikowiaceae(100);g__Metschnikowia(100);s__Metschnikowia_pulcherrima(100); | Metschnikowia | 13 | Saprotroph | \|Undefined Saprotroph\| | Yeast | NULL | Probable | As nectar-tap_saprotroph (Põlme et al. 2020); Decay sugar-rich substrates (Põlme et al. 2020) | Põlme S, et al. 2020. Fungal Diversity 105:1-16 ((https://doi.org/10.1007/s13225-020-00466-2)) |
| Otu46622 | 1 | k__Fungi(100);p__Ascomycota(100);c__Saccharomycetes(100);o__Saccharomycetales(100);f__Metschnikowiaceae(100);g__Metschnikowia(100);s__Metschnikowia_pulcherrima(100); | Metschnikowia | 13 | Saprotroph | \|Undefined Saprotroph\| | Yeast | NULL | Probable | As nectar-tap_saprotroph (Põlme et al. 2020); Decay sugar-rich substrates (Põlme et al. 2020) | Põlme S, et al. 2020. Fungal Diversity 105:1-16 ((https://doi.org/10.1007/s13225-020-00466-2)) |
| Otu46623 | 1 | k__Fungi(100);p__Ascomycota(100);c__Saccharomycetes(100);o__Saccharomycetales(100);f__Metschnikowiaceae(100);g__Metschnikowia(100);s__Metschnikowia_pulcherrima(100); | Metschnikowia | 13 | Saprotroph | \|Undefined Saprotroph\| | Yeast | NULL | Probable | As nectar-tap_saprotroph (Põlme et al. 2020); Decay sugar-rich substrates (Põlme et al. 2020) | Põlme S, et al. 2020. Fungal Diversity 105:1-16 ((https://doi.org/10.1007/s13225-020-00466-2)) |
| Otu46624 | 1 | k__Fungi(100);p__Ascomycota(100);c__Saccharomycetes(100);o__Saccharomycetales(100);f__Metschnikowiaceae(100);g__Metschnikowia(100);s__Metschnikowia_pulcherrima(100); | Metschnikowia | 13 | Saprotroph | \|Undefined Saprotroph\| | Yeast | NULL | Probable | As nectar-tap_saprotroph (Põlme et al. 2020); Decay sugar-rich substrates (Põlme et al. 2020) | Põlme S, et al. 2020. Fungal Diversity 105:1-16 ((https://doi.org/10.1007/s13225-020-00466-2)) |
| Otu46625 | 1 | k__Fungi(100);p__Ascomycota(100);c__Saccharomycetes(100);o__Saccharomycetales(100);f__Metschnikowiaceae(100);g__Metschnikowia(100);s__Metschnikowia_pulcherrima(100); | Metschnikowia | 13 | Saprotroph | \|Undefined Saprotroph\| | Yeast | NULL | Probable | As nectar-tap_saprotroph (Põlme et al. 2020); Decay sugar-rich substrates (Põlme et al. 2020) | Põlme S, et al. 2020. Fungal Diversity 105:1-16 ((https://doi.org/10.1007/s13225-020-00466-2)) |
| Otu46626 | 1 | k__Fungi(100);p__Ascomycota(100);c__Saccharomycetes(100);o__Saccharomycetales(100);f__Metschnikowiaceae(100);g__Metschnikowia(100);s__Metschnikowia_pulcherrima(100); | Metschnikowia | 13 | Saprotroph | \|Undefined Saprotroph\| | Yeast | NULL | Probable | As nectar-tap_saprotroph (Põlme et al. 2020); Decay sugar-rich substrates (Põlme et al. 2020) | Põlme S, et al. 2020. Fungal Diversity 105:1-16 ((https://doi.org/10.1007/s13225-020-00466-2)) |
| Otu46627 | 1 | k__Fungi(100);p__Ascomycota(100);c__Saccharomycetes(100);o__Saccharomycetales(100);f__Metschnikowiaceae(100);g__Metschnikowia(100);s__Metschnikowia_pulcherrima(100); | Metschnikowia | 13 | Saprotroph | \|Undefined Saprotroph\| | Yeast | NULL | Probable | As nectar-tap_saprotroph (Põlme et al. 2020); Decay sugar-rich substrates (Põlme et al. 2020) | Põlme S, et al. 2020. Fungal Diversity 105:1-16 ((https://doi.org/10.1007/s13225-020-00466-2)) |
| Otu46628 | 1 | k__Fungi(100);p__Ascomycota(100);c__Saccharomycetes(100);o__Saccharomycetales(100);f__Metschnikowiaceae(100);g__Metschnikowia(100);s__Metschnikowia_pulcherrima(100); | Metschnikowia | 13 | Saprotroph | \|Undefined Saprotroph\| | Yeast | NULL | Probable | As nectar-tap_saprotroph (Põlme et al. 2020); Decay sugar-rich substrates (Põlme et al. 2020) | Põlme S, et al. 2020. Fungal Diversity 105:1-16 ((https://doi.org/10.1007/s13225-020-00466-2)) |
| Otu46629 | 1 | k__Fungi(100);p__Ascomycota(100);c__Saccharomycetes(100);o__Saccharomycetales(100);f__Metschnikowiaceae(100);g__Metschnikowia(100);s__Metschnikowia_pulcherrima(100); | Metschnikowia | 13 | Saprotroph | \|Undefined Saprotroph\| | Yeast | NULL | Probable | As nectar-tap_saprotroph (Põlme et al. 2020); Decay sugar-rich substrates (Põlme et al. 2020) | Põlme S, et al. 2020. Fungal Diversity 105:1-16 ((https://doi.org/10.1007/s13225-020-00466-2)) |
| Otu46630 | 1 | k__Fungi(100);p__Ascomycota(100);c__Saccharomycetes(100);o__Saccharomycetales(100);f__Metschnikowiaceae(100);g__Metschnikowia(100);s__Metschnikowia_pulcherrima(100); | Metschnikowia | 13 | Saprotroph | \|Undefined Saprotroph\| | Yeast | NULL | Probable | As nectar-tap_saprotroph (Põlme et al. 2020); Decay sugar-rich substrates (Põlme et al. 2020) | Põlme S, et al. 2020. Fungal Diversity 105:1-16 ((https://doi.org/10.1007/s13225-020-00466-2)) |
| Otu46631 | 1 | k__Fungi(100);p__Ascomycota(100);c__Saccharomycetes(100);o__Saccharomycetales(100);f__Metschnikowiaceae(100);g__Metschnikowia(100);s__Metschnikowia_pulcherrima(100); | Metschnikowia | 13 | Saprotroph | \|Undefined Saprotroph\| | Yeast | NULL | Probable | As nectar-tap_saprotroph (Põlme et al. 2020); Decay sugar-rich substrates (Põlme et al. 2020) | Põlme S, et al. 2020. Fungal Diversity 105:1-16 ((https://doi.org/10.1007/s13225-020-00466-2)) |
| Otu46632 | 1 | k__Fungi(100);p__Ascomycota(100);c__Saccharomycetes(100);o__Saccharomycetales(100);f__Metschnikowiaceae(100);g__Metschnikowia(100);s__Metschnikowia_pulcherrima(100); | Metschnikowia | 13 | Saprotroph | \|Undefined Saprotroph\| | Yeast | NULL | Probable | As nectar-tap_saprotroph (Põlme et al. 2020); Decay sugar-rich substrates (Põlme et al. 2020) | Põlme S, et al. 2020. Fungal Diversity 105:1-16 ((https://doi.org/10.1007/s13225-020-00466-2)) |
| Otu46633 | 1 | k__Fungi(100);p__Ascomycota(100);c__Saccharomycetes(100);o__Saccharomycetales(100);f__Metschnikowiaceae(100);g__Metschnikowia(100);s__Metschnikowia_pulcherrima(100); | Metschnikowia | 13 | Saprotroph | \|Undefined Saprotroph\| | Yeast | NULL | Probable | As nectar-tap_saprotroph (Põlme et al. 2020); Decay sugar-rich substrates (Põlme et al. 2020) | Põlme S, et al. 2020. Fungal Diversity 105:1-16 ((https://doi.org/10.1007/s13225-020-00466-2)) |
| Otu46634 | 1 | k__Fungi(100);p__Ascomycota(100);c__Saccharomycetes(100);o__Saccharomycetales(100);f__Metschnikowiaceae(100);g__Metschnikowia(100);g__Metschnikowia_unclassified(100); | Metschnikowia | 13 | Saprotroph | \|Undefined Saprotroph\| | Yeast | NULL | Probable | As nectar-tap_saprotroph (Põlme et al. 2020); Decay sugar-rich substrates (Põlme et al. 2020) | Põlme S, et al. 2020. Fungal Diversity 105:1-16 ((https://doi.org/10.1007/s13225-020-00466-2)) |
| Otu46635 | 1 | k__Fungi(100);p__Ascomycota(100);c__Saccharomycetes(100);o__Saccharomycetales(100);f__Metschnikowiaceae(100);g__Metschnikowia(100);g__Metschnikowia_unclassified(100); | Metschnikowia | 13 | Saprotroph | \|Undefined Saprotroph\| | Yeast | NULL | Probable | As nectar-tap_saprotroph (Põlme et al. 2020); Decay sugar-rich substrates (Põlme et al. 2020) | Põlme S, et al. 2020. Fungal Diversity 105:1-16 ((https://doi.org/10.1007/s13225-020-00466-2)) |
| Otu46636 | 1 | k__Fungi(100);p__Ascomycota(100);c__Saccharomycetes(100);o__Saccharomycetales(100);f__Metschnikowiaceae(100);g__Metschnikowia(100);s__Metschnikowia_pulcherrima(100); | Metschnikowia | 13 | Saprotroph | \|Undefined Saprotroph\| | Yeast | NULL | Probable | As nectar-tap_saprotroph (Põlme et al. 2020); Decay sugar-rich substrates (Põlme et al. 2020) | Põlme S, et al. 2020. Fungal Diversity 105:1-16 ((https://doi.org/10.1007/s13225-020-00466-2)) |
| Otu46637 | 1 | k__Fungi(100);p__Ascomycota(100);c__Saccharomycetes(100);o__Saccharomycetales(100);f__Metschnikowiaceae(100);g__Metschnikowia(100);s__Metschnikowia_pulcherrima(100); | Metschnikowia | 13 | Saprotroph | \|Undefined Saprotroph\| | Yeast | NULL | Probable | As nectar-tap_saprotroph (Põlme et al. 2020); Decay sugar-rich substrates (Põlme et al. 2020) | Põlme S, et al. 2020. Fungal Diversity 105:1-16 ((https://doi.org/10.1007/s13225-020-00466-2)) |
| Otu46638 | 1 | k__Fungi(100);p__Ascomycota(100);c__Saccharomycetes(100);o__Saccharomycetales(100);f__Metschnikowiaceae(100);g__Metschnikowia(100);s__Metschnikowia_pulcherrima(100); | Metschnikowia | 13 | Saprotroph | \|Undefined Saprotroph\| | Yeast | NULL | Probable | As nectar-tap_saprotroph (Põlme et al. 2020); Decay sugar-rich substrates (Põlme et al. 2020) | Põlme S, et al. 2020. Fungal Diversity 105:1-16 ((https://doi.org/10.1007/s13225-020-00466-2)) |
| Otu46639 | 1 | k__Fungi(100);p__Ascomycota(100);c__Saccharomycetes(100);o__Saccharomycetales(100);f__Metschnikowiaceae(100);g__Metschnikowia(100);s__Metschnikowia_pulcherrima(100); | Metschnikowia | 13 | Saprotroph | \|Undefined Saprotroph\| | Yeast | NULL | Probable | As nectar-tap_saprotroph (Põlme et al. 2020); Decay sugar-rich substrates (Põlme et al. 2020) | Põlme S, et al. 2020. Fungal Diversity 105:1-16 ((https://doi.org/10.1007/s13225-020-00466-2)) |
| Otu46640 | 1 | k__Fungi(100);p__Ascomycota(100);c__Saccharomycetes(100);o__Saccharomycetales(100);f__Metschnikowiaceae(100);g__Metschnikowia(100);s__Metschnikowia_pulcherrima(100); | Metschnikowia | 13 | Saprotroph | \|Undefined Saprotroph\| | Yeast | NULL | Probable | As nectar-tap_saprotroph (Põlme et al. 2020); Decay sugar-rich substrates (Põlme et al. 2020) | Põlme S, et al. 2020. Fungal Diversity 105:1-16 ((https://doi.org/10.1007/s13225-020-00466-2)) |
| Otu46641 | 1 | k__Fungi(100);p__Ascomycota(100);c__Saccharomycetes(100);o__Saccharomycetales(100);f__Metschnikowiaceae(100);g__Metschnikowia(100);s__Metschnikowia_viticola(100); | Metschnikowia | 13 | Saprotroph | \|Undefined Saprotroph\| | Yeast | NULL | Probable | As nectar-tap_saprotroph (Põlme et al. 2020); Decay sugar-rich substrates (Põlme et al. 2020) | Põlme S, et al. 2020. Fungal Diversity 105:1-16 ((https://doi.org/10.1007/s13225-020-00466-2)) |
| Otu46642 | 1 | k__Fungi(100);p__Ascomycota(100);c__Saccharomycetes(100);o__Saccharomycetales(100);f__Metschnikowiaceae(100);g__Metschnikowia(100);s__Metschnikowia_pulcherrima(100); | Metschnikowia | 13 | Saprotroph | \|Undefined Saprotroph\| | Yeast | NULL | Probable | As nectar-tap_saprotroph (Põlme et al. 2020); Decay sugar-rich substrates (Põlme et al. 2020) | Põlme S, et al. 2020. Fungal Diversity 105:1-16 ((https://doi.org/10.1007/s13225-020-00466-2)) |
| Otu46643 | 1 | k__Fungi(100);p__Ascomycota(100);c__Saccharomycetes(100);o__Saccharomycetales(100);f__Metschnikowiaceae(100);g__Metschnikowia(100);s__Metschnikowia_pulcherrima(100); | Metschnikowia | 13 | Saprotroph | \|Undefined Saprotroph\| | Yeast | NULL | Probable | As nectar-tap_saprotroph (Põlme et al. 2020); Decay sugar-rich substrates (Põlme et al. 2020) | Põlme S, et al. 2020. Fungal Diversity 105:1-16 ((https://doi.org/10.1007/s13225-020-00466-2)) |
| Otu46644 | 1 | k__Fungi(100);p__Ascomycota(100);c__Saccharomycetes(100);o__Saccharomycetales(100);f__Metschnikowiaceae(100);g__Metschnikowia(100);s__Metschnikowia_pulcherrima(100); | Metschnikowia | 13 | Saprotroph | \|Undefined Saprotroph\| | Yeast | NULL | Probable | As nectar-tap_saprotroph (Põlme et al. 2020); Decay sugar-rich substrates (Põlme et al. 2020) | Põlme S, et al. 2020. Fungal Diversity 105:1-16 ((https://doi.org/10.1007/s13225-020-00466-2)) |
| Otu46645 | 1 | k__Fungi(100);p__Ascomycota(100);c__Saccharomycetes(100);o__Saccharomycetales(100);f__Metschnikowiaceae(100);g__Metschnikowia(100);s__Metschnikowia_pulcherrima(100); | Metschnikowia | 13 | Saprotroph | \|Undefined Saprotroph\| | Yeast | NULL | Probable | As nectar-tap_saprotroph (Põlme et al. 2020); Decay sugar-rich substrates (Põlme et al. 2020) | Põlme S, et al. 2020. Fungal Diversity 105:1-16 ((https://doi.org/10.1007/s13225-020-00466-2)) |
| Otu46646 | 1 | k__Fungi(100);p__Ascomycota(100);c__Saccharomycetes(100);o__Saccharomycetales(100);f__Saccharomycetales_family_Incertae_sedis(100);g__Candida(100);s__Candida_dosseyi(100); | Candida | 13 | Pathotroph-Saprotroph-Symbiotroph | Animal Pathogen-Endophyte-Endosymbiont-Epiphyte-\|Undefined Saprotroph\| | Dimorphic Yeast-Yeast | NULL | Probable | As nectar-tap_saprotroph (Põlme et al. 2020); Decay sugar-rich substrates (Põlme et al. 2020); This asexual genus is polyphyletic and contain species that function very differently depend on what they are. Caution should be taken when interpreting guilds for Candida species. | Manolakaki D, et al. 2010. Virulence. 1: 367-375 ((https://doi.org/10.4161/viru.1.5.12796)); Mekbib SB, et al. 2011. Tropical Plant Pathology 36:233-240 ((https://doi.org/10.1590/S1982-56762011000400004)); Põlme S, et al. 2020. Fungal Diversity 105:1-16 ((https://doi.org/10.1007/s13225-020-00466-2)) |
| Otu46647 | 1 | k__Fungi(100);p__Ascomycota(100);c__Saccharomycetes(100);o__Saccharomycetales(100);f__Metschnikowiaceae(100);g__Metschnikowia(100);g__Metschnikowia_unclassified(100); | Metschnikowia | 13 | Saprotroph | \|Undefined Saprotroph\| | Yeast | NULL | Probable | As nectar-tap_saprotroph (Põlme et al. 2020); Decay sugar-rich substrates (Põlme et al. 2020) | Põlme S, et al. 2020. Fungal Diversity 105:1-16 ((https://doi.org/10.1007/s13225-020-00466-2)) |
| Otu46648 | 1 | k__Fungi(100);p__Ascomycota(100);c__Saccharomycetes(100);o__Saccharomycetales(100);f__Metschnikowiaceae(100);g__Metschnikowia(100);s__Metschnikowia_pulcherrima(100); | Metschnikowia | 13 | Saprotroph | \|Undefined Saprotroph\| | Yeast | NULL | Probable | As nectar-tap_saprotroph (Põlme et al. 2020); Decay sugar-rich substrates (Põlme et al. 2020) | Põlme S, et al. 2020. Fungal Diversity 105:1-16 ((https://doi.org/10.1007/s13225-020-00466-2)) |
| Otu46649 | 1 | k__Fungi(100);p__Ascomycota(100);c__Saccharomycetes(100);o__Saccharomycetales(100);f__Metschnikowiaceae(100);g__Metschnikowia(100);s__Metschnikowia_pulcherrima(100); | Metschnikowia | 13 | Saprotroph | \|Undefined Saprotroph\| | Yeast | NULL | Probable | As nectar-tap_saprotroph (Põlme et al. 2020); Decay sugar-rich substrates (Põlme et al. 2020) | Põlme S, et al. 2020. Fungal Diversity 105:1-16 ((https://doi.org/10.1007/s13225-020-00466-2)) |
| Otu46650 | 1 | k__Fungi(100);p__Ascomycota(100);c__Saccharomycetes(100);o__Saccharomycetales(100);f__Metschnikowiaceae(100);g__Metschnikowia(100);s__Metschnikowia_pulcherrima(100); | Metschnikowia | 13 | Saprotroph | \|Undefined Saprotroph\| | Yeast | NULL | Probable | As nectar-tap_saprotroph (Põlme et al. 2020); Decay sugar-rich substrates (Põlme et al. 2020) | Põlme S, et al. 2020. Fungal Diversity 105:1-16 ((https://doi.org/10.1007/s13225-020-00466-2)) |
| Otu46651 | 1 | k__Fungi(100);p__Ascomycota(100);c__Saccharomycetes(100);o__Saccharomycetales(100);f__Metschnikowiaceae(100);g__Metschnikowia(100);s__Metschnikowia_pulcherrima(100); | Metschnikowia | 13 | Saprotroph | \|Undefined Saprotroph\| | Yeast | NULL | Probable | As nectar-tap_saprotroph (Põlme et al. 2020); Decay sugar-rich substrates (Põlme et al. 2020) | Põlme S, et al. 2020. Fungal Diversity 105:1-16 ((https://doi.org/10.1007/s13225-020-00466-2)) |
| Otu46652 | 1 | k__Fungi(100);p__Ascomycota(100);c__Saccharomycetes(100);o__Saccharomycetales(100);f__Metschnikowiaceae(100);g__Metschnikowia(100);s__Metschnikowia_pulcherrima(100); | Metschnikowia | 13 | Saprotroph | \|Undefined Saprotroph\| | Yeast | NULL | Probable | As nectar-tap_saprotroph (Põlme et al. 2020); Decay sugar-rich substrates (Põlme et al. 2020) | Põlme S, et al. 2020. Fungal Diversity 105:1-16 ((https://doi.org/10.1007/s13225-020-00466-2)) |
| Otu46655 | 1 | k__Fungi(100);p__Ascomycota(100);c__Saccharomycetes(100);o__Saccharomycetales(100);f__Metschnikowiaceae(100);g__Metschnikowia(100);s__Metschnikowia_pulcherrima(100); | Metschnikowia | 13 | Saprotroph | \|Undefined Saprotroph\| | Yeast | NULL | Probable | As nectar-tap_saprotroph (Põlme et al. 2020); Decay sugar-rich substrates (Põlme et al. 2020) | Põlme S, et al. 2020. Fungal Diversity 105:1-16 ((https://doi.org/10.1007/s13225-020-00466-2)) |
| Otu46656 | 1 | k__Fungi(100);p__Ascomycota(100);c__Saccharomycetes(100);o__Saccharomycetales(100);f__Metschnikowiaceae(100);g__Metschnikowia(100);s__Metschnikowia_pulcherrima(100); | Metschnikowia | 13 | Saprotroph | \|Undefined Saprotroph\| | Yeast | NULL | Probable | As nectar-tap_saprotroph (Põlme et al. 2020); Decay sugar-rich substrates (Põlme et al. 2020) | Põlme S, et al. 2020. Fungal Diversity 105:1-16 ((https://doi.org/10.1007/s13225-020-00466-2)) |
| Otu46657 | 1 | k__Fungi(100);p__Ascomycota(100);c__Saccharomycetes(100);o__Saccharomycetales(100);f__Metschnikowiaceae(100);g__Metschnikowia(100);s__Metschnikowia_pulcherrima(100); | Metschnikowia | 13 | Saprotroph | \|Undefined Saprotroph\| | Yeast | NULL | Probable | As nectar-tap_saprotroph (Põlme et al. 2020); Decay sugar-rich substrates (Põlme et al. 2020) | Põlme S, et al. 2020. Fungal Diversity 105:1-16 ((https://doi.org/10.1007/s13225-020-00466-2)) |
| Otu46658 | 1 | k__Fungi(100);p__Ascomycota(100);c__Saccharomycetes(100);o__Saccharomycetales(100);f__Metschnikowiaceae(100);g__Metschnikowia(100);g__Metschnikowia_unclassified(100); | Metschnikowia | 13 | Saprotroph | \|Undefined Saprotroph\| | Yeast | NULL | Probable | As nectar-tap_saprotroph (Põlme et al. 2020); Decay sugar-rich substrates (Põlme et al. 2020) | Põlme S, et al. 2020. Fungal Diversity 105:1-16 ((https://doi.org/10.1007/s13225-020-00466-2)) |
| Otu46660 | 1 | k__Fungi(100);p__Ascomycota(100);c__Saccharomycetes(100);o__Saccharomycetales(100);f__Metschnikowiaceae(100);g__Metschnikowia(100);s__Metschnikowia_pulcherrima(100); | Metschnikowia | 13 | Saprotroph | \|Undefined Saprotroph\| | Yeast | NULL | Probable | As nectar-tap_saprotroph (Põlme et al. 2020); Decay sugar-rich substrates (Põlme et al. 2020) | Põlme S, et al. 2020. Fungal Diversity 105:1-16 ((https://doi.org/10.1007/s13225-020-00466-2)) |
| Otu46661 | 1 | k__Fungi(100);p__Ascomycota(100);c__Saccharomycetes(100);o__Saccharomycetales(100);f__Metschnikowiaceae(100);g__Metschnikowia(100);s__Metschnikowia_pulcherrima(100); | Metschnikowia | 13 | Saprotroph | \|Undefined Saprotroph\| | Yeast | NULL | Probable | As nectar-tap_saprotroph (Põlme et al. 2020); Decay sugar-rich substrates (Põlme et al. 2020) | Põlme S, et al. 2020. Fungal Diversity 105:1-16 ((https://doi.org/10.1007/s13225-020-00466-2)) |
| Otu46662 | 1 | k__Fungi(100);p__Ascomycota(100);c__Saccharomycetes(100);o__Saccharomycetales(100);f__Metschnikowiaceae(100);g__Metschnikowia(100);g__Metschnikowia_unclassified(100); | Metschnikowia | 13 | Saprotroph | \|Undefined Saprotroph\| | Yeast | NULL | Probable | As nectar-tap_saprotroph (Põlme et al. 2020); Decay sugar-rich substrates (Põlme et al. 2020) | Põlme S, et al. 2020. Fungal Diversity 105:1-16 ((https://doi.org/10.1007/s13225-020-00466-2)) |
| Otu46663 | 1 | k__Fungi(100);p__Ascomycota(100);c__Saccharomycetes(100);o__Saccharomycetales(100);f__Metschnikowiaceae(100);g__Metschnikowia(100);s__Metschnikowia_pulcherrima(100); | Metschnikowia | 13 | Saprotroph | \|Undefined Saprotroph\| | Yeast | NULL | Probable | As nectar-tap_saprotroph (Põlme et al. 2020); Decay sugar-rich substrates (Põlme et al. 2020) | Põlme S, et al. 2020. Fungal Diversity 105:1-16 ((https://doi.org/10.1007/s13225-020-00466-2)) |
| Otu46666 | 1 | k__Fungi(100);p__Ascomycota(100);c__Saccharomycetes(100);o__Saccharomycetales(100);f__Metschnikowiaceae(100);g__Metschnikowia(100);s__Metschnikowia_pulcherrima(100); | Metschnikowia | 13 | Saprotroph | \|Undefined Saprotroph\| | Yeast | NULL | Probable | As nectar-tap_saprotroph (Põlme et al. 2020); Decay sugar-rich substrates (Põlme et al. 2020) | Põlme S, et al. 2020. Fungal Diversity 105:1-16 ((https://doi.org/10.1007/s13225-020-00466-2)) |
| Otu46668 | 1 | k__Fungi(100);p__Ascomycota(100);c__Saccharomycetes(100);o__Saccharomycetales(100);f__Metschnikowiaceae(100);g__Metschnikowia(100);s__Metschnikowia_pulcherrima(100); | Metschnikowia | 13 | Saprotroph | \|Undefined Saprotroph\| | Yeast | NULL | Probable | As nectar-tap_saprotroph (Põlme et al. 2020); Decay sugar-rich substrates (Põlme et al. 2020) | Põlme S, et al. 2020. Fungal Diversity 105:1-16 ((https://doi.org/10.1007/s13225-020-00466-2)) |
| Otu46669 | 1 | k__Fungi(100);p__Ascomycota(100);c__Saccharomycetes(100);o__Saccharomycetales(100);f__Metschnikowiaceae(100);g__Metschnikowia(100);g__Metschnikowia_unclassified(100); | Metschnikowia | 13 | Saprotroph | \|Undefined Saprotroph\| | Yeast | NULL | Probable | As nectar-tap_saprotroph (Põlme et al. 2020); Decay sugar-rich substrates (Põlme et al. 2020) | Põlme S, et al. 2020. Fungal Diversity 105:1-16 ((https://doi.org/10.1007/s13225-020-00466-2)) |
| Otu46670 | 1 | k__Fungi(100);p__Ascomycota(100);c__Saccharomycetes(100);o__Saccharomycetales(100);f__Metschnikowiaceae(100);g__Metschnikowia(100);s__Metschnikowia_pulcherrima(100); | Metschnikowia | 13 | Saprotroph | \|Undefined Saprotroph\| | Yeast | NULL | Probable | As nectar-tap_saprotroph (Põlme et al. 2020); Decay sugar-rich substrates (Põlme et al. 2020) | Põlme S, et al. 2020. Fungal Diversity 105:1-16 ((https://doi.org/10.1007/s13225-020-00466-2)) |
| Otu46671 | 1 | k__Fungi(100);p__Ascomycota(100);c__Saccharomycetes(100);o__Saccharomycetales(100);f__Metschnikowiaceae(100);g__Metschnikowia(100);g__Metschnikowia_unclassified(100); | Metschnikowia | 13 | Saprotroph | \|Undefined Saprotroph\| | Yeast | NULL | Probable | As nectar-tap_saprotroph (Põlme et al. 2020); Decay sugar-rich substrates (Põlme et al. 2020) | Põlme S, et al. 2020. Fungal Diversity 105:1-16 ((https://doi.org/10.1007/s13225-020-00466-2)) |
| Otu46672 | 1 | k__Fungi(100);p__Ascomycota(100);c__Saccharomycetes(100);o__Saccharomycetales(100);f__Metschnikowiaceae(100);g__Metschnikowia(100);g__Metschnikowia_unclassified(100); | Metschnikowia | 13 | Saprotroph | \|Undefined Saprotroph\| | Yeast | NULL | Probable | As nectar-tap_saprotroph (Põlme et al. 2020); Decay sugar-rich substrates (Põlme et al. 2020) | Põlme S, et al. 2020. Fungal Diversity 105:1-16 ((https://doi.org/10.1007/s13225-020-00466-2)) |
| Otu46673 | 1 | k__Fungi(100);p__Ascomycota(100);c__Saccharomycetes(100);o__Saccharomycetales(100);f__Metschnikowiaceae(100);g__Metschnikowia(100);s__Metschnikowia_pulcherrima(100); | Metschnikowia | 13 | Saprotroph | \|Undefined Saprotroph\| | Yeast | NULL | Probable | As nectar-tap_saprotroph (Põlme et al. 2020); Decay sugar-rich substrates (Põlme et al. 2020) | Põlme S, et al. 2020. Fungal Diversity 105:1-16 ((https://doi.org/10.1007/s13225-020-00466-2)) |
| Otu46675 | 1 | k__Fungi(100);p__Ascomycota(100);c__Saccharomycetes(100);o__Saccharomycetales(100);f__Metschnikowiaceae(100);g__Metschnikowia(100);s__Metschnikowia_pulcherrima(100); | Metschnikowia | 13 | Saprotroph | \|Undefined Saprotroph\| | Yeast | NULL | Probable | As nectar-tap_saprotroph (Põlme et al. 2020); Decay sugar-rich substrates (Põlme et al. 2020) | Põlme S, et al. 2020. Fungal Diversity 105:1-16 ((https://doi.org/10.1007/s13225-020-00466-2)) |
| Otu46676 | 1 | k__Fungi(100);p__Ascomycota(100);c__Saccharomycetes(100);o__Saccharomycetales(100);f__Metschnikowiaceae(100);g__Metschnikowia(100);g__Metschnikowia_unclassified(100); | Metschnikowia | 13 | Saprotroph | \|Undefined Saprotroph\| | Yeast | NULL | Probable | As nectar-tap_saprotroph (Põlme et al. 2020); Decay sugar-rich substrates (Põlme et al. 2020) | Põlme S, et al. 2020. Fungal Diversity 105:1-16 ((https://doi.org/10.1007/s13225-020-00466-2)) |
| Otu46678 | 1 | k__Fungi(100);p__Ascomycota(100);c__Saccharomycetes(100);o__Saccharomycetales(100);f__Metschnikowiaceae(100);g__Metschnikowia(100);s__Metschnikowia_pulcherrima(100); | Metschnikowia | 13 | Saprotroph | \|Undefined Saprotroph\| | Yeast | NULL | Probable | As nectar-tap_saprotroph (Põlme et al. 2020); Decay sugar-rich substrates (Põlme et al. 2020) | Põlme S, et al. 2020. Fungal Diversity 105:1-16 ((https://doi.org/10.1007/s13225-020-00466-2)) |
| Otu46679 | 1 | k__Fungi(100);p__Ascomycota(100);c__Saccharomycetes(100);o__Saccharomycetales(100);f__Metschnikowiaceae(100);g__Metschnikowia(100);s__Metschnikowia_pulcherrima(100); | Metschnikowia | 13 | Saprotroph | \|Undefined Saprotroph\| | Yeast | NULL | Probable | As nectar-tap_saprotroph (Põlme et al. 2020); Decay sugar-rich substrates (Põlme et al. 2020) | Põlme S, et al. 2020. Fungal Diversity 105:1-16 ((https://doi.org/10.1007/s13225-020-00466-2)) |
| Otu46680 | 1 | k__Fungi(100);p__Ascomycota(100);c__Saccharomycetes(100);o__Saccharomycetales(100);f__Metschnikowiaceae(100);g__Metschnikowia(100);s__Metschnikowia_pulcherrima(100); | Metschnikowia | 13 | Saprotroph | \|Undefined Saprotroph\| | Yeast | NULL | Probable | As nectar-tap_saprotroph (Põlme et al. 2020); Decay sugar-rich substrates (Põlme et al. 2020) | Põlme S, et al. 2020. Fungal Diversity 105:1-16 ((https://doi.org/10.1007/s13225-020-00466-2)) |
| Otu46681 | 1 | k__Fungi(100);p__Ascomycota(100);c__Saccharomycetes(100);o__Saccharomycetales(100);f__Metschnikowiaceae(100);g__Metschnikowia(100);s__Metschnikowia_pulcherrima(100); | Metschnikowia | 13 | Saprotroph | \|Undefined Saprotroph\| | Yeast | NULL | Probable | As nectar-tap_saprotroph (Põlme et al. 2020); Decay sugar-rich substrates (Põlme et al. 2020) | Põlme S, et al. 2020. Fungal Diversity 105:1-16 ((https://doi.org/10.1007/s13225-020-00466-2)) |
| Otu46682 | 1 | k__Fungi(100);p__Ascomycota(100);c__Saccharomycetes(100);o__Saccharomycetales(100);f__Metschnikowiaceae(100);g__Metschnikowia(100);s__Metschnikowia_pulcherrima(100); | Metschnikowia | 13 | Saprotroph | \|Undefined Saprotroph\| | Yeast | NULL | Probable | As nectar-tap_saprotroph (Põlme et al. 2020); Decay sugar-rich substrates (Põlme et al. 2020) | Põlme S, et al. 2020. Fungal Diversity 105:1-16 ((https://doi.org/10.1007/s13225-020-00466-2)) |
| Otu46683 | 1 | k__Fungi(100);p__Ascomycota(100);c__Saccharomycetes(100);o__Saccharomycetales(100);f__Metschnikowiaceae(100);g__Metschnikowia(100);s__Metschnikowia_pulcherrima(100); | Metschnikowia | 13 | Saprotroph | \|Undefined Saprotroph\| | Yeast | NULL | Probable | As nectar-tap_saprotroph (Põlme et al. 2020); Decay sugar-rich substrates (Põlme et al. 2020) | Põlme S, et al. 2020. Fungal Diversity 105:1-16 ((https://doi.org/10.1007/s13225-020-00466-2)) |
| Otu46685 | 1 | k__Fungi(100);p__Ascomycota(100);c__Saccharomycetes(100);o__Saccharomycetales(100);f__Metschnikowiaceae(100);g__Metschnikowia(100);s__Metschnikowia_pulcherrima(100); | Metschnikowia | 13 | Saprotroph | \|Undefined Saprotroph\| | Yeast | NULL | Probable | As nectar-tap_saprotroph (Põlme et al. 2020); Decay sugar-rich substrates (Põlme et al. 2020) | Põlme S, et al. 2020. Fungal Diversity 105:1-16 ((https://doi.org/10.1007/s13225-020-00466-2)) |
| Otu46686 | 1 | k__Fungi(100);p__Ascomycota(100);c__Saccharomycetes(100);o__Saccharomycetales(100);f__Metschnikowiaceae(100);g__Metschnikowia(100);s__Metschnikowia_pulcherrima(100); | Metschnikowia | 13 | Saprotroph | \|Undefined Saprotroph\| | Yeast | NULL | Probable | As nectar-tap_saprotroph (Põlme et al. 2020); Decay sugar-rich substrates (Põlme et al. 2020) | Põlme S, et al. 2020. Fungal Diversity 105:1-16 ((https://doi.org/10.1007/s13225-020-00466-2)) |
| Otu46687 | 1 | k__Fungi(100);p__Ascomycota(100);c__Saccharomycetes(100);o__Saccharomycetales(100);f__Metschnikowiaceae(100);g__Metschnikowia(100);s__Metschnikowia_pulcherrima(100); | Metschnikowia | 13 | Saprotroph | \|Undefined Saprotroph\| | Yeast | NULL | Probable | As nectar-tap_saprotroph (Põlme et al. 2020); Decay sugar-rich substrates (Põlme et al. 2020) | Põlme S, et al. 2020. Fungal Diversity 105:1-16 ((https://doi.org/10.1007/s13225-020-00466-2)) |
| Otu46688 | 1 | k__Fungi(100);p__Ascomycota(100);c__Saccharomycetes(100);o__Saccharomycetales(100);f__Metschnikowiaceae(100);g__Metschnikowia(100);s__Metschnikowia_pulcherrima(100); | Metschnikowia | 13 | Saprotroph | \|Undefined Saprotroph\| | Yeast | NULL | Probable | As nectar-tap_saprotroph (Põlme et al. 2020); Decay sugar-rich substrates (Põlme et al. 2020) | Põlme S, et al. 2020. Fungal Diversity 105:1-16 ((https://doi.org/10.1007/s13225-020-00466-2)) |
| Otu46689 | 1 | k__Fungi(100);p__Ascomycota(100);c__Saccharomycetes(100);o__Saccharomycetales(100);f__Metschnikowiaceae(100);g__Metschnikowia(100);s__Metschnikowia_pulcherrima(100); | Metschnikowia | 13 | Saprotroph | \|Undefined Saprotroph\| | Yeast | NULL | Probable | As nectar-tap_saprotroph (Põlme et al. 2020); Decay sugar-rich substrates (Põlme et al. 2020) | Põlme S, et al. 2020. Fungal Diversity 105:1-16 ((https://doi.org/10.1007/s13225-020-00466-2)) |
| Otu46690 | 1 | k__Fungi(100);p__Ascomycota(100);c__Saccharomycetes(100);o__Saccharomycetales(100);f__Metschnikowiaceae(100);g__Metschnikowia(100);s__Metschnikowia_pulcherrima(100); | Metschnikowia | 13 | Saprotroph | \|Undefined Saprotroph\| | Yeast | NULL | Probable | As nectar-tap_saprotroph (Põlme et al. 2020); Decay sugar-rich substrates (Põlme et al. 2020) | Põlme S, et al. 2020. Fungal Diversity 105:1-16 ((https://doi.org/10.1007/s13225-020-00466-2)) |
| Otu46691 | 1 | k__Fungi(100);p__Ascomycota(100);c__Saccharomycetes(100);o__Saccharomycetales(100);f__Metschnikowiaceae(100);g__Metschnikowia(100);s__Metschnikowia_pulcherrima(100); | Metschnikowia | 13 | Saprotroph | \|Undefined Saprotroph\| | Yeast | NULL | Probable | As nectar-tap_saprotroph (Põlme et al. 2020); Decay sugar-rich substrates (Põlme et al. 2020) | Põlme S, et al. 2020. Fungal Diversity 105:1-16 ((https://doi.org/10.1007/s13225-020-00466-2)) |
| Otu46692 | 1 | k__Fungi(100);p__Ascomycota(100);c__Saccharomycetes(100);o__Saccharomycetales(100);f__Metschnikowiaceae(100);g__Metschnikowia(100);s__Metschnikowia_pulcherrima(100); | Metschnikowia | 13 | Saprotroph | \|Undefined Saprotroph\| | Yeast | NULL | Probable | As nectar-tap_saprotroph (Põlme et al. 2020); Decay sugar-rich substrates (Põlme et al. 2020) | Põlme S, et al. 2020. Fungal Diversity 105:1-16 ((https://doi.org/10.1007/s13225-020-00466-2)) |
| Otu46693 | 1 | k__Fungi(100);p__Ascomycota(100);c__Saccharomycetes(100);o__Saccharomycetales(100);f__Metschnikowiaceae(100);g__Metschnikowia(100);s__Metschnikowia_pulcherrima(100); | Metschnikowia | 13 | Saprotroph | \|Undefined Saprotroph\| | Yeast | NULL | Probable | As nectar-tap_saprotroph (Põlme et al. 2020); Decay sugar-rich substrates (Põlme et al. 2020) | Põlme S, et al. 2020. Fungal Diversity 105:1-16 ((https://doi.org/10.1007/s13225-020-00466-2)) |
| Otu46694 | 1 | k__Fungi(100);p__Ascomycota(100);c__Saccharomycetes(100);o__Saccharomycetales(100);f__Metschnikowiaceae(100);g__Metschnikowia(100);g__Metschnikowia_unclassified(100); | Metschnikowia | 13 | Saprotroph | \|Undefined Saprotroph\| | Yeast | NULL | Probable | As nectar-tap_saprotroph (Põlme et al. 2020); Decay sugar-rich substrates (Põlme et al. 2020) | Põlme S, et al. 2020. Fungal Diversity 105:1-16 ((https://doi.org/10.1007/s13225-020-00466-2)) |
| Otu46695 | 1 | k__Fungi(100);p__Ascomycota(100);c__Saccharomycetes(100);o__Saccharomycetales(100);f__Metschnikowiaceae(100);g__Metschnikowia(100);s__Metschnikowia_pulcherrima(100); | Metschnikowia | 13 | Saprotroph | \|Undefined Saprotroph\| | Yeast | NULL | Probable | As nectar-tap_saprotroph (Põlme et al. 2020); Decay sugar-rich substrates (Põlme et al. 2020) | Põlme S, et al. 2020. Fungal Diversity 105:1-16 ((https://doi.org/10.1007/s13225-020-00466-2)) |
| Otu46696 | 1 | k__Fungi(100);p__Ascomycota(100);c__Saccharomycetes(100);o__Saccharomycetales(100);f__Metschnikowiaceae(100);g__Metschnikowia(100);s__Metschnikowia_pulcherrima(100); | Metschnikowia | 13 | Saprotroph | \|Undefined Saprotroph\| | Yeast | NULL | Probable | As nectar-tap_saprotroph (Põlme et al. 2020); Decay sugar-rich substrates (Põlme et al. 2020) | Põlme S, et al. 2020. Fungal Diversity 105:1-16 ((https://doi.org/10.1007/s13225-020-00466-2)) |
| Otu46697 | 1 | k__Fungi(100);p__Ascomycota(100);c__Saccharomycetes(100);o__Saccharomycetales(100);f__Metschnikowiaceae(100);g__Metschnikowia(100);g__Metschnikowia_unclassified(100); | Metschnikowia | 13 | Saprotroph | \|Undefined Saprotroph\| | Yeast | NULL | Probable | As nectar-tap_saprotroph (Põlme et al. 2020); Decay sugar-rich substrates (Põlme et al. 2020) | Põlme S, et al. 2020. Fungal Diversity 105:1-16 ((https://doi.org/10.1007/s13225-020-00466-2)) |
| Otu46698 | 1 | k__Fungi(100);p__Ascomycota(100);c__Saccharomycetes(100);o__Saccharomycetales(100);f__Metschnikowiaceae(100);g__Metschnikowia(100);s__Metschnikowia_pulcherrima(100); | Metschnikowia | 13 | Saprotroph | \|Undefined Saprotroph\| | Yeast | NULL | Probable | As nectar-tap_saprotroph (Põlme et al. 2020); Decay sugar-rich substrates (Põlme et al. 2020) | Põlme S, et al. 2020. Fungal Diversity 105:1-16 ((https://doi.org/10.1007/s13225-020-00466-2)) |
| Otu46699 | 1 | k__Fungi(100);p__Ascomycota(100);c__Saccharomycetes(100);o__Saccharomycetales(100);f__Metschnikowiaceae(100);g__Metschnikowia(100);s__Metschnikowia_pulcherrima(100); | Metschnikowia | 13 | Saprotroph | \|Undefined Saprotroph\| | Yeast | NULL | Probable | As nectar-tap_saprotroph (Põlme et al. 2020); Decay sugar-rich substrates (Põlme et al. 2020) | Põlme S, et al. 2020. Fungal Diversity 105:1-16 ((https://doi.org/10.1007/s13225-020-00466-2)) |
| Otu46700 | 1 | k__Fungi(100);p__Ascomycota(100);c__Saccharomycetes(100);o__Saccharomycetales(100);f__Metschnikowiaceae(100);g__Metschnikowia(100);s__Metschnikowia_pulcherrima(100); | Metschnikowia | 13 | Saprotroph | \|Undefined Saprotroph\| | Yeast | NULL | Probable | As nectar-tap_saprotroph (Põlme et al. 2020); Decay sugar-rich substrates (Põlme et al. 2020) | Põlme S, et al. 2020. Fungal Diversity 105:1-16 ((https://doi.org/10.1007/s13225-020-00466-2)) |
| Otu46701 | 1 | k__Fungi(100);p__Ascomycota(100);c__Saccharomycetes(100);o__Saccharomycetales(100);f__Metschnikowiaceae(100);g__Metschnikowia(100);s__Metschnikowia_pulcherrima(100); | Metschnikowia | 13 | Saprotroph | \|Undefined Saprotroph\| | Yeast | NULL | Probable | As nectar-tap_saprotroph (Põlme et al. 2020); Decay sugar-rich substrates (Põlme et al. 2020) | Põlme S, et al. 2020. Fungal Diversity 105:1-16 ((https://doi.org/10.1007/s13225-020-00466-2)) |
| Otu46702 | 1 | k__Fungi(100);p__Ascomycota(100);c__Saccharomycetes(100);o__Saccharomycetales(100);f__Metschnikowiaceae(100);g__Metschnikowia(100);s__Metschnikowia_pulcherrima(100); | Metschnikowia | 13 | Saprotroph | \|Undefined Saprotroph\| | Yeast | NULL | Probable | As nectar-tap_saprotroph (Põlme et al. 2020); Decay sugar-rich substrates (Põlme et al. 2020) | Põlme S, et al. 2020. Fungal Diversity 105:1-16 ((https://doi.org/10.1007/s13225-020-00466-2)) |
| Otu46703 | 1 | k__Fungi(100);p__Ascomycota(100);c__Saccharomycetes(100);o__Saccharomycetales(100);f__Metschnikowiaceae(100);g__Metschnikowia(100);s__Metschnikowia_pulcherrima(100); | Metschnikowia | 13 | Saprotroph | \|Undefined Saprotroph\| | Yeast | NULL | Probable | As nectar-tap_saprotroph (Põlme et al. 2020); Decay sugar-rich substrates (Põlme et al. 2020) | Põlme S, et al. 2020. Fungal Diversity 105:1-16 ((https://doi.org/10.1007/s13225-020-00466-2)) |
| Otu46704 | 1 | k__Fungi(100);p__Ascomycota(100);c__Saccharomycetes(100);o__Saccharomycetales(100);f__Metschnikowiaceae(100);g__Metschnikowia(100);s__Metschnikowia_pulcherrima(100); | Metschnikowia | 13 | Saprotroph | \|Undefined Saprotroph\| | Yeast | NULL | Probable | As nectar-tap_saprotroph (Põlme et al. 2020); Decay sugar-rich substrates (Põlme et al. 2020) | Põlme S, et al. 2020. Fungal Diversity 105:1-16 ((https://doi.org/10.1007/s13225-020-00466-2)) |
| Otu46705 | 1 | k__Fungi(100);p__Ascomycota(100);c__Saccharomycetes(100);o__Saccharomycetales(100);f__Metschnikowiaceae(100);g__Metschnikowia(100);g__Metschnikowia_unclassified(100); | Metschnikowia | 13 | Saprotroph | \|Undefined Saprotroph\| | Yeast | NULL | Probable | As nectar-tap_saprotroph (Põlme et al. 2020); Decay sugar-rich substrates (Põlme et al. 2020) | Põlme S, et al. 2020. Fungal Diversity 105:1-16 ((https://doi.org/10.1007/s13225-020-00466-2)) |
| Otu46706 | 1 | k__Fungi(100);p__Ascomycota(100);c__Saccharomycetes(100);o__Saccharomycetales(100);f__Metschnikowiaceae(100);g__Metschnikowia(100);g__Metschnikowia_unclassified(100); | Metschnikowia | 13 | Saprotroph | \|Undefined Saprotroph\| | Yeast | NULL | Probable | As nectar-tap_saprotroph (Põlme et al. 2020); Decay sugar-rich substrates (Põlme et al. 2020) | Põlme S, et al. 2020. Fungal Diversity 105:1-16 ((https://doi.org/10.1007/s13225-020-00466-2)) |
| Otu46707 | 1 | k__Fungi(100);p__Ascomycota(100);c__Saccharomycetes(100);o__Saccharomycetales(100);f__Metschnikowiaceae(100);g__Metschnikowia(100);g__Metschnikowia_unclassified(100); | Metschnikowia | 13 | Saprotroph | \|Undefined Saprotroph\| | Yeast | NULL | Probable | As nectar-tap_saprotroph (Põlme et al. 2020); Decay sugar-rich substrates (Põlme et al. 2020) | Põlme S, et al. 2020. Fungal Diversity 105:1-16 ((https://doi.org/10.1007/s13225-020-00466-2)) |
| Otu46708 | 1 | k__Fungi(100);p__Ascomycota(100);c__Saccharomycetes(100);o__Saccharomycetales(100);f__Metschnikowiaceae(100);g__Metschnikowia(100);g__Metschnikowia_unclassified(100); | Metschnikowia | 13 | Saprotroph | \|Undefined Saprotroph\| | Yeast | NULL | Probable | As nectar-tap_saprotroph (Põlme et al. 2020); Decay sugar-rich substrates (Põlme et al. 2020) | Põlme S, et al. 2020. Fungal Diversity 105:1-16 ((https://doi.org/10.1007/s13225-020-00466-2)) |
| Otu46709 | 1 | k__Fungi(100);p__Ascomycota(100);c__Saccharomycetes(100);o__Saccharomycetales(100);f__Metschnikowiaceae(100);g__Metschnikowia(100);g__Metschnikowia_unclassified(100); | Metschnikowia | 13 | Saprotroph | \|Undefined Saprotroph\| | Yeast | NULL | Probable | As nectar-tap_saprotroph (Põlme et al. 2020); Decay sugar-rich substrates (Põlme et al. 2020) | Põlme S, et al. 2020. Fungal Diversity 105:1-16 ((https://doi.org/10.1007/s13225-020-00466-2)) |
| Otu46711 | 1 | k__Fungi(100);p__Ascomycota(100);c__Saccharomycetes(100);o__Saccharomycetales(100);f__Metschnikowiaceae(100);g__Metschnikowia(100);g__Metschnikowia_unclassified(100); | Metschnikowia | 13 | Saprotroph | \|Undefined Saprotroph\| | Yeast | NULL | Probable | As nectar-tap_saprotroph (Põlme et al. 2020); Decay sugar-rich substrates (Põlme et al. 2020) | Põlme S, et al. 2020. Fungal Diversity 105:1-16 ((https://doi.org/10.1007/s13225-020-00466-2)) |
| Otu46713 | 1 | k__Fungi(100);p__Ascomycota(100);c__Saccharomycetes(100);o__Saccharomycetales(100);f__Metschnikowiaceae(100);g__Metschnikowia(100);s__Metschnikowia_pulcherrima(100); | Metschnikowia | 13 | Saprotroph | \|Undefined Saprotroph\| | Yeast | NULL | Probable | As nectar-tap_saprotroph (Põlme et al. 2020); Decay sugar-rich substrates (Põlme et al. 2020) | Põlme S, et al. 2020. Fungal Diversity 105:1-16 ((https://doi.org/10.1007/s13225-020-00466-2)) |
| Otu46715 | 1 | k__Fungi(100);p__Ascomycota(100);c__Saccharomycetes(100);o__Saccharomycetales(100);f__Metschnikowiaceae(100);g__Metschnikowia(100);s__Metschnikowia_pulcherrima(100); | Metschnikowia | 13 | Saprotroph | \|Undefined Saprotroph\| | Yeast | NULL | Probable | As nectar-tap_saprotroph (Põlme et al. 2020); Decay sugar-rich substrates (Põlme et al. 2020) | Põlme S, et al. 2020. Fungal Diversity 105:1-16 ((https://doi.org/10.1007/s13225-020-00466-2)) |
| Otu46716 | 1 | k__Fungi(100);p__Ascomycota(100);c__Saccharomycetes(100);o__Saccharomycetales(100);f__Metschnikowiaceae(100);g__Metschnikowia(100);s__Metschnikowia_pulcherrima(100); | Metschnikowia | 13 | Saprotroph | \|Undefined Saprotroph\| | Yeast | NULL | Probable | As nectar-tap_saprotroph (Põlme et al. 2020); Decay sugar-rich substrates (Põlme et al. 2020) | Põlme S, et al. 2020. Fungal Diversity 105:1-16 ((https://doi.org/10.1007/s13225-020-00466-2)) |
| Otu46717 | 1 | k__Fungi(100);p__Ascomycota(100);c__Saccharomycetes(100);o__Saccharomycetales(100);f__Metschnikowiaceae(100);g__Metschnikowia(100);s__Metschnikowia_pulcherrima(100); | Metschnikowia | 13 | Saprotroph | \|Undefined Saprotroph\| | Yeast | NULL | Probable | As nectar-tap_saprotroph (Põlme et al. 2020); Decay sugar-rich substrates (Põlme et al. 2020) | Põlme S, et al. 2020. Fungal Diversity 105:1-16 ((https://doi.org/10.1007/s13225-020-00466-2)) |
| Otu46718 | 1 | k__Fungi(100);p__Ascomycota(100);c__Saccharomycetes(100);o__Saccharomycetales(100);f__Metschnikowiaceae(100);g__Metschnikowia(100);s__Metschnikowia_pulcherrima(100); | Metschnikowia | 13 | Saprotroph | \|Undefined Saprotroph\| | Yeast | NULL | Probable | As nectar-tap_saprotroph (Põlme et al. 2020); Decay sugar-rich substrates (Põlme et al. 2020) | Põlme S, et al. 2020. Fungal Diversity 105:1-16 ((https://doi.org/10.1007/s13225-020-00466-2)) |
| Otu46720 | 1 | k__Fungi(100);p__Ascomycota(100);c__Saccharomycetes(100);o__Saccharomycetales(100);f__Metschnikowiaceae(100);g__Metschnikowia(100);s__Metschnikowia_pulcherrima(100); | Metschnikowia | 13 | Saprotroph | \|Undefined Saprotroph\| | Yeast | NULL | Probable | As nectar-tap_saprotroph (Põlme et al. 2020); Decay sugar-rich substrates (Põlme et al. 2020) | Põlme S, et al. 2020. Fungal Diversity 105:1-16 ((https://doi.org/10.1007/s13225-020-00466-2)) |
| Otu46721 | 1 | k__Fungi(100);p__Ascomycota(100);c__Saccharomycetes(100);o__Saccharomycetales(100);f__Metschnikowiaceae(100);g__Metschnikowia(100);g__Metschnikowia_unclassified(100); | Metschnikowia | 13 | Saprotroph | \|Undefined Saprotroph\| | Yeast | NULL | Probable | As nectar-tap_saprotroph (Põlme et al. 2020); Decay sugar-rich substrates (Põlme et al. 2020) | Põlme S, et al. 2020. Fungal Diversity 105:1-16 ((https://doi.org/10.1007/s13225-020-00466-2)) |
| Otu46722 | 1 | k__Fungi(100);p__Ascomycota(100);c__Saccharomycetes(100);o__Saccharomycetales(100);f__Metschnikowiaceae(100);g__Metschnikowia(100);s__Metschnikowia_pulcherrima(100); | Metschnikowia | 13 | Saprotroph | \|Undefined Saprotroph\| | Yeast | NULL | Probable | As nectar-tap_saprotroph (Põlme et al. 2020); Decay sugar-rich substrates (Põlme et al. 2020) | Põlme S, et al. 2020. Fungal Diversity 105:1-16 ((https://doi.org/10.1007/s13225-020-00466-2)) |
| Otu46724 | 1 | k__Fungi(100);p__Ascomycota(100);c__Saccharomycetes(100);o__Saccharomycetales(100);f__Metschnikowiaceae(100);g__Metschnikowia(100);s__Metschnikowia_pulcherrima(100); | Metschnikowia | 13 | Saprotroph | \|Undefined Saprotroph\| | Yeast | NULL | Probable | As nectar-tap_saprotroph (Põlme et al. 2020); Decay sugar-rich substrates (Põlme et al. 2020) | Põlme S, et al. 2020. Fungal Diversity 105:1-16 ((https://doi.org/10.1007/s13225-020-00466-2)) |
| Otu46725 | 1 | k__Fungi(100);p__Ascomycota(100);c__Saccharomycetes(100);o__Saccharomycetales(100);f__Metschnikowiaceae(100);g__Metschnikowia(100);g__Metschnikowia_unclassified(100); | Metschnikowia | 13 | Saprotroph | \|Undefined Saprotroph\| | Yeast | NULL | Probable | As nectar-tap_saprotroph (Põlme et al. 2020); Decay sugar-rich substrates (Põlme et al. 2020) | Põlme S, et al. 2020. Fungal Diversity 105:1-16 ((https://doi.org/10.1007/s13225-020-00466-2)) |
| Otu46726 | 1 | k__Fungi(100);p__Ascomycota(100);c__Saccharomycetes(100);o__Saccharomycetales(100);f__Metschnikowiaceae(100);g__Metschnikowia(100);g__Metschnikowia_unclassified(100); | Metschnikowia | 13 | Saprotroph | \|Undefined Saprotroph\| | Yeast | NULL | Probable | As nectar-tap_saprotroph (Põlme et al. 2020); Decay sugar-rich substrates (Põlme et al. 2020) | Põlme S, et al. 2020. Fungal Diversity 105:1-16 ((https://doi.org/10.1007/s13225-020-00466-2)) |
| Otu46727 | 1 | k__Fungi(100);p__Ascomycota(100);c__Saccharomycetes(100);o__Saccharomycetales(100);f__Metschnikowiaceae(100);g__Metschnikowia(100);s__Metschnikowia_pulcherrima(100); | Metschnikowia | 13 | Saprotroph | \|Undefined Saprotroph\| | Yeast | NULL | Probable | As nectar-tap_saprotroph (Põlme et al. 2020); Decay sugar-rich substrates (Põlme et al. 2020) | Põlme S, et al. 2020. Fungal Diversity 105:1-16 ((https://doi.org/10.1007/s13225-020-00466-2)) |
| Otu46728 | 1 | k__Fungi(100);p__Ascomycota(100);c__Saccharomycetes(100);o__Saccharomycetales(100);f__Metschnikowiaceae(100);g__Metschnikowia(100);s__Metschnikowia_pulcherrima(100); | Metschnikowia | 13 | Saprotroph | \|Undefined Saprotroph\| | Yeast | NULL | Probable | As nectar-tap_saprotroph (Põlme et al. 2020); Decay sugar-rich substrates (Põlme et al. 2020) | Põlme S, et al. 2020. Fungal Diversity 105:1-16 ((https://doi.org/10.1007/s13225-020-00466-2)) |
| Otu46729 | 1 | k__Fungi(100);p__Ascomycota(100);c__Saccharomycetes(100);o__Saccharomycetales(100);f__Metschnikowiaceae(100);g__Metschnikowia(100);s__Metschnikowia_pulcherrima(100); | Metschnikowia | 13 | Saprotroph | \|Undefined Saprotroph\| | Yeast | NULL | Probable | As nectar-tap_saprotroph (Põlme et al. 2020); Decay sugar-rich substrates (Põlme et al. 2020) | Põlme S, et al. 2020. Fungal Diversity 105:1-16 ((https://doi.org/10.1007/s13225-020-00466-2)) |
| Otu46730 | 1 | k__Fungi(100);p__Ascomycota(100);c__Saccharomycetes(100);o__Saccharomycetales(100);f__Metschnikowiaceae(100);g__Metschnikowia(100);s__Metschnikowia_pulcherrima(100); | Metschnikowia | 13 | Saprotroph | \|Undefined Saprotroph\| | Yeast | NULL | Probable | As nectar-tap_saprotroph (Põlme et al. 2020); Decay sugar-rich substrates (Põlme et al. 2020) | Põlme S, et al. 2020. Fungal Diversity 105:1-16 ((https://doi.org/10.1007/s13225-020-00466-2)) |
| Otu46731 | 1 | k__Fungi(100);p__Ascomycota(100);c__Saccharomycetes(100);o__Saccharomycetales(100);f__Metschnikowiaceae(100);g__Metschnikowia(100);s__Metschnikowia_pulcherrima(100); | Metschnikowia | 13 | Saprotroph | \|Undefined Saprotroph\| | Yeast | NULL | Probable | As nectar-tap_saprotroph (Põlme et al. 2020); Decay sugar-rich substrates (Põlme et al. 2020) | Põlme S, et al. 2020. Fungal Diversity 105:1-16 ((https://doi.org/10.1007/s13225-020-00466-2)) |
| Otu46732 | 1 | k__Fungi(100);p__Ascomycota(100);c__Saccharomycetes(100);o__Saccharomycetales(100);f__Metschnikowiaceae(100);g__Metschnikowia(100);g__Metschnikowia_unclassified(100); | Metschnikowia | 13 | Saprotroph | \|Undefined Saprotroph\| | Yeast | NULL | Probable | As nectar-tap_saprotroph (Põlme et al. 2020); Decay sugar-rich substrates (Põlme et al. 2020) | Põlme S, et al. 2020. Fungal Diversity 105:1-16 ((https://doi.org/10.1007/s13225-020-00466-2)) |
| Otu46733 | 1 | k__Fungi(100);p__Ascomycota(100);c__Saccharomycetes(100);o__Saccharomycetales(100);f__Metschnikowiaceae(100);g__Metschnikowia(100);g__Metschnikowia_unclassified(100); | Metschnikowia | 13 | Saprotroph | \|Undefined Saprotroph\| | Yeast | NULL | Probable | As nectar-tap_saprotroph (Põlme et al. 2020); Decay sugar-rich substrates (Põlme et al. 2020) | Põlme S, et al. 2020. Fungal Diversity 105:1-16 ((https://doi.org/10.1007/s13225-020-00466-2)) |
| Otu46734 | 1 | k__Fungi(100);p__Ascomycota(100);c__Saccharomycetes(100);o__Saccharomycetales(100);f__Metschnikowiaceae(100);g__Metschnikowia(100);s__Metschnikowia_pulcherrima(100); | Metschnikowia | 13 | Saprotroph | \|Undefined Saprotroph\| | Yeast | NULL | Probable | As nectar-tap_saprotroph (Põlme et al. 2020); Decay sugar-rich substrates (Põlme et al. 2020) | Põlme S, et al. 2020. Fungal Diversity 105:1-16 ((https://doi.org/10.1007/s13225-020-00466-2)) |
| Otu46735 | 1 | k__Fungi(100);p__Ascomycota(100);c__Saccharomycetes(100);o__Saccharomycetales(100);f__Metschnikowiaceae(100);g__Metschnikowia(100);g__Metschnikowia_unclassified(100); | Metschnikowia | 13 | Saprotroph | \|Undefined Saprotroph\| | Yeast | NULL | Probable | As nectar-tap_saprotroph (Põlme et al. 2020); Decay sugar-rich substrates (Põlme et al. 2020) | Põlme S, et al. 2020. Fungal Diversity 105:1-16 ((https://doi.org/10.1007/s13225-020-00466-2)) |
| Otu46736 | 1 | k__Fungi(100);p__Ascomycota(100);c__Saccharomycetes(100);o__Saccharomycetales(100);f__Metschnikowiaceae(100);g__Metschnikowia(100);s__Metschnikowia_pulcherrima(100); | Metschnikowia | 13 | Saprotroph | \|Undefined Saprotroph\| | Yeast | NULL | Probable | As nectar-tap_saprotroph (Põlme et al. 2020); Decay sugar-rich substrates (Põlme et al. 2020) | Põlme S, et al. 2020. Fungal Diversity 105:1-16 ((https://doi.org/10.1007/s13225-020-00466-2)) |
| Otu46737 | 1 | k__Fungi(100);p__Ascomycota(100);c__Saccharomycetes(100);o__Saccharomycetales(100);f__Metschnikowiaceae(100);g__Metschnikowia(100);s__Metschnikowia_pulcherrima(100); | Metschnikowia | 13 | Saprotroph | \|Undefined Saprotroph\| | Yeast | NULL | Probable | As nectar-tap_saprotroph (Põlme et al. 2020); Decay sugar-rich substrates (Põlme et al. 2020) | Põlme S, et al. 2020. Fungal Diversity 105:1-16 ((https://doi.org/10.1007/s13225-020-00466-2)) |
| Otu46738 | 1 | k__Fungi(100);p__Ascomycota(100);c__Saccharomycetes(100);o__Saccharomycetales(100);f__Metschnikowiaceae(100);g__Metschnikowia(100);g__Metschnikowia_unclassified(100); | Metschnikowia | 13 | Saprotroph | \|Undefined Saprotroph\| | Yeast | NULL | Probable | As nectar-tap_saprotroph (Põlme et al. 2020); Decay sugar-rich substrates (Põlme et al. 2020) | Põlme S, et al. 2020. Fungal Diversity 105:1-16 ((https://doi.org/10.1007/s13225-020-00466-2)) |
| Otu46739 | 1 | k__Fungi(100);p__Ascomycota(100);c__Saccharomycetes(100);o__Saccharomycetales(100);f__Metschnikowiaceae(100);g__Metschnikowia(100);g__Metschnikowia_unclassified(100); | Metschnikowia | 13 | Saprotroph | \|Undefined Saprotroph\| | Yeast | NULL | Probable | As nectar-tap_saprotroph (Põlme et al. 2020); Decay sugar-rich substrates (Põlme et al. 2020) | Põlme S, et al. 2020. Fungal Diversity 105:1-16 ((https://doi.org/10.1007/s13225-020-00466-2)) |
| Otu46740 | 1 | k__Fungi(100);p__Ascomycota(100);c__Saccharomycetes(100);o__Saccharomycetales(100);f__Metschnikowiaceae(100);g__Metschnikowia(100);s__Metschnikowia_pulcherrima(100); | Metschnikowia | 13 | Saprotroph | \|Undefined Saprotroph\| | Yeast | NULL | Probable | As nectar-tap_saprotroph (Põlme et al. 2020); Decay sugar-rich substrates (Põlme et al. 2020) | Põlme S, et al. 2020. Fungal Diversity 105:1-16 ((https://doi.org/10.1007/s13225-020-00466-2)) |
| Otu46741 | 1 | k__Fungi(100);p__Ascomycota(100);c__Saccharomycetes(100);o__Saccharomycetales(100);f__Metschnikowiaceae(100);g__Metschnikowia(100);s__Metschnikowia_pulcherrima(100); | Metschnikowia | 13 | Saprotroph | \|Undefined Saprotroph\| | Yeast | NULL | Probable | As nectar-tap_saprotroph (Põlme et al. 2020); Decay sugar-rich substrates (Põlme et al. 2020) | Põlme S, et al. 2020. Fungal Diversity 105:1-16 ((https://doi.org/10.1007/s13225-020-00466-2)) |
| Otu46742 | 1 | k__Fungi(100);p__Ascomycota(100);c__Saccharomycetes(100);o__Saccharomycetales(100);f__Metschnikowiaceae(100);g__Metschnikowia(100);s__Metschnikowia_pulcherrima(100); | Metschnikowia | 13 | Saprotroph | \|Undefined Saprotroph\| | Yeast | NULL | Probable | As nectar-tap_saprotroph (Põlme et al. 2020); Decay sugar-rich substrates (Põlme et al. 2020) | Põlme S, et al. 2020. Fungal Diversity 105:1-16 ((https://doi.org/10.1007/s13225-020-00466-2)) |
| Otu46743 | 1 | k__Fungi(100);p__Ascomycota(100);c__Saccharomycetes(100);o__Saccharomycetales(100);f__Metschnikowiaceae(100);g__Metschnikowia(100);s__Metschnikowia_pulcherrima(100); | Metschnikowia | 13 | Saprotroph | \|Undefined Saprotroph\| | Yeast | NULL | Probable | As nectar-tap_saprotroph (Põlme et al. 2020); Decay sugar-rich substrates (Põlme et al. 2020) | Põlme S, et al. 2020. Fungal Diversity 105:1-16 ((https://doi.org/10.1007/s13225-020-00466-2)) |
| Otu46744 | 1 | k__Fungi(100);p__Ascomycota(100);c__Saccharomycetes(100);o__Saccharomycetales(100);f__Metschnikowiaceae(100);g__Metschnikowia(100);s__Metschnikowia_pulcherrima(100); | Metschnikowia | 13 | Saprotroph | \|Undefined Saprotroph\| | Yeast | NULL | Probable | As nectar-tap_saprotroph (Põlme et al. 2020); Decay sugar-rich substrates (Põlme et al. 2020) | Põlme S, et al. 2020. Fungal Diversity 105:1-16 ((https://doi.org/10.1007/s13225-020-00466-2)) |
| Otu46745 | 1 | k__Fungi(100);p__Ascomycota(100);c__Saccharomycetes(100);o__Saccharomycetales(100);f__Metschnikowiaceae(100);g__Metschnikowia(100);s__Metschnikowia_pulcherrima(100); | Metschnikowia | 13 | Saprotroph | \|Undefined Saprotroph\| | Yeast | NULL | Probable | As nectar-tap_saprotroph (Põlme et al. 2020); Decay sugar-rich substrates (Põlme et al. 2020) | Põlme S, et al. 2020. Fungal Diversity 105:1-16 ((https://doi.org/10.1007/s13225-020-00466-2)) |
| Otu46746 | 1 | k__Fungi(100);p__Ascomycota(100);c__Saccharomycetes(100);o__Saccharomycetales(100);f__Metschnikowiaceae(100);g__Metschnikowia(100);s__Metschnikowia_pulcherrima(100); | Metschnikowia | 13 | Saprotroph | \|Undefined Saprotroph\| | Yeast | NULL | Probable | As nectar-tap_saprotroph (Põlme et al. 2020); Decay sugar-rich substrates (Põlme et al. 2020) | Põlme S, et al. 2020. Fungal Diversity 105:1-16 ((https://doi.org/10.1007/s13225-020-00466-2)) |
| Otu46748 | 1 | k__Fungi(100);p__Ascomycota(100);c__Saccharomycetes(100);o__Saccharomycetales(100);f__Metschnikowiaceae(100);g__Metschnikowia(100);s__Metschnikowia_pulcherrima(100); | Metschnikowia | 13 | Saprotroph | \|Undefined Saprotroph\| | Yeast | NULL | Probable | As nectar-tap_saprotroph (Põlme et al. 2020); Decay sugar-rich substrates (Põlme et al. 2020) | Põlme S, et al. 2020. Fungal Diversity 105:1-16 ((https://doi.org/10.1007/s13225-020-00466-2)) |
| Otu46749 | 1 | k__Fungi(100);p__Ascomycota(100);c__Saccharomycetes(100);o__Saccharomycetales(100);f__Metschnikowiaceae(100);g__Metschnikowia(100);s__Metschnikowia_pulcherrima(100); | Metschnikowia | 13 | Saprotroph | \|Undefined Saprotroph\| | Yeast | NULL | Probable | As nectar-tap_saprotroph (Põlme et al. 2020); Decay sugar-rich substrates (Põlme et al. 2020) | Põlme S, et al. 2020. Fungal Diversity 105:1-16 ((https://doi.org/10.1007/s13225-020-00466-2)) |
| Otu46750 | 1 | k__Fungi(100);p__Ascomycota(100);c__Saccharomycetes(100);o__Saccharomycetales(100);f__Metschnikowiaceae(100);g__Metschnikowia(100);s__Metschnikowia_pulcherrima(100); | Metschnikowia | 13 | Saprotroph | \|Undefined Saprotroph\| | Yeast | NULL | Probable | As nectar-tap_saprotroph (Põlme et al. 2020); Decay sugar-rich substrates (Põlme et al. 2020) | Põlme S, et al. 2020. Fungal Diversity 105:1-16 ((https://doi.org/10.1007/s13225-020-00466-2)) |
| Otu46751 | 1 | k__Fungi(100);p__Ascomycota(100);c__Saccharomycetes(100);o__Saccharomycetales(100);f__Metschnikowiaceae(100);g__Metschnikowia(100);s__Metschnikowia_pulcherrima(100); | Metschnikowia | 13 | Saprotroph | \|Undefined Saprotroph\| | Yeast | NULL | Probable | As nectar-tap_saprotroph (Põlme et al. 2020); Decay sugar-rich substrates (Põlme et al. 2020) | Põlme S, et al. 2020. Fungal Diversity 105:1-16 ((https://doi.org/10.1007/s13225-020-00466-2)) |
| Otu46752 | 1 | k__Fungi(100);p__Ascomycota(100);c__Saccharomycetes(100);o__Saccharomycetales(100);f__Metschnikowiaceae(100);g__Metschnikowia(100);s__Metschnikowia_pulcherrima(100); | Metschnikowia | 13 | Saprotroph | \|Undefined Saprotroph\| | Yeast | NULL | Probable | As nectar-tap_saprotroph (Põlme et al. 2020); Decay sugar-rich substrates (Põlme et al. 2020) | Põlme S, et al. 2020. Fungal Diversity 105:1-16 ((https://doi.org/10.1007/s13225-020-00466-2)) |
| Otu46753 | 1 | k__Fungi(100);p__Ascomycota(100);c__Saccharomycetes(100);o__Saccharomycetales(100);f__Metschnikowiaceae(100);g__Metschnikowia(100);s__Metschnikowia_pulcherrima(100); | Metschnikowia | 13 | Saprotroph | \|Undefined Saprotroph\| | Yeast | NULL | Probable | As nectar-tap_saprotroph (Põlme et al. 2020); Decay sugar-rich substrates (Põlme et al. 2020) | Põlme S, et al. 2020. Fungal Diversity 105:1-16 ((https://doi.org/10.1007/s13225-020-00466-2)) |
| Otu46754 | 1 | k__Fungi(100);p__Ascomycota(100);c__Saccharomycetes(100);o__Saccharomycetales(100);f__Metschnikowiaceae(100);g__Metschnikowia(100);s__Metschnikowia_pulcherrima(100); | Metschnikowia | 13 | Saprotroph | \|Undefined Saprotroph\| | Yeast | NULL | Probable | As nectar-tap_saprotroph (Põlme et al. 2020); Decay sugar-rich substrates (Põlme et al. 2020) | Põlme S, et al. 2020. Fungal Diversity 105:1-16 ((https://doi.org/10.1007/s13225-020-00466-2)) |
| Otu46755 | 1 | k__Fungi(100);p__Ascomycota(100);c__Saccharomycetes(100);o__Saccharomycetales(100);f__Metschnikowiaceae(100);g__Metschnikowia(100);s__Metschnikowia_pulcherrima(100); | Metschnikowia | 13 | Saprotroph | \|Undefined Saprotroph\| | Yeast | NULL | Probable | As nectar-tap_saprotroph (Põlme et al. 2020); Decay sugar-rich substrates (Põlme et al. 2020) | Põlme S, et al. 2020. Fungal Diversity 105:1-16 ((https://doi.org/10.1007/s13225-020-00466-2)) |
| Otu46756 | 1 | k__Fungi(100);p__Ascomycota(100);c__Saccharomycetes(100);o__Saccharomycetales(100);f__Metschnikowiaceae(100);g__Metschnikowia(100);s__Metschnikowia_pulcherrima(100); | Metschnikowia | 13 | Saprotroph | \|Undefined Saprotroph\| | Yeast | NULL | Probable | As nectar-tap_saprotroph (Põlme et al. 2020); Decay sugar-rich substrates (Põlme et al. 2020) | Põlme S, et al. 2020. Fungal Diversity 105:1-16 ((https://doi.org/10.1007/s13225-020-00466-2)) |
| Otu46757 | 1 | k__Fungi(100);p__Ascomycota(100);c__Saccharomycetes(100);o__Saccharomycetales(100);f__Metschnikowiaceae(100);g__Metschnikowia(100);s__Metschnikowia_pulcherrima(100); | Metschnikowia | 13 | Saprotroph | \|Undefined Saprotroph\| | Yeast | NULL | Probable | As nectar-tap_saprotroph (Põlme et al. 2020); Decay sugar-rich substrates (Põlme et al. 2020) | Põlme S, et al. 2020. Fungal Diversity 105:1-16 ((https://doi.org/10.1007/s13225-020-00466-2)) |
| Otu46758 | 1 | k__Fungi(100);p__Ascomycota(100);c__Saccharomycetes(100);o__Saccharomycetales(100);f__Metschnikowiaceae(100);g__Metschnikowia(100);s__Metschnikowia_pulcherrima(100); | Metschnikowia | 13 | Saprotroph | \|Undefined Saprotroph\| | Yeast | NULL | Probable | As nectar-tap_saprotroph (Põlme et al. 2020); Decay sugar-rich substrates (Põlme et al. 2020) | Põlme S, et al. 2020. Fungal Diversity 105:1-16 ((https://doi.org/10.1007/s13225-020-00466-2)) |
| Otu46759 | 1 | k__Fungi(100);p__Ascomycota(100);c__Saccharomycetes(100);o__Saccharomycetales(100);f__Metschnikowiaceae(100);g__Metschnikowia(100);s__Metschnikowia_pulcherrima(100); | Metschnikowia | 13 | Saprotroph | \|Undefined Saprotroph\| | Yeast | NULL | Probable | As nectar-tap_saprotroph (Põlme et al. 2020); Decay sugar-rich substrates (Põlme et al. 2020) | Põlme S, et al. 2020. Fungal Diversity 105:1-16 ((https://doi.org/10.1007/s13225-020-00466-2)) |
| Otu46760 | 1 | k__Fungi(100);p__Ascomycota(100);c__Saccharomycetes(100);o__Saccharomycetales(100);f__Metschnikowiaceae(100);g__Metschnikowia(100);g__Metschnikowia_unclassified(100); | Metschnikowia | 13 | Saprotroph | \|Undefined Saprotroph\| | Yeast | NULL | Probable | As nectar-tap_saprotroph (Põlme et al. 2020); Decay sugar-rich substrates (Põlme et al. 2020) | Põlme S, et al. 2020. Fungal Diversity 105:1-16 ((https://doi.org/10.1007/s13225-020-00466-2)) |
| Otu46761 | 1 | k__Fungi(100);p__Ascomycota(100);c__Saccharomycetes(100);o__Saccharomycetales(100);f__Metschnikowiaceae(100);g__Metschnikowia(100);g__Metschnikowia_unclassified(100); | Metschnikowia | 13 | Saprotroph | \|Undefined Saprotroph\| | Yeast | NULL | Probable | As nectar-tap_saprotroph (Põlme et al. 2020); Decay sugar-rich substrates (Põlme et al. 2020) | Põlme S, et al. 2020. Fungal Diversity 105:1-16 ((https://doi.org/10.1007/s13225-020-00466-2)) |
| Otu46762 | 1 | k__Fungi(100);p__Ascomycota(100);c__Saccharomycetes(100);o__Saccharomycetales(100);f__Metschnikowiaceae(100);g__Metschnikowia(100);g__Metschnikowia_unclassified(100); | Metschnikowia | 13 | Saprotroph | \|Undefined Saprotroph\| | Yeast | NULL | Probable | As nectar-tap_saprotroph (Põlme et al. 2020); Decay sugar-rich substrates (Põlme et al. 2020) | Põlme S, et al. 2020. Fungal Diversity 105:1-16 ((https://doi.org/10.1007/s13225-020-00466-2)) |
| Otu46763 | 1 | k__Fungi(100);p__Ascomycota(100);c__Saccharomycetes(100);o__Saccharomycetales(100);f__Metschnikowiaceae(100);g__Metschnikowia(100);s__Metschnikowia_pulcherrima(100); | Metschnikowia | 13 | Saprotroph | \|Undefined Saprotroph\| | Yeast | NULL | Probable | As nectar-tap_saprotroph (Põlme et al. 2020); Decay sugar-rich substrates (Põlme et al. 2020) | Põlme S, et al. 2020. Fungal Diversity 105:1-16 ((https://doi.org/10.1007/s13225-020-00466-2)) |
| Otu46764 | 1 | k__Fungi(100);p__Ascomycota(100);c__Saccharomycetes(100);o__Saccharomycetales(100);f__Metschnikowiaceae(100);g__Metschnikowia(100);g__Metschnikowia_unclassified(100); | Metschnikowia | 13 | Saprotroph | \|Undefined Saprotroph\| | Yeast | NULL | Probable | As nectar-tap_saprotroph (Põlme et al. 2020); Decay sugar-rich substrates (Põlme et al. 2020) | Põlme S, et al. 2020. Fungal Diversity 105:1-16 ((https://doi.org/10.1007/s13225-020-00466-2)) |
| Otu46765 | 1 | k__Fungi(100);p__Ascomycota(100);c__Saccharomycetes(100);o__Saccharomycetales(100);f__Metschnikowiaceae(100);g__Metschnikowia(100);s__Metschnikowia_pulcherrima(100); | Metschnikowia | 13 | Saprotroph | \|Undefined Saprotroph\| | Yeast | NULL | Probable | As nectar-tap_saprotroph (Põlme et al. 2020); Decay sugar-rich substrates (Põlme et al. 2020) | Põlme S, et al. 2020. Fungal Diversity 105:1-16 ((https://doi.org/10.1007/s13225-020-00466-2)) |
| Otu46766 | 1 | k__Fungi(100);p__Ascomycota(100);c__Saccharomycetes(100);o__Saccharomycetales(100);f__Metschnikowiaceae(100);g__Metschnikowia(100);g__Metschnikowia_unclassified(100); | Metschnikowia | 13 | Saprotroph | \|Undefined Saprotroph\| | Yeast | NULL | Probable | As nectar-tap_saprotroph (Põlme et al. 2020); Decay sugar-rich substrates (Põlme et al. 2020) | Põlme S, et al. 2020. Fungal Diversity 105:1-16 ((https://doi.org/10.1007/s13225-020-00466-2)) |
| Otu46767 | 1 | k__Fungi(100);p__Ascomycota(100);c__Saccharomycetes(100);o__Saccharomycetales(100);f__Metschnikowiaceae(100);g__Metschnikowia(100);s__Metschnikowia_pulcherrima(100); | Metschnikowia | 13 | Saprotroph | \|Undefined Saprotroph\| | Yeast | NULL | Probable | As nectar-tap_saprotroph (Põlme et al. 2020); Decay sugar-rich substrates (Põlme et al. 2020) | Põlme S, et al. 2020. Fungal Diversity 105:1-16 ((https://doi.org/10.1007/s13225-020-00466-2)) |
| Otu46768 | 1 | k__Fungi(100);p__Ascomycota(100);c__Saccharomycetes(100);o__Saccharomycetales(100);f__Metschnikowiaceae(100);g__Metschnikowia(100);s__Metschnikowia_pulcherrima(100); | Metschnikowia | 13 | Saprotroph | \|Undefined Saprotroph\| | Yeast | NULL | Probable | As nectar-tap_saprotroph (Põlme et al. 2020); Decay sugar-rich substrates (Põlme et al. 2020) | Põlme S, et al. 2020. Fungal Diversity 105:1-16 ((https://doi.org/10.1007/s13225-020-00466-2)) |
| Otu46770 | 1 | k__Fungi(100);p__Ascomycota(100);c__Saccharomycetes(100);o__Saccharomycetales(100);f__Metschnikowiaceae(100);g__Metschnikowia(100);s__Metschnikowia_pulcherrima(100); | Metschnikowia | 13 | Saprotroph | \|Undefined Saprotroph\| | Yeast | NULL | Probable | As nectar-tap_saprotroph (Põlme et al. 2020); Decay sugar-rich substrates (Põlme et al. 2020) | Põlme S, et al. 2020. Fungal Diversity 105:1-16 ((https://doi.org/10.1007/s13225-020-00466-2)) |
| Otu46771 | 1 | k__Fungi(100);p__Ascomycota(100);c__Saccharomycetes(100);o__Saccharomycetales(100);f__Metschnikowiaceae(100);g__Metschnikowia(100);s__Metschnikowia_pulcherrima(100); | Metschnikowia | 13 | Saprotroph | \|Undefined Saprotroph\| | Yeast | NULL | Probable | As nectar-tap_saprotroph (Põlme et al. 2020); Decay sugar-rich substrates (Põlme et al. 2020) | Põlme S, et al. 2020. Fungal Diversity 105:1-16 ((https://doi.org/10.1007/s13225-020-00466-2)) |
| Otu46772 | 1 | k__Fungi(100);p__Ascomycota(100);c__Saccharomycetes(100);o__Saccharomycetales(100);f__Metschnikowiaceae(100);g__Metschnikowia(100);s__Metschnikowia_pulcherrima(100); | Metschnikowia | 13 | Saprotroph | \|Undefined Saprotroph\| | Yeast | NULL | Probable | As nectar-tap_saprotroph (Põlme et al. 2020); Decay sugar-rich substrates (Põlme et al. 2020) | Põlme S, et al. 2020. Fungal Diversity 105:1-16 ((https://doi.org/10.1007/s13225-020-00466-2)) |
| Otu46773 | 1 | k__Fungi(100);p__Ascomycota(100);c__Saccharomycetes(100);o__Saccharomycetales(100);f__Metschnikowiaceae(100);g__Metschnikowia(100);s__Metschnikowia_pulcherrima(100); | Metschnikowia | 13 | Saprotroph | \|Undefined Saprotroph\| | Yeast | NULL | Probable | As nectar-tap_saprotroph (Põlme et al. 2020); Decay sugar-rich substrates (Põlme et al. 2020) | Põlme S, et al. 2020. Fungal Diversity 105:1-16 ((https://doi.org/10.1007/s13225-020-00466-2)) |
| Otu46774 | 1 | k__Fungi(100);p__Ascomycota(100);c__Saccharomycetes(100);o__Saccharomycetales(100);f__Metschnikowiaceae(100);g__Metschnikowia(100);s__Metschnikowia_pulcherrima(100); | Metschnikowia | 13 | Saprotroph | \|Undefined Saprotroph\| | Yeast | NULL | Probable | As nectar-tap_saprotroph (Põlme et al. 2020); Decay sugar-rich substrates (Põlme et al. 2020) | Põlme S, et al. 2020. Fungal Diversity 105:1-16 ((https://doi.org/10.1007/s13225-020-00466-2)) |
| Otu46775 | 1 | k__Fungi(100);p__Ascomycota(100);c__Saccharomycetes(100);o__Saccharomycetales(100);f__Metschnikowiaceae(100);g__Metschnikowia(100);s__Metschnikowia_pulcherrima(100); | Metschnikowia | 13 | Saprotroph | \|Undefined Saprotroph\| | Yeast | NULL | Probable | As nectar-tap_saprotroph (Põlme et al. 2020); Decay sugar-rich substrates (Põlme et al. 2020) | Põlme S, et al. 2020. Fungal Diversity 105:1-16 ((https://doi.org/10.1007/s13225-020-00466-2)) |
| Otu46776 | 1 | k__Fungi(100);p__Ascomycota(100);c__Saccharomycetes(100);o__Saccharomycetales(100);f__Metschnikowiaceae(100);g__Metschnikowia(100);s__Metschnikowia_pulcherrima(100); | Metschnikowia | 13 | Saprotroph | \|Undefined Saprotroph\| | Yeast | NULL | Probable | As nectar-tap_saprotroph (Põlme et al. 2020); Decay sugar-rich substrates (Põlme et al. 2020) | Põlme S, et al. 2020. Fungal Diversity 105:1-16 ((https://doi.org/10.1007/s13225-020-00466-2)) |
| Otu46777 | 1 | k__Fungi(100);p__Ascomycota(100);c__Saccharomycetes(100);o__Saccharomycetales(100);f__Metschnikowiaceae(100);g__Metschnikowia(100);s__Metschnikowia_pulcherrima(100); | Metschnikowia | 13 | Saprotroph | \|Undefined Saprotroph\| | Yeast | NULL | Probable | As nectar-tap_saprotroph (Põlme et al. 2020); Decay sugar-rich substrates (Põlme et al. 2020) | Põlme S, et al. 2020. Fungal Diversity 105:1-16 ((https://doi.org/10.1007/s13225-020-00466-2)) |
| Otu46778 | 1 | k__Fungi(100);p__Ascomycota(100);c__Saccharomycetes(100);o__Saccharomycetales(100);f__Metschnikowiaceae(100);g__Metschnikowia(100);s__Metschnikowia_pulcherrima(100); | Metschnikowia | 13 | Saprotroph | \|Undefined Saprotroph\| | Yeast | NULL | Probable | As nectar-tap_saprotroph (Põlme et al. 2020); Decay sugar-rich substrates (Põlme et al. 2020) | Põlme S, et al. 2020. Fungal Diversity 105:1-16 ((https://doi.org/10.1007/s13225-020-00466-2)) |
| Otu46780 | 1 | k__Fungi(100);p__Ascomycota(100);c__Saccharomycetes(100);o__Saccharomycetales(100);f__Metschnikowiaceae(100);g__Metschnikowia(100);g__Metschnikowia_unclassified(100); | Metschnikowia | 13 | Saprotroph | \|Undefined Saprotroph\| | Yeast | NULL | Probable | As nectar-tap_saprotroph (Põlme et al. 2020); Decay sugar-rich substrates (Põlme et al. 2020) | Põlme S, et al. 2020. Fungal Diversity 105:1-16 ((https://doi.org/10.1007/s13225-020-00466-2)) |
| Otu46781 | 1 | k__Fungi(100);p__Ascomycota(100);c__Saccharomycetes(100);o__Saccharomycetales(100);f__Metschnikowiaceae(100);g__Metschnikowia(100);s__Metschnikowia_pulcherrima(100); | Metschnikowia | 13 | Saprotroph | \|Undefined Saprotroph\| | Yeast | NULL | Probable | As nectar-tap_saprotroph (Põlme et al. 2020); Decay sugar-rich substrates (Põlme et al. 2020) | Põlme S, et al. 2020. Fungal Diversity 105:1-16 ((https://doi.org/10.1007/s13225-020-00466-2)) |
| Otu46782 | 1 | k__Fungi(100);p__Ascomycota(100);c__Saccharomycetes(100);o__Saccharomycetales(100);f__Metschnikowiaceae(100);g__Metschnikowia(100);s__Metschnikowia_pulcherrima(100); | Metschnikowia | 13 | Saprotroph | \|Undefined Saprotroph\| | Yeast | NULL | Probable | As nectar-tap_saprotroph (Põlme et al. 2020); Decay sugar-rich substrates (Põlme et al. 2020) | Põlme S, et al. 2020. Fungal Diversity 105:1-16 ((https://doi.org/10.1007/s13225-020-00466-2)) |
| Otu46783 | 1 | k__Fungi(100);p__Ascomycota(100);c__Saccharomycetes(100);o__Saccharomycetales(100);f__Metschnikowiaceae(100);g__Metschnikowia(100);g__Metschnikowia_unclassified(100); | Metschnikowia | 13 | Saprotroph | \|Undefined Saprotroph\| | Yeast | NULL | Probable | As nectar-tap_saprotroph (Põlme et al. 2020); Decay sugar-rich substrates (Põlme et al. 2020) | Põlme S, et al. 2020. Fungal Diversity 105:1-16 ((https://doi.org/10.1007/s13225-020-00466-2)) |
| Otu46785 | 1 | k__Fungi(100);p__Ascomycota(100);c__Saccharomycetes(100);o__Saccharomycetales(100);f__Metschnikowiaceae(100);g__Metschnikowia(100);g__Metschnikowia_unclassified(100); | Metschnikowia | 13 | Saprotroph | \|Undefined Saprotroph\| | Yeast | NULL | Probable | As nectar-tap_saprotroph (Põlme et al. 2020); Decay sugar-rich substrates (Põlme et al. 2020) | Põlme S, et al. 2020. Fungal Diversity 105:1-16 ((https://doi.org/10.1007/s13225-020-00466-2)) |
| Otu46788 | 1 | k__Fungi(100);p__Ascomycota(100);c__Saccharomycetes(100);o__Saccharomycetales(100);f__Metschnikowiaceae(100);g__Metschnikowia(100);s__Metschnikowia_pulcherrima(100); | Metschnikowia | 13 | Saprotroph | \|Undefined Saprotroph\| | Yeast | NULL | Probable | As nectar-tap_saprotroph (Põlme et al. 2020); Decay sugar-rich substrates (Põlme et al. 2020) | Põlme S, et al. 2020. Fungal Diversity 105:1-16 ((https://doi.org/10.1007/s13225-020-00466-2)) |
| Otu46789 | 1 | k__Fungi(100);p__Ascomycota(100);c__Saccharomycetes(100);o__Saccharomycetales(100);f__Metschnikowiaceae(100);g__Metschnikowia(100);g__Metschnikowia_unclassified(100); | Metschnikowia | 13 | Saprotroph | \|Undefined Saprotroph\| | Yeast | NULL | Probable | As nectar-tap_saprotroph (Põlme et al. 2020); Decay sugar-rich substrates (Põlme et al. 2020) | Põlme S, et al. 2020. Fungal Diversity 105:1-16 ((https://doi.org/10.1007/s13225-020-00466-2)) |
| Otu46790 | 1 | k__Fungi(100);p__Ascomycota(100);c__Saccharomycetes(100);o__Saccharomycetales(100);f__Metschnikowiaceae(100);g__Metschnikowia(100);s__Metschnikowia_pulcherrima(100); | Metschnikowia | 13 | Saprotroph | \|Undefined Saprotroph\| | Yeast | NULL | Probable | As nectar-tap_saprotroph (Põlme et al. 2020); Decay sugar-rich substrates (Põlme et al. 2020) | Põlme S, et al. 2020. Fungal Diversity 105:1-16 ((https://doi.org/10.1007/s13225-020-00466-2)) |
| Otu46791 | 1 | k__Fungi(100);p__Ascomycota(100);c__Saccharomycetes(100);o__Saccharomycetales(100);f__Metschnikowiaceae(100);g__Metschnikowia(100);g__Metschnikowia_unclassified(100); | Metschnikowia | 13 | Saprotroph | \|Undefined Saprotroph\| | Yeast | NULL | Probable | As nectar-tap_saprotroph (Põlme et al. 2020); Decay sugar-rich substrates (Põlme et al. 2020) | Põlme S, et al. 2020. Fungal Diversity 105:1-16 ((https://doi.org/10.1007/s13225-020-00466-2)) |
| Otu46792 | 1 | k__Fungi(100);p__Ascomycota(100);c__Saccharomycetes(100);o__Saccharomycetales(100);f__Metschnikowiaceae(100);g__Metschnikowia(100);s__Metschnikowia_pulcherrima(100); | Metschnikowia | 13 | Saprotroph | \|Undefined Saprotroph\| | Yeast | NULL | Probable | As nectar-tap_saprotroph (Põlme et al. 2020); Decay sugar-rich substrates (Põlme et al. 2020) | Põlme S, et al. 2020. Fungal Diversity 105:1-16 ((https://doi.org/10.1007/s13225-020-00466-2)) |
| Otu46793 | 1 | k__Fungi(100);p__Ascomycota(100);c__Saccharomycetes(100);o__Saccharomycetales(100);f__Metschnikowiaceae(100);g__Metschnikowia(100);s__Metschnikowia_pulcherrima(100); | Metschnikowia | 13 | Saprotroph | \|Undefined Saprotroph\| | Yeast | NULL | Probable | As nectar-tap_saprotroph (Põlme et al. 2020); Decay sugar-rich substrates (Põlme et al. 2020) | Põlme S, et al. 2020. Fungal Diversity 105:1-16 ((https://doi.org/10.1007/s13225-020-00466-2)) |
| Otu46794 | 1 | k__Fungi(100);p__Ascomycota(100);c__Saccharomycetes(100);o__Saccharomycetales(100);f__Metschnikowiaceae(100);g__Metschnikowia(100);s__Metschnikowia_pulcherrima(100); | Metschnikowia | 13 | Saprotroph | \|Undefined Saprotroph\| | Yeast | NULL | Probable | As nectar-tap_saprotroph (Põlme et al. 2020); Decay sugar-rich substrates (Põlme et al. 2020) | Põlme S, et al. 2020. Fungal Diversity 105:1-16 ((https://doi.org/10.1007/s13225-020-00466-2)) |
| Otu46795 | 1 | k__Fungi(100);p__Ascomycota(100);c__Saccharomycetes(100);o__Saccharomycetales(100);f__Metschnikowiaceae(100);g__Metschnikowia(100);s__Metschnikowia_pulcherrima(100); | Metschnikowia | 13 | Saprotroph | \|Undefined Saprotroph\| | Yeast | NULL | Probable | As nectar-tap_saprotroph (Põlme et al. 2020); Decay sugar-rich substrates (Põlme et al. 2020) | Põlme S, et al. 2020. Fungal Diversity 105:1-16 ((https://doi.org/10.1007/s13225-020-00466-2)) |

| | | | | | | | | | | | |
|---|---|---|---|---|---|---|---|---|---|---|---|
| Otu46796 | 1 | k__Fungi(100);p__Ascomycota(100);c__Saccharomycetes(100);o__Saccharomycetales(100);f__Metschnikowiaceae(100);g__Metschnikowia(100);s__Metschnikowia_pulcherrima(100); | Metschnikowia | 13 | Saprotroph | \|Undefined Saprotroph\| | Yeast | NULL | Probable | As nectar-tap_saprotroph (Põlme et al. 2020); Decay sugar-rich substrates (Põlme et al. 2020) | Põlme S, et al. 2020. Fungal Diversity 105:1-16 ((https://doi.org/10.1007/s13225-020-00466-2)) |
| Otu46797 | 1 | k__Fungi(100);p__Ascomycota(100);c__Saccharomycetes(100);o__Saccharomycetales(100);f__Metschnikowiaceae(100);g__Metschnikowia(100);s__Metschnikowia_pulcherrima(100); | Metschnikowia | 13 | Saprotroph | \|Undefined Saprotroph\| | Yeast | NULL | Probable | As nectar-tap_saprotroph (Põlme et al. 2020); Decay sugar-rich substrates (Põlme et al. 2020) | Põlme S, et al. 2020. Fungal Diversity 105:1-16 ((https://doi.org/10.1007/s13225-020-00466-2)) |
| Otu46798 | 1 | k__Fungi(100);p__Ascomycota(100);c__Saccharomycetes(100);o__Saccharomycetales(100);f__Metschnikowiaceae(100);g__Metschnikowia(100);s__Metschnikowia_pulcherrima(100); | Metschnikowia | 13 | Saprotroph | \|Undefined Saprotroph\| | Yeast | NULL | Probable | As nectar-tap_saprotroph (Põlme et al. 2020); Decay sugar-rich substrates (Põlme et al. 2020) | Põlme S, et al. 2020. Fungal Diversity 105:1-16 ((https://doi.org/10.1007/s13225-020-00466-2)) |
| Otu46799 | 1 | k__Fungi(100);p__Ascomycota(100);c__Saccharomycetes(100);o__Saccharomycetales(100);f__Metschnikowiaceae(100);g__Metschnikowia(100);g__Metschnikowia_unclassified(100); | Metschnikowia | 13 | Saprotroph | \|Undefined Saprotroph\| | Yeast | NULL | Probable | As nectar-tap_saprotroph (Põlme et al. 2020); Decay sugar-rich substrates (Põlme et al. 2020) | Põlme S, et al. 2020. Fungal Diversity 105:1-16 ((https://doi.org/10.1007/s13225-020-00466-2)) |
| Otu46800 | 1 | k__Fungi(100);p__Ascomycota(100);c__Saccharomycetes(100);o__Saccharomycetales(100);f__Metschnikowiaceae(100);g__Metschnikowia(100);s__Metschnikowia_pulcherrima(100); | Metschnikowia | 13 | Saprotroph | \|Undefined Saprotroph\| | Yeast | NULL | Probable | As nectar-tap_saprotroph (Põlme et al. 2020); Decay sugar-rich substrates (Põlme et al. 2020) | Põlme S, et al. 2020. Fungal Diversity 105:1-16 ((https://doi.org/10.1007/s13225-020-00466-2)) |
| Otu46801 | 1 | k__Fungi(100);p__Ascomycota(100);c__Saccharomycetes(100);o__Saccharomycetales(100);f__Metschnikowiaceae(100);g__Metschnikowia(100);g__Metschnikowia_unclassified(100); | Metschnikowia | 13 | Saprotroph | \|Undefined Saprotroph\| | Yeast | NULL | Probable | As nectar-tap_saprotroph (Põlme et al. 2020); Decay sugar-rich substrates (Põlme et al. 2020) | Põlme S, et al. 2020. Fungal Diversity 105:1-16 ((https://doi.org/10.1007/s13225-020-00466-2)) |
| Otu46802 | 1 | k__Fungi(100);p__Ascomycota(100);c__Saccharomycetes(100);o__Saccharomycetales(100);f__Metschnikowiaceae(100);g__Metschnikowia(100);s__Metschnikowia_pulcherrima(100); | Metschnikowia | 13 | Saprotroph | \|Undefined Saprotroph\| | Yeast | NULL | Probable | As nectar-tap_saprotroph (Põlme et al. 2020); Decay sugar-rich substrates (Põlme et al. 2020) | Põlme S, et al. 2020. Fungal Diversity 105:1-16 ((https://doi.org/10.1007/s13225-020-00466-2)) |
| Otu46803 | 1 | k__Fungi(100);p__Ascomycota(100);c__Saccharomycetes(100);o__Saccharomycetales(100);f__Metschnikowiaceae(100);g__Metschnikowia(100);s__Metschnikowia_pulcherrima(100); | Metschnikowia | 13 | Saprotroph | \|Undefined Saprotroph\| | Yeast | NULL | Probable | As nectar-tap_saprotroph (Põlme et al. 2020); Decay sugar-rich substrates (Põlme et al. 2020) | Põlme S, et al. 2020. Fungal Diversity 105:1-16 ((https://doi.org/10.1007/s13225-020-00466-2)) |
| Otu46804 | 1 | k__Fungi(100);p__Ascomycota(100);c__Saccharomycetes(100);o__Saccharomycetales(100);f__Metschnikowiaceae(100);g__Metschnikowia(100);s__Metschnikowia_pulcherrima(100); | Metschnikowia | 13 | Saprotroph | \|Undefined Saprotroph\| | Yeast | NULL | Probable | As nectar-tap_saprotroph (Põlme et al. 2020); Decay sugar-rich substrates (Põlme et al. 2020) | Põlme S, et al. 2020. Fungal Diversity 105:1-16 ((https://doi.org/10.1007/s13225-020-00466-2)) |
| Otu46805 | 1 | k__Fungi(100);p__Ascomycota(100);c__Saccharomycetes(100);o__Saccharomycetales(100);f__Metschnikowiaceae(100);g__Metschnikowia(100);s__Metschnikowia_pulcherrima(100); | Metschnikowia | 13 | Saprotroph | \|Undefined Saprotroph\| | Yeast | NULL | Probable | As nectar-tap_saprotroph (Põlme et al. 2020); Decay sugar-rich substrates (Põlme et al. 2020) | Põlme S, et al. 2020. Fungal Diversity 105:1-16 ((https://doi.org/10.1007/s13225-020-00466-2)) |
| Otu46806 | 1 | k__Fungi(100);p__Ascomycota(100);c__Saccharomycetes(100);o__Saccharomycetales(100);f__Metschnikowiaceae(100);g__Metschnikowia(100);s__Metschnikowia_pulcherrima(100); | Metschnikowia | 13 | Saprotroph | \|Undefined Saprotroph\| | Yeast | NULL | Probable | As nectar-tap_saprotroph (Põlme et al. 2020); Decay sugar-rich substrates (Põlme et al. 2020) | Põlme S, et al. 2020. Fungal Diversity 105:1-16 ((https://doi.org/10.1007/s13225-020-00466-2)) |
| Otu46808 | 1 | k__Fungi(100);p__Ascomycota(100);c__Saccharomycetes(100);o__Saccharomycetales(100);f__Metschnikowiaceae(100);g__Metschnikowia(100);s__Metschnikowia_pulcherrima(100); | Metschnikowia | 13 | Saprotroph | \|Undefined Saprotroph\| | Yeast | NULL | Probable | As nectar-tap_saprotroph (Põlme et al. 2020); Decay sugar-rich substrates (Põlme et al. 2020) | Põlme S, et al. 2020. Fungal Diversity 105:1-16 ((https://doi.org/10.1007/s13225-020-00466-2)) |
| Otu46809 | 1 | k__Fungi(100);p__Ascomycota(100);c__Saccharomycetes(100);o__Saccharomycetales(100);f__Metschnikowiaceae(100);g__Metschnikowia(100);s__Metschnikowia_pulcherrima(100); | Metschnikowia | 13 | Saprotroph | \|Undefined Saprotroph\| | Yeast | NULL | Probable | As nectar-tap_saprotroph (Põlme et al. 2020); Decay sugar-rich substrates (Põlme et al. 2020) | Põlme S, et al. 2020. Fungal Diversity 105:1-16 ((https://doi.org/10.1007/s13225-020-00466-2)) |
| Otu46810 | 1 | k__Fungi(100);p__Ascomycota(100);c__Saccharomycetes(100);o__Saccharomycetales(100);f__Metschnikowiaceae(100);g__Metschnikowia(100);s__Metschnikowia_pulcherrima(100); | Metschnikowia | 13 | Saprotroph | \|Undefined Saprotroph\| | Yeast | NULL | Probable | As nectar-tap_saprotroph (Põlme et al. 2020); Decay sugar-rich substrates (Põlme et al. 2020) | Põlme S, et al. 2020. Fungal Diversity 105:1-16 ((https://doi.org/10.1007/s13225-020-00466-2)) |
| Otu46811 | 1 | k__Fungi(100);p__Ascomycota(100);c__Saccharomycetes(100);o__Saccharomycetales(100);f__Metschnikowiaceae(100);g__Metschnikowia(100);g__Metschnikowia_unclassified(100); | Metschnikowia | 13 | Saprotroph | \|Undefined Saprotroph\| | Yeast | NULL | Probable | As nectar-tap_saprotroph (Põlme et al. 2020); Decay sugar-rich substrates (Põlme et al. 2020) | Põlme S, et al. 2020. Fungal Diversity 105:1-16 ((https://doi.org/10.1007/s13225-020-00466-2)) |
| Otu46812 | 1 | k__Fungi(100);p__Ascomycota(100);c__Saccharomycetes(100);o__Saccharomycetales(100);f__Metschnikowiaceae(100);g__Metschnikowia(100);s__Metschnikowia_pulcherrima(100); | Metschnikowia | 13 | Saprotroph | \|Undefined Saprotroph\| | Yeast | NULL | Probable | As nectar-tap_saprotroph (Põlme et al. 2020); Decay sugar-rich substrates (Põlme et al. 2020) | Põlme S, et al. 2020. Fungal Diversity 105:1-16 ((https://doi.org/10.1007/s13225-020-00466-2)) |
| Otu46813 | 1 | k__Fungi(100);p__Ascomycota(100);c__Saccharomycetes(100);o__Saccharomycetales(100);f__Metschnikowiaceae(100);g__Metschnikowia(100);s__Metschnikowia_pulcherrima(100); | Metschnikowia | 13 | Saprotroph | \|Undefined Saprotroph\| | Yeast | NULL | Probable | As nectar-tap_saprotroph (Põlme et al. 2020); Decay sugar-rich substrates (Põlme et al. 2020) | Põlme S, et al. 2020. Fungal Diversity 105:1-16 ((https://doi.org/10.1007/s13225-020-00466-2)) |
| Otu46814 | 1 | k__Fungi(100);p__Ascomycota(100);c__Saccharomycetes(100);o__Saccharomycetales(100);f__Metschnikowiaceae(100);g__Metschnikowia(100);s__Metschnikowia_pulcherrima(100); | Metschnikowia | 13 | Saprotroph | \|Undefined Saprotroph\| | Yeast | NULL | Probable | As nectar-tap_saprotroph (Põlme et al. 2020); Decay sugar-rich substrates (Põlme et al. 2020) | Põlme S, et al. 2020. Fungal Diversity 105:1-16 ((https://doi.org/10.1007/s13225-020-00466-2)) |
| Otu46815 | 1 | k__Fungi(100);p__Ascomycota(100);c__Saccharomycetes(100);o__Saccharomycetales(100);f__Metschnikowiaceae(100);g__Metschnikowia(100);s__Metschnikowia_pulcherrima(100); | Metschnikowia | 13 | Saprotroph | \|Undefined Saprotroph\| | Yeast | NULL | Probable | As nectar-tap_saprotroph (Põlme et al. 2020); Decay sugar-rich substrates (Põlme et al. 2020) | Põlme S, et al. 2020. Fungal Diversity 105:1-16 ((https://doi.org/10.1007/s13225-020-00466-2)) |
| Otu46816 | 1 | k__Fungi(100);p__Ascomycota(100);c__Saccharomycetes(100);o__Saccharomycetales(100);f__Metschnikowiaceae(100);g__Metschnikowia(100);s__Metschnikowia_pulcherrima(100); | Metschnikowia | 13 | Saprotroph | \|Undefined Saprotroph\| | Yeast | NULL | Probable | As nectar-tap_saprotroph (Põlme et al. 2020); Decay sugar-rich substrates (Põlme et al. 2020) | Põlme S, et al. 2020. Fungal Diversity 105:1-16 ((https://doi.org/10.1007/s13225-020-00466-2)) |
| Otu46817 | 1 | k__Fungi(100);p__Ascomycota(100);c__Saccharomycetes(100);o__Saccharomycetales(100);f__Metschnikowiaceae(100);g__Metschnikowia(100);s__Metschnikowia_pulcherrima(100); | Metschnikowia | 13 | Saprotroph | \|Undefined Saprotroph\| | Yeast | NULL | Probable | As nectar-tap_saprotroph (Põlme et al. 2020); Decay sugar-rich substrates (Põlme et al. 2020) | Põlme S, et al. 2020. Fungal Diversity 105:1-16 ((https://doi.org/10.1007/s13225-020-00466-2)) |
| Otu46818 | 1 | k__Fungi(100);p__Ascomycota(100);c__Saccharomycetes(100);o__Saccharomycetales(100);f__Metschnikowiaceae(100);g__Metschnikowia(100);s__Metschnikowia_pulcherrima(100); | Metschnikowia | 13 | Saprotroph | \|Undefined Saprotroph\| | Yeast | NULL | Probable | As nectar-tap_saprotroph (Põlme et al. 2020); Decay sugar-rich substrates (Põlme et al. 2020) | Põlme S, et al. 2020. Fungal Diversity 105:1-16 ((https://doi.org/10.1007/s13225-020-00466-2)) |
| Otu46819 | 1 | k__Fungi(100);p__Ascomycota(100);c__Saccharomycetes(100);o__Saccharomycetales(100);f__Metschnikowiaceae(100);g__Metschnikowia(100);s__Metschnikowia_pulcherrima(100); | Metschnikowia | 13 | Saprotroph | \|Undefined Saprotroph\| | Yeast | NULL | Probable | As nectar-tap_saprotroph (Põlme et al. 2020); Decay sugar-rich substrates (Põlme et al. 2020) | Põlme S, et al. 2020. Fungal Diversity 105:1-16 ((https://doi.org/10.1007/s13225-020-00466-2)) |
| Otu46821 | 1 | k__Fungi(100);p__Ascomycota(100);c__Saccharomycetes(100);o__Saccharomycetales(100);f__Metschnikowiaceae(100);g__Metschnikowia(100);s__Metschnikowia_pulcherrima(100); | Metschnikowia | 13 | Saprotroph | \|Undefined Saprotroph\| | Yeast | NULL | Probable | As nectar-tap_saprotroph (Põlme et al. 2020); Decay sugar-rich substrates (Põlme et al. 2020) | Põlme S, et al. 2020. Fungal Diversity 105:1-16 ((https://doi.org/10.1007/s13225-020-00466-2)) |
| Otu46822 | 1 | k__Fungi(100);p__Ascomycota(100);c__Saccharomycetes(100);o__Saccharomycetales(100);f__Metschnikowiaceae(100);g__Metschnikowia(100);s__Metschnikowia_pulcherrima(100); | Metschnikowia | 13 | Saprotroph | \|Undefined Saprotroph\| | Yeast | NULL | Probable | As nectar-tap_saprotroph (Põlme et al. 2020); Decay sugar-rich substrates (Põlme et al. 2020) | Põlme S, et al. 2020. Fungal Diversity 105:1-16 ((https://doi.org/10.1007/s13225-020-00466-2)) |
| Otu46823 | 1 | k__Fungi(100);p__Ascomycota(100);c__Saccharomycetes(100);o__Saccharomycetales(100);f__Metschnikowiaceae(100);g__Metschnikowia(100);s__Metschnikowia_pulcherrima(100); | Metschnikowia | 13 | Saprotroph | \|Undefined Saprotroph\| | Yeast | NULL | Probable | As nectar-tap_saprotroph (Põlme et al. 2020); Decay sugar-rich substrates (Põlme et al. 2020) | Põlme S, et al. 2020. Fungal Diversity 105:1-16 ((https://doi.org/10.1007/s13225-020-00466-2)) |
| Otu46826 | 1 | k__Fungi(100);p__Ascomycota(100);c__Saccharomycetes(100);o__Saccharomycetales(100);f__Metschnikowiaceae(100);g__Metschnikowia(100);s__Metschnikowia_pulcherrima(100); | Metschnikowia | 13 | Saprotroph | \|Undefined Saprotroph\| | Yeast | NULL | Probable | As nectar-tap_saprotroph (Põlme et al. 2020); Decay sugar-rich substrates (Põlme et al. 2020) | Põlme S, et al. 2020. Fungal Diversity 105:1-16 ((https://doi.org/10.1007/s13225-020-00466-2)) |
| Otu46827 | 1 | k__Fungi(100);p__Ascomycota(100);c__Saccharomycetes(100);o__Saccharomycetales(100);f__Metschnikowiaceae(100);g__Metschnikowia(100);s__Metschnikowia_pulcherrima(100); | Metschnikowia | 13 | Saprotroph | \|Undefined Saprotroph\| | Yeast | NULL | Probable | As nectar-tap_saprotroph (Põlme et al. 2020); Decay sugar-rich substrates (Põlme et al. 2020) | Põlme S, et al. 2020. Fungal Diversity 105:1-16 ((https://doi.org/10.1007/s13225-020-00466-2)) |
| Otu46828 | 1 | k__Fungi(100);p__Ascomycota(100);c__Saccharomycetes(100);o__Saccharomycetales(100);f__Metschnikowiaceae(100);g__Metschnikowia(100);s__Metschnikowia_pulcherrima(100); | Metschnikowia | 13 | Saprotroph | \|Undefined Saprotroph\| | Yeast | NULL | Probable | As nectar-tap_saprotroph (Põlme et al. 2020); Decay sugar-rich substrates (Põlme et al. 2020) | Põlme S, et al. 2020. Fungal Diversity 105:1-16 ((https://doi.org/10.1007/s13225-020-00466-2)) |
| Otu46829 | 1 | k__Fungi(100);p__Ascomycota(100);c__Saccharomycetes(100);o__Saccharomycetales(100);f__Metschnikowiaceae(100);g__Metschnikowia(100);g__Metschnikowia_unclassified(100); | Metschnikowia | 13 | Saprotroph | \|Undefined Saprotroph\| | Yeast | NULL | Probable | As nectar-tap_saprotroph (Põlme et al. 2020); Decay sugar-rich substrates (Põlme et al. 2020) | Põlme S, et al. 2020. Fungal Diversity 105:1-16 ((https://doi.org/10.1007/s13225-020-00466-2)) |
| Otu46830 | 1 | k__Fungi(100);p__Ascomycota(100);c__Saccharomycetes(100);o__Saccharomycetales(100);f__Metschnikowiaceae(100);g__Metschnikowia(100);g__Metschnikowia_unclassified(100); | Metschnikowia | 13 | Saprotroph | \|Undefined Saprotroph\| | Yeast | NULL | Probable | As nectar-tap_saprotroph (Põlme et al. 2020); Decay sugar-rich substrates (Põlme et al. 2020) | Põlme S, et al. 2020. Fungal Diversity 105:1-16 ((https://doi.org/10.1007/s13225-020-00466-2)) |
| Otu46831 | 1 | k__Fungi(100);p__Ascomycota(100);c__Saccharomycetes(100);o__Saccharomycetales(100);f__Metschnikowiaceae(100);g__Metschnikowia(100);s__Metschnikowia_pulcherrima(100); | Metschnikowia | 13 | Saprotroph | \|Undefined Saprotroph\| | Yeast | NULL | Probable | As nectar-tap_saprotroph (Põlme et al. 2020); Decay sugar-rich substrates (Põlme et al. 2020) | Põlme S, et al. 2020. Fungal Diversity 105:1-16 ((https://doi.org/10.1007/s13225-020-00466-2)) |
| Otu46834 | 1 | k__Fungi(100);p__Ascomycota(100);c__Saccharomycetes(100);o__Saccharomycetales(100);f__Metschnikowiaceae(100);g__Metschnikowia(100);s__Metschnikowia_pulcherrima(100); | Metschnikowia | 13 | Saprotroph | \|Undefined Saprotroph\| | Yeast | NULL | Probable | As nectar-tap_saprotroph (Põlme et al. 2020); Decay sugar-rich substrates (Põlme et al. 2020) | Põlme S, et al. 2020. Fungal Diversity 105:1-16 ((https://doi.org/10.1007/s13225-020-00466-2)) |
| Otu46835 | 1 | k__Fungi(100);p__Ascomycota(100);c__Saccharomycetes(100);o__Saccharomycetales(100);f__Metschnikowiaceae(100);g__Metschnikowia(100);s__Metschnikowia_pulcherrima(100); | Metschnikowia | 13 | Saprotroph | \|Undefined Saprotroph\| | Yeast | NULL | Probable | As nectar-tap_saprotroph (Põlme et al. 2020); Decay sugar-rich substrates (Põlme et al. 2020) | Põlme S, et al. 2020. Fungal Diversity 105:1-16 ((https://doi.org/10.1007/s13225-020-00466-2)) |
| Otu46836 | 1 | k__Fungi(100);p__Ascomycota(100);c__Saccharomycetes(100);o__Saccharomycetales(100);f__Metschnikowiaceae(100);g__Metschnikowia(100);s__Metschnikowia_pulcherrima(100); | Metschnikowia | 13 | Saprotroph | \|Undefined Saprotroph\| | Yeast | NULL | Probable | As nectar-tap_saprotroph (Põlme et al. 2020); Decay sugar-rich substrates (Põlme et al. 2020) | Põlme S, et al. 2020. Fungal Diversity 105:1-16 ((https://doi.org/10.1007/s13225-020-00466-2)) |
| Otu46837 | 1 | k__Fungi(100);p__Ascomycota(100);c__Saccharomycetes(100);o__Saccharomycetales(100);f__Metschnikowiaceae(100);g__Metschnikowia(100);s__Metschnikowia_pulcherrima(100); | Metschnikowia | 13 | Saprotroph | \|Undefined Saprotroph\| | Yeast | NULL | Probable | As nectar-tap_saprotroph (Põlme et al. 2020); Decay sugar-rich substrates (Põlme et al. 2020) | Põlme S, et al. 2020. Fungal Diversity 105:1-16 ((https://doi.org/10.1007/s13225-020-00466-2)) |
| Otu46839 | 1 | k__Fungi(100);p__Ascomycota(100);c__Saccharomycetes(100);o__Saccharomycetales(100);f__Metschnikowiaceae(100);g__Metschnikowia(100);g__Metschnikowia_unclassified(100); | Metschnikowia | 13 | Saprotroph | \|Undefined Saprotroph\| | Yeast | NULL | Probable | As nectar-tap_saprotroph (Põlme et al. 2020); Decay sugar-rich substrates (Põlme et al. 2020) | Põlme S, et al. 2020. Fungal Diversity 105:1-16 ((https://doi.org/10.1007/s13225-020-00466-2)) |
| Otu46840 | 1 | k__Fungi(100);p__Ascomycota(100);c__Saccharomycetes(100);o__Saccharomycetales(100);f__Metschnikowiaceae(100);g__Metschnikowia(100);s__Metschnikowia_pulcherrima(100); | Metschnikowia | 13 | Saprotroph | \|Undefined Saprotroph\| | Yeast | NULL | Probable | As nectar-tap_saprotroph (Põlme et al. 2020); Decay sugar-rich substrates (Põlme et al. 2020) | Põlme S, et al. 2020. Fungal Diversity 105:1-16 ((https://doi.org/10.1007/s13225-020-00466-2)) |
| Otu46841 | 1 | k__Fungi(100);p__Ascomycota(100);c__Saccharomycetes(100);o__Saccharomycetales(100);f__Metschnikowiaceae(100);g__Metschnikowia(100);g__Metschnikowia_unclassified(100); | Metschnikowia | 13 | Saprotroph | \|Undefined Saprotroph\| | Yeast | NULL | Probable | As nectar-tap_saprotroph (Põlme et al. 2020); Decay sugar-rich substrates (Põlme et al. 2020) | Põlme S, et al. 2020. Fungal Diversity 105:1-16 ((https://doi.org/10.1007/s13225-020-00466-2)) |
| Otu46842 | 1 | k__Fungi(100);p__Ascomycota(100);c__Saccharomycetes(100);o__Saccharomycetales(100);f__Metschnikowiaceae(100);g__Metschnikowia(100);s__Metschnikowia_pulcherrima(100); | Metschnikowia | 13 | Saprotroph | \|Undefined Saprotroph\| | Yeast | NULL | Probable | As nectar-tap_saprotroph (Põlme et al. 2020); Decay sugar-rich substrates (Põlme et al. 2020) | Põlme S, et al. 2020. Fungal Diversity 105:1-16 ((https://doi.org/10.1007/s13225-020-00466-2)) |
| Otu46843 | 1 | k__Fungi(100);p__Ascomycota(100);c__Saccharomycetes(100);o__Saccharomycetales(100);f__Metschnikowiaceae(100);g__Metschnikowia(100);s__Metschnikowia_pulcherrima(100); | Metschnikowia | 13 | Saprotroph | \|Undefined Saprotroph\| | Yeast | NULL | Probable | As nectar-tap_saprotroph (Põlme et al. 2020); Decay sugar-rich substrates (Põlme et al. 2020) | Põlme S, et al. 2020. Fungal Diversity 105:1-16 ((https://doi.org/10.1007/s13225-020-00466-2)) |
| Otu46844 | 1 | k__Fungi(100);p__Ascomycota(100);c__Saccharomycetes(100);o__Saccharomycetales(100);f__Metschnikowiaceae(100);g__Metschnikowia(100);s__Metschnikowia_pulcherrima(100); | Metschnikowia | 13 | Saprotroph | \|Undefined Saprotroph\| | Yeast | NULL | Probable | As nectar-tap_saprotroph (Põlme et al. 2020); Decay sugar-rich substrates (Põlme et al. 2020) | Põlme S, et al. 2020. Fungal Diversity 105:1-16 ((https://doi.org/10.1007/s13225-020-00466-2)) |
| Otu46845 | 1 | k__Fungi(100);p__Ascomycota(100);c__Saccharomycetes(100);o__Saccharomycetales(100);f__Metschnikowiaceae(100);g__Metschnikowia(100);s__Metschnikowia_pulcherrima(100); | Metschnikowia | 13 | Saprotroph | \|Undefined Saprotroph\| | Yeast | NULL | Probable | As nectar-tap_saprotroph (Põlme et al. 2020); Decay sugar-rich substrates (Põlme et al. 2020) | Põlme S, et al. 2020. Fungal Diversity 105:1-16 ((https://doi.org/10.1007/s13225-020-00466-2)) |
| Otu46846 | 1 | k__Fungi(100);p__Ascomycota(100);c__Saccharomycetes(100);o__Saccharomycetales(100);f__Metschnikowiaceae(100);g__Metschnikowia(100);s__Metschnikowia_pulcherrima(100); | Metschnikowia | 13 | Saprotroph | \|Undefined Saprotroph\| | Yeast | NULL | Probable | As nectar-tap_saprotroph (Põlme et al. 2020); Decay sugar-rich substrates (Põlme et al. 2020) | Põlme S, et al. 2020. Fungal Diversity 105:1-16 ((https://doi.org/10.1007/s13225-020-00466-2)) |
| Otu46847 | 1 | k__Fungi(100);p__Ascomycota(100);c__Saccharomycetes(100);o__Saccharomycetales(100);f__Metschnikowiaceae(100);g__Metschnikowia(100);s__Metschnikowia_pulcherrima(100); | Metschnikowia | 13 | Saprotroph | \|Undefined Saprotroph\| | Yeast | NULL | Probable | As nectar-tap_saprotroph (Põlme et al. 2020); Decay sugar-rich substrates (Põlme et al. 2020) | Põlme S, et al. 2020. Fungal Diversity 105:1-16 ((https://doi.org/10.1007/s13225-020-00466-2)) |
| Otu46848 | 1 | k__Fungi(100);p__Ascomycota(100);c__Saccharomycetes(100);o__Saccharomycetales(100);f__Metschnikowiaceae(100);g__Metschnikowia(100);s__Metschnikowia_pulcherrima(100); | Metschnikowia | 13 | Saprotroph | \|Undefined Saprotroph\| | Yeast | NULL | Probable | As nectar-tap_saprotroph (Põlme et al. 2020); Decay sugar-rich substrates (Põlme et al. 2020) | Põlme S, et al. 2020. Fungal Diversity 105:1-16 ((https://doi.org/10.1007/s13225-020-00466-2)) |
| Otu46849 | 1 | k__Fungi(100);p__Ascomycota(100);c__Saccharomycetes(100);o__Saccharomycetales(100);f__Metschnikowiaceae(100);g__Metschnikowia(100);s__Metschnikowia_pulcherrima(100); | Metschnikowia | 13 | Saprotroph | \|Undefined Saprotroph\| | Yeast | NULL | Probable | As nectar-tap_saprotroph (Põlme et al. 2020); Decay sugar-rich substrates (Põlme et al. 2020) | Põlme S, et al. 2020. Fungal Diversity 105:1-16 ((https://doi.org/10.1007/s13225-020-00466-2)) |
| Otu46850 | 1 | k__Fungi(100);p__Ascomycota(100);c__Saccharomycetes(100);o__Saccharomycetales(100);f__Metschnikowiaceae(100);g__Metschnikowia(100);s__Metschnikowia_pulcherrima(100); | Metschnikowia | 13 | Saprotroph | \|Undefined Saprotroph\| | Yeast | NULL | Probable | As nectar-tap_saprotroph (Põlme et al. 2020); Decay sugar-rich substrates (Põlme et al. 2020) | Põlme S, et al. 2020. Fungal Diversity 105:1-16 ((https://doi.org/10.1007/s13225-020-00466-2)) |
| Otu46851 | 1 | k__Fungi(100);p__Ascomycota(100);c__Saccharomycetes(100);o__Saccharomycetales(100);f__Metschnikowiaceae(100);g__Metschnikowia(100);g__Metschnikowia_unclassified(100); | Metschnikowia | 13 | Saprotroph | \|Undefined Saprotroph\| | Yeast | NULL | Probable | As nectar-tap_saprotroph (Põlme et al. 2020); Decay sugar-rich substrates (Põlme et al. 2020) | Põlme S, et al. 2020. Fungal Diversity 105:1-16 ((https://doi.org/10.1007/s13225-020-00466-2)) |
| Otu46852 | 1 | k__Fungi(100);p__Ascomycota(100);c__Saccharomycetes(100);o__Saccharomycetales(100);f__Metschnikowiaceae(100);g__Metschnikowia(100);s__Metschnikowia_pulcherrima(100); | Metschnikowia | 13 | Saprotroph | \|Undefined Saprotroph\| | Yeast | NULL | Probable | As nectar-tap_saprotroph (Põlme et al. 2020); Decay sugar-rich substrates (Põlme et al. 2020) | Põlme S, et al. 2020. Fungal Diversity 105:1-16 ((https://doi.org/10.1007/s13225-020-00466-2)) |
| Otu46853 | 1 | k__Fungi(100);p__Ascomycota(100);c__Saccharomycetes(100);o__Saccharomycetales(100);f__Metschnikowiaceae(100);g__Metschnikowia(100);s__Metschnikowia_pulcherrima(100); | Metschnikowia | 13 | Saprotroph | \|Undefined Saprotroph\| | Yeast | NULL | Probable | As nectar-tap_saprotroph (Põlme et al. 2020); Decay sugar-rich substrates (Põlme et al. 2020) | Põlme S, et al. 2020. Fungal Diversity 105:1-16 ((https://doi.org/10.1007/s13225-020-00466-2)) |
| Otu46854 | 1 | k__Fungi(100);p__Ascomycota(100);c__Saccharomycetes(100);o__Saccharomycetales(100);f__Metschnikowiaceae(100);g__Metschnikowia(100);s__Metschnikowia_pulcherrima(100); | Metschnikowia | 13 | Saprotroph | \|Undefined Saprotroph\| | Yeast | NULL | Probable | As nectar-tap_saprotroph (Põlme et al. 2020); Decay sugar-rich substrates (Põlme et al. 2020) | Põlme S, et al. 2020. Fungal Diversity 105:1-16 ((https://doi.org/10.1007/s13225-020-00466-2)) |
| Otu46855 | 1 | k__Fungi(100);p__Ascomycota(100);c__Saccharomycetes(100);o__Saccharomycetales(100);f__Metschnikowiaceae(100);g__Metschnikowia(100);s__Metschnikowia_pulcherrima(100); | Metschnikowia | 13 | Saprotroph | \|Undefined Saprotroph\| | Yeast | NULL | Probable | As nectar-tap_saprotroph (Põlme et al. 2020); Decay sugar-rich substrates (Põlme et al. 2020) | Põlme S, et al. 2020. Fungal Diversity 105:1-16 ((https://doi.org/10.1007/s13225-020-00466-2)) |
| Otu46856 | 1 | k__Fungi(100);p__Ascomycota(100);c__Sordariomycetes(100);o__Xylariales(100);f__Amphisphaeriaceae(100);f__Amphisphaeriaceae_unclassified(100);f__Amphisphaeriaceae_unclassified(100); | Amphisphaeriaceae | 9 | Saprotroph | Undefined Saprotroph | Microfungus | NULL | Possible | Saprobes or weak parasites of leaves, stems, etc., often appearing as secondary colonizers following damage to plants by other agents (Cannon & Kirk 2007) | Cannon PF, Kirk PM. 2007. Fungal Families of the World. CAB International, Cambridge (ISBN: 978-0851998275) |
| Otu46857 | 1 | k__Fungi(100);p__Ascomycota(100);c__Saccharomycetes(100);o__Saccharomycetales(100);f__Metschnikowiaceae(100);g__Metschnikowia(100);s__Metschnikowia_pulcherrima(100); | Metschnikowia | 13 | Saprotroph | \|Undefined Saprotroph\| | Yeast | NULL | Probable | As nectar-tap_saprotroph (Põlme et al. 2020); Decay sugar-rich substrates (Põlme et al. 2020) | Põlme S, et al. 2020. Fungal Diversity 105:1-16 ((https://doi.org/10.1007/s13225-020-00466-2)) |
| Otu46859 | 1 | k__Fungi(100);p__Ascomycota(100);c__Saccharomycetes(100);o__Saccharomycetales(100);f__Metschnikowiaceae(100);g__Metschnikowia(100);s__Metschnikowia_pulcherrima(100); | Metschnikowia | 13 | Saprotroph | \|Undefined Saprotroph\| | Yeast | NULL | Probable | As nectar-tap_saprotroph (Põlme et al. 2020); Decay sugar-rich substrates (Põlme et al. 2020) | Põlme S, et al. 2020. Fungal Diversity 105:1-16 ((https://doi.org/10.1007/s13225-020-00466-2)) |
| Otu46865 | 1 | k__Fungi(100);p__Ascomycota(100);c__Saccharomycetes(100);o__Saccharomycetales(100);f__Metschnikowiaceae(100);g__Metschnikowia(100);s__Metschnikowia_pulcherrima(100); | Metschnikowia | 13 | Saprotroph | \|Undefined Saprotroph\| | Yeast | NULL | Probable | As nectar-tap_saprotroph (Põlme et al. 2020); Decay sugar-rich substrates (Põlme et al. 2020) | Põlme S, et al. 2020. Fungal Diversity 105:1-16 ((https://doi.org/10.1007/s13225-020-00466-2)) |
| Otu46866 | 1 | k__Fungi(100);p__Ascomycota(100);c__Saccharomycetes(100);o__Saccharomycetales(100);f__Metschnikowiaceae(100);g__Metschnikowia(100);g__Metschnikowia_unclassified(100); | Metschnikowia | 13 | Saprotroph | \|Undefined Saprotroph\| | Yeast | NULL | Probable | As nectar-tap_saprotroph (Põlme et al. 2020); Decay sugar-rich substrates (Põlme et al. 2020) | Põlme S, et al. 2020. Fungal Diversity 105:1-16 ((https://doi.org/10.1007/s13225-020-00466-2)) |
| Otu46867 | 1 | k__Fungi(100);p__Ascomycota(100);c__Lecanoromycetes(100);o__Teloschistales(100);f__Caliciaceae(100);g__Calicium(100);g__Calicium_unclassified(100); | Calicium | 13 | Symbiotroph | \|Lichenized\| | Calicioid-Thallus | NULL | Highly Probable | NULL | Esslinger TL. 2014. North Dakota State University ((http://www.ndsu.edu/pubweb/~esslinge/chcklst/chcklst7.htm)); Põlme S, et al. 2020. Fungal Diversity 105:1-16 ((https://doi.org/10.1007/s13225-020-00466-2)) |
| Otu46870 | 1 | k__Fungi(100);p__Ascomycota(100);c__Saccharomycetes(100);o__Saccharomycetales(100);f__Metschnikowiaceae(100);g__Metschnikowia(100);g__Metschnikowia_unclassified(100); | Metschnikowia | 13 | Saprotroph | \|Undefined Saprotroph\| | Yeast | NULL | Probable | As nectar-tap_saprotroph (Põlme et al. 2020); Decay sugar-rich substrates (Põlme et al. 2020) | Põlme S, et al. 2020. Fungal Diversity 105:1-16 ((https://doi.org/10.1007/s13225-020-00466-2)) |
| Otu46871 | 1 | k__Fungi(100);p__Ascomycota(100);c__Saccharomycetes(100);o__Saccharomycetales(100);f__Metschnikowiaceae(100);g__Metschnikowia(100);s__Metschnikowia_pulcherrima(100); | Metschnikowia | 13 | Saprotroph | \|Undefined Saprotroph\| | Yeast | NULL | Probable | As nectar-tap_saprotroph (Põlme et al. 2020); Decay sugar-rich substrates (Põlme et al. 2020) | Põlme S, et al. 2020. Fungal Diversity 105:1-16 ((https://doi.org/10.1007/s13225-020-00466-2)) |
| Otu46872 | 1 | k__Fungi(100);p__Ascomycota(100);c__Saccharomycetes(100);o__Saccharomycetales(100);f__Metschnikowiaceae(100);g__Metschnikowia(100);s__Metschnikowia_pulcherrima(100); | Metschnikowia | 13 | Saprotroph | \|Undefined Saprotroph\| | Yeast | NULL | Probable | As nectar-tap_saprotroph (Põlme et al. 2020); Decay sugar-rich substrates (Põlme et al. 2020) | Põlme S, et al. 2020. Fungal Diversity 105:1-16 ((https://doi.org/10.1007/s13225-020-00466-2)) |
| Otu46873 | 1 | k__Fungi(100);p__Ascomycota(100);c__Saccharomycetes(100);o__Saccharomycetales(100);f__Metschnikowiaceae(100);g__Metschnikowia(100);s__Metschnikowia_pulcherrima(100); | Metschnikowia | 13 | Saprotroph | \|Undefined Saprotroph\| | Yeast | NULL | Probable | As nectar-tap_saprotroph (Põlme et al. 2020); Decay sugar-rich substrates (Põlme et al. 2020) | Põlme S, et al. 2020. Fungal Diversity 105:1-16 ((https://doi.org/10.1007/s13225-020-00466-2)) |
| Otu46874 | 1 | k__Fungi(100);p__Ascomycota(100);c__Saccharomycetes(100);o__Saccharomycetales(100);f__Metschnikowiaceae(100);g__Metschnikowia(100);s__Metschnikowia_pulcherrima(100); | Metschnikowia | 13 | Saprotroph | \|Undefined Saprotroph\| | Yeast | NULL | Probable | As nectar-tap_saprotroph (Põlme et al. 2020); Decay sugar-rich substrates (Põlme et al. 2020) | Põlme S, et al. 2020. Fungal Diversity 105:1-16 ((https://doi.org/10.1007/s13225-020-00466-2)) |
| Otu46875 | 1 | k__Fungi(100);p__Ascomycota(100);c__Saccharomycetes(100);o__Saccharomycetales(100);f__Metschnikowiaceae(100);g__Metschnikowia(100);s__Metschnikowia_pulcherrima(100); | Metschnikowia | 13 | Saprotroph | \|Undefined Saprotroph\| | Yeast | NULL | Probable | As nectar-tap_saprotroph (Põlme et al. 2020); Decay sugar-rich substrates (Põlme et al. 2020) | Põlme S, et al. 2020. Fungal Diversity 105:1-16 ((https://doi.org/10.1007/s13225-020-00466-2)) |
| Otu46876 | 1 | k__Fungi(100);p__Ascomycota(100);c__Saccharomycetes(100);o__Saccharomycetales(100);f__Metschnikowiaceae(100);g__Metschnikowia(100);s__Metschnikowia_pulcherrima(100); | Metschnikowia | 13 | Saprotroph | \|Undefined Saprotroph\| | Yeast | NULL | Probable | As nectar-tap_saprotroph (Põlme et al. 2020); Decay sugar-rich substrates (Põlme et al. 2020) | Põlme S, et al. 2020. Fungal Diversity 105:1-16 ((https://doi.org/10.1007/s13225-020-00466-2)) |
| Otu46877 | 1 | k__Fungi(100);p__Ascomycota(100);c__Saccharomycetes(100);o__Saccharomycetales(100);f__Metschnikowiaceae(100);g__Metschnikowia(100);s__Metschnikowia_pulcherrima(100); | Metschnikowia | 13 | Saprotroph | \|Undefined Saprotroph\| | Yeast | NULL | Probable | As nectar-tap_saprotroph (Põlme et al. 2020); Decay sugar-rich substrates (Põlme et al. 2020) | Põlme S, et al. 2020. Fungal Diversity 105:1-16 ((https://doi.org/10.1007/s13225-020-00466-2)) |
| Otu46879 | 1 | k__Fungi(100);p__Ascomycota(100);c__Saccharomycetes(100);o__Saccharomycetales(100);f__Metschnikowiaceae(100);g__Metschnikowia(100);s__Metschnikowia_pulcherrima(100); | Metschnikowia | 13 | Saprotroph | \|Undefined Saprotroph\| | Yeast | NULL | Probable | As nectar-tap_saprotroph (Põlme et al. 2020); Decay sugar-rich substrates (Põlme et al. 2020) | Põlme S, et al. 2020. Fungal Diversity 105:1-16 ((https://doi.org/10.1007/s13225-020-00466-2)) |
| Otu46881 | 1 | k__Fungi(100);p__Ascomycota(100);c__Saccharomycetes(100);o__Saccharomycetales(100);f__Metschnikowiaceae(100);g__Metschnikowia(100);s__Metschnikowia_pulcherrima(100); | Metschnikowia | 13 | Saprotroph | \|Undefined Saprotroph\| | Yeast | NULL | Probable | As nectar-tap_saprotroph (Põlme et al. 2020); Decay sugar-rich substrates (Põlme et al. 2020) | Põlme S, et al. 2020. Fungal Diversity 105:1-16 ((https://doi.org/10.1007/s13225-020-00466-2)) |
| Otu46882 | 1 | k__Fungi(100);p__Ascomycota(100);c__Saccharomycetes(100);o__Saccharomycetales(100);f__Metschnikowiaceae(100);g__Metschnikowia(100);s__Metschnikowia_pulcherrima(100); | Metschnikowia | 13 | Saprotroph | \|Undefined Saprotroph\| | Yeast | NULL | Probable | As nectar-tap_saprotroph (Põlme et al. 2020); Decay sugar-rich substrates (Põlme et al. 2020) | Põlme S, et al. 2020. Fungal Diversity 105:1-16 ((https://doi.org/10.1007/s13225-020-00466-2)) |
| Otu46883 | 1 | k__Fungi(100);p__Ascomycota(100);c__Saccharomycetes(100);o__Saccharomycetales(100);f__Metschnikowiaceae(100);g__Metschnikowia(100);g__Metschnikowia_unclassified(100); | Metschnikowia | 13 | Saprotroph | \|Undefined Saprotroph\| | Yeast | NULL | Probable | As nectar-tap_saprotroph (Põlme et al. 2020); Decay sugar-rich substrates (Põlme et al. 2020) | Põlme S, et al. 2020. Fungal Diversity 105:1-16 ((https://doi.org/10.1007/s13225-020-00466-2)) |
| Otu46884 | 1 | k__Fungi(100);p__Ascomycota(100);c__Saccharomycetes(100);o__Saccharomycetales(100);f__Metschnikowiaceae(100);g__Metschnikowia(100);s__Metschnikowia_pulcherrima(100); | Metschnikowia | 13 | Saprotroph | \|Undefined Saprotroph\| | Yeast | NULL | Probable | As nectar-tap_saprotroph (Põlme et al. 2020); Decay sugar-rich substrates (Põlme et al. 2020) | Põlme S, et al. 2020. Fungal Diversity 105:1-16 ((https://doi.org/10.1007/s13225-020-00466-2)) |
| Otu46885 | 1 | k__Fungi(100);p__Ascomycota(100);c__Saccharomycetes(100);o__Saccharomycetales(100);f__Metschnikowiaceae(100);g__Metschnikowia(100);s__Metschnikowia_pulcherrima(100); | Metschnikowia | 13 | Saprotroph | \|Undefined Saprotroph\| | Yeast | NULL | Probable | As nectar-tap_saprotroph (Põlme et al. 2020); Decay sugar-rich substrates (Põlme et al. 2020) | Põlme S, et al. 2020. Fungal Diversity 105:1-16 ((https://doi.org/10.1007/s13225-020-00466-2)) |
| Otu46886 | 1 | k__Fungi(100);p__Ascomycota(100);c__Saccharomycetes(100);o__Saccharomycetales(100);f__Metschnikowiaceae(100);g__Metschnikowia(100);s__Metschnikowia_pulcherrima(100); | Metschnikowia | 13 | Saprotroph | \|Undefined Saprotroph\| | Yeast | NULL | Probable | As nectar-tap_saprotroph (Põlme et al. 2020); Decay sugar-rich substrates (Põlme et al. 2020) | Põlme S, et al. 2020. Fungal Diversity 105:1-16 ((https://doi.org/10.1007/s13225-020-00466-2)) |
| Otu46887 | 1 | k__Fungi(100);p__Ascomycota(100);c__Saccharomycetes(100);o__Saccharomycetales(100);f__Metschnikowiaceae(100);g__Metschnikowia(100);s__Metschnikowia_pulcherrima(100); | Metschnikowia | 13 | Saprotroph | \|Undefined Saprotroph\| | Yeast | NULL | Probable | As nectar-tap_saprotroph (Põlme et al. 2020); Decay sugar-rich substrates (Põlme et al. 2020) | Põlme S, et al. 2020. Fungal Diversity 105:1-16 ((https://doi.org/10.1007/s13225-020-00466-2)) |
| Otu46888 | 1 | k__Fungi(100);p__Ascomycota(100);c__Saccharomycetes(100);o__Saccharomycetales(100);f__Metschnikowiaceae(100);g__Metschnikowia(100);s__Metschnikowia_pulcherrima(100); | Metschnikowia | 13 | Saprotroph | \|Undefined Saprotroph\| | Yeast | NULL | Probable | As nectar-tap_saprotroph (Põlme et al. 2020); Decay sugar-rich substrates (Põlme et al. 2020) | Põlme S, et al. 2020. Fungal Diversity 105:1-16 ((https://doi.org/10.1007/s13225-020-00466-2)) |
| Otu46889 | 1 | k__Fungi(100);p__Ascomycota(100);c__Saccharomycetes(100);o__Saccharomycetales(100);f__Metschnikowiaceae(100);g__Metschnikowia(100);s__Metschnikowia_pulcherrima(100); | Metschnikowia | 13 | Saprotroph | \|Undefined Saprotroph\| | Yeast | NULL | Probable | As nectar-tap_saprotroph (Põlme et al. 2020); Decay sugar-rich substrates (Põlme et al. 2020) | Põlme S, et al. 2020. Fungal Diversity 105:1-16 ((https://doi.org/10.1007/s13225-020-00466-2)) |
| Otu46890 | 1 | k__Fungi(100);p__Ascomycota(100);c__Saccharomycetes(100);o__Saccharomycetales(100);f__Metschnikowiaceae(100);g__Metschnikowia(100);g__Metschnikowia_unclassified(100); | Metschnikowia | 13 | Saprotroph | \|Undefined Saprotroph\| | Yeast | NULL | Probable | As nectar-tap_saprotroph (Põlme et al. 2020); Decay sugar-rich substrates (Põlme et al. 2020) | Põlme S, et al. 2020. Fungal Diversity 105:1-16 ((https://doi.org/10.1007/s13225-020-00466-2)) |
| Otu46891 | 1 | k__Fungi(100);p__Ascomycota(100);c__Saccharomycetes(100);o__Saccharomycetales(100);f__Metschnikowiaceae(100);g__Metschnikowia(100);s__Metschnikowia_pulcherrima(100); | Metschnikowia | 13 | Saprotroph | \|Undefined Saprotroph\| | Yeast | NULL | Probable | As nectar-tap_saprotroph (Põlme et al. 2020); Decay sugar-rich substrates (Põlme et al. 2020) | Põlme S, et al. 2020. Fungal Diversity 105:1-16 ((https://doi.org/10.1007/s13225-020-00466-2)) |
| Otu46892 | 1 | k__Fungi(100);p__Ascomycota(100);c__Saccharomycetes(100);o__Saccharomycetales(100);f__Metschnikowiaceae(100);g__Metschnikowia(100);s__Metschnikowia_pulcherrima(100); | Metschnikowia | 13 | Saprotroph | \|Undefined Saprotroph\| | Yeast | NULL | Probable | As nectar-tap_saprotroph (Põlme et al. 2020); Decay sugar-rich substrates (Põlme et al. 2020) | Põlme S, et al. 2020. Fungal Diversity 105:1-16 ((https://doi.org/10.1007/s13225-020-00466-2)) |
| Otu46893 | 1 | k__Fungi(100);p__Ascomycota(100);c__Saccharomycetes(100);o__Saccharomycetales(100);f__Metschnikowiaceae(100);g__Metschnikowia(100);s__Metschnikowia_pulcherrima(100); | Metschnikowia | 13 | Saprotroph | \|Undefined Saprotroph\| | Yeast | NULL | Probable | As nectar-tap_saprotroph (Põlme et al. 2020); Decay sugar-rich substrates (Põlme et al. 2020) | Põlme S, et al. 2020. Fungal Diversity 105:1-16 ((https://doi.org/10.1007/s13225-020-00466-2)) |
| Otu46894 | 1 | k__Fungi(100);p__Ascomycota(100);c__Saccharomycetes(100);o__Saccharomycetales(100);f__Metschnikowiaceae(100);g__Metschnikowia(100);s__Metschnikowia_pulcherrima(100); | Metschnikowia | 13 | Saprotroph | \|Undefined Saprotroph\| | Yeast | NULL | Probable | As nectar-tap_saprotroph (Põlme et al. 2020); Decay sugar-rich substrates (Põlme et al. 2020) | Põlme S, et al. 2020. Fungal Diversity 105:1-16 ((https://doi.org/10.1007/s13225-020-00466-2)) |
| Otu46895 | 1 | k__Fungi(100);p__Ascomycota(100);c__Saccharomycetes(100);o__Saccharomycetales(100);f__Metschnikowiaceae(100);g__Metschnikowia(100);s__Metschnikowia_pulcherrima(100); | Metschnikowia | 13 | Saprotroph | \|Undefined Saprotroph\| | Yeast | NULL | Probable | As nectar-tap_saprotroph (Põlme et al. 2020); Decay sugar-rich substrates (Põlme et al. 2020) | Põlme S, et al. 2020. Fungal Diversity 105:1-16 ((https://doi.org/10.1007/s13225-020-00466-2)) |
| Otu46896 | 1 | k__Fungi(100);p__Ascomycota(100);c__Saccharomycetes(100);o__Saccharomycetales(100);f__Metschnikowiaceae(100);g__Metschnikowia(100);s__Metschnikowia_pulcherrima(100); | Metschnikowia | 13 | Saprotroph | \|Undefined Saprotroph\| | Yeast | NULL | Probable | As nectar-tap_saprotroph (Põlme et al. 2020); Decay sugar-rich substrates (Põlme et al. 2020) | Põlme S, et al. 2020. Fungal Diversity 105:1-16 ((https://doi.org/10.1007/s13225-020-00466-2)) |
| Otu46897 | 1 | k__Fungi(100);p__Ascomycota(100);c__Saccharomycetes(100);o__Saccharomycetales(100);f__Metschnikowiaceae(100);g__Metschnikowia(100);s__Metschnikowia_pulcherrima(100); | Metschnikowia | 13 | Saprotroph | \|Undefined Saprotroph\| | Yeast | NULL | Probable | As nectar-tap_saprotroph (Põlme et al. 2020); Decay sugar-rich substrates (Põlme et al. 2020) | Põlme S, et al. 2020. Fungal Diversity 105:1-16 ((https://doi.org/10.1007/s13225-020-00466-2)) |
| Otu46898 | 1 | k__Fungi(100);p__Ascomycota(100);c__Saccharomycetes(100);o__Saccharomycetales(100);f__Metschnikowiaceae(100);g__Metschnikowia(100);s__Metschnikowia_pulcherrima(100); | Metschnikowia | 13 | Saprotroph | \|Undefined Saprotroph\| | Yeast | NULL | Probable | As nectar-tap_saprotroph (Põlme et al. 2020); Decay sugar-rich substrates (Põlme et al. 2020) | Põlme S, et al. 2020. Fungal Diversity 105:1-16 ((https://doi.org/10.1007/s13225-020-00466-2)) |
| Otu46899 | 1 | k__Fungi(100);p__Ascomycota(100);c__Saccharomycetes(100);o__Saccharomycetales(100);f__Metschnikowiaceae(100);g__Metschnikowia(100);s__Metschnikowia_pulcherrima(100); | Metschnikowia | 13 | Saprotroph | \|Undefined Saprotroph\| | Yeast | NULL | Probable | As nectar-tap_saprotroph (Põlme et al. 2020); Decay sugar-rich substrates (Põlme et al. 2020) | Põlme S, et al. 2020. Fungal Diversity 105:1-16 ((https://doi.org/10.1007/s13225-020-00466-2)) |
| Otu46900 | 1 | k__Fungi(100);p__Ascomycota(100);c__Saccharomycetes(100);o__Saccharomycetales(100);f__Metschnikowiaceae(100);g__Metschnikowia(100);s__Metschnikowia_pulcherrima(100); | Metschnikowia | 13 | Saprotroph | \|Undefined Saprotroph\| | Yeast | NULL | Probable | As nectar-tap_saprotroph (Põlme et al. 2020); Decay sugar-rich substrates (Põlme et al. 2020) | Põlme S, et al. 2020. Fungal Diversity 105:1-16 ((https://doi.org/10.1007/s13225-020-00466-2)) |
| Otu46901 | 1 | k__Fungi(100);p__Ascomycota(100);c__Saccharomycetes(100);o__Saccharomycetales(100);f__Metschnikowiaceae(100);g__Metschnikowia(100);s__Metschnikowia_pulcherrima(100); | Metschnikowia | 13 | Saprotroph | \|Undefined Saprotroph\| | Yeast | NULL | Probable | As nectar-tap_saprotroph (Põlme et al. 2020); Decay sugar-rich substrates (Põlme et al. 2020) | Põlme S, et al. 2020. Fungal Diversity 105:1-16 ((https://doi.org/10.1007/s13225-020-00466-2)) |
| Otu46902 | 1 | k__Fungi(100);p__Ascomycota(100);c__Saccharomycetes(100);o__Saccharomycetales(100);f__Metschnikowiaceae(100);g__Metschnikowia(100);g__Metschnikowia_unclassified(100); | Metschnikowia | 13 | Saprotroph | \|Undefined Saprotroph\| | Yeast | NULL | Probable | As nectar-tap_saprotroph (Põlme et al. 2020); Decay sugar-rich substrates (Põlme et al. 2020) | Põlme S, et al. 2020. Fungal Diversity 105:1-16 ((https://doi.org/10.1007/s13225-020-00466-2)) |
| Otu46903 | 1 | k__Fungi(100);p__Ascomycota(100);c__Saccharomycetes(100);o__Saccharomycetales(100);f__Metschnikowiaceae(100);g__Metschnikowia(100);s__Metschnikowia_pulcherrima(100); | Metschnikowia | 13 | Saprotroph | \|Undefined Saprotroph\| | Yeast | NULL | Probable | As nectar-tap_saprotroph (Põlme et al. 2020); Decay sugar-rich substrates (Põlme et al. 2020) | Põlme S, et al. 2020. Fungal Diversity 105:1-16 ((https://doi.org/10.1007/s13225-020-00466-2)) |
| Otu46904 | 1 | k__Fungi(100);p__Ascomycota(100);c__Saccharomycetes(100);o__Saccharomycetales(100);f__Metschnikowiaceae(100);g__Metschnikowia(100);s__Metschnikowia_pulcherrima(100); | Metschnikowia | 13 | Saprotroph | \|Undefined Saprotroph\| | Yeast | NULL | Probable | As nectar-tap_saprotroph (Põlme et al. 2020); Decay sugar-rich substrates (Põlme et al. 2020) | Põlme S, et al. 2020. Fungal Diversity 105:1-16 ((https://doi.org/10.1007/s13225-020-00466-2)) |
| Otu46905 | 1 | k__Fungi(100);p__Ascomycota(100);c__Saccharomycetes(100);o__Saccharomycetales(100);f__Metschnikowiaceae(100);g__Metschnikowia(100);s__Metschnikowia_pulcherrima(100); | Metschnikowia | 13 | Saprotroph | \|Undefined Saprotroph\| | Yeast | NULL | Probable | As nectar-tap_saprotroph (Põlme et al. 2020); Decay sugar-rich substrates (Põlme et al. 2020) | Põlme S, et al. 2020. Fungal Diversity 105:1-16 ((https://doi.org/10.1007/s13225-020-00466-2)) |
| Otu46906 | 1 | k__Fungi(100);p__Ascomycota(100);c__Saccharomycetes(100);o__Saccharomycetales(100);f__Metschnikowiaceae(100);g__Metschnikowia(100);s__Metschnikowia_pulcherrima(100); | Metschnikowia | 13 | Saprotroph | \|Undefined Saprotroph\| | Yeast | NULL | Probable | As nectar-tap_saprotroph (Põlme et al. 2020); Decay sugar-rich substrates (Põlme et al. 2020) | Põlme S, et al. 2020. Fungal Diversity 105:1-16 ((https://doi.org/10.1007/s13225-020-00466-2)) |
| Otu46907 | 1 | k__Fungi(100);p__Ascomycota(100);c__Saccharomycetes(100);o__Saccharomycetales(100);f__Metschnikowiaceae(100);g__Metschnikowia(100);g__Metschnikowia_unclassified(100); | Metschnikowia | 13 | Saprotroph | \|Undefined Saprotroph\| | Yeast | NULL | Probable | As nectar-tap_saprotroph (Põlme et al. 2020); Decay sugar-rich substrates (Põlme et al. 2020) | Põlme S, et al. 2020. Fungal Diversity 105:1-16 ((https://doi.org/10.1007/s13225-020-00466-2)) |
| Otu46908 | 1 | k__Fungi(100);p__Ascomycota(100);c__Saccharomycetes(100);o__Saccharomycetales(100);f__Metschnikowiaceae(100);g__Metschnikowia(100);s__Metschnikowia_pulcherrima(100); | Metschnikowia | 13 | Saprotroph | \|Undefined Saprotroph\| | Yeast | NULL | Probable | As nectar-tap_saprotroph (Põlme et al. 2020); Decay sugar-rich substrates (Põlme et al. 2020) | Põlme S, et al. 2020. Fungal Diversity 105:1-16 ((https://doi.org/10.1007/s13225-020-00466-2)) |
| Otu46909 | 1 | k__Fungi(100);p__Ascomycota(100);c__Saccharomycetes(100);o__Saccharomycetales(100);f__Metschnikowiaceae(100);g__Metschnikowia(100);s__Metschnikowia_pulcherrima(100); | Metschnikowia | 13 | Saprotroph | \|Undefined Saprotroph\| | Yeast | NULL | Probable | As nectar-tap_saprotroph (Põlme et al. 2020); Decay sugar-rich substrates (Põlme et al. 2020) | Põlme S, et al. 2020. Fungal Diversity 105:1-16 ((https://doi.org/10.1007/s13225-020-00466-2)) |
| Otu46910 | 1 | k__Fungi(100);p__Ascomycota(100);c__Saccharomycetes(100);o__Saccharomycetales(100);f__Metschnikowiaceae(100);g__Metschnikowia(100);s__Metschnikowia_pulcherrima(100); | Metschnikowia | 13 | Saprotroph | \|Undefined Saprotroph\| | Yeast | NULL | Probable | As nectar-tap_saprotroph (Põlme et al. 2020); Decay sugar-rich substrates (Põlme et al. 2020) | Põlme S, et al. 2020. Fungal Diversity 105:1-16 ((https://doi.org/10.1007/s13225-020-00466-2)) |
| Otu46911 | 1 | k__Fungi(100);p__Ascomycota(100);c__Saccharomycetes(100);o__Saccharomycetales(100);f__Metschnikowiaceae(100);g__Metschnikowia(100);s__Metschnikowia_pulcherrima(100); | Metschnikowia | 13 | Saprotroph | \|Undefined Saprotroph\| | Yeast | NULL | Probable | As nectar-tap_saprotroph (Põlme et al. 2020); Decay sugar-rich substrates (Põlme et al. 2020) | Põlme S, et al. 2020. Fungal Diversity 105:1-16 ((https://doi.org/10.1007/s13225-020-00466-2)) |
| Otu46912 | 1 | k__Fungi(100);p__Ascomycota(100);c__Saccharomycetes(100);o__Saccharomycetales(100);f__Metschnikowiaceae(100);g__Metschnikowia(100);s__Metschnikowia_pulcherrima(100); | Metschnikowia | 13 | Saprotroph | \|Undefined Saprotroph\| | Yeast | NULL | Probable | As nectar-tap_saprotroph (Põlme et al. 2020); Decay sugar-rich substrates (Põlme et al. 2020) | Põlme S, et al. 2020. Fungal Diversity 105:1-16 ((https://doi.org/10.1007/s13225-020-00466-2)) |
| Otu46913 | 1 | k__Fungi(100);p__Ascomycota(100);c__Saccharomycetes(100);o__Saccharomycetales(100);f__Metschnikowiaceae(100);g__Metschnikowia(100);s__Metschnikowia_pulcherrima(100); | Metschnikowia | 13 | Saprotroph | \|Undefined Saprotroph\| | Yeast | NULL | Probable | As nectar-tap_saprotroph (Põlme et al. 2020); Decay sugar-rich substrates (Põlme et al. 2020) | Põlme S, et al. 2020. Fungal Diversity 105:1-16 ((https://doi.org/10.1007/s13225-020-00466-2)) |
| Otu46914 | 1 | k__Fungi(100);p__Ascomycota(100);c__Saccharomycetes(100);o__Saccharomycetales(100);f__Metschnikowiaceae(100);g__Metschnikowia(100);s__Metschnikowia_pulcherrima(100); | Metschnikowia | 13 | Saprotroph | \|Undefined Saprotroph\| | Yeast | NULL | Probable | As nectar-tap_saprotroph (Põlme et al. 2020); Decay sugar-rich substrates (Põlme et al. 2020) | Põlme S, et al. 2020. Fungal Diversity 105:1-16 ((https://doi.org/10.1007/s13225-020-00466-2)) |
| Otu46915 | 1 | k__Fungi(100);p__Ascomycota(100);c__Saccharomycetes(100);o__Saccharomycetales(100);f__Metschnikowiaceae(100);g__Metschnikowia(100);g__Metschnikowia_unclassified(100); | Metschnikowia | 13 | Saprotroph | \|Undefined Saprotroph\| | Yeast | NULL | Probable | As nectar-tap_saprotroph (Põlme et al. 2020); Decay sugar-rich substrates (Põlme et al. 2020) | Põlme S, et al. 2020. Fungal Diversity 105:1-16 ((https://doi.org/10.1007/s13225-020-00466-2)) |
| Otu46916 | 1 | k__Fungi(100);p__Ascomycota(100);c__Saccharomycetes(100);o__Saccharomycetales(100);f__Metschnikowiaceae(100);g__Metschnikowia(100);g__Metschnikowia_unclassified(100); | Metschnikowia | 13 | Saprotroph | \|Undefined Saprotroph\| | Yeast | NULL | Probable | As nectar-tap_saprotroph (Põlme et al. 2020); Decay sugar-rich substrates (Põlme et al. 2020) | Põlme S, et al. 2020. Fungal Diversity 105:1-16 ((https://doi.org/10.1007/s13225-020-00466-2)) |
| Otu46917 | 1 | k__Fungi(100);p__Ascomycota(100);c__Saccharomycetes(100);o__Saccharomycetales(100);f__Metschnikowiaceae(100);g__Metschnikowia(100);g__Metschnikowia_unclassified(100); | Metschnikowia | 13 | Saprotroph | \|Undefined Saprotroph\| | Yeast | NULL | Probable | As nectar-tap_saprotroph (Põlme et al. 2020); Decay sugar-rich substrates (Põlme et al. 2020) | Põlme S, et al. 2020. Fungal Diversity 105:1-16 ((https://doi.org/10.1007/s13225-020-00466-2)) |
| Otu46918 | 1 | k__Fungi(100);p__Ascomycota(100);c__Saccharomycetes(100);o__Saccharomycetales(100);f__Metschnikowiaceae(100);g__Metschnikowia(100);g__Metschnikowia_unclassified(100); | Metschnikowia | 13 | Saprotroph | \|Undefined Saprotroph\| | Yeast | NULL | Probable | As nectar-tap_saprotroph (Põlme et al. 2020); Decay sugar-rich substrates (Põlme et al. 2020) | Põlme S, et al. 2020. Fungal Diversity 105:1-16 ((https://doi.org/10.1007/s13225-020-00466-2)) |
| Otu46919 | 1 | k__Fungi(100);p__Ascomycota(100);c__Saccharomycetes(100);o__Saccharomycetales(100);f__Metschnikowiaceae(100);g__Metschnikowia(100);g__Metschnikowia_unclassified(100); | Metschnikowia | 13 | Saprotroph | \|Undefined Saprotroph\| | Yeast | NULL | Probable | As nectar-tap_saprotroph (Põlme et al. 2020); Decay sugar-rich substrates (Põlme et al. 2020) | Põlme S, et al. 2020. Fungal Diversity 105:1-16 ((https://doi.org/10.1007/s13225-020-00466-2)) |
| Otu46920 | 1 | k__Fungi(100);p__Ascomycota(100);c__Saccharomycetes(100);o__Saccharomycetales(100);f__Metschnikowiaceae(100);g__Metschnikowia(100);s__Metschnikowia_pulcherrima(100); | Metschnikowia | 13 | Saprotroph | \|Undefined Saprotroph\| | Yeast | NULL | Probable | As nectar-tap_saprotroph (Põlme et al. 2020); Decay sugar-rich substrates (Põlme et al. 2020) | Põlme S, et al. 2020. Fungal Diversity 105:1-16 ((https://doi.org/10.1007/s13225-020-00466-2)) |
| Otu46921 | 1 | k__Fungi(100);p__Ascomycota(100);c__Saccharomycetes(100);o__Saccharomycetales(100);f__Metschnikowiaceae(100);g__Metschnikowia(100);s__Metschnikowia_pulcherrima(100); | Metschnikowia | 13 | Saprotroph | \|Undefined Saprotroph\| | Yeast | NULL | Probable | As nectar-tap_saprotroph (Põlme et al. 2020); Decay sugar-rich substrates (Põlme et al. 2020) | Põlme S, et al. 2020. Fungal Diversity 105:1-16 ((https://doi.org/10.1007/s13225-020-00466-2)) |
| Otu46922 | 1 | k__Fungi(100);p__Ascomycota(100);c__Saccharomycetes(100);o__Saccharomycetales(100);f__Metschnikowiaceae(100);g__Metschnikowia(100);s__Metschnikowia_pulcherrima(100); | Metschnikowia | 13 | Saprotroph | \|Undefined Saprotroph\| | Yeast | NULL | Probable | As nectar-tap_saprotroph (Põlme et al. 2020); Decay sugar-rich substrates (Põlme et al. 2020) | Põlme S, et al. 2020. Fungal Diversity 105:1-16 ((https://doi.org/10.1007/s13225-020-00466-2)) |
| Otu46923 | 1 | k__Fungi(100);p__Ascomycota(100);c__Saccharomycetes(100);o__Saccharomycetales(100);f__Metschnikowiaceae(100);g__Metschnikowia(100);s__Metschnikowia_pulcherrima(100); | Metschnikowia | 13 | Saprotroph | \|Undefined Saprotroph\| | Yeast | NULL | Probable | As nectar-tap_saprotroph (Põlme et al. 2020); Decay sugar-rich substrates (Põlme et al. 2020) | Põlme S, et al. 2020. Fungal Diversity 105:1-16 ((https://doi.org/10.1007/s13225-020-00466-2)) |
| Otu46924 | 1 | k__Fungi(100);p__Ascomycota(100);c__Saccharomycetes(100);o__Saccharomycetales(100);f__Metschnikowiaceae(100);g__Metschnikowia(100);s__Metschnikowia_pulcherrima(100); | Metschnikowia | 13 | Saprotroph | \|Undefined Saprotroph\| | Yeast | NULL | Probable | As nectar-tap_saprotroph (Põlme et al. 2020); Decay sugar-rich substrates (Põlme et al. 2020) | Põlme S, et al. 2020. Fungal Diversity 105:1-16 ((https://doi.org/10.1007/s13225-020-00466-2)) |
| Otu46925 | 1 | k__Fungi(100);p__Ascomycota(100);c__Saccharomycetes(100);o__Saccharomycetales(100);f__Metschnikowiaceae(100);g__Metschnikowia(100);s__Metschnikowia_pulcherrima(100); | Metschnikowia | 13 | Saprotroph | \|Undefined Saprotroph\| | Yeast | NULL | Probable | As nectar-tap_saprotroph (Põlme et al. 2020); Decay sugar-rich substrates (Põlme et al. 2020) | Põlme S, et al. 2020. Fungal Diversity 105:1-16 ((https://doi.org/10.1007/s13225-020-00466-2)) |
| Otu46926 | 1 | k__Fungi(100);p__Ascomycota(100);c__Saccharomycetes(100);o__Saccharomycetales(100);f__Metschnikowiaceae(100);g__Metschnikowia(100);g__Metschnikowia_unclassified(100); | Metschnikowia | 13 | Saprotroph | \|Undefined Saprotroph\| | Yeast | NULL | Probable | As nectar-tap_saprotroph (Põlme et al. 2020); Decay sugar-rich substrates (Põlme et al. 2020) | Põlme S, et al. 2020. Fungal Diversity 105:1-16 ((https://doi.org/10.1007/s13225-020-00466-2)) |
| Otu46927 | 1 | k__Fungi(100);p__Ascomycota(100);c__Saccharomycetes(100);o__Saccharomycetales(100);f__Metschnikowiaceae(100);g__Metschnikowia(100);s__Metschnikowia_pulcherrima(100); | Metschnikowia | 13 | Saprotroph | \|Undefined Saprotroph\| | Yeast | NULL | Probable | As nectar-tap_saprotroph (Põlme et al. 2020); Decay sugar-rich substrates (Põlme et al. 2020) | Põlme S, et al. 2020. Fungal Diversity 105:1-16 ((https://doi.org/10.1007/s13225-020-00466-2)) |
| Otu46928 | 1 | k__Fungi(100);p__Ascomycota(100);c__Saccharomycetes(100);o__Saccharomycetales(100);f__Metschnikowiaceae(100);g__Metschnikowia(100);g__Metschnikowia_unclassified(100); | Metschnikowia | 13 | Saprotroph | \|Undefined Saprotroph\| | Yeast | NULL | Probable | As nectar-tap_saprotroph (Põlme et al. 2020); Decay sugar-rich substrates (Põlme et al. 2020) | Põlme S, et al. 2020. Fungal Diversity 105:1-16 ((https://doi.org/10.1007/s13225-020-00466-2)) |
| Otu46929 | 1 | k__Fungi(100);p__Ascomycota(100);c__Saccharomycetes(100);o__Saccharomycetales(100);f__Metschnikowiaceae(100);g__Metschnikowia(100);s__Metschnikowia_pulcherrima(100); | Metschnikowia | 13 | Saprotroph | \|Undefined Saprotroph\| | Yeast | NULL | Probable | As nectar-tap_saprotroph (Põlme et al. 2020); Decay sugar-rich substrates (Põlme et al. 2020) | Põlme S, et al. 2020. Fungal Diversity 105:1-16 ((https://doi.org/10.1007/s13225-020-00466-2)) |
| Otu46930 | 1 | k__Fungi(100);p__Ascomycota(100);c__Saccharomycetes(100);o__Saccharomycetales(100);f__Metschnikowiaceae(100);g__Metschnikowia(100);s__Metschnikowia_pulcherrima(100); | Metschnikowia | 13 | Saprotroph | \|Undefined Saprotroph\| | Yeast | NULL | Probable | As nectar-tap_saprotroph (Põlme et al. 2020); Decay sugar-rich substrates (Põlme et al. 2020) | Põlme S, et al. 2020. Fungal Diversity 105:1-16 ((https://doi.org/10.1007/s13225-020-00466-2)) |
| Otu46931 | 1 | k__Fungi(100);p__Ascomycota(100);c__Saccharomycetes(100);o__Saccharomycetales(100);f__Metschnikowiaceae(100);g__Metschnikowia(100);s__Metschnikowia_pulcherrima(100); | Metschnikowia | 13 | Saprotroph | \|Undefined Saprotroph\| | Yeast | NULL | Probable | As nectar-tap_saprotroph (Põlme et al. 2020); Decay sugar-rich substrates (Põlme et al. 2020) | Põlme S, et al. 2020. Fungal Diversity 105:1-16 ((https://doi.org/10.1007/s13225-020-00466-2)) |
| Otu46932 | 1 | k__Fungi(100);p__Ascomycota(100);c__Saccharomycetes(100);o__Saccharomycetales(100);f__Metschnikowiaceae(100);g__Metschnikowia(100);s__Metschnikowia_pulcherrima(100); | Metschnikowia | 13 | Saprotroph | \|Undefined Saprotroph\| | Yeast | NULL | Probable | As nectar-tap_saprotroph (Põlme et al. 2020); Decay sugar-rich substrates (Põlme et al. 2020) | Põlme S, et al. 2020. Fungal Diversity 105:1-16 ((https://doi.org/10.1007/s13225-020-00466-2)) |
| Otu46933 | 1 | k__Fungi(100);p__Ascomycota(100);c__Saccharomycetes(100);o__Saccharomycetales(100);f__Metschnikowiaceae(100);g__Metschnikowia(100);s__Metschnikowia_pulcherrima(100); | Metschnikowia | 13 | Saprotroph | \|Undefined Saprotroph\| | Yeast | NULL | Probable | As nectar-tap_saprotroph (Põlme et al. 2020); Decay sugar-rich substrates (Põlme et al. 2020) | Põlme S, et al. 2020. Fungal Diversity 105:1-16 ((https://doi.org/10.1007/s13225-020-00466-2)) |
| Otu46934 | 1 | k__Fungi(100);p__Ascomycota(100);c__Saccharomycetes(100);o__Saccharomycetales(100);f__Metschnikowiaceae(100);g__Metschnikowia(100);s__Metschnikowia_pulcherrima(100); | Metschnikowia | 13 | Saprotroph | \|Undefined Saprotroph\| | Yeast | NULL | Probable | As nectar-tap_saprotroph (Põlme et al. 2020); Decay sugar-rich substrates (Põlme et al. 2020) | Põlme S, et al. 2020. Fungal Diversity 105:1-16 ((https://doi.org/10.1007/s13225-020-00466-2)) |
| Otu46935 | 1 | k__Fungi(100);p__Ascomycota(100);c__Saccharomycetes(100);o__Saccharomycetales(100);f__Metschnikowiaceae(100);g__Metschnikowia(100);s__Metschnikowia_pulcherrima(100); | Metschnikowia | 13 | Saprotroph | \|Undefined Saprotroph\| | Yeast | NULL | Probable | As nectar-tap_saprotroph (Põlme et al. 2020); Decay sugar-rich substrates (Põlme et al. 2020) | Põlme S, et al. 2020. Fungal Diversity 105:1-16 ((https://doi.org/10.1007/s13225-020-00466-2)) |
| Otu46936 | 1 | k__Fungi(100);p__Ascomycota(100);c__Saccharomycetes(100);o__Saccharomycetales(100);f__Metschnikowiaceae(100);g__Metschnikowia(100);g__Metschnikowia_unclassified(100); | Metschnikowia | 13 | Saprotroph | \|Undefined Saprotroph\| | Yeast | NULL | Probable | As nectar-tap_saprotroph (Põlme et al. 2020); Decay sugar-rich substrates (Põlme et al. 2020) | Põlme S, et al. 2020. Fungal Diversity 105:1-16 ((https://doi.org/10.1007/s13225-020-00466-2)) |
| Otu46937 | 1 | k__Fungi(100);p__Ascomycota(100);c__Saccharomycetes(100);o__Saccharomycetales(100);f__Metschnikowiaceae(100);g__Metschnikowia(100);s__Metschnikowia_pulcherrima(100); | Metschnikowia | 13 | Saprotroph | \|Undefined Saprotroph\| | Yeast | NULL | Probable | As nectar-tap_saprotroph (Põlme et al. 2020); Decay sugar-rich substrates (Põlme et al. 2020) | Põlme S, et al. 2020. Fungal Diversity 105:1-16 ((https://doi.org/10.1007/s13225-020-00466-2)) |
| Otu46938 | 1 | k__Fungi(100);p__Ascomycota(100);c__Saccharomycetes(100);o__Saccharomycetales(100);f__Metschnikowiaceae(100);g__Metschnikowia(100);g__Metschnikowia_unclassified(100); | Metschnikowia | 13 | Saprotroph | \|Undefined Saprotroph\| | Yeast | NULL | Probable | As nectar-tap_saprotroph (Põlme et al. 2020); Decay sugar-rich substrates (Põlme et al. 2020) | Põlme S, et al. 2020. Fungal Diversity 105:1-16 ((https://doi.org/10.1007/s13225-020-00466-2)) |
| Otu46939 | 1 | k__Fungi(100);p__Ascomycota(100);c__Saccharomycetes(100);o__Saccharomycetales(100);f__Metschnikowiaceae(100);g__Metschnikowia(100);s__Metschnikowia_pulcherrima(100); | Metschnikowia | 13 | Saprotroph | \|Undefined Saprotroph\| | Yeast | NULL | Probable | As nectar-tap_saprotroph (Põlme et al. 2020); Decay sugar-rich substrates (Põlme et al. 2020) | Põlme S, et al. 2020. Fungal Diversity 105:1-16 ((https://doi.org/10.1007/s13225-020-00466-2)) |
| Otu46941 | 1 | k__Fungi(100);p__Ascomycota(100);c__Saccharomycetes(100);o__Saccharomycetales(100);f__Metschnikowiaceae(100);g__Metschnikowia(100);s__Metschnikowia_pulcherrima(100); | Metschnikowia | 13 | Saprotroph | \|Undefined Saprotroph\| | Yeast | NULL | Probable | As nectar-tap_saprotroph (Põlme et al. 2020); Decay sugar-rich substrates (Põlme et al. 2020) | Põlme S, et al. 2020. Fungal Diversity 105:1-16 ((https://doi.org/10.1007/s13225-020-00466-2)) |
| Otu46942 | 1 | k__Fungi(100);p__Ascomycota(100);c__Saccharomycetes(100);o__Saccharomycetales(100);f__Metschnikowiaceae(100);g__Metschnikowia(100);g__Metschnikowia_unclassified(100); | Metschnikowia | 13 | Saprotroph | \|Undefined Saprotroph\| | Yeast | NULL | Probable | As nectar-tap_saprotroph (Põlme et al. 2020); Decay sugar-rich substrates (Põlme et al. 2020) | Põlme S, et al. 2020. Fungal Diversity 105:1-16 ((https://doi.org/10.1007/s13225-020-00466-2)) |
| Otu46943 | 1 | k__Fungi(100);p__Ascomycota(100);c__Saccharomycetes(100);o__Saccharomycetales(100);f__Metschnikowiaceae(100);g__Metschnikowia(100);g__Metschnikowia_unclassified(100); | Metschnikowia | 13 | Saprotroph | \|Undefined Saprotroph\| | Yeast | NULL | Probable | As nectar-tap_saprotroph (Põlme et al. 2020); Decay sugar-rich substrates (Põlme et al. 2020) | Põlme S, et al. 2020. Fungal Diversity 105:1-16 ((https://doi.org/10.1007/s13225-020-00466-2)) |
| Otu46944 | 1 | k__Fungi(100);p__Ascomycota(100);c__Saccharomycetes(100);o__Saccharomycetales(100);f__Metschnikowiaceae(100);g__Metschnikowia(100);s__Metschnikowia_pulcherrima(100); | Metschnikowia | 13 | Saprotroph | \|Undefined Saprotroph\| | Yeast | NULL | Probable | As nectar-tap_saprotroph (Põlme et al. 2020); Decay sugar-rich substrates (Põlme et al. 2020) | Põlme S, et al. 2020. Fungal Diversity 105:1-16 ((https://doi.org/10.1007/s13225-020-00466-2)) |
| Otu46945 | 1 | k__Fungi(100);p__Ascomycota(100);c__Saccharomycetes(100);o__Saccharomycetales(100);f__Metschnikowiaceae(100);g__Metschnikowia(100);g__Metschnikowia_unclassified(100); | Metschnikowia | 13 | Saprotroph | \|Undefined Saprotroph\| | Yeast | NULL | Probable | As nectar-tap_saprotroph (Põlme et al. 2020); Decay sugar-rich substrates (Põlme et al. 2020) | Põlme S, et al. 2020. Fungal Diversity 105:1-16 ((https://doi.org/10.1007/s13225-020-00466-2)) |
| Otu46946 | 1 | k__Fungi(100);p__Ascomycota(100);c__Saccharomycetes(100);o__Saccharomycetales(100);f__Metschnikowiaceae(100);g__Metschnikowia(100);g__Metschnikowia_unclassified(100); | Metschnikowia | 13 | Saprotroph | \|Undefined Saprotroph\| | Yeast | NULL | Probable | As nectar-tap_saprotroph (Põlme et al. 2020); Decay sugar-rich substrates (Põlme et al. 2020) | Põlme S, et al. 2020. Fungal Diversity 105:1-16 ((https://doi.org/10.1007/s13225-020-00466-2)) |
| Otu46947 | 1 | k__Fungi(100);p__Ascomycota(100);c__Saccharomycetes(100);o__Saccharomycetales(100);f__Metschnikowiaceae(100);g__Metschnikowia(100);s__Metschnikowia_pulcherrima(100); | Metschnikowia | 13 | Saprotroph | \|Undefined Saprotroph\| | Yeast | NULL | Probable | As nectar-tap_saprotroph (Põlme et al. 2020); Decay sugar-rich substrates (Põlme et al. 2020) | Põlme S, et al. 2020. Fungal Diversity 105:1-16 ((https://doi.org/10.1007/s13225-020-00466-2)) |
| Otu46948 | 1 | k__Fungi(100);p__Ascomycota(100);c__Saccharomycetes(100);o__Saccharomycetales(100);f__Metschnikowiaceae(100);g__Metschnikowia(100);g__Metschnikowia_unclassified(100); | Metschnikowia | 13 | Saprotroph | \|Undefined Saprotroph\| | Yeast | NULL | Probable | As nectar-tap_saprotroph (Põlme et al. 2020); Decay sugar-rich substrates (Põlme et al. 2020) | Põlme S, et al. 2020. Fungal Diversity 105:1-16 ((https://doi.org/10.1007/s13225-020-00466-2)) |
| Otu46949 | 1 | k__Fungi(100);p__Ascomycota(100);c__Saccharomycetes(100);o__Saccharomycetales(100);f__Metschnikowiaceae(100);g__Metschnikowia(100);g__Metschnikowia_unclassified(100); | Metschnikowia | 13 | Saprotroph | \|Undefined Saprotroph\| | Yeast | NULL | Probable | As nectar-tap_saprotroph (Põlme et al. 2020); Decay sugar-rich substrates (Põlme et al. 2020) | Põlme S, et al. 2020. Fungal Diversity 105:1-16 ((https://doi.org/10.1007/s13225-020-00466-2)) |
| Otu46950 | 1 | k__Fungi(100);p__Ascomycota(100);c__Saccharomycetes(100);o__Saccharomycetales(100);f__Metschnikowiaceae(100);g__Metschnikowia(100);s__Metschnikowia_pulcherrima(100); | Metschnikowia | 13 | Saprotroph | \|Undefined Saprotroph\| | Yeast | NULL | Probable | As nectar-tap_saprotroph (Põlme et al. 2020); Decay sugar-rich substrates (Põlme et al. 2020) | Põlme S, et al. 2020. Fungal Diversity 105:1-16 ((https://doi.org/10.1007/s13225-020-00466-2)) |
| Otu46951 | 1 | k__Fungi(100);p__Ascomycota(100);c__Saccharomycetes(100);o__Saccharomycetales(100);f__Metschnikowiaceae(100);g__Metschnikowia(100);s__Metschnikowia_pulcherrima(100); | Metschnikowia | 13 | Saprotroph | \|Undefined Saprotroph\| | Yeast | NULL | Probable | As nectar-tap_saprotroph (Põlme et al. 2020); Decay sugar-rich substrates (Põlme et al. 2020) | Põlme S, et al. 2020. Fungal Diversity 105:1-16 ((https://doi.org/10.1007/s13225-020-00466-2)) |
| Otu46952 | 1 | k__Fungi(100);p__Ascomycota(100);c__Saccharomycetes(100);o__Saccharomycetales(100);f__Metschnikowiaceae(100);g__Metschnikowia(100);g__Metschnikowia_unclassified(100); | Metschnikowia | 13 | Saprotroph | \|Undefined Saprotroph\| | Yeast | NULL | Probable | As nectar-tap_saprotroph (Põlme et al. 2020); Decay sugar-rich substrates (Põlme et al. 2020) | Põlme S, et al. 2020. Fungal Diversity 105:1-16 ((https://doi.org/10.1007/s13225-020-00466-2)) |
| Otu46953 | 1 | k__Fungi(100);p__Ascomycota(100);c__Saccharomycetes(100);o__Saccharomycetales(100);f__Metschnikowiaceae(100);g__Metschnikowia(100);s__Metschnikowia_pulcherrima(100); | Metschnikowia | 13 | Saprotroph | \|Undefined Saprotroph\| | Yeast | NULL | Probable | As nectar-tap_saprotroph (Põlme et al. 2020); Decay sugar-rich substrates (Põlme et al. 2020) | Põlme S, et al. 2020. Fungal Diversity 105:1-16 ((https://doi.org/10.1007/s13225-020-00466-2)) |
| Otu46954 | 1 | k__Fungi(100);p__Ascomycota(100);c__Saccharomycetes(100);o__Saccharomycetales(100);f__Metschnikowiaceae(100);g__Metschnikowia(100);s__Metschnikowia_pulcherrima(100); | Metschnikowia | 13 | Saprotroph | \|Undefined Saprotroph\| | Yeast | NULL | Probable | As nectar-tap_saprotroph (Põlme et al. 2020); Decay sugar-rich substrates (Põlme et al. 2020) | Põlme S, et al. 2020. Fungal Diversity 105:1-16 ((https://doi.org/10.1007/s13225-020-00466-2)) |
| Otu46955 | 1 | k__Fungi(100);p__Ascomycota(100);c__Saccharomycetes(100);o__Saccharomycetales(100);f__Metschnikowiaceae(100);g__Metschnikowia(100);s__Metschnikowia_pulcherrima(100); | Metschnikowia | 13 | Saprotroph | \|Undefined Saprotroph\| | Yeast | NULL | Probable | As nectar-tap_saprotroph (Põlme et al. 2020); Decay sugar-rich substrates (Põlme et al. 2020) | Põlme S, et al. 2020. Fungal Diversity 105:1-16 ((https://doi.org/10.1007/s13225-020-00466-2)) |
| Otu46956 | 1 | k__Fungi(100);p__Ascomycota(100);c__Saccharomycetes(100);o__Saccharomycetales(100);f__Metschnikowiaceae(100);g__Metschnikowia(100);s__Metschnikowia_pulcherrima(100); | Metschnikowia | 13 | Saprotroph | \|Undefined Saprotroph\| | Yeast | NULL | Probable | As nectar-tap_saprotroph (Põlme et al. 2020); Decay sugar-rich substrates (Põlme et al. 2020) | Põlme S, et al. 2020. Fungal Diversity 105:1-16 ((https://doi.org/10.1007/s13225-020-00466-2)) |
| Otu46957 | 1 | k__Fungi(100);p__Ascomycota(100);c__Saccharomycetes(100);o__Saccharomycetales(100);f__Metschnikowiaceae(100);g__Metschnikowia(100);s__Metschnikowia_pulcherrima(100); | Metschnikowia | 13 | Saprotroph | \|Undefined Saprotroph\| | Yeast | NULL | Probable | As nectar-tap_saprotroph (Põlme et al. 2020); Decay sugar-rich substrates (Põlme et al. 2020) | Põlme S, et al. 2020. Fungal Diversity 105:1-16 ((https://doi.org/10.1007/s13225-020-00466-2)) |
| Otu46958 | 1 | k__Fungi(100);p__Ascomycota(100);c__Saccharomycetes(100);o__Saccharomycetales(100);f__Metschnikowiaceae(100);g__Metschnikowia(100);s__Metschnikowia_pulcherrima(100); | Metschnikowia | 13 | Saprotroph | \|Undefined Saprotroph\| | Yeast | NULL | Probable | As nectar-tap_saprotroph (Põlme et al. 2020); Decay sugar-rich substrates (Põlme et al. 2020) | Põlme S, et al. 2020. Fungal Diversity 105:1-16 ((https://doi.org/10.1007/s13225-020-00466-2)) |
| Otu46959 | 1 | k__Fungi(100);p__Ascomycota(100);c__Saccharomycetes(100);o__Saccharomycetales(100);f__Metschnikowiaceae(100);g__Metschnikowia(100);s__Metschnikowia_pulcherrima(100); | Metschnikowia | 13 | Saprotroph | \|Undefined Saprotroph\| | Yeast | NULL | Probable | As nectar-tap_saprotroph (Põlme et al. 2020); Decay sugar-rich substrates (Põlme et al. 2020) | Põlme S, et al. 2020. Fungal Diversity 105:1-16 ((https://doi.org/10.1007/s13225-020-00466-2)) |
| Otu46960 | 1 | k__Fungi(100);p__Ascomycota(100);c__Saccharomycetes(100);o__Saccharomycetales(100);f__Metschnikowiaceae(100);g__Metschnikowia(100);s__Metschnikowia_pulcherrima(100); | Metschnikowia | 13 | Saprotroph | \|Undefined Saprotroph\| | Yeast | NULL | Probable | As nectar-tap_saprotroph (Põlme et al. 2020); Decay sugar-rich substrates (Põlme et al. 2020) | Põlme S, et al. 2020. Fungal Diversity 105:1-16 ((https://doi.org/10.1007/s13225-020-00466-2)) |
| Otu46961 | 1 | k__Fungi(100);p__Ascomycota(100);c__Saccharomycetes(100);o__Saccharomycetales(100);f__Metschnikowiaceae(100);g__Metschnikowia(100);s__Metschnikowia_pulcherrima(100); | Metschnikowia | 13 | Saprotroph | \|Undefined Saprotroph\| | Yeast | NULL | Probable | As nectar-tap_saprotroph (Põlme et al. 2020); Decay sugar-rich substrates (Põlme et al. 2020) | Põlme S, et al. 2020. Fungal Diversity 105:1-16 ((https://doi.org/10.1007/s13225-020-00466-2)) |
| Otu46962 | 1 | k__Fungi(100);p__Ascomycota(100);c__Saccharomycetes(100);o__Saccharomycetales(100);f__Metschnikowiaceae(100);g__Metschnikowia(100);s__Metschnikowia_pulcherrima(100); | Metschnikowia | 13 | Saprotroph | \|Undefined Saprotroph\| | Yeast | NULL | Probable | As nectar-tap_saprotroph (Põlme et al. 2020); Decay sugar-rich substrates (Põlme et al. 2020) | Põlme S, et al. 2020. Fungal Diversity 105:1-16 ((https://doi.org/10.1007/s13225-020-00466-2)) |
| Otu46963 | 1 | k__Fungi(100);p__Ascomycota(100);c__Saccharomycetes(100);o__Saccharomycetales(100);f__Metschnikowiaceae(100);g__Metschnikowia(100);s__Metschnikowia_pulcherrima(100); | Metschnikowia | 13 | Saprotroph | \|Undefined Saprotroph\| | Yeast | NULL | Probable | As nectar-tap_saprotroph (Põlme et al. 2020); Decay sugar-rich substrates (Põlme et al. 2020) | Põlme S, et al. 2020. Fungal Diversity 105:1-16 ((https://doi.org/10.1007/s13225-020-00466-2)) |
| Otu46964 | 1 | k__Fungi(100);p__Ascomycota(100);c__Saccharomycetes(100);o__Saccharomycetales(100);f__Metschnikowiaceae(100);g__Metschnikowia(100);s__Metschnikowia_pulcherrima(100); | Metschnikowia | 13 | Saprotroph | \|Undefined Saprotroph\| | Yeast | NULL | Probable | As nectar-tap_saprotroph (Põlme et al. 2020); Decay sugar-rich substrates (Põlme et al. 2020) | Põlme S, et al. 2020. Fungal Diversity 105:1-16 ((https://doi.org/10.1007/s13225-020-00466-2)) |
| Otu46965 | 1 | k__Fungi(100);p__Ascomycota(100);c__Saccharomycetes(100);o__Saccharomycetales(100);f__Metschnikowiaceae(100);g__Metschnikowia(100);s__Metschnikowia_pulcherrima(100); | Metschnikowia | 13 | Saprotroph | \|Undefined Saprotroph\| | Yeast | NULL | Probable | As nectar-tap_saprotroph (Põlme et al. 2020); Decay sugar-rich substrates (Põlme et al. 2020) | Põlme S, et al. 2020. Fungal Diversity 105:1-16 ((https://doi.org/10.1007/s13225-020-00466-2)) |
| Otu46966 | 1 | k__Fungi(100);p__Ascomycota(100);c__Saccharomycetes(100);o__Saccharomycetales(100);f__Metschnikowiaceae(100);g__Metschnikowia(100);g__Metschnikowia_unclassified(100); | Metschnikowia | 13 | Saprotroph | \|Undefined Saprotroph\| | Yeast | NULL | Probable | As nectar-tap_saprotroph (Põlme et al. 2020); Decay sugar-rich substrates (Põlme et al. 2020) | Põlme S, et al. 2020. Fungal Diversity 105:1-16 ((https://doi.org/10.1007/s13225-020-00466-2)) |
| Otu46967 | 1 | k__Fungi(100);p__Ascomycota(100);c__Saccharomycetes(100);o__Saccharomycetales(100);f__Metschnikowiaceae(100);g__Metschnikowia(100);s__Metschnikowia_pulcherrima(100); | Metschnikowia | 13 | Saprotroph | \|Undefined Saprotroph\| | Yeast | NULL | Probable | As nectar-tap_saprotroph (Põlme et al. 2020); Decay sugar-rich substrates (Põlme et al. 2020) | Põlme S, et al. 2020. Fungal Diversity 105:1-16 ((https://doi.org/10.1007/s13225-020-00466-2)) |
| Otu46968 | 1 | k__Fungi(100);p__Ascomycota(100);c__Saccharomycetes(100);o__Saccharomycetales(100);f__Metschnikowiaceae(100);g__Metschnikowia(100);s__Metschnikowia_pulcherrima(100); | Metschnikowia | 13 | Saprotroph | \|Undefined Saprotroph\| | Yeast | NULL | Probable | As nectar-tap_saprotroph (Põlme et al. 2020); Decay sugar-rich substrates (Põlme et al. 2020) | Põlme S, et al. 2020. Fungal Diversity 105:1-16 ((https://doi.org/10.1007/s13225-020-00466-2)) |
| Otu46969 | 1 | k__Fungi(100);p__Ascomycota(100);c__Saccharomycetes(100);o__Saccharomycetales(100);f__Metschnikowiaceae(100);g__Metschnikowia(100);s__Metschnikowia_pulcherrima(100); | Metschnikowia | 13 | Saprotroph | \|Undefined Saprotroph\| | Yeast | NULL | Probable | As nectar-tap_saprotroph (Põlme et al. 2020); Decay sugar-rich substrates (Põlme et al. 2020) | Põlme S, et al. 2020. Fungal Diversity 105:1-16 ((https://doi.org/10.1007/s13225-020-00466-2)) |
| Otu46970 | 1 | k__Fungi(100);p__Ascomycota(100);c__Saccharomycetes(100);o__Saccharomycetales(100);f__Metschnikowiaceae(100);g__Metschnikowia(100);s__Metschnikowia_pulcherrima(100); | Metschnikowia | 13 | Saprotroph | \|Undefined Saprotroph\| | Yeast | NULL | Probable | As nectar-tap_saprotroph (Põlme et al. 2020); Decay sugar-rich substrates (Põlme et al. 2020) | Põlme S, et al. 2020. Fungal Diversity 105:1-16 ((https://doi.org/10.1007/s13225-020-00466-2)) |
| Otu46971 | 1 | k__Fungi(100);p__Ascomycota(100);c__Saccharomycetes(100);o__Saccharomycetales(100);f__Metschnikowiaceae(100);g__Metschnikowia(100);s__Metschnikowia_pulcherrima(100); | Metschnikowia | 13 | Saprotroph | \|Undefined Saprotroph\| | Yeast | NULL | Probable | As nectar-tap_saprotroph (Põlme et al. 2020); Decay sugar-rich substrates (Põlme et al. 2020) | Põlme S, et al. 2020. Fungal Diversity 105:1-16 ((https://doi.org/10.1007/s13225-020-00466-2)) |
| Otu46972 | 1 | k__Fungi(100);p__Ascomycota(100);c__Saccharomycetes(100);o__Saccharomycetales(100);f__Metschnikowiaceae(100);g__Metschnikowia(100);s__Metschnikowia_pulcherrima(100); | Metschnikowia | 13 | Saprotroph | \|Undefined Saprotroph\| | Yeast | NULL | Probable | As nectar-tap_saprotroph (Põlme et al. 2020); Decay sugar-rich substrates (Põlme et al. 2020) | Põlme S, et al. 2020. Fungal Diversity 105:1-16 ((https://doi.org/10.1007/s13225-020-00466-2)) |
| Otu46973 | 1 | k__Fungi(100);p__Ascomycota(100);c__Saccharomycetes(100);o__Saccharomycetales(100);f__Metschnikowiaceae(100);g__Metschnikowia(100);s__Metschnikowia_pulcherrima(100); | Metschnikowia | 13 | Saprotroph | \|Undefined Saprotroph\| | Yeast | NULL | Probable | As nectar-tap_saprotroph (Põlme et al. 2020); Decay sugar-rich substrates (Põlme et al. 2020) | Põlme S, et al. 2020. Fungal Diversity 105:1-16 ((https://doi.org/10.1007/s13225-020-00466-2)) |
| Otu46974 | 1 | k__Fungi(100);p__Ascomycota(100);c__Saccharomycetes(100);o__Saccharomycetales(100);f__Metschnikowiaceae(100);g__Metschnikowia(100);g__Metschnikowia_unclassified(100); | Metschnikowia | 13 | Saprotroph | \|Undefined Saprotroph\| | Yeast | NULL | Probable | As nectar-tap_saprotroph (Põlme et al. 2020); Decay sugar-rich substrates (Põlme et al. 2020) | Põlme S, et al. 2020. Fungal Diversity 105:1-16 ((https://doi.org/10.1007/s13225-020-00466-2)) |
| Otu46975 | 1 | k__Fungi(100);p__Ascomycota(100);c__Saccharomycetes(100);o__Saccharomycetales(100);f__Metschnikowiaceae(100);g__Metschnikowia(100);s__Metschnikowia_pulcherrima(100); | Metschnikowia | 13 | Saprotroph | \|Undefined Saprotroph\| | Yeast | NULL | Probable | As nectar-tap_saprotroph (Põlme et al. 2020); Decay sugar-rich substrates (Põlme et al. 2020) | Põlme S, et al. 2020. Fungal Diversity 105:1-16 ((https://doi.org/10.1007/s13225-020-00466-2)) |
| Otu46977 | 1 | k__Fungi(100);p__Ascomycota(100);c__Saccharomycetes(100);o__Saccharomycetales(100);f__Metschnikowiaceae(100);g__Metschnikowia(100);s__Metschnikowia_pulcherrima(100); | Metschnikowia | 13 | Saprotroph | \|Undefined Saprotroph\| | Yeast | NULL | Probable | As nectar-tap_saprotroph (Põlme et al. 2020); Decay sugar-rich substrates (Põlme et al. 2020) | Põlme S, et al. 2020. Fungal Diversity 105:1-16 ((https://doi.org/10.1007/s13225-020-00466-2)) |
| Otu46978 | 1 | k__Fungi(100);p__Ascomycota(100);c__Saccharomycetes(100);o__Saccharomycetales(100);f__Metschnikowiaceae(100);g__Metschnikowia(100);s__Metschnikowia_pulcherrima(100); | Metschnikowia | 13 | Saprotroph | \|Undefined Saprotroph\| | Yeast | NULL | Probable | As nectar-tap_saprotroph (Põlme et al. 2020); Decay sugar-rich substrates (Põlme et al. 2020) | Põlme S, et al. 2020. Fungal Diversity 105:1-16 ((https://doi.org/10.1007/s13225-020-00466-2)) |
| Otu46979 | 1 | k__Fungi(100);p__Ascomycota(100);c__Saccharomycetes(100);o__Saccharomycetales(100);f__Metschnikowiaceae(100);g__Metschnikowia(100);g__Metschnikowia_unclassified(100); | Metschnikowia | 13 | Saprotroph | \|Undefined Saprotroph\| | Yeast | NULL | Probable | As nectar-tap_saprotroph (Põlme et al. 2020); Decay sugar-rich substrates (Põlme et al. 2020) | Põlme S, et al. 2020. Fungal Diversity 105:1-16 ((https://doi.org/10.1007/s13225-020-00466-2)) |
| Otu46980 | 1 | k__Fungi(100);p__Ascomycota(100);c__Saccharomycetes(100);o__Saccharomycetales(100);f__Metschnikowiaceae(100);g__Metschnikowia(100);s__Metschnikowia_pulcherrima(100); | Metschnikowia | 13 | Saprotroph | \|Undefined Saprotroph\| | Yeast | NULL | Probable | As nectar-tap_saprotroph (Põlme et al. 2020); Decay sugar-rich substrates (Põlme et al. 2020) | Põlme S, et al. 2020. Fungal Diversity 105:1-16 ((https://doi.org/10.1007/s13225-020-00466-2)) |
| Otu46981 | 1 | k__Fungi(100);p__Ascomycota(100);c__Saccharomycetes(100);o__Saccharomycetales(100);f__Metschnikowiaceae(100);g__Metschnikowia(100);g__Metschnikowia_unclassified(100); | Metschnikowia | 13 | Saprotroph | \|Undefined Saprotroph\| | Yeast | NULL | Probable | As nectar-tap_saprotroph (Põlme et al. 2020); Decay sugar-rich substrates (Põlme et al. 2020) | Põlme S, et al. 2020. Fungal Diversity 105:1-16 ((https://doi.org/10.1007/s13225-020-00466-2)) |
| Otu46982 | 1 | k__Fungi(100);p__Ascomycota(100);c__Saccharomycetes(100);o__Saccharomycetales(100);f__Metschnikowiaceae(100);g__Metschnikowia(100);g__Metschnikowia_unclassified(100); | Metschnikowia | 13 | Saprotroph | \|Undefined Saprotroph\| | Yeast | NULL | Probable | As nectar-tap_saprotroph (Põlme et al. 2020); Decay sugar-rich substrates (Põlme et al. 2020) | Põlme S, et al. 2020. Fungal Diversity 105:1-16 ((https://doi.org/10.1007/s13225-020-00466-2)) |
| Otu46983 | 1 | k__Fungi(100);p__Ascomycota(100);c__Saccharomycetes(100);o__Saccharomycetales(100);f__Metschnikowiaceae(100);g__Metschnikowia(100);s__Metschnikowia_pulcherrima(100); | Metschnikowia | 13 | Saprotroph | \|Undefined Saprotroph\| | Yeast | NULL | Probable | As nectar-tap_saprotroph (Põlme et al. 2020); Decay sugar-rich substrates (Põlme et al. 2020) | Põlme S, et al. 2020. Fungal Diversity 105:1-16 ((https://doi.org/10.1007/s13225-020-00466-2)) |
| Otu46984 | 1 | k__Fungi(100);p__Ascomycota(100);c__Saccharomycetes(100);o__Saccharomycetales(100);f__Metschnikowiaceae(100);g__Metschnikowia(100);s__Metschnikowia_pulcherrima(100); | Metschnikowia | 13 | Saprotroph | \|Undefined Saprotroph\| | Yeast | NULL | Probable | As nectar-tap_saprotroph (Põlme et al. 2020); Decay sugar-rich substrates (Põlme et al. 2020) | Põlme S, et al. 2020. Fungal Diversity 105:1-16 ((https://doi.org/10.1007/s13225-020-00466-2)) |
| Otu46985 | 1 | k__Fungi(100);p__Ascomycota(100);c__Saccharomycetes(100);o__Saccharomycetales(100);f__Metschnikowiaceae(100);g__Metschnikowia(100);g__Metschnikowia_unclassified(100); | Metschnikowia | 13 | Saprotroph | \|Undefined Saprotroph\| | Yeast | NULL | Probable | As nectar-tap_saprotroph (Põlme et al. 2020); Decay sugar-rich substrates (Põlme et al. 2020) | Põlme S, et al. 2020. Fungal Diversity 105:1-16 ((https://doi.org/10.1007/s13225-020-00466-2)) |
| Otu46986 | 1 | k__Fungi(100);p__Ascomycota(100);c__Saccharomycetes(100);o__Saccharomycetales(100);f__Metschnikowiaceae(100);g__Metschnikowia(100);s__Metschnikowia_pulcherrima(100); | Metschnikowia | 13 | Saprotroph | \|Undefined Saprotroph\| | Yeast | NULL | Probable | As nectar-tap_saprotroph (Põlme et al. 2020); Decay sugar-rich substrates (Põlme et al. 2020) | Põlme S, et al. 2020. Fungal Diversity 105:1-16 ((https://doi.org/10.1007/s13225-020-00466-2)) |
| Otu46987 | 1 | k__Fungi(100);p__Ascomycota(100);c__Saccharomycetes(100);o__Saccharomycetales(100);f__Metschnikowiaceae(100);g__Metschnikowia(100);s__Metschnikowia_pulcherrima(100); | Metschnikowia | 13 | Saprotroph | \|Undefined Saprotroph\| | Yeast | NULL | Probable | As nectar-tap_saprotroph (Põlme et al. 2020); Decay sugar-rich substrates (Põlme et al. 2020) | Põlme S, et al. 2020. Fungal Diversity 105:1-16 ((https://doi.org/10.1007/s13225-020-00466-2)) |
| Otu46988 | 1 | k__Fungi(100);p__Ascomycota(100);c__Saccharomycetes(100);o__Saccharomycetales(100);f__Metschnikowiaceae(100);g__Metschnikowia(100);g__Metschnikowia_unclassified(100); | Metschnikowia | 13 | Saprotroph | \|Undefined Saprotroph\| | Yeast | NULL | Probable | As nectar-tap_saprotroph (Põlme et al. 2020); Decay sugar-rich substrates (Põlme et al. 2020) | Põlme S, et al. 2020. Fungal Diversity 105:1-16 ((https://doi.org/10.1007/s13225-020-00466-2)) |
| Otu46989 | 1 | k__Fungi(100);p__Ascomycota(100);c__Saccharomycetes(100);o__Saccharomycetales(100);f__Metschnikowiaceae(100);g__Metschnikowia(100);s__Metschnikowia_pulcherrima(100); | Metschnikowia | 13 | Saprotroph | \|Undefined Saprotroph\| | Yeast | NULL | Probable | As nectar-tap_saprotroph (Põlme et al. 2020); Decay sugar-rich substrates (Põlme et al. 2020) | Põlme S, et al. 2020. Fungal Diversity 105:1-16 ((https://doi.org/10.1007/s13225-020-00466-2)) |
| Otu46990 | 1 | k__Fungi(100);p__Ascomycota(100);c__Saccharomycetes(100);o__Saccharomycetales(100);f__Metschnikowiaceae(100);g__Metschnikowia(100);s__Metschnikowia_pulcherrima(100); | Metschnikowia | 13 | Saprotroph | \|Undefined Saprotroph\| | Yeast | NULL | Probable | As nectar-tap_saprotroph (Põlme et al. 2020); Decay sugar-rich substrates (Põlme et al. 2020) | Põlme S, et al. 2020. Fungal Diversity 105:1-16 ((https://doi.org/10.1007/s13225-020-00466-2)) |
| Otu46991 | 1 | k__Fungi(100);p__Ascomycota(100);c__Saccharomycetes(100);o__Saccharomycetales(100);f__Metschnikowiaceae(100);g__Metschnikowia(100);g__Metschnikowia_unclassified(100); | Metschnikowia | 13 | Saprotroph | \|Undefined Saprotroph\| | Yeast | NULL | Probable | As nectar-tap_saprotroph (Põlme et al. 2020); Decay sugar-rich substrates (Põlme et al. 2020) | Põlme S, et al. 2020. Fungal Diversity 105:1-16 ((https://doi.org/10.1007/s13225-020-00466-2)) |
| Otu46993 | 1 | k__Fungi(100);p__Ascomycota(100);c__Saccharomycetes(100);o__Saccharomycetales(100);f__Metschnikowiaceae(100);g__Metschnikowia(100);s__Metschnikowia_pulcherrima(100); | Metschnikowia | 13 | Saprotroph | \|Undefined Saprotroph\| | Yeast | NULL | Probable | As nectar-tap_saprotroph (Põlme et al. 2020); Decay sugar-rich substrates (Põlme et al. 2020) | Põlme S, et al. 2020. Fungal Diversity 105:1-16 ((https://doi.org/10.1007/s13225-020-00466-2)) |
| Otu46994 | 1 | k__Fungi(100);p__Ascomycota(100);c__Saccharomycetes(100);o__Saccharomycetales(100);f__Metschnikowiaceae(100);g__Metschnikowia(100);s__Metschnikowia_pulcherrima(100); | Metschnikowia | 13 | Saprotroph | \|Undefined Saprotroph\| | Yeast | NULL | Probable | As nectar-tap_saprotroph (Põlme et al. 2020); Decay sugar-rich substrates (Põlme et al. 2020) | Põlme S, et al. 2020. Fungal Diversity 105:1-16 ((https://doi.org/10.1007/s13225-020-00466-2)) |
| Otu46995 | 1 | k__Fungi(100);p__Ascomycota(100);c__Saccharomycetes(100);o__Saccharomycetales(100);f__Metschnikowiaceae(100);g__Metschnikowia(100);s__Metschnikowia_pulcherrima(100); | Metschnikowia | 13 | Saprotroph | \|Undefined Saprotroph\| | Yeast | NULL | Probable | As nectar-tap_saprotroph (Põlme et al. 2020); Decay sugar-rich substrates (Põlme et al. 2020) | Põlme S, et al. 2020. Fungal Diversity 105:1-16 ((https://doi.org/10.1007/s13225-020-00466-2)) |
| Otu46997 | 1 | k__Fungi(100);p__Ascomycota(100);c__Saccharomycetes(100);o__Saccharomycetales(100);f__Metschnikowiaceae(100);g__Metschnikowia(100);s__Metschnikowia_pulcherrima(100); | Metschnikowia | 13 | Saprotroph | \|Undefined Saprotroph\| | Yeast | NULL | Probable | As nectar-tap_saprotroph (Põlme et al. 2020); Decay sugar-rich substrates (Põlme et al. 2020) | Põlme S, et al. 2020. Fungal Diversity 105:1-16 ((https://doi.org/10.1007/s13225-020-00466-2)) |
| Otu46998 | 1 | k__Fungi(100);p__Ascomycota(100);c__Saccharomycetes(100);o__Saccharomycetales(100);f__Metschnikowiaceae(100);g__Metschnikowia(100);s__Metschnikowia_pulcherrima(100); | Metschnikowia | 13 | Saprotroph | \|Undefined Saprotroph\| | Yeast | NULL | Probable | As nectar-tap_saprotroph (Põlme et al. 2020); Decay sugar-rich substrates (Põlme et al. 2020) | Põlme S, et al. 2020. Fungal Diversity 105:1-16 ((https://doi.org/10.1007/s13225-020-00466-2)) |
| Otu47000 | 1 | k__Fungi(100);p__Ascomycota(100);c__Saccharomycetes(100);o__Saccharomycetales(100);f__Metschnikowiaceae(100);g__Metschnikowia(100);s__Metschnikowia_pulcherrima(100); | Metschnikowia | 13 | Saprotroph | \|Undefined Saprotroph\| | Yeast | NULL | Probable | As nectar-tap_saprotroph (Põlme et al. 2020); Decay sugar-rich substrates (Põlme et al. 2020) | Põlme S, et al. 2020. Fungal Diversity 105:1-16 ((https://doi.org/10.1007/s13225-020-00466-2)) |
| Otu47001 | 1 | k__Fungi(100);p__Ascomycota(100);c__Saccharomycetes(100);o__Saccharomycetales(100);f__Metschnikowiaceae(100);g__Metschnikowia(100);g__Metschnikowia_unclassified(100); | Metschnikowia | 13 | Saprotroph | \|Undefined Saprotroph\| | Yeast | NULL | Probable | As nectar-tap_saprotroph (Põlme et al. 2020); Decay sugar-rich substrates (Põlme et al. 2020) | Põlme S, et al. 2020. Fungal Diversity 105:1-16 ((https://doi.org/10.1007/s13225-020-00466-2)) |
| Otu47002 | 1 | k__Fungi(100);p__Ascomycota(100);c__Saccharomycetes(100);o__Saccharomycetales(100);f__Metschnikowiaceae(100);g__Metschnikowia(100);s__Metschnikowia_pulcherrima(100); | Metschnikowia | 13 | Saprotroph | \|Undefined Saprotroph\| | Yeast | NULL | Probable | As nectar-tap_saprotroph (Põlme et al. 2020); Decay sugar-rich substrates (Põlme et al. 2020) | Põlme S, et al. 2020. Fungal Diversity 105:1-16 ((https://doi.org/10.1007/s13225-020-00466-2)) |
| Otu47003 | 1 | k__Fungi(100);p__Ascomycota(100);c__Saccharomycetes(100);o__Saccharomycetales(100);f__Metschnikowiaceae(100);g__Metschnikowia(100);s__Metschnikowia_pulcherrima(100); | Metschnikowia | 13 | Saprotroph | \|Undefined Saprotroph\| | Yeast | NULL | Probable | As nectar-tap_saprotroph (Põlme et al. 2020); Decay sugar-rich substrates (Põlme et al. 2020) | Põlme S, et al. 2020. Fungal Diversity 105:1-16 ((https://doi.org/10.1007/s13225-020-00466-2)) |
| Otu47004 | 1 | k__Fungi(100);p__Ascomycota(100);c__Saccharomycetes(100);o__Saccharomycetales(100);f__Metschnikowiaceae(100);g__Metschnikowia(100);g__Metschnikowia_unclassified(100); | Metschnikowia | 13 | Saprotroph | \|Undefined Saprotroph\| | Yeast | NULL | Probable | As nectar-tap_saprotroph (Põlme et al. 2020); Decay sugar-rich substrates (Põlme et al. 2020) | Põlme S, et al. 2020. Fungal Diversity 105:1-16 ((https://doi.org/10.1007/s13225-020-00466-2)) |
| Otu47005 | 1 | k__Fungi(100);p__Ascomycota(100);c__Saccharomycetes(100);o__Saccharomycetales(100);f__Metschnikowiaceae(100);g__Metschnikowia(100);g__Metschnikowia_unclassified(100); | Metschnikowia | 13 | Saprotroph | \|Undefined Saprotroph\| | Yeast | NULL | Probable | As nectar-tap_saprotroph (Põlme et al. 2020); Decay sugar-rich substrates (Põlme et al. 2020) | Põlme S, et al. 2020. Fungal Diversity 105:1-16 ((https://doi.org/10.1007/s13225-020-00466-2)) |
| Otu47006 | 1 | k__Fungi(100);p__Ascomycota(100);c__Saccharomycetes(100);o__Saccharomycetales(100);f__Metschnikowiaceae(100);g__Metschnikowia(100);g__Metschnikowia_unclassified(100); | Metschnikowia | 13 | Saprotroph | \|Undefined Saprotroph\| | Yeast | NULL | Probable | As nectar-tap_saprotroph (Põlme et al. 2020); Decay sugar-rich substrates (Põlme et al. 2020) | Põlme S, et al. 2020. Fungal Diversity 105:1-16 ((https://doi.org/10.1007/s13225-020-00466-2)) |
| Otu47007 | 1 | k__Fungi(100);p__Ascomycota(100);c__Saccharomycetes(100);o__Saccharomycetales(100);f__Metschnikowiaceae(100);g__Metschnikowia(100);s__Metschnikowia_pulcherrima(100); | Metschnikowia | 13 | Saprotroph | \|Undefined Saprotroph\| | Yeast | NULL | Probable | As nectar-tap_saprotroph (Põlme et al. 2020); Decay sugar-rich substrates (Põlme et al. 2020) | Põlme S, et al. 2020. Fungal Diversity 105:1-16 ((https://doi.org/10.1007/s13225-020-00466-2)) |
| Otu47008 | 1 | k__Fungi(100);p__Ascomycota(100);c__Saccharomycetes(100);o__Saccharomycetales(100);f__Metschnikowiaceae(100);g__Metschnikowia(100);s__Metschnikowia_pulcherrima(100); | Metschnikowia | 13 | Saprotroph | \|Undefined Saprotroph\| | Yeast | NULL | Probable | As nectar-tap_saprotroph (Põlme et al. 2020); Decay sugar-rich substrates (Põlme et al. 2020) | Põlme S, et al. 2020. Fungal Diversity 105:1-16 ((https://doi.org/10.1007/s13225-020-00466-2)) |
| Otu47009 | 1 | k__Fungi(100);p__Ascomycota(100);c__Saccharomycetes(100);o__Saccharomycetales(100);f__Metschnikowiaceae(100);g__Metschnikowia(100);s__Metschnikowia_pulcherrima(100); | Metschnikowia | 13 | Saprotroph | \|Undefined Saprotroph\| | Yeast | NULL | Probable | As nectar-tap_saprotroph (Põlme et al. 2020); Decay sugar-rich substrates (Põlme et al. 2020) | Põlme S, et al. 2020. Fungal Diversity 105:1-16 ((https://doi.org/10.1007/s13225-020-00466-2)) |
| Otu47010 | 1 | k__Fungi(100);p__Ascomycota(100);c__Saccharomycetes(100);o__Saccharomycetales(100);f__Metschnikowiaceae(100);g__Metschnikowia(100);s__Metschnikowia_pulcherrima(100); | Metschnikowia | 13 | Saprotroph | \|Undefined Saprotroph\| | Yeast | NULL | Probable | As nectar-tap_saprotroph (Põlme et al. 2020); Decay sugar-rich substrates (Põlme et al. 2020) | Põlme S, et al. 2020. Fungal Diversity 105:1-16 ((https://doi.org/10.1007/s13225-020-00466-2)) |
| Otu47011 | 1 | k__Fungi(100);p__Ascomycota(100);c__Saccharomycetes(100);o__Saccharomycetales(100);f__Metschnikowiaceae(100);g__Metschnikowia(100);s__Metschnikowia_pulcherrima(100); | Metschnikowia | 13 | Saprotroph | \|Undefined Saprotroph\| | Yeast | NULL | Probable | As nectar-tap_saprotroph (Põlme et al. 2020); Decay sugar-rich substrates (Põlme et al. 2020) | Põlme S, et al. 2020. Fungal Diversity 105:1-16 ((https://doi.org/10.1007/s13225-020-00466-2)) |
| Otu47012 | 1 | k__Fungi(100);p__Ascomycota(100);c__Saccharomycetes(100);o__Saccharomycetales(100);f__Metschnikowiaceae(100);g__Metschnikowia(100);s__Metschnikowia_pulcherrima(100); | Metschnikowia | 13 | Saprotroph | \|Undefined Saprotroph\| | Yeast | NULL | Probable | As nectar-tap_saprotroph (Põlme et al. 2020); Decay sugar-rich substrates (Põlme et al. 2020) | Põlme S, et al. 2020. Fungal Diversity 105:1-16 ((https://doi.org/10.1007/s13225-020-00466-2)) |
| Otu47013 | 1 | k__Fungi(100);p__Ascomycota(100);c__Saccharomycetes(100);o__Saccharomycetales(100);f__Metschnikowiaceae(100);g__Metschnikowia(100);s__Metschnikowia_pulcherrima(100); | Metschnikowia | 13 | Saprotroph | \|Undefined Saprotroph\| | Yeast | NULL | Probable | As nectar-tap_saprotroph (Põlme et al. 2020); Decay sugar-rich substrates (Põlme et al. 2020) | Põlme S, et al. 2020. Fungal Diversity 105:1-16 ((https://doi.org/10.1007/s13225-020-00466-2)) |
| Otu47014 | 1 | k__Fungi(100);p__Ascomycota(100);c__Saccharomycetes(100);o__Saccharomycetales(100);f__Metschnikowiaceae(100);g__Metschnikowia(100);g__Metschnikowia_unclassified(100); | Metschnikowia | 13 | Saprotroph | \|Undefined Saprotroph\| | Yeast | NULL | Probable | As nectar-tap_saprotroph (Põlme et al. 2020); Decay sugar-rich substrates (Põlme et al. 2020) | Põlme S, et al. 2020. Fungal Diversity 105:1-16 ((https://doi.org/10.1007/s13225-020-00466-2)) |
| Otu47015 | 1 | k__Fungi(100);p__Ascomycota(100);c__Saccharomycetes(100);o__Saccharomycetales(100);f__Metschnikowiaceae(100);g__Metschnikowia(100);s__Metschnikowia_pulcherrima(100); | Metschnikowia | 13 | Saprotroph | \|Undefined Saprotroph\| | Yeast | NULL | Probable | As nectar-tap_saprotroph (Põlme et al. 2020); Decay sugar-rich substrates (Põlme et al. 2020) | Põlme S, et al. 2020. Fungal Diversity 105:1-16 ((https://doi.org/10.1007/s13225-020-00466-2)) |
| Otu47016 | 1 | k__Fungi(100);p__Ascomycota(100);c__Saccharomycetes(100);o__Saccharomycetales(100);f__Metschnikowiaceae(100);g__Metschnikowia(100);s__Metschnikowia_pulcherrima(100); | Metschnikowia | 13 | Saprotroph | \|Undefined Saprotroph\| | Yeast | NULL | Probable | As nectar-tap_saprotroph (Põlme et al. 2020); Decay sugar-rich substrates (Põlme et al. 2020) | Põlme S, et al. 2020. Fungal Diversity 105:1-16 ((https://doi.org/10.1007/s13225-020-00466-2)) |
| Otu47017 | 1 | k__Fungi(100);p__Ascomycota(100);c__Saccharomycetes(100);o__Saccharomycetales(100);f__Metschnikowiaceae(100);g__Metschnikowia(100);s__Metschnikowia_pulcherrima(100); | Metschnikowia | 13 | Saprotroph | \|Undefined Saprotroph\| | Yeast | NULL | Probable | As nectar-tap_saprotroph (Põlme et al. 2020); Decay sugar-rich substrates (Põlme et al. 2020) | Põlme S, et al. 2020. Fungal Diversity 105:1-16 ((https://doi.org/10.1007/s13225-020-00466-2)) |
| Otu47018 | 1 | k__Fungi(100);p__Ascomycota(100);c__Saccharomycetes(100);o__Saccharomycetales(100);f__Metschnikowiaceae(100);g__Metschnikowia(100);s__Metschnikowia_pulcherrima(100); | Metschnikowia | 13 | Saprotroph | \|Undefined Saprotroph\| | Yeast | NULL | Probable | As nectar-tap_saprotroph (Põlme et al. 2020); Decay sugar-rich substrates (Põlme et al. 2020) | Põlme S, et al. 2020. Fungal Diversity 105:1-16 ((https://doi.org/10.1007/s13225-020-00466-2)) |
| Otu47019 | 1 | k__Fungi(100);p__Ascomycota(100);c__Saccharomycetes(100);o__Saccharomycetales(100);f__Metschnikowiaceae(100);g__Metschnikowia(100);s__Metschnikowia_pulcherrima(100); | Metschnikowia | 13 | Saprotroph | \|Undefined Saprotroph\| | Yeast | NULL | Probable | As nectar-tap_saprotroph (Põlme et al. 2020); Decay sugar-rich substrates (Põlme et al. 2020) | Põlme S, et al. 2020. Fungal Diversity 105:1-16 ((https://doi.org/10.1007/s13225-020-00466-2)) |
| Otu47020 | 1 | k__Fungi(100);p__Ascomycota(100);c__Saccharomycetes(100);o__Saccharomycetales(100);f__Metschnikowiaceae(100);g__Metschnikowia(100);s__Metschnikowia_pulcherrima(100); | Metschnikowia | 13 | Saprotroph | \|Undefined Saprotroph\| | Yeast | NULL | Probable | As nectar-tap_saprotroph (Põlme et al. 2020); Decay sugar-rich substrates (Põlme et al. 2020) | Põlme S, et al. 2020. Fungal Diversity 105:1-16 ((https://doi.org/10.1007/s13225-020-00466-2)) |
| Otu47021 | 1 | k__Fungi(100);p__Ascomycota(100);c__Saccharomycetes(100);o__Saccharomycetales(100);f__Metschnikowiaceae(100);g__Metschnikowia(100);s__Metschnikowia_pulcherrima(100); | Metschnikowia | 13 | Saprotroph | \|Undefined Saprotroph\| | Yeast | NULL | Probable | As nectar-tap_saprotroph (Põlme et al. 2020); Decay sugar-rich substrates (Põlme et al. 2020) | Põlme S, et al. 2020. Fungal Diversity 105:1-16 ((https://doi.org/10.1007/s13225-020-00466-2)) |
| Otu47022 | 1 | k__Fungi(100);p__Ascomycota(100);c__Saccharomycetes(100);o__Saccharomycetales(100);f__Metschnikowiaceae(100);g__Metschnikowia(100);g__Metschnikowia_unclassified(100); | Metschnikowia | 13 | Saprotroph | \|Undefined Saprotroph\| | Yeast | NULL | Probable | As nectar-tap_saprotroph (Põlme et al. 2020); Decay sugar-rich substrates (Põlme et al. 2020) | Põlme S, et al. 2020. Fungal Diversity 105:1-16 ((https://doi.org/10.1007/s13225-020-00466-2)) |
| Otu47023 | 1 | k__Fungi(100);p__Ascomycota(100);c__Saccharomycetes(100);o__Saccharomycetales(100);f__Metschnikowiaceae(100);g__Metschnikowia(100);s__Metschnikowia_pulcherrima(100); | Metschnikowia | 13 | Saprotroph | \|Undefined Saprotroph\| | Yeast | NULL | Probable | As nectar-tap_saprotroph (Põlme et al. 2020); Decay sugar-rich substrates (Põlme et al. 2020) | Põlme S, et al. 2020. Fungal Diversity 105:1-16 ((https://doi.org/10.1007/s13225-020-00466-2)) |
| Otu47024 | 1 | k__Fungi(100);p__Ascomycota(100);c__Saccharomycetes(100);o__Saccharomycetales(100);f__Metschnikowiaceae(100);g__Metschnikowia(100);g__Metschnikowia_unclassified(100); | Metschnikowia | 13 | Saprotroph | \|Undefined Saprotroph\| | Yeast | NULL | Probable | As nectar-tap_saprotroph (Põlme et al. 2020); Decay sugar-rich substrates (Põlme et al. 2020) | Põlme S, et al. 2020. Fungal Diversity 105:1-16 ((https://doi.org/10.1007/s13225-020-00466-2)) |
| Otu47025 | 1 | k__Fungi(100);p__Ascomycota(100);c__Saccharomycetes(100);o__Saccharomycetales(100);f__Metschnikowiaceae(100);g__Metschnikowia(100);s__Metschnikowia_pulcherrima(100); | Metschnikowia | 13 | Saprotroph | \|Undefined Saprotroph\| | Yeast | NULL | Probable | As nectar-tap_saprotroph (Põlme et al. 2020); Decay sugar-rich substrates (Põlme et al. 2020) | Põlme S, et al. 2020. Fungal Diversity 105:1-16 ((https://doi.org/10.1007/s13225-020-00466-2)) |
| Otu47026 | 1 | k__Fungi(100);p__Ascomycota(100);c__Saccharomycetes(100);o__Saccharomycetales(100);f__Metschnikowiaceae(100);g__Metschnikowia(100);s__Metschnikowia_pulcherrima(100); | Metschnikowia | 13 | Saprotroph | \|Undefined Saprotroph\| | Yeast | NULL | Probable | As nectar-tap_saprotroph (Põlme et al. 2020); Decay sugar-rich substrates (Põlme et al. 2020) | Põlme S, et al. 2020. Fungal Diversity 105:1-16 ((https://doi.org/10.1007/s13225-020-00466-2)) |
| Otu47027 | 1 | k__Fungi(100);p__Ascomycota(100);c__Saccharomycetes(100);o__Saccharomycetales(100);f__Metschnikowiaceae(100);g__Metschnikowia(100);s__Metschnikowia_pulcherrima(100); | Metschnikowia | 13 | Saprotroph | \|Undefined Saprotroph\| | Yeast | NULL | Probable | As nectar-tap_saprotroph (Põlme et al. 2020); Decay sugar-rich substrates (Põlme et al. 2020) | Põlme S, et al. 2020. Fungal Diversity 105:1-16 ((https://doi.org/10.1007/s13225-020-00466-2)) |
| Otu47028 | 1 | k__Fungi(100);p__Ascomycota(100);c__Saccharomycetes(100);o__Saccharomycetales(100);f__Metschnikowiaceae(100);g__Metschnikowia(100);s__Metschnikowia_pulcherrima(100); | Metschnikowia | 13 | Saprotroph | \|Undefined Saprotroph\| | Yeast | NULL | Probable | As nectar-tap_saprotroph (Põlme et al. 2020); Decay sugar-rich substrates (Põlme et al. 2020) | Põlme S, et al. 2020. Fungal Diversity 105:1-16 ((https://doi.org/10.1007/s13225-020-00466-2)) |
| Otu47029 | 1 | k__Fungi(100);p__Ascomycota(100);c__Saccharomycetes(100);o__Saccharomycetales(100);f__Metschnikowiaceae(100);g__Metschnikowia(100);s__Metschnikowia_pulcherrima(100); | Metschnikowia | 13 | Saprotroph | \|Undefined Saprotroph\| | Yeast | NULL | Probable | As nectar-tap_saprotroph (Põlme et al. 2020); Decay sugar-rich substrates (Põlme et al. 2020) | Põlme S, et al. 2020. Fungal Diversity 105:1-16 ((https://doi.org/10.1007/s13225-020-00466-2)) |
| Otu47030 | 1 | k__Fungi(100);p__Ascomycota(100);c__Saccharomycetes(100);o__Saccharomycetales(100);f__Metschnikowiaceae(100);g__Metschnikowia(100);g__Metschnikowia_unclassified(100); | Metschnikowia | 13 | Saprotroph | \|Undefined Saprotroph\| | Yeast | NULL | Probable | As nectar-tap_saprotroph (Põlme et al. 2020); Decay sugar-rich substrates (Põlme et al. 2020) | Põlme S, et al. 2020. Fungal Diversity 105:1-16 ((https://doi.org/10.1007/s13225-020-00466-2)) |
| Otu47031 | 1 | k__Fungi(100);p__Ascomycota(100);c__Saccharomycetes(100);o__Saccharomycetales(100);f__Metschnikowiaceae(100);g__Metschnikowia(100);s__Metschnikowia_pulcherrima(100); | Metschnikowia | 13 | Saprotroph | \|Undefined Saprotroph\| | Yeast | NULL | Probable | As nectar-tap_saprotroph (Põlme et al. 2020); Decay sugar-rich substrates (Põlme et al. 2020) | Põlme S, et al. 2020. Fungal Diversity 105:1-16 ((https://doi.org/10.1007/s13225-020-00466-2)) |
| Otu47032 | 1 | k__Fungi(100);p__Ascomycota(100);c__Saccharomycetes(100);o__Saccharomycetales(100);f__Metschnikowiaceae(100);g__Metschnikowia(100);g__Metschnikowia_unclassified(100); | Metschnikowia | 13 | Saprotroph | \|Undefined Saprotroph\| | Yeast | NULL | Probable | As nectar-tap_saprotroph (Põlme et al. 2020); Decay sugar-rich substrates (Põlme et al. 2020) | Põlme S, et al. 2020. Fungal Diversity 105:1-16 ((https://doi.org/10.1007/s13225-020-00466-2)) |
| Otu47033 | 1 | k__Fungi(100);p__Ascomycota(100);c__Saccharomycetes(100);o__Saccharomycetales(100);f__Metschnikowiaceae(100);g__Metschnikowia(100);g__Metschnikowia_unclassified(100); | Metschnikowia | 13 | Saprotroph | \|Undefined Saprotroph\| | Yeast | NULL | Probable | As nectar-tap_saprotroph (Põlme et al. 2020); Decay sugar-rich substrates (Põlme et al. 2020) | Põlme S, et al. 2020. Fungal Diversity 105:1-16 ((https://doi.org/10.1007/s13225-020-00466-2)) |

| | | | | | | | | | | | |
|---|---|---|---|---|---|---|---|---|---|---|---|
| Otu47034 | 1 | k__Fungi(100);p__Ascomycota(100);c__Saccharomycetes(100);o__Saccharomycetales(100);f__Metschnikowiaceae(100);g__Metschnikowia(100);s__Metschnikowia_pulcherrima(100); | Metschnikowia | 13 | Saprotroph | \|Undefined Saprotroph\| | Yeast | NULL | Probable | As nectar-tap_saprotroph (Põlme et al. 2020); Decay sugar-rich substrates (Põlme et al. 2020) | Põlme S, et al. 2020. Fungal Diversity 105:1-16 ((https://doi.org/10.1007/s13225-020-00466-2)) |
| Otu47035 | 1 | k__Fungi(100);p__Ascomycota(100);c__Saccharomycetes(100);o__Saccharomycetales(100);f__Metschnikowiaceae(100);g__Metschnikowia(100);s__Metschnikowia_pulcherrima(100); | Metschnikowia | 13 | Saprotroph | \|Undefined Saprotroph\| | Yeast | NULL | Probable | As nectar-tap_saprotroph (Põlme et al. 2020); Decay sugar-rich substrates (Põlme et al. 2020) | Põlme S, et al. 2020. Fungal Diversity 105:1-16 ((https://doi.org/10.1007/s13225-020-00466-2)) |
| Otu47036 | 1 | k__Fungi(100);p__Ascomycota(100);c__Saccharomycetes(100);o__Saccharomycetales(100);f__Metschnikowiaceae(100);g__Metschnikowia(100);g__Metschnikowia_unclassified(100); | Metschnikowia | 13 | Saprotroph | \|Undefined Saprotroph\| | Yeast | NULL | Probable | As nectar-tap_saprotroph (Põlme et al. 2020); Decay sugar-rich substrates (Põlme et al. 2020) | Põlme S, et al. 2020. Fungal Diversity 105:1-16 ((https://doi.org/10.1007/s13225-020-00466-2)) |
| Otu47037 | 1 | k__Fungi(100);p__Ascomycota(100);c__Saccharomycetes(100);o__Saccharomycetales(100);f__Metschnikowiaceae(100);g__Metschnikowia(100);s__Metschnikowia_pulcherrima(100); | Metschnikowia | 13 | Saprotroph | \|Undefined Saprotroph\| | Yeast | NULL | Probable | As nectar-tap_saprotroph (Põlme et al. 2020); Decay sugar-rich substrates (Põlme et al. 2020) | Põlme S, et al. 2020. Fungal Diversity 105:1-16 ((https://doi.org/10.1007/s13225-020-00466-2)) |
| Otu47038 | 1 | k__Fungi(100);p__Ascomycota(100);c__Saccharomycetes(100);o__Saccharomycetales(100);f__Metschnikowiaceae(100);g__Metschnikowia(100);g__Metschnikowia_unclassified(100); | Metschnikowia | 13 | Saprotroph | \|Undefined Saprotroph\| | Yeast | NULL | Probable | As nectar-tap_saprotroph (Põlme et al. 2020); Decay sugar-rich substrates (Põlme et al. 2020) | Põlme S, et al. 2020. Fungal Diversity 105:1-16 ((https://doi.org/10.1007/s13225-020-00466-2)) |
| Otu47039 | 1 | k__Fungi(100);p__Ascomycota(100);c__Saccharomycetes(100);o__Saccharomycetales(100);f__Metschnikowiaceae(100);g__Metschnikowia(100);s__Metschnikowia_pulcherrima(100); | Metschnikowia | 13 | Saprotroph | \|Undefined Saprotroph\| | Yeast | NULL | Probable | As nectar-tap_saprotroph (Põlme et al. 2020); Decay sugar-rich substrates (Põlme et al. 2020) | Põlme S, et al. 2020. Fungal Diversity 105:1-16 ((https://doi.org/10.1007/s13225-020-00466-2)) |
| Otu47040 | 1 | k__Fungi(100);p__Ascomycota(100);c__Saccharomycetes(100);o__Saccharomycetales(100);f__Metschnikowiaceae(100);g__Metschnikowia(100);s__Metschnikowia_pulcherrima(100); | Metschnikowia | 13 | Saprotroph | \|Undefined Saprotroph\| | Yeast | NULL | Probable | As nectar-tap_saprotroph (Põlme et al. 2020); Decay sugar-rich substrates (Põlme et al. 2020) | Põlme S, et al. 2020. Fungal Diversity 105:1-16 ((https://doi.org/10.1007/s13225-020-00466-2)) |
| Otu47041 | 1 | k__Fungi(100);p__Ascomycota(100);c__Saccharomycetes(100);o__Saccharomycetales(100);f__Metschnikowiaceae(100);g__Metschnikowia(100);s__Metschnikowia_pulcherrima(100); | Metschnikowia | 13 | Saprotroph | \|Undefined Saprotroph\| | Yeast | NULL | Probable | As nectar-tap_saprotroph (Põlme et al. 2020); Decay sugar-rich substrates (Põlme et al. 2020) | Põlme S, et al. 2020. Fungal Diversity 105:1-16 ((https://doi.org/10.1007/s13225-020-00466-2)) |
| Otu47042 | 1 | k__Fungi(100);p__Ascomycota(100);c__Saccharomycetes(100);o__Saccharomycetales(100);f__Metschnikowiaceae(100);g__Metschnikowia(100);s__Metschnikowia_pulcherrima(100); | Metschnikowia | 13 | Saprotroph | \|Undefined Saprotroph\| | Yeast | NULL | Probable | As nectar-tap_saprotroph (Põlme et al. 2020); Decay sugar-rich substrates (Põlme et al. 2020) | Põlme S, et al. 2020. Fungal Diversity 105:1-16 ((https://doi.org/10.1007/s13225-020-00466-2)) |
| Otu47043 | 1 | k__Fungi(100);p__Ascomycota(100);c__Saccharomycetes(100);o__Saccharomycetales(100);f__Metschnikowiaceae(100);g__Metschnikowia(100);s__Metschnikowia_pulcherrima(100); | Metschnikowia | 13 | Saprotroph | \|Undefined Saprotroph\| | Yeast | NULL | Probable | As nectar-tap_saprotroph (Põlme et al. 2020); Decay sugar-rich substrates (Põlme et al. 2020) | Põlme S, et al. 2020. Fungal Diversity 105:1-16 ((https://doi.org/10.1007/s13225-020-00466-2)) |
| Otu47044 | 1 | k__Fungi(100);p__Ascomycota(100);c__Saccharomycetes(100);o__Saccharomycetales(100);f__Metschnikowiaceae(100);g__Metschnikowia(100);s__Metschnikowia_pulcherrima(100); | Metschnikowia | 13 | Saprotroph | \|Undefined Saprotroph\| | Yeast | NULL | Probable | As nectar-tap_saprotroph (Põlme et al. 2020); Decay sugar-rich substrates (Põlme et al. 2020) | Põlme S, et al. 2020. Fungal Diversity 105:1-16 ((https://doi.org/10.1007/s13225-020-00466-2)) |
| Otu47046 | 1 | k__Fungi(100);p__Ascomycota(100);c__Saccharomycetes(100);o__Saccharomycetales(100);f__Metschnikowiaceae(100);g__Metschnikowia(100);s__Metschnikowia_pulcherrima(100); | Metschnikowia | 13 | Saprotroph | \|Undefined Saprotroph\| | Yeast | NULL | Probable | As nectar-tap_saprotroph (Põlme et al. 2020); Decay sugar-rich substrates (Põlme et al. 2020) | Põlme S, et al. 2020. Fungal Diversity 105:1-16 ((https://doi.org/10.1007/s13225-020-00466-2)) |
| Otu47047 | 1 | k__Fungi(100);p__Ascomycota(100);c__Saccharomycetes(100);o__Saccharomycetales(100);f__Metschnikowiaceae(100);g__Metschnikowia(100);s__Metschnikowia_pulcherrima(100); | Metschnikowia | 13 | Saprotroph | \|Undefined Saprotroph\| | Yeast | NULL | Probable | As nectar-tap_saprotroph (Põlme et al. 2020); Decay sugar-rich substrates (Põlme et al. 2020) | Põlme S, et al. 2020. Fungal Diversity 105:1-16 ((https://doi.org/10.1007/s13225-020-00466-2)) |
| Otu47048 | 1 | k__Fungi(100);p__Ascomycota(100);c__Saccharomycetes(100);o__Saccharomycetales(100);f__Metschnikowiaceae(100);g__Metschnikowia(100);g__Metschnikowia_unclassified(100); | Metschnikowia | 13 | Saprotroph | \|Undefined Saprotroph\| | Yeast | NULL | Probable | As nectar-tap_saprotroph (Põlme et al. 2020); Decay sugar-rich substrates (Põlme et al. 2020) | Põlme S, et al. 2020. Fungal Diversity 105:1-16 ((https://doi.org/10.1007/s13225-020-00466-2)) |
| Otu47049 | 1 | k__Fungi(100);p__Ascomycota(100);c__Saccharomycetes(100);o__Saccharomycetales(100);f__Metschnikowiaceae(100);g__Metschnikowia(100);g__Metschnikowia_unclassified(100); | Metschnikowia | 13 | Saprotroph | \|Undefined Saprotroph\| | Yeast | NULL | Probable | As nectar-tap_saprotroph (Põlme et al. 2020); Decay sugar-rich substrates (Põlme et al. 2020) | Põlme S, et al. 2020. Fungal Diversity 105:1-16 ((https://doi.org/10.1007/s13225-020-00466-2)) |
| Otu47050 | 1 | k__Fungi(100);p__Ascomycota(100);c__Saccharomycetes(100);o__Saccharomycetales(100);f__Metschnikowiaceae(100);g__Metschnikowia(100);s__Metschnikowia_pulcherrima(100); | Metschnikowia | 13 | Saprotroph | \|Undefined Saprotroph\| | Yeast | NULL | Probable | As nectar-tap_saprotroph (Põlme et al. 2020); Decay sugar-rich substrates (Põlme et al. 2020) | Põlme S, et al. 2020. Fungal Diversity 105:1-16 ((https://doi.org/10.1007/s13225-020-00466-2)) |
| Otu47051 | 1 | k__Fungi(100);p__Ascomycota(100);c__Saccharomycetes(100);o__Saccharomycetales(100);f__Metschnikowiaceae(100);g__Metschnikowia(100);g__Metschnikowia_unclassified(100); | Metschnikowia | 13 | Saprotroph | \|Undefined Saprotroph\| | Yeast | NULL | Probable | As nectar-tap_saprotroph (Põlme et al. 2020); Decay sugar-rich substrates (Põlme et al. 2020) | Põlme S, et al. 2020. Fungal Diversity 105:1-16 ((https://doi.org/10.1007/s13225-020-00466-2)) |
| Otu47052 | 1 | k__Fungi(100);p__Ascomycota(100);c__Saccharomycetes(100);o__Saccharomycetales(100);f__Metschnikowiaceae(100);g__Metschnikowia(100);s__Metschnikowia_pulcherrima(100); | Metschnikowia | 13 | Saprotroph | \|Undefined Saprotroph\| | Yeast | NULL | Probable | As nectar-tap_saprotroph (Põlme et al. 2020); Decay sugar-rich substrates (Põlme et al. 2020) | Põlme S, et al. 2020. Fungal Diversity 105:1-16 ((https://doi.org/10.1007/s13225-020-00466-2)) |
| Otu47053 | 1 | k__Fungi(100);p__Ascomycota(100);c__Saccharomycetes(100);o__Saccharomycetales(100);f__Metschnikowiaceae(100);g__Metschnikowia(100);s__Metschnikowia_pulcherrima(100); | Metschnikowia | 13 | Saprotroph | \|Undefined Saprotroph\| | Yeast | NULL | Probable | As nectar-tap_saprotroph (Põlme et al. 2020); Decay sugar-rich substrates (Põlme et al. 2020) | Põlme S, et al. 2020. Fungal Diversity 105:1-16 ((https://doi.org/10.1007/s13225-020-00466-2)) |
| Otu47055 | 1 | k__Fungi(100);p__Ascomycota(100);c__Saccharomycetes(100);o__Saccharomycetales(100);f__Metschnikowiaceae(100);g__Metschnikowia(100);s__Metschnikowia_pulcherrima(100); | Metschnikowia | 13 | Saprotroph | \|Undefined Saprotroph\| | Yeast | NULL | Probable | As nectar-tap_saprotroph (Põlme et al. 2020); Decay sugar-rich substrates (Põlme et al. 2020) | Põlme S, et al. 2020. Fungal Diversity 105:1-16 ((https://doi.org/10.1007/s13225-020-00466-2)) |
| Otu47056 | 1 | k__Fungi(100);p__Ascomycota(100);c__Saccharomycetes(100);o__Saccharomycetales(100);f__Metschnikowiaceae(100);g__Metschnikowia(100);s__Metschnikowia_pulcherrima(100); | Metschnikowia | 13 | Saprotroph | \|Undefined Saprotroph\| | Yeast | NULL | Probable | As nectar-tap_saprotroph (Põlme et al. 2020); Decay sugar-rich substrates (Põlme et al. 2020) | Põlme S, et al. 2020. Fungal Diversity 105:1-16 ((https://doi.org/10.1007/s13225-020-00466-2)) |
| Otu47057 | 1 | k__Fungi(100);p__Ascomycota(100);c__Saccharomycetes(100);o__Saccharomycetales(100);f__Metschnikowiaceae(100);g__Metschnikowia(100);g__Metschnikowia_unclassified(100); | Metschnikowia | 13 | Saprotroph | \|Undefined Saprotroph\| | Yeast | NULL | Probable | As nectar-tap_saprotroph (Põlme et al. 2020); Decay sugar-rich substrates (Põlme et al. 2020) | Põlme S, et al. 2020. Fungal Diversity 105:1-16 ((https://doi.org/10.1007/s13225-020-00466-2)) |
| Otu47058 | 1 | k__Fungi(100);p__Ascomycota(100);c__Saccharomycetes(100);o__Saccharomycetales(100);f__Metschnikowiaceae(100);g__Metschnikowia(100);s__Metschnikowia_pulcherrima(100); | Metschnikowia | 13 | Saprotroph | \|Undefined Saprotroph\| | Yeast | NULL | Probable | As nectar-tap_saprotroph (Põlme et al. 2020); Decay sugar-rich substrates (Põlme et al. 2020) | Põlme S, et al. 2020. Fungal Diversity 105:1-16 ((https://doi.org/10.1007/s13225-020-00466-2)) |
| Otu47059 | 1 | k__Fungi(100);p__Ascomycota(100);c__Saccharomycetes(100);o__Saccharomycetales(100);f__Metschnikowiaceae(100);g__Metschnikowia(100);g__Metschnikowia_unclassified(100); | Metschnikowia | 13 | Saprotroph | \|Undefined Saprotroph\| | Yeast | NULL | Probable | As nectar-tap_saprotroph (Põlme et al. 2020); Decay sugar-rich substrates (Põlme et al. 2020) | Põlme S, et al. 2020. Fungal Diversity 105:1-16 ((https://doi.org/10.1007/s13225-020-00466-2)) |
| Otu47060 | 1 | k__Fungi(100);p__Ascomycota(100);c__Saccharomycetes(100);o__Saccharomycetales(100);f__Metschnikowiaceae(100);g__Metschnikowia(100);s__Metschnikowia_pulcherrima(100); | Metschnikowia | 13 | Saprotroph | \|Undefined Saprotroph\| | Yeast | NULL | Probable | As nectar-tap_saprotroph (Põlme et al. 2020); Decay sugar-rich substrates (Põlme et al. 2020) | Põlme S, et al. 2020. Fungal Diversity 105:1-16 ((https://doi.org/10.1007/s13225-020-00466-2)) |
| Otu47061 | 1 | k__Fungi(100);p__Ascomycota(100);c__Saccharomycetes(100);o__Saccharomycetales(100);f__Metschnikowiaceae(100);g__Metschnikowia(100);g__Metschnikowia_unclassified(100); | Metschnikowia | 13 | Saprotroph | \|Undefined Saprotroph\| | Yeast | NULL | Probable | As nectar-tap_saprotroph (Põlme et al. 2020); Decay sugar-rich substrates (Põlme et al. 2020) | Põlme S, et al. 2020. Fungal Diversity 105:1-16 ((https://doi.org/10.1007/s13225-020-00466-2)) |
| Otu47062 | 1 | k__Fungi(100);p__Ascomycota(100);c__Saccharomycetes(100);o__Saccharomycetales(100);f__Metschnikowiaceae(100);g__Metschnikowia(100);s__Metschnikowia_pulcherrima(100); | Metschnikowia | 13 | Saprotroph | \|Undefined Saprotroph\| | Yeast | NULL | Probable | As nectar-tap_saprotroph (Põlme et al. 2020); Decay sugar-rich substrates (Põlme et al. 2020) | Põlme S, et al. 2020. Fungal Diversity 105:1-16 ((https://doi.org/10.1007/s13225-020-00466-2)) |
| Otu47063 | 1 | k__Fungi(100);p__Ascomycota(100);c__Saccharomycetes(100);o__Saccharomycetales(100);f__Metschnikowiaceae(100);g__Metschnikowia(100);g__Metschnikowia_unclassified(100); | Metschnikowia | 13 | Saprotroph | \|Undefined Saprotroph\| | Yeast | NULL | Probable | As nectar-tap_saprotroph (Põlme et al. 2020); Decay sugar-rich substrates (Põlme et al. 2020) | Põlme S, et al. 2020. Fungal Diversity 105:1-16 ((https://doi.org/10.1007/s13225-020-00466-2)) |
| Otu47064 | 1 | k__Fungi(100);p__Ascomycota(100);c__Saccharomycetes(100);o__Saccharomycetales(100);f__Metschnikowiaceae(100);g__Metschnikowia(100);g__Metschnikowia_unclassified(100); | Metschnikowia | 13 | Saprotroph | \|Undefined Saprotroph\| | Yeast | NULL | Probable | As nectar-tap_saprotroph (Põlme et al. 2020); Decay sugar-rich substrates (Põlme et al. 2020) | Põlme S, et al. 2020. Fungal Diversity 105:1-16 ((https://doi.org/10.1007/s13225-020-00466-2)) |
| Otu47065 | 1 | k__Fungi(100);p__Ascomycota(100);c__Saccharomycetes(100);o__Saccharomycetales(100);f__Metschnikowiaceae(100);g__Metschnikowia(100);g__Metschnikowia_unclassified(100); | Metschnikowia | 13 | Saprotroph | \|Undefined Saprotroph\| | Yeast | NULL | Probable | As nectar-tap_saprotroph (Põlme et al. 2020); Decay sugar-rich substrates (Põlme et al. 2020) | Põlme S, et al. 2020. Fungal Diversity 105:1-16 ((https://doi.org/10.1007/s13225-020-00466-2)) |
| Otu47066 | 1 | k__Fungi(100);p__Ascomycota(100);c__Saccharomycetes(100);o__Saccharomycetales(100);f__Metschnikowiaceae(100);g__Metschnikowia(100);s__Metschnikowia_pulcherrima(100); | Metschnikowia | 13 | Saprotroph | \|Undefined Saprotroph\| | Yeast | NULL | Probable | As nectar-tap_saprotroph (Põlme et al. 2020); Decay sugar-rich substrates (Põlme et al. 2020) | Põlme S, et al. 2020. Fungal Diversity 105:1-16 ((https://doi.org/10.1007/s13225-020-00466-2)) |
| Otu47067 | 1 | k__Fungi(100);p__Ascomycota(100);c__Saccharomycetes(100);o__Saccharomycetales(100);f__Metschnikowiaceae(100);g__Metschnikowia(100);s__Metschnikowia_pulcherrima(100); | Metschnikowia | 13 | Saprotroph | \|Undefined Saprotroph\| | Yeast | NULL | Probable | As nectar-tap_saprotroph (Põlme et al. 2020); Decay sugar-rich substrates (Põlme et al. 2020) | Põlme S, et al. 2020. Fungal Diversity 105:1-16 ((https://doi.org/10.1007/s13225-020-00466-2)) |
| Otu47068 | 1 | k__Fungi(100);p__Ascomycota(100);c__Saccharomycetes(100);o__Saccharomycetales(100);f__Metschnikowiaceae(100);g__Metschnikowia(100);s__Metschnikowia_pulcherrima(100); | Metschnikowia | 13 | Saprotroph | \|Undefined Saprotroph\| | Yeast | NULL | Probable | As nectar-tap_saprotroph (Põlme et al. 2020); Decay sugar-rich substrates (Põlme et al. 2020) | Põlme S, et al. 2020. Fungal Diversity 105:1-16 ((https://doi.org/10.1007/s13225-020-00466-2)) |
| Otu47069 | 1 | k__Fungi(100);p__Ascomycota(100);c__Saccharomycetes(100);o__Saccharomycetales(100);f__Metschnikowiaceae(100);g__Metschnikowia(100);g__Metschnikowia_unclassified(100); | Metschnikowia | 13 | Saprotroph | \|Undefined Saprotroph\| | Yeast | NULL | Probable | As nectar-tap_saprotroph (Põlme et al. 2020); Decay sugar-rich substrates (Põlme et al. 2020) | Põlme S, et al. 2020. Fungal Diversity 105:1-16 ((https://doi.org/10.1007/s13225-020-00466-2)) |
| Otu47070 | 1 | k__Fungi(100);p__Ascomycota(100);c__Saccharomycetes(100);o__Saccharomycetales(100);f__Metschnikowiaceae(100);g__Metschnikowia(100);g__Metschnikowia_unclassified(100); | Metschnikowia | 13 | Saprotroph | \|Undefined Saprotroph\| | Yeast | NULL | Probable | As nectar-tap_saprotroph (Põlme et al. 2020); Decay sugar-rich substrates (Põlme et al. 2020) | Põlme S, et al. 2020. Fungal Diversity 105:1-16 ((https://doi.org/10.1007/s13225-020-00466-2)) |
| Otu47071 | 1 | k__Fungi(100);p__Ascomycota(100);c__Saccharomycetes(100);o__Saccharomycetales(100);f__Metschnikowiaceae(100);g__Metschnikowia(100);s__Metschnikowia_pulcherrima(100); | Metschnikowia | 13 | Saprotroph | \|Undefined Saprotroph\| | Yeast | NULL | Probable | As nectar-tap_saprotroph (Põlme et al. 2020); Decay sugar-rich substrates (Põlme et al. 2020) | Põlme S, et al. 2020. Fungal Diversity 105:1-16 ((https://doi.org/10.1007/s13225-020-00466-2)) |
| Otu47072 | 1 | k__Fungi(100);p__Ascomycota(100);c__Saccharomycetes(100);o__Saccharomycetales(100);f__Metschnikowiaceae(100);g__Metschnikowia(100);g__Metschnikowia_unclassified(100); | Metschnikowia | 13 | Saprotroph | \|Undefined Saprotroph\| | Yeast | NULL | Probable | As nectar-tap_saprotroph (Põlme et al. 2020); Decay sugar-rich substrates (Põlme et al. 2020) | Põlme S, et al. 2020. Fungal Diversity 105:1-16 ((https://doi.org/10.1007/s13225-020-00466-2)) |
| Otu47073 | 1 | k__Fungi(100);p__Ascomycota(100);c__Saccharomycetes(100);o__Saccharomycetales(100);f__Metschnikowiaceae(100);g__Metschnikowia(100);s__Metschnikowia_pulcherrima(100); | Metschnikowia | 13 | Saprotroph | \|Undefined Saprotroph\| | Yeast | NULL | Probable | As nectar-tap_saprotroph (Põlme et al. 2020); Decay sugar-rich substrates (Põlme et al. 2020) | Põlme S, et al. 2020. Fungal Diversity 105:1-16 ((https://doi.org/10.1007/s13225-020-00466-2)) |
| Otu47074 | 1 | k__Fungi(100);p__Ascomycota(100);c__Saccharomycetes(100);o__Saccharomycetales(100);f__Metschnikowiaceae(100);g__Metschnikowia(100);g__Metschnikowia_unclassified(100); | Metschnikowia | 13 | Saprotroph | \|Undefined Saprotroph\| | Yeast | NULL | Probable | As nectar-tap_saprotroph (Põlme et al. 2020); Decay sugar-rich substrates (Põlme et al. 2020) | Põlme S, et al. 2020. Fungal Diversity 105:1-16 ((https://doi.org/10.1007/s13225-020-00466-2)) |
| Otu47075 | 1 | k__Fungi(100);p__Ascomycota(100);c__Saccharomycetes(100);o__Saccharomycetales(100);f__Metschnikowiaceae(100);g__Metschnikowia(100);s__Metschnikowia_pulcherrima(100); | Metschnikowia | 13 | Saprotroph | \|Undefined Saprotroph\| | Yeast | NULL | Probable | As nectar-tap_saprotroph (Põlme et al. 2020); Decay sugar-rich substrates (Põlme et al. 2020) | Põlme S, et al. 2020. Fungal Diversity 105:1-16 ((https://doi.org/10.1007/s13225-020-00466-2)) |
| Otu47076 | 1 | k__Fungi(100);p__Ascomycota(100);c__Saccharomycetes(100);o__Saccharomycetales(100);f__Metschnikowiaceae(100);g__Metschnikowia(100);s__Metschnikowia_pulcherrima(100); | Metschnikowia | 13 | Saprotroph | \|Undefined Saprotroph\| | Yeast | NULL | Probable | As nectar-tap_saprotroph (Põlme et al. 2020); Decay sugar-rich substrates (Põlme et al. 2020) | Põlme S, et al. 2020. Fungal Diversity 105:1-16 ((https://doi.org/10.1007/s13225-020-00466-2)) |
| Otu47077 | 1 | k__Fungi(100);p__Ascomycota(100);c__Saccharomycetes(100);o__Saccharomycetales(100);f__Metschnikowiaceae(100);g__Metschnikowia(100);s__Metschnikowia_pulcherrima(100); | Metschnikowia | 13 | Saprotroph | \|Undefined Saprotroph\| | Yeast | NULL | Probable | As nectar-tap_saprotroph (Põlme et al. 2020); Decay sugar-rich substrates (Põlme et al. 2020) | Põlme S, et al. 2020. Fungal Diversity 105:1-16 ((https://doi.org/10.1007/s13225-020-00466-2)) |
| Otu47078 | 1 | k__Fungi(100);p__Ascomycota(100);c__Saccharomycetes(100);o__Saccharomycetales(100);f__Metschnikowiaceae(100);g__Metschnikowia(100);g__Metschnikowia_unclassified(100); | Metschnikowia | 13 | Saprotroph | \|Undefined Saprotroph\| | Yeast | NULL | Probable | As nectar-tap_saprotroph (Põlme et al. 2020); Decay sugar-rich substrates (Põlme et al. 2020) | Põlme S, et al. 2020. Fungal Diversity 105:1-16 ((https://doi.org/10.1007/s13225-020-00466-2)) |
| Otu47079 | 1 | k__Fungi(100);p__Ascomycota(100);c__Saccharomycetes(100);o__Saccharomycetales(100);f__Metschnikowiaceae(100);g__Metschnikowia(100);g__Metschnikowia_unclassified(100); | Metschnikowia | 13 | Saprotroph | \|Undefined Saprotroph\| | Yeast | NULL | Probable | As nectar-tap_saprotroph (Põlme et al. 2020); Decay sugar-rich substrates (Põlme et al. 2020) | Põlme S, et al. 2020. Fungal Diversity 105:1-16 ((https://doi.org/10.1007/s13225-020-00466-2)) |
| Otu47080 | 1 | k__Fungi(100);p__Ascomycota(100);c__Saccharomycetes(100);o__Saccharomycetales(100);f__Metschnikowiaceae(100);g__Metschnikowia(100);s__Metschnikowia_pulcherrima(100); | Metschnikowia | 13 | Saprotroph | \|Undefined Saprotroph\| | Yeast | NULL | Probable | As nectar-tap_saprotroph (Põlme et al. 2020); Decay sugar-rich substrates (Põlme et al. 2020) | Põlme S, et al. 2020. Fungal Diversity 105:1-16 ((https://doi.org/10.1007/s13225-020-00466-2)) |
| Otu47081 | 1 | k__Fungi(100);p__Ascomycota(100);c__Saccharomycetes(100);o__Saccharomycetales(100);f__Metschnikowiaceae(100);g__Metschnikowia(100);g__Metschnikowia_unclassified(100); | Metschnikowia | 13 | Saprotroph | \|Undefined Saprotroph\| | Yeast | NULL | Probable | As nectar-tap_saprotroph (Põlme et al. 2020); Decay sugar-rich substrates (Põlme et al. 2020) | Põlme S, et al. 2020. Fungal Diversity 105:1-16 ((https://doi.org/10.1007/s13225-020-00466-2)) |
| Otu47082 | 1 | k__Fungi(100);p__Ascomycota(100);c__Saccharomycetes(100);o__Saccharomycetales(100);f__Metschnikowiaceae(100);g__Metschnikowia(100);s__Metschnikowia_pulcherrima(100); | Metschnikowia | 13 | Saprotroph | \|Undefined Saprotroph\| | Yeast | NULL | Probable | As nectar-tap_saprotroph (Põlme et al. 2020); Decay sugar-rich substrates (Põlme et al. 2020) | Põlme S, et al. 2020. Fungal Diversity 105:1-16 ((https://doi.org/10.1007/s13225-020-00466-2)) |
| Otu47083 | 1 | k__Fungi(100);p__Ascomycota(100);c__Saccharomycetes(100);o__Saccharomycetales(100);f__Metschnikowiaceae(100);g__Metschnikowia(100);s__Metschnikowia_pulcherrima(100); | Metschnikowia | 13 | Saprotroph | \|Undefined Saprotroph\| | Yeast | NULL | Probable | As nectar-tap_saprotroph (Põlme et al. 2020); Decay sugar-rich substrates (Põlme et al. 2020) | Põlme S, et al. 2020. Fungal Diversity 105:1-16 ((https://doi.org/10.1007/s13225-020-00466-2)) |
| Otu47084 | 1 | k__Fungi(100);p__Ascomycota(100);c__Saccharomycetes(100);o__Saccharomycetales(100);f__Metschnikowiaceae(100);g__Metschnikowia(100);s__Metschnikowia_pulcherrima(100); | Metschnikowia | 13 | Saprotroph | \|Undefined Saprotroph\| | Yeast | NULL | Probable | As nectar-tap_saprotroph (Põlme et al. 2020); Decay sugar-rich substrates (Põlme et al. 2020) | Põlme S, et al. 2020. Fungal Diversity 105:1-16 ((https://doi.org/10.1007/s13225-020-00466-2)) |
| Otu47085 | 1 | k__Fungi(100);p__Ascomycota(100);c__Saccharomycetes(100);o__Saccharomycetales(100);f__Metschnikowiaceae(100);g__Metschnikowia(100);g__Metschnikowia_unclassified(100); | Metschnikowia | 13 | Saprotroph | \|Undefined Saprotroph\| | Yeast | NULL | Probable | As nectar-tap_saprotroph (Põlme et al. 2020); Decay sugar-rich substrates (Põlme et al. 2020) | Põlme S, et al. 2020. Fungal Diversity 105:1-16 ((https://doi.org/10.1007/s13225-020-00466-2)) |
| Otu47086 | 1 | k__Fungi(100);p__Ascomycota(100);c__Saccharomycetes(100);o__Saccharomycetales(100);f__Metschnikowiaceae(100);g__Metschnikowia(100);s__Metschnikowia_pulcherrima(100); | Metschnikowia | 13 | Saprotroph | \|Undefined Saprotroph\| | Yeast | NULL | Probable | As nectar-tap_saprotroph (Põlme et al. 2020); Decay sugar-rich substrates (Põlme et al. 2020) | Põlme S, et al. 2020. Fungal Diversity 105:1-16 ((https://doi.org/10.1007/s13225-020-00466-2)) |
| Otu47087 | 1 | k__Fungi(100);p__Ascomycota(100);c__Saccharomycetes(100);o__Saccharomycetales(100);f__Metschnikowiaceae(100);g__Metschnikowia(100);s__Metschnikowia_pulcherrima(100); | Metschnikowia | 13 | Saprotroph | \|Undefined Saprotroph\| | Yeast | NULL | Probable | As nectar-tap_saprotroph (Põlme et al. 2020); Decay sugar-rich substrates (Põlme et al. 2020) | Põlme S, et al. 2020. Fungal Diversity 105:1-16 ((https://doi.org/10.1007/s13225-020-00466-2)) |
| Otu47088 | 1 | k__Fungi(100);p__Ascomycota(100);c__Saccharomycetes(100);o__Saccharomycetales(100);f__Metschnikowiaceae(100);g__Metschnikowia(100);s__Metschnikowia_pulcherrima(100); | Metschnikowia | 13 | Saprotroph | \|Undefined Saprotroph\| | Yeast | NULL | Probable | As nectar-tap_saprotroph (Põlme et al. 2020); Decay sugar-rich substrates (Põlme et al. 2020) | Põlme S, et al. 2020. Fungal Diversity 105:1-16 ((https://doi.org/10.1007/s13225-020-00466-2)) |
| Otu47089 | 1 | k__Fungi(100);p__Ascomycota(100);c__Saccharomycetes(100);o__Saccharomycetales(100);f__Metschnikowiaceae(100);g__Metschnikowia(100);s__Metschnikowia_pulcherrima(100); | Metschnikowia | 13 | Saprotroph | \|Undefined Saprotroph\| | Yeast | NULL | Probable | As nectar-tap_saprotroph (Põlme et al. 2020); Decay sugar-rich substrates (Põlme et al. 2020) | Põlme S, et al. 2020. Fungal Diversity 105:1-16 ((https://doi.org/10.1007/s13225-020-00466-2)) |
| Otu47090 | 1 | k__Fungi(100);p__Ascomycota(100);c__Saccharomycetes(100);o__Saccharomycetales(100);f__Metschnikowiaceae(100);g__Metschnikowia(100);s__Metschnikowia_pulcherrima(100); | Metschnikowia | 13 | Saprotroph | \|Undefined Saprotroph\| | Yeast | NULL | Probable | As nectar-tap_saprotroph (Põlme et al. 2020); Decay sugar-rich substrates (Põlme et al. 2020) | Põlme S, et al. 2020. Fungal Diversity 105:1-16 ((https://doi.org/10.1007/s13225-020-00466-2)) |
| Otu47091 | 1 | k__Fungi(100);p__Ascomycota(100);c__Saccharomycetes(100);o__Saccharomycetales(100);f__Metschnikowiaceae(100);g__Metschnikowia(100);s__Metschnikowia_pulcherrima(100); | Metschnikowia | 13 | Saprotroph | \|Undefined Saprotroph\| | Yeast | NULL | Probable | As nectar-tap_saprotroph (Põlme et al. 2020); Decay sugar-rich substrates (Põlme et al. 2020) | Põlme S, et al. 2020. Fungal Diversity 105:1-16 ((https://doi.org/10.1007/s13225-020-00466-2)) |
| Otu47092 | 1 | k__Fungi(100);p__Ascomycota(100);c__Saccharomycetes(100);o__Saccharomycetales(100);f__Metschnikowiaceae(100);g__Metschnikowia(100);s__Metschnikowia_pulcherrima(100); | Metschnikowia | 13 | Saprotroph | \|Undefined Saprotroph\| | Yeast | NULL | Probable | As nectar-tap_saprotroph (Põlme et al. 2020); Decay sugar-rich substrates (Põlme et al. 2020) | Põlme S, et al. 2020. Fungal Diversity 105:1-16 ((https://doi.org/10.1007/s13225-020-00466-2)) |
| Otu47093 | 1 | k__Fungi(100);p__Ascomycota(100);c__Saccharomycetes(100);o__Saccharomycetales(100);f__Metschnikowiaceae(100);g__Metschnikowia(100);s__Metschnikowia_pulcherrima(100); | Metschnikowia | 13 | Saprotroph | \|Undefined Saprotroph\| | Yeast | NULL | Probable | As nectar-tap_saprotroph (Põlme et al. 2020); Decay sugar-rich substrates (Põlme et al. 2020) | Põlme S, et al. 2020. Fungal Diversity 105:1-16 ((https://doi.org/10.1007/s13225-020-00466-2)) |
| Otu47095 | 1 | k__Fungi(100);p__Ascomycota(100);c__Saccharomycetes(100);o__Saccharomycetales(100);f__Metschnikowiaceae(100);g__Metschnikowia(100);s__Metschnikowia_pulcherrima(100); | Metschnikowia | 13 | Saprotroph | \|Undefined Saprotroph\| | Yeast | NULL | Probable | As nectar-tap_saprotroph (Põlme et al. 2020); Decay sugar-rich substrates (Põlme et al. 2020) | Põlme S, et al. 2020. Fungal Diversity 105:1-16 ((https://doi.org/10.1007/s13225-020-00466-2)) |
| Otu47096 | 1 | k__Fungi(100);p__Ascomycota(100);c__Saccharomycetes(100);o__Saccharomycetales(100);f__Metschnikowiaceae(100);g__Metschnikowia(100);s__Metschnikowia_pulcherrima(100); | Metschnikowia | 13 | Saprotroph | \|Undefined Saprotroph\| | Yeast | NULL | Probable | As nectar-tap_saprotroph (Põlme et al. 2020); Decay sugar-rich substrates (Põlme et al. 2020) | Põlme S, et al. 2020. Fungal Diversity 105:1-16 ((https://doi.org/10.1007/s13225-020-00466-2)) |
| Otu47097 | 1 | k__Fungi(100);p__Ascomycota(100);c__Saccharomycetes(100);o__Saccharomycetales(100);f__Metschnikowiaceae(100);g__Metschnikowia(100);s__Metschnikowia_pulcherrima(100); | Metschnikowia | 13 | Saprotroph | \|Undefined Saprotroph\| | Yeast | NULL | Probable | As nectar-tap_saprotroph (Põlme et al. 2020); Decay sugar-rich substrates (Põlme et al. 2020) | Põlme S, et al. 2020. Fungal Diversity 105:1-16 ((https://doi.org/10.1007/s13225-020-00466-2)) |
| Otu47098 | 1 | k__Fungi(100);p__Ascomycota(100);c__Saccharomycetes(100);o__Saccharomycetales(100);f__Metschnikowiaceae(100);g__Metschnikowia(100);s__Metschnikowia_pulcherrima(100); | Metschnikowia | 13 | Saprotroph | \|Undefined Saprotroph\| | Yeast | NULL | Probable | As nectar-tap_saprotroph (Põlme et al. 2020); Decay sugar-rich substrates (Põlme et al. 2020) | Põlme S, et al. 2020. Fungal Diversity 105:1-16 ((https://doi.org/10.1007/s13225-020-00466-2)) |
| Otu47103 | 1 | k__Fungi(100);p__Ascomycota(100);c__Saccharomycetes(100);o__Saccharomycetales(100);f__Metschnikowiaceae(100);g__Metschnikowia(100);s__Metschnikowia_pulcherrima(100); | Metschnikowia | 13 | Saprotroph | \|Undefined Saprotroph\| | Yeast | NULL | Probable | As nectar-tap_saprotroph (Põlme et al. 2020); Decay sugar-rich substrates (Põlme et al. 2020) | Põlme S, et al. 2020. Fungal Diversity 105:1-16 ((https://doi.org/10.1007/s13225-020-00466-2)) |
| Otu47104 | 1 | k__Fungi(100);p__Ascomycota(100);c__Saccharomycetes(100);o__Saccharomycetales(100);f__Metschnikowiaceae(100);g__Metschnikowia(100);s__Metschnikowia_pulcherrima(100); | Metschnikowia | 13 | Saprotroph | \|Undefined Saprotroph\| | Yeast | NULL | Probable | As nectar-tap_saprotroph (Põlme et al. 2020); Decay sugar-rich substrates (Põlme et al. 2020) | Põlme S, et al. 2020. Fungal Diversity 105:1-16 ((https://doi.org/10.1007/s13225-020-00466-2)) |
| Otu47105 | 1 | k__Fungi(100);p__Ascomycota(100);c__Saccharomycetes(100);o__Saccharomycetales(100);f__Metschnikowiaceae(100);g__Metschnikowia(100);g__Metschnikowia_unclassified(100); | Metschnikowia | 13 | Saprotroph | \|Undefined Saprotroph\| | Yeast | NULL | Probable | As nectar-tap_saprotroph (Põlme et al. 2020); Decay sugar-rich substrates (Põlme et al. 2020) | Põlme S, et al. 2020. Fungal Diversity 105:1-16 ((https://doi.org/10.1007/s13225-020-00466-2)) |
| Otu47106 | 1 | k__Fungi(100);p__Ascomycota(100);c__Saccharomycetes(100);o__Saccharomycetales(100);f__Metschnikowiaceae(100);g__Metschnikowia(100);s__Metschnikowia_pulcherrima(100); | Metschnikowia | 13 | Saprotroph | \|Undefined Saprotroph\| | Yeast | NULL | Probable | As nectar-tap_saprotroph (Põlme et al. 2020); Decay sugar-rich substrates (Põlme et al. 2020) | Põlme S, et al. 2020. Fungal Diversity 105:1-16 ((https://doi.org/10.1007/s13225-020-00466-2)) |
| Otu47107 | 1 | k__Fungi(100);p__Ascomycota(100);c__Saccharomycetes(100);o__Saccharomycetales(100);f__Metschnikowiaceae(100);g__Metschnikowia(100);s__Metschnikowia_pulcherrima(100); | Metschnikowia | 13 | Saprotroph | \|Undefined Saprotroph\| | Yeast | NULL | Probable | As nectar-tap_saprotroph (Põlme et al. 2020); Decay sugar-rich substrates (Põlme et al. 2020) | Põlme S, et al. 2020. Fungal Diversity 105:1-16 ((https://doi.org/10.1007/s13225-020-00466-2)) |
| Otu47108 | 1 | k__Fungi(100);p__Ascomycota(100);c__Saccharomycetes(100);o__Saccharomycetales(100);f__Metschnikowiaceae(100);g__Metschnikowia(100);s__Metschnikowia_pulcherrima(100); | Metschnikowia | 13 | Saprotroph | \|Undefined Saprotroph\| | Yeast | NULL | Probable | As nectar-tap_saprotroph (Põlme et al. 2020); Decay sugar-rich substrates (Põlme et al. 2020) | Põlme S, et al. 2020. Fungal Diversity 105:1-16 ((https://doi.org/10.1007/s13225-020-00466-2)) |
| Otu47109 | 1 | k__Fungi(100);p__Ascomycota(100);c__Saccharomycetes(100);o__Saccharomycetales(100);f__Metschnikowiaceae(100);g__Metschnikowia(100);s__Metschnikowia_pulcherrima(100); | Metschnikowia | 13 | Saprotroph | \|Undefined Saprotroph\| | Yeast | NULL | Probable | As nectar-tap_saprotroph (Põlme et al. 2020); Decay sugar-rich substrates (Põlme et al. 2020) | Põlme S, et al. 2020. Fungal Diversity 105:1-16 ((https://doi.org/10.1007/s13225-020-00466-2)) |
| Otu47110 | 1 | k__Fungi(100);p__Ascomycota(100);c__Saccharomycetes(100);o__Saccharomycetales(100);f__Metschnikowiaceae(100);g__Metschnikowia(100);g__Metschnikowia_unclassified(100); | Metschnikowia | 13 | Saprotroph | \|Undefined Saprotroph\| | Yeast | NULL | Probable | As nectar-tap_saprotroph (Põlme et al. 2020); Decay sugar-rich substrates (Põlme et al. 2020) | Põlme S, et al. 2020. Fungal Diversity 105:1-16 ((https://doi.org/10.1007/s13225-020-00466-2)) |
| Otu47112 | 1 | k__Fungi(100);p__Ascomycota(100);c__Saccharomycetes(100);o__Saccharomycetales(100);f__Metschnikowiaceae(100);g__Metschnikowia(100);s__Metschnikowia_pulcherrima(100); | Metschnikowia | 13 | Saprotroph | \|Undefined Saprotroph\| | Yeast | NULL | Probable | As nectar-tap_saprotroph (Põlme et al. 2020); Decay sugar-rich substrates (Põlme et al. 2020) | Põlme S, et al. 2020. Fungal Diversity 105:1-16 ((https://doi.org/10.1007/s13225-020-00466-2)) |
| Otu47113 | 1 | k__Fungi(100);p__Ascomycota(100);c__Saccharomycetes(100);o__Saccharomycetales(100);f__Metschnikowiaceae(100);g__Metschnikowia(100);s__Metschnikowia_pulcherrima(100); | Metschnikowia | 13 | Saprotroph | \|Undefined Saprotroph\| | Yeast | NULL | Probable | As nectar-tap_saprotroph (Põlme et al. 2020); Decay sugar-rich substrates (Põlme et al. 2020) | Põlme S, et al. 2020. Fungal Diversity 105:1-16 ((https://doi.org/10.1007/s13225-020-00466-2)) |
| Otu47114 | 1 | k__Fungi(100);p__Ascomycota(100);c__Saccharomycetes(100);o__Saccharomycetales(100);f__Metschnikowiaceae(100);g__Metschnikowia(100);s__Metschnikowia_pulcherrima(100); | Metschnikowia | 13 | Saprotroph | \|Undefined Saprotroph\| | Yeast | NULL | Probable | As nectar-tap_saprotroph (Põlme et al. 2020); Decay sugar-rich substrates (Põlme et al. 2020) | Põlme S, et al. 2020. Fungal Diversity 105:1-16 ((https://doi.org/10.1007/s13225-020-00466-2)) |
| Otu47115 | 1 | k__Fungi(100);p__Ascomycota(100);c__Saccharomycetes(100);o__Saccharomycetales(100);f__Metschnikowiaceae(100);g__Metschnikowia(100);g__Metschnikowia_unclassified(100); | Metschnikowia | 13 | Saprotroph | \|Undefined Saprotroph\| | Yeast | NULL | Probable | As nectar-tap_saprotroph (Põlme et al. 2020); Decay sugar-rich substrates (Põlme et al. 2020) | Põlme S, et al. 2020. Fungal Diversity 105:1-16 ((https://doi.org/10.1007/s13225-020-00466-2)) |
| Otu47116 | 1 | k__Fungi(100);p__Ascomycota(100);c__Saccharomycetes(100);o__Saccharomycetales(100);f__Metschnikowiaceae(100);g__Metschnikowia(100);s__Metschnikowia_pulcherrima(100); | Metschnikowia | 13 | Saprotroph | \|Undefined Saprotroph\| | Yeast | NULL | Probable | As nectar-tap_saprotroph (Põlme et al. 2020); Decay sugar-rich substrates (Põlme et al. 2020) | Põlme S, et al. 2020. Fungal Diversity 105:1-16 ((https://doi.org/10.1007/s13225-020-00466-2)) |
| Otu47117 | 1 | k__Fungi(100);p__Ascomycota(100);c__Saccharomycetes(100);o__Saccharomycetales(100);f__Metschnikowiaceae(100);g__Metschnikowia(100);g__Metschnikowia_unclassified(100); | Metschnikowia | 13 | Saprotroph | \|Undefined Saprotroph\| | Yeast | NULL | Probable | As nectar-tap_saprotroph (Põlme et al. 2020); Decay sugar-rich substrates (Põlme et al. 2020) | Põlme S, et al. 2020. Fungal Diversity 105:1-16 ((https://doi.org/10.1007/s13225-020-00466-2)) |
| Otu47118 | 1 | k__Fungi(100);p__Ascomycota(100);c__Saccharomycetes(100);o__Saccharomycetales(100);f__Metschnikowiaceae(100);g__Metschnikowia(100);g__Metschnikowia_unclassified(100); | Metschnikowia | 13 | Saprotroph | \|Undefined Saprotroph\| | Yeast | NULL | Probable | As nectar-tap_saprotroph (Põlme et al. 2020); Decay sugar-rich substrates (Põlme et al. 2020) | Põlme S, et al. 2020. Fungal Diversity 105:1-16 ((https://doi.org/10.1007/s13225-020-00466-2)) |
| Otu47119 | 1 | k__Fungi(100);p__Ascomycota(100);c__Saccharomycetes(100);o__Saccharomycetales(100);f__Metschnikowiaceae(100);g__Metschnikowia(100);s__Metschnikowia_pulcherrima(100); | Metschnikowia | 13 | Saprotroph | \|Undefined Saprotroph\| | Yeast | NULL | Probable | As nectar-tap_saprotroph (Põlme et al. 2020); Decay sugar-rich substrates (Põlme et al. 2020) | Põlme S, et al. 2020. Fungal Diversity 105:1-16 ((https://doi.org/10.1007/s13225-020-00466-2)) |
| Otu47120 | 1 | k__Fungi(100);p__Ascomycota(100);c__Saccharomycetes(100);o__Saccharomycetales(100);f__Metschnikowiaceae(100);g__Metschnikowia(100);s__Metschnikowia_pulcherrima(100); | Metschnikowia | 13 | Saprotroph | \|Undefined Saprotroph\| | Yeast | NULL | Probable | As nectar-tap_saprotroph (Põlme et al. 2020); Decay sugar-rich substrates (Põlme et al. 2020) | Põlme S, et al. 2020. Fungal Diversity 105:1-16 ((https://doi.org/10.1007/s13225-020-00466-2)) |
| Otu47122 | 1 | k__Fungi(100);p__Ascomycota(100);c__Saccharomycetes(100);o__Saccharomycetales(100);f__Metschnikowiaceae(100);g__Metschnikowia(100);s__Metschnikowia_pulcherrima(100); | Metschnikowia | 13 | Saprotroph | \|Undefined Saprotroph\| | Yeast | NULL | Probable | As nectar-tap_saprotroph (Põlme et al. 2020); Decay sugar-rich substrates (Põlme et al. 2020) | Põlme S, et al. 2020. Fungal Diversity 105:1-16 ((https://doi.org/10.1007/s13225-020-00466-2)) |
| Otu47123 | 1 | k__Fungi(100);p__Ascomycota(100);c__Saccharomycetes(100);o__Saccharomycetales(100);f__Metschnikowiaceae(100);g__Metschnikowia(100);s__Metschnikowia_pulcherrima(100); | Metschnikowia | 13 | Saprotroph | \|Undefined Saprotroph\| | Yeast | NULL | Probable | As nectar-tap_saprotroph (Põlme et al. 2020); Decay sugar-rich substrates (Põlme et al. 2020) | Põlme S, et al. 2020. Fungal Diversity 105:1-16 ((https://doi.org/10.1007/s13225-020-00466-2)) |
| Otu47124 | 1 | k__Fungi(100);p__Ascomycota(100);c__Saccharomycetes(100);o__Saccharomycetales(100);f__Metschnikowiaceae(100);g__Metschnikowia(100);s__Metschnikowia_pulcherrima(100); | Metschnikowia | 13 | Saprotroph | \|Undefined Saprotroph\| | Yeast | NULL | Probable | As nectar-tap_saprotroph (Põlme et al. 2020); Decay sugar-rich substrates (Põlme et al. 2020) | Põlme S, et al. 2020. Fungal Diversity 105:1-16 ((https://doi.org/10.1007/s13225-020-00466-2)) |
| Otu47125 | 1 | k__Fungi(100);p__Ascomycota(100);c__Saccharomycetes(100);o__Saccharomycetales(100);f__Metschnikowiaceae(100);g__Metschnikowia(100);s__Metschnikowia_pulcherrima(100); | Metschnikowia | 13 | Saprotroph | \|Undefined Saprotroph\| | Yeast | NULL | Probable | As nectar-tap_saprotroph (Põlme et al. 2020); Decay sugar-rich substrates (Põlme et al. 2020) | Põlme S, et al. 2020. Fungal Diversity 105:1-16 ((https://doi.org/10.1007/s13225-020-00466-2)) |
| Otu47126 | 1 | k__Fungi(100);p__Ascomycota(100);c__Saccharomycetes(100);o__Saccharomycetales(100);f__Metschnikowiaceae(100);g__Metschnikowia(100);s__Metschnikowia_pulcherrima(100); | Metschnikowia | 13 | Saprotroph | \|Undefined Saprotroph\| | Yeast | NULL | Probable | As nectar-tap_saprotroph (Põlme et al. 2020); Decay sugar-rich substrates (Põlme et al. 2020) | Põlme S, et al. 2020. Fungal Diversity 105:1-16 ((https://doi.org/10.1007/s13225-020-00466-2)) |
| Otu47127 | 1 | k__Fungi(100);p__Ascomycota(100);c__Saccharomycetes(100);o__Saccharomycetales(100);f__Metschnikowiaceae(100);g__Metschnikowia(100);s__Metschnikowia_pulcherrima(100); | Metschnikowia | 13 | Saprotroph | \|Undefined Saprotroph\| | Yeast | NULL | Probable | As nectar-tap_saprotroph (Põlme et al. 2020); Decay sugar-rich substrates (Põlme et al. 2020) | Põlme S, et al. 2020. Fungal Diversity 105:1-16 ((https://doi.org/10.1007/s13225-020-00466-2)) |
| Otu47128 | 1 | k__Fungi(100);p__Ascomycota(100);c__Saccharomycetes(100);o__Saccharomycetales(100);f__Metschnikowiaceae(100);g__Metschnikowia(100);s__Metschnikowia_pulcherrima(100); | Metschnikowia | 13 | Saprotroph | \|Undefined Saprotroph\| | Yeast | NULL | Probable | As nectar-tap_saprotroph (Põlme et al. 2020); Decay sugar-rich substrates (Põlme et al. 2020) | Põlme S, et al. 2020. Fungal Diversity 105:1-16 ((https://doi.org/10.1007/s13225-020-00466-2)) |
| Otu47129 | 1 | k__Fungi(100);p__Ascomycota(100);c__Saccharomycetes(100);o__Saccharomycetales(100);f__Metschnikowiaceae(100);g__Metschnikowia(100);s__Metschnikowia_pulcherrima(100); | Metschnikowia | 13 | Saprotroph | \|Undefined Saprotroph\| | Yeast | NULL | Probable | As nectar-tap_saprotroph (Põlme et al. 2020); Decay sugar-rich substrates (Põlme et al. 2020) | Põlme S, et al. 2020. Fungal Diversity 105:1-16 ((https://doi.org/10.1007/s13225-020-00466-2)) |
| Otu47130 | 1 | k__Fungi(100);p__Ascomycota(100);c__Saccharomycetes(100);o__Saccharomycetales(100);f__Metschnikowiaceae(100);g__Metschnikowia(100);s__Metschnikowia_pulcherrima(100); | Metschnikowia | 13 | Saprotroph | \|Undefined Saprotroph\| | Yeast | NULL | Probable | As nectar-tap_saprotroph (Põlme et al. 2020); Decay sugar-rich substrates (Põlme et al. 2020) | Põlme S, et al. 2020. Fungal Diversity 105:1-16 ((https://doi.org/10.1007/s13225-020-00466-2)) |
| Otu47131 | 1 | k__Fungi(100);p__Ascomycota(100);c__Saccharomycetes(100);o__Saccharomycetales(100);f__Metschnikowiaceae(100);g__Metschnikowia(100);s__Metschnikowia_pulcherrima(100); | Metschnikowia | 13 | Saprotroph | \|Undefined Saprotroph\| | Yeast | NULL | Probable | As nectar-tap_saprotroph (Põlme et al. 2020); Decay sugar-rich substrates (Põlme et al. 2020) | Põlme S, et al. 2020. Fungal Diversity 105:1-16 ((https://doi.org/10.1007/s13225-020-00466-2)) |
| Otu47132 | 1 | k__Fungi(100);p__Ascomycota(100);c__Saccharomycetes(100);o__Saccharomycetales(100);f__Metschnikowiaceae(100);g__Metschnikowia(100);s__Metschnikowia_pulcherrima(100); | Metschnikowia | 13 | Saprotroph | \|Undefined Saprotroph\| | Yeast | NULL | Probable | As nectar-tap_saprotroph (Põlme et al. 2020); Decay sugar-rich substrates (Põlme et al. 2020) | Põlme S, et al. 2020. Fungal Diversity 105:1-16 ((https://doi.org/10.1007/s13225-020-00466-2)) |
| Otu47133 | 1 | k__Fungi(100);p__Ascomycota(100);c__Saccharomycetes(100);o__Saccharomycetales(100);f__Metschnikowiaceae(100);g__Metschnikowia(100);s__Metschnikowia_pulcherrima(100); | Metschnikowia | 13 | Saprotroph | \|Undefined Saprotroph\| | Yeast | NULL | Probable | As nectar-tap_saprotroph (Põlme et al. 2020); Decay sugar-rich substrates (Põlme et al. 2020) | Põlme S, et al. 2020. Fungal Diversity 105:1-16 ((https://doi.org/10.1007/s13225-020-00466-2)) |
| Otu47134 | 1 | k__Fungi(100);p__Ascomycota(100);c__Saccharomycetes(100);o__Saccharomycetales(100);f__Metschnikowiaceae(100);g__Metschnikowia(100);g__Metschnikowia_unclassified(100); | Metschnikowia | 13 | Saprotroph | \|Undefined Saprotroph\| | Yeast | NULL | Probable | As nectar-tap_saprotroph (Põlme et al. 2020); Decay sugar-rich substrates (Põlme et al. 2020) | Põlme S, et al. 2020. Fungal Diversity 105:1-16 ((https://doi.org/10.1007/s13225-020-00466-2)) |
| Otu47135 | 1 | k__Fungi(100);p__Ascomycota(100);c__Saccharomycetes(100);o__Saccharomycetales(100);f__Metschnikowiaceae(100);g__Metschnikowia(100);s__Metschnikowia_pulcherrima(100); | Metschnikowia | 13 | Saprotroph | \|Undefined Saprotroph\| | Yeast | NULL | Probable | As nectar-tap_saprotroph (Põlme et al. 2020); Decay sugar-rich substrates (Põlme et al. 2020) | Põlme S, et al. 2020. Fungal Diversity 105:1-16 ((https://doi.org/10.1007/s13225-020-00466-2)) |
| Otu47136 | 1 | k__Fungi(100);p__Ascomycota(100);c__Saccharomycetes(100);o__Saccharomycetales(100);f__Metschnikowiaceae(100);g__Metschnikowia(100);s__Metschnikowia_pulcherrima(100); | Metschnikowia | 13 | Saprotroph | \|Undefined Saprotroph\| | Yeast | NULL | Probable | As nectar-tap_saprotroph (Põlme et al. 2020); Decay sugar-rich substrates (Põlme et al. 2020) | Põlme S, et al. 2020. Fungal Diversity 105:1-16 ((https://doi.org/10.1007/s13225-020-00466-2)) |
| Otu47138 | 1 | k__Fungi(100);p__Ascomycota(100);c__Saccharomycetes(100);o__Saccharomycetales(100);f__Metschnikowiaceae(100);g__Metschnikowia(100);s__Metschnikowia_pulcherrima(100); | Metschnikowia | 13 | Saprotroph | \|Undefined Saprotroph\| | Yeast | NULL | Probable | As nectar-tap_saprotroph (Põlme et al. 2020); Decay sugar-rich substrates (Põlme et al. 2020) | Põlme S, et al. 2020. Fungal Diversity 105:1-16 ((https://doi.org/10.1007/s13225-020-00466-2)) |
| Otu47139 | 1 | k__Fungi(100);p__Ascomycota(100);c__Saccharomycetes(100);o__Saccharomycetales(100);f__Metschnikowiaceae(100);g__Metschnikowia(100);s__Metschnikowia_pulcherrima(100); | Metschnikowia | 13 | Saprotroph | \|Undefined Saprotroph\| | Yeast | NULL | Probable | As nectar-tap_saprotroph (Põlme et al. 2020); Decay sugar-rich substrates (Põlme et al. 2020) | Põlme S, et al. 2020. Fungal Diversity 105:1-16 ((https://doi.org/10.1007/s13225-020-00466-2)) |
| Otu47140 | 1 | k__Fungi(100);p__Ascomycota(100);c__Saccharomycetes(100);o__Saccharomycetales(100);f__Metschnikowiaceae(100);g__Metschnikowia(100);s__Metschnikowia_pulcherrima(100); | Metschnikowia | 13 | Saprotroph | \|Undefined Saprotroph\| | Yeast | NULL | Probable | As nectar-tap_saprotroph (Põlme et al. 2020); Decay sugar-rich substrates (Põlme et al. 2020) | Põlme S, et al. 2020. Fungal Diversity 105:1-16 ((https://doi.org/10.1007/s13225-020-00466-2)) |
| Otu47141 | 1 | k__Fungi(100);p__Ascomycota(100);c__Saccharomycetes(100);o__Saccharomycetales(100);f__Metschnikowiaceae(100);g__Metschnikowia(100);g__Metschnikowia_unclassified(100); | Metschnikowia | 13 | Saprotroph | \|Undefined Saprotroph\| | Yeast | NULL | Probable | As nectar-tap_saprotroph (Põlme et al. 2020); Decay sugar-rich substrates (Põlme et al. 2020) | Põlme S, et al. 2020. Fungal Diversity 105:1-16 ((https://doi.org/10.1007/s13225-020-00466-2)) |
| Otu47144 | 1 | k__Fungi(100);p__Ascomycota(100);c__Saccharomycetes(100);o__Saccharomycetales(100);f__Metschnikowiaceae(100);g__Metschnikowia(100);s__Metschnikowia_pulcherrima(100); | Metschnikowia | 13 | Saprotroph | \|Undefined Saprotroph\| | Yeast | NULL | Probable | As nectar-tap_saprotroph (Põlme et al. 2020); Decay sugar-rich substrates (Põlme et al. 2020) | Põlme S, et al. 2020. Fungal Diversity 105:1-16 ((https://doi.org/10.1007/s13225-020-00466-2)) |
| Otu47146 | 1 | k__Fungi(100);p__Ascomycota(100);c__Saccharomycetes(100);o__Saccharomycetales(100);f__Metschnikowiaceae(100);g__Metschnikowia(100);s__Metschnikowia_pulcherrima(100); | Metschnikowia | 13 | Saprotroph | \|Undefined Saprotroph\| | Yeast | NULL | Probable | As nectar-tap_saprotroph (Põlme et al. 2020); Decay sugar-rich substrates (Põlme et al. 2020) | Põlme S, et al. 2020. Fungal Diversity 105:1-16 ((https://doi.org/10.1007/s13225-020-00466-2)) |
| Otu47147 | 1 | k__Fungi(100);p__Ascomycota(100);c__Saccharomycetes(100);o__Saccharomycetales(100);f__Metschnikowiaceae(100);g__Metschnikowia(100);s__Metschnikowia_pulcherrima(100); | Metschnikowia | 13 | Saprotroph | \|Undefined Saprotroph\| | Yeast | NULL | Probable | As nectar-tap_saprotroph (Põlme et al. 2020); Decay sugar-rich substrates (Põlme et al. 2020) | Põlme S, et al. 2020. Fungal Diversity 105:1-16 ((https://doi.org/10.1007/s13225-020-00466-2)) |
| Otu47148 | 1 | k__Fungi(100);p__Ascomycota(100);c__Saccharomycetes(100);o__Saccharomycetales(100);f__Metschnikowiaceae(100);g__Metschnikowia(100);s__Metschnikowia_pulcherrima(100); | Metschnikowia | 13 | Saprotroph | \|Undefined Saprotroph\| | Yeast | NULL | Probable | As nectar-tap_saprotroph (Põlme et al. 2020); Decay sugar-rich substrates (Põlme et al. 2020) | Põlme S, et al. 2020. Fungal Diversity 105:1-16 ((https://doi.org/10.1007/s13225-020-00466-2)) |
| Otu47149 | 1 | k__Fungi(100);p__Ascomycota(100);c__Saccharomycetes(100);o__Saccharomycetales(100);f__Metschnikowiaceae(100);g__Metschnikowia(100);s__Metschnikowia_pulcherrima(100); | Metschnikowia | 13 | Saprotroph | \|Undefined Saprotroph\| | Yeast | NULL | Probable | As nectar-tap_saprotroph (Põlme et al. 2020); Decay sugar-rich substrates (Põlme et al. 2020) | Põlme S, et al. 2020. Fungal Diversity 105:1-16 ((https://doi.org/10.1007/s13225-020-00466-2)) |
| Otu47150 | 1 | k__Fungi(100);p__Ascomycota(100);c__Saccharomycetes(100);o__Saccharomycetales(100);f__Metschnikowiaceae(100);g__Metschnikowia(100);g__Metschnikowia_unclassified(100); | Metschnikowia | 13 | Saprotroph | \|Undefined Saprotroph\| | Yeast | NULL | Probable | As nectar-tap_saprotroph (Põlme et al. 2020); Decay sugar-rich substrates (Põlme et al. 2020) | Põlme S, et al. 2020. Fungal Diversity 105:1-16 ((https://doi.org/10.1007/s13225-020-00466-2)) |
| Otu47151 | 1 | k__Fungi(100);p__Ascomycota(100);c__Saccharomycetes(100);o__Saccharomycetales(100);f__Metschnikowiaceae(100);g__Metschnikowia(100);s__Metschnikowia_pulcherrima(100); | Metschnikowia | 13 | Saprotroph | \|Undefined Saprotroph\| | Yeast | NULL | Probable | As nectar-tap_saprotroph (Põlme et al. 2020); Decay sugar-rich substrates (Põlme et al. 2020) | Põlme S, et al. 2020. Fungal Diversity 105:1-16 ((https://doi.org/10.1007/s13225-020-00466-2)) |
| Otu47152 | 1 | k__Fungi(100);p__Ascomycota(100);c__Saccharomycetes(100);o__Saccharomycetales(100);f__Metschnikowiaceae(100);g__Metschnikowia(100);s__Metschnikowia_pulcherrima(100); | Metschnikowia | 13 | Saprotroph | \|Undefined Saprotroph\| | Yeast | NULL | Probable | As nectar-tap_saprotroph (Põlme et al. 2020); Decay sugar-rich substrates (Põlme et al. 2020) | Põlme S, et al. 2020. Fungal Diversity 105:1-16 ((https://doi.org/10.1007/s13225-020-00466-2)) |
| Otu47153 | 1 | k__Fungi(100);p__Ascomycota(100);c__Saccharomycetes(100);o__Saccharomycetales(100);f__Metschnikowiaceae(100);g__Metschnikowia(100);g__Metschnikowia_unclassified(100); | Metschnikowia | 13 | Saprotroph | \|Undefined Saprotroph\| | Yeast | NULL | Probable | As nectar-tap_saprotroph (Põlme et al. 2020); Decay sugar-rich substrates (Põlme et al. 2020) | Põlme S, et al. 2020. Fungal Diversity 105:1-16 ((https://doi.org/10.1007/s13225-020-00466-2)) |
| Otu47155 | 1 | k__Fungi(100);p__Ascomycota(100);c__Saccharomycetes(100);o__Saccharomycetales(100);f__Metschnikowiaceae(100);g__Metschnikowia(100);s__Metschnikowia_pulcherrima(100); | Metschnikowia | 13 | Saprotroph | \|Undefined Saprotroph\| | Yeast | NULL | Probable | As nectar-tap_saprotroph (Põlme et al. 2020); Decay sugar-rich substrates (Põlme et al. 2020) | Põlme S, et al. 2020. Fungal Diversity 105:1-16 ((https://doi.org/10.1007/s13225-020-00466-2)) |
| Otu47156 | 1 | k__Fungi(100);p__Ascomycota(100);c__Saccharomycetes(100);o__Saccharomycetales(100);f__Metschnikowiaceae(100);g__Metschnikowia(100);s__Metschnikowia_pulcherrima(100); | Metschnikowia | 13 | Saprotroph | \|Undefined Saprotroph\| | Yeast | NULL | Probable | As nectar-tap_saprotroph (Põlme et al. 2020); Decay sugar-rich substrates (Põlme et al. 2020) | Põlme S, et al. 2020. Fungal Diversity 105:1-16 ((https://doi.org/10.1007/s13225-020-00466-2)) |
| Otu47157 | 1 | k__Fungi(100);p__Ascomycota(100);c__Saccharomycetes(100);o__Saccharomycetales(100);f__Metschnikowiaceae(100);g__Metschnikowia(100);s__Metschnikowia_pulcherrima(100); | Metschnikowia | 13 | Saprotroph | \|Undefined Saprotroph\| | Yeast | NULL | Probable | As nectar-tap_saprotroph (Põlme et al. 2020); Decay sugar-rich substrates (Põlme et al. 2020) | Põlme S, et al. 2020. Fungal Diversity 105:1-16 ((https://doi.org/10.1007/s13225-020-00466-2)) |
| Otu47158 | 1 | k__Fungi(100);p__Ascomycota(100);c__Saccharomycetes(100);o__Saccharomycetales(100);f__Metschnikowiaceae(100);g__Metschnikowia(100);s__Metschnikowia_pulcherrima(100); | Metschnikowia | 13 | Saprotroph | \|Undefined Saprotroph\| | Yeast | NULL | Probable | As nectar-tap_saprotroph (Põlme et al. 2020); Decay sugar-rich substrates (Põlme et al. 2020) | Põlme S, et al. 2020. Fungal Diversity 105:1-16 ((https://doi.org/10.1007/s13225-020-00466-2)) |
| Otu47159 | 1 | k__Fungi(100);p__Ascomycota(100);c__Saccharomycetes(100);o__Saccharomycetales(100);f__Metschnikowiaceae(100);g__Metschnikowia(100);s__Metschnikowia_pulcherrima(100); | Metschnikowia | 13 | Saprotroph | \|Undefined Saprotroph\| | Yeast | NULL | Probable | As nectar-tap_saprotroph (Põlme et al. 2020); Decay sugar-rich substrates (Põlme et al. 2020) | Põlme S, et al. 2020. Fungal Diversity 105:1-16 ((https://doi.org/10.1007/s13225-020-00466-2)) |
| Otu47160 | 1 | k__Fungi(100);p__Ascomycota(100);c__Saccharomycetes(100);o__Saccharomycetales(100);f__Metschnikowiaceae(100);g__Metschnikowia(100);s__Metschnikowia_pulcherrima(100); | Metschnikowia | 13 | Saprotroph | \|Undefined Saprotroph\| | Yeast | NULL | Probable | As nectar-tap_saprotroph (Põlme et al. 2020); Decay sugar-rich substrates (Põlme et al. 2020) | Põlme S, et al. 2020. Fungal Diversity 105:1-16 ((https://doi.org/10.1007/s13225-020-00466-2)) |
| Otu47161 | 1 | k__Fungi(100);p__Ascomycota(100);c__Saccharomycetes(100);o__Saccharomycetales(100);f__Metschnikowiaceae(100);g__Metschnikowia(100);s__Metschnikowia_pulcherrima(100); | Metschnikowia | 13 | Saprotroph | \|Undefined Saprotroph\| | Yeast | NULL | Probable | As nectar-tap_saprotroph (Põlme et al. 2020); Decay sugar-rich substrates (Põlme et al. 2020) | Põlme S, et al. 2020. Fungal Diversity 105:1-16 ((https://doi.org/10.1007/s13225-020-00466-2)) |
| Otu47162 | 1 | k__Fungi(100);p__Ascomycota(100);c__Saccharomycetes(100);o__Saccharomycetales(100);f__Metschnikowiaceae(100);g__Metschnikowia(100);s__Metschnikowia_pulcherrima(100); | Metschnikowia | 13 | Saprotroph | \|Undefined Saprotroph\| | Yeast | NULL | Probable | As nectar-tap_saprotroph (Põlme et al. 2020); Decay sugar-rich substrates (Põlme et al. 2020) | Põlme S, et al. 2020. Fungal Diversity 105:1-16 ((https://doi.org/10.1007/s13225-020-00466-2)) |
| Otu47163 | 1 | k__Fungi(100);p__Ascomycota(100);c__Saccharomycetes(100);o__Saccharomycetales(100);f__Metschnikowiaceae(100);g__Metschnikowia(100);s__Metschnikowia_pulcherrima(100); | Metschnikowia | 13 | Saprotroph | \|Undefined Saprotroph\| | Yeast | NULL | Probable | As nectar-tap_saprotroph (Põlme et al. 2020); Decay sugar-rich substrates (Põlme et al. 2020) | Põlme S, et al. 2020. Fungal Diversity 105:1-16 ((https://doi.org/10.1007/s13225-020-00466-2)) |
| Otu47164 | 1 | k__Fungi(100);p__Ascomycota(100);c__Saccharomycetes(100);o__Saccharomycetales(100);f__Metschnikowiaceae(100);g__Metschnikowia(100);s__Metschnikowia_pulcherrima(100); | Metschnikowia | 13 | Saprotroph | \|Undefined Saprotroph\| | Yeast | NULL | Probable | As nectar-tap_saprotroph (Põlme et al. 2020); Decay sugar-rich substrates (Põlme et al. 2020) | Põlme S, et al. 2020. Fungal Diversity 105:1-16 ((https://doi.org/10.1007/s13225-020-00466-2)) |
| Otu47165 | 1 | k__Fungi(100);p__Ascomycota(100);c__Saccharomycetes(100);o__Saccharomycetales(100);f__Metschnikowiaceae(100);g__Metschnikowia(100);s__Metschnikowia_pulcherrima(100); | Metschnikowia | 13 | Saprotroph | \|Undefined Saprotroph\| | Yeast | NULL | Probable | As nectar-tap_saprotroph (Põlme et al. 2020); Decay sugar-rich substrates (Põlme et al. 2020) | Põlme S, et al. 2020. Fungal Diversity 105:1-16 ((https://doi.org/10.1007/s13225-020-00466-2)) |
| Otu47166 | 1 | k__Fungi(100);p__Ascomycota(100);c__Saccharomycetes(100);o__Saccharomycetales(100);f__Metschnikowiaceae(100);g__Metschnikowia(100);g__Metschnikowia_unclassified(100); | Metschnikowia | 13 | Saprotroph | \|Undefined Saprotroph\| | Yeast | NULL | Probable | As nectar-tap_saprotroph (Põlme et al. 2020); Decay sugar-rich substrates (Põlme et al. 2020) | Põlme S, et al. 2020. Fungal Diversity 105:1-16 ((https://doi.org/10.1007/s13225-020-00466-2)) |
| Otu47167 | 1 | k__Fungi(100);p__Ascomycota(100);c__Saccharomycetes(100);o__Saccharomycetales(100);f__Metschnikowiaceae(100);g__Metschnikowia(100);s__Metschnikowia_pulcherrima(100); | Metschnikowia | 13 | Saprotroph | \|Undefined Saprotroph\| | Yeast | NULL | Probable | As nectar-tap_saprotroph (Põlme et al. 2020); Decay sugar-rich substrates (Põlme et al. 2020) | Põlme S, et al. 2020. Fungal Diversity 105:1-16 ((https://doi.org/10.1007/s13225-020-00466-2)) |
| Otu47168 | 1 | k__Fungi(100);p__Ascomycota(100);c__Saccharomycetes(100);o__Saccharomycetales(100);f__Metschnikowiaceae(100);g__Metschnikowia(100);s__Metschnikowia_pulcherrima(100); | Metschnikowia | 13 | Saprotroph | \|Undefined Saprotroph\| | Yeast | NULL | Probable | As nectar-tap_saprotroph (Põlme et al. 2020); Decay sugar-rich substrates (Põlme et al. 2020) | Põlme S, et al. 2020. Fungal Diversity 105:1-16 ((https://doi.org/10.1007/s13225-020-00466-2)) |
| Otu47169 | 1 | k__Fungi(100);p__Ascomycota(100);c__Saccharomycetes(100);o__Saccharomycetales(100);f__Metschnikowiaceae(100);g__Metschnikowia(100);s__Metschnikowia_pulcherrima(100); | Metschnikowia | 13 | Saprotroph | \|Undefined Saprotroph\| | Yeast | NULL | Probable | As nectar-tap_saprotroph (Põlme et al. 2020); Decay sugar-rich substrates (Põlme et al. 2020) | Põlme S, et al. 2020. Fungal Diversity 105:1-16 ((https://doi.org/10.1007/s13225-020-00466-2)) |
| Otu47170 | 1 | k__Fungi(100);p__Ascomycota(100);c__Saccharomycetes(100);o__Saccharomycetales(100);f__Metschnikowiaceae(100);g__Metschnikowia(100);g__Metschnikowia_unclassified(100); | Metschnikowia | 13 | Saprotroph | \|Undefined Saprotroph\| | Yeast | NULL | Probable | As nectar-tap_saprotroph (Põlme et al. 2020); Decay sugar-rich substrates (Põlme et al. 2020) | Põlme S, et al. 2020. Fungal Diversity 105:1-16 ((https://doi.org/10.1007/s13225-020-00466-2)) |
| Otu47171 | 1 | k__Fungi(100);p__Ascomycota(100);c__Saccharomycetes(100);o__Saccharomycetales(100);f__Metschnikowiaceae(100);g__Metschnikowia(100);s__Metschnikowia_pulcherrima(100); | Metschnikowia | 13 | Saprotroph | \|Undefined Saprotroph\| | Yeast | NULL | Probable | As nectar-tap_saprotroph (Põlme et al. 2020); Decay sugar-rich substrates (Põlme et al. 2020) | Põlme S, et al. 2020. Fungal Diversity 105:1-16 ((https://doi.org/10.1007/s13225-020-00466-2)) |
| Otu47172 | 1 | k__Fungi(100);p__Ascomycota(100);c__Saccharomycetes(100);o__Saccharomycetales(100);f__Metschnikowiaceae(100);g__Metschnikowia(100);s__Metschnikowia_pulcherrima(100); | Metschnikowia | 13 | Saprotroph | \|Undefined Saprotroph\| | Yeast | NULL | Probable | As nectar-tap_saprotroph (Põlme et al. 2020); Decay sugar-rich substrates (Põlme et al. 2020) | Põlme S, et al. 2020. Fungal Diversity 105:1-16 ((https://doi.org/10.1007/s13225-020-00466-2)) |
| Otu47173 | 1 | k__Fungi(100);p__Ascomycota(100);c__Saccharomycetes(100);o__Saccharomycetales(100);f__Metschnikowiaceae(100);g__Metschnikowia(100);s__Metschnikowia_pulcherrima(100); | Metschnikowia | 13 | Saprotroph | \|Undefined Saprotroph\| | Yeast | NULL | Probable | As nectar-tap_saprotroph (Põlme et al. 2020); Decay sugar-rich substrates (Põlme et al. 2020) | Põlme S, et al. 2020. Fungal Diversity 105:1-16 ((https://doi.org/10.1007/s13225-020-00466-2)) |
| Otu47174 | 1 | k__Fungi(100);p__Ascomycota(100);c__Saccharomycetes(100);o__Saccharomycetales(100);f__Metschnikowiaceae(100);g__Metschnikowia(100);s__Metschnikowia_pulcherrima(100); | Metschnikowia | 13 | Saprotroph | \|Undefined Saprotroph\| | Yeast | NULL | Probable | As nectar-tap_saprotroph (Põlme et al. 2020); Decay sugar-rich substrates (Põlme et al. 2020) | Põlme S, et al. 2020. Fungal Diversity 105:1-16 ((https://doi.org/10.1007/s13225-020-00466-2)) |
| Otu47175 | 1 | k__Fungi(100);p__Ascomycota(100);c__Saccharomycetes(100);o__Saccharomycetales(100);f__Metschnikowiaceae(100);g__Metschnikowia(100);g__Metschnikowia_unclassified(100); | Metschnikowia | 13 | Saprotroph | \|Undefined Saprotroph\| | Yeast | NULL | Probable | As nectar-tap_saprotroph (Põlme et al. 2020); Decay sugar-rich substrates (Põlme et al. 2020) | Põlme S, et al. 2020. Fungal Diversity 105:1-16 ((https://doi.org/10.1007/s13225-020-00466-2)) |
| Otu47176 | 1 | k__Fungi(100);p__Ascomycota(100);c__Saccharomycetes(100);o__Saccharomycetales(100);f__Metschnikowiaceae(100);g__Metschnikowia(100);s__Metschnikowia_pulcherrima(100); | Metschnikowia | 13 | Saprotroph | \|Undefined Saprotroph\| | Yeast | NULL | Probable | As nectar-tap_saprotroph (Põlme et al. 2020); Decay sugar-rich substrates (Põlme et al. 2020) | Põlme S, et al. 2020. Fungal Diversity 105:1-16 ((https://doi.org/10.1007/s13225-020-00466-2)) |
| Otu47177 | 1 | k__Fungi(100);p__Ascomycota(100);c__Saccharomycetes(100);o__Saccharomycetales(100);f__Metschnikowiaceae(100);g__Metschnikowia(100);g__Metschnikowia_unclassified(100); | Metschnikowia | 13 | Saprotroph | \|Undefined Saprotroph\| | Yeast | NULL | Probable | As nectar-tap_saprotroph (Põlme et al. 2020); Decay sugar-rich substrates (Põlme et al. 2020) | Põlme S, et al. 2020. Fungal Diversity 105:1-16 ((https://doi.org/10.1007/s13225-020-00466-2)) |
| Otu47178 | 1 | k__Fungi(100);p__Ascomycota(100);c__Saccharomycetes(100);o__Saccharomycetales(100);f__Metschnikowiaceae(100);g__Metschnikowia(100);s__Metschnikowia_pulcherrima(100); | Metschnikowia | 13 | Saprotroph | \|Undefined Saprotroph\| | Yeast | NULL | Probable | As nectar-tap_saprotroph (Põlme et al. 2020); Decay sugar-rich substrates (Põlme et al. 2020) | Põlme S, et al. 2020. Fungal Diversity 105:1-16 ((https://doi.org/10.1007/s13225-020-00466-2)) |
| Otu47179 | 1 | k__Fungi(100);p__Ascomycota(100);c__Saccharomycetes(100);o__Saccharomycetales(100);f__Metschnikowiaceae(100);g__Metschnikowia(100);s__Metschnikowia_pulcherrima(100); | Metschnikowia | 13 | Saprotroph | \|Undefined Saprotroph\| | Yeast | NULL | Probable | As nectar-tap_saprotroph (Põlme et al. 2020); Decay sugar-rich substrates (Põlme et al. 2020) | Põlme S, et al. 2020. Fungal Diversity 105:1-16 ((https://doi.org/10.1007/s13225-020-00466-2)) |
| Otu47180 | 1 | k__Fungi(100);p__Ascomycota(100);c__Saccharomycetes(100);o__Saccharomycetales(100);f__Metschnikowiaceae(100);g__Metschnikowia(100);s__Metschnikowia_pulcherrima(100); | Metschnikowia | 13 | Saprotroph | \|Undefined Saprotroph\| | Yeast | NULL | Probable | As nectar-tap_saprotroph (Põlme et al. 2020); Decay sugar-rich substrates (Põlme et al. 2020) | Põlme S, et al. 2020. Fungal Diversity 105:1-16 ((https://doi.org/10.1007/s13225-020-00466-2)) |
| Otu47182 | 1 | k__Fungi(100);p__Ascomycota(100);c__Saccharomycetes(100);o__Saccharomycetales(100);f__Metschnikowiaceae(100);g__Metschnikowia(100);s__Metschnikowia_pulcherrima(100); | Metschnikowia | 13 | Saprotroph | \|Undefined Saprotroph\| | Yeast | NULL | Probable | As nectar-tap_saprotroph (Põlme et al. 2020); Decay sugar-rich substrates (Põlme et al. 2020) | Põlme S, et al. 2020. Fungal Diversity 105:1-16 ((https://doi.org/10.1007/s13225-020-00466-2)) |
| Otu47183 | 1 | k__Fungi(100);p__Ascomycota(100);c__Saccharomycetes(100);o__Saccharomycetales(100);f__Metschnikowiaceae(100);g__Metschnikowia(100);s__Metschnikowia_pulcherrima(100); | Metschnikowia | 13 | Saprotroph | \|Undefined Saprotroph\| | Yeast | NULL | Probable | As nectar-tap_saprotroph (Põlme et al. 2020); Decay sugar-rich substrates (Põlme et al. 2020) | Põlme S, et al. 2020. Fungal Diversity 105:1-16 ((https://doi.org/10.1007/s13225-020-00466-2)) |
| Otu47184 | 1 | k__Fungi(100);p__Ascomycota(100);c__Saccharomycetes(100);o__Saccharomycetales(100);f__Metschnikowiaceae(100);g__Metschnikowia(100);g__Metschnikowia_unclassified(100); | Metschnikowia | 13 | Saprotroph | \|Undefined Saprotroph\| | Yeast | NULL | Probable | As nectar-tap_saprotroph (Põlme et al. 2020); Decay sugar-rich substrates (Põlme et al. 2020) | Põlme S, et al. 2020. Fungal Diversity 105:1-16 ((https://doi.org/10.1007/s13225-020-00466-2)) |
| Otu47185 | 1 | k__Fungi(100);p__Ascomycota(100);c__Saccharomycetes(100);o__Saccharomycetales(100);f__Metschnikowiaceae(100);g__Metschnikowia(100);s__Metschnikowia_pulcherrima(100); | Metschnikowia | 13 | Saprotroph | \|Undefined Saprotroph\| | Yeast | NULL | Probable | As nectar-tap_saprotroph (Põlme et al. 2020); Decay sugar-rich substrates (Põlme et al. 2020) | Põlme S, et al. 2020. Fungal Diversity 105:1-16 ((https://doi.org/10.1007/s13225-020-00466-2)) |
| Otu47186 | 1 | k__Fungi(100);p__Ascomycota(100);c__Saccharomycetes(100);o__Saccharomycetales(100);f__Metschnikowiaceae(100);g__Metschnikowia(100);s__Metschnikowia_pulcherrima(100); | Metschnikowia | 13 | Saprotroph | \|Undefined Saprotroph\| | Yeast | NULL | Probable | As nectar-tap_saprotroph (Põlme et al. 2020); Decay sugar-rich substrates (Põlme et al. 2020) | Põlme S, et al. 2020. Fungal Diversity 105:1-16 ((https://doi.org/10.1007/s13225-020-00466-2)) |
| Otu47189 | 1 | k__Fungi(100);p__Ascomycota(100);c__Saccharomycetes(100);o__Saccharomycetales(100);f__Metschnikowiaceae(100);g__Metschnikowia(100);s__Metschnikowia_pulcherrima(100); | Metschnikowia | 13 | Saprotroph | \|Undefined Saprotroph\| | Yeast | NULL | Probable | As nectar-tap_saprotroph (Põlme et al. 2020); Decay sugar-rich substrates (Põlme et al. 2020) | Põlme S, et al. 2020. Fungal Diversity 105:1-16 ((https://doi.org/10.1007/s13225-020-00466-2)) |
| Otu47190 | 1 | k__Fungi(100);p__Ascomycota(100);c__Saccharomycetes(100);o__Saccharomycetales(100);f__Metschnikowiaceae(100);g__Metschnikowia(100);g__Metschnikowia_unclassified(100); | Metschnikowia | 13 | Saprotroph | \|Undefined Saprotroph\| | Yeast | NULL | Probable | As nectar-tap_saprotroph (Põlme et al. 2020); Decay sugar-rich substrates (Põlme et al. 2020) | Põlme S, et al. 2020. Fungal Diversity 105:1-16 ((https://doi.org/10.1007/s13225-020-00466-2)) |
| Otu47191 | 1 | k__Fungi(100);p__Ascomycota(100);c__Saccharomycetes(100);o__Saccharomycetales(100);f__Metschnikowiaceae(100);g__Metschnikowia(100);s__Metschnikowia_pulcherrima(100); | Metschnikowia | 13 | Saprotroph | \|Undefined Saprotroph\| | Yeast | NULL | Probable | As nectar-tap_saprotroph (Põlme et al. 2020); Decay sugar-rich substrates (Põlme et al. 2020) | Põlme S, et al. 2020. Fungal Diversity 105:1-16 ((https://doi.org/10.1007/s13225-020-00466-2)) |
| Otu47192 | 1 | k__Fungi(100);p__Ascomycota(100);c__Saccharomycetes(100);o__Saccharomycetales(100);f__Metschnikowiaceae(100);g__Metschnikowia(100);g__Metschnikowia_unclassified(100); | Metschnikowia | 13 | Saprotroph | \|Undefined Saprotroph\| | Yeast | NULL | Probable | As nectar-tap_saprotroph (Põlme et al. 2020); Decay sugar-rich substrates (Põlme et al. 2020) | Põlme S, et al. 2020. Fungal Diversity 105:1-16 ((https://doi.org/10.1007/s13225-020-00466-2)) |
| Otu47193 | 1 | k__Fungi(100);p__Ascomycota(100);c__Saccharomycetes(100);o__Saccharomycetales(100);f__Metschnikowiaceae(100);g__Metschnikowia(100);s__Metschnikowia_pulcherrima(100); | Metschnikowia | 13 | Saprotroph | \|Undefined Saprotroph\| | Yeast | NULL | Probable | As nectar-tap_saprotroph (Põlme et al. 2020); Decay sugar-rich substrates (Põlme et al. 2020) | Põlme S, et al. 2020. Fungal Diversity 105:1-16 ((https://doi.org/10.1007/s13225-020-00466-2)) |
| Otu47195 | 1 | k__Fungi(100);p__Ascomycota(100);c__Saccharomycetes(100);o__Saccharomycetales(100);f__Metschnikowiaceae(100);g__Metschnikowia(100);g__Metschnikowia_unclassified(100); | Metschnikowia | 13 | Saprotroph | \|Undefined Saprotroph\| | Yeast | NULL | Probable | As nectar-tap_saprotroph (Põlme et al. 2020); Decay sugar-rich substrates (Põlme et al. 2020) | Põlme S, et al. 2020. Fungal Diversity 105:1-16 ((https://doi.org/10.1007/s13225-020-00466-2)) |
| Otu47196 | 1 | k__Fungi(100);p__Ascomycota(100);c__Saccharomycetes(100);o__Saccharomycetales(100);f__Metschnikowiaceae(100);g__Metschnikowia(100);g__Metschnikowia_unclassified(100); | Metschnikowia | 13 | Saprotroph | \|Undefined Saprotroph\| | Yeast | NULL | Probable | As nectar-tap_saprotroph (Põlme et al. 2020); Decay sugar-rich substrates (Põlme et al. 2020) | Põlme S, et al. 2020. Fungal Diversity 105:1-16 ((https://doi.org/10.1007/s13225-020-00466-2)) |
| Otu47197 | 1 | k__Fungi(100);p__Ascomycota(100);c__Saccharomycetes(100);o__Saccharomycetales(100);f__Metschnikowiaceae(100);g__Metschnikowia(100);s__Metschnikowia_pulcherrima(100); | Metschnikowia | 13 | Saprotroph | \|Undefined Saprotroph\| | Yeast | NULL | Probable | As nectar-tap_saprotroph (Põlme et al. 2020); Decay sugar-rich substrates (Põlme et al. 2020) | Põlme S, et al. 2020. Fungal Diversity 105:1-16 ((https://doi.org/10.1007/s13225-020-00466-2)) |
| Otu47198 | 1 | k__Fungi(100);p__Ascomycota(100);c__Saccharomycetes(100);o__Saccharomycetales(100);f__Metschnikowiaceae(100);g__Metschnikowia(100);g__Metschnikowia_unclassified(100); | Metschnikowia | 13 | Saprotroph | \|Undefined Saprotroph\| | Yeast | NULL | Probable | As nectar-tap_saprotroph (Põlme et al. 2020); Decay sugar-rich substrates (Põlme et al. 2020) | Põlme S, et al. 2020. Fungal Diversity 105:1-16 ((https://doi.org/10.1007/s13225-020-00466-2)) |
| Otu47199 | 1 | k__Fungi(100);p__Ascomycota(100);c__Saccharomycetes(100);o__Saccharomycetales(100);f__Metschnikowiaceae(100);g__Metschnikowia(100);g__Metschnikowia_unclassified(100); | Metschnikowia | 13 | Saprotroph | \|Undefined Saprotroph\| | Yeast | NULL | Probable | As nectar-tap_saprotroph (Põlme et al. 2020); Decay sugar-rich substrates (Põlme et al. 2020) | Põlme S, et al. 2020. Fungal Diversity 105:1-16 ((https://doi.org/10.1007/s13225-020-00466-2)) |
| Otu47200 | 1 | k__Fungi(100);p__Ascomycota(100);c__Saccharomycetes(100);o__Saccharomycetales(100);f__Metschnikowiaceae(100);g__Metschnikowia(100);s__Metschnikowia_pulcherrima(100); | Metschnikowia | 13 | Saprotroph | \|Undefined Saprotroph\| | Yeast | NULL | Probable | As nectar-tap_saprotroph (Põlme et al. 2020); Decay sugar-rich substrates (Põlme et al. 2020) | Põlme S, et al. 2020. Fungal Diversity 105:1-16 ((https://doi.org/10.1007/s13225-020-00466-2)) |
| Otu47201 | 1 | k__Fungi(100);p__Ascomycota(100);c__Saccharomycetes(100);o__Saccharomycetales(100);f__Metschnikowiaceae(100);g__Metschnikowia(100);s__Metschnikowia_pulcherrima(100); | Metschnikowia | 13 | Saprotroph | \|Undefined Saprotroph\| | Yeast | NULL | Probable | As nectar-tap_saprotroph (Põlme et al. 2020); Decay sugar-rich substrates (Põlme et al. 2020) | Põlme S, et al. 2020. Fungal Diversity 105:1-16 ((https://doi.org/10.1007/s13225-020-00466-2)) |
| Otu47202 | 1 | k__Fungi(100);p__Ascomycota(100);c__Saccharomycetes(100);o__Saccharomycetales(100);f__Metschnikowiaceae(100);g__Metschnikowia(100);s__Metschnikowia_pulcherrima(100); | Metschnikowia | 13 | Saprotroph | \|Undefined Saprotroph\| | Yeast | NULL | Probable | As nectar-tap_saprotroph (Põlme et al. 2020); Decay sugar-rich substrates (Põlme et al. 2020) | Põlme S, et al. 2020. Fungal Diversity 105:1-16 ((https://doi.org/10.1007/s13225-020-00466-2)) |
| Otu47203 | 1 | k__Fungi(100);p__Ascomycota(100);c__Saccharomycetes(100);o__Saccharomycetales(100);f__Metschnikowiaceae(100);g__Metschnikowia(100);s__Metschnikowia_pulcherrima(100); | Metschnikowia | 13 | Saprotroph | \|Undefined Saprotroph\| | Yeast | NULL | Probable | As nectar-tap_saprotroph (Põlme et al. 2020); Decay sugar-rich substrates (Põlme et al. 2020) | Põlme S, et al. 2020. Fungal Diversity 105:1-16 ((https://doi.org/10.1007/s13225-020-00466-2)) |
| Otu47204 | 1 | k__Fungi(100);p__Ascomycota(100);c__Saccharomycetes(100);o__Saccharomycetales(100);f__Metschnikowiaceae(100);g__Metschnikowia(100);s__Metschnikowia_pulcherrima(100); | Metschnikowia | 13 | Saprotroph | \|Undefined Saprotroph\| | Yeast | NULL | Probable | As nectar-tap_saprotroph (Põlme et al. 2020); Decay sugar-rich substrates (Põlme et al. 2020) | Põlme S, et al. 2020. Fungal Diversity 105:1-16 ((https://doi.org/10.1007/s13225-020-00466-2)) |
| Otu47205 | 1 | k__Fungi(100);p__Ascomycota(100);c__Saccharomycetes(100);o__Saccharomycetales(100);f__Metschnikowiaceae(100);g__Metschnikowia(100);s__Metschnikowia_pulcherrima(100); | Metschnikowia | 13 | Saprotroph | \|Undefined Saprotroph\| | Yeast | NULL | Probable | As nectar-tap_saprotroph (Põlme et al. 2020); Decay sugar-rich substrates (Põlme et al. 2020) | Põlme S, et al. 2020. Fungal Diversity 105:1-16 ((https://doi.org/10.1007/s13225-020-00466-2)) |
| Otu47206 | 1 | k__Fungi(100);p__Ascomycota(100);c__Saccharomycetes(100);o__Saccharomycetales(100);f__Metschnikowiaceae(100);g__Metschnikowia(100);g__Metschnikowia_unclassified(100); | Metschnikowia | 13 | Saprotroph | \|Undefined Saprotroph\| | Yeast | NULL | Probable | As nectar-tap_saprotroph (Põlme et al. 2020); Decay sugar-rich substrates (Põlme et al. 2020) | Põlme S, et al. 2020. Fungal Diversity 105:1-16 ((https://doi.org/10.1007/s13225-020-00466-2)) |
| Otu47207 | 1 | k__Fungi(100);p__Ascomycota(100);c__Saccharomycetes(100);o__Saccharomycetales(100);f__Metschnikowiaceae(100);g__Metschnikowia(100);s__Metschnikowia_pulcherrima(100); | Metschnikowia | 13 | Saprotroph | \|Undefined Saprotroph\| | Yeast | NULL | Probable | As nectar-tap_saprotroph (Põlme et al. 2020); Decay sugar-rich substrates (Põlme et al. 2020) | Põlme S, et al. 2020. Fungal Diversity 105:1-16 ((https://doi.org/10.1007/s13225-020-00466-2)) |
| Otu47208 | 1 | k__Fungi(100);p__Ascomycota(100);c__Saccharomycetes(100);o__Saccharomycetales(100);f__Metschnikowiaceae(100);g__Metschnikowia(100);s__Metschnikowia_pulcherrima(100); | Metschnikowia | 13 | Saprotroph | \|Undefined Saprotroph\| | Yeast | NULL | Probable | As nectar-tap_saprotroph (Põlme et al. 2020); Decay sugar-rich substrates (Põlme et al. 2020) | Põlme S, et al. 2020. Fungal Diversity 105:1-16 ((https://doi.org/10.1007/s13225-020-00466-2)) |
| Otu47209 | 1 | k__Fungi(100);p__Ascomycota(100);c__Saccharomycetes(100);o__Saccharomycetales(100);f__Metschnikowiaceae(100);g__Metschnikowia(100);s__Metschnikowia_pulcherrima(100); | Metschnikowia | 13 | Saprotroph | \|Undefined Saprotroph\| | Yeast | NULL | Probable | As nectar-tap_saprotroph (Põlme et al. 2020); Decay sugar-rich substrates (Põlme et al. 2020) | Põlme S, et al. 2020. Fungal Diversity 105:1-16 ((https://doi.org/10.1007/s13225-020-00466-2)) |
| Otu47210 | 1 | k__Fungi(100);p__Ascomycota(100);c__Saccharomycetes(100);o__Saccharomycetales(100);f__Metschnikowiaceae(100);g__Metschnikowia(100);g__Metschnikowia_unclassified(100); | Metschnikowia | 13 | Saprotroph | \|Undefined Saprotroph\| | Yeast | NULL | Probable | As nectar-tap_saprotroph (Põlme et al. 2020); Decay sugar-rich substrates (Põlme et al. 2020) | Põlme S, et al. 2020. Fungal Diversity 105:1-16 ((https://doi.org/10.1007/s13225-020-00466-2)) |
| Otu47211 | 1 | k__Fungi(100);p__Ascomycota(100);c__Saccharomycetes(100);o__Saccharomycetales(100);f__Metschnikowiaceae(100);g__Metschnikowia(100);s__Metschnikowia_pulcherrima(100); | Metschnikowia | 13 | Saprotroph | \|Undefined Saprotroph\| | Yeast | NULL | Probable | As nectar-tap_saprotroph (Põlme et al. 2020); Decay sugar-rich substrates (Põlme et al. 2020) | Põlme S, et al. 2020. Fungal Diversity 105:1-16 ((https://doi.org/10.1007/s13225-020-00466-2)) |
| Otu47212 | 1 | k__Fungi(100);p__Ascomycota(100);c__Saccharomycetes(100);o__Saccharomycetales(100);f__Metschnikowiaceae(100);g__Metschnikowia(100);s__Metschnikowia_pulcherrima(100); | Metschnikowia | 13 | Saprotroph | \|Undefined Saprotroph\| | Yeast | NULL | Probable | As nectar-tap_saprotroph (Põlme et al. 2020); Decay sugar-rich substrates (Põlme et al. 2020) | Põlme S, et al. 2020. Fungal Diversity 105:1-16 ((https://doi.org/10.1007/s13225-020-00466-2)) |
| Otu47213 | 1 | k__Fungi(100);p__Ascomycota(100);c__Saccharomycetes(100);o__Saccharomycetales(100);f__Metschnikowiaceae(100);g__Metschnikowia(100);g__Metschnikowia_unclassified(100); | Metschnikowia | 13 | Saprotroph | \|Undefined Saprotroph\| | Yeast | NULL | Probable | As nectar-tap_saprotroph (Põlme et al. 2020); Decay sugar-rich substrates (Põlme et al. 2020) | Põlme S, et al. 2020. Fungal Diversity 105:1-16 ((https://doi.org/10.1007/s13225-020-00466-2)) |
| Otu47214 | 1 | k__Fungi(100);p__Ascomycota(100);c__Saccharomycetes(100);o__Saccharomycetales(100);f__Metschnikowiaceae(100);g__Metschnikowia(100);s__Metschnikowia_pulcherrima(100); | Metschnikowia | 13 | Saprotroph | \|Undefined Saprotroph\| | Yeast | NULL | Probable | As nectar-tap_saprotroph (Põlme et al. 2020); Decay sugar-rich substrates (Põlme et al. 2020) | Põlme S, et al. 2020. Fungal Diversity 105:1-16 ((https://doi.org/10.1007/s13225-020-00466-2)) |
| Otu47215 | 1 | k__Fungi(100);p__Ascomycota(100);c__Saccharomycetes(100);o__Saccharomycetales(100);f__Metschnikowiaceae(100);g__Metschnikowia(100);s__Metschnikowia_pulcherrima(100); | Metschnikowia | 13 | Saprotroph | \|Undefined Saprotroph\| | Yeast | NULL | Probable | As nectar-tap_saprotroph (Põlme et al. 2020); Decay sugar-rich substrates (Põlme et al. 2020) | Põlme S, et al. 2020. Fungal Diversity 105:1-16 ((https://doi.org/10.1007/s13225-020-00466-2)) |
| Otu47216 | 1 | k__Fungi(100);p__Ascomycota(100);c__Saccharomycetes(100);o__Saccharomycetales(100);f__Metschnikowiaceae(100);g__Metschnikowia(100);s__Metschnikowia_pulcherrima(100); | Metschnikowia | 13 | Saprotroph | \|Undefined Saprotroph\| | Yeast | NULL | Probable | As nectar-tap_saprotroph (Põlme et al. 2020); Decay sugar-rich substrates (Põlme et al. 2020) | Põlme S, et al. 2020. Fungal Diversity 105:1-16 ((https://doi.org/10.1007/s13225-020-00466-2)) |
| Otu47217 | 1 | k__Fungi(100);p__Ascomycota(100);c__Saccharomycetes(100);o__Saccharomycetales(100);f__Metschnikowiaceae(100);g__Metschnikowia(100);s__Metschnikowia_pulcherrima(100); | Metschnikowia | 13 | Saprotroph | \|Undefined Saprotroph\| | Yeast | NULL | Probable | As nectar-tap_saprotroph (Põlme et al. 2020); Decay sugar-rich substrates (Põlme et al. 2020) | Põlme S, et al. 2020. Fungal Diversity 105:1-16 ((https://doi.org/10.1007/s13225-020-00466-2)) |
| Otu47218 | 1 | k__Fungi(100);p__Ascomycota(100);c__Saccharomycetes(100);o__Saccharomycetales(100);f__Metschnikowiaceae(100);g__Metschnikowia(100);s__Metschnikowia_pulcherrima(100); | Metschnikowia | 13 | Saprotroph | \|Undefined Saprotroph\| | Yeast | NULL | Probable | As nectar-tap_saprotroph (Põlme et al. 2020); Decay sugar-rich substrates (Põlme et al. 2020) | Põlme S, et al. 2020. Fungal Diversity 105:1-16 ((https://doi.org/10.1007/s13225-020-00466-2)) |
| Otu47219 | 1 | k__Fungi(100);p__Ascomycota(100);c__Saccharomycetes(100);o__Saccharomycetales(100);f__Metschnikowiaceae(100);g__Metschnikowia(100);s__Metschnikowia_pulcherrima(100); | Metschnikowia | 13 | Saprotroph | \|Undefined Saprotroph\| | Yeast | NULL | Probable | As nectar-tap_saprotroph (Põlme et al. 2020); Decay sugar-rich substrates (Põlme et al. 2020) | Põlme S, et al. 2020. Fungal Diversity 105:1-16 ((https://doi.org/10.1007/s13225-020-00466-2)) |
| Otu47220 | 1 | k__Fungi(100);p__Ascomycota(100);c__Saccharomycetes(100);o__Saccharomycetales(100);f__Metschnikowiaceae(100);g__Metschnikowia(100);s__Metschnikowia_pulcherrima(100); | Metschnikowia | 13 | Saprotroph | \|Undefined Saprotroph\| | Yeast | NULL | Probable | As nectar-tap_saprotroph (Põlme et al. 2020); Decay sugar-rich substrates (Põlme et al. 2020) | Põlme S, et al. 2020. Fungal Diversity 105:1-16 ((https://doi.org/10.1007/s13225-020-00466-2)) |
| Otu47221 | 1 | k__Fungi(100);p__Ascomycota(100);c__Saccharomycetes(100);o__Saccharomycetales(100);f__Metschnikowiaceae(100);g__Metschnikowia(100);s__Metschnikowia_pulcherrima(100); | Metschnikowia | 13 | Saprotroph | \|Undefined Saprotroph\| | Yeast | NULL | Probable | As nectar-tap_saprotroph (Põlme et al. 2020); Decay sugar-rich substrates (Põlme et al. 2020) | Põlme S, et al. 2020. Fungal Diversity 105:1-16 ((https://doi.org/10.1007/s13225-020-00466-2)) |
| Otu47222 | 1 | k__Fungi(100);p__Ascomycota(100);c__Saccharomycetes(100);o__Saccharomycetales(100);f__Metschnikowiaceae(100);g__Metschnikowia(100);s__Metschnikowia_pulcherrima(100); | Metschnikowia | 13 | Saprotroph | \|Undefined Saprotroph\| | Yeast | NULL | Probable | As nectar-tap_saprotroph (Põlme et al. 2020); Decay sugar-rich substrates (Põlme et al. 2020) | Põlme S, et al. 2020. Fungal Diversity 105:1-16 ((https://doi.org/10.1007/s13225-020-00466-2)) |
| Otu47223 | 1 | k__Fungi(100);p__Ascomycota(100);c__Saccharomycetes(100);o__Saccharomycetales(100);f__Metschnikowiaceae(100);g__Metschnikowia(100);g__Metschnikowia_unclassified(100); | Metschnikowia | 13 | Saprotroph | \|Undefined Saprotroph\| | Yeast | NULL | Probable | As nectar-tap_saprotroph (Põlme et al. 2020); Decay sugar-rich substrates (Põlme et al. 2020) | Põlme S, et al. 2020. Fungal Diversity 105:1-16 ((https://doi.org/10.1007/s13225-020-00466-2)) |
| Otu47224 | 1 | k__Fungi(100);p__Ascomycota(100);c__Saccharomycetes(100);o__Saccharomycetales(100);f__Metschnikowiaceae(100);g__Metschnikowia(100);s__Metschnikowia_pulcherrima(100); | Metschnikowia | 13 | Saprotroph | \|Undefined Saprotroph\| | Yeast | NULL | Probable | As nectar-tap_saprotroph (Põlme et al. 2020); Decay sugar-rich substrates (Põlme et al. 2020) | Põlme S, et al. 2020. Fungal Diversity 105:1-16 ((https://doi.org/10.1007/s13225-020-00466-2)) |
| Otu47225 | 1 | k__Fungi(100);p__Ascomycota(100);c__Saccharomycetes(100);o__Saccharomycetales(100);f__Metschnikowiaceae(100);g__Metschnikowia(100);s__Metschnikowia_pulcherrima(100); | Metschnikowia | 13 | Saprotroph | \|Undefined Saprotroph\| | Yeast | NULL | Probable | As nectar-tap_saprotroph (Põlme et al. 2020); Decay sugar-rich substrates (Põlme et al. 2020) | Põlme S, et al. 2020. Fungal Diversity 105:1-16 ((https://doi.org/10.1007/s13225-020-00466-2)) |
| Otu47227 | 1 | k__Fungi(100);p__Ascomycota(100);c__Saccharomycetes(100);o__Saccharomycetales(100);f__Metschnikowiaceae(100);g__Metschnikowia(100);s__Metschnikowia_pulcherrima(100); | Metschnikowia | 13 | Saprotroph | \|Undefined Saprotroph\| | Yeast | NULL | Probable | As nectar-tap_saprotroph (Põlme et al. 2020); Decay sugar-rich substrates (Põlme et al. 2020) | Põlme S, et al. 2020. Fungal Diversity 105:1-16 ((https://doi.org/10.1007/s13225-020-00466-2)) |
| Otu47228 | 1 | k__Fungi(100);p__Ascomycota(100);c__Saccharomycetes(100);o__Saccharomycetales(100);f__Metschnikowiaceae(100);g__Metschnikowia(100);s__Metschnikowia_pulcherrima(100); | Metschnikowia | 13 | Saprotroph | \|Undefined Saprotroph\| | Yeast | NULL | Probable | As nectar-tap_saprotroph (Põlme et al. 2020); Decay sugar-rich substrates (Põlme et al. 2020) | Põlme S, et al. 2020. Fungal Diversity 105:1-16 ((https://doi.org/10.1007/s13225-020-00466-2)) |
| Otu47230 | 1 | k__Fungi(100);p__Ascomycota(100);c__Saccharomycetes(100);o__Saccharomycetales(100);f__Metschnikowiaceae(100);g__Metschnikowia(100);s__Metschnikowia_pulcherrima(100); | Metschnikowia | 13 | Saprotroph | \|Undefined Saprotroph\| | Yeast | NULL | Probable | As nectar-tap_saprotroph (Põlme et al. 2020); Decay sugar-rich substrates (Põlme et al. 2020) | Põlme S, et al. 2020. Fungal Diversity 105:1-16 ((https://doi.org/10.1007/s13225-020-00466-2)) |
| Otu47231 | 1 | k__Fungi(100);p__Ascomycota(100);c__Saccharomycetes(100);o__Saccharomycetales(100);f__Metschnikowiaceae(100);g__Metschnikowia(100);s__Metschnikowia_pulcherrima(100); | Metschnikowia | 13 | Saprotroph | \|Undefined Saprotroph\| | Yeast | NULL | Probable | As nectar-tap_saprotroph (Põlme et al. 2020); Decay sugar-rich substrates (Põlme et al. 2020) | Põlme S, et al. 2020. Fungal Diversity 105:1-16 ((https://doi.org/10.1007/s13225-020-00466-2)) |
| Otu47232 | 1 | k__Fungi(100);p__Ascomycota(100);c__Saccharomycetes(100);o__Saccharomycetales(100);f__Metschnikowiaceae(100);g__Metschnikowia(100);s__Metschnikowia_pulcherrima(100); | Metschnikowia | 13 | Saprotroph | \|Undefined Saprotroph\| | Yeast | NULL | Probable | As nectar-tap_saprotroph (Põlme et al. 2020); Decay sugar-rich substrates (Põlme et al. 2020) | Põlme S, et al. 2020. Fungal Diversity 105:1-16 ((https://doi.org/10.1007/s13225-020-00466-2)) |
| Otu47233 | 1 | k__Fungi(100);p__Ascomycota(100);c__Saccharomycetes(100);o__Saccharomycetales(100);f__Metschnikowiaceae(100);g__Metschnikowia(100);s__Metschnikowia_pulcherrima(100); | Metschnikowia | 13 | Saprotroph | \|Undefined Saprotroph\| | Yeast | NULL | Probable | As nectar-tap_saprotroph (Põlme et al. 2020); Decay sugar-rich substrates (Põlme et al. 2020) | Põlme S, et al. 2020. Fungal Diversity 105:1-16 ((https://doi.org/10.1007/s13225-020-00466-2)) |
| Otu47234 | 1 | k__Fungi(100);p__Ascomycota(100);c__Saccharomycetes(100);o__Saccharomycetales(100);f__Metschnikowiaceae(100);g__Metschnikowia(100);g__Metschnikowia_unclassified(100); | Metschnikowia | 13 | Saprotroph | \|Undefined Saprotroph\| | Yeast | NULL | Probable | As nectar-tap_saprotroph (Põlme et al. 2020); Decay sugar-rich substrates (Põlme et al. 2020) | Põlme S, et al. 2020. Fungal Diversity 105:1-16 ((https://doi.org/10.1007/s13225-020-00466-2)) |
| Otu47235 | 1 | k__Fungi(100);p__Ascomycota(100);c__Saccharomycetes(100);o__Saccharomycetales(100);f__Metschnikowiaceae(100);g__Metschnikowia(100);g__Metschnikowia_unclassified(100); | Metschnikowia | 13 | Saprotroph | \|Undefined Saprotroph\| | Yeast | NULL | Probable | As nectar-tap_saprotroph (Põlme et al. 2020); Decay sugar-rich substrates (Põlme et al. 2020) | Põlme S, et al. 2020. Fungal Diversity 105:1-16 ((https://doi.org/10.1007/s13225-020-00466-2)) |
| Otu47236 | 1 | k__Fungi(100);p__Ascomycota(100);c__Saccharomycetes(100);o__Saccharomycetales(100);f__Metschnikowiaceae(100);g__Metschnikowia(100);s__Metschnikowia_pulcherrima(100); | Metschnikowia | 13 | Saprotroph | \|Undefined Saprotroph\| | Yeast | NULL | Probable | As nectar-tap_saprotroph (Põlme et al. 2020); Decay sugar-rich substrates (Põlme et al. 2020) | Põlme S, et al. 2020. Fungal Diversity 105:1-16 ((https://doi.org/10.1007/s13225-020-00466-2)) |
| Otu47237 | 1 | k__Fungi(100);p__Ascomycota(100);c__Saccharomycetes(100);o__Saccharomycetales(100);f__Metschnikowiaceae(100);g__Metschnikowia(100);g__Metschnikowia_unclassified(100); | Metschnikowia | 13 | Saprotroph | \|Undefined Saprotroph\| | Yeast | NULL | Probable | As nectar-tap_saprotroph (Põlme et al. 2020); Decay sugar-rich substrates (Põlme et al. 2020) | Põlme S, et al. 2020. Fungal Diversity 105:1-16 ((https://doi.org/10.1007/s13225-020-00466-2)) |
| Otu47238 | 1 | k__Fungi(100);p__Ascomycota(100);c__Saccharomycetes(100);o__Saccharomycetales(100);f__Metschnikowiaceae(100);g__Metschnikowia(100);s__Metschnikowia_pulcherrima(100); | Metschnikowia | 13 | Saprotroph | \|Undefined Saprotroph\| | Yeast | NULL | Probable | As nectar-tap_saprotroph (Põlme et al. 2020); Decay sugar-rich substrates (Põlme et al. 2020) | Põlme S, et al. 2020. Fungal Diversity 105:1-16 ((https://doi.org/10.1007/s13225-020-00466-2)) |
| Otu47239 | 1 | k__Fungi(100);p__Ascomycota(100);c__Saccharomycetes(100);o__Saccharomycetales(100);f__Metschnikowiaceae(100);g__Metschnikowia(100);s__Metschnikowia_pulcherrima(100); | Metschnikowia | 13 | Saprotroph | \|Undefined Saprotroph\| | Yeast | NULL | Probable | As nectar-tap_saprotroph (Põlme et al. 2020); Decay sugar-rich substrates (Põlme et al. 2020) | Põlme S, et al. 2020. Fungal Diversity 105:1-16 ((https://doi.org/10.1007/s13225-020-00466-2)) |
| Otu47240 | 1 | k__Fungi(100);p__Ascomycota(100);c__Saccharomycetes(100);o__Saccharomycetales(100);f__Metschnikowiaceae(100);g__Metschnikowia(100);s__Metschnikowia_pulcherrima(100); | Metschnikowia | 13 | Saprotroph | \|Undefined Saprotroph\| | Yeast | NULL | Probable | As nectar-tap_saprotroph (Põlme et al. 2020); Decay sugar-rich substrates (Põlme et al. 2020) | Põlme S, et al. 2020. Fungal Diversity 105:1-16 ((https://doi.org/10.1007/s13225-020-00466-2)) |
| Otu47241 | 1 | k__Fungi(100);p__Ascomycota(100);c__Saccharomycetes(100);o__Saccharomycetales(100);f__Metschnikowiaceae(100);g__Metschnikowia(100);s__Metschnikowia_pulcherrima(100); | Metschnikowia | 13 | Saprotroph | \|Undefined Saprotroph\| | Yeast | NULL | Probable | As nectar-tap_saprotroph (Põlme et al. 2020); Decay sugar-rich substrates (Põlme et al. 2020) | Põlme S, et al. 2020. Fungal Diversity 105:1-16 ((https://doi.org/10.1007/s13225-020-00466-2)) |
| Otu47243 | 1 | k__Fungi(100);p__Ascomycota(100);c__Saccharomycetes(100);o__Saccharomycetales(100);f__Metschnikowiaceae(100);g__Metschnikowia(100);g__Metschnikowia_unclassified(100); | Metschnikowia | 13 | Saprotroph | \|Undefined Saprotroph\| | Yeast | NULL | Probable | As nectar-tap_saprotroph (Põlme et al. 2020); Decay sugar-rich substrates (Põlme et al. 2020) | Põlme S, et al. 2020. Fungal Diversity 105:1-16 ((https://doi.org/10.1007/s13225-020-00466-2)) |
| Otu47244 | 1 | k__Fungi(100);p__Ascomycota(100);c__Saccharomycetes(100);o__Saccharomycetales(100);f__Metschnikowiaceae(100);g__Metschnikowia(100);s__Metschnikowia_pulcherrima(100); | Metschnikowia | 13 | Saprotroph | \|Undefined Saprotroph\| | Yeast | NULL | Probable | As nectar-tap_saprotroph (Põlme et al. 2020); Decay sugar-rich substrates (Põlme et al. 2020) | Põlme S, et al. 2020. Fungal Diversity 105:1-16 ((https://doi.org/10.1007/s13225-020-00466-2)) |
| Otu47245 | 1 | k__Fungi(100);p__Ascomycota(100);c__Saccharomycetes(100);o__Saccharomycetales(100);f__Metschnikowiaceae(100);g__Metschnikowia(100);s__Metschnikowia_pulcherrima(100); | Metschnikowia | 13 | Saprotroph | \|Undefined Saprotroph\| | Yeast | NULL | Probable | As nectar-tap_saprotroph (Põlme et al. 2020); Decay sugar-rich substrates (Põlme et al. 2020) | Põlme S, et al. 2020. Fungal Diversity 105:1-16 ((https://doi.org/10.1007/s13225-020-00466-2)) |
| Otu47246 | 1 | k__Fungi(100);p__Ascomycota(100);c__Saccharomycetes(100);o__Saccharomycetales(100);f__Metschnikowiaceae(100);g__Metschnikowia(100);s__Metschnikowia_pulcherrima(100); | Metschnikowia | 13 | Saprotroph | \|Undefined Saprotroph\| | Yeast | NULL | Probable | As nectar-tap_saprotroph (Põlme et al. 2020); Decay sugar-rich substrates (Põlme et al. 2020) | Põlme S, et al. 2020. Fungal Diversity 105:1-16 ((https://doi.org/10.1007/s13225-020-00466-2)) |
| Otu47247 | 1 | k__Fungi(100);p__Ascomycota(100);c__Saccharomycetes(100);o__Saccharomycetales(100);f__Metschnikowiaceae(100);g__Metschnikowia(100);s__Metschnikowia_pulcherrima(100); | Metschnikowia | 13 | Saprotroph | \|Undefined Saprotroph\| | Yeast | NULL | Probable | As nectar-tap_saprotroph (Põlme et al. 2020); Decay sugar-rich substrates (Põlme et al. 2020) | Põlme S, et al. 2020. Fungal Diversity 105:1-16 ((https://doi.org/10.1007/s13225-020-00466-2)) |
| Otu47248 | 1 | k__Fungi(100);p__Ascomycota(100);c__Saccharomycetes(100);o__Saccharomycetales(100);f__Metschnikowiaceae(100);g__Metschnikowia(100);s__Metschnikowia_pulcherrima(100); | Metschnikowia | 13 | Saprotroph | \|Undefined Saprotroph\| | Yeast | NULL | Probable | As nectar-tap_saprotroph (Põlme et al. 2020); Decay sugar-rich substrates (Põlme et al. 2020) | Põlme S, et al. 2020. Fungal Diversity 105:1-16 ((https://doi.org/10.1007/s13225-020-00466-2)) |
| Otu47250 | 1 | k__Fungi(100);p__Ascomycota(100);c__Saccharomycetes(100);o__Saccharomycetales(100);f__Metschnikowiaceae(100);g__Metschnikowia(100);s__Metschnikowia_pulcherrima(100); | Metschnikowia | 13 | Saprotroph | \|Undefined Saprotroph\| | Yeast | NULL | Probable | As nectar-tap_saprotroph (Põlme et al. 2020); Decay sugar-rich substrates (Põlme et al. 2020) | Põlme S, et al. 2020. Fungal Diversity 105:1-16 ((https://doi.org/10.1007/s13225-020-00466-2)) |
| Otu47251 | 1 | k__Fungi(100);p__Ascomycota(100);c__Saccharomycetes(100);o__Saccharomycetales(100);f__Metschnikowiaceae(100);g__Metschnikowia(100);g__Metschnikowia_unclassified(100); | Metschnikowia | 13 | Saprotroph | \|Undefined Saprotroph\| | Yeast | NULL | Probable | As nectar-tap_saprotroph (Põlme et al. 2020); Decay sugar-rich substrates (Põlme et al. 2020) | Põlme S, et al. 2020. Fungal Diversity 105:1-16 ((https://doi.org/10.1007/s13225-020-00466-2)) |
| Otu47252 | 1 | k__Fungi(100);p__Ascomycota(100);c__Saccharomycetes(100);o__Saccharomycetales(100);f__Metschnikowiaceae(100);g__Metschnikowia(100);s__Metschnikowia_pulcherrima(100); | Metschnikowia | 13 | Saprotroph | \|Undefined Saprotroph\| | Yeast | NULL | Probable | As nectar-tap_saprotroph (Põlme et al. 2020); Decay sugar-rich substrates (Põlme et al. 2020) | Põlme S, et al. 2020. Fungal Diversity 105:1-16 ((https://doi.org/10.1007/s13225-020-00466-2)) |
| Otu47253 | 1 | k__Fungi(100);p__Ascomycota(100);c__Saccharomycetes(100);o__Saccharomycetales(100);f__Metschnikowiaceae(100);g__Metschnikowia(100);s__Metschnikowia_pulcherrima(100); | Metschnikowia | 13 | Saprotroph | \|Undefined Saprotroph\| | Yeast | NULL | Probable | As nectar-tap_saprotroph (Põlme et al. 2020); Decay sugar-rich substrates (Põlme et al. 2020) | Põlme S, et al. 2020. Fungal Diversity 105:1-16 ((https://doi.org/10.1007/s13225-020-00466-2)) |
| Otu47254 | 1 | k__Fungi(100);p__Ascomycota(100);c__Saccharomycetes(100);o__Saccharomycetales(100);f__Metschnikowiaceae(100);g__Metschnikowia(100);s__Metschnikowia_pulcherrima(100); | Metschnikowia | 13 | Saprotroph | \|Undefined Saprotroph\| | Yeast | NULL | Probable | As nectar-tap_saprotroph (Põlme et al. 2020); Decay sugar-rich substrates (Põlme et al. 2020) | Põlme S, et al. 2020. Fungal Diversity 105:1-16 ((https://doi.org/10.1007/s13225-020-00466-2)) |
| Otu47255 | 1 | k__Fungi(100);p__Ascomycota(100);c__Saccharomycetes(100);o__Saccharomycetales(100);f__Metschnikowiaceae(100);g__Metschnikowia(100);s__Metschnikowia_pulcherrima(100); | Metschnikowia | 13 | Saprotroph | \|Undefined Saprotroph\| | Yeast | NULL | Probable | As nectar-tap_saprotroph (Põlme et al. 2020); Decay sugar-rich substrates (Põlme et al. 2020) | Põlme S, et al. 2020. Fungal Diversity 105:1-16 ((https://doi.org/10.1007/s13225-020-00466-2)) |
| Otu47256 | 1 | k__Fungi(100);p__Ascomycota(100);c__Saccharomycetes(100);o__Saccharomycetales(100);f__Metschnikowiaceae(100);g__Metschnikowia(100);s__Metschnikowia_pulcherrima(100); | Metschnikowia | 13 | Saprotroph | \|Undefined Saprotroph\| | Yeast | NULL | Probable | As nectar-tap_saprotroph (Põlme et al. 2020); Decay sugar-rich substrates (Põlme et al. 2020) | Põlme S, et al. 2020. Fungal Diversity 105:1-16 ((https://doi.org/10.1007/s13225-020-00466-2)) |
| Otu47257 | 1 | k__Fungi(100);p__Ascomycota(100);c__Saccharomycetes(100);o__Saccharomycetales(100);f__Metschnikowiaceae(100);g__Metschnikowia(100);g__Metschnikowia_unclassified(100); | Metschnikowia | 13 | Saprotroph | \|Undefined Saprotroph\| | Yeast | NULL | Probable | As nectar-tap_saprotroph (Põlme et al. 2020); Decay sugar-rich substrates (Põlme et al. 2020) | Põlme S, et al. 2020. Fungal Diversity 105:1-16 ((https://doi.org/10.1007/s13225-020-00466-2)) |
| Otu47258 | 1 | k__Fungi(100);p__Ascomycota(100);c__Saccharomycetes(100);o__Saccharomycetales(100);f__Metschnikowiaceae(100);g__Metschnikowia(100);s__Metschnikowia_pulcherrima(100); | Metschnikowia | 13 | Saprotroph | \|Undefined Saprotroph\| | Yeast | NULL | Probable | As nectar-tap_saprotroph (Põlme et al. 2020); Decay sugar-rich substrates (Põlme et al. 2020) | Põlme S, et al. 2020. Fungal Diversity 105:1-16 ((https://doi.org/10.1007/s13225-020-00466-2)) |
| Otu47259 | 1 | k__Fungi(100);p__Ascomycota(100);c__Saccharomycetes(100);o__Saccharomycetales(100);f__Metschnikowiaceae(100);g__Metschnikowia(100);s__Metschnikowia_pulcherrima(100); | Metschnikowia | 13 | Saprotroph | \|Undefined Saprotroph\| | Yeast | NULL | Probable | As nectar-tap_saprotroph (Põlme et al. 2020); Decay sugar-rich substrates (Põlme et al. 2020) | Põlme S, et al. 2020. Fungal Diversity 105:1-16 ((https://doi.org/10.1007/s13225-020-00466-2)) |
| Otu47260 | 1 | k__Fungi(100);p__Ascomycota(100);c__Saccharomycetes(100);o__Saccharomycetales(100);f__Metschnikowiaceae(100);g__Metschnikowia(100);s__Metschnikowia_pulcherrima(100); | Metschnikowia | 13 | Saprotroph | \|Undefined Saprotroph\| | Yeast | NULL | Probable | As nectar-tap_saprotroph (Põlme et al. 2020); Decay sugar-rich substrates (Põlme et al. 2020) | Põlme S, et al. 2020. Fungal Diversity 105:1-16 ((https://doi.org/10.1007/s13225-020-00466-2)) |
| Otu47261 | 1 | k__Fungi(100);p__Ascomycota(100);c__Saccharomycetes(100);o__Saccharomycetales(100);f__Metschnikowiaceae(100);g__Metschnikowia(100);g__Metschnikowia_unclassified(100); | Metschnikowia | 13 | Saprotroph | \|Undefined Saprotroph\| | Yeast | NULL | Probable | As nectar-tap_saprotroph (Põlme et al. 2020); Decay sugar-rich substrates (Põlme et al. 2020) | Põlme S, et al. 2020. Fungal Diversity 105:1-16 ((https://doi.org/10.1007/s13225-020-00466-2)) |
| Otu47262 | 1 | k__Fungi(100);p__Ascomycota(100);c__Saccharomycetes(100);o__Saccharomycetales(100);f__Metschnikowiaceae(100);g__Metschnikowia(100);g__Metschnikowia_unclassified(100); | Metschnikowia | 13 | Saprotroph | \|Undefined Saprotroph\| | Yeast | NULL | Probable | As nectar-tap_saprotroph (Põlme et al. 2020); Decay sugar-rich substrates (Põlme et al. 2020) | Põlme S, et al. 2020. Fungal Diversity 105:1-16 ((https://doi.org/10.1007/s13225-020-00466-2)) |
| Otu47263 | 1 | k__Fungi(100);p__Ascomycota(100);c__Saccharomycetes(100);o__Saccharomycetales(100);f__Metschnikowiaceae(100);g__Metschnikowia(100);s__Metschnikowia_pulcherrima(100); | Metschnikowia | 13 | Saprotroph | \|Undefined Saprotroph\| | Yeast | NULL | Probable | As nectar-tap_saprotroph (Põlme et al. 2020); Decay sugar-rich substrates (Põlme et al. 2020) | Põlme S, et al. 2020. Fungal Diversity 105:1-16 ((https://doi.org/10.1007/s13225-020-00466-2)) |
| Otu47264 | 1 | k__Fungi(100);p__Ascomycota(100);c__Saccharomycetes(100);o__Saccharomycetales(100);f__Metschnikowiaceae(100);g__Metschnikowia(100);s__Metschnikowia_pulcherrima(100); | Metschnikowia | 13 | Saprotroph | \|Undefined Saprotroph\| | Yeast | NULL | Probable | As nectar-tap_saprotroph (Põlme et al. 2020); Decay sugar-rich substrates (Põlme et al. 2020) | Põlme S, et al. 2020. Fungal Diversity 105:1-16 ((https://doi.org/10.1007/s13225-020-00466-2)) |
| Otu47265 | 1 | k__Fungi(100);p__Ascomycota(100);c__Saccharomycetes(100);o__Saccharomycetales(100);f__Metschnikowiaceae(100);g__Metschnikowia(100);s__Metschnikowia_pulcherrima(100); | Metschnikowia | 13 | Saprotroph | \|Undefined Saprotroph\| | Yeast | NULL | Probable | As nectar-tap_saprotroph (Põlme et al. 2020); Decay sugar-rich substrates (Põlme et al. 2020) | Põlme S, et al. 2020. Fungal Diversity 105:1-16 ((https://doi.org/10.1007/s13225-020-00466-2)) |
| Otu47266 | 1 | k__Fungi(100);p__Ascomycota(100);c__Saccharomycetes(100);o__Saccharomycetales(100);f__Metschnikowiaceae(100);g__Metschnikowia(100);s__Metschnikowia_pulcherrima(100); | Metschnikowia | 13 | Saprotroph | \|Undefined Saprotroph\| | Yeast | NULL | Probable | As nectar-tap_saprotroph (Põlme et al. 2020); Decay sugar-rich substrates (Põlme et al. 2020) | Põlme S, et al. 2020. Fungal Diversity 105:1-16 ((https://doi.org/10.1007/s13225-020-00466-2)) |
| Otu47267 | 1 | k__Fungi(100);p__Ascomycota(100);c__Saccharomycetes(100);o__Saccharomycetales(100);f__Metschnikowiaceae(100);g__Metschnikowia(100);s__Metschnikowia_pulcherrima(100); | Metschnikowia | 13 | Saprotroph | \|Undefined Saprotroph\| | Yeast | NULL | Probable | As nectar-tap_saprotroph (Põlme et al. 2020); Decay sugar-rich substrates (Põlme et al. 2020) | Põlme S, et al. 2020. Fungal Diversity 105:1-16 ((https://doi.org/10.1007/s13225-020-00466-2)) |
| Otu47268 | 1 | k__Fungi(100);p__Ascomycota(100);c__Saccharomycetes(100);o__Saccharomycetales(100);f__Metschnikowiaceae(100);g__Metschnikowia(100);s__Metschnikowia_pulcherrima(100); | Metschnikowia | 13 | Saprotroph | \|Undefined Saprotroph\| | Yeast | NULL | Probable | As nectar-tap_saprotroph (Põlme et al. 2020); Decay sugar-rich substrates (Põlme et al. 2020) | Põlme S, et al. 2020. Fungal Diversity 105:1-16 ((https://doi.org/10.1007/s13225-020-00466-2)) |
| Otu47269 | 1 | k__Fungi(100);p__Ascomycota(100);c__Saccharomycetes(100);o__Saccharomycetales(100);f__Metschnikowiaceae(100);g__Metschnikowia(100);g__Metschnikowia_unclassified(100); | Metschnikowia | 13 | Saprotroph | \|Undefined Saprotroph\| | Yeast | NULL | Probable | As nectar-tap_saprotroph (Põlme et al. 2020); Decay sugar-rich substrates (Põlme et al. 2020) | Põlme S, et al. 2020. Fungal Diversity 105:1-16 ((https://doi.org/10.1007/s13225-020-00466-2)) |
| Otu47270 | 1 | k__Fungi(100);p__Ascomycota(100);c__Saccharomycetes(100);o__Saccharomycetales(100);f__Metschnikowiaceae(100);g__Metschnikowia(100);g__Metschnikowia_unclassified(100); | Metschnikowia | 13 | Saprotroph | \|Undefined Saprotroph\| | Yeast | NULL | Probable | As nectar-tap_saprotroph (Põlme et al. 2020); Decay sugar-rich substrates (Põlme et al. 2020) | Põlme S, et al. 2020. Fungal Diversity 105:1-16 ((https://doi.org/10.1007/s13225-020-00466-2)) |
| Otu47271 | 1 | k__Fungi(100);p__Ascomycota(100);c__Saccharomycetes(100);o__Saccharomycetales(100);f__Metschnikowiaceae(100);g__Metschnikowia(100);g__Metschnikowia_unclassified(100); | Metschnikowia | 13 | Saprotroph | \|Undefined Saprotroph\| | Yeast | NULL | Probable | As nectar-tap_saprotroph (Põlme et al. 2020); Decay sugar-rich substrates (Põlme et al. 2020) | Põlme S, et al. 2020. Fungal Diversity 105:1-16 ((https://doi.org/10.1007/s13225-020-00466-2)) |

| | | | | | | | | | | | |
|---|---|---|---|---|---|---|---|---|---|---|---|
| Otu47272 | 1 | k__Fungi(100);p__Ascomycota(100);c__Saccharomycetes(100);o__Saccharomycetales(100);f__Metschnikowiaceae(100);g__Metschnikowia(100);s__Metschnikowia_pulcherrima(100); | Metschnikowia | 13 | Saprotroph | \|Undefined Saprotroph\| | Yeast | NULL | Probable | As nectar-tap_saprotroph (Põlme et al. 2020); Decay sugar-rich substrates (Põlme et al. 2020) | Põlme S, et al. 2020. Fungal Diversity 105:1-16 ((https://doi.org/10.1007/s13225-020-00466-2)) |
| Otu47273 | 1 | k__Fungi(100);p__Ascomycota(100);c__Saccharomycetes(100);o__Saccharomycetales(100);f__Metschnikowiaceae(100);g__Metschnikowia(100);s__Metschnikowia_pulcherrima(100); | Metschnikowia | 13 | Saprotroph | \|Undefined Saprotroph\| | Yeast | NULL | Probable | As nectar-tap_saprotroph (Põlme et al. 2020); Decay sugar-rich substrates (Põlme et al. 2020) | Põlme S, et al. 2020. Fungal Diversity 105:1-16 ((https://doi.org/10.1007/s13225-020-00466-2)) |
| Otu47274 | 1 | k__Fungi(100);p__Ascomycota(100);c__Saccharomycetes(100);o__Saccharomycetales(100);f__Metschnikowiaceae(100);g__Metschnikowia(100);s__Metschnikowia_pulcherrima(100); | Metschnikowia | 13 | Saprotroph | \|Undefined Saprotroph\| | Yeast | NULL | Probable | As nectar-tap_saprotroph (Põlme et al. 2020); Decay sugar-rich substrates (Põlme et al. 2020) | Põlme S, et al. 2020. Fungal Diversity 105:1-16 ((https://doi.org/10.1007/s13225-020-00466-2)) |
| Otu47275 | 1 | k__Fungi(100);p__Ascomycota(100);c__Saccharomycetes(100);o__Saccharomycetales(100);f__Metschnikowiaceae(100);g__Metschnikowia(100);g__Metschnikowia_unclassified(100); | Metschnikowia | 13 | Saprotroph | \|Undefined Saprotroph\| | Yeast | NULL | Probable | As nectar-tap_saprotroph (Põlme et al. 2020); Decay sugar-rich substrates (Põlme et al. 2020) | Põlme S, et al. 2020. Fungal Diversity 105:1-16 ((https://doi.org/10.1007/s13225-020-00466-2)) |
| Otu47276 | 1 | k__Fungi(100);p__Ascomycota(100);c__Saccharomycetes(100);o__Saccharomycetales(100);f__Metschnikowiaceae(100);g__Metschnikowia(100);s__Metschnikowia_pulcherrima(100); | Metschnikowia | 13 | Saprotroph | \|Undefined Saprotroph\| | Yeast | NULL | Probable | As nectar-tap_saprotroph (Põlme et al. 2020); Decay sugar-rich substrates (Põlme et al. 2020) | Põlme S, et al. 2020. Fungal Diversity 105:1-16 ((https://doi.org/10.1007/s13225-020-00466-2)) |
| Otu47277 | 1 | k__Fungi(100);p__Ascomycota(100);c__Saccharomycetes(100);o__Saccharomycetales(100);f__Metschnikowiaceae(100);g__Metschnikowia(100);g__Metschnikowia_unclassified(100); | Metschnikowia | 13 | Saprotroph | \|Undefined Saprotroph\| | Yeast | NULL | Probable | As nectar-tap_saprotroph (Põlme et al. 2020); Decay sugar-rich substrates (Põlme et al. 2020) | Põlme S, et al. 2020. Fungal Diversity 105:1-16 ((https://doi.org/10.1007/s13225-020-00466-2)) |
| Otu47278 | 1 | k__Fungi(100);p__Ascomycota(100);c__Saccharomycetes(100);o__Saccharomycetales(100);f__Metschnikowiaceae(100);g__Metschnikowia(100);g__Metschnikowia_unclassified(100); | Metschnikowia | 13 | Saprotroph | \|Undefined Saprotroph\| | Yeast | NULL | Probable | As nectar-tap_saprotroph (Põlme et al. 2020); Decay sugar-rich substrates (Põlme et al. 2020) | Põlme S, et al. 2020. Fungal Diversity 105:1-16 ((https://doi.org/10.1007/s13225-020-00466-2)) |
| Otu47279 | 1 | k__Fungi(100);p__Ascomycota(100);c__Saccharomycetes(100);o__Saccharomycetales(100);f__Metschnikowiaceae(100);g__Metschnikowia(100);g__Metschnikowia_unclassified(100); | Metschnikowia | 13 | Saprotroph | \|Undefined Saprotroph\| | Yeast | NULL | Probable | As nectar-tap_saprotroph (Põlme et al. 2020); Decay sugar-rich substrates (Põlme et al. 2020) | Põlme S, et al. 2020. Fungal Diversity 105:1-16 ((https://doi.org/10.1007/s13225-020-00466-2)) |
| Otu47280 | 1 | k__Fungi(100);p__Ascomycota(100);c__Saccharomycetes(100);o__Saccharomycetales(100);f__Metschnikowiaceae(100);g__Metschnikowia(100);g__Metschnikowia_unclassified(100); | Metschnikowia | 13 | Saprotroph | \|Undefined Saprotroph\| | Yeast | NULL | Probable | As nectar-tap_saprotroph (Põlme et al. 2020); Decay sugar-rich substrates (Põlme et al. 2020) | Põlme S, et al. 2020. Fungal Diversity 105:1-16 ((https://doi.org/10.1007/s13225-020-00466-2)) |
| Otu47281 | 1 | k__Fungi(100);p__Ascomycota(100);c__Saccharomycetes(100);o__Saccharomycetales(100);f__Metschnikowiaceae(100);g__Metschnikowia(100);s__Metschnikowia_pulcherrima(100); | Metschnikowia | 13 | Saprotroph | \|Undefined Saprotroph\| | Yeast | NULL | Probable | As nectar-tap_saprotroph (Põlme et al. 2020); Decay sugar-rich substrates (Põlme et al. 2020) | Põlme S, et al. 2020. Fungal Diversity 105:1-16 ((https://doi.org/10.1007/s13225-020-00466-2)) |
| Otu47282 | 1 | k__Fungi(100);p__Ascomycota(100);c__Saccharomycetes(100);o__Saccharomycetales(100);f__Metschnikowiaceae(100);g__Metschnikowia(100);s__Metschnikowia_pulcherrima(100); | Metschnikowia | 13 | Saprotroph | \|Undefined Saprotroph\| | Yeast | NULL | Probable | As nectar-tap_saprotroph (Põlme et al. 2020); Decay sugar-rich substrates (Põlme et al. 2020) | Põlme S, et al. 2020. Fungal Diversity 105:1-16 ((https://doi.org/10.1007/s13225-020-00466-2)) |
| Otu47283 | 1 | k__Fungi(100);p__Ascomycota(100);c__Saccharomycetes(100);o__Saccharomycetales(100);f__Metschnikowiaceae(100);g__Metschnikowia(100);s__Metschnikowia_pulcherrima(100); | Metschnikowia | 13 | Saprotroph | \|Undefined Saprotroph\| | Yeast | NULL | Probable | As nectar-tap_saprotroph (Põlme et al. 2020); Decay sugar-rich substrates (Põlme et al. 2020) | Põlme S, et al. 2020. Fungal Diversity 105:1-16 ((https://doi.org/10.1007/s13225-020-00466-2)) |
| Otu47284 | 1 | k__Fungi(100);p__Ascomycota(100);c__Saccharomycetes(100);o__Saccharomycetales(100);f__Metschnikowiaceae(100);g__Metschnikowia(100);s__Metschnikowia_pulcherrima(100); | Metschnikowia | 13 | Saprotroph | \|Undefined Saprotroph\| | Yeast | NULL | Probable | As nectar-tap_saprotroph (Põlme et al. 2020); Decay sugar-rich substrates (Põlme et al. 2020) | Põlme S, et al. 2020. Fungal Diversity 105:1-16 ((https://doi.org/10.1007/s13225-020-00466-2)) |
| Otu47285 | 1 | k__Fungi(100);p__Ascomycota(100);c__Saccharomycetes(100);o__Saccharomycetales(100);f__Metschnikowiaceae(100);g__Metschnikowia(100);s__Metschnikowia_pulcherrima(100); | Metschnikowia | 13 | Saprotroph | \|Undefined Saprotroph\| | Yeast | NULL | Probable | As nectar-tap_saprotroph (Põlme et al. 2020); Decay sugar-rich substrates (Põlme et al. 2020) | Põlme S, et al. 2020. Fungal Diversity 105:1-16 ((https://doi.org/10.1007/s13225-020-00466-2)) |
| Otu47286 | 1 | k__Fungi(100);p__Ascomycota(100);c__Saccharomycetes(100);o__Saccharomycetales(100);f__Metschnikowiaceae(100);g__Metschnikowia(100);g__Metschnikowia_unclassified(100); | Metschnikowia | 13 | Saprotroph | \|Undefined Saprotroph\| | Yeast | NULL | Probable | As nectar-tap_saprotroph (Põlme et al. 2020); Decay sugar-rich substrates (Põlme et al. 2020) | Põlme S, et al. 2020. Fungal Diversity 105:1-16 ((https://doi.org/10.1007/s13225-020-00466-2)) |
| Otu47287 | 1 | k__Fungi(100);p__Ascomycota(100);c__Saccharomycetes(100);o__Saccharomycetales(100);f__Metschnikowiaceae(100);g__Metschnikowia(100);s__Metschnikowia_pulcherrima(100); | Metschnikowia | 13 | Saprotroph | \|Undefined Saprotroph\| | Yeast | NULL | Probable | As nectar-tap_saprotroph (Põlme et al. 2020); Decay sugar-rich substrates (Põlme et al. 2020) | Põlme S, et al. 2020. Fungal Diversity 105:1-16 ((https://doi.org/10.1007/s13225-020-00466-2)) |
| Otu47288 | 1 | k__Fungi(100);p__Ascomycota(100);c__Saccharomycetes(100);o__Saccharomycetales(100);f__Metschnikowiaceae(100);g__Metschnikowia(100);s__Metschnikowia_pulcherrima(100); | Metschnikowia | 13 | Saprotroph | \|Undefined Saprotroph\| | Yeast | NULL | Probable | As nectar-tap_saprotroph (Põlme et al. 2020); Decay sugar-rich substrates (Põlme et al. 2020) | Põlme S, et al. 2020. Fungal Diversity 105:1-16 ((https://doi.org/10.1007/s13225-020-00466-2)) |
| Otu47289 | 1 | k__Fungi(100);p__Ascomycota(100);c__Saccharomycetes(100);o__Saccharomycetales(100);f__Metschnikowiaceae(100);g__Metschnikowia(100);s__Metschnikowia_pulcherrima(100); | Metschnikowia | 13 | Saprotroph | \|Undefined Saprotroph\| | Yeast | NULL | Probable | As nectar-tap_saprotroph (Põlme et al. 2020); Decay sugar-rich substrates (Põlme et al. 2020) | Põlme S, et al. 2020. Fungal Diversity 105:1-16 ((https://doi.org/10.1007/s13225-020-00466-2)) |
| Otu47290 | 1 | k__Fungi(100);p__Ascomycota(100);c__Saccharomycetes(100);o__Saccharomycetales(100);f__Metschnikowiaceae(100);g__Metschnikowia(100);s__Metschnikowia_pulcherrima(100); | Metschnikowia | 13 | Saprotroph | \|Undefined Saprotroph\| | Yeast | NULL | Probable | As nectar-tap_saprotroph (Põlme et al. 2020); Decay sugar-rich substrates (Põlme et al. 2020) | Põlme S, et al. 2020. Fungal Diversity 105:1-16 ((https://doi.org/10.1007/s13225-020-00466-2)) |
| Otu47291 | 1 | k__Fungi(100);p__Ascomycota(100);c__Saccharomycetes(100);o__Saccharomycetales(100);f__Metschnikowiaceae(100);g__Metschnikowia(100);g__Metschnikowia_unclassified(100); | Metschnikowia | 13 | Saprotroph | \|Undefined Saprotroph\| | Yeast | NULL | Probable | As nectar-tap_saprotroph (Põlme et al. 2020); Decay sugar-rich substrates (Põlme et al. 2020) | Põlme S, et al. 2020. Fungal Diversity 105:1-16 ((https://doi.org/10.1007/s13225-020-00466-2)) |
| Otu47292 | 1 | k__Fungi(100);p__Ascomycota(100);c__Saccharomycetes(100);o__Saccharomycetales(100);f__Metschnikowiaceae(100);g__Metschnikowia(100);s__Metschnikowia_pulcherrima(100); | Metschnikowia | 13 | Saprotroph | \|Undefined Saprotroph\| | Yeast | NULL | Probable | As nectar-tap_saprotroph (Põlme et al. 2020); Decay sugar-rich substrates (Põlme et al. 2020) | Põlme S, et al. 2020. Fungal Diversity 105:1-16 ((https://doi.org/10.1007/s13225-020-00466-2)) |
| Otu47293 | 1 | k__Fungi(100);p__Ascomycota(100);c__Saccharomycetes(100);o__Saccharomycetales(100);f__Metschnikowiaceae(100);g__Metschnikowia(100);s__Metschnikowia_pulcherrima(100); | Metschnikowia | 13 | Saprotroph | \|Undefined Saprotroph\| | Yeast | NULL | Probable | As nectar-tap_saprotroph (Põlme et al. 2020); Decay sugar-rich substrates (Põlme et al. 2020) | Põlme S, et al. 2020. Fungal Diversity 105:1-16 ((https://doi.org/10.1007/s13225-020-00466-2)) |
| Otu47294 | 1 | k__Fungi(100);p__Ascomycota(100);c__Saccharomycetes(100);o__Saccharomycetales(100);f__Metschnikowiaceae(100);g__Metschnikowia(100);s__Metschnikowia_pulcherrima(100); | Metschnikowia | 13 | Saprotroph | \|Undefined Saprotroph\| | Yeast | NULL | Probable | As nectar-tap_saprotroph (Põlme et al. 2020); Decay sugar-rich substrates (Põlme et al. 2020) | Põlme S, et al. 2020. Fungal Diversity 105:1-16 ((https://doi.org/10.1007/s13225-020-00466-2)) |
| Otu47295 | 1 | k__Fungi(100);p__Ascomycota(100);c__Saccharomycetes(100);o__Saccharomycetales(100);f__Metschnikowiaceae(100);g__Metschnikowia(100);s__Metschnikowia_pulcherrima(100); | Metschnikowia | 13 | Saprotroph | \|Undefined Saprotroph\| | Yeast | NULL | Probable | As nectar-tap_saprotroph (Põlme et al. 2020); Decay sugar-rich substrates (Põlme et al. 2020) | Põlme S, et al. 2020. Fungal Diversity 105:1-16 ((https://doi.org/10.1007/s13225-020-00466-2)) |
| Otu47296 | 1 | k__Fungi(100);p__Ascomycota(100);c__Saccharomycetes(100);o__Saccharomycetales(100);f__Metschnikowiaceae(100);g__Metschnikowia(100);s__Metschnikowia_pulcherrima(100); | Metschnikowia | 13 | Saprotroph | \|Undefined Saprotroph\| | Yeast | NULL | Probable | As nectar-tap_saprotroph (Põlme et al. 2020); Decay sugar-rich substrates (Põlme et al. 2020) | Põlme S, et al. 2020. Fungal Diversity 105:1-16 ((https://doi.org/10.1007/s13225-020-00466-2)) |
| Otu47297 | 1 | k__Fungi(100);p__Ascomycota(100);c__Saccharomycetes(100);o__Saccharomycetales(100);f__Metschnikowiaceae(100);g__Metschnikowia(100);s__Metschnikowia_pulcherrima(100); | Metschnikowia | 13 | Saprotroph | \|Undefined Saprotroph\| | Yeast | NULL | Probable | As nectar-tap_saprotroph (Põlme et al. 2020); Decay sugar-rich substrates (Põlme et al. 2020) | Põlme S, et al. 2020. Fungal Diversity 105:1-16 ((https://doi.org/10.1007/s13225-020-00466-2)) |
| Otu47298 | 1 | k__Fungi(100);p__Ascomycota(100);c__Saccharomycetes(100);o__Saccharomycetales(100);f__Metschnikowiaceae(100);g__Metschnikowia(100);s__Metschnikowia_pulcherrima(100); | Metschnikowia | 13 | Saprotroph | \|Undefined Saprotroph\| | Yeast | NULL | Probable | As nectar-tap_saprotroph (Põlme et al. 2020); Decay sugar-rich substrates (Põlme et al. 2020) | Põlme S, et al. 2020. Fungal Diversity 105:1-16 ((https://doi.org/10.1007/s13225-020-00466-2)) |
| Otu47299 | 1 | k__Fungi(100);p__Ascomycota(100);c__Saccharomycetes(100);o__Saccharomycetales(100);f__Metschnikowiaceae(100);g__Metschnikowia(100);s__Metschnikowia_pulcherrima(100); | Metschnikowia | 13 | Saprotroph | \|Undefined Saprotroph\| | Yeast | NULL | Probable | As nectar-tap_saprotroph (Põlme et al. 2020); Decay sugar-rich substrates (Põlme et al. 2020) | Põlme S, et al. 2020. Fungal Diversity 105:1-16 ((https://doi.org/10.1007/s13225-020-00466-2)) |
| Otu47300 | 1 | k__Fungi(100);p__Ascomycota(100);c__Saccharomycetes(100);o__Saccharomycetales(100);f__Metschnikowiaceae(100);g__Metschnikowia(100);s__Metschnikowia_pulcherrima(100); | Metschnikowia | 13 | Saprotroph | \|Undefined Saprotroph\| | Yeast | NULL | Probable | As nectar-tap_saprotroph (Põlme et al. 2020); Decay sugar-rich substrates (Põlme et al. 2020) | Põlme S, et al. 2020. Fungal Diversity 105:1-16 ((https://doi.org/10.1007/s13225-020-00466-2)) |
| Otu47301 | 1 | k__Fungi(100);p__Ascomycota(100);c__Saccharomycetes(100);o__Saccharomycetales(100);f__Metschnikowiaceae(100);g__Metschnikowia(100);s__Metschnikowia_pulcherrima(100); | Metschnikowia | 13 | Saprotroph | \|Undefined Saprotroph\| | Yeast | NULL | Probable | As nectar-tap_saprotroph (Põlme et al. 2020); Decay sugar-rich substrates (Põlme et al. 2020) | Põlme S, et al. 2020. Fungal Diversity 105:1-16 ((https://doi.org/10.1007/s13225-020-00466-2)) |
| Otu47302 | 1 | k__Fungi(100);p__Ascomycota(100);c__Saccharomycetes(100);o__Saccharomycetales(100);f__Metschnikowiaceae(100);g__Metschnikowia(100);s__Metschnikowia_pulcherrima(100); | Metschnikowia | 13 | Saprotroph | \|Undefined Saprotroph\| | Yeast | NULL | Probable | As nectar-tap_saprotroph (Põlme et al. 2020); Decay sugar-rich substrates (Põlme et al. 2020) | Põlme S, et al. 2020. Fungal Diversity 105:1-16 ((https://doi.org/10.1007/s13225-020-00466-2)) |
| Otu47303 | 1 | k__Fungi(100);p__Ascomycota(100);c__Saccharomycetes(100);o__Saccharomycetales(100);f__Metschnikowiaceae(100);g__Metschnikowia(100);s__Metschnikowia_pulcherrima(100); | Metschnikowia | 13 | Saprotroph | \|Undefined Saprotroph\| | Yeast | NULL | Probable | As nectar-tap_saprotroph (Põlme et al. 2020); Decay sugar-rich substrates (Põlme et al. 2020) | Põlme S, et al. 2020. Fungal Diversity 105:1-16 ((https://doi.org/10.1007/s13225-020-00466-2)) |
| Otu47304 | 1 | k__Fungi(100);p__Ascomycota(100);c__Saccharomycetes(100);o__Saccharomycetales(100);f__Metschnikowiaceae(100);g__Metschnikowia(100);s__Metschnikowia_pulcherrima(100); | Metschnikowia | 13 | Saprotroph | \|Undefined Saprotroph\| | Yeast | NULL | Probable | As nectar-tap_saprotroph (Põlme et al. 2020); Decay sugar-rich substrates (Põlme et al. 2020) | Põlme S, et al. 2020. Fungal Diversity 105:1-16 ((https://doi.org/10.1007/s13225-020-00466-2)) |
| Otu47305 | 1 | k__Fungi(100);p__Ascomycota(100);c__Saccharomycetes(100);o__Saccharomycetales(100);f__Metschnikowiaceae(100);g__Metschnikowia(100);s__Metschnikowia_pulcherrima(100); | Metschnikowia | 13 | Saprotroph | \|Undefined Saprotroph\| | Yeast | NULL | Probable | As nectar-tap_saprotroph (Põlme et al. 2020); Decay sugar-rich substrates (Põlme et al. 2020) | Põlme S, et al. 2020. Fungal Diversity 105:1-16 ((https://doi.org/10.1007/s13225-020-00466-2)) |
| Otu47306 | 1 | k__Fungi(100);p__Ascomycota(100);c__Saccharomycetes(100);o__Saccharomycetales(100);f__Metschnikowiaceae(100);g__Metschnikowia(100);s__Metschnikowia_pulcherrima(100); | Metschnikowia | 13 | Saprotroph | \|Undefined Saprotroph\| | Yeast | NULL | Probable | As nectar-tap_saprotroph (Põlme et al. 2020); Decay sugar-rich substrates (Põlme et al. 2020) | Põlme S, et al. 2020. Fungal Diversity 105:1-16 ((https://doi.org/10.1007/s13225-020-00466-2)) |
| Otu47307 | 1 | k__Fungi(100);p__Ascomycota(100);c__Saccharomycetes(100);o__Saccharomycetales(100);f__Metschnikowiaceae(100);g__Metschnikowia(100);s__Metschnikowia_pulcherrima(100); | Metschnikowia | 13 | Saprotroph | \|Undefined Saprotroph\| | Yeast | NULL | Probable | As nectar-tap_saprotroph (Põlme et al. 2020); Decay sugar-rich substrates (Põlme et al. 2020) | Põlme S, et al. 2020. Fungal Diversity 105:1-16 ((https://doi.org/10.1007/s13225-020-00466-2)) |
| Otu47308 | 1 | k__Fungi(100);p__Ascomycota(100);c__Saccharomycetes(100);o__Saccharomycetales(100);f__Metschnikowiaceae(100);g__Metschnikowia(100);s__Metschnikowia_pulcherrima(100); | Metschnikowia | 13 | Saprotroph | \|Undefined Saprotroph\| | Yeast | NULL | Probable | As nectar-tap_saprotroph (Põlme et al. 2020); Decay sugar-rich substrates (Põlme et al. 2020) | Põlme S, et al. 2020. Fungal Diversity 105:1-16 ((https://doi.org/10.1007/s13225-020-00466-2)) |
| Otu47309 | 1 | k__Fungi(100);p__Ascomycota(100);c__Saccharomycetes(100);o__Saccharomycetales(100);f__Metschnikowiaceae(100);g__Metschnikowia(100);s__Metschnikowia_pulcherrima(100); | Metschnikowia | 13 | Saprotroph | \|Undefined Saprotroph\| | Yeast | NULL | Probable | As nectar-tap_saprotroph (Põlme et al. 2020); Decay sugar-rich substrates (Põlme et al. 2020) | Põlme S, et al. 2020. Fungal Diversity 105:1-16 ((https://doi.org/10.1007/s13225-020-00466-2)) |
| Otu47310 | 1 | k__Fungi(100);p__Ascomycota(100);c__Saccharomycetes(100);o__Saccharomycetales(100);f__Metschnikowiaceae(100);g__Metschnikowia(100);s__Metschnikowia_pulcherrima(100); | Metschnikowia | 13 | Saprotroph | \|Undefined Saprotroph\| | Yeast | NULL | Probable | As nectar-tap_saprotroph (Põlme et al. 2020); Decay sugar-rich substrates (Põlme et al. 2020) | Põlme S, et al. 2020. Fungal Diversity 105:1-16 ((https://doi.org/10.1007/s13225-020-00466-2)) |
| Otu47312 | 1 | k__Fungi(100);p__Ascomycota(100);c__Saccharomycetes(100);o__Saccharomycetales(100);f__Metschnikowiaceae(100);g__Metschnikowia(100);s__Metschnikowia_pulcherrima(100); | Metschnikowia | 13 | Saprotroph | \|Undefined Saprotroph\| | Yeast | NULL | Probable | As nectar-tap_saprotroph (Põlme et al. 2020); Decay sugar-rich substrates (Põlme et al. 2020) | Põlme S, et al. 2020. Fungal Diversity 105:1-16 ((https://doi.org/10.1007/s13225-020-00466-2)) |
| Otu47313 | 1 | k__Fungi(100);p__Ascomycota(100);c__Saccharomycetes(100);o__Saccharomycetales(100);f__Metschnikowiaceae(100);g__Metschnikowia(100);g__Metschnikowia_unclassified(100); | Metschnikowia | 13 | Saprotroph | \|Undefined Saprotroph\| | Yeast | NULL | Probable | As nectar-tap_saprotroph (Põlme et al. 2020); Decay sugar-rich substrates (Põlme et al. 2020) | Põlme S, et al. 2020. Fungal Diversity 105:1-16 ((https://doi.org/10.1007/s13225-020-00466-2)) |
| Otu47314 | 1 | k__Fungi(100);p__Ascomycota(100);c__Saccharomycetes(100);o__Saccharomycetales(100);f__Metschnikowiaceae(100);g__Metschnikowia(100);s__Metschnikowia_pulcherrima(100); | Metschnikowia | 13 | Saprotroph | \|Undefined Saprotroph\| | Yeast | NULL | Probable | As nectar-tap_saprotroph (Põlme et al. 2020); Decay sugar-rich substrates (Põlme et al. 2020) | Põlme S, et al. 2020. Fungal Diversity 105:1-16 ((https://doi.org/10.1007/s13225-020-00466-2)) |
| Otu47316 | 1 | k__Fungi(100);p__Ascomycota(100);c__Saccharomycetes(100);o__Saccharomycetales(100);f__Metschnikowiaceae(100);g__Metschnikowia(100);s__Metschnikowia_pulcherrima(100); | Metschnikowia | 13 | Saprotroph | \|Undefined Saprotroph\| | Yeast | NULL | Probable | As nectar-tap_saprotroph (Põlme et al. 2020); Decay sugar-rich substrates (Põlme et al. 2020) | Põlme S, et al. 2020. Fungal Diversity 105:1-16 ((https://doi.org/10.1007/s13225-020-00466-2)) |
| Otu47317 | 1 | k__Fungi(100);p__Ascomycota(100);c__Saccharomycetes(100);o__Saccharomycetales(100);f__Metschnikowiaceae(100);g__Metschnikowia(100);s__Metschnikowia_pulcherrima(100); | Metschnikowia | 13 | Saprotroph | \|Undefined Saprotroph\| | Yeast | NULL | Probable | As nectar-tap_saprotroph (Põlme et al. 2020); Decay sugar-rich substrates (Põlme et al. 2020) | Põlme S, et al. 2020. Fungal Diversity 105:1-16 ((https://doi.org/10.1007/s13225-020-00466-2)) |
| Otu47318 | 1 | k__Fungi(100);p__Ascomycota(100);c__Saccharomycetes(100);o__Saccharomycetales(100);f__Metschnikowiaceae(100);g__Metschnikowia(100);s__Metschnikowia_pulcherrima(100); | Metschnikowia | 13 | Saprotroph | \|Undefined Saprotroph\| | Yeast | NULL | Probable | As nectar-tap_saprotroph (Põlme et al. 2020); Decay sugar-rich substrates (Põlme et al. 2020) | Põlme S, et al. 2020. Fungal Diversity 105:1-16 ((https://doi.org/10.1007/s13225-020-00466-2)) |
| Otu47319 | 1 | k__Fungi(100);p__Ascomycota(100);c__Saccharomycetes(100);o__Saccharomycetales(100);f__Metschnikowiaceae(100);g__Metschnikowia(100);s__Metschnikowia_pulcherrima(100); | Metschnikowia | 13 | Saprotroph | \|Undefined Saprotroph\| | Yeast | NULL | Probable | As nectar-tap_saprotroph (Põlme et al. 2020); Decay sugar-rich substrates (Põlme et al. 2020) | Põlme S, et al. 2020. Fungal Diversity 105:1-16 ((https://doi.org/10.1007/s13225-020-00466-2)) |
| Otu47320 | 1 | k__Fungi(100);p__Ascomycota(100);c__Saccharomycetes(100);o__Saccharomycetales(100);f__Metschnikowiaceae(100);g__Metschnikowia(100);s__Metschnikowia_pulcherrima(100); | Metschnikowia | 13 | Saprotroph | \|Undefined Saprotroph\| | Yeast | NULL | Probable | As nectar-tap_saprotroph (Põlme et al. 2020); Decay sugar-rich substrates (Põlme et al. 2020) | Põlme S, et al. 2020. Fungal Diversity 105:1-16 ((https://doi.org/10.1007/s13225-020-00466-2)) |
| Otu47321 | 1 | k__Fungi(100);p__Ascomycota(100);c__Saccharomycetes(100);o__Saccharomycetales(100);f__Metschnikowiaceae(100);g__Metschnikowia(100);s__Metschnikowia_pulcherrima(100); | Metschnikowia | 13 | Saprotroph | \|Undefined Saprotroph\| | Yeast | NULL | Probable | As nectar-tap_saprotroph (Põlme et al. 2020); Decay sugar-rich substrates (Põlme et al. 2020) | Põlme S, et al. 2020. Fungal Diversity 105:1-16 ((https://doi.org/10.1007/s13225-020-00466-2)) |
| Otu47322 | 1 | k__Fungi(100);p__Ascomycota(100);c__Saccharomycetes(100);o__Saccharomycetales(100);f__Metschnikowiaceae(100);g__Metschnikowia(100);s__Metschnikowia_pulcherrima(100); | Metschnikowia | 13 | Saprotroph | \|Undefined Saprotroph\| | Yeast | NULL | Probable | As nectar-tap_saprotroph (Põlme et al. 2020); Decay sugar-rich substrates (Põlme et al. 2020) | Põlme S, et al. 2020. Fungal Diversity 105:1-16 ((https://doi.org/10.1007/s13225-020-00466-2)) |
| Otu47323 | 1 | k__Fungi(100);p__Ascomycota(100);c__Saccharomycetes(100);o__Saccharomycetales(100);f__Metschnikowiaceae(100);g__Metschnikowia(100);g__Metschnikowia_unclassified(100); | Metschnikowia | 13 | Saprotroph | \|Undefined Saprotroph\| | Yeast | NULL | Probable | As nectar-tap_saprotroph (Põlme et al. 2020); Decay sugar-rich substrates (Põlme et al. 2020) | Põlme S, et al. 2020. Fungal Diversity 105:1-16 ((https://doi.org/10.1007/s13225-020-00466-2)) |
| Otu47324 | 1 | k__Fungi(100);p__Ascomycota(100);c__Saccharomycetes(100);o__Saccharomycetales(100);f__Metschnikowiaceae(100);g__Metschnikowia(100);s__Metschnikowia_pulcherrima(100); | Metschnikowia | 13 | Saprotroph | \|Undefined Saprotroph\| | Yeast | NULL | Probable | As nectar-tap_saprotroph (Põlme et al. 2020); Decay sugar-rich substrates (Põlme et al. 2020) | Põlme S, et al. 2020. Fungal Diversity 105:1-16 ((https://doi.org/10.1007/s13225-020-00466-2)) |
| Otu47325 | 1 | k__Fungi(100);p__Ascomycota(100);c__Saccharomycetes(100);o__Saccharomycetales(100);f__Metschnikowiaceae(100);g__Metschnikowia(100);s__Metschnikowia_pulcherrima(100); | Metschnikowia | 13 | Saprotroph | \|Undefined Saprotroph\| | Yeast | NULL | Probable | As nectar-tap_saprotroph (Põlme et al. 2020); Decay sugar-rich substrates (Põlme et al. 2020) | Põlme S, et al. 2020. Fungal Diversity 105:1-16 ((https://doi.org/10.1007/s13225-020-00466-2)) |
| Otu47327 | 1 | k__Fungi(100);p__Ascomycota(100);c__Saccharomycetes(100);o__Saccharomycetales(100);f__Metschnikowiaceae(100);g__Metschnikowia(100);s__Metschnikowia_pulcherrima(100); | Metschnikowia | 13 | Saprotroph | \|Undefined Saprotroph\| | Yeast | NULL | Probable | As nectar-tap_saprotroph (Põlme et al. 2020); Decay sugar-rich substrates (Põlme et al. 2020) | Põlme S, et al. 2020. Fungal Diversity 105:1-16 ((https://doi.org/10.1007/s13225-020-00466-2)) |
| Otu47328 | 1 | k__Fungi(100);p__Ascomycota(100);c__Saccharomycetes(100);o__Saccharomycetales(100);f__Metschnikowiaceae(100);g__Metschnikowia(100);s__Metschnikowia_pulcherrima(100); | Metschnikowia | 13 | Saprotroph | \|Undefined Saprotroph\| | Yeast | NULL | Probable | As nectar-tap_saprotroph (Põlme et al. 2020); Decay sugar-rich substrates (Põlme et al. 2020) | Põlme S, et al. 2020. Fungal Diversity 105:1-16 ((https://doi.org/10.1007/s13225-020-00466-2)) |
| Otu47329 | 1 | k__Fungi(100);p__Ascomycota(100);c__Saccharomycetes(100);o__Saccharomycetales(100);f__Metschnikowiaceae(100);g__Metschnikowia(100);g__Metschnikowia_unclassified(100); | Metschnikowia | 13 | Saprotroph | \|Undefined Saprotroph\| | Yeast | NULL | Probable | As nectar-tap_saprotroph (Põlme et al. 2020); Decay sugar-rich substrates (Põlme et al. 2020) | Põlme S, et al. 2020. Fungal Diversity 105:1-16 ((https://doi.org/10.1007/s13225-020-00466-2)) |
| Otu47330 | 1 | k__Fungi(100);p__Ascomycota(100);c__Saccharomycetes(100);o__Saccharomycetales(100);f__Metschnikowiaceae(100);g__Metschnikowia(100);s__Metschnikowia_pulcherrima(100); | Metschnikowia | 13 | Saprotroph | \|Undefined Saprotroph\| | Yeast | NULL | Probable | As nectar-tap_saprotroph (Põlme et al. 2020); Decay sugar-rich substrates (Põlme et al. 2020) | Põlme S, et al. 2020. Fungal Diversity 105:1-16 ((https://doi.org/10.1007/s13225-020-00466-2)) |
| Otu47331 | 1 | k__Fungi(100);p__Ascomycota(100);c__Saccharomycetes(100);o__Saccharomycetales(100);f__Metschnikowiaceae(100);g__Metschnikowia(100);s__Metschnikowia_pulcherrima(100); | Metschnikowia | 13 | Saprotroph | \|Undefined Saprotroph\| | Yeast | NULL | Probable | As nectar-tap_saprotroph (Põlme et al. 2020); Decay sugar-rich substrates (Põlme et al. 2020) | Põlme S, et al. 2020. Fungal Diversity 105:1-16 ((https://doi.org/10.1007/s13225-020-00466-2)) |
| Otu47332 | 1 | k__Fungi(100);p__Ascomycota(100);c__Saccharomycetes(100);o__Saccharomycetales(100);f__Metschnikowiaceae(100);g__Metschnikowia(100);g__Metschnikowia_unclassified(100); | Metschnikowia | 13 | Saprotroph | \|Undefined Saprotroph\| | Yeast | NULL | Probable | As nectar-tap_saprotroph (Põlme et al. 2020); Decay sugar-rich substrates (Põlme et al. 2020) | Põlme S, et al. 2020. Fungal Diversity 105:1-16 ((https://doi.org/10.1007/s13225-020-00466-2)) |
| Otu47333 | 1 | k__Fungi(100);p__Ascomycota(100);c__Saccharomycetes(100);o__Saccharomycetales(100);f__Metschnikowiaceae(100);g__Metschnikowia(100);s__Metschnikowia_pulcherrima(100); | Metschnikowia | 13 | Saprotroph | \|Undefined Saprotroph\| | Yeast | NULL | Probable | As nectar-tap_saprotroph (Põlme et al. 2020); Decay sugar-rich substrates (Põlme et al. 2020) | Põlme S, et al. 2020. Fungal Diversity 105:1-16 ((https://doi.org/10.1007/s13225-020-00466-2)) |
| Otu47334 | 1 | k__Fungi(100);p__Ascomycota(100);c__Saccharomycetes(100);o__Saccharomycetales(100);f__Metschnikowiaceae(100);g__Metschnikowia(100);g__Metschnikowia_unclassified(100); | Metschnikowia | 13 | Saprotroph | \|Undefined Saprotroph\| | Yeast | NULL | Probable | As nectar-tap_saprotroph (Põlme et al. 2020); Decay sugar-rich substrates (Põlme et al. 2020) | Põlme S, et al. 2020. Fungal Diversity 105:1-16 ((https://doi.org/10.1007/s13225-020-00466-2)) |
| Otu47335 | 1 | k__Fungi(100);p__Ascomycota(100);c__Saccharomycetes(100);o__Saccharomycetales(100);f__Metschnikowiaceae(100);g__Metschnikowia(100);s__Metschnikowia_pulcherrima(100); | Metschnikowia | 13 | Saprotroph | \|Undefined Saprotroph\| | Yeast | NULL | Probable | As nectar-tap_saprotroph (Põlme et al. 2020); Decay sugar-rich substrates (Põlme et al. 2020) | Põlme S, et al. 2020. Fungal Diversity 105:1-16 ((https://doi.org/10.1007/s13225-020-00466-2)) |
| Otu47336 | 1 | k__Fungi(100);p__Ascomycota(100);c__Saccharomycetes(100);o__Saccharomycetales(100);f__Metschnikowiaceae(100);g__Metschnikowia(100);g__Metschnikowia_unclassified(100); | Metschnikowia | 13 | Saprotroph | \|Undefined Saprotroph\| | Yeast | NULL | Probable | As nectar-tap_saprotroph (Põlme et al. 2020); Decay sugar-rich substrates (Põlme et al. 2020) | Põlme S, et al. 2020. Fungal Diversity 105:1-16 ((https://doi.org/10.1007/s13225-020-00466-2)) |
| Otu47337 | 1 | k__Fungi(100);p__Ascomycota(100);c__Saccharomycetes(100);o__Saccharomycetales(100);f__Metschnikowiaceae(100);g__Metschnikowia(100);s__Metschnikowia_pulcherrima(100); | Metschnikowia | 13 | Saprotroph | \|Undefined Saprotroph\| | Yeast | NULL | Probable | As nectar-tap_saprotroph (Põlme et al. 2020); Decay sugar-rich substrates (Põlme et al. 2020) | Põlme S, et al. 2020. Fungal Diversity 105:1-16 ((https://doi.org/10.1007/s13225-020-00466-2)) |
| Otu47338 | 1 | k__Fungi(100);p__Ascomycota(100);c__Saccharomycetes(100);o__Saccharomycetales(100);f__Metschnikowiaceae(100);g__Metschnikowia(100);s__Metschnikowia_pulcherrima(100); | Metschnikowia | 13 | Saprotroph | \|Undefined Saprotroph\| | Yeast | NULL | Probable | As nectar-tap_saprotroph (Põlme et al. 2020); Decay sugar-rich substrates (Põlme et al. 2020) | Põlme S, et al. 2020. Fungal Diversity 105:1-16 ((https://doi.org/10.1007/s13225-020-00466-2)) |
| Otu47339 | 1 | k__Fungi(100);p__Ascomycota(100);c__Saccharomycetes(100);o__Saccharomycetales(100);f__Metschnikowiaceae(100);g__Metschnikowia(100);s__Metschnikowia_pulcherrima(100); | Metschnikowia | 13 | Saprotroph | \|Undefined Saprotroph\| | Yeast | NULL | Probable | As nectar-tap_saprotroph (Põlme et al. 2020); Decay sugar-rich substrates (Põlme et al. 2020) | Põlme S, et al. 2020. Fungal Diversity 105:1-16 ((https://doi.org/10.1007/s13225-020-00466-2)) |
| Otu47340 | 1 | k__Fungi(100);p__Ascomycota(100);c__Saccharomycetes(100);o__Saccharomycetales(100);f__Metschnikowiaceae(100);g__Metschnikowia(100);s__Metschnikowia_pulcherrima(100); | Metschnikowia | 13 | Saprotroph | \|Undefined Saprotroph\| | Yeast | NULL | Probable | As nectar-tap_saprotroph (Põlme et al. 2020); Decay sugar-rich substrates (Põlme et al. 2020) | Põlme S, et al. 2020. Fungal Diversity 105:1-16 ((https://doi.org/10.1007/s13225-020-00466-2)) |
| Otu47341 | 1 | k__Fungi(100);p__Ascomycota(100);c__Saccharomycetes(100);o__Saccharomycetales(100);f__Metschnikowiaceae(100);g__Metschnikowia(100);s__Metschnikowia_pulcherrima(100); | Metschnikowia | 13 | Saprotroph | \|Undefined Saprotroph\| | Yeast | NULL | Probable | As nectar-tap_saprotroph (Põlme et al. 2020); Decay sugar-rich substrates (Põlme et al. 2020) | Põlme S, et al. 2020. Fungal Diversity 105:1-16 ((https://doi.org/10.1007/s13225-020-00466-2)) |
| Otu47342 | 1 | k__Fungi(100);p__Ascomycota(100);c__Saccharomycetes(100);o__Saccharomycetales(100);f__Metschnikowiaceae(100);g__Metschnikowia(100);s__Metschnikowia_pulcherrima(100); | Metschnikowia | 13 | Saprotroph | \|Undefined Saprotroph\| | Yeast | NULL | Probable | As nectar-tap_saprotroph (Põlme et al. 2020); Decay sugar-rich substrates (Põlme et al. 2020) | Põlme S, et al. 2020. Fungal Diversity 105:1-16 ((https://doi.org/10.1007/s13225-020-00466-2)) |
| Otu47343 | 1 | k__Fungi(100);p__Ascomycota(100);c__Saccharomycetes(100);o__Saccharomycetales(100);f__Metschnikowiaceae(100);g__Metschnikowia(100);g__Metschnikowia_unclassified(100); | Metschnikowia | 13 | Saprotroph | \|Undefined Saprotroph\| | Yeast | NULL | Probable | As nectar-tap_saprotroph (Põlme et al. 2020); Decay sugar-rich substrates (Põlme et al. 2020) | Põlme S, et al. 2020. Fungal Diversity 105:1-16 ((https://doi.org/10.1007/s13225-020-00466-2)) |
| Otu47344 | 1 | k__Fungi(100);p__Ascomycota(100);c__Saccharomycetes(100);o__Saccharomycetales(100);f__Metschnikowiaceae(100);g__Metschnikowia(100);g__Metschnikowia_unclassified(100); | Metschnikowia | 13 | Saprotroph | \|Undefined Saprotroph\| | Yeast | NULL | Probable | As nectar-tap_saprotroph (Põlme et al. 2020); Decay sugar-rich substrates (Põlme et al. 2020) | Põlme S, et al. 2020. Fungal Diversity 105:1-16 ((https://doi.org/10.1007/s13225-020-00466-2)) |
| Otu47345 | 1 | k__Fungi(100);p__Ascomycota(100);c__Saccharomycetes(100);o__Saccharomycetales(100);f__Metschnikowiaceae(100);g__Metschnikowia(100);s__Metschnikowia_pulcherrima(100); | Metschnikowia | 13 | Saprotroph | \|Undefined Saprotroph\| | Yeast | NULL | Probable | As nectar-tap_saprotroph (Põlme et al. 2020); Decay sugar-rich substrates (Põlme et al. 2020) | Põlme S, et al. 2020. Fungal Diversity 105:1-16 ((https://doi.org/10.1007/s13225-020-00466-2)) |
| Otu47346 | 1 | k__Fungi(100);p__Ascomycota(100);c__Saccharomycetes(100);o__Saccharomycetales(100);f__Metschnikowiaceae(100);g__Metschnikowia(100);s__Metschnikowia_pulcherrima(100); | Metschnikowia | 13 | Saprotroph | \|Undefined Saprotroph\| | Yeast | NULL | Probable | As nectar-tap_saprotroph (Põlme et al. 2020); Decay sugar-rich substrates (Põlme et al. 2020) | Põlme S, et al. 2020. Fungal Diversity 105:1-16 ((https://doi.org/10.1007/s13225-020-00466-2)) |
| Otu47347 | 1 | k__Fungi(100);p__Ascomycota(100);c__Saccharomycetes(100);o__Saccharomycetales(100);f__Metschnikowiaceae(100);g__Metschnikowia(100);g__Metschnikowia_unclassified(100); | Metschnikowia | 13 | Saprotroph | \|Undefined Saprotroph\| | Yeast | NULL | Probable | As nectar-tap_saprotroph (Põlme et al. 2020); Decay sugar-rich substrates (Põlme et al. 2020) | Põlme S, et al. 2020. Fungal Diversity 105:1-16 ((https://doi.org/10.1007/s13225-020-00466-2)) |
| Otu47348 | 1 | k__Fungi(100);p__Ascomycota(100);c__Saccharomycetes(100);o__Saccharomycetales(100);f__Metschnikowiaceae(100);g__Metschnikowia(100);g__Metschnikowia_unclassified(100); | Metschnikowia | 13 | Saprotroph | \|Undefined Saprotroph\| | Yeast | NULL | Probable | As nectar-tap_saprotroph (Põlme et al. 2020); Decay sugar-rich substrates (Põlme et al. 2020) | Põlme S, et al. 2020. Fungal Diversity 105:1-16 ((https://doi.org/10.1007/s13225-020-00466-2)) |
| Otu47349 | 1 | k__Fungi(100);p__Ascomycota(100);c__Saccharomycetes(100);o__Saccharomycetales(100);f__Metschnikowiaceae(100);g__Metschnikowia(100);g__Metschnikowia_unclassified(100); | Metschnikowia | 13 | Saprotroph | \|Undefined Saprotroph\| | Yeast | NULL | Probable | As nectar-tap_saprotroph (Põlme et al. 2020); Decay sugar-rich substrates (Põlme et al. 2020) | Põlme S, et al. 2020. Fungal Diversity 105:1-16 ((https://doi.org/10.1007/s13225-020-00466-2)) |
| Otu47350 | 1 | k__Fungi(100);p__Ascomycota(100);c__Saccharomycetes(100);o__Saccharomycetales(100);f__Metschnikowiaceae(100);g__Metschnikowia(100);s__Metschnikowia_pulcherrima(100); | Metschnikowia | 13 | Saprotroph | \|Undefined Saprotroph\| | Yeast | NULL | Probable | As nectar-tap_saprotroph (Põlme et al. 2020); Decay sugar-rich substrates (Põlme et al. 2020) | Põlme S, et al. 2020. Fungal Diversity 105:1-16 ((https://doi.org/10.1007/s13225-020-00466-2)) |
| Otu47351 | 1 | k__Fungi(100);p__Ascomycota(100);c__Saccharomycetes(100);o__Saccharomycetales(100);f__Metschnikowiaceae(100);g__Metschnikowia(100);s__Metschnikowia_pulcherrima(100); | Metschnikowia | 13 | Saprotroph | \|Undefined Saprotroph\| | Yeast | NULL | Probable | As nectar-tap_saprotroph (Põlme et al. 2020); Decay sugar-rich substrates (Põlme et al. 2020) | Põlme S, et al. 2020. Fungal Diversity 105:1-16 ((https://doi.org/10.1007/s13225-020-00466-2)) |
| Otu47352 | 1 | k__Fungi(100);p__Ascomycota(100);c__Saccharomycetes(100);o__Saccharomycetales(100);f__Metschnikowiaceae(100);g__Metschnikowia(100);s__Metschnikowia_pulcherrima(100); | Metschnikowia | 13 | Saprotroph | \|Undefined Saprotroph\| | Yeast | NULL | Probable | As nectar-tap_saprotroph (Põlme et al. 2020); Decay sugar-rich substrates (Põlme et al. 2020) | Põlme S, et al. 2020. Fungal Diversity 105:1-16 ((https://doi.org/10.1007/s13225-020-00466-2)) |
| Otu47353 | 1 | k__Fungi(100);p__Ascomycota(100);c__Saccharomycetes(100);o__Saccharomycetales(100);f__Metschnikowiaceae(100);g__Metschnikowia(100);g__Metschnikowia_unclassified(100); | Metschnikowia | 13 | Saprotroph | \|Undefined Saprotroph\| | Yeast | NULL | Probable | As nectar-tap_saprotroph (Põlme et al. 2020); Decay sugar-rich substrates (Põlme et al. 2020) | Põlme S, et al. 2020. Fungal Diversity 105:1-16 ((https://doi.org/10.1007/s13225-020-00466-2)) |
| Otu47354 | 1 | k__Fungi(100);p__Ascomycota(100);c__Saccharomycetes(100);o__Saccharomycetales(100);f__Metschnikowiaceae(100);g__Metschnikowia(100);s__Metschnikowia_pulcherrima(100); | Metschnikowia | 13 | Saprotroph | \|Undefined Saprotroph\| | Yeast | NULL | Probable | As nectar-tap_saprotroph (Põlme et al. 2020); Decay sugar-rich substrates (Põlme et al. 2020) | Põlme S, et al. 2020. Fungal Diversity 105:1-16 ((https://doi.org/10.1007/s13225-020-00466-2)) |
| Otu47356 | 1 | k__Fungi(100);p__Ascomycota(100);c__Saccharomycetes(100);o__Saccharomycetales(100);f__Metschnikowiaceae(100);g__Metschnikowia(100);s__Metschnikowia_pulcherrima(100); | Metschnikowia | 13 | Saprotroph | \|Undefined Saprotroph\| | Yeast | NULL | Probable | As nectar-tap_saprotroph (Põlme et al. 2020); Decay sugar-rich substrates (Põlme et al. 2020) | Põlme S, et al. 2020. Fungal Diversity 105:1-16 ((https://doi.org/10.1007/s13225-020-00466-2)) |
| Otu47357 | 1 | k__Fungi(100);p__Ascomycota(100);c__Saccharomycetes(100);o__Saccharomycetales(100);f__Metschnikowiaceae(100);g__Metschnikowia(100);s__Metschnikowia_pulcherrima(100); | Metschnikowia | 13 | Saprotroph | \|Undefined Saprotroph\| | Yeast | NULL | Probable | As nectar-tap_saprotroph (Põlme et al. 2020); Decay sugar-rich substrates (Põlme et al. 2020) | Põlme S, et al. 2020. Fungal Diversity 105:1-16 ((https://doi.org/10.1007/s13225-020-00466-2)) |
| Otu47358 | 1 | k__Fungi(100);p__Ascomycota(100);c__Saccharomycetes(100);o__Saccharomycetales(100);f__Metschnikowiaceae(100);g__Metschnikowia(100);s__Metschnikowia_pulcherrima(100); | Metschnikowia | 13 | Saprotroph | \|Undefined Saprotroph\| | Yeast | NULL | Probable | As nectar-tap_saprotroph (Põlme et al. 2020); Decay sugar-rich substrates (Põlme et al. 2020) | Põlme S, et al. 2020. Fungal Diversity 105:1-16 ((https://doi.org/10.1007/s13225-020-00466-2)) |
| Otu47360 | 1 | k__Fungi(100);p__Ascomycota(100);c__Dothideomycetes(100);o__Dothideomycetes_order_Incertae_sedis(100);f__Myxotrichaceae(100);g__Oidiodendron(100);s__Oidiodendron_maius(100); | Oidiodendron | 13 | Pathotroph-Saprotroph-Symbiotroph | Endophyte-Ericoid Mycorrhizal-\|Undefined Saprotroph\| | Microfungus | NULL | Probable | As root_endophyte (Põlme et al. 2020); As soil_saprotroph (Põlme et al. 2020); Dark Septate Endophyte; Decay soil (Põlme et al. 2020) | Newsham KK. 2011. New Phytologist 190:783-793 ((https://doi.org/10.1111/j.1469-8137.2010.03611.x)) (as dark septate root endophytes); Põlme S, et al. 2020. Fungal Diversity 105:1-16 ((https://doi.org/10.1007/s13225-020-00466-2)) |
| Otu47361 | 1 | k__Fungi(100);p__Ascomycota(100);c__Saccharomycetes(100);o__Saccharomycetales(100);f__Metschnikowiaceae(100);g__Metschnikowia(100);s__Metschnikowia_pulcherrima(100); | Metschnikowia | 13 | Saprotroph | \|Undefined Saprotroph\| | Yeast | NULL | Probable | As nectar-tap_saprotroph (Põlme et al. 2020); Decay sugar-rich substrates (Põlme et al. 2020) | Põlme S, et al. 2020. Fungal Diversity 105:1-16 ((https://doi.org/10.1007/s13225-020-00466-2)) |
| Otu47363 | 1 | k__Fungi(100);p__Ascomycota(100);c__Saccharomycetes(100);o__Saccharomycetales(100);f__Metschnikowiaceae(100);g__Metschnikowia(100);g__Metschnikowia_unclassified(100); | Metschnikowia | 13 | Saprotroph | \|Undefined Saprotroph\| | Yeast | NULL | Probable | As nectar-tap_saprotroph (Põlme et al. 2020); Decay sugar-rich substrates (Põlme et al. 2020) | Põlme S, et al. 2020. Fungal Diversity 105:1-16 ((https://doi.org/10.1007/s13225-020-00466-2)) |
| Otu47364 | 1 | k__Fungi(100);p__Ascomycota(100);c__Saccharomycetes(100);o__Saccharomycetales(100);f__Metschnikowiaceae(100);g__Metschnikowia(100);g__Metschnikowia_unclassified(100); | Metschnikowia | 13 | Saprotroph | \|Undefined Saprotroph\| | Yeast | NULL | Probable | As nectar-tap_saprotroph (Põlme et al. 2020); Decay sugar-rich substrates (Põlme et al. 2020) | Põlme S, et al. 2020. Fungal Diversity 105:1-16 ((https://doi.org/10.1007/s13225-020-00466-2)) |
| Otu47365 | 1 | k__Fungi(100);p__Ascomycota(100);c__Saccharomycetes(100);o__Saccharomycetales(100);f__Metschnikowiaceae(100);g__Metschnikowia(100);s__Metschnikowia_pulcherrima(100); | Metschnikowia | 13 | Saprotroph | \|Undefined Saprotroph\| | Yeast | NULL | Probable | As nectar-tap_saprotroph (Põlme et al. 2020); Decay sugar-rich substrates (Põlme et al. 2020) | Põlme S, et al. 2020. Fungal Diversity 105:1-16 ((https://doi.org/10.1007/s13225-020-00466-2)) |
| Otu47366 | 1 | k__Fungi(100);p__Ascomycota(100);c__Saccharomycetes(100);o__Saccharomycetales(100);f__Metschnikowiaceae(100);g__Metschnikowia(100);s__Metschnikowia_pulcherrima(100); | Metschnikowia | 13 | Saprotroph | \|Undefined Saprotroph\| | Yeast | NULL | Probable | As nectar-tap_saprotroph (Põlme et al. 2020); Decay sugar-rich substrates (Põlme et al. 2020) | Põlme S, et al. 2020. Fungal Diversity 105:1-16 ((https://doi.org/10.1007/s13225-020-00466-2)) |
| Otu47367 | 1 | k__Fungi(100);p__Ascomycota(100);c__Saccharomycetes(100);o__Saccharomycetales(100);f__Metschnikowiaceae(100);g__Metschnikowia(100);s__Metschnikowia_pulcherrima(100); | Metschnikowia | 13 | Saprotroph | \|Undefined Saprotroph\| | Yeast | NULL | Probable | As nectar-tap_saprotroph (Põlme et al. 2020); Decay sugar-rich substrates (Põlme et al. 2020) | Põlme S, et al. 2020. Fungal Diversity 105:1-16 ((https://doi.org/10.1007/s13225-020-00466-2)) |
| Otu47368 | 1 | k__Fungi(100);p__Ascomycota(100);c__Saccharomycetes(100);o__Saccharomycetales(100);f__Metschnikowiaceae(100);g__Metschnikowia(100);g__Metschnikowia_unclassified(100); | Metschnikowia | 13 | Saprotroph | \|Undefined Saprotroph\| | Yeast | NULL | Probable | As nectar-tap_saprotroph (Põlme et al. 2020); Decay sugar-rich substrates (Põlme et al. 2020) | Põlme S, et al. 2020. Fungal Diversity 105:1-16 ((https://doi.org/10.1007/s13225-020-00466-2)) |
| Otu47369 | 1 | k__Fungi(100);p__Ascomycota(100);c__Saccharomycetes(100);o__Saccharomycetales(100);f__Metschnikowiaceae(100);g__Metschnikowia(100);s__Metschnikowia_pulcherrima(100); | Metschnikowia | 13 | Saprotroph | \|Undefined Saprotroph\| | Yeast | NULL | Probable | As nectar-tap_saprotroph (Põlme et al. 2020); Decay sugar-rich substrates (Põlme et al. 2020) | Põlme S, et al. 2020. Fungal Diversity 105:1-16 ((https://doi.org/10.1007/s13225-020-00466-2)) |
| Otu47370 | 1 | k__Fungi(100);p__Ascomycota(100);c__Saccharomycetes(100);o__Saccharomycetales(100);f__Metschnikowiaceae(100);g__Metschnikowia(100);s__Metschnikowia_pulcherrima(100); | Metschnikowia | 13 | Saprotroph | \|Undefined Saprotroph\| | Yeast | NULL | Probable | As nectar-tap_saprotroph (Põlme et al. 2020); Decay sugar-rich substrates (Põlme et al. 2020) | Põlme S, et al. 2020. Fungal Diversity 105:1-16 ((https://doi.org/10.1007/s13225-020-00466-2)) |
| Otu47371 | 1 | k__Fungi(100);p__Ascomycota(100);c__Saccharomycetes(100);o__Saccharomycetales(100);f__Metschnikowiaceae(100);g__Metschnikowia(100);s__Metschnikowia_pulcherrima(100); | Metschnikowia | 13 | Saprotroph | \|Undefined Saprotroph\| | Yeast | NULL | Probable | As nectar-tap_saprotroph (Põlme et al. 2020); Decay sugar-rich substrates (Põlme et al. 2020) | Põlme S, et al. 2020. Fungal Diversity 105:1-16 ((https://doi.org/10.1007/s13225-020-00466-2)) |
| Otu47372 | 1 | k__Fungi(100);p__Ascomycota(100);c__Saccharomycetes(100);o__Saccharomycetales(100);f__Metschnikowiaceae(100);g__Metschnikowia(100);s__Metschnikowia_pulcherrima(100); | Metschnikowia | 13 | Saprotroph | \|Undefined Saprotroph\| | Yeast | NULL | Probable | As nectar-tap_saprotroph (Põlme et al. 2020); Decay sugar-rich substrates (Põlme et al. 2020) | Põlme S, et al. 2020. Fungal Diversity 105:1-16 ((https://doi.org/10.1007/s13225-020-00466-2)) |
| Otu47373 | 1 | k__Fungi(100);p__Ascomycota(100);c__Saccharomycetes(100);o__Saccharomycetales(100);f__Metschnikowiaceae(100);g__Metschnikowia(100);s__Metschnikowia_pulcherrima(100); | Metschnikowia | 13 | Saprotroph | \|Undefined Saprotroph\| | Yeast | NULL | Probable | As nectar-tap_saprotroph (Põlme et al. 2020); Decay sugar-rich substrates (Põlme et al. 2020) | Põlme S, et al. 2020. Fungal Diversity 105:1-16 ((https://doi.org/10.1007/s13225-020-00466-2)) |
| Otu47376 | 1 | k__Fungi(100);p__Ascomycota(100);c__Saccharomycetes(100);o__Saccharomycetales(100);f__Metschnikowiaceae(100);g__Metschnikowia(100);s__Metschnikowia_pulcherrima(100); | Metschnikowia | 13 | Saprotroph | \|Undefined Saprotroph\| | Yeast | NULL | Probable | As nectar-tap_saprotroph (Põlme et al. 2020); Decay sugar-rich substrates (Põlme et al. 2020) | Põlme S, et al. 2020. Fungal Diversity 105:1-16 ((https://doi.org/10.1007/s13225-020-00466-2)) |
| Otu47379 | 1 | k__Fungi(100);p__Ascomycota(100);c__Saccharomycetes(100);o__Saccharomycetales(100);f__Metschnikowiaceae(100);g__Metschnikowia(100);s__Metschnikowia_pulcherrima(100); | Metschnikowia | 13 | Saprotroph | \|Undefined Saprotroph\| | Yeast | NULL | Probable | As nectar-tap_saprotroph (Põlme et al. 2020); Decay sugar-rich substrates (Põlme et al. 2020) | Põlme S, et al. 2020. Fungal Diversity 105:1-16 ((https://doi.org/10.1007/s13225-020-00466-2)) |
| Otu47380 | 1 | k__Fungi(100);p__Ascomycota(100);c__Saccharomycetes(100);o__Saccharomycetales(100);f__Metschnikowiaceae(100);g__Metschnikowia(100);s__Metschnikowia_pulcherrima(100); | Metschnikowia | 13 | Saprotroph | \|Undefined Saprotroph\| | Yeast | NULL | Probable | As nectar-tap_saprotroph (Põlme et al. 2020); Decay sugar-rich substrates (Põlme et al. 2020) | Põlme S, et al. 2020. Fungal Diversity 105:1-16 ((https://doi.org/10.1007/s13225-020-00466-2)) |
| Otu47382 | 1 | k__Fungi(100);p__Ascomycota(100);c__Saccharomycetes(100);o__Saccharomycetales(100);f__Metschnikowiaceae(100);g__Metschnikowia(100);s__Metschnikowia_pulcherrima(100); | Metschnikowia | 13 | Saprotroph | \|Undefined Saprotroph\| | Yeast | NULL | Probable | As nectar-tap_saprotroph (Põlme et al. 2020); Decay sugar-rich substrates (Põlme et al. 2020) | Põlme S, et al. 2020. Fungal Diversity 105:1-16 ((https://doi.org/10.1007/s13225-020-00466-2)) |
| Otu47383 | 1 | k__Fungi(100);p__Ascomycota(100);c__Saccharomycetes(100);o__Saccharomycetales(100);f__Metschnikowiaceae(100);g__Metschnikowia(100);s__Metschnikowia_pulcherrima(100); | Metschnikowia | 13 | Saprotroph | \|Undefined Saprotroph\| | Yeast | NULL | Probable | As nectar-tap_saprotroph (Põlme et al. 2020); Decay sugar-rich substrates (Põlme et al. 2020) | Põlme S, et al. 2020. Fungal Diversity 105:1-16 ((https://doi.org/10.1007/s13225-020-00466-2)) |
| Otu47384 | 1 | k__Fungi(100);p__Ascomycota(100);c__Saccharomycetes(100);o__Saccharomycetales(100);f__Metschnikowiaceae(100);g__Metschnikowia(100);g__Metschnikowia_unclassified(100); | Metschnikowia | 13 | Saprotroph | \|Undefined Saprotroph\| | Yeast | NULL | Probable | As nectar-tap_saprotroph (Põlme et al. 2020); Decay sugar-rich substrates (Põlme et al. 2020) | Põlme S, et al. 2020. Fungal Diversity 105:1-16 ((https://doi.org/10.1007/s13225-020-00466-2)) |
| Otu47385 | 1 | k__Fungi(100);p__Ascomycota(100);c__Saccharomycetes(100);o__Saccharomycetales(100);f__Metschnikowiaceae(100);g__Metschnikowia(100);g__Metschnikowia_unclassified(100); | Metschnikowia | 13 | Saprotroph | \|Undefined Saprotroph\| | Yeast | NULL | Probable | As nectar-tap_saprotroph (Põlme et al. 2020); Decay sugar-rich substrates (Põlme et al. 2020) | Põlme S, et al. 2020. Fungal Diversity 105:1-16 ((https://doi.org/10.1007/s13225-020-00466-2)) |
| Otu47386 | 1 | k__Fungi(100);p__Ascomycota(100);c__Saccharomycetes(100);o__Saccharomycetales(100);f__Metschnikowiaceae(100);g__Metschnikowia(100);g__Metschnikowia_unclassified(100); | Metschnikowia | 13 | Saprotroph | \|Undefined Saprotroph\| | Yeast | NULL | Probable | As nectar-tap_saprotroph (Põlme et al. 2020); Decay sugar-rich substrates (Põlme et al. 2020) | Põlme S, et al. 2020. Fungal Diversity 105:1-16 ((https://doi.org/10.1007/s13225-020-00466-2)) |
| Otu47387 | 1 | k__Fungi(100);p__Ascomycota(100);c__Saccharomycetes(100);o__Saccharomycetales(100);f__Metschnikowiaceae(100);g__Metschnikowia(100);s__Metschnikowia_pulcherrima(100); | Metschnikowia | 13 | Saprotroph | \|Undefined Saprotroph\| | Yeast | NULL | Probable | As nectar-tap_saprotroph (Põlme et al. 2020); Decay sugar-rich substrates (Põlme et al. 2020) | Põlme S, et al. 2020. Fungal Diversity 105:1-16 ((https://doi.org/10.1007/s13225-020-00466-2)) |
| Otu47388 | 1 | k__Fungi(100);p__Ascomycota(100);c__Saccharomycetes(100);o__Saccharomycetales(100);f__Metschnikowiaceae(100);g__Metschnikowia(100);g__Metschnikowia_unclassified(100); | Metschnikowia | 13 | Saprotroph | \|Undefined Saprotroph\| | Yeast | NULL | Probable | As nectar-tap_saprotroph (Põlme et al. 2020); Decay sugar-rich substrates (Põlme et al. 2020) | Põlme S, et al. 2020. Fungal Diversity 105:1-16 ((https://doi.org/10.1007/s13225-020-00466-2)) |
| Otu47389 | 1 | k__Fungi(100);p__Ascomycota(100);c__Saccharomycetes(100);o__Saccharomycetales(100);f__Metschnikowiaceae(100);g__Metschnikowia(100);g__Metschnikowia_unclassified(100); | Metschnikowia | 13 | Saprotroph | \|Undefined Saprotroph\| | Yeast | NULL | Probable | As nectar-tap_saprotroph (Põlme et al. 2020); Decay sugar-rich substrates (Põlme et al. 2020) | Põlme S, et al. 2020. Fungal Diversity 105:1-16 ((https://doi.org/10.1007/s13225-020-00466-2)) |
| Otu47390 | 1 | k__Fungi(100);p__Ascomycota(100);c__Saccharomycetes(100);o__Saccharomycetales(100);f__Metschnikowiaceae(100);g__Metschnikowia(100);s__Metschnikowia_pulcherrima(100); | Metschnikowia | 13 | Saprotroph | \|Undefined Saprotroph\| | Yeast | NULL | Probable | As nectar-tap_saprotroph (Põlme et al. 2020); Decay sugar-rich substrates (Põlme et al. 2020) | Põlme S, et al. 2020. Fungal Diversity 105:1-16 ((https://doi.org/10.1007/s13225-020-00466-2)) |
| Otu47391 | 1 | k__Fungi(100);p__Ascomycota(100);c__Saccharomycetes(100);o__Saccharomycetales(100);f__Metschnikowiaceae(100);g__Metschnikowia(100);g__Metschnikowia_unclassified(100); | Metschnikowia | 13 | Saprotroph | \|Undefined Saprotroph\| | Yeast | NULL | Probable | As nectar-tap_saprotroph (Põlme et al. 2020); Decay sugar-rich substrates (Põlme et al. 2020) | Põlme S, et al. 2020. Fungal Diversity 105:1-16 ((https://doi.org/10.1007/s13225-020-00466-2)) |
| Otu47392 | 1 | k__Fungi(100);p__Ascomycota(100);c__Saccharomycetes(100);o__Saccharomycetales(100);f__Metschnikowiaceae(100);g__Metschnikowia(100);s__Metschnikowia_pulcherrima(100); | Metschnikowia | 13 | Saprotroph | \|Undefined Saprotroph\| | Yeast | NULL | Probable | As nectar-tap_saprotroph (Põlme et al. 2020); Decay sugar-rich substrates (Põlme et al. 2020) | Põlme S, et al. 2020. Fungal Diversity 105:1-16 ((https://doi.org/10.1007/s13225-020-00466-2)) |
| Otu47393 | 1 | k__Fungi(100);p__Ascomycota(100);c__Saccharomycetes(100);o__Saccharomycetales(100);f__Metschnikowiaceae(100);g__Metschnikowia(100);g__Metschnikowia_unclassified(100); | Metschnikowia | 13 | Saprotroph | \|Undefined Saprotroph\| | Yeast | NULL | Probable | As nectar-tap_saprotroph (Põlme et al. 2020); Decay sugar-rich substrates (Põlme et al. 2020) | Põlme S, et al. 2020. Fungal Diversity 105:1-16 ((https://doi.org/10.1007/s13225-020-00466-2)) |
| Otu47394 | 1 | k__Fungi(100);p__Ascomycota(100);c__Saccharomycetes(100);o__Saccharomycetales(100);f__Metschnikowiaceae(100);g__Metschnikowia(100);g__Metschnikowia_unclassified(100); | Metschnikowia | 13 | Saprotroph | \|Undefined Saprotroph\| | Yeast | NULL | Probable | As nectar-tap_saprotroph (Põlme et al. 2020); Decay sugar-rich substrates (Põlme et al. 2020) | Põlme S, et al. 2020. Fungal Diversity 105:1-16 ((https://doi.org/10.1007/s13225-020-00466-2)) |
| Otu47395 | 1 | k__Fungi(100);p__Ascomycota(100);c__Saccharomycetes(100);o__Saccharomycetales(100);f__Metschnikowiaceae(100);g__Metschnikowia(100);s__Metschnikowia_pulcherrima(100); | Metschnikowia | 13 | Saprotroph | \|Undefined Saprotroph\| | Yeast | NULL | Probable | As nectar-tap_saprotroph (Põlme et al. 2020); Decay sugar-rich substrates (Põlme et al. 2020) | Põlme S, et al. 2020. Fungal Diversity 105:1-16 ((https://doi.org/10.1007/s13225-020-00466-2)) |
| Otu47396 | 1 | k__Fungi(100);p__Ascomycota(100);c__Saccharomycetes(100);o__Saccharomycetales(100);f__Metschnikowiaceae(100);g__Metschnikowia(100);s__Metschnikowia_pulcherrima(100); | Metschnikowia | 13 | Saprotroph | \|Undefined Saprotroph\| | Yeast | NULL | Probable | As nectar-tap_saprotroph (Põlme et al. 2020); Decay sugar-rich substrates (Põlme et al. 2020) | Põlme S, et al. 2020. Fungal Diversity 105:1-16 ((https://doi.org/10.1007/s13225-020-00466-2)) |
| Otu47397 | 1 | k__Fungi(100);p__Ascomycota(100);c__Saccharomycetes(100);o__Saccharomycetales(100);f__Metschnikowiaceae(100);g__Metschnikowia(100);g__Metschnikowia_unclassified(100); | Metschnikowia | 13 | Saprotroph | \|Undefined Saprotroph\| | Yeast | NULL | Probable | As nectar-tap_saprotroph (Põlme et al. 2020); Decay sugar-rich substrates (Põlme et al. 2020) | Põlme S, et al. 2020. Fungal Diversity 105:1-16 ((https://doi.org/10.1007/s13225-020-00466-2)) |
| Otu47398 | 1 | k__Fungi(100);p__Ascomycota(100);c__Saccharomycetes(100);o__Saccharomycetales(100);f__Metschnikowiaceae(100);g__Metschnikowia(100);s__Metschnikowia_pulcherrima(100); | Metschnikowia | 13 | Saprotroph | \|Undefined Saprotroph\| | Yeast | NULL | Probable | As nectar-tap_saprotroph (Põlme et al. 2020); Decay sugar-rich substrates (Põlme et al. 2020) | Põlme S, et al. 2020. Fungal Diversity 105:1-16 ((https://doi.org/10.1007/s13225-020-00466-2)) |
| Otu47399 | 1 | k__Fungi(100);p__Ascomycota(100);c__Saccharomycetes(100);o__Saccharomycetales(100);f__Metschnikowiaceae(100);g__Metschnikowia(100);s__Metschnikowia_pulcherrima(100); | Metschnikowia | 13 | Saprotroph | \|Undefined Saprotroph\| | Yeast | NULL | Probable | As nectar-tap_saprotroph (Põlme et al. 2020); Decay sugar-rich substrates (Põlme et al. 2020) | Põlme S, et al. 2020. Fungal Diversity 105:1-16 ((https://doi.org/10.1007/s13225-020-00466-2)) |
| Otu47401 | 1 | k__Fungi(100);p__Ascomycota(100);c__Saccharomycetes(100);o__Saccharomycetales(100);f__Metschnikowiaceae(100);g__Metschnikowia(100);s__Metschnikowia_pulcherrima(100); | Metschnikowia | 13 | Saprotroph | \|Undefined Saprotroph\| | Yeast | NULL | Probable | As nectar-tap_saprotroph (Põlme et al. 2020); Decay sugar-rich substrates (Põlme et al. 2020) | Põlme S, et al. 2020. Fungal Diversity 105:1-16 ((https://doi.org/10.1007/s13225-020-00466-2)) |
| Otu47402 | 1 | k__Fungi(100);p__Ascomycota(100);c__Saccharomycetes(100);o__Saccharomycetales(100);f__Metschnikowiaceae(100);g__Metschnikowia(100);s__Metschnikowia_pulcherrima(100); | Metschnikowia | 13 | Saprotroph | \|Undefined Saprotroph\| | Yeast | NULL | Probable | As nectar-tap_saprotroph (Põlme et al. 2020); Decay sugar-rich substrates (Põlme et al. 2020) | Põlme S, et al. 2020. Fungal Diversity 105:1-16 ((https://doi.org/10.1007/s13225-020-00466-2)) |
| Otu47403 | 1 | k__Fungi(100);p__Ascomycota(100);c__Saccharomycetes(100);o__Saccharomycetales(100);f__Metschnikowiaceae(100);g__Metschnikowia(100);s__Metschnikowia_pulcherrima(100); | Metschnikowia | 13 | Saprotroph | \|Undefined Saprotroph\| | Yeast | NULL | Probable | As nectar-tap_saprotroph (Põlme et al. 2020); Decay sugar-rich substrates (Põlme et al. 2020) | Põlme S, et al. 2020. Fungal Diversity 105:1-16 ((https://doi.org/10.1007/s13225-020-00466-2)) |
| Otu47404 | 1 | k__Fungi(100);p__Ascomycota(100);c__Saccharomycetes(100);o__Saccharomycetales(100);f__Metschnikowiaceae(100);g__Metschnikowia(100);s__Metschnikowia_pulcherrima(100); | Metschnikowia | 13 | Saprotroph | \|Undefined Saprotroph\| | Yeast | NULL | Probable | As nectar-tap_saprotroph (Põlme et al. 2020); Decay sugar-rich substrates (Põlme et al. 2020) | Põlme S, et al. 2020. Fungal Diversity 105:1-16 ((https://doi.org/10.1007/s13225-020-00466-2)) |
| Otu47405 | 1 | k__Fungi(100);p__Ascomycota(100);c__Saccharomycetes(100);o__Saccharomycetales(100);f__Metschnikowiaceae(100);g__Metschnikowia(100);g__Metschnikowia_unclassified(100); | Metschnikowia | 13 | Saprotroph | \|Undefined Saprotroph\| | Yeast | NULL | Probable | As nectar-tap_saprotroph (Põlme et al. 2020); Decay sugar-rich substrates (Põlme et al. 2020) | Põlme S, et al. 2020. Fungal Diversity 105:1-16 ((https://doi.org/10.1007/s13225-020-00466-2)) |
| Otu47406 | 1 | k__Fungi(100);p__Ascomycota(100);c__Saccharomycetes(100);o__Saccharomycetales(100);f__Metschnikowiaceae(100);g__Metschnikowia(100);g__Metschnikowia_unclassified(100); | Metschnikowia | 13 | Saprotroph | \|Undefined Saprotroph\| | Yeast | NULL | Probable | As nectar-tap_saprotroph (Põlme et al. 2020); Decay sugar-rich substrates (Põlme et al. 2020) | Põlme S, et al. 2020. Fungal Diversity 105:1-16 ((https://doi.org/10.1007/s13225-020-00466-2)) |
| Otu47407 | 1 | k__Fungi(100);p__Ascomycota(100);c__Saccharomycetes(100);o__Saccharomycetales(100);f__Metschnikowiaceae(100);g__Metschnikowia(100);g__Metschnikowia_unclassified(100); | Metschnikowia | 13 | Saprotroph | \|Undefined Saprotroph\| | Yeast | NULL | Probable | As nectar-tap_saprotroph (Põlme et al. 2020); Decay sugar-rich substrates (Põlme et al. 2020) | Põlme S, et al. 2020. Fungal Diversity 105:1-16 ((https://doi.org/10.1007/s13225-020-00466-2)) |
| Otu47408 | 1 | k__Fungi(100);p__Ascomycota(100);c__Saccharomycetes(100);o__Saccharomycetales(100);f__Metschnikowiaceae(100);g__Metschnikowia(100);s__Metschnikowia_pulcherrima(100); | Metschnikowia | 13 | Saprotroph | \|Undefined Saprotroph\| | Yeast | NULL | Probable | As nectar-tap_saprotroph (Põlme et al. 2020); Decay sugar-rich substrates (Põlme et al. 2020) | Põlme S, et al. 2020. Fungal Diversity 105:1-16 ((https://doi.org/10.1007/s13225-020-00466-2)) |
| Otu47409 | 1 | k__Fungi(100);p__Ascomycota(100);c__Saccharomycetes(100);o__Saccharomycetales(100);f__Metschnikowiaceae(100);g__Metschnikowia(100);s__Metschnikowia_pulcherrima(100); | Metschnikowia | 13 | Saprotroph | \|Undefined Saprotroph\| | Yeast | NULL | Probable | As nectar-tap_saprotroph (Põlme et al. 2020); Decay sugar-rich substrates (Põlme et al. 2020) | Põlme S, et al. 2020. Fungal Diversity 105:1-16 ((https://doi.org/10.1007/s13225-020-00466-2)) |
| Otu47410 | 1 | k__Fungi(100);p__Ascomycota(100);c__Saccharomycetes(100);o__Saccharomycetales(100);f__Metschnikowiaceae(100);g__Metschnikowia(100);s__Metschnikowia_pulcherrima(100); | Metschnikowia | 13 | Saprotroph | \|Undefined Saprotroph\| | Yeast | NULL | Probable | As nectar-tap_saprotroph (Põlme et al. 2020); Decay sugar-rich substrates (Põlme et al. 2020) | Põlme S, et al. 2020. Fungal Diversity 105:1-16 ((https://doi.org/10.1007/s13225-020-00466-2)) |
| Otu47411 | 1 | k__Fungi(100);p__Ascomycota(100);c__Saccharomycetes(100);o__Saccharomycetales(100);f__Metschnikowiaceae(100);g__Metschnikowia(100);g__Metschnikowia_unclassified(100); | Metschnikowia | 13 | Saprotroph | \|Undefined Saprotroph\| | Yeast | NULL | Probable | As nectar-tap_saprotroph (Põlme et al. 2020); Decay sugar-rich substrates (Põlme et al. 2020) | Põlme S, et al. 2020. Fungal Diversity 105:1-16 ((https://doi.org/10.1007/s13225-020-00466-2)) |
| Otu47412 | 1 | k__Fungi(100);p__Ascomycota(100);c__Saccharomycetes(100);o__Saccharomycetales(100);f__Metschnikowiaceae(100);g__Metschnikowia(100);s__Metschnikowia_pulcherrima(100); | Metschnikowia | 13 | Saprotroph | \|Undefined Saprotroph\| | Yeast | NULL | Probable | As nectar-tap_saprotroph (Põlme et al. 2020); Decay sugar-rich substrates (Põlme et al. 2020) | Põlme S, et al. 2020. Fungal Diversity 105:1-16 ((https://doi.org/10.1007/s13225-020-00466-2)) |
| Otu47413 | 1 | k__Fungi(100);p__Ascomycota(100);c__Saccharomycetes(100);o__Saccharomycetales(100);f__Metschnikowiaceae(100);g__Metschnikowia(100);g__Metschnikowia_unclassified(100); | Metschnikowia | 13 | Saprotroph | \|Undefined Saprotroph\| | Yeast | NULL | Probable | As nectar-tap_saprotroph (Põlme et al. 2020); Decay sugar-rich substrates (Põlme et al. 2020) | Põlme S, et al. 2020. Fungal Diversity 105:1-16 ((https://doi.org/10.1007/s13225-020-00466-2)) |
| Otu47414 | 1 | k__Fungi(100);p__Ascomycota(100);c__Saccharomycetes(100);o__Saccharomycetales(100);f__Metschnikowiaceae(100);g__Metschnikowia(100);g__Metschnikowia_unclassified(100); | Metschnikowia | 13 | Saprotroph | \|Undefined Saprotroph\| | Yeast | NULL | Probable | As nectar-tap_saprotroph (Põlme et al. 2020); Decay sugar-rich substrates (Põlme et al. 2020) | Põlme S, et al. 2020. Fungal Diversity 105:1-16 ((https://doi.org/10.1007/s13225-020-00466-2)) |
| Otu47415 | 1 | k__Fungi(100);p__Ascomycota(100);c__Saccharomycetes(100);o__Saccharomycetales(100);f__Metschnikowiaceae(100);g__Metschnikowia(100);s__Metschnikowia_pulcherrima(100); | Metschnikowia | 13 | Saprotroph | \|Undefined Saprotroph\| | Yeast | NULL | Probable | As nectar-tap_saprotroph (Põlme et al. 2020); Decay sugar-rich substrates (Põlme et al. 2020) | Põlme S, et al. 2020. Fungal Diversity 105:1-16 ((https://doi.org/10.1007/s13225-020-00466-2)) |
| Otu47416 | 1 | k__Fungi(100);p__Ascomycota(100);c__Saccharomycetes(100);o__Saccharomycetales(100);f__Metschnikowiaceae(100);g__Metschnikowia(100);s__Metschnikowia_pulcherrima(100); | Metschnikowia | 13 | Saprotroph | \|Undefined Saprotroph\| | Yeast | NULL | Probable | As nectar-tap_saprotroph (Põlme et al. 2020); Decay sugar-rich substrates (Põlme et al. 2020) | Põlme S, et al. 2020. Fungal Diversity 105:1-16 ((https://doi.org/10.1007/s13225-020-00466-2)) |
| Otu47417 | 1 | k__Fungi(100);p__Ascomycota(100);c__Saccharomycetes(100);o__Saccharomycetales(100);f__Metschnikowiaceae(100);g__Metschnikowia(100);s__Metschnikowia_pulcherrima(100); | Metschnikowia | 13 | Saprotroph | \|Undefined Saprotroph\| | Yeast | NULL | Probable | As nectar-tap_saprotroph (Põlme et al. 2020); Decay sugar-rich substrates (Põlme et al. 2020) | Põlme S, et al. 2020. Fungal Diversity 105:1-16 ((https://doi.org/10.1007/s13225-020-00466-2)) |
| Otu47418 | 1 | k__Fungi(100);p__Ascomycota(100);c__Saccharomycetes(100);o__Saccharomycetales(100);f__Metschnikowiaceae(100);g__Metschnikowia(100);s__Metschnikowia_pulcherrima(100); | Metschnikowia | 13 | Saprotroph | \|Undefined Saprotroph\| | Yeast | NULL | Probable | As nectar-tap_saprotroph (Põlme et al. 2020); Decay sugar-rich substrates (Põlme et al. 2020) | Põlme S, et al. 2020. Fungal Diversity 105:1-16 ((https://doi.org/10.1007/s13225-020-00466-2)) |
| Otu47419 | 1 | k__Fungi(100);p__Ascomycota(100);c__Saccharomycetes(100);o__Saccharomycetales(100);f__Metschnikowiaceae(100);g__Metschnikowia(100);s__Metschnikowia_pulcherrima(100); | Metschnikowia | 13 | Saprotroph | \|Undefined Saprotroph\| | Yeast | NULL | Probable | As nectar-tap_saprotroph (Põlme et al. 2020); Decay sugar-rich substrates (Põlme et al. 2020) | Põlme S, et al. 2020. Fungal Diversity 105:1-16 ((https://doi.org/10.1007/s13225-020-00466-2)) |
| Otu47420 | 1 | k__Fungi(100);p__Ascomycota(100);c__Saccharomycetes(100);o__Saccharomycetales(100);f__Metschnikowiaceae(100);g__Metschnikowia(100);g__Metschnikowia_unclassified(100); | Metschnikowia | 13 | Saprotroph | \|Undefined Saprotroph\| | Yeast | NULL | Probable | As nectar-tap_saprotroph (Põlme et al. 2020); Decay sugar-rich substrates (Põlme et al. 2020) | Põlme S, et al. 2020. Fungal Diversity 105:1-16 ((https://doi.org/10.1007/s13225-020-00466-2)) |
| Otu47421 | 1 | k__Fungi(100);p__Ascomycota(100);c__Saccharomycetes(100);o__Saccharomycetales(100);f__Metschnikowiaceae(100);g__Metschnikowia(100);s__Metschnikowia_pulcherrima(100); | Metschnikowia | 13 | Saprotroph | \|Undefined Saprotroph\| | Yeast | NULL | Probable | As nectar-tap_saprotroph (Põlme et al. 2020); Decay sugar-rich substrates (Põlme et al. 2020) | Põlme S, et al. 2020. Fungal Diversity 105:1-16 ((https://doi.org/10.1007/s13225-020-00466-2)) |
| Otu47422 | 1 | k__Fungi(100);p__Ascomycota(100);c__Saccharomycetes(100);o__Saccharomycetales(100);f__Metschnikowiaceae(100);g__Metschnikowia(100);s__Metschnikowia_pulcherrima(100); | Metschnikowia | 13 | Saprotroph | \|Undefined Saprotroph\| | Yeast | NULL | Probable | As nectar-tap_saprotroph (Põlme et al. 2020); Decay sugar-rich substrates (Põlme et al. 2020) | Põlme S, et al. 2020. Fungal Diversity 105:1-16 ((https://doi.org/10.1007/s13225-020-00466-2)) |
| Otu47423 | 1 | k__Fungi(100);p__Ascomycota(100);c__Saccharomycetes(100);o__Saccharomycetales(100);f__Metschnikowiaceae(100);g__Metschnikowia(100);s__Metschnikowia_pulcherrima(100); | Metschnikowia | 13 | Saprotroph | \|Undefined Saprotroph\| | Yeast | NULL | Probable | As nectar-tap_saprotroph (Põlme et al. 2020); Decay sugar-rich substrates (Põlme et al. 2020) | Põlme S, et al. 2020. Fungal Diversity 105:1-16 ((https://doi.org/10.1007/s13225-020-00466-2)) |
| Otu47424 | 1 | k__Fungi(100);p__Ascomycota(100);c__Saccharomycetes(100);o__Saccharomycetales(100);f__Metschnikowiaceae(100);g__Metschnikowia(100);g__Metschnikowia_unclassified(100); | Metschnikowia | 13 | Saprotroph | \|Undefined Saprotroph\| | Yeast | NULL | Probable | As nectar-tap_saprotroph (Põlme et al. 2020); Decay sugar-rich substrates (Põlme et al. 2020) | Põlme S, et al. 2020. Fungal Diversity 105:1-16 ((https://doi.org/10.1007/s13225-020-00466-2)) |
| Otu47425 | 1 | k__Fungi(100);p__Ascomycota(100);c__Sordariomycetes(100);o__Hypocreales(100);f__Nectriaceae(100);g__Fusarium(100);g__Fusarium_unclassified(100); | Fusarium | 13 | Pathotroph-Saprotroph-Symbiotroph | Animal Pathogen-Endophyte-Lichen Parasite-\|Plant Pathogen\|-Plant Saprotroph-Undefined Saprotroph-Wood Saprotroph | Microfungus | Soft Rot | Possible | As foliar_endophyte (Põlme et al. 2020); As litter_saprotroph (Põlme et al. 2020); Causing damping-off disease (Põlme et al. 2020); Decay leaf, fruit, seed, roots, soil, animal material (Põlme et al. 2020); Likely opportunistic human pathogen (Irinyi et al. 2015) | Duncan CG, Eslyn WE. 1966. Mycologia 58:642-664 ((https://doi.org/10.2307/3757045)); Seehann G, et al. 1975. List of Fungi in Soft-Rot Tissue Tests. Institute fur Holzbiologie und Holzschutz, Bundesforschungsanstalt fur Forst - und Holzwirtshaft, Hamburg - Lohbrugge, Germany; Costa IPMW, et al. 2012. Mycotaxon 119:e26 ((http://mycotaxon.com/resources/checklists/costa_v119_checklist.pdf)); Tedersoo L, et al. 2014. Science 346:e1256688 ((https://doi.org/10.1126/science.1256688)); Irinyi L, et al. 2015. Medical Mycology 53:313-337 ((https://doi.org/10.1093/mmy/myv008)); Schroers, et al. 2016. Mycologia 108:806-819 ((https://doi.org/10.3852/15-255)); Põlme S, et al. 2020. Fungal Diversity 105:1-16 ((https://doi.org/10.1007/s13225-020-00466-2)); The American Phytopathological Society Common Names of Plant Diseases ((http://www.apsnet.org/publications/commonnames/Pages/default.aspx)); Lawrey JD, Diederich P. 2016. http://www.lichenicolous.net (accessed 24 May 2017) ((http://www.lichenicolous.net)) |
| Otu47426 | 1 | k__Fungi(100);p__Ascomycota(100);c__Saccharomycetes(100);o__Saccharomycetales(100);f__Metschnikowiaceae(100);g__Metschnikowia(100);g__Metschnikowia_unclassified(100); | Metschnikowia | 13 | Saprotroph | \|Undefined Saprotroph\| | Yeast | NULL | Probable | As nectar-tap_saprotroph (Põlme et al. 2020); Decay sugar-rich substrates (Põlme et al. 2020) | Põlme S, et al. 2020. Fungal Diversity 105:1-16 ((https://doi.org/10.1007/s13225-020-00466-2)) |
| Otu47427 | 1 | k__Fungi(100);p__Ascomycota(100);c__Saccharomycetes(100);o__Saccharomycetales(100);f__Metschnikowiaceae(100);g__Metschnikowia(100);s__Metschnikowia_pulcherrima(100); | Metschnikowia | 13 | Saprotroph | \|Undefined Saprotroph\| | Yeast | NULL | Probable | As nectar-tap_saprotroph (Põlme et al. 2020); Decay sugar-rich substrates (Põlme et al. 2020) | Põlme S, et al. 2020. Fungal Diversity 105:1-16 ((https://doi.org/10.1007/s13225-020-00466-2)) |
| Otu47428 | 1 | k__Fungi(100);p__Ascomycota(100);c__Saccharomycetes(100);o__Saccharomycetales(100);f__Metschnikowiaceae(100);g__Metschnikowia(100);g__Metschnikowia_unclassified(100); | Metschnikowia | 13 | Saprotroph | \|Undefined Saprotroph\| | Yeast | NULL | Probable | As nectar-tap_saprotroph (Põlme et al. 2020); Decay sugar-rich substrates (Põlme et al. 2020) | Põlme S, et al. 2020. Fungal Diversity 105:1-16 ((https://doi.org/10.1007/s13225-020-00466-2)) |
| Otu47429 | 1 | k__Fungi(100);p__Ascomycota(100);c__Saccharomycetes(100);o__Saccharomycetales(100);f__Metschnikowiaceae(100);g__Metschnikowia(100);s__Metschnikowia_pulcherrima(100); | Metschnikowia | 13 | Saprotroph | \|Undefined Saprotroph\| | Yeast | NULL | Probable | As nectar-tap_saprotroph (Põlme et al. 2020); Decay sugar-rich substrates (Põlme et al. 2020) | Põlme S, et al. 2020. Fungal Diversity 105:1-16 ((https://doi.org/10.1007/s13225-020-00466-2)) |
| Otu47430 | 1 | k__Fungi(100);p__Ascomycota(100);c__Saccharomycetes(100);o__Saccharomycetales(100);f__Metschnikowiaceae(100);g__Metschnikowia(100);s__Metschnikowia_pulcherrima(100); | Metschnikowia | 13 | Saprotroph | \|Undefined Saprotroph\| | Yeast | NULL | Probable | As nectar-tap_saprotroph (Põlme et al. 2020); Decay sugar-rich substrates (Põlme et al. 2020) | Põlme S, et al. 2020. Fungal Diversity 105:1-16 ((https://doi.org/10.1007/s13225-020-00466-2)) |
| Otu47431 | 1 | k__Fungi(100);p__Ascomycota(100);c__Saccharomycetes(100);o__Saccharomycetales(100);f__Metschnikowiaceae(100);g__Metschnikowia(100);s__Metschnikowia_pulcherrima(100); | Metschnikowia | 13 | Saprotroph | \|Undefined Saprotroph\| | Yeast | NULL | Probable | As nectar-tap_saprotroph (Põlme et al. 2020); Decay sugar-rich substrates (Põlme et al. 2020) | Põlme S, et al. 2020. Fungal Diversity 105:1-16 ((https://doi.org/10.1007/s13225-020-00466-2)) |
| Otu47432 | 1 | k__Fungi(100);p__Ascomycota(100);c__Saccharomycetes(100);o__Saccharomycetales(100);f__Metschnikowiaceae(100);g__Metschnikowia(100);s__Metschnikowia_pulcherrima(100); | Metschnikowia | 13 | Saprotroph | \|Undefined Saprotroph\| | Yeast | NULL | Probable | As nectar-tap_saprotroph (Põlme et al. 2020); Decay sugar-rich substrates (Põlme et al. 2020) | Põlme S, et al. 2020. Fungal Diversity 105:1-16 ((https://doi.org/10.1007/s13225-020-00466-2)) |
| Otu47433 | 1 | k__Fungi(100);p__Ascomycota(100);c__Saccharomycetes(100);o__Saccharomycetales(100);f__Metschnikowiaceae(100);g__Metschnikowia(100);s__Metschnikowia_pulcherrima(100); | Metschnikowia | 13 | Saprotroph | \|Undefined Saprotroph\| | Yeast | NULL | Probable | As nectar-tap_saprotroph (Põlme et al. 2020); Decay sugar-rich substrates (Põlme et al. 2020) | Põlme S, et al. 2020. Fungal Diversity 105:1-16 ((https://doi.org/10.1007/s13225-020-00466-2)) |
| Otu47434 | 1 | k__Fungi(100);p__Ascomycota(100);c__Saccharomycetes(100);o__Saccharomycetales(100);f__Metschnikowiaceae(100);g__Metschnikowia(100);s__Metschnikowia_pulcherrima(100); | Metschnikowia | 13 | Saprotroph | \|Undefined Saprotroph\| | Yeast | NULL | Probable | As nectar-tap_saprotroph (Põlme et al. 2020); Decay sugar-rich substrates (Põlme et al. 2020) | Põlme S, et al. 2020. Fungal Diversity 105:1-16 ((https://doi.org/10.1007/s13225-020-00466-2)) |
| Otu47435 | 1 | k__Fungi(100);p__Ascomycota(100);c__Saccharomycetes(100);o__Saccharomycetales(100);f__Metschnikowiaceae(100);g__Metschnikowia(100);g__Metschnikowia_unclassified(100); | Metschnikowia | 13 | Saprotroph | \|Undefined Saprotroph\| | Yeast | NULL | Probable | As nectar-tap_saprotroph (Põlme et al. 2020); Decay sugar-rich substrates (Põlme et al. 2020) | Põlme S, et al. 2020. Fungal Diversity 105:1-16 ((https://doi.org/10.1007/s13225-020-00466-2)) |
| Otu47436 | 1 | k__Fungi(100);p__Ascomycota(100);c__Saccharomycetes(100);o__Saccharomycetales(100);f__Metschnikowiaceae(100);g__Metschnikowia(100);s__Metschnikowia_pulcherrima(100); | Metschnikowia | 13 | Saprotroph | \|Undefined Saprotroph\| | Yeast | NULL | Probable | As nectar-tap_saprotroph (Põlme et al. 2020); Decay sugar-rich substrates (Põlme et al. 2020) | Põlme S, et al. 2020. Fungal Diversity 105:1-16 ((https://doi.org/10.1007/s13225-020-00466-2)) |
| Otu47437 | 1 | k__Fungi(100);p__Ascomycota(100);c__Saccharomycetes(100);o__Saccharomycetales(100);f__Metschnikowiaceae(100);g__Metschnikowia(100);g__Metschnikowia_unclassified(100); | Metschnikowia | 13 | Saprotroph | \|Undefined Saprotroph\| | Yeast | NULL | Probable | As nectar-tap_saprotroph (Põlme et al. 2020); Decay sugar-rich substrates (Põlme et al. 2020) | Põlme S, et al. 2020. Fungal Diversity 105:1-16 ((https://doi.org/10.1007/s13225-020-00466-2)) |
| Otu47438 | 1 | k__Fungi(100);p__Ascomycota(100);c__Saccharomycetes(100);o__Saccharomycetales(100);f__Metschnikowiaceae(100);g__Metschnikowia(100);s__Metschnikowia_pulcherrima(100); | Metschnikowia | 13 | Saprotroph | \|Undefined Saprotroph\| | Yeast | NULL | Probable | As nectar-tap_saprotroph (Põlme et al. 2020); Decay sugar-rich substrates (Põlme et al. 2020) | Põlme S, et al. 2020. Fungal Diversity 105:1-16 ((https://doi.org/10.1007/s13225-020-00466-2)) |
| Otu47439 | 1 | k__Fungi(100);p__Ascomycota(100);c__Saccharomycetes(100);o__Saccharomycetales(100);f__Metschnikowiaceae(100);g__Metschnikowia(100);g__Metschnikowia_unclassified(100); | Metschnikowia | 13 | Saprotroph | \|Undefined Saprotroph\| | Yeast | NULL | Probable | As nectar-tap_saprotroph (Põlme et al. 2020); Decay sugar-rich substrates (Põlme et al. 2020) | Põlme S, et al. 2020. Fungal Diversity 105:1-16 ((https://doi.org/10.1007/s13225-020-00466-2)) |
| Otu47440 | 1 | k__Fungi(100);p__Ascomycota(100);c__Saccharomycetes(100);o__Saccharomycetales(100);f__Metschnikowiaceae(100);g__Metschnikowia(100);s__Metschnikowia_pulcherrima(100); | Metschnikowia | 13 | Saprotroph | \|Undefined Saprotroph\| | Yeast | NULL | Probable | As nectar-tap_saprotroph (Põlme et al. 2020); Decay sugar-rich substrates (Põlme et al. 2020) | Põlme S, et al. 2020. Fungal Diversity 105:1-16 ((https://doi.org/10.1007/s13225-020-00466-2)) |
| Otu47441 | 1 | k__Fungi(100);p__Ascomycota(100);c__Saccharomycetes(100);o__Saccharomycetales(100);f__Metschnikowiaceae(100);g__Metschnikowia(100);g__Metschnikowia_unclassified(100); | Metschnikowia | 13 | Saprotroph | \|Undefined Saprotroph\| | Yeast | NULL | Probable | As nectar-tap_saprotroph (Põlme et al. 2020); Decay sugar-rich substrates (Põlme et al. 2020) | Põlme S, et al. 2020. Fungal Diversity 105:1-16 ((https://doi.org/10.1007/s13225-020-00466-2)) |
| Otu47442 | 1 | k__Fungi(100);p__Ascomycota(100);c__Saccharomycetes(100);o__Saccharomycetales(100);f__Metschnikowiaceae(100);g__Metschnikowia(100);g__Metschnikowia_unclassified(100); | Metschnikowia | 13 | Saprotroph | \|Undefined Saprotroph\| | Yeast | NULL | Probable | As nectar-tap_saprotroph (Põlme et al. 2020); Decay sugar-rich substrates (Põlme et al. 2020) | Põlme S, et al. 2020. Fungal Diversity 105:1-16 ((https://doi.org/10.1007/s13225-020-00466-2)) |
| Otu47443 | 1 | k__Fungi(100);p__Ascomycota(100);c__Saccharomycetes(100);o__Saccharomycetales(100);f__Metschnikowiaceae(100);g__Metschnikowia(100);s__Metschnikowia_pulcherrima(100); | Metschnikowia | 13 | Saprotroph | \|Undefined Saprotroph\| | Yeast | NULL | Probable | As nectar-tap_saprotroph (Põlme et al. 2020); Decay sugar-rich substrates (Põlme et al. 2020) | Põlme S, et al. 2020. Fungal Diversity 105:1-16 ((https://doi.org/10.1007/s13225-020-00466-2)) |
| Otu47444 | 1 | k__Fungi(100);p__Ascomycota(100);c__Saccharomycetes(100);o__Saccharomycetales(100);f__Metschnikowiaceae(100);g__Metschnikowia(100);s__Metschnikowia_pulcherrima(100); | Metschnikowia | 13 | Saprotroph | \|Undefined Saprotroph\| | Yeast | NULL | Probable | As nectar-tap_saprotroph (Põlme et al. 2020); Decay sugar-rich substrates (Põlme et al. 2020) | Põlme S, et al. 2020. Fungal Diversity 105:1-16 ((https://doi.org/10.1007/s13225-020-00466-2)) |
| Otu47445 | 1 | k__Fungi(100);p__Ascomycota(100);c__Saccharomycetes(100);o__Saccharomycetales(100);f__Metschnikowiaceae(100);g__Metschnikowia(100);g__Metschnikowia_unclassified(100); | Metschnikowia | 13 | Saprotroph | \|Undefined Saprotroph\| | Yeast | NULL | Probable | As nectar-tap_saprotroph (Põlme et al. 2020); Decay sugar-rich substrates (Põlme et al. 2020) | Põlme S, et al. 2020. Fungal Diversity 105:1-16 ((https://doi.org/10.1007/s13225-020-00466-2)) |
| Otu47446 | 1 | k__Fungi(100);p__Ascomycota(100);c__Saccharomycetes(100);o__Saccharomycetales(100);f__Metschnikowiaceae(100);g__Metschnikowia(100);s__Metschnikowia_pulcherrima(100); | Metschnikowia | 13 | Saprotroph | \|Undefined Saprotroph\| | Yeast | NULL | Probable | As nectar-tap_saprotroph (Põlme et al. 2020); Decay sugar-rich substrates (Põlme et al. 2020) | Põlme S, et al. 2020. Fungal Diversity 105:1-16 ((https://doi.org/10.1007/s13225-020-00466-2)) |
| Otu47447 | 1 | k__Fungi(100);p__Ascomycota(100);c__Saccharomycetes(100);o__Saccharomycetales(100);f__Metschnikowiaceae(100);g__Metschnikowia(100);s__Metschnikowia_pulcherrima(100); | Metschnikowia | 13 | Saprotroph | \|Undefined Saprotroph\| | Yeast | NULL | Probable | As nectar-tap_saprotroph (Põlme et al. 2020); Decay sugar-rich substrates (Põlme et al. 2020) | Põlme S, et al. 2020. Fungal Diversity 105:1-16 ((https://doi.org/10.1007/s13225-020-00466-2)) |
| Otu47448 | 1 | k__Fungi(100);p__Ascomycota(100);c__Saccharomycetes(100);o__Saccharomycetales(100);f__Metschnikowiaceae(100);g__Metschnikowia(100);s__Metschnikowia_pulcherrima(100); | Metschnikowia | 13 | Saprotroph | \|Undefined Saprotroph\| | Yeast | NULL | Probable | As nectar-tap_saprotroph (Põlme et al. 2020); Decay sugar-rich substrates (Põlme et al. 2020) | Põlme S, et al. 2020. Fungal Diversity 105:1-16 ((https://doi.org/10.1007/s13225-020-00466-2)) |
| Otu47449 | 1 | k__Fungi(100);p__Ascomycota(100);c__Saccharomycetes(100);o__Saccharomycetales(100);f__Metschnikowiaceae(100);g__Metschnikowia(100);s__Metschnikowia_pulcherrima(100); | Metschnikowia | 13 | Saprotroph | \|Undefined Saprotroph\| | Yeast | NULL | Probable | As nectar-tap_saprotroph (Põlme et al. 2020); Decay sugar-rich substrates (Põlme et al. 2020) | Põlme S, et al. 2020. Fungal Diversity 105:1-16 ((https://doi.org/10.1007/s13225-020-00466-2)) |
| Otu47450 | 1 | k__Fungi(100);p__Ascomycota(100);c__Saccharomycetes(100);o__Saccharomycetales(100);f__Metschnikowiaceae(100);g__Metschnikowia(100);g__Metschnikowia_unclassified(100); | Metschnikowia | 13 | Saprotroph | \|Undefined Saprotroph\| | Yeast | NULL | Probable | As nectar-tap_saprotroph (Põlme et al. 2020); Decay sugar-rich substrates (Põlme et al. 2020) | Põlme S, et al. 2020. Fungal Diversity 105:1-16 ((https://doi.org/10.1007/s13225-020-00466-2)) |
| Otu47451 | 1 | k__Fungi(100);p__Ascomycota(100);c__Saccharomycetes(100);o__Saccharomycetales(100);f__Metschnikowiaceae(100);g__Metschnikowia(100);g__Metschnikowia_unclassified(100); | Metschnikowia | 13 | Saprotroph | \|Undefined Saprotroph\| | Yeast | NULL | Probable | As nectar-tap_saprotroph (Põlme et al. 2020); Decay sugar-rich substrates (Põlme et al. 2020) | Põlme S, et al. 2020. Fungal Diversity 105:1-16 ((https://doi.org/10.1007/s13225-020-00466-2)) |
| Otu47452 | 1 | k__Fungi(100);p__Ascomycota(100);c__Saccharomycetes(100);o__Saccharomycetales(100);f__Metschnikowiaceae(100);g__Metschnikowia(100);s__Metschnikowia_pulcherrima(100); | Metschnikowia | 13 | Saprotroph | \|Undefined Saprotroph\| | Yeast | NULL | Probable | As nectar-tap_saprotroph (Põlme et al. 2020); Decay sugar-rich substrates (Põlme et al. 2020) | Põlme S, et al. 2020. Fungal Diversity 105:1-16 ((https://doi.org/10.1007/s13225-020-00466-2)) |
| Otu47453 | 1 | k__Fungi(100);p__Ascomycota(100);c__Saccharomycetes(100);o__Saccharomycetales(100);f__Metschnikowiaceae(100);g__Metschnikowia(100);g__Metschnikowia_unclassified(100); | Metschnikowia | 13 | Saprotroph | \|Undefined Saprotroph\| | Yeast | NULL | Probable | As nectar-tap_saprotroph (Põlme et al. 2020); Decay sugar-rich substrates (Põlme et al. 2020) | Põlme S, et al. 2020. Fungal Diversity 105:1-16 ((https://doi.org/10.1007/s13225-020-00466-2)) |
| Otu47454 | 1 | k__Fungi(100);p__Ascomycota(100);c__Saccharomycetes(100);o__Saccharomycetales(100);f__Metschnikowiaceae(100);g__Metschnikowia(100);s__Metschnikowia_pulcherrima(100); | Metschnikowia | 13 | Saprotroph | \|Undefined Saprotroph\| | Yeast | NULL | Probable | As nectar-tap_saprotroph (Põlme et al. 2020); Decay sugar-rich substrates (Põlme et al. 2020) | Põlme S, et al. 2020. Fungal Diversity 105:1-16 ((https://doi.org/10.1007/s13225-020-00466-2)) |
| Otu47455 | 1 | k__Fungi(100);p__Ascomycota(100);c__Saccharomycetes(100);o__Saccharomycetales(100);f__Metschnikowiaceae(100);g__Metschnikowia(100);s__Metschnikowia_pulcherrima(100); | Metschnikowia | 13 | Saprotroph | \|Undefined Saprotroph\| | Yeast | NULL | Probable | As nectar-tap_saprotroph (Põlme et al. 2020); Decay sugar-rich substrates (Põlme et al. 2020) | Põlme S, et al. 2020. Fungal Diversity 105:1-16 ((https://doi.org/10.1007/s13225-020-00466-2)) |
| Otu47456 | 1 | k__Fungi(100);p__Ascomycota(100);c__Saccharomycetes(100);o__Saccharomycetales(100);f__Metschnikowiaceae(100);g__Metschnikowia(100);g__Metschnikowia_unclassified(100); | Metschnikowia | 13 | Saprotroph | \|Undefined Saprotroph\| | Yeast | NULL | Probable | As nectar-tap_saprotroph (Põlme et al. 2020); Decay sugar-rich substrates (Põlme et al. 2020) | Põlme S, et al. 2020. Fungal Diversity 105:1-16 ((https://doi.org/10.1007/s13225-020-00466-2)) |
| Otu47457 | 1 | k__Fungi(100);p__Ascomycota(100);c__Saccharomycetes(100);o__Saccharomycetales(100);f__Metschnikowiaceae(100);g__Metschnikowia(100);s__Metschnikowia_pulcherrima(100); | Metschnikowia | 13 | Saprotroph | \|Undefined Saprotroph\| | Yeast | NULL | Probable | As nectar-tap_saprotroph (Põlme et al. 2020); Decay sugar-rich substrates (Põlme et al. 2020) | Põlme S, et al. 2020. Fungal Diversity 105:1-16 ((https://doi.org/10.1007/s13225-020-00466-2)) |
| Otu47458 | 1 | k__Fungi(100);p__Ascomycota(100);c__Saccharomycetes(100);o__Saccharomycetales(100);f__Metschnikowiaceae(100);g__Metschnikowia(100);s__Metschnikowia_pulcherrima(100); | Metschnikowia | 13 | Saprotroph | \|Undefined Saprotroph\| | Yeast | NULL | Probable | As nectar-tap_saprotroph (Põlme et al. 2020); Decay sugar-rich substrates (Põlme et al. 2020) | Põlme S, et al. 2020. Fungal Diversity 105:1-16 ((https://doi.org/10.1007/s13225-020-00466-2)) |
| Otu47459 | 1 | k__Fungi(100);p__Ascomycota(100);c__Saccharomycetes(100);o__Saccharomycetales(100);f__Metschnikowiaceae(100);g__Metschnikowia(100);s__Metschnikowia_pulcherrima(100); | Metschnikowia | 13 | Saprotroph | \|Undefined Saprotroph\| | Yeast | NULL | Probable | As nectar-tap_saprotroph (Põlme et al. 2020); Decay sugar-rich substrates (Põlme et al. 2020) | Põlme S, et al. 2020. Fungal Diversity 105:1-16 ((https://doi.org/10.1007/s13225-020-00466-2)) |
| Otu47460 | 1 | k__Fungi(100);p__Ascomycota(100);c__Saccharomycetes(100);o__Saccharomycetales(100);f__Metschnikowiaceae(100);g__Metschnikowia(100);s__Metschnikowia_pulcherrima(100); | Metschnikowia | 13 | Saprotroph | \|Undefined Saprotroph\| | Yeast | NULL | Probable | As nectar-tap_saprotroph (Põlme et al. 2020); Decay sugar-rich substrates (Põlme et al. 2020) | Põlme S, et al. 2020. Fungal Diversity 105:1-16 ((https://doi.org/10.1007/s13225-020-00466-2)) |
| Otu47461 | 1 | k__Fungi(100);p__Ascomycota(100);c__Saccharomycetes(100);o__Saccharomycetales(100);f__Saccharomycetales_family_Incertae_sedis(100);g__Candida(100);g__Candida_unclassified(100); | Candida | 13 | Pathotroph-Saprotroph-Symbiotroph | Animal Pathogen-Endophyte-Endosymbiont-Epiphyte-\|Undefined Saprotroph\| | Dimorphic Yeast-Yeast | NULL | Probable | As nectar-tap_saprotroph (Põlme et al. 2020); Decay sugar-rich substrates (Põlme et al. 2020); This asexual genus is polyphyletic and contain species that function very differently depend on what they are. Caution should be taken when interpreting guilds for Candida species. | Manolakaki D, et al. 2010. Virulence. 1: 367-375 ((https://doi.org/10.4161/viru.1.5.12796)); Mekbib SB, et al. 2011. Tropical Plant Pathology 36:233-240 ((http://dx.doi.org/10.1590/S1982-56762011000400004)); Põlme S, et al. 2020. Fungal Diversity 105:1-16 ((https://doi.org/10.1007/s13225-020-00466-2)) |
| Otu47462 | 1 | k__Fungi(100);p__Ascomycota(100);c__Saccharomycetes(100);o__Saccharomycetales(100);f__Metschnikowiaceae(100);g__Metschnikowia(100);s__Metschnikowia_pulcherrima(100); | Metschnikowia | 13 | Saprotroph | \|Undefined Saprotroph\| | Yeast | NULL | Probable | As nectar-tap_saprotroph (Põlme et al. 2020); Decay sugar-rich substrates (Põlme et al. 2020) | Põlme S, et al. 2020. Fungal Diversity 105:1-16 ((https://doi.org/10.1007/s13225-020-00466-2)) |
| Otu47463 | 1 | k__Fungi(100);p__Ascomycota(100);c__Saccharomycetes(100);o__Saccharomycetales(100);f__Metschnikowiaceae(100);g__Metschnikowia(100);s__Metschnikowia_pulcherrima(100); | Metschnikowia | 13 | Saprotroph | \|Undefined Saprotroph\| | Yeast | NULL | Probable | As nectar-tap_saprotroph (Põlme et al. 2020); Decay sugar-rich substrates (Põlme et al. 2020) | Põlme S, et al. 2020. Fungal Diversity 105:1-16 ((https://doi.org/10.1007/s13225-020-00466-2)) |
| Otu47464 | 1 | k__Fungi(100);p__Ascomycota(100);c__Saccharomycetes(100);o__Saccharomycetales(100);f__Metschnikowiaceae(100);g__Metschnikowia(100);s__Metschnikowia_pulcherrima(100); | Metschnikowia | 13 | Saprotroph | \|Undefined Saprotroph\| | Yeast | NULL | Probable | As nectar-tap_saprotroph (Põlme et al. 2020); Decay sugar-rich substrates (Põlme et al. 2020) | Põlme S, et al. 2020. Fungal Diversity 105:1-16 ((https://doi.org/10.1007/s13225-020-00466-2)) |
| Otu47465 | 1 | k__Fungi(100);p__Ascomycota(100);c__Saccharomycetes(100);o__Saccharomycetales(100);f__Metschnikowiaceae(100);g__Metschnikowia(100);s__Metschnikowia_pulcherrima(100); | Metschnikowia | 13 | Saprotroph | \|Undefined Saprotroph\| | Yeast | NULL | Probable | As nectar-tap_saprotroph (Põlme et al. 2020); Decay sugar-rich substrates (Põlme et al. 2020) | Põlme S, et al. 2020. Fungal Diversity 105:1-16 ((https://doi.org/10.1007/s13225-020-00466-2)) |
| Otu47466 | 1 | k__Fungi(100);p__Ascomycota(100);c__Saccharomycetes(100);o__Saccharomycetales(100);f__Metschnikowiaceae(100);g__Metschnikowia(100);g__Metschnikowia_unclassified(100); | Metschnikowia | 13 | Saprotroph | \|Undefined Saprotroph\| | Yeast | NULL | Probable | As nectar-tap_saprotroph (Põlme et al. 2020); Decay sugar-rich substrates (Põlme et al. 2020) | Põlme S, et al. 2020. Fungal Diversity 105:1-16 ((https://doi.org/10.1007/s13225-020-00466-2)) |
| Otu47467 | 1 | k__Fungi(100);p__Ascomycota(100);c__Saccharomycetes(100);o__Saccharomycetales(100);f__Metschnikowiaceae(100);g__Metschnikowia(100);s__Metschnikowia_pulcherrima(100); | Metschnikowia | 13 | Saprotroph | \|Undefined Saprotroph\| | Yeast | NULL | Probable | As nectar-tap_saprotroph (Põlme et al. 2020); Decay sugar-rich substrates (Põlme et al. 2020) | Põlme S, et al. 2020. Fungal Diversity 105:1-16 ((https://doi.org/10.1007/s13225-020-00466-2)) |
| Otu47468 | 1 | k__Fungi(100);p__Ascomycota(100);c__Saccharomycetes(100);o__Saccharomycetales(100);f__Metschnikowiaceae(100);g__Metschnikowia(100);s__Metschnikowia_pulcherrima(100); | Metschnikowia | 13 | Saprotroph | \|Undefined Saprotroph\| | Yeast | NULL | Probable | As nectar-tap_saprotroph (Põlme et al. 2020); Decay sugar-rich substrates (Põlme et al. 2020) | Põlme S, et al. 2020. Fungal Diversity 105:1-16 ((https://doi.org/10.1007/s13225-020-00466-2)) |
| Otu47469 | 1 | k__Fungi(100);p__Ascomycota(100);c__Saccharomycetes(100);o__Saccharomycetales(100);f__Metschnikowiaceae(100);g__Metschnikowia(100);g__Metschnikowia_unclassified(100); | Metschnikowia | 13 | Saprotroph | \|Undefined Saprotroph\| | Yeast | NULL | Probable | As nectar-tap_saprotroph (Põlme et al. 2020); Decay sugar-rich substrates (Põlme et al. 2020) | Põlme S, et al. 2020. Fungal Diversity 105:1-16 ((https://doi.org/10.1007/s13225-020-00466-2)) |
| Otu47470 | 1 | k__Fungi(100);p__Ascomycota(100);c__Saccharomycetes(100);o__Saccharomycetales(100);f__Metschnikowiaceae(100);g__Metschnikowia(100);g__Metschnikowia_unclassified(100); | Metschnikowia | 13 | Saprotroph | \|Undefined Saprotroph\| | Yeast | NULL | Probable | As nectar-tap_saprotroph (Põlme et al. 2020); Decay sugar-rich substrates (Põlme et al. 2020) | Põlme S, et al. 2020. Fungal Diversity 105:1-16 ((https://doi.org/10.1007/s13225-020-00466-2)) |
| Otu47472 | 1 | k__Fungi(100);p__Ascomycota(100);c__Saccharomycetes(100);o__Saccharomycetales(100);f__Metschnikowiaceae(100);g__Metschnikowia(100);g__Metschnikowia_unclassified(100); | Metschnikowia | 13 | Saprotroph | \|Undefined Saprotroph\| | Yeast | NULL | Probable | As nectar-tap_saprotroph (Põlme et al. 2020); Decay sugar-rich substrates (Põlme et al. 2020) | Põlme S, et al. 2020. Fungal Diversity 105:1-16 ((https://doi.org/10.1007/s13225-020-00466-2)) |
| Otu47473 | 1 | k__Fungi(100);p__Ascomycota(100);c__Saccharomycetes(100);o__Saccharomycetales(100);f__Metschnikowiaceae(100);g__Metschnikowia(100);s__Metschnikowia_pulcherrima(100); | Metschnikowia | 13 | Saprotroph | \|Undefined Saprotroph\| | Yeast | NULL | Probable | As nectar-tap_saprotroph (Põlme et al. 2020); Decay sugar-rich substrates (Põlme et al. 2020) | Põlme S, et al. 2020. Fungal Diversity 105:1-16 ((https://doi.org/10.1007/s13225-020-00466-2)) |
| Otu47474 | 1 | k__Fungi(100);p__Ascomycota(100);c__Saccharomycetes(100);o__Saccharomycetales(100);f__Metschnikowiaceae(100);g__Metschnikowia(100);s__Metschnikowia_pulcherrima(100); | Metschnikowia | 13 | Saprotroph | \|Undefined Saprotroph\| | Yeast | NULL | Probable | As nectar-tap_saprotroph (Põlme et al. 2020); Decay sugar-rich substrates (Põlme et al. 2020) | Põlme S, et al. 2020. Fungal Diversity 105:1-16 ((https://doi.org/10.1007/s13225-020-00466-2)) |
| Otu47475 | 1 | k__Fungi(100);p__Ascomycota(100);c__Saccharomycetes(100);o__Saccharomycetales(100);f__Metschnikowiaceae(100);g__Metschnikowia(100);s__Metschnikowia_pulcherrima(100); | Metschnikowia | 13 | Saprotroph | \|Undefined Saprotroph\| | Yeast | NULL | Probable | As nectar-tap_saprotroph (Põlme et al. 2020); Decay sugar-rich substrates (Põlme et al. 2020) | Põlme S, et al. 2020. Fungal Diversity 105:1-16 ((https://doi.org/10.1007/s13225-020-00466-2)) |
| Otu47476 | 1 | k__Fungi(100);p__Ascomycota(100);c__Saccharomycetes(100);o__Saccharomycetales(100);f__Metschnikowiaceae(100);g__Metschnikowia(100);s__Metschnikowia_pulcherrima(100); | Metschnikowia | 13 | Saprotroph | \|Undefined Saprotroph\| | Yeast | NULL | Probable | As nectar-tap_saprotroph (Põlme et al. 2020); Decay sugar-rich substrates (Põlme et al. 2020) | Põlme S, et al. 2020. Fungal Diversity 105:1-16 ((https://doi.org/10.1007/s13225-020-00466-2)) |
| Otu47477 | 1 | k__Fungi(100);p__Ascomycota(100);c__Saccharomycetes(100);o__Saccharomycetales(100);f__Metschnikowiaceae(100);g__Metschnikowia(100);g__Metschnikowia_unclassified(100); | Metschnikowia | 13 | Saprotroph | \|Undefined Saprotroph\| | Yeast | NULL | Probable | As nectar-tap_saprotroph (Põlme et al. 2020); Decay sugar-rich substrates (Põlme et al. 2020) | Põlme S, et al. 2020. Fungal Diversity 105:1-16 ((https://doi.org/10.1007/s13225-020-00466-2)) |
| Otu47478 | 1 | k__Fungi(100);p__Ascomycota(100);c__Saccharomycetes(100);o__Saccharomycetales(100);f__Metschnikowiaceae(100);g__Metschnikowia(100);g__Metschnikowia_unclassified(100); | Metschnikowia | 13 | Saprotroph | \|Undefined Saprotroph\| | Yeast | NULL | Probable | As nectar-tap_saprotroph (Põlme et al. 2020); Decay sugar-rich substrates (Põlme et al. 2020) | Põlme S, et al. 2020. Fungal Diversity 105:1-16 ((https://doi.org/10.1007/s13225-020-00466-2)) |
| Otu47479 | 1 | k__Fungi(100);p__Ascomycota(100);c__Saccharomycetes(100);o__Saccharomycetales(100);f__Metschnikowiaceae(100);g__Metschnikowia(100);g__Metschnikowia_unclassified(100); | Metschnikowia | 13 | Saprotroph | \|Undefined Saprotroph\| | Yeast | NULL | Probable | As nectar-tap_saprotroph (Põlme et al. 2020); Decay sugar-rich substrates (Põlme et al. 2020) | Põlme S, et al. 2020. Fungal Diversity 105:1-16 ((https://doi.org/10.1007/s13225-020-00466-2)) |
| Otu47480 | 1 | k__Fungi(100);p__Ascomycota(100);c__Saccharomycetes(100);o__Saccharomycetales(100);f__Metschnikowiaceae(100);g__Metschnikowia(100);s__Metschnikowia_pulcherrima(100); | Metschnikowia | 13 | Saprotroph | \|Undefined Saprotroph\| | Yeast | NULL | Probable | As nectar-tap_saprotroph (Põlme et al. 2020); Decay sugar-rich substrates (Põlme et al. 2020) | Põlme S, et al. 2020. Fungal Diversity 105:1-16 ((https://doi.org/10.1007/s13225-020-00466-2)) |
| Otu47481 | 1 | k__Fungi(100);p__Ascomycota(100);c__Saccharomycetes(100);o__Saccharomycetales(100);f__Metschnikowiaceae(100);g__Metschnikowia(100);s__Metschnikowia_pulcherrima(100); | Metschnikowia | 13 | Saprotroph | \|Undefined Saprotroph\| | Yeast | NULL | Probable | As nectar-tap_saprotroph (Põlme et al. 2020); Decay sugar-rich substrates (Põlme et al. 2020) | Põlme S, et al. 2020. Fungal Diversity 105:1-16 ((https://doi.org/10.1007/s13225-020-00466-2)) |
| Otu47482 | 1 | k__Fungi(100);p__Ascomycota(100);c__Saccharomycetes(100);o__Saccharomycetales(100);f__Metschnikowiaceae(100);g__Metschnikowia(100);s__Metschnikowia_pulcherrima(100); | Metschnikowia | 13 | Saprotroph | \|Undefined Saprotroph\| | Yeast | NULL | Probable | As nectar-tap_saprotroph (Põlme et al. 2020); Decay sugar-rich substrates (Põlme et al. 2020) | Põlme S, et al. 2020. Fungal Diversity 105:1-16 ((https://doi.org/10.1007/s13225-020-00466-2)) |
| Otu47483 | 1 | k__Fungi(100);p__Ascomycota(100);c__Saccharomycetes(100);o__Saccharomycetales(100);f__Metschnikowiaceae(100);g__Metschnikowia(100);g__Metschnikowia_unclassified(100); | Metschnikowia | 13 | Saprotroph | \|Undefined Saprotroph\| | Yeast | NULL | Probable | As nectar-tap_saprotroph (Põlme et al. 2020); Decay sugar-rich substrates (Põlme et al. 2020) | Põlme S, et al. 2020. Fungal Diversity 105:1-16 ((https://doi.org/10.1007/s13225-020-00466-2)) |
| Otu47484 | 1 | k__Fungi(100);p__Ascomycota(100);c__Saccharomycetes(100);o__Saccharomycetales(100);f__Metschnikowiaceae(100);g__Metschnikowia(100);s__Metschnikowia_pulcherrima(100); | Metschnikowia | 13 | Saprotroph | \|Undefined Saprotroph\| | Yeast | NULL | Probable | As nectar-tap_saprotroph (Põlme et al. 2020); Decay sugar-rich substrates (Põlme et al. 2020) | Põlme S, et al. 2020. Fungal Diversity 105:1-16 ((https://doi.org/10.1007/s13225-020-00466-2)) |
| Otu47485 | 1 | k__Fungi(100);p__Ascomycota(100);c__Saccharomycetes(100);o__Saccharomycetales(100);f__Metschnikowiaceae(100);g__Metschnikowia(100);s__Metschnikowia_pulcherrima(100); | Metschnikowia | 13 | Saprotroph | \|Undefined Saprotroph\| | Yeast | NULL | Probable | As nectar-tap_saprotroph (Põlme et al. 2020); Decay sugar-rich substrates (Põlme et al. 2020) | Põlme S, et al. 2020. Fungal Diversity 105:1-16 ((https://doi.org/10.1007/s13225-020-00466-2)) |
| Otu47486 | 1 | k__Fungi(100);p__Ascomycota(100);c__Saccharomycetes(100);o__Saccharomycetales(100);f__Metschnikowiaceae(100);g__Metschnikowia(100);s__Metschnikowia_pulcherrima(100); | Metschnikowia | 13 | Saprotroph | \|Undefined Saprotroph\| | Yeast | NULL | Probable | As nectar-tap_saprotroph (Põlme et al. 2020); Decay sugar-rich substrates (Põlme et al. 2020) | Põlme S, et al. 2020. Fungal Diversity 105:1-16 ((https://doi.org/10.1007/s13225-020-00466-2)) |
| Otu47487 | 1 | k__Fungi(100);p__Ascomycota(100);c__Saccharomycetes(100);o__Saccharomycetales(100);f__Metschnikowiaceae(100);g__Metschnikowia(100);s__Metschnikowia_pulcherrima(100); | Metschnikowia | 13 | Saprotroph | \|Undefined Saprotroph\| | Yeast | NULL | Probable | As nectar-tap_saprotroph (Põlme et al. 2020); Decay sugar-rich substrates (Põlme et al. 2020) | Põlme S, et al. 2020. Fungal Diversity 105:1-16 ((https://doi.org/10.1007/s13225-020-00466-2)) |
| Otu47488 | 1 | k__Fungi(100);p__Ascomycota(100);c__Saccharomycetes(100);o__Saccharomycetales(100);f__Metschnikowiaceae(100);g__Metschnikowia(100);s__Metschnikowia_pulcherrima(100); | Metschnikowia | 13 | Saprotroph | \|Undefined Saprotroph\| | Yeast | NULL | Probable | As nectar-tap_saprotroph (Põlme et al. 2020); Decay sugar-rich substrates (Põlme et al. 2020) | Põlme S, et al. 2020. Fungal Diversity 105:1-16 ((https://doi.org/10.1007/s13225-020-00466-2)) |
| Otu47489 | 1 | k__Fungi(100);p__Ascomycota(100);c__Saccharomycetes(100);o__Saccharomycetales(100);f__Metschnikowiaceae(100);g__Metschnikowia(100);s__Metschnikowia_pulcherrima(100); | Metschnikowia | 13 | Saprotroph | \|Undefined Saprotroph\| | Yeast | NULL | Probable | As nectar-tap_saprotroph (Põlme et al. 2020); Decay sugar-rich substrates (Põlme et al. 2020) | Põlme S, et al. 2020. Fungal Diversity 105:1-16 ((https://doi.org/10.1007/s13225-020-00466-2)) |
| Otu47490 | 1 | k__Fungi(100);p__Ascomycota(100);c__Saccharomycetes(100);o__Saccharomycetales(100);f__Metschnikowiaceae(100);g__Metschnikowia(100);s__Metschnikowia_pulcherrima(100); | Metschnikowia | 13 | Saprotroph | \|Undefined Saprotroph\| | Yeast | NULL | Probable | As nectar-tap_saprotroph (Põlme et al. 2020); Decay sugar-rich substrates (Põlme et al. 2020) | Põlme S, et al. 2020. Fungal Diversity 105:1-16 ((https://doi.org/10.1007/s13225-020-00466-2)) |
| Otu47491 | 1 | k__Fungi(100);p__Ascomycota(100);c__Saccharomycetes(100);o__Saccharomycetales(100);f__Metschnikowiaceae(100);g__Metschnikowia(100);g__Metschnikowia_unclassified(100); | Metschnikowia | 13 | Saprotroph | \|Undefined Saprotroph\| | Yeast | NULL | Probable | As nectar-tap_saprotroph (Põlme et al. 2020); Decay sugar-rich substrates (Põlme et al. 2020) | Põlme S, et al. 2020. Fungal Diversity 105:1-16 ((https://doi.org/10.1007/s13225-020-00466-2)) |
| Otu47492 | 1 | k__Fungi(100);p__Ascomycota(100);c__Saccharomycetes(100);o__Saccharomycetales(100);f__Metschnikowiaceae(100);g__Metschnikowia(100);s__Metschnikowia_pulcherrima(100); | Metschnikowia | 13 | Saprotroph | \|Undefined Saprotroph\| | Yeast | NULL | Probable | As nectar-tap_saprotroph (Põlme et al. 2020); Decay sugar-rich substrates (Põlme et al. 2020) | Põlme S, et al. 2020. Fungal Diversity 105:1-16 ((https://doi.org/10.1007/s13225-020-00466-2)) |
| Otu47493 | 1 | k__Fungi(100);p__Ascomycota(100);c__Saccharomycetes(100);o__Saccharomycetales(100);f__Metschnikowiaceae(100);g__Metschnikowia(100);s__Metschnikowia_pulcherrima(100); | Metschnikowia | 13 | Saprotroph | \|Undefined Saprotroph\| | Yeast | NULL | Probable | As nectar-tap_saprotroph (Põlme et al. 2020); Decay sugar-rich substrates (Põlme et al. 2020) | Põlme S, et al. 2020. Fungal Diversity 105:1-16 ((https://doi.org/10.1007/s13225-020-00466-2)) |

| | | | | | | | | | | | |
|---|---|---|---|---|---|---|---|---|---|---|---|
| Otu47494 | 1 | k__Fungi(100);p__Ascomycota(100);c__Saccharomycetes(100);o__Saccharomycetales(100);f__Metschnikowiaceae(100);g__Metschnikowia(100);s__Metschnikowia_pulcherrima(100); | Metschnikowia | 13 | Saprotroph | \|Undefined Saprotroph\| | Yeast | NULL | Probable | As nectar-tap_saprotroph (Põlme et al. 2020); Decay sugar-rich substrates (Põlme et al. 2020) | Põlme S, et al. 2020. Fungal Diversity 105:1-16 ((https://doi.org/10.1007/s13225-020-00466-2)) |
| Otu47495 | 1 | k__Fungi(100);p__Ascomycota(100);c__Saccharomycetes(100);o__Saccharomycetales(100);f__Metschnikowiaceae(100);g__Metschnikowia(100);g__Metschnikowia_unclassified(100); | Metschnikowia | 13 | Saprotroph | \|Undefined Saprotroph\| | Yeast | NULL | Probable | As nectar-tap_saprotroph (Põlme et al. 2020); Decay sugar-rich substrates (Põlme et al. 2020) | Põlme S, et al. 2020. Fungal Diversity 105:1-16 ((https://doi.org/10.1007/s13225-020-00466-2)) |
| Otu47496 | 1 | k__Fungi(100);p__Ascomycota(100);c__Saccharomycetes(100);o__Saccharomycetales(100);f__Metschnikowiaceae(100);g__Metschnikowia(100);s__Metschnikowia_pulcherrima(100); | Metschnikowia | 13 | Saprotroph | \|Undefined Saprotroph\| | Yeast | NULL | Probable | As nectar-tap_saprotroph (Põlme et al. 2020); Decay sugar-rich substrates (Põlme et al. 2020) | Põlme S, et al. 2020. Fungal Diversity 105:1-16 ((https://doi.org/10.1007/s13225-020-00466-2)) |
| Otu47497 | 1 | k__Fungi(100);p__Ascomycota(100);c__Saccharomycetes(100);o__Saccharomycetales(100);f__Metschnikowiaceae(100);g__Metschnikowia(100);s__Metschnikowia_pulcherrima(100); | Metschnikowia | 13 | Saprotroph | \|Undefined Saprotroph\| | Yeast | NULL | Probable | As nectar-tap_saprotroph (Põlme et al. 2020); Decay sugar-rich substrates (Põlme et al. 2020) | Põlme S, et al. 2020. Fungal Diversity 105:1-16 ((https://doi.org/10.1007/s13225-020-00466-2)) |
| Otu47498 | 1 | k__Fungi(100);p__Ascomycota(100);c__Saccharomycetes(100);o__Saccharomycetales(100);f__Metschnikowiaceae(100);g__Metschnikowia(100);s__Metschnikowia_pulcherrima(100); | Metschnikowia | 13 | Saprotroph | \|Undefined Saprotroph\| | Yeast | NULL | Probable | As nectar-tap_saprotroph (Põlme et al. 2020); Decay sugar-rich substrates (Põlme et al. 2020) | Põlme S, et al. 2020. Fungal Diversity 105:1-16 ((https://doi.org/10.1007/s13225-020-00466-2)) |
| Otu47499 | 1 | k__Fungi(100);p__Ascomycota(100);c__Saccharomycetes(100);o__Saccharomycetales(100);f__Metschnikowiaceae(100);g__Metschnikowia(100);s__Metschnikowia_pulcherrima(100); | Metschnikowia | 13 | Saprotroph | \|Undefined Saprotroph\| | Yeast | NULL | Probable | As nectar-tap_saprotroph (Põlme et al. 2020); Decay sugar-rich substrates (Põlme et al. 2020) | Põlme S, et al. 2020. Fungal Diversity 105:1-16 ((https://doi.org/10.1007/s13225-020-00466-2)) |
| Otu47500 | 1 | k__Fungi(100);p__Ascomycota(100);c__Saccharomycetes(100);o__Saccharomycetales(100);f__Metschnikowiaceae(100);g__Metschnikowia(100);s__Metschnikowia_pulcherrima(100); | Metschnikowia | 13 | Saprotroph | \|Undefined Saprotroph\| | Yeast | NULL | Probable | As nectar-tap_saprotroph (Põlme et al. 2020); Decay sugar-rich substrates (Põlme et al. 2020) | Põlme S, et al. 2020. Fungal Diversity 105:1-16 ((https://doi.org/10.1007/s13225-020-00466-2)) |
| Otu47501 | 1 | k__Fungi(100);p__Ascomycota(100);c__Saccharomycetes(100);o__Saccharomycetales(100);f__Metschnikowiaceae(100);g__Metschnikowia(100);s__Metschnikowia_pulcherrima(100); | Metschnikowia | 13 | Saprotroph | \|Undefined Saprotroph\| | Yeast | NULL | Probable | As nectar-tap_saprotroph (Põlme et al. 2020); Decay sugar-rich substrates (Põlme et al. 2020) | Põlme S, et al. 2020. Fungal Diversity 105:1-16 ((https://doi.org/10.1007/s13225-020-00466-2)) |
| Otu47502 | 1 | k__Fungi(100);p__Ascomycota(100);c__Saccharomycetes(100);o__Saccharomycetales(100);f__Metschnikowiaceae(100);g__Metschnikowia(100);s__Metschnikowia_pulcherrima(100); | Metschnikowia | 13 | Saprotroph | \|Undefined Saprotroph\| | Yeast | NULL | Probable | As nectar-tap_saprotroph (Põlme et al. 2020); Decay sugar-rich substrates (Põlme et al. 2020) | Põlme S, et al. 2020. Fungal Diversity 105:1-16 ((https://doi.org/10.1007/s13225-020-00466-2)) |
| Otu47503 | 1 | k__Fungi(100);p__Ascomycota(100);c__Saccharomycetes(100);o__Saccharomycetales(100);f__Metschnikowiaceae(100);g__Metschnikowia(100);s__Metschnikowia_pulcherrima(100); | Metschnikowia | 13 | Saprotroph | \|Undefined Saprotroph\| | Yeast | NULL | Probable | As nectar-tap_saprotroph (Põlme et al. 2020); Decay sugar-rich substrates (Põlme et al. 2020) | Põlme S, et al. 2020. Fungal Diversity 105:1-16 ((https://doi.org/10.1007/s13225-020-00466-2)) |
| Otu47504 | 1 | k__Fungi(100);p__Ascomycota(100);c__Saccharomycetes(100);o__Saccharomycetales(100);f__Metschnikowiaceae(100);g__Metschnikowia(100);s__Metschnikowia_pulcherrima(100); | Metschnikowia | 13 | Saprotroph | \|Undefined Saprotroph\| | Yeast | NULL | Probable | As nectar-tap_saprotroph (Põlme et al. 2020); Decay sugar-rich substrates (Põlme et al. 2020) | Põlme S, et al. 2020. Fungal Diversity 105:1-16 ((https://doi.org/10.1007/s13225-020-00466-2)) |
| Otu47505 | 1 | k__Fungi(100);p__Ascomycota(100);c__Saccharomycetes(100);o__Saccharomycetales(100);f__Metschnikowiaceae(100);g__Metschnikowia(100);s__Metschnikowia_pulcherrima(100); | Metschnikowia | 13 | Saprotroph | \|Undefined Saprotroph\| | Yeast | NULL | Probable | As nectar-tap_saprotroph (Põlme et al. 2020); Decay sugar-rich substrates (Põlme et al. 2020) | Põlme S, et al. 2020. Fungal Diversity 105:1-16 ((https://doi.org/10.1007/s13225-020-00466-2)) |
| Otu47506 | 1 | k__Fungi(100);p__Ascomycota(100);c__Saccharomycetes(100);o__Saccharomycetales(100);f__Metschnikowiaceae(100);g__Metschnikowia(100);s__Metschnikowia_pulcherrima(100); | Metschnikowia | 13 | Saprotroph | \|Undefined Saprotroph\| | Yeast | NULL | Probable | As nectar-tap_saprotroph (Põlme et al. 2020); Decay sugar-rich substrates (Põlme et al. 2020) | Põlme S, et al. 2020. Fungal Diversity 105:1-16 ((https://doi.org/10.1007/s13225-020-00466-2)) |
| Otu47507 | 1 | k__Fungi(100);p__Ascomycota(100);c__Saccharomycetes(100);o__Saccharomycetales(100);f__Metschnikowiaceae(100);g__Metschnikowia(100);s__Metschnikowia_pulcherrima(100); | Metschnikowia | 13 | Saprotroph | \|Undefined Saprotroph\| | Yeast | NULL | Probable | As nectar-tap_saprotroph (Põlme et al. 2020); Decay sugar-rich substrates (Põlme et al. 2020) | Põlme S, et al. 2020. Fungal Diversity 105:1-16 ((https://doi.org/10.1007/s13225-020-00466-2)) |
| Otu47508 | 1 | k__Fungi(100);p__Ascomycota(100);c__Saccharomycetes(100);o__Saccharomycetales(100);f__Metschnikowiaceae(100);g__Metschnikowia(100);s__Metschnikowia_pulcherrima(100); | Metschnikowia | 13 | Saprotroph | \|Undefined Saprotroph\| | Yeast | NULL | Probable | As nectar-tap_saprotroph (Põlme et al. 2020); Decay sugar-rich substrates (Põlme et al. 2020) | Põlme S, et al. 2020. Fungal Diversity 105:1-16 ((https://doi.org/10.1007/s13225-020-00466-2)) |
| Otu47509 | 1 | k__Fungi(100);p__Ascomycota(100);c__Saccharomycetes(100);o__Saccharomycetales(100);f__Metschnikowiaceae(100);g__Metschnikowia(100);s__Metschnikowia_pulcherrima(100); | Metschnikowia | 13 | Saprotroph | \|Undefined Saprotroph\| | Yeast | NULL | Probable | As nectar-tap_saprotroph (Põlme et al. 2020); Decay sugar-rich substrates (Põlme et al. 2020) | Põlme S, et al. 2020. Fungal Diversity 105:1-16 ((https://doi.org/10.1007/s13225-020-00466-2)) |
| Otu47510 | 1 | k__Fungi(100);p__Ascomycota(100);c__Saccharomycetes(100);o__Saccharomycetales(100);f__Metschnikowiaceae(100);g__Metschnikowia(100);s__Metschnikowia_pulcherrima(100); | Metschnikowia | 13 | Saprotroph | \|Undefined Saprotroph\| | Yeast | NULL | Probable | As nectar-tap_saprotroph (Põlme et al. 2020); Decay sugar-rich substrates (Põlme et al. 2020) | Põlme S, et al. 2020. Fungal Diversity 105:1-16 ((https://doi.org/10.1007/s13225-020-00466-2)) |
| Otu47511 | 1 | k__Fungi(100);p__Ascomycota(100);c__Saccharomycetes(100);o__Saccharomycetales(100);f__Metschnikowiaceae(100);g__Metschnikowia(100);s__Metschnikowia_pulcherrima(100); | Metschnikowia | 13 | Saprotroph | \|Undefined Saprotroph\| | Yeast | NULL | Probable | As nectar-tap_saprotroph (Põlme et al. 2020); Decay sugar-rich substrates (Põlme et al. 2020) | Põlme S, et al. 2020. Fungal Diversity 105:1-16 ((https://doi.org/10.1007/s13225-020-00466-2)) |
| Otu47512 | 1 | k__Fungi(100);p__Ascomycota(100);c__Saccharomycetes(100);o__Saccharomycetales(100);f__Metschnikowiaceae(100);g__Metschnikowia(100);s__Metschnikowia_pulcherrima(100); | Metschnikowia | 13 | Saprotroph | \|Undefined Saprotroph\| | Yeast | NULL | Probable | As nectar-tap_saprotroph (Põlme et al. 2020); Decay sugar-rich substrates (Põlme et al. 2020) | Põlme S, et al. 2020. Fungal Diversity 105:1-16 ((https://doi.org/10.1007/s13225-020-00466-2)) |
| Otu47513 | 1 | k__Fungi(100);p__Ascomycota(100);c__Saccharomycetes(100);o__Saccharomycetales(100);f__Metschnikowiaceae(100);g__Metschnikowia(100);g__Metschnikowia_unclassified(100); | Metschnikowia | 13 | Saprotroph | \|Undefined Saprotroph\| | Yeast | NULL | Probable | As nectar-tap_saprotroph (Põlme et al. 2020); Decay sugar-rich substrates (Põlme et al. 2020) | Põlme S, et al. 2020. Fungal Diversity 105:1-16 ((https://doi.org/10.1007/s13225-020-00466-2)) |
| Otu47514 | 1 | k__Fungi(100);p__Ascomycota(100);c__Saccharomycetes(100);o__Saccharomycetales(100);f__Metschnikowiaceae(100);g__Metschnikowia(100);s__Metschnikowia_pulcherrima(100); | Metschnikowia | 13 | Saprotroph | \|Undefined Saprotroph\| | Yeast | NULL | Probable | As nectar-tap_saprotroph (Põlme et al. 2020); Decay sugar-rich substrates (Põlme et al. 2020) | Põlme S, et al. 2020. Fungal Diversity 105:1-16 ((https://doi.org/10.1007/s13225-020-00466-2)) |
| Otu47515 | 1 | k__Fungi(100);p__Ascomycota(100);c__Saccharomycetes(100);o__Saccharomycetales(100);f__Metschnikowiaceae(100);g__Metschnikowia(100);s__Metschnikowia_pulcherrima(100); | Metschnikowia | 13 | Saprotroph | \|Undefined Saprotroph\| | Yeast | NULL | Probable | As nectar-tap_saprotroph (Põlme et al. 2020); Decay sugar-rich substrates (Põlme et al. 2020) | Põlme S, et al. 2020. Fungal Diversity 105:1-16 ((https://doi.org/10.1007/s13225-020-00466-2)) |
| Otu47517 | 1 | k__Fungi(100);p__Ascomycota(100);c__Saccharomycetes(100);o__Saccharomycetales(100);f__Metschnikowiaceae(100);g__Metschnikowia(100);s__Metschnikowia_pulcherrima(100); | Metschnikowia | 13 | Saprotroph | \|Undefined Saprotroph\| | Yeast | NULL | Probable | As nectar-tap_saprotroph (Põlme et al. 2020); Decay sugar-rich substrates (Põlme et al. 2020) | Põlme S, et al. 2020. Fungal Diversity 105:1-16 ((https://doi.org/10.1007/s13225-020-00466-2)) |
| Otu47518 | 1 | k__Fungi(100);p__Ascomycota(100);c__Saccharomycetes(100);o__Saccharomycetales(100);f__Metschnikowiaceae(100);g__Metschnikowia(100);s__Metschnikowia_pulcherrima(100); | Metschnikowia | 13 | Saprotroph | \|Undefined Saprotroph\| | Yeast | NULL | Probable | As nectar-tap_saprotroph (Põlme et al. 2020); Decay sugar-rich substrates (Põlme et al. 2020) | Põlme S, et al. 2020. Fungal Diversity 105:1-16 ((https://doi.org/10.1007/s13225-020-00466-2)) |
| Otu47519 | 1 | k__Fungi(100);p__Ascomycota(100);c__Saccharomycetes(100);o__Saccharomycetales(100);f__Metschnikowiaceae(100);g__Metschnikowia(100);s__Metschnikowia_pulcherrima(100); | Metschnikowia | 13 | Saprotroph | \|Undefined Saprotroph\| | Yeast | NULL | Probable | As nectar-tap_saprotroph (Põlme et al. 2020); Decay sugar-rich substrates (Põlme et al. 2020) | Põlme S, et al. 2020. Fungal Diversity 105:1-16 ((https://doi.org/10.1007/s13225-020-00466-2)) |
| Otu47520 | 1 | k__Fungi(100);p__Ascomycota(100);c__Saccharomycetes(100);o__Saccharomycetales(100);f__Metschnikowiaceae(100);g__Metschnikowia(100);s__Metschnikowia_pulcherrima(100); | Metschnikowia | 13 | Saprotroph | \|Undefined Saprotroph\| | Yeast | NULL | Probable | As nectar-tap_saprotroph (Põlme et al. 2020); Decay sugar-rich substrates (Põlme et al. 2020) | Põlme S, et al. 2020. Fungal Diversity 105:1-16 ((https://doi.org/10.1007/s13225-020-00466-2)) |
| Otu47521 | 1 | k__Fungi(100);p__Ascomycota(100);c__Saccharomycetes(100);o__Saccharomycetales(100);f__Metschnikowiaceae(100);g__Metschnikowia(100);s__Metschnikowia_pulcherrima(100); | Metschnikowia | 13 | Saprotroph | \|Undefined Saprotroph\| | Yeast | NULL | Probable | As nectar-tap_saprotroph (Põlme et al. 2020); Decay sugar-rich substrates (Põlme et al. 2020) | Põlme S, et al. 2020. Fungal Diversity 105:1-16 ((https://doi.org/10.1007/s13225-020-00466-2)) |
| Otu47522 | 1 | k__Fungi(100);p__Ascomycota(100);c__Saccharomycetes(100);o__Saccharomycetales(100);f__Metschnikowiaceae(100);g__Metschnikowia(100);s__Metschnikowia_pulcherrima(100); | Metschnikowia | 13 | Saprotroph | \|Undefined Saprotroph\| | Yeast | NULL | Probable | As nectar-tap_saprotroph (Põlme et al. 2020); Decay sugar-rich substrates (Põlme et al. 2020) | Põlme S, et al. 2020. Fungal Diversity 105:1-16 ((https://doi.org/10.1007/s13225-020-00466-2)) |
| Otu47523 | 1 | k__Fungi(100);p__Ascomycota(100);c__Saccharomycetes(100);o__Saccharomycetales(100);f__Metschnikowiaceae(100);g__Metschnikowia(100);s__Metschnikowia_pulcherrima(100); | Metschnikowia | 13 | Saprotroph | \|Undefined Saprotroph\| | Yeast | NULL | Probable | As nectar-tap_saprotroph (Põlme et al. 2020); Decay sugar-rich substrates (Põlme et al. 2020) | Põlme S, et al. 2020. Fungal Diversity 105:1-16 ((https://doi.org/10.1007/s13225-020-00466-2)) |
| Otu47524 | 1 | k__Fungi(100);p__Ascomycota(100);c__Saccharomycetes(100);o__Saccharomycetales(100);f__Metschnikowiaceae(100);g__Metschnikowia(100);s__Metschnikowia_pulcherrima(100); | Metschnikowia | 13 | Saprotroph | \|Undefined Saprotroph\| | Yeast | NULL | Probable | As nectar-tap_saprotroph (Põlme et al. 2020); Decay sugar-rich substrates (Põlme et al. 2020) | Põlme S, et al. 2020. Fungal Diversity 105:1-16 ((https://doi.org/10.1007/s13225-020-00466-2)) |
| Otu47525 | 1 | k__Fungi(100);p__Ascomycota(100);c__Saccharomycetes(100);o__Saccharomycetales(100);f__Metschnikowiaceae(100);g__Metschnikowia(100);s__Metschnikowia_pulcherrima(100); | Metschnikowia | 13 | Saprotroph | \|Undefined Saprotroph\| | Yeast | NULL | Probable | As nectar-tap_saprotroph (Põlme et al. 2020); Decay sugar-rich substrates (Põlme et al. 2020) | Põlme S, et al. 2020. Fungal Diversity 105:1-16 ((https://doi.org/10.1007/s13225-020-00466-2)) |
| Otu47526 | 1 | k__Fungi(100);p__Ascomycota(100);c__Saccharomycetes(100);o__Saccharomycetales(100);f__Metschnikowiaceae(100);g__Metschnikowia(100);g__Metschnikowia_unclassified(100); | Metschnikowia | 13 | Saprotroph | \|Undefined Saprotroph\| | Yeast | NULL | Probable | As nectar-tap_saprotroph (Põlme et al. 2020); Decay sugar-rich substrates (Põlme et al. 2020) | Põlme S, et al. 2020. Fungal Diversity 105:1-16 ((https://doi.org/10.1007/s13225-020-00466-2)) |
| Otu47527 | 1 | k__Fungi(100);p__Ascomycota(100);c__Saccharomycetes(100);o__Saccharomycetales(100);f__Metschnikowiaceae(100);g__Metschnikowia(100);s__Metschnikowia_pulcherrima(100); | Metschnikowia | 13 | Saprotroph | \|Undefined Saprotroph\| | Yeast | NULL | Probable | As nectar-tap_saprotroph (Põlme et al. 2020); Decay sugar-rich substrates (Põlme et al. 2020) | Põlme S, et al. 2020. Fungal Diversity 105:1-16 ((https://doi.org/10.1007/s13225-020-00466-2)) |
| Otu47528 | 1 | k__Fungi(100);p__Ascomycota(100);c__Saccharomycetes(100);o__Saccharomycetales(100);f__Metschnikowiaceae(100);g__Metschnikowia(100);s__Metschnikowia_pulcherrima(100); | Metschnikowia | 13 | Saprotroph | \|Undefined Saprotroph\| | Yeast | NULL | Probable | As nectar-tap_saprotroph (Põlme et al. 2020); Decay sugar-rich substrates (Põlme et al. 2020) | Põlme S, et al. 2020. Fungal Diversity 105:1-16 ((https://doi.org/10.1007/s13225-020-00466-2)) |
| Otu47529 | 1 | k__Fungi(100);p__Ascomycota(100);c__Saccharomycetes(100);o__Saccharomycetales(100);f__Metschnikowiaceae(100);g__Metschnikowia(100);s__Metschnikowia_pulcherrima(100); | Metschnikowia | 13 | Saprotroph | \|Undefined Saprotroph\| | Yeast | NULL | Probable | As nectar-tap_saprotroph (Põlme et al. 2020); Decay sugar-rich substrates (Põlme et al. 2020) | Põlme S, et al. 2020. Fungal Diversity 105:1-16 ((https://doi.org/10.1007/s13225-020-00466-2)) |
| Otu47530 | 1 | k__Fungi(100);p__Ascomycota(100);c__Saccharomycetes(100);o__Saccharomycetales(100);f__Metschnikowiaceae(100);g__Metschnikowia(100);s__Metschnikowia_pulcherrima(100); | Metschnikowia | 13 | Saprotroph | \|Undefined Saprotroph\| | Yeast | NULL | Probable | As nectar-tap_saprotroph (Põlme et al. 2020); Decay sugar-rich substrates (Põlme et al. 2020) | Põlme S, et al. 2020. Fungal Diversity 105:1-16 ((https://doi.org/10.1007/s13225-020-00466-2)) |
| Otu47531 | 1 | k__Fungi(100);p__Ascomycota(100);c__Saccharomycetes(100);o__Saccharomycetales(100);f__Metschnikowiaceae(100);g__Metschnikowia(100);s__Metschnikowia_pulcherrima(100); | Metschnikowia | 13 | Saprotroph | \|Undefined Saprotroph\| | Yeast | NULL | Probable | As nectar-tap_saprotroph (Põlme et al. 2020); Decay sugar-rich substrates (Põlme et al. 2020) | Põlme S, et al. 2020. Fungal Diversity 105:1-16 ((https://doi.org/10.1007/s13225-020-00466-2)) |
| Otu47532 | 1 | k__Fungi(100);p__Ascomycota(100);c__Saccharomycetes(100);o__Saccharomycetales(100);f__Metschnikowiaceae(100);g__Metschnikowia(100);s__Metschnikowia_pulcherrima(100); | Metschnikowia | 13 | Saprotroph | \|Undefined Saprotroph\| | Yeast | NULL | Probable | As nectar-tap_saprotroph (Põlme et al. 2020); Decay sugar-rich substrates (Põlme et al. 2020) | Põlme S, et al. 2020. Fungal Diversity 105:1-16 ((https://doi.org/10.1007/s13225-020-00466-2)) |
| Otu47533 | 1 | k__Fungi(100);p__Ascomycota(100);c__Saccharomycetes(100);o__Saccharomycetales(100);f__Metschnikowiaceae(100);g__Metschnikowia(100);s__Metschnikowia_pulcherrima(100); | Metschnikowia | 13 | Saprotroph | \|Undefined Saprotroph\| | Yeast | NULL | Probable | As nectar-tap_saprotroph (Põlme et al. 2020); Decay sugar-rich substrates (Põlme et al. 2020) | Põlme S, et al. 2020. Fungal Diversity 105:1-16 ((https://doi.org/10.1007/s13225-020-00466-2)) |
| Otu47534 | 1 | k__Fungi(100);p__Ascomycota(100);c__Saccharomycetes(100);o__Saccharomycetales(100);f__Metschnikowiaceae(100);g__Metschnikowia(100);s__Metschnikowia_pulcherrima(100); | Metschnikowia | 13 | Saprotroph | \|Undefined Saprotroph\| | Yeast | NULL | Probable | As nectar-tap_saprotroph (Põlme et al. 2020); Decay sugar-rich substrates (Põlme et al. 2020) | Põlme S, et al. 2020. Fungal Diversity 105:1-16 ((https://doi.org/10.1007/s13225-020-00466-2)) |
| Otu47535 | 1 | k__Fungi(100);p__Ascomycota(100);c__Saccharomycetes(100);o__Saccharomycetales(100);f__Metschnikowiaceae(100);g__Metschnikowia(100);s__Metschnikowia_pulcherrima(100); | Metschnikowia | 13 | Saprotroph | \|Undefined Saprotroph\| | Yeast | NULL | Probable | As nectar-tap_saprotroph (Põlme et al. 2020); Decay sugar-rich substrates (Põlme et al. 2020) | Põlme S, et al. 2020. Fungal Diversity 105:1-16 ((https://doi.org/10.1007/s13225-020-00466-2)) |
| Otu47536 | 1 | k__Fungi(100);p__Ascomycota(100);c__Saccharomycetes(100);o__Saccharomycetales(100);f__Metschnikowiaceae(100);g__Metschnikowia(100);s__Metschnikowia_pulcherrima(100); | Metschnikowia | 13 | Saprotroph | \|Undefined Saprotroph\| | Yeast | NULL | Probable | As nectar-tap_saprotroph (Põlme et al. 2020); Decay sugar-rich substrates (Põlme et al. 2020) | Põlme S, et al. 2020. Fungal Diversity 105:1-16 ((https://doi.org/10.1007/s13225-020-00466-2)) |
| Otu47537 | 1 | k__Fungi(100);p__Ascomycota(100);c__Saccharomycetes(100);o__Saccharomycetales(100);f__Metschnikowiaceae(100);g__Metschnikowia(100);s__Metschnikowia_pulcherrima(100); | Metschnikowia | 13 | Saprotroph | \|Undefined Saprotroph\| | Yeast | NULL | Probable | As nectar-tap_saprotroph (Põlme et al. 2020); Decay sugar-rich substrates (Põlme et al. 2020) | Põlme S, et al. 2020. Fungal Diversity 105:1-16 ((https://doi.org/10.1007/s13225-020-00466-2)) |
| Otu47544 | 1 | k__Fungi(100);p__Ascomycota(100);c__Saccharomycetes(100);o__Saccharomycetales(100);f__Metschnikowiaceae(100);g__Metschnikowia(100);s__Metschnikowia_pulcherrima(100); | Metschnikowia | 13 | Saprotroph | \|Undefined Saprotroph\| | Yeast | NULL | Probable | As nectar-tap_saprotroph (Põlme et al. 2020); Decay sugar-rich substrates (Põlme et al. 2020) | Põlme S, et al. 2020. Fungal Diversity 105:1-16 ((https://doi.org/10.1007/s13225-020-00466-2)) |
| Otu47545 | 1 | k__Fungi(100);p__Ascomycota(100);c__Saccharomycetes(100);o__Saccharomycetales(100);f__Metschnikowiaceae(100);g__Metschnikowia(100);s__Metschnikowia_pulcherrima(100); | Metschnikowia | 13 | Saprotroph | \|Undefined Saprotroph\| | Yeast | NULL | Probable | As nectar-tap_saprotroph (Põlme et al. 2020); Decay sugar-rich substrates (Põlme et al. 2020) | Põlme S, et al. 2020. Fungal Diversity 105:1-16 ((https://doi.org/10.1007/s13225-020-00466-2)) |
| Otu47546 | 1 | k__Fungi(100);p__Ascomycota(100);c__Saccharomycetes(100);o__Saccharomycetales(100);f__Metschnikowiaceae(100);g__Metschnikowia(100);s__Metschnikowia_pulcherrima(100); | Metschnikowia | 13 | Saprotroph | \|Undefined Saprotroph\| | Yeast | NULL | Probable | As nectar-tap_saprotroph (Põlme et al. 2020); Decay sugar-rich substrates (Põlme et al. 2020) | Põlme S, et al. 2020. Fungal Diversity 105:1-16 ((https://doi.org/10.1007/s13225-020-00466-2)) |
| Otu47547 | 1 | k__Fungi(100);p__Ascomycota(100);c__Saccharomycetes(100);o__Saccharomycetales(100);f__Metschnikowiaceae(100);g__Metschnikowia(100);s__Metschnikowia_pulcherrima(100); | Metschnikowia | 13 | Saprotroph | \|Undefined Saprotroph\| | Yeast | NULL | Probable | As nectar-tap_saprotroph (Põlme et al. 2020); Decay sugar-rich substrates (Põlme et al. 2020) | Põlme S, et al. 2020. Fungal Diversity 105:1-16 ((https://doi.org/10.1007/s13225-020-00466-2)) |
| Otu47548 | 1 | k__Fungi(100);p__Ascomycota(100);c__Saccharomycetes(100);o__Saccharomycetales(100);f__Metschnikowiaceae(100);g__Metschnikowia(100);s__Metschnikowia_pulcherrima(100); | Metschnikowia | 13 | Saprotroph | \|Undefined Saprotroph\| | Yeast | NULL | Probable | As nectar-tap_saprotroph (Põlme et al. 2020); Decay sugar-rich substrates (Põlme et al. 2020) | Põlme S, et al. 2020. Fungal Diversity 105:1-16 ((https://doi.org/10.1007/s13225-020-00466-2)) |
| Otu47550 | 1 | k__Fungi(100);p__Ascomycota(100);c__Saccharomycetes(100);o__Saccharomycetales(100);f__Metschnikowiaceae(100);g__Metschnikowia(100);s__Metschnikowia_pulcherrima(100); | Metschnikowia | 13 | Saprotroph | \|Undefined Saprotroph\| | Yeast | NULL | Probable | As nectar-tap_saprotroph (Põlme et al. 2020); Decay sugar-rich substrates (Põlme et al. 2020) | Põlme S, et al. 2020. Fungal Diversity 105:1-16 ((https://doi.org/10.1007/s13225-020-00466-2)) |
| Otu47555 | 1 | k__Fungi(100);p__Ascomycota(100);c__Saccharomycetes(100);o__Saccharomycetales(100);f__Metschnikowiaceae(100);g__Metschnikowia(100);s__Metschnikowia_pulcherrima(100); | Metschnikowia | 13 | Saprotroph | \|Undefined Saprotroph\| | Yeast | NULL | Probable | As nectar-tap_saprotroph (Põlme et al. 2020); Decay sugar-rich substrates (Põlme et al. 2020) | Põlme S, et al. 2020. Fungal Diversity 105:1-16 ((https://doi.org/10.1007/s13225-020-00466-2)) |
| Otu47556 | 1 | k__Fungi(100);p__Basidiomycota(100);c__Tremellomycetes(100);o__Tremellales(100);f__Tremellales_family_Incertae_sedis(100);g__Hannaella(100);s__Hannaella_luteola(100); | Hannaella | 13 | Pathotroph-Saprotroph | \|Animal Parasite\|-Undefined Saprotroph | Yeast | NULL | Probable | As animal_decomposer (Põlme et al. 2020); Decay animal material (Põlme et al. 2020) | Põlme S, et al. 2020. Fungal Diversity 105:1-16 ((https://doi.org/10.1007/s13225-020-00466-2)) |
| Otu47557 | 1 | k__Fungi(100);p__Ascomycota(100);c__Saccharomycetes(100);o__Saccharomycetales(100);f__Metschnikowiaceae(100);g__Metschnikowia(100);s__Metschnikowia_pulcherrima(100); | Metschnikowia | 13 | Saprotroph | \|Undefined Saprotroph\| | Yeast | NULL | Probable | As nectar-tap_saprotroph (Põlme et al. 2020); Decay sugar-rich substrates (Põlme et al. 2020) | Põlme S, et al. 2020. Fungal Diversity 105:1-16 ((https://doi.org/10.1007/s13225-020-00466-2)) |
| Otu47559 | 1 | k__Fungi(100);p__Ascomycota(100);c__Saccharomycetes(100);o__Saccharomycetales(100);f__Metschnikowiaceae(100);g__Metschnikowia(100);s__Metschnikowia_pulcherrima(100); | Metschnikowia | 13 | Saprotroph | \|Undefined Saprotroph\| | Yeast | NULL | Probable | As nectar-tap_saprotroph (Põlme et al. 2020); Decay sugar-rich substrates (Põlme et al. 2020) | Põlme S, et al. 2020. Fungal Diversity 105:1-16 ((https://doi.org/10.1007/s13225-020-00466-2)) |
| Otu47560 | 1 | k__Fungi(100);p__Ascomycota(100);c__Saccharomycetes(100);o__Saccharomycetales(100);f__Metschnikowiaceae(100);g__Metschnikowia(100);s__Metschnikowia_pulcherrima(100); | Metschnikowia | 13 | Saprotroph | \|Undefined Saprotroph\| | Yeast | NULL | Probable | As nectar-tap_saprotroph (Põlme et al. 2020); Decay sugar-rich substrates (Põlme et al. 2020) | Põlme S, et al. 2020. Fungal Diversity 105:1-16 ((https://doi.org/10.1007/s13225-020-00466-2)) |
| Otu47561 | 1 | k__Fungi(100);p__Ascomycota(100);c__Saccharomycetes(100);o__Saccharomycetales(100);f__Metschnikowiaceae(100);g__Metschnikowia(100);s__Metschnikowia_pulcherrima(100); | Metschnikowia | 13 | Saprotroph | \|Undefined Saprotroph\| | Yeast | NULL | Probable | As nectar-tap_saprotroph (Põlme et al. 2020); Decay sugar-rich substrates (Põlme et al. 2020) | Põlme S, et al. 2020. Fungal Diversity 105:1-16 ((https://doi.org/10.1007/s13225-020-00466-2)) |
| Otu47562 | 1 | k__Fungi(100);p__Ascomycota(100);c__Saccharomycetes(100);o__Saccharomycetales(100);f__Metschnikowiaceae(100);g__Metschnikowia(100);s__Metschnikowia_pulcherrima(100); | Metschnikowia | 13 | Saprotroph | \|Undefined Saprotroph\| | Yeast | NULL | Probable | As nectar-tap_saprotroph (Põlme et al. 2020); Decay sugar-rich substrates (Põlme et al. 2020) | Põlme S, et al. 2020. Fungal Diversity 105:1-16 ((https://doi.org/10.1007/s13225-020-00466-2)) |
| Otu47563 | 1 | k__Fungi(100);p__Ascomycota(100);c__Saccharomycetes(100);o__Saccharomycetales(100);f__Metschnikowiaceae(100);g__Metschnikowia(100);g__Metschnikowia_unclassified(100); | Metschnikowia | 13 | Saprotroph | \|Undefined Saprotroph\| | Yeast | NULL | Probable | As nectar-tap_saprotroph (Põlme et al. 2020); Decay sugar-rich substrates (Põlme et al. 2020) | Põlme S, et al. 2020. Fungal Diversity 105:1-16 ((https://doi.org/10.1007/s13225-020-00466-2)) |
| Otu47564 | 1 | k__Fungi(100);p__Ascomycota(100);c__Saccharomycetes(100);o__Saccharomycetales(100);f__Metschnikowiaceae(100);g__Metschnikowia(100);s__Metschnikowia_pulcherrima(100); | Metschnikowia | 13 | Saprotroph | \|Undefined Saprotroph\| | Yeast | NULL | Probable | As nectar-tap_saprotroph (Põlme et al. 2020); Decay sugar-rich substrates (Põlme et al. 2020) | Põlme S, et al. 2020. Fungal Diversity 105:1-16 ((https://doi.org/10.1007/s13225-020-00466-2)) |
| Otu47565 | 1 | k__Fungi(100);p__Ascomycota(100);c__Saccharomycetes(100);o__Saccharomycetales(100);f__Metschnikowiaceae(100);g__Metschnikowia(100);s__Metschnikowia_pulcherrima(100); | Metschnikowia | 13 | Saprotroph | \|Undefined Saprotroph\| | Yeast | NULL | Probable | As nectar-tap_saprotroph (Põlme et al. 2020); Decay sugar-rich substrates (Põlme et al. 2020) | Põlme S, et al. 2020. Fungal Diversity 105:1-16 ((https://doi.org/10.1007/s13225-020-00466-2)) |
| Otu47566 | 1 | k__Fungi(100);p__Ascomycota(100);c__Saccharomycetes(100);o__Saccharomycetales(100);f__Metschnikowiaceae(100);g__Metschnikowia(100);s__Metschnikowia_pulcherrima(100); | Metschnikowia | 13 | Saprotroph | \|Undefined Saprotroph\| | Yeast | NULL | Probable | As nectar-tap_saprotroph (Põlme et al. 2020); Decay sugar-rich substrates (Põlme et al. 2020) | Põlme S, et al. 2020. Fungal Diversity 105:1-16 ((https://doi.org/10.1007/s13225-020-00466-2)) |
| Otu47567 | 1 | k__Fungi(100);p__Ascomycota(100);c__Saccharomycetes(100);o__Saccharomycetales(100);f__Metschnikowiaceae(100);g__Metschnikowia(100);s__Metschnikowia_pulcherrima(100); | Metschnikowia | 13 | Saprotroph | \|Undefined Saprotroph\| | Yeast | NULL | Probable | As nectar-tap_saprotroph (Põlme et al. 2020); Decay sugar-rich substrates (Põlme et al. 2020) | Põlme S, et al. 2020. Fungal Diversity 105:1-16 ((https://doi.org/10.1007/s13225-020-00466-2)) |
| Otu47568 | 1 | k__Fungi(100);p__Ascomycota(100);c__Saccharomycetes(100);o__Saccharomycetales(100);f__Metschnikowiaceae(100);g__Metschnikowia(100);g__Metschnikowia_unclassified(100); | Metschnikowia | 13 | Saprotroph | \|Undefined Saprotroph\| | Yeast | NULL | Probable | As nectar-tap_saprotroph (Põlme et al. 2020); Decay sugar-rich substrates (Põlme et al. 2020) | Põlme S, et al. 2020. Fungal Diversity 105:1-16 ((https://doi.org/10.1007/s13225-020-00466-2)) |
| Otu47569 | 1 | k__Fungi(100);p__Ascomycota(100);c__Saccharomycetes(100);o__Saccharomycetales(100);f__Metschnikowiaceae(100);g__Metschnikowia(100);g__Metschnikowia_unclassified(100); | Metschnikowia | 13 | Saprotroph | \|Undefined Saprotroph\| | Yeast | NULL | Probable | As nectar-tap_saprotroph (Põlme et al. 2020); Decay sugar-rich substrates (Põlme et al. 2020) | Põlme S, et al. 2020. Fungal Diversity 105:1-16 ((https://doi.org/10.1007/s13225-020-00466-2)) |
| Otu47571 | 1 | k__Fungi(100);p__Ascomycota(100);c__Saccharomycetes(100);o__Saccharomycetales(100);f__Metschnikowiaceae(100);g__Metschnikowia(100);g__Metschnikowia_unclassified(100); | Metschnikowia | 13 | Saprotroph | \|Undefined Saprotroph\| | Yeast | NULL | Probable | As nectar-tap_saprotroph (Põlme et al. 2020); Decay sugar-rich substrates (Põlme et al. 2020) | Põlme S, et al. 2020. Fungal Diversity 105:1-16 ((https://doi.org/10.1007/s13225-020-00466-2)) |
| Otu47572 | 1 | k__Fungi(100);p__Ascomycota(100);c__Saccharomycetes(100);o__Saccharomycetales(100);f__Metschnikowiaceae(100);g__Metschnikowia(100);s__Metschnikowia_pulcherrima(100); | Metschnikowia | 13 | Saprotroph | \|Undefined Saprotroph\| | Yeast | NULL | Probable | As nectar-tap_saprotroph (Põlme et al. 2020); Decay sugar-rich substrates (Põlme et al. 2020) | Põlme S, et al. 2020. Fungal Diversity 105:1-16 ((https://doi.org/10.1007/s13225-020-00466-2)) |
| Otu47573 | 1 | k__Fungi(100);p__Ascomycota(100);c__Saccharomycetes(100);o__Saccharomycetales(100);f__Metschnikowiaceae(100);g__Metschnikowia(100);g__Metschnikowia_unclassified(100); | Metschnikowia | 13 | Saprotroph | \|Undefined Saprotroph\| | Yeast | NULL | Probable | As nectar-tap_saprotroph (Põlme et al. 2020); Decay sugar-rich substrates (Põlme et al. 2020) | Põlme S, et al. 2020. Fungal Diversity 105:1-16 ((https://doi.org/10.1007/s13225-020-00466-2)) |
| Otu47574 | 1 | k__Fungi(100);p__Ascomycota(100);c__Saccharomycetes(100);o__Saccharomycetales(100);f__Metschnikowiaceae(100);g__Metschnikowia(100);g__Metschnikowia_unclassified(100); | Metschnikowia | 13 | Saprotroph | \|Undefined Saprotroph\| | Yeast | NULL | Probable | As nectar-tap_saprotroph (Põlme et al. 2020); Decay sugar-rich substrates (Põlme et al. 2020) | Põlme S, et al. 2020. Fungal Diversity 105:1-16 ((https://doi.org/10.1007/s13225-020-00466-2)) |
| Otu47576 | 1 | k__Fungi(100);p__Ascomycota(100);c__Saccharomycetes(100);o__Saccharomycetales(100);f__Metschnikowiaceae(100);g__Metschnikowia(100);s__Metschnikowia_pulcherrima(100); | Metschnikowia | 13 | Saprotroph | \|Undefined Saprotroph\| | Yeast | NULL | Probable | As nectar-tap_saprotroph (Põlme et al. 2020); Decay sugar-rich substrates (Põlme et al. 2020) | Põlme S, et al. 2020. Fungal Diversity 105:1-16 ((https://doi.org/10.1007/s13225-020-00466-2)) |
| Otu47577 | 1 | k__Fungi(100);p__Ascomycota(100);c__Saccharomycetes(100);o__Saccharomycetales(100);f__Metschnikowiaceae(100);g__Metschnikowia(100);s__Metschnikowia_pulcherrima(100); | Metschnikowia | 13 | Saprotroph | \|Undefined Saprotroph\| | Yeast | NULL | Probable | As nectar-tap_saprotroph (Põlme et al. 2020); Decay sugar-rich substrates (Põlme et al. 2020) | Põlme S, et al. 2020. Fungal Diversity 105:1-16 ((https://doi.org/10.1007/s13225-020-00466-2)) |
| Otu47578 | 1 | k__Fungi(100);p__Ascomycota(100);c__Saccharomycetes(100);o__Saccharomycetales(100);f__Metschnikowiaceae(100);g__Metschnikowia(100);s__Metschnikowia_pulcherrima(100); | Metschnikowia | 13 | Saprotroph | \|Undefined Saprotroph\| | Yeast | NULL | Probable | As nectar-tap_saprotroph (Põlme et al. 2020); Decay sugar-rich substrates (Põlme et al. 2020) | Põlme S, et al. 2020. Fungal Diversity 105:1-16 ((https://doi.org/10.1007/s13225-020-00466-2)) |
| Otu47580 | 1 | k__Fungi(100);p__Ascomycota(100);c__Saccharomycetes(100);o__Saccharomycetales(100);f__Metschnikowiaceae(100);g__Metschnikowia(100);s__Metschnikowia_pulcherrima(100); | Metschnikowia | 13 | Saprotroph | \|Undefined Saprotroph\| | Yeast | NULL | Probable | As nectar-tap_saprotroph (Põlme et al. 2020); Decay sugar-rich substrates (Põlme et al. 2020) | Põlme S, et al. 2020. Fungal Diversity 105:1-16 ((https://doi.org/10.1007/s13225-020-00466-2)) |
| Otu47582 | 1 | k__Fungi(100);p__Ascomycota(100);c__Saccharomycetes(100);o__Saccharomycetales(100);f__Metschnikowiaceae(100);g__Metschnikowia(100);s__Metschnikowia_pulcherrima(100); | Metschnikowia | 13 | Saprotroph | \|Undefined Saprotroph\| | Yeast | NULL | Probable | As nectar-tap_saprotroph (Põlme et al. 2020); Decay sugar-rich substrates (Põlme et al. 2020) | Põlme S, et al. 2020. Fungal Diversity 105:1-16 ((https://doi.org/10.1007/s13225-020-00466-2)) |
| Otu47583 | 1 | k__Fungi(100);p__Ascomycota(100);c__Saccharomycetes(100);o__Saccharomycetales(100);f__Metschnikowiaceae(100);g__Metschnikowia(100);s__Metschnikowia_pulcherrima(100); | Metschnikowia | 13 | Saprotroph | \|Undefined Saprotroph\| | Yeast | NULL | Probable | As nectar-tap_saprotroph (Põlme et al. 2020); Decay sugar-rich substrates (Põlme et al. 2020) | Põlme S, et al. 2020. Fungal Diversity 105:1-16 ((https://doi.org/10.1007/s13225-020-00466-2)) |
| Otu47584 | 1 | k__Fungi(100);p__Ascomycota(100);c__Saccharomycetes(100);o__Saccharomycetales(100);f__Metschnikowiaceae(100);g__Metschnikowia(100);s__Metschnikowia_pulcherrima(100); | Metschnikowia | 13 | Saprotroph | \|Undefined Saprotroph\| | Yeast | NULL | Probable | As nectar-tap_saprotroph (Põlme et al. 2020); Decay sugar-rich substrates (Põlme et al. 2020) | Põlme S, et al. 2020. Fungal Diversity 105:1-16 ((https://doi.org/10.1007/s13225-020-00466-2)) |
| Otu47585 | 1 | k__Fungi(100);p__Ascomycota(100);c__Saccharomycetes(100);o__Saccharomycetales(100);f__Metschnikowiaceae(100);g__Metschnikowia(100);s__Metschnikowia_pulcherrima(100); | Metschnikowia | 13 | Saprotroph | \|Undefined Saprotroph\| | Yeast | NULL | Probable | As nectar-tap_saprotroph (Põlme et al. 2020); Decay sugar-rich substrates (Põlme et al. 2020) | Põlme S, et al. 2020. Fungal Diversity 105:1-16 ((https://doi.org/10.1007/s13225-020-00466-2)) |
| Otu47587 | 1 | k__Fungi(100);p__Ascomycota(100);c__Saccharomycetes(100);o__Saccharomycetales(100);f__Metschnikowiaceae(100);g__Metschnikowia(100);s__Metschnikowia_pulcherrima(100); | Metschnikowia | 13 | Saprotroph | \|Undefined Saprotroph\| | Yeast | NULL | Probable | As nectar-tap_saprotroph (Põlme et al. 2020); Decay sugar-rich substrates (Põlme et al. 2020) | Põlme S, et al. 2020. Fungal Diversity 105:1-16 ((https://doi.org/10.1007/s13225-020-00466-2)) |
| Otu47588 | 1 | k__Fungi(100);p__Ascomycota(100);c__Saccharomycetes(100);o__Saccharomycetales(100);f__Metschnikowiaceae(100);g__Metschnikowia(100);s__Metschnikowia_pulcherrima(100); | Metschnikowia | 13 | Saprotroph | \|Undefined Saprotroph\| | Yeast | NULL | Probable | As nectar-tap_saprotroph (Põlme et al. 2020); Decay sugar-rich substrates (Põlme et al. 2020) | Põlme S, et al. 2020. Fungal Diversity 105:1-16 ((https://doi.org/10.1007/s13225-020-00466-2)) |
| Otu47589 | 1 | k__Fungi(100);p__Ascomycota(100);c__Saccharomycetes(100);o__Saccharomycetales(100);f__Metschnikowiaceae(100);g__Metschnikowia(100);s__Metschnikowia_pulcherrima(100); | Metschnikowia | 13 | Saprotroph | \|Undefined Saprotroph\| | Yeast | NULL | Probable | As nectar-tap_saprotroph (Põlme et al. 2020); Decay sugar-rich substrates (Põlme et al. 2020) | Põlme S, et al. 2020. Fungal Diversity 105:1-16 ((https://doi.org/10.1007/s13225-020-00466-2)) |
| Otu47590 | 1 | k__Fungi(100);p__Ascomycota(100);c__Saccharomycetes(100);o__Saccharomycetales(100);f__Metschnikowiaceae(100);g__Metschnikowia(100);s__Metschnikowia_pulcherrima(100); | Metschnikowia | 13 | Saprotroph | \|Undefined Saprotroph\| | Yeast | NULL | Probable | As nectar-tap_saprotroph (Põlme et al. 2020); Decay sugar-rich substrates (Põlme et al. 2020) | Põlme S, et al. 2020. Fungal Diversity 105:1-16 ((https://doi.org/10.1007/s13225-020-00466-2)) |
| Otu47592 | 1 | k__Fungi(100);p__Ascomycota(100);c__Saccharomycetes(100);o__Saccharomycetales(100);f__Metschnikowiaceae(100);g__Metschnikowia(100);g__Metschnikowia_unclassified(100); | Metschnikowia | 13 | Saprotroph | \|Undefined Saprotroph\| | Yeast | NULL | Probable | As nectar-tap_saprotroph (Põlme et al. 2020); Decay sugar-rich substrates (Põlme et al. 2020) | Põlme S, et al. 2020. Fungal Diversity 105:1-16 ((https://doi.org/10.1007/s13225-020-00466-2)) |
| Otu47593 | 1 | k__Fungi(100);p__Ascomycota(100);c__Saccharomycetes(100);o__Saccharomycetales(100);f__Metschnikowiaceae(100);g__Metschnikowia(100);s__Metschnikowia_pulcherrima(100); | Metschnikowia | 13 | Saprotroph | \|Undefined Saprotroph\| | Yeast | NULL | Probable | As nectar-tap_saprotroph (Põlme et al. 2020); Decay sugar-rich substrates (Põlme et al. 2020) | Põlme S, et al. 2020. Fungal Diversity 105:1-16 ((https://doi.org/10.1007/s13225-020-00466-2)) |
| Otu47594 | 1 | k__Fungi(100);p__Ascomycota(100);c__Saccharomycetes(100);o__Saccharomycetales(100);f__Metschnikowiaceae(100);g__Metschnikowia(100);s__Metschnikowia_pulcherrima(100); | Metschnikowia | 13 | Saprotroph | \|Undefined Saprotroph\| | Yeast | NULL | Probable | As nectar-tap_saprotroph (Põlme et al. 2020); Decay sugar-rich substrates (Põlme et al. 2020) | Põlme S, et al. 2020. Fungal Diversity 105:1-16 ((https://doi.org/10.1007/s13225-020-00466-2)) |
| Otu47595 | 1 | k__Fungi(100);p__Ascomycota(100);c__Saccharomycetes(100);o__Saccharomycetales(100);f__Metschnikowiaceae(100);g__Metschnikowia(100);s__Metschnikowia_pulcherrima(100); | Metschnikowia | 13 | Saprotroph | \|Undefined Saprotroph\| | Yeast | NULL | Probable | As nectar-tap_saprotroph (Põlme et al. 2020); Decay sugar-rich substrates (Põlme et al. 2020) | Põlme S, et al. 2020. Fungal Diversity 105:1-16 ((https://doi.org/10.1007/s13225-020-00466-2)) |
| Otu47596 | 1 | k__Fungi(100);p__Ascomycota(100);c__Saccharomycetes(100);o__Saccharomycetales(100);f__Metschnikowiaceae(100);g__Metschnikowia(100);s__Metschnikowia_pulcherrima(100); | Metschnikowia | 13 | Saprotroph | \|Undefined Saprotroph\| | Yeast | NULL | Probable | As nectar-tap_saprotroph (Põlme et al. 2020); Decay sugar-rich substrates (Põlme et al. 2020) | Põlme S, et al. 2020. Fungal Diversity 105:1-16 ((https://doi.org/10.1007/s13225-020-00466-2)) |
| Otu47597 | 1 | k__Fungi(100);p__Ascomycota(100);c__Saccharomycetes(100);o__Saccharomycetales(100);f__Metschnikowiaceae(100);g__Metschnikowia(100);s__Metschnikowia_pulcherrima(100); | Metschnikowia | 13 | Saprotroph | \|Undefined Saprotroph\| | Yeast | NULL | Probable | As nectar-tap_saprotroph (Põlme et al. 2020); Decay sugar-rich substrates (Põlme et al. 2020) | Põlme S, et al. 2020. Fungal Diversity 105:1-16 ((https://doi.org/10.1007/s13225-020-00466-2)) |
| Otu47599 | 1 | k__Fungi(100);p__Ascomycota(100);c__Saccharomycetes(100);o__Saccharomycetales(100);f__Metschnikowiaceae(100);g__Metschnikowia(100);s__Metschnikowia_pulcherrima(100); | Metschnikowia | 13 | Saprotroph | \|Undefined Saprotroph\| | Yeast | NULL | Probable | As nectar-tap_saprotroph (Põlme et al. 2020); Decay sugar-rich substrates (Põlme et al. 2020) | Põlme S, et al. 2020. Fungal Diversity 105:1-16 ((https://doi.org/10.1007/s13225-020-00466-2)) |
| Otu47600 | 1 | k__Fungi(100);p__Ascomycota(100);c__Saccharomycetes(100);o__Saccharomycetales(100);f__Metschnikowiaceae(100);g__Metschnikowia(100);s__Metschnikowia_pulcherrima(100); | Metschnikowia | 13 | Saprotroph | \|Undefined Saprotroph\| | Yeast | NULL | Probable | As nectar-tap_saprotroph (Põlme et al. 2020); Decay sugar-rich substrates (Põlme et al. 2020) | Põlme S, et al. 2020. Fungal Diversity 105:1-16 ((https://doi.org/10.1007/s13225-020-00466-2)) |
| Otu47602 | 1 | k__Fungi(100);p__Ascomycota(100);c__Saccharomycetes(100);o__Saccharomycetales(100);f__Metschnikowiaceae(100);g__Metschnikowia(100);s__Metschnikowia_pulcherrima(100); | Metschnikowia | 13 | Saprotroph | \|Undefined Saprotroph\| | Yeast | NULL | Probable | As nectar-tap_saprotroph (Põlme et al. 2020); Decay sugar-rich substrates (Põlme et al. 2020) | Põlme S, et al. 2020. Fungal Diversity 105:1-16 ((https://doi.org/10.1007/s13225-020-00466-2)) |
| Otu47603 | 1 | k__Fungi(100);p__Ascomycota(100);c__Saccharomycetes(100);o__Saccharomycetales(100);f__Metschnikowiaceae(100);g__Metschnikowia(100);g__Metschnikowia_unclassified(100); | Metschnikowia | 13 | Saprotroph | \|Undefined Saprotroph\| | Yeast | NULL | Probable | As nectar-tap_saprotroph (Põlme et al. 2020); Decay sugar-rich substrates (Põlme et al. 2020) | Põlme S, et al. 2020. Fungal Diversity 105:1-16 ((https://doi.org/10.1007/s13225-020-00466-2)) |
| Otu47604 | 1 | k__Fungi(100);p__Ascomycota(100);c__Saccharomycetes(100);o__Saccharomycetales(100);f__Metschnikowiaceae(100);g__Metschnikowia(100);g__Metschnikowia_unclassified(100); | Metschnikowia | 13 | Saprotroph | \|Undefined Saprotroph\| | Yeast | NULL | Probable | As nectar-tap_saprotroph (Põlme et al. 2020); Decay sugar-rich substrates (Põlme et al. 2020) | Põlme S, et al. 2020. Fungal Diversity 105:1-16 ((https://doi.org/10.1007/s13225-020-00466-2)) |
| Otu47605 | 1 | k__Fungi(100);p__Ascomycota(100);c__Saccharomycetes(100);o__Saccharomycetales(100);f__Metschnikowiaceae(100);g__Metschnikowia(100);g__Metschnikowia_unclassified(100); | Metschnikowia | 13 | Saprotroph | \|Undefined Saprotroph\| | Yeast | NULL | Probable | As nectar-tap_saprotroph (Põlme et al. 2020); Decay sugar-rich substrates (Põlme et al. 2020) | Põlme S, et al. 2020. Fungal Diversity 105:1-16 ((https://doi.org/10.1007/s13225-020-00466-2)) |
| Otu47606 | 1 | k__Fungi(100);p__Ascomycota(100);c__Saccharomycetes(100);o__Saccharomycetales(100);f__Metschnikowiaceae(100);g__Metschnikowia(100);s__Metschnikowia_pulcherrima(100); | Metschnikowia | 13 | Saprotroph | \|Undefined Saprotroph\| | Yeast | NULL | Probable | As nectar-tap_saprotroph (Põlme et al. 2020); Decay sugar-rich substrates (Põlme et al. 2020) | Põlme S, et al. 2020. Fungal Diversity 105:1-16 ((https://doi.org/10.1007/s13225-020-00466-2)) |
| Otu47607 | 1 | k__Fungi(100);p__Ascomycota(100);c__Saccharomycetes(100);o__Saccharomycetales(100);f__Metschnikowiaceae(100);g__Metschnikowia(100);s__Metschnikowia_pulcherrima(100); | Metschnikowia | 13 | Saprotroph | \|Undefined Saprotroph\| | Yeast | NULL | Probable | As nectar-tap_saprotroph (Põlme et al. 2020); Decay sugar-rich substrates (Põlme et al. 2020) | Põlme S, et al. 2020. Fungal Diversity 105:1-16 ((https://doi.org/10.1007/s13225-020-00466-2)) |
| Otu47608 | 1 | k__Fungi(100);p__Ascomycota(100);c__Saccharomycetes(100);o__Saccharomycetales(100);f__Metschnikowiaceae(100);g__Metschnikowia(100);s__Metschnikowia_pulcherrima(100); | Metschnikowia | 13 | Saprotroph | \|Undefined Saprotroph\| | Yeast | NULL | Probable | As nectar-tap_saprotroph (Põlme et al. 2020); Decay sugar-rich substrates (Põlme et al. 2020) | Põlme S, et al. 2020. Fungal Diversity 105:1-16 ((https://doi.org/10.1007/s13225-020-00466-2)) |
| Otu47609 | 1 | k__Fungi(100);p__Ascomycota(100);c__Saccharomycetes(100);o__Saccharomycetales(100);f__Metschnikowiaceae(100);g__Metschnikowia(100);s__Metschnikowia_pulcherrima(100); | Metschnikowia | 13 | Saprotroph | \|Undefined Saprotroph\| | Yeast | NULL | Probable | As nectar-tap_saprotroph (Põlme et al. 2020); Decay sugar-rich substrates (Põlme et al. 2020) | Põlme S, et al. 2020. Fungal Diversity 105:1-16 ((https://doi.org/10.1007/s13225-020-00466-2)) |
| Otu47611 | 1 | k__Fungi(100);p__Ascomycota(100);c__Saccharomycetes(100);o__Saccharomycetales(100);f__Saccharomycetales_family_Incertae_sedis(100);g__Candida(100);s__Candida_cretensis(100); | Candida | 13 | Pathotroph-Saprotroph-Symbiotroph | Animal Pathogen-Endophyte-Endosymbiont-Epiphyte-\|Undefined Saprotroph\| | Dimorphic Yeast-Yeast | NULL | Probable | As nectar-tap_saprotroph (Põlme et al. 2020); Decay sugar-rich substrates (Põlme et al. 2020); This asexual genus is polyphyletic and contain species that function very differently depend on what they are. Caution should be taken when interpreting guilds for Candida species. | Manolakaki D, et al. 2010. Virulence. 1: 367-375 ((https://doi.org/10.4161/viru.1.5.12796)); Mekbib SB, et al. 2011. Tropical Plant Pathology 36:233-240 ((https://doi.org/10.1590/S1982-56762011000400004)); Põlme S, et al. 2020. Fungal Diversity 105:1-16 ((https://doi.org/10.1007/s13225-020-00466-2)) |
| Otu47612 | 1 | k__Fungi(100);p__Ascomycota(100);c__Saccharomycetes(100);o__Saccharomycetales(100);f__Metschnikowiaceae(100);g__Metschnikowia(100);s__Metschnikowia_pulcherrima(100); | Metschnikowia | 13 | Saprotroph | \|Undefined Saprotroph\| | Yeast | NULL | Probable | As nectar-tap_saprotroph (Põlme et al. 2020); Decay sugar-rich substrates (Põlme et al. 2020) | Põlme S, et al. 2020. Fungal Diversity 105:1-16 ((https://doi.org/10.1007/s13225-020-00466-2)) |
| Otu47614 | 1 | k__Fungi(100);p__Ascomycota(100);c__Saccharomycetes(100);o__Saccharomycetales(100);f__Metschnikowiaceae(100);g__Metschnikowia(100);g__Metschnikowia_unclassified(100); | Metschnikowia | 13 | Saprotroph | \|Undefined Saprotroph\| | Yeast | NULL | Probable | As nectar-tap_saprotroph (Põlme et al. 2020); Decay sugar-rich substrates (Põlme et al. 2020) | Põlme S, et al. 2020. Fungal Diversity 105:1-16 ((https://doi.org/10.1007/s13225-020-00466-2)) |
| Otu47615 | 1 | k__Fungi(100);p__Ascomycota(100);c__Saccharomycetes(100);o__Saccharomycetales(100);f__Metschnikowiaceae(100);g__Metschnikowia(100);s__Metschnikowia_pulcherrima(100); | Metschnikowia | 13 | Saprotroph | \|Undefined Saprotroph\| | Yeast | NULL | Probable | As nectar-tap_saprotroph (Põlme et al. 2020); Decay sugar-rich substrates (Põlme et al. 2020) | Põlme S, et al. 2020. Fungal Diversity 105:1-16 ((https://doi.org/10.1007/s13225-020-00466-2)) |
| Otu47616 | 1 | k__Fungi(100);p__Ascomycota(100);c__Saccharomycetes(100);o__Saccharomycetales(100);f__Metschnikowiaceae(100);g__Metschnikowia(100);s__Metschnikowia_pulcherrima(100); | Metschnikowia | 13 | Saprotroph | \|Undefined Saprotroph\| | Yeast | NULL | Probable | As nectar-tap_saprotroph (Põlme et al. 2020); Decay sugar-rich substrates (Põlme et al. 2020) | Põlme S, et al. 2020. Fungal Diversity 105:1-16 ((https://doi.org/10.1007/s13225-020-00466-2)) |
| Otu47617 | 1 | k__Fungi(100);p__Ascomycota(100);c__Saccharomycetes(100);o__Saccharomycetales(100);f__Metschnikowiaceae(100);g__Metschnikowia(100);s__Metschnikowia_pulcherrima(100); | Metschnikowia | 13 | Saprotroph | \|Undefined Saprotroph\| | Yeast | NULL | Probable | As nectar-tap_saprotroph (Põlme et al. 2020); Decay sugar-rich substrates (Põlme et al. 2020) | Põlme S, et al. 2020. Fungal Diversity 105:1-16 ((https://doi.org/10.1007/s13225-020-00466-2)) |
| Otu47618 | 1 | k__Fungi(100);p__Ascomycota(100);c__Saccharomycetes(100);o__Saccharomycetales(100);f__Metschnikowiaceae(100);g__Metschnikowia(100);s__Metschnikowia_pulcherrima(100); | Metschnikowia | 13 | Saprotroph | \|Undefined Saprotroph\| | Yeast | NULL | Probable | As nectar-tap_saprotroph (Põlme et al. 2020); Decay sugar-rich substrates (Põlme et al. 2020) | Põlme S, et al. 2020. Fungal Diversity 105:1-16 ((https://doi.org/10.1007/s13225-020-00466-2)) |
| Otu47619 | 1 | k__Fungi(100);p__Ascomycota(100);c__Saccharomycetes(100);o__Saccharomycetales(100);f__Metschnikowiaceae(100);g__Metschnikowia(100);s__Metschnikowia_pulcherrima(100); | Metschnikowia | 13 | Saprotroph | \|Undefined Saprotroph\| | Yeast | NULL | Probable | As nectar-tap_saprotroph (Põlme et al. 2020); Decay sugar-rich substrates (Põlme et al. 2020) | Põlme S, et al. 2020. Fungal Diversity 105:1-16 ((https://doi.org/10.1007/s13225-020-00466-2)) |
| Otu47620 | 1 | k__Fungi(100);p__Ascomycota(100);c__Saccharomycetes(100);o__Saccharomycetales(100);f__Metschnikowiaceae(100);g__Metschnikowia(100);s__Metschnikowia_pulcherrima(100); | Metschnikowia | 13 | Saprotroph | \|Undefined Saprotroph\| | Yeast | NULL | Probable | As nectar-tap_saprotroph (Põlme et al. 2020); Decay sugar-rich substrates (Põlme et al. 2020) | Põlme S, et al. 2020. Fungal Diversity 105:1-16 ((https://doi.org/10.1007/s13225-020-00466-2)) |
| Otu47621 | 1 | k__Fungi(100);p__Ascomycota(100);c__Saccharomycetes(100);o__Saccharomycetales(100);f__Metschnikowiaceae(100);g__Metschnikowia(100);s__Metschnikowia_pulcherrima(100); | Metschnikowia | 13 | Saprotroph | \|Undefined Saprotroph\| | Yeast | NULL | Probable | As nectar-tap_saprotroph (Põlme et al. 2020); Decay sugar-rich substrates (Põlme et al. 2020) | Põlme S, et al. 2020. Fungal Diversity 105:1-16 ((https://doi.org/10.1007/s13225-020-00466-2)) |
| Otu47622 | 1 | k__Fungi(100);p__Ascomycota(100);c__Saccharomycetes(100);o__Saccharomycetales(100);f__Metschnikowiaceae(100);g__Metschnikowia(100);s__Metschnikowia_pulcherrima(100); | Metschnikowia | 13 | Saprotroph | \|Undefined Saprotroph\| | Yeast | NULL | Probable | As nectar-tap_saprotroph (Põlme et al. 2020); Decay sugar-rich substrates (Põlme et al. 2020) | Põlme S, et al. 2020. Fungal Diversity 105:1-16 ((https://doi.org/10.1007/s13225-020-00466-2)) |
| Otu47625 | 1 | k__Fungi(100);p__Ascomycota(100);c__Saccharomycetes(100);o__Saccharomycetales(100);f__Metschnikowiaceae(100);g__Metschnikowia(100);s__Metschnikowia_pulcherrima(100); | Metschnikowia | 13 | Saprotroph | \|Undefined Saprotroph\| | Yeast | NULL | Probable | As nectar-tap_saprotroph (Põlme et al. 2020); Decay sugar-rich substrates (Põlme et al. 2020) | Põlme S, et al. 2020. Fungal Diversity 105:1-16 ((https://doi.org/10.1007/s13225-020-00466-2)) |
| Otu47626 | 1 | k__Fungi(100);p__Ascomycota(100);c__Saccharomycetes(100);o__Saccharomycetales(100);f__Metschnikowiaceae(100);g__Metschnikowia(100);s__Metschnikowia_pulcherrima(100); | Metschnikowia | 13 | Saprotroph | \|Undefined Saprotroph\| | Yeast | NULL | Probable | As nectar-tap_saprotroph (Põlme et al. 2020); Decay sugar-rich substrates (Põlme et al. 2020) | Põlme S, et al. 2020. Fungal Diversity 105:1-16 ((https://doi.org/10.1007/s13225-020-00466-2)) |
| Otu47627 | 1 | k__Fungi(100);p__Ascomycota(100);c__Saccharomycetes(100);o__Saccharomycetales(100);f__Metschnikowiaceae(100);g__Metschnikowia(100);s__Metschnikowia_pulcherrima(100); | Metschnikowia | 13 | Saprotroph | \|Undefined Saprotroph\| | Yeast | NULL | Probable | As nectar-tap_saprotroph (Põlme et al. 2020); Decay sugar-rich substrates (Põlme et al. 2020) | Põlme S, et al. 2020. Fungal Diversity 105:1-16 ((https://doi.org/10.1007/s13225-020-00466-2)) |
| Otu47628 | 1 | k__Fungi(100);p__Ascomycota(100);c__Saccharomycetes(100);o__Saccharomycetales(100);f__Metschnikowiaceae(100);g__Metschnikowia(100);s__Metschnikowia_pulcherrima(100); | Metschnikowia | 13 | Saprotroph | \|Undefined Saprotroph\| | Yeast | NULL | Probable | As nectar-tap_saprotroph (Põlme et al. 2020); Decay sugar-rich substrates (Põlme et al. 2020) | Põlme S, et al. 2020. Fungal Diversity 105:1-16 ((https://doi.org/10.1007/s13225-020-00466-2)) |
| Otu47629 | 1 | k__Fungi(100);p__Ascomycota(100);c__Saccharomycetes(100);o__Saccharomycetales(100);f__Metschnikowiaceae(100);g__Metschnikowia(100);s__Metschnikowia_pulcherrima(100); | Metschnikowia | 13 | Saprotroph | \|Undefined Saprotroph\| | Yeast | NULL | Probable | As nectar-tap_saprotroph (Põlme et al. 2020); Decay sugar-rich substrates (Põlme et al. 2020) | Põlme S, et al. 2020. Fungal Diversity 105:1-16 ((https://doi.org/10.1007/s13225-020-00466-2)) |
| Otu47630 | 1 | k__Fungi(100);p__Ascomycota(100);c__Saccharomycetes(100);o__Saccharomycetales(100);f__Metschnikowiaceae(100);g__Metschnikowia(100);s__Metschnikowia_pulcherrima(100); | Metschnikowia | 13 | Saprotroph | \|Undefined Saprotroph\| | Yeast | NULL | Probable | As nectar-tap_saprotroph (Põlme et al. 2020); Decay sugar-rich substrates (Põlme et al. 2020) | Põlme S, et al. 2020. Fungal Diversity 105:1-16 ((https://doi.org/10.1007/s13225-020-00466-2)) |
| Otu47631 | 1 | k__Fungi(100);p__Ascomycota(100);c__Saccharomycetes(100);o__Saccharomycetales(100);f__Metschnikowiaceae(100);g__Metschnikowia(100);s__Metschnikowia_pulcherrima(100); | Metschnikowia | 13 | Saprotroph | \|Undefined Saprotroph\| | Yeast | NULL | Probable | As nectar-tap_saprotroph (Põlme et al. 2020); Decay sugar-rich substrates (Põlme et al. 2020) | Põlme S, et al. 2020. Fungal Diversity 105:1-16 ((https://doi.org/10.1007/s13225-020-00466-2)) |
| Otu47632 | 1 | k__Fungi(100);p__Ascomycota(100);c__Saccharomycetes(100);o__Saccharomycetales(100);f__Metschnikowiaceae(100);g__Metschnikowia(100);s__Metschnikowia_pulcherrima(100); | Metschnikowia | 13 | Saprotroph | \|Undefined Saprotroph\| | Yeast | NULL | Probable | As nectar-tap_saprotroph (Põlme et al. 2020); Decay sugar-rich substrates (Põlme et al. 2020) | Põlme S, et al. 2020. Fungal Diversity 105:1-16 ((https://doi.org/10.1007/s13225-020-00466-2)) |
| Otu47633 | 1 | k__Fungi(100);p__Ascomycota(100);c__Saccharomycetes(100);o__Saccharomycetales(100);f__Metschnikowiaceae(100);g__Metschnikowia(100);s__Metschnikowia_pulcherrima(100); | Metschnikowia | 13 | Saprotroph | \|Undefined Saprotroph\| | Yeast | NULL | Probable | As nectar-tap_saprotroph (Põlme et al. 2020); Decay sugar-rich substrates (Põlme et al. 2020) | Põlme S, et al. 2020. Fungal Diversity 105:1-16 ((https://doi.org/10.1007/s13225-020-00466-2)) |
| Otu47634 | 1 | k__Fungi(100);p__Ascomycota(100);c__Saccharomycetes(100);o__Saccharomycetales(100);f__Metschnikowiaceae(100);g__Metschnikowia(100);s__Metschnikowia_pulcherrima(100); | Metschnikowia | 13 | Saprotroph | \|Undefined Saprotroph\| | Yeast | NULL | Probable | As nectar-tap_saprotroph (Põlme et al. 2020); Decay sugar-rich substrates (Põlme et al. 2020) | Põlme S, et al. 2020. Fungal Diversity 105:1-16 ((https://doi.org/10.1007/s13225-020-00466-2)) |
| Otu47635 | 1 | k__Fungi(100);p__Ascomycota(100);c__Saccharomycetes(100);o__Saccharomycetales(100);f__Metschnikowiaceae(100);g__Metschnikowia(100);g__Metschnikowia_unclassified(100); | Metschnikowia | 13 | Saprotroph | \|Undefined Saprotroph\| | Yeast | NULL | Probable | As nectar-tap_saprotroph (Põlme et al. 2020); Decay sugar-rich substrates (Põlme et al. 2020) | Põlme S, et al. 2020. Fungal Diversity 105:1-16 ((https://doi.org/10.1007/s13225-020-00466-2)) |
| Otu47637 | 1 | k__Fungi(100);p__Ascomycota(100);c__Saccharomycetes(100);o__Saccharomycetales(100);f__Metschnikowiaceae(100);g__Metschnikowia(100);s__Metschnikowia_pulcherrima(100); | Metschnikowia | 13 | Saprotroph | \|Undefined Saprotroph\| | Yeast | NULL | Probable | As nectar-tap_saprotroph (Põlme et al. 2020); Decay sugar-rich substrates (Põlme et al. 2020) | Põlme S, et al. 2020. Fungal Diversity 105:1-16 ((https://doi.org/10.1007/s13225-020-00466-2)) |
| Otu47638 | 1 | k__Fungi(100);p__Ascomycota(100);c__Saccharomycetes(100);o__Saccharomycetales(100);f__Metschnikowiaceae(100);g__Metschnikowia(100);s__Metschnikowia_pulcherrima(100); | Metschnikowia | 13 | Saprotroph | \|Undefined Saprotroph\| | Yeast | NULL | Probable | As nectar-tap_saprotroph (Põlme et al. 2020); Decay sugar-rich substrates (Põlme et al. 2020) | Põlme S, et al. 2020. Fungal Diversity 105:1-16 ((https://doi.org/10.1007/s13225-020-00466-2)) |
| Otu47639 | 1 | k__Fungi(100);p__Ascomycota(100);c__Saccharomycetes(100);o__Saccharomycetales(100);f__Metschnikowiaceae(100);g__Metschnikowia(100);s__Metschnikowia_pulcherrima(100); | Metschnikowia | 13 | Saprotroph | \|Undefined Saprotroph\| | Yeast | NULL | Probable | As nectar-tap_saprotroph (Põlme et al. 2020); Decay sugar-rich substrates (Põlme et al. 2020) | Põlme S, et al. 2020. Fungal Diversity 105:1-16 ((https://doi.org/10.1007/s13225-020-00466-2)) |
| Otu47640 | 1 | k__Fungi(100);p__Ascomycota(100);c__Saccharomycetes(100);o__Saccharomycetales(100);f__Metschnikowiaceae(100);g__Metschnikowia(100);s__Metschnikowia_pulcherrima(100); | Metschnikowia | 13 | Saprotroph | \|Undefined Saprotroph\| | Yeast | NULL | Probable | As nectar-tap_saprotroph (Põlme et al. 2020); Decay sugar-rich substrates (Põlme et al. 2020) | Põlme S, et al. 2020. Fungal Diversity 105:1-16 ((https://doi.org/10.1007/s13225-020-00466-2)) |
| Otu47641 | 1 | k__Fungi(100);p__Ascomycota(100);c__Saccharomycetes(100);o__Saccharomycetales(100);f__Metschnikowiaceae(100);g__Metschnikowia(100);s__Metschnikowia_pulcherrima(100); | Metschnikowia | 13 | Saprotroph | \|Undefined Saprotroph\| | Yeast | NULL | Probable | As nectar-tap_saprotroph (Põlme et al. 2020); Decay sugar-rich substrates (Põlme et al. 2020) | Põlme S, et al. 2020. Fungal Diversity 105:1-16 ((https://doi.org/10.1007/s13225-020-00466-2)) |
| Otu47642 | 1 | k__Fungi(100);p__Ascomycota(100);c__Saccharomycetes(100);o__Saccharomycetales(100);f__Metschnikowiaceae(100);g__Metschnikowia(100);s__Metschnikowia_pulcherrima(100); | Metschnikowia | 13 | Saprotroph | \|Undefined Saprotroph\| | Yeast | NULL | Probable | As nectar-tap_saprotroph (Põlme et al. 2020); Decay sugar-rich substrates (Põlme et al. 2020) | Põlme S, et al. 2020. Fungal Diversity 105:1-16 ((https://doi.org/10.1007/s13225-020-00466-2)) |
| Otu47643 | 1 | k__Fungi(100);p__Ascomycota(100);c__Saccharomycetes(100);o__Saccharomycetales(100);f__Metschnikowiaceae(100);g__Metschnikowia(100);g__Metschnikowia_unclassified(100); | Metschnikowia | 13 | Saprotroph | \|Undefined Saprotroph\| | Yeast | NULL | Probable | As nectar-tap_saprotroph (Põlme et al. 2020); Decay sugar-rich substrates (Põlme et al. 2020) | Põlme S, et al. 2020. Fungal Diversity 105:1-16 ((https://doi.org/10.1007/s13225-020-00466-2)) |
| Otu47644 | 1 | k__Fungi(100);p__Ascomycota(100);c__Saccharomycetes(100);o__Saccharomycetales(100);f__Metschnikowiaceae(100);g__Metschnikowia(100);s__Metschnikowia_pulcherrima(100); | Metschnikowia | 13 | Saprotroph | \|Undefined Saprotroph\| | Yeast | NULL | Probable | As nectar-tap_saprotroph (Põlme et al. 2020); Decay sugar-rich substrates (Põlme et al. 2020) | Põlme S, et al. 2020. Fungal Diversity 105:1-16 ((https://doi.org/10.1007/s13225-020-00466-2)) |
| Otu47645 | 1 | k__Fungi(100);p__Ascomycota(100);c__Saccharomycetes(100);o__Saccharomycetales(100);f__Metschnikowiaceae(100);g__Metschnikowia(100);g__Metschnikowia_unclassified(100); | Metschnikowia | 13 | Saprotroph | \|Undefined Saprotroph\| | Yeast | NULL | Probable | As nectar-tap_saprotroph (Põlme et al. 2020); Decay sugar-rich substrates (Põlme et al. 2020) | Põlme S, et al. 2020. Fungal Diversity 105:1-16 ((https://doi.org/10.1007/s13225-020-00466-2)) |
| Otu47646 | 1 | k__Fungi(100);p__Ascomycota(100);c__Saccharomycetes(100);o__Saccharomycetales(100);f__Metschnikowiaceae(100);g__Metschnikowia(100);s__Metschnikowia_pulcherrima(100); | Metschnikowia | 13 | Saprotroph | \|Undefined Saprotroph\| | Yeast | NULL | Probable | As nectar-tap_saprotroph (Põlme et al. 2020); Decay sugar-rich substrates (Põlme et al. 2020) | Põlme S, et al. 2020. Fungal Diversity 105:1-16 ((https://doi.org/10.1007/s13225-020-00466-2)) |
| Otu47647 | 1 | k__Fungi(100);p__Ascomycota(100);c__Saccharomycetes(100);o__Saccharomycetales(100);f__Metschnikowiaceae(100);g__Metschnikowia(100);s__Metschnikowia_pulcherrima(100); | Metschnikowia | 13 | Saprotroph | \|Undefined Saprotroph\| | Yeast | NULL | Probable | As nectar-tap_saprotroph (Põlme et al. 2020); Decay sugar-rich substrates (Põlme et al. 2020) | Põlme S, et al. 2020. Fungal Diversity 105:1-16 ((https://doi.org/10.1007/s13225-020-00466-2)) |
| Otu47648 | 1 | k__Fungi(100);p__Ascomycota(100);c__Saccharomycetes(100);o__Saccharomycetales(100);f__Metschnikowiaceae(100);g__Metschnikowia(100);s__Metschnikowia_pulcherrima(100); | Metschnikowia | 13 | Saprotroph | \|Undefined Saprotroph\| | Yeast | NULL | Probable | As nectar-tap_saprotroph (Põlme et al. 2020); Decay sugar-rich substrates (Põlme et al. 2020) | Põlme S, et al. 2020. Fungal Diversity 105:1-16 ((https://doi.org/10.1007/s13225-020-00466-2)) |
| Otu47649 | 1 | k__Fungi(100);p__Ascomycota(100);c__Saccharomycetes(100);o__Saccharomycetales(100);f__Metschnikowiaceae(100);g__Metschnikowia(100);s__Metschnikowia_pulcherrima(100); | Metschnikowia | 13 | Saprotroph | \|Undefined Saprotroph\| | Yeast | NULL | Probable | As nectar-tap_saprotroph (Põlme et al. 2020); Decay sugar-rich substrates (Põlme et al. 2020) | Põlme S, et al. 2020. Fungal Diversity 105:1-16 ((https://doi.org/10.1007/s13225-020-00466-2)) |
| Otu47650 | 1 | k__Fungi(100);p__Ascomycota(100);c__Saccharomycetes(100);o__Saccharomycetales(100);f__Metschnikowiaceae(100);g__Metschnikowia(100);g__Metschnikowia_unclassified(100); | Metschnikowia | 13 | Saprotroph | \|Undefined Saprotroph\| | Yeast | NULL | Probable | As nectar-tap_saprotroph (Põlme et al. 2020); Decay sugar-rich substrates (Põlme et al. 2020) | Põlme S, et al. 2020. Fungal Diversity 105:1-16 ((https://doi.org/10.1007/s13225-020-00466-2)) |
| Otu47651 | 1 | k__Fungi(100);p__Ascomycota(100);c__Saccharomycetes(100);o__Saccharomycetales(100);f__Metschnikowiaceae(100);g__Metschnikowia(100);s__Metschnikowia_pulcherrima(100); | Metschnikowia | 13 | Saprotroph | \|Undefined Saprotroph\| | Yeast | NULL | Probable | As nectar-tap_saprotroph (Põlme et al. 2020); Decay sugar-rich substrates (Põlme et al. 2020) | Põlme S, et al. 2020. Fungal Diversity 105:1-16 ((https://doi.org/10.1007/s13225-020-00466-2)) |
| Otu47652 | 1 | k__Fungi(100);p__Ascomycota(100);c__Saccharomycetes(100);o__Saccharomycetales(100);f__Metschnikowiaceae(100);g__Metschnikowia(100);s__Metschnikowia_pulcherrima(100); | Metschnikowia | 13 | Saprotroph | \|Undefined Saprotroph\| | Yeast | NULL | Probable | As nectar-tap_saprotroph (Põlme et al. 2020); Decay sugar-rich substrates (Põlme et al. 2020) | Põlme S, et al. 2020. Fungal Diversity 105:1-16 ((https://doi.org/10.1007/s13225-020-00466-2)) |
| Otu47653 | 1 | k__Fungi(100);p__Ascomycota(100);c__Saccharomycetes(100);o__Saccharomycetales(100);f__Metschnikowiaceae(100);g__Metschnikowia(100);s__Metschnikowia_pulcherrima(100); | Metschnikowia | 13 | Saprotroph | \|Undefined Saprotroph\| | Yeast | NULL | Probable | As nectar-tap_saprotroph (Põlme et al. 2020); Decay sugar-rich substrates (Põlme et al. 2020) | Põlme S, et al. 2020. Fungal Diversity 105:1-16 ((https://doi.org/10.1007/s13225-020-00466-2)) |
| Otu47654 | 1 | k__Fungi(100);p__Ascomycota(100);c__Saccharomycetes(100);o__Saccharomycetales(100);f__Metschnikowiaceae(100);g__Metschnikowia(100);s__Metschnikowia_pulcherrima(100); | Metschnikowia | 13 | Saprotroph | \|Undefined Saprotroph\| | Yeast | NULL | Probable | As nectar-tap_saprotroph (Põlme et al. 2020); Decay sugar-rich substrates (Põlme et al. 2020) | Põlme S, et al. 2020. Fungal Diversity 105:1-16 ((https://doi.org/10.1007/s13225-020-00466-2)) |
| Otu47655 | 1 | k__Fungi(100);p__Ascomycota(100);c__Saccharomycetes(100);o__Saccharomycetales(100);f__Metschnikowiaceae(100);g__Metschnikowia(100);s__Metschnikowia_pulcherrima(100); | Metschnikowia | 13 | Saprotroph | \|Undefined Saprotroph\| | Yeast | NULL | Probable | As nectar-tap_saprotroph (Põlme et al. 2020); Decay sugar-rich substrates (Põlme et al. 2020) | Põlme S, et al. 2020. Fungal Diversity 105:1-16 ((https://doi.org/10.1007/s13225-020-00466-2)) |
| Otu47656 | 1 | k__Fungi(100);p__Ascomycota(100);c__Saccharomycetes(100);o__Saccharomycetales(100);f__Metschnikowiaceae(100);g__Metschnikowia(100);s__Metschnikowia_pulcherrima(100); | Metschnikowia | 13 | Saprotroph | \|Undefined Saprotroph\| | Yeast | NULL | Probable | As nectar-tap_saprotroph (Põlme et al. 2020); Decay sugar-rich substrates (Põlme et al. 2020) | Põlme S, et al. 2020. Fungal Diversity 105:1-16 ((https://doi.org/10.1007/s13225-020-00466-2)) |
| Otu47657 | 1 | k__Fungi(100);p__Ascomycota(100);c__Saccharomycetes(100);o__Saccharomycetales(100);f__Metschnikowiaceae(100);g__Metschnikowia(100);s__Metschnikowia_pulcherrima(100); | Metschnikowia | 13 | Saprotroph | \|Undefined Saprotroph\| | Yeast | NULL | Probable | As nectar-tap_saprotroph (Põlme et al. 2020); Decay sugar-rich substrates (Põlme et al. 2020) | Põlme S, et al. 2020. Fungal Diversity 105:1-16 ((https://doi.org/10.1007/s13225-020-00466-2)) |
| Otu47658 | 1 | k__Fungi(100);p__Ascomycota(100);c__Saccharomycetes(100);o__Saccharomycetales(100);f__Metschnikowiaceae(100);g__Metschnikowia(100);s__Metschnikowia_pulcherrima(100); | Metschnikowia | 13 | Saprotroph | \|Undefined Saprotroph\| | Yeast | NULL | Probable | As nectar-tap_saprotroph (Põlme et al. 2020); Decay sugar-rich substrates (Põlme et al. 2020) | Põlme S, et al. 2020. Fungal Diversity 105:1-16 ((https://doi.org/10.1007/s13225-020-00466-2)) |
| Otu47660 | 1 | k__Fungi(100);p__Ascomycota(100);c__Saccharomycetes(100);o__Saccharomycetales(100);f__Metschnikowiaceae(100);g__Metschnikowia(100);s__Metschnikowia_pulcherrima(100); | Metschnikowia | 13 | Saprotroph | \|Undefined Saprotroph\| | Yeast | NULL | Probable | As nectar-tap_saprotroph (Põlme et al. 2020); Decay sugar-rich substrates (Põlme et al. 2020) | Põlme S, et al. 2020. Fungal Diversity 105:1-16 ((https://doi.org/10.1007/s13225-020-00466-2)) |
| Otu47661 | 1 | k__Fungi(100);p__Ascomycota(100);c__Saccharomycetes(100);o__Saccharomycetales(100);f__Metschnikowiaceae(100);g__Metschnikowia(100);g__Metschnikowia_unclassified(100); | Metschnikowia | 13 | Saprotroph | \|Undefined Saprotroph\| | Yeast | NULL | Probable | As nectar-tap_saprotroph (Põlme et al. 2020); Decay sugar-rich substrates (Põlme et al. 2020) | Põlme S, et al. 2020. Fungal Diversity 105:1-16 ((https://doi.org/10.1007/s13225-020-00466-2)) |
| Otu47662 | 1 | k__Fungi(100);p__Ascomycota(100);c__Saccharomycetes(100);o__Saccharomycetales(100);f__Metschnikowiaceae(100);g__Metschnikowia(100);s__Metschnikowia_pulcherrima(100); | Metschnikowia | 13 | Saprotroph | \|Undefined Saprotroph\| | Yeast | NULL | Probable | As nectar-tap_saprotroph (Põlme et al. 2020); Decay sugar-rich substrates (Põlme et al. 2020) | Põlme S, et al. 2020. Fungal Diversity 105:1-16 ((https://doi.org/10.1007/s13225-020-00466-2)) |
| Otu47663 | 1 | k__Fungi(100);p__Ascomycota(100);c__Saccharomycetes(100);o__Saccharomycetales(100);f__Metschnikowiaceae(100);g__Metschnikowia(100);s__Metschnikowia_pulcherrima(100); | Metschnikowia | 13 | Saprotroph | \|Undefined Saprotroph\| | Yeast | NULL | Probable | As nectar-tap_saprotroph (Põlme et al. 2020); Decay sugar-rich substrates (Põlme et al. 2020) | Põlme S, et al. 2020. Fungal Diversity 105:1-16 ((https://doi.org/10.1007/s13225-020-00466-2)) |
| Otu47664 | 1 | k__Fungi(100);p__Ascomycota(100);c__Saccharomycetes(100);o__Saccharomycetales(100);f__Metschnikowiaceae(100);g__Metschnikowia(100);s__Metschnikowia_pulcherrima(100); | Metschnikowia | 13 | Saprotroph | \|Undefined Saprotroph\| | Yeast | NULL | Probable | As nectar-tap_saprotroph (Põlme et al. 2020); Decay sugar-rich substrates (Põlme et al. 2020) | Põlme S, et al. 2020. Fungal Diversity 105:1-16 ((https://doi.org/10.1007/s13225-020-00466-2)) |
| Otu47665 | 1 | k__Fungi(100);p__Ascomycota(100);c__Saccharomycetes(100);o__Saccharomycetales(100);f__Metschnikowiaceae(100);g__Metschnikowia(100);s__Metschnikowia_pulcherrima(100); | Metschnikowia | 13 | Saprotroph | \|Undefined Saprotroph\| | Yeast | NULL | Probable | As nectar-tap_saprotroph (Põlme et al. 2020); Decay sugar-rich substrates (Põlme et al. 2020) | Põlme S, et al. 2020. Fungal Diversity 105:1-16 ((https://doi.org/10.1007/s13225-020-00466-2)) |
| Otu47666 | 1 | k__Fungi(100);p__Ascomycota(100);c__Saccharomycetes(100);o__Saccharomycetales(100);f__Metschnikowiaceae(100);g__Metschnikowia(100);g__Metschnikowia_unclassified(100); | Metschnikowia | 13 | Saprotroph | \|Undefined Saprotroph\| | Yeast | NULL | Probable | As nectar-tap_saprotroph (Põlme et al. 2020); Decay sugar-rich substrates (Põlme et al. 2020) | Põlme S, et al. 2020. Fungal Diversity 105:1-16 ((https://doi.org/10.1007/s13225-020-00466-2)) |
| Otu47667 | 1 | k__Fungi(100);p__Ascomycota(100);c__Saccharomycetes(100);o__Saccharomycetales(100);f__Metschnikowiaceae(100);g__Metschnikowia(100);g__Metschnikowia_unclassified(100); | Metschnikowia | 13 | Saprotroph | \|Undefined Saprotroph\| | Yeast | NULL | Probable | As nectar-tap_saprotroph (Põlme et al. 2020); Decay sugar-rich substrates (Põlme et al. 2020) | Põlme S, et al. 2020. Fungal Diversity 105:1-16 ((https://doi.org/10.1007/s13225-020-00466-2)) |
| Otu47668 | 1 | k__Fungi(100);p__Ascomycota(100);c__Saccharomycetes(100);o__Saccharomycetales(100);f__Metschnikowiaceae(100);g__Metschnikowia(100);s__Metschnikowia_pulcherrima(100); | Metschnikowia | 13 | Saprotroph | \|Undefined Saprotroph\| | Yeast | NULL | Probable | As nectar-tap_saprotroph (Põlme et al. 2020); Decay sugar-rich substrates (Põlme et al. 2020) | Põlme S, et al. 2020. Fungal Diversity 105:1-16 ((https://doi.org/10.1007/s13225-020-00466-2)) |
| Otu47669 | 1 | k__Fungi(100);p__Ascomycota(100);c__Saccharomycetes(100);o__Saccharomycetales(100);f__Metschnikowiaceae(100);g__Metschnikowia(100);s__Metschnikowia_pulcherrima(100); | Metschnikowia | 13 | Saprotroph | \|Undefined Saprotroph\| | Yeast | NULL | Probable | As nectar-tap_saprotroph (Põlme et al. 2020); Decay sugar-rich substrates (Põlme et al. 2020) | Põlme S, et al. 2020. Fungal Diversity 105:1-16 ((https://doi.org/10.1007/s13225-020-00466-2)) |
| Otu47670 | 1 | k__Fungi(100);p__Ascomycota(100);c__Saccharomycetes(100);o__Saccharomycetales(100);f__Metschnikowiaceae(100);g__Metschnikowia(100);s__Metschnikowia_pulcherrima(100); | Metschnikowia | 13 | Saprotroph | \|Undefined Saprotroph\| | Yeast | NULL | Probable | As nectar-tap_saprotroph (Põlme et al. 2020); Decay sugar-rich substrates (Põlme et al. 2020) | Põlme S, et al. 2020. Fungal Diversity 105:1-16 ((https://doi.org/10.1007/s13225-020-00466-2)) |
| Otu47671 | 1 | k__Fungi(100);p__Ascomycota(100);c__Saccharomycetes(100);o__Saccharomycetales(100);f__Metschnikowiaceae(100);g__Metschnikowia(100);s__Metschnikowia_pulcherrima(100); | Metschnikowia | 13 | Saprotroph | \|Undefined Saprotroph\| | Yeast | NULL | Probable | As nectar-tap_saprotroph (Põlme et al. 2020); Decay sugar-rich substrates (Põlme et al. 2020) | Põlme S, et al. 2020. Fungal Diversity 105:1-16 ((https://doi.org/10.1007/s13225-020-00466-2)) |
| Otu47672 | 1 | k__Fungi(100);p__Ascomycota(100);c__Saccharomycetes(100);o__Saccharomycetales(100);f__Metschnikowiaceae(100);g__Metschnikowia(100);s__Metschnikowia_pulcherrima(100); | Metschnikowia | 13 | Saprotroph | \|Undefined Saprotroph\| | Yeast | NULL | Probable | As nectar-tap_saprotroph (Põlme et al. 2020); Decay sugar-rich substrates (Põlme et al. 2020) | Põlme S, et al. 2020. Fungal Diversity 105:1-16 ((https://doi.org/10.1007/s13225-020-00466-2)) |
| Otu47673 | 1 | k__Fungi(100);p__Ascomycota(100);c__Saccharomycetes(100);o__Saccharomycetales(100);f__Metschnikowiaceae(100);g__Metschnikowia(100);s__Metschnikowia_pulcherrima(100); | Metschnikowia | 13 | Saprotroph | \|Undefined Saprotroph\| | Yeast | NULL | Probable | As nectar-tap_saprotroph (Põlme et al. 2020); Decay sugar-rich substrates (Põlme et al. 2020) | Põlme S, et al. 2020. Fungal Diversity 105:1-16 ((https://doi.org/10.1007/s13225-020-00466-2)) |
| Otu47674 | 1 | k__Fungi(100);p__Ascomycota(100);c__Saccharomycetes(100);o__Saccharomycetales(100);f__Metschnikowiaceae(100);g__Metschnikowia(100);s__Metschnikowia_pulcherrima(100); | Metschnikowia | 13 | Saprotroph | \|Undefined Saprotroph\| | Yeast | NULL | Probable | As nectar-tap_saprotroph (Põlme et al. 2020); Decay sugar-rich substrates (Põlme et al. 2020) | Põlme S, et al. 2020. Fungal Diversity 105:1-16 ((https://doi.org/10.1007/s13225-020-00466-2)) |
| Otu47675 | 1 | k__Fungi(100);p__Ascomycota(100);c__Saccharomycetes(100);o__Saccharomycetales(100);f__Metschnikowiaceae(100);g__Metschnikowia(100);s__Metschnikowia_pulcherrima(100); | Metschnikowia | 13 | Saprotroph | \|Undefined Saprotroph\| | Yeast | NULL | Probable | As nectar-tap_saprotroph (Põlme et al. 2020); Decay sugar-rich substrates (Põlme et al. 2020) | Põlme S, et al. 2020. Fungal Diversity 105:1-16 ((https://doi.org/10.1007/s13225-020-00466-2)) |
| Otu47676 | 1 | k__Fungi(100);p__Ascomycota(100);c__Saccharomycetes(100);o__Saccharomycetales(100);f__Metschnikowiaceae(100);g__Metschnikowia(100);g__Metschnikowia_unclassified(100); | Metschnikowia | 13 | Saprotroph | \|Undefined Saprotroph\| | Yeast | NULL | Probable | As nectar-tap_saprotroph (Põlme et al. 2020); Decay sugar-rich substrates (Põlme et al. 2020) | Põlme S, et al. 2020. Fungal Diversity 105:1-16 ((https://doi.org/10.1007/s13225-020-00466-2)) |
| Otu47677 | 1 | k__Fungi(100);p__Ascomycota(100);c__Saccharomycetes(100);o__Saccharomycetales(100);f__Metschnikowiaceae(100);g__Metschnikowia(100);s__Metschnikowia_pulcherrima(100); | Metschnikowia | 13 | Saprotroph | \|Undefined Saprotroph\| | Yeast | NULL | Probable | As nectar-tap_saprotroph (Põlme et al. 2020); Decay sugar-rich substrates (Põlme et al. 2020) | Põlme S, et al. 2020. Fungal Diversity 105:1-16 ((https://doi.org/10.1007/s13225-020-00466-2)) |
| Otu47678 | 1 | k__Fungi(100);p__Ascomycota(100);c__Saccharomycetes(100);o__Saccharomycetales(100);f__Metschnikowiaceae(100);g__Metschnikowia(100);s__Metschnikowia_pulcherrima(100); | Metschnikowia | 13 | Saprotroph | \|Undefined Saprotroph\| | Yeast | NULL | Probable | As nectar-tap_saprotroph (Põlme et al. 2020); Decay sugar-rich substrates (Põlme et al. 2020) | Põlme S, et al. 2020. Fungal Diversity 105:1-16 ((https://doi.org/10.1007/s13225-020-00466-2)) |
| Otu47679 | 1 | k__Fungi(100);p__Ascomycota(100);c__Saccharomycetes(100);o__Saccharomycetales(100);f__Metschnikowiaceae(100);g__Metschnikowia(100);s__Metschnikowia_pulcherrima(100); | Metschnikowia | 13 | Saprotroph | \|Undefined Saprotroph\| | Yeast | NULL | Probable | As nectar-tap_saprotroph (Põlme et al. 2020); Decay sugar-rich substrates (Põlme et al. 2020) | Põlme S, et al. 2020. Fungal Diversity 105:1-16 ((https://doi.org/10.1007/s13225-020-00466-2)) |
| Otu47680 | 1 | k__Fungi(100);p__Ascomycota(100);c__Saccharomycetes(100);o__Saccharomycetales(100);f__Metschnikowiaceae(100);g__Metschnikowia(100);s__Metschnikowia_pulcherrima(100); | Metschnikowia | 13 | Saprotroph | \|Undefined Saprotroph\| | Yeast | NULL | Probable | As nectar-tap_saprotroph (Põlme et al. 2020); Decay sugar-rich substrates (Põlme et al. 2020) | Põlme S, et al. 2020. Fungal Diversity 105:1-16 ((https://doi.org/10.1007/s13225-020-00466-2)) |
| Otu47681 | 1 | k__Fungi(100);p__Ascomycota(100);c__Saccharomycetes(100);o__Saccharomycetales(100);f__Metschnikowiaceae(100);g__Metschnikowia(100);s__Metschnikowia_pulcherrima(100); | Metschnikowia | 13 | Saprotroph | \|Undefined Saprotroph\| | Yeast | NULL | Probable | As nectar-tap_saprotroph (Põlme et al. 2020); Decay sugar-rich substrates (Põlme et al. 2020) | Põlme S, et al. 2020. Fungal Diversity 105:1-16 ((https://doi.org/10.1007/s13225-020-00466-2)) |
| Otu47682 | 1 | k__Fungi(100);p__Ascomycota(100);c__Saccharomycetes(100);o__Saccharomycetales(100);f__Metschnikowiaceae(100);g__Metschnikowia(100);s__Metschnikowia_pulcherrima(100); | Metschnikowia | 13 | Saprotroph | \|Undefined Saprotroph\| | Yeast | NULL | Probable | As nectar-tap_saprotroph (Põlme et al. 2020); Decay sugar-rich substrates (Põlme et al. 2020) | Põlme S, et al. 2020. Fungal Diversity 105:1-16 ((https://doi.org/10.1007/s13225-020-00466-2)) |
| Otu47683 | 1 | k__Fungi(100);p__Ascomycota(100);c__Saccharomycetes(100);o__Saccharomycetales(100);f__Metschnikowiaceae(100);g__Metschnikowia(100);s__Metschnikowia_pulcherrima(100); | Metschnikowia | 13 | Saprotroph | \|Undefined Saprotroph\| | Yeast | NULL | Probable | As nectar-tap_saprotroph (Põlme et al. 2020); Decay sugar-rich substrates (Põlme et al. 2020) | Põlme S, et al. 2020. Fungal Diversity 105:1-16 ((https://doi.org/10.1007/s13225-020-00466-2)) |
| Otu47684 | 1 | k__Fungi(100);p__Ascomycota(100);c__Saccharomycetes(100);o__Saccharomycetales(100);f__Metschnikowiaceae(100);g__Metschnikowia(100);s__Metschnikowia_pulcherrima(100); | Metschnikowia | 13 | Saprotroph | \|Undefined Saprotroph\| | Yeast | NULL | Probable | As nectar-tap_saprotroph (Põlme et al. 2020); Decay sugar-rich substrates (Põlme et al. 2020) | Põlme S, et al. 2020. Fungal Diversity 105:1-16 ((https://doi.org/10.1007/s13225-020-00466-2)) |
| Otu47685 | 1 | k__Fungi(100);p__Ascomycota(100);c__Saccharomycetes(100);o__Saccharomycetales(100);f__Metschnikowiaceae(100);g__Metschnikowia(100);s__Metschnikowia_pulcherrima(100); | Metschnikowia | 13 | Saprotroph | \|Undefined Saprotroph\| | Yeast | NULL | Probable | As nectar-tap_saprotroph (Põlme et al. 2020); Decay sugar-rich substrates (Põlme et al. 2020) | Põlme S, et al. 2020. Fungal Diversity 105:1-16 ((https://doi.org/10.1007/s13225-020-00466-2)) |
| Otu47686 | 1 | k__Fungi(100);p__Ascomycota(100);c__Saccharomycetes(100);o__Saccharomycetales(100);f__Metschnikowiaceae(100);g__Metschnikowia(100);s__Metschnikowia_pulcherrima(100); | Metschnikowia | 13 | Saprotroph | \|Undefined Saprotroph\| | Yeast | NULL | Probable | As nectar-tap_saprotroph (Põlme et al. 2020); Decay sugar-rich substrates (Põlme et al. 2020) | Põlme S, et al. 2020. Fungal Diversity 105:1-16 ((https://doi.org/10.1007/s13225-020-00466-2)) |
| Otu47687 | 1 | k__Fungi(100);p__Ascomycota(100);c__Saccharomycetes(100);o__Saccharomycetales(100);f__Metschnikowiaceae(100);g__Metschnikowia(100);s__Metschnikowia_pulcherrima(100); | Metschnikowia | 13 | Saprotroph | \|Undefined Saprotroph\| | Yeast | NULL | Probable | As nectar-tap_saprotroph (Põlme et al. 2020); Decay sugar-rich substrates (Põlme et al. 2020) | Põlme S, et al. 2020. Fungal Diversity 105:1-16 ((https://doi.org/10.1007/s13225-020-00466-2)) |
| Otu47688 | 1 | k__Fungi(100);p__Ascomycota(100);c__Saccharomycetes(100);o__Saccharomycetales(100);f__Metschnikowiaceae(100);g__Metschnikowia(100);s__Metschnikowia_pulcherrima(100); | Metschnikowia | 13 | Saprotroph | \|Undefined Saprotroph\| | Yeast | NULL | Probable | As nectar-tap_saprotroph (Põlme et al. 2020); Decay sugar-rich substrates (Põlme et al. 2020) | Põlme S, et al. 2020. Fungal Diversity 105:1-16 ((https://doi.org/10.1007/s13225-020-00466-2)) |
| Otu47689 | 1 | k__Fungi(100);p__Ascomycota(100);c__Saccharomycetes(100);o__Saccharomycetales(100);f__Metschnikowiaceae(100);g__Metschnikowia(100);s__Metschnikowia_pulcherrima(100); | Metschnikowia | 13 | Saprotroph | \|Undefined Saprotroph\| | Yeast | NULL | Probable | As nectar-tap_saprotroph (Põlme et al. 2020); Decay sugar-rich substrates (Põlme et al. 2020) | Põlme S, et al. 2020. Fungal Diversity 105:1-16 ((https://doi.org/10.1007/s13225-020-00466-2)) |
| Otu47690 | 1 | k__Fungi(100);p__Ascomycota(100);c__Saccharomycetes(100);o__Saccharomycetales(100);f__Metschnikowiaceae(100);g__Metschnikowia(100);s__Metschnikowia_pulcherrima(100); | Metschnikowia | 13 | Saprotroph | \|Undefined Saprotroph\| | Yeast | NULL | Probable | As nectar-tap_saprotroph (Põlme et al. 2020); Decay sugar-rich substrates (Põlme et al. 2020) | Põlme S, et al. 2020. Fungal Diversity 105:1-16 ((https://doi.org/10.1007/s13225-020-00466-2)) |
| Otu47691 | 1 | k__Fungi(100);p__Ascomycota(100);c__Saccharomycetes(100);o__Saccharomycetales(100);f__Metschnikowiaceae(100);g__Metschnikowia(100);s__Metschnikowia_pulcherrima(100); | Metschnikowia | 13 | Saprotroph | \|Undefined Saprotroph\| | Yeast | NULL | Probable | As nectar-tap_saprotroph (Põlme et al. 2020); Decay sugar-rich substrates (Põlme et al. 2020) | Põlme S, et al. 2020. Fungal Diversity 105:1-16 ((https://doi.org/10.1007/s13225-020-00466-2)) |
| Otu47692 | 1 | k__Fungi(100);p__Ascomycota(100);c__Saccharomycetes(100);o__Saccharomycetales(100);f__Metschnikowiaceae(100);g__Metschnikowia(100);s__Metschnikowia_pulcherrima(100); | Metschnikowia | 13 | Saprotroph | \|Undefined Saprotroph\| | Yeast | NULL | Probable | As nectar-tap_saprotroph (Põlme et al. 2020); Decay sugar-rich substrates (Põlme et al. 2020) | Põlme S, et al. 2020. Fungal Diversity 105:1-16 ((https://doi.org/10.1007/s13225-020-00466-2)) |
| Otu47694 | 1 | k__Fungi(100);p__Ascomycota(100);c__Saccharomycetes(100);o__Saccharomycetales(100);f__Metschnikowiaceae(100);g__Metschnikowia(100);s__Metschnikowia_pulcherrima(100); | Metschnikowia | 13 | Saprotroph | \|Undefined Saprotroph\| | Yeast | NULL | Probable | As nectar-tap_saprotroph (Põlme et al. 2020); Decay sugar-rich substrates (Põlme et al. 2020) | Põlme S, et al. 2020. Fungal Diversity 105:1-16 ((https://doi.org/10.1007/s13225-020-00466-2)) |
| Otu47695 | 1 | k__Fungi(100);p__Ascomycota(100);c__Saccharomycetes(100);o__Saccharomycetales(100);f__Metschnikowiaceae(100);g__Metschnikowia(100);g__Metschnikowia_unclassified(100); | Metschnikowia | 13 | Saprotroph | \|Undefined Saprotroph\| | Yeast | NULL | Probable | As nectar-tap_saprotroph (Põlme et al. 2020); Decay sugar-rich substrates (Põlme et al. 2020) | Põlme S, et al. 2020. Fungal Diversity 105:1-16 ((https://doi.org/10.1007/s13225-020-00466-2)) |
| Otu47696 | 1 | k__Fungi(100);p__Ascomycota(100);c__Saccharomycetes(100);o__Saccharomycetales(100);f__Metschnikowiaceae(100);g__Metschnikowia(100);s__Metschnikowia_pulcherrima(100); | Metschnikowia | 13 | Saprotroph | \|Undefined Saprotroph\| | Yeast | NULL | Probable | As nectar-tap_saprotroph (Põlme et al. 2020); Decay sugar-rich substrates (Põlme et al. 2020) | Põlme S, et al. 2020. Fungal Diversity 105:1-16 ((https://doi.org/10.1007/s13225-020-00466-2)) |
| Otu47697 | 1 | k__Fungi(100);p__Ascomycota(100);c__Saccharomycetes(100);o__Saccharomycetales(100);f__Metschnikowiaceae(100);g__Metschnikowia(100);s__Metschnikowia_pulcherrima(100); | Metschnikowia | 13 | Saprotroph | \|Undefined Saprotroph\| | Yeast | NULL | Probable | As nectar-tap_saprotroph (Põlme et al. 2020); Decay sugar-rich substrates (Põlme et al. 2020) | Põlme S, et al. 2020. Fungal Diversity 105:1-16 ((https://doi.org/10.1007/s13225-020-00466-2)) |
| Otu47698 | 1 | k__Fungi(100);p__Ascomycota(100);c__Saccharomycetes(100);o__Saccharomycetales(100);f__Metschnikowiaceae(100);g__Metschnikowia(100);s__Metschnikowia_pulcherrima(100); | Metschnikowia | 13 | Saprotroph | \|Undefined Saprotroph\| | Yeast | NULL | Probable | As nectar-tap_saprotroph (Põlme et al. 2020); Decay sugar-rich substrates (Põlme et al. 2020) | Põlme S, et al. 2020. Fungal Diversity 105:1-16 ((https://doi.org/10.1007/s13225-020-00466-2)) |
| Otu47699 | 1 | k__Fungi(100);p__Ascomycota(100);c__Saccharomycetes(100);o__Saccharomycetales(100);f__Metschnikowiaceae(100);g__Metschnikowia(100);s__Metschnikowia_pulcherrima(100); | Metschnikowia | 13 | Saprotroph | \|Undefined Saprotroph\| | Yeast | NULL | Probable | As nectar-tap_saprotroph (Põlme et al. 2020); Decay sugar-rich substrates (Põlme et al. 2020) | Põlme S, et al. 2020. Fungal Diversity 105:1-16 ((https://doi.org/10.1007/s13225-020-00466-2)) |
| Otu47700 | 1 | k__Fungi(100);p__Ascomycota(100);c__Saccharomycetes(100);o__Saccharomycetales(100);f__Metschnikowiaceae(100);g__Metschnikowia(100);s__Metschnikowia_pulcherrima(100); | Metschnikowia | 13 | Saprotroph | \|Undefined Saprotroph\| | Yeast | NULL | Probable | As nectar-tap_saprotroph (Põlme et al. 2020); Decay sugar-rich substrates (Põlme et al. 2020) | Põlme S, et al. 2020. Fungal Diversity 105:1-16 ((https://doi.org/10.1007/s13225-020-00466-2)) |
| Otu47701 | 1 | k__Fungi(100);p__Ascomycota(100);c__Saccharomycetes(100);o__Saccharomycetales(100);f__Metschnikowiaceae(100);g__Metschnikowia(100);s__Metschnikowia_pulcherrima(100); | Metschnikowia | 13 | Saprotroph | \|Undefined Saprotroph\| | Yeast | NULL | Probable | As nectar-tap_saprotroph (Põlme et al. 2020); Decay sugar-rich substrates (Põlme et al. 2020) | Põlme S, et al. 2020. Fungal Diversity 105:1-16 ((https://doi.org/10.1007/s13225-020-00466-2)) |
| Otu47702 | 1 | k__Fungi(100);p__Ascomycota(100);c__Saccharomycetes(100);o__Saccharomycetales(100);f__Metschnikowiaceae(100);g__Metschnikowia(100);g__Metschnikowia_unclassified(100); | Metschnikowia | 13 | Saprotroph | \|Undefined Saprotroph\| | Yeast | NULL | Probable | As nectar-tap_saprotroph (Põlme et al. 2020); Decay sugar-rich substrates (Põlme et al. 2020) | Põlme S, et al. 2020. Fungal Diversity 105:1-16 ((https://doi.org/10.1007/s13225-020-00466-2)) |
| Otu47703 | 1 | k__Fungi(100);p__Ascomycota(100);c__Saccharomycetes(100);o__Saccharomycetales(100);f__Metschnikowiaceae(100);g__Metschnikowia(100);s__Metschnikowia_pulcherrima(100); | Metschnikowia | 13 | Saprotroph | \|Undefined Saprotroph\| | Yeast | NULL | Probable | As nectar-tap_saprotroph (Põlme et al. 2020); Decay sugar-rich substrates (Põlme et al. 2020) | Põlme S, et al. 2020. Fungal Diversity 105:1-16 ((https://doi.org/10.1007/s13225-020-00466-2)) |
| Otu47704 | 1 | k__Fungi(100);p__Ascomycota(100);c__Saccharomycetes(100);o__Saccharomycetales(100);f__Metschnikowiaceae(100);g__Metschnikowia(100);g__Metschnikowia_unclassified(100); | Metschnikowia | 13 | Saprotroph | \|Undefined Saprotroph\| | Yeast | NULL | Probable | As nectar-tap_saprotroph (Põlme et al. 2020); Decay sugar-rich substrates (Põlme et al. 2020) | Põlme S, et al. 2020. Fungal Diversity 105:1-16 ((https://doi.org/10.1007/s13225-020-00466-2)) |
| Otu47705 | 1 | k__Fungi(100);p__Ascomycota(100);c__Saccharomycetes(100);o__Saccharomycetales(100);f__Metschnikowiaceae(100);g__Metschnikowia(100);s__Metschnikowia_noctiluminum(100); | Metschnikowia | 13 | Saprotroph | \|Undefined Saprotroph\| | Yeast | NULL | Probable | As nectar-tap_saprotroph (Põlme et al. 2020); Decay sugar-rich substrates (Põlme et al. 2020) | Põlme S, et al. 2020. Fungal Diversity 105:1-16 ((https://doi.org/10.1007/s13225-020-00466-2)) |
| Otu47706 | 1 | k__Fungi(100);p__Ascomycota(100);c__Saccharomycetes(100);o__Saccharomycetales(100);f__Metschnikowiaceae(100);g__Metschnikowia(100);g__Metschnikowia_unclassified(100); | Metschnikowia | 13 | Saprotroph | \|Undefined Saprotroph\| | Yeast | NULL | Probable | As nectar-tap_saprotroph (Põlme et al. 2020); Decay sugar-rich substrates (Põlme et al. 2020) | Põlme S, et al. 2020. Fungal Diversity 105:1-16 ((https://doi.org/10.1007/s13225-020-00466-2)) |
| Otu47707 | 1 | k__Fungi(100);p__Ascomycota(100);c__Saccharomycetes(100);o__Saccharomycetales(100);f__Metschnikowiaceae(100);g__Metschnikowia(100);s__Metschnikowia_pulcherrima(100); | Metschnikowia | 13 | Saprotroph | \|Undefined Saprotroph\| | Yeast | NULL | Probable | As nectar-tap_saprotroph (Põlme et al. 2020); Decay sugar-rich substrates (Põlme et al. 2020) | Põlme S, et al. 2020. Fungal Diversity 105:1-16 ((https://doi.org/10.1007/s13225-020-00466-2)) |
| Otu47708 | 1 | k__Fungi(100);p__Ascomycota(100);c__Saccharomycetes(100);o__Saccharomycetales(100);f__Metschnikowiaceae(100);g__Metschnikowia(100);s__Metschnikowia_pulcherrima(100); | Metschnikowia | 13 | Saprotroph | \|Undefined Saprotroph\| | Yeast | NULL | Probable | As nectar-tap_saprotroph (Põlme et al. 2020); Decay sugar-rich substrates (Põlme et al. 2020) | Põlme S, et al. 2020. Fungal Diversity 105:1-16 ((https://doi.org/10.1007/s13225-020-00466-2)) |
| Otu47709 | 1 | k__Fungi(100);p__Ascomycota(100);c__Saccharomycetes(100);o__Saccharomycetales(100);f__Metschnikowiaceae(100);g__Metschnikowia(100);s__Metschnikowia_pulcherrima(100); | Metschnikowia | 13 | Saprotroph | \|Undefined Saprotroph\| | Yeast | NULL | Probable | As nectar-tap_saprotroph (Põlme et al. 2020); Decay sugar-rich substrates (Põlme et al. 2020) | Põlme S, et al. 2020. Fungal Diversity 105:1-16 ((https://doi.org/10.1007/s13225-020-00466-2)) |
| Otu47710 | 1 | k__Fungi(100);p__Ascomycota(100);c__Saccharomycetes(100);o__Saccharomycetales(100);f__Metschnikowiaceae(100);g__Metschnikowia(100);s__Metschnikowia_pulcherrima(100); | Metschnikowia | 13 | Saprotroph | \|Undefined Saprotroph\| | Yeast | NULL | Probable | As nectar-tap_saprotroph (Põlme et al. 2020); Decay sugar-rich substrates (Põlme et al. 2020) | Põlme S, et al. 2020. Fungal Diversity 105:1-16 ((https://doi.org/10.1007/s13225-020-00466-2)) |
| Otu47711 | 1 | k__Fungi(100);p__Ascomycota(100);c__Saccharomycetes(100);o__Saccharomycetales(100);f__Metschnikowiaceae(100);g__Metschnikowia(100);g__Metschnikowia_unclassified(100); | Metschnikowia | 13 | Saprotroph | \|Undefined Saprotroph\| | Yeast | NULL | Probable | As nectar-tap_saprotroph (Põlme et al. 2020); Decay sugar-rich substrates (Põlme et al. 2020) | Põlme S, et al. 2020. Fungal Diversity 105:1-16 ((https://doi.org/10.1007/s13225-020-00466-2)) |
| Otu47712 | 1 | k__Fungi(100);p__Ascomycota(100);c__Saccharomycetes(100);o__Saccharomycetales(100);f__Metschnikowiaceae(100);g__Metschnikowia(100);s__Metschnikowia_pulcherrima(100); | Metschnikowia | 13 | Saprotroph | \|Undefined Saprotroph\| | Yeast | NULL | Probable | As nectar-tap_saprotroph (Põlme et al. 2020); Decay sugar-rich substrates (Põlme et al. 2020) | Põlme S, et al. 2020. Fungal Diversity 105:1-16 ((https://doi.org/10.1007/s13225-020-00466-2)) |
| Otu47713 | 1 | k__Fungi(100);p__Ascomycota(100);c__Saccharomycetes(100);o__Saccharomycetales(100);f__Metschnikowiaceae(100);g__Metschnikowia(100);s__Metschnikowia_pulcherrima(100); | Metschnikowia | 13 | Saprotroph | \|Undefined Saprotroph\| | Yeast | NULL | Probable | As nectar-tap_saprotroph (Põlme et al. 2020); Decay sugar-rich substrates (Põlme et al. 2020) | Põlme S, et al. 2020. Fungal Diversity 105:1-16 ((https://doi.org/10.1007/s13225-020-00466-2)) |
| Otu47714 | 1 | k__Fungi(100);p__Ascomycota(100);c__Saccharomycetes(100);o__Saccharomycetales(100);f__Metschnikowiaceae(100);g__Metschnikowia(100);s__Metschnikowia_pulcherrima(100); | Metschnikowia | 13 | Saprotroph | \|Undefined Saprotroph\| | Yeast | NULL | Probable | As nectar-tap_saprotroph (Põlme et al. 2020); Decay sugar-rich substrates (Põlme et al. 2020) | Põlme S, et al. 2020. Fungal Diversity 105:1-16 ((https://doi.org/10.1007/s13225-020-00466-2)) |
| Otu47715 | 1 | k__Fungi(100);p__Ascomycota(100);c__Saccharomycetes(100);o__Saccharomycetales(100);f__Metschnikowiaceae(100);g__Metschnikowia(100);g__Metschnikowia_unclassified(100); | Metschnikowia | 13 | Saprotroph | \|Undefined Saprotroph\| | Yeast | NULL | Probable | As nectar-tap_saprotroph (Põlme et al. 2020); Decay sugar-rich substrates (Põlme et al. 2020) | Põlme S, et al. 2020. Fungal Diversity 105:1-16 ((https://doi.org/10.1007/s13225-020-00466-2)) |
| Otu47716 | 1 | k__Fungi(100);p__Ascomycota(100);c__Saccharomycetes(100);o__Saccharomycetales(100);f__Metschnikowiaceae(100);g__Metschnikowia(100);s__Metschnikowia_pulcherrima(100); | Metschnikowia | 13 | Saprotroph | \|Undefined Saprotroph\| | Yeast | NULL | Probable | As nectar-tap_saprotroph (Põlme et al. 2020); Decay sugar-rich substrates (Põlme et al. 2020) | Põlme S, et al. 2020. Fungal Diversity 105:1-16 ((https://doi.org/10.1007/s13225-020-00466-2)) |
| Otu47717 | 1 | k__Fungi(100);p__Ascomycota(100);c__Saccharomycetes(100);o__Saccharomycetales(100);f__Metschnikowiaceae(100);g__Metschnikowia(100);s__Metschnikowia_pulcherrima(100); | Metschnikowia | 13 | Saprotroph | \|Undefined Saprotroph\| | Yeast | NULL | Probable | As nectar-tap_saprotroph (Põlme et al. 2020); Decay sugar-rich substrates (Põlme et al. 2020) | Põlme S, et al. 2020. Fungal Diversity 105:1-16 ((https://doi.org/10.1007/s13225-020-00466-2)) |
| Otu47718 | 1 | k__Fungi(100);p__Ascomycota(100);c__Saccharomycetes(100);o__Saccharomycetales(100);f__Metschnikowiaceae(100);g__Metschnikowia(100);s__Metschnikowia_pulcherrima(100); | Metschnikowia | 13 | Saprotroph | \|Undefined Saprotroph\| | Yeast | NULL | Probable | As nectar-tap_saprotroph (Põlme et al. 2020); Decay sugar-rich substrates (Põlme et al. 2020) | Põlme S, et al. 2020. Fungal Diversity 105:1-16 ((https://doi.org/10.1007/s13225-020-00466-2)) |
| Otu47719 | 1 | k__Fungi(100);p__Ascomycota(100);c__Saccharomycetes(100);o__Saccharomycetales(100);f__Metschnikowiaceae(100);g__Metschnikowia(100);s__Metschnikowia_pulcherrima(100); | Metschnikowia | 13 | Saprotroph | \|Undefined Saprotroph\| | Yeast | NULL | Probable | As nectar-tap_saprotroph (Põlme et al. 2020); Decay sugar-rich substrates (Põlme et al. 2020) | Põlme S, et al. 2020. Fungal Diversity 105:1-16 ((https://doi.org/10.1007/s13225-020-00466-2)) |
| Otu47721 | 1 | k__Fungi(100);p__Ascomycota(100);c__Saccharomycetes(100);o__Saccharomycetales(100);f__Metschnikowiaceae(100);g__Metschnikowia(100);s__Metschnikowia_pulcherrima(100); | Metschnikowia | 13 | Saprotroph | \|Undefined Saprotroph\| | Yeast | NULL | Probable | As nectar-tap_saprotroph (Põlme et al. 2020); Decay sugar-rich substrates (Põlme et al. 2020) | Põlme S, et al. 2020. Fungal Diversity 105:1-16 ((https://doi.org/10.1007/s13225-020-00466-2)) |
| Otu47724 | 1 | k__Fungi(100);p__Ascomycota(100);c__Saccharomycetes(100);o__Saccharomycetales(100);f__Metschnikowiaceae(100);g__Metschnikowia(100);s__Metschnikowia_pulcherrima(100); | Metschnikowia | 13 | Saprotroph | \|Undefined Saprotroph\| | Yeast | NULL | Probable | As nectar-tap_saprotroph (Põlme et al. 2020); Decay sugar-rich substrates (Põlme et al. 2020) | Põlme S, et al. 2020. Fungal Diversity 105:1-16 ((https://doi.org/10.1007/s13225-020-00466-2)) |
| Otu47725 | 1 | k__Fungi(100);p__Ascomycota(100);c__Saccharomycetes(100);o__Saccharomycetales(100);f__Metschnikowiaceae(100);g__Metschnikowia(100);g__Metschnikowia_unclassified(100); | Metschnikowia | 13 | Saprotroph | \|Undefined Saprotroph\| | Yeast | NULL | Probable | As nectar-tap_saprotroph (Põlme et al. 2020); Decay sugar-rich substrates (Põlme et al. 2020) | Põlme S, et al. 2020. Fungal Diversity 105:1-16 ((https://doi.org/10.1007/s13225-020-00466-2)) |
| Otu47726 | 1 | k__Fungi(100);p__Ascomycota(100);c__Saccharomycetes(100);o__Saccharomycetales(100);f__Metschnikowiaceae(100);g__Metschnikowia(100);s__Metschnikowia_pulcherrima(100); | Metschnikowia | 13 | Saprotroph | \|Undefined Saprotroph\| | Yeast | NULL | Probable | As nectar-tap_saprotroph (Põlme et al. 2020); Decay sugar-rich substrates (Põlme et al. 2020) | Põlme S, et al. 2020. Fungal Diversity 105:1-16 ((https://doi.org/10.1007/s13225-020-00466-2)) |
| Otu47727 | 1 | k__Fungi(100);p__Ascomycota(100);c__Saccharomycetes(100);o__Saccharomycetales(100);f__Metschnikowiaceae(100);g__Metschnikowia(100);s__Metschnikowia_pulcherrima(100); | Metschnikowia | 13 | Saprotroph | \|Undefined Saprotroph\| | Yeast | NULL | Probable | As nectar-tap_saprotroph (Põlme et al. 2020); Decay sugar-rich substrates (Põlme et al. 2020) | Põlme S, et al. 2020. Fungal Diversity 105:1-16 ((https://doi.org/10.1007/s13225-020-00466-2)) |
| Otu47728 | 1 | k__Fungi(100);p__Ascomycota(100);c__Saccharomycetes(100);o__Saccharomycetales(100);f__Metschnikowiaceae(100);g__Metschnikowia(100);s__Metschnikowia_pulcherrima(100); | Metschnikowia | 13 | Saprotroph | \|Undefined Saprotroph\| | Yeast | NULL | Probable | As nectar-tap_saprotroph (Põlme et al. 2020); Decay sugar-rich substrates (Põlme et al. 2020) | Põlme S, et al. 2020. Fungal Diversity 105:1-16 ((https://doi.org/10.1007/s13225-020-00466-2)) |
| Otu47729 | 1 | k__Fungi(100);p__Ascomycota(100);c__Saccharomycetes(100);o__Saccharomycetales(100);f__Metschnikowiaceae(100);g__Metschnikowia(100);s__Metschnikowia_pulcherrima(100); | Metschnikowia | 13 | Saprotroph | \|Undefined Saprotroph\| | Yeast | NULL | Probable | As nectar-tap_saprotroph (Põlme et al. 2020); Decay sugar-rich substrates (Põlme et al. 2020) | Põlme S, et al. 2020. Fungal Diversity 105:1-16 ((https://doi.org/10.1007/s13225-020-00466-2)) |
| Otu47732 | 1 | k__Fungi(100);p__Ascomycota(100);c__Saccharomycetes(100);o__Saccharomycetales(100);f__Metschnikowiaceae(100);g__Metschnikowia(100);g__Metschnikowia_unclassified(100); | Metschnikowia | 13 | Saprotroph | \|Undefined Saprotroph\| | Yeast | NULL | Probable | As nectar-tap_saprotroph (Põlme et al. 2020); Decay sugar-rich substrates (Põlme et al. 2020) | Põlme S, et al. 2020. Fungal Diversity 105:1-16 ((https://doi.org/10.1007/s13225-020-00466-2)) |
| Otu47733 | 1 | k__Fungi(100);p__Ascomycota(100);c__Saccharomycetes(100);o__Saccharomycetales(100);f__Metschnikowiaceae(100);g__Metschnikowia(100);s__Metschnikowia_pulcherrima(100); | Metschnikowia | 13 | Saprotroph | \|Undefined Saprotroph\| | Yeast | NULL | Probable | As nectar-tap_saprotroph (Põlme et al. 2020); Decay sugar-rich substrates (Põlme et al. 2020) | Põlme S, et al. 2020. Fungal Diversity 105:1-16 ((https://doi.org/10.1007/s13225-020-00466-2)) |
| Otu47734 | 1 | k__Fungi(100);p__Ascomycota(100);c__Saccharomycetes(100);o__Saccharomycetales(100);f__Metschnikowiaceae(100);g__Metschnikowia(100);s__Metschnikowia_pulcherrima(100); | Metschnikowia | 13 | Saprotroph | \|Undefined Saprotroph\| | Yeast | NULL | Probable | As nectar-tap_saprotroph (Põlme et al. 2020); Decay sugar-rich substrates (Põlme et al. 2020) | Põlme S, et al. 2020. Fungal Diversity 105:1-16 ((https://doi.org/10.1007/s13225-020-00466-2)) |
| Otu47735 | 1 | k__Fungi(100);p__Ascomycota(100);c__Saccharomycetes(100);o__Saccharomycetales(100);f__Metschnikowiaceae(100);g__Metschnikowia(100);s__Metschnikowia_pulcherrima(100); | Metschnikowia | 13 | Saprotroph | \|Undefined Saprotroph\| | Yeast | NULL | Probable | As nectar-tap_saprotroph (Põlme et al. 2020); Decay sugar-rich substrates (Põlme et al. 2020) | Põlme S, et al. 2020. Fungal Diversity 105:1-16 ((https://doi.org/10.1007/s13225-020-00466-2)) |
| Otu47736 | 1 | k__Fungi(100);p__Ascomycota(100);c__Saccharomycetes(100);o__Saccharomycetales(100);f__Metschnikowiaceae(100);g__Metschnikowia(100);g__Metschnikowia_unclassified(100); | Metschnikowia | 13 | Saprotroph | \|Undefined Saprotroph\| | Yeast | NULL | Probable | As nectar-tap_saprotroph (Põlme et al. 2020); Decay sugar-rich substrates (Põlme et al. 2020) | Põlme S, et al. 2020. Fungal Diversity 105:1-16 ((https://doi.org/10.1007/s13225-020-00466-2)) |
| Otu47737 | 1 | k__Fungi(100);p__Ascomycota(100);c__Saccharomycetes(100);o__Saccharomycetales(100);f__Metschnikowiaceae(100);g__Metschnikowia(100);s__Metschnikowia_pulcherrima(100); | Metschnikowia | 13 | Saprotroph | \|Undefined Saprotroph\| | Yeast | NULL | Probable | As nectar-tap_saprotroph (Põlme et al. 2020); Decay sugar-rich substrates (Põlme et al. 2020) | Põlme S, et al. 2020. Fungal Diversity 105:1-16 ((https://doi.org/10.1007/s13225-020-00466-2)) |
| Otu47738 | 1 | k__Fungi(100);p__Ascomycota(100);c__Saccharomycetes(100);o__Saccharomycetales(100);f__Metschnikowiaceae(100);g__Metschnikowia(100);s__Metschnikowia_pulcherrima(100); | Metschnikowia | 13 | Saprotroph | \|Undefined Saprotroph\| | Yeast | NULL | Probable | As nectar-tap_saprotroph (Põlme et al. 2020); Decay sugar-rich substrates (Põlme et al. 2020) | Põlme S, et al. 2020. Fungal Diversity 105:1-16 ((https://doi.org/10.1007/s13225-020-00466-2)) |
| Otu47739 | 1 | k__Fungi(100);p__Ascomycota(100);c__Saccharomycetes(100);o__Saccharomycetales(100);f__Metschnikowiaceae(100);g__Metschnikowia(100);s__Metschnikowia_pulcherrima(100); | Metschnikowia | 13 | Saprotroph | \|Undefined Saprotroph\| | Yeast | NULL | Probable | As nectar-tap_saprotroph (Põlme et al. 2020); Decay sugar-rich substrates (Põlme et al. 2020) | Põlme S, et al. 2020. Fungal Diversity 105:1-16 ((https://doi.org/10.1007/s13225-020-00466-2)) |
| Otu47740 | 1 | k__Fungi(100);p__Ascomycota(100);c__Saccharomycetes(100);o__Saccharomycetales(100);f__Metschnikowiaceae(100);g__Metschnikowia(100);s__Metschnikowia_pulcherrima(100); | Metschnikowia | 13 | Saprotroph | \|Undefined Saprotroph\| | Yeast | NULL | Probable | As nectar-tap_saprotroph (Põlme et al. 2020); Decay sugar-rich substrates (Põlme et al. 2020) | Põlme S, et al. 2020. Fungal Diversity 105:1-16 ((https://doi.org/10.1007/s13225-020-00466-2)) |
| Otu47741 | 1 | k__Fungi(100);p__Ascomycota(100);c__Saccharomycetes(100);o__Saccharomycetales(100);f__Metschnikowiaceae(100);g__Metschnikowia(100);g__Metschnikowia_unclassified(100); | Metschnikowia | 13 | Saprotroph | \|Undefined Saprotroph\| | Yeast | NULL | Probable | As nectar-tap_saprotroph (Põlme et al. 2020); Decay sugar-rich substrates (Põlme et al. 2020) | Põlme S, et al. 2020. Fungal Diversity 105:1-16 ((https://doi.org/10.1007/s13225-020-00466-2)) |

| | | | | | | | | | | | |
|---|---|---|---|---|---|---|---|---|---|---|---|
| Otu47742 | 1 | k__Fungi(100);p__Ascomycota(100);c__Saccharomycetes(100);o__Saccharomycetales(100);f__Metschnikowiaceae(100);g__Metschnikowia(100);s__Metschnikowia_pulcherrima(100); | Metschnikowia | 13 | Saprotroph | \|Undefined Saprotroph\| | Yeast | NULL | Probable | As nectar-tap_saprotroph (Põlme et al. 2020); Decay sugar-rich substrates (Põlme et al. 2020) | Põlme S, et al. 2020. Fungal Diversity 105:1-16 ((https://doi.org/10.1007/s13225-020-00466-2)) |
| Otu47743 | 1 | k__Fungi(100);p__Ascomycota(100);c__Saccharomycetes(100);o__Saccharomycetales(100);f__Metschnikowiaceae(100);g__Metschnikowia(100);g__Metschnikowia_unclassified(100); | Metschnikowia | 13 | Saprotroph | \|Undefined Saprotroph\| | Yeast | NULL | Probable | As nectar-tap_saprotroph (Põlme et al. 2020); Decay sugar-rich substrates (Põlme et al. 2020) | Põlme S, et al. 2020. Fungal Diversity 105:1-16 ((https://doi.org/10.1007/s13225-020-00466-2)) |
| Otu47744 | 1 | k__Fungi(100);p__Ascomycota(100);c__Saccharomycetes(100);o__Saccharomycetales(100);f__Metschnikowiaceae(100);g__Metschnikowia(100);g__Metschnikowia_unclassified(100); | Metschnikowia | 13 | Saprotroph | \|Undefined Saprotroph\| | Yeast | NULL | Probable | As nectar-tap_saprotroph (Põlme et al. 2020); Decay sugar-rich substrates (Põlme et al. 2020) | Põlme S, et al. 2020. Fungal Diversity 105:1-16 ((https://doi.org/10.1007/s13225-020-00466-2)) |
| Otu47745 | 1 | k__Fungi(100);p__Ascomycota(100);c__Saccharomycetes(100);o__Saccharomycetales(100);f__Metschnikowiaceae(100);g__Metschnikowia(100);g__Metschnikowia_unclassified(100); | Metschnikowia | 13 | Saprotroph | \|Undefined Saprotroph\| | Yeast | NULL | Probable | As nectar-tap_saprotroph (Põlme et al. 2020); Decay sugar-rich substrates (Põlme et al. 2020) | Põlme S, et al. 2020. Fungal Diversity 105:1-16 ((https://doi.org/10.1007/s13225-020-00466-2)) |
| Otu47746 | 1 | k__Fungi(100);p__Ascomycota(100);c__Saccharomycetes(100);o__Saccharomycetales(100);f__Metschnikowiaceae(100);g__Metschnikowia(100);s__Metschnikowia_pulcherrima(100); | Metschnikowia | 13 | Saprotroph | \|Undefined Saprotroph\| | Yeast | NULL | Probable | As nectar-tap_saprotroph (Põlme et al. 2020); Decay sugar-rich substrates (Põlme et al. 2020) | Põlme S, et al. 2020. Fungal Diversity 105:1-16 ((https://doi.org/10.1007/s13225-020-00466-2)) |
| Otu47747 | 1 | k__Fungi(100);p__Ascomycota(100);c__Saccharomycetes(100);o__Saccharomycetales(100);f__Metschnikowiaceae(100);g__Metschnikowia(100);s__Metschnikowia_pulcherrima(100); | Metschnikowia | 13 | Saprotroph | \|Undefined Saprotroph\| | Yeast | NULL | Probable | As nectar-tap_saprotroph (Põlme et al. 2020); Decay sugar-rich substrates (Põlme et al. 2020) | Põlme S, et al. 2020. Fungal Diversity 105:1-16 ((https://doi.org/10.1007/s13225-020-00466-2)) |
| Otu47748 | 1 | k__Fungi(100);p__Ascomycota(100);c__Saccharomycetes(100);o__Saccharomycetales(100);f__Metschnikowiaceae(100);g__Metschnikowia(100);g__Metschnikowia_unclassified(100); | Metschnikowia | 13 | Saprotroph | \|Undefined Saprotroph\| | Yeast | NULL | Probable | As nectar-tap_saprotroph (Põlme et al. 2020); Decay sugar-rich substrates (Põlme et al. 2020) | Põlme S, et al. 2020. Fungal Diversity 105:1-16 ((https://doi.org/10.1007/s13225-020-00466-2)) |
| Otu47749 | 1 | k__Fungi(100);p__Ascomycota(100);c__Saccharomycetes(100);o__Saccharomycetales(100);f__Metschnikowiaceae(100);g__Metschnikowia(100);s__Metschnikowia_pulcherrima(100); | Metschnikowia | 13 | Saprotroph | \|Undefined Saprotroph\| | Yeast | NULL | Probable | As nectar-tap_saprotroph (Põlme et al. 2020); Decay sugar-rich substrates (Põlme et al. 2020) | Põlme S, et al. 2020. Fungal Diversity 105:1-16 ((https://doi.org/10.1007/s13225-020-00466-2)) |
| Otu47750 | 1 | k__Fungi(100);p__Ascomycota(100);c__Saccharomycetes(100);o__Saccharomycetales(100);f__Metschnikowiaceae(100);g__Metschnikowia(100);g__Metschnikowia_unclassified(100); | Metschnikowia | 13 | Saprotroph | \|Undefined Saprotroph\| | Yeast | NULL | Probable | As nectar-tap_saprotroph (Põlme et al. 2020); Decay sugar-rich substrates (Põlme et al. 2020) | Põlme S, et al. 2020. Fungal Diversity 105:1-16 ((https://doi.org/10.1007/s13225-020-00466-2)) |
| Otu47751 | 1 | k__Fungi(100);p__Ascomycota(100);c__Saccharomycetes(100);o__Saccharomycetales(100);f__Metschnikowiaceae(100);g__Metschnikowia(100);g__Metschnikowia_unclassified(100); | Metschnikowia | 13 | Saprotroph | \|Undefined Saprotroph\| | Yeast | NULL | Probable | As nectar-tap_saprotroph (Põlme et al. 2020); Decay sugar-rich substrates (Põlme et al. 2020) | Põlme S, et al. 2020. Fungal Diversity 105:1-16 ((https://doi.org/10.1007/s13225-020-00466-2)) |
| Otu47752 | 1 | k__Fungi(100);p__Ascomycota(100);c__Saccharomycetes(100);o__Saccharomycetales(100);f__Metschnikowiaceae(100);g__Metschnikowia(100);g__Metschnikowia_unclassified(100); | Metschnikowia | 13 | Saprotroph | \|Undefined Saprotroph\| | Yeast | NULL | Probable | As nectar-tap_saprotroph (Põlme et al. 2020); Decay sugar-rich substrates (Põlme et al. 2020) | Põlme S, et al. 2020. Fungal Diversity 105:1-16 ((https://doi.org/10.1007/s13225-020-00466-2)) |
| Otu47753 | 1 | k__Fungi(100);p__Ascomycota(100);c__Saccharomycetes(100);o__Saccharomycetales(100);f__Metschnikowiaceae(100);g__Metschnikowia(100);s__Metschnikowia_pulcherrima(100); | Metschnikowia | 13 | Saprotroph | \|Undefined Saprotroph\| | Yeast | NULL | Probable | As nectar-tap_saprotroph (Põlme et al. 2020); Decay sugar-rich substrates (Põlme et al. 2020) | Põlme S, et al. 2020. Fungal Diversity 105:1-16 ((https://doi.org/10.1007/s13225-020-00466-2)) |
| Otu47754 | 1 | k__Fungi(100);p__Ascomycota(100);c__Saccharomycetes(100);o__Saccharomycetales(100);f__Metschnikowiaceae(100);g__Metschnikowia(100);g__Metschnikowia_unclassified(100); | Metschnikowia | 13 | Saprotroph | \|Undefined Saprotroph\| | Yeast | NULL | Probable | As nectar-tap_saprotroph (Põlme et al. 2020); Decay sugar-rich substrates (Põlme et al. 2020) | Põlme S, et al. 2020. Fungal Diversity 105:1-16 ((https://doi.org/10.1007/s13225-020-00466-2)) |
| Otu47755 | 1 | k__Fungi(100);p__Ascomycota(100);c__Saccharomycetes(100);o__Saccharomycetales(100);f__Metschnikowiaceae(100);g__Metschnikowia(100);s__Metschnikowia_pulcherrima(100); | Metschnikowia | 13 | Saprotroph | \|Undefined Saprotroph\| | Yeast | NULL | Probable | As nectar-tap_saprotroph (Põlme et al. 2020); Decay sugar-rich substrates (Põlme et al. 2020) | Põlme S, et al. 2020. Fungal Diversity 105:1-16 ((https://doi.org/10.1007/s13225-020-00466-2)) |
| Otu47756 | 1 | k__Fungi(100);p__Ascomycota(100);c__Saccharomycetes(100);o__Saccharomycetales(100);f__Metschnikowiaceae(100);g__Metschnikowia(100);s__Metschnikowia_pulcherrima(100); | Metschnikowia | 13 | Saprotroph | \|Undefined Saprotroph\| | Yeast | NULL | Probable | As nectar-tap_saprotroph (Põlme et al. 2020); Decay sugar-rich substrates (Põlme et al. 2020) | Põlme S, et al. 2020. Fungal Diversity 105:1-16 ((https://doi.org/10.1007/s13225-020-00466-2)) |
| Otu47757 | 1 | k__Fungi(100);p__Ascomycota(100);c__Saccharomycetes(100);o__Saccharomycetales(100);f__Metschnikowiaceae(100);g__Metschnikowia(100);g__Metschnikowia_unclassified(100); | Metschnikowia | 13 | Saprotroph | \|Undefined Saprotroph\| | Yeast | NULL | Probable | As nectar-tap_saprotroph (Põlme et al. 2020); Decay sugar-rich substrates (Põlme et al. 2020) | Põlme S, et al. 2020. Fungal Diversity 105:1-16 ((https://doi.org/10.1007/s13225-020-00466-2)) |
| Otu47758 | 1 | k__Fungi(100);p__Ascomycota(100);c__Saccharomycetes(100);o__Saccharomycetales(100);f__Metschnikowiaceae(100);g__Metschnikowia(100);s__Metschnikowia_pulcherrima(100); | Metschnikowia | 13 | Saprotroph | \|Undefined Saprotroph\| | Yeast | NULL | Probable | As nectar-tap_saprotroph (Põlme et al. 2020); Decay sugar-rich substrates (Põlme et al. 2020) | Põlme S, et al. 2020. Fungal Diversity 105:1-16 ((https://doi.org/10.1007/s13225-020-00466-2)) |
| Otu47759 | 1 | k__Fungi(100);p__Ascomycota(100);c__Saccharomycetes(100);o__Saccharomycetales(100);f__Metschnikowiaceae(100);g__Metschnikowia(100);s__Metschnikowia_pulcherrima(100); | Metschnikowia | 13 | Saprotroph | \|Undefined Saprotroph\| | Yeast | NULL | Probable | As nectar-tap_saprotroph (Põlme et al. 2020); Decay sugar-rich substrates (Põlme et al. 2020) | Põlme S, et al. 2020. Fungal Diversity 105:1-16 ((https://doi.org/10.1007/s13225-020-00466-2)) |
| Otu47760 | 1 | k__Fungi(100);p__Ascomycota(100);c__Saccharomycetes(100);o__Saccharomycetales(100);f__Metschnikowiaceae(100);g__Metschnikowia(100);s__Metschnikowia_pulcherrima(100); | Metschnikowia | 13 | Saprotroph | \|Undefined Saprotroph\| | Yeast | NULL | Probable | As nectar-tap_saprotroph (Põlme et al. 2020); Decay sugar-rich substrates (Põlme et al. 2020) | Põlme S, et al. 2020. Fungal Diversity 105:1-16 ((https://doi.org/10.1007/s13225-020-00466-2)) |
| Otu47761 | 1 | k__Fungi(100);p__Ascomycota(100);c__Saccharomycetes(100);o__Saccharomycetales(100);f__Metschnikowiaceae(100);g__Metschnikowia(100);s__Metschnikowia_pulcherrima(100); | Metschnikowia | 13 | Saprotroph | \|Undefined Saprotroph\| | Yeast | NULL | Probable | As nectar-tap_saprotroph (Põlme et al. 2020); Decay sugar-rich substrates (Põlme et al. 2020) | Põlme S, et al. 2020. Fungal Diversity 105:1-16 ((https://doi.org/10.1007/s13225-020-00466-2)) |
| Otu47762 | 1 | k__Fungi(100);p__Ascomycota(100);c__Saccharomycetes(100);o__Saccharomycetales(100);f__Metschnikowiaceae(100);g__Metschnikowia(100);s__Metschnikowia_pulcherrima(100); | Metschnikowia | 13 | Saprotroph | \|Undefined Saprotroph\| | Yeast | NULL | Probable | As nectar-tap_saprotroph (Põlme et al. 2020); Decay sugar-rich substrates (Põlme et al. 2020) | Põlme S, et al. 2020. Fungal Diversity 105:1-16 ((https://doi.org/10.1007/s13225-020-00466-2)) |
| Otu47763 | 1 | k__Fungi(100);p__Ascomycota(100);c__Saccharomycetes(100);o__Saccharomycetales(100);f__Metschnikowiaceae(100);g__Metschnikowia(100);s__Metschnikowia_pulcherrima(100); | Metschnikowia | 13 | Saprotroph | \|Undefined Saprotroph\| | Yeast | NULL | Probable | As nectar-tap_saprotroph (Põlme et al. 2020); Decay sugar-rich substrates (Põlme et al. 2020) | Põlme S, et al. 2020. Fungal Diversity 105:1-16 ((https://doi.org/10.1007/s13225-020-00466-2)) |
| Otu47764 | 1 | k__Fungi(100);p__Ascomycota(100);c__Saccharomycetes(100);o__Saccharomycetales(100);f__Metschnikowiaceae(100);g__Metschnikowia(100);s__Metschnikowia_pulcherrima(100); | Metschnikowia | 13 | Saprotroph | \|Undefined Saprotroph\| | Yeast | NULL | Probable | As nectar-tap_saprotroph (Põlme et al. 2020); Decay sugar-rich substrates (Põlme et al. 2020) | Põlme S, et al. 2020. Fungal Diversity 105:1-16 ((https://doi.org/10.1007/s13225-020-00466-2)) |
| Otu47765 | 1 | k__Fungi(100);p__Ascomycota(100);c__Saccharomycetes(100);o__Saccharomycetales(100);f__Metschnikowiaceae(100);g__Metschnikowia(100);s__Metschnikowia_pulcherrima(100); | Metschnikowia | 13 | Saprotroph | \|Undefined Saprotroph\| | Yeast | NULL | Probable | As nectar-tap_saprotroph (Põlme et al. 2020); Decay sugar-rich substrates (Põlme et al. 2020) | Põlme S, et al. 2020. Fungal Diversity 105:1-16 ((https://doi.org/10.1007/s13225-020-00466-2)) |
| Otu47766 | 1 | k__Fungi(100);p__Ascomycota(100);c__Saccharomycetes(100);o__Saccharomycetales(100);f__Metschnikowiaceae(100);g__Metschnikowia(100);s__Metschnikowia_pulcherrima(100); | Metschnikowia | 13 | Saprotroph | \|Undefined Saprotroph\| | Yeast | NULL | Probable | As nectar-tap_saprotroph (Põlme et al. 2020); Decay sugar-rich substrates (Põlme et al. 2020) | Põlme S, et al. 2020. Fungal Diversity 105:1-16 ((https://doi.org/10.1007/s13225-020-00466-2)) |
| Otu47767 | 1 | k__Fungi(100);p__Ascomycota(100);c__Saccharomycetes(100);o__Saccharomycetales(100);f__Metschnikowiaceae(100);g__Metschnikowia(100);s__Metschnikowia_pulcherrima(100); | Metschnikowia | 13 | Saprotroph | \|Undefined Saprotroph\| | Yeast | NULL | Probable | As nectar-tap_saprotroph (Põlme et al. 2020); Decay sugar-rich substrates (Põlme et al. 2020) | Põlme S, et al. 2020. Fungal Diversity 105:1-16 ((https://doi.org/10.1007/s13225-020-00466-2)) |
| Otu47768 | 1 | k__Fungi(100);p__Ascomycota(100);c__Saccharomycetes(100);o__Saccharomycetales(100);f__Metschnikowiaceae(100);g__Metschnikowia(100);s__Metschnikowia_pulcherrima(100); | Metschnikowia | 13 | Saprotroph | \|Undefined Saprotroph\| | Yeast | NULL | Probable | As nectar-tap_saprotroph (Põlme et al. 2020); Decay sugar-rich substrates (Põlme et al. 2020) | Põlme S, et al. 2020. Fungal Diversity 105:1-16 ((https://doi.org/10.1007/s13225-020-00466-2)) |
| Otu47769 | 1 | k__Fungi(100);p__Ascomycota(100);c__Saccharomycetes(100);o__Saccharomycetales(100);f__Metschnikowiaceae(100);g__Metschnikowia(100);s__Metschnikowia_pulcherrima(100); | Metschnikowia | 13 | Saprotroph | \|Undefined Saprotroph\| | Yeast | NULL | Probable | As nectar-tap_saprotroph (Põlme et al. 2020); Decay sugar-rich substrates (Põlme et al. 2020) | Põlme S, et al. 2020. Fungal Diversity 105:1-16 ((https://doi.org/10.1007/s13225-020-00466-2)) |
| Otu47770 | 1 | k__Fungi(100);p__Ascomycota(100);c__Saccharomycetes(100);o__Saccharomycetales(100);f__Metschnikowiaceae(100);g__Metschnikowia(100);s__Metschnikowia_pulcherrima(100); | Metschnikowia | 13 | Saprotroph | \|Undefined Saprotroph\| | Yeast | NULL | Probable | As nectar-tap_saprotroph (Põlme et al. 2020); Decay sugar-rich substrates (Põlme et al. 2020) | Põlme S, et al. 2020. Fungal Diversity 105:1-16 ((https://doi.org/10.1007/s13225-020-00466-2)) |
| Otu47772 | 1 | k__Fungi(100);p__Ascomycota(100);c__Saccharomycetes(100);o__Saccharomycetales(100);f__Metschnikowiaceae(100);g__Metschnikowia(100);g__Metschnikowia_unclassified(100); | Metschnikowia | 13 | Saprotroph | \|Undefined Saprotroph\| | Yeast | NULL | Probable | As nectar-tap_saprotroph (Põlme et al. 2020); Decay sugar-rich substrates (Põlme et al. 2020) | Põlme S, et al. 2020. Fungal Diversity 105:1-16 ((https://doi.org/10.1007/s13225-020-00466-2)) |
| Otu47773 | 1 | k__Fungi(100);p__Ascomycota(100);c__Saccharomycetes(100);o__Saccharomycetales(100);f__Metschnikowiaceae(100);g__Metschnikowia(100);s__Metschnikowia_pulcherrima(100); | Metschnikowia | 13 | Saprotroph | \|Undefined Saprotroph\| | Yeast | NULL | Probable | As nectar-tap_saprotroph (Põlme et al. 2020); Decay sugar-rich substrates (Põlme et al. 2020) | Põlme S, et al. 2020. Fungal Diversity 105:1-16 ((https://doi.org/10.1007/s13225-020-00466-2)) |
| Otu47774 | 1 | k__Fungi(100);p__Ascomycota(100);c__Saccharomycetes(100);o__Saccharomycetales(100);f__Metschnikowiaceae(100);g__Metschnikowia(100);s__Metschnikowia_pulcherrima(100); | Metschnikowia | 13 | Saprotroph | \|Undefined Saprotroph\| | Yeast | NULL | Probable | As nectar-tap_saprotroph (Põlme et al. 2020); Decay sugar-rich substrates (Põlme et al. 2020) | Põlme S, et al. 2020. Fungal Diversity 105:1-16 ((https://doi.org/10.1007/s13225-020-00466-2)) |
| Otu47775 | 1 | k__Fungi(100);p__Ascomycota(100);c__Saccharomycetes(100);o__Saccharomycetales(100);f__Metschnikowiaceae(100);g__Metschnikowia(100);s__Metschnikowia_pulcherrima(100); | Metschnikowia | 13 | Saprotroph | \|Undefined Saprotroph\| | Yeast | NULL | Probable | As nectar-tap_saprotroph (Põlme et al. 2020); Decay sugar-rich substrates (Põlme et al. 2020) | Põlme S, et al. 2020. Fungal Diversity 105:1-16 ((https://doi.org/10.1007/s13225-020-00466-2)) |
| Otu47776 | 1 | k__Fungi(100);p__Ascomycota(100);c__Saccharomycetes(100);o__Saccharomycetales(100);f__Metschnikowiaceae(100);g__Metschnikowia(100);g__Metschnikowia_unclassified(100); | Metschnikowia | 13 | Saprotroph | \|Undefined Saprotroph\| | Yeast | NULL | Probable | As nectar-tap_saprotroph (Põlme et al. 2020); Decay sugar-rich substrates (Põlme et al. 2020) | Põlme S, et al. 2020. Fungal Diversity 105:1-16 ((https://doi.org/10.1007/s13225-020-00466-2)) |
| Otu47777 | 1 | k__Fungi(100);p__Ascomycota(100);c__Saccharomycetes(100);o__Saccharomycetales(100);f__Metschnikowiaceae(100);g__Metschnikowia(100);s__Metschnikowia_pulcherrima(100); | Metschnikowia | 13 | Saprotroph | \|Undefined Saprotroph\| | Yeast | NULL | Probable | As nectar-tap_saprotroph (Põlme et al. 2020); Decay sugar-rich substrates (Põlme et al. 2020) | Põlme S, et al. 2020. Fungal Diversity 105:1-16 ((https://doi.org/10.1007/s13225-020-00466-2)) |
| Otu47778 | 1 | k__Fungi(100);p__Ascomycota(100);c__Saccharomycetes(100);o__Saccharomycetales(100);f__Metschnikowiaceae(100);g__Metschnikowia(100);g__Metschnikowia_unclassified(100); | Metschnikowia | 13 | Saprotroph | \|Undefined Saprotroph\| | Yeast | NULL | Probable | As nectar-tap_saprotroph (Põlme et al. 2020); Decay sugar-rich substrates (Põlme et al. 2020) | Põlme S, et al. 2020. Fungal Diversity 105:1-16 ((https://doi.org/10.1007/s13225-020-00466-2)) |
| Otu47779 | 1 | k__Fungi(100);p__Ascomycota(100);c__Saccharomycetes(100);o__Saccharomycetales(100);f__Metschnikowiaceae(100);g__Metschnikowia(100);s__Metschnikowia_pulcherrima(100); | Metschnikowia | 13 | Saprotroph | \|Undefined Saprotroph\| | Yeast | NULL | Probable | As nectar-tap_saprotroph (Põlme et al. 2020); Decay sugar-rich substrates (Põlme et al. 2020) | Põlme S, et al. 2020. Fungal Diversity 105:1-16 ((https://doi.org/10.1007/s13225-020-00466-2)) |
| Otu47780 | 1 | k__Fungi(100);p__Ascomycota(100);c__Saccharomycetes(100);o__Saccharomycetales(100);f__Saccharomycetales_family_Incertae_sedis(100);g__Candida(100);s__Candida_sp_AS23486(100); | Candida | 13 | Pathotroph-Saprotroph-Symbiotroph | Animal Pathogen-Endophyte-Endosymbiont-Epiphyte-\|Undefined Saprotroph\| | Dimorphic Yeast-Yeast | NULL | Probable | As nectar-tap_saprotroph (Põlme et al. 2020); Decay sugar-rich substrates (Põlme et al. 2020); This asexual genus is polyphyletic and contain species that function very differently depend on what they are. Caution should be taken when interpreting guilds for Candida species. | Manolakaki D, et al. 2010. Virulence. 1: 367-375 ((https://doi.org/10.4161/viru.1.5.12796)); Mekbib SB, et al. 2011. Tropical Plant Pathology 36:233-240 ((https://doi.org/10.1590/S1982-56762011000400004)); Põlme S, et al. 2020. Fungal Diversity 105:1-16 ((https://doi.org/10.1007/s13225-020-00466-2)) |
| Otu47781 | 1 | k__Fungi(100);p__Ascomycota(100);c__Saccharomycetes(100);o__Saccharomycetales(100);f__Metschnikowiaceae(100);g__Metschnikowia(100);g__Metschnikowia_unclassified(100); | Metschnikowia | 13 | Saprotroph | \|Undefined Saprotroph\| | Yeast | NULL | Probable | As nectar-tap_saprotroph (Põlme et al. 2020); Decay sugar-rich substrates (Põlme et al. 2020) | Põlme S, et al. 2020. Fungal Diversity 105:1-16 ((https://doi.org/10.1007/s13225-020-00466-2)) |
| Otu47782 | 1 | k__Fungi(100);p__Ascomycota(100);c__Saccharomycetes(100);o__Saccharomycetales(100);f__Metschnikowiaceae(100);g__Metschnikowia(100);g__Metschnikowia_unclassified(100); | Metschnikowia | 13 | Saprotroph | \|Undefined Saprotroph\| | Yeast | NULL | Probable | As nectar-tap_saprotroph (Põlme et al. 2020); Decay sugar-rich substrates (Põlme et al. 2020) | Põlme S, et al. 2020. Fungal Diversity 105:1-16 ((https://doi.org/10.1007/s13225-020-00466-2)) |
| Otu47783 | 1 | k__Fungi(100);p__Ascomycota(100);c__Saccharomycetes(100);o__Saccharomycetales(100);f__Metschnikowiaceae(100);g__Metschnikowia(100);s__Metschnikowia_pulcherrima(100); | Metschnikowia | 13 | Saprotroph | \|Undefined Saprotroph\| | Yeast | NULL | Probable | As nectar-tap_saprotroph (Põlme et al. 2020); Decay sugar-rich substrates (Põlme et al. 2020) | Põlme S, et al. 2020. Fungal Diversity 105:1-16 ((https://doi.org/10.1007/s13225-020-00466-2)) |
| Otu47784 | 1 | k__Fungi(100);p__Ascomycota(100);c__Saccharomycetes(100);o__Saccharomycetales(100);f__Metschnikowiaceae(100);g__Metschnikowia(100);s__Metschnikowia_pulcherrima(100); | Metschnikowia | 13 | Saprotroph | \|Undefined Saprotroph\| | Yeast | NULL | Probable | As nectar-tap_saprotroph (Põlme et al. 2020); Decay sugar-rich substrates (Põlme et al. 2020) | Põlme S, et al. 2020. Fungal Diversity 105:1-16 ((https://doi.org/10.1007/s13225-020-00466-2)) |
| Otu47785 | 1 | k__Fungi(100);p__Ascomycota(100);c__Saccharomycetes(100);o__Saccharomycetales(100);f__Metschnikowiaceae(100);g__Metschnikowia(100);s__Metschnikowia_pulcherrima(100); | Metschnikowia | 13 | Saprotroph | \|Undefined Saprotroph\| | Yeast | NULL | Probable | As nectar-tap_saprotroph (Põlme et al. 2020); Decay sugar-rich substrates (Põlme et al. 2020) | Põlme S, et al. 2020. Fungal Diversity 105:1-16 ((https://doi.org/10.1007/s13225-020-00466-2)) |
| Otu47786 | 1 | k__Fungi(100);p__Ascomycota(100);c__Saccharomycetes(100);o__Saccharomycetales(100);f__Metschnikowiaceae(100);g__Metschnikowia(100);g__Metschnikowia_unclassified(100); | Metschnikowia | 13 | Saprotroph | \|Undefined Saprotroph\| | Yeast | NULL | Probable | As nectar-tap_saprotroph (Põlme et al. 2020); Decay sugar-rich substrates (Põlme et al. 2020) | Põlme S, et al. 2020. Fungal Diversity 105:1-16 ((https://doi.org/10.1007/s13225-020-00466-2)) |
| Otu47787 | 1 | k__Fungi(100);p__Ascomycota(100);c__Saccharomycetes(100);o__Saccharomycetales(100);f__Metschnikowiaceae(100);g__Metschnikowia(100);s__Metschnikowia_pulcherrima(100); | Metschnikowia | 13 | Saprotroph | \|Undefined Saprotroph\| | Yeast | NULL | Probable | As nectar-tap_saprotroph (Põlme et al. 2020); Decay sugar-rich substrates (Põlme et al. 2020) | Põlme S, et al. 2020. Fungal Diversity 105:1-16 ((https://doi.org/10.1007/s13225-020-00466-2)) |
| Otu47788 | 1 | k__Fungi(100);p__Ascomycota(100);c__Saccharomycetes(100);o__Saccharomycetales(100);f__Metschnikowiaceae(100);g__Metschnikowia(100);g__Metschnikowia_unclassified(100); | Metschnikowia | 13 | Saprotroph | \|Undefined Saprotroph\| | Yeast | NULL | Probable | As nectar-tap_saprotroph (Põlme et al. 2020); Decay sugar-rich substrates (Põlme et al. 2020) | Põlme S, et al. 2020. Fungal Diversity 105:1-16 ((https://doi.org/10.1007/s13225-020-00466-2)) |
| Otu47789 | 1 | k__Fungi(100);p__Ascomycota(100);c__Saccharomycetes(100);o__Saccharomycetales(100);f__Metschnikowiaceae(100);g__Metschnikowia(100);s__Metschnikowia_pulcherrima(100); | Metschnikowia | 13 | Saprotroph | \|Undefined Saprotroph\| | Yeast | NULL | Probable | As nectar-tap_saprotroph (Põlme et al. 2020); Decay sugar-rich substrates (Põlme et al. 2020) | Põlme S, et al. 2020. Fungal Diversity 105:1-16 ((https://doi.org/10.1007/s13225-020-00466-2)) |
| Otu47791 | 1 | k__Fungi(100);p__Ascomycota(100);c__Saccharomycetes(100);o__Saccharomycetales(100);f__Metschnikowiaceae(100);g__Metschnikowia(100);s__Metschnikowia_pulcherrima(100); | Metschnikowia | 13 | Saprotroph | \|Undefined Saprotroph\| | Yeast | NULL | Probable | As nectar-tap_saprotroph (Põlme et al. 2020); Decay sugar-rich substrates (Põlme et al. 2020) | Põlme S, et al. 2020. Fungal Diversity 105:1-16 ((https://doi.org/10.1007/s13225-020-00466-2)) |
| Otu47792 | 1 | k__Fungi(100);p__Ascomycota(100);c__Saccharomycetes(100);o__Saccharomycetales(100);f__Metschnikowiaceae(100);g__Metschnikowia(100);s__Metschnikowia_pulcherrima(100); | Metschnikowia | 13 | Saprotroph | \|Undefined Saprotroph\| | Yeast | NULL | Probable | As nectar-tap_saprotroph (Põlme et al. 2020); Decay sugar-rich substrates (Põlme et al. 2020) | Põlme S, et al. 2020. Fungal Diversity 105:1-16 ((https://doi.org/10.1007/s13225-020-00466-2)) |
| Otu47793 | 1 | k__Fungi(100);p__Ascomycota(100);c__Saccharomycetes(100);o__Saccharomycetales(100);f__Metschnikowiaceae(100);g__Metschnikowia(100);s__Metschnikowia_pulcherrima(100); | Metschnikowia | 13 | Saprotroph | \|Undefined Saprotroph\| | Yeast | NULL | Probable | As nectar-tap_saprotroph (Põlme et al. 2020); Decay sugar-rich substrates (Põlme et al. 2020) | Põlme S, et al. 2020. Fungal Diversity 105:1-16 ((https://doi.org/10.1007/s13225-020-00466-2)) |
| Otu47794 | 1 | k__Fungi(100);p__Ascomycota(100);c__Saccharomycetes(100);o__Saccharomycetales(100);f__Metschnikowiaceae(100);g__Metschnikowia(100);s__Metschnikowia_pulcherrima(100); | Metschnikowia | 13 | Saprotroph | \|Undefined Saprotroph\| | Yeast | NULL | Probable | As nectar-tap_saprotroph (Põlme et al. 2020); Decay sugar-rich substrates (Põlme et al. 2020) | Põlme S, et al. 2020. Fungal Diversity 105:1-16 ((https://doi.org/10.1007/s13225-020-00466-2)) |
| Otu47795 | 1 | k__Fungi(100);p__Ascomycota(100);c__Saccharomycetes(100);o__Saccharomycetales(100);f__Metschnikowiaceae(100);g__Metschnikowia(100);s__Metschnikowia_pulcherrima(100); | Metschnikowia | 13 | Saprotroph | \|Undefined Saprotroph\| | Yeast | NULL | Probable | As nectar-tap_saprotroph (Põlme et al. 2020); Decay sugar-rich substrates (Põlme et al. 2020) | Põlme S, et al. 2020. Fungal Diversity 105:1-16 ((https://doi.org/10.1007/s13225-020-00466-2)) |
| Otu47796 | 1 | k__Fungi(100);p__Ascomycota(100);c__Saccharomycetes(100);o__Saccharomycetales(100);f__Metschnikowiaceae(100);g__Metschnikowia(100);s__Metschnikowia_pulcherrima(100); | Metschnikowia | 13 | Saprotroph | \|Undefined Saprotroph\| | Yeast | NULL | Probable | As nectar-tap_saprotroph (Põlme et al. 2020); Decay sugar-rich substrates (Põlme et al. 2020) | Põlme S, et al. 2020. Fungal Diversity 105:1-16 ((https://doi.org/10.1007/s13225-020-00466-2)) |
| Otu47797 | 1 | k__Fungi(100);p__Ascomycota(100);c__Saccharomycetes(100);o__Saccharomycetales(100);f__Metschnikowiaceae(100);g__Metschnikowia(100);s__Metschnikowia_pulcherrima(100); | Metschnikowia | 13 | Saprotroph | \|Undefined Saprotroph\| | Yeast | NULL | Probable | As nectar-tap_saprotroph (Põlme et al. 2020); Decay sugar-rich substrates (Põlme et al. 2020) | Põlme S, et al. 2020. Fungal Diversity 105:1-16 ((https://doi.org/10.1007/s13225-020-00466-2)) |
| Otu47798 | 1 | k__Fungi(100);p__Ascomycota(100);c__Saccharomycetes(100);o__Saccharomycetales(100);f__Metschnikowiaceae(100);g__Metschnikowia(100);s__Metschnikowia_pulcherrima(100); | Metschnikowia | 13 | Saprotroph | \|Undefined Saprotroph\| | Yeast | NULL | Probable | As nectar-tap_saprotroph (Põlme et al. 2020); Decay sugar-rich substrates (Põlme et al. 2020) | Põlme S, et al. 2020. Fungal Diversity 105:1-16 ((https://doi.org/10.1007/s13225-020-00466-2)) |
| Otu47800 | 1 | k__Fungi(100);p__Ascomycota(100);c__Saccharomycetes(100);o__Saccharomycetales(100);f__Metschnikowiaceae(100);g__Metschnikowia(100);s__Metschnikowia_pulcherrima(100); | Metschnikowia | 13 | Saprotroph | \|Undefined Saprotroph\| | Yeast | NULL | Probable | As nectar-tap_saprotroph (Põlme et al. 2020); Decay sugar-rich substrates (Põlme et al. 2020) | Põlme S, et al. 2020. Fungal Diversity 105:1-16 ((https://doi.org/10.1007/s13225-020-00466-2)) |
| Otu47801 | 1 | k__Fungi(100);p__Ascomycota(100);c__Saccharomycetes(100);o__Saccharomycetales(100);f__Metschnikowiaceae(100);g__Metschnikowia(100);g__Metschnikowia_unclassified(100); | Metschnikowia | 13 | Saprotroph | \|Undefined Saprotroph\| | Yeast | NULL | Probable | As nectar-tap_saprotroph (Põlme et al. 2020); Decay sugar-rich substrates (Põlme et al. 2020) | Põlme S, et al. 2020. Fungal Diversity 105:1-16 ((https://doi.org/10.1007/s13225-020-00466-2)) |
| Otu47805 | 1 | k__Fungi(100);p__Ascomycota(100);c__Saccharomycetes(100);o__Saccharomycetales(100);f__Metschnikowiaceae(100);g__Metschnikowia(100);s__Metschnikowia_pulcherrima(100); | Metschnikowia | 13 | Saprotroph | \|Undefined Saprotroph\| | Yeast | NULL | Probable | As nectar-tap_saprotroph (Põlme et al. 2020); Decay sugar-rich substrates (Põlme et al. 2020) | Põlme S, et al. 2020. Fungal Diversity 105:1-16 ((https://doi.org/10.1007/s13225-020-00466-2)) |
| Otu47806 | 1 | k__Fungi(100);p__Ascomycota(100);c__Saccharomycetes(100);o__Saccharomycetales(100);f__Metschnikowiaceae(100);g__Metschnikowia(100);s__Metschnikowia_pulcherrima(100); | Metschnikowia | 13 | Saprotroph | \|Undefined Saprotroph\| | Yeast | NULL | Probable | As nectar-tap_saprotroph (Põlme et al. 2020); Decay sugar-rich substrates (Põlme et al. 2020) | Põlme S, et al. 2020. Fungal Diversity 105:1-16 ((https://doi.org/10.1007/s13225-020-00466-2)) |
| Otu47807 | 1 | k__Fungi(100);p__Ascomycota(100);c__Saccharomycetes(100);o__Saccharomycetales(100);f__Metschnikowiaceae(100);g__Metschnikowia(100);g__Metschnikowia_unclassified(100); | Metschnikowia | 13 | Saprotroph | \|Undefined Saprotroph\| | Yeast | NULL | Probable | As nectar-tap_saprotroph (Põlme et al. 2020); Decay sugar-rich substrates (Põlme et al. 2020) | Põlme S, et al. 2020. Fungal Diversity 105:1-16 ((https://doi.org/10.1007/s13225-020-00466-2)) |
| Otu47808 | 1 | k__Fungi(100);p__Ascomycota(100);c__Saccharomycetes(100);o__Saccharomycetales(100);f__Metschnikowiaceae(100);g__Metschnikowia(100);s__Metschnikowia_pulcherrima(100); | Metschnikowia | 13 | Saprotroph | \|Undefined Saprotroph\| | Yeast | NULL | Probable | As nectar-tap_saprotroph (Põlme et al. 2020); Decay sugar-rich substrates (Põlme et al. 2020) | Põlme S, et al. 2020. Fungal Diversity 105:1-16 ((https://doi.org/10.1007/s13225-020-00466-2)) |
| Otu47809 | 1 | k__Fungi(100);p__Ascomycota(100);c__Saccharomycetes(100);o__Saccharomycetales(100);f__Metschnikowiaceae(100);g__Metschnikowia(100);s__Metschnikowia_pulcherrima(100); | Metschnikowia | 13 | Saprotroph | \|Undefined Saprotroph\| | Yeast | NULL | Probable | As nectar-tap_saprotroph (Põlme et al. 2020); Decay sugar-rich substrates (Põlme et al. 2020) | Põlme S, et al. 2020. Fungal Diversity 105:1-16 ((https://doi.org/10.1007/s13225-020-00466-2)) |
| Otu47810 | 1 | k__Fungi(100);p__Ascomycota(100);c__Saccharomycetes(100);o__Saccharomycetales(100);f__Metschnikowiaceae(100);g__Metschnikowia(100);s__Metschnikowia_pulcherrima(100); | Metschnikowia | 13 | Saprotroph | \|Undefined Saprotroph\| | Yeast | NULL | Probable | As nectar-tap_saprotroph (Põlme et al. 2020); Decay sugar-rich substrates (Põlme et al. 2020) | Põlme S, et al. 2020. Fungal Diversity 105:1-16 ((https://doi.org/10.1007/s13225-020-00466-2)) |
| Otu47811 | 1 | k__Fungi(100);p__Ascomycota(100);c__Saccharomycetes(100);o__Saccharomycetales(100);f__Metschnikowiaceae(100);g__Metschnikowia(100);s__Metschnikowia_pulcherrima(100); | Metschnikowia | 13 | Saprotroph | \|Undefined Saprotroph\| | Yeast | NULL | Probable | As nectar-tap_saprotroph (Põlme et al. 2020); Decay sugar-rich substrates (Põlme et al. 2020) | Põlme S, et al. 2020. Fungal Diversity 105:1-16 ((https://doi.org/10.1007/s13225-020-00466-2)) |
| Otu47812 | 1 | k__Fungi(100);p__Ascomycota(100);c__Saccharomycetes(100);o__Saccharomycetales(100);f__Metschnikowiaceae(100);g__Metschnikowia(100);s__Metschnikowia_pulcherrima(100); | Metschnikowia | 13 | Saprotroph | \|Undefined Saprotroph\| | Yeast | NULL | Probable | As nectar-tap_saprotroph (Põlme et al. 2020); Decay sugar-rich substrates (Põlme et al. 2020) | Põlme S, et al. 2020. Fungal Diversity 105:1-16 ((https://doi.org/10.1007/s13225-020-00466-2)) |
| Otu47813 | 1 | k__Fungi(100);p__Ascomycota(100);c__Saccharomycetes(100);o__Saccharomycetales(100);f__Metschnikowiaceae(100);g__Metschnikowia(100);s__Metschnikowia_pulcherrima(100); | Metschnikowia | 13 | Saprotroph | \|Undefined Saprotroph\| | Yeast | NULL | Probable | As nectar-tap_saprotroph (Põlme et al. 2020); Decay sugar-rich substrates (Põlme et al. 2020) | Põlme S, et al. 2020. Fungal Diversity 105:1-16 ((https://doi.org/10.1007/s13225-020-00466-2)) |
| Otu47814 | 1 | k__Fungi(100);p__Ascomycota(100);c__Saccharomycetes(100);o__Saccharomycetales(100);f__Metschnikowiaceae(100);g__Metschnikowia(100);s__Metschnikowia_pulcherrima(100); | Metschnikowia | 13 | Saprotroph | \|Undefined Saprotroph\| | Yeast | NULL | Probable | As nectar-tap_saprotroph (Põlme et al. 2020); Decay sugar-rich substrates (Põlme et al. 2020) | Põlme S, et al. 2020. Fungal Diversity 105:1-16 ((https://doi.org/10.1007/s13225-020-00466-2)) |
| Otu47815 | 1 | k__Fungi(100);p__Ascomycota(100);c__Saccharomycetes(100);o__Saccharomycetales(100);f__Metschnikowiaceae(100);g__Metschnikowia(100);s__Metschnikowia_pulcherrima(100); | Metschnikowia | 13 | Saprotroph | \|Undefined Saprotroph\| | Yeast | NULL | Probable | As nectar-tap_saprotroph (Põlme et al. 2020); Decay sugar-rich substrates (Põlme et al. 2020) | Põlme S, et al. 2020. Fungal Diversity 105:1-16 ((https://doi.org/10.1007/s13225-020-00466-2)) |
| Otu47816 | 1 | k__Fungi(100);p__Ascomycota(100);c__Saccharomycetes(100);o__Saccharomycetales(100);f__Metschnikowiaceae(100);g__Metschnikowia(100);s__Metschnikowia_pulcherrima(100); | Metschnikowia | 13 | Saprotroph | \|Undefined Saprotroph\| | Yeast | NULL | Probable | As nectar-tap_saprotroph (Põlme et al. 2020); Decay sugar-rich substrates (Põlme et al. 2020) | Põlme S, et al. 2020. Fungal Diversity 105:1-16 ((https://doi.org/10.1007/s13225-020-00466-2)) |
| Otu47817 | 1 | k__Fungi(100);p__Ascomycota(100);c__Saccharomycetes(100);o__Saccharomycetales(100);f__Metschnikowiaceae(100);g__Metschnikowia(100);s__Metschnikowia_pulcherrima(100); | Metschnikowia | 13 | Saprotroph | \|Undefined Saprotroph\| | Yeast | NULL | Probable | As nectar-tap_saprotroph (Põlme et al. 2020); Decay sugar-rich substrates (Põlme et al. 2020) | Põlme S, et al. 2020. Fungal Diversity 105:1-16 ((https://doi.org/10.1007/s13225-020-00466-2)) |
| Otu47818 | 1 | k__Fungi(100);p__Ascomycota(100);c__Saccharomycetes(100);o__Saccharomycetales(100);f__Metschnikowiaceae(100);g__Metschnikowia(100);s__Metschnikowia_pulcherrima(100); | Metschnikowia | 13 | Saprotroph | \|Undefined Saprotroph\| | Yeast | NULL | Probable | As nectar-tap_saprotroph (Põlme et al. 2020); Decay sugar-rich substrates (Põlme et al. 2020) | Põlme S, et al. 2020. Fungal Diversity 105:1-16 ((https://doi.org/10.1007/s13225-020-00466-2)) |
| Otu47819 | 1 | k__Fungi(100);p__Ascomycota(100);c__Saccharomycetes(100);o__Saccharomycetales(100);f__Metschnikowiaceae(100);g__Metschnikowia(100);s__Metschnikowia_pulcherrima(100); | Metschnikowia | 13 | Saprotroph | \|Undefined Saprotroph\| | Yeast | NULL | Probable | As nectar-tap_saprotroph (Põlme et al. 2020); Decay sugar-rich substrates (Põlme et al. 2020) | Põlme S, et al. 2020. Fungal Diversity 105:1-16 ((https://doi.org/10.1007/s13225-020-00466-2)) |
| Otu47820 | 1 | k__Fungi(100);p__Ascomycota(100);c__Saccharomycetes(100);o__Saccharomycetales(100);f__Metschnikowiaceae(100);g__Metschnikowia(100);s__Metschnikowia_pulcherrima(100); | Metschnikowia | 13 | Saprotroph | \|Undefined Saprotroph\| | Yeast | NULL | Probable | As nectar-tap_saprotroph (Põlme et al. 2020); Decay sugar-rich substrates (Põlme et al. 2020) | Põlme S, et al. 2020. Fungal Diversity 105:1-16 ((https://doi.org/10.1007/s13225-020-00466-2)) |
| Otu47821 | 1 | k__Fungi(100);p__Ascomycota(100);c__Saccharomycetes(100);o__Saccharomycetales(100);f__Metschnikowiaceae(100);g__Metschnikowia(100);s__Metschnikowia_pulcherrima(100); | Metschnikowia | 13 | Saprotroph | \|Undefined Saprotroph\| | Yeast | NULL | Probable | As nectar-tap_saprotroph (Põlme et al. 2020); Decay sugar-rich substrates (Põlme et al. 2020) | Põlme S, et al. 2020. Fungal Diversity 105:1-16 ((https://doi.org/10.1007/s13225-020-00466-2)) |
| Otu47822 | 1 | k__Fungi(100);p__Ascomycota(100);c__Saccharomycetes(100);o__Saccharomycetales(100);f__Metschnikowiaceae(100);g__Metschnikowia(100);s__Metschnikowia_pulcherrima(100); | Metschnikowia | 13 | Saprotroph | \|Undefined Saprotroph\| | Yeast | NULL | Probable | As nectar-tap_saprotroph (Põlme et al. 2020); Decay sugar-rich substrates (Põlme et al. 2020) | Põlme S, et al. 2020. Fungal Diversity 105:1-16 ((https://doi.org/10.1007/s13225-020-00466-2)) |
| Otu47823 | 1 | k__Fungi(100);p__Ascomycota(100);c__Saccharomycetes(100);o__Saccharomycetales(100);f__Metschnikowiaceae(100);g__Metschnikowia(100);s__Metschnikowia_pulcherrima(100); | Metschnikowia | 13 | Saprotroph | \|Undefined Saprotroph\| | Yeast | NULL | Probable | As nectar-tap_saprotroph (Põlme et al. 2020); Decay sugar-rich substrates (Põlme et al. 2020) | Põlme S, et al. 2020. Fungal Diversity 105:1-16 ((https://doi.org/10.1007/s13225-020-00466-2)) |
| Otu47824 | 1 | k__Fungi(100);p__Ascomycota(100);c__Saccharomycetes(100);o__Saccharomycetales(100);f__Metschnikowiaceae(100);g__Metschnikowia(100);g__Metschnikowia_unclassified(100); | Metschnikowia | 13 | Saprotroph | \|Undefined Saprotroph\| | Yeast | NULL | Probable | As nectar-tap_saprotroph (Põlme et al. 2020); Decay sugar-rich substrates (Põlme et al. 2020) | Põlme S, et al. 2020. Fungal Diversity 105:1-16 ((https://doi.org/10.1007/s13225-020-00466-2)) |
| Otu47826 | 1 | k__Fungi(100);p__Ascomycota(100);c__Saccharomycetes(100);o__Saccharomycetales(100);f__Metschnikowiaceae(100);g__Metschnikowia(100);s__Metschnikowia_pulcherrima(100); | Metschnikowia | 13 | Saprotroph | \|Undefined Saprotroph\| | Yeast | NULL | Probable | As nectar-tap_saprotroph (Põlme et al. 2020); Decay sugar-rich substrates (Põlme et al. 2020) | Põlme S, et al. 2020. Fungal Diversity 105:1-16 ((https://doi.org/10.1007/s13225-020-00466-2)) |
| Otu47827 | 1 | k__Fungi(100);p__Ascomycota(100);c__Saccharomycetes(100);o__Saccharomycetales(100);f__Metschnikowiaceae(100);g__Metschnikowia(100);s__Metschnikowia_pulcherrima(100); | Metschnikowia | 13 | Saprotroph | \|Undefined Saprotroph\| | Yeast | NULL | Probable | As nectar-tap_saprotroph (Põlme et al. 2020); Decay sugar-rich substrates (Põlme et al. 2020) | Põlme S, et al. 2020. Fungal Diversity 105:1-16 ((https://doi.org/10.1007/s13225-020-00466-2)) |
| Otu47828 | 1 | k__Fungi(100);p__Ascomycota(100);c__Saccharomycetes(100);o__Saccharomycetales(100);f__Metschnikowiaceae(100);g__Metschnikowia(100);s__Metschnikowia_pulcherrima(100); | Metschnikowia | 13 | Saprotroph | \|Undefined Saprotroph\| | Yeast | NULL | Probable | As nectar-tap_saprotroph (Põlme et al. 2020); Decay sugar-rich substrates (Põlme et al. 2020) | Põlme S, et al. 2020. Fungal Diversity 105:1-16 ((https://doi.org/10.1007/s13225-020-00466-2)) |
| Otu47829 | 1 | k__Fungi(100);p__Ascomycota(100);c__Saccharomycetes(100);o__Saccharomycetales(100);f__Metschnikowiaceae(100);g__Metschnikowia(100);g__Metschnikowia_unclassified(100); | Metschnikowia | 13 | Saprotroph | \|Undefined Saprotroph\| | Yeast | NULL | Probable | As nectar-tap_saprotroph (Põlme et al. 2020); Decay sugar-rich substrates (Põlme et al. 2020) | Põlme S, et al. 2020. Fungal Diversity 105:1-16 ((https://doi.org/10.1007/s13225-020-00466-2)) |
| Otu47830 | 1 | k__Fungi(100);p__Ascomycota(100);c__Saccharomycetes(100);o__Saccharomycetales(100);f__Metschnikowiaceae(100);g__Metschnikowia(100);s__Metschnikowia_pulcherrima(100); | Metschnikowia | 13 | Saprotroph | \|Undefined Saprotroph\| | Yeast | NULL | Probable | As nectar-tap_saprotroph (Põlme et al. 2020); Decay sugar-rich substrates (Põlme et al. 2020) | Põlme S, et al. 2020. Fungal Diversity 105:1-16 ((https://doi.org/10.1007/s13225-020-00466-2)) |
| Otu47831 | 1 | k__Fungi(100);p__Ascomycota(100);c__Saccharomycetes(100);o__Saccharomycetales(100);f__Metschnikowiaceae(100);g__Metschnikowia(100);s__Metschnikowia_pulcherrima(100); | Metschnikowia | 13 | Saprotroph | \|Undefined Saprotroph\| | Yeast | NULL | Probable | As nectar-tap_saprotroph (Põlme et al. 2020); Decay sugar-rich substrates (Põlme et al. 2020) | Põlme S, et al. 2020. Fungal Diversity 105:1-16 ((https://doi.org/10.1007/s13225-020-00466-2)) |
| Otu47832 | 1 | k__Fungi(100);p__Ascomycota(100);c__Saccharomycetes(100);o__Saccharomycetales(100);f__Metschnikowiaceae(100);g__Metschnikowia(100);s__Metschnikowia_pulcherrima(100); | Metschnikowia | 13 | Saprotroph | \|Undefined Saprotroph\| | Yeast | NULL | Probable | As nectar-tap_saprotroph (Põlme et al. 2020); Decay sugar-rich substrates (Põlme et al. 2020) | Põlme S, et al. 2020. Fungal Diversity 105:1-16 ((https://doi.org/10.1007/s13225-020-00466-2)) |
| Otu47833 | 1 | k__Fungi(100);p__Ascomycota(100);c__Saccharomycetes(100);o__Saccharomycetales(100);f__Metschnikowiaceae(100);g__Metschnikowia(100);g__Metschnikowia_unclassified(100); | Metschnikowia | 13 | Saprotroph | \|Undefined Saprotroph\| | Yeast | NULL | Probable | As nectar-tap_saprotroph (Põlme et al. 2020); Decay sugar-rich substrates (Põlme et al. 2020) | Põlme S, et al. 2020. Fungal Diversity 105:1-16 ((https://doi.org/10.1007/s13225-020-00466-2)) |
| Otu47834 | 1 | k__Fungi(100);p__Ascomycota(100);c__Saccharomycetes(100);o__Saccharomycetales(100);f__Metschnikowiaceae(100);g__Metschnikowia(100);g__Metschnikowia_unclassified(100); | Metschnikowia | 13 | Saprotroph | \|Undefined Saprotroph\| | Yeast | NULL | Probable | As nectar-tap_saprotroph (Põlme et al. 2020); Decay sugar-rich substrates (Põlme et al. 2020) | Põlme S, et al. 2020. Fungal Diversity 105:1-16 ((https://doi.org/10.1007/s13225-020-00466-2)) |
| Otu47835 | 1 | k__Fungi(100);p__Ascomycota(100);c__Saccharomycetes(100);o__Saccharomycetales(100);f__Metschnikowiaceae(100);g__Metschnikowia(100);s__Metschnikowia_pulcherrima(100); | Metschnikowia | 13 | Saprotroph | \|Undefined Saprotroph\| | Yeast | NULL | Probable | As nectar-tap_saprotroph (Põlme et al. 2020); Decay sugar-rich substrates (Põlme et al. 2020) | Põlme S, et al. 2020. Fungal Diversity 105:1-16 ((https://doi.org/10.1007/s13225-020-00466-2)) |
| Otu47836 | 1 | k__Fungi(100);p__Ascomycota(100);c__Saccharomycetes(100);o__Saccharomycetales(100);f__Metschnikowiaceae(100);g__Metschnikowia(100);g__Metschnikowia_unclassified(100); | Metschnikowia | 13 | Saprotroph | \|Undefined Saprotroph\| | Yeast | NULL | Probable | As nectar-tap_saprotroph (Põlme et al. 2020); Decay sugar-rich substrates (Põlme et al. 2020) | Põlme S, et al. 2020. Fungal Diversity 105:1-16 ((https://doi.org/10.1007/s13225-020-00466-2)) |
| Otu47837 | 1 | k__Fungi(100);p__Ascomycota(100);c__Saccharomycetes(100);o__Saccharomycetales(100);f__Metschnikowiaceae(100);g__Metschnikowia(100);s__Metschnikowia_pulcherrima(100); | Metschnikowia | 13 | Saprotroph | \|Undefined Saprotroph\| | Yeast | NULL | Probable | As nectar-tap_saprotroph (Põlme et al. 2020); Decay sugar-rich substrates (Põlme et al. 2020) | Põlme S, et al. 2020. Fungal Diversity 105:1-16 ((https://doi.org/10.1007/s13225-020-00466-2)) |
| Otu47838 | 1 | k__Fungi(100);p__Ascomycota(100);c__Saccharomycetes(100);o__Saccharomycetales(100);f__Metschnikowiaceae(100);g__Metschnikowia(100);s__Metschnikowia_pulcherrima(100); | Metschnikowia | 13 | Saprotroph | \|Undefined Saprotroph\| | Yeast | NULL | Probable | As nectar-tap_saprotroph (Põlme et al. 2020); Decay sugar-rich substrates (Põlme et al. 2020) | Põlme S, et al. 2020. Fungal Diversity 105:1-16 ((https://doi.org/10.1007/s13225-020-00466-2)) |
| Otu47839 | 1 | k__Fungi(100);p__Ascomycota(100);c__Saccharomycetes(100);o__Saccharomycetales(100);f__Metschnikowiaceae(100);g__Metschnikowia(100);s__Metschnikowia_pulcherrima(100); | Metschnikowia | 13 | Saprotroph | \|Undefined Saprotroph\| | Yeast | NULL | Probable | As nectar-tap_saprotroph (Põlme et al. 2020); Decay sugar-rich substrates (Põlme et al. 2020) | Põlme S, et al. 2020. Fungal Diversity 105:1-16 ((https://doi.org/10.1007/s13225-020-00466-2)) |
| Otu47842 | 1 | k__Fungi(100);p__Ascomycota(100);c__Saccharomycetes(100);o__Saccharomycetales(100);f__Metschnikowiaceae(100);g__Metschnikowia(100);s__Metschnikowia_pulcherrima(100); | Metschnikowia | 13 | Saprotroph | \|Undefined Saprotroph\| | Yeast | NULL | Probable | As nectar-tap_saprotroph (Põlme et al. 2020); Decay sugar-rich substrates (Põlme et al. 2020) | Põlme S, et al. 2020. Fungal Diversity 105:1-16 ((https://doi.org/10.1007/s13225-020-00466-2)) |
| Otu47843 | 1 | k__Fungi(100);p__Ascomycota(100);c__Saccharomycetes(100);o__Saccharomycetales(100);f__Metschnikowiaceae(100);g__Metschnikowia(100);s__Metschnikowia_pulcherrima(100); | Metschnikowia | 13 | Saprotroph | \|Undefined Saprotroph\| | Yeast | NULL | Probable | As nectar-tap_saprotroph (Põlme et al. 2020); Decay sugar-rich substrates (Põlme et al. 2020) | Põlme S, et al. 2020. Fungal Diversity 105:1-16 ((https://doi.org/10.1007/s13225-020-00466-2)) |
| Otu47844 | 1 | k__Fungi(100);p__Ascomycota(100);c__Saccharomycetes(100);o__Saccharomycetales(100);f__Metschnikowiaceae(100);g__Metschnikowia(100);s__Metschnikowia_pulcherrima(100); | Metschnikowia | 13 | Saprotroph | \|Undefined Saprotroph\| | Yeast | NULL | Probable | As nectar-tap_saprotroph (Põlme et al. 2020); Decay sugar-rich substrates (Põlme et al. 2020) | Põlme S, et al. 2020. Fungal Diversity 105:1-16 ((https://doi.org/10.1007/s13225-020-00466-2)) |
| Otu47845 | 1 | k__Fungi(100);p__Ascomycota(100);c__Saccharomycetes(100);o__Saccharomycetales(100);f__Metschnikowiaceae(100);g__Metschnikowia(100);s__Metschnikowia_pulcherrima(100); | Metschnikowia | 13 | Saprotroph | \|Undefined Saprotroph\| | Yeast | NULL | Probable | As nectar-tap_saprotroph (Põlme et al. 2020); Decay sugar-rich substrates (Põlme et al. 2020) | Põlme S, et al. 2020. Fungal Diversity 105:1-16 ((https://doi.org/10.1007/s13225-020-00466-2)) |
| Otu47846 | 1 | k__Fungi(100);p__Ascomycota(100);c__Saccharomycetes(100);o__Saccharomycetales(100);f__Metschnikowiaceae(100);g__Metschnikowia(100);s__Metschnikowia_pulcherrima(100); | Metschnikowia | 13 | Saprotroph | \|Undefined Saprotroph\| | Yeast | NULL | Probable | As nectar-tap_saprotroph (Põlme et al. 2020); Decay sugar-rich substrates (Põlme et al. 2020) | Põlme S, et al. 2020. Fungal Diversity 105:1-16 ((https://doi.org/10.1007/s13225-020-00466-2)) |
| Otu47847 | 1 | k__Fungi(100);p__Ascomycota(100);c__Saccharomycetes(100);o__Saccharomycetales(100);f__Metschnikowiaceae(100);g__Metschnikowia(100);s__Metschnikowia_pulcherrima(100); | Metschnikowia | 13 | Saprotroph | \|Undefined Saprotroph\| | Yeast | NULL | Probable | As nectar-tap_saprotroph (Põlme et al. 2020); Decay sugar-rich substrates (Põlme et al. 2020) | Põlme S, et al. 2020. Fungal Diversity 105:1-16 ((https://doi.org/10.1007/s13225-020-00466-2)) |
| Otu47848 | 1 | k__Fungi(100);p__Ascomycota(100);c__Saccharomycetes(100);o__Saccharomycetales(100);f__Metschnikowiaceae(100);g__Metschnikowia(100);s__Metschnikowia_pulcherrima(100); | Metschnikowia | 13 | Saprotroph | \|Undefined Saprotroph\| | Yeast | NULL | Probable | As nectar-tap_saprotroph (Põlme et al. 2020); Decay sugar-rich substrates (Põlme et al. 2020) | Põlme S, et al. 2020. Fungal Diversity 105:1-16 ((https://doi.org/10.1007/s13225-020-00466-2)) |
| Otu47850 | 1 | k__Fungi(100);p__Ascomycota(100);c__Saccharomycetes(100);o__Saccharomycetales(100);f__Metschnikowiaceae(100);g__Metschnikowia(100);s__Metschnikowia_pulcherrima(100); | Metschnikowia | 13 | Saprotroph | \|Undefined Saprotroph\| | Yeast | NULL | Probable | As nectar-tap_saprotroph (Põlme et al. 2020); Decay sugar-rich substrates (Põlme et al. 2020) | Põlme S, et al. 2020. Fungal Diversity 105:1-16 ((https://doi.org/10.1007/s13225-020-00466-2)) |
| Otu47851 | 1 | k__Fungi(100);p__Ascomycota(100);c__Saccharomycetes(100);o__Saccharomycetales(100);f__Metschnikowiaceae(100);g__Metschnikowia(100);s__Metschnikowia_pulcherrima(100); | Metschnikowia | 13 | Saprotroph | \|Undefined Saprotroph\| | Yeast | NULL | Probable | As nectar-tap_saprotroph (Põlme et al. 2020); Decay sugar-rich substrates (Põlme et al. 2020) | Põlme S, et al. 2020. Fungal Diversity 105:1-16 ((https://doi.org/10.1007/s13225-020-00466-2)) |
| Otu47852 | 1 | k__Fungi(100);p__Ascomycota(100);c__Saccharomycetes(100);o__Saccharomycetales(100);f__Metschnikowiaceae(100);g__Metschnikowia(100);s__Metschnikowia_pulcherrima(100); | Metschnikowia | 13 | Saprotroph | \|Undefined Saprotroph\| | Yeast | NULL | Probable | As nectar-tap_saprotroph (Põlme et al. 2020); Decay sugar-rich substrates (Põlme et al. 2020) | Põlme S, et al. 2020. Fungal Diversity 105:1-16 ((https://doi.org/10.1007/s13225-020-00466-2)) |
| Otu47853 | 1 | k__Fungi(100);p__Ascomycota(100);c__Saccharomycetes(100);o__Saccharomycetales(100);f__Metschnikowiaceae(100);g__Metschnikowia(100);g__Metschnikowia_unclassified(100); | Metschnikowia | 13 | Saprotroph | \|Undefined Saprotroph\| | Yeast | NULL | Probable | As nectar-tap_saprotroph (Põlme et al. 2020); Decay sugar-rich substrates (Põlme et al. 2020) | Põlme S, et al. 2020. Fungal Diversity 105:1-16 ((https://doi.org/10.1007/s13225-020-00466-2)) |
| Otu47854 | 1 | k__Fungi(100);p__Ascomycota(100);c__Saccharomycetes(100);o__Saccharomycetales(100);f__Metschnikowiaceae(100);g__Metschnikowia(100);s__Metschnikowia_pulcherrima(100); | Metschnikowia | 13 | Saprotroph | \|Undefined Saprotroph\| | Yeast | NULL | Probable | As nectar-tap_saprotroph (Põlme et al. 2020); Decay sugar-rich substrates (Põlme et al. 2020) | Põlme S, et al. 2020. Fungal Diversity 105:1-16 ((https://doi.org/10.1007/s13225-020-00466-2)) |
| Otu47855 | 1 | k__Fungi(100);p__Ascomycota(100);c__Saccharomycetes(100);o__Saccharomycetales(100);f__Metschnikowiaceae(100);g__Metschnikowia(100);s__Metschnikowia_pulcherrima(100); | Metschnikowia | 13 | Saprotroph | \|Undefined Saprotroph\| | Yeast | NULL | Probable | As nectar-tap_saprotroph (Põlme et al. 2020); Decay sugar-rich substrates (Põlme et al. 2020) | Põlme S, et al. 2020. Fungal Diversity 105:1-16 ((https://doi.org/10.1007/s13225-020-00466-2)) |
| Otu47856 | 1 | k__Fungi(100);p__Ascomycota(100);c__Saccharomycetes(100);o__Saccharomycetales(100);f__Metschnikowiaceae(100);g__Metschnikowia(100);g__Metschnikowia_unclassified(100); | Metschnikowia | 13 | Saprotroph | \|Undefined Saprotroph\| | Yeast | NULL | Probable | As nectar-tap_saprotroph (Põlme et al. 2020); Decay sugar-rich substrates (Põlme et al. 2020) | Põlme S, et al. 2020. Fungal Diversity 105:1-16 ((https://doi.org/10.1007/s13225-020-00466-2)) |
| Otu47857 | 1 | k__Fungi(100);p__Ascomycota(100);c__Saccharomycetes(100);o__Saccharomycetales(100);f__Metschnikowiaceae(100);g__Metschnikowia(100);s__Metschnikowia_pulcherrima(100); | Metschnikowia | 13 | Saprotroph | \|Undefined Saprotroph\| | Yeast | NULL | Probable | As nectar-tap_saprotroph (Põlme et al. 2020); Decay sugar-rich substrates (Põlme et al. 2020) | Põlme S, et al. 2020. Fungal Diversity 105:1-16 ((https://doi.org/10.1007/s13225-020-00466-2)) |
| Otu47858 | 1 | k__Fungi(100);p__Ascomycota(100);c__Saccharomycetes(100);o__Saccharomycetales(100);f__Metschnikowiaceae(100);g__Metschnikowia(100);s__Metschnikowia_pulcherrima(100); | Metschnikowia | 13 | Saprotroph | \|Undefined Saprotroph\| | Yeast | NULL | Probable | As nectar-tap_saprotroph (Põlme et al. 2020); Decay sugar-rich substrates (Põlme et al. 2020) | Põlme S, et al. 2020. Fungal Diversity 105:1-16 ((https://doi.org/10.1007/s13225-020-00466-2)) |
| Otu47859 | 1 | k__Fungi(100);p__Ascomycota(100);c__Saccharomycetes(100);o__Saccharomycetales(100);f__Metschnikowiaceae(100);g__Metschnikowia(100);s__Metschnikowia_pulcherrima(100); | Metschnikowia | 13 | Saprotroph | \|Undefined Saprotroph\| | Yeast | NULL | Probable | As nectar-tap_saprotroph (Põlme et al. 2020); Decay sugar-rich substrates (Põlme et al. 2020) | Põlme S, et al. 2020. Fungal Diversity 105:1-16 ((https://doi.org/10.1007/s13225-020-00466-2)) |
| Otu47860 | 1 | k__Fungi(100);p__Ascomycota(100);c__Saccharomycetes(100);o__Saccharomycetales(100);f__Metschnikowiaceae(100);g__Metschnikowia(100);s__Metschnikowia_pulcherrima(100); | Metschnikowia | 13 | Saprotroph | \|Undefined Saprotroph\| | Yeast | NULL | Probable | As nectar-tap_saprotroph (Põlme et al. 2020); Decay sugar-rich substrates (Põlme et al. 2020) | Põlme S, et al. 2020. Fungal Diversity 105:1-16 ((https://doi.org/10.1007/s13225-020-00466-2)) |
| Otu47861 | 1 | k__Fungi(100);p__Ascomycota(100);c__Saccharomycetes(100);o__Saccharomycetales(100);f__Metschnikowiaceae(100);g__Metschnikowia(100);s__Metschnikowia_pulcherrima(100); | Metschnikowia | 13 | Saprotroph | \|Undefined Saprotroph\| | Yeast | NULL | Probable | As nectar-tap_saprotroph (Põlme et al. 2020); Decay sugar-rich substrates (Põlme et al. 2020) | Põlme S, et al. 2020. Fungal Diversity 105:1-16 ((https://doi.org/10.1007/s13225-020-00466-2)) |
| Otu47862 | 1 | k__Fungi(100);p__Ascomycota(100);c__Saccharomycetes(100);o__Saccharomycetales(100);f__Metschnikowiaceae(100);g__Metschnikowia(100);g__Metschnikowia_unclassified(100); | Metschnikowia | 13 | Saprotroph | \|Undefined Saprotroph\| | Yeast | NULL | Probable | As nectar-tap_saprotroph (Põlme et al. 2020); Decay sugar-rich substrates (Põlme et al. 2020) | Põlme S, et al. 2020. Fungal Diversity 105:1-16 ((https://doi.org/10.1007/s13225-020-00466-2)) |
| Otu47863 | 1 | k__Fungi(100);p__Ascomycota(100);c__Saccharomycetes(100);o__Saccharomycetales(100);f__Metschnikowiaceae(100);g__Metschnikowia(100);s__Metschnikowia_pulcherrima(100); | Metschnikowia | 13 | Saprotroph | \|Undefined Saprotroph\| | Yeast | NULL | Probable | As nectar-tap_saprotroph (Põlme et al. 2020); Decay sugar-rich substrates (Põlme et al. 2020) | Põlme S, et al. 2020. Fungal Diversity 105:1-16 ((https://doi.org/10.1007/s13225-020-00466-2)) |
| Otu47864 | 1 | k__Fungi(100);p__Ascomycota(100);c__Saccharomycetes(100);o__Saccharomycetales(100);f__Metschnikowiaceae(100);g__Metschnikowia(100);s__Metschnikowia_pulcherrima(100); | Metschnikowia | 13 | Saprotroph | \|Undefined Saprotroph\| | Yeast | NULL | Probable | As nectar-tap_saprotroph (Põlme et al. 2020); Decay sugar-rich substrates (Põlme et al. 2020) | Põlme S, et al. 2020. Fungal Diversity 105:1-16 ((https://doi.org/10.1007/s13225-020-00466-2)) |
| Otu47865 | 1 | k__Fungi(100);p__Ascomycota(100);c__Saccharomycetes(100);o__Saccharomycetales(100);f__Metschnikowiaceae(100);g__Metschnikowia(100);s__Metschnikowia_pulcherrima(100); | Metschnikowia | 13 | Saprotroph | \|Undefined Saprotroph\| | Yeast | NULL | Probable | As nectar-tap_saprotroph (Põlme et al. 2020); Decay sugar-rich substrates (Põlme et al. 2020) | Põlme S, et al. 2020. Fungal Diversity 105:1-16 ((https://doi.org/10.1007/s13225-020-00466-2)) |
| Otu47867 | 1 | k__Fungi(100);p__Ascomycota(100);c__Saccharomycetes(100);o__Saccharomycetales(100);f__Metschnikowiaceae(100);g__Metschnikowia(100);g__Metschnikowia_unclassified(100); | Metschnikowia | 13 | Saprotroph | \|Undefined Saprotroph\| | Yeast | NULL | Probable | As nectar-tap_saprotroph (Põlme et al. 2020); Decay sugar-rich substrates (Põlme et al. 2020) | Põlme S, et al. 2020. Fungal Diversity 105:1-16 ((https://doi.org/10.1007/s13225-020-00466-2)) |
| Otu47868 | 1 | k__Fungi(100);p__Ascomycota(100);c__Saccharomycetes(100);o__Saccharomycetales(100);f__Metschnikowiaceae(100);g__Metschnikowia(100);s__Metschnikowia_pulcherrima(100); | Metschnikowia | 13 | Saprotroph | \|Undefined Saprotroph\| | Yeast | NULL | Probable | As nectar-tap_saprotroph (Põlme et al. 2020); Decay sugar-rich substrates (Põlme et al. 2020) | Põlme S, et al. 2020. Fungal Diversity 105:1-16 ((https://doi.org/10.1007/s13225-020-00466-2)) |
| Otu47871 | 1 | k__Fungi(100);p__Ascomycota(100);c__Saccharomycetes(100);o__Saccharomycetales(100);f__Metschnikowiaceae(100);g__Metschnikowia(100);g__Metschnikowia_unclassified(100); | Metschnikowia | 13 | Saprotroph | \|Undefined Saprotroph\| | Yeast | NULL | Probable | As nectar-tap_saprotroph (Põlme et al. 2020); Decay sugar-rich substrates (Põlme et al. 2020) | Põlme S, et al. 2020. Fungal Diversity 105:1-16 ((https://doi.org/10.1007/s13225-020-00466-2)) |
| Otu47873 | 1 | k__Fungi(100);p__Ascomycota(100);c__Saccharomycetes(100);o__Saccharomycetales(100);f__Metschnikowiaceae(100);g__Metschnikowia(100);s__Metschnikowia_pulcherrima(100); | Metschnikowia | 13 | Saprotroph | \|Undefined Saprotroph\| | Yeast | NULL | Probable | As nectar-tap_saprotroph (Põlme et al. 2020); Decay sugar-rich substrates (Põlme et al. 2020) | Põlme S, et al. 2020. Fungal Diversity 105:1-16 ((https://doi.org/10.1007/s13225-020-00466-2)) |
| Otu47874 | 1 | k__Fungi(100);p__Ascomycota(100);c__Saccharomycetes(100);o__Saccharomycetales(100);f__Metschnikowiaceae(100);g__Metschnikowia(100);s__Metschnikowia_pulcherrima(100); | Metschnikowia | 13 | Saprotroph | \|Undefined Saprotroph\| | Yeast | NULL | Probable | As nectar-tap_saprotroph (Põlme et al. 2020); Decay sugar-rich substrates (Põlme et al. 2020) | Põlme S, et al. 2020. Fungal Diversity 105:1-16 ((https://doi.org/10.1007/s13225-020-00466-2)) |
| Otu47875 | 1 | k__Fungi(100);p__Ascomycota(100);c__Saccharomycetes(100);o__Saccharomycetales(100);f__Metschnikowiaceae(100);g__Metschnikowia(100);s__Metschnikowia_pulcherrima(100); | Metschnikowia | 13 | Saprotroph | \|Undefined Saprotroph\| | Yeast | NULL | Probable | As nectar-tap_saprotroph (Põlme et al. 2020); Decay sugar-rich substrates (Põlme et al. 2020) | Põlme S, et al. 2020. Fungal Diversity 105:1-16 ((https://doi.org/10.1007/s13225-020-00466-2)) |
| Otu47877 | 1 | k__Fungi(100);p__Ascomycota(100);c__Saccharomycetes(100);o__Saccharomycetales(100);f__Metschnikowiaceae(100);g__Metschnikowia(100);s__Metschnikowia_pulcherrima(100); | Metschnikowia | 13 | Saprotroph | \|Undefined Saprotroph\| | Yeast | NULL | Probable | As nectar-tap_saprotroph (Põlme et al. 2020); Decay sugar-rich substrates (Põlme et al. 2020) | Põlme S, et al. 2020. Fungal Diversity 105:1-16 ((https://doi.org/10.1007/s13225-020-00466-2)) |
| Otu47878 | 1 | k__Fungi(100);p__Ascomycota(100);c__Saccharomycetes(100);o__Saccharomycetales(100);f__Metschnikowiaceae(100);g__Metschnikowia(100);s__Metschnikowia_pulcherrima(100); | Metschnikowia | 13 | Saprotroph | \|Undefined Saprotroph\| | Yeast | NULL | Probable | As nectar-tap_saprotroph (Põlme et al. 2020); Decay sugar-rich substrates (Põlme et al. 2020) | Põlme S, et al. 2020. Fungal Diversity 105:1-16 ((https://doi.org/10.1007/s13225-020-00466-2)) |
| Otu47879 | 1 | k__Fungi(100);p__Ascomycota(100);c__Saccharomycetes(100);o__Saccharomycetales(100);f__Metschnikowiaceae(100);g__Metschnikowia(100);s__Metschnikowia_pulcherrima(100); | Metschnikowia | 13 | Saprotroph | \|Undefined Saprotroph\| | Yeast | NULL | Probable | As nectar-tap_saprotroph (Põlme et al. 2020); Decay sugar-rich substrates (Põlme et al. 2020) | Põlme S, et al. 2020. Fungal Diversity 105:1-16 ((https://doi.org/10.1007/s13225-020-00466-2)) |
| Otu47880 | 1 | k__Fungi(100);p__Ascomycota(100);c__Saccharomycetes(100);o__Saccharomycetales(100);f__Metschnikowiaceae(100);g__Metschnikowia(100);s__Metschnikowia_pulcherrima(100); | Metschnikowia | 13 | Saprotroph | \|Undefined Saprotroph\| | Yeast | NULL | Probable | As nectar-tap_saprotroph (Põlme et al. 2020); Decay sugar-rich substrates (Põlme et al. 2020) | Põlme S, et al. 2020. Fungal Diversity 105:1-16 ((https://doi.org/10.1007/s13225-020-00466-2)) |
| Otu47881 | 1 | k__Fungi(100);p__Ascomycota(100);c__Saccharomycetes(100);o__Saccharomycetales(100);f__Metschnikowiaceae(100);g__Metschnikowia(100);s__Metschnikowia_pulcherrima(100); | Metschnikowia | 13 | Saprotroph | \|Undefined Saprotroph\| | Yeast | NULL | Probable | As nectar-tap_saprotroph (Põlme et al. 2020); Decay sugar-rich substrates (Põlme et al. 2020) | Põlme S, et al. 2020. Fungal Diversity 105:1-16 ((https://doi.org/10.1007/s13225-020-00466-2)) |
| Otu47882 | 1 | k__Fungi(100);p__Ascomycota(100);c__Saccharomycetes(100);o__Saccharomycetales(100);f__Metschnikowiaceae(100);g__Metschnikowia(100);s__Metschnikowia_pulcherrima(100); | Metschnikowia | 13 | Saprotroph | \|Undefined Saprotroph\| | Yeast | NULL | Probable | As nectar-tap_saprotroph (Põlme et al. 2020); Decay sugar-rich substrates (Põlme et al. 2020) | Põlme S, et al. 2020. Fungal Diversity 105:1-16 ((https://doi.org/10.1007/s13225-020-00466-2)) |
| Otu47883 | 1 | k__Fungi(100);p__Ascomycota(100);c__Dothideomycetes(100);o__Pleosporales(100);f__Montagnulaceae(100);g__Paraphaeosphaeria(100);g__Paraphaeosphaeria_unclassified(100); | Paraphaeosphaeria | 13 | Saprotroph | Undefined Saprotroph-\|Wood Saprotroph\| | Microfungus | NULL | Probable | Decay wood (Põlme et al. 2020) | Tedersoo L, et al. 2014. Science 346:e1256688 ((https://doi.org/10.1126/science.1256688)); Põlme S, et al. 2020. Fungal Diversity 105:1-16 ((https://doi.org/10.1007/s13225-020-00466-2)) |
| Otu47884 | 1 | k__Fungi(100);p__Ascomycota(100);c__Eurotiomycetes(100);o__Eurotiomycetes_order_Incertae_sedis(100);f__Eremascaceae(100);g__Eremascus(100);s__Eremascus_sp(100); | Eremascus | 13 | Saprotroph | \|Undefined Saprotroph\| | Microfungus | NULL | Probable | NULL | Tedersoo L, et al. 2014. Science 346:e1256688 ((https://doi.org/10.1126/science.1256688)); Põlme S, et al. 2020. Fungal Diversity 105:1-16 ((https://doi.org/10.1007/s13225-020-00466-2)) |
| Otu47885 | 1 | k__Fungi(100);p__Ascomycota(100);c__Saccharomycetes(100);o__Saccharomycetales(100);f__Metschnikowiaceae(100);g__Metschnikowia(100);g__Metschnikowia_unclassified(100); | Metschnikowia | 13 | Saprotroph | \|Undefined Saprotroph\| | Yeast | NULL | Probable | As nectar-tap_saprotroph (Põlme et al. 2020); Decay sugar-rich substrates (Põlme et al. 2020) | Põlme S, et al. 2020. Fungal Diversity 105:1-16 ((https://doi.org/10.1007/s13225-020-00466-2)) |
| Otu47886 | 1 | k__Fungi(100);p__Ascomycota(100);c__Saccharomycetes(100);o__Saccharomycetales(100);f__Metschnikowiaceae(100);g__Metschnikowia(100);s__Metschnikowia_pulcherrima(100); | Metschnikowia | 13 | Saprotroph | \|Undefined Saprotroph\| | Yeast | NULL | Probable | As nectar-tap_saprotroph (Põlme et al. 2020); Decay sugar-rich substrates (Põlme et al. 2020) | Põlme S, et al. 2020. Fungal Diversity 105:1-16 ((https://doi.org/10.1007/s13225-020-00466-2)) |
| Otu47887 | 1 | k__Fungi(100);p__Ascomycota(100);c__Saccharomycetes(100);o__Saccharomycetales(100);f__Metschnikowiaceae(100);g__Metschnikowia(100);s__Metschnikowia_pulcherrima(100); | Metschnikowia | 13 | Saprotroph | \|Undefined Saprotroph\| | Yeast | NULL | Probable | As nectar-tap_saprotroph (Põlme et al. 2020); Decay sugar-rich substrates (Põlme et al. 2020) | Põlme S, et al. 2020. Fungal Diversity 105:1-16 ((https://doi.org/10.1007/s13225-020-00466-2)) |
| Otu47888 | 1 | k__Fungi(100);p__Ascomycota(100);c__Saccharomycetes(100);o__Saccharomycetales(100);f__Metschnikowiaceae(100);g__Metschnikowia(100);s__Metschnikowia_pulcherrima(100); | Metschnikowia | 13 | Saprotroph | \|Undefined Saprotroph\| | Yeast | NULL | Probable | As nectar-tap_saprotroph (Põlme et al. 2020); Decay sugar-rich substrates (Põlme et al. 2020) | Põlme S, et al. 2020. Fungal Diversity 105:1-16 ((https://doi.org/10.1007/s13225-020-00466-2)) |
| Otu47889 | 1 | k__Fungi(100);p__Ascomycota(100);c__Saccharomycetes(100);o__Saccharomycetales(100);f__Metschnikowiaceae(100);g__Metschnikowia(100);g__Metschnikowia_unclassified(100); | Metschnikowia | 13 | Saprotroph | \|Undefined Saprotroph\| | Yeast | NULL | Probable | As nectar-tap_saprotroph (Põlme et al. 2020); Decay sugar-rich substrates (Põlme et al. 2020) | Põlme S, et al. 2020. Fungal Diversity 105:1-16 ((https://doi.org/10.1007/s13225-020-00466-2)) |
| Otu47890 | 1 | k__Fungi(100);p__Ascomycota(100);c__Saccharomycetes(100);o__Saccharomycetales(100);f__Metschnikowiaceae(100);g__Metschnikowia(100);s__Metschnikowia_pulcherrima(100); | Metschnikowia | 13 | Saprotroph | \|Undefined Saprotroph\| | Yeast | NULL | Probable | As nectar-tap_saprotroph (Põlme et al. 2020); Decay sugar-rich substrates (Põlme et al. 2020) | Põlme S, et al. 2020. Fungal Diversity 105:1-16 ((https://doi.org/10.1007/s13225-020-00466-2)) |
| Otu47891 | 1 | k__Fungi(100);p__Ascomycota(100);c__Saccharomycetes(100);o__Saccharomycetales(100);f__Metschnikowiaceae(100);g__Metschnikowia(100);s__Metschnikowia_pulcherrima(100); | Metschnikowia | 13 | Saprotroph | \|Undefined Saprotroph\| | Yeast | NULL | Probable | As nectar-tap_saprotroph (Põlme et al. 2020); Decay sugar-rich substrates (Põlme et al. 2020) | Põlme S, et al. 2020. Fungal Diversity 105:1-16 ((https://doi.org/10.1007/s13225-020-00466-2)) |
| Otu47892 | 1 | k__Fungi(100);p__Ascomycota(100);c__Saccharomycetes(100);o__Saccharomycetales(100);f__Metschnikowiaceae(100);g__Metschnikowia(100);s__Metschnikowia_pulcherrima(100); | Metschnikowia | 13 | Saprotroph | \|Undefined Saprotroph\| | Yeast | NULL | Probable | As nectar-tap_saprotroph (Põlme et al. 2020); Decay sugar-rich substrates (Põlme et al. 2020) | Põlme S, et al. 2020. Fungal Diversity 105:1-16 ((https://doi.org/10.1007/s13225-020-00466-2)) |
| Otu47893 | 1 | k__Fungi(100);p__Ascomycota(100);c__Saccharomycetes(100);o__Saccharomycetales(100);f__Metschnikowiaceae(100);g__Metschnikowia(100);s__Metschnikowia_pulcherrima(100); | Metschnikowia | 13 | Saprotroph | \|Undefined Saprotroph\| | Yeast | NULL | Probable | As nectar-tap_saprotroph (Põlme et al. 2020); Decay sugar-rich substrates (Põlme et al. 2020) | Põlme S, et al. 2020. Fungal Diversity 105:1-16 ((https://doi.org/10.1007/s13225-020-00466-2)) |
| Otu47894 | 1 | k__Fungi(100);p__Ascomycota(100);c__Saccharomycetes(100);o__Saccharomycetales(100);f__Metschnikowiaceae(100);g__Metschnikowia(100);s__Metschnikowia_pulcherrima(100); | Metschnikowia | 13 | Saprotroph | \|Undefined Saprotroph\| | Yeast | NULL | Probable | As nectar-tap_saprotroph (Põlme et al. 2020); Decay sugar-rich substrates (Põlme et al. 2020) | Põlme S, et al. 2020. Fungal Diversity 105:1-16 ((https://doi.org/10.1007/s13225-020-00466-2)) |
| Otu47895 | 1 | k__Fungi(100);p__Ascomycota(100);c__Saccharomycetes(100);o__Saccharomycetales(100);f__Metschnikowiaceae(100);g__Metschnikowia(100);g__Metschnikowia_unclassified(100); | Metschnikowia | 13 | Saprotroph | \|Undefined Saprotroph\| | Yeast | NULL | Probable | As nectar-tap_saprotroph (Põlme et al. 2020); Decay sugar-rich substrates (Põlme et al. 2020) | Põlme S, et al. 2020. Fungal Diversity 105:1-16 ((https://doi.org/10.1007/s13225-020-00466-2)) |
| Otu47896 | 1 | k__Fungi(100);p__Ascomycota(100);c__Saccharomycetes(100);o__Saccharomycetales(100);f__Metschnikowiaceae(100);g__Metschnikowia(100);s__Metschnikowia_pulcherrima(100); | Metschnikowia | 13 | Saprotroph | \|Undefined Saprotroph\| | Yeast | NULL | Probable | As nectar-tap_saprotroph (Põlme et al. 2020); Decay sugar-rich substrates (Põlme et al. 2020) | Põlme S, et al. 2020. Fungal Diversity 105:1-16 ((https://doi.org/10.1007/s13225-020-00466-2)) |
| Otu47897 | 1 | k__Fungi(100);p__Ascomycota(100);c__Saccharomycetes(100);o__Saccharomycetales(100);f__Metschnikowiaceae(100);g__Metschnikowia(100);s__Metschnikowia_pulcherrima(100); | Metschnikowia | 13 | Saprotroph | \|Undefined Saprotroph\| | Yeast | NULL | Probable | As nectar-tap_saprotroph (Põlme et al. 2020); Decay sugar-rich substrates (Põlme et al. 2020) | Põlme S, et al. 2020. Fungal Diversity 105:1-16 ((https://doi.org/10.1007/s13225-020-00466-2)) |
| Otu47898 | 1 | k__Fungi(100);p__Ascomycota(100);c__Saccharomycetes(100);o__Saccharomycetales(100);f__Metschnikowiaceae(100);g__Metschnikowia(100);s__Metschnikowia_pulcherrima(100); | Metschnikowia | 13 | Saprotroph | \|Undefined Saprotroph\| | Yeast | NULL | Probable | As nectar-tap_saprotroph (Põlme et al. 2020); Decay sugar-rich substrates (Põlme et al. 2020) | Põlme S, et al. 2020. Fungal Diversity 105:1-16 ((https://doi.org/10.1007/s13225-020-00466-2)) |
| Otu47899 | 1 | k__Fungi(100);p__Ascomycota(100);c__Saccharomycetes(100);o__Saccharomycetales(100);f__Metschnikowiaceae(100);g__Metschnikowia(100);g__Metschnikowia_unclassified(100); | Metschnikowia | 13 | Saprotroph | \|Undefined Saprotroph\| | Yeast | NULL | Probable | As nectar-tap_saprotroph (Põlme et al. 2020); Decay sugar-rich substrates (Põlme et al. 2020) | Põlme S, et al. 2020. Fungal Diversity 105:1-16 ((https://doi.org/10.1007/s13225-020-00466-2)) |
| Otu47900 | 1 | k__Fungi(100);p__Ascomycota(100);c__Saccharomycetes(100);o__Saccharomycetales(100);f__Metschnikowiaceae(100);g__Metschnikowia(100);s__Metschnikowia_pulcherrima(100); | Metschnikowia | 13 | Saprotroph | \|Undefined Saprotroph\| | Yeast | NULL | Probable | As nectar-tap_saprotroph (Põlme et al. 2020); Decay sugar-rich substrates (Põlme et al. 2020) | Põlme S, et al. 2020. Fungal Diversity 105:1-16 ((https://doi.org/10.1007/s13225-020-00466-2)) |
| Otu47901 | 1 | k__Fungi(100);p__Ascomycota(100);c__Saccharomycetes(100);o__Saccharomycetales(100);f__Metschnikowiaceae(100);g__Metschnikowia(100);s__Metschnikowia_pulcherrima(100); | Metschnikowia | 13 | Saprotroph | \|Undefined Saprotroph\| | Yeast | NULL | Probable | As nectar-tap_saprotroph (Põlme et al. 2020); Decay sugar-rich substrates (Põlme et al. 2020) | Põlme S, et al. 2020. Fungal Diversity 105:1-16 ((https://doi.org/10.1007/s13225-020-00466-2)) |
| Otu47902 | 1 | k__Fungi(100);p__Ascomycota(100);c__Saccharomycetes(100);o__Saccharomycetales(100);f__Metschnikowiaceae(100);g__Metschnikowia(100);g__Metschnikowia_unclassified(100); | Metschnikowia | 13 | Saprotroph | \|Undefined Saprotroph\| | Yeast | NULL | Probable | As nectar-tap_saprotroph (Põlme et al. 2020); Decay sugar-rich substrates (Põlme et al. 2020) | Põlme S, et al. 2020. Fungal Diversity 105:1-16 ((https://doi.org/10.1007/s13225-020-00466-2)) |
| Otu47903 | 1 | k__Fungi(100);p__Ascomycota(100);c__Saccharomycetes(100);o__Saccharomycetales(100);f__Metschnikowiaceae(100);g__Metschnikowia(100);g__Metschnikowia_unclassified(100); | Metschnikowia | 13 | Saprotroph | \|Undefined Saprotroph\| | Yeast | NULL | Probable | As nectar-tap_saprotroph (Põlme et al. 2020); Decay sugar-rich substrates (Põlme et al. 2020) | Põlme S, et al. 2020. Fungal Diversity 105:1-16 ((https://doi.org/10.1007/s13225-020-00466-2)) |
| Otu47904 | 1 | k__Fungi(100);p__Ascomycota(100);c__Saccharomycetes(100);o__Saccharomycetales(100);f__Metschnikowiaceae(100);g__Metschnikowia(100);s__Metschnikowia_pulcherrima(100); | Metschnikowia | 13 | Saprotroph | \|Undefined Saprotroph\| | Yeast | NULL | Probable | As nectar-tap_saprotroph (Põlme et al. 2020); Decay sugar-rich substrates (Põlme et al. 2020) | Põlme S, et al. 2020. Fungal Diversity 105:1-16 ((https://doi.org/10.1007/s13225-020-00466-2)) |
| Otu47905 | 1 | k__Fungi(100);p__Ascomycota(100);c__Saccharomycetes(100);o__Saccharomycetales(100);f__Metschnikowiaceae(100);g__Metschnikowia(100);s__Metschnikowia_pulcherrima(100); | Metschnikowia | 13 | Saprotroph | \|Undefined Saprotroph\| | Yeast | NULL | Probable | As nectar-tap_saprotroph (Põlme et al. 2020); Decay sugar-rich substrates (Põlme et al. 2020) | Põlme S, et al. 2020. Fungal Diversity 105:1-16 ((https://doi.org/10.1007/s13225-020-00466-2)) |
| Otu47906 | 1 | k__Fungi(100);p__Ascomycota(100);c__Saccharomycetes(100);o__Saccharomycetales(100);f__Metschnikowiaceae(100);g__Metschnikowia(100);g__Metschnikowia_unclassified(100); | Metschnikowia | 13 | Saprotroph | \|Undefined Saprotroph\| | Yeast | NULL | Probable | As nectar-tap_saprotroph (Põlme et al. 2020); Decay sugar-rich substrates (Põlme et al. 2020) | Põlme S, et al. 2020. Fungal Diversity 105:1-16 ((https://doi.org/10.1007/s13225-020-00466-2)) |
| Otu47907 | 1 | k__Fungi(100);p__Ascomycota(100);c__Saccharomycetes(100);o__Saccharomycetales(100);f__Metschnikowiaceae(100);g__Metschnikowia(100);s__Metschnikowia_pulcherrima(100); | Metschnikowia | 13 | Saprotroph | \|Undefined Saprotroph\| | Yeast | NULL | Probable | As nectar-tap_saprotroph (Põlme et al. 2020); Decay sugar-rich substrates (Põlme et al. 2020) | Põlme S, et al. 2020. Fungal Diversity 105:1-16 ((https://doi.org/10.1007/s13225-020-00466-2)) |
| Otu47908 | 1 | k__Fungi(100);p__Ascomycota(100);c__Saccharomycetes(100);o__Saccharomycetales(100);f__Metschnikowiaceae(100);g__Metschnikowia(100);g__Metschnikowia_unclassified(100); | Metschnikowia | 13 | Saprotroph | \|Undefined Saprotroph\| | Yeast | NULL | Probable | As nectar-tap_saprotroph (Põlme et al. 2020); Decay sugar-rich substrates (Põlme et al. 2020) | Põlme S, et al. 2020. Fungal Diversity 105:1-16 ((https://doi.org/10.1007/s13225-020-00466-2)) |
| Otu47909 | 1 | k__Fungi(100);p__Ascomycota(100);c__Saccharomycetes(100);o__Saccharomycetales(100);f__Metschnikowiaceae(100);g__Metschnikowia(100);s__Metschnikowia_pulcherrima(100); | Metschnikowia | 13 | Saprotroph | \|Undefined Saprotroph\| | Yeast | NULL | Probable | As nectar-tap_saprotroph (Põlme et al. 2020); Decay sugar-rich substrates (Põlme et al. 2020) | Põlme S, et al. 2020. Fungal Diversity 105:1-16 ((https://doi.org/10.1007/s13225-020-00466-2)) |
| Otu47910 | 1 | k__Fungi(100);p__Ascomycota(100);c__Saccharomycetes(100);o__Saccharomycetales(100);f__Metschnikowiaceae(100);g__Metschnikowia(100);g__Metschnikowia_unclassified(100); | Metschnikowia | 13 | Saprotroph | \|Undefined Saprotroph\| | Yeast | NULL | Probable | As nectar-tap_saprotroph (Põlme et al. 2020); Decay sugar-rich substrates (Põlme et al. 2020) | Põlme S, et al. 2020. Fungal Diversity 105:1-16 ((https://doi.org/10.1007/s13225-020-00466-2)) |
| Otu47911 | 1 | k__Fungi(100);p__Ascomycota(100);c__Saccharomycetes(100);o__Saccharomycetales(100);f__Metschnikowiaceae(100);g__Metschnikowia(100);g__Metschnikowia_unclassified(100); | Metschnikowia | 13 | Saprotroph | \|Undefined Saprotroph\| | Yeast | NULL | Probable | As nectar-tap_saprotroph (Põlme et al. 2020); Decay sugar-rich substrates (Põlme et al. 2020) | Põlme S, et al. 2020. Fungal Diversity 105:1-16 ((https://doi.org/10.1007/s13225-020-00466-2)) |
| Otu47912 | 1 | k__Fungi(100);p__Ascomycota(100);c__Saccharomycetes(100);o__Saccharomycetales(100);f__Metschnikowiaceae(100);g__Metschnikowia(100);s__Metschnikowia_pulcherrima(100); | Metschnikowia | 13 | Saprotroph | \|Undefined Saprotroph\| | Yeast | NULL | Probable | As nectar-tap_saprotroph (Põlme et al. 2020); Decay sugar-rich substrates (Põlme et al. 2020) | Põlme S, et al. 2020. Fungal Diversity 105:1-16 ((https://doi.org/10.1007/s13225-020-00466-2)) |
| Otu47913 | 1 | k__Fungi(100);p__Ascomycota(100);c__Saccharomycetes(100);o__Saccharomycetales(100);f__Metschnikowiaceae(100);g__Metschnikowia(100);g__Metschnikowia_unclassified(100); | Metschnikowia | 13 | Saprotroph | \|Undefined Saprotroph\| | Yeast | NULL | Probable | As nectar-tap_saprotroph (Põlme et al. 2020); Decay sugar-rich substrates (Põlme et al. 2020) | Põlme S, et al. 2020. Fungal Diversity 105:1-16 ((https://doi.org/10.1007/s13225-020-00466-2)) |
| Otu47914 | 1 | k__Fungi(100);p__Ascomycota(100);c__Saccharomycetes(100);o__Saccharomycetales(100);f__Metschnikowiaceae(100);g__Metschnikowia(100);g__Metschnikowia_unclassified(100); | Metschnikowia | 13 | Saprotroph | \|Undefined Saprotroph\| | Yeast | NULL | Probable | As nectar-tap_saprotroph (Põlme et al. 2020); Decay sugar-rich substrates (Põlme et al. 2020) | Põlme S, et al. 2020. Fungal Diversity 105:1-16 ((https://doi.org/10.1007/s13225-020-00466-2)) |
| Otu47915 | 1 | k__Fungi(100);p__Ascomycota(100);c__Saccharomycetes(100);o__Saccharomycetales(100);f__Metschnikowiaceae(100);g__Metschnikowia(100);s__Metschnikowia_pulcherrima(100); | Metschnikowia | 13 | Saprotroph | \|Undefined Saprotroph\| | Yeast | NULL | Probable | As nectar-tap_saprotroph (Põlme et al. 2020); Decay sugar-rich substrates (Põlme et al. 2020) | Põlme S, et al. 2020. Fungal Diversity 105:1-16 ((https://doi.org/10.1007/s13225-020-00466-2)) |
| Otu47916 | 1 | k__Fungi(100);p__Ascomycota(100);c__Saccharomycetes(100);o__Saccharomycetales(100);f__Metschnikowiaceae(100);g__Metschnikowia(100);s__Metschnikowia_pulcherrima(100); | Metschnikowia | 13 | Saprotroph | \|Undefined Saprotroph\| | Yeast | NULL | Probable | As nectar-tap_saprotroph (Põlme et al. 2020); Decay sugar-rich substrates (Põlme et al. 2020) | Põlme S, et al. 2020. Fungal Diversity 105:1-16 ((https://doi.org/10.1007/s13225-020-00466-2)) |
| Otu47917 | 1 | k__Fungi(100);p__Ascomycota(100);c__Saccharomycetes(100);o__Saccharomycetales(100);f__Metschnikowiaceae(100);g__Metschnikowia(100);s__Metschnikowia_pulcherrima(100); | Metschnikowia | 13 | Saprotroph | \|Undefined Saprotroph\| | Yeast | NULL | Probable | As nectar-tap_saprotroph (Põlme et al. 2020); Decay sugar-rich substrates (Põlme et al. 2020) | Põlme S, et al. 2020. Fungal Diversity 105:1-16 ((https://doi.org/10.1007/s13225-020-00466-2)) |
| Otu47918 | 1 | k__Fungi(100);p__Ascomycota(100);c__Saccharomycetes(100);o__Saccharomycetales(100);f__Metschnikowiaceae(100);g__Metschnikowia(100);s__Metschnikowia_pulcherrima(100); | Metschnikowia | 13 | Saprotroph | \|Undefined Saprotroph\| | Yeast | NULL | Probable | As nectar-tap_saprotroph (Põlme et al. 2020); Decay sugar-rich substrates (Põlme et al. 2020) | Põlme S, et al. 2020. Fungal Diversity 105:1-16 ((https://doi.org/10.1007/s13225-020-00466-2)) |
| Otu47919 | 1 | k__Fungi(100);p__Ascomycota(100);c__Saccharomycetes(100);o__Saccharomycetales(100);f__Metschnikowiaceae(100);g__Metschnikowia(100);s__Metschnikowia_pulcherrima(100); | Metschnikowia | 13 | Saprotroph | \|Undefined Saprotroph\| | Yeast | NULL | Probable | As nectar-tap_saprotroph (Põlme et al. 2020); Decay sugar-rich substrates (Põlme et al. 2020) | Põlme S, et al. 2020. Fungal Diversity 105:1-16 ((https://doi.org/10.1007/s13225-020-00466-2)) |
| Otu47920 | 1 | k__Fungi(100);p__Ascomycota(100);c__Saccharomycetes(100);o__Saccharomycetales(100);f__Metschnikowiaceae(100);g__Metschnikowia(100);s__Metschnikowia_pulcherrima(100); | Metschnikowia | 13 | Saprotroph | \|Undefined Saprotroph\| | Yeast | NULL | Probable | As nectar-tap_saprotroph (Põlme et al. 2020); Decay sugar-rich substrates (Põlme et al. 2020) | Põlme S, et al. 2020. Fungal Diversity 105:1-16 ((https://doi.org/10.1007/s13225-020-00466-2)) |
| Otu47921 | 1 | k__Fungi(100);p__Ascomycota(100);c__Saccharomycetes(100);o__Saccharomycetales(100);f__Metschnikowiaceae(100);g__Metschnikowia(100);s__Metschnikowia_pulcherrima(100); | Metschnikowia | 13 | Saprotroph | \|Undefined Saprotroph\| | Yeast | NULL | Probable | As nectar-tap_saprotroph (Põlme et al. 2020); Decay sugar-rich substrates (Põlme et al. 2020) | Põlme S, et al. 2020. Fungal Diversity 105:1-16 ((https://doi.org/10.1007/s13225-020-00466-2)) |
| Otu47922 | 1 | k__Fungi(100);p__Ascomycota(100);c__Saccharomycetes(100);o__Saccharomycetales(100);f__Metschnikowiaceae(100);g__Metschnikowia(100);s__Metschnikowia_pulcherrima(100); | Metschnikowia | 13 | Saprotroph | \|Undefined Saprotroph\| | Yeast | NULL | Probable | As nectar-tap_saprotroph (Põlme et al. 2020); Decay sugar-rich substrates (Põlme et al. 2020) | Põlme S, et al. 2020. Fungal Diversity 105:1-16 ((https://doi.org/10.1007/s13225-020-00466-2)) |
| Otu47923 | 1 | k__Fungi(100);p__Ascomycota(100);c__Saccharomycetes(100);o__Saccharomycetales(100);f__Metschnikowiaceae(100);g__Metschnikowia(100);g__Metschnikowia_unclassified(100); | Metschnikowia | 13 | Saprotroph | \|Undefined Saprotroph\| | Yeast | NULL | Probable | As nectar-tap_saprotroph (Põlme et al. 2020); Decay sugar-rich substrates (Põlme et al. 2020) | Põlme S, et al. 2020. Fungal Diversity 105:1-16 ((https://doi.org/10.1007/s13225-020-00466-2)) |
| Otu47924 | 1 | k__Fungi(100);p__Ascomycota(100);c__Saccharomycetes(100);o__Saccharomycetales(100);f__Metschnikowiaceae(100);g__Metschnikowia(100);s__Metschnikowia_pulcherrima(100); | Metschnikowia | 13 | Saprotroph | \|Undefined Saprotroph\| | Yeast | NULL | Probable | As nectar-tap_saprotroph (Põlme et al. 2020); Decay sugar-rich substrates (Põlme et al. 2020) | Põlme S, et al. 2020. Fungal Diversity 105:1-16 ((https://doi.org/10.1007/s13225-020-00466-2)) |
| Otu47925 | 1 | k__Fungi(100);p__Ascomycota(100);c__Saccharomycetes(100);o__Saccharomycetales(100);f__Metschnikowiaceae(100);g__Metschnikowia(100);s__Metschnikowia_pulcherrima(100); | Metschnikowia | 13 | Saprotroph | \|Undefined Saprotroph\| | Yeast | NULL | Probable | As nectar-tap_saprotroph (Põlme et al. 2020); Decay sugar-rich substrates (Põlme et al. 2020) | Põlme S, et al. 2020. Fungal Diversity 105:1-16 ((https://doi.org/10.1007/s13225-020-00466-2)) |
| Otu47926 | 1 | k__Fungi(100);p__Ascomycota(100);c__Saccharomycetes(100);o__Saccharomycetales(100);f__Metschnikowiaceae(100);g__Metschnikowia(100);s__Metschnikowia_pulcherrima(100); | Metschnikowia | 13 | Saprotroph | \|Undefined Saprotroph\| | Yeast | NULL | Probable | As nectar-tap_saprotroph (Põlme et al. 2020); Decay sugar-rich substrates (Põlme et al. 2020) | Põlme S, et al. 2020. Fungal Diversity 105:1-16 ((https://doi.org/10.1007/s13225-020-00466-2)) |
| Otu47927 | 1 | k__Fungi(100);p__Ascomycota(100);c__Saccharomycetes(100);o__Saccharomycetales(100);f__Metschnikowiaceae(100);g__Metschnikowia(100);s__Metschnikowia_pulcherrima(100); | Metschnikowia | 13 | Saprotroph | \|Undefined Saprotroph\| | Yeast | NULL | Probable | As nectar-tap_saprotroph (Põlme et al. 2020); Decay sugar-rich substrates (Põlme et al. 2020) | Põlme S, et al. 2020. Fungal Diversity 105:1-16 ((https://doi.org/10.1007/s13225-020-00466-2)) |
| Otu47928 | 1 | k__Fungi(100);p__Ascomycota(100);c__Saccharomycetes(100);o__Saccharomycetales(100);f__Metschnikowiaceae(100);g__Metschnikowia(100);s__Metschnikowia_pulcherrima(100); | Metschnikowia | 13 | Saprotroph | \|Undefined Saprotroph\| | Yeast | NULL | Probable | As nectar-tap_saprotroph (Põlme et al. 2020); Decay sugar-rich substrates (Põlme et al. 2020) | Põlme S, et al. 2020. Fungal Diversity 105:1-16 ((https://doi.org/10.1007/s13225-020-00466-2)) |
| Otu47929 | 1 | k__Fungi(100);p__Ascomycota(100);c__Saccharomycetes(100);o__Saccharomycetales(100);f__Metschnikowiaceae(100);g__Metschnikowia(100);s__Metschnikowia_pulcherrima(100); | Metschnikowia | 13 | Saprotroph | \|Undefined Saprotroph\| | Yeast | NULL | Probable | As nectar-tap_saprotroph (Põlme et al. 2020); Decay sugar-rich substrates (Põlme et al. 2020) | Põlme S, et al. 2020. Fungal Diversity 105:1-16 ((https://doi.org/10.1007/s13225-020-00466-2)) |
| Otu47930 | 1 | k__Fungi(100);p__Ascomycota(100);c__Saccharomycetes(100);o__Saccharomycetales(100);f__Metschnikowiaceae(100);g__Metschnikowia(100);s__Metschnikowia_pulcherrima(100); | Metschnikowia | 13 | Saprotroph | \|Undefined Saprotroph\| | Yeast | NULL | Probable | As nectar-tap_saprotroph (Põlme et al. 2020); Decay sugar-rich substrates (Põlme et al. 2020) | Põlme S, et al. 2020. Fungal Diversity 105:1-16 ((https://doi.org/10.1007/s13225-020-00466-2)) |
| Otu47931 | 1 | k__Fungi(100);p__Ascomycota(100);c__Saccharomycetes(100);o__Saccharomycetales(100);f__Metschnikowiaceae(100);g__Metschnikowia(100);s__Metschnikowia_pulcherrima(100); | Metschnikowia | 13 | Saprotroph | \|Undefined Saprotroph\| | Yeast | NULL | Probable | As nectar-tap_saprotroph (Põlme et al. 2020); Decay sugar-rich substrates (Põlme et al. 2020) | Põlme S, et al. 2020. Fungal Diversity 105:1-16 ((https://doi.org/10.1007/s13225-020-00466-2)) |
| Otu47932 | 1 | k__Fungi(100);p__Ascomycota(100);c__Saccharomycetes(100);o__Saccharomycetales(100);f__Metschnikowiaceae(100);g__Metschnikowia(100);s__Metschnikowia_pulcherrima(100); | Metschnikowia | 13 | Saprotroph | \|Undefined Saprotroph\| | Yeast | NULL | Probable | As nectar-tap_saprotroph (Põlme et al. 2020); Decay sugar-rich substrates (Põlme et al. 2020) | Põlme S, et al. 2020. Fungal Diversity 105:1-16 ((https://doi.org/10.1007/s13225-020-00466-2)) |
| Otu47933 | 1 | k__Fungi(100);p__Ascomycota(100);c__Saccharomycetes(100);o__Saccharomycetales(100);f__Metschnikowiaceae(100);g__Metschnikowia(100);s__Metschnikowia_pulcherrima(100); | Metschnikowia | 13 | Saprotroph | \|Undefined Saprotroph\| | Yeast | NULL | Probable | As nectar-tap_saprotroph (Põlme et al. 2020); Decay sugar-rich substrates (Põlme et al. 2020) | Põlme S, et al. 2020. Fungal Diversity 105:1-16 ((https://doi.org/10.1007/s13225-020-00466-2)) |
| Otu47934 | 1 | k__Fungi(100);p__Ascomycota(100);c__Saccharomycetes(100);o__Saccharomycetales(100);f__Metschnikowiaceae(100);g__Metschnikowia(100);s__Metschnikowia_pulcherrima(100); | Metschnikowia | 13 | Saprotroph | \|Undefined Saprotroph\| | Yeast | NULL | Probable | As nectar-tap_saprotroph (Põlme et al. 2020); Decay sugar-rich substrates (Põlme et al. 2020) | Põlme S, et al. 2020. Fungal Diversity 105:1-16 ((https://doi.org/10.1007/s13225-020-00466-2)) |
| Otu47935 | 1 | k__Fungi(100);p__Ascomycota(100);c__Saccharomycetes(100);o__Saccharomycetales(100);f__Metschnikowiaceae(100);g__Metschnikowia(100);g__Metschnikowia_unclassified(100); | Metschnikowia | 13 | Saprotroph | \|Undefined Saprotroph\| | Yeast | NULL | Probable | As nectar-tap_saprotroph (Põlme et al. 2020); Decay sugar-rich substrates (Põlme et al. 2020) | Põlme S, et al. 2020. Fungal Diversity 105:1-16 ((https://doi.org/10.1007/s13225-020-00466-2)) |
| Otu47936 | 1 | k__Fungi(100);p__Ascomycota(100);c__Saccharomycetes(100);o__Saccharomycetales(100);f__Metschnikowiaceae(100);g__Metschnikowia(100);s__Metschnikowia_pulcherrima(100); | Metschnikowia | 13 | Saprotroph | \|Undefined Saprotroph\| | Yeast | NULL | Probable | As nectar-tap_saprotroph (Põlme et al. 2020); Decay sugar-rich substrates (Põlme et al. 2020) | Põlme S, et al. 2020. Fungal Diversity 105:1-16 ((https://doi.org/10.1007/s13225-020-00466-2)) |
| Otu47937 | 1 | k__Fungi(100);p__Ascomycota(100);c__Saccharomycetes(100);o__Saccharomycetales(100);f__Metschnikowiaceae(100);g__Metschnikowia(100);g__Metschnikowia_unclassified(100); | Metschnikowia | 13 | Saprotroph | \|Undefined Saprotroph\| | Yeast | NULL | Probable | As nectar-tap_saprotroph (Põlme et al. 2020); Decay sugar-rich substrates (Põlme et al. 2020) | Põlme S, et al. 2020. Fungal Diversity 105:1-16 ((https://doi.org/10.1007/s13225-020-00466-2)) |
| Otu47938 | 1 | k__Fungi(100);p__Ascomycota(100);c__Saccharomycetes(100);o__Saccharomycetales(100);f__Metschnikowiaceae(100);g__Metschnikowia(100);s__Metschnikowia_pulcherrima(100); | Metschnikowia | 13 | Saprotroph | \|Undefined Saprotroph\| | Yeast | NULL | Probable | As nectar-tap_saprotroph (Põlme et al. 2020); Decay sugar-rich substrates (Põlme et al. 2020) | Põlme S, et al. 2020. Fungal Diversity 105:1-16 ((https://doi.org/10.1007/s13225-020-00466-2)) |
| Otu47939 | 1 | k__Fungi(100);p__Ascomycota(100);c__Saccharomycetes(100);o__Saccharomycetales(100);f__Metschnikowiaceae(100);g__Metschnikowia(100);s__Metschnikowia_pulcherrima(100); | Metschnikowia | 13 | Saprotroph | \|Undefined Saprotroph\| | Yeast | NULL | Probable | As nectar-tap_saprotroph (Põlme et al. 2020); Decay sugar-rich substrates (Põlme et al. 2020) | Põlme S, et al. 2020. Fungal Diversity 105:1-16 ((https://doi.org/10.1007/s13225-020-00466-2)) |
| Otu47940 | 1 | k__Fungi(100);p__Ascomycota(100);c__Saccharomycetes(100);o__Saccharomycetales(100);f__Metschnikowiaceae(100);g__Metschnikowia(100);g__Metschnikowia_unclassified(100); | Metschnikowia | 13 | Saprotroph | \|Undefined Saprotroph\| | Yeast | NULL | Probable | As nectar-tap_saprotroph (Põlme et al. 2020); Decay sugar-rich substrates (Põlme et al. 2020) | Põlme S, et al. 2020. Fungal Diversity 105:1-16 ((https://doi.org/10.1007/s13225-020-00466-2)) |
| Otu47941 | 1 | k__Fungi(100);p__Ascomycota(100);c__Saccharomycetes(100);o__Saccharomycetales(100);f__Metschnikowiaceae(100);g__Metschnikowia(100);s__Metschnikowia_pulcherrima(100); | Metschnikowia | 13 | Saprotroph | \|Undefined Saprotroph\| | Yeast | NULL | Probable | As nectar-tap_saprotroph (Põlme et al. 2020); Decay sugar-rich substrates (Põlme et al. 2020) | Põlme S, et al. 2020. Fungal Diversity 105:1-16 ((https://doi.org/10.1007/s13225-020-00466-2)) |
| Otu47942 | 1 | k__Fungi(100);p__Ascomycota(100);c__Saccharomycetes(100);o__Saccharomycetales(100);f__Metschnikowiaceae(100);g__Metschnikowia(100);g__Metschnikowia_unclassified(100); | Metschnikowia | 13 | Saprotroph | \|Undefined Saprotroph\| | Yeast | NULL | Probable | As nectar-tap_saprotroph (Põlme et al. 2020); Decay sugar-rich substrates (Põlme et al. 2020) | Põlme S, et al. 2020. Fungal Diversity 105:1-16 ((https://doi.org/10.1007/s13225-020-00466-2)) |
| Otu47943 | 1 | k__Fungi(100);p__Ascomycota(100);c__Saccharomycetes(100);o__Saccharomycetales(100);f__Metschnikowiaceae(100);g__Metschnikowia(100);g__Metschnikowia_unclassified(100); | Metschnikowia | 13 | Saprotroph | \|Undefined Saprotroph\| | Yeast | NULL | Probable | As nectar-tap_saprotroph (Põlme et al. 2020); Decay sugar-rich substrates (Põlme et al. 2020) | Põlme S, et al. 2020. Fungal Diversity 105:1-16 ((https://doi.org/10.1007/s13225-020-00466-2)) |
| Otu47944 | 1 | k__Fungi(100);p__Ascomycota(100);c__Saccharomycetes(100);o__Saccharomycetales(100);f__Metschnikowiaceae(100);g__Metschnikowia(100);s__Metschnikowia_pulcherrima(100); | Metschnikowia | 13 | Saprotroph | \|Undefined Saprotroph\| | Yeast | NULL | Probable | As nectar-tap_saprotroph (Põlme et al. 2020); Decay sugar-rich substrates (Põlme et al. 2020) | Põlme S, et al. 2020. Fungal Diversity 105:1-16 ((https://doi.org/10.1007/s13225-020-00466-2)) |
| Otu47945 | 1 | k__Fungi(100);p__Ascomycota(100);c__Saccharomycetes(100);o__Saccharomycetales(100);f__Metschnikowiaceae(100);g__Metschnikowia(100);g__Metschnikowia_unclassified(100); | Metschnikowia | 13 | Saprotroph | \|Undefined Saprotroph\| | Yeast | NULL | Probable | As nectar-tap_saprotroph (Põlme et al. 2020); Decay sugar-rich substrates (Põlme et al. 2020) | Põlme S, et al. 2020. Fungal Diversity 105:1-16 ((https://doi.org/10.1007/s13225-020-00466-2)) |
| Otu47946 | 1 | k__Fungi(100);p__Ascomycota(100);c__Saccharomycetes(100);o__Saccharomycetales(100);f__Metschnikowiaceae(100);g__Metschnikowia(100);g__Metschnikowia_unclassified(100); | Metschnikowia | 13 | Saprotroph | \|Undefined Saprotroph\| | Yeast | NULL | Probable | As nectar-tap_saprotroph (Põlme et al. 2020); Decay sugar-rich substrates (Põlme et al. 2020) | Põlme S, et al. 2020. Fungal Diversity 105:1-16 ((https://doi.org/10.1007/s13225-020-00466-2)) |
| Otu47947 | 1 | k__Fungi(100);p__Ascomycota(100);c__Saccharomycetes(100);o__Saccharomycetales(100);f__Metschnikowiaceae(100);g__Metschnikowia(100);s__Metschnikowia_pulcherrima(100); | Metschnikowia | 13 | Saprotroph | \|Undefined Saprotroph\| | Yeast | NULL | Probable | As nectar-tap_saprotroph (Põlme et al. 2020); Decay sugar-rich substrates (Põlme et al. 2020) | Põlme S, et al. 2020. Fungal Diversity 105:1-16 ((https://doi.org/10.1007/s13225-020-00466-2)) |
| Otu47948 | 1 | k__Fungi(100);p__Ascomycota(100);c__Saccharomycetes(100);o__Saccharomycetales(100);f__Metschnikowiaceae(100);g__Metschnikowia(100);s__Metschnikowia_pulcherrima(100); | Metschnikowia | 13 | Saprotroph | \|Undefined Saprotroph\| | Yeast | NULL | Probable | As nectar-tap_saprotroph (Põlme et al. 2020); Decay sugar-rich substrates (Põlme et al. 2020) | Põlme S, et al. 2020. Fungal Diversity 105:1-16 ((https://doi.org/10.1007/s13225-020-00466-2)) |
| Otu47949 | 1 | k__Fungi(100);p__Ascomycota(100);c__Saccharomycetes(100);o__Saccharomycetales(100);f__Metschnikowiaceae(100);g__Metschnikowia(100);s__Metschnikowia_pulcherrima(100); | Metschnikowia | 13 | Saprotroph | \|Undefined Saprotroph\| | Yeast | NULL | Probable | As nectar-tap_saprotroph (Põlme et al. 2020); Decay sugar-rich substrates (Põlme et al. 2020) | Põlme S, et al. 2020. Fungal Diversity 105:1-16 ((https://doi.org/10.1007/s13225-020-00466-2)) |
| Otu47950 | 1 | k__Fungi(100);p__Ascomycota(100);c__Saccharomycetes(100);o__Saccharomycetales(100);f__Metschnikowiaceae(100);g__Metschnikowia(100);s__Metschnikowia_pulcherrima(100); | Metschnikowia | 13 | Saprotroph | \|Undefined Saprotroph\| | Yeast | NULL | Probable | As nectar-tap_saprotroph (Põlme et al. 2020); Decay sugar-rich substrates (Põlme et al. 2020) | Põlme S, et al. 2020. Fungal Diversity 105:1-16 ((https://doi.org/10.1007/s13225-020-00466-2)) |
| Otu47951 | 1 | k__Fungi(100);p__Ascomycota(100);c__Saccharomycetes(100);o__Saccharomycetales(100);f__Metschnikowiaceae(100);g__Metschnikowia(100);s__Metschnikowia_pulcherrima(100); | Metschnikowia | 13 | Saprotroph | \|Undefined Saprotroph\| | Yeast | NULL | Probable | As nectar-tap_saprotroph (Põlme et al. 2020); Decay sugar-rich substrates (Põlme et al. 2020) | Põlme S, et al. 2020. Fungal Diversity 105:1-16 ((https://doi.org/10.1007/s13225-020-00466-2)) |
| Otu47952 | 1 | k__Fungi(100);p__Ascomycota(100);c__Saccharomycetes(100);o__Saccharomycetales(100);f__Metschnikowiaceae(100);g__Metschnikowia(100);s__Metschnikowia_pulcherrima(100); | Metschnikowia | 13 | Saprotroph | \|Undefined Saprotroph\| | Yeast | NULL | Probable | As nectar-tap_saprotroph (Põlme et al. 2020); Decay sugar-rich substrates (Põlme et al. 2020) | Põlme S, et al. 2020. Fungal Diversity 105:1-16 ((https://doi.org/10.1007/s13225-020-00466-2)) |
| Otu47953 | 1 | k__Fungi(100);p__Ascomycota(100);c__Saccharomycetes(100);o__Saccharomycetales(100);f__Metschnikowiaceae(100);g__Metschnikowia(100);g__Metschnikowia_unclassified(100); | Metschnikowia | 13 | Saprotroph | \|Undefined Saprotroph\| | Yeast | NULL | Probable | As nectar-tap_saprotroph (Põlme et al. 2020); Decay sugar-rich substrates (Põlme et al. 2020) | Põlme S, et al. 2020. Fungal Diversity 105:1-16 ((https://doi.org/10.1007/s13225-020-00466-2)) |
| Otu47954 | 1 | k__Fungi(100);p__Ascomycota(100);c__Saccharomycetes(100);o__Saccharomycetales(100);f__Metschnikowiaceae(100);g__Metschnikowia(100);g__Metschnikowia_unclassified(100); | Metschnikowia | 13 | Saprotroph | \|Undefined Saprotroph\| | Yeast | NULL | Probable | As nectar-tap_saprotroph (Põlme et al. 2020); Decay sugar-rich substrates (Põlme et al. 2020) | Põlme S, et al. 2020. Fungal Diversity 105:1-16 ((https://doi.org/10.1007/s13225-020-00466-2)) |
| Otu47955 | 1 | k__Fungi(100);p__Ascomycota(100);c__Saccharomycetes(100);o__Saccharomycetales(100);f__Metschnikowiaceae(100);g__Metschnikowia(100);s__Metschnikowia_pulcherrima(100); | Metschnikowia | 13 | Saprotroph | \|Undefined Saprotroph\| | Yeast | NULL | Probable | As nectar-tap_saprotroph (Põlme et al. 2020); Decay sugar-rich substrates (Põlme et al. 2020) | Põlme S, et al. 2020. Fungal Diversity 105:1-16 ((https://doi.org/10.1007/s13225-020-00466-2)) |
| Otu47956 | 1 | k__Fungi(100);p__Ascomycota(100);c__Saccharomycetes(100);o__Saccharomycetales(100);f__Metschnikowiaceae(100);g__Metschnikowia(100);s__Metschnikowia_pulcherrima(100); | Metschnikowia | 13 | Saprotroph | \|Undefined Saprotroph\| | Yeast | NULL | Probable | As nectar-tap_saprotroph (Põlme et al. 2020); Decay sugar-rich substrates (Põlme et al. 2020) | Põlme S, et al. 2020. Fungal Diversity 105:1-16 ((https://doi.org/10.1007/s13225-020-00466-2)) |
| Otu47957 | 1 | k__Fungi(100);p__Ascomycota(100);c__Saccharomycetes(100);o__Saccharomycetales(100);f__Metschnikowiaceae(100);g__Metschnikowia(100);g__Metschnikowia_unclassified(100); | Metschnikowia | 13 | Saprotroph | \|Undefined Saprotroph\| | Yeast | NULL | Probable | As nectar-tap_saprotroph (Põlme et al. 2020); Decay sugar-rich substrates (Põlme et al. 2020) | Põlme S, et al. 2020. Fungal Diversity 105:1-16 ((https://doi.org/10.1007/s13225-020-00466-2)) |
| Otu47958 | 1 | k__Fungi(100);p__Ascomycota(100);c__Saccharomycetes(100);o__Saccharomycetales(100);f__Metschnikowiaceae(100);g__Metschnikowia(100);s__Metschnikowia_pulcherrima(100); | Metschnikowia | 13 | Saprotroph | \|Undefined Saprotroph\| | Yeast | NULL | Probable | As nectar-tap_saprotroph (Põlme et al. 2020); Decay sugar-rich substrates (Põlme et al. 2020) | Põlme S, et al. 2020. Fungal Diversity 105:1-16 ((https://doi.org/10.1007/s13225-020-00466-2)) |
| Otu47959 | 1 | k__Fungi(100);p__Ascomycota(100);c__Saccharomycetes(100);o__Saccharomycetales(100);f__Metschnikowiaceae(100);g__Metschnikowia(100);s__Metschnikowia_pulcherrima(100); | Metschnikowia | 13 | Saprotroph | \|Undefined Saprotroph\| | Yeast | NULL | Probable | As nectar-tap_saprotroph (Põlme et al. 2020); Decay sugar-rich substrates (Põlme et al. 2020) | Põlme S, et al. 2020. Fungal Diversity 105:1-16 ((https://doi.org/10.1007/s13225-020-00466-2)) |
| Otu47960 | 1 | k__Fungi(100);p__Ascomycota(100);c__Saccharomycetes(100);o__Saccharomycetales(100);f__Metschnikowiaceae(100);g__Metschnikowia(100);s__Metschnikowia_pulcherrima(100); | Metschnikowia | 13 | Saprotroph | \|Undefined Saprotroph\| | Yeast | NULL | Probable | As nectar-tap_saprotroph (Põlme et al. 2020); Decay sugar-rich substrates (Põlme et al. 2020) | Põlme S, et al. 2020. Fungal Diversity 105:1-16 ((https://doi.org/10.1007/s13225-020-00466-2)) |
| Otu47961 | 1 | k__Fungi(100);p__Ascomycota(100);c__Saccharomycetes(100);o__Saccharomycetales(100);f__Metschnikowiaceae(100);g__Metschnikowia(100);s__Metschnikowia_pulcherrima(100); | Metschnikowia | 13 | Saprotroph | \|Undefined Saprotroph\| | Yeast | NULL | Probable | As nectar-tap_saprotroph (Põlme et al. 2020); Decay sugar-rich substrates (Põlme et al. 2020) | Põlme S, et al. 2020. Fungal Diversity 105:1-16 ((https://doi.org/10.1007/s13225-020-00466-2)) |
| Otu47962 | 1 | k__Fungi(100);p__Ascomycota(100);c__Saccharomycetes(100);o__Saccharomycetales(100);f__Metschnikowiaceae(100);g__Metschnikowia(100);s__Metschnikowia_pulcherrima(100); | Metschnikowia | 13 | Saprotroph | \|Undefined Saprotroph\| | Yeast | NULL | Probable | As nectar-tap_saprotroph (Põlme et al. 2020); Decay sugar-rich substrates (Põlme et al. 2020) | Põlme S, et al. 2020. Fungal Diversity 105:1-16 ((https://doi.org/10.1007/s13225-020-00466-2)) |
| Otu47963 | 1 | k__Fungi(100);p__Ascomycota(100);c__Saccharomycetes(100);o__Saccharomycetales(100);f__Metschnikowiaceae(100);g__Metschnikowia(100);s__Metschnikowia_pulcherrima(100); | Metschnikowia | 13 | Saprotroph | \|Undefined Saprotroph\| | Yeast | NULL | Probable | As nectar-tap_saprotroph (Põlme et al. 2020); Decay sugar-rich substrates (Põlme et al. 2020) | Põlme S, et al. 2020. Fungal Diversity 105:1-16 ((https://doi.org/10.1007/s13225-020-00466-2)) |
| Otu47964 | 1 | k__Fungi(100);p__Ascomycota(100);c__Saccharomycetes(100);o__Saccharomycetales(100);f__Metschnikowiaceae(100);g__Metschnikowia(100);s__Metschnikowia_pulcherrima(100); | Metschnikowia | 13 | Saprotroph | \|Undefined Saprotroph\| | Yeast | NULL | Probable | As nectar-tap_saprotroph (Põlme et al. 2020); Decay sugar-rich substrates (Põlme et al. 2020) | Põlme S, et al. 2020. Fungal Diversity 105:1-16 ((https://doi.org/10.1007/s13225-020-00466-2)) |
| Otu47965 | 1 | k__Fungi(100);p__Ascomycota(100);c__Saccharomycetes(100);o__Saccharomycetales(100);f__Metschnikowiaceae(100);g__Metschnikowia(100);g__Metschnikowia_unclassified(100); | Metschnikowia | 13 | Saprotroph | \|Undefined Saprotroph\| | Yeast | NULL | Probable | As nectar-tap_saprotroph (Põlme et al. 2020); Decay sugar-rich substrates (Põlme et al. 2020) | Põlme S, et al. 2020. Fungal Diversity 105:1-16 ((https://doi.org/10.1007/s13225-020-00466-2)) |
| Otu47966 | 1 | k__Fungi(100);p__Ascomycota(100);c__Saccharomycetes(100);o__Saccharomycetales(100);f__Metschnikowiaceae(100);g__Metschnikowia(100);s__Metschnikowia_pulcherrima(100); | Metschnikowia | 13 | Saprotroph | \|Undefined Saprotroph\| | Yeast | NULL | Probable | As nectar-tap_saprotroph (Põlme et al. 2020); Decay sugar-rich substrates (Põlme et al. 2020) | Põlme S, et al. 2020. Fungal Diversity 105:1-16 ((https://doi.org/10.1007/s13225-020-00466-2)) |
| Otu47967 | 1 | k__Fungi(100);p__Ascomycota(100);c__Saccharomycetes(100);o__Saccharomycetales(100);f__Metschnikowiaceae(100);g__Metschnikowia(100);s__Metschnikowia_pulcherrima(100); | Metschnikowia | 13 | Saprotroph | \|Undefined Saprotroph\| | Yeast | NULL | Probable | As nectar-tap_saprotroph (Põlme et al. 2020); Decay sugar-rich substrates (Põlme et al. 2020) | Põlme S, et al. 2020. Fungal Diversity 105:1-16 ((https://doi.org/10.1007/s13225-020-00466-2)) |
| Otu47968 | 1 | k__Fungi(100);p__Ascomycota(100);c__Saccharomycetes(100);o__Saccharomycetales(100);f__Metschnikowiaceae(100);g__Metschnikowia(100);s__Metschnikowia_pulcherrima(100); | Metschnikowia | 13 | Saprotroph | \|Undefined Saprotroph\| | Yeast | NULL | Probable | As nectar-tap_saprotroph (Põlme et al. 2020); Decay sugar-rich substrates (Põlme et al. 2020) | Põlme S, et al. 2020. Fungal Diversity 105:1-16 ((https://doi.org/10.1007/s13225-020-00466-2)) |
| Otu47969 | 1 | k__Fungi(100);p__Ascomycota(100);c__Saccharomycetes(100);o__Saccharomycetales(100);f__Metschnikowiaceae(100);g__Metschnikowia(100);g__Metschnikowia_unclassified(100); | Metschnikowia | 13 | Saprotroph | \|Undefined Saprotroph\| | Yeast | NULL | Probable | As nectar-tap_saprotroph (Põlme et al. 2020); Decay sugar-rich substrates (Põlme et al. 2020) | Põlme S, et al. 2020. Fungal Diversity 105:1-16 ((https://doi.org/10.1007/s13225-020-00466-2)) |
| Otu47971 | 1 | k__Fungi(100);p__Ascomycota(100);c__Saccharomycetes(100);o__Saccharomycetales(100);f__Metschnikowiaceae(100);g__Metschnikowia(100);s__Metschnikowia_pulcherrima(100); | Metschnikowia | 13 | Saprotroph | \|Undefined Saprotroph\| | Yeast | NULL | Probable | As nectar-tap_saprotroph (Põlme et al. 2020); Decay sugar-rich substrates (Põlme et al. 2020) | Põlme S, et al. 2020. Fungal Diversity 105:1-16 ((https://doi.org/10.1007/s13225-020-00466-2)) |

| | | | | | | | | | | | |
|---|---|---|---|---|---|---|---|---|---|---|---|
| Otu47972 | 1 | k__Fungi(100);p__Ascomycota(100);c__Saccharomycetes(100);o__Saccharomycetales(100);f__Metschnikowiaceae(100);g__Metschnikowia(100);s__Metschnikowia_pulcherrima(100); | Metschnikowia | 13 | Saprotroph | \|Undefined Saprotroph\| | Yeast | NULL | Probable | As nectar-tap_saprotroph (Põlme et al. 2020); Decay sugar-rich substrates (Põlme et al. 2020) | Põlme S, et al. 2020. Fungal Diversity 105:1-16 ((https://doi.org/10.1007/s13225-020-00466-2)) |
| Otu47973 | 1 | k__Fungi(100);p__Ascomycota(100);c__Saccharomycetes(100);o__Saccharomycetales(100);f__Metschnikowiaceae(100);g__Metschnikowia(100);s__Metschnikowia_pulcherrima(100); | Metschnikowia | 13 | Saprotroph | \|Undefined Saprotroph\| | Yeast | NULL | Probable | As nectar-tap_saprotroph (Põlme et al. 2020); Decay sugar-rich substrates (Põlme et al. 2020) | Põlme S, et al. 2020. Fungal Diversity 105:1-16 ((https://doi.org/10.1007/s13225-020-00466-2)) |
| Otu47974 | 1 | k__Fungi(100);p__Ascomycota(100);c__Saccharomycetes(100);o__Saccharomycetales(100);f__Metschnikowiaceae(100);g__Metschnikowia(100);s__Metschnikowia_pulcherrima(100); | Metschnikowia | 13 | Saprotroph | \|Undefined Saprotroph\| | Yeast | NULL | Probable | As nectar-tap_saprotroph (Põlme et al. 2020); Decay sugar-rich substrates (Põlme et al. 2020) | Põlme S, et al. 2020. Fungal Diversity 105:1-16 ((https://doi.org/10.1007/s13225-020-00466-2)) |
| Otu47975 | 1 | k__Fungi(100);p__Ascomycota(100);c__Saccharomycetes(100);o__Saccharomycetales(100);f__Metschnikowiaceae(100);g__Metschnikowia(100);s__Metschnikowia_pulcherrima(100); | Metschnikowia | 13 | Saprotroph | \|Undefined Saprotroph\| | Yeast | NULL | Probable | As nectar-tap_saprotroph (Põlme et al. 2020); Decay sugar-rich substrates (Põlme et al. 2020) | Põlme S, et al. 2020. Fungal Diversity 105:1-16 ((https://doi.org/10.1007/s13225-020-00466-2)) |
| Otu47977 | 1 | k__Fungi(100);p__Ascomycota(100);c__Saccharomycetes(100);o__Saccharomycetales(100);f__Metschnikowiaceae(100);g__Metschnikowia(100);s__Metschnikowia_pulcherrima(100); | Metschnikowia | 13 | Saprotroph | \|Undefined Saprotroph\| | Yeast | NULL | Probable | As nectar-tap_saprotroph (Põlme et al. 2020); Decay sugar-rich substrates (Põlme et al. 2020) | Põlme S, et al. 2020. Fungal Diversity 105:1-16 ((https://doi.org/10.1007/s13225-020-00466-2)) |
| Otu47978 | 1 | k__Fungi(100);p__Ascomycota(100);c__Saccharomycetes(100);o__Saccharomycetales(100);f__Metschnikowiaceae(100);g__Metschnikowia(100);s__Metschnikowia_pulcherrima(100); | Metschnikowia | 13 | Saprotroph | \|Undefined Saprotroph\| | Yeast | NULL | Probable | As nectar-tap_saprotroph (Põlme et al. 2020); Decay sugar-rich substrates (Põlme et al. 2020) | Põlme S, et al. 2020. Fungal Diversity 105:1-16 ((https://doi.org/10.1007/s13225-020-00466-2)) |
| Otu47979 | 1 | k__Fungi(100);p__Ascomycota(100);c__Saccharomycetes(100);o__Saccharomycetales(100);f__Metschnikowiaceae(100);g__Metschnikowia(100);s__Metschnikowia_pulcherrima(100); | Metschnikowia | 13 | Saprotroph | \|Undefined Saprotroph\| | Yeast | NULL | Probable | As nectar-tap_saprotroph (Põlme et al. 2020); Decay sugar-rich substrates (Põlme et al. 2020) | Põlme S, et al. 2020. Fungal Diversity 105:1-16 ((https://doi.org/10.1007/s13225-020-00466-2)) |
| Otu47980 | 1 | k__Fungi(100);p__Ascomycota(100);c__Saccharomycetes(100);o__Saccharomycetales(100);f__Metschnikowiaceae(100);g__Metschnikowia(100);s__Metschnikowia_pulcherrima(100); | Metschnikowia | 13 | Saprotroph | \|Undefined Saprotroph\| | Yeast | NULL | Probable | As nectar-tap_saprotroph (Põlme et al. 2020); Decay sugar-rich substrates (Põlme et al. 2020) | Põlme S, et al. 2020. Fungal Diversity 105:1-16 ((https://doi.org/10.1007/s13225-020-00466-2)) |
| Otu47981 | 1 | k__Fungi(100);p__Ascomycota(100);c__Saccharomycetes(100);o__Saccharomycetales(100);f__Metschnikowiaceae(100);g__Metschnikowia(100);s__Metschnikowia_pulcherrima(100); | Metschnikowia | 13 | Saprotroph | \|Undefined Saprotroph\| | Yeast | NULL | Probable | As nectar-tap_saprotroph (Põlme et al. 2020); Decay sugar-rich substrates (Põlme et al. 2020) | Põlme S, et al. 2020. Fungal Diversity 105:1-16 ((https://doi.org/10.1007/s13225-020-00466-2)) |
| Otu47982 | 1 | k__Fungi(100);p__Ascomycota(100);c__Saccharomycetes(100);o__Saccharomycetales(100);f__Metschnikowiaceae(100);g__Metschnikowia(100);s__Metschnikowia_pulcherrima(100); | Metschnikowia | 13 | Saprotroph | \|Undefined Saprotroph\| | Yeast | NULL | Probable | As nectar-tap_saprotroph (Põlme et al. 2020); Decay sugar-rich substrates (Põlme et al. 2020) | Põlme S, et al. 2020. Fungal Diversity 105:1-16 ((https://doi.org/10.1007/s13225-020-00466-2)) |
| Otu47983 | 1 | k__Fungi(100);p__Ascomycota(100);c__Saccharomycetes(100);o__Saccharomycetales(100);f__Metschnikowiaceae(100);g__Metschnikowia(100);s__Metschnikowia_pulcherrima(100); | Metschnikowia | 13 | Saprotroph | \|Undefined Saprotroph\| | Yeast | NULL | Probable | As nectar-tap_saprotroph (Põlme et al. 2020); Decay sugar-rich substrates (Põlme et al. 2020) | Põlme S, et al. 2020. Fungal Diversity 105:1-16 ((https://doi.org/10.1007/s13225-020-00466-2)) |
| Otu47984 | 1 | k__Fungi(100);p__Ascomycota(100);c__Saccharomycetes(100);o__Saccharomycetales(100);f__Metschnikowiaceae(100);g__Metschnikowia(100);s__Metschnikowia_pulcherrima(100); | Metschnikowia | 13 | Saprotroph | \|Undefined Saprotroph\| | Yeast | NULL | Probable | As nectar-tap_saprotroph (Põlme et al. 2020); Decay sugar-rich substrates (Põlme et al. 2020) | Põlme S, et al. 2020. Fungal Diversity 105:1-16 ((https://doi.org/10.1007/s13225-020-00466-2)) |
| Otu47985 | 1 | k__Fungi(100);p__Ascomycota(100);c__Saccharomycetes(100);o__Saccharomycetales(100);f__Metschnikowiaceae(100);g__Metschnikowia(100);g__Metschnikowia_unclassified(100); | Metschnikowia | 13 | Saprotroph | \|Undefined Saprotroph\| | Yeast | NULL | Probable | As nectar-tap_saprotroph (Põlme et al. 2020); Decay sugar-rich substrates (Põlme et al. 2020) | Põlme S, et al. 2020. Fungal Diversity 105:1-16 ((https://doi.org/10.1007/s13225-020-00466-2)) |
| Otu47986 | 1 | k__Fungi(100);p__Ascomycota(100);c__Saccharomycetes(100);o__Saccharomycetales(100);f__Metschnikowiaceae(100);g__Metschnikowia(100);s__Metschnikowia_pulcherrima(100); | Metschnikowia | 13 | Saprotroph | \|Undefined Saprotroph\| | Yeast | NULL | Probable | As nectar-tap_saprotroph (Põlme et al. 2020); Decay sugar-rich substrates (Põlme et al. 2020) | Põlme S, et al. 2020. Fungal Diversity 105:1-16 ((https://doi.org/10.1007/s13225-020-00466-2)) |
| Otu47987 | 1 | k__Fungi(100);p__Ascomycota(100);c__Saccharomycetes(100);o__Saccharomycetales(100);f__Metschnikowiaceae(100);g__Metschnikowia(100);g__Metschnikowia_unclassified(100); | Metschnikowia | 13 | Saprotroph | \|Undefined Saprotroph\| | Yeast | NULL | Probable | As nectar-tap_saprotroph (Põlme et al. 2020); Decay sugar-rich substrates (Põlme et al. 2020) | Põlme S, et al. 2020. Fungal Diversity 105:1-16 ((https://doi.org/10.1007/s13225-020-00466-2)) |
| Otu47989 | 1 | k__Fungi(100);p__Ascomycota(100);c__Saccharomycetes(100);o__Saccharomycetales(100);f__Metschnikowiaceae(100);g__Metschnikowia(100);s__Metschnikowia_pulcherrima(100); | Metschnikowia | 13 | Saprotroph | \|Undefined Saprotroph\| | Yeast | NULL | Probable | As nectar-tap_saprotroph (Põlme et al. 2020); Decay sugar-rich substrates (Põlme et al. 2020) | Põlme S, et al. 2020. Fungal Diversity 105:1-16 ((https://doi.org/10.1007/s13225-020-00466-2)) |
| Otu47990 | 1 | k__Fungi(100);p__Ascomycota(100);c__Saccharomycetes(100);o__Saccharomycetales(100);f__Metschnikowiaceae(100);g__Metschnikowia(100);s__Metschnikowia_pulcherrima(100); | Metschnikowia | 13 | Saprotroph | \|Undefined Saprotroph\| | Yeast | NULL | Probable | As nectar-tap_saprotroph (Põlme et al. 2020); Decay sugar-rich substrates (Põlme et al. 2020) | Põlme S, et al. 2020. Fungal Diversity 105:1-16 ((https://doi.org/10.1007/s13225-020-00466-2)) |
| Otu47991 | 1 | k__Fungi(100);p__Ascomycota(100);c__Saccharomycetes(100);o__Saccharomycetales(100);f__Metschnikowiaceae(100);g__Metschnikowia(100);g__Metschnikowia_unclassified(100); | Metschnikowia | 13 | Saprotroph | \|Undefined Saprotroph\| | Yeast | NULL | Probable | As nectar-tap_saprotroph (Põlme et al. 2020); Decay sugar-rich substrates (Põlme et al. 2020) | Põlme S, et al. 2020. Fungal Diversity 105:1-16 ((https://doi.org/10.1007/s13225-020-00466-2)) |
| Otu47992 | 1 | k__Fungi(100);p__Ascomycota(100);c__Saccharomycetes(100);o__Saccharomycetales(100);f__Metschnikowiaceae(100);g__Metschnikowia(100);s__Metschnikowia_pulcherrima(100); | Metschnikowia | 13 | Saprotroph | \|Undefined Saprotroph\| | Yeast | NULL | Probable | As nectar-tap_saprotroph (Põlme et al. 2020); Decay sugar-rich substrates (Põlme et al. 2020) | Põlme S, et al. 2020. Fungal Diversity 105:1-16 ((https://doi.org/10.1007/s13225-020-00466-2)) |
| Otu47993 | 1 | k__Fungi(100);p__Ascomycota(100);c__Saccharomycetes(100);o__Saccharomycetales(100);f__Metschnikowiaceae(100);g__Metschnikowia(100);g__Metschnikowia_unclassified(100); | Metschnikowia | 13 | Saprotroph | \|Undefined Saprotroph\| | Yeast | NULL | Probable | As nectar-tap_saprotroph (Põlme et al. 2020); Decay sugar-rich substrates (Põlme et al. 2020) | Põlme S, et al. 2020. Fungal Diversity 105:1-16 ((https://doi.org/10.1007/s13225-020-00466-2)) |
| Otu47994 | 1 | k__Fungi(100);p__Ascomycota(100);c__Saccharomycetes(100);o__Saccharomycetales(100);f__Metschnikowiaceae(100);g__Metschnikowia(100);s__Metschnikowia_pulcherrima(100); | Metschnikowia | 13 | Saprotroph | \|Undefined Saprotroph\| | Yeast | NULL | Probable | As nectar-tap_saprotroph (Põlme et al. 2020); Decay sugar-rich substrates (Põlme et al. 2020) | Põlme S, et al. 2020. Fungal Diversity 105:1-16 ((https://doi.org/10.1007/s13225-020-00466-2)) |
| Otu47995 | 1 | k__Fungi(100);p__Ascomycota(100);c__Saccharomycetes(100);o__Saccharomycetales(100);f__Metschnikowiaceae(100);g__Metschnikowia(100);s__Metschnikowia_pulcherrima(100); | Metschnikowia | 13 | Saprotroph | \|Undefined Saprotroph\| | Yeast | NULL | Probable | As nectar-tap_saprotroph (Põlme et al. 2020); Decay sugar-rich substrates (Põlme et al. 2020) | Põlme S, et al. 2020. Fungal Diversity 105:1-16 ((https://doi.org/10.1007/s13225-020-00466-2)) |
| Otu47996 | 1 | k__Fungi(100);p__Ascomycota(100);c__Saccharomycetes(100);o__Saccharomycetales(100);f__Metschnikowiaceae(100);g__Metschnikowia(100);g__Metschnikowia_unclassified(100); | Metschnikowia | 13 | Saprotroph | \|Undefined Saprotroph\| | Yeast | NULL | Probable | As nectar-tap_saprotroph (Põlme et al. 2020); Decay sugar-rich substrates (Põlme et al. 2020) | Põlme S, et al. 2020. Fungal Diversity 105:1-16 ((https://doi.org/10.1007/s13225-020-00466-2)) |
| Otu47997 | 1 | k__Fungi(100);p__Ascomycota(100);c__Saccharomycetes(100);o__Saccharomycetales(100);f__Metschnikowiaceae(100);g__Metschnikowia(100);s__Metschnikowia_pulcherrima(100); | Metschnikowia | 13 | Saprotroph | \|Undefined Saprotroph\| | Yeast | NULL | Probable | As nectar-tap_saprotroph (Põlme et al. 2020); Decay sugar-rich substrates (Põlme et al. 2020) | Põlme S, et al. 2020. Fungal Diversity 105:1-16 ((https://doi.org/10.1007/s13225-020-00466-2)) |
| Otu47998 | 1 | k__Fungi(100);p__Ascomycota(100);c__Saccharomycetes(100);o__Saccharomycetales(100);f__Metschnikowiaceae(100);g__Metschnikowia(100);s__Metschnikowia_pulcherrima(100); | Metschnikowia | 13 | Saprotroph | \|Undefined Saprotroph\| | Yeast | NULL | Probable | As nectar-tap_saprotroph (Põlme et al. 2020); Decay sugar-rich substrates (Põlme et al. 2020) | Põlme S, et al. 2020. Fungal Diversity 105:1-16 ((https://doi.org/10.1007/s13225-020-00466-2)) |
| Otu47999 | 1 | k__Fungi(100);p__Ascomycota(100);c__Saccharomycetes(100);o__Saccharomycetales(100);f__Metschnikowiaceae(100);g__Metschnikowia(100);s__Metschnikowia_pulcherrima(100); | Metschnikowia | 13 | Saprotroph | \|Undefined Saprotroph\| | Yeast | NULL | Probable | As nectar-tap_saprotroph (Põlme et al. 2020); Decay sugar-rich substrates (Põlme et al. 2020) | Põlme S, et al. 2020. Fungal Diversity 105:1-16 ((https://doi.org/10.1007/s13225-020-00466-2)) |
| Otu48000 | 1 | k__Fungi(100);p__Ascomycota(100);c__Saccharomycetes(100);o__Saccharomycetales(100);f__Metschnikowiaceae(100);g__Metschnikowia(100);s__Metschnikowia_pulcherrima(100); | Metschnikowia | 13 | Saprotroph | \|Undefined Saprotroph\| | Yeast | NULL | Probable | As nectar-tap_saprotroph (Põlme et al. 2020); Decay sugar-rich substrates (Põlme et al. 2020) | Põlme S, et al. 2020. Fungal Diversity 105:1-16 ((https://doi.org/10.1007/s13225-020-00466-2)) |
| Otu48001 | 1 | k__Fungi(100);p__Ascomycota(100);c__Saccharomycetes(100);o__Saccharomycetales(100);f__Metschnikowiaceae(100);g__Metschnikowia(100);s__Metschnikowia_pulcherrima(100); | Metschnikowia | 13 | Saprotroph | \|Undefined Saprotroph\| | Yeast | NULL | Probable | As nectar-tap_saprotroph (Põlme et al. 2020); Decay sugar-rich substrates (Põlme et al. 2020) | Põlme S, et al. 2020. Fungal Diversity 105:1-16 ((https://doi.org/10.1007/s13225-020-00466-2)) |
| Otu48002 | 1 | k__Fungi(100);p__Ascomycota(100);c__Saccharomycetes(100);o__Saccharomycetales(100);f__Metschnikowiaceae(100);g__Metschnikowia(100);s__Metschnikowia_pulcherrima(100); | Metschnikowia | 13 | Saprotroph | \|Undefined Saprotroph\| | Yeast | NULL | Probable | As nectar-tap_saprotroph (Põlme et al. 2020); Decay sugar-rich substrates (Põlme et al. 2020) | Põlme S, et al. 2020. Fungal Diversity 105:1-16 ((https://doi.org/10.1007/s13225-020-00466-2)) |
| Otu48003 | 1 | k__Fungi(100);p__Ascomycota(100);c__Saccharomycetes(100);o__Saccharomycetales(100);f__Metschnikowiaceae(100);g__Metschnikowia(100);s__Metschnikowia_pulcherrima(100); | Metschnikowia | 13 | Saprotroph | \|Undefined Saprotroph\| | Yeast | NULL | Probable | As nectar-tap_saprotroph (Põlme et al. 2020); Decay sugar-rich substrates (Põlme et al. 2020) | Põlme S, et al. 2020. Fungal Diversity 105:1-16 ((https://doi.org/10.1007/s13225-020-00466-2)) |
| Otu48004 | 1 | k__Fungi(100);p__Ascomycota(100);c__Saccharomycetes(100);o__Saccharomycetales(100);f__Metschnikowiaceae(100);g__Metschnikowia(100);s__Metschnikowia_pulcherrima(100); | Metschnikowia | 13 | Saprotroph | \|Undefined Saprotroph\| | Yeast | NULL | Probable | As nectar-tap_saprotroph (Põlme et al. 2020); Decay sugar-rich substrates (Põlme et al. 2020) | Põlme S, et al. 2020. Fungal Diversity 105:1-16 ((https://doi.org/10.1007/s13225-020-00466-2)) |
| Otu48005 | 1 | k__Fungi(100);p__Ascomycota(100);c__Saccharomycetes(100);o__Saccharomycetales(100);f__Metschnikowiaceae(100);g__Metschnikowia(100);s__Metschnikowia_pulcherrima(100); | Metschnikowia | 13 | Saprotroph | \|Undefined Saprotroph\| | Yeast | NULL | Probable | As nectar-tap_saprotroph (Põlme et al. 2020); Decay sugar-rich substrates (Põlme et al. 2020) | Põlme S, et al. 2020. Fungal Diversity 105:1-16 ((https://doi.org/10.1007/s13225-020-00466-2)) |
| Otu48006 | 1 | k__Fungi(100);p__Ascomycota(100);c__Saccharomycetes(100);o__Saccharomycetales(100);f__Metschnikowiaceae(100);g__Metschnikowia(100);s__Metschnikowia_pulcherrima(100); | Metschnikowia | 13 | Saprotroph | \|Undefined Saprotroph\| | Yeast | NULL | Probable | As nectar-tap_saprotroph (Põlme et al. 2020); Decay sugar-rich substrates (Põlme et al. 2020) | Põlme S, et al. 2020. Fungal Diversity 105:1-16 ((https://doi.org/10.1007/s13225-020-00466-2)) |
| Otu48007 | 1 | k__Fungi(100);p__Ascomycota(100);c__Saccharomycetes(100);o__Saccharomycetales(100);f__Metschnikowiaceae(100);g__Metschnikowia(100);s__Metschnikowia_pulcherrima(100); | Metschnikowia | 13 | Saprotroph | \|Undefined Saprotroph\| | Yeast | NULL | Probable | As nectar-tap_saprotroph (Põlme et al. 2020); Decay sugar-rich substrates (Põlme et al. 2020) | Põlme S, et al. 2020. Fungal Diversity 105:1-16 ((https://doi.org/10.1007/s13225-020-00466-2)) |
| Otu48008 | 1 | k__Fungi(100);p__Ascomycota(100);c__Saccharomycetes(100);o__Saccharomycetales(100);f__Metschnikowiaceae(100);g__Metschnikowia(100);s__Metschnikowia_pulcherrima(100); | Metschnikowia | 13 | Saprotroph | \|Undefined Saprotroph\| | Yeast | NULL | Probable | As nectar-tap_saprotroph (Põlme et al. 2020); Decay sugar-rich substrates (Põlme et al. 2020) | Põlme S, et al. 2020. Fungal Diversity 105:1-16 ((https://doi.org/10.1007/s13225-020-00466-2)) |
| Otu48009 | 1 | k__Fungi(100);p__Ascomycota(100);c__Saccharomycetes(100);o__Saccharomycetales(100);f__Metschnikowiaceae(100);g__Metschnikowia(100);s__Metschnikowia_pulcherrima(100); | Metschnikowia | 13 | Saprotroph | \|Undefined Saprotroph\| | Yeast | NULL | Probable | As nectar-tap_saprotroph (Põlme et al. 2020); Decay sugar-rich substrates (Põlme et al. 2020) | Põlme S, et al. 2020. Fungal Diversity 105:1-16 ((https://doi.org/10.1007/s13225-020-00466-2)) |
| Otu48010 | 1 | k__Fungi(100);p__Ascomycota(100);c__Saccharomycetes(100);o__Saccharomycetales(100);f__Metschnikowiaceae(100);g__Metschnikowia(100);s__Metschnikowia_pulcherrima(100); | Metschnikowia | 13 | Saprotroph | \|Undefined Saprotroph\| | Yeast | NULL | Probable | As nectar-tap_saprotroph (Põlme et al. 2020); Decay sugar-rich substrates (Põlme et al. 2020) | Põlme S, et al. 2020. Fungal Diversity 105:1-16 ((https://doi.org/10.1007/s13225-020-00466-2)) |
| Otu48012 | 1 | k__Fungi(100);p__Ascomycota(100);c__Saccharomycetes(100);o__Saccharomycetales(100);f__Metschnikowiaceae(100);g__Metschnikowia(100);s__Metschnikowia_pulcherrima(100); | Metschnikowia | 13 | Saprotroph | \|Undefined Saprotroph\| | Yeast | NULL | Probable | As nectar-tap_saprotroph (Põlme et al. 2020); Decay sugar-rich substrates (Põlme et al. 2020) | Põlme S, et al. 2020. Fungal Diversity 105:1-16 ((https://doi.org/10.1007/s13225-020-00466-2)) |
| Otu48014 | 1 | k__Fungi(100);p__Ascomycota(100);c__Saccharomycetes(100);o__Saccharomycetales(100);f__Metschnikowiaceae(100);g__Metschnikowia(100);s__Metschnikowia_pulcherrima(100); | Metschnikowia | 13 | Saprotroph | \|Undefined Saprotroph\| | Yeast | NULL | Probable | As nectar-tap_saprotroph (Põlme et al. 2020); Decay sugar-rich substrates (Põlme et al. 2020) | Põlme S, et al. 2020. Fungal Diversity 105:1-16 ((https://doi.org/10.1007/s13225-020-00466-2)) |
| Otu48015 | 1 | k__Fungi(100);p__Ascomycota(100);c__Saccharomycetes(100);o__Saccharomycetales(100);f__Metschnikowiaceae(100);g__Metschnikowia(100);s__Metschnikowia_pulcherrima(100); | Metschnikowia | 13 | Saprotroph | \|Undefined Saprotroph\| | Yeast | NULL | Probable | As nectar-tap_saprotroph (Põlme et al. 2020); Decay sugar-rich substrates (Põlme et al. 2020) | Põlme S, et al. 2020. Fungal Diversity 105:1-16 ((https://doi.org/10.1007/s13225-020-00466-2)) |
| Otu48016 | 1 | k__Fungi(100);p__Ascomycota(100);c__Saccharomycetes(100);o__Saccharomycetales(100);f__Metschnikowiaceae(100);g__Metschnikowia(100);s__Metschnikowia_pulcherrima(100); | Metschnikowia | 13 | Saprotroph | \|Undefined Saprotroph\| | Yeast | NULL | Probable | As nectar-tap_saprotroph (Põlme et al. 2020); Decay sugar-rich substrates (Põlme et al. 2020) | Põlme S, et al. 2020. Fungal Diversity 105:1-16 ((https://doi.org/10.1007/s13225-020-00466-2)) |
| Otu48017 | 1 | k__Fungi(100);p__Ascomycota(100);c__Saccharomycetes(100);o__Saccharomycetales(100);f__Metschnikowiaceae(100);g__Metschnikowia(100);s__Metschnikowia_pulcherrima(100); | Metschnikowia | 13 | Saprotroph | \|Undefined Saprotroph\| | Yeast | NULL | Probable | As nectar-tap_saprotroph (Põlme et al. 2020); Decay sugar-rich substrates (Põlme et al. 2020) | Põlme S, et al. 2020. Fungal Diversity 105:1-16 ((https://doi.org/10.1007/s13225-020-00466-2)) |
| Otu48018 | 1 | k__Fungi(100);p__Ascomycota(100);c__Saccharomycetes(100);o__Saccharomycetales(100);f__Metschnikowiaceae(100);g__Metschnikowia(100);g__Metschnikowia_unclassified(100); | Metschnikowia | 13 | Saprotroph | \|Undefined Saprotroph\| | Yeast | NULL | Probable | As nectar-tap_saprotroph (Põlme et al. 2020); Decay sugar-rich substrates (Põlme et al. 2020) | Põlme S, et al. 2020. Fungal Diversity 105:1-16 ((https://doi.org/10.1007/s13225-020-00466-2)) |
| Otu48019 | 1 | k__Fungi(100);p__Ascomycota(100);c__Saccharomycetes(100);o__Saccharomycetales(100);f__Metschnikowiaceae(100);g__Metschnikowia(100);s__Metschnikowia_pulcherrima(100); | Metschnikowia | 13 | Saprotroph | \|Undefined Saprotroph\| | Yeast | NULL | Probable | As nectar-tap_saprotroph (Põlme et al. 2020); Decay sugar-rich substrates (Põlme et al. 2020) | Põlme S, et al. 2020. Fungal Diversity 105:1-16 ((https://doi.org/10.1007/s13225-020-00466-2)) |
| Otu48021 | 1 | k__Fungi(100);p__Ascomycota(100);c__Saccharomycetes(100);o__Saccharomycetales(100);f__Metschnikowiaceae(100);g__Metschnikowia(100);s__Metschnikowia_pulcherrima(100); | Metschnikowia | 13 | Saprotroph | \|Undefined Saprotroph\| | Yeast | NULL | Probable | As nectar-tap_saprotroph (Põlme et al. 2020); Decay sugar-rich substrates (Põlme et al. 2020) | Põlme S, et al. 2020. Fungal Diversity 105:1-16 ((https://doi.org/10.1007/s13225-020-00466-2)) |
| Otu48022 | 1 | k__Fungi(100);p__Ascomycota(100);c__Saccharomycetes(100);o__Saccharomycetales(100);f__Metschnikowiaceae(100);g__Metschnikowia(100);s__Metschnikowia_pulcherrima(100); | Metschnikowia | 13 | Saprotroph | \|Undefined Saprotroph\| | Yeast | NULL | Probable | As nectar-tap_saprotroph (Põlme et al. 2020); Decay sugar-rich substrates (Põlme et al. 2020) | Põlme S, et al. 2020. Fungal Diversity 105:1-16 ((https://doi.org/10.1007/s13225-020-00466-2)) |
| Otu48024 | 1 | k__Fungi(100);p__Ascomycota(100);c__Saccharomycetes(100);o__Saccharomycetales(100);f__Metschnikowiaceae(100);g__Metschnikowia(100);s__Metschnikowia_pulcherrima(100); | Metschnikowia | 13 | Saprotroph | \|Undefined Saprotroph\| | Yeast | NULL | Probable | As nectar-tap_saprotroph (Põlme et al. 2020); Decay sugar-rich substrates (Põlme et al. 2020) | Põlme S, et al. 2020. Fungal Diversity 105:1-16 ((https://doi.org/10.1007/s13225-020-00466-2)) |
| Otu48025 | 1 | k__Fungi(100);p__Ascomycota(100);c__Saccharomycetes(100);o__Saccharomycetales(100);f__Metschnikowiaceae(100);g__Metschnikowia(100);g__Metschnikowia_unclassified(100); | Metschnikowia | 13 | Saprotroph | \|Undefined Saprotroph\| | Yeast | NULL | Probable | As nectar-tap_saprotroph (Põlme et al. 2020); Decay sugar-rich substrates (Põlme et al. 2020) | Põlme S, et al. 2020. Fungal Diversity 105:1-16 ((https://doi.org/10.1007/s13225-020-00466-2)) |
| Otu48026 | 1 | k__Fungi(100);p__Ascomycota(100);c__Saccharomycetes(100);o__Saccharomycetales(100);f__Metschnikowiaceae(100);g__Metschnikowia(100);s__Metschnikowia_pulcherrima(100); | Metschnikowia | 13 | Saprotroph | \|Undefined Saprotroph\| | Yeast | NULL | Probable | As nectar-tap_saprotroph (Põlme et al. 2020); Decay sugar-rich substrates (Põlme et al. 2020) | Põlme S, et al. 2020. Fungal Diversity 105:1-16 ((https://doi.org/10.1007/s13225-020-00466-2)) |
| Otu48027 | 1 | k__Fungi(100);p__Ascomycota(100);c__Saccharomycetes(100);o__Saccharomycetales(100);f__Metschnikowiaceae(100);g__Metschnikowia(100);s__Metschnikowia_pulcherrima(100); | Metschnikowia | 13 | Saprotroph | \|Undefined Saprotroph\| | Yeast | NULL | Probable | As nectar-tap_saprotroph (Põlme et al. 2020); Decay sugar-rich substrates (Põlme et al. 2020) | Põlme S, et al. 2020. Fungal Diversity 105:1-16 ((https://doi.org/10.1007/s13225-020-00466-2)) |
| Otu48028 | 1 | k__Fungi(100);p__Ascomycota(100);c__Saccharomycetes(100);o__Saccharomycetales(100);f__Metschnikowiaceae(100);g__Metschnikowia(100);g__Metschnikowia_unclassified(100); | Metschnikowia | 13 | Saprotroph | \|Undefined Saprotroph\| | Yeast | NULL | Probable | As nectar-tap_saprotroph (Põlme et al. 2020); Decay sugar-rich substrates (Põlme et al. 2020) | Põlme S, et al. 2020. Fungal Diversity 105:1-16 ((https://doi.org/10.1007/s13225-020-00466-2)) |
| Otu48029 | 1 | k__Fungi(100);p__Ascomycota(100);c__Saccharomycetes(100);o__Saccharomycetales(100);f__Metschnikowiaceae(100);g__Metschnikowia(100);s__Metschnikowia_pulcherrima(100); | Metschnikowia | 13 | Saprotroph | \|Undefined Saprotroph\| | Yeast | NULL | Probable | As nectar-tap_saprotroph (Põlme et al. 2020); Decay sugar-rich substrates (Põlme et al. 2020) | Põlme S, et al. 2020. Fungal Diversity 105:1-16 ((https://doi.org/10.1007/s13225-020-00466-2)) |
| Otu48030 | 1 | k__Fungi(100);p__Ascomycota(100);c__Saccharomycetes(100);o__Saccharomycetales(100);f__Metschnikowiaceae(100);g__Metschnikowia(100);g__Metschnikowia_unclassified(100); | Metschnikowia | 13 | Saprotroph | \|Undefined Saprotroph\| | Yeast | NULL | Probable | As nectar-tap_saprotroph (Põlme et al. 2020); Decay sugar-rich substrates (Põlme et al. 2020) | Põlme S, et al. 2020. Fungal Diversity 105:1-16 ((https://doi.org/10.1007/s13225-020-00466-2)) |
| Otu48031 | 1 | k__Fungi(100);p__Ascomycota(100);c__Saccharomycetes(100);o__Saccharomycetales(100);f__Metschnikowiaceae(100);g__Metschnikowia(100);s__Metschnikowia_pulcherrima(100); | Metschnikowia | 13 | Saprotroph | \|Undefined Saprotroph\| | Yeast | NULL | Probable | As nectar-tap_saprotroph (Põlme et al. 2020); Decay sugar-rich substrates (Põlme et al. 2020) | Põlme S, et al. 2020. Fungal Diversity 105:1-16 ((https://doi.org/10.1007/s13225-020-00466-2)) |
| Otu48032 | 1 | k__Fungi(100);p__Ascomycota(100);c__Saccharomycetes(100);o__Saccharomycetales(100);f__Metschnikowiaceae(100);g__Metschnikowia(100);g__Metschnikowia_unclassified(100); | Metschnikowia | 13 | Saprotroph | \|Undefined Saprotroph\| | Yeast | NULL | Probable | As nectar-tap_saprotroph (Põlme et al. 2020); Decay sugar-rich substrates (Põlme et al. 2020) | Põlme S, et al. 2020. Fungal Diversity 105:1-16 ((https://doi.org/10.1007/s13225-020-00466-2)) |
| Otu48033 | 1 | k__Fungi(100);p__Ascomycota(100);c__Saccharomycetes(100);o__Saccharomycetales(100);f__Metschnikowiaceae(100);g__Metschnikowia(100);s__Metschnikowia_pulcherrima(100); | Metschnikowia | 13 | Saprotroph | \|Undefined Saprotroph\| | Yeast | NULL | Probable | As nectar-tap_saprotroph (Põlme et al. 2020); Decay sugar-rich substrates (Põlme et al. 2020) | Põlme S, et al. 2020. Fungal Diversity 105:1-16 ((https://doi.org/10.1007/s13225-020-00466-2)) |
| Otu48034 | 1 | k__Fungi(100);p__Ascomycota(100);c__Saccharomycetes(100);o__Saccharomycetales(100);f__Metschnikowiaceae(100);g__Metschnikowia(100);s__Metschnikowia_pulcherrima(100); | Metschnikowia | 13 | Saprotroph | \|Undefined Saprotroph\| | Yeast | NULL | Probable | As nectar-tap_saprotroph (Põlme et al. 2020); Decay sugar-rich substrates (Põlme et al. 2020) | Põlme S, et al. 2020. Fungal Diversity 105:1-16 ((https://doi.org/10.1007/s13225-020-00466-2)) |
| Otu48035 | 1 | k__Fungi(100);p__Ascomycota(100);c__Saccharomycetes(100);o__Saccharomycetales(100);f__Metschnikowiaceae(100);g__Metschnikowia(100);s__Metschnikowia_pulcherrima(100); | Metschnikowia | 13 | Saprotroph | \|Undefined Saprotroph\| | Yeast | NULL | Probable | As nectar-tap_saprotroph (Põlme et al. 2020); Decay sugar-rich substrates (Põlme et al. 2020) | Põlme S, et al. 2020. Fungal Diversity 105:1-16 ((https://doi.org/10.1007/s13225-020-00466-2)) |
| Otu48036 | 1 | k__Fungi(100);p__Ascomycota(100);c__Saccharomycetes(100);o__Saccharomycetales(100);f__Metschnikowiaceae(100);g__Metschnikowia(100);g__Metschnikowia_unclassified(100); | Metschnikowia | 13 | Saprotroph | \|Undefined Saprotroph\| | Yeast | NULL | Probable | As nectar-tap_saprotroph (Põlme et al. 2020); Decay sugar-rich substrates (Põlme et al. 2020) | Põlme S, et al. 2020. Fungal Diversity 105:1-16 ((https://doi.org/10.1007/s13225-020-00466-2)) |
| Otu48037 | 1 | k__Fungi(100);p__Ascomycota(100);c__Saccharomycetes(100);o__Saccharomycetales(100);f__Metschnikowiaceae(100);g__Metschnikowia(100);g__Metschnikowia_unclassified(100); | Metschnikowia | 13 | Saprotroph | \|Undefined Saprotroph\| | Yeast | NULL | Probable | As nectar-tap_saprotroph (Põlme et al. 2020); Decay sugar-rich substrates (Põlme et al. 2020) | Põlme S, et al. 2020. Fungal Diversity 105:1-16 ((https://doi.org/10.1007/s13225-020-00466-2)) |
| Otu48038 | 1 | k__Fungi(100);p__Ascomycota(100);c__Saccharomycetes(100);o__Saccharomycetales(100);f__Metschnikowiaceae(100);g__Metschnikowia(100);s__Metschnikowia_pulcherrima(100); | Metschnikowia | 13 | Saprotroph | \|Undefined Saprotroph\| | Yeast | NULL | Probable | As nectar-tap_saprotroph (Põlme et al. 2020); Decay sugar-rich substrates (Põlme et al. 2020) | Põlme S, et al. 2020. Fungal Diversity 105:1-16 ((https://doi.org/10.1007/s13225-020-00466-2)) |
| Otu48039 | 1 | k__Fungi(100);p__Ascomycota(100);c__Saccharomycetes(100);o__Saccharomycetales(100);f__Metschnikowiaceae(100);g__Metschnikowia(100);s__Metschnikowia_pulcherrima(100); | Metschnikowia | 13 | Saprotroph | \|Undefined Saprotroph\| | Yeast | NULL | Probable | As nectar-tap_saprotroph (Põlme et al. 2020); Decay sugar-rich substrates (Põlme et al. 2020) | Põlme S, et al. 2020. Fungal Diversity 105:1-16 ((https://doi.org/10.1007/s13225-020-00466-2)) |
| Otu48040 | 1 | k__Fungi(100);p__Ascomycota(100);c__Saccharomycetes(100);o__Saccharomycetales(100);f__Metschnikowiaceae(100);g__Metschnikowia(100);s__Metschnikowia_pulcherrima(100); | Metschnikowia | 13 | Saprotroph | \|Undefined Saprotroph\| | Yeast | NULL | Probable | As nectar-tap_saprotroph (Põlme et al. 2020); Decay sugar-rich substrates (Põlme et al. 2020) | Põlme S, et al. 2020. Fungal Diversity 105:1-16 ((https://doi.org/10.1007/s13225-020-00466-2)) |
| Otu48041 | 1 | k__Fungi(100);p__Ascomycota(100);c__Saccharomycetes(100);o__Saccharomycetales(100);f__Metschnikowiaceae(100);g__Metschnikowia(100);s__Metschnikowia_pulcherrima(100); | Metschnikowia | 13 | Saprotroph | \|Undefined Saprotroph\| | Yeast | NULL | Probable | As nectar-tap_saprotroph (Põlme et al. 2020); Decay sugar-rich substrates (Põlme et al. 2020) | Põlme S, et al. 2020. Fungal Diversity 105:1-16 ((https://doi.org/10.1007/s13225-020-00466-2)) |
| Otu48042 | 1 | k__Fungi(100);p__Ascomycota(100);c__Saccharomycetes(100);o__Saccharomycetales(100);f__Metschnikowiaceae(100);g__Metschnikowia(100);s__Metschnikowia_pulcherrima(100); | Metschnikowia | 13 | Saprotroph | \|Undefined Saprotroph\| | Yeast | NULL | Probable | As nectar-tap_saprotroph (Põlme et al. 2020); Decay sugar-rich substrates (Põlme et al. 2020) | Põlme S, et al. 2020. Fungal Diversity 105:1-16 ((https://doi.org/10.1007/s13225-020-00466-2)) |
| Otu48043 | 1 | k__Fungi(100);p__Ascomycota(100);c__Saccharomycetes(100);o__Saccharomycetales(100);f__Metschnikowiaceae(100);g__Metschnikowia(100);s__Metschnikowia_pulcherrima(100); | Metschnikowia | 13 | Saprotroph | \|Undefined Saprotroph\| | Yeast | NULL | Probable | As nectar-tap_saprotroph (Põlme et al. 2020); Decay sugar-rich substrates (Põlme et al. 2020) | Põlme S, et al. 2020. Fungal Diversity 105:1-16 ((https://doi.org/10.1007/s13225-020-00466-2)) |
| Otu48045 | 1 | k__Fungi(100);p__Ascomycota(100);c__Saccharomycetes(100);o__Saccharomycetales(100);f__Metschnikowiaceae(100);g__Metschnikowia(100);g__Metschnikowia_unclassified(100); | Metschnikowia | 13 | Saprotroph | \|Undefined Saprotroph\| | Yeast | NULL | Probable | As nectar-tap_saprotroph (Põlme et al. 2020); Decay sugar-rich substrates (Põlme et al. 2020) | Põlme S, et al. 2020. Fungal Diversity 105:1-16 ((https://doi.org/10.1007/s13225-020-00466-2)) |
| Otu48046 | 1 | k__Fungi(100);p__Ascomycota(100);c__Saccharomycetes(100);o__Saccharomycetales(100);f__Metschnikowiaceae(100);g__Metschnikowia(100);s__Metschnikowia_pulcherrima(100); | Metschnikowia | 13 | Saprotroph | \|Undefined Saprotroph\| | Yeast | NULL | Probable | As nectar-tap_saprotroph (Põlme et al. 2020); Decay sugar-rich substrates (Põlme et al. 2020) | Põlme S, et al. 2020. Fungal Diversity 105:1-16 ((https://doi.org/10.1007/s13225-020-00466-2)) |
| Otu48047 | 1 | k__Fungi(100);p__Ascomycota(100);c__Saccharomycetes(100);o__Saccharomycetales(100);f__Metschnikowiaceae(100);g__Metschnikowia(100);g__Metschnikowia_unclassified(100); | Metschnikowia | 13 | Saprotroph | \|Undefined Saprotroph\| | Yeast | NULL | Probable | As nectar-tap_saprotroph (Põlme et al. 2020); Decay sugar-rich substrates (Põlme et al. 2020) | Põlme S, et al. 2020. Fungal Diversity 105:1-16 ((https://doi.org/10.1007/s13225-020-00466-2)) |
| Otu48048 | 1 | k__Fungi(100);p__Ascomycota(100);c__Saccharomycetes(100);o__Saccharomycetales(100);f__Metschnikowiaceae(100);g__Metschnikowia(100);s__Metschnikowia_pulcherrima(100); | Metschnikowia | 13 | Saprotroph | \|Undefined Saprotroph\| | Yeast | NULL | Probable | As nectar-tap_saprotroph (Põlme et al. 2020); Decay sugar-rich substrates (Põlme et al. 2020) | Põlme S, et al. 2020. Fungal Diversity 105:1-16 ((https://doi.org/10.1007/s13225-020-00466-2)) |
| Otu48049 | 1 | k__Fungi(100);p__Ascomycota(100);c__Saccharomycetes(100);o__Saccharomycetales(100);f__Metschnikowiaceae(100);g__Metschnikowia(100);s__Metschnikowia_pulcherrima(100); | Metschnikowia | 13 | Saprotroph | \|Undefined Saprotroph\| | Yeast | NULL | Probable | As nectar-tap_saprotroph (Põlme et al. 2020); Decay sugar-rich substrates (Põlme et al. 2020) | Põlme S, et al. 2020. Fungal Diversity 105:1-16 ((https://doi.org/10.1007/s13225-020-00466-2)) |
| Otu48050 | 1 | k__Fungi(100);p__Ascomycota(100);c__Saccharomycetes(100);o__Saccharomycetales(100);f__Metschnikowiaceae(100);g__Metschnikowia(100);s__Metschnikowia_pulcherrima(100); | Metschnikowia | 13 | Saprotroph | \|Undefined Saprotroph\| | Yeast | NULL | Probable | As nectar-tap_saprotroph (Põlme et al. 2020); Decay sugar-rich substrates (Põlme et al. 2020) | Põlme S, et al. 2020. Fungal Diversity 105:1-16 ((https://doi.org/10.1007/s13225-020-00466-2)) |
| Otu48051 | 1 | k__Fungi(100);p__Ascomycota(100);c__Saccharomycetes(100);o__Saccharomycetales(100);f__Metschnikowiaceae(100);g__Metschnikowia(100);s__Metschnikowia_pulcherrima(100); | Metschnikowia | 13 | Saprotroph | \|Undefined Saprotroph\| | Yeast | NULL | Probable | As nectar-tap_saprotroph (Põlme et al. 2020); Decay sugar-rich substrates (Põlme et al. 2020) | Põlme S, et al. 2020. Fungal Diversity 105:1-16 ((https://doi.org/10.1007/s13225-020-00466-2)) |
| Otu48052 | 1 | k__Fungi(100);p__Ascomycota(100);c__Saccharomycetes(100);o__Saccharomycetales(100);f__Metschnikowiaceae(100);g__Metschnikowia(100);s__Metschnikowia_pulcherrima(100); | Metschnikowia | 13 | Saprotroph | \|Undefined Saprotroph\| | Yeast | NULL | Probable | As nectar-tap_saprotroph (Põlme et al. 2020); Decay sugar-rich substrates (Põlme et al. 2020) | Põlme S, et al. 2020. Fungal Diversity 105:1-16 ((https://doi.org/10.1007/s13225-020-00466-2)) |
| Otu48053 | 1 | k__Fungi(100);p__Ascomycota(100);c__Saccharomycetes(100);o__Saccharomycetales(100);f__Metschnikowiaceae(100);g__Metschnikowia(100);s__Metschnikowia_pulcherrima(100); | Metschnikowia | 13 | Saprotroph | \|Undefined Saprotroph\| | Yeast | NULL | Probable | As nectar-tap_saprotroph (Põlme et al. 2020); Decay sugar-rich substrates (Põlme et al. 2020) | Põlme S, et al. 2020. Fungal Diversity 105:1-16 ((https://doi.org/10.1007/s13225-020-00466-2)) |
| Otu48055 | 1 | k__Fungi(100);p__Ascomycota(100);c__Saccharomycetes(100);o__Saccharomycetales(100);f__Metschnikowiaceae(100);g__Metschnikowia(100);s__Metschnikowia_pulcherrima(100); | Metschnikowia | 13 | Saprotroph | \|Undefined Saprotroph\| | Yeast | NULL | Probable | As nectar-tap_saprotroph (Põlme et al. 2020); Decay sugar-rich substrates (Põlme et al. 2020) | Põlme S, et al. 2020. Fungal Diversity 105:1-16 ((https://doi.org/10.1007/s13225-020-00466-2)) |
| Otu48056 | 1 | k__Fungi(100);p__Ascomycota(100);c__Saccharomycetes(100);o__Saccharomycetales(100);f__Metschnikowiaceae(100);g__Metschnikowia(100);s__Metschnikowia_pulcherrima(100); | Metschnikowia | 13 | Saprotroph | \|Undefined Saprotroph\| | Yeast | NULL | Probable | As nectar-tap_saprotroph (Põlme et al. 2020); Decay sugar-rich substrates (Põlme et al. 2020) | Põlme S, et al. 2020. Fungal Diversity 105:1-16 ((https://doi.org/10.1007/s13225-020-00466-2)) |
| Otu48057 | 1 | k__Fungi(100);p__Ascomycota(100);c__Saccharomycetes(100);o__Saccharomycetales(100);f__Metschnikowiaceae(100);g__Metschnikowia(100);s__Metschnikowia_pulcherrima(100); | Metschnikowia | 13 | Saprotroph | \|Undefined Saprotroph\| | Yeast | NULL | Probable | As nectar-tap_saprotroph (Põlme et al. 2020); Decay sugar-rich substrates (Põlme et al. 2020) | Põlme S, et al. 2020. Fungal Diversity 105:1-16 ((https://doi.org/10.1007/s13225-020-00466-2)) |
| Otu48058 | 1 | k__Fungi(100);p__Ascomycota(100);c__Saccharomycetes(100);o__Saccharomycetales(100);f__Metschnikowiaceae(100);g__Metschnikowia(100);s__Metschnikowia_pulcherrima(100); | Metschnikowia | 13 | Saprotroph | \|Undefined Saprotroph\| | Yeast | NULL | Probable | As nectar-tap_saprotroph (Põlme et al. 2020); Decay sugar-rich substrates (Põlme et al. 2020) | Põlme S, et al. 2020. Fungal Diversity 105:1-16 ((https://doi.org/10.1007/s13225-020-00466-2)) |
| Otu48059 | 1 | k__Fungi(100);p__Ascomycota(100);c__Saccharomycetes(100);o__Saccharomycetales(100);f__Metschnikowiaceae(100);g__Metschnikowia(100);s__Metschnikowia_pulcherrima(100); | Metschnikowia | 13 | Saprotroph | \|Undefined Saprotroph\| | Yeast | NULL | Probable | As nectar-tap_saprotroph (Põlme et al. 2020); Decay sugar-rich substrates (Põlme et al. 2020) | Põlme S, et al. 2020. Fungal Diversity 105:1-16 ((https://doi.org/10.1007/s13225-020-00466-2)) |
| Otu48060 | 1 | k__Fungi(100);p__Ascomycota(100);c__Saccharomycetes(100);o__Saccharomycetales(100);f__Metschnikowiaceae(100);g__Metschnikowia(100);s__Metschnikowia_pulcherrima(100); | Metschnikowia | 13 | Saprotroph | \|Undefined Saprotroph\| | Yeast | NULL | Probable | As nectar-tap_saprotroph (Põlme et al. 2020); Decay sugar-rich substrates (Põlme et al. 2020) | Põlme S, et al. 2020. Fungal Diversity 105:1-16 ((https://doi.org/10.1007/s13225-020-00466-2)) |
| Otu48061 | 1 | k__Fungi(100);p__Ascomycota(100);c__Saccharomycetes(100);o__Saccharomycetales(100);f__Metschnikowiaceae(100);g__Metschnikowia(100);s__Metschnikowia_pulcherrima(100); | Metschnikowia | 13 | Saprotroph | \|Undefined Saprotroph\| | Yeast | NULL | Probable | As nectar-tap_saprotroph (Põlme et al. 2020); Decay sugar-rich substrates (Põlme et al. 2020) | Põlme S, et al. 2020. Fungal Diversity 105:1-16 ((https://doi.org/10.1007/s13225-020-00466-2)) |
| Otu48062 | 1 | k__Fungi(100);p__Ascomycota(100);c__Saccharomycetes(100);o__Saccharomycetales(100);f__Metschnikowiaceae(100);g__Metschnikowia(100);s__Metschnikowia_pulcherrima(100); | Metschnikowia | 13 | Saprotroph | \|Undefined Saprotroph\| | Yeast | NULL | Probable | As nectar-tap_saprotroph (Põlme et al. 2020); Decay sugar-rich substrates (Põlme et al. 2020) | Põlme S, et al. 2020. Fungal Diversity 105:1-16 ((https://doi.org/10.1007/s13225-020-00466-2)) |
| Otu48064 | 1 | k__Fungi(100);p__Ascomycota(100);c__Saccharomycetes(100);o__Saccharomycetales(100);f__Metschnikowiaceae(100);g__Metschnikowia(100);s__Metschnikowia_pulcherrima(100); | Metschnikowia | 13 | Saprotroph | \|Undefined Saprotroph\| | Yeast | NULL | Probable | As nectar-tap_saprotroph (Põlme et al. 2020); Decay sugar-rich substrates (Põlme et al. 2020) | Põlme S, et al. 2020. Fungal Diversity 105:1-16 ((https://doi.org/10.1007/s13225-020-00466-2)) |
| Otu48065 | 1 | k__Fungi(100);p__Ascomycota(100);c__Saccharomycetes(100);o__Saccharomycetales(100);f__Metschnikowiaceae(100);g__Metschnikowia(100);s__Metschnikowia_pulcherrima(100); | Metschnikowia | 13 | Saprotroph | \|Undefined Saprotroph\| | Yeast | NULL | Probable | As nectar-tap_saprotroph (Põlme et al. 2020); Decay sugar-rich substrates (Põlme et al. 2020) | Põlme S, et al. 2020. Fungal Diversity 105:1-16 ((https://doi.org/10.1007/s13225-020-00466-2)) |
| Otu48066 | 1 | k__Fungi(100);p__Ascomycota(100);c__Saccharomycetes(100);o__Saccharomycetales(100);f__Metschnikowiaceae(100);g__Metschnikowia(100);s__Metschnikowia_pulcherrima(100); | Metschnikowia | 13 | Saprotroph | \|Undefined Saprotroph\| | Yeast | NULL | Probable | As nectar-tap_saprotroph (Põlme et al. 2020); Decay sugar-rich substrates (Põlme et al. 2020) | Põlme S, et al. 2020. Fungal Diversity 105:1-16 ((https://doi.org/10.1007/s13225-020-00466-2)) |
| Otu48067 | 1 | k__Fungi(100);p__Ascomycota(100);c__Saccharomycetes(100);o__Saccharomycetales(100);f__Metschnikowiaceae(100);g__Metschnikowia(100);s__Metschnikowia_pulcherrima(100); | Metschnikowia | 13 | Saprotroph | \|Undefined Saprotroph\| | Yeast | NULL | Probable | As nectar-tap_saprotroph (Põlme et al. 2020); Decay sugar-rich substrates (Põlme et al. 2020) | Põlme S, et al. 2020. Fungal Diversity 105:1-16 ((https://doi.org/10.1007/s13225-020-00466-2)) |
| Otu48068 | 1 | k__Fungi(100);p__Ascomycota(100);c__Saccharomycetes(100);o__Saccharomycetales(100);f__Metschnikowiaceae(100);g__Metschnikowia(100);g__Metschnikowia_unclassified(100); | Metschnikowia | 13 | Saprotroph | \|Undefined Saprotroph\| | Yeast | NULL | Probable | As nectar-tap_saprotroph (Põlme et al. 2020); Decay sugar-rich substrates (Põlme et al. 2020) | Põlme S, et al. 2020. Fungal Diversity 105:1-16 ((https://doi.org/10.1007/s13225-020-00466-2)) |
| Otu48069 | 1 | k__Fungi(100);p__Ascomycota(100);c__Saccharomycetes(100);o__Saccharomycetales(100);f__Metschnikowiaceae(100);g__Metschnikowia(100);s__Metschnikowia_pulcherrima(100); | Metschnikowia | 13 | Saprotroph | \|Undefined Saprotroph\| | Yeast | NULL | Probable | As nectar-tap_saprotroph (Põlme et al. 2020); Decay sugar-rich substrates (Põlme et al. 2020) | Põlme S, et al. 2020. Fungal Diversity 105:1-16 ((https://doi.org/10.1007/s13225-020-00466-2)) |
| Otu48070 | 1 | k__Fungi(100);p__Ascomycota(100);c__Saccharomycetes(100);o__Saccharomycetales(100);f__Metschnikowiaceae(100);g__Metschnikowia(100);s__Metschnikowia_pulcherrima(100); | Metschnikowia | 13 | Saprotroph | \|Undefined Saprotroph\| | Yeast | NULL | Probable | As nectar-tap_saprotroph (Põlme et al. 2020); Decay sugar-rich substrates (Põlme et al. 2020) | Põlme S, et al. 2020. Fungal Diversity 105:1-16 ((https://doi.org/10.1007/s13225-020-00466-2)) |
| Otu48071 | 1 | k__Fungi(100);p__Ascomycota(100);c__Saccharomycetes(100);o__Saccharomycetales(100);f__Metschnikowiaceae(100);g__Metschnikowia(100);s__Metschnikowia_pulcherrima(100); | Metschnikowia | 13 | Saprotroph | \|Undefined Saprotroph\| | Yeast | NULL | Probable | As nectar-tap_saprotroph (Põlme et al. 2020); Decay sugar-rich substrates (Põlme et al. 2020) | Põlme S, et al. 2020. Fungal Diversity 105:1-16 ((https://doi.org/10.1007/s13225-020-00466-2)) |
| Otu48072 | 1 | k__Fungi(100);p__Ascomycota(100);c__Saccharomycetes(100);o__Saccharomycetales(100);f__Metschnikowiaceae(100);g__Metschnikowia(100);g__Metschnikowia_unclassified(100); | Metschnikowia | 13 | Saprotroph | \|Undefined Saprotroph\| | Yeast | NULL | Probable | As nectar-tap_saprotroph (Põlme et al. 2020); Decay sugar-rich substrates (Põlme et al. 2020) | Põlme S, et al. 2020. Fungal Diversity 105:1-16 ((https://doi.org/10.1007/s13225-020-00466-2)) |
| Otu48073 | 1 | k__Fungi(100);p__Ascomycota(100);c__Saccharomycetes(100);o__Saccharomycetales(100);f__Metschnikowiaceae(100);g__Metschnikowia(100);s__Metschnikowia_pulcherrima(100); | Metschnikowia | 13 | Saprotroph | \|Undefined Saprotroph\| | Yeast | NULL | Probable | As nectar-tap_saprotroph (Põlme et al. 2020); Decay sugar-rich substrates (Põlme et al. 2020) | Põlme S, et al. 2020. Fungal Diversity 105:1-16 ((https://doi.org/10.1007/s13225-020-00466-2)) |
| Otu48074 | 1 | k__Fungi(100);p__Ascomycota(100);c__Saccharomycetes(100);o__Saccharomycetales(100);f__Metschnikowiaceae(100);g__Metschnikowia(100);s__Metschnikowia_pulcherrima(100); | Metschnikowia | 13 | Saprotroph | \|Undefined Saprotroph\| | Yeast | NULL | Probable | As nectar-tap_saprotroph (Põlme et al. 2020); Decay sugar-rich substrates (Põlme et al. 2020) | Põlme S, et al. 2020. Fungal Diversity 105:1-16 ((https://doi.org/10.1007/s13225-020-00466-2)) |
| Otu48075 | 1 | k__Fungi(100);p__Ascomycota(100);c__Saccharomycetes(100);o__Saccharomycetales(100);f__Metschnikowiaceae(100);g__Metschnikowia(100);s__Metschnikowia_pulcherrima(100); | Metschnikowia | 13 | Saprotroph | \|Undefined Saprotroph\| | Yeast | NULL | Probable | As nectar-tap_saprotroph (Põlme et al. 2020); Decay sugar-rich substrates (Põlme et al. 2020) | Põlme S, et al. 2020. Fungal Diversity 105:1-16 ((https://doi.org/10.1007/s13225-020-00466-2)) |
| Otu48076 | 1 | k__Fungi(100);p__Ascomycota(100);c__Saccharomycetes(100);o__Saccharomycetales(100);f__Metschnikowiaceae(100);g__Metschnikowia(100);s__Metschnikowia_pulcherrima(100); | Metschnikowia | 13 | Saprotroph | \|Undefined Saprotroph\| | Yeast | NULL | Probable | As nectar-tap_saprotroph (Põlme et al. 2020); Decay sugar-rich substrates (Põlme et al. 2020) | Põlme S, et al. 2020. Fungal Diversity 105:1-16 ((https://doi.org/10.1007/s13225-020-00466-2)) |
| Otu48077 | 1 | k__Fungi(100);p__Ascomycota(100);c__Saccharomycetes(100);o__Saccharomycetales(100);f__Metschnikowiaceae(100);g__Metschnikowia(100);s__Metschnikowia_pulcherrima(100); | Metschnikowia | 13 | Saprotroph | \|Undefined Saprotroph\| | Yeast | NULL | Probable | As nectar-tap_saprotroph (Põlme et al. 2020); Decay sugar-rich substrates (Põlme et al. 2020) | Põlme S, et al. 2020. Fungal Diversity 105:1-16 ((https://doi.org/10.1007/s13225-020-00466-2)) |
| Otu48078 | 1 | k__Fungi(100);p__Ascomycota(100);c__Saccharomycetes(100);o__Saccharomycetales(100);f__Metschnikowiaceae(100);g__Metschnikowia(100);s__Metschnikowia_pulcherrima(100); | Metschnikowia | 13 | Saprotroph | \|Undefined Saprotroph\| | Yeast | NULL | Probable | As nectar-tap_saprotroph (Põlme et al. 2020); Decay sugar-rich substrates (Põlme et al. 2020) | Põlme S, et al. 2020. Fungal Diversity 105:1-16 ((https://doi.org/10.1007/s13225-020-00466-2)) |
| Otu48079 | 1 | k__Fungi(100);p__Ascomycota(100);c__Saccharomycetes(100);o__Saccharomycetales(100);f__Metschnikowiaceae(100);g__Metschnikowia(100);s__Metschnikowia_pulcherrima(100); | Metschnikowia | 13 | Saprotroph | \|Undefined Saprotroph\| | Yeast | NULL | Probable | As nectar-tap_saprotroph (Põlme et al. 2020); Decay sugar-rich substrates (Põlme et al. 2020) | Põlme S, et al. 2020. Fungal Diversity 105:1-16 ((https://doi.org/10.1007/s13225-020-00466-2)) |
| Otu48080 | 1 | k__Fungi(100);p__Ascomycota(100);c__Saccharomycetes(100);o__Saccharomycetales(100);f__Metschnikowiaceae(100);g__Metschnikowia(100);s__Metschnikowia_pulcherrima(100); | Metschnikowia | 13 | Saprotroph | \|Undefined Saprotroph\| | Yeast | NULL | Probable | As nectar-tap_saprotroph (Põlme et al. 2020); Decay sugar-rich substrates (Põlme et al. 2020) | Põlme S, et al. 2020. Fungal Diversity 105:1-16 ((https://doi.org/10.1007/s13225-020-00466-2)) |
| Otu48081 | 1 | k__Fungi(100);p__Ascomycota(100);c__Saccharomycetes(100);o__Saccharomycetales(100);f__Metschnikowiaceae(100);g__Metschnikowia(100);s__Metschnikowia_pulcherrima(100); | Metschnikowia | 13 | Saprotroph | \|Undefined Saprotroph\| | Yeast | NULL | Probable | As nectar-tap_saprotroph (Põlme et al. 2020); Decay sugar-rich substrates (Põlme et al. 2020) | Põlme S, et al. 2020. Fungal Diversity 105:1-16 ((https://doi.org/10.1007/s13225-020-00466-2)) |
| Otu48082 | 1 | k__Fungi(100);p__Ascomycota(100);c__Saccharomycetes(100);o__Saccharomycetales(100);f__Metschnikowiaceae(100);g__Metschnikowia(100);s__Metschnikowia_pulcherrima(100); | Metschnikowia | 13 | Saprotroph | \|Undefined Saprotroph\| | Yeast | NULL | Probable | As nectar-tap_saprotroph (Põlme et al. 2020); Decay sugar-rich substrates (Põlme et al. 2020) | Põlme S, et al. 2020. Fungal Diversity 105:1-16 ((https://doi.org/10.1007/s13225-020-00466-2)) |
| Otu48083 | 1 | k__Fungi(100);p__Ascomycota(100);c__Saccharomycetes(100);o__Saccharomycetales(100);f__Metschnikowiaceae(100);g__Metschnikowia(100);g__Metschnikowia_unclassified(100); | Metschnikowia | 13 | Saprotroph | \|Undefined Saprotroph\| | Yeast | NULL | Probable | As nectar-tap_saprotroph (Põlme et al. 2020); Decay sugar-rich substrates (Põlme et al. 2020) | Põlme S, et al. 2020. Fungal Diversity 105:1-16 ((https://doi.org/10.1007/s13225-020-00466-2)) |
| Otu48084 | 1 | k__Fungi(100);p__Ascomycota(100);c__Saccharomycetes(100);o__Saccharomycetales(100);f__Metschnikowiaceae(100);g__Metschnikowia(100);s__Metschnikowia_pulcherrima(100); | Metschnikowia | 13 | Saprotroph | \|Undefined Saprotroph\| | Yeast | NULL | Probable | As nectar-tap_saprotroph (Põlme et al. 2020); Decay sugar-rich substrates (Põlme et al. 2020) | Põlme S, et al. 2020. Fungal Diversity 105:1-16 ((https://doi.org/10.1007/s13225-020-00466-2)) |
| Otu48085 | 1 | k__Fungi(100);p__Ascomycota(100);c__Saccharomycetes(100);o__Saccharomycetales(100);f__Metschnikowiaceae(100);g__Metschnikowia(100);s__Metschnikowia_pulcherrima(100); | Metschnikowia | 13 | Saprotroph | \|Undefined Saprotroph\| | Yeast | NULL | Probable | As nectar-tap_saprotroph (Põlme et al. 2020); Decay sugar-rich substrates (Põlme et al. 2020) | Põlme S, et al. 2020. Fungal Diversity 105:1-16 ((https://doi.org/10.1007/s13225-020-00466-2)) |
| Otu48086 | 1 | k__Fungi(100);p__Ascomycota(100);c__Saccharomycetes(100);o__Saccharomycetales(100);f__Metschnikowiaceae(100);g__Metschnikowia(100);s__Metschnikowia_pulcherrima(100); | Metschnikowia | 13 | Saprotroph | \|Undefined Saprotroph\| | Yeast | NULL | Probable | As nectar-tap_saprotroph (Põlme et al. 2020); Decay sugar-rich substrates (Põlme et al. 2020) | Põlme S, et al. 2020. Fungal Diversity 105:1-16 ((https://doi.org/10.1007/s13225-020-00466-2)) |
| Otu48087 | 1 | k__Fungi(100);p__Ascomycota(100);c__Saccharomycetes(100);o__Saccharomycetales(100);f__Metschnikowiaceae(100);g__Metschnikowia(100);s__Metschnikowia_pulcherrima(100); | Metschnikowia | 13 | Saprotroph | \|Undefined Saprotroph\| | Yeast | NULL | Probable | As nectar-tap_saprotroph (Põlme et al. 2020); Decay sugar-rich substrates (Põlme et al. 2020) | Põlme S, et al. 2020. Fungal Diversity 105:1-16 ((https://doi.org/10.1007/s13225-020-00466-2)) |
| Otu48088 | 1 | k__Fungi(100);p__Ascomycota(100);c__Saccharomycetes(100);o__Saccharomycetales(100);f__Metschnikowiaceae(100);g__Metschnikowia(100);g__Metschnikowia_unclassified(100); | Metschnikowia | 13 | Saprotroph | \|Undefined Saprotroph\| | Yeast | NULL | Probable | As nectar-tap_saprotroph (Põlme et al. 2020); Decay sugar-rich substrates (Põlme et al. 2020) | Põlme S, et al. 2020. Fungal Diversity 105:1-16 ((https://doi.org/10.1007/s13225-020-00466-2)) |
| Otu48089 | 1 | k__Fungi(100);p__Ascomycota(100);c__Saccharomycetes(100);o__Saccharomycetales(100);f__Metschnikowiaceae(100);g__Metschnikowia(100);s__Metschnikowia_pulcherrima(100); | Metschnikowia | 13 | Saprotroph | \|Undefined Saprotroph\| | Yeast | NULL | Probable | As nectar-tap_saprotroph (Põlme et al. 2020); Decay sugar-rich substrates (Põlme et al. 2020) | Põlme S, et al. 2020. Fungal Diversity 105:1-16 ((https://doi.org/10.1007/s13225-020-00466-2)) |
| Otu48090 | 1 | k__Fungi(100);p__Ascomycota(100);c__Saccharomycetes(100);o__Saccharomycetales(100);f__Metschnikowiaceae(100);g__Metschnikowia(100);s__Metschnikowia_pulcherrima(100); | Metschnikowia | 13 | Saprotroph | \|Undefined Saprotroph\| | Yeast | NULL | Probable | As nectar-tap_saprotroph (Põlme et al. 2020); Decay sugar-rich substrates (Põlme et al. 2020) | Põlme S, et al. 2020. Fungal Diversity 105:1-16 ((https://doi.org/10.1007/s13225-020-00466-2)) |
| Otu48091 | 1 | k__Fungi(100);p__Ascomycota(100);c__Saccharomycetes(100);o__Saccharomycetales(100);f__Metschnikowiaceae(100);g__Metschnikowia(100);g__Metschnikowia_unclassified(100); | Metschnikowia | 13 | Saprotroph | \|Undefined Saprotroph\| | Yeast | NULL | Probable | As nectar-tap_saprotroph (Põlme et al. 2020); Decay sugar-rich substrates (Põlme et al. 2020) | Põlme S, et al. 2020. Fungal Diversity 105:1-16 ((https://doi.org/10.1007/s13225-020-00466-2)) |
| Otu48092 | 1 | k__Fungi(100);p__Ascomycota(100);c__Saccharomycetes(100);o__Saccharomycetales(100);f__Metschnikowiaceae(100);g__Metschnikowia(100);g__Metschnikowia_unclassified(100); | Metschnikowia | 13 | Saprotroph | \|Undefined Saprotroph\| | Yeast | NULL | Probable | As nectar-tap_saprotroph (Põlme et al. 2020); Decay sugar-rich substrates (Põlme et al. 2020) | Põlme S, et al. 2020. Fungal Diversity 105:1-16 ((https://doi.org/10.1007/s13225-020-00466-2)) |
| Otu48093 | 1 | k__Fungi(100);p__Ascomycota(100);c__Saccharomycetes(100);o__Saccharomycetales(100);f__Metschnikowiaceae(100);g__Metschnikowia(100);s__Metschnikowia_pulcherrima(100); | Metschnikowia | 13 | Saprotroph | \|Undefined Saprotroph\| | Yeast | NULL | Probable | As nectar-tap_saprotroph (Põlme et al. 2020); Decay sugar-rich substrates (Põlme et al. 2020) | Põlme S, et al. 2020. Fungal Diversity 105:1-16 ((https://doi.org/10.1007/s13225-020-00466-2)) |
| Otu48095 | 1 | k__Fungi(100);p__Ascomycota(100);c__Saccharomycetes(100);o__Saccharomycetales(100);f__Metschnikowiaceae(100);g__Metschnikowia(100);s__Metschnikowia_pulcherrima(100); | Metschnikowia | 13 | Saprotroph | \|Undefined Saprotroph\| | Yeast | NULL | Probable | As nectar-tap_saprotroph (Põlme et al. 2020); Decay sugar-rich substrates (Põlme et al. 2020) | Põlme S, et al. 2020. Fungal Diversity 105:1-16 ((https://doi.org/10.1007/s13225-020-00466-2)) |
| Otu48096 | 1 | k__Fungi(100);p__Ascomycota(100);c__Saccharomycetes(100);o__Saccharomycetales(100);f__Metschnikowiaceae(100);g__Metschnikowia(100);s__Metschnikowia_pulcherrima(100); | Metschnikowia | 13 | Saprotroph | \|Undefined Saprotroph\| | Yeast | NULL | Probable | As nectar-tap_saprotroph (Põlme et al. 2020); Decay sugar-rich substrates (Põlme et al. 2020) | Põlme S, et al. 2020. Fungal Diversity 105:1-16 ((https://doi.org/10.1007/s13225-020-00466-2)) |
| Otu48097 | 1 | k__Fungi(100);p__Ascomycota(100);c__Saccharomycetes(100);o__Saccharomycetales(100);f__Metschnikowiaceae(100);g__Metschnikowia(100);s__Metschnikowia_pulcherrima(100); | Metschnikowia | 13 | Saprotroph | \|Undefined Saprotroph\| | Yeast | NULL | Probable | As nectar-tap_saprotroph (Põlme et al. 2020); Decay sugar-rich substrates (Põlme et al. 2020) | Põlme S, et al. 2020. Fungal Diversity 105:1-16 ((https://doi.org/10.1007/s13225-020-00466-2)) |
| Otu48098 | 1 | k__Fungi(100);p__Ascomycota(100);c__Saccharomycetes(100);o__Saccharomycetales(100);f__Metschnikowiaceae(100);g__Metschnikowia(100);s__Metschnikowia_pulcherrima(100); | Metschnikowia | 13 | Saprotroph | \|Undefined Saprotroph\| | Yeast | NULL | Probable | As nectar-tap_saprotroph (Põlme et al. 2020); Decay sugar-rich substrates (Põlme et al. 2020) | Põlme S, et al. 2020. Fungal Diversity 105:1-16 ((https://doi.org/10.1007/s13225-020-00466-2)) |
| Otu48099 | 1 | k__Fungi(100);p__Ascomycota(100);c__Saccharomycetes(100);o__Saccharomycetales(100);f__Metschnikowiaceae(100);g__Metschnikowia(100);g__Metschnikowia_unclassified(100); | Metschnikowia | 13 | Saprotroph | \|Undefined Saprotroph\| | Yeast | NULL | Probable | As nectar-tap_saprotroph (Põlme et al. 2020); Decay sugar-rich substrates (Põlme et al. 2020) | Põlme S, et al. 2020. Fungal Diversity 105:1-16 ((https://doi.org/10.1007/s13225-020-00466-2)) |
| Otu48100 | 1 | k__Fungi(100);p__Ascomycota(100);c__Saccharomycetes(100);o__Saccharomycetales(100);f__Metschnikowiaceae(100);g__Metschnikowia(100);g__Metschnikowia_unclassified(100); | Metschnikowia | 13 | Saprotroph | \|Undefined Saprotroph\| | Yeast | NULL | Probable | As nectar-tap_saprotroph (Põlme et al. 2020); Decay sugar-rich substrates (Põlme et al. 2020) | Põlme S, et al. 2020. Fungal Diversity 105:1-16 ((https://doi.org/10.1007/s13225-020-00466-2)) |
| Otu48101 | 1 | k__Fungi(100);p__Ascomycota(100);c__Saccharomycetes(100);o__Saccharomycetales(100);f__Metschnikowiaceae(100);g__Metschnikowia(100);s__Metschnikowia_pulcherrima(100); | Metschnikowia | 13 | Saprotroph | \|Undefined Saprotroph\| | Yeast | NULL | Probable | As nectar-tap_saprotroph (Põlme et al. 2020); Decay sugar-rich substrates (Põlme et al. 2020) | Põlme S, et al. 2020. Fungal Diversity 105:1-16 ((https://doi.org/10.1007/s13225-020-00466-2)) |
| Otu48102 | 1 | k__Fungi(100);p__Ascomycota(100);c__Saccharomycetes(100);o__Saccharomycetales(100);f__Metschnikowiaceae(100);g__Metschnikowia(100);s__Metschnikowia_pulcherrima(100); | Metschnikowia | 13 | Saprotroph | \|Undefined Saprotroph\| | Yeast | NULL | Probable | As nectar-tap_saprotroph (Põlme et al. 2020); Decay sugar-rich substrates (Põlme et al. 2020) | Põlme S, et al. 2020. Fungal Diversity 105:1-16 ((https://doi.org/10.1007/s13225-020-00466-2)) |
| Otu48103 | 1 | k__Fungi(100);p__Ascomycota(100);c__Saccharomycetes(100);o__Saccharomycetales(100);f__Metschnikowiaceae(100);g__Metschnikowia(100);s__Metschnikowia_pulcherrima(100); | Metschnikowia | 13 | Saprotroph | \|Undefined Saprotroph\| | Yeast | NULL | Probable | As nectar-tap_saprotroph (Põlme et al. 2020); Decay sugar-rich substrates (Põlme et al. 2020) | Põlme S, et al. 2020. Fungal Diversity 105:1-16 ((https://doi.org/10.1007/s13225-020-00466-2)) |
| Otu48104 | 1 | k__Fungi(100);p__Ascomycota(100);c__Saccharomycetes(100);o__Saccharomycetales(100);f__Metschnikowiaceae(100);g__Metschnikowia(100);g__Metschnikowia_unclassified(100); | Metschnikowia | 13 | Saprotroph | \|Undefined Saprotroph\| | Yeast | NULL | Probable | As nectar-tap_saprotroph (Põlme et al. 2020); Decay sugar-rich substrates (Põlme et al. 2020) | Põlme S, et al. 2020. Fungal Diversity 105:1-16 ((https://doi.org/10.1007/s13225-020-00466-2)) |
| Otu48105 | 1 | k__Fungi(100);p__Ascomycota(100);c__Saccharomycetes(100);o__Saccharomycetales(100);f__Metschnikowiaceae(100);g__Metschnikowia(100);g__Metschnikowia_unclassified(100); | Metschnikowia | 13 | Saprotroph | \|Undefined Saprotroph\| | Yeast | NULL | Probable | As nectar-tap_saprotroph (Põlme et al. 2020); Decay sugar-rich substrates (Põlme et al. 2020) | Põlme S, et al. 2020. Fungal Diversity 105:1-16 ((https://doi.org/10.1007/s13225-020-00466-2)) |
| Otu48106 | 1 | k__Fungi(100);p__Ascomycota(100);c__Saccharomycetes(100);o__Saccharomycetales(100);f__Metschnikowiaceae(100);g__Metschnikowia(100);s__Metschnikowia_pulcherrima(100); | Metschnikowia | 13 | Saprotroph | \|Undefined Saprotroph\| | Yeast | NULL | Probable | As nectar-tap_saprotroph (Põlme et al. 2020); Decay sugar-rich substrates (Põlme et al. 2020) | Põlme S, et al. 2020. Fungal Diversity 105:1-16 ((https://doi.org/10.1007/s13225-020-00466-2)) |
| Otu48108 | 1 | k__Fungi(100);p__Ascomycota(100);c__Dothideomycetes(100);o__Capnodiales(100);f__Mycosphaerellaceae(100);f__Mycosphaerellaceae_unclassified(100);f__Mycosphaerellaceae_unclassified(100); | Mycosphaerellaceae | 9 | Pathotroph-Saprotroph | Plant Pathogen-Undefined Saprotroph | Microfungus | NULL | Probable | Biotrophic, necrotrophic or saprobic on various plant tissues (Cannon & Kirk 2007) | Cannon PF, Kirk PM. 2007. Fungal Families of the World. CAB International, Cambridge (ISBN: 978-0851998275) |
| Otu48109 | 1 | k__Fungi(100);p__Ascomycota(100);c__Eurotiomycetes(100);o__Chaetothyriales(100);f__Herpotrichiellaceae(100);g__Rhinocladiella(100);s__Rhinocladiella_sp_VL271(100); | Rhinocladiella | 13 | Pathotroph-Saprotroph-Symbiotroph | Animal Parasite-Endophyte-\|Undefined Saprotroph\|-Wood Saprotroph | Dimorphic-Facultative Yeast | Soft Rot | Probable | As root_endophyte_dark_septate (Põlme et al. 2020); As soil_saprotroph (Põlme et al. 2020); Decay soil, animal material (Põlme et al. 2020) | Nilsson T. 1973. Studia Forestalia Suecica 104:1-40 ((https://pub.epsilon.slu.se/5711/)); Seehann G, et al. 1975. List of Fungi in Soft-Rot Tissue Tests. Institute fur Holzbiologie und Holzschutz, Bundesforschungsanstalt fur Forst - und Holzwirtshaft, Hamburg - Lohbrugge, Germany; Costa IPMW, et al. 2012. Mycotaxon 119:e26 ((http://mycotaxon.com/resources/checklists/costa_v119_checklist.pdf)); Tedersoo L, et al. 2014. Science 346:e1256688 ((https://doi.org/10.1126/science.1256688)); Põlme S, et al. 2020. Fungal Diversity 105:1-16 ((https://doi.org/10.1007/s13225-020-00466-2)) |
| Otu48110 | 1 | k__Fungi(100);p__Ascomycota(100);c__Saccharomycetes(100);o__Saccharomycetales(100);f__Metschnikowiaceae(100);g__Metschnikowia(100);s__Metschnikowia_pulcherrima(100); | Metschnikowia | 13 | Saprotroph | \|Undefined Saprotroph\| | Yeast | NULL | Probable | As nectar-tap_saprotroph (Põlme et al. 2020); Decay sugar-rich substrates (Põlme et al. 2020) | Põlme S, et al. 2020. Fungal Diversity 105:1-16 ((https://doi.org/10.1007/s13225-020-00466-2)) |
| Otu48111 | 1 | k__Fungi(100);p__Ascomycota(100);c__Saccharomycetes(100);o__Saccharomycetales(100);f__Metschnikowiaceae(100);g__Metschnikowia(100);s__Metschnikowia_pulcherrima(100); | Metschnikowia | 13 | Saprotroph | \|Undefined Saprotroph\| | Yeast | NULL | Probable | As nectar-tap_saprotroph (Põlme et al. 2020); Decay sugar-rich substrates (Põlme et al. 2020) | Põlme S, et al. 2020. Fungal Diversity 105:1-16 ((https://doi.org/10.1007/s13225-020-00466-2)) |
| Otu48112 | 1 | k__Fungi(100);p__Ascomycota(100);c__Saccharomycetes(100);o__Saccharomycetales(100);f__Metschnikowiaceae(100);g__Metschnikowia(100);s__Metschnikowia_pulcherrima(100); | Metschnikowia | 13 | Saprotroph | \|Undefined Saprotroph\| | Yeast | NULL | Probable | As nectar-tap_saprotroph (Põlme et al. 2020); Decay sugar-rich substrates (Põlme et al. 2020) | Põlme S, et al. 2020. Fungal Diversity 105:1-16 ((https://doi.org/10.1007/s13225-020-00466-2)) |
| Otu48113 | 1 | k__Fungi(100);p__Ascomycota(100);c__Saccharomycetes(100);o__Saccharomycetales(100);f__Metschnikowiaceae(100);g__Metschnikowia(100);s__Metschnikowia_pulcherrima(100); | Metschnikowia | 13 | Saprotroph | \|Undefined Saprotroph\| | Yeast | NULL | Probable | As nectar-tap_saprotroph (Põlme et al. 2020); Decay sugar-rich substrates (Põlme et al. 2020) | Põlme S, et al. 2020. Fungal Diversity 105:1-16 ((https://doi.org/10.1007/s13225-020-00466-2)) |
| Otu48114 | 1 | k__Fungi(100);p__Ascomycota(100);c__Saccharomycetes(100);o__Saccharomycetales(100);f__Metschnikowiaceae(100);g__Metschnikowia(100);g__Metschnikowia_unclassified(100); | Metschnikowia | 13 | Saprotroph | \|Undefined Saprotroph\| | Yeast | NULL | Probable | As nectar-tap_saprotroph (Põlme et al. 2020); Decay sugar-rich substrates (Põlme et al. 2020) | Põlme S, et al. 2020. Fungal Diversity 105:1-16 ((https://doi.org/10.1007/s13225-020-00466-2)) |
| Otu48115 | 1 | k__Fungi(100);p__Ascomycota(100);c__Saccharomycetes(100);o__Saccharomycetales(100);f__Metschnikowiaceae(100);g__Metschnikowia(100);g__Metschnikowia_unclassified(100); | Metschnikowia | 13 | Saprotroph | \|Undefined Saprotroph\| | Yeast | NULL | Probable | As nectar-tap_saprotroph (Põlme et al. 2020); Decay sugar-rich substrates (Põlme et al. 2020) | Põlme S, et al. 2020. Fungal Diversity 105:1-16 ((https://doi.org/10.1007/s13225-020-00466-2)) |
| Otu48116 | 1 | k__Fungi(100);p__Ascomycota(100);c__Saccharomycetes(100);o__Saccharomycetales(100);f__Metschnikowiaceae(100);g__Metschnikowia(100);g__Metschnikowia_unclassified(100); | Metschnikowia | 13 | Saprotroph | \|Undefined Saprotroph\| | Yeast | NULL | Probable | As nectar-tap_saprotroph (Põlme et al. 2020); Decay sugar-rich substrates (Põlme et al. 2020) | Põlme S, et al. 2020. Fungal Diversity 105:1-16 ((https://doi.org/10.1007/s13225-020-00466-2)) |
| Otu48117 | 1 | k__Fungi(100);p__Ascomycota(100);c__Saccharomycetes(100);o__Saccharomycetales(100);f__Metschnikowiaceae(100);g__Metschnikowia(100);g__Metschnikowia_unclassified(100); | Metschnikowia | 13 | Saprotroph | \|Undefined Saprotroph\| | Yeast | NULL | Probable | As nectar-tap_saprotroph (Põlme et al. 2020); Decay sugar-rich substrates (Põlme et al. 2020) | Põlme S, et al. 2020. Fungal Diversity 105:1-16 ((https://doi.org/10.1007/s13225-020-00466-2)) |
| Otu48118 | 1 | k__Fungi(100);p__Ascomycota(100);c__Saccharomycetes(100);o__Saccharomycetales(100);f__Metschnikowiaceae(100);g__Metschnikowia(100);s__Metschnikowia_pulcherrima(100); | Metschnikowia | 13 | Saprotroph | \|Undefined Saprotroph\| | Yeast | NULL | Probable | As nectar-tap_saprotroph (Põlme et al. 2020); Decay sugar-rich substrates (Põlme et al. 2020) | Põlme S, et al. 2020. Fungal Diversity 105:1-16 ((https://doi.org/10.1007/s13225-020-00466-2)) |
| Otu48119 | 1 | k__Fungi(100);p__Ascomycota(100);c__Saccharomycetes(100);o__Saccharomycetales(100);f__Metschnikowiaceae(100);g__Metschnikowia(100);g__Metschnikowia_unclassified(100); | Metschnikowia | 13 | Saprotroph | \|Undefined Saprotroph\| | Yeast | NULL | Probable | As nectar-tap_saprotroph (Põlme et al. 2020); Decay sugar-rich substrates (Põlme et al. 2020) | Põlme S, et al. 2020. Fungal Diversity 105:1-16 ((https://doi.org/10.1007/s13225-020-00466-2)) |
| Otu48120 | 1 | k__Fungi(100);p__Ascomycota(100);c__Saccharomycetes(100);o__Saccharomycetales(100);f__Metschnikowiaceae(100);g__Metschnikowia(100);s__Metschnikowia_pulcherrima(100); | Metschnikowia | 13 | Saprotroph | \|Undefined Saprotroph\| | Yeast | NULL | Probable | As nectar-tap_saprotroph (Põlme et al. 2020); Decay sugar-rich substrates (Põlme et al. 2020) | Põlme S, et al. 2020. Fungal Diversity 105:1-16 ((https://doi.org/10.1007/s13225-020-00466-2)) |
| Otu48121 | 1 | k__Fungi(100);p__Ascomycota(100);c__Saccharomycetes(100);o__Saccharomycetales(100);f__Metschnikowiaceae(100);g__Metschnikowia(100);s__Metschnikowia_pulcherrima(100); | Metschnikowia | 13 | Saprotroph | \|Undefined Saprotroph\| | Yeast | NULL | Probable | As nectar-tap_saprotroph (Põlme et al. 2020); Decay sugar-rich substrates (Põlme et al. 2020) | Põlme S, et al. 2020. Fungal Diversity 105:1-16 ((https://doi.org/10.1007/s13225-020-00466-2)) |
| Otu48122 | 1 | k__Fungi(100);p__Ascomycota(100);c__Saccharomycetes(100);o__Saccharomycetales(100);f__Metschnikowiaceae(100);g__Metschnikowia(100);s__Metschnikowia_pulcherrima(100); | Metschnikowia | 13 | Saprotroph | \|Undefined Saprotroph\| | Yeast | NULL | Probable | As nectar-tap_saprotroph (Põlme et al. 2020); Decay sugar-rich substrates (Põlme et al. 2020) | Põlme S, et al. 2020. Fungal Diversity 105:1-16 ((https://doi.org/10.1007/s13225-020-00466-2)) |
| Otu48123 | 1 | k__Fungi(100);p__Ascomycota(100);c__Saccharomycetes(100);o__Saccharomycetales(100);f__Metschnikowiaceae(100);g__Metschnikowia(100);s__Metschnikowia_pulcherrima(100); | Metschnikowia | 13 | Saprotroph | \|Undefined Saprotroph\| | Yeast | NULL | Probable | As nectar-tap_saprotroph (Põlme et al. 2020); Decay sugar-rich substrates (Põlme et al. 2020) | Põlme S, et al. 2020. Fungal Diversity 105:1-16 ((https://doi.org/10.1007/s13225-020-00466-2)) |
| Otu48124 | 1 | k__Fungi(100);p__Ascomycota(100);c__Saccharomycetes(100);o__Saccharomycetales(100);f__Metschnikowiaceae(100);g__Metschnikowia(100);s__Metschnikowia_pulcherrima(100); | Metschnikowia | 13 | Saprotroph | \|Undefined Saprotroph\| | Yeast | NULL | Probable | As nectar-tap_saprotroph (Põlme et al. 2020); Decay sugar-rich substrates (Põlme et al. 2020) | Põlme S, et al. 2020. Fungal Diversity 105:1-16 ((https://doi.org/10.1007/s13225-020-00466-2)) |
| Otu48125 | 1 | k__Fungi(100);p__Ascomycota(100);c__Saccharomycetes(100);o__Saccharomycetales(100);f__Metschnikowiaceae(100);g__Metschnikowia(100);s__Metschnikowia_pulcherrima(100); | Metschnikowia | 13 | Saprotroph | \|Undefined Saprotroph\| | Yeast | NULL | Probable | As nectar-tap_saprotroph (Põlme et al. 2020); Decay sugar-rich substrates (Põlme et al. 2020) | Põlme S, et al. 2020. Fungal Diversity 105:1-16 ((https://doi.org/10.1007/s13225-020-00466-2)) |
| Otu48126 | 1 | k__Fungi(100);p__Ascomycota(100);c__Saccharomycetes(100);o__Saccharomycetales(100);f__Metschnikowiaceae(100);g__Metschnikowia(100);g__Metschnikowia_unclassified(100); | Metschnikowia | 13 | Saprotroph | \|Undefined Saprotroph\| | Yeast | NULL | Probable | As nectar-tap_saprotroph (Põlme et al. 2020); Decay sugar-rich substrates (Põlme et al. 2020) | Põlme S, et al. 2020. Fungal Diversity 105:1-16 ((https://doi.org/10.1007/s13225-020-00466-2)) |
| Otu48127 | 1 | k__Fungi(100);p__Ascomycota(100);c__Saccharomycetes(100);o__Saccharomycetales(100);f__Metschnikowiaceae(100);g__Metschnikowia(100);s__Metschnikowia_pulcherrima(100); | Metschnikowia | 13 | Saprotroph | \|Undefined Saprotroph\| | Yeast | NULL | Probable | As nectar-tap_saprotroph (Põlme et al. 2020); Decay sugar-rich substrates (Põlme et al. 2020) | Põlme S, et al. 2020. Fungal Diversity 105:1-16 ((https://doi.org/10.1007/s13225-020-00466-2)) |
| Otu48128 | 1 | k__Fungi(100);p__Ascomycota(100);c__Saccharomycetes(100);o__Saccharomycetales(100);f__Metschnikowiaceae(100);g__Metschnikowia(100);s__Metschnikowia_pulcherrima(100); | Metschnikowia | 13 | Saprotroph | \|Undefined Saprotroph\| | Yeast | NULL | Probable | As nectar-tap_saprotroph (Põlme et al. 2020); Decay sugar-rich substrates (Põlme et al. 2020) | Põlme S, et al. 2020. Fungal Diversity 105:1-16 ((https://doi.org/10.1007/s13225-020-00466-2)) |
| Otu48129 | 1 | k__Fungi(100);p__Ascomycota(100);c__Saccharomycetes(100);o__Saccharomycetales(100);f__Metschnikowiaceae(100);g__Metschnikowia(100);s__Metschnikowia_pulcherrima(100); | Metschnikowia | 13 | Saprotroph | \|Undefined Saprotroph\| | Yeast | NULL | Probable | As nectar-tap_saprotroph (Põlme et al. 2020); Decay sugar-rich substrates (Põlme et al. 2020) | Põlme S, et al. 2020. Fungal Diversity 105:1-16 ((https://doi.org/10.1007/s13225-020-00466-2)) |
| Otu48130 | 1 | k__Fungi(100);p__Ascomycota(100);c__Saccharomycetes(100);o__Saccharomycetales(100);f__Metschnikowiaceae(100);g__Metschnikowia(100);s__Metschnikowia_pulcherrima(100); | Metschnikowia | 13 | Saprotroph | \|Undefined Saprotroph\| | Yeast | NULL | Probable | As nectar-tap_saprotroph (Põlme et al. 2020); Decay sugar-rich substrates (Põlme et al. 2020) | Põlme S, et al. 2020. Fungal Diversity 105:1-16 ((https://doi.org/10.1007/s13225-020-00466-2)) |
| Otu48131 | 1 | k__Fungi(100);p__Ascomycota(100);c__Saccharomycetes(100);o__Saccharomycetales(100);f__Metschnikowiaceae(100);g__Metschnikowia(100);s__Metschnikowia_pulcherrima(100); | Metschnikowia | 13 | Saprotroph | \|Undefined Saprotroph\| | Yeast | NULL | Probable | As nectar-tap_saprotroph (Põlme et al. 2020); Decay sugar-rich substrates (Põlme et al. 2020) | Põlme S, et al. 2020. Fungal Diversity 105:1-16 ((https://doi.org/10.1007/s13225-020-00466-2)) |
| Otu48132 | 1 | k__Fungi(100);p__Ascomycota(100);c__Saccharomycetes(100);o__Saccharomycetales(100);f__Metschnikowiaceae(100);g__Metschnikowia(100);s__Metschnikowia_pulcherrima(100); | Metschnikowia | 13 | Saprotroph | \|Undefined Saprotroph\| | Yeast | NULL | Probable | As nectar-tap_saprotroph (Põlme et al. 2020); Decay sugar-rich substrates (Põlme et al. 2020) | Põlme S, et al. 2020. Fungal Diversity 105:1-16 ((https://doi.org/10.1007/s13225-020-00466-2)) |
| Otu48133 | 1 | k__Fungi(100);p__Ascomycota(100);c__Saccharomycetes(100);o__Saccharomycetales(100);f__Metschnikowiaceae(100);g__Metschnikowia(100);s__Metschnikowia_pulcherrima(100); | Metschnikowia | 13 | Saprotroph | \|Undefined Saprotroph\| | Yeast | NULL | Probable | As nectar-tap_saprotroph (Põlme et al. 2020); Decay sugar-rich substrates (Põlme et al. 2020) | Põlme S, et al. 2020. Fungal Diversity 105:1-16 ((https://doi.org/10.1007/s13225-020-00466-2)) |
| Otu48134 | 1 | k__Fungi(100);p__Ascomycota(100);c__Saccharomycetes(100);o__Saccharomycetales(100);f__Metschnikowiaceae(100);g__Metschnikowia(100);s__Metschnikowia_pulcherrima(100); | Metschnikowia | 13 | Saprotroph | \|Undefined Saprotroph\| | Yeast | NULL | Probable | As nectar-tap_saprotroph (Põlme et al. 2020); Decay sugar-rich substrates (Põlme et al. 2020) | Põlme S, et al. 2020. Fungal Diversity 105:1-16 ((https://doi.org/10.1007/s13225-020-00466-2)) |
| Otu48135 | 1 | k__Fungi(100);p__Ascomycota(100);c__Saccharomycetes(100);o__Saccharomycetales(100);f__Metschnikowiaceae(100);g__Metschnikowia(100);s__Metschnikowia_pulcherrima(100); | Metschnikowia | 13 | Saprotroph | \|Undefined Saprotroph\| | Yeast | NULL | Probable | As nectar-tap_saprotroph (Põlme et al. 2020); Decay sugar-rich substrates (Põlme et al. 2020) | Põlme S, et al. 2020. Fungal Diversity 105:1-16 ((https://doi.org/10.1007/s13225-020-00466-2)) |
| Otu48136 | 1 | k__Fungi(100);p__Ascomycota(100);c__Saccharomycetes(100);o__Saccharomycetales(100);f__Metschnikowiaceae(100);g__Metschnikowia(100);s__Metschnikowia_pulcherrima(100); | Metschnikowia | 13 | Saprotroph | \|Undefined Saprotroph\| | Yeast | NULL | Probable | As nectar-tap_saprotroph (Põlme et al. 2020); Decay sugar-rich substrates (Põlme et al. 2020) | Põlme S, et al. 2020. Fungal Diversity 105:1-16 ((https://doi.org/10.1007/s13225-020-00466-2)) |
| Otu48137 | 1 | k__Fungi(100);p__Ascomycota(100);c__Saccharomycetes(100);o__Saccharomycetales(100);f__Metschnikowiaceae(100);g__Metschnikowia(100);s__Metschnikowia_pulcherrima(100); | Metschnikowia | 13 | Saprotroph | \|Undefined Saprotroph\| | Yeast | NULL | Probable | As nectar-tap_saprotroph (Põlme et al. 2020); Decay sugar-rich substrates (Põlme et al. 2020) | Põlme S, et al. 2020. Fungal Diversity 105:1-16 ((https://doi.org/10.1007/s13225-020-00466-2)) |
| Otu48138 | 1 | k__Fungi(100);p__Ascomycota(100);c__Saccharomycetes(100);o__Saccharomycetales(100);f__Metschnikowiaceae(100);g__Metschnikowia(100);s__Metschnikowia_pulcherrima(100); | Metschnikowia | 13 | Saprotroph | \|Undefined Saprotroph\| | Yeast | NULL | Probable | As nectar-tap_saprotroph (Põlme et al. 2020); Decay sugar-rich substrates (Põlme et al. 2020) | Põlme S, et al. 2020. Fungal Diversity 105:1-16 ((https://doi.org/10.1007/s13225-020-00466-2)) |
| Otu48139 | 1 | k__Fungi(100);p__Ascomycota(100);c__Saccharomycetes(100);o__Saccharomycetales(100);f__Metschnikowiaceae(100);g__Metschnikowia(100);g__Metschnikowia_unclassified(100); | Metschnikowia | 13 | Saprotroph | \|Undefined Saprotroph\| | Yeast | NULL | Probable | As nectar-tap_saprotroph (Põlme et al. 2020); Decay sugar-rich substrates (Põlme et al. 2020) | Põlme S, et al. 2020. Fungal Diversity 105:1-16 ((https://doi.org/10.1007/s13225-020-00466-2)) |
| Otu48140 | 1 | k__Fungi(100);p__Ascomycota(100);c__Saccharomycetes(100);o__Saccharomycetales(100);f__Metschnikowiaceae(100);g__Metschnikowia(100);s__Metschnikowia_pulcherrima(100); | Metschnikowia | 13 | Saprotroph | \|Undefined Saprotroph\| | Yeast | NULL | Probable | As nectar-tap_saprotroph (Põlme et al. 2020); Decay sugar-rich substrates (Põlme et al. 2020) | Põlme S, et al. 2020. Fungal Diversity 105:1-16 ((https://doi.org/10.1007/s13225-020-00466-2)) |
| Otu48141 | 1 | k__Fungi(100);p__Ascomycota(100);c__Saccharomycetes(100);o__Saccharomycetales(100);f__Metschnikowiaceae(100);g__Metschnikowia(100);g__Metschnikowia_unclassified(100); | Metschnikowia | 13 | Saprotroph | \|Undefined Saprotroph\| | Yeast | NULL | Probable | As nectar-tap_saprotroph (Põlme et al. 2020); Decay sugar-rich substrates (Põlme et al. 2020) | Põlme S, et al. 2020. Fungal Diversity 105:1-16 ((https://doi.org/10.1007/s13225-020-00466-2)) |
| Otu48142 | 1 | k__Fungi(100);p__Ascomycota(100);c__Saccharomycetes(100);o__Saccharomycetales(100);f__Metschnikowiaceae(100);g__Metschnikowia(100);s__Metschnikowia_pulcherrima(100); | Metschnikowia | 13 | Saprotroph | \|Undefined Saprotroph\| | Yeast | NULL | Probable | As nectar-tap_saprotroph (Põlme et al. 2020); Decay sugar-rich substrates (Põlme et al. 2020) | Põlme S, et al. 2020. Fungal Diversity 105:1-16 ((https://doi.org/10.1007/s13225-020-00466-2)) |
| Otu48143 | 1 | k__Fungi(100);p__Ascomycota(100);c__Saccharomycetes(100);o__Saccharomycetales(100);f__Metschnikowiaceae(100);g__Metschnikowia(100);s__Metschnikowia_pulcherrima(100); | Metschnikowia | 13 | Saprotroph | \|Undefined Saprotroph\| | Yeast | NULL | Probable | As nectar-tap_saprotroph (Põlme et al. 2020); Decay sugar-rich substrates (Põlme et al. 2020) | Põlme S, et al. 2020. Fungal Diversity 105:1-16 ((https://doi.org/10.1007/s13225-020-00466-2)) |
| Otu48144 | 1 | k__Fungi(100);p__Ascomycota(100);c__Saccharomycetes(100);o__Saccharomycetales(100);f__Metschnikowiaceae(100);g__Metschnikowia(100);s__Metschnikowia_pulcherrima(100); | Metschnikowia | 13 | Saprotroph | \|Undefined Saprotroph\| | Yeast | NULL | Probable | As nectar-tap_saprotroph (Põlme et al. 2020); Decay sugar-rich substrates (Põlme et al. 2020) | Põlme S, et al. 2020. Fungal Diversity 105:1-16 ((https://doi.org/10.1007/s13225-020-00466-2)) |
| Otu48145 | 1 | k__Fungi(100);p__Ascomycota(100);c__Saccharomycetes(100);o__Saccharomycetales(100);f__Metschnikowiaceae(100);g__Metschnikowia(100);s__Metschnikowia_pulcherrima(100); | Metschnikowia | 13 | Saprotroph | \|Undefined Saprotroph\| | Yeast | NULL | Probable | As nectar-tap_saprotroph (Põlme et al. 2020); Decay sugar-rich substrates (Põlme et al. 2020) | Põlme S, et al. 2020. Fungal Diversity 105:1-16 ((https://doi.org/10.1007/s13225-020-00466-2)) |
| Otu48146 | 1 | k__Fungi(100);p__Ascomycota(100);c__Saccharomycetes(100);o__Saccharomycetales(100);f__Metschnikowiaceae(100);g__Metschnikowia(100);s__Metschnikowia_pulcherrima(100); | Metschnikowia | 13 | Saprotroph | \|Undefined Saprotroph\| | Yeast | NULL | Probable | As nectar-tap_saprotroph (Põlme et al. 2020); Decay sugar-rich substrates (Põlme et al. 2020) | Põlme S, et al. 2020. Fungal Diversity 105:1-16 ((https://doi.org/10.1007/s13225-020-00466-2)) |
| Otu48147 | 1 | k__Fungi(100);p__Ascomycota(100);c__Saccharomycetes(100);o__Saccharomycetales(100);f__Metschnikowiaceae(100);g__Metschnikowia(100);s__Metschnikowia_pulcherrima(100); | Metschnikowia | 13 | Saprotroph | \|Undefined Saprotroph\| | Yeast | NULL | Probable | As nectar-tap_saprotroph (Põlme et al. 2020); Decay sugar-rich substrates (Põlme et al. 2020) | Põlme S, et al. 2020. Fungal Diversity 105:1-16 ((https://doi.org/10.1007/s13225-020-00466-2)) |
| Otu48148 | 1 | k__Fungi(100);p__Ascomycota(100);c__Saccharomycetes(100);o__Saccharomycetales(100);f__Metschnikowiaceae(100);g__Metschnikowia(100);s__Metschnikowia_pulcherrima(100); | Metschnikowia | 13 | Saprotroph | \|Undefined Saprotroph\| | Yeast | NULL | Probable | As nectar-tap_saprotroph (Põlme et al. 2020); Decay sugar-rich substrates (Põlme et al. 2020) | Põlme S, et al. 2020. Fungal Diversity 105:1-16 ((https://doi.org/10.1007/s13225-020-00466-2)) |
| Otu48149 | 1 | k__Fungi(100);p__Ascomycota(100);c__Saccharomycetes(100);o__Saccharomycetales(100);f__Metschnikowiaceae(100);g__Metschnikowia(100);s__Metschnikowia_pulcherrima(100); | Metschnikowia | 13 | Saprotroph | \|Undefined Saprotroph\| | Yeast | NULL | Probable | As nectar-tap_saprotroph (Põlme et al. 2020); Decay sugar-rich substrates (Põlme et al. 2020) | Põlme S, et al. 2020. Fungal Diversity 105:1-16 ((https://doi.org/10.1007/s13225-020-00466-2)) |
| Otu48150 | 1 | k__Fungi(100);p__Ascomycota(100);c__Saccharomycetes(100);o__Saccharomycetales(100);f__Metschnikowiaceae(100);g__Metschnikowia(100);s__Metschnikowia_pulcherrima(100); | Metschnikowia | 13 | Saprotroph | \|Undefined Saprotroph\| | Yeast | NULL | Probable | As nectar-tap_saprotroph (Põlme et al. 2020); Decay sugar-rich substrates (Põlme et al. 2020) | Põlme S, et al. 2020. Fungal Diversity 105:1-16 ((https://doi.org/10.1007/s13225-020-00466-2)) |
| Otu48151 | 1 | k__Fungi(100);p__Ascomycota(100);c__Saccharomycetes(100);o__Saccharomycetales(100);f__Metschnikowiaceae(100);g__Metschnikowia(100);g__Metschnikowia_unclassified(100); | Metschnikowia | 13 | Saprotroph | \|Undefined Saprotroph\| | Yeast | NULL | Probable | As nectar-tap_saprotroph (Põlme et al. 2020); Decay sugar-rich substrates (Põlme et al. 2020) | Põlme S, et al. 2020. Fungal Diversity 105:1-16 ((https://doi.org/10.1007/s13225-020-00466-2)) |
| Otu48152 | 1 | k__Fungi(100);p__Ascomycota(100);c__Saccharomycetes(100);o__Saccharomycetales(100);f__Metschnikowiaceae(100);g__Metschnikowia(100);g__Metschnikowia_unclassified(100); | Metschnikowia | 13 | Saprotroph | \|Undefined Saprotroph\| | Yeast | NULL | Probable | As nectar-tap_saprotroph (Põlme et al. 2020); Decay sugar-rich substrates (Põlme et al. 2020) | Põlme S, et al. 2020. Fungal Diversity 105:1-16 ((https://doi.org/10.1007/s13225-020-00466-2)) |
| Otu48153 | 1 | k__Fungi(100);p__Ascomycota(100);c__Saccharomycetes(100);o__Saccharomycetales(100);f__Metschnikowiaceae(100);g__Metschnikowia(100);g__Metschnikowia_unclassified(100); | Metschnikowia | 13 | Saprotroph | \|Undefined Saprotroph\| | Yeast | NULL | Probable | As nectar-tap_saprotroph (Põlme et al. 2020); Decay sugar-rich substrates (Põlme et al. 2020) | Põlme S, et al. 2020. Fungal Diversity 105:1-16 ((https://doi.org/10.1007/s13225-020-00466-2)) |
| Otu48154 | 1 | k__Fungi(100);p__Ascomycota(100);c__Saccharomycetes(100);o__Saccharomycetales(100);f__Metschnikowiaceae(100);g__Metschnikowia(100);s__Metschnikowia_pulcherrima(100); | Metschnikowia | 13 | Saprotroph | \|Undefined Saprotroph\| | Yeast | NULL | Probable | As nectar-tap_saprotroph (Põlme et al. 2020); Decay sugar-rich substrates (Põlme et al. 2020) | Põlme S, et al. 2020. Fungal Diversity 105:1-16 ((https://doi.org/10.1007/s13225-020-00466-2)) |
| Otu48155 | 1 | k__Fungi(100);p__Ascomycota(100);c__Saccharomycetes(100);o__Saccharomycetales(100);f__Metschnikowiaceae(100);g__Metschnikowia(100);g__Metschnikowia_unclassified(100); | Metschnikowia | 13 | Saprotroph | \|Undefined Saprotroph\| | Yeast | NULL | Probable | As nectar-tap_saprotroph (Põlme et al. 2020); Decay sugar-rich substrates (Põlme et al. 2020) | Põlme S, et al. 2020. Fungal Diversity 105:1-16 ((https://doi.org/10.1007/s13225-020-00466-2)) |
| Otu48156 | 1 | k__Fungi(100);p__Ascomycota(100);c__Saccharomycetes(100);o__Saccharomycetales(100);f__Metschnikowiaceae(100);g__Metschnikowia(100);g__Metschnikowia_unclassified(100); | Metschnikowia | 13 | Saprotroph | \|Undefined Saprotroph\| | Yeast | NULL | Probable | As nectar-tap_saprotroph (Põlme et al. 2020); Decay sugar-rich substrates (Põlme et al. 2020) | Põlme S, et al. 2020. Fungal Diversity 105:1-16 ((https://doi.org/10.1007/s13225-020-00466-2)) |
| Otu48157 | 1 | k__Fungi(100);p__Ascomycota(100);c__Saccharomycetes(100);o__Saccharomycetales(100);f__Metschnikowiaceae(100);g__Metschnikowia(100);g__Metschnikowia_unclassified(100); | Metschnikowia | 13 | Saprotroph | \|Undefined Saprotroph\| | Yeast | NULL | Probable | As nectar-tap_saprotroph (Põlme et al. 2020); Decay sugar-rich substrates (Põlme et al. 2020) | Põlme S, et al. 2020. Fungal Diversity 105:1-16 ((https://doi.org/10.1007/s13225-020-00466-2)) |
| Otu48158 | 1 | k__Fungi(100);p__Ascomycota(100);c__Saccharomycetes(100);o__Saccharomycetales(100);f__Metschnikowiaceae(100);g__Metschnikowia(100);s__Metschnikowia_pulcherrima(100); | Metschnikowia | 13 | Saprotroph | \|Undefined Saprotroph\| | Yeast | NULL | Probable | As nectar-tap_saprotroph (Põlme et al. 2020); Decay sugar-rich substrates (Põlme et al. 2020) | Põlme S, et al. 2020. Fungal Diversity 105:1-16 ((https://doi.org/10.1007/s13225-020-00466-2)) |
| Otu48159 | 1 | k__Fungi(100);p__Ascomycota(100);c__Saccharomycetes(100);o__Saccharomycetales(100);f__Metschnikowiaceae(100);g__Metschnikowia(100);s__Metschnikowia_pulcherrima(100); | Metschnikowia | 13 | Saprotroph | \|Undefined Saprotroph\| | Yeast | NULL | Probable | As nectar-tap_saprotroph (Põlme et al. 2020); Decay sugar-rich substrates (Põlme et al. 2020) | Põlme S, et al. 2020. Fungal Diversity 105:1-16 ((https://doi.org/10.1007/s13225-020-00466-2)) |
| Otu48160 | 1 | k__Fungi(100);p__Ascomycota(100);c__Saccharomycetes(100);o__Saccharomycetales(100);f__Metschnikowiaceae(100);g__Metschnikowia(100);s__Metschnikowia_pulcherrima(100); | Metschnikowia | 13 | Saprotroph | \|Undefined Saprotroph\| | Yeast | NULL | Probable | As nectar-tap_saprotroph (Põlme et al. 2020); Decay sugar-rich substrates (Põlme et al. 2020) | Põlme S, et al. 2020. Fungal Diversity 105:1-16 ((https://doi.org/10.1007/s13225-020-00466-2)) |
| Otu48161 | 1 | k__Fungi(100);p__Ascomycota(100);c__Saccharomycetes(100);o__Saccharomycetales(100);f__Metschnikowiaceae(100);g__Metschnikowia(100);s__Metschnikowia_pulcherrima(100); | Metschnikowia | 13 | Saprotroph | \|Undefined Saprotroph\| | Yeast | NULL | Probable | As nectar-tap_saprotroph (Põlme et al. 2020); Decay sugar-rich substrates (Põlme et al. 2020) | Põlme S, et al. 2020. Fungal Diversity 105:1-16 ((https://doi.org/10.1007/s13225-020-00466-2)) |
| Otu48162 | 1 | k__Fungi(100);p__Ascomycota(100);c__Saccharomycetes(100);o__Saccharomycetales(100);f__Metschnikowiaceae(100);g__Metschnikowia(100);s__Metschnikowia_pulcherrima(100); | Metschnikowia | 13 | Saprotroph | \|Undefined Saprotroph\| | Yeast | NULL | Probable | As nectar-tap_saprotroph (Põlme et al. 2020); Decay sugar-rich substrates (Põlme et al. 2020) | Põlme S, et al. 2020. Fungal Diversity 105:1-16 ((https://doi.org/10.1007/s13225-020-00466-2)) |
| Otu48163 | 1 | k__Fungi(100);p__Ascomycota(100);c__Saccharomycetes(100);o__Saccharomycetales(100);f__Metschnikowiaceae(100);g__Metschnikowia(100);s__Metschnikowia_pulcherrima(100); | Metschnikowia | 13 | Saprotroph | \|Undefined Saprotroph\| | Yeast | NULL | Probable | As nectar-tap_saprotroph (Põlme et al. 2020); Decay sugar-rich substrates (Põlme et al. 2020) | Põlme S, et al. 2020. Fungal Diversity 105:1-16 ((https://doi.org/10.1007/s13225-020-00466-2)) |
| Otu48164 | 1 | k__Fungi(100);p__Ascomycota(100);c__Saccharomycetes(100);o__Saccharomycetales(100);f__Metschnikowiaceae(100);g__Metschnikowia(100);s__Metschnikowia_pulcherrima(100); | Metschnikowia | 13 | Saprotroph | \|Undefined Saprotroph\| | Yeast | NULL | Probable | As nectar-tap_saprotroph (Põlme et al. 2020); Decay sugar-rich substrates (Põlme et al. 2020) | Põlme S, et al. 2020. Fungal Diversity 105:1-16 ((https://doi.org/10.1007/s13225-020-00466-2)) |
| Otu48165 | 1 | k__Fungi(100);p__Ascomycota(100);c__Saccharomycetes(100);o__Saccharomycetales(100);f__Metschnikowiaceae(100);g__Metschnikowia(100);g__Metschnikowia_unclassified(100); | Metschnikowia | 13 | Saprotroph | \|Undefined Saprotroph\| | Yeast | NULL | Probable | As nectar-tap_saprotroph (Põlme et al. 2020); Decay sugar-rich substrates (Põlme et al. 2020) | Põlme S, et al. 2020. Fungal Diversity 105:1-16 ((https://doi.org/10.1007/s13225-020-00466-2)) |
| Otu48166 | 1 | k__Fungi(100);p__Ascomycota(100);c__Saccharomycetes(100);o__Saccharomycetales(100);f__Metschnikowiaceae(100);g__Metschnikowia(100);s__Metschnikowia_pulcherrima(100); | Metschnikowia | 13 | Saprotroph | \|Undefined Saprotroph\| | Yeast | NULL | Probable | As nectar-tap_saprotroph (Põlme et al. 2020); Decay sugar-rich substrates (Põlme et al. 2020) | Põlme S, et al. 2020. Fungal Diversity 105:1-16 ((https://doi.org/10.1007/s13225-020-00466-2)) |
| Otu48167 | 1 | k__Fungi(100);p__Ascomycota(100);c__Saccharomycetes(100);o__Saccharomycetales(100);f__Saccharomycetales_family_Incertae_sedis(100);g__Candida(100);s__Candida_dosseyi(100); | Candida | 13 | Pathotroph-Saprotroph-Symbiotroph | Animal Pathogen-Endophyte-Endosymbiont-Epiphyte-\|Undefined Saprotroph\| | Dimorphic Yeast-Yeast | NULL | Probable | As nectar-tap_saprotroph (Põlme et al. 2020); Decay sugar-rich substrates (Põlme et al. 2020); This asexual genus is polyphyletic and contain species that function very differently depend on what they are. Caution should be taken when interpreting guilds for Candida species. | Manolakaki D, et al. 2010. Virulence. 1: 367-375 ((https://doi.org/10.4161/viru.1.5.12796)); Mekbib SB, et al. 2011. Tropical Plant Pathology 36:233-240 ((https://doi.org/10.1590/S1982-56762011000400004)); Põlme S, et al. 2020. Fungal Diversity 105:1-16 ((https://doi.org/10.1007/s13225-020-00466-2)) |
| Otu48168 | 1 | k__Fungi(100);p__Ascomycota(100);c__Saccharomycetes(100);o__Saccharomycetales(100);f__Metschnikowiaceae(100);g__Metschnikowia(100);s__Metschnikowia_pulcherrima(100); | Metschnikowia | 13 | Saprotroph | \|Undefined Saprotroph\| | Yeast | NULL | Probable | As nectar-tap_saprotroph (Põlme et al. 2020); Decay sugar-rich substrates (Põlme et al. 2020) | Põlme S, et al. 2020. Fungal Diversity 105:1-16 ((https://doi.org/10.1007/s13225-020-00466-2)) |
| Otu48169 | 1 | k__Fungi(100);p__Ascomycota(100);c__Saccharomycetes(100);o__Saccharomycetales(100);f__Metschnikowiaceae(100);g__Metschnikowia(100);s__Metschnikowia_pulcherrima(100); | Metschnikowia | 13 | Saprotroph | \|Undefined Saprotroph\| | Yeast | NULL | Probable | As nectar-tap_saprotroph (Põlme et al. 2020); Decay sugar-rich substrates (Põlme et al. 2020) | Põlme S, et al. 2020. Fungal Diversity 105:1-16 ((https://doi.org/10.1007/s13225-020-00466-2)) |
| Otu48170 | 1 | k__Fungi(100);p__Ascomycota(100);c__Saccharomycetes(100);o__Saccharomycetales(100);f__Metschnikowiaceae(100);g__Metschnikowia(100);g__Metschnikowia_unclassified(100); | Metschnikowia | 13 | Saprotroph | \|Undefined Saprotroph\| | Yeast | NULL | Probable | As nectar-tap_saprotroph (Põlme et al. 2020); Decay sugar-rich substrates (Põlme et al. 2020) | Põlme S, et al. 2020. Fungal Diversity 105:1-16 ((https://doi.org/10.1007/s13225-020-00466-2)) |
| Otu48171 | 1 | k__Fungi(100);p__Ascomycota(100);c__Saccharomycetes(100);o__Saccharomycetales(100);f__Metschnikowiaceae(100);g__Metschnikowia(100);s__Metschnikowia_pulcherrima(100); | Metschnikowia | 13 | Saprotroph | \|Undefined Saprotroph\| | Yeast | NULL | Probable | As nectar-tap_saprotroph (Põlme et al. 2020); Decay sugar-rich substrates (Põlme et al. 2020) | Põlme S, et al. 2020. Fungal Diversity 105:1-16 ((https://doi.org/10.1007/s13225-020-00466-2)) |
| Otu48172 | 1 | k__Fungi(100);p__Ascomycota(100);c__Saccharomycetes(100);o__Saccharomycetales(100);f__Metschnikowiaceae(100);g__Metschnikowia(100);s__Metschnikowia_pulcherrima(100); | Metschnikowia | 13 | Saprotroph | \|Undefined Saprotroph\| | Yeast | NULL | Probable | As nectar-tap_saprotroph (Põlme et al. 2020); Decay sugar-rich substrates (Põlme et al. 2020) | Põlme S, et al. 2020. Fungal Diversity 105:1-16 ((https://doi.org/10.1007/s13225-020-00466-2)) |
| Otu48173 | 1 | k__Fungi(100);p__Ascomycota(100);c__Saccharomycetes(100);o__Saccharomycetales(100);f__Metschnikowiaceae(100);g__Metschnikowia(100);s__Metschnikowia_pulcherrima(100); | Metschnikowia | 13 | Saprotroph | \|Undefined Saprotroph\| | Yeast | NULL | Probable | As nectar-tap_saprotroph (Põlme et al. 2020); Decay sugar-rich substrates (Põlme et al. 2020) | Põlme S, et al. 2020. Fungal Diversity 105:1-16 ((https://doi.org/10.1007/s13225-020-00466-2)) |
| Otu48174 | 1 | k__Fungi(100);p__Ascomycota(100);c__Saccharomycetes(100);o__Saccharomycetales(100);f__Metschnikowiaceae(100);g__Metschnikowia(100);s__Metschnikowia_pulcherrima(100); | Metschnikowia | 13 | Saprotroph | \|Undefined Saprotroph\| | Yeast | NULL | Probable | As nectar-tap_saprotroph (Põlme et al. 2020); Decay sugar-rich substrates (Põlme et al. 2020) | Põlme S, et al. 2020. Fungal Diversity 105:1-16 ((https://doi.org/10.1007/s13225-020-00466-2)) |
| Otu48175 | 1 | k__Fungi(100);p__Ascomycota(100);c__Saccharomycetes(100);o__Saccharomycetales(100);f__Metschnikowiaceae(100);g__Metschnikowia(100);s__Metschnikowia_pulcherrima(100); | Metschnikowia | 13 | Saprotroph | \|Undefined Saprotroph\| | Yeast | NULL | Probable | As nectar-tap_saprotroph (Põlme et al. 2020); Decay sugar-rich substrates (Põlme et al. 2020) | Põlme S, et al. 2020. Fungal Diversity 105:1-16 ((https://doi.org/10.1007/s13225-020-00466-2)) |
| Otu48176 | 1 | k__Fungi(100);p__Ascomycota(100);c__Saccharomycetes(100);o__Saccharomycetales(100);f__Metschnikowiaceae(100);g__Metschnikowia(100);s__Metschnikowia_pulcherrima(100); | Metschnikowia | 13 | Saprotroph | \|Undefined Saprotroph\| | Yeast | NULL | Probable | As nectar-tap_saprotroph (Põlme et al. 2020); Decay sugar-rich substrates (Põlme et al. 2020) | Põlme S, et al. 2020. Fungal Diversity 105:1-16 ((https://doi.org/10.1007/s13225-020-00466-2)) |
| Otu48177 | 1 | k__Fungi(100);p__Ascomycota(100);c__Saccharomycetes(100);o__Saccharomycetales(100);f__Metschnikowiaceae(100);g__Metschnikowia(100);s__Metschnikowia_pulcherrima(100); | Metschnikowia | 13 | Saprotroph | \|Undefined Saprotroph\| | Yeast | NULL | Probable | As nectar-tap_saprotroph (Põlme et al. 2020); Decay sugar-rich substrates (Põlme et al. 2020) | Põlme S, et al. 2020. Fungal Diversity 105:1-16 ((https://doi.org/10.1007/s13225-020-00466-2)) |
| Otu48178 | 1 | k__Fungi(100);p__Ascomycota(100);c__Saccharomycetes(100);o__Saccharomycetales(100);f__Metschnikowiaceae(100);g__Metschnikowia(100);s__Metschnikowia_pulcherrima(100); | Metschnikowia | 13 | Saprotroph | \|Undefined Saprotroph\| | Yeast | NULL | Probable | As nectar-tap_saprotroph (Põlme et al. 2020); Decay sugar-rich substrates (Põlme et al. 2020) | Põlme S, et al. 2020. Fungal Diversity 105:1-16 ((https://doi.org/10.1007/s13225-020-00466-2)) |
| Otu48179 | 1 | k__Fungi(100);p__Ascomycota(100);c__Saccharomycetes(100);o__Saccharomycetales(100);f__Metschnikowiaceae(100);g__Metschnikowia(100);s__Metschnikowia_pulcherrima(100); | Metschnikowia | 13 | Saprotroph | \|Undefined Saprotroph\| | Yeast | NULL | Probable | As nectar-tap_saprotroph (Põlme et al. 2020); Decay sugar-rich substrates (Põlme et al. 2020) | Põlme S, et al. 2020. Fungal Diversity 105:1-16 ((https://doi.org/10.1007/s13225-020-00466-2)) |
| Otu48180 | 1 | k__Fungi(100);p__Ascomycota(100);c__Saccharomycetes(100);o__Saccharomycetales(100);f__Metschnikowiaceae(100);g__Metschnikowia(100);s__Metschnikowia_pulcherrima(100); | Metschnikowia | 13 | Saprotroph | \|Undefined Saprotroph\| | Yeast | NULL | Probable | As nectar-tap_saprotroph (Põlme et al. 2020); Decay sugar-rich substrates (Põlme et al. 2020) | Põlme S, et al. 2020. Fungal Diversity 105:1-16 ((https://doi.org/10.1007/s13225-020-00466-2)) |
| Otu48181 | 1 | k__Fungi(100);p__Ascomycota(100);c__Saccharomycetes(100);o__Saccharomycetales(100);f__Metschnikowiaceae(100);g__Metschnikowia(100);s__Metschnikowia_pulcherrima(100); | Metschnikowia | 13 | Saprotroph | \|Undefined Saprotroph\| | Yeast | NULL | Probable | As nectar-tap_saprotroph (Põlme et al. 2020); Decay sugar-rich substrates (Põlme et al. 2020) | Põlme S, et al. 2020. Fungal Diversity 105:1-16 ((https://doi.org/10.1007/s13225-020-00466-2)) |
| Otu48182 | 1 | k__Fungi(100);p__Ascomycota(100);c__Saccharomycetes(100);o__Saccharomycetales(100);f__Metschnikowiaceae(100);g__Metschnikowia(100);g__Metschnikowia_unclassified(100); | Metschnikowia | 13 | Saprotroph | \|Undefined Saprotroph\| | Yeast | NULL | Probable | As nectar-tap_saprotroph (Põlme et al. 2020); Decay sugar-rich substrates (Põlme et al. 2020) | Põlme S, et al. 2020. Fungal Diversity 105:1-16 ((https://doi.org/10.1007/s13225-020-00466-2)) |
| Otu48183 | 1 | k__Fungi(100);p__Ascomycota(100);c__Saccharomycetes(100);o__Saccharomycetales(100);f__Metschnikowiaceae(100);g__Metschnikowia(100);s__Metschnikowia_pulcherrima(100); | Metschnikowia | 13 | Saprotroph | \|Undefined Saprotroph\| | Yeast | NULL | Probable | As nectar-tap_saprotroph (Põlme et al. 2020); Decay sugar-rich substrates (Põlme et al. 2020) | Põlme S, et al. 2020. Fungal Diversity 105:1-16 ((https://doi.org/10.1007/s13225-020-00466-2)) |
| Otu48184 | 1 | k__Fungi(100);p__Ascomycota(100);c__Saccharomycetes(100);o__Saccharomycetales(100);f__Metschnikowiaceae(100);g__Metschnikowia(100);s__Metschnikowia_pulcherrima(100); | Metschnikowia | 13 | Saprotroph | \|Undefined Saprotroph\| | Yeast | NULL | Probable | As nectar-tap_saprotroph (Põlme et al. 2020); Decay sugar-rich substrates (Põlme et al. 2020) | Põlme S, et al. 2020. Fungal Diversity 105:1-16 ((https://doi.org/10.1007/s13225-020-00466-2)) |
| Otu48185 | 1 | k__Fungi(100);p__Ascomycota(100);c__Saccharomycetes(100);o__Saccharomycetales(100);f__Metschnikowiaceae(100);g__Metschnikowia(100);s__Metschnikowia_pulcherrima(100); | Metschnikowia | 13 | Saprotroph | \|Undefined Saprotroph\| | Yeast | NULL | Probable | As nectar-tap_saprotroph (Põlme et al. 2020); Decay sugar-rich substrates (Põlme et al. 2020) | Põlme S, et al. 2020. Fungal Diversity 105:1-16 ((https://doi.org/10.1007/s13225-020-00466-2)) |
| Otu48186 | 1 | k__Fungi(100);p__Ascomycota(100);c__Saccharomycetes(100);o__Saccharomycetales(100);f__Metschnikowiaceae(100);g__Metschnikowia(100);s__Metschnikowia_pulcherrima(100); | Metschnikowia | 13 | Saprotroph | \|Undefined Saprotroph\| | Yeast | NULL | Probable | As nectar-tap_saprotroph (Põlme et al. 2020); Decay sugar-rich substrates (Põlme et al. 2020) | Põlme S, et al. 2020. Fungal Diversity 105:1-16 ((https://doi.org/10.1007/s13225-020-00466-2)) |
| Otu48187 | 1 | k__Fungi(100);p__Ascomycota(100);c__Saccharomycetes(100);o__Saccharomycetales(100);f__Metschnikowiaceae(100);g__Metschnikowia(100);g__Metschnikowia_unclassified(100); | Metschnikowia | 13 | Saprotroph | \|Undefined Saprotroph\| | Yeast | NULL | Probable | As nectar-tap_saprotroph (Põlme et al. 2020); Decay sugar-rich substrates (Põlme et al. 2020) | Põlme S, et al. 2020. Fungal Diversity 105:1-16 ((https://doi.org/10.1007/s13225-020-00466-2)) |
| Otu48188 | 1 | k__Fungi(100);p__Ascomycota(100);c__Saccharomycetes(100);o__Saccharomycetales(100);f__Metschnikowiaceae(100);g__Metschnikowia(100);s__Metschnikowia_pulcherrima(100); | Metschnikowia | 13 | Saprotroph | \|Undefined Saprotroph\| | Yeast | NULL | Probable | As nectar-tap_saprotroph (Põlme et al. 2020); Decay sugar-rich substrates (Põlme et al. 2020) | Põlme S, et al. 2020. Fungal Diversity 105:1-16 ((https://doi.org/10.1007/s13225-020-00466-2)) |
| Otu48189 | 1 | k__Fungi(100);p__Ascomycota(100);c__Saccharomycetes(100);o__Saccharomycetales(100);f__Metschnikowiaceae(100);g__Metschnikowia(100);g__Metschnikowia_unclassified(100); | Metschnikowia | 13 | Saprotroph | \|Undefined Saprotroph\| | Yeast | NULL | Probable | As nectar-tap_saprotroph (Põlme et al. 2020); Decay sugar-rich substrates (Põlme et al. 2020) | Põlme S, et al. 2020. Fungal Diversity 105:1-16 ((https://doi.org/10.1007/s13225-020-00466-2)) |
| Otu48190 | 1 | k__Fungi(100);p__Ascomycota(100);c__Saccharomycetes(100);o__Saccharomycetales(100);f__Metschnikowiaceae(100);g__Metschnikowia(100);s__Metschnikowia_pulcherrima(100); | Metschnikowia | 13 | Saprotroph | \|Undefined Saprotroph\| | Yeast | NULL | Probable | As nectar-tap_saprotroph (Põlme et al. 2020); Decay sugar-rich substrates (Põlme et al. 2020) | Põlme S, et al. 2020. Fungal Diversity 105:1-16 ((https://doi.org/10.1007/s13225-020-00466-2)) |
| Otu48191 | 1 | k__Fungi(100);p__Ascomycota(100);c__Saccharomycetes(100);o__Saccharomycetales(100);f__Metschnikowiaceae(100);g__Metschnikowia(100);s__Metschnikowia_pulcherrima(100); | Metschnikowia | 13 | Saprotroph | \|Undefined Saprotroph\| | Yeast | NULL | Probable | As nectar-tap_saprotroph (Põlme et al. 2020); Decay sugar-rich substrates (Põlme et al. 2020) | Põlme S, et al. 2020. Fungal Diversity 105:1-16 ((https://doi.org/10.1007/s13225-020-00466-2)) |
| Otu48192 | 1 | k__Fungi(100);p__Ascomycota(100);c__Saccharomycetes(100);o__Saccharomycetales(100);f__Metschnikowiaceae(100);g__Metschnikowia(100);s__Metschnikowia_pulcherrima(100); | Metschnikowia | 13 | Saprotroph | \|Undefined Saprotroph\| | Yeast | NULL | Probable | As nectar-tap_saprotroph (Põlme et al. 2020); Decay sugar-rich substrates (Põlme et al. 2020) | Põlme S, et al. 2020. Fungal Diversity 105:1-16 ((https://doi.org/10.1007/s13225-020-00466-2)) |
| Otu48193 | 1 | k__Fungi(100);p__Ascomycota(100);c__Saccharomycetes(100);o__Saccharomycetales(100);f__Metschnikowiaceae(100);g__Metschnikowia(100);s__Metschnikowia_pulcherrima(100); | Metschnikowia | 13 | Saprotroph | \|Undefined Saprotroph\| | Yeast | NULL | Probable | As nectar-tap_saprotroph (Põlme et al. 2020); Decay sugar-rich substrates (Põlme et al. 2020) | Põlme S, et al. 2020. Fungal Diversity 105:1-16 ((https://doi.org/10.1007/s13225-020-00466-2)) |
| Otu48194 | 1 | k__Fungi(100);p__Ascomycota(100);c__Saccharomycetes(100);o__Saccharomycetales(100);f__Metschnikowiaceae(100);g__Metschnikowia(100);s__Metschnikowia_pulcherrima(100); | Metschnikowia | 13 | Saprotroph | \|Undefined Saprotroph\| | Yeast | NULL | Probable | As nectar-tap_saprotroph (Põlme et al. 2020); Decay sugar-rich substrates (Põlme et al. 2020) | Põlme S, et al. 2020. Fungal Diversity 105:1-16 ((https://doi.org/10.1007/s13225-020-00466-2)) |

| | | | | | | | | | | | |
|---|---|---|---|---|---|---|---|---|---|---|---|
| Otu48195 | 1 | k__Fungi(100);p__Ascomycota(100);c__Saccharomycetes(100);o__Saccharomycetales(100);f__Metschnikowiaceae(100);g__Metschnikowia(100);s__Metschnikowia_pulcherrima(100); | Metschnikowia | 13 | Saprotroph | \|Undefined Saprotroph\| | Yeast | NULL | Probable | As nectar-tap_saprotroph (Põlme et al. 2020); Decay sugar-rich substrates (Põlme et al. 2020) | Põlme S, et al. 2020. Fungal Diversity 105:1-16 ((https://doi.org/10.1007/s13225-020-00466-2)) |
| Otu48196 | 1 | k__Fungi(100);p__Ascomycota(100);c__Saccharomycetes(100);o__Saccharomycetales(100);f__Metschnikowiaceae(100);g__Metschnikowia(100);s__Metschnikowia_pulcherrima(100); | Metschnikowia | 13 | Saprotroph | \|Undefined Saprotroph\| | Yeast | NULL | Probable | As nectar-tap_saprotroph (Põlme et al. 2020); Decay sugar-rich substrates (Põlme et al. 2020) | Põlme S, et al. 2020. Fungal Diversity 105:1-16 ((https://doi.org/10.1007/s13225-020-00466-2)) |
| Otu48197 | 1 | k__Fungi(100);p__Ascomycota(100);c__Saccharomycetes(100);o__Saccharomycetales(100);f__Metschnikowiaceae(100);g__Metschnikowia(100);s__Metschnikowia_pulcherrima(100); | Metschnikowia | 13 | Saprotroph | \|Undefined Saprotroph\| | Yeast | NULL | Probable | As nectar-tap_saprotroph (Põlme et al. 2020); Decay sugar-rich substrates (Põlme et al. 2020) | Põlme S, et al. 2020. Fungal Diversity 105:1-16 ((https://doi.org/10.1007/s13225-020-00466-2)) |
| Otu48198 | 1 | k__Fungi(100);p__Ascomycota(100);c__Saccharomycetes(100);o__Saccharomycetales(100);f__Metschnikowiaceae(100);g__Metschnikowia(100);s__Metschnikowia_pulcherrima(100); | Metschnikowia | 13 | Saprotroph | \|Undefined Saprotroph\| | Yeast | NULL | Probable | As nectar-tap_saprotroph (Põlme et al. 2020); Decay sugar-rich substrates (Põlme et al. 2020) | Põlme S, et al. 2020. Fungal Diversity 105:1-16 ((https://doi.org/10.1007/s13225-020-00466-2)) |
| Otu48199 | 1 | k__Fungi(100);p__Ascomycota(100);c__Saccharomycetes(100);o__Saccharomycetales(100);f__Metschnikowiaceae(100);g__Metschnikowia(100);s__Metschnikowia_pulcherrima(100); | Metschnikowia | 13 | Saprotroph | \|Undefined Saprotroph\| | Yeast | NULL | Probable | As nectar-tap_saprotroph (Põlme et al. 2020); Decay sugar-rich substrates (Põlme et al. 2020) | Põlme S, et al. 2020. Fungal Diversity 105:1-16 ((https://doi.org/10.1007/s13225-020-00466-2)) |
| Otu48200 | 1 | k__Fungi(100);p__Ascomycota(100);c__Saccharomycetes(100);o__Saccharomycetales(100);f__Metschnikowiaceae(100);g__Metschnikowia(100);s__Metschnikowia_pulcherrima(100); | Metschnikowia | 13 | Saprotroph | \|Undefined Saprotroph\| | Yeast | NULL | Probable | As nectar-tap_saprotroph (Põlme et al. 2020); Decay sugar-rich substrates (Põlme et al. 2020) | Põlme S, et al. 2020. Fungal Diversity 105:1-16 ((https://doi.org/10.1007/s13225-020-00466-2)) |
| Otu48201 | 1 | k__Fungi(100);p__Ascomycota(100);c__Saccharomycetes(100);o__Saccharomycetales(100);f__Metschnikowiaceae(100);g__Metschnikowia(100);s__Metschnikowia_pulcherrima(100); | Metschnikowia | 13 | Saprotroph | \|Undefined Saprotroph\| | Yeast | NULL | Probable | As nectar-tap_saprotroph (Põlme et al. 2020); Decay sugar-rich substrates (Põlme et al. 2020) | Põlme S, et al. 2020. Fungal Diversity 105:1-16 ((https://doi.org/10.1007/s13225-020-00466-2)) |
| Otu48202 | 1 | k__Fungi(100);p__Ascomycota(100);c__Saccharomycetes(100);o__Saccharomycetales(100);f__Metschnikowiaceae(100);g__Metschnikowia(100);s__Metschnikowia_pulcherrima(100); | Metschnikowia | 13 | Saprotroph | \|Undefined Saprotroph\| | Yeast | NULL | Probable | As nectar-tap_saprotroph (Põlme et al. 2020); Decay sugar-rich substrates (Põlme et al. 2020) | Põlme S, et al. 2020. Fungal Diversity 105:1-16 ((https://doi.org/10.1007/s13225-020-00466-2)) |
| Otu48203 | 1 | k__Fungi(100);p__Ascomycota(100);c__Saccharomycetes(100);o__Saccharomycetales(100);f__Metschnikowiaceae(100);g__Metschnikowia(100);s__Metschnikowia_pulcherrima(100); | Metschnikowia | 13 | Saprotroph | \|Undefined Saprotroph\| | Yeast | NULL | Probable | As nectar-tap_saprotroph (Põlme et al. 2020); Decay sugar-rich substrates (Põlme et al. 2020) | Põlme S, et al. 2020. Fungal Diversity 105:1-16 ((https://doi.org/10.1007/s13225-020-00466-2)) |
| Otu48204 | 1 | k__Fungi(100);p__Ascomycota(100);c__Saccharomycetes(100);o__Saccharomycetales(100);f__Metschnikowiaceae(100);g__Metschnikowia(100);s__Metschnikowia_pulcherrima(100); | Metschnikowia | 13 | Saprotroph | \|Undefined Saprotroph\| | Yeast | NULL | Probable | As nectar-tap_saprotroph (Põlme et al. 2020); Decay sugar-rich substrates (Põlme et al. 2020) | Põlme S, et al. 2020. Fungal Diversity 105:1-16 ((https://doi.org/10.1007/s13225-020-00466-2)) |
| Otu48205 | 1 | k__Fungi(100);p__Ascomycota(100);c__Saccharomycetes(100);o__Saccharomycetales(100);f__Metschnikowiaceae(100);g__Metschnikowia(100);s__Metschnikowia_pulcherrima(100); | Metschnikowia | 13 | Saprotroph | \|Undefined Saprotroph\| | Yeast | NULL | Probable | As nectar-tap_saprotroph (Põlme et al. 2020); Decay sugar-rich substrates (Põlme et al. 2020) | Põlme S, et al. 2020. Fungal Diversity 105:1-16 ((https://doi.org/10.1007/s13225-020-00466-2)) |
| Otu48206 | 1 | k__Fungi(100);p__Ascomycota(100);c__Saccharomycetes(100);o__Saccharomycetales(100);f__Metschnikowiaceae(100);g__Metschnikowia(100);s__Metschnikowia_pulcherrima(100); | Metschnikowia | 13 | Saprotroph | \|Undefined Saprotroph\| | Yeast | NULL | Probable | As nectar-tap_saprotroph (Põlme et al. 2020); Decay sugar-rich substrates (Põlme et al. 2020) | Põlme S, et al. 2020. Fungal Diversity 105:1-16 ((https://doi.org/10.1007/s13225-020-00466-2)) |
| Otu48207 | 1 | k__Fungi(100);p__Ascomycota(100);c__Saccharomycetes(100);o__Saccharomycetales(100);f__Metschnikowiaceae(100);g__Metschnikowia(100);s__Metschnikowia_pulcherrima(100); | Metschnikowia | 13 | Saprotroph | \|Undefined Saprotroph\| | Yeast | NULL | Probable | As nectar-tap_saprotroph (Põlme et al. 2020); Decay sugar-rich substrates (Põlme et al. 2020) | Põlme S, et al. 2020. Fungal Diversity 105:1-16 ((https://doi.org/10.1007/s13225-020-00466-2)) |
| Otu48208 | 1 | k__Fungi(100);p__Ascomycota(100);c__Saccharomycetes(100);o__Saccharomycetales(100);f__Metschnikowiaceae(100);g__Metschnikowia(100);s__Metschnikowia_pulcherrima(100); | Metschnikowia | 13 | Saprotroph | \|Undefined Saprotroph\| | Yeast | NULL | Probable | As nectar-tap_saprotroph (Põlme et al. 2020); Decay sugar-rich substrates (Põlme et al. 2020) | Põlme S, et al. 2020. Fungal Diversity 105:1-16 ((https://doi.org/10.1007/s13225-020-00466-2)) |
| Otu48209 | 1 | k__Fungi(100);p__Ascomycota(100);c__Saccharomycetes(100);o__Saccharomycetales(100);f__Metschnikowiaceae(100);g__Metschnikowia(100);g__Metschnikowia_unclassified(100); | Metschnikowia | 13 | Saprotroph | \|Undefined Saprotroph\| | Yeast | NULL | Probable | As nectar-tap_saprotroph (Põlme et al. 2020); Decay sugar-rich substrates (Põlme et al. 2020) | Põlme S, et al. 2020. Fungal Diversity 105:1-16 ((https://doi.org/10.1007/s13225-020-00466-2)) |
| Otu48210 | 1 | k__Fungi(100);p__Ascomycota(100);c__Saccharomycetes(100);o__Saccharomycetales(100);f__Metschnikowiaceae(100);g__Metschnikowia(100);g__Metschnikowia_unclassified(100); | Metschnikowia | 13 | Saprotroph | \|Undefined Saprotroph\| | Yeast | NULL | Probable | As nectar-tap_saprotroph (Põlme et al. 2020); Decay sugar-rich substrates (Põlme et al. 2020) | Põlme S, et al. 2020. Fungal Diversity 105:1-16 ((https://doi.org/10.1007/s13225-020-00466-2)) |
| Otu48211 | 1 | k__Fungi(100);p__Ascomycota(100);c__Saccharomycetes(100);o__Saccharomycetales(100);f__Metschnikowiaceae(100);g__Metschnikowia(100);s__Metschnikowia_pulcherrima(100); | Metschnikowia | 13 | Saprotroph | \|Undefined Saprotroph\| | Yeast | NULL | Probable | As nectar-tap_saprotroph (Põlme et al. 2020); Decay sugar-rich substrates (Põlme et al. 2020) | Põlme S, et al. 2020. Fungal Diversity 105:1-16 ((https://doi.org/10.1007/s13225-020-00466-2)) |
| Otu48212 | 1 | k__Fungi(100);p__Ascomycota(100);c__Saccharomycetes(100);o__Saccharomycetales(100);f__Metschnikowiaceae(100);g__Metschnikowia(100);g__Metschnikowia_unclassified(100); | Metschnikowia | 13 | Saprotroph | \|Undefined Saprotroph\| | Yeast | NULL | Probable | As nectar-tap_saprotroph (Põlme et al. 2020); Decay sugar-rich substrates (Põlme et al. 2020) | Põlme S, et al. 2020. Fungal Diversity 105:1-16 ((https://doi.org/10.1007/s13225-020-00466-2)) |
| Otu48213 | 1 | k__Fungi(100);p__Ascomycota(100);c__Saccharomycetes(100);o__Saccharomycetales(100);f__Metschnikowiaceae(100);g__Metschnikowia(100);s__Metschnikowia_pulcherrima(100); | Metschnikowia | 13 | Saprotroph | \|Undefined Saprotroph\| | Yeast | NULL | Probable | As nectar-tap_saprotroph (Põlme et al. 2020); Decay sugar-rich substrates (Põlme et al. 2020) | Põlme S, et al. 2020. Fungal Diversity 105:1-16 ((https://doi.org/10.1007/s13225-020-00466-2)) |
| Otu48214 | 1 | k__Fungi(100);p__Ascomycota(100);c__Saccharomycetes(100);o__Saccharomycetales(100);f__Metschnikowiaceae(100);g__Metschnikowia(100);s__Metschnikowia_pulcherrima(100); | Metschnikowia | 13 | Saprotroph | \|Undefined Saprotroph\| | Yeast | NULL | Probable | As nectar-tap_saprotroph (Põlme et al. 2020); Decay sugar-rich substrates (Põlme et al. 2020) | Põlme S, et al. 2020. Fungal Diversity 105:1-16 ((https://doi.org/10.1007/s13225-020-00466-2)) |
| Otu48215 | 1 | k__Fungi(100);p__Ascomycota(100);c__Saccharomycetes(100);o__Saccharomycetales(100);f__Metschnikowiaceae(100);g__Metschnikowia(100);g__Metschnikowia_unclassified(100); | Metschnikowia | 13 | Saprotroph | \|Undefined Saprotroph\| | Yeast | NULL | Probable | As nectar-tap_saprotroph (Põlme et al. 2020); Decay sugar-rich substrates (Põlme et al. 2020) | Põlme S, et al. 2020. Fungal Diversity 105:1-16 ((https://doi.org/10.1007/s13225-020-00466-2)) |
| Otu48216 | 1 | k__Fungi(100);p__Ascomycota(100);c__Saccharomycetes(100);o__Saccharomycetales(100);f__Metschnikowiaceae(100);g__Metschnikowia(100);g__Metschnikowia_unclassified(100); | Metschnikowia | 13 | Saprotroph | \|Undefined Saprotroph\| | Yeast | NULL | Probable | As nectar-tap_saprotroph (Põlme et al. 2020); Decay sugar-rich substrates (Põlme et al. 2020) | Põlme S, et al. 2020. Fungal Diversity 105:1-16 ((https://doi.org/10.1007/s13225-020-00466-2)) |
| Otu48217 | 1 | k__Fungi(100);p__Ascomycota(100);c__Saccharomycetes(100);o__Saccharomycetales(100);f__Metschnikowiaceae(100);g__Metschnikowia(100);s__Metschnikowia_pulcherrima(100); | Metschnikowia | 13 | Saprotroph | \|Undefined Saprotroph\| | Yeast | NULL | Probable | As nectar-tap_saprotroph (Põlme et al. 2020); Decay sugar-rich substrates (Põlme et al. 2020) | Põlme S, et al. 2020. Fungal Diversity 105:1-16 ((https://doi.org/10.1007/s13225-020-00466-2)) |
| Otu48218 | 1 | k__Fungi(100);p__Ascomycota(100);c__Saccharomycetes(100);o__Saccharomycetales(100);f__Metschnikowiaceae(100);g__Metschnikowia(100);s__Metschnikowia_pulcherrima(100); | Metschnikowia | 13 | Saprotroph | \|Undefined Saprotroph\| | Yeast | NULL | Probable | As nectar-tap_saprotroph (Põlme et al. 2020); Decay sugar-rich substrates (Põlme et al. 2020) | Põlme S, et al. 2020. Fungal Diversity 105:1-16 ((https://doi.org/10.1007/s13225-020-00466-2)) |
| Otu48219 | 1 | k__Fungi(100);p__Ascomycota(100);c__Saccharomycetes(100);o__Saccharomycetales(100);f__Metschnikowiaceae(100);g__Metschnikowia(100);s__Metschnikowia_pulcherrima(100); | Metschnikowia | 13 | Saprotroph | \|Undefined Saprotroph\| | Yeast | NULL | Probable | As nectar-tap_saprotroph (Põlme et al. 2020); Decay sugar-rich substrates (Põlme et al. 2020) | Põlme S, et al. 2020. Fungal Diversity 105:1-16 ((https://doi.org/10.1007/s13225-020-00466-2)) |
| Otu48220 | 1 | k__Fungi(100);p__Ascomycota(100);c__Saccharomycetes(100);o__Saccharomycetales(100);f__Metschnikowiaceae(100);g__Metschnikowia(100);g__Metschnikowia_unclassified(100); | Metschnikowia | 13 | Saprotroph | \|Undefined Saprotroph\| | Yeast | NULL | Probable | As nectar-tap_saprotroph (Põlme et al. 2020); Decay sugar-rich substrates (Põlme et al. 2020) | Põlme S, et al. 2020. Fungal Diversity 105:1-16 ((https://doi.org/10.1007/s13225-020-00466-2)) |
| Otu48221 | 1 | k__Fungi(100);p__Ascomycota(100);c__Saccharomycetes(100);o__Saccharomycetales(100);f__Metschnikowiaceae(100);g__Metschnikowia(100);s__Metschnikowia_pulcherrima(100); | Metschnikowia | 13 | Saprotroph | \|Undefined Saprotroph\| | Yeast | NULL | Probable | As nectar-tap_saprotroph (Põlme et al. 2020); Decay sugar-rich substrates (Põlme et al. 2020) | Põlme S, et al. 2020. Fungal Diversity 105:1-16 ((https://doi.org/10.1007/s13225-020-00466-2)) |
| Otu48222 | 1 | k__Fungi(100);p__Ascomycota(100);c__Saccharomycetes(100);o__Saccharomycetales(100);f__Metschnikowiaceae(100);g__Metschnikowia(100);s__Metschnikowia_pulcherrima(100); | Metschnikowia | 13 | Saprotroph | \|Undefined Saprotroph\| | Yeast | NULL | Probable | As nectar-tap_saprotroph (Põlme et al. 2020); Decay sugar-rich substrates (Põlme et al. 2020) | Põlme S, et al. 2020. Fungal Diversity 105:1-16 ((https://doi.org/10.1007/s13225-020-00466-2)) |
| Otu48223 | 1 | k__Fungi(100);p__Ascomycota(100);c__Saccharomycetes(100);o__Saccharomycetales(100);f__Metschnikowiaceae(100);g__Metschnikowia(100);s__Metschnikowia_pulcherrima(100); | Metschnikowia | 13 | Saprotroph | \|Undefined Saprotroph\| | Yeast | NULL | Probable | As nectar-tap_saprotroph (Põlme et al. 2020); Decay sugar-rich substrates (Põlme et al. 2020) | Põlme S, et al. 2020. Fungal Diversity 105:1-16 ((https://doi.org/10.1007/s13225-020-00466-2)) |
| Otu48224 | 1 | k__Fungi(100);p__Ascomycota(100);c__Saccharomycetes(100);o__Saccharomycetales(100);f__Metschnikowiaceae(100);g__Metschnikowia(100);g__Metschnikowia_unclassified(100); | Metschnikowia | 13 | Saprotroph | \|Undefined Saprotroph\| | Yeast | NULL | Probable | As nectar-tap_saprotroph (Põlme et al. 2020); Decay sugar-rich substrates (Põlme et al. 2020) | Põlme S, et al. 2020. Fungal Diversity 105:1-16 ((https://doi.org/10.1007/s13225-020-00466-2)) |
| Otu48225 | 1 | k__Fungi(100);p__Ascomycota(100);c__Saccharomycetes(100);o__Saccharomycetales(100);f__Metschnikowiaceae(100);g__Metschnikowia(100);g__Metschnikowia_unclassified(100); | Metschnikowia | 13 | Saprotroph | \|Undefined Saprotroph\| | Yeast | NULL | Probable | As nectar-tap_saprotroph (Põlme et al. 2020); Decay sugar-rich substrates (Põlme et al. 2020) | Põlme S, et al. 2020. Fungal Diversity 105:1-16 ((https://doi.org/10.1007/s13225-020-00466-2)) |
| Otu48226 | 1 | k__Fungi(100);p__Ascomycota(100);c__Saccharomycetes(100);o__Saccharomycetales(100);f__Metschnikowiaceae(100);g__Metschnikowia(100);g__Metschnikowia_unclassified(100); | Metschnikowia | 13 | Saprotroph | \|Undefined Saprotroph\| | Yeast | NULL | Probable | As nectar-tap_saprotroph (Põlme et al. 2020); Decay sugar-rich substrates (Põlme et al. 2020) | Põlme S, et al. 2020. Fungal Diversity 105:1-16 ((https://doi.org/10.1007/s13225-020-00466-2)) |
| Otu48227 | 1 | k__Fungi(100);p__Ascomycota(100);c__Saccharomycetes(100);o__Saccharomycetales(100);f__Metschnikowiaceae(100);g__Metschnikowia(100);s__Metschnikowia_pulcherrima(100); | Metschnikowia | 13 | Saprotroph | \|Undefined Saprotroph\| | Yeast | NULL | Probable | As nectar-tap_saprotroph (Põlme et al. 2020); Decay sugar-rich substrates (Põlme et al. 2020) | Põlme S, et al. 2020. Fungal Diversity 105:1-16 ((https://doi.org/10.1007/s13225-020-00466-2)) |
| Otu48228 | 1 | k__Fungi(100);p__Ascomycota(100);c__Saccharomycetes(100);o__Saccharomycetales(100);f__Metschnikowiaceae(100);g__Metschnikowia(100);s__Metschnikowia_pulcherrima(100); | Metschnikowia | 13 | Saprotroph | \|Undefined Saprotroph\| | Yeast | NULL | Probable | As nectar-tap_saprotroph (Põlme et al. 2020); Decay sugar-rich substrates (Põlme et al. 2020) | Põlme S, et al. 2020. Fungal Diversity 105:1-16 ((https://doi.org/10.1007/s13225-020-00466-2)) |
| Otu48229 | 1 | k__Fungi(100);p__Ascomycota(100);c__Saccharomycetes(100);o__Saccharomycetales(100);f__Metschnikowiaceae(100);g__Metschnikowia(100);g__Metschnikowia_unclassified(100); | Metschnikowia | 13 | Saprotroph | \|Undefined Saprotroph\| | Yeast | NULL | Probable | As nectar-tap_saprotroph (Põlme et al. 2020); Decay sugar-rich substrates (Põlme et al. 2020) | Põlme S, et al. 2020. Fungal Diversity 105:1-16 ((https://doi.org/10.1007/s13225-020-00466-2)) |
| Otu48230 | 1 | k__Fungi(100);p__Ascomycota(100);c__Saccharomycetes(100);o__Saccharomycetales(100);f__Metschnikowiaceae(100);g__Metschnikowia(100);s__Metschnikowia_pulcherrima(100); | Metschnikowia | 13 | Saprotroph | \|Undefined Saprotroph\| | Yeast | NULL | Probable | As nectar-tap_saprotroph (Põlme et al. 2020); Decay sugar-rich substrates (Põlme et al. 2020) | Põlme S, et al. 2020. Fungal Diversity 105:1-16 ((https://doi.org/10.1007/s13225-020-00466-2)) |
| Otu48231 | 1 | k__Fungi(100);p__Ascomycota(100);c__Saccharomycetes(100);o__Saccharomycetales(100);f__Metschnikowiaceae(100);g__Metschnikowia(100);s__Metschnikowia_pulcherrima(100); | Metschnikowia | 13 | Saprotroph | \|Undefined Saprotroph\| | Yeast | NULL | Probable | As nectar-tap_saprotroph (Põlme et al. 2020); Decay sugar-rich substrates (Põlme et al. 2020) | Põlme S, et al. 2020. Fungal Diversity 105:1-16 ((https://doi.org/10.1007/s13225-020-00466-2)) |
| Otu48232 | 1 | k__Fungi(100);p__Ascomycota(100);c__Saccharomycetes(100);o__Saccharomycetales(100);f__Metschnikowiaceae(100);g__Metschnikowia(100);g__Metschnikowia_unclassified(100); | Metschnikowia | 13 | Saprotroph | \|Undefined Saprotroph\| | Yeast | NULL | Probable | As nectar-tap_saprotroph (Põlme et al. 2020); Decay sugar-rich substrates (Põlme et al. 2020) | Põlme S, et al. 2020. Fungal Diversity 105:1-16 ((https://doi.org/10.1007/s13225-020-00466-2)) |
| Otu48233 | 1 | k__Fungi(100);p__Ascomycota(100);c__Saccharomycetes(100);o__Saccharomycetales(100);f__Metschnikowiaceae(100);g__Metschnikowia(100);g__Metschnikowia_unclassified(100); | Metschnikowia | 13 | Saprotroph | \|Undefined Saprotroph\| | Yeast | NULL | Probable | As nectar-tap_saprotroph (Põlme et al. 2020); Decay sugar-rich substrates (Põlme et al. 2020) | Põlme S, et al. 2020. Fungal Diversity 105:1-16 ((https://doi.org/10.1007/s13225-020-00466-2)) |
| Otu48234 | 1 | k__Fungi(100);p__Ascomycota(100);c__Saccharomycetes(100);o__Saccharomycetales(100);f__Metschnikowiaceae(100);g__Metschnikowia(100);s__Metschnikowia_pulcherrima(100); | Metschnikowia | 13 | Saprotroph | \|Undefined Saprotroph\| | Yeast | NULL | Probable | As nectar-tap_saprotroph (Põlme et al. 2020); Decay sugar-rich substrates (Põlme et al. 2020) | Põlme S, et al. 2020. Fungal Diversity 105:1-16 ((https://doi.org/10.1007/s13225-020-00466-2)) |
| Otu48235 | 1 | k__Fungi(100);p__Ascomycota(100);c__Saccharomycetes(100);o__Saccharomycetales(100);f__Metschnikowiaceae(100);g__Metschnikowia(100);s__Metschnikowia_pulcherrima(100); | Metschnikowia | 13 | Saprotroph | \|Undefined Saprotroph\| | Yeast | NULL | Probable | As nectar-tap_saprotroph (Põlme et al. 2020); Decay sugar-rich substrates (Põlme et al. 2020) | Põlme S, et al. 2020. Fungal Diversity 105:1-16 ((https://doi.org/10.1007/s13225-020-00466-2)) |
| Otu48236 | 1 | k__Fungi(100);p__Ascomycota(100);c__Saccharomycetes(100);o__Saccharomycetales(100);f__Metschnikowiaceae(100);g__Metschnikowia(100);g__Metschnikowia_unclassified(100); | Metschnikowia | 13 | Saprotroph | \|Undefined Saprotroph\| | Yeast | NULL | Probable | As nectar-tap_saprotroph (Põlme et al. 2020); Decay sugar-rich substrates (Põlme et al. 2020) | Põlme S, et al. 2020. Fungal Diversity 105:1-16 ((https://doi.org/10.1007/s13225-020-00466-2)) |
| Otu48238 | 1 | k__Fungi(100);p__Ascomycota(100);c__Saccharomycetes(100);o__Saccharomycetales(100);f__Metschnikowiaceae(100);g__Metschnikowia(100);g__Metschnikowia_unclassified(100); | Metschnikowia | 13 | Saprotroph | \|Undefined Saprotroph\| | Yeast | NULL | Probable | As nectar-tap_saprotroph (Põlme et al. 2020); Decay sugar-rich substrates (Põlme et al. 2020) | Põlme S, et al. 2020. Fungal Diversity 105:1-16 ((https://doi.org/10.1007/s13225-020-00466-2)) |
| Otu48239 | 1 | k__Fungi(100);p__Ascomycota(100);c__Saccharomycetes(100);o__Saccharomycetales(100);f__Metschnikowiaceae(100);g__Metschnikowia(100);s__Metschnikowia_pulcherrima(100); | Metschnikowia | 13 | Saprotroph | \|Undefined Saprotroph\| | Yeast | NULL | Probable | As nectar-tap_saprotroph (Põlme et al. 2020); Decay sugar-rich substrates (Põlme et al. 2020) | Põlme S, et al. 2020. Fungal Diversity 105:1-16 ((https://doi.org/10.1007/s13225-020-00466-2)) |
| Otu48240 | 1 | k__Fungi(100);p__Ascomycota(100);c__Saccharomycetes(100);o__Saccharomycetales(100);f__Metschnikowiaceae(100);g__Metschnikowia(100);s__Metschnikowia_pulcherrima(100); | Metschnikowia | 13 | Saprotroph | \|Undefined Saprotroph\| | Yeast | NULL | Probable | As nectar-tap_saprotroph (Põlme et al. 2020); Decay sugar-rich substrates (Põlme et al. 2020) | Põlme S, et al. 2020. Fungal Diversity 105:1-16 ((https://doi.org/10.1007/s13225-020-00466-2)) |
| Otu48241 | 1 | k__Fungi(100);p__Ascomycota(100);c__Saccharomycetes(100);o__Saccharomycetales(100);f__Metschnikowiaceae(100);g__Metschnikowia(100);s__Metschnikowia_pulcherrima(100); | Metschnikowia | 13 | Saprotroph | \|Undefined Saprotroph\| | Yeast | NULL | Probable | As nectar-tap_saprotroph (Põlme et al. 2020); Decay sugar-rich substrates (Põlme et al. 2020) | Põlme S, et al. 2020. Fungal Diversity 105:1-16 ((https://doi.org/10.1007/s13225-020-00466-2)) |
| Otu48242 | 1 | k__Fungi(100);p__Ascomycota(100);c__Saccharomycetes(100);o__Saccharomycetales(100);f__Metschnikowiaceae(100);g__Metschnikowia(100);s__Metschnikowia_pulcherrima(100); | Metschnikowia | 13 | Saprotroph | \|Undefined Saprotroph\| | Yeast | NULL | Probable | As nectar-tap_saprotroph (Põlme et al. 2020); Decay sugar-rich substrates (Põlme et al. 2020) | Põlme S, et al. 2020. Fungal Diversity 105:1-16 ((https://doi.org/10.1007/s13225-020-00466-2)) |
| Otu48243 | 1 | k__Fungi(100);p__Ascomycota(100);c__Saccharomycetes(100);o__Saccharomycetales(100);f__Metschnikowiaceae(100);g__Metschnikowia(100);s__Metschnikowia_pulcherrima(100); | Metschnikowia | 13 | Saprotroph | \|Undefined Saprotroph\| | Yeast | NULL | Probable | As nectar-tap_saprotroph (Põlme et al. 2020); Decay sugar-rich substrates (Põlme et al. 2020) | Põlme S, et al. 2020. Fungal Diversity 105:1-16 ((https://doi.org/10.1007/s13225-020-00466-2)) |
| Otu48244 | 1 | k__Fungi(100);p__Ascomycota(100);c__Saccharomycetes(100);o__Saccharomycetales(100);f__Metschnikowiaceae(100);g__Metschnikowia(100);s__Metschnikowia_pulcherrima(100); | Metschnikowia | 13 | Saprotroph | \|Undefined Saprotroph\| | Yeast | NULL | Probable | As nectar-tap_saprotroph (Põlme et al. 2020); Decay sugar-rich substrates (Põlme et al. 2020) | Põlme S, et al. 2020. Fungal Diversity 105:1-16 ((https://doi.org/10.1007/s13225-020-00466-2)) |
| Otu48245 | 1 | k__Fungi(100);p__Ascomycota(100);c__Saccharomycetes(100);o__Saccharomycetales(100);f__Metschnikowiaceae(100);g__Metschnikowia(100);s__Metschnikowia_pulcherrima(100); | Metschnikowia | 13 | Saprotroph | \|Undefined Saprotroph\| | Yeast | NULL | Probable | As nectar-tap_saprotroph (Põlme et al. 2020); Decay sugar-rich substrates (Põlme et al. 2020) | Põlme S, et al. 2020. Fungal Diversity 105:1-16 ((https://doi.org/10.1007/s13225-020-00466-2)) |
| Otu48246 | 1 | k__Fungi(100);p__Ascomycota(100);c__Saccharomycetes(100);o__Saccharomycetales(100);f__Metschnikowiaceae(100);g__Metschnikowia(100);s__Metschnikowia_pulcherrima(100); | Metschnikowia | 13 | Saprotroph | \|Undefined Saprotroph\| | Yeast | NULL | Probable | As nectar-tap_saprotroph (Põlme et al. 2020); Decay sugar-rich substrates (Põlme et al. 2020) | Põlme S, et al. 2020. Fungal Diversity 105:1-16 ((https://doi.org/10.1007/s13225-020-00466-2)) |
| Otu48247 | 1 | k__Fungi(100);p__Ascomycota(100);c__Saccharomycetes(100);o__Saccharomycetales(100);f__Metschnikowiaceae(100);g__Metschnikowia(100);s__Metschnikowia_pulcherrima(100); | Metschnikowia | 13 | Saprotroph | \|Undefined Saprotroph\| | Yeast | NULL | Probable | As nectar-tap_saprotroph (Põlme et al. 2020); Decay sugar-rich substrates (Põlme et al. 2020) | Põlme S, et al. 2020. Fungal Diversity 105:1-16 ((https://doi.org/10.1007/s13225-020-00466-2)) |
| Otu48248 | 1 | k__Fungi(100);p__Ascomycota(100);c__Saccharomycetes(100);o__Saccharomycetales(100);f__Metschnikowiaceae(100);g__Metschnikowia(100);g__Metschnikowia_unclassified(100); | Metschnikowia | 13 | Saprotroph | \|Undefined Saprotroph\| | Yeast | NULL | Probable | As nectar-tap_saprotroph (Põlme et al. 2020); Decay sugar-rich substrates (Põlme et al. 2020) | Põlme S, et al. 2020. Fungal Diversity 105:1-16 ((https://doi.org/10.1007/s13225-020-00466-2)) |
| Otu48249 | 1 | k__Fungi(100);p__Ascomycota(100);c__Saccharomycetes(100);o__Saccharomycetales(100);f__Metschnikowiaceae(100);g__Metschnikowia(100);s__Metschnikowia_pulcherrima(100); | Metschnikowia | 13 | Saprotroph | \|Undefined Saprotroph\| | Yeast | NULL | Probable | As nectar-tap_saprotroph (Põlme et al. 2020); Decay sugar-rich substrates (Põlme et al. 2020) | Põlme S, et al. 2020. Fungal Diversity 105:1-16 ((https://doi.org/10.1007/s13225-020-00466-2)) |
| Otu48250 | 1 | k__Fungi(100);p__Ascomycota(100);c__Saccharomycetes(100);o__Saccharomycetales(100);f__Metschnikowiaceae(100);g__Metschnikowia(100);s__Metschnikowia_pulcherrima(100); | Metschnikowia | 13 | Saprotroph | \|Undefined Saprotroph\| | Yeast | NULL | Probable | As nectar-tap_saprotroph (Põlme et al. 2020); Decay sugar-rich substrates (Põlme et al. 2020) | Põlme S, et al. 2020. Fungal Diversity 105:1-16 ((https://doi.org/10.1007/s13225-020-00466-2)) |
| Otu48251 | 1 | k__Fungi(100);p__Ascomycota(100);c__Saccharomycetes(100);o__Saccharomycetales(100);f__Metschnikowiaceae(100);g__Metschnikowia(100);s__Metschnikowia_pulcherrima(100); | Metschnikowia | 13 | Saprotroph | \|Undefined Saprotroph\| | Yeast | NULL | Probable | As nectar-tap_saprotroph (Põlme et al. 2020); Decay sugar-rich substrates (Põlme et al. 2020) | Põlme S, et al. 2020. Fungal Diversity 105:1-16 ((https://doi.org/10.1007/s13225-020-00466-2)) |
| Otu48252 | 1 | k__Fungi(100);p__Ascomycota(100);c__Saccharomycetes(100);o__Saccharomycetales(100);f__Metschnikowiaceae(100);g__Metschnikowia(100);g__Metschnikowia_unclassified(100); | Metschnikowia | 13 | Saprotroph | \|Undefined Saprotroph\| | Yeast | NULL | Probable | As nectar-tap_saprotroph (Põlme et al. 2020); Decay sugar-rich substrates (Põlme et al. 2020) | Põlme S, et al. 2020. Fungal Diversity 105:1-16 ((https://doi.org/10.1007/s13225-020-00466-2)) |
| Otu48253 | 1 | k__Fungi(100);p__Ascomycota(100);c__Saccharomycetes(100);o__Saccharomycetales(100);f__Metschnikowiaceae(100);g__Metschnikowia(100);s__Metschnikowia_pulcherrima(100); | Metschnikowia | 13 | Saprotroph | \|Undefined Saprotroph\| | Yeast | NULL | Probable | As nectar-tap_saprotroph (Põlme et al. 2020); Decay sugar-rich substrates (Põlme et al. 2020) | Põlme S, et al. 2020. Fungal Diversity 105:1-16 ((https://doi.org/10.1007/s13225-020-00466-2)) |
| Otu48254 | 1 | k__Fungi(100);p__Ascomycota(100);c__Saccharomycetes(100);o__Saccharomycetales(100);f__Metschnikowiaceae(100);g__Metschnikowia(100);s__Metschnikowia_pulcherrima(100); | Metschnikowia | 13 | Saprotroph | \|Undefined Saprotroph\| | Yeast | NULL | Probable | As nectar-tap_saprotroph (Põlme et al. 2020); Decay sugar-rich substrates (Põlme et al. 2020) | Põlme S, et al. 2020. Fungal Diversity 105:1-16 ((https://doi.org/10.1007/s13225-020-00466-2)) |
| Otu48255 | 1 | k__Fungi(100);p__Ascomycota(100);c__Saccharomycetes(100);o__Saccharomycetales(100);f__Metschnikowiaceae(100);g__Metschnikowia(100);s__Metschnikowia_pulcherrima(100); | Metschnikowia | 13 | Saprotroph | \|Undefined Saprotroph\| | Yeast | NULL | Probable | As nectar-tap_saprotroph (Põlme et al. 2020); Decay sugar-rich substrates (Põlme et al. 2020) | Põlme S, et al. 2020. Fungal Diversity 105:1-16 ((https://doi.org/10.1007/s13225-020-00466-2)) |
| Otu48256 | 1 | k__Fungi(100);p__Ascomycota(100);c__Saccharomycetes(100);o__Saccharomycetales(100);f__Metschnikowiaceae(100);g__Metschnikowia(100);g__Metschnikowia_unclassified(100); | Metschnikowia | 13 | Saprotroph | \|Undefined Saprotroph\| | Yeast | NULL | Probable | As nectar-tap_saprotroph (Põlme et al. 2020); Decay sugar-rich substrates (Põlme et al. 2020) | Põlme S, et al. 2020. Fungal Diversity 105:1-16 ((https://doi.org/10.1007/s13225-020-00466-2)) |
| Otu48257 | 1 | k__Fungi(100);p__Ascomycota(100);c__Saccharomycetes(100);o__Saccharomycetales(100);f__Metschnikowiaceae(100);g__Metschnikowia(100);s__Metschnikowia_pulcherrima(100); | Metschnikowia | 13 | Saprotroph | \|Undefined Saprotroph\| | Yeast | NULL | Probable | As nectar-tap_saprotroph (Põlme et al. 2020); Decay sugar-rich substrates (Põlme et al. 2020) | Põlme S, et al. 2020. Fungal Diversity 105:1-16 ((https://doi.org/10.1007/s13225-020-00466-2)) |
| Otu48258 | 1 | k__Fungi(100);p__Ascomycota(100);c__Saccharomycetes(100);o__Saccharomycetales(100);f__Metschnikowiaceae(100);g__Metschnikowia(100);s__Metschnikowia_pulcherrima(100); | Metschnikowia | 13 | Saprotroph | \|Undefined Saprotroph\| | Yeast | NULL | Probable | As nectar-tap_saprotroph (Põlme et al. 2020); Decay sugar-rich substrates (Põlme et al. 2020) | Põlme S, et al. 2020. Fungal Diversity 105:1-16 ((https://doi.org/10.1007/s13225-020-00466-2)) |
| Otu48259 | 1 | k__Fungi(100);p__Ascomycota(100);c__Saccharomycetes(100);o__Saccharomycetales(100);f__Metschnikowiaceae(100);g__Metschnikowia(100);g__Metschnikowia_unclassified(100); | Metschnikowia | 13 | Saprotroph | \|Undefined Saprotroph\| | Yeast | NULL | Probable | As nectar-tap_saprotroph (Põlme et al. 2020); Decay sugar-rich substrates (Põlme et al. 2020) | Põlme S, et al. 2020. Fungal Diversity 105:1-16 ((https://doi.org/10.1007/s13225-020-00466-2)) |
| Otu48260 | 1 | k__Fungi(100);p__Ascomycota(100);c__Saccharomycetes(100);o__Saccharomycetales(100);f__Metschnikowiaceae(100);g__Metschnikowia(100);s__Metschnikowia_pulcherrima(100); | Metschnikowia | 13 | Saprotroph | \|Undefined Saprotroph\| | Yeast | NULL | Probable | As nectar-tap_saprotroph (Põlme et al. 2020); Decay sugar-rich substrates (Põlme et al. 2020) | Põlme S, et al. 2020. Fungal Diversity 105:1-16 ((https://doi.org/10.1007/s13225-020-00466-2)) |
| Otu48262 | 1 | k__Fungi(100);p__Ascomycota(100);c__Saccharomycetes(100);o__Saccharomycetales(100);f__Metschnikowiaceae(100);g__Metschnikowia(100);s__Metschnikowia_pulcherrima(100); | Metschnikowia | 13 | Saprotroph | \|Undefined Saprotroph\| | Yeast | NULL | Probable | As nectar-tap_saprotroph (Põlme et al. 2020); Decay sugar-rich substrates (Põlme et al. 2020) | Põlme S, et al. 2020. Fungal Diversity 105:1-16 ((https://doi.org/10.1007/s13225-020-00466-2)) |
| Otu48263 | 1 | k__Fungi(100);p__Ascomycota(100);c__Saccharomycetes(100);o__Saccharomycetales(100);f__Metschnikowiaceae(100);g__Metschnikowia(100);s__Metschnikowia_pulcherrima(100); | Metschnikowia | 13 | Saprotroph | \|Undefined Saprotroph\| | Yeast | NULL | Probable | As nectar-tap_saprotroph (Põlme et al. 2020); Decay sugar-rich substrates (Põlme et al. 2020) | Põlme S, et al. 2020. Fungal Diversity 105:1-16 ((https://doi.org/10.1007/s13225-020-00466-2)) |
| Otu48264 | 1 | k__Fungi(100);p__Ascomycota(100);c__Saccharomycetes(100);o__Saccharomycetales(100);f__Metschnikowiaceae(100);g__Metschnikowia(100);s__Metschnikowia_pulcherrima(100); | Metschnikowia | 13 | Saprotroph | \|Undefined Saprotroph\| | Yeast | NULL | Probable | As nectar-tap_saprotroph (Põlme et al. 2020); Decay sugar-rich substrates (Põlme et al. 2020) | Põlme S, et al. 2020. Fungal Diversity 105:1-16 ((https://doi.org/10.1007/s13225-020-00466-2)) |
| Otu48266 | 1 | k__Fungi(100);p__Ascomycota(100);c__Saccharomycetes(100);o__Saccharomycetales(100);f__Metschnikowiaceae(100);g__Metschnikowia(100);s__Metschnikowia_pulcherrima(100); | Metschnikowia | 13 | Saprotroph | \|Undefined Saprotroph\| | Yeast | NULL | Probable | As nectar-tap_saprotroph (Põlme et al. 2020); Decay sugar-rich substrates (Põlme et al. 2020) | Põlme S, et al. 2020. Fungal Diversity 105:1-16 ((https://doi.org/10.1007/s13225-020-00466-2)) |
| Otu48267 | 1 | k__Fungi(100);p__Ascomycota(100);c__Saccharomycetes(100);o__Saccharomycetales(100);f__Metschnikowiaceae(100);g__Metschnikowia(100);s__Metschnikowia_pulcherrima(100); | Metschnikowia | 13 | Saprotroph | \|Undefined Saprotroph\| | Yeast | NULL | Probable | As nectar-tap_saprotroph (Põlme et al. 2020); Decay sugar-rich substrates (Põlme et al. 2020) | Põlme S, et al. 2020. Fungal Diversity 105:1-16 ((https://doi.org/10.1007/s13225-020-00466-2)) |
| Otu48268 | 1 | k__Fungi(100);p__Ascomycota(100);c__Saccharomycetes(100);o__Saccharomycetales(100);f__Metschnikowiaceae(100);g__Metschnikowia(100);s__Metschnikowia_pulcherrima(100); | Metschnikowia | 13 | Saprotroph | \|Undefined Saprotroph\| | Yeast | NULL | Probable | As nectar-tap_saprotroph (Põlme et al. 2020); Decay sugar-rich substrates (Põlme et al. 2020) | Põlme S, et al. 2020. Fungal Diversity 105:1-16 ((https://doi.org/10.1007/s13225-020-00466-2)) |
| Otu48269 | 1 | k__Fungi(100);p__Ascomycota(100);c__Saccharomycetes(100);o__Saccharomycetales(100);f__Metschnikowiaceae(100);g__Metschnikowia(100);s__Metschnikowia_pulcherrima(100); | Metschnikowia | 13 | Saprotroph | \|Undefined Saprotroph\| | Yeast | NULL | Probable | As nectar-tap_saprotroph (Põlme et al. 2020); Decay sugar-rich substrates (Põlme et al. 2020) | Põlme S, et al. 2020. Fungal Diversity 105:1-16 ((https://doi.org/10.1007/s13225-020-00466-2)) |
| Otu48270 | 1 | k__Fungi(100);p__Ascomycota(100);c__Saccharomycetes(100);o__Saccharomycetales(100);f__Metschnikowiaceae(100);g__Metschnikowia(100);s__Metschnikowia_pulcherrima(100); | Metschnikowia | 13 | Saprotroph | \|Undefined Saprotroph\| | Yeast | NULL | Probable | As nectar-tap_saprotroph (Põlme et al. 2020); Decay sugar-rich substrates (Põlme et al. 2020) | Põlme S, et al. 2020. Fungal Diversity 105:1-16 ((https://doi.org/10.1007/s13225-020-00466-2)) |
| Otu48271 | 1 | k__Fungi(100);p__Ascomycota(100);c__Saccharomycetes(100);o__Saccharomycetales(100);f__Metschnikowiaceae(100);g__Metschnikowia(100);s__Metschnikowia_pulcherrima(100); | Metschnikowia | 13 | Saprotroph | \|Undefined Saprotroph\| | Yeast | NULL | Probable | As nectar-tap_saprotroph (Põlme et al. 2020); Decay sugar-rich substrates (Põlme et al. 2020) | Põlme S, et al. 2020. Fungal Diversity 105:1-16 ((https://doi.org/10.1007/s13225-020-00466-2)) |
| Otu48272 | 1 | k__Fungi(100);p__Ascomycota(100);c__Saccharomycetes(100);o__Saccharomycetales(100);f__Metschnikowiaceae(100);g__Metschnikowia(100);g__Metschnikowia_unclassified(100); | Metschnikowia | 13 | Saprotroph | \|Undefined Saprotroph\| | Yeast | NULL | Probable | As nectar-tap_saprotroph (Põlme et al. 2020); Decay sugar-rich substrates (Põlme et al. 2020) | Põlme S, et al. 2020. Fungal Diversity 105:1-16 ((https://doi.org/10.1007/s13225-020-00466-2)) |
| Otu48273 | 1 | k__Fungi(100);p__Ascomycota(100);c__Saccharomycetes(100);o__Saccharomycetales(100);f__Metschnikowiaceae(100);g__Metschnikowia(100);s__Metschnikowia_pulcherrima(100); | Metschnikowia | 13 | Saprotroph | \|Undefined Saprotroph\| | Yeast | NULL | Probable | As nectar-tap_saprotroph (Põlme et al. 2020); Decay sugar-rich substrates (Põlme et al. 2020) | Põlme S, et al. 2020. Fungal Diversity 105:1-16 ((https://doi.org/10.1007/s13225-020-00466-2)) |
| Otu48274 | 1 | k__Fungi(100);p__Ascomycota(100);c__Saccharomycetes(100);o__Saccharomycetales(100);f__Metschnikowiaceae(100);g__Metschnikowia(100);s__Metschnikowia_pulcherrima(100); | Metschnikowia | 13 | Saprotroph | \|Undefined Saprotroph\| | Yeast | NULL | Probable | As nectar-tap_saprotroph (Põlme et al. 2020); Decay sugar-rich substrates (Põlme et al. 2020) | Põlme S, et al. 2020. Fungal Diversity 105:1-16 ((https://doi.org/10.1007/s13225-020-00466-2)) |
| Otu48275 | 1 | k__Fungi(100);p__Ascomycota(100);c__Saccharomycetes(100);o__Saccharomycetales(100);f__Metschnikowiaceae(100);g__Metschnikowia(100);s__Metschnikowia_pulcherrima(100); | Metschnikowia | 13 | Saprotroph | \|Undefined Saprotroph\| | Yeast | NULL | Probable | As nectar-tap_saprotroph (Põlme et al. 2020); Decay sugar-rich substrates (Põlme et al. 2020) | Põlme S, et al. 2020. Fungal Diversity 105:1-16 ((https://doi.org/10.1007/s13225-020-00466-2)) |
| Otu48276 | 1 | k__Fungi(100);p__Ascomycota(100);c__Saccharomycetes(100);o__Saccharomycetales(100);f__Metschnikowiaceae(100);g__Metschnikowia(100);s__Metschnikowia_pulcherrima(100); | Metschnikowia | 13 | Saprotroph | \|Undefined Saprotroph\| | Yeast | NULL | Probable | As nectar-tap_saprotroph (Põlme et al. 2020); Decay sugar-rich substrates (Põlme et al. 2020) | Põlme S, et al. 2020. Fungal Diversity 105:1-16 ((https://doi.org/10.1007/s13225-020-00466-2)) |
| Otu48277 | 1 | k__Fungi(100);p__Ascomycota(100);c__Saccharomycetes(100);o__Saccharomycetales(100);f__Metschnikowiaceae(100);g__Metschnikowia(100);s__Metschnikowia_pulcherrima(100); | Metschnikowia | 13 | Saprotroph | \|Undefined Saprotroph\| | Yeast | NULL | Probable | As nectar-tap_saprotroph (Põlme et al. 2020); Decay sugar-rich substrates (Põlme et al. 2020) | Põlme S, et al. 2020. Fungal Diversity 105:1-16 ((https://doi.org/10.1007/s13225-020-00466-2)) |
| Otu48278 | 1 | k__Fungi(100);p__Ascomycota(100);c__Saccharomycetes(100);o__Saccharomycetales(100);f__Metschnikowiaceae(100);g__Metschnikowia(100);s__Metschnikowia_pulcherrima(100); | Metschnikowia | 13 | Saprotroph | \|Undefined Saprotroph\| | Yeast | NULL | Probable | As nectar-tap_saprotroph (Põlme et al. 2020); Decay sugar-rich substrates (Põlme et al. 2020) | Põlme S, et al. 2020. Fungal Diversity 105:1-16 ((https://doi.org/10.1007/s13225-020-00466-2)) |
| Otu48279 | 1 | k__Fungi(100);p__Ascomycota(100);c__Saccharomycetes(100);o__Saccharomycetales(100);f__Metschnikowiaceae(100);g__Metschnikowia(100);s__Metschnikowia_pulcherrima(100); | Metschnikowia | 13 | Saprotroph | \|Undefined Saprotroph\| | Yeast | NULL | Probable | As nectar-tap_saprotroph (Põlme et al. 2020); Decay sugar-rich substrates (Põlme et al. 2020) | Põlme S, et al. 2020. Fungal Diversity 105:1-16 ((https://doi.org/10.1007/s13225-020-00466-2)) |
| Otu48280 | 1 | k__Fungi(100);p__Ascomycota(100);c__Saccharomycetes(100);o__Saccharomycetales(100);f__Metschnikowiaceae(100);g__Metschnikowia(100);s__Metschnikowia_pulcherrima(100); | Metschnikowia | 13 | Saprotroph | \|Undefined Saprotroph\| | Yeast | NULL | Probable | As nectar-tap_saprotroph (Põlme et al. 2020); Decay sugar-rich substrates (Põlme et al. 2020) | Põlme S, et al. 2020. Fungal Diversity 105:1-16 ((https://doi.org/10.1007/s13225-020-00466-2)) |
| Otu48281 | 1 | k__Fungi(100);p__Ascomycota(100);c__Saccharomycetes(100);o__Saccharomycetales(100);f__Metschnikowiaceae(100);g__Metschnikowia(100);s__Metschnikowia_pulcherrima(100); | Metschnikowia | 13 | Saprotroph | \|Undefined Saprotroph\| | Yeast | NULL | Probable | As nectar-tap_saprotroph (Põlme et al. 2020); Decay sugar-rich substrates (Põlme et al. 2020) | Põlme S, et al. 2020. Fungal Diversity 105:1-16 ((https://doi.org/10.1007/s13225-020-00466-2)) |
| Otu48282 | 1 | k__Fungi(100);p__Ascomycota(100);c__Saccharomycetes(100);o__Saccharomycetales(100);f__Metschnikowiaceae(100);g__Metschnikowia(100);s__Metschnikowia_pulcherrima(100); | Metschnikowia | 13 | Saprotroph | \|Undefined Saprotroph\| | Yeast | NULL | Probable | As nectar-tap_saprotroph (Põlme et al. 2020); Decay sugar-rich substrates (Põlme et al. 2020) | Põlme S, et al. 2020. Fungal Diversity 105:1-16 ((https://doi.org/10.1007/s13225-020-00466-2)) |
| Otu48283 | 1 | k__Fungi(100);p__Ascomycota(100);c__Saccharomycetes(100);o__Saccharomycetales(100);f__Metschnikowiaceae(100);g__Metschnikowia(100);s__Metschnikowia_pulcherrima(100); | Metschnikowia | 13 | Saprotroph | \|Undefined Saprotroph\| | Yeast | NULL | Probable | As nectar-tap_saprotroph (Põlme et al. 2020); Decay sugar-rich substrates (Põlme et al. 2020) | Põlme S, et al. 2020. Fungal Diversity 105:1-16 ((https://doi.org/10.1007/s13225-020-00466-2)) |
| Otu48284 | 1 | k__Fungi(100);p__Ascomycota(100);c__Saccharomycetes(100);o__Saccharomycetales(100);f__Metschnikowiaceae(100);g__Metschnikowia(100);s__Metschnikowia_pulcherrima(100); | Metschnikowia | 13 | Saprotroph | \|Undefined Saprotroph\| | Yeast | NULL | Probable | As nectar-tap_saprotroph (Põlme et al. 2020); Decay sugar-rich substrates (Põlme et al. 2020) | Põlme S, et al. 2020. Fungal Diversity 105:1-16 ((https://doi.org/10.1007/s13225-020-00466-2)) |
| Otu48285 | 1 | k__Fungi(100);p__Ascomycota(100);c__Saccharomycetes(100);o__Saccharomycetales(100);f__Metschnikowiaceae(100);g__Metschnikowia(100);s__Metschnikowia_pulcherrima(100); | Metschnikowia | 13 | Saprotroph | \|Undefined Saprotroph\| | Yeast | NULL | Probable | As nectar-tap_saprotroph (Põlme et al. 2020); Decay sugar-rich substrates (Põlme et al. 2020) | Põlme S, et al. 2020. Fungal Diversity 105:1-16 ((https://doi.org/10.1007/s13225-020-00466-2)) |
| Otu48286 | 1 | k__Fungi(100);p__Ascomycota(100);c__Saccharomycetes(100);o__Saccharomycetales(100);f__Metschnikowiaceae(100);g__Metschnikowia(100);g__Metschnikowia_unclassified(100); | Metschnikowia | 13 | Saprotroph | \|Undefined Saprotroph\| | Yeast | NULL | Probable | As nectar-tap_saprotroph (Põlme et al. 2020); Decay sugar-rich substrates (Põlme et al. 2020) | Põlme S, et al. 2020. Fungal Diversity 105:1-16 ((https://doi.org/10.1007/s13225-020-00466-2)) |
| Otu48287 | 1 | k__Fungi(100);p__Ascomycota(100);c__Saccharomycetes(100);o__Saccharomycetales(100);f__Metschnikowiaceae(100);g__Metschnikowia(100);s__Metschnikowia_pulcherrima(100); | Metschnikowia | 13 | Saprotroph | \|Undefined Saprotroph\| | Yeast | NULL | Probable | As nectar-tap_saprotroph (Põlme et al. 2020); Decay sugar-rich substrates (Põlme et al. 2020) | Põlme S, et al. 2020. Fungal Diversity 105:1-16 ((https://doi.org/10.1007/s13225-020-00466-2)) |
| Otu48288 | 1 | k__Fungi(100);p__Ascomycota(100);c__Saccharomycetes(100);o__Saccharomycetales(100);f__Metschnikowiaceae(100);g__Metschnikowia(100);s__Metschnikowia_pulcherrima(100); | Metschnikowia | 13 | Saprotroph | \|Undefined Saprotroph\| | Yeast | NULL | Probable | As nectar-tap_saprotroph (Põlme et al. 2020); Decay sugar-rich substrates (Põlme et al. 2020) | Põlme S, et al. 2020. Fungal Diversity 105:1-16 ((https://doi.org/10.1007/s13225-020-00466-2)) |
| Otu48289 | 1 | k__Fungi(100);p__Ascomycota(100);c__Saccharomycetes(100);o__Saccharomycetales(100);f__Metschnikowiaceae(100);g__Metschnikowia(100);g__Metschnikowia_unclassified(100); | Metschnikowia | 13 | Saprotroph | \|Undefined Saprotroph\| | Yeast | NULL | Probable | As nectar-tap_saprotroph (Põlme et al. 2020); Decay sugar-rich substrates (Põlme et al. 2020) | Põlme S, et al. 2020. Fungal Diversity 105:1-16 ((https://doi.org/10.1007/s13225-020-00466-2)) |
| Otu48290 | 1 | k__Fungi(100);p__Ascomycota(100);c__Saccharomycetes(100);o__Saccharomycetales(100);f__Metschnikowiaceae(100);g__Metschnikowia(100);g__Metschnikowia_unclassified(100); | Metschnikowia | 13 | Saprotroph | \|Undefined Saprotroph\| | Yeast | NULL | Probable | As nectar-tap_saprotroph (Põlme et al. 2020); Decay sugar-rich substrates (Põlme et al. 2020) | Põlme S, et al. 2020. Fungal Diversity 105:1-16 ((https://doi.org/10.1007/s13225-020-00466-2)) |
| Otu48291 | 1 | k__Fungi(100);p__Ascomycota(100);c__Saccharomycetes(100);o__Saccharomycetales(100);f__Metschnikowiaceae(100);g__Metschnikowia(100);s__Metschnikowia_pulcherrima(100); | Metschnikowia | 13 | Saprotroph | \|Undefined Saprotroph\| | Yeast | NULL | Probable | As nectar-tap_saprotroph (Põlme et al. 2020); Decay sugar-rich substrates (Põlme et al. 2020) | Põlme S, et al. 2020. Fungal Diversity 105:1-16 ((https://doi.org/10.1007/s13225-020-00466-2)) |
| Otu48292 | 1 | k__Fungi(100);p__Ascomycota(100);c__Saccharomycetes(100);o__Saccharomycetales(100);f__Metschnikowiaceae(100);g__Metschnikowia(100);s__Metschnikowia_pulcherrima(100); | Metschnikowia | 13 | Saprotroph | \|Undefined Saprotroph\| | Yeast | NULL | Probable | As nectar-tap_saprotroph (Põlme et al. 2020); Decay sugar-rich substrates (Põlme et al. 2020) | Põlme S, et al. 2020. Fungal Diversity 105:1-16 ((https://doi.org/10.1007/s13225-020-00466-2)) |
| Otu48293 | 1 | k__Fungi(100);p__Ascomycota(100);c__Saccharomycetes(100);o__Saccharomycetales(100);f__Metschnikowiaceae(100);g__Metschnikowia(100);s__Metschnikowia_pulcherrima(100); | Metschnikowia | 13 | Saprotroph | \|Undefined Saprotroph\| | Yeast | NULL | Probable | As nectar-tap_saprotroph (Põlme et al. 2020); Decay sugar-rich substrates (Põlme et al. 2020) | Põlme S, et al. 2020. Fungal Diversity 105:1-16 ((https://doi.org/10.1007/s13225-020-00466-2)) |
| Otu48294 | 1 | k__Fungi(100);p__Ascomycota(100);c__Saccharomycetes(100);o__Saccharomycetales(100);f__Metschnikowiaceae(100);g__Metschnikowia(100);s__Metschnikowia_pulcherrima(100); | Metschnikowia | 13 | Saprotroph | \|Undefined Saprotroph\| | Yeast | NULL | Probable | As nectar-tap_saprotroph (Põlme et al. 2020); Decay sugar-rich substrates (Põlme et al. 2020) | Põlme S, et al. 2020. Fungal Diversity 105:1-16 ((https://doi.org/10.1007/s13225-020-00466-2)) |
| Otu48295 | 1 | k__Fungi(100);p__Ascomycota(100);c__Saccharomycetes(100);o__Saccharomycetales(100);f__Metschnikowiaceae(100);g__Metschnikowia(100);s__Metschnikowia_pulcherrima(100); | Metschnikowia | 13 | Saprotroph | \|Undefined Saprotroph\| | Yeast | NULL | Probable | As nectar-tap_saprotroph (Põlme et al. 2020); Decay sugar-rich substrates (Põlme et al. 2020) | Põlme S, et al. 2020. Fungal Diversity 105:1-16 ((https://doi.org/10.1007/s13225-020-00466-2)) |
| Otu48296 | 1 | k__Fungi(100);p__Ascomycota(100);c__Saccharomycetes(100);o__Saccharomycetales(100);f__Metschnikowiaceae(100);g__Metschnikowia(100);s__Metschnikowia_pulcherrima(100); | Metschnikowia | 13 | Saprotroph | \|Undefined Saprotroph\| | Yeast | NULL | Probable | As nectar-tap_saprotroph (Põlme et al. 2020); Decay sugar-rich substrates (Põlme et al. 2020) | Põlme S, et al. 2020. Fungal Diversity 105:1-16 ((https://doi.org/10.1007/s13225-020-00466-2)) |
| Otu48297 | 1 | k__Fungi(100);p__Ascomycota(100);c__Saccharomycetes(100);o__Saccharomycetales(100);f__Metschnikowiaceae(100);g__Metschnikowia(100);s__Metschnikowia_pulcherrima(100); | Metschnikowia | 13 | Saprotroph | \|Undefined Saprotroph\| | Yeast | NULL | Probable | As nectar-tap_saprotroph (Põlme et al. 2020); Decay sugar-rich substrates (Põlme et al. 2020) | Põlme S, et al. 2020. Fungal Diversity 105:1-16 ((https://doi.org/10.1007/s13225-020-00466-2)) |
| Otu48298 | 1 | k__Fungi(100);p__Ascomycota(100);c__Saccharomycetes(100);o__Saccharomycetales(100);f__Metschnikowiaceae(100);g__Metschnikowia(100);s__Metschnikowia_pulcherrima(100); | Metschnikowia | 13 | Saprotroph | \|Undefined Saprotroph\| | Yeast | NULL | Probable | As nectar-tap_saprotroph (Põlme et al. 2020); Decay sugar-rich substrates (Põlme et al. 2020) | Põlme S, et al. 2020. Fungal Diversity 105:1-16 ((https://doi.org/10.1007/s13225-020-00466-2)) |
| Otu48299 | 1 | k__Fungi(100);p__Ascomycota(100);c__Saccharomycetes(100);o__Saccharomycetales(100);f__Metschnikowiaceae(100);g__Metschnikowia(100);s__Metschnikowia_pulcherrima(100); | Metschnikowia | 13 | Saprotroph | \|Undefined Saprotroph\| | Yeast | NULL | Probable | As nectar-tap_saprotroph (Põlme et al. 2020); Decay sugar-rich substrates (Põlme et al. 2020) | Põlme S, et al. 2020. Fungal Diversity 105:1-16 ((https://doi.org/10.1007/s13225-020-00466-2)) |
| Otu48300 | 1 | k__Fungi(100);p__Ascomycota(100);c__Saccharomycetes(100);o__Saccharomycetales(100);f__Metschnikowiaceae(100);g__Metschnikowia(100);s__Metschnikowia_pulcherrima(100); | Metschnikowia | 13 | Saprotroph | \|Undefined Saprotroph\| | Yeast | NULL | Probable | As nectar-tap_saprotroph (Põlme et al. 2020); Decay sugar-rich substrates (Põlme et al. 2020) | Põlme S, et al. 2020. Fungal Diversity 105:1-16 ((https://doi.org/10.1007/s13225-020-00466-2)) |
| Otu48301 | 1 | k__Fungi(100);p__Ascomycota(100);c__Saccharomycetes(100);o__Saccharomycetales(100);f__Metschnikowiaceae(100);g__Metschnikowia(100);s__Metschnikowia_pulcherrima(100); | Metschnikowia | 13 | Saprotroph | \|Undefined Saprotroph\| | Yeast | NULL | Probable | As nectar-tap_saprotroph (Põlme et al. 2020); Decay sugar-rich substrates (Põlme et al. 2020) | Põlme S, et al. 2020. Fungal Diversity 105:1-16 ((https://doi.org/10.1007/s13225-020-00466-2)) |
| Otu48302 | 1 | k__Fungi(100);p__Ascomycota(100);c__Saccharomycetes(100);o__Saccharomycetales(100);f__Metschnikowiaceae(100);g__Metschnikowia(100);g__Metschnikowia_unclassified(100); | Metschnikowia | 13 | Saprotroph | \|Undefined Saprotroph\| | Yeast | NULL | Probable | As nectar-tap_saprotroph (Põlme et al. 2020); Decay sugar-rich substrates (Põlme et al. 2020) | Põlme S, et al. 2020. Fungal Diversity 105:1-16 ((https://doi.org/10.1007/s13225-020-00466-2)) |
| Otu48303 | 1 | k__Fungi(100);p__Ascomycota(100);c__Saccharomycetes(100);o__Saccharomycetales(100);f__Metschnikowiaceae(100);g__Metschnikowia(100);s__Metschnikowia_pulcherrima(100); | Metschnikowia | 13 | Saprotroph | \|Undefined Saprotroph\| | Yeast | NULL | Probable | As nectar-tap_saprotroph (Põlme et al. 2020); Decay sugar-rich substrates (Põlme et al. 2020) | Põlme S, et al. 2020. Fungal Diversity 105:1-16 ((https://doi.org/10.1007/s13225-020-00466-2)) |
| Otu48304 | 1 | k__Fungi(100);p__Ascomycota(100);c__Saccharomycetes(100);o__Saccharomycetales(100);f__Metschnikowiaceae(100);g__Metschnikowia(100);s__Metschnikowia_pulcherrima(100); | Metschnikowia | 13 | Saprotroph | \|Undefined Saprotroph\| | Yeast | NULL | Probable | As nectar-tap_saprotroph (Põlme et al. 2020); Decay sugar-rich substrates (Põlme et al. 2020) | Põlme S, et al. 2020. Fungal Diversity 105:1-16 ((https://doi.org/10.1007/s13225-020-00466-2)) |
| Otu48305 | 1 | k__Fungi(100);p__Ascomycota(100);c__Saccharomycetes(100);o__Saccharomycetales(100);f__Metschnikowiaceae(100);g__Metschnikowia(100);s__Metschnikowia_pulcherrima(100); | Metschnikowia | 13 | Saprotroph | \|Undefined Saprotroph\| | Yeast | NULL | Probable | As nectar-tap_saprotroph (Põlme et al. 2020); Decay sugar-rich substrates (Põlme et al. 2020) | Põlme S, et al. 2020. Fungal Diversity 105:1-16 ((https://doi.org/10.1007/s13225-020-00466-2)) |
| Otu48306 | 1 | k__Fungi(100);p__Ascomycota(100);c__Saccharomycetes(100);o__Saccharomycetales(100);f__Metschnikowiaceae(100);g__Metschnikowia(100);s__Metschnikowia_pulcherrima(100); | Metschnikowia | 13 | Saprotroph | \|Undefined Saprotroph\| | Yeast | NULL | Probable | As nectar-tap_saprotroph (Põlme et al. 2020); Decay sugar-rich substrates (Põlme et al. 2020) | Põlme S, et al. 2020. Fungal Diversity 105:1-16 ((https://doi.org/10.1007/s13225-020-00466-2)) |
| Otu48307 | 1 | k__Fungi(100);p__Ascomycota(100);c__Saccharomycetes(100);o__Saccharomycetales(100);f__Metschnikowiaceae(100);g__Metschnikowia(100);s__Metschnikowia_pulcherrima(100); | Metschnikowia | 13 | Saprotroph | \|Undefined Saprotroph\| | Yeast | NULL | Probable | As nectar-tap_saprotroph (Põlme et al. 2020); Decay sugar-rich substrates (Põlme et al. 2020) | Põlme S, et al. 2020. Fungal Diversity 105:1-16 ((https://doi.org/10.1007/s13225-020-00466-2)) |
| Otu48308 | 1 | k__Fungi(100);p__Ascomycota(100);c__Saccharomycetes(100);o__Saccharomycetales(100);f__Metschnikowiaceae(100);g__Metschnikowia(100);s__Metschnikowia_pulcherrima(100); | Metschnikowia | 13 | Saprotroph | \|Undefined Saprotroph\| | Yeast | NULL | Probable | As nectar-tap_saprotroph (Põlme et al. 2020); Decay sugar-rich substrates (Põlme et al. 2020) | Põlme S, et al. 2020. Fungal Diversity 105:1-16 ((https://doi.org/10.1007/s13225-020-00466-2)) |
| Otu48309 | 1 | k__Fungi(100);p__Ascomycota(100);c__Saccharomycetes(100);o__Saccharomycetales(100);f__Metschnikowiaceae(100);g__Metschnikowia(100);s__Metschnikowia_pulcherrima(100); | Metschnikowia | 13 | Saprotroph | \|Undefined Saprotroph\| | Yeast | NULL | Probable | As nectar-tap_saprotroph (Põlme et al. 2020); Decay sugar-rich substrates (Põlme et al. 2020) | Põlme S, et al. 2020. Fungal Diversity 105:1-16 ((https://doi.org/10.1007/s13225-020-00466-2)) |
| Otu48310 | 1 | k__Fungi(100);p__Ascomycota(100);c__Saccharomycetes(100);o__Saccharomycetales(100);f__Metschnikowiaceae(100);g__Metschnikowia(100);s__Metschnikowia_pulcherrima(100); | Metschnikowia | 13 | Saprotroph | \|Undefined Saprotroph\| | Yeast | NULL | Probable | As nectar-tap_saprotroph (Põlme et al. 2020); Decay sugar-rich substrates (Põlme et al. 2020) | Põlme S, et al. 2020. Fungal Diversity 105:1-16 ((https://doi.org/10.1007/s13225-020-00466-2)) |
| Otu48311 | 1 | k__Fungi(100);p__Ascomycota(100);c__Saccharomycetes(100);o__Saccharomycetales(100);f__Metschnikowiaceae(100);g__Metschnikowia(100);g__Metschnikowia_unclassified(100); | Metschnikowia | 13 | Saprotroph | \|Undefined Saprotroph\| | Yeast | NULL | Probable | As nectar-tap_saprotroph (Põlme et al. 2020); Decay sugar-rich substrates (Põlme et al. 2020) | Põlme S, et al. 2020. Fungal Diversity 105:1-16 ((https://doi.org/10.1007/s13225-020-00466-2)) |
| Otu48312 | 1 | k__Fungi(100);p__Ascomycota(100);c__Saccharomycetes(100);o__Saccharomycetales(100);f__Metschnikowiaceae(100);g__Metschnikowia(100);s__Metschnikowia_pulcherrima(100); | Metschnikowia | 13 | Saprotroph | \|Undefined Saprotroph\| | Yeast | NULL | Probable | As nectar-tap_saprotroph (Põlme et al. 2020); Decay sugar-rich substrates (Põlme et al. 2020) | Põlme S, et al. 2020. Fungal Diversity 105:1-16 ((https://doi.org/10.1007/s13225-020-00466-2)) |
| Otu48313 | 1 | k__Fungi(100);p__Ascomycota(100);c__Saccharomycetes(100);o__Saccharomycetales(100);f__Metschnikowiaceae(100);g__Metschnikowia(100);s__Metschnikowia_pulcherrima(100); | Metschnikowia | 13 | Saprotroph | \|Undefined Saprotroph\| | Yeast | NULL | Probable | As nectar-tap_saprotroph (Põlme et al. 2020); Decay sugar-rich substrates (Põlme et al. 2020) | Põlme S, et al. 2020. Fungal Diversity 105:1-16 ((https://doi.org/10.1007/s13225-020-00466-2)) |
| Otu48314 | 1 | k__Fungi(100);p__Ascomycota(100);c__Saccharomycetes(100);o__Saccharomycetales(100);f__Metschnikowiaceae(100);g__Metschnikowia(100);s__Metschnikowia_pulcherrima(100); | Metschnikowia | 13 | Saprotroph | \|Undefined Saprotroph\| | Yeast | NULL | Probable | As nectar-tap_saprotroph (Põlme et al. 2020); Decay sugar-rich substrates (Põlme et al. 2020) | Põlme S, et al. 2020. Fungal Diversity 105:1-16 ((https://doi.org/10.1007/s13225-020-00466-2)) |
| Otu48315 | 1 | k__Fungi(100);p__Ascomycota(100);c__Saccharomycetes(100);o__Saccharomycetales(100);f__Metschnikowiaceae(100);g__Metschnikowia(100);g__Metschnikowia_unclassified(100); | Metschnikowia | 13 | Saprotroph | \|Undefined Saprotroph\| | Yeast | NULL | Probable | As nectar-tap_saprotroph (Põlme et al. 2020); Decay sugar-rich substrates (Põlme et al. 2020) | Põlme S, et al. 2020. Fungal Diversity 105:1-16 ((https://doi.org/10.1007/s13225-020-00466-2)) |
| Otu48316 | 1 | k__Fungi(100);p__Ascomycota(100);c__Saccharomycetes(100);o__Saccharomycetales(100);f__Metschnikowiaceae(100);g__Metschnikowia(100);s__Metschnikowia_pulcherrima(100); | Metschnikowia | 13 | Saprotroph | \|Undefined Saprotroph\| | Yeast | NULL | Probable | As nectar-tap_saprotroph (Põlme et al. 2020); Decay sugar-rich substrates (Põlme et al. 2020) | Põlme S, et al. 2020. Fungal Diversity 105:1-16 ((https://doi.org/10.1007/s13225-020-00466-2)) |
| Otu48317 | 1 | k__Fungi(100);p__Ascomycota(100);c__Saccharomycetes(100);o__Saccharomycetales(100);f__Metschnikowiaceae(100);g__Metschnikowia(100);s__Metschnikowia_pulcherrima(100); | Metschnikowia | 13 | Saprotroph | \|Undefined Saprotroph\| | Yeast | NULL | Probable | As nectar-tap_saprotroph (Põlme et al. 2020); Decay sugar-rich substrates (Põlme et al. 2020) | Põlme S, et al. 2020. Fungal Diversity 105:1-16 ((https://doi.org/10.1007/s13225-020-00466-2)) |
| Otu48318 | 1 | k__Fungi(100);p__Ascomycota(100);c__Saccharomycetes(100);o__Saccharomycetales(100);f__Metschnikowiaceae(100);g__Metschnikowia(100);s__Metschnikowia_pulcherrima(100); | Metschnikowia | 13 | Saprotroph | \|Undefined Saprotroph\| | Yeast | NULL | Probable | As nectar-tap_saprotroph (Põlme et al. 2020); Decay sugar-rich substrates (Põlme et al. 2020) | Põlme S, et al. 2020. Fungal Diversity 105:1-16 ((https://doi.org/10.1007/s13225-020-00466-2)) |
| Otu48319 | 1 | k__Fungi(100);p__Ascomycota(100);c__Saccharomycetes(100);o__Saccharomycetales(100);f__Metschnikowiaceae(100);g__Metschnikowia(100);s__Metschnikowia_pulcherrima(100); | Metschnikowia | 13 | Saprotroph | \|Undefined Saprotroph\| | Yeast | NULL | Probable | As nectar-tap_saprotroph (Põlme et al. 2020); Decay sugar-rich substrates (Põlme et al. 2020) | Põlme S, et al. 2020. Fungal Diversity 105:1-16 ((https://doi.org/10.1007/s13225-020-00466-2)) |
| Otu48320 | 1 | k__Fungi(100);p__Ascomycota(100);c__Saccharomycetes(100);o__Saccharomycetales(100);f__Metschnikowiaceae(100);g__Metschnikowia(100);g__Metschnikowia_unclassified(100); | Metschnikowia | 13 | Saprotroph | \|Undefined Saprotroph\| | Yeast | NULL | Probable | As nectar-tap_saprotroph (Põlme et al. 2020); Decay sugar-rich substrates (Põlme et al. 2020) | Põlme S, et al. 2020. Fungal Diversity 105:1-16 ((https://doi.org/10.1007/s13225-020-00466-2)) |
| Otu48321 | 1 | k__Fungi(100);p__Ascomycota(100);c__Saccharomycetes(100);o__Saccharomycetales(100);f__Metschnikowiaceae(100);g__Metschnikowia(100);s__Metschnikowia_pulcherrima(100); | Metschnikowia | 13 | Saprotroph | \|Undefined Saprotroph\| | Yeast | NULL | Probable | As nectar-tap_saprotroph (Põlme et al. 2020); Decay sugar-rich substrates (Põlme et al. 2020) | Põlme S, et al. 2020. Fungal Diversity 105:1-16 ((https://doi.org/10.1007/s13225-020-00466-2)) |
| Otu48322 | 1 | k__Fungi(100);p__Ascomycota(100);c__Saccharomycetes(100);o__Saccharomycetales(100);f__Metschnikowiaceae(100);g__Metschnikowia(100);s__Metschnikowia_pulcherrima(100); | Metschnikowia | 13 | Saprotroph | \|Undefined Saprotroph\| | Yeast | NULL | Probable | As nectar-tap_saprotroph (Põlme et al. 2020); Decay sugar-rich substrates (Põlme et al. 2020) | Põlme S, et al. 2020. Fungal Diversity 105:1-16 ((https://doi.org/10.1007/s13225-020-00466-2)) |
| Otu48323 | 1 | k__Fungi(100);p__Ascomycota(100);c__Saccharomycetes(100);o__Saccharomycetales(100);f__Metschnikowiaceae(100);g__Metschnikowia(100);s__Metschnikowia_pulcherrima(100); | Metschnikowia | 13 | Saprotroph | \|Undefined Saprotroph\| | Yeast | NULL | Probable | As nectar-tap_saprotroph (Põlme et al. 2020); Decay sugar-rich substrates (Põlme et al. 2020) | Põlme S, et al. 2020. Fungal Diversity 105:1-16 ((https://doi.org/10.1007/s13225-020-00466-2)) |
| Otu48324 | 1 | k__Fungi(100);p__Ascomycota(100);c__Saccharomycetes(100);o__Saccharomycetales(100);f__Metschnikowiaceae(100);g__Metschnikowia(100);s__Metschnikowia_pulcherrima(100); | Metschnikowia | 13 | Saprotroph | \|Undefined Saprotroph\| | Yeast | NULL | Probable | As nectar-tap_saprotroph (Põlme et al. 2020); Decay sugar-rich substrates (Põlme et al. 2020) | Põlme S, et al. 2020. Fungal Diversity 105:1-16 ((https://doi.org/10.1007/s13225-020-00466-2)) |
| Otu48325 | 1 | k__Fungi(100);p__Ascomycota(100);c__Saccharomycetes(100);o__Saccharomycetales(100);f__Metschnikowiaceae(100);g__Metschnikowia(100);s__Metschnikowia_pulcherrima(100); | Metschnikowia | 13 | Saprotroph | \|Undefined Saprotroph\| | Yeast | NULL | Probable | As nectar-tap_saprotroph (Põlme et al. 2020); Decay sugar-rich substrates (Põlme et al. 2020) | Põlme S, et al. 2020. Fungal Diversity 105:1-16 ((https://doi.org/10.1007/s13225-020-00466-2)) |
| Otu48326 | 1 | k__Fungi(100);p__Ascomycota(100);c__Saccharomycetes(100);o__Saccharomycetales(100);f__Metschnikowiaceae(100);g__Metschnikowia(100);s__Metschnikowia_pulcherrima(100); | Metschnikowia | 13 | Saprotroph | \|Undefined Saprotroph\| | Yeast | NULL | Probable | As nectar-tap_saprotroph (Põlme et al. 2020); Decay sugar-rich substrates (Põlme et al. 2020) | Põlme S, et al. 2020. Fungal Diversity 105:1-16 ((https://doi.org/10.1007/s13225-020-00466-2)) |
| Otu48327 | 1 | k__Fungi(100);p__Ascomycota(100);c__Saccharomycetes(100);o__Saccharomycetales(100);f__Metschnikowiaceae(100);g__Metschnikowia(100);s__Metschnikowia_pulcherrima(100); | Metschnikowia | 13 | Saprotroph | \|Undefined Saprotroph\| | Yeast | NULL | Probable | As nectar-tap_saprotroph (Põlme et al. 2020); Decay sugar-rich substrates (Põlme et al. 2020) | Põlme S, et al. 2020. Fungal Diversity 105:1-16 ((https://doi.org/10.1007/s13225-020-00466-2)) |
| Otu48328 | 1 | k__Fungi(100);p__Ascomycota(100);c__Saccharomycetes(100);o__Saccharomycetales(100);f__Metschnikowiaceae(100);g__Metschnikowia(100);s__Metschnikowia_pulcherrima(100); | Metschnikowia | 13 | Saprotroph | \|Undefined Saprotroph\| | Yeast | NULL | Probable | As nectar-tap_saprotroph (Põlme et al. 2020); Decay sugar-rich substrates (Põlme et al. 2020) | Põlme S, et al. 2020. Fungal Diversity 105:1-16 ((https://doi.org/10.1007/s13225-020-00466-2)) |
| Otu48329 | 1 | k__Fungi(100);p__Ascomycota(100);c__Saccharomycetes(100);o__Saccharomycetales(100);f__Metschnikowiaceae(100);g__Metschnikowia(100);s__Metschnikowia_pulcherrima(100); | Metschnikowia | 13 | Saprotroph | \|Undefined Saprotroph\| | Yeast | NULL | Probable | As nectar-tap_saprotroph (Põlme et al. 2020); Decay sugar-rich substrates (Põlme et al. 2020) | Põlme S, et al. 2020. Fungal Diversity 105:1-16 ((https://doi.org/10.1007/s13225-020-00466-2)) |
| Otu48330 | 1 | k__Fungi(100);p__Ascomycota(100);c__Saccharomycetes(100);o__Saccharomycetales(100);f__Metschnikowiaceae(100);g__Metschnikowia(100);s__Metschnikowia_pulcherrima(100); | Metschnikowia | 13 | Saprotroph | \|Undefined Saprotroph\| | Yeast | NULL | Probable | As nectar-tap_saprotroph (Põlme et al. 2020); Decay sugar-rich substrates (Põlme et al. 2020) | Põlme S, et al. 2020. Fungal Diversity 105:1-16 ((https://doi.org/10.1007/s13225-020-00466-2)) |
| Otu48331 | 1 | k__Fungi(100);p__Ascomycota(100);c__Saccharomycetes(100);o__Saccharomycetales(100);f__Metschnikowiaceae(100);g__Metschnikowia(100);g__Metschnikowia_unclassified(100); | Metschnikowia | 13 | Saprotroph | \|Undefined Saprotroph\| | Yeast | NULL | Probable | As nectar-tap_saprotroph (Põlme et al. 2020); Decay sugar-rich substrates (Põlme et al. 2020) | Põlme S, et al. 2020. Fungal Diversity 105:1-16 ((https://doi.org/10.1007/s13225-020-00466-2)) |
| Otu48332 | 1 | k__Fungi(100);p__Ascomycota(100);c__Saccharomycetes(100);o__Saccharomycetales(100);f__Metschnikowiaceae(100);g__Metschnikowia(100);s__Metschnikowia_pulcherrima(100); | Metschnikowia | 13 | Saprotroph | \|Undefined Saprotroph\| | Yeast | NULL | Probable | As nectar-tap_saprotroph (Põlme et al. 2020); Decay sugar-rich substrates (Põlme et al. 2020) | Põlme S, et al. 2020. Fungal Diversity 105:1-16 ((https://doi.org/10.1007/s13225-020-00466-2)) |
| Otu48333 | 1 | k__Fungi(100);p__Ascomycota(100);c__Saccharomycetes(100);o__Saccharomycetales(100);f__Metschnikowiaceae(100);g__Metschnikowia(100);s__Metschnikowia_pulcherrima(100); | Metschnikowia | 13 | Saprotroph | \|Undefined Saprotroph\| | Yeast | NULL | Probable | As nectar-tap_saprotroph (Põlme et al. 2020); Decay sugar-rich substrates (Põlme et al. 2020) | Põlme S, et al. 2020. Fungal Diversity 105:1-16 ((https://doi.org/10.1007/s13225-020-00466-2)) |
| Otu48334 | 1 | k__Fungi(100);p__Ascomycota(100);c__Saccharomycetes(100);o__Saccharomycetales(100);f__Metschnikowiaceae(100);g__Metschnikowia(100);s__Metschnikowia_pulcherrima(100); | Metschnikowia | 13 | Saprotroph | \|Undefined Saprotroph\| | Yeast | NULL | Probable | As nectar-tap_saprotroph (Põlme et al. 2020); Decay sugar-rich substrates (Põlme et al. 2020) | Põlme S, et al. 2020. Fungal Diversity 105:1-16 ((https://doi.org/10.1007/s13225-020-00466-2)) |
| Otu48335 | 1 | k__Fungi(100);p__Ascomycota(100);c__Saccharomycetes(100);o__Saccharomycetales(100);f__Metschnikowiaceae(100);g__Metschnikowia(100);s__Metschnikowia_pulcherrima(100); | Metschnikowia | 13 | Saprotroph | \|Undefined Saprotroph\| | Yeast | NULL | Probable | As nectar-tap_saprotroph (Põlme et al. 2020); Decay sugar-rich substrates (Põlme et al. 2020) | Põlme S, et al. 2020. Fungal Diversity 105:1-16 ((https://doi.org/10.1007/s13225-020-00466-2)) |
| Otu48336 | 1 | k__Fungi(100);p__Ascomycota(100);c__Saccharomycetes(100);o__Saccharomycetales(100);f__Metschnikowiaceae(100);g__Metschnikowia(100);s__Metschnikowia_pulcherrima(100); | Metschnikowia | 13 | Saprotroph | \|Undefined Saprotroph\| | Yeast | NULL | Probable | As nectar-tap_saprotroph (Põlme et al. 2020); Decay sugar-rich substrates (Põlme et al. 2020) | Põlme S, et al. 2020. Fungal Diversity 105:1-16 ((https://doi.org/10.1007/s13225-020-00466-2)) |
| Otu48337 | 1 | k__Fungi(100);p__Ascomycota(100);c__Saccharomycetes(100);o__Saccharomycetales(100);f__Metschnikowiaceae(100);g__Metschnikowia(100);g__Metschnikowia_unclassified(100); | Metschnikowia | 13 | Saprotroph | \|Undefined Saprotroph\| | Yeast | NULL | Probable | As nectar-tap_saprotroph (Põlme et al. 2020); Decay sugar-rich substrates (Põlme et al. 2020) | Põlme S, et al. 2020. Fungal Diversity 105:1-16 ((https://doi.org/10.1007/s13225-020-00466-2)) |
| Otu48338 | 1 | k__Fungi(100);p__Ascomycota(100);c__Saccharomycetes(100);o__Saccharomycetales(100);f__Metschnikowiaceae(100);g__Metschnikowia(100);g__Metschnikowia_unclassified(100); | Metschnikowia | 13 | Saprotroph | \|Undefined Saprotroph\| | Yeast | NULL | Probable | As nectar-tap_saprotroph (Põlme et al. 2020); Decay sugar-rich substrates (Põlme et al. 2020) | Põlme S, et al. 2020. Fungal Diversity 105:1-16 ((https://doi.org/10.1007/s13225-020-00466-2)) |
| Otu48339 | 1 | k__Fungi(100);p__Ascomycota(100);c__Saccharomycetes(100);o__Saccharomycetales(100);f__Metschnikowiaceae(100);g__Metschnikowia(100);s__Metschnikowia_pulcherrima(100); | Metschnikowia | 13 | Saprotroph | \|Undefined Saprotroph\| | Yeast | NULL | Probable | As nectar-tap_saprotroph (Põlme et al. 2020); Decay sugar-rich substrates (Põlme et al. 2020) | Põlme S, et al. 2020. Fungal Diversity 105:1-16 ((https://doi.org/10.1007/s13225-020-00466-2)) |
| Otu48340 | 1 | k__Fungi(100);p__Ascomycota(100);c__Saccharomycetes(100);o__Saccharomycetales(100);f__Trichomonascaceae(100);g__Blastobotrys(100);s__Blastobotrys_nivea(100); | Blastobotrys | 13 | Saprotroph | \|Undefined Saprotroph\| | Yeast | Its unusual metal | Highly Probable | As nectar-tap_saprotroph (Põlme et al. 2020); Decay sugar-rich substrates (Põlme et al. 2020); In the course of identifying alternative microorganisms for biotechnological interest, A. adeninivorans has already proved its strengthened competitiveness as a promising cell factory for many more applications. | Kunze G, et al. 2014. Biotechnology for Biofuels 7:66 ((https://doi.org/10.1186/1754-6834-7-66)); Põlme S, et al. 2020. Fungal Diversity 105:1-16 ((https://doi.org/10.1007/s13225-020-00466-2)) |
| Otu48341 | 1 | k__Fungi(100);p__Ascomycota(100);c__Saccharomycetes(100);o__Saccharomycetales(100);f__Metschnikowiaceae(100);g__Metschnikowia(100);g__Metschnikowia_unclassified(100); | Metschnikowia | 13 | Saprotroph | \|Undefined Saprotroph\| | Yeast | NULL | Probable | As nectar-tap_saprotroph (Põlme et al. 2020); Decay sugar-rich substrates (Põlme et al. 2020) | Põlme S, et al. 2020. Fungal Diversity 105:1-16 ((https://doi.org/10.1007/s13225-020-00466-2)) |
| Otu48342 | 1 | k__Fungi(100);p__Ascomycota(100);c__Saccharomycetes(100);o__Saccharomycetales(100);f__Metschnikowiaceae(100);g__Metschnikowia(100);s__Metschnikowia_pulcherrima(100); | Metschnikowia | 13 | Saprotroph | \|Undefined Saprotroph\| | Yeast | NULL | Probable | As nectar-tap_saprotroph (Põlme et al. 2020); Decay sugar-rich substrates (Põlme et al. 2020) | Põlme S, et al. 2020. Fungal Diversity 105:1-16 ((https://doi.org/10.1007/s13225-020-00466-2)) |
| Otu48343 | 1 | k__Fungi(100);p__Ascomycota(100);c__Saccharomycetes(100);o__Saccharomycetales(100);f__Metschnikowiaceae(100);g__Metschnikowia(100);g__Metschnikowia_unclassified(100); | Metschnikowia | 13 | Saprotroph | \|Undefined Saprotroph\| | Yeast | NULL | Probable | As nectar-tap_saprotroph (Põlme et al. 2020); Decay sugar-rich substrates (Põlme et al. 2020) | Põlme S, et al. 2020. Fungal Diversity 105:1-16 ((https://doi.org/10.1007/s13225-020-00466-2)) |
| Otu48344 | 1 | k__Fungi(100);p__Ascomycota(100);c__Saccharomycetes(100);o__Saccharomycetales(100);f__Metschnikowiaceae(100);g__Metschnikowia(100);s__Metschnikowia_pulcherrima(100); | Metschnikowia | 13 | Saprotroph | \|Undefined Saprotroph\| | Yeast | NULL | Probable | As nectar-tap_saprotroph (Põlme et al. 2020); Decay sugar-rich substrates (Põlme et al. 2020) | Põlme S, et al. 2020. Fungal Diversity 105:1-16 ((https://doi.org/10.1007/s13225-020-00466-2)) |
| Otu48345 | 1 | k__Fungi(100);p__Ascomycota(100);c__Saccharomycetes(100);o__Saccharomycetales(100);f__Metschnikowiaceae(100);g__Metschnikowia(100);s__Metschnikowia_pulcherrima(100); | Metschnikowia | 13 | Saprotroph | \|Undefined Saprotroph\| | Yeast | NULL | Probable | As nectar-tap_saprotroph (Põlme et al. 2020); Decay sugar-rich substrates (Põlme et al. 2020) | Põlme S, et al. 2020. Fungal Diversity 105:1-16 ((https://doi.org/10.1007/s13225-020-00466-2)) |
| Otu48346 | 1 | k__Fungi(100);p__Ascomycota(100);c__Saccharomycetes(100);o__Saccharomycetales(100);f__Metschnikowiaceae(100);g__Metschnikowia(100);s__Metschnikowia_pulcherrima(100); | Metschnikowia | 13 | Saprotroph | \|Undefined Saprotroph\| | Yeast | NULL | Probable | As nectar-tap_saprotroph (Põlme et al. 2020); Decay sugar-rich substrates (Põlme et al. 2020) | Põlme S, et al. 2020. Fungal Diversity 105:1-16 ((https://doi.org/10.1007/s13225-020-00466-2)) |
| Otu48347 | 1 | k__Fungi(100);p__Ascomycota(100);c__Saccharomycetes(100);o__Saccharomycetales(100);f__Metschnikowiaceae(100);g__Metschnikowia(100);s__Metschnikowia_pulcherrima(100); | Metschnikowia | 13 | Saprotroph | \|Undefined Saprotroph\| | Yeast | NULL | Probable | As nectar-tap_saprotroph (Põlme et al. 2020); Decay sugar-rich substrates (Põlme et al. 2020) | Põlme S, et al. 2020. Fungal Diversity 105:1-16 ((https://doi.org/10.1007/s13225-020-00466-2)) |
| Otu48348 | 1 | k__Fungi(100);p__Ascomycota(100);c__Saccharomycetes(100);o__Saccharomycetales(100);f__Metschnikowiaceae(100);g__Metschnikowia(100);s__Metschnikowia_pulcherrima(100); | Metschnikowia | 13 | Saprotroph | \|Undefined Saprotroph\| | Yeast | NULL | Probable | As nectar-tap_saprotroph (Põlme et al. 2020); Decay sugar-rich substrates (Põlme et al. 2020) | Põlme S, et al. 2020. Fungal Diversity 105:1-16 ((https://doi.org/10.1007/s13225-020-00466-2)) |
| Otu48349 | 1 | k__Fungi(100);p__Ascomycota(100);c__Saccharomycetes(100);o__Saccharomycetales(100);f__Metschnikowiaceae(100);g__Metschnikowia(100);s__Metschnikowia_pulcherrima(100); | Metschnikowia | 13 | Saprotroph | \|Undefined Saprotroph\| | Yeast | NULL | Probable | As nectar-tap_saprotroph (Põlme et al. 2020); Decay sugar-rich substrates (Põlme et al. 2020) | Põlme S, et al. 2020. Fungal Diversity 105:1-16 ((https://doi.org/10.1007/s13225-020-00466-2)) |
| Otu48350 | 1 | k__Fungi(100);p__Ascomycota(100);c__Saccharomycetes(100);o__Saccharomycetales(100);f__Metschnikowiaceae(100);g__Metschnikowia(100);s__Metschnikowia_pulcherrima(100); | Metschnikowia | 13 | Saprotroph | \|Undefined Saprotroph\| | Yeast | NULL | Probable | As nectar-tap_saprotroph (Põlme et al. 2020); Decay sugar-rich substrates (Põlme et al. 2020) | Põlme S, et al. 2020. Fungal Diversity 105:1-16 ((https://doi.org/10.1007/s13225-020-00466-2)) |
| Otu48351 | 1 | k__Fungi(100);p__Ascomycota(100);c__Saccharomycetes(100);o__Saccharomycetales(100);f__Metschnikowiaceae(100);g__Metschnikowia(100);s__Metschnikowia_pulcherrima(100); | Metschnikowia | 13 | Saprotroph | \|Undefined Saprotroph\| | Yeast | NULL | Probable | As nectar-tap_saprotroph (Põlme et al. 2020); Decay sugar-rich substrates (Põlme et al. 2020) | Põlme S, et al. 2020. Fungal Diversity 105:1-16 ((https://doi.org/10.1007/s13225-020-00466-2)) |
| Otu48352 | 1 | k__Fungi(100);p__Ascomycota(100);c__Saccharomycetes(100);o__Saccharomycetales(100);f__Metschnikowiaceae(100);g__Metschnikowia(100);g__Metschnikowia_unclassified(100); | Metschnikowia | 13 | Saprotroph | \|Undefined Saprotroph\| | Yeast | NULL | Probable | As nectar-tap_saprotroph (Põlme et al. 2020); Decay sugar-rich substrates (Põlme et al. 2020) | Põlme S, et al. 2020. Fungal Diversity 105:1-16 ((https://doi.org/10.1007/s13225-020-00466-2)) |
| Otu48353 | 1 | k__Fungi(100);p__Ascomycota(100);c__Saccharomycetes(100);o__Saccharomycetales(100);f__Metschnikowiaceae(100);g__Metschnikowia(100);g__Metschnikowia_unclassified(100); | Metschnikowia | 13 | Saprotroph | \|Undefined Saprotroph\| | Yeast | NULL | Probable | As nectar-tap_saprotroph (Põlme et al. 2020); Decay sugar-rich substrates (Põlme et al. 2020) | Põlme S, et al. 2020. Fungal Diversity 105:1-16 ((https://doi.org/10.1007/s13225-020-00466-2)) |
| Otu48354 | 1 | k__Fungi(100);p__Ascomycota(100);c__Saccharomycetes(100);o__Saccharomycetales(100);f__Metschnikowiaceae(100);g__Metschnikowia(100);g__Metschnikowia_unclassified(100); | Metschnikowia | 13 | Saprotroph | \|Undefined Saprotroph\| | Yeast | NULL | Probable | As nectar-tap_saprotroph (Põlme et al. 2020); Decay sugar-rich substrates (Põlme et al. 2020) | Põlme S, et al. 2020. Fungal Diversity 105:1-16 ((https://doi.org/10.1007/s13225-020-00466-2)) |
| Otu48355 | 1 | k__Fungi(100);p__Ascomycota(100);c__Saccharomycetes(100);o__Saccharomycetales(100);f__Metschnikowiaceae(100);g__Metschnikowia(100);g__Metschnikowia_unclassified(100); | Metschnikowia | 13 | Saprotroph | \|Undefined Saprotroph\| | Yeast | NULL | Probable | As nectar-tap_saprotroph (Põlme et al. 2020); Decay sugar-rich substrates (Põlme et al. 2020) | Põlme S, et al. 2020. Fungal Diversity 105:1-16 ((https://doi.org/10.1007/s13225-020-00466-2)) |
| Otu48356 | 1 | k__Fungi(100);p__Ascomycota(100);c__Saccharomycetes(100);o__Saccharomycetales(100);f__Metschnikowiaceae(100);g__Metschnikowia(100);g__Metschnikowia_unclassified(100); | Metschnikowia | 13 | Saprotroph | \|Undefined Saprotroph\| | Yeast | NULL | Probable | As nectar-tap_saprotroph (Põlme et al. 2020); Decay sugar-rich substrates (Põlme et al. 2020) | Põlme S, et al. 2020. Fungal Diversity 105:1-16 ((https://doi.org/10.1007/s13225-020-00466-2)) |
| Otu48357 | 1 | k__Fungi(100);p__Ascomycota(100);c__Saccharomycetes(100);o__Saccharomycetales(100);f__Metschnikowiaceae(100);g__Metschnikowia(100);g__Metschnikowia_unclassified(100); | Metschnikowia | 13 | Saprotroph | \|Undefined Saprotroph\| | Yeast | NULL | Probable | As nectar-tap_saprotroph (Põlme et al. 2020); Decay sugar-rich substrates (Põlme et al. 2020) | Põlme S, et al. 2020. Fungal Diversity 105:1-16 ((https://doi.org/10.1007/s13225-020-00466-2)) |
| Otu48358 | 1 | k__Fungi(100);p__Ascomycota(100);c__Saccharomycetes(100);o__Saccharomycetales(100);f__Metschnikowiaceae(100);g__Metschnikowia(100);g__Metschnikowia_unclassified(100); | Metschnikowia | 13 | Saprotroph | \|Undefined Saprotroph\| | Yeast | NULL | Probable | As nectar-tap_saprotroph (Põlme et al. 2020); Decay sugar-rich substrates (Põlme et al. 2020) | Põlme S, et al. 2020. Fungal Diversity 105:1-16 ((https://doi.org/10.1007/s13225-020-00466-2)) |
| Otu48359 | 1 | k__Fungi(100);p__Ascomycota(100);c__Saccharomycetes(100);o__Saccharomycetales(100);f__Metschnikowiaceae(100);g__Metschnikowia(100);g__Metschnikowia_unclassified(100); | Metschnikowia | 13 | Saprotroph | \|Undefined Saprotroph\| | Yeast | NULL | Probable | As nectar-tap_saprotroph (Põlme et al. 2020); Decay sugar-rich substrates (Põlme et al. 2020) | Põlme S, et al. 2020. Fungal Diversity 105:1-16 ((https://doi.org/10.1007/s13225-020-00466-2)) |
| Otu48360 | 1 | k__Fungi(100);p__Ascomycota(100);c__Saccharomycetes(100);o__Saccharomycetales(100);f__Metschnikowiaceae(100);g__Metschnikowia(100);g__Metschnikowia_unclassified(100); | Metschnikowia | 13 | Saprotroph | \|Undefined Saprotroph\| | Yeast | NULL | Probable | As nectar-tap_saprotroph (Põlme et al. 2020); Decay sugar-rich substrates (Põlme et al. 2020) | Põlme S, et al. 2020. Fungal Diversity 105:1-16 ((https://doi.org/10.1007/s13225-020-00466-2)) |
| Otu48361 | 1 | k__Fungi(100);p__Ascomycota(100);c__Saccharomycetes(100);o__Saccharomycetales(100);f__Metschnikowiaceae(100);g__Metschnikowia(100);g__Metschnikowia_unclassified(100); | Metschnikowia | 13 | Saprotroph | \|Undefined Saprotroph\| | Yeast | NULL | Probable | As nectar-tap_saprotroph (Põlme et al. 2020); Decay sugar-rich substrates (Põlme et al. 2020) | Põlme S, et al. 2020. Fungal Diversity 105:1-16 ((https://doi.org/10.1007/s13225-020-00466-2)) |
| Otu48362 | 1 | k__Fungi(100);p__Ascomycota(100);c__Saccharomycetes(100);o__Saccharomycetales(100);f__Metschnikowiaceae(100);g__Metschnikowia(100);g__Metschnikowia_unclassified(100); | Metschnikowia | 13 | Saprotroph | \|Undefined Saprotroph\| | Yeast | NULL | Probable | As nectar-tap_saprotroph (Põlme et al. 2020); Decay sugar-rich substrates (Põlme et al. 2020) | Põlme S, et al. 2020. Fungal Diversity 105:1-16 ((https://doi.org/10.1007/s13225-020-00466-2)) |
| Otu48363 | 1 | k__Fungi(100);p__Ascomycota(100);c__Saccharomycetes(100);o__Saccharomycetales(100);f__Metschnikowiaceae(100);g__Metschnikowia(100);g__Metschnikowia_unclassified(100); | Metschnikowia | 13 | Saprotroph | \|Undefined Saprotroph\| | Yeast | NULL | Probable | As nectar-tap_saprotroph (Põlme et al. 2020); Decay sugar-rich substrates (Põlme et al. 2020) | Põlme S, et al. 2020. Fungal Diversity 105:1-16 ((https://doi.org/10.1007/s13225-020-00466-2)) |
| Otu48364 | 1 | k__Fungi(100);p__Ascomycota(100);c__Saccharomycetes(100);o__Saccharomycetales(100);f__Metschnikowiaceae(100);g__Metschnikowia(100);g__Metschnikowia_unclassified(100); | Metschnikowia | 13 | Saprotroph | \|Undefined Saprotroph\| | Yeast | NULL | Probable | As nectar-tap_saprotroph (Põlme et al. 2020); Decay sugar-rich substrates (Põlme et al. 2020) | Põlme S, et al. 2020. Fungal Diversity 105:1-16 ((https://doi.org/10.1007/s13225-020-00466-2)) |
| Otu48365 | 1 | k__Fungi(100);p__Ascomycota(100);c__Saccharomycetes(100);o__Saccharomycetales(100);f__Metschnikowiaceae(100);g__Metschnikowia(100);g__Metschnikowia_unclassified(100); | Metschnikowia | 13 | Saprotroph | \|Undefined Saprotroph\| | Yeast | NULL | Probable | As nectar-tap_saprotroph (Põlme et al. 2020); Decay sugar-rich substrates (Põlme et al. 2020) | Põlme S, et al. 2020. Fungal Diversity 105:1-16 ((https://doi.org/10.1007/s13225-020-00466-2)) |
| Otu48366 | 1 | k__Fungi(100);p__Ascomycota(100);c__Saccharomycetes(100);o__Saccharomycetales(100);f__Metschnikowiaceae(100);g__Metschnikowia(100);g__Metschnikowia_unclassified(100); | Metschnikowia | 13 | Saprotroph | \|Undefined Saprotroph\| | Yeast | NULL | Probable | As nectar-tap_saprotroph (Põlme et al. 2020); Decay sugar-rich substrates (Põlme et al. 2020) | Põlme S, et al. 2020. Fungal Diversity 105:1-16 ((https://doi.org/10.1007/s13225-020-00466-2)) |
| Otu48367 | 1 | k__Fungi(100);p__Ascomycota(100);c__Saccharomycetes(100);o__Saccharomycetales(100);f__Metschnikowiaceae(100);g__Metschnikowia(100);s__Metschnikowia_pulcherrima(100); | Metschnikowia | 13 | Saprotroph | \|Undefined Saprotroph\| | Yeast | NULL | Probable | As nectar-tap_saprotroph (Põlme et al. 2020); Decay sugar-rich substrates (Põlme et al. 2020) | Põlme S, et al. 2020. Fungal Diversity 105:1-16 ((https://doi.org/10.1007/s13225-020-00466-2)) |
| Otu48368 | 1 | k__Fungi(100);p__Ascomycota(100);c__Saccharomycetes(100);o__Saccharomycetales(100);f__Metschnikowiaceae(100);g__Metschnikowia(100);s__Metschnikowia_pulcherrima(100); | Metschnikowia | 13 | Saprotroph | \|Undefined Saprotroph\| | Yeast | NULL | Probable | As nectar-tap_saprotroph (Põlme et al. 2020); Decay sugar-rich substrates (Põlme et al. 2020) | Põlme S, et al. 2020. Fungal Diversity 105:1-16 ((https://doi.org/10.1007/s13225-020-00466-2)) |
| Otu48369 | 1 | k__Fungi(100);p__Ascomycota(100);c__Saccharomycetes(100);o__Saccharomycetales(100);f__Metschnikowiaceae(100);g__Metschnikowia(100);s__Metschnikowia_pulcherrima(100); | Metschnikowia | 13 | Saprotroph | \|Undefined Saprotroph\| | Yeast | NULL | Probable | As nectar-tap_saprotroph (Põlme et al. 2020); Decay sugar-rich substrates (Põlme et al. 2020) | Põlme S, et al. 2020. Fungal Diversity 105:1-16 ((https://doi.org/10.1007/s13225-020-00466-2)) |
| Otu48370 | 1 | k__Fungi(100);p__Ascomycota(100);c__Saccharomycetes(100);o__Saccharomycetales(100);f__Metschnikowiaceae(100);g__Metschnikowia(100);s__Metschnikowia_pulcherrima(100); | Metschnikowia | 13 | Saprotroph | \|Undefined Saprotroph\| | Yeast | NULL | Probable | As nectar-tap_saprotroph (Põlme et al. 2020); Decay sugar-rich substrates (Põlme et al. 2020) | Põlme S, et al. 2020. Fungal Diversity 105:1-16 ((https://doi.org/10.1007/s13225-020-00466-2)) |
| Otu48371 | 1 | k__Fungi(100);p__Ascomycota(100);c__Saccharomycetes(100);o__Saccharomycetales(100);f__Metschnikowiaceae(100);g__Metschnikowia(100);s__Metschnikowia_pulcherrima(100); | Metschnikowia | 13 | Saprotroph | \|Undefined Saprotroph\| | Yeast | NULL | Probable | As nectar-tap_saprotroph (Põlme et al. 2020); Decay sugar-rich substrates (Põlme et al. 2020) | Põlme S, et al. 2020. Fungal Diversity 105:1-16 ((https://doi.org/10.1007/s13225-020-00466-2)) |
| Otu48372 | 1 | k__Fungi(100);p__Ascomycota(100);c__Saccharomycetes(100);o__Saccharomycetales(100);f__Metschnikowiaceae(100);g__Metschnikowia(100);s__Metschnikowia_pulcherrima(100); | Metschnikowia | 13 | Saprotroph | \|Undefined Saprotroph\| | Yeast | NULL | Probable | As nectar-tap_saprotroph (Põlme et al. 2020); Decay sugar-rich substrates (Põlme et al. 2020) | Põlme S, et al. 2020. Fungal Diversity 105:1-16 ((https://doi.org/10.1007/s13225-020-00466-2)) |
| Otu48373 | 1 | k__Fungi(100);p__Ascomycota(100);c__Saccharomycetes(100);o__Saccharomycetales(100);f__Metschnikowiaceae(100);g__Metschnikowia(100);s__Metschnikowia_pulcherrima(100); | Metschnikowia | 13 | Saprotroph | \|Undefined Saprotroph\| | Yeast | NULL | Probable | As nectar-tap_saprotroph (Põlme et al. 2020); Decay sugar-rich substrates (Põlme et al. 2020) | Põlme S, et al. 2020. Fungal Diversity 105:1-16 ((https://doi.org/10.1007/s13225-020-00466-2)) |
| Otu48374 | 1 | k__Fungi(100);p__Ascomycota(100);c__Saccharomycetes(100);o__Saccharomycetales(100);f__Metschnikowiaceae(100);g__Metschnikowia(100);s__Metschnikowia_pulcherrima(100); | Metschnikowia | 13 | Saprotroph | \|Undefined Saprotroph\| | Yeast | NULL | Probable | As nectar-tap_saprotroph (Põlme et al. 2020); Decay sugar-rich substrates (Põlme et al. 2020) | Põlme S, et al. 2020. Fungal Diversity 105:1-16 ((https://doi.org/10.1007/s13225-020-00466-2)) |
| Otu48375 | 1 | k__Fungi(100);p__Ascomycota(100);c__Saccharomycetes(100);o__Saccharomycetales(100);f__Metschnikowiaceae(100);g__Metschnikowia(100);s__Metschnikowia_pulcherrima(100); | Metschnikowia | 13 | Saprotroph | \|Undefined Saprotroph\| | Yeast | NULL | Probable | As nectar-tap_saprotroph (Põlme et al. 2020); Decay sugar-rich substrates (Põlme et al. 2020) | Põlme S, et al. 2020. Fungal Diversity 105:1-16 ((https://doi.org/10.1007/s13225-020-00466-2)) |
| Otu48376 | 1 | k__Fungi(100);p__Ascomycota(100);c__Saccharomycetes(100);o__Saccharomycetales(100);f__Metschnikowiaceae(100);g__Metschnikowia(100);s__Metschnikowia_pulcherrima(100); | Metschnikowia | 13 | Saprotroph | \|Undefined Saprotroph\| | Yeast | NULL | Probable | As nectar-tap_saprotroph (Põlme et al. 2020); Decay sugar-rich substrates (Põlme et al. 2020) | Põlme S, et al. 2020. Fungal Diversity 105:1-16 ((https://doi.org/10.1007/s13225-020-00466-2)) |
| Otu48377 | 1 | k__Fungi(100);p__Ascomycota(100);c__Saccharomycetes(100);o__Saccharomycetales(100);f__Metschnikowiaceae(100);g__Metschnikowia(100);g__Metschnikowia_unclassified(100); | Metschnikowia | 13 | Saprotroph | \|Undefined Saprotroph\| | Yeast | NULL | Probable | As nectar-tap_saprotroph (Põlme et al. 2020); Decay sugar-rich substrates (Põlme et al. 2020) | Põlme S, et al. 2020. Fungal Diversity 105:1-16 ((https://doi.org/10.1007/s13225-020-00466-2)) |
| Otu48378 | 1 | k__Fungi(100);p__Ascomycota(100);c__Saccharomycetes(100);o__Saccharomycetales(100);f__Metschnikowiaceae(100);g__Metschnikowia(100);s__Metschnikowia_pulcherrima(100); | Metschnikowia | 13 | Saprotroph | \|Undefined Saprotroph\| | Yeast | NULL | Probable | As nectar-tap_saprotroph (Põlme et al. 2020); Decay sugar-rich substrates (Põlme et al. 2020) | Põlme S, et al. 2020. Fungal Diversity 105:1-16 ((https://doi.org/10.1007/s13225-020-00466-2)) |
| Otu48379 | 1 | k__Fungi(100);p__Ascomycota(100);c__Saccharomycetes(100);o__Saccharomycetales(100);f__Metschnikowiaceae(100);g__Metschnikowia(100);s__Metschnikowia_pulcherrima(100); | Metschnikowia | 13 | Saprotroph | \|Undefined Saprotroph\| | Yeast | NULL | Probable | As nectar-tap_saprotroph (Põlme et al. 2020); Decay sugar-rich substrates (Põlme et al. 2020) | Põlme S, et al. 2020. Fungal Diversity 105:1-16 ((https://doi.org/10.1007/s13225-020-00466-2)) |
| Otu48380 | 1 | k__Fungi(100);p__Ascomycota(100);c__Saccharomycetes(100);o__Saccharomycetales(100);f__Metschnikowiaceae(100);g__Metschnikowia(100);s__Metschnikowia_pulcherrima(100); | Metschnikowia | 13 | Saprotroph | \|Undefined Saprotroph\| | Yeast | NULL | Probable | As nectar-tap_saprotroph (Põlme et al. 2020); Decay sugar-rich substrates (Põlme et al. 2020) | Põlme S, et al. 2020. Fungal Diversity 105:1-16 ((https://doi.org/10.1007/s13225-020-00466-2)) |
| Otu48381 | 1 | k__Fungi(100);p__Ascomycota(100);c__Saccharomycetes(100);o__Saccharomycetales(100);f__Metschnikowiaceae(100);g__Metschnikowia(100);s__Metschnikowia_pulcherrima(100); | Metschnikowia | 13 | Saprotroph | \|Undefined Saprotroph\| | Yeast | NULL | Probable | As nectar-tap_saprotroph (Põlme et al. 2020); Decay sugar-rich substrates (Põlme et al. 2020) | Põlme S, et al. 2020. Fungal Diversity 105:1-16 ((https://doi.org/10.1007/s13225-020-00466-2)) |
| Otu48382 | 1 | k__Fungi(100);p__Ascomycota(100);c__Saccharomycetes(100);o__Saccharomycetales(100);f__Metschnikowiaceae(100);g__Metschnikowia(100);g__Metschnikowia_unclassified(100); | Metschnikowia | 13 | Saprotroph | \|Undefined Saprotroph\| | Yeast | NULL | Probable | As nectar-tap_saprotroph (Põlme et al. 2020); Decay sugar-rich substrates (Põlme et al. 2020) | Põlme S, et al. 2020. Fungal Diversity 105:1-16 ((https://doi.org/10.1007/s13225-020-00466-2)) |
| Otu48383 | 1 | k__Fungi(100);p__Ascomycota(100);c__Saccharomycetes(100);o__Saccharomycetales(100);f__Metschnikowiaceae(100);g__Metschnikowia(100);s__Metschnikowia_pulcherrima(100); | Metschnikowia | 13 | Saprotroph | \|Undefined Saprotroph\| | Yeast | NULL | Probable | As nectar-tap_saprotroph (Põlme et al. 2020); Decay sugar-rich substrates (Põlme et al. 2020) | Põlme S, et al. 2020. Fungal Diversity 105:1-16 ((https://doi.org/10.1007/s13225-020-00466-2)) |
| Otu48384 | 1 | k__Fungi(100);p__Ascomycota(100);c__Saccharomycetes(100);o__Saccharomycetales(100);f__Metschnikowiaceae(100);g__Metschnikowia(100);s__Metschnikowia_pulcherrima(100); | Metschnikowia | 13 | Saprotroph | \|Undefined Saprotroph\| | Yeast | NULL | Probable | As nectar-tap_saprotroph (Põlme et al. 2020); Decay sugar-rich substrates (Põlme et al. 2020) | Põlme S, et al. 2020. Fungal Diversity 105:1-16 ((https://doi.org/10.1007/s13225-020-00466-2)) |
| Otu48385 | 1 | k__Fungi(100);p__Ascomycota(100);c__Saccharomycetes(100);o__Saccharomycetales(100);f__Metschnikowiaceae(100);g__Metschnikowia(100);g__Metschnikowia_unclassified(100); | Metschnikowia | 13 | Saprotroph | \|Undefined Saprotroph\| | Yeast | NULL | Probable | As nectar-tap_saprotroph (Põlme et al. 2020); Decay sugar-rich substrates (Põlme et al. 2020) | Põlme S, et al. 2020. Fungal Diversity 105:1-16 ((https://doi.org/10.1007/s13225-020-00466-2)) |
| Otu48386 | 1 | k__Fungi(100);p__Ascomycota(100);c__Saccharomycetes(100);o__Saccharomycetales(100);f__Metschnikowiaceae(100);g__Metschnikowia(100);s__Metschnikowia_pulcherrima(100); | Metschnikowia | 13 | Saprotroph | \|Undefined Saprotroph\| | Yeast | NULL | Probable | As nectar-tap_saprotroph (Põlme et al. 2020); Decay sugar-rich substrates (Põlme et al. 2020) | Põlme S, et al. 2020. Fungal Diversity 105:1-16 ((https://doi.org/10.1007/s13225-020-00466-2)) |
| Otu48387 | 1 | k__Fungi(100);p__Ascomycota(100);c__Saccharomycetes(100);o__Saccharomycetales(100);f__Metschnikowiaceae(100);g__Metschnikowia(100);s__Metschnikowia_pulcherrima(100); | Metschnikowia | 13 | Saprotroph | \|Undefined Saprotroph\| | Yeast | NULL | Probable | As nectar-tap_saprotroph (Põlme et al. 2020); Decay sugar-rich substrates (Põlme et al. 2020) | Põlme S, et al. 2020. Fungal Diversity 105:1-16 ((https://doi.org/10.1007/s13225-020-00466-2)) |
| Otu48388 | 1 | k__Fungi(100);p__Ascomycota(100);c__Saccharomycetes(100);o__Saccharomycetales(100);f__Metschnikowiaceae(100);g__Metschnikowia(100);s__Metschnikowia_pulcherrima(100); | Metschnikowia | 13 | Saprotroph | \|Undefined Saprotroph\| | Yeast | NULL | Probable | As nectar-tap_saprotroph (Põlme et al. 2020); Decay sugar-rich substrates (Põlme et al. 2020) | Põlme S, et al. 2020. Fungal Diversity 105:1-16 ((https://doi.org/10.1007/s13225-020-00466-2)) |
| Otu48389 | 1 | k__Fungi(100);p__Ascomycota(100);c__Saccharomycetes(100);o__Saccharomycetales(100);f__Metschnikowiaceae(100);g__Metschnikowia(100);s__Metschnikowia_pulcherrima(100); | Metschnikowia | 13 | Saprotroph | \|Undefined Saprotroph\| | Yeast | NULL | Probable | As nectar-tap_saprotroph (Põlme et al. 2020); Decay sugar-rich substrates (Põlme et al. 2020) | Põlme S, et al. 2020. Fungal Diversity 105:1-16 ((https://doi.org/10.1007/s13225-020-00466-2)) |
| Otu48390 | 1 | k__Fungi(100);p__Ascomycota(100);c__Saccharomycetes(100);o__Saccharomycetales(100);f__Metschnikowiaceae(100);g__Metschnikowia(100);s__Metschnikowia_pulcherrima(100); | Metschnikowia | 13 | Saprotroph | \|Undefined Saprotroph\| | Yeast | NULL | Probable | As nectar-tap_saprotroph (Põlme et al. 2020); Decay sugar-rich substrates (Põlme et al. 2020) | Põlme S, et al. 2020. Fungal Diversity 105:1-16 ((https://doi.org/10.1007/s13225-020-00466-2)) |
| Otu48391 | 1 | k__Fungi(100);p__Ascomycota(100);c__Saccharomycetes(100);o__Saccharomycetales(100);f__Metschnikowiaceae(100);g__Metschnikowia(100);s__Metschnikowia_pulcherrima(100); | Metschnikowia | 13 | Saprotroph | \|Undefined Saprotroph\| | Yeast | NULL | Probable | As nectar-tap_saprotroph (Põlme et al. 2020); Decay sugar-rich substrates (Põlme et al. 2020) | Põlme S, et al. 2020. Fungal Diversity 105:1-16 ((https://doi.org/10.1007/s13225-020-00466-2)) |
| Otu48392 | 1 | k__Fungi(100);p__Ascomycota(100);c__Saccharomycetes(100);o__Saccharomycetales(100);f__Metschnikowiaceae(100);g__Metschnikowia(100);g__Metschnikowia_unclassified(100); | Metschnikowia | 13 | Saprotroph | \|Undefined Saprotroph\| | Yeast | NULL | Probable | As nectar-tap_saprotroph (Põlme et al. 2020); Decay sugar-rich substrates (Põlme et al. 2020) | Põlme S, et al. 2020. Fungal Diversity 105:1-16 ((https://doi.org/10.1007/s13225-020-00466-2)) |
| Otu48393 | 1 | k__Fungi(100);p__Ascomycota(100);c__Saccharomycetes(100);o__Saccharomycetales(100);f__Metschnikowiaceae(100);g__Metschnikowia(100);g__Metschnikowia_unclassified(100); | Metschnikowia | 13 | Saprotroph | \|Undefined Saprotroph\| | Yeast | NULL | Probable | As nectar-tap_saprotroph (Põlme et al. 2020); Decay sugar-rich substrates (Põlme et al. 2020) | Põlme S, et al. 2020. Fungal Diversity 105:1-16 ((https://doi.org/10.1007/s13225-020-00466-2)) |
| Otu48394 | 1 | k__Fungi(100);p__Ascomycota(100);c__Saccharomycetes(100);o__Saccharomycetales(100);f__Metschnikowiaceae(100);g__Metschnikowia(100);s__Metschnikowia_pulcherrima(100); | Metschnikowia | 13 | Saprotroph | \|Undefined Saprotroph\| | Yeast | NULL | Probable | As nectar-tap_saprotroph (Põlme et al. 2020); Decay sugar-rich substrates (Põlme et al. 2020) | Põlme S, et al. 2020. Fungal Diversity 105:1-16 ((https://doi.org/10.1007/s13225-020-00466-2)) |
| Otu48395 | 1 | k__Fungi(100);p__Ascomycota(100);c__Saccharomycetes(100);o__Saccharomycetales(100);f__Metschnikowiaceae(100);g__Metschnikowia(100);g__Metschnikowia_unclassified(100); | Metschnikowia | 13 | Saprotroph | \|Undefined Saprotroph\| | Yeast | NULL | Probable | As nectar-tap_saprotroph (Põlme et al. 2020); Decay sugar-rich substrates (Põlme et al. 2020) | Põlme S, et al. 2020. Fungal Diversity 105:1-16 ((https://doi.org/10.1007/s13225-020-00466-2)) |
| Otu48396 | 1 | k__Fungi(100);p__Ascomycota(100);c__Saccharomycetes(100);o__Saccharomycetales(100);f__Metschnikowiaceae(100);g__Metschnikowia(100);s__Metschnikowia_pulcherrima(100); | Metschnikowia | 13 | Saprotroph | \|Undefined Saprotroph\| | Yeast | NULL | Probable | As nectar-tap_saprotroph (Põlme et al. 2020); Decay sugar-rich substrates (Põlme et al. 2020) | Põlme S, et al. 2020. Fungal Diversity 105:1-16 ((https://doi.org/10.1007/s13225-020-00466-2)) |
| Otu48397 | 1 | k__Fungi(100);p__Ascomycota(100);c__Saccharomycetes(100);o__Saccharomycetales(100);f__Metschnikowiaceae(100);g__Metschnikowia(100);g__Metschnikowia_unclassified(100); | Metschnikowia | 13 | Saprotroph | \|Undefined Saprotroph\| | Yeast | NULL | Probable | As nectar-tap_saprotroph (Põlme et al. 2020); Decay sugar-rich substrates (Põlme et al. 2020) | Põlme S, et al. 2020. Fungal Diversity 105:1-16 ((https://doi.org/10.1007/s13225-020-00466-2)) |
| Otu48398 | 1 | k__Fungi(100);p__Ascomycota(100);c__Saccharomycetes(100);o__Saccharomycetales(100);f__Metschnikowiaceae(100);g__Metschnikowia(100);s__Metschnikowia_pulcherrima(100); | Metschnikowia | 13 | Saprotroph | \|Undefined Saprotroph\| | Yeast | NULL | Probable | As nectar-tap_saprotroph (Põlme et al. 2020); Decay sugar-rich substrates (Põlme et al. 2020) | Põlme S, et al. 2020. Fungal Diversity 105:1-16 ((https://doi.org/10.1007/s13225-020-00466-2)) |
| Otu48399 | 1 | k__Fungi(100);p__Ascomycota(100);c__Saccharomycetes(100);o__Saccharomycetales(100);f__Metschnikowiaceae(100);g__Metschnikowia(100);s__Metschnikowia_pulcherrima(100); | Metschnikowia | 13 | Saprotroph | \|Undefined Saprotroph\| | Yeast | NULL | Probable | As nectar-tap_saprotroph (Põlme et al. 2020); Decay sugar-rich substrates (Põlme et al. 2020) | Põlme S, et al. 2020. Fungal Diversity 105:1-16 ((https://doi.org/10.1007/s13225-020-00466-2)) |
| Otu48400 | 1 | k__Fungi(100);p__Ascomycota(100);c__Saccharomycetes(100);o__Saccharomycetales(100);f__Metschnikowiaceae(100);g__Metschnikowia(100);s__Metschnikowia_pulcherrima(100); | Metschnikowia | 13 | Saprotroph | \|Undefined Saprotroph\| | Yeast | NULL | Probable | As nectar-tap_saprotroph (Põlme et al. 2020); Decay sugar-rich substrates (Põlme et al. 2020) | Põlme S, et al. 2020. Fungal Diversity 105:1-16 ((https://doi.org/10.1007/s13225-020-00466-2)) |
| Otu48401 | 1 | k__Fungi(100);p__Ascomycota(100);c__Saccharomycetes(100);o__Saccharomycetales(100);f__Metschnikowiaceae(100);g__Metschnikowia(100);s__Metschnikowia_pulcherrima(100); | Metschnikowia | 13 | Saprotroph | \|Undefined Saprotroph\| | Yeast | NULL | Probable | As nectar-tap_saprotroph (Põlme et al. 2020); Decay sugar-rich substrates (Põlme et al. 2020) | Põlme S, et al. 2020. Fungal Diversity 105:1-16 ((https://doi.org/10.1007/s13225-020-00466-2)) |
| Otu48402 | 1 | k__Fungi(100);p__Ascomycota(100);c__Saccharomycetes(100);o__Saccharomycetales(100);f__Metschnikowiaceae(100);g__Metschnikowia(100);s__Metschnikowia_pulcherrima(100); | Metschnikowia | 13 | Saprotroph | \|Undefined Saprotroph\| | Yeast | NULL | Probable | As nectar-tap_saprotroph (Põlme et al. 2020); Decay sugar-rich substrates (Põlme et al. 2020) | Põlme S, et al. 2020. Fungal Diversity 105:1-16 ((https://doi.org/10.1007/s13225-020-00466-2)) |
| Otu48403 | 1 | k__Fungi(100);p__Ascomycota(100);c__Saccharomycetes(100);o__Saccharomycetales(100);f__Metschnikowiaceae(100);g__Metschnikowia(100);s__Metschnikowia_pulcherrima(100); | Metschnikowia | 13 | Saprotroph | \|Undefined Saprotroph\| | Yeast | NULL | Probable | As nectar-tap_saprotroph (Põlme et al. 2020); Decay sugar-rich substrates (Põlme et al. 2020) | Põlme S, et al. 2020. Fungal Diversity 105:1-16 ((https://doi.org/10.1007/s13225-020-00466-2)) |
| Otu48404 | 1 | k__Fungi(100);p__Ascomycota(100);c__Saccharomycetes(100);o__Saccharomycetales(100);f__Metschnikowiaceae(100);g__Metschnikowia(100);g__Metschnikowia_unclassified(100); | Metschnikowia | 13 | Saprotroph | \|Undefined Saprotroph\| | Yeast | NULL | Probable | As nectar-tap_saprotroph (Põlme et al. 2020); Decay sugar-rich substrates (Põlme et al. 2020) | Põlme S, et al. 2020. Fungal Diversity 105:1-16 ((https://doi.org/10.1007/s13225-020-00466-2)) |
| Otu48405 | 1 | k__Fungi(100);p__Ascomycota(100);c__Saccharomycetes(100);o__Saccharomycetales(100);f__Metschnikowiaceae(100);g__Metschnikowia(100);s__Metschnikowia_pulcherrima(100); | Metschnikowia | 13 | Saprotroph | \|Undefined Saprotroph\| | Yeast | NULL | Probable | As nectar-tap_saprotroph (Põlme et al. 2020); Decay sugar-rich substrates (Põlme et al. 2020) | Põlme S, et al. 2020. Fungal Diversity 105:1-16 ((https://doi.org/10.1007/s13225-020-00466-2)) |
| Otu48406 | 1 | k__Fungi(100);p__Ascomycota(100);c__Saccharomycetes(100);o__Saccharomycetales(100);f__Metschnikowiaceae(100);g__Metschnikowia(100);s__Metschnikowia_pulcherrima(100); | Metschnikowia | 13 | Saprotroph | \|Undefined Saprotroph\| | Yeast | NULL | Probable | As nectar-tap_saprotroph (Põlme et al. 2020); Decay sugar-rich substrates (Põlme et al. 2020) | Põlme S, et al. 2020. Fungal Diversity 105:1-16 ((https://doi.org/10.1007/s13225-020-00466-2)) |
| Otu48407 | 1 | k__Fungi(100);p__Ascomycota(100);c__Saccharomycetes(100);o__Saccharomycetales(100);f__Metschnikowiaceae(100);g__Metschnikowia(100);g__Metschnikowia_unclassified(100); | Metschnikowia | 13 | Saprotroph | \|Undefined Saprotroph\| | Yeast | NULL | Probable | As nectar-tap_saprotroph (Põlme et al. 2020); Decay sugar-rich substrates (Põlme et al. 2020) | Põlme S, et al. 2020. Fungal Diversity 105:1-16 ((https://doi.org/10.1007/s13225-020-00466-2)) |
| Otu48408 | 1 | k__Fungi(100);p__Ascomycota(100);c__Saccharomycetes(100);o__Saccharomycetales(100);f__Metschnikowiaceae(100);g__Metschnikowia(100);g__Metschnikowia_unclassified(100); | Metschnikowia | 13 | Saprotroph | \|Undefined Saprotroph\| | Yeast | NULL | Probable | As nectar-tap_saprotroph (Põlme et al. 2020); Decay sugar-rich substrates (Põlme et al. 2020) | Põlme S, et al. 2020. Fungal Diversity 105:1-16 ((https://doi.org/10.1007/s13225-020-00466-2)) |
| Otu48409 | 1 | k__Fungi(100);p__Ascomycota(100);c__Saccharomycetes(100);o__Saccharomycetales(100);f__Metschnikowiaceae(100);g__Metschnikowia(100);s__Metschnikowia_pulcherrima(100); | Metschnikowia | 13 | Saprotroph | \|Undefined Saprotroph\| | Yeast | NULL | Probable | As nectar-tap_saprotroph (Põlme et al. 2020); Decay sugar-rich substrates (Põlme et al. 2020) | Põlme S, et al. 2020. Fungal Diversity 105:1-16 ((https://doi.org/10.1007/s13225-020-00466-2)) |
| Otu48410 | 1 | k__Fungi(100);p__Ascomycota(100);c__Saccharomycetes(100);o__Saccharomycetales(100);f__Metschnikowiaceae(100);g__Metschnikowia(100);s__Metschnikowia_pulcherrima(100); | Metschnikowia | 13 | Saprotroph | \|Undefined Saprotroph\| | Yeast | NULL | Probable | As nectar-tap_saprotroph (Põlme et al. 2020); Decay sugar-rich substrates (Põlme et al. 2020) | Põlme S, et al. 2020. Fungal Diversity 105:1-16 ((https://doi.org/10.1007/s13225-020-00466-2)) |
| Otu48411 | 1 | k__Fungi(100);p__Ascomycota(100);c__Saccharomycetes(100);o__Saccharomycetales(100);f__Metschnikowiaceae(100);g__Metschnikowia(100);s__Metschnikowia_pulcherrima(100); | Metschnikowia | 13 | Saprotroph | \|Undefined Saprotroph\| | Yeast | NULL | Probable | As nectar-tap_saprotroph (Põlme et al. 2020); Decay sugar-rich substrates (Põlme et al. 2020) | Põlme S, et al. 2020. Fungal Diversity 105:1-16 ((https://doi.org/10.1007/s13225-020-00466-2)) |
| Otu48413 | 1 | k__Fungi(100);p__Ascomycota(100);c__Saccharomycetes(100);o__Saccharomycetales(100);f__Metschnikowiaceae(100);g__Metschnikowia(100);s__Metschnikowia_pulcherrima(100); | Metschnikowia | 13 | Saprotroph | \|Undefined Saprotroph\| | Yeast | NULL | Probable | As nectar-tap_saprotroph (Põlme et al. 2020); Decay sugar-rich substrates (Põlme et al. 2020) | Põlme S, et al. 2020. Fungal Diversity 105:1-16 ((https://doi.org/10.1007/s13225-020-00466-2)) |
| Otu48414 | 1 | k__Fungi(100);p__Ascomycota(100);c__Saccharomycetes(100);o__Saccharomycetales(100);f__Metschnikowiaceae(100);g__Metschnikowia(100);g__Metschnikowia_unclassified(100); | Metschnikowia | 13 | Saprotroph | \|Undefined Saprotroph\| | Yeast | NULL | Probable | As nectar-tap_saprotroph (Põlme et al. 2020); Decay sugar-rich substrates (Põlme et al. 2020) | Põlme S, et al. 2020. Fungal Diversity 105:1-16 ((https://doi.org/10.1007/s13225-020-00466-2)) |

| | | | | | | | | | | | |
|---|---|---|---|---|---|---|---|---|---|---|---|
| Otu48648 | 1 | k__Fungi(100);p__Ascomycota(100);c__Saccharomycetes(100);o__Saccharomycetales(100);f__Metschnikowiaceae(100);g__Metschnikowia(100);s__Metschnikowia_pulcherrima(100); | Metschnikowia | 13 | Saprotroph | \|Undefined Saprotroph\| | Yeast | NULL | Probable | As nectar-tap_saprotroph (Põlme et al. 2020); Decay sugar-rich substrates (Põlme et al. 2020) | Põlme S, et al. 2020. Fungal Diversity 105:1-16 ((https://doi.org/10.1007/s13225-020-00466-2)) |
| Otu48649 | 1 | k__Fungi(100);p__Ascomycota(100);c__Saccharomycetes(100);o__Saccharomycetales(100);f__Metschnikowiaceae(100);g__Metschnikowia(100);g__Metschnikowia_unclassified(100); | Metschnikowia | 13 | Saprotroph | \|Undefined Saprotroph\| | Yeast | NULL | Probable | As nectar-tap_saprotroph (Põlme et al. 2020); Decay sugar-rich substrates (Põlme et al. 2020) | Põlme S, et al. 2020. Fungal Diversity 105:1-16 ((https://doi.org/10.1007/s13225-020-00466-2)) |
| Otu48650 | 1 | k__Fungi(100);p__Ascomycota(100);c__Saccharomycetes(100);o__Saccharomycetales(100);f__Metschnikowiaceae(100);g__Metschnikowia(100);s__Metschnikowia_pulcherrima(100); | Metschnikowia | 13 | Saprotroph | \|Undefined Saprotroph\| | Yeast | NULL | Probable | As nectar-tap_saprotroph (Põlme et al. 2020); Decay sugar-rich substrates (Põlme et al. 2020) | Põlme S, et al. 2020. Fungal Diversity 105:1-16 ((https://doi.org/10.1007/s13225-020-00466-2)) |
| Otu48651 | 1 | k__Fungi(100);p__Ascomycota(100);c__Saccharomycetes(100);o__Saccharomycetales(100);f__Metschnikowiaceae(100);g__Metschnikowia(100);g__Metschnikowia_unclassified(100); | Metschnikowia | 13 | Saprotroph | \|Undefined Saprotroph\| | Yeast | NULL | Probable | As nectar-tap_saprotroph (Põlme et al. 2020); Decay sugar-rich substrates (Põlme et al. 2020) | Põlme S, et al. 2020. Fungal Diversity 105:1-16 ((https://doi.org/10.1007/s13225-020-00466-2)) |
| Otu48652 | 1 | k__Fungi(100);p__Ascomycota(100);c__Saccharomycetes(100);o__Saccharomycetales(100);f__Metschnikowiaceae(100);g__Metschnikowia(100);s__Metschnikowia_pulcherrima(100); | Metschnikowia | 13 | Saprotroph | \|Undefined Saprotroph\| | Yeast | NULL | Probable | As nectar-tap_saprotroph (Põlme et al. 2020); Decay sugar-rich substrates (Põlme et al. 2020) | Põlme S, et al. 2020. Fungal Diversity 105:1-16 ((https://doi.org/10.1007/s13225-020-00466-2)) |
| Otu48653 | 1 | k__Fungi(100);p__Ascomycota(100);c__Saccharomycetes(100);o__Saccharomycetales(100);f__Metschnikowiaceae(100);g__Metschnikowia(100);s__Metschnikowia_pulcherrima(100); | Metschnikowia | 13 | Saprotroph | \|Undefined Saprotroph\| | Yeast | NULL | Probable | As nectar-tap_saprotroph (Põlme et al. 2020); Decay sugar-rich substrates (Põlme et al. 2020) | Põlme S, et al. 2020. Fungal Diversity 105:1-16 ((https://doi.org/10.1007/s13225-020-00466-2)) |
| Otu48654 | 1 | k__Fungi(100);p__Ascomycota(100);c__Saccharomycetes(100);o__Saccharomycetales(100);f__Metschnikowiaceae(100);g__Metschnikowia(100);s__Metschnikowia_pulcherrima(100); | Metschnikowia | 13 | Saprotroph | \|Undefined Saprotroph\| | Yeast | NULL | Probable | As nectar-tap_saprotroph (Põlme et al. 2020); Decay sugar-rich substrates (Põlme et al. 2020) | Põlme S, et al. 2020. Fungal Diversity 105:1-16 ((https://doi.org/10.1007/s13225-020-00466-2)) |
| Otu48655 | 1 | k__Fungi(100);p__Ascomycota(100);c__Saccharomycetes(100);o__Saccharomycetales(100);f__Metschnikowiaceae(100);g__Metschnikowia(100);s__Metschnikowia_pulcherrima(100); | Metschnikowia | 13 | Saprotroph | \|Undefined Saprotroph\| | Yeast | NULL | Probable | As nectar-tap_saprotroph (Põlme et al. 2020); Decay sugar-rich substrates (Põlme et al. 2020) | Põlme S, et al. 2020. Fungal Diversity 105:1-16 ((https://doi.org/10.1007/s13225-020-00466-2)) |
| Otu48656 | 1 | k__Fungi(100);p__Ascomycota(100);c__Saccharomycetes(100);o__Saccharomycetales(100);f__Metschnikowiaceae(100);g__Metschnikowia(100);s__Metschnikowia_pulcherrima(100); | Metschnikowia | 13 | Saprotroph | \|Undefined Saprotroph\| | Yeast | NULL | Probable | As nectar-tap_saprotroph (Põlme et al. 2020); Decay sugar-rich substrates (Põlme et al. 2020) | Põlme S, et al. 2020. Fungal Diversity 105:1-16 ((https://doi.org/10.1007/s13225-020-00466-2)) |
| Otu48657 | 1 | k__Fungi(100);p__Ascomycota(100);c__Saccharomycetes(100);o__Saccharomycetales(100);f__Metschnikowiaceae(100);g__Metschnikowia(100);s__Metschnikowia_pulcherrima(100); | Metschnikowia | 13 | Saprotroph | \|Undefined Saprotroph\| | Yeast | NULL | Probable | As nectar-tap_saprotroph (Põlme et al. 2020); Decay sugar-rich substrates (Põlme et al. 2020) | Põlme S, et al. 2020. Fungal Diversity 105:1-16 ((https://doi.org/10.1007/s13225-020-00466-2)) |
| Otu48658 | 1 | k__Fungi(100);p__Ascomycota(100);c__Saccharomycetes(100);o__Saccharomycetales(100);f__Metschnikowiaceae(100);g__Metschnikowia(100);s__Metschnikowia_pulcherrima(100); | Metschnikowia | 13 | Saprotroph | \|Undefined Saprotroph\| | Yeast | NULL | Probable | As nectar-tap_saprotroph (Põlme et al. 2020); Decay sugar-rich substrates (Põlme et al. 2020) | Põlme S, et al. 2020. Fungal Diversity 105:1-16 ((https://doi.org/10.1007/s13225-020-00466-2)) |
| Otu48659 | 1 | k__Fungi(100);p__Ascomycota(100);c__Saccharomycetes(100);o__Saccharomycetales(100);f__Metschnikowiaceae(100);g__Metschnikowia(100);s__Metschnikowia_pulcherrima(100); | Metschnikowia | 13 | Saprotroph | \|Undefined Saprotroph\| | Yeast | NULL | Probable | As nectar-tap_saprotroph (Põlme et al. 2020); Decay sugar-rich substrates (Põlme et al. 2020) | Põlme S, et al. 2020. Fungal Diversity 105:1-16 ((https://doi.org/10.1007/s13225-020-00466-2)) |
| Otu48660 | 1 | k__Fungi(100);p__Ascomycota(100);c__Saccharomycetes(100);o__Saccharomycetales(100);f__Metschnikowiaceae(100);g__Metschnikowia(100);s__Metschnikowia_pulcherrima(100); | Metschnikowia | 13 | Saprotroph | \|Undefined Saprotroph\| | Yeast | NULL | Probable | As nectar-tap_saprotroph (Põlme et al. 2020); Decay sugar-rich substrates (Põlme et al. 2020) | Põlme S, et al. 2020. Fungal Diversity 105:1-16 ((https://doi.org/10.1007/s13225-020-00466-2)) |
| Otu48661 | 1 | k__Fungi(100);p__Ascomycota(100);c__Saccharomycetes(100);o__Saccharomycetales(100);f__Metschnikowiaceae(100);g__Metschnikowia(100);s__Metschnikowia_pulcherrima(100); | Metschnikowia | 13 | Saprotroph | \|Undefined Saprotroph\| | Yeast | NULL | Probable | As nectar-tap_saprotroph (Põlme et al. 2020); Decay sugar-rich substrates (Põlme et al. 2020) | Põlme S, et al. 2020. Fungal Diversity 105:1-16 ((https://doi.org/10.1007/s13225-020-00466-2)) |
| Otu48662 | 1 | k__Fungi(100);p__Ascomycota(100);c__Saccharomycetes(100);o__Saccharomycetales(100);f__Metschnikowiaceae(100);g__Metschnikowia(100);s__Metschnikowia_pulcherrima(100); | Metschnikowia | 13 | Saprotroph | \|Undefined Saprotroph\| | Yeast | NULL | Probable | As nectar-tap_saprotroph (Põlme et al. 2020); Decay sugar-rich substrates (Põlme et al. 2020) | Põlme S, et al. 2020. Fungal Diversity 105:1-16 ((https://doi.org/10.1007/s13225-020-00466-2)) |
| Otu48663 | 1 | k__Fungi(100);p__Ascomycota(100);c__Saccharomycetes(100);o__Saccharomycetales(100);f__Metschnikowiaceae(100);g__Metschnikowia(100);s__Metschnikowia_pulcherrima(100); | Metschnikowia | 13 | Saprotroph | \|Undefined Saprotroph\| | Yeast | NULL | Probable | As nectar-tap_saprotroph (Põlme et al. 2020); Decay sugar-rich substrates (Põlme et al. 2020) | Põlme S, et al. 2020. Fungal Diversity 105:1-16 ((https://doi.org/10.1007/s13225-020-00466-2)) |
| Otu48664 | 1 | k__Fungi(100);p__Ascomycota(100);c__Saccharomycetes(100);o__Saccharomycetales(100);f__Metschnikowiaceae(100);g__Metschnikowia(100);g__Metschnikowia_unclassified(100); | Metschnikowia | 13 | Saprotroph | \|Undefined Saprotroph\| | Yeast | NULL | Probable | As nectar-tap_saprotroph (Põlme et al. 2020); Decay sugar-rich substrates (Põlme et al. 2020) | Põlme S, et al. 2020. Fungal Diversity 105:1-16 ((https://doi.org/10.1007/s13225-020-00466-2)) |
| Otu48665 | 1 | k__Fungi(100);p__Ascomycota(100);c__Saccharomycetes(100);o__Saccharomycetales(100);f__Metschnikowiaceae(100);g__Metschnikowia(100);g__Metschnikowia_unclassified(100); | Metschnikowia | 13 | Saprotroph | \|Undefined Saprotroph\| | Yeast | NULL | Probable | As nectar-tap_saprotroph (Põlme et al. 2020); Decay sugar-rich substrates (Põlme et al. 2020) | Põlme S, et al. 2020. Fungal Diversity 105:1-16 ((https://doi.org/10.1007/s13225-020-00466-2)) |
| Otu48666 | 1 | k__Fungi(100);p__Ascomycota(100);c__Saccharomycetes(100);o__Saccharomycetales(100);f__Metschnikowiaceae(100);g__Metschnikowia(100);s__Metschnikowia_pulcherrima(100); | Metschnikowia | 13 | Saprotroph | \|Undefined Saprotroph\| | Yeast | NULL | Probable | As nectar-tap_saprotroph (Põlme et al. 2020); Decay sugar-rich substrates (Põlme et al. 2020) | Põlme S, et al. 2020. Fungal Diversity 105:1-16 ((https://doi.org/10.1007/s13225-020-00466-2)) |
| Otu48667 | 1 | k__Fungi(100);p__Ascomycota(100);c__Saccharomycetes(100);o__Saccharomycetales(100);f__Metschnikowiaceae(100);g__Metschnikowia(100);s__Metschnikowia_pulcherrima(100); | Metschnikowia | 13 | Saprotroph | \|Undefined Saprotroph\| | Yeast | NULL | Probable | As nectar-tap_saprotroph (Põlme et al. 2020); Decay sugar-rich substrates (Põlme et al. 2020) | Põlme S, et al. 2020. Fungal Diversity 105:1-16 ((https://doi.org/10.1007/s13225-020-00466-2)) |
| Otu48668 | 1 | k__Fungi(100);p__Ascomycota(100);c__Saccharomycetes(100);o__Saccharomycetales(100);f__Metschnikowiaceae(100);g__Metschnikowia(100);g__Metschnikowia_unclassified(100); | Metschnikowia | 13 | Saprotroph | \|Undefined Saprotroph\| | Yeast | NULL | Probable | As nectar-tap_saprotroph (Põlme et al. 2020); Decay sugar-rich substrates (Põlme et al. 2020) | Põlme S, et al. 2020. Fungal Diversity 105:1-16 ((https://doi.org/10.1007/s13225-020-00466-2)) |
| Otu48669 | 1 | k__Fungi(100);p__Ascomycota(100);c__Saccharomycetes(100);o__Saccharomycetales(100);f__Metschnikowiaceae(100);g__Metschnikowia(100);s__Metschnikowia_pulcherrima(100); | Metschnikowia | 13 | Saprotroph | \|Undefined Saprotroph\| | Yeast | NULL | Probable | As nectar-tap_saprotroph (Põlme et al. 2020); Decay sugar-rich substrates (Põlme et al. 2020) | Põlme S, et al. 2020. Fungal Diversity 105:1-16 ((https://doi.org/10.1007/s13225-020-00466-2)) |
| Otu48670 | 1 | k__Fungi(100);p__Ascomycota(100);c__Saccharomycetes(100);o__Saccharomycetales(100);f__Metschnikowiaceae(100);g__Metschnikowia(100);s__Metschnikowia_pulcherrima(100); | Metschnikowia | 13 | Saprotroph | \|Undefined Saprotroph\| | Yeast | NULL | Probable | As nectar-tap_saprotroph (Põlme et al. 2020); Decay sugar-rich substrates (Põlme et al. 2020) | Põlme S, et al. 2020. Fungal Diversity 105:1-16 ((https://doi.org/10.1007/s13225-020-00466-2)) |
| Otu48671 | 1 | k__Fungi(100);p__Ascomycota(100);c__Saccharomycetes(100);o__Saccharomycetales(100);f__Metschnikowiaceae(100);g__Metschnikowia(100);s__Metschnikowia_pulcherrima(100); | Metschnikowia | 13 | Saprotroph | \|Undefined Saprotroph\| | Yeast | NULL | Probable | As nectar-tap_saprotroph (Põlme et al. 2020); Decay sugar-rich substrates (Põlme et al. 2020) | Põlme S, et al. 2020. Fungal Diversity 105:1-16 ((https://doi.org/10.1007/s13225-020-00466-2)) |
| Otu48672 | 1 | k__Fungi(100);p__Ascomycota(100);c__Saccharomycetes(100);o__Saccharomycetales(100);f__Metschnikowiaceae(100);g__Metschnikowia(100);g__Metschnikowia_unclassified(100); | Metschnikowia | 13 | Saprotroph | \|Undefined Saprotroph\| | Yeast | NULL | Probable | As nectar-tap_saprotroph (Põlme et al. 2020); Decay sugar-rich substrates (Põlme et al. 2020) | Põlme S, et al. 2020. Fungal Diversity 105:1-16 ((https://doi.org/10.1007/s13225-020-00466-2)) |
| Otu48673 | 1 | k__Fungi(100);p__Ascomycota(100);c__Saccharomycetes(100);o__Saccharomycetales(100);f__Metschnikowiaceae(100);g__Metschnikowia(100);s__Metschnikowia_pulcherrima(100); | Metschnikowia | 13 | Saprotroph | \|Undefined Saprotroph\| | Yeast | NULL | Probable | As nectar-tap_saprotroph (Põlme et al. 2020); Decay sugar-rich substrates (Põlme et al. 2020) | Põlme S, et al. 2020. Fungal Diversity 105:1-16 ((https://doi.org/10.1007/s13225-020-00466-2)) |
| Otu48674 | 1 | k__Fungi(100);p__Ascomycota(100);c__Saccharomycetes(100);o__Saccharomycetales(100);f__Metschnikowiaceae(100);g__Metschnikowia(100);s__Metschnikowia_pulcherrima(100); | Metschnikowia | 13 | Saprotroph | \|Undefined Saprotroph\| | Yeast | NULL | Probable | As nectar-tap_saprotroph (Põlme et al. 2020); Decay sugar-rich substrates (Põlme et al. 2020) | Põlme S, et al. 2020. Fungal Diversity 105:1-16 ((https://doi.org/10.1007/s13225-020-00466-2)) |
| Otu48675 | 1 | k__Fungi(100);p__Ascomycota(100);c__Saccharomycetes(100);o__Saccharomycetales(100);f__Metschnikowiaceae(100);g__Metschnikowia(100);s__Metschnikowia_pulcherrima(100); | Metschnikowia | 13 | Saprotroph | \|Undefined Saprotroph\| | Yeast | NULL | Probable | As nectar-tap_saprotroph (Põlme et al. 2020); Decay sugar-rich substrates (Põlme et al. 2020) | Põlme S, et al. 2020. Fungal Diversity 105:1-16 ((https://doi.org/10.1007/s13225-020-00466-2)) |
| Otu48676 | 1 | k__Fungi(100);p__Ascomycota(100);c__Saccharomycetes(100);o__Saccharomycetales(100);f__Metschnikowiaceae(100);g__Metschnikowia(100);g__Metschnikowia_unclassified(100); | Metschnikowia | 13 | Saprotroph | \|Undefined Saprotroph\| | Yeast | NULL | Probable | As nectar-tap_saprotroph (Põlme et al. 2020); Decay sugar-rich substrates (Põlme et al. 2020) | Põlme S, et al. 2020. Fungal Diversity 105:1-16 ((https://doi.org/10.1007/s13225-020-00466-2)) |
| Otu48677 | 1 | k__Fungi(100);p__Ascomycota(100);c__Saccharomycetes(100);o__Saccharomycetales(100);f__Metschnikowiaceae(100);g__Metschnikowia(100);s__Metschnikowia_pulcherrima(100); | Metschnikowia | 13 | Saprotroph | \|Undefined Saprotroph\| | Yeast | NULL | Probable | As nectar-tap_saprotroph (Põlme et al. 2020); Decay sugar-rich substrates (Põlme et al. 2020) | Põlme S, et al. 2020. Fungal Diversity 105:1-16 ((https://doi.org/10.1007/s13225-020-00466-2)) |
| Otu48678 | 1 | k__Fungi(100);p__Ascomycota(100);c__Saccharomycetes(100);o__Saccharomycetales(100);f__Metschnikowiaceae(100);g__Metschnikowia(100);s__Metschnikowia_pulcherrima(100); | Metschnikowia | 13 | Saprotroph | \|Undefined Saprotroph\| | Yeast | NULL | Probable | As nectar-tap_saprotroph (Põlme et al. 2020); Decay sugar-rich substrates (Põlme et al. 2020) | Põlme S, et al. 2020. Fungal Diversity 105:1-16 ((https://doi.org/10.1007/s13225-020-00466-2)) |
| Otu48679 | 1 | k__Fungi(100);p__Ascomycota(100);c__Saccharomycetes(100);o__Saccharomycetales(100);f__Metschnikowiaceae(100);g__Metschnikowia(100);s__Metschnikowia_pulcherrima(100); | Metschnikowia | 13 | Saprotroph | \|Undefined Saprotroph\| | Yeast | NULL | Probable | As nectar-tap_saprotroph (Põlme et al. 2020); Decay sugar-rich substrates (Põlme et al. 2020) | Põlme S, et al. 2020. Fungal Diversity 105:1-16 ((https://doi.org/10.1007/s13225-020-00466-2)) |
| Otu48680 | 1 | k__Fungi(100);p__Ascomycota(100);c__Saccharomycetes(100);o__Saccharomycetales(100);f__Metschnikowiaceae(100);g__Metschnikowia(100);s__Metschnikowia_pulcherrima(100); | Metschnikowia | 13 | Saprotroph | \|Undefined Saprotroph\| | Yeast | NULL | Probable | As nectar-tap_saprotroph (Põlme et al. 2020); Decay sugar-rich substrates (Põlme et al. 2020) | Põlme S, et al. 2020. Fungal Diversity 105:1-16 ((https://doi.org/10.1007/s13225-020-00466-2)) |
| Otu48681 | 1 | k__Fungi(100);p__Ascomycota(100);c__Saccharomycetes(100);o__Saccharomycetales(100);f__Metschnikowiaceae(100);g__Metschnikowia(100);s__Metschnikowia_pulcherrima(100); | Metschnikowia | 13 | Saprotroph | \|Undefined Saprotroph\| | Yeast | NULL | Probable | As nectar-tap_saprotroph (Põlme et al. 2020); Decay sugar-rich substrates (Põlme et al. 2020) | Põlme S, et al. 2020. Fungal Diversity 105:1-16 ((https://doi.org/10.1007/s13225-020-00466-2)) |
| Otu48682 | 1 | k__Fungi(100);p__Ascomycota(100);c__Saccharomycetes(100);o__Saccharomycetales(100);f__Metschnikowiaceae(100);g__Metschnikowia(100);s__Metschnikowia_pulcherrima(100); | Metschnikowia | 13 | Saprotroph | \|Undefined Saprotroph\| | Yeast | NULL | Probable | As nectar-tap_saprotroph (Põlme et al. 2020); Decay sugar-rich substrates (Põlme et al. 2020) | Põlme S, et al. 2020. Fungal Diversity 105:1-16 ((https://doi.org/10.1007/s13225-020-00466-2)) |
| Otu48683 | 1 | k__Fungi(100);p__Ascomycota(100);c__Saccharomycetes(100);o__Saccharomycetales(100);f__Metschnikowiaceae(100);g__Metschnikowia(100);s__Metschnikowia_pulcherrima(100); | Metschnikowia | 13 | Saprotroph | \|Undefined Saprotroph\| | Yeast | NULL | Probable | As nectar-tap_saprotroph (Põlme et al. 2020); Decay sugar-rich substrates (Põlme et al. 2020) | Põlme S, et al. 2020. Fungal Diversity 105:1-16 ((https://doi.org/10.1007/s13225-020-00466-2)) |
| Otu48684 | 1 | k__Fungi(100);p__Ascomycota(100);c__Saccharomycetes(100);o__Saccharomycetales(100);f__Metschnikowiaceae(100);g__Metschnikowia(100);s__Metschnikowia_pulcherrima(100); | Metschnikowia | 13 | Saprotroph | \|Undefined Saprotroph\| | Yeast | NULL | Probable | As nectar-tap_saprotroph (Põlme et al. 2020); Decay sugar-rich substrates (Põlme et al. 2020) | Põlme S, et al. 2020. Fungal Diversity 105:1-16 ((https://doi.org/10.1007/s13225-020-00466-2)) |
| Otu48685 | 1 | k__Fungi(100);p__Ascomycota(100);c__Saccharomycetes(100);o__Saccharomycetales(100);f__Metschnikowiaceae(100);g__Metschnikowia(100);s__Metschnikowia_sp(100); | Metschnikowia | 13 | Saprotroph | \|Undefined Saprotroph\| | Yeast | NULL | Probable | As nectar-tap_saprotroph (Põlme et al. 2020); Decay sugar-rich substrates (Põlme et al. 2020) | Põlme S, et al. 2020. Fungal Diversity 105:1-16 ((https://doi.org/10.1007/s13225-020-00466-2)) |
| Otu48686 | 1 | k__Fungi(100);p__Ascomycota(100);c__Saccharomycetes(100);o__Saccharomycetales(100);f__Metschnikowiaceae(100);g__Metschnikowia(100);s__Metschnikowia_pulcherrima(100); | Metschnikowia | 13 | Saprotroph | \|Undefined Saprotroph\| | Yeast | NULL | Probable | As nectar-tap_saprotroph (Põlme et al. 2020); Decay sugar-rich substrates (Põlme et al. 2020) | Põlme S, et al. 2020. Fungal Diversity 105:1-16 ((https://doi.org/10.1007/s13225-020-00466-2)) |
| Otu48687 | 1 | k__Fungi(100);p__Ascomycota(100);c__Saccharomycetes(100);o__Saccharomycetales(100);f__Metschnikowiaceae(100);g__Metschnikowia(100);s__Metschnikowia_pulcherrima(100); | Metschnikowia | 13 | Saprotroph | \|Undefined Saprotroph\| | Yeast | NULL | Probable | As nectar-tap_saprotroph (Põlme et al. 2020); Decay sugar-rich substrates (Põlme et al. 2020) | Põlme S, et al. 2020. Fungal Diversity 105:1-16 ((https://doi.org/10.1007/s13225-020-00466-2)) |
| Otu48688 | 1 | k__Fungi(100);p__Ascomycota(100);c__Saccharomycetes(100);o__Saccharomycetales(100);f__Metschnikowiaceae(100);g__Metschnikowia(100);s__Metschnikowia_pulcherrima(100); | Metschnikowia | 13 | Saprotroph | \|Undefined Saprotroph\| | Yeast | NULL | Probable | As nectar-tap_saprotroph (Põlme et al. 2020); Decay sugar-rich substrates (Põlme et al. 2020) | Põlme S, et al. 2020. Fungal Diversity 105:1-16 ((https://doi.org/10.1007/s13225-020-00466-2)) |
| Otu48689 | 1 | k__Fungi(100);p__Ascomycota(100);c__Saccharomycetes(100);o__Saccharomycetales(100);f__Metschnikowiaceae(100);g__Metschnikowia(100);s__Metschnikowia_pulcherrima(100); | Metschnikowia | 13 | Saprotroph | \|Undefined Saprotroph\| | Yeast | NULL | Probable | As nectar-tap_saprotroph (Põlme et al. 2020); Decay sugar-rich substrates (Põlme et al. 2020) | Põlme S, et al. 2020. Fungal Diversity 105:1-16 ((https://doi.org/10.1007/s13225-020-00466-2)) |
| Otu48690 | 1 | k__Fungi(100);p__Ascomycota(100);c__Saccharomycetes(100);o__Saccharomycetales(100);f__Metschnikowiaceae(100);g__Metschnikowia(100);s__Metschnikowia_pulcherrima(100); | Metschnikowia | 13 | Saprotroph | \|Undefined Saprotroph\| | Yeast | NULL | Probable | As nectar-tap_saprotroph (Põlme et al. 2020); Decay sugar-rich substrates (Põlme et al. 2020) | Põlme S, et al. 2020. Fungal Diversity 105:1-16 ((https://doi.org/10.1007/s13225-020-00466-2)) |
| Otu48691 | 1 | k__Fungi(100);p__Ascomycota(100);c__Saccharomycetes(100);o__Saccharomycetales(100);f__Metschnikowiaceae(100);g__Metschnikowia(100);g__Metschnikowia_unclassified(100); | Metschnikowia | 13 | Saprotroph | \|Undefined Saprotroph\| | Yeast | NULL | Probable | As nectar-tap_saprotroph (Põlme et al. 2020); Decay sugar-rich substrates (Põlme et al. 2020) | Põlme S, et al. 2020. Fungal Diversity 105:1-16 ((https://doi.org/10.1007/s13225-020-00466-2)) |
| Otu48692 | 1 | k__Fungi(100);p__Ascomycota(100);c__Saccharomycetes(100);o__Saccharomycetales(100);f__Metschnikowiaceae(100);g__Metschnikowia(100);s__Metschnikowia_pulcherrima(100); | Metschnikowia | 13 | Saprotroph | \|Undefined Saprotroph\| | Yeast | NULL | Probable | As nectar-tap_saprotroph (Põlme et al. 2020); Decay sugar-rich substrates (Põlme et al. 2020) | Põlme S, et al. 2020. Fungal Diversity 105:1-16 ((https://doi.org/10.1007/s13225-020-00466-2)) |
| Otu48693 | 1 | k__Fungi(100);p__Ascomycota(100);c__Saccharomycetes(100);o__Saccharomycetales(100);f__Metschnikowiaceae(100);g__Metschnikowia(100);s__Metschnikowia_pulcherrima(100); | Metschnikowia | 13 | Saprotroph | \|Undefined Saprotroph\| | Yeast | NULL | Probable | As nectar-tap_saprotroph (Põlme et al. 2020); Decay sugar-rich substrates (Põlme et al. 2020) | Põlme S, et al. 2020. Fungal Diversity 105:1-16 ((https://doi.org/10.1007/s13225-020-00466-2)) |
| Otu48694 | 1 | k__Fungi(100);p__Ascomycota(100);c__Saccharomycetes(100);o__Saccharomycetales(100);f__Metschnikowiaceae(100);g__Metschnikowia(100);s__Metschnikowia_pulcherrima(100); | Metschnikowia | 13 | Saprotroph | \|Undefined Saprotroph\| | Yeast | NULL | Probable | As nectar-tap_saprotroph (Põlme et al. 2020); Decay sugar-rich substrates (Põlme et al. 2020) | Põlme S, et al. 2020. Fungal Diversity 105:1-16 ((https://doi.org/10.1007/s13225-020-00466-2)) |
| Otu48695 | 1 | k__Fungi(100);p__Ascomycota(100);c__Saccharomycetes(100);o__Saccharomycetales(100);f__Metschnikowiaceae(100);g__Metschnikowia(100);g__Metschnikowia_unclassified(100); | Metschnikowia | 13 | Saprotroph | \|Undefined Saprotroph\| | Yeast | NULL | Probable | As nectar-tap_saprotroph (Põlme et al. 2020); Decay sugar-rich substrates (Põlme et al. 2020) | Põlme S, et al. 2020. Fungal Diversity 105:1-16 ((https://doi.org/10.1007/s13225-020-00466-2)) |
| Otu48696 | 1 | k__Fungi(100);p__Ascomycota(100);c__Saccharomycetes(100);o__Saccharomycetales(100);f__Metschnikowiaceae(100);g__Metschnikowia(100);s__Metschnikowia_pulcherrima(100); | Metschnikowia | 13 | Saprotroph | \|Undefined Saprotroph\| | Yeast | NULL | Probable | As nectar-tap_saprotroph (Põlme et al. 2020); Decay sugar-rich substrates (Põlme et al. 2020) | Põlme S, et al. 2020. Fungal Diversity 105:1-16 ((https://doi.org/10.1007/s13225-020-00466-2)) |
| Otu48697 | 1 | k__Fungi(100);p__Ascomycota(100);c__Saccharomycetes(100);o__Saccharomycetales(100);f__Metschnikowiaceae(100);g__Metschnikowia(100);g__Metschnikowia_unclassified(100); | Metschnikowia | 13 | Saprotroph | \|Undefined Saprotroph\| | Yeast | NULL | Probable | As nectar-tap_saprotroph (Põlme et al. 2020); Decay sugar-rich substrates (Põlme et al. 2020) | Põlme S, et al. 2020. Fungal Diversity 105:1-16 ((https://doi.org/10.1007/s13225-020-00466-2)) |
| Otu48698 | 1 | k__Fungi(100);p__Ascomycota(100);c__Saccharomycetes(100);o__Saccharomycetales(100);f__Metschnikowiaceae(100);g__Metschnikowia(100);s__Metschnikowia_pulcherrima(100); | Metschnikowia | 13 | Saprotroph | \|Undefined Saprotroph\| | Yeast | NULL | Probable | As nectar-tap_saprotroph (Põlme et al. 2020); Decay sugar-rich substrates (Põlme et al. 2020) | Põlme S, et al. 2020. Fungal Diversity 105:1-16 ((https://doi.org/10.1007/s13225-020-00466-2)) |
| Otu48699 | 1 | k__Fungi(100);p__Ascomycota(100);c__Saccharomycetes(100);o__Saccharomycetales(100);f__Metschnikowiaceae(100);g__Metschnikowia(100);s__Metschnikowia_pulcherrima(100); | Metschnikowia | 13 | Saprotroph | \|Undefined Saprotroph\| | Yeast | NULL | Probable | As nectar-tap_saprotroph (Põlme et al. 2020); Decay sugar-rich substrates (Põlme et al. 2020) | Põlme S, et al. 2020. Fungal Diversity 105:1-16 ((https://doi.org/10.1007/s13225-020-00466-2)) |
| Otu48700 | 1 | k__Fungi(100);p__Ascomycota(100);c__Saccharomycetes(100);o__Saccharomycetales(100);f__Metschnikowiaceae(100);g__Metschnikowia(100);s__Metschnikowia_viticola(100); | Metschnikowia | 13 | Saprotroph | \|Undefined Saprotroph\| | Yeast | NULL | Probable | As nectar-tap_saprotroph (Põlme et al. 2020); Decay sugar-rich substrates (Põlme et al. 2020) | Põlme S, et al. 2020. Fungal Diversity 105:1-16 ((https://doi.org/10.1007/s13225-020-00466-2)) |
| Otu48701 | 1 | k__Fungi(100);p__Ascomycota(100);c__Saccharomycetes(100);o__Saccharomycetales(100);f__Metschnikowiaceae(100);g__Metschnikowia(100);s__Metschnikowia_pulcherrima(100); | Metschnikowia | 13 | Saprotroph | \|Undefined Saprotroph\| | Yeast | NULL | Probable | As nectar-tap_saprotroph (Põlme et al. 2020); Decay sugar-rich substrates (Põlme et al. 2020) | Põlme S, et al. 2020. Fungal Diversity 105:1-16 ((https://doi.org/10.1007/s13225-020-00466-2)) |
| Otu48702 | 1 | k__Fungi(100);p__Ascomycota(100);c__Saccharomycetes(100);o__Saccharomycetales(100);f__Metschnikowiaceae(100);g__Metschnikowia(100);s__Metschnikowia_pulcherrima(100); | Metschnikowia | 13 | Saprotroph | \|Undefined Saprotroph\| | Yeast | NULL | Probable | As nectar-tap_saprotroph (Põlme et al. 2020); Decay sugar-rich substrates (Põlme et al. 2020) | Põlme S, et al. 2020. Fungal Diversity 105:1-16 ((https://doi.org/10.1007/s13225-020-00466-2)) |
| Otu48703 | 1 | k__Fungi(100);p__Ascomycota(100);c__Saccharomycetes(100);o__Saccharomycetales(100);f__Metschnikowiaceae(100);g__Metschnikowia(100);g__Metschnikowia_unclassified(100); | Metschnikowia | 13 | Saprotroph | \|Undefined Saprotroph\| | Yeast | NULL | Probable | As nectar-tap_saprotroph (Põlme et al. 2020); Decay sugar-rich substrates (Põlme et al. 2020) | Põlme S, et al. 2020. Fungal Diversity 105:1-16 ((https://doi.org/10.1007/s13225-020-00466-2)) |
| Otu48704 | 1 | k__Fungi(100);p__Ascomycota(100);c__Saccharomycetes(100);o__Saccharomycetales(100);f__Metschnikowiaceae(100);g__Metschnikowia(100);g__Metschnikowia_unclassified(100); | Metschnikowia | 13 | Saprotroph | \|Undefined Saprotroph\| | Yeast | NULL | Probable | As nectar-tap_saprotroph (Põlme et al. 2020); Decay sugar-rich substrates (Põlme et al. 2020) | Põlme S, et al. 2020. Fungal Diversity 105:1-16 ((https://doi.org/10.1007/s13225-020-00466-2)) |
| Otu48705 | 1 | k__Fungi(100);p__Ascomycota(100);c__Saccharomycetes(100);o__Saccharomycetales(100);f__Metschnikowiaceae(100);g__Metschnikowia(100);s__Metschnikowia_pulcherrima(100); | Metschnikowia | 13 | Saprotroph | \|Undefined Saprotroph\| | Yeast | NULL | Probable | As nectar-tap_saprotroph (Põlme et al. 2020); Decay sugar-rich substrates (Põlme et al. 2020) | Põlme S, et al. 2020. Fungal Diversity 105:1-16 ((https://doi.org/10.1007/s13225-020-00466-2)) |
| Otu48706 | 1 | k__Fungi(100);p__Ascomycota(100);c__Saccharomycetes(100);o__Saccharomycetales(100);f__Metschnikowiaceae(100);g__Metschnikowia(100);g__Metschnikowia_unclassified(100); | Metschnikowia | 13 | Saprotroph | \|Undefined Saprotroph\| | Yeast | NULL | Probable | As nectar-tap_saprotroph (Põlme et al. 2020); Decay sugar-rich substrates (Põlme et al. 2020) | Põlme S, et al. 2020. Fungal Diversity 105:1-16 ((https://doi.org/10.1007/s13225-020-00466-2)) |
| Otu48707 | 1 | k__Fungi(100);p__Ascomycota(100);c__Saccharomycetes(100);o__Saccharomycetales(100);f__Metschnikowiaceae(100);g__Metschnikowia(100);s__Metschnikowia_pulcherrima(100); | Metschnikowia | 13 | Saprotroph | \|Undefined Saprotroph\| | Yeast | NULL | Probable | As nectar-tap_saprotroph (Põlme et al. 2020); Decay sugar-rich substrates (Põlme et al. 2020) | Põlme S, et al. 2020. Fungal Diversity 105:1-16 ((https://doi.org/10.1007/s13225-020-00466-2)) |
| Otu48708 | 1 | k__Fungi(100);p__Ascomycota(100);c__Saccharomycetes(100);o__Saccharomycetales(100);f__Metschnikowiaceae(100);g__Metschnikowia(100);s__Metschnikowia_pulcherrima(100); | Metschnikowia | 13 | Saprotroph | \|Undefined Saprotroph\| | Yeast | NULL | Probable | As nectar-tap_saprotroph (Põlme et al. 2020); Decay sugar-rich substrates (Põlme et al. 2020) | Põlme S, et al. 2020. Fungal Diversity 105:1-16 ((https://doi.org/10.1007/s13225-020-00466-2)) |
| Otu48709 | 1 | k__Fungi(100);p__Ascomycota(100);c__Saccharomycetes(100);o__Saccharomycetales(100);f__Metschnikowiaceae(100);g__Metschnikowia(100);g__Metschnikowia_unclassified(100); | Metschnikowia | 13 | Saprotroph | \|Undefined Saprotroph\| | Yeast | NULL | Probable | As nectar-tap_saprotroph (Põlme et al. 2020); Decay sugar-rich substrates (Põlme et al. 2020) | Põlme S, et al. 2020. Fungal Diversity 105:1-16 ((https://doi.org/10.1007/s13225-020-00466-2)) |
| Otu48710 | 1 | k__Fungi(100);p__Ascomycota(100);c__Saccharomycetes(100);o__Saccharomycetales(100);f__Metschnikowiaceae(100);g__Metschnikowia(100);s__Metschnikowia_pulcherrima(100); | Metschnikowia | 13 | Saprotroph | \|Undefined Saprotroph\| | Yeast | NULL | Probable | As nectar-tap_saprotroph (Põlme et al. 2020); Decay sugar-rich substrates (Põlme et al. 2020) | Põlme S, et al. 2020. Fungal Diversity 105:1-16 ((https://doi.org/10.1007/s13225-020-00466-2)) |
| Otu48711 | 1 | k__Fungi(100);p__Ascomycota(100);c__Saccharomycetes(100);o__Saccharomycetales(100);f__Metschnikowiaceae(100);g__Metschnikowia(100);s__Metschnikowia_pulcherrima(100); | Metschnikowia | 13 | Saprotroph | \|Undefined Saprotroph\| | Yeast | NULL | Probable | As nectar-tap_saprotroph (Põlme et al. 2020); Decay sugar-rich substrates (Põlme et al. 2020) | Põlme S, et al. 2020. Fungal Diversity 105:1-16 ((https://doi.org/10.1007/s13225-020-00466-2)) |
| Otu48712 | 1 | k__Fungi(100);p__Ascomycota(100);c__Saccharomycetes(100);o__Saccharomycetales(100);f__Metschnikowiaceae(100);g__Metschnikowia(100);s__Metschnikowia_pulcherrima(100); | Metschnikowia | 13 | Saprotroph | \|Undefined Saprotroph\| | Yeast | NULL | Probable | As nectar-tap_saprotroph (Põlme et al. 2020); Decay sugar-rich substrates (Põlme et al. 2020) | Põlme S, et al. 2020. Fungal Diversity 105:1-16 ((https://doi.org/10.1007/s13225-020-00466-2)) |
| Otu48713 | 1 | k__Fungi(100);p__Ascomycota(100);c__Saccharomycetes(100);o__Saccharomycetales(100);f__Metschnikowiaceae(100);g__Metschnikowia(100);s__Metschnikowia_pulcherrima(100); | Metschnikowia | 13 | Saprotroph | \|Undefined Saprotroph\| | Yeast | NULL | Probable | As nectar-tap_saprotroph (Põlme et al. 2020); Decay sugar-rich substrates (Põlme et al. 2020) | Põlme S, et al. 2020. Fungal Diversity 105:1-16 ((https://doi.org/10.1007/s13225-020-00466-2)) |
| Otu48714 | 1 | k__Fungi(100);p__Ascomycota(100);c__Saccharomycetes(100);o__Saccharomycetales(100);f__Metschnikowiaceae(100);g__Metschnikowia(100);s__Metschnikowia_pulcherrima(100); | Metschnikowia | 13 | Saprotroph | \|Undefined Saprotroph\| | Yeast | NULL | Probable | As nectar-tap_saprotroph (Põlme et al. 2020); Decay sugar-rich substrates (Põlme et al. 2020) | Põlme S, et al. 2020. Fungal Diversity 105:1-16 ((https://doi.org/10.1007/s13225-020-00466-2)) |
| Otu48715 | 1 | k__Fungi(100);p__Ascomycota(100);c__Saccharomycetes(100);o__Saccharomycetales(100);f__Metschnikowiaceae(100);g__Metschnikowia(100);s__Metschnikowia_pulcherrima(100); | Metschnikowia | 13 | Saprotroph | \|Undefined Saprotroph\| | Yeast | NULL | Probable | As nectar-tap_saprotroph (Põlme et al. 2020); Decay sugar-rich substrates (Põlme et al. 2020) | Põlme S, et al. 2020. Fungal Diversity 105:1-16 ((https://doi.org/10.1007/s13225-020-00466-2)) |
| Otu48716 | 1 | k__Fungi(100);p__Ascomycota(100);c__Saccharomycetes(100);o__Saccharomycetales(100);f__Metschnikowiaceae(100);g__Metschnikowia(100);s__Metschnikowia_pulcherrima(100); | Metschnikowia | 13 | Saprotroph | \|Undefined Saprotroph\| | Yeast | NULL | Probable | As nectar-tap_saprotroph (Põlme et al. 2020); Decay sugar-rich substrates (Põlme et al. 2020) | Põlme S, et al. 2020. Fungal Diversity 105:1-16 ((https://doi.org/10.1007/s13225-020-00466-2)) |
| Otu48717 | 1 | k__Fungi(100);p__Ascomycota(100);c__Saccharomycetes(100);o__Saccharomycetales(100);f__Metschnikowiaceae(100);g__Metschnikowia(100);s__Metschnikowia_pulcherrima(100); | Metschnikowia | 13 | Saprotroph | \|Undefined Saprotroph\| | Yeast | NULL | Probable | As nectar-tap_saprotroph (Põlme et al. 2020); Decay sugar-rich substrates (Põlme et al. 2020) | Põlme S, et al. 2020. Fungal Diversity 105:1-16 ((https://doi.org/10.1007/s13225-020-00466-2)) |
| Otu48718 | 1 | k__Fungi(100);p__Ascomycota(100);c__Saccharomycetes(100);o__Saccharomycetales(100);f__Metschnikowiaceae(100);g__Metschnikowia(100);s__Metschnikowia_pulcherrima(100); | Metschnikowia | 13 | Saprotroph | \|Undefined Saprotroph\| | Yeast | NULL | Probable | As nectar-tap_saprotroph (Põlme et al. 2020); Decay sugar-rich substrates (Põlme et al. 2020) | Põlme S, et al. 2020. Fungal Diversity 105:1-16 ((https://doi.org/10.1007/s13225-020-00466-2)) |
| Otu48719 | 1 | k__Fungi(100);p__Ascomycota(100);c__Saccharomycetes(100);o__Saccharomycetales(100);f__Metschnikowiaceae(100);g__Metschnikowia(100);s__Metschnikowia_pulcherrima(100); | Metschnikowia | 13 | Saprotroph | \|Undefined Saprotroph\| | Yeast | NULL | Probable | As nectar-tap_saprotroph (Põlme et al. 2020); Decay sugar-rich substrates (Põlme et al. 2020) | Põlme S, et al. 2020. Fungal Diversity 105:1-16 ((https://doi.org/10.1007/s13225-020-00466-2)) |
| Otu48720 | 1 | k__Fungi(100);p__Ascomycota(100);c__Saccharomycetes(100);o__Saccharomycetales(100);f__Metschnikowiaceae(100);g__Metschnikowia(100);s__Metschnikowia_pulcherrima(100); | Metschnikowia | 13 | Saprotroph | \|Undefined Saprotroph\| | Yeast | NULL | Probable | As nectar-tap_saprotroph (Põlme et al. 2020); Decay sugar-rich substrates (Põlme et al. 2020) | Põlme S, et al. 2020. Fungal Diversity 105:1-16 ((https://doi.org/10.1007/s13225-020-00466-2)) |
| Otu48721 | 1 | k__Fungi(100);p__Ascomycota(100);c__Saccharomycetes(100);o__Saccharomycetales(100);f__Metschnikowiaceae(100);g__Metschnikowia(100);s__Metschnikowia_pulcherrima(100); | Metschnikowia | 13 | Saprotroph | \|Undefined Saprotroph\| | Yeast | NULL | Probable | As nectar-tap_saprotroph (Põlme et al. 2020); Decay sugar-rich substrates (Põlme et al. 2020) | Põlme S, et al. 2020. Fungal Diversity 105:1-16 ((https://doi.org/10.1007/s13225-020-00466-2)) |
| Otu48722 | 1 | k__Fungi(100);p__Ascomycota(100);c__Saccharomycetes(100);o__Saccharomycetales(100);f__Metschnikowiaceae(100);g__Metschnikowia(100);s__Metschnikowia_pulcherrima(100); | Metschnikowia | 13 | Saprotroph | \|Undefined Saprotroph\| | Yeast | NULL | Probable | As nectar-tap_saprotroph (Põlme et al. 2020); Decay sugar-rich substrates (Põlme et al. 2020) | Põlme S, et al. 2020. Fungal Diversity 105:1-16 ((https://doi.org/10.1007/s13225-020-00466-2)) |
| Otu48723 | 1 | k__Fungi(100);p__Ascomycota(100);c__Saccharomycetes(100);o__Saccharomycetales(100);f__Metschnikowiaceae(100);g__Metschnikowia(100);s__Metschnikowia_pulcherrima(100); | Metschnikowia | 13 | Saprotroph | \|Undefined Saprotroph\| | Yeast | NULL | Probable | As nectar-tap_saprotroph (Põlme et al. 2020); Decay sugar-rich substrates (Põlme et al. 2020) | Põlme S, et al. 2020. Fungal Diversity 105:1-16 ((https://doi.org/10.1007/s13225-020-00466-2)) |
| Otu48724 | 1 | k__Fungi(100);p__Ascomycota(100);c__Saccharomycetes(100);o__Saccharomycetales(100);f__Metschnikowiaceae(100);g__Metschnikowia(100);g__Metschnikowia_unclassified(100); | Metschnikowia | 13 | Saprotroph | \|Undefined Saprotroph\| | Yeast | NULL | Probable | As nectar-tap_saprotroph (Põlme et al. 2020); Decay sugar-rich substrates (Põlme et al. 2020) | Põlme S, et al. 2020. Fungal Diversity 105:1-16 ((https://doi.org/10.1007/s13225-020-00466-2)) |
| Otu48725 | 1 | k__Fungi(100);p__Ascomycota(100);c__Saccharomycetes(100);o__Saccharomycetales(100);f__Metschnikowiaceae(100);g__Metschnikowia(100);s__Metschnikowia_pulcherrima(100); | Metschnikowia | 13 | Saprotroph | \|Undefined Saprotroph\| | Yeast | NULL | Probable | As nectar-tap_saprotroph (Põlme et al. 2020); Decay sugar-rich substrates (Põlme et al. 2020) | Põlme S, et al. 2020. Fungal Diversity 105:1-16 ((https://doi.org/10.1007/s13225-020-00466-2)) |
| Otu48726 | 1 | k__Fungi(100);p__Ascomycota(100);c__Saccharomycetes(100);o__Saccharomycetales(100);f__Metschnikowiaceae(100);g__Metschnikowia(100);s__Metschnikowia_pulcherrima(100); | Metschnikowia | 13 | Saprotroph | \|Undefined Saprotroph\| | Yeast | NULL | Probable | As nectar-tap_saprotroph (Põlme et al. 2020); Decay sugar-rich substrates (Põlme et al. 2020) | Põlme S, et al. 2020. Fungal Diversity 105:1-16 ((https://doi.org/10.1007/s13225-020-00466-2)) |
| Otu48727 | 1 | k__Fungi(100);p__Ascomycota(100);c__Saccharomycetes(100);o__Saccharomycetales(100);f__Metschnikowiaceae(100);g__Metschnikowia(100);s__Metschnikowia_pulcherrima(100); | Metschnikowia | 13 | Saprotroph | \|Undefined Saprotroph\| | Yeast | NULL | Probable | As nectar-tap_saprotroph (Põlme et al. 2020); Decay sugar-rich substrates (Põlme et al. 2020) | Põlme S, et al. 2020. Fungal Diversity 105:1-16 ((https://doi.org/10.1007/s13225-020-00466-2)) |
| Otu48728 | 1 | k__Fungi(100);p__Ascomycota(100);c__Saccharomycetes(100);o__Saccharomycetales(100);f__Metschnikowiaceae(100);g__Metschnikowia(100);s__Metschnikowia_pulcherrima(100); | Metschnikowia | 13 | Saprotroph | \|Undefined Saprotroph\| | Yeast | NULL | Probable | As nectar-tap_saprotroph (Põlme et al. 2020); Decay sugar-rich substrates (Põlme et al. 2020) | Põlme S, et al. 2020. Fungal Diversity 105:1-16 ((https://doi.org/10.1007/s13225-020-00466-2)) |
| Otu48729 | 1 | k__Fungi(100);p__Ascomycota(100);c__Saccharomycetes(100);o__Saccharomycetales(100);f__Metschnikowiaceae(100);g__Metschnikowia(100);s__Metschnikowia_pulcherrima(100); | Metschnikowia | 13 | Saprotroph | \|Undefined Saprotroph\| | Yeast | NULL | Probable | As nectar-tap_saprotroph (Põlme et al. 2020); Decay sugar-rich substrates (Põlme et al. 2020) | Põlme S, et al. 2020. Fungal Diversity 105:1-16 ((https://doi.org/10.1007/s13225-020-00466-2)) |
| Otu48730 | 1 | k__Fungi(100);p__Ascomycota(100);c__Saccharomycetes(100);o__Saccharomycetales(100);f__Metschnikowiaceae(100);g__Metschnikowia(100);g__Metschnikowia_unclassified(100); | Metschnikowia | 13 | Saprotroph | \|Undefined Saprotroph\| | Yeast | NULL | Probable | As nectar-tap_saprotroph (Põlme et al. 2020); Decay sugar-rich substrates (Põlme et al. 2020) | Põlme S, et al. 2020. Fungal Diversity 105:1-16 ((https://doi.org/10.1007/s13225-020-00466-2)) |
| Otu48731 | 1 | k__Fungi(100);p__Ascomycota(100);c__Saccharomycetes(100);o__Saccharomycetales(100);f__Metschnikowiaceae(100);g__Metschnikowia(100);s__Metschnikowia_pulcherrima(100); | Metschnikowia | 13 | Saprotroph | \|Undefined Saprotroph\| | Yeast | NULL | Probable | As nectar-tap_saprotroph (Põlme et al. 2020); Decay sugar-rich substrates (Põlme et al. 2020) | Põlme S, et al. 2020. Fungal Diversity 105:1-16 ((https://doi.org/10.1007/s13225-020-00466-2)) |
| Otu48732 | 1 | k__Fungi(100);p__Ascomycota(100);c__Saccharomycetes(100);o__Saccharomycetales(100);f__Metschnikowiaceae(100);g__Metschnikowia(100);s__Metschnikowia_pulcherrima(100); | Metschnikowia | 13 | Saprotroph | \|Undefined Saprotroph\| | Yeast | NULL | Probable | As nectar-tap_saprotroph (Põlme et al. 2020); Decay sugar-rich substrates (Põlme et al. 2020) | Põlme S, et al. 2020. Fungal Diversity 105:1-16 ((https://doi.org/10.1007/s13225-020-00466-2)) |
| Otu48733 | 1 | k__Fungi(100);p__Ascomycota(100);c__Saccharomycetes(100);o__Saccharomycetales(100);f__Metschnikowiaceae(100);g__Metschnikowia(100);s__Metschnikowia_pulcherrima(100); | Metschnikowia | 13 | Saprotroph | \|Undefined Saprotroph\| | Yeast | NULL | Probable | As nectar-tap_saprotroph (Põlme et al. 2020); Decay sugar-rich substrates (Põlme et al. 2020) | Põlme S, et al. 2020. Fungal Diversity 105:1-16 ((https://doi.org/10.1007/s13225-020-00466-2)) |
| Otu48734 | 1 | k__Fungi(100);p__Ascomycota(100);c__Saccharomycetes(100);o__Saccharomycetales(100);f__Metschnikowiaceae(100);g__Metschnikowia(100);s__Metschnikowia_pulcherrima(100); | Metschnikowia | 13 | Saprotroph | \|Undefined Saprotroph\| | Yeast | NULL | Probable | As nectar-tap_saprotroph (Põlme et al. 2020); Decay sugar-rich substrates (Põlme et al. 2020) | Põlme S, et al. 2020. Fungal Diversity 105:1-16 ((https://doi.org/10.1007/s13225-020-00466-2)) |
| Otu48735 | 1 | k__Fungi(100);p__Ascomycota(100);c__Saccharomycetes(100);o__Saccharomycetales(100);f__Metschnikowiaceae(100);g__Metschnikowia(100);s__Metschnikowia_pulcherrima(100); | Metschnikowia | 13 | Saprotroph | \|Undefined Saprotroph\| | Yeast | NULL | Probable | As nectar-tap_saprotroph (Põlme et al. 2020); Decay sugar-rich substrates (Põlme et al. 2020) | Põlme S, et al. 2020. Fungal Diversity 105:1-16 ((https://doi.org/10.1007/s13225-020-00466-2)) |
| Otu48736 | 1 | k__Fungi(100);p__Ascomycota(100);c__Saccharomycetes(100);o__Saccharomycetales(100);f__Metschnikowiaceae(100);g__Metschnikowia(100);s__Metschnikowia_pulcherrima(100); | Metschnikowia | 13 | Saprotroph | \|Undefined Saprotroph\| | Yeast | NULL | Probable | As nectar-tap_saprotroph (Põlme et al. 2020); Decay sugar-rich substrates (Põlme et al. 2020) | Põlme S, et al. 2020. Fungal Diversity 105:1-16 ((https://doi.org/10.1007/s13225-020-00466-2)) |
| Otu48737 | 1 | k__Fungi(100);p__Ascomycota(100);c__Saccharomycetes(100);o__Saccharomycetales(100);f__Metschnikowiaceae(100);g__Metschnikowia(100);s__Metschnikowia_pulcherrima(100); | Metschnikowia | 13 | Saprotroph | \|Undefined Saprotroph\| | Yeast | NULL | Probable | As nectar-tap_saprotroph (Põlme et al. 2020); Decay sugar-rich substrates (Põlme et al. 2020) | Põlme S, et al. 2020. Fungal Diversity 105:1-16 ((https://doi.org/10.1007/s13225-020-00466-2)) |
| Otu48738 | 1 | k__Fungi(100);p__Ascomycota(100);c__Saccharomycetes(100);o__Saccharomycetales(100);f__Metschnikowiaceae(100);g__Metschnikowia(100);g__Metschnikowia_unclassified(100); | Metschnikowia | 13 | Saprotroph | \|Undefined Saprotroph\| | Yeast | NULL | Probable | As nectar-tap_saprotroph (Põlme et al. 2020); Decay sugar-rich substrates (Põlme et al. 2020) | Põlme S, et al. 2020. Fungal Diversity 105:1-16 ((https://doi.org/10.1007/s13225-020-00466-2)) |
| Otu48739 | 1 | k__Fungi(100);p__Ascomycota(100);c__Saccharomycetes(100);o__Saccharomycetales(100);f__Metschnikowiaceae(100);g__Metschnikowia(100);s__Metschnikowia_pulcherrima(100); | Metschnikowia | 13 | Saprotroph | \|Undefined Saprotroph\| | Yeast | NULL | Probable | As nectar-tap_saprotroph (Põlme et al. 2020); Decay sugar-rich substrates (Põlme et al. 2020) | Põlme S, et al. 2020. Fungal Diversity 105:1-16 ((https://doi.org/10.1007/s13225-020-00466-2)) |
| Otu48740 | 1 | k__Fungi(100);p__Ascomycota(100);c__Saccharomycetes(100);o__Saccharomycetales(100);f__Metschnikowiaceae(100);g__Metschnikowia(100);s__Metschnikowia_pulcherrima(100); | Metschnikowia | 13 | Saprotroph | \|Undefined Saprotroph\| | Yeast | NULL | Probable | As nectar-tap_saprotroph (Põlme et al. 2020); Decay sugar-rich substrates (Põlme et al. 2020) | Põlme S, et al. 2020. Fungal Diversity 105:1-16 ((https://doi.org/10.1007/s13225-020-00466-2)) |
| Otu48741 | 1 | k__Fungi(100);p__Ascomycota(100);c__Saccharomycetes(100);o__Saccharomycetales(100);f__Metschnikowiaceae(100);g__Metschnikowia(100);s__Metschnikowia_pulcherrima(100); | Metschnikowia | 13 | Saprotroph | \|Undefined Saprotroph\| | Yeast | NULL | Probable | As nectar-tap_saprotroph (Põlme et al. 2020); Decay sugar-rich substrates (Põlme et al. 2020) | Põlme S, et al. 2020. Fungal Diversity 105:1-16 ((https://doi.org/10.1007/s13225-020-00466-2)) |
| Otu48742 | 1 | k__Fungi(100);p__Ascomycota(100);c__Saccharomycetes(100);o__Saccharomycetales(100);f__Metschnikowiaceae(100);g__Metschnikowia(100);s__Metschnikowia_pulcherrima(100); | Metschnikowia | 13 | Saprotroph | \|Undefined Saprotroph\| | Yeast | NULL | Probable | As nectar-tap_saprotroph (Põlme et al. 2020); Decay sugar-rich substrates (Põlme et al. 2020) | Põlme S, et al. 2020. Fungal Diversity 105:1-16 ((https://doi.org/10.1007/s13225-020-00466-2)) |
| Otu48743 | 1 | k__Fungi(100);p__Ascomycota(100);c__Saccharomycetes(100);o__Saccharomycetales(100);f__Metschnikowiaceae(100);g__Metschnikowia(100);g__Metschnikowia_unclassified(100); | Metschnikowia | 13 | Saprotroph | \|Undefined Saprotroph\| | Yeast | NULL | Probable | As nectar-tap_saprotroph (Põlme et al. 2020); Decay sugar-rich substrates (Põlme et al. 2020) | Põlme S, et al. 2020. Fungal Diversity 105:1-16 ((https://doi.org/10.1007/s13225-020-00466-2)) |
| Otu48744 | 1 | k__Fungi(100);p__Ascomycota(100);c__Saccharomycetes(100);o__Saccharomycetales(100);f__Metschnikowiaceae(100);g__Metschnikowia(100);g__Metschnikowia_unclassified(100); | Metschnikowia | 13 | Saprotroph | \|Undefined Saprotroph\| | Yeast | NULL | Probable | As nectar-tap_saprotroph (Põlme et al. 2020); Decay sugar-rich substrates (Põlme et al. 2020) | Põlme S, et al. 2020. Fungal Diversity 105:1-16 ((https://doi.org/10.1007/s13225-020-00466-2)) |
| Otu48745 | 1 | k__Fungi(100);p__Ascomycota(100);c__Saccharomycetes(100);o__Saccharomycetales(100);f__Metschnikowiaceae(100);g__Metschnikowia(100);s__Metschnikowia_pulcherrima(100); | Metschnikowia | 13 | Saprotroph | \|Undefined Saprotroph\| | Yeast | NULL | Probable | As nectar-tap_saprotroph (Põlme et al. 2020); Decay sugar-rich substrates (Põlme et al. 2020) | Põlme S, et al. 2020. Fungal Diversity 105:1-16 ((https://doi.org/10.1007/s13225-020-00466-2)) |
| Otu48746 | 1 | k__Fungi(100);p__Ascomycota(100);c__Saccharomycetes(100);o__Saccharomycetales(100);f__Metschnikowiaceae(100);g__Metschnikowia(100);s__Metschnikowia_pulcherrima(100); | Metschnikowia | 13 | Saprotroph | \|Undefined Saprotroph\| | Yeast | NULL | Probable | As nectar-tap_saprotroph (Põlme et al. 2020); Decay sugar-rich substrates (Põlme et al. 2020) | Põlme S, et al. 2020. Fungal Diversity 105:1-16 ((https://doi.org/10.1007/s13225-020-00466-2)) |
| Otu48747 | 1 | k__Fungi(100);p__Ascomycota(100);c__Saccharomycetes(100);o__Saccharomycetales(100);f__Metschnikowiaceae(100);g__Metschnikowia(100);s__Metschnikowia_pulcherrima(100); | Metschnikowia | 13 | Saprotroph | \|Undefined Saprotroph\| | Yeast | NULL | Probable | As nectar-tap_saprotroph (Põlme et al. 2020); Decay sugar-rich substrates (Põlme et al. 2020) | Põlme S, et al. 2020. Fungal Diversity 105:1-16 ((https://doi.org/10.1007/s13225-020-00466-2)) |
| Otu48748 | 1 | k__Fungi(100);p__Ascomycota(100);c__Saccharomycetes(100);o__Saccharomycetales(100);f__Metschnikowiaceae(100);g__Metschnikowia(100);s__Metschnikowia_pulcherrima(100); | Metschnikowia | 13 | Saprotroph | \|Undefined Saprotroph\| | Yeast | NULL | Probable | As nectar-tap_saprotroph (Põlme et al. 2020); Decay sugar-rich substrates (Põlme et al. 2020) | Põlme S, et al. 2020. Fungal Diversity 105:1-16 ((https://doi.org/10.1007/s13225-020-00466-2)) |
| Otu48749 | 1 | k__Fungi(100);p__Ascomycota(100);c__Saccharomycetes(100);o__Saccharomycetales(100);f__Metschnikowiaceae(100);g__Metschnikowia(100);g__Metschnikowia_unclassified(100); | Metschnikowia | 13 | Saprotroph | \|Undefined Saprotroph\| | Yeast | NULL | Probable | As nectar-tap_saprotroph (Põlme et al. 2020); Decay sugar-rich substrates (Põlme et al. 2020) | Põlme S, et al. 2020. Fungal Diversity 105:1-16 ((https://doi.org/10.1007/s13225-020-00466-2)) |
| Otu48750 | 1 | k__Fungi(100);p__Ascomycota(100);c__Saccharomycetes(100);o__Saccharomycetales(100);f__Metschnikowiaceae(100);g__Metschnikowia(100);s__Metschnikowia_pulcherrima(100); | Metschnikowia | 13 | Saprotroph | \|Undefined Saprotroph\| | Yeast | NULL | Probable | As nectar-tap_saprotroph (Põlme et al. 2020); Decay sugar-rich substrates (Põlme et al. 2020) | Põlme S, et al. 2020. Fungal Diversity 105:1-16 ((https://doi.org/10.1007/s13225-020-00466-2)) |
| Otu48751 | 1 | k__Fungi(100);p__Ascomycota(100);c__Saccharomycetes(100);o__Saccharomycetales(100);f__Metschnikowiaceae(100);g__Metschnikowia(100);g__Metschnikowia_unclassified(100); | Metschnikowia | 13 | Saprotroph | \|Undefined Saprotroph\| | Yeast | NULL | Probable | As nectar-tap_saprotroph (Põlme et al. 2020); Decay sugar-rich substrates (Põlme et al. 2020) | Põlme S, et al. 2020. Fungal Diversity 105:1-16 ((https://doi.org/10.1007/s13225-020-00466-2)) |
| Otu48752 | 1 | k__Fungi(100);p__Ascomycota(100);c__Saccharomycetes(100);o__Saccharomycetales(100);f__Metschnikowiaceae(100);g__Metschnikowia(100);s__Metschnikowia_pulcherrima(100); | Metschnikowia | 13 | Saprotroph | \|Undefined Saprotroph\| | Yeast | NULL | Probable | As nectar-tap_saprotroph (Põlme et al. 2020); Decay sugar-rich substrates (Põlme et al. 2020) | Põlme S, et al. 2020. Fungal Diversity 105:1-16 ((https://doi.org/10.1007/s13225-020-00466-2)) |
| Otu48753 | 1 | k__Fungi(100);p__Ascomycota(100);c__Saccharomycetes(100);o__Saccharomycetales(100);f__Metschnikowiaceae(100);g__Metschnikowia(100);g__Metschnikowia_unclassified(100); | Metschnikowia | 13 | Saprotroph | \|Undefined Saprotroph\| | Yeast | NULL | Probable | As nectar-tap_saprotroph (Põlme et al. 2020); Decay sugar-rich substrates (Põlme et al. 2020) | Põlme S, et al. 2020. Fungal Diversity 105:1-16 ((https://doi.org/10.1007/s13225-020-00466-2)) |
| Otu48754 | 1 | k__Fungi(100);p__Ascomycota(100);c__Saccharomycetes(100);o__Saccharomycetales(100);f__Metschnikowiaceae(100);g__Metschnikowia(100);s__Metschnikowia_pulcherrima(100); | Metschnikowia | 13 | Saprotroph | \|Undefined Saprotroph\| | Yeast | NULL | Probable | As nectar-tap_saprotroph (Põlme et al. 2020); Decay sugar-rich substrates (Põlme et al. 2020) | Põlme S, et al. 2020. Fungal Diversity 105:1-16 ((https://doi.org/10.1007/s13225-020-00466-2)) |
| Otu48755 | 1 | k__Fungi(100);p__Ascomycota(100);c__Saccharomycetes(100);o__Saccharomycetales(100);f__Metschnikowiaceae(100);g__Metschnikowia(100);g__Metschnikowia_unclassified(100); | Metschnikowia | 13 | Saprotroph | \|Undefined Saprotroph\| | Yeast | NULL | Probable | As nectar-tap_saprotroph (Põlme et al. 2020); Decay sugar-rich substrates (Põlme et al. 2020) | Põlme S, et al. 2020. Fungal Diversity 105:1-16 ((https://doi.org/10.1007/s13225-020-00466-2)) |
| Otu48756 | 1 | k__Fungi(100);p__Ascomycota(100);c__Saccharomycetes(100);o__Saccharomycetales(100);f__Metschnikowiaceae(100);g__Metschnikowia(100);s__Metschnikowia_pulcherrima(100); | Metschnikowia | 13 | Saprotroph | \|Undefined Saprotroph\| | Yeast | NULL | Probable | As nectar-tap_saprotroph (Põlme et al. 2020); Decay sugar-rich substrates (Põlme et al. 2020) | Põlme S, et al. 2020. Fungal Diversity 105:1-16 ((https://doi.org/10.1007/s13225-020-00466-2)) |
| Otu48757 | 1 | k__Fungi(100);p__Ascomycota(100);c__Saccharomycetes(100);o__Saccharomycetales(100);f__Metschnikowiaceae(100);g__Metschnikowia(100);g__Metschnikowia_unclassified(100); | Metschnikowia | 13 | Saprotroph | \|Undefined Saprotroph\| | Yeast | NULL | Probable | As nectar-tap_saprotroph (Põlme et al. 2020); Decay sugar-rich substrates (Põlme et al. 2020) | Põlme S, et al. 2020. Fungal Diversity 105:1-16 ((https://doi.org/10.1007/s13225-020-00466-2)) |
| Otu48758 | 1 | k__Fungi(100);p__Ascomycota(100);c__Saccharomycetes(100);o__Saccharomycetales(100);f__Metschnikowiaceae(100);g__Metschnikowia(100);g__Metschnikowia_unclassified(100); | Metschnikowia | 13 | Saprotroph | \|Undefined Saprotroph\| | Yeast | NULL | Probable | As nectar-tap_saprotroph (Põlme et al. 2020); Decay sugar-rich substrates (Põlme et al. 2020) | Põlme S, et al. 2020. Fungal Diversity 105:1-16 ((https://doi.org/10.1007/s13225-020-00466-2)) |
| Otu48759 | 1 | k__Fungi(100);p__Ascomycota(100);c__Saccharomycetes(100);o__Saccharomycetales(100);f__Metschnikowiaceae(100);g__Metschnikowia(100);g__Metschnikowia_unclassified(100); | Metschnikowia | 13 | Saprotroph | \|Undefined Saprotroph\| | Yeast | NULL | Probable | As nectar-tap_saprotroph (Põlme et al. 2020); Decay sugar-rich substrates (Põlme et al. 2020) | Põlme S, et al. 2020. Fungal Diversity 105:1-16 ((https://doi.org/10.1007/s13225-020-00466-2)) |
| Otu48760 | 1 | k__Fungi(100);p__Ascomycota(100);c__Saccharomycetes(100);o__Saccharomycetales(100);f__Metschnikowiaceae(100);g__Metschnikowia(100);g__Metschnikowia_unclassified(100); | Metschnikowia | 13 | Saprotroph | \|Undefined Saprotroph\| | Yeast | NULL | Probable | As nectar-tap_saprotroph (Põlme et al. 2020); Decay sugar-rich substrates (Põlme et al. 2020) | Põlme S, et al. 2020. Fungal Diversity 105:1-16 ((https://doi.org/10.1007/s13225-020-00466-2)) |
| Otu48761 | 1 | k__Fungi(100);p__Ascomycota(100);c__Saccharomycetes(100);o__Saccharomycetales(100);f__Metschnikowiaceae(100);g__Metschnikowia(100);g__Metschnikowia_unclassified(100); | Metschnikowia | 13 | Saprotroph | \|Undefined Saprotroph\| | Yeast | NULL | Probable | As nectar-tap_saprotroph (Põlme et al. 2020); Decay sugar-rich substrates (Põlme et al. 2020) | Põlme S, et al. 2020. Fungal Diversity 105:1-16 ((https://doi.org/10.1007/s13225-020-00466-2)) |
| Otu48762 | 1 | k__Fungi(100);p__Ascomycota(100);c__Saccharomycetes(100);o__Saccharomycetales(100);f__Metschnikowiaceae(100);g__Metschnikowia(100);g__Metschnikowia_unclassified(100); | Metschnikowia | 13 | Saprotroph | \|Undefined Saprotroph\| | Yeast | NULL | Probable | As nectar-tap_saprotroph (Põlme et al. 2020); Decay sugar-rich substrates (Põlme et al. 2020) | Põlme S, et al. 2020. Fungal Diversity 105:1-16 ((https://doi.org/10.1007/s13225-020-00466-2)) |
| Otu48763 | 1 | k__Fungi(100);p__Ascomycota(100);c__Saccharomycetes(100);o__Saccharomycetales(100);f__Metschnikowiaceae(100);g__Metschnikowia(100);g__Metschnikowia_unclassified(100); | Metschnikowia | 13 | Saprotroph | \|Undefined Saprotroph\| | Yeast | NULL | Probable | As nectar-tap_saprotroph (Põlme et al. 2020); Decay sugar-rich substrates (Põlme et al. 2020) | Põlme S, et al. 2020. Fungal Diversity 105:1-16 ((https://doi.org/10.1007/s13225-020-00466-2)) |
| Otu48764 | 1 | k__Fungi(100);p__Ascomycota(100);c__Saccharomycetes(100);o__Saccharomycetales(100);f__Metschnikowiaceae(100);g__Metschnikowia(100);g__Metschnikowia_unclassified(100); | Metschnikowia | 13 | Saprotroph | \|Undefined Saprotroph\| | Yeast | NULL | Probable | As nectar-tap_saprotroph (Põlme et al. 2020); Decay sugar-rich substrates (Põlme et al. 2020) | Põlme S, et al. 2020. Fungal Diversity 105:1-16 ((https://doi.org/10.1007/s13225-020-00466-2)) |
| Otu48765 | 1 | k__Fungi(100);p__Ascomycota(100);c__Saccharomycetes(100);o__Saccharomycetales(100);f__Metschnikowiaceae(100);g__Metschnikowia(100);g__Metschnikowia_unclassified(100); | Metschnikowia | 13 | Saprotroph | \|Undefined Saprotroph\| | Yeast | NULL | Probable | As nectar-tap_saprotroph (Põlme et al. 2020); Decay sugar-rich substrates (Põlme et al. 2020) | Põlme S, et al. 2020. Fungal Diversity 105:1-16 ((https://doi.org/10.1007/s13225-020-00466-2)) |
| Otu48766 | 1 | k__Fungi(100);p__Ascomycota(100);c__Saccharomycetes(100);o__Saccharomycetales(100);f__Metschnikowiaceae(100);g__Metschnikowia(100);s__Metschnikowia_pulcherrima(100); | Metschnikowia | 13 | Saprotroph | \|Undefined Saprotroph\| | Yeast | NULL | Probable | As nectar-tap_saprotroph (Põlme et al. 2020); Decay sugar-rich substrates (Põlme et al. 2020) | Põlme S, et al. 2020. Fungal Diversity 105:1-16 ((https://doi.org/10.1007/s13225-020-00466-2)) |
| Otu48767 | 1 | k__Fungi(100);p__Ascomycota(100);c__Saccharomycetes(100);o__Saccharomycetales(100);f__Metschnikowiaceae(100);g__Metschnikowia(100);s__Metschnikowia_pulcherrima(100); | Metschnikowia | 13 | Saprotroph | \|Undefined Saprotroph\| | Yeast | NULL | Probable | As nectar-tap_saprotroph (Põlme et al. 2020); Decay sugar-rich substrates (Põlme et al. 2020) | Põlme S, et al. 2020. Fungal Diversity 105:1-16 ((https://doi.org/10.1007/s13225-020-00466-2)) |
| Otu48768 | 1 | k__Fungi(100);p__Ascomycota(100);c__Saccharomycetes(100);o__Saccharomycetales(100);f__Metschnikowiaceae(100);g__Metschnikowia(100);g__Metschnikowia_unclassified(100); | Metschnikowia | 13 | Saprotroph | \|Undefined Saprotroph\| | Yeast | NULL | Probable | As nectar-tap_saprotroph (Põlme et al. 2020); Decay sugar-rich substrates (Põlme et al. 2020) | Põlme S, et al. 2020. Fungal Diversity 105:1-16 ((https://doi.org/10.1007/s13225-020-00466-2)) |
| Otu48769 | 1 | k__Fungi(100);p__Ascomycota(100);c__Saccharomycetes(100);o__Saccharomycetales(100);f__Metschnikowiaceae(100);g__Metschnikowia(100);s__Metschnikowia_pulcherrima(100); | Metschnikowia | 13 | Saprotroph | \|Undefined Saprotroph\| | Yeast | NULL | Probable | As nectar-tap_saprotroph (Põlme et al. 2020); Decay sugar-rich substrates (Põlme et al. 2020) | Põlme S, et al. 2020. Fungal Diversity 105:1-16 ((https://doi.org/10.1007/s13225-020-00466-2)) |
| Otu48770 | 1 | k__Fungi(100);p__Ascomycota(100);c__Saccharomycetes(100);o__Saccharomycetales(100);f__Metschnikowiaceae(100);g__Metschnikowia(100);g__Metschnikowia_unclassified(100); | Metschnikowia | 13 | Saprotroph | \|Undefined Saprotroph\| | Yeast | NULL | Probable | As nectar-tap_saprotroph (Põlme et al. 2020); Decay sugar-rich substrates (Põlme et al. 2020) | Põlme S, et al. 2020. Fungal Diversity 105:1-16 ((https://doi.org/10.1007/s13225-020-00466-2)) |
| Otu48771 | 1 | k__Fungi(100);p__Ascomycota(100);c__Saccharomycetes(100);o__Saccharomycetales(100);f__Metschnikowiaceae(100);g__Metschnikowia(100);s__Metschnikowia_pulcherrima(100); | Metschnikowia | 13 | Saprotroph | \|Undefined Saprotroph\| | Yeast | NULL | Probable | As nectar-tap_saprotroph (Põlme et al. 2020); Decay sugar-rich substrates (Põlme et al. 2020) | Põlme S, et al. 2020. Fungal Diversity 105:1-16 ((https://doi.org/10.1007/s13225-020-00466-2)) |
| Otu48772 | 1 | k__Fungi(100);p__Ascomycota(100);c__Saccharomycetes(100);o__Saccharomycetales(100);f__Metschnikowiaceae(100);g__Metschnikowia(100);s__Metschnikowia_pulcherrima(100); | Metschnikowia | 13 | Saprotroph | \|Undefined Saprotroph\| | Yeast | NULL | Probable | As nectar-tap_saprotroph (Põlme et al. 2020); Decay sugar-rich substrates (Põlme et al. 2020) | Põlme S, et al. 2020. Fungal Diversity 105:1-16 ((https://doi.org/10.1007/s13225-020-00466-2)) |
| Otu48773 | 1 | k__Fungi(100);p__Ascomycota(100);c__Saccharomycetes(100);o__Saccharomycetales(100);f__Metschnikowiaceae(100);g__Metschnikowia(100);s__Metschnikowia_pulcherrima(100); | Metschnikowia | 13 | Saprotroph | \|Undefined Saprotroph\| | Yeast | NULL | Probable | As nectar-tap_saprotroph (Põlme et al. 2020); Decay sugar-rich substrates (Põlme et al. 2020) | Põlme S, et al. 2020. Fungal Diversity 105:1-16 ((https://doi.org/10.1007/s13225-020-00466-2)) |
| Otu48774 | 1 | k__Fungi(100);p__Ascomycota(100);c__Saccharomycetes(100);o__Saccharomycetales(100);f__Metschnikowiaceae(100);g__Metschnikowia(100);g__Metschnikowia_unclassified(100); | Metschnikowia | 13 | Saprotroph | \|Undefined Saprotroph\| | Yeast | NULL | Probable | As nectar-tap_saprotroph (Põlme et al. 2020); Decay sugar-rich substrates (Põlme et al. 2020) | Põlme S, et al. 2020. Fungal Diversity 105:1-16 ((https://doi.org/10.1007/s13225-020-00466-2)) |
| Otu48776 | 1 | k__Fungi(100);p__Ascomycota(100);c__Saccharomycetes(100);o__Saccharomycetales(100);f__Metschnikowiaceae(100);g__Metschnikowia(100);s__Metschnikowia_pulcherrima(100); | Metschnikowia | 13 | Saprotroph | \|Undefined Saprotroph\| | Yeast | NULL | Probable | As nectar-tap_saprotroph (Põlme et al. 2020); Decay sugar-rich substrates (Põlme et al. 2020) | Põlme S, et al. 2020. Fungal Diversity 105:1-16 ((https://doi.org/10.1007/s13225-020-00466-2)) |
| Otu48777 | 1 | k__Fungi(100);p__Ascomycota(100);c__Saccharomycetes(100);o__Saccharomycetales(100);f__Metschnikowiaceae(100);g__Metschnikowia(100);s__Metschnikowia_pulcherrima(100); | Metschnikowia | 13 | Saprotroph | \|Undefined Saprotroph\| | Yeast | NULL | Probable | As nectar-tap_saprotroph (Põlme et al. 2020); Decay sugar-rich substrates (Põlme et al. 2020) | Põlme S, et al. 2020. Fungal Diversity 105:1-16 ((https://doi.org/10.1007/s13225-020-00466-2)) |
| Otu48778 | 1 | k__Fungi(100);p__Ascomycota(100);c__Saccharomycetes(100);o__Saccharomycetales(100);f__Metschnikowiaceae(100);g__Metschnikowia(100);s__Metschnikowia_pulcherrima(100); | Metschnikowia | 13 | Saprotroph | \|Undefined Saprotroph\| | Yeast | NULL | Probable | As nectar-tap_saprotroph (Põlme et al. 2020); Decay sugar-rich substrates (Põlme et al. 2020) | Põlme S, et al. 2020. Fungal Diversity 105:1-16 ((https://doi.org/10.1007/s13225-020-00466-2)) |
| Otu48779 | 1 | k__Fungi(100);p__Ascomycota(100);c__Saccharomycetes(100);o__Saccharomycetales(100);f__Metschnikowiaceae(100);g__Metschnikowia(100);s__Metschnikowia_pulcherrima(100); | Metschnikowia | 13 | Saprotroph | \|Undefined Saprotroph\| | Yeast | NULL | Probable | As nectar-tap_saprotroph (Põlme et al. 2020); Decay sugar-rich substrates (Põlme et al. 2020) | Põlme S, et al. 2020. Fungal Diversity 105:1-16 ((https://doi.org/10.1007/s13225-020-00466-2)) |
| Otu48780 | 1 | k__Fungi(100);p__Ascomycota(100);c__Saccharomycetes(100);o__Saccharomycetales(100);f__Metschnikowiaceae(100);g__Metschnikowia(100);s__Metschnikowia_pulcherrima(100); | Metschnikowia | 13 | Saprotroph | \|Undefined Saprotroph\| | Yeast | NULL | Probable | As nectar-tap_saprotroph (Põlme et al. 2020); Decay sugar-rich substrates (Põlme et al. 2020) | Põlme S, et al. 2020. Fungal Diversity 105:1-16 ((https://doi.org/10.1007/s13225-020-00466-2)) |
| Otu48781 | 1 | k__Fungi(100);p__Ascomycota(100);c__Saccharomycetes(100);o__Saccharomycetales(100);f__Metschnikowiaceae(100);g__Metschnikowia(100);s__Metschnikowia_pulcherrima(100); | Metschnikowia | 13 | Saprotroph | \|Undefined Saprotroph\| | Yeast | NULL | Probable | As nectar-tap_saprotroph (Põlme et al. 2020); Decay sugar-rich substrates (Põlme et al. 2020) | Põlme S, et al. 2020. Fungal Diversity 105:1-16 ((https://doi.org/10.1007/s13225-020-00466-2)) |
| Otu48782 | 1 | k__Fungi(100);p__Ascomycota(100);c__Saccharomycetes(100);o__Saccharomycetales(100);f__Metschnikowiaceae(100);g__Metschnikowia(100);s__Metschnikowia_pulcherrima(100); | Metschnikowia | 13 | Saprotroph | \|Undefined Saprotroph\| | Yeast | NULL | Probable | As nectar-tap_saprotroph (Põlme et al. 2020); Decay sugar-rich substrates (Põlme et al. 2020) | Põlme S, et al. 2020. Fungal Diversity 105:1-16 ((https://doi.org/10.1007/s13225-020-00466-2)) |
| Otu48783 | 1 | k__Fungi(100);p__Ascomycota(100);c__Saccharomycetes(100);o__Saccharomycetales(100);f__Metschnikowiaceae(100);g__Metschnikowia(100);g__Metschnikowia_unclassified(100); | Metschnikowia | 13 | Saprotroph | \|Undefined Saprotroph\| | Yeast | NULL | Probable | As nectar-tap_saprotroph (Põlme et al. 2020); Decay sugar-rich substrates (Põlme et al. 2020) | Põlme S, et al. 2020. Fungal Diversity 105:1-16 ((https://doi.org/10.1007/s13225-020-00466-2)) |
| Otu48784 | 1 | k__Fungi(100);p__Ascomycota(100);c__Saccharomycetes(100);o__Saccharomycetales(100);f__Metschnikowiaceae(100);g__Metschnikowia(100);g__Metschnikowia_unclassified(100); | Metschnikowia | 13 | Saprotroph | \|Undefined Saprotroph\| | Yeast | NULL | Probable | As nectar-tap_saprotroph (Põlme et al. 2020); Decay sugar-rich substrates (Põlme et al. 2020) | Põlme S, et al. 2020. Fungal Diversity 105:1-16 ((https://doi.org/10.1007/s13225-020-00466-2)) |
| Otu48785 | 1 | k__Fungi(100);p__Ascomycota(100);c__Saccharomycetes(100);o__Saccharomycetales(100);f__Metschnikowiaceae(100);g__Metschnikowia(100);s__Metschnikowia_pulcherrima(100); | Metschnikowia | 13 | Saprotroph | \|Undefined Saprotroph\| | Yeast | NULL | Probable | As nectar-tap_saprotroph (Põlme et al. 2020); Decay sugar-rich substrates (Põlme et al. 2020) | Põlme S, et al. 2020. Fungal Diversity 105:1-16 ((https://doi.org/10.1007/s13225-020-00466-2)) |
| Otu48786 | 1 | k__Fungi(100);p__Ascomycota(100);c__Saccharomycetes(100);o__Saccharomycetales(100);f__Metschnikowiaceae(100);g__Metschnikowia(100);g__Metschnikowia_unclassified(100); | Metschnikowia | 13 | Saprotroph | \|Undefined Saprotroph\| | Yeast | NULL | Probable | As nectar-tap_saprotroph (Põlme et al. 2020); Decay sugar-rich substrates (Põlme et al. 2020) | Põlme S, et al. 2020. Fungal Diversity 105:1-16 ((https://doi.org/10.1007/s13225-020-00466-2)) |
| Otu48787 | 1 | k__Fungi(100);p__Ascomycota(100);c__Saccharomycetes(100);o__Saccharomycetales(100);f__Metschnikowiaceae(100);g__Metschnikowia(100);s__Metschnikowia_pulcherrima(100); | Metschnikowia | 13 | Saprotroph | \|Undefined Saprotroph\| | Yeast | NULL | Probable | As nectar-tap_saprotroph (Põlme et al. 2020); Decay sugar-rich substrates (Põlme et al. 2020) | Põlme S, et al. 2020. Fungal Diversity 105:1-16 ((https://doi.org/10.1007/s13225-020-00466-2)) |
| Otu48788 | 1 | k__Fungi(100);p__Ascomycota(100);c__Saccharomycetes(100);o__Saccharomycetales(100);f__Metschnikowiaceae(100);g__Metschnikowia(100);g__Metschnikowia_unclassified(100); | Metschnikowia | 13 | Saprotroph | \|Undefined Saprotroph\| | Yeast | NULL | Probable | As nectar-tap_saprotroph (Põlme et al. 2020); Decay sugar-rich substrates (Põlme et al. 2020) | Põlme S, et al. 2020. Fungal Diversity 105:1-16 ((https://doi.org/10.1007/s13225-020-00466-2)) |
| Otu48789 | 1 | k__Fungi(100);p__Ascomycota(100);c__Saccharomycetes(100);o__Saccharomycetales(100);f__Metschnikowiaceae(100);g__Metschnikowia(100);g__Metschnikowia_unclassified(100); | Metschnikowia | 13 | Saprotroph | \|Undefined Saprotroph\| | Yeast | NULL | Probable | As nectar-tap_saprotroph (Põlme et al. 2020); Decay sugar-rich substrates (Põlme et al. 2020) | Põlme S, et al. 2020. Fungal Diversity 105:1-16 ((https://doi.org/10.1007/s13225-020-00466-2)) |
| Otu48790 | 1 | k__Fungi(100);p__Ascomycota(100);c__Saccharomycetes(100);o__Saccharomycetales(100);f__Metschnikowiaceae(100);g__Metschnikowia(100);s__Metschnikowia_pulcherrima(100); | Metschnikowia | 13 | Saprotroph | \|Undefined Saprotroph\| | Yeast | NULL | Probable | As nectar-tap_saprotroph (Põlme et al. 2020); Decay sugar-rich substrates (Põlme et al. 2020) | Põlme S, et al. 2020. Fungal Diversity 105:1-16 ((https://doi.org/10.1007/s13225-020-00466-2)) |
| Otu48791 | 1 | k__Fungi(100);p__Ascomycota(100);c__Saccharomycetes(100);o__Saccharomycetales(100);f__Metschnikowiaceae(100);g__Metschnikowia(100);s__Metschnikowia_pulcherrima(100); | Metschnikowia | 13 | Saprotroph | \|Undefined Saprotroph\| | Yeast | NULL | Probable | As nectar-tap_saprotroph (Põlme et al. 2020); Decay sugar-rich substrates (Põlme et al. 2020) | Põlme S, et al. 2020. Fungal Diversity 105:1-16 ((https://doi.org/10.1007/s13225-020-00466-2)) |
| Otu48792 | 1 | k__Fungi(100);p__Ascomycota(100);c__Saccharomycetes(100);o__Saccharomycetales(100);f__Metschnikowiaceae(100);g__Metschnikowia(100);g__Metschnikowia_unclassified(100); | Metschnikowia | 13 | Saprotroph | \|Undefined Saprotroph\| | Yeast | NULL | Probable | As nectar-tap_saprotroph (Põlme et al. 2020); Decay sugar-rich substrates (Põlme et al. 2020) | Põlme S, et al. 2020. Fungal Diversity 105:1-16 ((https://doi.org/10.1007/s13225-020-00466-2)) |
| Otu48793 | 1 | k__Fungi(100);p__Ascomycota(100);c__Saccharomycetes(100);o__Saccharomycetales(100);f__Metschnikowiaceae(100);g__Metschnikowia(100);s__Metschnikowia_pulcherrima(100); | Metschnikowia | 13 | Saprotroph | \|Undefined Saprotroph\| | Yeast | NULL | Probable | As nectar-tap_saprotroph (Põlme et al. 2020); Decay sugar-rich substrates (Põlme et al. 2020) | Põlme S, et al. 2020. Fungal Diversity 105:1-16 ((https://doi.org/10.1007/s13225-020-00466-2)) |
| Otu48794 | 1 | k__Fungi(100);p__Ascomycota(100);c__Saccharomycetes(100);o__Saccharomycetales(100);f__Metschnikowiaceae(100);g__Metschnikowia(100);g__Metschnikowia_unclassified(100); | Metschnikowia | 13 | Saprotroph | \|Undefined Saprotroph\| | Yeast | NULL | Probable | As nectar-tap_saprotroph (Põlme et al. 2020); Decay sugar-rich substrates (Põlme et al. 2020) | Põlme S, et al. 2020. Fungal Diversity 105:1-16 ((https://doi.org/10.1007/s13225-020-00466-2)) |
| Otu48795 | 1 | k__Fungi(100);p__Ascomycota(100);c__Saccharomycetes(100);o__Saccharomycetales(100);f__Metschnikowiaceae(100);g__Metschnikowia(100);s__Metschnikowia_pulcherrima(100); | Metschnikowia | 13 | Saprotroph | \|Undefined Saprotroph\| | Yeast | NULL | Probable | As nectar-tap_saprotroph (Põlme et al. 2020); Decay sugar-rich substrates (Põlme et al. 2020) | Põlme S, et al. 2020. Fungal Diversity 105:1-16 ((https://doi.org/10.1007/s13225-020-00466-2)) |
| Otu48796 | 1 | k__Fungi(100);p__Ascomycota(100);c__Saccharomycetes(100);o__Saccharomycetales(100);f__Metschnikowiaceae(100);g__Metschnikowia(100);g__Metschnikowia_unclassified(100); | Metschnikowia | 13 | Saprotroph | \|Undefined Saprotroph\| | Yeast | NULL | Probable | As nectar-tap_saprotroph (Põlme et al. 2020); Decay sugar-rich substrates (Põlme et al. 2020) | Põlme S, et al. 2020. Fungal Diversity 105:1-16 ((https://doi.org/10.1007/s13225-020-00466-2)) |
| Otu48797 | 1 | k__Fungi(100);p__Ascomycota(100);c__Saccharomycetes(100);o__Saccharomycetales(100);f__Metschnikowiaceae(100);g__Metschnikowia(100);s__Metschnikowia_pulcherrima(100); | Metschnikowia | 13 | Saprotroph | \|Undefined Saprotroph\| | Yeast | NULL | Probable | As nectar-tap_saprotroph (Põlme et al. 2020); Decay sugar-rich substrates (Põlme et al. 2020) | Põlme S, et al. 2020. Fungal Diversity 105:1-16 ((https://doi.org/10.1007/s13225-020-00466-2)) |
| Otu48798 | 1 | k__Fungi(100);p__Ascomycota(100);c__Saccharomycetes(100);o__Saccharomycetales(100);f__Metschnikowiaceae(100);g__Metschnikowia(100);g__Metschnikowia_unclassified(100); | Metschnikowia | 13 | Saprotroph | \|Undefined Saprotroph\| | Yeast | NULL | Probable | As nectar-tap_saprotroph (Põlme et al. 2020); Decay sugar-rich substrates (Põlme et al. 2020) | Põlme S, et al. 2020. Fungal Diversity 105:1-16 ((https://doi.org/10.1007/s13225-020-00466-2)) |
| Otu48799 | 1 | k__Fungi(100);p__Ascomycota(100);c__Saccharomycetes(100);o__Saccharomycetales(100);f__Metschnikowiaceae(100);g__Metschnikowia(100);s__Metschnikowia_pulcherrima(100); | Metschnikowia | 13 | Saprotroph | \|Undefined Saprotroph\| | Yeast | NULL | Probable | As nectar-tap_saprotroph (Põlme et al. 2020); Decay sugar-rich substrates (Põlme et al. 2020) | Põlme S, et al. 2020. Fungal Diversity 105:1-16 ((https://doi.org/10.1007/s13225-020-00466-2)) |
| Otu48800 | 1 | k__Fungi(100);p__Ascomycota(100);c__Saccharomycetes(100);o__Saccharomycetales(100);f__Metschnikowiaceae(100);g__Metschnikowia(100);s__Metschnikowia_pulcherrima(100); | Metschnikowia | 13 | Saprotroph | \|Undefined Saprotroph\| | Yeast | NULL | Probable | As nectar-tap_saprotroph (Põlme et al. 2020); Decay sugar-rich substrates (Põlme et al. 2020) | Põlme S, et al. 2020. Fungal Diversity 105:1-16 ((https://doi.org/10.1007/s13225-020-00466-2)) |
| Otu48801 | 1 | k__Fungi(100);p__Ascomycota(100);c__Saccharomycetes(100);o__Saccharomycetales(100);f__Metschnikowiaceae(100);g__Metschnikowia(100);s__Metschnikowia_pulcherrima(100); | Metschnikowia | 13 | Saprotroph | \|Undefined Saprotroph\| | Yeast | NULL | Probable | As nectar-tap_saprotroph (Põlme et al. 2020); Decay sugar-rich substrates (Põlme et al. 2020) | Põlme S, et al. 2020. Fungal Diversity 105:1-16 ((https://doi.org/10.1007/s13225-020-00466-2)) |
| Otu48802 | 1 | k__Fungi(100);p__Ascomycota(100);c__Saccharomycetes(100);o__Saccharomycetales(100);f__Metschnikowiaceae(100);g__Metschnikowia(100);s__Metschnikowia_pulcherrima(100); | Metschnikowia | 13 | Saprotroph | \|Undefined Saprotroph\| | Yeast | NULL | Probable | As nectar-tap_saprotroph (Põlme et al. 2020); Decay sugar-rich substrates (Põlme et al. 2020) | Põlme S, et al. 2020. Fungal Diversity 105:1-16 ((https://doi.org/10.1007/s13225-020-00466-2)) |
| Otu48804 | 1 | k__Fungi(100);p__Ascomycota(100);c__Saccharomycetes(100);o__Saccharomycetales(100);f__Metschnikowiaceae(100);g__Metschnikowia(100);s__Metschnikowia_pulcherrima(100); | Metschnikowia | 13 | Saprotroph | \|Undefined Saprotroph\| | Yeast | NULL | Probable | As nectar-tap_saprotroph (Põlme et al. 2020); Decay sugar-rich substrates (Põlme et al. 2020) | Põlme S, et al. 2020. Fungal Diversity 105:1-16 ((https://doi.org/10.1007/s13225-020-00466-2)) |
| Otu48805 | 1 | k__Fungi(100);p__Ascomycota(100);c__Saccharomycetes(100);o__Saccharomycetales(100);f__Metschnikowiaceae(100);g__Metschnikowia(100);g__Metschnikowia_unclassified(100); | Metschnikowia | 13 | Saprotroph | \|Undefined Saprotroph\| | Yeast | NULL | Probable | As nectar-tap_saprotroph (Põlme et al. 2020); Decay sugar-rich substrates (Põlme et al. 2020) | Põlme S, et al. 2020. Fungal Diversity 105:1-16 ((https://doi.org/10.1007/s13225-020-00466-2)) |
| Otu48806 | 1 | k__Fungi(100);p__Ascomycota(100);c__Saccharomycetes(100);o__Saccharomycetales(100);f__Metschnikowiaceae(100);g__Metschnikowia(100);s__Metschnikowia_pulcherrima(100); | Metschnikowia | 13 | Saprotroph | \|Undefined Saprotroph\| | Yeast | NULL | Probable | As nectar-tap_saprotroph (Põlme et al. 2020); Decay sugar-rich substrates (Põlme et al. 2020) | Põlme S, et al. 2020. Fungal Diversity 105:1-16 ((https://doi.org/10.1007/s13225-020-00466-2)) |
| Otu48807 | 1 | k__Fungi(100);p__Ascomycota(100);c__Saccharomycetes(100);o__Saccharomycetales(100);f__Metschnikowiaceae(100);g__Metschnikowia(100);s__Metschnikowia_pulcherrima(100); | Metschnikowia | 13 | Saprotroph | \|Undefined Saprotroph\| | Yeast | NULL | Probable | As nectar-tap_saprotroph (Põlme et al. 2020); Decay sugar-rich substrates (Põlme et al. 2020) | Põlme S, et al. 2020. Fungal Diversity 105:1-16 ((https://doi.org/10.1007/s13225-020-00466-2)) |
| Otu48808 | 1 | k__Fungi(100);p__Ascomycota(100);c__Saccharomycetes(100);o__Saccharomycetales(100);f__Metschnikowiaceae(100);g__Metschnikowia(100);g__Metschnikowia_unclassified(100); | Metschnikowia | 13 | Saprotroph | \|Undefined Saprotroph\| | Yeast | NULL | Probable | As nectar-tap_saprotroph (Põlme et al. 2020); Decay sugar-rich substrates (Põlme et al. 2020) | Põlme S, et al. 2020. Fungal Diversity 105:1-16 ((https://doi.org/10.1007/s13225-020-00466-2)) |
| Otu48809 | 1 | k__Fungi(100);p__Ascomycota(100);c__Saccharomycetes(100);o__Saccharomycetales(100);f__Metschnikowiaceae(100);g__Metschnikowia(100);s__Metschnikowia_pulcherrima(100); | Metschnikowia | 13 | Saprotroph | \|Undefined Saprotroph\| | Yeast | NULL | Probable | As nectar-tap_saprotroph (Põlme et al. 2020); Decay sugar-rich substrates (Põlme et al. 2020) | Põlme S, et al. 2020. Fungal Diversity 105:1-16 ((https://doi.org/10.1007/s13225-020-00466-2)) |
| Otu48810 | 1 | k__Fungi(100);p__Ascomycota(100);c__Saccharomycetes(100);o__Saccharomycetales(100);f__Metschnikowiaceae(100);g__Metschnikowia(100);g__Metschnikowia_unclassified(100); | Metschnikowia | 13 | Saprotroph | \|Undefined Saprotroph\| | Yeast | NULL | Probable | As nectar-tap_saprotroph (Põlme et al. 2020); Decay sugar-rich substrates (Põlme et al. 2020) | Põlme S, et al. 2020. Fungal Diversity 105:1-16 ((https://doi.org/10.1007/s13225-020-00466-2)) |
| Otu48811 | 1 | k__Fungi(100);p__Ascomycota(100);c__Saccharomycetes(100);o__Saccharomycetales(100);f__Metschnikowiaceae(100);g__Metschnikowia(100);s__Metschnikowia_pulcherrima(100); | Metschnikowia | 13 | Saprotroph | \|Undefined Saprotroph\| | Yeast | NULL | Probable | As nectar-tap_saprotroph (Põlme et al. 2020); Decay sugar-rich substrates (Põlme et al. 2020) | Põlme S, et al. 2020. Fungal Diversity 105:1-16 ((https://doi.org/10.1007/s13225-020-00466-2)) |
| Otu48812 | 1 | k__Fungi(100);p__Ascomycota(100);c__Saccharomycetes(100);o__Saccharomycetales(100);f__Metschnikowiaceae(100);g__Metschnikowia(100);s__Metschnikowia_pulcherrima(100); | Metschnikowia | 13 | Saprotroph | \|Undefined Saprotroph\| | Yeast | NULL | Probable | As nectar-tap_saprotroph (Põlme et al. 2020); Decay sugar-rich substrates (Põlme et al. 2020) | Põlme S, et al. 2020. Fungal Diversity 105:1-16 ((https://doi.org/10.1007/s13225-020-00466-2)) |
| Otu48813 | 1 | k__Fungi(100);p__Ascomycota(100);c__Saccharomycetes(100);o__Saccharomycetales(100);f__Metschnikowiaceae(100);g__Metschnikowia(100);s__Metschnikowia_pulcherrima(100); | Metschnikowia | 13 | Saprotroph | \|Undefined Saprotroph\| | Yeast | NULL | Probable | As nectar-tap_saprotroph (Põlme et al. 2020); Decay sugar-rich substrates (Põlme et al. 2020) | Põlme S, et al. 2020. Fungal Diversity 105:1-16 ((https://doi.org/10.1007/s13225-020-00466-2)) |
| Otu48814 | 1 | k__Fungi(100);p__Ascomycota(100);c__Saccharomycetes(100);o__Saccharomycetales(100);f__Metschnikowiaceae(100);g__Metschnikowia(100);s__Metschnikowia_pulcherrima(100); | Metschnikowia | 13 | Saprotroph | \|Undefined Saprotroph\| | Yeast | NULL | Probable | As nectar-tap_saprotroph (Põlme et al. 2020); Decay sugar-rich substrates (Põlme et al. 2020) | Põlme S, et al. 2020. Fungal Diversity 105:1-16 ((https://doi.org/10.1007/s13225-020-00466-2)) |
| Otu48815 | 1 | k__Fungi(100);p__Ascomycota(100);c__Saccharomycetes(100);o__Saccharomycetales(100);f__Metschnikowiaceae(100);g__Metschnikowia(100);g__Metschnikowia_unclassified(100); | Metschnikowia | 13 | Saprotroph | \|Undefined Saprotroph\| | Yeast | NULL | Probable | As nectar-tap_saprotroph (Põlme et al. 2020); Decay sugar-rich substrates (Põlme et al. 2020) | Põlme S, et al. 2020. Fungal Diversity 105:1-16 ((https://doi.org/10.1007/s13225-020-00466-2)) |
| Otu48816 | 1 | k__Fungi(100);p__Ascomycota(100);c__Saccharomycetes(100);o__Saccharomycetales(100);f__Metschnikowiaceae(100);g__Metschnikowia(100);g__Metschnikowia_unclassified(100); | Metschnikowia | 13 | Saprotroph | \|Undefined Saprotroph\| | Yeast | NULL | Probable | As nectar-tap_saprotroph (Põlme et al. 2020); Decay sugar-rich substrates (Põlme et al. 2020) | Põlme S, et al. 2020. Fungal Diversity 105:1-16 ((https://doi.org/10.1007/s13225-020-00466-2)) |
| Otu48818 | 1 | k__Fungi(100);p__Ascomycota(100);c__Saccharomycetes(100);o__Saccharomycetales(100);f__Metschnikowiaceae(100);g__Metschnikowia(100);g__Metschnikowia_unclassified(100); | Metschnikowia | 13 | Saprotroph | \|Undefined Saprotroph\| | Yeast | NULL | Probable | As nectar-tap_saprotroph (Põlme et al. 2020); Decay sugar-rich substrates (Põlme et al. 2020) | Põlme S, et al. 2020. Fungal Diversity 105:1-16 ((https://doi.org/10.1007/s13225-020-00466-2)) |
| Otu48819 | 1 | k__Fungi(100);p__Ascomycota(100);c__Saccharomycetes(100);o__Saccharomycetales(100);f__Metschnikowiaceae(100);g__Metschnikowia(100);s__Metschnikowia_pulcherrima(100); | Metschnikowia | 13 | Saprotroph | \|Undefined Saprotroph\| | Yeast | NULL | Probable | As nectar-tap_saprotroph (Põlme et al. 2020); Decay sugar-rich substrates (Põlme et al. 2020) | Põlme S, et al. 2020. Fungal Diversity 105:1-16 ((https://doi.org/10.1007/s13225-020-00466-2)) |
| Otu48820 | 1 | k__Fungi(100);p__Ascomycota(100);c__Saccharomycetes(100);o__Saccharomycetales(100);f__Metschnikowiaceae(100);g__Metschnikowia(100);s__Metschnikowia_pulcherrima(100); | Metschnikowia | 13 | Saprotroph | \|Undefined Saprotroph\| | Yeast | NULL | Probable | As nectar-tap_saprotroph (Põlme et al. 2020); Decay sugar-rich substrates (Põlme et al. 2020) | Põlme S, et al. 2020. Fungal Diversity 105:1-16 ((https://doi.org/10.1007/s13225-020-00466-2)) |
| Otu48821 | 1 | k__Fungi(100);p__Ascomycota(100);c__Saccharomycetes(100);o__Saccharomycetales(100);f__Metschnikowiaceae(100);g__Metschnikowia(100);s__Metschnikowia_pulcherrima(100); | Metschnikowia | 13 | Saprotroph | \|Undefined Saprotroph\| | Yeast | NULL | Probable | As nectar-tap_saprotroph (Põlme et al. 2020); Decay sugar-rich substrates (Põlme et al. 2020) | Põlme S, et al. 2020. Fungal Diversity 105:1-16 ((https://doi.org/10.1007/s13225-020-00466-2)) |
| Otu48822 | 1 | k__Fungi(100);p__Ascomycota(100);c__Saccharomycetes(100);o__Saccharomycetales(100);f__Metschnikowiaceae(100);g__Metschnikowia(100);g__Metschnikowia_unclassified(100); | Metschnikowia | 13 | Saprotroph | \|Undefined Saprotroph\| | Yeast | NULL | Probable | As nectar-tap_saprotroph (Põlme et al. 2020); Decay sugar-rich substrates (Põlme et al. 2020) | Põlme S, et al. 2020. Fungal Diversity 105:1-16 ((https://doi.org/10.1007/s13225-020-00466-2)) |
| Otu48823 | 1 | k__Fungi(100);p__Ascomycota(100);c__Saccharomycetes(100);o__Saccharomycetales(100);f__Metschnikowiaceae(100);g__Metschnikowia(100);g__Metschnikowia_unclassified(100); | Metschnikowia | 13 | Saprotroph | \|Undefined Saprotroph\| | Yeast | NULL | Probable | As nectar-tap_saprotroph (Põlme et al. 2020); Decay sugar-rich substrates (Põlme et al. 2020) | Põlme S, et al. 2020. Fungal Diversity 105:1-16 ((https://doi.org/10.1007/s13225-020-00466-2)) |
| Otu48824 | 1 | k__Fungi(100);p__Ascomycota(100);c__Saccharomycetes(100);o__Saccharomycetales(100);f__Metschnikowiaceae(100);g__Metschnikowia(100);s__Metschnikowia_pulcherrima(100); | Metschnikowia | 13 | Saprotroph | \|Undefined Saprotroph\| | Yeast | NULL | Probable | As nectar-tap_saprotroph (Põlme et al. 2020); Decay sugar-rich substrates (Põlme et al. 2020) | Põlme S, et al. 2020. Fungal Diversity 105:1-16 ((https://doi.org/10.1007/s13225-020-00466-2)) |
| Otu48825 | 1 | k__Fungi(100);p__Ascomycota(100);c__Saccharomycetes(100);o__Saccharomycetales(100);f__Saccharomycetales_family_Incertae_sedis(100);g__Candida(100);g__Candida_unclassified(100); | Candida | 13 | Pathotroph-Saprotroph-Symbiotroph | Animal Pathogen-Endophyte-Endosymbiont-Epiphyte-\|Undefined Saprotroph\| | Dimorphic Yeast-Yeast | NULL | Probable | As nectar-tap_saprotroph (Põlme et al. 2020); Decay sugar-rich substrates (Põlme et al. 2020); This asexual genus is polyphyletic and contain species that function very differently depend on what they are. Caution should be taken when interpreting guilds for Candida species. | Manolakaki D, et al. 2010. Virulence. 1: 367-375 ((https://doi.org/10.4161/viru.1.5.12796)); Mekbib SB, et al. 2011. Tropical Plant Pathology 36:233-240 ((https://doi.org/10.1590/S1982-56762011000400004)); Põlme S, et al. 2020. Fungal Diversity 105:1-16 ((https://doi.org/10.1007/s13225-020-00466-2)) |
| Otu48826 | 1 | k__Fungi(100);p__Ascomycota(100);c__Dothideomycetes(100);o__Pleosporales(100);f__Pleosporaceae(100);g__Alternaria(100);g__Alternaria_unclassified(100); | Alternaria | 13 | Pathotroph-Saprotroph-Symbiotroph | Animal Pathogen-Endophyte-\|Plant Pathogen\|-Plant Saprotroph-Wood Saprotroph | Microfungus | Soft Rot | Possible | As foliar_endophyte (Põlme et al. 2020); As litter_saprotroph (Põlme et al. 2020); Decay leaf, fruit, seed (Põlme et al. 2020) | Duncan CG, Eslyn WE. 1966. Mycologia 58:642-664 ((https://doi.org/10.2307/3757045)); Seehann G, et al. 1975. List of Fungi in Soft-Rot Tissue Tests. Institute fur Holzbiologie und Holzschutz, Bundesforschungsanstalt fur Forst - und Holzwirtshaft, Hamburg - Lohbrugge, Germany ((https://www.irg-wp.com/irgdocs/details.php?d6493033-c9ea-40f3-8e3e-5697f32d640d)); Tedersoo L, et al. 2014. Science 346: e1256688 ((https://doi.org/10.1126/science.1256688)); Põlme S, et al. 2020. Fungal Diversity 105:1-16 ((https://doi.org/10.1007/s13225-020-00466-2)) |
| Otu48827 | 1 | k__Fungi(100);p__Ascomycota(100);c__Saccharomycetes(100);o__Saccharomycetales(100);f__Metschnikowiaceae(100);g__Metschnikowia(100);g__Metschnikowia_unclassified(100); | Metschnikowia | 13 | Saprotroph | \|Undefined Saprotroph\| | Yeast | NULL | Probable | As nectar-tap_saprotroph (Põlme et al. 2020); Decay sugar-rich substrates (Põlme et al. 2020) | Põlme S, et al. 2020. Fungal Diversity 105:1-16 ((https://doi.org/10.1007/s13225-020-00466-2)) |
| Otu48828 | 1 | k__Fungi(100);p__Ascomycota(100);c__Saccharomycetes(100);o__Saccharomycetales(100);f__Metschnikowiaceae(100);g__Metschnikowia(100);s__Metschnikowia_pulcherrima(100); | Metschnikowia | 13 | Saprotroph | \|Undefined Saprotroph\| | Yeast | NULL | Probable | As nectar-tap_saprotroph (Põlme et al. 2020); Decay sugar-rich substrates (Põlme et al. 2020) | Põlme S, et al. 2020. Fungal Diversity 105:1-16 ((https://doi.org/10.1007/s13225-020-00466-2)) |
| Otu48829 | 1 | k__Fungi(100);p__Ascomycota(100);c__Saccharomycetes(100);o__Saccharomycetales(100);f__Metschnikowiaceae(100);g__Metschnikowia(100);s__Metschnikowia_pulcherrima(100); | Metschnikowia | 13 | Saprotroph | \|Undefined Saprotroph\| | Yeast | NULL | Probable | As nectar-tap_saprotroph (Põlme et al. 2020); Decay sugar-rich substrates (Põlme et al. 2020) | Põlme S, et al. 2020. Fungal Diversity 105:1-16 ((https://doi.org/10.1007/s13225-020-00466-2)) |
| Otu48830 | 1 | k__Fungi(100);p__Ascomycota(100);c__Saccharomycetes(100);o__Saccharomycetales(100);f__Metschnikowiaceae(100);g__Metschnikowia(100);s__Metschnikowia_pulcherrima(100); | Metschnikowia | 13 | Saprotroph | \|Undefined Saprotroph\| | Yeast | NULL | Probable | As nectar-tap_saprotroph (Põlme et al. 2020); Decay sugar-rich substrates (Põlme et al. 2020) | Põlme S, et al. 2020. Fungal Diversity 105:1-16 ((https://doi.org/10.1007/s13225-020-00466-2)) |
| Otu48831 | 1 | k__Fungi(100);p__Ascomycota(100);c__Saccharomycetes(100);o__Saccharomycetales(100);f__Metschnikowiaceae(100);g__Metschnikowia(100);s__Metschnikowia_pulcherrima(100); | Metschnikowia | 13 | Saprotroph | \|Undefined Saprotroph\| | Yeast | NULL | Probable | As nectar-tap_saprotroph (Põlme et al. 2020); Decay sugar-rich substrates (Põlme et al. 2020) | Põlme S, et al. 2020. Fungal Diversity 105:1-16 ((https://doi.org/10.1007/s13225-020-00466-2)) |
| Otu48832 | 1 | k__Fungi(100);p__Ascomycota(100);c__Saccharomycetes(100);o__Saccharomycetales(100);f__Metschnikowiaceae(100);g__Metschnikowia(100);s__Metschnikowia_pulcherrima(100); | Metschnikowia | 13 | Saprotroph | \|Undefined Saprotroph\| | Yeast | NULL | Probable | As nectar-tap_saprotroph (Põlme et al. 2020); Decay sugar-rich substrates (Põlme et al. 2020) | Põlme S, et al. 2020. Fungal Diversity 105:1-16 ((https://doi.org/10.1007/s13225-020-00466-2)) |
| Otu48833 | 1 | k__Fungi(100);p__Ascomycota(100);c__Saccharomycetes(100);o__Saccharomycetales(100);f__Metschnikowiaceae(100);g__Metschnikowia(100);s__Metschnikowia_pulcherrima(100); | Metschnikowia | 13 | Saprotroph | \|Undefined Saprotroph\| | Yeast | NULL | Probable | As nectar-tap_saprotroph (Põlme et al. 2020); Decay sugar-rich substrates (Põlme et al. 2020) | Põlme S, et al. 2020. Fungal Diversity 105:1-16 ((https://doi.org/10.1007/s13225-020-00466-2)) |
| Otu48834 | 1 | k__Fungi(100);p__Ascomycota(100);c__Saccharomycetes(100);o__Saccharomycetales(100);f__Metschnikowiaceae(100);g__Metschnikowia(100);g__Metschnikowia_unclassified(100); | Metschnikowia | 13 | Saprotroph | \|Undefined Saprotroph\| | Yeast | NULL | Probable | As nectar-tap_saprotroph (Põlme et al. 2020); Decay sugar-rich substrates (Põlme et al. 2020) | Põlme S, et al. 2020. Fungal Diversity 105:1-16 ((https://doi.org/10.1007/s13225-020-00466-2)) |
| Otu48835 | 1 | k__Fungi(100);p__Ascomycota(100);c__Saccharomycetes(100);o__Saccharomycetales(100);f__Metschnikowiaceae(100);g__Metschnikowia(100);s__Metschnikowia_pulcherrima(100); | Metschnikowia | 13 | Saprotroph | \|Undefined Saprotroph\| | Yeast | NULL | Probable | As nectar-tap_saprotroph (Põlme et al. 2020); Decay sugar-rich substrates (Põlme et al. 2020) | Põlme S, et al. 2020. Fungal Diversity 105:1-16 ((https://doi.org/10.1007/s13225-020-00466-2)) |
| Otu48836 | 1 | k__Fungi(100);p__Ascomycota(100);c__Saccharomycetes(100);o__Saccharomycetales(100);f__Metschnikowiaceae(100);g__Metschnikowia(100);s__Metschnikowia_pulcherrima(100); | Metschnikowia | 13 | Saprotroph | \|Undefined Saprotroph\| | Yeast | NULL | Probable | As nectar-tap_saprotroph (Põlme et al. 2020); Decay sugar-rich substrates (Põlme et al. 2020) | Põlme S, et al. 2020. Fungal Diversity 105:1-16 ((https://doi.org/10.1007/s13225-020-00466-2)) |
| Otu48837 | 1 | k__Fungi(100);p__Ascomycota(100);c__Saccharomycetes(100);o__Saccharomycetales(100);f__Metschnikowiaceae(100);g__Metschnikowia(100);s__Metschnikowia_pulcherrima(100); | Metschnikowia | 13 | Saprotroph | \|Undefined Saprotroph\| | Yeast | NULL | Probable | As nectar-tap_saprotroph (Põlme et al. 2020); Decay sugar-rich substrates (Põlme et al. 2020) | Põlme S, et al. 2020. Fungal Diversity 105:1-16 ((https://doi.org/10.1007/s13225-020-00466-2)) |
| Otu48838 | 1 | k__Fungi(100);p__Ascomycota(100);c__Saccharomycetes(100);o__Saccharomycetales(100);f__Metschnikowiaceae(100);g__Metschnikowia(100);g__Metschnikowia_unclassified(100); | Metschnikowia | 13 | Saprotroph | \|Undefined Saprotroph\| | Yeast | NULL | Probable | As nectar-tap_saprotroph (Põlme et al. 2020); Decay sugar-rich substrates (Põlme et al. 2020) | Põlme S, et al. 2020. Fungal Diversity 105:1-16 ((https://doi.org/10.1007/s13225-020-00466-2)) |
| Otu48839 | 1 | k__Fungi(100);p__Ascomycota(100);c__Saccharomycetes(100);o__Saccharomycetales(100);f__Metschnikowiaceae(100);g__Metschnikowia(100);s__Metschnikowia_pulcherrima(100); | Metschnikowia | 13 | Saprotroph | \|Undefined Saprotroph\| | Yeast | NULL | Probable | As nectar-tap_saprotroph (Põlme et al. 2020); Decay sugar-rich substrates (Põlme et al. 2020) | Põlme S, et al. 2020. Fungal Diversity 105:1-16 ((https://doi.org/10.1007/s13225-020-00466-2)) |
| Otu48840 | 1 | k__Fungi(100);p__Ascomycota(100);c__Saccharomycetes(100);o__Saccharomycetales(100);f__Metschnikowiaceae(100);g__Metschnikowia(100);s__Metschnikowia_pulcherrima(100); | Metschnikowia | 13 | Saprotroph | \|Undefined Saprotroph\| | Yeast | NULL | Probable | As nectar-tap_saprotroph (Põlme et al. 2020); Decay sugar-rich substrates (Põlme et al. 2020) | Põlme S, et al. 2020. Fungal Diversity 105:1-16 ((https://doi.org/10.1007/s13225-020-00466-2)) |
| Otu48841 | 1 | k__Fungi(100);p__Ascomycota(100);c__Saccharomycetes(100);o__Saccharomycetales(100);f__Metschnikowiaceae(100);g__Metschnikowia(100);g__Metschnikowia_unclassified(100); | Metschnikowia | 13 | Saprotroph | \|Undefined Saprotroph\| | Yeast | NULL | Probable | As nectar-tap_saprotroph (Põlme et al. 2020); Decay sugar-rich substrates (Põlme et al. 2020) | Põlme S, et al. 2020. Fungal Diversity 105:1-16 ((https://doi.org/10.1007/s13225-020-00466-2)) |
| Otu48842 | 1 | k__Fungi(100);p__Ascomycota(100);c__Saccharomycetes(100);o__Saccharomycetales(100);f__Metschnikowiaceae(100);g__Metschnikowia(100);s__Metschnikowia_pulcherrima(100); | Metschnikowia | 13 | Saprotroph | \|Undefined Saprotroph\| | Yeast | NULL | Probable | As nectar-tap_saprotroph (Põlme et al. 2020); Decay sugar-rich substrates (Põlme et al. 2020) | Põlme S, et al. 2020. Fungal Diversity 105:1-16 ((https://doi.org/10.1007/s13225-020-00466-2)) |
| Otu48843 | 1 | k__Fungi(100);p__Ascomycota(100);c__Saccharomycetes(100);o__Saccharomycetales(100);f__Metschnikowiaceae(100);g__Metschnikowia(100);s__Metschnikowia_pulcherrima(100); | Metschnikowia | 13 | Saprotroph | \|Undefined Saprotroph\| | Yeast | NULL | Probable | As nectar-tap_saprotroph (Põlme et al. 2020); Decay sugar-rich substrates (Põlme et al. 2020) | Põlme S, et al. 2020. Fungal Diversity 105:1-16 ((https://doi.org/10.1007/s13225-020-00466-2)) |
| Otu48845 | 1 | k__Fungi(100);p__Ascomycota(100);c__Saccharomycetes(100);o__Saccharomycetales(100);f__Metschnikowiaceae(100);g__Metschnikowia(100);s__Metschnikowia_pulcherrima(100); | Metschnikowia | 13 | Saprotroph | \|Undefined Saprotroph\| | Yeast | NULL | Probable | As nectar-tap_saprotroph (Põlme et al. 2020); Decay sugar-rich substrates (Põlme et al. 2020) | Põlme S, et al. 2020. Fungal Diversity 105:1-16 ((https://doi.org/10.1007/s13225-020-00466-2)) |
| Otu48846 | 1 | k__Fungi(100);p__Ascomycota(100);c__Saccharomycetes(100);o__Saccharomycetales(100);f__Metschnikowiaceae(100);g__Metschnikowia(100);s__Metschnikowia_pulcherrima(100); | Metschnikowia | 13 | Saprotroph | \|Undefined Saprotroph\| | Yeast | NULL | Probable | As nectar-tap_saprotroph (Põlme et al. 2020); Decay sugar-rich substrates (Põlme et al. 2020) | Põlme S, et al. 2020. Fungal Diversity 105:1-16 ((https://doi.org/10.1007/s13225-020-00466-2)) |
| Otu48847 | 1 | k__Fungi(100);p__Ascomycota(100);c__Saccharomycetes(100);o__Saccharomycetales(100);f__Metschnikowiaceae(100);g__Metschnikowia(100);s__Metschnikowia_pulcherrima(100); | Metschnikowia | 13 | Saprotroph | \|Undefined Saprotroph\| | Yeast | NULL | Probable | As nectar-tap_saprotroph (Põlme et al. 2020); Decay sugar-rich substrates (Põlme et al. 2020) | Põlme S, et al. 2020. Fungal Diversity 105:1-16 ((https://doi.org/10.1007/s13225-020-00466-2)) |
| Otu48848 | 1 | k__Fungi(100);p__Ascomycota(100);c__Saccharomycetes(100);o__Saccharomycetales(100);f__Metschnikowiaceae(100);g__Metschnikowia(100);s__Metschnikowia_pulcherrima(100); | Metschnikowia | 13 | Saprotroph | \|Undefined Saprotroph\| | Yeast | NULL | Probable | As nectar-tap_saprotroph (Põlme et al. 2020); Decay sugar-rich substrates (Põlme et al. 2020) | Põlme S, et al. 2020. Fungal Diversity 105:1-16 ((https://doi.org/10.1007/s13225-020-00466-2)) |
| Otu48849 | 1 | k__Fungi(100);p__Ascomycota(100);c__Saccharomycetes(100);o__Saccharomycetales(100);f__Metschnikowiaceae(100);g__Metschnikowia(100);s__Metschnikowia_pulcherrima(100); | Metschnikowia | 13 | Saprotroph | \|Undefined Saprotroph\| | Yeast | NULL | Probable | As nectar-tap_saprotroph (Põlme et al. 2020); Decay sugar-rich substrates (Põlme et al. 2020) | Põlme S, et al. 2020. Fungal Diversity 105:1-16 ((https://doi.org/10.1007/s13225-020-00466-2)) |
| Otu48850 | 1 | k__Fungi(100);p__Ascomycota(100);c__Saccharomycetes(100);o__Saccharomycetales(100);f__Metschnikowiaceae(100);g__Metschnikowia(100);s__Metschnikowia_pulcherrima(100); | Metschnikowia | 13 | Saprotroph | \|Undefined Saprotroph\| | Yeast | NULL | Probable | As nectar-tap_saprotroph (Põlme et al. 2020); Decay sugar-rich substrates (Põlme et al. 2020) | Põlme S, et al. 2020. Fungal Diversity 105:1-16 ((https://doi.org/10.1007/s13225-020-00466-2)) |
| Otu48851 | 1 | k__Fungi(100);p__Ascomycota(100);c__Saccharomycetes(100);o__Saccharomycetales(100);f__Metschnikowiaceae(100);g__Metschnikowia(100);s__Metschnikowia_pulcherrima(100); | Metschnikowia | 13 | Saprotroph | \|Undefined Saprotroph\| | Yeast | NULL | Probable | As nectar-tap_saprotroph (Põlme et al. 2020); Decay sugar-rich substrates (Põlme et al. 2020) | Põlme S, et al. 2020. Fungal Diversity 105:1-16 ((https://doi.org/10.1007/s13225-020-00466-2)) |
| Otu48852 | 1 | k__Fungi(100);p__Ascomycota(100);c__Saccharomycetes(100);o__Saccharomycetales(100);f__Metschnikowiaceae(100);g__Metschnikowia(100);g__Metschnikowia_unclassified(100); | Metschnikowia | 13 | Saprotroph | \|Undefined Saprotroph\| | Yeast | NULL | Probable | As nectar-tap_saprotroph (Põlme et al. 2020); Decay sugar-rich substrates (Põlme et al. 2020) | Põlme S, et al. 2020. Fungal Diversity 105:1-16 ((https://doi.org/10.1007/s13225-020-00466-2)) |
| Otu48853 | 1 | k__Fungi(100);p__Ascomycota(100);c__Saccharomycetes(100);o__Saccharomycetales(100);f__Metschnikowiaceae(100);g__Metschnikowia(100);s__Metschnikowia_pulcherrima(100); | Metschnikowia | 13 | Saprotroph | \|Undefined Saprotroph\| | Yeast | NULL | Probable | As nectar-tap_saprotroph (Põlme et al. 2020); Decay sugar-rich substrates (Põlme et al. 2020) | Põlme S, et al. 2020. Fungal Diversity 105:1-16 ((https://doi.org/10.1007/s13225-020-00466-2)) |
| Otu48854 | 1 | k__Fungi(100);p__Ascomycota(100);c__Saccharomycetes(100);o__Saccharomycetales(100);f__Metschnikowiaceae(100);g__Metschnikowia(100);s__Metschnikowia_pulcherrima(100); | Metschnikowia | 13 | Saprotroph | \|Undefined Saprotroph\| | Yeast | NULL | Probable | As nectar-tap_saprotroph (Põlme et al. 2020); Decay sugar-rich substrates (Põlme et al. 2020) | Põlme S, et al. 2020. Fungal Diversity 105:1-16 ((https://doi.org/10.1007/s13225-020-00466-2)) |
| Otu48855 | 1 | k__Fungi(100);p__Ascomycota(100);c__Saccharomycetes(100);o__Saccharomycetales(100);f__Metschnikowiaceae(100);g__Metschnikowia(100);g__Metschnikowia_unclassified(100); | Metschnikowia | 13 | Saprotroph | \|Undefined Saprotroph\| | Yeast | NULL | Probable | As nectar-tap_saprotroph (Põlme et al. 2020); Decay sugar-rich substrates (Põlme et al. 2020) | Põlme S, et al. 2020. Fungal Diversity 105:1-16 ((https://doi.org/10.1007/s13225-020-00466-2)) |
| Otu48856 | 1 | k__Fungi(100);p__Ascomycota(100);c__Saccharomycetes(100);o__Saccharomycetales(100);f__Metschnikowiaceae(100);g__Metschnikowia(100);s__Metschnikowia_pulcherrima(100); | Metschnikowia | 13 | Saprotroph | \|Undefined Saprotroph\| | Yeast | NULL | Probable | As nectar-tap_saprotroph (Põlme et al. 2020); Decay sugar-rich substrates (Põlme et al. 2020) | Põlme S, et al. 2020. Fungal Diversity 105:1-16 ((https://doi.org/10.1007/s13225-020-00466-2)) |
| Otu48857 | 1 | k__Fungi(100);p__Ascomycota(100);c__Saccharomycetes(100);o__Saccharomycetales(100);f__Metschnikowiaceae(100);g__Metschnikowia(100);g__Metschnikowia_unclassified(100); | Metschnikowia | 13 | Saprotroph | \|Undefined Saprotroph\| | Yeast | NULL | Probable | As nectar-tap_saprotroph (Põlme et al. 2020); Decay sugar-rich substrates (Põlme et al. 2020) | Põlme S, et al. 2020. Fungal Diversity 105:1-16 ((https://doi.org/10.1007/s13225-020-00466-2)) |
| Otu48858 | 1 | k__Fungi(100);p__Ascomycota(100);c__Saccharomycetes(100);o__Saccharomycetales(100);f__Metschnikowiaceae(100);g__Metschnikowia(100);s__Metschnikowia_pulcherrima(100); | Metschnikowia | 13 | Saprotroph | \|Undefined Saprotroph\| | Yeast | NULL | Probable | As nectar-tap_saprotroph (Põlme et al. 2020); Decay sugar-rich substrates (Põlme et al. 2020) | Põlme S, et al. 2020. Fungal Diversity 105:1-16 ((https://doi.org/10.1007/s13225-020-00466-2)) |
| Otu48859 | 1 | k__Fungi(100);p__Ascomycota(100);c__Saccharomycetes(100);o__Saccharomycetales(100);f__Metschnikowiaceae(100);g__Metschnikowia(100);g__Metschnikowia_unclassified(100); | Metschnikowia | 13 | Saprotroph | \|Undefined Saprotroph\| | Yeast | NULL | Probable | As nectar-tap_saprotroph (Põlme et al. 2020); Decay sugar-rich substrates (Põlme et al. 2020) | Põlme S, et al. 2020. Fungal Diversity 105:1-16 ((https://doi.org/10.1007/s13225-020-00466-2)) |
| Otu48860 | 1 | k__Fungi(100);p__Ascomycota(100);c__Saccharomycetes(100);o__Saccharomycetales(100);f__Metschnikowiaceae(100);g__Metschnikowia(100);s__Metschnikowia_pulcherrima(100); | Metschnikowia | 13 | Saprotroph | \|Undefined Saprotroph\| | Yeast | NULL | Probable | As nectar-tap_saprotroph (Põlme et al. 2020); Decay sugar-rich substrates (Põlme et al. 2020) | Põlme S, et al. 2020. Fungal Diversity 105:1-16 ((https://doi.org/10.1007/s13225-020-00466-2)) |
| Otu48861 | 1 | k__Fungi(100);p__Ascomycota(100);c__Saccharomycetes(100);o__Saccharomycetales(100);f__Metschnikowiaceae(100);g__Metschnikowia(100);s__Metschnikowia_pulcherrima(100); | Metschnikowia | 13 | Saprotroph | \|Undefined Saprotroph\| | Yeast | NULL | Probable | As nectar-tap_saprotroph (Põlme et al. 2020); Decay sugar-rich substrates (Põlme et al. 2020) | Põlme S, et al. 2020. Fungal Diversity 105:1-16 ((https://doi.org/10.1007/s13225-020-00466-2)) |
| Otu48862 | 1 | k__Fungi(100);p__Ascomycota(100);c__Saccharomycetes(100);o__Saccharomycetales(100);f__Metschnikowiaceae(100);g__Metschnikowia(100);g__Metschnikowia_unclassified(100); | Metschnikowia | 13 | Saprotroph | \|Undefined Saprotroph\| | Yeast | NULL | Probable | As nectar-tap_saprotroph (Põlme et al. 2020); Decay sugar-rich substrates (Põlme et al. 2020) | Põlme S, et al. 2020. Fungal Diversity 105:1-16 ((https://doi.org/10.1007/s13225-020-00466-2)) |
| Otu48863 | 1 | k__Fungi(100);p__Ascomycota(100);c__Saccharomycetes(100);o__Saccharomycetales(100);f__Metschnikowiaceae(100);g__Metschnikowia(100);s__Metschnikowia_pulcherrima(100); | Metschnikowia | 13 | Saprotroph | \|Undefined Saprotroph\| | Yeast | NULL | Probable | As nectar-tap_saprotroph (Põlme et al. 2020); Decay sugar-rich substrates (Põlme et al. 2020) | Põlme S, et al. 2020. Fungal Diversity 105:1-16 ((https://doi.org/10.1007/s13225-020-00466-2)) |

| | | | | | | | | | | | |
|---|---|---|---|---|---|---|---|---|---|---|---|
| Otu48864 | 1 | k__Fungi(100);p__Ascomycota(100);c__Saccharomycetes(100);o__Saccharomycetales(100);f__Metschnikowiaceae(100);g__Metschnikowia(100);s__Metschnikowia_pulcherrima(100); | Metschnikowia | 13 | Saprotroph | \|Undefined Saprotroph\| | Yeast | NULL | Probable | As nectar-tap_saprotroph (Põlme et al. 2020); Decay sugar-rich substrates (Põlme et al. 2020) | Põlme S, et al. 2020. Fungal Diversity 105:1-16 ((https://doi.org/10.1007/s13225-020-00466-2)) |
| Otu48865 | 1 | k__Fungi(100);p__Ascomycota(100);c__Saccharomycetes(100);o__Saccharomycetales(100);f__Metschnikowiaceae(100);g__Metschnikowia(100);g__Metschnikowia_unclassified(100); | Metschnikowia | 13 | Saprotroph | \|Undefined Saprotroph\| | Yeast | NULL | Probable | As nectar-tap_saprotroph (Põlme et al. 2020); Decay sugar-rich substrates (Põlme et al. 2020) | Põlme S, et al. 2020. Fungal Diversity 105:1-16 ((https://doi.org/10.1007/s13225-020-00466-2)) |
| Otu48866 | 1 | k__Fungi(100);p__Ascomycota(100);c__Saccharomycetes(100);o__Saccharomycetales(100);f__Metschnikowiaceae(100);g__Metschnikowia(100);s__Metschnikowia_pulcherrima(100); | Metschnikowia | 13 | Saprotroph | \|Undefined Saprotroph\| | Yeast | NULL | Probable | As nectar-tap_saprotroph (Põlme et al. 2020); Decay sugar-rich substrates (Põlme et al. 2020) | Põlme S, et al. 2020. Fungal Diversity 105:1-16 ((https://doi.org/10.1007/s13225-020-00466-2)) |
| Otu48867 | 1 | k__Fungi(100);p__Ascomycota(100);c__Saccharomycetes(100);o__Saccharomycetales(100);f__Metschnikowiaceae(100);g__Metschnikowia(100);s__Metschnikowia_pulcherrima(100); | Metschnikowia | 13 | Saprotroph | \|Undefined Saprotroph\| | Yeast | NULL | Probable | As nectar-tap_saprotroph (Põlme et al. 2020); Decay sugar-rich substrates (Põlme et al. 2020) | Põlme S, et al. 2020. Fungal Diversity 105:1-16 ((https://doi.org/10.1007/s13225-020-00466-2)) |
| Otu48868 | 1 | k__Fungi(100);p__Ascomycota(100);c__Saccharomycetes(100);o__Saccharomycetales(100);f__Metschnikowiaceae(100);g__Metschnikowia(100);s__Metschnikowia_pulcherrima(100); | Metschnikowia | 13 | Saprotroph | \|Undefined Saprotroph\| | Yeast | NULL | Probable | As nectar-tap_saprotroph (Põlme et al. 2020); Decay sugar-rich substrates (Põlme et al. 2020) | Põlme S, et al. 2020. Fungal Diversity 105:1-16 ((https://doi.org/10.1007/s13225-020-00466-2)) |
| Otu48869 | 1 | k__Fungi(100);p__Ascomycota(100);c__Saccharomycetes(100);o__Saccharomycetales(100);f__Metschnikowiaceae(100);g__Metschnikowia(100);g__Metschnikowia_unclassified(100); | Metschnikowia | 13 | Saprotroph | \|Undefined Saprotroph\| | Yeast | NULL | Probable | As nectar-tap_saprotroph (Põlme et al. 2020); Decay sugar-rich substrates (Põlme et al. 2020) | Põlme S, et al. 2020. Fungal Diversity 105:1-16 ((https://doi.org/10.1007/s13225-020-00466-2)) |
| Otu48870 | 1 | k__Fungi(100);p__Ascomycota(100);c__Saccharomycetes(100);o__Saccharomycetales(100);f__Metschnikowiaceae(100);g__Metschnikowia(100);s__Metschnikowia_pulcherrima(100); | Metschnikowia | 13 | Saprotroph | \|Undefined Saprotroph\| | Yeast | NULL | Probable | As nectar-tap_saprotroph (Põlme et al. 2020); Decay sugar-rich substrates (Põlme et al. 2020) | Põlme S, et al. 2020. Fungal Diversity 105:1-16 ((https://doi.org/10.1007/s13225-020-00466-2)) |
| Otu48871 | 1 | k__Fungi(100);p__Ascomycota(100);c__Saccharomycetes(100);o__Saccharomycetales(100);f__Metschnikowiaceae(100);g__Metschnikowia(100);g__Metschnikowia_unclassified(100); | Metschnikowia | 13 | Saprotroph | \|Undefined Saprotroph\| | Yeast | NULL | Probable | As nectar-tap_saprotroph (Põlme et al. 2020); Decay sugar-rich substrates (Põlme et al. 2020) | Põlme S, et al. 2020. Fungal Diversity 105:1-16 ((https://doi.org/10.1007/s13225-020-00466-2)) |
| Otu48872 | 1 | k__Fungi(100);p__Ascomycota(100);c__Saccharomycetes(100);o__Saccharomycetales(100);f__Metschnikowiaceae(100);g__Metschnikowia(100);s__Metschnikowia_pulcherrima(100); | Metschnikowia | 13 | Saprotroph | \|Undefined Saprotroph\| | Yeast | NULL | Probable | As nectar-tap_saprotroph (Põlme et al. 2020); Decay sugar-rich substrates (Põlme et al. 2020) | Põlme S, et al. 2020. Fungal Diversity 105:1-16 ((https://doi.org/10.1007/s13225-020-00466-2)) |
| Otu48873 | 1 | k__Fungi(100);p__Ascomycota(100);c__Saccharomycetes(100);o__Saccharomycetales(100);f__Metschnikowiaceae(100);g__Metschnikowia(100);s__Metschnikowia_pulcherrima(100); | Metschnikowia | 13 | Saprotroph | \|Undefined Saprotroph\| | Yeast | NULL | Probable | As nectar-tap_saprotroph (Põlme et al. 2020); Decay sugar-rich substrates (Põlme et al. 2020) | Põlme S, et al. 2020. Fungal Diversity 105:1-16 ((https://doi.org/10.1007/s13225-020-00466-2)) |
| Otu48874 | 1 | k__Fungi(100);p__Ascomycota(100);c__Saccharomycetes(100);o__Saccharomycetales(100);f__Metschnikowiaceae(100);g__Metschnikowia(100);s__Metschnikowia_pulcherrima(100); | Metschnikowia | 13 | Saprotroph | \|Undefined Saprotroph\| | Yeast | NULL | Probable | As nectar-tap_saprotroph (Põlme et al. 2020); Decay sugar-rich substrates (Põlme et al. 2020) | Põlme S, et al. 2020. Fungal Diversity 105:1-16 ((https://doi.org/10.1007/s13225-020-00466-2)) |
| Otu48875 | 1 | k__Fungi(100);p__Ascomycota(100);c__Saccharomycetes(100);o__Saccharomycetales(100);f__Metschnikowiaceae(100);g__Metschnikowia(100);s__Metschnikowia_pulcherrima(100); | Metschnikowia | 13 | Saprotroph | \|Undefined Saprotroph\| | Yeast | NULL | Probable | As nectar-tap_saprotroph (Põlme et al. 2020); Decay sugar-rich substrates (Põlme et al. 2020) | Põlme S, et al. 2020. Fungal Diversity 105:1-16 ((https://doi.org/10.1007/s13225-020-00466-2)) |
| Otu48876 | 1 | k__Fungi(100);p__Ascomycota(100);c__Saccharomycetes(100);o__Saccharomycetales(100);f__Metschnikowiaceae(100);g__Metschnikowia(100);s__Metschnikowia_pulcherrima(100); | Metschnikowia | 13 | Saprotroph | \|Undefined Saprotroph\| | Yeast | NULL | Probable | As nectar-tap_saprotroph (Põlme et al. 2020); Decay sugar-rich substrates (Põlme et al. 2020) | Põlme S, et al. 2020. Fungal Diversity 105:1-16 ((https://doi.org/10.1007/s13225-020-00466-2)) |
| Otu48877 | 1 | k__Fungi(100);p__Ascomycota(100);c__Saccharomycetes(100);o__Saccharomycetales(100);f__Metschnikowiaceae(100);g__Metschnikowia(100);s__Metschnikowia_pulcherrima(100); | Metschnikowia | 13 | Saprotroph | \|Undefined Saprotroph\| | Yeast | NULL | Probable | As nectar-tap_saprotroph (Põlme et al. 2020); Decay sugar-rich substrates (Põlme et al. 2020) | Põlme S, et al. 2020. Fungal Diversity 105:1-16 ((https://doi.org/10.1007/s13225-020-00466-2)) |
| Otu48878 | 1 | k__Fungi(100);p__Ascomycota(100);c__Saccharomycetes(100);o__Saccharomycetales(100);f__Metschnikowiaceae(100);g__Metschnikowia(100);s__Metschnikowia_pulcherrima(100); | Metschnikowia | 13 | Saprotroph | \|Undefined Saprotroph\| | Yeast | NULL | Probable | As nectar-tap_saprotroph (Põlme et al. 2020); Decay sugar-rich substrates (Põlme et al. 2020) | Põlme S, et al. 2020. Fungal Diversity 105:1-16 ((https://doi.org/10.1007/s13225-020-00466-2)) |
| Otu48879 | 1 | k__Fungi(100);p__Ascomycota(100);c__Saccharomycetes(100);o__Saccharomycetales(100);f__Metschnikowiaceae(100);g__Metschnikowia(100);s__Metschnikowia_pulcherrima(100); | Metschnikowia | 13 | Saprotroph | \|Undefined Saprotroph\| | Yeast | NULL | Probable | As nectar-tap_saprotroph (Põlme et al. 2020); Decay sugar-rich substrates (Põlme et al. 2020) | Põlme S, et al. 2020. Fungal Diversity 105:1-16 ((https://doi.org/10.1007/s13225-020-00466-2)) |
| Otu48880 | 1 | k__Fungi(100);p__Ascomycota(100);c__Saccharomycetes(100);o__Saccharomycetales(100);f__Metschnikowiaceae(100);g__Metschnikowia(100);s__Metschnikowia_pulcherrima(100); | Metschnikowia | 13 | Saprotroph | \|Undefined Saprotroph\| | Yeast | NULL | Probable | As nectar-tap_saprotroph (Põlme et al. 2020); Decay sugar-rich substrates (Põlme et al. 2020) | Põlme S, et al. 2020. Fungal Diversity 105:1-16 ((https://doi.org/10.1007/s13225-020-00466-2)) |
| Otu48881 | 1 | k__Fungi(100);p__Ascomycota(100);c__Saccharomycetes(100);o__Saccharomycetales(100);f__Metschnikowiaceae(100);g__Metschnikowia(100);g__Metschnikowia_unclassified(100); | Metschnikowia | 13 | Saprotroph | \|Undefined Saprotroph\| | Yeast | NULL | Probable | As nectar-tap_saprotroph (Põlme et al. 2020); Decay sugar-rich substrates (Põlme et al. 2020) | Põlme S, et al. 2020. Fungal Diversity 105:1-16 ((https://doi.org/10.1007/s13225-020-00466-2)) |
| Otu48882 | 1 | k__Fungi(100);p__Ascomycota(100);c__Saccharomycetes(100);o__Saccharomycetales(100);f__Metschnikowiaceae(100);g__Metschnikowia(100);s__Metschnikowia_pulcherrima(100); | Metschnikowia | 13 | Saprotroph | \|Undefined Saprotroph\| | Yeast | NULL | Probable | As nectar-tap_saprotroph (Põlme et al. 2020); Decay sugar-rich substrates (Põlme et al. 2020) | Põlme S, et al. 2020. Fungal Diversity 105:1-16 ((https://doi.org/10.1007/s13225-020-00466-2)) |
| Otu48883 | 1 | k__Fungi(100);p__Ascomycota(100);c__Saccharomycetes(100);o__Saccharomycetales(100);f__Metschnikowiaceae(100);g__Metschnikowia(100);s__Metschnikowia_pulcherrima(100); | Metschnikowia | 13 | Saprotroph | \|Undefined Saprotroph\| | Yeast | NULL | Probable | As nectar-tap_saprotroph (Põlme et al. 2020); Decay sugar-rich substrates (Põlme et al. 2020) | Põlme S, et al. 2020. Fungal Diversity 105:1-16 ((https://doi.org/10.1007/s13225-020-00466-2)) |
| Otu48884 | 1 | k__Fungi(100);p__Ascomycota(100);c__Saccharomycetes(100);o__Saccharomycetales(100);f__Metschnikowiaceae(100);g__Metschnikowia(100);g__Metschnikowia_unclassified(100); | Metschnikowia | 13 | Saprotroph | \|Undefined Saprotroph\| | Yeast | NULL | Probable | As nectar-tap_saprotroph (Põlme et al. 2020); Decay sugar-rich substrates (Põlme et al. 2020) | Põlme S, et al. 2020. Fungal Diversity 105:1-16 ((https://doi.org/10.1007/s13225-020-00466-2)) |
| Otu48885 | 1 | k__Fungi(100);p__Ascomycota(100);c__Saccharomycetes(100);o__Saccharomycetales(100);f__Metschnikowiaceae(100);g__Metschnikowia(100);g__Metschnikowia_unclassified(100); | Metschnikowia | 13 | Saprotroph | \|Undefined Saprotroph\| | Yeast | NULL | Probable | As nectar-tap_saprotroph (Põlme et al. 2020); Decay sugar-rich substrates (Põlme et al. 2020) | Põlme S, et al. 2020. Fungal Diversity 105:1-16 ((https://doi.org/10.1007/s13225-020-00466-2)) |
| Otu48886 | 1 | k__Fungi(100);p__Ascomycota(100);c__Saccharomycetes(100);o__Saccharomycetales(100);f__Metschnikowiaceae(100);g__Metschnikowia(100);s__Metschnikowia_pulcherrima(100); | Metschnikowia | 13 | Saprotroph | \|Undefined Saprotroph\| | Yeast | NULL | Probable | As nectar-tap_saprotroph (Põlme et al. 2020); Decay sugar-rich substrates (Põlme et al. 2020) | Põlme S, et al. 2020. Fungal Diversity 105:1-16 ((https://doi.org/10.1007/s13225-020-00466-2)) |
| Otu48887 | 1 | k__Fungi(100);p__Ascomycota(100);c__Saccharomycetes(100);o__Saccharomycetales(100);f__Metschnikowiaceae(100);g__Metschnikowia(100);s__Metschnikowia_pulcherrima(100); | Metschnikowia | 13 | Saprotroph | \|Undefined Saprotroph\| | Yeast | NULL | Probable | As nectar-tap_saprotroph (Põlme et al. 2020); Decay sugar-rich substrates (Põlme et al. 2020) | Põlme S, et al. 2020. Fungal Diversity 105:1-16 ((https://doi.org/10.1007/s13225-020-00466-2)) |
| Otu48888 | 1 | k__Fungi(100);p__Ascomycota(100);c__Saccharomycetes(100);o__Saccharomycetales(100);f__Metschnikowiaceae(100);g__Metschnikowia(100);s__Metschnikowia_pulcherrima(100); | Metschnikowia | 13 | Saprotroph | \|Undefined Saprotroph\| | Yeast | NULL | Probable | As nectar-tap_saprotroph (Põlme et al. 2020); Decay sugar-rich substrates (Põlme et al. 2020) | Põlme S, et al. 2020. Fungal Diversity 105:1-16 ((https://doi.org/10.1007/s13225-020-00466-2)) |
| Otu48889 | 1 | k__Fungi(100);p__Ascomycota(100);c__Saccharomycetes(100);o__Saccharomycetales(100);f__Metschnikowiaceae(100);g__Metschnikowia(100);s__Metschnikowia_pulcherrima(100); | Metschnikowia | 13 | Saprotroph | \|Undefined Saprotroph\| | Yeast | NULL | Probable | As nectar-tap_saprotroph (Põlme et al. 2020); Decay sugar-rich substrates (Põlme et al. 2020) | Põlme S, et al. 2020. Fungal Diversity 105:1-16 ((https://doi.org/10.1007/s13225-020-00466-2)) |
| Otu48890 | 1 | k__Fungi(100);p__Ascomycota(100);c__Saccharomycetes(100);o__Saccharomycetales(100);f__Metschnikowiaceae(100);g__Metschnikowia(100);s__Metschnikowia_pulcherrima(100); | Metschnikowia | 13 | Saprotroph | \|Undefined Saprotroph\| | Yeast | NULL | Probable | As nectar-tap_saprotroph (Põlme et al. 2020); Decay sugar-rich substrates (Põlme et al. 2020) | Põlme S, et al. 2020. Fungal Diversity 105:1-16 ((https://doi.org/10.1007/s13225-020-00466-2)) |
| Otu48891 | 1 | k__Fungi(100);p__Ascomycota(100);c__Saccharomycetes(100);o__Saccharomycetales(100);f__Metschnikowiaceae(100);g__Metschnikowia(100);s__Metschnikowia_pulcherrima(100); | Metschnikowia | 13 | Saprotroph | \|Undefined Saprotroph\| | Yeast | NULL | Probable | As nectar-tap_saprotroph (Põlme et al. 2020); Decay sugar-rich substrates (Põlme et al. 2020) | Põlme S, et al. 2020. Fungal Diversity 105:1-16 ((https://doi.org/10.1007/s13225-020-00466-2)) |
| Otu48892 | 1 | k__Fungi(100);p__Ascomycota(100);c__Saccharomycetes(100);o__Saccharomycetales(100);f__Metschnikowiaceae(100);g__Metschnikowia(100);g__Metschnikowia_unclassified(100); | Metschnikowia | 13 | Saprotroph | \|Undefined Saprotroph\| | Yeast | NULL | Probable | As nectar-tap_saprotroph (Põlme et al. 2020); Decay sugar-rich substrates (Põlme et al. 2020) | Põlme S, et al. 2020. Fungal Diversity 105:1-16 ((https://doi.org/10.1007/s13225-020-00466-2)) |
| Otu48893 | 1 | k__Fungi(100);p__Ascomycota(100);c__Saccharomycetes(100);o__Saccharomycetales(100);f__Metschnikowiaceae(100);g__Metschnikowia(100);s__Metschnikowia_pulcherrima(100); | Metschnikowia | 13 | Saprotroph | \|Undefined Saprotroph\| | Yeast | NULL | Probable | As nectar-tap_saprotroph (Põlme et al. 2020); Decay sugar-rich substrates (Põlme et al. 2020) | Põlme S, et al. 2020. Fungal Diversity 105:1-16 ((https://doi.org/10.1007/s13225-020-00466-2)) |
| Otu48894 | 1 | k__Fungi(100);p__Ascomycota(100);c__Saccharomycetes(100);o__Saccharomycetales(100);f__Metschnikowiaceae(100);g__Metschnikowia(100);g__Metschnikowia_unclassified(100); | Metschnikowia | 13 | Saprotroph | \|Undefined Saprotroph\| | Yeast | NULL | Probable | As nectar-tap_saprotroph (Põlme et al. 2020); Decay sugar-rich substrates (Põlme et al. 2020) | Põlme S, et al. 2020. Fungal Diversity 105:1-16 ((https://doi.org/10.1007/s13225-020-00466-2)) |
| Otu48895 | 1 | k__Fungi(100);p__Ascomycota(100);c__Saccharomycetes(100);o__Saccharomycetales(100);f__Metschnikowiaceae(100);g__Metschnikowia(100);g__Metschnikowia_unclassified(100); | Metschnikowia | 13 | Saprotroph | \|Undefined Saprotroph\| | Yeast | NULL | Probable | As nectar-tap_saprotroph (Põlme et al. 2020); Decay sugar-rich substrates (Põlme et al. 2020) | Põlme S, et al. 2020. Fungal Diversity 105:1-16 ((https://doi.org/10.1007/s13225-020-00466-2)) |
| Otu48896 | 1 | k__Fungi(100);p__Ascomycota(100);c__Saccharomycetes(100);o__Saccharomycetales(100);f__Metschnikowiaceae(100);g__Metschnikowia(100);s__Metschnikowia_pulcherrima(100); | Metschnikowia | 13 | Saprotroph | \|Undefined Saprotroph\| | Yeast | NULL | Probable | As nectar-tap_saprotroph (Põlme et al. 2020); Decay sugar-rich substrates (Põlme et al. 2020) | Põlme S, et al. 2020. Fungal Diversity 105:1-16 ((https://doi.org/10.1007/s13225-020-00466-2)) |
| Otu48897 | 1 | k__Fungi(100);p__Ascomycota(100);c__Saccharomycetes(100);o__Saccharomycetales(100);f__Metschnikowiaceae(100);g__Metschnikowia(100);g__Metschnikowia_unclassified(100); | Metschnikowia | 13 | Saprotroph | \|Undefined Saprotroph\| | Yeast | NULL | Probable | As nectar-tap_saprotroph (Põlme et al. 2020); Decay sugar-rich substrates (Põlme et al. 2020) | Põlme S, et al. 2020. Fungal Diversity 105:1-16 ((https://doi.org/10.1007/s13225-020-00466-2)) |
| Otu48898 | 1 | k__Fungi(100);p__Ascomycota(100);c__Saccharomycetes(100);o__Saccharomycetales(100);f__Metschnikowiaceae(100);g__Metschnikowia(100);s__Metschnikowia_pulcherrima(100); | Metschnikowia | 13 | Saprotroph | \|Undefined Saprotroph\| | Yeast | NULL | Probable | As nectar-tap_saprotroph (Põlme et al. 2020); Decay sugar-rich substrates (Põlme et al. 2020) | Põlme S, et al. 2020. Fungal Diversity 105:1-16 ((https://doi.org/10.1007/s13225-020-00466-2)) |
| Otu48899 | 1 | k__Fungi(100);p__Ascomycota(100);c__Saccharomycetes(100);o__Saccharomycetales(100);f__Metschnikowiaceae(100);g__Metschnikowia(100);s__Metschnikowia_pulcherrima(100); | Metschnikowia | 13 | Saprotroph | \|Undefined Saprotroph\| | Yeast | NULL | Probable | As nectar-tap_saprotroph (Põlme et al. 2020); Decay sugar-rich substrates (Põlme et al. 2020) | Põlme S, et al. 2020. Fungal Diversity 105:1-16 ((https://doi.org/10.1007/s13225-020-00466-2)) |
| Otu48900 | 1 | k__Fungi(100);p__Ascomycota(100);c__Saccharomycetes(100);o__Saccharomycetales(100);f__Metschnikowiaceae(100);g__Metschnikowia(100);s__Metschnikowia_pulcherrima(100); | Metschnikowia | 13 | Saprotroph | \|Undefined Saprotroph\| | Yeast | NULL | Probable | As nectar-tap_saprotroph (Põlme et al. 2020); Decay sugar-rich substrates (Põlme et al. 2020) | Põlme S, et al. 2020. Fungal Diversity 105:1-16 ((https://doi.org/10.1007/s13225-020-00466-2)) |
| Otu48901 | 1 | k__Fungi(100);p__Ascomycota(100);c__Saccharomycetes(100);o__Saccharomycetales(100);f__Metschnikowiaceae(100);g__Metschnikowia(100);s__Metschnikowia_pulcherrima(100); | Metschnikowia | 13 | Saprotroph | \|Undefined Saprotroph\| | Yeast | NULL | Probable | As nectar-tap_saprotroph (Põlme et al. 2020); Decay sugar-rich substrates (Põlme et al. 2020) | Põlme S, et al. 2020. Fungal Diversity 105:1-16 ((https://doi.org/10.1007/s13225-020-00466-2)) |
| Otu48902 | 1 | k__Fungi(100);p__Ascomycota(100);c__Saccharomycetes(100);o__Saccharomycetales(100);f__Metschnikowiaceae(100);g__Metschnikowia(100);s__Metschnikowia_pulcherrima(100); | Metschnikowia | 13 | Saprotroph | \|Undefined Saprotroph\| | Yeast | NULL | Probable | As nectar-tap_saprotroph (Põlme et al. 2020); Decay sugar-rich substrates (Põlme et al. 2020) | Põlme S, et al. 2020. Fungal Diversity 105:1-16 ((https://doi.org/10.1007/s13225-020-00466-2)) |
| Otu48903 | 1 | k__Fungi(100);p__Ascomycota(100);c__Saccharomycetes(100);o__Saccharomycetales(100);f__Metschnikowiaceae(100);g__Metschnikowia(100);s__Metschnikowia_pulcherrima(100); | Metschnikowia | 13 | Saprotroph | \|Undefined Saprotroph\| | Yeast | NULL | Probable | As nectar-tap_saprotroph (Põlme et al. 2020); Decay sugar-rich substrates (Põlme et al. 2020) | Põlme S, et al. 2020. Fungal Diversity 105:1-16 ((https://doi.org/10.1007/s13225-020-00466-2)) |
| Otu48904 | 1 | k__Fungi(100);p__Ascomycota(100);c__Saccharomycetes(100);o__Saccharomycetales(100);f__Metschnikowiaceae(100);g__Metschnikowia(100);g__Metschnikowia_unclassified(100); | Metschnikowia | 13 | Saprotroph | \|Undefined Saprotroph\| | Yeast | NULL | Probable | As nectar-tap_saprotroph (Põlme et al. 2020); Decay sugar-rich substrates (Põlme et al. 2020) | Põlme S, et al. 2020. Fungal Diversity 105:1-16 ((https://doi.org/10.1007/s13225-020-00466-2)) |
| Otu48905 | 1 | k__Fungi(100);p__Ascomycota(100);c__Saccharomycetes(100);o__Saccharomycetales(100);f__Metschnikowiaceae(100);g__Metschnikowia(100);g__Metschnikowia_unclassified(100); | Metschnikowia | 13 | Saprotroph | \|Undefined Saprotroph\| | Yeast | NULL | Probable | As nectar-tap_saprotroph (Põlme et al. 2020); Decay sugar-rich substrates (Põlme et al. 2020) | Põlme S, et al. 2020. Fungal Diversity 105:1-16 ((https://doi.org/10.1007/s13225-020-00466-2)) |
| Otu48906 | 1 | k__Fungi(100);p__Ascomycota(100);c__Saccharomycetes(100);o__Saccharomycetales(100);f__Metschnikowiaceae(100);g__Metschnikowia(100);s__Metschnikowia_pulcherrima(100); | Metschnikowia | 13 | Saprotroph | \|Undefined Saprotroph\| | Yeast | NULL | Probable | As nectar-tap_saprotroph (Põlme et al. 2020); Decay sugar-rich substrates (Põlme et al. 2020) | Põlme S, et al. 2020. Fungal Diversity 105:1-16 ((https://doi.org/10.1007/s13225-020-00466-2)) |
| Otu48907 | 1 | k__Fungi(100);p__Ascomycota(100);c__Saccharomycetes(100);o__Saccharomycetales(100);f__Metschnikowiaceae(100);g__Metschnikowia(100);g__Metschnikowia_unclassified(100); | Metschnikowia | 13 | Saprotroph | \|Undefined Saprotroph\| | Yeast | NULL | Probable | As nectar-tap_saprotroph (Põlme et al. 2020); Decay sugar-rich substrates (Põlme et al. 2020) | Põlme S, et al. 2020. Fungal Diversity 105:1-16 ((https://doi.org/10.1007/s13225-020-00466-2)) |
| Otu48908 | 1 | k__Fungi(100);p__Ascomycota(100);c__Saccharomycetes(100);o__Saccharomycetales(100);f__Metschnikowiaceae(100);g__Metschnikowia(100);s__Metschnikowia_pulcherrima(100); | Metschnikowia | 13 | Saprotroph | \|Undefined Saprotroph\| | Yeast | NULL | Probable | As nectar-tap_saprotroph (Põlme et al. 2020); Decay sugar-rich substrates (Põlme et al. 2020) | Põlme S, et al. 2020. Fungal Diversity 105:1-16 ((https://doi.org/10.1007/s13225-020-00466-2)) |
| Otu48909 | 1 | k__Fungi(100);p__Ascomycota(100);c__Saccharomycetes(100);o__Saccharomycetales(100);f__Metschnikowiaceae(100);g__Metschnikowia(100);g__Metschnikowia_unclassified(100); | Metschnikowia | 13 | Saprotroph | \|Undefined Saprotroph\| | Yeast | NULL | Probable | As nectar-tap_saprotroph (Põlme et al. 2020); Decay sugar-rich substrates (Põlme et al. 2020) | Põlme S, et al. 2020. Fungal Diversity 105:1-16 ((https://doi.org/10.1007/s13225-020-00466-2)) |
| Otu48910 | 1 | k__Fungi(100);p__Ascomycota(100);c__Saccharomycetes(100);o__Saccharomycetales(100);f__Metschnikowiaceae(100);g__Metschnikowia(100);s__Metschnikowia_pulcherrima(100); | Metschnikowia | 13 | Saprotroph | \|Undefined Saprotroph\| | Yeast | NULL | Probable | As nectar-tap_saprotroph (Põlme et al. 2020); Decay sugar-rich substrates (Põlme et al. 2020) | Põlme S, et al. 2020. Fungal Diversity 105:1-16 ((https://doi.org/10.1007/s13225-020-00466-2)) |
| Otu48911 | 1 | k__Fungi(100);p__Ascomycota(100);c__Saccharomycetes(100);o__Saccharomycetales(100);f__Saccharomycetales_family_Incertae_sedis(100);f__Saccharomycetales_family_Incertae_sedis_unclassified(100);f__Saccharomycetales_family_Incertae_sedis_unclassified(100); | Saccharomycetales | 7 | Saprotroph | Undefined Saprotroph | Yeast | NULL | Possible | NULL | Sterkenburg E, et al. 2015. New Phytologist 207:1145-1158 ((https://doi.org/10.1111/nph.13426)) |
| Otu48912 | 1 | k__Fungi(100);p__Ascomycota(100);c__Saccharomycetes(100);o__Saccharomycetales(100);f__Metschnikowiaceae(100);g__Metschnikowia(100);s__Metschnikowia_pulcherrima(100); | Metschnikowia | 13 | Saprotroph | \|Undefined Saprotroph\| | Yeast | NULL | Probable | As nectar-tap_saprotroph (Põlme et al. 2020); Decay sugar-rich substrates (Põlme et al. 2020) | Põlme S, et al. 2020. Fungal Diversity 105:1-16 ((https://doi.org/10.1007/s13225-020-00466-2)) |
| Otu48913 | 1 | k__Fungi(100);p__Ascomycota(100);c__Saccharomycetes(100);o__Saccharomycetales(100);f__Metschnikowiaceae(100);g__Metschnikowia(100);g__Metschnikowia_unclassified(100); | Metschnikowia | 13 | Saprotroph | \|Undefined Saprotroph\| | Yeast | NULL | Probable | As nectar-tap_saprotroph (Põlme et al. 2020); Decay sugar-rich substrates (Põlme et al. 2020) | Põlme S, et al. 2020. Fungal Diversity 105:1-16 ((https://doi.org/10.1007/s13225-020-00466-2)) |
| Otu48914 | 1 | k__Fungi(100);p__Ascomycota(100);c__Saccharomycetes(100);o__Saccharomycetales(100);f__Metschnikowiaceae(100);g__Metschnikowia(100);g__Metschnikowia_unclassified(100); | Metschnikowia | 13 | Saprotroph | \|Undefined Saprotroph\| | Yeast | NULL | Probable | As nectar-tap_saprotroph (Põlme et al. 2020); Decay sugar-rich substrates (Põlme et al. 2020) | Põlme S, et al. 2020. Fungal Diversity 105:1-16 ((https://doi.org/10.1007/s13225-020-00466-2)) |
| Otu48915 | 1 | k__Fungi(100);p__Ascomycota(100);c__Saccharomycetes(100);o__Saccharomycetales(100);f__Metschnikowiaceae(100);g__Metschnikowia(100);s__Metschnikowia_pulcherrima(100); | Metschnikowia | 13 | Saprotroph | \|Undefined Saprotroph\| | Yeast | NULL | Probable | As nectar-tap_saprotroph (Põlme et al. 2020); Decay sugar-rich substrates (Põlme et al. 2020) | Põlme S, et al. 2020. Fungal Diversity 105:1-16 ((https://doi.org/10.1007/s13225-020-00466-2)) |
| Otu48916 | 1 | k__Fungi(100);p__Ascomycota(100);c__Saccharomycetes(100);o__Saccharomycetales(100);f__Metschnikowiaceae(100);g__Metschnikowia(100);s__Metschnikowia_pulcherrima(100); | Metschnikowia | 13 | Saprotroph | \|Undefined Saprotroph\| | Yeast | NULL | Probable | As nectar-tap_saprotroph (Põlme et al. 2020); Decay sugar-rich substrates (Põlme et al. 2020) | Põlme S, et al. 2020. Fungal Diversity 105:1-16 ((https://doi.org/10.1007/s13225-020-00466-2)) |
| Otu48917 | 1 | k__Fungi(100);p__Ascomycota(100);c__Saccharomycetes(100);o__Saccharomycetales(100);f__Metschnikowiaceae(100);g__Metschnikowia(100);s__Metschnikowia_pulcherrima(100); | Metschnikowia | 13 | Saprotroph | \|Undefined Saprotroph\| | Yeast | NULL | Probable | As nectar-tap_saprotroph (Põlme et al. 2020); Decay sugar-rich substrates (Põlme et al. 2020) | Põlme S, et al. 2020. Fungal Diversity 105:1-16 ((https://doi.org/10.1007/s13225-020-00466-2)) |
| Otu48918 | 1 | k__Fungi(100);p__Ascomycota(100);c__Saccharomycetes(100);o__Saccharomycetales(100);f__Metschnikowiaceae(100);g__Metschnikowia(100);g__Metschnikowia_unclassified(100); | Metschnikowia | 13 | Saprotroph | \|Undefined Saprotroph\| | Yeast | NULL | Probable | As nectar-tap_saprotroph (Põlme et al. 2020); Decay sugar-rich substrates (Põlme et al. 2020) | Põlme S, et al. 2020. Fungal Diversity 105:1-16 ((https://doi.org/10.1007/s13225-020-00466-2)) |
| Otu48919 | 1 | k__Fungi(100);p__Ascomycota(100);c__Saccharomycetes(100);o__Saccharomycetales(100);f__Metschnikowiaceae(100);g__Metschnikowia(100);s__Metschnikowia_pulcherrima(100); | Metschnikowia | 13 | Saprotroph | \|Undefined Saprotroph\| | Yeast | NULL | Probable | As nectar-tap_saprotroph (Põlme et al. 2020); Decay sugar-rich substrates (Põlme et al. 2020) | Põlme S, et al. 2020. Fungal Diversity 105:1-16 ((https://doi.org/10.1007/s13225-020-00466-2)) |
| Otu48920 | 1 | k__Fungi(100);p__Ascomycota(100);c__Saccharomycetes(100);o__Saccharomycetales(100);f__Metschnikowiaceae(100);g__Metschnikowia(100);s__Metschnikowia_pulcherrima(100); | Metschnikowia | 13 | Saprotroph | \|Undefined Saprotroph\| | Yeast | NULL | Probable | As nectar-tap_saprotroph (Põlme et al. 2020); Decay sugar-rich substrates (Põlme et al. 2020) | Põlme S, et al. 2020. Fungal Diversity 105:1-16 ((https://doi.org/10.1007/s13225-020-00466-2)) |
| Otu48921 | 1 | k__Fungi(100);p__Ascomycota(100);c__Saccharomycetes(100);o__Saccharomycetales(100);f__Metschnikowiaceae(100);g__Metschnikowia(100);s__Metschnikowia_pulcherrima(100); | Metschnikowia | 13 | Saprotroph | \|Undefined Saprotroph\| | Yeast | NULL | Probable | As nectar-tap_saprotroph (Põlme et al. 2020); Decay sugar-rich substrates (Põlme et al. 2020) | Põlme S, et al. 2020. Fungal Diversity 105:1-16 ((https://doi.org/10.1007/s13225-020-00466-2)) |
| Otu48922 | 1 | k__Fungi(100);p__Ascomycota(100);c__Saccharomycetes(100);o__Saccharomycetales(100);f__Metschnikowiaceae(100);g__Metschnikowia(100);s__Metschnikowia_pulcherrima(100); | Metschnikowia | 13 | Saprotroph | \|Undefined Saprotroph\| | Yeast | NULL | Probable | As nectar-tap_saprotroph (Põlme et al. 2020); Decay sugar-rich substrates (Põlme et al. 2020) | Põlme S, et al. 2020. Fungal Diversity 105:1-16 ((https://doi.org/10.1007/s13225-020-00466-2)) |
| Otu48923 | 1 | k__Fungi(100);p__Ascomycota(100);c__Saccharomycetes(100);o__Saccharomycetales(100);f__Metschnikowiaceae(100);g__Metschnikowia(100);s__Metschnikowia_pulcherrima(100); | Metschnikowia | 13 | Saprotroph | \|Undefined Saprotroph\| | Yeast | NULL | Probable | As nectar-tap_saprotroph (Põlme et al. 2020); Decay sugar-rich substrates (Põlme et al. 2020) | Põlme S, et al. 2020. Fungal Diversity 105:1-16 ((https://doi.org/10.1007/s13225-020-00466-2)) |
| Otu48924 | 1 | k__Fungi(100);p__Ascomycota(100);c__Saccharomycetes(100);o__Saccharomycetales(100);f__Metschnikowiaceae(100);g__Metschnikowia(100);s__Metschnikowia_pulcherrima(100); | Metschnikowia | 13 | Saprotroph | \|Undefined Saprotroph\| | Yeast | NULL | Probable | As nectar-tap_saprotroph (Põlme et al. 2020); Decay sugar-rich substrates (Põlme et al. 2020) | Põlme S, et al. 2020. Fungal Diversity 105:1-16 ((https://doi.org/10.1007/s13225-020-00466-2)) |
| Otu48925 | 1 | k__Fungi(100);p__Ascomycota(100);c__Saccharomycetes(100);o__Saccharomycetales(100);f__Metschnikowiaceae(100);g__Metschnikowia(100);s__Metschnikowia_pulcherrima(100); | Metschnikowia | 13 | Saprotroph | \|Undefined Saprotroph\| | Yeast | NULL | Probable | As nectar-tap_saprotroph (Põlme et al. 2020); Decay sugar-rich substrates (Põlme et al. 2020) | Põlme S, et al. 2020. Fungal Diversity 105:1-16 ((https://doi.org/10.1007/s13225-020-00466-2)) |
| Otu48926 | 1 | k__Fungi(100);p__Ascomycota(100);c__Saccharomycetes(100);o__Saccharomycetales(100);f__Metschnikowiaceae(100);g__Metschnikowia(100);s__Metschnikowia_pulcherrima(100); | Metschnikowia | 13 | Saprotroph | \|Undefined Saprotroph\| | Yeast | NULL | Probable | As nectar-tap_saprotroph (Põlme et al. 2020); Decay sugar-rich substrates (Põlme et al. 2020) | Põlme S, et al. 2020. Fungal Diversity 105:1-16 ((https://doi.org/10.1007/s13225-020-00466-2)) |
| Otu48927 | 1 | k__Fungi(100);p__Ascomycota(100);c__Saccharomycetes(100);o__Saccharomycetales(100);f__Metschnikowiaceae(100);g__Metschnikowia(100);s__Metschnikowia_pulcherrima(100); | Metschnikowia | 13 | Saprotroph | \|Undefined Saprotroph\| | Yeast | NULL | Probable | As nectar-tap_saprotroph (Põlme et al. 2020); Decay sugar-rich substrates (Põlme et al. 2020) | Põlme S, et al. 2020. Fungal Diversity 105:1-16 ((https://doi.org/10.1007/s13225-020-00466-2)) |
| Otu48928 | 1 | k__Fungi(100);p__Ascomycota(100);c__Saccharomycetes(100);o__Saccharomycetales(100);f__Metschnikowiaceae(100);g__Metschnikowia(100);s__Metschnikowia_pulcherrima(100); | Metschnikowia | 13 | Saprotroph | \|Undefined Saprotroph\| | Yeast | NULL | Probable | As nectar-tap_saprotroph (Põlme et al. 2020); Decay sugar-rich substrates (Põlme et al. 2020) | Põlme S, et al. 2020. Fungal Diversity 105:1-16 ((https://doi.org/10.1007/s13225-020-00466-2)) |
| Otu48929 | 1 | k__Fungi(100);p__Ascomycota(100);c__Saccharomycetes(100);o__Saccharomycetales(100);f__Metschnikowiaceae(100);g__Metschnikowia(100);s__Metschnikowia_pulcherrima(100); | Metschnikowia | 13 | Saprotroph | \|Undefined Saprotroph\| | Yeast | NULL | Probable | As nectar-tap_saprotroph (Põlme et al. 2020); Decay sugar-rich substrates (Põlme et al. 2020) | Põlme S, et al. 2020. Fungal Diversity 105:1-16 ((https://doi.org/10.1007/s13225-020-00466-2)) |
| Otu48930 | 1 | k__Fungi(100);p__Ascomycota(100);c__Saccharomycetes(100);o__Saccharomycetales(100);f__Metschnikowiaceae(100);g__Metschnikowia(100);s__Metschnikowia_pulcherrima(100); | Metschnikowia | 13 | Saprotroph | \|Undefined Saprotroph\| | Yeast | NULL | Probable | As nectar-tap_saprotroph (Põlme et al. 2020); Decay sugar-rich substrates (Põlme et al. 2020) | Põlme S, et al. 2020. Fungal Diversity 105:1-16 ((https://doi.org/10.1007/s13225-020-00466-2)) |
| Otu48931 | 1 | k__Fungi(100);p__Ascomycota(100);c__Saccharomycetes(100);o__Saccharomycetales(100);f__Metschnikowiaceae(100);g__Metschnikowia(100);g__Metschnikowia_unclassified(100); | Metschnikowia | 13 | Saprotroph | \|Undefined Saprotroph\| | Yeast | NULL | Probable | As nectar-tap_saprotroph (Põlme et al. 2020); Decay sugar-rich substrates (Põlme et al. 2020) | Põlme S, et al. 2020. Fungal Diversity 105:1-16 ((https://doi.org/10.1007/s13225-020-00466-2)) |
| Otu48932 | 1 | k__Fungi(100);p__Ascomycota(100);c__Saccharomycetes(100);o__Saccharomycetales(100);f__Metschnikowiaceae(100);g__Metschnikowia(100);s__Metschnikowia_pulcherrima(100); | Metschnikowia | 13 | Saprotroph | \|Undefined Saprotroph\| | Yeast | NULL | Probable | As nectar-tap_saprotroph (Põlme et al. 2020); Decay sugar-rich substrates (Põlme et al. 2020) | Põlme S, et al. 2020. Fungal Diversity 105:1-16 ((https://doi.org/10.1007/s13225-020-00466-2)) |
| Otu48933 | 1 | k__Fungi(100);p__Ascomycota(100);c__Saccharomycetes(100);o__Saccharomycetales(100);f__Metschnikowiaceae(100);g__Metschnikowia(100);s__Metschnikowia_pulcherrima(100); | Metschnikowia | 13 | Saprotroph | \|Undefined Saprotroph\| | Yeast | NULL | Probable | As nectar-tap_saprotroph (Põlme et al. 2020); Decay sugar-rich substrates (Põlme et al. 2020) | Põlme S, et al. 2020. Fungal Diversity 105:1-16 ((https://doi.org/10.1007/s13225-020-00466-2)) |
| Otu48934 | 1 | k__Fungi(100);p__Ascomycota(100);c__Saccharomycetes(100);o__Saccharomycetales(100);f__Metschnikowiaceae(100);g__Metschnikowia(100);s__Metschnikowia_pulcherrima(100); | Metschnikowia | 13 | Saprotroph | \|Undefined Saprotroph\| | Yeast | NULL | Probable | As nectar-tap_saprotroph (Põlme et al. 2020); Decay sugar-rich substrates (Põlme et al. 2020) | Põlme S, et al. 2020. Fungal Diversity 105:1-16 ((https://doi.org/10.1007/s13225-020-00466-2)) |
| Otu48935 | 1 | k__Fungi(100);p__Ascomycota(100);c__Saccharomycetes(100);o__Saccharomycetales(100);f__Metschnikowiaceae(100);g__Metschnikowia(100);s__Metschnikowia_pulcherrima(100); | Metschnikowia | 13 | Saprotroph | \|Undefined Saprotroph\| | Yeast | NULL | Probable | As nectar-tap_saprotroph (Põlme et al. 2020); Decay sugar-rich substrates (Põlme et al. 2020) | Põlme S, et al. 2020. Fungal Diversity 105:1-16 ((https://doi.org/10.1007/s13225-020-00466-2)) |
| Otu48936 | 1 | k__Fungi(100);p__Ascomycota(100);c__Saccharomycetes(100);o__Saccharomycetales(100);f__Metschnikowiaceae(100);g__Metschnikowia(100);s__Metschnikowia_pulcherrima(100); | Metschnikowia | 13 | Saprotroph | \|Undefined Saprotroph\| | Yeast | NULL | Probable | As nectar-tap_saprotroph (Põlme et al. 2020); Decay sugar-rich substrates (Põlme et al. 2020) | Põlme S, et al. 2020. Fungal Diversity 105:1-16 ((https://doi.org/10.1007/s13225-020-00466-2)) |
| Otu48938 | 1 | k__Fungi(100);p__Ascomycota(100);c__Saccharomycetes(100);o__Saccharomycetales(100);f__Metschnikowiaceae(100);g__Metschnikowia(100);g__Metschnikowia_unclassified(100); | Metschnikowia | 13 | Saprotroph | \|Undefined Saprotroph\| | Yeast | NULL | Probable | As nectar-tap_saprotroph (Põlme et al. 2020); Decay sugar-rich substrates (Põlme et al. 2020) | Põlme S, et al. 2020. Fungal Diversity 105:1-16 ((https://doi.org/10.1007/s13225-020-00466-2)) |
| Otu48940 | 1 | k__Fungi(100);p__Ascomycota(100);c__Saccharomycetes(100);o__Saccharomycetales(100);f__Metschnikowiaceae(100);g__Metschnikowia(100);g__Metschnikowia_unclassified(100); | Metschnikowia | 13 | Saprotroph | \|Undefined Saprotroph\| | Yeast | NULL | Probable | As nectar-tap_saprotroph (Põlme et al. 2020); Decay sugar-rich substrates (Põlme et al. 2020) | Põlme S, et al. 2020. Fungal Diversity 105:1-16 ((https://doi.org/10.1007/s13225-020-00466-2)) |
| Otu48941 | 1 | k__Fungi(100);p__Ascomycota(100);c__Saccharomycetes(100);o__Saccharomycetales(100);f__Metschnikowiaceae(100);g__Metschnikowia(100);g__Metschnikowia_unclassified(100); | Metschnikowia | 13 | Saprotroph | \|Undefined Saprotroph\| | Yeast | NULL | Probable | As nectar-tap_saprotroph (Põlme et al. 2020); Decay sugar-rich substrates (Põlme et al. 2020) | Põlme S, et al. 2020. Fungal Diversity 105:1-16 ((https://doi.org/10.1007/s13225-020-00466-2)) |
| Otu48942 | 1 | k__Fungi(100);p__Ascomycota(100);c__Saccharomycetes(100);o__Saccharomycetales(100);f__Metschnikowiaceae(100);g__Metschnikowia(100);s__Metschnikowia_pulcherrima(100); | Metschnikowia | 13 | Saprotroph | \|Undefined Saprotroph\| | Yeast | NULL | Probable | As nectar-tap_saprotroph (Põlme et al. 2020); Decay sugar-rich substrates (Põlme et al. 2020) | Põlme S, et al. 2020. Fungal Diversity 105:1-16 ((https://doi.org/10.1007/s13225-020-00466-2)) |
| Otu48943 | 1 | k__Fungi(100);p__Ascomycota(100);c__Saccharomycetes(100);o__Saccharomycetales(100);f__Metschnikowiaceae(100);g__Metschnikowia(100);s__Metschnikowia_pulcherrima(100); | Metschnikowia | 13 | Saprotroph | \|Undefined Saprotroph\| | Yeast | NULL | Probable | As nectar-tap_saprotroph (Põlme et al. 2020); Decay sugar-rich substrates (Põlme et al. 2020) | Põlme S, et al. 2020. Fungal Diversity 105:1-16 ((https://doi.org/10.1007/s13225-020-00466-2)) |
| Otu48944 | 1 | k__Fungi(100);p__Ascomycota(100);c__Saccharomycetes(100);o__Saccharomycetales(100);f__Metschnikowiaceae(100);g__Metschnikowia(100);s__Metschnikowia_pulcherrima(100); | Metschnikowia | 13 | Saprotroph | \|Undefined Saprotroph\| | Yeast | NULL | Probable | As nectar-tap_saprotroph (Põlme et al. 2020); Decay sugar-rich substrates (Põlme et al. 2020) | Põlme S, et al. 2020. Fungal Diversity 105:1-16 ((https://doi.org/10.1007/s13225-020-00466-2)) |
| Otu48945 | 1 | k__Fungi(100);p__Ascomycota(100);c__Saccharomycetes(100);o__Saccharomycetales(100);f__Metschnikowiaceae(100);g__Metschnikowia(100);s__Metschnikowia_pulcherrima(100); | Metschnikowia | 13 | Saprotroph | \|Undefined Saprotroph\| | Yeast | NULL | Probable | As nectar-tap_saprotroph (Põlme et al. 2020); Decay sugar-rich substrates (Põlme et al. 2020) | Põlme S, et al. 2020. Fungal Diversity 105:1-16 ((https://doi.org/10.1007/s13225-020-00466-2)) |
| Otu48946 | 1 | k__Fungi(100);p__Ascomycota(100);c__Saccharomycetes(100);o__Saccharomycetales(100);f__Metschnikowiaceae(100);g__Metschnikowia(100);s__Metschnikowia_pulcherrima(100); | Metschnikowia | 13 | Saprotroph | \|Undefined Saprotroph\| | Yeast | NULL | Probable | As nectar-tap_saprotroph (Põlme et al. 2020); Decay sugar-rich substrates (Põlme et al. 2020) | Põlme S, et al. 2020. Fungal Diversity 105:1-16 ((https://doi.org/10.1007/s13225-020-00466-2)) |
| Otu48947 | 1 | k__Fungi(100);p__Ascomycota(100);c__Saccharomycetes(100);o__Saccharomycetales(100);f__Metschnikowiaceae(100);g__Metschnikowia(100);s__Metschnikowia_pulcherrima(100); | Metschnikowia | 13 | Saprotroph | \|Undefined Saprotroph\| | Yeast | NULL | Probable | As nectar-tap_saprotroph (Põlme et al. 2020); Decay sugar-rich substrates (Põlme et al. 2020) | Põlme S, et al. 2020. Fungal Diversity 105:1-16 ((https://doi.org/10.1007/s13225-020-00466-2)) |
| Otu48948 | 1 | k__Fungi(100);p__Ascomycota(100);c__Saccharomycetes(100);o__Saccharomycetales(100);f__Metschnikowiaceae(100);g__Metschnikowia(100);s__Metschnikowia_pulcherrima(100); | Metschnikowia | 13 | Saprotroph | \|Undefined Saprotroph\| | Yeast | NULL | Probable | As nectar-tap_saprotroph (Põlme et al. 2020); Decay sugar-rich substrates (Põlme et al. 2020) | Põlme S, et al. 2020. Fungal Diversity 105:1-16 ((https://doi.org/10.1007/s13225-020-00466-2)) |
| Otu48949 | 1 | k__Fungi(100);p__Ascomycota(100);c__Saccharomycetes(100);o__Saccharomycetales(100);f__Metschnikowiaceae(100);g__Metschnikowia(100);g__Metschnikowia_unclassified(100); | Metschnikowia | 13 | Saprotroph | \|Undefined Saprotroph\| | Yeast | NULL | Probable | As nectar-tap_saprotroph (Põlme et al. 2020); Decay sugar-rich substrates (Põlme et al. 2020) | Põlme S, et al. 2020. Fungal Diversity 105:1-16 ((https://doi.org/10.1007/s13225-020-00466-2)) |
| Otu48950 | 1 | k__Fungi(100);p__Ascomycota(100);c__Saccharomycetes(100);o__Saccharomycetales(100);f__Metschnikowiaceae(100);g__Metschnikowia(100);s__Metschnikowia_pulcherrima(100); | Metschnikowia | 13 | Saprotroph | \|Undefined Saprotroph\| | Yeast | NULL | Probable | As nectar-tap_saprotroph (Põlme et al. 2020); Decay sugar-rich substrates (Põlme et al. 2020) | Põlme S, et al. 2020. Fungal Diversity 105:1-16 ((https://doi.org/10.1007/s13225-020-00466-2)) |
| Otu48951 | 1 | k__Fungi(100);p__Ascomycota(100);c__Saccharomycetes(100);o__Saccharomycetales(100);f__Metschnikowiaceae(100);g__Metschnikowia(100);s__Metschnikowia_pulcherrima(100); | Metschnikowia | 13 | Saprotroph | \|Undefined Saprotroph\| | Yeast | NULL | Probable | As nectar-tap_saprotroph (Põlme et al. 2020); Decay sugar-rich substrates (Põlme et al. 2020) | Põlme S, et al. 2020. Fungal Diversity 105:1-16 ((https://doi.org/10.1007/s13225-020-00466-2)) |
| Otu48952 | 1 | k__Fungi(100);p__Ascomycota(100);c__Saccharomycetes(100);o__Saccharomycetales(100);f__Metschnikowiaceae(100);g__Metschnikowia(100);s__Metschnikowia_pulcherrima(100); | Metschnikowia | 13 | Saprotroph | \|Undefined Saprotroph\| | Yeast | NULL | Probable | As nectar-tap_saprotroph (Põlme et al. 2020); Decay sugar-rich substrates (Põlme et al. 2020) | Põlme S, et al. 2020. Fungal Diversity 105:1-16 ((https://doi.org/10.1007/s13225-020-00466-2)) |
| Otu48953 | 1 | k__Fungi(100);p__Ascomycota(100);c__Saccharomycetes(100);o__Saccharomycetales(100);f__Metschnikowiaceae(100);g__Metschnikowia(100);s__Metschnikowia_pulcherrima(100); | Metschnikowia | 13 | Saprotroph | \|Undefined Saprotroph\| | Yeast | NULL | Probable | As nectar-tap_saprotroph (Põlme et al. 2020); Decay sugar-rich substrates (Põlme et al. 2020) | Põlme S, et al. 2020. Fungal Diversity 105:1-16 ((https://doi.org/10.1007/s13225-020-00466-2)) |
| Otu48954 | 1 | k__Fungi(100);p__Ascomycota(100);c__Saccharomycetes(100);o__Saccharomycetales(100);f__Metschnikowiaceae(100);g__Metschnikowia(100);s__Metschnikowia_pulcherrima(100); | Metschnikowia | 13 | Saprotroph | \|Undefined Saprotroph\| | Yeast | NULL | Probable | As nectar-tap_saprotroph (Põlme et al. 2020); Decay sugar-rich substrates (Põlme et al. 2020) | Põlme S, et al. 2020. Fungal Diversity 105:1-16 ((https://doi.org/10.1007/s13225-020-00466-2)) |
| Otu48955 | 1 | k__Fungi(100);p__Ascomycota(100);c__Saccharomycetes(100);o__Saccharomycetales(100);f__Metschnikowiaceae(100);g__Metschnikowia(100);s__Metschnikowia_pulcherrima(100); | Metschnikowia | 13 | Saprotroph | \|Undefined Saprotroph\| | Yeast | NULL | Probable | As nectar-tap_saprotroph (Põlme et al. 2020); Decay sugar-rich substrates (Põlme et al. 2020) | Põlme S, et al. 2020. Fungal Diversity 105:1-16 ((https://doi.org/10.1007/s13225-020-00466-2)) |
| Otu48956 | 1 | k__Fungi(100);p__Ascomycota(100);c__Saccharomycetes(100);o__Saccharomycetales(100);f__Metschnikowiaceae(100);g__Metschnikowia(100);s__Metschnikowia_pulcherrima(100); | Metschnikowia | 13 | Saprotroph | \|Undefined Saprotroph\| | Yeast | NULL | Probable | As nectar-tap_saprotroph (Põlme et al. 2020); Decay sugar-rich substrates (Põlme et al. 2020) | Põlme S, et al. 2020. Fungal Diversity 105:1-16 ((https://doi.org/10.1007/s13225-020-00466-2)) |
| Otu48957 | 1 | k__Fungi(100);p__Ascomycota(100);c__Saccharomycetes(100);o__Saccharomycetales(100);f__Metschnikowiaceae(100);g__Metschnikowia(100);g__Metschnikowia_unclassified(100); | Metschnikowia | 13 | Saprotroph | \|Undefined Saprotroph\| | Yeast | NULL | Probable | As nectar-tap_saprotroph (Põlme et al. 2020); Decay sugar-rich substrates (Põlme et al. 2020) | Põlme S, et al. 2020. Fungal Diversity 105:1-16 ((https://doi.org/10.1007/s13225-020-00466-2)) |
| Otu48958 | 1 | k__Fungi(100);p__Ascomycota(100);c__Saccharomycetes(100);o__Saccharomycetales(100);f__Metschnikowiaceae(100);g__Metschnikowia(100);s__Metschnikowia_pulcherrima(100); | Metschnikowia | 13 | Saprotroph | \|Undefined Saprotroph\| | Yeast | NULL | Probable | As nectar-tap_saprotroph (Põlme et al. 2020); Decay sugar-rich substrates (Põlme et al. 2020) | Põlme S, et al. 2020. Fungal Diversity 105:1-16 ((https://doi.org/10.1007/s13225-020-00466-2)) |
| Otu48961 | 1 | k__Fungi(100);p__Ascomycota(100);c__Saccharomycetes(100);o__Saccharomycetales(100);f__Metschnikowiaceae(100);g__Metschnikowia(100);g__Metschnikowia_unclassified(100); | Metschnikowia | 13 | Saprotroph | \|Undefined Saprotroph\| | Yeast | NULL | Probable | As nectar-tap_saprotroph (Põlme et al. 2020); Decay sugar-rich substrates (Põlme et al. 2020) | Põlme S, et al. 2020. Fungal Diversity 105:1-16 ((https://doi.org/10.1007/s13225-020-00466-2)) |
| Otu48962 | 1 | k__Fungi(100);p__Ascomycota(100);c__Saccharomycetes(100);o__Saccharomycetales(100);f__Metschnikowiaceae(100);g__Metschnikowia(100);s__Metschnikowia_pulcherrima(100); | Metschnikowia | 13 | Saprotroph | \|Undefined Saprotroph\| | Yeast | NULL | Probable | As nectar-tap_saprotroph (Põlme et al. 2020); Decay sugar-rich substrates (Põlme et al. 2020) | Põlme S, et al. 2020. Fungal Diversity 105:1-16 ((https://doi.org/10.1007/s13225-020-00466-2)) |
| Otu48963 | 1 | k__Fungi(100);p__Ascomycota(100);c__Saccharomycetes(100);o__Saccharomycetales(100);f__Metschnikowiaceae(100);g__Metschnikowia(100);s__Metschnikowia_pulcherrima(100); | Metschnikowia | 13 | Saprotroph | \|Undefined Saprotroph\| | Yeast | NULL | Probable | As nectar-tap_saprotroph (Põlme et al. 2020); Decay sugar-rich substrates (Põlme et al. 2020) | Põlme S, et al. 2020. Fungal Diversity 105:1-16 ((https://doi.org/10.1007/s13225-020-00466-2)) |
| Otu48965 | 1 | k__Fungi(100);p__Ascomycota(100);c__Saccharomycetes(100);o__Saccharomycetales(100);f__Metschnikowiaceae(100);g__Metschnikowia(100);s__Metschnikowia_pulcherrima(100); | Metschnikowia | 13 | Saprotroph | \|Undefined Saprotroph\| | Yeast | NULL | Probable | As nectar-tap_saprotroph (Põlme et al. 2020); Decay sugar-rich substrates (Põlme et al. 2020) | Põlme S, et al. 2020. Fungal Diversity 105:1-16 ((https://doi.org/10.1007/s13225-020-00466-2)) |
| Otu48966 | 1 | k__Fungi(100);p__Ascomycota(100);c__Saccharomycetes(100);o__Saccharomycetales(100);f__Metschnikowiaceae(100);g__Metschnikowia(100);s__Metschnikowia_pulcherrima(100); | Metschnikowia | 13 | Saprotroph | \|Undefined Saprotroph\| | Yeast | NULL | Probable | As nectar-tap_saprotroph (Põlme et al. 2020); Decay sugar-rich substrates (Põlme et al. 2020) | Põlme S, et al. 2020. Fungal Diversity 105:1-16 ((https://doi.org/10.1007/s13225-020-00466-2)) |
| Otu48967 | 1 | k__Fungi(100);p__Ascomycota(100);c__Saccharomycetes(100);o__Saccharomycetales(100);f__Metschnikowiaceae(100);g__Metschnikowia(100);s__Metschnikowia_pulcherrima(100); | Metschnikowia | 13 | Saprotroph | \|Undefined Saprotroph\| | Yeast | NULL | Probable | As nectar-tap_saprotroph (Põlme et al. 2020); Decay sugar-rich substrates (Põlme et al. 2020) | Põlme S, et al. 2020. Fungal Diversity 105:1-16 ((https://doi.org/10.1007/s13225-020-00466-2)) |
| Otu48968 | 1 | k__Fungi(100);p__Ascomycota(100);c__Saccharomycetes(100);o__Saccharomycetales(100);f__Metschnikowiaceae(100);g__Metschnikowia(100);s__Metschnikowia_pulcherrima(100); | Metschnikowia | 13 | Saprotroph | \|Undefined Saprotroph\| | Yeast | NULL | Probable | As nectar-tap_saprotroph (Põlme et al. 2020); Decay sugar-rich substrates (Põlme et al. 2020) | Põlme S, et al. 2020. Fungal Diversity 105:1-16 ((https://doi.org/10.1007/s13225-020-00466-2)) |
| Otu48969 | 1 | k__Fungi(100);p__Ascomycota(100);c__Saccharomycetes(100);o__Saccharomycetales(100);f__Metschnikowiaceae(100);g__Metschnikowia(100);s__Metschnikowia_pulcherrima(100); | Metschnikowia | 13 | Saprotroph | \|Undefined Saprotroph\| | Yeast | NULL | Probable | As nectar-tap_saprotroph (Põlme et al. 2020); Decay sugar-rich substrates (Põlme et al. 2020) | Põlme S, et al. 2020. Fungal Diversity 105:1-16 ((https://doi.org/10.1007/s13225-020-00466-2)) |
| Otu48970 | 1 | k__Fungi(100);p__Ascomycota(100);c__Saccharomycetes(100);o__Saccharomycetales(100);f__Metschnikowiaceae(100);g__Metschnikowia(100);s__Metschnikowia_pulcherrima(100); | Metschnikowia | 13 | Saprotroph | \|Undefined Saprotroph\| | Yeast | NULL | Probable | As nectar-tap_saprotroph (Põlme et al. 2020); Decay sugar-rich substrates (Põlme et al. 2020) | Põlme S, et al. 2020. Fungal Diversity 105:1-16 ((https://doi.org/10.1007/s13225-020-00466-2)) |
| Otu48971 | 1 | k__Fungi(100);p__Ascomycota(100);c__Saccharomycetes(100);o__Saccharomycetales(100);f__Metschnikowiaceae(100);g__Metschnikowia(100);s__Metschnikowia_pulcherrima(100); | Metschnikowia | 13 | Saprotroph | \|Undefined Saprotroph\| | Yeast | NULL | Probable | As nectar-tap_saprotroph (Põlme et al. 2020); Decay sugar-rich substrates (Põlme et al. 2020) | Põlme S, et al. 2020. Fungal Diversity 105:1-16 ((https://doi.org/10.1007/s13225-020-00466-2)) |
| Otu48972 | 1 | k__Fungi(100);p__Ascomycota(100);c__Saccharomycetes(100);o__Saccharomycetales(100);f__Metschnikowiaceae(100);g__Metschnikowia(100);s__Metschnikowia_pulcherrima(100); | Metschnikowia | 13 | Saprotroph | \|Undefined Saprotroph\| | Yeast | NULL | Probable | As nectar-tap_saprotroph (Põlme et al. 2020); Decay sugar-rich substrates (Põlme et al. 2020) | Põlme S, et al. 2020. Fungal Diversity 105:1-16 ((https://doi.org/10.1007/s13225-020-00466-2)) |
| Otu48973 | 1 | k__Fungi(100);p__Ascomycota(100);c__Saccharomycetes(100);o__Saccharomycetales(100);f__Metschnikowiaceae(100);g__Metschnikowia(100);g__Metschnikowia_unclassified(100); | Metschnikowia | 13 | Saprotroph | \|Undefined Saprotroph\| | Yeast | NULL | Probable | As nectar-tap_saprotroph (Põlme et al. 2020); Decay sugar-rich substrates (Põlme et al. 2020) | Põlme S, et al. 2020. Fungal Diversity 105:1-16 ((https://doi.org/10.1007/s13225-020-00466-2)) |
| Otu48974 | 1 | k__Fungi(100);p__Ascomycota(100);c__Saccharomycetes(100);o__Saccharomycetales(100);f__Metschnikowiaceae(100);g__Metschnikowia(100);s__Metschnikowia_pulcherrima(100); | Metschnikowia | 13 | Saprotroph | \|Undefined Saprotroph\| | Yeast | NULL | Probable | As nectar-tap_saprotroph (Põlme et al. 2020); Decay sugar-rich substrates (Põlme et al. 2020) | Põlme S, et al. 2020. Fungal Diversity 105:1-16 ((https://doi.org/10.1007/s13225-020-00466-2)) |
| Otu48975 | 1 | k__Fungi(100);p__Ascomycota(100);c__Saccharomycetes(100);o__Saccharomycetales(100);f__Metschnikowiaceae(100);g__Metschnikowia(100);s__Metschnikowia_pulcherrima(100); | Metschnikowia | 13 | Saprotroph | \|Undefined Saprotroph\| | Yeast | NULL | Probable | As nectar-tap_saprotroph (Põlme et al. 2020); Decay sugar-rich substrates (Põlme et al. 2020) | Põlme S, et al. 2020. Fungal Diversity 105:1-16 ((https://doi.org/10.1007/s13225-020-00466-2)) |
| Otu48977 | 1 | k__Fungi(100);p__Ascomycota(100);c__Saccharomycetes(100);o__Saccharomycetales(100);f__Metschnikowiaceae(100);g__Metschnikowia(100);s__Metschnikowia_pulcherrima(100); | Metschnikowia | 13 | Saprotroph | \|Undefined Saprotroph\| | Yeast | NULL | Probable | As nectar-tap_saprotroph (Põlme et al. 2020); Decay sugar-rich substrates (Põlme et al. 2020) | Põlme S, et al. 2020. Fungal Diversity 105:1-16 ((https://doi.org/10.1007/s13225-020-00466-2)) |
| Otu48978 | 1 | k__Fungi(100);p__Ascomycota(100);c__Saccharomycetes(100);o__Saccharomycetales(100);f__Metschnikowiaceae(100);g__Metschnikowia(100);s__Metschnikowia_pulcherrima(100); | Metschnikowia | 13 | Saprotroph | \|Undefined Saprotroph\| | Yeast | NULL | Probable | As nectar-tap_saprotroph (Põlme et al. 2020); Decay sugar-rich substrates (Põlme et al. 2020) | Põlme S, et al. 2020. Fungal Diversity 105:1-16 ((https://doi.org/10.1007/s13225-020-00466-2)) |
| Otu48979 | 1 | k__Fungi(100);p__Ascomycota(100);c__Saccharomycetes(100);o__Saccharomycetales(100);f__Metschnikowiaceae(100);g__Metschnikowia(100);s__Metschnikowia_pulcherrima(100); | Metschnikowia | 13 | Saprotroph | \|Undefined Saprotroph\| | Yeast | NULL | Probable | As nectar-tap_saprotroph (Põlme et al. 2020); Decay sugar-rich substrates (Põlme et al. 2020) | Põlme S, et al. 2020. Fungal Diversity 105:1-16 ((https://doi.org/10.1007/s13225-020-00466-2)) |
| Otu48980 | 1 | k__Fungi(100);p__Ascomycota(100);c__Saccharomycetes(100);o__Saccharomycetales(100);f__Metschnikowiaceae(100);g__Metschnikowia(100);s__Metschnikowia_pulcherrima(100); | Metschnikowia | 13 | Saprotroph | \|Undefined Saprotroph\| | Yeast | NULL | Probable | As nectar-tap_saprotroph (Põlme et al. 2020); Decay sugar-rich substrates (Põlme et al. 2020) | Põlme S, et al. 2020. Fungal Diversity 105:1-16 ((https://doi.org/10.1007/s13225-020-00466-2)) |
| Otu48981 | 1 | k__Fungi(100);p__Ascomycota(100);c__Saccharomycetes(100);o__Saccharomycetales(100);f__Metschnikowiaceae(100);g__Metschnikowia(100);s__Metschnikowia_pulcherrima(100); | Metschnikowia | 13 | Saprotroph | \|Undefined Saprotroph\| | Yeast | NULL | Probable | As nectar-tap_saprotroph (Põlme et al. 2020); Decay sugar-rich substrates (Põlme et al. 2020) | Põlme S, et al. 2020. Fungal Diversity 105:1-16 ((https://doi.org/10.1007/s13225-020-00466-2)) |
| Otu48982 | 1 | k__Fungi(100);p__Ascomycota(100);c__Saccharomycetes(100);o__Saccharomycetales(100);f__Metschnikowiaceae(100);g__Metschnikowia(100);s__Metschnikowia_pulcherrima(100); | Metschnikowia | 13 | Saprotroph | \|Undefined Saprotroph\| | Yeast | NULL | Probable | As nectar-tap_saprotroph (Põlme et al. 2020); Decay sugar-rich substrates (Põlme et al. 2020) | Põlme S, et al. 2020. Fungal Diversity 105:1-16 ((https://doi.org/10.1007/s13225-020-00466-2)) |
| Otu48983 | 1 | k__Fungi(100);p__Ascomycota(100);c__Saccharomycetes(100);o__Saccharomycetales(100);f__Metschnikowiaceae(100);g__Metschnikowia(100);s__Metschnikowia_pulcherrima(100); | Metschnikowia | 13 | Saprotroph | \|Undefined Saprotroph\| | Yeast | NULL | Probable | As nectar-tap_saprotroph (Põlme et al. 2020); Decay sugar-rich substrates (Põlme et al. 2020) | Põlme S, et al. 2020. Fungal Diversity 105:1-16 ((https://doi.org/10.1007/s13225-020-00466-2)) |
| Otu48984 | 1 | k__Fungi(100);p__Ascomycota(100);c__Saccharomycetes(100);o__Saccharomycetales(100);f__Metschnikowiaceae(100);g__Metschnikowia(100);s__Metschnikowia_pulcherrima(100); | Metschnikowia | 13 | Saprotroph | \|Undefined Saprotroph\| | Yeast | NULL | Probable | As nectar-tap_saprotroph (Põlme et al. 2020); Decay sugar-rich substrates (Põlme et al. 2020) | Põlme S, et al. 2020. Fungal Diversity 105:1-16 ((https://doi.org/10.1007/s13225-020-00466-2)) |
| Otu48985 | 1 | k__Fungi(100);p__Ascomycota(100);c__Saccharomycetes(100);o__Saccharomycetales(100);f__Metschnikowiaceae(100);g__Metschnikowia(100);s__Metschnikowia_pulcherrima(100); | Metschnikowia | 13 | Saprotroph | \|Undefined Saprotroph\| | Yeast | NULL | Probable | As nectar-tap_saprotroph (Põlme et al. 2020); Decay sugar-rich substrates (Põlme et al. 2020) | Põlme S, et al. 2020. Fungal Diversity 105:1-16 ((https://doi.org/10.1007/s13225-020-00466-2)) |
| Otu48986 | 1 | k__Fungi(100);p__Ascomycota(100);c__Saccharomycetes(100);o__Saccharomycetales(100);f__Metschnikowiaceae(100);g__Metschnikowia(100);g__Metschnikowia_unclassified(100); | Metschnikowia | 13 | Saprotroph | \|Undefined Saprotroph\| | Yeast | NULL | Probable | As nectar-tap_saprotroph (Põlme et al. 2020); Decay sugar-rich substrates (Põlme et al. 2020) | Põlme S, et al. 2020. Fungal Diversity 105:1-16 ((https://doi.org/10.1007/s13225-020-00466-2)) |
| Otu48987 | 1 | k__Fungi(100);p__Ascomycota(100);c__Saccharomycetes(100);o__Saccharomycetales(100);f__Metschnikowiaceae(100);g__Metschnikowia(100);s__Metschnikowia_pulcherrima(100); | Metschnikowia | 13 | Saprotroph | \|Undefined Saprotroph\| | Yeast | NULL | Probable | As nectar-tap_saprotroph (Põlme et al. 2020); Decay sugar-rich substrates (Põlme et al. 2020) | Põlme S, et al. 2020. Fungal Diversity 105:1-16 ((https://doi.org/10.1007/s13225-020-00466-2)) |
| Otu48989 | 1 | k__Fungi(100);p__Ascomycota(100);c__Saccharomycetes(100);o__Saccharomycetales(100);f__Metschnikowiaceae(100);g__Metschnikowia(100);s__Metschnikowia_pulcherrima(100); | Metschnikowia | 13 | Saprotroph | \|Undefined Saprotroph\| | Yeast | NULL | Probable | As nectar-tap_saprotroph (Põlme et al. 2020); Decay sugar-rich substrates (Põlme et al. 2020) | Põlme S, et al. 2020. Fungal Diversity 105:1-16 ((https://doi.org/10.1007/s13225-020-00466-2)) |
| Otu48990 | 1 | k__Fungi(100);p__Ascomycota(100);c__Saccharomycetes(100);o__Saccharomycetales(100);f__Metschnikowiaceae(100);g__Metschnikowia(100);s__Metschnikowia_pulcherrima(100); | Metschnikowia | 13 | Saprotroph | \|Undefined Saprotroph\| | Yeast | NULL | Probable | As nectar-tap_saprotroph (Põlme et al. 2020); Decay sugar-rich substrates (Põlme et al. 2020) | Põlme S, et al. 2020. Fungal Diversity 105:1-16 ((https://doi.org/10.1007/s13225-020-00466-2)) |
| Otu48991 | 1 | k__Fungi(100);p__Ascomycota(100);c__Saccharomycetes(100);o__Saccharomycetales(100);f__Metschnikowiaceae(100);g__Metschnikowia(100);s__Metschnikowia_pulcherrima(100); | Metschnikowia | 13 | Saprotroph | \|Undefined Saprotroph\| | Yeast | NULL | Probable | As nectar-tap_saprotroph (Põlme et al. 2020); Decay sugar-rich substrates (Põlme et al. 2020) | Põlme S, et al. 2020. Fungal Diversity 105:1-16 ((https://doi.org/10.1007/s13225-020-00466-2)) |
| Otu48992 | 1 | k__Fungi(100);p__Ascomycota(100);c__Saccharomycetes(100);o__Saccharomycetales(100);f__Metschnikowiaceae(100);g__Metschnikowia(100);s__Metschnikowia_pulcherrima(100); | Metschnikowia | 13 | Saprotroph | \|Undefined Saprotroph\| | Yeast | NULL | Probable | As nectar-tap_saprotroph (Põlme et al. 2020); Decay sugar-rich substrates (Põlme et al. 2020) | Põlme S, et al. 2020. Fungal Diversity 105:1-16 ((https://doi.org/10.1007/s13225-020-00466-2)) |
| Otu48993 | 1 | k__Fungi(100);p__Ascomycota(100);c__Saccharomycetes(100);o__Saccharomycetales(100);f__Metschnikowiaceae(100);g__Metschnikowia(100);s__Metschnikowia_pulcherrima(100); | Metschnikowia | 13 | Saprotroph | \|Undefined Saprotroph\| | Yeast | NULL | Probable | As nectar-tap_saprotroph (Põlme et al. 2020); Decay sugar-rich substrates (Põlme et al. 2020) | Põlme S, et al. 2020. Fungal Diversity 105:1-16 ((https://doi.org/10.1007/s13225-020-00466-2)) |
| Otu48994 | 1 | k__Fungi(100);p__Ascomycota(100);c__Saccharomycetes(100);o__Saccharomycetales(100);f__Metschnikowiaceae(100);g__Metschnikowia(100);s__Metschnikowia_pulcherrima(100); | Metschnikowia | 13 | Saprotroph | \|Undefined Saprotroph\| | Yeast | NULL | Probable | As nectar-tap_saprotroph (Põlme et al. 2020); Decay sugar-rich substrates (Põlme et al. 2020) | Põlme S, et al. 2020. Fungal Diversity 105:1-16 ((https://doi.org/10.1007/s13225-020-00466-2)) |
| Otu48995 | 1 | k__Fungi(100);p__Ascomycota(100);c__Saccharomycetes(100);o__Saccharomycetales(100);f__Metschnikowiaceae(100);g__Metschnikowia(100);s__Metschnikowia_pulcherrima(100); | Metschnikowia | 13 | Saprotroph | \|Undefined Saprotroph\| | Yeast | NULL | Probable | As nectar-tap_saprotroph (Põlme et al. 2020); Decay sugar-rich substrates (Põlme et al. 2020) | Põlme S, et al. 2020. Fungal Diversity 105:1-16 ((https://doi.org/10.1007/s13225-020-00466-2)) |
| Otu48996 | 1 | k__Fungi(100);p__Ascomycota(100);c__Saccharomycetes(100);o__Saccharomycetales(100);f__Metschnikowiaceae(100);g__Metschnikowia(100);s__Metschnikowia_pulcherrima(100); | Metschnikowia | 13 | Saprotroph | \|Undefined Saprotroph\| | Yeast | NULL | Probable | As nectar-tap_saprotroph (Põlme et al. 2020); Decay sugar-rich substrates (Põlme et al. 2020) | Põlme S, et al. 2020. Fungal Diversity 105:1-16 ((https://doi.org/10.1007/s13225-020-00466-2)) |
| Otu48997 | 1 | k__Fungi(100);p__Ascomycota(100);c__Saccharomycetes(100);o__Saccharomycetales(100);f__Metschnikowiaceae(100);g__Metschnikowia(100);g__Metschnikowia_unclassified(100); | Metschnikowia | 13 | Saprotroph | \|Undefined Saprotroph\| | Yeast | NULL | Probable | As nectar-tap_saprotroph (Põlme et al. 2020); Decay sugar-rich substrates (Põlme et al. 2020) | Põlme S, et al. 2020. Fungal Diversity 105:1-16 ((https://doi.org/10.1007/s13225-020-00466-2)) |
| Otu48998 | 1 | k__Fungi(100);p__Ascomycota(100);c__Saccharomycetes(100);o__Saccharomycetales(100);f__Metschnikowiaceae(100);g__Metschnikowia(100);s__Metschnikowia_pulcherrima(100); | Metschnikowia | 13 | Saprotroph | \|Undefined Saprotroph\| | Yeast | NULL | Probable | As nectar-tap_saprotroph (Põlme et al. 2020); Decay sugar-rich substrates (Põlme et al. 2020) | Põlme S, et al. 2020. Fungal Diversity 105:1-16 ((https://doi.org/10.1007/s13225-020-00466-2)) |
| Otu48999 | 1 | k__Fungi(100);p__Ascomycota(100);c__Saccharomycetes(100);o__Saccharomycetales(100);f__Metschnikowiaceae(100);g__Metschnikowia(100);s__Metschnikowia_pulcherrima(100); | Metschnikowia | 13 | Saprotroph | \|Undefined Saprotroph\| | Yeast | NULL | Probable | As nectar-tap_saprotroph (Põlme et al. 2020); Decay sugar-rich substrates (Põlme et al. 2020) | Põlme S, et al. 2020. Fungal Diversity 105:1-16 ((https://doi.org/10.1007/s13225-020-00466-2)) |
| Otu49000 | 1 | k__Fungi(100);p__Ascomycota(100);c__Saccharomycetes(100);o__Saccharomycetales(100);f__Metschnikowiaceae(100);g__Metschnikowia(100);g__Metschnikowia_unclassified(100); | Metschnikowia | 13 | Saprotroph | \|Undefined Saprotroph\| | Yeast | NULL | Probable | As nectar-tap_saprotroph (Põlme et al. 2020); Decay sugar-rich substrates (Põlme et al. 2020) | Põlme S, et al. 2020. Fungal Diversity 105:1-16 ((https://doi.org/10.1007/s13225-020-00466-2)) |
| Otu49001 | 1 | k__Fungi(100);p__Ascomycota(100);c__Saccharomycetes(100);o__Saccharomycetales(100);f__Metschnikowiaceae(100);g__Metschnikowia(100);s__Metschnikowia_pulcherrima(100); | Metschnikowia | 13 | Saprotroph | \|Undefined Saprotroph\| | Yeast | NULL | Probable | As nectar-tap_saprotroph (Põlme et al. 2020); Decay sugar-rich substrates (Põlme et al. 2020) | Põlme S, et al. 2020. Fungal Diversity 105:1-16 ((https://doi.org/10.1007/s13225-020-00466-2)) |
| Otu49002 | 1 | k__Fungi(100);p__Ascomycota(100);c__Saccharomycetes(100);o__Saccharomycetales(100);f__Metschnikowiaceae(100);g__Metschnikowia(100);g__Metschnikowia_unclassified(100); | Metschnikowia | 13 | Saprotroph | \|Undefined Saprotroph\| | Yeast | NULL | Probable | As nectar-tap_saprotroph (Põlme et al. 2020); Decay sugar-rich substrates (Põlme et al. 2020) | Põlme S, et al. 2020. Fungal Diversity 105:1-16 ((https://doi.org/10.1007/s13225-020-00466-2)) |
| Otu49003 | 1 | k__Fungi(100);p__Ascomycota(100);c__Saccharomycetes(100);o__Saccharomycetales(100);f__Metschnikowiaceae(100);g__Metschnikowia(100);s__Metschnikowia_pulcherrima(100); | Metschnikowia | 13 | Saprotroph | \|Undefined Saprotroph\| | Yeast | NULL | Probable | As nectar-tap_saprotroph (Põlme et al. 2020); Decay sugar-rich substrates (Põlme et al. 2020) | Põlme S, et al. 2020. Fungal Diversity 105:1-16 ((https://doi.org/10.1007/s13225-020-00466-2)) |
| Otu49004 | 1 | k__Fungi(100);p__Ascomycota(100);c__Saccharomycetes(100);o__Saccharomycetales(100);f__Metschnikowiaceae(100);g__Metschnikowia(100);s__Metschnikowia_pulcherrima(100); | Metschnikowia | 13 | Saprotroph | \|Undefined Saprotroph\| | Yeast | NULL | Probable | As nectar-tap_saprotroph (Põlme et al. 2020); Decay sugar-rich substrates (Põlme et al. 2020) | Põlme S, et al. 2020. Fungal Diversity 105:1-16 ((https://doi.org/10.1007/s13225-020-00466-2)) |
| Otu49005 | 1 | k__Fungi(100);p__Ascomycota(100);c__Saccharomycetes(100);o__Saccharomycetales(100);f__Metschnikowiaceae(100);g__Metschnikowia(100);s__Metschnikowia_pulcherrima(100); | Metschnikowia | 13 | Saprotroph | \|Undefined Saprotroph\| | Yeast | NULL | Probable | As nectar-tap_saprotroph (Põlme et al. 2020); Decay sugar-rich substrates (Põlme et al. 2020) | Põlme S, et al. 2020. Fungal Diversity 105:1-16 ((https://doi.org/10.1007/s13225-020-00466-2)) |
| Otu49006 | 1 | k__Fungi(100);p__Ascomycota(100);c__Saccharomycetes(100);o__Saccharomycetales(100);f__Metschnikowiaceae(100);g__Metschnikowia(100);g__Metschnikowia_unclassified(100); | Metschnikowia | 13 | Saprotroph | \|Undefined Saprotroph\| | Yeast | NULL | Probable | As nectar-tap_saprotroph (Põlme et al. 2020); Decay sugar-rich substrates (Põlme et al. 2020) | Põlme S, et al. 2020. Fungal Diversity 105:1-16 ((https://doi.org/10.1007/s13225-020-00466-2)) |
| Otu49007 | 1 | k__Fungi(100);p__Ascomycota(100);c__Saccharomycetes(100);o__Saccharomycetales(100);f__Metschnikowiaceae(100);g__Metschnikowia(100);s__Metschnikowia_pulcherrima(100); | Metschnikowia | 13 | Saprotroph | \|Undefined Saprotroph\| | Yeast | NULL | Probable | As nectar-tap_saprotroph (Põlme et al. 2020); Decay sugar-rich substrates (Põlme et al. 2020) | Põlme S, et al. 2020. Fungal Diversity 105:1-16 ((https://doi.org/10.1007/s13225-020-00466-2)) |
| Otu49008 | 1 | k__Fungi(100);p__Ascomycota(100);c__Saccharomycetes(100);o__Saccharomycetales(100);f__Metschnikowiaceae(100);g__Metschnikowia(100);s__Metschnikowia_pulcherrima(100); | Metschnikowia | 13 | Saprotroph | \|Undefined Saprotroph\| | Yeast | NULL | Probable | As nectar-tap_saprotroph (Põlme et al. 2020); Decay sugar-rich substrates (Põlme et al. 2020) | Põlme S, et al. 2020. Fungal Diversity 105:1-16 ((https://doi.org/10.1007/s13225-020-00466-2)) |
| Otu49009 | 1 | k__Fungi(100);p__Ascomycota(100);c__Saccharomycetes(100);o__Saccharomycetales(100);f__Metschnikowiaceae(100);g__Metschnikowia(100);g__Metschnikowia_unclassified(100); | Metschnikowia | 13 | Saprotroph | \|Undefined Saprotroph\| | Yeast | NULL | Probable | As nectar-tap_saprotroph (Põlme et al. 2020); Decay sugar-rich substrates (Põlme et al. 2020) | Põlme S, et al. 2020. Fungal Diversity 105:1-16 ((https://doi.org/10.1007/s13225-020-00466-2)) |
| Otu49010 | 1 | k__Fungi(100);p__Ascomycota(100);c__Saccharomycetes(100);o__Saccharomycetales(100);f__Metschnikowiaceae(100);g__Metschnikowia(100);g__Metschnikowia_unclassified(100); | Metschnikowia | 13 | Saprotroph | \|Undefined Saprotroph\| | Yeast | NULL | Probable | As nectar-tap_saprotroph (Põlme et al. 2020); Decay sugar-rich substrates (Põlme et al. 2020) | Põlme S, et al. 2020. Fungal Diversity 105:1-16 ((https://doi.org/10.1007/s13225-020-00466-2)) |
| Otu49011 | 1 | k__Fungi(100);p__Ascomycota(100);c__Saccharomycetes(100);o__Saccharomycetales(100);f__Metschnikowiaceae(100);g__Metschnikowia(100);s__Metschnikowia_pulcherrima(100); | Metschnikowia | 13 | Saprotroph | \|Undefined Saprotroph\| | Yeast | NULL | Probable | As nectar-tap_saprotroph (Põlme et al. 2020); Decay sugar-rich substrates (Põlme et al. 2020) | Põlme S, et al. 2020. Fungal Diversity 105:1-16 ((https://doi.org/10.1007/s13225-020-00466-2)) |
| Otu49012 | 1 | k__Fungi(100);p__Ascomycota(100);c__Saccharomycetes(100);o__Saccharomycetales(100);f__Metschnikowiaceae(100);g__Metschnikowia(100);s__Metschnikowia_pulcherrima(100); | Metschnikowia | 13 | Saprotroph | \|Undefined Saprotroph\| | Yeast | NULL | Probable | As nectar-tap_saprotroph (Põlme et al. 2020); Decay sugar-rich substrates (Põlme et al. 2020) | Põlme S, et al. 2020. Fungal Diversity 105:1-16 ((https://doi.org/10.1007/s13225-020-00466-2)) |
| Otu49013 | 1 | k__Fungi(100);p__Ascomycota(100);c__Saccharomycetes(100);o__Saccharomycetales(100);f__Metschnikowiaceae(100);g__Metschnikowia(100);s__Metschnikowia_pulcherrima(100); | Metschnikowia | 13 | Saprotroph | \|Undefined Saprotroph\| | Yeast | NULL | Probable | As nectar-tap_saprotroph (Põlme et al. 2020); Decay sugar-rich substrates (Põlme et al. 2020) | Põlme S, et al. 2020. Fungal Diversity 105:1-16 ((https://doi.org/10.1007/s13225-020-00466-2)) |
| Otu49014 | 1 | k__Fungi(100);p__Ascomycota(100);c__Saccharomycetes(100);o__Saccharomycetales(100);f__Metschnikowiaceae(100);g__Metschnikowia(100);s__Metschnikowia_pulcherrima(100); | Metschnikowia | 13 | Saprotroph | \|Undefined Saprotroph\| | Yeast | NULL | Probable | As nectar-tap_saprotroph (Põlme et al. 2020); Decay sugar-rich substrates (Põlme et al. 2020) | Põlme S, et al. 2020. Fungal Diversity 105:1-16 ((https://doi.org/10.1007/s13225-020-00466-2)) |
| Otu49015 | 1 | k__Fungi(100);p__Ascomycota(100);c__Saccharomycetes(100);o__Saccharomycetales(100);f__Metschnikowiaceae(100);g__Metschnikowia(100);g__Metschnikowia_unclassified(100); | Metschnikowia | 13 | Saprotroph | \|Undefined Saprotroph\| | Yeast | NULL | Probable | As nectar-tap_saprotroph (Põlme et al. 2020); Decay sugar-rich substrates (Põlme et al. 2020) | Põlme S, et al. 2020. Fungal Diversity 105:1-16 ((https://doi.org/10.1007/s13225-020-00466-2)) |
| Otu49016 | 1 | k__Fungi(100);p__Ascomycota(100);c__Saccharomycetes(100);o__Saccharomycetales(100);f__Metschnikowiaceae(100);g__Metschnikowia(100);s__Metschnikowia_pulcherrima(100); | Metschnikowia | 13 | Saprotroph | \|Undefined Saprotroph\| | Yeast | NULL | Probable | As nectar-tap_saprotroph (Põlme et al. 2020); Decay sugar-rich substrates (Põlme et al. 2020) | Põlme S, et al. 2020. Fungal Diversity 105:1-16 ((https://doi.org/10.1007/s13225-020-00466-2)) |
| Otu49017 | 1 | k__Fungi(100);p__Ascomycota(100);c__Saccharomycetes(100);o__Saccharomycetales(100);f__Metschnikowiaceae(100);g__Metschnikowia(100);s__Metschnikowia_pulcherrima(100); | Metschnikowia | 13 | Saprotroph | \|Undefined Saprotroph\| | Yeast | NULL | Probable | As nectar-tap_saprotroph (Põlme et al. 2020); Decay sugar-rich substrates (Põlme et al. 2020) | Põlme S, et al. 2020. Fungal Diversity 105:1-16 ((https://doi.org/10.1007/s13225-020-00466-2)) |
| Otu49018 | 1 | k__Fungi(100);p__Ascomycota(100);c__Saccharomycetes(100);o__Saccharomycetales(100);f__Metschnikowiaceae(100);g__Metschnikowia(100);s__Metschnikowia_pulcherrima(100); | Metschnikowia | 13 | Saprotroph | \|Undefined Saprotroph\| | Yeast | NULL | Probable | As nectar-tap_saprotroph (Põlme et al. 2020); Decay sugar-rich substrates (Põlme et al. 2020) | Põlme S, et al. 2020. Fungal Diversity 105:1-16 ((https://doi.org/10.1007/s13225-020-00466-2)) |
| Otu49019 | 1 | k__Fungi(100);p__Ascomycota(100);c__Saccharomycetes(100);o__Saccharomycetales(100);f__Metschnikowiaceae(100);g__Metschnikowia(100);s__Metschnikowia_pulcherrima(100); | Metschnikowia | 13 | Saprotroph | \|Undefined Saprotroph\| | Yeast | NULL | Probable | As nectar-tap_saprotroph (Põlme et al. 2020); Decay sugar-rich substrates (Põlme et al. 2020) | Põlme S, et al. 2020. Fungal Diversity 105:1-16 ((https://doi.org/10.1007/s13225-020-00466-2)) |
| Otu49020 | 1 | k__Fungi(100);p__Ascomycota(100);c__Saccharomycetes(100);o__Saccharomycetales(100);f__Metschnikowiaceae(100);g__Metschnikowia(100);s__Metschnikowia_pulcherrima(100); | Metschnikowia | 13 | Saprotroph | \|Undefined Saprotroph\| | Yeast | NULL | Probable | As nectar-tap_saprotroph (Põlme et al. 2020); Decay sugar-rich substrates (Põlme et al. 2020) | Põlme S, et al. 2020. Fungal Diversity 105:1-16 ((https://doi.org/10.1007/s13225-020-00466-2)) |
| Otu49021 | 1 | k__Fungi(100);p__Ascomycota(100);c__Saccharomycetes(100);o__Saccharomycetales(100);f__Metschnikowiaceae(100);g__Metschnikowia(100);s__Metschnikowia_pulcherrima(100); | Metschnikowia | 13 | Saprotroph | \|Undefined Saprotroph\| | Yeast | NULL | Probable | As nectar-tap_saprotroph (Põlme et al. 2020); Decay sugar-rich substrates (Põlme et al. 2020) | Põlme S, et al. 2020. Fungal Diversity 105:1-16 ((https://doi.org/10.1007/s13225-020-00466-2)) |
| Otu49022 | 1 | k__Fungi(100);p__Ascomycota(100);c__Saccharomycetes(100);o__Saccharomycetales(100);f__Metschnikowiaceae(100);g__Metschnikowia(100);s__Metschnikowia_pulcherrima(100); | Metschnikowia | 13 | Saprotroph | \|Undefined Saprotroph\| | Yeast | NULL | Probable | As nectar-tap_saprotroph (Põlme et al. 2020); Decay sugar-rich substrates (Põlme et al. 2020) | Põlme S, et al. 2020. Fungal Diversity 105:1-16 ((https://doi.org/10.1007/s13225-020-00466-2)) |
| Otu49023 | 1 | k__Fungi(100);p__Ascomycota(100);c__Saccharomycetes(100);o__Saccharomycetales(100);f__Metschnikowiaceae(100);g__Metschnikowia(100);s__Metschnikowia_pulcherrima(100); | Metschnikowia | 13 | Saprotroph | \|Undefined Saprotroph\| | Yeast | NULL | Probable | As nectar-tap_saprotroph (Põlme et al. 2020); Decay sugar-rich substrates (Põlme et al. 2020) | Põlme S, et al. 2020. Fungal Diversity 105:1-16 ((https://doi.org/10.1007/s13225-020-00466-2)) |
| Otu49024 | 1 | k__Fungi(100);p__Ascomycota(100);c__Saccharomycetes(100);o__Saccharomycetales(100);f__Metschnikowiaceae(100);g__Metschnikowia(100);s__Metschnikowia_pulcherrima(100); | Metschnikowia | 13 | Saprotroph | \|Undefined Saprotroph\| | Yeast | NULL | Probable | As nectar-tap_saprotroph (Põlme et al. 2020); Decay sugar-rich substrates (Põlme et al. 2020) | Põlme S, et al. 2020. Fungal Diversity 105:1-16 ((https://doi.org/10.1007/s13225-020-00466-2)) |
| Otu49025 | 1 | k__Fungi(100);p__Ascomycota(100);c__Saccharomycetes(100);o__Saccharomycetales(100);f__Metschnikowiaceae(100);g__Metschnikowia(100);s__Metschnikowia_pulcherrima(100); | Metschnikowia | 13 | Saprotroph | \|Undefined Saprotroph\| | Yeast | NULL | Probable | As nectar-tap_saprotroph (Põlme et al. 2020); Decay sugar-rich substrates (Põlme et al. 2020) | Põlme S, et al. 2020. Fungal Diversity 105:1-16 ((https://doi.org/10.1007/s13225-020-00466-2)) |
| Otu49026 | 1 | k__Fungi(100);p__Ascomycota(100);c__Saccharomycetes(100);o__Saccharomycetales(100);f__Metschnikowiaceae(100);g__Metschnikowia(100);s__Metschnikowia_pulcherrima(100); | Metschnikowia | 13 | Saprotroph | \|Undefined Saprotroph\| | Yeast | NULL | Probable | As nectar-tap_saprotroph (Põlme et al. 2020); Decay sugar-rich substrates (Põlme et al. 2020) | Põlme S, et al. 2020. Fungal Diversity 105:1-16 ((https://doi.org/10.1007/s13225-020-00466-2)) |
| Otu49027 | 1 | k__Fungi(100);p__Ascomycota(100);c__Saccharomycetes(100);o__Saccharomycetales(100);f__Metschnikowiaceae(100);g__Metschnikowia(100);s__Metschnikowia_pulcherrima(100); | Metschnikowia | 13 | Saprotroph | \|Undefined Saprotroph\| | Yeast | NULL | Probable | As nectar-tap_saprotroph (Põlme et al. 2020); Decay sugar-rich substrates (Põlme et al. 2020) | Põlme S, et al. 2020. Fungal Diversity 105:1-16 ((https://doi.org/10.1007/s13225-020-00466-2)) |
| Otu49028 | 1 | k__Fungi(100);p__Ascomycota(100);c__Saccharomycetes(100);o__Saccharomycetales(100);f__Metschnikowiaceae(100);g__Metschnikowia(100);s__Metschnikowia_pulcherrima(100); | Metschnikowia | 13 | Saprotroph | \|Undefined Saprotroph\| | Yeast | NULL | Probable | As nectar-tap_saprotroph (Põlme et al. 2020); Decay sugar-rich substrates (Põlme et al. 2020) | Põlme S, et al. 2020. Fungal Diversity 105:1-16 ((https://doi.org/10.1007/s13225-020-00466-2)) |
| Otu49029 | 1 | k__Fungi(100);p__Ascomycota(100);c__Saccharomycetes(100);o__Saccharomycetales(100);f__Metschnikowiaceae(100);g__Metschnikowia(100);g__Metschnikowia_unclassified(100); | Metschnikowia | 13 | Saprotroph | \|Undefined Saprotroph\| | Yeast | NULL | Probable | As nectar-tap_saprotroph (Põlme et al. 2020); Decay sugar-rich substrates (Põlme et al. 2020) | Põlme S, et al. 2020. Fungal Diversity 105:1-16 ((https://doi.org/10.1007/s13225-020-00466-2)) |
| Otu49030 | 1 | k__Fungi(100);p__Ascomycota(100);c__Saccharomycetes(100);o__Saccharomycetales(100);f__Metschnikowiaceae(100);g__Metschnikowia(100);s__Metschnikowia_pulcherrima(100); | Metschnikowia | 13 | Saprotroph | \|Undefined Saprotroph\| | Yeast | NULL | Probable | As nectar-tap_saprotroph (Põlme et al. 2020); Decay sugar-rich substrates (Põlme et al. 2020) | Põlme S, et al. 2020. Fungal Diversity 105:1-16 ((https://doi.org/10.1007/s13225-020-00466-2)) |
| Otu49031 | 1 | k__Fungi(100);p__Ascomycota(100);c__Saccharomycetes(100);o__Saccharomycetales(100);f__Metschnikowiaceae(100);g__Metschnikowia(100);s__Metschnikowia_pulcherrima(100); | Metschnikowia | 13 | Saprotroph | \|Undefined Saprotroph\| | Yeast | NULL | Probable | As nectar-tap_saprotroph (Põlme et al. 2020); Decay sugar-rich substrates (Põlme et al. 2020) | Põlme S, et al. 2020. Fungal Diversity 105:1-16 ((https://doi.org/10.1007/s13225-020-00466-2)) |
| Otu49032 | 1 | k__Fungi(100);p__Ascomycota(100);c__Saccharomycetes(100);o__Saccharomycetales(100);f__Metschnikowiaceae(100);g__Metschnikowia(100);g__Metschnikowia_unclassified(100); | Metschnikowia | 13 | Saprotroph | \|Undefined Saprotroph\| | Yeast | NULL | Probable | As nectar-tap_saprotroph (Põlme et al. 2020); Decay sugar-rich substrates (Põlme et al. 2020) | Põlme S, et al. 2020. Fungal Diversity 105:1-16 ((https://doi.org/10.1007/s13225-020-00466-2)) |
| Otu49033 | 1 | k__Fungi(100);p__Ascomycota(100);c__Saccharomycetes(100);o__Saccharomycetales(100);f__Metschnikowiaceae(100);g__Metschnikowia(100);s__Metschnikowia_pulcherrima(100); | Metschnikowia | 13 | Saprotroph | \|Undefined Saprotroph\| | Yeast | NULL | Probable | As nectar-tap_saprotroph (Põlme et al. 2020); Decay sugar-rich substrates (Põlme et al. 2020) | Põlme S, et al. 2020. Fungal Diversity 105:1-16 ((https://doi.org/10.1007/s13225-020-00466-2)) |
| Otu49036 | 1 | k__Fungi(100);p__Ascomycota(100);c__Saccharomycetes(100);o__Saccharomycetales(100);f__Metschnikowiaceae(100);g__Metschnikowia(100);s__Metschnikowia_pulcherrima(100); | Metschnikowia | 13 | Saprotroph | \|Undefined Saprotroph\| | Yeast | NULL | Probable | As nectar-tap_saprotroph (Põlme et al. 2020); Decay sugar-rich substrates (Põlme et al. 2020) | Põlme S, et al. 2020. Fungal Diversity 105:1-16 ((https://doi.org/10.1007/s13225-020-00466-2)) |
| Otu49037 | 1 | k__Fungi(100);p__Ascomycota(100);c__Saccharomycetes(100);o__Saccharomycetales(100);f__Metschnikowiaceae(100);g__Metschnikowia(100);s__Metschnikowia_pulcherrima(100); | Metschnikowia | 13 | Saprotroph | \|Undefined Saprotroph\| | Yeast | NULL | Probable | As nectar-tap_saprotroph (Põlme et al. 2020); Decay sugar-rich substrates (Põlme et al. 2020) | Põlme S, et al. 2020. Fungal Diversity 105:1-16 ((https://doi.org/10.1007/s13225-020-00466-2)) |
| Otu49038 | 1 | k__Fungi(100);p__Ascomycota(100);c__Saccharomycetes(100);o__Saccharomycetales(100);f__Metschnikowiaceae(100);g__Metschnikowia(100);g__Metschnikowia_unclassified(100); | Metschnikowia | 13 | Saprotroph | \|Undefined Saprotroph\| | Yeast | NULL | Probable | As nectar-tap_saprotroph (Põlme et al. 2020); Decay sugar-rich substrates (Põlme et al. 2020) | Põlme S, et al. 2020. Fungal Diversity 105:1-16 ((https://doi.org/10.1007/s13225-020-00466-2)) |
| Otu49039 | 1 | k__Fungi(100);p__Ascomycota(100);c__Saccharomycetes(100);o__Saccharomycetales(100);f__Metschnikowiaceae(100);g__Metschnikowia(100);s__Metschnikowia_pulcherrima(100); | Metschnikowia | 13 | Saprotroph | \|Undefined Saprotroph\| | Yeast | NULL | Probable | As nectar-tap_saprotroph (Põlme et al. 2020); Decay sugar-rich substrates (Põlme et al. 2020) | Põlme S, et al. 2020. Fungal Diversity 105:1-16 ((https://doi.org/10.1007/s13225-020-00466-2)) |
| Otu49040 | 1 | k__Fungi(100);p__Ascomycota(100);c__Saccharomycetes(100);o__Saccharomycetales(100);f__Metschnikowiaceae(100);g__Metschnikowia(100);s__Metschnikowia_pulcherrima(100); | Metschnikowia | 13 | Saprotroph | \|Undefined Saprotroph\| | Yeast | NULL | Probable | As nectar-tap_saprotroph (Põlme et al. 2020); Decay sugar-rich substrates (Põlme et al. 2020) | Põlme S, et al. 2020. Fungal Diversity 105:1-16 ((https://doi.org/10.1007/s13225-020-00466-2)) |
| Otu49041 | 1 | k__Fungi(100);p__Ascomycota(100);c__Saccharomycetes(100);o__Saccharomycetales(100);f__Metschnikowiaceae(100);g__Metschnikowia(100);s__Metschnikowia_pulcherrima(100); | Metschnikowia | 13 | Saprotroph | \|Undefined Saprotroph\| | Yeast | NULL | Probable | As nectar-tap_saprotroph (Põlme et al. 2020); Decay sugar-rich substrates (Põlme et al. 2020) | Põlme S, et al. 2020. Fungal Diversity 105:1-16 ((https://doi.org/10.1007/s13225-020-00466-2)) |
| Otu49042 | 1 | k__Fungi(100);p__Ascomycota(100);c__Saccharomycetes(100);o__Saccharomycetales(100);f__Metschnikowiaceae(100);g__Metschnikowia(100);s__Metschnikowia_pulcherrima(100); | Metschnikowia | 13 | Saprotroph | \|Undefined Saprotroph\| | Yeast | NULL | Probable | As nectar-tap_saprotroph (Põlme et al. 2020); Decay sugar-rich substrates (Põlme et al. 2020) | Põlme S, et al. 2020. Fungal Diversity 105:1-16 ((https://doi.org/10.1007/s13225-020-00466-2)) |
| Otu49044 | 1 | k__Fungi(100);p__Ascomycota(100);c__Saccharomycetes(100);o__Saccharomycetales(100);f__Metschnikowiaceae(100);g__Metschnikowia(100);s__Metschnikowia_pulcherrima(100); | Metschnikowia | 13 | Saprotroph | \|Undefined Saprotroph\| | Yeast | NULL | Probable | As nectar-tap_saprotroph (Põlme et al. 2020); Decay sugar-rich substrates (Põlme et al. 2020) | Põlme S, et al. 2020. Fungal Diversity 105:1-16 ((https://doi.org/10.1007/s13225-020-00466-2)) |
| Otu49045 | 1 | k__Fungi(100);p__Ascomycota(100);c__Saccharomycetes(100);o__Saccharomycetales(100);f__Metschnikowiaceae(100);g__Metschnikowia(100);s__Metschnikowia_pulcherrima(100); | Metschnikowia | 13 | Saprotroph | \|Undefined Saprotroph\| | Yeast | NULL | Probable | As nectar-tap_saprotroph (Põlme et al. 2020); Decay sugar-rich substrates (Põlme et al. 2020) | Põlme S, et al. 2020. Fungal Diversity 105:1-16 ((https://doi.org/10.1007/s13225-020-00466-2)) |
| Otu49046 | 1 | k__Fungi(100);p__Ascomycota(100);c__Saccharomycetes(100);o__Saccharomycetales(100);f__Metschnikowiaceae(100);g__Metschnikowia(100);s__Metschnikowia_pulcherrima(100); | Metschnikowia | 13 | Saprotroph | \|Undefined Saprotroph\| | Yeast | NULL | Probable | As nectar-tap_saprotroph (Põlme et al. 2020); Decay sugar-rich substrates (Põlme et al. 2020) | Põlme S, et al. 2020. Fungal Diversity 105:1-16 ((https://doi.org/10.1007/s13225-020-00466-2)) |
| Otu49047 | 1 | k__Fungi(100);p__Ascomycota(100);c__Saccharomycetes(100);o__Saccharomycetales(100);f__Metschnikowiaceae(100);g__Metschnikowia(100);s__Metschnikowia_pulcherrima(100); | Metschnikowia | 13 | Saprotroph | \|Undefined Saprotroph\| | Yeast | NULL | Probable | As nectar-tap_saprotroph (Põlme et al. 2020); Decay sugar-rich substrates (Põlme et al. 2020) | Põlme S, et al. 2020. Fungal Diversity 105:1-16 ((https://doi.org/10.1007/s13225-020-00466-2)) |
| Otu49048 | 1 | k__Fungi(100);p__Ascomycota(100);c__Saccharomycetes(100);o__Saccharomycetales(100);f__Metschnikowiaceae(100);g__Metschnikowia(100);s__Metschnikowia_pulcherrima(100); | Metschnikowia | 13 | Saprotroph | \|Undefined Saprotroph\| | Yeast | NULL | Probable | As nectar-tap_saprotroph (Põlme et al. 2020); Decay sugar-rich substrates (Põlme et al. 2020) | Põlme S, et al. 2020. Fungal Diversity 105:1-16 ((https://doi.org/10.1007/s13225-020-00466-2)) |
| Otu49049 | 1 | k__Fungi(100);p__Ascomycota(100);c__Saccharomycetes(100);o__Saccharomycetales(100);f__Metschnikowiaceae(100);g__Metschnikowia(100);s__Metschnikowia_pulcherrima(100); | Metschnikowia | 13 | Saprotroph | \|Undefined Saprotroph\| | Yeast | NULL | Probable | As nectar-tap_saprotroph (Põlme et al. 2020); Decay sugar-rich substrates (Põlme et al. 2020) | Põlme S, et al. 2020. Fungal Diversity 105:1-16 ((https://doi.org/10.1007/s13225-020-00466-2)) |
| Otu49050 | 1 | k__Fungi(100);p__Ascomycota(100);c__Saccharomycetes(100);o__Saccharomycetales(100);f__Metschnikowiaceae(100);g__Metschnikowia(100);s__Metschnikowia_pulcherrima(100); | Metschnikowia | 13 | Saprotroph | \|Undefined Saprotroph\| | Yeast | NULL | Probable | As nectar-tap_saprotroph (Põlme et al. 2020); Decay sugar-rich substrates (Põlme et al. 2020) | Põlme S, et al. 2020. Fungal Diversity 105:1-16 ((https://doi.org/10.1007/s13225-020-00466-2)) |
| Otu49051 | 1 | k__Fungi(100);p__Ascomycota(100);c__Saccharomycetes(100);o__Saccharomycetales(100);f__Metschnikowiaceae(100);g__Metschnikowia(100);s__Metschnikowia_pulcherrima(100); | Metschnikowia | 13 | Saprotroph | \|Undefined Saprotroph\| | Yeast | NULL | Probable | As nectar-tap_saprotroph (Põlme et al. 2020); Decay sugar-rich substrates (Põlme et al. 2020) | Põlme S, et al. 2020. Fungal Diversity 105:1-16 ((https://doi.org/10.1007/s13225-020-00466-2)) |
| Otu49052 | 1 | k__Fungi(100);p__Ascomycota(100);c__Saccharomycetes(100);o__Saccharomycetales(100);f__Metschnikowiaceae(100);g__Metschnikowia(100);g__Metschnikowia_unclassified(100); | Metschnikowia | 13 | Saprotroph | \|Undefined Saprotroph\| | Yeast | NULL | Probable | As nectar-tap_saprotroph (Põlme et al. 2020); Decay sugar-rich substrates (Põlme et al. 2020) | Põlme S, et al. 2020. Fungal Diversity 105:1-16 ((https://doi.org/10.1007/s13225-020-00466-2)) |
| Otu49053 | 1 | k__Fungi(100);p__Ascomycota(100);c__Saccharomycetes(100);o__Saccharomycetales(100);f__Metschnikowiaceae(100);g__Metschnikowia(100);s__Metschnikowia_pulcherrima(100); | Metschnikowia | 13 | Saprotroph | \|Undefined Saprotroph\| | Yeast | NULL | Probable | As nectar-tap_saprotroph (Põlme et al. 2020); Decay sugar-rich substrates (Põlme et al. 2020) | Põlme S, et al. 2020. Fungal Diversity 105:1-16 ((https://doi.org/10.1007/s13225-020-00466-2)) |
| Otu49054 | 1 | k__Fungi(100);p__Ascomycota(100);c__Saccharomycetes(100);o__Saccharomycetales(100);f__Metschnikowiaceae(100);g__Metschnikowia(100);s__Metschnikowia_pulcherrima(100); | Metschnikowia | 13 | Saprotroph | \|Undefined Saprotroph\| | Yeast | NULL | Probable | As nectar-tap_saprotroph (Põlme et al. 2020); Decay sugar-rich substrates (Põlme et al. 2020) | Põlme S, et al. 2020. Fungal Diversity 105:1-16 ((https://doi.org/10.1007/s13225-020-00466-2)) |
| Otu49055 | 1 | k__Fungi(100);p__Ascomycota(100);c__Saccharomycetes(100);o__Saccharomycetales(100);f__Metschnikowiaceae(100);g__Metschnikowia(100);g__Metschnikowia_unclassified(100); | Metschnikowia | 13 | Saprotroph | \|Undefined Saprotroph\| | Yeast | NULL | Probable | As nectar-tap_saprotroph (Põlme et al. 2020); Decay sugar-rich substrates (Põlme et al. 2020) | Põlme S, et al. 2020. Fungal Diversity 105:1-16 ((https://doi.org/10.1007/s13225-020-00466-2)) |
| Otu49056 | 1 | k__Fungi(100);p__Ascomycota(100);c__Saccharomycetes(100);o__Saccharomycetales(100);f__Metschnikowiaceae(100);g__Metschnikowia(100);s__Metschnikowia_pulcherrima(100); | Metschnikowia | 13 | Saprotroph | \|Undefined Saprotroph\| | Yeast | NULL | Probable | As nectar-tap_saprotroph (Põlme et al. 2020); Decay sugar-rich substrates (Põlme et al. 2020) | Põlme S, et al. 2020. Fungal Diversity 105:1-16 ((https://doi.org/10.1007/s13225-020-00466-2)) |
| Otu49057 | 1 | k__Fungi(100);p__Ascomycota(100);c__Saccharomycetes(100);o__Saccharomycetales(100);f__Metschnikowiaceae(100);g__Metschnikowia(100);s__Metschnikowia_pulcherrima(100); | Metschnikowia | 13 | Saprotroph | \|Undefined Saprotroph\| | Yeast | NULL | Probable | As nectar-tap_saprotroph (Põlme et al. 2020); Decay sugar-rich substrates (Põlme et al. 2020) | Põlme S, et al. 2020. Fungal Diversity 105:1-16 ((https://doi.org/10.1007/s13225-020-00466-2)) |
| Otu49058 | 1 | k__Fungi(100);p__Ascomycota(100);c__Saccharomycetes(100);o__Saccharomycetales(100);f__Metschnikowiaceae(100);g__Metschnikowia(100);s__Metschnikowia_pulcherrima(100); | Metschnikowia | 13 | Saprotroph | \|Undefined Saprotroph\| | Yeast | NULL | Probable | As nectar-tap_saprotroph (Põlme et al. 2020); Decay sugar-rich substrates (Põlme et al. 2020) | Põlme S, et al. 2020. Fungal Diversity 105:1-16 ((https://doi.org/10.1007/s13225-020-00466-2)) |
| Otu49060 | 1 | k__Fungi(100);p__Ascomycota(100);c__Saccharomycetes(100);o__Saccharomycetales(100);f__Metschnikowiaceae(100);g__Metschnikowia(100);g__Metschnikowia_unclassified(100); | Metschnikowia | 13 | Saprotroph | \|Undefined Saprotroph\| | Yeast | NULL | Probable | As nectar-tap_saprotroph (Põlme et al. 2020); Decay sugar-rich substrates (Põlme et al. 2020) | Põlme S, et al. 2020. Fungal Diversity 105:1-16 ((https://doi.org/10.1007/s13225-020-00466-2)) |
| Otu49061 | 1 | k__Fungi(100);p__Ascomycota(100);c__Saccharomycetes(100);o__Saccharomycetales(100);f__Metschnikowiaceae(100);g__Metschnikowia(100);g__Metschnikowia_unclassified(100); | Metschnikowia | 13 | Saprotroph | \|Undefined Saprotroph\| | Yeast | NULL | Probable | As nectar-tap_saprotroph (Põlme et al. 2020); Decay sugar-rich substrates (Põlme et al. 2020) | Põlme S, et al. 2020. Fungal Diversity 105:1-16 ((https://doi.org/10.1007/s13225-020-00466-2)) |
| Otu49062 | 1 | k__Fungi(100);p__Ascomycota(100);c__Saccharomycetes(100);o__Saccharomycetales(100);f__Metschnikowiaceae(100);g__Metschnikowia(100);g__Metschnikowia_unclassified(100); | Metschnikowia | 13 | Saprotroph | \|Undefined Saprotroph\| | Yeast | NULL | Probable | As nectar-tap_saprotroph (Põlme et al. 2020); Decay sugar-rich substrates (Põlme et al. 2020) | Põlme S, et al. 2020. Fungal Diversity 105:1-16 ((https://doi.org/10.1007/s13225-020-00466-2)) |
| Otu49063 | 1 | k__Fungi(100);p__Ascomycota(100);c__Saccharomycetes(100);o__Saccharomycetales(100);f__Metschnikowiaceae(100);g__Metschnikowia(100);g__Metschnikowia_unclassified(100); | Metschnikowia | 13 | Saprotroph | \|Undefined Saprotroph\| | Yeast | NULL | Probable | As nectar-tap_saprotroph (Põlme et al. 2020); Decay sugar-rich substrates (Põlme et al. 2020) | Põlme S, et al. 2020. Fungal Diversity 105:1-16 ((https://doi.org/10.1007/s13225-020-00466-2)) |
| Otu49064 | 1 | k__Fungi(100);p__Ascomycota(100);c__Saccharomycetes(100);o__Saccharomycetales(100);f__Metschnikowiaceae(100);g__Metschnikowia(100);g__Metschnikowia_unclassified(100); | Metschnikowia | 13 | Saprotroph | \|Undefined Saprotroph\| | Yeast | NULL | Probable | As nectar-tap_saprotroph (Põlme et al. 2020); Decay sugar-rich substrates (Põlme et al. 2020) | Põlme S, et al. 2020. Fungal Diversity 105:1-16 ((https://doi.org/10.1007/s13225-020-00466-2)) |
| Otu49065 | 1 | k__Fungi(100);p__Ascomycota(100);c__Saccharomycetes(100);o__Saccharomycetales(100);f__Metschnikowiaceae(100);g__Metschnikowia(100);s__Metschnikowia_pulcherrima(100); | Metschnikowia | 13 | Saprotroph | \|Undefined Saprotroph\| | Yeast | NULL | Probable | As nectar-tap_saprotroph (Põlme et al. 2020); Decay sugar-rich substrates (Põlme et al. 2020) | Põlme S, et al. 2020. Fungal Diversity 105:1-16 ((https://doi.org/10.1007/s13225-020-00466-2)) |
| Otu49066 | 1 | k__Fungi(100);p__Ascomycota(100);c__Saccharomycetes(100);o__Saccharomycetales(100);f__Metschnikowiaceae(100);g__Metschnikowia(100);s__Metschnikowia_pulcherrima(100); | Metschnikowia | 13 | Saprotroph | \|Undefined Saprotroph\| | Yeast | NULL | Probable | As nectar-tap_saprotroph (Põlme et al. 2020); Decay sugar-rich substrates (Põlme et al. 2020) | Põlme S, et al. 2020. Fungal Diversity 105:1-16 ((https://doi.org/10.1007/s13225-020-00466-2)) |
| Otu49067 | 1 | k__Fungi(100);p__Ascomycota(100);c__Saccharomycetes(100);o__Saccharomycetales(100);f__Metschnikowiaceae(100);g__Metschnikowia(100);s__Metschnikowia_pulcherrima(100); | Metschnikowia | 13 | Saprotroph | \|Undefined Saprotroph\| | Yeast | NULL | Probable | As nectar-tap_saprotroph (Põlme et al. 2020); Decay sugar-rich substrates (Põlme et al. 2020) | Põlme S, et al. 2020. Fungal Diversity 105:1-16 ((https://doi.org/10.1007/s13225-020-00466-2)) |
| Otu49068 | 1 | k__Fungi(100);p__Ascomycota(100);c__Saccharomycetes(100);o__Saccharomycetales(100);f__Metschnikowiaceae(100);g__Metschnikowia(100);s__Metschnikowia_pulcherrima(100); | Metschnikowia | 13 | Saprotroph | \|Undefined Saprotroph\| | Yeast | NULL | Probable | As nectar-tap_saprotroph (Põlme et al. 2020); Decay sugar-rich substrates (Põlme et al. 2020) | Põlme S, et al. 2020. Fungal Diversity 105:1-16 ((https://doi.org/10.1007/s13225-020-00466-2)) |
| Otu49069 | 1 | k__Fungi(100);p__Ascomycota(100);c__Saccharomycetes(100);o__Saccharomycetales(100);f__Metschnikowiaceae(100);g__Metschnikowia(100);s__Metschnikowia_pulcherrima(100); | Metschnikowia | 13 | Saprotroph | \|Undefined Saprotroph\| | Yeast | NULL | Probable | As nectar-tap_saprotroph (Põlme et al. 2020); Decay sugar-rich substrates (Põlme et al. 2020) | Põlme S, et al. 2020. Fungal Diversity 105:1-16 ((https://doi.org/10.1007/s13225-020-00466-2)) |
| Otu49070 | 1 | k__Fungi(100);p__Ascomycota(100);c__Saccharomycetes(100);o__Saccharomycetales(100);f__Metschnikowiaceae(100);g__Metschnikowia(100);s__Metschnikowia_pulcherrima(100); | Metschnikowia | 13 | Saprotroph | \|Undefined Saprotroph\| | Yeast | NULL | Probable | As nectar-tap_saprotroph (Põlme et al. 2020); Decay sugar-rich substrates (Põlme et al. 2020) | Põlme S, et al. 2020. Fungal Diversity 105:1-16 ((https://doi.org/10.1007/s13225-020-00466-2)) |
| Otu49071 | 1 | k__Fungi(100);p__Ascomycota(100);c__Saccharomycetes(100);o__Saccharomycetales(100);f__Metschnikowiaceae(100);g__Metschnikowia(100);g__Metschnikowia_unclassified(100); | Metschnikowia | 13 | Saprotroph | \|Undefined Saprotroph\| | Yeast | NULL | Probable | As nectar-tap_saprotroph (Põlme et al. 2020); Decay sugar-rich substrates (Põlme et al. 2020) | Põlme S, et al. 2020. Fungal Diversity 105:1-16 ((https://doi.org/10.1007/s13225-020-00466-2)) |
| Otu49072 | 1 | k__Fungi(100);p__Ascomycota(100);c__Saccharomycetes(100);o__Saccharomycetales(100);f__Metschnikowiaceae(100);g__Metschnikowia(100);s__Metschnikowia_pulcherrima(100); | Metschnikowia | 13 | Saprotroph | \|Undefined Saprotroph\| | Yeast | NULL | Probable | As nectar-tap_saprotroph (Põlme et al. 2020); Decay sugar-rich substrates (Põlme et al. 2020) | Põlme S, et al. 2020. Fungal Diversity 105:1-16 ((https://doi.org/10.1007/s13225-020-00466-2)) |
| Otu49073 | 1 | k__Fungi(100);p__Ascomycota(100);c__Saccharomycetes(100);o__Saccharomycetales(100);f__Metschnikowiaceae(100);g__Metschnikowia(100);s__Metschnikowia_pulcherrima(100); | Metschnikowia | 13 | Saprotroph | \|Undefined Saprotroph\| | Yeast | NULL | Probable | As nectar-tap_saprotroph (Põlme et al. 2020); Decay sugar-rich substrates (Põlme et al. 2020) | Põlme S, et al. 2020. Fungal Diversity 105:1-16 ((https://doi.org/10.1007/s13225-020-00466-2)) |
| Otu49074 | 1 | k__Fungi(100);p__Ascomycota(100);c__Saccharomycetes(100);o__Saccharomycetales(100);f__Metschnikowiaceae(100);g__Metschnikowia(100);s__Metschnikowia_viticola(100); | Metschnikowia | 13 | Saprotroph | \|Undefined Saprotroph\| | Yeast | NULL | Probable | As nectar-tap_saprotroph (Põlme et al. 2020); Decay sugar-rich substrates (Põlme et al. 2020) | Põlme S, et al. 2020. Fungal Diversity 105:1-16 ((https://doi.org/10.1007/s13225-020-00466-2)) |
| Otu49075 | 1 | k__Fungi(100);p__Ascomycota(100);c__Saccharomycetes(100);o__Saccharomycetales(100);f__Metschnikowiaceae(100);g__Metschnikowia(100);s__Metschnikowia_pulcherrima(100); | Metschnikowia | 13 | Saprotroph | \|Undefined Saprotroph\| | Yeast | NULL | Probable | As nectar-tap_saprotroph (Põlme et al. 2020); Decay sugar-rich substrates (Põlme et al. 2020) | Põlme S, et al. 2020. Fungal Diversity 105:1-16 ((https://doi.org/10.1007/s13225-020-00466-2)) |
| Otu49076 | 1 | k__Fungi(100);p__Ascomycota(100);c__Saccharomycetes(100);o__Saccharomycetales(100);f__Metschnikowiaceae(100);g__Metschnikowia(100);s__Metschnikowia_pulcherrima(100); | Metschnikowia | 13 | Saprotroph | \|Undefined Saprotroph\| | Yeast | NULL | Probable | As nectar-tap_saprotroph (Põlme et al. 2020); Decay sugar-rich substrates (Põlme et al. 2020) | Põlme S, et al. 2020. Fungal Diversity 105:1-16 ((https://doi.org/10.1007/s13225-020-00466-2)) |
| Otu49077 | 1 | k__Fungi(100);p__Ascomycota(100);c__Saccharomycetes(100);o__Saccharomycetales(100);f__Metschnikowiaceae(100);g__Metschnikowia(100);s__Metschnikowia_pulcherrima(100); | Metschnikowia | 13 | Saprotroph | \|Undefined Saprotroph\| | Yeast | NULL | Probable | As nectar-tap_saprotroph (Põlme et al. 2020); Decay sugar-rich substrates (Põlme et al. 2020) | Põlme S, et al. 2020. Fungal Diversity 105:1-16 ((https://doi.org/10.1007/s13225-020-00466-2)) |
| Otu49078 | 1 | k__Fungi(100);p__Ascomycota(100);c__Saccharomycetes(100);o__Saccharomycetales(100);f__Metschnikowiaceae(100);g__Metschnikowia(100);s__Metschnikowia_pulcherrima(100); | Metschnikowia | 13 | Saprotroph | \|Undefined Saprotroph\| | Yeast | NULL | Probable | As nectar-tap_saprotroph (Põlme et al. 2020); Decay sugar-rich substrates (Põlme et al. 2020) | Põlme S, et al. 2020. Fungal Diversity 105:1-16 ((https://doi.org/10.1007/s13225-020-00466-2)) |
| Otu49079 | 1 | k__Fungi(100);p__Ascomycota(100);c__Saccharomycetes(100);o__Saccharomycetales(100);f__Metschnikowiaceae(100);g__Metschnikowia(100);s__Metschnikowia_pulcherrima(100); | Metschnikowia | 13 | Saprotroph | \|Undefined Saprotroph\| | Yeast | NULL | Probable | As nectar-tap_saprotroph (Põlme et al. 2020); Decay sugar-rich substrates (Põlme et al. 2020) | Põlme S, et al. 2020. Fungal Diversity 105:1-16 ((https://doi.org/10.1007/s13225-020-00466-2)) |
| Otu49080 | 1 | k__Fungi(100);p__Ascomycota(100);c__Saccharomycetes(100);o__Saccharomycetales(100);f__Metschnikowiaceae(100);g__Metschnikowia(100);s__Metschnikowia_pulcherrima(100); | Metschnikowia | 13 | Saprotroph | \|Undefined Saprotroph\| | Yeast | NULL | Probable | As nectar-tap_saprotroph (Põlme et al. 2020); Decay sugar-rich substrates (Põlme et al. 2020) | Põlme S, et al. 2020. Fungal Diversity 105:1-16 ((https://doi.org/10.1007/s13225-020-00466-2)) |
| Otu49081 | 1 | k__Fungi(100);p__Ascomycota(100);c__Saccharomycetes(100);o__Saccharomycetales(100);f__Metschnikowiaceae(100);g__Metschnikowia(100);s__Metschnikowia_pulcherrima(100); | Metschnikowia | 13 | Saprotroph | \|Undefined Saprotroph\| | Yeast | NULL | Probable | As nectar-tap_saprotroph (Põlme et al. 2020); Decay sugar-rich substrates (Põlme et al. 2020) | Põlme S, et al. 2020. Fungal Diversity 105:1-16 ((https://doi.org/10.1007/s13225-020-00466-2)) |
| Otu49082 | 1 | k__Fungi(100);p__Ascomycota(100);c__Saccharomycetes(100);o__Saccharomycetales(100);f__Metschnikowiaceae(100);g__Metschnikowia(100);s__Metschnikowia_pulcherrima(100); | Metschnikowia | 13 | Saprotroph | \|Undefined Saprotroph\| | Yeast | NULL | Probable | As nectar-tap_saprotroph (Põlme et al. 2020); Decay sugar-rich substrates (Põlme et al. 2020) | Põlme S, et al. 2020. Fungal Diversity 105:1-16 ((https://doi.org/10.1007/s13225-020-00466-2)) |
| Otu49083 | 1 | k__Fungi(100);p__Ascomycota(100);c__Saccharomycetes(100);o__Saccharomycetales(100);f__Metschnikowiaceae(100);g__Metschnikowia(100);s__Metschnikowia_pulcherrima(100); | Metschnikowia | 13 | Saprotroph | \|Undefined Saprotroph\| | Yeast | NULL | Probable | As nectar-tap_saprotroph (Põlme et al. 2020); Decay sugar-rich substrates (Põlme et al. 2020) | Põlme S, et al. 2020. Fungal Diversity 105:1-16 ((https://doi.org/10.1007/s13225-020-00466-2)) |
| Otu49084 | 1 | k__Fungi(100);p__Ascomycota(100);c__Saccharomycetes(100);o__Saccharomycetales(100);f__Metschnikowiaceae(100);g__Metschnikowia(100);s__Metschnikowia_pulcherrima(100); | Metschnikowia | 13 | Saprotroph | \|Undefined Saprotroph\| | Yeast | NULL | Probable | As nectar-tap_saprotroph (Põlme et al. 2020); Decay sugar-rich substrates (Põlme et al. 2020) | Põlme S, et al. 2020. Fungal Diversity 105:1-16 ((https://doi.org/10.1007/s13225-020-00466-2)) |
| Otu49085 | 1 | k__Fungi(100);p__Ascomycota(100);c__Saccharomycetes(100);o__Saccharomycetales(100);f__Metschnikowiaceae(100);g__Metschnikowia(100);g__Metschnikowia_unclassified(100); | Metschnikowia | 13 | Saprotroph | \|Undefined Saprotroph\| | Yeast | NULL | Probable | As nectar-tap_saprotroph (Põlme et al. 2020); Decay sugar-rich substrates (Põlme et al. 2020) | Põlme S, et al. 2020. Fungal Diversity 105:1-16 ((https://doi.org/10.1007/s13225-020-00466-2)) |
| Otu49086 | 1 | k__Fungi(100);p__Ascomycota(100);c__Saccharomycetes(100);o__Saccharomycetales(100);f__Metschnikowiaceae(100);g__Metschnikowia(100);g__Metschnikowia_unclassified(100); | Metschnikowia | 13 | Saprotroph | \|Undefined Saprotroph\| | Yeast | NULL | Probable | As nectar-tap_saprotroph (Põlme et al. 2020); Decay sugar-rich substrates (Põlme et al. 2020) | Põlme S, et al. 2020. Fungal Diversity 105:1-16 ((https://doi.org/10.1007/s13225-020-00466-2)) |
| Otu49087 | 1 | k__Fungi(100);p__Ascomycota(100);c__Saccharomycetes(100);o__Saccharomycetales(100);f__Metschnikowiaceae(100);g__Metschnikowia(100);g__Metschnikowia_unclassified(100); | Metschnikowia | 13 | Saprotroph | \|Undefined Saprotroph\| | Yeast | NULL | Probable | As nectar-tap_saprotroph (Põlme et al. 2020); Decay sugar-rich substrates (Põlme et al. 2020) | Põlme S, et al. 2020. Fungal Diversity 105:1-16 ((https://doi.org/10.1007/s13225-020-00466-2)) |
| Otu49088 | 1 | k__Fungi(100);p__Ascomycota(100);c__Saccharomycetes(100);o__Saccharomycetales(100);f__Metschnikowiaceae(100);g__Metschnikowia(100);s__Metschnikowia_pulcherrima(100); | Metschnikowia | 13 | Saprotroph | \|Undefined Saprotroph\| | Yeast | NULL | Probable | As nectar-tap_saprotroph (Põlme et al. 2020); Decay sugar-rich substrates (Põlme et al. 2020) | Põlme S, et al. 2020. Fungal Diversity 105:1-16 ((https://doi.org/10.1007/s13225-020-00466-2)) |
| Otu49089 | 1 | k__Fungi(100);p__Ascomycota(100);c__Saccharomycetes(100);o__Saccharomycetales(100);f__Metschnikowiaceae(100);g__Metschnikowia(100);s__Metschnikowia_pulcherrima(100); | Metschnikowia | 13 | Saprotroph | \|Undefined Saprotroph\| | Yeast | NULL | Probable | As nectar-tap_saprotroph (Põlme et al. 2020); Decay sugar-rich substrates (Põlme et al. 2020) | Põlme S, et al. 2020. Fungal Diversity 105:1-16 ((https://doi.org/10.1007/s13225-020-00466-2)) |
| Otu49090 | 1 | k__Fungi(100);p__Ascomycota(100);c__Saccharomycetes(100);o__Saccharomycetales(100);f__Metschnikowiaceae(100);g__Metschnikowia(100);s__Metschnikowia_pulcherrima(100); | Metschnikowia | 13 | Saprotroph | \|Undefined Saprotroph\| | Yeast | NULL | Probable | As nectar-tap_saprotroph (Põlme et al. 2020); Decay sugar-rich substrates (Põlme et al. 2020) | Põlme S, et al. 2020. Fungal Diversity 105:1-16 ((https://doi.org/10.1007/s13225-020-00466-2)) |

| | | | | | | | | | | | |
|---|---|---|---|---|---|---|---|---|---|---|---|
| Otu49091 | 1 | k__Fungi(100);p__Ascomycota(100);c__Saccharomycetes(100);o__Saccharomycetales(100);f__Metschnikowiaceae(100);g__Metschnikowia(100);s__Metschnikowia_pulcherrima(100); | Metschnikowia | 13 | Saprotroph | \|Undefined Saprotroph\| | Yeast | NULL | Probable | As nectar-tap_saprotroph (Põlme et al. 2020); Decay sugar-rich substrates (Põlme et al. 2020) | Põlme S, et al. 2020. Fungal Diversity 105:1-16 ((https://doi.org/10.1007/s13225-020-00466-2)) |
| Otu49092 | 1 | k__Fungi(100);p__Ascomycota(100);c__Saccharomycetes(100);o__Saccharomycetales(100);f__Metschnikowiaceae(100);g__Metschnikowia(100);g__Metschnikowia_unclassified(100); | Metschnikowia | 13 | Saprotroph | \|Undefined Saprotroph\| | Yeast | NULL | Probable | As nectar-tap_saprotroph (Põlme et al. 2020); Decay sugar-rich substrates (Põlme et al. 2020) | Põlme S, et al. 2020. Fungal Diversity 105:1-16 ((https://doi.org/10.1007/s13225-020-00466-2)) |
| Otu49093 | 1 | k__Fungi(100);p__Ascomycota(100);c__Saccharomycetes(100);o__Saccharomycetales(100);f__Metschnikowiaceae(100);g__Metschnikowia(100);g__Metschnikowia_unclassified(100); | Metschnikowia | 13 | Saprotroph | \|Undefined Saprotroph\| | Yeast | NULL | Probable | As nectar-tap_saprotroph (Põlme et al. 2020); Decay sugar-rich substrates (Põlme et al. 2020) | Põlme S, et al. 2020. Fungal Diversity 105:1-16 ((https://doi.org/10.1007/s13225-020-00466-2)) |
| Otu49094 | 1 | k__Fungi(100);p__Ascomycota(100);c__Saccharomycetes(100);o__Saccharomycetales(100);f__Metschnikowiaceae(100);g__Metschnikowia(100);s__Metschnikowia_pulcherrima(100); | Metschnikowia | 13 | Saprotroph | \|Undefined Saprotroph\| | Yeast | NULL | Probable | As nectar-tap_saprotroph (Põlme et al. 2020); Decay sugar-rich substrates (Põlme et al. 2020) | Põlme S, et al. 2020. Fungal Diversity 105:1-16 ((https://doi.org/10.1007/s13225-020-00466-2)) |
| Otu49095 | 1 | k__Fungi(100);p__Ascomycota(100);c__Saccharomycetes(100);o__Saccharomycetales(100);f__Metschnikowiaceae(100);g__Metschnikowia(100);g__Metschnikowia_unclassified(100); | Metschnikowia | 13 | Saprotroph | \|Undefined Saprotroph\| | Yeast | NULL | Probable | As nectar-tap_saprotroph (Põlme et al. 2020); Decay sugar-rich substrates (Põlme et al. 2020) | Põlme S, et al. 2020. Fungal Diversity 105:1-16 ((https://doi.org/10.1007/s13225-020-00466-2)) |
| Otu49096 | 1 | k__Fungi(100);p__Ascomycota(100);c__Saccharomycetes(100);o__Saccharomycetales(100);f__Metschnikowiaceae(100);g__Metschnikowia(100);s__Metschnikowia_pulcherrima(100); | Metschnikowia | 13 | Saprotroph | \|Undefined Saprotroph\| | Yeast | NULL | Probable | As nectar-tap_saprotroph (Põlme et al. 2020); Decay sugar-rich substrates (Põlme et al. 2020) | Põlme S, et al. 2020. Fungal Diversity 105:1-16 ((https://doi.org/10.1007/s13225-020-00466-2)) |
| Otu49097 | 1 | k__Fungi(100);p__Ascomycota(100);c__Saccharomycetes(100);o__Saccharomycetales(100);f__Metschnikowiaceae(100);g__Metschnikowia(100);s__Metschnikowia_pulcherrima(100); | Metschnikowia | 13 | Saprotroph | \|Undefined Saprotroph\| | Yeast | NULL | Probable | As nectar-tap_saprotroph (Põlme et al. 2020); Decay sugar-rich substrates (Põlme et al. 2020) | Põlme S, et al. 2020. Fungal Diversity 105:1-16 ((https://doi.org/10.1007/s13225-020-00466-2)) |
| Otu49098 | 1 | k__Fungi(100);p__Ascomycota(100);c__Saccharomycetes(100);o__Saccharomycetales(100);f__Metschnikowiaceae(100);g__Metschnikowia(100);s__Metschnikowia_pulcherrima(100); | Metschnikowia | 13 | Saprotroph | \|Undefined Saprotroph\| | Yeast | NULL | Probable | As nectar-tap_saprotroph (Põlme et al. 2020); Decay sugar-rich substrates (Põlme et al. 2020) | Põlme S, et al. 2020. Fungal Diversity 105:1-16 ((https://doi.org/10.1007/s13225-020-00466-2)) |
| Otu49099 | 1 | k__Fungi(100);p__Ascomycota(100);c__Saccharomycetes(100);o__Saccharomycetales(100);f__Metschnikowiaceae(100);g__Metschnikowia(100);g__Metschnikowia_unclassified(100); | Metschnikowia | 13 | Saprotroph | \|Undefined Saprotroph\| | Yeast | NULL | Probable | As nectar-tap_saprotroph (Põlme et al. 2020); Decay sugar-rich substrates (Põlme et al. 2020) | Põlme S, et al. 2020. Fungal Diversity 105:1-16 ((https://doi.org/10.1007/s13225-020-00466-2)) |
| Otu49100 | 1 | k__Fungi(100);p__Ascomycota(100);c__Saccharomycetes(100);o__Saccharomycetales(100);f__Metschnikowiaceae(100);g__Metschnikowia(100);s__Metschnikowia_pulcherrima(100); | Metschnikowia | 13 | Saprotroph | \|Undefined Saprotroph\| | Yeast | NULL | Probable | As nectar-tap_saprotroph (Põlme et al. 2020); Decay sugar-rich substrates (Põlme et al. 2020) | Põlme S, et al. 2020. Fungal Diversity 105:1-16 ((https://doi.org/10.1007/s13225-020-00466-2)) |
| Otu49101 | 1 | k__Fungi(100);p__Ascomycota(100);c__Saccharomycetes(100);o__Saccharomycetales(100);f__Metschnikowiaceae(100);g__Metschnikowia(100);s__Metschnikowia_pulcherrima(100); | Metschnikowia | 13 | Saprotroph | \|Undefined Saprotroph\| | Yeast | NULL | Probable | As nectar-tap_saprotroph (Põlme et al. 2020); Decay sugar-rich substrates (Põlme et al. 2020) | Põlme S, et al. 2020. Fungal Diversity 105:1-16 ((https://doi.org/10.1007/s13225-020-00466-2)) |
| Otu49102 | 1 | k__Fungi(100);p__Ascomycota(100);c__Saccharomycetes(100);o__Saccharomycetales(100);f__Metschnikowiaceae(100);g__Metschnikowia(100);s__Metschnikowia_pulcherrima(100); | Metschnikowia | 13 | Saprotroph | \|Undefined Saprotroph\| | Yeast | NULL | Probable | As nectar-tap_saprotroph (Põlme et al. 2020); Decay sugar-rich substrates (Põlme et al. 2020) | Põlme S, et al. 2020. Fungal Diversity 105:1-16 ((https://doi.org/10.1007/s13225-020-00466-2)) |
| Otu49103 | 1 | k__Fungi(100);p__Ascomycota(100);c__Saccharomycetes(100);o__Saccharomycetales(100);f__Metschnikowiaceae(100);g__Metschnikowia(100);s__Metschnikowia_pulcherrima(100); | Metschnikowia | 13 | Saprotroph | \|Undefined Saprotroph\| | Yeast | NULL | Probable | As nectar-tap_saprotroph (Põlme et al. 2020); Decay sugar-rich substrates (Põlme et al. 2020) | Põlme S, et al. 2020. Fungal Diversity 105:1-16 ((https://doi.org/10.1007/s13225-020-00466-2)) |
| Otu49104 | 1 | k__Fungi(100);p__Ascomycota(100);c__Saccharomycetes(100);o__Saccharomycetales(100);f__Metschnikowiaceae(100);g__Metschnikowia(100);s__Metschnikowia_pulcherrima(100); | Metschnikowia | 13 | Saprotroph | \|Undefined Saprotroph\| | Yeast | NULL | Probable | As nectar-tap_saprotroph (Põlme et al. 2020); Decay sugar-rich substrates (Põlme et al. 2020) | Põlme S, et al. 2020. Fungal Diversity 105:1-16 ((https://doi.org/10.1007/s13225-020-00466-2)) |
| Otu49105 | 1 | k__Fungi(100);p__Ascomycota(100);c__Saccharomycetes(100);o__Saccharomycetales(100);f__Metschnikowiaceae(100);g__Metschnikowia(100);s__Metschnikowia_pulcherrima(100); | Metschnikowia | 13 | Saprotroph | \|Undefined Saprotroph\| | Yeast | NULL | Probable | As nectar-tap_saprotroph (Põlme et al. 2020); Decay sugar-rich substrates (Põlme et al. 2020) | Põlme S, et al. 2020. Fungal Diversity 105:1-16 ((https://doi.org/10.1007/s13225-020-00466-2)) |
| Otu49106 | 1 | k__Fungi(100);p__Ascomycota(100);c__Saccharomycetes(100);o__Saccharomycetales(100);f__Metschnikowiaceae(100);g__Metschnikowia(100);s__Metschnikowia_pulcherrima(100); | Metschnikowia | 13 | Saprotroph | \|Undefined Saprotroph\| | Yeast | NULL | Probable | As nectar-tap_saprotroph (Põlme et al. 2020); Decay sugar-rich substrates (Põlme et al. 2020) | Põlme S, et al. 2020. Fungal Diversity 105:1-16 ((https://doi.org/10.1007/s13225-020-00466-2)) |
| Otu49107 | 1 | k__Fungi(100);p__Ascomycota(100);c__Saccharomycetes(100);o__Saccharomycetales(100);f__Metschnikowiaceae(100);g__Metschnikowia(100);s__Metschnikowia_pulcherrima(100); | Metschnikowia | 13 | Saprotroph | \|Undefined Saprotroph\| | Yeast | NULL | Probable | As nectar-tap_saprotroph (Põlme et al. 2020); Decay sugar-rich substrates (Põlme et al. 2020) | Põlme S, et al. 2020. Fungal Diversity 105:1-16 ((https://doi.org/10.1007/s13225-020-00466-2)) |
| Otu49108 | 1 | k__Fungi(100);p__Ascomycota(100);c__Saccharomycetes(100);o__Saccharomycetales(100);f__Metschnikowiaceae(100);g__Metschnikowia(100);s__Metschnikowia_pulcherrima(100); | Metschnikowia | 13 | Saprotroph | \|Undefined Saprotroph\| | Yeast | NULL | Probable | As nectar-tap_saprotroph (Põlme et al. 2020); Decay sugar-rich substrates (Põlme et al. 2020) | Põlme S, et al. 2020. Fungal Diversity 105:1-16 ((https://doi.org/10.1007/s13225-020-00466-2)) |
| Otu49109 | 1 | k__Fungi(100);p__Ascomycota(100);c__Saccharomycetes(100);o__Saccharomycetales(100);f__Metschnikowiaceae(100);g__Metschnikowia(100);s__Metschnikowia_pulcherrima(100); | Metschnikowia | 13 | Saprotroph | \|Undefined Saprotroph\| | Yeast | NULL | Probable | As nectar-tap_saprotroph (Põlme et al. 2020); Decay sugar-rich substrates (Põlme et al. 2020) | Põlme S, et al. 2020. Fungal Diversity 105:1-16 ((https://doi.org/10.1007/s13225-020-00466-2)) |
| Otu49110 | 1 | k__Fungi(100);p__Ascomycota(100);c__Saccharomycetes(100);o__Saccharomycetales(100);f__Metschnikowiaceae(100);g__Metschnikowia(100);s__Metschnikowia_pulcherrima(100); | Metschnikowia | 13 | Saprotroph | \|Undefined Saprotroph\| | Yeast | NULL | Probable | As nectar-tap_saprotroph (Põlme et al. 2020); Decay sugar-rich substrates (Põlme et al. 2020) | Põlme S, et al. 2020. Fungal Diversity 105:1-16 ((https://doi.org/10.1007/s13225-020-00466-2)) |
| Otu49111 | 1 | k__Fungi(100);p__Ascomycota(100);c__Saccharomycetes(100);o__Saccharomycetales(100);f__Metschnikowiaceae(100);g__Metschnikowia(100);s__Metschnikowia_pulcherrima(100); | Metschnikowia | 13 | Saprotroph | \|Undefined Saprotroph\| | Yeast | NULL | Probable | As nectar-tap_saprotroph (Põlme et al. 2020); Decay sugar-rich substrates (Põlme et al. 2020) | Põlme S, et al. 2020. Fungal Diversity 105:1-16 ((https://doi.org/10.1007/s13225-020-00466-2)) |
| Otu49112 | 1 | k__Fungi(100);p__Ascomycota(100);c__Saccharomycetes(100);o__Saccharomycetales(100);f__Metschnikowiaceae(100);g__Metschnikowia(100);s__Metschnikowia_pulcherrima(100); | Metschnikowia | 13 | Saprotroph | \|Undefined Saprotroph\| | Yeast | NULL | Probable | As nectar-tap_saprotroph (Põlme et al. 2020); Decay sugar-rich substrates (Põlme et al. 2020) | Põlme S, et al. 2020. Fungal Diversity 105:1-16 ((https://doi.org/10.1007/s13225-020-00466-2)) |
| Otu49113 | 1 | k__Fungi(100);p__Ascomycota(100);c__Saccharomycetes(100);o__Saccharomycetales(100);f__Metschnikowiaceae(100);g__Metschnikowia(100);s__Metschnikowia_pulcherrima(100); | Metschnikowia | 13 | Saprotroph | \|Undefined Saprotroph\| | Yeast | NULL | Probable | As nectar-tap_saprotroph (Põlme et al. 2020); Decay sugar-rich substrates (Põlme et al. 2020) | Põlme S, et al. 2020. Fungal Diversity 105:1-16 ((https://doi.org/10.1007/s13225-020-00466-2)) |
| Otu49114 | 1 | k__Fungi(100);p__Ascomycota(100);c__Saccharomycetes(100);o__Saccharomycetales(100);f__Metschnikowiaceae(100);g__Metschnikowia(100);g__Metschnikowia_unclassified(100); | Metschnikowia | 13 | Saprotroph | \|Undefined Saprotroph\| | Yeast | NULL | Probable | As nectar-tap_saprotroph (Põlme et al. 2020); Decay sugar-rich substrates (Põlme et al. 2020) | Põlme S, et al. 2020. Fungal Diversity 105:1-16 ((https://doi.org/10.1007/s13225-020-00466-2)) |
| Otu49115 | 1 | k__Fungi(100);p__Ascomycota(100);c__Saccharomycetes(100);o__Saccharomycetales(100);f__Metschnikowiaceae(100);g__Metschnikowia(100);g__Metschnikowia_unclassified(100); | Metschnikowia | 13 | Saprotroph | \|Undefined Saprotroph\| | Yeast | NULL | Probable | As nectar-tap_saprotroph (Põlme et al. 2020); Decay sugar-rich substrates (Põlme et al. 2020) | Põlme S, et al. 2020. Fungal Diversity 105:1-16 ((https://doi.org/10.1007/s13225-020-00466-2)) |
| Otu49116 | 1 | k__Fungi(100);p__Ascomycota(100);c__Saccharomycetes(100);o__Saccharomycetales(100);f__Metschnikowiaceae(100);g__Metschnikowia(100);g__Metschnikowia_unclassified(100); | Metschnikowia | 13 | Saprotroph | \|Undefined Saprotroph\| | Yeast | NULL | Probable | As nectar-tap_saprotroph (Põlme et al. 2020); Decay sugar-rich substrates (Põlme et al. 2020) | Põlme S, et al. 2020. Fungal Diversity 105:1-16 ((https://doi.org/10.1007/s13225-020-00466-2)) |
| Otu49117 | 1 | k__Fungi(100);p__Ascomycota(100);c__Saccharomycetes(100);o__Saccharomycetales(100);f__Metschnikowiaceae(100);g__Metschnikowia(100);g__Metschnikowia_unclassified(100); | Metschnikowia | 13 | Saprotroph | \|Undefined Saprotroph\| | Yeast | NULL | Probable | As nectar-tap_saprotroph (Põlme et al. 2020); Decay sugar-rich substrates (Põlme et al. 2020) | Põlme S, et al. 2020. Fungal Diversity 105:1-16 ((https://doi.org/10.1007/s13225-020-00466-2)) |
| Otu49118 | 1 | k__Fungi(100);p__Ascomycota(100);c__Saccharomycetes(100);o__Saccharomycetales(100);f__Metschnikowiaceae(100);g__Metschnikowia(100);g__Metschnikowia_unclassified(100); | Metschnikowia | 13 | Saprotroph | \|Undefined Saprotroph\| | Yeast | NULL | Probable | As nectar-tap_saprotroph (Põlme et al. 2020); Decay sugar-rich substrates (Põlme et al. 2020) | Põlme S, et al. 2020. Fungal Diversity 105:1-16 ((https://doi.org/10.1007/s13225-020-00466-2)) |
| Otu49119 | 1 | k__Fungi(100);p__Ascomycota(100);c__Saccharomycetes(100);o__Saccharomycetales(100);f__Metschnikowiaceae(100);g__Metschnikowia(100);s__Metschnikowia_pulcherrima(100); | Metschnikowia | 13 | Saprotroph | \|Undefined Saprotroph\| | Yeast | NULL | Probable | As nectar-tap_saprotroph (Põlme et al. 2020); Decay sugar-rich substrates (Põlme et al. 2020) | Põlme S, et al. 2020. Fungal Diversity 105:1-16 ((https://doi.org/10.1007/s13225-020-00466-2)) |
| Otu49120 | 1 | k__Fungi(100);p__Ascomycota(100);c__Saccharomycetes(100);o__Saccharomycetales(100);f__Metschnikowiaceae(100);g__Metschnikowia(100);g__Metschnikowia_unclassified(100); | Metschnikowia | 13 | Saprotroph | \|Undefined Saprotroph\| | Yeast | NULL | Probable | As nectar-tap_saprotroph (Põlme et al. 2020); Decay sugar-rich substrates (Põlme et al. 2020) | Põlme S, et al. 2020. Fungal Diversity 105:1-16 ((https://doi.org/10.1007/s13225-020-00466-2)) |
| Otu49121 | 1 | k__Fungi(100);p__Ascomycota(100);c__Saccharomycetes(100);o__Saccharomycetales(100);f__Metschnikowiaceae(100);g__Metschnikowia(100);s__Metschnikowia_pulcherrima(100); | Metschnikowia | 13 | Saprotroph | \|Undefined Saprotroph\| | Yeast | NULL | Probable | As nectar-tap_saprotroph (Põlme et al. 2020); Decay sugar-rich substrates (Põlme et al. 2020) | Põlme S, et al. 2020. Fungal Diversity 105:1-16 ((https://doi.org/10.1007/s13225-020-00466-2)) |
| Otu49122 | 1 | k__Fungi(100);p__Ascomycota(100);c__Saccharomycetes(100);o__Saccharomycetales(100);f__Metschnikowiaceae(100);g__Metschnikowia(100);s__Metschnikowia_pulcherrima(100); | Metschnikowia | 13 | Saprotroph | \|Undefined Saprotroph\| | Yeast | NULL | Probable | As nectar-tap_saprotroph (Põlme et al. 2020); Decay sugar-rich substrates (Põlme et al. 2020) | Põlme S, et al. 2020. Fungal Diversity 105:1-16 ((https://doi.org/10.1007/s13225-020-00466-2)) |
| Otu49123 | 1 | k__Fungi(100);p__Ascomycota(100);c__Saccharomycetes(100);o__Saccharomycetales(100);f__Metschnikowiaceae(100);g__Metschnikowia(100);s__Metschnikowia_pulcherrima(100); | Metschnikowia | 13 | Saprotroph | \|Undefined Saprotroph\| | Yeast | NULL | Probable | As nectar-tap_saprotroph (Põlme et al. 2020); Decay sugar-rich substrates (Põlme et al. 2020) | Põlme S, et al. 2020. Fungal Diversity 105:1-16 ((https://doi.org/10.1007/s13225-020-00466-2)) |
| Otu49124 | 1 | k__Fungi(100);p__Ascomycota(100);c__Saccharomycetes(100);o__Saccharomycetales(100);f__Metschnikowiaceae(100);g__Metschnikowia(100);s__Metschnikowia_pulcherrima(100); | Metschnikowia | 13 | Saprotroph | \|Undefined Saprotroph\| | Yeast | NULL | Probable | As nectar-tap_saprotroph (Põlme et al. 2020); Decay sugar-rich substrates (Põlme et al. 2020) | Põlme S, et al. 2020. Fungal Diversity 105:1-16 ((https://doi.org/10.1007/s13225-020-00466-2)) |
| Otu49125 | 1 | k__Fungi(100);p__Ascomycota(100);c__Saccharomycetes(100);o__Saccharomycetales(100);f__Metschnikowiaceae(100);g__Metschnikowia(100);g__Metschnikowia_unclassified(100); | Metschnikowia | 13 | Saprotroph | \|Undefined Saprotroph\| | Yeast | NULL | Probable | As nectar-tap_saprotroph (Põlme et al. 2020); Decay sugar-rich substrates (Põlme et al. 2020) | Põlme S, et al. 2020. Fungal Diversity 105:1-16 ((https://doi.org/10.1007/s13225-020-00466-2)) |
| Otu49126 | 1 | k__Fungi(100);p__Ascomycota(100);c__Saccharomycetes(100);o__Saccharomycetales(100);f__Metschnikowiaceae(100);g__Metschnikowia(100);s__Metschnikowia_pulcherrima(100); | Metschnikowia | 13 | Saprotroph | \|Undefined Saprotroph\| | Yeast | NULL | Probable | As nectar-tap_saprotroph (Põlme et al. 2020); Decay sugar-rich substrates (Põlme et al. 2020) | Põlme S, et al. 2020. Fungal Diversity 105:1-16 ((https://doi.org/10.1007/s13225-020-00466-2)) |
| Otu49127 | 1 | k__Fungi(100);p__Ascomycota(100);c__Saccharomycetes(100);o__Saccharomycetales(100);f__Metschnikowiaceae(100);g__Metschnikowia(100);s__Metschnikowia_pulcherrima(100); | Metschnikowia | 13 | Saprotroph | \|Undefined Saprotroph\| | Yeast | NULL | Probable | As nectar-tap_saprotroph (Põlme et al. 2020); Decay sugar-rich substrates (Põlme et al. 2020) | Põlme S, et al. 2020. Fungal Diversity 105:1-16 ((https://doi.org/10.1007/s13225-020-00466-2)) |
| Otu49128 | 1 | k__Fungi(100);p__Ascomycota(100);c__Saccharomycetes(100);o__Saccharomycetales(100);f__Metschnikowiaceae(100);g__Metschnikowia(100);s__Metschnikowia_pulcherrima(100); | Metschnikowia | 13 | Saprotroph | \|Undefined Saprotroph\| | Yeast | NULL | Probable | As nectar-tap_saprotroph (Põlme et al. 2020); Decay sugar-rich substrates (Põlme et al. 2020) | Põlme S, et al. 2020. Fungal Diversity 105:1-16 ((https://doi.org/10.1007/s13225-020-00466-2)) |
| Otu49129 | 1 | k__Fungi(100);p__Ascomycota(100);c__Saccharomycetes(100);o__Saccharomycetales(100);f__Metschnikowiaceae(100);g__Metschnikowia(100);g__Metschnikowia_unclassified(100); | Metschnikowia | 13 | Saprotroph | \|Undefined Saprotroph\| | Yeast | NULL | Probable | As nectar-tap_saprotroph (Põlme et al. 2020); Decay sugar-rich substrates (Põlme et al. 2020) | Põlme S, et al. 2020. Fungal Diversity 105:1-16 ((https://doi.org/10.1007/s13225-020-00466-2)) |
| Otu49130 | 1 | k__Fungi(100);p__Ascomycota(100);c__Saccharomycetes(100);o__Saccharomycetales(100);f__Metschnikowiaceae(100);g__Metschnikowia(100);s__Metschnikowia_pulcherrima(100); | Metschnikowia | 13 | Saprotroph | \|Undefined Saprotroph\| | Yeast | NULL | Probable | As nectar-tap_saprotroph (Põlme et al. 2020); Decay sugar-rich substrates (Põlme et al. 2020) | Põlme S, et al. 2020. Fungal Diversity 105:1-16 ((https://doi.org/10.1007/s13225-020-00466-2)) |
| Otu49131 | 1 | k__Fungi(100);p__Ascomycota(100);c__Saccharomycetes(100);o__Saccharomycetales(100);f__Metschnikowiaceae(100);g__Metschnikowia(100);s__Metschnikowia_pulcherrima(100); | Metschnikowia | 13 | Saprotroph | \|Undefined Saprotroph\| | Yeast | NULL | Probable | As nectar-tap_saprotroph (Põlme et al. 2020); Decay sugar-rich substrates (Põlme et al. 2020) | Põlme S, et al. 2020. Fungal Diversity 105:1-16 ((https://doi.org/10.1007/s13225-020-00466-2)) |
| Otu49132 | 1 | k__Fungi(100);p__Ascomycota(100);c__Saccharomycetes(100);o__Saccharomycetales(100);f__Metschnikowiaceae(100);g__Metschnikowia(100);g__Metschnikowia_unclassified(100); | Metschnikowia | 13 | Saprotroph | \|Undefined Saprotroph\| | Yeast | NULL | Probable | As nectar-tap_saprotroph (Põlme et al. 2020); Decay sugar-rich substrates (Põlme et al. 2020) | Põlme S, et al. 2020. Fungal Diversity 105:1-16 ((https://doi.org/10.1007/s13225-020-00466-2)) |
| Otu49133 | 1 | k__Fungi(100);p__Ascomycota(100);c__Saccharomycetes(100);o__Saccharomycetales(100);f__Metschnikowiaceae(100);g__Metschnikowia(100);s__Metschnikowia_pulcherrima(100); | Metschnikowia | 13 | Saprotroph | \|Undefined Saprotroph\| | Yeast | NULL | Probable | As nectar-tap_saprotroph (Põlme et al. 2020); Decay sugar-rich substrates (Põlme et al. 2020) | Põlme S, et al. 2020. Fungal Diversity 105:1-16 ((https://doi.org/10.1007/s13225-020-00466-2)) |
| Otu49134 | 1 | k__Fungi(100);p__Ascomycota(100);c__Saccharomycetes(100);o__Saccharomycetales(100);f__Metschnikowiaceae(100);g__Metschnikowia(100);s__Metschnikowia_pulcherrima(100); | Metschnikowia | 13 | Saprotroph | \|Undefined Saprotroph\| | Yeast | NULL | Probable | As nectar-tap_saprotroph (Põlme et al. 2020); Decay sugar-rich substrates (Põlme et al. 2020) | Põlme S, et al. 2020. Fungal Diversity 105:1-16 ((https://doi.org/10.1007/s13225-020-00466-2)) |
| Otu49135 | 1 | k__Fungi(100);p__Ascomycota(100);c__Saccharomycetes(100);o__Saccharomycetales(100);f__Metschnikowiaceae(100);g__Metschnikowia(100);s__Metschnikowia_pulcherrima(100); | Metschnikowia | 13 | Saprotroph | \|Undefined Saprotroph\| | Yeast | NULL | Probable | As nectar-tap_saprotroph (Põlme et al. 2020); Decay sugar-rich substrates (Põlme et al. 2020) | Põlme S, et al. 2020. Fungal Diversity 105:1-16 ((https://doi.org/10.1007/s13225-020-00466-2)) |
| Otu49136 | 1 | k__Fungi(100);p__Ascomycota(100);c__Saccharomycetes(100);o__Saccharomycetales(100);f__Metschnikowiaceae(100);g__Metschnikowia(100);s__Metschnikowia_pulcherrima(100); | Metschnikowia | 13 | Saprotroph | \|Undefined Saprotroph\| | Yeast | NULL | Probable | As nectar-tap_saprotroph (Põlme et al. 2020); Decay sugar-rich substrates (Põlme et al. 2020) | Põlme S, et al. 2020. Fungal Diversity 105:1-16 ((https://doi.org/10.1007/s13225-020-00466-2)) |
| Otu49137 | 1 | k__Fungi(100);p__Ascomycota(100);c__Saccharomycetes(100);o__Saccharomycetales(100);f__Metschnikowiaceae(100);g__Metschnikowia(100);g__Metschnikowia_unclassified(100); | Metschnikowia | 13 | Saprotroph | \|Undefined Saprotroph\| | Yeast | NULL | Probable | As nectar-tap_saprotroph (Põlme et al. 2020); Decay sugar-rich substrates (Põlme et al. 2020) | Põlme S, et al. 2020. Fungal Diversity 105:1-16 ((https://doi.org/10.1007/s13225-020-00466-2)) |
| Otu49138 | 1 | k__Fungi(100);p__Ascomycota(100);c__Saccharomycetes(100);o__Saccharomycetales(100);f__Metschnikowiaceae(100);g__Metschnikowia(100);s__Metschnikowia_pulcherrima(100); | Metschnikowia | 13 | Saprotroph | \|Undefined Saprotroph\| | Yeast | NULL | Probable | As nectar-tap_saprotroph (Põlme et al. 2020); Decay sugar-rich substrates (Põlme et al. 2020) | Põlme S, et al. 2020. Fungal Diversity 105:1-16 ((https://doi.org/10.1007/s13225-020-00466-2)) |
| Otu49139 | 1 | k__Fungi(100);p__Ascomycota(100);c__Saccharomycetes(100);o__Saccharomycetales(100);f__Metschnikowiaceae(100);g__Metschnikowia(100);g__Metschnikowia_unclassified(100); | Metschnikowia | 13 | Saprotroph | \|Undefined Saprotroph\| | Yeast | NULL | Probable | As nectar-tap_saprotroph (Põlme et al. 2020); Decay sugar-rich substrates (Põlme et al. 2020) | Põlme S, et al. 2020. Fungal Diversity 105:1-16 ((https://doi.org/10.1007/s13225-020-00466-2)) |
| Otu49140 | 1 | k__Fungi(100);p__Ascomycota(100);c__Saccharomycetes(100);o__Saccharomycetales(100);f__Metschnikowiaceae(100);g__Metschnikowia(100);g__Metschnikowia_unclassified(100); | Metschnikowia | 13 | Saprotroph | \|Undefined Saprotroph\| | Yeast | NULL | Probable | As nectar-tap_saprotroph (Põlme et al. 2020); Decay sugar-rich substrates (Põlme et al. 2020) | Põlme S, et al. 2020. Fungal Diversity 105:1-16 ((https://doi.org/10.1007/s13225-020-00466-2)) |
| Otu49141 | 1 | k__Fungi(100);p__Ascomycota(100);c__Saccharomycetes(100);o__Saccharomycetales(100);f__Metschnikowiaceae(100);g__Metschnikowia(100);s__Metschnikowia_pulcherrima(100); | Metschnikowia | 13 | Saprotroph | \|Undefined Saprotroph\| | Yeast | NULL | Probable | As nectar-tap_saprotroph (Põlme et al. 2020); Decay sugar-rich substrates (Põlme et al. 2020) | Põlme S, et al. 2020. Fungal Diversity 105:1-16 ((https://doi.org/10.1007/s13225-020-00466-2)) |
| Otu49142 | 1 | k__Fungi(100);p__Ascomycota(100);c__Saccharomycetes(100);o__Saccharomycetales(100);f__Metschnikowiaceae(100);g__Metschnikowia(100);s__Metschnikowia_pulcherrima(100); | Metschnikowia | 13 | Saprotroph | \|Undefined Saprotroph\| | Yeast | NULL | Probable | As nectar-tap_saprotroph (Põlme et al. 2020); Decay sugar-rich substrates (Põlme et al. 2020) | Põlme S, et al. 2020. Fungal Diversity 105:1-16 ((https://doi.org/10.1007/s13225-020-00466-2)) |
| Otu49143 | 1 | k__Fungi(100);p__Ascomycota(100);c__Saccharomycetes(100);o__Saccharomycetales(100);f__Metschnikowiaceae(100);g__Metschnikowia(100);s__Metschnikowia_pulcherrima(100); | Metschnikowia | 13 | Saprotroph | \|Undefined Saprotroph\| | Yeast | NULL | Probable | As nectar-tap_saprotroph (Põlme et al. 2020); Decay sugar-rich substrates (Põlme et al. 2020) | Põlme S, et al. 2020. Fungal Diversity 105:1-16 ((https://doi.org/10.1007/s13225-020-00466-2)) |
| Otu49144 | 1 | k__Fungi(100);p__Ascomycota(100);c__Saccharomycetes(100);o__Saccharomycetales(100);f__Metschnikowiaceae(100);g__Metschnikowia(100);s__Metschnikowia_pulcherrima(100); | Metschnikowia | 13 | Saprotroph | \|Undefined Saprotroph\| | Yeast | NULL | Probable | As nectar-tap_saprotroph (Põlme et al. 2020); Decay sugar-rich substrates (Põlme et al. 2020) | Põlme S, et al. 2020. Fungal Diversity 105:1-16 ((https://doi.org/10.1007/s13225-020-00466-2)) |
| Otu49145 | 1 | k__Fungi(100);p__Ascomycota(100);c__Saccharomycetes(100);o__Saccharomycetales(100);f__Metschnikowiaceae(100);g__Metschnikowia(100);s__Metschnikowia_pulcherrima(100); | Metschnikowia | 13 | Saprotroph | \|Undefined Saprotroph\| | Yeast | NULL | Probable | As nectar-tap_saprotroph (Põlme et al. 2020); Decay sugar-rich substrates (Põlme et al. 2020) | Põlme S, et al. 2020. Fungal Diversity 105:1-16 ((https://doi.org/10.1007/s13225-020-00466-2)) |
| Otu49146 | 1 | k__Fungi(100);p__Ascomycota(100);c__Saccharomycetes(100);o__Saccharomycetales(100);f__Metschnikowiaceae(100);g__Metschnikowia(100);s__Metschnikowia_pulcherrima(100); | Metschnikowia | 13 | Saprotroph | \|Undefined Saprotroph\| | Yeast | NULL | Probable | As nectar-tap_saprotroph (Põlme et al. 2020); Decay sugar-rich substrates (Põlme et al. 2020) | Põlme S, et al. 2020. Fungal Diversity 105:1-16 ((https://doi.org/10.1007/s13225-020-00466-2)) |
| Otu49147 | 1 | k__Fungi(100);p__Ascomycota(100);c__Saccharomycetes(100);o__Saccharomycetales(100);f__Metschnikowiaceae(100);g__Metschnikowia(100);s__Metschnikowia_pulcherrima(100); | Metschnikowia | 13 | Saprotroph | \|Undefined Saprotroph\| | Yeast | NULL | Probable | As nectar-tap_saprotroph (Põlme et al. 2020); Decay sugar-rich substrates (Põlme et al. 2020) | Põlme S, et al. 2020. Fungal Diversity 105:1-16 ((https://doi.org/10.1007/s13225-020-00466-2)) |
| Otu49148 | 1 | k__Fungi(100);p__Ascomycota(100);c__Saccharomycetes(100);o__Saccharomycetales(100);f__Metschnikowiaceae(100);g__Metschnikowia(100);s__Metschnikowia_pulcherrima(100); | Metschnikowia | 13 | Saprotroph | \|Undefined Saprotroph\| | Yeast | NULL | Probable | As nectar-tap_saprotroph (Põlme et al. 2020); Decay sugar-rich substrates (Põlme et al. 2020) | Põlme S, et al. 2020. Fungal Diversity 105:1-16 ((https://doi.org/10.1007/s13225-020-00466-2)) |
| Otu49149 | 1 | k__Fungi(100);p__Ascomycota(100);c__Saccharomycetes(100);o__Saccharomycetales(100);f__Metschnikowiaceae(100);g__Metschnikowia(100);s__Metschnikowia_pulcherrima(100); | Metschnikowia | 13 | Saprotroph | \|Undefined Saprotroph\| | Yeast | NULL | Probable | As nectar-tap_saprotroph (Põlme et al. 2020); Decay sugar-rich substrates (Põlme et al. 2020) | Põlme S, et al. 2020. Fungal Diversity 105:1-16 ((https://doi.org/10.1007/s13225-020-00466-2)) |
| Otu49150 | 1 | k__Fungi(100);p__Ascomycota(100);c__Saccharomycetes(100);o__Saccharomycetales(100);f__Metschnikowiaceae(100);g__Metschnikowia(100);s__Metschnikowia_pulcherrima(100); | Metschnikowia | 13 | Saprotroph | \|Undefined Saprotroph\| | Yeast | NULL | Probable | As nectar-tap_saprotroph (Põlme et al. 2020); Decay sugar-rich substrates (Põlme et al. 2020) | Põlme S, et al. 2020. Fungal Diversity 105:1-16 ((https://doi.org/10.1007/s13225-020-00466-2)) |
| Otu49151 | 1 | k__Fungi(100);p__Ascomycota(100);c__Saccharomycetes(100);o__Saccharomycetales(100);f__Metschnikowiaceae(100);g__Metschnikowia(100);s__Metschnikowia_pulcherrima(100); | Metschnikowia | 13 | Saprotroph | \|Undefined Saprotroph\| | Yeast | NULL | Probable | As nectar-tap_saprotroph (Põlme et al. 2020); Decay sugar-rich substrates (Põlme et al. 2020) | Põlme S, et al. 2020. Fungal Diversity 105:1-16 ((https://doi.org/10.1007/s13225-020-00466-2)) |
| Otu49152 | 1 | k__Fungi(100);p__Ascomycota(100);c__Saccharomycetes(100);o__Saccharomycetales(100);f__Metschnikowiaceae(100);g__Metschnikowia(100);s__Metschnikowia_pulcherrima(100); | Metschnikowia | 13 | Saprotroph | \|Undefined Saprotroph\| | Yeast | NULL | Probable | As nectar-tap_saprotroph (Põlme et al. 2020); Decay sugar-rich substrates (Põlme et al. 2020) | Põlme S, et al. 2020. Fungal Diversity 105:1-16 ((https://doi.org/10.1007/s13225-020-00466-2)) |
| Otu49153 | 1 | k__Fungi(100);p__Ascomycota(100);c__Saccharomycetes(100);o__Saccharomycetales(100);f__Metschnikowiaceae(100);g__Metschnikowia(100);s__Metschnikowia_pulcherrima(100); | Metschnikowia | 13 | Saprotroph | \|Undefined Saprotroph\| | Yeast | NULL | Probable | As nectar-tap_saprotroph (Põlme et al. 2020); Decay sugar-rich substrates (Põlme et al. 2020) | Põlme S, et al. 2020. Fungal Diversity 105:1-16 ((https://doi.org/10.1007/s13225-020-00466-2)) |
| Otu49154 | 1 | k__Fungi(100);p__Ascomycota(100);c__Saccharomycetes(100);o__Saccharomycetales(100);f__Metschnikowiaceae(100);g__Metschnikowia(100);s__Metschnikowia_pulcherrima(100); | Metschnikowia | 13 | Saprotroph | \|Undefined Saprotroph\| | Yeast | NULL | Probable | As nectar-tap_saprotroph (Põlme et al. 2020); Decay sugar-rich substrates (Põlme et al. 2020) | Põlme S, et al. 2020. Fungal Diversity 105:1-16 ((https://doi.org/10.1007/s13225-020-00466-2)) |
| Otu49155 | 1 | k__Fungi(100);p__Ascomycota(100);c__Saccharomycetes(100);o__Saccharomycetales(100);f__Metschnikowiaceae(100);g__Metschnikowia(100);g__Metschnikowia_unclassified(100); | Metschnikowia | 13 | Saprotroph | \|Undefined Saprotroph\| | Yeast | NULL | Probable | As nectar-tap_saprotroph (Põlme et al. 2020); Decay sugar-rich substrates (Põlme et al. 2020) | Põlme S, et al. 2020. Fungal Diversity 105:1-16 ((https://doi.org/10.1007/s13225-020-00466-2)) |
| Otu49157 | 1 | k__Fungi(100);p__Ascomycota(100);c__Saccharomycetes(100);o__Saccharomycetales(100);f__Metschnikowiaceae(100);g__Metschnikowia(100);s__Metschnikowia_pulcherrima(100); | Metschnikowia | 13 | Saprotroph | \|Undefined Saprotroph\| | Yeast | NULL | Probable | As nectar-tap_saprotroph (Põlme et al. 2020); Decay sugar-rich substrates (Põlme et al. 2020) | Põlme S, et al. 2020. Fungal Diversity 105:1-16 ((https://doi.org/10.1007/s13225-020-00466-2)) |
| Otu49158 | 1 | k__Fungi(100);p__Ascomycota(100);c__Saccharomycetes(100);o__Saccharomycetales(100);f__Metschnikowiaceae(100);g__Metschnikowia(100);s__Metschnikowia_pulcherrima(100); | Metschnikowia | 13 | Saprotroph | \|Undefined Saprotroph\| | Yeast | NULL | Probable | As nectar-tap_saprotroph (Põlme et al. 2020); Decay sugar-rich substrates (Põlme et al. 2020) | Põlme S, et al. 2020. Fungal Diversity 105:1-16 ((https://doi.org/10.1007/s13225-020-00466-2)) |
| Otu49159 | 1 | k__Fungi(100);p__Ascomycota(100);c__Saccharomycetes(100);o__Saccharomycetales(100);f__Metschnikowiaceae(100);g__Metschnikowia(100);s__Metschnikowia_pulcherrima(100); | Metschnikowia | 13 | Saprotroph | \|Undefined Saprotroph\| | Yeast | NULL | Probable | As nectar-tap_saprotroph (Põlme et al. 2020); Decay sugar-rich substrates (Põlme et al. 2020) | Põlme S, et al. 2020. Fungal Diversity 105:1-16 ((https://doi.org/10.1007/s13225-020-00466-2)) |
| Otu49160 | 1 | k__Fungi(100);p__Ascomycota(100);c__Saccharomycetes(100);o__Saccharomycetales(100);f__Metschnikowiaceae(100);g__Metschnikowia(100);s__Metschnikowia_pulcherrima(100); | Metschnikowia | 13 | Saprotroph | \|Undefined Saprotroph\| | Yeast | NULL | Probable | As nectar-tap_saprotroph (Põlme et al. 2020); Decay sugar-rich substrates (Põlme et al. 2020) | Põlme S, et al. 2020. Fungal Diversity 105:1-16 ((https://doi.org/10.1007/s13225-020-00466-2)) |
| Otu49161 | 1 | k__Fungi(100);p__Ascomycota(100);c__Saccharomycetes(100);o__Saccharomycetales(100);f__Metschnikowiaceae(100);g__Metschnikowia(100);s__Metschnikowia_pulcherrima(100); | Metschnikowia | 13 | Saprotroph | \|Undefined Saprotroph\| | Yeast | NULL | Probable | As nectar-tap_saprotroph (Põlme et al. 2020); Decay sugar-rich substrates (Põlme et al. 2020) | Põlme S, et al. 2020. Fungal Diversity 105:1-16 ((https://doi.org/10.1007/s13225-020-00466-2)) |
| Otu49162 | 1 | k__Fungi(100);p__Ascomycota(100);c__Saccharomycetes(100);o__Saccharomycetales(100);f__Metschnikowiaceae(100);g__Metschnikowia(100);s__Metschnikowia_pulcherrima(100); | Metschnikowia | 13 | Saprotroph | \|Undefined Saprotroph\| | Yeast | NULL | Probable | As nectar-tap_saprotroph (Põlme et al. 2020); Decay sugar-rich substrates (Põlme et al. 2020) | Põlme S, et al. 2020. Fungal Diversity 105:1-16 ((https://doi.org/10.1007/s13225-020-00466-2)) |
| Otu49163 | 1 | k__Fungi(100);p__Ascomycota(100);c__Saccharomycetes(100);o__Saccharomycetales(100);f__Metschnikowiaceae(100);g__Metschnikowia(100);s__Metschnikowia_pulcherrima(100); | Metschnikowia | 13 | Saprotroph | \|Undefined Saprotroph\| | Yeast | NULL | Probable | As nectar-tap_saprotroph (Põlme et al. 2020); Decay sugar-rich substrates (Põlme et al. 2020) | Põlme S, et al. 2020. Fungal Diversity 105:1-16 ((https://doi.org/10.1007/s13225-020-00466-2)) |
| Otu49164 | 1 | k__Fungi(100);p__Ascomycota(100);c__Saccharomycetes(100);o__Saccharomycetales(100);f__Metschnikowiaceae(100);g__Metschnikowia(100);g__Metschnikowia_unclassified(100); | Metschnikowia | 13 | Saprotroph | \|Undefined Saprotroph\| | Yeast | NULL | Probable | As nectar-tap_saprotroph (Põlme et al. 2020); Decay sugar-rich substrates (Põlme et al. 2020) | Põlme S, et al. 2020. Fungal Diversity 105:1-16 ((https://doi.org/10.1007/s13225-020-00466-2)) |
| Otu49165 | 1 | k__Fungi(100);p__Ascomycota(100);c__Saccharomycetes(100);o__Saccharomycetales(100);f__Metschnikowiaceae(100);g__Metschnikowia(100);s__Metschnikowia_pulcherrima(100); | Metschnikowia | 13 | Saprotroph | \|Undefined Saprotroph\| | Yeast | NULL | Probable | As nectar-tap_saprotroph (Põlme et al. 2020); Decay sugar-rich substrates (Põlme et al. 2020) | Põlme S, et al. 2020. Fungal Diversity 105:1-16 ((https://doi.org/10.1007/s13225-020-00466-2)) |
| Otu49166 | 1 | k__Fungi(100);p__Ascomycota(100);c__Saccharomycetes(100);o__Saccharomycetales(100);f__Metschnikowiaceae(100);g__Metschnikowia(100);s__Metschnikowia_pulcherrima(100); | Metschnikowia | 13 | Saprotroph | \|Undefined Saprotroph\| | Yeast | NULL | Probable | As nectar-tap_saprotroph (Põlme et al. 2020); Decay sugar-rich substrates (Põlme et al. 2020) | Põlme S, et al. 2020. Fungal Diversity 105:1-16 ((https://doi.org/10.1007/s13225-020-00466-2)) |
| Otu49167 | 1 | k__Fungi(100);p__Ascomycota(100);c__Saccharomycetes(100);o__Saccharomycetales(100);f__Metschnikowiaceae(100);g__Metschnikowia(100);s__Metschnikowia_pulcherrima(100); | Metschnikowia | 13 | Saprotroph | \|Undefined Saprotroph\| | Yeast | NULL | Probable | As nectar-tap_saprotroph (Põlme et al. 2020); Decay sugar-rich substrates (Põlme et al. 2020) | Põlme S, et al. 2020. Fungal Diversity 105:1-16 ((https://doi.org/10.1007/s13225-020-00466-2)) |
| Otu49168 | 1 | k__Fungi(100);p__Ascomycota(100);c__Saccharomycetes(100);o__Saccharomycetales(100);f__Metschnikowiaceae(100);g__Metschnikowia(100);s__Metschnikowia_pulcherrima(100); | Metschnikowia | 13 | Saprotroph | \|Undefined Saprotroph\| | Yeast | NULL | Probable | As nectar-tap_saprotroph (Põlme et al. 2020); Decay sugar-rich substrates (Põlme et al. 2020) | Põlme S, et al. 2020. Fungal Diversity 105:1-16 ((https://doi.org/10.1007/s13225-020-00466-2)) |
| Otu49169 | 1 | k__Fungi(100);p__Ascomycota(100);c__Saccharomycetes(100);o__Saccharomycetales(100);f__Metschnikowiaceae(100);g__Metschnikowia(100);g__Metschnikowia_unclassified(100); | Metschnikowia | 13 | Saprotroph | \|Undefined Saprotroph\| | Yeast | NULL | Probable | As nectar-tap_saprotroph (Põlme et al. 2020); Decay sugar-rich substrates (Põlme et al. 2020) | Põlme S, et al. 2020. Fungal Diversity 105:1-16 ((https://doi.org/10.1007/s13225-020-00466-2)) |
| Otu49170 | 1 | k__Fungi(100);p__Ascomycota(100);c__Saccharomycetes(100);o__Saccharomycetales(100);f__Metschnikowiaceae(100);g__Metschnikowia(100);s__Metschnikowia_pulcherrima(100); | Metschnikowia | 13 | Saprotroph | \|Undefined Saprotroph\| | Yeast | NULL | Probable | As nectar-tap_saprotroph (Põlme et al. 2020); Decay sugar-rich substrates (Põlme et al. 2020) | Põlme S, et al. 2020. Fungal Diversity 105:1-16 ((https://doi.org/10.1007/s13225-020-00466-2)) |
| Otu49171 | 1 | k__Fungi(100);p__Ascomycota(100);c__Saccharomycetes(100);o__Saccharomycetales(100);f__Metschnikowiaceae(100);g__Metschnikowia(100);s__Metschnikowia_pulcherrima(100); | Metschnikowia | 13 | Saprotroph | \|Undefined Saprotroph\| | Yeast | NULL | Probable | As nectar-tap_saprotroph (Põlme et al. 2020); Decay sugar-rich substrates (Põlme et al. 2020) | Põlme S, et al. 2020. Fungal Diversity 105:1-16 ((https://doi.org/10.1007/s13225-020-00466-2)) |
| Otu49172 | 1 | k__Fungi(100);p__Ascomycota(100);c__Saccharomycetes(100);o__Saccharomycetales(100);f__Metschnikowiaceae(100);g__Metschnikowia(100);g__Metschnikowia_unclassified(100); | Metschnikowia | 13 | Saprotroph | \|Undefined Saprotroph\| | Yeast | NULL | Probable | As nectar-tap_saprotroph (Põlme et al. 2020); Decay sugar-rich substrates (Põlme et al. 2020) | Põlme S, et al. 2020. Fungal Diversity 105:1-16 ((https://doi.org/10.1007/s13225-020-00466-2)) |
| Otu49173 | 1 | k__Fungi(100);p__Ascomycota(100);c__Saccharomycetes(100);o__Saccharomycetales(100);f__Metschnikowiaceae(100);g__Metschnikowia(100);s__Metschnikowia_pulcherrima(100); | Metschnikowia | 13 | Saprotroph | \|Undefined Saprotroph\| | Yeast | NULL | Probable | As nectar-tap_saprotroph (Põlme et al. 2020); Decay sugar-rich substrates (Põlme et al. 2020) | Põlme S, et al. 2020. Fungal Diversity 105:1-16 ((https://doi.org/10.1007/s13225-020-00466-2)) |
| Otu49174 | 1 | k__Fungi(100);p__Ascomycota(100);c__Saccharomycetes(100);o__Saccharomycetales(100);f__Metschnikowiaceae(100);g__Metschnikowia(100);s__Metschnikowia_pulcherrima(100); | Metschnikowia | 13 | Saprotroph | \|Undefined Saprotroph\| | Yeast | NULL | Probable | As nectar-tap_saprotroph (Põlme et al. 2020); Decay sugar-rich substrates (Põlme et al. 2020) | Põlme S, et al. 2020. Fungal Diversity 105:1-16 ((https://doi.org/10.1007/s13225-020-00466-2)) |
| Otu49175 | 1 | k__Fungi(100);p__Ascomycota(100);c__Saccharomycetes(100);o__Saccharomycetales(100);f__Metschnikowiaceae(100);g__Metschnikowia(100);s__Metschnikowia_pulcherrima(100); | Metschnikowia | 13 | Saprotroph | \|Undefined Saprotroph\| | Yeast | NULL | Probable | As nectar-tap_saprotroph (Põlme et al. 2020); Decay sugar-rich substrates (Põlme et al. 2020) | Põlme S, et al. 2020. Fungal Diversity 105:1-16 ((https://doi.org/10.1007/s13225-020-00466-2)) |
| Otu49176 | 1 | k__Fungi(100);p__Ascomycota(100);c__Saccharomycetes(100);o__Saccharomycetales(100);f__Metschnikowiaceae(100);g__Metschnikowia(100);g__Metschnikowia_unclassified(100); | Metschnikowia | 13 | Saprotroph | \|Undefined Saprotroph\| | Yeast | NULL | Probable | As nectar-tap_saprotroph (Põlme et al. 2020); Decay sugar-rich substrates (Põlme et al. 2020) | Põlme S, et al. 2020. Fungal Diversity 105:1-16 ((https://doi.org/10.1007/s13225-020-00466-2)) |
| Otu49177 | 1 | k__Fungi(100);p__Ascomycota(100);c__Saccharomycetes(100);o__Saccharomycetales(100);f__Metschnikowiaceae(100);g__Metschnikowia(100);s__Metschnikowia_pulcherrima(100); | Metschnikowia | 13 | Saprotroph | \|Undefined Saprotroph\| | Yeast | NULL | Probable | As nectar-tap_saprotroph (Põlme et al. 2020); Decay sugar-rich substrates (Põlme et al. 2020) | Põlme S, et al. 2020. Fungal Diversity 105:1-16 ((https://doi.org/10.1007/s13225-020-00466-2)) |
| Otu49178 | 1 | k__Fungi(100);p__Ascomycota(100);c__Saccharomycetes(100);o__Saccharomycetales(100);f__Metschnikowiaceae(100);g__Metschnikowia(100);s__Metschnikowia_pulcherrima(100); | Metschnikowia | 13 | Saprotroph | \|Undefined Saprotroph\| | Yeast | NULL | Probable | As nectar-tap_saprotroph (Põlme et al. 2020); Decay sugar-rich substrates (Põlme et al. 2020) | Põlme S, et al. 2020. Fungal Diversity 105:1-16 ((https://doi.org/10.1007/s13225-020-00466-2)) |
| Otu49179 | 1 | k__Fungi(100);p__Ascomycota(100);c__Saccharomycetes(100);o__Saccharomycetales(100);f__Metschnikowiaceae(100);g__Metschnikowia(100);g__Metschnikowia_unclassified(100); | Metschnikowia | 13 | Saprotroph | \|Undefined Saprotroph\| | Yeast | NULL | Probable | As nectar-tap_saprotroph (Põlme et al. 2020); Decay sugar-rich substrates (Põlme et al. 2020) | Põlme S, et al. 2020. Fungal Diversity 105:1-16 ((https://doi.org/10.1007/s13225-020-00466-2)) |
| Otu49181 | 1 | k__Fungi(100);p__Ascomycota(100);c__Saccharomycetes(100);o__Saccharomycetales(100);f__Metschnikowiaceae(100);g__Metschnikowia(100);s__Metschnikowia_pulcherrima(100); | Metschnikowia | 13 | Saprotroph | \|Undefined Saprotroph\| | Yeast | NULL | Probable | As nectar-tap_saprotroph (Põlme et al. 2020); Decay sugar-rich substrates (Põlme et al. 2020) | Põlme S, et al. 2020. Fungal Diversity 105:1-16 ((https://doi.org/10.1007/s13225-020-00466-2)) |
| Otu49182 | 1 | k__Fungi(100);p__Ascomycota(100);c__Saccharomycetes(100);o__Saccharomycetales(100);f__Metschnikowiaceae(100);g__Metschnikowia(100);s__Metschnikowia_pulcherrima(100); | Metschnikowia | 13 | Saprotroph | \|Undefined Saprotroph\| | Yeast | NULL | Probable | As nectar-tap_saprotroph (Põlme et al. 2020); Decay sugar-rich substrates (Põlme et al. 2020) | Põlme S, et al. 2020. Fungal Diversity 105:1-16 ((https://doi.org/10.1007/s13225-020-00466-2)) |
| Otu49183 | 1 | k__Fungi(100);p__Ascomycota(100);c__Saccharomycetes(100);o__Saccharomycetales(100);f__Metschnikowiaceae(100);g__Metschnikowia(100);s__Metschnikowia_pulcherrima(100); | Metschnikowia | 13 | Saprotroph | \|Undefined Saprotroph\| | Yeast | NULL | Probable | As nectar-tap_saprotroph (Põlme et al. 2020); Decay sugar-rich substrates (Põlme et al. 2020) | Põlme S, et al. 2020. Fungal Diversity 105:1-16 ((https://doi.org/10.1007/s13225-020-00466-2)) |
| Otu49184 | 1 | k__Fungi(100);p__Ascomycota(100);c__Saccharomycetes(100);o__Saccharomycetales(100);f__Metschnikowiaceae(100);g__Metschnikowia(100);s__Metschnikowia_pulcherrima(100); | Metschnikowia | 13 | Saprotroph | \|Undefined Saprotroph\| | Yeast | NULL | Probable | As nectar-tap_saprotroph (Põlme et al. 2020); Decay sugar-rich substrates (Põlme et al. 2020) | Põlme S, et al. 2020. Fungal Diversity 105:1-16 ((https://doi.org/10.1007/s13225-020-00466-2)) |
| Otu49185 | 1 | k__Fungi(100);p__Ascomycota(100);c__Saccharomycetes(100);o__Saccharomycetales(100);f__Metschnikowiaceae(100);g__Metschnikowia(100);s__Metschnikowia_pulcherrima(100); | Metschnikowia | 13 | Saprotroph | \|Undefined Saprotroph\| | Yeast | NULL | Probable | As nectar-tap_saprotroph (Põlme et al. 2020); Decay sugar-rich substrates (Põlme et al. 2020) | Põlme S, et al. 2020. Fungal Diversity 105:1-16 ((https://doi.org/10.1007/s13225-020-00466-2)) |
| Otu49186 | 1 | k__Fungi(100);p__Ascomycota(100);c__Saccharomycetes(100);o__Saccharomycetales(100);f__Metschnikowiaceae(100);g__Metschnikowia(100);s__Metschnikowia_pulcherrima(100); | Metschnikowia | 13 | Saprotroph | \|Undefined Saprotroph\| | Yeast | NULL | Probable | As nectar-tap_saprotroph (Põlme et al. 2020); Decay sugar-rich substrates (Põlme et al. 2020) | Põlme S, et al. 2020. Fungal Diversity 105:1-16 ((https://doi.org/10.1007/s13225-020-00466-2)) |
| Otu49187 | 1 | k__Fungi(100);p__Ascomycota(100);c__Saccharomycetes(100);o__Saccharomycetales(100);f__Metschnikowiaceae(100);g__Metschnikowia(100);g__Metschnikowia_unclassified(100); | Metschnikowia | 13 | Saprotroph | \|Undefined Saprotroph\| | Yeast | NULL | Probable | As nectar-tap_saprotroph (Põlme et al. 2020); Decay sugar-rich substrates (Põlme et al. 2020) | Põlme S, et al. 2020. Fungal Diversity 105:1-16 ((https://doi.org/10.1007/s13225-020-00466-2)) |
| Otu49188 | 1 | k__Fungi(100);p__Ascomycota(100);c__Saccharomycetes(100);o__Saccharomycetales(100);f__Metschnikowiaceae(100);g__Metschnikowia(100);s__Metschnikowia_pulcherrima(100); | Metschnikowia | 13 | Saprotroph | \|Undefined Saprotroph\| | Yeast | NULL | Probable | As nectar-tap_saprotroph (Põlme et al. 2020); Decay sugar-rich substrates (Põlme et al. 2020) | Põlme S, et al. 2020. Fungal Diversity 105:1-16 ((https://doi.org/10.1007/s13225-020-00466-2)) |
| Otu49189 | 1 | k__Fungi(100);p__Ascomycota(100);c__Saccharomycetes(100);o__Saccharomycetales(100);f__Metschnikowiaceae(100);g__Metschnikowia(100);s__Metschnikowia_pulcherrima(100); | Metschnikowia | 13 | Saprotroph | \|Undefined Saprotroph\| | Yeast | NULL | Probable | As nectar-tap_saprotroph (Põlme et al. 2020); Decay sugar-rich substrates (Põlme et al. 2020) | Põlme S, et al. 2020. Fungal Diversity 105:1-16 ((https://doi.org/10.1007/s13225-020-00466-2)) |
| Otu49190 | 1 | k__Fungi(100);p__Ascomycota(100);c__Saccharomycetes(100);o__Saccharomycetales(100);f__Metschnikowiaceae(100);g__Metschnikowia(100);s__Metschnikowia_pulcherrima(100); | Metschnikowia | 13 | Saprotroph | \|Undefined Saprotroph\| | Yeast | NULL | Probable | As nectar-tap_saprotroph (Põlme et al. 2020); Decay sugar-rich substrates (Põlme et al. 2020) | Põlme S, et al. 2020. Fungal Diversity 105:1-16 ((https://doi.org/10.1007/s13225-020-00466-2)) |
| Otu49191 | 1 | k__Fungi(100);p__Ascomycota(100);c__Saccharomycetes(100);o__Saccharomycetales(100);f__Metschnikowiaceae(100);g__Metschnikowia(100);s__Metschnikowia_pulcherrima(100); | Metschnikowia | 13 | Saprotroph | \|Undefined Saprotroph\| | Yeast | NULL | Probable | As nectar-tap_saprotroph (Põlme et al. 2020); Decay sugar-rich substrates (Põlme et al. 2020) | Põlme S, et al. 2020. Fungal Diversity 105:1-16 ((https://doi.org/10.1007/s13225-020-00466-2)) |
| Otu49192 | 1 | k__Fungi(100);p__Ascomycota(100);c__Saccharomycetes(100);o__Saccharomycetales(100);f__Metschnikowiaceae(100);g__Metschnikowia(100);s__Metschnikowia_pulcherrima(100); | Metschnikowia | 13 | Saprotroph | \|Undefined Saprotroph\| | Yeast | NULL | Probable | As nectar-tap_saprotroph (Põlme et al. 2020); Decay sugar-rich substrates (Põlme et al. 2020) | Põlme S, et al. 2020. Fungal Diversity 105:1-16 ((https://doi.org/10.1007/s13225-020-00466-2)) |
| Otu49193 | 1 | k__Fungi(100);p__Ascomycota(100);c__Saccharomycetes(100);o__Saccharomycetales(100);f__Metschnikowiaceae(100);g__Metschnikowia(100);s__Metschnikowia_pulcherrima(100); | Metschnikowia | 13 | Saprotroph | \|Undefined Saprotroph\| | Yeast | NULL | Probable | As nectar-tap_saprotroph (Põlme et al. 2020); Decay sugar-rich substrates (Põlme et al. 2020) | Põlme S, et al. 2020. Fungal Diversity 105:1-16 ((https://doi.org/10.1007/s13225-020-00466-2)) |
| Otu49194 | 1 | k__Fungi(100);p__Ascomycota(100);c__Saccharomycetes(100);o__Saccharomycetales(100);f__Metschnikowiaceae(100);g__Metschnikowia(100);g__Metschnikowia_unclassified(100); | Metschnikowia | 13 | Saprotroph | \|Undefined Saprotroph\| | Yeast | NULL | Probable | As nectar-tap_saprotroph (Põlme et al. 2020); Decay sugar-rich substrates (Põlme et al. 2020) | Põlme S, et al. 2020. Fungal Diversity 105:1-16 ((https://doi.org/10.1007/s13225-020-00466-2)) |
| Otu49195 | 1 | k__Fungi(100);p__Ascomycota(100);c__Saccharomycetes(100);o__Saccharomycetales(100);f__Metschnikowiaceae(100);g__Metschnikowia(100);s__Metschnikowia_pulcherrima(100); | Metschnikowia | 13 | Saprotroph | \|Undefined Saprotroph\| | Yeast | NULL | Probable | As nectar-tap_saprotroph (Põlme et al. 2020); Decay sugar-rich substrates (Põlme et al. 2020) | Põlme S, et al. 2020. Fungal Diversity 105:1-16 ((https://doi.org/10.1007/s13225-020-00466-2)) |
| Otu49196 | 1 | k__Fungi(100);p__Ascomycota(100);c__Saccharomycetes(100);o__Saccharomycetales(100);f__Metschnikowiaceae(100);g__Metschnikowia(100);g__Metschnikowia_unclassified(100); | Metschnikowia | 13 | Saprotroph | \|Undefined Saprotroph\| | Yeast | NULL | Probable | As nectar-tap_saprotroph (Põlme et al. 2020); Decay sugar-rich substrates (Põlme et al. 2020) | Põlme S, et al. 2020. Fungal Diversity 105:1-16 ((https://doi.org/10.1007/s13225-020-00466-2)) |
| Otu49197 | 1 | k__Fungi(100);p__Ascomycota(100);c__Saccharomycetes(100);o__Saccharomycetales(100);f__Metschnikowiaceae(100);g__Metschnikowia(100);s__Metschnikowia_pulcherrima(100); | Metschnikowia | 13 | Saprotroph | \|Undefined Saprotroph\| | Yeast | NULL | Probable | As nectar-tap_saprotroph (Põlme et al. 2020); Decay sugar-rich substrates (Põlme et al. 2020) | Põlme S, et al. 2020. Fungal Diversity 105:1-16 ((https://doi.org/10.1007/s13225-020-00466-2)) |
| Otu49198 | 1 | k__Fungi(100);p__Ascomycota(100);c__Saccharomycetes(100);o__Saccharomycetales(100);f__Metschnikowiaceae(100);g__Metschnikowia(100);g__Metschnikowia_unclassified(100); | Metschnikowia | 13 | Saprotroph | \|Undefined Saprotroph\| | Yeast | NULL | Probable | As nectar-tap_saprotroph (Põlme et al. 2020); Decay sugar-rich substrates (Põlme et al. 2020) | Põlme S, et al. 2020. Fungal Diversity 105:1-16 ((https://doi.org/10.1007/s13225-020-00466-2)) |
| Otu49199 | 1 | k__Fungi(100);p__Ascomycota(100);c__Saccharomycetes(100);o__Saccharomycetales(100);f__Metschnikowiaceae(100);g__Metschnikowia(100);s__Metschnikowia_pulcherrima(100); | Metschnikowia | 13 | Saprotroph | \|Undefined Saprotroph\| | Yeast | NULL | Probable | As nectar-tap_saprotroph (Põlme et al. 2020); Decay sugar-rich substrates (Põlme et al. 2020) | Põlme S, et al. 2020. Fungal Diversity 105:1-16 ((https://doi.org/10.1007/s13225-020-00466-2)) |
| Otu49200 | 1 | k__Fungi(100);p__Ascomycota(100);c__Saccharomycetes(100);o__Saccharomycetales(100);f__Metschnikowiaceae(100);g__Metschnikowia(100);s__Metschnikowia_pulcherrima(100); | Metschnikowia | 13 | Saprotroph | \|Undefined Saprotroph\| | Yeast | NULL | Probable | As nectar-tap_saprotroph (Põlme et al. 2020); Decay sugar-rich substrates (Põlme et al. 2020) | Põlme S, et al. 2020. Fungal Diversity 105:1-16 ((https://doi.org/10.1007/s13225-020-00466-2)) |
| Otu49201 | 1 | k__Fungi(100);p__Ascomycota(100);c__Saccharomycetes(100);o__Saccharomycetales(100);f__Metschnikowiaceae(100);g__Metschnikowia(100);g__Metschnikowia_unclassified(100); | Metschnikowia | 13 | Saprotroph | \|Undefined Saprotroph\| | Yeast | NULL | Probable | As nectar-tap_saprotroph (Põlme et al. 2020); Decay sugar-rich substrates (Põlme et al. 2020) | Põlme S, et al. 2020. Fungal Diversity 105:1-16 ((https://doi.org/10.1007/s13225-020-00466-2)) |
| Otu49202 | 1 | k__Fungi(100);p__Ascomycota(100);c__Saccharomycetes(100);o__Saccharomycetales(100);f__Metschnikowiaceae(100);g__Metschnikowia(100);s__Metschnikowia_pulcherrima(100); | Metschnikowia | 13 | Saprotroph | \|Undefined Saprotroph\| | Yeast | NULL | Probable | As nectar-tap_saprotroph (Põlme et al. 2020); Decay sugar-rich substrates (Põlme et al. 2020) | Põlme S, et al. 2020. Fungal Diversity 105:1-16 ((https://doi.org/10.1007/s13225-020-00466-2)) |
| Otu49203 | 1 | k__Fungi(100);p__Ascomycota(100);c__Saccharomycetes(100);o__Saccharomycetales(100);f__Metschnikowiaceae(100);g__Metschnikowia(100);s__Metschnikowia_pulcherrima(100); | Metschnikowia | 13 | Saprotroph | \|Undefined Saprotroph\| | Yeast | NULL | Probable | As nectar-tap_saprotroph (Põlme et al. 2020); Decay sugar-rich substrates (Põlme et al. 2020) | Põlme S, et al. 2020. Fungal Diversity 105:1-16 ((https://doi.org/10.1007/s13225-020-00466-2)) |
| Otu49204 | 1 | k__Fungi(100);p__Ascomycota(100);c__Saccharomycetes(100);o__Saccharomycetales(100);f__Metschnikowiaceae(100);g__Metschnikowia(100);s__Metschnikowia_pulcherrima(100); | Metschnikowia | 13 | Saprotroph | \|Undefined Saprotroph\| | Yeast | NULL | Probable | As nectar-tap_saprotroph (Põlme et al. 2020); Decay sugar-rich substrates (Põlme et al. 2020) | Põlme S, et al. 2020. Fungal Diversity 105:1-16 ((https://doi.org/10.1007/s13225-020-00466-2)) |
| Otu49205 | 1 | k__Fungi(100);p__Ascomycota(100);c__Saccharomycetes(100);o__Saccharomycetales(100);f__Metschnikowiaceae(100);g__Metschnikowia(100);s__Metschnikowia_pulcherrima(100); | Metschnikowia | 13 | Saprotroph | \|Undefined Saprotroph\| | Yeast | NULL | Probable | As nectar-tap_saprotroph (Põlme et al. 2020); Decay sugar-rich substrates (Põlme et al. 2020) | Põlme S, et al. 2020. Fungal Diversity 105:1-16 ((https://doi.org/10.1007/s13225-020-00466-2)) |
| Otu49206 | 1 | k__Fungi(100);p__Ascomycota(100);c__Saccharomycetes(100);o__Saccharomycetales(100);f__Metschnikowiaceae(100);g__Metschnikowia(100);g__Metschnikowia_unclassified(100); | Metschnikowia | 13 | Saprotroph | \|Undefined Saprotroph\| | Yeast | NULL | Probable | As nectar-tap_saprotroph (Põlme et al. 2020); Decay sugar-rich substrates (Põlme et al. 2020) | Põlme S, et al. 2020. Fungal Diversity 105:1-16 ((https://doi.org/10.1007/s13225-020-00466-2)) |
| Otu49207 | 1 | k__Fungi(100);p__Ascomycota(100);c__Saccharomycetes(100);o__Saccharomycetales(100);f__Metschnikowiaceae(100);g__Metschnikowia(100);s__Metschnikowia_pulcherrima(100); | Metschnikowia | 13 | Saprotroph | \|Undefined Saprotroph\| | Yeast | NULL | Probable | As nectar-tap_saprotroph (Põlme et al. 2020); Decay sugar-rich substrates (Põlme et al. 2020) | Põlme S, et al. 2020. Fungal Diversity 105:1-16 ((https://doi.org/10.1007/s13225-020-00466-2)) |
| Otu49208 | 1 | k__Fungi(100);p__Ascomycota(100);c__Saccharomycetes(100);o__Saccharomycetales(100);f__Metschnikowiaceae(100);g__Metschnikowia(100);s__Metschnikowia_pulcherrima(100); | Metschnikowia | 13 | Saprotroph | \|Undefined Saprotroph\| | Yeast | NULL | Probable | As nectar-tap_saprotroph (Põlme et al. 2020); Decay sugar-rich substrates (Põlme et al. 2020) | Põlme S, et al. 2020. Fungal Diversity 105:1-16 ((https://doi.org/10.1007/s13225-020-00466-2)) |
| Otu49209 | 1 | k__Fungi(100);p__Ascomycota(100);c__Saccharomycetes(100);o__Saccharomycetales(100);f__Metschnikowiaceae(100);g__Metschnikowia(100);s__Metschnikowia_pulcherrima(100); | Metschnikowia | 13 | Saprotroph | \|Undefined Saprotroph\| | Yeast | NULL | Probable | As nectar-tap_saprotroph (Põlme et al. 2020); Decay sugar-rich substrates (Põlme et al. 2020) | Põlme S, et al. 2020. Fungal Diversity 105:1-16 ((https://doi.org/10.1007/s13225-020-00466-2)) |
| Otu49210 | 1 | k__Fungi(100);p__Ascomycota(100);c__Saccharomycetes(100);o__Saccharomycetales(100);f__Metschnikowiaceae(100);g__Metschnikowia(100);s__Metschnikowia_pulcherrima(100); | Metschnikowia | 13 | Saprotroph | \|Undefined Saprotroph\| | Yeast | NULL | Probable | As nectar-tap_saprotroph (Põlme et al. 2020); Decay sugar-rich substrates (Põlme et al. 2020) | Põlme S, et al. 2020. Fungal Diversity 105:1-16 ((https://doi.org/10.1007/s13225-020-00466-2)) |
| Otu49211 | 1 | k__Fungi(100);p__Ascomycota(100);c__Saccharomycetes(100);o__Saccharomycetales(100);f__Metschnikowiaceae(100);g__Metschnikowia(100);s__Metschnikowia_pulcherrima(100); | Metschnikowia | 13 | Saprotroph | \|Undefined Saprotroph\| | Yeast | NULL | Probable | As nectar-tap_saprotroph (Põlme et al. 2020); Decay sugar-rich substrates (Põlme et al. 2020) | Põlme S, et al. 2020. Fungal Diversity 105:1-16 ((https://doi.org/10.1007/s13225-020-00466-2)) |
| Otu49212 | 1 | k__Fungi(100);p__Ascomycota(100);c__Saccharomycetes(100);o__Saccharomycetales(100);f__Metschnikowiaceae(100);g__Metschnikowia(100);s__Metschnikowia_pulcherrima(100); | Metschnikowia | 13 | Saprotroph | \|Undefined Saprotroph\| | Yeast | NULL | Probable | As nectar-tap_saprotroph (Põlme et al. 2020); Decay sugar-rich substrates (Põlme et al. 2020) | Põlme S, et al. 2020. Fungal Diversity 105:1-16 ((https://doi.org/10.1007/s13225-020-00466-2)) |
| Otu49213 | 1 | k__Fungi(100);p__Ascomycota(100);c__Saccharomycetes(100);o__Saccharomycetales(100);f__Metschnikowiaceae(100);g__Metschnikowia(100);s__Metschnikowia_pulcherrima(100); | Metschnikowia | 13 | Saprotroph | \|Undefined Saprotroph\| | Yeast | NULL | Probable | As nectar-tap_saprotroph (Põlme et al. 2020); Decay sugar-rich substrates (Põlme et al. 2020) | Põlme S, et al. 2020. Fungal Diversity 105:1-16 ((https://doi.org/10.1007/s13225-020-00466-2)) |
| Otu49214 | 1 | k__Fungi(100);p__Ascomycota(100);c__Saccharomycetes(100);o__Saccharomycetales(100);f__Metschnikowiaceae(100);g__Metschnikowia(100);s__Metschnikowia_pulcherrima(100); | Metschnikowia | 13 | Saprotroph | \|Undefined Saprotroph\| | Yeast | NULL | Probable | As nectar-tap_saprotroph (Põlme et al. 2020); Decay sugar-rich substrates (Põlme et al. 2020) | Põlme S, et al. 2020. Fungal Diversity 105:1-16 ((https://doi.org/10.1007/s13225-020-00466-2)) |
| Otu49215 | 1 | k__Fungi(100);p__Ascomycota(100);c__Saccharomycetes(100);o__Saccharomycetales(100);f__Metschnikowiaceae(100);g__Metschnikowia(100);s__Metschnikowia_pulcherrima(100); | Metschnikowia | 13 | Saprotroph | \|Undefined Saprotroph\| | Yeast | NULL | Probable | As nectar-tap_saprotroph (Põlme et al. 2020); Decay sugar-rich substrates (Põlme et al. 2020) | Põlme S, et al. 2020. Fungal Diversity 105:1-16 ((https://doi.org/10.1007/s13225-020-00466-2)) |
| Otu49216 | 1 | k__Fungi(100);p__Ascomycota(100);c__Saccharomycetes(100);o__Saccharomycetales(100);f__Metschnikowiaceae(100);g__Metschnikowia(100);s__Metschnikowia_pulcherrima(100); | Metschnikowia | 13 | Saprotroph | \|Undefined Saprotroph\| | Yeast | NULL | Probable | As nectar-tap_saprotroph (Põlme et al. 2020); Decay sugar-rich substrates (Põlme et al. 2020) | Põlme S, et al. 2020. Fungal Diversity 105:1-16 ((https://doi.org/10.1007/s13225-020-00466-2)) |
| Otu49217 | 1 | k__Fungi(100);p__Ascomycota(100);c__Saccharomycetes(100);o__Saccharomycetales(100);f__Metschnikowiaceae(100);g__Metschnikowia(100);s__Metschnikowia_pulcherrima(100); | Metschnikowia | 13 | Saprotroph | \|Undefined Saprotroph\| | Yeast | NULL | Probable | As nectar-tap_saprotroph (Põlme et al. 2020); Decay sugar-rich substrates (Põlme et al. 2020) | Põlme S, et al. 2020. Fungal Diversity 105:1-16 ((https://doi.org/10.1007/s13225-020-00466-2)) |
| Otu49218 | 1 | k__Fungi(100);p__Ascomycota(100);c__Saccharomycetes(100);o__Saccharomycetales(100);f__Metschnikowiaceae(100);g__Metschnikowia(100);s__Metschnikowia_pulcherrima(100); | Metschnikowia | 13 | Saprotroph | \|Undefined Saprotroph\| | Yeast | NULL | Probable | As nectar-tap_saprotroph (Põlme et al. 2020); Decay sugar-rich substrates (Põlme et al. 2020) | Põlme S, et al. 2020. Fungal Diversity 105:1-16 ((https://doi.org/10.1007/s13225-020-00466-2)) |
| Otu49220 | 1 | k__Fungi(100);p__Ascomycota(100);c__Saccharomycetes(100);o__Saccharomycetales(100);f__Metschnikowiaceae(100);g__Metschnikowia(100);s__Metschnikowia_pulcherrima(100); | Metschnikowia | 13 | Saprotroph | \|Undefined Saprotroph\| | Yeast | NULL | Probable | As nectar-tap_saprotroph (Põlme et al. 2020); Decay sugar-rich substrates (Põlme et al. 2020) | Põlme S, et al. 2020. Fungal Diversity 105:1-16 ((https://doi.org/10.1007/s13225-020-00466-2)) |
| Otu49221 | 1 | k__Fungi(100);p__Ascomycota(100);c__Saccharomycetes(100);o__Saccharomycetales(100);f__Metschnikowiaceae(100);g__Metschnikowia(100);s__Metschnikowia_pulcherrima(100); | Metschnikowia | 13 | Saprotroph | \|Undefined Saprotroph\| | Yeast | NULL | Probable | As nectar-tap_saprotroph (Põlme et al. 2020); Decay sugar-rich substrates (Põlme et al. 2020) | Põlme S, et al. 2020. Fungal Diversity 105:1-16 ((https://doi.org/10.1007/s13225-020-00466-2)) |
| Otu49222 | 1 | k__Fungi(100);p__Ascomycota(100);c__Saccharomycetes(100);o__Saccharomycetales(100);f__Metschnikowiaceae(100);g__Metschnikowia(100);g__Metschnikowia_unclassified(100); | Metschnikowia | 13 | Saprotroph | \|Undefined Saprotroph\| | Yeast | NULL | Probable | As nectar-tap_saprotroph (Põlme et al. 2020); Decay sugar-rich substrates (Põlme et al. 2020) | Põlme S, et al. 2020. Fungal Diversity 105:1-16 ((https://doi.org/10.1007/s13225-020-00466-2)) |
| Otu49223 | 1 | k__Fungi(100);p__Ascomycota(100);c__Saccharomycetes(100);o__Saccharomycetales(100);f__Metschnikowiaceae(100);g__Metschnikowia(100);s__Metschnikowia_pulcherrima(100); | Metschnikowia | 13 | Saprotroph | \|Undefined Saprotroph\| | Yeast | NULL | Probable | As nectar-tap_saprotroph (Põlme et al. 2020); Decay sugar-rich substrates (Põlme et al. 2020) | Põlme S, et al. 2020. Fungal Diversity 105:1-16 ((https://doi.org/10.1007/s13225-020-00466-2)) |
| Otu49224 | 1 | k__Fungi(100);p__Ascomycota(100);c__Saccharomycetes(100);o__Saccharomycetales(100);f__Metschnikowiaceae(100);g__Metschnikowia(100);s__Metschnikowia_pulcherrima(100); | Metschnikowia | 13 | Saprotroph | \|Undefined Saprotroph\| | Yeast | NULL | Probable | As nectar-tap_saprotroph (Põlme et al. 2020); Decay sugar-rich substrates (Põlme et al. 2020) | Põlme S, et al. 2020. Fungal Diversity 105:1-16 ((https://doi.org/10.1007/s13225-020-00466-2)) |
| Otu49225 | 1 | k__Fungi(100);p__Ascomycota(100);c__Saccharomycetes(100);o__Saccharomycetales(100);f__Metschnikowiaceae(100);g__Metschnikowia(100);s__Metschnikowia_pulcherrima(100); | Metschnikowia | 13 | Saprotroph | \|Undefined Saprotroph\| | Yeast | NULL | Probable | As nectar-tap_saprotroph (Põlme et al. 2020); Decay sugar-rich substrates (Põlme et al. 2020) | Põlme S, et al. 2020. Fungal Diversity 105:1-16 ((https://doi.org/10.1007/s13225-020-00466-2)) |
| Otu49226 | 1 | k__Fungi(100);p__Ascomycota(100);c__Saccharomycetes(100);o__Saccharomycetales(100);f__Metschnikowiaceae(100);g__Metschnikowia(100);s__Metschnikowia_pulcherrima(100); | Metschnikowia | 13 | Saprotroph | \|Undefined Saprotroph\| | Yeast | NULL | Probable | As nectar-tap_saprotroph (Põlme et al. 2020); Decay sugar-rich substrates (Põlme et al. 2020) | Põlme S, et al. 2020. Fungal Diversity 105:1-16 ((https://doi.org/10.1007/s13225-020-00466-2)) |
| Otu49227 | 1 | k__Fungi(100);p__Ascomycota(100);c__Saccharomycetes(100);o__Saccharomycetales(100);f__Metschnikowiaceae(100);g__Metschnikowia(100);s__Metschnikowia_pulcherrima(100); | Metschnikowia | 13 | Saprotroph | \|Undefined Saprotroph\| | Yeast | NULL | Probable | As nectar-tap_saprotroph (Põlme et al. 2020); Decay sugar-rich substrates (Põlme et al. 2020) | Põlme S, et al. 2020. Fungal Diversity 105:1-16 ((https://doi.org/10.1007/s13225-020-00466-2)) |
| Otu49228 | 1 | k__Fungi(100);p__Ascomycota(100);c__Saccharomycetes(100);o__Saccharomycetales(100);f__Metschnikowiaceae(100);g__Metschnikowia(100);g__Metschnikowia_unclassified(100); | Metschnikowia | 13 | Saprotroph | \|Undefined Saprotroph\| | Yeast | NULL | Probable | As nectar-tap_saprotroph (Põlme et al. 2020); Decay sugar-rich substrates (Põlme et al. 2020) | Põlme S, et al. 2020. Fungal Diversity 105:1-16 ((https://doi.org/10.1007/s13225-020-00466-2)) |
| Otu49229 | 1 | k__Fungi(100);p__Ascomycota(100);c__Saccharomycetes(100);o__Saccharomycetales(100);f__Metschnikowiaceae(100);g__Metschnikowia(100);s__Metschnikowia_pulcherrima(100); | Metschnikowia | 13 | Saprotroph | \|Undefined Saprotroph\| | Yeast | NULL | Probable | As nectar-tap_saprotroph (Põlme et al. 2020); Decay sugar-rich substrates (Põlme et al. 2020) | Põlme S, et al. 2020. Fungal Diversity 105:1-16 ((https://doi.org/10.1007/s13225-020-00466-2)) |
| Otu49230 | 1 | k__Fungi(100);p__Ascomycota(100);c__Saccharomycetes(100);o__Saccharomycetales(100);f__Metschnikowiaceae(100);g__Metschnikowia(100);s__Metschnikowia_pulcherrima(100); | Metschnikowia | 13 | Saprotroph | \|Undefined Saprotroph\| | Yeast | NULL | Probable | As nectar-tap_saprotroph (Põlme et al. 2020); Decay sugar-rich substrates (Põlme et al. 2020) | Põlme S, et al. 2020. Fungal Diversity 105:1-16 ((https://doi.org/10.1007/s13225-020-00466-2)) |
| Otu49231 | 1 | k__Fungi(100);p__Ascomycota(100);c__Saccharomycetes(100);o__Saccharomycetales(100);f__Metschnikowiaceae(100);g__Metschnikowia(100);s__Metschnikowia_pulcherrima(100); | Metschnikowia | 13 | Saprotroph | \|Undefined Saprotroph\| | Yeast | NULL | Probable | As nectar-tap_saprotroph (Põlme et al. 2020); Decay sugar-rich substrates (Põlme et al. 2020) | Põlme S, et al. 2020. Fungal Diversity 105:1-16 ((https://doi.org/10.1007/s13225-020-00466-2)) |
| Otu49232 | 1 | k__Fungi(100);p__Ascomycota(100);c__Saccharomycetes(100);o__Saccharomycetales(100);f__Metschnikowiaceae(100);g__Metschnikowia(100);s__Metschnikowia_pulcherrima(100); | Metschnikowia | 13 | Saprotroph | \|Undefined Saprotroph\| | Yeast | NULL | Probable | As nectar-tap_saprotroph (Põlme et al. 2020); Decay sugar-rich substrates (Põlme et al. 2020) | Põlme S, et al. 2020. Fungal Diversity 105:1-16 ((https://doi.org/10.1007/s13225-020-00466-2)) |
| Otu49233 | 1 | k__Fungi(100);p__Ascomycota(100);c__Saccharomycetes(100);o__Saccharomycetales(100);f__Metschnikowiaceae(100);g__Metschnikowia(100);s__Metschnikowia_pulcherrima(100); | Metschnikowia | 13 | Saprotroph | \|Undefined Saprotroph\| | Yeast | NULL | Probable | As nectar-tap_saprotroph (Põlme et al. 2020); Decay sugar-rich substrates (Põlme et al. 2020) | Põlme S, et al. 2020. Fungal Diversity 105:1-16 ((https://doi.org/10.1007/s13225-020-00466-2)) |
| Otu49235 | 1 | k__Fungi(100);p__Ascomycota(100);c__Saccharomycetes(100);o__Saccharomycetales(100);f__Metschnikowiaceae(100);g__Metschnikowia(100);s__Metschnikowia_pulcherrima(100); | Metschnikowia | 13 | Saprotroph | \|Undefined Saprotroph\| | Yeast | NULL | Probable | As nectar-tap_saprotroph (Põlme et al. 2020); Decay sugar-rich substrates (Põlme et al. 2020) | Põlme S, et al. 2020. Fungal Diversity 105:1-16 ((https://doi.org/10.1007/s13225-020-00466-2)) |
| Otu49236 | 1 | k__Fungi(100);p__Ascomycota(100);c__Saccharomycetes(100);o__Saccharomycetales(100);f__Metschnikowiaceae(100);g__Metschnikowia(100);s__Metschnikowia_pulcherrima(100); | Metschnikowia | 13 | Saprotroph | \|Undefined Saprotroph\| | Yeast | NULL | Probable | As nectar-tap_saprotroph (Põlme et al. 2020); Decay sugar-rich substrates (Põlme et al. 2020) | Põlme S, et al. 2020. Fungal Diversity 105:1-16 ((https://doi.org/10.1007/s13225-020-00466-2)) |
| Otu49237 | 1 | k__Fungi(100);p__Ascomycota(100);c__Saccharomycetes(100);o__Saccharomycetales(100);f__Metschnikowiaceae(100);g__Metschnikowia(100);s__Metschnikowia_pulcherrima(100); | Metschnikowia | 13 | Saprotroph | \|Undefined Saprotroph\| | Yeast | NULL | Probable | As nectar-tap_saprotroph (Põlme et al. 2020); Decay sugar-rich substrates (Põlme et al. 2020) | Põlme S, et al. 2020. Fungal Diversity 105:1-16 ((https://doi.org/10.1007/s13225-020-00466-2)) |
| Otu49238 | 1 | k__Fungi(100);p__Ascomycota(100);c__Saccharomycetes(100);o__Saccharomycetales(100);f__Metschnikowiaceae(100);g__Metschnikowia(100);s__Metschnikowia_pulcherrima(100); | Metschnikowia | 13 | Saprotroph | \|Undefined Saprotroph\| | Yeast | NULL | Probable | As nectar-tap_saprotroph (Põlme et al. 2020); Decay sugar-rich substrates (Põlme et al. 2020) | Põlme S, et al. 2020. Fungal Diversity 105:1-16 ((https://doi.org/10.1007/s13225-020-00466-2)) |
| Otu49241 | 1 | k__Fungi(100);p__Ascomycota(100);c__Saccharomycetes(100);o__Saccharomycetales(100);f__Metschnikowiaceae(100);g__Metschnikowia(100);s__Metschnikowia_pulcherrima(100); | Metschnikowia | 13 | Saprotroph | \|Undefined Saprotroph\| | Yeast | NULL | Probable | As nectar-tap_saprotroph (Põlme et al. 2020); Decay sugar-rich substrates (Põlme et al. 2020) | Põlme S, et al. 2020. Fungal Diversity 105:1-16 ((https://doi.org/10.1007/s13225-020-00466-2)) |
| Otu49242 | 1 | k__Fungi(100);p__Ascomycota(100);c__Saccharomycetes(100);o__Saccharomycetales(100);f__Metschnikowiaceae(100);g__Metschnikowia(100);s__Metschnikowia_pulcherrima(100); | Metschnikowia | 13 | Saprotroph | \|Undefined Saprotroph\| | Yeast | NULL | Probable | As nectar-tap_saprotroph (Põlme et al. 2020); Decay sugar-rich substrates (Põlme et al. 2020) | Põlme S, et al. 2020. Fungal Diversity 105:1-16 ((https://doi.org/10.1007/s13225-020-00466-2)) |
| Otu49243 | 1 | k__Fungi(100);p__Ascomycota(100);c__Saccharomycetes(100);o__Saccharomycetales(100);f__Metschnikowiaceae(100);g__Metschnikowia(100);s__Metschnikowia_pulcherrima(100); | Metschnikowia | 13 | Saprotroph | \|Undefined Saprotroph\| | Yeast | NULL | Probable | As nectar-tap_saprotroph (Põlme et al. 2020); Decay sugar-rich substrates (Põlme et al. 2020) | Põlme S, et al. 2020. Fungal Diversity 105:1-16 ((https://doi.org/10.1007/s13225-020-00466-2)) |
| Otu49245 | 1 | k__Fungi(100);p__Ascomycota(100);c__Saccharomycetes(100);o__Saccharomycetales(100);f__Metschnikowiaceae(100);g__Metschnikowia(100);s__Metschnikowia_pulcherrima(100); | Metschnikowia | 13 | Saprotroph | \|Undefined Saprotroph\| | Yeast | NULL | Probable | As nectar-tap_saprotroph (Põlme et al. 2020); Decay sugar-rich substrates (Põlme et al. 2020) | Põlme S, et al. 2020. Fungal Diversity 105:1-16 ((https://doi.org/10.1007/s13225-020-00466-2)) |
| Otu49246 | 1 | k__Fungi(100);p__Ascomycota(100);c__Saccharomycetes(100);o__Saccharomycetales(100);f__Metschnikowiaceae(100);g__Metschnikowia(100);g__Metschnikowia_unclassified(100); | Metschnikowia | 13 | Saprotroph | \|Undefined Saprotroph\| | Yeast | NULL | Probable | As nectar-tap_saprotroph (Põlme et al. 2020); Decay sugar-rich substrates (Põlme et al. 2020) | Põlme S, et al. 2020. Fungal Diversity 105:1-16 ((https://doi.org/10.1007/s13225-020-00466-2)) |
| Otu49247 | 1 | k__Fungi(100);p__Ascomycota(100);c__Saccharomycetes(100);o__Saccharomycetales(100);f__Metschnikowiaceae(100);g__Metschnikowia(100);s__Metschnikowia_pulcherrima(100); | Metschnikowia | 13 | Saprotroph | \|Undefined Saprotroph\| | Yeast | NULL | Probable | As nectar-tap_saprotroph (Põlme et al. 2020); Decay sugar-rich substrates (Põlme et al. 2020) | Põlme S, et al. 2020. Fungal Diversity 105:1-16 ((https://doi.org/10.1007/s13225-020-00466-2)) |
| Otu49248 | 1 | k__Fungi(100);p__Ascomycota(100);c__Saccharomycetes(100);o__Saccharomycetales(100);f__Metschnikowiaceae(100);g__Metschnikowia(100);s__Metschnikowia_pulcherrima(100); | Metschnikowia | 13 | Saprotroph | \|Undefined Saprotroph\| | Yeast | NULL | Probable | As nectar-tap_saprotroph (Põlme et al. 2020); Decay sugar-rich substrates (Põlme et al. 2020) | Põlme S, et al. 2020. Fungal Diversity 105:1-16 ((https://doi.org/10.1007/s13225-020-00466-2)) |
| Otu49249 | 1 | k__Fungi(100);p__Ascomycota(100);c__Saccharomycetes(100);o__Saccharomycetales(100);f__Metschnikowiaceae(100);g__Metschnikowia(100);s__Metschnikowia_pulcherrima(100); | Metschnikowia | 13 | Saprotroph | \|Undefined Saprotroph\| | Yeast | NULL | Probable | As nectar-tap_saprotroph (Põlme et al. 2020); Decay sugar-rich substrates (Põlme et al. 2020) | Põlme S, et al. 2020. Fungal Diversity 105:1-16 ((https://doi.org/10.1007/s13225-020-00466-2)) |
| Otu49250 | 1 | k__Fungi(100);p__Ascomycota(100);c__Saccharomycetes(100);o__Saccharomycetales(100);f__Metschnikowiaceae(100);g__Metschnikowia(100);s__Metschnikowia_pulcherrima(100); | Metschnikowia | 13 | Saprotroph | \|Undefined Saprotroph\| | Yeast | NULL | Probable | As nectar-tap_saprotroph (Põlme et al. 2020); Decay sugar-rich substrates (Põlme et al. 2020) | Põlme S, et al. 2020. Fungal Diversity 105:1-16 ((https://doi.org/10.1007/s13225-020-00466-2)) |
| Otu49251 | 1 | k__Fungi(100);p__Ascomycota(100);c__Saccharomycetes(100);o__Saccharomycetales(100);f__Metschnikowiaceae(100);g__Metschnikowia(100);s__Metschnikowia_pulcherrima(100); | Metschnikowia | 13 | Saprotroph | \|Undefined Saprotroph\| | Yeast | NULL | Probable | As nectar-tap_saprotroph (Põlme et al. 2020); Decay sugar-rich substrates (Põlme et al. 2020) | Põlme S, et al. 2020. Fungal Diversity 105:1-16 ((https://doi.org/10.1007/s13225-020-00466-2)) |
| Otu49252 | 1 | k__Fungi(100);p__Ascomycota(100);c__Saccharomycetes(100);o__Saccharomycetales(100);f__Metschnikowiaceae(100);g__Metschnikowia(100);s__Metschnikowia_pulcherrima(100); | Metschnikowia | 13 | Saprotroph | \|Undefined Saprotroph\| | Yeast | NULL | Probable | As nectar-tap_saprotroph (Põlme et al. 2020); Decay sugar-rich substrates (Põlme et al. 2020) | Põlme S, et al. 2020. Fungal Diversity 105:1-16 ((https://doi.org/10.1007/s13225-020-00466-2)) |
| Otu49253 | 1 | k__Fungi(100);p__Ascomycota(100);c__Saccharomycetes(100);o__Saccharomycetales(100);f__Metschnikowiaceae(100);g__Metschnikowia(100);g__Metschnikowia_unclassified(100); | Metschnikowia | 13 | Saprotroph | \|Undefined Saprotroph\| | Yeast | NULL | Probable | As nectar-tap_saprotroph (Põlme et al. 2020); Decay sugar-rich substrates (Põlme et al. 2020) | Põlme S, et al. 2020. Fungal Diversity 105:1-16 ((https://doi.org/10.1007/s13225-020-00466-2)) |
| Otu49254 | 1 | k__Fungi(100);p__Ascomycota(100);c__Saccharomycetes(100);o__Saccharomycetales(100);f__Metschnikowiaceae(100);g__Metschnikowia(100);s__Metschnikowia_pulcherrima(100); | Metschnikowia | 13 | Saprotroph | \|Undefined Saprotroph\| | Yeast | NULL | Probable | As nectar-tap_saprotroph (Põlme et al. 2020); Decay sugar-rich substrates (Põlme et al. 2020) | Põlme S, et al. 2020. Fungal Diversity 105:1-16 ((https://doi.org/10.1007/s13225-020-00466-2)) |
| Otu49255 | 1 | k__Fungi(100);p__Ascomycota(100);c__Saccharomycetes(100);o__Saccharomycetales(100);f__Metschnikowiaceae(100);g__Metschnikowia(100);s__Metschnikowia_pulcherrima(100); | Metschnikowia | 13 | Saprotroph | \|Undefined Saprotroph\| | Yeast | NULL | Probable | As nectar-tap_saprotroph (Põlme et al. 2020); Decay sugar-rich substrates (Põlme et al. 2020) | Põlme S, et al. 2020. Fungal Diversity 105:1-16 ((https://doi.org/10.1007/s13225-020-00466-2)) |
| Otu49256 | 1 | k__Fungi(100);p__Ascomycota(100);c__Saccharomycetes(100);o__Saccharomycetales(100);f__Metschnikowiaceae(100);g__Metschnikowia(100);s__Metschnikowia_pulcherrima(100); | Metschnikowia | 13 | Saprotroph | \|Undefined Saprotroph\| | Yeast | NULL | Probable | As nectar-tap_saprotroph (Põlme et al. 2020); Decay sugar-rich substrates (Põlme et al. 2020) | Põlme S, et al. 2020. Fungal Diversity 105:1-16 ((https://doi.org/10.1007/s13225-020-00466-2)) |
| Otu49257 | 1 | k__Fungi(100);p__Ascomycota(100);c__Saccharomycetes(100);o__Saccharomycetales(100);f__Metschnikowiaceae(100);g__Metschnikowia(100);s__Metschnikowia_pulcherrima(100); | Metschnikowia | 13 | Saprotroph | \|Undefined Saprotroph\| | Yeast | NULL | Probable | As nectar-tap_saprotroph (Põlme et al. 2020); Decay sugar-rich substrates (Põlme et al. 2020) | Põlme S, et al. 2020. Fungal Diversity 105:1-16 ((https://doi.org/10.1007/s13225-020-00466-2)) |
| Otu49258 | 1 | k__Fungi(100);p__Ascomycota(100);c__Saccharomycetes(100);o__Saccharomycetales(100);f__Metschnikowiaceae(100);g__Metschnikowia(100);s__Metschnikowia_pulcherrima(100); | Metschnikowia | 13 | Saprotroph | \|Undefined Saprotroph\| | Yeast | NULL | Probable | As nectar-tap_saprotroph (Põlme et al. 2020); Decay sugar-rich substrates (Põlme et al. 2020) | Põlme S, et al. 2020. Fungal Diversity 105:1-16 ((https://doi.org/10.1007/s13225-020-00466-2)) |
| Otu49259 | 1 | k__Fungi(100);p__Ascomycota(100);c__Saccharomycetes(100);o__Saccharomycetales(100);f__Metschnikowiaceae(100);g__Metschnikowia(100);s__Metschnikowia_pulcherrima(100); | Metschnikowia | 13 | Saprotroph | \|Undefined Saprotroph\| | Yeast | NULL | Probable | As nectar-tap_saprotroph (Põlme et al. 2020); Decay sugar-rich substrates (Põlme et al. 2020) | Põlme S, et al. 2020. Fungal Diversity 105:1-16 ((https://doi.org/10.1007/s13225-020-00466-2)) |
| Otu49260 | 1 | k__Fungi(100);p__Ascomycota(100);c__Saccharomycetes(100);o__Saccharomycetales(100);f__Metschnikowiaceae(100);g__Metschnikowia(100);s__Metschnikowia_pulcherrima(100); | Metschnikowia | 13 | Saprotroph | \|Undefined Saprotroph\| | Yeast | NULL | Probable | As nectar-tap_saprotroph (Põlme et al. 2020); Decay sugar-rich substrates (Põlme et al. 2020) | Põlme S, et al. 2020. Fungal Diversity 105:1-16 ((https://doi.org/10.1007/s13225-020-00466-2)) |
| Otu49261 | 1 | k__Fungi(100);p__Ascomycota(100);c__Saccharomycetes(100);o__Saccharomycetales(100);f__Metschnikowiaceae(100);g__Metschnikowia(100);s__Metschnikowia_pulcherrima(100); | Metschnikowia | 13 | Saprotroph | \|Undefined Saprotroph\| | Yeast | NULL | Probable | As nectar-tap_saprotroph (Põlme et al. 2020); Decay sugar-rich substrates (Põlme et al. 2020) | Põlme S, et al. 2020. Fungal Diversity 105:1-16 ((https://doi.org/10.1007/s13225-020-00466-2)) |
| Otu49262 | 1 | k__Fungi(100);p__Ascomycota(100);c__Saccharomycetes(100);o__Saccharomycetales(100);f__Metschnikowiaceae(100);g__Metschnikowia(100);s__Metschnikowia_pulcherrima(100); | Metschnikowia | 13 | Saprotroph | \|Undefined Saprotroph\| | Yeast | NULL | Probable | As nectar-tap_saprotroph (Põlme et al. 2020); Decay sugar-rich substrates (Põlme et al. 2020) | Põlme S, et al. 2020. Fungal Diversity 105:1-16 ((https://doi.org/10.1007/s13225-020-00466-2)) |
| Otu49263 | 1 | k__Fungi(100);p__Ascomycota(100);c__Saccharomycetes(100);o__Saccharomycetales(100);f__Metschnikowiaceae(100);g__Metschnikowia(100);g__Metschnikowia_unclassified(100); | Metschnikowia | 13 | Saprotroph | \|Undefined Saprotroph\| | Yeast | NULL | Probable | As nectar-tap_saprotroph (Põlme et al. 2020); Decay sugar-rich substrates (Põlme et al. 2020) | Põlme S, et al. 2020. Fungal Diversity 105:1-16 ((https://doi.org/10.1007/s13225-020-00466-2)) |
| Otu49264 | 1 | k__Fungi(100);p__Ascomycota(100);c__Saccharomycetes(100);o__Saccharomycetales(100);f__Metschnikowiaceae(100);g__Metschnikowia(100);s__Metschnikowia_pulcherrima(100); | Metschnikowia | 13 | Saprotroph | \|Undefined Saprotroph\| | Yeast | NULL | Probable | As nectar-tap_saprotroph (Põlme et al. 2020); Decay sugar-rich substrates (Põlme et al. 2020) | Põlme S, et al. 2020. Fungal Diversity 105:1-16 ((https://doi.org/10.1007/s13225-020-00466-2)) |
| Otu49266 | 1 | k__Fungi(100);p__Ascomycota(100);c__Saccharomycetes(100);o__Saccharomycetales(100);f__Metschnikowiaceae(100);g__Metschnikowia(100);g__Metschnikowia_unclassified(100); | Metschnikowia | 13 | Saprotroph | \|Undefined Saprotroph\| | Yeast | NULL | Probable | As nectar-tap_saprotroph (Põlme et al. 2020); Decay sugar-rich substrates (Põlme et al. 2020) | Põlme S, et al. 2020. Fungal Diversity 105:1-16 ((https://doi.org/10.1007/s13225-020-00466-2)) |
| Otu49268 | 1 | k__Fungi(100);p__Ascomycota(100);c__Saccharomycetes(100);o__Saccharomycetales(100);f__Metschnikowiaceae(100);g__Metschnikowia(100);s__Metschnikowia_pulcherrima(100); | Metschnikowia | 13 | Saprotroph | \|Undefined Saprotroph\| | Yeast | NULL | Probable | As nectar-tap_saprotroph (Põlme et al. 2020); Decay sugar-rich substrates (Põlme et al. 2020) | Põlme S, et al. 2020. Fungal Diversity 105:1-16 ((https://doi.org/10.1007/s13225-020-00466-2)) |
| Otu49269 | 1 | k__Fungi(100);p__Ascomycota(100);c__Saccharomycetes(100);o__Saccharomycetales(100);f__Metschnikowiaceae(100);g__Metschnikowia(100);s__Metschnikowia_pulcherrima(100); | Metschnikowia | 13 | Saprotroph | \|Undefined Saprotroph\| | Yeast | NULL | Probable | As nectar-tap_saprotroph (Põlme et al. 2020); Decay sugar-rich substrates (Põlme et al. 2020) | Põlme S, et al. 2020. Fungal Diversity 105:1-16 ((https://doi.org/10.1007/s13225-020-00466-2)) |
| Otu49270 | 1 | k__Fungi(100);p__Ascomycota(100);c__Saccharomycetes(100);o__Saccharomycetales(100);f__Metschnikowiaceae(100);g__Metschnikowia(100);s__Metschnikowia_pulcherrima(100); | Metschnikowia | 13 | Saprotroph | \|Undefined Saprotroph\| | Yeast | NULL | Probable | As nectar-tap_saprotroph (Põlme et al. 2020); Decay sugar-rich substrates (Põlme et al. 2020) | Põlme S, et al. 2020. Fungal Diversity 105:1-16 ((https://doi.org/10.1007/s13225-020-00466-2)) |
| Otu49271 | 1 | k__Fungi(100);p__Ascomycota(100);c__Saccharomycetes(100);o__Saccharomycetales(100);f__Metschnikowiaceae(100);g__Metschnikowia(100);s__Metschnikowia_pulcherrima(100); | Metschnikowia | 13 | Saprotroph | \|Undefined Saprotroph\| | Yeast | NULL | Probable | As nectar-tap_saprotroph (Põlme et al. 2020); Decay sugar-rich substrates (Põlme et al. 2020) | Põlme S, et al. 2020. Fungal Diversity 105:1-16 ((https://doi.org/10.1007/s13225-020-00466-2)) |
| Otu49272 | 1 | k__Fungi(100);p__Ascomycota(100);c__Saccharomycetes(100);o__Saccharomycetales(100);f__Metschnikowiaceae(100);g__Metschnikowia(100);s__Metschnikowia_pulcherrima(100); | Metschnikowia | 13 | Saprotroph | \|Undefined Saprotroph\| | Yeast | NULL | Probable | As nectar-tap_saprotroph (Põlme et al. 2020); Decay sugar-rich substrates (Põlme et al. 2020) | Põlme S, et al. 2020. Fungal Diversity 105:1-16 ((https://doi.org/10.1007/s13225-020-00466-2)) |
| Otu49273 | 1 | k__Fungi(100);p__Ascomycota(100);c__Saccharomycetes(100);o__Saccharomycetales(100);f__Metschnikowiaceae(100);g__Metschnikowia(100);g__Metschnikowia_unclassified(100); | Metschnikowia | 13 | Saprotroph | \|Undefined Saprotroph\| | Yeast | NULL | Probable | As nectar-tap_saprotroph (Põlme et al. 2020); Decay sugar-rich substrates (Põlme et al. 2020) | Põlme S, et al. 2020. Fungal Diversity 105:1-16 ((https://doi.org/10.1007/s13225-020-00466-2)) |
| Otu49274 | 1 | k__Fungi(100);p__Ascomycota(100);c__Saccharomycetes(100);o__Saccharomycetales(100);f__Metschnikowiaceae(100);g__Metschnikowia(100);s__Metschnikowia_pulcherrima(100); | Metschnikowia | 13 | Saprotroph | \|Undefined Saprotroph\| | Yeast | NULL | Probable | As nectar-tap_saprotroph (Põlme et al. 2020); Decay sugar-rich substrates (Põlme et al. 2020) | Põlme S, et al. 2020. Fungal Diversity 105:1-16 ((https://doi.org/10.1007/s13225-020-00466-2)) |
| Otu49275 | 1 | k__Fungi(100);p__Ascomycota(100);c__Saccharomycetes(100);o__Saccharomycetales(100);f__Metschnikowiaceae(100);g__Metschnikowia(100);g__Metschnikowia_unclassified(100); | Metschnikowia | 13 | Saprotroph | \|Undefined Saprotroph\| | Yeast | NULL | Probable | As nectar-tap_saprotroph (Põlme et al. 2020); Decay sugar-rich substrates (Põlme et al. 2020) | Põlme S, et al. 2020. Fungal Diversity 105:1-16 ((https://doi.org/10.1007/s13225-020-00466-2)) |
| Otu49276 | 1 | k__Fungi(100);p__Ascomycota(100);c__Saccharomycetes(100);o__Saccharomycetales(100);f__Metschnikowiaceae(100);g__Metschnikowia(100);s__Metschnikowia_pulcherrima(100); | Metschnikowia | 13 | Saprotroph | \|Undefined Saprotroph\| | Yeast | NULL | Probable | As nectar-tap_saprotroph (Põlme et al. 2020); Decay sugar-rich substrates (Põlme et al. 2020) | Põlme S, et al. 2020. Fungal Diversity 105:1-16 ((https://doi.org/10.1007/s13225-020-00466-2)) |
| Otu49277 | 1 | k__Fungi(100);p__Ascomycota(100);c__Saccharomycetes(100);o__Saccharomycetales(100);f__Metschnikowiaceae(100);g__Metschnikowia(100);g__Metschnikowia_unclassified(100); | Metschnikowia | 13 | Saprotroph | \|Undefined Saprotroph\| | Yeast | NULL | Probable | As nectar-tap_saprotroph (Põlme et al. 2020); Decay sugar-rich substrates (Põlme et al. 2020) | Põlme S, et al. 2020. Fungal Diversity 105:1-16 ((https://doi.org/10.1007/s13225-020-00466-2)) |
| Otu49278 | 1 | k__Fungi(100);p__Ascomycota(100);c__Saccharomycetes(100);o__Saccharomycetales(100);f__Metschnikowiaceae(100);g__Metschnikowia(100);s__Metschnikowia_pulcherrima(100); | Metschnikowia | 13 | Saprotroph | \|Undefined Saprotroph\| | Yeast | NULL | Probable | As nectar-tap_saprotroph (Põlme et al. 2020); Decay sugar-rich substrates (Põlme et al. 2020) | Põlme S, et al. 2020. Fungal Diversity 105:1-16 ((https://doi.org/10.1007/s13225-020-00466-2)) |
| Otu49279 | 1 | k__Fungi(100);p__Ascomycota(100);c__Saccharomycetes(100);o__Saccharomycetales(100);f__Metschnikowiaceae(100);g__Metschnikowia(100);s__Metschnikowia_pulcherrima(100); | Metschnikowia | 13 | Saprotroph | \|Undefined Saprotroph\| | Yeast | NULL | Probable | As nectar-tap_saprotroph (Põlme et al. 2020); Decay sugar-rich substrates (Põlme et al. 2020) | Põlme S, et al. 2020. Fungal Diversity 105:1-16 ((https://doi.org/10.1007/s13225-020-00466-2)) |
| Otu49280 | 1 | k__Fungi(100);p__Ascomycota(100);c__Saccharomycetes(100);o__Saccharomycetales(100);f__Metschnikowiaceae(100);g__Metschnikowia(100);s__Metschnikowia_pulcherrima(100); | Metschnikowia | 13 | Saprotroph | \|Undefined Saprotroph\| | Yeast | NULL | Probable | As nectar-tap_saprotroph (Põlme et al. 2020); Decay sugar-rich substrates (Põlme et al. 2020) | Põlme S, et al. 2020. Fungal Diversity 105:1-16 ((https://doi.org/10.1007/s13225-020-00466-2)) |
| Otu49281 | 1 | k__Fungi(100);p__Ascomycota(100);c__Saccharomycetes(100);o__Saccharomycetales(100);f__Metschnikowiaceae(100);g__Metschnikowia(100);s__Metschnikowia_pulcherrima(100); | Metschnikowia | 13 | Saprotroph | \|Undefined Saprotroph\| | Yeast | NULL | Probable | As nectar-tap_saprotroph (Põlme et al. 2020); Decay sugar-rich substrates (Põlme et al. 2020) | Põlme S, et al. 2020. Fungal Diversity 105:1-16 ((https://doi.org/10.1007/s13225-020-00466-2)) |
| Otu49282 | 1 | k__Fungi(100);p__Ascomycota(100);c__Saccharomycetes(100);o__Saccharomycetales(100);f__Metschnikowiaceae(100);g__Metschnikowia(100);s__Metschnikowia_pulcherrima(100); | Metschnikowia | 13 | Saprotroph | \|Undefined Saprotroph\| | Yeast | NULL | Probable | As nectar-tap_saprotroph (Põlme et al. 2020); Decay sugar-rich substrates (Põlme et al. 2020) | Põlme S, et al. 2020. Fungal Diversity 105:1-16 ((https://doi.org/10.1007/s13225-020-00466-2)) |
| Otu49283 | 1 | k__Fungi(100);p__Ascomycota(100);c__Saccharomycetes(100);o__Saccharomycetales(100);f__Metschnikowiaceae(100);g__Metschnikowia(100);s__Metschnikowia_pulcherrima(100); | Metschnikowia | 13 | Saprotroph | \|Undefined Saprotroph\| | Yeast | NULL | Probable | As nectar-tap_saprotroph (Põlme et al. 2020); Decay sugar-rich substrates (Põlme et al. 2020) | Põlme S, et al. 2020. Fungal Diversity 105:1-16 ((https://doi.org/10.1007/s13225-020-00466-2)) |
| Otu49284 | 1 | k__Fungi(100);p__Ascomycota(100);c__Saccharomycetes(100);o__Saccharomycetales(100);f__Metschnikowiaceae(100);g__Metschnikowia(100);g__Metschnikowia_unclassified(100); | Metschnikowia | 13 | Saprotroph | \|Undefined Saprotroph\| | Yeast | NULL | Probable | As nectar-tap_saprotroph (Põlme et al. 2020); Decay sugar-rich substrates (Põlme et al. 2020) | Põlme S, et al. 2020. Fungal Diversity 105:1-16 ((https://doi.org/10.1007/s13225-020-00466-2)) |
| Otu49285 | 1 | k__Fungi(100);p__Ascomycota(100);c__Saccharomycetes(100);o__Saccharomycetales(100);f__Metschnikowiaceae(100);g__Metschnikowia(100);s__Metschnikowia_pulcherrima(100); | Metschnikowia | 13 | Saprotroph | \|Undefined Saprotroph\| | Yeast | NULL | Probable | As nectar-tap_saprotroph (Põlme et al. 2020); Decay sugar-rich substrates (Põlme et al. 2020) | Põlme S, et al. 2020. Fungal Diversity 105:1-16 ((https://doi.org/10.1007/s13225-020-00466-2)) |
| Otu49286 | 1 | k__Fungi(100);p__Ascomycota(100);c__Saccharomycetes(100);o__Saccharomycetales(100);f__Metschnikowiaceae(100);g__Metschnikowia(100);g__Metschnikowia_unclassified(100); | Metschnikowia | 13 | Saprotroph | \|Undefined Saprotroph\| | Yeast | NULL | Probable | As nectar-tap_saprotroph (Põlme et al. 2020); Decay sugar-rich substrates (Põlme et al. 2020) | Põlme S, et al. 2020. Fungal Diversity 105:1-16 ((https://doi.org/10.1007/s13225-020-00466-2)) |
| Otu49287 | 1 | k__Fungi(100);p__Ascomycota(100);c__Saccharomycetes(100);o__Saccharomycetales(100);f__Metschnikowiaceae(100);g__Metschnikowia(100);g__Metschnikowia_unclassified(100); | Metschnikowia | 13 | Saprotroph | \|Undefined Saprotroph\| | Yeast | NULL | Probable | As nectar-tap_saprotroph (Põlme et al. 2020); Decay sugar-rich substrates (Põlme et al. 2020) | Põlme S, et al. 2020. Fungal Diversity 105:1-16 ((https://doi.org/10.1007/s13225-020-00466-2)) |
| Otu49288 | 1 | k__Fungi(100);p__Ascomycota(100);c__Saccharomycetes(100);o__Saccharomycetales(100);f__Metschnikowiaceae(100);g__Metschnikowia(100);g__Metschnikowia_unclassified(100); | Metschnikowia | 13 | Saprotroph | \|Undefined Saprotroph\| | Yeast | NULL | Probable | As nectar-tap_saprotroph (Põlme et al. 2020); Decay sugar-rich substrates (Põlme et al. 2020) | Põlme S, et al. 2020. Fungal Diversity 105:1-16 ((https://doi.org/10.1007/s13225-020-00466-2)) |
| Otu49289 | 1 | k__Fungi(100);p__Ascomycota(100);c__Saccharomycetes(100);o__Saccharomycetales(100);f__Metschnikowiaceae(100);g__Metschnikowia(100);g__Metschnikowia_unclassified(100); | Metschnikowia | 13 | Saprotroph | \|Undefined Saprotroph\| | Yeast | NULL | Probable | As nectar-tap_saprotroph (Põlme et al. 2020); Decay sugar-rich substrates (Põlme et al. 2020) | Põlme S, et al. 2020. Fungal Diversity 105:1-16 ((https://doi.org/10.1007/s13225-020-00466-2)) |
| Otu49290 | 1 | k__Fungi(100);p__Ascomycota(100);c__Saccharomycetes(100);o__Saccharomycetales(100);f__Metschnikowiaceae(100);g__Metschnikowia(100);g__Metschnikowia_unclassified(100); | Metschnikowia | 13 | Saprotroph | \|Undefined Saprotroph\| | Yeast | NULL | Probable | As nectar-tap_saprotroph (Põlme et al. 2020); Decay sugar-rich substrates (Põlme et al. 2020) | Põlme S, et al. 2020. Fungal Diversity 105:1-16 ((https://doi.org/10.1007/s13225-020-00466-2)) |
| Otu49291 | 1 | k__Fungi(100);p__Ascomycota(100);c__Saccharomycetes(100);o__Saccharomycetales(100);f__Metschnikowiaceae(100);g__Metschnikowia(100);g__Metschnikowia_unclassified(100); | Metschnikowia | 13 | Saprotroph | \|Undefined Saprotroph\| | Yeast | NULL | Probable | As nectar-tap_saprotroph (Põlme et al. 2020); Decay sugar-rich substrates (Põlme et al. 2020) | Põlme S, et al. 2020. Fungal Diversity 105:1-16 ((https://doi.org/10.1007/s13225-020-00466-2)) |
| Otu49292 | 1 | k__Fungi(100);p__Ascomycota(100);c__Saccharomycetes(100);o__Saccharomycetales(100);f__Metschnikowiaceae(100);g__Metschnikowia(100);g__Metschnikowia_unclassified(100); | Metschnikowia | 13 | Saprotroph | \|Undefined Saprotroph\| | Yeast | NULL | Probable | As nectar-tap_saprotroph (Põlme et al. 2020); Decay sugar-rich substrates (Põlme et al. 2020) | Põlme S, et al. 2020. Fungal Diversity 105:1-16 ((https://doi.org/10.1007/s13225-020-00466-2)) |
| Otu49293 | 1 | k__Fungi(100);p__Ascomycota(100);c__Saccharomycetes(100);o__Saccharomycetales(100);f__Metschnikowiaceae(100);g__Metschnikowia(100);s__Metschnikowia_pulcherrima(100); | Metschnikowia | 13 | Saprotroph | \|Undefined Saprotroph\| | Yeast | NULL | Probable | As nectar-tap_saprotroph (Põlme et al. 2020); Decay sugar-rich substrates (Põlme et al. 2020) | Põlme S, et al. 2020. Fungal Diversity 105:1-16 ((https://doi.org/10.1007/s13225-020-00466-2)) |
| Otu49294 | 1 | k__Fungi(100);p__Ascomycota(100);c__Saccharomycetes(100);o__Saccharomycetales(100);f__Metschnikowiaceae(100);g__Metschnikowia(100);s__Metschnikowia_pulcherrima(100); | Metschnikowia | 13 | Saprotroph | \|Undefined Saprotroph\| | Yeast | NULL | Probable | As nectar-tap_saprotroph (Põlme et al. 2020); Decay sugar-rich substrates (Põlme et al. 2020) | Põlme S, et al. 2020. Fungal Diversity 105:1-16 ((https://doi.org/10.1007/s13225-020-00466-2)) |
| Otu49295 | 1 | k__Fungi(100);p__Ascomycota(100);c__Saccharomycetes(100);o__Saccharomycetales(100);f__Metschnikowiaceae(100);g__Metschnikowia(100);g__Metschnikowia_unclassified(100); | Metschnikowia | 13 | Saprotroph | \|Undefined Saprotroph\| | Yeast | NULL | Probable | As nectar-tap_saprotroph (Põlme et al. 2020); Decay sugar-rich substrates (Põlme et al. 2020) | Põlme S, et al. 2020. Fungal Diversity 105:1-16 ((https://doi.org/10.1007/s13225-020-00466-2)) |
| Otu49296 | 1 | k__Fungi(100);p__Ascomycota(100);c__Saccharomycetes(100);o__Saccharomycetales(100);f__Metschnikowiaceae(100);g__Metschnikowia(100);s__Metschnikowia_pulcherrima(100); | Metschnikowia | 13 | Saprotroph | \|Undefined Saprotroph\| | Yeast | NULL | Probable | As nectar-tap_saprotroph (Põlme et al. 2020); Decay sugar-rich substrates (Põlme et al. 2020) | Põlme S, et al. 2020. Fungal Diversity 105:1-16 ((https://doi.org/10.1007/s13225-020-00466-2)) |
| Otu49297 | 1 | k__Fungi(100);p__Ascomycota(100);c__Saccharomycetes(100);o__Saccharomycetales(100);f__Metschnikowiaceae(100);g__Metschnikowia(100);s__Metschnikowia_pulcherrima(100); | Metschnikowia | 13 | Saprotroph | \|Undefined Saprotroph\| | Yeast | NULL | Probable | As nectar-tap_saprotroph (Põlme et al. 2020); Decay sugar-rich substrates (Põlme et al. 2020) | Põlme S, et al. 2020. Fungal Diversity 105:1-16 ((https://doi.org/10.1007/s13225-020-00466-2)) |
| Otu49298 | 1 | k__Fungi(100);p__Ascomycota(100);c__Saccharomycetes(100);o__Saccharomycetales(100);f__Metschnikowiaceae(100);g__Metschnikowia(100);g__Metschnikowia_unclassified(100); | Metschnikowia | 13 | Saprotroph | \|Undefined Saprotroph\| | Yeast | NULL | Probable | As nectar-tap_saprotroph (Põlme et al. 2020); Decay sugar-rich substrates (Põlme et al. 2020) | Põlme S, et al. 2020. Fungal Diversity 105:1-16 ((https://doi.org/10.1007/s13225-020-00466-2)) |
| Otu49299 | 1 | k__Fungi(100);p__Ascomycota(100);c__Saccharomycetes(100);o__Saccharomycetales(100);f__Metschnikowiaceae(100);g__Metschnikowia(100);s__Metschnikowia_pulcherrima(100); | Metschnikowia | 13 | Saprotroph | \|Undefined Saprotroph\| | Yeast | NULL | Probable | As nectar-tap_saprotroph (Põlme et al. 2020); Decay sugar-rich substrates (Põlme et al. 2020) | Põlme S, et al. 2020. Fungal Diversity 105:1-16 ((https://doi.org/10.1007/s13225-020-00466-2)) |
| Otu49300 | 1 | k__Fungi(100);p__Ascomycota(100);c__Saccharomycetes(100);o__Saccharomycetales(100);f__Metschnikowiaceae(100);g__Metschnikowia(100);s__Metschnikowia_pulcherrima(100); | Metschnikowia | 13 | Saprotroph | \|Undefined Saprotroph\| | Yeast | NULL | Probable | As nectar-tap_saprotroph (Põlme et al. 2020); Decay sugar-rich substrates (Põlme et al. 2020) | Põlme S, et al. 2020. Fungal Diversity 105:1-16 ((https://doi.org/10.1007/s13225-020-00466-2)) |
| Otu49302 | 1 | k__Fungi(100);p__Ascomycota(100);c__Saccharomycetes(100);o__Saccharomycetales(100);f__Metschnikowiaceae(100);g__Metschnikowia(100);s__Metschnikowia_pulcherrima(100); | Metschnikowia | 13 | Saprotroph | \|Undefined Saprotroph\| | Yeast | NULL | Probable | As nectar-tap_saprotroph (Põlme et al. 2020); Decay sugar-rich substrates (Põlme et al. 2020) | Põlme S, et al. 2020. Fungal Diversity 105:1-16 ((https://doi.org/10.1007/s13225-020-00466-2)) |
| Otu49303 | 1 | k__Fungi(100);p__Ascomycota(100);c__Saccharomycetes(100);o__Saccharomycetales(100);f__Metschnikowiaceae(100);g__Metschnikowia(100);s__Metschnikowia_pulcherrima(100); | Metschnikowia | 13 | Saprotroph | \|Undefined Saprotroph\| | Yeast | NULL | Probable | As nectar-tap_saprotroph (Põlme et al. 2020); Decay sugar-rich substrates (Põlme et al. 2020) | Põlme S, et al. 2020. Fungal Diversity 105:1-16 ((https://doi.org/10.1007/s13225-020-00466-2)) |
| Otu49304 | 1 | k__Fungi(100);p__Ascomycota(100);c__Saccharomycetes(100);o__Saccharomycetales(100);f__Metschnikowiaceae(100);g__Metschnikowia(100);s__Metschnikowia_pulcherrima(100); | Metschnikowia | 13 | Saprotroph | \|Undefined Saprotroph\| | Yeast | NULL | Probable | As nectar-tap_saprotroph (Põlme et al. 2020); Decay sugar-rich substrates (Põlme et al. 2020) | Põlme S, et al. 2020. Fungal Diversity 105:1-16 ((https://doi.org/10.1007/s13225-020-00466-2)) |
| Otu49305 | 1 | k__Fungi(100);p__Ascomycota(100);c__Saccharomycetes(100);o__Saccharomycetales(100);f__Metschnikowiaceae(100);g__Metschnikowia(100);s__Metschnikowia_pulcherrima(100); | Metschnikowia | 13 | Saprotroph | \|Undefined Saprotroph\| | Yeast | NULL | Probable | As nectar-tap_saprotroph (Põlme et al. 2020); Decay sugar-rich substrates (Põlme et al. 2020) | Põlme S, et al. 2020. Fungal Diversity 105:1-16 ((https://doi.org/10.1007/s13225-020-00466-2)) |
| Otu49307 | 1 | k__Fungi(100);p__Ascomycota(100);c__Saccharomycetes(100);o__Saccharomycetales(100);f__Metschnikowiaceae(100);g__Metschnikowia(100);s__Metschnikowia_pulcherrima(100); | Metschnikowia | 13 | Saprotroph | \|Undefined Saprotroph\| | Yeast | NULL | Probable | As nectar-tap_saprotroph (Põlme et al. 2020); Decay sugar-rich substrates (Põlme et al. 2020) | Põlme S, et al. 2020. Fungal Diversity 105:1-16 ((https://doi.org/10.1007/s13225-020-00466-2)) |
| Otu49308 | 1 | k__Fungi(100);p__Ascomycota(100);c__Saccharomycetes(100);o__Saccharomycetales(100);f__Metschnikowiaceae(100);g__Metschnikowia(100);s__Metschnikowia_pulcherrima(100); | Metschnikowia | 13 | Saprotroph | \|Undefined Saprotroph\| | Yeast | NULL | Probable | As nectar-tap_saprotroph (Põlme et al. 2020); Decay sugar-rich substrates (Põlme et al. 2020) | Põlme S, et al. 2020. Fungal Diversity 105:1-16 ((https://doi.org/10.1007/s13225-020-00466-2)) |
| Otu49309 | 1 | k__Fungi(100);p__Ascomycota(100);c__Saccharomycetes(100);o__Saccharomycetales(100);f__Metschnikowiaceae(100);g__Metschnikowia(100);s__Metschnikowia_pulcherrima(100); | Metschnikowia | 13 | Saprotroph | \|Undefined Saprotroph\| | Yeast | NULL | Probable | As nectar-tap_saprotroph (Põlme et al. 2020); Decay sugar-rich substrates (Põlme et al. 2020) | Põlme S, et al. 2020. Fungal Diversity 105:1-16 ((https://doi.org/10.1007/s13225-020-00466-2)) |
| Otu49310 | 1 | k__Fungi(100);p__Ascomycota(100);c__Saccharomycetes(100);o__Saccharomycetales(100);f__Metschnikowiaceae(100);g__Metschnikowia(100);s__Metschnikowia_pulcherrima(100); | Metschnikowia | 13 | Saprotroph | \|Undefined Saprotroph\| | Yeast | NULL | Probable | As nectar-tap_saprotroph (Põlme et al. 2020); Decay sugar-rich substrates (Põlme et al. 2020) | Põlme S, et al. 2020. Fungal Diversity 105:1-16 ((https://doi.org/10.1007/s13225-020-00466-2)) |
| Otu49311 | 1 | k__Fungi(100);p__Ascomycota(100);c__Saccharomycetes(100);o__Saccharomycetales(100);f__Metschnikowiaceae(100);g__Metschnikowia(100);g__Metschnikowia_unclassified(100); | Metschnikowia | 13 | Saprotroph | \|Undefined Saprotroph\| | Yeast | NULL | Probable | As nectar-tap_saprotroph (Põlme et al. 2020); Decay sugar-rich substrates (Põlme et al. 2020) | Põlme S, et al. 2020. Fungal Diversity 105:1-16 ((https://doi.org/10.1007/s13225-020-00466-2)) |
| Otu49312 | 1 | k__Fungi(100);p__Ascomycota(100);c__Saccharomycetes(100);o__Saccharomycetales(100);f__Metschnikowiaceae(100);g__Metschnikowia(100);s__Metschnikowia_pulcherrima(100); | Metschnikowia | 13 | Saprotroph | \|Undefined Saprotroph\| | Yeast | NULL | Probable | As nectar-tap_saprotroph (Põlme et al. 2020); Decay sugar-rich substrates (Põlme et al. 2020) | Põlme S, et al. 2020. Fungal Diversity 105:1-16 ((https://doi.org/10.1007/s13225-020-00466-2)) |
| Otu49313 | 1 | k__Fungi(100);p__Ascomycota(100);c__Saccharomycetes(100);o__Saccharomycetales(100);f__Metschnikowiaceae(100);g__Metschnikowia(100);g__Metschnikowia_unclassified(100); | Metschnikowia | 13 | Saprotroph | \|Undefined Saprotroph\| | Yeast | NULL | Probable | As nectar-tap_saprotroph (Põlme et al. 2020); Decay sugar-rich substrates (Põlme et al. 2020) | Põlme S, et al. 2020. Fungal Diversity 105:1-16 ((https://doi.org/10.1007/s13225-020-00466-2)) |
| Otu49314 | 1 | k__Fungi(100);p__Ascomycota(100);c__Saccharomycetes(100);o__Saccharomycetales(100);f__Metschnikowiaceae(100);g__Metschnikowia(100);g__Metschnikowia_unclassified(100); | Metschnikowia | 13 | Saprotroph | \|Undefined Saprotroph\| | Yeast | NULL | Probable | As nectar-tap_saprotroph (Põlme et al. 2020); Decay sugar-rich substrates (Põlme et al. 2020) | Põlme S, et al. 2020. Fungal Diversity 105:1-16 ((https://doi.org/10.1007/s13225-020-00466-2)) |
| Otu49315 | 1 | k__Fungi(100);p__Ascomycota(100);c__Saccharomycetes(100);o__Saccharomycetales(100);f__Metschnikowiaceae(100);g__Metschnikowia(100);s__Metschnikowia_pulcherrima(100); | Metschnikowia | 13 | Saprotroph | \|Undefined Saprotroph\| | Yeast | NULL | Probable | As nectar-tap_saprotroph (Põlme et al. 2020); Decay sugar-rich substrates (Põlme et al. 2020) | Põlme S, et al. 2020. Fungal Diversity 105:1-16 ((https://doi.org/10.1007/s13225-020-00466-2)) |
| Otu49316 | 1 | k__Fungi(100);p__Ascomycota(100);c__Saccharomycetes(100);o__Saccharomycetales(100);f__Metschnikowiaceae(100);g__Metschnikowia(100);s__Metschnikowia_pulcherrima(100); | Metschnikowia | 13 | Saprotroph | \|Undefined Saprotroph\| | Yeast | NULL | Probable | As nectar-tap_saprotroph (Põlme et al. 2020); Decay sugar-rich substrates (Põlme et al. 2020) | Põlme S, et al. 2020. Fungal Diversity 105:1-16 ((https://doi.org/10.1007/s13225-020-00466-2)) |
| Otu49317 | 1 | k__Fungi(100);p__Ascomycota(100);c__Saccharomycetes(100);o__Saccharomycetales(100);f__Metschnikowiaceae(100);g__Metschnikowia(100);s__Metschnikowia_pulcherrima(100); | Metschnikowia | 13 | Saprotroph | \|Undefined Saprotroph\| | Yeast | NULL | Probable | As nectar-tap_saprotroph (Põlme et al. 2020); Decay sugar-rich substrates (Põlme et al. 2020) | Põlme S, et al. 2020. Fungal Diversity 105:1-16 ((https://doi.org/10.1007/s13225-020-00466-2)) |

| | | | | | | | | | | | |
|---|---|---|---|---|---|---|---|---|---|---|---|
| Otu49318 | 1 | k__Fungi(100);p__Ascomycota(100);c__Saccharomycetes(100);o__Saccharomycetales(100);f__Metschnikowiaceae(100);g__Metschnikowia(100);s__Metschnikowia_pulcherrima(100); | Metschnikowia | 13 | Saprotroph | \|Undefined Saprotroph\| | Yeast | NULL | Probable | As nectar-tap_saprotroph (Põlme et al. 2020); Decay sugar-rich substrates (Põlme et al. 2020) | Põlme S, et al. 2020. Fungal Diversity 105:1-16 ((https://doi.org/10.1007/s13225-020-00466-2)) |
| Otu49319 | 1 | k__Fungi(100);p__Ascomycota(100);c__Saccharomycetes(100);o__Saccharomycetales(100);f__Metschnikowiaceae(100);g__Metschnikowia(100);s__Metschnikowia_pulcherrima(100); | Metschnikowia | 13 | Saprotroph | \|Undefined Saprotroph\| | Yeast | NULL | Probable | As nectar-tap_saprotroph (Põlme et al. 2020); Decay sugar-rich substrates (Põlme et al. 2020) | Põlme S, et al. 2020. Fungal Diversity 105:1-16 ((https://doi.org/10.1007/s13225-020-00466-2)) |
| Otu49320 | 1 | k__Fungi(100);p__Ascomycota(100);c__Saccharomycetes(100);o__Saccharomycetales(100);f__Metschnikowiaceae(100);g__Metschnikowia(100);s__Metschnikowia_pulcherrima(100); | Metschnikowia | 13 | Saprotroph | \|Undefined Saprotroph\| | Yeast | NULL | Probable | As nectar-tap_saprotroph (Põlme et al. 2020); Decay sugar-rich substrates (Põlme et al. 2020) | Põlme S, et al. 2020. Fungal Diversity 105:1-16 ((https://doi.org/10.1007/s13225-020-00466-2)) |
| Otu49321 | 1 | k__Fungi(100);p__Ascomycota(100);c__Saccharomycetes(100);o__Saccharomycetales(100);f__Metschnikowiaceae(100);g__Metschnikowia(100);s__Metschnikowia_pulcherrima(100); | Metschnikowia | 13 | Saprotroph | \|Undefined Saprotroph\| | Yeast | NULL | Probable | As nectar-tap_saprotroph (Põlme et al. 2020); Decay sugar-rich substrates (Põlme et al. 2020) | Põlme S, et al. 2020. Fungal Diversity 105:1-16 ((https://doi.org/10.1007/s13225-020-00466-2)) |
| Otu49322 | 1 | k__Fungi(100);p__Ascomycota(100);c__Saccharomycetes(100);o__Saccharomycetales(100);f__Metschnikowiaceae(100);g__Metschnikowia(100);s__Metschnikowia_pulcherrima(100); | Metschnikowia | 13 | Saprotroph | \|Undefined Saprotroph\| | Yeast | NULL | Probable | As nectar-tap_saprotroph (Põlme et al. 2020); Decay sugar-rich substrates (Põlme et al. 2020) | Põlme S, et al. 2020. Fungal Diversity 105:1-16 ((https://doi.org/10.1007/s13225-020-00466-2)) |
| Otu49323 | 1 | k__Fungi(100);p__Ascomycota(100);c__Saccharomycetes(100);o__Saccharomycetales(100);f__Metschnikowiaceae(100);g__Metschnikowia(100);g__Metschnikowia_unclassified(100); | Metschnikowia | 13 | Saprotroph | \|Undefined Saprotroph\| | Yeast | NULL | Probable | As nectar-tap_saprotroph (Põlme et al. 2020); Decay sugar-rich substrates (Põlme et al. 2020) | Põlme S, et al. 2020. Fungal Diversity 105:1-16 ((https://doi.org/10.1007/s13225-020-00466-2)) |
| Otu49324 | 1 | k__Fungi(100);p__Ascomycota(100);c__Saccharomycetes(100);o__Saccharomycetales(100);f__Metschnikowiaceae(100);g__Metschnikowia(100);s__Metschnikowia_pulcherrima(100); | Metschnikowia | 13 | Saprotroph | \|Undefined Saprotroph\| | Yeast | NULL | Probable | As nectar-tap_saprotroph (Põlme et al. 2020); Decay sugar-rich substrates (Põlme et al. 2020) | Põlme S, et al. 2020. Fungal Diversity 105:1-16 ((https://doi.org/10.1007/s13225-020-00466-2)) |
| Otu49325 | 1 | k__Fungi(100);p__Ascomycota(100);c__Saccharomycetes(100);o__Saccharomycetales(100);f__Metschnikowiaceae(100);g__Metschnikowia(100);s__Metschnikowia_pulcherrima(100); | Metschnikowia | 13 | Saprotroph | \|Undefined Saprotroph\| | Yeast | NULL | Probable | As nectar-tap_saprotroph (Põlme et al. 2020); Decay sugar-rich substrates (Põlme et al. 2020) | Põlme S, et al. 2020. Fungal Diversity 105:1-16 ((https://doi.org/10.1007/s13225-020-00466-2)) |
| Otu49327 | 1 | k__Fungi(100);p__Ascomycota(100);c__Saccharomycetes(100);o__Saccharomycetales(100);f__Metschnikowiaceae(100);g__Metschnikowia(100);s__Metschnikowia_pulcherrima(100); | Metschnikowia | 13 | Saprotroph | \|Undefined Saprotroph\| | Yeast | NULL | Probable | As nectar-tap_saprotroph (Põlme et al. 2020); Decay sugar-rich substrates (Põlme et al. 2020) | Põlme S, et al. 2020. Fungal Diversity 105:1-16 ((https://doi.org/10.1007/s13225-020-00466-2)) |
| Otu49328 | 1 | k__Fungi(100);p__Ascomycota(100);c__Saccharomycetes(100);o__Saccharomycetales(100);f__Metschnikowiaceae(100);g__Metschnikowia(100);s__Metschnikowia_pulcherrima(100); | Metschnikowia | 13 | Saprotroph | \|Undefined Saprotroph\| | Yeast | NULL | Probable | As nectar-tap_saprotroph (Põlme et al. 2020); Decay sugar-rich substrates (Põlme et al. 2020) | Põlme S, et al. 2020. Fungal Diversity 105:1-16 ((https://doi.org/10.1007/s13225-020-00466-2)) |
| Otu49329 | 1 | k__Fungi(100);p__Ascomycota(100);c__Saccharomycetes(100);o__Saccharomycetales(100);f__Metschnikowiaceae(100);g__Metschnikowia(100);s__Metschnikowia_pulcherrima(100); | Metschnikowia | 13 | Saprotroph | \|Undefined Saprotroph\| | Yeast | NULL | Probable | As nectar-tap_saprotroph (Põlme et al. 2020); Decay sugar-rich substrates (Põlme et al. 2020) | Põlme S, et al. 2020. Fungal Diversity 105:1-16 ((https://doi.org/10.1007/s13225-020-00466-2)) |
| Otu49330 | 1 | k__Fungi(100);p__Ascomycota(100);c__Saccharomycetes(100);o__Saccharomycetales(100);f__Metschnikowiaceae(100);g__Metschnikowia(100);s__Metschnikowia_pulcherrima(100); | Metschnikowia | 13 | Saprotroph | \|Undefined Saprotroph\| | Yeast | NULL | Probable | As nectar-tap_saprotroph (Põlme et al. 2020); Decay sugar-rich substrates (Põlme et al. 2020) | Põlme S, et al. 2020. Fungal Diversity 105:1-16 ((https://doi.org/10.1007/s13225-020-00466-2)) |
| Otu49331 | 1 | k__Fungi(100);p__Ascomycota(100);c__Saccharomycetes(100);o__Saccharomycetales(100);f__Metschnikowiaceae(100);g__Metschnikowia(100);g__Metschnikowia_unclassified(100); | Metschnikowia | 13 | Saprotroph | \|Undefined Saprotroph\| | Yeast | NULL | Probable | As nectar-tap_saprotroph (Põlme et al. 2020); Decay sugar-rich substrates (Põlme et al. 2020) | Põlme S, et al. 2020. Fungal Diversity 105:1-16 ((https://doi.org/10.1007/s13225-020-00466-2)) |
| Otu49332 | 1 | k__Fungi(100);p__Ascomycota(100);c__Saccharomycetes(100);o__Saccharomycetales(100);f__Metschnikowiaceae(100);g__Metschnikowia(100);s__Metschnikowia_pulcherrima(100); | Metschnikowia | 13 | Saprotroph | \|Undefined Saprotroph\| | Yeast | NULL | Probable | As nectar-tap_saprotroph (Põlme et al. 2020); Decay sugar-rich substrates (Põlme et al. 2020) | Põlme S, et al. 2020. Fungal Diversity 105:1-16 ((https://doi.org/10.1007/s13225-020-00466-2)) |
| Otu49333 | 1 | k__Fungi(100);p__Ascomycota(100);c__Saccharomycetes(100);o__Saccharomycetales(100);f__Metschnikowiaceae(100);g__Metschnikowia(100);g__Metschnikowia_unclassified(100); | Metschnikowia | 13 | Saprotroph | \|Undefined Saprotroph\| | Yeast | NULL | Probable | As nectar-tap_saprotroph (Põlme et al. 2020); Decay sugar-rich substrates (Põlme et al. 2020) | Põlme S, et al. 2020. Fungal Diversity 105:1-16 ((https://doi.org/10.1007/s13225-020-00466-2)) |
| Otu49334 | 1 | k__Fungi(100);p__Ascomycota(100);c__Saccharomycetes(100);o__Saccharomycetales(100);f__Metschnikowiaceae(100);g__Metschnikowia(100);s__Metschnikowia_pulcherrima(100); | Metschnikowia | 13 | Saprotroph | \|Undefined Saprotroph\| | Yeast | NULL | Probable | As nectar-tap_saprotroph (Põlme et al. 2020); Decay sugar-rich substrates (Põlme et al. 2020) | Põlme S, et al. 2020. Fungal Diversity 105:1-16 ((https://doi.org/10.1007/s13225-020-00466-2)) |
| Otu49336 | 1 | k__Fungi(100);p__Ascomycota(100);c__Saccharomycetes(100);o__Saccharomycetales(100);f__Metschnikowiaceae(100);g__Metschnikowia(100);g__Metschnikowia_unclassified(100); | Metschnikowia | 13 | Saprotroph | \|Undefined Saprotroph\| | Yeast | NULL | Probable | As nectar-tap_saprotroph (Põlme et al. 2020); Decay sugar-rich substrates (Põlme et al. 2020) | Põlme S, et al. 2020. Fungal Diversity 105:1-16 ((https://doi.org/10.1007/s13225-020-00466-2)) |
| Otu49337 | 1 | k__Fungi(100);p__Ascomycota(100);c__Saccharomycetes(100);o__Saccharomycetales(100);f__Metschnikowiaceae(100);g__Metschnikowia(100);g__Metschnikowia_unclassified(100); | Metschnikowia | 13 | Saprotroph | \|Undefined Saprotroph\| | Yeast | NULL | Probable | As nectar-tap_saprotroph (Põlme et al. 2020); Decay sugar-rich substrates (Põlme et al. 2020) | Põlme S, et al. 2020. Fungal Diversity 105:1-16 ((https://doi.org/10.1007/s13225-020-00466-2)) |
| Otu49338 | 1 | k__Fungi(100);p__Ascomycota(100);c__Saccharomycetes(100);o__Saccharomycetales(100);f__Metschnikowiaceae(100);g__Metschnikowia(100);g__Metschnikowia_unclassified(100); | Metschnikowia | 13 | Saprotroph | \|Undefined Saprotroph\| | Yeast | NULL | Probable | As nectar-tap_saprotroph (Põlme et al. 2020); Decay sugar-rich substrates (Põlme et al. 2020) | Põlme S, et al. 2020. Fungal Diversity 105:1-16 ((https://doi.org/10.1007/s13225-020-00466-2)) |
| Otu49339 | 1 | k__Fungi(100);p__Ascomycota(100);c__Saccharomycetes(100);o__Saccharomycetales(100);f__Metschnikowiaceae(100);g__Metschnikowia(100);g__Metschnikowia_unclassified(100); | Metschnikowia | 13 | Saprotroph | \|Undefined Saprotroph\| | Yeast | NULL | Probable | As nectar-tap_saprotroph (Põlme et al. 2020); Decay sugar-rich substrates (Põlme et al. 2020) | Põlme S, et al. 2020. Fungal Diversity 105:1-16 ((https://doi.org/10.1007/s13225-020-00466-2)) |
| Otu49340 | 1 | k__Fungi(100);p__Ascomycota(100);c__Saccharomycetes(100);o__Saccharomycetales(100);f__Metschnikowiaceae(100);g__Metschnikowia(100);s__Metschnikowia_pulcherrima(100); | Metschnikowia | 13 | Saprotroph | \|Undefined Saprotroph\| | Yeast | NULL | Probable | As nectar-tap_saprotroph (Põlme et al. 2020); Decay sugar-rich substrates (Põlme et al. 2020) | Põlme S, et al. 2020. Fungal Diversity 105:1-16 ((https://doi.org/10.1007/s13225-020-00466-2)) |
| Otu49341 | 1 | k__Fungi(100);p__Ascomycota(100);c__Saccharomycetes(100);o__Saccharomycetales(100);f__Metschnikowiaceae(100);g__Metschnikowia(100);s__Metschnikowia_pulcherrima(100); | Metschnikowia | 13 | Saprotroph | \|Undefined Saprotroph\| | Yeast | NULL | Probable | As nectar-tap_saprotroph (Põlme et al. 2020); Decay sugar-rich substrates (Põlme et al. 2020) | Põlme S, et al. 2020. Fungal Diversity 105:1-16 ((https://doi.org/10.1007/s13225-020-00466-2)) |
| Otu49342 | 1 | k__Fungi(100);p__Ascomycota(100);c__Saccharomycetes(100);o__Saccharomycetales(100);f__Metschnikowiaceae(100);g__Metschnikowia(100);s__Metschnikowia_pulcherrima(100); | Metschnikowia | 13 | Saprotroph | \|Undefined Saprotroph\| | Yeast | NULL | Probable | As nectar-tap_saprotroph (Põlme et al. 2020); Decay sugar-rich substrates (Põlme et al. 2020) | Põlme S, et al. 2020. Fungal Diversity 105:1-16 ((https://doi.org/10.1007/s13225-020-00466-2)) |
| Otu49343 | 1 | k__Fungi(100);p__Ascomycota(100);c__Saccharomycetes(100);o__Saccharomycetales(100);f__Metschnikowiaceae(100);g__Metschnikowia(100);s__Metschnikowia_pulcherrima(100); | Metschnikowia | 13 | Saprotroph | \|Undefined Saprotroph\| | Yeast | NULL | Probable | As nectar-tap_saprotroph (Põlme et al. 2020); Decay sugar-rich substrates (Põlme et al. 2020) | Põlme S, et al. 2020. Fungal Diversity 105:1-16 ((https://doi.org/10.1007/s13225-020-00466-2)) |
| Otu49344 | 1 | k__Fungi(100);p__Ascomycota(100);c__Saccharomycetes(100);o__Saccharomycetales(100);f__Metschnikowiaceae(100);g__Metschnikowia(100);s__Metschnikowia_pulcherrima(100); | Metschnikowia | 13 | Saprotroph | \|Undefined Saprotroph\| | Yeast | NULL | Probable | As nectar-tap_saprotroph (Põlme et al. 2020); Decay sugar-rich substrates (Põlme et al. 2020) | Põlme S, et al. 2020. Fungal Diversity 105:1-16 ((https://doi.org/10.1007/s13225-020-00466-2)) |
| Otu49345 | 1 | k__Fungi(100);p__Ascomycota(100);c__Saccharomycetes(100);o__Saccharomycetales(100);f__Metschnikowiaceae(100);g__Metschnikowia(100);g__Metschnikowia_unclassified(100); | Metschnikowia | 13 | Saprotroph | \|Undefined Saprotroph\| | Yeast | NULL | Probable | As nectar-tap_saprotroph (Põlme et al. 2020); Decay sugar-rich substrates (Põlme et al. 2020) | Põlme S, et al. 2020. Fungal Diversity 105:1-16 ((https://doi.org/10.1007/s13225-020-00466-2)) |
| Otu49346 | 1 | k__Fungi(100);p__Ascomycota(100);c__Saccharomycetes(100);o__Saccharomycetales(100);f__Metschnikowiaceae(100);g__Metschnikowia(100);s__Metschnikowia_pulcherrima(100); | Metschnikowia | 13 | Saprotroph | \|Undefined Saprotroph\| | Yeast | NULL | Probable | As nectar-tap_saprotroph (Põlme et al. 2020); Decay sugar-rich substrates (Põlme et al. 2020) | Põlme S, et al. 2020. Fungal Diversity 105:1-16 ((https://doi.org/10.1007/s13225-020-00466-2)) |
| Otu49347 | 1 | k__Fungi(100);p__Ascomycota(100);c__Saccharomycetes(100);o__Saccharomycetales(100);f__Metschnikowiaceae(100);g__Metschnikowia(100);s__Metschnikowia_pulcherrima(100); | Metschnikowia | 13 | Saprotroph | \|Undefined Saprotroph\| | Yeast | NULL | Probable | As nectar-tap_saprotroph (Põlme et al. 2020); Decay sugar-rich substrates (Põlme et al. 2020) | Põlme S, et al. 2020. Fungal Diversity 105:1-16 ((https://doi.org/10.1007/s13225-020-00466-2)) |
| Otu49348 | 1 | k__Fungi(100);p__Ascomycota(100);c__Saccharomycetes(100);o__Saccharomycetales(100);f__Metschnikowiaceae(100);g__Metschnikowia(100);s__Metschnikowia_pulcherrima(100); | Metschnikowia | 13 | Saprotroph | \|Undefined Saprotroph\| | Yeast | NULL | Probable | As nectar-tap_saprotroph (Põlme et al. 2020); Decay sugar-rich substrates (Põlme et al. 2020) | Põlme S, et al. 2020. Fungal Diversity 105:1-16 ((https://doi.org/10.1007/s13225-020-00466-2)) |
| Otu49349 | 1 | k__Fungi(100);p__Ascomycota(100);c__Saccharomycetes(100);o__Saccharomycetales(100);f__Metschnikowiaceae(100);g__Metschnikowia(100);s__Metschnikowia_pulcherrima(100); | Metschnikowia | 13 | Saprotroph | \|Undefined Saprotroph\| | Yeast | NULL | Probable | As nectar-tap_saprotroph (Põlme et al. 2020); Decay sugar-rich substrates (Põlme et al. 2020) | Põlme S, et al. 2020. Fungal Diversity 105:1-16 ((https://doi.org/10.1007/s13225-020-00466-2)) |
| Otu49350 | 1 | k__Fungi(100);p__Ascomycota(100);c__Saccharomycetes(100);o__Saccharomycetales(100);f__Metschnikowiaceae(100);g__Metschnikowia(100);s__Metschnikowia_pulcherrima(100); | Metschnikowia | 13 | Saprotroph | \|Undefined Saprotroph\| | Yeast | NULL | Probable | As nectar-tap_saprotroph (Põlme et al. 2020); Decay sugar-rich substrates (Põlme et al. 2020) | Põlme S, et al. 2020. Fungal Diversity 105:1-16 ((https://doi.org/10.1007/s13225-020-00466-2)) |
| Otu49351 | 1 | k__Fungi(100);p__Ascomycota(100);c__Saccharomycetes(100);o__Saccharomycetales(100);f__Metschnikowiaceae(100);g__Metschnikowia(100);g__Metschnikowia_unclassified(100); | Metschnikowia | 13 | Saprotroph | \|Undefined Saprotroph\| | Yeast | NULL | Probable | As nectar-tap_saprotroph (Põlme et al. 2020); Decay sugar-rich substrates (Põlme et al. 2020) | Põlme S, et al. 2020. Fungal Diversity 105:1-16 ((https://doi.org/10.1007/s13225-020-00466-2)) |
| Otu49352 | 1 | k__Fungi(100);p__Ascomycota(100);c__Saccharomycetes(100);o__Saccharomycetales(100);f__Metschnikowiaceae(100);g__Metschnikowia(100);s__Metschnikowia_pulcherrima(100); | Metschnikowia | 13 | Saprotroph | \|Undefined Saprotroph\| | Yeast | NULL | Probable | As nectar-tap_saprotroph (Põlme et al. 2020); Decay sugar-rich substrates (Põlme et al. 2020) | Põlme S, et al. 2020. Fungal Diversity 105:1-16 ((https://doi.org/10.1007/s13225-020-00466-2)) |
| Otu49353 | 1 | k__Fungi(100);p__Ascomycota(100);c__Saccharomycetes(100);o__Saccharomycetales(100);f__Metschnikowiaceae(100);g__Metschnikowia(100);s__Metschnikowia_pulcherrima(100); | Metschnikowia | 13 | Saprotroph | \|Undefined Saprotroph\| | Yeast | NULL | Probable | As nectar-tap_saprotroph (Põlme et al. 2020); Decay sugar-rich substrates (Põlme et al. 2020) | Põlme S, et al. 2020. Fungal Diversity 105:1-16 ((https://doi.org/10.1007/s13225-020-00466-2)) |
| Otu49354 | 1 | k__Fungi(100);p__Ascomycota(100);c__Saccharomycetes(100);o__Saccharomycetales(100);f__Metschnikowiaceae(100);g__Metschnikowia(100);s__Metschnikowia_pulcherrima(100); | Metschnikowia | 13 | Saprotroph | \|Undefined Saprotroph\| | Yeast | NULL | Probable | As nectar-tap_saprotroph (Põlme et al. 2020); Decay sugar-rich substrates (Põlme et al. 2020) | Põlme S, et al. 2020. Fungal Diversity 105:1-16 ((https://doi.org/10.1007/s13225-020-00466-2)) |
| Otu49355 | 1 | k__Fungi(100);p__Ascomycota(100);c__Saccharomycetes(100);o__Saccharomycetales(100);f__Metschnikowiaceae(100);g__Metschnikowia(100);s__Metschnikowia_pulcherrima(100); | Metschnikowia | 13 | Saprotroph | \|Undefined Saprotroph\| | Yeast | NULL | Probable | As nectar-tap_saprotroph (Põlme et al. 2020); Decay sugar-rich substrates (Põlme et al. 2020) | Põlme S, et al. 2020. Fungal Diversity 105:1-16 ((https://doi.org/10.1007/s13225-020-00466-2)) |
| Otu49356 | 1 | k__Fungi(100);p__Ascomycota(100);c__Saccharomycetes(100);o__Saccharomycetales(100);f__Metschnikowiaceae(100);g__Metschnikowia(100);s__Metschnikowia_pulcherrima(100); | Metschnikowia | 13 | Saprotroph | \|Undefined Saprotroph\| | Yeast | NULL | Probable | As nectar-tap_saprotroph (Põlme et al. 2020); Decay sugar-rich substrates (Põlme et al. 2020) | Põlme S, et al. 2020. Fungal Diversity 105:1-16 ((https://doi.org/10.1007/s13225-020-00466-2)) |
| Otu49357 | 1 | k__Fungi(100);p__Ascomycota(100);c__Saccharomycetes(100);o__Saccharomycetales(100);f__Metschnikowiaceae(100);g__Metschnikowia(100);s__Metschnikowia_pulcherrima(100); | Metschnikowia | 13 | Saprotroph | \|Undefined Saprotroph\| | Yeast | NULL | Probable | As nectar-tap_saprotroph (Põlme et al. 2020); Decay sugar-rich substrates (Põlme et al. 2020) | Põlme S, et al. 2020. Fungal Diversity 105:1-16 ((https://doi.org/10.1007/s13225-020-00466-2)) |
| Otu49358 | 1 | k__Fungi(100);p__Ascomycota(100);c__Saccharomycetes(100);o__Saccharomycetales(100);f__Metschnikowiaceae(100);g__Metschnikowia(100);s__Metschnikowia_pulcherrima(100); | Metschnikowia | 13 | Saprotroph | \|Undefined Saprotroph\| | Yeast | NULL | Probable | As nectar-tap_saprotroph (Põlme et al. 2020); Decay sugar-rich substrates (Põlme et al. 2020) | Põlme S, et al. 2020. Fungal Diversity 105:1-16 ((https://doi.org/10.1007/s13225-020-00466-2)) |
| Otu49359 | 1 | k__Fungi(100);p__Ascomycota(100);c__Saccharomycetes(100);o__Saccharomycetales(100);f__Metschnikowiaceae(100);g__Metschnikowia(100);g__Metschnikowia_unclassified(100); | Metschnikowia | 13 | Saprotroph | \|Undefined Saprotroph\| | Yeast | NULL | Probable | As nectar-tap_saprotroph (Põlme et al. 2020); Decay sugar-rich substrates (Põlme et al. 2020) | Põlme S, et al. 2020. Fungal Diversity 105:1-16 ((https://doi.org/10.1007/s13225-020-00466-2)) |
| Otu49360 | 1 | k__Fungi(100);p__Ascomycota(100);c__Saccharomycetes(100);o__Saccharomycetales(100);f__Metschnikowiaceae(100);g__Metschnikowia(100);g__Metschnikowia_unclassified(100); | Metschnikowia | 13 | Saprotroph | \|Undefined Saprotroph\| | Yeast | NULL | Probable | As nectar-tap_saprotroph (Põlme et al. 2020); Decay sugar-rich substrates (Põlme et al. 2020) | Põlme S, et al. 2020. Fungal Diversity 105:1-16 ((https://doi.org/10.1007/s13225-020-00466-2)) |
| Otu49361 | 1 | k__Fungi(100);p__Ascomycota(100);c__Saccharomycetes(100);o__Saccharomycetales(100);f__Metschnikowiaceae(100);g__Metschnikowia(100);g__Metschnikowia_unclassified(100); | Metschnikowia | 13 | Saprotroph | \|Undefined Saprotroph\| | Yeast | NULL | Probable | As nectar-tap_saprotroph (Põlme et al. 2020); Decay sugar-rich substrates (Põlme et al. 2020) | Põlme S, et al. 2020. Fungal Diversity 105:1-16 ((https://doi.org/10.1007/s13225-020-00466-2)) |
| Otu49362 | 1 | k__Fungi(100);p__Ascomycota(100);c__Saccharomycetes(100);o__Saccharomycetales(100);f__Metschnikowiaceae(100);g__Metschnikowia(100);s__Metschnikowia_pulcherrima(100); | Metschnikowia | 13 | Saprotroph | \|Undefined Saprotroph\| | Yeast | NULL | Probable | As nectar-tap_saprotroph (Põlme et al. 2020); Decay sugar-rich substrates (Põlme et al. 2020) | Põlme S, et al. 2020. Fungal Diversity 105:1-16 ((https://doi.org/10.1007/s13225-020-00466-2)) |
| Otu49363 | 1 | k__Fungi(100);p__Ascomycota(100);c__Saccharomycetes(100);o__Saccharomycetales(100);f__Metschnikowiaceae(100);g__Metschnikowia(100);s__Metschnikowia_pulcherrima(100); | Metschnikowia | 13 | Saprotroph | \|Undefined Saprotroph\| | Yeast | NULL | Probable | As nectar-tap_saprotroph (Põlme et al. 2020); Decay sugar-rich substrates (Põlme et al. 2020) | Põlme S, et al. 2020. Fungal Diversity 105:1-16 ((https://doi.org/10.1007/s13225-020-00466-2)) |
| Otu49364 | 1 | k__Fungi(100);p__Ascomycota(100);c__Saccharomycetes(100);o__Saccharomycetales(100);f__Metschnikowiaceae(100);g__Metschnikowia(100);s__Metschnikowia_pulcherrima(100); | Metschnikowia | 13 | Saprotroph | \|Undefined Saprotroph\| | Yeast | NULL | Probable | As nectar-tap_saprotroph (Põlme et al. 2020); Decay sugar-rich substrates (Põlme et al. 2020) | Põlme S, et al. 2020. Fungal Diversity 105:1-16 ((https://doi.org/10.1007/s13225-020-00466-2)) |
| Otu49365 | 1 | k__Fungi(100);p__Ascomycota(100);c__Saccharomycetes(100);o__Saccharomycetales(100);f__Saccharomycetaceae(100);g__Kazachstania(100);s__Kazachstania_intestinalis(100); | Kazachstania | 13 | Saprotroph | \|Undefined Saprotroph\| | Yeast | NULL | Probable | As nectar-tap_saprotroph (Põlme et al. 2020); Decay sugar-rich substrates (Põlme et al. 2020) | Põlme S, et al. 2020. Fungal Diversity 105:1-16 ((https://doi.org/10.1007/s13225-020-00466-2)) |
| Otu49366 | 1 | k__Fungi(100);p__Ascomycota(100);c__Saccharomycetes(100);o__Saccharomycetales(100);f__Metschnikowiaceae(100);g__Metschnikowia(100);s__Metschnikowia_pulcherrima(100); | Metschnikowia | 13 | Saprotroph | \|Undefined Saprotroph\| | Yeast | NULL | Probable | As nectar-tap_saprotroph (Põlme et al. 2020); Decay sugar-rich substrates (Põlme et al. 2020) | Põlme S, et al. 2020. Fungal Diversity 105:1-16 ((https://doi.org/10.1007/s13225-020-00466-2)) |
| Otu49367 | 1 | k__Fungi(100);p__Ascomycota(100);c__Saccharomycetes(100);o__Saccharomycetales(100);f__Metschnikowiaceae(100);g__Metschnikowia(100);s__Metschnikowia_pulcherrima(100); | Metschnikowia | 13 | Saprotroph | \|Undefined Saprotroph\| | Yeast | NULL | Probable | As nectar-tap_saprotroph (Põlme et al. 2020); Decay sugar-rich substrates (Põlme et al. 2020) | Põlme S, et al. 2020. Fungal Diversity 105:1-16 ((https://doi.org/10.1007/s13225-020-00466-2)) |
| Otu49368 | 1 | k__Fungi(100);p__Ascomycota(100);c__Saccharomycetes(100);o__Saccharomycetales(100);f__Metschnikowiaceae(100);g__Metschnikowia(100);s__Metschnikowia_pulcherrima(100); | Metschnikowia | 13 | Saprotroph | \|Undefined Saprotroph\| | Yeast | NULL | Probable | As nectar-tap_saprotroph (Põlme et al. 2020); Decay sugar-rich substrates (Põlme et al. 2020) | Põlme S, et al. 2020. Fungal Diversity 105:1-16 ((https://doi.org/10.1007/s13225-020-00466-2)) |
| Otu49369 | 1 | k__Fungi(100);p__Ascomycota(100);c__Saccharomycetes(100);o__Saccharomycetales(100);f__Metschnikowiaceae(100);g__Metschnikowia(100);s__Metschnikowia_pulcherrima(100); | Metschnikowia | 13 | Saprotroph | \|Undefined Saprotroph\| | Yeast | NULL | Probable | As nectar-tap_saprotroph (Põlme et al. 2020); Decay sugar-rich substrates (Põlme et al. 2020) | Põlme S, et al. 2020. Fungal Diversity 105:1-16 ((https://doi.org/10.1007/s13225-020-00466-2)) |
| Otu49370 | 1 | k__Fungi(100);p__Ascomycota(100);c__Saccharomycetes(100);o__Saccharomycetales(100);f__Metschnikowiaceae(100);g__Metschnikowia(100);g__Metschnikowia_unclassified(100); | Metschnikowia | 13 | Saprotroph | \|Undefined Saprotroph\| | Yeast | NULL | Probable | As nectar-tap_saprotroph (Põlme et al. 2020); Decay sugar-rich substrates (Põlme et al. 2020) | Põlme S, et al. 2020. Fungal Diversity 105:1-16 ((https://doi.org/10.1007/s13225-020-00466-2)) |
| Otu49371 | 1 | k__Fungi(100);p__Ascomycota(100);c__Saccharomycetes(100);o__Saccharomycetales(100);f__Metschnikowiaceae(100);g__Metschnikowia(100);s__Metschnikowia_pulcherrima(100); | Metschnikowia | 13 | Saprotroph | \|Undefined Saprotroph\| | Yeast | NULL | Probable | As nectar-tap_saprotroph (Põlme et al. 2020); Decay sugar-rich substrates (Põlme et al. 2020) | Põlme S, et al. 2020. Fungal Diversity 105:1-16 ((https://doi.org/10.1007/s13225-020-00466-2)) |
| Otu49372 | 1 | k__Fungi(100);p__Ascomycota(100);c__Saccharomycetes(100);o__Saccharomycetales(100);f__Metschnikowiaceae(100);g__Metschnikowia(100);s__Metschnikowia_pulcherrima(100); | Metschnikowia | 13 | Saprotroph | \|Undefined Saprotroph\| | Yeast | NULL | Probable | As nectar-tap_saprotroph (Põlme et al. 2020); Decay sugar-rich substrates (Põlme et al. 2020) | Põlme S, et al. 2020. Fungal Diversity 105:1-16 ((https://doi.org/10.1007/s13225-020-00466-2)) |
| Otu49373 | 1 | k__Fungi(100);p__Ascomycota(100);c__Saccharomycetes(100);o__Saccharomycetales(100);f__Metschnikowiaceae(100);g__Metschnikowia(100);s__Metschnikowia_pulcherrima(100); | Metschnikowia | 13 | Saprotroph | \|Undefined Saprotroph\| | Yeast | NULL | Probable | As nectar-tap_saprotroph (Põlme et al. 2020); Decay sugar-rich substrates (Põlme et al. 2020) | Põlme S, et al. 2020. Fungal Diversity 105:1-16 ((https://doi.org/10.1007/s13225-020-00466-2)) |
| Otu49374 | 1 | k__Fungi(100);p__Ascomycota(100);c__Saccharomycetes(100);o__Saccharomycetales(100);f__Metschnikowiaceae(100);g__Metschnikowia(100);s__Metschnikowia_pulcherrima(100); | Metschnikowia | 13 | Saprotroph | \|Undefined Saprotroph\| | Yeast | NULL | Probable | As nectar-tap_saprotroph (Põlme et al. 2020); Decay sugar-rich substrates (Põlme et al. 2020) | Põlme S, et al. 2020. Fungal Diversity 105:1-16 ((https://doi.org/10.1007/s13225-020-00466-2)) |
| Otu49375 | 1 | k__Fungi(100);p__Ascomycota(100);c__Saccharomycetes(100);o__Saccharomycetales(100);f__Metschnikowiaceae(100);g__Metschnikowia(100);s__Metschnikowia_pulcherrima(100); | Metschnikowia | 13 | Saprotroph | \|Undefined Saprotroph\| | Yeast | NULL | Probable | As nectar-tap_saprotroph (Põlme et al. 2020); Decay sugar-rich substrates (Põlme et al. 2020) | Põlme S, et al. 2020. Fungal Diversity 105:1-16 ((https://doi.org/10.1007/s13225-020-00466-2)) |
| Otu49376 | 1 | k__Fungi(100);p__Ascomycota(100);c__Saccharomycetes(100);o__Saccharomycetales(100);f__Metschnikowiaceae(100);g__Metschnikowia(100);s__Metschnikowia_pulcherrima(100); | Metschnikowia | 13 | Saprotroph | \|Undefined Saprotroph\| | Yeast | NULL | Probable | As nectar-tap_saprotroph (Põlme et al. 2020); Decay sugar-rich substrates (Põlme et al. 2020) | Põlme S, et al. 2020. Fungal Diversity 105:1-16 ((https://doi.org/10.1007/s13225-020-00466-2)) |
| Otu49377 | 1 | k__Fungi(100);p__Ascomycota(100);c__Saccharomycetes(100);o__Saccharomycetales(100);f__Metschnikowiaceae(100);g__Metschnikowia(100);s__Metschnikowia_pulcherrima(100); | Metschnikowia | 13 | Saprotroph | \|Undefined Saprotroph\| | Yeast | NULL | Probable | As nectar-tap_saprotroph (Põlme et al. 2020); Decay sugar-rich substrates (Põlme et al. 2020) | Põlme S, et al. 2020. Fungal Diversity 105:1-16 ((https://doi.org/10.1007/s13225-020-00466-2)) |
| Otu49378 | 1 | k__Fungi(100);p__Ascomycota(100);c__Saccharomycetes(100);o__Saccharomycetales(100);f__Metschnikowiaceae(100);g__Metschnikowia(100);s__Metschnikowia_pulcherrima(100); | Metschnikowia | 13 | Saprotroph | \|Undefined Saprotroph\| | Yeast | NULL | Probable | As nectar-tap_saprotroph (Põlme et al. 2020); Decay sugar-rich substrates (Põlme et al. 2020) | Põlme S, et al. 2020. Fungal Diversity 105:1-16 ((https://doi.org/10.1007/s13225-020-00466-2)) |
| Otu49379 | 1 | k__Fungi(100);p__Ascomycota(100);c__Saccharomycetes(100);o__Saccharomycetales(100);f__Metschnikowiaceae(100);g__Metschnikowia(100);g__Metschnikowia_unclassified(100); | Metschnikowia | 13 | Saprotroph | \|Undefined Saprotroph\| | Yeast | NULL | Probable | As nectar-tap_saprotroph (Põlme et al. 2020); Decay sugar-rich substrates (Põlme et al. 2020) | Põlme S, et al. 2020. Fungal Diversity 105:1-16 ((https://doi.org/10.1007/s13225-020-00466-2)) |
| Otu49380 | 1 | k__Fungi(100);p__Ascomycota(100);c__Saccharomycetes(100);o__Saccharomycetales(100);f__Metschnikowiaceae(100);g__Metschnikowia(100);s__Metschnikowia_pulcherrima(100); | Metschnikowia | 13 | Saprotroph | \|Undefined Saprotroph\| | Yeast | NULL | Probable | As nectar-tap_saprotroph (Põlme et al. 2020); Decay sugar-rich substrates (Põlme et al. 2020) | Põlme S, et al. 2020. Fungal Diversity 105:1-16 ((https://doi.org/10.1007/s13225-020-00466-2)) |
| Otu49381 | 1 | k__Fungi(100);p__Ascomycota(100);c__Saccharomycetes(100);o__Saccharomycetales(100);f__Metschnikowiaceae(100);g__Metschnikowia(100);g__Metschnikowia_unclassified(100); | Metschnikowia | 13 | Saprotroph | \|Undefined Saprotroph\| | Yeast | NULL | Probable | As nectar-tap_saprotroph (Põlme et al. 2020); Decay sugar-rich substrates (Põlme et al. 2020) | Põlme S, et al. 2020. Fungal Diversity 105:1-16 ((https://doi.org/10.1007/s13225-020-00466-2)) |
| Otu49382 | 1 | k__Fungi(100);p__Ascomycota(100);c__Saccharomycetes(100);o__Saccharomycetales(100);f__Metschnikowiaceae(100);g__Metschnikowia(100);s__Metschnikowia_pulcherrima(100); | Metschnikowia | 13 | Saprotroph | \|Undefined Saprotroph\| | Yeast | NULL | Probable | As nectar-tap_saprotroph (Põlme et al. 2020); Decay sugar-rich substrates (Põlme et al. 2020) | Põlme S, et al. 2020. Fungal Diversity 105:1-16 ((https://doi.org/10.1007/s13225-020-00466-2)) |
| Otu49383 | 1 | k__Fungi(100);p__Ascomycota(100);c__Saccharomycetes(100);o__Saccharomycetales(100);f__Metschnikowiaceae(100);g__Metschnikowia(100);s__Metschnikowia_pulcherrima(100); | Metschnikowia | 13 | Saprotroph | \|Undefined Saprotroph\| | Yeast | NULL | Probable | As nectar-tap_saprotroph (Põlme et al. 2020); Decay sugar-rich substrates (Põlme et al. 2020) | Põlme S, et al. 2020. Fungal Diversity 105:1-16 ((https://doi.org/10.1007/s13225-020-00466-2)) |
| Otu49384 | 1 | k__Fungi(100);p__Ascomycota(100);c__Saccharomycetes(100);o__Saccharomycetales(100);f__Metschnikowiaceae(100);g__Metschnikowia(100);g__Metschnikowia_unclassified(100); | Metschnikowia | 13 | Saprotroph | \|Undefined Saprotroph\| | Yeast | NULL | Probable | As nectar-tap_saprotroph (Põlme et al. 2020); Decay sugar-rich substrates (Põlme et al. 2020) | Põlme S, et al. 2020. Fungal Diversity 105:1-16 ((https://doi.org/10.1007/s13225-020-00466-2)) |
| Otu49385 | 1 | k__Fungi(100);p__Ascomycota(100);c__Saccharomycetes(100);o__Saccharomycetales(100);f__Metschnikowiaceae(100);g__Metschnikowia(100);g__Metschnikowia_unclassified(100); | Metschnikowia | 13 | Saprotroph | \|Undefined Saprotroph\| | Yeast | NULL | Probable | As nectar-tap_saprotroph (Põlme et al. 2020); Decay sugar-rich substrates (Põlme et al. 2020) | Põlme S, et al. 2020. Fungal Diversity 105:1-16 ((https://doi.org/10.1007/s13225-020-00466-2)) |
| Otu49386 | 1 | k__Fungi(100);p__Ascomycota(100);c__Saccharomycetes(100);o__Saccharomycetales(100);f__Metschnikowiaceae(100);g__Metschnikowia(100);g__Metschnikowia_unclassified(100); | Metschnikowia | 13 | Saprotroph | \|Undefined Saprotroph\| | Yeast | NULL | Probable | As nectar-tap_saprotroph (Põlme et al. 2020); Decay sugar-rich substrates (Põlme et al. 2020) | Põlme S, et al. 2020. Fungal Diversity 105:1-16 ((https://doi.org/10.1007/s13225-020-00466-2)) |
| Otu49387 | 1 | k__Fungi(100);p__Ascomycota(100);c__Saccharomycetes(100);o__Saccharomycetales(100);f__Metschnikowiaceae(100);g__Metschnikowia(100);s__Metschnikowia_pulcherrima(100); | Metschnikowia | 13 | Saprotroph | \|Undefined Saprotroph\| | Yeast | NULL | Probable | As nectar-tap_saprotroph (Põlme et al. 2020); Decay sugar-rich substrates (Põlme et al. 2020) | Põlme S, et al. 2020. Fungal Diversity 105:1-16 ((https://doi.org/10.1007/s13225-020-00466-2)) |
| Otu49388 | 1 | k__Fungi(100);p__Ascomycota(100);c__Saccharomycetes(100);o__Saccharomycetales(100);f__Metschnikowiaceae(100);g__Metschnikowia(100);s__Metschnikowia_pulcherrima(100); | Metschnikowia | 13 | Saprotroph | \|Undefined Saprotroph\| | Yeast | NULL | Probable | As nectar-tap_saprotroph (Põlme et al. 2020); Decay sugar-rich substrates (Põlme et al. 2020) | Põlme S, et al. 2020. Fungal Diversity 105:1-16 ((https://doi.org/10.1007/s13225-020-00466-2)) |
| Otu49389 | 1 | k__Fungi(100);p__Ascomycota(100);c__Saccharomycetes(100);o__Saccharomycetales(100);f__Metschnikowiaceae(100);g__Metschnikowia(100);s__Metschnikowia_pulcherrima(100); | Metschnikowia | 13 | Saprotroph | \|Undefined Saprotroph\| | Yeast | NULL | Probable | As nectar-tap_saprotroph (Põlme et al. 2020); Decay sugar-rich substrates (Põlme et al. 2020) | Põlme S, et al. 2020. Fungal Diversity 105:1-16 ((https://doi.org/10.1007/s13225-020-00466-2)) |
| Otu49390 | 1 | k__Fungi(100);p__Ascomycota(100);c__Saccharomycetes(100);o__Saccharomycetales(100);f__Metschnikowiaceae(100);g__Metschnikowia(100);s__Metschnikowia_pulcherrima(100); | Metschnikowia | 13 | Saprotroph | \|Undefined Saprotroph\| | Yeast | NULL | Probable | As nectar-tap_saprotroph (Põlme et al. 2020); Decay sugar-rich substrates (Põlme et al. 2020) | Põlme S, et al. 2020. Fungal Diversity 105:1-16 ((https://doi.org/10.1007/s13225-020-00466-2)) |
| Otu49391 | 1 | k__Fungi(100);p__Ascomycota(100);c__Saccharomycetes(100);o__Saccharomycetales(100);f__Metschnikowiaceae(100);g__Metschnikowia(100);s__Metschnikowia_pulcherrima(100); | Metschnikowia | 13 | Saprotroph | \|Undefined Saprotroph\| | Yeast | NULL | Probable | As nectar-tap_saprotroph (Põlme et al. 2020); Decay sugar-rich substrates (Põlme et al. 2020) | Põlme S, et al. 2020. Fungal Diversity 105:1-16 ((https://doi.org/10.1007/s13225-020-00466-2)) |
| Otu49392 | 1 | k__Fungi(100);p__Ascomycota(100);c__Saccharomycetes(100);o__Saccharomycetales(100);f__Metschnikowiaceae(100);g__Metschnikowia(100);s__Metschnikowia_pulcherrima(100); | Metschnikowia | 13 | Saprotroph | \|Undefined Saprotroph\| | Yeast | NULL | Probable | As nectar-tap_saprotroph (Põlme et al. 2020); Decay sugar-rich substrates (Põlme et al. 2020) | Põlme S, et al. 2020. Fungal Diversity 105:1-16 ((https://doi.org/10.1007/s13225-020-00466-2)) |
| Otu49393 | 1 | k__Fungi(100);p__Ascomycota(100);c__Saccharomycetes(100);o__Saccharomycetales(100);f__Metschnikowiaceae(100);g__Metschnikowia(100);s__Metschnikowia_pulcherrima(100); | Metschnikowia | 13 | Saprotroph | \|Undefined Saprotroph\| | Yeast | NULL | Probable | As nectar-tap_saprotroph (Põlme et al. 2020); Decay sugar-rich substrates (Põlme et al. 2020) | Põlme S, et al. 2020. Fungal Diversity 105:1-16 ((https://doi.org/10.1007/s13225-020-00466-2)) |
| Otu49394 | 1 | k__Fungi(100);p__Ascomycota(100);c__Saccharomycetes(100);o__Saccharomycetales(100);f__Metschnikowiaceae(100);g__Metschnikowia(100);s__Metschnikowia_pulcherrima(100); | Metschnikowia | 13 | Saprotroph | \|Undefined Saprotroph\| | Yeast | NULL | Probable | As nectar-tap_saprotroph (Põlme et al. 2020); Decay sugar-rich substrates (Põlme et al. 2020) | Põlme S, et al. 2020. Fungal Diversity 105:1-16 ((https://doi.org/10.1007/s13225-020-00466-2)) |
| Otu49395 | 1 | k__Fungi(100);p__Ascomycota(100);c__Saccharomycetes(100);o__Saccharomycetales(100);f__Metschnikowiaceae(100);g__Metschnikowia(100);s__Metschnikowia_pulcherrima(100); | Metschnikowia | 13 | Saprotroph | \|Undefined Saprotroph\| | Yeast | NULL | Probable | As nectar-tap_saprotroph (Põlme et al. 2020); Decay sugar-rich substrates (Põlme et al. 2020) | Põlme S, et al. 2020. Fungal Diversity 105:1-16 ((https://doi.org/10.1007/s13225-020-00466-2)) |
| Otu49396 | 1 | k__Fungi(100);p__Ascomycota(100);c__Saccharomycetes(100);o__Saccharomycetales(100);f__Metschnikowiaceae(100);g__Metschnikowia(100);s__Metschnikowia_pulcherrima(100); | Metschnikowia | 13 | Saprotroph | \|Undefined Saprotroph\| | Yeast | NULL | Probable | As nectar-tap_saprotroph (Põlme et al. 2020); Decay sugar-rich substrates (Põlme et al. 2020) | Põlme S, et al. 2020. Fungal Diversity 105:1-16 ((https://doi.org/10.1007/s13225-020-00466-2)) |
| Otu49397 | 1 | k__Fungi(100);p__Ascomycota(100);c__Saccharomycetes(100);o__Saccharomycetales(100);f__Metschnikowiaceae(100);g__Metschnikowia(100);s__Metschnikowia_pulcherrima(100); | Metschnikowia | 13 | Saprotroph | \|Undefined Saprotroph\| | Yeast | NULL | Probable | As nectar-tap_saprotroph (Põlme et al. 2020); Decay sugar-rich substrates (Põlme et al. 2020) | Põlme S, et al. 2020. Fungal Diversity 105:1-16 ((https://doi.org/10.1007/s13225-020-00466-2)) |
| Otu49398 | 1 | k__Fungi(100);p__Ascomycota(100);c__Saccharomycetes(100);o__Saccharomycetales(100);f__Metschnikowiaceae(100);g__Metschnikowia(100);g__Metschnikowia_unclassified(100); | Metschnikowia | 13 | Saprotroph | \|Undefined Saprotroph\| | Yeast | NULL | Probable | As nectar-tap_saprotroph (Põlme et al. 2020); Decay sugar-rich substrates (Põlme et al. 2020) | Põlme S, et al. 2020. Fungal Diversity 105:1-16 ((https://doi.org/10.1007/s13225-020-00466-2)) |
| Otu49399 | 1 | k__Fungi(100);p__Ascomycota(100);c__Saccharomycetes(100);o__Saccharomycetales(100);f__Metschnikowiaceae(100);g__Metschnikowia(100);g__Metschnikowia_unclassified(100); | Metschnikowia | 13 | Saprotroph | \|Undefined Saprotroph\| | Yeast | NULL | Probable | As nectar-tap_saprotroph (Põlme et al. 2020); Decay sugar-rich substrates (Põlme et al. 2020) | Põlme S, et al. 2020. Fungal Diversity 105:1-16 ((https://doi.org/10.1007/s13225-020-00466-2)) |
| Otu49400 | 1 | k__Fungi(100);p__Ascomycota(100);c__Saccharomycetes(100);o__Saccharomycetales(100);f__Metschnikowiaceae(100);g__Metschnikowia(100);s__Metschnikowia_pulcherrima(100); | Metschnikowia | 13 | Saprotroph | \|Undefined Saprotroph\| | Yeast | NULL | Probable | As nectar-tap_saprotroph (Põlme et al. 2020); Decay sugar-rich substrates (Põlme et al. 2020) | Põlme S, et al. 2020. Fungal Diversity 105:1-16 ((https://doi.org/10.1007/s13225-020-00466-2)) |
| Otu49401 | 1 | k__Fungi(100);p__Ascomycota(100);c__Saccharomycetes(100);o__Saccharomycetales(100);f__Metschnikowiaceae(100);g__Metschnikowia(100);s__Metschnikowia_pulcherrima(100); | Metschnikowia | 13 | Saprotroph | \|Undefined Saprotroph\| | Yeast | NULL | Probable | As nectar-tap_saprotroph (Põlme et al. 2020); Decay sugar-rich substrates (Põlme et al. 2020) | Põlme S, et al. 2020. Fungal Diversity 105:1-16 ((https://doi.org/10.1007/s13225-020-00466-2)) |
| Otu49402 | 1 | k__Fungi(100);p__Ascomycota(100);c__Saccharomycetes(100);o__Saccharomycetales(100);f__Metschnikowiaceae(100);g__Metschnikowia(100);s__Metschnikowia_pulcherrima(100); | Metschnikowia | 13 | Saprotroph | \|Undefined Saprotroph\| | Yeast | NULL | Probable | As nectar-tap_saprotroph (Põlme et al. 2020); Decay sugar-rich substrates (Põlme et al. 2020) | Põlme S, et al. 2020. Fungal Diversity 105:1-16 ((https://doi.org/10.1007/s13225-020-00466-2)) |
| Otu49403 | 1 | k__Fungi(100);p__Ascomycota(100);c__Saccharomycetes(100);o__Saccharomycetales(100);f__Metschnikowiaceae(100);g__Metschnikowia(100);g__Metschnikowia_unclassified(100); | Metschnikowia | 13 | Saprotroph | \|Undefined Saprotroph\| | Yeast | NULL | Probable | As nectar-tap_saprotroph (Põlme et al. 2020); Decay sugar-rich substrates (Põlme et al. 2020) | Põlme S, et al. 2020. Fungal Diversity 105:1-16 ((https://doi.org/10.1007/s13225-020-00466-2)) |
| Otu49404 | 1 | k__Fungi(100);p__Ascomycota(100);c__Saccharomycetes(100);o__Saccharomycetales(100);f__Metschnikowiaceae(100);g__Metschnikowia(100);g__Metschnikowia_unclassified(100); | Metschnikowia | 13 | Saprotroph | \|Undefined Saprotroph\| | Yeast | NULL | Probable | As nectar-tap_saprotroph (Põlme et al. 2020); Decay sugar-rich substrates (Põlme et al. 2020) | Põlme S, et al. 2020. Fungal Diversity 105:1-16 ((https://doi.org/10.1007/s13225-020-00466-2)) |
| Otu49405 | 1 | k__Fungi(100);p__Ascomycota(100);c__Saccharomycetes(100);o__Saccharomycetales(100);f__Metschnikowiaceae(100);g__Metschnikowia(100);s__Metschnikowia_pulcherrima(100); | Metschnikowia | 13 | Saprotroph | \|Undefined Saprotroph\| | Yeast | NULL | Probable | As nectar-tap_saprotroph (Põlme et al. 2020); Decay sugar-rich substrates (Põlme et al. 2020) | Põlme S, et al. 2020. Fungal Diversity 105:1-16 ((https://doi.org/10.1007/s13225-020-00466-2)) |
| Otu49406 | 1 | k__Fungi(100);p__Ascomycota(100);c__Saccharomycetes(100);o__Saccharomycetales(100);f__Metschnikowiaceae(100);g__Metschnikowia(100);g__Metschnikowia_unclassified(100); | Metschnikowia | 13 | Saprotroph | \|Undefined Saprotroph\| | Yeast | NULL | Probable | As nectar-tap_saprotroph (Põlme et al. 2020); Decay sugar-rich substrates (Põlme et al. 2020) | Põlme S, et al. 2020. Fungal Diversity 105:1-16 ((https://doi.org/10.1007/s13225-020-00466-2)) |
| Otu49407 | 1 | k__Fungi(100);p__Ascomycota(100);c__Saccharomycetes(100);o__Saccharomycetales(100);f__Metschnikowiaceae(100);g__Metschnikowia(100);s__Metschnikowia_pulcherrima(100); | Metschnikowia | 13 | Saprotroph | \|Undefined Saprotroph\| | Yeast | NULL | Probable | As nectar-tap_saprotroph (Põlme et al. 2020); Decay sugar-rich substrates (Põlme et al. 2020) | Põlme S, et al. 2020. Fungal Diversity 105:1-16 ((https://doi.org/10.1007/s13225-020-00466-2)) |
| Otu49408 | 1 | k__Fungi(100);p__Ascomycota(100);c__Saccharomycetes(100);o__Saccharomycetales(100);f__Metschnikowiaceae(100);g__Metschnikowia(100);g__Metschnikowia_unclassified(100); | Metschnikowia | 13 | Saprotroph | \|Undefined Saprotroph\| | Yeast | NULL | Probable | As nectar-tap_saprotroph (Põlme et al. 2020); Decay sugar-rich substrates (Põlme et al. 2020) | Põlme S, et al. 2020. Fungal Diversity 105:1-16 ((https://doi.org/10.1007/s13225-020-00466-2)) |
| Otu49409 | 1 | k__Fungi(100);p__Ascomycota(100);c__Saccharomycetes(100);o__Saccharomycetales(100);f__Metschnikowiaceae(100);g__Metschnikowia(100);g__Metschnikowia_unclassified(100); | Metschnikowia | 13 | Saprotroph | \|Undefined Saprotroph\| | Yeast | NULL | Probable | As nectar-tap_saprotroph (Põlme et al. 2020); Decay sugar-rich substrates (Põlme et al. 2020) | Põlme S, et al. 2020. Fungal Diversity 105:1-16 ((https://doi.org/10.1007/s13225-020-00466-2)) |
| Otu49410 | 1 | k__Fungi(100);p__Ascomycota(100);c__Saccharomycetes(100);o__Saccharomycetales(100);f__Metschnikowiaceae(100);g__Metschnikowia(100);g__Metschnikowia_unclassified(100); | Metschnikowia | 13 | Saprotroph | \|Undefined Saprotroph\| | Yeast | NULL | Probable | As nectar-tap_saprotroph (Põlme et al. 2020); Decay sugar-rich substrates (Põlme et al. 2020) | Põlme S, et al. 2020. Fungal Diversity 105:1-16 ((https://doi.org/10.1007/s13225-020-00466-2)) |
| Otu49411 | 1 | k__Fungi(100);p__Ascomycota(100);c__Saccharomycetes(100);o__Saccharomycetales(100);f__Metschnikowiaceae(100);g__Metschnikowia(100);s__Metschnikowia_pulcherrima(100); | Metschnikowia | 13 | Saprotroph | \|Undefined Saprotroph\| | Yeast | NULL | Probable | As nectar-tap_saprotroph (Põlme et al. 2020); Decay sugar-rich substrates (Põlme et al. 2020) | Põlme S, et al. 2020. Fungal Diversity 105:1-16 ((https://doi.org/10.1007/s13225-020-00466-2)) |
| Otu49412 | 1 | k__Fungi(100);p__Ascomycota(100);c__Saccharomycetes(100);o__Saccharomycetales(100);f__Metschnikowiaceae(100);g__Metschnikowia(100);s__Metschnikowia_pulcherrima(100); | Metschnikowia | 13 | Saprotroph | \|Undefined Saprotroph\| | Yeast | NULL | Probable | As nectar-tap_saprotroph (Põlme et al. 2020); Decay sugar-rich substrates (Põlme et al. 2020) | Põlme S, et al. 2020. Fungal Diversity 105:1-16 ((https://doi.org/10.1007/s13225-020-00466-2)) |
| Otu49413 | 1 | k__Fungi(100);p__Ascomycota(100);c__Saccharomycetes(100);o__Saccharomycetales(100);f__Metschnikowiaceae(100);g__Metschnikowia(100);s__Metschnikowia_pulcherrima(100); | Metschnikowia | 13 | Saprotroph | \|Undefined Saprotroph\| | Yeast | NULL | Probable | As nectar-tap_saprotroph (Põlme et al. 2020); Decay sugar-rich substrates (Põlme et al. 2020) | Põlme S, et al. 2020. Fungal Diversity 105:1-16 ((https://doi.org/10.1007/s13225-020-00466-2)) |
| Otu49414 | 1 | k__Fungi(100);p__Ascomycota(100);c__Saccharomycetes(100);o__Saccharomycetales(100);f__Metschnikowiaceae(100);g__Metschnikowia(100);s__Metschnikowia_pulcherrima(100); | Metschnikowia | 13 | Saprotroph | \|Undefined Saprotroph\| | Yeast | NULL | Probable | As nectar-tap_saprotroph (Põlme et al. 2020); Decay sugar-rich substrates (Põlme et al. 2020) | Põlme S, et al. 2020. Fungal Diversity 105:1-16 ((https://doi.org/10.1007/s13225-020-00466-2)) |
| Otu49415 | 1 | k__Fungi(100);p__Ascomycota(100);c__Saccharomycetes(100);o__Saccharomycetales(100);f__Metschnikowiaceae(100);g__Metschnikowia(100);s__Metschnikowia_pulcherrima(100); | Metschnikowia | 13 | Saprotroph | \|Undefined Saprotroph\| | Yeast | NULL | Probable | As nectar-tap_saprotroph (Põlme et al. 2020); Decay sugar-rich substrates (Põlme et al. 2020) | Põlme S, et al. 2020. Fungal Diversity 105:1-16 ((https://doi.org/10.1007/s13225-020-00466-2)) |
| Otu49416 | 1 | k__Fungi(100);p__Ascomycota(100);c__Saccharomycetes(100);o__Saccharomycetales(100);f__Metschnikowiaceae(100);g__Metschnikowia(100);s__Metschnikowia_pulcherrima(100); | Metschnikowia | 13 | Saprotroph | \|Undefined Saprotroph\| | Yeast | NULL | Probable | As nectar-tap_saprotroph (Põlme et al. 2020); Decay sugar-rich substrates (Põlme et al. 2020) | Põlme S, et al. 2020. Fungal Diversity 105:1-16 ((https://doi.org/10.1007/s13225-020-00466-2)) |
| Otu49417 | 1 | k__Fungi(100);p__Ascomycota(100);c__Saccharomycetes(100);o__Saccharomycetales(100);f__Metschnikowiaceae(100);g__Metschnikowia(100);s__Metschnikowia_pulcherrima(100); | Metschnikowia | 13 | Saprotroph | \|Undefined Saprotroph\| | Yeast | NULL | Probable | As nectar-tap_saprotroph (Põlme et al. 2020); Decay sugar-rich substrates (Põlme et al. 2020) | Põlme S, et al. 2020. Fungal Diversity 105:1-16 ((https://doi.org/10.1007/s13225-020-00466-2)) |
| Otu49418 | 1 | k__Fungi(100);p__Ascomycota(100);c__Saccharomycetes(100);o__Saccharomycetales(100);f__Metschnikowiaceae(100);g__Metschnikowia(100);s__Metschnikowia_pulcherrima(100); | Metschnikowia | 13 | Saprotroph | \|Undefined Saprotroph\| | Yeast | NULL | Probable | As nectar-tap_saprotroph (Põlme et al. 2020); Decay sugar-rich substrates (Põlme et al. 2020) | Põlme S, et al. 2020. Fungal Diversity 105:1-16 ((https://doi.org/10.1007/s13225-020-00466-2)) |
| Otu49419 | 1 | k__Fungi(100);p__Ascomycota(100);c__Saccharomycetes(100);o__Saccharomycetales(100);f__Metschnikowiaceae(100);g__Metschnikowia(100);g__Metschnikowia_unclassified(100); | Metschnikowia | 13 | Saprotroph | \|Undefined Saprotroph\| | Yeast | NULL | Probable | As nectar-tap_saprotroph (Põlme et al. 2020); Decay sugar-rich substrates (Põlme et al. 2020) | Põlme S, et al. 2020. Fungal Diversity 105:1-16 ((https://doi.org/10.1007/s13225-020-00466-2)) |
| Otu49420 | 1 | k__Fungi(100);p__Ascomycota(100);c__Saccharomycetes(100);o__Saccharomycetales(100);f__Metschnikowiaceae(100);g__Metschnikowia(100);s__Metschnikowia_pulcherrima(100); | Metschnikowia | 13 | Saprotroph | \|Undefined Saprotroph\| | Yeast | NULL | Probable | As nectar-tap_saprotroph (Põlme et al. 2020); Decay sugar-rich substrates (Põlme et al. 2020) | Põlme S, et al. 2020. Fungal Diversity 105:1-16 ((https://doi.org/10.1007/s13225-020-00466-2)) |
| Otu49421 | 1 | k__Fungi(100);p__Ascomycota(100);c__Saccharomycetes(100);o__Saccharomycetales(100);f__Metschnikowiaceae(100);g__Metschnikowia(100);s__Metschnikowia_pulcherrima(100); | Metschnikowia | 13 | Saprotroph | \|Undefined Saprotroph\| | Yeast | NULL | Probable | As nectar-tap_saprotroph (Põlme et al. 2020); Decay sugar-rich substrates (Põlme et al. 2020) | Põlme S, et al. 2020. Fungal Diversity 105:1-16 ((https://doi.org/10.1007/s13225-020-00466-2)) |
| Otu49422 | 1 | k__Fungi(100);p__Ascomycota(100);c__Saccharomycetes(100);o__Saccharomycetales(100);f__Metschnikowiaceae(100);g__Metschnikowia(100);s__Metschnikowia_pulcherrima(100); | Metschnikowia | 13 | Saprotroph | \|Undefined Saprotroph\| | Yeast | NULL | Probable | As nectar-tap_saprotroph (Põlme et al. 2020); Decay sugar-rich substrates (Põlme et al. 2020) | Põlme S, et al. 2020. Fungal Diversity 105:1-16 ((https://doi.org/10.1007/s13225-020-00466-2)) |
| Otu49423 | 1 | k__Fungi(100);p__Ascomycota(100);c__Saccharomycetes(100);o__Saccharomycetales(100);f__Metschnikowiaceae(100);g__Metschnikowia(100);s__Metschnikowia_pulcherrima(100); | Metschnikowia | 13 | Saprotroph | \|Undefined Saprotroph\| | Yeast | NULL | Probable | As nectar-tap_saprotroph (Põlme et al. 2020); Decay sugar-rich substrates (Põlme et al. 2020) | Põlme S, et al. 2020. Fungal Diversity 105:1-16 ((https://doi.org/10.1007/s13225-020-00466-2)) |
| Otu49424 | 1 | k__Fungi(100);p__Ascomycota(100);c__Saccharomycetes(100);o__Saccharomycetales(100);f__Metschnikowiaceae(100);g__Metschnikowia(100);g__Metschnikowia_unclassified(100); | Metschnikowia | 13 | Saprotroph | \|Undefined Saprotroph\| | Yeast | NULL | Probable | As nectar-tap_saprotroph (Põlme et al. 2020); Decay sugar-rich substrates (Põlme et al. 2020) | Põlme S, et al. 2020. Fungal Diversity 105:1-16 ((https://doi.org/10.1007/s13225-020-00466-2)) |
| Otu49425 | 1 | k__Fungi(100);p__Ascomycota(100);c__Saccharomycetes(100);o__Saccharomycetales(100);f__Metschnikowiaceae(100);g__Metschnikowia(100);s__Metschnikowia_pulcherrima(100); | Metschnikowia | 13 | Saprotroph | \|Undefined Saprotroph\| | Yeast | NULL | Probable | As nectar-tap_saprotroph (Põlme et al. 2020); Decay sugar-rich substrates (Põlme et al. 2020) | Põlme S, et al. 2020. Fungal Diversity 105:1-16 ((https://doi.org/10.1007/s13225-020-00466-2)) |
| Otu49426 | 1 | k__Fungi(100);p__Ascomycota(100);c__Saccharomycetes(100);o__Saccharomycetales(100);f__Metschnikowiaceae(100);g__Metschnikowia(100);s__Metschnikowia_pulcherrima(100); | Metschnikowia | 13 | Saprotroph | \|Undefined Saprotroph\| | Yeast | NULL | Probable | As nectar-tap_saprotroph (Põlme et al. 2020); Decay sugar-rich substrates (Põlme et al. 2020) | Põlme S, et al. 2020. Fungal Diversity 105:1-16 ((https://doi.org/10.1007/s13225-020-00466-2)) |
| Otu49427 | 1 | k__Fungi(100);p__Ascomycota(100);c__Saccharomycetes(100);o__Saccharomycetales(100);f__Metschnikowiaceae(100);g__Metschnikowia(100);g__Metschnikowia_unclassified(100); | Metschnikowia | 13 | Saprotroph | \|Undefined Saprotroph\| | Yeast | NULL | Probable | As nectar-tap_saprotroph (Põlme et al. 2020); Decay sugar-rich substrates (Põlme et al. 2020) | Põlme S, et al. 2020. Fungal Diversity 105:1-16 ((https://doi.org/10.1007/s13225-020-00466-2)) |
| Otu49428 | 1 | k__Fungi(100);p__Ascomycota(100);c__Saccharomycetes(100);o__Saccharomycetales(100);f__Metschnikowiaceae(100);g__Metschnikowia(100);s__Metschnikowia_pulcherrima(100); | Metschnikowia | 13 | Saprotroph | \|Undefined Saprotroph\| | Yeast | NULL | Probable | As nectar-tap_saprotroph (Põlme et al. 2020); Decay sugar-rich substrates (Põlme et al. 2020) | Põlme S, et al. 2020. Fungal Diversity 105:1-16 ((https://doi.org/10.1007/s13225-020-00466-2)) |
| Otu49429 | 1 | k__Fungi(100);p__Ascomycota(100);c__Saccharomycetes(100);o__Saccharomycetales(100);f__Metschnikowiaceae(100);g__Metschnikowia(100);s__Metschnikowia_pulcherrima(100); | Metschnikowia | 13 | Saprotroph | \|Undefined Saprotroph\| | Yeast | NULL | Probable | As nectar-tap_saprotroph (Põlme et al. 2020); Decay sugar-rich substrates (Põlme et al. 2020) | Põlme S, et al. 2020. Fungal Diversity 105:1-16 ((https://doi.org/10.1007/s13225-020-00466-2)) |
| Otu49430 | 1 | k__Fungi(100);p__Ascomycota(100);c__Saccharomycetes(100);o__Saccharomycetales(100);f__Metschnikowiaceae(100);g__Metschnikowia(100);g__Metschnikowia_unclassified(100); | Metschnikowia | 13 | Saprotroph | \|Undefined Saprotroph\| | Yeast | NULL | Probable | As nectar-tap_saprotroph (Põlme et al. 2020); Decay sugar-rich substrates (Põlme et al. 2020) | Põlme S, et al. 2020. Fungal Diversity 105:1-16 ((https://doi.org/10.1007/s13225-020-00466-2)) |
| Otu49431 | 1 | k__Fungi(100);p__Ascomycota(100);c__Saccharomycetes(100);o__Saccharomycetales(100);f__Metschnikowiaceae(100);g__Metschnikowia(100);g__Metschnikowia_unclassified(100); | Metschnikowia | 13 | Saprotroph | \|Undefined Saprotroph\| | Yeast | NULL | Probable | As nectar-tap_saprotroph (Põlme et al. 2020); Decay sugar-rich substrates (Põlme et al. 2020) | Põlme S, et al. 2020. Fungal Diversity 105:1-16 ((https://doi.org/10.1007/s13225-020-00466-2)) |
| Otu49432 | 1 | k__Fungi(100);p__Ascomycota(100);c__Saccharomycetes(100);o__Saccharomycetales(100);f__Metschnikowiaceae(100);g__Metschnikowia(100);s__Metschnikowia_pulcherrima(100); | Metschnikowia | 13 | Saprotroph | \|Undefined Saprotroph\| | Yeast | NULL | Probable | As nectar-tap_saprotroph (Põlme et al. 2020); Decay sugar-rich substrates (Põlme et al. 2020) | Põlme S, et al. 2020. Fungal Diversity 105:1-16 ((https://doi.org/10.1007/s13225-020-00466-2)) |
| Otu49433 | 1 | k__Fungi(100);p__Ascomycota(100);c__Saccharomycetes(100);o__Saccharomycetales(100);f__Metschnikowiaceae(100);g__Metschnikowia(100);g__Metschnikowia_unclassified(100); | Metschnikowia | 13 | Saprotroph | \|Undefined Saprotroph\| | Yeast | NULL | Probable | As nectar-tap_saprotroph (Põlme et al. 2020); Decay sugar-rich substrates (Põlme et al. 2020) | Põlme S, et al. 2020. Fungal Diversity 105:1-16 ((https://doi.org/10.1007/s13225-020-00466-2)) |
| Otu49434 | 1 | k__Fungi(100);p__Ascomycota(100);c__Saccharomycetes(100);o__Saccharomycetales(100);f__Metschnikowiaceae(100);g__Metschnikowia(100);s__Metschnikowia_pulcherrima(100); | Metschnikowia | 13 | Saprotroph | \|Undefined Saprotroph\| | Yeast | NULL | Probable | As nectar-tap_saprotroph (Põlme et al. 2020); Decay sugar-rich substrates (Põlme et al. 2020) | Põlme S, et al. 2020. Fungal Diversity 105:1-16 ((https://doi.org/10.1007/s13225-020-00466-2)) |
| Otu49435 | 1 | k__Fungi(100);p__Ascomycota(100);c__Saccharomycetes(100);o__Saccharomycetales(100);f__Metschnikowiaceae(100);g__Metschnikowia(100);s__Metschnikowia_pulcherrima(100); | Metschnikowia | 13 | Saprotroph | \|Undefined Saprotroph\| | Yeast | NULL | Probable | As nectar-tap_saprotroph (Põlme et al. 2020); Decay sugar-rich substrates (Põlme et al. 2020) | Põlme S, et al. 2020. Fungal Diversity 105:1-16 ((https://doi.org/10.1007/s13225-020-00466-2)) |
| Otu49436 | 1 | k__Fungi(100);p__Ascomycota(100);c__Saccharomycetes(100);o__Saccharomycetales(100);f__Metschnikowiaceae(100);g__Metschnikowia(100);s__Metschnikowia_pulcherrima(100); | Metschnikowia | 13 | Saprotroph | \|Undefined Saprotroph\| | Yeast | NULL | Probable | As nectar-tap_saprotroph (Põlme et al. 2020); Decay sugar-rich substrates (Põlme et al. 2020) | Põlme S, et al. 2020. Fungal Diversity 105:1-16 ((https://doi.org/10.1007/s13225-020-00466-2)) |
| Otu49437 | 1 | k__Fungi(100);p__Ascomycota(100);c__Saccharomycetes(100);o__Saccharomycetales(100);f__Metschnikowiaceae(100);g__Metschnikowia(100);s__Metschnikowia_pulcherrima(100); | Metschnikowia | 13 | Saprotroph | \|Undefined Saprotroph\| | Yeast | NULL | Probable | As nectar-tap_saprotroph (Põlme et al. 2020); Decay sugar-rich substrates (Põlme et al. 2020) | Põlme S, et al. 2020. Fungal Diversity 105:1-16 ((https://doi.org/10.1007/s13225-020-00466-2)) |
| Otu49438 | 1 | k__Fungi(100);p__Ascomycota(100);c__Saccharomycetes(100);o__Saccharomycetales(100);f__Metschnikowiaceae(100);g__Metschnikowia(100);g__Metschnikowia_unclassified(100); | Metschnikowia | 13 | Saprotroph | \|Undefined Saprotroph\| | Yeast | NULL | Probable | As nectar-tap_saprotroph (Põlme et al. 2020); Decay sugar-rich substrates (Põlme et al. 2020) | Põlme S, et al. 2020. Fungal Diversity 105:1-16 ((https://doi.org/10.1007/s13225-020-00466-2)) |
| Otu49439 | 1 | k__Fungi(100);p__Ascomycota(100);c__Saccharomycetes(100);o__Saccharomycetales(100);f__Metschnikowiaceae(100);g__Metschnikowia(100);s__Metschnikowia_pulcherrima(100); | Metschnikowia | 13 | Saprotroph | \|Undefined Saprotroph\| | Yeast | NULL | Probable | As nectar-tap_saprotroph (Põlme et al. 2020); Decay sugar-rich substrates (Põlme et al. 2020) | Põlme S, et al. 2020. Fungal Diversity 105:1-16 ((https://doi.org/10.1007/s13225-020-00466-2)) |
| Otu49440 | 1 | k__Fungi(100);p__Ascomycota(100);c__Saccharomycetes(100);o__Saccharomycetales(100);f__Metschnikowiaceae(100);g__Metschnikowia(100);s__Metschnikowia_pulcherrima(100); | Metschnikowia | 13 | Saprotroph | \|Undefined Saprotroph\| | Yeast | NULL | Probable | As nectar-tap_saprotroph (Põlme et al. 2020); Decay sugar-rich substrates (Põlme et al. 2020) | Põlme S, et al. 2020. Fungal Diversity 105:1-16 ((https://doi.org/10.1007/s13225-020-00466-2)) |
| Otu49441 | 1 | k__Fungi(100);p__Ascomycota(100);c__Saccharomycetes(100);o__Saccharomycetales(100);f__Metschnikowiaceae(100);g__Metschnikowia(100);s__Metschnikowia_pulcherrima(100); | Metschnikowia | 13 | Saprotroph | \|Undefined Saprotroph\| | Yeast | NULL | Probable | As nectar-tap_saprotroph (Põlme et al. 2020); Decay sugar-rich substrates (Põlme et al. 2020) | Põlme S, et al. 2020. Fungal Diversity 105:1-16 ((https://doi.org/10.1007/s13225-020-00466-2)) |
| Otu49442 | 1 | k__Fungi(100);p__Ascomycota(100);c__Saccharomycetes(100);o__Saccharomycetales(100);f__Metschnikowiaceae(100);g__Metschnikowia(100);s__Metschnikowia_pulcherrima(100); | Metschnikowia | 13 | Saprotroph | \|Undefined Saprotroph\| | Yeast | NULL | Probable | As nectar-tap_saprotroph (Põlme et al. 2020); Decay sugar-rich substrates (Põlme et al. 2020) | Põlme S, et al. 2020. Fungal Diversity 105:1-16 ((https://doi.org/10.1007/s13225-020-00466-2)) |
| Otu49443 | 1 | k__Fungi(100);p__Ascomycota(100);c__Saccharomycetes(100);o__Saccharomycetales(100);f__Metschnikowiaceae(100);g__Metschnikowia(100);g__Metschnikowia_unclassified(100); | Metschnikowia | 13 | Saprotroph | \|Undefined Saprotroph\| | Yeast | NULL | Probable | As nectar-tap_saprotroph (Põlme et al. 2020); Decay sugar-rich substrates (Põlme et al. 2020) | Põlme S, et al. 2020. Fungal Diversity 105:1-16 ((https://doi.org/10.1007/s13225-020-00466-2)) |
| Otu49444 | 1 | k__Fungi(100);p__Ascomycota(100);c__Saccharomycetes(100);o__Saccharomycetales(100);f__Metschnikowiaceae(100);g__Metschnikowia(100);s__Metschnikowia_pulcherrima(100); | Metschnikowia | 13 | Saprotroph | \|Undefined Saprotroph\| | Yeast | NULL | Probable | As nectar-tap_saprotroph (Põlme et al. 2020); Decay sugar-rich substrates (Põlme et al. 2020) | Põlme S, et al. 2020. Fungal Diversity 105:1-16 ((https://doi.org/10.1007/s13225-020-00466-2)) |
| Otu49445 | 1 | k__Fungi(100);p__Ascomycota(100);c__Saccharomycetes(100);o__Saccharomycetales(100);f__Metschnikowiaceae(100);g__Metschnikowia(100);s__Metschnikowia_pulcherrima(100); | Metschnikowia | 13 | Saprotroph | \|Undefined Saprotroph\| | Yeast | NULL | Probable | As nectar-tap_saprotroph (Põlme et al. 2020); Decay sugar-rich substrates (Põlme et al. 2020) | Põlme S, et al. 2020. Fungal Diversity 105:1-16 ((https://doi.org/10.1007/s13225-020-00466-2)) |
| Otu49446 | 1 | k__Fungi(100);p__Ascomycota(100);c__Saccharomycetes(100);o__Saccharomycetales(100);f__Metschnikowiaceae(100);g__Metschnikowia(100);s__Metschnikowia_pulcherrima(100); | Metschnikowia | 13 | Saprotroph | \|Undefined Saprotroph\| | Yeast | NULL | Probable | As nectar-tap_saprotroph (Põlme et al. 2020); Decay sugar-rich substrates (Põlme et al. 2020) | Põlme S, et al. 2020. Fungal Diversity 105:1-16 ((https://doi.org/10.1007/s13225-020-00466-2)) |
| Otu49447 | 1 | k__Fungi(100);p__Ascomycota(100);c__Saccharomycetes(100);o__Saccharomycetales(100);f__Metschnikowiaceae(100);g__Metschnikowia(100);s__Metschnikowia_pulcherrima(100); | Metschnikowia | 13 | Saprotroph | \|Undefined Saprotroph\| | Yeast | NULL | Probable | As nectar-tap_saprotroph (Põlme et al. 2020); Decay sugar-rich substrates (Põlme et al. 2020) | Põlme S, et al. 2020. Fungal Diversity 105:1-16 ((https://doi.org/10.1007/s13225-020-00466-2)) |
| Otu49448 | 1 | k__Fungi(100);p__Ascomycota(100);c__Saccharomycetes(100);o__Saccharomycetales(100);f__Metschnikowiaceae(100);g__Metschnikowia(100);s__Metschnikowia_pulcherrima(100); | Metschnikowia | 13 | Saprotroph | \|Undefined Saprotroph\| | Yeast | NULL | Probable | As nectar-tap_saprotroph (Põlme et al. 2020); Decay sugar-rich substrates (Põlme et al. 2020) | Põlme S, et al. 2020. Fungal Diversity 105:1-16 ((https://doi.org/10.1007/s13225-020-00466-2)) |
| Otu49449 | 1 | k__Fungi(100);p__Ascomycota(100);c__Saccharomycetes(100);o__Saccharomycetales(100);f__Metschnikowiaceae(100);g__Metschnikowia(100);s__Metschnikowia_pulcherrima(100); | Metschnikowia | 13 | Saprotroph | \|Undefined Saprotroph\| | Yeast | NULL | Probable | As nectar-tap_saprotroph (Põlme et al. 2020); Decay sugar-rich substrates (Põlme et al. 2020) | Põlme S, et al. 2020. Fungal Diversity 105:1-16 ((https://doi.org/10.1007/s13225-020-00466-2)) |
| Otu49450 | 1 | k__Fungi(100);p__Ascomycota(100);c__Saccharomycetes(100);o__Saccharomycetales(100);f__Metschnikowiaceae(100);g__Metschnikowia(100);g__Metschnikowia_unclassified(100); | Metschnikowia | 13 | Saprotroph | \|Undefined Saprotroph\| | Yeast | NULL | Probable | As nectar-tap_saprotroph (Põlme et al. 2020); Decay sugar-rich substrates (Põlme et al. 2020) | Põlme S, et al. 2020. Fungal Diversity 105:1-16 ((https://doi.org/10.1007/s13225-020-00466-2)) |
| Otu49451 | 1 | k__Fungi(100);p__Ascomycota(100);c__Saccharomycetes(100);o__Saccharomycetales(100);f__Metschnikowiaceae(100);g__Metschnikowia(100);g__Metschnikowia_unclassified(100); | Metschnikowia | 13 | Saprotroph | \|Undefined Saprotroph\| | Yeast | NULL | Probable | As nectar-tap_saprotroph (Põlme et al. 2020); Decay sugar-rich substrates (Põlme et al. 2020) | Põlme S, et al. 2020. Fungal Diversity 105:1-16 ((https://doi.org/10.1007/s13225-020-00466-2)) |
| Otu49452 | 1 | k__Fungi(100);p__Ascomycota(100);c__Saccharomycetes(100);o__Saccharomycetales(100);f__Metschnikowiaceae(100);g__Metschnikowia(100);s__Metschnikowia_pulcherrima(100); | Metschnikowia | 13 | Saprotroph | \|Undefined Saprotroph\| | Yeast | NULL | Probable | As nectar-tap_saprotroph (Põlme et al. 2020); Decay sugar-rich substrates (Põlme et al. 2020) | Põlme S, et al. 2020. Fungal Diversity 105:1-16 ((https://doi.org/10.1007/s13225-020-00466-2)) |
| Otu49453 | 1 | k__Fungi(100);p__Ascomycota(100);c__Saccharomycetes(100);o__Saccharomycetales(100);f__Metschnikowiaceae(100);g__Metschnikowia(100);s__Metschnikowia_pulcherrima(100); | Metschnikowia | 13 | Saprotroph | \|Undefined Saprotroph\| | Yeast | NULL | Probable | As nectar-tap_saprotroph (Põlme et al. 2020); Decay sugar-rich substrates (Põlme et al. 2020) | Põlme S, et al. 2020. Fungal Diversity 105:1-16 ((https://doi.org/10.1007/s13225-020-00466-2)) |
| Otu49454 | 1 | k__Fungi(100);p__Ascomycota(100);c__Saccharomycetes(100);o__Saccharomycetales(100);f__Metschnikowiaceae(100);g__Metschnikowia(100);s__Metschnikowia_pulcherrima(100); | Metschnikowia | 13 | Saprotroph | \|Undefined Saprotroph\| | Yeast | NULL | Probable | As nectar-tap_saprotroph (Põlme et al. 2020); Decay sugar-rich substrates (Põlme et al. 2020) | Põlme S, et al. 2020. Fungal Diversity 105:1-16 ((https://doi.org/10.1007/s13225-020-00466-2)) |
| Otu49455 | 1 | k__Fungi(100);p__Ascomycota(100);c__Saccharomycetes(100);o__Saccharomycetales(100);f__Metschnikowiaceae(100);g__Metschnikowia(100);s__Metschnikowia_pulcherrima(100); | Metschnikowia | 13 | Saprotroph | \|Undefined Saprotroph\| | Yeast | NULL | Probable | As nectar-tap_saprotroph (Põlme et al. 2020); Decay sugar-rich substrates (Põlme et al. 2020) | Põlme S, et al. 2020. Fungal Diversity 105:1-16 ((https://doi.org/10.1007/s13225-020-00466-2)) |
| Otu49456 | 1 | k__Fungi(100);p__Ascomycota(100);c__Saccharomycetes(100);o__Saccharomycetales(100);f__Metschnikowiaceae(100);g__Metschnikowia(100);s__Metschnikowia_pulcherrima(100); | Metschnikowia | 13 | Saprotroph | \|Undefined Saprotroph\| | Yeast | NULL | Probable | As nectar-tap_saprotroph (Põlme et al. 2020); Decay sugar-rich substrates (Põlme et al. 2020) | Põlme S, et al. 2020. Fungal Diversity 105:1-16 ((https://doi.org/10.1007/s13225-020-00466-2)) |
| Otu49457 | 1 | k__Fungi(100);p__Ascomycota(100);c__Saccharomycetes(100);o__Saccharomycetales(100);f__Metschnikowiaceae(100);g__Metschnikowia(100);s__Metschnikowia_pulcherrima(100); | Metschnikowia | 13 | Saprotroph | \|Undefined Saprotroph\| | Yeast | NULL | Probable | As nectar-tap_saprotroph (Põlme et al. 2020); Decay sugar-rich substrates (Põlme et al. 2020) | Põlme S, et al. 2020. Fungal Diversity 105:1-16 ((https://doi.org/10.1007/s13225-020-00466-2)) |
| Otu49458 | 1 | k__Fungi(100);p__Ascomycota(100);c__Saccharomycetes(100);o__Saccharomycetales(100);f__Metschnikowiaceae(100);g__Metschnikowia(100);g__Metschnikowia_unclassified(100); | Metschnikowia | 13 | Saprotroph | \|Undefined Saprotroph\| | Yeast | NULL | Probable | As nectar-tap_saprotroph (Põlme et al. 2020); Decay sugar-rich substrates (Põlme et al. 2020) | Põlme S, et al. 2020. Fungal Diversity 105:1-16 ((https://doi.org/10.1007/s13225-020-00466-2)) |
| Otu49459 | 1 | k__Fungi(100);p__Ascomycota(100);c__Saccharomycetes(100);o__Saccharomycetales(100);f__Metschnikowiaceae(100);g__Metschnikowia(100);s__Metschnikowia_pulcherrima(100); | Metschnikowia | 13 | Saprotroph | \|Undefined Saprotroph\| | Yeast | NULL | Probable | As nectar-tap_saprotroph (Põlme et al. 2020); Decay sugar-rich substrates (Põlme et al. 2020) | Põlme S, et al. 2020. Fungal Diversity 105:1-16 ((https://doi.org/10.1007/s13225-020-00466-2)) |
| Otu49460 | 1 | k__Fungi(100);p__Ascomycota(100);c__Saccharomycetes(100);o__Saccharomycetales(100);f__Metschnikowiaceae(100);g__Metschnikowia(100);s__Metschnikowia_pulcherrima(100); | Metschnikowia | 13 | Saprotroph | \|Undefined Saprotroph\| | Yeast | NULL | Probable | As nectar-tap_saprotroph (Põlme et al. 2020); Decay sugar-rich substrates (Põlme et al. 2020) | Põlme S, et al. 2020. Fungal Diversity 105:1-16 ((https://doi.org/10.1007/s13225-020-00466-2)) |
| Otu49461 | 1 | k__Fungi(100);p__Ascomycota(100);c__Saccharomycetes(100);o__Saccharomycetales(100);f__Metschnikowiaceae(100);g__Metschnikowia(100);s__Metschnikowia_pulcherrima(100); | Metschnikowia | 13 | Saprotroph | \|Undefined Saprotroph\| | Yeast | NULL | Probable | As nectar-tap_saprotroph (Põlme et al. 2020); Decay sugar-rich substrates (Põlme et al. 2020) | Põlme S, et al. 2020. Fungal Diversity 105:1-16 ((https://doi.org/10.1007/s13225-020-00466-2)) |
| Otu49462 | 1 | k__Fungi(100);p__Ascomycota(100);c__Saccharomycetes(100);o__Saccharomycetales(100);f__Metschnikowiaceae(100);g__Metschnikowia(100);g__Metschnikowia_unclassified(100); | Metschnikowia | 13 | Saprotroph | \|Undefined Saprotroph\| | Yeast | NULL | Probable | As nectar-tap_saprotroph (Põlme et al. 2020); Decay sugar-rich substrates (Põlme et al. 2020) | Põlme S, et al. 2020. Fungal Diversity 105:1-16 ((https://doi.org/10.1007/s13225-020-00466-2)) |
| Otu49463 | 1 | k__Fungi(100);p__Ascomycota(100);c__Saccharomycetes(100);o__Saccharomycetales(100);f__Metschnikowiaceae(100);g__Metschnikowia(100);s__Metschnikowia_pulcherrima(100); | Metschnikowia | 13 | Saprotroph | \|Undefined Saprotroph\| | Yeast | NULL | Probable | As nectar-tap_saprotroph (Põlme et al. 2020); Decay sugar-rich substrates (Põlme et al. 2020) | Põlme S, et al. 2020. Fungal Diversity 105:1-16 ((https://doi.org/10.1007/s13225-020-00466-2)) |
| Otu49464 | 1 | k__Fungi(100);p__Ascomycota(100);c__Saccharomycetes(100);o__Saccharomycetales(100);f__Metschnikowiaceae(100);g__Metschnikowia(100);s__Metschnikowia_pulcherrima(100); | Metschnikowia | 13 | Saprotroph | \|Undefined Saprotroph\| | Yeast | NULL | Probable | As nectar-tap_saprotroph (Põlme et al. 2020); Decay sugar-rich substrates (Põlme et al. 2020) | Põlme S, et al. 2020. Fungal Diversity 105:1-16 ((https://doi.org/10.1007/s13225-020-00466-2)) |
| Otu49465 | 1 | k__Fungi(100);p__Ascomycota(100);c__Saccharomycetes(100);o__Saccharomycetales(100);f__Metschnikowiaceae(100);g__Metschnikowia(100);s__Metschnikowia_pulcherrima(100); | Metschnikowia | 13 | Saprotroph | \|Undefined Saprotroph\| | Yeast | NULL | Probable | As nectar-tap_saprotroph (Põlme et al. 2020); Decay sugar-rich substrates (Põlme et al. 2020) | Põlme S, et al. 2020. Fungal Diversity 105:1-16 ((https://doi.org/10.1007/s13225-020-00466-2)) |
| Otu49466 | 1 | k__Fungi(100);p__Ascomycota(100);c__Saccharomycetes(100);o__Saccharomycetales(100);f__Metschnikowiaceae(100);g__Metschnikowia(100);s__Metschnikowia_pulcherrima(100); | Metschnikowia | 13 | Saprotroph | \|Undefined Saprotroph\| | Yeast | NULL | Probable | As nectar-tap_saprotroph (Põlme et al. 2020); Decay sugar-rich substrates (Põlme et al. 2020) | Põlme S, et al. 2020. Fungal Diversity 105:1-16 ((https://doi.org/10.1007/s13225-020-00466-2)) |
| Otu49467 | 1 | k__Fungi(100);p__Ascomycota(100);c__Saccharomycetes(100);o__Saccharomycetales(100);f__Metschnikowiaceae(100);g__Metschnikowia(100);s__Metschnikowia_pulcherrima(100); | Metschnikowia | 13 | Saprotroph | \|Undefined Saprotroph\| | Yeast | NULL | Probable | As nectar-tap_saprotroph (Põlme et al. 2020); Decay sugar-rich substrates (Põlme et al. 2020) | Põlme S, et al. 2020. Fungal Diversity 105:1-16 ((https://doi.org/10.1007/s13225-020-00466-2)) |
| Otu49468 | 1 | k__Fungi(100);p__Ascomycota(100);c__Saccharomycetes(100);o__Saccharomycetales(100);f__Metschnikowiaceae(100);g__Metschnikowia(100);s__Metschnikowia_pulcherrima(100); | Metschnikowia | 13 | Saprotroph | \|Undefined Saprotroph\| | Yeast | NULL | Probable | As nectar-tap_saprotroph (Põlme et al. 2020); Decay sugar-rich substrates (Põlme et al. 2020) | Põlme S, et al. 2020. Fungal Diversity 105:1-16 ((https://doi.org/10.1007/s13225-020-00466-2)) |
| Otu49469 | 1 | k__Fungi(100);p__Ascomycota(100);c__Saccharomycetes(100);o__Saccharomycetales(100);f__Metschnikowiaceae(100);g__Metschnikowia(100);s__Metschnikowia_pulcherrima(100); | Metschnikowia | 13 | Saprotroph | \|Undefined Saprotroph\| | Yeast | NULL | Probable | As nectar-tap_saprotroph (Põlme et al. 2020); Decay sugar-rich substrates (Põlme et al. 2020) | Põlme S, et al. 2020. Fungal Diversity 105:1-16 ((https://doi.org/10.1007/s13225-020-00466-2)) |
| Otu49470 | 1 | k__Fungi(100);p__Ascomycota(100);c__Saccharomycetes(100);o__Saccharomycetales(100);f__Metschnikowiaceae(100);g__Metschnikowia(100);s__Metschnikowia_pulcherrima(100); | Metschnikowia | 13 | Saprotroph | \|Undefined Saprotroph\| | Yeast | NULL | Probable | As nectar-tap_saprotroph (Põlme et al. 2020); Decay sugar-rich substrates (Põlme et al. 2020) | Põlme S, et al. 2020. Fungal Diversity 105:1-16 ((https://doi.org/10.1007/s13225-020-00466-2)) |
| Otu49471 | 1 | k__Fungi(100);p__Ascomycota(100);c__Saccharomycetes(100);o__Saccharomycetales(100);f__Metschnikowiaceae(100);g__Metschnikowia(100);s__Metschnikowia_pulcherrima(100); | Metschnikowia | 13 | Saprotroph | \|Undefined Saprotroph\| | Yeast | NULL | Probable | As nectar-tap_saprotroph (Põlme et al. 2020); Decay sugar-rich substrates (Põlme et al. 2020) | Põlme S, et al. 2020. Fungal Diversity 105:1-16 ((https://doi.org/10.1007/s13225-020-00466-2)) |
| Otu49472 | 1 | k__Fungi(100);p__Ascomycota(100);c__Saccharomycetes(100);o__Saccharomycetales(100);f__Metschnikowiaceae(100);g__Metschnikowia(100);s__Metschnikowia_pulcherrima(100); | Metschnikowia | 13 | Saprotroph | \|Undefined Saprotroph\| | Yeast | NULL | Probable | As nectar-tap_saprotroph (Põlme et al. 2020); Decay sugar-rich substrates (Põlme et al. 2020) | Põlme S, et al. 2020. Fungal Diversity 105:1-16 ((https://doi.org/10.1007/s13225-020-00466-2)) |
| Otu49473 | 1 | k__Fungi(100);p__Ascomycota(100);c__Saccharomycetes(100);o__Saccharomycetales(100);f__Metschnikowiaceae(100);g__Metschnikowia(100);s__Metschnikowia_pulcherrima(100); | Metschnikowia | 13 | Saprotroph | \|Undefined Saprotroph\| | Yeast | NULL | Probable | As nectar-tap_saprotroph (Põlme et al. 2020); Decay sugar-rich substrates (Põlme et al. 2020) | Põlme S, et al. 2020. Fungal Diversity 105:1-16 ((https://doi.org/10.1007/s13225-020-00466-2)) |
| Otu49474 | 1 | k__Fungi(100);p__Ascomycota(100);c__Saccharomycetes(100);o__Saccharomycetales(100);f__Metschnikowiaceae(100);g__Metschnikowia(100);g__Metschnikowia_unclassified(100); | Metschnikowia | 13 | Saprotroph | \|Undefined Saprotroph\| | Yeast | NULL | Probable | As nectar-tap_saprotroph (Põlme et al. 2020); Decay sugar-rich substrates (Põlme et al. 2020) | Põlme S, et al. 2020. Fungal Diversity 105:1-16 ((https://doi.org/10.1007/s13225-020-00466-2)) |
| Otu49475 | 1 | k__Fungi(100);p__Ascomycota(100);c__Saccharomycetes(100);o__Saccharomycetales(100);f__Metschnikowiaceae(100);g__Metschnikowia(100);g__Metschnikowia_unclassified(100); | Metschnikowia | 13 | Saprotroph | \|Undefined Saprotroph\| | Yeast | NULL | Probable | As nectar-tap_saprotroph (Põlme et al. 2020); Decay sugar-rich substrates (Põlme et al. 2020) | Põlme S, et al. 2020. Fungal Diversity 105:1-16 ((https://doi.org/10.1007/s13225-020-00466-2)) |
| Otu49476 | 1 | k__Fungi(100);p__Ascomycota(100);c__Saccharomycetes(100);o__Saccharomycetales(100);f__Metschnikowiaceae(100);g__Metschnikowia(100);s__Metschnikowia_pulcherrima(100); | Metschnikowia | 13 | Saprotroph | \|Undefined Saprotroph\| | Yeast | NULL | Probable | As nectar-tap_saprotroph (Põlme et al. 2020); Decay sugar-rich substrates (Põlme et al. 2020) | Põlme S, et al. 2020. Fungal Diversity 105:1-16 ((https://doi.org/10.1007/s13225-020-00466-2)) |
| Otu49477 | 1 | k__Fungi(100);p__Ascomycota(100);c__Saccharomycetes(100);o__Saccharomycetales(100);f__Metschnikowiaceae(100);g__Metschnikowia(100);s__Metschnikowia_pulcherrima(100); | Metschnikowia | 13 | Saprotroph | \|Undefined Saprotroph\| | Yeast | NULL | Probable | As nectar-tap_saprotroph (Põlme et al. 2020); Decay sugar-rich substrates (Põlme et al. 2020) | Põlme S, et al. 2020. Fungal Diversity 105:1-16 ((https://doi.org/10.1007/s13225-020-00466-2)) |
| Otu49478 | 1 | k__Fungi(100);p__Ascomycota(100);c__Saccharomycetes(100);o__Saccharomycetales(100);f__Metschnikowiaceae(100);g__Metschnikowia(100);g__Metschnikowia_unclassified(100); | Metschnikowia | 13 | Saprotroph | \|Undefined Saprotroph\| | Yeast | NULL | Probable | As nectar-tap_saprotroph (Põlme et al. 2020); Decay sugar-rich substrates (Põlme et al. 2020) | Põlme S, et al. 2020. Fungal Diversity 105:1-16 ((https://doi.org/10.1007/s13225-020-00466-2)) |
| Otu49479 | 1 | k__Fungi(100);p__Ascomycota(100);c__Saccharomycetes(100);o__Saccharomycetales(100);f__Metschnikowiaceae(100);g__Metschnikowia(100);s__Metschnikowia_pulcherrima(100); | Metschnikowia | 13 | Saprotroph | \|Undefined Saprotroph\| | Yeast | NULL | Probable | As nectar-tap_saprotroph (Põlme et al. 2020); Decay sugar-rich substrates (Põlme et al. 2020) | Põlme S, et al. 2020. Fungal Diversity 105:1-16 ((https://doi.org/10.1007/s13225-020-00466-2)) |
| Otu49480 | 1 | k__Fungi(100);p__Ascomycota(100);c__Saccharomycetes(100);o__Saccharomycetales(100);f__Metschnikowiaceae(100);g__Metschnikowia(100);g__Metschnikowia_unclassified(100); | Metschnikowia | 13 | Saprotroph | \|Undefined Saprotroph\| | Yeast | NULL | Probable | As nectar-tap_saprotroph (Põlme et al. 2020); Decay sugar-rich substrates (Põlme et al. 2020) | Põlme S, et al. 2020. Fungal Diversity 105:1-16 ((https://doi.org/10.1007/s13225-020-00466-2)) |
| Otu49481 | 1 | k__Fungi(100);p__Ascomycota(100);c__Saccharomycetes(100);o__Saccharomycetales(100);f__Metschnikowiaceae(100);g__Metschnikowia(100);s__Metschnikowia_pulcherrima(100); | Metschnikowia | 13 | Saprotroph | \|Undefined Saprotroph\| | Yeast | NULL | Probable | As nectar-tap_saprotroph (Põlme et al. 2020); Decay sugar-rich substrates (Põlme et al. 2020) | Põlme S, et al. 2020. Fungal Diversity 105:1-16 ((https://doi.org/10.1007/s13225-020-00466-2)) |
| Otu49483 | 1 | k__Fungi(100);p__Ascomycota(100);c__Saccharomycetes(100);o__Saccharomycetales(100);f__Metschnikowiaceae(100);g__Metschnikowia(100);g__Metschnikowia_unclassified(100); | Metschnikowia | 13 | Saprotroph | \|Undefined Saprotroph\| | Yeast | NULL | Probable | As nectar-tap_saprotroph (Põlme et al. 2020); Decay sugar-rich substrates (Põlme et al. 2020) | Põlme S, et al. 2020. Fungal Diversity 105:1-16 ((https://doi.org/10.1007/s13225-020-00466-2)) |
| Otu49484 | 1 | k__Fungi(100);p__Ascomycota(100);c__Saccharomycetes(100);o__Saccharomycetales(100);f__Metschnikowiaceae(100);g__Metschnikowia(100);s__Metschnikowia_pulcherrima(100); | Metschnikowia | 13 | Saprotroph | \|Undefined Saprotroph\| | Yeast | NULL | Probable | As nectar-tap_saprotroph (Põlme et al. 2020); Decay sugar-rich substrates (Põlme et al. 2020) | Põlme S, et al. 2020. Fungal Diversity 105:1-16 ((https://doi.org/10.1007/s13225-020-00466-2)) |
| Otu49485 | 1 | k__Fungi(100);p__Ascomycota(100);c__Saccharomycetes(100);o__Saccharomycetales(100);f__Metschnikowiaceae(100);g__Metschnikowia(100);s__Metschnikowia_pulcherrima(100); | Metschnikowia | 13 | Saprotroph | \|Undefined Saprotroph\| | Yeast | NULL | Probable | As nectar-tap_saprotroph (Põlme et al. 2020); Decay sugar-rich substrates (Põlme et al. 2020) | Põlme S, et al. 2020. Fungal Diversity 105:1-16 ((https://doi.org/10.1007/s13225-020-00466-2)) |
| Otu49486 | 1 | k__Fungi(100);p__Ascomycota(100);c__Saccharomycetes(100);o__Saccharomycetales(100);f__Metschnikowiaceae(100);g__Metschnikowia(100);g__Metschnikowia_unclassified(100); | Metschnikowia | 13 | Saprotroph | \|Undefined Saprotroph\| | Yeast | NULL | Probable | As nectar-tap_saprotroph (Põlme et al. 2020); Decay sugar-rich substrates (Põlme et al. 2020) | Põlme S, et al. 2020. Fungal Diversity 105:1-16 ((https://doi.org/10.1007/s13225-020-00466-2)) |
| Otu49487 | 1 | k__Fungi(100);p__Ascomycota(100);c__Saccharomycetes(100);o__Saccharomycetales(100);f__Metschnikowiaceae(100);g__Metschnikowia(100);s__Metschnikowia_pulcherrima(100); | Metschnikowia | 13 | Saprotroph | \|Undefined Saprotroph\| | Yeast | NULL | Probable | As nectar-tap_saprotroph (Põlme et al. 2020); Decay sugar-rich substrates (Põlme et al. 2020) | Põlme S, et al. 2020. Fungal Diversity 105:1-16 ((https://doi.org/10.1007/s13225-020-00466-2)) |
| Otu49488 | 1 | k__Fungi(100);p__Ascomycota(100);c__Saccharomycetes(100);o__Saccharomycetales(100);f__Metschnikowiaceae(100);g__Metschnikowia(100);s__Metschnikowia_pulcherrima(100); | Metschnikowia | 13 | Saprotroph | \|Undefined Saprotroph\| | Yeast | NULL | Probable | As nectar-tap_saprotroph (Põlme et al. 2020); Decay sugar-rich substrates (Põlme et al. 2020) | Põlme S, et al. 2020. Fungal Diversity 105:1-16 ((https://doi.org/10.1007/s13225-020-00466-2)) |
| Otu49489 | 1 | k__Fungi(100);p__Ascomycota(100);c__Saccharomycetes(100);o__Saccharomycetales(100);f__Metschnikowiaceae(100);g__Metschnikowia(100);s__Metschnikowia_pulcherrima(100); | Metschnikowia | 13 | Saprotroph | \|Undefined Saprotroph\| | Yeast | NULL | Probable | As nectar-tap_saprotroph (Põlme et al. 2020); Decay sugar-rich substrates (Põlme et al. 2020) | Põlme S, et al. 2020. Fungal Diversity 105:1-16 ((https://doi.org/10.1007/s13225-020-00466-2)) |
| Otu49490 | 1 | k__Fungi(100);p__Ascomycota(100);c__Saccharomycetes(100);o__Saccharomycetales(100);f__Metschnikowiaceae(100);g__Metschnikowia(100);s__Metschnikowia_pulcherrima(100); | Metschnikowia | 13 | Saprotroph | \|Undefined Saprotroph\| | Yeast | NULL | Probable | As nectar-tap_saprotroph (Põlme et al. 2020); Decay sugar-rich substrates (Põlme et al. 2020) | Põlme S, et al. 2020. Fungal Diversity 105:1-16 ((https://doi.org/10.1007/s13225-020-00466-2)) |
| Otu49491 | 1 | k__Fungi(100);p__Ascomycota(100);c__Saccharomycetes(100);o__Saccharomycetales(100);f__Metschnikowiaceae(100);g__Metschnikowia(100);s__Metschnikowia_pulcherrima(100); | Metschnikowia | 13 | Saprotroph | \|Undefined Saprotroph\| | Yeast | NULL | Probable | As nectar-tap_saprotroph (Põlme et al. 2020); Decay sugar-rich substrates (Põlme et al. 2020) | Põlme S, et al. 2020. Fungal Diversity 105:1-16 ((https://doi.org/10.1007/s13225-020-00466-2)) |
| Otu49492 | 1 | k__Fungi(100);p__Ascomycota(100);c__Saccharomycetes(100);o__Saccharomycetales(100);f__Metschnikowiaceae(100);g__Metschnikowia(100);g__Metschnikowia_unclassified(100); | Metschnikowia | 13 | Saprotroph | \|Undefined Saprotroph\| | Yeast | NULL | Probable | As nectar-tap_saprotroph (Põlme et al. 2020); Decay sugar-rich substrates (Põlme et al. 2020) | Põlme S, et al. 2020. Fungal Diversity 105:1-16 ((https://doi.org/10.1007/s13225-020-00466-2)) |
| Otu49493 | 1 | k__Fungi(100);p__Ascomycota(100);c__Saccharomycetes(100);o__Saccharomycetales(100);f__Metschnikowiaceae(100);g__Metschnikowia(100);s__Metschnikowia_pulcherrima(100); | Metschnikowia | 13 | Saprotroph | \|Undefined Saprotroph\| | Yeast | NULL | Probable | As nectar-tap_saprotroph (Põlme et al. 2020); Decay sugar-rich substrates (Põlme et al. 2020) | Põlme S, et al. 2020. Fungal Diversity 105:1-16 ((https://doi.org/10.1007/s13225-020-00466-2)) |
| Otu49494 | 1 | k__Fungi(100);p__Ascomycota(100);c__Saccharomycetes(100);o__Saccharomycetales(100);f__Metschnikowiaceae(100);g__Metschnikowia(100);s__Metschnikowia_pulcherrima(100); | Metschnikowia | 13 | Saprotroph | \|Undefined Saprotroph\| | Yeast | NULL | Probable | As nectar-tap_saprotroph (Põlme et al. 2020); Decay sugar-rich substrates (Põlme et al. 2020) | Põlme S, et al. 2020. Fungal Diversity 105:1-16 ((https://doi.org/10.1007/s13225-020-00466-2)) |
| Otu49495 | 1 | k__Fungi(100);p__Ascomycota(100);c__Saccharomycetes(100);o__Saccharomycetales(100);f__Metschnikowiaceae(100);g__Metschnikowia(100);g__Metschnikowia_unclassified(100); | Metschnikowia | 13 | Saprotroph | \|Undefined Saprotroph\| | Yeast | NULL | Probable | As nectar-tap_saprotroph (Põlme et al. 2020); Decay sugar-rich substrates (Põlme et al. 2020) | Põlme S, et al. 2020. Fungal Diversity 105:1-16 ((https://doi.org/10.1007/s13225-020-00466-2)) |
| Otu49496 | 1 | k__Fungi(100);p__Ascomycota(100);c__Saccharomycetes(100);o__Saccharomycetales(100);f__Metschnikowiaceae(100);g__Metschnikowia(100);s__Metschnikowia_pulcherrima(100); | Metschnikowia | 13 | Saprotroph | \|Undefined Saprotroph\| | Yeast | NULL | Probable | As nectar-tap_saprotroph (Põlme et al. 2020); Decay sugar-rich substrates (Põlme et al. 2020) | Põlme S, et al. 2020. Fungal Diversity 105:1-16 ((https://doi.org/10.1007/s13225-020-00466-2)) |
| Otu49497 | 1 | k__Fungi(100);p__Ascomycota(100);c__Saccharomycetes(100);o__Saccharomycetales(100);f__Metschnikowiaceae(100);g__Metschnikowia(100);s__Metschnikowia_pulcherrima(100); | Metschnikowia | 13 | Saprotroph | \|Undefined Saprotroph\| | Yeast | NULL | Probable | As nectar-tap_saprotroph (Põlme et al. 2020); Decay sugar-rich substrates (Põlme et al. 2020) | Põlme S, et al. 2020. Fungal Diversity 105:1-16 ((https://doi.org/10.1007/s13225-020-00466-2)) |
| Otu49498 | 1 | k__Fungi(100);p__Ascomycota(100);c__Saccharomycetes(100);o__Saccharomycetales(100);f__Metschnikowiaceae(100);g__Metschnikowia(100);s__Metschnikowia_pulcherrima(100); | Metschnikowia | 13 | Saprotroph | \|Undefined Saprotroph\| | Yeast | NULL | Probable | As nectar-tap_saprotroph (Põlme et al. 2020); Decay sugar-rich substrates (Põlme et al. 2020) | Põlme S, et al. 2020. Fungal Diversity 105:1-16 ((https://doi.org/10.1007/s13225-020-00466-2)) |
| Otu49499 | 1 | k__Fungi(100);p__Ascomycota(100);c__Saccharomycetes(100);o__Saccharomycetales(100);f__Metschnikowiaceae(100);g__Metschnikowia(100);s__Metschnikowia_pulcherrima(100); | Metschnikowia | 13 | Saprotroph | \|Undefined Saprotroph\| | Yeast | NULL | Probable | As nectar-tap_saprotroph (Põlme et al. 2020); Decay sugar-rich substrates (Põlme et al. 2020) | Põlme S, et al. 2020. Fungal Diversity 105:1-16 ((https://doi.org/10.1007/s13225-020-00466-2)) |
| Otu49500 | 1 | k__Fungi(100);p__Ascomycota(100);c__Saccharomycetes(100);o__Saccharomycetales(100);f__Metschnikowiaceae(100);g__Metschnikowia(100);s__Metschnikowia_pulcherrima(100); | Metschnikowia | 13 | Saprotroph | \|Undefined Saprotroph\| | Yeast | NULL | Probable | As nectar-tap_saprotroph (Põlme et al. 2020); Decay sugar-rich substrates (Põlme et al. 2020) | Põlme S, et al. 2020. Fungal Diversity 105:1-16 ((https://doi.org/10.1007/s13225-020-00466-2)) |
| Otu49501 | 1 | k__Fungi(100);p__Ascomycota(100);c__Saccharomycetes(100);o__Saccharomycetales(100);f__Metschnikowiaceae(100);g__Metschnikowia(100);s__Metschnikowia_pulcherrima(100); | Metschnikowia | 13 | Saprotroph | \|Undefined Saprotroph\| | Yeast | NULL | Probable | As nectar-tap_saprotroph (Põlme et al. 2020); Decay sugar-rich substrates (Põlme et al. 2020) | Põlme S, et al. 2020. Fungal Diversity 105:1-16 ((https://doi.org/10.1007/s13225-020-00466-2)) |
| Otu49504 | 1 | k__Fungi(100);p__Ascomycota(100);c__Saccharomycetes(100);o__Saccharomycetales(100);f__Metschnikowiaceae(100);g__Metschnikowia(100);s__Metschnikowia_pulcherrima(100); | Metschnikowia | 13 | Saprotroph | \|Undefined Saprotroph\| | Yeast | NULL | Probable | As nectar-tap_saprotroph (Põlme et al. 2020); Decay sugar-rich substrates (Põlme et al. 2020) | Põlme S, et al. 2020. Fungal Diversity 105:1-16 ((https://doi.org/10.1007/s13225-020-00466-2)) |
| Otu49505 | 1 | k__Fungi(100);p__Ascomycota(100);c__Saccharomycetes(100);o__Saccharomycetales(100);f__Metschnikowiaceae(100);g__Metschnikowia(100);s__Metschnikowia_pulcherrima(100); | Metschnikowia | 13 | Saprotroph | \|Undefined Saprotroph\| | Yeast | NULL | Probable | As nectar-tap_saprotroph (Põlme et al. 2020); Decay sugar-rich substrates (Põlme et al. 2020) | Põlme S, et al. 2020. Fungal Diversity 105:1-16 ((https://doi.org/10.1007/s13225-020-00466-2)) |
| Otu49506 | 1 | k__Fungi(100);p__Ascomycota(100);c__Saccharomycetes(100);o__Saccharomycetales(100);f__Metschnikowiaceae(100);g__Metschnikowia(100);s__Metschnikowia_pulcherrima(100); | Metschnikowia | 13 | Saprotroph | \|Undefined Saprotroph\| | Yeast | NULL | Probable | As nectar-tap_saprotroph (Põlme et al. 2020); Decay sugar-rich substrates (Põlme et al. 2020) | Põlme S, et al. 2020. Fungal Diversity 105:1-16 ((https://doi.org/10.1007/s13225-020-00466-2)) |
| Otu49507 | 1 | k__Fungi(100);p__Ascomycota(100);c__Saccharomycetes(100);o__Saccharomycetales(100);f__Metschnikowiaceae(100);g__Metschnikowia(100);g__Metschnikowia_unclassified(100); | Metschnikowia | 13 | Saprotroph | \|Undefined Saprotroph\| | Yeast | NULL | Probable | As nectar-tap_saprotroph (Põlme et al. 2020); Decay sugar-rich substrates (Põlme et al. 2020) | Põlme S, et al. 2020. Fungal Diversity 105:1-16 ((https://doi.org/10.1007/s13225-020-00466-2)) |
| Otu49508 | 1 | k__Fungi(100);p__Ascomycota(100);c__Saccharomycetes(100);o__Saccharomycetales(100);f__Metschnikowiaceae(100);g__Metschnikowia(100);s__Metschnikowia_pulcherrima(100); | Metschnikowia | 13 | Saprotroph | \|Undefined Saprotroph\| | Yeast | NULL | Probable | As nectar-tap_saprotroph (Põlme et al. 2020); Decay sugar-rich substrates (Põlme et al. 2020) | Põlme S, et al. 2020. Fungal Diversity 105:1-16 ((https://doi.org/10.1007/s13225-020-00466-2)) |
| Otu49509 | 1 | k__Fungi(100);p__Ascomycota(100);c__Saccharomycetes(100);o__Saccharomycetales(100);f__Metschnikowiaceae(100);g__Metschnikowia(100);s__Metschnikowia_pulcherrima(100); | Metschnikowia | 13 | Saprotroph | \|Undefined Saprotroph\| | Yeast | NULL | Probable | As nectar-tap_saprotroph (Põlme et al. 2020); Decay sugar-rich substrates (Põlme et al. 2020) | Põlme S, et al. 2020. Fungal Diversity 105:1-16 ((https://doi.org/10.1007/s13225-020-00466-2)) |
| Otu49510 | 1 | k__Fungi(100);p__Ascomycota(100);c__Saccharomycetes(100);o__Saccharomycetales(100);f__Metschnikowiaceae(100);g__Metschnikowia(100);s__Metschnikowia_pulcherrima(100); | Metschnikowia | 13 | Saprotroph | \|Undefined Saprotroph\| | Yeast | NULL | Probable | As nectar-tap_saprotroph (Põlme et al. 2020); Decay sugar-rich substrates (Põlme et al. 2020) | Põlme S, et al. 2020. Fungal Diversity 105:1-16 ((https://doi.org/10.1007/s13225-020-00466-2)) |
| Otu49511 | 1 | k__Fungi(100);p__Ascomycota(100);c__Saccharomycetes(100);o__Saccharomycetales(100);f__Metschnikowiaceae(100);g__Metschnikowia(100);s__Metschnikowia_pulcherrima(100); | Metschnikowia | 13 | Saprotroph | \|Undefined Saprotroph\| | Yeast | NULL | Probable | As nectar-tap_saprotroph (Põlme et al. 2020); Decay sugar-rich substrates (Põlme et al. 2020) | Põlme S, et al. 2020. Fungal Diversity 105:1-16 ((https://doi.org/10.1007/s13225-020-00466-2)) |
| Otu49512 | 1 | k__Fungi(100);p__Ascomycota(100);c__Saccharomycetes(100);o__Saccharomycetales(100);f__Metschnikowiaceae(100);g__Metschnikowia(100);s__Metschnikowia_pulcherrima(100); | Metschnikowia | 13 | Saprotroph | \|Undefined Saprotroph\| | Yeast | NULL | Probable | As nectar-tap_saprotroph (Põlme et al. 2020); Decay sugar-rich substrates (Põlme et al. 2020) | Põlme S, et al. 2020. Fungal Diversity 105:1-16 ((https://doi.org/10.1007/s13225-020-00466-2)) |
| Otu49513 | 1 | k__Fungi(100);p__Ascomycota(100);c__Saccharomycetes(100);o__Saccharomycetales(100);f__Metschnikowiaceae(100);g__Metschnikowia(100);g__Metschnikowia_unclassified(100); | Metschnikowia | 13 | Saprotroph | \|Undefined Saprotroph\| | Yeast | NULL | Probable | As nectar-tap_saprotroph (Põlme et al. 2020); Decay sugar-rich substrates (Põlme et al. 2020) | Põlme S, et al. 2020. Fungal Diversity 105:1-16 ((https://doi.org/10.1007/s13225-020-00466-2)) |
| Otu49514 | 1 | k__Fungi(100);p__Ascomycota(100);c__Saccharomycetes(100);o__Saccharomycetales(100);f__Metschnikowiaceae(100);g__Metschnikowia(100);s__Metschnikowia_pulcherrima(100); | Metschnikowia | 13 | Saprotroph | \|Undefined Saprotroph\| | Yeast | NULL | Probable | As nectar-tap_saprotroph (Põlme et al. 2020); Decay sugar-rich substrates (Põlme et al. 2020) | Põlme S, et al. 2020. Fungal Diversity 105:1-16 ((https://doi.org/10.1007/s13225-020-00466-2)) |
| Otu49515 | 1 | k__Fungi(100);p__Ascomycota(100);c__Saccharomycetes(100);o__Saccharomycetales(100);f__Metschnikowiaceae(100);g__Metschnikowia(100);g__Metschnikowia_unclassified(100); | Metschnikowia | 13 | Saprotroph | \|Undefined Saprotroph\| | Yeast | NULL | Probable | As nectar-tap_saprotroph (Põlme et al. 2020); Decay sugar-rich substrates (Põlme et al. 2020) | Põlme S, et al. 2020. Fungal Diversity 105:1-16 ((https://doi.org/10.1007/s13225-020-00466-2)) |
| Otu49516 | 1 | k__Fungi(100);p__Ascomycota(100);c__Saccharomycetes(100);o__Saccharomycetales(100);f__Metschnikowiaceae(100);g__Metschnikowia(100);s__Metschnikowia_pulcherrima(100); | Metschnikowia | 13 | Saprotroph | \|Undefined Saprotroph\| | Yeast | NULL | Probable | As nectar-tap_saprotroph (Põlme et al. 2020); Decay sugar-rich substrates (Põlme et al. 2020) | Põlme S, et al. 2020. Fungal Diversity 105:1-16 ((https://doi.org/10.1007/s13225-020-00466-2)) |
| Otu49517 | 1 | k__Fungi(100);p__Ascomycota(100);c__Saccharomycetes(100);o__Saccharomycetales(100);f__Metschnikowiaceae(100);g__Metschnikowia(100);g__Metschnikowia_unclassified(100); | Metschnikowia | 13 | Saprotroph | \|Undefined Saprotroph\| | Yeast | NULL | Probable | As nectar-tap_saprotroph (Põlme et al. 2020); Decay sugar-rich substrates (Põlme et al. 2020) | Põlme S, et al. 2020. Fungal Diversity 105:1-16 ((https://doi.org/10.1007/s13225-020-00466-2)) |
| Otu49518 | 1 | k__Fungi(100);p__Ascomycota(100);c__Saccharomycetes(100);o__Saccharomycetales(100);f__Metschnikowiaceae(100);g__Metschnikowia(100);s__Metschnikowia_pulcherrima(100); | Metschnikowia | 13 | Saprotroph | \|Undefined Saprotroph\| | Yeast | NULL | Probable | As nectar-tap_saprotroph (Põlme et al. 2020); Decay sugar-rich substrates (Põlme et al. 2020) | Põlme S, et al. 2020. Fungal Diversity 105:1-16 ((https://doi.org/10.1007/s13225-020-00466-2)) |
| Otu49519 | 1 | k__Fungi(100);p__Ascomycota(100);c__Saccharomycetes(100);o__Saccharomycetales(100);f__Metschnikowiaceae(100);g__Metschnikowia(100);s__Metschnikowia_pulcherrima(100); | Metschnikowia | 13 | Saprotroph | \|Undefined Saprotroph\| | Yeast | NULL | Probable | As nectar-tap_saprotroph (Põlme et al. 2020); Decay sugar-rich substrates (Põlme et al. 2020) | Põlme S, et al. 2020. Fungal Diversity 105:1-16 ((https://doi.org/10.1007/s13225-020-00466-2)) |
| Otu49520 | 1 | k__Fungi(100);p__Ascomycota(100);c__Saccharomycetes(100);o__Saccharomycetales(100);f__Metschnikowiaceae(100);g__Metschnikowia(100);s__Metschnikowia_pulcherrima(100); | Metschnikowia | 13 | Saprotroph | \|Undefined Saprotroph\| | Yeast | NULL | Probable | As nectar-tap_saprotroph (Põlme et al. 2020); Decay sugar-rich substrates (Põlme et al. 2020) | Põlme S, et al. 2020. Fungal Diversity 105:1-16 ((https://doi.org/10.1007/s13225-020-00466-2)) |
| Otu49521 | 1 | k__Fungi(100);p__Ascomycota(100);c__Saccharomycetes(100);o__Saccharomycetales(100);f__Metschnikowiaceae(100);g__Metschnikowia(100);g__Metschnikowia_unclassified(100); | Metschnikowia | 13 | Saprotroph | \|Undefined Saprotroph\| | Yeast | NULL | Probable | As nectar-tap_saprotroph (Põlme et al. 2020); Decay sugar-rich substrates (Põlme et al. 2020) | Põlme S, et al. 2020. Fungal Diversity 105:1-16 ((https://doi.org/10.1007/s13225-020-00466-2)) |
| Otu49522 | 1 | k__Fungi(100);p__Ascomycota(100);c__Saccharomycetes(100);o__Saccharomycetales(100);f__Metschnikowiaceae(100);g__Metschnikowia(100);s__Metschnikowia_pulcherrima(100); | Metschnikowia | 13 | Saprotroph | \|Undefined Saprotroph\| | Yeast | NULL | Probable | As nectar-tap_saprotroph (Põlme et al. 2020); Decay sugar-rich substrates (Põlme et al. 2020) | Põlme S, et al. 2020. Fungal Diversity 105:1-16 ((https://doi.org/10.1007/s13225-020-00466-2)) |
| Otu49523 | 1 | k__Fungi(100);p__Ascomycota(100);c__Saccharomycetes(100);o__Saccharomycetales(100);f__Metschnikowiaceae(100);g__Metschnikowia(100);s__Metschnikowia_pulcherrima(100); | Metschnikowia | 13 | Saprotroph | \|Undefined Saprotroph\| | Yeast | NULL | Probable | As nectar-tap_saprotroph (Põlme et al. 2020); Decay sugar-rich substrates (Põlme et al. 2020) | Põlme S, et al. 2020. Fungal Diversity 105:1-16 ((https://doi.org/10.1007/s13225-020-00466-2)) |
| Otu49524 | 1 | k__Fungi(100);p__Ascomycota(100);c__Saccharomycetes(100);o__Saccharomycetales(100);f__Metschnikowiaceae(100);g__Metschnikowia(100);s__Metschnikowia_pulcherrima(100); | Metschnikowia | 13 | Saprotroph | \|Undefined Saprotroph\| | Yeast | NULL | Probable | As nectar-tap_saprotroph (Põlme et al. 2020); Decay sugar-rich substrates (Põlme et al. 2020) | Põlme S, et al. 2020. Fungal Diversity 105:1-16 ((https://doi.org/10.1007/s13225-020-00466-2)) |
| Otu49525 | 1 | k__Fungi(100);p__Ascomycota(100);c__Saccharomycetes(100);o__Saccharomycetales(100);f__Metschnikowiaceae(100);g__Metschnikowia(100);s__Metschnikowia_pulcherrima(100); | Metschnikowia | 13 | Saprotroph | \|Undefined Saprotroph\| | Yeast | NULL | Probable | As nectar-tap_saprotroph (Põlme et al. 2020); Decay sugar-rich substrates (Põlme et al. 2020) | Põlme S, et al. 2020. Fungal Diversity 105:1-16 ((https://doi.org/10.1007/s13225-020-00466-2)) |
| Otu49526 | 1 | k__Fungi(100);p__Ascomycota(100);c__Saccharomycetes(100);o__Saccharomycetales(100);f__Metschnikowiaceae(100);g__Metschnikowia(100);s__Metschnikowia_pulcherrima(100); | Metschnikowia | 13 | Saprotroph | \|Undefined Saprotroph\| | Yeast | NULL | Probable | As nectar-tap_saprotroph (Põlme et al. 2020); Decay sugar-rich substrates (Põlme et al. 2020) | Põlme S, et al. 2020. Fungal Diversity 105:1-16 ((https://doi.org/10.1007/s13225-020-00466-2)) |
| Otu49527 | 1 | k__Fungi(100);p__Ascomycota(100);c__Saccharomycetes(100);o__Saccharomycetales(100);f__Metschnikowiaceae(100);g__Metschnikowia(100);s__Metschnikowia_pulcherrima(100); | Metschnikowia | 13 | Saprotroph | \|Undefined Saprotroph\| | Yeast | NULL | Probable | As nectar-tap_saprotroph (Põlme et al. 2020); Decay sugar-rich substrates (Põlme et al. 2020) | Põlme S, et al. 2020. Fungal Diversity 105:1-16 ((https://doi.org/10.1007/s13225-020-00466-2)) |
| Otu49528 | 1 | k__Fungi(100);p__Ascomycota(100);c__Saccharomycetes(100);o__Saccharomycetales(100);f__Metschnikowiaceae(100);g__Metschnikowia(100);s__Metschnikowia_pulcherrima(100); | Metschnikowia | 13 | Saprotroph | \|Undefined Saprotroph\| | Yeast | NULL | Probable | As nectar-tap_saprotroph (Põlme et al. 2020); Decay sugar-rich substrates (Põlme et al. 2020) | Põlme S, et al. 2020. Fungal Diversity 105:1-16 ((https://doi.org/10.1007/s13225-020-00466-2)) |
| Otu49529 | 1 | k__Fungi(100);p__Ascomycota(100);c__Saccharomycetes(100);o__Saccharomycetales(100);f__Metschnikowiaceae(100);g__Metschnikowia(100);s__Metschnikowia_pulcherrima(100); | Metschnikowia | 13 | Saprotroph | \|Undefined Saprotroph\| | Yeast | NULL | Probable | As nectar-tap_saprotroph (Põlme et al. 2020); Decay sugar-rich substrates (Põlme et al. 2020) | Põlme S, et al. 2020. Fungal Diversity 105:1-16 ((https://doi.org/10.1007/s13225-020-00466-2)) |
| Otu49530 | 1 | k__Fungi(100);p__Ascomycota(100);c__Saccharomycetes(100);o__Saccharomycetales(100);f__Metschnikowiaceae(100);g__Metschnikowia(100);g__Metschnikowia_unclassified(100); | Metschnikowia | 13 | Saprotroph | \|Undefined Saprotroph\| | Yeast | NULL | Probable | As nectar-tap_saprotroph (Põlme et al. 2020); Decay sugar-rich substrates (Põlme et al. 2020) | Põlme S, et al. 2020. Fungal Diversity 105:1-16 ((https://doi.org/10.1007/s13225-020-00466-2)) |
| Otu49531 | 1 | k__Fungi(100);p__Ascomycota(100);c__Saccharomycetes(100);o__Saccharomycetales(100);f__Metschnikowiaceae(100);g__Metschnikowia(100);s__Metschnikowia_pulcherrima(100); | Metschnikowia | 13 | Saprotroph | \|Undefined Saprotroph\| | Yeast | NULL | Probable | As nectar-tap_saprotroph (Põlme et al. 2020); Decay sugar-rich substrates (Põlme et al. 2020) | Põlme S, et al. 2020. Fungal Diversity 105:1-16 ((https://doi.org/10.1007/s13225-020-00466-2)) |
| Otu49532 | 1 | k__Fungi(100);p__Ascomycota(100);c__Saccharomycetes(100);o__Saccharomycetales(100);f__Metschnikowiaceae(100);g__Metschnikowia(100);s__Metschnikowia_pulcherrima(100); | Metschnikowia | 13 | Saprotroph | \|Undefined Saprotroph\| | Yeast | NULL | Probable | As nectar-tap_saprotroph (Põlme et al. 2020); Decay sugar-rich substrates (Põlme et al. 2020) | Põlme S, et al. 2020. Fungal Diversity 105:1-16 ((https://doi.org/10.1007/s13225-020-00466-2)) |
| Otu49533 | 1 | k__Fungi(100);p__Ascomycota(100);c__Saccharomycetes(100);o__Saccharomycetales(100);f__Metschnikowiaceae(100);g__Metschnikowia(100);s__Metschnikowia_pulcherrima(100); | Metschnikowia | 13 | Saprotroph | \|Undefined Saprotroph\| | Yeast | NULL | Probable | As nectar-tap_saprotroph (Põlme et al. 2020); Decay sugar-rich substrates (Põlme et al. 2020) | Põlme S, et al. 2020. Fungal Diversity 105:1-16 ((https://doi.org/10.1007/s13225-020-00466-2)) |
| Otu49534 | 1 | k__Fungi(100);p__Ascomycota(100);c__Saccharomycetes(100);o__Saccharomycetales(100);f__Metschnikowiaceae(100);g__Metschnikowia(100);g__Metschnikowia_unclassified(100); | Metschnikowia | 13 | Saprotroph | \|Undefined Saprotroph\| | Yeast | NULL | Probable | As nectar-tap_saprotroph (Põlme et al. 2020); Decay sugar-rich substrates (Põlme et al. 2020) | Põlme S, et al. 2020. Fungal Diversity 105:1-16 ((https://doi.org/10.1007/s13225-020-00466-2)) |
| Otu49535 | 1 | k__Fungi(100);p__Ascomycota(100);c__Saccharomycetes(100);o__Saccharomycetales(100);f__Metschnikowiaceae(100);g__Metschnikowia(100);g__Metschnikowia_unclassified(100); | Metschnikowia | 13 | Saprotroph | \|Undefined Saprotroph\| | Yeast | NULL | Probable | As nectar-tap_saprotroph (Põlme et al. 2020); Decay sugar-rich substrates (Põlme et al. 2020) | Põlme S, et al. 2020. Fungal Diversity 105:1-16 ((https://doi.org/10.1007/s13225-020-00466-2)) |
| Otu49536 | 1 | k__Fungi(100);p__Ascomycota(100);c__Saccharomycetes(100);o__Saccharomycetales(100);f__Metschnikowiaceae(100);g__Metschnikowia(100);g__Metschnikowia_unclassified(100); | Metschnikowia | 13 | Saprotroph | \|Undefined Saprotroph\| | Yeast | NULL | Probable | As nectar-tap_saprotroph (Põlme et al. 2020); Decay sugar-rich substrates (Põlme et al. 2020) | Põlme S, et al. 2020. Fungal Diversity 105:1-16 ((https://doi.org/10.1007/s13225-020-00466-2)) |
| Otu49537 | 1 | k__Fungi(100);p__Ascomycota(100);c__Saccharomycetes(100);o__Saccharomycetales(100);f__Metschnikowiaceae(100);g__Metschnikowia(100);g__Metschnikowia_unclassified(100); | Metschnikowia | 13 | Saprotroph | \|Undefined Saprotroph\| | Yeast | NULL | Probable | As nectar-tap_saprotroph (Põlme et al. 2020); Decay sugar-rich substrates (Põlme et al. 2020) | Põlme S, et al. 2020. Fungal Diversity 105:1-16 ((https://doi.org/10.1007/s13225-020-00466-2)) |
| Otu49538 | 1 | k__Fungi(100);p__Ascomycota(100);c__Saccharomycetes(100);o__Saccharomycetales(100);f__Metschnikowiaceae(100);g__Metschnikowia(100);g__Metschnikowia_unclassified(100); | Metschnikowia | 13 | Saprotroph | \|Undefined Saprotroph\| | Yeast | NULL | Probable | As nectar-tap_saprotroph (Põlme et al. 2020); Decay sugar-rich substrates (Põlme et al. 2020) | Põlme S, et al. 2020. Fungal Diversity 105:1-16 ((https://doi.org/10.1007/s13225-020-00466-2)) |

| | | | | | | | | | | | | |
|---|---|---|---|---|---|---|---|---|---|---|---|---|
| Otu49540 | 1 | k__Fungi(100);p__Ascomycota(100);c__Saccharomycetes(100);o__Saccharomycetales(100);f__Metschnikowiaceae(100);g__Metschnikowia(100);g__Metschnikowia_unclassified(100); | Metschnikowia | | 13 | Saprotroph | \|Undefined Saprotroph\| | Yeast | NULL | Probable | As nectar-tap_saprotroph (Põlme et al. 2020); Decay sugar-rich substrates (Põlme et al. 2020) | Põlme S, et al. 2020. Fungal Diversity 105:1-16 ((https://doi.org/10.1007/s13225-020-00466-2)) |
| Otu49541 | 1 | k__Fungi(100);p__Ascomycota(100);c__Saccharomycetes(100);o__Saccharomycetales(100);f__Metschnikowiaceae(100);g__Metschnikowia(100);g__Metschnikowia_unclassified(100); | Metschnikowia | | 13 | Saprotroph | \|Undefined Saprotroph\| | Yeast | NULL | Probable | As nectar-tap_saprotroph (Põlme et al. 2020); Decay sugar-rich substrates (Põlme et al. 2020) | Põlme S, et al. 2020. Fungal Diversity 105:1-16 ((https://doi.org/10.1007/s13225-020-00466-2)) |
| Otu49542 | 1 | k__Fungi(100);p__Ascomycota(100);c__Saccharomycetes(100);o__Saccharomycetales(100);f__Metschnikowiaceae(100);g__Metschnikowia(100);g__Metschnikowia_unclassified(100); | Metschnikowia | | 13 | Saprotroph | \|Undefined Saprotroph\| | Yeast | NULL | Probable | As nectar-tap_saprotroph (Põlme et al. 2020); Decay sugar-rich substrates (Põlme et al. 2020) | Põlme S, et al. 2020. Fungal Diversity 105:1-16 ((https://doi.org/10.1007/s13225-020-00466-2)) |
| Otu49544 | 1 | k__Fungi(100);p__Ascomycota(100);c__Saccharomycetes(100);o__Saccharomycetales(100);f__Metschnikowiaceae(100);g__Metschnikowia(100);s__Metschnikowia_pulcherrima(100); | Metschnikowia | | 13 | Saprotroph | \|Undefined Saprotroph\| | Yeast | NULL | Probable | As nectar-tap_saprotroph (Põlme et al. 2020); Decay sugar-rich substrates (Põlme et al. 2020) | Põlme S, et al. 2020. Fungal Diversity 105:1-16 ((https://doi.org/10.1007/s13225-020-00466-2)) |
| Otu49545 | 1 | k__Fungi(100);p__Ascomycota(100);c__Saccharomycetes(100);o__Saccharomycetales(100);f__Metschnikowiaceae(100);g__Metschnikowia(100);s__Metschnikowia_pulcherrima(100); | Metschnikowia | | 13 | Saprotroph | \|Undefined Saprotroph\| | Yeast | NULL | Probable | As nectar-tap_saprotroph (Põlme et al. 2020); Decay sugar-rich substrates (Põlme et al. 2020) | Põlme S, et al. 2020. Fungal Diversity 105:1-16 ((https://doi.org/10.1007/s13225-020-00466-2)) |
| Otu49546 | 1 | k__Fungi(100);p__Ascomycota(100);c__Saccharomycetes(100);o__Saccharomycetales(100);f__Metschnikowiaceae(100);g__Metschnikowia(100);s__Metschnikowia_pulcherrima(100); | Metschnikowia | | 13 | Saprotroph | \|Undefined Saprotroph\| | Yeast | NULL | Probable | As nectar-tap_saprotroph (Põlme et al. 2020); Decay sugar-rich substrates (Põlme et al. 2020) | Põlme S, et al. 2020. Fungal Diversity 105:1-16 ((https://doi.org/10.1007/s13225-020-00466-2)) |
| Otu49547 | 1 | k__Fungi(100);p__Ascomycota(100);c__Saccharomycetes(100);o__Saccharomycetales(100);f__Metschnikowiaceae(100);g__Metschnikowia(100);s__Metschnikowia_pulcherrima(100); | Metschnikowia | | 13 | Saprotroph | \|Undefined Saprotroph\| | Yeast | NULL | Probable | As nectar-tap_saprotroph (Põlme et al. 2020); Decay sugar-rich substrates (Põlme et al. 2020) | Põlme S, et al. 2020. Fungal Diversity 105:1-16 ((https://doi.org/10.1007/s13225-020-00466-2)) |
| Otu49548 | 1 | k__Fungi(100);p__Ascomycota(100);c__Saccharomycetes(100);o__Saccharomycetales(100);f__Metschnikowiaceae(100);g__Metschnikowia(100);s__Metschnikowia_pulcherrima(100); | Metschnikowia | | 13 | Saprotroph | \|Undefined Saprotroph\| | Yeast | NULL | Probable | As nectar-tap_saprotroph (Põlme et al. 2020); Decay sugar-rich substrates (Põlme et al. 2020) | Põlme S, et al. 2020. Fungal Diversity 105:1-16 ((https://doi.org/10.1007/s13225-020-00466-2)) |
| Otu49549 | 1 | k__Fungi(100);p__Ascomycota(100);c__Saccharomycetes(100);o__Saccharomycetales(100);f__Metschnikowiaceae(100);g__Metschnikowia(100);g__Metschnikowia_unclassified(100); | Metschnikowia | | 13 | Saprotroph | \|Undefined Saprotroph\| | Yeast | NULL | Probable | As nectar-tap_saprotroph (Põlme et al. 2020); Decay sugar-rich substrates (Põlme et al. 2020) | Põlme S, et al. 2020. Fungal Diversity 105:1-16 ((https://doi.org/10.1007/s13225-020-00466-2)) |
| Otu49550 | 1 | k__Fungi(100);p__Ascomycota(100);c__Saccharomycetes(100);o__Saccharomycetales(100);f__Metschnikowiaceae(100);g__Metschnikowia(100);s__Metschnikowia_pulcherrima(100); | Metschnikowia | | 13 | Saprotroph | \|Undefined Saprotroph\| | Yeast | NULL | Probable | As nectar-tap_saprotroph (Põlme et al. 2020); Decay sugar-rich substrates (Põlme et al. 2020) | Põlme S, et al. 2020. Fungal Diversity 105:1-16 ((https://doi.org/10.1007/s13225-020-00466-2)) |
| Otu49551 | 1 | k__Fungi(100);p__Ascomycota(100);c__Saccharomycetes(100);o__Saccharomycetales(100);f__Metschnikowiaceae(100);g__Metschnikowia(100);g__Metschnikowia_unclassified(100); | Metschnikowia | | 13 | Saprotroph | \|Undefined Saprotroph\| | Yeast | NULL | Probable | As nectar-tap_saprotroph (Põlme et al. 2020); Decay sugar-rich substrates (Põlme et al. 2020) | Põlme S, et al. 2020. Fungal Diversity 105:1-16 ((https://doi.org/10.1007/s13225-020-00466-2)) |
| Otu49552 | 1 | k__Fungi(100);p__Ascomycota(100);c__Saccharomycetes(100);o__Saccharomycetales(100);f__Metschnikowiaceae(100);g__Metschnikowia(100);s__Metschnikowia_pulcherrima(100); | Metschnikowia | | 13 | Saprotroph | \|Undefined Saprotroph\| | Yeast | NULL | Probable | As nectar-tap_saprotroph (Põlme et al. 2020); Decay sugar-rich substrates (Põlme et al. 2020) | Põlme S, et al. 2020. Fungal Diversity 105:1-16 ((https://doi.org/10.1007/s13225-020-00466-2)) |
| Otu49553 | 1 | k__Fungi(100);p__Ascomycota(100);c__Saccharomycetes(100);o__Saccharomycetales(100);f__Metschnikowiaceae(100);g__Metschnikowia(100);s__Metschnikowia_pulcherrima(100); | Metschnikowia | | 13 | Saprotroph | \|Undefined Saprotroph\| | Yeast | NULL | Probable | As nectar-tap_saprotroph (Põlme et al. 2020); Decay sugar-rich substrates (Põlme et al. 2020) | Põlme S, et al. 2020. Fungal Diversity 105:1-16 ((https://doi.org/10.1007/s13225-020-00466-2)) |
| Otu49554 | 1 | k__Fungi(100);p__Ascomycota(100);c__Saccharomycetes(100);o__Saccharomycetales(100);f__Metschnikowiaceae(100);g__Metschnikowia(100);s__Metschnikowia_pulcherrima(100); | Metschnikowia | | 13 | Saprotroph | \|Undefined Saprotroph\| | Yeast | NULL | Probable | As nectar-tap_saprotroph (Põlme et al. 2020); Decay sugar-rich substrates (Põlme et al. 2020) | Põlme S, et al. 2020. Fungal Diversity 105:1-16 ((https://doi.org/10.1007/s13225-020-00466-2)) |
| Otu49555 | 1 | k__Fungi(100);p__Ascomycota(100);c__Saccharomycetes(100);o__Saccharomycetales(100);f__Metschnikowiaceae(100);g__Metschnikowia(100);s__Metschnikowia_pulcherrima(100); | Metschnikowia | | 13 | Saprotroph | \|Undefined Saprotroph\| | Yeast | NULL | Probable | As nectar-tap_saprotroph (Põlme et al. 2020); Decay sugar-rich substrates (Põlme et al. 2020) | Põlme S, et al. 2020. Fungal Diversity 105:1-16 ((https://doi.org/10.1007/s13225-020-00466-2)) |
| Otu49556 | 1 | k__Fungi(100);p__Ascomycota(100);c__Saccharomycetes(100);o__Saccharomycetales(100);f__Metschnikowiaceae(100);g__Metschnikowia(100);s__Metschnikowia_pulcherrima(100); | Metschnikowia | | 13 | Saprotroph | \|Undefined Saprotroph\| | Yeast | NULL | Probable | As nectar-tap_saprotroph (Põlme et al. 2020); Decay sugar-rich substrates (Põlme et al. 2020) | Põlme S, et al. 2020. Fungal Diversity 105:1-16 ((https://doi.org/10.1007/s13225-020-00466-2)) |
| Otu49557 | 1 | k__Fungi(100);p__Ascomycota(100);c__Saccharomycetes(100);o__Saccharomycetales(100);f__Metschnikowiaceae(100);g__Metschnikowia(100);s__Metschnikowia_pulcherrima(100); | Metschnikowia | | 13 | Saprotroph | \|Undefined Saprotroph\| | Yeast | NULL | Probable | As nectar-tap_saprotroph (Põlme et al. 2020); Decay sugar-rich substrates (Põlme et al. 2020) | Põlme S, et al. 2020. Fungal Diversity 105:1-16 ((https://doi.org/10.1007/s13225-020-00466-2)) |
| Otu49558 | 1 | k__Fungi(100);p__Ascomycota(100);c__Saccharomycetes(100);o__Saccharomycetales(100);f__Metschnikowiaceae(100);g__Metschnikowia(100);s__Metschnikowia_pulcherrima(100); | Metschnikowia | | 13 | Saprotroph | \|Undefined Saprotroph\| | Yeast | NULL | Probable | As nectar-tap_saprotroph (Põlme et al. 2020); Decay sugar-rich substrates (Põlme et al. 2020) | Põlme S, et al. 2020. Fungal Diversity 105:1-16 ((https://doi.org/10.1007/s13225-020-00466-2)) |
| Otu49559 | 1 | k__Fungi(100);p__Ascomycota(100);c__Saccharomycetes(100);o__Saccharomycetales(100);f__Metschnikowiaceae(100);g__Metschnikowia(100);s__Metschnikowia_pulcherrima(100); | Metschnikowia | | 13 | Saprotroph | \|Undefined Saprotroph\| | Yeast | NULL | Probable | As nectar-tap_saprotroph (Põlme et al. 2020); Decay sugar-rich substrates (Põlme et al. 2020) | Põlme S, et al. 2020. Fungal Diversity 105:1-16 ((https://doi.org/10.1007/s13225-020-00466-2)) |
| Otu49560 | 1 | k__Fungi(100);p__Ascomycota(100);c__Saccharomycetes(100);o__Saccharomycetales(100);f__Metschnikowiaceae(100);g__Metschnikowia(100);s__Metschnikowia_pulcherrima(100); | Metschnikowia | | 13 | Saprotroph | \|Undefined Saprotroph\| | Yeast | NULL | Probable | As nectar-tap_saprotroph (Põlme et al. 2020); Decay sugar-rich substrates (Põlme et al. 2020) | Põlme S, et al. 2020. Fungal Diversity 105:1-16 ((https://doi.org/10.1007/s13225-020-00466-2)) |
| Otu49561 | 1 | k__Fungi(100);p__Ascomycota(100);c__Saccharomycetes(100);o__Saccharomycetales(100);f__Metschnikowiaceae(100);g__Metschnikowia(100);s__Metschnikowia_pulcherrima(100); | Metschnikowia | | 13 | Saprotroph | \|Undefined Saprotroph\| | Yeast | NULL | Probable | As nectar-tap_saprotroph (Põlme et al. 2020); Decay sugar-rich substrates (Põlme et al. 2020) | Põlme S, et al. 2020. Fungal Diversity 105:1-16 ((https://doi.org/10.1007/s13225-020-00466-2)) |
| Otu49562 | 1 | k__Fungi(100);p__Ascomycota(100);c__Saccharomycetes(100);o__Saccharomycetales(100);f__Metschnikowiaceae(100);g__Metschnikowia(100);s__Metschnikowia_pulcherrima(100); | Metschnikowia | | 13 | Saprotroph | \|Undefined Saprotroph\| | Yeast | NULL | Probable | As nectar-tap_saprotroph (Põlme et al. 2020); Decay sugar-rich substrates (Põlme et al. 2020) | Põlme S, et al. 2020. Fungal Diversity 105:1-16 ((https://doi.org/10.1007/s13225-020-00466-2)) |
| Otu49563 | 1 | k__Fungi(100);p__Ascomycota(100);c__Saccharomycetes(100);o__Saccharomycetales(100);f__Metschnikowiaceae(100);g__Metschnikowia(100);s__Metschnikowia_pulcherrima(100); | Metschnikowia | | 13 | Saprotroph | \|Undefined Saprotroph\| | Yeast | NULL | Probable | As nectar-tap_saprotroph (Põlme et al. 2020); Decay sugar-rich substrates (Põlme et al. 2020) | Põlme S, et al. 2020. Fungal Diversity 105:1-16 ((https://doi.org/10.1007/s13225-020-00466-2)) |
| Otu49564 | 1 | k__Fungi(100);p__Ascomycota(100);c__Saccharomycetes(100);o__Saccharomycetales(100);f__Metschnikowiaceae(100);g__Metschnikowia(100);s__Metschnikowia_pulcherrima(100); | Metschnikowia | | 13 | Saprotroph | \|Undefined Saprotroph\| | Yeast | NULL | Probable | As nectar-tap_saprotroph (Põlme et al. 2020); Decay sugar-rich substrates (Põlme et al. 2020) | Põlme S, et al. 2020. Fungal Diversity 105:1-16 ((https://doi.org/10.1007/s13225-020-00466-2)) |
| Otu49565 | 1 | k__Fungi(100);p__Ascomycota(100);c__Saccharomycetes(100);o__Saccharomycetales(100);f__Metschnikowiaceae(100);g__Metschnikowia(100);g__Metschnikowia_unclassified(100); | Metschnikowia | | 13 | Saprotroph | \|Undefined Saprotroph\| | Yeast | NULL | Probable | As nectar-tap_saprotroph (Põlme et al. 2020); Decay sugar-rich substrates (Põlme et al. 2020) | Põlme S, et al. 2020. Fungal Diversity 105:1-16 ((https://doi.org/10.1007/s13225-020-00466-2)) |
| Otu49567 | 1 | k__Fungi(100);p__Ascomycota(100);c__Saccharomycetes(100);o__Saccharomycetales(100);f__Metschnikowiaceae(100);g__Metschnikowia(100);s__Metschnikowia_pulcherrima(100); | Metschnikowia | | 13 | Saprotroph | \|Undefined Saprotroph\| | Yeast | NULL | Probable | As nectar-tap_saprotroph (Põlme et al. 2020); Decay sugar-rich substrates (Põlme et al. 2020) | Põlme S, et al. 2020. Fungal Diversity 105:1-16 ((https://doi.org/10.1007/s13225-020-00466-2)) |
| Otu49568 | 1 | k__Fungi(100);p__Ascomycota(100);c__Saccharomycetes(100);o__Saccharomycetales(100);f__Metschnikowiaceae(100);g__Metschnikowia(100);g__Metschnikowia_unclassified(100); | Metschnikowia | | 13 | Saprotroph | \|Undefined Saprotroph\| | Yeast | NULL | Probable | As nectar-tap_saprotroph (Põlme et al. 2020); Decay sugar-rich substrates (Põlme et al. 2020) | Põlme S, et al. 2020. Fungal Diversity 105:1-16 ((https://doi.org/10.1007/s13225-020-00466-2)) |
| Otu49569 | 1 | k__Fungi(100);p__Ascomycota(100);c__Saccharomycetes(100);o__Saccharomycetales(100);f__Metschnikowiaceae(100);g__Metschnikowia(100);s__Metschnikowia_pulcherrima(100); | Metschnikowia | | 13 | Saprotroph | \|Undefined Saprotroph\| | Yeast | NULL | Probable | As nectar-tap_saprotroph (Põlme et al. 2020); Decay sugar-rich substrates (Põlme et al. 2020) | Põlme S, et al. 2020. Fungal Diversity 105:1-16 ((https://doi.org/10.1007/s13225-020-00466-2)) |
| Otu49570 | 1 | k__Fungi(100);p__Ascomycota(100);c__Saccharomycetes(100);o__Saccharomycetales(100);f__Metschnikowiaceae(100);g__Metschnikowia(100);s__Metschnikowia_pulcherrima(100); | Metschnikowia | | 13 | Saprotroph | \|Undefined Saprotroph\| | Yeast | NULL | Probable | As nectar-tap_saprotroph (Põlme et al. 2020); Decay sugar-rich substrates (Põlme et al. 2020) | Põlme S, et al. 2020. Fungal Diversity 105:1-16 ((https://doi.org/10.1007/s13225-020-00466-2)) |
| Otu49571 | 1 | k__Fungi(100);p__Ascomycota(100);c__Saccharomycetes(100);o__Saccharomycetales(100);f__Metschnikowiaceae(100);g__Metschnikowia(100);s__Metschnikowia_pulcherrima(100); | Metschnikowia | | 13 | Saprotroph | \|Undefined Saprotroph\| | Yeast | NULL | Probable | As nectar-tap_saprotroph (Põlme et al. 2020); Decay sugar-rich substrates (Põlme et al. 2020) | Põlme S, et al. 2020. Fungal Diversity 105:1-16 ((https://doi.org/10.1007/s13225-020-00466-2)) |
| Otu49572 | 1 | k__Fungi(100);p__Ascomycota(100);c__Saccharomycetes(100);o__Saccharomycetales(100);f__Metschnikowiaceae(100);g__Metschnikowia(100);s__Metschnikowia_pulcherrima(100); | Metschnikowia | | 13 | Saprotroph | \|Undefined Saprotroph\| | Yeast | NULL | Probable | As nectar-tap_saprotroph (Põlme et al. 2020); Decay sugar-rich substrates (Põlme et al. 2020) | Põlme S, et al. 2020. Fungal Diversity 105:1-16 ((https://doi.org/10.1007/s13225-020-00466-2)) |
| Otu49573 | 1 | k__Fungi(100);p__Ascomycota(100);c__Saccharomycetes(100);o__Saccharomycetales(100);f__Metschnikowiaceae(100);g__Metschnikowia(100);g__Metschnikowia_unclassified(100); | Metschnikowia | | 13 | Saprotroph | \|Undefined Saprotroph\| | Yeast | NULL | Probable | As nectar-tap_saprotroph (Põlme et al. 2020); Decay sugar-rich substrates (Põlme et al. 2020) | Põlme S, et al. 2020. Fungal Diversity 105:1-16 ((https://doi.org/10.1007/s13225-020-00466-2)) |
| Otu49574 | 1 | k__Fungi(100);p__Ascomycota(100);c__Saccharomycetes(100);o__Saccharomycetales(100);f__Metschnikowiaceae(100);g__Metschnikowia(100);s__Metschnikowia_pulcherrima(100); | Metschnikowia | | 13 | Saprotroph | \|Undefined Saprotroph\| | Yeast | NULL | Probable | As nectar-tap_saprotroph (Põlme et al. 2020); Decay sugar-rich substrates (Põlme et al. 2020) | Põlme S, et al. 2020. Fungal Diversity 105:1-16 ((https://doi.org/10.1007/s13225-020-00466-2)) |
| Otu49575 | 1 | k__Fungi(100);p__Ascomycota(100);c__Saccharomycetes(100);o__Saccharomycetales(100);f__Metschnikowiaceae(100);g__Metschnikowia(100);g__Metschnikowia_unclassified(100); | Metschnikowia | | 13 | Saprotroph | \|Undefined Saprotroph\| | Yeast | NULL | Probable | As nectar-tap_saprotroph (Põlme et al. 2020); Decay sugar-rich substrates (Põlme et al. 2020) | Põlme S, et al. 2020. Fungal Diversity 105:1-16 ((https://doi.org/10.1007/s13225-020-00466-2)) |
| Otu49576 | 1 | k__Fungi(100);p__Ascomycota(100);c__Saccharomycetes(100);o__Saccharomycetales(100);f__Metschnikowiaceae(100);g__Metschnikowia(100);s__Metschnikowia_pulcherrima(100); | Metschnikowia | | 13 | Saprotroph | \|Undefined Saprotroph\| | Yeast | NULL | Probable | As nectar-tap_saprotroph (Põlme et al. 2020); Decay sugar-rich substrates (Põlme et al. 2020) | Põlme S, et al. 2020. Fungal Diversity 105:1-16 ((https://doi.org/10.1007/s13225-020-00466-2)) |
| Otu49577 | 1 | k__Fungi(100);p__Ascomycota(100);c__Saccharomycetes(100);o__Saccharomycetales(100);f__Metschnikowiaceae(100);g__Metschnikowia(100);s__Metschnikowia_pulcherrima(100); | Metschnikowia | | 13 | Saprotroph | \|Undefined Saprotroph\| | Yeast | NULL | Probable | As nectar-tap_saprotroph (Põlme et al. 2020); Decay sugar-rich substrates (Põlme et al. 2020) | Põlme S, et al. 2020. Fungal Diversity 105:1-16 ((https://doi.org/10.1007/s13225-020-00466-2)) |
| Otu49578 | 1 | k__Fungi(100);p__Ascomycota(100);c__Saccharomycetes(100);o__Saccharomycetales(100);f__Metschnikowiaceae(100);g__Metschnikowia(100);g__Metschnikowia_unclassified(100); | Metschnikowia | | 13 | Saprotroph | \|Undefined Saprotroph\| | Yeast | NULL | Probable | As nectar-tap_saprotroph (Põlme et al. 2020); Decay sugar-rich substrates (Põlme et al. 2020) | Põlme S, et al. 2020. Fungal Diversity 105:1-16 ((https://doi.org/10.1007/s13225-020-00466-2)) |
| Otu49580 | 1 | k__Fungi(100);p__Ascomycota(100);c__Saccharomycetes(100);o__Saccharomycetales(100);f__Metschnikowiaceae(100);g__Metschnikowia(100);s__Metschnikowia_pulcherrima(100); | Metschnikowia | | 13 | Saprotroph | \|Undefined Saprotroph\| | Yeast | NULL | Probable | As nectar-tap_saprotroph (Põlme et al. 2020); Decay sugar-rich substrates (Põlme et al. 2020) | Põlme S, et al. 2020. Fungal Diversity 105:1-16 ((https://doi.org/10.1007/s13225-020-00466-2)) |
| Otu49581 | 1 | k__Fungi(100);p__Ascomycota(100);c__Sordariomycetes(100);o__Sordariales(100);f__Chaetomiaceae(100);g__Humicola(100);g__Humicola_unclassified(100); | Humicola | | 13 | Pathotroph-Saprotroph | Plant Pathogen-Undefined Saprotroph-\|Wood Saprotroph\| | Microfungus | Soft Rot | Probable | Decay wood (Põlme et al. 2020) | Seehann G, et al. 1975. List of Fungi in Soft-Rot Tissue Tests. Institute fur Holzbiologie und Holzschutz, Bundesforschungsanstalt fur Forst - und Holzwirtshaft, Hamburg - Lohbrugge, Germany; Costa IPMW, et al. 2012. Mycotaxon 119:e26 ((http://mycotaxon.com/resources/checklists/costa_v119_checklist.pdf)); Tedersoo L, et al. 2014. Science 346:e1256688 ((https://doi.org/10.1126/science.1256688)); Põlme S, et al. 2020. Fungal Diversity 105:1-16 ((https://doi.org/10.1007/s13225-020-00466-2)) |
| Otu49582 | 1 | k__Fungi(100);p__Ascomycota(100);c__Saccharomycetes(100);o__Saccharomycetales(100);f__Metschnikowiaceae(100);g__Metschnikowia(100);s__Metschnikowia_pulcherrima(100); | Metschnikowia | | 13 | Saprotroph | \|Undefined Saprotroph\| | Yeast | NULL | Probable | As nectar-tap_saprotroph (Põlme et al. 2020); Decay sugar-rich substrates (Põlme et al. 2020) | Põlme S, et al. 2020. Fungal Diversity 105:1-16 ((https://doi.org/10.1007/s13225-020-00466-2)) |
| Otu49583 | 1 | k__Fungi(100);p__Ascomycota(100);c__Saccharomycetes(100);o__Saccharomycetales(100);f__Metschnikowiaceae(100);g__Metschnikowia(100);s__Metschnikowia_pulcherrima(100); | Metschnikowia | | 13 | Saprotroph | \|Undefined Saprotroph\| | Yeast | NULL | Probable | As nectar-tap_saprotroph (Põlme et al. 2020); Decay sugar-rich substrates (Põlme et al. 2020) | Põlme S, et al. 2020. Fungal Diversity 105:1-16 ((https://doi.org/10.1007/s13225-020-00466-2)) |
| Otu49584 | 1 | k__Fungi(100);p__Ascomycota(100);c__Saccharomycetes(100);o__Saccharomycetales(100);f__Metschnikowiaceae(100);g__Metschnikowia(100);g__Metschnikowia_unclassified(100); | Metschnikowia | | 13 | Saprotroph | \|Undefined Saprotroph\| | Yeast | NULL | Probable | As nectar-tap_saprotroph (Põlme et al. 2020); Decay sugar-rich substrates (Põlme et al. 2020) | Põlme S, et al. 2020. Fungal Diversity 105:1-16 ((https://doi.org/10.1007/s13225-020-00466-2)) |
| Otu49585 | 1 | k__Fungi(100);p__Ascomycota(100);c__Saccharomycetes(100);o__Saccharomycetales(100);f__Metschnikowiaceae(100);g__Metschnikowia(100);s__Metschnikowia_pulcherrima(100); | Metschnikowia | | 13 | Saprotroph | \|Undefined Saprotroph\| | Yeast | NULL | Probable | As nectar-tap_saprotroph (Põlme et al. 2020); Decay sugar-rich substrates (Põlme et al. 2020) | Põlme S, et al. 2020. Fungal Diversity 105:1-16 ((https://doi.org/10.1007/s13225-020-00466-2)) |
| Otu49586 | 1 | k__Fungi(100);p__Ascomycota(100);c__Saccharomycetes(100);o__Saccharomycetales(100);f__Metschnikowiaceae(100);g__Metschnikowia(100);s__Metschnikowia_pulcherrima(100); | Metschnikowia | | 13 | Saprotroph | \|Undefined Saprotroph\| | Yeast | NULL | Probable | As nectar-tap_saprotroph (Põlme et al. 2020); Decay sugar-rich substrates (Põlme et al. 2020) | Põlme S, et al. 2020. Fungal Diversity 105:1-16 ((https://doi.org/10.1007/s13225-020-00466-2)) |
| Otu49587 | 1 | k__Fungi(100);p__Ascomycota(100);c__Saccharomycetes(100);o__Saccharomycetales(100);f__Metschnikowiaceae(100);g__Metschnikowia(100);g__Metschnikowia_unclassified(100); | Metschnikowia | | 13 | Saprotroph | \|Undefined Saprotroph\| | Yeast | NULL | Probable | As nectar-tap_saprotroph (Põlme et al. 2020); Decay sugar-rich substrates (Põlme et al. 2020) | Põlme S, et al. 2020. Fungal Diversity 105:1-16 ((https://doi.org/10.1007/s13225-020-00466-2)) |
| Otu49588 | 1 | k__Fungi(100);p__Ascomycota(100);c__Saccharomycetes(100);o__Saccharomycetales(100);f__Metschnikowiaceae(100);g__Metschnikowia(100);s__Metschnikowia_pulcherrima(100); | Metschnikowia | | 13 | Saprotroph | \|Undefined Saprotroph\| | Yeast | NULL | Probable | As nectar-tap_saprotroph (Põlme et al. 2020); Decay sugar-rich substrates (Põlme et al. 2020) | Põlme S, et al. 2020. Fungal Diversity 105:1-16 ((https://doi.org/10.1007/s13225-020-00466-2)) |
| Otu49589 | 1 | k__Fungi(100);p__Ascomycota(100);c__Saccharomycetes(100);o__Saccharomycetales(100);f__Metschnikowiaceae(100);g__Metschnikowia(100);g__Metschnikowia_unclassified(100); | Metschnikowia | | 13 | Saprotroph | \|Undefined Saprotroph\| | Yeast | NULL | Probable | As nectar-tap_saprotroph (Põlme et al. 2020); Decay sugar-rich substrates (Põlme et al. 2020) | Põlme S, et al. 2020. Fungal Diversity 105:1-16 ((https://doi.org/10.1007/s13225-020-00466-2)) |
| Otu49590 | 1 | k__Fungi(100);p__Ascomycota(100);c__Saccharomycetes(100);o__Saccharomycetales(100);f__Metschnikowiaceae(100);g__Metschnikowia(100);s__Metschnikowia_pulcherrima(100); | Metschnikowia | | 13 | Saprotroph | \|Undefined Saprotroph\| | Yeast | NULL | Probable | As nectar-tap_saprotroph (Põlme et al. 2020); Decay sugar-rich substrates (Põlme et al. 2020) | Põlme S, et al. 2020. Fungal Diversity 105:1-16 ((https://doi.org/10.1007/s13225-020-00466-2)) |
| Otu49591 | 1 | k__Fungi(100);p__Ascomycota(100);c__Saccharomycetes(100);o__Saccharomycetales(100);f__Metschnikowiaceae(100);g__Metschnikowia(100);s__Metschnikowia_pulcherrima(100); | Metschnikowia | | 13 | Saprotroph | \|Undefined Saprotroph\| | Yeast | NULL | Probable | As nectar-tap_saprotroph (Põlme et al. 2020); Decay sugar-rich substrates (Põlme et al. 2020) | Põlme S, et al. 2020. Fungal Diversity 105:1-16 ((https://doi.org/10.1007/s13225-020-00466-2)) |
| Otu49592 | 1 | k__Fungi(100);p__Ascomycota(100);c__Saccharomycetes(100);o__Saccharomycetales(100);f__Metschnikowiaceae(100);g__Metschnikowia(100);g__Metschnikowia_unclassified(100); | Metschnikowia | | 13 | Saprotroph | \|Undefined Saprotroph\| | Yeast | NULL | Probable | As nectar-tap_saprotroph (Põlme et al. 2020); Decay sugar-rich substrates (Põlme et al. 2020) | Põlme S, et al. 2020. Fungal Diversity 105:1-16 ((https://doi.org/10.1007/s13225-020-00466-2)) |
| Otu49593 | 1 | k__Fungi(100);p__Ascomycota(100);c__Saccharomycetes(100);o__Saccharomycetales(100);f__Metschnikowiaceae(100);g__Metschnikowia(100);g__Metschnikowia_unclassified(100); | Metschnikowia | | 13 | Saprotroph | \|Undefined Saprotroph\| | Yeast | NULL | Probable | As nectar-tap_saprotroph (Põlme et al. 2020); Decay sugar-rich substrates (Põlme et al. 2020) | Põlme S, et al. 2020. Fungal Diversity 105:1-16 ((https://doi.org/10.1007/s13225-020-00466-2)) |
| Otu49594 | 1 | k__Fungi(100);p__Ascomycota(100);c__Saccharomycetes(100);o__Saccharomycetales(100);f__Metschnikowiaceae(100);g__Metschnikowia(100);s__Metschnikowia_pulcherrima(100); | Metschnikowia | | 13 | Saprotroph | \|Undefined Saprotroph\| | Yeast | NULL | Probable | As nectar-tap_saprotroph (Põlme et al. 2020); Decay sugar-rich substrates (Põlme et al. 2020) | Põlme S, et al. 2020. Fungal Diversity 105:1-16 ((https://doi.org/10.1007/s13225-020-00466-2)) |
| Otu49595 | 1 | k__Fungi(100);p__Ascomycota(100);c__Saccharomycetes(100);o__Saccharomycetales(100);f__Metschnikowiaceae(100);g__Metschnikowia(100);g__Metschnikowia_unclassified(100); | Metschnikowia | | 13 | Saprotroph | \|Undefined Saprotroph\| | Yeast | NULL | Probable | As nectar-tap_saprotroph (Põlme et al. 2020); Decay sugar-rich substrates (Põlme et al. 2020) | Põlme S, et al. 2020. Fungal Diversity 105:1-16 ((https://doi.org/10.1007/s13225-020-00466-2)) |
| Otu49596 | 1 | k__Fungi(100);p__Ascomycota(100);c__Saccharomycetes(100);o__Saccharomycetales(100);f__Metschnikowiaceae(100);g__Metschnikowia(100);s__Metschnikowia_pulcherrima(100); | Metschnikowia | | 13 | Saprotroph | \|Undefined Saprotroph\| | Yeast | NULL | Probable | As nectar-tap_saprotroph (Põlme et al. 2020); Decay sugar-rich substrates (Põlme et al. 2020) | Põlme S, et al. 2020. Fungal Diversity 105:1-16 ((https://doi.org/10.1007/s13225-020-00466-2)) |
| Otu49597 | 1 | k__Fungi(100);p__Ascomycota(100);c__Saccharomycetes(100);o__Saccharomycetales(100);f__Metschnikowiaceae(100);g__Metschnikowia(100);s__Metschnikowia_pulcherrima(100); | Metschnikowia | | 13 | Saprotroph | \|Undefined Saprotroph\| | Yeast | NULL | Probable | As nectar-tap_saprotroph (Põlme et al. 2020); Decay sugar-rich substrates (Põlme et al. 2020) | Põlme S, et al. 2020. Fungal Diversity 105:1-16 ((https://doi.org/10.1007/s13225-020-00466-2)) |
| Otu49598 | 1 | k__Fungi(100);p__Ascomycota(100);c__Saccharomycetes(100);o__Saccharomycetales(100);f__Metschnikowiaceae(100);g__Metschnikowia(100);s__Metschnikowia_pulcherrima(100); | Metschnikowia | | 13 | Saprotroph | \|Undefined Saprotroph\| | Yeast | NULL | Probable | As nectar-tap_saprotroph (Põlme et al. 2020); Decay sugar-rich substrates (Põlme et al. 2020) | Põlme S, et al. 2020. Fungal Diversity 105:1-16 ((https://doi.org/10.1007/s13225-020-00466-2)) |
| Otu49599 | 1 | k__Fungi(100);p__Ascomycota(100);c__Saccharomycetes(100);o__Saccharomycetales(100);f__Metschnikowiaceae(100);g__Metschnikowia(100);s__Metschnikowia_pulcherrima(100); | Metschnikowia | | 13 | Saprotroph | \|Undefined Saprotroph\| | Yeast | NULL | Probable | As nectar-tap_saprotroph (Põlme et al. 2020); Decay sugar-rich substrates (Põlme et al. 2020) | Põlme S, et al. 2020. Fungal Diversity 105:1-16 ((https://doi.org/10.1007/s13225-020-00466-2)) |
| Otu49600 | 1 | k__Fungi(100);p__Ascomycota(100);c__Saccharomycetes(100);o__Saccharomycetales(100);f__Metschnikowiaceae(100);g__Metschnikowia(100);s__Metschnikowia_pulcherrima(100); | Metschnikowia | | 13 | Saprotroph | \|Undefined Saprotroph\| | Yeast | NULL | Probable | As nectar-tap_saprotroph (Põlme et al. 2020); Decay sugar-rich substrates (Põlme et al. 2020) | Põlme S, et al. 2020. Fungal Diversity 105:1-16 ((https://doi.org/10.1007/s13225-020-00466-2)) |
| Otu49601 | 1 | k__Fungi(100);p__Ascomycota(100);c__Saccharomycetes(100);o__Saccharomycetales(100);f__Metschnikowiaceae(100);g__Metschnikowia(100);s__Metschnikowia_pulcherrima(100); | Metschnikowia | | 13 | Saprotroph | \|Undefined Saprotroph\| | Yeast | NULL | Probable | As nectar-tap_saprotroph (Põlme et al. 2020); Decay sugar-rich substrates (Põlme et al. 2020) | Põlme S, et al. 2020. Fungal Diversity 105:1-16 ((https://doi.org/10.1007/s13225-020-00466-2)) |
| Otu49602 | 1 | k__Fungi(100);p__Ascomycota(100);c__Sordariomycetes(100);o__Hypocreales(100);f__Nectriaceae(100);g__Fusarium(100);g__Fusarium_unclassified(100); | Fusarium | | 13 | Pathotroph-Saprotroph-Symbiotroph | Animal Pathogen-Endophyte-Lichen Parasite-\|Plant Pathogen\|-Plant Saprotroph-Undefined Saprotroph-Wood Saprotroph | Microfungus | Soft Rot | Possible | As foliar_endophyte (Põlme et al. 2020); As litter_saprotroph (Põlme et al. 2020); Causing damping-off disease (Põlme et al. 2020); Decay leaf, fruit, seed, roots, soil, animal material (Põlme et al. 2020); Likely opportunistic human pathogen (Irinyi et al. 2015) | Duncan CG, Eslyn WE. 1966. Mycologia 58:642-664 ((https://doi.org/10.2307/3757045)); Seehann G, et al. 1975. List of Fungi in Soft-Rot Tissue Tests. Institute fur Holzbiologie und Holzschutz, Bundesforschungsanstalt fur Forst - und Holzwirtshaft, Hamburg - Lohbrugge, Germany; Costa IPMW, et al. 2012. Mycotaxon 119:e26 ((http://mycotaxon.com/resources/checklists/costa_v119_checklist.pdf)); Tedersoo L, et al. 2014. Science 346:e1256688 ((https://doi.org/10.1126/science.1256688)); Irinyi L, et al. 2015. Medical Mycology 53:313-337 ((https://doi.org/10.1093/mmy/myv008)); Schroers, et al. 2016. Mycologia 108:806-819 ((https://doi.org/10.3852/15-255)); Põlme S, et al. 2020. Fungal Diversity 105:1-16 ((https://doi.org/10.1007/s13225-020-00466-2)); The American Phytopathological Society Common Names of Plant Diseases ((http://www.apsnet.org/publications/commonnames/Pages/default.aspx)); Lawrey JD, Diederich P. 2016. http://www.lichenicolous.net (accessed 24 May 2017) ((http://www.lichenicolous.net)) |
| Otu49603 | 1 | k__Fungi(100);p__Ascomycota(100);c__Saccharomycetes(100);o__Saccharomycetales(100);f__Metschnikowiaceae(100);g__Metschnikowia(100);g__Metschnikowia_unclassified(100); | Metschnikowia | | 13 | Saprotroph | \|Undefined Saprotroph\| | Yeast | NULL | Probable | As nectar-tap_saprotroph (Põlme et al. 2020); Decay sugar-rich substrates (Põlme et al. 2020) | Põlme S, et al. 2020. Fungal Diversity 105:1-16 ((https://doi.org/10.1007/s13225-020-00466-2)) |
| Otu49604 | 1 | k__Fungi(100);p__Ascomycota(100);c__Saccharomycetes(100);o__Saccharomycetales(100);f__Metschnikowiaceae(100);g__Metschnikowia(100);s__Metschnikowia_pulcherrima(100); | Metschnikowia | | 13 | Saprotroph | \|Undefined Saprotroph\| | Yeast | NULL | Probable | As nectar-tap_saprotroph (Põlme et al. 2020); Decay sugar-rich substrates (Põlme et al. 2020) | Põlme S, et al. 2020. Fungal Diversity 105:1-16 ((https://doi.org/10.1007/s13225-020-00466-2)) |
| Otu49605 | 1 | k__Fungi(100);p__Ascomycota(100);c__Saccharomycetes(100);o__Saccharomycetales(100);f__Metschnikowiaceae(100);g__Metschnikowia(100);g__Metschnikowia_unclassified(100); | Metschnikowia | | 13 | Saprotroph | \|Undefined Saprotroph\| | Yeast | NULL | Probable | As nectar-tap_saprotroph (Põlme et al. 2020); Decay sugar-rich substrates (Põlme et al. 2020) | Põlme S, et al. 2020. Fungal Diversity 105:1-16 ((https://doi.org/10.1007/s13225-020-00466-2)) |
| Otu49606 | 1 | k__Fungi(100);p__Ascomycota(100);c__Saccharomycetes(100);o__Saccharomycetales(100);f__Metschnikowiaceae(100);g__Metschnikowia(100);s__Metschnikowia_pulcherrima(100); | Metschnikowia | | 13 | Saprotroph | \|Undefined Saprotroph\| | Yeast | NULL | Probable | As nectar-tap_saprotroph (Põlme et al. 2020); Decay sugar-rich substrates (Põlme et al. 2020) | Põlme S, et al. 2020. Fungal Diversity 105:1-16 ((https://doi.org/10.1007/s13225-020-00466-2)) |
| Otu49607 | 1 | k__Fungi(100);p__Ascomycota(100);c__Saccharomycetes(100);o__Saccharomycetales(100);f__Metschnikowiaceae(100);g__Metschnikowia(100);s__Metschnikowia_pulcherrima(100); | Metschnikowia | | 13 | Saprotroph | \|Undefined Saprotroph\| | Yeast | NULL | Probable | As nectar-tap_saprotroph (Põlme et al. 2020); Decay sugar-rich substrates (Põlme et al. 2020) | Põlme S, et al. 2020. Fungal Diversity 105:1-16 ((https://doi.org/10.1007/s13225-020-00466-2)) |
| Otu49608 | 1 | k__Fungi(100);p__Ascomycota(100);c__Saccharomycetes(100);o__Saccharomycetales(100);f__Metschnikowiaceae(100);g__Metschnikowia(100);g__Metschnikowia_unclassified(100); | Metschnikowia | | 13 | Saprotroph | \|Undefined Saprotroph\| | Yeast | NULL | Probable | As nectar-tap_saprotroph (Põlme et al. 2020); Decay sugar-rich substrates (Põlme et al. 2020) | Põlme S, et al. 2020. Fungal Diversity 105:1-16 ((https://doi.org/10.1007/s13225-020-00466-2)) |
| Otu49610 | 1 | k__Fungi(100);p__Ascomycota(100);c__Saccharomycetes(100);o__Saccharomycetales(100);f__Metschnikowiaceae(100);g__Metschnikowia(100);s__Metschnikowia_pulcherrima(100); | Metschnikowia | | 13 | Saprotroph | \|Undefined Saprotroph\| | Yeast | NULL | Probable | As nectar-tap_saprotroph (Põlme et al. 2020); Decay sugar-rich substrates (Põlme et al. 2020) | Põlme S, et al. 2020. Fungal Diversity 105:1-16 ((https://doi.org/10.1007/s13225-020-00466-2)) |
| Otu49611 | 1 | k__Fungi(100);p__Ascomycota(100);c__Saccharomycetes(100);o__Saccharomycetales(100);f__Metschnikowiaceae(100);g__Metschnikowia(100);s__Metschnikowia_pulcherrima(100); | Metschnikowia | | 13 | Saprotroph | \|Undefined Saprotroph\| | Yeast | NULL | Probable | As nectar-tap_saprotroph (Põlme et al. 2020); Decay sugar-rich substrates (Põlme et al. 2020) | Põlme S, et al. 2020. Fungal Diversity 105:1-16 ((https://doi.org/10.1007/s13225-020-00466-2)) |
| Otu49612 | 1 | k__Fungi(100);p__Ascomycota(100);c__Saccharomycetes(100);o__Saccharomycetales(100);f__Metschnikowiaceae(100);g__Metschnikowia(100);s__Metschnikowia_pulcherrima(100); | Metschnikowia | | 13 | Saprotroph | \|Undefined Saprotroph\| | Yeast | NULL | Probable | As nectar-tap_saprotroph (Põlme et al. 2020); Decay sugar-rich substrates (Põlme et al. 2020) | Põlme S, et al. 2020. Fungal Diversity 105:1-16 ((https://doi.org/10.1007/s13225-020-00466-2)) |
| Otu49613 | 1 | k__Fungi(100);p__Ascomycota(100);c__Saccharomycetes(100);o__Saccharomycetales(100);f__Metschnikowiaceae(100);g__Metschnikowia(100);s__Metschnikowia_pulcherrima(100); | Metschnikowia | | 13 | Saprotroph | \|Undefined Saprotroph\| | Yeast | NULL | Probable | As nectar-tap_saprotroph (Põlme et al. 2020); Decay sugar-rich substrates (Põlme et al. 2020) | Põlme S, et al. 2020. Fungal Diversity 105:1-16 ((https://doi.org/10.1007/s13225-020-00466-2)) |
| Otu49614 | 1 | k__Fungi(100);p__Ascomycota(100);c__Saccharomycetes(100);o__Saccharomycetales(100);f__Metschnikowiaceae(100);g__Metschnikowia(100);s__Metschnikowia_pulcherrima(100); | Metschnikowia | | 13 | Saprotroph | \|Undefined Saprotroph\| | Yeast | NULL | Probable | As nectar-tap_saprotroph (Põlme et al. 2020); Decay sugar-rich substrates (Põlme et al. 2020) | Põlme S, et al. 2020. Fungal Diversity 105:1-16 ((https://doi.org/10.1007/s13225-020-00466-2)) |
| Otu49615 | 1 | k__Fungi(100);p__Ascomycota(100);c__Saccharomycetes(100);o__Saccharomycetales(100);f__Metschnikowiaceae(100);g__Metschnikowia(100);s__Metschnikowia_pulcherrima(100); | Metschnikowia | | 13 | Saprotroph | \|Undefined Saprotroph\| | Yeast | NULL | Probable | As nectar-tap_saprotroph (Põlme et al. 2020); Decay sugar-rich substrates (Põlme et al. 2020) | Põlme S, et al. 2020. Fungal Diversity 105:1-16 ((https://doi.org/10.1007/s13225-020-00466-2)) |
| Otu49616 | 1 | k__Fungi(100);p__Ascomycota(100);c__Saccharomycetes(100);o__Saccharomycetales(100);f__Metschnikowiaceae(100);g__Metschnikowia(100);g__Metschnikowia_unclassified(100); | Metschnikowia | | 13 | Saprotroph | \|Undefined Saprotroph\| | Yeast | NULL | Probable | As nectar-tap_saprotroph (Põlme et al. 2020); Decay sugar-rich substrates (Põlme et al. 2020) | Põlme S, et al. 2020. Fungal Diversity 105:1-16 ((https://doi.org/10.1007/s13225-020-00466-2)) |
| Otu49617 | 1 | k__Fungi(100);p__Ascomycota(100);c__Saccharomycetes(100);o__Saccharomycetales(100);f__Metschnikowiaceae(100);g__Metschnikowia(100);s__Metschnikowia_pulcherrima(100); | Metschnikowia | | 13 | Saprotroph | \|Undefined Saprotroph\| | Yeast | NULL | Probable | As nectar-tap_saprotroph (Põlme et al. 2020); Decay sugar-rich substrates (Põlme et al. 2020) | Põlme S, et al. 2020. Fungal Diversity 105:1-16 ((https://doi.org/10.1007/s13225-020-00466-2)) |
| Otu49618 | 1 | k__Fungi(100);p__Ascomycota(100);c__Saccharomycetes(100);o__Saccharomycetales(100);f__Metschnikowiaceae(100);g__Metschnikowia(100);s__Metschnikowia_pulcherrima(100); | Metschnikowia | | 13 | Saprotroph | \|Undefined Saprotroph\| | Yeast | NULL | Probable | As nectar-tap_saprotroph (Põlme et al. 2020); Decay sugar-rich substrates (Põlme et al. 2020) | Põlme S, et al. 2020. Fungal Diversity 105:1-16 ((https://doi.org/10.1007/s13225-020-00466-2)) |
| Otu49619 | 1 | k__Fungi(100);p__Ascomycota(100);c__Saccharomycetes(100);o__Saccharomycetales(100);f__Metschnikowiaceae(100);g__Metschnikowia(100);s__Metschnikowia_pulcherrima(100); | Metschnikowia | | 13 | Saprotroph | \|Undefined Saprotroph\| | Yeast | NULL | Probable | As nectar-tap_saprotroph (Põlme et al. 2020); Decay sugar-rich substrates (Põlme et al. 2020) | Põlme S, et al. 2020. Fungal Diversity 105:1-16 ((https://doi.org/10.1007/s13225-020-00466-2)) |
| Otu49620 | 1 | k__Fungi(100);p__Ascomycota(100);c__Saccharomycetes(100);o__Saccharomycetales(100);f__Metschnikowiaceae(100);g__Metschnikowia(100);s__Metschnikowia_pulcherrima(100); | Metschnikowia | | 13 | Saprotroph | \|Undefined Saprotroph\| | Yeast | NULL | Probable | As nectar-tap_saprotroph (Põlme et al. 2020); Decay sugar-rich substrates (Põlme et al. 2020) | Põlme S, et al. 2020. Fungal Diversity 105:1-16 ((https://doi.org/10.1007/s13225-020-00466-2)) |
| Otu49621 | 1 | k__Fungi(100);p__Ascomycota(100);c__Saccharomycetes(100);o__Saccharomycetales(100);f__Metschnikowiaceae(100);g__Metschnikowia(100);s__Metschnikowia_pulcherrima(100); | Metschnikowia | | 13 | Saprotroph | \|Undefined Saprotroph\| | Yeast | NULL | Probable | As nectar-tap_saprotroph (Põlme et al. 2020); Decay sugar-rich substrates (Põlme et al. 2020) | Põlme S, et al. 2020. Fungal Diversity 105:1-16 ((https://doi.org/10.1007/s13225-020-00466-2)) |
| Otu49622 | 1 | k__Fungi(100);p__Ascomycota(100);c__Saccharomycetes(100);o__Saccharomycetales(100);f__Metschnikowiaceae(100);g__Metschnikowia(100);g__Metschnikowia_unclassified(100); | Metschnikowia | | 13 | Saprotroph | \|Undefined Saprotroph\| | Yeast | NULL | Probable | As nectar-tap_saprotroph (Põlme et al. 2020); Decay sugar-rich substrates (Põlme et al. 2020) | Põlme S, et al. 2020. Fungal Diversity 105:1-16 ((https://doi.org/10.1007/s13225-020-00466-2)) |
| Otu49623 | 1 | k__Fungi(100);p__Ascomycota(100);c__Saccharomycetes(100);o__Saccharomycetales(100);f__Metschnikowiaceae(100);g__Metschnikowia(100);s__Metschnikowia_pulcherrima(100); | Metschnikowia | | 13 | Saprotroph | \|Undefined Saprotroph\| | Yeast | NULL | Probable | As nectar-tap_saprotroph (Põlme et al. 2020); Decay sugar-rich substrates (Põlme et al. 2020) | Põlme S, et al. 2020. Fungal Diversity 105:1-16 ((https://doi.org/10.1007/s13225-020-00466-2)) |
| Otu49624 | 1 | k__Fungi(100);p__Ascomycota(100);c__Saccharomycetes(100);o__Saccharomycetales(100);f__Metschnikowiaceae(100);g__Metschnikowia(100);s__Metschnikowia_pulcherrima(100); | Metschnikowia | | 13 | Saprotroph | \|Undefined Saprotroph\| | Yeast | NULL | Probable | As nectar-tap_saprotroph (Põlme et al. 2020); Decay sugar-rich substrates (Põlme et al. 2020) | Põlme S, et al. 2020. Fungal Diversity 105:1-16 ((https://doi.org/10.1007/s13225-020-00466-2)) |
| Otu49625 | 1 | k__Fungi(100);p__Ascomycota(100);c__Saccharomycetes(100);o__Saccharomycetales(100);f__Metschnikowiaceae(100);g__Metschnikowia(100);s__Metschnikowia_pulcherrima(100); | Metschnikowia | | 13 | Saprotroph | \|Undefined Saprotroph\| | Yeast | NULL | Probable | As nectar-tap_saprotroph (Põlme et al. 2020); Decay sugar-rich substrates (Põlme et al. 2020) | Põlme S, et al. 2020. Fungal Diversity 105:1-16 ((https://doi.org/10.1007/s13225-020-00466-2)) |
| Otu49626 | 1 | k__Fungi(100);p__Ascomycota(100);c__Saccharomycetes(100);o__Saccharomycetales(100);f__Metschnikowiaceae(100);g__Metschnikowia(100);g__Metschnikowia_unclassified(100); | Metschnikowia | | 13 | Saprotroph | \|Undefined Saprotroph\| | Yeast | NULL | Probable | As nectar-tap_saprotroph (Põlme et al. 2020); Decay sugar-rich substrates (Põlme et al. 2020) | Põlme S, et al. 2020. Fungal Diversity 105:1-16 ((https://doi.org/10.1007/s13225-020-00466-2)) |
| Otu49627 | 1 | k__Fungi(100);p__Ascomycota(100);c__Saccharomycetes(100);o__Saccharomycetales(100);f__Metschnikowiaceae(100);g__Metschnikowia(100);s__Metschnikowia_pulcherrima(100); | Metschnikowia | | 13 | Saprotroph | \|Undefined Saprotroph\| | Yeast | NULL | Probable | As nectar-tap_saprotroph (Põlme et al. 2020); Decay sugar-rich substrates (Põlme et al. 2020) | Põlme S, et al. 2020. Fungal Diversity 105:1-16 ((https://doi.org/10.1007/s13225-020-00466-2)) |
| Otu49628 | 1 | k__Fungi(100);p__Ascomycota(100);c__Saccharomycetes(100);o__Saccharomycetales(100);f__Metschnikowiaceae(100);g__Metschnikowia(100);g__Metschnikowia_unclassified(100); | Metschnikowia | | 13 | Saprotroph | \|Undefined Saprotroph\| | Yeast | NULL | Probable | As nectar-tap_saprotroph (Põlme et al. 2020); Decay sugar-rich substrates (Põlme et al. 2020) | Põlme S, et al. 2020. Fungal Diversity 105:1-16 ((https://doi.org/10.1007/s13225-020-00466-2)) |
| Otu49629 | 1 | k__Fungi(100);p__Ascomycota(100);c__Saccharomycetes(100);o__Saccharomycetales(100);f__Metschnikowiaceae(100);g__Metschnikowia(100);s__Metschnikowia_pulcherrima(100); | Metschnikowia | | 13 | Saprotroph | \|Undefined Saprotroph\| | Yeast | NULL | Probable | As nectar-tap_saprotroph (Põlme et al. 2020); Decay sugar-rich substrates (Põlme et al. 2020) | Põlme S, et al. 2020. Fungal Diversity 105:1-16 ((https://doi.org/10.1007/s13225-020-00466-2)) |
| Otu49630 | 1 | k__Fungi(100);p__Ascomycota(100);c__Saccharomycetes(100);o__Saccharomycetales(100);f__Metschnikowiaceae(100);g__Metschnikowia(100);s__Metschnikowia_pulcherrima(100); | Metschnikowia | | 13 | Saprotroph | \|Undefined Saprotroph\| | Yeast | NULL | Probable | As nectar-tap_saprotroph (Põlme et al. 2020); Decay sugar-rich substrates (Põlme et al. 2020) | Põlme S, et al. 2020. Fungal Diversity 105:1-16 ((https://doi.org/10.1007/s13225-020-00466-2)) |
| Otu49631 | 1 | k__Fungi(100);p__Ascomycota(100);c__Saccharomycetes(100);o__Saccharomycetales(100);f__Metschnikowiaceae(100);g__Metschnikowia(100);g__Metschnikowia_unclassified(100); | Metschnikowia | | 13 | Saprotroph | \|Undefined Saprotroph\| | Yeast | NULL | Probable | As nectar-tap_saprotroph (Põlme et al. 2020); Decay sugar-rich substrates (Põlme et al. 2020) | Põlme S, et al. 2020. Fungal Diversity 105:1-16 ((https://doi.org/10.1007/s13225-020-00466-2)) |
| Otu49632 | 1 | k__Fungi(100);p__Ascomycota(100);c__Saccharomycetes(100);o__Saccharomycetales(100);f__Metschnikowiaceae(100);g__Metschnikowia(100);s__Metschnikowia_pulcherrima(100); | Metschnikowia | | 13 | Saprotroph | \|Undefined Saprotroph\| | Yeast | NULL | Probable | As nectar-tap_saprotroph (Põlme et al. 2020); Decay sugar-rich substrates (Põlme et al. 2020) | Põlme S, et al. 2020. Fungal Diversity 105:1-16 ((https://doi.org/10.1007/s13225-020-00466-2)) |
| Otu49633 | 1 | k__Fungi(100);p__Ascomycota(100);c__Saccharomycetes(100);o__Saccharomycetales(100);f__Metschnikowiaceae(100);g__Metschnikowia(100);g__Metschnikowia_unclassified(100); | Metschnikowia | | 13 | Saprotroph | \|Undefined Saprotroph\| | Yeast | NULL | Probable | As nectar-tap_saprotroph (Põlme et al. 2020); Decay sugar-rich substrates (Põlme et al. 2020) | Põlme S, et al. 2020. Fungal Diversity 105:1-16 ((https://doi.org/10.1007/s13225-020-00466-2)) |
| Otu49634 | 1 | k__Fungi(100);p__Ascomycota(100);c__Saccharomycetes(100);o__Saccharomycetales(100);f__Metschnikowiaceae(100);g__Metschnikowia(100);g__Metschnikowia_unclassified(100); | Metschnikowia | | 13 | Saprotroph | \|Undefined Saprotroph\| | Yeast | NULL | Probable | As nectar-tap_saprotroph (Põlme et al. 2020); Decay sugar-rich substrates (Põlme et al. 2020) | Põlme S, et al. 2020. Fungal Diversity 105:1-16 ((https://doi.org/10.1007/s13225-020-00466-2)) |
| Otu49635 | 1 | k__Fungi(100);p__Ascomycota(100);c__Saccharomycetes(100);o__Saccharomycetales(100);f__Metschnikowiaceae(100);g__Metschnikowia(100);s__Metschnikowia_pulcherrima(100); | Metschnikowia | | 13 | Saprotroph | \|Undefined Saprotroph\| | Yeast | NULL | Probable | As nectar-tap_saprotroph (Põlme et al. 2020); Decay sugar-rich substrates (Põlme et al. 2020) | Põlme S, et al. 2020. Fungal Diversity 105:1-16 ((https://doi.org/10.1007/s13225-020-00466-2)) |
| Otu49636 | 1 | k__Fungi(100);p__Ascomycota(100);c__Saccharomycetes(100);o__Saccharomycetales(100);f__Metschnikowiaceae(100);g__Metschnikowia(100);s__Metschnikowia_pulcherrima(100); | Metschnikowia | | 13 | Saprotroph | \|Undefined Saprotroph\| | Yeast | NULL | Probable | As nectar-tap_saprotroph (Põlme et al. 2020); Decay sugar-rich substrates (Põlme et al. 2020) | Põlme S, et al. 2020. Fungal Diversity 105:1-16 ((https://doi.org/10.1007/s13225-020-00466-2)) |
| Otu49637 | 1 | k__Fungi(100);p__Ascomycota(100);c__Saccharomycetes(100);o__Saccharomycetales(100);f__Metschnikowiaceae(100);g__Metschnikowia(100);s__Metschnikowia_pulcherrima(100); | Metschnikowia | | 13 | Saprotroph | \|Undefined Saprotroph\| | Yeast | NULL | Probable | As nectar-tap_saprotroph (Põlme et al. 2020); Decay sugar-rich substrates (Põlme et al. 2020) | Põlme S, et al. 2020. Fungal Diversity 105:1-16 ((https://doi.org/10.1007/s13225-020-00466-2)) |
| Otu49638 | 1 | k__Fungi(100);p__Ascomycota(100);c__Saccharomycetes(100);o__Saccharomycetales(100);f__Metschnikowiaceae(100);g__Metschnikowia(100);s__Metschnikowia_pulcherrima(100); | Metschnikowia | | 13 | Saprotroph | \|Undefined Saprotroph\| | Yeast | NULL | Probable | As nectar-tap_saprotroph (Põlme et al. 2020); Decay sugar-rich substrates (Põlme et al. 2020) | Põlme S, et al. 2020. Fungal Diversity 105:1-16 ((https://doi.org/10.1007/s13225-020-00466-2)) |
| Otu49639 | 1 | k__Fungi(100);p__Ascomycota(100);c__Saccharomycetes(100);o__Saccharomycetales(100);f__Metschnikowiaceae(100);g__Metschnikowia(100);s__Metschnikowia_pulcherrima(100); | Metschnikowia | | 13 | Saprotroph | \|Undefined Saprotroph\| | Yeast | NULL | Probable | As nectar-tap_saprotroph (Põlme et al. 2020); Decay sugar-rich substrates (Põlme et al. 2020) | Põlme S, et al. 2020. Fungal Diversity 105:1-16 ((https://doi.org/10.1007/s13225-020-00466-2)) |
| Otu49641 | 1 | k__Fungi(100);p__Ascomycota(100);c__Saccharomycetes(100);o__Saccharomycetales(100);f__Metschnikowiaceae(100);g__Metschnikowia(100);s__Metschnikowia_pulcherrima(100); | Metschnikowia | | 13 | Saprotroph | \|Undefined Saprotroph\| | Yeast | NULL | Probable | As nectar-tap_saprotroph (Põlme et al. 2020); Decay sugar-rich substrates (Põlme et al. 2020) | Põlme S, et al. 2020. Fungal Diversity 105:1-16 ((https://doi.org/10.1007/s13225-020-00466-2)) |
| Otu49642 | 1 | k__Fungi(100);p__Ascomycota(100);c__Saccharomycetes(100);o__Saccharomycetales(100);f__Metschnikowiaceae(100);g__Metschnikowia(100);s__Metschnikowia_pulcherrima(100); | Metschnikowia | | 13 | Saprotroph | \|Undefined Saprotroph\| | Yeast | NULL | Probable | As nectar-tap_saprotroph (Põlme et al. 2020); Decay sugar-rich substrates (Põlme et al. 2020) | Põlme S, et al. 2020. Fungal Diversity 105:1-16 ((https://doi.org/10.1007/s13225-020-00466-2)) |
| Otu49643 | 1 | k__Fungi(100);p__Ascomycota(100);c__Saccharomycetes(100);o__Saccharomycetales(100);f__Metschnikowiaceae(100);g__Metschnikowia(100);s__Metschnikowia_pulcherrima(100); | Metschnikowia | | 13 | Saprotroph | \|Undefined Saprotroph\| | Yeast | NULL | Probable | As nectar-tap_saprotroph (Põlme et al. 2020); Decay sugar-rich substrates (Põlme et al. 2020) | Põlme S, et al. 2020. Fungal Diversity 105:1-16 ((https://doi.org/10.1007/s13225-020-00466-2)) |
| Otu49644 | 1 | k__Fungi(100);p__Ascomycota(100);c__Saccharomycetes(100);o__Saccharomycetales(100);f__Metschnikowiaceae(100);g__Metschnikowia(100);g__Metschnikowia_unclassified(100); | Metschnikowia | | 13 | Saprotroph | \|Undefined Saprotroph\| | Yeast | NULL | Probable | As nectar-tap_saprotroph (Põlme et al. 2020); Decay sugar-rich substrates (Põlme et al. 2020) | Põlme S, et al. 2020. Fungal Diversity 105:1-16 ((https://doi.org/10.1007/s13225-020-00466-2)) |
| Otu49645 | 1 | k__Fungi(100);p__Ascomycota(100);c__Saccharomycetes(100);o__Saccharomycetales(100);f__Metschnikowiaceae(100);g__Metschnikowia(100);g__Metschnikowia_unclassified(100); | Metschnikowia | | 13 | Saprotroph | \|Undefined Saprotroph\| | Yeast | NULL | Probable | As nectar-tap_saprotroph (Põlme et al. 2020); Decay sugar-rich substrates (Põlme et al. 2020) | Põlme S, et al. 2020. Fungal Diversity 105:1-16 ((https://doi.org/10.1007/s13225-020-00466-2)) |
| Otu49646 | 1 | k__Fungi(100);p__Ascomycota(100);c__Saccharomycetes(100);o__Saccharomycetales(100);f__Metschnikowiaceae(100);g__Metschnikowia(100);g__Metschnikowia_unclassified(100); | Metschnikowia | | 13 | Saprotroph | \|Undefined Saprotroph\| | Yeast | NULL | Probable | As nectar-tap_saprotroph (Põlme et al. 2020); Decay sugar-rich substrates (Põlme et al. 2020) | Põlme S, et al. 2020. Fungal Diversity 105:1-16 ((https://doi.org/10.1007/s13225-020-00466-2)) |
| Otu49647 | 1 | k__Fungi(100);p__Ascomycota(100);c__Saccharomycetes(100);o__Saccharomycetales(100);f__Metschnikowiaceae(100);g__Metschnikowia(100);s__Metschnikowia_pulcherrima(100); | Metschnikowia | | 13 | Saprotroph | \|Undefined Saprotroph\| | Yeast | NULL | Probable | As nectar-tap_saprotroph (Põlme et al. 2020); Decay sugar-rich substrates (Põlme et al. 2020) | Põlme S, et al. 2020. Fungal Diversity 105:1-16 ((https://doi.org/10.1007/s13225-020-00466-2)) |
| Otu49648 | 1 | k__Fungi(100);p__Ascomycota(100);c__Saccharomycetes(100);o__Saccharomycetales(100);f__Metschnikowiaceae(100);g__Metschnikowia(100);g__Metschnikowia_unclassified(100); | Metschnikowia | | 13 | Saprotroph | \|Undefined Saprotroph\| | Yeast | NULL | Probable | As nectar-tap_saprotroph (Põlme et al. 2020); Decay sugar-rich substrates (Põlme et al. 2020) | Põlme S, et al. 2020. Fungal Diversity 105:1-16 ((https://doi.org/10.1007/s13225-020-00466-2)) |
| Otu49649 | 1 | k__Fungi(100);p__Ascomycota(100);c__Saccharomycetes(100);o__Saccharomycetales(100);f__Metschnikowiaceae(100);g__Metschnikowia(100);g__Metschnikowia_unclassified(100); | Metschnikowia | | 13 | Saprotroph | \|Undefined Saprotroph\| | Yeast | NULL | Probable | As nectar-tap_saprotroph (Põlme et al. 2020); Decay sugar-rich substrates (Põlme et al. 2020) | Põlme S, et al. 2020. Fungal Diversity 105:1-16 ((https://doi.org/10.1007/s13225-020-00466-2)) |
| Otu49650 | 1 | k__Fungi(100);p__Ascomycota(100);c__Saccharomycetes(100);o__Saccharomycetales(100);f__Metschnikowiaceae(100);g__Metschnikowia(100);g__Metschnikowia_unclassified(100); | Metschnikowia | | 13 | Saprotroph | \|Undefined Saprotroph\| | Yeast | NULL | Probable | As nectar-tap_saprotroph (Põlme et al. 2020); Decay sugar-rich substrates (Põlme et al. 2020) | Põlme S, et al. 2020. Fungal Diversity 105:1-16 ((https://doi.org/10.1007/s13225-020-00466-2)) |
| Otu49651 | 1 | k__Fungi(100);p__Ascomycota(100);c__Saccharomycetes(100);o__Saccharomycetales(100);f__Metschnikowiaceae(100);g__Metschnikowia(100);g__Metschnikowia_unclassified(100); | Metschnikowia | | 13 | Saprotroph | \|Undefined Saprotroph\| | Yeast | NULL | Probable | As nectar-tap_saprotroph (Põlme et al. 2020); Decay sugar-rich substrates (Põlme et al. 2020) | Põlme S, et al. 2020. Fungal Diversity 105:1-16 ((https://doi.org/10.1007/s13225-020-00466-2)) |
| Otu49652 | 1 | k__Fungi(100);p__Ascomycota(100);c__Saccharomycetes(100);o__Saccharomycetales(100);f__Metschnikowiaceae(100);g__Metschnikowia(100);s__Metschnikowia_pulcherrima(100); | Metschnikowia | | 13 | Saprotroph | \|Undefined Saprotroph\| | Yeast | NULL | Probable | As nectar-tap_saprotroph (Põlme et al. 2020); Decay sugar-rich substrates (Põlme et al. 2020) | Põlme S, et al. 2020. Fungal Diversity 105:1-16 ((https://doi.org/10.1007/s13225-020-00466-2)) |
| Otu49653 | 1 | k__Fungi(100);p__Ascomycota(100);c__Saccharomycetes(100);o__Saccharomycetales(100);f__Metschnikowiaceae(100);g__Metschnikowia(100);s__Metschnikowia_pulcherrima(100); | Metschnikowia | | 13 | Saprotroph | \|Undefined Saprotroph\| | Yeast | NULL | Probable | As nectar-tap_saprotroph (Põlme et al. 2020); Decay sugar-rich substrates (Põlme et al. 2020) | Põlme S, et al. 2020. Fungal Diversity 105:1-16 ((https://doi.org/10.1007/s13225-020-00466-2)) |
| Otu49654 | 1 | k__Fungi(100);p__Ascomycota(100);c__Saccharomycetes(100);o__Saccharomycetales(100);f__Metschnikowiaceae(100);g__Metschnikowia(100);s__Metschnikowia_pulcherrima(100); | Metschnikowia | | 13 | Saprotroph | \|Undefined Saprotroph\| | Yeast | NULL | Probable | As nectar-tap_saprotroph (Põlme et al. 2020); Decay sugar-rich substrates (Põlme et al. 2020) | Põlme S, et al. 2020. Fungal Diversity 105:1-16 ((https://doi.org/10.1007/s13225-020-00466-2)) |
| Otu49655 | 1 | k__Fungi(100);p__Ascomycota(100);c__Saccharomycetes(100);o__Saccharomycetales(100);f__Metschnikowiaceae(100);g__Metschnikowia(100);s__Metschnikowia_pulcherrima(100); | Metschnikowia | | 13 | Saprotroph | \|Undefined Saprotroph\| | Yeast | NULL | Probable | As nectar-tap_saprotroph (Põlme et al. 2020); Decay sugar-rich substrates (Põlme et al. 2020) | Põlme S, et al. 2020. Fungal Diversity 105:1-16 ((https://doi.org/10.1007/s13225-020-00466-2)) |
| Otu49656 | 1 | k__Fungi(100);p__Ascomycota(100);c__Saccharomycetes(100);o__Saccharomycetales(100);f__Metschnikowiaceae(100);g__Metschnikowia(100);g__Metschnikowia_unclassified(100); | Metschnikowia | | 13 | Saprotroph | \|Undefined Saprotroph\| | Yeast | NULL | Probable | As nectar-tap_saprotroph (Põlme et al. 2020); Decay sugar-rich substrates (Põlme et al. 2020) | Põlme S, et al. 2020. Fungal Diversity 105:1-16 ((https://doi.org/10.1007/s13225-020-00466-2)) |
| Otu49657 | 1 | k__Fungi(100);p__Ascomycota(100);c__Saccharomycetes(100);o__Saccharomycetales(100);f__Metschnikowiaceae(100);g__Metschnikowia(100);s__Metschnikowia_pulcherrima(100); | Metschnikowia | | 13 | Saprotroph | \|Undefined Saprotroph\| | Yeast | NULL | Probable | As nectar-tap_saprotroph (Põlme et al. 2020); Decay sugar-rich substrates (Põlme et al. 2020) | Põlme S, et al. 2020. Fungal Diversity 105:1-16 ((https://doi.org/10.1007/s13225-020-00466-2)) |
| Otu49658 | 1 | k__Fungi(100);p__Ascomycota(100);c__Saccharomycetes(100);o__Saccharomycetales(100);f__Metschnikowiaceae(100);g__Metschnikowia(100);s__Metschnikowia_pulcherrima(100); | Metschnikowia | | 13 | Saprotroph | \|Undefined Saprotroph\| | Yeast | NULL | Probable | As nectar-tap_saprotroph (Põlme et al. 2020); Decay sugar-rich substrates (Põlme et al. 2020) | Põlme S, et al. 2020. Fungal Diversity 105:1-16 ((https://doi.org/10.1007/s13225-020-00466-2)) |
| Otu49659 | 1 | k__Fungi(100);p__Ascomycota(100);c__Saccharomycetes(100);o__Saccharomycetales(100);f__Metschnikowiaceae(100);g__Metschnikowia(100);s__Metschnikowia_pulcherrima(100); | Metschnikowia | | 13 | Saprotroph | \|Undefined Saprotroph\| | Yeast | NULL | Probable | As nectar-tap_saprotroph (Põlme et al. 2020); Decay sugar-rich substrates (Põlme et al. 2020) | Põlme S, et al. 2020. Fungal Diversity 105:1-16 ((https://doi.org/10.1007/s13225-020-00466-2)) |
| Otu49660 | 1 | k__Fungi(100);p__Ascomycota(100);c__Saccharomycetes(100);o__Saccharomycetales(100);f__Metschnikowiaceae(100);g__Metschnikowia(100);g__Metschnikowia_unclassified(100); | Metschnikowia | | 13 | Saprotroph | \|Undefined Saprotroph\| | Yeast | NULL | Probable | As nectar-tap_saprotroph (Põlme et al. 2020); Decay sugar-rich substrates (Põlme et al. 2020) | Põlme S, et al. 2020. Fungal Diversity 105:1-16 ((https://doi.org/10.1007/s13225-020-00466-2)) |
| Otu49661 | 1 | k__Fungi(100);p__Ascomycota(100);c__Saccharomycetes(100);o__Saccharomycetales(100);f__Metschnikowiaceae(100);g__Metschnikowia(100);s__Metschnikowia_pulcherrima(100); | Metschnikowia | | 13 | Saprotroph | \|Undefined Saprotroph\| | Yeast | NULL | Probable | As nectar-tap_saprotroph (Põlme et al. 2020); Decay sugar-rich substrates (Põlme et al. 2020) | Põlme S, et al. 2020. Fungal Diversity 105:1-16 ((https://doi.org/10.1007/s13225-020-00466-2)) |
| Otu49662 | 1 | k__Fungi(100);p__Ascomycota(100);c__Saccharomycetes(100);o__Saccharomycetales(100);f__Metschnikowiaceae(100);g__Metschnikowia(100);s__Metschnikowia_pulcherrima(100); | Metschnikowia | | 13 | Saprotroph | \|Undefined Saprotroph\| | Yeast | NULL | Probable | As nectar-tap_saprotroph (Põlme et al. 2020); Decay sugar-rich substrates (Põlme et al. 2020) | Põlme S, et al. 2020. Fungal Diversity 105:1-16 ((https://doi.org/10.1007/s13225-020-00466-2)) |
| Otu49663 | 1 | k__Fungi(100);p__Ascomycota(100);c__Saccharomycetes(100);o__Saccharomycetales(100);f__Metschnikowiaceae(100);g__Metschnikowia(100);s__Metschnikowia_pulcherrima(100); | Metschnikowia | | 13 | Saprotroph | \|Undefined Saprotroph\| | Yeast | NULL | Probable | As nectar-tap_saprotroph (Põlme et al. 2020); Decay sugar-rich substrates (Põlme et al. 2020) | Põlme S, et al. 2020. Fungal Diversity 105:1-16 ((https://doi.org/10.1007/s13225-020-00466-2)) |
| Otu49664 | 1 | k__Fungi(100);p__Ascomycota(100);c__Saccharomycetes(100);o__Saccharomycetales(100);f__Metschnikowiaceae(100);g__Metschnikowia(100);s__Metschnikowia_pulcherrima(100); | Metschnikowia | | 13 | Saprotroph | \|Undefined Saprotroph\| | Yeast | NULL | Probable | As nectar-tap_saprotroph (Põlme et al. 2020); Decay sugar-rich substrates (Põlme et al. 2020) | Põlme S, et al. 2020. Fungal Diversity 105:1-16 ((https://doi.org/10.1007/s13225-020-00466-2)) |
| Otu49665 | 1 | k__Fungi(100);p__Ascomycota(100);c__Saccharomycetes(100);o__Saccharomycetales(100);f__Metschnikowiaceae(100);g__Metschnikowia(100);g__Metschnikowia_unclassified(100); | Metschnikowia | | 13 | Saprotroph | \|Undefined Saprotroph\| | Yeast | NULL | Probable | As nectar-tap_saprotroph (Põlme et al. 2020); Decay sugar-rich substrates (Põlme et al. 2020) | Põlme S, et al. 2020. Fungal Diversity 105:1-16 ((https://doi.org/10.1007/s13225-020-00466-2)) |
| Otu49666 | 1 | k__Fungi(100);p__Ascomycota(100);c__Saccharomycetes(100);o__Saccharomycetales(100);f__Metschnikowiaceae(100);g__Metschnikowia(100);s__Metschnikowia_pulcherrima(100); | Metschnikowia | | 13 | Saprotroph | \|Undefined Saprotroph\| | Yeast | NULL | Probable | As nectar-tap_saprotroph (Põlme et al. 2020); Decay sugar-rich substrates (Põlme et al. 2020) | Põlme S, et al. 2020. Fungal Diversity 105:1-16 ((https://doi.org/10.1007/s13225-020-00466-2)) |
| Otu49667 | 1 | k__Fungi(100);p__Ascomycota(100);c__Saccharomycetes(100);o__Saccharomycetales(100);f__Metschnikowiaceae(100);g__Metschnikowia(100);s__Metschnikowia_pulcherrima(100); | Metschnikowia | | 13 | Saprotroph | \|Undefined Saprotroph\| | Yeast | NULL | Probable | As nectar-tap_saprotroph (Põlme et al. 2020); Decay sugar-rich substrates (Põlme et al. 2020) | Põlme S, et al. 2020. Fungal Diversity 105:1-16 ((https://doi.org/10.1007/s13225-020-00466-2)) |
| Otu49668 | 1 | k__Fungi(100);p__Ascomycota(100);c__Saccharomycetes(100);o__Saccharomycetales(100);f__Metschnikowiaceae(100);g__Metschnikowia(100);s__Metschnikowia_pulcherrima(100); | Metschnikowia | | 13 | Saprotroph | \|Undefined Saprotroph\| | Yeast | NULL | Probable | As nectar-tap_saprotroph (Põlme et al. 2020); Decay sugar-rich substrates (Põlme et al. 2020) | Põlme S, et al. 2020. Fungal Diversity 105:1-16 ((https://doi.org/10.1007/s13225-020-00466-2)) |
| Otu49669 | 1 | k__Fungi(100);p__Ascomycota(100);c__Saccharomycetes(100);o__Saccharomycetales(100);f__Metschnikowiaceae(100);g__Metschnikowia(100);s__Metschnikowia_pulcherrima(100); | Metschnikowia | | 13 | Saprotroph | \|Undefined Saprotroph\| | Yeast | NULL | Probable | As nectar-tap_saprotroph (Põlme et al. 2020); Decay sugar-rich substrates (Põlme et al. 2020) | Põlme S, et al. 2020. Fungal Diversity 105:1-16 ((https://doi.org/10.1007/s13225-020-00466-2)) |
| Otu49670 | 1 | k__Fungi(100);p__Ascomycota(100);c__Saccharomycetes(100);o__Saccharomycetales(100);f__Metschnikowiaceae(100);g__Metschnikowia(100);s__Metschnikowia_pulcherrima(100); | Metschnikowia | | 13 | Saprotroph | \|Undefined Saprotroph\| | Yeast | NULL | Probable | As nectar-tap_saprotroph (Põlme et al. 2020); Decay sugar-rich substrates (Põlme et al. 2020) | Põlme S, et al. 2020. Fungal Diversity 105:1-16 ((https://doi.org/10.1007/s13225-020-00466-2)) |
| Otu49671 | 1 | k__Fungi(100);p__Ascomycota(100);c__Saccharomycetes(100);o__Saccharomycetales(100);f__Metschnikowiaceae(100);g__Metschnikowia(100);g__Metschnikowia_unclassified(100); | Metschnikowia | | 13 | Saprotroph | \|Undefined Saprotroph\| | Yeast | NULL | Probable | As nectar-tap_saprotroph (Põlme et al. 2020); Decay sugar-rich substrates (Põlme et al. 2020) | Põlme S, et al. 2020. Fungal Diversity 105:1-16 ((https://doi.org/10.1007/s13225-020-00466-2)) |
| Otu49672 | 1 | k__Fungi(100);p__Ascomycota(100);c__Saccharomycetes(100);o__Saccharomycetales(100);f__Metschnikowiaceae(100);g__Metschnikowia(100);g__Metschnikowia_unclassified(100); | Metschnikowia | | 13 | Saprotroph | \|Undefined Saprotroph\| | Yeast | NULL | Probable | As nectar-tap_saprotroph (Põlme et al. 2020); Decay sugar-rich substrates (Põlme et al. 2020) | Põlme S, et al. 2020. Fungal Diversity 105:1-16 ((https://doi.org/10.1007/s13225-020-00466-2)) |
| Otu49673 | 1 | k__Fungi(100);p__Ascomycota(100);c__Saccharomycetes(100);o__Saccharomycetales(100);f__Metschnikowiaceae(100);g__Metschnikowia(100);g__Metschnikowia_unclassified(100); | Metschnikowia | | 13 | Saprotroph | \|Undefined Saprotroph\| | Yeast | NULL | Probable | As nectar-tap_saprotroph (Põlme et al. 2020); Decay sugar-rich substrates (Põlme et al. 2020) | Põlme S, et al. 2020. Fungal Diversity 105:1-16 ((https://doi.org/10.1007/s13225-020-00466-2)) |
| Otu49674 | 1 | k__Fungi(100);p__Ascomycota(100);c__Saccharomycetes(100);o__Saccharomycetales(100);f__Metschnikowiaceae(100);g__Metschnikowia(100);g__Metschnikowia_unclassified(100); | Metschnikowia | | 13 | Saprotroph | \|Undefined Saprotroph\| | Yeast | NULL | Probable | As nectar-tap_saprotroph (Põlme et al. 2020); Decay sugar-rich substrates (Põlme et al. 2020) | Põlme S, et al. 2020. Fungal Diversity 105:1-16 ((https://doi.org/10.1007/s13225-020-00466-2)) |
| Otu49675 | 1 | k__Fungi(100);p__Ascomycota(100);c__Saccharomycetes(100);o__Saccharomycetales(100);f__Metschnikowiaceae(100);g__Metschnikowia(100);g__Metschnikowia_unclassified(100); | Metschnikowia | | 13 | Saprotroph | \|Undefined Saprotroph\| | Yeast | NULL | Probable | As nectar-tap_saprotroph (Põlme et al. 2020); Decay sugar-rich substrates (Põlme et al. 2020) | Põlme S, et al. 2020. Fungal Diversity 105:1-16 ((https://doi.org/10.1007/s13225-020-00466-2)) |
| Otu49676 | 1 | k__Fungi(100);p__Ascomycota(100);c__Saccharomycetes(100);o__Saccharomycetales(100);f__Metschnikowiaceae(100);g__Metschnikowia(100);s__Metschnikowia_pulcherrima(100); | Metschnikowia | | 13 | Saprotroph | \|Undefined Saprotroph\| | Yeast | NULL | Probable | As nectar-tap_saprotroph (Põlme et al. 2020); Decay sugar-rich substrates (Põlme et al. 2020) | Põlme S, et al. 2020. Fungal Diversity 105:1-16 ((https://doi.org/10.1007/s13225-020-00466-2)) |
| Otu49677 | 1 | k__Fungi(100);p__Ascomycota(100);c__Saccharomycetes(100);o__Saccharomycetales(100);f__Metschnikowiaceae(100);g__Metschnikowia(100);s__Metschnikowia_pulcherrima(100); | Metschnikowia | | 13 | Saprotroph | \|Undefined Saprotroph\| | Yeast | NULL | Probable | As nectar-tap_saprotroph (Põlme et al. 2020); Decay sugar-rich substrates (Põlme et al. 2020) | Põlme S, et al. 2020. Fungal Diversity 105:1-16 ((https://doi.org/10.1007/s13225-020-00466-2)) |
| Otu49678 | 1 | k__Fungi(100);p__Ascomycota(100);c__Saccharomycetes(100);o__Saccharomycetales(100);f__Metschnikowiaceae(100);g__Metschnikowia(100);s__Metschnikowia_pulcherrima(100); | Metschnikowia | | 13 | Saprotroph | \|Undefined Saprotroph\| | Yeast | NULL | Probable | As nectar-tap_saprotroph (Põlme et al. 2020); Decay sugar-rich substrates (Põlme et al. 2020) | Põlme S, et al. 2020. Fungal Diversity 105:1-16 ((https://doi.org/10.1007/s13225-020-00466-2)) |
| Otu49679 | 1 | k__Fungi(100);p__Ascomycota(100);c__Saccharomycetes(100);o__Saccharomycetales(100);f__Metschnikowiaceae(100);g__Metschnikowia(100);s__Metschnikowia_pulcherrima(100); | Metschnikowia | | 13 | Saprotroph | \|Undefined Saprotroph\| | Yeast | NULL | Probable | As nectar-tap_saprotroph (Põlme et al. 2020); Decay sugar-rich substrates (Põlme et al. 2020) | Põlme S, et al. 2020. Fungal Diversity 105:1-16 ((https://doi.org/10.1007/s13225-020-00466-2)) |
| Otu49680 | 1 | k__Fungi(100);p__Ascomycota(100);c__Saccharomycetes(100);o__Saccharomycetales(100);f__Metschnikowiaceae(100);g__Metschnikowia(100);g__Metschnikowia_unclassified(100); | Metschnikowia | | 13 | Saprotroph | \|Undefined Saprotroph\| | Yeast | NULL | Probable | As nectar-tap_saprotroph (Põlme et al. 2020); Decay sugar-rich substrates (Põlme et al. 2020) | Põlme S, et al. 2020. Fungal Diversity 105:1-16 ((https://doi.org/10.1007/s13225-020-00466-2)) |
| Otu49681 | 1 | k__Fungi(100);p__Ascomycota(100);c__Saccharomycetes(100);o__Saccharomycetales(100);f__Metschnikowiaceae(100);g__Metschnikowia(100);s__Metschnikowia_pulcherrima(100); | Metschnikowia | | 13 | Saprotroph | \|Undefined Saprotroph\| | Yeast | NULL | Probable | As nectar-tap_saprotroph (Põlme et al. 2020); Decay sugar-rich substrates (Põlme et al. 2020) | Põlme S, et al. 2020. Fungal Diversity 105:1-16 ((https://doi.org/10.1007/s13225-020-00466-2)) |
| Otu49682 | 1 | k__Fungi(100);p__Ascomycota(100);c__Saccharomycetes(100);o__Saccharomycetales(100);f__Metschnikowiaceae(100);g__Metschnikowia(100);s__Metschnikowia_pulcherrima(100); | Metschnikowia | | 13 | Saprotroph | \|Undefined Saprotroph\| | Yeast | NULL | Probable | As nectar-tap_saprotroph (Põlme et al. 2020); Decay sugar-rich substrates (Põlme et al. 2020) | Põlme S, et al. 2020. Fungal Diversity 105:1-16 ((https://doi.org/10.1007/s13225-020-00466-2)) |
| Otu49683 | 1 | k__Fungi(100);p__Ascomycota(100);c__Saccharomycetes(100);o__Saccharomycetales(100);f__Metschnikowiaceae(100);g__Metschnikowia(100);s__Metschnikowia_pulcherrima(100); | Metschnikowia | | 13 | Saprotroph | \|Undefined Saprotroph\| | Yeast | NULL | Probable | As nectar-tap_saprotroph (Põlme et al. 2020); Decay sugar-rich substrates (Põlme et al. 2020) | Põlme S, et al. 2020. Fungal Diversity 105:1-16 ((https://doi.org/10.1007/s13225-020-00466-2)) |
| Otu49684 | 1 | k__Fungi(100);p__Ascomycota(100);c__Saccharomycetes(100);o__Saccharomycetales(100);f__Metschnikowiaceae(100);g__Metschnikowia(100);g__Metschnikowia_unclassified(100); | Metschnikowia | | 13 | Saprotroph | \|Undefined Saprotroph\| | Yeast | NULL | Probable | As nectar-tap_saprotroph (Põlme et al. 2020); Decay sugar-rich substrates (Põlme et al. 2020) | Põlme S, et al. 2020. Fungal Diversity 105:1-16 ((https://doi.org/10.1007/s13225-020-00466-2)) |
| Otu49685 | 1 | k__Fungi(100);p__Ascomycota(100);c__Saccharomycetes(100);o__Saccharomycetales(100);f__Metschnikowiaceae(100);g__Metschnikowia(100);g__Metschnikowia_unclassified(100); | Metschnikowia | | 13 | Saprotroph | \|Undefined Saprotroph\| | Yeast | NULL | Probable | As nectar-tap_saprotroph (Põlme et al. 2020); Decay sugar-rich substrates (Põlme et al. 2020) | Põlme S, et al. 2020. Fungal Diversity 105:1-16 ((https://doi.org/10.1007/s13225-020-00466-2)) |
| Otu49686 | 1 | k__Fungi(100);p__Ascomycota(100);c__Saccharomycetes(100);o__Saccharomycetales(100);f__Metschnikowiaceae(100);g__Metschnikowia(100);s__Metschnikowia_pulcherrima(100); | Metschnikowia | | 13 | Saprotroph | \|Undefined Saprotroph\| | Yeast | NULL | Probable | As nectar-tap_saprotroph (Põlme et al. 2020); Decay sugar-rich substrates (Põlme et al. 2020) | Põlme S, et al. 2020. Fungal Diversity 105:1-16 ((https://doi.org/10.1007/s13225-020-00466-2)) |
| Otu49687 | 1 | k__Fungi(100);p__Ascomycota(100);c__Saccharomycetes(100);o__Saccharomycetales(100);f__Metschnikowiaceae(100);g__Metschnikowia(100);g__Metschnikowia_unclassified(100); | Metschnikowia | | 13 | Saprotroph | \|Undefined Saprotroph\| | Yeast | NULL | Probable | As nectar-tap_saprotroph (Põlme et al. 2020); Decay sugar-rich substrates (Põlme et al. 2020) | Põlme S, et al. 2020. Fungal Diversity 105:1-16 ((https://doi.org/10.1007/s13225-020-00466-2)) |
| Otu49688 | 1 | k__Fungi(100);p__Ascomycota(100);c__Saccharomycetes(100);o__Saccharomycetales(100);f__Metschnikowiaceae(100);g__Metschnikowia(100);s__Metschnikowia_pulcherrima(100); | Metschnikowia | | 13 | Saprotroph | \|Undefined Saprotroph\| | Yeast | NULL | Probable | As nectar-tap_saprotroph (Põlme et al. 2020); Decay sugar-rich substrates (Põlme et al. 2020) | Põlme S, et al. 2020. Fungal Diversity 105:1-16 ((https://doi.org/10.1007/s13225-020-00466-2)) |
| Otu49689 | 1 | k__Fungi(100);p__Ascomycota(100);c__Saccharomycetes(100);o__Saccharomycetales(100);f__Metschnikowiaceae(100);g__Metschnikowia(100);s__Metschnikowia_pulcherrima(100); | Metschnikowia | | 13 | Saprotroph | \|Undefined Saprotroph\| | Yeast | NULL | Probable | As nectar-tap_saprotroph (Põlme et al. 2020); Decay sugar-rich substrates (Põlme et al. 2020) | Põlme S, et al. 2020. Fungal Diversity 105:1-16 ((https://doi.org/10.1007/s13225-020-00466-2)) |
| Otu49690 | 1 | k__Fungi(100);p__Ascomycota(100);c__Saccharomycetes(100);o__Saccharomycetales(100);f__Metschnikowiaceae(100);g__Metschnikowia(100);s__Metschnikowia_pulcherrima(100); | Metschnikowia | | 13 | Saprotroph | \|Undefined Saprotroph\| | Yeast | NULL | Probable | As nectar-tap_saprotroph (Põlme et al. 2020); Decay sugar-rich substrates (Põlme et al. 2020) | Põlme S, et al. 2020. Fungal Diversity 105:1-16 ((https://doi.org/10.1007/s13225-020-00466-2)) |
| Otu49691 | 1 | k__Fungi(100);p__Ascomycota(100);c__Saccharomycetes(100);o__Saccharomycetales(100);f__Metschnikowiaceae(100);g__Metschnikowia(100);s__Metschnikowia_pulcherrima(100); | Metschnikowia | | 13 | Saprotroph | \|Undefined Saprotroph\| | Yeast | NULL | Probable | As nectar-tap_saprotroph (Põlme et al. 2020); Decay sugar-rich substrates (Põlme et al. 2020) | Põlme S, et al. 2020. Fungal Diversity 105:1-16 ((https://doi.org/10.1007/s13225-020-00466-2)) |
| Otu49692 | 1 | k__Fungi(100);p__Ascomycota(100);c__Saccharomycetes(100);o__Saccharomycetales(100);f__Metschnikowiaceae(100);g__Metschnikowia(100);s__Metschnikowia_pulcherrima(100); | Metschnikowia | | 13 | Saprotroph | \|Undefined Saprotroph\| | Yeast | NULL | Probable | As nectar-tap_saprotroph (Põlme et al. 2020); Decay sugar-rich substrates (Põlme et al. 2020) | Põlme S, et al. 2020. Fungal Diversity 105:1-16 ((https://doi.org/10.1007/s13225-020-00466-2)) |
| Otu49693 | 1 | k__Fungi(100);p__Ascomycota(100);c__Saccharomycetes(100);o__Saccharomycetales(100);f__Metschnikowiaceae(100);g__Metschnikowia(100);s__Metschnikowia_pulcherrima(100); | Metschnikowia | | 13 | Saprotroph | \|Undefined Saprotroph\| | Yeast | NULL | Probable | As nectar-tap_saprotroph (Põlme et al. 2020); Decay sugar-rich substrates (Põlme et al. 2020) | Põlme S, et al. 2020. Fungal Diversity 105:1-16 ((https://doi.org/10.1007/s13225-020-00466-2)) |
| Otu49694 | 1 | k__Fungi(100);p__Ascomycota(100);c__Saccharomycetes(100);o__Saccharomycetales(100);f__Metschnikowiaceae(100);g__Metschnikowia(100);s__Metschnikowia_pulcherrima(100); | Metschnikowia | | 13 | Saprotroph | \|Undefined Saprotroph\| | Yeast | NULL | Probable | As nectar-tap_saprotroph (Põlme et al. 2020); Decay sugar-rich substrates (Põlme et al. 2020) | Põlme S, et al. 2020. Fungal Diversity 105:1-16 ((https://doi.org/10.1007/s13225-020-00466-2)) |
| Otu49695 | 1 | k__Fungi(100);p__Ascomycota(100);c__Saccharomycetes(100);o__Saccharomycetales(100);f__Metschnikowiaceae(100);g__Metschnikowia(100);s__Metschnikowia_pulcherrima(100); | Metschnikowia | | 13 | Saprotroph | \|Undefined Saprotroph\| | Yeast | NULL | Probable | As nectar-tap_saprotroph (Põlme et al. 2020); Decay sugar-rich substrates (Põlme et al. 2020) | Põlme S, et al. 2020. Fungal Diversity 105:1-16 ((https://doi.org/10.1007/s13225-020-00466-2)) |
| Otu49696 | 1 | k__Fungi(100);p__Ascomycota(100);c__Saccharomycetes(100);o__Saccharomycetales(100);f__Metschnikowiaceae(100);g__Metschnikowia(100);s__Metschnikowia_pulcherrima(100); | Metschnikowia | | 13 | Saprotroph | \|Undefined Saprotroph\| | Yeast | NULL | Probable | As nectar-tap_saprotroph (Põlme et al. 2020); Decay sugar-rich substrates (Põlme et al. 2020) | Põlme S, et al. 2020. Fungal Diversity 105:1-16 ((https://doi.org/10.1007/s13225-020-00466-2)) |
| Otu49697 | 1 | k__Fungi(100);p__Ascomycota(100);c__Saccharomycetes(100);o__Saccharomycetales(100);f__Metschnikowiaceae(100);g__Metschnikowia(100);s__Metschnikowia_pulcherrima(100); | Metschnikowia | | 13 | Saprotroph | \|Undefined Saprotroph\| | Yeast | NULL | Probable | As nectar-tap_saprotroph (Põlme et al. 2020); Decay sugar-rich substrates (Põlme et al. 2020) | Põlme S, et al. 2020. Fungal Diversity 105:1-16 ((https://doi.org/10.1007/s13225-020-00466-2)) |
| Otu49698 | 1 | k__Fungi(100);p__Ascomycota(100);c__Saccharomycetes(100);o__Saccharomycetales(100);f__Metschnikowiaceae(100);g__Metschnikowia(100);s__Metschnikowia_pulcherrima(100); | Metschnikowia | | 13 | Saprotroph | \|Undefined Saprotroph\| | Yeast | NULL | Probable | As nectar-tap_saprotroph (Põlme et al. 2020); Decay sugar-rich substrates (Põlme et al. 2020) | Põlme S, et al. 2020. Fungal Diversity 105:1-16 ((https://doi.org/10.1007/s13225-020-00466-2)) |
| Otu49699 | 1 | k__Fungi(100);p__Ascomycota(100);c__Saccharomycetes(100);o__Saccharomycetales(100);f__Metschnikowiaceae(100);g__Metschnikowia(100);s__Metschnikowia_pulcherrima(100); | Metschnikowia | | 13 | Saprotroph | \|Undefined Saprotroph\| | Yeast | NULL | Probable | As nectar-tap_saprotroph (Põlme et al. 2020); Decay sugar-rich substrates (Põlme et al. 2020) | Põlme S, et al. 2020. Fungal Diversity 105:1-16 ((https://doi.org/10.1007/s13225-020-00466-2)) |
| Otu49700 | 1 | k__Fungi(100);p__Ascomycota(100);c__Saccharomycetes(100);o__Saccharomycetales(100);f__Metschnikowiaceae(100);g__Metschnikowia(100);s__Metschnikowia_pulcherrima(100); | Metschnikowia | | 13 | Saprotroph | \|Undefined Saprotroph\| | Yeast | NULL | Probable | As nectar-tap_saprotroph (Põlme et al. 2020); Decay sugar-rich substrates (Põlme et al. 2020) | Põlme S, et al. 2020. Fungal Diversity 105:1-16 ((https://doi.org/10.1007/s13225-020-00466-2)) |
| Otu49701 | 1 | k__Fungi(100);p__Ascomycota(100);c__Saccharomycetes(100);o__Saccharomycetales(100);f__Metschnikowiaceae(100);g__Metschnikowia(100);g__Metschnikowia_unclassified(100); | Metschnikowia | | 13 | Saprotroph | \|Undefined Saprotroph\| | Yeast | NULL | Probable | As nectar-tap_saprotroph (Põlme et al. 2020); Decay sugar-rich substrates (Põlme et al. 2020) | Põlme S, et al. 2020. Fungal Diversity 105:1-16 ((https://doi.org/10.1007/s13225-020-00466-2)) |
| Otu49702 | 1 | k__Fungi(100);p__Ascomycota(100);c__Saccharomycetes(100);o__Saccharomycetales(100);f__Metschnikowiaceae(100);g__Metschnikowia(100);s__Metschnikowia_pulcherrima(100); | Metschnikowia | | 13 | Saprotroph | \|Undefined Saprotroph\| | Yeast | NULL | Probable | As nectar-tap_saprotroph (Põlme et al. 2020); Decay sugar-rich substrates (Põlme et al. 2020) | Põlme S, et al. 2020. Fungal Diversity 105:1-16 ((https://doi.org/10.1007/s13225-020-00466-2)) |
| Otu49703 | 1 | k__Fungi(100);p__Ascomycota(100);c__Saccharomycetes(100);o__Saccharomycetales(100);f__Metschnikowiaceae(100);g__Metschnikowia(100);s__Metschnikowia_pulcherrima(100); | Metschnikowia | | 13 | Saprotroph | \|Undefined Saprotroph\| | Yeast | NULL | Probable | As nectar-tap_saprotroph (Põlme et al. 2020); Decay sugar-rich substrates (Põlme et al. 2020) | Põlme S, et al. 2020. Fungal Diversity 105:1-16 ((https://doi.org/10.1007/s13225-020-00466-2)) |
| Otu49704 | 1 | k__Fungi(100);p__Ascomycota(100);c__Saccharomycetes(100);o__Saccharomycetales(100);f__Metschnikowiaceae(100);g__Metschnikowia(100);s__Metschnikowia_pulcherrima(100); | Metschnikowia | | 13 | Saprotroph | \|Undefined Saprotroph\| | Yeast | NULL | Probable | As nectar-tap_saprotroph (Põlme et al. 2020); Decay sugar-rich substrates (Põlme et al. 2020) | Põlme S, et al. 2020. Fungal Diversity 105:1-16 ((https://doi.org/10.1007/s13225-020-00466-2)) |
| Otu49705 | 1 | k__Fungi(100);p__Ascomycota(100);c__Saccharomycetes(100);o__Saccharomycetales(100);f__Metschnikowiaceae(100);g__Metschnikowia(100);s__Metschnikowia_pulcherrima(100); | Metschnikowia | | 13 | Saprotroph | \|Undefined Saprotroph\| | Yeast | NULL | Probable | As nectar-tap_saprotroph (Põlme et al. 2020); Decay sugar-rich substrates (Põlme et al. 2020) | Põlme S, et al. 2020. Fungal Diversity 105:1-16 ((https://doi.org/10.1007/s13225-020-00466-2)) |
| Otu49706 | 1 | k__Fungi(100);p__Ascomycota(100);c__Saccharomycetes(100);o__Saccharomycetales(100);f__Metschnikowiaceae(100);g__Metschnikowia(100);s__Metschnikowia_pulcherrima(100); | Metschnikowia | | 13 | Saprotroph | \|Undefined Saprotroph\| | Yeast | NULL | Probable | As nectar-tap_saprotroph (Põlme et al. 2020); Decay sugar-rich substrates (Põlme et al. 2020) | Põlme S, et al. 2020. Fungal Diversity 105:1-16 ((https://doi.org/10.1007/s13225-020-00466-2)) |
| Otu49707 | 1 | k__Fungi(100);p__Ascomycota(100);c__Saccharomycetes(100);o__Saccharomycetales(100);f__Metschnikowiaceae(100);g__Metschnikowia(100);s__Metschnikowia_pulcherrima(100); | Metschnikowia | | 13 | Saprotroph | \|Undefined Saprotroph\| | Yeast | NULL | Probable | As nectar-tap_saprotroph (Põlme et al. 2020); Decay sugar-rich substrates (Põlme et al. 2020) | Põlme S, et al. 2020. Fungal Diversity 105:1-16 ((https://doi.org/10.1007/s13225-020-00466-2)) |
| Otu49708 | 1 | k__Fungi(100);p__Ascomycota(100);c__Saccharomycetes(100);o__Saccharomycetales(100);f__Metschnikowiaceae(100);g__Metschnikowia(100);s__Metschnikowia_pulcherrima(100); | Metschnikowia | | 13 | Saprotroph | \|Undefined Saprotroph\| | Yeast | NULL | Probable | As nectar-tap_saprotroph (Põlme et al. 2020); Decay sugar-rich substrates (Põlme et al. 2020) | Põlme S, et al. 2020. Fungal Diversity 105:1-16 ((https://doi.org/10.1007/s13225-020-00466-2)) |
| Otu49709 | 1 | k__Fungi(100);p__Ascomycota(100);c__Saccharomycetes(100);o__Saccharomycetales(100);f__Metschnikowiaceae(100);g__Metschnikowia(100);s__Metschnikowia_pulcherrima(100); | Metschnikowia | | 13 | Saprotroph | \|Undefined Saprotroph\| | Yeast | NULL | Probable | As nectar-tap_saprotroph (Põlme et al. 2020); Decay sugar-rich substrates (Põlme et al. 2020) | Põlme S, et al. 2020. Fungal Diversity 105:1-16 ((https://doi.org/10.1007/s13225-020-00466-2)) |
| Otu49710 | 1 | k__Fungi(100);p__Ascomycota(100);c__Saccharomycetes(100);o__Saccharomycetales(100);f__Metschnikowiaceae(100);g__Metschnikowia(100);g__Metschnikowia_unclassified(100); | Metschnikowia | | 13 | Saprotroph | \|Undefined Saprotroph\| | Yeast | NULL | Probable | As nectar-tap_saprotroph (Põlme et al. 2020); Decay sugar-rich substrates (Põlme et al. 2020) | Põlme S, et al. 2020. Fungal Diversity 105:1-16 ((https://doi.org/10.1007/s13225-020-00466-2)) |
| Otu49711 | 1 | k__Fungi(100);p__Ascomycota(100);c__Saccharomycetes(100);o__Saccharomycetales(100);f__Metschnikowiaceae(100);g__Metschnikowia(100);s__Metschnikowia_pulcherrima(100); | Metschnikowia | | 13 | Saprotroph | \|Undefined Saprotroph\| | Yeast | NULL | Probable | As nectar-tap_saprotroph (Põlme et al. 2020); Decay sugar-rich substrates (Põlme et al. 2020) | Põlme S, et al. 2020. Fungal Diversity 105:1-16 ((https://doi.org/10.1007/s13225-020-00466-2)) |
| Otu49712 | 1 | k__Fungi(100);p__Ascomycota(100);c__Saccharomycetes(100);o__Saccharomycetales(100);f__Metschnikowiaceae(100);g__Metschnikowia(100);g__Metschnikowia_unclassified(100); | Metschnikowia | | 13 | Saprotroph | \|Undefined Saprotroph\| | Yeast | NULL | Probable | As nectar-tap_saprotroph (Põlme et al. 2020); Decay sugar-rich substrates (Põlme et al. 2020) | Põlme S, et al. 2020. Fungal Diversity 105:1-16 ((https://doi.org/10.1007/s13225-020-00466-2)) |
| Otu49713 | 1 | k__Fungi(100);p__Ascomycota(100);c__Saccharomycetes(100);o__Saccharomycetales(100);f__Metschnikowiaceae(100);g__Metschnikowia(100);s__Metschnikowia_pulcherrima(100); | Metschnikowia | | 13 | Saprotroph | \|Undefined Saprotroph\| | Yeast | NULL | Probable | As nectar-tap_saprotroph (Põlme et al. 2020); Decay sugar-rich substrates (Põlme et al. 2020) | Põlme S, et al. 2020. Fungal Diversity 105:1-16 ((https://doi.org/10.1007/s13225-020-00466-2)) |
| Otu49714 | 1 | k__Fungi(100);p__Ascomycota(100);c__Saccharomycetes(100);o__Saccharomycetales(100);f__Metschnikowiaceae(100);g__Metschnikowia(100);g__Metschnikowia_unclassified(100); | Metschnikowia | | 13 | Saprotroph | \|Undefined Saprotroph\| | Yeast | NULL | Probable | As nectar-tap_saprotroph (Põlme et al. 2020); Decay sugar-rich substrates (Põlme et al. 2020) | Põlme S, et al. 2020. Fungal Diversity 105:1-16 ((https://doi.org/10.1007/s13225-020-00466-2)) |
| Otu49715 | 1 | k__Fungi(100);p__Ascomycota(100);c__Saccharomycetes(100);o__Saccharomycetales(100);f__Metschnikowiaceae(100);g__Metschnikowia(100);s__Metschnikowia_pulcherrima(100); | Metschnikowia | | 13 | Saprotroph | \|Undefined Saprotroph\| | Yeast | NULL | Probable | As nectar-tap_saprotroph (Põlme et al. 2020); Decay sugar-rich substrates (Põlme et al. 2020) | Põlme S, et al. 2020. Fungal Diversity 105:1-16 ((https://doi.org/10.1007/s13225-020-00466-2)) |
| Otu49716 | 1 | k__Fungi(100);p__Ascomycota(100);c__Saccharomycetes(100);o__Saccharomycetales(100);f__Metschnikowiaceae(100);g__Metschnikowia(100);g__Metschnikowia_unclassified(100); | Metschnikowia | | 13 | Saprotroph | \|Undefined Saprotroph\| | Yeast | NULL | Probable | As nectar-tap_saprotroph (Põlme et al. 2020); Decay sugar-rich substrates (Põlme et al. 2020) | Põlme S, et al. 2020. Fungal Diversity 105:1-16 ((https://doi.org/10.1007/s13225-020-00466-2)) |
| Otu49717 | 1 | k__Fungi(100);p__Ascomycota(100);c__Saccharomycetes(100);o__Saccharomycetales(100);f__Metschnikowiaceae(100);g__Metschnikowia(100);s__Metschnikowia_pulcherrima(100); | Metschnikowia | | 13 | Saprotroph | \|Undefined Saprotroph\| | Yeast | NULL | Probable | As nectar-tap_saprotroph (Põlme et al. 2020); Decay sugar-rich substrates (Põlme et al. 2020) | Põlme S, et al. 2020. Fungal Diversity 105:1-16 ((https://doi.org/10.1007/s13225-020-00466-2)) |
| Otu49718 | 1 | k__Fungi(100);p__Ascomycota(100);c__Saccharomycetes(100);o__Saccharomycetales(100);f__Metschnikowiaceae(100);g__Metschnikowia(100);s__Metschnikowia_pulcherrima(100); | Metschnikowia | | 13 | Saprotroph | \|Undefined Saprotroph\| | Yeast | NULL | Probable | As nectar-tap_saprotroph (Põlme et al. 2020); Decay sugar-rich substrates (Põlme et al. 2020) | Põlme S, et al. 2020. Fungal Diversity 105:1-16 ((https://doi.org/10.1007/s13225-020-00466-2)) |
| Otu49719 | 1 | k__Fungi(100);p__Ascomycota(100);c__Saccharomycetes(100);o__Saccharomycetales(100);f__Metschnikowiaceae(100);g__Metschnikowia(100);s__Metschnikowia_pulcherrima(100); | Metschnikowia | | 13 | Saprotroph | \|Undefined Saprotroph\| | Yeast | NULL | Probable | As nectar-tap_saprotroph (Põlme et al. 2020); Decay sugar-rich substrates (Põlme et al. 2020) | Põlme S, et al. 2020. Fungal Diversity 105:1-16 ((https://doi.org/10.1007/s13225-020-00466-2)) |
| Otu49720 | 1 | k__Fungi(100);p__Ascomycota(100);c__Saccharomycetes(100);o__Saccharomycetales(100);f__Metschnikowiaceae(100);g__Metschnikowia(100);s__Metschnikowia_pulcherrima(100); | Metschnikowia | | 13 | Saprotroph | \|Undefined Saprotroph\| | Yeast | NULL | Probable | As nectar-tap_saprotroph (Põlme et al. 2020); Decay sugar-rich substrates (Põlme et al. 2020) | Põlme S, et al. 2020. Fungal Diversity 105:1-16 ((https://doi.org/10.1007/s13225-020-00466-2)) |
| Otu49721 | 1 | k__Fungi(100);p__Ascomycota(100);c__Saccharomycetes(100);o__Saccharomycetales(100);f__Metschnikowiaceae(100);g__Metschnikowia(100);s__Metschnikowia_pulcherrima(100); | Metschnikowia | | 13 | Saprotroph | \|Undefined Saprotroph\| | Yeast | NULL | Probable | As nectar-tap_saprotroph (Põlme et al. 2020); Decay sugar-rich substrates (Põlme et al. 2020) | Põlme S, et al. 2020. Fungal Diversity 105:1-16 ((https://doi.org/10.1007/s13225-020-00466-2)) |
| Otu49722 | 1 | k__Fungi(100);p__Ascomycota(100);c__Saccharomycetes(100);o__Saccharomycetales(100);f__Metschnikowiaceae(100);g__Metschnikowia(100);s__Metschnikowia_pulcherrima(100); | Metschnikowia | | 13 | Saprotroph | \|Undefined Saprotroph\| | Yeast | NULL | Probable | As nectar-tap_saprotroph (Põlme et al. 2020); Decay sugar-rich substrates (Põlme et al. 2020) | Põlme S, et al. 2020. Fungal Diversity 105:1-16 ((https://doi.org/10.1007/s13225-020-00466-2)) |
| Otu49723 | 1 | k__Fungi(100);p__Ascomycota(100);c__Saccharomycetes(100);o__Saccharomycetales(100);f__Metschnikowiaceae(100);g__Metschnikowia(100);s__Metschnikowia_pulcherrima(100); | Metschnikowia | | 13 | Saprotroph | \|Undefined Saprotroph\| | Yeast | NULL | Probable | As nectar-tap_saprotroph (Põlme et al. 2020); Decay sugar-rich substrates (Põlme et al. 2020) | Põlme S, et al. 2020. Fungal Diversity 105:1-16 ((https://doi.org/10.1007/s13225-020-00466-2)) |
| Otu49724 | 1 | k__Fungi(100);p__Ascomycota(100);c__Saccharomycetes(100);o__Saccharomycetales(100);f__Metschnikowiaceae(100);g__Metschnikowia(100);g__Metschnikowia_unclassified(100); | Metschnikowia | | 13 | Saprotroph | \|Undefined Saprotroph\| | Yeast | NULL | Probable | As nectar-tap_saprotroph (Põlme et al. 2020); Decay sugar-rich substrates (Põlme et al. 2020) | Põlme S, et al. 2020. Fungal Diversity 105:1-16 ((https://doi.org/10.1007/s13225-020-00466-2)) |
| Otu49726 | 1 | k__Fungi(100);p__Ascomycota(100);c__Saccharomycetes(100);o__Saccharomycetales(100);f__Metschnikowiaceae(100);g__Metschnikowia(100);s__Metschnikowia_pulcherrima(100); | Metschnikowia | | 13 | Saprotroph | \|Undefined Saprotroph\| | Yeast | NULL | Probable | As nectar-tap_saprotroph (Põlme et al. 2020); Decay sugar-rich substrates (Põlme et al. 2020) | Põlme S, et al. 2020. Fungal Diversity 105:1-16 ((https://doi.org/10.1007/s13225-020-00466-2)) |
| Otu49727 | 1 | k__Fungi(100);p__Ascomycota(100);c__Saccharomycetes(100);o__Saccharomycetales(100);f__Metschnikowiaceae(100);g__Metschnikowia(100);g__Metschnikowia_unclassified(100); | Metschnikowia | | 13 | Saprotroph | \|Undefined Saprotroph\| | Yeast | NULL | Probable | As nectar-tap_saprotroph (Põlme et al. 2020); Decay sugar-rich substrates (Põlme et al. 2020) | Põlme S, et al. 2020. Fungal Diversity 105:1-16 ((https://doi.org/10.1007/s13225-020-00466-2)) |
| Otu49728 | 1 | k__Fungi(100);p__Ascomycota(100);c__Saccharomycetes(100);o__Saccharomycetales(100);f__Metschnikowiaceae(100);g__Metschnikowia(100);s__Metschnikowia_pulcherrima(100); | Metschnikowia | | 13 | Saprotroph | \|Undefined Saprotroph\| | Yeast | NULL | Probable | As nectar-tap_saprotroph (Põlme et al. 2020); Decay sugar-rich substrates (Põlme et al. 2020) | Põlme S, et al. 2020. Fungal Diversity 105:1-16 ((https://doi.org/10.1007/s13225-020-00466-2)) |
| Otu49729 | 1 | k__Fungi(100);p__Ascomycota(100);c__Saccharomycetes(100);o__Saccharomycetales(100);f__Metschnikowiaceae(100);g__Metschnikowia(100);s__Metschnikowia_pulcherrima(100); | Metschnikowia | | 13 | Saprotroph | \|Undefined Saprotroph\| | Yeast | NULL | Probable | As nectar-tap_saprotroph (Põlme et al. 2020); Decay sugar-rich substrates (Põlme et al. 2020) | Põlme S, et al. 2020. Fungal Diversity 105:1-16 ((https://doi.org/10.1007/s13225-020-00466-2)) |
| Otu49730 | 1 | k__Fungi(100);p__Ascomycota(100);c__Saccharomycetes(100);o__Saccharomycetales(100);f__Metschnikowiaceae(100);g__Metschnikowia(100);s__Metschnikowia_pulcherrima(100); | Metschnikowia | | 13 | Saprotroph | \|Undefined Saprotroph\| | Yeast | NULL | Probable | As nectar-tap_saprotroph (Põlme et al. 2020); Decay sugar-rich substrates (Põlme et al. 2020) | Põlme S, et al. 2020. Fungal Diversity 105:1-16 ((https://doi.org/10.1007/s13225-020-00466-2)) |
| Otu49731 | 1 | k__Fungi(100);p__Ascomycota(100);c__Saccharomycetes(100);o__Saccharomycetales(100);f__Metschnikowiaceae(100);g__Metschnikowia(100);s__Metschnikowia_pulcherrima(100); | Metschnikowia | | 13 | Saprotroph | \|Undefined Saprotroph\| | Yeast | NULL | Probable | As nectar-tap_saprotroph (Põlme et al. 2020); Decay sugar-rich substrates (Põlme et al. 2020) | Põlme S, et al. 2020. Fungal Diversity 105:1-16 ((https://doi.org/10.1007/s13225-020-00466-2)) |
| Otu49732 | 1 | k__Fungi(100);p__Ascomycota(100);c__Saccharomycetes(100);o__Saccharomycetales(100);f__Metschnikowiaceae(100);g__Metschnikowia(100);s__Metschnikowia_pulcherrima(100); | Metschnikowia | | 13 | Saprotroph | \|Undefined Saprotroph\| | Yeast | NULL | Probable | As nectar-tap_saprotroph (Põlme et al. 2020); Decay sugar-rich substrates (Põlme et al. 2020) | Põlme S, et al. 2020. Fungal Diversity 105:1-16 ((https://doi.org/10.1007/s13225-020-00466-2)) |
| Otu49733 | 1 | k__Fungi(100);p__Ascomycota(100);c__Saccharomycetes(100);o__Saccharomycetales(100);f__Metschnikowiaceae(100);g__Metschnikowia(100);g__Metschnikowia_unclassified(100); | Metschnikowia | | 13 | Saprotroph | \|Undefined Saprotroph\| | Yeast | NULL | Probable | As nectar-tap_saprotroph (Põlme et al. 2020); Decay sugar-rich substrates (Põlme et al. 2020) | Põlme S, et al. 2020. Fungal Diversity 105:1-16 ((https://doi.org/10.1007/s13225-020-00466-2)) |
| Otu49734 | 1 | k__Fungi(100);p__Ascomycota(100);c__Saccharomycetes(100);o__Saccharomycetales(100);f__Metschnikowiaceae(100);g__Metschnikowia(100);s__Metschnikowia_pulcherrima(100); | Metschnikowia | | 13 | Saprotroph | \|Undefined Saprotroph\| | Yeast | NULL | Probable | As nectar-tap_saprotroph (Põlme et al. 2020); Decay sugar-rich substrates (Põlme et al. 2020) | Põlme S, et al. 2020. Fungal Diversity 105:1-16 ((https://doi.org/10.1007/s13225-020-00466-2)) |
| Otu49735 | 1 | k__Fungi(100);p__Ascomycota(100);c__Saccharomycetes(100);o__Saccharomycetales(100);f__Metschnikowiaceae(100);g__Metschnikowia(100);s__Metschnikowia_pulcherrima(100); | Metschnikowia | | 13 | Saprotroph | \|Undefined Saprotroph\| | Yeast | NULL | Probable | As nectar-tap_saprotroph (Põlme et al. 2020); Decay sugar-rich substrates (Põlme et al. 2020) | Põlme S, et al. 2020. Fungal Diversity 105:1-16 ((https://doi.org/10.1007/s13225-020-00466-2)) |
| Otu49736 | 1 | k__Fungi(100);p__Ascomycota(100);c__Saccharomycetes(100);o__Saccharomycetales(100);f__Metschnikowiaceae(100);g__Metschnikowia(100);s__Metschnikowia_pulcherrima(100); | Metschnikowia | | 13 | Saprotroph | \|Undefined Saprotroph\| | Yeast | NULL | Probable | As nectar-tap_saprotroph (Põlme et al. 2020); Decay sugar-rich substrates (Põlme et al. 2020) | Põlme S, et al. 2020. Fungal Diversity 105:1-16 ((https://doi.org/10.1007/s13225-020-00466-2)) |
| Otu49737 | 1 | k__Fungi(100);p__Ascomycota(100);c__Saccharomycetes(100);o__Saccharomycetales(100);f__Metschnikowiaceae(100);g__Metschnikowia(100);s__Metschnikowia_pulcherrima(100); | Metschnikowia | | 13 | Saprotroph | \|Undefined Saprotroph\| | Yeast | NULL | Probable | As nectar-tap_saprotroph (Põlme et al. 2020); Decay sugar-rich substrates (Põlme et al. 2020) | Põlme S, et al. 2020. Fungal Diversity 105:1-16 ((https://doi.org/10.1007/s13225-020-00466-2)) |
| Otu49738 | 1 | k__Fungi(100);p__Ascomycota(100);c__Saccharomycetes(100);o__Saccharomycetales(100);f__Metschnikowiaceae(100);g__Metschnikowia(100);g__Metschnikowia_unclassified(100); | Metschnikowia | | 13 | Saprotroph | \|Undefined Saprotroph\| | Yeast | NULL | Probable | As nectar-tap_saprotroph (Põlme et al. 2020); Decay sugar-rich substrates (Põlme et al. 2020) | Põlme S, et al. 2020. Fungal Diversity 105:1-16 ((https://doi.org/10.1007/s13225-020-00466-2)) |
| Otu49739 | 1 | k__Fungi(100);p__Ascomycota(100);c__Saccharomycetes(100);o__Saccharomycetales(100);f__Metschnikowiaceae(100);g__Metschnikowia(100);s__Metschnikowia_pulcherrima(100); | Metschnikowia | | 13 | Saprotroph | \|Undefined Saprotroph\| | Yeast | NULL | Probable | As nectar-tap_saprotroph (Põlme et al. 2020); Decay sugar-rich substrates (Põlme et al. 2020) | Põlme S, et al. 2020. Fungal Diversity 105:1-16 ((https://doi.org/10.1007/s13225-020-00466-2)) |
| Otu49740 | 1 | k__Fungi(100);p__Ascomycota(100);c__Saccharomycetes(100);o__Saccharomycetales(100);f__Metschnikowiaceae(100);g__Metschnikowia(100);s__Metschnikowia_pulcherrima(100); | Metschnikowia | | 13 | Saprotroph | \|Undefined Saprotroph\| | Yeast | NULL | Probable | As nectar-tap_saprotroph (Põlme et al. 2020); Decay sugar-rich substrates (Põlme et al. 2020) | Põlme S, et al. 2020. Fungal Diversity 105:1-16 ((https://doi.org/10.1007/s13225-020-00466-2)) |
| Otu49741 | 1 | k__Fungi(100);p__Ascomycota(100);c__Saccharomycetes(100);o__Saccharomycetales(100);f__Metschnikowiaceae(100);g__Metschnikowia(100);g__Metschnikowia_unclassified(100); | Metschnikowia | | 13 | Saprotroph | \|Undefined Saprotroph\| | Yeast | NULL | Probable | As nectar-tap_saprotroph (Põlme et al. 2020); Decay sugar-rich substrates (Põlme et al. 2020) | Põlme S, et al. 2020. Fungal Diversity 105:1-16 ((https://doi.org/10.1007/s13225-020-00466-2)) |
| Otu49742 | 1 | k__Fungi(100);p__Ascomycota(100);c__Saccharomycetes(100);o__Saccharomycetales(100);f__Metschnikowiaceae(100);g__Metschnikowia(100);s__Metschnikowia_pulcherrima(100); | Metschnikowia | | 13 | Saprotroph | \|Undefined Saprotroph\| | Yeast | NULL | Probable | As nectar-tap_saprotroph (Põlme et al. 2020); Decay sugar-rich substrates (Põlme et al. 2020) | Põlme S, et al. 2020. Fungal Diversity 105:1-16 ((https://doi.org/10.1007/s13225-020-00466-2)) |
| Otu49743 | 1 | k__Fungi(100);p__Ascomycota(100);c__Saccharomycetes(100);o__Saccharomycetales(100);f__Metschnikowiaceae(100);g__Metschnikowia(100);s__Metschnikowia_pulcherrima(100); | Metschnikowia | | 13 | Saprotroph | \|Undefined Saprotroph\| | Yeast | NULL | Probable | As nectar-tap_saprotroph (Põlme et al. 2020); Decay sugar-rich substrates (Põlme et al. 2020) | Põlme S, et al. 2020. Fungal Diversity 105:1-16 ((https://doi.org/10.1007/s13225-020-00466-2)) |
| Otu49744 | 1 | k__Fungi(100);p__Ascomycota(100);c__Saccharomycetes(100);o__Saccharomycetales(100);f__Metschnikowiaceae(100);g__Metschnikowia(100);s__Metschnikowia_pulcherrima(100); | Metschnikowia | | 13 | Saprotroph | \|Undefined Saprotroph\| | Yeast | NULL | Probable | As nectar-tap_saprotroph (Põlme et al. 2020); Decay sugar-rich substrates (Põlme et al. 2020) | Põlme S, et al. 2020. Fungal Diversity 105:1-16 ((https://doi.org/10.1007/s13225-020-00466-2)) |
| Otu49745 | 1 | k__Fungi(100);p__Ascomycota(100);c__Saccharomycetes(100);o__Saccharomycetales(100);f__Metschnikowiaceae(100);g__Metschnikowia(100);g__Metschnikowia_unclassified(100); | Metschnikowia | | 13 | Saprotroph | \|Undefined Saprotroph\| | Yeast | NULL | Probable | As nectar-tap_saprotroph (Põlme et al. 2020); Decay sugar-rich substrates (Põlme et al. 2020) | Põlme S, et al. 2020. Fungal Diversity 105:1-16 ((https://doi.org/10.1007/s13225-020-00466-2)) |
| Otu49746 | 1 | k__Fungi(100);p__Ascomycota(100);c__Saccharomycetes(100);o__Saccharomycetales(100);f__Metschnikowiaceae(100);g__Metschnikowia(100);s__Metschnikowia_pulcherrima(100); | Metschnikowia | | 13 | Saprotroph | \|Undefined Saprotroph\| | Yeast | NULL | Probable | As nectar-tap_saprotroph (Põlme et al. 2020); Decay sugar-rich substrates (Põlme et al. 2020) | Põlme S, et al. 2020. Fungal Diversity 105:1-16 ((https://doi.org/10.1007/s13225-020-00466-2)) |
| Otu49747 | 1 | k__Fungi(100);p__Ascomycota(100);c__Saccharomycetes(100);o__Saccharomycetales(100);f__Metschnikowiaceae(100);g__Metschnikowia(100);s__Metschnikowia_pulcherrima(100); | Metschnikowia | | 13 | Saprotroph | \|Undefined Saprotroph\| | Yeast | NULL | Probable | As nectar-tap_saprotroph (Põlme et al. 2020); Decay sugar-rich substrates (Põlme et al. 2020) | Põlme S, et al. 2020. Fungal Diversity 105:1-16 ((https://doi.org/10.1007/s13225-020-00466-2)) |
| Otu49748 | 1 | k__Fungi(100);p__Ascomycota(100);c__Saccharomycetes(100);o__Saccharomycetales(100);f__Metschnikowiaceae(100);g__Metschnikowia(100);g__Metschnikowia_unclassified(100); | Metschnikowia | | 13 | Saprotroph | \|Undefined Saprotroph\| | Yeast | NULL | Probable | As nectar-tap_saprotroph (Põlme et al. 2020); Decay sugar-rich substrates (Põlme et al. 2020) | Põlme S, et al. 2020. Fungal Diversity 105:1-16 ((https://doi.org/10.1007/s13225-020-00466-2)) |
| Otu49749 | 1 | k__Fungi(100);p__Ascomycota(100);c__Saccharomycetes(100);o__Saccharomycetales(100);f__Metschnikowiaceae(100);g__Metschnikowia(100);s__Metschnikowia_pulcherrima(100); | Metschnikowia | | 13 | Saprotroph | \|Undefined Saprotroph\| | Yeast | NULL | Probable | As nectar-tap_saprotroph (Põlme et al. 2020); Decay sugar-rich substrates (Põlme et al. 2020) | Põlme S, et al. 2020. Fungal Diversity 105:1-16 ((https://doi.org/10.1007/s13225-020-00466-2)) |
| Otu49750 | 1 | k__Fungi(100);p__Ascomycota(100);c__Saccharomycetes(100);o__Saccharomycetales(100);f__Metschnikowiaceae(100);g__Metschnikowia(100);s__Metschnikowia_pulcherrima(100); | Metschnikowia | | 13 | Saprotroph | \|Undefined Saprotroph\| | Yeast | NULL | Probable | As nectar-tap_saprotroph (Põlme et al. 2020); Decay sugar-rich substrates (Põlme et al. 2020) | Põlme S, et al. 2020. Fungal Diversity 105:1-16 ((https://doi.org/10.1007/s13225-020-00466-2)) |
| Otu49751 | 1 | k__Fungi(100);p__Ascomycota(100);c__Saccharomycetes(100);o__Saccharomycetales(100);f__Metschnikowiaceae(100);g__Metschnikowia(100);s__Metschnikowia_pulcherrima(100); | Metschnikowia | | 13 | Saprotroph | \|Undefined Saprotroph\| | Yeast | NULL | Probable | As nectar-tap_saprotroph (Põlme et al. 2020); Decay sugar-rich substrates (Põlme et al. 2020) | Põlme S, et al. 2020. Fungal Diversity 105:1-16 ((https://doi.org/10.1007/s13225-020-00466-2)) |
| Otu49752 | 1 | k__Fungi(100);p__Ascomycota(100);c__Saccharomycetes(100);o__Saccharomycetales(100);f__Metschnikowiaceae(100);g__Metschnikowia(100);s__Metschnikowia_pulcherrima(100); | Metschnikowia | | 13 | Saprotroph | \|Undefined Saprotroph\| | Yeast | NULL | Probable | As nectar-tap_saprotroph (Põlme et al. 2020); Decay sugar-rich substrates (Põlme et al. 2020) | Põlme S, et al. 2020. Fungal Diversity 105:1-16 ((https://doi.org/10.1007/s13225-020-00466-2)) |
| Otu49753 | 1 | k__Fungi(100);p__Ascomycota(100);c__Saccharomycetes(100);o__Saccharomycetales(100);f__Metschnikowiaceae(100);g__Metschnikowia(100);g__Metschnikowia_unclassified(100); | Metschnikowia | | 13 | Saprotroph | \|Undefined Saprotroph\| | Yeast | NULL | Probable | As nectar-tap_saprotroph (Põlme et al. 2020); Decay sugar-rich substrates (Põlme et al. 2020) | Põlme S, et al. 2020. Fungal Diversity 105:1-16 ((https://doi.org/10.1007/s13225-020-00466-2)) |
| Otu49755 | 1 | k__Fungi(100);p__Ascomycota(100);c__Saccharomycetes(100);o__Saccharomycetales(100);f__Metschnikowiaceae(100);g__Metschnikowia(100);s__Metschnikowia_pulcherrima(100); | Metschnikowia | | 13 | Saprotroph | \|Undefined Saprotroph\| | Yeast | NULL | Probable | As nectar-tap_saprotroph (Põlme et al. 2020); Decay sugar-rich substrates (Põlme et al. 2020) | Põlme S, et al. 2020. Fungal Diversity 105:1-16 ((https://doi.org/10.1007/s13225-020-00466-2)) |

| | | | | | | | | | | | |
|---|---|---|---|---|---|---|---|---|---|---|---|
| Otu49756 | 1 | k__Fungi(100);p__Ascomycota(100);c__Saccharomycetes(100);o__Saccharomycetales(100);f__Metschnikowiaceae(100);g__Metschnikowia(100);s__Metschnikowia_pulcherrima(100); | Metschnikowia | 13 | Saprotroph | \|Undefined Saprotroph\| | Yeast | NULL | Probable | As nectar-tap_saprotroph (Põlme et al. 2020); Decay sugar-rich substrates (Põlme et al. 2020) | Põlme S, et al. 2020. Fungal Diversity 105:1-16 ((https://doi.org/10.1007/s13225-020-00466-2)) |
| Otu49757 | 1 | k__Fungi(100);p__Ascomycota(100);c__Saccharomycetes(100);o__Saccharomycetales(100);f__Metschnikowiaceae(100);g__Metschnikowia(100);s__Metschnikowia_noctiluminum(100); | Metschnikowia | 13 | Saprotroph | \|Undefined Saprotroph\| | Yeast | NULL | Probable | As nectar-tap_saprotroph (Põlme et al. 2020); Decay sugar-rich substrates (Põlme et al. 2020) | Põlme S, et al. 2020. Fungal Diversity 105:1-16 ((https://doi.org/10.1007/s13225-020-00466-2)) |
| Otu49758 | 1 | k__Fungi(100);p__Ascomycota(100);c__Saccharomycetes(100);o__Saccharomycetales(100);f__Metschnikowiaceae(100);g__Metschnikowia(100);s__Metschnikowia_pulcherrima(100); | Metschnikowia | 13 | Saprotroph | \|Undefined Saprotroph\| | Yeast | NULL | Probable | As nectar-tap_saprotroph (Põlme et al. 2020); Decay sugar-rich substrates (Põlme et al. 2020) | Põlme S, et al. 2020. Fungal Diversity 105:1-16 ((https://doi.org/10.1007/s13225-020-00466-2)) |
| Otu49759 | 1 | k__Fungi(100);p__Ascomycota(100);c__Saccharomycetes(100);o__Saccharomycetales(100);f__Metschnikowiaceae(100);g__Metschnikowia(100);s__Metschnikowia_pulcherrima(100); | Metschnikowia | 13 | Saprotroph | \|Undefined Saprotroph\| | Yeast | NULL | Probable | As nectar-tap_saprotroph (Põlme et al. 2020); Decay sugar-rich substrates (Põlme et al. 2020) | Põlme S, et al. 2020. Fungal Diversity 105:1-16 ((https://doi.org/10.1007/s13225-020-00466-2)) |
| Otu49760 | 1 | k__Fungi(100);p__Ascomycota(100);c__Saccharomycetes(100);o__Saccharomycetales(100);f__Metschnikowiaceae(100);g__Metschnikowia(100);s__Metschnikowia_pulcherrima(100); | Metschnikowia | 13 | Saprotroph | \|Undefined Saprotroph\| | Yeast | NULL | Probable | As nectar-tap_saprotroph (Põlme et al. 2020); Decay sugar-rich substrates (Põlme et al. 2020) | Põlme S, et al. 2020. Fungal Diversity 105:1-16 ((https://doi.org/10.1007/s13225-020-00466-2)) |
| Otu49761 | 1 | k__Fungi(100);p__Ascomycota(100);c__Saccharomycetes(100);o__Saccharomycetales(100);f__Metschnikowiaceae(100);g__Metschnikowia(100);s__Metschnikowia_pulcherrima(100); | Metschnikowia | 13 | Saprotroph | \|Undefined Saprotroph\| | Yeast | NULL | Probable | As nectar-tap_saprotroph (Põlme et al. 2020); Decay sugar-rich substrates (Põlme et al. 2020) | Põlme S, et al. 2020. Fungal Diversity 105:1-16 ((https://doi.org/10.1007/s13225-020-00466-2)) |
| Otu49762 | 1 | k__Fungi(100);p__Ascomycota(100);c__Saccharomycetes(100);o__Saccharomycetales(100);f__Metschnikowiaceae(100);g__Metschnikowia(100);s__Metschnikowia_pulcherrima(100); | Metschnikowia | 13 | Saprotroph | \|Undefined Saprotroph\| | Yeast | NULL | Probable | As nectar-tap_saprotroph (Põlme et al. 2020); Decay sugar-rich substrates (Põlme et al. 2020) | Põlme S, et al. 2020. Fungal Diversity 105:1-16 ((https://doi.org/10.1007/s13225-020-00466-2)) |
| Otu49763 | 1 | k__Fungi(100);p__Ascomycota(100);c__Saccharomycetes(100);o__Saccharomycetales(100);f__Metschnikowiaceae(100);g__Metschnikowia(100);g__Metschnikowia_unclassified(100); | Metschnikowia | 13 | Saprotroph | \|Undefined Saprotroph\| | Yeast | NULL | Probable | As nectar-tap_saprotroph (Põlme et al. 2020); Decay sugar-rich substrates (Põlme et al. 2020) | Põlme S, et al. 2020. Fungal Diversity 105:1-16 ((https://doi.org/10.1007/s13225-020-00466-2)) |
| Otu49764 | 1 | k__Fungi(100);p__Ascomycota(100);c__Saccharomycetes(100);o__Saccharomycetales(100);f__Metschnikowiaceae(100);g__Metschnikowia(100);g__Metschnikowia_unclassified(100); | Metschnikowia | 13 | Saprotroph | \|Undefined Saprotroph\| | Yeast | NULL | Probable | As nectar-tap_saprotroph (Põlme et al. 2020); Decay sugar-rich substrates (Põlme et al. 2020) | Põlme S, et al. 2020. Fungal Diversity 105:1-16 ((https://doi.org/10.1007/s13225-020-00466-2)) |
| Otu49766 | 1 | k__Fungi(100);p__Ascomycota(100);c__Saccharomycetes(100);o__Saccharomycetales(100);f__Metschnikowiaceae(100);g__Metschnikowia(100);s__Metschnikowia_pulcherrima(100); | Metschnikowia | 13 | Saprotroph | \|Undefined Saprotroph\| | Yeast | NULL | Probable | As nectar-tap_saprotroph (Põlme et al. 2020); Decay sugar-rich substrates (Põlme et al. 2020) | Põlme S, et al. 2020. Fungal Diversity 105:1-16 ((https://doi.org/10.1007/s13225-020-00466-2)) |
| Otu49767 | 1 | k__Fungi(100);p__Ascomycota(100);c__Saccharomycetes(100);o__Saccharomycetales(100);f__Metschnikowiaceae(100);g__Metschnikowia(100);s__Metschnikowia_pulcherrima(100); | Metschnikowia | 13 | Saprotroph | \|Undefined Saprotroph\| | Yeast | NULL | Probable | As nectar-tap_saprotroph (Põlme et al. 2020); Decay sugar-rich substrates (Põlme et al. 2020) | Põlme S, et al. 2020. Fungal Diversity 105:1-16 ((https://doi.org/10.1007/s13225-020-00466-2)) |
| Otu49768 | 1 | k__Fungi(100);p__Ascomycota(100);c__Saccharomycetes(100);o__Saccharomycetales(100);f__Metschnikowiaceae(100);g__Metschnikowia(100);s__Metschnikowia_pulcherrima(100); | Metschnikowia | 13 | Saprotroph | \|Undefined Saprotroph\| | Yeast | NULL | Probable | As nectar-tap_saprotroph (Põlme et al. 2020); Decay sugar-rich substrates (Põlme et al. 2020) | Põlme S, et al. 2020. Fungal Diversity 105:1-16 ((https://doi.org/10.1007/s13225-020-00466-2)) |
| Otu49769 | 1 | k__Fungi(100);p__Ascomycota(100);c__Saccharomycetes(100);o__Saccharomycetales(100);f__Metschnikowiaceae(100);g__Metschnikowia(100);s__Metschnikowia_pulcherrima(100); | Metschnikowia | 13 | Saprotroph | \|Undefined Saprotroph\| | Yeast | NULL | Probable | As nectar-tap_saprotroph (Põlme et al. 2020); Decay sugar-rich substrates (Põlme et al. 2020) | Põlme S, et al. 2020. Fungal Diversity 105:1-16 ((https://doi.org/10.1007/s13225-020-00466-2)) |
| Otu49770 | 1 | k__Fungi(100);p__Ascomycota(100);c__Saccharomycetes(100);o__Saccharomycetales(100);f__Metschnikowiaceae(100);g__Metschnikowia(100);s__Metschnikowia_pulcherrima(100); | Metschnikowia | 13 | Saprotroph | \|Undefined Saprotroph\| | Yeast | NULL | Probable | As nectar-tap_saprotroph (Põlme et al. 2020); Decay sugar-rich substrates (Põlme et al. 2020) | Põlme S, et al. 2020. Fungal Diversity 105:1-16 ((https://doi.org/10.1007/s13225-020-00466-2)) |
| Otu49771 | 1 | k__Fungi(100);p__Ascomycota(100);c__Saccharomycetes(100);o__Saccharomycetales(100);f__Metschnikowiaceae(100);g__Metschnikowia(100);s__Metschnikowia_pulcherrima(100); | Metschnikowia | 13 | Saprotroph | \|Undefined Saprotroph\| | Yeast | NULL | Probable | As nectar-tap_saprotroph (Põlme et al. 2020); Decay sugar-rich substrates (Põlme et al. 2020) | Põlme S, et al. 2020. Fungal Diversity 105:1-16 ((https://doi.org/10.1007/s13225-020-00466-2)) |
| Otu49772 | 1 | k__Fungi(100);p__Ascomycota(100);c__Saccharomycetes(100);o__Saccharomycetales(100);f__Metschnikowiaceae(100);g__Metschnikowia(100);g__Metschnikowia_unclassified(100); | Metschnikowia | 13 | Saprotroph | \|Undefined Saprotroph\| | Yeast | NULL | Probable | As nectar-tap_saprotroph (Põlme et al. 2020); Decay sugar-rich substrates (Põlme et al. 2020) | Põlme S, et al. 2020. Fungal Diversity 105:1-16 ((https://doi.org/10.1007/s13225-020-00466-2)) |
| Otu49773 | 1 | k__Fungi(100);p__Ascomycota(100);c__Saccharomycetes(100);o__Saccharomycetales(100);f__Metschnikowiaceae(100);g__Metschnikowia(100);s__Metschnikowia_pulcherrima(100); | Metschnikowia | 13 | Saprotroph | \|Undefined Saprotroph\| | Yeast | NULL | Probable | As nectar-tap_saprotroph (Põlme et al. 2020); Decay sugar-rich substrates (Põlme et al. 2020) | Põlme S, et al. 2020. Fungal Diversity 105:1-16 ((https://doi.org/10.1007/s13225-020-00466-2)) |
| Otu49774 | 1 | k__Fungi(100);p__Ascomycota(100);c__Saccharomycetes(100);o__Saccharomycetales(100);f__Metschnikowiaceae(100);g__Metschnikowia(100);s__Metschnikowia_pulcherrima(100); | Metschnikowia | 13 | Saprotroph | \|Undefined Saprotroph\| | Yeast | NULL | Probable | As nectar-tap_saprotroph (Põlme et al. 2020); Decay sugar-rich substrates (Põlme et al. 2020) | Põlme S, et al. 2020. Fungal Diversity 105:1-16 ((https://doi.org/10.1007/s13225-020-00466-2)) |
| Otu49775 | 1 | k__Fungi(100);p__Ascomycota(100);c__Saccharomycetes(100);o__Saccharomycetales(100);f__Metschnikowiaceae(100);g__Metschnikowia(100);g__Metschnikowia_unclassified(100); | Metschnikowia | 13 | Saprotroph | \|Undefined Saprotroph\| | Yeast | NULL | Probable | As nectar-tap_saprotroph (Põlme et al. 2020); Decay sugar-rich substrates (Põlme et al. 2020) | Põlme S, et al. 2020. Fungal Diversity 105:1-16 ((https://doi.org/10.1007/s13225-020-00466-2)) |
| Otu49776 | 1 | k__Fungi(100);p__Ascomycota(100);c__Saccharomycetes(100);o__Saccharomycetales(100);f__Metschnikowiaceae(100);g__Metschnikowia(100);g__Metschnikowia_unclassified(100); | Metschnikowia | 13 | Saprotroph | \|Undefined Saprotroph\| | Yeast | NULL | Probable | As nectar-tap_saprotroph (Põlme et al. 2020); Decay sugar-rich substrates (Põlme et al. 2020) | Põlme S, et al. 2020. Fungal Diversity 105:1-16 ((https://doi.org/10.1007/s13225-020-00466-2)) |
| Otu49777 | 1 | k__Fungi(100);p__Ascomycota(100);c__Saccharomycetes(100);o__Saccharomycetales(100);f__Metschnikowiaceae(100);g__Metschnikowia(100);s__Metschnikowia_pulcherrima(100); | Metschnikowia | 13 | Saprotroph | \|Undefined Saprotroph\| | Yeast | NULL | Probable | As nectar-tap_saprotroph (Põlme et al. 2020); Decay sugar-rich substrates (Põlme et al. 2020) | Põlme S, et al. 2020. Fungal Diversity 105:1-16 ((https://doi.org/10.1007/s13225-020-00466-2)) |
| Otu49778 | 1 | k__Fungi(100);p__Ascomycota(100);c__Saccharomycetes(100);o__Saccharomycetales(100);f__Metschnikowiaceae(100);g__Metschnikowia(100);g__Metschnikowia_unclassified(100); | Metschnikowia | 13 | Saprotroph | \|Undefined Saprotroph\| | Yeast | NULL | Probable | As nectar-tap_saprotroph (Põlme et al. 2020); Decay sugar-rich substrates (Põlme et al. 2020) | Põlme S, et al. 2020. Fungal Diversity 105:1-16 ((https://doi.org/10.1007/s13225-020-00466-2)) |
| Otu49779 | 1 | k__Fungi(100);p__Ascomycota(100);c__Saccharomycetes(100);o__Saccharomycetales(100);f__Metschnikowiaceae(100);g__Metschnikowia(100);s__Metschnikowia_pulcherrima(100); | Metschnikowia | 13 | Saprotroph | \|Undefined Saprotroph\| | Yeast | NULL | Probable | As nectar-tap_saprotroph (Põlme et al. 2020); Decay sugar-rich substrates (Põlme et al. 2020) | Põlme S, et al. 2020. Fungal Diversity 105:1-16 ((https://doi.org/10.1007/s13225-020-00466-2)) |
| Otu49780 | 1 | k__Fungi(100);p__Ascomycota(100);c__Saccharomycetes(100);o__Saccharomycetales(100);f__Metschnikowiaceae(100);g__Metschnikowia(100);s__Metschnikowia_pulcherrima(100); | Metschnikowia | 13 | Saprotroph | \|Undefined Saprotroph\| | Yeast | NULL | Probable | As nectar-tap_saprotroph (Põlme et al. 2020); Decay sugar-rich substrates (Põlme et al. 2020) | Põlme S, et al. 2020. Fungal Diversity 105:1-16 ((https://doi.org/10.1007/s13225-020-00466-2)) |
| Otu49781 | 1 | k__Fungi(100);p__Ascomycota(100);c__Saccharomycetes(100);o__Saccharomycetales(100);f__Metschnikowiaceae(100);g__Metschnikowia(100);s__Metschnikowia_pulcherrima(100); | Metschnikowia | 13 | Saprotroph | \|Undefined Saprotroph\| | Yeast | NULL | Probable | As nectar-tap_saprotroph (Põlme et al. 2020); Decay sugar-rich substrates (Põlme et al. 2020) | Põlme S, et al. 2020. Fungal Diversity 105:1-16 ((https://doi.org/10.1007/s13225-020-00466-2)) |
| Otu49782 | 1 | k__Fungi(100);p__Ascomycota(100);c__Saccharomycetes(100);o__Saccharomycetales(100);f__Metschnikowiaceae(100);g__Metschnikowia(100);s__Metschnikowia_pulcherrima(100); | Metschnikowia | 13 | Saprotroph | \|Undefined Saprotroph\| | Yeast | NULL | Probable | As nectar-tap_saprotroph (Põlme et al. 2020); Decay sugar-rich substrates (Põlme et al. 2020) | Põlme S, et al. 2020. Fungal Diversity 105:1-16 ((https://doi.org/10.1007/s13225-020-00466-2)) |
| Otu49783 | 1 | k__Fungi(100);p__Ascomycota(100);c__Saccharomycetes(100);o__Saccharomycetales(100);f__Metschnikowiaceae(100);g__Metschnikowia(100);s__Metschnikowia_pulcherrima(100); | Metschnikowia | 13 | Saprotroph | \|Undefined Saprotroph\| | Yeast | NULL | Probable | As nectar-tap_saprotroph (Põlme et al. 2020); Decay sugar-rich substrates (Põlme et al. 2020) | Põlme S, et al. 2020. Fungal Diversity 105:1-16 ((https://doi.org/10.1007/s13225-020-00466-2)) |
| Otu49784 | 1 | k__Fungi(100);p__Ascomycota(100);c__Saccharomycetes(100);o__Saccharomycetales(100);f__Metschnikowiaceae(100);g__Metschnikowia(100);g__Metschnikowia_unclassified(100); | Metschnikowia | 13 | Saprotroph | \|Undefined Saprotroph\| | Yeast | NULL | Probable | As nectar-tap_saprotroph (Põlme et al. 2020); Decay sugar-rich substrates (Põlme et al. 2020) | Põlme S, et al. 2020. Fungal Diversity 105:1-16 ((https://doi.org/10.1007/s13225-020-00466-2)) |
| Otu49785 | 1 | k__Fungi(100);p__Ascomycota(100);c__Saccharomycetes(100);o__Saccharomycetales(100);f__Metschnikowiaceae(100);g__Metschnikowia(100);s__Metschnikowia_pulcherrima(100); | Metschnikowia | 13 | Saprotroph | \|Undefined Saprotroph\| | Yeast | NULL | Probable | As nectar-tap_saprotroph (Põlme et al. 2020); Decay sugar-rich substrates (Põlme et al. 2020) | Põlme S, et al. 2020. Fungal Diversity 105:1-16 ((https://doi.org/10.1007/s13225-020-00466-2)) |
| Otu49786 | 1 | k__Fungi(100);p__Ascomycota(100);c__Saccharomycetes(100);o__Saccharomycetales(100);f__Metschnikowiaceae(100);g__Metschnikowia(100);s__Metschnikowia_pulcherrima(100); | Metschnikowia | 13 | Saprotroph | \|Undefined Saprotroph\| | Yeast | NULL | Probable | As nectar-tap_saprotroph (Põlme et al. 2020); Decay sugar-rich substrates (Põlme et al. 2020) | Põlme S, et al. 2020. Fungal Diversity 105:1-16 ((https://doi.org/10.1007/s13225-020-00466-2)) |
| Otu49787 | 1 | k__Fungi(100);p__Ascomycota(100);c__Saccharomycetes(100);o__Saccharomycetales(100);f__Metschnikowiaceae(100);g__Metschnikowia(100);g__Metschnikowia_unclassified(100); | Metschnikowia | 13 | Saprotroph | \|Undefined Saprotroph\| | Yeast | NULL | Probable | As nectar-tap_saprotroph (Põlme et al. 2020); Decay sugar-rich substrates (Põlme et al. 2020) | Põlme S, et al. 2020. Fungal Diversity 105:1-16 ((https://doi.org/10.1007/s13225-020-00466-2)) |
| Otu49788 | 1 | k__Fungi(100);p__Ascomycota(100);c__Saccharomycetes(100);o__Saccharomycetales(100);f__Metschnikowiaceae(100);g__Metschnikowia(100);s__Metschnikowia_pulcherrima(100); | Metschnikowia | 13 | Saprotroph | \|Undefined Saprotroph\| | Yeast | NULL | Probable | As nectar-tap_saprotroph (Põlme et al. 2020); Decay sugar-rich substrates (Põlme et al. 2020) | Põlme S, et al. 2020. Fungal Diversity 105:1-16 ((https://doi.org/10.1007/s13225-020-00466-2)) |
| Otu49790 | 1 | k__Fungi(100);p__Ascomycota(100);c__Saccharomycetes(100);o__Saccharomycetales(100);f__Metschnikowiaceae(100);g__Metschnikowia(100);s__Metschnikowia_pulcherrima(100); | Metschnikowia | 13 | Saprotroph | \|Undefined Saprotroph\| | Yeast | NULL | Probable | As nectar-tap_saprotroph (Põlme et al. 2020); Decay sugar-rich substrates (Põlme et al. 2020) | Põlme S, et al. 2020. Fungal Diversity 105:1-16 ((https://doi.org/10.1007/s13225-020-00466-2)) |
| Otu49791 | 1 | k__Fungi(100);p__Ascomycota(100);c__Saccharomycetes(100);o__Saccharomycetales(100);f__Metschnikowiaceae(100);g__Metschnikowia(100);s__Metschnikowia_pulcherrima(100); | Metschnikowia | 13 | Saprotroph | \|Undefined Saprotroph\| | Yeast | NULL | Probable | As nectar-tap_saprotroph (Põlme et al. 2020); Decay sugar-rich substrates (Põlme et al. 2020) | Põlme S, et al. 2020. Fungal Diversity 105:1-16 ((https://doi.org/10.1007/s13225-020-00466-2)) |
| Otu49792 | 1 | k__Fungi(100);p__Ascomycota(100);c__Saccharomycetes(100);o__Saccharomycetales(100);f__Metschnikowiaceae(100);g__Metschnikowia(100);s__Metschnikowia_pulcherrima(100); | Metschnikowia | 13 | Saprotroph | \|Undefined Saprotroph\| | Yeast | NULL | Probable | As nectar-tap_saprotroph (Põlme et al. 2020); Decay sugar-rich substrates (Põlme et al. 2020) | Põlme S, et al. 2020. Fungal Diversity 105:1-16 ((https://doi.org/10.1007/s13225-020-00466-2)) |
| Otu49793 | 1 | k__Fungi(100);p__Ascomycota(100);c__Saccharomycetes(100);o__Saccharomycetales(100);f__Metschnikowiaceae(100);g__Metschnikowia(100);s__Metschnikowia_pulcherrima(100); | Metschnikowia | 13 | Saprotroph | \|Undefined Saprotroph\| | Yeast | NULL | Probable | As nectar-tap_saprotroph (Põlme et al. 2020); Decay sugar-rich substrates (Põlme et al. 2020) | Põlme S, et al. 2020. Fungal Diversity 105:1-16 ((https://doi.org/10.1007/s13225-020-00466-2)) |
| Otu49794 | 1 | k__Fungi(100);p__Ascomycota(100);c__Saccharomycetes(100);o__Saccharomycetales(100);f__Metschnikowiaceae(100);g__Metschnikowia(100);s__Metschnikowia_pulcherrima(100); | Metschnikowia | 13 | Saprotroph | \|Undefined Saprotroph\| | Yeast | NULL | Probable | As nectar-tap_saprotroph (Põlme et al. 2020); Decay sugar-rich substrates (Põlme et al. 2020) | Põlme S, et al. 2020. Fungal Diversity 105:1-16 ((https://doi.org/10.1007/s13225-020-00466-2)) |
| Otu49795 | 1 | k__Fungi(100);p__Ascomycota(100);c__Saccharomycetes(100);o__Saccharomycetales(100);f__Metschnikowiaceae(100);g__Metschnikowia(100);s__Metschnikowia_pulcherrima(100); | Metschnikowia | 13 | Saprotroph | \|Undefined Saprotroph\| | Yeast | NULL | Probable | As nectar-tap_saprotroph (Põlme et al. 2020); Decay sugar-rich substrates (Põlme et al. 2020) | Põlme S, et al. 2020. Fungal Diversity 105:1-16 ((https://doi.org/10.1007/s13225-020-00466-2)) |
| Otu49796 | 1 | k__Fungi(100);p__Ascomycota(100);c__Saccharomycetes(100);o__Saccharomycetales(100);f__Metschnikowiaceae(100);g__Metschnikowia(100);s__Metschnikowia_pulcherrima(100); | Metschnikowia | 13 | Saprotroph | \|Undefined Saprotroph\| | Yeast | NULL | Probable | As nectar-tap_saprotroph (Põlme et al. 2020); Decay sugar-rich substrates (Põlme et al. 2020) | Põlme S, et al. 2020. Fungal Diversity 105:1-16 ((https://doi.org/10.1007/s13225-020-00466-2)) |
| Otu49797 | 1 | k__Fungi(100);p__Ascomycota(100);c__Saccharomycetes(100);o__Saccharomycetales(100);f__Metschnikowiaceae(100);g__Metschnikowia(100);g__Metschnikowia_unclassified(100); | Metschnikowia | 13 | Saprotroph | \|Undefined Saprotroph\| | Yeast | NULL | Probable | As nectar-tap_saprotroph (Põlme et al. 2020); Decay sugar-rich substrates (Põlme et al. 2020) | Põlme S, et al. 2020. Fungal Diversity 105:1-16 ((https://doi.org/10.1007/s13225-020-00466-2)) |
| Otu49798 | 1 | k__Fungi(100);p__Ascomycota(100);c__Saccharomycetes(100);o__Saccharomycetales(100);f__Metschnikowiaceae(100);g__Metschnikowia(100);g__Metschnikowia_unclassified(100); | Metschnikowia | 13 | Saprotroph | \|Undefined Saprotroph\| | Yeast | NULL | Probable | As nectar-tap_saprotroph (Põlme et al. 2020); Decay sugar-rich substrates (Põlme et al. 2020) | Põlme S, et al. 2020. Fungal Diversity 105:1-16 ((https://doi.org/10.1007/s13225-020-00466-2)) |
| Otu49799 | 1 | k__Fungi(100);p__Ascomycota(100);c__Saccharomycetes(100);o__Saccharomycetales(100);f__Metschnikowiaceae(100);g__Metschnikowia(100);s__Metschnikowia_pulcherrima(100); | Metschnikowia | 13 | Saprotroph | \|Undefined Saprotroph\| | Yeast | NULL | Probable | As nectar-tap_saprotroph (Põlme et al. 2020); Decay sugar-rich substrates (Põlme et al. 2020) | Põlme S, et al. 2020. Fungal Diversity 105:1-16 ((https://doi.org/10.1007/s13225-020-00466-2)) |
| Otu49800 | 1 | k__Fungi(100);p__Ascomycota(100);c__Saccharomycetes(100);o__Saccharomycetales(100);f__Metschnikowiaceae(100);g__Metschnikowia(100);s__Metschnikowia_pulcherrima(100); | Metschnikowia | 13 | Saprotroph | \|Undefined Saprotroph\| | Yeast | NULL | Probable | As nectar-tap_saprotroph (Põlme et al. 2020); Decay sugar-rich substrates (Põlme et al. 2020) | Põlme S, et al. 2020. Fungal Diversity 105:1-16 ((https://doi.org/10.1007/s13225-020-00466-2)) |
| Otu49801 | 1 | k__Fungi(100);p__Ascomycota(100);c__Saccharomycetes(100);o__Saccharomycetales(100);f__Metschnikowiaceae(100);g__Metschnikowia(100);s__Metschnikowia_pulcherrima(100); | Metschnikowia | 13 | Saprotroph | \|Undefined Saprotroph\| | Yeast | NULL | Probable | As nectar-tap_saprotroph (Põlme et al. 2020); Decay sugar-rich substrates (Põlme et al. 2020) | Põlme S, et al. 2020. Fungal Diversity 105:1-16 ((https://doi.org/10.1007/s13225-020-00466-2)) |
| Otu49802 | 1 | k__Fungi(100);p__Ascomycota(100);c__Saccharomycetes(100);o__Saccharomycetales(100);f__Metschnikowiaceae(100);g__Metschnikowia(100);s__Metschnikowia_pulcherrima(100); | Metschnikowia | 13 | Saprotroph | \|Undefined Saprotroph\| | Yeast | NULL | Probable | As nectar-tap_saprotroph (Põlme et al. 2020); Decay sugar-rich substrates (Põlme et al. 2020) | Põlme S, et al. 2020. Fungal Diversity 105:1-16 ((https://doi.org/10.1007/s13225-020-00466-2)) |
| Otu49803 | 1 | k__Fungi(100);p__Ascomycota(100);c__Saccharomycetes(100);o__Saccharomycetales(100);f__Metschnikowiaceae(100);g__Metschnikowia(100);s__Metschnikowia_pulcherrima(100); | Metschnikowia | 13 | Saprotroph | \|Undefined Saprotroph\| | Yeast | NULL | Probable | As nectar-tap_saprotroph (Põlme et al. 2020); Decay sugar-rich substrates (Põlme et al. 2020) | Põlme S, et al. 2020. Fungal Diversity 105:1-16 ((https://doi.org/10.1007/s13225-020-00466-2)) |
| Otu49804 | 1 | k__Fungi(100);p__Ascomycota(100);c__Saccharomycetes(100);o__Saccharomycetales(100);f__Metschnikowiaceae(100);g__Metschnikowia(100);g__Metschnikowia_unclassified(100); | Metschnikowia | 13 | Saprotroph | \|Undefined Saprotroph\| | Yeast | NULL | Probable | As nectar-tap_saprotroph (Põlme et al. 2020); Decay sugar-rich substrates (Põlme et al. 2020) | Põlme S, et al. 2020. Fungal Diversity 105:1-16 ((https://doi.org/10.1007/s13225-020-00466-2)) |
| Otu49805 | 1 | k__Fungi(100);p__Ascomycota(100);c__Saccharomycetes(100);o__Saccharomycetales(100);f__Metschnikowiaceae(100);g__Metschnikowia(100);s__Metschnikowia_pulcherrima(100); | Metschnikowia | 13 | Saprotroph | \|Undefined Saprotroph\| | Yeast | NULL | Probable | As nectar-tap_saprotroph (Põlme et al. 2020); Decay sugar-rich substrates (Põlme et al. 2020) | Põlme S, et al. 2020. Fungal Diversity 105:1-16 ((https://doi.org/10.1007/s13225-020-00466-2)) |
| Otu49806 | 1 | k__Fungi(100);p__Ascomycota(100);c__Saccharomycetes(100);o__Saccharomycetales(100);f__Metschnikowiaceae(100);g__Metschnikowia(100);g__Metschnikowia_unclassified(100); | Metschnikowia | 13 | Saprotroph | \|Undefined Saprotroph\| | Yeast | NULL | Probable | As nectar-tap_saprotroph (Põlme et al. 2020); Decay sugar-rich substrates (Põlme et al. 2020) | Põlme S, et al. 2020. Fungal Diversity 105:1-16 ((https://doi.org/10.1007/s13225-020-00466-2)) |
| Otu49807 | 1 | k__Fungi(100);p__Ascomycota(100);c__Saccharomycetes(100);o__Saccharomycetales(100);f__Metschnikowiaceae(100);g__Metschnikowia(100);g__Metschnikowia_unclassified(100); | Metschnikowia | 13 | Saprotroph | \|Undefined Saprotroph\| | Yeast | NULL | Probable | As nectar-tap_saprotroph (Põlme et al. 2020); Decay sugar-rich substrates (Põlme et al. 2020) | Põlme S, et al. 2020. Fungal Diversity 105:1-16 ((https://doi.org/10.1007/s13225-020-00466-2)) |
| Otu49808 | 1 | k__Fungi(100);p__Ascomycota(100);c__Saccharomycetes(100);o__Saccharomycetales(100);f__Metschnikowiaceae(100);g__Metschnikowia(100);s__Metschnikowia_pulcherrima(100); | Metschnikowia | 13 | Saprotroph | \|Undefined Saprotroph\| | Yeast | NULL | Probable | As nectar-tap_saprotroph (Põlme et al. 2020); Decay sugar-rich substrates (Põlme et al. 2020) | Põlme S, et al. 2020. Fungal Diversity 105:1-16 ((https://doi.org/10.1007/s13225-020-00466-2)) |
| Otu49809 | 1 | k__Fungi(100);p__Ascomycota(100);c__Saccharomycetes(100);o__Saccharomycetales(100);f__Metschnikowiaceae(100);g__Metschnikowia(100);s__Metschnikowia_pulcherrima(100); | Metschnikowia | 13 | Saprotroph | \|Undefined Saprotroph\| | Yeast | NULL | Probable | As nectar-tap_saprotroph (Põlme et al. 2020); Decay sugar-rich substrates (Põlme et al. 2020) | Põlme S, et al. 2020. Fungal Diversity 105:1-16 ((https://doi.org/10.1007/s13225-020-00466-2)) |
| Otu49810 | 1 | k__Fungi(100);p__Ascomycota(100);c__Saccharomycetes(100);o__Saccharomycetales(100);f__Metschnikowiaceae(100);g__Metschnikowia(100);s__Metschnikowia_pulcherrima(100); | Metschnikowia | 13 | Saprotroph | \|Undefined Saprotroph\| | Yeast | NULL | Probable | As nectar-tap_saprotroph (Põlme et al. 2020); Decay sugar-rich substrates (Põlme et al. 2020) | Põlme S, et al. 2020. Fungal Diversity 105:1-16 ((https://doi.org/10.1007/s13225-020-00466-2)) |
| Otu49811 | 1 | k__Fungi(100);p__Ascomycota(100);c__Saccharomycetes(100);o__Saccharomycetales(100);f__Metschnikowiaceae(100);g__Metschnikowia(100);s__Metschnikowia_pulcherrima(100); | Metschnikowia | 13 | Saprotroph | \|Undefined Saprotroph\| | Yeast | NULL | Probable | As nectar-tap_saprotroph (Põlme et al. 2020); Decay sugar-rich substrates (Põlme et al. 2020) | Põlme S, et al. 2020. Fungal Diversity 105:1-16 ((https://doi.org/10.1007/s13225-020-00466-2)) |
| Otu49812 | 1 | k__Fungi(100);p__Ascomycota(100);c__Saccharomycetes(100);o__Saccharomycetales(100);f__Metschnikowiaceae(100);g__Metschnikowia(100);s__Metschnikowia_pulcherrima(100); | Metschnikowia | 13 | Saprotroph | \|Undefined Saprotroph\| | Yeast | NULL | Probable | As nectar-tap_saprotroph (Põlme et al. 2020); Decay sugar-rich substrates (Põlme et al. 2020) | Põlme S, et al. 2020. Fungal Diversity 105:1-16 ((https://doi.org/10.1007/s13225-020-00466-2)) |
| Otu49813 | 1 | k__Fungi(100);p__Ascomycota(100);c__Saccharomycetes(100);o__Saccharomycetales(100);f__Metschnikowiaceae(100);g__Metschnikowia(100);s__Metschnikowia_pulcherrima(100); | Metschnikowia | 13 | Saprotroph | \|Undefined Saprotroph\| | Yeast | NULL | Probable | As nectar-tap_saprotroph (Põlme et al. 2020); Decay sugar-rich substrates (Põlme et al. 2020) | Põlme S, et al. 2020. Fungal Diversity 105:1-16 ((https://doi.org/10.1007/s13225-020-00466-2)) |
| Otu49814 | 1 | k__Fungi(100);p__Ascomycota(100);c__Saccharomycetes(100);o__Saccharomycetales(100);f__Metschnikowiaceae(100);g__Metschnikowia(100);s__Metschnikowia_pulcherrima(100); | Metschnikowia | 13 | Saprotroph | \|Undefined Saprotroph\| | Yeast | NULL | Probable | As nectar-tap_saprotroph (Põlme et al. 2020); Decay sugar-rich substrates (Põlme et al. 2020) | Põlme S, et al. 2020. Fungal Diversity 105:1-16 ((https://doi.org/10.1007/s13225-020-00466-2)) |
| Otu49815 | 1 | k__Fungi(100);p__Ascomycota(100);c__Saccharomycetes(100);o__Saccharomycetales(100);f__Metschnikowiaceae(100);g__Metschnikowia(100);s__Metschnikowia_pulcherrima(100); | Metschnikowia | 13 | Saprotroph | \|Undefined Saprotroph\| | Yeast | NULL | Probable | As nectar-tap_saprotroph (Põlme et al. 2020); Decay sugar-rich substrates (Põlme et al. 2020) | Põlme S, et al. 2020. Fungal Diversity 105:1-16 ((https://doi.org/10.1007/s13225-020-00466-2)) |
| Otu49816 | 1 | k__Fungi(100);p__Ascomycota(100);c__Saccharomycetes(100);o__Saccharomycetales(100);f__Metschnikowiaceae(100);g__Metschnikowia(100);s__Metschnikowia_pulcherrima(100); | Metschnikowia | 13 | Saprotroph | \|Undefined Saprotroph\| | Yeast | NULL | Probable | As nectar-tap_saprotroph (Põlme et al. 2020); Decay sugar-rich substrates (Põlme et al. 2020) | Põlme S, et al. 2020. Fungal Diversity 105:1-16 ((https://doi.org/10.1007/s13225-020-00466-2)) |
| Otu49817 | 1 | k__Fungi(100);p__Ascomycota(100);c__Saccharomycetes(100);o__Saccharomycetales(100);f__Metschnikowiaceae(100);g__Metschnikowia(100);s__Metschnikowia_pulcherrima(100); | Metschnikowia | 13 | Saprotroph | \|Undefined Saprotroph\| | Yeast | NULL | Probable | As nectar-tap_saprotroph (Põlme et al. 2020); Decay sugar-rich substrates (Põlme et al. 2020) | Põlme S, et al. 2020. Fungal Diversity 105:1-16 ((https://doi.org/10.1007/s13225-020-00466-2)) |
| Otu49818 | 1 | k__Fungi(100);p__Ascomycota(100);c__Saccharomycetes(100);o__Saccharomycetales(100);f__Metschnikowiaceae(100);g__Metschnikowia(100);s__Metschnikowia_pulcherrima(100); | Metschnikowia | 13 | Saprotroph | \|Undefined Saprotroph\| | Yeast | NULL | Probable | As nectar-tap_saprotroph (Põlme et al. 2020); Decay sugar-rich substrates (Põlme et al. 2020) | Põlme S, et al. 2020. Fungal Diversity 105:1-16 ((https://doi.org/10.1007/s13225-020-00466-2)) |
| Otu49819 | 1 | k__Fungi(100);p__Ascomycota(100);c__Saccharomycetes(100);o__Saccharomycetales(100);f__Metschnikowiaceae(100);g__Metschnikowia(100);s__Metschnikowia_pulcherrima(100); | Metschnikowia | 13 | Saprotroph | \|Undefined Saprotroph\| | Yeast | NULL | Probable | As nectar-tap_saprotroph (Põlme et al. 2020); Decay sugar-rich substrates (Põlme et al. 2020) | Põlme S, et al. 2020. Fungal Diversity 105:1-16 ((https://doi.org/10.1007/s13225-020-00466-2)) |
| Otu49820 | 1 | k__Fungi(100);p__Ascomycota(100);c__Saccharomycetes(100);o__Saccharomycetales(100);f__Metschnikowiaceae(100);g__Metschnikowia(100);s__Metschnikowia_pulcherrima(100); | Metschnikowia | 13 | Saprotroph | \|Undefined Saprotroph\| | Yeast | NULL | Probable | As nectar-tap_saprotroph (Põlme et al. 2020); Decay sugar-rich substrates (Põlme et al. 2020) | Põlme S, et al. 2020. Fungal Diversity 105:1-16 ((https://doi.org/10.1007/s13225-020-00466-2)) |
| Otu49821 | 1 | k__Fungi(100);p__Ascomycota(100);c__Saccharomycetes(100);o__Saccharomycetales(100);f__Metschnikowiaceae(100);g__Metschnikowia(100);g__Metschnikowia_unclassified(100); | Metschnikowia | 13 | Saprotroph | \|Undefined Saprotroph\| | Yeast | NULL | Probable | As nectar-tap_saprotroph (Põlme et al. 2020); Decay sugar-rich substrates (Põlme et al. 2020) | Põlme S, et al. 2020. Fungal Diversity 105:1-16 ((https://doi.org/10.1007/s13225-020-00466-2)) |
| Otu49822 | 1 | k__Fungi(100);p__Ascomycota(100);c__Saccharomycetes(100);o__Saccharomycetales(100);f__Metschnikowiaceae(100);g__Metschnikowia(100);g__Metschnikowia_unclassified(100); | Metschnikowia | 13 | Saprotroph | \|Undefined Saprotroph\| | Yeast | NULL | Probable | As nectar-tap_saprotroph (Põlme et al. 2020); Decay sugar-rich substrates (Põlme et al. 2020) | Põlme S, et al. 2020. Fungal Diversity 105:1-16 ((https://doi.org/10.1007/s13225-020-00466-2)) |
| Otu49823 | 1 | k__Fungi(100);p__Ascomycota(100);c__Saccharomycetes(100);o__Saccharomycetales(100);f__Metschnikowiaceae(100);g__Metschnikowia(100);s__Metschnikowia_pulcherrima(100); | Metschnikowia | 13 | Saprotroph | \|Undefined Saprotroph\| | Yeast | NULL | Probable | As nectar-tap_saprotroph (Põlme et al. 2020); Decay sugar-rich substrates (Põlme et al. 2020) | Põlme S, et al. 2020. Fungal Diversity 105:1-16 ((https://doi.org/10.1007/s13225-020-00466-2)) |
| Otu49824 | 1 | k__Fungi(100);p__Ascomycota(100);c__Saccharomycetes(100);o__Saccharomycetales(100);f__Metschnikowiaceae(100);g__Metschnikowia(100);s__Metschnikowia_pulcherrima(100); | Metschnikowia | 13 | Saprotroph | \|Undefined Saprotroph\| | Yeast | NULL | Probable | As nectar-tap_saprotroph (Põlme et al. 2020); Decay sugar-rich substrates (Põlme et al. 2020) | Põlme S, et al. 2020. Fungal Diversity 105:1-16 ((https://doi.org/10.1007/s13225-020-00466-2)) |
| Otu49825 | 1 | k__Fungi(100);p__Ascomycota(100);c__Saccharomycetes(100);o__Saccharomycetales(100);f__Metschnikowiaceae(100);g__Metschnikowia(100);s__Metschnikowia_pulcherrima(100); | Metschnikowia | 13 | Saprotroph | \|Undefined Saprotroph\| | Yeast | NULL | Probable | As nectar-tap_saprotroph (Põlme et al. 2020); Decay sugar-rich substrates (Põlme et al. 2020) | Põlme S, et al. 2020. Fungal Diversity 105:1-16 ((https://doi.org/10.1007/s13225-020-00466-2)) |
| Otu49826 | 1 | k__Fungi(100);p__Ascomycota(100);c__Saccharomycetes(100);o__Saccharomycetales(100);f__Metschnikowiaceae(100);g__Metschnikowia(100);g__Metschnikowia_unclassified(100); | Metschnikowia | 13 | Saprotroph | \|Undefined Saprotroph\| | Yeast | NULL | Probable | As nectar-tap_saprotroph (Põlme et al. 2020); Decay sugar-rich substrates (Põlme et al. 2020) | Põlme S, et al. 2020. Fungal Diversity 105:1-16 ((https://doi.org/10.1007/s13225-020-00466-2)) |
| Otu49827 | 1 | k__Fungi(100);p__Ascomycota(100);c__Saccharomycetes(100);o__Saccharomycetales(100);f__Metschnikowiaceae(100);g__Metschnikowia(100);s__Metschnikowia_pulcherrima(100); | Metschnikowia | 13 | Saprotroph | \|Undefined Saprotroph\| | Yeast | NULL | Probable | As nectar-tap_saprotroph (Põlme et al. 2020); Decay sugar-rich substrates (Põlme et al. 2020) | Põlme S, et al. 2020. Fungal Diversity 105:1-16 ((https://doi.org/10.1007/s13225-020-00466-2)) |
| Otu49828 | 1 | k__Fungi(100);p__Ascomycota(100);c__Saccharomycetes(100);o__Saccharomycetales(100);f__Metschnikowiaceae(100);g__Metschnikowia(100);g__Metschnikowia_unclassified(100); | Metschnikowia | 13 | Saprotroph | \|Undefined Saprotroph\| | Yeast | NULL | Probable | As nectar-tap_saprotroph (Põlme et al. 2020); Decay sugar-rich substrates (Põlme et al. 2020) | Põlme S, et al. 2020. Fungal Diversity 105:1-16 ((https://doi.org/10.1007/s13225-020-00466-2)) |
| Otu49829 | 1 | k__Fungi(100);p__Ascomycota(100);c__Saccharomycetes(100);o__Saccharomycetales(100);f__Metschnikowiaceae(100);g__Metschnikowia(100);s__Metschnikowia_pulcherrima(100); | Metschnikowia | 13 | Saprotroph | \|Undefined Saprotroph\| | Yeast | NULL | Probable | As nectar-tap_saprotroph (Põlme et al. 2020); Decay sugar-rich substrates (Põlme et al. 2020) | Põlme S, et al. 2020. Fungal Diversity 105:1-16 ((https://doi.org/10.1007/s13225-020-00466-2)) |
| Otu49830 | 1 | k__Fungi(100);p__Ascomycota(100);c__Saccharomycetes(100);o__Saccharomycetales(100);f__Metschnikowiaceae(100);g__Metschnikowia(100);s__Metschnikowia_pulcherrima(100); | Metschnikowia | 13 | Saprotroph | \|Undefined Saprotroph\| | Yeast | NULL | Probable | As nectar-tap_saprotroph (Põlme et al. 2020); Decay sugar-rich substrates (Põlme et al. 2020) | Põlme S, et al. 2020. Fungal Diversity 105:1-16 ((https://doi.org/10.1007/s13225-020-00466-2)) |
| Otu49831 | 1 | k__Fungi(100);p__Ascomycota(100);c__Saccharomycetes(100);o__Saccharomycetales(100);f__Metschnikowiaceae(100);g__Metschnikowia(100);s__Metschnikowia_pulcherrima(100); | Metschnikowia | 13 | Saprotroph | \|Undefined Saprotroph\| | Yeast | NULL | Probable | As nectar-tap_saprotroph (Põlme et al. 2020); Decay sugar-rich substrates (Põlme et al. 2020) | Põlme S, et al. 2020. Fungal Diversity 105:1-16 ((https://doi.org/10.1007/s13225-020-00466-2)) |
| Otu49832 | 1 | k__Fungi(100);p__Ascomycota(100);c__Saccharomycetes(100);o__Saccharomycetales(100);f__Metschnikowiaceae(100);g__Metschnikowia(100);s__Metschnikowia_pulcherrima(100); | Metschnikowia | 13 | Saprotroph | \|Undefined Saprotroph\| | Yeast | NULL | Probable | As nectar-tap_saprotroph (Põlme et al. 2020); Decay sugar-rich substrates (Põlme et al. 2020) | Põlme S, et al. 2020. Fungal Diversity 105:1-16 ((https://doi.org/10.1007/s13225-020-00466-2)) |
| Otu49833 | 1 | k__Fungi(100);p__Ascomycota(100);c__Saccharomycetes(100);o__Saccharomycetales(100);f__Metschnikowiaceae(100);g__Metschnikowia(100);s__Metschnikowia_pulcherrima(100); | Metschnikowia | 13 | Saprotroph | \|Undefined Saprotroph\| | Yeast | NULL | Probable | As nectar-tap_saprotroph (Põlme et al. 2020); Decay sugar-rich substrates (Põlme et al. 2020) | Põlme S, et al. 2020. Fungal Diversity 105:1-16 ((https://doi.org/10.1007/s13225-020-00466-2)) |
| Otu49834 | 1 | k__Fungi(100);p__Ascomycota(100);c__Saccharomycetes(100);o__Saccharomycetales(100);f__Metschnikowiaceae(100);g__Metschnikowia(100);s__Metschnikowia_pulcherrima(100); | Metschnikowia | 13 | Saprotroph | \|Undefined Saprotroph\| | Yeast | NULL | Probable | As nectar-tap_saprotroph (Põlme et al. 2020); Decay sugar-rich substrates (Põlme et al. 2020) | Põlme S, et al. 2020. Fungal Diversity 105:1-16 ((https://doi.org/10.1007/s13225-020-00466-2)) |
| Otu49835 | 1 | k__Fungi(100);p__Ascomycota(100);c__Saccharomycetes(100);o__Saccharomycetales(100);f__Metschnikowiaceae(100);g__Metschnikowia(100);g__Metschnikowia_unclassified(100); | Metschnikowia | 13 | Saprotroph | \|Undefined Saprotroph\| | Yeast | NULL | Probable | As nectar-tap_saprotroph (Põlme et al. 2020); Decay sugar-rich substrates (Põlme et al. 2020) | Põlme S, et al. 2020. Fungal Diversity 105:1-16 ((https://doi.org/10.1007/s13225-020-00466-2)) |
| Otu49836 | 1 | k__Fungi(100);p__Ascomycota(100);c__Saccharomycetes(100);o__Saccharomycetales(100);f__Metschnikowiaceae(100);g__Metschnikowia(100);s__Metschnikowia_pulcherrima(100); | Metschnikowia | 13 | Saprotroph | \|Undefined Saprotroph\| | Yeast | NULL | Probable | As nectar-tap_saprotroph (Põlme et al. 2020); Decay sugar-rich substrates (Põlme et al. 2020) | Põlme S, et al. 2020. Fungal Diversity 105:1-16 ((https://doi.org/10.1007/s13225-020-00466-2)) |
| Otu49837 | 1 | k__Fungi(100);p__Ascomycota(100);c__Saccharomycetes(100);o__Saccharomycetales(100);f__Metschnikowiaceae(100);g__Metschnikowia(100);g__Metschnikowia_unclassified(100); | Metschnikowia | 13 | Saprotroph | \|Undefined Saprotroph\| | Yeast | NULL | Probable | As nectar-tap_saprotroph (Põlme et al. 2020); Decay sugar-rich substrates (Põlme et al. 2020) | Põlme S, et al. 2020. Fungal Diversity 105:1-16 ((https://doi.org/10.1007/s13225-020-00466-2)) |
| Otu49838 | 1 | k__Fungi(100);p__Ascomycota(100);c__Saccharomycetes(100);o__Saccharomycetales(100);f__Metschnikowiaceae(100);g__Metschnikowia(100);s__Metschnikowia_pulcherrima(100); | Metschnikowia | 13 | Saprotroph | \|Undefined Saprotroph\| | Yeast | NULL | Probable | As nectar-tap_saprotroph (Põlme et al. 2020); Decay sugar-rich substrates (Põlme et al. 2020) | Põlme S, et al. 2020. Fungal Diversity 105:1-16 ((https://doi.org/10.1007/s13225-020-00466-2)) |
| Otu49839 | 1 | k__Fungi(100);p__Ascomycota(100);c__Saccharomycetes(100);o__Saccharomycetales(100);f__Metschnikowiaceae(100);g__Metschnikowia(100);g__Metschnikowia_unclassified(100); | Metschnikowia | 13 | Saprotroph | \|Undefined Saprotroph\| | Yeast | NULL | Probable | As nectar-tap_saprotroph (Põlme et al. 2020); Decay sugar-rich substrates (Põlme et al. 2020) | Põlme S, et al. 2020. Fungal Diversity 105:1-16 ((https://doi.org/10.1007/s13225-020-00466-2)) |
| Otu49840 | 1 | k__Fungi(100);p__Ascomycota(100);c__Saccharomycetes(100);o__Saccharomycetales(100);f__Metschnikowiaceae(100);g__Metschnikowia(100);g__Metschnikowia_unclassified(100); | Metschnikowia | 13 | Saprotroph | \|Undefined Saprotroph\| | Yeast | NULL | Probable | As nectar-tap_saprotroph (Põlme et al. 2020); Decay sugar-rich substrates (Põlme et al. 2020) | Põlme S, et al. 2020. Fungal Diversity 105:1-16 ((https://doi.org/10.1007/s13225-020-00466-2)) |
| Otu49841 | 1 | k__Fungi(100);p__Ascomycota(100);c__Saccharomycetes(100);o__Saccharomycetales(100);f__Metschnikowiaceae(100);g__Metschnikowia(100);s__Metschnikowia_pulcherrima(100); | Metschnikowia | 13 | Saprotroph | \|Undefined Saprotroph\| | Yeast | NULL | Probable | As nectar-tap_saprotroph (Põlme et al. 2020); Decay sugar-rich substrates (Põlme et al. 2020) | Põlme S, et al. 2020. Fungal Diversity 105:1-16 ((https://doi.org/10.1007/s13225-020-00466-2)) |
| Otu49842 | 1 | k__Fungi(100);p__Ascomycota(100);c__Saccharomycetes(100);o__Saccharomycetales(100);f__Metschnikowiaceae(100);g__Metschnikowia(100);s__Metschnikowia_pulcherrima(100); | Metschnikowia | 13 | Saprotroph | \|Undefined Saprotroph\| | Yeast | NULL | Probable | As nectar-tap_saprotroph (Põlme et al. 2020); Decay sugar-rich substrates (Põlme et al. 2020) | Põlme S, et al. 2020. Fungal Diversity 105:1-16 ((https://doi.org/10.1007/s13225-020-00466-2)) |
| Otu49843 | 1 | k__Fungi(100);p__Ascomycota(100);c__Saccharomycetes(100);o__Saccharomycetales(100);f__Metschnikowiaceae(100);g__Metschnikowia(100);s__Metschnikowia_pulcherrima(100); | Metschnikowia | 13 | Saprotroph | \|Undefined Saprotroph\| | Yeast | NULL | Probable | As nectar-tap_saprotroph (Põlme et al. 2020); Decay sugar-rich substrates (Põlme et al. 2020) | Põlme S, et al. 2020. Fungal Diversity 105:1-16 ((https://doi.org/10.1007/s13225-020-00466-2)) |
| Otu49844 | 1 | k__Fungi(100);p__Ascomycota(100);c__Saccharomycetes(100);o__Saccharomycetales(100);f__Metschnikowiaceae(100);g__Metschnikowia(100);s__Metschnikowia_pulcherrima(100); | Metschnikowia | 13 | Saprotroph | \|Undefined Saprotroph\| | Yeast | NULL | Probable | As nectar-tap_saprotroph (Põlme et al. 2020); Decay sugar-rich substrates (Põlme et al. 2020) | Põlme S, et al. 2020. Fungal Diversity 105:1-16 ((https://doi.org/10.1007/s13225-020-00466-2)) |
| Otu49845 | 1 | k__Fungi(100);p__Ascomycota(100);c__Saccharomycetes(100);o__Saccharomycetales(100);f__Metschnikowiaceae(100);g__Metschnikowia(100);s__Metschnikowia_pulcherrima(100); | Metschnikowia | 13 | Saprotroph | \|Undefined Saprotroph\| | Yeast | NULL | Probable | As nectar-tap_saprotroph (Põlme et al. 2020); Decay sugar-rich substrates (Põlme et al. 2020) | Põlme S, et al. 2020. Fungal Diversity 105:1-16 ((https://doi.org/10.1007/s13225-020-00466-2)) |
| Otu49846 | 1 | k__Fungi(100);p__Ascomycota(100);c__Saccharomycetes(100);o__Saccharomycetales(100);f__Metschnikowiaceae(100);g__Metschnikowia(100);s__Metschnikowia_pulcherrima(100); | Metschnikowia | 13 | Saprotroph | \|Undefined Saprotroph\| | Yeast | NULL | Probable | As nectar-tap_saprotroph (Põlme et al. 2020); Decay sugar-rich substrates (Põlme et al. 2020) | Põlme S, et al. 2020. Fungal Diversity 105:1-16 ((https://doi.org/10.1007/s13225-020-00466-2)) |
| Otu49847 | 1 | k__Fungi(100);p__Ascomycota(100);c__Saccharomycetes(100);o__Saccharomycetales(100);f__Metschnikowiaceae(100);g__Metschnikowia(100);s__Metschnikowia_pulcherrima(100); | Metschnikowia | 13 | Saprotroph | \|Undefined Saprotroph\| | Yeast | NULL | Probable | As nectar-tap_saprotroph (Põlme et al. 2020); Decay sugar-rich substrates (Põlme et al. 2020) | Põlme S, et al. 2020. Fungal Diversity 105:1-16 ((https://doi.org/10.1007/s13225-020-00466-2)) |
| Otu49848 | 1 | k__Fungi(100);p__Ascomycota(100);c__Saccharomycetes(100);o__Saccharomycetales(100);f__Metschnikowiaceae(100);g__Metschnikowia(100);s__Metschnikowia_pulcherrima(100); | Metschnikowia | 13 | Saprotroph | \|Undefined Saprotroph\| | Yeast | NULL | Probable | As nectar-tap_saprotroph (Põlme et al. 2020); Decay sugar-rich substrates (Põlme et al. 2020) | Põlme S, et al. 2020. Fungal Diversity 105:1-16 ((https://doi.org/10.1007/s13225-020-00466-2)) |
| Otu49849 | 1 | k__Fungi(100);p__Ascomycota(100);c__Saccharomycetes(100);o__Saccharomycetales(100);f__Metschnikowiaceae(100);g__Metschnikowia(100);s__Metschnikowia_pulcherrima(100); | Metschnikowia | 13 | Saprotroph | \|Undefined Saprotroph\| | Yeast | NULL | Probable | As nectar-tap_saprotroph (Põlme et al. 2020); Decay sugar-rich substrates (Põlme et al. 2020) | Põlme S, et al. 2020. Fungal Diversity 105:1-16 ((https://doi.org/10.1007/s13225-020-00466-2)) |
| Otu49850 | 1 | k__Fungi(100);p__Ascomycota(100);c__Saccharomycetes(100);o__Saccharomycetales(100);f__Metschnikowiaceae(100);g__Metschnikowia(100);g__Metschnikowia_unclassified(100); | Metschnikowia | 13 | Saprotroph | \|Undefined Saprotroph\| | Yeast | NULL | Probable | As nectar-tap_saprotroph (Põlme et al. 2020); Decay sugar-rich substrates (Põlme et al. 2020) | Põlme S, et al. 2020. Fungal Diversity 105:1-16 ((https://doi.org/10.1007/s13225-020-00466-2)) |
| Otu49851 | 1 | k__Fungi(100);p__Ascomycota(100);c__Saccharomycetes(100);o__Saccharomycetales(100);f__Metschnikowiaceae(100);g__Metschnikowia(100);s__Metschnikowia_pulcherrima(100); | Metschnikowia | 13 | Saprotroph | \|Undefined Saprotroph\| | Yeast | NULL | Probable | As nectar-tap_saprotroph (Põlme et al. 2020); Decay sugar-rich substrates (Põlme et al. 2020) | Põlme S, et al. 2020. Fungal Diversity 105:1-16 ((https://doi.org/10.1007/s13225-020-00466-2)) |
| Otu49852 | 1 | k__Fungi(100);p__Ascomycota(100);c__Saccharomycetes(100);o__Saccharomycetales(100);f__Metschnikowiaceae(100);g__Metschnikowia(100);g__Metschnikowia_unclassified(100); | Metschnikowia | 13 | Saprotroph | \|Undefined Saprotroph\| | Yeast | NULL | Probable | As nectar-tap_saprotroph (Põlme et al. 2020); Decay sugar-rich substrates (Põlme et al. 2020) | Põlme S, et al. 2020. Fungal Diversity 105:1-16 ((https://doi.org/10.1007/s13225-020-00466-2)) |
| Otu49853 | 1 | k__Fungi(100);p__Ascomycota(100);c__Saccharomycetes(100);o__Saccharomycetales(100);f__Metschnikowiaceae(100);g__Metschnikowia(100);s__Metschnikowia_pulcherrima(100); | Metschnikowia | 13 | Saprotroph | \|Undefined Saprotroph\| | Yeast | NULL | Probable | As nectar-tap_saprotroph (Põlme et al. 2020); Decay sugar-rich substrates (Põlme et al. 2020) | Põlme S, et al. 2020. Fungal Diversity 105:1-16 ((https://doi.org/10.1007/s13225-020-00466-2)) |
| Otu49854 | 1 | k__Fungi(100);p__Ascomycota(100);c__Saccharomycetes(100);o__Saccharomycetales(100);f__Metschnikowiaceae(100);g__Metschnikowia(100);s__Metschnikowia_pulcherrima(100); | Metschnikowia | 13 | Saprotroph | \|Undefined Saprotroph\| | Yeast | NULL | Probable | As nectar-tap_saprotroph (Põlme et al. 2020); Decay sugar-rich substrates (Põlme et al. 2020) | Põlme S, et al. 2020. Fungal Diversity 105:1-16 ((https://doi.org/10.1007/s13225-020-00466-2)) |
| Otu49855 | 1 | k__Fungi(100);p__Ascomycota(100);c__Saccharomycetes(100);o__Saccharomycetales(100);f__Metschnikowiaceae(100);g__Metschnikowia(100);g__Metschnikowia_unclassified(100); | Metschnikowia | 13 | Saprotroph | \|Undefined Saprotroph\| | Yeast | NULL | Probable | As nectar-tap_saprotroph (Põlme et al. 2020); Decay sugar-rich substrates (Põlme et al. 2020) | Põlme S, et al. 2020. Fungal Diversity 105:1-16 ((https://doi.org/10.1007/s13225-020-00466-2)) |
| Otu49856 | 1 | k__Fungi(100);p__Ascomycota(100);c__Saccharomycetes(100);o__Saccharomycetales(100);f__Metschnikowiaceae(100);g__Metschnikowia(100);s__Metschnikowia_pulcherrima(100); | Metschnikowia | 13 | Saprotroph | \|Undefined Saprotroph\| | Yeast | NULL | Probable | As nectar-tap_saprotroph (Põlme et al. 2020); Decay sugar-rich substrates (Põlme et al. 2020) | Põlme S, et al. 2020. Fungal Diversity 105:1-16 ((https://doi.org/10.1007/s13225-020-00466-2)) |
| Otu49858 | 1 | k__Fungi(100);p__Ascomycota(100);c__Saccharomycetes(100);o__Saccharomycetales(100);f__Metschnikowiaceae(100);g__Metschnikowia(100);s__Metschnikowia_pulcherrima(100); | Metschnikowia | 13 | Saprotroph | \|Undefined Saprotroph\| | Yeast | NULL | Probable | As nectar-tap_saprotroph (Põlme et al. 2020); Decay sugar-rich substrates (Põlme et al. 2020) | Põlme S, et al. 2020. Fungal Diversity 105:1-16 ((https://doi.org/10.1007/s13225-020-00466-2)) |
| Otu49859 | 1 | k__Fungi(100);p__Ascomycota(100);c__Saccharomycetes(100);o__Saccharomycetales(100);f__Metschnikowiaceae(100);g__Metschnikowia(100);g__Metschnikowia_unclassified(100); | Metschnikowia | 13 | Saprotroph | \|Undefined Saprotroph\| | Yeast | NULL | Probable | As nectar-tap_saprotroph (Põlme et al. 2020); Decay sugar-rich substrates (Põlme et al. 2020) | Põlme S, et al. 2020. Fungal Diversity 105:1-16 ((https://doi.org/10.1007/s13225-020-00466-2)) |
| Otu49860 | 1 | k__Fungi(100);p__Ascomycota(100);c__Saccharomycetes(100);o__Saccharomycetales(100);f__Metschnikowiaceae(100);g__Metschnikowia(100);g__Metschnikowia_unclassified(100); | Metschnikowia | 13 | Saprotroph | \|Undefined Saprotroph\| | Yeast | NULL | Probable | As nectar-tap_saprotroph (Põlme et al. 2020); Decay sugar-rich substrates (Põlme et al. 2020) | Põlme S, et al. 2020. Fungal Diversity 105:1-16 ((https://doi.org/10.1007/s13225-020-00466-2)) |
| Otu49861 | 1 | k__Fungi(100);p__Ascomycota(100);c__Saccharomycetes(100);o__Saccharomycetales(100);f__Metschnikowiaceae(100);g__Metschnikowia(100);g__Metschnikowia_unclassified(100); | Metschnikowia | 13 | Saprotroph | \|Undefined Saprotroph\| | Yeast | NULL | Probable | As nectar-tap_saprotroph (Põlme et al. 2020); Decay sugar-rich substrates (Põlme et al. 2020) | Põlme S, et al. 2020. Fungal Diversity 105:1-16 ((https://doi.org/10.1007/s13225-020-00466-2)) |
| Otu49862 | 1 | k__Fungi(100);p__Ascomycota(100);c__Saccharomycetes(100);o__Saccharomycetales(100);f__Metschnikowiaceae(100);g__Metschnikowia(100);s__Metschnikowia_pulcherrima(100); | Metschnikowia | 13 | Saprotroph | \|Undefined Saprotroph\| | Yeast | NULL | Probable | As nectar-tap_saprotroph (Põlme et al. 2020); Decay sugar-rich substrates (Põlme et al. 2020) | Põlme S, et al. 2020. Fungal Diversity 105:1-16 ((https://doi.org/10.1007/s13225-020-00466-2)) |
| Otu49863 | 1 | k__Fungi(100);p__Ascomycota(100);c__Saccharomycetes(100);o__Saccharomycetales(100);f__Metschnikowiaceae(100);g__Metschnikowia(100);s__Metschnikowia_pulcherrima(100); | Metschnikowia | 13 | Saprotroph | \|Undefined Saprotroph\| | Yeast | NULL | Probable | As nectar-tap_saprotroph (Põlme et al. 2020); Decay sugar-rich substrates (Põlme et al. 2020) | Põlme S, et al. 2020. Fungal Diversity 105:1-16 ((https://doi.org/10.1007/s13225-020-00466-2)) |
| Otu49864 | 1 | k__Fungi(100);p__Ascomycota(100);c__Saccharomycetes(100);o__Saccharomycetales(100);f__Metschnikowiaceae(100);g__Metschnikowia(100);g__Metschnikowia_unclassified(100); | Metschnikowia | 13 | Saprotroph | \|Undefined Saprotroph\| | Yeast | NULL | Probable | As nectar-tap_saprotroph (Põlme et al. 2020); Decay sugar-rich substrates (Põlme et al. 2020) | Põlme S, et al. 2020. Fungal Diversity 105:1-16 ((https://doi.org/10.1007/s13225-020-00466-2)) |
| Otu49865 | 1 | k__Fungi(100);p__Ascomycota(100);c__Saccharomycetes(100);o__Saccharomycetales(100);f__Metschnikowiaceae(100);g__Metschnikowia(100);s__Metschnikowia_pulcherrima(100); | Metschnikowia | 13 | Saprotroph | \|Undefined Saprotroph\| | Yeast | NULL | Probable | As nectar-tap_saprotroph (Põlme et al. 2020); Decay sugar-rich substrates (Põlme et al. 2020) | Põlme S, et al. 2020. Fungal Diversity 105:1-16 ((https://doi.org/10.1007/s13225-020-00466-2)) |
| Otu49866 | 1 | k__Fungi(100);p__Ascomycota(100);c__Saccharomycetes(100);o__Saccharomycetales(100);f__Metschnikowiaceae(100);g__Metschnikowia(100);g__Metschnikowia_unclassified(100); | Metschnikowia | 13 | Saprotroph | \|Undefined Saprotroph\| | Yeast | NULL | Probable | As nectar-tap_saprotroph (Põlme et al. 2020); Decay sugar-rich substrates (Põlme et al. 2020) | Põlme S, et al. 2020. Fungal Diversity 105:1-16 ((https://doi.org/10.1007/s13225-020-00466-2)) |
| Otu49867 | 1 | k__Fungi(100);p__Ascomycota(100);c__Saccharomycetes(100);o__Saccharomycetales(100);f__Metschnikowiaceae(100);g__Metschnikowia(100);s__Metschnikowia_pulcherrima(100); | Metschnikowia | 13 | Saprotroph | \|Undefined Saprotroph\| | Yeast | NULL | Probable | As nectar-tap_saprotroph (Põlme et al. 2020); Decay sugar-rich substrates (Põlme et al. 2020) | Põlme S, et al. 2020. Fungal Diversity 105:1-16 ((https://doi.org/10.1007/s13225-020-00466-2)) |
| Otu49868 | 1 | k__Fungi(100);p__Ascomycota(100);c__Saccharomycetes(100);o__Saccharomycetales(100);f__Metschnikowiaceae(100);g__Metschnikowia(100);s__Metschnikowia_pulcherrima(100); | Metschnikowia | 13 | Saprotroph | \|Undefined Saprotroph\| | Yeast | NULL | Probable | As nectar-tap_saprotroph (Põlme et al. 2020); Decay sugar-rich substrates (Põlme et al. 2020) | Põlme S, et al. 2020. Fungal Diversity 105:1-16 ((https://doi.org/10.1007/s13225-020-00466-2)) |
| Otu49869 | 1 | k__Fungi(100);p__Ascomycota(100);c__Saccharomycetes(100);o__Saccharomycetales(100);f__Metschnikowiaceae(100);g__Metschnikowia(100);s__Metschnikowia_pulcherrima(100); | Metschnikowia | 13 | Saprotroph | \|Undefined Saprotroph\| | Yeast | NULL | Probable | As nectar-tap_saprotroph (Põlme et al. 2020); Decay sugar-rich substrates (Põlme et al. 2020) | Põlme S, et al. 2020. Fungal Diversity 105:1-16 ((https://doi.org/10.1007/s13225-020-00466-2)) |
| Otu49870 | 1 | k__Fungi(100);p__Ascomycota(100);c__Saccharomycetes(100);o__Saccharomycetales(100);f__Metschnikowiaceae(100);g__Metschnikowia(100);s__Metschnikowia_pulcherrima(100); | Metschnikowia | 13 | Saprotroph | \|Undefined Saprotroph\| | Yeast | NULL | Probable | As nectar-tap_saprotroph (Põlme et al. 2020); Decay sugar-rich substrates (Põlme et al. 2020) | Põlme S, et al. 2020. Fungal Diversity 105:1-16 ((https://doi.org/10.1007/s13225-020-00466-2)) |
| Otu49871 | 1 | k__Fungi(100);p__Ascomycota(100);c__Saccharomycetes(100);o__Saccharomycetales(100);f__Metschnikowiaceae(100);g__Metschnikowia(100);s__Metschnikowia_pulcherrima(100); | Metschnikowia | 13 | Saprotroph | \|Undefined Saprotroph\| | Yeast | NULL | Probable | As nectar-tap_saprotroph (Põlme et al. 2020); Decay sugar-rich substrates (Põlme et al. 2020) | Põlme S, et al. 2020. Fungal Diversity 105:1-16 ((https://doi.org/10.1007/s13225-020-00466-2)) |
| Otu49872 | 1 | k__Fungi(100);p__Ascomycota(100);c__Saccharomycetes(100);o__Saccharomycetales(100);f__Metschnikowiaceae(100);g__Metschnikowia(100);s__Metschnikowia_pulcherrima(100); | Metschnikowia | 13 | Saprotroph | \|Undefined Saprotroph\| | Yeast | NULL | Probable | As nectar-tap_saprotroph (Põlme et al. 2020); Decay sugar-rich substrates (Põlme et al. 2020) | Põlme S, et al. 2020. Fungal Diversity 105:1-16 ((https://doi.org/10.1007/s13225-020-00466-2)) |
| Otu49874 | 1 | k__Fungi(100);p__Ascomycota(100);c__Saccharomycetes(100);o__Saccharomycetales(100);f__Metschnikowiaceae(100);g__Metschnikowia(100);s__Metschnikowia_pulcherrima(100); | Metschnikowia | 13 | Saprotroph | \|Undefined Saprotroph\| | Yeast | NULL | Probable | As nectar-tap_saprotroph (Põlme et al. 2020); Decay sugar-rich substrates (Põlme et al. 2020) | Põlme S, et al. 2020. Fungal Diversity 105:1-16 ((https://doi.org/10.1007/s13225-020-00466-2)) |
| Otu49875 | 1 | k__Fungi(100);p__Ascomycota(100);c__Saccharomycetes(100);o__Saccharomycetales(100);f__Metschnikowiaceae(100);g__Metschnikowia(100);s__Metschnikowia_pulcherrima(100); | Metschnikowia | 13 | Saprotroph | \|Undefined Saprotroph\| | Yeast | NULL | Probable | As nectar-tap_saprotroph (Põlme et al. 2020); Decay sugar-rich substrates (Põlme et al. 2020) | Põlme S, et al. 2020. Fungal Diversity 105:1-16 ((https://doi.org/10.1007/s13225-020-00466-2)) |
| Otu49876 | 1 | k__Fungi(100);p__Ascomycota(100);c__Saccharomycetes(100);o__Saccharomycetales(100);f__Metschnikowiaceae(100);g__Metschnikowia(100);s__Metschnikowia_pulcherrima(100); | Metschnikowia | 13 | Saprotroph | \|Undefined Saprotroph\| | Yeast | NULL | Probable | As nectar-tap_saprotroph (Põlme et al. 2020); Decay sugar-rich substrates (Põlme et al. 2020) | Põlme S, et al. 2020. Fungal Diversity 105:1-16 ((https://doi.org/10.1007/s13225-020-00466-2)) |
| Otu49877 | 1 | k__Fungi(100);p__Ascomycota(100);c__Saccharomycetes(100);o__Saccharomycetales(100);f__Metschnikowiaceae(100);g__Metschnikowia(100);s__Metschnikowia_pulcherrima(100); | Metschnikowia | 13 | Saprotroph | \|Undefined Saprotroph\| | Yeast | NULL | Probable | As nectar-tap_saprotroph (Põlme et al. 2020); Decay sugar-rich substrates (Põlme et al. 2020) | Põlme S, et al. 2020. Fungal Diversity 105:1-16 ((https://doi.org/10.1007/s13225-020-00466-2)) |
| Otu49878 | 1 | k__Fungi(100);p__Ascomycota(100);c__Saccharomycetes(100);o__Saccharomycetales(100);f__Metschnikowiaceae(100);g__Metschnikowia(100);s__Metschnikowia_pulcherrima(100); | Metschnikowia | 13 | Saprotroph | \|Undefined Saprotroph\| | Yeast | NULL | Probable | As nectar-tap_saprotroph (Põlme et al. 2020); Decay sugar-rich substrates (Põlme et al. 2020) | Põlme S, et al. 2020. Fungal Diversity 105:1-16 ((https://doi.org/10.1007/s13225-020-00466-2)) |
| Otu49879 | 1 | k__Fungi(100);p__Ascomycota(100);c__Saccharomycetes(100);o__Saccharomycetales(100);f__Metschnikowiaceae(100);g__Metschnikowia(100);s__Metschnikowia_pulcherrima(100); | Metschnikowia | 13 | Saprotroph | \|Undefined Saprotroph\| | Yeast | NULL | Probable | As nectar-tap_saprotroph (Põlme et al. 2020); Decay sugar-rich substrates (Põlme et al. 2020) | Põlme S, et al. 2020. Fungal Diversity 105:1-16 ((https://doi.org/10.1007/s13225-020-00466-2)) |
| Otu49880 | 1 | k__Fungi(100);p__Ascomycota(100);c__Saccharomycetes(100);o__Saccharomycetales(100);f__Metschnikowiaceae(100);g__Metschnikowia(100);s__Metschnikowia_pulcherrima(100); | Metschnikowia | 13 | Saprotroph | \|Undefined Saprotroph\| | Yeast | NULL | Probable | As nectar-tap_saprotroph (Põlme et al. 2020); Decay sugar-rich substrates (Põlme et al. 2020) | Põlme S, et al. 2020. Fungal Diversity 105:1-16 ((https://doi.org/10.1007/s13225-020-00466-2)) |
| Otu49881 | 1 | k__Fungi(100);p__Ascomycota(100);c__Saccharomycetes(100);o__Saccharomycetales(100);f__Metschnikowiaceae(100);g__Metschnikowia(100);s__Metschnikowia_pulcherrima(100); | Metschnikowia | 13 | Saprotroph | \|Undefined Saprotroph\| | Yeast | NULL | Probable | As nectar-tap_saprotroph (Põlme et al. 2020); Decay sugar-rich substrates (Põlme et al. 2020) | Põlme S, et al. 2020. Fungal Diversity 105:1-16 ((https://doi.org/10.1007/s13225-020-00466-2)) |
| Otu49882 | 1 | k__Fungi(100);p__Ascomycota(100);c__Saccharomycetes(100);o__Saccharomycetales(100);f__Metschnikowiaceae(100);g__Metschnikowia(100);s__Metschnikowia_pulcherrima(100); | Metschnikowia | 13 | Saprotroph | \|Undefined Saprotroph\| | Yeast | NULL | Probable | As nectar-tap_saprotroph (Põlme et al. 2020); Decay sugar-rich substrates (Põlme et al. 2020) | Põlme S, et al. 2020. Fungal Diversity 105:1-16 ((https://doi.org/10.1007/s13225-020-00466-2)) |
| Otu49883 | 1 | k__Fungi(100);p__Ascomycota(100);c__Saccharomycetes(100);o__Saccharomycetales(100);f__Metschnikowiaceae(100);g__Metschnikowia(100);s__Metschnikowia_pulcherrima(100); | Metschnikowia | 13 | Saprotroph | \|Undefined Saprotroph\| | Yeast | NULL | Probable | As nectar-tap_saprotroph (Põlme et al. 2020); Decay sugar-rich substrates (Põlme et al. 2020) | Põlme S, et al. 2020. Fungal Diversity 105:1-16 ((https://doi.org/10.1007/s13225-020-00466-2)) |
| Otu49884 | 1 | k__Fungi(100);p__Ascomycota(100);c__Saccharomycetes(100);o__Saccharomycetales(100);f__Metschnikowiaceae(100);g__Metschnikowia(100);s__Metschnikowia_pulcherrima(100); | Metschnikowia | 13 | Saprotroph | \|Undefined Saprotroph\| | Yeast | NULL | Probable | As nectar-tap_saprotroph (Põlme et al. 2020); Decay sugar-rich substrates (Põlme et al. 2020) | Põlme S, et al. 2020. Fungal Diversity 105:1-16 ((https://doi.org/10.1007/s13225-020-00466-2)) |
| Otu49885 | 1 | k__Fungi(100);p__Ascomycota(100);c__Saccharomycetes(100);o__Saccharomycetales(100);f__Metschnikowiaceae(100);g__Metschnikowia(100);s__Metschnikowia_pulcherrima(100); | Metschnikowia | 13 | Saprotroph | \|Undefined Saprotroph\| | Yeast | NULL | Probable | As nectar-tap_saprotroph (Põlme et al. 2020); Decay sugar-rich substrates (Põlme et al. 2020) | Põlme S, et al. 2020. Fungal Diversity 105:1-16 ((https://doi.org/10.1007/s13225-020-00466-2)) |
| Otu49886 | 1 | k__Fungi(100);p__Ascomycota(100);c__Saccharomycetes(100);o__Saccharomycetales(100);f__Metschnikowiaceae(100);g__Metschnikowia(100);s__Metschnikowia_pulcherrima(100); | Metschnikowia | 13 | Saprotroph | \|Undefined Saprotroph\| | Yeast | NULL | Probable | As nectar-tap_saprotroph (Põlme et al. 2020); Decay sugar-rich substrates (Põlme et al. 2020) | Põlme S, et al. 2020. Fungal Diversity 105:1-16 ((https://doi.org/10.1007/s13225-020-00466-2)) |
| Otu49887 | 1 | k__Fungi(100);p__Ascomycota(100);c__Saccharomycetes(100);o__Saccharomycetales(100);f__Metschnikowiaceae(100);g__Metschnikowia(100);g__Metschnikowia_unclassified(100); | Metschnikowia | 13 | Saprotroph | \|Undefined Saprotroph\| | Yeast | NULL | Probable | As nectar-tap_saprotroph (Põlme et al. 2020); Decay sugar-rich substrates (Põlme et al. 2020) | Põlme S, et al. 2020. Fungal Diversity 105:1-16 ((https://doi.org/10.1007/s13225-020-00466-2)) |
| Otu49889 | 1 | k__Fungi(100);p__Ascomycota(100);c__Saccharomycetes(100);o__Saccharomycetales(100);f__Metschnikowiaceae(100);g__Metschnikowia(100);s__Metschnikowia_pulcherrima(100); | Metschnikowia | 13 | Saprotroph | \|Undefined Saprotroph\| | Yeast | NULL | Probable | As nectar-tap_saprotroph (Põlme et al. 2020); Decay sugar-rich substrates (Põlme et al. 2020) | Põlme S, et al. 2020. Fungal Diversity 105:1-16 ((https://doi.org/10.1007/s13225-020-00466-2)) |
| Otu49890 | 1 | k__Fungi(100);p__Ascomycota(100);c__Saccharomycetes(100);o__Saccharomycetales(100);f__Metschnikowiaceae(100);g__Metschnikowia(100);s__Metschnikowia_pulcherrima(100); | Metschnikowia | 13 | Saprotroph | \|Undefined Saprotroph\| | Yeast | NULL | Probable | As nectar-tap_saprotroph (Põlme et al. 2020); Decay sugar-rich substrates (Põlme et al. 2020) | Põlme S, et al. 2020. Fungal Diversity 105:1-16 ((https://doi.org/10.1007/s13225-020-00466-2)) |
| Otu49891 | 1 | k__Fungi(100);p__Ascomycota(100);c__Saccharomycetes(100);o__Saccharomycetales(100);f__Metschnikowiaceae(100);g__Metschnikowia(100);g__Metschnikowia_unclassified(100); | Metschnikowia | 13 | Saprotroph | \|Undefined Saprotroph\| | Yeast | NULL | Probable | As nectar-tap_saprotroph (Põlme et al. 2020); Decay sugar-rich substrates (Põlme et al. 2020) | Põlme S, et al. 2020. Fungal Diversity 105:1-16 ((https://doi.org/10.1007/s13225-020-00466-2)) |
| Otu49892 | 1 | k__Fungi(100);p__Ascomycota(100);c__Saccharomycetes(100);o__Saccharomycetales(100);f__Metschnikowiaceae(100);g__Metschnikowia(100);g__Metschnikowia_unclassified(100); | Metschnikowia | 13 | Saprotroph | \|Undefined Saprotroph\| | Yeast | NULL | Probable | As nectar-tap_saprotroph (Põlme et al. 2020); Decay sugar-rich substrates (Põlme et al. 2020) | Põlme S, et al. 2020. Fungal Diversity 105:1-16 ((https://doi.org/10.1007/s13225-020-00466-2)) |
| Otu49893 | 1 | k__Fungi(100);p__Ascomycota(100);c__Saccharomycetes(100);o__Saccharomycetales(100);f__Metschnikowiaceae(100);g__Metschnikowia(100);g__Metschnikowia_unclassified(100); | Metschnikowia | 13 | Saprotroph | \|Undefined Saprotroph\| | Yeast | NULL | Probable | As nectar-tap_saprotroph (Põlme et al. 2020); Decay sugar-rich substrates (Põlme et al. 2020) | Põlme S, et al. 2020. Fungal Diversity 105:1-16 ((https://doi.org/10.1007/s13225-020-00466-2)) |
| Otu49894 | 1 | k__Fungi(100);p__Ascomycota(100);c__Saccharomycetes(100);o__Saccharomycetales(100);f__Metschnikowiaceae(100);g__Metschnikowia(100);s__Metschnikowia_pulcherrima(100); | Metschnikowia | 13 | Saprotroph | \|Undefined Saprotroph\| | Yeast | NULL | Probable | As nectar-tap_saprotroph (Põlme et al. 2020); Decay sugar-rich substrates (Põlme et al. 2020) | Põlme S, et al. 2020. Fungal Diversity 105:1-16 ((https://doi.org/10.1007/s13225-020-00466-2)) |
| Otu49895 | 1 | k__Fungi(100);p__Ascomycota(100);c__Saccharomycetes(100);o__Saccharomycetales(100);f__Metschnikowiaceae(100);g__Metschnikowia(100);s__Metschnikowia_pulcherrima(100); | Metschnikowia | 13 | Saprotroph | \|Undefined Saprotroph\| | Yeast | NULL | Probable | As nectar-tap_saprotroph (Põlme et al. 2020); Decay sugar-rich substrates (Põlme et al. 2020) | Põlme S, et al. 2020. Fungal Diversity 105:1-16 ((https://doi.org/10.1007/s13225-020-00466-2)) |
| Otu49896 | 1 | k__Fungi(100);p__Ascomycota(100);c__Saccharomycetes(100);o__Saccharomycetales(100);f__Metschnikowiaceae(100);g__Metschnikowia(100);s__Metschnikowia_pulcherrima(100); | Metschnikowia | 13 | Saprotroph | \|Undefined Saprotroph\| | Yeast | NULL | Probable | As nectar-tap_saprotroph (Põlme et al. 2020); Decay sugar-rich substrates (Põlme et al. 2020) | Põlme S, et al. 2020. Fungal Diversity 105:1-16 ((https://doi.org/10.1007/s13225-020-00466-2)) |
| Otu49897 | 1 | k__Fungi(100);p__Ascomycota(100);c__Saccharomycetes(100);o__Saccharomycetales(100);f__Metschnikowiaceae(100);g__Metschnikowia(100);s__Metschnikowia_pulcherrima(100); | Metschnikowia | 13 | Saprotroph | \|Undefined Saprotroph\| | Yeast | NULL | Probable | As nectar-tap_saprotroph (Põlme et al. 2020); Decay sugar-rich substrates (Põlme et al. 2020) | Põlme S, et al. 2020. Fungal Diversity 105:1-16 ((https://doi.org/10.1007/s13225-020-00466-2)) |
| Otu49898 | 1 | k__Fungi(100);p__Ascomycota(100);c__Saccharomycetes(100);o__Saccharomycetales(100);f__Metschnikowiaceae(100);g__Metschnikowia(100);s__Metschnikowia_pulcherrima(100); | Metschnikowia | 13 | Saprotroph | \|Undefined Saprotroph\| | Yeast | NULL | Probable | As nectar-tap_saprotroph (Põlme et al. 2020); Decay sugar-rich substrates (Põlme et al. 2020) | Põlme S, et al. 2020. Fungal Diversity 105:1-16 ((https://doi.org/10.1007/s13225-020-00466-2)) |
| Otu49899 | 1 | k__Fungi(100);p__Ascomycota(100);c__Saccharomycetes(100);o__Saccharomycetales(100);f__Metschnikowiaceae(100);g__Metschnikowia(100);s__Metschnikowia_pulcherrima(100); | Metschnikowia | 13 | Saprotroph | \|Undefined Saprotroph\| | Yeast | NULL | Probable | As nectar-tap_saprotroph (Põlme et al. 2020); Decay sugar-rich substrates (Põlme et al. 2020) | Põlme S, et al. 2020. Fungal Diversity 105:1-16 ((https://doi.org/10.1007/s13225-020-00466-2)) |
| Otu49900 | 1 | k__Fungi(100);p__Ascomycota(100);c__Saccharomycetes(100);o__Saccharomycetales(100);f__Metschnikowiaceae(100);g__Metschnikowia(100);s__Metschnikowia_pulcherrima(100); | Metschnikowia | 13 | Saprotroph | \|Undefined Saprotroph\| | Yeast | NULL | Probable | As nectar-tap_saprotroph (Põlme et al. 2020); Decay sugar-rich substrates (Põlme et al. 2020) | Põlme S, et al. 2020. Fungal Diversity 105:1-16 ((https://doi.org/10.1007/s13225-020-00466-2)) |
| Otu49901 | 1 | k__Fungi(100);p__Ascomycota(100);c__Saccharomycetes(100);o__Saccharomycetales(100);f__Metschnikowiaceae(100);g__Metschnikowia(100);s__Metschnikowia_pulcherrima(100); | Metschnikowia | 13 | Saprotroph | \|Undefined Saprotroph\| | Yeast | NULL | Probable | As nectar-tap_saprotroph (Põlme et al. 2020); Decay sugar-rich substrates (Põlme et al. 2020) | Põlme S, et al. 2020. Fungal Diversity 105:1-16 ((https://doi.org/10.1007/s13225-020-00466-2)) |
| Otu49902 | 1 | k__Fungi(100);p__Ascomycota(100);c__Saccharomycetes(100);o__Saccharomycetales(100);f__Metschnikowiaceae(100);g__Metschnikowia(100);s__Metschnikowia_pulcherrima(100); | Metschnikowia | 13 | Saprotroph | \|Undefined Saprotroph\| | Yeast | NULL | Probable | As nectar-tap_saprotroph (Põlme et al. 2020); Decay sugar-rich substrates (Põlme et al. 2020) | Põlme S, et al. 2020. Fungal Diversity 105:1-16 ((https://doi.org/10.1007/s13225-020-00466-2)) |
| Otu49903 | 1 | k__Fungi(100);p__Ascomycota(100);c__Saccharomycetes(100);o__Saccharomycetales(100);f__Metschnikowiaceae(100);g__Metschnikowia(100);s__Metschnikowia_pulcherrima(100); | Metschnikowia | 13 | Saprotroph | \|Undefined Saprotroph\| | Yeast | NULL | Probable | As nectar-tap_saprotroph (Põlme et al. 2020); Decay sugar-rich substrates (Põlme et al. 2020) | Põlme S, et al. 2020. Fungal Diversity 105:1-16 ((https://doi.org/10.1007/s13225-020-00466-2)) |
| Otu49904 | 1 | k__Fungi(100);p__Ascomycota(100);c__Saccharomycetes(100);o__Saccharomycetales(100);f__Metschnikowiaceae(100);g__Metschnikowia(100);s__Metschnikowia_pulcherrima(100); | Metschnikowia | 13 | Saprotroph | \|Undefined Saprotroph\| | Yeast | NULL | Probable | As nectar-tap_saprotroph (Põlme et al. 2020); Decay sugar-rich substrates (Põlme et al. 2020) | Põlme S, et al. 2020. Fungal Diversity 105:1-16 ((https://doi.org/10.1007/s13225-020-00466-2)) |
| Otu49905 | 1 | k__Fungi(100);p__Ascomycota(100);c__Saccharomycetes(100);o__Saccharomycetales(100);f__Metschnikowiaceae(100);g__Metschnikowia(100);s__Metschnikowia_pulcherrima(100); | Metschnikowia | 13 | Saprotroph | \|Undefined Saprotroph\| | Yeast | NULL | Probable | As nectar-tap_saprotroph (Põlme et al. 2020); Decay sugar-rich substrates (Põlme et al. 2020) | Põlme S, et al. 2020. Fungal Diversity 105:1-16 ((https://doi.org/10.1007/s13225-020-00466-2)) |
| Otu49906 | 1 | k__Fungi(100);p__Ascomycota(100);c__Saccharomycetes(100);o__Saccharomycetales(100);f__Metschnikowiaceae(100);g__Metschnikowia(100);g__Metschnikowia_unclassified(100); | Metschnikowia | 13 | Saprotroph | \|Undefined Saprotroph\| | Yeast | NULL | Probable | As nectar-tap_saprotroph (Põlme et al. 2020); Decay sugar-rich substrates (Põlme et al. 2020) | Põlme S, et al. 2020. Fungal Diversity 105:1-16 ((https://doi.org/10.1007/s13225-020-00466-2)) |
| Otu49907 | 1 | k__Fungi(100);p__Ascomycota(100);c__Saccharomycetes(100);o__Saccharomycetales(100);f__Metschnikowiaceae(100);g__Metschnikowia(100);s__Metschnikowia_pulcherrima(100); | Metschnikowia | 13 | Saprotroph | \|Undefined Saprotroph\| | Yeast | NULL | Probable | As nectar-tap_saprotroph (Põlme et al. 2020); Decay sugar-rich substrates (Põlme et al. 2020) | Põlme S, et al. 2020. Fungal Diversity 105:1-16 ((https://doi.org/10.1007/s13225-020-00466-2)) |
| Otu49908 | 1 | k__Fungi(100);p__Ascomycota(100);c__Saccharomycetes(100);o__Saccharomycetales(100);f__Metschnikowiaceae(100);g__Metschnikowia(100);s__Metschnikowia_pulcherrima(100); | Metschnikowia | 13 | Saprotroph | \|Undefined Saprotroph\| | Yeast | NULL | Probable | As nectar-tap_saprotroph (Põlme et al. 2020); Decay sugar-rich substrates (Põlme et al. 2020) | Põlme S, et al. 2020. Fungal Diversity 105:1-16 ((https://doi.org/10.1007/s13225-020-00466-2)) |
| Otu49909 | 1 | k__Fungi(100);p__Ascomycota(100);c__Saccharomycetes(100);o__Saccharomycetales(100);f__Metschnikowiaceae(100);g__Metschnikowia(100);s__Metschnikowia_pulcherrima(100); | Metschnikowia | 13 | Saprotroph | \|Undefined Saprotroph\| | Yeast | NULL | Probable | As nectar-tap_saprotroph (Põlme et al. 2020); Decay sugar-rich substrates (Põlme et al. 2020) | Põlme S, et al. 2020. Fungal Diversity 105:1-16 ((https://doi.org/10.1007/s13225-020-00466-2)) |
| Otu49910 | 1 | k__Fungi(100);p__Ascomycota(100);c__Saccharomycetes(100);o__Saccharomycetales(100);f__Metschnikowiaceae(100);g__Metschnikowia(100);s__Metschnikowia_pulcherrima(100); | Metschnikowia | 13 | Saprotroph | \|Undefined Saprotroph\| | Yeast | NULL | Probable | As nectar-tap_saprotroph (Põlme et al. 2020); Decay sugar-rich substrates (Põlme et al. 2020) | Põlme S, et al. 2020. Fungal Diversity 105:1-16 ((https://doi.org/10.1007/s13225-020-00466-2)) |
| Otu49911 | 1 | k__Fungi(100);p__Ascomycota(100);c__Saccharomycetes(100);o__Saccharomycetales(100);f__Metschnikowiaceae(100);g__Metschnikowia(100);s__Metschnikowia_pulcherrima(100); | Metschnikowia | 13 | Saprotroph | \|Undefined Saprotroph\| | Yeast | NULL | Probable | As nectar-tap_saprotroph (Põlme et al. 2020); Decay sugar-rich substrates (Põlme et al. 2020) | Põlme S, et al. 2020. Fungal Diversity 105:1-16 ((https://doi.org/10.1007/s13225-020-00466-2)) |
| Otu49912 | 1 | k__Fungi(100);p__Ascomycota(100);c__Saccharomycetes(100);o__Saccharomycetales(100);f__Metschnikowiaceae(100);g__Metschnikowia(100);s__Metschnikowia_pulcherrima(100); | Metschnikowia | 13 | Saprotroph | \|Undefined Saprotroph\| | Yeast | NULL | Probable | As nectar-tap_saprotroph (Põlme et al. 2020); Decay sugar-rich substrates (Põlme et al. 2020) | Põlme S, et al. 2020. Fungal Diversity 105:1-16 ((https://doi.org/10.1007/s13225-020-00466-2)) |
| Otu49913 | 1 | k__Fungi(100);p__Ascomycota(100);c__Saccharomycetes(100);o__Saccharomycetales(100);f__Metschnikowiaceae(100);g__Metschnikowia(100);s__Metschnikowia_pulcherrima(100); | Metschnikowia | 13 | Saprotroph | \|Undefined Saprotroph\| | Yeast | NULL | Probable | As nectar-tap_saprotroph (Põlme et al. 2020); Decay sugar-rich substrates (Põlme et al. 2020) | Põlme S, et al. 2020. Fungal Diversity 105:1-16 ((https://doi.org/10.1007/s13225-020-00466-2)) |
| Otu49914 | 1 | k__Fungi(100);p__Ascomycota(100);c__Saccharomycetes(100);o__Saccharomycetales(100);f__Metschnikowiaceae(100);g__Metschnikowia(100);g__Metschnikowia_unclassified(100); | Metschnikowia | 13 | Saprotroph | \|Undefined Saprotroph\| | Yeast | NULL | Probable | As nectar-tap_saprotroph (Põlme et al. 2020); Decay sugar-rich substrates (Põlme et al. 2020) | Põlme S, et al. 2020. Fungal Diversity 105:1-16 ((https://doi.org/10.1007/s13225-020-00466-2)) |
| Otu49915 | 1 | k__Fungi(100);p__Ascomycota(100);c__Saccharomycetes(100);o__Saccharomycetales(100);f__Metschnikowiaceae(100);g__Metschnikowia(100);s__Metschnikowia_pulcherrima(100); | Metschnikowia | 13 | Saprotroph | \|Undefined Saprotroph\| | Yeast | NULL | Probable | As nectar-tap_saprotroph (Põlme et al. 2020); Decay sugar-rich substrates (Põlme et al. 2020) | Põlme S, et al. 2020. Fungal Diversity 105:1-16 ((https://doi.org/10.1007/s13225-020-00466-2)) |
| Otu49916 | 1 | k__Fungi(100);p__Ascomycota(100);c__Saccharomycetes(100);o__Saccharomycetales(100);f__Metschnikowiaceae(100);g__Metschnikowia(100);g__Metschnikowia_unclassified(100); | Metschnikowia | 13 | Saprotroph | \|Undefined Saprotroph\| | Yeast | NULL | Probable | As nectar-tap_saprotroph (Põlme et al. 2020); Decay sugar-rich substrates (Põlme et al. 2020) | Põlme S, et al. 2020. Fungal Diversity 105:1-16 ((https://doi.org/10.1007/s13225-020-00466-2)) |
| Otu49917 | 1 | k__Fungi(100);p__Ascomycota(100);c__Saccharomycetes(100);o__Saccharomycetales(100);f__Metschnikowiaceae(100);g__Metschnikowia(100);g__Metschnikowia_unclassified(100); | Metschnikowia | 13 | Saprotroph | \|Undefined Saprotroph\| | Yeast | NULL | Probable | As nectar-tap_saprotroph (Põlme et al. 2020); Decay sugar-rich substrates (Põlme et al. 2020) | Põlme S, et al. 2020. Fungal Diversity 105:1-16 ((https://doi.org/10.1007/s13225-020-00466-2)) |
| Otu49918 | 1 | k__Fungi(100);p__Ascomycota(100);c__Saccharomycetes(100);o__Saccharomycetales(100);f__Metschnikowiaceae(100);g__Metschnikowia(100);s__Metschnikowia_pulcherrima(100); | Metschnikowia | 13 | Saprotroph | \|Undefined Saprotroph\| | Yeast | NULL | Probable | As nectar-tap_saprotroph (Põlme et al. 2020); Decay sugar-rich substrates (Põlme et al. 2020) | Põlme S, et al. 2020. Fungal Diversity 105:1-16 ((https://doi.org/10.1007/s13225-020-00466-2)) |
| Otu49919 | 1 | k__Fungi(100);p__Ascomycota(100);c__Saccharomycetes(100);o__Saccharomycetales(100);f__Metschnikowiaceae(100);g__Metschnikowia(100);s__Metschnikowia_pulcherrima(100); | Metschnikowia | 13 | Saprotroph | \|Undefined Saprotroph\| | Yeast | NULL | Probable | As nectar-tap_saprotroph (Põlme et al. 2020); Decay sugar-rich substrates (Põlme et al. 2020) | Põlme S, et al. 2020. Fungal Diversity 105:1-16 ((https://doi.org/10.1007/s13225-020-00466-2)) |
| Otu49920 | 1 | k__Fungi(100);p__Ascomycota(100);c__Saccharomycetes(100);o__Saccharomycetales(100);f__Metschnikowiaceae(100);g__Metschnikowia(100);s__Metschnikowia_pulcherrima(100); | Metschnikowia | 13 | Saprotroph | \|Undefined Saprotroph\| | Yeast | NULL | Probable | As nectar-tap_saprotroph (Põlme et al. 2020); Decay sugar-rich substrates (Põlme et al. 2020) | Põlme S, et al. 2020. Fungal Diversity 105:1-16 ((https://doi.org/10.1007/s13225-020-00466-2)) |
| Otu49921 | 1 | k__Fungi(100);p__Ascomycota(100);c__Saccharomycetes(100);o__Saccharomycetales(100);f__Metschnikowiaceae(100);g__Metschnikowia(100);s__Metschnikowia_pulcherrima(100); | Metschnikowia | 13 | Saprotroph | \|Undefined Saprotroph\| | Yeast | NULL | Probable | As nectar-tap_saprotroph (Põlme et al. 2020); Decay sugar-rich substrates (Põlme et al. 2020) | Põlme S, et al. 2020. Fungal Diversity 105:1-16 ((https://doi.org/10.1007/s13225-020-00466-2)) |
| Otu49922 | 1 | k__Fungi(100);p__Ascomycota(100);c__Saccharomycetes(100);o__Saccharomycetales(100);f__Metschnikowiaceae(100);g__Metschnikowia(100);s__Metschnikowia_pulcherrima(100); | Metschnikowia | 13 | Saprotroph | \|Undefined Saprotroph\| | Yeast | NULL | Probable | As nectar-tap_saprotroph (Põlme et al. 2020); Decay sugar-rich substrates (Põlme et al. 2020) | Põlme S, et al. 2020. Fungal Diversity 105:1-16 ((https://doi.org/10.1007/s13225-020-00466-2)) |
| Otu49923 | 1 | k__Fungi(100);p__Ascomycota(100);c__Saccharomycetes(100);o__Saccharomycetales(100);f__Metschnikowiaceae(100);g__Metschnikowia(100);s__Metschnikowia_pulcherrima(100); | Metschnikowia | 13 | Saprotroph | \|Undefined Saprotroph\| | Yeast | NULL | Probable | As nectar-tap_saprotroph (Põlme et al. 2020); Decay sugar-rich substrates (Põlme et al. 2020) | Põlme S, et al. 2020. Fungal Diversity 105:1-16 ((https://doi.org/10.1007/s13225-020-00466-2)) |
| Otu49924 | 1 | k__Fungi(100);p__Ascomycota(100);c__Saccharomycetes(100);o__Saccharomycetales(100);f__Metschnikowiaceae(100);g__Metschnikowia(100);s__Metschnikowia_pulcherrima(100); | Metschnikowia | 13 | Saprotroph | \|Undefined Saprotroph\| | Yeast | NULL | Probable | As nectar-tap_saprotroph (Põlme et al. 2020); Decay sugar-rich substrates (Põlme et al. 2020) | Põlme S, et al. 2020. Fungal Diversity 105:1-16 ((https://doi.org/10.1007/s13225-020-00466-2)) |
| Otu49925 | 1 | k__Fungi(100);p__Ascomycota(100);c__Saccharomycetes(100);o__Saccharomycetales(100);f__Metschnikowiaceae(100);g__Metschnikowia(100);g__Metschnikowia_unclassified(100); | Metschnikowia | 13 | Saprotroph | \|Undefined Saprotroph\| | Yeast | NULL | Probable | As nectar-tap_saprotroph (Põlme et al. 2020); Decay sugar-rich substrates (Põlme et al. 2020) | Põlme S, et al. 2020. Fungal Diversity 105:1-16 ((https://doi.org/10.1007/s13225-020-00466-2)) |
| Otu49926 | 1 | k__Fungi(100);p__Ascomycota(100);c__Saccharomycetes(100);o__Saccharomycetales(100);f__Metschnikowiaceae(100);g__Metschnikowia(100);g__Metschnikowia_unclassified(100); | Metschnikowia | 13 | Saprotroph | \|Undefined Saprotroph\| | Yeast | NULL | Probable | As nectar-tap_saprotroph (Põlme et al. 2020); Decay sugar-rich substrates (Põlme et al. 2020) | Põlme S, et al. 2020. Fungal Diversity 105:1-16 ((https://doi.org/10.1007/s13225-020-00466-2)) |
| Otu49927 | 1 | k__Fungi(100);p__Ascomycota(100);c__Saccharomycetes(100);o__Saccharomycetales(100);f__Metschnikowiaceae(100);g__Metschnikowia(100);g__Metschnikowia_unclassified(100); | Metschnikowia | 13 | Saprotroph | \|Undefined Saprotroph\| | Yeast | NULL | Probable | As nectar-tap_saprotroph (Põlme et al. 2020); Decay sugar-rich substrates (Põlme et al. 2020) | Põlme S, et al. 2020. Fungal Diversity 105:1-16 ((https://doi.org/10.1007/s13225-020-00466-2)) |
| Otu49928 | 1 | k__Fungi(100);p__Ascomycota(100);c__Saccharomycetes(100);o__Saccharomycetales(100);f__Metschnikowiaceae(100);g__Metschnikowia(100);g__Metschnikowia_unclassified(100); | Metschnikowia | 13 | Saprotroph | \|Undefined Saprotroph\| | Yeast | NULL | Probable | As nectar-tap_saprotroph (Põlme et al. 2020); Decay sugar-rich substrates (Põlme et al. 2020) | Põlme S, et al. 2020. Fungal Diversity 105:1-16 ((https://doi.org/10.1007/s13225-020-00466-2)) |
| Otu49929 | 1 | k__Fungi(100);p__Ascomycota(100);c__Saccharomycetes(100);o__Saccharomycetales(100);f__Metschnikowiaceae(100);g__Metschnikowia(100);s__Metschnikowia_pulcherrima(100); | Metschnikowia | 13 | Saprotroph | \|Undefined Saprotroph\| | Yeast | NULL | Probable | As nectar-tap_saprotroph (Põlme et al. 2020); Decay sugar-rich substrates (Põlme et al. 2020) | Põlme S, et al. 2020. Fungal Diversity 105:1-16 ((https://doi.org/10.1007/s13225-020-00466-2)) |
| Otu49930 | 1 | k__Fungi(100);p__Ascomycota(100);c__Saccharomycetes(100);o__Saccharomycetales(100);f__Metschnikowiaceae(100);g__Metschnikowia(100);s__Metschnikowia_pulcherrima(100); | Metschnikowia | 13 | Saprotroph | \|Undefined Saprotroph\| | Yeast | NULL | Probable | As nectar-tap_saprotroph (Põlme et al. 2020); Decay sugar-rich substrates (Põlme et al. 2020) | Põlme S, et al. 2020. Fungal Diversity 105:1-16 ((https://doi.org/10.1007/s13225-020-00466-2)) |
| Otu49931 | 1 | k__Fungi(100);p__Ascomycota(100);c__Saccharomycetes(100);o__Saccharomycetales(100);f__Metschnikowiaceae(100);g__Metschnikowia(100);s__Metschnikowia_pulcherrima(100); | Metschnikowia | 13 | Saprotroph | \|Undefined Saprotroph\| | Yeast | NULL | Probable | As nectar-tap_saprotroph (Põlme et al. 2020); Decay sugar-rich substrates (Põlme et al. 2020) | Põlme S, et al. 2020. Fungal Diversity 105:1-16 ((https://doi.org/10.1007/s13225-020-00466-2)) |
| Otu49932 | 1 | k__Fungi(100);p__Ascomycota(100);c__Saccharomycetes(100);o__Saccharomycetales(100);f__Metschnikowiaceae(100);g__Metschnikowia(100);s__Metschnikowia_pulcherrima(100); | Metschnikowia | 13 | Saprotroph | \|Undefined Saprotroph\| | Yeast | NULL | Probable | As nectar-tap_saprotroph (Põlme et al. 2020); Decay sugar-rich substrates (Põlme et al. 2020) | Põlme S, et al. 2020. Fungal Diversity 105:1-16 ((https://doi.org/10.1007/s13225-020-00466-2)) |
| Otu49933 | 1 | k__Fungi(100);p__Ascomycota(100);c__Saccharomycetes(100);o__Saccharomycetales(100);f__Metschnikowiaceae(100);g__Metschnikowia(100);s__Metschnikowia_pulcherrima(100); | Metschnikowia | 13 | Saprotroph | \|Undefined Saprotroph\| | Yeast | NULL | Probable | As nectar-tap_saprotroph (Põlme et al. 2020); Decay sugar-rich substrates (Põlme et al. 2020) | Põlme S, et al. 2020. Fungal Diversity 105:1-16 ((https://doi.org/10.1007/s13225-020-00466-2)) |
| Otu49934 | 1 | k__Fungi(100);p__Ascomycota(100);c__Saccharomycetes(100);o__Saccharomycetales(100);f__Metschnikowiaceae(100);g__Metschnikowia(100);s__Metschnikowia_pulcherrima(100); | Metschnikowia | 13 | Saprotroph | \|Undefined Saprotroph\| | Yeast | NULL | Probable | As nectar-tap_saprotroph (Põlme et al. 2020); Decay sugar-rich substrates (Põlme et al. 2020) | Põlme S, et al. 2020. Fungal Diversity 105:1-16 ((https://doi.org/10.1007/s13225-020-00466-2)) |
| Otu49935 | 1 | k__Fungi(100);p__Ascomycota(100);c__Saccharomycetes(100);o__Saccharomycetales(100);f__Metschnikowiaceae(100);g__Metschnikowia(100);g__Metschnikowia_unclassified(100); | Metschnikowia | 13 | Saprotroph | \|Undefined Saprotroph\| | Yeast | NULL | Probable | As nectar-tap_saprotroph (Põlme et al. 2020); Decay sugar-rich substrates (Põlme et al. 2020) | Põlme S, et al. 2020. Fungal Diversity 105:1-16 ((https://doi.org/10.1007/s13225-020-00466-2)) |
| Otu49936 | 1 | k__Fungi(100);p__Ascomycota(100);c__Saccharomycetes(100);o__Saccharomycetales(100);f__Metschnikowiaceae(100);g__Metschnikowia(100);g__Metschnikowia_unclassified(100); | Metschnikowia | 13 | Saprotroph | \|Undefined Saprotroph\| | Yeast | NULL | Probable | As nectar-tap_saprotroph (Põlme et al. 2020); Decay sugar-rich substrates (Põlme et al. 2020) | Põlme S, et al. 2020. Fungal Diversity 105:1-16 ((https://doi.org/10.1007/s13225-020-00466-2)) |
| Otu49937 | 1 | k__Fungi(100);p__Ascomycota(100);c__Saccharomycetes(100);o__Saccharomycetales(100);f__Metschnikowiaceae(100);g__Metschnikowia(100);s__Metschnikowia_pulcherrima(100); | Metschnikowia | 13 | Saprotroph | \|Undefined Saprotroph\| | Yeast | NULL | Probable | As nectar-tap_saprotroph (Põlme et al. 2020); Decay sugar-rich substrates (Põlme et al. 2020) | Põlme S, et al. 2020. Fungal Diversity 105:1-16 ((https://doi.org/10.1007/s13225-020-00466-2)) |
| Otu49938 | 1 | k__Fungi(100);p__Ascomycota(100);c__Saccharomycetes(100);o__Saccharomycetales(100);f__Metschnikowiaceae(100);g__Metschnikowia(100);g__Metschnikowia_unclassified(100); | Metschnikowia | 13 | Saprotroph | \|Undefined Saprotroph\| | Yeast | NULL | Probable | As nectar-tap_saprotroph (Põlme et al. 2020); Decay sugar-rich substrates (Põlme et al. 2020) | Põlme S, et al. 2020. Fungal Diversity 105:1-16 ((https://doi.org/10.1007/s13225-020-00466-2)) |
| Otu49939 | 1 | k__Fungi(100);p__Ascomycota(100);c__Saccharomycetes(100);o__Saccharomycetales(100);f__Metschnikowiaceae(100);g__Metschnikowia(100);s__Metschnikowia_pulcherrima(100); | Metschnikowia | 13 | Saprotroph | \|Undefined Saprotroph\| | Yeast | NULL | Probable | As nectar-tap_saprotroph (Põlme et al. 2020); Decay sugar-rich substrates (Põlme et al. 2020) | Põlme S, et al. 2020. Fungal Diversity 105:1-16 ((https://doi.org/10.1007/s13225-020-00466-2)) |
| Otu49940 | 1 | k__Fungi(100);p__Ascomycota(100);c__Saccharomycetes(100);o__Saccharomycetales(100);f__Metschnikowiaceae(100);g__Metschnikowia(100);s__Metschnikowia_pulcherrima(100); | Metschnikowia | 13 | Saprotroph | \|Undefined Saprotroph\| | Yeast | NULL | Probable | As nectar-tap_saprotroph (Põlme et al. 2020); Decay sugar-rich substrates (Põlme et al. 2020) | Põlme S, et al. 2020. Fungal Diversity 105:1-16 ((https://doi.org/10.1007/s13225-020-00466-2)) |
| Otu49941 | 1 | k__Fungi(100);p__Ascomycota(100);c__Saccharomycetes(100);o__Saccharomycetales(100);f__Metschnikowiaceae(100);g__Metschnikowia(100);g__Metschnikowia_unclassified(100); | Metschnikowia | 13 | Saprotroph | \|Undefined Saprotroph\| | Yeast | NULL | Probable | As nectar-tap_saprotroph (Põlme et al. 2020); Decay sugar-rich substrates (Põlme et al. 2020) | Põlme S, et al. 2020. Fungal Diversity 105:1-16 ((https://doi.org/10.1007/s13225-020-00466-2)) |
| Otu49942 | 1 | k__Fungi(100);p__Ascomycota(100);c__Saccharomycetes(100);o__Saccharomycetales(100);f__Metschnikowiaceae(100);g__Metschnikowia(100);s__Metschnikowia_pulcherrima(100); | Metschnikowia | 13 | Saprotroph | \|Undefined Saprotroph\| | Yeast | NULL | Probable | As nectar-tap_saprotroph (Põlme et al. 2020); Decay sugar-rich substrates (Põlme et al. 2020) | Põlme S, et al. 2020. Fungal Diversity 105:1-16 ((https://doi.org/10.1007/s13225-020-00466-2)) |
| Otu49943 | 1 | k__Fungi(100);p__Ascomycota(100);c__Saccharomycetes(100);o__Saccharomycetales(100);f__Metschnikowiaceae(100);g__Metschnikowia(100);s__Metschnikowia_pulcherrima(100); | Metschnikowia | 13 | Saprotroph | \|Undefined Saprotroph\| | Yeast | NULL | Probable | As nectar-tap_saprotroph (Põlme et al. 2020); Decay sugar-rich substrates (Põlme et al. 2020) | Põlme S, et al. 2020. Fungal Diversity 105:1-16 ((https://doi.org/10.1007/s13225-020-00466-2)) |
| Otu49944 | 1 | k__Fungi(100);p__Ascomycota(100);c__Saccharomycetes(100);o__Saccharomycetales(100);f__Metschnikowiaceae(100);g__Metschnikowia(100);s__Metschnikowia_pulcherrima(100); | Metschnikowia | 13 | Saprotroph | \|Undefined Saprotroph\| | Yeast | NULL | Probable | As nectar-tap_saprotroph (Põlme et al. 2020); Decay sugar-rich substrates (Põlme et al. 2020) | Põlme S, et al. 2020. Fungal Diversity 105:1-16 ((https://doi.org/10.1007/s13225-020-00466-2)) |
| Otu49945 | 1 | k__Fungi(100);p__Ascomycota(100);c__Saccharomycetes(100);o__Saccharomycetales(100);f__Metschnikowiaceae(100);g__Metschnikowia(100);s__Metschnikowia_pulcherrima(100); | Metschnikowia | 13 | Saprotroph | \|Undefined Saprotroph\| | Yeast | NULL | Probable | As nectar-tap_saprotroph (Põlme et al. 2020); Decay sugar-rich substrates (Põlme et al. 2020) | Põlme S, et al. 2020. Fungal Diversity 105:1-16 ((https://doi.org/10.1007/s13225-020-00466-2)) |
| Otu49946 | 1 | k__Fungi(100);p__Ascomycota(100);c__Saccharomycetes(100);o__Saccharomycetales(100);f__Metschnikowiaceae(100);g__Metschnikowia(100);s__Metschnikowia_pulcherrima(100); | Metschnikowia | 13 | Saprotroph | \|Undefined Saprotroph\| | Yeast | NULL | Probable | As nectar-tap_saprotroph (Põlme et al. 2020); Decay sugar-rich substrates (Põlme et al. 2020) | Põlme S, et al. 2020. Fungal Diversity 105:1-16 ((https://doi.org/10.1007/s13225-020-00466-2)) |
| Otu49947 | 1 | k__Fungi(100);p__Ascomycota(100);c__Saccharomycetes(100);o__Saccharomycetales(100);f__Metschnikowiaceae(100);g__Metschnikowia(100);s__Metschnikowia_pulcherrima(100); | Metschnikowia | 13 | Saprotroph | \|Undefined Saprotroph\| | Yeast | NULL | Probable | As nectar-tap_saprotroph (Põlme et al. 2020); Decay sugar-rich substrates (Põlme et al. 2020) | Põlme S, et al. 2020. Fungal Diversity 105:1-16 ((https://doi.org/10.1007/s13225-020-00466-2)) |
| Otu49948 | 1 | k__Fungi(100);p__Ascomycota(100);c__Saccharomycetes(100);o__Saccharomycetales(100);f__Metschnikowiaceae(100);g__Metschnikowia(100);g__Metschnikowia_unclassified(100); | Metschnikowia | 13 | Saprotroph | \|Undefined Saprotroph\| | Yeast | NULL | Probable | As nectar-tap_saprotroph (Põlme et al. 2020); Decay sugar-rich substrates (Põlme et al. 2020) | Põlme S, et al. 2020. Fungal Diversity 105:1-16 ((https://doi.org/10.1007/s13225-020-00466-2)) |
| Otu49949 | 1 | k__Fungi(100);p__Ascomycota(100);c__Saccharomycetes(100);o__Saccharomycetales(100);f__Metschnikowiaceae(100);g__Metschnikowia(100);s__Metschnikowia_pulcherrima(100); | Metschnikowia | 13 | Saprotroph | \|Undefined Saprotroph\| | Yeast | NULL | Probable | As nectar-tap_saprotroph (Põlme et al. 2020); Decay sugar-rich substrates (Põlme et al. 2020) | Põlme S, et al. 2020. Fungal Diversity 105:1-16 ((https://doi.org/10.1007/s13225-020-00466-2)) |
| Otu49950 | 1 | k__Fungi(100);p__Ascomycota(100);c__Saccharomycetes(100);o__Saccharomycetales(100);f__Metschnikowiaceae(100);g__Metschnikowia(100);g__Metschnikowia_unclassified(100); | Metschnikowia | 13 | Saprotroph | \|Undefined Saprotroph\| | Yeast | NULL | Probable | As nectar-tap_saprotroph (Põlme et al. 2020); Decay sugar-rich substrates (Põlme et al. 2020) | Põlme S, et al. 2020. Fungal Diversity 105:1-16 ((https://doi.org/10.1007/s13225-020-00466-2)) |
| Otu49951 | 1 | k__Fungi(100);p__Ascomycota(100);c__Saccharomycetes(100);o__Saccharomycetales(100);f__Metschnikowiaceae(100);g__Metschnikowia(100);g__Metschnikowia_unclassified(100); | Metschnikowia | 13 | Saprotroph | \|Undefined Saprotroph\| | Yeast | NULL | Probable | As nectar-tap_saprotroph (Põlme et al. 2020); Decay sugar-rich substrates (Põlme et al. 2020) | Põlme S, et al. 2020. Fungal Diversity 105:1-16 ((https://doi.org/10.1007/s13225-020-00466-2)) |
| Otu49952 | 1 | k__Fungi(100);p__Ascomycota(100);c__Saccharomycetes(100);o__Saccharomycetales(100);f__Metschnikowiaceae(100);g__Metschnikowia(100);s__Metschnikowia_pulcherrima(100); | Metschnikowia | 13 | Saprotroph | \|Undefined Saprotroph\| | Yeast | NULL | Probable | As nectar-tap_saprotroph (Põlme et al. 2020); Decay sugar-rich substrates (Põlme et al. 2020) | Põlme S, et al. 2020. Fungal Diversity 105:1-16 ((https://doi.org/10.1007/s13225-020-00466-2)) |
| Otu49953 | 1 | k__Fungi(100);p__Ascomycota(100);c__Saccharomycetes(100);o__Saccharomycetales(100);f__Metschnikowiaceae(100);g__Metschnikowia(100);g__Metschnikowia_unclassified(100); | Metschnikowia | 13 | Saprotroph | \|Undefined Saprotroph\| | Yeast | NULL | Probable | As nectar-tap_saprotroph (Põlme et al. 2020); Decay sugar-rich substrates (Põlme et al. 2020) | Põlme S, et al. 2020. Fungal Diversity 105:1-16 ((https://doi.org/10.1007/s13225-020-00466-2)) |
| Otu49954 | 1 | k__Fungi(100);p__Ascomycota(100);c__Saccharomycetes(100);o__Saccharomycetales(100);f__Metschnikowiaceae(100);g__Metschnikowia(100);s__Metschnikowia_pulcherrima(100); | Metschnikowia | 13 | Saprotroph | \|Undefined Saprotroph\| | Yeast | NULL | Probable | As nectar-tap_saprotroph (Põlme et al. 2020); Decay sugar-rich substrates (Põlme et al. 2020) | Põlme S, et al. 2020. Fungal Diversity 105:1-16 ((https://doi.org/10.1007/s13225-020-00466-2)) |
| Otu49955 | 1 | k__Fungi(100);p__Ascomycota(100);c__Saccharomycetes(100);o__Saccharomycetales(100);f__Metschnikowiaceae(100);g__Metschnikowia(100);s__Metschnikowia_pulcherrima(100); | Metschnikowia | 13 | Saprotroph | \|Undefined Saprotroph\| | Yeast | NULL | Probable | As nectar-tap_saprotroph (Põlme et al. 2020); Decay sugar-rich substrates (Põlme et al. 2020) | Põlme S, et al. 2020. Fungal Diversity 105:1-16 ((https://doi.org/10.1007/s13225-020-00466-2)) |
| Otu49956 | 1 | k__Fungi(100);p__Ascomycota(100);c__Saccharomycetes(100);o__Saccharomycetales(100);f__Metschnikowiaceae(100);g__Metschnikowia(100);s__Metschnikowia_pulcherrima(100); | Metschnikowia | 13 | Saprotroph | \|Undefined Saprotroph\| | Yeast | NULL | Probable | As nectar-tap_saprotroph (Põlme et al. 2020); Decay sugar-rich substrates (Põlme et al. 2020) | Põlme S, et al. 2020. Fungal Diversity 105:1-16 ((https://doi.org/10.1007/s13225-020-00466-2)) |
| Otu49957 | 1 | k__Fungi(100);p__Ascomycota(100);c__Saccharomycetes(100);o__Saccharomycetales(100);f__Metschnikowiaceae(100);g__Metschnikowia(100);g__Metschnikowia_unclassified(100); | Metschnikowia | 13 | Saprotroph | \|Undefined Saprotroph\| | Yeast | NULL | Probable | As nectar-tap_saprotroph (Põlme et al. 2020); Decay sugar-rich substrates (Põlme et al. 2020) | Põlme S, et al. 2020. Fungal Diversity 105:1-16 ((https://doi.org/10.1007/s13225-020-00466-2)) |
| Otu49958 | 1 | k__Fungi(100);p__Ascomycota(100);c__Saccharomycetes(100);o__Saccharomycetales(100);f__Metschnikowiaceae(100);g__Metschnikowia(100);g__Metschnikowia_unclassified(100); | Metschnikowia | 13 | Saprotroph | \|Undefined Saprotroph\| | Yeast | NULL | Probable | As nectar-tap_saprotroph (Põlme et al. 2020); Decay sugar-rich substrates (Põlme et al. 2020) | Põlme S, et al. 2020. Fungal Diversity 105:1-16 ((https://doi.org/10.1007/s13225-020-00466-2)) |
| Otu49959 | 1 | k__Fungi(100);p__Ascomycota(100);c__Saccharomycetes(100);o__Saccharomycetales(100);f__Metschnikowiaceae(100);g__Metschnikowia(100);s__Metschnikowia_pulcherrima(100); | Metschnikowia | 13 | Saprotroph | \|Undefined Saprotroph\| | Yeast | NULL | Probable | As nectar-tap_saprotroph (Põlme et al. 2020); Decay sugar-rich substrates (Põlme et al. 2020) | Põlme S, et al. 2020. Fungal Diversity 105:1-16 ((https://doi.org/10.1007/s13225-020-00466-2)) |
| Otu49960 | 1 | k__Fungi(100);p__Ascomycota(100);c__Saccharomycetes(100);o__Saccharomycetales(100);f__Metschnikowiaceae(100);g__Metschnikowia(100);s__Metschnikowia_pulcherrima(100); | Metschnikowia | 13 | Saprotroph | \|Undefined Saprotroph\| | Yeast | NULL | Probable | As nectar-tap_saprotroph (Põlme et al. 2020); Decay sugar-rich substrates (Põlme et al. 2020) | Põlme S, et al. 2020. Fungal Diversity 105:1-16 ((https://doi.org/10.1007/s13225-020-00466-2)) |
| Otu49961 | 1 | k__Fungi(100);p__Ascomycota(100);c__Saccharomycetes(100);o__Saccharomycetales(100);f__Metschnikowiaceae(100);g__Metschnikowia(100);s__Metschnikowia_pulcherrima(100); | Metschnikowia | 13 | Saprotroph | \|Undefined Saprotroph\| | Yeast | NULL | Probable | As nectar-tap_saprotroph (Põlme et al. 2020); Decay sugar-rich substrates (Põlme et al. 2020) | Põlme S, et al. 2020. Fungal Diversity 105:1-16 ((https://doi.org/10.1007/s13225-020-00466-2)) |
| Otu49962 | 1 | k__Fungi(100);p__Ascomycota(100);c__Saccharomycetes(100);o__Saccharomycetales(100);f__Metschnikowiaceae(100);g__Metschnikowia(100);s__Metschnikowia_pulcherrima(100); | Metschnikowia | 13 | Saprotroph | \|Undefined Saprotroph\| | Yeast | NULL | Probable | As nectar-tap_saprotroph (Põlme et al. 2020); Decay sugar-rich substrates (Põlme et al. 2020) | Põlme S, et al. 2020. Fungal Diversity 105:1-16 ((https://doi.org/10.1007/s13225-020-00466-2)) |
| Otu49963 | 1 | k__Fungi(100);p__Ascomycota(100);c__Saccharomycetes(100);o__Saccharomycetales(100);f__Metschnikowiaceae(100);g__Metschnikowia(100);s__Metschnikowia_pulcherrima(100); | Metschnikowia | 13 | Saprotroph | \|Undefined Saprotroph\| | Yeast | NULL | Probable | As nectar-tap_saprotroph (Põlme et al. 2020); Decay sugar-rich substrates (Põlme et al. 2020) | Põlme S, et al. 2020. Fungal Diversity 105:1-16 ((https://doi.org/10.1007/s13225-020-00466-2)) |
| Otu49964 | 1 | k__Fungi(100);p__Ascomycota(100);c__Saccharomycetes(100);o__Saccharomycetales(100);f__Metschnikowiaceae(100);g__Metschnikowia(100);s__Metschnikowia_pulcherrima(100); | Metschnikowia | 13 | Saprotroph | \|Undefined Saprotroph\| | Yeast | NULL | Probable | As nectar-tap_saprotroph (Põlme et al. 2020); Decay sugar-rich substrates (Põlme et al. 2020) | Põlme S, et al. 2020. Fungal Diversity 105:1-16 ((https://doi.org/10.1007/s13225-020-00466-2)) |
| Otu49965 | 1 | k__Fungi(100);p__Ascomycota(100);c__Saccharomycetes(100);o__Saccharomycetales(100);f__Metschnikowiaceae(100);g__Metschnikowia(100);s__Metschnikowia_pulcherrima(100); | Metschnikowia | 13 | Saprotroph | \|Undefined Saprotroph\| | Yeast | NULL | Probable | As nectar-tap_saprotroph (Põlme et al. 2020); Decay sugar-rich substrates (Põlme et al. 2020) | Põlme S, et al. 2020. Fungal Diversity 105:1-16 ((https://doi.org/10.1007/s13225-020-00466-2)) |
| Otu49966 | 1 | k__Fungi(100);p__Ascomycota(100);c__Saccharomycetes(100);o__Saccharomycetales(100);f__Metschnikowiaceae(100);g__Metschnikowia(100);s__Metschnikowia_pulcherrima(100); | Metschnikowia | 13 | Saprotroph | \|Undefined Saprotroph\| | Yeast | NULL | Probable | As nectar-tap_saprotroph (Põlme et al. 2020); Decay sugar-rich substrates (Põlme et al. 2020) | Põlme S, et al. 2020. Fungal Diversity 105:1-16 ((https://doi.org/10.1007/s13225-020-00466-2)) |
| Otu49967 | 1 | k__Fungi(100);p__Ascomycota(100);c__Saccharomycetes(100);o__Saccharomycetales(100);f__Metschnikowiaceae(100);g__Metschnikowia(100);s__Metschnikowia_pulcherrima(100); | Metschnikowia | 13 | Saprotroph | \|Undefined Saprotroph\| | Yeast | NULL | Probable | As nectar-tap_saprotroph (Põlme et al. 2020); Decay sugar-rich substrates (Põlme et al. 2020) | Põlme S, et al. 2020. Fungal Diversity 105:1-16 ((https://doi.org/10.1007/s13225-020-00466-2)) |
| Otu49968 | 1 | k__Fungi(100);p__Ascomycota(100);c__Saccharomycetes(100);o__Saccharomycetales(100);f__Metschnikowiaceae(100);g__Metschnikowia(100);s__Metschnikowia_pulcherrima(100); | Metschnikowia | 13 | Saprotroph | \|Undefined Saprotroph\| | Yeast | NULL | Probable | As nectar-tap_saprotroph (Põlme et al. 2020); Decay sugar-rich substrates (Põlme et al. 2020) | Põlme S, et al. 2020. Fungal Diversity 105:1-16 ((https://doi.org/10.1007/s13225-020-00466-2)) |
| Otu49969 | 1 | k__Fungi(100);p__Ascomycota(100);c__Saccharomycetes(100);o__Saccharomycetales(100);f__Metschnikowiaceae(100);g__Metschnikowia(100);s__Metschnikowia_pulcherrima(100); | Metschnikowia | 13 | Saprotroph | \|Undefined Saprotroph\| | Yeast | NULL | Probable | As nectar-tap_saprotroph (Põlme et al. 2020); Decay sugar-rich substrates (Põlme et al. 2020) | Põlme S, et al. 2020. Fungal Diversity 105:1-16 ((https://doi.org/10.1007/s13225-020-00466-2)) |
| Otu49970 | 1 | k__Fungi(100);p__Ascomycota(100);c__Saccharomycetes(100);o__Saccharomycetales(100);f__Metschnikowiaceae(100);g__Metschnikowia(100);s__Metschnikowia_pulcherrima(100); | Metschnikowia | 13 | Saprotroph | \|Undefined Saprotroph\| | Yeast | NULL | Probable | As nectar-tap_saprotroph (Põlme et al. 2020); Decay sugar-rich substrates (Põlme et al. 2020) | Põlme S, et al. 2020. Fungal Diversity 105:1-16 ((https://doi.org/10.1007/s13225-020-00466-2)) |
| Otu49971 | 1 | k__Fungi(100);p__Ascomycota(100);c__Saccharomycetes(100);o__Saccharomycetales(100);f__Metschnikowiaceae(100);g__Metschnikowia(100);g__Metschnikowia_unclassified(100); | Metschnikowia | 13 | Saprotroph | \|Undefined Saprotroph\| | Yeast | NULL | Probable | As nectar-tap_saprotroph (Põlme et al. 2020); Decay sugar-rich substrates (Põlme et al. 2020) | Põlme S, et al. 2020. Fungal Diversity 105:1-16 ((https://doi.org/10.1007/s13225-020-00466-2)) |
| Otu49972 | 1 | k__Fungi(100);p__Ascomycota(100);c__Saccharomycetes(100);o__Saccharomycetales(100);f__Metschnikowiaceae(100);g__Metschnikowia(100);g__Metschnikowia_unclassified(100); | Metschnikowia | 13 | Saprotroph | \|Undefined Saprotroph\| | Yeast | NULL | Probable | As nectar-tap_saprotroph (Põlme et al. 2020); Decay sugar-rich substrates (Põlme et al. 2020) | Põlme S, et al. 2020. Fungal Diversity 105:1-16 ((https://doi.org/10.1007/s13225-020-00466-2)) |
| Otu49973 | 1 | k__Fungi(100);p__Ascomycota(100);c__Saccharomycetes(100);o__Saccharomycetales(100);f__Metschnikowiaceae(100);g__Metschnikowia(100);s__Metschnikowia_pulcherrima(100); | Metschnikowia | 13 | Saprotroph | \|Undefined Saprotroph\| | Yeast | NULL | Probable | As nectar-tap_saprotroph (Põlme et al. 2020); Decay sugar-rich substrates (Põlme et al. 2020) | Põlme S, et al. 2020. Fungal Diversity 105:1-16 ((https://doi.org/10.1007/s13225-020-00466-2)) |
| Otu49974 | 1 | k__Fungi(100);p__Ascomycota(100);c__Saccharomycetes(100);o__Saccharomycetales(100);f__Metschnikowiaceae(100);g__Metschnikowia(100);g__Metschnikowia_unclassified(100); | Metschnikowia | 13 | Saprotroph | \|Undefined Saprotroph\| | Yeast | NULL | Probable | As nectar-tap_saprotroph (Põlme et al. 2020); Decay sugar-rich substrates (Põlme et al. 2020) | Põlme S, et al. 2020. Fungal Diversity 105:1-16 ((https://doi.org/10.1007/s13225-020-00466-2)) |
| Otu49975 | 1 | k__Fungi(100);p__Ascomycota(100);c__Saccharomycetes(100);o__Saccharomycetales(100);f__Metschnikowiaceae(100);g__Metschnikowia(100);g__Metschnikowia_unclassified(100); | Metschnikowia | 13 | Saprotroph | \|Undefined Saprotroph\| | Yeast | NULL | Probable | As nectar-tap_saprotroph (Põlme et al. 2020); Decay sugar-rich substrates (Põlme et al. 2020) | Põlme S, et al. 2020. Fungal Diversity 105:1-16 ((https://doi.org/10.1007/s13225-020-00466-2)) |
| Otu49976 | 1 | k__Fungi(100);p__Ascomycota(100);c__Saccharomycetes(100);o__Saccharomycetales(100);f__Metschnikowiaceae(100);g__Metschnikowia(100);g__Metschnikowia_unclassified(100); | Metschnikowia | 13 | Saprotroph | \|Undefined Saprotroph\| | Yeast | NULL | Probable | As nectar-tap_saprotroph (Põlme et al. 2020); Decay sugar-rich substrates (Põlme et al. 2020) | Põlme S, et al. 2020. Fungal Diversity 105:1-16 ((https://doi.org/10.1007/s13225-020-00466-2)) |

| | | | | | | | | | | | |
|---|---|---|---|---|---|---|---|---|---|---|---|
| Otu49977 | 1 | k__Fungi(100);p__Ascomycota(100);c__Saccharomycetes(100);o__Saccharomycetales(100);f__Metschnikowiaceae(100);g__Metschnikowia(100);s__Metschnikowia_pulcherrima(100); | Metschnikowia | 13 | Saprotroph | \|Undefined Saprotroph\| | Yeast | NULL | Probable | As nectar-tap_saprotroph (Põlme et al. 2020); Decay sugar-rich substrates (Põlme et al. 2020) | Põlme S, et al. 2020. Fungal Diversity 105:1-16 ((https://doi.org/10.1007/s13225-020-00466-2)) |
| Otu49978 | 1 | k__Fungi(100);p__Ascomycota(100);c__Saccharomycetes(100);o__Saccharomycetales(100);f__Metschnikowiaceae(100);g__Metschnikowia(100);s__Metschnikowia_pulcherrima(100); | Metschnikowia | 13 | Saprotroph | \|Undefined Saprotroph\| | Yeast | NULL | Probable | As nectar-tap_saprotroph (Põlme et al. 2020); Decay sugar-rich substrates (Põlme et al. 2020) | Põlme S, et al. 2020. Fungal Diversity 105:1-16 ((https://doi.org/10.1007/s13225-020-00466-2)) |
| Otu49979 | 1 | k__Fungi(100);p__Ascomycota(100);c__Saccharomycetes(100);o__Saccharomycetales(100);f__Metschnikowiaceae(100);g__Metschnikowia(100);s__Metschnikowia_pulcherrima(100); | Metschnikowia | 13 | Saprotroph | \|Undefined Saprotroph\| | Yeast | NULL | Probable | As nectar-tap_saprotroph (Põlme et al. 2020); Decay sugar-rich substrates (Põlme et al. 2020) | Põlme S, et al. 2020. Fungal Diversity 105:1-16 ((https://doi.org/10.1007/s13225-020-00466-2)) |
| Otu49980 | 1 | k__Fungi(100);p__Ascomycota(100);c__Saccharomycetes(100);o__Saccharomycetales(100);f__Metschnikowiaceae(100);g__Metschnikowia(100);s__Metschnikowia_pulcherrima(100); | Metschnikowia | 13 | Saprotroph | \|Undefined Saprotroph\| | Yeast | NULL | Probable | As nectar-tap_saprotroph (Põlme et al. 2020); Decay sugar-rich substrates (Põlme et al. 2020) | Põlme S, et al. 2020. Fungal Diversity 105:1-16 ((https://doi.org/10.1007/s13225-020-00466-2)) |
| Otu49981 | 1 | k__Fungi(100);p__Ascomycota(100);c__Saccharomycetes(100);o__Saccharomycetales(100);f__Metschnikowiaceae(100);g__Metschnikowia(100);s__Metschnikowia_pulcherrima(100); | Metschnikowia | 13 | Saprotroph | \|Undefined Saprotroph\| | Yeast | NULL | Probable | As nectar-tap_saprotroph (Põlme et al. 2020); Decay sugar-rich substrates (Põlme et al. 2020) | Põlme S, et al. 2020. Fungal Diversity 105:1-16 ((https://doi.org/10.1007/s13225-020-00466-2)) |
| Otu49982 | 1 | k__Fungi(100);p__Ascomycota(100);c__Saccharomycetes(100);o__Saccharomycetales(100);f__Metschnikowiaceae(100);g__Metschnikowia(100);s__Metschnikowia_pulcherrima(100); | Metschnikowia | 13 | Saprotroph | \|Undefined Saprotroph\| | Yeast | NULL | Probable | As nectar-tap_saprotroph (Põlme et al. 2020); Decay sugar-rich substrates (Põlme et al. 2020) | Põlme S, et al. 2020. Fungal Diversity 105:1-16 ((https://doi.org/10.1007/s13225-020-00466-2)) |
| Otu49983 | 1 | k__Fungi(100);p__Ascomycota(100);c__Saccharomycetes(100);o__Saccharomycetales(100);f__Metschnikowiaceae(100);g__Metschnikowia(100);s__Metschnikowia_pulcherrima(100); | Metschnikowia | 13 | Saprotroph | \|Undefined Saprotroph\| | Yeast | NULL | Probable | As nectar-tap_saprotroph (Põlme et al. 2020); Decay sugar-rich substrates (Põlme et al. 2020) | Põlme S, et al. 2020. Fungal Diversity 105:1-16 ((https://doi.org/10.1007/s13225-020-00466-2)) |
| Otu49984 | 1 | k__Fungi(100);p__Ascomycota(100);c__Saccharomycetes(100);o__Saccharomycetales(100);f__Metschnikowiaceae(100);g__Metschnikowia(100);s__Metschnikowia_pulcherrima(100); | Metschnikowia | 13 | Saprotroph | \|Undefined Saprotroph\| | Yeast | NULL | Probable | As nectar-tap_saprotroph (Põlme et al. 2020); Decay sugar-rich substrates (Põlme et al. 2020) | Põlme S, et al. 2020. Fungal Diversity 105:1-16 ((https://doi.org/10.1007/s13225-020-00466-2)) |
| Otu49985 | 1 | k__Fungi(100);p__Ascomycota(100);c__Saccharomycetes(100);o__Saccharomycetales(100);f__Metschnikowiaceae(100);g__Metschnikowia(100);s__Metschnikowia_pulcherrima(100); | Metschnikowia | 13 | Saprotroph | \|Undefined Saprotroph\| | Yeast | NULL | Probable | As nectar-tap_saprotroph (Põlme et al. 2020); Decay sugar-rich substrates (Põlme et al. 2020) | Põlme S, et al. 2020. Fungal Diversity 105:1-16 ((https://doi.org/10.1007/s13225-020-00466-2)) |
| Otu49986 | 1 | k__Fungi(100);p__Ascomycota(100);c__Saccharomycetes(100);o__Saccharomycetales(100);f__Metschnikowiaceae(100);g__Metschnikowia(100);s__Metschnikowia_pulcherrima(100); | Metschnikowia | 13 | Saprotroph | \|Undefined Saprotroph\| | Yeast | NULL | Probable | As nectar-tap_saprotroph (Põlme et al. 2020); Decay sugar-rich substrates (Põlme et al. 2020) | Põlme S, et al. 2020. Fungal Diversity 105:1-16 ((https://doi.org/10.1007/s13225-020-00466-2)) |
| Otu49987 | 1 | k__Fungi(100);p__Ascomycota(100);c__Saccharomycetes(100);o__Saccharomycetales(100);f__Metschnikowiaceae(100);g__Metschnikowia(100);s__Metschnikowia_pulcherrima(100); | Metschnikowia | 13 | Saprotroph | \|Undefined Saprotroph\| | Yeast | NULL | Probable | As nectar-tap_saprotroph (Põlme et al. 2020); Decay sugar-rich substrates (Põlme et al. 2020) | Põlme S, et al. 2020. Fungal Diversity 105:1-16 ((https://doi.org/10.1007/s13225-020-00466-2)) |
| Otu49988 | 1 | k__Fungi(100);p__Ascomycota(100);c__Saccharomycetes(100);o__Saccharomycetales(100);f__Metschnikowiaceae(100);g__Metschnikowia(100);s__Metschnikowia_pulcherrima(100); | Metschnikowia | 13 | Saprotroph | \|Undefined Saprotroph\| | Yeast | NULL | Probable | As nectar-tap_saprotroph (Põlme et al. 2020); Decay sugar-rich substrates (Põlme et al. 2020) | Põlme S, et al. 2020. Fungal Diversity 105:1-16 ((https://doi.org/10.1007/s13225-020-00466-2)) |
| Otu49989 | 1 | k__Fungi(100);p__Ascomycota(100);c__Saccharomycetes(100);o__Saccharomycetales(100);f__Metschnikowiaceae(100);g__Metschnikowia(100);s__Metschnikowia_pulcherrima(100); | Metschnikowia | 13 | Saprotroph | \|Undefined Saprotroph\| | Yeast | NULL | Probable | As nectar-tap_saprotroph (Põlme et al. 2020); Decay sugar-rich substrates (Põlme et al. 2020) | Põlme S, et al. 2020. Fungal Diversity 105:1-16 ((https://doi.org/10.1007/s13225-020-00466-2)) |
| Otu49990 | 1 | k__Fungi(100);p__Ascomycota(100);c__Saccharomycetes(100);o__Saccharomycetales(100);f__Metschnikowiaceae(100);g__Metschnikowia(100);g__Metschnikowia_unclassified(100); | Metschnikowia | 13 | Saprotroph | \|Undefined Saprotroph\| | Yeast | NULL | Probable | As nectar-tap_saprotroph (Põlme et al. 2020); Decay sugar-rich substrates (Põlme et al. 2020) | Põlme S, et al. 2020. Fungal Diversity 105:1-16 ((https://doi.org/10.1007/s13225-020-00466-2)) |
| Otu49991 | 1 | k__Fungi(100);p__Ascomycota(100);c__Saccharomycetes(100);o__Saccharomycetales(100);f__Metschnikowiaceae(100);g__Metschnikowia(100);g__Metschnikowia_unclassified(100); | Metschnikowia | 13 | Saprotroph | \|Undefined Saprotroph\| | Yeast | NULL | Probable | As nectar-tap_saprotroph (Põlme et al. 2020); Decay sugar-rich substrates (Põlme et al. 2020) | Põlme S, et al. 2020. Fungal Diversity 105:1-16 ((https://doi.org/10.1007/s13225-020-00466-2)) |
| Otu49992 | 1 | k__Fungi(100);p__Ascomycota(100);c__Saccharomycetes(100);o__Saccharomycetales(100);f__Metschnikowiaceae(100);g__Metschnikowia(100);s__Metschnikowia_pulcherrima(100); | Metschnikowia | 13 | Saprotroph | \|Undefined Saprotroph\| | Yeast | NULL | Probable | As nectar-tap_saprotroph (Põlme et al. 2020); Decay sugar-rich substrates (Põlme et al. 2020) | Põlme S, et al. 2020. Fungal Diversity 105:1-16 ((https://doi.org/10.1007/s13225-020-00466-2)) |
| Otu49993 | 1 | k__Fungi(100);p__Ascomycota(100);c__Saccharomycetes(100);o__Saccharomycetales(100);f__Metschnikowiaceae(100);g__Metschnikowia(100);s__Metschnikowia_pulcherrima(100); | Metschnikowia | 13 | Saprotroph | \|Undefined Saprotroph\| | Yeast | NULL | Probable | As nectar-tap_saprotroph (Põlme et al. 2020); Decay sugar-rich substrates (Põlme et al. 2020) | Põlme S, et al. 2020. Fungal Diversity 105:1-16 ((https://doi.org/10.1007/s13225-020-00466-2)) |
| Otu49994 | 1 | k__Fungi(100);p__Ascomycota(100);c__Saccharomycetes(100);o__Saccharomycetales(100);f__Metschnikowiaceae(100);g__Metschnikowia(100);s__Metschnikowia_pulcherrima(100); | Metschnikowia | 13 | Saprotroph | \|Undefined Saprotroph\| | Yeast | NULL | Probable | As nectar-tap_saprotroph (Põlme et al. 2020); Decay sugar-rich substrates (Põlme et al. 2020) | Põlme S, et al. 2020. Fungal Diversity 105:1-16 ((https://doi.org/10.1007/s13225-020-00466-2)) |
| Otu49995 | 1 | k__Fungi(100);p__Ascomycota(100);c__Saccharomycetes(100);o__Saccharomycetales(100);f__Metschnikowiaceae(100);g__Metschnikowia(100);g__Metschnikowia_unclassified(100); | Metschnikowia | 13 | Saprotroph | \|Undefined Saprotroph\| | Yeast | NULL | Probable | As nectar-tap_saprotroph (Põlme et al. 2020); Decay sugar-rich substrates (Põlme et al. 2020) | Põlme S, et al. 2020. Fungal Diversity 105:1-16 ((https://doi.org/10.1007/s13225-020-00466-2)) |
| Otu49996 | 1 | k__Fungi(100);p__Ascomycota(100);c__Saccharomycetes(100);o__Saccharomycetales(100);f__Metschnikowiaceae(100);g__Metschnikowia(100);g__Metschnikowia_unclassified(100); | Metschnikowia | 13 | Saprotroph | \|Undefined Saprotroph\| | Yeast | NULL | Probable | As nectar-tap_saprotroph (Põlme et al. 2020); Decay sugar-rich substrates (Põlme et al. 2020) | Põlme S, et al. 2020. Fungal Diversity 105:1-16 ((https://doi.org/10.1007/s13225-020-00466-2)) |
| Otu49997 | 1 | k__Fungi(100);p__Ascomycota(100);c__Saccharomycetes(100);o__Saccharomycetales(100);f__Metschnikowiaceae(100);g__Metschnikowia(100);g__Metschnikowia_unclassified(100); | Metschnikowia | 13 | Saprotroph | \|Undefined Saprotroph\| | Yeast | NULL | Probable | As nectar-tap_saprotroph (Põlme et al. 2020); Decay sugar-rich substrates (Põlme et al. 2020) | Põlme S, et al. 2020. Fungal Diversity 105:1-16 ((https://doi.org/10.1007/s13225-020-00466-2)) |
| Otu49998 | 1 | k__Fungi(100);p__Ascomycota(100);c__Saccharomycetes(100);o__Saccharomycetales(100);f__Metschnikowiaceae(100);g__Metschnikowia(100);s__Metschnikowia_pulcherrima(100); | Metschnikowia | 13 | Saprotroph | \|Undefined Saprotroph\| | Yeast | NULL | Probable | As nectar-tap_saprotroph (Põlme et al. 2020); Decay sugar-rich substrates (Põlme et al. 2020) | Põlme S, et al. 2020. Fungal Diversity 105:1-16 ((https://doi.org/10.1007/s13225-020-00466-2)) |
| Otu49999 | 1 | k__Fungi(100);p__Ascomycota(100);c__Saccharomycetes(100);o__Saccharomycetales(100);f__Metschnikowiaceae(100);g__Metschnikowia(100);s__Metschnikowia_pulcherrima(100); | Metschnikowia | 13 | Saprotroph | \|Undefined Saprotroph\| | Yeast | NULL | Probable | As nectar-tap_saprotroph (Põlme et al. 2020); Decay sugar-rich substrates (Põlme et al. 2020) | Põlme S, et al. 2020. Fungal Diversity 105:1-16 ((https://doi.org/10.1007/s13225-020-00466-2)) |
| Otu50000 | 1 | k__Fungi(100);p__Ascomycota(100);c__Saccharomycetes(100);o__Saccharomycetales(100);f__Metschnikowiaceae(100);g__Metschnikowia(100);s__Metschnikowia_pulcherrima(100); | Metschnikowia | 13 | Saprotroph | \|Undefined Saprotroph\| | Yeast | NULL | Probable | As nectar-tap_saprotroph (Põlme et al. 2020); Decay sugar-rich substrates (Põlme et al. 2020) | Põlme S, et al. 2020. Fungal Diversity 105:1-16 ((https://doi.org/10.1007/s13225-020-00466-2)) |
| Otu50001 | 1 | k__Fungi(100);p__Ascomycota(100);c__Saccharomycetes(100);o__Saccharomycetales(100);f__Metschnikowiaceae(100);g__Metschnikowia(100);s__Metschnikowia_pulcherrima(100); | Metschnikowia | 13 | Saprotroph | \|Undefined Saprotroph\| | Yeast | NULL | Probable | As nectar-tap_saprotroph (Põlme et al. 2020); Decay sugar-rich substrates (Põlme et al. 2020) | Põlme S, et al. 2020. Fungal Diversity 105:1-16 ((https://doi.org/10.1007/s13225-020-00466-2)) |
| Otu50002 | 1 | k__Fungi(100);p__Ascomycota(100);c__Saccharomycetes(100);o__Saccharomycetales(100);f__Metschnikowiaceae(100);g__Metschnikowia(100);s__Metschnikowia_pulcherrima(100); | Metschnikowia | 13 | Saprotroph | \|Undefined Saprotroph\| | Yeast | NULL | Probable | As nectar-tap_saprotroph (Põlme et al. 2020); Decay sugar-rich substrates (Põlme et al. 2020) | Põlme S, et al. 2020. Fungal Diversity 105:1-16 ((https://doi.org/10.1007/s13225-020-00466-2)) |
| Otu50003 | 1 | k__Fungi(100);p__Ascomycota(100);c__Saccharomycetes(100);o__Saccharomycetales(100);f__Metschnikowiaceae(100);g__Metschnikowia(100);g__Metschnikowia_unclassified(100); | Metschnikowia | 13 | Saprotroph | \|Undefined Saprotroph\| | Yeast | NULL | Probable | As nectar-tap_saprotroph (Põlme et al. 2020); Decay sugar-rich substrates (Põlme et al. 2020) | Põlme S, et al. 2020. Fungal Diversity 105:1-16 ((https://doi.org/10.1007/s13225-020-00466-2)) |
| Otu50004 | 1 | k__Fungi(100);p__Ascomycota(100);c__Saccharomycetes(100);o__Saccharomycetales(100);f__Metschnikowiaceae(100);g__Metschnikowia(100);s__Metschnikowia_pulcherrima(100); | Metschnikowia | 13 | Saprotroph | \|Undefined Saprotroph\| | Yeast | NULL | Probable | As nectar-tap_saprotroph (Põlme et al. 2020); Decay sugar-rich substrates (Põlme et al. 2020) | Põlme S, et al. 2020. Fungal Diversity 105:1-16 ((https://doi.org/10.1007/s13225-020-00466-2)) |
| Otu50005 | 1 | k__Fungi(100);p__Ascomycota(100);c__Saccharomycetes(100);o__Saccharomycetales(100);f__Metschnikowiaceae(100);g__Metschnikowia(100);s__Metschnikowia_pulcherrima(100); | Metschnikowia | 13 | Saprotroph | \|Undefined Saprotroph\| | Yeast | NULL | Probable | As nectar-tap_saprotroph (Põlme et al. 2020); Decay sugar-rich substrates (Põlme et al. 2020) | Põlme S, et al. 2020. Fungal Diversity 105:1-16 ((https://doi.org/10.1007/s13225-020-00466-2)) |
| Otu50006 | 1 | k__Fungi(100);p__Ascomycota(100);c__Saccharomycetes(100);o__Saccharomycetales(100);f__Metschnikowiaceae(100);g__Metschnikowia(100);s__Metschnikowia_pulcherrima(100); | Metschnikowia | 13 | Saprotroph | \|Undefined Saprotroph\| | Yeast | NULL | Probable | As nectar-tap_saprotroph (Põlme et al. 2020); Decay sugar-rich substrates (Põlme et al. 2020) | Põlme S, et al. 2020. Fungal Diversity 105:1-16 ((https://doi.org/10.1007/s13225-020-00466-2)) |
| Otu50007 | 1 | k__Fungi(100);p__Ascomycota(100);c__Saccharomycetes(100);o__Saccharomycetales(100);f__Metschnikowiaceae(100);g__Metschnikowia(100);s__Metschnikowia_pulcherrima(100); | Metschnikowia | 13 | Saprotroph | \|Undefined Saprotroph\| | Yeast | NULL | Probable | As nectar-tap_saprotroph (Põlme et al. 2020); Decay sugar-rich substrates (Põlme et al. 2020) | Põlme S, et al. 2020. Fungal Diversity 105:1-16 ((https://doi.org/10.1007/s13225-020-00466-2)) |
| Otu50008 | 1 | k__Fungi(100);p__Ascomycota(100);c__Saccharomycetes(100);o__Saccharomycetales(100);f__Metschnikowiaceae(100);g__Metschnikowia(100);s__Metschnikowia_pulcherrima(100); | Metschnikowia | 13 | Saprotroph | \|Undefined Saprotroph\| | Yeast | NULL | Probable | As nectar-tap_saprotroph (Põlme et al. 2020); Decay sugar-rich substrates (Põlme et al. 2020) | Põlme S, et al. 2020. Fungal Diversity 105:1-16 ((https://doi.org/10.1007/s13225-020-00466-2)) |
| Otu50009 | 1 | k__Fungi(100);p__Ascomycota(100);c__Saccharomycetes(100);o__Saccharomycetales(100);f__Metschnikowiaceae(100);g__Metschnikowia(100);s__Metschnikowia_pulcherrima(100); | Metschnikowia | 13 | Saprotroph | \|Undefined Saprotroph\| | Yeast | NULL | Probable | As nectar-tap_saprotroph (Põlme et al. 2020); Decay sugar-rich substrates (Põlme et al. 2020) | Põlme S, et al. 2020. Fungal Diversity 105:1-16 ((https://doi.org/10.1007/s13225-020-00466-2)) |
| Otu50010 | 1 | k__Fungi(100);p__Ascomycota(100);c__Saccharomycetes(100);o__Saccharomycetales(100);f__Metschnikowiaceae(100);g__Metschnikowia(100);s__Metschnikowia_pulcherrima(100); | Metschnikowia | 13 | Saprotroph | \|Undefined Saprotroph\| | Yeast | NULL | Probable | As nectar-tap_saprotroph (Põlme et al. 2020); Decay sugar-rich substrates (Põlme et al. 2020) | Põlme S, et al. 2020. Fungal Diversity 105:1-16 ((https://doi.org/10.1007/s13225-020-00466-2)) |
| Otu50011 | 1 | k__Fungi(100);p__Ascomycota(100);c__Saccharomycetes(100);o__Saccharomycetales(100);f__Metschnikowiaceae(100);g__Metschnikowia(100);s__Metschnikowia_pulcherrima(100); | Metschnikowia | 13 | Saprotroph | \|Undefined Saprotroph\| | Yeast | NULL | Probable | As nectar-tap_saprotroph (Põlme et al. 2020); Decay sugar-rich substrates (Põlme et al. 2020) | Põlme S, et al. 2020. Fungal Diversity 105:1-16 ((https://doi.org/10.1007/s13225-020-00466-2)) |
| Otu50012 | 1 | k__Fungi(100);p__Ascomycota(100);c__Saccharomycetes(100);o__Saccharomycetales(100);f__Metschnikowiaceae(100);g__Metschnikowia(100);s__Metschnikowia_pulcherrima(100); | Metschnikowia | 13 | Saprotroph | \|Undefined Saprotroph\| | Yeast | NULL | Probable | As nectar-tap_saprotroph (Põlme et al. 2020); Decay sugar-rich substrates (Põlme et al. 2020) | Põlme S, et al. 2020. Fungal Diversity 105:1-16 ((https://doi.org/10.1007/s13225-020-00466-2)) |
| Otu50013 | 1 | k__Fungi(100);p__Ascomycota(100);c__Saccharomycetes(100);o__Saccharomycetales(100);f__Metschnikowiaceae(100);g__Metschnikowia(100);s__Metschnikowia_pulcherrima(100); | Metschnikowia | 13 | Saprotroph | \|Undefined Saprotroph\| | Yeast | NULL | Probable | As nectar-tap_saprotroph (Põlme et al. 2020); Decay sugar-rich substrates (Põlme et al. 2020) | Põlme S, et al. 2020. Fungal Diversity 105:1-16 ((https://doi.org/10.1007/s13225-020-00466-2)) |
| Otu50014 | 1 | k__Fungi(100);p__Ascomycota(100);c__Saccharomycetes(100);o__Saccharomycetales(100);f__Metschnikowiaceae(100);g__Metschnikowia(100);s__Metschnikowia_pulcherrima(100); | Metschnikowia | 13 | Saprotroph | \|Undefined Saprotroph\| | Yeast | NULL | Probable | As nectar-tap_saprotroph (Põlme et al. 2020); Decay sugar-rich substrates (Põlme et al. 2020) | Põlme S, et al. 2020. Fungal Diversity 105:1-16 ((https://doi.org/10.1007/s13225-020-00466-2)) |
| Otu50016 | 1 | k__Fungi(100);p__Ascomycota(100);c__Saccharomycetes(100);o__Saccharomycetales(100);f__Metschnikowiaceae(100);g__Metschnikowia(100);s__Metschnikowia_pulcherrima(100); | Metschnikowia | 13 | Saprotroph | \|Undefined Saprotroph\| | Yeast | NULL | Probable | As nectar-tap_saprotroph (Põlme et al. 2020); Decay sugar-rich substrates (Põlme et al. 2020) | Põlme S, et al. 2020. Fungal Diversity 105:1-16 ((https://doi.org/10.1007/s13225-020-00466-2)) |
| Otu50017 | 1 | k__Fungi(100);p__Ascomycota(100);c__Saccharomycetes(100);o__Saccharomycetales(100);f__Metschnikowiaceae(100);g__Metschnikowia(100);s__Metschnikowia_pulcherrima(100); | Metschnikowia | 13 | Saprotroph | \|Undefined Saprotroph\| | Yeast | NULL | Probable | As nectar-tap_saprotroph (Põlme et al. 2020); Decay sugar-rich substrates (Põlme et al. 2020) | Põlme S, et al. 2020. Fungal Diversity 105:1-16 ((https://doi.org/10.1007/s13225-020-00466-2)) |
| Otu50018 | 1 | k__Fungi(100);p__Ascomycota(100);c__Saccharomycetes(100);o__Saccharomycetales(100);f__Metschnikowiaceae(100);g__Metschnikowia(100);g__Metschnikowia_unclassified(100); | Metschnikowia | 13 | Saprotroph | \|Undefined Saprotroph\| | Yeast | NULL | Probable | As nectar-tap_saprotroph (Põlme et al. 2020); Decay sugar-rich substrates (Põlme et al. 2020) | Põlme S, et al. 2020. Fungal Diversity 105:1-16 ((https://doi.org/10.1007/s13225-020-00466-2)) |
| Otu50020 | 1 | k__Fungi(100);p__Ascomycota(100);c__Saccharomycetes(100);o__Saccharomycetales(100);f__Metschnikowiaceae(100);g__Metschnikowia(100);s__Metschnikowia_pulcherrima(100); | Metschnikowia | 13 | Saprotroph | \|Undefined Saprotroph\| | Yeast | NULL | Probable | As nectar-tap_saprotroph (Põlme et al. 2020); Decay sugar-rich substrates (Põlme et al. 2020) | Põlme S, et al. 2020. Fungal Diversity 105:1-16 ((https://doi.org/10.1007/s13225-020-00466-2)) |
| Otu50021 | 1 | k__Fungi(100);p__Ascomycota(100);c__Saccharomycetes(100);o__Saccharomycetales(100);f__Metschnikowiaceae(100);g__Metschnikowia(100);s__Metschnikowia_pulcherrima(100); | Metschnikowia | 13 | Saprotroph | \|Undefined Saprotroph\| | Yeast | NULL | Probable | As nectar-tap_saprotroph (Põlme et al. 2020); Decay sugar-rich substrates (Põlme et al. 2020) | Põlme S, et al. 2020. Fungal Diversity 105:1-16 ((https://doi.org/10.1007/s13225-020-00466-2)) |
| Otu50022 | 1 | k__Fungi(100);p__Ascomycota(100);c__Saccharomycetes(100);o__Saccharomycetales(100);f__Metschnikowiaceae(100);g__Metschnikowia(100);s__Metschnikowia_pulcherrima(100); | Metschnikowia | 13 | Saprotroph | \|Undefined Saprotroph\| | Yeast | NULL | Probable | As nectar-tap_saprotroph (Põlme et al. 2020); Decay sugar-rich substrates (Põlme et al. 2020) | Põlme S, et al. 2020. Fungal Diversity 105:1-16 ((https://doi.org/10.1007/s13225-020-00466-2)) |
| Otu50023 | 1 | k__Fungi(100);p__Ascomycota(100);c__Saccharomycetes(100);o__Saccharomycetales(100);f__Metschnikowiaceae(100);g__Metschnikowia(100);s__Metschnikowia_pulcherrima(100); | Metschnikowia | 13 | Saprotroph | \|Undefined Saprotroph\| | Yeast | NULL | Probable | As nectar-tap_saprotroph (Põlme et al. 2020); Decay sugar-rich substrates (Põlme et al. 2020) | Põlme S, et al. 2020. Fungal Diversity 105:1-16 ((https://doi.org/10.1007/s13225-020-00466-2)) |
| Otu50024 | 1 | k__Fungi(100);p__Ascomycota(100);c__Saccharomycetes(100);o__Saccharomycetales(100);f__Metschnikowiaceae(100);g__Metschnikowia(100);g__Metschnikowia_unclassified(100); | Metschnikowia | 13 | Saprotroph | \|Undefined Saprotroph\| | Yeast | NULL | Probable | As nectar-tap_saprotroph (Põlme et al. 2020); Decay sugar-rich substrates (Põlme et al. 2020) | Põlme S, et al. 2020. Fungal Diversity 105:1-16 ((https://doi.org/10.1007/s13225-020-00466-2)) |
| Otu50025 | 1 | k__Fungi(100);p__Ascomycota(100);c__Saccharomycetes(100);o__Saccharomycetales(100);f__Metschnikowiaceae(100);g__Metschnikowia(100);s__Metschnikowia_pulcherrima(100); | Metschnikowia | 13 | Saprotroph | \|Undefined Saprotroph\| | Yeast | NULL | Probable | As nectar-tap_saprotroph (Põlme et al. 2020); Decay sugar-rich substrates (Põlme et al. 2020) | Põlme S, et al. 2020. Fungal Diversity 105:1-16 ((https://doi.org/10.1007/s13225-020-00466-2)) |
| Otu50026 | 1 | k__Fungi(100);p__Ascomycota(100);c__Saccharomycetes(100);o__Saccharomycetales(100);f__Metschnikowiaceae(100);g__Metschnikowia(100);s__Metschnikowia_pulcherrima(100); | Metschnikowia | 13 | Saprotroph | \|Undefined Saprotroph\| | Yeast | NULL | Probable | As nectar-tap_saprotroph (Põlme et al. 2020); Decay sugar-rich substrates (Põlme et al. 2020) | Põlme S, et al. 2020. Fungal Diversity 105:1-16 ((https://doi.org/10.1007/s13225-020-00466-2)) |
| Otu50027 | 1 | k__Fungi(100);p__Ascomycota(100);c__Saccharomycetes(100);o__Saccharomycetales(100);f__Metschnikowiaceae(100);g__Metschnikowia(100);s__Metschnikowia_pulcherrima(100); | Metschnikowia | 13 | Saprotroph | \|Undefined Saprotroph\| | Yeast | NULL | Probable | As nectar-tap_saprotroph (Põlme et al. 2020); Decay sugar-rich substrates (Põlme et al. 2020) | Põlme S, et al. 2020. Fungal Diversity 105:1-16 ((https://doi.org/10.1007/s13225-020-00466-2)) |
| Otu50028 | 1 | k__Fungi(100);p__Ascomycota(100);c__Saccharomycetes(100);o__Saccharomycetales(100);f__Metschnikowiaceae(100);g__Metschnikowia(100);s__Metschnikowia_pulcherrima(100); | Metschnikowia | 13 | Saprotroph | \|Undefined Saprotroph\| | Yeast | NULL | Probable | As nectar-tap_saprotroph (Põlme et al. 2020); Decay sugar-rich substrates (Põlme et al. 2020) | Põlme S, et al. 2020. Fungal Diversity 105:1-16 ((https://doi.org/10.1007/s13225-020-00466-2)) |
| Otu50029 | 1 | k__Fungi(100);p__Ascomycota(100);c__Saccharomycetes(100);o__Saccharomycetales(100);f__Metschnikowiaceae(100);g__Metschnikowia(100);s__Metschnikowia_pulcherrima(100); | Metschnikowia | 13 | Saprotroph | \|Undefined Saprotroph\| | Yeast | NULL | Probable | As nectar-tap_saprotroph (Põlme et al. 2020); Decay sugar-rich substrates (Põlme et al. 2020) | Põlme S, et al. 2020. Fungal Diversity 105:1-16 ((https://doi.org/10.1007/s13225-020-00466-2)) |
| Otu50030 | 1 | k__Fungi(100);p__Ascomycota(100);c__Saccharomycetes(100);o__Saccharomycetales(100);f__Metschnikowiaceae(100);g__Metschnikowia(100);s__Metschnikowia_pulcherrima(100); | Metschnikowia | 13 | Saprotroph | \|Undefined Saprotroph\| | Yeast | NULL | Probable | As nectar-tap_saprotroph (Põlme et al. 2020); Decay sugar-rich substrates (Põlme et al. 2020) | Põlme S, et al. 2020. Fungal Diversity 105:1-16 ((https://doi.org/10.1007/s13225-020-00466-2)) |
| Otu50031 | 1 | k__Fungi(100);p__Ascomycota(100);c__Saccharomycetes(100);o__Saccharomycetales(100);f__Metschnikowiaceae(100);g__Metschnikowia(100);s__Metschnikowia_pulcherrima(100); | Metschnikowia | 13 | Saprotroph | \|Undefined Saprotroph\| | Yeast | NULL | Probable | As nectar-tap_saprotroph (Põlme et al. 2020); Decay sugar-rich substrates (Põlme et al. 2020) | Põlme S, et al. 2020. Fungal Diversity 105:1-16 ((https://doi.org/10.1007/s13225-020-00466-2)) |
| Otu50032 | 1 | k__Fungi(100);p__Ascomycota(100);c__Saccharomycetes(100);o__Saccharomycetales(100);f__Metschnikowiaceae(100);g__Metschnikowia(100);g__Metschnikowia_unclassified(100); | Metschnikowia | 13 | Saprotroph | \|Undefined Saprotroph\| | Yeast | NULL | Probable | As nectar-tap_saprotroph (Põlme et al. 2020); Decay sugar-rich substrates (Põlme et al. 2020) | Põlme S, et al. 2020. Fungal Diversity 105:1-16 ((https://doi.org/10.1007/s13225-020-00466-2)) |
| Otu50033 | 1 | k__Fungi(100);p__Ascomycota(100);c__Saccharomycetes(100);o__Saccharomycetales(100);f__Metschnikowiaceae(100);g__Metschnikowia(100);s__Metschnikowia_pulcherrima(100); | Metschnikowia | 13 | Saprotroph | \|Undefined Saprotroph\| | Yeast | NULL | Probable | As nectar-tap_saprotroph (Põlme et al. 2020); Decay sugar-rich substrates (Põlme et al. 2020) | Põlme S, et al. 2020. Fungal Diversity 105:1-16 ((https://doi.org/10.1007/s13225-020-00466-2)) |
| Otu50034 | 1 | k__Fungi(100);p__Ascomycota(100);c__Saccharomycetes(100);o__Saccharomycetales(100);f__Metschnikowiaceae(100);g__Metschnikowia(100);s__Metschnikowia_pulcherrima(100); | Metschnikowia | 13 | Saprotroph | \|Undefined Saprotroph\| | Yeast | NULL | Probable | As nectar-tap_saprotroph (Põlme et al. 2020); Decay sugar-rich substrates (Põlme et al. 2020) | Põlme S, et al. 2020. Fungal Diversity 105:1-16 ((https://doi.org/10.1007/s13225-020-00466-2)) |
| Otu50035 | 1 | k__Fungi(100);p__Ascomycota(100);c__Saccharomycetes(100);o__Saccharomycetales(100);f__Metschnikowiaceae(100);g__Metschnikowia(100);g__Metschnikowia_unclassified(100); | Metschnikowia | 13 | Saprotroph | \|Undefined Saprotroph\| | Yeast | NULL | Probable | As nectar-tap_saprotroph (Põlme et al. 2020); Decay sugar-rich substrates (Põlme et al. 2020) | Põlme S, et al. 2020. Fungal Diversity 105:1-16 ((https://doi.org/10.1007/s13225-020-00466-2)) |
| Otu50036 | 1 | k__Fungi(100);p__Ascomycota(100);c__Saccharomycetes(100);o__Saccharomycetales(100);f__Metschnikowiaceae(100);g__Metschnikowia(100);s__Metschnikowia_pulcherrima(100); | Metschnikowia | 13 | Saprotroph | \|Undefined Saprotroph\| | Yeast | NULL | Probable | As nectar-tap_saprotroph (Põlme et al. 2020); Decay sugar-rich substrates (Põlme et al. 2020) | Põlme S, et al. 2020. Fungal Diversity 105:1-16 ((https://doi.org/10.1007/s13225-020-00466-2)) |
| Otu50037 | 1 | k__Fungi(100);p__Ascomycota(100);c__Saccharomycetes(100);o__Saccharomycetales(100);f__Metschnikowiaceae(100);g__Metschnikowia(100);g__Metschnikowia_unclassified(100); | Metschnikowia | 13 | Saprotroph | \|Undefined Saprotroph\| | Yeast | NULL | Probable | As nectar-tap_saprotroph (Põlme et al. 2020); Decay sugar-rich substrates (Põlme et al. 2020) | Põlme S, et al. 2020. Fungal Diversity 105:1-16 ((https://doi.org/10.1007/s13225-020-00466-2)) |
| Otu50038 | 1 | k__Fungi(100);p__Ascomycota(100);c__Saccharomycetes(100);o__Saccharomycetales(100);f__Metschnikowiaceae(100);g__Metschnikowia(100);s__Metschnikowia_pulcherrima(100); | Metschnikowia | 13 | Saprotroph | \|Undefined Saprotroph\| | Yeast | NULL | Probable | As nectar-tap_saprotroph (Põlme et al. 2020); Decay sugar-rich substrates (Põlme et al. 2020) | Põlme S, et al. 2020. Fungal Diversity 105:1-16 ((https://doi.org/10.1007/s13225-020-00466-2)) |
| Otu50039 | 1 | k__Fungi(100);p__Ascomycota(100);c__Saccharomycetes(100);o__Saccharomycetales(100);f__Metschnikowiaceae(100);g__Metschnikowia(100);s__Metschnikowia_pulcherrima(100); | Metschnikowia | 13 | Saprotroph | \|Undefined Saprotroph\| | Yeast | NULL | Probable | As nectar-tap_saprotroph (Põlme et al. 2020); Decay sugar-rich substrates (Põlme et al. 2020) | Põlme S, et al. 2020. Fungal Diversity 105:1-16 ((https://doi.org/10.1007/s13225-020-00466-2)) |
| Otu50040 | 1 | k__Fungi(100);p__Ascomycota(100);c__Saccharomycetes(100);o__Saccharomycetales(100);f__Metschnikowiaceae(100);g__Metschnikowia(100);s__Metschnikowia_pulcherrima(100); | Metschnikowia | 13 | Saprotroph | \|Undefined Saprotroph\| | Yeast | NULL | Probable | As nectar-tap_saprotroph (Põlme et al. 2020); Decay sugar-rich substrates (Põlme et al. 2020) | Põlme S, et al. 2020. Fungal Diversity 105:1-16 ((https://doi.org/10.1007/s13225-020-00466-2)) |
| Otu50041 | 1 | k__Fungi(100);p__Ascomycota(100);c__Saccharomycetes(100);o__Saccharomycetales(100);f__Metschnikowiaceae(100);g__Metschnikowia(100);s__Metschnikowia_pulcherrima(100); | Metschnikowia | 13 | Saprotroph | \|Undefined Saprotroph\| | Yeast | NULL | Probable | As nectar-tap_saprotroph (Põlme et al. 2020); Decay sugar-rich substrates (Põlme et al. 2020) | Põlme S, et al. 2020. Fungal Diversity 105:1-16 ((https://doi.org/10.1007/s13225-020-00466-2)) |
| Otu50042 | 1 | k__Fungi(100);p__Ascomycota(100);c__Saccharomycetes(100);o__Saccharomycetales(100);f__Metschnikowiaceae(100);g__Metschnikowia(100);s__Metschnikowia_pulcherrima(100); | Metschnikowia | 13 | Saprotroph | \|Undefined Saprotroph\| | Yeast | NULL | Probable | As nectar-tap_saprotroph (Põlme et al. 2020); Decay sugar-rich substrates (Põlme et al. 2020) | Põlme S, et al. 2020. Fungal Diversity 105:1-16 ((https://doi.org/10.1007/s13225-020-00466-2)) |
| Otu50043 | 1 | k__Fungi(100);p__Ascomycota(100);c__Saccharomycetes(100);o__Saccharomycetales(100);f__Metschnikowiaceae(100);g__Metschnikowia(100);s__Metschnikowia_pulcherrima(100); | Metschnikowia | 13 | Saprotroph | \|Undefined Saprotroph\| | Yeast | NULL | Probable | As nectar-tap_saprotroph (Põlme et al. 2020); Decay sugar-rich substrates (Põlme et al. 2020) | Põlme S, et al. 2020. Fungal Diversity 105:1-16 ((https://doi.org/10.1007/s13225-020-00466-2)) |
| Otu50044 | 1 | k__Fungi(100);p__Ascomycota(100);c__Saccharomycetes(100);o__Saccharomycetales(100);f__Metschnikowiaceae(100);g__Metschnikowia(100);s__Metschnikowia_pulcherrima(100); | Metschnikowia | 13 | Saprotroph | \|Undefined Saprotroph\| | Yeast | NULL | Probable | As nectar-tap_saprotroph (Põlme et al. 2020); Decay sugar-rich substrates (Põlme et al. 2020) | Põlme S, et al. 2020. Fungal Diversity 105:1-16 ((https://doi.org/10.1007/s13225-020-00466-2)) |
| Otu50045 | 1 | k__Fungi(100);p__Ascomycota(100);c__Saccharomycetes(100);o__Saccharomycetales(100);f__Metschnikowiaceae(100);g__Metschnikowia(100);g__Metschnikowia_unclassified(100); | Metschnikowia | 13 | Saprotroph | \|Undefined Saprotroph\| | Yeast | NULL | Probable | As nectar-tap_saprotroph (Põlme et al. 2020); Decay sugar-rich substrates (Põlme et al. 2020) | Põlme S, et al. 2020. Fungal Diversity 105:1-16 ((https://doi.org/10.1007/s13225-020-00466-2)) |
| Otu50046 | 1 | k__Fungi(100);p__Ascomycota(100);c__Saccharomycetes(100);o__Saccharomycetales(100);f__Metschnikowiaceae(100);g__Metschnikowia(100);s__Metschnikowia_pulcherrima(100); | Metschnikowia | 13 | Saprotroph | \|Undefined Saprotroph\| | Yeast | NULL | Probable | As nectar-tap_saprotroph (Põlme et al. 2020); Decay sugar-rich substrates (Põlme et al. 2020) | Põlme S, et al. 2020. Fungal Diversity 105:1-16 ((https://doi.org/10.1007/s13225-020-00466-2)) |
| Otu50047 | 1 | k__Fungi(100);p__Ascomycota(100);c__Saccharomycetes(100);o__Saccharomycetales(100);f__Metschnikowiaceae(100);g__Metschnikowia(100);s__Metschnikowia_pulcherrima(100); | Metschnikowia | 13 | Saprotroph | \|Undefined Saprotroph\| | Yeast | NULL | Probable | As nectar-tap_saprotroph (Põlme et al. 2020); Decay sugar-rich substrates (Põlme et al. 2020) | Põlme S, et al. 2020. Fungal Diversity 105:1-16 ((https://doi.org/10.1007/s13225-020-00466-2)) |
| Otu50048 | 1 | k__Fungi(100);p__Ascomycota(100);c__Saccharomycetes(100);o__Saccharomycetales(100);f__Metschnikowiaceae(100);g__Metschnikowia(100);s__Metschnikowia_pulcherrima(100); | Metschnikowia | 13 | Saprotroph | \|Undefined Saprotroph\| | Yeast | NULL | Probable | As nectar-tap_saprotroph (Põlme et al. 2020); Decay sugar-rich substrates (Põlme et al. 2020) | Põlme S, et al. 2020. Fungal Diversity 105:1-16 ((https://doi.org/10.1007/s13225-020-00466-2)) |
| Otu50049 | 1 | k__Fungi(100);p__Ascomycota(100);c__Saccharomycetes(100);o__Saccharomycetales(100);f__Metschnikowiaceae(100);g__Metschnikowia(100);s__Metschnikowia_pulcherrima(100); | Metschnikowia | 13 | Saprotroph | \|Undefined Saprotroph\| | Yeast | NULL | Probable | As nectar-tap_saprotroph (Põlme et al. 2020); Decay sugar-rich substrates (Põlme et al. 2020) | Põlme S, et al. 2020. Fungal Diversity 105:1-16 ((https://doi.org/10.1007/s13225-020-00466-2)) |
| Otu50050 | 1 | k__Fungi(100);p__Ascomycota(100);c__Saccharomycetes(100);o__Saccharomycetales(100);f__Metschnikowiaceae(100);g__Metschnikowia(100);s__Metschnikowia_pulcherrima(100); | Metschnikowia | 13 | Saprotroph | \|Undefined Saprotroph\| | Yeast | NULL | Probable | As nectar-tap_saprotroph (Põlme et al. 2020); Decay sugar-rich substrates (Põlme et al. 2020) | Põlme S, et al. 2020. Fungal Diversity 105:1-16 ((https://doi.org/10.1007/s13225-020-00466-2)) |
| Otu50051 | 1 | k__Fungi(100);p__Ascomycota(100);c__Saccharomycetes(100);o__Saccharomycetales(100);f__Metschnikowiaceae(100);g__Metschnikowia(100);s__Metschnikowia_pulcherrima(100); | Metschnikowia | 13 | Saprotroph | \|Undefined Saprotroph\| | Yeast | NULL | Probable | As nectar-tap_saprotroph (Põlme et al. 2020); Decay sugar-rich substrates (Põlme et al. 2020) | Põlme S, et al. 2020. Fungal Diversity 105:1-16 ((https://doi.org/10.1007/s13225-020-00466-2)) |
| Otu50052 | 1 | k__Fungi(100);p__Ascomycota(100);c__Saccharomycetes(100);o__Saccharomycetales(100);f__Metschnikowiaceae(100);g__Metschnikowia(100);s__Metschnikowia_pulcherrima(100); | Metschnikowia | 13 | Saprotroph | \|Undefined Saprotroph\| | Yeast | NULL | Probable | As nectar-tap_saprotroph (Põlme et al. 2020); Decay sugar-rich substrates (Põlme et al. 2020) | Põlme S, et al. 2020. Fungal Diversity 105:1-16 ((https://doi.org/10.1007/s13225-020-00466-2)) |
| Otu50053 | 1 | k__Fungi(100);p__Ascomycota(100);c__Saccharomycetes(100);o__Saccharomycetales(100);f__Metschnikowiaceae(100);g__Metschnikowia(100);s__Metschnikowia_pulcherrima(100); | Metschnikowia | 13 | Saprotroph | \|Undefined Saprotroph\| | Yeast | NULL | Probable | As nectar-tap_saprotroph (Põlme et al. 2020); Decay sugar-rich substrates (Põlme et al. 2020) | Põlme S, et al. 2020. Fungal Diversity 105:1-16 ((https://doi.org/10.1007/s13225-020-00466-2)) |
| Otu50054 | 1 | k__Fungi(100);p__Ascomycota(100);c__Saccharomycetes(100);o__Saccharomycetales(100);f__Metschnikowiaceae(100);g__Metschnikowia(100);s__Metschnikowia_pulcherrima(100); | Metschnikowia | 13 | Saprotroph | \|Undefined Saprotroph\| | Yeast | NULL | Probable | As nectar-tap_saprotroph (Põlme et al. 2020); Decay sugar-rich substrates (Põlme et al. 2020) | Põlme S, et al. 2020. Fungal Diversity 105:1-16 ((https://doi.org/10.1007/s13225-020-00466-2)) |
| Otu50055 | 1 | k__Fungi(100);p__Ascomycota(100);c__Saccharomycetes(100);o__Saccharomycetales(100);f__Metschnikowiaceae(100);g__Metschnikowia(100);g__Metschnikowia_unclassified(100); | Metschnikowia | 13 | Saprotroph | \|Undefined Saprotroph\| | Yeast | NULL | Probable | As nectar-tap_saprotroph (Põlme et al. 2020); Decay sugar-rich substrates (Põlme et al. 2020) | Põlme S, et al. 2020. Fungal Diversity 105:1-16 ((https://doi.org/10.1007/s13225-020-00466-2)) |
| Otu50056 | 1 | k__Fungi(100);p__Ascomycota(100);c__Saccharomycetes(100);o__Saccharomycetales(100);f__Metschnikowiaceae(100);g__Metschnikowia(100);g__Metschnikowia_unclassified(100); | Metschnikowia | 13 | Saprotroph | \|Undefined Saprotroph\| | Yeast | NULL | Probable | As nectar-tap_saprotroph (Põlme et al. 2020); Decay sugar-rich substrates (Põlme et al. 2020) | Põlme S, et al. 2020. Fungal Diversity 105:1-16 ((https://doi.org/10.1007/s13225-020-00466-2)) |
| Otu50057 | 1 | k__Fungi(100);p__Ascomycota(100);c__Saccharomycetes(100);o__Saccharomycetales(100);f__Metschnikowiaceae(100);g__Metschnikowia(100);s__Metschnikowia_pulcherrima(100); | Metschnikowia | 13 | Saprotroph | \|Undefined Saprotroph\| | Yeast | NULL | Probable | As nectar-tap_saprotroph (Põlme et al. 2020); Decay sugar-rich substrates (Põlme et al. 2020) | Põlme S, et al. 2020. Fungal Diversity 105:1-16 ((https://doi.org/10.1007/s13225-020-00466-2)) |
| Otu50058 | 1 | k__Fungi(100);p__Ascomycota(100);c__Saccharomycetes(100);o__Saccharomycetales(100);f__Metschnikowiaceae(100);g__Metschnikowia(100);s__Metschnikowia_pulcherrima(100); | Metschnikowia | 13 | Saprotroph | \|Undefined Saprotroph\| | Yeast | NULL | Probable | As nectar-tap_saprotroph (Põlme et al. 2020); Decay sugar-rich substrates (Põlme et al. 2020) | Põlme S, et al. 2020. Fungal Diversity 105:1-16 ((https://doi.org/10.1007/s13225-020-00466-2)) |
| Otu50059 | 1 | k__Fungi(100);p__Ascomycota(100);c__Saccharomycetes(100);o__Saccharomycetales(100);f__Metschnikowiaceae(100);g__Metschnikowia(100);g__Metschnikowia_unclassified(100); | Metschnikowia | 13 | Saprotroph | \|Undefined Saprotroph\| | Yeast | NULL | Probable | As nectar-tap_saprotroph (Põlme et al. 2020); Decay sugar-rich substrates (Põlme et al. 2020) | Põlme S, et al. 2020. Fungal Diversity 105:1-16 ((https://doi.org/10.1007/s13225-020-00466-2)) |
| Otu50060 | 1 | k__Fungi(100);p__Ascomycota(100);c__Saccharomycetes(100);o__Saccharomycetales(100);f__Metschnikowiaceae(100);g__Metschnikowia(100);s__Metschnikowia_pulcherrima(100); | Metschnikowia | 13 | Saprotroph | \|Undefined Saprotroph\| | Yeast | NULL | Probable | As nectar-tap_saprotroph (Põlme et al. 2020); Decay sugar-rich substrates (Põlme et al. 2020) | Põlme S, et al. 2020. Fungal Diversity 105:1-16 ((https://doi.org/10.1007/s13225-020-00466-2)) |
| Otu50061 | 1 | k__Fungi(100);p__Ascomycota(100);c__Saccharomycetes(100);o__Saccharomycetales(100);f__Metschnikowiaceae(100);g__Metschnikowia(100);s__Metschnikowia_pulcherrima(100); | Metschnikowia | 13 | Saprotroph | \|Undefined Saprotroph\| | Yeast | NULL | Probable | As nectar-tap_saprotroph (Põlme et al. 2020); Decay sugar-rich substrates (Põlme et al. 2020) | Põlme S, et al. 2020. Fungal Diversity 105:1-16 ((https://doi.org/10.1007/s13225-020-00466-2)) |
| Otu50062 | 1 | k__Fungi(100);p__Ascomycota(100);c__Saccharomycetes(100);o__Saccharomycetales(100);f__Metschnikowiaceae(100);g__Metschnikowia(100);g__Metschnikowia_unclassified(100); | Metschnikowia | 13 | Saprotroph | \|Undefined Saprotroph\| | Yeast | NULL | Probable | As nectar-tap_saprotroph (Põlme et al. 2020); Decay sugar-rich substrates (Põlme et al. 2020) | Põlme S, et al. 2020. Fungal Diversity 105:1-16 ((https://doi.org/10.1007/s13225-020-00466-2)) |
| Otu50063 | 1 | k__Fungi(100);p__Ascomycota(100);c__Saccharomycetes(100);o__Saccharomycetales(100);f__Metschnikowiaceae(100);g__Metschnikowia(100);g__Metschnikowia_unclassified(100); | Metschnikowia | 13 | Saprotroph | \|Undefined Saprotroph\| | Yeast | NULL | Probable | As nectar-tap_saprotroph (Põlme et al. 2020); Decay sugar-rich substrates (Põlme et al. 2020) | Põlme S, et al. 2020. Fungal Diversity 105:1-16 ((https://doi.org/10.1007/s13225-020-00466-2)) |
| Otu50064 | 1 | k__Fungi(100);p__Ascomycota(100);c__Saccharomycetes(100);o__Saccharomycetales(100);f__Metschnikowiaceae(100);g__Metschnikowia(100);s__Metschnikowia_pulcherrima(100); | Metschnikowia | 13 | Saprotroph | \|Undefined Saprotroph\| | Yeast | NULL | Probable | As nectar-tap_saprotroph (Põlme et al. 2020); Decay sugar-rich substrates (Põlme et al. 2020) | Põlme S, et al. 2020. Fungal Diversity 105:1-16 ((https://doi.org/10.1007/s13225-020-00466-2)) |
| Otu50065 | 1 | k__Fungi(100);p__Ascomycota(100);c__Saccharomycetes(100);o__Saccharomycetales(100);f__Metschnikowiaceae(100);g__Metschnikowia(100);s__Metschnikowia_pulcherrima(100); | Metschnikowia | 13 | Saprotroph | \|Undefined Saprotroph\| | Yeast | NULL | Probable | As nectar-tap_saprotroph (Põlme et al. 2020); Decay sugar-rich substrates (Põlme et al. 2020) | Põlme S, et al. 2020. Fungal Diversity 105:1-16 ((https://doi.org/10.1007/s13225-020-00466-2)) |
| Otu50066 | 1 | k__Fungi(100);p__Ascomycota(100);c__Saccharomycetes(100);o__Saccharomycetales(100);f__Metschnikowiaceae(100);g__Metschnikowia(100);s__Metschnikowia_pulcherrima(100); | Metschnikowia | 13 | Saprotroph | \|Undefined Saprotroph\| | Yeast | NULL | Probable | As nectar-tap_saprotroph (Põlme et al. 2020); Decay sugar-rich substrates (Põlme et al. 2020) | Põlme S, et al. 2020. Fungal Diversity 105:1-16 ((https://doi.org/10.1007/s13225-020-00466-2)) |
| Otu50067 | 1 | k__Fungi(100);p__Ascomycota(100);c__Saccharomycetes(100);o__Saccharomycetales(100);f__Metschnikowiaceae(100);g__Metschnikowia(100);s__Metschnikowia_pulcherrima(100); | Metschnikowia | 13 | Saprotroph | \|Undefined Saprotroph\| | Yeast | NULL | Probable | As nectar-tap_saprotroph (Põlme et al. 2020); Decay sugar-rich substrates (Põlme et al. 2020) | Põlme S, et al. 2020. Fungal Diversity 105:1-16 ((https://doi.org/10.1007/s13225-020-00466-2)) |
| Otu50068 | 1 | k__Fungi(100);p__Ascomycota(100);c__Saccharomycetes(100);o__Saccharomycetales(100);f__Metschnikowiaceae(100);g__Metschnikowia(100);s__Metschnikowia_pulcherrima(100); | Metschnikowia | 13 | Saprotroph | \|Undefined Saprotroph\| | Yeast | NULL | Probable | As nectar-tap_saprotroph (Põlme et al. 2020); Decay sugar-rich substrates (Põlme et al. 2020) | Põlme S, et al. 2020. Fungal Diversity 105:1-16 ((https://doi.org/10.1007/s13225-020-00466-2)) |
| Otu50069 | 1 | k__Fungi(100);p__Ascomycota(100);c__Saccharomycetes(100);o__Saccharomycetales(100);f__Metschnikowiaceae(100);g__Metschnikowia(100);s__Metschnikowia_pulcherrima(100); | Metschnikowia | 13 | Saprotroph | \|Undefined Saprotroph\| | Yeast | NULL | Probable | As nectar-tap_saprotroph (Põlme et al. 2020); Decay sugar-rich substrates (Põlme et al. 2020) | Põlme S, et al. 2020. Fungal Diversity 105:1-16 ((https://doi.org/10.1007/s13225-020-00466-2)) |
| Otu50070 | 1 | k__Fungi(100);p__Ascomycota(100);c__Saccharomycetes(100);o__Saccharomycetales(100);f__Metschnikowiaceae(100);g__Metschnikowia(100);g__Metschnikowia_unclassified(100); | Metschnikowia | 13 | Saprotroph | \|Undefined Saprotroph\| | Yeast | NULL | Probable | As nectar-tap_saprotroph (Põlme et al. 2020); Decay sugar-rich substrates (Põlme et al. 2020) | Põlme S, et al. 2020. Fungal Diversity 105:1-16 ((https://doi.org/10.1007/s13225-020-00466-2)) |
| Otu50071 | 1 | k__Fungi(100);p__Ascomycota(100);c__Saccharomycetes(100);o__Saccharomycetales(100);f__Metschnikowiaceae(100);g__Metschnikowia(100);s__Metschnikowia_pulcherrima(100); | Metschnikowia | 13 | Saprotroph | \|Undefined Saprotroph\| | Yeast | NULL | Probable | As nectar-tap_saprotroph (Põlme et al. 2020); Decay sugar-rich substrates (Põlme et al. 2020) | Põlme S, et al. 2020. Fungal Diversity 105:1-16 ((https://doi.org/10.1007/s13225-020-00466-2)) |
| Otu50072 | 1 | k__Fungi(100);p__Ascomycota(100);c__Saccharomycetes(100);o__Saccharomycetales(100);f__Metschnikowiaceae(100);g__Metschnikowia(100);s__Metschnikowia_pulcherrima(100); | Metschnikowia | 13 | Saprotroph | \|Undefined Saprotroph\| | Yeast | NULL | Probable | As nectar-tap_saprotroph (Põlme et al. 2020); Decay sugar-rich substrates (Põlme et al. 2020) | Põlme S, et al. 2020. Fungal Diversity 105:1-16 ((https://doi.org/10.1007/s13225-020-00466-2)) |
| Otu50073 | 1 | k__Fungi(100);p__Ascomycota(100);c__Saccharomycetes(100);o__Saccharomycetales(100);f__Metschnikowiaceae(100);g__Metschnikowia(100);g__Metschnikowia_unclassified(100); | Metschnikowia | 13 | Saprotroph | \|Undefined Saprotroph\| | Yeast | NULL | Probable | As nectar-tap_saprotroph (Põlme et al. 2020); Decay sugar-rich substrates (Põlme et al. 2020) | Põlme S, et al. 2020. Fungal Diversity 105:1-16 ((https://doi.org/10.1007/s13225-020-00466-2)) |
| Otu50074 | 1 | k__Fungi(100);p__Ascomycota(100);c__Saccharomycetes(100);o__Saccharomycetales(100);f__Metschnikowiaceae(100);g__Metschnikowia(100);s__Metschnikowia_pulcherrima(100); | Metschnikowia | 13 | Saprotroph | \|Undefined Saprotroph\| | Yeast | NULL | Probable | As nectar-tap_saprotroph (Põlme et al. 2020); Decay sugar-rich substrates (Põlme et al. 2020) | Põlme S, et al. 2020. Fungal Diversity 105:1-16 ((https://doi.org/10.1007/s13225-020-00466-2)) |
| Otu50075 | 1 | k__Fungi(100);p__Ascomycota(100);c__Saccharomycetes(100);o__Saccharomycetales(100);f__Metschnikowiaceae(100);g__Metschnikowia(100);s__Metschnikowia_pulcherrima(100); | Metschnikowia | 13 | Saprotroph | \|Undefined Saprotroph\| | Yeast | NULL | Probable | As nectar-tap_saprotroph (Põlme et al. 2020); Decay sugar-rich substrates (Põlme et al. 2020) | Põlme S, et al. 2020. Fungal Diversity 105:1-16 ((https://doi.org/10.1007/s13225-020-00466-2)) |
| Otu50077 | 1 | k__Fungi(100);p__Ascomycota(100);c__Saccharomycetes(100);o__Saccharomycetales(100);f__Metschnikowiaceae(100);g__Metschnikowia(100);s__Metschnikowia_pulcherrima(100); | Metschnikowia | 13 | Saprotroph | \|Undefined Saprotroph\| | Yeast | NULL | Probable | As nectar-tap_saprotroph (Põlme et al. 2020); Decay sugar-rich substrates (Põlme et al. 2020) | Põlme S, et al. 2020. Fungal Diversity 105:1-16 ((https://doi.org/10.1007/s13225-020-00466-2)) |
| Otu50078 | 1 | k__Fungi(100);p__Ascomycota(100);c__Saccharomycetes(100);o__Saccharomycetales(100);f__Metschnikowiaceae(100);g__Metschnikowia(100);s__Metschnikowia_pulcherrima(100); | Metschnikowia | 13 | Saprotroph | \|Undefined Saprotroph\| | Yeast | NULL | Probable | As nectar-tap_saprotroph (Põlme et al. 2020); Decay sugar-rich substrates (Põlme et al. 2020) | Põlme S, et al. 2020. Fungal Diversity 105:1-16 ((https://doi.org/10.1007/s13225-020-00466-2)) |
| Otu50080 | 1 | k__Fungi(100);p__Ascomycota(100);c__Saccharomycetes(100);o__Saccharomycetales(100);f__Metschnikowiaceae(100);g__Metschnikowia(100);g__Metschnikowia_unclassified(100); | Metschnikowia | 13 | Saprotroph | \|Undefined Saprotroph\| | Yeast | NULL | Probable | As nectar-tap_saprotroph (Põlme et al. 2020); Decay sugar-rich substrates (Põlme et al. 2020) | Põlme S, et al. 2020. Fungal Diversity 105:1-16 ((https://doi.org/10.1007/s13225-020-00466-2)) |
| Otu50081 | 1 | k__Fungi(100);p__Ascomycota(100);c__Saccharomycetes(100);o__Saccharomycetales(100);f__Metschnikowiaceae(100);g__Metschnikowia(100);s__Metschnikowia_pulcherrima(100); | Metschnikowia | 13 | Saprotroph | \|Undefined Saprotroph\| | Yeast | NULL | Probable | As nectar-tap_saprotroph (Põlme et al. 2020); Decay sugar-rich substrates (Põlme et al. 2020) | Põlme S, et al. 2020. Fungal Diversity 105:1-16 ((https://doi.org/10.1007/s13225-020-00466-2)) |
| Otu50082 | 1 | k__Fungi(100);p__Ascomycota(100);c__Saccharomycetes(100);o__Saccharomycetales(100);f__Metschnikowiaceae(100);g__Metschnikowia(100);s__Metschnikowia_pulcherrima(100); | Metschnikowia | 13 | Saprotroph | \|Undefined Saprotroph\| | Yeast | NULL | Probable | As nectar-tap_saprotroph (Põlme et al. 2020); Decay sugar-rich substrates (Põlme et al. 2020) | Põlme S, et al. 2020. Fungal Diversity 105:1-16 ((https://doi.org/10.1007/s13225-020-00466-2)) |
| Otu50083 | 1 | k__Fungi(100);p__Ascomycota(100);c__Saccharomycetes(100);o__Saccharomycetales(100);f__Metschnikowiaceae(100);g__Metschnikowia(100);g__Metschnikowia_unclassified(100); | Metschnikowia | 13 | Saprotroph | \|Undefined Saprotroph\| | Yeast | NULL | Probable | As nectar-tap_saprotroph (Põlme et al. 2020); Decay sugar-rich substrates (Põlme et al. 2020) | Põlme S, et al. 2020. Fungal Diversity 105:1-16 ((https://doi.org/10.1007/s13225-020-00466-2)) |
| Otu50084 | 1 | k__Fungi(100);p__Ascomycota(100);c__Saccharomycetes(100);o__Saccharomycetales(100);f__Metschnikowiaceae(100);g__Metschnikowia(100);g__Metschnikowia_unclassified(100); | Metschnikowia | 13 | Saprotroph | \|Undefined Saprotroph\| | Yeast | NULL | Probable | As nectar-tap_saprotroph (Põlme et al. 2020); Decay sugar-rich substrates (Põlme et al. 2020) | Põlme S, et al. 2020. Fungal Diversity 105:1-16 ((https://doi.org/10.1007/s13225-020-00466-2)) |
| Otu50085 | 1 | k__Fungi(100);p__Ascomycota(100);c__Saccharomycetes(100);o__Saccharomycetales(100);f__Metschnikowiaceae(100);g__Metschnikowia(100);g__Metschnikowia_unclassified(100); | Metschnikowia | 13 | Saprotroph | \|Undefined Saprotroph\| | Yeast | NULL | Probable | As nectar-tap_saprotroph (Põlme et al. 2020); Decay sugar-rich substrates (Põlme et al. 2020) | Põlme S, et al. 2020. Fungal Diversity 105:1-16 ((https://doi.org/10.1007/s13225-020-00466-2)) |
| Otu50086 | 1 | k__Fungi(100);p__Ascomycota(100);c__Saccharomycetes(100);o__Saccharomycetales(100);f__Metschnikowiaceae(100);g__Metschnikowia(100);s__Metschnikowia_pulcherrima(100); | Metschnikowia | 13 | Saprotroph | \|Undefined Saprotroph\| | Yeast | NULL | Probable | As nectar-tap_saprotroph (Põlme et al. 2020); Decay sugar-rich substrates (Põlme et al. 2020) | Põlme S, et al. 2020. Fungal Diversity 105:1-16 ((https://doi.org/10.1007/s13225-020-00466-2)) |
| Otu50088 | 1 | k__Fungi(100);p__Ascomycota(100);c__Saccharomycetes(100);o__Saccharomycetales(100);f__Metschnikowiaceae(100);g__Metschnikowia(100);g__Metschnikowia_unclassified(100); | Metschnikowia | 13 | Saprotroph | \|Undefined Saprotroph\| | Yeast | NULL | Probable | As nectar-tap_saprotroph (Põlme et al. 2020); Decay sugar-rich substrates (Põlme et al. 2020) | Põlme S, et al. 2020. Fungal Diversity 105:1-16 ((https://doi.org/10.1007/s13225-020-00466-2)) |
| Otu50089 | 1 | k__Fungi(100);p__Ascomycota(100);c__Saccharomycetes(100);o__Saccharomycetales(100);f__Metschnikowiaceae(100);g__Metschnikowia(100);g__Metschnikowia_unclassified(100); | Metschnikowia | 13 | Saprotroph | \|Undefined Saprotroph\| | Yeast | NULL | Probable | As nectar-tap_saprotroph (Põlme et al. 2020); Decay sugar-rich substrates (Põlme et al. 2020) | Põlme S, et al. 2020. Fungal Diversity 105:1-16 ((https://doi.org/10.1007/s13225-020-00466-2)) |
| Otu50090 | 1 | k__Fungi(100);p__Ascomycota(100);c__Saccharomycetes(100);o__Saccharomycetales(100);f__Metschnikowiaceae(100);g__Metschnikowia(100);s__Metschnikowia_pulcherrima(100); | Metschnikowia | 13 | Saprotroph | \|Undefined Saprotroph\| | Yeast | NULL | Probable | As nectar-tap_saprotroph (Põlme et al. 2020); Decay sugar-rich substrates (Põlme et al. 2020) | Põlme S, et al. 2020. Fungal Diversity 105:1-16 ((https://doi.org/10.1007/s13225-020-00466-2)) |
| Otu50091 | 1 | k__Fungi(100);p__Ascomycota(100);c__Saccharomycetes(100);o__Saccharomycetales(100);f__Metschnikowiaceae(100);g__Metschnikowia(100);g__Metschnikowia_unclassified(100); | Metschnikowia | 13 | Saprotroph | \|Undefined Saprotroph\| | Yeast | NULL | Probable | As nectar-tap_saprotroph (Põlme et al. 2020); Decay sugar-rich substrates (Põlme et al. 2020) | Põlme S, et al. 2020. Fungal Diversity 105:1-16 ((https://doi.org/10.1007/s13225-020-00466-2)) |
| Otu50092 | 1 | k__Fungi(100);p__Ascomycota(100);c__Saccharomycetes(100);o__Saccharomycetales(100);f__Metschnikowiaceae(100);g__Metschnikowia(100);s__Metschnikowia_pulcherrima(100); | Metschnikowia | 13 | Saprotroph | \|Undefined Saprotroph\| | Yeast | NULL | Probable | As nectar-tap_saprotroph (Põlme et al. 2020); Decay sugar-rich substrates (Põlme et al. 2020) | Põlme S, et al. 2020. Fungal Diversity 105:1-16 ((https://doi.org/10.1007/s13225-020-00466-2)) |
| Otu50093 | 1 | k__Fungi(100);p__Ascomycota(100);c__Saccharomycetes(100);o__Saccharomycetales(100);f__Metschnikowiaceae(100);g__Metschnikowia(100);s__Metschnikowia_pulcherrima(100); | Metschnikowia | 13 | Saprotroph | \|Undefined Saprotroph\| | Yeast | NULL | Probable | As nectar-tap_saprotroph (Põlme et al. 2020); Decay sugar-rich substrates (Põlme et al. 2020) | Põlme S, et al. 2020. Fungal Diversity 105:1-16 ((https://doi.org/10.1007/s13225-020-00466-2)) |
| Otu50094 | 1 | k__Fungi(100);p__Ascomycota(100);c__Saccharomycetes(100);o__Saccharomycetales(100);f__Metschnikowiaceae(100);g__Metschnikowia(100);g__Metschnikowia_unclassified(100); | Metschnikowia | 13 | Saprotroph | \|Undefined Saprotroph\| | Yeast | NULL | Probable | As nectar-tap_saprotroph (Põlme et al. 2020); Decay sugar-rich substrates (Põlme et al. 2020) | Põlme S, et al. 2020. Fungal Diversity 105:1-16 ((https://doi.org/10.1007/s13225-020-00466-2)) |
| Otu50095 | 1 | k__Fungi(100);p__Ascomycota(100);c__Saccharomycetes(100);o__Saccharomycetales(100);f__Metschnikowiaceae(100);g__Metschnikowia(100);s__Metschnikowia_pulcherrima(100); | Metschnikowia | 13 | Saprotroph | \|Undefined Saprotroph\| | Yeast | NULL | Probable | As nectar-tap_saprotroph (Põlme et al. 2020); Decay sugar-rich substrates (Põlme et al. 2020) | Põlme S, et al. 2020. Fungal Diversity 105:1-16 ((https://doi.org/10.1007/s13225-020-00466-2)) |
| Otu50096 | 1 | k__Fungi(100);p__Ascomycota(100);c__Saccharomycetes(100);o__Saccharomycetales(100);f__Metschnikowiaceae(100);g__Metschnikowia(100);g__Metschnikowia_unclassified(100); | Metschnikowia | 13 | Saprotroph | \|Undefined Saprotroph\| | Yeast | NULL | Probable | As nectar-tap_saprotroph (Põlme et al. 2020); Decay sugar-rich substrates (Põlme et al. 2020) | Põlme S, et al. 2020. Fungal Diversity 105:1-16 ((https://doi.org/10.1007/s13225-020-00466-2)) |
| Otu50097 | 1 | k__Fungi(100);p__Ascomycota(100);c__Saccharomycetes(100);o__Saccharomycetales(100);f__Metschnikowiaceae(100);g__Metschnikowia(100);s__Metschnikowia_pulcherrima(100); | Metschnikowia | 13 | Saprotroph | \|Undefined Saprotroph\| | Yeast | NULL | Probable | As nectar-tap_saprotroph (Põlme et al. 2020); Decay sugar-rich substrates (Põlme et al. 2020) | Põlme S, et al. 2020. Fungal Diversity 105:1-16 ((https://doi.org/10.1007/s13225-020-00466-2)) |
| Otu50098 | 1 | k__Fungi(100);p__Ascomycota(100);c__Saccharomycetes(100);o__Saccharomycetales(100);f__Metschnikowiaceae(100);g__Metschnikowia(100);s__Metschnikowia_pulcherrima(100); | Metschnikowia | 13 | Saprotroph | \|Undefined Saprotroph\| | Yeast | NULL | Probable | As nectar-tap_saprotroph (Põlme et al. 2020); Decay sugar-rich substrates (Põlme et al. 2020) | Põlme S, et al. 2020. Fungal Diversity 105:1-16 ((https://doi.org/10.1007/s13225-020-00466-2)) |
| Otu50099 | 1 | k__Fungi(100);p__Ascomycota(100);c__Saccharomycetes(100);o__Saccharomycetales(100);f__Metschnikowiaceae(100);g__Metschnikowia(100);s__Metschnikowia_pulcherrima(100); | Metschnikowia | 13 | Saprotroph | \|Undefined Saprotroph\| | Yeast | NULL | Probable | As nectar-tap_saprotroph (Põlme et al. 2020); Decay sugar-rich substrates (Põlme et al. 2020) | Põlme S, et al. 2020. Fungal Diversity 105:1-16 ((https://doi.org/10.1007/s13225-020-00466-2)) |
| Otu50100 | 1 | k__Fungi(100);p__Ascomycota(100);c__Saccharomycetes(100);o__Saccharomycetales(100);f__Metschnikowiaceae(100);g__Metschnikowia(100);s__Metschnikowia_pulcherrima(100); | Metschnikowia | 13 | Saprotroph | \|Undefined Saprotroph\| | Yeast | NULL | Probable | As nectar-tap_saprotroph (Põlme et al. 2020); Decay sugar-rich substrates (Põlme et al. 2020) | Põlme S, et al. 2020. Fungal Diversity 105:1-16 ((https://doi.org/10.1007/s13225-020-00466-2)) |
| Otu50101 | 1 | k__Fungi(100);p__Ascomycota(100);c__Saccharomycetes(100);o__Saccharomycetales(100);f__Metschnikowiaceae(100);g__Metschnikowia(100);s__Metschnikowia_pulcherrima(100); | Metschnikowia | 13 | Saprotroph | \|Undefined Saprotroph\| | Yeast | NULL | Probable | As nectar-tap_saprotroph (Põlme et al. 2020); Decay sugar-rich substrates (Põlme et al. 2020) | Põlme S, et al. 2020. Fungal Diversity 105:1-16 ((https://doi.org/10.1007/s13225-020-00466-2)) |
| Otu50102 | 1 | k__Fungi(100);p__Ascomycota(100);c__Saccharomycetes(100);o__Saccharomycetales(100);f__Metschnikowiaceae(100);g__Metschnikowia(100);s__Metschnikowia_pulcherrima(100); | Metschnikowia | 13 | Saprotroph | \|Undefined Saprotroph\| | Yeast | NULL | Probable | As nectar-tap_saprotroph (Põlme et al. 2020); Decay sugar-rich substrates (Põlme et al. 2020) | Põlme S, et al. 2020. Fungal Diversity 105:1-16 ((https://doi.org/10.1007/s13225-020-00466-2)) |
| Otu50103 | 1 | k__Fungi(100);p__Ascomycota(100);c__Saccharomycetes(100);o__Saccharomycetales(100);f__Metschnikowiaceae(100);g__Metschnikowia(100);s__Metschnikowia_pulcherrima(100); | Metschnikowia | 13 | Saprotroph | \|Undefined Saprotroph\| | Yeast | NULL | Probable | As nectar-tap_saprotroph (Põlme et al. 2020); Decay sugar-rich substrates (Põlme et al. 2020) | Põlme S, et al. 2020. Fungal Diversity 105:1-16 ((https://doi.org/10.1007/s13225-020-00466-2)) |
| Otu50104 | 1 | k__Fungi(100);p__Ascomycota(100);c__Saccharomycetes(100);o__Saccharomycetales(100);f__Metschnikowiaceae(100);g__Metschnikowia(100);s__Metschnikowia_pulcherrima(100); | Metschnikowia | 13 | Saprotroph | \|Undefined Saprotroph\| | Yeast | NULL | Probable | As nectar-tap_saprotroph (Põlme et al. 2020); Decay sugar-rich substrates (Põlme et al. 2020) | Põlme S, et al. 2020. Fungal Diversity 105:1-16 ((https://doi.org/10.1007/s13225-020-00466-2)) |
| Otu50105 | 1 | k__Fungi(100);p__Ascomycota(100);c__Saccharomycetes(100);o__Saccharomycetales(100);f__Metschnikowiaceae(100);g__Metschnikowia(100);s__Metschnikowia_pulcherrima(100); | Metschnikowia | 13 | Saprotroph | \|Undefined Saprotroph\| | Yeast | NULL | Probable | As nectar-tap_saprotroph (Põlme et al. 2020); Decay sugar-rich substrates (Põlme et al. 2020) | Põlme S, et al. 2020. Fungal Diversity 105:1-16 ((https://doi.org/10.1007/s13225-020-00466-2)) |
| Otu50107 | 1 | k__Fungi(100);p__Ascomycota(100);c__Saccharomycetes(100);o__Saccharomycetales(100);f__Metschnikowiaceae(100);g__Metschnikowia(100);s__Metschnikowia_pulcherrima(100); | Metschnikowia | 13 | Saprotroph | \|Undefined Saprotroph\| | Yeast | NULL | Probable | As nectar-tap_saprotroph (Põlme et al. 2020); Decay sugar-rich substrates (Põlme et al. 2020) | Põlme S, et al. 2020. Fungal Diversity 105:1-16 ((https://doi.org/10.1007/s13225-020-00466-2)) |
| Otu50108 | 1 | k__Fungi(100);p__Ascomycota(100);c__Saccharomycetes(100);o__Saccharomycetales(100);f__Metschnikowiaceae(100);g__Metschnikowia(100);s__Metschnikowia_pulcherrima(100); | Metschnikowia | 13 | Saprotroph | \|Undefined Saprotroph\| | Yeast | NULL | Probable | As nectar-tap_saprotroph (Põlme et al. 2020); Decay sugar-rich substrates (Põlme et al. 2020) | Põlme S, et al. 2020. Fungal Diversity 105:1-16 ((https://doi.org/10.1007/s13225-020-00466-2)) |
| Otu50109 | 1 | k__Fungi(100);p__Ascomycota(100);c__Saccharomycetes(100);o__Saccharomycetales(100);f__Metschnikowiaceae(100);g__Metschnikowia(100);g__Metschnikowia_unclassified(100); | Metschnikowia | 13 | Saprotroph | \|Undefined Saprotroph\| | Yeast | NULL | Probable | As nectar-tap_saprotroph (Põlme et al. 2020); Decay sugar-rich substrates (Põlme et al. 2020) | Põlme S, et al. 2020. Fungal Diversity 105:1-16 ((https://doi.org/10.1007/s13225-020-00466-2)) |
| Otu50110 | 1 | k__Fungi(100);p__Ascomycota(100);c__Saccharomycetes(100);o__Saccharomycetales(100);f__Saccharomycetaceae(100);g__Kazachstania(100);s__Kazachstania_intestinalis(100); | Kazachstania | 13 | Saprotroph | \|Undefined Saprotroph\| | Yeast | NULL | Probable | As nectar-tap_saprotroph (Põlme et al. 2020); Decay sugar-rich substrates (Põlme et al. 2020) | Põlme S, et al. 2020. Fungal Diversity 105:1-16 ((https://doi.org/10.1007/s13225-020-00466-2)) |
| Otu50111 | 1 | k__Fungi(100);p__Ascomycota(100);c__Saccharomycetes(100);o__Saccharomycetales(100);f__Metschnikowiaceae(100);g__Metschnikowia(100);s__Metschnikowia_pulcherrima(100); | Metschnikowia | 13 | Saprotroph | \|Undefined Saprotroph\| | Yeast | NULL | Probable | As nectar-tap_saprotroph (Põlme et al. 2020); Decay sugar-rich substrates (Põlme et al. 2020) | Põlme S, et al. 2020. Fungal Diversity 105:1-16 ((https://doi.org/10.1007/s13225-020-00466-2)) |
| Otu50112 | 1 | k__Fungi(100);p__Ascomycota(100);c__Saccharomycetes(100);o__Saccharomycetales(100);f__Metschnikowiaceae(100);g__Metschnikowia(100);g__Metschnikowia_unclassified(100); | Metschnikowia | 13 | Saprotroph | \|Undefined Saprotroph\| | Yeast | NULL | Probable | As nectar-tap_saprotroph (Põlme et al. 2020); Decay sugar-rich substrates (Põlme et al. 2020) | Põlme S, et al. 2020. Fungal Diversity 105:1-16 ((https://doi.org/10.1007/s13225-020-00466-2)) |
| Otu50113 | 1 | k__Fungi(100);p__Ascomycota(100);c__Saccharomycetes(100);o__Saccharomycetales(100);f__Metschnikowiaceae(100);g__Metschnikowia(100);s__Metschnikowia_pulcherrima(100); | Metschnikowia | 13 | Saprotroph | \|Undefined Saprotroph\| | Yeast | NULL | Probable | As nectar-tap_saprotroph (Põlme et al. 2020); Decay sugar-rich substrates (Põlme et al. 2020) | Põlme S, et al. 2020. Fungal Diversity 105:1-16 ((https://doi.org/10.1007/s13225-020-00466-2)) |
| Otu50114 | 1 | k__Fungi(100);p__Ascomycota(100);c__Saccharomycetes(100);o__Saccharomycetales(100);f__Metschnikowiaceae(100);g__Metschnikowia(100);s__Metschnikowia_pulcherrima(100); | Metschnikowia | 13 | Saprotroph | \|Undefined Saprotroph\| | Yeast | NULL | Probable | As nectar-tap_saprotroph (Põlme et al. 2020); Decay sugar-rich substrates (Põlme et al. 2020) | Põlme S, et al. 2020. Fungal Diversity 105:1-16 ((https://doi.org/10.1007/s13225-020-00466-2)) |
| Otu50115 | 1 | k__Fungi(100);p__Ascomycota(100);c__Saccharomycetes(100);o__Saccharomycetales(100);f__Metschnikowiaceae(100);g__Metschnikowia(100);s__Metschnikowia_pulcherrima(100); | Metschnikowia | 13 | Saprotroph | \|Undefined Saprotroph\| | Yeast | NULL | Probable | As nectar-tap_saprotroph (Põlme et al. 2020); Decay sugar-rich substrates (Põlme et al. 2020) | Põlme S, et al. 2020. Fungal Diversity 105:1-16 ((https://doi.org/10.1007/s13225-020-00466-2)) |
| Otu50116 | 1 | k__Fungi(100);p__Ascomycota(100);c__Saccharomycetes(100);o__Saccharomycetales(100);f__Metschnikowiaceae(100);g__Metschnikowia(100);s__Metschnikowia_pulcherrima(100); | Metschnikowia | 13 | Saprotroph | \|Undefined Saprotroph\| | Yeast | NULL | Probable | As nectar-tap_saprotroph (Põlme et al. 2020); Decay sugar-rich substrates (Põlme et al. 2020) | Põlme S, et al. 2020. Fungal Diversity 105:1-16 ((https://doi.org/10.1007/s13225-020-00466-2)) |
| Otu50117 | 1 | k__Fungi(100);p__Ascomycota(100);c__Saccharomycetes(100);o__Saccharomycetales(100);f__Metschnikowiaceae(100);g__Metschnikowia(100);s__Metschnikowia_pulcherrima(100); | Metschnikowia | 13 | Saprotroph | \|Undefined Saprotroph\| | Yeast | NULL | Probable | As nectar-tap_saprotroph (Põlme et al. 2020); Decay sugar-rich substrates (Põlme et al. 2020) | Põlme S, et al. 2020. Fungal Diversity 105:1-16 ((https://doi.org/10.1007/s13225-020-00466-2)) |
| Otu50118 | 1 | k__Fungi(100);p__Ascomycota(100);c__Saccharomycetes(100);o__Saccharomycetales(100);f__Metschnikowiaceae(100);g__Metschnikowia(100);s__Metschnikowia_pulcherrima(100); | Metschnikowia | 13 | Saprotroph | \|Undefined Saprotroph\| | Yeast | NULL | Probable | As nectar-tap_saprotroph (Põlme et al. 2020); Decay sugar-rich substrates (Põlme et al. 2020) | Põlme S, et al. 2020. Fungal Diversity 105:1-16 ((https://doi.org/10.1007/s13225-020-00466-2)) |
| Otu50119 | 1 | k__Fungi(100);p__Ascomycota(100);c__Saccharomycetes(100);o__Saccharomycetales(100);f__Metschnikowiaceae(100);g__Metschnikowia(100);g__Metschnikowia_unclassified(100); | Metschnikowia | 13 | Saprotroph | \|Undefined Saprotroph\| | Yeast | NULL | Probable | As nectar-tap_saprotroph (Põlme et al. 2020); Decay sugar-rich substrates (Põlme et al. 2020) | Põlme S, et al. 2020. Fungal Diversity 105:1-16 ((https://doi.org/10.1007/s13225-020-00466-2)) |
| Otu50120 | 1 | k__Fungi(100);p__Ascomycota(100);c__Saccharomycetes(100);o__Saccharomycetales(100);f__Metschnikowiaceae(100);g__Metschnikowia(100);s__Metschnikowia_pulcherrima(100); | Metschnikowia | 13 | Saprotroph | \|Undefined Saprotroph\| | Yeast | NULL | Probable | As nectar-tap_saprotroph (Põlme et al. 2020); Decay sugar-rich substrates (Põlme et al. 2020) | Põlme S, et al. 2020. Fungal Diversity 105:1-16 ((https://doi.org/10.1007/s13225-020-00466-2)) |
| Otu50121 | 1 | k__Fungi(100);p__Ascomycota(100);c__Saccharomycetes(100);o__Saccharomycetales(100);f__Metschnikowiaceae(100);g__Metschnikowia(100);s__Metschnikowia_pulcherrima(100); | Metschnikowia | 13 | Saprotroph | \|Undefined Saprotroph\| | Yeast | NULL | Probable | As nectar-tap_saprotroph (Põlme et al. 2020); Decay sugar-rich substrates (Põlme et al. 2020) | Põlme S, et al. 2020. Fungal Diversity 105:1-16 ((https://doi.org/10.1007/s13225-020-00466-2)) |
| Otu50122 | 1 | k__Fungi(100);p__Ascomycota(100);c__Saccharomycetes(100);o__Saccharomycetales(100);f__Metschnikowiaceae(100);g__Metschnikowia(100);s__Metschnikowia_pulcherrima(100); | Metschnikowia | 13 | Saprotroph | \|Undefined Saprotroph\| | Yeast | NULL | Probable | As nectar-tap_saprotroph (Põlme et al. 2020); Decay sugar-rich substrates (Põlme et al. 2020) | Põlme S, et al. 2020. Fungal Diversity 105:1-16 ((https://doi.org/10.1007/s13225-020-00466-2)) |
| Otu50123 | 1 | k__Fungi(100);p__Ascomycota(100);c__Saccharomycetes(100);o__Saccharomycetales(100);f__Metschnikowiaceae(100);g__Metschnikowia(100);s__Metschnikowia_pulcherrima(100); | Metschnikowia | 13 | Saprotroph | \|Undefined Saprotroph\| | Yeast | NULL | Probable | As nectar-tap_saprotroph (Põlme et al. 2020); Decay sugar-rich substrates (Põlme et al. 2020) | Põlme S, et al. 2020. Fungal Diversity 105:1-16 ((https://doi.org/10.1007/s13225-020-00466-2)) |
| Otu50124 | 1 | k__Fungi(100);p__Ascomycota(100);c__Saccharomycetes(100);o__Saccharomycetales(100);f__Metschnikowiaceae(100);g__Metschnikowia(100);s__Metschnikowia_pulcherrima(100); | Metschnikowia | 13 | Saprotroph | \|Undefined Saprotroph\| | Yeast | NULL | Probable | As nectar-tap_saprotroph (Põlme et al. 2020); Decay sugar-rich substrates (Põlme et al. 2020) | Põlme S, et al. 2020. Fungal Diversity 105:1-16 ((https://doi.org/10.1007/s13225-020-00466-2)) |
| Otu50126 | 1 | k__Fungi(100);p__Ascomycota(100);c__Saccharomycetes(100);o__Saccharomycetales(100);f__Metschnikowiaceae(100);g__Metschnikowia(100);g__Metschnikowia_unclassified(100); | Metschnikowia | 13 | Saprotroph | \|Undefined Saprotroph\| | Yeast | NULL | Probable | As nectar-tap_saprotroph (Põlme et al. 2020); Decay sugar-rich substrates (Põlme et al. 2020) | Põlme S, et al. 2020. Fungal Diversity 105:1-16 ((https://doi.org/10.1007/s13225-020-00466-2)) |
| Otu50127 | 1 | k__Fungi(100);p__Ascomycota(100);c__Saccharomycetes(100);o__Saccharomycetales(100);f__Metschnikowiaceae(100);g__Metschnikowia(100);s__Metschnikowia_pulcherrima(100); | Metschnikowia | 13 | Saprotroph | \|Undefined Saprotroph\| | Yeast | NULL | Probable | As nectar-tap_saprotroph (Põlme et al. 2020); Decay sugar-rich substrates (Põlme et al. 2020) | Põlme S, et al. 2020. Fungal Diversity 105:1-16 ((https://doi.org/10.1007/s13225-020-00466-2)) |
| Otu50128 | 1 | k__Fungi(100);p__Ascomycota(100);c__Saccharomycetes(100);o__Saccharomycetales(100);f__Metschnikowiaceae(100);g__Metschnikowia(100);s__Metschnikowia_pulcherrima(100); | Metschnikowia | 13 | Saprotroph | \|Undefined Saprotroph\| | Yeast | NULL | Probable | As nectar-tap_saprotroph (Põlme et al. 2020); Decay sugar-rich substrates (Põlme et al. 2020) | Põlme S, et al. 2020. Fungal Diversity 105:1-16 ((https://doi.org/10.1007/s13225-020-00466-2)) |
| Otu50129 | 1 | k__Fungi(100);p__Ascomycota(100);c__Saccharomycetes(100);o__Saccharomycetales(100);f__Metschnikowiaceae(100);g__Metschnikowia(100);g__Metschnikowia_unclassified(100); | Metschnikowia | 13 | Saprotroph | \|Undefined Saprotroph\| | Yeast | NULL | Probable | As nectar-tap_saprotroph (Põlme et al. 2020); Decay sugar-rich substrates (Põlme et al. 2020) | Põlme S, et al. 2020. Fungal Diversity 105:1-16 ((https://doi.org/10.1007/s13225-020-00466-2)) |
| Otu50130 | 1 | k__Fungi(100);p__Ascomycota(100);c__Saccharomycetes(100);o__Saccharomycetales(100);f__Metschnikowiaceae(100);g__Metschnikowia(100);g__Metschnikowia_unclassified(100); | Metschnikowia | 13 | Saprotroph | \|Undefined Saprotroph\| | Yeast | NULL | Probable | As nectar-tap_saprotroph (Põlme et al. 2020); Decay sugar-rich substrates (Põlme et al. 2020) | Põlme S, et al. 2020. Fungal Diversity 105:1-16 ((https://doi.org/10.1007/s13225-020-00466-2)) |
| Otu50131 | 1 | k__Fungi(100);p__Ascomycota(100);c__Saccharomycetes(100);o__Saccharomycetales(100);f__Metschnikowiaceae(100);g__Metschnikowia(100);g__Metschnikowia_unclassified(100); | Metschnikowia | 13 | Saprotroph | \|Undefined Saprotroph\| | Yeast | NULL | Probable | As nectar-tap_saprotroph (Põlme et al. 2020); Decay sugar-rich substrates (Põlme et al. 2020) | Põlme S, et al. 2020. Fungal Diversity 105:1-16 ((https://doi.org/10.1007/s13225-020-00466-2)) |
| Otu50132 | 1 | k__Fungi(100);p__Ascomycota(100);c__Saccharomycetes(100);o__Saccharomycetales(100);f__Metschnikowiaceae(100);g__Metschnikowia(100);s__Metschnikowia_pulcherrima(100); | Metschnikowia | 13 | Saprotroph | \|Undefined Saprotroph\| | Yeast | NULL | Probable | As nectar-tap_saprotroph (Põlme et al. 2020); Decay sugar-rich substrates (Põlme et al. 2020) | Põlme S, et al. 2020. Fungal Diversity 105:1-16 ((https://doi.org/10.1007/s13225-020-00466-2)) |
| Otu50133 | 1 | k__Fungi(100);p__Ascomycota(100);c__Saccharomycetes(100);o__Saccharomycetales(100);f__Metschnikowiaceae(100);g__Metschnikowia(100);s__Metschnikowia_pulcherrima(100); | Metschnikowia | 13 | Saprotroph | \|Undefined Saprotroph\| | Yeast | NULL | Probable | As nectar-tap_saprotroph (Põlme et al. 2020); Decay sugar-rich substrates (Põlme et al. 2020) | Põlme S, et al. 2020. Fungal Diversity 105:1-16 ((https://doi.org/10.1007/s13225-020-00466-2)) |
| Otu50134 | 1 | k__Fungi(100);p__Ascomycota(100);c__Saccharomycetes(100);o__Saccharomycetales(100);f__Metschnikowiaceae(100);g__Metschnikowia(100);g__Metschnikowia_unclassified(100); | Metschnikowia | 13 | Saprotroph | \|Undefined Saprotroph\| | Yeast | NULL | Probable | As nectar-tap_saprotroph (Põlme et al. 2020); Decay sugar-rich substrates (Põlme et al. 2020) | Põlme S, et al. 2020. Fungal Diversity 105:1-16 ((https://doi.org/10.1007/s13225-020-00466-2)) |
| Otu50135 | 1 | k__Fungi(100);p__Ascomycota(100);c__Saccharomycetes(100);o__Saccharomycetales(100);f__Metschnikowiaceae(100);g__Metschnikowia(100);g__Metschnikowia_unclassified(100); | Metschnikowia | 13 | Saprotroph | \|Undefined Saprotroph\| | Yeast | NULL | Probable | As nectar-tap_saprotroph (Põlme et al. 2020); Decay sugar-rich substrates (Põlme et al. 2020) | Põlme S, et al. 2020. Fungal Diversity 105:1-16 ((https://doi.org/10.1007/s13225-020-00466-2)) |
| Otu50136 | 1 | k__Fungi(100);p__Ascomycota(100);c__Saccharomycetes(100);o__Saccharomycetales(100);f__Metschnikowiaceae(100);g__Metschnikowia(100);g__Metschnikowia_unclassified(100); | Metschnikowia | 13 | Saprotroph | \|Undefined Saprotroph\| | Yeast | NULL | Probable | As nectar-tap_saprotroph (Põlme et al. 2020); Decay sugar-rich substrates (Põlme et al. 2020) | Põlme S, et al. 2020. Fungal Diversity 105:1-16 ((https://doi.org/10.1007/s13225-020-00466-2)) |
| Otu50137 | 1 | k__Fungi(100);p__Ascomycota(100);c__Saccharomycetes(100);o__Saccharomycetales(100);f__Metschnikowiaceae(100);g__Metschnikowia(100);s__Metschnikowia_pulcherrima(100); | Metschnikowia | 13 | Saprotroph | \|Undefined Saprotroph\| | Yeast | NULL | Probable | As nectar-tap_saprotroph (Põlme et al. 2020); Decay sugar-rich substrates (Põlme et al. 2020) | Põlme S, et al. 2020. Fungal Diversity 105:1-16 ((https://doi.org/10.1007/s13225-020-00466-2)) |
| Otu50138 | 1 | k__Fungi(100);p__Ascomycota(100);c__Saccharomycetes(100);o__Saccharomycetales(100);f__Metschnikowiaceae(100);g__Metschnikowia(100);s__Metschnikowia_pulcherrima(100); | Metschnikowia | 13 | Saprotroph | \|Undefined Saprotroph\| | Yeast | NULL | Probable | As nectar-tap_saprotroph (Põlme et al. 2020); Decay sugar-rich substrates (Põlme et al. 2020) | Põlme S, et al. 2020. Fungal Diversity 105:1-16 ((https://doi.org/10.1007/s13225-020-00466-2)) |
| Otu50139 | 1 | k__Fungi(100);p__Ascomycota(100);c__Saccharomycetes(100);o__Saccharomycetales(100);o__Saccharomycetales_unclassified(100);o__Saccharomycetales_unclassified(100);o__Saccharomycetales_unclassified(100); | Saccharomycetales | 7 | Saprotroph | Undefined Saprotroph | Yeast | NULL | Possible | NULL | Sterkenburg E, et al. 2015. New Phytologist 207:1145-1158 ((https://doi.org/10.1111/nph.13426)) |
| Otu50140 | 1 | k__Fungi(100);p__Ascomycota(100);c__Saccharomycetes(100);o__Saccharomycetales(100);f__Metschnikowiaceae(100);g__Metschnikowia(100);g__Metschnikowia_unclassified(100); | Metschnikowia | 13 | Saprotroph | \|Undefined Saprotroph\| | Yeast | NULL | Probable | As nectar-tap_saprotroph (Põlme et al. 2020); Decay sugar-rich substrates (Põlme et al. 2020) | Põlme S, et al. 2020. Fungal Diversity 105:1-16 ((https://doi.org/10.1007/s13225-020-00466-2)) |
| Otu50141 | 1 | k__Fungi(100);p__Ascomycota(100);c__Saccharomycetes(100);o__Saccharomycetales(100);f__Metschnikowiaceae(100);g__Metschnikowia(100);s__Metschnikowia_pulcherrima(100); | Metschnikowia | 13 | Saprotroph | \|Undefined Saprotroph\| | Yeast | NULL | Probable | As nectar-tap_saprotroph (Põlme et al. 2020); Decay sugar-rich substrates (Põlme et al. 2020) | Põlme S, et al. 2020. Fungal Diversity 105:1-16 ((https://doi.org/10.1007/s13225-020-00466-2)) |
| Otu50142 | 1 | k__Fungi(100);p__Ascomycota(100);c__Saccharomycetes(100);o__Saccharomycetales(100);f__Metschnikowiaceae(100);g__Metschnikowia(100);s__Metschnikowia_pulcherrima(100); | Metschnikowia | 13 | Saprotroph | \|Undefined Saprotroph\| | Yeast | NULL | Probable | As nectar-tap_saprotroph (Põlme et al. 2020); Decay sugar-rich substrates (Põlme et al. 2020) | Põlme S, et al. 2020. Fungal Diversity 105:1-16 ((https://doi.org/10.1007/s13225-020-00466-2)) |
| Otu50145 | 1 | k__Fungi(100);p__Ascomycota(100);c__Saccharomycetes(100);o__Saccharomycetales(100);f__Metschnikowiaceae(100);g__Metschnikowia(100);s__Metschnikowia_pulcherrima(100); | Metschnikowia | 13 | Saprotroph | \|Undefined Saprotroph\| | Yeast | NULL | Probable | As nectar-tap_saprotroph (Põlme et al. 2020); Decay sugar-rich substrates (Põlme et al. 2020) | Põlme S, et al. 2020. Fungal Diversity 105:1-16 ((https://doi.org/10.1007/s13225-020-00466-2)) |
| Otu50146 | 1 | k__Fungi(100);p__Ascomycota(100);c__Saccharomycetes(100);o__Saccharomycetales(100);f__Metschnikowiaceae(100);g__Metschnikowia(100);g__Metschnikowia_unclassified(100); | Metschnikowia | 13 | Saprotroph | \|Undefined Saprotroph\| | Yeast | NULL | Probable | As nectar-tap_saprotroph (Põlme et al. 2020); Decay sugar-rich substrates (Põlme et al. 2020) | Põlme S, et al. 2020. Fungal Diversity 105:1-16 ((https://doi.org/10.1007/s13225-020-00466-2)) |
| Otu50147 | 1 | k__Fungi(100);p__Ascomycota(100);c__Saccharomycetes(100);o__Saccharomycetales(100);f__Metschnikowiaceae(100);g__Metschnikowia(100);s__Metschnikowia_pulcherrima(100); | Metschnikowia | 13 | Saprotroph | \|Undefined Saprotroph\| | Yeast | NULL | Probable | As nectar-tap_saprotroph (Põlme et al. 2020); Decay sugar-rich substrates (Põlme et al. 2020) | Põlme S, et al. 2020. Fungal Diversity 105:1-16 ((https://doi.org/10.1007/s13225-020-00466-2)) |
| Otu50148 | 1 | k__Fungi(100);p__Ascomycota(100);c__Saccharomycetes(100);o__Saccharomycetales(100);f__Metschnikowiaceae(100);g__Metschnikowia(100);g__Metschnikowia_unclassified(100); | Metschnikowia | 13 | Saprotroph | \|Undefined Saprotroph\| | Yeast | NULL | Probable | As nectar-tap_saprotroph (Põlme et al. 2020); Decay sugar-rich substrates (Põlme et al. 2020) | Põlme S, et al. 2020. Fungal Diversity 105:1-16 ((https://doi.org/10.1007/s13225-020-00466-2)) |
| Otu50149 | 1 | k__Fungi(100);p__Ascomycota(100);c__Saccharomycetes(100);o__Saccharomycetales(100);f__Metschnikowiaceae(100);g__Metschnikowia(100);g__Metschnikowia_unclassified(100); | Metschnikowia | 13 | Saprotroph | \|Undefined Saprotroph\| | Yeast | NULL | Probable | As nectar-tap_saprotroph (Põlme et al. 2020); Decay sugar-rich substrates (Põlme et al. 2020) | Põlme S, et al. 2020. Fungal Diversity 105:1-16 ((https://doi.org/10.1007/s13225-020-00466-2)) |
| Otu50151 | 1 | k__Fungi(100);p__Ascomycota(100);c__Saccharomycetes(100);o__Saccharomycetales(100);f__Metschnikowiaceae(100);g__Metschnikowia(100);s__Metschnikowia_pulcherrima(100); | Metschnikowia | 13 | Saprotroph | \|Undefined Saprotroph\| | Yeast | NULL | Probable | As nectar-tap_saprotroph (Põlme et al. 2020); Decay sugar-rich substrates (Põlme et al. 2020) | Põlme S, et al. 2020. Fungal Diversity 105:1-16 ((https://doi.org/10.1007/s13225-020-00466-2)) |
| Otu50152 | 1 | k__Fungi(100);p__Ascomycota(100);c__Saccharomycetes(100);o__Saccharomycetales(100);f__Metschnikowiaceae(100);g__Metschnikowia(100);g__Metschnikowia_unclassified(100); | Metschnikowia | 13 | Saprotroph | \|Undefined Saprotroph\| | Yeast | NULL | Probable | As nectar-tap_saprotroph (Põlme et al. 2020); Decay sugar-rich substrates (Põlme et al. 2020) | Põlme S, et al. 2020. Fungal Diversity 105:1-16 ((https://doi.org/10.1007/s13225-020-00466-2)) |
| Otu50153 | 1 | k__Fungi(100);p__Ascomycota(100);c__Saccharomycetes(100);o__Saccharomycetales(100);f__Metschnikowiaceae(100);g__Metschnikowia(100);s__Metschnikowia_pulcherrima(100); | Metschnikowia | 13 | Saprotroph | \|Undefined Saprotroph\| | Yeast | NULL | Probable | As nectar-tap_saprotroph (Põlme et al. 2020); Decay sugar-rich substrates (Põlme et al. 2020) | Põlme S, et al. 2020. Fungal Diversity 105:1-16 ((https://doi.org/10.1007/s13225-020-00466-2)) |
| Otu50154 | 1 | k__Fungi(100);p__Ascomycota(100);c__Saccharomycetes(100);o__Saccharomycetales(100);f__Metschnikowiaceae(100);g__Metschnikowia(100);s__Metschnikowia_pulcherrima(100); | Metschnikowia | 13 | Saprotroph | \|Undefined Saprotroph\| | Yeast | NULL | Probable | As nectar-tap_saprotroph (Põlme et al. 2020); Decay sugar-rich substrates (Põlme et al. 2020) | Põlme S, et al. 2020. Fungal Diversity 105:1-16 ((https://doi.org/10.1007/s13225-020-00466-2)) |
| Otu50155 | 1 | k__Fungi(100);p__Ascomycota(100);c__Saccharomycetes(100);o__Saccharomycetales(100);f__Metschnikowiaceae(100);g__Metschnikowia(100);s__Metschnikowia_pulcherrima(100); | Metschnikowia | 13 | Saprotroph | \|Undefined Saprotroph\| | Yeast | NULL | Probable | As nectar-tap_saprotroph (Põlme et al. 2020); Decay sugar-rich substrates (Põlme et al. 2020) | Põlme S, et al. 2020. Fungal Diversity 105:1-16 ((https://doi.org/10.1007/s13225-020-00466-2)) |
| Otu50156 | 1 | k__Fungi(100);p__Ascomycota(100);c__Saccharomycetes(100);o__Saccharomycetales(100);f__Metschnikowiaceae(100);g__Metschnikowia(100);g__Metschnikowia_unclassified(100); | Metschnikowia | 13 | Saprotroph | \|Undefined Saprotroph\| | Yeast | NULL | Probable | As nectar-tap_saprotroph (Põlme et al. 2020); Decay sugar-rich substrates (Põlme et al. 2020) | Põlme S, et al. 2020. Fungal Diversity 105:1-16 ((https://doi.org/10.1007/s13225-020-00466-2)) |
| Otu50157 | 1 | k__Fungi(100);p__Ascomycota(100);c__Saccharomycetes(100);o__Saccharomycetales(100);f__Metschnikowiaceae(100);g__Metschnikowia(100);s__Metschnikowia_pulcherrima(100); | Metschnikowia | 13 | Saprotroph | \|Undefined Saprotroph\| | Yeast | NULL | Probable | As nectar-tap_saprotroph (Põlme et al. 2020); Decay sugar-rich substrates (Põlme et al. 2020) | Põlme S, et al. 2020. Fungal Diversity 105:1-16 ((https://doi.org/10.1007/s13225-020-00466-2)) |
| Otu50158 | 1 | k__Fungi(100);p__Ascomycota(100);c__Saccharomycetes(100);o__Saccharomycetales(100);f__Metschnikowiaceae(100);g__Metschnikowia(100);g__Metschnikowia_unclassified(100); | Metschnikowia | 13 | Saprotroph | \|Undefined Saprotroph\| | Yeast | NULL | Probable | As nectar-tap_saprotroph (Põlme et al. 2020); Decay sugar-rich substrates (Põlme et al. 2020) | Põlme S, et al. 2020. Fungal Diversity 105:1-16 ((https://doi.org/10.1007/s13225-020-00466-2)) |
| Otu50159 | 1 | k__Fungi(100);p__Ascomycota(100);c__Saccharomycetes(100);o__Saccharomycetales(100);f__Metschnikowiaceae(100);g__Metschnikowia(100);s__Metschnikowia_pulcherrima(100); | Metschnikowia | 13 | Saprotroph | \|Undefined Saprotroph\| | Yeast | NULL | Probable | As nectar-tap_saprotroph (Põlme et al. 2020); Decay sugar-rich substrates (Põlme et al. 2020) | Põlme S, et al. 2020. Fungal Diversity 105:1-16 ((https://doi.org/10.1007/s13225-020-00466-2)) |
| Otu50160 | 1 | k__Fungi(100);p__Ascomycota(100);c__Saccharomycetes(100);o__Saccharomycetales(100);f__Metschnikowiaceae(100);g__Metschnikowia(100);g__Metschnikowia_unclassified(100); | Metschnikowia | 13 | Saprotroph | \|Undefined Saprotroph\| | Yeast | NULL | Probable | As nectar-tap_saprotroph (Põlme et al. 2020); Decay sugar-rich substrates (Põlme et al. 2020) | Põlme S, et al. 2020. Fungal Diversity 105:1-16 ((https://doi.org/10.1007/s13225-020-00466-2)) |
| Otu50161 | 1 | k__Fungi(100);p__Ascomycota(100);c__Saccharomycetes(100);o__Saccharomycetales(100);f__Metschnikowiaceae(100);g__Metschnikowia(100);g__Metschnikowia_unclassified(100); | Metschnikowia | 13 | Saprotroph | \|Undefined Saprotroph\| | Yeast | NULL | Probable | As nectar-tap_saprotroph (Põlme et al. 2020); Decay sugar-rich substrates (Põlme et al. 2020) | Põlme S, et al. 2020. Fungal Diversity 105:1-16 ((https://doi.org/10.1007/s13225-020-00466-2)) |
| Otu50162 | 1 | k__Fungi(100);p__Ascomycota(100);c__Saccharomycetes(100);o__Saccharomycetales(100);f__Metschnikowiaceae(100);g__Metschnikowia(100);g__Metschnikowia_unclassified(100); | Metschnikowia | 13 | Saprotroph | \|Undefined Saprotroph\| | Yeast | NULL | Probable | As nectar-tap_saprotroph (Põlme et al. 2020); Decay sugar-rich substrates (Põlme et al. 2020) | Põlme S, et al. 2020. Fungal Diversity 105:1-16 ((https://doi.org/10.1007/s13225-020-00466-2)) |
| Otu50163 | 1 | k__Fungi(100);p__Ascomycota(100);c__Saccharomycetes(100);o__Saccharomycetales(100);f__Metschnikowiaceae(100);g__Metschnikowia(100);s__Metschnikowia_pulcherrima(100); | Metschnikowia | 13 | Saprotroph | \|Undefined Saprotroph\| | Yeast | NULL | Probable | As nectar-tap_saprotroph (Põlme et al. 2020); Decay sugar-rich substrates (Põlme et al. 2020) | Põlme S, et al. 2020. Fungal Diversity 105:1-16 ((https://doi.org/10.1007/s13225-020-00466-2)) |
| Otu50164 | 1 | k__Fungi(100);p__Ascomycota(100);c__Saccharomycetes(100);o__Saccharomycetales(100);f__Metschnikowiaceae(100);g__Metschnikowia(100);s__Metschnikowia_pulcherrima(100); | Metschnikowia | 13 | Saprotroph | \|Undefined Saprotroph\| | Yeast | NULL | Probable | As nectar-tap_saprotroph (Põlme et al. 2020); Decay sugar-rich substrates (Põlme et al. 2020) | Põlme S, et al. 2020. Fungal Diversity 105:1-16 ((https://doi.org/10.1007/s13225-020-00466-2)) |
| Otu50165 | 1 | k__Fungi(100);p__Ascomycota(100);c__Saccharomycetes(100);o__Saccharomycetales(100);f__Metschnikowiaceae(100);g__Metschnikowia(100);g__Metschnikowia_unclassified(100); | Metschnikowia | 13 | Saprotroph | \|Undefined Saprotroph\| | Yeast | NULL | Probable | As nectar-tap_saprotroph (Põlme et al. 2020); Decay sugar-rich substrates (Põlme et al. 2020) | Põlme S, et al. 2020. Fungal Diversity 105:1-16 ((https://doi.org/10.1007/s13225-020-00466-2)) |
| Otu50166 | 1 | k__Fungi(100);p__Ascomycota(100);c__Saccharomycetes(100);o__Saccharomycetales(100);f__Metschnikowiaceae(100);g__Metschnikowia(100);s__Metschnikowia_pulcherrima(100); | Metschnikowia | 13 | Saprotroph | \|Undefined Saprotroph\| | Yeast | NULL | Probable | As nectar-tap_saprotroph (Põlme et al. 2020); Decay sugar-rich substrates (Põlme et al. 2020) | Põlme S, et al. 2020. Fungal Diversity 105:1-16 ((https://doi.org/10.1007/s13225-020-00466-2)) |
| Otu50167 | 1 | k__Fungi(100);p__Ascomycota(100);c__Saccharomycetes(100);o__Saccharomycetales(100);f__Metschnikowiaceae(100);g__Metschnikowia(100);s__Metschnikowia_pulcherrima(100); | Metschnikowia | 13 | Saprotroph | \|Undefined Saprotroph\| | Yeast | NULL | Probable | As nectar-tap_saprotroph (Põlme et al. 2020); Decay sugar-rich substrates (Põlme et al. 2020) | Põlme S, et al. 2020. Fungal Diversity 105:1-16 ((https://doi.org/10.1007/s13225-020-00466-2)) |
| Otu50168 | 1 | k__Fungi(100);p__Ascomycota(100);c__Saccharomycetes(100);o__Saccharomycetales(100);f__Metschnikowiaceae(100);g__Metschnikowia(100);s__Metschnikowia_pulcherrima(100); | Metschnikowia | 13 | Saprotroph | \|Undefined Saprotroph\| | Yeast | NULL | Probable | As nectar-tap_saprotroph (Põlme et al. 2020); Decay sugar-rich substrates (Põlme et al. 2020) | Põlme S, et al. 2020. Fungal Diversity 105:1-16 ((https://doi.org/10.1007/s13225-020-00466-2)) |
| Otu50169 | 1 | k__Fungi(100);p__Ascomycota(100);c__Saccharomycetes(100);o__Saccharomycetales(100);f__Metschnikowiaceae(100);g__Metschnikowia(100);s__Metschnikowia_pulcherrima(100); | Metschnikowia | 13 | Saprotroph | \|Undefined Saprotroph\| | Yeast | NULL | Probable | As nectar-tap_saprotroph (Põlme et al. 2020); Decay sugar-rich substrates (Põlme et al. 2020) | Põlme S, et al. 2020. Fungal Diversity 105:1-16 ((https://doi.org/10.1007/s13225-020-00466-2)) |
| Otu50170 | 1 | k__Fungi(100);p__Ascomycota(100);c__Saccharomycetes(100);o__Saccharomycetales(100);f__Metschnikowiaceae(100);g__Metschnikowia(100);g__Metschnikowia_unclassified(100); | Metschnikowia | 13 | Saprotroph | \|Undefined Saprotroph\| | Yeast | NULL | Probable | As nectar-tap_saprotroph (Põlme et al. 2020); Decay sugar-rich substrates (Põlme et al. 2020) | Põlme S, et al. 2020. Fungal Diversity 105:1-16 ((https://doi.org/10.1007/s13225-020-00466-2)) |
| Otu50171 | 1 | k__Fungi(100);p__Ascomycota(100);c__Saccharomycetes(100);o__Saccharomycetales(100);f__Metschnikowiaceae(100);g__Metschnikowia(100);s__Metschnikowia_pulcherrima(100); | Metschnikowia | 13 | Saprotroph | \|Undefined Saprotroph\| | Yeast | NULL | Probable | As nectar-tap_saprotroph (Põlme et al. 2020); Decay sugar-rich substrates (Põlme et al. 2020) | Põlme S, et al. 2020. Fungal Diversity 105:1-16 ((https://doi.org/10.1007/s13225-020-00466-2)) |
| Otu50172 | 1 | k__Fungi(100);p__Ascomycota(100);c__Saccharomycetes(100);o__Saccharomycetales(100);f__Metschnikowiaceae(100);g__Metschnikowia(100);s__Metschnikowia_pulcherrima(100); | Metschnikowia | 13 | Saprotroph | \|Undefined Saprotroph\| | Yeast | NULL | Probable | As nectar-tap_saprotroph (Põlme et al. 2020); Decay sugar-rich substrates (Põlme et al. 2020) | Põlme S, et al. 2020. Fungal Diversity 105:1-16 ((https://doi.org/10.1007/s13225-020-00466-2)) |
| Otu50173 | 1 | k__Fungi(100);p__Ascomycota(100);c__Saccharomycetes(100);o__Saccharomycetales(100);f__Metschnikowiaceae(100);g__Metschnikowia(100);s__Metschnikowia_pulcherrima(100); | Metschnikowia | 13 | Saprotroph | \|Undefined Saprotroph\| | Yeast | NULL | Probable | As nectar-tap_saprotroph (Põlme et al. 2020); Decay sugar-rich substrates (Põlme et al. 2020) | Põlme S, et al. 2020. Fungal Diversity 105:1-16 ((https://doi.org/10.1007/s13225-020-00466-2)) |
| Otu50174 | 1 | k__Fungi(100);p__Ascomycota(100);c__Saccharomycetes(100);o__Saccharomycetales(100);f__Metschnikowiaceae(100);g__Metschnikowia(100);g__Metschnikowia_unclassified(100); | Metschnikowia | 13 | Saprotroph | \|Undefined Saprotroph\| | Yeast | NULL | Probable | As nectar-tap_saprotroph (Põlme et al. 2020); Decay sugar-rich substrates (Põlme et al. 2020) | Põlme S, et al. 2020. Fungal Diversity 105:1-16 ((https://doi.org/10.1007/s13225-020-00466-2)) |
| Otu50175 | 1 | k__Fungi(100);p__Ascomycota(100);c__Saccharomycetes(100);o__Saccharomycetales(100);f__Metschnikowiaceae(100);g__Metschnikowia(100);s__Metschnikowia_pulcherrima(100); | Metschnikowia | 13 | Saprotroph | \|Undefined Saprotroph\| | Yeast | NULL | Probable | As nectar-tap_saprotroph (Põlme et al. 2020); Decay sugar-rich substrates (Põlme et al. 2020) | Põlme S, et al. 2020. Fungal Diversity 105:1-16 ((https://doi.org/10.1007/s13225-020-00466-2)) |
| Otu50176 | 1 | k__Fungi(100);p__Ascomycota(100);c__Saccharomycetes(100);o__Saccharomycetales(100);f__Metschnikowiaceae(100);g__Metschnikowia(100);s__Metschnikowia_pulcherrima(100); | Metschnikowia | 13 | Saprotroph | \|Undefined Saprotroph\| | Yeast | NULL | Probable | As nectar-tap_saprotroph (Põlme et al. 2020); Decay sugar-rich substrates (Põlme et al. 2020) | Põlme S, et al. 2020. Fungal Diversity 105:1-16 ((https://doi.org/10.1007/s13225-020-00466-2)) |
| Otu50177 | 1 | k__Fungi(100);p__Ascomycota(100);c__Saccharomycetes(100);o__Saccharomycetales(100);f__Metschnikowiaceae(100);g__Metschnikowia(100);s__Metschnikowia_pulcherrima(100); | Metschnikowia | 13 | Saprotroph | \|Undefined Saprotroph\| | Yeast | NULL | Probable | As nectar-tap_saprotroph (Põlme et al. 2020); Decay sugar-rich substrates (Põlme et al. 2020) | Põlme S, et al. 2020. Fungal Diversity 105:1-16 ((https://doi.org/10.1007/s13225-020-00466-2)) |
| Otu50178 | 1 | k__Fungi(100);p__Ascomycota(100);c__Saccharomycetes(100);o__Saccharomycetales(100);f__Metschnikowiaceae(100);g__Metschnikowia(100);s__Metschnikowia_pulcherrima(100); | Metschnikowia | 13 | Saprotroph | \|Undefined Saprotroph\| | Yeast | NULL | Probable | As nectar-tap_saprotroph (Põlme et al. 2020); Decay sugar-rich substrates (Põlme et al. 2020) | Põlme S, et al. 2020. Fungal Diversity 105:1-16 ((https://doi.org/10.1007/s13225-020-00466-2)) |
| Otu50179 | 1 | k__Fungi(100);p__Ascomycota(100);c__Saccharomycetes(100);o__Saccharomycetales(100);f__Metschnikowiaceae(100);g__Metschnikowia(100);g__Metschnikowia_unclassified(100); | Metschnikowia | 13 | Saprotroph | \|Undefined Saprotroph\| | Yeast | NULL | Probable | As nectar-tap_saprotroph (Põlme et al. 2020); Decay sugar-rich substrates (Põlme et al. 2020) | Põlme S, et al. 2020. Fungal Diversity 105:1-16 ((https://doi.org/10.1007/s13225-020-00466-2)) |
| Otu50180 | 1 | k__Fungi(100);p__Ascomycota(100);c__Saccharomycetes(100);o__Saccharomycetales(100);f__Metschnikowiaceae(100);g__Metschnikowia(100);g__Metschnikowia_unclassified(100); | Metschnikowia | 13 | Saprotroph | \|Undefined Saprotroph\| | Yeast | NULL | Probable | As nectar-tap_saprotroph (Põlme et al. 2020); Decay sugar-rich substrates (Põlme et al. 2020) | Põlme S, et al. 2020. Fungal Diversity 105:1-16 ((https://doi.org/10.1007/s13225-020-00466-2)) |
| Otu50181 | 1 | k__Fungi(100);p__Ascomycota(100);c__Saccharomycetes(100);o__Saccharomycetales(100);f__Metschnikowiaceae(100);g__Metschnikowia(100);s__Metschnikowia_pulcherrima(100); | Metschnikowia | 13 | Saprotroph | \|Undefined Saprotroph\| | Yeast | NULL | Probable | As nectar-tap_saprotroph (Põlme et al. 2020); Decay sugar-rich substrates (Põlme et al. 2020) | Põlme S, et al. 2020. Fungal Diversity 105:1-16 ((https://doi.org/10.1007/s13225-020-00466-2)) |
| Otu50182 | 1 | k__Fungi(100);p__Ascomycota(100);c__Saccharomycetes(100);o__Saccharomycetales(100);f__Metschnikowiaceae(100);g__Metschnikowia(100);s__Metschnikowia_pulcherrima(100); | Metschnikowia | 13 | Saprotroph | \|Undefined Saprotroph\| | Yeast | NULL | Probable | As nectar-tap_saprotroph (Põlme et al. 2020); Decay sugar-rich substrates (Põlme et al. 2020) | Põlme S, et al. 2020. Fungal Diversity 105:1-16 ((https://doi.org/10.1007/s13225-020-00466-2)) |
| Otu50183 | 1 | k__Fungi(100);p__Ascomycota(100);c__Saccharomycetes(100);o__Saccharomycetales(100);f__Metschnikowiaceae(100);g__Metschnikowia(100);s__Metschnikowia_pulcherrima(100); | Metschnikowia | 13 | Saprotroph | \|Undefined Saprotroph\| | Yeast | NULL | Probable | As nectar-tap_saprotroph (Põlme et al. 2020); Decay sugar-rich substrates (Põlme et al. 2020) | Põlme S, et al. 2020. Fungal Diversity 105:1-16 ((https://doi.org/10.1007/s13225-020-00466-2)) |
| Otu50184 | 1 | k__Fungi(100);p__Ascomycota(100);c__Saccharomycetes(100);o__Saccharomycetales(100);f__Metschnikowiaceae(100);g__Metschnikowia(100);s__Metschnikowia_pulcherrima(100); | Metschnikowia | 13 | Saprotroph | \|Undefined Saprotroph\| | Yeast | NULL | Probable | As nectar-tap_saprotroph (Põlme et al. 2020); Decay sugar-rich substrates (Põlme et al. 2020) | Põlme S, et al. 2020. Fungal Diversity 105:1-16 ((https://doi.org/10.1007/s13225-020-00466-2)) |
| Otu50185 | 1 | k__Fungi(100);p__Ascomycota(100);c__Saccharomycetes(100);o__Saccharomycetales(100);f__Metschnikowiaceae(100);g__Metschnikowia(100);s__Metschnikowia_pulcherrima(100); | Metschnikowia | 13 | Saprotroph | \|Undefined Saprotroph\| | Yeast | NULL | Probable | As nectar-tap_saprotroph (Põlme et al. 2020); Decay sugar-rich substrates (Põlme et al. 2020) | Põlme S, et al. 2020. Fungal Diversity 105:1-16 ((https://doi.org/10.1007/s13225-020-00466-2)) |
| Otu50187 | 1 | k__Fungi(100);p__Ascomycota(100);c__Saccharomycetes(100);o__Saccharomycetales(100);f__Metschnikowiaceae(100);g__Metschnikowia(100);g__Metschnikowia_unclassified(100); | Metschnikowia | 13 | Saprotroph | \|Undefined Saprotroph\| | Yeast | NULL | Probable | As nectar-tap_saprotroph (Põlme et al. 2020); Decay sugar-rich substrates (Põlme et al. 2020) | Põlme S, et al. 2020. Fungal Diversity 105:1-16 ((https://doi.org/10.1007/s13225-020-00466-2)) |
| Otu50188 | 1 | k__Fungi(100);p__Ascomycota(100);c__Saccharomycetes(100);o__Saccharomycetales(100);f__Metschnikowiaceae(100);g__Metschnikowia(100);s__Metschnikowia_pulcherrima(100); | Metschnikowia | 13 | Saprotroph | \|Undefined Saprotroph\| | Yeast | NULL | Probable | As nectar-tap_saprotroph (Põlme et al. 2020); Decay sugar-rich substrates (Põlme et al. 2020) | Põlme S, et al. 2020. Fungal Diversity 105:1-16 ((https://doi.org/10.1007/s13225-020-00466-2)) |
| Otu50189 | 1 | k__Fungi(100);p__Ascomycota(100);c__Saccharomycetes(100);o__Saccharomycetales(100);f__Metschnikowiaceae(100);g__Metschnikowia(100);s__Metschnikowia_pulcherrima(100); | Metschnikowia | 13 | Saprotroph | \|Undefined Saprotroph\| | Yeast | NULL | Probable | As nectar-tap_saprotroph (Põlme et al. 2020); Decay sugar-rich substrates (Põlme et al. 2020) | Põlme S, et al. 2020. Fungal Diversity 105:1-16 ((https://doi.org/10.1007/s13225-020-00466-2)) |
| Otu50190 | 1 | k__Fungi(100);p__Ascomycota(100);c__Saccharomycetes(100);o__Saccharomycetales(100);f__Metschnikowiaceae(100);g__Metschnikowia(100);s__Metschnikowia_pulcherrima(100); | Metschnikowia | 13 | Saprotroph | \|Undefined Saprotroph\| | Yeast | NULL | Probable | As nectar-tap_saprotroph (Põlme et al. 2020); Decay sugar-rich substrates (Põlme et al. 2020) | Põlme S, et al. 2020. Fungal Diversity 105:1-16 ((https://doi.org/10.1007/s13225-020-00466-2)) |
| Otu50191 | 1 | k__Fungi(100);p__Ascomycota(100);c__Saccharomycetes(100);o__Saccharomycetales(100);f__Metschnikowiaceae(100);g__Metschnikowia(100);s__Metschnikowia_pulcherrima(100); | Metschnikowia | 13 | Saprotroph | \|Undefined Saprotroph\| | Yeast | NULL | Probable | As nectar-tap_saprotroph (Põlme et al. 2020); Decay sugar-rich substrates (Põlme et al. 2020) | Põlme S, et al. 2020. Fungal Diversity 105:1-16 ((https://doi.org/10.1007/s13225-020-00466-2)) |
| Otu50192 | 1 | k__Fungi(100);p__Ascomycota(100);c__Saccharomycetes(100);o__Saccharomycetales(100);f__Metschnikowiaceae(100);g__Metschnikowia(100);s__Metschnikowia_pulcherrima(100); | Metschnikowia | 13 | Saprotroph | \|Undefined Saprotroph\| | Yeast | NULL | Probable | As nectar-tap_saprotroph (Põlme et al. 2020); Decay sugar-rich substrates (Põlme et al. 2020) | Põlme S, et al. 2020. Fungal Diversity 105:1-16 ((https://doi.org/10.1007/s13225-020-00466-2)) |
| Otu50193 | 1 | k__Fungi(100);p__Ascomycota(100);c__Saccharomycetes(100);o__Saccharomycetales(100);f__Metschnikowiaceae(100);g__Metschnikowia(100);s__Metschnikowia_pulcherrima(100); | Metschnikowia | 13 | Saprotroph | \|Undefined Saprotroph\| | Yeast | NULL | Probable | As nectar-tap_saprotroph (Põlme et al. 2020); Decay sugar-rich substrates (Põlme et al. 2020) | Põlme S, et al. 2020. Fungal Diversity 105:1-16 ((https://doi.org/10.1007/s13225-020-00466-2)) |
| Otu50194 | 1 | k__Fungi(100);p__Ascomycota(100);c__Saccharomycetes(100);o__Saccharomycetales(100);f__Metschnikowiaceae(100);g__Metschnikowia(100);s__Metschnikowia_pulcherrima(100); | Metschnikowia | 13 | Saprotroph | \|Undefined Saprotroph\| | Yeast | NULL | Probable | As nectar-tap_saprotroph (Põlme et al. 2020); Decay sugar-rich substrates (Põlme et al. 2020) | Põlme S, et al. 2020. Fungal Diversity 105:1-16 ((https://doi.org/10.1007/s13225-020-00466-2)) |
| Otu50195 | 1 | k__Fungi(100);p__Ascomycota(100);c__Saccharomycetes(100);o__Saccharomycetales(100);f__Metschnikowiaceae(100);g__Metschnikowia(100);s__Metschnikowia_pulcherrima(100); | Metschnikowia | 13 | Saprotroph | \|Undefined Saprotroph\| | Yeast | NULL | Probable | As nectar-tap_saprotroph (Põlme et al. 2020); Decay sugar-rich substrates (Põlme et al. 2020) | Põlme S, et al. 2020. Fungal Diversity 105:1-16 ((https://doi.org/10.1007/s13225-020-00466-2)) |
| Otu50196 | 1 | k__Fungi(100);p__Ascomycota(100);c__Saccharomycetes(100);o__Saccharomycetales(100);f__Metschnikowiaceae(100);g__Metschnikowia(100);g__Metschnikowia_unclassified(100); | Metschnikowia | 13 | Saprotroph | \|Undefined Saprotroph\| | Yeast | NULL | Probable | As nectar-tap_saprotroph (Põlme et al. 2020); Decay sugar-rich substrates (Põlme et al. 2020) | Põlme S, et al. 2020. Fungal Diversity 105:1-16 ((https://doi.org/10.1007/s13225-020-00466-2)) |
| Otu50197 | 1 | k__Fungi(100);p__Ascomycota(100);c__Saccharomycetes(100);o__Saccharomycetales(100);f__Metschnikowiaceae(100);g__Metschnikowia(100);s__Metschnikowia_pulcherrima(100); | Metschnikowia | 13 | Saprotroph | \|Undefined Saprotroph\| | Yeast | NULL | Probable | As nectar-tap_saprotroph (Põlme et al. 2020); Decay sugar-rich substrates (Põlme et al. 2020) | Põlme S, et al. 2020. Fungal Diversity 105:1-16 ((https://doi.org/10.1007/s13225-020-00466-2)) |
| Otu50198 | 1 | k__Fungi(100);p__Ascomycota(100);c__Saccharomycetes(100);o__Saccharomycetales(100);f__Metschnikowiaceae(100);g__Metschnikowia(100);g__Metschnikowia_unclassified(100); | Metschnikowia | 13 | Saprotroph | \|Undefined Saprotroph\| | Yeast | NULL | Probable | As nectar-tap_saprotroph (Põlme et al. 2020); Decay sugar-rich substrates (Põlme et al. 2020) | Põlme S, et al. 2020. Fungal Diversity 105:1-16 ((https://doi.org/10.1007/s13225-020-00466-2)) |
| Otu50199 | 1 | k__Fungi(100);p__Ascomycota(100);c__Saccharomycetes(100);o__Saccharomycetales(100);f__Metschnikowiaceae(100);g__Metschnikowia(100);g__Metschnikowia_unclassified(100); | Metschnikowia | 13 | Saprotroph | \|Undefined Saprotroph\| | Yeast | NULL | Probable | As nectar-tap_saprotroph (Põlme et al. 2020); Decay sugar-rich substrates (Põlme et al. 2020) | Põlme S, et al. 2020. Fungal Diversity 105:1-16 ((https://doi.org/10.1007/s13225-020-00466-2)) |
| Otu50200 | 1 | k__Fungi(100);p__Ascomycota(100);c__Saccharomycetes(100);o__Saccharomycetales(100);f__Metschnikowiaceae(100);g__Metschnikowia(100);g__Metschnikowia_unclassified(100); | Metschnikowia | 13 | Saprotroph | \|Undefined Saprotroph\| | Yeast | NULL | Probable | As nectar-tap_saprotroph (Põlme et al. 2020); Decay sugar-rich substrates (Põlme et al. 2020) | Põlme S, et al. 2020. Fungal Diversity 105:1-16 ((https://doi.org/10.1007/s13225-020-00466-2)) |
| Otu50201 | 1 | k__Fungi(100);p__Ascomycota(100);c__Saccharomycetes(100);o__Saccharomycetales(100);f__Metschnikowiaceae(100);g__Metschnikowia(100);g__Metschnikowia_unclassified(100); | Metschnikowia | 13 | Saprotroph | \|Undefined Saprotroph\| | Yeast | NULL | Probable | As nectar-tap_saprotroph (Põlme et al. 2020); Decay sugar-rich substrates (Põlme et al. 2020) | Põlme S, et al. 2020. Fungal Diversity 105:1-16 ((https://doi.org/10.1007/s13225-020-00466-2)) |
| Otu50202 | 1 | k__Fungi(100);p__Ascomycota(100);c__Saccharomycetes(100);o__Saccharomycetales(100);f__Metschnikowiaceae(100);g__Metschnikowia(100);s__Metschnikowia_pulcherrima(100); | Metschnikowia | 13 | Saprotroph | \|Undefined Saprotroph\| | Yeast | NULL | Probable | As nectar-tap_saprotroph (Põlme et al. 2020); Decay sugar-rich substrates (Põlme et al. 2020) | Põlme S, et al. 2020. Fungal Diversity 105:1-16 ((https://doi.org/10.1007/s13225-020-00466-2)) |
| Otu50203 | 1 | k__Fungi(100);p__Ascomycota(100);c__Saccharomycetes(100);o__Saccharomycetales(100);f__Metschnikowiaceae(100);g__Metschnikowia(100);s__Metschnikowia_pulcherrima(100); | Metschnikowia | 13 | Saprotroph | \|Undefined Saprotroph\| | Yeast | NULL | Probable | As nectar-tap_saprotroph (Põlme et al. 2020); Decay sugar-rich substrates (Põlme et al. 2020) | Põlme S, et al. 2020. Fungal Diversity 105:1-16 ((https://doi.org/10.1007/s13225-020-00466-2)) |

| | | | | | | | | | | | |
|---|---|---|---|---|---|---|---|---|---|---|---|
| Otu50204 | 1 | k__Fungi(100);p__Ascomycota(100);c__Saccharomycetes(100);o__Saccharomycetales(100);f__Metschnikowiaceae(100);g__Metschnikowia(100);g__Metschnikowia_unclassified(100); | Metschnikowia | 13 | Saprotroph | \|Undefined Saprotroph\| | Yeast | NULL | Probable | As nectar-tap_saprotroph (Põlme et al. 2020); Decay sugar-rich substrates (Põlme et al. 2020) | Põlme S, et al. 2020. Fungal Diversity 105:1-16 ((https://doi.org/10.1007/s13225-020-00466-2)) |
| Otu50205 | 1 | k__Fungi(100);p__Ascomycota(100);c__Saccharomycetes(100);o__Saccharomycetales(100);f__Metschnikowiaceae(100);g__Metschnikowia(100);s__Metschnikowia_pulcherrima(100); | Metschnikowia | 13 | Saprotroph | \|Undefined Saprotroph\| | Yeast | NULL | Probable | As nectar-tap_saprotroph (Põlme et al. 2020); Decay sugar-rich substrates (Põlme et al. 2020) | Põlme S, et al. 2020. Fungal Diversity 105:1-16 ((https://doi.org/10.1007/s13225-020-00466-2)) |
| Otu50206 | 1 | k__Fungi(100);p__Ascomycota(100);c__Saccharomycetes(100);o__Saccharomycetales(100);f__Metschnikowiaceae(100);g__Metschnikowia(100);s__Metschnikowia_pulcherrima(100); | Metschnikowia | 13 | Saprotroph | \|Undefined Saprotroph\| | Yeast | NULL | Probable | As nectar-tap_saprotroph (Põlme et al. 2020); Decay sugar-rich substrates (Põlme et al. 2020) | Põlme S, et al. 2020. Fungal Diversity 105:1-16 ((https://doi.org/10.1007/s13225-020-00466-2)) |
| Otu50207 | 1 | k__Fungi(100);p__Ascomycota(100);c__Saccharomycetes(100);o__Saccharomycetales(100);f__Metschnikowiaceae(100);g__Metschnikowia(100);s__Metschnikowia_pulcherrima(100); | Metschnikowia | 13 | Saprotroph | \|Undefined Saprotroph\| | Yeast | NULL | Probable | As nectar-tap_saprotroph (Põlme et al. 2020); Decay sugar-rich substrates (Põlme et al. 2020) | Põlme S, et al. 2020. Fungal Diversity 105:1-16 ((https://doi.org/10.1007/s13225-020-00466-2)) |
| Otu50208 | 1 | k__Fungi(100);p__Ascomycota(100);c__Saccharomycetes(100);o__Saccharomycetales(100);f__Metschnikowiaceae(100);g__Metschnikowia(100);s__Metschnikowia_pulcherrima(100); | Metschnikowia | 13 | Saprotroph | \|Undefined Saprotroph\| | Yeast | NULL | Probable | As nectar-tap_saprotroph (Põlme et al. 2020); Decay sugar-rich substrates (Põlme et al. 2020) | Põlme S, et al. 2020. Fungal Diversity 105:1-16 ((https://doi.org/10.1007/s13225-020-00466-2)) |
| Otu50209 | 1 | k__Fungi(100);p__Ascomycota(100);c__Saccharomycetes(100);o__Saccharomycetales(100);f__Metschnikowiaceae(100);g__Metschnikowia(100);s__Metschnikowia_pulcherrima(100); | Metschnikowia | 13 | Saprotroph | \|Undefined Saprotroph\| | Yeast | NULL | Probable | As nectar-tap_saprotroph (Põlme et al. 2020); Decay sugar-rich substrates (Põlme et al. 2020) | Põlme S, et al. 2020. Fungal Diversity 105:1-16 ((https://doi.org/10.1007/s13225-020-00466-2)) |
| Otu50210 | 1 | k__Fungi(100);p__Ascomycota(100);c__Saccharomycetes(100);o__Saccharomycetales(100);f__Metschnikowiaceae(100);g__Metschnikowia(100);g__Metschnikowia_unclassified(100); | Metschnikowia | 13 | Saprotroph | \|Undefined Saprotroph\| | Yeast | NULL | Probable | As nectar-tap_saprotroph (Põlme et al. 2020); Decay sugar-rich substrates (Põlme et al. 2020) | Põlme S, et al. 2020. Fungal Diversity 105:1-16 ((https://doi.org/10.1007/s13225-020-00466-2)) |
| Otu50211 | 1 | k__Fungi(100);p__Ascomycota(100);c__Saccharomycetes(100);o__Saccharomycetales(100);f__Metschnikowiaceae(100);g__Metschnikowia(100);s__Metschnikowia_pulcherrima(100); | Metschnikowia | 13 | Saprotroph | \|Undefined Saprotroph\| | Yeast | NULL | Probable | As nectar-tap_saprotroph (Põlme et al. 2020); Decay sugar-rich substrates (Põlme et al. 2020) | Põlme S, et al. 2020. Fungal Diversity 105:1-16 ((https://doi.org/10.1007/s13225-020-00466-2)) |
| Otu50212 | 1 | k__Fungi(100);p__Ascomycota(100);c__Saccharomycetes(100);o__Saccharomycetales(100);f__Metschnikowiaceae(100);g__Metschnikowia(100);s__Metschnikowia_pulcherrima(100); | Metschnikowia | 13 | Saprotroph | \|Undefined Saprotroph\| | Yeast | NULL | Probable | As nectar-tap_saprotroph (Põlme et al. 2020); Decay sugar-rich substrates (Põlme et al. 2020) | Põlme S, et al. 2020. Fungal Diversity 105:1-16 ((https://doi.org/10.1007/s13225-020-00466-2)) |
| Otu50213 | 1 | k__Fungi(100);p__Ascomycota(100);c__Saccharomycetes(100);o__Saccharomycetales(100);f__Metschnikowiaceae(100);g__Metschnikowia(100);s__Metschnikowia_pulcherrima(100); | Metschnikowia | 13 | Saprotroph | \|Undefined Saprotroph\| | Yeast | NULL | Probable | As nectar-tap_saprotroph (Põlme et al. 2020); Decay sugar-rich substrates (Põlme et al. 2020) | Põlme S, et al. 2020. Fungal Diversity 105:1-16 ((https://doi.org/10.1007/s13225-020-00466-2)) |
| Otu50214 | 1 | k__Fungi(100);p__Ascomycota(100);c__Saccharomycetes(100);o__Saccharomycetales(100);f__Metschnikowiaceae(100);g__Metschnikowia(100);s__Metschnikowia_pulcherrima(100); | Metschnikowia | 13 | Saprotroph | \|Undefined Saprotroph\| | Yeast | NULL | Probable | As nectar-tap_saprotroph (Põlme et al. 2020); Decay sugar-rich substrates (Põlme et al. 2020) | Põlme S, et al. 2020. Fungal Diversity 105:1-16 ((https://doi.org/10.1007/s13225-020-00466-2)) |
| Otu50215 | 1 | k__Fungi(100);p__Ascomycota(100);c__Saccharomycetes(100);o__Saccharomycetales(100);f__Metschnikowiaceae(100);g__Metschnikowia(100);s__Metschnikowia_pulcherrima(100); | Metschnikowia | 13 | Saprotroph | \|Undefined Saprotroph\| | Yeast | NULL | Probable | As nectar-tap_saprotroph (Põlme et al. 2020); Decay sugar-rich substrates (Põlme et al. 2020) | Põlme S, et al. 2020. Fungal Diversity 105:1-16 ((https://doi.org/10.1007/s13225-020-00466-2)) |
| Otu50216 | 1 | k__Fungi(100);p__Ascomycota(100);c__Saccharomycetes(100);o__Saccharomycetales(100);f__Metschnikowiaceae(100);g__Metschnikowia(100);g__Metschnikowia_unclassified(100); | Metschnikowia | 13 | Saprotroph | \|Undefined Saprotroph\| | Yeast | NULL | Probable | As nectar-tap_saprotroph (Põlme et al. 2020); Decay sugar-rich substrates (Põlme et al. 2020) | Põlme S, et al. 2020. Fungal Diversity 105:1-16 ((https://doi.org/10.1007/s13225-020-00466-2)) |
| Otu50217 | 1 | k__Fungi(100);p__Ascomycota(100);c__Saccharomycetes(100);o__Saccharomycetales(100);f__Metschnikowiaceae(100);g__Metschnikowia(100);g__Metschnikowia_unclassified(100); | Metschnikowia | 13 | Saprotroph | \|Undefined Saprotroph\| | Yeast | NULL | Probable | As nectar-tap_saprotroph (Põlme et al. 2020); Decay sugar-rich substrates (Põlme et al. 2020) | Põlme S, et al. 2020. Fungal Diversity 105:1-16 ((https://doi.org/10.1007/s13225-020-00466-2)) |
| Otu50218 | 1 | k__Fungi(100);p__Ascomycota(100);c__Saccharomycetes(100);o__Saccharomycetales(100);f__Metschnikowiaceae(100);g__Metschnikowia(100);s__Metschnikowia_pulcherrima(100); | Metschnikowia | 13 | Saprotroph | \|Undefined Saprotroph\| | Yeast | NULL | Probable | As nectar-tap_saprotroph (Põlme et al. 2020); Decay sugar-rich substrates (Põlme et al. 2020) | Põlme S, et al. 2020. Fungal Diversity 105:1-16 ((https://doi.org/10.1007/s13225-020-00466-2)) |
| Otu50219 | 1 | k__Fungi(100);p__Ascomycota(100);c__Saccharomycetes(100);o__Saccharomycetales(100);f__Metschnikowiaceae(100);g__Metschnikowia(100);s__Metschnikowia_pulcherrima(100); | Metschnikowia | 13 | Saprotroph | \|Undefined Saprotroph\| | Yeast | NULL | Probable | As nectar-tap_saprotroph (Põlme et al. 2020); Decay sugar-rich substrates (Põlme et al. 2020) | Põlme S, et al. 2020. Fungal Diversity 105:1-16 ((https://doi.org/10.1007/s13225-020-00466-2)) |
| Otu50220 | 1 | k__Fungi(100);p__Ascomycota(100);c__Saccharomycetes(100);o__Saccharomycetales(100);f__Metschnikowiaceae(100);g__Metschnikowia(100);g__Metschnikowia_unclassified(100); | Metschnikowia | 13 | Saprotroph | \|Undefined Saprotroph\| | Yeast | NULL | Probable | As nectar-tap_saprotroph (Põlme et al. 2020); Decay sugar-rich substrates (Põlme et al. 2020) | Põlme S, et al. 2020. Fungal Diversity 105:1-16 ((https://doi.org/10.1007/s13225-020-00466-2)) |
| Otu50221 | 1 | k__Fungi(100);p__Ascomycota(100);c__Saccharomycetes(100);o__Saccharomycetales(100);f__Metschnikowiaceae(100);g__Metschnikowia(100);s__Metschnikowia_pulcherrima(100); | Metschnikowia | 13 | Saprotroph | \|Undefined Saprotroph\| | Yeast | NULL | Probable | As nectar-tap_saprotroph (Põlme et al. 2020); Decay sugar-rich substrates (Põlme et al. 2020) | Põlme S, et al. 2020. Fungal Diversity 105:1-16 ((https://doi.org/10.1007/s13225-020-00466-2)) |
| Otu50222 | 1 | k__Fungi(100);p__Ascomycota(100);c__Saccharomycetes(100);o__Saccharomycetales(100);f__Metschnikowiaceae(100);g__Metschnikowia(100);s__Metschnikowia_pulcherrima(100); | Metschnikowia | 13 | Saprotroph | \|Undefined Saprotroph\| | Yeast | NULL | Probable | As nectar-tap_saprotroph (Põlme et al. 2020); Decay sugar-rich substrates (Põlme et al. 2020) | Põlme S, et al. 2020. Fungal Diversity 105:1-16 ((https://doi.org/10.1007/s13225-020-00466-2)) |
| Otu50223 | 1 | k__Fungi(100);p__Ascomycota(100);c__Saccharomycetes(100);o__Saccharomycetales(100);f__Metschnikowiaceae(100);g__Metschnikowia(100);s__Metschnikowia_pulcherrima(100); | Metschnikowia | 13 | Saprotroph | \|Undefined Saprotroph\| | Yeast | NULL | Probable | As nectar-tap_saprotroph (Põlme et al. 2020); Decay sugar-rich substrates (Põlme et al. 2020) | Põlme S, et al. 2020. Fungal Diversity 105:1-16 ((https://doi.org/10.1007/s13225-020-00466-2)) |
| Otu50224 | 1 | k__Fungi(100);p__Ascomycota(100);c__Saccharomycetes(100);o__Saccharomycetales(100);f__Metschnikowiaceae(100);g__Metschnikowia(100);s__Metschnikowia_pulcherrima(100); | Metschnikowia | 13 | Saprotroph | \|Undefined Saprotroph\| | Yeast | NULL | Probable | As nectar-tap_saprotroph (Põlme et al. 2020); Decay sugar-rich substrates (Põlme et al. 2020) | Põlme S, et al. 2020. Fungal Diversity 105:1-16 ((https://doi.org/10.1007/s13225-020-00466-2)) |
| Otu50225 | 1 | k__Fungi(100);p__Ascomycota(100);c__Saccharomycetes(100);o__Saccharomycetales(100);f__Metschnikowiaceae(100);g__Metschnikowia(100);s__Metschnikowia_pulcherrima(100); | Metschnikowia | 13 | Saprotroph | \|Undefined Saprotroph\| | Yeast | NULL | Probable | As nectar-tap_saprotroph (Põlme et al. 2020); Decay sugar-rich substrates (Põlme et al. 2020) | Põlme S, et al. 2020. Fungal Diversity 105:1-16 ((https://doi.org/10.1007/s13225-020-00466-2)) |
| Otu50226 | 1 | k__Fungi(100);p__Ascomycota(100);c__Saccharomycetes(100);o__Saccharomycetales(100);f__Metschnikowiaceae(100);g__Metschnikowia(100);g__Metschnikowia_unclassified(100); | Metschnikowia | 13 | Saprotroph | \|Undefined Saprotroph\| | Yeast | NULL | Probable | As nectar-tap_saprotroph (Põlme et al. 2020); Decay sugar-rich substrates (Põlme et al. 2020) | Põlme S, et al. 2020. Fungal Diversity 105:1-16 ((https://doi.org/10.1007/s13225-020-00466-2)) |
| Otu50227 | 1 | k__Fungi(100);p__Ascomycota(100);c__Saccharomycetes(100);o__Saccharomycetales(100);f__Metschnikowiaceae(100);g__Metschnikowia(100);s__Metschnikowia_pulcherrima(100); | Metschnikowia | 13 | Saprotroph | \|Undefined Saprotroph\| | Yeast | NULL | Probable | As nectar-tap_saprotroph (Põlme et al. 2020); Decay sugar-rich substrates (Põlme et al. 2020) | Põlme S, et al. 2020. Fungal Diversity 105:1-16 ((https://doi.org/10.1007/s13225-020-00466-2)) |
| Otu50228 | 1 | k__Fungi(100);p__Ascomycota(100);c__Saccharomycetes(100);o__Saccharomycetales(100);f__Metschnikowiaceae(100);g__Metschnikowia(100);s__Metschnikowia_pulcherrima(100); | Metschnikowia | 13 | Saprotroph | \|Undefined Saprotroph\| | Yeast | NULL | Probable | As nectar-tap_saprotroph (Põlme et al. 2020); Decay sugar-rich substrates (Põlme et al. 2020) | Põlme S, et al. 2020. Fungal Diversity 105:1-16 ((https://doi.org/10.1007/s13225-020-00466-2)) |
| Otu50229 | 1 | k__Fungi(100);p__Ascomycota(100);c__Saccharomycetes(100);o__Saccharomycetales(100);f__Metschnikowiaceae(100);g__Metschnikowia(100);s__Metschnikowia_pulcherrima(100); | Metschnikowia | 13 | Saprotroph | \|Undefined Saprotroph\| | Yeast | NULL | Probable | As nectar-tap_saprotroph (Põlme et al. 2020); Decay sugar-rich substrates (Põlme et al. 2020) | Põlme S, et al. 2020. Fungal Diversity 105:1-16 ((https://doi.org/10.1007/s13225-020-00466-2)) |
| Otu50230 | 1 | k__Fungi(100);p__Ascomycota(100);c__Saccharomycetes(100);o__Saccharomycetales(100);f__Metschnikowiaceae(100);g__Metschnikowia(100);s__Metschnikowia_pulcherrima(100); | Metschnikowia | 13 | Saprotroph | \|Undefined Saprotroph\| | Yeast | NULL | Probable | As nectar-tap_saprotroph (Põlme et al. 2020); Decay sugar-rich substrates (Põlme et al. 2020) | Põlme S, et al. 2020. Fungal Diversity 105:1-16 ((https://doi.org/10.1007/s13225-020-00466-2)) |
| Otu50231 | 1 | k__Fungi(100);p__Ascomycota(100);c__Saccharomycetes(100);o__Saccharomycetales(100);f__Metschnikowiaceae(100);g__Metschnikowia(100);s__Metschnikowia_pulcherrima(100); | Metschnikowia | 13 | Saprotroph | \|Undefined Saprotroph\| | Yeast | NULL | Probable | As nectar-tap_saprotroph (Põlme et al. 2020); Decay sugar-rich substrates (Põlme et al. 2020) | Põlme S, et al. 2020. Fungal Diversity 105:1-16 ((https://doi.org/10.1007/s13225-020-00466-2)) |
| Otu50232 | 1 | k__Fungi(100);p__Ascomycota(100);c__Saccharomycetes(100);o__Saccharomycetales(100);f__Metschnikowiaceae(100);g__Metschnikowia(100);s__Metschnikowia_pulcherrima(100); | Metschnikowia | 13 | Saprotroph | \|Undefined Saprotroph\| | Yeast | NULL | Probable | As nectar-tap_saprotroph (Põlme et al. 2020); Decay sugar-rich substrates (Põlme et al. 2020) | Põlme S, et al. 2020. Fungal Diversity 105:1-16 ((https://doi.org/10.1007/s13225-020-00466-2)) |
| Otu50233 | 1 | k__Fungi(100);p__Ascomycota(100);c__Saccharomycetes(100);o__Saccharomycetales(100);f__Metschnikowiaceae(100);g__Metschnikowia(100);s__Metschnikowia_pulcherrima(100); | Metschnikowia | 13 | Saprotroph | \|Undefined Saprotroph\| | Yeast | NULL | Probable | As nectar-tap_saprotroph (Põlme et al. 2020); Decay sugar-rich substrates (Põlme et al. 2020) | Põlme S, et al. 2020. Fungal Diversity 105:1-16 ((https://doi.org/10.1007/s13225-020-00466-2)) |
| Otu50234 | 1 | k__Fungi(100);p__Ascomycota(100);c__Saccharomycetes(100);o__Saccharomycetales(100);f__Metschnikowiaceae(100);g__Metschnikowia(100);g__Metschnikowia_unclassified(100); | Metschnikowia | 13 | Saprotroph | \|Undefined Saprotroph\| | Yeast | NULL | Probable | As nectar-tap_saprotroph (Põlme et al. 2020); Decay sugar-rich substrates (Põlme et al. 2020) | Põlme S, et al. 2020. Fungal Diversity 105:1-16 ((https://doi.org/10.1007/s13225-020-00466-2)) |
| Otu50235 | 1 | k__Fungi(100);p__Ascomycota(100);c__Saccharomycetes(100);o__Saccharomycetales(100);f__Metschnikowiaceae(100);g__Metschnikowia(100);s__Metschnikowia_pulcherrima(100); | Metschnikowia | 13 | Saprotroph | \|Undefined Saprotroph\| | Yeast | NULL | Probable | As nectar-tap_saprotroph (Põlme et al. 2020); Decay sugar-rich substrates (Põlme et al. 2020) | Põlme S, et al. 2020. Fungal Diversity 105:1-16 ((https://doi.org/10.1007/s13225-020-00466-2)) |
| Otu50236 | 1 | k__Fungi(100);p__Ascomycota(100);c__Saccharomycetes(100);o__Saccharomycetales(100);f__Metschnikowiaceae(100);g__Metschnikowia(100);s__Metschnikowia_pulcherrima(100); | Metschnikowia | 13 | Saprotroph | \|Undefined Saprotroph\| | Yeast | NULL | Probable | As nectar-tap_saprotroph (Põlme et al. 2020); Decay sugar-rich substrates (Põlme et al. 2020) | Põlme S, et al. 2020. Fungal Diversity 105:1-16 ((https://doi.org/10.1007/s13225-020-00466-2)) |
| Otu50237 | 1 | k__Fungi(100);p__Ascomycota(100);c__Saccharomycetes(100);o__Saccharomycetales(100);f__Metschnikowiaceae(100);g__Metschnikowia(100);s__Metschnikowia_pulcherrima(100); | Metschnikowia | 13 | Saprotroph | \|Undefined Saprotroph\| | Yeast | NULL | Probable | As nectar-tap_saprotroph (Põlme et al. 2020); Decay sugar-rich substrates (Põlme et al. 2020) | Põlme S, et al. 2020. Fungal Diversity 105:1-16 ((https://doi.org/10.1007/s13225-020-00466-2)) |
| Otu50238 | 1 | k__Fungi(100);p__Ascomycota(100);c__Saccharomycetes(100);o__Saccharomycetales(100);f__Metschnikowiaceae(100);g__Metschnikowia(100);s__Metschnikowia_pulcherrima(100); | Metschnikowia | 13 | Saprotroph | \|Undefined Saprotroph\| | Yeast | NULL | Probable | As nectar-tap_saprotroph (Põlme et al. 2020); Decay sugar-rich substrates (Põlme et al. 2020) | Põlme S, et al. 2020. Fungal Diversity 105:1-16 ((https://doi.org/10.1007/s13225-020-00466-2)) |
| Otu50239 | 1 | k__Fungi(100);p__Ascomycota(100);c__Saccharomycetes(100);o__Saccharomycetales(100);f__Metschnikowiaceae(100);g__Metschnikowia(100);g__Metschnikowia_unclassified(100); | Metschnikowia | 13 | Saprotroph | \|Undefined Saprotroph\| | Yeast | NULL | Probable | As nectar-tap_saprotroph (Põlme et al. 2020); Decay sugar-rich substrates (Põlme et al. 2020) | Põlme S, et al. 2020. Fungal Diversity 105:1-16 ((https://doi.org/10.1007/s13225-020-00466-2)) |
| Otu50240 | 1 | k__Fungi(100);p__Ascomycota(100);c__Saccharomycetes(100);o__Saccharomycetales(100);f__Metschnikowiaceae(100);g__Metschnikowia(100);s__Metschnikowia_pulcherrima(100); | Metschnikowia | 13 | Saprotroph | \|Undefined Saprotroph\| | Yeast | NULL | Probable | As nectar-tap_saprotroph (Põlme et al. 2020); Decay sugar-rich substrates (Põlme et al. 2020) | Põlme S, et al. 2020. Fungal Diversity 105:1-16 ((https://doi.org/10.1007/s13225-020-00466-2)) |
| Otu50243 | 1 | k__Fungi(100);p__Ascomycota(100);c__Saccharomycetes(100);o__Saccharomycetales(100);f__Metschnikowiaceae(100);g__Metschnikowia(100);g__Metschnikowia_unclassified(100); | Metschnikowia | 13 | Saprotroph | \|Undefined Saprotroph\| | Yeast | NULL | Probable | As nectar-tap_saprotroph (Põlme et al. 2020); Decay sugar-rich substrates (Põlme et al. 2020) | Põlme S, et al. 2020. Fungal Diversity 105:1-16 ((https://doi.org/10.1007/s13225-020-00466-2)) |
| Otu50244 | 1 | k__Fungi(100);p__Ascomycota(100);c__Saccharomycetes(100);o__Saccharomycetales(100);f__Metschnikowiaceae(100);g__Metschnikowia(100);s__Metschnikowia_pulcherrima(100); | Metschnikowia | 13 | Saprotroph | \|Undefined Saprotroph\| | Yeast | NULL | Probable | As nectar-tap_saprotroph (Põlme et al. 2020); Decay sugar-rich substrates (Põlme et al. 2020) | Põlme S, et al. 2020. Fungal Diversity 105:1-16 ((https://doi.org/10.1007/s13225-020-00466-2)) |
| Otu50245 | 1 | k__Fungi(100);p__Ascomycota(100);c__Saccharomycetes(100);o__Saccharomycetales(100);f__Metschnikowiaceae(100);g__Metschnikowia(100);s__Metschnikowia_pulcherrima(100); | Metschnikowia | 13 | Saprotroph | \|Undefined Saprotroph\| | Yeast | NULL | Probable | As nectar-tap_saprotroph (Põlme et al. 2020); Decay sugar-rich substrates (Põlme et al. 2020) | Põlme S, et al. 2020. Fungal Diversity 105:1-16 ((https://doi.org/10.1007/s13225-020-00466-2)) |
| Otu50246 | 1 | k__Fungi(100);p__Ascomycota(100);c__Saccharomycetes(100);o__Saccharomycetales(100);f__Metschnikowiaceae(100);g__Metschnikowia(100);s__Metschnikowia_pulcherrima(100); | Metschnikowia | 13 | Saprotroph | \|Undefined Saprotroph\| | Yeast | NULL | Probable | As nectar-tap_saprotroph (Põlme et al. 2020); Decay sugar-rich substrates (Põlme et al. 2020) | Põlme S, et al. 2020. Fungal Diversity 105:1-16 ((https://doi.org/10.1007/s13225-020-00466-2)) |
| Otu50247 | 1 | k__Fungi(100);p__Ascomycota(100);c__Saccharomycetes(100);o__Saccharomycetales(100);f__Metschnikowiaceae(100);g__Metschnikowia(100);s__Metschnikowia_pulcherrima(100); | Metschnikowia | 13 | Saprotroph | \|Undefined Saprotroph\| | Yeast | NULL | Probable | As nectar-tap_saprotroph (Põlme et al. 2020); Decay sugar-rich substrates (Põlme et al. 2020) | Põlme S, et al. 2020. Fungal Diversity 105:1-16 ((https://doi.org/10.1007/s13225-020-00466-2)) |
| Otu50248 | 1 | k__Fungi(100);p__Ascomycota(100);c__Saccharomycetes(100);o__Saccharomycetales(100);f__Metschnikowiaceae(100);g__Metschnikowia(100);g__Metschnikowia_unclassified(100); | Metschnikowia | 13 | Saprotroph | \|Undefined Saprotroph\| | Yeast | NULL | Probable | As nectar-tap_saprotroph (Põlme et al. 2020); Decay sugar-rich substrates (Põlme et al. 2020) | Põlme S, et al. 2020. Fungal Diversity 105:1-16 ((https://doi.org/10.1007/s13225-020-00466-2)) |
| Otu50249 | 1 | k__Fungi(100);p__Ascomycota(100);c__Saccharomycetes(100);o__Saccharomycetales(100);f__Metschnikowiaceae(100);g__Metschnikowia(100);s__Metschnikowia_pulcherrima(100); | Metschnikowia | 13 | Saprotroph | \|Undefined Saprotroph\| | Yeast | NULL | Probable | As nectar-tap_saprotroph (Põlme et al. 2020); Decay sugar-rich substrates (Põlme et al. 2020) | Põlme S, et al. 2020. Fungal Diversity 105:1-16 ((https://doi.org/10.1007/s13225-020-00466-2)) |
| Otu50250 | 1 | k__Fungi(100);p__Ascomycota(100);c__Saccharomycetes(100);o__Saccharomycetales(100);f__Metschnikowiaceae(100);g__Metschnikowia(100);s__Metschnikowia_pulcherrima(100); | Metschnikowia | 13 | Saprotroph | \|Undefined Saprotroph\| | Yeast | NULL | Probable | As nectar-tap_saprotroph (Põlme et al. 2020); Decay sugar-rich substrates (Põlme et al. 2020) | Põlme S, et al. 2020. Fungal Diversity 105:1-16 ((https://doi.org/10.1007/s13225-020-00466-2)) |
| Otu50251 | 1 | k__Fungi(100);p__Ascomycota(100);c__Saccharomycetes(100);o__Saccharomycetales(100);f__Metschnikowiaceae(100);g__Metschnikowia(100);g__Metschnikowia_unclassified(100); | Metschnikowia | 13 | Saprotroph | \|Undefined Saprotroph\| | Yeast | NULL | Probable | As nectar-tap_saprotroph (Põlme et al. 2020); Decay sugar-rich substrates (Põlme et al. 2020) | Põlme S, et al. 2020. Fungal Diversity 105:1-16 ((https://doi.org/10.1007/s13225-020-00466-2)) |
| Otu50252 | 1 | k__Fungi(100);p__Ascomycota(100);c__Saccharomycetes(100);o__Saccharomycetales(100);f__Metschnikowiaceae(100);g__Metschnikowia(100);s__Metschnikowia_pulcherrima(100); | Metschnikowia | 13 | Saprotroph | \|Undefined Saprotroph\| | Yeast | NULL | Probable | As nectar-tap_saprotroph (Põlme et al. 2020); Decay sugar-rich substrates (Põlme et al. 2020) | Põlme S, et al. 2020. Fungal Diversity 105:1-16 ((https://doi.org/10.1007/s13225-020-00466-2)) |
| Otu50253 | 1 | k__Fungi(100);p__Ascomycota(100);c__Saccharomycetes(100);o__Saccharomycetales(100);f__Metschnikowiaceae(100);g__Metschnikowia(100);s__Metschnikowia_pulcherrima(100); | Metschnikowia | 13 | Saprotroph | \|Undefined Saprotroph\| | Yeast | NULL | Probable | As nectar-tap_saprotroph (Põlme et al. 2020); Decay sugar-rich substrates (Põlme et al. 2020) | Põlme S, et al. 2020. Fungal Diversity 105:1-16 ((https://doi.org/10.1007/s13225-020-00466-2)) |
| Otu50255 | 1 | k__Fungi(100);p__Ascomycota(100);c__Saccharomycetes(100);o__Saccharomycetales(100);f__Metschnikowiaceae(100);g__Metschnikowia(100);s__Metschnikowia_pulcherrima(100); | Metschnikowia | 13 | Saprotroph | \|Undefined Saprotroph\| | Yeast | NULL | Probable | As nectar-tap_saprotroph (Põlme et al. 2020); Decay sugar-rich substrates (Põlme et al. 2020) | Põlme S, et al. 2020. Fungal Diversity 105:1-16 ((https://doi.org/10.1007/s13225-020-00466-2)) |
| Otu50256 | 1 | k__Fungi(100);p__Ascomycota(100);c__Saccharomycetes(100);o__Saccharomycetales(100);f__Metschnikowiaceae(100);g__Metschnikowia(100);s__Metschnikowia_pulcherrima(100); | Metschnikowia | 13 | Saprotroph | \|Undefined Saprotroph\| | Yeast | NULL | Probable | As nectar-tap_saprotroph (Põlme et al. 2020); Decay sugar-rich substrates (Põlme et al. 2020) | Põlme S, et al. 2020. Fungal Diversity 105:1-16 ((https://doi.org/10.1007/s13225-020-00466-2)) |
| Otu50257 | 1 | k__Fungi(100);p__Ascomycota(100);c__Saccharomycetes(100);o__Saccharomycetales(100);f__Metschnikowiaceae(100);g__Metschnikowia(100);s__Metschnikowia_pulcherrima(100); | Metschnikowia | 13 | Saprotroph | \|Undefined Saprotroph\| | Yeast | NULL | Probable | As nectar-tap_saprotroph (Põlme et al. 2020); Decay sugar-rich substrates (Põlme et al. 2020) | Põlme S, et al. 2020. Fungal Diversity 105:1-16 ((https://doi.org/10.1007/s13225-020-00466-2)) |
| Otu50258 | 1 | k__Fungi(100);p__Ascomycota(100);c__Saccharomycetes(100);o__Saccharomycetales(100);f__Metschnikowiaceae(100);g__Metschnikowia(100);s__Metschnikowia_pulcherrima(100); | Metschnikowia | 13 | Saprotroph | \|Undefined Saprotroph\| | Yeast | NULL | Probable | As nectar-tap_saprotroph (Põlme et al. 2020); Decay sugar-rich substrates (Põlme et al. 2020) | Põlme S, et al. 2020. Fungal Diversity 105:1-16 ((https://doi.org/10.1007/s13225-020-00466-2)) |
| Otu50259 | 1 | k__Fungi(100);p__Ascomycota(100);c__Saccharomycetes(100);o__Saccharomycetales(100);f__Metschnikowiaceae(100);g__Metschnikowia(100);s__Metschnikowia_pulcherrima(100); | Metschnikowia | 13 | Saprotroph | \|Undefined Saprotroph\| | Yeast | NULL | Probable | As nectar-tap_saprotroph (Põlme et al. 2020); Decay sugar-rich substrates (Põlme et al. 2020) | Põlme S, et al. 2020. Fungal Diversity 105:1-16 ((https://doi.org/10.1007/s13225-020-00466-2)) |
| Otu50260 | 1 | k__Fungi(100);p__Ascomycota(100);c__Saccharomycetes(100);o__Saccharomycetales(100);f__Metschnikowiaceae(100);g__Metschnikowia(100);s__Metschnikowia_pulcherrima(100); | Metschnikowia | 13 | Saprotroph | \|Undefined Saprotroph\| | Yeast | NULL | Probable | As nectar-tap_saprotroph (Põlme et al. 2020); Decay sugar-rich substrates (Põlme et al. 2020) | Põlme S, et al. 2020. Fungal Diversity 105:1-16 ((https://doi.org/10.1007/s13225-020-00466-2)) |
| Otu50261 | 1 | k__Fungi(100);p__Ascomycota(100);c__Saccharomycetes(100);o__Saccharomycetales(100);f__Metschnikowiaceae(100);g__Metschnikowia(100);s__Metschnikowia_pulcherrima(100); | Metschnikowia | 13 | Saprotroph | \|Undefined Saprotroph\| | Yeast | NULL | Probable | As nectar-tap_saprotroph (Põlme et al. 2020); Decay sugar-rich substrates (Põlme et al. 2020) | Põlme S, et al. 2020. Fungal Diversity 105:1-16 ((https://doi.org/10.1007/s13225-020-00466-2)) |
| Otu50262 | 1 | k__Fungi(100);p__Ascomycota(100);c__Saccharomycetes(100);o__Saccharomycetales(100);f__Metschnikowiaceae(100);g__Metschnikowia(100);s__Metschnikowia_pulcherrima(100); | Metschnikowia | 13 | Saprotroph | \|Undefined Saprotroph\| | Yeast | NULL | Probable | As nectar-tap_saprotroph (Põlme et al. 2020); Decay sugar-rich substrates (Põlme et al. 2020) | Põlme S, et al. 2020. Fungal Diversity 105:1-16 ((https://doi.org/10.1007/s13225-020-00466-2)) |
| Otu50263 | 1 | k__Fungi(100);p__Ascomycota(100);c__Saccharomycetes(100);o__Saccharomycetales(100);f__Metschnikowiaceae(100);g__Metschnikowia(100);g__Metschnikowia_unclassified(100); | Metschnikowia | 13 | Saprotroph | \|Undefined Saprotroph\| | Yeast | NULL | Probable | As nectar-tap_saprotroph (Põlme et al. 2020); Decay sugar-rich substrates (Põlme et al. 2020) | Põlme S, et al. 2020. Fungal Diversity 105:1-16 ((https://doi.org/10.1007/s13225-020-00466-2)) |
| Otu50264 | 1 | k__Fungi(100);p__Ascomycota(100);c__Saccharomycetes(100);o__Saccharomycetales(100);f__Metschnikowiaceae(100);g__Metschnikowia(100);g__Metschnikowia_unclassified(100); | Metschnikowia | 13 | Saprotroph | \|Undefined Saprotroph\| | Yeast | NULL | Probable | As nectar-tap_saprotroph (Põlme et al. 2020); Decay sugar-rich substrates (Põlme et al. 2020) | Põlme S, et al. 2020. Fungal Diversity 105:1-16 ((https://doi.org/10.1007/s13225-020-00466-2)) |
| Otu50265 | 1 | k__Fungi(100);p__Ascomycota(100);c__Saccharomycetes(100);o__Saccharomycetales(100);f__Metschnikowiaceae(100);g__Metschnikowia(100);g__Metschnikowia_unclassified(100); | Metschnikowia | 13 | Saprotroph | \|Undefined Saprotroph\| | Yeast | NULL | Probable | As nectar-tap_saprotroph (Põlme et al. 2020); Decay sugar-rich substrates (Põlme et al. 2020) | Põlme S, et al. 2020. Fungal Diversity 105:1-16 ((https://doi.org/10.1007/s13225-020-00466-2)) |
| Otu50266 | 1 | k__Fungi(100);p__Ascomycota(100);c__Saccharomycetes(100);o__Saccharomycetales(100);f__Metschnikowiaceae(100);g__Metschnikowia(100);g__Metschnikowia_unclassified(100); | Metschnikowia | 13 | Saprotroph | \|Undefined Saprotroph\| | Yeast | NULL | Probable | As nectar-tap_saprotroph (Põlme et al. 2020); Decay sugar-rich substrates (Põlme et al. 2020) | Põlme S, et al. 2020. Fungal Diversity 105:1-16 ((https://doi.org/10.1007/s13225-020-00466-2)) |
| Otu50267 | 1 | k__Fungi(100);p__Ascomycota(100);c__Saccharomycetes(100);o__Saccharomycetales(100);f__Metschnikowiaceae(100);g__Metschnikowia(100);g__Metschnikowia_unclassified(100); | Metschnikowia | 13 | Saprotroph | \|Undefined Saprotroph\| | Yeast | NULL | Probable | As nectar-tap_saprotroph (Põlme et al. 2020); Decay sugar-rich substrates (Põlme et al. 2020) | Põlme S, et al. 2020. Fungal Diversity 105:1-16 ((https://doi.org/10.1007/s13225-020-00466-2)) |
| Otu50268 | 1 | k__Fungi(100);p__Ascomycota(100);c__Saccharomycetes(100);o__Saccharomycetales(100);f__Metschnikowiaceae(100);g__Metschnikowia(100);s__Metschnikowia_pulcherrima(100); | Metschnikowia | 13 | Saprotroph | \|Undefined Saprotroph\| | Yeast | NULL | Probable | As nectar-tap_saprotroph (Põlme et al. 2020); Decay sugar-rich substrates (Põlme et al. 2020) | Põlme S, et al. 2020. Fungal Diversity 105:1-16 ((https://doi.org/10.1007/s13225-020-00466-2)) |
| Otu50269 | 1 | k__Fungi(100);p__Ascomycota(100);c__Saccharomycetes(100);o__Saccharomycetales(100);f__Metschnikowiaceae(100);g__Metschnikowia(100);g__Metschnikowia_unclassified(100); | Metschnikowia | 13 | Saprotroph | \|Undefined Saprotroph\| | Yeast | NULL | Probable | As nectar-tap_saprotroph (Põlme et al. 2020); Decay sugar-rich substrates (Põlme et al. 2020) | Põlme S, et al. 2020. Fungal Diversity 105:1-16 ((https://doi.org/10.1007/s13225-020-00466-2)) |
| Otu50270 | 1 | k__Fungi(100);p__Ascomycota(100);c__Saccharomycetes(100);o__Saccharomycetales(100);f__Metschnikowiaceae(100);g__Metschnikowia(100);s__Metschnikowia_pulcherrima(100); | Metschnikowia | 13 | Saprotroph | \|Undefined Saprotroph\| | Yeast | NULL | Probable | As nectar-tap_saprotroph (Põlme et al. 2020); Decay sugar-rich substrates (Põlme et al. 2020) | Põlme S, et al. 2020. Fungal Diversity 105:1-16 ((https://doi.org/10.1007/s13225-020-00466-2)) |
| Otu50271 | 1 | k__Fungi(100);p__Ascomycota(100);c__Saccharomycetes(100);o__Saccharomycetales(100);f__Metschnikowiaceae(100);g__Metschnikowia(100);s__Metschnikowia_pulcherrima(100); | Metschnikowia | 13 | Saprotroph | \|Undefined Saprotroph\| | Yeast | NULL | Probable | As nectar-tap_saprotroph (Põlme et al. 2020); Decay sugar-rich substrates (Põlme et al. 2020) | Põlme S, et al. 2020. Fungal Diversity 105:1-16 ((https://doi.org/10.1007/s13225-020-00466-2)) |
| Otu50272 | 1 | k__Fungi(100);p__Ascomycota(100);c__Saccharomycetes(100);o__Saccharomycetales(100);f__Metschnikowiaceae(100);g__Metschnikowia(100);g__Metschnikowia_unclassified(100); | Metschnikowia | 13 | Saprotroph | \|Undefined Saprotroph\| | Yeast | NULL | Probable | As nectar-tap_saprotroph (Põlme et al. 2020); Decay sugar-rich substrates (Põlme et al. 2020) | Põlme S, et al. 2020. Fungal Diversity 105:1-16 ((https://doi.org/10.1007/s13225-020-00466-2)) |
| Otu50273 | 1 | k__Fungi(100);p__Ascomycota(100);c__Saccharomycetes(100);o__Saccharomycetales(100);f__Metschnikowiaceae(100);g__Metschnikowia(100);g__Metschnikowia_unclassified(100); | Metschnikowia | 13 | Saprotroph | \|Undefined Saprotroph\| | Yeast | NULL | Probable | As nectar-tap_saprotroph (Põlme et al. 2020); Decay sugar-rich substrates (Põlme et al. 2020) | Põlme S, et al. 2020. Fungal Diversity 105:1-16 ((https://doi.org/10.1007/s13225-020-00466-2)) |
| Otu50275 | 1 | k__Fungi(100);p__Ascomycota(100);c__Saccharomycetes(100);o__Saccharomycetales(100);f__Metschnikowiaceae(100);g__Metschnikowia(100);s__Metschnikowia_pulcherrima(100); | Metschnikowia | 13 | Saprotroph | \|Undefined Saprotroph\| | Yeast | NULL | Probable | As nectar-tap_saprotroph (Põlme et al. 2020); Decay sugar-rich substrates (Põlme et al. 2020) | Põlme S, et al. 2020. Fungal Diversity 105:1-16 ((https://doi.org/10.1007/s13225-020-00466-2)) |
| Otu50276 | 1 | k__Fungi(100);p__Ascomycota(100);c__Saccharomycetes(100);o__Saccharomycetales(100);f__Metschnikowiaceae(100);g__Metschnikowia(100);s__Metschnikowia_pulcherrima(100); | Metschnikowia | 13 | Saprotroph | \|Undefined Saprotroph\| | Yeast | NULL | Probable | As nectar-tap_saprotroph (Põlme et al. 2020); Decay sugar-rich substrates (Põlme et al. 2020) | Põlme S, et al. 2020. Fungal Diversity 105:1-16 ((https://doi.org/10.1007/s13225-020-00466-2)) |
| Otu50277 | 1 | k__Fungi(100);p__Ascomycota(100);c__Saccharomycetes(100);o__Saccharomycetales(100);f__Metschnikowiaceae(100);g__Metschnikowia(100);g__Metschnikowia_unclassified(100); | Metschnikowia | 13 | Saprotroph | \|Undefined Saprotroph\| | Yeast | NULL | Probable | As nectar-tap_saprotroph (Põlme et al. 2020); Decay sugar-rich substrates (Põlme et al. 2020) | Põlme S, et al. 2020. Fungal Diversity 105:1-16 ((https://doi.org/10.1007/s13225-020-00466-2)) |
| Otu50278 | 1 | k__Fungi(100);p__Ascomycota(100);c__Saccharomycetes(100);o__Saccharomycetales(100);f__Metschnikowiaceae(100);g__Metschnikowia(100);s__Metschnikowia_pulcherrima(100); | Metschnikowia | 13 | Saprotroph | \|Undefined Saprotroph\| | Yeast | NULL | Probable | As nectar-tap_saprotroph (Põlme et al. 2020); Decay sugar-rich substrates (Põlme et al. 2020) | Põlme S, et al. 2020. Fungal Diversity 105:1-16 ((https://doi.org/10.1007/s13225-020-00466-2)) |
| Otu50279 | 1 | k__Fungi(100);p__Ascomycota(100);c__Saccharomycetes(100);o__Saccharomycetales(100);f__Metschnikowiaceae(100);g__Metschnikowia(100);s__Metschnikowia_pulcherrima(100); | Metschnikowia | 13 | Saprotroph | \|Undefined Saprotroph\| | Yeast | NULL | Probable | As nectar-tap_saprotroph (Põlme et al. 2020); Decay sugar-rich substrates (Põlme et al. 2020) | Põlme S, et al. 2020. Fungal Diversity 105:1-16 ((https://doi.org/10.1007/s13225-020-00466-2)) |
| Otu50280 | 1 | k__Fungi(100);p__Ascomycota(100);c__Saccharomycetes(100);o__Saccharomycetales(100);f__Metschnikowiaceae(100);g__Metschnikowia(100);g__Metschnikowia_unclassified(100); | Metschnikowia | 13 | Saprotroph | \|Undefined Saprotroph\| | Yeast | NULL | Probable | As nectar-tap_saprotroph (Põlme et al. 2020); Decay sugar-rich substrates (Põlme et al. 2020) | Põlme S, et al. 2020. Fungal Diversity 105:1-16 ((https://doi.org/10.1007/s13225-020-00466-2)) |
| Otu50281 | 1 | k__Fungi(100);p__Ascomycota(100);c__Saccharomycetes(100);o__Saccharomycetales(100);f__Metschnikowiaceae(100);g__Metschnikowia(100);s__Metschnikowia_pulcherrima(100); | Metschnikowia | 13 | Saprotroph | \|Undefined Saprotroph\| | Yeast | NULL | Probable | As nectar-tap_saprotroph (Põlme et al. 2020); Decay sugar-rich substrates (Põlme et al. 2020) | Põlme S, et al. 2020. Fungal Diversity 105:1-16 ((https://doi.org/10.1007/s13225-020-00466-2)) |
| Otu50282 | 1 | k__Fungi(100);p__Ascomycota(100);c__Saccharomycetes(100);o__Saccharomycetales(100);f__Metschnikowiaceae(100);g__Metschnikowia(100);g__Metschnikowia_unclassified(100); | Metschnikowia | 13 | Saprotroph | \|Undefined Saprotroph\| | Yeast | NULL | Probable | As nectar-tap_saprotroph (Põlme et al. 2020); Decay sugar-rich substrates (Põlme et al. 2020) | Põlme S, et al. 2020. Fungal Diversity 105:1-16 ((https://doi.org/10.1007/s13225-020-00466-2)) |
| Otu50283 | 1 | k__Fungi(100);p__Ascomycota(100);c__Saccharomycetes(100);o__Saccharomycetales(100);f__Metschnikowiaceae(100);g__Metschnikowia(100);g__Metschnikowia_unclassified(100); | Metschnikowia | 13 | Saprotroph | \|Undefined Saprotroph\| | Yeast | NULL | Probable | As nectar-tap_saprotroph (Põlme et al. 2020); Decay sugar-rich substrates (Põlme et al. 2020) | Põlme S, et al. 2020. Fungal Diversity 105:1-16 ((https://doi.org/10.1007/s13225-020-00466-2)) |
| Otu50284 | 1 | k__Fungi(100);p__Ascomycota(100);c__Saccharomycetes(100);o__Saccharomycetales(100);f__Metschnikowiaceae(100);g__Metschnikowia(100);g__Metschnikowia_unclassified(100); | Metschnikowia | 13 | Saprotroph | \|Undefined Saprotroph\| | Yeast | NULL | Probable | As nectar-tap_saprotroph (Põlme et al. 2020); Decay sugar-rich substrates (Põlme et al. 2020) | Põlme S, et al. 2020. Fungal Diversity 105:1-16 ((https://doi.org/10.1007/s13225-020-00466-2)) |
| Otu50285 | 1 | k__Fungi(100);p__Ascomycota(100);c__Saccharomycetes(100);o__Saccharomycetales(100);f__Metschnikowiaceae(100);g__Metschnikowia(100);s__Metschnikowia_pulcherrima(100); | Metschnikowia | 13 | Saprotroph | \|Undefined Saprotroph\| | Yeast | NULL | Probable | As nectar-tap_saprotroph (Põlme et al. 2020); Decay sugar-rich substrates (Põlme et al. 2020) | Põlme S, et al. 2020. Fungal Diversity 105:1-16 ((https://doi.org/10.1007/s13225-020-00466-2)) |
| Otu50286 | 1 | k__Fungi(100);p__Ascomycota(100);c__Saccharomycetes(100);o__Saccharomycetales(100);f__Metschnikowiaceae(100);g__Metschnikowia(100);s__Metschnikowia_pulcherrima(100); | Metschnikowia | 13 | Saprotroph | \|Undefined Saprotroph\| | Yeast | NULL | Probable | As nectar-tap_saprotroph (Põlme et al. 2020); Decay sugar-rich substrates (Põlme et al. 2020) | Põlme S, et al. 2020. Fungal Diversity 105:1-16 ((https://doi.org/10.1007/s13225-020-00466-2)) |
| Otu50287 | 1 | k__Fungi(100);p__Ascomycota(100);c__Saccharomycetes(100);o__Saccharomycetales(100);f__Metschnikowiaceae(100);g__Metschnikowia(100);s__Metschnikowia_pulcherrima(100); | Metschnikowia | 13 | Saprotroph | \|Undefined Saprotroph\| | Yeast | NULL | Probable | As nectar-tap_saprotroph (Põlme et al. 2020); Decay sugar-rich substrates (Põlme et al. 2020) | Põlme S, et al. 2020. Fungal Diversity 105:1-16 ((https://doi.org/10.1007/s13225-020-00466-2)) |
| Otu50288 | 1 | k__Fungi(100);p__Ascomycota(100);c__Saccharomycetes(100);o__Saccharomycetales(100);f__Metschnikowiaceae(100);g__Metschnikowia(100);s__Metschnikowia_pulcherrima(100); | Metschnikowia | 13 | Saprotroph | \|Undefined Saprotroph\| | Yeast | NULL | Probable | As nectar-tap_saprotroph (Põlme et al. 2020); Decay sugar-rich substrates (Põlme et al. 2020) | Põlme S, et al. 2020. Fungal Diversity 105:1-16 ((https://doi.org/10.1007/s13225-020-00466-2)) |
| Otu50289 | 1 | k__Fungi(100);p__Ascomycota(100);c__Saccharomycetes(100);o__Saccharomycetales(100);f__Metschnikowiaceae(100);g__Metschnikowia(100);s__Metschnikowia_pulcherrima(100); | Metschnikowia | 13 | Saprotroph | \|Undefined Saprotroph\| | Yeast | NULL | Probable | As nectar-tap_saprotroph (Põlme et al. 2020); Decay sugar-rich substrates (Põlme et al. 2020) | Põlme S, et al. 2020. Fungal Diversity 105:1-16 ((https://doi.org/10.1007/s13225-020-00466-2)) |
| Otu50290 | 1 | k__Fungi(100);p__Ascomycota(100);c__Saccharomycetes(100);o__Saccharomycetales(100);f__Metschnikowiaceae(100);g__Metschnikowia(100);s__Metschnikowia_pulcherrima(100); | Metschnikowia | 13 | Saprotroph | \|Undefined Saprotroph\| | Yeast | NULL | Probable | As nectar-tap_saprotroph (Põlme et al. 2020); Decay sugar-rich substrates (Põlme et al. 2020) | Põlme S, et al. 2020. Fungal Diversity 105:1-16 ((https://doi.org/10.1007/s13225-020-00466-2)) |
| Otu50291 | 1 | k__Fungi(100);p__Ascomycota(100);c__Saccharomycetes(100);o__Saccharomycetales(100);f__Metschnikowiaceae(100);g__Metschnikowia(100);s__Metschnikowia_pulcherrima(100); | Metschnikowia | 13 | Saprotroph | \|Undefined Saprotroph\| | Yeast | NULL | Probable | As nectar-tap_saprotroph (Põlme et al. 2020); Decay sugar-rich substrates (Põlme et al. 2020) | Põlme S, et al. 2020. Fungal Diversity 105:1-16 ((https://doi.org/10.1007/s13225-020-00466-2)) |
| Otu50292 | 1 | k__Fungi(100);p__Ascomycota(100);c__Saccharomycetes(100);o__Saccharomycetales(100);f__Metschnikowiaceae(100);g__Metschnikowia(100);s__Metschnikowia_pulcherrima(100); | Metschnikowia | 13 | Saprotroph | \|Undefined Saprotroph\| | Yeast | NULL | Probable | As nectar-tap_saprotroph (Põlme et al. 2020); Decay sugar-rich substrates (Põlme et al. 2020) | Põlme S, et al. 2020. Fungal Diversity 105:1-16 ((https://doi.org/10.1007/s13225-020-00466-2)) |
| Otu50293 | 1 | k__Fungi(100);p__Ascomycota(100);c__Saccharomycetes(100);o__Saccharomycetales(100);f__Metschnikowiaceae(100);g__Metschnikowia(100);g__Metschnikowia_unclassified(100); | Metschnikowia | 13 | Saprotroph | \|Undefined Saprotroph\| | Yeast | NULL | Probable | As nectar-tap_saprotroph (Põlme et al. 2020); Decay sugar-rich substrates (Põlme et al. 2020) | Põlme S, et al. 2020. Fungal Diversity 105:1-16 ((https://doi.org/10.1007/s13225-020-00466-2)) |
| Otu50294 | 1 | k__Fungi(100);p__Ascomycota(100);c__Saccharomycetes(100);o__Saccharomycetales(100);f__Metschnikowiaceae(100);g__Metschnikowia(100);s__Metschnikowia_pulcherrima(100); | Metschnikowia | 13 | Saprotroph | \|Undefined Saprotroph\| | Yeast | NULL | Probable | As nectar-tap_saprotroph (Põlme et al. 2020); Decay sugar-rich substrates (Põlme et al. 2020) | Põlme S, et al. 2020. Fungal Diversity 105:1-16 ((https://doi.org/10.1007/s13225-020-00466-2)) |
| Otu50295 | 1 | k__Fungi(100);p__Ascomycota(100);c__Saccharomycetes(100);o__Saccharomycetales(100);f__Metschnikowiaceae(100);g__Metschnikowia(100);s__Metschnikowia_pulcherrima(100); | Metschnikowia | 13 | Saprotroph | \|Undefined Saprotroph\| | Yeast | NULL | Probable | As nectar-tap_saprotroph (Põlme et al. 2020); Decay sugar-rich substrates (Põlme et al. 2020) | Põlme S, et al. 2020. Fungal Diversity 105:1-16 ((https://doi.org/10.1007/s13225-020-00466-2)) |
| Otu50296 | 1 | k__Fungi(100);p__Ascomycota(100);c__Saccharomycetes(100);o__Saccharomycetales(100);f__Metschnikowiaceae(100);g__Metschnikowia(100);g__Metschnikowia_unclassified(100); | Metschnikowia | 13 | Saprotroph | \|Undefined Saprotroph\| | Yeast | NULL | Probable | As nectar-tap_saprotroph (Põlme et al. 2020); Decay sugar-rich substrates (Põlme et al. 2020) | Põlme S, et al. 2020. Fungal Diversity 105:1-16 ((https://doi.org/10.1007/s13225-020-00466-2)) |
| Otu50297 | 1 | k__Fungi(100);p__Ascomycota(100);c__Saccharomycetes(100);o__Saccharomycetales(100);f__Metschnikowiaceae(100);g__Metschnikowia(100);s__Metschnikowia_pulcherrima(100); | Metschnikowia | 13 | Saprotroph | \|Undefined Saprotroph\| | Yeast | NULL | Probable | As nectar-tap_saprotroph (Põlme et al. 2020); Decay sugar-rich substrates (Põlme et al. 2020) | Põlme S, et al. 2020. Fungal Diversity 105:1-16 ((https://doi.org/10.1007/s13225-020-00466-2)) |
| Otu50298 | 1 | k__Fungi(100);p__Ascomycota(100);c__Saccharomycetes(100);o__Saccharomycetales(100);f__Metschnikowiaceae(100);g__Metschnikowia(100);s__Metschnikowia_pulcherrima(100); | Metschnikowia | 13 | Saprotroph | \|Undefined Saprotroph\| | Yeast | NULL | Probable | As nectar-tap_saprotroph (Põlme et al. 2020); Decay sugar-rich substrates (Põlme et al. 2020) | Põlme S, et al. 2020. Fungal Diversity 105:1-16 ((https://doi.org/10.1007/s13225-020-00466-2)) |
| Otu50299 | 1 | k__Fungi(100);p__Ascomycota(100);c__Saccharomycetes(100);o__Saccharomycetales(100);f__Metschnikowiaceae(100);g__Metschnikowia(100);s__Metschnikowia_pulcherrima(100); | Metschnikowia | 13 | Saprotroph | \|Undefined Saprotroph\| | Yeast | NULL | Probable | As nectar-tap_saprotroph (Põlme et al. 2020); Decay sugar-rich substrates (Põlme et al. 2020) | Põlme S, et al. 2020. Fungal Diversity 105:1-16 ((https://doi.org/10.1007/s13225-020-00466-2)) |
| Otu50300 | 1 | k__Fungi(100);p__Ascomycota(100);c__Saccharomycetes(100);o__Saccharomycetales(100);f__Metschnikowiaceae(100);g__Metschnikowia(100);s__Metschnikowia_pulcherrima(100); | Metschnikowia | 13 | Saprotroph | \|Undefined Saprotroph\| | Yeast | NULL | Probable | As nectar-tap_saprotroph (Põlme et al. 2020); Decay sugar-rich substrates (Põlme et al. 2020) | Põlme S, et al. 2020. Fungal Diversity 105:1-16 ((https://doi.org/10.1007/s13225-020-00466-2)) |
| Otu50301 | 1 | k__Fungi(100);p__Ascomycota(100);c__Saccharomycetes(100);o__Saccharomycetales(100);f__Metschnikowiaceae(100);g__Metschnikowia(100);s__Metschnikowia_pulcherrima(100); | Metschnikowia | 13 | Saprotroph | \|Undefined Saprotroph\| | Yeast | NULL | Probable | As nectar-tap_saprotroph (Põlme et al. 2020); Decay sugar-rich substrates (Põlme et al. 2020) | Põlme S, et al. 2020. Fungal Diversity 105:1-16 ((https://doi.org/10.1007/s13225-020-00466-2)) |
| Otu50302 | 1 | k__Fungi(100);p__Ascomycota(100);c__Saccharomycetes(100);o__Saccharomycetales(100);f__Metschnikowiaceae(100);g__Metschnikowia(100);s__Metschnikowia_pulcherrima(100); | Metschnikowia | 13 | Saprotroph | \|Undefined Saprotroph\| | Yeast | NULL | Probable | As nectar-tap_saprotroph (Põlme et al. 2020); Decay sugar-rich substrates (Põlme et al. 2020) | Põlme S, et al. 2020. Fungal Diversity 105:1-16 ((https://doi.org/10.1007/s13225-020-00466-2)) |
| Otu50303 | 1 | k__Fungi(100);p__Ascomycota(100);c__Saccharomycetes(100);o__Saccharomycetales(100);f__Metschnikowiaceae(100);g__Metschnikowia(100);g__Metschnikowia_unclassified(100); | Metschnikowia | 13 | Saprotroph | \|Undefined Saprotroph\| | Yeast | NULL | Probable | As nectar-tap_saprotroph (Põlme et al. 2020); Decay sugar-rich substrates (Põlme et al. 2020) | Põlme S, et al. 2020. Fungal Diversity 105:1-16 ((https://doi.org/10.1007/s13225-020-00466-2)) |
| Otu50304 | 1 | k__Fungi(100);p__Ascomycota(100);c__Saccharomycetes(100);o__Saccharomycetales(100);f__Metschnikowiaceae(100);g__Metschnikowia(100);s__Metschnikowia_pulcherrima(100); | Metschnikowia | 13 | Saprotroph | \|Undefined Saprotroph\| | Yeast | NULL | Probable | As nectar-tap_saprotroph (Põlme et al. 2020); Decay sugar-rich substrates (Põlme et al. 2020) | Põlme S, et al. 2020. Fungal Diversity 105:1-16 ((https://doi.org/10.1007/s13225-020-00466-2)) |
| Otu50305 | 1 | k__Fungi(100);p__Ascomycota(100);c__Saccharomycetes(100);o__Saccharomycetales(100);f__Metschnikowiaceae(100);g__Metschnikowia(100);s__Metschnikowia_pulcherrima(100); | Metschnikowia | 13 | Saprotroph | \|Undefined Saprotroph\| | Yeast | NULL | Probable | As nectar-tap_saprotroph (Põlme et al. 2020); Decay sugar-rich substrates (Põlme et al. 2020) | Põlme S, et al. 2020. Fungal Diversity 105:1-16 ((https://doi.org/10.1007/s13225-020-00466-2)) |
| Otu50306 | 1 | k__Fungi(100);p__Ascomycota(100);c__Saccharomycetes(100);o__Saccharomycetales(100);f__Metschnikowiaceae(100);g__Metschnikowia(100);s__Metschnikowia_pulcherrima(100); | Metschnikowia | 13 | Saprotroph | \|Undefined Saprotroph\| | Yeast | NULL | Probable | As nectar-tap_saprotroph (Põlme et al. 2020); Decay sugar-rich substrates (Põlme et al. 2020) | Põlme S, et al. 2020. Fungal Diversity 105:1-16 ((https://doi.org/10.1007/s13225-020-00466-2)) |
| Otu50307 | 1 | k__Fungi(100);p__Ascomycota(100);c__Saccharomycetes(100);o__Saccharomycetales(100);f__Metschnikowiaceae(100);g__Metschnikowia(100);s__Metschnikowia_pulcherrima(100); | Metschnikowia | 13 | Saprotroph | \|Undefined Saprotroph\| | Yeast | NULL | Probable | As nectar-tap_saprotroph (Põlme et al. 2020); Decay sugar-rich substrates (Põlme et al. 2020) | Põlme S, et al. 2020. Fungal Diversity 105:1-16 ((https://doi.org/10.1007/s13225-020-00466-2)) |
| Otu50308 | 1 | k__Fungi(100);p__Ascomycota(100);c__Saccharomycetes(100);o__Saccharomycetales(100);f__Metschnikowiaceae(100);g__Metschnikowia(100);s__Metschnikowia_pulcherrima(100); | Metschnikowia | 13 | Saprotroph | \|Undefined Saprotroph\| | Yeast | NULL | Probable | As nectar-tap_saprotroph (Põlme et al. 2020); Decay sugar-rich substrates (Põlme et al. 2020) | Põlme S, et al. 2020. Fungal Diversity 105:1-16 ((https://doi.org/10.1007/s13225-020-00466-2)) |
| Otu50309 | 1 | k__Fungi(100);p__Ascomycota(100);c__Saccharomycetes(100);o__Saccharomycetales(100);f__Metschnikowiaceae(100);g__Metschnikowia(100);s__Metschnikowia_pulcherrima(100); | Metschnikowia | 13 | Saprotroph | \|Undefined Saprotroph\| | Yeast | NULL | Probable | As nectar-tap_saprotroph (Põlme et al. 2020); Decay sugar-rich substrates (Põlme et al. 2020) | Põlme S, et al. 2020. Fungal Diversity 105:1-16 ((https://doi.org/10.1007/s13225-020-00466-2)) |
| Otu50310 | 1 | k__Fungi(100);p__Ascomycota(100);c__Saccharomycetes(100);o__Saccharomycetales(100);f__Metschnikowiaceae(100);g__Metschnikowia(100);s__Metschnikowia_pulcherrima(100); | Metschnikowia | 13 | Saprotroph | \|Undefined Saprotroph\| | Yeast | NULL | Probable | As nectar-tap_saprotroph (Põlme et al. 2020); Decay sugar-rich substrates (Põlme et al. 2020) | Põlme S, et al. 2020. Fungal Diversity 105:1-16 ((https://doi.org/10.1007/s13225-020-00466-2)) |
| Otu50311 | 1 | k__Fungi(100);p__Ascomycota(100);c__Saccharomycetes(100);o__Saccharomycetales(100);f__Metschnikowiaceae(100);g__Metschnikowia(100);s__Metschnikowia_pulcherrima(100); | Metschnikowia | 13 | Saprotroph | \|Undefined Saprotroph\| | Yeast | NULL | Probable | As nectar-tap_saprotroph (Põlme et al. 2020); Decay sugar-rich substrates (Põlme et al. 2020) | Põlme S, et al. 2020. Fungal Diversity 105:1-16 ((https://doi.org/10.1007/s13225-020-00466-2)) |
| Otu50312 | 1 | k__Fungi(100);p__Ascomycota(100);c__Saccharomycetes(100);o__Saccharomycetales(100);f__Metschnikowiaceae(100);g__Metschnikowia(100);s__Metschnikowia_pulcherrima(100); | Metschnikowia | 13 | Saprotroph | \|Undefined Saprotroph\| | Yeast | NULL | Probable | As nectar-tap_saprotroph (Põlme et al. 2020); Decay sugar-rich substrates (Põlme et al. 2020) | Põlme S, et al. 2020. Fungal Diversity 105:1-16 ((https://doi.org/10.1007/s13225-020-00466-2)) |
| Otu50313 | 1 | k__Fungi(100);p__Ascomycota(100);c__Saccharomycetes(100);o__Saccharomycetales(100);f__Metschnikowiaceae(100);g__Metschnikowia(100);s__Metschnikowia_pulcherrima(100); | Metschnikowia | 13 | Saprotroph | \|Undefined Saprotroph\| | Yeast | NULL | Probable | As nectar-tap_saprotroph (Põlme et al. 2020); Decay sugar-rich substrates (Põlme et al. 2020) | Põlme S, et al. 2020. Fungal Diversity 105:1-16 ((https://doi.org/10.1007/s13225-020-00466-2)) |
| Otu50314 | 1 | k__Fungi(100);p__Ascomycota(100);c__Saccharomycetes(100);o__Saccharomycetales(100);f__Metschnikowiaceae(100);g__Metschnikowia(100);s__Metschnikowia_pulcherrima(100); | Metschnikowia | 13 | Saprotroph | \|Undefined Saprotroph\| | Yeast | NULL | Probable | As nectar-tap_saprotroph (Põlme et al. 2020); Decay sugar-rich substrates (Põlme et al. 2020) | Põlme S, et al. 2020. Fungal Diversity 105:1-16 ((https://doi.org/10.1007/s13225-020-00466-2)) |
| Otu50315 | 1 | k__Fungi(100);p__Ascomycota(100);c__Saccharomycetes(100);o__Saccharomycetales(100);f__Metschnikowiaceae(100);g__Metschnikowia(100);s__Metschnikowia_pulcherrima(100); | Metschnikowia | 13 | Saprotroph | \|Undefined Saprotroph\| | Yeast | NULL | Probable | As nectar-tap_saprotroph (Põlme et al. 2020); Decay sugar-rich substrates (Põlme et al. 2020) | Põlme S, et al. 2020. Fungal Diversity 105:1-16 ((https://doi.org/10.1007/s13225-020-00466-2)) |
| Otu50316 | 1 | k__Fungi(100);p__Ascomycota(100);c__Saccharomycetes(100);o__Saccharomycetales(100);f__Metschnikowiaceae(100);g__Metschnikowia(100);g__Metschnikowia_unclassified(100); | Metschnikowia | 13 | Saprotroph | \|Undefined Saprotroph\| | Yeast | NULL | Probable | As nectar-tap_saprotroph (Põlme et al. 2020); Decay sugar-rich substrates (Põlme et al. 2020) | Põlme S, et al. 2020. Fungal Diversity 105:1-16 ((https://doi.org/10.1007/s13225-020-00466-2)) |
| Otu50317 | 1 | k__Fungi(100);p__Ascomycota(100);c__Saccharomycetes(100);o__Saccharomycetales(100);f__Metschnikowiaceae(100);g__Metschnikowia(100);s__Metschnikowia_pulcherrima(100); | Metschnikowia | 13 | Saprotroph | \|Undefined Saprotroph\| | Yeast | NULL | Probable | As nectar-tap_saprotroph (Põlme et al. 2020); Decay sugar-rich substrates (Põlme et al. 2020) | Põlme S, et al. 2020. Fungal Diversity 105:1-16 ((https://doi.org/10.1007/s13225-020-00466-2)) |
| Otu50318 | 1 | k__Fungi(100);p__Ascomycota(100);c__Saccharomycetes(100);o__Saccharomycetales(100);f__Metschnikowiaceae(100);g__Metschnikowia(100);s__Metschnikowia_pulcherrima(100); | Metschnikowia | 13 | Saprotroph | \|Undefined Saprotroph\| | Yeast | NULL | Probable | As nectar-tap_saprotroph (Põlme et al. 2020); Decay sugar-rich substrates (Põlme et al. 2020) | Põlme S, et al. 2020. Fungal Diversity 105:1-16 ((https://doi.org/10.1007/s13225-020-00466-2)) |
| Otu50319 | 1 | k__Fungi(100);p__Ascomycota(100);c__Saccharomycetes(100);o__Saccharomycetales(100);f__Metschnikowiaceae(100);g__Metschnikowia(100);s__Metschnikowia_pulcherrima(100); | Metschnikowia | 13 | Saprotroph | \|Undefined Saprotroph\| | Yeast | NULL | Probable | As nectar-tap_saprotroph (Põlme et al. 2020); Decay sugar-rich substrates (Põlme et al. 2020) | Põlme S, et al. 2020. Fungal Diversity 105:1-16 ((https://doi.org/10.1007/s13225-020-00466-2)) |
| Otu50320 | 1 | k__Fungi(100);p__Ascomycota(100);c__Saccharomycetes(100);o__Saccharomycetales(100);f__Metschnikowiaceae(100);g__Metschnikowia(100);s__Metschnikowia_pulcherrima(100); | Metschnikowia | 13 | Saprotroph | \|Undefined Saprotroph\| | Yeast | NULL | Probable | As nectar-tap_saprotroph (Põlme et al. 2020); Decay sugar-rich substrates (Põlme et al. 2020) | Põlme S, et al. 2020. Fungal Diversity 105:1-16 ((https://doi.org/10.1007/s13225-020-00466-2)) |
| Otu50321 | 1 | k__Fungi(100);p__Ascomycota(100);c__Saccharomycetes(100);o__Saccharomycetales(100);f__Metschnikowiaceae(100);g__Metschnikowia(100);s__Metschnikowia_pulcherrima(100); | Metschnikowia | 13 | Saprotroph | \|Undefined Saprotroph\| | Yeast | NULL | Probable | As nectar-tap_saprotroph (Põlme et al. 2020); Decay sugar-rich substrates (Põlme et al. 2020) | Põlme S, et al. 2020. Fungal Diversity 105:1-16 ((https://doi.org/10.1007/s13225-020-00466-2)) |
| Otu50322 | 1 | k__Fungi(100);p__Ascomycota(100);c__Saccharomycetes(100);o__Saccharomycetales(100);f__Metschnikowiaceae(100);g__Metschnikowia(100);s__Metschnikowia_pulcherrima(100); | Metschnikowia | 13 | Saprotroph | \|Undefined Saprotroph\| | Yeast | NULL | Probable | As nectar-tap_saprotroph (Põlme et al. 2020); Decay sugar-rich substrates (Põlme et al. 2020) | Põlme S, et al. 2020. Fungal Diversity 105:1-16 ((https://doi.org/10.1007/s13225-020-00466-2)) |
| Otu50323 | 1 | k__Fungi(100);p__Ascomycota(100);c__Saccharomycetes(100);o__Saccharomycetales(100);f__Metschnikowiaceae(100);g__Metschnikowia(100);g__Metschnikowia_unclassified(100); | Metschnikowia | 13 | Saprotroph | \|Undefined Saprotroph\| | Yeast | NULL | Probable | As nectar-tap_saprotroph (Põlme et al. 2020); Decay sugar-rich substrates (Põlme et al. 2020) | Põlme S, et al. 2020. Fungal Diversity 105:1-16 ((https://doi.org/10.1007/s13225-020-00466-2)) |
| Otu50324 | 1 | k__Fungi(100);p__Ascomycota(100);c__Saccharomycetes(100);o__Saccharomycetales(100);f__Metschnikowiaceae(100);g__Metschnikowia(100);s__Metschnikowia_pulcherrima(100); | Metschnikowia | 13 | Saprotroph | \|Undefined Saprotroph\| | Yeast | NULL | Probable | As nectar-tap_saprotroph (Põlme et al. 2020); Decay sugar-rich substrates (Põlme et al. 2020) | Põlme S, et al. 2020. Fungal Diversity 105:1-16 ((https://doi.org/10.1007/s13225-020-00466-2)) |
| Otu50328 | 1 | k__Fungi(100);p__Ascomycota(100);c__Saccharomycetes(100);o__Saccharomycetales(100);f__Metschnikowiaceae(100);g__Metschnikowia(100);s__Metschnikowia_pulcherrima(100); | Metschnikowia | 13 | Saprotroph | \|Undefined Saprotroph\| | Yeast | NULL | Probable | As nectar-tap_saprotroph (Põlme et al. 2020); Decay sugar-rich substrates (Põlme et al. 2020) | Põlme S, et al. 2020. Fungal Diversity 105:1-16 ((https://doi.org/10.1007/s13225-020-00466-2)) |
| Otu50329 | 1 | k__Fungi(100);p__Ascomycota(100);c__Saccharomycetes(100);o__Saccharomycetales(100);f__Metschnikowiaceae(100);g__Metschnikowia(100);s__Metschnikowia_pulcherrima(100); | Metschnikowia | 13 | Saprotroph | \|Undefined Saprotroph\| | Yeast | NULL | Probable | As nectar-tap_saprotroph (Põlme et al. 2020); Decay sugar-rich substrates (Põlme et al. 2020) | Põlme S, et al. 2020. Fungal Diversity 105:1-16 ((https://doi.org/10.1007/s13225-020-00466-2)) |
| Otu50330 | 1 | k__Fungi(100);p__Ascomycota(100);c__Saccharomycetes(100);o__Saccharomycetales(100);f__Metschnikowiaceae(100);g__Metschnikowia(100);s__Metschnikowia_pulcherrima(100); | Metschnikowia | 13 | Saprotroph | \|Undefined Saprotroph\| | Yeast | NULL | Probable | As nectar-tap_saprotroph (Põlme et al. 2020); Decay sugar-rich substrates (Põlme et al. 2020) | Põlme S, et al. 2020. Fungal Diversity 105:1-16 ((https://doi.org/10.1007/s13225-020-00466-2)) |
| Otu50333 | 1 | k__Fungi(100);p__Ascomycota(100);c__Saccharomycetes(100);o__Saccharomycetales(100);f__Metschnikowiaceae(100);g__Metschnikowia(100);s__Metschnikowia_pulcherrima(100); | Metschnikowia | 13 | Saprotroph | \|Undefined Saprotroph\| | Yeast | NULL | Probable | As nectar-tap_saprotroph (Põlme et al. 2020); Decay sugar-rich substrates (Põlme et al. 2020) | Põlme S, et al. 2020. Fungal Diversity 105:1-16 ((https://doi.org/10.1007/s13225-020-00466-2)) |
| Otu50334 | 1 | k__Fungi(100);p__Ascomycota(100);c__Saccharomycetes(100);o__Saccharomycetales(100);f__Metschnikowiaceae(100);g__Metschnikowia(100);g__Metschnikowia_unclassified(100); | Metschnikowia | 13 | Saprotroph | \|Undefined Saprotroph\| | Yeast | NULL | Probable | As nectar-tap_saprotroph (Põlme et al. 2020); Decay sugar-rich substrates (Põlme et al. 2020) | Põlme S, et al. 2020. Fungal Diversity 105:1-16 ((https://doi.org/10.1007/s13225-020-00466-2)) |
| Otu50335 | 1 | k__Fungi(100);p__Ascomycota(100);c__Saccharomycetes(100);o__Saccharomycetales(100);f__Metschnikowiaceae(100);g__Metschnikowia(100);s__Metschnikowia_pulcherrima(100); | Metschnikowia | 13 | Saprotroph | \|Undefined Saprotroph\| | Yeast | NULL | Probable | As nectar-tap_saprotroph (Põlme et al. 2020); Decay sugar-rich substrates (Põlme et al. 2020) | Põlme S, et al. 2020. Fungal Diversity 105:1-16 ((https://doi.org/10.1007/s13225-020-00466-2)) |
| Otu50336 | 1 | k__Fungi(100);p__Ascomycota(100);c__Saccharomycetes(100);o__Saccharomycetales(100);f__Metschnikowiaceae(100);g__Metschnikowia(100);s__Metschnikowia_pulcherrima(100); | Metschnikowia | 13 | Saprotroph | \|Undefined Saprotroph\| | Yeast | NULL | Probable | As nectar-tap_saprotroph (Põlme et al. 2020); Decay sugar-rich substrates (Põlme et al. 2020) | Põlme S, et al. 2020. Fungal Diversity 105:1-16 ((https://doi.org/10.1007/s13225-020-00466-2)) |
| Otu50337 | 1 | k__Fungi(100);p__Ascomycota(100);c__Saccharomycetes(100);o__Saccharomycetales(100);f__Metschnikowiaceae(100);g__Metschnikowia(100);s__Metschnikowia_pulcherrima(100); | Metschnikowia | 13 | Saprotroph | \|Undefined Saprotroph\| | Yeast | NULL | Probable | As nectar-tap_saprotroph (Põlme et al. 2020); Decay sugar-rich substrates (Põlme et al. 2020) | Põlme S, et al. 2020. Fungal Diversity 105:1-16 ((https://doi.org/10.1007/s13225-020-00466-2)) |
| Otu50338 | 1 | k__Fungi(100);p__Ascomycota(100);c__Saccharomycetes(100);o__Saccharomycetales(100);f__Metschnikowiaceae(100);g__Metschnikowia(100);s__Metschnikowia_pulcherrima(100); | Metschnikowia | 13 | Saprotroph | \|Undefined Saprotroph\| | Yeast | NULL | Probable | As nectar-tap_saprotroph (Põlme et al. 2020); Decay sugar-rich substrates (Põlme et al. 2020) | Põlme S, et al. 2020. Fungal Diversity 105:1-16 ((https://doi.org/10.1007/s13225-020-00466-2)) |
| Otu50339 | 1 | k__Fungi(100);p__Ascomycota(100);c__Saccharomycetes(100);o__Saccharomycetales(100);f__Metschnikowiaceae(100);g__Metschnikowia(100);g__Metschnikowia_unclassified(100); | Metschnikowia | 13 | Saprotroph | \|Undefined Saprotroph\| | Yeast | NULL | Probable | As nectar-tap_saprotroph (Põlme et al. 2020); Decay sugar-rich substrates (Põlme et al. 2020) | Põlme S, et al. 2020. Fungal Diversity 105:1-16 ((https://doi.org/10.1007/s13225-020-00466-2)) |
| Otu50340 | 1 | k__Fungi(100);p__Ascomycota(100);c__Saccharomycetes(100);o__Saccharomycetales(100);f__Metschnikowiaceae(100);g__Metschnikowia(100);s__Metschnikowia_pulcherrima(100); | Metschnikowia | 13 | Saprotroph | \|Undefined Saprotroph\| | Yeast | NULL | Probable | As nectar-tap_saprotroph (Põlme et al. 2020); Decay sugar-rich substrates (Põlme et al. 2020) | Põlme S, et al. 2020. Fungal Diversity 105:1-16 ((https://doi.org/10.1007/s13225-020-00466-2)) |
| Otu50341 | 1 | k__Fungi(100);p__Ascomycota(100);c__Saccharomycetes(100);o__Saccharomycetales(100);f__Metschnikowiaceae(100);g__Metschnikowia(100);s__Metschnikowia_pulcherrima(100); | Metschnikowia | 13 | Saprotroph | \|Undefined Saprotroph\| | Yeast | NULL | Probable | As nectar-tap_saprotroph (Põlme et al. 2020); Decay sugar-rich substrates (Põlme et al. 2020) | Põlme S, et al. 2020. Fungal Diversity 105:1-16 ((https://doi.org/10.1007/s13225-020-00466-2)) |
| Otu50342 | 1 | k__Fungi(100);p__Ascomycota(100);c__Saccharomycetes(100);o__Saccharomycetales(100);f__Metschnikowiaceae(100);g__Metschnikowia(100);s__Metschnikowia_pulcherrima(100); | Metschnikowia | 13 | Saprotroph | \|Undefined Saprotroph\| | Yeast | NULL | Probable | As nectar-tap_saprotroph (Põlme et al. 2020); Decay sugar-rich substrates (Põlme et al. 2020) | Põlme S, et al. 2020. Fungal Diversity 105:1-16 ((https://doi.org/10.1007/s13225-020-00466-2)) |
| Otu50343 | 1 | k__Fungi(100);p__Ascomycota(100);c__Saccharomycetes(100);o__Saccharomycetales(100);f__Metschnikowiaceae(100);g__Metschnikowia(100);s__Metschnikowia_pulcherrima(100); | Metschnikowia | 13 | Saprotroph | \|Undefined Saprotroph\| | Yeast | NULL | Probable | As nectar-tap_saprotroph (Põlme et al. 2020); Decay sugar-rich substrates (Põlme et al. 2020) | Põlme S, et al. 2020. Fungal Diversity 105:1-16 ((https://doi.org/10.1007/s13225-020-00466-2)) |
| Otu50344 | 1 | k__Fungi(100);p__Ascomycota(100);c__Saccharomycetes(100);o__Saccharomycetales(100);f__Metschnikowiaceae(100);g__Metschnikowia(100);s__Metschnikowia_pulcherrima(100); | Metschnikowia | 13 | Saprotroph | \|Undefined Saprotroph\| | Yeast | NULL | Probable | As nectar-tap_saprotroph (Põlme et al. 2020); Decay sugar-rich substrates (Põlme et al. 2020) | Põlme S, et al. 2020. Fungal Diversity 105:1-16 ((https://doi.org/10.1007/s13225-020-00466-2)) |
| Otu50345 | 1 | k__Fungi(100);p__Ascomycota(100);c__Saccharomycetes(100);o__Saccharomycetales(100);f__Metschnikowiaceae(100);g__Metschnikowia(100);s__Metschnikowia_pulcherrima(100); | Metschnikowia | 13 | Saprotroph | \|Undefined Saprotroph\| | Yeast | NULL | Probable | As nectar-tap_saprotroph (Põlme et al. 2020); Decay sugar-rich substrates (Põlme et al. 2020) | Põlme S, et al. 2020. Fungal Diversity 105:1-16 ((https://doi.org/10.1007/s13225-020-00466-2)) |
| Otu50346 | 1 | k__Fungi(100);p__Ascomycota(100);c__Saccharomycetes(100);o__Saccharomycetales(100);f__Metschnikowiaceae(100);g__Metschnikowia(100);s__Metschnikowia_pulcherrima(100); | Metschnikowia | 13 | Saprotroph | \|Undefined Saprotroph\| | Yeast | NULL | Probable | As nectar-tap_saprotroph (Põlme et al. 2020); Decay sugar-rich substrates (Põlme et al. 2020) | Põlme S, et al. 2020. Fungal Diversity 105:1-16 ((https://doi.org/10.1007/s13225-020-00466-2)) |
| Otu50347 | 1 | k__Fungi(100);p__Ascomycota(100);c__Saccharomycetes(100);o__Saccharomycetales(100);f__Metschnikowiaceae(100);g__Metschnikowia(100);s__Metschnikowia_pulcherrima(100); | Metschnikowia | 13 | Saprotroph | \|Undefined Saprotroph\| | Yeast | NULL | Probable | As nectar-tap_saprotroph (Põlme et al. 2020); Decay sugar-rich substrates (Põlme et al. 2020) | Põlme S, et al. 2020. Fungal Diversity 105:1-16 ((https://doi.org/10.1007/s13225-020-00466-2)) |
| Otu50348 | 1 | k__Fungi(100);p__Ascomycota(100);c__Saccharomycetes(100);o__Saccharomycetales(100);f__Metschnikowiaceae(100);g__Metschnikowia(100);s__Metschnikowia_pulcherrima(100); | Metschnikowia | 13 | Saprotroph | \|Undefined Saprotroph\| | Yeast | NULL | Probable | As nectar-tap_saprotroph (Põlme et al. 2020); Decay sugar-rich substrates (Põlme et al. 2020) | Põlme S, et al. 2020. Fungal Diversity 105:1-16 ((https://doi.org/10.1007/s13225-020-00466-2)) |
| Otu50349 | 1 | k__Fungi(100);p__Ascomycota(100);c__Saccharomycetes(100);o__Saccharomycetales(100);f__Metschnikowiaceae(100);g__Metschnikowia(100);s__Metschnikowia_pulcherrima(100); | Metschnikowia | 13 | Saprotroph | \|Undefined Saprotroph\| | Yeast | NULL | Probable | As nectar-tap_saprotroph (Põlme et al. 2020); Decay sugar-rich substrates (Põlme et al. 2020) | Põlme S, et al. 2020. Fungal Diversity 105:1-16 ((https://doi.org/10.1007/s13225-020-00466-2)) |
| Otu50350 | 1 | k__Fungi(100);p__Ascomycota(100);c__Saccharomycetes(100);o__Saccharomycetales(100);f__Metschnikowiaceae(100);g__Metschnikowia(100);g__Metschnikowia_unclassified(100); | Metschnikowia | 13 | Saprotroph | \|Undefined Saprotroph\| | Yeast | NULL | Probable | As nectar-tap_saprotroph (Põlme et al. 2020); Decay sugar-rich substrates (Põlme et al. 2020) | Põlme S, et al. 2020. Fungal Diversity 105:1-16 ((https://doi.org/10.1007/s13225-020-00466-2)) |
| Otu50351 | 1 | k__Fungi(100);p__Ascomycota(100);c__Saccharomycetes(100);o__Saccharomycetales(100);f__Metschnikowiaceae(100);g__Metschnikowia(100);s__Metschnikowia_pulcherrima(100); | Metschnikowia | 13 | Saprotroph | \|Undefined Saprotroph\| | Yeast | NULL | Probable | As nectar-tap_saprotroph (Põlme et al. 2020); Decay sugar-rich substrates (Põlme et al. 2020) | Põlme S, et al. 2020. Fungal Diversity 105:1-16 ((https://doi.org/10.1007/s13225-020-00466-2)) |
| Otu50352 | 1 | k__Fungi(100);p__Ascomycota(100);c__Saccharomycetes(100);o__Saccharomycetales(100);f__Metschnikowiaceae(100);g__Metschnikowia(100);g__Metschnikowia_unclassified(100); | Metschnikowia | 13 | Saprotroph | \|Undefined Saprotroph\| | Yeast | NULL | Probable | As nectar-tap_saprotroph (Põlme et al. 2020); Decay sugar-rich substrates (Põlme et al. 2020) | Põlme S, et al. 2020. Fungal Diversity 105:1-16 ((https://doi.org/10.1007/s13225-020-00466-2)) |
| Otu50353 | 1 | k__Fungi(100);p__Ascomycota(100);c__Saccharomycetes(100);o__Saccharomycetales(100);f__Metschnikowiaceae(100);g__Metschnikowia(100);g__Metschnikowia_unclassified(100); | Metschnikowia | 13 | Saprotroph | \|Undefined Saprotroph\| | Yeast | NULL | Probable | As nectar-tap_saprotroph (Põlme et al. 2020); Decay sugar-rich substrates (Põlme et al. 2020) | Põlme S, et al. 2020. Fungal Diversity 105:1-16 ((https://doi.org/10.1007/s13225-020-00466-2)) |
| Otu50354 | 1 | k__Fungi(100);p__Ascomycota(100);c__Saccharomycetes(100);o__Saccharomycetales(100);f__Metschnikowiaceae(100);g__Metschnikowia(100);g__Metschnikowia_unclassified(100); | Metschnikowia | 13 | Saprotroph | \|Undefined Saprotroph\| | Yeast | NULL | Probable | As nectar-tap_saprotroph (Põlme et al. 2020); Decay sugar-rich substrates (Põlme et al. 2020) | Põlme S, et al. 2020. Fungal Diversity 105:1-16 ((https://doi.org/10.1007/s13225-020-00466-2)) |
| Otu50355 | 1 | k__Fungi(100);p__Ascomycota(100);c__Saccharomycetes(100);o__Saccharomycetales(100);f__Metschnikowiaceae(100);g__Metschnikowia(100);s__Metschnikowia_pulcherrima(100); | Metschnikowia | 13 | Saprotroph | \|Undefined Saprotroph\| | Yeast | NULL | Probable | As nectar-tap_saprotroph (Põlme et al. 2020); Decay sugar-rich substrates (Põlme et al. 2020) | Põlme S, et al. 2020. Fungal Diversity 105:1-16 ((https://doi.org/10.1007/s13225-020-00466-2)) |
| Otu50356 | 1 | k__Fungi(100);p__Ascomycota(100);c__Saccharomycetes(100);o__Saccharomycetales(100);f__Metschnikowiaceae(100);g__Metschnikowia(100);s__Metschnikowia_pulcherrima(100); | Metschnikowia | 13 | Saprotroph | \|Undefined Saprotroph\| | Yeast | NULL | Probable | As nectar-tap_saprotroph (Põlme et al. 2020); Decay sugar-rich substrates (Põlme et al. 2020) | Põlme S, et al. 2020. Fungal Diversity 105:1-16 ((https://doi.org/10.1007/s13225-020-00466-2)) |
| Otu50357 | 1 | k__Fungi(100);p__Ascomycota(100);c__Saccharomycetes(100);o__Saccharomycetales(100);f__Metschnikowiaceae(100);g__Metschnikowia(100);s__Metschnikowia_pulcherrima(100); | Metschnikowia | 13 | Saprotroph | \|Undefined Saprotroph\| | Yeast | NULL | Probable | As nectar-tap_saprotroph (Põlme et al. 2020); Decay sugar-rich substrates (Põlme et al. 2020) | Põlme S, et al. 2020. Fungal Diversity 105:1-16 ((https://doi.org/10.1007/s13225-020-00466-2)) |
| Otu50358 | 1 | k__Fungi(100);p__Ascomycota(100);c__Saccharomycetes(100);o__Saccharomycetales(100);f__Metschnikowiaceae(100);g__Metschnikowia(100);g__Metschnikowia_unclassified(100); | Metschnikowia | 13 | Saprotroph | \|Undefined Saprotroph\| | Yeast | NULL | Probable | As nectar-tap_saprotroph (Põlme et al. 2020); Decay sugar-rich substrates (Põlme et al. 2020) | Põlme S, et al. 2020. Fungal Diversity 105:1-16 ((https://doi.org/10.1007/s13225-020-00466-2)) |
| Otu50359 | 1 | k__Fungi(100);p__Ascomycota(100);c__Saccharomycetes(100);o__Saccharomycetales(100);f__Metschnikowiaceae(100);g__Metschnikowia(100);s__Metschnikowia_pulcherrima(100); | Metschnikowia | 13 | Saprotroph | \|Undefined Saprotroph\| | Yeast | NULL | Probable | As nectar-tap_saprotroph (Põlme et al. 2020); Decay sugar-rich substrates (Põlme et al. 2020) | Põlme S, et al. 2020. Fungal Diversity 105:1-16 ((https://doi.org/10.1007/s13225-020-00466-2)) |
| Otu50360 | 1 | k__Fungi(100);p__Ascomycota(100);c__Saccharomycetes(100);o__Saccharomycetales(100);f__Metschnikowiaceae(100);g__Metschnikowia(100);s__Metschnikowia_pulcherrima(100); | Metschnikowia | 13 | Saprotroph | \|Undefined Saprotroph\| | Yeast | NULL | Probable | As nectar-tap_saprotroph (Põlme et al. 2020); Decay sugar-rich substrates (Põlme et al. 2020) | Põlme S, et al. 2020. Fungal Diversity 105:1-16 ((https://doi.org/10.1007/s13225-020-00466-2)) |
| Otu50361 | 1 | k__Fungi(100);p__Ascomycota(100);c__Saccharomycetes(100);o__Saccharomycetales(100);f__Metschnikowiaceae(100);g__Metschnikowia(100);g__Metschnikowia_unclassified(100); | Metschnikowia | 13 | Saprotroph | \|Undefined Saprotroph\| | Yeast | NULL | Probable | As nectar-tap_saprotroph (Põlme et al. 2020); Decay sugar-rich substrates (Põlme et al. 2020) | Põlme S, et al. 2020. Fungal Diversity 105:1-16 ((https://doi.org/10.1007/s13225-020-00466-2)) |
| Otu50362 | 1 | k__Fungi(100);p__Ascomycota(100);c__Saccharomycetes(100);o__Saccharomycetales(100);f__Metschnikowiaceae(100);g__Metschnikowia(100);s__Metschnikowia_pulcherrima(100); | Metschnikowia | 13 | Saprotroph | \|Undefined Saprotroph\| | Yeast | NULL | Probable | As nectar-tap_saprotroph (Põlme et al. 2020); Decay sugar-rich substrates (Põlme et al. 2020) | Põlme S, et al. 2020. Fungal Diversity 105:1-16 ((https://doi.org/10.1007/s13225-020-00466-2)) |
| Otu50363 | 1 | k__Fungi(100);p__Ascomycota(100);c__Saccharomycetes(100);o__Saccharomycetales(100);f__Metschnikowiaceae(100);g__Metschnikowia(100);s__Metschnikowia_pulcherrima(100); | Metschnikowia | 13 | Saprotroph | \|Undefined Saprotroph\| | Yeast | NULL | Probable | As nectar-tap_saprotroph (Põlme et al. 2020); Decay sugar-rich substrates (Põlme et al. 2020) | Põlme S, et al. 2020. Fungal Diversity 105:1-16 ((https://doi.org/10.1007/s13225-020-00466-2)) |
| Otu50364 | 1 | k__Fungi(100);p__Ascomycota(100);c__Saccharomycetes(100);o__Saccharomycetales(100);f__Metschnikowiaceae(100);g__Metschnikowia(100);g__Metschnikowia_unclassified(100); | Metschnikowia | 13 | Saprotroph | \|Undefined Saprotroph\| | Yeast | NULL | Probable | As nectar-tap_saprotroph (Põlme et al. 2020); Decay sugar-rich substrates (Põlme et al. 2020) | Põlme S, et al. 2020. Fungal Diversity 105:1-16 ((https://doi.org/10.1007/s13225-020-00466-2)) |
| Otu50365 | 1 | k__Fungi(100);p__Ascomycota(100);c__Saccharomycetes(100);o__Saccharomycetales(100);f__Metschnikowiaceae(100);g__Metschnikowia(100);s__Metschnikowia_pulcherrima(100); | Metschnikowia | 13 | Saprotroph | \|Undefined Saprotroph\| | Yeast | NULL | Probable | As nectar-tap_saprotroph (Põlme et al. 2020); Decay sugar-rich substrates (Põlme et al. 2020) | Põlme S, et al. 2020. Fungal Diversity 105:1-16 ((https://doi.org/10.1007/s13225-020-00466-2)) |
| Otu50368 | 1 | k__Fungi(100);p__Ascomycota(100);c__Saccharomycetes(100);o__Saccharomycetales(100);f__Metschnikowiaceae(100);g__Metschnikowia(100);s__Metschnikowia_pulcherrima(100); | Metschnikowia | 13 | Saprotroph | \|Undefined Saprotroph\| | Yeast | NULL | Probable | As nectar-tap_saprotroph (Põlme et al. 2020); Decay sugar-rich substrates (Põlme et al. 2020) | Põlme S, et al. 2020. Fungal Diversity 105:1-16 ((https://doi.org/10.1007/s13225-020-00466-2)) |
| Otu50369 | 1 | k__Fungi(100);p__Ascomycota(100);c__Saccharomycetes(100);o__Saccharomycetales(100);f__Metschnikowiaceae(100);g__Metschnikowia(100);s__Metschnikowia_pulcherrima(100); | Metschnikowia | 13 | Saprotroph | \|Undefined Saprotroph\| | Yeast | NULL | Probable | As nectar-tap_saprotroph (Põlme et al. 2020); Decay sugar-rich substrates (Põlme et al. 2020) | Põlme S, et al. 2020. Fungal Diversity 105:1-16 ((https://doi.org/10.1007/s13225-020-00466-2)) |
| Otu50370 | 1 | k__Fungi(100);p__Ascomycota(100);c__Saccharomycetes(100);o__Saccharomycetales(100);f__Metschnikowiaceae(100);g__Metschnikowia(100);g__Metschnikowia_unclassified(100); | Metschnikowia | 13 | Saprotroph | \|Undefined Saprotroph\| | Yeast | NULL | Probable | As nectar-tap_saprotroph (Põlme et al. 2020); Decay sugar-rich substrates (Põlme et al. 2020) | Põlme S, et al. 2020. Fungal Diversity 105:1-16 ((https://doi.org/10.1007/s13225-020-00466-2)) |
| Otu50371 | 1 | k__Fungi(100);p__Ascomycota(100);c__Saccharomycetes(100);o__Saccharomycetales(100);f__Metschnikowiaceae(100);g__Metschnikowia(100);s__Metschnikowia_pulcherrima(100); | Metschnikowia | 13 | Saprotroph | \|Undefined Saprotroph\| | Yeast | NULL | Probable | As nectar-tap_saprotroph (Põlme et al. 2020); Decay sugar-rich substrates (Põlme et al. 2020) | Põlme S, et al. 2020. Fungal Diversity 105:1-16 ((https://doi.org/10.1007/s13225-020-00466-2)) |
| Otu50372 | 1 | k__Fungi(100);p__Ascomycota(100);c__Saccharomycetes(100);o__Saccharomycetales(100);f__Metschnikowiaceae(100);g__Metschnikowia(100);s__Metschnikowia_pulcherrima(100); | Metschnikowia | 13 | Saprotroph | \|Undefined Saprotroph\| | Yeast | NULL | Probable | As nectar-tap_saprotroph (Põlme et al. 2020); Decay sugar-rich substrates (Põlme et al. 2020) | Põlme S, et al. 2020. Fungal Diversity 105:1-16 ((https://doi.org/10.1007/s13225-020-00466-2)) |
| Otu50373 | 1 | k__Fungi(100);p__Ascomycota(100);c__Saccharomycetes(100);o__Saccharomycetales(100);f__Metschnikowiaceae(100);g__Metschnikowia(100);s__Metschnikowia_pulcherrima(100); | Metschnikowia | 13 | Saprotroph | \|Undefined Saprotroph\| | Yeast | NULL | Probable | As nectar-tap_saprotroph (Põlme et al. 2020); Decay sugar-rich substrates (Põlme et al. 2020) | Põlme S, et al. 2020. Fungal Diversity 105:1-16 ((https://doi.org/10.1007/s13225-020-00466-2)) |
| Otu50375 | 1 | k__Fungi(100);p__Ascomycota(100);c__Saccharomycetes(100);o__Saccharomycetales(100);o__Saccharomycetales_unclassified(100);o__Saccharomycetales_unclassified(100);o__Saccharomycetales_unclassified(100); | Saccharomycetales | 7 | Saprotroph | Undefined Saprotroph | Yeast | NULL | Possible | NULL | Sterkenburg E, et al. 2015. New Phytologist 207:1145-1158 ((https://doi.org/10.1111/nph.13426)) |
| Otu50376 | 1 | k__Fungi(100);p__Ascomycota(100);c__Saccharomycetes(100);o__Saccharomycetales(100);f__Metschnikowiaceae(100);g__Metschnikowia(100);s__Metschnikowia_pulcherrima(100); | Metschnikowia | 13 | Saprotroph | \|Undefined Saprotroph\| | Yeast | NULL | Probable | As nectar-tap_saprotroph (Põlme et al. 2020); Decay sugar-rich substrates (Põlme et al. 2020) | Põlme S, et al. 2020. Fungal Diversity 105:1-16 ((https://doi.org/10.1007/s13225-020-00466-2)) |
| Otu50377 | 1 | k__Fungi(100);p__Ascomycota(100);c__Saccharomycetes(100);o__Saccharomycetales(100);f__Metschnikowiaceae(100);g__Metschnikowia(100);s__Metschnikowia_pulcherrima(100); | Metschnikowia | 13 | Saprotroph | \|Undefined Saprotroph\| | Yeast | NULL | Probable | As nectar-tap_saprotroph (Põlme et al. 2020); Decay sugar-rich substrates (Põlme et al. 2020) | Põlme S, et al. 2020. Fungal Diversity 105:1-16 ((https://doi.org/10.1007/s13225-020-00466-2)) |
| Otu50378 | 1 | k__Fungi(100);p__Ascomycota(100);c__Saccharomycetes(100);o__Saccharomycetales(100);f__Metschnikowiaceae(100);g__Metschnikowia(100);s__Metschnikowia_pulcherrima(100); | Metschnikowia | 13 | Saprotroph | \|Undefined Saprotroph\| | Yeast | NULL | Probable | As nectar-tap_saprotroph (Põlme et al. 2020); Decay sugar-rich substrates (Põlme et al. 2020) | Põlme S, et al. 2020. Fungal Diversity 105:1-16 ((https://doi.org/10.1007/s13225-020-00466-2)) |
| Otu50379 | 1 | k__Fungi(100);p__Ascomycota(100);c__Saccharomycetes(100);o__Saccharomycetales(100);o__Saccharomycetales_unclassified(100);o__Saccharomycetales_unclassified(100);o__Saccharomycetales_unclassified(100); | Saccharomycetales | 7 | Saprotroph | Undefined Saprotroph | Yeast | NULL | Possible | NULL | Sterkenburg E, et al. 2015. New Phytologist 207:1145-1158 ((https://doi.org/10.1111/nph.13426)) |
| Otu50381 | 1 | k__Fungi(100);p__Ascomycota(100);c__Saccharomycetes(100);o__Saccharomycetales(100);f__Metschnikowiaceae(100);g__Metschnikowia(100);s__Metschnikowia_pulcherrima(100); | Metschnikowia | 13 | Saprotroph | \|Undefined Saprotroph\| | Yeast | NULL | Probable | As nectar-tap_saprotroph (Põlme et al. 2020); Decay sugar-rich substrates (Põlme et al. 2020) | Põlme S, et al. 2020. Fungal Diversity 105:1-16 ((https://doi.org/10.1007/s13225-020-00466-2)) |
| Otu50382 | 1 | k__Fungi(100);p__Ascomycota(100);c__Saccharomycetes(100);o__Saccharomycetales(100);f__Metschnikowiaceae(100);g__Metschnikowia(100);g__Metschnikowia_unclassified(100); | Metschnikowia | 13 | Saprotroph | \|Undefined Saprotroph\| | Yeast | NULL | Probable | As nectar-tap_saprotroph (Põlme et al. 2020); Decay sugar-rich substrates (Põlme et al. 2020) | Põlme S, et al. 2020. Fungal Diversity 105:1-16 ((https://doi.org/10.1007/s13225-020-00466-2)) |
| Otu50383 | 1 | k__Fungi(100);p__Ascomycota(100);c__Saccharomycetes(100);o__Saccharomycetales(100);f__Metschnikowiaceae(100);g__Metschnikowia(100);g__Metschnikowia_unclassified(100); | Metschnikowia | 13 | Saprotroph | \|Undefined Saprotroph\| | Yeast | NULL | Probable | As nectar-tap_saprotroph (Põlme et al. 2020); Decay sugar-rich substrates (Põlme et al. 2020) | Põlme S, et al. 2020. Fungal Diversity 105:1-16 ((https://doi.org/10.1007/s13225-020-00466-2)) |
| Otu50384 | 1 | k__Fungi(100);p__Ascomycota(100);c__Saccharomycetes(100);o__Saccharomycetales(100);f__Metschnikowiaceae(100);g__Metschnikowia(100);s__Metschnikowia_pulcherrima(100); | Metschnikowia | 13 | Saprotroph | \|Undefined Saprotroph\| | Yeast | NULL | Probable | As nectar-tap_saprotroph (Põlme et al. 2020); Decay sugar-rich substrates (Põlme et al. 2020) | Põlme S, et al. 2020. Fungal Diversity 105:1-16 ((https://doi.org/10.1007/s13225-020-00466-2)) |
| Otu50385 | 1 | k__Fungi(100);p__Ascomycota(100);c__Saccharomycetes(100);o__Saccharomycetales(100);f__Metschnikowiaceae(100);g__Metschnikowia(100);s__Metschnikowia_pulcherrima(100); | Metschnikowia | 13 | Saprotroph | \|Undefined Saprotroph\| | Yeast | NULL | Probable | As nectar-tap_saprotroph (Põlme et al. 2020); Decay sugar-rich substrates (Põlme et al. 2020) | Põlme S, et al. 2020. Fungal Diversity 105:1-16 ((https://doi.org/10.1007/s13225-020-00466-2)) |
| Otu50386 | 1 | k__Fungi(100);p__Ascomycota(100);c__Saccharomycetes(100);o__Saccharomycetales(100);f__Metschnikowiaceae(100);g__Metschnikowia(100);s__Metschnikowia_pulcherrima(100); | Metschnikowia | 13 | Saprotroph | \|Undefined Saprotroph\| | Yeast | NULL | Probable | As nectar-tap_saprotroph (Põlme et al. 2020); Decay sugar-rich substrates (Põlme et al. 2020) | Põlme S, et al. 2020. Fungal Diversity 105:1-16 ((https://doi.org/10.1007/s13225-020-00466-2)) |
| Otu50387 | 1 | k__Fungi(100);p__Ascomycota(100);c__Saccharomycetes(100);o__Saccharomycetales(100);f__Metschnikowiaceae(100);g__Metschnikowia(100);s__Metschnikowia_pulcherrima(100); | Metschnikowia | 13 | Saprotroph | \|Undefined Saprotroph\| | Yeast | NULL | Probable | As nectar-tap_saprotroph (Põlme et al. 2020); Decay sugar-rich substrates (Põlme et al. 2020) | Põlme S, et al. 2020. Fungal Diversity 105:1-16 ((https://doi.org/10.1007/s13225-020-00466-2)) |
| Otu50388 | 1 | k__Fungi(100);p__Ascomycota(100);c__Saccharomycetes(100);o__Saccharomycetales(100);f__Metschnikowiaceae(100);g__Metschnikowia(100);s__Metschnikowia_pulcherrima(100); | Metschnikowia | 13 | Saprotroph | \|Undefined Saprotroph\| | Yeast | NULL | Probable | As nectar-tap_saprotroph (Põlme et al. 2020); Decay sugar-rich substrates (Põlme et al. 2020) | Põlme S, et al. 2020. Fungal Diversity 105:1-16 ((https://doi.org/10.1007/s13225-020-00466-2)) |
| Otu50390 | 1 | k__Fungi(100);p__Ascomycota(100);c__Saccharomycetes(100);o__Saccharomycetales(100);f__Metschnikowiaceae(100);g__Metschnikowia(100);g__Metschnikowia_unclassified(100); | Metschnikowia | 13 | Saprotroph | \|Undefined Saprotroph\| | Yeast | NULL | Probable | As nectar-tap_saprotroph (Põlme et al. 2020); Decay sugar-rich substrates (Põlme et al. 2020) | Põlme S, et al. 2020. Fungal Diversity 105:1-16 ((https://doi.org/10.1007/s13225-020-00466-2)) |
| Otu50391 | 1 | k__Fungi(100);p__Ascomycota(100);c__Saccharomycetes(100);o__Saccharomycetales(100);f__Metschnikowiaceae(100);g__Metschnikowia(100);s__Metschnikowia_pulcherrima(100); | Metschnikowia | 13 | Saprotroph | \|Undefined Saprotroph\| | Yeast | NULL | Probable | As nectar-tap_saprotroph (Põlme et al. 2020); Decay sugar-rich substrates (Põlme et al. 2020) | Põlme S, et al. 2020. Fungal Diversity 105:1-16 ((https://doi.org/10.1007/s13225-020-00466-2)) |
| Otu50392 | 1 | k__Fungi(100);p__Ascomycota(100);c__Saccharomycetes(100);o__Saccharomycetales(100);f__Metschnikowiaceae(100);g__Metschnikowia(100);s__Metschnikowia_pulcherrima(100); | Metschnikowia | 13 | Saprotroph | \|Undefined Saprotroph\| | Yeast | NULL | Probable | As nectar-tap_saprotroph (Põlme et al. 2020); Decay sugar-rich substrates (Põlme et al. 2020) | Põlme S, et al. 2020. Fungal Diversity 105:1-16 ((https://doi.org/10.1007/s13225-020-00466-2)) |
| Otu50393 | 1 | k__Fungi(100);p__Ascomycota(100);c__Saccharomycetes(100);o__Saccharomycetales(100);f__Metschnikowiaceae(100);g__Metschnikowia(100);s__Metschnikowia_pulcherrima(100); | Metschnikowia | 13 | Saprotroph | \|Undefined Saprotroph\| | Yeast | NULL | Probable | As nectar-tap_saprotroph (Põlme et al. 2020); Decay sugar-rich substrates (Põlme et al. 2020) | Põlme S, et al. 2020. Fungal Diversity 105:1-16 ((https://doi.org/10.1007/s13225-020-00466-2)) |
| Otu50394 | 1 | k__Fungi(100);p__Ascomycota(100);c__Saccharomycetes(100);o__Saccharomycetales(100);f__Metschnikowiaceae(100);g__Metschnikowia(100);s__Metschnikowia_pulcherrima(100); | Metschnikowia | 13 | Saprotroph | \|Undefined Saprotroph\| | Yeast | NULL | Probable | As nectar-tap_saprotroph (Põlme et al. 2020); Decay sugar-rich substrates (Põlme et al. 2020) | Põlme S, et al. 2020. Fungal Diversity 105:1-16 ((https://doi.org/10.1007/s13225-020-00466-2)) |
| Otu50395 | 1 | k__Fungi(100);p__Ascomycota(100);c__Saccharomycetes(100);o__Saccharomycetales(100);f__Metschnikowiaceae(100);g__Metschnikowia(100);s__Metschnikowia_pulcherrima(100); | Metschnikowia | 13 | Saprotroph | \|Undefined Saprotroph\| | Yeast | NULL | Probable | As nectar-tap_saprotroph (Põlme et al. 2020); Decay sugar-rich substrates (Põlme et al. 2020) | Põlme S, et al. 2020. Fungal Diversity 105:1-16 ((https://doi.org/10.1007/s13225-020-00466-2)) |
| Otu50396 | 1 | k__Fungi(100);p__Ascomycota(100);c__Saccharomycetes(100);o__Saccharomycetales(100);f__Metschnikowiaceae(100);g__Metschnikowia(100);g__Metschnikowia_unclassified(100); | Metschnikowia | 13 | Saprotroph | \|Undefined Saprotroph\| | Yeast | NULL | Probable | As nectar-tap_saprotroph (Põlme et al. 2020); Decay sugar-rich substrates (Põlme et al. 2020) | Põlme S, et al. 2020. Fungal Diversity 105:1-16 ((https://doi.org/10.1007/s13225-020-00466-2)) |
| Otu50397 | 1 | k__Fungi(100);p__Ascomycota(100);c__Saccharomycetes(100);o__Saccharomycetales(100);f__Metschnikowiaceae(100);g__Metschnikowia(100);s__Metschnikowia_pulcherrima(100); | Metschnikowia | 13 | Saprotroph | \|Undefined Saprotroph\| | Yeast | NULL | Probable | As nectar-tap_saprotroph (Põlme et al. 2020); Decay sugar-rich substrates (Põlme et al. 2020) | Põlme S, et al. 2020. Fungal Diversity 105:1-16 ((https://doi.org/10.1007/s13225-020-00466-2)) |
| Otu50398 | 1 | k__Fungi(100);p__Ascomycota(100);c__Saccharomycetes(100);o__Saccharomycetales(100);f__Metschnikowiaceae(100);g__Metschnikowia(100);s__Metschnikowia_pulcherrima(100); | Metschnikowia | 13 | Saprotroph | \|Undefined Saprotroph\| | Yeast | NULL | Probable | As nectar-tap_saprotroph (Põlme et al. 2020); Decay sugar-rich substrates (Põlme et al. 2020) | Põlme S, et al. 2020. Fungal Diversity 105:1-16 ((https://doi.org/10.1007/s13225-020-00466-2)) |
| Otu50399 | 1 | k__Fungi(100);p__Ascomycota(100);c__Saccharomycetes(100);o__Saccharomycetales(100);f__Metschnikowiaceae(100);g__Metschnikowia(100);g__Metschnikowia_unclassified(100); | Metschnikowia | 13 | Saprotroph | \|Undefined Saprotroph\| | Yeast | NULL | Probable | As nectar-tap_saprotroph (Põlme et al. 2020); Decay sugar-rich substrates (Põlme et al. 2020) | Põlme S, et al. 2020. Fungal Diversity 105:1-16 ((https://doi.org/10.1007/s13225-020-00466-2)) |
| Otu50400 | 1 | k__Fungi(100);p__Ascomycota(100);c__Saccharomycetes(100);o__Saccharomycetales(100);f__Metschnikowiaceae(100);g__Metschnikowia(100);s__Metschnikowia_pulcherrima(100); | Metschnikowia | 13 | Saprotroph | \|Undefined Saprotroph\| | Yeast | NULL | Probable | As nectar-tap_saprotroph (Põlme et al. 2020); Decay sugar-rich substrates (Põlme et al. 2020) | Põlme S, et al. 2020. Fungal Diversity 105:1-16 ((https://doi.org/10.1007/s13225-020-00466-2)) |
| Otu50401 | 1 | k__Fungi(100);p__Ascomycota(100);c__Saccharomycetes(100);o__Saccharomycetales(100);f__Metschnikowiaceae(100);g__Metschnikowia(100);g__Metschnikowia_unclassified(100); | Metschnikowia | 13 | Saprotroph | \|Undefined Saprotroph\| | Yeast | NULL | Probable | As nectar-tap_saprotroph (Põlme et al. 2020); Decay sugar-rich substrates (Põlme et al. 2020) | Põlme S, et al. 2020. Fungal Diversity 105:1-16 ((https://doi.org/10.1007/s13225-020-00466-2)) |
| Otu50403 | 1 | k__Fungi(100);p__Ascomycota(100);c__Saccharomycetes(100);o__Saccharomycetales(100);f__Metschnikowiaceae(100);g__Metschnikowia(100);s__Metschnikowia_pulcherrima(100); | Metschnikowia | 13 | Saprotroph | \|Undefined Saprotroph\| | Yeast | NULL | Probable | As nectar-tap_saprotroph (Põlme et al. 2020); Decay sugar-rich substrates (Põlme et al. 2020) | Põlme S, et al. 2020. Fungal Diversity 105:1-16 ((https://doi.org/10.1007/s13225-020-00466-2)) |
| Otu50404 | 1 | k__Fungi(100);p__Ascomycota(100);c__Saccharomycetes(100);o__Saccharomycetales(100);f__Metschnikowiaceae(100);g__Metschnikowia(100);g__Metschnikowia_unclassified(100); | Metschnikowia | 13 | Saprotroph | \|Undefined Saprotroph\| | Yeast | NULL | Probable | As nectar-tap_saprotroph (Põlme et al. 2020); Decay sugar-rich substrates (Põlme et al. 2020) | Põlme S, et al. 2020. Fungal Diversity 105:1-16 ((https://doi.org/10.1007/s13225-020-00466-2)) |
| Otu50405 | 1 | k__Fungi(100);p__Ascomycota(100);c__Saccharomycetes(100);o__Saccharomycetales(100);f__Metschnikowiaceae(100);g__Metschnikowia(100);s__Metschnikowia_pulcherrima(100); | Metschnikowia | 13 | Saprotroph | \|Undefined Saprotroph\| | Yeast | NULL | Probable | As nectar-tap_saprotroph (Põlme et al. 2020); Decay sugar-rich substrates (Põlme et al. 2020) | Põlme S, et al. 2020. Fungal Diversity 105:1-16 ((https://doi.org/10.1007/s13225-020-00466-2)) |
| Otu50406 | 1 | k__Fungi(100);p__Ascomycota(100);c__Saccharomycetes(100);o__Saccharomycetales(100);f__Metschnikowiaceae(100);g__Metschnikowia(100);s__Metschnikowia_pulcherrima(100); | Metschnikowia | 13 | Saprotroph | \|Undefined Saprotroph\| | Yeast | NULL | Probable | As nectar-tap_saprotroph (Põlme et al. 2020); Decay sugar-rich substrates (Põlme et al. 2020) | Põlme S, et al. 2020. Fungal Diversity 105:1-16 ((https://doi.org/10.1007/s13225-020-00466-2)) |
| Otu50407 | 1 | k__Fungi(100);p__Ascomycota(100);c__Saccharomycetes(100);o__Saccharomycetales(100);f__Metschnikowiaceae(100);g__Metschnikowia(100);s__Metschnikowia_pulcherrima(100); | Metschnikowia | 13 | Saprotroph | \|Undefined Saprotroph\| | Yeast | NULL | Probable | As nectar-tap_saprotroph (Põlme et al. 2020); Decay sugar-rich substrates (Põlme et al. 2020) | Põlme S, et al. 2020. Fungal Diversity 105:1-16 ((https://doi.org/10.1007/s13225-020-00466-2)) |
| Otu50408 | 1 | k__Fungi(100);p__Ascomycota(100);c__Saccharomycetes(100);o__Saccharomycetales(100);f__Metschnikowiaceae(100);g__Metschnikowia(100);g__Metschnikowia_unclassified(100); | Metschnikowia | 13 | Saprotroph | \|Undefined Saprotroph\| | Yeast | NULL | Probable | As nectar-tap_saprotroph (Põlme et al. 2020); Decay sugar-rich substrates (Põlme et al. 2020) | Põlme S, et al. 2020. Fungal Diversity 105:1-16 ((https://doi.org/10.1007/s13225-020-00466-2)) |
| Otu50410 | 1 | k__Fungi(100);p__Ascomycota(100);c__Saccharomycetes(100);o__Saccharomycetales(100);f__Metschnikowiaceae(100);g__Metschnikowia(100);s__Metschnikowia_pulcherrima(100); | Metschnikowia | 13 | Saprotroph | \|Undefined Saprotroph\| | Yeast | NULL | Probable | As nectar-tap_saprotroph (Põlme et al. 2020); Decay sugar-rich substrates (Põlme et al. 2020) | Põlme S, et al. 2020. Fungal Diversity 105:1-16 ((https://doi.org/10.1007/s13225-020-00466-2)) |
| Otu50412 | 1 | k__Fungi(100);p__Ascomycota(100);c__Saccharomycetes(100);o__Saccharomycetales(100);f__Metschnikowiaceae(100);g__Metschnikowia(100);s__Metschnikowia_pulcherrima(100); | Metschnikowia | 13 | Saprotroph | \|Undefined Saprotroph\| | Yeast | NULL | Probable | As nectar-tap_saprotroph (Põlme et al. 2020); Decay sugar-rich substrates (Põlme et al. 2020) | Põlme S, et al. 2020. Fungal Diversity 105:1-16 ((https://doi.org/10.1007/s13225-020-00466-2)) |
| Otu50413 | 1 | k__Fungi(100);p__Ascomycota(100);c__Saccharomycetes(100);o__Saccharomycetales(100);f__Metschnikowiaceae(100);g__Metschnikowia(100);s__Metschnikowia_pulcherrima(100); | Metschnikowia | 13 | Saprotroph | \|Undefined Saprotroph\| | Yeast | NULL | Probable | As nectar-tap_saprotroph (Põlme et al. 2020); Decay sugar-rich substrates (Põlme et al. 2020) | Põlme S, et al. 2020. Fungal Diversity 105:1-16 ((https://doi.org/10.1007/s13225-020-00466-2)) |
| Otu50414 | 1 | k__Fungi(100);p__Ascomycota(100);c__Saccharomycetes(100);o__Saccharomycetales(100);f__Metschnikowiaceae(100);g__Metschnikowia(100);s__Metschnikowia_pulcherrima(100); | Metschnikowia | 13 | Saprotroph | \|Undefined Saprotroph\| | Yeast | NULL | Probable | As nectar-tap_saprotroph (Põlme et al. 2020); Decay sugar-rich substrates (Põlme et al. 2020) | Põlme S, et al. 2020. Fungal Diversity 105:1-16 ((https://doi.org/10.1007/s13225-020-00466-2)) |
| Otu50415 | 1 | k__Fungi(100);p__Ascomycota(100);c__Saccharomycetes(100);o__Saccharomycetales(100);f__Metschnikowiaceae(100);g__Metschnikowia(100);s__Metschnikowia_pulcherrima(100); | Metschnikowia | 13 | Saprotroph | \|Undefined Saprotroph\| | Yeast | NULL | Probable | As nectar-tap_saprotroph (Põlme et al. 2020); Decay sugar-rich substrates (Põlme et al. 2020) | Põlme S, et al. 2020. Fungal Diversity 105:1-16 ((https://doi.org/10.1007/s13225-020-00466-2)) |
| Otu50417 | 1 | k__Fungi(100);p__Ascomycota(100);c__Saccharomycetes(100);o__Saccharomycetales(100);f__Metschnikowiaceae(100);g__Metschnikowia(100);g__Metschnikowia_unclassified(100); | Metschnikowia | 13 | Saprotroph | \|Undefined Saprotroph\| | Yeast | NULL | Probable | As nectar-tap_saprotroph (Põlme et al. 2020); Decay sugar-rich substrates (Põlme et al. 2020) | Põlme S, et al. 2020. Fungal Diversity 105:1-16 ((https://doi.org/10.1007/s13225-020-00466-2)) |
| Otu50418 | 1 | k__Fungi(100);p__Ascomycota(100);c__Saccharomycetes(100);o__Saccharomycetales(100);f__Metschnikowiaceae(100);g__Metschnikowia(100);g__Metschnikowia_unclassified(100); | Metschnikowia | 13 | Saprotroph | \|Undefined Saprotroph\| | Yeast | NULL | Probable | As nectar-tap_saprotroph (Põlme et al. 2020); Decay sugar-rich substrates (Põlme et al. 2020) | Põlme S, et al. 2020. Fungal Diversity 105:1-16 ((https://doi.org/10.1007/s13225-020-00466-2)) |
| Otu50419 | 1 | k__Fungi(100);p__Ascomycota(100);c__Saccharomycetes(100);o__Saccharomycetales(100);f__Metschnikowiaceae(100);g__Metschnikowia(100);s__Metschnikowia_pulcherrima(100); | Metschnikowia | 13 | Saprotroph | \|Undefined Saprotroph\| | Yeast | NULL | Probable | As nectar-tap_saprotroph (Põlme et al. 2020); Decay sugar-rich substrates (Põlme et al. 2020) | Põlme S, et al. 2020. Fungal Diversity 105:1-16 ((https://doi.org/10.1007/s13225-020-00466-2)) |
| Otu50420 | 1 | k__Fungi(100);p__Ascomycota(100);c__Saccharomycetes(100);o__Saccharomycetales(100);f__Metschnikowiaceae(100);g__Metschnikowia(100);g__Metschnikowia_unclassified(100); | Metschnikowia | 13 | Saprotroph | \|Undefined Saprotroph\| | Yeast | NULL | Probable | As nectar-tap_saprotroph (Põlme et al. 2020); Decay sugar-rich substrates (Põlme et al. 2020) | Põlme S, et al. 2020. Fungal Diversity 105:1-16 ((https://doi.org/10.1007/s13225-020-00466-2)) |
| Otu50421 | 1 | k__Fungi(100);p__Ascomycota(100);c__Saccharomycetes(100);o__Saccharomycetales(100);f__Metschnikowiaceae(100);g__Metschnikowia(100);s__Metschnikowia_pulcherrima(100); | Metschnikowia | 13 | Saprotroph | \|Undefined Saprotroph\| | Yeast | NULL | Probable | As nectar-tap_saprotroph (Põlme et al. 2020); Decay sugar-rich substrates (Põlme et al. 2020) | Põlme S, et al. 2020. Fungal Diversity 105:1-16 ((https://doi.org/10.1007/s13225-020-00466-2)) |
| Otu50422 | 1 | k__Fungi(100);p__Ascomycota(100);c__Saccharomycetes(100);o__Saccharomycetales(100);f__Metschnikowiaceae(100);g__Metschnikowia(100);s__Metschnikowia_pulcherrima(100); | Metschnikowia | 13 | Saprotroph | \|Undefined Saprotroph\| | Yeast | NULL | Probable | As nectar-tap_saprotroph (Põlme et al. 2020); Decay sugar-rich substrates (Põlme et al. 2020) | Põlme S, et al. 2020. Fungal Diversity 105:1-16 ((https://doi.org/10.1007/s13225-020-00466-2)) |
| Otu50423 | 1 | k__Fungi(100);p__Ascomycota(100);c__Saccharomycetes(100);o__Saccharomycetales(100);f__Metschnikowiaceae(100);g__Metschnikowia(100);s__Metschnikowia_pulcherrima(100); | Metschnikowia | 13 | Saprotroph | \|Undefined Saprotroph\| | Yeast | NULL | Probable | As nectar-tap_saprotroph (Põlme et al. 2020); Decay sugar-rich substrates (Põlme et al. 2020) | Põlme S, et al. 2020. Fungal Diversity 105:1-16 ((https://doi.org/10.1007/s13225-020-00466-2)) |
| Otu50424 | 1 | k__Fungi(100);p__Ascomycota(100);c__Saccharomycetes(100);o__Saccharomycetales(100);f__Metschnikowiaceae(100);g__Metschnikowia(100);s__Metschnikowia_pulcherrima(100); | Metschnikowia | 13 | Saprotroph | \|Undefined Saprotroph\| | Yeast | NULL | Probable | As nectar-tap_saprotroph (Põlme et al. 2020); Decay sugar-rich substrates (Põlme et al. 2020) | Põlme S, et al. 2020. Fungal Diversity 105:1-16 ((https://doi.org/10.1007/s13225-020-00466-2)) |
| Otu50425 | 1 | k__Fungi(100);p__Ascomycota(100);c__Saccharomycetes(100);o__Saccharomycetales(100);f__Metschnikowiaceae(100);g__Metschnikowia(100);s__Metschnikowia_pulcherrima(100); | Metschnikowia | 13 | Saprotroph | \|Undefined Saprotroph\| | Yeast | NULL | Probable | As nectar-tap_saprotroph (Põlme et al. 2020); Decay sugar-rich substrates (Põlme et al. 2020) | Põlme S, et al. 2020. Fungal Diversity 105:1-16 ((https://doi.org/10.1007/s13225-020-00466-2)) |
| Otu50426 | 1 | k__Fungi(100);p__Ascomycota(100);c__Saccharomycetes(100);o__Saccharomycetales(100);f__Metschnikowiaceae(100);g__Metschnikowia(100);s__Metschnikowia_pulcherrima(100); | Metschnikowia | 13 | Saprotroph | \|Undefined Saprotroph\| | Yeast | NULL | Probable | As nectar-tap_saprotroph (Põlme et al. 2020); Decay sugar-rich substrates (Põlme et al. 2020) | Põlme S, et al. 2020. Fungal Diversity 105:1-16 ((https://doi.org/10.1007/s13225-020-00466-2)) |
| Otu50427 | 1 | k__Fungi(100);p__Ascomycota(100);c__Saccharomycetes(100);o__Saccharomycetales(100);f__Metschnikowiaceae(100);g__Metschnikowia(100);g__Metschnikowia_unclassified(100); | Metschnikowia | 13 | Saprotroph | \|Undefined Saprotroph\| | Yeast | NULL | Probable | As nectar-tap_saprotroph (Põlme et al. 2020); Decay sugar-rich substrates (Põlme et al. 2020) | Põlme S, et al. 2020. Fungal Diversity 105:1-16 ((https://doi.org/10.1007/s13225-020-00466-2)) |
| Otu50428 | 1 | k__Fungi(100);p__Ascomycota(100);c__Saccharomycetes(100);o__Saccharomycetales(100);f__Metschnikowiaceae(100);g__Metschnikowia(100);s__Metschnikowia_pulcherrima(100); | Metschnikowia | 13 | Saprotroph | \|Undefined Saprotroph\| | Yeast | NULL | Probable | As nectar-tap_saprotroph (Põlme et al. 2020); Decay sugar-rich substrates (Põlme et al. 2020) | Põlme S, et al. 2020. Fungal Diversity 105:1-16 ((https://doi.org/10.1007/s13225-020-00466-2)) |
| Otu50429 | 1 | k__Fungi(100);p__Ascomycota(100);c__Saccharomycetes(100);o__Saccharomycetales(100);f__Metschnikowiaceae(100);g__Metschnikowia(100);s__Metschnikowia_pulcherrima(100); | Metschnikowia | 13 | Saprotroph | \|Undefined Saprotroph\| | Yeast | NULL | Probable | As nectar-tap_saprotroph (Põlme et al. 2020); Decay sugar-rich substrates (Põlme et al. 2020) | Põlme S, et al. 2020. Fungal Diversity 105:1-16 ((https://doi.org/10.1007/s13225-020-00466-2)) |
| Otu50430 | 1 | k__Fungi(100);p__Ascomycota(100);c__Saccharomycetes(100);o__Saccharomycetales(100);f__Metschnikowiaceae(100);g__Metschnikowia(100);s__Metschnikowia_pulcherrima(100); | Metschnikowia | 13 | Saprotroph | \|Undefined Saprotroph\| | Yeast | NULL | Probable | As nectar-tap_saprotroph (Põlme et al. 2020); Decay sugar-rich substrates (Põlme et al. 2020) | Põlme S, et al. 2020. Fungal Diversity 105:1-16 ((https://doi.org/10.1007/s13225-020-00466-2)) |
| Otu50431 | 1 | k__Fungi(100);p__Ascomycota(100);c__Saccharomycetes(100);o__Saccharomycetales(100);f__Metschnikowiaceae(100);g__Metschnikowia(100);s__Metschnikowia_pulcherrima(100); | Metschnikowia | 13 | Saprotroph | \|Undefined Saprotroph\| | Yeast | NULL | Probable | As nectar-tap_saprotroph (Põlme et al. 2020); Decay sugar-rich substrates (Põlme et al. 2020) | Põlme S, et al. 2020. Fungal Diversity 105:1-16 ((https://doi.org/10.1007/s13225-020-00466-2)) |
| Otu50432 | 1 | k__Fungi(100);p__Ascomycota(100);c__Saccharomycetes(100);o__Saccharomycetales(100);f__Metschnikowiaceae(100);g__Metschnikowia(100);s__Metschnikowia_pulcherrima(100); | Metschnikowia | 13 | Saprotroph | \|Undefined Saprotroph\| | Yeast | NULL | Probable | As nectar-tap_saprotroph (Põlme et al. 2020); Decay sugar-rich substrates (Põlme et al. 2020) | Põlme S, et al. 2020. Fungal Diversity 105:1-16 ((https://doi.org/10.1007/s13225-020-00466-2)) |
| Otu50433 | 1 | k__Fungi(100);p__Ascomycota(100);c__Saccharomycetes(100);o__Saccharomycetales(100);f__Metschnikowiaceae(100);g__Metschnikowia(100);g__Metschnikowia_unclassified(100); | Metschnikowia | 13 | Saprotroph | \|Undefined Saprotroph\| | Yeast | NULL | Probable | As nectar-tap_saprotroph (Põlme et al. 2020); Decay sugar-rich substrates (Põlme et al. 2020) | Põlme S, et al. 2020. Fungal Diversity 105:1-16 ((https://doi.org/10.1007/s13225-020-00466-2)) |
| Otu50434 | 1 | k__Fungi(100);p__Ascomycota(100);c__Saccharomycetes(100);o__Saccharomycetales(100);f__Metschnikowiaceae(100);g__Metschnikowia(100);g__Metschnikowia_unclassified(100); | Metschnikowia | 13 | Saprotroph | \|Undefined Saprotroph\| | Yeast | NULL | Probable | As nectar-tap_saprotroph (Põlme et al. 2020); Decay sugar-rich substrates (Põlme et al. 2020) | Põlme S, et al. 2020. Fungal Diversity 105:1-16 ((https://doi.org/10.1007/s13225-020-00466-2)) |
| Otu50435 | 1 | k__Fungi(100);p__Ascomycota(100);c__Saccharomycetes(100);o__Saccharomycetales(100);f__Metschnikowiaceae(100);g__Metschnikowia(100);s__Metschnikowia_pulcherrima(100); | Metschnikowia | 13 | Saprotroph | \|Undefined Saprotroph\| | Yeast | NULL | Probable | As nectar-tap_saprotroph (Põlme et al. 2020); Decay sugar-rich substrates (Põlme et al. 2020) | Põlme S, et al. 2020. Fungal Diversity 105:1-16 ((https://doi.org/10.1007/s13225-020-00466-2)) |
| Otu50436 | 1 | k__Fungi(100);p__Ascomycota(100);c__Saccharomycetes(100);o__Saccharomycetales(100);f__Metschnikowiaceae(100);g__Metschnikowia(100);s__Metschnikowia_pulcherrima(100); | Metschnikowia | 13 | Saprotroph | \|Undefined Saprotroph\| | Yeast | NULL | Probable | As nectar-tap_saprotroph (Põlme et al. 2020); Decay sugar-rich substrates (Põlme et al. 2020) | Põlme S, et al. 2020. Fungal Diversity 105:1-16 ((https://doi.org/10.1007/s13225-020-00466-2)) |
| Otu50437 | 1 | k__Fungi(100);p__Ascomycota(100);c__Saccharomycetes(100);o__Saccharomycetales(100);f__Metschnikowiaceae(100);g__Metschnikowia(100);s__Metschnikowia_pulcherrima(100); | Metschnikowia | 13 | Saprotroph | \|Undefined Saprotroph\| | Yeast | NULL | Probable | As nectar-tap_saprotroph (Põlme et al. 2020); Decay sugar-rich substrates (Põlme et al. 2020) | Põlme S, et al. 2020. Fungal Diversity 105:1-16 ((https://doi.org/10.1007/s13225-020-00466-2)) |

| | | | | | | | | | | | |
|---|---|---|---|---|---|---|---|---|---|---|---|
| Otu50438 | 1 | k__Fungi(100);p__Ascomycota(100);c__Saccharomycetes(100);o__Saccharomycetales(100);f__Metschnikowiaceae(100);g__Metschnikowia(100);s__Metschnikowia_pulcherrima(100); | Metschnikowia | 13 | Saprotroph | \|Undefined Saprotroph\| | Yeast | NULL | Probable | As nectar-tap_saprotroph (Põlme et al. 2020); Decay sugar-rich substrates (Põlme et al. 2020) | Põlme S, et al. 2020. Fungal Diversity 105:1-16 ((https://doi.org/10.1007/s13225-020-00466-2)) |
| Otu50439 | 1 | k__Fungi(100);p__Ascomycota(100);c__Saccharomycetes(100);o__Saccharomycetales(100);f__Metschnikowiaceae(100);g__Metschnikowia(100);s__Metschnikowia_pulcherrima(100); | Metschnikowia | 13 | Saprotroph | \|Undefined Saprotroph\| | Yeast | NULL | Probable | As nectar-tap_saprotroph (Põlme et al. 2020); Decay sugar-rich substrates (Põlme et al. 2020) | Põlme S, et al. 2020. Fungal Diversity 105:1-16 ((https://doi.org/10.1007/s13225-020-00466-2)) |
| Otu50440 | 1 | k__Fungi(100);p__Ascomycota(100);c__Saccharomycetes(100);o__Saccharomycetales(100);f__Metschnikowiaceae(100);g__Metschnikowia(100);s__Metschnikowia_pulcherrima(100); | Metschnikowia | 13 | Saprotroph | \|Undefined Saprotroph\| | Yeast | NULL | Probable | As nectar-tap_saprotroph (Põlme et al. 2020); Decay sugar-rich substrates (Põlme et al. 2020) | Põlme S, et al. 2020. Fungal Diversity 105:1-16 ((https://doi.org/10.1007/s13225-020-00466-2)) |
| Otu50441 | 1 | k__Fungi(100);p__Ascomycota(100);c__Saccharomycetes(100);o__Saccharomycetales(100);f__Metschnikowiaceae(100);g__Metschnikowia(100);s__Metschnikowia_pulcherrima(100); | Metschnikowia | 13 | Saprotroph | \|Undefined Saprotroph\| | Yeast | NULL | Probable | As nectar-tap_saprotroph (Põlme et al. 2020); Decay sugar-rich substrates (Põlme et al. 2020) | Põlme S, et al. 2020. Fungal Diversity 105:1-16 ((https://doi.org/10.1007/s13225-020-00466-2)) |
| Otu50442 | 1 | k__Fungi(100);p__Ascomycota(100);c__Saccharomycetes(100);o__Saccharomycetales(100);f__Metschnikowiaceae(100);g__Metschnikowia(100);g__Metschnikowia_unclassified(100); | Metschnikowia | 13 | Saprotroph | \|Undefined Saprotroph\| | Yeast | NULL | Probable | As nectar-tap_saprotroph (Põlme et al. 2020); Decay sugar-rich substrates (Põlme et al. 2020) | Põlme S, et al. 2020. Fungal Diversity 105:1-16 ((https://doi.org/10.1007/s13225-020-00466-2)) |
| Otu50443 | 1 | k__Fungi(100);p__Ascomycota(100);c__Saccharomycetes(100);o__Saccharomycetales(100);f__Metschnikowiaceae(100);g__Metschnikowia(100);g__Metschnikowia_unclassified(100); | Metschnikowia | 13 | Saprotroph | \|Undefined Saprotroph\| | Yeast | NULL | Probable | As nectar-tap_saprotroph (Põlme et al. 2020); Decay sugar-rich substrates (Põlme et al. 2020) | Põlme S, et al. 2020. Fungal Diversity 105:1-16 ((https://doi.org/10.1007/s13225-020-00466-2)) |
| Otu50444 | 1 | k__Fungi(100);p__Ascomycota(100);c__Saccharomycetes(100);o__Saccharomycetales(100);f__Metschnikowiaceae(100);g__Metschnikowia(100);s__Metschnikowia_pulcherrima(100); | Metschnikowia | 13 | Saprotroph | \|Undefined Saprotroph\| | Yeast | NULL | Probable | As nectar-tap_saprotroph (Põlme et al. 2020); Decay sugar-rich substrates (Põlme et al. 2020) | Põlme S, et al. 2020. Fungal Diversity 105:1-16 ((https://doi.org/10.1007/s13225-020-00466-2)) |
| Otu50445 | 1 | k__Fungi(100);p__Ascomycota(100);c__Saccharomycetes(100);o__Saccharomycetales(100);f__Metschnikowiaceae(100);g__Metschnikowia(100);g__Metschnikowia_unclassified(100); | Metschnikowia | 13 | Saprotroph | \|Undefined Saprotroph\| | Yeast | NULL | Probable | As nectar-tap_saprotroph (Põlme et al. 2020); Decay sugar-rich substrates (Põlme et al. 2020) | Põlme S, et al. 2020. Fungal Diversity 105:1-16 ((https://doi.org/10.1007/s13225-020-00466-2)) |
| Otu50446 | 1 | k__Fungi(100);p__Ascomycota(100);c__Saccharomycetes(100);o__Saccharomycetales(100);f__Metschnikowiaceae(100);g__Metschnikowia(100);s__Metschnikowia_pulcherrima(100); | Metschnikowia | 13 | Saprotroph | \|Undefined Saprotroph\| | Yeast | NULL | Probable | As nectar-tap_saprotroph (Põlme et al. 2020); Decay sugar-rich substrates (Põlme et al. 2020) | Põlme S, et al. 2020. Fungal Diversity 105:1-16 ((https://doi.org/10.1007/s13225-020-00466-2)) |
| Otu50448 | 1 | k__Fungi(100);p__Ascomycota(100);c__Saccharomycetes(100);o__Saccharomycetales(100);f__Metschnikowiaceae(100);g__Metschnikowia(100);s__Metschnikowia_pulcherrima(100); | Metschnikowia | 13 | Saprotroph | \|Undefined Saprotroph\| | Yeast | NULL | Probable | As nectar-tap_saprotroph (Põlme et al. 2020); Decay sugar-rich substrates (Põlme et al. 2020) | Põlme S, et al. 2020. Fungal Diversity 105:1-16 ((https://doi.org/10.1007/s13225-020-00466-2)) |
| Otu50450 | 1 | k__Fungi(100);p__Ascomycota(100);c__Saccharomycetes(100);o__Saccharomycetales(100);f__Metschnikowiaceae(100);g__Metschnikowia(100);s__Metschnikowia_pulcherrima(100); | Metschnikowia | 13 | Saprotroph | \|Undefined Saprotroph\| | Yeast | NULL | Probable | As nectar-tap_saprotroph (Põlme et al. 2020); Decay sugar-rich substrates (Põlme et al. 2020) | Põlme S, et al. 2020. Fungal Diversity 105:1-16 ((https://doi.org/10.1007/s13225-020-00466-2)) |
| Otu50451 | 1 | k__Fungi(100);p__Ascomycota(100);c__Saccharomycetes(100);o__Saccharomycetales(100);f__Metschnikowiaceae(100);g__Metschnikowia(100);s__Metschnikowia_pulcherrima(100); | Metschnikowia | 13 | Saprotroph | \|Undefined Saprotroph\| | Yeast | NULL | Probable | As nectar-tap_saprotroph (Põlme et al. 2020); Decay sugar-rich substrates (Põlme et al. 2020) | Põlme S, et al. 2020. Fungal Diversity 105:1-16 ((https://doi.org/10.1007/s13225-020-00466-2)) |
| Otu50452 | 1 | k__Fungi(100);p__Ascomycota(100);c__Saccharomycetes(100);o__Saccharomycetales(100);f__Metschnikowiaceae(100);g__Metschnikowia(100);s__Metschnikowia_pulcherrima(100); | Metschnikowia | 13 | Saprotroph | \|Undefined Saprotroph\| | Yeast | NULL | Probable | As nectar-tap_saprotroph (Põlme et al. 2020); Decay sugar-rich substrates (Põlme et al. 2020) | Põlme S, et al. 2020. Fungal Diversity 105:1-16 ((https://doi.org/10.1007/s13225-020-00466-2)) |
| Otu50453 | 1 | k__Fungi(100);p__Ascomycota(100);c__Saccharomycetes(100);o__Saccharomycetales(100);f__Metschnikowiaceae(100);g__Metschnikowia(100);g__Metschnikowia_unclassified(100); | Metschnikowia | 13 | Saprotroph | \|Undefined Saprotroph\| | Yeast | NULL | Probable | As nectar-tap_saprotroph (Põlme et al. 2020); Decay sugar-rich substrates (Põlme et al. 2020) | Põlme S, et al. 2020. Fungal Diversity 105:1-16 ((https://doi.org/10.1007/s13225-020-00466-2)) |
| Otu50454 | 1 | k__Fungi(100);p__Ascomycota(100);c__Saccharomycetes(100);o__Saccharomycetales(100);f__Metschnikowiaceae(100);g__Metschnikowia(100);s__Metschnikowia_pulcherrima(100); | Metschnikowia | 13 | Saprotroph | \|Undefined Saprotroph\| | Yeast | NULL | Probable | As nectar-tap_saprotroph (Põlme et al. 2020); Decay sugar-rich substrates (Põlme et al. 2020) | Põlme S, et al. 2020. Fungal Diversity 105:1-16 ((https://doi.org/10.1007/s13225-020-00466-2)) |
| Otu50455 | 1 | k__Fungi(100);p__Ascomycota(100);c__Saccharomycetes(100);o__Saccharomycetales(100);f__Metschnikowiaceae(100);g__Metschnikowia(100);g__Metschnikowia_unclassified(100); | Metschnikowia | 13 | Saprotroph | \|Undefined Saprotroph\| | Yeast | NULL | Probable | As nectar-tap_saprotroph (Põlme et al. 2020); Decay sugar-rich substrates (Põlme et al. 2020) | Põlme S, et al. 2020. Fungal Diversity 105:1-16 ((https://doi.org/10.1007/s13225-020-00466-2)) |
| Otu50456 | 1 | k__Fungi(100);p__Ascomycota(100);c__Saccharomycetes(100);o__Saccharomycetales(100);f__Metschnikowiaceae(100);g__Metschnikowia(100);g__Metschnikowia_unclassified(100); | Metschnikowia | 13 | Saprotroph | \|Undefined Saprotroph\| | Yeast | NULL | Probable | As nectar-tap_saprotroph (Põlme et al. 2020); Decay sugar-rich substrates (Põlme et al. 2020) | Põlme S, et al. 2020. Fungal Diversity 105:1-16 ((https://doi.org/10.1007/s13225-020-00466-2)) |
| Otu50457 | 1 | k__Fungi(100);p__Ascomycota(100);c__Saccharomycetes(100);o__Saccharomycetales(100);f__Metschnikowiaceae(100);g__Metschnikowia(100);g__Metschnikowia_unclassified(100); | Metschnikowia | 13 | Saprotroph | \|Undefined Saprotroph\| | Yeast | NULL | Probable | As nectar-tap_saprotroph (Põlme et al. 2020); Decay sugar-rich substrates (Põlme et al. 2020) | Põlme S, et al. 2020. Fungal Diversity 105:1-16 ((https://doi.org/10.1007/s13225-020-00466-2)) |
| Otu50458 | 1 | k__Fungi(100);p__Ascomycota(100);c__Saccharomycetes(100);o__Saccharomycetales(100);f__Metschnikowiaceae(100);g__Metschnikowia(100);s__Metschnikowia_pulcherrima(100); | Metschnikowia | 13 | Saprotroph | \|Undefined Saprotroph\| | Yeast | NULL | Probable | As nectar-tap_saprotroph (Põlme et al. 2020); Decay sugar-rich substrates (Põlme et al. 2020) | Põlme S, et al. 2020. Fungal Diversity 105:1-16 ((https://doi.org/10.1007/s13225-020-00466-2)) |
| Otu50459 | 1 | k__Fungi(100);p__Ascomycota(100);c__Saccharomycetes(100);o__Saccharomycetales(100);f__Metschnikowiaceae(100);g__Metschnikowia(100);s__Metschnikowia_pulcherrima(100); | Metschnikowia | 13 | Saprotroph | \|Undefined Saprotroph\| | Yeast | NULL | Probable | As nectar-tap_saprotroph (Põlme et al. 2020); Decay sugar-rich substrates (Põlme et al. 2020) | Põlme S, et al. 2020. Fungal Diversity 105:1-16 ((https://doi.org/10.1007/s13225-020-00466-2)) |
| Otu50460 | 1 | k__Fungi(100);p__Ascomycota(100);c__Saccharomycetes(100);o__Saccharomycetales(100);f__Metschnikowiaceae(100);g__Metschnikowia(100);s__Metschnikowia_pulcherrima(100); | Metschnikowia | 13 | Saprotroph | \|Undefined Saprotroph\| | Yeast | NULL | Probable | As nectar-tap_saprotroph (Põlme et al. 2020); Decay sugar-rich substrates (Põlme et al. 2020) | Põlme S, et al. 2020. Fungal Diversity 105:1-16 ((https://doi.org/10.1007/s13225-020-00466-2)) |
| Otu50461 | 1 | k__Fungi(100);p__Ascomycota(100);c__Saccharomycetes(100);o__Saccharomycetales(100);f__Metschnikowiaceae(100);g__Metschnikowia(100);s__Metschnikowia_pulcherrima(100); | Metschnikowia | 13 | Saprotroph | \|Undefined Saprotroph\| | Yeast | NULL | Probable | As nectar-tap_saprotroph (Põlme et al. 2020); Decay sugar-rich substrates (Põlme et al. 2020) | Põlme S, et al. 2020. Fungal Diversity 105:1-16 ((https://doi.org/10.1007/s13225-020-00466-2)) |
| Otu50462 | 1 | k__Fungi(100);p__Ascomycota(100);c__Saccharomycetes(100);o__Saccharomycetales(100);f__Metschnikowiaceae(100);g__Metschnikowia(100);s__Metschnikowia_pulcherrima(100); | Metschnikowia | 13 | Saprotroph | \|Undefined Saprotroph\| | Yeast | NULL | Probable | As nectar-tap_saprotroph (Põlme et al. 2020); Decay sugar-rich substrates (Põlme et al. 2020) | Põlme S, et al. 2020. Fungal Diversity 105:1-16 ((https://doi.org/10.1007/s13225-020-00466-2)) |
| Otu50463 | 1 | k__Fungi(100);p__Ascomycota(100);c__Saccharomycetes(100);o__Saccharomycetales(100);f__Metschnikowiaceae(100);g__Metschnikowia(100);s__Metschnikowia_pulcherrima(100); | Metschnikowia | 13 | Saprotroph | \|Undefined Saprotroph\| | Yeast | NULL | Probable | As nectar-tap_saprotroph (Põlme et al. 2020); Decay sugar-rich substrates (Põlme et al. 2020) | Põlme S, et al. 2020. Fungal Diversity 105:1-16 ((https://doi.org/10.1007/s13225-020-00466-2)) |
| Otu50464 | 1 | k__Fungi(100);p__Ascomycota(100);c__Saccharomycetes(100);o__Saccharomycetales(100);f__Metschnikowiaceae(100);g__Metschnikowia(100);s__Metschnikowia_pulcherrima(100); | Metschnikowia | 13 | Saprotroph | \|Undefined Saprotroph\| | Yeast | NULL | Probable | As nectar-tap_saprotroph (Põlme et al. 2020); Decay sugar-rich substrates (Põlme et al. 2020) | Põlme S, et al. 2020. Fungal Diversity 105:1-16 ((https://doi.org/10.1007/s13225-020-00466-2)) |
| Otu50465 | 1 | k__Fungi(100);p__Ascomycota(100);c__Saccharomycetes(100);o__Saccharomycetales(100);f__Metschnikowiaceae(100);g__Metschnikowia(100);s__Metschnikowia_pulcherrima(100); | Metschnikowia | 13 | Saprotroph | \|Undefined Saprotroph\| | Yeast | NULL | Probable | As nectar-tap_saprotroph (Põlme et al. 2020); Decay sugar-rich substrates (Põlme et al. 2020) | Põlme S, et al. 2020. Fungal Diversity 105:1-16 ((https://doi.org/10.1007/s13225-020-00466-2)) |
| Otu50466 | 1 | k__Fungi(100);p__Ascomycota(100);c__Saccharomycetes(100);o__Saccharomycetales(100);f__Metschnikowiaceae(100);g__Metschnikowia(100);s__Metschnikowia_pulcherrima(100); | Metschnikowia | 13 | Saprotroph | \|Undefined Saprotroph\| | Yeast | NULL | Probable | As nectar-tap_saprotroph (Põlme et al. 2020); Decay sugar-rich substrates (Põlme et al. 2020) | Põlme S, et al. 2020. Fungal Diversity 105:1-16 ((https://doi.org/10.1007/s13225-020-00466-2)) |
| Otu50467 | 1 | k__Fungi(100);p__Ascomycota(100);c__Saccharomycetes(100);o__Saccharomycetales(100);f__Metschnikowiaceae(100);g__Metschnikowia(100);g__Metschnikowia_unclassified(100); | Metschnikowia | 13 | Saprotroph | \|Undefined Saprotroph\| | Yeast | NULL | Probable | As nectar-tap_saprotroph (Põlme et al. 2020); Decay sugar-rich substrates (Põlme et al. 2020) | Põlme S, et al. 2020. Fungal Diversity 105:1-16 ((https://doi.org/10.1007/s13225-020-00466-2)) |
| Otu50468 | 1 | k__Fungi(100);p__Ascomycota(100);c__Saccharomycetes(100);o__Saccharomycetales(100);f__Metschnikowiaceae(100);g__Metschnikowia(100);s__Metschnikowia_pulcherrima(100); | Metschnikowia | 13 | Saprotroph | \|Undefined Saprotroph\| | Yeast | NULL | Probable | As nectar-tap_saprotroph (Põlme et al. 2020); Decay sugar-rich substrates (Põlme et al. 2020) | Põlme S, et al. 2020. Fungal Diversity 105:1-16 ((https://doi.org/10.1007/s13225-020-00466-2)) |
| Otu50469 | 1 | k__Fungi(100);p__Ascomycota(100);c__Saccharomycetes(100);o__Saccharomycetales(100);f__Saccharomycetaceae(100);g__Kazachstania(100);s__Kazachstania_intestinalis(100); | Kazachstania | 13 | Saprotroph | \|Undefined Saprotroph\| | Yeast | NULL | Probable | As nectar-tap_saprotroph (Põlme et al. 2020); Decay sugar-rich substrates (Põlme et al. 2020) | Põlme S, et al. 2020. Fungal Diversity 105:1-16 ((https://doi.org/10.1007/s13225-020-00466-2)) |
| Otu50470 | 1 | k__Fungi(100);p__Ascomycota(100);c__Saccharomycetes(100);o__Saccharomycetales(100);f__Metschnikowiaceae(100);g__Metschnikowia(100);s__Metschnikowia_pulcherrima(100); | Metschnikowia | 13 | Saprotroph | \|Undefined Saprotroph\| | Yeast | NULL | Probable | As nectar-tap_saprotroph (Põlme et al. 2020); Decay sugar-rich substrates (Põlme et al. 2020) | Põlme S, et al. 2020. Fungal Diversity 105:1-16 ((https://doi.org/10.1007/s13225-020-00466-2)) |
| Otu50471 | 1 | k__Fungi(100);p__Ascomycota(100);c__Saccharomycetes(100);o__Saccharomycetales(100);f__Metschnikowiaceae(100);g__Metschnikowia(100);s__Metschnikowia_pulcherrima(100); | Metschnikowia | 13 | Saprotroph | \|Undefined Saprotroph\| | Yeast | NULL | Probable | As nectar-tap_saprotroph (Põlme et al. 2020); Decay sugar-rich substrates (Põlme et al. 2020) | Põlme S, et al. 2020. Fungal Diversity 105:1-16 ((https://doi.org/10.1007/s13225-020-00466-2)) |
| Otu50472 | 1 | k__Fungi(100);p__Ascomycota(100);c__Saccharomycetes(100);o__Saccharomycetales(100);f__Metschnikowiaceae(100);g__Metschnikowia(100);s__Metschnikowia_pulcherrima(100); | Metschnikowia | 13 | Saprotroph | \|Undefined Saprotroph\| | Yeast | NULL | Probable | As nectar-tap_saprotroph (Põlme et al. 2020); Decay sugar-rich substrates (Põlme et al. 2020) | Põlme S, et al. 2020. Fungal Diversity 105:1-16 ((https://doi.org/10.1007/s13225-020-00466-2)) |
| Otu50473 | 1 | k__Fungi(100);p__Ascomycota(100);c__Saccharomycetes(100);o__Saccharomycetales(100);f__Metschnikowiaceae(100);g__Metschnikowia(100);g__Metschnikowia_unclassified(100); | Metschnikowia | 13 | Saprotroph | \|Undefined Saprotroph\| | Yeast | NULL | Probable | As nectar-tap_saprotroph (Põlme et al. 2020); Decay sugar-rich substrates (Põlme et al. 2020) | Põlme S, et al. 2020. Fungal Diversity 105:1-16 ((https://doi.org/10.1007/s13225-020-00466-2)) |
| Otu50475 | 1 | k__Fungi(100);p__Ascomycota(100);c__Saccharomycetes(100);o__Saccharomycetales(100);f__Metschnikowiaceae(100);g__Metschnikowia(100);s__Metschnikowia_pulcherrima(100); | Metschnikowia | 13 | Saprotroph | \|Undefined Saprotroph\| | Yeast | NULL | Probable | As nectar-tap_saprotroph (Põlme et al. 2020); Decay sugar-rich substrates (Põlme et al. 2020) | Põlme S, et al. 2020. Fungal Diversity 105:1-16 ((https://doi.org/10.1007/s13225-020-00466-2)) |
| Otu50477 | 1 | k__Fungi(100);p__Ascomycota(100);c__Saccharomycetes(100);o__Saccharomycetales(100);f__Metschnikowiaceae(100);g__Metschnikowia(100);s__Metschnikowia_pulcherrima(100); | Metschnikowia | 13 | Saprotroph | \|Undefined Saprotroph\| | Yeast | NULL | Probable | As nectar-tap_saprotroph (Põlme et al. 2020); Decay sugar-rich substrates (Põlme et al. 2020) | Põlme S, et al. 2020. Fungal Diversity 105:1-16 ((https://doi.org/10.1007/s13225-020-00466-2)) |
| Otu50478 | 1 | k__Fungi(100);p__Ascomycota(100);c__Saccharomycetes(100);o__Saccharomycetales(100);f__Metschnikowiaceae(100);g__Metschnikowia(100);s__Metschnikowia_pulcherrima(100); | Metschnikowia | 13 | Saprotroph | \|Undefined Saprotroph\| | Yeast | NULL | Probable | As nectar-tap_saprotroph (Põlme et al. 2020); Decay sugar-rich substrates (Põlme et al. 2020) | Põlme S, et al. 2020. Fungal Diversity 105:1-16 ((https://doi.org/10.1007/s13225-020-00466-2)) |
| Otu50479 | 1 | k__Fungi(100);p__Ascomycota(100);c__Saccharomycetes(100);o__Saccharomycetales(100);f__Metschnikowiaceae(100);g__Metschnikowia(100);s__Metschnikowia_pulcherrima(100); | Metschnikowia | 13 | Saprotroph | \|Undefined Saprotroph\| | Yeast | NULL | Probable | As nectar-tap_saprotroph (Põlme et al. 2020); Decay sugar-rich substrates (Põlme et al. 2020) | Põlme S, et al. 2020. Fungal Diversity 105:1-16 ((https://doi.org/10.1007/s13225-020-00466-2)) |
| Otu50480 | 1 | k__Fungi(100);p__Ascomycota(100);c__Saccharomycetes(100);o__Saccharomycetales(100);f__Metschnikowiaceae(100);g__Metschnikowia(100);s__Metschnikowia_pulcherrima(100); | Metschnikowia | 13 | Saprotroph | \|Undefined Saprotroph\| | Yeast | NULL | Probable | As nectar-tap_saprotroph (Põlme et al. 2020); Decay sugar-rich substrates (Põlme et al. 2020) | Põlme S, et al. 2020. Fungal Diversity 105:1-16 ((https://doi.org/10.1007/s13225-020-00466-2)) |
| Otu50481 | 1 | k__Fungi(100);p__Ascomycota(100);c__Saccharomycetes(100);o__Saccharomycetales(100);f__Metschnikowiaceae(100);g__Metschnikowia(100);s__Metschnikowia_pulcherrima(100); | Metschnikowia | 13 | Saprotroph | \|Undefined Saprotroph\| | Yeast | NULL | Probable | As nectar-tap_saprotroph (Põlme et al. 2020); Decay sugar-rich substrates (Põlme et al. 2020) | Põlme S, et al. 2020. Fungal Diversity 105:1-16 ((https://doi.org/10.1007/s13225-020-00466-2)) |
| Otu50482 | 1 | k__Fungi(100);p__Ascomycota(100);c__Saccharomycetes(100);o__Saccharomycetales(100);f__Metschnikowiaceae(100);g__Metschnikowia(100);g__Metschnikowia_unclassified(100); | Metschnikowia | 13 | Saprotroph | \|Undefined Saprotroph\| | Yeast | NULL | Probable | As nectar-tap_saprotroph (Põlme et al. 2020); Decay sugar-rich substrates (Põlme et al. 2020) | Põlme S, et al. 2020. Fungal Diversity 105:1-16 ((https://doi.org/10.1007/s13225-020-00466-2)) |
| Otu50484 | 1 | k__Fungi(100);p__Ascomycota(100);c__Saccharomycetes(100);o__Saccharomycetales(100);f__Metschnikowiaceae(100);g__Metschnikowia(100);g__Metschnikowia_unclassified(100); | Metschnikowia | 13 | Saprotroph | \|Undefined Saprotroph\| | Yeast | NULL | Probable | As nectar-tap_saprotroph (Põlme et al. 2020); Decay sugar-rich substrates (Põlme et al. 2020) | Põlme S, et al. 2020. Fungal Diversity 105:1-16 ((https://doi.org/10.1007/s13225-020-00466-2)) |
| Otu50490 | 1 | k__Fungi(100);p__Ascomycota(100);c__Saccharomycetes(100);o__Saccharomycetales(100);f__Metschnikowiaceae(100);g__Metschnikowia(100);g__Metschnikowia_unclassified(100); | Metschnikowia | 13 | Saprotroph | \|Undefined Saprotroph\| | Yeast | NULL | Probable | As nectar-tap_saprotroph (Põlme et al. 2020); Decay sugar-rich substrates (Põlme et al. 2020) | Põlme S, et al. 2020. Fungal Diversity 105:1-16 ((https://doi.org/10.1007/s13225-020-00466-2)) |
| Otu50491 | 1 | k__Fungi(100);p__Ascomycota(100);c__Saccharomycetes(100);o__Saccharomycetales(100);f__Metschnikowiaceae(100);g__Metschnikowia(100);s__Metschnikowia_pulcherrima(100); | Metschnikowia | 13 | Saprotroph | \|Undefined Saprotroph\| | Yeast | NULL | Probable | As nectar-tap_saprotroph (Põlme et al. 2020); Decay sugar-rich substrates (Põlme et al. 2020) | Põlme S, et al. 2020. Fungal Diversity 105:1-16 ((https://doi.org/10.1007/s13225-020-00466-2)) |
| Otu50492 | 1 | k__Fungi(100);p__Ascomycota(100);c__Saccharomycetes(100);o__Saccharomycetales(100);f__Metschnikowiaceae(100);g__Metschnikowia(100);s__Metschnikowia_pulcherrima(100); | Metschnikowia | 13 | Saprotroph | \|Undefined Saprotroph\| | Yeast | NULL | Probable | As nectar-tap_saprotroph (Põlme et al. 2020); Decay sugar-rich substrates (Põlme et al. 2020) | Põlme S, et al. 2020. Fungal Diversity 105:1-16 ((https://doi.org/10.1007/s13225-020-00466-2)) |
| Otu50493 | 1 | k__Fungi(100);p__Ascomycota(100);c__Saccharomycetes(100);o__Saccharomycetales(100);f__Metschnikowiaceae(100);g__Metschnikowia(100);s__Metschnikowia_pulcherrima(100); | Metschnikowia | 13 | Saprotroph | \|Undefined Saprotroph\| | Yeast | NULL | Probable | As nectar-tap_saprotroph (Põlme et al. 2020); Decay sugar-rich substrates (Põlme et al. 2020) | Põlme S, et al. 2020. Fungal Diversity 105:1-16 ((https://doi.org/10.1007/s13225-020-00466-2)) |
| Otu50495 | 1 | k__Fungi(100);p__Ascomycota(100);c__Saccharomycetes(100);o__Saccharomycetales(100);f__Metschnikowiaceae(100);g__Metschnikowia(100);g__Metschnikowia_unclassified(100); | Metschnikowia | 13 | Saprotroph | \|Undefined Saprotroph\| | Yeast | NULL | Probable | As nectar-tap_saprotroph (Põlme et al. 2020); Decay sugar-rich substrates (Põlme et al. 2020) | Põlme S, et al. 2020. Fungal Diversity 105:1-16 ((https://doi.org/10.1007/s13225-020-00466-2)) |
| Otu50496 | 1 | k__Fungi(100);p__Ascomycota(100);c__Saccharomycetes(100);o__Saccharomycetales(100);f__Metschnikowiaceae(100);g__Metschnikowia(100);s__Metschnikowia_pulcherrima(100); | Metschnikowia | 13 | Saprotroph | \|Undefined Saprotroph\| | Yeast | NULL | Probable | As nectar-tap_saprotroph (Põlme et al. 2020); Decay sugar-rich substrates (Põlme et al. 2020) | Põlme S, et al. 2020. Fungal Diversity 105:1-16 ((https://doi.org/10.1007/s13225-020-00466-2)) |
| Otu50497 | 1 | k__Fungi(100);p__Ascomycota(100);c__Saccharomycetes(100);o__Saccharomycetales(100);f__Metschnikowiaceae(100);g__Metschnikowia(100);s__Metschnikowia_pulcherrima(100); | Metschnikowia | 13 | Saprotroph | \|Undefined Saprotroph\| | Yeast | NULL | Probable | As nectar-tap_saprotroph (Põlme et al. 2020); Decay sugar-rich substrates (Põlme et al. 2020) | Põlme S, et al. 2020. Fungal Diversity 105:1-16 ((https://doi.org/10.1007/s13225-020-00466-2)) |
| Otu50498 | 1 | k__Fungi(100);p__Ascomycota(100);c__Saccharomycetes(100);o__Saccharomycetales(100);f__Metschnikowiaceae(100);g__Metschnikowia(100);s__Metschnikowia_pulcherrima(100); | Metschnikowia | 13 | Saprotroph | \|Undefined Saprotroph\| | Yeast | NULL | Probable | As nectar-tap_saprotroph (Põlme et al. 2020); Decay sugar-rich substrates (Põlme et al. 2020) | Põlme S, et al. 2020. Fungal Diversity 105:1-16 ((https://doi.org/10.1007/s13225-020-00466-2)) |
| Otu50499 | 1 | k__Fungi(100);p__Ascomycota(100);c__Saccharomycetes(100);o__Saccharomycetales(100);f__Metschnikowiaceae(100);g__Metschnikowia(100);s__Metschnikowia_pulcherrima(100); | Metschnikowia | 13 | Saprotroph | \|Undefined Saprotroph\| | Yeast | NULL | Probable | As nectar-tap_saprotroph (Põlme et al. 2020); Decay sugar-rich substrates (Põlme et al. 2020) | Põlme S, et al. 2020. Fungal Diversity 105:1-16 ((https://doi.org/10.1007/s13225-020-00466-2)) |
| Otu50500 | 1 | k__Fungi(100);p__Ascomycota(100);c__Saccharomycetes(100);o__Saccharomycetales(100);f__Metschnikowiaceae(100);g__Metschnikowia(100);s__Metschnikowia_pulcherrima(100); | Metschnikowia | 13 | Saprotroph | \|Undefined Saprotroph\| | Yeast | NULL | Probable | As nectar-tap_saprotroph (Põlme et al. 2020); Decay sugar-rich substrates (Põlme et al. 2020) | Põlme S, et al. 2020. Fungal Diversity 105:1-16 ((https://doi.org/10.1007/s13225-020-00466-2)) |
| Otu50501 | 1 | k__Fungi(100);p__Ascomycota(100);c__Saccharomycetes(100);o__Saccharomycetales(100);f__Metschnikowiaceae(100);g__Metschnikowia(100);s__Metschnikowia_pulcherrima(100); | Metschnikowia | 13 | Saprotroph | \|Undefined Saprotroph\| | Yeast | NULL | Probable | As nectar-tap_saprotroph (Põlme et al. 2020); Decay sugar-rich substrates (Põlme et al. 2020) | Põlme S, et al. 2020. Fungal Diversity 105:1-16 ((https://doi.org/10.1007/s13225-020-00466-2)) |
| Otu50502 | 1 | k__Fungi(100);p__Ascomycota(100);c__Saccharomycetes(100);o__Saccharomycetales(100);f__Metschnikowiaceae(100);g__Metschnikowia(100);g__Metschnikowia_unclassified(100); | Metschnikowia | 13 | Saprotroph | \|Undefined Saprotroph\| | Yeast | NULL | Probable | As nectar-tap_saprotroph (Põlme et al. 2020); Decay sugar-rich substrates (Põlme et al. 2020) | Põlme S, et al. 2020. Fungal Diversity 105:1-16 ((https://doi.org/10.1007/s13225-020-00466-2)) |
| Otu50503 | 1 | k__Fungi(100);p__Ascomycota(100);c__Saccharomycetes(100);o__Saccharomycetales(100);f__Metschnikowiaceae(100);g__Metschnikowia(100);s__Metschnikowia_pulcherrima(100); | Metschnikowia | 13 | Saprotroph | \|Undefined Saprotroph\| | Yeast | NULL | Probable | As nectar-tap_saprotroph (Põlme et al. 2020); Decay sugar-rich substrates (Põlme et al. 2020) | Põlme S, et al. 2020. Fungal Diversity 105:1-16 ((https://doi.org/10.1007/s13225-020-00466-2)) |
| Otu50504 | 1 | k__Fungi(100);p__Ascomycota(100);c__Saccharomycetes(100);o__Saccharomycetales(100);f__Metschnikowiaceae(100);g__Metschnikowia(100);s__Metschnikowia_pulcherrima(100); | Metschnikowia | 13 | Saprotroph | \|Undefined Saprotroph\| | Yeast | NULL | Probable | As nectar-tap_saprotroph (Põlme et al. 2020); Decay sugar-rich substrates (Põlme et al. 2020) | Põlme S, et al. 2020. Fungal Diversity 105:1-16 ((https://doi.org/10.1007/s13225-020-00466-2)) |
| Otu50505 | 1 | k__Fungi(100);p__Ascomycota(100);c__Saccharomycetes(100);o__Saccharomycetales(100);f__Metschnikowiaceae(100);g__Metschnikowia(100);s__Metschnikowia_pulcherrima(100); | Metschnikowia | 13 | Saprotroph | \|Undefined Saprotroph\| | Yeast | NULL | Probable | As nectar-tap_saprotroph (Põlme et al. 2020); Decay sugar-rich substrates (Põlme et al. 2020) | Põlme S, et al. 2020. Fungal Diversity 105:1-16 ((https://doi.org/10.1007/s13225-020-00466-2)) |
| Otu50506 | 1 | k__Fungi(100);p__Ascomycota(100);c__Saccharomycetes(100);o__Saccharomycetales(100);f__Metschnikowiaceae(100);g__Metschnikowia(100);s__Metschnikowia_pulcherrima(100); | Metschnikowia | 13 | Saprotroph | \|Undefined Saprotroph\| | Yeast | NULL | Probable | As nectar-tap_saprotroph (Põlme et al. 2020); Decay sugar-rich substrates (Põlme et al. 2020) | Põlme S, et al. 2020. Fungal Diversity 105:1-16 ((https://doi.org/10.1007/s13225-020-00466-2)) |
| Otu50507 | 1 | k__Fungi(100);p__Ascomycota(100);c__Saccharomycetes(100);o__Saccharomycetales(100);f__Metschnikowiaceae(100);g__Metschnikowia(100);s__Metschnikowia_pulcherrima(100); | Metschnikowia | 13 | Saprotroph | \|Undefined Saprotroph\| | Yeast | NULL | Probable | As nectar-tap_saprotroph (Põlme et al. 2020); Decay sugar-rich substrates (Põlme et al. 2020) | Põlme S, et al. 2020. Fungal Diversity 105:1-16 ((https://doi.org/10.1007/s13225-020-00466-2)) |
| Otu50508 | 1 | k__Fungi(100);p__Ascomycota(100);c__Saccharomycetes(100);o__Saccharomycetales(100);f__Metschnikowiaceae(100);g__Metschnikowia(100);s__Metschnikowia_pulcherrima(100); | Metschnikowia | 13 | Saprotroph | \|Undefined Saprotroph\| | Yeast | NULL | Probable | As nectar-tap_saprotroph (Põlme et al. 2020); Decay sugar-rich substrates (Põlme et al. 2020) | Põlme S, et al. 2020. Fungal Diversity 105:1-16 ((https://doi.org/10.1007/s13225-020-00466-2)) |
| Otu50509 | 1 | k__Fungi(100);p__Ascomycota(100);c__Saccharomycetes(100);o__Saccharomycetales(100);f__Metschnikowiaceae(100);g__Metschnikowia(100);s__Metschnikowia_pulcherrima(100); | Metschnikowia | 13 | Saprotroph | \|Undefined Saprotroph\| | Yeast | NULL | Probable | As nectar-tap_saprotroph (Põlme et al. 2020); Decay sugar-rich substrates (Põlme et al. 2020) | Põlme S, et al. 2020. Fungal Diversity 105:1-16 ((https://doi.org/10.1007/s13225-020-00466-2)) |
| Otu50510 | 1 | k__Fungi(100);p__Ascomycota(100);c__Saccharomycetes(100);o__Saccharomycetales(100);f__Metschnikowiaceae(100);g__Metschnikowia(100);g__Metschnikowia_unclassified(100); | Metschnikowia | 13 | Saprotroph | \|Undefined Saprotroph\| | Yeast | NULL | Probable | As nectar-tap_saprotroph (Põlme et al. 2020); Decay sugar-rich substrates (Põlme et al. 2020) | Põlme S, et al. 2020. Fungal Diversity 105:1-16 ((https://doi.org/10.1007/s13225-020-00466-2)) |
| Otu50512 | 1 | k__Fungi(100);p__Ascomycota(100);c__Saccharomycetes(100);o__Saccharomycetales(100);f__Metschnikowiaceae(100);g__Metschnikowia(100);g__Metschnikowia_unclassified(100); | Metschnikowia | 13 | Saprotroph | \|Undefined Saprotroph\| | Yeast | NULL | Probable | As nectar-tap_saprotroph (Põlme et al. 2020); Decay sugar-rich substrates (Põlme et al. 2020) | Põlme S, et al. 2020. Fungal Diversity 105:1-16 ((https://doi.org/10.1007/s13225-020-00466-2)) |
| Otu50513 | 1 | k__Fungi(100);p__Ascomycota(100);c__Saccharomycetes(100);o__Saccharomycetales(100);f__Metschnikowiaceae(100);g__Metschnikowia(100);s__Metschnikowia_pulcherrima(100); | Metschnikowia | 13 | Saprotroph | \|Undefined Saprotroph\| | Yeast | NULL | Probable | As nectar-tap_saprotroph (Põlme et al. 2020); Decay sugar-rich substrates (Põlme et al. 2020) | Põlme S, et al. 2020. Fungal Diversity 105:1-16 ((https://doi.org/10.1007/s13225-020-00466-2)) |
| Otu50514 | 1 | k__Fungi(100);p__Ascomycota(100);c__Saccharomycetes(100);o__Saccharomycetales(100);f__Metschnikowiaceae(100);g__Metschnikowia(100);s__Metschnikowia_pulcherrima(100); | Metschnikowia | 13 | Saprotroph | \|Undefined Saprotroph\| | Yeast | NULL | Probable | As nectar-tap_saprotroph (Põlme et al. 2020); Decay sugar-rich substrates (Põlme et al. 2020) | Põlme S, et al. 2020. Fungal Diversity 105:1-16 ((https://doi.org/10.1007/s13225-020-00466-2)) |
| Otu50515 | 1 | k__Fungi(100);p__Ascomycota(100);c__Saccharomycetes(100);o__Saccharomycetales(100);f__Metschnikowiaceae(100);g__Metschnikowia(100);s__Metschnikowia_pulcherrima(100); | Metschnikowia | 13 | Saprotroph | \|Undefined Saprotroph\| | Yeast | NULL | Probable | As nectar-tap_saprotroph (Põlme et al. 2020); Decay sugar-rich substrates (Põlme et al. 2020) | Põlme S, et al. 2020. Fungal Diversity 105:1-16 ((https://doi.org/10.1007/s13225-020-00466-2)) |
| Otu50516 | 1 | k__Fungi(100);p__Ascomycota(100);c__Saccharomycetes(100);o__Saccharomycetales(100);f__Metschnikowiaceae(100);g__Metschnikowia(100);s__Metschnikowia_pulcherrima(100); | Metschnikowia | 13 | Saprotroph | \|Undefined Saprotroph\| | Yeast | NULL | Probable | As nectar-tap_saprotroph (Põlme et al. 2020); Decay sugar-rich substrates (Põlme et al. 2020) | Põlme S, et al. 2020. Fungal Diversity 105:1-16 ((https://doi.org/10.1007/s13225-020-00466-2)) |
| Otu50517 | 1 | k__Fungi(100);p__Ascomycota(100);c__Saccharomycetes(100);o__Saccharomycetales(100);f__Metschnikowiaceae(100);g__Metschnikowia(100);g__Metschnikowia_unclassified(100); | Metschnikowia | 13 | Saprotroph | \|Undefined Saprotroph\| | Yeast | NULL | Probable | As nectar-tap_saprotroph (Põlme et al. 2020); Decay sugar-rich substrates (Põlme et al. 2020) | Põlme S, et al. 2020. Fungal Diversity 105:1-16 ((https://doi.org/10.1007/s13225-020-00466-2)) |
| Otu50518 | 1 | k__Fungi(100);p__Ascomycota(100);c__Saccharomycetes(100);o__Saccharomycetales(100);f__Metschnikowiaceae(100);g__Metschnikowia(100);s__Metschnikowia_pulcherrima(100); | Metschnikowia | 13 | Saprotroph | \|Undefined Saprotroph\| | Yeast | NULL | Probable | As nectar-tap_saprotroph (Põlme et al. 2020); Decay sugar-rich substrates (Põlme et al. 2020) | Põlme S, et al. 2020. Fungal Diversity 105:1-16 ((https://doi.org/10.1007/s13225-020-00466-2)) |
| Otu50519 | 1 | k__Fungi(100);p__Ascomycota(100);c__Saccharomycetes(100);o__Saccharomycetales(100);f__Metschnikowiaceae(100);g__Metschnikowia(100);s__Metschnikowia_pulcherrima(100); | Metschnikowia | 13 | Saprotroph | \|Undefined Saprotroph\| | Yeast | NULL | Probable | As nectar-tap_saprotroph (Põlme et al. 2020); Decay sugar-rich substrates (Põlme et al. 2020) | Põlme S, et al. 2020. Fungal Diversity 105:1-16 ((https://doi.org/10.1007/s13225-020-00466-2)) |
| Otu50520 | 1 | k__Fungi(100);p__Ascomycota(100);c__Saccharomycetes(100);o__Saccharomycetales(100);f__Metschnikowiaceae(100);g__Metschnikowia(100);s__Metschnikowia_pulcherrima(100); | Metschnikowia | 13 | Saprotroph | \|Undefined Saprotroph\| | Yeast | NULL | Probable | As nectar-tap_saprotroph (Põlme et al. 2020); Decay sugar-rich substrates (Põlme et al. 2020) | Põlme S, et al. 2020. Fungal Diversity 105:1-16 ((https://doi.org/10.1007/s13225-020-00466-2)) |
| Otu50521 | 1 | k__Fungi(100);p__Ascomycota(100);c__Saccharomycetes(100);o__Saccharomycetales(100);f__Metschnikowiaceae(100);g__Metschnikowia(100);s__Metschnikowia_pulcherrima(100); | Metschnikowia | 13 | Saprotroph | \|Undefined Saprotroph\| | Yeast | NULL | Probable | As nectar-tap_saprotroph (Põlme et al. 2020); Decay sugar-rich substrates (Põlme et al. 2020) | Põlme S, et al. 2020. Fungal Diversity 105:1-16 ((https://doi.org/10.1007/s13225-020-00466-2)) |
| Otu50522 | 1 | k__Fungi(100);p__Ascomycota(100);c__Saccharomycetes(100);o__Saccharomycetales(100);f__Metschnikowiaceae(100);g__Metschnikowia(100);g__Metschnikowia_unclassified(100); | Metschnikowia | 13 | Saprotroph | \|Undefined Saprotroph\| | Yeast | NULL | Probable | As nectar-tap_saprotroph (Põlme et al. 2020); Decay sugar-rich substrates (Põlme et al. 2020) | Põlme S, et al. 2020. Fungal Diversity 105:1-16 ((https://doi.org/10.1007/s13225-020-00466-2)) |
| Otu50523 | 1 | k__Fungi(100);p__Ascomycota(100);c__Saccharomycetes(100);o__Saccharomycetales(100);f__Metschnikowiaceae(100);g__Metschnikowia(100);s__Metschnikowia_pulcherrima(100); | Metschnikowia | 13 | Saprotroph | \|Undefined Saprotroph\| | Yeast | NULL | Probable | As nectar-tap_saprotroph (Põlme et al. 2020); Decay sugar-rich substrates (Põlme et al. 2020) | Põlme S, et al. 2020. Fungal Diversity 105:1-16 ((https://doi.org/10.1007/s13225-020-00466-2)) |
| Otu50524 | 1 | k__Fungi(100);p__Ascomycota(100);c__Saccharomycetes(100);o__Saccharomycetales(100);f__Metschnikowiaceae(100);g__Metschnikowia(100);g__Metschnikowia_unclassified(100); | Metschnikowia | 13 | Saprotroph | \|Undefined Saprotroph\| | Yeast | NULL | Probable | As nectar-tap_saprotroph (Põlme et al. 2020); Decay sugar-rich substrates (Põlme et al. 2020) | Põlme S, et al. 2020. Fungal Diversity 105:1-16 ((https://doi.org/10.1007/s13225-020-00466-2)) |
| Otu50525 | 1 | k__Fungi(100);p__Ascomycota(100);c__Saccharomycetes(100);o__Saccharomycetales(100);f__Metschnikowiaceae(100);g__Metschnikowia(100);g__Metschnikowia_unclassified(100); | Metschnikowia | 13 | Saprotroph | \|Undefined Saprotroph\| | Yeast | NULL | Probable | As nectar-tap_saprotroph (Põlme et al. 2020); Decay sugar-rich substrates (Põlme et al. 2020) | Põlme S, et al. 2020. Fungal Diversity 105:1-16 ((https://doi.org/10.1007/s13225-020-00466-2)) |
| Otu50526 | 1 | k__Fungi(100);p__Ascomycota(100);c__Saccharomycetes(100);o__Saccharomycetales(100);f__Metschnikowiaceae(100);g__Metschnikowia(100);s__Metschnikowia_pulcherrima(100); | Metschnikowia | 13 | Saprotroph | \|Undefined Saprotroph\| | Yeast | NULL | Probable | As nectar-tap_saprotroph (Põlme et al. 2020); Decay sugar-rich substrates (Põlme et al. 2020) | Põlme S, et al. 2020. Fungal Diversity 105:1-16 ((https://doi.org/10.1007/s13225-020-00466-2)) |
| Otu50527 | 1 | k__Fungi(100);p__Ascomycota(100);c__Saccharomycetes(100);o__Saccharomycetales(100);f__Metschnikowiaceae(100);g__Metschnikowia(100);g__Metschnikowia_unclassified(100); | Metschnikowia | 13 | Saprotroph | \|Undefined Saprotroph\| | Yeast | NULL | Probable | As nectar-tap_saprotroph (Põlme et al. 2020); Decay sugar-rich substrates (Põlme et al. 2020) | Põlme S, et al. 2020. Fungal Diversity 105:1-16 ((https://doi.org/10.1007/s13225-020-00466-2)) |
| Otu50528 | 1 | k__Fungi(100);p__Ascomycota(100);c__Saccharomycetes(100);o__Saccharomycetales(100);f__Metschnikowiaceae(100);g__Metschnikowia(100);s__Metschnikowia_pulcherrima(100); | Metschnikowia | 13 | Saprotroph | \|Undefined Saprotroph\| | Yeast | NULL | Probable | As nectar-tap_saprotroph (Põlme et al. 2020); Decay sugar-rich substrates (Põlme et al. 2020) | Põlme S, et al. 2020. Fungal Diversity 105:1-16 ((https://doi.org/10.1007/s13225-020-00466-2)) |
| Otu50529 | 1 | k__Fungi(100);p__Ascomycota(100);c__Saccharomycetes(100);o__Saccharomycetales(100);f__Metschnikowiaceae(100);g__Metschnikowia(100);s__Metschnikowia_pulcherrima(100); | Metschnikowia | 13 | Saprotroph | \|Undefined Saprotroph\| | Yeast | NULL | Probable | As nectar-tap_saprotroph (Põlme et al. 2020); Decay sugar-rich substrates (Põlme et al. 2020) | Põlme S, et al. 2020. Fungal Diversity 105:1-16 ((https://doi.org/10.1007/s13225-020-00466-2)) |
| Otu50530 | 1 | k__Fungi(100);p__Ascomycota(100);c__Saccharomycetes(100);o__Saccharomycetales(100);f__Metschnikowiaceae(100);g__Metschnikowia(100);s__Metschnikowia_pulcherrima(100); | Metschnikowia | 13 | Saprotroph | \|Undefined Saprotroph\| | Yeast | NULL | Probable | As nectar-tap_saprotroph (Põlme et al. 2020); Decay sugar-rich substrates (Põlme et al. 2020) | Põlme S, et al. 2020. Fungal Diversity 105:1-16 ((https://doi.org/10.1007/s13225-020-00466-2)) |
| Otu50531 | 1 | k__Fungi(100);p__Ascomycota(100);c__Saccharomycetes(100);o__Saccharomycetales(100);f__Metschnikowiaceae(100);g__Metschnikowia(100);s__Metschnikowia_pulcherrima(100); | Metschnikowia | 13 | Saprotroph | \|Undefined Saprotroph\| | Yeast | NULL | Probable | As nectar-tap_saprotroph (Põlme et al. 2020); Decay sugar-rich substrates (Põlme et al. 2020) | Põlme S, et al. 2020. Fungal Diversity 105:1-16 ((https://doi.org/10.1007/s13225-020-00466-2)) |
| Otu50532 | 1 | k__Fungi(100);p__Ascomycota(100);c__Saccharomycetes(100);o__Saccharomycetales(100);f__Metschnikowiaceae(100);g__Metschnikowia(100);g__Metschnikowia_unclassified(100); | Metschnikowia | 13 | Saprotroph | \|Undefined Saprotroph\| | Yeast | NULL | Probable | As nectar-tap_saprotroph (Põlme et al. 2020); Decay sugar-rich substrates (Põlme et al. 2020) | Põlme S, et al. 2020. Fungal Diversity 105:1-16 ((https://doi.org/10.1007/s13225-020-00466-2)) |
| Otu50533 | 1 | k__Fungi(100);p__Ascomycota(100);c__Saccharomycetes(100);o__Saccharomycetales(100);f__Metschnikowiaceae(100);g__Metschnikowia(100);s__Metschnikowia_pulcherrima(100); | Metschnikowia | 13 | Saprotroph | \|Undefined Saprotroph\| | Yeast | NULL | Probable | As nectar-tap_saprotroph (Põlme et al. 2020); Decay sugar-rich substrates (Põlme et al. 2020) | Põlme S, et al. 2020. Fungal Diversity 105:1-16 ((https://doi.org/10.1007/s13225-020-00466-2)) |
| Otu50534 | 1 | k__Fungi(100);p__Ascomycota(100);c__Saccharomycetes(100);o__Saccharomycetales(100);f__Metschnikowiaceae(100);g__Metschnikowia(100);g__Metschnikowia_unclassified(100); | Metschnikowia | 13 | Saprotroph | \|Undefined Saprotroph\| | Yeast | NULL | Probable | As nectar-tap_saprotroph (Põlme et al. 2020); Decay sugar-rich substrates (Põlme et al. 2020) | Põlme S, et al. 2020. Fungal Diversity 105:1-16 ((https://doi.org/10.1007/s13225-020-00466-2)) |
| Otu50535 | 1 | k__Fungi(100);p__Ascomycota(100);c__Saccharomycetes(100);o__Saccharomycetales(100);f__Metschnikowiaceae(100);g__Metschnikowia(100);s__Metschnikowia_pulcherrima(100); | Metschnikowia | 13 | Saprotroph | \|Undefined Saprotroph\| | Yeast | NULL | Probable | As nectar-tap_saprotroph (Põlme et al. 2020); Decay sugar-rich substrates (Põlme et al. 2020) | Põlme S, et al. 2020. Fungal Diversity 105:1-16 ((https://doi.org/10.1007/s13225-020-00466-2)) |
| Otu50536 | 1 | k__Fungi(100);p__Ascomycota(100);c__Saccharomycetes(100);o__Saccharomycetales(100);f__Metschnikowiaceae(100);g__Metschnikowia(100);s__Metschnikowia_pulcherrima(100); | Metschnikowia | 13 | Saprotroph | \|Undefined Saprotroph\| | Yeast | NULL | Probable | As nectar-tap_saprotroph (Põlme et al. 2020); Decay sugar-rich substrates (Põlme et al. 2020) | Põlme S, et al. 2020. Fungal Diversity 105:1-16 ((https://doi.org/10.1007/s13225-020-00466-2)) |
| Otu50537 | 1 | k__Fungi(100);p__Ascomycota(100);c__Saccharomycetes(100);o__Saccharomycetales(100);f__Metschnikowiaceae(100);g__Metschnikowia(100);s__Metschnikowia_pulcherrima(100); | Metschnikowia | 13 | Saprotroph | \|Undefined Saprotroph\| | Yeast | NULL | Probable | As nectar-tap_saprotroph (Põlme et al. 2020); Decay sugar-rich substrates (Põlme et al. 2020) | Põlme S, et al. 2020. Fungal Diversity 105:1-16 ((https://doi.org/10.1007/s13225-020-00466-2)) |
| Otu50538 | 1 | k__Fungi(100);p__Ascomycota(100);c__Saccharomycetes(100);o__Saccharomycetales(100);f__Metschnikowiaceae(100);g__Metschnikowia(100);s__Metschnikowia_pulcherrima(100); | Metschnikowia | 13 | Saprotroph | \|Undefined Saprotroph\| | Yeast | NULL | Probable | As nectar-tap_saprotroph (Põlme et al. 2020); Decay sugar-rich substrates (Põlme et al. 2020) | Põlme S, et al. 2020. Fungal Diversity 105:1-16 ((https://doi.org/10.1007/s13225-020-00466-2)) |
| Otu50539 | 1 | k__Fungi(100);p__Ascomycota(100);c__Saccharomycetes(100);o__Saccharomycetales(100);f__Metschnikowiaceae(100);g__Metschnikowia(100);s__Metschnikowia_pulcherrima(100); | Metschnikowia | 13 | Saprotroph | \|Undefined Saprotroph\| | Yeast | NULL | Probable | As nectar-tap_saprotroph (Põlme et al. 2020); Decay sugar-rich substrates (Põlme et al. 2020) | Põlme S, et al. 2020. Fungal Diversity 105:1-16 ((https://doi.org/10.1007/s13225-020-00466-2)) |
| Otu50540 | 1 | k__Fungi(100);p__Ascomycota(100);c__Saccharomycetes(100);o__Saccharomycetales(100);f__Metschnikowiaceae(100);g__Metschnikowia(100);s__Metschnikowia_pulcherrima(100); | Metschnikowia | 13 | Saprotroph | \|Undefined Saprotroph\| | Yeast | NULL | Probable | As nectar-tap_saprotroph (Põlme et al. 2020); Decay sugar-rich substrates (Põlme et al. 2020) | Põlme S, et al. 2020. Fungal Diversity 105:1-16 ((https://doi.org/10.1007/s13225-020-00466-2)) |
| Otu50541 | 1 | k__Fungi(100);p__Ascomycota(100);c__Saccharomycetes(100);o__Saccharomycetales(100);f__Metschnikowiaceae(100);g__Metschnikowia(100);s__Metschnikowia_pulcherrima(100); | Metschnikowia | 13 | Saprotroph | \|Undefined Saprotroph\| | Yeast | NULL | Probable | As nectar-tap_saprotroph (Põlme et al. 2020); Decay sugar-rich substrates (Põlme et al. 2020) | Põlme S, et al. 2020. Fungal Diversity 105:1-16 ((https://doi.org/10.1007/s13225-020-00466-2)) |
| Otu50542 | 1 | k__Fungi(100);p__Ascomycota(100);c__Saccharomycetes(100);o__Saccharomycetales(100);f__Metschnikowiaceae(100);g__Metschnikowia(100);s__Metschnikowia_pulcherrima(100); | Metschnikowia | 13 | Saprotroph | \|Undefined Saprotroph\| | Yeast | NULL | Probable | As nectar-tap_saprotroph (Põlme et al. 2020); Decay sugar-rich substrates (Põlme et al. 2020) | Põlme S, et al. 2020. Fungal Diversity 105:1-16 ((https://doi.org/10.1007/s13225-020-00466-2)) |
| Otu50543 | 1 | k__Fungi(100);p__Ascomycota(100);c__Saccharomycetes(100);o__Saccharomycetales(100);f__Metschnikowiaceae(100);g__Metschnikowia(100);g__Metschnikowia_unclassified(100); | Metschnikowia | 13 | Saprotroph | \|Undefined Saprotroph\| | Yeast | NULL | Probable | As nectar-tap_saprotroph (Põlme et al. 2020); Decay sugar-rich substrates (Põlme et al. 2020) | Põlme S, et al. 2020. Fungal Diversity 105:1-16 ((https://doi.org/10.1007/s13225-020-00466-2)) |
| Otu50545 | 1 | k__Fungi(100);p__Ascomycota(100);c__Saccharomycetes(100);o__Saccharomycetales(100);f__Metschnikowiaceae(100);g__Metschnikowia(100);s__Metschnikowia_pulcherrima(100); | Metschnikowia | 13 | Saprotroph | \|Undefined Saprotroph\| | Yeast | NULL | Probable | As nectar-tap_saprotroph (Põlme et al. 2020); Decay sugar-rich substrates (Põlme et al. 2020) | Põlme S, et al. 2020. Fungal Diversity 105:1-16 ((https://doi.org/10.1007/s13225-020-00466-2)) |
| Otu50546 | 1 | k__Fungi(100);p__Ascomycota(100);c__Saccharomycetes(100);o__Saccharomycetales(100);f__Metschnikowiaceae(100);g__Metschnikowia(100);g__Metschnikowia_unclassified(100); | Metschnikowia | 13 | Saprotroph | \|Undefined Saprotroph\| | Yeast | NULL | Probable | As nectar-tap_saprotroph (Põlme et al. 2020); Decay sugar-rich substrates (Põlme et al. 2020) | Põlme S, et al. 2020. Fungal Diversity 105:1-16 ((https://doi.org/10.1007/s13225-020-00466-2)) |
| Otu50547 | 1 | k__Fungi(100);p__Ascomycota(100);c__Saccharomycetes(100);o__Saccharomycetales(100);f__Metschnikowiaceae(100);g__Metschnikowia(100);s__Metschnikowia_pulcherrima(100); | Metschnikowia | 13 | Saprotroph | \|Undefined Saprotroph\| | Yeast | NULL | Probable | As nectar-tap_saprotroph (Põlme et al. 2020); Decay sugar-rich substrates (Põlme et al. 2020) | Põlme S, et al. 2020. Fungal Diversity 105:1-16 ((https://doi.org/10.1007/s13225-020-00466-2)) |
| Otu50548 | 1 | k__Fungi(100);p__Ascomycota(100);c__Saccharomycetes(100);o__Saccharomycetales(100);f__Metschnikowiaceae(100);g__Metschnikowia(100);s__Metschnikowia_pulcherrima(100); | Metschnikowia | 13 | Saprotroph | \|Undefined Saprotroph\| | Yeast | NULL | Probable | As nectar-tap_saprotroph (Põlme et al. 2020); Decay sugar-rich substrates (Põlme et al. 2020) | Põlme S, et al. 2020. Fungal Diversity 105:1-16 ((https://doi.org/10.1007/s13225-020-00466-2)) |
| Otu50549 | 1 | k__Fungi(100);p__Ascomycota(100);c__Saccharomycetes(100);o__Saccharomycetales(100);f__Metschnikowiaceae(100);g__Metschnikowia(100);s__Metschnikowia_pulcherrima(100); | Metschnikowia | 13 | Saprotroph | \|Undefined Saprotroph\| | Yeast | NULL | Probable | As nectar-tap_saprotroph (Põlme et al. 2020); Decay sugar-rich substrates (Põlme et al. 2020) | Põlme S, et al. 2020. Fungal Diversity 105:1-16 ((https://doi.org/10.1007/s13225-020-00466-2)) |
| Otu50550 | 1 | k__Fungi(100);p__Ascomycota(100);c__Saccharomycetes(100);o__Saccharomycetales(100);f__Metschnikowiaceae(100);g__Metschnikowia(100);s__Metschnikowia_pulcherrima(100); | Metschnikowia | 13 | Saprotroph | \|Undefined Saprotroph\| | Yeast | NULL | Probable | As nectar-tap_saprotroph (Põlme et al. 2020); Decay sugar-rich substrates (Põlme et al. 2020) | Põlme S, et al. 2020. Fungal Diversity 105:1-16 ((https://doi.org/10.1007/s13225-020-00466-2)) |
| Otu50551 | 1 | k__Fungi(100);p__Ascomycota(100);c__Saccharomycetes(100);o__Saccharomycetales(100);f__Metschnikowiaceae(100);g__Metschnikowia(100);s__Metschnikowia_pulcherrima(100); | Metschnikowia | 13 | Saprotroph | \|Undefined Saprotroph\| | Yeast | NULL | Probable | As nectar-tap_saprotroph (Põlme et al. 2020); Decay sugar-rich substrates (Põlme et al. 2020) | Põlme S, et al. 2020. Fungal Diversity 105:1-16 ((https://doi.org/10.1007/s13225-020-00466-2)) |
| Otu50552 | 1 | k__Fungi(100);p__Ascomycota(100);c__Saccharomycetes(100);o__Saccharomycetales(100);f__Metschnikowiaceae(100);g__Metschnikowia(100);s__Metschnikowia_pulcherrima(100); | Metschnikowia | 13 | Saprotroph | \|Undefined Saprotroph\| | Yeast | NULL | Probable | As nectar-tap_saprotroph (Põlme et al. 2020); Decay sugar-rich substrates (Põlme et al. 2020) | Põlme S, et al. 2020. Fungal Diversity 105:1-16 ((https://doi.org/10.1007/s13225-020-00466-2)) |
| Otu50553 | 1 | k__Fungi(100);p__Ascomycota(100);c__Saccharomycetes(100);o__Saccharomycetales(100);f__Metschnikowiaceae(100);g__Metschnikowia(100);s__Metschnikowia_pulcherrima(100); | Metschnikowia | 13 | Saprotroph | \|Undefined Saprotroph\| | Yeast | NULL | Probable | As nectar-tap_saprotroph (Põlme et al. 2020); Decay sugar-rich substrates (Põlme et al. 2020) | Põlme S, et al. 2020. Fungal Diversity 105:1-16 ((https://doi.org/10.1007/s13225-020-00466-2)) |
| Otu50554 | 1 | k__Fungi(100);p__Ascomycota(100);c__Saccharomycetes(100);o__Saccharomycetales(100);f__Metschnikowiaceae(100);g__Metschnikowia(100);s__Metschnikowia_pulcherrima(100); | Metschnikowia | 13 | Saprotroph | \|Undefined Saprotroph\| | Yeast | NULL | Probable | As nectar-tap_saprotroph (Põlme et al. 2020); Decay sugar-rich substrates (Põlme et al. 2020) | Põlme S, et al. 2020. Fungal Diversity 105:1-16 ((https://doi.org/10.1007/s13225-020-00466-2)) |
| Otu50555 | 1 | k__Fungi(100);p__Ascomycota(100);c__Saccharomycetes(100);o__Saccharomycetales(100);f__Metschnikowiaceae(100);g__Metschnikowia(100);s__Metschnikowia_pulcherrima(100); | Metschnikowia | 13 | Saprotroph | \|Undefined Saprotroph\| | Yeast | NULL | Probable | As nectar-tap_saprotroph (Põlme et al. 2020); Decay sugar-rich substrates (Põlme et al. 2020) | Põlme S, et al. 2020. Fungal Diversity 105:1-16 ((https://doi.org/10.1007/s13225-020-00466-2)) |
| Otu50556 | 1 | k__Fungi(100);p__Ascomycota(100);c__Saccharomycetes(100);o__Saccharomycetales(100);f__Metschnikowiaceae(100);g__Metschnikowia(100);g__Metschnikowia_unclassified(100); | Metschnikowia | 13 | Saprotroph | \|Undefined Saprotroph\| | Yeast | NULL | Probable | As nectar-tap_saprotroph (Põlme et al. 2020); Decay sugar-rich substrates (Põlme et al. 2020) | Põlme S, et al. 2020. Fungal Diversity 105:1-16 ((https://doi.org/10.1007/s13225-020-00466-2)) |
| Otu50557 | 1 | k__Fungi(100);p__Ascomycota(100);c__Saccharomycetes(100);o__Saccharomycetales(100);f__Metschnikowiaceae(100);g__Metschnikowia(100);s__Metschnikowia_pulcherrima(100); | Metschnikowia | 13 | Saprotroph | \|Undefined Saprotroph\| | Yeast | NULL | Probable | As nectar-tap_saprotroph (Põlme et al. 2020); Decay sugar-rich substrates (Põlme et al. 2020) | Põlme S, et al. 2020. Fungal Diversity 105:1-16 ((https://doi.org/10.1007/s13225-020-00466-2)) |
| Otu50558 | 1 | k__Fungi(100);p__Ascomycota(100);c__Saccharomycetes(100);o__Saccharomycetales(100);f__Metschnikowiaceae(100);g__Metschnikowia(100);g__Metschnikowia_unclassified(100); | Metschnikowia | 13 | Saprotroph | \|Undefined Saprotroph\| | Yeast | NULL | Probable | As nectar-tap_saprotroph (Põlme et al. 2020); Decay sugar-rich substrates (Põlme et al. 2020) | Põlme S, et al. 2020. Fungal Diversity 105:1-16 ((https://doi.org/10.1007/s13225-020-00466-2)) |
| Otu50559 | 1 | k__Fungi(100);p__Ascomycota(100);c__Saccharomycetes(100);o__Saccharomycetales(100);f__Metschnikowiaceae(100);g__Metschnikowia(100);s__Metschnikowia_pulcherrima(100); | Metschnikowia | 13 | Saprotroph | \|Undefined Saprotroph\| | Yeast | NULL | Probable | As nectar-tap_saprotroph (Põlme et al. 2020); Decay sugar-rich substrates (Põlme et al. 2020) | Põlme S, et al. 2020. Fungal Diversity 105:1-16 ((https://doi.org/10.1007/s13225-020-00466-2)) |
| Otu50560 | 1 | k__Fungi(100);p__Ascomycota(100);c__Saccharomycetes(100);o__Saccharomycetales(100);f__Metschnikowiaceae(100);g__Metschnikowia(100);s__Metschnikowia_pulcherrima(100); | Metschnikowia | 13 | Saprotroph | \|Undefined Saprotroph\| | Yeast | NULL | Probable | As nectar-tap_saprotroph (Põlme et al. 2020); Decay sugar-rich substrates (Põlme et al. 2020) | Põlme S, et al. 2020. Fungal Diversity 105:1-16 ((https://doi.org/10.1007/s13225-020-00466-2)) |
| Otu50561 | 1 | k__Fungi(100);p__Ascomycota(100);c__Saccharomycetes(100);o__Saccharomycetales(100);f__Metschnikowiaceae(100);g__Metschnikowia(100);s__Metschnikowia_pulcherrima(100); | Metschnikowia | 13 | Saprotroph | \|Undefined Saprotroph\| | Yeast | NULL | Probable | As nectar-tap_saprotroph (Põlme et al. 2020); Decay sugar-rich substrates (Põlme et al. 2020) | Põlme S, et al. 2020. Fungal Diversity 105:1-16 ((https://doi.org/10.1007/s13225-020-00466-2)) |
| Otu50562 | 1 | k__Fungi(100);p__Ascomycota(100);c__Saccharomycetes(100);o__Saccharomycetales(100);f__Metschnikowiaceae(100);g__Metschnikowia(100);s__Metschnikowia_pulcherrima(100); | Metschnikowia | 13 | Saprotroph | \|Undefined Saprotroph\| | Yeast | NULL | Probable | As nectar-tap_saprotroph (Põlme et al. 2020); Decay sugar-rich substrates (Põlme et al. 2020) | Põlme S, et al. 2020. Fungal Diversity 105:1-16 ((https://doi.org/10.1007/s13225-020-00466-2)) |
| Otu50563 | 1 | k__Fungi(100);p__Ascomycota(100);c__Saccharomycetes(100);o__Saccharomycetales(100);f__Metschnikowiaceae(100);g__Metschnikowia(100);s__Metschnikowia_pulcherrima(100); | Metschnikowia | 13 | Saprotroph | \|Undefined Saprotroph\| | Yeast | NULL | Probable | As nectar-tap_saprotroph (Põlme et al. 2020); Decay sugar-rich substrates (Põlme et al. 2020) | Põlme S, et al. 2020. Fungal Diversity 105:1-16 ((https://doi.org/10.1007/s13225-020-00466-2)) |
| Otu50564 | 1 | k__Fungi(100);p__Ascomycota(100);c__Saccharomycetes(100);o__Saccharomycetales(100);f__Metschnikowiaceae(100);g__Metschnikowia(100);s__Metschnikowia_pulcherrima(100); | Metschnikowia | 13 | Saprotroph | \|Undefined Saprotroph\| | Yeast | NULL | Probable | As nectar-tap_saprotroph (Põlme et al. 2020); Decay sugar-rich substrates (Põlme et al. 2020) | Põlme S, et al. 2020. Fungal Diversity 105:1-16 ((https://doi.org/10.1007/s13225-020-00466-2)) |
| Otu50565 | 1 | k__Fungi(100);p__Ascomycota(100);c__Saccharomycetes(100);o__Saccharomycetales(100);f__Metschnikowiaceae(100);g__Metschnikowia(100);s__Metschnikowia_pulcherrima(100); | Metschnikowia | 13 | Saprotroph | \|Undefined Saprotroph\| | Yeast | NULL | Probable | As nectar-tap_saprotroph (Põlme et al. 2020); Decay sugar-rich substrates (Põlme et al. 2020) | Põlme S, et al. 2020. Fungal Diversity 105:1-16 ((https://doi.org/10.1007/s13225-020-00466-2)) |
| Otu50566 | 1 | k__Fungi(100);p__Ascomycota(100);c__Saccharomycetes(100);o__Saccharomycetales(100);f__Metschnikowiaceae(100);g__Metschnikowia(100);g__Metschnikowia_unclassified(100); | Metschnikowia | 13 | Saprotroph | \|Undefined Saprotroph\| | Yeast | NULL | Probable | As nectar-tap_saprotroph (Põlme et al. 2020); Decay sugar-rich substrates (Põlme et al. 2020) | Põlme S, et al. 2020. Fungal Diversity 105:1-16 ((https://doi.org/10.1007/s13225-020-00466-2)) |
| Otu50567 | 1 | k__Fungi(100);p__Ascomycota(100);c__Saccharomycetes(100);o__Saccharomycetales(100);f__Metschnikowiaceae(100);g__Metschnikowia(100);g__Metschnikowia_unclassified(100); | Metschnikowia | 13 | Saprotroph | \|Undefined Saprotroph\| | Yeast | NULL | Probable | As nectar-tap_saprotroph (Põlme et al. 2020); Decay sugar-rich substrates (Põlme et al. 2020) | Põlme S, et al. 2020. Fungal Diversity 105:1-16 ((https://doi.org/10.1007/s13225-020-00466-2)) |
| Otu50568 | 1 | k__Fungi(100);p__Ascomycota(100);c__Saccharomycetes(100);o__Saccharomycetales(100);f__Metschnikowiaceae(100);g__Metschnikowia(100);s__Metschnikowia_pulcherrima(100); | Metschnikowia | 13 | Saprotroph | \|Undefined Saprotroph\| | Yeast | NULL | Probable | As nectar-tap_saprotroph (Põlme et al. 2020); Decay sugar-rich substrates (Põlme et al. 2020) | Põlme S, et al. 2020. Fungal Diversity 105:1-16 ((https://doi.org/10.1007/s13225-020-00466-2)) |
| Otu50569 | 1 | k__Fungi(100);p__Ascomycota(100);c__Saccharomycetes(100);o__Saccharomycetales(100);f__Metschnikowiaceae(100);g__Metschnikowia(100);s__Metschnikowia_pulcherrima(100); | Metschnikowia | 13 | Saprotroph | \|Undefined Saprotroph\| | Yeast | NULL | Probable | As nectar-tap_saprotroph (Põlme et al. 2020); Decay sugar-rich substrates (Põlme et al. 2020) | Põlme S, et al. 2020. Fungal Diversity 105:1-16 ((https://doi.org/10.1007/s13225-020-00466-2)) |
| Otu50570 | 1 | k__Fungi(100);p__Ascomycota(100);c__Saccharomycetes(100);o__Saccharomycetales(100);f__Metschnikowiaceae(100);g__Metschnikowia(100);s__Metschnikowia_pulcherrima(100); | Metschnikowia | 13 | Saprotroph | \|Undefined Saprotroph\| | Yeast | NULL | Probable | As nectar-tap_saprotroph (Põlme et al. 2020); Decay sugar-rich substrates (Põlme et al. 2020) | Põlme S, et al. 2020. Fungal Diversity 105:1-16 ((https://doi.org/10.1007/s13225-020-00466-2)) |
| Otu50571 | 1 | k__Fungi(100);p__Ascomycota(100);c__Saccharomycetes(100);o__Saccharomycetales(100);f__Metschnikowiaceae(100);g__Metschnikowia(100);g__Metschnikowia_unclassified(100); | Metschnikowia | 13 | Saprotroph | \|Undefined Saprotroph\| | Yeast | NULL | Probable | As nectar-tap_saprotroph (Põlme et al. 2020); Decay sugar-rich substrates (Põlme et al. 2020) | Põlme S, et al. 2020. Fungal Diversity 105:1-16 ((https://doi.org/10.1007/s13225-020-00466-2)) |
| Otu50572 | 1 | k__Fungi(100);p__Ascomycota(100);c__Saccharomycetes(100);o__Saccharomycetales(100);f__Metschnikowiaceae(100);g__Metschnikowia(100);g__Metschnikowia_unclassified(100); | Metschnikowia | 13 | Saprotroph | \|Undefined Saprotroph\| | Yeast | NULL | Probable | As nectar-tap_saprotroph (Põlme et al. 2020); Decay sugar-rich substrates (Põlme et al. 2020) | Põlme S, et al. 2020. Fungal Diversity 105:1-16 ((https://doi.org/10.1007/s13225-020-00466-2)) |
| Otu50573 | 1 | k__Fungi(100);p__Ascomycota(100);c__Saccharomycetes(100);o__Saccharomycetales(100);f__Metschnikowiaceae(100);g__Metschnikowia(100);s__Metschnikowia_pulcherrima(100); | Metschnikowia | 13 | Saprotroph | \|Undefined Saprotroph\| | Yeast | NULL | Probable | As nectar-tap_saprotroph (Põlme et al. 2020); Decay sugar-rich substrates (Põlme et al. 2020) | Põlme S, et al. 2020. Fungal Diversity 105:1-16 ((https://doi.org/10.1007/s13225-020-00466-2)) |
| Otu50574 | 1 | k__Fungi(100);p__Ascomycota(100);c__Saccharomycetes(100);o__Saccharomycetales(100);f__Metschnikowiaceae(100);g__Metschnikowia(100);g__Metschnikowia_unclassified(100); | Metschnikowia | 13 | Saprotroph | \|Undefined Saprotroph\| | Yeast | NULL | Probable | As nectar-tap_saprotroph (Põlme et al. 2020); Decay sugar-rich substrates (Põlme et al. 2020) | Põlme S, et al. 2020. Fungal Diversity 105:1-16 ((https://doi.org/10.1007/s13225-020-00466-2)) |
| Otu50575 | 1 | k__Fungi(100);p__Ascomycota(100);c__Saccharomycetes(100);o__Saccharomycetales(100);f__Metschnikowiaceae(100);g__Metschnikowia(100);g__Metschnikowia_unclassified(100); | Metschnikowia | 13 | Saprotroph | \|Undefined Saprotroph\| | Yeast | NULL | Probable | As nectar-tap_saprotroph (Põlme et al. 2020); Decay sugar-rich substrates (Põlme et al. 2020) | Põlme S, et al. 2020. Fungal Diversity 105:1-16 ((https://doi.org/10.1007/s13225-020-00466-2)) |
| Otu50576 | 1 | k__Fungi(100);p__Ascomycota(100);c__Saccharomycetes(100);o__Saccharomycetales(100);f__Metschnikowiaceae(100);g__Metschnikowia(100);g__Metschnikowia_unclassified(100); | Metschnikowia | 13 | Saprotroph | \|Undefined Saprotroph\| | Yeast | NULL | Probable | As nectar-tap_saprotroph (Põlme et al. 2020); Decay sugar-rich substrates (Põlme et al. 2020) | Põlme S, et al. 2020. Fungal Diversity 105:1-16 ((https://doi.org/10.1007/s13225-020-00466-2)) |
| Otu50577 | 1 | k__Fungi(100);p__Ascomycota(100);c__Saccharomycetes(100);o__Saccharomycetales(100);f__Metschnikowiaceae(100);g__Metschnikowia(100);s__Metschnikowia_pulcherrima(100); | Metschnikowia | 13 | Saprotroph | \|Undefined Saprotroph\| | Yeast | NULL | Probable | As nectar-tap_saprotroph (Põlme et al. 2020); Decay sugar-rich substrates (Põlme et al. 2020) | Põlme S, et al. 2020. Fungal Diversity 105:1-16 ((https://doi.org/10.1007/s13225-020-00466-2)) |
| Otu50578 | 1 | k__Fungi(100);p__Ascomycota(100);c__Saccharomycetes(100);o__Saccharomycetales(100);f__Metschnikowiaceae(100);g__Metschnikowia(100);s__Metschnikowia_pulcherrima(100); | Metschnikowia | 13 | Saprotroph | \|Undefined Saprotroph\| | Yeast | NULL | Probable | As nectar-tap_saprotroph (Põlme et al. 2020); Decay sugar-rich substrates (Põlme et al. 2020) | Põlme S, et al. 2020. Fungal Diversity 105:1-16 ((https://doi.org/10.1007/s13225-020-00466-2)) |
| Otu50579 | 1 | k__Fungi(100);p__Ascomycota(100);c__Saccharomycetes(100);o__Saccharomycetales(100);f__Metschnikowiaceae(100);g__Metschnikowia(100);s__Metschnikowia_pulcherrima(100); | Metschnikowia | 13 | Saprotroph | \|Undefined Saprotroph\| | Yeast | NULL | Probable | As nectar-tap_saprotroph (Põlme et al. 2020); Decay sugar-rich substrates (Põlme et al. 2020) | Põlme S, et al. 2020. Fungal Diversity 105:1-16 ((https://doi.org/10.1007/s13225-020-00466-2)) |
| Otu50580 | 1 | k__Fungi(100);p__Ascomycota(100);c__Saccharomycetes(100);o__Saccharomycetales(100);f__Metschnikowiaceae(100);g__Metschnikowia(100);s__Metschnikowia_pulcherrima(100); | Metschnikowia | 13 | Saprotroph | \|Undefined Saprotroph\| | Yeast | NULL | Probable | As nectar-tap_saprotroph (Põlme et al. 2020); Decay sugar-rich substrates (Põlme et al. 2020) | Põlme S, et al. 2020. Fungal Diversity 105:1-16 ((https://doi.org/10.1007/s13225-020-00466-2)) |
| Otu50581 | 1 | k__Fungi(100);p__Ascomycota(100);c__Saccharomycetes(100);o__Saccharomycetales(100);f__Metschnikowiaceae(100);g__Metschnikowia(100);s__Metschnikowia_pulcherrima(100); | Metschnikowia | 13 | Saprotroph | \|Undefined Saprotroph\| | Yeast | NULL | Probable | As nectar-tap_saprotroph (Põlme et al. 2020); Decay sugar-rich substrates (Põlme et al. 2020) | Põlme S, et al. 2020. Fungal Diversity 105:1-16 ((https://doi.org/10.1007/s13225-020-00466-2)) |
| Otu50582 | 1 | k__Fungi(100);p__Ascomycota(100);c__Saccharomycetes(100);o__Saccharomycetales(100);f__Metschnikowiaceae(100);g__Metschnikowia(100);s__Metschnikowia_pulcherrima(100); | Metschnikowia | 13 | Saprotroph | \|Undefined Saprotroph\| | Yeast | NULL | Probable | As nectar-tap_saprotroph (Põlme et al. 2020); Decay sugar-rich substrates (Põlme et al. 2020) | Põlme S, et al. 2020. Fungal Diversity 105:1-16 ((https://doi.org/10.1007/s13225-020-00466-2)) |
| Otu50583 | 1 | k__Fungi(100);p__Ascomycota(100);c__Saccharomycetes(100);o__Saccharomycetales(100);f__Metschnikowiaceae(100);g__Metschnikowia(100);g__Metschnikowia_unclassified(100); | Metschnikowia | 13 | Saprotroph | \|Undefined Saprotroph\| | Yeast | NULL | Probable | As nectar-tap_saprotroph (Põlme et al. 2020); Decay sugar-rich substrates (Põlme et al. 2020) | Põlme S, et al. 2020. Fungal Diversity 105:1-16 ((https://doi.org/10.1007/s13225-020-00466-2)) |
| Otu50584 | 1 | k__Fungi(100);p__Ascomycota(100);c__Saccharomycetes(100);o__Saccharomycetales(100);f__Metschnikowiaceae(100);g__Metschnikowia(100);s__Metschnikowia_pulcherrima(100); | Metschnikowia | 13 | Saprotroph | \|Undefined Saprotroph\| | Yeast | NULL | Probable | As nectar-tap_saprotroph (Põlme et al. 2020); Decay sugar-rich substrates (Põlme et al. 2020) | Põlme S, et al. 2020. Fungal Diversity 105:1-16 ((https://doi.org/10.1007/s13225-020-00466-2)) |
| Otu50585 | 1 | k__Fungi(100);p__Ascomycota(100);c__Saccharomycetes(100);o__Saccharomycetales(100);f__Metschnikowiaceae(100);g__Metschnikowia(100);g__Metschnikowia_unclassified(100); | Metschnikowia | 13 | Saprotroph | \|Undefined Saprotroph\| | Yeast | NULL | Probable | As nectar-tap_saprotroph (Põlme et al. 2020); Decay sugar-rich substrates (Põlme et al. 2020) | Põlme S, et al. 2020. Fungal Diversity 105:1-16 ((https://doi.org/10.1007/s13225-020-00466-2)) |
| Otu50586 | 1 | k__Fungi(100);p__Ascomycota(100);c__Saccharomycetes(100);o__Saccharomycetales(100);f__Metschnikowiaceae(100);g__Metschnikowia(100);s__Metschnikowia_pulcherrima(100); | Metschnikowia | 13 | Saprotroph | \|Undefined Saprotroph\| | Yeast | NULL | Probable | As nectar-tap_saprotroph (Põlme et al. 2020); Decay sugar-rich substrates (Põlme et al. 2020) | Põlme S, et al. 2020. Fungal Diversity 105:1-16 ((https://doi.org/10.1007/s13225-020-00466-2)) |
| Otu50587 | 1 | k__Fungi(100);p__Ascomycota(100);c__Saccharomycetes(100);o__Saccharomycetales(100);f__Metschnikowiaceae(100);g__Metschnikowia(100);s__Metschnikowia_pulcherrima(100); | Metschnikowia | 13 | Saprotroph | \|Undefined Saprotroph\| | Yeast | NULL | Probable | As nectar-tap_saprotroph (Põlme et al. 2020); Decay sugar-rich substrates (Põlme et al. 2020) | Põlme S, et al. 2020. Fungal Diversity 105:1-16 ((https://doi.org/10.1007/s13225-020-00466-2)) |
| Otu50588 | 1 | k__Fungi(100);p__Ascomycota(100);c__Saccharomycetes(100);o__Saccharomycetales(100);f__Metschnikowiaceae(100);g__Metschnikowia(100);s__Metschnikowia_pulcherrima(100); | Metschnikowia | 13 | Saprotroph | \|Undefined Saprotroph\| | Yeast | NULL | Probable | As nectar-tap_saprotroph (Põlme et al. 2020); Decay sugar-rich substrates (Põlme et al. 2020) | Põlme S, et al. 2020. Fungal Diversity 105:1-16 ((https://doi.org/10.1007/s13225-020-00466-2)) |
| Otu50589 | 1 | k__Fungi(100);p__Ascomycota(100);c__Saccharomycetes(100);o__Saccharomycetales(100);f__Metschnikowiaceae(100);g__Metschnikowia(100);s__Metschnikowia_pulcherrima(100); | Metschnikowia | 13 | Saprotroph | \|Undefined Saprotroph\| | Yeast | NULL | Probable | As nectar-tap_saprotroph (Põlme et al. 2020); Decay sugar-rich substrates (Põlme et al. 2020) | Põlme S, et al. 2020. Fungal Diversity 105:1-16 ((https://doi.org/10.1007/s13225-020-00466-2)) |
| Otu50590 | 1 | k__Fungi(100);p__Ascomycota(100);c__Saccharomycetes(100);o__Saccharomycetales(100);f__Metschnikowiaceae(100);g__Metschnikowia(100);s__Metschnikowia_pulcherrima(100); | Metschnikowia | 13 | Saprotroph | \|Undefined Saprotroph\| | Yeast | NULL | Probable | As nectar-tap_saprotroph (Põlme et al. 2020); Decay sugar-rich substrates (Põlme et al. 2020) | Põlme S, et al. 2020. Fungal Diversity 105:1-16 ((https://doi.org/10.1007/s13225-020-00466-2)) |
| Otu50591 | 1 | k__Fungi(100);p__Ascomycota(100);c__Saccharomycetes(100);o__Saccharomycetales(100);f__Metschnikowiaceae(100);g__Metschnikowia(100);g__Metschnikowia_unclassified(100); | Metschnikowia | 13 | Saprotroph | \|Undefined Saprotroph\| | Yeast | NULL | Probable | As nectar-tap_saprotroph (Põlme et al. 2020); Decay sugar-rich substrates (Põlme et al. 2020) | Põlme S, et al. 2020. Fungal Diversity 105:1-16 ((https://doi.org/10.1007/s13225-020-00466-2)) |
| Otu50592 | 1 | k__Fungi(100);p__Ascomycota(100);c__Saccharomycetes(100);o__Saccharomycetales(100);f__Metschnikowiaceae(100);g__Metschnikowia(100);g__Metschnikowia_unclassified(100); | Metschnikowia | 13 | Saprotroph | \|Undefined Saprotroph\| | Yeast | NULL | Probable | As nectar-tap_saprotroph (Põlme et al. 2020); Decay sugar-rich substrates (Põlme et al. 2020) | Põlme S, et al. 2020. Fungal Diversity 105:1-16 ((https://doi.org/10.1007/s13225-020-00466-2)) |
| Otu50593 | 1 | k__Fungi(100);p__Ascomycota(100);c__Saccharomycetes(100);o__Saccharomycetales(100);f__Metschnikowiaceae(100);g__Metschnikowia(100);g__Metschnikowia_unclassified(100); | Metschnikowia | 13 | Saprotroph | \|Undefined Saprotroph\| | Yeast | NULL | Probable | As nectar-tap_saprotroph (Põlme et al. 2020); Decay sugar-rich substrates (Põlme et al. 2020) | Põlme S, et al. 2020. Fungal Diversity 105:1-16 ((https://doi.org/10.1007/s13225-020-00466-2)) |
| Otu50594 | 1 | k__Fungi(100);p__Ascomycota(100);c__Saccharomycetes(100);o__Saccharomycetales(100);f__Metschnikowiaceae(100);g__Metschnikowia(100);s__Metschnikowia_pulcherrima(100); | Metschnikowia | 13 | Saprotroph | \|Undefined Saprotroph\| | Yeast | NULL | Probable | As nectar-tap_saprotroph (Põlme et al. 2020); Decay sugar-rich substrates (Põlme et al. 2020) | Põlme S, et al. 2020. Fungal Diversity 105:1-16 ((https://doi.org/10.1007/s13225-020-00466-2)) |
| Otu50595 | 1 | k__Fungi(100);p__Ascomycota(100);c__Saccharomycetes(100);o__Saccharomycetales(100);f__Metschnikowiaceae(100);g__Metschnikowia(100);g__Metschnikowia_unclassified(100); | Metschnikowia | 13 | Saprotroph | \|Undefined Saprotroph\| | Yeast | NULL | Probable | As nectar-tap_saprotroph (Põlme et al. 2020); Decay sugar-rich substrates (Põlme et al. 2020) | Põlme S, et al. 2020. Fungal Diversity 105:1-16 ((https://doi.org/10.1007/s13225-020-00466-2)) |
| Otu50596 | 1 | k__Fungi(100);p__Ascomycota(100);c__Saccharomycetes(100);o__Saccharomycetales(100);f__Metschnikowiaceae(100);g__Metschnikowia(100);s__Metschnikowia_pulcherrima(100); | Metschnikowia | 13 | Saprotroph | \|Undefined Saprotroph\| | Yeast | NULL | Probable | As nectar-tap_saprotroph (Põlme et al. 2020); Decay sugar-rich substrates (Põlme et al. 2020) | Põlme S, et al. 2020. Fungal Diversity 105:1-16 ((https://doi.org/10.1007/s13225-020-00466-2)) |
| Otu50597 | 1 | k__Fungi(100);p__Ascomycota(100);c__Saccharomycetes(100);o__Saccharomycetales(100);f__Metschnikowiaceae(100);g__Metschnikowia(100);g__Metschnikowia_unclassified(100); | Metschnikowia | 13 | Saprotroph | \|Undefined Saprotroph\| | Yeast | NULL | Probable | As nectar-tap_saprotroph (Põlme et al. 2020); Decay sugar-rich substrates (Põlme et al. 2020) | Põlme S, et al. 2020. Fungal Diversity 105:1-16 ((https://doi.org/10.1007/s13225-020-00466-2)) |
| Otu50598 | 1 | k__Fungi(100);p__Ascomycota(100);c__Saccharomycetes(100);o__Saccharomycetales(100);f__Metschnikowiaceae(100);g__Metschnikowia(100);s__Metschnikowia_pulcherrima(100); | Metschnikowia | 13 | Saprotroph | \|Undefined Saprotroph\| | Yeast | NULL | Probable | As nectar-tap_saprotroph (Põlme et al. 2020); Decay sugar-rich substrates (Põlme et al. 2020) | Põlme S, et al. 2020. Fungal Diversity 105:1-16 ((https://doi.org/10.1007/s13225-020-00466-2)) |
| Otu50599 | 1 | k__Fungi(100);p__Ascomycota(100);c__Saccharomycetes(100);o__Saccharomycetales(100);f__Metschnikowiaceae(100);g__Metschnikowia(100);s__Metschnikowia_pulcherrima(100); | Metschnikowia | 13 | Saprotroph | \|Undefined Saprotroph\| | Yeast | NULL | Probable | As nectar-tap_saprotroph (Põlme et al. 2020); Decay sugar-rich substrates (Põlme et al. 2020) | Põlme S, et al. 2020. Fungal Diversity 105:1-16 ((https://doi.org/10.1007/s13225-020-00466-2)) |
| Otu50600 | 1 | k__Fungi(100);p__Ascomycota(100);c__Saccharomycetes(100);o__Saccharomycetales(100);f__Metschnikowiaceae(100);g__Metschnikowia(100);s__Metschnikowia_pulcherrima(100); | Metschnikowia | 13 | Saprotroph | \|Undefined Saprotroph\| | Yeast | NULL | Probable | As nectar-tap_saprotroph (Põlme et al. 2020); Decay sugar-rich substrates (Põlme et al. 2020) | Põlme S, et al. 2020. Fungal Diversity 105:1-16 ((https://doi.org/10.1007/s13225-020-00466-2)) |
| Otu50601 | 1 | k__Fungi(100);p__Ascomycota(100);c__Saccharomycetes(100);o__Saccharomycetales(100);f__Metschnikowiaceae(100);g__Metschnikowia(100);g__Metschnikowia_unclassified(100); | Metschnikowia | 13 | Saprotroph | \|Undefined Saprotroph\| | Yeast | NULL | Probable | As nectar-tap_saprotroph (Põlme et al. 2020); Decay sugar-rich substrates (Põlme et al. 2020) | Põlme S, et al. 2020. Fungal Diversity 105:1-16 ((https://doi.org/10.1007/s13225-020-00466-2)) |
| Otu50602 | 1 | k__Fungi(100);p__Ascomycota(100);c__Saccharomycetes(100);o__Saccharomycetales(100);f__Metschnikowiaceae(100);g__Metschnikowia(100);s__Metschnikowia_pulcherrima(100); | Metschnikowia | 13 | Saprotroph | \|Undefined Saprotroph\| | Yeast | NULL | Probable | As nectar-tap_saprotroph (Põlme et al. 2020); Decay sugar-rich substrates (Põlme et al. 2020) | Põlme S, et al. 2020. Fungal Diversity 105:1-16 ((https://doi.org/10.1007/s13225-020-00466-2)) |
| Otu50603 | 1 | k__Fungi(100);p__Ascomycota(100);c__Saccharomycetes(100);o__Saccharomycetales(100);f__Metschnikowiaceae(100);g__Metschnikowia(100);s__Metschnikowia_pulcherrima(100); | Metschnikowia | 13 | Saprotroph | \|Undefined Saprotroph\| | Yeast | NULL | Probable | As nectar-tap_saprotroph (Põlme et al. 2020); Decay sugar-rich substrates (Põlme et al. 2020) | Põlme S, et al. 2020. Fungal Diversity 105:1-16 ((https://doi.org/10.1007/s13225-020-00466-2)) |
| Otu50604 | 1 | k__Fungi(100);p__Ascomycota(100);c__Saccharomycetes(100);o__Saccharomycetales(100);f__Metschnikowiaceae(100);g__Metschnikowia(100);s__Metschnikowia_pulcherrima(100); | Metschnikowia | 13 | Saprotroph | \|Undefined Saprotroph\| | Yeast | NULL | Probable | As nectar-tap_saprotroph (Põlme et al. 2020); Decay sugar-rich substrates (Põlme et al. 2020) | Põlme S, et al. 2020. Fungal Diversity 105:1-16 ((https://doi.org/10.1007/s13225-020-00466-2)) |
| Otu50605 | 1 | k__Fungi(100);p__Ascomycota(100);c__Saccharomycetes(100);o__Saccharomycetales(100);f__Metschnikowiaceae(100);g__Metschnikowia(100);s__Metschnikowia_pulcherrima(100); | Metschnikowia | 13 | Saprotroph | \|Undefined Saprotroph\| | Yeast | NULL | Probable | As nectar-tap_saprotroph (Põlme et al. 2020); Decay sugar-rich substrates (Põlme et al. 2020) | Põlme S, et al. 2020. Fungal Diversity 105:1-16 ((https://doi.org/10.1007/s13225-020-00466-2)) |
| Otu50606 | 1 | k__Fungi(100);p__Ascomycota(100);c__Saccharomycetes(100);o__Saccharomycetales(100);f__Metschnikowiaceae(100);g__Metschnikowia(100);s__Metschnikowia_pulcherrima(100); | Metschnikowia | 13 | Saprotroph | \|Undefined Saprotroph\| | Yeast | NULL | Probable | As nectar-tap_saprotroph (Põlme et al. 2020); Decay sugar-rich substrates (Põlme et al. 2020) | Põlme S, et al. 2020. Fungal Diversity 105:1-16 ((https://doi.org/10.1007/s13225-020-00466-2)) |
| Otu50608 | 1 | k__Fungi(100);p__Ascomycota(100);c__Saccharomycetes(100);o__Saccharomycetales(100);f__Metschnikowiaceae(100);g__Metschnikowia(100);s__Metschnikowia_pulcherrima(100); | Metschnikowia | 13 | Saprotroph | \|Undefined Saprotroph\| | Yeast | NULL | Probable | As nectar-tap_saprotroph (Põlme et al. 2020); Decay sugar-rich substrates (Põlme et al. 2020) | Põlme S, et al. 2020. Fungal Diversity 105:1-16 ((https://doi.org/10.1007/s13225-020-00466-2)) |
| Otu50609 | 1 | k__Fungi(100);p__Ascomycota(100);c__Saccharomycetes(100);o__Saccharomycetales(100);f__Metschnikowiaceae(100);g__Metschnikowia(100);s__Metschnikowia_pulcherrima(100); | Metschnikowia | 13 | Saprotroph | \|Undefined Saprotroph\| | Yeast | NULL | Probable | As nectar-tap_saprotroph (Põlme et al. 2020); Decay sugar-rich substrates (Põlme et al. 2020) | Põlme S, et al. 2020. Fungal Diversity 105:1-16 ((https://doi.org/10.1007/s13225-020-00466-2)) |
| Otu50610 | 1 | k__Fungi(100);p__Ascomycota(100);c__Saccharomycetes(100);o__Saccharomycetales(100);f__Metschnikowiaceae(100);g__Metschnikowia(100);s__Metschnikowia_pulcherrima(100); | Metschnikowia | 13 | Saprotroph | \|Undefined Saprotroph\| | Yeast | NULL | Probable | As nectar-tap_saprotroph (Põlme et al. 2020); Decay sugar-rich substrates (Põlme et al. 2020) | Põlme S, et al. 2020. Fungal Diversity 105:1-16 ((https://doi.org/10.1007/s13225-020-00466-2)) |
| Otu50611 | 1 | k__Fungi(100);p__Ascomycota(100);c__Saccharomycetes(100);o__Saccharomycetales(100);f__Metschnikowiaceae(100);g__Metschnikowia(100);s__Metschnikowia_pulcherrima(100); | Metschnikowia | 13 | Saprotroph | \|Undefined Saprotroph\| | Yeast | NULL | Probable | As nectar-tap_saprotroph (Põlme et al. 2020); Decay sugar-rich substrates (Põlme et al. 2020) | Põlme S, et al. 2020. Fungal Diversity 105:1-16 ((https://doi.org/10.1007/s13225-020-00466-2)) |
| Otu50612 | 1 | k__Fungi(100);p__Ascomycota(100);c__Saccharomycetes(100);o__Saccharomycetales(100);f__Metschnikowiaceae(100);g__Metschnikowia(100);s__Metschnikowia_pulcherrima(100); | Metschnikowia | 13 | Saprotroph | \|Undefined Saprotroph\| | Yeast | NULL | Probable | As nectar-tap_saprotroph (Põlme et al. 2020); Decay sugar-rich substrates (Põlme et al. 2020) | Põlme S, et al. 2020. Fungal Diversity 105:1-16 ((https://doi.org/10.1007/s13225-020-00466-2)) |
| Otu50613 | 1 | k__Fungi(100);p__Ascomycota(100);c__Saccharomycetes(100);o__Saccharomycetales(100);f__Metschnikowiaceae(100);g__Metschnikowia(100);s__Metschnikowia_pulcherrima(100); | Metschnikowia | 13 | Saprotroph | \|Undefined Saprotroph\| | Yeast | NULL | Probable | As nectar-tap_saprotroph (Põlme et al. 2020); Decay sugar-rich substrates (Põlme et al. 2020) | Põlme S, et al. 2020. Fungal Diversity 105:1-16 ((https://doi.org/10.1007/s13225-020-00466-2)) |
| Otu50614 | 1 | k__Fungi(100);p__Ascomycota(100);c__Saccharomycetes(100);o__Saccharomycetales(100);f__Metschnikowiaceae(100);g__Metschnikowia(100);s__Metschnikowia_pulcherrima(100); | Metschnikowia | 13 | Saprotroph | \|Undefined Saprotroph\| | Yeast | NULL | Probable | As nectar-tap_saprotroph (Põlme et al. 2020); Decay sugar-rich substrates (Põlme et al. 2020) | Põlme S, et al. 2020. Fungal Diversity 105:1-16 ((https://doi.org/10.1007/s13225-020-00466-2)) |
| Otu50615 | 1 | k__Fungi(100);p__Ascomycota(100);c__Saccharomycetes(100);o__Saccharomycetales(100);f__Metschnikowiaceae(100);g__Metschnikowia(100);s__Metschnikowia_pulcherrima(100); | Metschnikowia | 13 | Saprotroph | \|Undefined Saprotroph\| | Yeast | NULL | Probable | As nectar-tap_saprotroph (Põlme et al. 2020); Decay sugar-rich substrates (Põlme et al. 2020) | Põlme S, et al. 2020. Fungal Diversity 105:1-16 ((https://doi.org/10.1007/s13225-020-00466-2)) |
| Otu50616 | 1 | k__Fungi(100);p__Ascomycota(100);c__Saccharomycetes(100);o__Saccharomycetales(100);f__Metschnikowiaceae(100);g__Metschnikowia(100);s__Metschnikowia_pulcherrima(100); | Metschnikowia | 13 | Saprotroph | \|Undefined Saprotroph\| | Yeast | NULL | Probable | As nectar-tap_saprotroph (Põlme et al. 2020); Decay sugar-rich substrates (Põlme et al. 2020) | Põlme S, et al. 2020. Fungal Diversity 105:1-16 ((https://doi.org/10.1007/s13225-020-00466-2)) |
| Otu50617 | 1 | k__Fungi(100);p__Ascomycota(100);c__Saccharomycetes(100);o__Saccharomycetales(100);f__Metschnikowiaceae(100);g__Metschnikowia(100);s__Metschnikowia_pulcherrima(100); | Metschnikowia | 13 | Saprotroph | \|Undefined Saprotroph\| | Yeast | NULL | Probable | As nectar-tap_saprotroph (Põlme et al. 2020); Decay sugar-rich substrates (Põlme et al. 2020) | Põlme S, et al. 2020. Fungal Diversity 105:1-16 ((https://doi.org/10.1007/s13225-020-00466-2)) |
| Otu50618 | 1 | k__Fungi(100);p__Ascomycota(100);c__Saccharomycetes(100);o__Saccharomycetales(100);f__Metschnikowiaceae(100);g__Metschnikowia(100);s__Metschnikowia_pulcherrima(100); | Metschnikowia | 13 | Saprotroph | \|Undefined Saprotroph\| | Yeast | NULL | Probable | As nectar-tap_saprotroph (Põlme et al. 2020); Decay sugar-rich substrates (Põlme et al. 2020) | Põlme S, et al. 2020. Fungal Diversity 105:1-16 ((https://doi.org/10.1007/s13225-020-00466-2)) |
| Otu50619 | 1 | k__Fungi(100);p__Ascomycota(100);c__Saccharomycetes(100);o__Saccharomycetales(100);f__Metschnikowiaceae(100);g__Metschnikowia(100);g__Metschnikowia_unclassified(100); | Metschnikowia | 13 | Saprotroph | \|Undefined Saprotroph\| | Yeast | NULL | Probable | As nectar-tap_saprotroph (Põlme et al. 2020); Decay sugar-rich substrates (Põlme et al. 2020) | Põlme S, et al. 2020. Fungal Diversity 105:1-16 ((https://doi.org/10.1007/s13225-020-00466-2)) |
| Otu50620 | 1 | k__Fungi(100);p__Ascomycota(100);c__Saccharomycetes(100);o__Saccharomycetales(100);f__Metschnikowiaceae(100);g__Metschnikowia(100);s__Metschnikowia_pulcherrima(100); | Metschnikowia | 13 | Saprotroph | \|Undefined Saprotroph\| | Yeast | NULL | Probable | As nectar-tap_saprotroph (Põlme et al. 2020); Decay sugar-rich substrates (Põlme et al. 2020) | Põlme S, et al. 2020. Fungal Diversity 105:1-16 ((https://doi.org/10.1007/s13225-020-00466-2)) |
| Otu50621 | 1 | k__Fungi(100);p__Ascomycota(100);c__Saccharomycetes(100);o__Saccharomycetales(100);f__Metschnikowiaceae(100);g__Metschnikowia(100);s__Metschnikowia_pulcherrima(100); | Metschnikowia | 13 | Saprotroph | \|Undefined Saprotroph\| | Yeast | NULL | Probable | As nectar-tap_saprotroph (Põlme et al. 2020); Decay sugar-rich substrates (Põlme et al. 2020) | Põlme S, et al. 2020. Fungal Diversity 105:1-16 ((https://doi.org/10.1007/s13225-020-00466-2)) |
| Otu50622 | 1 | k__Fungi(100);p__Ascomycota(100);c__Saccharomycetes(100);o__Saccharomycetales(100);f__Metschnikowiaceae(100);g__Metschnikowia(100);s__Metschnikowia_pulcherrima(100); | Metschnikowia | 13 | Saprotroph | \|Undefined Saprotroph\| | Yeast | NULL | Probable | As nectar-tap_saprotroph (Põlme et al. 2020); Decay sugar-rich substrates (Põlme et al. 2020) | Põlme S, et al. 2020. Fungal Diversity 105:1-16 ((https://doi.org/10.1007/s13225-020-00466-2)) |
| Otu50623 | 1 | k__Fungi(100);p__Ascomycota(100);c__Saccharomycetes(100);o__Saccharomycetales(100);f__Metschnikowiaceae(100);g__Metschnikowia(100);s__Metschnikowia_pulcherrima(100); | Metschnikowia | 13 | Saprotroph | \|Undefined Saprotroph\| | Yeast | NULL | Probable | As nectar-tap_saprotroph (Põlme et al. 2020); Decay sugar-rich substrates (Põlme et al. 2020) | Põlme S, et al. 2020. Fungal Diversity 105:1-16 ((https://doi.org/10.1007/s13225-020-00466-2)) |
| Otu50624 | 1 | k__Fungi(100);p__Ascomycota(100);c__Saccharomycetes(100);o__Saccharomycetales(100);f__Metschnikowiaceae(100);g__Metschnikowia(100);s__Metschnikowia_pulcherrima(100); | Metschnikowia | 13 | Saprotroph | \|Undefined Saprotroph\| | Yeast | NULL | Probable | As nectar-tap_saprotroph (Põlme et al. 2020); Decay sugar-rich substrates (Põlme et al. 2020) | Põlme S, et al. 2020. Fungal Diversity 105:1-16 ((https://doi.org/10.1007/s13225-020-00466-2)) |
| Otu50625 | 1 | k__Fungi(100);p__Ascomycota(100);c__Saccharomycetes(100);o__Saccharomycetales(100);f__Metschnikowiaceae(100);g__Metschnikowia(100);g__Metschnikowia_unclassified(100); | Metschnikowia | 13 | Saprotroph | \|Undefined Saprotroph\| | Yeast | NULL | Probable | As nectar-tap_saprotroph (Põlme et al. 2020); Decay sugar-rich substrates (Põlme et al. 2020) | Põlme S, et al. 2020. Fungal Diversity 105:1-16 ((https://doi.org/10.1007/s13225-020-00466-2)) |
| Otu50626 | 1 | k__Fungi(100);p__Ascomycota(100);c__Saccharomycetes(100);o__Saccharomycetales(100);f__Metschnikowiaceae(100);g__Metschnikowia(100);g__Metschnikowia_unclassified(100); | Metschnikowia | 13 | Saprotroph | \|Undefined Saprotroph\| | Yeast | NULL | Probable | As nectar-tap_saprotroph (Põlme et al. 2020); Decay sugar-rich substrates (Põlme et al. 2020) | Põlme S, et al. 2020. Fungal Diversity 105:1-16 ((https://doi.org/10.1007/s13225-020-00466-2)) |
| Otu50627 | 1 | k__Fungi(100);p__Ascomycota(100);c__Saccharomycetes(100);o__Saccharomycetales(100);f__Metschnikowiaceae(100);g__Metschnikowia(100);s__Metschnikowia_pulcherrima(100); | Metschnikowia | 13 | Saprotroph | \|Undefined Saprotroph\| | Yeast | NULL | Probable | As nectar-tap_saprotroph (Põlme et al. 2020); Decay sugar-rich substrates (Põlme et al. 2020) | Põlme S, et al. 2020. Fungal Diversity 105:1-16 ((https://doi.org/10.1007/s13225-020-00466-2)) |
| Otu50628 | 1 | k__Fungi(100);p__Ascomycota(100);c__Saccharomycetes(100);o__Saccharomycetales(100);f__Metschnikowiaceae(100);g__Metschnikowia(100);s__Metschnikowia_pulcherrima(100); | Metschnikowia | 13 | Saprotroph | \|Undefined Saprotroph\| | Yeast | NULL | Probable | As nectar-tap_saprotroph (Põlme et al. 2020); Decay sugar-rich substrates (Põlme et al. 2020) | Põlme S, et al. 2020. Fungal Diversity 105:1-16 ((https://doi.org/10.1007/s13225-020-00466-2)) |
| Otu50629 | 1 | k__Fungi(100);p__Ascomycota(100);c__Saccharomycetes(100);o__Saccharomycetales(100);f__Metschnikowiaceae(100);g__Metschnikowia(100);s__Metschnikowia_pulcherrima(100); | Metschnikowia | 13 | Saprotroph | \|Undefined Saprotroph\| | Yeast | NULL | Probable | As nectar-tap_saprotroph (Põlme et al. 2020); Decay sugar-rich substrates (Põlme et al. 2020) | Põlme S, et al. 2020. Fungal Diversity 105:1-16 ((https://doi.org/10.1007/s13225-020-00466-2)) |
| Otu50630 | 1 | k__Fungi(100);p__Ascomycota(100);c__Saccharomycetes(100);o__Saccharomycetales(100);f__Metschnikowiaceae(100);g__Metschnikowia(100);s__Metschnikowia_pulcherrima(100); | Metschnikowia | 13 | Saprotroph | \|Undefined Saprotroph\| | Yeast | NULL | Probable | As nectar-tap_saprotroph (Põlme et al. 2020); Decay sugar-rich substrates (Põlme et al. 2020) | Põlme S, et al. 2020. Fungal Diversity 105:1-16 ((https://doi.org/10.1007/s13225-020-00466-2)) |
| Otu50631 | 1 | k__Fungi(100);p__Ascomycota(100);c__Saccharomycetes(100);o__Saccharomycetales(100);f__Metschnikowiaceae(100);g__Metschnikowia(100);s__Metschnikowia_pulcherrima(100); | Metschnikowia | 13 | Saprotroph | \|Undefined Saprotroph\| | Yeast | NULL | Probable | As nectar-tap_saprotroph (Põlme et al. 2020); Decay sugar-rich substrates (Põlme et al. 2020) | Põlme S, et al. 2020. Fungal Diversity 105:1-16 ((https://doi.org/10.1007/s13225-020-00466-2)) |
| Otu50632 | 1 | k__Fungi(100);p__Ascomycota(100);c__Saccharomycetes(100);o__Saccharomycetales(100);f__Metschnikowiaceae(100);g__Metschnikowia(100);s__Metschnikowia_pulcherrima(100); | Metschnikowia | 13 | Saprotroph | \|Undefined Saprotroph\| | Yeast | NULL | Probable | As nectar-tap_saprotroph (Põlme et al. 2020); Decay sugar-rich substrates (Põlme et al. 2020) | Põlme S, et al. 2020. Fungal Diversity 105:1-16 ((https://doi.org/10.1007/s13225-020-00466-2)) |
| Otu50633 | 1 | k__Fungi(100);p__Ascomycota(100);c__Saccharomycetes(100);o__Saccharomycetales(100);f__Metschnikowiaceae(100);g__Metschnikowia(100);s__Metschnikowia_pulcherrima(100); | Metschnikowia | 13 | Saprotroph | \|Undefined Saprotroph\| | Yeast | NULL | Probable | As nectar-tap_saprotroph (Põlme et al. 2020); Decay sugar-rich substrates (Põlme et al. 2020) | Põlme S, et al. 2020. Fungal Diversity 105:1-16 ((https://doi.org/10.1007/s13225-020-00466-2)) |
| Otu50634 | 1 | k__Fungi(100);p__Ascomycota(100);c__Saccharomycetes(100);o__Saccharomycetales(100);f__Metschnikowiaceae(100);g__Metschnikowia(100);g__Metschnikowia_unclassified(100); | Metschnikowia | 13 | Saprotroph | \|Undefined Saprotroph\| | Yeast | NULL | Probable | As nectar-tap_saprotroph (Põlme et al. 2020); Decay sugar-rich substrates (Põlme et al. 2020) | Põlme S, et al. 2020. Fungal Diversity 105:1-16 ((https://doi.org/10.1007/s13225-020-00466-2)) |
| Otu50635 | 1 | k__Fungi(100);p__Ascomycota(100);c__Saccharomycetes(100);o__Saccharomycetales(100);f__Metschnikowiaceae(100);g__Metschnikowia(100);s__Metschnikowia_pulcherrima(100); | Metschnikowia | 13 | Saprotroph | \|Undefined Saprotroph\| | Yeast | NULL | Probable | As nectar-tap_saprotroph (Põlme et al. 2020); Decay sugar-rich substrates (Põlme et al. 2020) | Põlme S, et al. 2020. Fungal Diversity 105:1-16 ((https://doi.org/10.1007/s13225-020-00466-2)) |
| Otu50636 | 1 | k__Fungi(100);p__Ascomycota(100);c__Saccharomycetes(100);o__Saccharomycetales(100);f__Metschnikowiaceae(100);g__Metschnikowia(100);s__Metschnikowia_pulcherrima(100); | Metschnikowia | 13 | Saprotroph | \|Undefined Saprotroph\| | Yeast | NULL | Probable | As nectar-tap_saprotroph (Põlme et al. 2020); Decay sugar-rich substrates (Põlme et al. 2020) | Põlme S, et al. 2020. Fungal Diversity 105:1-16 ((https://doi.org/10.1007/s13225-020-00466-2)) |
| Otu50637 | 1 | k__Fungi(100);p__Ascomycota(100);c__Saccharomycetes(100);o__Saccharomycetales(100);f__Metschnikowiaceae(100);g__Metschnikowia(100);s__Metschnikowia_pulcherrima(100); | Metschnikowia | 13 | Saprotroph | \|Undefined Saprotroph\| | Yeast | NULL | Probable | As nectar-tap_saprotroph (Põlme et al. 2020); Decay sugar-rich substrates (Põlme et al. 2020) | Põlme S, et al. 2020. Fungal Diversity 105:1-16 ((https://doi.org/10.1007/s13225-020-00466-2)) |
| Otu50638 | 1 | k__Fungi(100);p__Ascomycota(100);c__Saccharomycetes(100);o__Saccharomycetales(100);f__Metschnikowiaceae(100);g__Metschnikowia(100);s__Metschnikowia_pulcherrima(100); | Metschnikowia | 13 | Saprotroph | \|Undefined Saprotroph\| | Yeast | NULL | Probable | As nectar-tap_saprotroph (Põlme et al. 2020); Decay sugar-rich substrates (Põlme et al. 2020) | Põlme S, et al. 2020. Fungal Diversity 105:1-16 ((https://doi.org/10.1007/s13225-020-00466-2)) |
| Otu50639 | 1 | k__Fungi(100);p__Ascomycota(100);c__Saccharomycetes(100);o__Saccharomycetales(100);f__Metschnikowiaceae(100);g__Metschnikowia(100);g__Metschnikowia_unclassified(100); | Metschnikowia | 13 | Saprotroph | \|Undefined Saprotroph\| | Yeast | NULL | Probable | As nectar-tap_saprotroph (Põlme et al. 2020); Decay sugar-rich substrates (Põlme et al. 2020) | Põlme S, et al. 2020. Fungal Diversity 105:1-16 ((https://doi.org/10.1007/s13225-020-00466-2)) |
| Otu50640 | 1 | k__Fungi(100);p__Ascomycota(100);c__Saccharomycetes(100);o__Saccharomycetales(100);f__Metschnikowiaceae(100);g__Metschnikowia(100);g__Metschnikowia_unclassified(100); | Metschnikowia | 13 | Saprotroph | \|Undefined Saprotroph\| | Yeast | NULL | Probable | As nectar-tap_saprotroph (Põlme et al. 2020); Decay sugar-rich substrates (Põlme et al. 2020) | Põlme S, et al. 2020. Fungal Diversity 105:1-16 ((https://doi.org/10.1007/s13225-020-00466-2)) |
| Otu50641 | 1 | k__Fungi(100);p__Ascomycota(100);c__Saccharomycetes(100);o__Saccharomycetales(100);f__Metschnikowiaceae(100);g__Metschnikowia(100);s__Metschnikowia_pulcherrima(100); | Metschnikowia | 13 | Saprotroph | \|Undefined Saprotroph\| | Yeast | NULL | Probable | As nectar-tap_saprotroph (Põlme et al. 2020); Decay sugar-rich substrates (Põlme et al. 2020) | Põlme S, et al. 2020. Fungal Diversity 105:1-16 ((https://doi.org/10.1007/s13225-020-00466-2)) |
| Otu50643 | 1 | k__Fungi(100);p__Ascomycota(100);c__Saccharomycetes(100);o__Saccharomycetales(100);f__Metschnikowiaceae(100);g__Metschnikowia(100);s__Metschnikowia_pulcherrima(100); | Metschnikowia | 13 | Saprotroph | \|Undefined Saprotroph\| | Yeast | NULL | Probable | As nectar-tap_saprotroph (Põlme et al. 2020); Decay sugar-rich substrates (Põlme et al. 2020) | Põlme S, et al. 2020. Fungal Diversity 105:1-16 ((https://doi.org/10.1007/s13225-020-00466-2)) |
| Otu50644 | 1 | k__Fungi(100);p__Ascomycota(100);c__Saccharomycetes(100);o__Saccharomycetales(100);f__Metschnikowiaceae(100);g__Metschnikowia(100);s__Metschnikowia_pulcherrima(100); | Metschnikowia | 13 | Saprotroph | \|Undefined Saprotroph\| | Yeast | NULL | Probable | As nectar-tap_saprotroph (Põlme et al. 2020); Decay sugar-rich substrates (Põlme et al. 2020) | Põlme S, et al. 2020. Fungal Diversity 105:1-16 ((https://doi.org/10.1007/s13225-020-00466-2)) |
| Otu50645 | 1 | k__Fungi(100);p__Ascomycota(100);c__Saccharomycetes(100);o__Saccharomycetales(100);f__Saccharomycetales_family_Incertae_sedis(100);f__Saccharomycetales_family_Incertae_sedis_unclassified(100);f__Saccharomycetales_family_Incertae_sedis_unclassified(100); | Saccharomycetales | 7 | Saprotroph | Undefined Saprotroph | Yeast | NULL | Possible | NULL | Sterkenburg E, et al. 2015. New Phytologist 207:1145-1158 ((https://doi.org/10.1111/nph.13426)) |
| Otu50648 | 1 | k__Fungi(100);p__Ascomycota(100);c__Saccharomycetes(100);o__Saccharomycetales(100);f__Metschnikowiaceae(100);g__Metschnikowia(100);s__Metschnikowia_pulcherrima(100); | Metschnikowia | 13 | Saprotroph | \|Undefined Saprotroph\| | Yeast | NULL | Probable | As nectar-tap_saprotroph (Põlme et al. 2020); Decay sugar-rich substrates (Põlme et al. 2020) | Põlme S, et al. 2020. Fungal Diversity 105:1-16 ((https://doi.org/10.1007/s13225-020-00466-2)) |
| Otu50650 | 1 | k__Fungi(100);p__Ascomycota(100);c__Saccharomycetes(100);o__Saccharomycetales(100);f__Metschnikowiaceae(100);g__Metschnikowia(100);s__Metschnikowia_pulcherrima(100); | Metschnikowia | 13 | Saprotroph | \|Undefined Saprotroph\| | Yeast | NULL | Probable | As nectar-tap_saprotroph (Põlme et al. 2020); Decay sugar-rich substrates (Põlme et al. 2020) | Põlme S, et al. 2020. Fungal Diversity 105:1-16 ((https://doi.org/10.1007/s13225-020-00466-2)) |
| Otu50653 | 1 | k__Fungi(100);p__Ascomycota(100);c__Saccharomycetes(100);o__Saccharomycetales(100);f__Metschnikowiaceae(100);g__Metschnikowia(100);s__Metschnikowia_pulcherrima(100); | Metschnikowia | 13 | Saprotroph | \|Undefined Saprotroph\| | Yeast | NULL | Probable | As nectar-tap_saprotroph (Põlme et al. 2020); Decay sugar-rich substrates (Põlme et al. 2020) | Põlme S, et al. 2020. Fungal Diversity 105:1-16 ((https://doi.org/10.1007/s13225-020-00466-2)) |
| Otu50654 | 1 | k__Fungi(100);p__Ascomycota(100);c__Saccharomycetes(100);o__Saccharomycetales(100);f__Metschnikowiaceae(100);g__Metschnikowia(100);s__Metschnikowia_pulcherrima(100); | Metschnikowia | 13 | Saprotroph | \|Undefined Saprotroph\| | Yeast | NULL | Probable | As nectar-tap_saprotroph (Põlme et al. 2020); Decay sugar-rich substrates (Põlme et al. 2020) | Põlme S, et al. 2020. Fungal Diversity 105:1-16 ((https://doi.org/10.1007/s13225-020-00466-2)) |
| Otu50655 | 1 | k__Fungi(100);p__Ascomycota(100);c__Saccharomycetes(100);o__Saccharomycetales(100);f__Metschnikowiaceae(100);g__Metschnikowia(100);s__Metschnikowia_pulcherrima(100); | Metschnikowia | 13 | Saprotroph | \|Undefined Saprotroph\| | Yeast | NULL | Probable | As nectar-tap_saprotroph (Põlme et al. 2020); Decay sugar-rich substrates (Põlme et al. 2020) | Põlme S, et al. 2020. Fungal Diversity 105:1-16 ((https://doi.org/10.1007/s13225-020-00466-2)) |
| Otu50657 | 1 | k__Fungi(100);p__Ascomycota(100);c__Saccharomycetes(100);o__Saccharomycetales(100);o__Saccharomycetales_unclassified(100);o__Saccharomycetales_unclassified(100);o__Saccharomycetales_unclassified(100); | Saccharomycetales | 7 | Saprotroph | Undefined Saprotroph | Yeast | NULL | Possible | NULL | Sterkenburg E, et al. 2015. New Phytologist 207:1145-1158 ((https://doi.org/10.1111/nph.13426)) |
| Otu50658 | 1 | k__Fungi(100);p__Ascomycota(100);c__Saccharomycetes(100);o__Saccharomycetales(100);f__Metschnikowiaceae(100);g__Metschnikowia(100);s__Metschnikowia_pulcherrima(100); | Metschnikowia | 13 | Saprotroph | \|Undefined Saprotroph\| | Yeast | NULL | Probable | As nectar-tap_saprotroph (Põlme et al. 2020); Decay sugar-rich substrates (Põlme et al. 2020) | Põlme S, et al. 2020. Fungal Diversity 105:1-16 ((https://doi.org/10.1007/s13225-020-00466-2)) |
| Otu50660 | 1 | k__Fungi(100);p__Ascomycota(100);c__Dothideomycetes(100);o__Capnodiales(100);f__Capnodiales_family_Incertae_sedis(100);g__Toxicocladosporium(100);s__Toxicocladosporium_sp(100); | Toxicocladosporium | 13 | Pathotroph-Symbiotroph | Endophyte-\|Plant Pathogen\| | Microfungus | NULL | Probable | As foliar_endophyte (Põlme et al. 2020); Decay leaf, fruit, seed (Põlme et al. 2020); Isolated from leaves of Posoqueria and Eucalyptus camaldulensis (occurring within a single small lesion) as well as moldy paint | Crous PW, et al. 2009. Persoonia 22:83-94 ((https://doi.org/10.3767/003158509X439364)); Põlme S, et al. 2020. Fungal Diversity 105:1-16 ((https://doi.org/10.1007/s13225-020-00466-2)) |
| Otu50661 | 1 | k__Fungi(100);p__Ascomycota(100);c__Saccharomycetes(100);o__Saccharomycetales(100);f__Metschnikowiaceae(100);g__Metschnikowia(100);g__Metschnikowia_unclassified(100); | Metschnikowia | 13 | Saprotroph | \|Undefined Saprotroph\| | Yeast | NULL | Probable | As nectar-tap_saprotroph (Põlme et al. 2020); Decay sugar-rich substrates (Põlme et al. 2020) | Põlme S, et al. 2020. Fungal Diversity 105:1-16 ((https://doi.org/10.1007/s13225-020-00466-2)) |
| Otu50662 | 1 | k__Fungi(100);p__Ascomycota(100);c__Saccharomycetes(100);o__Saccharomycetales(100);f__Metschnikowiaceae(100);g__Metschnikowia(100);g__Metschnikowia_unclassified(100); | Metschnikowia | 13 | Saprotroph | \|Undefined Saprotroph\| | Yeast | NULL | Probable | As nectar-tap_saprotroph (Põlme et al. 2020); Decay sugar-rich substrates (Põlme et al. 2020) | Põlme S, et al. 2020. Fungal Diversity 105:1-16 ((https://doi.org/10.1007/s13225-020-00466-2)) |
| Otu50663 | 1 | k__Fungi(100);p__Ascomycota(100);c__Saccharomycetes(100);o__Saccharomycetales(100);f__Metschnikowiaceae(100);g__Metschnikowia(100);s__Metschnikowia_pulcherrima(100); | Metschnikowia | 13 | Saprotroph | \|Undefined Saprotroph\| | Yeast | NULL | Probable | As nectar-tap_saprotroph (Põlme et al. 2020); Decay sugar-rich substrates (Põlme et al. 2020) | Põlme S, et al. 2020. Fungal Diversity 105:1-16 ((https://doi.org/10.1007/s13225-020-00466-2)) |
| Otu50664 | 1 | k__Fungi(100);p__Ascomycota(100);c__Saccharomycetes(100);o__Saccharomycetales(100);f__Metschnikowiaceae(100);g__Metschnikowia(100);s__Metschnikowia_pulcherrima(100); | Metschnikowia | 13 | Saprotroph | \|Undefined Saprotroph\| | Yeast | NULL | Probable | As nectar-tap_saprotroph (Põlme et al. 2020); Decay sugar-rich substrates (Põlme et al. 2020) | Põlme S, et al. 2020. Fungal Diversity 105:1-16 ((https://doi.org/10.1007/s13225-020-00466-2)) |
| Otu50665 | 1 | k__Fungi(100);p__Ascomycota(100);c__Saccharomycetes(100);o__Saccharomycetales(100);f__Metschnikowiaceae(100);g__Metschnikowia(100);s__Metschnikowia_pulcherrima(100); | Metschnikowia | 13 | Saprotroph | \|Undefined Saprotroph\| | Yeast | NULL | Probable | As nectar-tap_saprotroph (Põlme et al. 2020); Decay sugar-rich substrates (Põlme et al. 2020) | Põlme S, et al. 2020. Fungal Diversity 105:1-16 ((https://doi.org/10.1007/s13225-020-00466-2)) |
| Otu50666 | 1 | k__Fungi(100);p__Ascomycota(100);c__Saccharomycetes(100);o__Saccharomycetales(100);f__Metschnikowiaceae(100);g__Metschnikowia(100);s__Metschnikowia_pulcherrima(100); | Metschnikowia | 13 | Saprotroph | \|Undefined Saprotroph\| | Yeast | NULL | Probable | As nectar-tap_saprotroph (Põlme et al. 2020); Decay sugar-rich substrates (Põlme et al. 2020) | Põlme S, et al. 2020. Fungal Diversity 105:1-16 ((https://doi.org/10.1007/s13225-020-00466-2)) |
| Otu50667 | 1 | k__Fungi(100);p__Ascomycota(100);c__Saccharomycetes(100);o__Saccharomycetales(100);f__Metschnikowiaceae(100);g__Metschnikowia(100);s__Metschnikowia_pulcherrima(100); | Metschnikowia | 13 | Saprotroph | \|Undefined Saprotroph\| | Yeast | NULL | Probable | As nectar-tap_saprotroph (Põlme et al. 2020); Decay sugar-rich substrates (Põlme et al. 2020) | Põlme S, et al. 2020. Fungal Diversity 105:1-16 ((https://doi.org/10.1007/s13225-020-00466-2)) |
| Otu50668 | 1 | k__Fungi(100);p__Ascomycota(100);c__Saccharomycetes(100);o__Saccharomycetales(100);f__Metschnikowiaceae(100);g__Metschnikowia(100);g__Metschnikowia_unclassified(100); | Metschnikowia | 13 | Saprotroph | \|Undefined Saprotroph\| | Yeast | NULL | Probable | As nectar-tap_saprotroph (Põlme et al. 2020); Decay sugar-rich substrates (Põlme et al. 2020) | Põlme S, et al. 2020. Fungal Diversity 105:1-16 ((https://doi.org/10.1007/s13225-020-00466-2)) |
| Otu50669 | 1 | k__Fungi(100);p__Ascomycota(100);c__Saccharomycetes(100);o__Saccharomycetales(100);f__Metschnikowiaceae(100);g__Metschnikowia(100);g__Metschnikowia_unclassified(100); | Metschnikowia | 13 | Saprotroph | \|Undefined Saprotroph\| | Yeast | NULL | Probable | As nectar-tap_saprotroph (Põlme et al. 2020); Decay sugar-rich substrates (Põlme et al. 2020) | Põlme S, et al. 2020. Fungal Diversity 105:1-16 ((https://doi.org/10.1007/s13225-020-00466-2)) |
| Otu50670 | 1 | k__Fungi(100);p__Ascomycota(100);c__Saccharomycetes(100);o__Saccharomycetales(100);f__Metschnikowiaceae(100);g__Metschnikowia(100);s__Metschnikowia_pulcherrima(100); | Metschnikowia | 13 | Saprotroph | \|Undefined Saprotroph\| | Yeast | NULL | Probable | As nectar-tap_saprotroph (Põlme et al. 2020); Decay sugar-rich substrates (Põlme et al. 2020) | Põlme S, et al. 2020. Fungal Diversity 105:1-16 ((https://doi.org/10.1007/s13225-020-00466-2)) |
| Otu50671 | 1 | k__Fungi(100);p__Ascomycota(100);c__Saccharomycetes(100);o__Saccharomycetales(100);f__Metschnikowiaceae(100);g__Metschnikowia(100);s__Metschnikowia_pulcherrima(100); | Metschnikowia | 13 | Saprotroph | \|Undefined Saprotroph\| | Yeast | NULL | Probable | As nectar-tap_saprotroph (Põlme et al. 2020); Decay sugar-rich substrates (Põlme et al. 2020) | Põlme S, et al. 2020. Fungal Diversity 105:1-16 ((https://doi.org/10.1007/s13225-020-00466-2)) |
| Otu50672 | 1 | k__Fungi(100);p__Ascomycota(100);c__Saccharomycetes(100);o__Saccharomycetales(100);f__Metschnikowiaceae(100);g__Metschnikowia(100);g__Metschnikowia_unclassified(100); | Metschnikowia | 13 | Saprotroph | \|Undefined Saprotroph\| | Yeast | NULL | Probable | As nectar-tap_saprotroph (Põlme et al. 2020); Decay sugar-rich substrates (Põlme et al. 2020) | Põlme S, et al. 2020. Fungal Diversity 105:1-16 ((https://doi.org/10.1007/s13225-020-00466-2)) |
| Otu50673 | 1 | k__Fungi(100);p__Ascomycota(100);c__Saccharomycetes(100);o__Saccharomycetales(100);f__Metschnikowiaceae(100);g__Metschnikowia(100);g__Metschnikowia_unclassified(100); | Metschnikowia | 13 | Saprotroph | \|Undefined Saprotroph\| | Yeast | NULL | Probable | As nectar-tap_saprotroph (Põlme et al. 2020); Decay sugar-rich substrates (Põlme et al. 2020) | Põlme S, et al. 2020. Fungal Diversity 105:1-16 ((https://doi.org/10.1007/s13225-020-00466-2)) |
| Otu50674 | 1 | k__Fungi(100);p__Ascomycota(100);c__Saccharomycetes(100);o__Saccharomycetales(100);f__Metschnikowiaceae(100);g__Metschnikowia(100);s__Metschnikowia_pulcherrima(100); | Metschnikowia | 13 | Saprotroph | \|Undefined Saprotroph\| | Yeast | NULL | Probable | As nectar-tap_saprotroph (Põlme et al. 2020); Decay sugar-rich substrates (Põlme et al. 2020) | Põlme S, et al. 2020. Fungal Diversity 105:1-16 ((https://doi.org/10.1007/s13225-020-00466-2)) |
| Otu50675 | 1 | k__Fungi(100);p__Ascomycota(100);c__Saccharomycetes(100);o__Saccharomycetales(100);f__Metschnikowiaceae(100);g__Metschnikowia(100);s__Metschnikowia_pulcherrima(100); | Metschnikowia | 13 | Saprotroph | \|Undefined Saprotroph\| | Yeast | NULL | Probable | As nectar-tap_saprotroph (Põlme et al. 2020); Decay sugar-rich substrates (Põlme et al. 2020) | Põlme S, et al. 2020. Fungal Diversity 105:1-16 ((https://doi.org/10.1007/s13225-020-00466-2)) |

| | | | | | | | | | | | |
|---|---|---|---|---|---|---|---|---|---|---|---|
| Otu50676 | 1 | k__Fungi(100);p__Ascomycota(100);c__Saccharomycetes(100);o__Saccharomycetales(100);f__Metschnikowiaceae(100);g__Metschnikowia(100);s__Metschnikowia_pulcherrima(100); | Metschnikowia | 13 | Saprotroph | \|Undefined Saprotroph\| | Yeast | NULL | Probable | As nectar-tap_saprotroph (Põlme et al. 2020); Decay sugar-rich substrates (Põlme et al. 2020) | Põlme S, et al. 2020. Fungal Diversity 105:1-16 ((https://doi.org/10.1007/s13225-020-00466-2)) |
| Otu50677 | 1 | k__Fungi(100);p__Ascomycota(100);c__Saccharomycetes(100);o__Saccharomycetales(100);f__Metschnikowiaceae(100);g__Metschnikowia(100);s__Metschnikowia_pulcherrima(100); | Metschnikowia | 13 | Saprotroph | \|Undefined Saprotroph\| | Yeast | NULL | Probable | As nectar-tap_saprotroph (Põlme et al. 2020); Decay sugar-rich substrates (Põlme et al. 2020) | Põlme S, et al. 2020. Fungal Diversity 105:1-16 ((https://doi.org/10.1007/s13225-020-00466-2)) |
| Otu50678 | 1 | k__Fungi(100);p__Ascomycota(100);c__Saccharomycetes(100);o__Saccharomycetales(100);f__Metschnikowiaceae(100);g__Metschnikowia(100);s__Metschnikowia_pulcherrima(100); | Metschnikowia | 13 | Saprotroph | \|Undefined Saprotroph\| | Yeast | NULL | Probable | As nectar-tap_saprotroph (Põlme et al. 2020); Decay sugar-rich substrates (Põlme et al. 2020) | Põlme S, et al. 2020. Fungal Diversity 105:1-16 ((https://doi.org/10.1007/s13225-020-00466-2)) |
| Otu50679 | 1 | k__Fungi(100);p__Ascomycota(100);c__Saccharomycetes(100);o__Saccharomycetales(100);f__Metschnikowiaceae(100);g__Metschnikowia(100);s__Metschnikowia_pulcherrima(100); | Metschnikowia | 13 | Saprotroph | \|Undefined Saprotroph\| | Yeast | NULL | Probable | As nectar-tap_saprotroph (Põlme et al. 2020); Decay sugar-rich substrates (Põlme et al. 2020) | Põlme S, et al. 2020. Fungal Diversity 105:1-16 ((https://doi.org/10.1007/s13225-020-00466-2)) |
| Otu50680 | 1 | k__Fungi(100);p__Ascomycota(100);c__Saccharomycetes(100);o__Saccharomycetales(100);f__Metschnikowiaceae(100);g__Metschnikowia(100);s__Metschnikowia_pulcherrima(100); | Metschnikowia | 13 | Saprotroph | \|Undefined Saprotroph\| | Yeast | NULL | Probable | As nectar-tap_saprotroph (Põlme et al. 2020); Decay sugar-rich substrates (Põlme et al. 2020) | Põlme S, et al. 2020. Fungal Diversity 105:1-16 ((https://doi.org/10.1007/s13225-020-00466-2)) |
| Otu50681 | 1 | k__Fungi(100);p__Ascomycota(100);c__Saccharomycetes(100);o__Saccharomycetales(100);f__Metschnikowiaceae(100);g__Metschnikowia(100);s__Metschnikowia_pulcherrima(100); | Metschnikowia | 13 | Saprotroph | \|Undefined Saprotroph\| | Yeast | NULL | Probable | As nectar-tap_saprotroph (Põlme et al. 2020); Decay sugar-rich substrates (Põlme et al. 2020) | Põlme S, et al. 2020. Fungal Diversity 105:1-16 ((https://doi.org/10.1007/s13225-020-00466-2)) |
| Otu50682 | 1 | k__Fungi(100);p__Ascomycota(100);c__Saccharomycetes(100);o__Saccharomycetales(100);f__Metschnikowiaceae(100);g__Metschnikowia(100);s__Metschnikowia_pulcherrima(100); | Metschnikowia | 13 | Saprotroph | \|Undefined Saprotroph\| | Yeast | NULL | Probable | As nectar-tap_saprotroph (Põlme et al. 2020); Decay sugar-rich substrates (Põlme et al. 2020) | Põlme S, et al. 2020. Fungal Diversity 105:1-16 ((https://doi.org/10.1007/s13225-020-00466-2)) |
| Otu50683 | 1 | k__Fungi(100);p__Ascomycota(100);c__Saccharomycetes(100);o__Saccharomycetales(100);f__Metschnikowiaceae(100);g__Metschnikowia(100);s__Metschnikowia_pulcherrima(100); | Metschnikowia | 13 | Saprotroph | \|Undefined Saprotroph\| | Yeast | NULL | Probable | As nectar-tap_saprotroph (Põlme et al. 2020); Decay sugar-rich substrates (Põlme et al. 2020) | Põlme S, et al. 2020. Fungal Diversity 105:1-16 ((https://doi.org/10.1007/s13225-020-00466-2)) |
| Otu50684 | 1 | k__Fungi(100);p__Ascomycota(100);c__Saccharomycetes(100);o__Saccharomycetales(100);f__Metschnikowiaceae(100);g__Metschnikowia(100);s__Metschnikowia_pulcherrima(100); | Metschnikowia | 13 | Saprotroph | \|Undefined Saprotroph\| | Yeast | NULL | Probable | As nectar-tap_saprotroph (Põlme et al. 2020); Decay sugar-rich substrates (Põlme et al. 2020) | Põlme S, et al. 2020. Fungal Diversity 105:1-16 ((https://doi.org/10.1007/s13225-020-00466-2)) |
| Otu50685 | 1 | k__Fungi(100);p__Ascomycota(100);c__Saccharomycetes(100);o__Saccharomycetales(100);f__Metschnikowiaceae(100);g__Metschnikowia(100);s__Metschnikowia_pulcherrima(100); | Metschnikowia | 13 | Saprotroph | \|Undefined Saprotroph\| | Yeast | NULL | Probable | As nectar-tap_saprotroph (Põlme et al. 2020); Decay sugar-rich substrates (Põlme et al. 2020) | Põlme S, et al. 2020. Fungal Diversity 105:1-16 ((https://doi.org/10.1007/s13225-020-00466-2)) |
| Otu50686 | 1 | k__Fungi(100);p__Ascomycota(100);c__Saccharomycetes(100);o__Saccharomycetales(100);f__Metschnikowiaceae(100);g__Metschnikowia(100);s__Metschnikowia_pulcherrima(100); | Metschnikowia | 13 | Saprotroph | \|Undefined Saprotroph\| | Yeast | NULL | Probable | As nectar-tap_saprotroph (Põlme et al. 2020); Decay sugar-rich substrates (Põlme et al. 2020) | Põlme S, et al. 2020. Fungal Diversity 105:1-16 ((https://doi.org/10.1007/s13225-020-00466-2)) |
| Otu50687 | 1 | k__Fungi(100);p__Ascomycota(100);c__Saccharomycetes(100);o__Saccharomycetales(100);f__Metschnikowiaceae(100);g__Metschnikowia(100);g__Metschnikowia_unclassified(100); | Metschnikowia | 13 | Saprotroph | \|Undefined Saprotroph\| | Yeast | NULL | Probable | As nectar-tap_saprotroph (Põlme et al. 2020); Decay sugar-rich substrates (Põlme et al. 2020) | Põlme S, et al. 2020. Fungal Diversity 105:1-16 ((https://doi.org/10.1007/s13225-020-00466-2)) |
| Otu50688 | 1 | k__Fungi(100);p__Ascomycota(100);c__Saccharomycetes(100);o__Saccharomycetales(100);f__Metschnikowiaceae(100);g__Metschnikowia(100);s__Metschnikowia_pulcherrima(100); | Metschnikowia | 13 | Saprotroph | \|Undefined Saprotroph\| | Yeast | NULL | Probable | As nectar-tap_saprotroph (Põlme et al. 2020); Decay sugar-rich substrates (Põlme et al. 2020) | Põlme S, et al. 2020. Fungal Diversity 105:1-16 ((https://doi.org/10.1007/s13225-020-00466-2)) |
| Otu50689 | 1 | k__Fungi(100);p__Ascomycota(100);c__Saccharomycetes(100);o__Saccharomycetales(100);f__Metschnikowiaceae(100);g__Metschnikowia(100);s__Metschnikowia_pulcherrima(100); | Metschnikowia | 13 | Saprotroph | \|Undefined Saprotroph\| | Yeast | NULL | Probable | As nectar-tap_saprotroph (Põlme et al. 2020); Decay sugar-rich substrates (Põlme et al. 2020) | Põlme S, et al. 2020. Fungal Diversity 105:1-16 ((https://doi.org/10.1007/s13225-020-00466-2)) |
| Otu50690 | 1 | k__Fungi(100);p__Ascomycota(100);c__Saccharomycetes(100);o__Saccharomycetales(100);f__Metschnikowiaceae(100);g__Metschnikowia(100);g__Metschnikowia_unclassified(100); | Metschnikowia | 13 | Saprotroph | \|Undefined Saprotroph\| | Yeast | NULL | Probable | As nectar-tap_saprotroph (Põlme et al. 2020); Decay sugar-rich substrates (Põlme et al. 2020) | Põlme S, et al. 2020. Fungal Diversity 105:1-16 ((https://doi.org/10.1007/s13225-020-00466-2)) |
| Otu50691 | 1 | k__Fungi(100);p__Ascomycota(100);c__Saccharomycetes(100);o__Saccharomycetales(100);f__Metschnikowiaceae(100);g__Metschnikowia(100);s__Metschnikowia_pulcherrima(100); | Metschnikowia | 13 | Saprotroph | \|Undefined Saprotroph\| | Yeast | NULL | Probable | As nectar-tap_saprotroph (Põlme et al. 2020); Decay sugar-rich substrates (Põlme et al. 2020) | Põlme S, et al. 2020. Fungal Diversity 105:1-16 ((https://doi.org/10.1007/s13225-020-00466-2)) |
| Otu50692 | 1 | k__Fungi(100);p__Ascomycota(100);c__Saccharomycetes(100);o__Saccharomycetales(100);f__Metschnikowiaceae(100);g__Metschnikowia(100);g__Metschnikowia_unclassified(100); | Metschnikowia | 13 | Saprotroph | \|Undefined Saprotroph\| | Yeast | NULL | Probable | As nectar-tap_saprotroph (Põlme et al. 2020); Decay sugar-rich substrates (Põlme et al. 2020) | Põlme S, et al. 2020. Fungal Diversity 105:1-16 ((https://doi.org/10.1007/s13225-020-00466-2)) |
| Otu50693 | 1 | k__Fungi(100);p__Ascomycota(100);c__Saccharomycetes(100);o__Saccharomycetales(100);f__Metschnikowiaceae(100);g__Metschnikowia(100);s__Metschnikowia_pulcherrima(100); | Metschnikowia | 13 | Saprotroph | \|Undefined Saprotroph\| | Yeast | NULL | Probable | As nectar-tap_saprotroph (Põlme et al. 2020); Decay sugar-rich substrates (Põlme et al. 2020) | Põlme S, et al. 2020. Fungal Diversity 105:1-16 ((https://doi.org/10.1007/s13225-020-00466-2)) |
| Otu50694 | 1 | k__Fungi(100);p__Ascomycota(100);c__Saccharomycetes(100);o__Saccharomycetales(100);f__Metschnikowiaceae(100);g__Metschnikowia(100);s__Metschnikowia_pulcherrima(100); | Metschnikowia | 13 | Saprotroph | \|Undefined Saprotroph\| | Yeast | NULL | Probable | As nectar-tap_saprotroph (Põlme et al. 2020); Decay sugar-rich substrates (Põlme et al. 2020) | Põlme S, et al. 2020. Fungal Diversity 105:1-16 ((https://doi.org/10.1007/s13225-020-00466-2)) |
| Otu50696 | 1 | k__Fungi(100);p__Ascomycota(100);c__Saccharomycetes(100);o__Saccharomycetales(100);f__Metschnikowiaceae(100);g__Metschnikowia(100);s__Metschnikowia_pulcherrima(100); | Metschnikowia | 13 | Saprotroph | \|Undefined Saprotroph\| | Yeast | NULL | Probable | As nectar-tap_saprotroph (Põlme et al. 2020); Decay sugar-rich substrates (Põlme et al. 2020) | Põlme S, et al. 2020. Fungal Diversity 105:1-16 ((https://doi.org/10.1007/s13225-020-00466-2)) |
| Otu50697 | 1 | k__Fungi(100);p__Ascomycota(100);c__Saccharomycetes(100);o__Saccharomycetales(100);f__Metschnikowiaceae(100);g__Metschnikowia(100);g__Metschnikowia_unclassified(100); | Metschnikowia | 13 | Saprotroph | \|Undefined Saprotroph\| | Yeast | NULL | Probable | As nectar-tap_saprotroph (Põlme et al. 2020); Decay sugar-rich substrates (Põlme et al. 2020) | Põlme S, et al. 2020. Fungal Diversity 105:1-16 ((https://doi.org/10.1007/s13225-020-00466-2)) |
| Otu50699 | 1 | k__Fungi(100);p__Ascomycota(100);c__Saccharomycetes(100);o__Saccharomycetales(100);f__Metschnikowiaceae(100);g__Metschnikowia(100);g__Metschnikowia_unclassified(100); | Metschnikowia | 13 | Saprotroph | \|Undefined Saprotroph\| | Yeast | NULL | Probable | As nectar-tap_saprotroph (Põlme et al. 2020); Decay sugar-rich substrates (Põlme et al. 2020) | Põlme S, et al. 2020. Fungal Diversity 105:1-16 ((https://doi.org/10.1007/s13225-020-00466-2)) |
| Otu50700 | 1 | k__Fungi(100);p__Ascomycota(100);c__Saccharomycetes(100);o__Saccharomycetales(100);f__Metschnikowiaceae(100);g__Metschnikowia(100);s__Metschnikowia_pulcherrima(100); | Metschnikowia | 13 | Saprotroph | \|Undefined Saprotroph\| | Yeast | NULL | Probable | As nectar-tap_saprotroph (Põlme et al. 2020); Decay sugar-rich substrates (Põlme et al. 2020) | Põlme S, et al. 2020. Fungal Diversity 105:1-16 ((https://doi.org/10.1007/s13225-020-00466-2)) |
| Otu50701 | 1 | k__Fungi(100);p__Ascomycota(100);c__Saccharomycetes(100);o__Saccharomycetales(100);f__Metschnikowiaceae(100);g__Metschnikowia(100);s__Metschnikowia_pulcherrima(100); | Metschnikowia | 13 | Saprotroph | \|Undefined Saprotroph\| | Yeast | NULL | Probable | As nectar-tap_saprotroph (Põlme et al. 2020); Decay sugar-rich substrates (Põlme et al. 2020) | Põlme S, et al. 2020. Fungal Diversity 105:1-16 ((https://doi.org/10.1007/s13225-020-00466-2)) |
| Otu50702 | 1 | k__Fungi(100);p__Ascomycota(100);c__Saccharomycetes(100);o__Saccharomycetales(100);f__Metschnikowiaceae(100);g__Metschnikowia(100);s__Metschnikowia_pulcherrima(100); | Metschnikowia | 13 | Saprotroph | \|Undefined Saprotroph\| | Yeast | NULL | Probable | As nectar-tap_saprotroph (Põlme et al. 2020); Decay sugar-rich substrates (Põlme et al. 2020) | Põlme S, et al. 2020. Fungal Diversity 105:1-16 ((https://doi.org/10.1007/s13225-020-00466-2)) |
| Otu50703 | 1 | k__Fungi(100);p__Ascomycota(100);c__Saccharomycetes(100);o__Saccharomycetales(100);f__Metschnikowiaceae(100);g__Metschnikowia(100);s__Metschnikowia_pulcherrima(100); | Metschnikowia | 13 | Saprotroph | \|Undefined Saprotroph\| | Yeast | NULL | Probable | As nectar-tap_saprotroph (Põlme et al. 2020); Decay sugar-rich substrates (Põlme et al. 2020) | Põlme S, et al. 2020. Fungal Diversity 105:1-16 ((https://doi.org/10.1007/s13225-020-00466-2)) |
| Otu50704 | 1 | k__Fungi(100);p__Ascomycota(100);c__Saccharomycetes(100);o__Saccharomycetales(100);f__Metschnikowiaceae(100);g__Metschnikowia(100);s__Metschnikowia_pulcherrima(100); | Metschnikowia | 13 | Saprotroph | \|Undefined Saprotroph\| | Yeast | NULL | Probable | As nectar-tap_saprotroph (Põlme et al. 2020); Decay sugar-rich substrates (Põlme et al. 2020) | Põlme S, et al. 2020. Fungal Diversity 105:1-16 ((https://doi.org/10.1007/s13225-020-00466-2)) |
| Otu50705 | 1 | k__Fungi(100);p__Ascomycota(100);c__Saccharomycetes(100);o__Saccharomycetales(100);f__Metschnikowiaceae(100);g__Metschnikowia(100);s__Metschnikowia_pulcherrima(100); | Metschnikowia | 13 | Saprotroph | \|Undefined Saprotroph\| | Yeast | NULL | Probable | As nectar-tap_saprotroph (Põlme et al. 2020); Decay sugar-rich substrates (Põlme et al. 2020) | Põlme S, et al. 2020. Fungal Diversity 105:1-16 ((https://doi.org/10.1007/s13225-020-00466-2)) |
| Otu50706 | 1 | k__Fungi(100);p__Ascomycota(100);c__Saccharomycetes(100);o__Saccharomycetales(100);f__Metschnikowiaceae(100);g__Metschnikowia(100);s__Metschnikowia_pulcherrima(100); | Metschnikowia | 13 | Saprotroph | \|Undefined Saprotroph\| | Yeast | NULL | Probable | As nectar-tap_saprotroph (Põlme et al. 2020); Decay sugar-rich substrates (Põlme et al. 2020) | Põlme S, et al. 2020. Fungal Diversity 105:1-16 ((https://doi.org/10.1007/s13225-020-00466-2)) |
| Otu50707 | 1 | k__Fungi(100);p__Ascomycota(100);c__Saccharomycetes(100);o__Saccharomycetales(100);f__Metschnikowiaceae(100);g__Metschnikowia(100);s__Metschnikowia_pulcherrima(100); | Metschnikowia | 13 | Saprotroph | \|Undefined Saprotroph\| | Yeast | NULL | Probable | As nectar-tap_saprotroph (Põlme et al. 2020); Decay sugar-rich substrates (Põlme et al. 2020) | Põlme S, et al. 2020. Fungal Diversity 105:1-16 ((https://doi.org/10.1007/s13225-020-00466-2)) |
| Otu50709 | 1 | k__Fungi(100);p__Ascomycota(100);c__Saccharomycetes(100);o__Saccharomycetales(100);f__Saccharomycetales_family_Incertae_sedis(100);g__Candida(100);g__Candida_unclassified(100); | Candida | 13 | Pathotroph-Saprotroph-Symbiotroph | Animal Pathogen-Endophyte-Endosymbiont-Epiphyte-\|Undefined Saprotroph\| | Dimorphic Yeast-Yeast | NULL | Probable | As nectar-tap_saprotroph (Põlme et al. 2020); Decay sugar-rich substrates (Põlme et al. 2020); This asexual genus is polyphyletic and contain species that function very differently depend on what they are. Caution should be taken when interpreting guilds for Candida species. | Manolakaki D, et al. 2010. Virulence. 1: 367-375 ((https://doi.org/10.4161/viru.1.5.12796)); Mekbib SB, et al. 2011. Tropical Plant Pathology 36:233-240 ((https://doi.org/10.1590/S1982-56762011000400004)); Põlme S, et al. 2020. Fungal Diversity 105:1-16 ((https://doi.org/10.1007/s13225-020-00466-2)) |
| Otu50711 | 1 | k__Fungi(100);p__Ascomycota(100);c__Saccharomycetes(100);o__Saccharomycetales(100);f__Metschnikowiaceae(100);g__Metschnikowia(100);s__Metschnikowia_pulcherrima(100); | Metschnikowia | 13 | Saprotroph | \|Undefined Saprotroph\| | Yeast | NULL | Probable | As nectar-tap_saprotroph (Põlme et al. 2020); Decay sugar-rich substrates (Põlme et al. 2020) | Põlme S, et al. 2020. Fungal Diversity 105:1-16 ((https://doi.org/10.1007/s13225-020-00466-2)) |
| Otu50712 | 1 | k__Fungi(100);p__Ascomycota(100);c__Saccharomycetes(100);o__Saccharomycetales(100);f__Saccharomycetales_family_Incertae_sedis(100);g__Candida(100);g__Candida_unclassified(100); | Candida | 13 | Pathotroph-Saprotroph-Symbiotroph | Animal Pathogen-Endophyte-Endosymbiont-Epiphyte-\|Undefined Saprotroph\| | Dimorphic Yeast-Yeast | NULL | Probable | As nectar-tap_saprotroph (Põlme et al. 2020); Decay sugar-rich substrates (Põlme et al. 2020); This asexual genus is polyphyletic and contain species that function very differently depend on what they are. Caution should be taken when interpreting guilds for Candida species. | Manolakaki D, et al. 2010. Virulence. 1: 367-375 ((https://doi.org/10.4161/viru.1.5.12796)); Mekbib SB, et al. 2011. Tropical Plant Pathology 36:233-240 ((https://doi.org/10.1590/S1982-56762011000400004)); Põlme S, et al. 2020. Fungal Diversity 105:1-16 ((https://doi.org/10.1007/s13225-020-00466-2)) |
| Otu50714 | 1 | k__Fungi(100);p__Ascomycota(100);c__Saccharomycetes(100);o__Saccharomycetales(100);f__Metschnikowiaceae(100);g__Metschnikowia(100);s__Metschnikowia_pulcherrima(100); | Metschnikowia | 13 | Saprotroph | \|Undefined Saprotroph\| | Yeast | NULL | Probable | As nectar-tap_saprotroph (Põlme et al. 2020); Decay sugar-rich substrates (Põlme et al. 2020) | Põlme S, et al. 2020. Fungal Diversity 105:1-16 ((https://doi.org/10.1007/s13225-020-00466-2)) |
| Otu50715 | 1 | k__Fungi(100);p__Ascomycota(100);c__Saccharomycetes(100);o__Saccharomycetales(100);f__Metschnikowiaceae(100);g__Metschnikowia(100);g__Metschnikowia_unclassified(100); | Metschnikowia | 13 | Saprotroph | \|Undefined Saprotroph\| | Yeast | NULL | Probable | As nectar-tap_saprotroph (Põlme et al. 2020); Decay sugar-rich substrates (Põlme et al. 2020) | Põlme S, et al. 2020. Fungal Diversity 105:1-16 ((https://doi.org/10.1007/s13225-020-00466-2)) |
| Otu50716 | 1 | k__Fungi(100);p__Ascomycota(100);c__Saccharomycetes(100);o__Saccharomycetales(100);f__Metschnikowiaceae(100);g__Metschnikowia(100);s__Metschnikowia_pulcherrima(100); | Metschnikowia | 13 | Saprotroph | \|Undefined Saprotroph\| | Yeast | NULL | Probable | As nectar-tap_saprotroph (Põlme et al. 2020); Decay sugar-rich substrates (Põlme et al. 2020) | Põlme S, et al. 2020. Fungal Diversity 105:1-16 ((https://doi.org/10.1007/s13225-020-00466-2)) |
| Otu50717 | 1 | k__Fungi(100);p__Ascomycota(100);c__Saccharomycetes(100);o__Saccharomycetales(100);f__Metschnikowiaceae(100);g__Metschnikowia(100);g__Metschnikowia_unclassified(100); | Metschnikowia | 13 | Saprotroph | \|Undefined Saprotroph\| | Yeast | NULL | Probable | As nectar-tap_saprotroph (Põlme et al. 2020); Decay sugar-rich substrates (Põlme et al. 2020) | Põlme S, et al. 2020. Fungal Diversity 105:1-16 ((https://doi.org/10.1007/s13225-020-00466-2)) |
| Otu50719 | 1 | k__Fungi(100);p__Ascomycota(100);c__Saccharomycetes(100);o__Saccharomycetales(100);f__Metschnikowiaceae(100);g__Metschnikowia(100);s__Metschnikowia_pulcherrima(100); | Metschnikowia | 13 | Saprotroph | \|Undefined Saprotroph\| | Yeast | NULL | Probable | As nectar-tap_saprotroph (Põlme et al. 2020); Decay sugar-rich substrates (Põlme et al. 2020) | Põlme S, et al. 2020. Fungal Diversity 105:1-16 ((https://doi.org/10.1007/s13225-020-00466-2)) |
| Otu50720 | 1 | k__Fungi(100);p__Ascomycota(100);c__Saccharomycetes(100);o__Saccharomycetales(100);f__Metschnikowiaceae(100);g__Metschnikowia(100);s__Metschnikowia_pulcherrima(100); | Metschnikowia | 13 | Saprotroph | \|Undefined Saprotroph\| | Yeast | NULL | Probable | As nectar-tap_saprotroph (Põlme et al. 2020); Decay sugar-rich substrates (Põlme et al. 2020) | Põlme S, et al. 2020. Fungal Diversity 105:1-16 ((https://doi.org/10.1007/s13225-020-00466-2)) |
| Otu50721 | 1 | k__Fungi(100);p__Ascomycota(100);c__Saccharomycetes(100);o__Saccharomycetales(100);f__Metschnikowiaceae(100);g__Metschnikowia(100);s__Metschnikowia_pulcherrima(100); | Metschnikowia | 13 | Saprotroph | \|Undefined Saprotroph\| | Yeast | NULL | Probable | As nectar-tap_saprotroph (Põlme et al. 2020); Decay sugar-rich substrates (Põlme et al. 2020) | Põlme S, et al. 2020. Fungal Diversity 105:1-16 ((https://doi.org/10.1007/s13225-020-00466-2)) |
| Otu50722 | 1 | k__Fungi(100);p__Ascomycota(100);c__Saccharomycetes(100);o__Saccharomycetales(100);f__Metschnikowiaceae(100);g__Metschnikowia(100);g__Metschnikowia_unclassified(100); | Metschnikowia | 13 | Saprotroph | \|Undefined Saprotroph\| | Yeast | NULL | Probable | As nectar-tap_saprotroph (Põlme et al. 2020); Decay sugar-rich substrates (Põlme et al. 2020) | Põlme S, et al. 2020. Fungal Diversity 105:1-16 ((https://doi.org/10.1007/s13225-020-00466-2)) |
| Otu50724 | 1 | k__Fungi(100);p__Ascomycota(100);c__Saccharomycetes(100);o__Saccharomycetales(100);f__Metschnikowiaceae(100);g__Metschnikowia(100);s__Metschnikowia_pulcherrima(100); | Metschnikowia | 13 | Saprotroph | \|Undefined Saprotroph\| | Yeast | NULL | Probable | As nectar-tap_saprotroph (Põlme et al. 2020); Decay sugar-rich substrates (Põlme et al. 2020) | Põlme S, et al. 2020. Fungal Diversity 105:1-16 ((https://doi.org/10.1007/s13225-020-00466-2)) |
| Otu50725 | 1 | k__Fungi(100);p__Ascomycota(100);c__Saccharomycetes(100);o__Saccharomycetales(100);f__Metschnikowiaceae(100);g__Metschnikowia(100);s__Metschnikowia_pulcherrima(100); | Metschnikowia | 13 | Saprotroph | \|Undefined Saprotroph\| | Yeast | NULL | Probable | As nectar-tap_saprotroph (Põlme et al. 2020); Decay sugar-rich substrates (Põlme et al. 2020) | Põlme S, et al. 2020. Fungal Diversity 105:1-16 ((https://doi.org/10.1007/s13225-020-00466-2)) |
| Otu50726 | 1 | k__Fungi(100);p__Ascomycota(100);c__Saccharomycetes(100);o__Saccharomycetales(100);f__Metschnikowiaceae(100);g__Metschnikowia(100);s__Metschnikowia_pulcherrima(100); | Metschnikowia | 13 | Saprotroph | \|Undefined Saprotroph\| | Yeast | NULL | Probable | As nectar-tap_saprotroph (Põlme et al. 2020); Decay sugar-rich substrates (Põlme et al. 2020) | Põlme S, et al. 2020. Fungal Diversity 105:1-16 ((https://doi.org/10.1007/s13225-020-00466-2)) |
| Otu50727 | 1 | k__Fungi(100);p__Ascomycota(100);c__Saccharomycetes(100);o__Saccharomycetales(100);f__Metschnikowiaceae(100);g__Metschnikowia(100);s__Metschnikowia_pulcherrima(100); | Metschnikowia | 13 | Saprotroph | \|Undefined Saprotroph\| | Yeast | NULL | Probable | As nectar-tap_saprotroph (Põlme et al. 2020); Decay sugar-rich substrates (Põlme et al. 2020) | Põlme S, et al. 2020. Fungal Diversity 105:1-16 ((https://doi.org/10.1007/s13225-020-00466-2)) |
| Otu50728 | 1 | k__Fungi(100);p__Ascomycota(100);c__Saccharomycetes(100);o__Saccharomycetales(100);f__Metschnikowiaceae(100);g__Metschnikowia(100);g__Metschnikowia_unclassified(100); | Metschnikowia | 13 | Saprotroph | \|Undefined Saprotroph\| | Yeast | NULL | Probable | As nectar-tap_saprotroph (Põlme et al. 2020); Decay sugar-rich substrates (Põlme et al. 2020) | Põlme S, et al. 2020. Fungal Diversity 105:1-16 ((https://doi.org/10.1007/s13225-020-00466-2)) |
| Otu50729 | 1 | k__Fungi(100);p__Ascomycota(100);c__Saccharomycetes(100);o__Saccharomycetales(100);f__Metschnikowiaceae(100);g__Metschnikowia(100);s__Metschnikowia_pulcherrima(100); | Metschnikowia | 13 | Saprotroph | \|Undefined Saprotroph\| | Yeast | NULL | Probable | As nectar-tap_saprotroph (Põlme et al. 2020); Decay sugar-rich substrates (Põlme et al. 2020) | Põlme S, et al. 2020. Fungal Diversity 105:1-16 ((https://doi.org/10.1007/s13225-020-00466-2)) |
| Otu50730 | 1 | k__Fungi(100);p__Ascomycota(100);c__Saccharomycetes(100);o__Saccharomycetales(100);f__Metschnikowiaceae(100);g__Metschnikowia(100);s__Metschnikowia_pulcherrima(100); | Metschnikowia | 13 | Saprotroph | \|Undefined Saprotroph\| | Yeast | NULL | Probable | As nectar-tap_saprotroph (Põlme et al. 2020); Decay sugar-rich substrates (Põlme et al. 2020) | Põlme S, et al. 2020. Fungal Diversity 105:1-16 ((https://doi.org/10.1007/s13225-020-00466-2)) |
| Otu50731 | 1 | k__Fungi(100);p__Ascomycota(100);c__Saccharomycetes(100);o__Saccharomycetales(100);f__Metschnikowiaceae(100);g__Metschnikowia(100);s__Metschnikowia_pulcherrima(100); | Metschnikowia | 13 | Saprotroph | \|Undefined Saprotroph\| | Yeast | NULL | Probable | As nectar-tap_saprotroph (Põlme et al. 2020); Decay sugar-rich substrates (Põlme et al. 2020) | Põlme S, et al. 2020. Fungal Diversity 105:1-16 ((https://doi.org/10.1007/s13225-020-00466-2)) |
| Otu50732 | 1 | k__Fungi(100);p__Ascomycota(100);c__Saccharomycetes(100);o__Saccharomycetales(100);f__Metschnikowiaceae(100);g__Metschnikowia(100);s__Metschnikowia_pulcherrima(100); | Metschnikowia | 13 | Saprotroph | \|Undefined Saprotroph\| | Yeast | NULL | Probable | As nectar-tap_saprotroph (Põlme et al. 2020); Decay sugar-rich substrates (Põlme et al. 2020) | Põlme S, et al. 2020. Fungal Diversity 105:1-16 ((https://doi.org/10.1007/s13225-020-00466-2)) |
| Otu50733 | 1 | k__Fungi(100);p__Ascomycota(100);c__Saccharomycetes(100);o__Saccharomycetales(100);f__Metschnikowiaceae(100);g__Metschnikowia(100);g__Metschnikowia_unclassified(100); | Metschnikowia | 13 | Saprotroph | \|Undefined Saprotroph\| | Yeast | NULL | Probable | As nectar-tap_saprotroph (Põlme et al. 2020); Decay sugar-rich substrates (Põlme et al. 2020) | Põlme S, et al. 2020. Fungal Diversity 105:1-16 ((https://doi.org/10.1007/s13225-020-00466-2)) |
| Otu50734 | 1 | k__Fungi(100);p__Ascomycota(100);c__Saccharomycetes(100);o__Saccharomycetales(100);f__Metschnikowiaceae(100);g__Metschnikowia(100);s__Metschnikowia_pulcherrima(100); | Metschnikowia | 13 | Saprotroph | \|Undefined Saprotroph\| | Yeast | NULL | Probable | As nectar-tap_saprotroph (Põlme et al. 2020); Decay sugar-rich substrates (Põlme et al. 2020) | Põlme S, et al. 2020. Fungal Diversity 105:1-16 ((https://doi.org/10.1007/s13225-020-00466-2)) |
| Otu50735 | 1 | k__Fungi(100);p__Ascomycota(100);c__Saccharomycetes(100);o__Saccharomycetales(100);f__Metschnikowiaceae(100);g__Metschnikowia(100);g__Metschnikowia_unclassified(100); | Metschnikowia | 13 | Saprotroph | \|Undefined Saprotroph\| | Yeast | NULL | Probable | As nectar-tap_saprotroph (Põlme et al. 2020); Decay sugar-rich substrates (Põlme et al. 2020) | Põlme S, et al. 2020. Fungal Diversity 105:1-16 ((https://doi.org/10.1007/s13225-020-00466-2)) |
| Otu50736 | 1 | k__Fungi(100);p__Ascomycota(100);c__Saccharomycetes(100);o__Saccharomycetales(100);f__Metschnikowiaceae(100);g__Metschnikowia(100);g__Metschnikowia_unclassified(100); | Metschnikowia | 13 | Saprotroph | \|Undefined Saprotroph\| | Yeast | NULL | Probable | As nectar-tap_saprotroph (Põlme et al. 2020); Decay sugar-rich substrates (Põlme et al. 2020) | Põlme S, et al. 2020. Fungal Diversity 105:1-16 ((https://doi.org/10.1007/s13225-020-00466-2)) |
| Otu50737 | 1 | k__Fungi(100);p__Ascomycota(100);c__Saccharomycetes(100);o__Saccharomycetales(100);f__Metschnikowiaceae(100);g__Metschnikowia(100);s__Metschnikowia_pulcherrima(100); | Metschnikowia | 13 | Saprotroph | \|Undefined Saprotroph\| | Yeast | NULL | Probable | As nectar-tap_saprotroph (Põlme et al. 2020); Decay sugar-rich substrates (Põlme et al. 2020) | Põlme S, et al. 2020. Fungal Diversity 105:1-16 ((https://doi.org/10.1007/s13225-020-00466-2)) |
| Otu50738 | 1 | k__Fungi(100);p__Ascomycota(100);c__Saccharomycetes(100);o__Saccharomycetales(100);f__Metschnikowiaceae(100);g__Metschnikowia(100);g__Metschnikowia_unclassified(100); | Metschnikowia | 13 | Saprotroph | \|Undefined Saprotroph\| | Yeast | NULL | Probable | As nectar-tap_saprotroph (Põlme et al. 2020); Decay sugar-rich substrates (Põlme et al. 2020) | Põlme S, et al. 2020. Fungal Diversity 105:1-16 ((https://doi.org/10.1007/s13225-020-00466-2)) |
| Otu50739 | 1 | k__Fungi(100);p__Ascomycota(100);c__Saccharomycetes(100);o__Saccharomycetales(100);f__Metschnikowiaceae(100);g__Metschnikowia(100);s__Metschnikowia_pulcherrima(100); | Metschnikowia | 13 | Saprotroph | \|Undefined Saprotroph\| | Yeast | NULL | Probable | As nectar-tap_saprotroph (Põlme et al. 2020); Decay sugar-rich substrates (Põlme et al. 2020) | Põlme S, et al. 2020. Fungal Diversity 105:1-16 ((https://doi.org/10.1007/s13225-020-00466-2)) |
| Otu50740 | 1 | k__Fungi(100);p__Ascomycota(100);c__Saccharomycetes(100);o__Saccharomycetales(100);f__Metschnikowiaceae(100);g__Metschnikowia(100);s__Metschnikowia_pulcherrima(100); | Metschnikowia | 13 | Saprotroph | \|Undefined Saprotroph\| | Yeast | NULL | Probable | As nectar-tap_saprotroph (Põlme et al. 2020); Decay sugar-rich substrates (Põlme et al. 2020) | Põlme S, et al. 2020. Fungal Diversity 105:1-16 ((https://doi.org/10.1007/s13225-020-00466-2)) |
| Otu50741 | 1 | k__Fungi(100);p__Ascomycota(100);c__Saccharomycetes(100);o__Saccharomycetales(100);f__Metschnikowiaceae(100);g__Metschnikowia(100);g__Metschnikowia_unclassified(100); | Metschnikowia | 13 | Saprotroph | \|Undefined Saprotroph\| | Yeast | NULL | Probable | As nectar-tap_saprotroph (Põlme et al. 2020); Decay sugar-rich substrates (Põlme et al. 2020) | Põlme S, et al. 2020. Fungal Diversity 105:1-16 ((https://doi.org/10.1007/s13225-020-00466-2)) |
| Otu50742 | 1 | k__Fungi(100);p__Ascomycota(100);c__Saccharomycetes(100);o__Saccharomycetales(100);f__Metschnikowiaceae(100);g__Metschnikowia(100);s__Metschnikowia_pulcherrima(100); | Metschnikowia | 13 | Saprotroph | \|Undefined Saprotroph\| | Yeast | NULL | Probable | As nectar-tap_saprotroph (Põlme et al. 2020); Decay sugar-rich substrates (Põlme et al. 2020) | Põlme S, et al. 2020. Fungal Diversity 105:1-16 ((https://doi.org/10.1007/s13225-020-00466-2)) |
| Otu50743 | 1 | k__Fungi(100);p__Ascomycota(100);c__Saccharomycetes(100);o__Saccharomycetales(100);f__Metschnikowiaceae(100);g__Metschnikowia(100);s__Metschnikowia_pulcherrima(100); | Metschnikowia | 13 | Saprotroph | \|Undefined Saprotroph\| | Yeast | NULL | Probable | As nectar-tap_saprotroph (Põlme et al. 2020); Decay sugar-rich substrates (Põlme et al. 2020) | Põlme S, et al. 2020. Fungal Diversity 105:1-16 ((https://doi.org/10.1007/s13225-020-00466-2)) |
| Otu50744 | 1 | k__Fungi(100);p__Ascomycota(100);c__Saccharomycetes(100);o__Saccharomycetales(100);f__Metschnikowiaceae(100);g__Metschnikowia(100);g__Metschnikowia_unclassified(100); | Metschnikowia | 13 | Saprotroph | \|Undefined Saprotroph\| | Yeast | NULL | Probable | As nectar-tap_saprotroph (Põlme et al. 2020); Decay sugar-rich substrates (Põlme et al. 2020) | Põlme S, et al. 2020. Fungal Diversity 105:1-16 ((https://doi.org/10.1007/s13225-020-00466-2)) |
| Otu50745 | 1 | k__Fungi(100);p__Ascomycota(100);c__Saccharomycetes(100);o__Saccharomycetales(100);f__Metschnikowiaceae(100);g__Metschnikowia(100);s__Metschnikowia_pulcherrima(100); | Metschnikowia | 13 | Saprotroph | \|Undefined Saprotroph\| | Yeast | NULL | Probable | As nectar-tap_saprotroph (Põlme et al. 2020); Decay sugar-rich substrates (Põlme et al. 2020) | Põlme S, et al. 2020. Fungal Diversity 105:1-16 ((https://doi.org/10.1007/s13225-020-00466-2)) |
| Otu50746 | 1 | k__Fungi(100);p__Ascomycota(100);c__Saccharomycetes(100);o__Saccharomycetales(100);f__Metschnikowiaceae(100);g__Metschnikowia(100);g__Metschnikowia_unclassified(100); | Metschnikowia | 13 | Saprotroph | \|Undefined Saprotroph\| | Yeast | NULL | Probable | As nectar-tap_saprotroph (Põlme et al. 2020); Decay sugar-rich substrates (Põlme et al. 2020) | Põlme S, et al. 2020. Fungal Diversity 105:1-16 ((https://doi.org/10.1007/s13225-020-00466-2)) |
| Otu50747 | 1 | k__Fungi(100);p__Ascomycota(100);c__Saccharomycetes(100);o__Saccharomycetales(100);f__Metschnikowiaceae(100);g__Metschnikowia(100);s__Metschnikowia_pulcherrima(100); | Metschnikowia | 13 | Saprotroph | \|Undefined Saprotroph\| | Yeast | NULL | Probable | As nectar-tap_saprotroph (Põlme et al. 2020); Decay sugar-rich substrates (Põlme et al. 2020) | Põlme S, et al. 2020. Fungal Diversity 105:1-16 ((https://doi.org/10.1007/s13225-020-00466-2)) |
| Otu50748 | 1 | k__Fungi(100);p__Ascomycota(100);c__Saccharomycetes(100);o__Saccharomycetales(100);f__Metschnikowiaceae(100);g__Metschnikowia(100);s__Metschnikowia_pulcherrima(100); | Metschnikowia | 13 | Saprotroph | \|Undefined Saprotroph\| | Yeast | NULL | Probable | As nectar-tap_saprotroph (Põlme et al. 2020); Decay sugar-rich substrates (Põlme et al. 2020) | Põlme S, et al. 2020. Fungal Diversity 105:1-16 ((https://doi.org/10.1007/s13225-020-00466-2)) |
| Otu50749 | 1 | k__Fungi(100);p__Ascomycota(100);c__Saccharomycetes(100);o__Saccharomycetales(100);f__Metschnikowiaceae(100);g__Metschnikowia(100);s__Metschnikowia_pulcherrima(100); | Metschnikowia | 13 | Saprotroph | \|Undefined Saprotroph\| | Yeast | NULL | Probable | As nectar-tap_saprotroph (Põlme et al. 2020); Decay sugar-rich substrates (Põlme et al. 2020) | Põlme S, et al. 2020. Fungal Diversity 105:1-16 ((https://doi.org/10.1007/s13225-020-00466-2)) |
| Otu50750 | 1 | k__Fungi(100);p__Ascomycota(100);c__Saccharomycetes(100);o__Saccharomycetales(100);f__Metschnikowiaceae(100);g__Metschnikowia(100);s__Metschnikowia_pulcherrima(100); | Metschnikowia | 13 | Saprotroph | \|Undefined Saprotroph\| | Yeast | NULL | Probable | As nectar-tap_saprotroph (Põlme et al. 2020); Decay sugar-rich substrates (Põlme et al. 2020) | Põlme S, et al. 2020. Fungal Diversity 105:1-16 ((https://doi.org/10.1007/s13225-020-00466-2)) |
| Otu50755 | 1 | k__Fungi(100);p__Ascomycota(100);c__Saccharomycetes(100);o__Saccharomycetales(100);f__Metschnikowiaceae(100);g__Metschnikowia(100);g__Metschnikowia_unclassified(100); | Metschnikowia | 13 | Saprotroph | \|Undefined Saprotroph\| | Yeast | NULL | Probable | As nectar-tap_saprotroph (Põlme et al. 2020); Decay sugar-rich substrates (Põlme et al. 2020) | Põlme S, et al. 2020. Fungal Diversity 105:1-16 ((https://doi.org/10.1007/s13225-020-00466-2)) |
| Otu50756 | 1 | k__Fungi(100);p__Ascomycota(100);c__Saccharomycetes(100);o__Saccharomycetales(100);f__Metschnikowiaceae(100);g__Metschnikowia(100);s__Metschnikowia_pulcherrima(100); | Metschnikowia | 13 | Saprotroph | \|Undefined Saprotroph\| | Yeast | NULL | Probable | As nectar-tap_saprotroph (Põlme et al. 2020); Decay sugar-rich substrates (Põlme et al. 2020) | Põlme S, et al. 2020. Fungal Diversity 105:1-16 ((https://doi.org/10.1007/s13225-020-00466-2)) |
| Otu50757 | 1 | k__Fungi(100);p__Ascomycota(100);c__Saccharomycetes(100);o__Saccharomycetales(100);f__Metschnikowiaceae(100);g__Metschnikowia(100);s__Metschnikowia_pulcherrima(100); | Metschnikowia | 13 | Saprotroph | \|Undefined Saprotroph\| | Yeast | NULL | Probable | As nectar-tap_saprotroph (Põlme et al. 2020); Decay sugar-rich substrates (Põlme et al. 2020) | Põlme S, et al. 2020. Fungal Diversity 105:1-16 ((https://doi.org/10.1007/s13225-020-00466-2)) |
| Otu50758 | 1 | k__Fungi(100);p__Ascomycota(100);c__Saccharomycetes(100);o__Saccharomycetales(100);f__Metschnikowiaceae(100);g__Metschnikowia(100);s__Metschnikowia_pulcherrima(100); | Metschnikowia | 13 | Saprotroph | \|Undefined Saprotroph\| | Yeast | NULL | Probable | As nectar-tap_saprotroph (Põlme et al. 2020); Decay sugar-rich substrates (Põlme et al. 2020) | Põlme S, et al. 2020. Fungal Diversity 105:1-16 ((https://doi.org/10.1007/s13225-020-00466-2)) |
| Otu50760 | 1 | k__Fungi(100);p__Ascomycota(100);c__Saccharomycetes(100);o__Saccharomycetales(100);f__Metschnikowiaceae(100);g__Metschnikowia(100);g__Metschnikowia_unclassified(100); | Metschnikowia | 13 | Saprotroph | \|Undefined Saprotroph\| | Yeast | NULL | Probable | As nectar-tap_saprotroph (Põlme et al. 2020); Decay sugar-rich substrates (Põlme et al. 2020) | Põlme S, et al. 2020. Fungal Diversity 105:1-16 ((https://doi.org/10.1007/s13225-020-00466-2)) |
| Otu50761 | 1 | k__Fungi(100);p__Ascomycota(100);c__Saccharomycetes(100);o__Saccharomycetales(100);f__Metschnikowiaceae(100);g__Metschnikowia(100);s__Metschnikowia_pulcherrima(100); | Metschnikowia | 13 | Saprotroph | \|Undefined Saprotroph\| | Yeast | NULL | Probable | As nectar-tap_saprotroph (Põlme et al. 2020); Decay sugar-rich substrates (Põlme et al. 2020) | Põlme S, et al. 2020. Fungal Diversity 105:1-16 ((https://doi.org/10.1007/s13225-020-00466-2)) |
| Otu50762 | 1 | k__Fungi(100);p__Ascomycota(100);c__Saccharomycetes(100);o__Saccharomycetales(100);f__Metschnikowiaceae(100);g__Metschnikowia(100);s__Metschnikowia_pulcherrima(100); | Metschnikowia | 13 | Saprotroph | \|Undefined Saprotroph\| | Yeast | NULL | Probable | As nectar-tap_saprotroph (Põlme et al. 2020); Decay sugar-rich substrates (Põlme et al. 2020) | Põlme S, et al. 2020. Fungal Diversity 105:1-16 ((https://doi.org/10.1007/s13225-020-00466-2)) |
| Otu50763 | 1 | k__Fungi(100);p__Ascomycota(100);c__Saccharomycetes(100);o__Saccharomycetales(100);f__Metschnikowiaceae(100);g__Metschnikowia(100);g__Metschnikowia_unclassified(100); | Metschnikowia | 13 | Saprotroph | \|Undefined Saprotroph\| | Yeast | NULL | Probable | As nectar-tap_saprotroph (Põlme et al. 2020); Decay sugar-rich substrates (Põlme et al. 2020) | Põlme S, et al. 2020. Fungal Diversity 105:1-16 ((https://doi.org/10.1007/s13225-020-00466-2)) |
| Otu50764 | 1 | k__Fungi(100);p__Ascomycota(100);c__Saccharomycetes(100);o__Saccharomycetales(100);f__Metschnikowiaceae(100);g__Metschnikowia(100);s__Metschnikowia_pulcherrima(100); | Metschnikowia | 13 | Saprotroph | \|Undefined Saprotroph\| | Yeast | NULL | Probable | As nectar-tap_saprotroph (Põlme et al. 2020); Decay sugar-rich substrates (Põlme et al. 2020) | Põlme S, et al. 2020. Fungal Diversity 105:1-16 ((https://doi.org/10.1007/s13225-020-00466-2)) |
| Otu50765 | 1 | k__Fungi(100);p__Ascomycota(100);c__Saccharomycetes(100);o__Saccharomycetales(100);f__Metschnikowiaceae(100);g__Metschnikowia(100);s__Metschnikowia_pulcherrima(100); | Metschnikowia | 13 | Saprotroph | \|Undefined Saprotroph\| | Yeast | NULL | Probable | As nectar-tap_saprotroph (Põlme et al. 2020); Decay sugar-rich substrates (Põlme et al. 2020) | Põlme S, et al. 2020. Fungal Diversity 105:1-16 ((https://doi.org/10.1007/s13225-020-00466-2)) |
| Otu50766 | 1 | k__Fungi(100);p__Ascomycota(100);c__Saccharomycetes(100);o__Saccharomycetales(100);f__Metschnikowiaceae(100);g__Metschnikowia(100);s__Metschnikowia_pulcherrima(100); | Metschnikowia | 13 | Saprotroph | \|Undefined Saprotroph\| | Yeast | NULL | Probable | As nectar-tap_saprotroph (Põlme et al. 2020); Decay sugar-rich substrates (Põlme et al. 2020) | Põlme S, et al. 2020. Fungal Diversity 105:1-16 ((https://doi.org/10.1007/s13225-020-00466-2)) |
| Otu50768 | 1 | k__Fungi(100);p__Ascomycota(100);c__Saccharomycetes(100);o__Saccharomycetales(100);f__Metschnikowiaceae(100);g__Metschnikowia(100);g__Metschnikowia_unclassified(100); | Metschnikowia | 13 | Saprotroph | \|Undefined Saprotroph\| | Yeast | NULL | Probable | As nectar-tap_saprotroph (Põlme et al. 2020); Decay sugar-rich substrates (Põlme et al. 2020) | Põlme S, et al. 2020. Fungal Diversity 105:1-16 ((https://doi.org/10.1007/s13225-020-00466-2)) |
| Otu50769 | 1 | k__Fungi(100);p__Ascomycota(100);c__Saccharomycetes(100);o__Saccharomycetales(100);f__Metschnikowiaceae(100);g__Metschnikowia(100);g__Metschnikowia_unclassified(100); | Metschnikowia | 13 | Saprotroph | \|Undefined Saprotroph\| | Yeast | NULL | Probable | As nectar-tap_saprotroph (Põlme et al. 2020); Decay sugar-rich substrates (Põlme et al. 2020) | Põlme S, et al. 2020. Fungal Diversity 105:1-16 ((https://doi.org/10.1007/s13225-020-00466-2)) |
| Otu50770 | 1 | k__Fungi(100);p__Ascomycota(100);c__Saccharomycetes(100);o__Saccharomycetales(100);f__Metschnikowiaceae(100);g__Metschnikowia(100);s__Metschnikowia_pulcherrima(100); | Metschnikowia | 13 | Saprotroph | \|Undefined Saprotroph\| | Yeast | NULL | Probable | As nectar-tap_saprotroph (Põlme et al. 2020); Decay sugar-rich substrates (Põlme et al. 2020) | Põlme S, et al. 2020. Fungal Diversity 105:1-16 ((https://doi.org/10.1007/s13225-020-00466-2)) |
| Otu50771 | 1 | k__Fungi(100);p__Ascomycota(100);c__Saccharomycetes(100);o__Saccharomycetales(100);f__Metschnikowiaceae(100);g__Metschnikowia(100);g__Metschnikowia_unclassified(100); | Metschnikowia | 13 | Saprotroph | \|Undefined Saprotroph\| | Yeast | NULL | Probable | As nectar-tap_saprotroph (Põlme et al. 2020); Decay sugar-rich substrates (Põlme et al. 2020) | Põlme S, et al. 2020. Fungal Diversity 105:1-16 ((https://doi.org/10.1007/s13225-020-00466-2)) |
| Otu50772 | 1 | k__Fungi(100);p__Ascomycota(100);c__Saccharomycetes(100);o__Saccharomycetales(100);f__Metschnikowiaceae(100);g__Metschnikowia(100);g__Metschnikowia_unclassified(100); | Metschnikowia | 13 | Saprotroph | \|Undefined Saprotroph\| | Yeast | NULL | Probable | As nectar-tap_saprotroph (Põlme et al. 2020); Decay sugar-rich substrates (Põlme et al. 2020) | Põlme S, et al. 2020. Fungal Diversity 105:1-16 ((https://doi.org/10.1007/s13225-020-00466-2)) |
| Otu50773 | 1 | k__Fungi(100);p__Ascomycota(100);c__Saccharomycetes(100);o__Saccharomycetales(100);f__Metschnikowiaceae(100);g__Metschnikowia(100);s__Metschnikowia_pulcherrima(100); | Metschnikowia | 13 | Saprotroph | \|Undefined Saprotroph\| | Yeast | NULL | Probable | As nectar-tap_saprotroph (Põlme et al. 2020); Decay sugar-rich substrates (Põlme et al. 2020) | Põlme S, et al. 2020. Fungal Diversity 105:1-16 ((https://doi.org/10.1007/s13225-020-00466-2)) |
| Otu50774 | 1 | k__Fungi(100);p__Ascomycota(100);c__Saccharomycetes(100);o__Saccharomycetales(100);f__Metschnikowiaceae(100);g__Metschnikowia(100);s__Metschnikowia_pulcherrima(100); | Metschnikowia | 13 | Saprotroph | \|Undefined Saprotroph\| | Yeast | NULL | Probable | As nectar-tap_saprotroph (Põlme et al. 2020); Decay sugar-rich substrates (Põlme et al. 2020) | Põlme S, et al. 2020. Fungal Diversity 105:1-16 ((https://doi.org/10.1007/s13225-020-00466-2)) |
| Otu50775 | 1 | k__Fungi(100);p__Ascomycota(100);c__Saccharomycetes(100);o__Saccharomycetales(100);f__Metschnikowiaceae(100);g__Metschnikowia(100);s__Metschnikowia_pulcherrima(100); | Metschnikowia | 13 | Saprotroph | \|Undefined Saprotroph\| | Yeast | NULL | Probable | As nectar-tap_saprotroph (Põlme et al. 2020); Decay sugar-rich substrates (Põlme et al. 2020) | Põlme S, et al. 2020. Fungal Diversity 105:1-16 ((https://doi.org/10.1007/s13225-020-00466-2)) |
| Otu50776 | 1 | k__Fungi(100);p__Ascomycota(100);c__Saccharomycetes(100);o__Saccharomycetales(100);f__Metschnikowiaceae(100);g__Metschnikowia(100);g__Metschnikowia_unclassified(100); | Metschnikowia | 13 | Saprotroph | \|Undefined Saprotroph\| | Yeast | NULL | Probable | As nectar-tap_saprotroph (Põlme et al. 2020); Decay sugar-rich substrates (Põlme et al. 2020) | Põlme S, et al. 2020. Fungal Diversity 105:1-16 ((https://doi.org/10.1007/s13225-020-00466-2)) |
| Otu50777 | 1 | k__Fungi(100);p__Ascomycota(100);c__Saccharomycetes(100);o__Saccharomycetales(100);f__Metschnikowiaceae(100);g__Metschnikowia(100);s__Metschnikowia_pulcherrima(100); | Metschnikowia | 13 | Saprotroph | \|Undefined Saprotroph\| | Yeast | NULL | Probable | As nectar-tap_saprotroph (Põlme et al. 2020); Decay sugar-rich substrates (Põlme et al. 2020) | Põlme S, et al. 2020. Fungal Diversity 105:1-16 ((https://doi.org/10.1007/s13225-020-00466-2)) |
| Otu50779 | 1 | k__Fungi(100);p__Ascomycota(100);c__Saccharomycetes(100);o__Saccharomycetales(100);f__Metschnikowiaceae(100);g__Metschnikowia(100);s__Metschnikowia_pulcherrima(100); | Metschnikowia | 13 | Saprotroph | \|Undefined Saprotroph\| | Yeast | NULL | Probable | As nectar-tap_saprotroph (Põlme et al. 2020); Decay sugar-rich substrates (Põlme et al. 2020) | Põlme S, et al. 2020. Fungal Diversity 105:1-16 ((https://doi.org/10.1007/s13225-020-00466-2)) |
| Otu50780 | 1 | k__Fungi(100);p__Ascomycota(100);c__Saccharomycetes(100);o__Saccharomycetales(100);f__Metschnikowiaceae(100);g__Metschnikowia(100);s__Metschnikowia_pulcherrima(100); | Metschnikowia | 13 | Saprotroph | \|Undefined Saprotroph\| | Yeast | NULL | Probable | As nectar-tap_saprotroph (Põlme et al. 2020); Decay sugar-rich substrates (Põlme et al. 2020) | Põlme S, et al. 2020. Fungal Diversity 105:1-16 ((https://doi.org/10.1007/s13225-020-00466-2)) |
| Otu50781 | 1 | k__Fungi(100);p__Ascomycota(100);c__Saccharomycetes(100);o__Saccharomycetales(100);f__Metschnikowiaceae(100);g__Metschnikowia(100);s__Metschnikowia_pulcherrima(100); | Metschnikowia | 13 | Saprotroph | \|Undefined Saprotroph\| | Yeast | NULL | Probable | As nectar-tap_saprotroph (Põlme et al. 2020); Decay sugar-rich substrates (Põlme et al. 2020) | Põlme S, et al. 2020. Fungal Diversity 105:1-16 ((https://doi.org/10.1007/s13225-020-00466-2)) |
| Otu50782 | 1 | k__Fungi(100);p__Ascomycota(100);c__Saccharomycetes(100);o__Saccharomycetales(100);f__Metschnikowiaceae(100);g__Metschnikowia(100);s__Metschnikowia_pulcherrima(100); | Metschnikowia | 13 | Saprotroph | \|Undefined Saprotroph\| | Yeast | NULL | Probable | As nectar-tap_saprotroph (Põlme et al. 2020); Decay sugar-rich substrates (Põlme et al. 2020) | Põlme S, et al. 2020. Fungal Diversity 105:1-16 ((https://doi.org/10.1007/s13225-020-00466-2)) |
| Otu50783 | 1 | k__Fungi(100);p__Ascomycota(100);c__Saccharomycetes(100);o__Saccharomycetales(100);f__Metschnikowiaceae(100);g__Metschnikowia(100);g__Metschnikowia_unclassified(100); | Metschnikowia | 13 | Saprotroph | \|Undefined Saprotroph\| | Yeast | NULL | Probable | As nectar-tap_saprotroph (Põlme et al. 2020); Decay sugar-rich substrates (Põlme et al. 2020) | Põlme S, et al. 2020. Fungal Diversity 105:1-16 ((https://doi.org/10.1007/s13225-020-00466-2)) |
| Otu50784 | 1 | k__Fungi(100);p__Ascomycota(100);c__Saccharomycetes(100);o__Saccharomycetales(100);f__Metschnikowiaceae(100);g__Metschnikowia(100);s__Metschnikowia_pulcherrima(100); | Metschnikowia | 13 | Saprotroph | \|Undefined Saprotroph\| | Yeast | NULL | Probable | As nectar-tap_saprotroph (Põlme et al. 2020); Decay sugar-rich substrates (Põlme et al. 2020) | Põlme S, et al. 2020. Fungal Diversity 105:1-16 ((https://doi.org/10.1007/s13225-020-00466-2)) |
| Otu50785 | 1 | k__Fungi(100);p__Ascomycota(100);c__Saccharomycetes(100);o__Saccharomycetales(100);f__Metschnikowiaceae(100);g__Metschnikowia(100);s__Metschnikowia_pulcherrima(100); | Metschnikowia | 13 | Saprotroph | \|Undefined Saprotroph\| | Yeast | NULL | Probable | As nectar-tap_saprotroph (Põlme et al. 2020); Decay sugar-rich substrates (Põlme et al. 2020) | Põlme S, et al. 2020. Fungal Diversity 105:1-16 ((https://doi.org/10.1007/s13225-020-00466-2)) |
| Otu50786 | 1 | k__Fungi(100);p__Ascomycota(100);c__Saccharomycetes(100);o__Saccharomycetales(100);f__Metschnikowiaceae(100);g__Metschnikowia(100);g__Metschnikowia_unclassified(100); | Metschnikowia | 13 | Saprotroph | \|Undefined Saprotroph\| | Yeast | NULL | Probable | As nectar-tap_saprotroph (Põlme et al. 2020); Decay sugar-rich substrates (Põlme et al. 2020) | Põlme S, et al. 2020. Fungal Diversity 105:1-16 ((https://doi.org/10.1007/s13225-020-00466-2)) |
| Otu50787 | 1 | k__Fungi(100);p__Ascomycota(100);c__Saccharomycetes(100);o__Saccharomycetales(100);f__Metschnikowiaceae(100);g__Metschnikowia(100);g__Metschnikowia_unclassified(100); | Metschnikowia | 13 | Saprotroph | \|Undefined Saprotroph\| | Yeast | NULL | Probable | As nectar-tap_saprotroph (Põlme et al. 2020); Decay sugar-rich substrates (Põlme et al. 2020) | Põlme S, et al. 2020. Fungal Diversity 105:1-16 ((https://doi.org/10.1007/s13225-020-00466-2)) |
| Otu50788 | 1 | k__Fungi(100);p__Ascomycota(100);c__Saccharomycetes(100);o__Saccharomycetales(100);f__Metschnikowiaceae(100);g__Metschnikowia(100);s__Metschnikowia_pulcherrima(100); | Metschnikowia | 13 | Saprotroph | \|Undefined Saprotroph\| | Yeast | NULL | Probable | As nectar-tap_saprotroph (Põlme et al. 2020); Decay sugar-rich substrates (Põlme et al. 2020) | Põlme S, et al. 2020. Fungal Diversity 105:1-16 ((https://doi.org/10.1007/s13225-020-00466-2)) |
| Otu50790 | 1 | k__Fungi(100);p__Ascomycota(100);c__Saccharomycetes(100);o__Saccharomycetales(100);f__Metschnikowiaceae(100);g__Metschnikowia(100);s__Metschnikowia_pulcherrima(100); | Metschnikowia | 13 | Saprotroph | \|Undefined Saprotroph\| | Yeast | NULL | Probable | As nectar-tap_saprotroph (Põlme et al. 2020); Decay sugar-rich substrates (Põlme et al. 2020) | Põlme S, et al. 2020. Fungal Diversity 105:1-16 ((https://doi.org/10.1007/s13225-020-00466-2)) |
| Otu50791 | 1 | k__Fungi(100);p__Ascomycota(100);c__Saccharomycetes(100);o__Saccharomycetales(100);f__Metschnikowiaceae(100);g__Metschnikowia(100);g__Metschnikowia_unclassified(100); | Metschnikowia | 13 | Saprotroph | \|Undefined Saprotroph\| | Yeast | NULL | Probable | As nectar-tap_saprotroph (Põlme et al. 2020); Decay sugar-rich substrates (Põlme et al. 2020) | Põlme S, et al. 2020. Fungal Diversity 105:1-16 ((https://doi.org/10.1007/s13225-020-00466-2)) |
| Otu50792 | 1 | k__Fungi(100);p__Ascomycota(100);c__Saccharomycetes(100);o__Saccharomycetales(100);f__Metschnikowiaceae(100);g__Metschnikowia(100);s__Metschnikowia_pulcherrima(100); | Metschnikowia | 13 | Saprotroph | \|Undefined Saprotroph\| | Yeast | NULL | Probable | As nectar-tap_saprotroph (Põlme et al. 2020); Decay sugar-rich substrates (Põlme et al. 2020) | Põlme S, et al. 2020. Fungal Diversity 105:1-16 ((https://doi.org/10.1007/s13225-020-00466-2)) |
| Otu50793 | 1 | k__Fungi(100);p__Ascomycota(100);c__Saccharomycetes(100);o__Saccharomycetales(100);f__Metschnikowiaceae(100);g__Metschnikowia(100);s__Metschnikowia_pulcherrima(100); | Metschnikowia | 13 | Saprotroph | \|Undefined Saprotroph\| | Yeast | NULL | Probable | As nectar-tap_saprotroph (Põlme et al. 2020); Decay sugar-rich substrates (Põlme et al. 2020) | Põlme S, et al. 2020. Fungal Diversity 105:1-16 ((https://doi.org/10.1007/s13225-020-00466-2)) |
| Otu50794 | 1 | k__Fungi(100);p__Ascomycota(100);c__Saccharomycetes(100);o__Saccharomycetales(100);f__Metschnikowiaceae(100);g__Metschnikowia(100);s__Metschnikowia_pulcherrima(100); | Metschnikowia | 13 | Saprotroph | \|Undefined Saprotroph\| | Yeast | NULL | Probable | As nectar-tap_saprotroph (Põlme et al. 2020); Decay sugar-rich substrates (Põlme et al. 2020) | Põlme S, et al. 2020. Fungal Diversity 105:1-16 ((https://doi.org/10.1007/s13225-020-00466-2)) |
| Otu50795 | 1 | k__Fungi(100);p__Ascomycota(100);c__Saccharomycetes(100);o__Saccharomycetales(100);f__Metschnikowiaceae(100);g__Metschnikowia(100);s__Metschnikowia_pulcherrima(100); | Metschnikowia | 13 | Saprotroph | \|Undefined Saprotroph\| | Yeast | NULL | Probable | As nectar-tap_saprotroph (Põlme et al. 2020); Decay sugar-rich substrates (Põlme et al. 2020) | Põlme S, et al. 2020. Fungal Diversity 105:1-16 ((https://doi.org/10.1007/s13225-020-00466-2)) |
| Otu50796 | 1 | k__Fungi(100);p__Ascomycota(100);c__Saccharomycetes(100);o__Saccharomycetales(100);f__Metschnikowiaceae(100);g__Metschnikowia(100);s__Metschnikowia_pulcherrima(100); | Metschnikowia | 13 | Saprotroph | \|Undefined Saprotroph\| | Yeast | NULL | Probable | As nectar-tap_saprotroph (Põlme et al. 2020); Decay sugar-rich substrates (Põlme et al. 2020) | Põlme S, et al. 2020. Fungal Diversity 105:1-16 ((https://doi.org/10.1007/s13225-020-00466-2)) |
| Otu50797 | 1 | k__Fungi(100);p__Ascomycota(100);c__Saccharomycetes(100);o__Saccharomycetales(100);f__Metschnikowiaceae(100);g__Metschnikowia(100);s__Metschnikowia_pulcherrima(100); | Metschnikowia | 13 | Saprotroph | \|Undefined Saprotroph\| | Yeast | NULL | Probable | As nectar-tap_saprotroph (Põlme et al. 2020); Decay sugar-rich substrates (Põlme et al. 2020) | Põlme S, et al. 2020. Fungal Diversity 105:1-16 ((https://doi.org/10.1007/s13225-020-00466-2)) |
| Otu50798 | 1 | k__Fungi(100);p__Ascomycota(100);c__Saccharomycetes(100);o__Saccharomycetales(100);f__Metschnikowiaceae(100);g__Metschnikowia(100);s__Metschnikowia_pulcherrima(100); | Metschnikowia | 13 | Saprotroph | \|Undefined Saprotroph\| | Yeast | NULL | Probable | As nectar-tap_saprotroph (Põlme et al. 2020); Decay sugar-rich substrates (Põlme et al. 2020) | Põlme S, et al. 2020. Fungal Diversity 105:1-16 ((https://doi.org/10.1007/s13225-020-00466-2)) |
| Otu50799 | 1 | k__Fungi(100);p__Ascomycota(100);c__Saccharomycetes(100);o__Saccharomycetales(100);f__Metschnikowiaceae(100);g__Metschnikowia(100);s__Metschnikowia_pulcherrima(100); | Metschnikowia | 13 | Saprotroph | \|Undefined Saprotroph\| | Yeast | NULL | Probable | As nectar-tap_saprotroph (Põlme et al. 2020); Decay sugar-rich substrates (Põlme et al. 2020) | Põlme S, et al. 2020. Fungal Diversity 105:1-16 ((https://doi.org/10.1007/s13225-020-00466-2)) |
| Otu50800 | 1 | k__Fungi(100);p__Ascomycota(100);c__Saccharomycetes(100);o__Saccharomycetales(100);f__Metschnikowiaceae(100);g__Metschnikowia(100);g__Metschnikowia_unclassified(100); | Metschnikowia | 13 | Saprotroph | \|Undefined Saprotroph\| | Yeast | NULL | Probable | As nectar-tap_saprotroph (Põlme et al. 2020); Decay sugar-rich substrates (Põlme et al. 2020) | Põlme S, et al. 2020. Fungal Diversity 105:1-16 ((https://doi.org/10.1007/s13225-020-00466-2)) |
| Otu50802 | 1 | k__Fungi(100);p__Ascomycota(100);c__Saccharomycetes(100);o__Saccharomycetales(100);f__Metschnikowiaceae(100);g__Metschnikowia(100);g__Metschnikowia_unclassified(100); | Metschnikowia | 13 | Saprotroph | \|Undefined Saprotroph\| | Yeast | NULL | Probable | As nectar-tap_saprotroph (Põlme et al. 2020); Decay sugar-rich substrates (Põlme et al. 2020) | Põlme S, et al. 2020. Fungal Diversity 105:1-16 ((https://doi.org/10.1007/s13225-020-00466-2)) |
| Otu50803 | 1 | k__Fungi(100);p__Ascomycota(100);c__Saccharomycetes(100);o__Saccharomycetales(100);f__Metschnikowiaceae(100);g__Metschnikowia(100);s__Metschnikowia_pulcherrima(100); | Metschnikowia | 13 | Saprotroph | \|Undefined Saprotroph\| | Yeast | NULL | Probable | As nectar-tap_saprotroph (Põlme et al. 2020); Decay sugar-rich substrates (Põlme et al. 2020) | Põlme S, et al. 2020. Fungal Diversity 105:1-16 ((https://doi.org/10.1007/s13225-020-00466-2)) |
| Otu50804 | 1 | k__Fungi(100);p__Ascomycota(100);c__Saccharomycetes(100);o__Saccharomycetales(100);f__Metschnikowiaceae(100);g__Metschnikowia(100);g__Metschnikowia_unclassified(100); | Metschnikowia | 13 | Saprotroph | \|Undefined Saprotroph\| | Yeast | NULL | Probable | As nectar-tap_saprotroph (Põlme et al. 2020); Decay sugar-rich substrates (Põlme et al. 2020) | Põlme S, et al. 2020. Fungal Diversity 105:1-16 ((https://doi.org/10.1007/s13225-020-00466-2)) |
| Otu50805 | 1 | k__Fungi(100);p__Ascomycota(100);c__Saccharomycetes(100);o__Saccharomycetales(100);f__Metschnikowiaceae(100);g__Metschnikowia(100);g__Metschnikowia_unclassified(100); | Metschnikowia | 13 | Saprotroph | \|Undefined Saprotroph\| | Yeast | NULL | Probable | As nectar-tap_saprotroph (Põlme et al. 2020); Decay sugar-rich substrates (Põlme et al. 2020) | Põlme S, et al. 2020. Fungal Diversity 105:1-16 ((https://doi.org/10.1007/s13225-020-00466-2)) |
| Otu50807 | 1 | k__Fungi(100);p__Ascomycota(100);c__Saccharomycetes(100);o__Saccharomycetales(100);f__Metschnikowiaceae(100);g__Metschnikowia(100);s__Metschnikowia_pulcherrima(100); | Metschnikowia | 13 | Saprotroph | \|Undefined Saprotroph\| | Yeast | NULL | Probable | As nectar-tap_saprotroph (Põlme et al. 2020); Decay sugar-rich substrates (Põlme et al. 2020) | Põlme S, et al. 2020. Fungal Diversity 105:1-16 ((https://doi.org/10.1007/s13225-020-00466-2)) |
| Otu50808 | 1 | k__Fungi(100);p__Ascomycota(100);c__Saccharomycetes(100);o__Saccharomycetales(100);f__Metschnikowiaceae(100);g__Metschnikowia(100);s__Metschnikowia_pulcherrima(100); | Metschnikowia | 13 | Saprotroph | \|Undefined Saprotroph\| | Yeast | NULL | Probable | As nectar-tap_saprotroph (Põlme et al. 2020); Decay sugar-rich substrates (Põlme et al. 2020) | Põlme S, et al. 2020. Fungal Diversity 105:1-16 ((https://doi.org/10.1007/s13225-020-00466-2)) |
| Otu50809 | 1 | k__Fungi(100);p__Ascomycota(100);c__Saccharomycetes(100);o__Saccharomycetales(100);f__Metschnikowiaceae(100);g__Metschnikowia(100);g__Metschnikowia_unclassified(100); | Metschnikowia | 13 | Saprotroph | \|Undefined Saprotroph\| | Yeast | NULL | Probable | As nectar-tap_saprotroph (Põlme et al. 2020); Decay sugar-rich substrates (Põlme et al. 2020) | Põlme S, et al. 2020. Fungal Diversity 105:1-16 ((https://doi.org/10.1007/s13225-020-00466-2)) |
| Otu50810 | 1 | k__Fungi(100);p__Ascomycota(100);c__Saccharomycetes(100);o__Saccharomycetales(100);f__Metschnikowiaceae(100);g__Metschnikowia(100);s__Metschnikowia_pulcherrima(100); | Metschnikowia | 13 | Saprotroph | \|Undefined Saprotroph\| | Yeast | NULL | Probable | As nectar-tap_saprotroph (Põlme et al. 2020); Decay sugar-rich substrates (Põlme et al. 2020) | Põlme S, et al. 2020. Fungal Diversity 105:1-16 ((https://doi.org/10.1007/s13225-020-00466-2)) |
| Otu50811 | 1 | k__Fungi(100);p__Ascomycota(100);c__Saccharomycetes(100);o__Saccharomycetales(100);f__Metschnikowiaceae(100);g__Metschnikowia(100);g__Metschnikowia_unclassified(100); | Metschnikowia | 13 | Saprotroph | \|Undefined Saprotroph\| | Yeast | NULL | Probable | As nectar-tap_saprotroph (Põlme et al. 2020); Decay sugar-rich substrates (Põlme et al. 2020) | Põlme S, et al. 2020. Fungal Diversity 105:1-16 ((https://doi.org/10.1007/s13225-020-00466-2)) |
| Otu50812 | 1 | k__Fungi(100);p__Ascomycota(100);c__Saccharomycetes(100);o__Saccharomycetales(100);f__Metschnikowiaceae(100);g__Metschnikowia(100);s__Metschnikowia_pulcherrima(100); | Metschnikowia | 13 | Saprotroph | \|Undefined Saprotroph\| | Yeast | NULL | Probable | As nectar-tap_saprotroph (Põlme et al. 2020); Decay sugar-rich substrates (Põlme et al. 2020) | Põlme S, et al. 2020. Fungal Diversity 105:1-16 ((https://doi.org/10.1007/s13225-020-00466-2)) |
| Otu50813 | 1 | k__Fungi(100);p__Ascomycota(100);c__Saccharomycetes(100);o__Saccharomycetales(100);f__Metschnikowiaceae(100);g__Metschnikowia(100);s__Metschnikowia_pulcherrima(100); | Metschnikowia | 13 | Saprotroph | \|Undefined Saprotroph\| | Yeast | NULL | Probable | As nectar-tap_saprotroph (Põlme et al. 2020); Decay sugar-rich substrates (Põlme et al. 2020) | Põlme S, et al. 2020. Fungal Diversity 105:1-16 ((https://doi.org/10.1007/s13225-020-00466-2)) |
| Otu50814 | 1 | k__Fungi(100);p__Ascomycota(100);c__Saccharomycetes(100);o__Saccharomycetales(100);f__Metschnikowiaceae(100);g__Metschnikowia(100);g__Metschnikowia_unclassified(100); | Metschnikowia | 13 | Saprotroph | \|Undefined Saprotroph\| | Yeast | NULL | Probable | As nectar-tap_saprotroph (Põlme et al. 2020); Decay sugar-rich substrates (Põlme et al. 2020) | Põlme S, et al. 2020. Fungal Diversity 105:1-16 ((https://doi.org/10.1007/s13225-020-00466-2)) |
| Otu50815 | 1 | k__Fungi(100);p__Ascomycota(100);c__Saccharomycetes(100);o__Saccharomycetales(100);f__Metschnikowiaceae(100);g__Metschnikowia(100);s__Metschnikowia_pulcherrima(100); | Metschnikowia | 13 | Saprotroph | \|Undefined Saprotroph\| | Yeast | NULL | Probable | As nectar-tap_saprotroph (Põlme et al. 2020); Decay sugar-rich substrates (Põlme et al. 2020) | Põlme S, et al. 2020. Fungal Diversity 105:1-16 ((https://doi.org/10.1007/s13225-020-00466-2)) |
| Otu50816 | 1 | k__Fungi(100);p__Ascomycota(100);c__Saccharomycetes(100);o__Saccharomycetales(100);f__Metschnikowiaceae(100);g__Metschnikowia(100);s__Metschnikowia_pulcherrima(100); | Metschnikowia | 13 | Saprotroph | \|Undefined Saprotroph\| | Yeast | NULL | Probable | As nectar-tap_saprotroph (Põlme et al. 2020); Decay sugar-rich substrates (Põlme et al. 2020) | Põlme S, et al. 2020. Fungal Diversity 105:1-16 ((https://doi.org/10.1007/s13225-020-00466-2)) |
| Otu50817 | 1 | k__Fungi(100);p__Ascomycota(100);c__Saccharomycetes(100);o__Saccharomycetales(100);f__Metschnikowiaceae(100);g__Metschnikowia(100);s__Metschnikowia_pulcherrima(100); | Metschnikowia | 13 | Saprotroph | \|Undefined Saprotroph\| | Yeast | NULL | Probable | As nectar-tap_saprotroph (Põlme et al. 2020); Decay sugar-rich substrates (Põlme et al. 2020) | Põlme S, et al. 2020. Fungal Diversity 105:1-16 ((https://doi.org/10.1007/s13225-020-00466-2)) |
| Otu50818 | 1 | k__Fungi(100);p__Ascomycota(100);c__Saccharomycetes(100);o__Saccharomycetales(100);f__Metschnikowiaceae(100);g__Metschnikowia(100);s__Metschnikowia_pulcherrima(100); | Metschnikowia | 13 | Saprotroph | \|Undefined Saprotroph\| | Yeast | NULL | Probable | As nectar-tap_saprotroph (Põlme et al. 2020); Decay sugar-rich substrates (Põlme et al. 2020) | Põlme S, et al. 2020. Fungal Diversity 105:1-16 ((https://doi.org/10.1007/s13225-020-00466-2)) |
| Otu50819 | 1 | k__Fungi(100);p__Ascomycota(100);c__Saccharomycetes(100);o__Saccharomycetales(100);f__Metschnikowiaceae(100);g__Metschnikowia(100);s__Metschnikowia_pulcherrima(100); | Metschnikowia | 13 | Saprotroph | \|Undefined Saprotroph\| | Yeast | NULL | Probable | As nectar-tap_saprotroph (Põlme et al. 2020); Decay sugar-rich substrates (Põlme et al. 2020) | Põlme S, et al. 2020. Fungal Diversity 105:1-16 ((https://doi.org/10.1007/s13225-020-00466-2)) |
| Otu50820 | 1 | k__Fungi(100);p__Ascomycota(100);c__Saccharomycetes(100);o__Saccharomycetales(100);f__Metschnikowiaceae(100);g__Metschnikowia(100);s__Metschnikowia_pulcherrima(100); | Metschnikowia | 13 | Saprotroph | \|Undefined Saprotroph\| | Yeast | NULL | Probable | As nectar-tap_saprotroph (Põlme et al. 2020); Decay sugar-rich substrates (Põlme et al. 2020) | Põlme S, et al. 2020. Fungal Diversity 105:1-16 ((https://doi.org/10.1007/s13225-020-00466-2)) |
| Otu50821 | 1 | k__Fungi(100);p__Ascomycota(100);c__Saccharomycetes(100);o__Saccharomycetales(100);f__Metschnikowiaceae(100);g__Metschnikowia(100);s__Metschnikowia_pulcherrima(100); | Metschnikowia | 13 | Saprotroph | \|Undefined Saprotroph\| | Yeast | NULL | Probable | As nectar-tap_saprotroph (Põlme et al. 2020); Decay sugar-rich substrates (Põlme et al. 2020) | Põlme S, et al. 2020. Fungal Diversity 105:1-16 ((https://doi.org/10.1007/s13225-020-00466-2)) |
| Otu50822 | 1 | k__Fungi(100);p__Ascomycota(100);c__Saccharomycetes(100);o__Saccharomycetales(100);f__Metschnikowiaceae(100);g__Metschnikowia(100);s__Metschnikowia_pulcherrima(100); | Metschnikowia | 13 | Saprotroph | \|Undefined Saprotroph\| | Yeast | NULL | Probable | As nectar-tap_saprotroph (Põlme et al. 2020); Decay sugar-rich substrates (Põlme et al. 2020) | Põlme S, et al. 2020. Fungal Diversity 105:1-16 ((https://doi.org/10.1007/s13225-020-00466-2)) |
| Otu50823 | 1 | k__Fungi(100);p__Ascomycota(100);c__Saccharomycetes(100);o__Saccharomycetales(100);f__Metschnikowiaceae(100);g__Metschnikowia(100);s__Metschnikowia_pulcherrima(100); | Metschnikowia | 13 | Saprotroph | \|Undefined Saprotroph\| | Yeast | NULL | Probable | As nectar-tap_saprotroph (Põlme et al. 2020); Decay sugar-rich substrates (Põlme et al. 2020) | Põlme S, et al. 2020. Fungal Diversity 105:1-16 ((https://doi.org/10.1007/s13225-020-00466-2)) |
| Otu50824 | 1 | k__Fungi(100);p__Ascomycota(100);c__Saccharomycetes(100);o__Saccharomycetales(100);f__Metschnikowiaceae(100);g__Metschnikowia(100);s__Metschnikowia_pulcherrima(100); | Metschnikowia | 13 | Saprotroph | \|Undefined Saprotroph\| | Yeast | NULL | Probable | As nectar-tap_saprotroph (Põlme et al. 2020); Decay sugar-rich substrates (Põlme et al. 2020) | Põlme S, et al. 2020. Fungal Diversity 105:1-16 ((https://doi.org/10.1007/s13225-020-00466-2)) |
| Otu50825 | 1 | k__Fungi(100);p__Ascomycota(100);c__Saccharomycetes(100);o__Saccharomycetales(100);o__Saccharomycetales_unclassified(100);o__Saccharomycetales_unclassified(100);o__Saccharomycetales_unclassified(100); | Saccharomycetales | 7 | Saprotroph | Undefined Saprotroph | Yeast | NULL | Possible | NULL | Sterkenburg E, et al. 2015. New Phytologist 207:1145-1158 ((https://doi.org/10.1111/nph.13426)) |
| Otu50826 | 1 | k__Fungi(100);p__Ascomycota(100);c__Saccharomycetes(100);o__Saccharomycetales(100);f__Metschnikowiaceae(100);g__Metschnikowia(100);s__Metschnikowia_pulcherrima(100); | Metschnikowia | 13 | Saprotroph | \|Undefined Saprotroph\| | Yeast | NULL | Probable | As nectar-tap_saprotroph (Põlme et al. 2020); Decay sugar-rich substrates (Põlme et al. 2020) | Põlme S, et al. 2020. Fungal Diversity 105:1-16 ((https://doi.org/10.1007/s13225-020-00466-2)) |
| Otu50827 | 1 | k__Fungi(100);p__Ascomycota(100);c__Saccharomycetes(100);o__Saccharomycetales(100);f__Metschnikowiaceae(100);g__Metschnikowia(100);s__Metschnikowia_pulcherrima(100); | Metschnikowia | 13 | Saprotroph | \|Undefined Saprotroph\| | Yeast | NULL | Probable | As nectar-tap_saprotroph (Põlme et al. 2020); Decay sugar-rich substrates (Põlme et al. 2020) | Põlme S, et al. 2020. Fungal Diversity 105:1-16 ((https://doi.org/10.1007/s13225-020-00466-2)) |
| Otu50828 | 1 | k__Fungi(100);p__Ascomycota(100);c__Saccharomycetes(100);o__Saccharomycetales(100);f__Saccharomycetales_family_Incertae_sedis(100);g__Candida(100);g__Candida_unclassified(100); | Candida | 13 | Pathotroph-Saprotroph-Symbiotroph | Animal Pathogen-Endophyte-Endosymbiont-Epiphyte-\|Undefined Saprotroph\| | Dimorphic Yeast-Yeast | NULL | Probable | As nectar-tap_saprotroph (Põlme et al. 2020); Decay sugar-rich substrates (Põlme et al. 2020); This asexual genus is polyphyletic and contain species that function very differently depend on what they are. Caution should be taken when interpreting guilds for Candida species. | Manolakaki D, et al. 2010. Virulence. 1: 367-375 ((https://doi.org/10.4161/viru.1.5.12796)); Mekbib SB, et al. 2011. Tropical Plant Pathology 36:233-240 ((https://doi.org/10.1590/S1982-56762011000400004)); Põlme S, et al. 2020. Fungal Diversity 105:1-16 ((https://doi.org/10.1007/s13225-020-00466-2)) |
| Otu50829 | 1 | k__Fungi(100);p__Ascomycota(100);c__Saccharomycetes(100);o__Saccharomycetales(100);f__Metschnikowiaceae(100);g__Metschnikowia(100);s__Metschnikowia_pulcherrima(100); | Metschnikowia | 13 | Saprotroph | \|Undefined Saprotroph\| | Yeast | NULL | Probable | As nectar-tap_saprotroph (Põlme et al. 2020); Decay sugar-rich substrates (Põlme et al. 2020) | Põlme S, et al. 2020. Fungal Diversity 105:1-16 ((https://doi.org/10.1007/s13225-020-00466-2)) |
| Otu50830 | 1 | k__Fungi(100);p__Ascomycota(100);c__Saccharomycetes(100);o__Saccharomycetales(100);f__Metschnikowiaceae(100);g__Metschnikowia(100);s__Metschnikowia_pulcherrima(100); | Metschnikowia | 13 | Saprotroph | \|Undefined Saprotroph\| | Yeast | NULL | Probable | As nectar-tap_saprotroph (Põlme et al. 2020); Decay sugar-rich substrates (Põlme et al. 2020) | Põlme S, et al. 2020. Fungal Diversity 105:1-16 ((https://doi.org/10.1007/s13225-020-00466-2)) |
| Otu50831 | 1 | k__Fungi(100);p__Ascomycota(100);c__Saccharomycetes(100);o__Saccharomycetales(100);f__Metschnikowiaceae(100);g__Metschnikowia(100);s__Metschnikowia_pulcherrima(100); | Metschnikowia | 13 | Saprotroph | \|Undefined Saprotroph\| | Yeast | NULL | Probable | As nectar-tap_saprotroph (Põlme et al. 2020); Decay sugar-rich substrates (Põlme et al. 2020) | Põlme S, et al. 2020. Fungal Diversity 105:1-16 ((https://doi.org/10.1007/s13225-020-00466-2)) |
| Otu50833 | 1 | k__Fungi(100);p__Ascomycota(100);c__Saccharomycetes(100);o__Saccharomycetales(100);f__Saccharomycetales_family_Incertae_sedis(100);g__Candida(100);g__Candida_unclassified(100); | Candida | 13 | Pathotroph-Saprotroph-Symbiotroph | Animal Pathogen-Endophyte-Endosymbiont-Epiphyte-\|Undefined Saprotroph\| | Dimorphic Yeast-Yeast | NULL | Probable | As nectar-tap_saprotroph (Põlme et al. 2020); Decay sugar-rich substrates (Põlme et al. 2020); This asexual genus is polyphyletic and contain species that function very differently depend on what they are. Caution should be taken when interpreting guilds for Candida species. | Manolakaki D, et al. 2010. Virulence. 1: 367-375 ((https://doi.org/10.4161/viru.1.5.12796)); Mekbib SB, et al. 2011. Tropical Plant Pathology 36:233-240 ((https://doi.org/10.1590/S1982-56762011000400004)); Põlme S, et al. 2020. Fungal Diversity 105:1-16 ((https://doi.org/10.1007/s13225-020-00466-2)) |
| Otu50839 | 1 | k__Fungi(100);p__Ascomycota(100);c__Saccharomycetes(100);o__Saccharomycetales(100);f__Metschnikowiaceae(100);g__Metschnikowia(100);g__Metschnikowia_unclassified(100); | Metschnikowia | 13 | Saprotroph | \|Undefined Saprotroph\| | Yeast | NULL | Probable | As nectar-tap_saprotroph (Põlme et al. 2020); Decay sugar-rich substrates (Põlme et al. 2020) | Põlme S, et al. 2020. Fungal Diversity 105:1-16 ((https://doi.org/10.1007/s13225-020-00466-2)) |
| Otu50840 | 1 | k__Fungi(100);p__Ascomycota(100);c__Saccharomycetes(100);o__Saccharomycetales(100);f__Saccharomycetales_family_Incertae_sedis(100);g__Candida(100);g__Candida_unclassified(100); | Candida | 13 | Pathotroph-Saprotroph-Symbiotroph | Animal Pathogen-Endophyte-Endosymbiont-Epiphyte-\|Undefined Saprotroph\| | Dimorphic Yeast-Yeast | NULL | Probable | As nectar-tap_saprotroph (Põlme et al. 2020); Decay sugar-rich substrates (Põlme et al. 2020); This asexual genus is polyphyletic and contain species that function very differently depend on what they are. Caution should be taken when interpreting guilds for Candida species. | Manolakaki D, et al. 2010. Virulence. 1: 367-375 ((https://doi.org/10.4161/viru.1.5.12796)); Mekbib SB, et al. 2011. Tropical Plant Pathology 36:233-240 ((https://doi.org/10.1590/S1982-56762011000400004)); Põlme S, et al. 2020. Fungal Diversity 105:1-16 ((https://doi.org/10.1007/s13225-020-00466-2)) |
| Otu50841 | 1 | k__Fungi(100);p__Ascomycota(100);c__Saccharomycetes(100);o__Saccharomycetales(100);f__Metschnikowiaceae(100);g__Metschnikowia(100);s__Metschnikowia_pulcherrima(100); | Metschnikowia | 13 | Saprotroph | \|Undefined Saprotroph\| | Yeast | NULL | Probable | As nectar-tap_saprotroph (Põlme et al. 2020); Decay sugar-rich substrates (Põlme et al. 2020) | Põlme S, et al. 2020. Fungal Diversity 105:1-16 ((https://doi.org/10.1007/s13225-020-00466-2)) |
| Otu50842 | 1 | k__Fungi(100);p__Ascomycota(100);c__Saccharomycetes(100);o__Saccharomycetales(100);f__Metschnikowiaceae(100);g__Metschnikowia(100);g__Metschnikowia_unclassified(100); | Metschnikowia | 13 | Saprotroph | \|Undefined Saprotroph\| | Yeast | NULL | Probable | As nectar-tap_saprotroph (Põlme et al. 2020); Decay sugar-rich substrates (Põlme et al. 2020) | Põlme S, et al. 2020. Fungal Diversity 105:1-16 ((https://doi.org/10.1007/s13225-020-00466-2)) |
| Otu50843 | 1 | k__Fungi(100);p__Ascomycota(100);c__Saccharomycetes(100);o__Saccharomycetales(100);f__Metschnikowiaceae(100);g__Metschnikowia(100);s__Metschnikowia_pulcherrima(100); | Metschnikowia | 13 | Saprotroph | \|Undefined Saprotroph\| | Yeast | NULL | Probable | As nectar-tap_saprotroph (Põlme et al. 2020); Decay sugar-rich substrates (Põlme et al. 2020) | Põlme S, et al. 2020. Fungal Diversity 105:1-16 ((https://doi.org/10.1007/s13225-020-00466-2)) |
| Otu50844 | 1 | k__Fungi(100);p__Ascomycota(100);c__Saccharomycetes(100);o__Saccharomycetales(100);f__Metschnikowiaceae(100);g__Metschnikowia(100);g__Metschnikowia_unclassified(100); | Metschnikowia | 13 | Saprotroph | \|Undefined Saprotroph\| | Yeast | NULL | Probable | As nectar-tap_saprotroph (Põlme et al. 2020); Decay sugar-rich substrates (Põlme et al. 2020) | Põlme S, et al. 2020. Fungal Diversity 105:1-16 ((https://doi.org/10.1007/s13225-020-00466-2)) |
| Otu50846 | 1 | k__Fungi(100);p__Ascomycota(100);c__Saccharomycetes(100);o__Saccharomycetales(100);f__Metschnikowiaceae(100);g__Metschnikowia(100);s__Metschnikowia_pulcherrima(100); | Metschnikowia | 13 | Saprotroph | \|Undefined Saprotroph\| | Yeast | NULL | Probable | As nectar-tap_saprotroph (Põlme et al. 2020); Decay sugar-rich substrates (Põlme et al. 2020) | Põlme S, et al. 2020. Fungal Diversity 105:1-16 ((https://doi.org/10.1007/s13225-020-00466-2)) |
| Otu50851 | 1 | k__Fungi(100);p__Ascomycota(100);c__Saccharomycetes(100);o__Saccharomycetales(100);f__Metschnikowiaceae(100);g__Metschnikowia(100);g__Metschnikowia_unclassified(100); | Metschnikowia | 13 | Saprotroph | \|Undefined Saprotroph\| | Yeast | NULL | Probable | As nectar-tap_saprotroph (Põlme et al. 2020); Decay sugar-rich substrates (Põlme et al. 2020) | Põlme S, et al. 2020. Fungal Diversity 105:1-16 ((https://doi.org/10.1007/s13225-020-00466-2)) |
| Otu50852 | 1 | k__Fungi(100);p__Ascomycota(100);c__Saccharomycetes(100);o__Saccharomycetales(100);f__Metschnikowiaceae(100);g__Metschnikowia(100);s__Metschnikowia_pulcherrima(100); | Metschnikowia | 13 | Saprotroph | \|Undefined Saprotroph\| | Yeast | NULL | Probable | As nectar-tap_saprotroph (Põlme et al. 2020); Decay sugar-rich substrates (Põlme et al. 2020) | Põlme S, et al. 2020. Fungal Diversity 105:1-16 ((https://doi.org/10.1007/s13225-020-00466-2)) |
| Otu50861 | 1 | k__Fungi(100);p__Ascomycota(100);c__Saccharomycetes(100);o__Saccharomycetales(100);f__Metschnikowiaceae(100);g__Metschnikowia(100);g__Metschnikowia_unclassified(100); | Metschnikowia | 13 | Saprotroph | \|Undefined Saprotroph\| | Yeast | NULL | Probable | As nectar-tap_saprotroph (Põlme et al. 2020); Decay sugar-rich substrates (Põlme et al. 2020) | Põlme S, et al. 2020. Fungal Diversity 105:1-16 ((https://doi.org/10.1007/s13225-020-00466-2)) |
| Otu50862 | 1 | k__Fungi(100);p__Ascomycota(100);c__Saccharomycetes(100);o__Saccharomycetales(100);f__Saccharomycetales_family_Incertae_sedis(100);g__Candida(100);s__Candida_cretensis(100); | Candida | 13 | Pathotroph-Saprotroph-Symbiotroph | Animal Pathogen-Endophyte-Endosymbiont-Epiphyte-\|Undefined Saprotroph\| | Dimorphic Yeast-Yeast | NULL | Probable | As nectar-tap_saprotroph (Põlme et al. 2020); Decay sugar-rich substrates (Põlme et al. 2020); This asexual genus is polyphyletic and contain species that function very differently depend on what they are. Caution should be taken when interpreting guilds for Candida species. | Manolakaki D, et al. 2010. Virulence. 1: 367-375 ((https://doi.org/10.4161/viru.1.5.12796)); Mekbib SB, et al. 2011. Tropical Plant Pathology 36:233-240 ((https://doi.org/10.1590/S1982-56762011000400004)); Põlme S, et al. 2020. Fungal Diversity 105:1-16 ((https://doi.org/10.1007/s13225-020-00466-2)) |
| Otu50863 | 1 | k__Fungi(100);p__Ascomycota(100);c__Saccharomycetes(100);o__Saccharomycetales(100);f__Metschnikowiaceae(100);g__Metschnikowia(100);s__Metschnikowia_pulcherrima(100); | Metschnikowia | 13 | Saprotroph | \|Undefined Saprotroph\| | Yeast | NULL | Probable | As nectar-tap_saprotroph (Põlme et al. 2020); Decay sugar-rich substrates (Põlme et al. 2020) | Põlme S, et al. 2020. Fungal Diversity 105:1-16 ((https://doi.org/10.1007/s13225-020-00466-2)) |
| Otu50864 | 1 | k__Fungi(100);p__Ascomycota(100);c__Saccharomycetes(100);o__Saccharomycetales(100);f__Metschnikowiaceae(100);g__Metschnikowia(100);s__Metschnikowia_pulcherrima(100); | Metschnikowia | 13 | Saprotroph | \|Undefined Saprotroph\| | Yeast | NULL | Probable | As nectar-tap_saprotroph (Põlme et al. 2020); Decay sugar-rich substrates (Põlme et al. 2020) | Põlme S, et al. 2020. Fungal Diversity 105:1-16 ((https://doi.org/10.1007/s13225-020-00466-2)) |
| Otu50865 | 1 | k__Fungi(100);p__Ascomycota(100);c__Saccharomycetes(100);o__Saccharomycetales(100);f__Metschnikowiaceae(100);g__Metschnikowia(100);s__Metschnikowia_pulcherrima(100); | Metschnikowia | 13 | Saprotroph | \|Undefined Saprotroph\| | Yeast | NULL | Probable | As nectar-tap_saprotroph (Põlme et al. 2020); Decay sugar-rich substrates (Põlme et al. 2020) | Põlme S, et al. 2020. Fungal Diversity 105:1-16 ((https://doi.org/10.1007/s13225-020-00466-2)) |
| Otu50866 | 1 | k__Fungi(100);p__Ascomycota(100);c__Saccharomycetes(100);o__Saccharomycetales(100);f__Metschnikowiaceae(100);g__Metschnikowia(100);g__Metschnikowia_unclassified(100); | Metschnikowia | 13 | Saprotroph | \|Undefined Saprotroph\| | Yeast | NULL | Probable | As nectar-tap_saprotroph (Põlme et al. 2020); Decay sugar-rich substrates (Põlme et al. 2020) | Põlme S, et al. 2020. Fungal Diversity 105:1-16 ((https://doi.org/10.1007/s13225-020-00466-2)) |
| Otu50867 | 1 | k__Fungi(100);p__Ascomycota(100);c__Saccharomycetes(100);o__Saccharomycetales(100);f__Metschnikowiaceae(100);g__Metschnikowia(100);s__Metschnikowia_pulcherrima(100); | Metschnikowia | 13 | Saprotroph | \|Undefined Saprotroph\| | Yeast | NULL | Probable | As nectar-tap_saprotroph (Põlme et al. 2020); Decay sugar-rich substrates (Põlme et al. 2020) | Põlme S, et al. 2020. Fungal Diversity 105:1-16 ((https://doi.org/10.1007/s13225-020-00466-2)) |
| Otu50870 | 1 | k__Fungi(100);p__Ascomycota(100);c__Saccharomycetes(100);o__Saccharomycetales(100);f__Metschnikowiaceae(100);g__Metschnikowia(100);s__Metschnikowia_pulcherrima(100); | Metschnikowia | 13 | Saprotroph | \|Undefined Saprotroph\| | Yeast | NULL | Probable | As nectar-tap_saprotroph (Põlme et al. 2020); Decay sugar-rich substrates (Põlme et al. 2020) | Põlme S, et al. 2020. Fungal Diversity 105:1-16 ((https://doi.org/10.1007/s13225-020-00466-2)) |
| Otu50871 | 1 | k__Fungi(100);p__Ascomycota(100);c__Saccharomycetes(100);o__Saccharomycetales(100);f__Metschnikowiaceae(100);g__Metschnikowia(100);s__Metschnikowia_pulcherrima(100); | Metschnikowia | 13 | Saprotroph | \|Undefined Saprotroph\| | Yeast | NULL | Probable | As nectar-tap_saprotroph (Põlme et al. 2020); Decay sugar-rich substrates (Põlme et al. 2020) | Põlme S, et al. 2020. Fungal Diversity 105:1-16 ((https://doi.org/10.1007/s13225-020-00466-2)) |
| Otu50872 | 1 | k__Fungi(100);p__Ascomycota(100);c__Saccharomycetes(100);o__Saccharomycetales(100);f__Metschnikowiaceae(100);g__Metschnikowia(100);s__Metschnikowia_pulcherrima(100); | Metschnikowia | 13 | Saprotroph | \|Undefined Saprotroph\| | Yeast | NULL | Probable | As nectar-tap_saprotroph (Põlme et al. 2020); Decay sugar-rich substrates (Põlme et al. 2020) | Põlme S, et al. 2020. Fungal Diversity 105:1-16 ((https://doi.org/10.1007/s13225-020-00466-2)) |
| Otu50874 | 1 | k__Fungi(100);p__Ascomycota(100);c__Saccharomycetes(100);o__Saccharomycetales(100);f__Saccharomycetales_family_Incertae_sedis(100);f__Saccharomycetales_family_Incertae_sedis_unclassified(100);f__Saccharomycetales_family_Incertae_sedis_unclassified(100); | Saccharomycetales | 7 | Saprotroph | Undefined Saprotroph | Yeast | NULL | Possible | NULL | Sterkenburg E, et al. 2015. New Phytologist 207:1145-1158 ((https://doi.org/10.1111/nph.13426)) |
| Otu50879 | 1 | k__Fungi(100);p__Ascomycota(100);c__Saccharomycetes(100);o__Saccharomycetales(100);f__Metschnikowiaceae(100);g__Metschnikowia(100);g__Metschnikowia_unclassified(100); | Metschnikowia | 13 | Saprotroph | \|Undefined Saprotroph\| | Yeast | NULL | Probable | As nectar-tap_saprotroph (Põlme et al. 2020); Decay sugar-rich substrates (Põlme et al. 2020) | Põlme S, et al. 2020. Fungal Diversity 105:1-16 ((https://doi.org/10.1007/s13225-020-00466-2)) |
| Otu50880 | 1 | k__Fungi(100);p__Ascomycota(100);c__Saccharomycetes(100);o__Saccharomycetales(100);f__Metschnikowiaceae(100);g__Metschnikowia(100);s__Metschnikowia_pulcherrima(100); | Metschnikowia | 13 | Saprotroph | \|Undefined Saprotroph\| | Yeast | NULL | Probable | As nectar-tap_saprotroph (Põlme et al. 2020); Decay sugar-rich substrates (Põlme et al. 2020) | Põlme S, et al. 2020. Fungal Diversity 105:1-16 ((https://doi.org/10.1007/s13225-020-00466-2)) |
| Otu50881 | 1 | k__Fungi(100);p__Ascomycota(100);c__Saccharomycetes(100);o__Saccharomycetales(100);f__Metschnikowiaceae(100);g__Metschnikowia(100);s__Metschnikowia_pulcherrima(100); | Metschnikowia | 13 | Saprotroph | \|Undefined Saprotroph\| | Yeast | NULL | Probable | As nectar-tap_saprotroph (Põlme et al. 2020); Decay sugar-rich substrates (Põlme et al. 2020) | Põlme S, et al. 2020. Fungal Diversity 105:1-16 ((https://doi.org/10.1007/s13225-020-00466-2)) |
| Otu50882 | 1 | k__Fungi(100);p__Basidiomycota(100);c__Microbotryomycetes(100);o__Sporidiobolales(100);f__Sporidiobolales_family_Incertae_sedis(100);g__Sporobolomyces(100);s__Sporobolomyces_lactosus(100); | Sporobolomyces | 13 | Pathotroph-Saprotroph | \|Fungal Parasite\|-Plant Saprotroph | Yeast | NULL | Probable | As litter_saprotroph (Põlme et al. 2020); As mycoparasite (Põlme et al. 2020); Decay fungal material (Põlme et al. 2020) | Tedersoo L, et al. 2014. Science 346:e1256688 ((https://doi.org/10.1126/science.1256688)); Purahong W, et al. 2016. Molecular Ecology 25:4059-4074 ((https://doi.org/10.1111/mec.13739)); Põlme S, et al. 2020. Fungal Diversity 105:1-16 ((https://doi.org/10.1007/s13225-020-00466-2)) |
| Otu50884 | 1 | k__Fungi(100);p__Ascomycota(100);c__Saccharomycetes(100);o__Saccharomycetales(100);f__Metschnikowiaceae(100);g__Metschnikowia(100);s__Metschnikowia_pulcherrima(100); | Metschnikowia | 13 | Saprotroph | \|Undefined Saprotroph\| | Yeast | NULL | Probable | As nectar-tap_saprotroph (Põlme et al. 2020); Decay sugar-rich substrates (Põlme et al. 2020) | Põlme S, et al. 2020. Fungal Diversity 105:1-16 ((https://doi.org/10.1007/s13225-020-00466-2)) |
| Otu50885 | 1 | k__Fungi(100);p__Ascomycota(100);c__Saccharomycetes(100);o__Saccharomycetales(100);f__Metschnikowiaceae(100);g__Metschnikowia(100);s__Metschnikowia_pulcherrima(100); | Metschnikowia | 13 | Saprotroph | \|Undefined Saprotroph\| | Yeast | NULL | Probable | As nectar-tap_saprotroph (Põlme et al. 2020); Decay sugar-rich substrates (Põlme et al. 2020) | Põlme S, et al. 2020. Fungal Diversity 105:1-16 ((https://doi.org/10.1007/s13225-020-00466-2)) |
| Otu50886 | 1 | k__Fungi(100);p__Ascomycota(100);c__Saccharomycetes(100);o__Saccharomycetales(100);f__Metschnikowiaceae(100);g__Metschnikowia(100);s__Metschnikowia_pulcherrima(100); | Metschnikowia | 13 | Saprotroph | \|Undefined Saprotroph\| | Yeast | NULL | Probable | As nectar-tap_saprotroph (Põlme et al. 2020); Decay sugar-rich substrates (Põlme et al. 2020) | Põlme S, et al. 2020. Fungal Diversity 105:1-16 ((https://doi.org/10.1007/s13225-020-00466-2)) |
| Otu50887 | 1 | k__Fungi(100);p__Ascomycota(100);c__Saccharomycetes(100);o__Saccharomycetales(100);f__Metschnikowiaceae(100);g__Metschnikowia(100);g__Metschnikowia_unclassified(100); | Metschnikowia | 13 | Saprotroph | \|Undefined Saprotroph\| | Yeast | NULL | Probable | As nectar-tap_saprotroph (Põlme et al. 2020); Decay sugar-rich substrates (Põlme et al. 2020) | Põlme S, et al. 2020. Fungal Diversity 105:1-16 ((https://doi.org/10.1007/s13225-020-00466-2)) |
| Otu50888 | 1 | k__Fungi(100);p__Ascomycota(100);c__Saccharomycetes(100);o__Saccharomycetales(100);f__Metschnikowiaceae(100);g__Metschnikowia(100);g__Metschnikowia_unclassified(100); | Metschnikowia | 13 | Saprotroph | \|Undefined Saprotroph\| | Yeast | NULL | Probable | As nectar-tap_saprotroph (Põlme et al. 2020); Decay sugar-rich substrates (Põlme et al. 2020) | Põlme S, et al. 2020. Fungal Diversity 105:1-16 ((https://doi.org/10.1007/s13225-020-00466-2)) |
| Otu50889 | 1 | k__Fungi(100);p__Ascomycota(100);c__Saccharomycetes(100);o__Saccharomycetales(100);f__Metschnikowiaceae(100);g__Metschnikowia(100);s__Metschnikowia_pulcherrima(100); | Metschnikowia | 13 | Saprotroph | \|Undefined Saprotroph\| | Yeast | NULL | Probable | As nectar-tap_saprotroph (Põlme et al. 2020); Decay sugar-rich substrates (Põlme et al. 2020) | Põlme S, et al. 2020. Fungal Diversity 105:1-16 ((https://doi.org/10.1007/s13225-020-00466-2)) |
| Otu50890 | 1 | k__Fungi(100);p__Ascomycota(100);c__Saccharomycetes(100);o__Saccharomycetales(100);f__Metschnikowiaceae(100);g__Metschnikowia(100);s__Metschnikowia_pulcherrima(100); | Metschnikowia | 13 | Saprotroph | \|Undefined Saprotroph\| | Yeast | NULL | Probable | As nectar-tap_saprotroph (Põlme et al. 2020); Decay sugar-rich substrates (Põlme et al. 2020) | Põlme S, et al. 2020. Fungal Diversity 105:1-16 ((https://doi.org/10.1007/s13225-020-00466-2)) |
| Otu50891 | 1 | k__Fungi(100);p__Ascomycota(100);c__Saccharomycetes(100);o__Saccharomycetales(100);f__Metschnikowiaceae(100);g__Metschnikowia(100);s__Metschnikowia_pulcherrima(100); | Metschnikowia | 13 | Saprotroph | \|Undefined Saprotroph\| | Yeast | NULL | Probable | As nectar-tap_saprotroph (Põlme et al. 2020); Decay sugar-rich substrates (Põlme et al. 2020) | Põlme S, et al. 2020. Fungal Diversity 105:1-16 ((https://doi.org/10.1007/s13225-020-00466-2)) |
| Otu50892 | 1 | k__Fungi(100);p__Ascomycota(100);c__Saccharomycetes(100);o__Saccharomycetales(100);f__Metschnikowiaceae(100);g__Metschnikowia(100);g__Metschnikowia_unclassified(100); | Metschnikowia | 13 | Saprotroph | \|Undefined Saprotroph\| | Yeast | NULL | Probable | As nectar-tap_saprotroph (Põlme et al. 2020); Decay sugar-rich substrates (Põlme et al. 2020) | Põlme S, et al. 2020. Fungal Diversity 105:1-16 ((https://doi.org/10.1007/s13225-020-00466-2)) |
| Otu50893 | 1 | k__Fungi(100);p__Ascomycota(100);c__Saccharomycetes(100);o__Saccharomycetales(100);f__Metschnikowiaceae(100);g__Metschnikowia(100);s__Metschnikowia_pulcherrima(100); | Metschnikowia | 13 | Saprotroph | \|Undefined Saprotroph\| | Yeast | NULL | Probable | As nectar-tap_saprotroph (Põlme et al. 2020); Decay sugar-rich substrates (Põlme et al. 2020) | Põlme S, et al. 2020. Fungal Diversity 105:1-16 ((https://doi.org/10.1007/s13225-020-00466-2)) |
| Otu50894 | 1 | k__Fungi(100);p__Ascomycota(100);c__Saccharomycetes(100);o__Saccharomycetales(100);f__Metschnikowiaceae(100);g__Metschnikowia(100);s__Metschnikowia_pulcherrima(100); | Metschnikowia | 13 | Saprotroph | \|Undefined Saprotroph\| | Yeast | NULL | Probable | As nectar-tap_saprotroph (Põlme et al. 2020); Decay sugar-rich substrates (Põlme et al. 2020) | Põlme S, et al. 2020. Fungal Diversity 105:1-16 ((https://doi.org/10.1007/s13225-020-00466-2)) |
| Otu50895 | 1 | k__Fungi(100);p__Ascomycota(100);c__Saccharomycetes(100);o__Saccharomycetales(100);f__Metschnikowiaceae(100);g__Metschnikowia(100);s__Metschnikowia_pulcherrima(100); | Metschnikowia | 13 | Saprotroph | \|Undefined Saprotroph\| | Yeast | NULL | Probable | As nectar-tap_saprotroph (Põlme et al. 2020); Decay sugar-rich substrates (Põlme et al. 2020) | Põlme S, et al. 2020. Fungal Diversity 105:1-16 ((https://doi.org/10.1007/s13225-020-00466-2)) |
| Otu50896 | 1 | k__Fungi(100);p__Ascomycota(100);c__Saccharomycetes(100);o__Saccharomycetales(100);f__Metschnikowiaceae(100);g__Metschnikowia(100);s__Metschnikowia_pulcherrima(100); | Metschnikowia | 13 | Saprotroph | \|Undefined Saprotroph\| | Yeast | NULL | Probable | As nectar-tap_saprotroph (Põlme et al. 2020); Decay sugar-rich substrates (Põlme et al. 2020) | Põlme S, et al. 2020. Fungal Diversity 105:1-16 ((https://doi.org/10.1007/s13225-020-00466-2)) |
| Otu50897 | 1 | k__Fungi(100);p__Ascomycota(100);c__Saccharomycetes(100);o__Saccharomycetales(100);f__Metschnikowiaceae(100);g__Metschnikowia(100);s__Metschnikowia_pulcherrima(100); | Metschnikowia | 13 | Saprotroph | \|Undefined Saprotroph\| | Yeast | NULL | Probable | As nectar-tap_saprotroph (Põlme et al. 2020); Decay sugar-rich substrates (Põlme et al. 2020) | Põlme S, et al. 2020. Fungal Diversity 105:1-16 ((https://doi.org/10.1007/s13225-020-00466-2)) |
| Otu50898 | 1 | k__Fungi(100);p__Ascomycota(100);c__Saccharomycetes(100);o__Saccharomycetales(100);f__Metschnikowiaceae(100);g__Metschnikowia(100);g__Metschnikowia_unclassified(100); | Metschnikowia | 13 | Saprotroph | \|Undefined Saprotroph\| | Yeast | NULL | Probable | As nectar-tap_saprotroph (Põlme et al. 2020); Decay sugar-rich substrates (Põlme et al. 2020) | Põlme S, et al. 2020. Fungal Diversity 105:1-16 ((https://doi.org/10.1007/s13225-020-00466-2)) |
| Otu50899 | 1 | k__Fungi(100);p__Ascomycota(100);c__Saccharomycetes(100);o__Saccharomycetales(100);f__Metschnikowiaceae(100);g__Metschnikowia(100);s__Metschnikowia_pulcherrima(100); | Metschnikowia | 13 | Saprotroph | \|Undefined Saprotroph\| | Yeast | NULL | Probable | As nectar-tap_saprotroph (Põlme et al. 2020); Decay sugar-rich substrates (Põlme et al. 2020) | Põlme S, et al. 2020. Fungal Diversity 105:1-16 ((https://doi.org/10.1007/s13225-020-00466-2)) |
| Otu50900 | 1 | k__Fungi(100);p__Ascomycota(100);c__Saccharomycetes(100);o__Saccharomycetales(100);f__Metschnikowiaceae(100);g__Metschnikowia(100);s__Metschnikowia_pulcherrima(100); | Metschnikowia | 13 | Saprotroph | \|Undefined Saprotroph\| | Yeast | NULL | Probable | As nectar-tap_saprotroph (Põlme et al. 2020); Decay sugar-rich substrates (Põlme et al. 2020) | Põlme S, et al. 2020. Fungal Diversity 105:1-16 ((https://doi.org/10.1007/s13225-020-00466-2)) |
| Otu50901 | 1 | k__Fungi(100);p__Ascomycota(100);c__Saccharomycetes(100);o__Saccharomycetales(100);f__Metschnikowiaceae(100);g__Metschnikowia(100);s__Metschnikowia_pulcherrima(100); | Metschnikowia | 13 | Saprotroph | \|Undefined Saprotroph\| | Yeast | NULL | Probable | As nectar-tap_saprotroph (Põlme et al. 2020); Decay sugar-rich substrates (Põlme et al. 2020) | Põlme S, et al. 2020. Fungal Diversity 105:1-16 ((https://doi.org/10.1007/s13225-020-00466-2)) |
| Otu50902 | 1 | k__Fungi(100);p__Ascomycota(100);c__Saccharomycetes(100);o__Saccharomycetales(100);f__Metschnikowiaceae(100);g__Metschnikowia(100);g__Metschnikowia_unclassified(100); | Metschnikowia | 13 | Saprotroph | \|Undefined Saprotroph\| | Yeast | NULL | Probable | As nectar-tap_saprotroph (Põlme et al. 2020); Decay sugar-rich substrates (Põlme et al. 2020) | Põlme S, et al. 2020. Fungal Diversity 105:1-16 ((https://doi.org/10.1007/s13225-020-00466-2)) |
| Otu50903 | 1 | k__Fungi(100);p__Ascomycota(100);c__Saccharomycetes(100);o__Saccharomycetales(100);f__Metschnikowiaceae(100);g__Metschnikowia(100);g__Metschnikowia_unclassified(100); | Metschnikowia | 13 | Saprotroph | \|Undefined Saprotroph\| | Yeast | NULL | Probable | As nectar-tap_saprotroph (Põlme et al. 2020); Decay sugar-rich substrates (Põlme et al. 2020) | Põlme S, et al. 2020. Fungal Diversity 105:1-16 ((https://doi.org/10.1007/s13225-020-00466-2)) |
| Otu50904 | 1 | k__Fungi(100);p__Ascomycota(100);c__Saccharomycetes(100);o__Saccharomycetales(100);f__Metschnikowiaceae(100);g__Metschnikowia(100);g__Metschnikowia_unclassified(100); | Metschnikowia | 13 | Saprotroph | \|Undefined Saprotroph\| | Yeast | NULL | Probable | As nectar-tap_saprotroph (Põlme et al. 2020); Decay sugar-rich substrates (Põlme et al. 2020) | Põlme S, et al. 2020. Fungal Diversity 105:1-16 ((https://doi.org/10.1007/s13225-020-00466-2)) |
| Otu50907 | 1 | k__Fungi(100);p__Ascomycota(100);c__Saccharomycetes(100);o__Saccharomycetales(100);f__Metschnikowiaceae(100);g__Metschnikowia(100);s__Metschnikowia_pulcherrima(100); | Metschnikowia | 13 | Saprotroph | \|Undefined Saprotroph\| | Yeast | NULL | Probable | As nectar-tap_saprotroph (Põlme et al. 2020); Decay sugar-rich substrates (Põlme et al. 2020) | Põlme S, et al. 2020. Fungal Diversity 105:1-16 ((https://doi.org/10.1007/s13225-020-00466-2)) |
| Otu50913 | 1 | k__Fungi(100);p__Ascomycota(100);c__Saccharomycetes(100);o__Saccharomycetales(100);f__Saccharomycetales_family_Incertae_sedis(100);g__Candida(100);g__Candida_unclassified(100); | Candida | 13 | Pathotroph-Saprotroph-Symbiotroph | Animal Pathogen-Endophyte-Endosymbiont-Epiphyte-\|Undefined Saprotroph\| | Dimorphic Yeast-Yeast | NULL | Probable | As nectar-tap_saprotroph (Põlme et al. 2020); Decay sugar-rich substrates (Põlme et al. 2020); This asexual genus is polyphyletic and contain species that function very differently depend on what they are. Caution should be taken when interpreting guilds for Candida species. | Manolakaki D, et al. 2010. Virulence. 1: 367-375 ((https://doi.org/10.4161/viru.1.5.12796)); Mekbib SB, et al. 2011. Tropical Plant Pathology 36:233-240 ((https://doi.org/10.1590/S1982-56762011000400004)); Põlme S, et al. 2020. Fungal Diversity 105:1-16 ((https://doi.org/10.1007/s13225-020-00466-2)) |
| Otu50914 | 1 | k__Fungi(100);p__Ascomycota(100);c__Saccharomycetes(100);o__Saccharomycetales(100);f__Saccharomycetales_family_Incertae_sedis(100);g__Candida(100);g__Candida_unclassified(100); | Candida | 13 | Pathotroph-Saprotroph-Symbiotroph | Animal Pathogen-Endophyte-Endosymbiont-Epiphyte-\|Undefined Saprotroph\| | Dimorphic Yeast-Yeast | NULL | Probable | As nectar-tap_saprotroph (Põlme et al. 2020); Decay sugar-rich substrates (Põlme et al. 2020); This asexual genus is polyphyletic and contain species that function very differently depend on what they are. Caution should be taken when interpreting guilds for Candida species. | Manolakaki D, et al. 2010. Virulence. 1: 367-375 ((https://doi.org/10.4161/viru.1.5.12796)); Mekbib SB, et al. 2011. Tropical Plant Pathology 36:233-240 ((https://doi.org/10.1590/S1982-56762011000400004)); Põlme S, et al. 2020. Fungal Diversity 105:1-16 ((https://doi.org/10.1007/s13225-020-00466-2)) |
| Otu50915 | 1 | k__Fungi(100);p__Ascomycota(100);c__Saccharomycetes(100);o__Saccharomycetales(100);o__Saccharomycetales_unclassified(100);o__Saccharomycetales_unclassified(100);o__Saccharomycetales_unclassified(100); | Saccharomycetales | 7 | Saprotroph | Undefined Saprotroph | Yeast | NULL | Possible | NULL | Sterkenburg E, et al. 2015. New Phytologist 207:1145-1158 ((https://doi.org/10.1111/nph.13426)) |
| Otu50917 | 1 | k__Fungi(100);p__Ascomycota(100);c__Saccharomycetes(100);o__Saccharomycetales(100);f__Metschnikowiaceae(100);g__Metschnikowia(100);s__Metschnikowia_pulcherrima(100); | Metschnikowia | 13 | Saprotroph | \|Undefined Saprotroph\| | Yeast | NULL | Probable | As nectar-tap_saprotroph (Põlme et al. 2020); Decay sugar-rich substrates (Põlme et al. 2020) | Põlme S, et al. 2020. Fungal Diversity 105:1-16 ((https://doi.org/10.1007/s13225-020-00466-2)) |
| Otu50918 | 1 | k__Fungi(100);p__Ascomycota(100);c__Saccharomycetes(100);o__Saccharomycetales(100);f__Metschnikowiaceae(100);g__Metschnikowia(100);g__Metschnikowia_unclassified(100); | Metschnikowia | 13 | Saprotroph | \|Undefined Saprotroph\| | Yeast | NULL | Probable | As nectar-tap_saprotroph (Põlme et al. 2020); Decay sugar-rich substrates (Põlme et al. 2020) | Põlme S, et al. 2020. Fungal Diversity 105:1-16 ((https://doi.org/10.1007/s13225-020-00466-2)) |
| Otu50919 | 1 | k__Fungi(100);p__Ascomycota(100);c__Saccharomycetes(100);o__Saccharomycetales(100);f__Saccharomycetales_family_Incertae_sedis(100);g__Candida(100);g__Candida_unclassified(100); | Candida | 13 | Pathotroph-Saprotroph-Symbiotroph | Animal Pathogen-Endophyte-Endosymbiont-Epiphyte-\|Undefined Saprotroph\| | Dimorphic Yeast-Yeast | NULL | Probable | As nectar-tap_saprotroph (Põlme et al. 2020); Decay sugar-rich substrates (Põlme et al. 2020); This asexual genus is polyphyletic and contain species that function very differently depend on what they are. Caution should be taken when interpreting guilds for Candida species. | Manolakaki D, et al. 2010. Virulence. 1: 367-375 ((https://doi.org/10.4161/viru.1.5.12796)); Mekbib SB, et al. 2011. Tropical Plant Pathology 36:233-240 ((https://doi.org/10.1590/S1982-56762011000400004)); Põlme S, et al. 2020. Fungal Diversity 105:1-16 ((https://doi.org/10.1007/s13225-020-00466-2)) |
| Otu50920 | 1 | k__Fungi(100);p__Ascomycota(100);c__Saccharomycetes(100);o__Saccharomycetales(100);f__Metschnikowiaceae(100);g__Metschnikowia(100);s__Metschnikowia_pulcherrima(100); | Metschnikowia | 13 | Saprotroph | \|Undefined Saprotroph\| | Yeast | NULL | Probable | As nectar-tap_saprotroph (Põlme et al. 2020); Decay sugar-rich substrates (Põlme et al. 2020) | Põlme S, et al. 2020. Fungal Diversity 105:1-16 ((https://doi.org/10.1007/s13225-020-00466-2)) |
| Otu50921 | 1 | k__Fungi(100);p__Ascomycota(100);c__Saccharomycetes(100);o__Saccharomycetales(100);f__Metschnikowiaceae(100);g__Metschnikowia(100);s__Metschnikowia_pulcherrima(100); | Metschnikowia | 13 | Saprotroph | \|Undefined Saprotroph\| | Yeast | NULL | Probable | As nectar-tap_saprotroph (Põlme et al. 2020); Decay sugar-rich substrates (Põlme et al. 2020) | Põlme S, et al. 2020. Fungal Diversity 105:1-16 ((https://doi.org/10.1007/s13225-020-00466-2)) |
| Otu50922 | 1 | k__Fungi(100);p__Ascomycota(100);c__Saccharomycetes(100);o__Saccharomycetales(100);f__Metschnikowiaceae(100);g__Metschnikowia(100);s__Metschnikowia_pulcherrima(100); | Metschnikowia | 13 | Saprotroph | \|Undefined Saprotroph\| | Yeast | NULL | Probable | As nectar-tap_saprotroph (Põlme et al. 2020); Decay sugar-rich substrates (Põlme et al. 2020) | Põlme S, et al. 2020. Fungal Diversity 105:1-16 ((https://doi.org/10.1007/s13225-020-00466-2)) |
| Otu50924 | 1 | k__Fungi(100);p__Ascomycota(100);c__Saccharomycetes(100);o__Saccharomycetales(100);f__Saccharomycetales_family_Incertae_sedis(100);g__Candida(100);g__Candida_unclassified(100); | Candida | 13 | Pathotroph-Saprotroph-Symbiotroph | Animal Pathogen-Endophyte-Endosymbiont-Epiphyte-\|Undefined Saprotroph\| | Dimorphic Yeast-Yeast | NULL | Probable | As nectar-tap_saprotroph (Põlme et al. 2020); Decay sugar-rich substrates (Põlme et al. 2020); This asexual genus is polyphyletic and contain species that function very differently depend on what they are. Caution should be taken when interpreting guilds for Candida species. | Manolakaki D, et al. 2010. Virulence. 1: 367-375 ((https://doi.org/10.4161/viru.1.5.12796)); Mekbib SB, et al. 2011. Tropical Plant Pathology 36:233-240 ((https://doi.org/10.1590/S1982-56762011000400004)); Põlme S, et al. 2020. Fungal Diversity 105:1-16 ((https://doi.org/10.1007/s13225-020-00466-2)) |
| Otu50927 | 1 | k__Fungi(100);p__Ascomycota(100);c__Saccharomycetes(100);o__Saccharomycetales(100);f__Metschnikowiaceae(100);g__Metschnikowia(100);s__Metschnikowia_pulcherrima(100); | Metschnikowia | 13 | Saprotroph | \|Undefined Saprotroph\| | Yeast | NULL | Probable | As nectar-tap_saprotroph (Põlme et al. 2020); Decay sugar-rich substrates (Põlme et al. 2020) | Põlme S, et al. 2020. Fungal Diversity 105:1-16 ((https://doi.org/10.1007/s13225-020-00466-2)) |
| Otu50928 | 1 | k__Fungi(100);p__Ascomycota(100);c__Saccharomycetes(100);o__Saccharomycetales(100);f__Metschnikowiaceae(100);g__Metschnikowia(100);s__Metschnikowia_pulcherrima(100); | Metschnikowia | 13 | Saprotroph | \|Undefined Saprotroph\| | Yeast | NULL | Probable | As nectar-tap_saprotroph (Põlme et al. 2020); Decay sugar-rich substrates (Põlme et al. 2020) | Põlme S, et al. 2020. Fungal Diversity 105:1-16 ((https://doi.org/10.1007/s13225-020-00466-2)) |
| Otu50929 | 1 | k__Fungi(100);p__Ascomycota(100);c__Saccharomycetes(100);o__Saccharomycetales(100);f__Metschnikowiaceae(100);g__Metschnikowia(100);g__Metschnikowia_unclassified(100); | Metschnikowia | 13 | Saprotroph | \|Undefined Saprotroph\| | Yeast | NULL | Probable | As nectar-tap_saprotroph (Põlme et al. 2020); Decay sugar-rich substrates (Põlme et al. 2020) | Põlme S, et al. 2020. Fungal Diversity 105:1-16 ((https://doi.org/10.1007/s13225-020-00466-2)) |
| Otu50930 | 1 | k__Fungi(100);p__Ascomycota(100);c__Saccharomycetes(100);o__Saccharomycetales(100);f__Metschnikowiaceae(100);g__Metschnikowia(100);g__Metschnikowia_unclassified(100); | Metschnikowia | 13 | Saprotroph | \|Undefined Saprotroph\| | Yeast | NULL | Probable | As nectar-tap_saprotroph (Põlme et al. 2020); Decay sugar-rich substrates (Põlme et al. 2020) | Põlme S, et al. 2020. Fungal Diversity 105:1-16 ((https://doi.org/10.1007/s13225-020-00466-2)) |
| Otu50931 | 1 | k__Fungi(100);p__Ascomycota(100);c__Saccharomycetes(100);o__Saccharomycetales(100);f__Metschnikowiaceae(100);g__Metschnikowia(100);s__Metschnikowia_pulcherrima(100); | Metschnikowia | 13 | Saprotroph | \|Undefined Saprotroph\| | Yeast | NULL | Probable | As nectar-tap_saprotroph (Põlme et al. 2020); Decay sugar-rich substrates (Põlme et al. 2020) | Põlme S, et al. 2020. Fungal Diversity 105:1-16 ((https://doi.org/10.1007/s13225-020-00466-2)) |

| | | | | | | | | | | | |
|---|---|---|---|---|---|---|---|---|---|---|---|
| Otu50932 | 1 | k__Fungi(100);p__Ascomycota(100);c__Saccharomycetes(100);o__Saccharomycetales(100);f__Metschnikowiaceae(100);g__Metschnikowia(100);s__Metschnikowia_pulcherrima(100); | Metschnikowia | 13 | Saprotroph | \|Undefined Saprotroph\| | Yeast | NULL | Probable | As nectar-tap_saprotroph (Põlme et al. 2020); Decay sugar-rich substrates (Põlme et al. 2020) | Põlme S, et al. 2020. Fungal Diversity 105:1-16 ((https://doi.org/10.1007/s13225-020-00466-2)) |
| Otu50933 | 1 | k__Fungi(100);p__Ascomycota(100);c__Saccharomycetes(100);o__Saccharomycetales(100);f__Metschnikowiaceae(100);g__Metschnikowia(100);s__Metschnikowia_pulcherrima(100); | Metschnikowia | 13 | Saprotroph | \|Undefined Saprotroph\| | Yeast | NULL | Probable | As nectar-tap_saprotroph (Põlme et al. 2020); Decay sugar-rich substrates (Põlme et al. 2020) | Põlme S, et al. 2020. Fungal Diversity 105:1-16 ((https://doi.org/10.1007/s13225-020-00466-2)) |
| Otu50934 | 1 | k__Fungi(100);p__Ascomycota(100);c__Saccharomycetes(100);o__Saccharomycetales(100);f__Metschnikowiaceae(100);g__Metschnikowia(100);s__Metschnikowia_pulcherrima(100); | Metschnikowia | 13 | Saprotroph | \|Undefined Saprotroph\| | Yeast | NULL | Probable | As nectar-tap_saprotroph (Põlme et al. 2020); Decay sugar-rich substrates (Põlme et al. 2020) | Põlme S, et al. 2020. Fungal Diversity 105:1-16 ((https://doi.org/10.1007/s13225-020-00466-2)) |
| Otu50935 | 1 | k__Fungi(100);p__Ascomycota(100);c__Saccharomycetes(100);o__Saccharomycetales(100);f__Metschnikowiaceae(100);g__Metschnikowia(100);s__Metschnikowia_pulcherrima(100); | Metschnikowia | 13 | Saprotroph | \|Undefined Saprotroph\| | Yeast | NULL | Probable | As nectar-tap_saprotroph (Põlme et al. 2020); Decay sugar-rich substrates (Põlme et al. 2020) | Põlme S, et al. 2020. Fungal Diversity 105:1-16 ((https://doi.org/10.1007/s13225-020-00466-2)) |
| Otu50936 | 1 | k__Fungi(100);p__Ascomycota(100);c__Saccharomycetes(100);o__Saccharomycetales(100);f__Metschnikowiaceae(100);g__Metschnikowia(100);g__Metschnikowia_unclassified(100); | Metschnikowia | 13 | Saprotroph | \|Undefined Saprotroph\| | Yeast | NULL | Probable | As nectar-tap_saprotroph (Põlme et al. 2020); Decay sugar-rich substrates (Põlme et al. 2020) | Põlme S, et al. 2020. Fungal Diversity 105:1-16 ((https://doi.org/10.1007/s13225-020-00466-2)) |
| Otu50937 | 1 | k__Fungi(100);p__Ascomycota(100);c__Saccharomycetes(100);o__Saccharomycetales(100);f__Metschnikowiaceae(100);g__Metschnikowia(100);g__Metschnikowia_unclassified(100); | Metschnikowia | 13 | Saprotroph | \|Undefined Saprotroph\| | Yeast | NULL | Probable | As nectar-tap_saprotroph (Põlme et al. 2020); Decay sugar-rich substrates (Põlme et al. 2020) | Põlme S, et al. 2020. Fungal Diversity 105:1-16 ((https://doi.org/10.1007/s13225-020-00466-2)) |
| Otu50938 | 1 | k__Fungi(100);p__Ascomycota(100);c__Saccharomycetes(100);o__Saccharomycetales(100);f__Metschnikowiaceae(100);g__Metschnikowia(100);s__Metschnikowia_pulcherrima(100); | Metschnikowia | 13 | Saprotroph | \|Undefined Saprotroph\| | Yeast | NULL | Probable | As nectar-tap_saprotroph (Põlme et al. 2020); Decay sugar-rich substrates (Põlme et al. 2020) | Põlme S, et al. 2020. Fungal Diversity 105:1-16 ((https://doi.org/10.1007/s13225-020-00466-2)) |
| Otu50940 | 1 | k__Fungi(100);p__Ascomycota(100);c__Saccharomycetes(100);o__Saccharomycetales(100);f__Metschnikowiaceae(100);g__Metschnikowia(100);s__Metschnikowia_pulcherrima(100); | Metschnikowia | 13 | Saprotroph | \|Undefined Saprotroph\| | Yeast | NULL | Probable | As nectar-tap_saprotroph (Põlme et al. 2020); Decay sugar-rich substrates (Põlme et al. 2020) | Põlme S, et al. 2020. Fungal Diversity 105:1-16 ((https://doi.org/10.1007/s13225-020-00466-2)) |
| Otu50942 | 1 | k__Fungi(100);p__Ascomycota(100);c__Saccharomycetes(100);o__Saccharomycetales(100);f__Metschnikowiaceae(100);g__Metschnikowia(100);s__Metschnikowia_pulcherrima(100); | Metschnikowia | 13 | Saprotroph | \|Undefined Saprotroph\| | Yeast | NULL | Probable | As nectar-tap_saprotroph (Põlme et al. 2020); Decay sugar-rich substrates (Põlme et al. 2020) | Põlme S, et al. 2020. Fungal Diversity 105:1-16 ((https://doi.org/10.1007/s13225-020-00466-2)) |
| Otu50943 | 1 | k__Fungi(100);p__Ascomycota(100);c__Saccharomycetes(100);o__Saccharomycetales(100);f__Metschnikowiaceae(100);g__Metschnikowia(100);s__Metschnikowia_pulcherrima(100); | Metschnikowia | 13 | Saprotroph | \|Undefined Saprotroph\| | Yeast | NULL | Probable | As nectar-tap_saprotroph (Põlme et al. 2020); Decay sugar-rich substrates (Põlme et al. 2020) | Põlme S, et al. 2020. Fungal Diversity 105:1-16 ((https://doi.org/10.1007/s13225-020-00466-2)) |
| Otu50944 | 1 | k__Fungi(100);p__Ascomycota(100);c__Saccharomycetes(100);o__Saccharomycetales(100);f__Metschnikowiaceae(100);g__Metschnikowia(100);g__Metschnikowia_unclassified(100); | Metschnikowia | 13 | Saprotroph | \|Undefined Saprotroph\| | Yeast | NULL | Probable | As nectar-tap_saprotroph (Põlme et al. 2020); Decay sugar-rich substrates (Põlme et al. 2020) | Põlme S, et al. 2020. Fungal Diversity 105:1-16 ((https://doi.org/10.1007/s13225-020-00466-2)) |
| Otu50945 | 1 | k__Fungi(100);p__Ascomycota(100);c__Saccharomycetes(100);o__Saccharomycetales(100);f__Metschnikowiaceae(100);g__Metschnikowia(100);s__Metschnikowia_pulcherrima(100); | Metschnikowia | 13 | Saprotroph | \|Undefined Saprotroph\| | Yeast | NULL | Probable | As nectar-tap_saprotroph (Põlme et al. 2020); Decay sugar-rich substrates (Põlme et al. 2020) | Põlme S, et al. 2020. Fungal Diversity 105:1-16 ((https://doi.org/10.1007/s13225-020-00466-2)) |
| Otu50946 | 1 | k__Fungi(100);p__Ascomycota(100);c__Saccharomycetes(100);o__Saccharomycetales(100);f__Metschnikowiaceae(100);g__Metschnikowia(100);s__Metschnikowia_pulcherrima(100); | Metschnikowia | 13 | Saprotroph | \|Undefined Saprotroph\| | Yeast | NULL | Probable | As nectar-tap_saprotroph (Põlme et al. 2020); Decay sugar-rich substrates (Põlme et al. 2020) | Põlme S, et al. 2020. Fungal Diversity 105:1-16 ((https://doi.org/10.1007/s13225-020-00466-2)) |
| Otu50947 | 1 | k__Fungi(100);p__Ascomycota(100);c__Saccharomycetes(100);o__Saccharomycetales(100);f__Metschnikowiaceae(100);g__Metschnikowia(100);s__Metschnikowia_pulcherrima(100); | Metschnikowia | 13 | Saprotroph | \|Undefined Saprotroph\| | Yeast | NULL | Probable | As nectar-tap_saprotroph (Põlme et al. 2020); Decay sugar-rich substrates (Põlme et al. 2020) | Põlme S, et al. 2020. Fungal Diversity 105:1-16 ((https://doi.org/10.1007/s13225-020-00466-2)) |
| Otu50948 | 1 | k__Fungi(100);p__Ascomycota(100);c__Saccharomycetes(100);o__Saccharomycetales(100);f__Metschnikowiaceae(100);g__Metschnikowia(100);s__Metschnikowia_pulcherrima(100); | Metschnikowia | 13 | Saprotroph | \|Undefined Saprotroph\| | Yeast | NULL | Probable | As nectar-tap_saprotroph (Põlme et al. 2020); Decay sugar-rich substrates (Põlme et al. 2020) | Põlme S, et al. 2020. Fungal Diversity 105:1-16 ((https://doi.org/10.1007/s13225-020-00466-2)) |
| Otu50949 | 1 | k__Fungi(100);p__Ascomycota(100);c__Saccharomycetes(100);o__Saccharomycetales(100);f__Metschnikowiaceae(100);g__Metschnikowia(100);s__Metschnikowia_pulcherrima(100); | Metschnikowia | 13 | Saprotroph | \|Undefined Saprotroph\| | Yeast | NULL | Probable | As nectar-tap_saprotroph (Põlme et al. 2020); Decay sugar-rich substrates (Põlme et al. 2020) | Põlme S, et al. 2020. Fungal Diversity 105:1-16 ((https://doi.org/10.1007/s13225-020-00466-2)) |
| Otu50950 | 1 | k__Fungi(100);p__Ascomycota(100);c__Saccharomycetes(100);o__Saccharomycetales(100);f__Saccharomycetales_family_Incertae_sedis(100);f__Saccharomycetales_family_Incertae_sedis_unclassified(100);f__Saccharomycetales_family_Incertae_sedis_unclassified(100); | Saccharomycetales | 7 | Saprotroph | Undefined Saprotroph | Yeast | NULL | Possible | NULL | Sterkenburg E, et al. 2015. New Phytologist 207:1145-1158 ((https://doi.org/10.1111/nph.13426)) |
| Otu50951 | 1 | k__Fungi(100);p__Ascomycota(100);c__Saccharomycetes(100);o__Saccharomycetales(100);f__Metschnikowiaceae(100);g__Metschnikowia(100);s__Metschnikowia_pulcherrima(100); | Metschnikowia | 13 | Saprotroph | \|Undefined Saprotroph\| | Yeast | NULL | Probable | As nectar-tap_saprotroph (Põlme et al. 2020); Decay sugar-rich substrates (Põlme et al. 2020) | Põlme S, et al. 2020. Fungal Diversity 105:1-16 ((https://doi.org/10.1007/s13225-020-00466-2)) |
| Otu50952 | 1 | k__Fungi(100);p__Ascomycota(100);c__Saccharomycetes(100);o__Saccharomycetales(100);f__Metschnikowiaceae(100);g__Metschnikowia(100);s__Metschnikowia_pulcherrima(100); | Metschnikowia | 13 | Saprotroph | \|Undefined Saprotroph\| | Yeast | NULL | Probable | As nectar-tap_saprotroph (Põlme et al. 2020); Decay sugar-rich substrates (Põlme et al. 2020) | Põlme S, et al. 2020. Fungal Diversity 105:1-16 ((https://doi.org/10.1007/s13225-020-00466-2)) |
| Otu50953 | 1 | k__Fungi(100);p__Ascomycota(100);c__Saccharomycetes(100);o__Saccharomycetales(100);f__Metschnikowiaceae(100);g__Metschnikowia(100);s__Metschnikowia_pulcherrima(100); | Metschnikowia | 13 | Saprotroph | \|Undefined Saprotroph\| | Yeast | NULL | Probable | As nectar-tap_saprotroph (Põlme et al. 2020); Decay sugar-rich substrates (Põlme et al. 2020) | Põlme S, et al. 2020. Fungal Diversity 105:1-16 ((https://doi.org/10.1007/s13225-020-00466-2)) |
| Otu50954 | 1 | k__Fungi(100);p__Ascomycota(100);c__Saccharomycetes(100);o__Saccharomycetales(100);f__Metschnikowiaceae(100);g__Metschnikowia(100);g__Metschnikowia_unclassified(100); | Metschnikowia | 13 | Saprotroph | \|Undefined Saprotroph\| | Yeast | NULL | Probable | As nectar-tap_saprotroph (Põlme et al. 2020); Decay sugar-rich substrates (Põlme et al. 2020) | Põlme S, et al. 2020. Fungal Diversity 105:1-16 ((https://doi.org/10.1007/s13225-020-00466-2)) |
| Otu50955 | 1 | k__Fungi(100);p__Ascomycota(100);c__Saccharomycetes(100);o__Saccharomycetales(100);f__Metschnikowiaceae(100);g__Metschnikowia(100);s__Metschnikowia_pulcherrima(100); | Metschnikowia | 13 | Saprotroph | \|Undefined Saprotroph\| | Yeast | NULL | Probable | As nectar-tap_saprotroph (Põlme et al. 2020); Decay sugar-rich substrates (Põlme et al. 2020) | Põlme S, et al. 2020. Fungal Diversity 105:1-16 ((https://doi.org/10.1007/s13225-020-00466-2)) |
| Otu50956 | 1 | k__Fungi(100);p__Ascomycota(100);c__Saccharomycetes(100);o__Saccharomycetales(100);f__Metschnikowiaceae(100);g__Metschnikowia(100);g__Metschnikowia_unclassified(100); | Metschnikowia | 13 | Saprotroph | \|Undefined Saprotroph\| | Yeast | NULL | Probable | As nectar-tap_saprotroph (Põlme et al. 2020); Decay sugar-rich substrates (Põlme et al. 2020) | Põlme S, et al. 2020. Fungal Diversity 105:1-16 ((https://doi.org/10.1007/s13225-020-00466-2)) |
| Otu50957 | 1 | k__Fungi(100);p__Ascomycota(100);c__Saccharomycetes(100);o__Saccharomycetales(100);f__Metschnikowiaceae(100);g__Metschnikowia(100);g__Metschnikowia_unclassified(100); | Metschnikowia | 13 | Saprotroph | \|Undefined Saprotroph\| | Yeast | NULL | Probable | As nectar-tap_saprotroph (Põlme et al. 2020); Decay sugar-rich substrates (Põlme et al. 2020) | Põlme S, et al. 2020. Fungal Diversity 105:1-16 ((https://doi.org/10.1007/s13225-020-00466-2)) |
| Otu50959 | 1 | k__Fungi(100);p__Ascomycota(100);c__Saccharomycetes(100);o__Saccharomycetales(100);f__Metschnikowiaceae(100);g__Metschnikowia(100);s__Metschnikowia_pulcherrima(100); | Metschnikowia | 13 | Saprotroph | \|Undefined Saprotroph\| | Yeast | NULL | Probable | As nectar-tap_saprotroph (Põlme et al. 2020); Decay sugar-rich substrates (Põlme et al. 2020) | Põlme S, et al. 2020. Fungal Diversity 105:1-16 ((https://doi.org/10.1007/s13225-020-00466-2)) |
| Otu50960 | 1 | k__Fungi(100);p__Ascomycota(100);c__Saccharomycetes(100);o__Saccharomycetales(100);f__Metschnikowiaceae(100);g__Metschnikowia(100);s__Metschnikowia_pulcherrima(100); | Metschnikowia | 13 | Saprotroph | \|Undefined Saprotroph\| | Yeast | NULL | Probable | As nectar-tap_saprotroph (Põlme et al. 2020); Decay sugar-rich substrates (Põlme et al. 2020) | Põlme S, et al. 2020. Fungal Diversity 105:1-16 ((https://doi.org/10.1007/s13225-020-00466-2)) |
| Otu50961 | 1 | k__Fungi(100);p__Ascomycota(100);c__Saccharomycetes(100);o__Saccharomycetales(100);f__Metschnikowiaceae(100);g__Metschnikowia(100);s__Metschnikowia_pulcherrima(100); | Metschnikowia | 13 | Saprotroph | \|Undefined Saprotroph\| | Yeast | NULL | Probable | As nectar-tap_saprotroph (Põlme et al. 2020); Decay sugar-rich substrates (Põlme et al. 2020) | Põlme S, et al. 2020. Fungal Diversity 105:1-16 ((https://doi.org/10.1007/s13225-020-00466-2)) |
| Otu50962 | 1 | k__Fungi(100);p__Ascomycota(100);c__Saccharomycetes(100);o__Saccharomycetales(100);f__Metschnikowiaceae(100);g__Metschnikowia(100);s__Metschnikowia_pulcherrima(100); | Metschnikowia | 13 | Saprotroph | \|Undefined Saprotroph\| | Yeast | NULL | Probable | As nectar-tap_saprotroph (Põlme et al. 2020); Decay sugar-rich substrates (Põlme et al. 2020) | Põlme S, et al. 2020. Fungal Diversity 105:1-16 ((https://doi.org/10.1007/s13225-020-00466-2)) |
| Otu50964 | 1 | k__Fungi(100);p__Ascomycota(100);c__Saccharomycetes(100);o__Saccharomycetales(100);f__Metschnikowiaceae(100);g__Metschnikowia(100);s__Metschnikowia_pulcherrima(100); | Metschnikowia | 13 | Saprotroph | \|Undefined Saprotroph\| | Yeast | NULL | Probable | As nectar-tap_saprotroph (Põlme et al. 2020); Decay sugar-rich substrates (Põlme et al. 2020) | Põlme S, et al. 2020. Fungal Diversity 105:1-16 ((https://doi.org/10.1007/s13225-020-00466-2)) |
| Otu50966 | 1 | k__Fungi(100);p__Ascomycota(100);c__Saccharomycetes(100);o__Saccharomycetales(100);f__Metschnikowiaceae(100);g__Metschnikowia(100);s__Metschnikowia_pulcherrima(100); | Metschnikowia | 13 | Saprotroph | \|Undefined Saprotroph\| | Yeast | NULL | Probable | As nectar-tap_saprotroph (Põlme et al. 2020); Decay sugar-rich substrates (Põlme et al. 2020) | Põlme S, et al. 2020. Fungal Diversity 105:1-16 ((https://doi.org/10.1007/s13225-020-00466-2)) |
| Otu50967 | 1 | k__Fungi(100);p__Ascomycota(100);c__Saccharomycetes(100);o__Saccharomycetales(100);f__Metschnikowiaceae(100);g__Metschnikowia(100);g__Metschnikowia_unclassified(100); | Metschnikowia | 13 | Saprotroph | \|Undefined Saprotroph\| | Yeast | NULL | Probable | As nectar-tap_saprotroph (Põlme et al. 2020); Decay sugar-rich substrates (Põlme et al. 2020) | Põlme S, et al. 2020. Fungal Diversity 105:1-16 ((https://doi.org/10.1007/s13225-020-00466-2)) |
| Otu50968 | 1 | k__Fungi(100);p__Ascomycota(100);c__Saccharomycetes(100);o__Saccharomycetales(100);f__Metschnikowiaceae(100);g__Metschnikowia(100);s__Metschnikowia_pulcherrima(100); | Metschnikowia | 13 | Saprotroph | \|Undefined Saprotroph\| | Yeast | NULL | Probable | As nectar-tap_saprotroph (Põlme et al. 2020); Decay sugar-rich substrates (Põlme et al. 2020) | Põlme S, et al. 2020. Fungal Diversity 105:1-16 ((https://doi.org/10.1007/s13225-020-00466-2)) |
| Otu50970 | 1 | k__Fungi(100);p__Ascomycota(100);c__Saccharomycetes(100);o__Saccharomycetales(100);f__Metschnikowiaceae(100);g__Metschnikowia(100);s__Metschnikowia_pulcherrima(100); | Metschnikowia | 13 | Saprotroph | \|Undefined Saprotroph\| | Yeast | NULL | Probable | As nectar-tap_saprotroph (Põlme et al. 2020); Decay sugar-rich substrates (Põlme et al. 2020) | Põlme S, et al. 2020. Fungal Diversity 105:1-16 ((https://doi.org/10.1007/s13225-020-00466-2)) |
| Otu50974 | 1 | k__Fungi(100);p__Ascomycota(100);c__Saccharomycetes(100);o__Saccharomycetales(100);f__Metschnikowiaceae(100);g__Metschnikowia(100);s__Metschnikowia_pulcherrima(100); | Metschnikowia | 13 | Saprotroph | \|Undefined Saprotroph\| | Yeast | NULL | Probable | As nectar-tap_saprotroph (Põlme et al. 2020); Decay sugar-rich substrates (Põlme et al. 2020) | Põlme S, et al. 2020. Fungal Diversity 105:1-16 ((https://doi.org/10.1007/s13225-020-00466-2)) |
| Otu50977 | 1 | k__Fungi(100);p__Ascomycota(100);c__Saccharomycetes(100);o__Saccharomycetales(100);f__Metschnikowiaceae(100);g__Metschnikowia(100);g__Metschnikowia_unclassified(100); | Metschnikowia | 13 | Saprotroph | \|Undefined Saprotroph\| | Yeast | NULL | Probable | As nectar-tap_saprotroph (Põlme et al. 2020); Decay sugar-rich substrates (Põlme et al. 2020) | Põlme S, et al. 2020. Fungal Diversity 105:1-16 ((https://doi.org/10.1007/s13225-020-00466-2)) |
| Otu50978 | 1 | k__Fungi(100);p__Ascomycota(100);c__Saccharomycetes(100);o__Saccharomycetales(100);f__Metschnikowiaceae(100);g__Metschnikowia(100);g__Metschnikowia_unclassified(100); | Metschnikowia | 13 | Saprotroph | \|Undefined Saprotroph\| | Yeast | NULL | Probable | As nectar-tap_saprotroph (Põlme et al. 2020); Decay sugar-rich substrates (Põlme et al. 2020) | Põlme S, et al. 2020. Fungal Diversity 105:1-16 ((https://doi.org/10.1007/s13225-020-00466-2)) |
| Otu50979 | 1 | k__Fungi(100);p__Ascomycota(100);c__Saccharomycetes(100);o__Saccharomycetales(100);f__Metschnikowiaceae(100);g__Metschnikowia(100);s__Metschnikowia_pulcherrima(100); | Metschnikowia | 13 | Saprotroph | \|Undefined Saprotroph\| | Yeast | NULL | Probable | As nectar-tap_saprotroph (Põlme et al. 2020); Decay sugar-rich substrates (Põlme et al. 2020) | Põlme S, et al. 2020. Fungal Diversity 105:1-16 ((https://doi.org/10.1007/s13225-020-00466-2)) |
| Otu50980 | 1 | k__Fungi(100);p__Ascomycota(100);c__Saccharomycetes(100);o__Saccharomycetales(100);f__Metschnikowiaceae(100);g__Metschnikowia(100);s__Metschnikowia_pulcherrima(100); | Metschnikowia | 13 | Saprotroph | \|Undefined Saprotroph\| | Yeast | NULL | Probable | As nectar-tap_saprotroph (Põlme et al. 2020); Decay sugar-rich substrates (Põlme et al. 2020) | Põlme S, et al. 2020. Fungal Diversity 105:1-16 ((https://doi.org/10.1007/s13225-020-00466-2)) |
| Otu50981 | 1 | k__Fungi(100);p__Ascomycota(100);c__Saccharomycetes(100);o__Saccharomycetales(100);f__Metschnikowiaceae(100);g__Metschnikowia(100);s__Metschnikowia_pulcherrima(100); | Metschnikowia | 13 | Saprotroph | \|Undefined Saprotroph\| | Yeast | NULL | Probable | As nectar-tap_saprotroph (Põlme et al. 2020); Decay sugar-rich substrates (Põlme et al. 2020) | Põlme S, et al. 2020. Fungal Diversity 105:1-16 ((https://doi.org/10.1007/s13225-020-00466-2)) |
| Otu50982 | 1 | k__Fungi(100);p__Ascomycota(100);c__Saccharomycetes(100);o__Saccharomycetales(100);f__Metschnikowiaceae(100);g__Metschnikowia(100);s__Metschnikowia_pulcherrima(100); | Metschnikowia | 13 | Saprotroph | \|Undefined Saprotroph\| | Yeast | NULL | Probable | As nectar-tap_saprotroph (Põlme et al. 2020); Decay sugar-rich substrates (Põlme et al. 2020) | Põlme S, et al. 2020. Fungal Diversity 105:1-16 ((https://doi.org/10.1007/s13225-020-00466-2)) |
| Otu50983 | 1 | k__Fungi(100);p__Ascomycota(100);c__Saccharomycetes(100);o__Saccharomycetales(100);f__Metschnikowiaceae(100);g__Metschnikowia(100);s__Metschnikowia_pulcherrima(100); | Metschnikowia | 13 | Saprotroph | \|Undefined Saprotroph\| | Yeast | NULL | Probable | As nectar-tap_saprotroph (Põlme et al. 2020); Decay sugar-rich substrates (Põlme et al. 2020) | Põlme S, et al. 2020. Fungal Diversity 105:1-16 ((https://doi.org/10.1007/s13225-020-00466-2)) |
| Otu50986 | 1 | k__Fungi(100);p__Ascomycota(100);c__Saccharomycetes(100);o__Saccharomycetales(100);f__Saccharomycetales_family_Incertae_sedis(100);g__Candida(100);s__Candida_cretensis(100); | Candida | 13 | Pathotroph-Saprotroph-Symbiotroph | Animal Pathogen-Endophyte-Endosymbiont-Epiphyte-\|Undefined Saprotroph\| | Dimorphic Yeast-Yeast | NULL | Probable | As nectar-tap_saprotroph (Põlme et al. 2020); Decay sugar-rich substrates (Põlme et al. 2020); This asexual genus is polyphyletic and contain species that function very differently depend on what they are. Caution should be taken when interpreting guilds for Candida species. | Manolakaki D, et al. 2010. Virulence. 1: 367-375 ((https://doi.org/10.4161/viru.1.5.12796)); Mekbib SB, et al. 2011. Tropical Plant Pathology 36:233-240 ((https://doi.org/10.1590/S1982-56762011000400004)); Põlme S, et al. 2020. Fungal Diversity 105:1-16 ((https://doi.org/10.1007/s13225-020-00466-2)) |
| Otu50990 | 1 | k__Fungi(100);p__Ascomycota(100);c__Saccharomycetes(100);o__Saccharomycetales(100);f__Metschnikowiaceae(100);g__Metschnikowia(100);s__Metschnikowia_pulcherrima(100); | Metschnikowia | 13 | Saprotroph | \|Undefined Saprotroph\| | Yeast | NULL | Probable | As nectar-tap_saprotroph (Põlme et al. 2020); Decay sugar-rich substrates (Põlme et al. 2020) | Põlme S, et al. 2020. Fungal Diversity 105:1-16 ((https://doi.org/10.1007/s13225-020-00466-2)) |
| Otu50994 | 1 | k__Fungi(100);p__Ascomycota(100);c__Saccharomycetes(100);o__Saccharomycetales(100);f__Metschnikowiaceae(100);g__Metschnikowia(100);g__Metschnikowia_unclassified(100); | Metschnikowia | 13 | Saprotroph | \|Undefined Saprotroph\| | Yeast | NULL | Probable | As nectar-tap_saprotroph (Põlme et al. 2020); Decay sugar-rich substrates (Põlme et al. 2020) | Põlme S, et al. 2020. Fungal Diversity 105:1-16 ((https://doi.org/10.1007/s13225-020-00466-2)) |
| Otu50995 | 1 | k__Fungi(100);p__Ascomycota(100);c__Saccharomycetes(100);o__Saccharomycetales(100);f__Metschnikowiaceae(100);g__Metschnikowia(100);s__Metschnikowia_pulcherrima(100); | Metschnikowia | 13 | Saprotroph | \|Undefined Saprotroph\| | Yeast | NULL | Probable | As nectar-tap_saprotroph (Põlme et al. 2020); Decay sugar-rich substrates (Põlme et al. 2020) | Põlme S, et al. 2020. Fungal Diversity 105:1-16 ((https://doi.org/10.1007/s13225-020-00466-2)) |
| Otu50997 | 1 | k__Fungi(100);p__Ascomycota(100);c__Saccharomycetes(100);o__Saccharomycetales(100);f__Metschnikowiaceae(100);g__Metschnikowia(100);s__Metschnikowia_pulcherrima(100); | Metschnikowia | 13 | Saprotroph | \|Undefined Saprotroph\| | Yeast | NULL | Probable | As nectar-tap_saprotroph (Põlme et al. 2020); Decay sugar-rich substrates (Põlme et al. 2020) | Põlme S, et al. 2020. Fungal Diversity 105:1-16 ((https://doi.org/10.1007/s13225-020-00466-2)) |
| Otu50998 | 1 | k__Fungi(100);p__Ascomycota(100);c__Saccharomycetes(100);o__Saccharomycetales(100);f__Metschnikowiaceae(100);g__Metschnikowia(100);s__Metschnikowia_pulcherrima(100); | Metschnikowia | 13 | Saprotroph | \|Undefined Saprotroph\| | Yeast | NULL | Probable | As nectar-tap_saprotroph (Põlme et al. 2020); Decay sugar-rich substrates (Põlme et al. 2020) | Põlme S, et al. 2020. Fungal Diversity 105:1-16 ((https://doi.org/10.1007/s13225-020-00466-2)) |
| Otu50999 | 1 | k__Fungi(100);p__Ascomycota(100);c__Saccharomycetes(100);o__Saccharomycetales(100);f__Metschnikowiaceae(100);g__Metschnikowia(100);s__Metschnikowia_pulcherrima(100); | Metschnikowia | 13 | Saprotroph | \|Undefined Saprotroph\| | Yeast | NULL | Probable | As nectar-tap_saprotroph (Põlme et al. 2020); Decay sugar-rich substrates (Põlme et al. 2020) | Põlme S, et al. 2020. Fungal Diversity 105:1-16 ((https://doi.org/10.1007/s13225-020-00466-2)) |
| Otu51000 | 1 | k__Fungi(100);p__Ascomycota(100);c__Saccharomycetes(100);o__Saccharomycetales(100);f__Metschnikowiaceae(100);g__Metschnikowia(100);s__Metschnikowia_pulcherrima(100); | Metschnikowia | 13 | Saprotroph | \|Undefined Saprotroph\| | Yeast | NULL | Probable | As nectar-tap_saprotroph (Põlme et al. 2020); Decay sugar-rich substrates (Põlme et al. 2020) | Põlme S, et al. 2020. Fungal Diversity 105:1-16 ((https://doi.org/10.1007/s13225-020-00466-2)) |
| Otu51001 | 1 | k__Fungi(100);p__Ascomycota(100);c__Saccharomycetes(100);o__Saccharomycetales(100);f__Metschnikowiaceae(100);g__Metschnikowia(100);s__Metschnikowia_pulcherrima(100); | Metschnikowia | 13 | Saprotroph | \|Undefined Saprotroph\| | Yeast | NULL | Probable | As nectar-tap_saprotroph (Põlme et al. 2020); Decay sugar-rich substrates (Põlme et al. 2020) | Põlme S, et al. 2020. Fungal Diversity 105:1-16 ((https://doi.org/10.1007/s13225-020-00466-2)) |
| Otu51002 | 1 | k__Fungi(100);p__Ascomycota(100);c__Saccharomycetes(100);o__Saccharomycetales(100);f__Metschnikowiaceae(100);g__Metschnikowia(100);s__Metschnikowia_pulcherrima(100); | Metschnikowia | 13 | Saprotroph | \|Undefined Saprotroph\| | Yeast | NULL | Probable | As nectar-tap_saprotroph (Põlme et al. 2020); Decay sugar-rich substrates (Põlme et al. 2020) | Põlme S, et al. 2020. Fungal Diversity 105:1-16 ((https://doi.org/10.1007/s13225-020-00466-2)) |
| Otu51004 | 1 | k__Fungi(100);p__Ascomycota(100);c__Saccharomycetes(100);o__Saccharomycetales(100);f__Saccharomycetales_family_Incertae_sedis(100);g__Candida(100);g__Candida_unclassified(100); | Candida | 13 | Pathotroph-Saprotroph-Symbiotroph | Animal Pathogen-Endophyte-Endosymbiont-Epiphyte-\|Undefined Saprotroph\| | Dimorphic Yeast-Yeast | NULL | Probable | As nectar-tap_saprotroph (Põlme et al. 2020); Decay sugar-rich substrates (Põlme et al. 2020); This asexual genus is polyphyletic and contain species that function very differently depend on what they are. Caution should be taken when interpreting guilds for Candida species. | Manolakaki D, et al. 2010. Virulence. 1: 367-375 ((https://doi.org/10.4161/viru.1.5.12796)); Mekbib SB, et al. 2011. Tropical Plant Pathology 36:233-240 ((https://doi.org/10.1590/S1982-56762011000400004)); Põlme S, et al. 2020. Fungal Diversity 105:1-16 ((https://doi.org/10.1007/s13225-020-00466-2)) |
| Otu51005 | 1 | k__Fungi(100);p__Ascomycota(100);c__Saccharomycetes(100);o__Saccharomycetales(100);f__Metschnikowiaceae(100);g__Metschnikowia(100);s__Metschnikowia_pulcherrima(100); | Metschnikowia | 13 | Saprotroph | \|Undefined Saprotroph\| | Yeast | NULL | Probable | As nectar-tap_saprotroph (Põlme et al. 2020); Decay sugar-rich substrates (Põlme et al. 2020) | Põlme S, et al. 2020. Fungal Diversity 105:1-16 ((https://doi.org/10.1007/s13225-020-00466-2)) |
| Otu51006 | 1 | k__Fungi(100);p__Ascomycota(100);c__Saccharomycetes(100);o__Saccharomycetales(100);f__Metschnikowiaceae(100);g__Metschnikowia(100);s__Metschnikowia_pulcherrima(100); | Metschnikowia | 13 | Saprotroph | \|Undefined Saprotroph\| | Yeast | NULL | Probable | As nectar-tap_saprotroph (Põlme et al. 2020); Decay sugar-rich substrates (Põlme et al. 2020) | Põlme S, et al. 2020. Fungal Diversity 105:1-16 ((https://doi.org/10.1007/s13225-020-00466-2)) |
| Otu51007 | 1 | k__Fungi(100);p__Ascomycota(100);c__Saccharomycetes(100);o__Saccharomycetales(100);f__Metschnikowiaceae(100);g__Metschnikowia(100);s__Metschnikowia_pulcherrima(100); | Metschnikowia | 13 | Saprotroph | \|Undefined Saprotroph\| | Yeast | NULL | Probable | As nectar-tap_saprotroph (Põlme et al. 2020); Decay sugar-rich substrates (Põlme et al. 2020) | Põlme S, et al. 2020. Fungal Diversity 105:1-16 ((https://doi.org/10.1007/s13225-020-00466-2)) |
| Otu51008 | 1 | k__Fungi(100);p__Ascomycota(100);c__Saccharomycetes(100);o__Saccharomycetales(100);f__Metschnikowiaceae(100);g__Metschnikowia(100);s__Metschnikowia_pulcherrima(100); | Metschnikowia | 13 | Saprotroph | \|Undefined Saprotroph\| | Yeast | NULL | Probable | As nectar-tap_saprotroph (Põlme et al. 2020); Decay sugar-rich substrates (Põlme et al. 2020) | Põlme S, et al. 2020. Fungal Diversity 105:1-16 ((https://doi.org/10.1007/s13225-020-00466-2)) |
| Otu51009 | 1 | k__Fungi(100);p__Ascomycota(100);c__Saccharomycetes(100);o__Saccharomycetales(100);f__Metschnikowiaceae(100);g__Metschnikowia(100);g__Metschnikowia_unclassified(100); | Metschnikowia | 13 | Saprotroph | \|Undefined Saprotroph\| | Yeast | NULL | Probable | As nectar-tap_saprotroph (Põlme et al. 2020); Decay sugar-rich substrates (Põlme et al. 2020) | Põlme S, et al. 2020. Fungal Diversity 105:1-16 ((https://doi.org/10.1007/s13225-020-00466-2)) |
| Otu51010 | 1 | k__Fungi(100);p__Ascomycota(100);c__Saccharomycetes(100);o__Saccharomycetales(100);f__Metschnikowiaceae(100);g__Metschnikowia(100);s__Metschnikowia_pulcherrima(100); | Metschnikowia | 13 | Saprotroph | \|Undefined Saprotroph\| | Yeast | NULL | Probable | As nectar-tap_saprotroph (Põlme et al. 2020); Decay sugar-rich substrates (Põlme et al. 2020) | Põlme S, et al. 2020. Fungal Diversity 105:1-16 ((https://doi.org/10.1007/s13225-020-00466-2)) |
| Otu51011 | 1 | k__Fungi(100);p__Ascomycota(100);c__Saccharomycetes(100);o__Saccharomycetales(100);f__Metschnikowiaceae(100);g__Metschnikowia(100);s__Metschnikowia_pulcherrima(100); | Metschnikowia | 13 | Saprotroph | \|Undefined Saprotroph\| | Yeast | NULL | Probable | As nectar-tap_saprotroph (Põlme et al. 2020); Decay sugar-rich substrates (Põlme et al. 2020) | Põlme S, et al. 2020. Fungal Diversity 105:1-16 ((https://doi.org/10.1007/s13225-020-00466-2)) |
| Otu51012 | 1 | k__Fungi(100);p__Ascomycota(100);c__Saccharomycetes(100);o__Saccharomycetales(100);f__Metschnikowiaceae(100);g__Metschnikowia(100);s__Metschnikowia_pulcherrima(100); | Metschnikowia | 13 | Saprotroph | \|Undefined Saprotroph\| | Yeast | NULL | Probable | As nectar-tap_saprotroph (Põlme et al. 2020); Decay sugar-rich substrates (Põlme et al. 2020) | Põlme S, et al. 2020. Fungal Diversity 105:1-16 ((https://doi.org/10.1007/s13225-020-00466-2)) |
| Otu51013 | 1 | k__Fungi(100);p__Ascomycota(100);c__Saccharomycetes(100);o__Saccharomycetales(100);f__Metschnikowiaceae(100);g__Metschnikowia(100);s__Metschnikowia_pulcherrima(100); | Metschnikowia | 13 | Saprotroph | \|Undefined Saprotroph\| | Yeast | NULL | Probable | As nectar-tap_saprotroph (Põlme et al. 2020); Decay sugar-rich substrates (Põlme et al. 2020) | Põlme S, et al. 2020. Fungal Diversity 105:1-16 ((https://doi.org/10.1007/s13225-020-00466-2)) |
| Otu51014 | 1 | k__Fungi(100);p__Ascomycota(100);c__Saccharomycetes(100);o__Saccharomycetales(100);f__Metschnikowiaceae(100);g__Metschnikowia(100);s__Metschnikowia_pulcherrima(100); | Metschnikowia | 13 | Saprotroph | \|Undefined Saprotroph\| | Yeast | NULL | Probable | As nectar-tap_saprotroph (Põlme et al. 2020); Decay sugar-rich substrates (Põlme et al. 2020) | Põlme S, et al. 2020. Fungal Diversity 105:1-16 ((https://doi.org/10.1007/s13225-020-00466-2)) |
| Otu51015 | 1 | k__Fungi(100);p__Ascomycota(100);c__Saccharomycetes(100);o__Saccharomycetales(100);f__Metschnikowiaceae(100);g__Metschnikowia(100);s__Metschnikowia_pulcherrima(100); | Metschnikowia | 13 | Saprotroph | \|Undefined Saprotroph\| | Yeast | NULL | Probable | As nectar-tap_saprotroph (Põlme et al. 2020); Decay sugar-rich substrates (Põlme et al. 2020) | Põlme S, et al. 2020. Fungal Diversity 105:1-16 ((https://doi.org/10.1007/s13225-020-00466-2)) |
| Otu51016 | 1 | k__Fungi(100);p__Ascomycota(100);c__Saccharomycetes(100);o__Saccharomycetales(100);f__Metschnikowiaceae(100);g__Metschnikowia(100);g__Metschnikowia_unclassified(100); | Metschnikowia | 13 | Saprotroph | \|Undefined Saprotroph\| | Yeast | NULL | Probable | As nectar-tap_saprotroph (Põlme et al. 2020); Decay sugar-rich substrates (Põlme et al. 2020) | Põlme S, et al. 2020. Fungal Diversity 105:1-16 ((https://doi.org/10.1007/s13225-020-00466-2)) |
| Otu51017 | 1 | k__Fungi(100);p__Ascomycota(100);c__Saccharomycetes(100);o__Saccharomycetales(100);f__Metschnikowiaceae(100);g__Metschnikowia(100);s__Metschnikowia_pulcherrima(100); | Metschnikowia | 13 | Saprotroph | \|Undefined Saprotroph\| | Yeast | NULL | Probable | As nectar-tap_saprotroph (Põlme et al. 2020); Decay sugar-rich substrates (Põlme et al. 2020) | Põlme S, et al. 2020. Fungal Diversity 105:1-16 ((https://doi.org/10.1007/s13225-020-00466-2)) |
| Otu51018 | 1 | k__Fungi(100);p__Ascomycota(100);c__Saccharomycetes(100);o__Saccharomycetales(100);f__Metschnikowiaceae(100);g__Metschnikowia(100);s__Metschnikowia_pulcherrima(100); | Metschnikowia | 13 | Saprotroph | \|Undefined Saprotroph\| | Yeast | NULL | Probable | As nectar-tap_saprotroph (Põlme et al. 2020); Decay sugar-rich substrates (Põlme et al. 2020) | Põlme S, et al. 2020. Fungal Diversity 105:1-16 ((https://doi.org/10.1007/s13225-020-00466-2)) |
| Otu51019 | 1 | k__Fungi(100);p__Ascomycota(100);c__Saccharomycetes(100);o__Saccharomycetales(100);f__Metschnikowiaceae(100);g__Metschnikowia(100);s__Metschnikowia_pulcherrima(100); | Metschnikowia | 13 | Saprotroph | \|Undefined Saprotroph\| | Yeast | NULL | Probable | As nectar-tap_saprotroph (Põlme et al. 2020); Decay sugar-rich substrates (Põlme et al. 2020) | Põlme S, et al. 2020. Fungal Diversity 105:1-16 ((https://doi.org/10.1007/s13225-020-00466-2)) |
| Otu51020 | 1 | k__Fungi(100);p__Ascomycota(100);c__Saccharomycetes(100);o__Saccharomycetales(100);f__Metschnikowiaceae(100);g__Metschnikowia(100);g__Metschnikowia_unclassified(100); | Metschnikowia | 13 | Saprotroph | \|Undefined Saprotroph\| | Yeast | NULL | Probable | As nectar-tap_saprotroph (Põlme et al. 2020); Decay sugar-rich substrates (Põlme et al. 2020) | Põlme S, et al. 2020. Fungal Diversity 105:1-16 ((https://doi.org/10.1007/s13225-020-00466-2)) |
| Otu51021 | 1 | k__Fungi(100);p__Ascomycota(100);c__Saccharomycetes(100);o__Saccharomycetales(100);f__Metschnikowiaceae(100);g__Metschnikowia(100);g__Metschnikowia_unclassified(100); | Metschnikowia | 13 | Saprotroph | \|Undefined Saprotroph\| | Yeast | NULL | Probable | As nectar-tap_saprotroph (Põlme et al. 2020); Decay sugar-rich substrates (Põlme et al. 2020) | Põlme S, et al. 2020. Fungal Diversity 105:1-16 ((https://doi.org/10.1007/s13225-020-00466-2)) |
| Otu51022 | 1 | k__Fungi(100);p__Ascomycota(100);c__Saccharomycetes(100);o__Saccharomycetales(100);f__Metschnikowiaceae(100);g__Metschnikowia(100);s__Metschnikowia_pulcherrima(100); | Metschnikowia | 13 | Saprotroph | \|Undefined Saprotroph\| | Yeast | NULL | Probable | As nectar-tap_saprotroph (Põlme et al. 2020); Decay sugar-rich substrates (Põlme et al. 2020) | Põlme S, et al. 2020. Fungal Diversity 105:1-16 ((https://doi.org/10.1007/s13225-020-00466-2)) |
| Otu51023 | 1 | k__Fungi(100);p__Ascomycota(100);c__Saccharomycetes(100);o__Saccharomycetales(100);f__Metschnikowiaceae(100);g__Metschnikowia(100);s__Metschnikowia_pulcherrima(100); | Metschnikowia | 13 | Saprotroph | \|Undefined Saprotroph\| | Yeast | NULL | Probable | As nectar-tap_saprotroph (Põlme et al. 2020); Decay sugar-rich substrates (Põlme et al. 2020) | Põlme S, et al. 2020. Fungal Diversity 105:1-16 ((https://doi.org/10.1007/s13225-020-00466-2)) |
| Otu51024 | 1 | k__Fungi(100);p__Ascomycota(100);c__Saccharomycetes(100);o__Saccharomycetales(100);f__Metschnikowiaceae(100);g__Metschnikowia(100);s__Metschnikowia_pulcherrima(100); | Metschnikowia | 13 | Saprotroph | \|Undefined Saprotroph\| | Yeast | NULL | Probable | As nectar-tap_saprotroph (Põlme et al. 2020); Decay sugar-rich substrates (Põlme et al. 2020) | Põlme S, et al. 2020. Fungal Diversity 105:1-16 ((https://doi.org/10.1007/s13225-020-00466-2)) |
| Otu51025 | 1 | k__Fungi(100);p__Ascomycota(100);c__Saccharomycetes(100);o__Saccharomycetales(100);f__Metschnikowiaceae(100);g__Metschnikowia(100);g__Metschnikowia_unclassified(100); | Metschnikowia | 13 | Saprotroph | \|Undefined Saprotroph\| | Yeast | NULL | Probable | As nectar-tap_saprotroph (Põlme et al. 2020); Decay sugar-rich substrates (Põlme et al. 2020) | Põlme S, et al. 2020. Fungal Diversity 105:1-16 ((https://doi.org/10.1007/s13225-020-00466-2)) |
| Otu51031 | 1 | k__Fungi(100);p__Ascomycota(100);c__Saccharomycetes(100);o__Saccharomycetales(100);f__Metschnikowiaceae(100);g__Metschnikowia(100);s__Metschnikowia_pulcherrima(100); | Metschnikowia | 13 | Saprotroph | \|Undefined Saprotroph\| | Yeast | NULL | Probable | As nectar-tap_saprotroph (Põlme et al. 2020); Decay sugar-rich substrates (Põlme et al. 2020) | Põlme S, et al. 2020. Fungal Diversity 105:1-16 ((https://doi.org/10.1007/s13225-020-00466-2)) |
| Otu51035 | 1 | k__Fungi(100);p__Ascomycota(100);c__Saccharomycetes(100);o__Saccharomycetales(100);f__Metschnikowiaceae(100);g__Metschnikowia(100);s__Metschnikowia_pulcherrima(100); | Metschnikowia | 13 | Saprotroph | \|Undefined Saprotroph\| | Yeast | NULL | Probable | As nectar-tap_saprotroph (Põlme et al. 2020); Decay sugar-rich substrates (Põlme et al. 2020) | Põlme S, et al. 2020. Fungal Diversity 105:1-16 ((https://doi.org/10.1007/s13225-020-00466-2)) |
| Otu51037 | 1 | k__Fungi(100);p__Ascomycota(100);c__Saccharomycetes(100);o__Saccharomycetales(100);f__Metschnikowiaceae(100);g__Metschnikowia(100);s__Metschnikowia_pulcherrima(100); | Metschnikowia | 13 | Saprotroph | \|Undefined Saprotroph\| | Yeast | NULL | Probable | As nectar-tap_saprotroph (Põlme et al. 2020); Decay sugar-rich substrates (Põlme et al. 2020) | Põlme S, et al. 2020. Fungal Diversity 105:1-16 ((https://doi.org/10.1007/s13225-020-00466-2)) |
| Otu51038 | 1 | k__Fungi(100);p__Ascomycota(100);c__Saccharomycetes(100);o__Saccharomycetales(100);f__Metschnikowiaceae(100);g__Metschnikowia(100);g__Metschnikowia_unclassified(100); | Metschnikowia | 13 | Saprotroph | \|Undefined Saprotroph\| | Yeast | NULL | Probable | As nectar-tap_saprotroph (Põlme et al. 2020); Decay sugar-rich substrates (Põlme et al. 2020) | Põlme S, et al. 2020. Fungal Diversity 105:1-16 ((https://doi.org/10.1007/s13225-020-00466-2)) |
| Otu51039 | 1 | k__Fungi(100);p__Ascomycota(100);c__Saccharomycetes(100);o__Saccharomycetales(100);f__Metschnikowiaceae(100);g__Metschnikowia(100);g__Metschnikowia_unclassified(100); | Metschnikowia | 13 | Saprotroph | \|Undefined Saprotroph\| | Yeast | NULL | Probable | As nectar-tap_saprotroph (Põlme et al. 2020); Decay sugar-rich substrates (Põlme et al. 2020) | Põlme S, et al. 2020. Fungal Diversity 105:1-16 ((https://doi.org/10.1007/s13225-020-00466-2)) |
| Otu51040 | 1 | k__Fungi(100);p__Ascomycota(100);c__Saccharomycetes(100);o__Saccharomycetales(100);f__Metschnikowiaceae(100);g__Metschnikowia(100);g__Metschnikowia_unclassified(100); | Metschnikowia | 13 | Saprotroph | \|Undefined Saprotroph\| | Yeast | NULL | Probable | As nectar-tap_saprotroph (Põlme et al. 2020); Decay sugar-rich substrates (Põlme et al. 2020) | Põlme S, et al. 2020. Fungal Diversity 105:1-16 ((https://doi.org/10.1007/s13225-020-00466-2)) |
| Otu51041 | 1 | k__Fungi(100);p__Ascomycota(100);c__Saccharomycetes(100);o__Saccharomycetales(100);f__Metschnikowiaceae(100);g__Metschnikowia(100);s__Metschnikowia_pulcherrima(100); | Metschnikowia | 13 | Saprotroph | \|Undefined Saprotroph\| | Yeast | NULL | Probable | As nectar-tap_saprotroph (Põlme et al. 2020); Decay sugar-rich substrates (Põlme et al. 2020) | Põlme S, et al. 2020. Fungal Diversity 105:1-16 ((https://doi.org/10.1007/s13225-020-00466-2)) |
| Otu51042 | 1 | k__Fungi(100);p__Ascomycota(100);c__Saccharomycetes(100);o__Saccharomycetales(100);f__Metschnikowiaceae(100);g__Metschnikowia(100);s__Metschnikowia_pulcherrima(100); | Metschnikowia | 13 | Saprotroph | \|Undefined Saprotroph\| | Yeast | NULL | Probable | As nectar-tap_saprotroph (Põlme et al. 2020); Decay sugar-rich substrates (Põlme et al. 2020) | Põlme S, et al. 2020. Fungal Diversity 105:1-16 ((https://doi.org/10.1007/s13225-020-00466-2)) |
| Otu51043 | 1 | k__Fungi(100);p__Ascomycota(100);c__Saccharomycetes(100);o__Saccharomycetales(100);f__Metschnikowiaceae(100);g__Metschnikowia(100);s__Metschnikowia_pulcherrima(100); | Metschnikowia | 13 | Saprotroph | \|Undefined Saprotroph\| | Yeast | NULL | Probable | As nectar-tap_saprotroph (Põlme et al. 2020); Decay sugar-rich substrates (Põlme et al. 2020) | Põlme S, et al. 2020. Fungal Diversity 105:1-16 ((https://doi.org/10.1007/s13225-020-00466-2)) |
| Otu51044 | 1 | k__Fungi(100);p__Ascomycota(100);c__Saccharomycetes(100);o__Saccharomycetales(100);f__Metschnikowiaceae(100);g__Metschnikowia(100);s__Metschnikowia_pulcherrima(100); | Metschnikowia | 13 | Saprotroph | \|Undefined Saprotroph\| | Yeast | NULL | Probable | As nectar-tap_saprotroph (Põlme et al. 2020); Decay sugar-rich substrates (Põlme et al. 2020) | Põlme S, et al. 2020. Fungal Diversity 105:1-16 ((https://doi.org/10.1007/s13225-020-00466-2)) |
| Otu51046 | 1 | k__Fungi(100);p__Ascomycota(100);c__Saccharomycetes(100);o__Saccharomycetales(100);f__Metschnikowiaceae(100);g__Metschnikowia(100);s__Metschnikowia_pulcherrima(100); | Metschnikowia | 13 | Saprotroph | \|Undefined Saprotroph\| | Yeast | NULL | Probable | As nectar-tap_saprotroph (Põlme et al. 2020); Decay sugar-rich substrates (Põlme et al. 2020) | Põlme S, et al. 2020. Fungal Diversity 105:1-16 ((https://doi.org/10.1007/s13225-020-00466-2)) |
| Otu51047 | 1 | k__Fungi(100);p__Ascomycota(100);c__Saccharomycetes(100);o__Saccharomycetales(100);f__Metschnikowiaceae(100);g__Metschnikowia(100);s__Metschnikowia_pulcherrima(100); | Metschnikowia | 13 | Saprotroph | \|Undefined Saprotroph\| | Yeast | NULL | Probable | As nectar-tap_saprotroph (Põlme et al. 2020); Decay sugar-rich substrates (Põlme et al. 2020) | Põlme S, et al. 2020. Fungal Diversity 105:1-16 ((https://doi.org/10.1007/s13225-020-00466-2)) |
| Otu51048 | 1 | k__Fungi(100);p__Ascomycota(100);c__Saccharomycetes(100);o__Saccharomycetales(100);f__Metschnikowiaceae(100);g__Metschnikowia(100);s__Metschnikowia_pulcherrima(100); | Metschnikowia | 13 | Saprotroph | \|Undefined Saprotroph\| | Yeast | NULL | Probable | As nectar-tap_saprotroph (Põlme et al. 2020); Decay sugar-rich substrates (Põlme et al. 2020) | Põlme S, et al. 2020. Fungal Diversity 105:1-16 ((https://doi.org/10.1007/s13225-020-00466-2)) |
| Otu51049 | 1 | k__Fungi(100);p__Ascomycota(100);c__Saccharomycetes(100);o__Saccharomycetales(100);f__Metschnikowiaceae(100);g__Metschnikowia(100);s__Metschnikowia_pulcherrima(100); | Metschnikowia | 13 | Saprotroph | \|Undefined Saprotroph\| | Yeast | NULL | Probable | As nectar-tap_saprotroph (Põlme et al. 2020); Decay sugar-rich substrates (Põlme et al. 2020) | Põlme S, et al. 2020. Fungal Diversity 105:1-16 ((https://doi.org/10.1007/s13225-020-00466-2)) |
| Otu51050 | 1 | k__Fungi(100);p__Ascomycota(100);c__Saccharomycetes(100);o__Saccharomycetales(100);f__Metschnikowiaceae(100);g__Metschnikowia(100);s__Metschnikowia_pulcherrima(100); | Metschnikowia | 13 | Saprotroph | \|Undefined Saprotroph\| | Yeast | NULL | Probable | As nectar-tap_saprotroph (Põlme et al. 2020); Decay sugar-rich substrates (Põlme et al. 2020) | Põlme S, et al. 2020. Fungal Diversity 105:1-16 ((https://doi.org/10.1007/s13225-020-00466-2)) |
| Otu51051 | 1 | k__Fungi(100);p__Ascomycota(100);c__Saccharomycetes(100);o__Saccharomycetales(100);f__Metschnikowiaceae(100);g__Metschnikowia(100);s__Metschnikowia_pulcherrima(100); | Metschnikowia | 13 | Saprotroph | \|Undefined Saprotroph\| | Yeast | NULL | Probable | As nectar-tap_saprotroph (Põlme et al. 2020); Decay sugar-rich substrates (Põlme et al. 2020) | Põlme S, et al. 2020. Fungal Diversity 105:1-16 ((https://doi.org/10.1007/s13225-020-00466-2)) |
| Otu51052 | 1 | k__Fungi(100);p__Ascomycota(100);c__Saccharomycetes(100);o__Saccharomycetales(100);f__Metschnikowiaceae(100);g__Metschnikowia(100);s__Metschnikowia_pulcherrima(100); | Metschnikowia | 13 | Saprotroph | \|Undefined Saprotroph\| | Yeast | NULL | Probable | As nectar-tap_saprotroph (Põlme et al. 2020); Decay sugar-rich substrates (Põlme et al. 2020) | Põlme S, et al. 2020. Fungal Diversity 105:1-16 ((https://doi.org/10.1007/s13225-020-00466-2)) |
| Otu51053 | 1 | k__Fungi(100);p__Ascomycota(100);c__Saccharomycetes(100);o__Saccharomycetales(100);f__Metschnikowiaceae(100);g__Metschnikowia(100);s__Metschnikowia_pulcherrima(100); | Metschnikowia | 13 | Saprotroph | \|Undefined Saprotroph\| | Yeast | NULL | Probable | As nectar-tap_saprotroph (Põlme et al. 2020); Decay sugar-rich substrates (Põlme et al. 2020) | Põlme S, et al. 2020. Fungal Diversity 105:1-16 ((https://doi.org/10.1007/s13225-020-00466-2)) |
| Otu51054 | 1 | k__Fungi(100);p__Ascomycota(100);c__Saccharomycetes(100);o__Saccharomycetales(100);f__Metschnikowiaceae(100);g__Metschnikowia(100);s__Metschnikowia_pulcherrima(100); | Metschnikowia | 13 | Saprotroph | \|Undefined Saprotroph\| | Yeast | NULL | Probable | As nectar-tap_saprotroph (Põlme et al. 2020); Decay sugar-rich substrates (Põlme et al. 2020) | Põlme S, et al. 2020. Fungal Diversity 105:1-16 ((https://doi.org/10.1007/s13225-020-00466-2)) |
| Otu51055 | 1 | k__Fungi(100);p__Ascomycota(100);c__Saccharomycetes(100);o__Saccharomycetales(100);f__Metschnikowiaceae(100);g__Metschnikowia(100);s__Metschnikowia_pulcherrima(100); | Metschnikowia | 13 | Saprotroph | \|Undefined Saprotroph\| | Yeast | NULL | Probable | As nectar-tap_saprotroph (Põlme et al. 2020); Decay sugar-rich substrates (Põlme et al. 2020) | Põlme S, et al. 2020. Fungal Diversity 105:1-16 ((https://doi.org/10.1007/s13225-020-00466-2)) |
| Otu51056 | 1 | k__Fungi(100);p__Ascomycota(100);c__Saccharomycetes(100);o__Saccharomycetales(100);f__Metschnikowiaceae(100);g__Metschnikowia(100);s__Metschnikowia_pulcherrima(100); | Metschnikowia | 13 | Saprotroph | \|Undefined Saprotroph\| | Yeast | NULL | Probable | As nectar-tap_saprotroph (Põlme et al. 2020); Decay sugar-rich substrates (Põlme et al. 2020) | Põlme S, et al. 2020. Fungal Diversity 105:1-16 ((https://doi.org/10.1007/s13225-020-00466-2)) |
| Otu51057 | 1 | k__Fungi(100);p__Ascomycota(100);c__Saccharomycetes(100);o__Saccharomycetales(100);f__Metschnikowiaceae(100);g__Metschnikowia(100);s__Metschnikowia_pulcherrima(100); | Metschnikowia | 13 | Saprotroph | \|Undefined Saprotroph\| | Yeast | NULL | Probable | As nectar-tap_saprotroph (Põlme et al. 2020); Decay sugar-rich substrates (Põlme et al. 2020) | Põlme S, et al. 2020. Fungal Diversity 105:1-16 ((https://doi.org/10.1007/s13225-020-00466-2)) |
| Otu51058 | 1 | k__Fungi(100);p__Ascomycota(100);c__Saccharomycetes(100);o__Saccharomycetales(100);f__Metschnikowiaceae(100);g__Metschnikowia(100);s__Metschnikowia_pulcherrima(100); | Metschnikowia | 13 | Saprotroph | \|Undefined Saprotroph\| | Yeast | NULL | Probable | As nectar-tap_saprotroph (Põlme et al. 2020); Decay sugar-rich substrates (Põlme et al. 2020) | Põlme S, et al. 2020. Fungal Diversity 105:1-16 ((https://doi.org/10.1007/s13225-020-00466-2)) |
| Otu51059 | 1 | k__Fungi(100);p__Ascomycota(100);c__Saccharomycetes(100);o__Saccharomycetales(100);f__Metschnikowiaceae(100);g__Metschnikowia(100);s__Metschnikowia_pulcherrima(100); | Metschnikowia | 13 | Saprotroph | \|Undefined Saprotroph\| | Yeast | NULL | Probable | As nectar-tap_saprotroph (Põlme et al. 2020); Decay sugar-rich substrates (Põlme et al. 2020) | Põlme S, et al. 2020. Fungal Diversity 105:1-16 ((https://doi.org/10.1007/s13225-020-00466-2)) |
| Otu51060 | 1 | k__Fungi(100);p__Ascomycota(100);c__Saccharomycetes(100);o__Saccharomycetales(100);f__Metschnikowiaceae(100);g__Metschnikowia(100);g__Metschnikowia_unclassified(100); | Metschnikowia | 13 | Saprotroph | \|Undefined Saprotroph\| | Yeast | NULL | Probable | As nectar-tap_saprotroph (Põlme et al. 2020); Decay sugar-rich substrates (Põlme et al. 2020) | Põlme S, et al. 2020. Fungal Diversity 105:1-16 ((https://doi.org/10.1007/s13225-020-00466-2)) |
| Otu51061 | 1 | k__Fungi(100);p__Ascomycota(100);c__Saccharomycetes(100);o__Saccharomycetales(100);f__Metschnikowiaceae(100);g__Metschnikowia(100);g__Metschnikowia_unclassified(100); | Metschnikowia | 13 | Saprotroph | \|Undefined Saprotroph\| | Yeast | NULL | Probable | As nectar-tap_saprotroph (Põlme et al. 2020); Decay sugar-rich substrates (Põlme et al. 2020) | Põlme S, et al. 2020. Fungal Diversity 105:1-16 ((https://doi.org/10.1007/s13225-020-00466-2)) |
| Otu51062 | 1 | k__Fungi(100);p__Ascomycota(100);c__Saccharomycetes(100);o__Saccharomycetales(100);f__Metschnikowiaceae(100);g__Metschnikowia(100);s__Metschnikowia_pulcherrima(100); | Metschnikowia | 13 | Saprotroph | \|Undefined Saprotroph\| | Yeast | NULL | Probable | As nectar-tap_saprotroph (Põlme et al. 2020); Decay sugar-rich substrates (Põlme et al. 2020) | Põlme S, et al. 2020. Fungal Diversity 105:1-16 ((https://doi.org/10.1007/s13225-020-00466-2)) |
| Otu51064 | 1 | k__Fungi(100);p__Ascomycota(100);c__Saccharomycetes(100);o__Saccharomycetales(100);f__Metschnikowiaceae(100);g__Metschnikowia(100);g__Metschnikowia_unclassified(100); | Metschnikowia | 13 | Saprotroph | \|Undefined Saprotroph\| | Yeast | NULL | Probable | As nectar-tap_saprotroph (Põlme et al. 2020); Decay sugar-rich substrates (Põlme et al. 2020) | Põlme S, et al. 2020. Fungal Diversity 105:1-16 ((https://doi.org/10.1007/s13225-020-00466-2)) |
| Otu51065 | 1 | k__Fungi(100);p__Ascomycota(100);c__Saccharomycetes(100);o__Saccharomycetales(100);f__Metschnikowiaceae(100);g__Metschnikowia(100);s__Metschnikowia_pulcherrima(100); | Metschnikowia | 13 | Saprotroph | \|Undefined Saprotroph\| | Yeast | NULL | Probable | As nectar-tap_saprotroph (Põlme et al. 2020); Decay sugar-rich substrates (Põlme et al. 2020) | Põlme S, et al. 2020. Fungal Diversity 105:1-16 ((https://doi.org/10.1007/s13225-020-00466-2)) |
| Otu51067 | 1 | k__Fungi(100);p__Ascomycota(100);c__Saccharomycetes(100);o__Saccharomycetales(100);f__Metschnikowiaceae(100);g__Metschnikowia(100);s__Metschnikowia_pulcherrima(100); | Metschnikowia | 13 | Saprotroph | \|Undefined Saprotroph\| | Yeast | NULL | Probable | As nectar-tap_saprotroph (Põlme et al. 2020); Decay sugar-rich substrates (Põlme et al. 2020) | Põlme S, et al. 2020. Fungal Diversity 105:1-16 ((https://doi.org/10.1007/s13225-020-00466-2)) |
| Otu51068 | 1 | k__Fungi(100);p__Ascomycota(100);c__Saccharomycetes(100);o__Saccharomycetales(100);f__Metschnikowiaceae(100);g__Metschnikowia(100);s__Metschnikowia_pulcherrima(100); | Metschnikowia | 13 | Saprotroph | \|Undefined Saprotroph\| | Yeast | NULL | Probable | As nectar-tap_saprotroph (Põlme et al. 2020); Decay sugar-rich substrates (Põlme et al. 2020) | Põlme S, et al. 2020. Fungal Diversity 105:1-16 ((https://doi.org/10.1007/s13225-020-00466-2)) |
| Otu51069 | 1 | k__Fungi(100);p__Ascomycota(100);c__Saccharomycetes(100);o__Saccharomycetales(100);f__Metschnikowiaceae(100);g__Metschnikowia(100);s__Metschnikowia_pulcherrima(100); | Metschnikowia | 13 | Saprotroph | \|Undefined Saprotroph\| | Yeast | NULL | Probable | As nectar-tap_saprotroph (Põlme et al. 2020); Decay sugar-rich substrates (Põlme et al. 2020) | Põlme S, et al. 2020. Fungal Diversity 105:1-16 ((https://doi.org/10.1007/s13225-020-00466-2)) |
| Otu51070 | 1 | k__Fungi(100);p__Ascomycota(100);c__Saccharomycetes(100);o__Saccharomycetales(100);f__Metschnikowiaceae(100);g__Metschnikowia(100);g__Metschnikowia_unclassified(100); | Metschnikowia | 13 | Saprotroph | \|Undefined Saprotroph\| | Yeast | NULL | Probable | As nectar-tap_saprotroph (Põlme et al. 2020); Decay sugar-rich substrates (Põlme et al. 2020) | Põlme S, et al. 2020. Fungal Diversity 105:1-16 ((https://doi.org/10.1007/s13225-020-00466-2)) |
| Otu51071 | 1 | k__Fungi(100);p__Ascomycota(100);c__Saccharomycetes(100);o__Saccharomycetales(100);f__Metschnikowiaceae(100);g__Metschnikowia(100);s__Metschnikowia_pulcherrima(100); | Metschnikowia | 13 | Saprotroph | \|Undefined Saprotroph\| | Yeast | NULL | Probable | As nectar-tap_saprotroph (Põlme et al. 2020); Decay sugar-rich substrates (Põlme et al. 2020) | Põlme S, et al. 2020. Fungal Diversity 105:1-16 ((https://doi.org/10.1007/s13225-020-00466-2)) |
| Otu51072 | 1 | k__Fungi(100);p__Ascomycota(100);c__Saccharomycetes(100);o__Saccharomycetales(100);f__Metschnikowiaceae(100);g__Metschnikowia(100);s__Metschnikowia_pulcherrima(100); | Metschnikowia | 13 | Saprotroph | \|Undefined Saprotroph\| | Yeast | NULL | Probable | As nectar-tap_saprotroph (Põlme et al. 2020); Decay sugar-rich substrates (Põlme et al. 2020) | Põlme S, et al. 2020. Fungal Diversity 105:1-16 ((https://doi.org/10.1007/s13225-020-00466-2)) |
| Otu51073 | 1 | k__Fungi(100);p__Ascomycota(100);c__Saccharomycetes(100);o__Saccharomycetales(100);f__Metschnikowiaceae(100);g__Metschnikowia(100);s__Metschnikowia_pulcherrima(100); | Metschnikowia | 13 | Saprotroph | \|Undefined Saprotroph\| | Yeast | NULL | Probable | As nectar-tap_saprotroph (Põlme et al. 2020); Decay sugar-rich substrates (Põlme et al. 2020) | Põlme S, et al. 2020. Fungal Diversity 105:1-16 ((https://doi.org/10.1007/s13225-020-00466-2)) |
| Otu51074 | 1 | k__Fungi(100);p__Ascomycota(100);c__Saccharomycetes(100);o__Saccharomycetales(100);f__Metschnikowiaceae(100);g__Metschnikowia(100);g__Metschnikowia_unclassified(100); | Metschnikowia | 13 | Saprotroph | \|Undefined Saprotroph\| | Yeast | NULL | Probable | As nectar-tap_saprotroph (Põlme et al. 2020); Decay sugar-rich substrates (Põlme et al. 2020) | Põlme S, et al. 2020. Fungal Diversity 105:1-16 ((https://doi.org/10.1007/s13225-020-00466-2)) |
| Otu51075 | 1 | k__Fungi(100);p__Ascomycota(100);c__Saccharomycetes(100);o__Saccharomycetales(100);f__Metschnikowiaceae(100);g__Metschnikowia(100);s__Metschnikowia_pulcherrima(100); | Metschnikowia | 13 | Saprotroph | \|Undefined Saprotroph\| | Yeast | NULL | Probable | As nectar-tap_saprotroph (Põlme et al. 2020); Decay sugar-rich substrates (Põlme et al. 2020) | Põlme S, et al. 2020. Fungal Diversity 105:1-16 ((https://doi.org/10.1007/s13225-020-00466-2)) |
| Otu51076 | 1 | k__Fungi(100);p__Ascomycota(100);c__Saccharomycetes(100);o__Saccharomycetales(100);f__Metschnikowiaceae(100);g__Metschnikowia(100);g__Metschnikowia_unclassified(100); | Metschnikowia | 13 | Saprotroph | \|Undefined Saprotroph\| | Yeast | NULL | Probable | As nectar-tap_saprotroph (Põlme et al. 2020); Decay sugar-rich substrates (Põlme et al. 2020) | Põlme S, et al. 2020. Fungal Diversity 105:1-16 ((https://doi.org/10.1007/s13225-020-00466-2)) |
| Otu51077 | 1 | k__Fungi(100);p__Ascomycota(100);c__Saccharomycetes(100);o__Saccharomycetales(100);f__Metschnikowiaceae(100);g__Metschnikowia(100);g__Metschnikowia_unclassified(100); | Metschnikowia | 13 | Saprotroph | \|Undefined Saprotroph\| | Yeast | NULL | Probable | As nectar-tap_saprotroph (Põlme et al. 2020); Decay sugar-rich substrates (Põlme et al. 2020) | Põlme S, et al. 2020. Fungal Diversity 105:1-16 ((https://doi.org/10.1007/s13225-020-00466-2)) |
| Otu51078 | 1 | k__Fungi(100);p__Ascomycota(100);c__Saccharomycetes(100);o__Saccharomycetales(100);f__Metschnikowiaceae(100);g__Metschnikowia(100);s__Metschnikowia_pulcherrima(100); | Metschnikowia | 13 | Saprotroph | \|Undefined Saprotroph\| | Yeast | NULL | Probable | As nectar-tap_saprotroph (Põlme et al. 2020); Decay sugar-rich substrates (Põlme et al. 2020) | Põlme S, et al. 2020. Fungal Diversity 105:1-16 ((https://doi.org/10.1007/s13225-020-00466-2)) |
| Otu51079 | 1 | k__Fungi(100);p__Ascomycota(100);c__Saccharomycetes(100);o__Saccharomycetales(100);f__Metschnikowiaceae(100);g__Metschnikowia(100);g__Metschnikowia_unclassified(100); | Metschnikowia | 13 | Saprotroph | \|Undefined Saprotroph\| | Yeast | NULL | Probable | As nectar-tap_saprotroph (Põlme et al. 2020); Decay sugar-rich substrates (Põlme et al. 2020) | Põlme S, et al. 2020. Fungal Diversity 105:1-16 ((https://doi.org/10.1007/s13225-020-00466-2)) |
| Otu51080 | 1 | k__Fungi(100);p__Ascomycota(100);c__Saccharomycetes(100);o__Saccharomycetales(100);f__Metschnikowiaceae(100);g__Metschnikowia(100);g__Metschnikowia_unclassified(100); | Metschnikowia | 13 | Saprotroph | \|Undefined Saprotroph\| | Yeast | NULL | Probable | As nectar-tap_saprotroph (Põlme et al. 2020); Decay sugar-rich substrates (Põlme et al. 2020) | Põlme S, et al. 2020. Fungal Diversity 105:1-16 ((https://doi.org/10.1007/s13225-020-00466-2)) |
| Otu51081 | 1 | k__Fungi(100);p__Ascomycota(100);c__Saccharomycetes(100);o__Saccharomycetales(100);f__Metschnikowiaceae(100);g__Metschnikowia(100);s__Metschnikowia_pulcherrima(100); | Metschnikowia | 13 | Saprotroph | \|Undefined Saprotroph\| | Yeast | NULL | Probable | As nectar-tap_saprotroph (Põlme et al. 2020); Decay sugar-rich substrates (Põlme et al. 2020) | Põlme S, et al. 2020. Fungal Diversity 105:1-16 ((https://doi.org/10.1007/s13225-020-00466-2)) |
| Otu51082 | 1 | k__Fungi(100);p__Ascomycota(100);c__Saccharomycetes(100);o__Saccharomycetales(100);f__Metschnikowiaceae(100);g__Metschnikowia(100);s__Metschnikowia_pulcherrima(100); | Metschnikowia | 13 | Saprotroph | \|Undefined Saprotroph\| | Yeast | NULL | Probable | As nectar-tap_saprotroph (Põlme et al. 2020); Decay sugar-rich substrates (Põlme et al. 2020) | Põlme S, et al. 2020. Fungal Diversity 105:1-16 ((https://doi.org/10.1007/s13225-020-00466-2)) |
| Otu51083 | 1 | k__Fungi(100);p__Ascomycota(100);c__Saccharomycetes(100);o__Saccharomycetales(100);f__Metschnikowiaceae(100);g__Metschnikowia(100);s__Metschnikowia_pulcherrima(100); | Metschnikowia | 13 | Saprotroph | \|Undefined Saprotroph\| | Yeast | NULL | Probable | As nectar-tap_saprotroph (Põlme et al. 2020); Decay sugar-rich substrates (Põlme et al. 2020) | Põlme S, et al. 2020. Fungal Diversity 105:1-16 ((https://doi.org/10.1007/s13225-020-00466-2)) |
| Otu51084 | 1 | k__Fungi(100);p__Ascomycota(100);c__Saccharomycetes(100);o__Saccharomycetales(100);f__Metschnikowiaceae(100);g__Metschnikowia(100);s__Metschnikowia_pulcherrima(100); | Metschnikowia | 13 | Saprotroph | \|Undefined Saprotroph\| | Yeast | NULL | Probable | As nectar-tap_saprotroph (Põlme et al. 2020); Decay sugar-rich substrates (Põlme et al. 2020) | Põlme S, et al. 2020. Fungal Diversity 105:1-16 ((https://doi.org/10.1007/s13225-020-00466-2)) |
| Otu51085 | 1 | k__Fungi(100);p__Ascomycota(100);c__Saccharomycetes(100);o__Saccharomycetales(100);f__Metschnikowiaceae(100);g__Metschnikowia(100);s__Metschnikowia_pulcherrima(100); | Metschnikowia | 13 | Saprotroph | \|Undefined Saprotroph\| | Yeast | NULL | Probable | As nectar-tap_saprotroph (Põlme et al. 2020); Decay sugar-rich substrates (Põlme et al. 2020) | Põlme S, et al. 2020. Fungal Diversity 105:1-16 ((https://doi.org/10.1007/s13225-020-00466-2)) |
| Otu51086 | 1 | k__Fungi(100);p__Ascomycota(100);c__Saccharomycetes(100);o__Saccharomycetales(100);f__Metschnikowiaceae(100);g__Metschnikowia(100);g__Metschnikowia_unclassified(100); | Metschnikowia | 13 | Saprotroph | \|Undefined Saprotroph\| | Yeast | NULL | Probable | As nectar-tap_saprotroph (Põlme et al. 2020); Decay sugar-rich substrates (Põlme et al. 2020) | Põlme S, et al. 2020. Fungal Diversity 105:1-16 ((https://doi.org/10.1007/s13225-020-00466-2)) |
| Otu51087 | 1 | k__Fungi(100);p__Ascomycota(100);c__Saccharomycetes(100);o__Saccharomycetales(100);f__Metschnikowiaceae(100);g__Metschnikowia(100);s__Metschnikowia_viticola(100); | Metschnikowia | 13 | Saprotroph | \|Undefined Saprotroph\| | Yeast | NULL | Probable | As nectar-tap_saprotroph (Põlme et al. 2020); Decay sugar-rich substrates (Põlme et al. 2020) | Põlme S, et al. 2020. Fungal Diversity 105:1-16 ((https://doi.org/10.1007/s13225-020-00466-2)) |
| Otu51088 | 1 | k__Fungi(100);p__Ascomycota(100);c__Saccharomycetes(100);o__Saccharomycetales(100);f__Metschnikowiaceae(100);g__Metschnikowia(100);g__Metschnikowia_unclassified(100); | Metschnikowia | 13 | Saprotroph | \|Undefined Saprotroph\| | Yeast | NULL | Probable | As nectar-tap_saprotroph (Põlme et al. 2020); Decay sugar-rich substrates (Põlme et al. 2020) | Põlme S, et al. 2020. Fungal Diversity 105:1-16 ((https://doi.org/10.1007/s13225-020-00466-2)) |
| Otu51089 | 1 | k__Fungi(100);p__Ascomycota(100);c__Saccharomycetes(100);o__Saccharomycetales(100);f__Metschnikowiaceae(100);g__Metschnikowia(100);s__Metschnikowia_pulcherrima(100); | Metschnikowia | 13 | Saprotroph | \|Undefined Saprotroph\| | Yeast | NULL | Probable | As nectar-tap_saprotroph (Põlme et al. 2020); Decay sugar-rich substrates (Põlme et al. 2020) | Põlme S, et al. 2020. Fungal Diversity 105:1-16 ((https://doi.org/10.1007/s13225-020-00466-2)) |
| Otu51090 | 1 | k__Fungi(100);p__Ascomycota(100);c__Saccharomycetes(100);o__Saccharomycetales(100);f__Metschnikowiaceae(100);g__Metschnikowia(100);s__Metschnikowia_pulcherrima(100); | Metschnikowia | 13 | Saprotroph | \|Undefined Saprotroph\| | Yeast | NULL | Probable | As nectar-tap_saprotroph (Põlme et al. 2020); Decay sugar-rich substrates (Põlme et al. 2020) | Põlme S, et al. 2020. Fungal Diversity 105:1-16 ((https://doi.org/10.1007/s13225-020-00466-2)) |
| Otu51092 | 1 | k__Fungi(100);p__Ascomycota(100);c__Saccharomycetes(100);o__Saccharomycetales(100);f__Metschnikowiaceae(100);g__Metschnikowia(100);s__Metschnikowia_pulcherrima(100); | Metschnikowia | 13 | Saprotroph | \|Undefined Saprotroph\| | Yeast | NULL | Probable | As nectar-tap_saprotroph (Põlme et al. 2020); Decay sugar-rich substrates (Põlme et al. 2020) | Põlme S, et al. 2020. Fungal Diversity 105:1-16 ((https://doi.org/10.1007/s13225-020-00466-2)) |
| Otu51093 | 1 | k__Fungi(100);p__Ascomycota(100);c__Saccharomycetes(100);o__Saccharomycetales(100);f__Metschnikowiaceae(100);g__Metschnikowia(100);g__Metschnikowia_unclassified(100); | Metschnikowia | 13 | Saprotroph | \|Undefined Saprotroph\| | Yeast | NULL | Probable | As nectar-tap_saprotroph (Põlme et al. 2020); Decay sugar-rich substrates (Põlme et al. 2020) | Põlme S, et al. 2020. Fungal Diversity 105:1-16 ((https://doi.org/10.1007/s13225-020-00466-2)) |
| Otu51095 | 1 | k__Fungi(100);p__Ascomycota(100);c__Saccharomycetes(100);o__Saccharomycetales(100);f__Metschnikowiaceae(100);g__Metschnikowia(100);s__Metschnikowia_pulcherrima(100); | Metschnikowia | 13 | Saprotroph | \|Undefined Saprotroph\| | Yeast | NULL | Probable | As nectar-tap_saprotroph (Põlme et al. 2020); Decay sugar-rich substrates (Põlme et al. 2020) | Põlme S, et al. 2020. Fungal Diversity 105:1-16 ((https://doi.org/10.1007/s13225-020-00466-2)) |
| Otu51096 | 1 | k__Fungi(100);p__Ascomycota(100);c__Saccharomycetes(100);o__Saccharomycetales(100);f__Metschnikowiaceae(100);g__Metschnikowia(100);s__Metschnikowia_pulcherrima(100); | Metschnikowia | 13 | Saprotroph | \|Undefined Saprotroph\| | Yeast | NULL | Probable | As nectar-tap_saprotroph (Põlme et al. 2020); Decay sugar-rich substrates (Põlme et al. 2020) | Põlme S, et al. 2020. Fungal Diversity 105:1-16 ((https://doi.org/10.1007/s13225-020-00466-2)) |
| Otu51097 | 1 | k__Fungi(100);p__Ascomycota(100);c__Saccharomycetes(100);o__Saccharomycetales(100);f__Metschnikowiaceae(100);g__Metschnikowia(100);s__Metschnikowia_pulcherrima(100); | Metschnikowia | 13 | Saprotroph | \|Undefined Saprotroph\| | Yeast | NULL | Probable | As nectar-tap_saprotroph (Põlme et al. 2020); Decay sugar-rich substrates (Põlme et al. 2020) | Põlme S, et al. 2020. Fungal Diversity 105:1-16 ((https://doi.org/10.1007/s13225-020-00466-2)) |
| Otu51098 | 1 | k__Fungi(100);p__Ascomycota(100);c__Saccharomycetes(100);o__Saccharomycetales(100);f__Metschnikowiaceae(100);g__Metschnikowia(100);s__Metschnikowia_pulcherrima(100); | Metschnikowia | 13 | Saprotroph | \|Undefined Saprotroph\| | Yeast | NULL | Probable | As nectar-tap_saprotroph (Põlme et al. 2020); Decay sugar-rich substrates (Põlme et al. 2020) | Põlme S, et al. 2020. Fungal Diversity 105:1-16 ((https://doi.org/10.1007/s13225-020-00466-2)) |
| Otu51099 | 1 | k__Fungi(100);p__Ascomycota(100);c__Saccharomycetes(100);o__Saccharomycetales(100);f__Saccharomycetales_family_Incertae_sedis(100);g__Candida(100);g__Candida_unclassified(100); | Candida | 13 | Pathotroph-Saprotroph-Symbiotroph | Animal Pathogen-Endophyte-Endosymbiont-Epiphyte-\|Undefined Saprotroph\| | Dimorphic Yeast-Yeast | NULL | Probable | As nectar-tap_saprotroph (Põlme et al. 2020); Decay sugar-rich substrates (Põlme et al. 2020); This asexual genus is polyphyletic and contain species that function very differently depend on what they are. Caution should be taken when interpreting guilds for Candida species. | Manolakaki D, et al. 2010. Virulence. 1: 367-375 ((https://doi.org/10.4161/viru.1.5.12796)); Mekbib SB, et al. 2011. Tropical Plant Pathology 36:233-240 ((https://doi.org/10.1590/S1982-56762011000400004)); Põlme S, et al. 2020. Fungal Diversity 105:1-16 ((https://doi.org/10.1007/s13225-020-00466-2)) |
| Otu51100 | 1 | k__Fungi(100);p__Ascomycota(100);c__Saccharomycetes(100);o__Saccharomycetales(100);f__Metschnikowiaceae(100);g__Metschnikowia(100);s__Metschnikowia_pulcherrima(100); | Metschnikowia | 13 | Saprotroph | \|Undefined Saprotroph\| | Yeast | NULL | Probable | As nectar-tap_saprotroph (Põlme et al. 2020); Decay sugar-rich substrates (Põlme et al. 2020) | Põlme S, et al. 2020. Fungal Diversity 105:1-16 ((https://doi.org/10.1007/s13225-020-00466-2)) |
| Otu51103 | 1 | k__Fungi(100);p__Ascomycota(100);c__Saccharomycetes(100);o__Saccharomycetales(100);f__Metschnikowiaceae(100);g__Metschnikowia(100);s__Metschnikowia_pulcherrima(100); | Metschnikowia | 13 | Saprotroph | \|Undefined Saprotroph\| | Yeast | NULL | Probable | As nectar-tap_saprotroph (Põlme et al. 2020); Decay sugar-rich substrates (Põlme et al. 2020) | Põlme S, et al. 2020. Fungal Diversity 105:1-16 ((https://doi.org/10.1007/s13225-020-00466-2)) |
| Otu51107 | 1 | k__Fungi(100);p__Ascomycota(100);c__Saccharomycetes(100);o__Saccharomycetales(100);f__Metschnikowiaceae(100);g__Metschnikowia(100);g__Metschnikowia_unclassified(100); | Metschnikowia | 13 | Saprotroph | \|Undefined Saprotroph\| | Yeast | NULL | Probable | As nectar-tap_saprotroph (Põlme et al. 2020); Decay sugar-rich substrates (Põlme et al. 2020) | Põlme S, et al. 2020. Fungal Diversity 105:1-16 ((https://doi.org/10.1007/s13225-020-00466-2)) |
| Otu51108 | 1 | k__Fungi(100);p__Ascomycota(100);c__Saccharomycetes(100);o__Saccharomycetales(100);f__Metschnikowiaceae(100);g__Metschnikowia(100);g__Metschnikowia_unclassified(100); | Metschnikowia | 13 | Saprotroph | \|Undefined Saprotroph\| | Yeast | NULL | Probable | As nectar-tap_saprotroph (Põlme et al. 2020); Decay sugar-rich substrates (Põlme et al. 2020) | Põlme S, et al. 2020. Fungal Diversity 105:1-16 ((https://doi.org/10.1007/s13225-020-00466-2)) |
| Otu51109 | 1 | k__Fungi(100);p__Ascomycota(100);c__Saccharomycetes(100);o__Saccharomycetales(100);f__Metschnikowiaceae(100);g__Metschnikowia(100);s__Metschnikowia_pulcherrima(100); | Metschnikowia | 13 | Saprotroph | \|Undefined Saprotroph\| | Yeast | NULL | Probable | As nectar-tap_saprotroph (Põlme et al. 2020); Decay sugar-rich substrates (Põlme et al. 2020) | Põlme S, et al. 2020. Fungal Diversity 105:1-16 ((https://doi.org/10.1007/s13225-020-00466-2)) |
| Otu51112 | 1 | k__Fungi(100);p__Ascomycota(100);c__Saccharomycetes(100);o__Saccharomycetales(100);f__Metschnikowiaceae(100);g__Metschnikowia(100);s__Metschnikowia_pulcherrima(100); | Metschnikowia | 13 | Saprotroph | \|Undefined Saprotroph\| | Yeast | NULL | Probable | As nectar-tap_saprotroph (Põlme et al. 2020); Decay sugar-rich substrates (Põlme et al. 2020) | Põlme S, et al. 2020. Fungal Diversity 105:1-16 ((https://doi.org/10.1007/s13225-020-00466-2)) |
| Otu51113 | 1 | k__Fungi(100);p__Ascomycota(100);c__Saccharomycetes(100);o__Saccharomycetales(100);f__Metschnikowiaceae(100);g__Metschnikowia(100);s__Metschnikowia_pulcherrima(100); | Metschnikowia | 13 | Saprotroph | \|Undefined Saprotroph\| | Yeast | NULL | Probable | As nectar-tap_saprotroph (Põlme et al. 2020); Decay sugar-rich substrates (Põlme et al. 2020) | Põlme S, et al. 2020. Fungal Diversity 105:1-16 ((https://doi.org/10.1007/s13225-020-00466-2)) |
| Otu51114 | 1 | k__Fungi(100);p__Ascomycota(100);c__Saccharomycetes(100);o__Saccharomycetales(100);f__Metschnikowiaceae(100);g__Metschnikowia(100);s__Metschnikowia_pulcherrima(100); | Metschnikowia | 13 | Saprotroph | \|Undefined Saprotroph\| | Yeast | NULL | Probable | As nectar-tap_saprotroph (Põlme et al. 2020); Decay sugar-rich substrates (Põlme et al. 2020) | Põlme S, et al. 2020. Fungal Diversity 105:1-16 ((https://doi.org/10.1007/s13225-020-00466-2)) |
| Otu51115 | 1 | k__Fungi(100);p__Ascomycota(100);c__Saccharomycetes(100);o__Saccharomycetales(100);f__Metschnikowiaceae(100);g__Metschnikowia(100);s__Metschnikowia_pulcherrima(100); | Metschnikowia | 13 | Saprotroph | \|Undefined Saprotroph\| | Yeast | NULL | Probable | As nectar-tap_saprotroph (Põlme et al. 2020); Decay sugar-rich substrates (Põlme et al. 2020) | Põlme S, et al. 2020. Fungal Diversity 105:1-16 ((https://doi.org/10.1007/s13225-020-00466-2)) |
| Otu51116 | 1 | k__Fungi(100);p__Basidiomycota(100);c__Microbotryomycetes(100);o__Sporidiobolales(100);f__Sporidiobolales_family_Incertae_sedis(100);g__Sporobolomyces(100);s__Sporobolomyces_lactosus(100); | Sporobolomyces | 13 | Pathotroph-Saprotroph | \|Fungal Parasite\|-Plant Saprotroph | Yeast | NULL | Probable | As litter_saprotroph (Põlme et al. 2020); As mycoparasite (Põlme et al. 2020); Decay fungal material (Põlme et al. 2020) | Tedersoo L, et al. 2014. Science 346:e1256688 ((https://doi.org/10.1126/science.1256688)); Purahong W, et al. 2016. Molecular Ecology 25:4059-4074 ((https://doi.org/10.1111/mec.13739)); Põlme S, et al. 2020. Fungal Diversity 105:1-16 ((https://doi.org/10.1007/s13225-020-00466-2)) |
| Otu51117 | 1 | k__Fungi(100);p__Ascomycota(100);c__Saccharomycetes(100);o__Saccharomycetales(100);f__Metschnikowiaceae(100);g__Metschnikowia(100);g__Metschnikowia_unclassified(100); | Metschnikowia | 13 | Saprotroph | \|Undefined Saprotroph\| | Yeast | NULL | Probable | As nectar-tap_saprotroph (Põlme et al. 2020); Decay sugar-rich substrates (Põlme et al. 2020) | Põlme S, et al. 2020. Fungal Diversity 105:1-16 ((https://doi.org/10.1007/s13225-020-00466-2)) |
| Otu51118 | 1 | k__Fungi(100);p__Ascomycota(100);c__Saccharomycetes(100);o__Saccharomycetales(100);f__Metschnikowiaceae(100);g__Metschnikowia(100);s__Metschnikowia_pulcherrima(100); | Metschnikowia | 13 | Saprotroph | \|Undefined Saprotroph\| | Yeast | NULL | Probable | As nectar-tap_saprotroph (Põlme et al. 2020); Decay sugar-rich substrates (Põlme et al. 2020) | Põlme S, et al. 2020. Fungal Diversity 105:1-16 ((https://doi.org/10.1007/s13225-020-00466-2)) |
| Otu51119 | 1 | k__Fungi(100);p__Ascomycota(100);c__Saccharomycetes(100);o__Saccharomycetales(100);f__Metschnikowiaceae(100);g__Metschnikowia(100);s__Metschnikowia_pulcherrima(100); | Metschnikowia | 13 | Saprotroph | \|Undefined Saprotroph\| | Yeast | NULL | Probable | As nectar-tap_saprotroph (Põlme et al. 2020); Decay sugar-rich substrates (Põlme et al. 2020) | Põlme S, et al. 2020. Fungal Diversity 105:1-16 ((https://doi.org/10.1007/s13225-020-00466-2)) |
| Otu51120 | 1 | k__Fungi(100);p__Ascomycota(100);c__Saccharomycetes(100);o__Saccharomycetales(100);f__Metschnikowiaceae(100);g__Metschnikowia(100);s__Metschnikowia_pulcherrima(100); | Metschnikowia | 13 | Saprotroph | \|Undefined Saprotroph\| | Yeast | NULL | Probable | As nectar-tap_saprotroph (Põlme et al. 2020); Decay sugar-rich substrates (Põlme et al. 2020) | Põlme S, et al. 2020. Fungal Diversity 105:1-16 ((https://doi.org/10.1007/s13225-020-00466-2)) |
| Otu51121 | 1 | k__Fungi(100);p__Ascomycota(100);c__Saccharomycetes(100);o__Saccharomycetales(100);f__Metschnikowiaceae(100);g__Metschnikowia(100);g__Metschnikowia_unclassified(100); | Metschnikowia | 13 | Saprotroph | \|Undefined Saprotroph\| | Yeast | NULL | Probable | As nectar-tap_saprotroph (Põlme et al. 2020); Decay sugar-rich substrates (Põlme et al. 2020) | Põlme S, et al. 2020. Fungal Diversity 105:1-16 ((https://doi.org/10.1007/s13225-020-00466-2)) |
| Otu51122 | 1 | k__Fungi(100);p__Ascomycota(100);c__Saccharomycetes(100);o__Saccharomycetales(100);f__Metschnikowiaceae(100);g__Metschnikowia(100);s__Metschnikowia_pulcherrima(100); | Metschnikowia | 13 | Saprotroph | \|Undefined Saprotroph\| | Yeast | NULL | Probable | As nectar-tap_saprotroph (Põlme et al. 2020); Decay sugar-rich substrates (Põlme et al. 2020) | Põlme S, et al. 2020. Fungal Diversity 105:1-16 ((https://doi.org/10.1007/s13225-020-00466-2)) |
| Otu51124 | 1 | k__Fungi(100);p__Ascomycota(100);c__Saccharomycetes(100);o__Saccharomycetales(100);f__Metschnikowiaceae(100);g__Metschnikowia(100);g__Metschnikowia_unclassified(100); | Metschnikowia | 13 | Saprotroph | \|Undefined Saprotroph\| | Yeast | NULL | Probable | As nectar-tap_saprotroph (Põlme et al. 2020); Decay sugar-rich substrates (Põlme et al. 2020) | Põlme S, et al. 2020. Fungal Diversity 105:1-16 ((https://doi.org/10.1007/s13225-020-00466-2)) |
| Otu51125 | 1 | k__Fungi(100);p__Ascomycota(100);c__Saccharomycetes(100);o__Saccharomycetales(100);f__Metschnikowiaceae(100);g__Metschnikowia(100);s__Metschnikowia_pulcherrima(100); | Metschnikowia | 13 | Saprotroph | \|Undefined Saprotroph\| | Yeast | NULL | Probable | As nectar-tap_saprotroph (Põlme et al. 2020); Decay sugar-rich substrates (Põlme et al. 2020) | Põlme S, et al. 2020. Fungal Diversity 105:1-16 ((https://doi.org/10.1007/s13225-020-00466-2)) |
| Otu51126 | 1 | k__Fungi(100);p__Ascomycota(100);c__Saccharomycetes(100);o__Saccharomycetales(100);f__Metschnikowiaceae(100);g__Metschnikowia(100);s__Metschnikowia_pulcherrima(100); | Metschnikowia | 13 | Saprotroph | \|Undefined Saprotroph\| | Yeast | NULL | Probable | As nectar-tap_saprotroph (Põlme et al. 2020); Decay sugar-rich substrates (Põlme et al. 2020) | Põlme S, et al. 2020. Fungal Diversity 105:1-16 ((https://doi.org/10.1007/s13225-020-00466-2)) |
| Otu51127 | 1 | k__Fungi(100);p__Ascomycota(100);c__Saccharomycetes(100);o__Saccharomycetales(100);f__Metschnikowiaceae(100);g__Metschnikowia(100);g__Metschnikowia_unclassified(100); | Metschnikowia | 13 | Saprotroph | \|Undefined Saprotroph\| | Yeast | NULL | Probable | As nectar-tap_saprotroph (Põlme et al. 2020); Decay sugar-rich substrates (Põlme et al. 2020) | Põlme S, et al. 2020. Fungal Diversity 105:1-16 ((https://doi.org/10.1007/s13225-020-00466-2)) |
| Otu51128 | 1 | k__Fungi(100);p__Ascomycota(100);c__Saccharomycetes(100);o__Saccharomycetales(100);f__Metschnikowiaceae(100);g__Metschnikowia(100);s__Metschnikowia_pulcherrima(100); | Metschnikowia | 13 | Saprotroph | \|Undefined Saprotroph\| | Yeast | NULL | Probable | As nectar-tap_saprotroph (Põlme et al. 2020); Decay sugar-rich substrates (Põlme et al. 2020) | Põlme S, et al. 2020. Fungal Diversity 105:1-16 ((https://doi.org/10.1007/s13225-020-00466-2)) |
| Otu51129 | 1 | k__Fungi(100);p__Ascomycota(100);c__Saccharomycetes(100);o__Saccharomycetales(100);f__Metschnikowiaceae(100);g__Metschnikowia(100);s__Metschnikowia_pulcherrima(100); | Metschnikowia | 13 | Saprotroph | \|Undefined Saprotroph\| | Yeast | NULL | Probable | As nectar-tap_saprotroph (Põlme et al. 2020); Decay sugar-rich substrates (Põlme et al. 2020) | Põlme S, et al. 2020. Fungal Diversity 105:1-16 ((https://doi.org/10.1007/s13225-020-00466-2)) |
| Otu51130 | 1 | k__Fungi(100);p__Ascomycota(100);c__Saccharomycetes(100);o__Saccharomycetales(100);f__Metschnikowiaceae(100);g__Metschnikowia(100);s__Metschnikowia_pulcherrima(100); | Metschnikowia | 13 | Saprotroph | \|Undefined Saprotroph\| | Yeast | NULL | Probable | As nectar-tap_saprotroph (Põlme et al. 2020); Decay sugar-rich substrates (Põlme et al. 2020) | Põlme S, et al. 2020. Fungal Diversity 105:1-16 ((https://doi.org/10.1007/s13225-020-00466-2)) |
| Otu51131 | 1 | k__Fungi(100);p__Ascomycota(100);c__Saccharomycetes(100);o__Saccharomycetales(100);f__Metschnikowiaceae(100);g__Metschnikowia(100);g__Metschnikowia_unclassified(100); | Metschnikowia | 13 | Saprotroph | \|Undefined Saprotroph\| | Yeast | NULL | Probable | As nectar-tap_saprotroph (Põlme et al. 2020); Decay sugar-rich substrates (Põlme et al. 2020) | Põlme S, et al. 2020. Fungal Diversity 105:1-16 ((https://doi.org/10.1007/s13225-020-00466-2)) |
| Otu51132 | 1 | k__Fungi(100);p__Ascomycota(100);c__Saccharomycetes(100);o__Saccharomycetales(100);f__Metschnikowiaceae(100);g__Metschnikowia(100);s__Metschnikowia_pulcherrima(100); | Metschnikowia | 13 | Saprotroph | \|Undefined Saprotroph\| | Yeast | NULL | Probable | As nectar-tap_saprotroph (Põlme et al. 2020); Decay sugar-rich substrates (Põlme et al. 2020) | Põlme S, et al. 2020. Fungal Diversity 105:1-16 ((https://doi.org/10.1007/s13225-020-00466-2)) |
| Otu51133 | 1 | k__Fungi(100);p__Ascomycota(100);c__Saccharomycetes(100);o__Saccharomycetales(100);f__Metschnikowiaceae(100);g__Metschnikowia(100);s__Metschnikowia_pulcherrima(100); | Metschnikowia | 13 | Saprotroph | \|Undefined Saprotroph\| | Yeast | NULL | Probable | As nectar-tap_saprotroph (Põlme et al. 2020); Decay sugar-rich substrates (Põlme et al. 2020) | Põlme S, et al. 2020. Fungal Diversity 105:1-16 ((https://doi.org/10.1007/s13225-020-00466-2)) |
| Otu51134 | 1 | k__Fungi(100);p__Ascomycota(100);c__Saccharomycetes(100);o__Saccharomycetales(100);f__Metschnikowiaceae(100);g__Metschnikowia(100);s__Metschnikowia_pulcherrima(100); | Metschnikowia | 13 | Saprotroph | \|Undefined Saprotroph\| | Yeast | NULL | Probable | As nectar-tap_saprotroph (Põlme et al. 2020); Decay sugar-rich substrates (Põlme et al. 2020) | Põlme S, et al. 2020. Fungal Diversity 105:1-16 ((https://doi.org/10.1007/s13225-020-00466-2)) |
| Otu51135 | 1 | k__Fungi(100);p__Ascomycota(100);c__Saccharomycetes(100);o__Saccharomycetales(100);f__Metschnikowiaceae(100);g__Metschnikowia(100);s__Metschnikowia_pulcherrima(100); | Metschnikowia | 13 | Saprotroph | \|Undefined Saprotroph\| | Yeast | NULL | Probable | As nectar-tap_saprotroph (Põlme et al. 2020); Decay sugar-rich substrates (Põlme et al. 2020) | Põlme S, et al. 2020. Fungal Diversity 105:1-16 ((https://doi.org/10.1007/s13225-020-00466-2)) |
| Otu51136 | 1 | k__Fungi(100);p__Ascomycota(100);c__Saccharomycetes(100);o__Saccharomycetales(100);f__Metschnikowiaceae(100);g__Metschnikowia(100);s__Metschnikowia_pulcherrima(100); | Metschnikowia | 13 | Saprotroph | \|Undefined Saprotroph\| | Yeast | NULL | Probable | As nectar-tap_saprotroph (Põlme et al. 2020); Decay sugar-rich substrates (Põlme et al. 2020) | Põlme S, et al. 2020. Fungal Diversity 105:1-16 ((https://doi.org/10.1007/s13225-020-00466-2)) |
| Otu51137 | 1 | k__Fungi(100);p__Ascomycota(100);c__Saccharomycetes(100);o__Saccharomycetales(100);f__Metschnikowiaceae(100);g__Metschnikowia(100);s__Metschnikowia_pulcherrima(100); | Metschnikowia | 13 | Saprotroph | \|Undefined Saprotroph\| | Yeast | NULL | Probable | As nectar-tap_saprotroph (Põlme et al. 2020); Decay sugar-rich substrates (Põlme et al. 2020) | Põlme S, et al. 2020. Fungal Diversity 105:1-16 ((https://doi.org/10.1007/s13225-020-00466-2)) |
| Otu51138 | 1 | k__Fungi(100);p__Ascomycota(100);c__Saccharomycetes(100);o__Saccharomycetales(100);f__Metschnikowiaceae(100);g__Metschnikowia(100);s__Metschnikowia_pulcherrima(100); | Metschnikowia | 13 | Saprotroph | \|Undefined Saprotroph\| | Yeast | NULL | Probable | As nectar-tap_saprotroph (Põlme et al. 2020); Decay sugar-rich substrates (Põlme et al. 2020) | Põlme S, et al. 2020. Fungal Diversity 105:1-16 ((https://doi.org/10.1007/s13225-020-00466-2)) |
| Otu51139 | 1 | k__Fungi(100);p__Ascomycota(100);c__Saccharomycetes(100);o__Saccharomycetales(100);f__Metschnikowiaceae(100);g__Metschnikowia(100);s__Metschnikowia_pulcherrima(100); | Metschnikowia | 13 | Saprotroph | \|Undefined Saprotroph\| | Yeast | NULL | Probable | As nectar-tap_saprotroph (Põlme et al. 2020); Decay sugar-rich substrates (Põlme et al. 2020) | Põlme S, et al. 2020. Fungal Diversity 105:1-16 ((https://doi.org/10.1007/s13225-020-00466-2)) |
| Otu51140 | 1 | k__Fungi(100);p__Ascomycota(100);c__Saccharomycetes(100);o__Saccharomycetales(100);f__Metschnikowiaceae(100);g__Metschnikowia(100);g__Metschnikowia_unclassified(100); | Metschnikowia | 13 | Saprotroph | \|Undefined Saprotroph\| | Yeast | NULL | Probable | As nectar-tap_saprotroph (Põlme et al. 2020); Decay sugar-rich substrates (Põlme et al. 2020) | Põlme S, et al. 2020. Fungal Diversity 105:1-16 ((https://doi.org/10.1007/s13225-020-00466-2)) |
| Otu51141 | 1 | k__Fungi(100);p__Ascomycota(100);c__Saccharomycetes(100);o__Saccharomycetales(100);f__Metschnikowiaceae(100);g__Metschnikowia(100);s__Metschnikowia_pulcherrima(100); | Metschnikowia | 13 | Saprotroph | \|Undefined Saprotroph\| | Yeast | NULL | Probable | As nectar-tap_saprotroph (Põlme et al. 2020); Decay sugar-rich substrates (Põlme et al. 2020) | Põlme S, et al. 2020. Fungal Diversity 105:1-16 ((https://doi.org/10.1007/s13225-020-00466-2)) |
| Otu51142 | 1 | k__Fungi(100);p__Ascomycota(100);c__Saccharomycetes(100);o__Saccharomycetales(100);f__Metschnikowiaceae(100);g__Metschnikowia(100);g__Metschnikowia_unclassified(100); | Metschnikowia | 13 | Saprotroph | \|Undefined Saprotroph\| | Yeast | NULL | Probable | As nectar-tap_saprotroph (Põlme et al. 2020); Decay sugar-rich substrates (Põlme et al. 2020) | Põlme S, et al. 2020. Fungal Diversity 105:1-16 ((https://doi.org/10.1007/s13225-020-00466-2)) |
| Otu51143 | 1 | k__Fungi(100);p__Ascomycota(100);c__Saccharomycetes(100);o__Saccharomycetales(100);f__Metschnikowiaceae(100);g__Metschnikowia(100);s__Metschnikowia_pulcherrima(100); | Metschnikowia | 13 | Saprotroph | \|Undefined Saprotroph\| | Yeast | NULL | Probable | As nectar-tap_saprotroph (Põlme et al. 2020); Decay sugar-rich substrates (Põlme et al. 2020) | Põlme S, et al. 2020. Fungal Diversity 105:1-16 ((https://doi.org/10.1007/s13225-020-00466-2)) |
| Otu51144 | 1 | k__Fungi(100);p__Ascomycota(100);c__Saccharomycetes(100);o__Saccharomycetales(100);f__Metschnikowiaceae(100);g__Metschnikowia(100);s__Metschnikowia_pulcherrima(100); | Metschnikowia | 13 | Saprotroph | \|Undefined Saprotroph\| | Yeast | NULL | Probable | As nectar-tap_saprotroph (Põlme et al. 2020); Decay sugar-rich substrates (Põlme et al. 2020) | Põlme S, et al. 2020. Fungal Diversity 105:1-16 ((https://doi.org/10.1007/s13225-020-00466-2)) |
| Otu51146 | 1 | k__Fungi(100);p__Ascomycota(100);c__Saccharomycetes(100);o__Saccharomycetales(100);f__Metschnikowiaceae(100);g__Metschnikowia(100);s__Metschnikowia_pulcherrima(100); | Metschnikowia | 13 | Saprotroph | \|Undefined Saprotroph\| | Yeast | NULL | Probable | As nectar-tap_saprotroph (Põlme et al. 2020); Decay sugar-rich substrates (Põlme et al. 2020) | Põlme S, et al. 2020. Fungal Diversity 105:1-16 ((https://doi.org/10.1007/s13225-020-00466-2)) |
| Otu51147 | 1 | k__Fungi(100);p__Ascomycota(100);c__Saccharomycetes(100);o__Saccharomycetales(100);f__Metschnikowiaceae(100);g__Metschnikowia(100);s__Metschnikowia_pulcherrima(100); | Metschnikowia | 13 | Saprotroph | \|Undefined Saprotroph\| | Yeast | NULL | Probable | As nectar-tap_saprotroph (Põlme et al. 2020); Decay sugar-rich substrates (Põlme et al. 2020) | Põlme S, et al. 2020. Fungal Diversity 105:1-16 ((https://doi.org/10.1007/s13225-020-00466-2)) |
| Otu51148 | 1 | k__Fungi(100);p__Ascomycota(100);c__Saccharomycetes(100);o__Saccharomycetales(100);f__Metschnikowiaceae(100);g__Metschnikowia(100);s__Metschnikowia_pulcherrima(100); | Metschnikowia | 13 | Saprotroph | \|Undefined Saprotroph\| | Yeast | NULL | Probable | As nectar-tap_saprotroph (Põlme et al. 2020); Decay sugar-rich substrates (Põlme et al. 2020) | Põlme S, et al. 2020. Fungal Diversity 105:1-16 ((https://doi.org/10.1007/s13225-020-00466-2)) |
| Otu51149 | 1 | k__Fungi(100);p__Ascomycota(100);c__Saccharomycetes(100);o__Saccharomycetales(100);f__Metschnikowiaceae(100);g__Metschnikowia(100);s__Metschnikowia_pulcherrima(100); | Metschnikowia | 13 | Saprotroph | \|Undefined Saprotroph\| | Yeast | NULL | Probable | As nectar-tap_saprotroph (Põlme et al. 2020); Decay sugar-rich substrates (Põlme et al. 2020) | Põlme S, et al. 2020. Fungal Diversity 105:1-16 ((https://doi.org/10.1007/s13225-020-00466-2)) |
| Otu51151 | 1 | k__Fungi(100);p__Ascomycota(100);c__Saccharomycetes(100);o__Saccharomycetales(100);f__Metschnikowiaceae(100);g__Metschnikowia(100);s__Metschnikowia_pulcherrima(100); | Metschnikowia | 13 | Saprotroph | \|Undefined Saprotroph\| | Yeast | NULL | Probable | As nectar-tap_saprotroph (Põlme et al. 2020); Decay sugar-rich substrates (Põlme et al. 2020) | Põlme S, et al. 2020. Fungal Diversity 105:1-16 ((https://doi.org/10.1007/s13225-020-00466-2)) |
| Otu51152 | 1 | k__Fungi(100);p__Ascomycota(100);c__Saccharomycetes(100);o__Saccharomycetales(100);f__Metschnikowiaceae(100);g__Metschnikowia(100);s__Metschnikowia_pulcherrima(100); | Metschnikowia | 13 | Saprotroph | \|Undefined Saprotroph\| | Yeast | NULL | Probable | As nectar-tap_saprotroph (Põlme et al. 2020); Decay sugar-rich substrates (Põlme et al. 2020) | Põlme S, et al. 2020. Fungal Diversity 105:1-16 ((https://doi.org/10.1007/s13225-020-00466-2)) |
| Otu51153 | 1 | k__Fungi(100);p__Ascomycota(100);c__Saccharomycetes(100);o__Saccharomycetales(100);f__Metschnikowiaceae(100);g__Metschnikowia(100);s__Metschnikowia_pulcherrima(100); | Metschnikowia | 13 | Saprotroph | \|Undefined Saprotroph\| | Yeast | NULL | Probable | As nectar-tap_saprotroph (Põlme et al. 2020); Decay sugar-rich substrates (Põlme et al. 2020) | Põlme S, et al. 2020. Fungal Diversity 105:1-16 ((https://doi.org/10.1007/s13225-020-00466-2)) |
| Otu51155 | 1 | k__Fungi(100);p__Ascomycota(100);c__Saccharomycetes(100);o__Saccharomycetales(100);f__Metschnikowiaceae(100);g__Metschnikowia(100);s__Metschnikowia_pulcherrima(100); | Metschnikowia | 13 | Saprotroph | \|Undefined Saprotroph\| | Yeast | NULL | Probable | As nectar-tap_saprotroph (Põlme et al. 2020); Decay sugar-rich substrates (Põlme et al. 2020) | Põlme S, et al. 2020. Fungal Diversity 105:1-16 ((https://doi.org/10.1007/s13225-020-00466-2)) |
| Otu51160 | 1 | k__Fungi(100);p__Ascomycota(100);c__Saccharomycetes(100);o__Saccharomycetales(100);f__Metschnikowiaceae(100);g__Metschnikowia(100);s__Metschnikowia_pulcherrima(100); | Metschnikowia | 13 | Saprotroph | \|Undefined Saprotroph\| | Yeast | NULL | Probable | As nectar-tap_saprotroph (Põlme et al. 2020); Decay sugar-rich substrates (Põlme et al. 2020) | Põlme S, et al. 2020. Fungal Diversity 105:1-16 ((https://doi.org/10.1007/s13225-020-00466-2)) |
| Otu51161 | 1 | k__Fungi(100);p__Ascomycota(100);c__Saccharomycetes(100);o__Saccharomycetales(100);f__Metschnikowiaceae(100);g__Metschnikowia(100);s__Metschnikowia_pulcherrima(100); | Metschnikowia | 13 | Saprotroph | \|Undefined Saprotroph\| | Yeast | NULL | Probable | As nectar-tap_saprotroph (Põlme et al. 2020); Decay sugar-rich substrates (Põlme et al. 2020) | Põlme S, et al. 2020. Fungal Diversity 105:1-16 ((https://doi.org/10.1007/s13225-020-00466-2)) |
| Otu51162 | 1 | k__Fungi(100);p__Ascomycota(100);c__Saccharomycetes(100);o__Saccharomycetales(100);f__Metschnikowiaceae(100);g__Metschnikowia(100);g__Metschnikowia_unclassified(100); | Metschnikowia | 13 | Saprotroph | \|Undefined Saprotroph\| | Yeast | NULL | Probable | As nectar-tap_saprotroph (Põlme et al. 2020); Decay sugar-rich substrates (Põlme et al. 2020) | Põlme S, et al. 2020. Fungal Diversity 105:1-16 ((https://doi.org/10.1007/s13225-020-00466-2)) |
| Otu51163 | 1 | k__Fungi(100);p__Ascomycota(100);c__Saccharomycetes(100);o__Saccharomycetales(100);f__Metschnikowiaceae(100);g__Metschnikowia(100);s__Metschnikowia_pulcherrima(100); | Metschnikowia | 13 | Saprotroph | \|Undefined Saprotroph\| | Yeast | NULL | Probable | As nectar-tap_saprotroph (Põlme et al. 2020); Decay sugar-rich substrates (Põlme et al. 2020) | Põlme S, et al. 2020. Fungal Diversity 105:1-16 ((https://doi.org/10.1007/s13225-020-00466-2)) |
| Otu51165 | 1 | k__Fungi(100);p__Ascomycota(100);c__Saccharomycetes(100);o__Saccharomycetales(100);f__Metschnikowiaceae(100);g__Metschnikowia(100);s__Metschnikowia_pulcherrima(100); | Metschnikowia | 13 | Saprotroph | \|Undefined Saprotroph\| | Yeast | NULL | Probable | As nectar-tap_saprotroph (Põlme et al. 2020); Decay sugar-rich substrates (Põlme et al. 2020) | Põlme S, et al. 2020. Fungal Diversity 105:1-16 ((https://doi.org/10.1007/s13225-020-00466-2)) |
| Otu51166 | 1 | k__Fungi(100);p__Ascomycota(100);c__Saccharomycetes(100);o__Saccharomycetales(100);f__Metschnikowiaceae(100);g__Metschnikowia(100);s__Metschnikowia_pulcherrima(100); | Metschnikowia | 13 | Saprotroph | \|Undefined Saprotroph\| | Yeast | NULL | Probable | As nectar-tap_saprotroph (Põlme et al. 2020); Decay sugar-rich substrates (Põlme et al. 2020) | Põlme S, et al. 2020. Fungal Diversity 105:1-16 ((https://doi.org/10.1007/s13225-020-00466-2)) |
| Otu51167 | 1 | k__Fungi(100);p__Ascomycota(100);c__Saccharomycetes(100);o__Saccharomycetales(100);f__Metschnikowiaceae(100);g__Metschnikowia(100);g__Metschnikowia_unclassified(100); | Metschnikowia | 13 | Saprotroph | \|Undefined Saprotroph\| | Yeast | NULL | Probable | As nectar-tap_saprotroph (Põlme et al. 2020); Decay sugar-rich substrates (Põlme et al. 2020) | Põlme S, et al. 2020. Fungal Diversity 105:1-16 ((https://doi.org/10.1007/s13225-020-00466-2)) |
| Otu51168 | 1 | k__Fungi(100);p__Ascomycota(100);c__Saccharomycetes(100);o__Saccharomycetales(100);f__Metschnikowiaceae(100);g__Metschnikowia(100);s__Metschnikowia_pulcherrima(100); | Metschnikowia | 13 | Saprotroph | \|Undefined Saprotroph\| | Yeast | NULL | Probable | As nectar-tap_saprotroph (Põlme et al. 2020); Decay sugar-rich substrates (Põlme et al. 2020) | Põlme S, et al. 2020. Fungal Diversity 105:1-16 ((https://doi.org/10.1007/s13225-020-00466-2)) |
| Otu51169 | 1 | k__Fungi(100);p__Ascomycota(100);c__Saccharomycetes(100);o__Saccharomycetales(100);f__Metschnikowiaceae(100);g__Metschnikowia(100);g__Metschnikowia_unclassified(100); | Metschnikowia | 13 | Saprotroph | \|Undefined Saprotroph\| | Yeast | NULL | Probable | As nectar-tap_saprotroph (Põlme et al. 2020); Decay sugar-rich substrates (Põlme et al. 2020) | Põlme S, et al. 2020. Fungal Diversity 105:1-16 ((https://doi.org/10.1007/s13225-020-00466-2)) |
| Otu51170 | 1 | k__Fungi(100);p__Ascomycota(100);c__Saccharomycetes(100);o__Saccharomycetales(100);f__Metschnikowiaceae(100);g__Metschnikowia(100);s__Metschnikowia_pulcherrima(100); | Metschnikowia | 13 | Saprotroph | \|Undefined Saprotroph\| | Yeast | NULL | Probable | As nectar-tap_saprotroph (Põlme et al. 2020); Decay sugar-rich substrates (Põlme et al. 2020) | Põlme S, et al. 2020. Fungal Diversity 105:1-16 ((https://doi.org/10.1007/s13225-020-00466-2)) |
| Otu51171 | 1 | k__Fungi(100);p__Ascomycota(100);c__Saccharomycetes(100);o__Saccharomycetales(100);f__Metschnikowiaceae(100);g__Metschnikowia(100);s__Metschnikowia_pulcherrima(100); | Metschnikowia | 13 | Saprotroph | \|Undefined Saprotroph\| | Yeast | NULL | Probable | As nectar-tap_saprotroph (Põlme et al. 2020); Decay sugar-rich substrates (Põlme et al. 2020) | Põlme S, et al. 2020. Fungal Diversity 105:1-16 ((https://doi.org/10.1007/s13225-020-00466-2)) |
| Otu51172 | 1 | k__Fungi(100);p__Ascomycota(100);c__Saccharomycetes(100);o__Saccharomycetales(100);f__Metschnikowiaceae(100);g__Metschnikowia(100);s__Metschnikowia_pulcherrima(100); | Metschnikowia | 13 | Saprotroph | \|Undefined Saprotroph\| | Yeast | NULL | Probable | As nectar-tap_saprotroph (Põlme et al. 2020); Decay sugar-rich substrates (Põlme et al. 2020) | Põlme S, et al. 2020. Fungal Diversity 105:1-16 ((https://doi.org/10.1007/s13225-020-00466-2)) |
| Otu51173 | 1 | k__Fungi(100);p__Ascomycota(100);c__Saccharomycetes(100);o__Saccharomycetales(100);f__Metschnikowiaceae(100);g__Metschnikowia(100);s__Metschnikowia_pulcherrima(100); | Metschnikowia | 13 | Saprotroph | \|Undefined Saprotroph\| | Yeast | NULL | Probable | As nectar-tap_saprotroph (Põlme et al. 2020); Decay sugar-rich substrates (Põlme et al. 2020) | Põlme S, et al. 2020. Fungal Diversity 105:1-16 ((https://doi.org/10.1007/s13225-020-00466-2)) |
| Otu51175 | 1 | k__Fungi(100);p__Ascomycota(100);c__Saccharomycetes(100);o__Saccharomycetales(100);f__Metschnikowiaceae(100);g__Metschnikowia(100);g__Metschnikowia_unclassified(100); | Metschnikowia | 13 | Saprotroph | \|Undefined Saprotroph\| | Yeast | NULL | Probable | As nectar-tap_saprotroph (Põlme et al. 2020); Decay sugar-rich substrates (Põlme et al. 2020) | Põlme S, et al. 2020. Fungal Diversity 105:1-16 ((https://doi.org/10.1007/s13225-020-00466-2)) |
| Otu51176 | 1 | k__Fungi(100);p__Ascomycota(100);c__Saccharomycetes(100);o__Saccharomycetales(100);f__Metschnikowiaceae(100);g__Metschnikowia(100);s__Metschnikowia_pulcherrima(100); | Metschnikowia | 13 | Saprotroph | \|Undefined Saprotroph\| | Yeast | NULL | Probable | As nectar-tap_saprotroph (Põlme et al. 2020); Decay sugar-rich substrates (Põlme et al. 2020) | Põlme S, et al. 2020. Fungal Diversity 105:1-16 ((https://doi.org/10.1007/s13225-020-00466-2)) |
| Otu51177 | 1 | k__Fungi(100);p__Ascomycota(100);c__Saccharomycetes(100);o__Saccharomycetales(100);f__Metschnikowiaceae(100);g__Metschnikowia(100);s__Metschnikowia_pulcherrima(100); | Metschnikowia | 13 | Saprotroph | \|Undefined Saprotroph\| | Yeast | NULL | Probable | As nectar-tap_saprotroph (Põlme et al. 2020); Decay sugar-rich substrates (Põlme et al. 2020) | Põlme S, et al. 2020. Fungal Diversity 105:1-16 ((https://doi.org/10.1007/s13225-020-00466-2)) |
| Otu51178 | 1 | k__Fungi(100);p__Ascomycota(100);c__Saccharomycetes(100);o__Saccharomycetales(100);f__Metschnikowiaceae(100);g__Metschnikowia(100);s__Metschnikowia_pulcherrima(100); | Metschnikowia | 13 | Saprotroph | \|Undefined Saprotroph\| | Yeast | NULL | Probable | As nectar-tap_saprotroph (Põlme et al. 2020); Decay sugar-rich substrates (Põlme et al. 2020) | Põlme S, et al. 2020. Fungal Diversity 105:1-16 ((https://doi.org/10.1007/s13225-020-00466-2)) |
| Otu51179 | 1 | k__Fungi(100);p__Ascomycota(100);c__Saccharomycetes(100);o__Saccharomycetales(100);f__Metschnikowiaceae(100);g__Metschnikowia(100);s__Metschnikowia_pulcherrima(100); | Metschnikowia | 13 | Saprotroph | \|Undefined Saprotroph\| | Yeast | NULL | Probable | As nectar-tap_saprotroph (Põlme et al. 2020); Decay sugar-rich substrates (Põlme et al. 2020) | Põlme S, et al. 2020. Fungal Diversity 105:1-16 ((https://doi.org/10.1007/s13225-020-00466-2)) |
| Otu51180 | 1 | k__Fungi(100);p__Ascomycota(100);c__Saccharomycetes(100);o__Saccharomycetales(100);f__Metschnikowiaceae(100);g__Metschnikowia(100);g__Metschnikowia_unclassified(100); | Metschnikowia | 13 | Saprotroph | \|Undefined Saprotroph\| | Yeast | NULL | Probable | As nectar-tap_saprotroph (Põlme et al. 2020); Decay sugar-rich substrates (Põlme et al. 2020) | Põlme S, et al. 2020. Fungal Diversity 105:1-16 ((https://doi.org/10.1007/s13225-020-00466-2)) |
| Otu51181 | 1 | k__Fungi(100);p__Ascomycota(100);c__Saccharomycetes(100);o__Saccharomycetales(100);f__Metschnikowiaceae(100);g__Metschnikowia(100);s__Metschnikowia_pulcherrima(100); | Metschnikowia | 13 | Saprotroph | \|Undefined Saprotroph\| | Yeast | NULL | Probable | As nectar-tap_saprotroph (Põlme et al. 2020); Decay sugar-rich substrates (Põlme et al. 2020) | Põlme S, et al. 2020. Fungal Diversity 105:1-16 ((https://doi.org/10.1007/s13225-020-00466-2)) |
| Otu51182 | 1 | k__Fungi(100);p__Ascomycota(100);c__Saccharomycetes(100);o__Saccharomycetales(100);f__Metschnikowiaceae(100);g__Metschnikowia(100);s__Metschnikowia_pulcherrima(100); | Metschnikowia | 13 | Saprotroph | \|Undefined Saprotroph\| | Yeast | NULL | Probable | As nectar-tap_saprotroph (Põlme et al. 2020); Decay sugar-rich substrates (Põlme et al. 2020) | Põlme S, et al. 2020. Fungal Diversity 105:1-16 ((https://doi.org/10.1007/s13225-020-00466-2)) |
| Otu51183 | 1 | k__Fungi(100);p__Ascomycota(100);c__Saccharomycetes(100);o__Saccharomycetales(100);f__Metschnikowiaceae(100);g__Metschnikowia(100);s__Metschnikowia_pulcherrima(100); | Metschnikowia | 13 | Saprotroph | \|Undefined Saprotroph\| | Yeast | NULL | Probable | As nectar-tap_saprotroph (Põlme et al. 2020); Decay sugar-rich substrates (Põlme et al. 2020) | Põlme S, et al. 2020. Fungal Diversity 105:1-16 ((https://doi.org/10.1007/s13225-020-00466-2)) |
| Otu51185 | 1 | k__Fungi(100);p__Ascomycota(100);c__Saccharomycetes(100);o__Saccharomycetales(100);f__Metschnikowiaceae(100);g__Metschnikowia(100);g__Metschnikowia_unclassified(100); | Metschnikowia | 13 | Saprotroph | \|Undefined Saprotroph\| | Yeast | NULL | Probable | As nectar-tap_saprotroph (Põlme et al. 2020); Decay sugar-rich substrates (Põlme et al. 2020) | Põlme S, et al. 2020. Fungal Diversity 105:1-16 ((https://doi.org/10.1007/s13225-020-00466-2)) |
| Otu51186 | 1 | k__Fungi(100);p__Ascomycota(100);c__Saccharomycetes(100);o__Saccharomycetales(100);f__Metschnikowiaceae(100);g__Metschnikowia(100);s__Metschnikowia_pulcherrima(100); | Metschnikowia | 13 | Saprotroph | \|Undefined Saprotroph\| | Yeast | NULL | Probable | As nectar-tap_saprotroph (Põlme et al. 2020); Decay sugar-rich substrates (Põlme et al. 2020) | Põlme S, et al. 2020. Fungal Diversity 105:1-16 ((https://doi.org/10.1007/s13225-020-00466-2)) |
| Otu51187 | 1 | k__Fungi(100);p__Ascomycota(100);c__Saccharomycetes(100);o__Saccharomycetales(100);f__Metschnikowiaceae(100);g__Metschnikowia(100);s__Metschnikowia_pulcherrima(100); | Metschnikowia | 13 | Saprotroph | \|Undefined Saprotroph\| | Yeast | NULL | Probable | As nectar-tap_saprotroph (Põlme et al. 2020); Decay sugar-rich substrates (Põlme et al. 2020) | Põlme S, et al. 2020. Fungal Diversity 105:1-16 ((https://doi.org/10.1007/s13225-020-00466-2)) |
| Otu51188 | 1 | k__Fungi(100);p__Ascomycota(100);c__Saccharomycetes(100);o__Saccharomycetales(100);f__Metschnikowiaceae(100);g__Metschnikowia(100);s__Metschnikowia_pulcherrima(100); | Metschnikowia | 13 | Saprotroph | \|Undefined Saprotroph\| | Yeast | NULL | Probable | As nectar-tap_saprotroph (Põlme et al. 2020); Decay sugar-rich substrates (Põlme et al. 2020) | Põlme S, et al. 2020. Fungal Diversity 105:1-16 ((https://doi.org/10.1007/s13225-020-00466-2)) |
| Otu51189 | 1 | k__Fungi(100);p__Ascomycota(100);c__Saccharomycetes(100);o__Saccharomycetales(100);f__Metschnikowiaceae(100);g__Metschnikowia(100);s__Metschnikowia_pulcherrima(100); | Metschnikowia | 13 | Saprotroph | \|Undefined Saprotroph\| | Yeast | NULL | Probable | As nectar-tap_saprotroph (Põlme et al. 2020); Decay sugar-rich substrates (Põlme et al. 2020) | Põlme S, et al. 2020. Fungal Diversity 105:1-16 ((https://doi.org/10.1007/s13225-020-00466-2)) |
| Otu51190 | 1 | k__Fungi(100);p__Ascomycota(100);c__Saccharomycetes(100);o__Saccharomycetales(100);f__Metschnikowiaceae(100);g__Metschnikowia(100);s__Metschnikowia_pulcherrima(100); | Metschnikowia | 13 | Saprotroph | \|Undefined Saprotroph\| | Yeast | NULL | Probable | As nectar-tap_saprotroph (Põlme et al. 2020); Decay sugar-rich substrates (Põlme et al. 2020) | Põlme S, et al. 2020. Fungal Diversity 105:1-16 ((https://doi.org/10.1007/s13225-020-00466-2)) |
| Otu51191 | 1 | k__Fungi(100);p__Ascomycota(100);c__Saccharomycetes(100);o__Saccharomycetales(100);f__Metschnikowiaceae(100);g__Metschnikowia(100);g__Metschnikowia_unclassified(100); | Metschnikowia | 13 | Saprotroph | \|Undefined Saprotroph\| | Yeast | NULL | Probable | As nectar-tap_saprotroph (Põlme et al. 2020); Decay sugar-rich substrates (Põlme et al. 2020) | Põlme S, et al. 2020. Fungal Diversity 105:1-16 ((https://doi.org/10.1007/s13225-020-00466-2)) |
| Otu51192 | 1 | k__Fungi(100);p__Ascomycota(100);c__Saccharomycetes(100);o__Saccharomycetales(100);f__Metschnikowiaceae(100);g__Metschnikowia(100);g__Metschnikowia_unclassified(100); | Metschnikowia | 13 | Saprotroph | \|Undefined Saprotroph\| | Yeast | NULL | Probable | As nectar-tap_saprotroph (Põlme et al. 2020); Decay sugar-rich substrates (Põlme et al. 2020) | Põlme S, et al. 2020. Fungal Diversity 105:1-16 ((https://doi.org/10.1007/s13225-020-00466-2)) |
| Otu51194 | 1 | k__Fungi(100);p__Ascomycota(100);c__Saccharomycetes(100);o__Saccharomycetales(100);f__Metschnikowiaceae(100);g__Metschnikowia(100);g__Metschnikowia_unclassified(100); | Metschnikowia | 13 | Saprotroph | \|Undefined Saprotroph\| | Yeast | NULL | Probable | As nectar-tap_saprotroph (Põlme et al. 2020); Decay sugar-rich substrates (Põlme et al. 2020) | Põlme S, et al. 2020. Fungal Diversity 105:1-16 ((https://doi.org/10.1007/s13225-020-00466-2)) |
| Otu51195 | 1 | k__Fungi(100);p__Ascomycota(100);c__Saccharomycetes(100);o__Saccharomycetales(100);f__Metschnikowiaceae(100);g__Metschnikowia(100);g__Metschnikowia_unclassified(100); | Metschnikowia | 13 | Saprotroph | \|Undefined Saprotroph\| | Yeast | NULL | Probable | As nectar-tap_saprotroph (Põlme et al. 2020); Decay sugar-rich substrates (Põlme et al. 2020) | Põlme S, et al. 2020. Fungal Diversity 105:1-16 ((https://doi.org/10.1007/s13225-020-00466-2)) |
| Otu51196 | 1 | k__Fungi(100);p__Ascomycota(100);c__Saccharomycetes(100);o__Saccharomycetales(100);f__Metschnikowiaceae(100);g__Metschnikowia(100);s__Metschnikowia_pulcherrima(100); | Metschnikowia | 13 | Saprotroph | \|Undefined Saprotroph\| | Yeast | NULL | Probable | As nectar-tap_saprotroph (Põlme et al. 2020); Decay sugar-rich substrates (Põlme et al. 2020) | Põlme S, et al. 2020. Fungal Diversity 105:1-16 ((https://doi.org/10.1007/s13225-020-00466-2)) |

| | | | | | | | | | | | |
|---|---|---|---|---|---|---|---|---|---|---|---|
| Otu51197 | 1 | k__Fungi(100);p__Ascomycota(100);c__Saccharomycetes(100);o__Saccharomycetales(100);f__Metschnikowiaceae(100);g__Metschnikowia(100);s__Metschnikowia_pulcherrima(100); | Metschnikowia | 13 | Saprotroph | \|Undefined Saprotroph\| | Yeast | NULL | Probable | As nectar-tap_saprotroph (Põlme et al. 2020); Decay sugar-rich substrates (Põlme et al. 2020) | Põlme S, et al. 2020. Fungal Diversity 105:1-16 ((https://doi.org/10.1007/s13225-020-00466-2)) |
| Otu51198 | 1 | k__Fungi(100);p__Ascomycota(100);c__Saccharomycetes(100);o__Saccharomycetales(100);f__Metschnikowiaceae(100);g__Metschnikowia(100);s__Metschnikowia_pulcherrima(100); | Metschnikowia | 13 | Saprotroph | \|Undefined Saprotroph\| | Yeast | NULL | Probable | As nectar-tap_saprotroph (Põlme et al. 2020); Decay sugar-rich substrates (Põlme et al. 2020) | Põlme S, et al. 2020. Fungal Diversity 105:1-16 ((https://doi.org/10.1007/s13225-020-00466-2)) |
| Otu51199 | 1 | k__Fungi(100);p__Ascomycota(100);c__Saccharomycetes(100);o__Saccharomycetales(100);f__Metschnikowiaceae(100);g__Metschnikowia(100);s__Metschnikowia_pulcherrima(100); | Metschnikowia | 13 | Saprotroph | \|Undefined Saprotroph\| | Yeast | NULL | Probable | As nectar-tap_saprotroph (Põlme et al. 2020); Decay sugar-rich substrates (Põlme et al. 2020) | Põlme S, et al. 2020. Fungal Diversity 105:1-16 ((https://doi.org/10.1007/s13225-020-00466-2)) |
| Otu51200 | 1 | k__Fungi(100);p__Ascomycota(100);c__Saccharomycetes(100);o__Saccharomycetales(100);f__Metschnikowiaceae(100);g__Metschnikowia(100);g__Metschnikowia_unclassified(100); | Metschnikowia | 13 | Saprotroph | \|Undefined Saprotroph\| | Yeast | NULL | Probable | As nectar-tap_saprotroph (Põlme et al. 2020); Decay sugar-rich substrates (Põlme et al. 2020) | Põlme S, et al. 2020. Fungal Diversity 105:1-16 ((https://doi.org/10.1007/s13225-020-00466-2)) |
| Otu51201 | 1 | k__Fungi(100);p__Ascomycota(100);c__Saccharomycetes(100);o__Saccharomycetales(100);f__Metschnikowiaceae(100);g__Metschnikowia(100);s__Metschnikowia_pulcherrima(100); | Metschnikowia | 13 | Saprotroph | \|Undefined Saprotroph\| | Yeast | NULL | Probable | As nectar-tap_saprotroph (Põlme et al. 2020); Decay sugar-rich substrates (Põlme et al. 2020) | Põlme S, et al. 2020. Fungal Diversity 105:1-16 ((https://doi.org/10.1007/s13225-020-00466-2)) |
| Otu51202 | 1 | k__Fungi(100);p__Ascomycota(100);c__Saccharomycetes(100);o__Saccharomycetales(100);f__Metschnikowiaceae(100);g__Metschnikowia(100);s__Metschnikowia_pulcherrima(100); | Metschnikowia | 13 | Saprotroph | \|Undefined Saprotroph\| | Yeast | NULL | Probable | As nectar-tap_saprotroph (Põlme et al. 2020); Decay sugar-rich substrates (Põlme et al. 2020) | Põlme S, et al. 2020. Fungal Diversity 105:1-16 ((https://doi.org/10.1007/s13225-020-00466-2)) |
| Otu51203 | 1 | k__Fungi(100);p__Ascomycota(100);c__Saccharomycetes(100);o__Saccharomycetales(100);f__Metschnikowiaceae(100);g__Metschnikowia(100);s__Metschnikowia_pulcherrima(100); | Metschnikowia | 13 | Saprotroph | \|Undefined Saprotroph\| | Yeast | NULL | Probable | As nectar-tap_saprotroph (Põlme et al. 2020); Decay sugar-rich substrates (Põlme et al. 2020) | Põlme S, et al. 2020. Fungal Diversity 105:1-16 ((https://doi.org/10.1007/s13225-020-00466-2)) |
| Otu51204 | 1 | k__Fungi(100);p__Ascomycota(100);c__Saccharomycetes(100);o__Saccharomycetales(100);f__Metschnikowiaceae(100);g__Metschnikowia(100);g__Metschnikowia_unclassified(100); | Metschnikowia | 13 | Saprotroph | \|Undefined Saprotroph\| | Yeast | NULL | Probable | As nectar-tap_saprotroph (Põlme et al. 2020); Decay sugar-rich substrates (Põlme et al. 2020) | Põlme S, et al. 2020. Fungal Diversity 105:1-16 ((https://doi.org/10.1007/s13225-020-00466-2)) |
| Otu51205 | 1 | k__Fungi(100);p__Ascomycota(100);c__Saccharomycetes(100);o__Saccharomycetales(100);f__Saccharomycetales_family_Incertae_sedis(100);g__Candida(100);g__Candida_unclassified(100); | Candida | 13 | Pathotroph-Saprotroph-Symbiotroph | Animal Pathogen-Endophyte-Endosymbiont-Epiphyte-\|Undefined Saprotroph\| | Dimorphic Yeast-Yeast | NULL | Probable | As nectar-tap_saprotroph (Põlme et al. 2020); Decay sugar-rich substrates (Põlme et al. 2020); This asexual genus is polyphyletic and contain species that function very differently depend on what they are. Caution should be taken when interpreting guilds for Candida species. | Manolakaki D, et al. 2010. Virulence. 1: 367-375 ((https://doi.org/10.4161/viru.1.5.12796)); Mekbib SB, et al. 2011. Tropical Plant Pathology 36:233-240 ((https://doi.org/10.1590/S1982-56762011000400004)); Põlme S, et al. 2020. Fungal Diversity 105:1-16 ((https://doi.org/10.1007/s13225-020-00466-2)) |
| Otu51206 | 1 | k__Fungi(100);p__Ascomycota(100);c__Saccharomycetes(100);o__Saccharomycetales(100);f__Saccharomycetales_family_Incertae_sedis(100);g__Candida(100);s__Candida_cretensis(100); | Candida | 13 | Pathotroph-Saprotroph-Symbiotroph | Animal Pathogen-Endophyte-Endosymbiont-Epiphyte-\|Undefined Saprotroph\| | Dimorphic Yeast-Yeast | NULL | Probable | As nectar-tap_saprotroph (Põlme et al. 2020); Decay sugar-rich substrates (Põlme et al. 2020); This asexual genus is polyphyletic and contain species that function very differently depend on what they are. Caution should be taken when interpreting guilds for Candida species. | Manolakaki D, et al. 2010. Virulence. 1: 367-375 ((https://doi.org/10.4161/viru.1.5.12796)); Mekbib SB, et al. 2011. Tropical Plant Pathology 36:233-240 ((https://doi.org/10.1590/S1982-56762011000400004)); Põlme S, et al. 2020. Fungal Diversity 105:1-16 ((https://doi.org/10.1007/s13225-020-00466-2)) |
| Otu51208 | 1 | k__Fungi(100);p__Ascomycota(100);c__Saccharomycetes(100);o__Saccharomycetales(100);f__Metschnikowiaceae(100);g__Metschnikowia(100);s__Metschnikowia_pulcherrima(100); | Metschnikowia | 13 | Saprotroph | \|Undefined Saprotroph\| | Yeast | NULL | Probable | As nectar-tap_saprotroph (Põlme et al. 2020); Decay sugar-rich substrates (Põlme et al. 2020) | Põlme S, et al. 2020. Fungal Diversity 105:1-16 ((https://doi.org/10.1007/s13225-020-00466-2)) |
| Otu51209 | 1 | k__Fungi(100);p__Ascomycota(100);c__Saccharomycetes(100);o__Saccharomycetales(100);f__Metschnikowiaceae(100);g__Metschnikowia(100);s__Metschnikowia_pulcherrima(100); | Metschnikowia | 13 | Saprotroph | \|Undefined Saprotroph\| | Yeast | NULL | Probable | As nectar-tap_saprotroph (Põlme et al. 2020); Decay sugar-rich substrates (Põlme et al. 2020) | Põlme S, et al. 2020. Fungal Diversity 105:1-16 ((https://doi.org/10.1007/s13225-020-00466-2)) |
| Otu51210 | 1 | k__Fungi(100);p__Ascomycota(100);c__Saccharomycetes(100);o__Saccharomycetales(100);f__Metschnikowiaceae(100);g__Metschnikowia(100);s__Metschnikowia_pulcherrima(100); | Metschnikowia | 13 | Saprotroph | \|Undefined Saprotroph\| | Yeast | NULL | Probable | As nectar-tap_saprotroph (Põlme et al. 2020); Decay sugar-rich substrates (Põlme et al. 2020) | Põlme S, et al. 2020. Fungal Diversity 105:1-16 ((https://doi.org/10.1007/s13225-020-00466-2)) |
| Otu51211 | 1 | k__Fungi(100);p__Ascomycota(100);c__Saccharomycetes(100);o__Saccharomycetales(100);f__Metschnikowiaceae(100);g__Metschnikowia(100);s__Metschnikowia_pulcherrima(100); | Metschnikowia | 13 | Saprotroph | \|Undefined Saprotroph\| | Yeast | NULL | Probable | As nectar-tap_saprotroph (Põlme et al. 2020); Decay sugar-rich substrates (Põlme et al. 2020) | Põlme S, et al. 2020. Fungal Diversity 105:1-16 ((https://doi.org/10.1007/s13225-020-00466-2)) |
| Otu51212 | 1 | k__Fungi(100);p__Ascomycota(100);c__Saccharomycetes(100);o__Saccharomycetales(100);f__Metschnikowiaceae(100);g__Metschnikowia(100);s__Metschnikowia_pulcherrima(100); | Metschnikowia | 13 | Saprotroph | \|Undefined Saprotroph\| | Yeast | NULL | Probable | As nectar-tap_saprotroph (Põlme et al. 2020); Decay sugar-rich substrates (Põlme et al. 2020) | Põlme S, et al. 2020. Fungal Diversity 105:1-16 ((https://doi.org/10.1007/s13225-020-00466-2)) |
| Otu51213 | 1 | k__Fungi(100);p__Ascomycota(100);c__Saccharomycetes(100);o__Saccharomycetales(100);f__Metschnikowiaceae(100);g__Metschnikowia(100);s__Metschnikowia_pulcherrima(100); | Metschnikowia | 13 | Saprotroph | \|Undefined Saprotroph\| | Yeast | NULL | Probable | As nectar-tap_saprotroph (Põlme et al. 2020); Decay sugar-rich substrates (Põlme et al. 2020) | Põlme S, et al. 2020. Fungal Diversity 105:1-16 ((https://doi.org/10.1007/s13225-020-00466-2)) |
| Otu51214 | 1 | k__Fungi(100);p__Ascomycota(100);c__Saccharomycetes(100);o__Saccharomycetales(100);f__Metschnikowiaceae(100);g__Metschnikowia(100);s__Metschnikowia_pulcherrima(100); | Metschnikowia | 13 | Saprotroph | \|Undefined Saprotroph\| | Yeast | NULL | Probable | As nectar-tap_saprotroph (Põlme et al. 2020); Decay sugar-rich substrates (Põlme et al. 2020) | Põlme S, et al. 2020. Fungal Diversity 105:1-16 ((https://doi.org/10.1007/s13225-020-00466-2)) |
| Otu51215 | 1 | k__Fungi(100);p__Ascomycota(100);c__Saccharomycetes(100);o__Saccharomycetales(100);f__Metschnikowiaceae(100);g__Metschnikowia(100);g__Metschnikowia_unclassified(100); | Metschnikowia | 13 | Saprotroph | \|Undefined Saprotroph\| | Yeast | NULL | Probable | As nectar-tap_saprotroph (Põlme et al. 2020); Decay sugar-rich substrates (Põlme et al. 2020) | Põlme S, et al. 2020. Fungal Diversity 105:1-16 ((https://doi.org/10.1007/s13225-020-00466-2)) |
| Otu51216 | 1 | k__Fungi(100);p__Ascomycota(100);c__Saccharomycetes(100);o__Saccharomycetales(100);f__Metschnikowiaceae(100);g__Metschnikowia(100);g__Metschnikowia_unclassified(100); | Metschnikowia | 13 | Saprotroph | \|Undefined Saprotroph\| | Yeast | NULL | Probable | As nectar-tap_saprotroph (Põlme et al. 2020); Decay sugar-rich substrates (Põlme et al. 2020) | Põlme S, et al. 2020. Fungal Diversity 105:1-16 ((https://doi.org/10.1007/s13225-020-00466-2)) |
| Otu51217 | 1 | k__Fungi(100);p__Ascomycota(100);c__Saccharomycetes(100);o__Saccharomycetales(100);f__Metschnikowiaceae(100);g__Metschnikowia(100);s__Metschnikowia_pulcherrima(100); | Metschnikowia | 13 | Saprotroph | \|Undefined Saprotroph\| | Yeast | NULL | Probable | As nectar-tap_saprotroph (Põlme et al. 2020); Decay sugar-rich substrates (Põlme et al. 2020) | Põlme S, et al. 2020. Fungal Diversity 105:1-16 ((https://doi.org/10.1007/s13225-020-00466-2)) |
| Otu51218 | 1 | k__Fungi(100);p__Ascomycota(100);c__Saccharomycetes(100);o__Saccharomycetales(100);f__Metschnikowiaceae(100);g__Metschnikowia(100);s__Metschnikowia_pulcherrima(100); | Metschnikowia | 13 | Saprotroph | \|Undefined Saprotroph\| | Yeast | NULL | Probable | As nectar-tap_saprotroph (Põlme et al. 2020); Decay sugar-rich substrates (Põlme et al. 2020) | Põlme S, et al. 2020. Fungal Diversity 105:1-16 ((https://doi.org/10.1007/s13225-020-00466-2)) |
| Otu51220 | 1 | k__Fungi(100);p__Ascomycota(100);c__Saccharomycetes(100);o__Saccharomycetales(100);f__Metschnikowiaceae(100);g__Metschnikowia(100);g__Metschnikowia_unclassified(100); | Metschnikowia | 13 | Saprotroph | \|Undefined Saprotroph\| | Yeast | NULL | Probable | As nectar-tap_saprotroph (Põlme et al. 2020); Decay sugar-rich substrates (Põlme et al. 2020) | Põlme S, et al. 2020. Fungal Diversity 105:1-16 ((https://doi.org/10.1007/s13225-020-00466-2)) |
| Otu51221 | 1 | k__Fungi(100);p__Ascomycota(100);c__Saccharomycetes(100);o__Saccharomycetales(100);f__Metschnikowiaceae(100);g__Metschnikowia(100);g__Metschnikowia_unclassified(100); | Metschnikowia | 13 | Saprotroph | \|Undefined Saprotroph\| | Yeast | NULL | Probable | As nectar-tap_saprotroph (Põlme et al. 2020); Decay sugar-rich substrates (Põlme et al. 2020) | Põlme S, et al. 2020. Fungal Diversity 105:1-16 ((https://doi.org/10.1007/s13225-020-00466-2)) |
| Otu51222 | 1 | k__Fungi(100);p__Ascomycota(100);c__Saccharomycetes(100);o__Saccharomycetales(100);f__Metschnikowiaceae(100);g__Metschnikowia(100);g__Metschnikowia_unclassified(100); | Metschnikowia | 13 | Saprotroph | \|Undefined Saprotroph\| | Yeast | NULL | Probable | As nectar-tap_saprotroph (Põlme et al. 2020); Decay sugar-rich substrates (Põlme et al. 2020) | Põlme S, et al. 2020. Fungal Diversity 105:1-16 ((https://doi.org/10.1007/s13225-020-00466-2)) |
| Otu51223 | 1 | k__Fungi(100);p__Ascomycota(100);c__Saccharomycetes(100);o__Saccharomycetales(100);f__Metschnikowiaceae(100);g__Metschnikowia(100);g__Metschnikowia_unclassified(100); | Metschnikowia | 13 | Saprotroph | \|Undefined Saprotroph\| | Yeast | NULL | Probable | As nectar-tap_saprotroph (Põlme et al. 2020); Decay sugar-rich substrates (Põlme et al. 2020) | Põlme S, et al. 2020. Fungal Diversity 105:1-16 ((https://doi.org/10.1007/s13225-020-00466-2)) |
| Otu51224 | 1 | k__Fungi(100);p__Ascomycota(100);c__Saccharomycetes(100);o__Saccharomycetales(100);f__Metschnikowiaceae(100);g__Metschnikowia(100);g__Metschnikowia_unclassified(100); | Metschnikowia | 13 | Saprotroph | \|Undefined Saprotroph\| | Yeast | NULL | Probable | As nectar-tap_saprotroph (Põlme et al. 2020); Decay sugar-rich substrates (Põlme et al. 2020) | Põlme S, et al. 2020. Fungal Diversity 105:1-16 ((https://doi.org/10.1007/s13225-020-00466-2)) |
| Otu51225 | 1 | k__Fungi(100);p__Ascomycota(100);c__Saccharomycetes(100);o__Saccharomycetales(100);f__Metschnikowiaceae(100);g__Metschnikowia(100);s__Metschnikowia_pulcherrima(100); | Metschnikowia | 13 | Saprotroph | \|Undefined Saprotroph\| | Yeast | NULL | Probable | As nectar-tap_saprotroph (Põlme et al. 2020); Decay sugar-rich substrates (Põlme et al. 2020) | Põlme S, et al. 2020. Fungal Diversity 105:1-16 ((https://doi.org/10.1007/s13225-020-00466-2)) |
| Otu51226 | 1 | k__Fungi(100);p__Ascomycota(100);c__Saccharomycetes(100);o__Saccharomycetales(100);f__Metschnikowiaceae(100);g__Metschnikowia(100);s__Metschnikowia_pulcherrima(100); | Metschnikowia | 13 | Saprotroph | \|Undefined Saprotroph\| | Yeast | NULL | Probable | As nectar-tap_saprotroph (Põlme et al. 2020); Decay sugar-rich substrates (Põlme et al. 2020) | Põlme S, et al. 2020. Fungal Diversity 105:1-16 ((https://doi.org/10.1007/s13225-020-00466-2)) |
| Otu51227 | 1 | k__Fungi(100);p__Ascomycota(100);c__Saccharomycetes(100);o__Saccharomycetales(100);f__Metschnikowiaceae(100);g__Metschnikowia(100);s__Metschnikowia_pulcherrima(100); | Metschnikowia | 13 | Saprotroph | \|Undefined Saprotroph\| | Yeast | NULL | Probable | As nectar-tap_saprotroph (Põlme et al. 2020); Decay sugar-rich substrates (Põlme et al. 2020) | Põlme S, et al. 2020. Fungal Diversity 105:1-16 ((https://doi.org/10.1007/s13225-020-00466-2)) |
| Otu51228 | 1 | k__Fungi(100);p__Ascomycota(100);c__Saccharomycetes(100);o__Saccharomycetales(100);f__Metschnikowiaceae(100);g__Metschnikowia(100);s__Metschnikowia_pulcherrima(100); | Metschnikowia | 13 | Saprotroph | \|Undefined Saprotroph\| | Yeast | NULL | Probable | As nectar-tap_saprotroph (Põlme et al. 2020); Decay sugar-rich substrates (Põlme et al. 2020) | Põlme S, et al. 2020. Fungal Diversity 105:1-16 ((https://doi.org/10.1007/s13225-020-00466-2)) |
| Otu51229 | 1 | k__Fungi(100);p__Ascomycota(100);c__Saccharomycetes(100);o__Saccharomycetales(100);f__Metschnikowiaceae(100);g__Metschnikowia(100);s__Metschnikowia_pulcherrima(100); | Metschnikowia | 13 | Saprotroph | \|Undefined Saprotroph\| | Yeast | NULL | Probable | As nectar-tap_saprotroph (Põlme et al. 2020); Decay sugar-rich substrates (Põlme et al. 2020) | Põlme S, et al. 2020. Fungal Diversity 105:1-16 ((https://doi.org/10.1007/s13225-020-00466-2)) |
| Otu51230 | 1 | k__Fungi(100);p__Ascomycota(100);c__Saccharomycetes(100);o__Saccharomycetales(100);f__Metschnikowiaceae(100);g__Metschnikowia(100);s__Metschnikowia_pulcherrima(100); | Metschnikowia | 13 | Saprotroph | \|Undefined Saprotroph\| | Yeast | NULL | Probable | As nectar-tap_saprotroph (Põlme et al. 2020); Decay sugar-rich substrates (Põlme et al. 2020) | Põlme S, et al. 2020. Fungal Diversity 105:1-16 ((https://doi.org/10.1007/s13225-020-00466-2)) |
| Otu51231 | 1 | k__Fungi(100);p__Ascomycota(100);c__Saccharomycetes(100);o__Saccharomycetales(100);f__Metschnikowiaceae(100);g__Metschnikowia(100);s__Metschnikowia_pulcherrima(100); | Metschnikowia | 13 | Saprotroph | \|Undefined Saprotroph\| | Yeast | NULL | Probable | As nectar-tap_saprotroph (Põlme et al. 2020); Decay sugar-rich substrates (Põlme et al. 2020) | Põlme S, et al. 2020. Fungal Diversity 105:1-16 ((https://doi.org/10.1007/s13225-020-00466-2)) |
| Otu51232 | 1 | k__Fungi(100);p__Ascomycota(100);c__Saccharomycetes(100);o__Saccharomycetales(100);f__Metschnikowiaceae(100);g__Metschnikowia(100);s__Metschnikowia_pulcherrima(100); | Metschnikowia | 13 | Saprotroph | \|Undefined Saprotroph\| | Yeast | NULL | Probable | As nectar-tap_saprotroph (Põlme et al. 2020); Decay sugar-rich substrates (Põlme et al. 2020) | Põlme S, et al. 2020. Fungal Diversity 105:1-16 ((https://doi.org/10.1007/s13225-020-00466-2)) |
| Otu51233 | 1 | k__Fungi(100);p__Ascomycota(100);c__Saccharomycetes(100);o__Saccharomycetales(100);f__Metschnikowiaceae(100);g__Metschnikowia(100);g__Metschnikowia_unclassified(100); | Metschnikowia | 13 | Saprotroph | \|Undefined Saprotroph\| | Yeast | NULL | Probable | As nectar-tap_saprotroph (Põlme et al. 2020); Decay sugar-rich substrates (Põlme et al. 2020) | Põlme S, et al. 2020. Fungal Diversity 105:1-16 ((https://doi.org/10.1007/s13225-020-00466-2)) |
| Otu51234 | 1 | k__Fungi(100);p__Ascomycota(100);c__Saccharomycetes(100);o__Saccharomycetales(100);f__Metschnikowiaceae(100);g__Metschnikowia(100);g__Metschnikowia_unclassified(100); | Metschnikowia | 13 | Saprotroph | \|Undefined Saprotroph\| | Yeast | NULL | Probable | As nectar-tap_saprotroph (Põlme et al. 2020); Decay sugar-rich substrates (Põlme et al. 2020) | Põlme S, et al. 2020. Fungal Diversity 105:1-16 ((https://doi.org/10.1007/s13225-020-00466-2)) |
| Otu51235 | 1 | k__Fungi(100);p__Ascomycota(100);c__Saccharomycetes(100);o__Saccharomycetales(100);f__Metschnikowiaceae(100);g__Metschnikowia(100);s__Metschnikowia_pulcherrima(100); | Metschnikowia | 13 | Saprotroph | \|Undefined Saprotroph\| | Yeast | NULL | Probable | As nectar-tap_saprotroph (Põlme et al. 2020); Decay sugar-rich substrates (Põlme et al. 2020) | Põlme S, et al. 2020. Fungal Diversity 105:1-16 ((https://doi.org/10.1007/s13225-020-00466-2)) |
| Otu51237 | 1 | k__Fungi(100);p__Ascomycota(100);c__Saccharomycetes(100);o__Saccharomycetales(100);f__Metschnikowiaceae(100);g__Metschnikowia(100);g__Metschnikowia_unclassified(100); | Metschnikowia | 13 | Saprotroph | \|Undefined Saprotroph\| | Yeast | NULL | Probable | As nectar-tap_saprotroph (Põlme et al. 2020); Decay sugar-rich substrates (Põlme et al. 2020) | Põlme S, et al. 2020. Fungal Diversity 105:1-16 ((https://doi.org/10.1007/s13225-020-00466-2)) |
| Otu51238 | 1 | k__Fungi(100);p__Ascomycota(100);c__Saccharomycetes(100);o__Saccharomycetales(100);f__Metschnikowiaceae(100);g__Metschnikowia(100);g__Metschnikowia_unclassified(100); | Metschnikowia | 13 | Saprotroph | \|Undefined Saprotroph\| | Yeast | NULL | Probable | As nectar-tap_saprotroph (Põlme et al. 2020); Decay sugar-rich substrates (Põlme et al. 2020) | Põlme S, et al. 2020. Fungal Diversity 105:1-16 ((https://doi.org/10.1007/s13225-020-00466-2)) |
| Otu51239 | 1 | k__Fungi(100);p__Ascomycota(100);c__Saccharomycetes(100);o__Saccharomycetales(100);f__Metschnikowiaceae(100);g__Metschnikowia(100);g__Metschnikowia_unclassified(100); | Metschnikowia | 13 | Saprotroph | \|Undefined Saprotroph\| | Yeast | NULL | Probable | As nectar-tap_saprotroph (Põlme et al. 2020); Decay sugar-rich substrates (Põlme et al. 2020) | Põlme S, et al. 2020. Fungal Diversity 105:1-16 ((https://doi.org/10.1007/s13225-020-00466-2)) |
| Otu51240 | 1 | k__Fungi(100);p__Ascomycota(100);c__Saccharomycetes(100);o__Saccharomycetales(100);f__Metschnikowiaceae(100);g__Metschnikowia(100);s__Metschnikowia_pulcherrima(100); | Metschnikowia | 13 | Saprotroph | \|Undefined Saprotroph\| | Yeast | NULL | Probable | As nectar-tap_saprotroph (Põlme et al. 2020); Decay sugar-rich substrates (Põlme et al. 2020) | Põlme S, et al. 2020. Fungal Diversity 105:1-16 ((https://doi.org/10.1007/s13225-020-00466-2)) |
| Otu51241 | 1 | k__Fungi(100);p__Ascomycota(100);c__Saccharomycetes(100);o__Saccharomycetales(100);f__Metschnikowiaceae(100);g__Metschnikowia(100);s__Metschnikowia_pulcherrima(100); | Metschnikowia | 13 | Saprotroph | \|Undefined Saprotroph\| | Yeast | NULL | Probable | As nectar-tap_saprotroph (Põlme et al. 2020); Decay sugar-rich substrates (Põlme et al. 2020) | Põlme S, et al. 2020. Fungal Diversity 105:1-16 ((https://doi.org/10.1007/s13225-020-00466-2)) |
| Otu51242 | 1 | k__Fungi(100);p__Ascomycota(100);c__Saccharomycetes(100);o__Saccharomycetales(100);f__Metschnikowiaceae(100);g__Metschnikowia(100);s__Metschnikowia_pulcherrima(100); | Metschnikowia | 13 | Saprotroph | \|Undefined Saprotroph\| | Yeast | NULL | Probable | As nectar-tap_saprotroph (Põlme et al. 2020); Decay sugar-rich substrates (Põlme et al. 2020) | Põlme S, et al. 2020. Fungal Diversity 105:1-16 ((https://doi.org/10.1007/s13225-020-00466-2)) |
| Otu51243 | 1 | k__Fungi(100);p__Ascomycota(100);c__Saccharomycetes(100);o__Saccharomycetales(100);f__Metschnikowiaceae(100);g__Metschnikowia(100);s__Metschnikowia_pulcherrima(100); | Metschnikowia | 13 | Saprotroph | \|Undefined Saprotroph\| | Yeast | NULL | Probable | As nectar-tap_saprotroph (Põlme et al. 2020); Decay sugar-rich substrates (Põlme et al. 2020) | Põlme S, et al. 2020. Fungal Diversity 105:1-16 ((https://doi.org/10.1007/s13225-020-00466-2)) |
| Otu51244 | 1 | k__Fungi(100);p__Ascomycota(100);c__Saccharomycetes(100);o__Saccharomycetales(100);f__Metschnikowiaceae(100);g__Metschnikowia(100);s__Metschnikowia_pulcherrima(100); | Metschnikowia | 13 | Saprotroph | \|Undefined Saprotroph\| | Yeast | NULL | Probable | As nectar-tap_saprotroph (Põlme et al. 2020); Decay sugar-rich substrates (Põlme et al. 2020) | Põlme S, et al. 2020. Fungal Diversity 105:1-16 ((https://doi.org/10.1007/s13225-020-00466-2)) |
| Otu51245 | 1 | k__Fungi(100);p__Ascomycota(100);c__Saccharomycetes(100);o__Saccharomycetales(100);f__Metschnikowiaceae(100);g__Metschnikowia(100);s__Metschnikowia_pulcherrima(100); | Metschnikowia | 13 | Saprotroph | \|Undefined Saprotroph\| | Yeast | NULL | Probable | As nectar-tap_saprotroph (Põlme et al. 2020); Decay sugar-rich substrates (Põlme et al. 2020) | Põlme S, et al. 2020. Fungal Diversity 105:1-16 ((https://doi.org/10.1007/s13225-020-00466-2)) |
| Otu51246 | 1 | k__Fungi(100);p__Ascomycota(100);c__Saccharomycetes(100);o__Saccharomycetales(100);f__Metschnikowiaceae(100);g__Metschnikowia(100);s__Metschnikowia_pulcherrima(100); | Metschnikowia | 13 | Saprotroph | \|Undefined Saprotroph\| | Yeast | NULL | Probable | As nectar-tap_saprotroph (Põlme et al. 2020); Decay sugar-rich substrates (Põlme et al. 2020) | Põlme S, et al. 2020. Fungal Diversity 105:1-16 ((https://doi.org/10.1007/s13225-020-00466-2)) |
| Otu51247 | 1 | k__Fungi(100);p__Ascomycota(100);c__Saccharomycetes(100);o__Saccharomycetales(100);f__Metschnikowiaceae(100);g__Metschnikowia(100);s__Metschnikowia_pulcherrima(100); | Metschnikowia | 13 | Saprotroph | \|Undefined Saprotroph\| | Yeast | NULL | Probable | As nectar-tap_saprotroph (Põlme et al. 2020); Decay sugar-rich substrates (Põlme et al. 2020) | Põlme S, et al. 2020. Fungal Diversity 105:1-16 ((https://doi.org/10.1007/s13225-020-00466-2)) |
| Otu51248 | 1 | k__Fungi(100);p__Ascomycota(100);c__Saccharomycetes(100);o__Saccharomycetales(100);f__Metschnikowiaceae(100);g__Metschnikowia(100);g__Metschnikowia_unclassified(100); | Metschnikowia | 13 | Saprotroph | \|Undefined Saprotroph\| | Yeast | NULL | Probable | As nectar-tap_saprotroph (Põlme et al. 2020); Decay sugar-rich substrates (Põlme et al. 2020) | Põlme S, et al. 2020. Fungal Diversity 105:1-16 ((https://doi.org/10.1007/s13225-020-00466-2)) |
| Otu51249 | 1 | k__Fungi(100);p__Ascomycota(100);c__Saccharomycetes(100);o__Saccharomycetales(100);f__Metschnikowiaceae(100);g__Metschnikowia(100);s__Metschnikowia_pulcherrima(100); | Metschnikowia | 13 | Saprotroph | \|Undefined Saprotroph\| | Yeast | NULL | Probable | As nectar-tap_saprotroph (Põlme et al. 2020); Decay sugar-rich substrates (Põlme et al. 2020) | Põlme S, et al. 2020. Fungal Diversity 105:1-16 ((https://doi.org/10.1007/s13225-020-00466-2)) |
| Otu51251 | 1 | k__Fungi(100);p__Ascomycota(100);c__Saccharomycetes(100);o__Saccharomycetales(100);f__Metschnikowiaceae(100);g__Metschnikowia(100);s__Metschnikowia_pulcherrima(100); | Metschnikowia | 13 | Saprotroph | \|Undefined Saprotroph\| | Yeast | NULL | Probable | As nectar-tap_saprotroph (Põlme et al. 2020); Decay sugar-rich substrates (Põlme et al. 2020) | Põlme S, et al. 2020. Fungal Diversity 105:1-16 ((https://doi.org/10.1007/s13225-020-00466-2)) |
| Otu51252 | 1 | k__Fungi(100);p__Ascomycota(100);c__Saccharomycetes(100);o__Saccharomycetales(100);f__Metschnikowiaceae(100);g__Metschnikowia(100);g__Metschnikowia_unclassified(100); | Metschnikowia | 13 | Saprotroph | \|Undefined Saprotroph\| | Yeast | NULL | Probable | As nectar-tap_saprotroph (Põlme et al. 2020); Decay sugar-rich substrates (Põlme et al. 2020) | Põlme S, et al. 2020. Fungal Diversity 105:1-16 ((https://doi.org/10.1007/s13225-020-00466-2)) |
| Otu51253 | 1 | k__Fungi(100);p__Ascomycota(100);c__Saccharomycetes(100);o__Saccharomycetales(100);f__Metschnikowiaceae(100);g__Metschnikowia(100);g__Metschnikowia_unclassified(100); | Metschnikowia | 13 | Saprotroph | \|Undefined Saprotroph\| | Yeast | NULL | Probable | As nectar-tap_saprotroph (Põlme et al. 2020); Decay sugar-rich substrates (Põlme et al. 2020) | Põlme S, et al. 2020. Fungal Diversity 105:1-16 ((https://doi.org/10.1007/s13225-020-00466-2)) |
| Otu51254 | 1 | k__Fungi(100);p__Ascomycota(100);c__Saccharomycetes(100);o__Saccharomycetales(100);f__Metschnikowiaceae(100);g__Metschnikowia(100);s__Metschnikowia_pulcherrima(100); | Metschnikowia | 13 | Saprotroph | \|Undefined Saprotroph\| | Yeast | NULL | Probable | As nectar-tap_saprotroph (Põlme et al. 2020); Decay sugar-rich substrates (Põlme et al. 2020) | Põlme S, et al. 2020. Fungal Diversity 105:1-16 ((https://doi.org/10.1007/s13225-020-00466-2)) |
| Otu51255 | 1 | k__Fungi(100);p__Ascomycota(100);c__Saccharomycetes(100);o__Saccharomycetales(100);f__Metschnikowiaceae(100);g__Metschnikowia(100);s__Metschnikowia_pulcherrima(100); | Metschnikowia | 13 | Saprotroph | \|Undefined Saprotroph\| | Yeast | NULL | Probable | As nectar-tap_saprotroph (Põlme et al. 2020); Decay sugar-rich substrates (Põlme et al. 2020) | Põlme S, et al. 2020. Fungal Diversity 105:1-16 ((https://doi.org/10.1007/s13225-020-00466-2)) |
| Otu51256 | 1 | k__Fungi(100);p__Ascomycota(100);c__Saccharomycetes(100);o__Saccharomycetales(100);f__Metschnikowiaceae(100);g__Metschnikowia(100);s__Metschnikowia_pulcherrima(100); | Metschnikowia | 13 | Saprotroph | \|Undefined Saprotroph\| | Yeast | NULL | Probable | As nectar-tap_saprotroph (Põlme et al. 2020); Decay sugar-rich substrates (Põlme et al. 2020) | Põlme S, et al. 2020. Fungal Diversity 105:1-16 ((https://doi.org/10.1007/s13225-020-00466-2)) |
| Otu51257 | 1 | k__Fungi(100);p__Ascomycota(100);c__Saccharomycetes(100);o__Saccharomycetales(100);f__Metschnikowiaceae(100);g__Metschnikowia(100);g__Metschnikowia_unclassified(100); | Metschnikowia | 13 | Saprotroph | \|Undefined Saprotroph\| | Yeast | NULL | Probable | As nectar-tap_saprotroph (Põlme et al. 2020); Decay sugar-rich substrates (Põlme et al. 2020) | Põlme S, et al. 2020. Fungal Diversity 105:1-16 ((https://doi.org/10.1007/s13225-020-00466-2)) |
| Otu51258 | 1 | k__Fungi(100);p__Ascomycota(100);c__Saccharomycetes(100);o__Saccharomycetales(100);f__Metschnikowiaceae(100);g__Metschnikowia(100);g__Metschnikowia_unclassified(100); | Metschnikowia | 13 | Saprotroph | \|Undefined Saprotroph\| | Yeast | NULL | Probable | As nectar-tap_saprotroph (Põlme et al. 2020); Decay sugar-rich substrates (Põlme et al. 2020) | Põlme S, et al. 2020. Fungal Diversity 105:1-16 ((https://doi.org/10.1007/s13225-020-00466-2)) |
| Otu51259 | 1 | k__Fungi(100);p__Ascomycota(100);c__Saccharomycetes(100);o__Saccharomycetales(100);f__Metschnikowiaceae(100);g__Metschnikowia(100);s__Metschnikowia_pulcherrima(100); | Metschnikowia | 13 | Saprotroph | \|Undefined Saprotroph\| | Yeast | NULL | Probable | As nectar-tap_saprotroph (Põlme et al. 2020); Decay sugar-rich substrates (Põlme et al. 2020) | Põlme S, et al. 2020. Fungal Diversity 105:1-16 ((https://doi.org/10.1007/s13225-020-00466-2)) |
| Otu51260 | 1 | k__Fungi(100);p__Ascomycota(100);c__Saccharomycetes(100);o__Saccharomycetales(100);f__Metschnikowiaceae(100);g__Metschnikowia(100);g__Metschnikowia_unclassified(100); | Metschnikowia | 13 | Saprotroph | \|Undefined Saprotroph\| | Yeast | NULL | Probable | As nectar-tap_saprotroph (Põlme et al. 2020); Decay sugar-rich substrates (Põlme et al. 2020) | Põlme S, et al. 2020. Fungal Diversity 105:1-16 ((https://doi.org/10.1007/s13225-020-00466-2)) |
| Otu51261 | 1 | k__Fungi(100);p__Ascomycota(100);c__Saccharomycetes(100);o__Saccharomycetales(100);f__Metschnikowiaceae(100);g__Metschnikowia(100);g__Metschnikowia_unclassified(100); | Metschnikowia | 13 | Saprotroph | \|Undefined Saprotroph\| | Yeast | NULL | Probable | As nectar-tap_saprotroph (Põlme et al. 2020); Decay sugar-rich substrates (Põlme et al. 2020) | Põlme S, et al. 2020. Fungal Diversity 105:1-16 ((https://doi.org/10.1007/s13225-020-00466-2)) |
| Otu51262 | 1 | k__Fungi(100);p__Ascomycota(100);c__Saccharomycetes(100);o__Saccharomycetales(100);f__Metschnikowiaceae(100);g__Metschnikowia(100);s__Metschnikowia_pulcherrima(100); | Metschnikowia | 13 | Saprotroph | \|Undefined Saprotroph\| | Yeast | NULL | Probable | As nectar-tap_saprotroph (Põlme et al. 2020); Decay sugar-rich substrates (Põlme et al. 2020) | Põlme S, et al. 2020. Fungal Diversity 105:1-16 ((https://doi.org/10.1007/s13225-020-00466-2)) |
| Otu51263 | 1 | k__Fungi(100);p__Ascomycota(100);c__Saccharomycetes(100);o__Saccharomycetales(100);f__Metschnikowiaceae(100);g__Metschnikowia(100);s__Metschnikowia_pulcherrima(100); | Metschnikowia | 13 | Saprotroph | \|Undefined Saprotroph\| | Yeast | NULL | Probable | As nectar-tap_saprotroph (Põlme et al. 2020); Decay sugar-rich substrates (Põlme et al. 2020) | Põlme S, et al. 2020. Fungal Diversity 105:1-16 ((https://doi.org/10.1007/s13225-020-00466-2)) |
| Otu51264 | 1 | k__Fungi(100);p__Ascomycota(100);c__Saccharomycetes(100);o__Saccharomycetales(100);f__Metschnikowiaceae(100);g__Metschnikowia(100);s__Metschnikowia_pulcherrima(100); | Metschnikowia | 13 | Saprotroph | \|Undefined Saprotroph\| | Yeast | NULL | Probable | As nectar-tap_saprotroph (Põlme et al. 2020); Decay sugar-rich substrates (Põlme et al. 2020) | Põlme S, et al. 2020. Fungal Diversity 105:1-16 ((https://doi.org/10.1007/s13225-020-00466-2)) |
| Otu51265 | 1 | k__Fungi(100);p__Ascomycota(100);c__Saccharomycetes(100);o__Saccharomycetales(100);f__Metschnikowiaceae(100);g__Metschnikowia(100);s__Metschnikowia_pulcherrima(100); | Metschnikowia | 13 | Saprotroph | \|Undefined Saprotroph\| | Yeast | NULL | Probable | As nectar-tap_saprotroph (Põlme et al. 2020); Decay sugar-rich substrates (Põlme et al. 2020) | Põlme S, et al. 2020. Fungal Diversity 105:1-16 ((https://doi.org/10.1007/s13225-020-00466-2)) |
| Otu51266 | 1 | k__Fungi(100);p__Ascomycota(100);c__Saccharomycetes(100);o__Saccharomycetales(100);f__Metschnikowiaceae(100);g__Metschnikowia(100);s__Metschnikowia_pulcherrima(100); | Metschnikowia | 13 | Saprotroph | \|Undefined Saprotroph\| | Yeast | NULL | Probable | As nectar-tap_saprotroph (Põlme et al. 2020); Decay sugar-rich substrates (Põlme et al. 2020) | Põlme S, et al. 2020. Fungal Diversity 105:1-16 ((https://doi.org/10.1007/s13225-020-00466-2)) |
| Otu51267 | 1 | k__Fungi(100);p__Ascomycota(100);c__Saccharomycetes(100);o__Saccharomycetales(100);f__Metschnikowiaceae(100);g__Metschnikowia(100);s__Metschnikowia_pulcherrima(100); | Metschnikowia | 13 | Saprotroph | \|Undefined Saprotroph\| | Yeast | NULL | Probable | As nectar-tap_saprotroph (Põlme et al. 2020); Decay sugar-rich substrates (Põlme et al. 2020) | Põlme S, et al. 2020. Fungal Diversity 105:1-16 ((https://doi.org/10.1007/s13225-020-00466-2)) |
| Otu51268 | 1 | k__Fungi(100);p__Ascomycota(100);c__Saccharomycetes(100);o__Saccharomycetales(100);f__Metschnikowiaceae(100);g__Metschnikowia(100);s__Metschnikowia_pulcherrima(100); | Metschnikowia | 13 | Saprotroph | \|Undefined Saprotroph\| | Yeast | NULL | Probable | As nectar-tap_saprotroph (Põlme et al. 2020); Decay sugar-rich substrates (Põlme et al. 2020) | Põlme S, et al. 2020. Fungal Diversity 105:1-16 ((https://doi.org/10.1007/s13225-020-00466-2)) |
| Otu51270 | 1 | k__Fungi(100);p__Ascomycota(100);c__Saccharomycetes(100);o__Saccharomycetales(100);f__Metschnikowiaceae(100);g__Metschnikowia(100);s__Metschnikowia_pulcherrima(100); | Metschnikowia | 13 | Saprotroph | \|Undefined Saprotroph\| | Yeast | NULL | Probable | As nectar-tap_saprotroph (Põlme et al. 2020); Decay sugar-rich substrates (Põlme et al. 2020) | Põlme S, et al. 2020. Fungal Diversity 105:1-16 ((https://doi.org/10.1007/s13225-020-00466-2)) |
| Otu51271 | 1 | k__Fungi(100);p__Ascomycota(100);c__Saccharomycetes(100);o__Saccharomycetales(100);f__Metschnikowiaceae(100);g__Metschnikowia(100);s__Metschnikowia_pulcherrima(100); | Metschnikowia | 13 | Saprotroph | \|Undefined Saprotroph\| | Yeast | NULL | Probable | As nectar-tap_saprotroph (Põlme et al. 2020); Decay sugar-rich substrates (Põlme et al. 2020) | Põlme S, et al. 2020. Fungal Diversity 105:1-16 ((https://doi.org/10.1007/s13225-020-00466-2)) |
| Otu51272 | 1 | k__Fungi(100);p__Ascomycota(100);c__Saccharomycetes(100);o__Saccharomycetales(100);f__Metschnikowiaceae(100);g__Metschnikowia(100);s__Metschnikowia_pulcherrima(100); | Metschnikowia | 13 | Saprotroph | \|Undefined Saprotroph\| | Yeast | NULL | Probable | As nectar-tap_saprotroph (Põlme et al. 2020); Decay sugar-rich substrates (Põlme et al. 2020) | Põlme S, et al. 2020. Fungal Diversity 105:1-16 ((https://doi.org/10.1007/s13225-020-00466-2)) |
| Otu51273 | 1 | k__Fungi(100);p__Ascomycota(100);c__Saccharomycetes(100);o__Saccharomycetales(100);f__Metschnikowiaceae(100);g__Metschnikowia(100);s__Metschnikowia_pulcherrima(100); | Metschnikowia | 13 | Saprotroph | \|Undefined Saprotroph\| | Yeast | NULL | Probable | As nectar-tap_saprotroph (Põlme et al. 2020); Decay sugar-rich substrates (Põlme et al. 2020) | Põlme S, et al. 2020. Fungal Diversity 105:1-16 ((https://doi.org/10.1007/s13225-020-00466-2)) |
| Otu51274 | 1 | k__Fungi(100);p__Ascomycota(100);c__Saccharomycetes(100);o__Saccharomycetales(100);f__Metschnikowiaceae(100);g__Metschnikowia(100);s__Metschnikowia_pulcherrima(100); | Metschnikowia | 13 | Saprotroph | \|Undefined Saprotroph\| | Yeast | NULL | Probable | As nectar-tap_saprotroph (Põlme et al. 2020); Decay sugar-rich substrates (Põlme et al. 2020) | Põlme S, et al. 2020. Fungal Diversity 105:1-16 ((https://doi.org/10.1007/s13225-020-00466-2)) |
| Otu51275 | 1 | k__Fungi(100);p__Ascomycota(100);c__Saccharomycetes(100);o__Saccharomycetales(100);f__Metschnikowiaceae(100);g__Metschnikowia(100);g__Metschnikowia_unclassified(100); | Metschnikowia | 13 | Saprotroph | \|Undefined Saprotroph\| | Yeast | NULL | Probable | As nectar-tap_saprotroph (Põlme et al. 2020); Decay sugar-rich substrates (Põlme et al. 2020) | Põlme S, et al. 2020. Fungal Diversity 105:1-16 ((https://doi.org/10.1007/s13225-020-00466-2)) |
| Otu51276 | 1 | k__Fungi(100);p__Ascomycota(100);c__Saccharomycetes(100);o__Saccharomycetales(100);f__Metschnikowiaceae(100);g__Metschnikowia(100);s__Metschnikowia_pulcherrima(100); | Metschnikowia | 13 | Saprotroph | \|Undefined Saprotroph\| | Yeast | NULL | Probable | As nectar-tap_saprotroph (Põlme et al. 2020); Decay sugar-rich substrates (Põlme et al. 2020) | Põlme S, et al. 2020. Fungal Diversity 105:1-16 ((https://doi.org/10.1007/s13225-020-00466-2)) |
| Otu51277 | 1 | k__Fungi(100);p__Ascomycota(100);c__Saccharomycetes(100);o__Saccharomycetales(100);f__Metschnikowiaceae(100);g__Metschnikowia(100);s__Metschnikowia_pulcherrima(100); | Metschnikowia | 13 | Saprotroph | \|Undefined Saprotroph\| | Yeast | NULL | Probable | As nectar-tap_saprotroph (Põlme et al. 2020); Decay sugar-rich substrates (Põlme et al. 2020) | Põlme S, et al. 2020. Fungal Diversity 105:1-16 ((https://doi.org/10.1007/s13225-020-00466-2)) |
| Otu51278 | 1 | k__Fungi(100);p__Ascomycota(100);c__Saccharomycetes(100);o__Saccharomycetales(100);f__Metschnikowiaceae(100);g__Metschnikowia(100);s__Metschnikowia_pulcherrima(100); | Metschnikowia | 13 | Saprotroph | \|Undefined Saprotroph\| | Yeast | NULL | Probable | As nectar-tap_saprotroph (Põlme et al. 2020); Decay sugar-rich substrates (Põlme et al. 2020) | Põlme S, et al. 2020. Fungal Diversity 105:1-16 ((https://doi.org/10.1007/s13225-020-00466-2)) |
| Otu51279 | 1 | k__Fungi(100);p__Ascomycota(100);c__Saccharomycetes(100);o__Saccharomycetales(100);f__Metschnikowiaceae(100);g__Metschnikowia(100);s__Metschnikowia_pulcherrima(100); | Metschnikowia | 13 | Saprotroph | \|Undefined Saprotroph\| | Yeast | NULL | Probable | As nectar-tap_saprotroph (Põlme et al. 2020); Decay sugar-rich substrates (Põlme et al. 2020) | Põlme S, et al. 2020. Fungal Diversity 105:1-16 ((https://doi.org/10.1007/s13225-020-00466-2)) |
| Otu51280 | 1 | k__Fungi(100);p__Ascomycota(100);c__Saccharomycetes(100);o__Saccharomycetales(100);f__Metschnikowiaceae(100);g__Metschnikowia(100);s__Metschnikowia_pulcherrima(100); | Metschnikowia | 13 | Saprotroph | \|Undefined Saprotroph\| | Yeast | NULL | Probable | As nectar-tap_saprotroph (Põlme et al. 2020); Decay sugar-rich substrates (Põlme et al. 2020) | Põlme S, et al. 2020. Fungal Diversity 105:1-16 ((https://doi.org/10.1007/s13225-020-00466-2)) |
| Otu51282 | 1 | k__Fungi(100);p__Ascomycota(100);c__Saccharomycetes(100);o__Saccharomycetales(100);f__Metschnikowiaceae(100);g__Metschnikowia(100);s__Metschnikowia_pulcherrima(100); | Metschnikowia | 13 | Saprotroph | \|Undefined Saprotroph\| | Yeast | NULL | Probable | As nectar-tap_saprotroph (Põlme et al. 2020); Decay sugar-rich substrates (Põlme et al. 2020) | Põlme S, et al. 2020. Fungal Diversity 105:1-16 ((https://doi.org/10.1007/s13225-020-00466-2)) |
| Otu51286 | 1 | k__Fungi(100);p__Ascomycota(100);c__Saccharomycetes(100);o__Saccharomycetales(100);f__Metschnikowiaceae(100);g__Metschnikowia(100);s__Metschnikowia_pulcherrima(100); | Metschnikowia | 13 | Saprotroph | \|Undefined Saprotroph\| | Yeast | NULL | Probable | As nectar-tap_saprotroph (Põlme et al. 2020); Decay sugar-rich substrates (Põlme et al. 2020) | Põlme S, et al. 2020. Fungal Diversity 105:1-16 ((https://doi.org/10.1007/s13225-020-00466-2)) |
| Otu51287 | 1 | k__Fungi(100);p__Ascomycota(100);c__Saccharomycetes(100);o__Saccharomycetales(100);f__Metschnikowiaceae(100);g__Metschnikowia(100);s__Metschnikowia_pulcherrima(100); | Metschnikowia | 13 | Saprotroph | \|Undefined Saprotroph\| | Yeast | NULL | Probable | As nectar-tap_saprotroph (Põlme et al. 2020); Decay sugar-rich substrates (Põlme et al. 2020) | Põlme S, et al. 2020. Fungal Diversity 105:1-16 ((https://doi.org/10.1007/s13225-020-00466-2)) |
| Otu51288 | 1 | k__Fungi(100);p__Ascomycota(100);c__Saccharomycetes(100);o__Saccharomycetales(100);f__Metschnikowiaceae(100);g__Metschnikowia(100);s__Metschnikowia_pulcherrima(100); | Metschnikowia | 13 | Saprotroph | \|Undefined Saprotroph\| | Yeast | NULL | Probable | As nectar-tap_saprotroph (Põlme et al. 2020); Decay sugar-rich substrates (Põlme et al. 2020) | Põlme S, et al. 2020. Fungal Diversity 105:1-16 ((https://doi.org/10.1007/s13225-020-00466-2)) |
| Otu51290 | 1 | k__Fungi(100);p__Ascomycota(100);c__Saccharomycetes(100);o__Saccharomycetales(100);f__Metschnikowiaceae(100);g__Metschnikowia(100);s__Metschnikowia_pulcherrima(100); | Metschnikowia | 13 | Saprotroph | \|Undefined Saprotroph\| | Yeast | NULL | Probable | As nectar-tap_saprotroph (Põlme et al. 2020); Decay sugar-rich substrates (Põlme et al. 2020) | Põlme S, et al. 2020. Fungal Diversity 105:1-16 ((https://doi.org/10.1007/s13225-020-00466-2)) |
| Otu51292 | 1 | k__Fungi(100);p__Ascomycota(100);c__Saccharomycetes(100);o__Saccharomycetales(100);f__Metschnikowiaceae(100);g__Metschnikowia(100);s__Metschnikowia_pulcherrima(100); | Metschnikowia | 13 | Saprotroph | \|Undefined Saprotroph\| | Yeast | NULL | Probable | As nectar-tap_saprotroph (Põlme et al. 2020); Decay sugar-rich substrates (Põlme et al. 2020) | Põlme S, et al. 2020. Fungal Diversity 105:1-16 ((https://doi.org/10.1007/s13225-020-00466-2)) |
| Otu51293 | 1 | k__Fungi(100);p__Ascomycota(100);c__Saccharomycetes(100);o__Saccharomycetales(100);f__Metschnikowiaceae(100);g__Metschnikowia(100);s__Metschnikowia_pulcherrima(100); | Metschnikowia | 13 | Saprotroph | \|Undefined Saprotroph\| | Yeast | NULL | Probable | As nectar-tap_saprotroph (Põlme et al. 2020); Decay sugar-rich substrates (Põlme et al. 2020) | Põlme S, et al. 2020. Fungal Diversity 105:1-16 ((https://doi.org/10.1007/s13225-020-00466-2)) |
| Otu51294 | 1 | k__Fungi(100);p__Ascomycota(100);c__Saccharomycetes(100);o__Saccharomycetales(100);f__Metschnikowiaceae(100);g__Metschnikowia(100);s__Metschnikowia_pulcherrima(100); | Metschnikowia | 13 | Saprotroph | \|Undefined Saprotroph\| | Yeast | NULL | Probable | As nectar-tap_saprotroph (Põlme et al. 2020); Decay sugar-rich substrates (Põlme et al. 2020) | Põlme S, et al. 2020. Fungal Diversity 105:1-16 ((https://doi.org/10.1007/s13225-020-00466-2)) |
| Otu51295 | 1 | k__Fungi(100);p__Ascomycota(100);c__Saccharomycetes(100);o__Saccharomycetales(100);f__Metschnikowiaceae(100);g__Metschnikowia(100);s__Metschnikowia_pulcherrima(100); | Metschnikowia | 13 | Saprotroph | \|Undefined Saprotroph\| | Yeast | NULL | Probable | As nectar-tap_saprotroph (Põlme et al. 2020); Decay sugar-rich substrates (Põlme et al. 2020) | Põlme S, et al. 2020. Fungal Diversity 105:1-16 ((https://doi.org/10.1007/s13225-020-00466-2)) |
| Otu51296 | 1 | k__Fungi(100);p__Ascomycota(100);c__Saccharomycetes(100);o__Saccharomycetales(100);f__Metschnikowiaceae(100);g__Metschnikowia(100);s__Metschnikowia_pulcherrima(100); | Metschnikowia | 13 | Saprotroph | \|Undefined Saprotroph\| | Yeast | NULL | Probable | As nectar-tap_saprotroph (Põlme et al. 2020); Decay sugar-rich substrates (Põlme et al. 2020) | Põlme S, et al. 2020. Fungal Diversity 105:1-16 ((https://doi.org/10.1007/s13225-020-00466-2)) |
| Otu51297 | 1 | k__Fungi(100);p__Ascomycota(100);c__Saccharomycetes(100);o__Saccharomycetales(100);f__Metschnikowiaceae(100);g__Metschnikowia(100);s__Metschnikowia_pulcherrima(100); | Metschnikowia | 13 | Saprotroph | \|Undefined Saprotroph\| | Yeast | NULL | Probable | As nectar-tap_saprotroph (Põlme et al. 2020); Decay sugar-rich substrates (Põlme et al. 2020) | Põlme S, et al. 2020. Fungal Diversity 105:1-16 ((https://doi.org/10.1007/s13225-020-00466-2)) |
| Otu51298 | 1 | k__Fungi(100);p__Ascomycota(100);c__Saccharomycetes(100);o__Saccharomycetales(100);f__Metschnikowiaceae(100);g__Metschnikowia(100);g__Metschnikowia_unclassified(100); | Metschnikowia | 13 | Saprotroph | \|Undefined Saprotroph\| | Yeast | NULL | Probable | As nectar-tap_saprotroph (Põlme et al. 2020); Decay sugar-rich substrates (Põlme et al. 2020) | Põlme S, et al. 2020. Fungal Diversity 105:1-16 ((https://doi.org/10.1007/s13225-020-00466-2)) |
| Otu51299 | 1 | k__Fungi(100);p__Ascomycota(100);c__Saccharomycetes(100);o__Saccharomycetales(100);f__Metschnikowiaceae(100);g__Metschnikowia(100);s__Metschnikowia_pulcherrima(100); | Metschnikowia | 13 | Saprotroph | \|Undefined Saprotroph\| | Yeast | NULL | Probable | As nectar-tap_saprotroph (Põlme et al. 2020); Decay sugar-rich substrates (Põlme et al. 2020) | Põlme S, et al. 2020. Fungal Diversity 105:1-16 ((https://doi.org/10.1007/s13225-020-00466-2)) |
| Otu51300 | 1 | k__Fungi(100);p__Ascomycota(100);c__Saccharomycetes(100);o__Saccharomycetales(100);f__Metschnikowiaceae(100);g__Metschnikowia(100);s__Metschnikowia_pulcherrima(100); | Metschnikowia | 13 | Saprotroph | \|Undefined Saprotroph\| | Yeast | NULL | Probable | As nectar-tap_saprotroph (Põlme et al. 2020); Decay sugar-rich substrates (Põlme et al. 2020) | Põlme S, et al. 2020. Fungal Diversity 105:1-16 ((https://doi.org/10.1007/s13225-020-00466-2)) |
| Otu51301 | 1 | k__Fungi(100);p__Ascomycota(100);c__Saccharomycetes(100);o__Saccharomycetales(100);f__Metschnikowiaceae(100);g__Metschnikowia(100);s__Metschnikowia_pulcherrima(100); | Metschnikowia | 13 | Saprotroph | \|Undefined Saprotroph\| | Yeast | NULL | Probable | As nectar-tap_saprotroph (Põlme et al. 2020); Decay sugar-rich substrates (Põlme et al. 2020) | Põlme S, et al. 2020. Fungal Diversity 105:1-16 ((https://doi.org/10.1007/s13225-020-00466-2)) |
| Otu51302 | 1 | k__Fungi(100);p__Ascomycota(100);c__Saccharomycetes(100);o__Saccharomycetales(100);f__Metschnikowiaceae(100);g__Metschnikowia(100);s__Metschnikowia_pulcherrima(100); | Metschnikowia | 13 | Saprotroph | \|Undefined Saprotroph\| | Yeast | NULL | Probable | As nectar-tap_saprotroph (Põlme et al. 2020); Decay sugar-rich substrates (Põlme et al. 2020) | Põlme S, et al. 2020. Fungal Diversity 105:1-16 ((https://doi.org/10.1007/s13225-020-00466-2)) |
| Otu51303 | 1 | k__Fungi(100);p__Ascomycota(100);c__Saccharomycetes(100);o__Saccharomycetales(100);f__Metschnikowiaceae(100);g__Metschnikowia(100);g__Metschnikowia_unclassified(100); | Metschnikowia | 13 | Saprotroph | \|Undefined Saprotroph\| | Yeast | NULL | Probable | As nectar-tap_saprotroph (Põlme et al. 2020); Decay sugar-rich substrates (Põlme et al. 2020) | Põlme S, et al. 2020. Fungal Diversity 105:1-16 ((https://doi.org/10.1007/s13225-020-00466-2)) |
| Otu51304 | 1 | k__Fungi(100);p__Ascomycota(100);c__Saccharomycetes(100);o__Saccharomycetales(100);f__Metschnikowiaceae(100);g__Metschnikowia(100);s__Metschnikowia_pulcherrima(100); | Metschnikowia | 13 | Saprotroph | \|Undefined Saprotroph\| | Yeast | NULL | Probable | As nectar-tap_saprotroph (Põlme et al. 2020); Decay sugar-rich substrates (Põlme et al. 2020) | Põlme S, et al. 2020. Fungal Diversity 105:1-16 ((https://doi.org/10.1007/s13225-020-00466-2)) |
| Otu51305 | 1 | k__Fungi(100);p__Ascomycota(100);c__Saccharomycetes(100);o__Saccharomycetales(100);f__Metschnikowiaceae(100);g__Metschnikowia(100);g__Metschnikowia_unclassified(100); | Metschnikowia | 13 | Saprotroph | \|Undefined Saprotroph\| | Yeast | NULL | Probable | As nectar-tap_saprotroph (Põlme et al. 2020); Decay sugar-rich substrates (Põlme et al. 2020) | Põlme S, et al. 2020. Fungal Diversity 105:1-16 ((https://doi.org/10.1007/s13225-020-00466-2)) |
| Otu51306 | 1 | k__Fungi(100);p__Ascomycota(100);c__Saccharomycetes(100);o__Saccharomycetales(100);f__Metschnikowiaceae(100);g__Metschnikowia(100);s__Metschnikowia_pulcherrima(100); | Metschnikowia | 13 | Saprotroph | \|Undefined Saprotroph\| | Yeast | NULL | Probable | As nectar-tap_saprotroph (Põlme et al. 2020); Decay sugar-rich substrates (Põlme et al. 2020) | Põlme S, et al. 2020. Fungal Diversity 105:1-16 ((https://doi.org/10.1007/s13225-020-00466-2)) |
| Otu51307 | 1 | k__Fungi(100);p__Ascomycota(100);c__Saccharomycetes(100);o__Saccharomycetales(100);f__Metschnikowiaceae(100);g__Metschnikowia(100);s__Metschnikowia_pulcherrima(100); | Metschnikowia | 13 | Saprotroph | \|Undefined Saprotroph\| | Yeast | NULL | Probable | As nectar-tap_saprotroph (Põlme et al. 2020); Decay sugar-rich substrates (Põlme et al. 2020) | Põlme S, et al. 2020. Fungal Diversity 105:1-16 ((https://doi.org/10.1007/s13225-020-00466-2)) |
| Otu51308 | 1 | k__Fungi(100);p__Ascomycota(100);c__Saccharomycetes(100);o__Saccharomycetales(100);f__Metschnikowiaceae(100);g__Metschnikowia(100);s__Metschnikowia_pulcherrima(100); | Metschnikowia | 13 | Saprotroph | \|Undefined Saprotroph\| | Yeast | NULL | Probable | As nectar-tap_saprotroph (Põlme et al. 2020); Decay sugar-rich substrates (Põlme et al. 2020) | Põlme S, et al. 2020. Fungal Diversity 105:1-16 ((https://doi.org/10.1007/s13225-020-00466-2)) |
| Otu51309 | 1 | k__Fungi(100);p__Ascomycota(100);c__Saccharomycetes(100);o__Saccharomycetales(100);f__Metschnikowiaceae(100);g__Metschnikowia(100);s__Metschnikowia_pulcherrima(100); | Metschnikowia | 13 | Saprotroph | \|Undefined Saprotroph\| | Yeast | NULL | Probable | As nectar-tap_saprotroph (Põlme et al. 2020); Decay sugar-rich substrates (Põlme et al. 2020) | Põlme S, et al. 2020. Fungal Diversity 105:1-16 ((https://doi.org/10.1007/s13225-020-00466-2)) |
| Otu51310 | 1 | k__Fungi(100);p__Ascomycota(100);c__Saccharomycetes(100);o__Saccharomycetales(100);f__Metschnikowiaceae(100);g__Metschnikowia(100);s__Metschnikowia_pulcherrima(100); | Metschnikowia | 13 | Saprotroph | \|Undefined Saprotroph\| | Yeast | NULL | Probable | As nectar-tap_saprotroph (Põlme et al. 2020); Decay sugar-rich substrates (Põlme et al. 2020) | Põlme S, et al. 2020. Fungal Diversity 105:1-16 ((https://doi.org/10.1007/s13225-020-00466-2)) |
| Otu51311 | 1 | k__Fungi(100);p__Ascomycota(100);c__Saccharomycetes(100);o__Saccharomycetales(100);f__Metschnikowiaceae(100);g__Metschnikowia(100);s__Metschnikowia_pulcherrima(100); | Metschnikowia | 13 | Saprotroph | \|Undefined Saprotroph\| | Yeast | NULL | Probable | As nectar-tap_saprotroph (Põlme et al. 2020); Decay sugar-rich substrates (Põlme et al. 2020) | Põlme S, et al. 2020. Fungal Diversity 105:1-16 ((https://doi.org/10.1007/s13225-020-00466-2)) |
| Otu51312 | 1 | k__Fungi(100);p__Ascomycota(100);c__Saccharomycetes(100);o__Saccharomycetales(100);f__Metschnikowiaceae(100);g__Metschnikowia(100);g__Metschnikowia_unclassified(100); | Metschnikowia | 13 | Saprotroph | \|Undefined Saprotroph\| | Yeast | NULL | Probable | As nectar-tap_saprotroph (Põlme et al. 2020); Decay sugar-rich substrates (Põlme et al. 2020) | Põlme S, et al. 2020. Fungal Diversity 105:1-16 ((https://doi.org/10.1007/s13225-020-00466-2)) |
| Otu51313 | 1 | k__Fungi(100);p__Ascomycota(100);c__Saccharomycetes(100);o__Saccharomycetales(100);f__Metschnikowiaceae(100);g__Metschnikowia(100);g__Metschnikowia_unclassified(100); | Metschnikowia | 13 | Saprotroph | \|Undefined Saprotroph\| | Yeast | NULL | Probable | As nectar-tap_saprotroph (Põlme et al. 2020); Decay sugar-rich substrates (Põlme et al. 2020) | Põlme S, et al. 2020. Fungal Diversity 105:1-16 ((https://doi.org/10.1007/s13225-020-00466-2)) |
| Otu51314 | 1 | k__Fungi(100);p__Ascomycota(100);c__Saccharomycetes(100);o__Saccharomycetales(100);f__Metschnikowiaceae(100);g__Metschnikowia(100);s__Metschnikowia_pulcherrima(100); | Metschnikowia | 13 | Saprotroph | \|Undefined Saprotroph\| | Yeast | NULL | Probable | As nectar-tap_saprotroph (Põlme et al. 2020); Decay sugar-rich substrates (Põlme et al. 2020) | Põlme S, et al. 2020. Fungal Diversity 105:1-16 ((https://doi.org/10.1007/s13225-020-00466-2)) |
| Otu51315 | 1 | k__Fungi(100);p__Ascomycota(100);c__Saccharomycetes(100);o__Saccharomycetales(100);f__Metschnikowiaceae(100);g__Metschnikowia(100);g__Metschnikowia_unclassified(100); | Metschnikowia | 13 | Saprotroph | \|Undefined Saprotroph\| | Yeast | NULL | Probable | As nectar-tap_saprotroph (Põlme et al. 2020); Decay sugar-rich substrates (Põlme et al. 2020) | Põlme S, et al. 2020. Fungal Diversity 105:1-16 ((https://doi.org/10.1007/s13225-020-00466-2)) |
| Otu51316 | 1 | k__Fungi(100);p__Ascomycota(100);c__Saccharomycetes(100);o__Saccharomycetales(100);f__Metschnikowiaceae(100);g__Metschnikowia(100);s__Metschnikowia_pulcherrima(100); | Metschnikowia | 13 | Saprotroph | \|Undefined Saprotroph\| | Yeast | NULL | Probable | As nectar-tap_saprotroph (Põlme et al. 2020); Decay sugar-rich substrates (Põlme et al. 2020) | Põlme S, et al. 2020. Fungal Diversity 105:1-16 ((https://doi.org/10.1007/s13225-020-00466-2)) |
| Otu51317 | 1 | k__Fungi(100);p__Ascomycota(100);c__Saccharomycetes(100);o__Saccharomycetales(100);f__Metschnikowiaceae(100);g__Metschnikowia(100);s__Metschnikowia_pulcherrima(100); | Metschnikowia | 13 | Saprotroph | \|Undefined Saprotroph\| | Yeast | NULL | Probable | As nectar-tap_saprotroph (Põlme et al. 2020); Decay sugar-rich substrates (Põlme et al. 2020) | Põlme S, et al. 2020. Fungal Diversity 105:1-16 ((https://doi.org/10.1007/s13225-020-00466-2)) |
| Otu51318 | 1 | k__Fungi(100);p__Ascomycota(100);c__Saccharomycetes(100);o__Saccharomycetales(100);f__Metschnikowiaceae(100);g__Metschnikowia(100);g__Metschnikowia_unclassified(100); | Metschnikowia | 13 | Saprotroph | \|Undefined Saprotroph\| | Yeast | NULL | Probable | As nectar-tap_saprotroph (Põlme et al. 2020); Decay sugar-rich substrates (Põlme et al. 2020) | Põlme S, et al. 2020. Fungal Diversity 105:1-16 ((https://doi.org/10.1007/s13225-020-00466-2)) |
| Otu51319 | 1 | k__Fungi(100);p__Ascomycota(100);c__Saccharomycetes(100);o__Saccharomycetales(100);f__Metschnikowiaceae(100);g__Metschnikowia(100);s__Metschnikowia_pulcherrima(100); | Metschnikowia | 13 | Saprotroph | \|Undefined Saprotroph\| | Yeast | NULL | Probable | As nectar-tap_saprotroph (Põlme et al. 2020); Decay sugar-rich substrates (Põlme et al. 2020) | Põlme S, et al. 2020. Fungal Diversity 105:1-16 ((https://doi.org/10.1007/s13225-020-00466-2)) |
| Otu51320 | 1 | k__Fungi(100);p__Ascomycota(100);c__Saccharomycetes(100);o__Saccharomycetales(100);f__Metschnikowiaceae(100);g__Metschnikowia(100);s__Metschnikowia_pulcherrima(100); | Metschnikowia | 13 | Saprotroph | \|Undefined Saprotroph\| | Yeast | NULL | Probable | As nectar-tap_saprotroph (Põlme et al. 2020); Decay sugar-rich substrates (Põlme et al. 2020) | Põlme S, et al. 2020. Fungal Diversity 105:1-16 ((https://doi.org/10.1007/s13225-020-00466-2)) |
| Otu51321 | 1 | k__Fungi(100);p__Ascomycota(100);c__Saccharomycetes(100);o__Saccharomycetales(100);f__Metschnikowiaceae(100);g__Metschnikowia(100);s__Metschnikowia_pulcherrima(100); | Metschnikowia | 13 | Saprotroph | \|Undefined Saprotroph\| | Yeast | NULL | Probable | As nectar-tap_saprotroph (Põlme et al. 2020); Decay sugar-rich substrates (Põlme et al. 2020) | Põlme S, et al. 2020. Fungal Diversity 105:1-16 ((https://doi.org/10.1007/s13225-020-00466-2)) |
| Otu51322 | 1 | k__Fungi(100);p__Ascomycota(100);c__Saccharomycetes(100);o__Saccharomycetales(100);f__Metschnikowiaceae(100);g__Metschnikowia(100);s__Metschnikowia_pulcherrima(100); | Metschnikowia | 13 | Saprotroph | \|Undefined Saprotroph\| | Yeast | NULL | Probable | As nectar-tap_saprotroph (Põlme et al. 2020); Decay sugar-rich substrates (Põlme et al. 2020) | Põlme S, et al. 2020. Fungal Diversity 105:1-16 ((https://doi.org/10.1007/s13225-020-00466-2)) |
| Otu51323 | 1 | k__Fungi(100);p__Ascomycota(100);c__Saccharomycetes(100);o__Saccharomycetales(100);f__Metschnikowiaceae(100);g__Metschnikowia(100);g__Metschnikowia_unclassified(100); | Metschnikowia | 13 | Saprotroph | \|Undefined Saprotroph\| | Yeast | NULL | Probable | As nectar-tap_saprotroph (Põlme et al. 2020); Decay sugar-rich substrates (Põlme et al. 2020) | Põlme S, et al. 2020. Fungal Diversity 105:1-16 ((https://doi.org/10.1007/s13225-020-00466-2)) |
| Otu51324 | 1 | k__Fungi(100);p__Ascomycota(100);c__Saccharomycetes(100);o__Saccharomycetales(100);f__Metschnikowiaceae(100);g__Metschnikowia(100);g__Metschnikowia_unclassified(100); | Metschnikowia | 13 | Saprotroph | \|Undefined Saprotroph\| | Yeast | NULL | Probable | As nectar-tap_saprotroph (Põlme et al. 2020); Decay sugar-rich substrates (Põlme et al. 2020) | Põlme S, et al. 2020. Fungal Diversity 105:1-16 ((https://doi.org/10.1007/s13225-020-00466-2)) |
| Otu51325 | 1 | k__Fungi(100);p__Ascomycota(100);c__Saccharomycetes(100);o__Saccharomycetales(100);f__Metschnikowiaceae(100);g__Metschnikowia(100);s__Metschnikowia_pulcherrima(100); | Metschnikowia | 13 | Saprotroph | \|Undefined Saprotroph\| | Yeast | NULL | Probable | As nectar-tap_saprotroph (Põlme et al. 2020); Decay sugar-rich substrates (Põlme et al. 2020) | Põlme S, et al. 2020. Fungal Diversity 105:1-16 ((https://doi.org/10.1007/s13225-020-00466-2)) |
| Otu51326 | 1 | k__Fungi(100);p__Ascomycota(100);c__Saccharomycetes(100);o__Saccharomycetales(100);f__Metschnikowiaceae(100);g__Metschnikowia(100);s__Metschnikowia_pulcherrima(100); | Metschnikowia | 13 | Saprotroph | \|Undefined Saprotroph\| | Yeast | NULL | Probable | As nectar-tap_saprotroph (Põlme et al. 2020); Decay sugar-rich substrates (Põlme et al. 2020) | Põlme S, et al. 2020. Fungal Diversity 105:1-16 ((https://doi.org/10.1007/s13225-020-00466-2)) |
| Otu51327 | 1 | k__Fungi(100);p__Ascomycota(100);c__Saccharomycetes(100);o__Saccharomycetales(100);f__Metschnikowiaceae(100);g__Metschnikowia(100);s__Metschnikowia_pulcherrima(100); | Metschnikowia | 13 | Saprotroph | \|Undefined Saprotroph\| | Yeast | NULL | Probable | As nectar-tap_saprotroph (Põlme et al. 2020); Decay sugar-rich substrates (Põlme et al. 2020) | Põlme S, et al. 2020. Fungal Diversity 105:1-16 ((https://doi.org/10.1007/s13225-020-00466-2)) |
| Otu51328 | 1 | k__Fungi(100);p__Ascomycota(100);c__Saccharomycetes(100);o__Saccharomycetales(100);f__Metschnikowiaceae(100);g__Metschnikowia(100);s__Metschnikowia_pulcherrima(100); | Metschnikowia | 13 | Saprotroph | \|Undefined Saprotroph\| | Yeast | NULL | Probable | As nectar-tap_saprotroph (Põlme et al. 2020); Decay sugar-rich substrates (Põlme et al. 2020) | Põlme S, et al. 2020. Fungal Diversity 105:1-16 ((https://doi.org/10.1007/s13225-020-00466-2)) |
| Otu51329 | 1 | k__Fungi(100);p__Ascomycota(100);c__Saccharomycetes(100);o__Saccharomycetales(100);f__Metschnikowiaceae(100);g__Metschnikowia(100);s__Metschnikowia_pulcherrima(100); | Metschnikowia | 13 | Saprotroph | \|Undefined Saprotroph\| | Yeast | NULL | Probable | As nectar-tap_saprotroph (Põlme et al. 2020); Decay sugar-rich substrates (Põlme et al. 2020) | Põlme S, et al. 2020. Fungal Diversity 105:1-16 ((https://doi.org/10.1007/s13225-020-00466-2)) |
| Otu51330 | 1 | k__Fungi(100);p__Ascomycota(100);c__Saccharomycetes(100);o__Saccharomycetales(100);f__Metschnikowiaceae(100);g__Metschnikowia(100);s__Metschnikowia_pulcherrima(100); | Metschnikowia | 13 | Saprotroph | \|Undefined Saprotroph\| | Yeast | NULL | Probable | As nectar-tap_saprotroph (Põlme et al. 2020); Decay sugar-rich substrates (Põlme et al. 2020) | Põlme S, et al. 2020. Fungal Diversity 105:1-16 ((https://doi.org/10.1007/s13225-020-00466-2)) |
| Otu51331 | 1 | k__Fungi(100);p__Ascomycota(100);c__Saccharomycetes(100);o__Saccharomycetales(100);f__Metschnikowiaceae(100);g__Metschnikowia(100);g__Metschnikowia_unclassified(100); | Metschnikowia | 13 | Saprotroph | \|Undefined Saprotroph\| | Yeast | NULL | Probable | As nectar-tap_saprotroph (Põlme et al. 2020); Decay sugar-rich substrates (Põlme et al. 2020) | Põlme S, et al. 2020. Fungal Diversity 105:1-16 ((https://doi.org/10.1007/s13225-020-00466-2)) |
| Otu51332 | 1 | k__Fungi(100);p__Ascomycota(100);c__Saccharomycetes(100);o__Saccharomycetales(100);f__Metschnikowiaceae(100);g__Metschnikowia(100);s__Metschnikowia_pulcherrima(100); | Metschnikowia | 13 | Saprotroph | \|Undefined Saprotroph\| | Yeast | NULL | Probable | As nectar-tap_saprotroph (Põlme et al. 2020); Decay sugar-rich substrates (Põlme et al. 2020) | Põlme S, et al. 2020. Fungal Diversity 105:1-16 ((https://doi.org/10.1007/s13225-020-00466-2)) |
| Otu51333 | 1 | k__Fungi(100);p__Ascomycota(100);c__Saccharomycetes(100);o__Saccharomycetales(100);f__Metschnikowiaceae(100);g__Metschnikowia(100);s__Metschnikowia_pulcherrima(100); | Metschnikowia | 13 | Saprotroph | \|Undefined Saprotroph\| | Yeast | NULL | Probable | As nectar-tap_saprotroph (Põlme et al. 2020); Decay sugar-rich substrates (Põlme et al. 2020) | Põlme S, et al. 2020. Fungal Diversity 105:1-16 ((https://doi.org/10.1007/s13225-020-00466-2)) |
| Otu51334 | 1 | k__Fungi(100);p__Ascomycota(100);c__Saccharomycetes(100);o__Saccharomycetales(100);f__Metschnikowiaceae(100);g__Metschnikowia(100);s__Metschnikowia_pulcherrima(100); | Metschnikowia | 13 | Saprotroph | \|Undefined Saprotroph\| | Yeast | NULL | Probable | As nectar-tap_saprotroph (Põlme et al. 2020); Decay sugar-rich substrates (Põlme et al. 2020) | Põlme S, et al. 2020. Fungal Diversity 105:1-16 ((https://doi.org/10.1007/s13225-020-00466-2)) |
| Otu51335 | 1 | k__Fungi(100);p__Ascomycota(100);c__Saccharomycetes(100);o__Saccharomycetales(100);f__Metschnikowiaceae(100);g__Metschnikowia(100);s__Metschnikowia_pulcherrima(100); | Metschnikowia | 13 | Saprotroph | \|Undefined Saprotroph\| | Yeast | NULL | Probable | As nectar-tap_saprotroph (Põlme et al. 2020); Decay sugar-rich substrates (Põlme et al. 2020) | Põlme S, et al. 2020. Fungal Diversity 105:1-16 ((https://doi.org/10.1007/s13225-020-00466-2)) |
| Otu51336 | 1 | k__Fungi(100);p__Ascomycota(100);c__Saccharomycetes(100);o__Saccharomycetales(100);f__Metschnikowiaceae(100);g__Metschnikowia(100);s__Metschnikowia_pulcherrima(100); | Metschnikowia | 13 | Saprotroph | \|Undefined Saprotroph\| | Yeast | NULL | Probable | As nectar-tap_saprotroph (Põlme et al. 2020); Decay sugar-rich substrates (Põlme et al. 2020) | Põlme S, et al. 2020. Fungal Diversity 105:1-16 ((https://doi.org/10.1007/s13225-020-00466-2)) |
| Otu51337 | 1 | k__Fungi(100);p__Ascomycota(100);c__Saccharomycetes(100);o__Saccharomycetales(100);f__Metschnikowiaceae(100);g__Metschnikowia(100);g__Metschnikowia_unclassified(100); | Metschnikowia | 13 | Saprotroph | \|Undefined Saprotroph\| | Yeast | NULL | Probable | As nectar-tap_saprotroph (Põlme et al. 2020); Decay sugar-rich substrates (Põlme et al. 2020) | Põlme S, et al. 2020. Fungal Diversity 105:1-16 ((https://doi.org/10.1007/s13225-020-00466-2)) |
| Otu51338 | 1 | k__Fungi(100);p__Ascomycota(100);c__Saccharomycetes(100);o__Saccharomycetales(100);f__Metschnikowiaceae(100);g__Metschnikowia(100);g__Metschnikowia_unclassified(100); | Metschnikowia | 13 | Saprotroph | \|Undefined Saprotroph\| | Yeast | NULL | Probable | As nectar-tap_saprotroph (Põlme et al. 2020); Decay sugar-rich substrates (Põlme et al. 2020) | Põlme S, et al. 2020. Fungal Diversity 105:1-16 ((https://doi.org/10.1007/s13225-020-00466-2)) |
| Otu51339 | 1 | k__Fungi(100);p__Ascomycota(100);c__Saccharomycetes(100);o__Saccharomycetales(100);f__Metschnikowiaceae(100);g__Metschnikowia(100);g__Metschnikowia_unclassified(100); | Metschnikowia | 13 | Saprotroph | \|Undefined Saprotroph\| | Yeast | NULL | Probable | As nectar-tap_saprotroph (Põlme et al. 2020); Decay sugar-rich substrates (Põlme et al. 2020) | Põlme S, et al. 2020. Fungal Diversity 105:1-16 ((https://doi.org/10.1007/s13225-020-00466-2)) |
| Otu51340 | 1 | k__Fungi(100);p__Ascomycota(100);c__Saccharomycetes(100);o__Saccharomycetales(100);f__Metschnikowiaceae(100);g__Metschnikowia(100);g__Metschnikowia_unclassified(100); | Metschnikowia | 13 | Saprotroph | \|Undefined Saprotroph\| | Yeast | NULL | Probable | As nectar-tap_saprotroph (Põlme et al. 2020); Decay sugar-rich substrates (Põlme et al. 2020) | Põlme S, et al. 2020. Fungal Diversity 105:1-16 ((https://doi.org/10.1007/s13225-020-00466-2)) |
| Otu51342 | 1 | k__Fungi(100);p__Ascomycota(100);c__Saccharomycetes(100);o__Saccharomycetales(100);f__Metschnikowiaceae(100);g__Metschnikowia(100);s__Metschnikowia_pulcherrima(100); | Metschnikowia | 13 | Saprotroph | \|Undefined Saprotroph\| | Yeast | NULL | Probable | As nectar-tap_saprotroph (Põlme et al. 2020); Decay sugar-rich substrates (Põlme et al. 2020) | Põlme S, et al. 2020. Fungal Diversity 105:1-16 ((https://doi.org/10.1007/s13225-020-00466-2)) |
| Otu51343 | 1 | k__Fungi(100);p__Ascomycota(100);c__Saccharomycetes(100);o__Saccharomycetales(100);f__Metschnikowiaceae(100);g__Metschnikowia(100);s__Metschnikowia_pulcherrima(100); | Metschnikowia | 13 | Saprotroph | \|Undefined Saprotroph\| | Yeast | NULL | Probable | As nectar-tap_saprotroph (Põlme et al. 2020); Decay sugar-rich substrates (Põlme et al. 2020) | Põlme S, et al. 2020. Fungal Diversity 105:1-16 ((https://doi.org/10.1007/s13225-020-00466-2)) |
| Otu51344 | 1 | k__Fungi(100);p__Ascomycota(100);c__Saccharomycetes(100);o__Saccharomycetales(100);f__Metschnikowiaceae(100);g__Metschnikowia(100);g__Metschnikowia_unclassified(100); | Metschnikowia | 13 | Saprotroph | \|Undefined Saprotroph\| | Yeast | NULL | Probable | As nectar-tap_saprotroph (Põlme et al. 2020); Decay sugar-rich substrates (Põlme et al. 2020) | Põlme S, et al. 2020. Fungal Diversity 105:1-16 ((https://doi.org/10.1007/s13225-020-00466-2)) |
| Otu51345 | 1 | k__Fungi(100);p__Ascomycota(100);c__Saccharomycetes(100);o__Saccharomycetales(100);f__Metschnikowiaceae(100);g__Metschnikowia(100);g__Metschnikowia_unclassified(100); | Metschnikowia | 13 | Saprotroph | \|Undefined Saprotroph\| | Yeast | NULL | Probable | As nectar-tap_saprotroph (Põlme et al. 2020); Decay sugar-rich substrates (Põlme et al. 2020) | Põlme S, et al. 2020. Fungal Diversity 105:1-16 ((https://doi.org/10.1007/s13225-020-00466-2)) |
| Otu51346 | 1 | k__Fungi(100);p__Ascomycota(100);c__Saccharomycetes(100);o__Saccharomycetales(100);f__Metschnikowiaceae(100);g__Metschnikowia(100);g__Metschnikowia_unclassified(100); | Metschnikowia | 13 | Saprotroph | \|Undefined Saprotroph\| | Yeast | NULL | Probable | As nectar-tap_saprotroph (Põlme et al. 2020); Decay sugar-rich substrates (Põlme et al. 2020) | Põlme S, et al. 2020. Fungal Diversity 105:1-16 ((https://doi.org/10.1007/s13225-020-00466-2)) |
| Otu51347 | 1 | k__Fungi(100);p__Ascomycota(100);c__Saccharomycetes(100);o__Saccharomycetales(100);f__Metschnikowiaceae(100);g__Metschnikowia(100);g__Metschnikowia_unclassified(100); | Metschnikowia | 13 | Saprotroph | \|Undefined Saprotroph\| | Yeast | NULL | Probable | As nectar-tap_saprotroph (Põlme et al. 2020); Decay sugar-rich substrates (Põlme et al. 2020) | Põlme S, et al. 2020. Fungal Diversity 105:1-16 ((https://doi.org/10.1007/s13225-020-00466-2)) |
| Otu51348 | 1 | k__Fungi(100);p__Ascomycota(100);c__Saccharomycetes(100);o__Saccharomycetales(100);f__Metschnikowiaceae(100);g__Metschnikowia(100);s__Metschnikowia_pulcherrima(100); | Metschnikowia | 13 | Saprotroph | \|Undefined Saprotroph\| | Yeast | NULL | Probable | As nectar-tap_saprotroph (Põlme et al. 2020); Decay sugar-rich substrates (Põlme et al. 2020) | Põlme S, et al. 2020. Fungal Diversity 105:1-16 ((https://doi.org/10.1007/s13225-020-00466-2)) |
| Otu51349 | 1 | k__Fungi(100);p__Ascomycota(100);c__Saccharomycetes(100);o__Saccharomycetales(100);f__Metschnikowiaceae(100);g__Metschnikowia(100);s__Metschnikowia_pulcherrima(100); | Metschnikowia | 13 | Saprotroph | \|Undefined Saprotroph\| | Yeast | NULL | Probable | As nectar-tap_saprotroph (Põlme et al. 2020); Decay sugar-rich substrates (Põlme et al. 2020) | Põlme S, et al. 2020. Fungal Diversity 105:1-16 ((https://doi.org/10.1007/s13225-020-00466-2)) |
| Otu51350 | 1 | k__Fungi(100);p__Ascomycota(100);c__Saccharomycetes(100);o__Saccharomycetales(100);f__Metschnikowiaceae(100);g__Metschnikowia(100);s__Metschnikowia_pulcherrima(100); | Metschnikowia | 13 | Saprotroph | \|Undefined Saprotroph\| | Yeast | NULL | Probable | As nectar-tap_saprotroph (Põlme et al. 2020); Decay sugar-rich substrates (Põlme et al. 2020) | Põlme S, et al. 2020. Fungal Diversity 105:1-16 ((https://doi.org/10.1007/s13225-020-00466-2)) |
| Otu51351 | 1 | k__Fungi(100);p__Ascomycota(100);c__Saccharomycetes(100);o__Saccharomycetales(100);f__Metschnikowiaceae(100);g__Metschnikowia(100);g__Metschnikowia_unclassified(100); | Metschnikowia | 13 | Saprotroph | \|Undefined Saprotroph\| | Yeast | NULL | Probable | As nectar-tap_saprotroph (Põlme et al. 2020); Decay sugar-rich substrates (Põlme et al. 2020) | Põlme S, et al. 2020. Fungal Diversity 105:1-16 ((https://doi.org/10.1007/s13225-020-00466-2)) |
| Otu51352 | 1 | k__Fungi(100);p__Ascomycota(100);c__Saccharomycetes(100);o__Saccharomycetales(100);f__Metschnikowiaceae(100);g__Metschnikowia(100);s__Metschnikowia_pulcherrima(100); | Metschnikowia | 13 | Saprotroph | \|Undefined Saprotroph\| | Yeast | NULL | Probable | As nectar-tap_saprotroph (Põlme et al. 2020); Decay sugar-rich substrates (Põlme et al. 2020) | Põlme S, et al. 2020. Fungal Diversity 105:1-16 ((https://doi.org/10.1007/s13225-020-00466-2)) |
| Otu51353 | 1 | k__Fungi(100);p__Ascomycota(100);c__Saccharomycetes(100);o__Saccharomycetales(100);f__Metschnikowiaceae(100);g__Metschnikowia(100);g__Metschnikowia_unclassified(100); | Metschnikowia | 13 | Saprotroph | \|Undefined Saprotroph\| | Yeast | NULL | Probable | As nectar-tap_saprotroph (Põlme et al. 2020); Decay sugar-rich substrates (Põlme et al. 2020) | Põlme S, et al. 2020. Fungal Diversity 105:1-16 ((https://doi.org/10.1007/s13225-020-00466-2)) |
| Otu51354 | 1 | k__Fungi(100);p__Ascomycota(100);c__Saccharomycetes(100);o__Saccharomycetales(100);f__Metschnikowiaceae(100);g__Metschnikowia(100);s__Metschnikowia_pulcherrima(100); | Metschnikowia | 13 | Saprotroph | \|Undefined Saprotroph\| | Yeast | NULL | Probable | As nectar-tap_saprotroph (Põlme et al. 2020); Decay sugar-rich substrates (Põlme et al. 2020) | Põlme S, et al. 2020. Fungal Diversity 105:1-16 ((https://doi.org/10.1007/s13225-020-00466-2)) |
| Otu51355 | 1 | k__Fungi(100);p__Ascomycota(100);c__Saccharomycetes(100);o__Saccharomycetales(100);f__Metschnikowiaceae(100);g__Metschnikowia(100);s__Metschnikowia_pulcherrima(100); | Metschnikowia | 13 | Saprotroph | \|Undefined Saprotroph\| | Yeast | NULL | Probable | As nectar-tap_saprotroph (Põlme et al. 2020); Decay sugar-rich substrates (Põlme et al. 2020) | Põlme S, et al. 2020. Fungal Diversity 105:1-16 ((https://doi.org/10.1007/s13225-020-00466-2)) |
| Otu51356 | 1 | k__Fungi(100);p__Ascomycota(100);c__Saccharomycetes(100);o__Saccharomycetales(100);f__Metschnikowiaceae(100);g__Metschnikowia(100);s__Metschnikowia_pulcherrima(100); | Metschnikowia | 13 | Saprotroph | \|Undefined Saprotroph\| | Yeast | NULL | Probable | As nectar-tap_saprotroph (Põlme et al. 2020); Decay sugar-rich substrates (Põlme et al. 2020) | Põlme S, et al. 2020. Fungal Diversity 105:1-16 ((https://doi.org/10.1007/s13225-020-00466-2)) |
| Otu51357 | 1 | k__Fungi(100);p__Ascomycota(100);c__Saccharomycetes(100);o__Saccharomycetales(100);f__Metschnikowiaceae(100);g__Metschnikowia(100);g__Metschnikowia_unclassified(100); | Metschnikowia | 13 | Saprotroph | \|Undefined Saprotroph\| | Yeast | NULL | Probable | As nectar-tap_saprotroph (Põlme et al. 2020); Decay sugar-rich substrates (Põlme et al. 2020) | Põlme S, et al. 2020. Fungal Diversity 105:1-16 ((https://doi.org/10.1007/s13225-020-00466-2)) |
| Otu51358 | 1 | k__Fungi(100);p__Ascomycota(100);c__Saccharomycetes(100);o__Saccharomycetales(100);f__Metschnikowiaceae(100);g__Metschnikowia(100);s__Metschnikowia_pulcherrima(100); | Metschnikowia | 13 | Saprotroph | \|Undefined Saprotroph\| | Yeast | NULL | Probable | As nectar-tap_saprotroph (Põlme et al. 2020); Decay sugar-rich substrates (Põlme et al. 2020) | Põlme S, et al. 2020. Fungal Diversity 105:1-16 ((https://doi.org/10.1007/s13225-020-00466-2)) |
| Otu51359 | 1 | k__Fungi(100);p__Ascomycota(100);c__Saccharomycetes(100);o__Saccharomycetales(100);f__Metschnikowiaceae(100);g__Metschnikowia(100);s__Metschnikowia_pulcherrima(100); | Metschnikowia | 13 | Saprotroph | \|Undefined Saprotroph\| | Yeast | NULL | Probable | As nectar-tap_saprotroph (Põlme et al. 2020); Decay sugar-rich substrates (Põlme et al. 2020) | Põlme S, et al. 2020. Fungal Diversity 105:1-16 ((https://doi.org/10.1007/s13225-020-00466-2)) |
| Otu51360 | 1 | k__Fungi(100);p__Ascomycota(100);c__Saccharomycetes(100);o__Saccharomycetales(100);f__Metschnikowiaceae(100);g__Metschnikowia(100);s__Metschnikowia_pulcherrima(100); | Metschnikowia | 13 | Saprotroph | \|Undefined Saprotroph\| | Yeast | NULL | Probable | As nectar-tap_saprotroph (Põlme et al. 2020); Decay sugar-rich substrates (Põlme et al. 2020) | Põlme S, et al. 2020. Fungal Diversity 105:1-16 ((https://doi.org/10.1007/s13225-020-00466-2)) |
| Otu51361 | 1 | k__Fungi(100);p__Ascomycota(100);c__Saccharomycetes(100);o__Saccharomycetales(100);f__Metschnikowiaceae(100);g__Metschnikowia(100);s__Metschnikowia_pulcherrima(100); | Metschnikowia | 13 | Saprotroph | \|Undefined Saprotroph\| | Yeast | NULL | Probable | As nectar-tap_saprotroph (Põlme et al. 2020); Decay sugar-rich substrates (Põlme et al. 2020) | Põlme S, et al. 2020. Fungal Diversity 105:1-16 ((https://doi.org/10.1007/s13225-020-00466-2)) |
| Otu51362 | 1 | k__Fungi(100);p__Ascomycota(100);c__Saccharomycetes(100);o__Saccharomycetales(100);f__Metschnikowiaceae(100);g__Metschnikowia(100);s__Metschnikowia_pulcherrima(100); | Metschnikowia | 13 | Saprotroph | \|Undefined Saprotroph\| | Yeast | NULL | Probable | As nectar-tap_saprotroph (Põlme et al. 2020); Decay sugar-rich substrates (Põlme et al. 2020) | Põlme S, et al. 2020. Fungal Diversity 105:1-16 ((https://doi.org/10.1007/s13225-020-00466-2)) |
| Otu51363 | 1 | k__Fungi(100);p__Ascomycota(100);c__Saccharomycetes(100);o__Saccharomycetales(100);f__Metschnikowiaceae(100);g__Metschnikowia(100);g__Metschnikowia_unclassified(100); | Metschnikowia | 13 | Saprotroph | \|Undefined Saprotroph\| | Yeast | NULL | Probable | As nectar-tap_saprotroph (Põlme et al. 2020); Decay sugar-rich substrates (Põlme et al. 2020) | Põlme S, et al. 2020. Fungal Diversity 105:1-16 ((https://doi.org/10.1007/s13225-020-00466-2)) |
| Otu51364 | 1 | k__Fungi(100);p__Ascomycota(100);c__Saccharomycetes(100);o__Saccharomycetales(100);f__Metschnikowiaceae(100);g__Metschnikowia(100);s__Metschnikowia_pulcherrima(100); | Metschnikowia | 13 | Saprotroph | \|Undefined Saprotroph\| | Yeast | NULL | Probable | As nectar-tap_saprotroph (Põlme et al. 2020); Decay sugar-rich substrates (Põlme et al. 2020) | Põlme S, et al. 2020. Fungal Diversity 105:1-16 ((https://doi.org/10.1007/s13225-020-00466-2)) |
| Otu51365 | 1 | k__Fungi(100);p__Ascomycota(100);c__Saccharomycetes(100);o__Saccharomycetales(100);f__Metschnikowiaceae(100);g__Metschnikowia(100);s__Metschnikowia_pulcherrima(100); | Metschnikowia | 13 | Saprotroph | \|Undefined Saprotroph\| | Yeast | NULL | Probable | As nectar-tap_saprotroph (Põlme et al. 2020); Decay sugar-rich substrates (Põlme et al. 2020) | Põlme S, et al. 2020. Fungal Diversity 105:1-16 ((https://doi.org/10.1007/s13225-020-00466-2)) |
| Otu51366 | 1 | k__Fungi(100);p__Ascomycota(100);c__Saccharomycetes(100);o__Saccharomycetales(100);f__Metschnikowiaceae(100);g__Metschnikowia(100);s__Metschnikowia_pulcherrima(100); | Metschnikowia | 13 | Saprotroph | \|Undefined Saprotroph\| | Yeast | NULL | Probable | As nectar-tap_saprotroph (Põlme et al. 2020); Decay sugar-rich substrates (Põlme et al. 2020) | Põlme S, et al. 2020. Fungal Diversity 105:1-16 ((https://doi.org/10.1007/s13225-020-00466-2)) |
| Otu51367 | 1 | k__Fungi(100);p__Ascomycota(100);c__Saccharomycetes(100);o__Saccharomycetales(100);f__Metschnikowiaceae(100);g__Metschnikowia(100);s__Metschnikowia_pulcherrima(100); | Metschnikowia | 13 | Saprotroph | \|Undefined Saprotroph\| | Yeast | NULL | Probable | As nectar-tap_saprotroph (Põlme et al. 2020); Decay sugar-rich substrates (Põlme et al. 2020) | Põlme S, et al. 2020. Fungal Diversity 105:1-16 ((https://doi.org/10.1007/s13225-020-00466-2)) |
| Otu51368 | 1 | k__Fungi(100);p__Ascomycota(100);c__Saccharomycetes(100);o__Saccharomycetales(100);f__Metschnikowiaceae(100);g__Metschnikowia(100);s__Metschnikowia_pulcherrima(100); | Metschnikowia | 13 | Saprotroph | \|Undefined Saprotroph\| | Yeast | NULL | Probable | As nectar-tap_saprotroph (Põlme et al. 2020); Decay sugar-rich substrates (Põlme et al. 2020) | Põlme S, et al. 2020. Fungal Diversity 105:1-16 ((https://doi.org/10.1007/s13225-020-00466-2)) |
| Otu51369 | 1 | k__Fungi(100);p__Ascomycota(100);c__Saccharomycetes(100);o__Saccharomycetales(100);f__Metschnikowiaceae(100);g__Metschnikowia(100);s__Metschnikowia_pulcherrima(100); | Metschnikowia | 13 | Saprotroph | \|Undefined Saprotroph\| | Yeast | NULL | Probable | As nectar-tap_saprotroph (Põlme et al. 2020); Decay sugar-rich substrates (Põlme et al. 2020) | Põlme S, et al. 2020. Fungal Diversity 105:1-16 ((https://doi.org/10.1007/s13225-020-00466-2)) |
| Otu51370 | 1 | k__Fungi(100);p__Ascomycota(100);c__Saccharomycetes(100);o__Saccharomycetales(100);f__Metschnikowiaceae(100);g__Metschnikowia(100);s__Metschnikowia_pulcherrima(100); | Metschnikowia | 13 | Saprotroph | \|Undefined Saprotroph\| | Yeast | NULL | Probable | As nectar-tap_saprotroph (Põlme et al. 2020); Decay sugar-rich substrates (Põlme et al. 2020) | Põlme S, et al. 2020. Fungal Diversity 105:1-16 ((https://doi.org/10.1007/s13225-020-00466-2)) |
| Otu51371 | 1 | k__Fungi(100);p__Ascomycota(100);c__Saccharomycetes(100);o__Saccharomycetales(100);f__Metschnikowiaceae(100);g__Metschnikowia(100);g__Metschnikowia_unclassified(100); | Metschnikowia | 13 | Saprotroph | \|Undefined Saprotroph\| | Yeast | NULL | Probable | As nectar-tap_saprotroph (Põlme et al. 2020); Decay sugar-rich substrates (Põlme et al. 2020) | Põlme S, et al. 2020. Fungal Diversity 105:1-16 ((https://doi.org/10.1007/s13225-020-00466-2)) |
| Otu51372 | 1 | k__Fungi(100);p__Ascomycota(100);c__Saccharomycetes(100);o__Saccharomycetales(100);f__Metschnikowiaceae(100);g__Metschnikowia(100);g__Metschnikowia_unclassified(100); | Metschnikowia | 13 | Saprotroph | \|Undefined Saprotroph\| | Yeast | NULL | Probable | As nectar-tap_saprotroph (Põlme et al. 2020); Decay sugar-rich substrates (Põlme et al. 2020) | Põlme S, et al. 2020. Fungal Diversity 105:1-16 ((https://doi.org/10.1007/s13225-020-00466-2)) |
| Otu51373 | 1 | k__Fungi(100);p__Ascomycota(100);c__Saccharomycetes(100);o__Saccharomycetales(100);f__Metschnikowiaceae(100);g__Metschnikowia(100);g__Metschnikowia_unclassified(100); | Metschnikowia | 13 | Saprotroph | \|Undefined Saprotroph\| | Yeast | NULL | Probable | As nectar-tap_saprotroph (Põlme et al. 2020); Decay sugar-rich substrates (Põlme et al. 2020) | Põlme S, et al. 2020. Fungal Diversity 105:1-16 ((https://doi.org/10.1007/s13225-020-00466-2)) |
| Otu51374 | 1 | k__Fungi(100);p__Ascomycota(100);c__Saccharomycetes(100);o__Saccharomycetales(100);f__Metschnikowiaceae(100);g__Metschnikowia(100);s__Metschnikowia_pulcherrima(100); | Metschnikowia | 13 | Saprotroph | \|Undefined Saprotroph\| | Yeast | NULL | Probable | As nectar-tap_saprotroph (Põlme et al. 2020); Decay sugar-rich substrates (Põlme et al. 2020) | Põlme S, et al. 2020. Fungal Diversity 105:1-16 ((https://doi.org/10.1007/s13225-020-00466-2)) |
| Otu51375 | 1 | k__Fungi(100);p__Ascomycota(100);c__Saccharomycetes(100);o__Saccharomycetales(100);f__Metschnikowiaceae(100);g__Metschnikowia(100);g__Metschnikowia_unclassified(100); | Metschnikowia | 13 | Saprotroph | \|Undefined Saprotroph\| | Yeast | NULL | Probable | As nectar-tap_saprotroph (Põlme et al. 2020); Decay sugar-rich substrates (Põlme et al. 2020) | Põlme S, et al. 2020. Fungal Diversity 105:1-16 ((https://doi.org/10.1007/s13225-020-00466-2)) |
| Otu51376 | 1 | k__Fungi(100);p__Ascomycota(100);c__Saccharomycetes(100);o__Saccharomycetales(100);f__Metschnikowiaceae(100);g__Metschnikowia(100);s__Metschnikowia_pulcherrima(100); | Metschnikowia | 13 | Saprotroph | \|Undefined Saprotroph\| | Yeast | NULL | Probable | As nectar-tap_saprotroph (Põlme et al. 2020); Decay sugar-rich substrates (Põlme et al. 2020) | Põlme S, et al. 2020. Fungal Diversity 105:1-16 ((https://doi.org/10.1007/s13225-020-00466-2)) |
| Otu51377 | 1 | k__Fungi(100);p__Ascomycota(100);c__Saccharomycetes(100);o__Saccharomycetales(100);f__Metschnikowiaceae(100);g__Metschnikowia(100);g__Metschnikowia_unclassified(100); | Metschnikowia | 13 | Saprotroph | \|Undefined Saprotroph\| | Yeast | NULL | Probable | As nectar-tap_saprotroph (Põlme et al. 2020); Decay sugar-rich substrates (Põlme et al. 2020) | Põlme S, et al. 2020. Fungal Diversity 105:1-16 ((https://doi.org/10.1007/s13225-020-00466-2)) |
| Otu51378 | 1 | k__Fungi(100);p__Ascomycota(100);c__Saccharomycetes(100);o__Saccharomycetales(100);f__Metschnikowiaceae(100);g__Metschnikowia(100);s__Metschnikowia_pulcherrima(100); | Metschnikowia | 13 | Saprotroph | \|Undefined Saprotroph\| | Yeast | NULL | Probable | As nectar-tap_saprotroph (Põlme et al. 2020); Decay sugar-rich substrates (Põlme et al. 2020) | Põlme S, et al. 2020. Fungal Diversity 105:1-16 ((https://doi.org/10.1007/s13225-020-00466-2)) |
| Otu51379 | 1 | k__Fungi(100);p__Ascomycota(100);c__Saccharomycetes(100);o__Saccharomycetales(100);f__Metschnikowiaceae(100);g__Metschnikowia(100);g__Metschnikowia_unclassified(100); | Metschnikowia | 13 | Saprotroph | \|Undefined Saprotroph\| | Yeast | NULL | Probable | As nectar-tap_saprotroph (Põlme et al. 2020); Decay sugar-rich substrates (Põlme et al. 2020) | Põlme S, et al. 2020. Fungal Diversity 105:1-16 ((https://doi.org/10.1007/s13225-020-00466-2)) |
| Otu51380 | 1 | k__Fungi(100);p__Ascomycota(100);c__Saccharomycetes(100);o__Saccharomycetales(100);f__Metschnikowiaceae(100);g__Metschnikowia(100);s__Metschnikowia_pulcherrima(100); | Metschnikowia | 13 | Saprotroph | \|Undefined Saprotroph\| | Yeast | NULL | Probable | As nectar-tap_saprotroph (Põlme et al. 2020); Decay sugar-rich substrates (Põlme et al. 2020) | Põlme S, et al. 2020. Fungal Diversity 105:1-16 ((https://doi.org/10.1007/s13225-020-00466-2)) |
| Otu51381 | 1 | k__Fungi(100);p__Ascomycota(100);c__Saccharomycetes(100);o__Saccharomycetales(100);f__Metschnikowiaceae(100);g__Metschnikowia(100);s__Metschnikowia_pulcherrima(100); | Metschnikowia | 13 | Saprotroph | \|Undefined Saprotroph\| | Yeast | NULL | Probable | As nectar-tap_saprotroph (Põlme et al. 2020); Decay sugar-rich substrates (Põlme et al. 2020) | Põlme S, et al. 2020. Fungal Diversity 105:1-16 ((https://doi.org/10.1007/s13225-020-00466-2)) |
| Otu51382 | 1 | k__Fungi(100);p__Ascomycota(100);c__Saccharomycetes(100);o__Saccharomycetales(100);f__Metschnikowiaceae(100);g__Metschnikowia(100);s__Metschnikowia_pulcherrima(100); | Metschnikowia | 13 | Saprotroph | \|Undefined Saprotroph\| | Yeast | NULL | Probable | As nectar-tap_saprotroph (Põlme et al. 2020); Decay sugar-rich substrates (Põlme et al. 2020) | Põlme S, et al. 2020. Fungal Diversity 105:1-16 ((https://doi.org/10.1007/s13225-020-00466-2)) |
| Otu51383 | 1 | k__Fungi(100);p__Ascomycota(100);c__Saccharomycetes(100);o__Saccharomycetales(100);f__Metschnikowiaceae(100);g__Metschnikowia(100);s__Metschnikowia_pulcherrima(100); | Metschnikowia | 13 | Saprotroph | \|Undefined Saprotroph\| | Yeast | NULL | Probable | As nectar-tap_saprotroph (Põlme et al. 2020); Decay sugar-rich substrates (Põlme et al. 2020) | Põlme S, et al. 2020. Fungal Diversity 105:1-16 ((https://doi.org/10.1007/s13225-020-00466-2)) |
| Otu51384 | 1 | k__Fungi(100);p__Ascomycota(100);c__Saccharomycetes(100);o__Saccharomycetales(100);f__Metschnikowiaceae(100);g__Metschnikowia(100);s__Metschnikowia_pulcherrima(100); | Metschnikowia | 13 | Saprotroph | \|Undefined Saprotroph\| | Yeast | NULL | Probable | As nectar-tap_saprotroph (Põlme et al. 2020); Decay sugar-rich substrates (Põlme et al. 2020) | Põlme S, et al. 2020. Fungal Diversity 105:1-16 ((https://doi.org/10.1007/s13225-020-00466-2)) |
| Otu51385 | 1 | k__Fungi(100);p__Ascomycota(100);c__Saccharomycetes(100);o__Saccharomycetales(100);f__Metschnikowiaceae(100);g__Metschnikowia(100);g__Metschnikowia_unclassified(100); | Metschnikowia | 13 | Saprotroph | \|Undefined Saprotroph\| | Yeast | NULL | Probable | As nectar-tap_saprotroph (Põlme et al. 2020); Decay sugar-rich substrates (Põlme et al. 2020) | Põlme S, et al. 2020. Fungal Diversity 105:1-16 ((https://doi.org/10.1007/s13225-020-00466-2)) |
| Otu51386 | 1 | k__Fungi(100);p__Ascomycota(100);c__Saccharomycetes(100);o__Saccharomycetales(100);f__Metschnikowiaceae(100);g__Metschnikowia(100);s__Metschnikowia_pulcherrima(100); | Metschnikowia | 13 | Saprotroph | \|Undefined Saprotroph\| | Yeast | NULL | Probable | As nectar-tap_saprotroph (Põlme et al. 2020); Decay sugar-rich substrates (Põlme et al. 2020) | Põlme S, et al. 2020. Fungal Diversity 105:1-16 ((https://doi.org/10.1007/s13225-020-00466-2)) |
| Otu51387 | 1 | k__Fungi(100);p__Ascomycota(100);c__Saccharomycetes(100);o__Saccharomycetales(100);f__Metschnikowiaceae(100);g__Metschnikowia(100);s__Metschnikowia_pulcherrima(100); | Metschnikowia | 13 | Saprotroph | \|Undefined Saprotroph\| | Yeast | NULL | Probable | As nectar-tap_saprotroph (Põlme et al. 2020); Decay sugar-rich substrates (Põlme et al. 2020) | Põlme S, et al. 2020. Fungal Diversity 105:1-16 ((https://doi.org/10.1007/s13225-020-00466-2)) |
| Otu51388 | 1 | k__Fungi(100);p__Ascomycota(100);c__Saccharomycetes(100);o__Saccharomycetales(100);f__Metschnikowiaceae(100);g__Metschnikowia(100);g__Metschnikowia_unclassified(100); | Metschnikowia | 13 | Saprotroph | \|Undefined Saprotroph\| | Yeast | NULL | Probable | As nectar-tap_saprotroph (Põlme et al. 2020); Decay sugar-rich substrates (Põlme et al. 2020) | Põlme S, et al. 2020. Fungal Diversity 105:1-16 ((https://doi.org/10.1007/s13225-020-00466-2)) |
| Otu51389 | 1 | k__Fungi(100);p__Ascomycota(100);c__Saccharomycetes(100);o__Saccharomycetales(100);f__Metschnikowiaceae(100);g__Metschnikowia(100);s__Metschnikowia_pulcherrima(100); | Metschnikowia | 13 | Saprotroph | \|Undefined Saprotroph\| | Yeast | NULL | Probable | As nectar-tap_saprotroph (Põlme et al. 2020); Decay sugar-rich substrates (Põlme et al. 2020) | Põlme S, et al. 2020. Fungal Diversity 105:1-16 ((https://doi.org/10.1007/s13225-020-00466-2)) |
| Otu51390 | 1 | k__Fungi(100);p__Ascomycota(100);c__Saccharomycetes(100);o__Saccharomycetales(100);f__Metschnikowiaceae(100);g__Metschnikowia(100);s__Metschnikowia_pulcherrima(100); | Metschnikowia | 13 | Saprotroph | \|Undefined Saprotroph\| | Yeast | NULL | Probable | As nectar-tap_saprotroph (Põlme et al. 2020); Decay sugar-rich substrates (Põlme et al. 2020) | Põlme S, et al. 2020. Fungal Diversity 105:1-16 ((https://doi.org/10.1007/s13225-020-00466-2)) |
| Otu51391 | 1 | k__Fungi(100);p__Ascomycota(100);c__Saccharomycetes(100);o__Saccharomycetales(100);f__Metschnikowiaceae(100);g__Metschnikowia(100);s__Metschnikowia_pulcherrima(100); | Metschnikowia | 13 | Saprotroph | \|Undefined Saprotroph\| | Yeast | NULL | Probable | As nectar-tap_saprotroph (Põlme et al. 2020); Decay sugar-rich substrates (Põlme et al. 2020) | Põlme S, et al. 2020. Fungal Diversity 105:1-16 ((https://doi.org/10.1007/s13225-020-00466-2)) |
| Otu51392 | 1 | k__Fungi(100);p__Ascomycota(100);c__Saccharomycetes(100);o__Saccharomycetales(100);f__Metschnikowiaceae(100);g__Metschnikowia(100);s__Metschnikowia_pulcherrima(100); | Metschnikowia | 13 | Saprotroph | \|Undefined Saprotroph\| | Yeast | NULL | Probable | As nectar-tap_saprotroph (Põlme et al. 2020); Decay sugar-rich substrates (Põlme et al. 2020) | Põlme S, et al. 2020. Fungal Diversity 105:1-16 ((https://doi.org/10.1007/s13225-020-00466-2)) |
| Otu51393 | 1 | k__Fungi(100);p__Ascomycota(100);c__Saccharomycetes(100);o__Saccharomycetales(100);f__Metschnikowiaceae(100);g__Metschnikowia(100);g__Metschnikowia_unclassified(100); | Metschnikowia | 13 | Saprotroph | \|Undefined Saprotroph\| | Yeast | NULL | Probable | As nectar-tap_saprotroph (Põlme et al. 2020); Decay sugar-rich substrates (Põlme et al. 2020) | Põlme S, et al. 2020. Fungal Diversity 105:1-16 ((https://doi.org/10.1007/s13225-020-00466-2)) |
| Otu51394 | 1 | k__Fungi(100);p__Ascomycota(100);c__Saccharomycetes(100);o__Saccharomycetales(100);f__Metschnikowiaceae(100);g__Metschnikowia(100);s__Metschnikowia_pulcherrima(100); | Metschnikowia | 13 | Saprotroph | \|Undefined Saprotroph\| | Yeast | NULL | Probable | As nectar-tap_saprotroph (Põlme et al. 2020); Decay sugar-rich substrates (Põlme et al. 2020) | Põlme S, et al. 2020. Fungal Diversity 105:1-16 ((https://doi.org/10.1007/s13225-020-00466-2)) |
| Otu51395 | 1 | k__Fungi(100);p__Ascomycota(100);c__Saccharomycetes(100);o__Saccharomycetales(100);f__Metschnikowiaceae(100);g__Metschnikowia(100);g__Metschnikowia_unclassified(100); | Metschnikowia | 13 | Saprotroph | \|Undefined Saprotroph\| | Yeast | NULL | Probable | As nectar-tap_saprotroph (Põlme et al. 2020); Decay sugar-rich substrates (Põlme et al. 2020) | Põlme S, et al. 2020. Fungal Diversity 105:1-16 ((https://doi.org/10.1007/s13225-020-00466-2)) |
| Otu51396 | 1 | k__Fungi(100);p__Ascomycota(100);c__Saccharomycetes(100);o__Saccharomycetales(100);f__Metschnikowiaceae(100);g__Metschnikowia(100);s__Metschnikowia_pulcherrima(100); | Metschnikowia | 13 | Saprotroph | \|Undefined Saprotroph\| | Yeast | NULL | Probable | As nectar-tap_saprotroph (Põlme et al. 2020); Decay sugar-rich substrates (Põlme et al. 2020) | Põlme S, et al. 2020. Fungal Diversity 105:1-16 ((https://doi.org/10.1007/s13225-020-00466-2)) |
| Otu51397 | 1 | k__Fungi(100);p__Ascomycota(100);c__Saccharomycetes(100);o__Saccharomycetales(100);f__Metschnikowiaceae(100);g__Metschnikowia(100);s__Metschnikowia_pulcherrima(100); | Metschnikowia | 13 | Saprotroph | \|Undefined Saprotroph\| | Yeast | NULL | Probable | As nectar-tap_saprotroph (Põlme et al. 2020); Decay sugar-rich substrates (Põlme et al. 2020) | Põlme S, et al. 2020. Fungal Diversity 105:1-16 ((https://doi.org/10.1007/s13225-020-00466-2)) |
| Otu51399 | 1 | k__Fungi(100);p__Ascomycota(100);c__Saccharomycetes(100);o__Saccharomycetales(100);f__Metschnikowiaceae(100);g__Metschnikowia(100);s__Metschnikowia_pulcherrima(100); | Metschnikowia | 13 | Saprotroph | \|Undefined Saprotroph\| | Yeast | NULL | Probable | As nectar-tap_saprotroph (Põlme et al. 2020); Decay sugar-rich substrates (Põlme et al. 2020) | Põlme S, et al. 2020. Fungal Diversity 105:1-16 ((https://doi.org/10.1007/s13225-020-00466-2)) |
| Otu51400 | 1 | k__Fungi(100);p__Ascomycota(100);c__Saccharomycetes(100);o__Saccharomycetales(100);f__Metschnikowiaceae(100);g__Metschnikowia(100);s__Metschnikowia_pulcherrima(100); | Metschnikowia | 13 | Saprotroph | \|Undefined Saprotroph\| | Yeast | NULL | Probable | As nectar-tap_saprotroph (Põlme et al. 2020); Decay sugar-rich substrates (Põlme et al. 2020) | Põlme S, et al. 2020. Fungal Diversity 105:1-16 ((https://doi.org/10.1007/s13225-020-00466-2)) |
| Otu51401 | 1 | k__Fungi(100);p__Ascomycota(100);c__Saccharomycetes(100);o__Saccharomycetales(100);f__Metschnikowiaceae(100);g__Metschnikowia(100);g__Metschnikowia_unclassified(100); | Metschnikowia | 13 | Saprotroph | \|Undefined Saprotroph\| | Yeast | NULL | Probable | As nectar-tap_saprotroph (Põlme et al. 2020); Decay sugar-rich substrates (Põlme et al. 2020) | Põlme S, et al. 2020. Fungal Diversity 105:1-16 ((https://doi.org/10.1007/s13225-020-00466-2)) |
| Otu51402 | 1 | k__Fungi(100);p__Ascomycota(100);c__Saccharomycetes(100);o__Saccharomycetales(100);f__Metschnikowiaceae(100);g__Metschnikowia(100);s__Metschnikowia_pulcherrima(100); | Metschnikowia | 13 | Saprotroph | \|Undefined Saprotroph\| | Yeast | NULL | Probable | As nectar-tap_saprotroph (Põlme et al. 2020); Decay sugar-rich substrates (Põlme et al. 2020) | Põlme S, et al. 2020. Fungal Diversity 105:1-16 ((https://doi.org/10.1007/s13225-020-00466-2)) |
| Otu51403 | 1 | k__Fungi(100);p__Ascomycota(100);c__Saccharomycetes(100);o__Saccharomycetales(100);f__Metschnikowiaceae(100);g__Metschnikowia(100);s__Metschnikowia_pulcherrima(100); | Metschnikowia | 13 | Saprotroph | \|Undefined Saprotroph\| | Yeast | NULL | Probable | As nectar-tap_saprotroph (Põlme et al. 2020); Decay sugar-rich substrates (Põlme et al. 2020) | Põlme S, et al. 2020. Fungal Diversity 105:1-16 ((https://doi.org/10.1007/s13225-020-00466-2)) |
| Otu51404 | 1 | k__Fungi(100);p__Ascomycota(100);c__Saccharomycetes(100);o__Saccharomycetales(100);f__Metschnikowiaceae(100);g__Metschnikowia(100);s__Metschnikowia_pulcherrima(100); | Metschnikowia | 13 | Saprotroph | \|Undefined Saprotroph\| | Yeast | NULL | Probable | As nectar-tap_saprotroph (Põlme et al. 2020); Decay sugar-rich substrates (Põlme et al. 2020) | Põlme S, et al. 2020. Fungal Diversity 105:1-16 ((https://doi.org/10.1007/s13225-020-00466-2)) |
| Otu51405 | 1 | k__Fungi(100);p__Ascomycota(100);c__Saccharomycetes(100);o__Saccharomycetales(100);f__Metschnikowiaceae(100);g__Metschnikowia(100);s__Metschnikowia_pulcherrima(100); | Metschnikowia | 13 | Saprotroph | \|Undefined Saprotroph\| | Yeast | NULL | Probable | As nectar-tap_saprotroph (Põlme et al. 2020); Decay sugar-rich substrates (Põlme et al. 2020) | Põlme S, et al. 2020. Fungal Diversity 105:1-16 ((https://doi.org/10.1007/s13225-020-00466-2)) |
| Otu51406 | 1 | k__Fungi(100);p__Ascomycota(100);c__Saccharomycetes(100);o__Saccharomycetales(100);f__Metschnikowiaceae(100);g__Metschnikowia(100);s__Metschnikowia_pulcherrima(100); | Metschnikowia | 13 | Saprotroph | \|Undefined Saprotroph\| | Yeast | NULL | Probable | As nectar-tap_saprotroph (Põlme et al. 2020); Decay sugar-rich substrates (Põlme et al. 2020) | Põlme S, et al. 2020. Fungal Diversity 105:1-16 ((https://doi.org/10.1007/s13225-020-00466-2)) |
| Otu51409 | 1 | k__Fungi(100);p__Ascomycota(100);c__Saccharomycetes(100);o__Saccharomycetales(100);f__Metschnikowiaceae(100);g__Metschnikowia(100);s__Metschnikowia_pulcherrima(100); | Metschnikowia | 13 | Saprotroph | \|Undefined Saprotroph\| | Yeast | NULL | Probable | As nectar-tap_saprotroph (Põlme et al. 2020); Decay sugar-rich substrates (Põlme et al. 2020) | Põlme S, et al. 2020. Fungal Diversity 105:1-16 ((https://doi.org/10.1007/s13225-020-00466-2)) |
| Otu51411 | 1 | k__Fungi(100);p__Ascomycota(100);c__Saccharomycetes(100);o__Saccharomycetales(100);f__Metschnikowiaceae(100);g__Metschnikowia(100);s__Metschnikowia_pulcherrima(100); | Metschnikowia | 13 | Saprotroph | \|Undefined Saprotroph\| | Yeast | NULL | Probable | As nectar-tap_saprotroph (Põlme et al. 2020); Decay sugar-rich substrates (Põlme et al. 2020) | Põlme S, et al. 2020. Fungal Diversity 105:1-16 ((https://doi.org/10.1007/s13225-020-00466-2)) |
| Otu51412 | 1 | k__Fungi(100);p__Ascomycota(100);c__Saccharomycetes(100);o__Saccharomycetales(100);f__Metschnikowiaceae(100);g__Metschnikowia(100);s__Metschnikowia_pulcherrima(100); | Metschnikowia | 13 | Saprotroph | \|Undefined Saprotroph\| | Yeast | NULL | Probable | As nectar-tap_saprotroph (Põlme et al. 2020); Decay sugar-rich substrates (Põlme et al. 2020) | Põlme S, et al. 2020. Fungal Diversity 105:1-16 ((https://doi.org/10.1007/s13225-020-00466-2)) |
| Otu51413 | 1 | k__Fungi(100);p__Ascomycota(100);c__Saccharomycetes(100);o__Saccharomycetales(100);f__Metschnikowiaceae(100);g__Metschnikowia(100);g__Metschnikowia_unclassified(100); | Metschnikowia | 13 | Saprotroph | \|Undefined Saprotroph\| | Yeast | NULL | Probable | As nectar-tap_saprotroph (Põlme et al. 2020); Decay sugar-rich substrates (Põlme et al. 2020) | Põlme S, et al. 2020. Fungal Diversity 105:1-16 ((https://doi.org/10.1007/s13225-020-00466-2)) |
| Otu51414 | 1 | k__Fungi(100);p__Ascomycota(100);c__Saccharomycetes(100);o__Saccharomycetales(100);f__Metschnikowiaceae(100);g__Metschnikowia(100);s__Metschnikowia_pulcherrima(100); | Metschnikowia | 13 | Saprotroph | \|Undefined Saprotroph\| | Yeast | NULL | Probable | As nectar-tap_saprotroph (Põlme et al. 2020); Decay sugar-rich substrates (Põlme et al. 2020) | Põlme S, et al. 2020. Fungal Diversity 105:1-16 ((https://doi.org/10.1007/s13225-020-00466-2)) |
| Otu51415 | 1 | k__Fungi(100);p__Ascomycota(100);c__Saccharomycetes(100);o__Saccharomycetales(100);f__Metschnikowiaceae(100);g__Metschnikowia(100);s__Metschnikowia_pulcherrima(100); | Metschnikowia | 13 | Saprotroph | \|Undefined Saprotroph\| | Yeast | NULL | Probable | As nectar-tap_saprotroph (Põlme et al. 2020); Decay sugar-rich substrates (Põlme et al. 2020) | Põlme S, et al. 2020. Fungal Diversity 105:1-16 ((https://doi.org/10.1007/s13225-020-00466-2)) |
| Otu51416 | 1 | k__Fungi(100);p__Ascomycota(100);c__Saccharomycetes(100);o__Saccharomycetales(100);f__Metschnikowiaceae(100);g__Metschnikowia(100);s__Metschnikowia_pulcherrima(100); | Metschnikowia | 13 | Saprotroph | \|Undefined Saprotroph\| | Yeast | NULL | Probable | As nectar-tap_saprotroph (Põlme et al. 2020); Decay sugar-rich substrates (Põlme et al. 2020) | Põlme S, et al. 2020. Fungal Diversity 105:1-16 ((https://doi.org/10.1007/s13225-020-00466-2)) |
| Otu51417 | 1 | k__Fungi(100);p__Ascomycota(100);c__Saccharomycetes(100);o__Saccharomycetales(100);f__Metschnikowiaceae(100);g__Metschnikowia(100);s__Metschnikowia_pulcherrima(100); | Metschnikowia | 13 | Saprotroph | \|Undefined Saprotroph\| | Yeast | NULL | Probable | As nectar-tap_saprotroph (Põlme et al. 2020); Decay sugar-rich substrates (Põlme et al. 2020) | Põlme S, et al. 2020. Fungal Diversity 105:1-16 ((https://doi.org/10.1007/s13225-020-00466-2)) |
| Otu51418 | 1 | k__Fungi(100);p__Ascomycota(100);c__Saccharomycetes(100);o__Saccharomycetales(100);f__Metschnikowiaceae(100);g__Metschnikowia(100);s__Metschnikowia_pulcherrima(100); | Metschnikowia | 13 | Saprotroph | \|Undefined Saprotroph\| | Yeast | NULL | Probable | As nectar-tap_saprotroph (Põlme et al. 2020); Decay sugar-rich substrates (Põlme et al. 2020) | Põlme S, et al. 2020. Fungal Diversity 105:1-16 ((https://doi.org/10.1007/s13225-020-00466-2)) |
| Otu51419 | 1 | k__Fungi(100);p__Ascomycota(100);c__Saccharomycetes(100);o__Saccharomycetales(100);f__Metschnikowiaceae(100);g__Metschnikowia(100);s__Metschnikowia_pulcherrima(100); | Metschnikowia | 13 | Saprotroph | \|Undefined Saprotroph\| | Yeast | NULL | Probable | As nectar-tap_saprotroph (Põlme et al. 2020); Decay sugar-rich substrates (Põlme et al. 2020) | Põlme S, et al. 2020. Fungal Diversity 105:1-16 ((https://doi.org/10.1007/s13225-020-00466-2)) |
| Otu51420 | 1 | k__Fungi(100);p__Ascomycota(100);c__Saccharomycetes(100);o__Saccharomycetales(100);f__Metschnikowiaceae(100);g__Metschnikowia(100);s__Metschnikowia_pulcherrima(100); | Metschnikowia | 13 | Saprotroph | \|Undefined Saprotroph\| | Yeast | NULL | Probable | As nectar-tap_saprotroph (Põlme et al. 2020); Decay sugar-rich substrates (Põlme et al. 2020) | Põlme S, et al. 2020. Fungal Diversity 105:1-16 ((https://doi.org/10.1007/s13225-020-00466-2)) |
| Otu51421 | 1 | k__Fungi(100);p__Ascomycota(100);c__Saccharomycetes(100);o__Saccharomycetales(100);f__Metschnikowiaceae(100);g__Metschnikowia(100);g__Metschnikowia_unclassified(100); | Metschnikowia | 13 | Saprotroph | \|Undefined Saprotroph\| | Yeast | NULL | Probable | As nectar-tap_saprotroph (Põlme et al. 2020); Decay sugar-rich substrates (Põlme et al. 2020) | Põlme S, et al. 2020. Fungal Diversity 105:1-16 ((https://doi.org/10.1007/s13225-020-00466-2)) |
| Otu51422 | 1 | k__Fungi(100);p__Ascomycota(100);c__Saccharomycetes(100);o__Saccharomycetales(100);f__Metschnikowiaceae(100);g__Metschnikowia(100);s__Metschnikowia_pulcherrima(100); | Metschnikowia | 13 | Saprotroph | \|Undefined Saprotroph\| | Yeast | NULL | Probable | As nectar-tap_saprotroph (Põlme et al. 2020); Decay sugar-rich substrates (Põlme et al. 2020) | Põlme S, et al. 2020. Fungal Diversity 105:1-16 ((https://doi.org/10.1007/s13225-020-00466-2)) |
| Otu51423 | 1 | k__Fungi(100);p__Ascomycota(100);c__Saccharomycetes(100);o__Saccharomycetales(100);f__Metschnikowiaceae(100);g__Metschnikowia(100);g__Metschnikowia_unclassified(100); | Metschnikowia | 13 | Saprotroph | \|Undefined Saprotroph\| | Yeast | NULL | Probable | As nectar-tap_saprotroph (Põlme et al. 2020); Decay sugar-rich substrates (Põlme et al. 2020) | Põlme S, et al. 2020. Fungal Diversity 105:1-16 ((https://doi.org/10.1007/s13225-020-00466-2)) |
| Otu51424 | 1 | k__Fungi(100);p__Ascomycota(100);c__Saccharomycetes(100);o__Saccharomycetales(100);f__Metschnikowiaceae(100);g__Metschnikowia(100);s__Metschnikowia_pulcherrima(100); | Metschnikowia | 13 | Saprotroph | \|Undefined Saprotroph\| | Yeast | NULL | Probable | As nectar-tap_saprotroph (Põlme et al. 2020); Decay sugar-rich substrates (Põlme et al. 2020) | Põlme S, et al. 2020. Fungal Diversity 105:1-16 ((https://doi.org/10.1007/s13225-020-00466-2)) |
| Otu51425 | 1 | k__Fungi(100);p__Ascomycota(100);c__Saccharomycetes(100);o__Saccharomycetales(100);f__Metschnikowiaceae(100);g__Metschnikowia(100);s__Metschnikowia_pulcherrima(100); | Metschnikowia | 13 | Saprotroph | \|Undefined Saprotroph\| | Yeast | NULL | Probable | As nectar-tap_saprotroph (Põlme et al. 2020); Decay sugar-rich substrates (Põlme et al. 2020) | Põlme S, et al. 2020. Fungal Diversity 105:1-16 ((https://doi.org/10.1007/s13225-020-00466-2)) |
| Otu51426 | 1 | k__Fungi(100);p__Ascomycota(100);c__Saccharomycetes(100);o__Saccharomycetales(100);f__Metschnikowiaceae(100);g__Metschnikowia(100);s__Metschnikowia_pulcherrima(100); | Metschnikowia | 13 | Saprotroph | \|Undefined Saprotroph\| | Yeast | NULL | Probable | As nectar-tap_saprotroph (Põlme et al. 2020); Decay sugar-rich substrates (Põlme et al. 2020) | Põlme S, et al. 2020. Fungal Diversity 105:1-16 ((https://doi.org/10.1007/s13225-020-00466-2)) |

| | | | | | | | | | | | |
|---|---|---|---|---|---|---|---|---|---|---|---|
| Otu51427 | 1 | k__Fungi(100);p__Ascomycota(100);c__Saccharomycetes(100);o__Saccharomycetales(100);f__Metschnikowiaceae(100);g__Metschnikowia(100);s__Metschnikowia_pulcherrima(100); | Metschnikowia | 13 | Saprotroph | \|Undefined Saprotroph\| | Yeast | NULL | Probable | As nectar-tap_saprotroph (Põlme et al. 2020); Decay sugar-rich substrates (Põlme et al. 2020) | Põlme S, et al. 2020. Fungal Diversity 105:1-16 ((https://doi.org/10.1007/s13225-020-00466-2)) |
| Otu51428 | 1 | k__Fungi(100);p__Ascomycota(100);c__Saccharomycetes(100);o__Saccharomycetales(100);f__Metschnikowiaceae(100);g__Metschnikowia(100);s__Metschnikowia_pulcherrima(100); | Metschnikowia | 13 | Saprotroph | \|Undefined Saprotroph\| | Yeast | NULL | Probable | As nectar-tap_saprotroph (Põlme et al. 2020); Decay sugar-rich substrates (Põlme et al. 2020) | Põlme S, et al. 2020. Fungal Diversity 105:1-16 ((https://doi.org/10.1007/s13225-020-00466-2)) |
| Otu51429 | 1 | k__Fungi(100);p__Ascomycota(100);c__Saccharomycetes(100);o__Saccharomycetales(100);f__Metschnikowiaceae(100);g__Metschnikowia(100);s__Metschnikowia_pulcherrima(100); | Metschnikowia | 13 | Saprotroph | \|Undefined Saprotroph\| | Yeast | NULL | Probable | As nectar-tap_saprotroph (Põlme et al. 2020); Decay sugar-rich substrates (Põlme et al. 2020) | Põlme S, et al. 2020. Fungal Diversity 105:1-16 ((https://doi.org/10.1007/s13225-020-00466-2)) |
| Otu51430 | 1 | k__Fungi(100);p__Ascomycota(100);c__Saccharomycetes(100);o__Saccharomycetales(100);f__Metschnikowiaceae(100);g__Metschnikowia(100);g__Metschnikowia_unclassified(100); | Metschnikowia | 13 | Saprotroph | \|Undefined Saprotroph\| | Yeast | NULL | Probable | As nectar-tap_saprotroph (Põlme et al. 2020); Decay sugar-rich substrates (Põlme et al. 2020) | Põlme S, et al. 2020. Fungal Diversity 105:1-16 ((https://doi.org/10.1007/s13225-020-00466-2)) |
| Otu51431 | 1 | k__Fungi(100);p__Ascomycota(100);c__Saccharomycetes(100);o__Saccharomycetales(100);f__Metschnikowiaceae(100);g__Metschnikowia(100);s__Metschnikowia_pulcherrima(100); | Metschnikowia | 13 | Saprotroph | \|Undefined Saprotroph\| | Yeast | NULL | Probable | As nectar-tap_saprotroph (Põlme et al. 2020); Decay sugar-rich substrates (Põlme et al. 2020) | Põlme S, et al. 2020. Fungal Diversity 105:1-16 ((https://doi.org/10.1007/s13225-020-00466-2)) |
| Otu51432 | 1 | k__Fungi(100);p__Ascomycota(100);c__Saccharomycetes(100);o__Saccharomycetales(100);f__Metschnikowiaceae(100);g__Metschnikowia(100);s__Metschnikowia_pulcherrima(100); | Metschnikowia | 13 | Saprotroph | \|Undefined Saprotroph\| | Yeast | NULL | Probable | As nectar-tap_saprotroph (Põlme et al. 2020); Decay sugar-rich substrates (Põlme et al. 2020) | Põlme S, et al. 2020. Fungal Diversity 105:1-16 ((https://doi.org/10.1007/s13225-020-00466-2)) |
| Otu51433 | 1 | k__Fungi(100);p__Ascomycota(100);c__Saccharomycetes(100);o__Saccharomycetales(100);f__Metschnikowiaceae(100);g__Metschnikowia(100);s__Metschnikowia_pulcherrima(100); | Metschnikowia | 13 | Saprotroph | \|Undefined Saprotroph\| | Yeast | NULL | Probable | As nectar-tap_saprotroph (Põlme et al. 2020); Decay sugar-rich substrates (Põlme et al. 2020) | Põlme S, et al. 2020. Fungal Diversity 105:1-16 ((https://doi.org/10.1007/s13225-020-00466-2)) |
| Otu51434 | 1 | k__Fungi(100);p__Ascomycota(100);c__Saccharomycetes(100);o__Saccharomycetales(100);f__Metschnikowiaceae(100);g__Metschnikowia(100);g__Metschnikowia_unclassified(100); | Metschnikowia | 13 | Saprotroph | \|Undefined Saprotroph\| | Yeast | NULL | Probable | As nectar-tap_saprotroph (Põlme et al. 2020); Decay sugar-rich substrates (Põlme et al. 2020) | Põlme S, et al. 2020. Fungal Diversity 105:1-16 ((https://doi.org/10.1007/s13225-020-00466-2)) |
| Otu51435 | 1 | k__Fungi(100);p__Ascomycota(100);c__Saccharomycetes(100);o__Saccharomycetales(100);f__Metschnikowiaceae(100);g__Metschnikowia(100);s__Metschnikowia_pulcherrima(100); | Metschnikowia | 13 | Saprotroph | \|Undefined Saprotroph\| | Yeast | NULL | Probable | As nectar-tap_saprotroph (Põlme et al. 2020); Decay sugar-rich substrates (Põlme et al. 2020) | Põlme S, et al. 2020. Fungal Diversity 105:1-16 ((https://doi.org/10.1007/s13225-020-00466-2)) |
| Otu51436 | 1 | k__Fungi(100);p__Ascomycota(100);c__Saccharomycetes(100);o__Saccharomycetales(100);f__Metschnikowiaceae(100);g__Metschnikowia(100);g__Metschnikowia_unclassified(100); | Metschnikowia | 13 | Saprotroph | \|Undefined Saprotroph\| | Yeast | NULL | Probable | As nectar-tap_saprotroph (Põlme et al. 2020); Decay sugar-rich substrates (Põlme et al. 2020) | Põlme S, et al. 2020. Fungal Diversity 105:1-16 ((https://doi.org/10.1007/s13225-020-00466-2)) |
| Otu51437 | 1 | k__Fungi(100);p__Ascomycota(100);c__Saccharomycetes(100);o__Saccharomycetales(100);f__Metschnikowiaceae(100);g__Metschnikowia(100);s__Metschnikowia_pulcherrima(100); | Metschnikowia | 13 | Saprotroph | \|Undefined Saprotroph\| | Yeast | NULL | Probable | As nectar-tap_saprotroph (Põlme et al. 2020); Decay sugar-rich substrates (Põlme et al. 2020) | Põlme S, et al. 2020. Fungal Diversity 105:1-16 ((https://doi.org/10.1007/s13225-020-00466-2)) |
| Otu51439 | 1 | k__Fungi(100);p__Ascomycota(100);c__Saccharomycetes(100);o__Saccharomycetales(100);f__Metschnikowiaceae(100);g__Metschnikowia(100);s__Metschnikowia_pulcherrima(100); | Metschnikowia | 13 | Saprotroph | \|Undefined Saprotroph\| | Yeast | NULL | Probable | As nectar-tap_saprotroph (Põlme et al. 2020); Decay sugar-rich substrates (Põlme et al. 2020) | Põlme S, et al. 2020. Fungal Diversity 105:1-16 ((https://doi.org/10.1007/s13225-020-00466-2)) |
| Otu51440 | 1 | k__Fungi(100);p__Ascomycota(100);c__Saccharomycetes(100);o__Saccharomycetales(100);f__Metschnikowiaceae(100);g__Metschnikowia(100);s__Metschnikowia_pulcherrima(100); | Metschnikowia | 13 | Saprotroph | \|Undefined Saprotroph\| | Yeast | NULL | Probable | As nectar-tap_saprotroph (Põlme et al. 2020); Decay sugar-rich substrates (Põlme et al. 2020) | Põlme S, et al. 2020. Fungal Diversity 105:1-16 ((https://doi.org/10.1007/s13225-020-00466-2)) |
| Otu51441 | 1 | k__Fungi(100);p__Ascomycota(100);c__Saccharomycetes(100);o__Saccharomycetales(100);f__Metschnikowiaceae(100);g__Metschnikowia(100);s__Metschnikowia_pulcherrima(100); | Metschnikowia | 13 | Saprotroph | \|Undefined Saprotroph\| | Yeast | NULL | Probable | As nectar-tap_saprotroph (Põlme et al. 2020); Decay sugar-rich substrates (Põlme et al. 2020) | Põlme S, et al. 2020. Fungal Diversity 105:1-16 ((https://doi.org/10.1007/s13225-020-00466-2)) |
| Otu51442 | 1 | k__Fungi(100);p__Ascomycota(100);c__Saccharomycetes(100);o__Saccharomycetales(100);f__Metschnikowiaceae(100);g__Metschnikowia(100);s__Metschnikowia_pulcherrima(100); | Metschnikowia | 13 | Saprotroph | \|Undefined Saprotroph\| | Yeast | NULL | Probable | As nectar-tap_saprotroph (Põlme et al. 2020); Decay sugar-rich substrates (Põlme et al. 2020) | Põlme S, et al. 2020. Fungal Diversity 105:1-16 ((https://doi.org/10.1007/s13225-020-00466-2)) |
| Otu51443 | 1 | k__Fungi(100);p__Ascomycota(100);c__Saccharomycetes(100);o__Saccharomycetales(100);f__Metschnikowiaceae(100);g__Metschnikowia(100);g__Metschnikowia_unclassified(100); | Metschnikowia | 13 | Saprotroph | \|Undefined Saprotroph\| | Yeast | NULL | Probable | As nectar-tap_saprotroph (Põlme et al. 2020); Decay sugar-rich substrates (Põlme et al. 2020) | Põlme S, et al. 2020. Fungal Diversity 105:1-16 ((https://doi.org/10.1007/s13225-020-00466-2)) |
| Otu51444 | 1 | k__Fungi(100);p__Ascomycota(100);c__Saccharomycetes(100);o__Saccharomycetales(100);f__Metschnikowiaceae(100);g__Metschnikowia(100);s__Metschnikowia_pulcherrima(100); | Metschnikowia | 13 | Saprotroph | \|Undefined Saprotroph\| | Yeast | NULL | Probable | As nectar-tap_saprotroph (Põlme et al. 2020); Decay sugar-rich substrates (Põlme et al. 2020) | Põlme S, et al. 2020. Fungal Diversity 105:1-16 ((https://doi.org/10.1007/s13225-020-00466-2)) |
| Otu51445 | 1 | k__Fungi(100);p__Ascomycota(100);c__Saccharomycetes(100);o__Saccharomycetales(100);f__Metschnikowiaceae(100);g__Metschnikowia(100);s__Metschnikowia_pulcherrima(100); | Metschnikowia | 13 | Saprotroph | \|Undefined Saprotroph\| | Yeast | NULL | Probable | As nectar-tap_saprotroph (Põlme et al. 2020); Decay sugar-rich substrates (Põlme et al. 2020) | Põlme S, et al. 2020. Fungal Diversity 105:1-16 ((https://doi.org/10.1007/s13225-020-00466-2)) |
| Otu51446 | 1 | k__Fungi(100);p__Ascomycota(100);c__Saccharomycetes(100);o__Saccharomycetales(100);f__Metschnikowiaceae(100);g__Metschnikowia(100);g__Metschnikowia_unclassified(100); | Metschnikowia | 13 | Saprotroph | \|Undefined Saprotroph\| | Yeast | NULL | Probable | As nectar-tap_saprotroph (Põlme et al. 2020); Decay sugar-rich substrates (Põlme et al. 2020) | Põlme S, et al. 2020. Fungal Diversity 105:1-16 ((https://doi.org/10.1007/s13225-020-00466-2)) |
| Otu51447 | 1 | k__Fungi(100);p__Ascomycota(100);c__Saccharomycetes(100);o__Saccharomycetales(100);f__Metschnikowiaceae(100);g__Metschnikowia(100);g__Metschnikowia_unclassified(100); | Metschnikowia | 13 | Saprotroph | \|Undefined Saprotroph\| | Yeast | NULL | Probable | As nectar-tap_saprotroph (Põlme et al. 2020); Decay sugar-rich substrates (Põlme et al. 2020) | Põlme S, et al. 2020. Fungal Diversity 105:1-16 ((https://doi.org/10.1007/s13225-020-00466-2)) |
| Otu51448 | 1 | k__Fungi(100);p__Ascomycota(100);c__Saccharomycetes(100);o__Saccharomycetales(100);f__Metschnikowiaceae(100);g__Metschnikowia(100);s__Metschnikowia_pulcherrima(100); | Metschnikowia | 13 | Saprotroph | \|Undefined Saprotroph\| | Yeast | NULL | Probable | As nectar-tap_saprotroph (Põlme et al. 2020); Decay sugar-rich substrates (Põlme et al. 2020) | Põlme S, et al. 2020. Fungal Diversity 105:1-16 ((https://doi.org/10.1007/s13225-020-00466-2)) |
| Otu51449 | 1 | k__Fungi(100);p__Ascomycota(100);c__Saccharomycetes(100);o__Saccharomycetales(100);f__Metschnikowiaceae(100);g__Metschnikowia(100);s__Metschnikowia_pulcherrima(100); | Metschnikowia | 13 | Saprotroph | \|Undefined Saprotroph\| | Yeast | NULL | Probable | As nectar-tap_saprotroph (Põlme et al. 2020); Decay sugar-rich substrates (Põlme et al. 2020) | Põlme S, et al. 2020. Fungal Diversity 105:1-16 ((https://doi.org/10.1007/s13225-020-00466-2)) |
| Otu51450 | 1 | k__Fungi(100);p__Ascomycota(100);c__Saccharomycetes(100);o__Saccharomycetales(100);f__Metschnikowiaceae(100);g__Metschnikowia(100);s__Metschnikowia_pulcherrima(100); | Metschnikowia | 13 | Saprotroph | \|Undefined Saprotroph\| | Yeast | NULL | Probable | As nectar-tap_saprotroph (Põlme et al. 2020); Decay sugar-rich substrates (Põlme et al. 2020) | Põlme S, et al. 2020. Fungal Diversity 105:1-16 ((https://doi.org/10.1007/s13225-020-00466-2)) |
| Otu51451 | 1 | k__Fungi(100);p__Ascomycota(100);c__Saccharomycetes(100);o__Saccharomycetales(100);f__Metschnikowiaceae(100);g__Metschnikowia(100);s__Metschnikowia_pulcherrima(100); | Metschnikowia | 13 | Saprotroph | \|Undefined Saprotroph\| | Yeast | NULL | Probable | As nectar-tap_saprotroph (Põlme et al. 2020); Decay sugar-rich substrates (Põlme et al. 2020) | Põlme S, et al. 2020. Fungal Diversity 105:1-16 ((https://doi.org/10.1007/s13225-020-00466-2)) |
| Otu51453 | 1 | k__Fungi(100);p__Ascomycota(100);c__Saccharomycetes(100);o__Saccharomycetales(100);f__Metschnikowiaceae(100);g__Metschnikowia(100);s__Metschnikowia_pulcherrima(100); | Metschnikowia | 13 | Saprotroph | \|Undefined Saprotroph\| | Yeast | NULL | Probable | As nectar-tap_saprotroph (Põlme et al. 2020); Decay sugar-rich substrates (Põlme et al. 2020) | Põlme S, et al. 2020. Fungal Diversity 105:1-16 ((https://doi.org/10.1007/s13225-020-00466-2)) |
| Otu51454 | 1 | k__Fungi(100);p__Ascomycota(100);c__Saccharomycetes(100);o__Saccharomycetales(100);f__Metschnikowiaceae(100);g__Metschnikowia(100);g__Metschnikowia_unclassified(100); | Metschnikowia | 13 | Saprotroph | \|Undefined Saprotroph\| | Yeast | NULL | Probable | As nectar-tap_saprotroph (Põlme et al. 2020); Decay sugar-rich substrates (Põlme et al. 2020) | Põlme S, et al. 2020. Fungal Diversity 105:1-16 ((https://doi.org/10.1007/s13225-020-00466-2)) |
| Otu51455 | 1 | k__Fungi(100);p__Ascomycota(100);c__Saccharomycetes(100);o__Saccharomycetales(100);f__Metschnikowiaceae(100);g__Metschnikowia(100);s__Metschnikowia_pulcherrima(100); | Metschnikowia | 13 | Saprotroph | \|Undefined Saprotroph\| | Yeast | NULL | Probable | As nectar-tap_saprotroph (Põlme et al. 2020); Decay sugar-rich substrates (Põlme et al. 2020) | Põlme S, et al. 2020. Fungal Diversity 105:1-16 ((https://doi.org/10.1007/s13225-020-00466-2)) |
| Otu51456 | 1 | k__Fungi(100);p__Ascomycota(100);c__Saccharomycetes(100);o__Saccharomycetales(100);f__Metschnikowiaceae(100);g__Metschnikowia(100);s__Metschnikowia_pulcherrima(100); | Metschnikowia | 13 | Saprotroph | \|Undefined Saprotroph\| | Yeast | NULL | Probable | As nectar-tap_saprotroph (Põlme et al. 2020); Decay sugar-rich substrates (Põlme et al. 2020) | Põlme S, et al. 2020. Fungal Diversity 105:1-16 ((https://doi.org/10.1007/s13225-020-00466-2)) |
| Otu51457 | 1 | k__Fungi(100);p__Ascomycota(100);c__Saccharomycetes(100);o__Saccharomycetales(100);f__Metschnikowiaceae(100);g__Metschnikowia(100);g__Metschnikowia_unclassified(100); | Metschnikowia | 13 | Saprotroph | \|Undefined Saprotroph\| | Yeast | NULL | Probable | As nectar-tap_saprotroph (Põlme et al. 2020); Decay sugar-rich substrates (Põlme et al. 2020) | Põlme S, et al. 2020. Fungal Diversity 105:1-16 ((https://doi.org/10.1007/s13225-020-00466-2)) |
| Otu51458 | 1 | k__Fungi(100);p__Ascomycota(100);c__Saccharomycetes(100);o__Saccharomycetales(100);f__Metschnikowiaceae(100);g__Metschnikowia(100);s__Metschnikowia_pulcherrima(100); | Metschnikowia | 13 | Saprotroph | \|Undefined Saprotroph\| | Yeast | NULL | Probable | As nectar-tap_saprotroph (Põlme et al. 2020); Decay sugar-rich substrates (Põlme et al. 2020) | Põlme S, et al. 2020. Fungal Diversity 105:1-16 ((https://doi.org/10.1007/s13225-020-00466-2)) |
| Otu51459 | 1 | k__Fungi(100);p__Ascomycota(100);c__Saccharomycetes(100);o__Saccharomycetales(100);f__Metschnikowiaceae(100);g__Metschnikowia(100);g__Metschnikowia_unclassified(100); | Metschnikowia | 13 | Saprotroph | \|Undefined Saprotroph\| | Yeast | NULL | Probable | As nectar-tap_saprotroph (Põlme et al. 2020); Decay sugar-rich substrates (Põlme et al. 2020) | Põlme S, et al. 2020. Fungal Diversity 105:1-16 ((https://doi.org/10.1007/s13225-020-00466-2)) |
| Otu51460 | 1 | k__Fungi(100);p__Ascomycota(100);c__Saccharomycetes(100);o__Saccharomycetales(100);f__Metschnikowiaceae(100);g__Metschnikowia(100);g__Metschnikowia_unclassified(100); | Metschnikowia | 13 | Saprotroph | \|Undefined Saprotroph\| | Yeast | NULL | Probable | As nectar-tap_saprotroph (Põlme et al. 2020); Decay sugar-rich substrates (Põlme et al. 2020) | Põlme S, et al. 2020. Fungal Diversity 105:1-16 ((https://doi.org/10.1007/s13225-020-00466-2)) |
| Otu51461 | 1 | k__Fungi(100);p__Ascomycota(100);c__Saccharomycetes(100);o__Saccharomycetales(100);f__Metschnikowiaceae(100);g__Metschnikowia(100);s__Metschnikowia_pulcherrima(100); | Metschnikowia | 13 | Saprotroph | \|Undefined Saprotroph\| | Yeast | NULL | Probable | As nectar-tap_saprotroph (Põlme et al. 2020); Decay sugar-rich substrates (Põlme et al. 2020) | Põlme S, et al. 2020. Fungal Diversity 105:1-16 ((https://doi.org/10.1007/s13225-020-00466-2)) |
| Otu51462 | 1 | k__Fungi(100);p__Ascomycota(100);c__Saccharomycetes(100);o__Saccharomycetales(100);f__Metschnikowiaceae(100);g__Metschnikowia(100);s__Metschnikowia_pulcherrima(100); | Metschnikowia | 13 | Saprotroph | \|Undefined Saprotroph\| | Yeast | NULL | Probable | As nectar-tap_saprotroph (Põlme et al. 2020); Decay sugar-rich substrates (Põlme et al. 2020) | Põlme S, et al. 2020. Fungal Diversity 105:1-16 ((https://doi.org/10.1007/s13225-020-00466-2)) |
| Otu51463 | 1 | k__Fungi(100);p__Ascomycota(100);c__Saccharomycetes(100);o__Saccharomycetales(100);f__Metschnikowiaceae(100);g__Metschnikowia(100);s__Metschnikowia_pulcherrima(100); | Metschnikowia | 13 | Saprotroph | \|Undefined Saprotroph\| | Yeast | NULL | Probable | As nectar-tap_saprotroph (Põlme et al. 2020); Decay sugar-rich substrates (Põlme et al. 2020) | Põlme S, et al. 2020. Fungal Diversity 105:1-16 ((https://doi.org/10.1007/s13225-020-00466-2)) |
| Otu51464 | 1 | k__Fungi(100);p__Ascomycota(100);c__Saccharomycetes(100);o__Saccharomycetales(100);f__Metschnikowiaceae(100);g__Metschnikowia(100);s__Metschnikowia_pulcherrima(100); | Metschnikowia | 13 | Saprotroph | \|Undefined Saprotroph\| | Yeast | NULL | Probable | As nectar-tap_saprotroph (Põlme et al. 2020); Decay sugar-rich substrates (Põlme et al. 2020) | Põlme S, et al. 2020. Fungal Diversity 105:1-16 ((https://doi.org/10.1007/s13225-020-00466-2)) |
| Otu51465 | 1 | k__Fungi(100);p__Ascomycota(100);c__Saccharomycetes(100);o__Saccharomycetales(100);f__Metschnikowiaceae(100);g__Metschnikowia(100);s__Metschnikowia_pulcherrima(100); | Metschnikowia | 13 | Saprotroph | \|Undefined Saprotroph\| | Yeast | NULL | Probable | As nectar-tap_saprotroph (Põlme et al. 2020); Decay sugar-rich substrates (Põlme et al. 2020) | Põlme S, et al. 2020. Fungal Diversity 105:1-16 ((https://doi.org/10.1007/s13225-020-00466-2)) |
| Otu51466 | 1 | k__Fungi(100);p__Ascomycota(100);c__Saccharomycetes(100);o__Saccharomycetales(100);f__Metschnikowiaceae(100);g__Metschnikowia(100);s__Metschnikowia_pulcherrima(100); | Metschnikowia | 13 | Saprotroph | \|Undefined Saprotroph\| | Yeast | NULL | Probable | As nectar-tap_saprotroph (Põlme et al. 2020); Decay sugar-rich substrates (Põlme et al. 2020) | Põlme S, et al. 2020. Fungal Diversity 105:1-16 ((https://doi.org/10.1007/s13225-020-00466-2)) |
| Otu51467 | 1 | k__Fungi(100);p__Ascomycota(100);c__Saccharomycetes(100);o__Saccharomycetales(100);f__Metschnikowiaceae(100);g__Metschnikowia(100);g__Metschnikowia_unclassified(100); | Metschnikowia | 13 | Saprotroph | \|Undefined Saprotroph\| | Yeast | NULL | Probable | As nectar-tap_saprotroph (Põlme et al. 2020); Decay sugar-rich substrates (Põlme et al. 2020) | Põlme S, et al. 2020. Fungal Diversity 105:1-16 ((https://doi.org/10.1007/s13225-020-00466-2)) |
| Otu51468 | 1 | k__Fungi(100);p__Ascomycota(100);c__Saccharomycetes(100);o__Saccharomycetales(100);f__Metschnikowiaceae(100);g__Metschnikowia(100);s__Metschnikowia_pulcherrima(100); | Metschnikowia | 13 | Saprotroph | \|Undefined Saprotroph\| | Yeast | NULL | Probable | As nectar-tap_saprotroph (Põlme et al. 2020); Decay sugar-rich substrates (Põlme et al. 2020) | Põlme S, et al. 2020. Fungal Diversity 105:1-16 ((https://doi.org/10.1007/s13225-020-00466-2)) |
| Otu51470 | 1 | k__Fungi(100);p__Ascomycota(100);c__Saccharomycetes(100);o__Saccharomycetales(100);f__Metschnikowiaceae(100);g__Metschnikowia(100);s__Metschnikowia_pulcherrima(100); | Metschnikowia | 13 | Saprotroph | \|Undefined Saprotroph\| | Yeast | NULL | Probable | As nectar-tap_saprotroph (Põlme et al. 2020); Decay sugar-rich substrates (Põlme et al. 2020) | Põlme S, et al. 2020. Fungal Diversity 105:1-16 ((https://doi.org/10.1007/s13225-020-00466-2)) |
| Otu51472 | 1 | k__Fungi(100);p__Ascomycota(100);c__Saccharomycetes(100);o__Saccharomycetales(100);f__Metschnikowiaceae(100);g__Metschnikowia(100);s__Metschnikowia_pulcherrima(100); | Metschnikowia | 13 | Saprotroph | \|Undefined Saprotroph\| | Yeast | NULL | Probable | As nectar-tap_saprotroph (Põlme et al. 2020); Decay sugar-rich substrates (Põlme et al. 2020) | Põlme S, et al. 2020. Fungal Diversity 105:1-16 ((https://doi.org/10.1007/s13225-020-00466-2)) |
| Otu51473 | 1 | k__Fungi(100);p__Ascomycota(100);c__Saccharomycetes(100);o__Saccharomycetales(100);f__Metschnikowiaceae(100);g__Metschnikowia(100);s__Metschnikowia_pulcherrima(100); | Metschnikowia | 13 | Saprotroph | \|Undefined Saprotroph\| | Yeast | NULL | Probable | As nectar-tap_saprotroph (Põlme et al. 2020); Decay sugar-rich substrates (Põlme et al. 2020) | Põlme S, et al. 2020. Fungal Diversity 105:1-16 ((https://doi.org/10.1007/s13225-020-00466-2)) |
| Otu51474 | 1 | k__Fungi(100);p__Ascomycota(100);c__Saccharomycetes(100);o__Saccharomycetales(100);f__Metschnikowiaceae(100);g__Metschnikowia(100);s__Metschnikowia_pulcherrima(100); | Metschnikowia | 13 | Saprotroph | \|Undefined Saprotroph\| | Yeast | NULL | Probable | As nectar-tap_saprotroph (Põlme et al. 2020); Decay sugar-rich substrates (Põlme et al. 2020) | Põlme S, et al. 2020. Fungal Diversity 105:1-16 ((https://doi.org/10.1007/s13225-020-00466-2)) |
| Otu51475 | 1 | k__Fungi(100);p__Ascomycota(100);c__Saccharomycetes(100);o__Saccharomycetales(100);f__Metschnikowiaceae(100);g__Metschnikowia(100);s__Metschnikowia_pulcherrima(100); | Metschnikowia | 13 | Saprotroph | \|Undefined Saprotroph\| | Yeast | NULL | Probable | As nectar-tap_saprotroph (Põlme et al. 2020); Decay sugar-rich substrates (Põlme et al. 2020) | Põlme S, et al. 2020. Fungal Diversity 105:1-16 ((https://doi.org/10.1007/s13225-020-00466-2)) |
| Otu51476 | 1 | k__Fungi(100);p__Ascomycota(100);c__Saccharomycetes(100);o__Saccharomycetales(100);f__Metschnikowiaceae(100);g__Metschnikowia(100);g__Metschnikowia_unclassified(100); | Metschnikowia | 13 | Saprotroph | \|Undefined Saprotroph\| | Yeast | NULL | Probable | As nectar-tap_saprotroph (Põlme et al. 2020); Decay sugar-rich substrates (Põlme et al. 2020) | Põlme S, et al. 2020. Fungal Diversity 105:1-16 ((https://doi.org/10.1007/s13225-020-00466-2)) |
| Otu51477 | 1 | k__Fungi(100);p__Ascomycota(100);c__Saccharomycetes(100);o__Saccharomycetales(100);f__Metschnikowiaceae(100);g__Metschnikowia(100);s__Metschnikowia_pulcherrima(100); | Metschnikowia | 13 | Saprotroph | \|Undefined Saprotroph\| | Yeast | NULL | Probable | As nectar-tap_saprotroph (Põlme et al. 2020); Decay sugar-rich substrates (Põlme et al. 2020) | Põlme S, et al. 2020. Fungal Diversity 105:1-16 ((https://doi.org/10.1007/s13225-020-00466-2)) |
| Otu51479 | 1 | k__Fungi(100);p__Ascomycota(100);c__Saccharomycetes(100);o__Saccharomycetales(100);f__Metschnikowiaceae(100);g__Metschnikowia(100);s__Metschnikowia_pulcherrima(100); | Metschnikowia | 13 | Saprotroph | \|Undefined Saprotroph\| | Yeast | NULL | Probable | As nectar-tap_saprotroph (Põlme et al. 2020); Decay sugar-rich substrates (Põlme et al. 2020) | Põlme S, et al. 2020. Fungal Diversity 105:1-16 ((https://doi.org/10.1007/s13225-020-00466-2)) |
| Otu51480 | 1 | k__Fungi(100);p__Ascomycota(100);c__Saccharomycetes(100);o__Saccharomycetales(100);f__Metschnikowiaceae(100);g__Metschnikowia(100);g__Metschnikowia_unclassified(100); | Metschnikowia | 13 | Saprotroph | \|Undefined Saprotroph\| | Yeast | NULL | Probable | As nectar-tap_saprotroph (Põlme et al. 2020); Decay sugar-rich substrates (Põlme et al. 2020) | Põlme S, et al. 2020. Fungal Diversity 105:1-16 ((https://doi.org/10.1007/s13225-020-00466-2)) |
| Otu51481 | 1 | k__Fungi(100);p__Ascomycota(100);c__Saccharomycetes(100);o__Saccharomycetales(100);f__Metschnikowiaceae(100);g__Metschnikowia(100);s__Metschnikowia_pulcherrima(100); | Metschnikowia | 13 | Saprotroph | \|Undefined Saprotroph\| | Yeast | NULL | Probable | As nectar-tap_saprotroph (Põlme et al. 2020); Decay sugar-rich substrates (Põlme et al. 2020) | Põlme S, et al. 2020. Fungal Diversity 105:1-16 ((https://doi.org/10.1007/s13225-020-00466-2)) |
| Otu51482 | 1 | k__Fungi(100);p__Ascomycota(100);c__Saccharomycetes(100);o__Saccharomycetales(100);f__Metschnikowiaceae(100);g__Metschnikowia(100);g__Metschnikowia_unclassified(100); | Metschnikowia | 13 | Saprotroph | \|Undefined Saprotroph\| | Yeast | NULL | Probable | As nectar-tap_saprotroph (Põlme et al. 2020); Decay sugar-rich substrates (Põlme et al. 2020) | Põlme S, et al. 2020. Fungal Diversity 105:1-16 ((https://doi.org/10.1007/s13225-020-00466-2)) |
| Otu51483 | 1 | k__Fungi(100);p__Ascomycota(100);c__Saccharomycetes(100);o__Saccharomycetales(100);f__Metschnikowiaceae(100);g__Metschnikowia(100);s__Metschnikowia_pulcherrima(100); | Metschnikowia | 13 | Saprotroph | \|Undefined Saprotroph\| | Yeast | NULL | Probable | As nectar-tap_saprotroph (Põlme et al. 2020); Decay sugar-rich substrates (Põlme et al. 2020) | Põlme S, et al. 2020. Fungal Diversity 105:1-16 ((https://doi.org/10.1007/s13225-020-00466-2)) |
| Otu51485 | 1 | k__Fungi(100);p__Ascomycota(100);c__Saccharomycetes(100);o__Saccharomycetales(100);f__Metschnikowiaceae(100);g__Metschnikowia(100);g__Metschnikowia_unclassified(100); | Metschnikowia | 13 | Saprotroph | \|Undefined Saprotroph\| | Yeast | NULL | Probable | As nectar-tap_saprotroph (Põlme et al. 2020); Decay sugar-rich substrates (Põlme et al. 2020) | Põlme S, et al. 2020. Fungal Diversity 105:1-16 ((https://doi.org/10.1007/s13225-020-00466-2)) |
| Otu51486 | 1 | k__Fungi(100);p__Ascomycota(100);c__Saccharomycetes(100);o__Saccharomycetales(100);f__Metschnikowiaceae(100);g__Metschnikowia(100);g__Metschnikowia_unclassified(100); | Metschnikowia | 13 | Saprotroph | \|Undefined Saprotroph\| | Yeast | NULL | Probable | As nectar-tap_saprotroph (Põlme et al. 2020); Decay sugar-rich substrates (Põlme et al. 2020) | Põlme S, et al. 2020. Fungal Diversity 105:1-16 ((https://doi.org/10.1007/s13225-020-00466-2)) |
| Otu51487 | 1 | k__Fungi(100);p__Ascomycota(100);c__Saccharomycetes(100);o__Saccharomycetales(100);f__Metschnikowiaceae(100);g__Metschnikowia(100);s__Metschnikowia_pulcherrima(100); | Metschnikowia | 13 | Saprotroph | \|Undefined Saprotroph\| | Yeast | NULL | Probable | As nectar-tap_saprotroph (Põlme et al. 2020); Decay sugar-rich substrates (Põlme et al. 2020) | Põlme S, et al. 2020. Fungal Diversity 105:1-16 ((https://doi.org/10.1007/s13225-020-00466-2)) |
| Otu51488 | 1 | k__Fungi(100);p__Ascomycota(100);c__Saccharomycetes(100);o__Saccharomycetales(100);f__Metschnikowiaceae(100);g__Metschnikowia(100);g__Metschnikowia_unclassified(100); | Metschnikowia | 13 | Saprotroph | \|Undefined Saprotroph\| | Yeast | NULL | Probable | As nectar-tap_saprotroph (Põlme et al. 2020); Decay sugar-rich substrates (Põlme et al. 2020) | Põlme S, et al. 2020. Fungal Diversity 105:1-16 ((https://doi.org/10.1007/s13225-020-00466-2)) |
| Otu51489 | 1 | k__Fungi(100);p__Ascomycota(100);c__Saccharomycetes(100);o__Saccharomycetales(100);f__Metschnikowiaceae(100);g__Metschnikowia(100);s__Metschnikowia_pulcherrima(100); | Metschnikowia | 13 | Saprotroph | \|Undefined Saprotroph\| | Yeast | NULL | Probable | As nectar-tap_saprotroph (Põlme et al. 2020); Decay sugar-rich substrates (Põlme et al. 2020) | Põlme S, et al. 2020. Fungal Diversity 105:1-16 ((https://doi.org/10.1007/s13225-020-00466-2)) |
| Otu51490 | 1 | k__Fungi(100);p__Ascomycota(100);c__Saccharomycetes(100);o__Saccharomycetales(100);f__Metschnikowiaceae(100);g__Metschnikowia(100);s__Metschnikowia_pulcherrima(100); | Metschnikowia | 13 | Saprotroph | \|Undefined Saprotroph\| | Yeast | NULL | Probable | As nectar-tap_saprotroph (Põlme et al. 2020); Decay sugar-rich substrates (Põlme et al. 2020) | Põlme S, et al. 2020. Fungal Diversity 105:1-16 ((https://doi.org/10.1007/s13225-020-00466-2)) |
| Otu51491 | 1 | k__Fungi(100);p__Ascomycota(100);c__Saccharomycetes(100);o__Saccharomycetales(100);f__Metschnikowiaceae(100);g__Metschnikowia(100);s__Metschnikowia_pulcherrima(100); | Metschnikowia | 13 | Saprotroph | \|Undefined Saprotroph\| | Yeast | NULL | Probable | As nectar-tap_saprotroph (Põlme et al. 2020); Decay sugar-rich substrates (Põlme et al. 2020) | Põlme S, et al. 2020. Fungal Diversity 105:1-16 ((https://doi.org/10.1007/s13225-020-00466-2)) |
| Otu51492 | 1 | k__Fungi(100);p__Ascomycota(100);c__Saccharomycetes(100);o__Saccharomycetales(100);f__Metschnikowiaceae(100);g__Metschnikowia(100);s__Metschnikowia_pulcherrima(100); | Metschnikowia | 13 | Saprotroph | \|Undefined Saprotroph\| | Yeast | NULL | Probable | As nectar-tap_saprotroph (Põlme et al. 2020); Decay sugar-rich substrates (Põlme et al. 2020) | Põlme S, et al. 2020. Fungal Diversity 105:1-16 ((https://doi.org/10.1007/s13225-020-00466-2)) |
| Otu51493 | 1 | k__Fungi(100);p__Ascomycota(100);c__Saccharomycetes(100);o__Saccharomycetales(100);f__Metschnikowiaceae(100);g__Metschnikowia(100);s__Metschnikowia_pulcherrima(100); | Metschnikowia | 13 | Saprotroph | \|Undefined Saprotroph\| | Yeast | NULL | Probable | As nectar-tap_saprotroph (Põlme et al. 2020); Decay sugar-rich substrates (Põlme et al. 2020) | Põlme S, et al. 2020. Fungal Diversity 105:1-16 ((https://doi.org/10.1007/s13225-020-00466-2)) |
| Otu51494 | 1 | k__Fungi(100);p__Ascomycota(100);c__Saccharomycetes(100);o__Saccharomycetales(100);f__Metschnikowiaceae(100);g__Metschnikowia(100);s__Metschnikowia_pulcherrima(100); | Metschnikowia | 13 | Saprotroph | \|Undefined Saprotroph\| | Yeast | NULL | Probable | As nectar-tap_saprotroph (Põlme et al. 2020); Decay sugar-rich substrates (Põlme et al. 2020) | Põlme S, et al. 2020. Fungal Diversity 105:1-16 ((https://doi.org/10.1007/s13225-020-00466-2)) |
| Otu51495 | 1 | k__Fungi(100);p__Ascomycota(100);c__Saccharomycetes(100);o__Saccharomycetales(100);f__Metschnikowiaceae(100);g__Metschnikowia(100);s__Metschnikowia_pulcherrima(100); | Metschnikowia | 13 | Saprotroph | \|Undefined Saprotroph\| | Yeast | NULL | Probable | As nectar-tap_saprotroph (Põlme et al. 2020); Decay sugar-rich substrates (Põlme et al. 2020) | Põlme S, et al. 2020. Fungal Diversity 105:1-16 ((https://doi.org/10.1007/s13225-020-00466-2)) |
| Otu51496 | 1 | k__Fungi(100);p__Ascomycota(100);c__Saccharomycetes(100);o__Saccharomycetales(100);f__Metschnikowiaceae(100);g__Metschnikowia(100);s__Metschnikowia_pulcherrima(100); | Metschnikowia | 13 | Saprotroph | \|Undefined Saprotroph\| | Yeast | NULL | Probable | As nectar-tap_saprotroph (Põlme et al. 2020); Decay sugar-rich substrates (Põlme et al. 2020) | Põlme S, et al. 2020. Fungal Diversity 105:1-16 ((https://doi.org/10.1007/s13225-020-00466-2)) |
| Otu51497 | 1 | k__Fungi(100);p__Ascomycota(100);c__Saccharomycetes(100);o__Saccharomycetales(100);f__Metschnikowiaceae(100);g__Metschnikowia(100);s__Metschnikowia_pulcherrima(100); | Metschnikowia | 13 | Saprotroph | \|Undefined Saprotroph\| | Yeast | NULL | Probable | As nectar-tap_saprotroph (Põlme et al. 2020); Decay sugar-rich substrates (Põlme et al. 2020) | Põlme S, et al. 2020. Fungal Diversity 105:1-16 ((https://doi.org/10.1007/s13225-020-00466-2)) |
| Otu51498 | 1 | k__Fungi(100);p__Ascomycota(100);c__Saccharomycetes(100);o__Saccharomycetales(100);f__Metschnikowiaceae(100);g__Metschnikowia(100);s__Metschnikowia_pulcherrima(100); | Metschnikowia | 13 | Saprotroph | \|Undefined Saprotroph\| | Yeast | NULL | Probable | As nectar-tap_saprotroph (Põlme et al. 2020); Decay sugar-rich substrates (Põlme et al. 2020) | Põlme S, et al. 2020. Fungal Diversity 105:1-16 ((https://doi.org/10.1007/s13225-020-00466-2)) |
| Otu51499 | 1 | k__Fungi(100);p__Ascomycota(100);c__Saccharomycetes(100);o__Saccharomycetales(100);f__Metschnikowiaceae(100);g__Metschnikowia(100);g__Metschnikowia_unclassified(100); | Metschnikowia | 13 | Saprotroph | \|Undefined Saprotroph\| | Yeast | NULL | Probable | As nectar-tap_saprotroph (Põlme et al. 2020); Decay sugar-rich substrates (Põlme et al. 2020) | Põlme S, et al. 2020. Fungal Diversity 105:1-16 ((https://doi.org/10.1007/s13225-020-00466-2)) |
| Otu51500 | 1 | k__Fungi(100);p__Ascomycota(100);c__Saccharomycetes(100);o__Saccharomycetales(100);f__Metschnikowiaceae(100);g__Metschnikowia(100);s__Metschnikowia_pulcherrima(100); | Metschnikowia | 13 | Saprotroph | \|Undefined Saprotroph\| | Yeast | NULL | Probable | As nectar-tap_saprotroph (Põlme et al. 2020); Decay sugar-rich substrates (Põlme et al. 2020) | Põlme S, et al. 2020. Fungal Diversity 105:1-16 ((https://doi.org/10.1007/s13225-020-00466-2)) |
| Otu51501 | 1 | k__Fungi(100);p__Ascomycota(100);c__Saccharomycetes(100);o__Saccharomycetales(100);f__Metschnikowiaceae(100);g__Metschnikowia(100);s__Metschnikowia_pulcherrima(100); | Metschnikowia | 13 | Saprotroph | \|Undefined Saprotroph\| | Yeast | NULL | Probable | As nectar-tap_saprotroph (Põlme et al. 2020); Decay sugar-rich substrates (Põlme et al. 2020) | Põlme S, et al. 2020. Fungal Diversity 105:1-16 ((https://doi.org/10.1007/s13225-020-00466-2)) |
| Otu51506 | 1 | k__Fungi(100);p__Ascomycota(100);c__Saccharomycetes(100);o__Saccharomycetales(100);f__Metschnikowiaceae(100);g__Metschnikowia(100);g__Metschnikowia_unclassified(100); | Metschnikowia | 13 | Saprotroph | \|Undefined Saprotroph\| | Yeast | NULL | Probable | As nectar-tap_saprotroph (Põlme et al. 2020); Decay sugar-rich substrates (Põlme et al. 2020) | Põlme S, et al. 2020. Fungal Diversity 105:1-16 ((https://doi.org/10.1007/s13225-020-00466-2)) |
| Otu51507 | 1 | k__Fungi(100);p__Ascomycota(100);c__Saccharomycetes(100);o__Saccharomycetales(100);f__Metschnikowiaceae(100);g__Metschnikowia(100);s__Metschnikowia_pulcherrima(100); | Metschnikowia | 13 | Saprotroph | \|Undefined Saprotroph\| | Yeast | NULL | Probable | As nectar-tap_saprotroph (Põlme et al. 2020); Decay sugar-rich substrates (Põlme et al. 2020) | Põlme S, et al. 2020. Fungal Diversity 105:1-16 ((https://doi.org/10.1007/s13225-020-00466-2)) |
| Otu51510 | 1 | k__Fungi(100);p__Ascomycota(100);c__Saccharomycetes(100);o__Saccharomycetales(100);f__Metschnikowiaceae(100);g__Metschnikowia(100);g__Metschnikowia_unclassified(100); | Metschnikowia | 13 | Saprotroph | \|Undefined Saprotroph\| | Yeast | NULL | Probable | As nectar-tap_saprotroph (Põlme et al. 2020); Decay sugar-rich substrates (Põlme et al. 2020) | Põlme S, et al. 2020. Fungal Diversity 105:1-16 ((https://doi.org/10.1007/s13225-020-00466-2)) |
| Otu51511 | 1 | k__Fungi(100);p__Ascomycota(100);c__Saccharomycetes(100);o__Saccharomycetales(100);f__Metschnikowiaceae(100);g__Metschnikowia(100);s__Metschnikowia_pulcherrima(100); | Metschnikowia | 13 | Saprotroph | \|Undefined Saprotroph\| | Yeast | NULL | Probable | As nectar-tap_saprotroph (Põlme et al. 2020); Decay sugar-rich substrates (Põlme et al. 2020) | Põlme S, et al. 2020. Fungal Diversity 105:1-16 ((https://doi.org/10.1007/s13225-020-00466-2)) |
| Otu51514 | 1 | k__Fungi(100);p__Ascomycota(100);c__Saccharomycetes(100);o__Saccharomycetales(100);f__Metschnikowiaceae(100);g__Metschnikowia(100);s__Metschnikowia_pulcherrima(100); | Metschnikowia | 13 | Saprotroph | \|Undefined Saprotroph\| | Yeast | NULL | Probable | As nectar-tap_saprotroph (Põlme et al. 2020); Decay sugar-rich substrates (Põlme et al. 2020) | Põlme S, et al. 2020. Fungal Diversity 105:1-16 ((https://doi.org/10.1007/s13225-020-00466-2)) |
| Otu51516 | 1 | k__Fungi(100);p__Ascomycota(100);c__Saccharomycetes(100);o__Saccharomycetales(100);f__Metschnikowiaceae(100);g__Metschnikowia(100);s__Metschnikowia_pulcherrima(100); | Metschnikowia | 13 | Saprotroph | \|Undefined Saprotroph\| | Yeast | NULL | Probable | As nectar-tap_saprotroph (Põlme et al. 2020); Decay sugar-rich substrates (Põlme et al. 2020) | Põlme S, et al. 2020. Fungal Diversity 105:1-16 ((https://doi.org/10.1007/s13225-020-00466-2)) |
| Otu51525 | 1 | k__Fungi(100);p__Ascomycota(100);c__Saccharomycetes(100);o__Saccharomycetales(100);f__Metschnikowiaceae(100);g__Metschnikowia(100);s__Metschnikowia_pulcherrima(100); | Metschnikowia | 13 | Saprotroph | \|Undefined Saprotroph\| | Yeast | NULL | Probable | As nectar-tap_saprotroph (Põlme et al. 2020); Decay sugar-rich substrates (Põlme et al. 2020) | Põlme S, et al. 2020. Fungal Diversity 105:1-16 ((https://doi.org/10.1007/s13225-020-00466-2)) |
| Otu51527 | 1 | k__Fungi(100);p__Ascomycota(100);c__Saccharomycetes(100);o__Saccharomycetales(100);f__Metschnikowiaceae(100);g__Metschnikowia(100);s__Metschnikowia_pulcherrima(100); | Metschnikowia | 13 | Saprotroph | \|Undefined Saprotroph\| | Yeast | NULL | Probable | As nectar-tap_saprotroph (Põlme et al. 2020); Decay sugar-rich substrates (Põlme et al. 2020) | Põlme S, et al. 2020. Fungal Diversity 105:1-16 ((https://doi.org/10.1007/s13225-020-00466-2)) |
| Otu51528 | 1 | k__Fungi(100);p__Ascomycota(100);c__Saccharomycetes(100);o__Saccharomycetales(100);f__Metschnikowiaceae(100);g__Metschnikowia(100);s__Metschnikowia_pulcherrima(100); | Metschnikowia | 13 | Saprotroph | \|Undefined Saprotroph\| | Yeast | NULL | Probable | As nectar-tap_saprotroph (Põlme et al. 2020); Decay sugar-rich substrates (Põlme et al. 2020) | Põlme S, et al. 2020. Fungal Diversity 105:1-16 ((https://doi.org/10.1007/s13225-020-00466-2)) |
| Otu51531 | 1 | k__Fungi(100);p__Ascomycota(100);c__Saccharomycetes(100);o__Saccharomycetales(100);f__Metschnikowiaceae(100);g__Metschnikowia(100);s__Metschnikowia_pulcherrima(100); | Metschnikowia | 13 | Saprotroph | \|Undefined Saprotroph\| | Yeast | NULL | Probable | As nectar-tap_saprotroph (Põlme et al. 2020); Decay sugar-rich substrates (Põlme et al. 2020) | Põlme S, et al. 2020. Fungal Diversity 105:1-16 ((https://doi.org/10.1007/s13225-020-00466-2)) |
| Otu51532 | 1 | k__Fungi(100);p__Ascomycota(100);c__Saccharomycetes(100);o__Saccharomycetales(100);f__Metschnikowiaceae(100);g__Metschnikowia(100);g__Metschnikowia_unclassified(100); | Metschnikowia | 13 | Saprotroph | \|Undefined Saprotroph\| | Yeast | NULL | Probable | As nectar-tap_saprotroph (Põlme et al. 2020); Decay sugar-rich substrates (Põlme et al. 2020) | Põlme S, et al. 2020. Fungal Diversity 105:1-16 ((https://doi.org/10.1007/s13225-020-00466-2)) |
| Otu51533 | 1 | k__Fungi(100);p__Ascomycota(100);c__Saccharomycetes(100);o__Saccharomycetales(100);f__Metschnikowiaceae(100);g__Metschnikowia(100);s__Metschnikowia_pulcherrima(100); | Metschnikowia | 13 | Saprotroph | \|Undefined Saprotroph\| | Yeast | NULL | Probable | As nectar-tap_saprotroph (Põlme et al. 2020); Decay sugar-rich substrates (Põlme et al. 2020) | Põlme S, et al. 2020. Fungal Diversity 105:1-16 ((https://doi.org/10.1007/s13225-020-00466-2)) |
| Otu51534 | 1 | k__Fungi(100);p__Ascomycota(100);c__Saccharomycetes(100);o__Saccharomycetales(100);f__Metschnikowiaceae(100);g__Metschnikowia(100);s__Metschnikowia_pulcherrima(100); | Metschnikowia | 13 | Saprotroph | \|Undefined Saprotroph\| | Yeast | NULL | Probable | As nectar-tap_saprotroph (Põlme et al. 2020); Decay sugar-rich substrates (Põlme et al. 2020) | Põlme S, et al. 2020. Fungal Diversity 105:1-16 ((https://doi.org/10.1007/s13225-020-00466-2)) |
| Otu51535 | 1 | k__Fungi(100);p__Ascomycota(100);c__Saccharomycetes(100);o__Saccharomycetales(100);f__Metschnikowiaceae(100);g__Metschnikowia(100);s__Metschnikowia_pulcherrima(100); | Metschnikowia | 13 | Saprotroph | \|Undefined Saprotroph\| | Yeast | NULL | Probable | As nectar-tap_saprotroph (Põlme et al. 2020); Decay sugar-rich substrates (Põlme et al. 2020) | Põlme S, et al. 2020. Fungal Diversity 105:1-16 ((https://doi.org/10.1007/s13225-020-00466-2)) |
| Otu51536 | 1 | k__Fungi(100);p__Ascomycota(100);c__Saccharomycetes(100);o__Saccharomycetales(100);f__Metschnikowiaceae(100);g__Metschnikowia(100);g__Metschnikowia_unclassified(100); | Metschnikowia | 13 | Saprotroph | \|Undefined Saprotroph\| | Yeast | NULL | Probable | As nectar-tap_saprotroph (Põlme et al. 2020); Decay sugar-rich substrates (Põlme et al. 2020) | Põlme S, et al. 2020. Fungal Diversity 105:1-16 ((https://doi.org/10.1007/s13225-020-00466-2)) |
| Otu51537 | 1 | k__Fungi(100);p__Ascomycota(100);c__Saccharomycetes(100);o__Saccharomycetales(100);f__Metschnikowiaceae(100);g__Metschnikowia(100);g__Metschnikowia_unclassified(100); | Metschnikowia | 13 | Saprotroph | \|Undefined Saprotroph\| | Yeast | NULL | Probable | As nectar-tap_saprotroph (Põlme et al. 2020); Decay sugar-rich substrates (Põlme et al. 2020) | Põlme S, et al. 2020. Fungal Diversity 105:1-16 ((https://doi.org/10.1007/s13225-020-00466-2)) |
| Otu51538 | 1 | k__Fungi(100);p__Ascomycota(100);c__Saccharomycetes(100);o__Saccharomycetales(100);f__Metschnikowiaceae(100);g__Metschnikowia(100);s__Metschnikowia_pulcherrima(100); | Metschnikowia | 13 | Saprotroph | \|Undefined Saprotroph\| | Yeast | NULL | Probable | As nectar-tap_saprotroph (Põlme et al. 2020); Decay sugar-rich substrates (Põlme et al. 2020) | Põlme S, et al. 2020. Fungal Diversity 105:1-16 ((https://doi.org/10.1007/s13225-020-00466-2)) |
| Otu51539 | 1 | k__Fungi(100);p__Ascomycota(100);c__Saccharomycetes(100);o__Saccharomycetales(100);f__Metschnikowiaceae(100);g__Metschnikowia(100);s__Metschnikowia_pulcherrima(100); | Metschnikowia | 13 | Saprotroph | \|Undefined Saprotroph\| | Yeast | NULL | Probable | As nectar-tap_saprotroph (Põlme et al. 2020); Decay sugar-rich substrates (Põlme et al. 2020) | Põlme S, et al. 2020. Fungal Diversity 105:1-16 ((https://doi.org/10.1007/s13225-020-00466-2)) |
| Otu51541 | 1 | k__Fungi(100);p__Ascomycota(100);c__Saccharomycetes(100);o__Saccharomycetales(100);f__Metschnikowiaceae(100);g__Metschnikowia(100);s__Metschnikowia_pulcherrima(100); | Metschnikowia | 13 | Saprotroph | \|Undefined Saprotroph\| | Yeast | NULL | Probable | As nectar-tap_saprotroph (Põlme et al. 2020); Decay sugar-rich substrates (Põlme et al. 2020) | Põlme S, et al. 2020. Fungal Diversity 105:1-16 ((https://doi.org/10.1007/s13225-020-00466-2)) |
| Otu51542 | 1 | k__Fungi(100);p__Ascomycota(100);c__Saccharomycetes(100);o__Saccharomycetales(100);f__Metschnikowiaceae(100);g__Metschnikowia(100);s__Metschnikowia_pulcherrima(100); | Metschnikowia | 13 | Saprotroph | \|Undefined Saprotroph\| | Yeast | NULL | Probable | As nectar-tap_saprotroph (Põlme et al. 2020); Decay sugar-rich substrates (Põlme et al. 2020) | Põlme S, et al. 2020. Fungal Diversity 105:1-16 ((https://doi.org/10.1007/s13225-020-00466-2)) |
| Otu51543 | 1 | k__Fungi(100);p__Ascomycota(100);c__Saccharomycetes(100);o__Saccharomycetales(100);f__Metschnikowiaceae(100);g__Metschnikowia(100);s__Metschnikowia_pulcherrima(100); | Metschnikowia | 13 | Saprotroph | \|Undefined Saprotroph\| | Yeast | NULL | Probable | As nectar-tap_saprotroph (Põlme et al. 2020); Decay sugar-rich substrates (Põlme et al. 2020) | Põlme S, et al. 2020. Fungal Diversity 105:1-16 ((https://doi.org/10.1007/s13225-020-00466-2)) |
| Otu51544 | 1 | k__Fungi(100);p__Ascomycota(100);c__Saccharomycetes(100);o__Saccharomycetales(100);f__Metschnikowiaceae(100);g__Metschnikowia(100);s__Metschnikowia_pulcherrima(100); | Metschnikowia | 13 | Saprotroph | \|Undefined Saprotroph\| | Yeast | NULL | Probable | As nectar-tap_saprotroph (Põlme et al. 2020); Decay sugar-rich substrates (Põlme et al. 2020) | Põlme S, et al. 2020. Fungal Diversity 105:1-16 ((https://doi.org/10.1007/s13225-020-00466-2)) |
| Otu51545 | 1 | k__Fungi(100);p__Ascomycota(100);c__Saccharomycetes(100);o__Saccharomycetales(100);f__Metschnikowiaceae(100);g__Metschnikowia(100);s__Metschnikowia_pulcherrima(100); | Metschnikowia | 13 | Saprotroph | \|Undefined Saprotroph\| | Yeast | NULL | Probable | As nectar-tap_saprotroph (Põlme et al. 2020); Decay sugar-rich substrates (Põlme et al. 2020) | Põlme S, et al. 2020. Fungal Diversity 105:1-16 ((https://doi.org/10.1007/s13225-020-00466-2)) |
| Otu51546 | 1 | k__Fungi(100);p__Ascomycota(100);c__Saccharomycetes(100);o__Saccharomycetales(100);f__Metschnikowiaceae(100);g__Metschnikowia(100);s__Metschnikowia_pulcherrima(100); | Metschnikowia | 13 | Saprotroph | \|Undefined Saprotroph\| | Yeast | NULL | Probable | As nectar-tap_saprotroph (Põlme et al. 2020); Decay sugar-rich substrates (Põlme et al. 2020) | Põlme S, et al. 2020. Fungal Diversity 105:1-16 ((https://doi.org/10.1007/s13225-020-00466-2)) |
| Otu51547 | 1 | k__Fungi(100);p__Ascomycota(100);c__Saccharomycetes(100);o__Saccharomycetales(100);f__Metschnikowiaceae(100);g__Metschnikowia(100);g__Metschnikowia_unclassified(100); | Metschnikowia | 13 | Saprotroph | \|Undefined Saprotroph\| | Yeast | NULL | Probable | As nectar-tap_saprotroph (Põlme et al. 2020); Decay sugar-rich substrates (Põlme et al. 2020) | Põlme S, et al. 2020. Fungal Diversity 105:1-16 ((https://doi.org/10.1007/s13225-020-00466-2)) |
| Otu51548 | 1 | k__Fungi(100);p__Ascomycota(100);c__Saccharomycetes(100);o__Saccharomycetales(100);f__Metschnikowiaceae(100);g__Metschnikowia(100);s__Metschnikowia_pulcherrima(100); | Metschnikowia | 13 | Saprotroph | \|Undefined Saprotroph\| | Yeast | NULL | Probable | As nectar-tap_saprotroph (Põlme et al. 2020); Decay sugar-rich substrates (Põlme et al. 2020) | Põlme S, et al. 2020. Fungal Diversity 105:1-16 ((https://doi.org/10.1007/s13225-020-00466-2)) |
| Otu51549 | 1 | k__Fungi(100);p__Ascomycota(100);c__Saccharomycetes(100);o__Saccharomycetales(100);f__Metschnikowiaceae(100);g__Metschnikowia(100);s__Metschnikowia_pulcherrima(100); | Metschnikowia | 13 | Saprotroph | \|Undefined Saprotroph\| | Yeast | NULL | Probable | As nectar-tap_saprotroph (Põlme et al. 2020); Decay sugar-rich substrates (Põlme et al. 2020) | Põlme S, et al. 2020. Fungal Diversity 105:1-16 ((https://doi.org/10.1007/s13225-020-00466-2)) |
| Otu51554 | 1 | k__Fungi(100);p__Ascomycota(100);c__Saccharomycetes(100);o__Saccharomycetales(100);f__Metschnikowiaceae(100);g__Metschnikowia(100);g__Metschnikowia_unclassified(100); | Metschnikowia | 13 | Saprotroph | \|Undefined Saprotroph\| | Yeast | NULL | Probable | As nectar-tap_saprotroph (Põlme et al. 2020); Decay sugar-rich substrates (Põlme et al. 2020) | Põlme S, et al. 2020. Fungal Diversity 105:1-16 ((https://doi.org/10.1007/s13225-020-00466-2)) |
| Otu51555 | 1 | k__Fungi(100);p__Ascomycota(100);c__Saccharomycetes(100);o__Saccharomycetales(100);f__Metschnikowiaceae(100);g__Metschnikowia(100);s__Metschnikowia_pulcherrima(100); | Metschnikowia | 13 | Saprotroph | \|Undefined Saprotroph\| | Yeast | NULL | Probable | As nectar-tap_saprotroph (Põlme et al. 2020); Decay sugar-rich substrates (Põlme et al. 2020) | Põlme S, et al. 2020. Fungal Diversity 105:1-16 ((https://doi.org/10.1007/s13225-020-00466-2)) |
| Otu51556 | 1 | k__Fungi(100);p__Ascomycota(100);c__Saccharomycetes(100);o__Saccharomycetales(100);f__Metschnikowiaceae(100);g__Metschnikowia(100);s__Metschnikowia_pulcherrima(100); | Metschnikowia | 13 | Saprotroph | \|Undefined Saprotroph\| | Yeast | NULL | Probable | As nectar-tap_saprotroph (Põlme et al. 2020); Decay sugar-rich substrates (Põlme et al. 2020) | Põlme S, et al. 2020. Fungal Diversity 105:1-16 ((https://doi.org/10.1007/s13225-020-00466-2)) |
| Otu51557 | 1 | k__Fungi(100);p__Ascomycota(100);c__Saccharomycetes(100);o__Saccharomycetales(100);f__Metschnikowiaceae(100);g__Metschnikowia(100);s__Metschnikowia_pulcherrima(100); | Metschnikowia | 13 | Saprotroph | \|Undefined Saprotroph\| | Yeast | NULL | Probable | As nectar-tap_saprotroph (Põlme et al. 2020); Decay sugar-rich substrates (Põlme et al. 2020) | Põlme S, et al. 2020. Fungal Diversity 105:1-16 ((https://doi.org/10.1007/s13225-020-00466-2)) |
| Otu51558 | 1 | k__Fungi(100);p__Ascomycota(100);c__Saccharomycetes(100);o__Saccharomycetales(100);f__Metschnikowiaceae(100);g__Metschnikowia(100);s__Metschnikowia_pulcherrima(100); | Metschnikowia | 13 | Saprotroph | \|Undefined Saprotroph\| | Yeast | NULL | Probable | As nectar-tap_saprotroph (Põlme et al. 2020); Decay sugar-rich substrates (Põlme et al. 2020) | Põlme S, et al. 2020. Fungal Diversity 105:1-16 ((https://doi.org/10.1007/s13225-020-00466-2)) |
| Otu51559 | 1 | k__Fungi(100);p__Ascomycota(100);c__Saccharomycetes(100);o__Saccharomycetales(100);f__Metschnikowiaceae(100);g__Metschnikowia(100);s__Metschnikowia_pulcherrima(100); | Metschnikowia | 13 | Saprotroph | \|Undefined Saprotroph\| | Yeast | NULL | Probable | As nectar-tap_saprotroph (Põlme et al. 2020); Decay sugar-rich substrates (Põlme et al. 2020) | Põlme S, et al. 2020. Fungal Diversity 105:1-16 ((https://doi.org/10.1007/s13225-020-00466-2)) |
| Otu51560 | 1 | k__Fungi(100);p__Ascomycota(100);c__Saccharomycetes(100);o__Saccharomycetales(100);f__Metschnikowiaceae(100);g__Metschnikowia(100);s__Metschnikowia_pulcherrima(100); | Metschnikowia | 13 | Saprotroph | \|Undefined Saprotroph\| | Yeast | NULL | Probable | As nectar-tap_saprotroph (Põlme et al. 2020); Decay sugar-rich substrates (Põlme et al. 2020) | Põlme S, et al. 2020. Fungal Diversity 105:1-16 ((https://doi.org/10.1007/s13225-020-00466-2)) |
| Otu51561 | 1 | k__Fungi(100);p__Ascomycota(100);c__Saccharomycetes(100);o__Saccharomycetales(100);f__Metschnikowiaceae(100);g__Metschnikowia(100);s__Metschnikowia_pulcherrima(100); | Metschnikowia | 13 | Saprotroph | \|Undefined Saprotroph\| | Yeast | NULL | Probable | As nectar-tap_saprotroph (Põlme et al. 2020); Decay sugar-rich substrates (Põlme et al. 2020) | Põlme S, et al. 2020. Fungal Diversity 105:1-16 ((https://doi.org/10.1007/s13225-020-00466-2)) |
| Otu51563 | 1 | k__Fungi(100);p__Ascomycota(100);c__Saccharomycetes(100);o__Saccharomycetales(100);f__Metschnikowiaceae(100);g__Metschnikowia(100);s__Metschnikowia_pulcherrima(100); | Metschnikowia | 13 | Saprotroph | \|Undefined Saprotroph\| | Yeast | NULL | Probable | As nectar-tap_saprotroph (Põlme et al. 2020); Decay sugar-rich substrates (Põlme et al. 2020) | Põlme S, et al. 2020. Fungal Diversity 105:1-16 ((https://doi.org/10.1007/s13225-020-00466-2)) |
| Otu51564 | 1 | k__Fungi(100);p__Ascomycota(100);c__Saccharomycetes(100);o__Saccharomycetales(100);f__Metschnikowiaceae(100);g__Metschnikowia(100);s__Metschnikowia_pulcherrima(100); | Metschnikowia | 13 | Saprotroph | \|Undefined Saprotroph\| | Yeast | NULL | Probable | As nectar-tap_saprotroph (Põlme et al. 2020); Decay sugar-rich substrates (Põlme et al. 2020) | Põlme S, et al. 2020. Fungal Diversity 105:1-16 ((https://doi.org/10.1007/s13225-020-00466-2)) |
| Otu51565 | 1 | k__Fungi(100);p__Ascomycota(100);c__Saccharomycetes(100);o__Saccharomycetales(100);f__Metschnikowiaceae(100);g__Metschnikowia(100);s__Metschnikowia_pulcherrima(100); | Metschnikowia | 13 | Saprotroph | \|Undefined Saprotroph\| | Yeast | NULL | Probable | As nectar-tap_saprotroph (Põlme et al. 2020); Decay sugar-rich substrates (Põlme et al. 2020) | Põlme S, et al. 2020. Fungal Diversity 105:1-16 ((https://doi.org/10.1007/s13225-020-00466-2)) |
| Otu51566 | 1 | k__Fungi(100);p__Ascomycota(100);c__Saccharomycetes(100);o__Saccharomycetales(100);f__Metschnikowiaceae(100);g__Metschnikowia(100);s__Metschnikowia_pulcherrima(100); | Metschnikowia | 13 | Saprotroph | \|Undefined Saprotroph\| | Yeast | NULL | Probable | As nectar-tap_saprotroph (Põlme et al. 2020); Decay sugar-rich substrates (Põlme et al. 2020) | Põlme S, et al. 2020. Fungal Diversity 105:1-16 ((https://doi.org/10.1007/s13225-020-00466-2)) |
| Otu51567 | 1 | k__Fungi(100);p__Ascomycota(100);c__Saccharomycetes(100);o__Saccharomycetales(100);f__Metschnikowiaceae(100);g__Metschnikowia(100);s__Metschnikowia_pulcherrima(100); | Metschnikowia | 13 | Saprotroph | \|Undefined Saprotroph\| | Yeast | NULL | Probable | As nectar-tap_saprotroph (Põlme et al. 2020); Decay sugar-rich substrates (Põlme et al. 2020) | Põlme S, et al. 2020. Fungal Diversity 105:1-16 ((https://doi.org/10.1007/s13225-020-00466-2)) |
| Otu51568 | 1 | k__Fungi(100);p__Ascomycota(100);c__Saccharomycetes(100);o__Saccharomycetales(100);f__Metschnikowiaceae(100);g__Metschnikowia(100);s__Metschnikowia_pulcherrima(100); | Metschnikowia | 13 | Saprotroph | \|Undefined Saprotroph\| | Yeast | NULL | Probable | As nectar-tap_saprotroph (Põlme et al. 2020); Decay sugar-rich substrates (Põlme et al. 2020) | Põlme S, et al. 2020. Fungal Diversity 105:1-16 ((https://doi.org/10.1007/s13225-020-00466-2)) |
| Otu51571 | 1 | k__Fungi(100);p__Ascomycota(100);c__Saccharomycetes(100);o__Saccharomycetales(100);f__Metschnikowiaceae(100);g__Metschnikowia(100);s__Metschnikowia_pulcherrima(100); | Metschnikowia | 13 | Saprotroph | \|Undefined Saprotroph\| | Yeast | NULL | Probable | As nectar-tap_saprotroph (Põlme et al. 2020); Decay sugar-rich substrates (Põlme et al. 2020) | Põlme S, et al. 2020. Fungal Diversity 105:1-16 ((https://doi.org/10.1007/s13225-020-00466-2)) |
| Otu51573 | 1 | k__Fungi(100);p__Ascomycota(100);c__Saccharomycetes(100);o__Saccharomycetales(100);f__Metschnikowiaceae(100);g__Metschnikowia(100);s__Metschnikowia_pulcherrima(100); | Metschnikowia | 13 | Saprotroph | \|Undefined Saprotroph\| | Yeast | NULL | Probable | As nectar-tap_saprotroph (Põlme et al. 2020); Decay sugar-rich substrates (Põlme et al. 2020) | Põlme S, et al. 2020. Fungal Diversity 105:1-16 ((https://doi.org/10.1007/s13225-020-00466-2)) |
| Otu51576 | 1 | k__Fungi(100);p__Ascomycota(100);c__Saccharomycetes(100);o__Saccharomycetales(100);f__Metschnikowiaceae(100);g__Metschnikowia(100);s__Metschnikowia_pulcherrima(100); | Metschnikowia | 13 | Saprotroph | \|Undefined Saprotroph\| | Yeast | NULL | Probable | As nectar-tap_saprotroph (Põlme et al. 2020); Decay sugar-rich substrates (Põlme et al. 2020) | Põlme S, et al. 2020. Fungal Diversity 105:1-16 ((https://doi.org/10.1007/s13225-020-00466-2)) |
| Otu51577 | 1 | k__Fungi(100);p__Ascomycota(100);c__Saccharomycetes(100);o__Saccharomycetales(100);f__Metschnikowiaceae(100);g__Metschnikowia(100);s__Metschnikowia_pulcherrima(100); | Metschnikowia | 13 | Saprotroph | \|Undefined Saprotroph\| | Yeast | NULL | Probable | As nectar-tap_saprotroph (Põlme et al. 2020); Decay sugar-rich substrates (Põlme et al. 2020) | Põlme S, et al. 2020. Fungal Diversity 105:1-16 ((https://doi.org/10.1007/s13225-020-00466-2)) |
| Otu51578 | 1 | k__Fungi(100);p__Ascomycota(100);c__Saccharomycetes(100);o__Saccharomycetales(100);f__Metschnikowiaceae(100);g__Metschnikowia(100);g__Metschnikowia_unclassified(100); | Metschnikowia | 13 | Saprotroph | \|Undefined Saprotroph\| | Yeast | NULL | Probable | As nectar-tap_saprotroph (Põlme et al. 2020); Decay sugar-rich substrates (Põlme et al. 2020) | Põlme S, et al. 2020. Fungal Diversity 105:1-16 ((https://doi.org/10.1007/s13225-020-00466-2)) |
| Otu51579 | 1 | k__Fungi(100);p__Ascomycota(100);c__Saccharomycetes(100);o__Saccharomycetales(100);f__Metschnikowiaceae(100);g__Metschnikowia(100);g__Metschnikowia_unclassified(100); | Metschnikowia | 13 | Saprotroph | \|Undefined Saprotroph\| | Yeast | NULL | Probable | As nectar-tap_saprotroph (Põlme et al. 2020); Decay sugar-rich substrates (Põlme et al. 2020) | Põlme S, et al. 2020. Fungal Diversity 105:1-16 ((https://doi.org/10.1007/s13225-020-00466-2)) |
| Otu51580 | 1 | k__Fungi(100);p__Ascomycota(100);c__Saccharomycetes(100);o__Saccharomycetales(100);f__Metschnikowiaceae(100);g__Metschnikowia(100);s__Metschnikowia_pulcherrima(100); | Metschnikowia | 13 | Saprotroph | \|Undefined Saprotroph\| | Yeast | NULL | Probable | As nectar-tap_saprotroph (Põlme et al. 2020); Decay sugar-rich substrates (Põlme et al. 2020) | Põlme S, et al. 2020. Fungal Diversity 105:1-16 ((https://doi.org/10.1007/s13225-020-00466-2)) |
| Otu51581 | 1 | k__Fungi(100);p__Ascomycota(100);c__Saccharomycetes(100);o__Saccharomycetales(100);f__Metschnikowiaceae(100);g__Metschnikowia(100);s__Metschnikowia_pulcherrima(100); | Metschnikowia | 13 | Saprotroph | \|Undefined Saprotroph\| | Yeast | NULL | Probable | As nectar-tap_saprotroph (Põlme et al. 2020); Decay sugar-rich substrates (Põlme et al. 2020) | Põlme S, et al. 2020. Fungal Diversity 105:1-16 ((https://doi.org/10.1007/s13225-020-00466-2)) |
| Otu51582 | 1 | k__Fungi(100);p__Ascomycota(100);c__Saccharomycetes(100);o__Saccharomycetales(100);f__Metschnikowiaceae(100);g__Metschnikowia(100);s__Metschnikowia_pulcherrima(100); | Metschnikowia | 13 | Saprotroph | \|Undefined Saprotroph\| | Yeast | NULL | Probable | As nectar-tap_saprotroph (Põlme et al. 2020); Decay sugar-rich substrates (Põlme et al. 2020) | Põlme S, et al. 2020. Fungal Diversity 105:1-16 ((https://doi.org/10.1007/s13225-020-00466-2)) |
| Otu51583 | 1 | k__Fungi(100);p__Ascomycota(100);c__Saccharomycetes(100);o__Saccharomycetales(100);f__Metschnikowiaceae(100);g__Metschnikowia(100);s__Metschnikowia_pulcherrima(100); | Metschnikowia | 13 | Saprotroph | \|Undefined Saprotroph\| | Yeast | NULL | Probable | As nectar-tap_saprotroph (Põlme et al. 2020); Decay sugar-rich substrates (Põlme et al. 2020) | Põlme S, et al. 2020. Fungal Diversity 105:1-16 ((https://doi.org/10.1007/s13225-020-00466-2)) |
| Otu51584 | 1 | k__Fungi(100);p__Ascomycota(100);c__Saccharomycetes(100);o__Saccharomycetales(100);f__Metschnikowiaceae(100);g__Metschnikowia(100);s__Metschnikowia_pulcherrima(100); | Metschnikowia | 13 | Saprotroph | \|Undefined Saprotroph\| | Yeast | NULL | Probable | As nectar-tap_saprotroph (Põlme et al. 2020); Decay sugar-rich substrates (Põlme et al. 2020) | Põlme S, et al. 2020. Fungal Diversity 105:1-16 ((https://doi.org/10.1007/s13225-020-00466-2)) |
| Otu51585 | 1 | k__Fungi(100);p__Ascomycota(100);c__Saccharomycetes(100);o__Saccharomycetales(100);f__Metschnikowiaceae(100);g__Metschnikowia(100);s__Metschnikowia_pulcherrima(100); | Metschnikowia | 13 | Saprotroph | \|Undefined Saprotroph\| | Yeast | NULL | Probable | As nectar-tap_saprotroph (Põlme et al. 2020); Decay sugar-rich substrates (Põlme et al. 2020) | Põlme S, et al. 2020. Fungal Diversity 105:1-16 ((https://doi.org/10.1007/s13225-020-00466-2)) |
| Otu51586 | 1 | k__Fungi(100);p__Ascomycota(100);c__Saccharomycetes(100);o__Saccharomycetales(100);f__Metschnikowiaceae(100);g__Metschnikowia(100);s__Metschnikowia_pulcherrima(100); | Metschnikowia | 13 | Saprotroph | \|Undefined Saprotroph\| | Yeast | NULL | Probable | As nectar-tap_saprotroph (Põlme et al. 2020); Decay sugar-rich substrates (Põlme et al. 2020) | Põlme S, et al. 2020. Fungal Diversity 105:1-16 ((https://doi.org/10.1007/s13225-020-00466-2)) |
| Otu51587 | 1 | k__Fungi(100);p__Ascomycota(100);c__Saccharomycetes(100);o__Saccharomycetales(100);f__Metschnikowiaceae(100);g__Metschnikowia(100);s__Metschnikowia_pulcherrima(100); | Metschnikowia | 13 | Saprotroph | \|Undefined Saprotroph\| | Yeast | NULL | Probable | As nectar-tap_saprotroph (Põlme et al. 2020); Decay sugar-rich substrates (Põlme et al. 2020) | Põlme S, et al. 2020. Fungal Diversity 105:1-16 ((https://doi.org/10.1007/s13225-020-00466-2)) |
| Otu51588 | 1 | k__Fungi(100);p__Ascomycota(100);c__Saccharomycetes(100);o__Saccharomycetales(100);f__Metschnikowiaceae(100);g__Metschnikowia(100);s__Metschnikowia_pulcherrima(100); | Metschnikowia | 13 | Saprotroph | \|Undefined Saprotroph\| | Yeast | NULL | Probable | As nectar-tap_saprotroph (Põlme et al. 2020); Decay sugar-rich substrates (Põlme et al. 2020) | Põlme S, et al. 2020. Fungal Diversity 105:1-16 ((https://doi.org/10.1007/s13225-020-00466-2)) |
| Otu51589 | 1 | k__Fungi(100);p__Ascomycota(100);c__Saccharomycetes(100);o__Saccharomycetales(100);f__Metschnikowiaceae(100);g__Metschnikowia(100);s__Metschnikowia_pulcherrima(100); | Metschnikowia | 13 | Saprotroph | \|Undefined Saprotroph\| | Yeast | NULL | Probable | As nectar-tap_saprotroph (Põlme et al. 2020); Decay sugar-rich substrates (Põlme et al. 2020) | Põlme S, et al. 2020. Fungal Diversity 105:1-16 ((https://doi.org/10.1007/s13225-020-00466-2)) |
| Otu51590 | 1 | k__Fungi(100);p__Ascomycota(100);c__Saccharomycetes(100);o__Saccharomycetales(100);f__Metschnikowiaceae(100);g__Metschnikowia(100);s__Metschnikowia_pulcherrima(100); | Metschnikowia | 13 | Saprotroph | \|Undefined Saprotroph\| | Yeast | NULL | Probable | As nectar-tap_saprotroph (Põlme et al. 2020); Decay sugar-rich substrates (Põlme et al. 2020) | Põlme S, et al. 2020. Fungal Diversity 105:1-16 ((https://doi.org/10.1007/s13225-020-00466-2)) |
| Otu51591 | 1 | k__Fungi(100);p__Ascomycota(100);c__Saccharomycetes(100);o__Saccharomycetales(100);f__Metschnikowiaceae(100);g__Metschnikowia(100);g__Metschnikowia_unclassified(100); | Metschnikowia | 13 | Saprotroph | \|Undefined Saprotroph\| | Yeast | NULL | Probable | As nectar-tap_saprotroph (Põlme et al. 2020); Decay sugar-rich substrates (Põlme et al. 2020) | Põlme S, et al. 2020. Fungal Diversity 105:1-16 ((https://doi.org/10.1007/s13225-020-00466-2)) |
| Otu51592 | 1 | k__Fungi(100);p__Ascomycota(100);c__Saccharomycetes(100);o__Saccharomycetales(100);f__Metschnikowiaceae(100);g__Metschnikowia(100);g__Metschnikowia_unclassified(100); | Metschnikowia | 13 | Saprotroph | \|Undefined Saprotroph\| | Yeast | NULL | Probable | As nectar-tap_saprotroph (Põlme et al. 2020); Decay sugar-rich substrates (Põlme et al. 2020) | Põlme S, et al. 2020. Fungal Diversity 105:1-16 ((https://doi.org/10.1007/s13225-020-00466-2)) |
| Otu51593 | 1 | k__Fungi(100);p__Ascomycota(100);c__Saccharomycetes(100);o__Saccharomycetales(100);f__Metschnikowiaceae(100);g__Metschnikowia(100);g__Metschnikowia_unclassified(100); | Metschnikowia | 13 | Saprotroph | \|Undefined Saprotroph\| | Yeast | NULL | Probable | As nectar-tap_saprotroph (Põlme et al. 2020); Decay sugar-rich substrates (Põlme et al. 2020) | Põlme S, et al. 2020. Fungal Diversity 105:1-16 ((https://doi.org/10.1007/s13225-020-00466-2)) |
| Otu51594 | 1 | k__Fungi(100);p__Ascomycota(100);c__Saccharomycetes(100);o__Saccharomycetales(100);f__Metschnikowiaceae(100);g__Metschnikowia(100);s__Metschnikowia_pulcherrima(100); | Metschnikowia | 13 | Saprotroph | \|Undefined Saprotroph\| | Yeast | NULL | Probable | As nectar-tap_saprotroph (Põlme et al. 2020); Decay sugar-rich substrates (Põlme et al. 2020) | Põlme S, et al. 2020. Fungal Diversity 105:1-16 ((https://doi.org/10.1007/s13225-020-00466-2)) |
| Otu51595 | 1 | k__Fungi(100);p__Ascomycota(100);c__Saccharomycetes(100);o__Saccharomycetales(100);f__Metschnikowiaceae(100);g__Metschnikowia(100);s__Metschnikowia_pulcherrima(100); | Metschnikowia | 13 | Saprotroph | \|Undefined Saprotroph\| | Yeast | NULL | Probable | As nectar-tap_saprotroph (Põlme et al. 2020); Decay sugar-rich substrates (Põlme et al. 2020) | Põlme S, et al. 2020. Fungal Diversity 105:1-16 ((https://doi.org/10.1007/s13225-020-00466-2)) |
| Otu51596 | 1 | k__Fungi(100);p__Ascomycota(100);c__Saccharomycetes(100);o__Saccharomycetales(100);f__Metschnikowiaceae(100);g__Metschnikowia(100);g__Metschnikowia_unclassified(100); | Metschnikowia | 13 | Saprotroph | \|Undefined Saprotroph\| | Yeast | NULL | Probable | As nectar-tap_saprotroph (Põlme et al. 2020); Decay sugar-rich substrates (Põlme et al. 2020) | Põlme S, et al. 2020. Fungal Diversity 105:1-16 ((https://doi.org/10.1007/s13225-020-00466-2)) |
| Otu51597 | 1 | k__Fungi(100);p__Ascomycota(100);c__Saccharomycetes(100);o__Saccharomycetales(100);f__Metschnikowiaceae(100);g__Metschnikowia(100);g__Metschnikowia_unclassified(100); | Metschnikowia | 13 | Saprotroph | \|Undefined Saprotroph\| | Yeast | NULL | Probable | As nectar-tap_saprotroph (Põlme et al. 2020); Decay sugar-rich substrates (Põlme et al. 2020) | Põlme S, et al. 2020. Fungal Diversity 105:1-16 ((https://doi.org/10.1007/s13225-020-00466-2)) |
| Otu51598 | 1 | k__Fungi(100);p__Ascomycota(100);c__Saccharomycetes(100);o__Saccharomycetales(100);f__Metschnikowiaceae(100);g__Metschnikowia(100);g__Metschnikowia_unclassified(100); | Metschnikowia | 13 | Saprotroph | \|Undefined Saprotroph\| | Yeast | NULL | Probable | As nectar-tap_saprotroph (Põlme et al. 2020); Decay sugar-rich substrates (Põlme et al. 2020) | Põlme S, et al. 2020. Fungal Diversity 105:1-16 ((https://doi.org/10.1007/s13225-020-00466-2)) |
| Otu51599 | 1 | k__Fungi(100);p__Ascomycota(100);c__Saccharomycetes(100);o__Saccharomycetales(100);f__Metschnikowiaceae(100);g__Metschnikowia(100);s__Metschnikowia_pulcherrima(100); | Metschnikowia | 13 | Saprotroph | \|Undefined Saprotroph\| | Yeast | NULL | Probable | As nectar-tap_saprotroph (Põlme et al. 2020); Decay sugar-rich substrates (Põlme et al. 2020) | Põlme S, et al. 2020. Fungal Diversity 105:1-16 ((https://doi.org/10.1007/s13225-020-00466-2)) |
| Otu51600 | 1 | k__Fungi(100);p__Ascomycota(100);c__Saccharomycetes(100);o__Saccharomycetales(100);f__Metschnikowiaceae(100);g__Metschnikowia(100);s__Metschnikowia_pulcherrima(100); | Metschnikowia | 13 | Saprotroph | \|Undefined Saprotroph\| | Yeast | NULL | Probable | As nectar-tap_saprotroph (Põlme et al. 2020); Decay sugar-rich substrates (Põlme et al. 2020) | Põlme S, et al. 2020. Fungal Diversity 105:1-16 ((https://doi.org/10.1007/s13225-020-00466-2)) |
| Otu51601 | 1 | k__Fungi(100);p__Ascomycota(100);c__Saccharomycetes(100);o__Saccharomycetales(100);f__Metschnikowiaceae(100);g__Metschnikowia(100);s__Metschnikowia_pulcherrima(100); | Metschnikowia | 13 | Saprotroph | \|Undefined Saprotroph\| | Yeast | NULL | Probable | As nectar-tap_saprotroph (Põlme et al. 2020); Decay sugar-rich substrates (Põlme et al. 2020) | Põlme S, et al. 2020. Fungal Diversity 105:1-16 ((https://doi.org/10.1007/s13225-020-00466-2)) |
| Otu51602 | 1 | k__Fungi(100);p__Ascomycota(100);c__Saccharomycetes(100);o__Saccharomycetales(100);f__Metschnikowiaceae(100);g__Metschnikowia(100);s__Metschnikowia_pulcherrima(100); | Metschnikowia | 13 | Saprotroph | \|Undefined Saprotroph\| | Yeast | NULL | Probable | As nectar-tap_saprotroph (Põlme et al. 2020); Decay sugar-rich substrates (Põlme et al. 2020) | Põlme S, et al. 2020. Fungal Diversity 105:1-16 ((https://doi.org/10.1007/s13225-020-00466-2)) |
| Otu51603 | 1 | k__Fungi(100);p__Ascomycota(100);c__Saccharomycetes(100);o__Saccharomycetales(100);f__Metschnikowiaceae(100);g__Metschnikowia(100);s__Metschnikowia_pulcherrima(100); | Metschnikowia | 13 | Saprotroph | \|Undefined Saprotroph\| | Yeast | NULL | Probable | As nectar-tap_saprotroph (Põlme et al. 2020); Decay sugar-rich substrates (Põlme et al. 2020) | Põlme S, et al. 2020. Fungal Diversity 105:1-16 ((https://doi.org/10.1007/s13225-020-00466-2)) |
| Otu51604 | 1 | k__Fungi(100);p__Ascomycota(100);c__Saccharomycetes(100);o__Saccharomycetales(100);f__Metschnikowiaceae(100);g__Metschnikowia(100);s__Metschnikowia_pulcherrima(100); | Metschnikowia | 13 | Saprotroph | \|Undefined Saprotroph\| | Yeast | NULL | Probable | As nectar-tap_saprotroph (Põlme et al. 2020); Decay sugar-rich substrates (Põlme et al. 2020) | Põlme S, et al. 2020. Fungal Diversity 105:1-16 ((https://doi.org/10.1007/s13225-020-00466-2)) |
| Otu51605 | 1 | k__Fungi(100);p__Ascomycota(100);c__Saccharomycetes(100);o__Saccharomycetales(100);f__Metschnikowiaceae(100);g__Metschnikowia(100);g__Metschnikowia_unclassified(100); | Metschnikowia | 13 | Saprotroph | \|Undefined Saprotroph\| | Yeast | NULL | Probable | As nectar-tap_saprotroph (Põlme et al. 2020); Decay sugar-rich substrates (Põlme et al. 2020) | Põlme S, et al. 2020. Fungal Diversity 105:1-16 ((https://doi.org/10.1007/s13225-020-00466-2)) |
| Otu51606 | 1 | k__Fungi(100);p__Ascomycota(100);c__Saccharomycetes(100);o__Saccharomycetales(100);f__Metschnikowiaceae(100);g__Metschnikowia(100);g__Metschnikowia_unclassified(100); | Metschnikowia | 13 | Saprotroph | \|Undefined Saprotroph\| | Yeast | NULL | Probable | As nectar-tap_saprotroph (Põlme et al. 2020); Decay sugar-rich substrates (Põlme et al. 2020) | Põlme S, et al. 2020. Fungal Diversity 105:1-16 ((https://doi.org/10.1007/s13225-020-00466-2)) |
| Otu51608 | 1 | k__Fungi(100);p__Ascomycota(100);c__Saccharomycetes(100);o__Saccharomycetales(100);f__Metschnikowiaceae(100);g__Metschnikowia(100);s__Metschnikowia_pulcherrima(100); | Metschnikowia | 13 | Saprotroph | \|Undefined Saprotroph\| | Yeast | NULL | Probable | As nectar-tap_saprotroph (Põlme et al. 2020); Decay sugar-rich substrates (Põlme et al. 2020) | Põlme S, et al. 2020. Fungal Diversity 105:1-16 ((https://doi.org/10.1007/s13225-020-00466-2)) |
| Otu51609 | 1 | k__Fungi(100);p__Ascomycota(100);c__Saccharomycetes(100);o__Saccharomycetales(100);f__Metschnikowiaceae(100);g__Metschnikowia(100);s__Metschnikowia_pulcherrima(100); | Metschnikowia | 13 | Saprotroph | \|Undefined Saprotroph\| | Yeast | NULL | Probable | As nectar-tap_saprotroph (Põlme et al. 2020); Decay sugar-rich substrates (Põlme et al. 2020) | Põlme S, et al. 2020. Fungal Diversity 105:1-16 ((https://doi.org/10.1007/s13225-020-00466-2)) |
| Otu51610 | 1 | k__Fungi(100);p__Ascomycota(100);c__Saccharomycetes(100);o__Saccharomycetales(100);f__Metschnikowiaceae(100);g__Metschnikowia(100);s__Metschnikowia_pulcherrima(100); | Metschnikowia | 13 | Saprotroph | \|Undefined Saprotroph\| | Yeast | NULL | Probable | As nectar-tap_saprotroph (Põlme et al. 2020); Decay sugar-rich substrates (Põlme et al. 2020) | Põlme S, et al. 2020. Fungal Diversity 105:1-16 ((https://doi.org/10.1007/s13225-020-00466-2)) |
| Otu51611 | 1 | k__Fungi(100);p__Ascomycota(100);c__Saccharomycetes(100);o__Saccharomycetales(100);f__Metschnikowiaceae(100);g__Metschnikowia(100);s__Metschnikowia_pulcherrima(100); | Metschnikowia | 13 | Saprotroph | \|Undefined Saprotroph\| | Yeast | NULL | Probable | As nectar-tap_saprotroph (Põlme et al. 2020); Decay sugar-rich substrates (Põlme et al. 2020) | Põlme S, et al. 2020. Fungal Diversity 105:1-16 ((https://doi.org/10.1007/s13225-020-00466-2)) |
| Otu51612 | 1 | k__Fungi(100);p__Ascomycota(100);c__Saccharomycetes(100);o__Saccharomycetales(100);f__Metschnikowiaceae(100);g__Metschnikowia(100);s__Metschnikowia_pulcherrima(100); | Metschnikowia | 13 | Saprotroph | \|Undefined Saprotroph\| | Yeast | NULL | Probable | As nectar-tap_saprotroph (Põlme et al. 2020); Decay sugar-rich substrates (Põlme et al. 2020) | Põlme S, et al. 2020. Fungal Diversity 105:1-16 ((https://doi.org/10.1007/s13225-020-00466-2)) |
| Otu51613 | 1 | k__Fungi(100);p__Ascomycota(100);c__Saccharomycetes(100);o__Saccharomycetales(100);f__Metschnikowiaceae(100);g__Metschnikowia(100);g__Metschnikowia_unclassified(100); | Metschnikowia | 13 | Saprotroph | \|Undefined Saprotroph\| | Yeast | NULL | Probable | As nectar-tap_saprotroph (Põlme et al. 2020); Decay sugar-rich substrates (Põlme et al. 2020) | Põlme S, et al. 2020. Fungal Diversity 105:1-16 ((https://doi.org/10.1007/s13225-020-00466-2)) |
| Otu51614 | 1 | k__Fungi(100);p__Ascomycota(100);c__Saccharomycetes(100);o__Saccharomycetales(100);f__Metschnikowiaceae(100);g__Metschnikowia(100);s__Metschnikowia_pulcherrima(100); | Metschnikowia | 13 | Saprotroph | \|Undefined Saprotroph\| | Yeast | NULL | Probable | As nectar-tap_saprotroph (Põlme et al. 2020); Decay sugar-rich substrates (Põlme et al. 2020) | Põlme S, et al. 2020. Fungal Diversity 105:1-16 ((https://doi.org/10.1007/s13225-020-00466-2)) |
| Otu51615 | 1 | k__Fungi(100);p__Ascomycota(100);c__Saccharomycetes(100);o__Saccharomycetales(100);f__Metschnikowiaceae(100);g__Metschnikowia(100);g__Metschnikowia_unclassified(100); | Metschnikowia | 13 | Saprotroph | \|Undefined Saprotroph\| | Yeast | NULL | Probable | As nectar-tap_saprotroph (Põlme et al. 2020); Decay sugar-rich substrates (Põlme et al. 2020) | Põlme S, et al. 2020. Fungal Diversity 105:1-16 ((https://doi.org/10.1007/s13225-020-00466-2)) |
| Otu51616 | 1 | k__Fungi(100);p__Ascomycota(100);c__Saccharomycetes(100);o__Saccharomycetales(100);f__Metschnikowiaceae(100);g__Metschnikowia(100);s__Metschnikowia_pulcherrima(100); | Metschnikowia | 13 | Saprotroph | \|Undefined Saprotroph\| | Yeast | NULL | Probable | As nectar-tap_saprotroph (Põlme et al. 2020); Decay sugar-rich substrates (Põlme et al. 2020) | Põlme S, et al. 2020. Fungal Diversity 105:1-16 ((https://doi.org/10.1007/s13225-020-00466-2)) |
| Otu51617 | 1 | k__Fungi(100);p__Ascomycota(100);c__Saccharomycetes(100);o__Saccharomycetales(100);f__Metschnikowiaceae(100);g__Metschnikowia(100);g__Metschnikowia_unclassified(100); | Metschnikowia | 13 | Saprotroph | \|Undefined Saprotroph\| | Yeast | NULL | Probable | As nectar-tap_saprotroph (Põlme et al. 2020); Decay sugar-rich substrates (Põlme et al. 2020) | Põlme S, et al. 2020. Fungal Diversity 105:1-16 ((https://doi.org/10.1007/s13225-020-00466-2)) |
| Otu51618 | 1 | k__Fungi(100);p__Ascomycota(100);c__Saccharomycetes(100);o__Saccharomycetales(100);f__Metschnikowiaceae(100);g__Metschnikowia(100);s__Metschnikowia_pulcherrima(100); | Metschnikowia | 13 | Saprotroph | \|Undefined Saprotroph\| | Yeast | NULL | Probable | As nectar-tap_saprotroph (Põlme et al. 2020); Decay sugar-rich substrates (Põlme et al. 2020) | Põlme S, et al. 2020. Fungal Diversity 105:1-16 ((https://doi.org/10.1007/s13225-020-00466-2)) |
| Otu51619 | 1 | k__Fungi(100);p__Ascomycota(100);c__Saccharomycetes(100);o__Saccharomycetales(100);f__Metschnikowiaceae(100);g__Metschnikowia(100);s__Metschnikowia_pulcherrima(100); | Metschnikowia | 13 | Saprotroph | \|Undefined Saprotroph\| | Yeast | NULL | Probable | As nectar-tap_saprotroph (Põlme et al. 2020); Decay sugar-rich substrates (Põlme et al. 2020) | Põlme S, et al. 2020. Fungal Diversity 105:1-16 ((https://doi.org/10.1007/s13225-020-00466-2)) |
| Otu51620 | 1 | k__Fungi(100);p__Ascomycota(100);c__Saccharomycetes(100);o__Saccharomycetales(100);f__Metschnikowiaceae(100);g__Metschnikowia(100);s__Metschnikowia_pulcherrima(100); | Metschnikowia | 13 | Saprotroph | \|Undefined Saprotroph\| | Yeast | NULL | Probable | As nectar-tap_saprotroph (Põlme et al. 2020); Decay sugar-rich substrates (Põlme et al. 2020) | Põlme S, et al. 2020. Fungal Diversity 105:1-16 ((https://doi.org/10.1007/s13225-020-00466-2)) |
| Otu51621 | 1 | k__Fungi(100);p__Ascomycota(100);c__Saccharomycetes(100);o__Saccharomycetales(100);f__Metschnikowiaceae(100);g__Metschnikowia(100);s__Metschnikowia_pulcherrima(100); | Metschnikowia | 13 | Saprotroph | \|Undefined Saprotroph\| | Yeast | NULL | Probable | As nectar-tap_saprotroph (Põlme et al. 2020); Decay sugar-rich substrates (Põlme et al. 2020) | Põlme S, et al. 2020. Fungal Diversity 105:1-16 ((https://doi.org/10.1007/s13225-020-00466-2)) |
| Otu51622 | 1 | k__Fungi(100);p__Ascomycota(100);c__Saccharomycetes(100);o__Saccharomycetales(100);f__Metschnikowiaceae(100);g__Metschnikowia(100);s__Metschnikowia_pulcherrima(100); | Metschnikowia | 13 | Saprotroph | \|Undefined Saprotroph\| | Yeast | NULL | Probable | As nectar-tap_saprotroph (Põlme et al. 2020); Decay sugar-rich substrates (Põlme et al. 2020) | Põlme S, et al. 2020. Fungal Diversity 105:1-16 ((https://doi.org/10.1007/s13225-020-00466-2)) |
| Otu51623 | 1 | k__Fungi(100);p__Ascomycota(100);c__Saccharomycetes(100);o__Saccharomycetales(100);f__Metschnikowiaceae(100);g__Metschnikowia(100);g__Metschnikowia_unclassified(100); | Metschnikowia | 13 | Saprotroph | \|Undefined Saprotroph\| | Yeast | NULL | Probable | As nectar-tap_saprotroph (Põlme et al. 2020); Decay sugar-rich substrates (Põlme et al. 2020) | Põlme S, et al. 2020. Fungal Diversity 105:1-16 ((https://doi.org/10.1007/s13225-020-00466-2)) |
| Otu51624 | 1 | k__Fungi(100);p__Ascomycota(100);c__Saccharomycetes(100);o__Saccharomycetales(100);f__Metschnikowiaceae(100);g__Metschnikowia(100);s__Metschnikowia_pulcherrima(100); | Metschnikowia | 13 | Saprotroph | \|Undefined Saprotroph\| | Yeast | NULL | Probable | As nectar-tap_saprotroph (Põlme et al. 2020); Decay sugar-rich substrates (Põlme et al. 2020) | Põlme S, et al. 2020. Fungal Diversity 105:1-16 ((https://doi.org/10.1007/s13225-020-00466-2)) |
| Otu51625 | 1 | k__Fungi(100);p__Ascomycota(100);c__Saccharomycetes(100);o__Saccharomycetales(100);f__Metschnikowiaceae(100);g__Metschnikowia(100);g__Metschnikowia_unclassified(100); | Metschnikowia | 13 | Saprotroph | \|Undefined Saprotroph\| | Yeast | NULL | Probable | As nectar-tap_saprotroph (Põlme et al. 2020); Decay sugar-rich substrates (Põlme et al. 2020) | Põlme S, et al. 2020. Fungal Diversity 105:1-16 ((https://doi.org/10.1007/s13225-020-00466-2)) |
| Otu51626 | 1 | k__Fungi(100);p__Ascomycota(100);c__Saccharomycetes(100);o__Saccharomycetales(100);f__Metschnikowiaceae(100);g__Metschnikowia(100);s__Metschnikowia_pulcherrima(100); | Metschnikowia | 13 | Saprotroph | \|Undefined Saprotroph\| | Yeast | NULL | Probable | As nectar-tap_saprotroph (Põlme et al. 2020); Decay sugar-rich substrates (Põlme et al. 2020) | Põlme S, et al. 2020. Fungal Diversity 105:1-16 ((https://doi.org/10.1007/s13225-020-00466-2)) |
| Otu51627 | 1 | k__Fungi(100);p__Ascomycota(100);c__Saccharomycetes(100);o__Saccharomycetales(100);f__Metschnikowiaceae(100);g__Metschnikowia(100);s__Metschnikowia_pulcherrima(100); | Metschnikowia | 13 | Saprotroph | \|Undefined Saprotroph\| | Yeast | NULL | Probable | As nectar-tap_saprotroph (Põlme et al. 2020); Decay sugar-rich substrates (Põlme et al. 2020) | Põlme S, et al. 2020. Fungal Diversity 105:1-16 ((https://doi.org/10.1007/s13225-020-00466-2)) |
| Otu51628 | 1 | k__Fungi(100);p__Ascomycota(100);c__Saccharomycetes(100);o__Saccharomycetales(100);f__Metschnikowiaceae(100);g__Metschnikowia(100);s__Metschnikowia_pulcherrima(100); | Metschnikowia | 13 | Saprotroph | \|Undefined Saprotroph\| | Yeast | NULL | Probable | As nectar-tap_saprotroph (Põlme et al. 2020); Decay sugar-rich substrates (Põlme et al. 2020) | Põlme S, et al. 2020. Fungal Diversity 105:1-16 ((https://doi.org/10.1007/s13225-020-00466-2)) |
| Otu51629 | 1 | k__Fungi(100);p__Ascomycota(100);c__Saccharomycetes(100);o__Saccharomycetales(100);f__Metschnikowiaceae(100);g__Metschnikowia(100);g__Metschnikowia_unclassified(100); | Metschnikowia | 13 | Saprotroph | \|Undefined Saprotroph\| | Yeast | NULL | Probable | As nectar-tap_saprotroph (Põlme et al. 2020); Decay sugar-rich substrates (Põlme et al. 2020) | Põlme S, et al. 2020. Fungal Diversity 105:1-16 ((https://doi.org/10.1007/s13225-020-00466-2)) |
| Otu51630 | 1 | k__Fungi(100);p__Ascomycota(100);c__Saccharomycetes(100);o__Saccharomycetales(100);f__Metschnikowiaceae(100);g__Metschnikowia(100);s__Metschnikowia_pulcherrima(100); | Metschnikowia | 13 | Saprotroph | \|Undefined Saprotroph\| | Yeast | NULL | Probable | As nectar-tap_saprotroph (Põlme et al. 2020); Decay sugar-rich substrates (Põlme et al. 2020) | Põlme S, et al. 2020. Fungal Diversity 105:1-16 ((https://doi.org/10.1007/s13225-020-00466-2)) |
| Otu51631 | 1 | k__Fungi(100);p__Ascomycota(100);c__Saccharomycetes(100);o__Saccharomycetales(100);f__Metschnikowiaceae(100);g__Metschnikowia(100);g__Metschnikowia_unclassified(100); | Metschnikowia | 13 | Saprotroph | \|Undefined Saprotroph\| | Yeast | NULL | Probable | As nectar-tap_saprotroph (Põlme et al. 2020); Decay sugar-rich substrates (Põlme et al. 2020) | Põlme S, et al. 2020. Fungal Diversity 105:1-16 ((https://doi.org/10.1007/s13225-020-00466-2)) |
| Otu51632 | 1 | k__Fungi(100);p__Ascomycota(100);c__Saccharomycetes(100);o__Saccharomycetales(100);f__Metschnikowiaceae(100);g__Metschnikowia(100);s__Metschnikowia_pulcherrima(100); | Metschnikowia | 13 | Saprotroph | \|Undefined Saprotroph\| | Yeast | NULL | Probable | As nectar-tap_saprotroph (Põlme et al. 2020); Decay sugar-rich substrates (Põlme et al. 2020) | Põlme S, et al. 2020. Fungal Diversity 105:1-16 ((https://doi.org/10.1007/s13225-020-00466-2)) |
| Otu51633 | 1 | k__Fungi(100);p__Ascomycota(100);c__Saccharomycetes(100);o__Saccharomycetales(100);f__Metschnikowiaceae(100);g__Metschnikowia(100);s__Metschnikowia_pulcherrima(100); | Metschnikowia | 13 | Saprotroph | \|Undefined Saprotroph\| | Yeast | NULL | Probable | As nectar-tap_saprotroph (Põlme et al. 2020); Decay sugar-rich substrates (Põlme et al. 2020) | Põlme S, et al. 2020. Fungal Diversity 105:1-16 ((https://doi.org/10.1007/s13225-020-00466-2)) |
| Otu51634 | 1 | k__Fungi(100);p__Ascomycota(100);c__Saccharomycetes(100);o__Saccharomycetales(100);f__Metschnikowiaceae(100);g__Metschnikowia(100);s__Metschnikowia_pulcherrima(100); | Metschnikowia | 13 | Saprotroph | \|Undefined Saprotroph\| | Yeast | NULL | Probable | As nectar-tap_saprotroph (Põlme et al. 2020); Decay sugar-rich substrates (Põlme et al. 2020) | Põlme S, et al. 2020. Fungal Diversity 105:1-16 ((https://doi.org/10.1007/s13225-020-00466-2)) |
| Otu51635 | 1 | k__Fungi(100);p__Ascomycota(100);c__Saccharomycetes(100);o__Saccharomycetales(100);f__Metschnikowiaceae(100);g__Metschnikowia(100);s__Metschnikowia_pulcherrima(100); | Metschnikowia | 13 | Saprotroph | \|Undefined Saprotroph\| | Yeast | NULL | Probable | As nectar-tap_saprotroph (Põlme et al. 2020); Decay sugar-rich substrates (Põlme et al. 2020) | Põlme S, et al. 2020. Fungal Diversity 105:1-16 ((https://doi.org/10.1007/s13225-020-00466-2)) |
| Otu51636 | 1 | k__Fungi(100);p__Ascomycota(100);c__Saccharomycetes(100);o__Saccharomycetales(100);f__Metschnikowiaceae(100);g__Metschnikowia(100);s__Metschnikowia_pulcherrima(100); | Metschnikowia | 13 | Saprotroph | \|Undefined Saprotroph\| | Yeast | NULL | Probable | As nectar-tap_saprotroph (Põlme et al. 2020); Decay sugar-rich substrates (Põlme et al. 2020) | Põlme S, et al. 2020. Fungal Diversity 105:1-16 ((https://doi.org/10.1007/s13225-020-00466-2)) |
| Otu51637 | 1 | k__Fungi(100);p__Ascomycota(100);c__Saccharomycetes(100);o__Saccharomycetales(100);f__Metschnikowiaceae(100);g__Metschnikowia(100);s__Metschnikowia_pulcherrima(100); | Metschnikowia | 13 | Saprotroph | \|Undefined Saprotroph\| | Yeast | NULL | Probable | As nectar-tap_saprotroph (Põlme et al. 2020); Decay sugar-rich substrates (Põlme et al. 2020) | Põlme S, et al. 2020. Fungal Diversity 105:1-16 ((https://doi.org/10.1007/s13225-020-00466-2)) |
| Otu51638 | 1 | k__Fungi(100);p__Ascomycota(100);c__Saccharomycetes(100);o__Saccharomycetales(100);f__Metschnikowiaceae(100);g__Metschnikowia(100);s__Metschnikowia_pulcherrima(100); | Metschnikowia | 13 | Saprotroph | \|Undefined Saprotroph\| | Yeast | NULL | Probable | As nectar-tap_saprotroph (Põlme et al. 2020); Decay sugar-rich substrates (Põlme et al. 2020) | Põlme S, et al. 2020. Fungal Diversity 105:1-16 ((https://doi.org/10.1007/s13225-020-00466-2)) |
| Otu51639 | 1 | k__Fungi(100);p__Ascomycota(100);c__Saccharomycetes(100);o__Saccharomycetales(100);f__Metschnikowiaceae(100);g__Metschnikowia(100);s__Metschnikowia_pulcherrima(100); | Metschnikowia | 13 | Saprotroph | \|Undefined Saprotroph\| | Yeast | NULL | Probable | As nectar-tap_saprotroph (Põlme et al. 2020); Decay sugar-rich substrates (Põlme et al. 2020) | Põlme S, et al. 2020. Fungal Diversity 105:1-16 ((https://doi.org/10.1007/s13225-020-00466-2)) |
| Otu51640 | 1 | k__Fungi(100);p__Ascomycota(100);c__Saccharomycetes(100);o__Saccharomycetales(100);f__Metschnikowiaceae(100);g__Metschnikowia(100);g__Metschnikowia_unclassified(100); | Metschnikowia | 13 | Saprotroph | \|Undefined Saprotroph\| | Yeast | NULL | Probable | As nectar-tap_saprotroph (Põlme et al. 2020); Decay sugar-rich substrates (Põlme et al. 2020) | Põlme S, et al. 2020. Fungal Diversity 105:1-16 ((https://doi.org/10.1007/s13225-020-00466-2)) |
| Otu51641 | 1 | k__Fungi(100);p__Ascomycota(100);c__Saccharomycetes(100);o__Saccharomycetales(100);f__Metschnikowiaceae(100);g__Metschnikowia(100);g__Metschnikowia_unclassified(100); | Metschnikowia | 13 | Saprotroph | \|Undefined Saprotroph\| | Yeast | NULL | Probable | As nectar-tap_saprotroph (Põlme et al. 2020); Decay sugar-rich substrates (Põlme et al. 2020) | Põlme S, et al. 2020. Fungal Diversity 105:1-16 ((https://doi.org/10.1007/s13225-020-00466-2)) |
| Otu51642 | 1 | k__Fungi(100);p__Ascomycota(100);c__Saccharomycetes(100);o__Saccharomycetales(100);f__Metschnikowiaceae(100);g__Metschnikowia(100);s__Metschnikowia_pulcherrima(100); | Metschnikowia | 13 | Saprotroph | \|Undefined Saprotroph\| | Yeast | NULL | Probable | As nectar-tap_saprotroph (Põlme et al. 2020); Decay sugar-rich substrates (Põlme et al. 2020) | Põlme S, et al. 2020. Fungal Diversity 105:1-16 ((https://doi.org/10.1007/s13225-020-00466-2)) |
| Otu51643 | 1 | k__Fungi(100);p__Ascomycota(100);c__Saccharomycetes(100);o__Saccharomycetales(100);f__Metschnikowiaceae(100);g__Metschnikowia(100);g__Metschnikowia_unclassified(100); | Metschnikowia | 13 | Saprotroph | \|Undefined Saprotroph\| | Yeast | NULL | Probable | As nectar-tap_saprotroph (Põlme et al. 2020); Decay sugar-rich substrates (Põlme et al. 2020) | Põlme S, et al. 2020. Fungal Diversity 105:1-16 ((https://doi.org/10.1007/s13225-020-00466-2)) |
| Otu51644 | 1 | k__Fungi(100);p__Ascomycota(100);c__Saccharomycetes(100);o__Saccharomycetales(100);f__Metschnikowiaceae(100);g__Metschnikowia(100);s__Metschnikowia_pulcherrima(100); | Metschnikowia | 13 | Saprotroph | \|Undefined Saprotroph\| | Yeast | NULL | Probable | As nectar-tap_saprotroph (Põlme et al. 2020); Decay sugar-rich substrates (Põlme et al. 2020) | Põlme S, et al. 2020. Fungal Diversity 105:1-16 ((https://doi.org/10.1007/s13225-020-00466-2)) |
| Otu51645 | 1 | k__Fungi(100);p__Ascomycota(100);c__Saccharomycetes(100);o__Saccharomycetales(100);f__Metschnikowiaceae(100);g__Metschnikowia(100);s__Metschnikowia_pulcherrima(100); | Metschnikowia | 13 | Saprotroph | \|Undefined Saprotroph\| | Yeast | NULL | Probable | As nectar-tap_saprotroph (Põlme et al. 2020); Decay sugar-rich substrates (Põlme et al. 2020) | Põlme S, et al. 2020. Fungal Diversity 105:1-16 ((https://doi.org/10.1007/s13225-020-00466-2)) |
| Otu51646 | 1 | k__Fungi(100);p__Ascomycota(100);c__Saccharomycetes(100);o__Saccharomycetales(100);f__Metschnikowiaceae(100);g__Metschnikowia(100);g__Metschnikowia_unclassified(100); | Metschnikowia | 13 | Saprotroph | \|Undefined Saprotroph\| | Yeast | NULL | Probable | As nectar-tap_saprotroph (Põlme et al. 2020); Decay sugar-rich substrates (Põlme et al. 2020) | Põlme S, et al. 2020. Fungal Diversity 105:1-16 ((https://doi.org/10.1007/s13225-020-00466-2)) |
| Otu51647 | 1 | k__Fungi(100);p__Ascomycota(100);c__Saccharomycetes(100);o__Saccharomycetales(100);f__Metschnikowiaceae(100);g__Metschnikowia(100);s__Metschnikowia_pulcherrima(100); | Metschnikowia | 13 | Saprotroph | \|Undefined Saprotroph\| | Yeast | NULL | Probable | As nectar-tap_saprotroph (Põlme et al. 2020); Decay sugar-rich substrates (Põlme et al. 2020) | Põlme S, et al. 2020. Fungal Diversity 105:1-16 ((https://doi.org/10.1007/s13225-020-00466-2)) |
| Otu51648 | 1 | k__Fungi(100);p__Ascomycota(100);c__Saccharomycetes(100);o__Saccharomycetales(100);f__Metschnikowiaceae(100);g__Metschnikowia(100);s__Metschnikowia_pulcherrima(100); | Metschnikowia | 13 | Saprotroph | \|Undefined Saprotroph\| | Yeast | NULL | Probable | As nectar-tap_saprotroph (Põlme et al. 2020); Decay sugar-rich substrates (Põlme et al. 2020) | Põlme S, et al. 2020. Fungal Diversity 105:1-16 ((https://doi.org/10.1007/s13225-020-00466-2)) |
| Otu51649 | 1 | k__Fungi(100);p__Ascomycota(100);c__Saccharomycetes(100);o__Saccharomycetales(100);f__Metschnikowiaceae(100);g__Metschnikowia(100);g__Metschnikowia_unclassified(100); | Metschnikowia | 13 | Saprotroph | \|Undefined Saprotroph\| | Yeast | NULL | Probable | As nectar-tap_saprotroph (Põlme et al. 2020); Decay sugar-rich substrates (Põlme et al. 2020) | Põlme S, et al. 2020. Fungal Diversity 105:1-16 ((https://doi.org/10.1007/s13225-020-00466-2)) |
| Otu51650 | 1 | k__Fungi(100);p__Ascomycota(100);c__Saccharomycetes(100);o__Saccharomycetales(100);f__Metschnikowiaceae(100);g__Metschnikowia(100);s__Metschnikowia_pulcherrima(100); | Metschnikowia | 13 | Saprotroph | \|Undefined Saprotroph\| | Yeast | NULL | Probable | As nectar-tap_saprotroph (Põlme et al. 2020); Decay sugar-rich substrates (Põlme et al. 2020) | Põlme S, et al. 2020. Fungal Diversity 105:1-16 ((https://doi.org/10.1007/s13225-020-00466-2)) |
| Otu51651 | 1 | k__Fungi(100);p__Ascomycota(100);c__Saccharomycetes(100);o__Saccharomycetales(100);f__Metschnikowiaceae(100);g__Metschnikowia(100);g__Metschnikowia_unclassified(100); | Metschnikowia | 13 | Saprotroph | \|Undefined Saprotroph\| | Yeast | NULL | Probable | As nectar-tap_saprotroph (Põlme et al. 2020); Decay sugar-rich substrates (Põlme et al. 2020) | Põlme S, et al. 2020. Fungal Diversity 105:1-16 ((https://doi.org/10.1007/s13225-020-00466-2)) |
| Otu51652 | 1 | k__Fungi(100);p__Ascomycota(100);c__Saccharomycetes(100);o__Saccharomycetales(100);f__Metschnikowiaceae(100);g__Metschnikowia(100);g__Metschnikowia_unclassified(100); | Metschnikowia | 13 | Saprotroph | \|Undefined Saprotroph\| | Yeast | NULL | Probable | As nectar-tap_saprotroph (Põlme et al. 2020); Decay sugar-rich substrates (Põlme et al. 2020) | Põlme S, et al. 2020. Fungal Diversity 105:1-16 ((https://doi.org/10.1007/s13225-020-00466-2)) |
| Otu51653 | 1 | k__Fungi(100);p__Ascomycota(100);c__Saccharomycetes(100);o__Saccharomycetales(100);f__Metschnikowiaceae(100);g__Metschnikowia(100);s__Metschnikowia_pulcherrima(100); | Metschnikowia | 13 | Saprotroph | \|Undefined Saprotroph\| | Yeast | NULL | Probable | As nectar-tap_saprotroph (Põlme et al. 2020); Decay sugar-rich substrates (Põlme et al. 2020) | Põlme S, et al. 2020. Fungal Diversity 105:1-16 ((https://doi.org/10.1007/s13225-020-00466-2)) |
| Otu51654 | 1 | k__Fungi(100);p__Ascomycota(100);c__Saccharomycetes(100);o__Saccharomycetales(100);f__Metschnikowiaceae(100);g__Metschnikowia(100);s__Metschnikowia_pulcherrima(100); | Metschnikowia | 13 | Saprotroph | \|Undefined Saprotroph\| | Yeast | NULL | Probable | As nectar-tap_saprotroph (Põlme et al. 2020); Decay sugar-rich substrates (Põlme et al. 2020) | Põlme S, et al. 2020. Fungal Diversity 105:1-16 ((https://doi.org/10.1007/s13225-020-00466-2)) |
| Otu51655 | 1 | k__Fungi(100);p__Ascomycota(100);c__Saccharomycetes(100);o__Saccharomycetales(100);f__Metschnikowiaceae(100);g__Metschnikowia(100);g__Metschnikowia_unclassified(100); | Metschnikowia | 13 | Saprotroph | \|Undefined Saprotroph\| | Yeast | NULL | Probable | As nectar-tap_saprotroph (Põlme et al. 2020); Decay sugar-rich substrates (Põlme et al. 2020) | Põlme S, et al. 2020. Fungal Diversity 105:1-16 ((https://doi.org/10.1007/s13225-020-00466-2)) |
| Otu51656 | 1 | k__Fungi(100);p__Ascomycota(100);c__Saccharomycetes(100);o__Saccharomycetales(100);f__Metschnikowiaceae(100);g__Metschnikowia(100);s__Metschnikowia_pulcherrima(100); | Metschnikowia | 13 | Saprotroph | \|Undefined Saprotroph\| | Yeast | NULL | Probable | As nectar-tap_saprotroph (Põlme et al. 2020); Decay sugar-rich substrates (Põlme et al. 2020) | Põlme S, et al. 2020. Fungal Diversity 105:1-16 ((https://doi.org/10.1007/s13225-020-00466-2)) |
| Otu51657 | 1 | k__Fungi(100);p__Ascomycota(100);c__Saccharomycetes(100);o__Saccharomycetales(100);f__Metschnikowiaceae(100);g__Metschnikowia(100);s__Metschnikowia_pulcherrima(100); | Metschnikowia | 13 | Saprotroph | \|Undefined Saprotroph\| | Yeast | NULL | Probable | As nectar-tap_saprotroph (Põlme et al. 2020); Decay sugar-rich substrates (Põlme et al. 2020) | Põlme S, et al. 2020. Fungal Diversity 105:1-16 ((https://doi.org/10.1007/s13225-020-00466-2)) |
| Otu51658 | 1 | k__Fungi(100);p__Ascomycota(100);c__Saccharomycetes(100);o__Saccharomycetales(100);f__Metschnikowiaceae(100);g__Metschnikowia(100);s__Metschnikowia_pulcherrima(100); | Metschnikowia | 13 | Saprotroph | \|Undefined Saprotroph\| | Yeast | NULL | Probable | As nectar-tap_saprotroph (Põlme et al. 2020); Decay sugar-rich substrates (Põlme et al. 2020) | Põlme S, et al. 2020. Fungal Diversity 105:1-16 ((https://doi.org/10.1007/s13225-020-00466-2)) |
| Otu51659 | 1 | k__Fungi(100);p__Ascomycota(100);c__Saccharomycetes(100);o__Saccharomycetales(100);f__Metschnikowiaceae(100);g__Metschnikowia(100);s__Metschnikowia_pulcherrima(100); | Metschnikowia | 13 | Saprotroph | \|Undefined Saprotroph\| | Yeast | NULL | Probable | As nectar-tap_saprotroph (Põlme et al. 2020); Decay sugar-rich substrates (Põlme et al. 2020) | Põlme S, et al. 2020. Fungal Diversity 105:1-16 ((https://doi.org/10.1007/s13225-020-00466-2)) |
| Otu51660 | 1 | k__Fungi(100);p__Ascomycota(100);c__Saccharomycetes(100);o__Saccharomycetales(100);f__Metschnikowiaceae(100);g__Metschnikowia(100);s__Metschnikowia_pulcherrima(100); | Metschnikowia | 13 | Saprotroph | \|Undefined Saprotroph\| | Yeast | NULL | Probable | As nectar-tap_saprotroph (Põlme et al. 2020); Decay sugar-rich substrates (Põlme et al. 2020) | Põlme S, et al. 2020. Fungal Diversity 105:1-16 ((https://doi.org/10.1007/s13225-020-00466-2)) |
| Otu51661 | 1 | k__Fungi(100);p__Ascomycota(100);c__Saccharomycetes(100);o__Saccharomycetales(100);f__Metschnikowiaceae(100);g__Metschnikowia(100);s__Metschnikowia_pulcherrima(100); | Metschnikowia | 13 | Saprotroph | \|Undefined Saprotroph\| | Yeast | NULL | Probable | As nectar-tap_saprotroph (Põlme et al. 2020); Decay sugar-rich substrates (Põlme et al. 2020) | Põlme S, et al. 2020. Fungal Diversity 105:1-16 ((https://doi.org/10.1007/s13225-020-00466-2)) |
| Otu51662 | 1 | k__Fungi(100);p__Ascomycota(100);c__Saccharomycetes(100);o__Saccharomycetales(100);f__Metschnikowiaceae(100);g__Metschnikowia(100);s__Metschnikowia_pulcherrima(100); | Metschnikowia | 13 | Saprotroph | \|Undefined Saprotroph\| | Yeast | NULL | Probable | As nectar-tap_saprotroph (Põlme et al. 2020); Decay sugar-rich substrates (Põlme et al. 2020) | Põlme S, et al. 2020. Fungal Diversity 105:1-16 ((https://doi.org/10.1007/s13225-020-00466-2)) |
| Otu51663 | 1 | k__Fungi(100);p__Ascomycota(100);c__Saccharomycetes(100);o__Saccharomycetales(100);f__Metschnikowiaceae(100);g__Metschnikowia(100);g__Metschnikowia_unclassified(100); | Metschnikowia | 13 | Saprotroph | \|Undefined Saprotroph\| | Yeast | NULL | Probable | As nectar-tap_saprotroph (Põlme et al. 2020); Decay sugar-rich substrates (Põlme et al. 2020) | Põlme S, et al. 2020. Fungal Diversity 105:1-16 ((https://doi.org/10.1007/s13225-020-00466-2)) |
| Otu51664 | 1 | k__Fungi(100);p__Ascomycota(100);c__Saccharomycetes(100);o__Saccharomycetales(100);f__Metschnikowiaceae(100);g__Metschnikowia(100);s__Metschnikowia_pulcherrima(100); | Metschnikowia | 13 | Saprotroph | \|Undefined Saprotroph\| | Yeast | NULL | Probable | As nectar-tap_saprotroph (Põlme et al. 2020); Decay sugar-rich substrates (Põlme et al. 2020) | Põlme S, et al. 2020. Fungal Diversity 105:1-16 ((https://doi.org/10.1007/s13225-020-00466-2)) |
| Otu51665 | 1 | k__Fungi(100);p__Ascomycota(100);c__Saccharomycetes(100);o__Saccharomycetales(100);f__Metschnikowiaceae(100);g__Metschnikowia(100);s__Metschnikowia_pulcherrima(100); | Metschnikowia | 13 | Saprotroph | \|Undefined Saprotroph\| | Yeast | NULL | Probable | As nectar-tap_saprotroph (Põlme et al. 2020); Decay sugar-rich substrates (Põlme et al. 2020) | Põlme S, et al. 2020. Fungal Diversity 105:1-16 ((https://doi.org/10.1007/s13225-020-00466-2)) |
| Otu51666 | 1 | k__Fungi(100);p__Ascomycota(100);c__Saccharomycetes(100);o__Saccharomycetales(100);f__Metschnikowiaceae(100);g__Metschnikowia(100);s__Metschnikowia_pulcherrima(100); | Metschnikowia | 13 | Saprotroph | \|Undefined Saprotroph\| | Yeast | NULL | Probable | As nectar-tap_saprotroph (Põlme et al. 2020); Decay sugar-rich substrates (Põlme et al. 2020) | Põlme S, et al. 2020. Fungal Diversity 105:1-16 ((https://doi.org/10.1007/s13225-020-00466-2)) |
| Otu51667 | 1 | k__Fungi(100);p__Ascomycota(100);c__Saccharomycetes(100);o__Saccharomycetales(100);f__Metschnikowiaceae(100);g__Metschnikowia(100);s__Metschnikowia_pulcherrima(100); | Metschnikowia | 13 | Saprotroph | \|Undefined Saprotroph\| | Yeast | NULL | Probable | As nectar-tap_saprotroph (Põlme et al. 2020); Decay sugar-rich substrates (Põlme et al. 2020) | Põlme S, et al. 2020. Fungal Diversity 105:1-16 ((https://doi.org/10.1007/s13225-020-00466-2)) |
| Otu51668 | 1 | k__Fungi(100);p__Ascomycota(100);c__Saccharomycetes(100);o__Saccharomycetales(100);f__Metschnikowiaceae(100);g__Metschnikowia(100);s__Metschnikowia_pulcherrima(100); | Metschnikowia | 13 | Saprotroph | \|Undefined Saprotroph\| | Yeast | NULL | Probable | As nectar-tap_saprotroph (Põlme et al. 2020); Decay sugar-rich substrates (Põlme et al. 2020) | Põlme S, et al. 2020. Fungal Diversity 105:1-16 ((https://doi.org/10.1007/s13225-020-00466-2)) |
| Otu51669 | 1 | k__Fungi(100);p__Ascomycota(100);c__Saccharomycetes(100);o__Saccharomycetales(100);f__Metschnikowiaceae(100);g__Metschnikowia(100);s__Metschnikowia_pulcherrima(100); | Metschnikowia | 13 | Saprotroph | \|Undefined Saprotroph\| | Yeast | NULL | Probable | As nectar-tap_saprotroph (Põlme et al. 2020); Decay sugar-rich substrates (Põlme et al. 2020) | Põlme S, et al. 2020. Fungal Diversity 105:1-16 ((https://doi.org/10.1007/s13225-020-00466-2)) |
| Otu51670 | 1 | k__Fungi(100);p__Ascomycota(100);c__Saccharomycetes(100);o__Saccharomycetales(100);f__Metschnikowiaceae(100);g__Metschnikowia(100);s__Metschnikowia_pulcherrima(100); | Metschnikowia | 13 | Saprotroph | \|Undefined Saprotroph\| | Yeast | NULL | Probable | As nectar-tap_saprotroph (Põlme et al. 2020); Decay sugar-rich substrates (Põlme et al. 2020) | Põlme S, et al. 2020. Fungal Diversity 105:1-16 ((https://doi.org/10.1007/s13225-020-00466-2)) |
| Otu51671 | 1 | k__Fungi(100);p__Ascomycota(100);c__Saccharomycetes(100);o__Saccharomycetales(100);f__Metschnikowiaceae(100);g__Metschnikowia(100);s__Metschnikowia_pulcherrima(100); | Metschnikowia | 13 | Saprotroph | \|Undefined Saprotroph\| | Yeast | NULL | Probable | As nectar-tap_saprotroph (Põlme et al. 2020); Decay sugar-rich substrates (Põlme et al. 2020) | Põlme S, et al. 2020. Fungal Diversity 105:1-16 ((https://doi.org/10.1007/s13225-020-00466-2)) |
| Otu51672 | 1 | k__Fungi(100);p__Ascomycota(100);c__Saccharomycetes(100);o__Saccharomycetales(100);f__Metschnikowiaceae(100);g__Metschnikowia(100);s__Metschnikowia_pulcherrima(100); | Metschnikowia | 13 | Saprotroph | \|Undefined Saprotroph\| | Yeast | NULL | Probable | As nectar-tap_saprotroph (Põlme et al. 2020); Decay sugar-rich substrates (Põlme et al. 2020) | Põlme S, et al. 2020. Fungal Diversity 105:1-16 ((https://doi.org/10.1007/s13225-020-00466-2)) |
| Otu51673 | 1 | k__Fungi(100);p__Ascomycota(100);c__Saccharomycetes(100);o__Saccharomycetales(100);f__Metschnikowiaceae(100);g__Metschnikowia(100);s__Metschnikowia_pulcherrima(100); | Metschnikowia | 13 | Saprotroph | \|Undefined Saprotroph\| | Yeast | NULL | Probable | As nectar-tap_saprotroph (Põlme et al. 2020); Decay sugar-rich substrates (Põlme et al. 2020) | Põlme S, et al. 2020. Fungal Diversity 105:1-16 ((https://doi.org/10.1007/s13225-020-00466-2)) |
| Otu51674 | 1 | k__Fungi(100);p__Ascomycota(100);c__Saccharomycetes(100);o__Saccharomycetales(100);f__Metschnikowiaceae(100);g__Metschnikowia(100);s__Metschnikowia_pulcherrima(100); | Metschnikowia | 13 | Saprotroph | \|Undefined Saprotroph\| | Yeast | NULL | Probable | As nectar-tap_saprotroph (Põlme et al. 2020); Decay sugar-rich substrates (Põlme et al. 2020) | Põlme S, et al. 2020. Fungal Diversity 105:1-16 ((https://doi.org/10.1007/s13225-020-00466-2)) |
| Otu51675 | 1 | k__Fungi(100);p__Ascomycota(100);c__Saccharomycetes(100);o__Saccharomycetales(100);f__Metschnikowiaceae(100);g__Metschnikowia(100);s__Metschnikowia_pulcherrima(100); | Metschnikowia | 13 | Saprotroph | \|Undefined Saprotroph\| | Yeast | NULL | Probable | As nectar-tap_saprotroph (Põlme et al. 2020); Decay sugar-rich substrates (Põlme et al. 2020) | Põlme S, et al. 2020. Fungal Diversity 105:1-16 ((https://doi.org/10.1007/s13225-020-00466-2)) |
| Otu51676 | 1 | k__Fungi(100);p__Ascomycota(100);c__Saccharomycetes(100);o__Saccharomycetales(100);f__Metschnikowiaceae(100);g__Metschnikowia(100);g__Metschnikowia_unclassified(100); | Metschnikowia | 13 | Saprotroph | \|Undefined Saprotroph\| | Yeast | NULL | Probable | As nectar-tap_saprotroph (Põlme et al. 2020); Decay sugar-rich substrates (Põlme et al. 2020) | Põlme S, et al. 2020. Fungal Diversity 105:1-16 ((https://doi.org/10.1007/s13225-020-00466-2)) |
| Otu51677 | 1 | k__Fungi(100);p__Ascomycota(100);c__Saccharomycetes(100);o__Saccharomycetales(100);f__Metschnikowiaceae(100);g__Metschnikowia(100);g__Metschnikowia_unclassified(100); | Metschnikowia | 13 | Saprotroph | \|Undefined Saprotroph\| | Yeast | NULL | Probable | As nectar-tap_saprotroph (Põlme et al. 2020); Decay sugar-rich substrates (Põlme et al. 2020) | Põlme S, et al. 2020. Fungal Diversity 105:1-16 ((https://doi.org/10.1007/s13225-020-00466-2)) |
| Otu51678 | 1 | k__Fungi(100);p__Ascomycota(100);c__Saccharomycetes(100);o__Saccharomycetales(100);f__Metschnikowiaceae(100);g__Metschnikowia(100);s__Metschnikowia_pulcherrima(100); | Metschnikowia | 13 | Saprotroph | \|Undefined Saprotroph\| | Yeast | NULL | Probable | As nectar-tap_saprotroph (Põlme et al. 2020); Decay sugar-rich substrates (Põlme et al. 2020) | Põlme S, et al. 2020. Fungal Diversity 105:1-16 ((https://doi.org/10.1007/s13225-020-00466-2)) |
| Otu51679 | 1 | k__Fungi(100);p__Ascomycota(100);c__Saccharomycetes(100);o__Saccharomycetales(100);f__Metschnikowiaceae(100);g__Metschnikowia(100);s__Metschnikowia_pulcherrima(100); | Metschnikowia | 13 | Saprotroph | \|Undefined Saprotroph\| | Yeast | NULL | Probable | As nectar-tap_saprotroph (Põlme et al. 2020); Decay sugar-rich substrates (Põlme et al. 2020) | Põlme S, et al. 2020. Fungal Diversity 105:1-16 ((https://doi.org/10.1007/s13225-020-00466-2)) |
| Otu51680 | 1 | k__Fungi(100);p__Ascomycota(100);c__Saccharomycetes(100);o__Saccharomycetales(100);f__Metschnikowiaceae(100);g__Metschnikowia(100);s__Metschnikowia_pulcherrima(100); | Metschnikowia | 13 | Saprotroph | \|Undefined Saprotroph\| | Yeast | NULL | Probable | As nectar-tap_saprotroph (Põlme et al. 2020); Decay sugar-rich substrates (Põlme et al. 2020) | Põlme S, et al. 2020. Fungal Diversity 105:1-16 ((https://doi.org/10.1007/s13225-020-00466-2)) |

| | | | | | | | | | | | |
|---|---|---|---|---|---|---|---|---|---|---|---|
| Otu51681 | 1 | k__Fungi(100);p__Ascomycota(100);c__Saccharomycetes(100);o__Saccharomycetales(100);f__Metschnikowiaceae(100);g__Metschnikowia(100);g__Metschnikowia_unclassified(100); | Metschnikowia | 13 | Saprotroph | \|Undefined Saprotroph\| | Yeast | NULL | Probable | As nectar-tap_saprotroph (Põlme et al. 2020); Decay sugar-rich substrates (Põlme et al. 2020) | Põlme S, et al. 2020. Fungal Diversity 105:1-16 ((https://doi.org/10.1007/s13225-020-00466-2)) |
| Otu51683 | 1 | k__Fungi(100);p__Ascomycota(100);c__Saccharomycetes(100);o__Saccharomycetales(100);f__Metschnikowiaceae(100);g__Metschnikowia(100);s__Metschnikowia_pulcherrima(100); | Metschnikowia | 13 | Saprotroph | \|Undefined Saprotroph\| | Yeast | NULL | Probable | As nectar-tap_saprotroph (Põlme et al. 2020); Decay sugar-rich substrates (Põlme et al. 2020) | Põlme S, et al. 2020. Fungal Diversity 105:1-16 ((https://doi.org/10.1007/s13225-020-00466-2)) |
| Otu51684 | 1 | k__Fungi(100);p__Ascomycota(100);c__Saccharomycetes(100);o__Saccharomycetales(100);f__Metschnikowiaceae(100);g__Metschnikowia(100);s__Metschnikowia_pulcherrima(100); | Metschnikowia | 13 | Saprotroph | \|Undefined Saprotroph\| | Yeast | NULL | Probable | As nectar-tap_saprotroph (Põlme et al. 2020); Decay sugar-rich substrates (Põlme et al. 2020) | Põlme S, et al. 2020. Fungal Diversity 105:1-16 ((https://doi.org/10.1007/s13225-020-00466-2)) |
| Otu51685 | 1 | k__Fungi(100);p__Ascomycota(100);c__Saccharomycetes(100);o__Saccharomycetales(100);f__Metschnikowiaceae(100);g__Metschnikowia(100);s__Metschnikowia_pulcherrima(100); | Metschnikowia | 13 | Saprotroph | \|Undefined Saprotroph\| | Yeast | NULL | Probable | As nectar-tap_saprotroph (Põlme et al. 2020); Decay sugar-rich substrates (Põlme et al. 2020) | Põlme S, et al. 2020. Fungal Diversity 105:1-16 ((https://doi.org/10.1007/s13225-020-00466-2)) |
| Otu51686 | 1 | k__Fungi(100);p__Ascomycota(100);c__Saccharomycetes(100);o__Saccharomycetales(100);f__Metschnikowiaceae(100);g__Metschnikowia(100);s__Metschnikowia_pulcherrima(100); | Metschnikowia | 13 | Saprotroph | \|Undefined Saprotroph\| | Yeast | NULL | Probable | As nectar-tap_saprotroph (Põlme et al. 2020); Decay sugar-rich substrates (Põlme et al. 2020) | Põlme S, et al. 2020. Fungal Diversity 105:1-16 ((https://doi.org/10.1007/s13225-020-00466-2)) |
| Otu51687 | 1 | k__Fungi(100);p__Ascomycota(100);c__Saccharomycetes(100);o__Saccharomycetales(100);f__Metschnikowiaceae(100);g__Metschnikowia(100);s__Metschnikowia_pulcherrima(100); | Metschnikowia | 13 | Saprotroph | \|Undefined Saprotroph\| | Yeast | NULL | Probable | As nectar-tap_saprotroph (Põlme et al. 2020); Decay sugar-rich substrates (Põlme et al. 2020) | Põlme S, et al. 2020. Fungal Diversity 105:1-16 ((https://doi.org/10.1007/s13225-020-00466-2)) |
| Otu51688 | 1 | k__Fungi(100);p__Ascomycota(100);c__Saccharomycetes(100);o__Saccharomycetales(100);f__Metschnikowiaceae(100);g__Metschnikowia(100);s__Metschnikowia_pulcherrima(100); | Metschnikowia | 13 | Saprotroph | \|Undefined Saprotroph\| | Yeast | NULL | Probable | As nectar-tap_saprotroph (Põlme et al. 2020); Decay sugar-rich substrates (Põlme et al. 2020) | Põlme S, et al. 2020. Fungal Diversity 105:1-16 ((https://doi.org/10.1007/s13225-020-00466-2)) |
| Otu51689 | 1 | k__Fungi(100);p__Ascomycota(100);c__Saccharomycetes(100);o__Saccharomycetales(100);f__Metschnikowiaceae(100);g__Metschnikowia(100);s__Metschnikowia_pulcherrima(100); | Metschnikowia | 13 | Saprotroph | \|Undefined Saprotroph\| | Yeast | NULL | Probable | As nectar-tap_saprotroph (Põlme et al. 2020); Decay sugar-rich substrates (Põlme et al. 2020) | Põlme S, et al. 2020. Fungal Diversity 105:1-16 ((https://doi.org/10.1007/s13225-020-00466-2)) |
| Otu51691 | 1 | k__Fungi(100);p__Ascomycota(100);c__Saccharomycetes(100);o__Saccharomycetales(100);f__Metschnikowiaceae(100);g__Metschnikowia(100);g__Metschnikowia_unclassified(100); | Metschnikowia | 13 | Saprotroph | \|Undefined Saprotroph\| | Yeast | NULL | Probable | As nectar-tap_saprotroph (Põlme et al. 2020); Decay sugar-rich substrates (Põlme et al. 2020) | Põlme S, et al. 2020. Fungal Diversity 105:1-16 ((https://doi.org/10.1007/s13225-020-00466-2)) |
| Otu51692 | 1 | k__Fungi(100);p__Ascomycota(100);c__Saccharomycetes(100);o__Saccharomycetales(100);f__Metschnikowiaceae(100);g__Metschnikowia(100);s__Metschnikowia_pulcherrima(100); | Metschnikowia | 13 | Saprotroph | \|Undefined Saprotroph\| | Yeast | NULL | Probable | As nectar-tap_saprotroph (Põlme et al. 2020); Decay sugar-rich substrates (Põlme et al. 2020) | Põlme S, et al. 2020. Fungal Diversity 105:1-16 ((https://doi.org/10.1007/s13225-020-00466-2)) |
| Otu51693 | 1 | k__Fungi(100);p__Ascomycota(100);c__Saccharomycetes(100);o__Saccharomycetales(100);f__Metschnikowiaceae(100);g__Metschnikowia(100);s__Metschnikowia_pulcherrima(100); | Metschnikowia | 13 | Saprotroph | \|Undefined Saprotroph\| | Yeast | NULL | Probable | As nectar-tap_saprotroph (Põlme et al. 2020); Decay sugar-rich substrates (Põlme et al. 2020) | Põlme S, et al. 2020. Fungal Diversity 105:1-16 ((https://doi.org/10.1007/s13225-020-00466-2)) |
| Otu51694 | 1 | k__Fungi(100);p__Ascomycota(100);c__Saccharomycetes(100);o__Saccharomycetales(100);f__Metschnikowiaceae(100);g__Metschnikowia(100);s__Metschnikowia_pulcherrima(100); | Metschnikowia | 13 | Saprotroph | \|Undefined Saprotroph\| | Yeast | NULL | Probable | As nectar-tap_saprotroph (Põlme et al. 2020); Decay sugar-rich substrates (Põlme et al. 2020) | Põlme S, et al. 2020. Fungal Diversity 105:1-16 ((https://doi.org/10.1007/s13225-020-00466-2)) |
| Otu51695 | 1 | k__Fungi(100);p__Ascomycota(100);c__Saccharomycetes(100);o__Saccharomycetales(100);f__Metschnikowiaceae(100);g__Metschnikowia(100);s__Metschnikowia_pulcherrima(100); | Metschnikowia | 13 | Saprotroph | \|Undefined Saprotroph\| | Yeast | NULL | Probable | As nectar-tap_saprotroph (Põlme et al. 2020); Decay sugar-rich substrates (Põlme et al. 2020) | Põlme S, et al. 2020. Fungal Diversity 105:1-16 ((https://doi.org/10.1007/s13225-020-00466-2)) |
| Otu51696 | 1 | k__Fungi(100);p__Ascomycota(100);c__Saccharomycetes(100);o__Saccharomycetales(100);f__Metschnikowiaceae(100);g__Metschnikowia(100);s__Metschnikowia_pulcherrima(100); | Metschnikowia | 13 | Saprotroph | \|Undefined Saprotroph\| | Yeast | NULL | Probable | As nectar-tap_saprotroph (Põlme et al. 2020); Decay sugar-rich substrates (Põlme et al. 2020) | Põlme S, et al. 2020. Fungal Diversity 105:1-16 ((https://doi.org/10.1007/s13225-020-00466-2)) |
| Otu51697 | 1 | k__Fungi(100);p__Ascomycota(100);c__Saccharomycetes(100);o__Saccharomycetales(100);f__Metschnikowiaceae(100);g__Metschnikowia(100);g__Metschnikowia_unclassified(100); | Metschnikowia | 13 | Saprotroph | \|Undefined Saprotroph\| | Yeast | NULL | Probable | As nectar-tap_saprotroph (Põlme et al. 2020); Decay sugar-rich substrates (Põlme et al. 2020) | Põlme S, et al. 2020. Fungal Diversity 105:1-16 ((https://doi.org/10.1007/s13225-020-00466-2)) |
| Otu51698 | 1 | k__Fungi(100);p__Ascomycota(100);c__Saccharomycetes(100);o__Saccharomycetales(100);f__Metschnikowiaceae(100);g__Metschnikowia(100);g__Metschnikowia_unclassified(100); | Metschnikowia | 13 | Saprotroph | \|Undefined Saprotroph\| | Yeast | NULL | Probable | As nectar-tap_saprotroph (Põlme et al. 2020); Decay sugar-rich substrates (Põlme et al. 2020) | Põlme S, et al. 2020. Fungal Diversity 105:1-16 ((https://doi.org/10.1007/s13225-020-00466-2)) |
| Otu51699 | 1 | k__Fungi(100);p__Ascomycota(100);c__Saccharomycetes(100);o__Saccharomycetales(100);f__Metschnikowiaceae(100);g__Metschnikowia(100);s__Metschnikowia_pulcherrima(100); | Metschnikowia | 13 | Saprotroph | \|Undefined Saprotroph\| | Yeast | NULL | Probable | As nectar-tap_saprotroph (Põlme et al. 2020); Decay sugar-rich substrates (Põlme et al. 2020) | Põlme S, et al. 2020. Fungal Diversity 105:1-16 ((https://doi.org/10.1007/s13225-020-00466-2)) |
| Otu51700 | 1 | k__Fungi(100);p__Ascomycota(100);c__Saccharomycetes(100);o__Saccharomycetales(100);f__Metschnikowiaceae(100);g__Metschnikowia(100);s__Metschnikowia_pulcherrima(100); | Metschnikowia | 13 | Saprotroph | \|Undefined Saprotroph\| | Yeast | NULL | Probable | As nectar-tap_saprotroph (Põlme et al. 2020); Decay sugar-rich substrates (Põlme et al. 2020) | Põlme S, et al. 2020. Fungal Diversity 105:1-16 ((https://doi.org/10.1007/s13225-020-00466-2)) |
| Otu51701 | 1 | k__Fungi(100);p__Ascomycota(100);c__Saccharomycetes(100);o__Saccharomycetales(100);f__Metschnikowiaceae(100);g__Metschnikowia(100);s__Metschnikowia_pulcherrima(100); | Metschnikowia | 13 | Saprotroph | \|Undefined Saprotroph\| | Yeast | NULL | Probable | As nectar-tap_saprotroph (Põlme et al. 2020); Decay sugar-rich substrates (Põlme et al. 2020) | Põlme S, et al. 2020. Fungal Diversity 105:1-16 ((https://doi.org/10.1007/s13225-020-00466-2)) |
| Otu51702 | 1 | k__Fungi(100);p__Ascomycota(100);c__Saccharomycetes(100);o__Saccharomycetales(100);f__Metschnikowiaceae(100);g__Metschnikowia(100);s__Metschnikowia_pulcherrima(100); | Metschnikowia | 13 | Saprotroph | \|Undefined Saprotroph\| | Yeast | NULL | Probable | As nectar-tap_saprotroph (Põlme et al. 2020); Decay sugar-rich substrates (Põlme et al. 2020) | Põlme S, et al. 2020. Fungal Diversity 105:1-16 ((https://doi.org/10.1007/s13225-020-00466-2)) |
| Otu51703 | 1 | k__Fungi(100);p__Ascomycota(100);c__Saccharomycetes(100);o__Saccharomycetales(100);f__Metschnikowiaceae(100);g__Metschnikowia(100);s__Metschnikowia_pulcherrima(100); | Metschnikowia | 13 | Saprotroph | \|Undefined Saprotroph\| | Yeast | NULL | Probable | As nectar-tap_saprotroph (Põlme et al. 2020); Decay sugar-rich substrates (Põlme et al. 2020) | Põlme S, et al. 2020. Fungal Diversity 105:1-16 ((https://doi.org/10.1007/s13225-020-00466-2)) |
| Otu51704 | 1 | k__Fungi(100);p__Ascomycota(100);c__Saccharomycetes(100);o__Saccharomycetales(100);f__Metschnikowiaceae(100);g__Metschnikowia(100);g__Metschnikowia_unclassified(100); | Metschnikowia | 13 | Saprotroph | \|Undefined Saprotroph\| | Yeast | NULL | Probable | As nectar-tap_saprotroph (Põlme et al. 2020); Decay sugar-rich substrates (Põlme et al. 2020) | Põlme S, et al. 2020. Fungal Diversity 105:1-16 ((https://doi.org/10.1007/s13225-020-00466-2)) |
| Otu51705 | 1 | k__Fungi(100);p__Ascomycota(100);c__Saccharomycetes(100);o__Saccharomycetales(100);f__Metschnikowiaceae(100);g__Metschnikowia(100);s__Metschnikowia_pulcherrima(100); | Metschnikowia | 13 | Saprotroph | \|Undefined Saprotroph\| | Yeast | NULL | Probable | As nectar-tap_saprotroph (Põlme et al. 2020); Decay sugar-rich substrates (Põlme et al. 2020) | Põlme S, et al. 2020. Fungal Diversity 105:1-16 ((https://doi.org/10.1007/s13225-020-00466-2)) |
| Otu51706 | 1 | k__Fungi(100);p__Ascomycota(100);c__Saccharomycetes(100);o__Saccharomycetales(100);f__Metschnikowiaceae(100);g__Metschnikowia(100);s__Metschnikowia_pulcherrima(100); | Metschnikowia | 13 | Saprotroph | \|Undefined Saprotroph\| | Yeast | NULL | Probable | As nectar-tap_saprotroph (Põlme et al. 2020); Decay sugar-rich substrates (Põlme et al. 2020) | Põlme S, et al. 2020. Fungal Diversity 105:1-16 ((https://doi.org/10.1007/s13225-020-00466-2)) |
| Otu51707 | 1 | k__Fungi(100);p__Ascomycota(100);c__Saccharomycetes(100);o__Saccharomycetales(100);f__Metschnikowiaceae(100);g__Metschnikowia(100);s__Metschnikowia_pulcherrima(100); | Metschnikowia | 13 | Saprotroph | \|Undefined Saprotroph\| | Yeast | NULL | Probable | As nectar-tap_saprotroph (Põlme et al. 2020); Decay sugar-rich substrates (Põlme et al. 2020) | Põlme S, et al. 2020. Fungal Diversity 105:1-16 ((https://doi.org/10.1007/s13225-020-00466-2)) |
| Otu51708 | 1 | k__Fungi(100);p__Ascomycota(100);c__Saccharomycetes(100);o__Saccharomycetales(100);f__Metschnikowiaceae(100);g__Metschnikowia(100);g__Metschnikowia_unclassified(100); | Metschnikowia | 13 | Saprotroph | \|Undefined Saprotroph\| | Yeast | NULL | Probable | As nectar-tap_saprotroph (Põlme et al. 2020); Decay sugar-rich substrates (Põlme et al. 2020) | Põlme S, et al. 2020. Fungal Diversity 105:1-16 ((https://doi.org/10.1007/s13225-020-00466-2)) |
| Otu51709 | 1 | k__Fungi(100);p__Ascomycota(100);c__Saccharomycetes(100);o__Saccharomycetales(100);f__Metschnikowiaceae(100);g__Metschnikowia(100);g__Metschnikowia_unclassified(100); | Metschnikowia | 13 | Saprotroph | \|Undefined Saprotroph\| | Yeast | NULL | Probable | As nectar-tap_saprotroph (Põlme et al. 2020); Decay sugar-rich substrates (Põlme et al. 2020) | Põlme S, et al. 2020. Fungal Diversity 105:1-16 ((https://doi.org/10.1007/s13225-020-00466-2)) |
| Otu51710 | 1 | k__Fungi(100);p__Ascomycota(100);c__Saccharomycetes(100);o__Saccharomycetales(100);f__Metschnikowiaceae(100);g__Metschnikowia(100);s__Metschnikowia_pulcherrima(100); | Metschnikowia | 13 | Saprotroph | \|Undefined Saprotroph\| | Yeast | NULL | Probable | As nectar-tap_saprotroph (Põlme et al. 2020); Decay sugar-rich substrates (Põlme et al. 2020) | Põlme S, et al. 2020. Fungal Diversity 105:1-16 ((https://doi.org/10.1007/s13225-020-00466-2)) |
| Otu51711 | 1 | k__Fungi(100);p__Ascomycota(100);c__Saccharomycetes(100);o__Saccharomycetales(100);f__Metschnikowiaceae(100);g__Metschnikowia(100);g__Metschnikowia_unclassified(100); | Metschnikowia | 13 | Saprotroph | \|Undefined Saprotroph\| | Yeast | NULL | Probable | As nectar-tap_saprotroph (Põlme et al. 2020); Decay sugar-rich substrates (Põlme et al. 2020) | Põlme S, et al. 2020. Fungal Diversity 105:1-16 ((https://doi.org/10.1007/s13225-020-00466-2)) |
| Otu51712 | 1 | k__Fungi(100);p__Ascomycota(100);c__Saccharomycetes(100);o__Saccharomycetales(100);f__Metschnikowiaceae(100);g__Metschnikowia(100);g__Metschnikowia_unclassified(100); | Metschnikowia | 13 | Saprotroph | \|Undefined Saprotroph\| | Yeast | NULL | Probable | As nectar-tap_saprotroph (Põlme et al. 2020); Decay sugar-rich substrates (Põlme et al. 2020) | Põlme S, et al. 2020. Fungal Diversity 105:1-16 ((https://doi.org/10.1007/s13225-020-00466-2)) |
| Otu51713 | 1 | k__Fungi(100);p__Ascomycota(100);c__Saccharomycetes(100);o__Saccharomycetales(100);f__Metschnikowiaceae(100);g__Metschnikowia(100);g__Metschnikowia_unclassified(100); | Metschnikowia | 13 | Saprotroph | \|Undefined Saprotroph\| | Yeast | NULL | Probable | As nectar-tap_saprotroph (Põlme et al. 2020); Decay sugar-rich substrates (Põlme et al. 2020) | Põlme S, et al. 2020. Fungal Diversity 105:1-16 ((https://doi.org/10.1007/s13225-020-00466-2)) |
| Otu51714 | 1 | k__Fungi(100);p__Ascomycota(100);c__Saccharomycetes(100);o__Saccharomycetales(100);f__Metschnikowiaceae(100);g__Metschnikowia(100);s__Metschnikowia_pulcherrima(100); | Metschnikowia | 13 | Saprotroph | \|Undefined Saprotroph\| | Yeast | NULL | Probable | As nectar-tap_saprotroph (Põlme et al. 2020); Decay sugar-rich substrates (Põlme et al. 2020) | Põlme S, et al. 2020. Fungal Diversity 105:1-16 ((https://doi.org/10.1007/s13225-020-00466-2)) |
| Otu51715 | 1 | k__Fungi(100);p__Ascomycota(100);c__Saccharomycetes(100);o__Saccharomycetales(100);f__Metschnikowiaceae(100);g__Metschnikowia(100);s__Metschnikowia_pulcherrima(100); | Metschnikowia | 13 | Saprotroph | \|Undefined Saprotroph\| | Yeast | NULL | Probable | As nectar-tap_saprotroph (Põlme et al. 2020); Decay sugar-rich substrates (Põlme et al. 2020) | Põlme S, et al. 2020. Fungal Diversity 105:1-16 ((https://doi.org/10.1007/s13225-020-00466-2)) |
| Otu51716 | 1 | k__Fungi(100);p__Ascomycota(100);c__Saccharomycetes(100);o__Saccharomycetales(100);f__Metschnikowiaceae(100);g__Metschnikowia(100);s__Metschnikowia_pulcherrima(100); | Metschnikowia | 13 | Saprotroph | \|Undefined Saprotroph\| | Yeast | NULL | Probable | As nectar-tap_saprotroph (Põlme et al. 2020); Decay sugar-rich substrates (Põlme et al. 2020) | Põlme S, et al. 2020. Fungal Diversity 105:1-16 ((https://doi.org/10.1007/s13225-020-00466-2)) |
| Otu51717 | 1 | k__Fungi(100);p__Ascomycota(100);c__Saccharomycetes(100);o__Saccharomycetales(100);f__Metschnikowiaceae(100);g__Metschnikowia(100);g__Metschnikowia_unclassified(100); | Metschnikowia | 13 | Saprotroph | \|Undefined Saprotroph\| | Yeast | NULL | Probable | As nectar-tap_saprotroph (Põlme et al. 2020); Decay sugar-rich substrates (Põlme et al. 2020) | Põlme S, et al. 2020. Fungal Diversity 105:1-16 ((https://doi.org/10.1007/s13225-020-00466-2)) |
| Otu51718 | 1 | k__Fungi(100);p__Ascomycota(100);c__Saccharomycetes(100);o__Saccharomycetales(100);f__Metschnikowiaceae(100);g__Metschnikowia(100);s__Metschnikowia_pulcherrima(100); | Metschnikowia | 13 | Saprotroph | \|Undefined Saprotroph\| | Yeast | NULL | Probable | As nectar-tap_saprotroph (Põlme et al. 2020); Decay sugar-rich substrates (Põlme et al. 2020) | Põlme S, et al. 2020. Fungal Diversity 105:1-16 ((https://doi.org/10.1007/s13225-020-00466-2)) |
| Otu51719 | 1 | k__Fungi(100);p__Ascomycota(100);c__Saccharomycetes(100);o__Saccharomycetales(100);f__Metschnikowiaceae(100);g__Metschnikowia(100);s__Metschnikowia_pulcherrima(100); | Metschnikowia | 13 | Saprotroph | \|Undefined Saprotroph\| | Yeast | NULL | Probable | As nectar-tap_saprotroph (Põlme et al. 2020); Decay sugar-rich substrates (Põlme et al. 2020) | Põlme S, et al. 2020. Fungal Diversity 105:1-16 ((https://doi.org/10.1007/s13225-020-00466-2)) |
| Otu51720 | 1 | k__Fungi(100);p__Ascomycota(100);c__Saccharomycetes(100);o__Saccharomycetales(100);f__Metschnikowiaceae(100);g__Metschnikowia(100);g__Metschnikowia_unclassified(100); | Metschnikowia | 13 | Saprotroph | \|Undefined Saprotroph\| | Yeast | NULL | Probable | As nectar-tap_saprotroph (Põlme et al. 2020); Decay sugar-rich substrates (Põlme et al. 2020) | Põlme S, et al. 2020. Fungal Diversity 105:1-16 ((https://doi.org/10.1007/s13225-020-00466-2)) |
| Otu51721 | 1 | k__Fungi(100);p__Ascomycota(100);c__Saccharomycetes(100);o__Saccharomycetales(100);f__Metschnikowiaceae(100);g__Metschnikowia(100);g__Metschnikowia_unclassified(100); | Metschnikowia | 13 | Saprotroph | \|Undefined Saprotroph\| | Yeast | NULL | Probable | As nectar-tap_saprotroph (Põlme et al. 2020); Decay sugar-rich substrates (Põlme et al. 2020) | Põlme S, et al. 2020. Fungal Diversity 105:1-16 ((https://doi.org/10.1007/s13225-020-00466-2)) |
| Otu51722 | 1 | k__Fungi(100);p__Ascomycota(100);c__Saccharomycetes(100);o__Saccharomycetales(100);f__Metschnikowiaceae(100);g__Metschnikowia(100);g__Metschnikowia_unclassified(100); | Metschnikowia | 13 | Saprotroph | \|Undefined Saprotroph\| | Yeast | NULL | Probable | As nectar-tap_saprotroph (Põlme et al. 2020); Decay sugar-rich substrates (Põlme et al. 2020) | Põlme S, et al. 2020. Fungal Diversity 105:1-16 ((https://doi.org/10.1007/s13225-020-00466-2)) |
| Otu51723 | 1 | k__Fungi(100);p__Ascomycota(100);c__Saccharomycetes(100);o__Saccharomycetales(100);f__Metschnikowiaceae(100);g__Metschnikowia(100);s__Metschnikowia_pulcherrima(100); | Metschnikowia | 13 | Saprotroph | \|Undefined Saprotroph\| | Yeast | NULL | Probable | As nectar-tap_saprotroph (Põlme et al. 2020); Decay sugar-rich substrates (Põlme et al. 2020) | Põlme S, et al. 2020. Fungal Diversity 105:1-16 ((https://doi.org/10.1007/s13225-020-00466-2)) |
| Otu51724 | 1 | k__Fungi(100);p__Ascomycota(100);c__Saccharomycetes(100);o__Saccharomycetales(100);f__Metschnikowiaceae(100);g__Metschnikowia(100);s__Metschnikowia_pulcherrima(100); | Metschnikowia | 13 | Saprotroph | \|Undefined Saprotroph\| | Yeast | NULL | Probable | As nectar-tap_saprotroph (Põlme et al. 2020); Decay sugar-rich substrates (Põlme et al. 2020) | Põlme S, et al. 2020. Fungal Diversity 105:1-16 ((https://doi.org/10.1007/s13225-020-00466-2)) |
| Otu51725 | 1 | k__Fungi(100);p__Ascomycota(100);c__Saccharomycetes(100);o__Saccharomycetales(100);f__Metschnikowiaceae(100);g__Metschnikowia(100);g__Metschnikowia_unclassified(100); | Metschnikowia | 13 | Saprotroph | \|Undefined Saprotroph\| | Yeast | NULL | Probable | As nectar-tap_saprotroph (Põlme et al. 2020); Decay sugar-rich substrates (Põlme et al. 2020) | Põlme S, et al. 2020. Fungal Diversity 105:1-16 ((https://doi.org/10.1007/s13225-020-00466-2)) |
| Otu51726 | 1 | k__Fungi(100);p__Ascomycota(100);c__Saccharomycetes(100);o__Saccharomycetales(100);f__Metschnikowiaceae(100);g__Metschnikowia(100);g__Metschnikowia_unclassified(100); | Metschnikowia | 13 | Saprotroph | \|Undefined Saprotroph\| | Yeast | NULL | Probable | As nectar-tap_saprotroph (Põlme et al. 2020); Decay sugar-rich substrates (Põlme et al. 2020) | Põlme S, et al. 2020. Fungal Diversity 105:1-16 ((https://doi.org/10.1007/s13225-020-00466-2)) |
| Otu51727 | 1 | k__Fungi(100);p__Ascomycota(100);c__Saccharomycetes(100);o__Saccharomycetales(100);f__Metschnikowiaceae(100);g__Metschnikowia(100);s__Metschnikowia_pulcherrima(100); | Metschnikowia | 13 | Saprotroph | \|Undefined Saprotroph\| | Yeast | NULL | Probable | As nectar-tap_saprotroph (Põlme et al. 2020); Decay sugar-rich substrates (Põlme et al. 2020) | Põlme S, et al. 2020. Fungal Diversity 105:1-16 ((https://doi.org/10.1007/s13225-020-00466-2)) |
| Otu51728 | 1 | k__Fungi(100);p__Ascomycota(100);c__Saccharomycetes(100);o__Saccharomycetales(100);f__Metschnikowiaceae(100);g__Metschnikowia(100);s__Metschnikowia_pulcherrima(100); | Metschnikowia | 13 | Saprotroph | \|Undefined Saprotroph\| | Yeast | NULL | Probable | As nectar-tap_saprotroph (Põlme et al. 2020); Decay sugar-rich substrates (Põlme et al. 2020) | Põlme S, et al. 2020. Fungal Diversity 105:1-16 ((https://doi.org/10.1007/s13225-020-00466-2)) |
| Otu51729 | 1 | k__Fungi(100);p__Ascomycota(100);c__Saccharomycetes(100);o__Saccharomycetales(100);f__Metschnikowiaceae(100);g__Metschnikowia(100);s__Metschnikowia_pulcherrima(100); | Metschnikowia | 13 | Saprotroph | \|Undefined Saprotroph\| | Yeast | NULL | Probable | As nectar-tap_saprotroph (Põlme et al. 2020); Decay sugar-rich substrates (Põlme et al. 2020) | Põlme S, et al. 2020. Fungal Diversity 105:1-16 ((https://doi.org/10.1007/s13225-020-00466-2)) |
| Otu51730 | 1 | k__Fungi(100);p__Ascomycota(100);c__Saccharomycetes(100);o__Saccharomycetales(100);f__Metschnikowiaceae(100);g__Metschnikowia(100);s__Metschnikowia_pulcherrima(100); | Metschnikowia | 13 | Saprotroph | \|Undefined Saprotroph\| | Yeast | NULL | Probable | As nectar-tap_saprotroph (Põlme et al. 2020); Decay sugar-rich substrates (Põlme et al. 2020) | Põlme S, et al. 2020. Fungal Diversity 105:1-16 ((https://doi.org/10.1007/s13225-020-00466-2)) |
| Otu51731 | 1 | k__Fungi(100);p__Ascomycota(100);c__Saccharomycetes(100);o__Saccharomycetales(100);f__Metschnikowiaceae(100);g__Metschnikowia(100);s__Metschnikowia_pulcherrima(100); | Metschnikowia | 13 | Saprotroph | \|Undefined Saprotroph\| | Yeast | NULL | Probable | As nectar-tap_saprotroph (Põlme et al. 2020); Decay sugar-rich substrates (Põlme et al. 2020) | Põlme S, et al. 2020. Fungal Diversity 105:1-16 ((https://doi.org/10.1007/s13225-020-00466-2)) |
| Otu51732 | 1 | k__Fungi(100);p__Ascomycota(100);c__Saccharomycetes(100);o__Saccharomycetales(100);f__Metschnikowiaceae(100);g__Metschnikowia(100);s__Metschnikowia_pulcherrima(100); | Metschnikowia | 13 | Saprotroph | \|Undefined Saprotroph\| | Yeast | NULL | Probable | As nectar-tap_saprotroph (Põlme et al. 2020); Decay sugar-rich substrates (Põlme et al. 2020) | Põlme S, et al. 2020. Fungal Diversity 105:1-16 ((https://doi.org/10.1007/s13225-020-00466-2)) |
| Otu51733 | 1 | k__Fungi(100);p__Ascomycota(100);c__Saccharomycetes(100);o__Saccharomycetales(100);f__Metschnikowiaceae(100);g__Metschnikowia(100);s__Metschnikowia_pulcherrima(100); | Metschnikowia | 13 | Saprotroph | \|Undefined Saprotroph\| | Yeast | NULL | Probable | As nectar-tap_saprotroph (Põlme et al. 2020); Decay sugar-rich substrates (Põlme et al. 2020) | Põlme S, et al. 2020. Fungal Diversity 105:1-16 ((https://doi.org/10.1007/s13225-020-00466-2)) |
| Otu51734 | 1 | k__Fungi(100);p__Ascomycota(100);c__Saccharomycetes(100);o__Saccharomycetales(100);f__Metschnikowiaceae(100);g__Metschnikowia(100);s__Metschnikowia_pulcherrima(100); | Metschnikowia | 13 | Saprotroph | \|Undefined Saprotroph\| | Yeast | NULL | Probable | As nectar-tap_saprotroph (Põlme et al. 2020); Decay sugar-rich substrates (Põlme et al. 2020) | Põlme S, et al. 2020. Fungal Diversity 105:1-16 ((https://doi.org/10.1007/s13225-020-00466-2)) |
| Otu51735 | 1 | k__Fungi(100);p__Ascomycota(100);c__Saccharomycetes(100);o__Saccharomycetales(100);f__Metschnikowiaceae(100);g__Metschnikowia(100);s__Metschnikowia_pulcherrima(100); | Metschnikowia | 13 | Saprotroph | \|Undefined Saprotroph\| | Yeast | NULL | Probable | As nectar-tap_saprotroph (Põlme et al. 2020); Decay sugar-rich substrates (Põlme et al. 2020) | Põlme S, et al. 2020. Fungal Diversity 105:1-16 ((https://doi.org/10.1007/s13225-020-00466-2)) |
| Otu51736 | 1 | k__Fungi(100);p__Ascomycota(100);c__Saccharomycetes(100);o__Saccharomycetales(100);f__Metschnikowiaceae(100);g__Metschnikowia(100);s__Metschnikowia_pulcherrima(100); | Metschnikowia | 13 | Saprotroph | \|Undefined Saprotroph\| | Yeast | NULL | Probable | As nectar-tap_saprotroph (Põlme et al. 2020); Decay sugar-rich substrates (Põlme et al. 2020) | Põlme S, et al. 2020. Fungal Diversity 105:1-16 ((https://doi.org/10.1007/s13225-020-00466-2)) |
| Otu51738 | 1 | k__Fungi(100);p__Ascomycota(100);c__Saccharomycetes(100);o__Saccharomycetales(100);f__Metschnikowiaceae(100);g__Metschnikowia(100);s__Metschnikowia_pulcherrima(100); | Metschnikowia | 13 | Saprotroph | \|Undefined Saprotroph\| | Yeast | NULL | Probable | As nectar-tap_saprotroph (Põlme et al. 2020); Decay sugar-rich substrates (Põlme et al. 2020) | Põlme S, et al. 2020. Fungal Diversity 105:1-16 ((https://doi.org/10.1007/s13225-020-00466-2)) |
| Otu51739 | 1 | k__Fungi(100);p__Ascomycota(100);c__Saccharomycetes(100);o__Saccharomycetales(100);f__Metschnikowiaceae(100);g__Metschnikowia(100);s__Metschnikowia_pulcherrima(100); | Metschnikowia | 13 | Saprotroph | \|Undefined Saprotroph\| | Yeast | NULL | Probable | As nectar-tap_saprotroph (Põlme et al. 2020); Decay sugar-rich substrates (Põlme et al. 2020) | Põlme S, et al. 2020. Fungal Diversity 105:1-16 ((https://doi.org/10.1007/s13225-020-00466-2)) |
| Otu51740 | 1 | k__Fungi(100);p__Ascomycota(100);c__Saccharomycetes(100);o__Saccharomycetales(100);f__Metschnikowiaceae(100);g__Metschnikowia(100);s__Metschnikowia_pulcherrima(100); | Metschnikowia | 13 | Saprotroph | \|Undefined Saprotroph\| | Yeast | NULL | Probable | As nectar-tap_saprotroph (Põlme et al. 2020); Decay sugar-rich substrates (Põlme et al. 2020) | Põlme S, et al. 2020. Fungal Diversity 105:1-16 ((https://doi.org/10.1007/s13225-020-00466-2)) |
| Otu51741 | 1 | k__Fungi(100);p__Ascomycota(100);c__Saccharomycetes(100);o__Saccharomycetales(100);f__Metschnikowiaceae(100);g__Metschnikowia(100);s__Metschnikowia_pulcherrima(100); | Metschnikowia | 13 | Saprotroph | \|Undefined Saprotroph\| | Yeast | NULL | Probable | As nectar-tap_saprotroph (Põlme et al. 2020); Decay sugar-rich substrates (Põlme et al. 2020) | Põlme S, et al. 2020. Fungal Diversity 105:1-16 ((https://doi.org/10.1007/s13225-020-00466-2)) |
| Otu51742 | 1 | k__Fungi(100);p__Ascomycota(100);c__Saccharomycetes(100);o__Saccharomycetales(100);f__Metschnikowiaceae(100);g__Metschnikowia(100);s__Metschnikowia_pulcherrima(100); | Metschnikowia | 13 | Saprotroph | \|Undefined Saprotroph\| | Yeast | NULL | Probable | As nectar-tap_saprotroph (Põlme et al. 2020); Decay sugar-rich substrates (Põlme et al. 2020) | Põlme S, et al. 2020. Fungal Diversity 105:1-16 ((https://doi.org/10.1007/s13225-020-00466-2)) |
| Otu51743 | 1 | k__Fungi(100);p__Ascomycota(100);c__Saccharomycetes(100);o__Saccharomycetales(100);f__Metschnikowiaceae(100);g__Metschnikowia(100);s__Metschnikowia_pulcherrima(100); | Metschnikowia | 13 | Saprotroph | \|Undefined Saprotroph\| | Yeast | NULL | Probable | As nectar-tap_saprotroph (Põlme et al. 2020); Decay sugar-rich substrates (Põlme et al. 2020) | Põlme S, et al. 2020. Fungal Diversity 105:1-16 ((https://doi.org/10.1007/s13225-020-00466-2)) |
| Otu51744 | 1 | k__Fungi(100);p__Ascomycota(100);c__Saccharomycetes(100);o__Saccharomycetales(100);f__Metschnikowiaceae(100);g__Metschnikowia(100);s__Metschnikowia_pulcherrima(100); | Metschnikowia | 13 | Saprotroph | \|Undefined Saprotroph\| | Yeast | NULL | Probable | As nectar-tap_saprotroph (Põlme et al. 2020); Decay sugar-rich substrates (Põlme et al. 2020) | Põlme S, et al. 2020. Fungal Diversity 105:1-16 ((https://doi.org/10.1007/s13225-020-00466-2)) |
| Otu51745 | 1 | k__Fungi(100);p__Ascomycota(100);c__Saccharomycetes(100);o__Saccharomycetales(100);f__Metschnikowiaceae(100);g__Metschnikowia(100);g__Metschnikowia_unclassified(100); | Metschnikowia | 13 | Saprotroph | \|Undefined Saprotroph\| | Yeast | NULL | Probable | As nectar-tap_saprotroph (Põlme et al. 2020); Decay sugar-rich substrates (Põlme et al. 2020) | Põlme S, et al. 2020. Fungal Diversity 105:1-16 ((https://doi.org/10.1007/s13225-020-00466-2)) |
| Otu51747 | 1 | k__Fungi(100);p__Ascomycota(100);c__Saccharomycetes(100);o__Saccharomycetales(100);f__Metschnikowiaceae(100);g__Metschnikowia(100);g__Metschnikowia_unclassified(100); | Metschnikowia | 13 | Saprotroph | \|Undefined Saprotroph\| | Yeast | NULL | Probable | As nectar-tap_saprotroph (Põlme et al. 2020); Decay sugar-rich substrates (Põlme et al. 2020) | Põlme S, et al. 2020. Fungal Diversity 105:1-16 ((https://doi.org/10.1007/s13225-020-00466-2)) |
| Otu51748 | 1 | k__Fungi(100);p__Ascomycota(100);c__Saccharomycetes(100);o__Saccharomycetales(100);f__Metschnikowiaceae(100);g__Metschnikowia(100);s__Metschnikowia_pulcherrima(100); | Metschnikowia | 13 | Saprotroph | \|Undefined Saprotroph\| | Yeast | NULL | Probable | As nectar-tap_saprotroph (Põlme et al. 2020); Decay sugar-rich substrates (Põlme et al. 2020) | Põlme S, et al. 2020. Fungal Diversity 105:1-16 ((https://doi.org/10.1007/s13225-020-00466-2)) |
| Otu51749 | 1 | k__Fungi(100);p__Ascomycota(100);c__Saccharomycetes(100);o__Saccharomycetales(100);f__Metschnikowiaceae(100);g__Metschnikowia(100);s__Metschnikowia_pulcherrima(100); | Metschnikowia | 13 | Saprotroph | \|Undefined Saprotroph\| | Yeast | NULL | Probable | As nectar-tap_saprotroph (Põlme et al. 2020); Decay sugar-rich substrates (Põlme et al. 2020) | Põlme S, et al. 2020. Fungal Diversity 105:1-16 ((https://doi.org/10.1007/s13225-020-00466-2)) |
| Otu51750 | 1 | k__Fungi(100);p__Ascomycota(100);c__Saccharomycetes(100);o__Saccharomycetales(100);f__Metschnikowiaceae(100);g__Metschnikowia(100);s__Metschnikowia_pulcherrima(100); | Metschnikowia | 13 | Saprotroph | \|Undefined Saprotroph\| | Yeast | NULL | Probable | As nectar-tap_saprotroph (Põlme et al. 2020); Decay sugar-rich substrates (Põlme et al. 2020) | Põlme S, et al. 2020. Fungal Diversity 105:1-16 ((https://doi.org/10.1007/s13225-020-00466-2)) |
| Otu51751 | 1 | k__Fungi(100);p__Ascomycota(100);c__Saccharomycetes(100);o__Saccharomycetales(100);f__Metschnikowiaceae(100);g__Metschnikowia(100);s__Metschnikowia_pulcherrima(100); | Metschnikowia | 13 | Saprotroph | \|Undefined Saprotroph\| | Yeast | NULL | Probable | As nectar-tap_saprotroph (Põlme et al. 2020); Decay sugar-rich substrates (Põlme et al. 2020) | Põlme S, et al. 2020. Fungal Diversity 105:1-16 ((https://doi.org/10.1007/s13225-020-00466-2)) |
| Otu51752 | 1 | k__Fungi(100);p__Ascomycota(100);c__Saccharomycetes(100);o__Saccharomycetales(100);f__Metschnikowiaceae(100);g__Metschnikowia(100);s__Metschnikowia_pulcherrima(100); | Metschnikowia | 13 | Saprotroph | \|Undefined Saprotroph\| | Yeast | NULL | Probable | As nectar-tap_saprotroph (Põlme et al. 2020); Decay sugar-rich substrates (Põlme et al. 2020) | Põlme S, et al. 2020. Fungal Diversity 105:1-16 ((https://doi.org/10.1007/s13225-020-00466-2)) |
| Otu51753 | 1 | k__Fungi(100);p__Ascomycota(100);c__Saccharomycetes(100);o__Saccharomycetales(100);f__Metschnikowiaceae(100);g__Metschnikowia(100);s__Metschnikowia_pulcherrima(100); | Metschnikowia | 13 | Saprotroph | \|Undefined Saprotroph\| | Yeast | NULL | Probable | As nectar-tap_saprotroph (Põlme et al. 2020); Decay sugar-rich substrates (Põlme et al. 2020) | Põlme S, et al. 2020. Fungal Diversity 105:1-16 ((https://doi.org/10.1007/s13225-020-00466-2)) |
| Otu51754 | 1 | k__Fungi(100);p__Ascomycota(100);c__Saccharomycetes(100);o__Saccharomycetales(100);f__Metschnikowiaceae(100);g__Metschnikowia(100);s__Metschnikowia_pulcherrima(100); | Metschnikowia | 13 | Saprotroph | \|Undefined Saprotroph\| | Yeast | NULL | Probable | As nectar-tap_saprotroph (Põlme et al. 2020); Decay sugar-rich substrates (Põlme et al. 2020) | Põlme S, et al. 2020. Fungal Diversity 105:1-16 ((https://doi.org/10.1007/s13225-020-00466-2)) |
| Otu51755 | 1 | k__Fungi(100);p__Ascomycota(100);c__Saccharomycetes(100);o__Saccharomycetales(100);f__Metschnikowiaceae(100);g__Metschnikowia(100);s__Metschnikowia_pulcherrima(100); | Metschnikowia | 13 | Saprotroph | \|Undefined Saprotroph\| | Yeast | NULL | Probable | As nectar-tap_saprotroph (Põlme et al. 2020); Decay sugar-rich substrates (Põlme et al. 2020) | Põlme S, et al. 2020. Fungal Diversity 105:1-16 ((https://doi.org/10.1007/s13225-020-00466-2)) |
| Otu51756 | 1 | k__Fungi(100);p__Ascomycota(100);c__Saccharomycetes(100);o__Saccharomycetales(100);f__Metschnikowiaceae(100);g__Metschnikowia(100);s__Metschnikowia_pulcherrima(100); | Metschnikowia | 13 | Saprotroph | \|Undefined Saprotroph\| | Yeast | NULL | Probable | As nectar-tap_saprotroph (Põlme et al. 2020); Decay sugar-rich substrates (Põlme et al. 2020) | Põlme S, et al. 2020. Fungal Diversity 105:1-16 ((https://doi.org/10.1007/s13225-020-00466-2)) |
| Otu51757 | 1 | k__Fungi(100);p__Ascomycota(100);c__Saccharomycetes(100);o__Saccharomycetales(100);f__Metschnikowiaceae(100);g__Metschnikowia(100);s__Metschnikowia_pulcherrima(100); | Metschnikowia | 13 | Saprotroph | \|Undefined Saprotroph\| | Yeast | NULL | Probable | As nectar-tap_saprotroph (Põlme et al. 2020); Decay sugar-rich substrates (Põlme et al. 2020) | Põlme S, et al. 2020. Fungal Diversity 105:1-16 ((https://doi.org/10.1007/s13225-020-00466-2)) |
| Otu51758 | 1 | k__Fungi(100);p__Ascomycota(100);c__Saccharomycetes(100);o__Saccharomycetales(100);f__Metschnikowiaceae(100);g__Metschnikowia(100);g__Metschnikowia_unclassified(100); | Metschnikowia | 13 | Saprotroph | \|Undefined Saprotroph\| | Yeast | NULL | Probable | As nectar-tap_saprotroph (Põlme et al. 2020); Decay sugar-rich substrates (Põlme et al. 2020) | Põlme S, et al. 2020. Fungal Diversity 105:1-16 ((https://doi.org/10.1007/s13225-020-00466-2)) |
| Otu51759 | 1 | k__Fungi(100);p__Ascomycota(100);c__Saccharomycetes(100);o__Saccharomycetales(100);f__Metschnikowiaceae(100);g__Metschnikowia(100);s__Metschnikowia_pulcherrima(100); | Metschnikowia | 13 | Saprotroph | \|Undefined Saprotroph\| | Yeast | NULL | Probable | As nectar-tap_saprotroph (Põlme et al. 2020); Decay sugar-rich substrates (Põlme et al. 2020) | Põlme S, et al. 2020. Fungal Diversity 105:1-16 ((https://doi.org/10.1007/s13225-020-00466-2)) |
| Otu51760 | 1 | k__Fungi(100);p__Ascomycota(100);c__Saccharomycetes(100);o__Saccharomycetales(100);f__Metschnikowiaceae(100);g__Metschnikowia(100);s__Metschnikowia_pulcherrima(100); | Metschnikowia | 13 | Saprotroph | \|Undefined Saprotroph\| | Yeast | NULL | Probable | As nectar-tap_saprotroph (Põlme et al. 2020); Decay sugar-rich substrates (Põlme et al. 2020) | Põlme S, et al. 2020. Fungal Diversity 105:1-16 ((https://doi.org/10.1007/s13225-020-00466-2)) |
| Otu51761 | 1 | k__Fungi(100);p__Ascomycota(100);c__Saccharomycetes(100);o__Saccharomycetales(100);f__Metschnikowiaceae(100);g__Metschnikowia(100);s__Metschnikowia_pulcherrima(100); | Metschnikowia | 13 | Saprotroph | \|Undefined Saprotroph\| | Yeast | NULL | Probable | As nectar-tap_saprotroph (Põlme et al. 2020); Decay sugar-rich substrates (Põlme et al. 2020) | Põlme S, et al. 2020. Fungal Diversity 105:1-16 ((https://doi.org/10.1007/s13225-020-00466-2)) |
| Otu51765 | 1 | k__Fungi(100);p__Ascomycota(100);c__Saccharomycetes(100);o__Saccharomycetales(100);o__Saccharomycetales_unclassified(100);o__Saccharomycetales_unclassified(100);o__Saccharomycetales_unclassified(100); | Saccharomycetales | 7 | Saprotroph | Undefined Saprotroph | Yeast | NULL | Possible | NULL | Sterkenburg E, et al. 2015. New Phytologist 207:1145-1158 ((https://doi.org/10.1111/nph.13426)) |
| Otu51767 | 1 | k__Fungi(100);p__Ascomycota(100);c__Saccharomycetes(100);o__Saccharomycetales(100);f__Metschnikowiaceae(100);g__Metschnikowia(100);s__Metschnikowia_pulcherrima(100); | Metschnikowia | 13 | Saprotroph | \|Undefined Saprotroph\| | Yeast | NULL | Probable | As nectar-tap_saprotroph (Põlme et al. 2020); Decay sugar-rich substrates (Põlme et al. 2020) | Põlme S, et al. 2020. Fungal Diversity 105:1-16 ((https://doi.org/10.1007/s13225-020-00466-2)) |
| Otu51771 | 1 | k__Fungi(100);p__Ascomycota(100);c__Saccharomycetes(100);o__Saccharomycetales(100);f__Metschnikowiaceae(100);g__Metschnikowia(100);s__Metschnikowia_pulcherrima(100); | Metschnikowia | 13 | Saprotroph | \|Undefined Saprotroph\| | Yeast | NULL | Probable | As nectar-tap_saprotroph (Põlme et al. 2020); Decay sugar-rich substrates (Põlme et al. 2020) | Põlme S, et al. 2020. Fungal Diversity 105:1-16 ((https://doi.org/10.1007/s13225-020-00466-2)) |
| Otu51772 | 1 | k__Fungi(100);p__Ascomycota(100);c__Saccharomycetes(100);o__Saccharomycetales(100);f__Metschnikowiaceae(100);g__Metschnikowia(100);s__Metschnikowia_pulcherrima(100); | Metschnikowia | 13 | Saprotroph | \|Undefined Saprotroph\| | Yeast | NULL | Probable | As nectar-tap_saprotroph (Põlme et al. 2020); Decay sugar-rich substrates (Põlme et al. 2020) | Põlme S, et al. 2020. Fungal Diversity 105:1-16 ((https://doi.org/10.1007/s13225-020-00466-2)) |
| Otu51773 | 1 | k__Fungi(100);p__Ascomycota(100);c__Saccharomycetes(100);o__Saccharomycetales(100);f__Metschnikowiaceae(100);g__Metschnikowia(100);s__Metschnikowia_pulcherrima(100); | Metschnikowia | 13 | Saprotroph | \|Undefined Saprotroph\| | Yeast | NULL | Probable | As nectar-tap_saprotroph (Põlme et al. 2020); Decay sugar-rich substrates (Põlme et al. 2020) | Põlme S, et al. 2020. Fungal Diversity 105:1-16 ((https://doi.org/10.1007/s13225-020-00466-2)) |
| Otu51774 | 1 | k__Fungi(100);p__Ascomycota(100);c__Saccharomycetes(100);o__Saccharomycetales(100);f__Metschnikowiaceae(100);g__Metschnikowia(100);s__Metschnikowia_pulcherrima(100); | Metschnikowia | 13 | Saprotroph | \|Undefined Saprotroph\| | Yeast | NULL | Probable | As nectar-tap_saprotroph (Põlme et al. 2020); Decay sugar-rich substrates (Põlme et al. 2020) | Põlme S, et al. 2020. Fungal Diversity 105:1-16 ((https://doi.org/10.1007/s13225-020-00466-2)) |
| Otu51775 | 1 | k__Fungi(100);p__Ascomycota(100);c__Saccharomycetes(100);o__Saccharomycetales(100);f__Metschnikowiaceae(100);g__Metschnikowia(100);s__Metschnikowia_pulcherrima(100); | Metschnikowia | 13 | Saprotroph | \|Undefined Saprotroph\| | Yeast | NULL | Probable | As nectar-tap_saprotroph (Põlme et al. 2020); Decay sugar-rich substrates (Põlme et al. 2020) | Põlme S, et al. 2020. Fungal Diversity 105:1-16 ((https://doi.org/10.1007/s13225-020-00466-2)) |
| Otu51777 | 1 | k__Fungi(100);p__Ascomycota(100);c__Saccharomycetes(100);o__Saccharomycetales(100);f__Metschnikowiaceae(100);g__Metschnikowia(100);s__Metschnikowia_pulcherrima(100); | Metschnikowia | 13 | Saprotroph | \|Undefined Saprotroph\| | Yeast | NULL | Probable | As nectar-tap_saprotroph (Põlme et al. 2020); Decay sugar-rich substrates (Põlme et al. 2020) | Põlme S, et al. 2020. Fungal Diversity 105:1-16 ((https://doi.org/10.1007/s13225-020-00466-2)) |
| Otu51778 | 1 | k__Fungi(100);p__Ascomycota(100);c__Saccharomycetes(100);o__Saccharomycetales(100);f__Metschnikowiaceae(100);g__Metschnikowia(100);g__Metschnikowia_unclassified(100); | Metschnikowia | 13 | Saprotroph | \|Undefined Saprotroph\| | Yeast | NULL | Probable | As nectar-tap_saprotroph (Põlme et al. 2020); Decay sugar-rich substrates (Põlme et al. 2020) | Põlme S, et al. 2020. Fungal Diversity 105:1-16 ((https://doi.org/10.1007/s13225-020-00466-2)) |
| Otu51779 | 1 | k__Fungi(100);p__Ascomycota(100);c__Saccharomycetes(100);o__Saccharomycetales(100);f__Metschnikowiaceae(100);g__Metschnikowia(100);s__Metschnikowia_pulcherrima(100); | Metschnikowia | 13 | Saprotroph | \|Undefined Saprotroph\| | Yeast | NULL | Probable | As nectar-tap_saprotroph (Põlme et al. 2020); Decay sugar-rich substrates (Põlme et al. 2020) | Põlme S, et al. 2020. Fungal Diversity 105:1-16 ((https://doi.org/10.1007/s13225-020-00466-2)) |
| Otu51780 | 1 | k__Fungi(100);p__Ascomycota(100);c__Saccharomycetes(100);o__Saccharomycetales(100);f__Metschnikowiaceae(100);g__Metschnikowia(100);g__Metschnikowia_unclassified(100); | Metschnikowia | 13 | Saprotroph | \|Undefined Saprotroph\| | Yeast | NULL | Probable | As nectar-tap_saprotroph (Põlme et al. 2020); Decay sugar-rich substrates (Põlme et al. 2020) | Põlme S, et al. 2020. Fungal Diversity 105:1-16 ((https://doi.org/10.1007/s13225-020-00466-2)) |
| Otu51785 | 1 | k__Fungi(100);p__Ascomycota(100);c__Saccharomycetes(100);o__Saccharomycetales(100);f__Metschnikowiaceae(100);g__Metschnikowia(100);g__Metschnikowia_unclassified(100); | Metschnikowia | 13 | Saprotroph | \|Undefined Saprotroph\| | Yeast | NULL | Probable | As nectar-tap_saprotroph (Põlme et al. 2020); Decay sugar-rich substrates (Põlme et al. 2020) | Põlme S, et al. 2020. Fungal Diversity 105:1-16 ((https://doi.org/10.1007/s13225-020-00466-2)) |
| Otu51786 | 1 | k__Fungi(100);p__Ascomycota(100);c__Saccharomycetes(100);o__Saccharomycetales(100);f__Metschnikowiaceae(100);g__Metschnikowia(100);g__Metschnikowia_unclassified(100); | Metschnikowia | 13 | Saprotroph | \|Undefined Saprotroph\| | Yeast | NULL | Probable | As nectar-tap_saprotroph (Põlme et al. 2020); Decay sugar-rich substrates (Põlme et al. 2020) | Põlme S, et al. 2020. Fungal Diversity 105:1-16 ((https://doi.org/10.1007/s13225-020-00466-2)) |
| Otu51787 | 1 | k__Fungi(100);p__Ascomycota(100);c__Saccharomycetes(100);o__Saccharomycetales(100);f__Metschnikowiaceae(100);g__Metschnikowia(100);s__Metschnikowia_pulcherrima(100); | Metschnikowia | 13 | Saprotroph | \|Undefined Saprotroph\| | Yeast | NULL | Probable | As nectar-tap_saprotroph (Põlme et al. 2020); Decay sugar-rich substrates (Põlme et al. 2020) | Põlme S, et al. 2020. Fungal Diversity 105:1-16 ((https://doi.org/10.1007/s13225-020-00466-2)) |
| Otu51788 | 1 | k__Fungi(100);p__Ascomycota(100);c__Saccharomycetes(100);o__Saccharomycetales(100);f__Metschnikowiaceae(100);g__Metschnikowia(100);s__Metschnikowia_pulcherrima(100); | Metschnikowia | 13 | Saprotroph | \|Undefined Saprotroph\| | Yeast | NULL | Probable | As nectar-tap_saprotroph (Põlme et al. 2020); Decay sugar-rich substrates (Põlme et al. 2020) | Põlme S, et al. 2020. Fungal Diversity 105:1-16 ((https://doi.org/10.1007/s13225-020-00466-2)) |
| Otu51789 | 1 | k__Fungi(100);p__Ascomycota(100);c__Saccharomycetes(100);o__Saccharomycetales(100);f__Metschnikowiaceae(100);g__Metschnikowia(100);g__Metschnikowia_unclassified(100); | Metschnikowia | 13 | Saprotroph | \|Undefined Saprotroph\| | Yeast | NULL | Probable | As nectar-tap_saprotroph (Põlme et al. 2020); Decay sugar-rich substrates (Põlme et al. 2020) | Põlme S, et al. 2020. Fungal Diversity 105:1-16 ((https://doi.org/10.1007/s13225-020-00466-2)) |
| Otu51790 | 1 | k__Fungi(100);p__Ascomycota(100);c__Saccharomycetes(100);o__Saccharomycetales(100);f__Metschnikowiaceae(100);g__Metschnikowia(100);s__Metschnikowia_pulcherrima(100); | Metschnikowia | 13 | Saprotroph | \|Undefined Saprotroph\| | Yeast | NULL | Probable | As nectar-tap_saprotroph (Põlme et al. 2020); Decay sugar-rich substrates (Põlme et al. 2020) | Põlme S, et al. 2020. Fungal Diversity 105:1-16 ((https://doi.org/10.1007/s13225-020-00466-2)) |
| Otu51791 | 1 | k__Fungi(100);p__Ascomycota(100);c__Saccharomycetes(100);o__Saccharomycetales(100);f__Metschnikowiaceae(100);g__Metschnikowia(100);s__Metschnikowia_pulcherrima(100); | Metschnikowia | 13 | Saprotroph | \|Undefined Saprotroph\| | Yeast | NULL | Probable | As nectar-tap_saprotroph (Põlme et al. 2020); Decay sugar-rich substrates (Põlme et al. 2020) | Põlme S, et al. 2020. Fungal Diversity 105:1-16 ((https://doi.org/10.1007/s13225-020-00466-2)) |
| Otu51792 | 1 | k__Fungi(100);p__Ascomycota(100);c__Saccharomycetes(100);o__Saccharomycetales(100);f__Metschnikowiaceae(100);g__Metschnikowia(100);s__Metschnikowia_pulcherrima(100); | Metschnikowia | 13 | Saprotroph | \|Undefined Saprotroph\| | Yeast | NULL | Probable | As nectar-tap_saprotroph (Põlme et al. 2020); Decay sugar-rich substrates (Põlme et al. 2020) | Põlme S, et al. 2020. Fungal Diversity 105:1-16 ((https://doi.org/10.1007/s13225-020-00466-2)) |
| Otu51793 | 1 | k__Fungi(100);p__Ascomycota(100);c__Saccharomycetes(100);o__Saccharomycetales(100);f__Metschnikowiaceae(100);g__Metschnikowia(100);s__Metschnikowia_pulcherrima(100); | Metschnikowia | 13 | Saprotroph | \|Undefined Saprotroph\| | Yeast | NULL | Probable | As nectar-tap_saprotroph (Põlme et al. 2020); Decay sugar-rich substrates (Põlme et al. 2020) | Põlme S, et al. 2020. Fungal Diversity 105:1-16 ((https://doi.org/10.1007/s13225-020-00466-2)) |
| Otu51794 | 1 | k__Fungi(100);p__Ascomycota(100);c__Saccharomycetes(100);o__Saccharomycetales(100);f__Metschnikowiaceae(100);g__Metschnikowia(100);g__Metschnikowia_unclassified(100); | Metschnikowia | 13 | Saprotroph | \|Undefined Saprotroph\| | Yeast | NULL | Probable | As nectar-tap_saprotroph (Põlme et al. 2020); Decay sugar-rich substrates (Põlme et al. 2020) | Põlme S, et al. 2020. Fungal Diversity 105:1-16 ((https://doi.org/10.1007/s13225-020-00466-2)) |
| Otu51795 | 1 | k__Fungi(100);p__Ascomycota(100);c__Saccharomycetes(100);o__Saccharomycetales(100);f__Metschnikowiaceae(100);g__Metschnikowia(100);s__Metschnikowia_pulcherrima(100); | Metschnikowia | 13 | Saprotroph | \|Undefined Saprotroph\| | Yeast | NULL | Probable | As nectar-tap_saprotroph (Põlme et al. 2020); Decay sugar-rich substrates (Põlme et al. 2020) | Põlme S, et al. 2020. Fungal Diversity 105:1-16 ((https://doi.org/10.1007/s13225-020-00466-2)) |
| Otu51796 | 1 | k__Fungi(100);p__Ascomycota(100);c__Saccharomycetes(100);o__Saccharomycetales(100);f__Metschnikowiaceae(100);g__Metschnikowia(100);s__Metschnikowia_pulcherrima(100); | Metschnikowia | 13 | Saprotroph | \|Undefined Saprotroph\| | Yeast | NULL | Probable | As nectar-tap_saprotroph (Põlme et al. 2020); Decay sugar-rich substrates (Põlme et al. 2020) | Põlme S, et al. 2020. Fungal Diversity 105:1-16 ((https://doi.org/10.1007/s13225-020-00466-2)) |
| Otu51797 | 1 | k__Fungi(100);p__Ascomycota(100);c__Saccharomycetes(100);o__Saccharomycetales(100);f__Metschnikowiaceae(100);g__Metschnikowia(100);s__Metschnikowia_pulcherrima(100); | Metschnikowia | 13 | Saprotroph | \|Undefined Saprotroph\| | Yeast | NULL | Probable | As nectar-tap_saprotroph (Põlme et al. 2020); Decay sugar-rich substrates (Põlme et al. 2020) | Põlme S, et al. 2020. Fungal Diversity 105:1-16 ((https://doi.org/10.1007/s13225-020-00466-2)) |
| Otu51798 | 1 | k__Fungi(100);p__Ascomycota(100);c__Saccharomycetes(100);o__Saccharomycetales(100);f__Metschnikowiaceae(100);g__Metschnikowia(100);s__Metschnikowia_pulcherrima(100); | Metschnikowia | 13 | Saprotroph | \|Undefined Saprotroph\| | Yeast | NULL | Probable | As nectar-tap_saprotroph (Põlme et al. 2020); Decay sugar-rich substrates (Põlme et al. 2020) | Põlme S, et al. 2020. Fungal Diversity 105:1-16 ((https://doi.org/10.1007/s13225-020-00466-2)) |
| Otu51799 | 1 | k__Fungi(100);p__Ascomycota(100);c__Saccharomycetes(100);o__Saccharomycetales(100);f__Metschnikowiaceae(100);g__Metschnikowia(100);s__Metschnikowia_pulcherrima(100); | Metschnikowia | 13 | Saprotroph | \|Undefined Saprotroph\| | Yeast | NULL | Probable | As nectar-tap_saprotroph (Põlme et al. 2020); Decay sugar-rich substrates (Põlme et al. 2020) | Põlme S, et al. 2020. Fungal Diversity 105:1-16 ((https://doi.org/10.1007/s13225-020-00466-2)) |
| Otu51801 | 1 | k__Fungi(100);p__Ascomycota(100);c__Saccharomycetes(100);o__Saccharomycetales(100);f__Metschnikowiaceae(100);g__Metschnikowia(100);s__Metschnikowia_pulcherrima(100); | Metschnikowia | 13 | Saprotroph | \|Undefined Saprotroph\| | Yeast | NULL | Probable | As nectar-tap_saprotroph (Põlme et al. 2020); Decay sugar-rich substrates (Põlme et al. 2020) | Põlme S, et al. 2020. Fungal Diversity 105:1-16 ((https://doi.org/10.1007/s13225-020-00466-2)) |
| Otu51802 | 1 | k__Fungi(100);p__Ascomycota(100);c__Saccharomycetes(100);o__Saccharomycetales(100);f__Metschnikowiaceae(100);g__Metschnikowia(100);s__Metschnikowia_pulcherrima(100); | Metschnikowia | 13 | Saprotroph | \|Undefined Saprotroph\| | Yeast | NULL | Probable | As nectar-tap_saprotroph (Põlme et al. 2020); Decay sugar-rich substrates (Põlme et al. 2020) | Põlme S, et al. 2020. Fungal Diversity 105:1-16 ((https://doi.org/10.1007/s13225-020-00466-2)) |
| Otu51803 | 1 | k__Fungi(100);p__Ascomycota(100);c__Saccharomycetes(100);o__Saccharomycetales(100);f__Metschnikowiaceae(100);g__Metschnikowia(100);s__Metschnikowia_pulcherrima(100); | Metschnikowia | 13 | Saprotroph | \|Undefined Saprotroph\| | Yeast | NULL | Probable | As nectar-tap_saprotroph (Põlme et al. 2020); Decay sugar-rich substrates (Põlme et al. 2020) | Põlme S, et al. 2020. Fungal Diversity 105:1-16 ((https://doi.org/10.1007/s13225-020-00466-2)) |
| Otu51804 | 1 | k__Fungi(100);p__Ascomycota(100);c__Saccharomycetes(100);o__Saccharomycetales(100);f__Metschnikowiaceae(100);g__Metschnikowia(100);g__Metschnikowia_unclassified(100); | Metschnikowia | 13 | Saprotroph | \|Undefined Saprotroph\| | Yeast | NULL | Probable | As nectar-tap_saprotroph (Põlme et al. 2020); Decay sugar-rich substrates (Põlme et al. 2020) | Põlme S, et al. 2020. Fungal Diversity 105:1-16 ((https://doi.org/10.1007/s13225-020-00466-2)) |
| Otu51805 | 1 | k__Fungi(100);p__Ascomycota(100);c__Saccharomycetes(100);o__Saccharomycetales(100);f__Metschnikowiaceae(100);g__Metschnikowia(100);s__Metschnikowia_pulcherrima(100); | Metschnikowia | 13 | Saprotroph | \|Undefined Saprotroph\| | Yeast | NULL | Probable | As nectar-tap_saprotroph (Põlme et al. 2020); Decay sugar-rich substrates (Põlme et al. 2020) | Põlme S, et al. 2020. Fungal Diversity 105:1-16 ((https://doi.org/10.1007/s13225-020-00466-2)) |
| Otu51806 | 1 | k__Fungi(100);p__Ascomycota(100);c__Saccharomycetes(100);o__Saccharomycetales(100);f__Metschnikowiaceae(100);g__Metschnikowia(100);s__Metschnikowia_pulcherrima(100); | Metschnikowia | 13 | Saprotroph | \|Undefined Saprotroph\| | Yeast | NULL | Probable | As nectar-tap_saprotroph (Põlme et al. 2020); Decay sugar-rich substrates (Põlme et al. 2020) | Põlme S, et al. 2020. Fungal Diversity 105:1-16 ((https://doi.org/10.1007/s13225-020-00466-2)) |
| Otu51807 | 1 | k__Fungi(100);p__Ascomycota(100);c__Saccharomycetes(100);o__Saccharomycetales(100);f__Metschnikowiaceae(100);g__Metschnikowia(100);s__Metschnikowia_pulcherrima(100); | Metschnikowia | 13 | Saprotroph | \|Undefined Saprotroph\| | Yeast | NULL | Probable | As nectar-tap_saprotroph (Põlme et al. 2020); Decay sugar-rich substrates (Põlme et al. 2020) | Põlme S, et al. 2020. Fungal Diversity 105:1-16 ((https://doi.org/10.1007/s13225-020-00466-2)) |
| Otu51808 | 1 | k__Fungi(100);p__Ascomycota(100);c__Saccharomycetes(100);o__Saccharomycetales(100);f__Metschnikowiaceae(100);g__Metschnikowia(100);s__Metschnikowia_pulcherrima(100); | Metschnikowia | 13 | Saprotroph | \|Undefined Saprotroph\| | Yeast | NULL | Probable | As nectar-tap_saprotroph (Põlme et al. 2020); Decay sugar-rich substrates (Põlme et al. 2020) | Põlme S, et al. 2020. Fungal Diversity 105:1-16 ((https://doi.org/10.1007/s13225-020-00466-2)) |
| Otu51809 | 1 | k__Fungi(100);p__Ascomycota(100);c__Saccharomycetes(100);o__Saccharomycetales(100);f__Metschnikowiaceae(100);g__Metschnikowia(100);s__Metschnikowia_pulcherrima(100); | Metschnikowia | 13 | Saprotroph | \|Undefined Saprotroph\| | Yeast | NULL | Probable | As nectar-tap_saprotroph (Põlme et al. 2020); Decay sugar-rich substrates (Põlme et al. 2020) | Põlme S, et al. 2020. Fungal Diversity 105:1-16 ((https://doi.org/10.1007/s13225-020-00466-2)) |
| Otu51813 | 1 | k__Fungi(100);p__Ascomycota(100);c__Saccharomycetes(100);o__Saccharomycetales(100);f__Metschnikowiaceae(100);g__Metschnikowia(100);s__Metschnikowia_pulcherrima(100); | Metschnikowia | 13 | Saprotroph | \|Undefined Saprotroph\| | Yeast | NULL | Probable | As nectar-tap_saprotroph (Põlme et al. 2020); Decay sugar-rich substrates (Põlme et al. 2020) | Põlme S, et al. 2020. Fungal Diversity 105:1-16 ((https://doi.org/10.1007/s13225-020-00466-2)) |
| Otu51814 | 1 | k__Fungi(100);p__Ascomycota(100);c__Saccharomycetes(100);o__Saccharomycetales(100);f__Metschnikowiaceae(100);g__Metschnikowia(100);s__Metschnikowia_pulcherrima(100); | Metschnikowia | 13 | Saprotroph | \|Undefined Saprotroph\| | Yeast | NULL | Probable | As nectar-tap_saprotroph (Põlme et al. 2020); Decay sugar-rich substrates (Põlme et al. 2020) | Põlme S, et al. 2020. Fungal Diversity 105:1-16 ((https://doi.org/10.1007/s13225-020-00466-2)) |
| Otu51815 | 1 | k__Fungi(100);p__Ascomycota(100);c__Saccharomycetes(100);o__Saccharomycetales(100);f__Metschnikowiaceae(100);g__Metschnikowia(100);g__Metschnikowia_unclassified(100); | Metschnikowia | 13 | Saprotroph | \|Undefined Saprotroph\| | Yeast | NULL | Probable | As nectar-tap_saprotroph (Põlme et al. 2020); Decay sugar-rich substrates (Põlme et al. 2020) | Põlme S, et al. 2020. Fungal Diversity 105:1-16 ((https://doi.org/10.1007/s13225-020-00466-2)) |
| Otu51816 | 1 | k__Fungi(100);p__Ascomycota(100);c__Saccharomycetes(100);o__Saccharomycetales(100);f__Metschnikowiaceae(100);g__Metschnikowia(100);s__Metschnikowia_pulcherrima(100); | Metschnikowia | 13 | Saprotroph | \|Undefined Saprotroph\| | Yeast | NULL | Probable | As nectar-tap_saprotroph (Põlme et al. 2020); Decay sugar-rich substrates (Põlme et al. 2020) | Põlme S, et al. 2020. Fungal Diversity 105:1-16 ((https://doi.org/10.1007/s13225-020-00466-2)) |
| Otu51817 | 1 | k__Fungi(100);p__Ascomycota(100);c__Saccharomycetes(100);o__Saccharomycetales(100);f__Metschnikowiaceae(100);g__Metschnikowia(100);s__Metschnikowia_pulcherrima(100); | Metschnikowia | 13 | Saprotroph | \|Undefined Saprotroph\| | Yeast | NULL | Probable | As nectar-tap_saprotroph (Põlme et al. 2020); Decay sugar-rich substrates (Põlme et al. 2020) | Põlme S, et al. 2020. Fungal Diversity 105:1-16 ((https://doi.org/10.1007/s13225-020-00466-2)) |
| Otu51819 | 1 | k__Fungi(100);p__Ascomycota(100);c__Saccharomycetes(100);o__Saccharomycetales(100);f__Metschnikowiaceae(100);g__Metschnikowia(100);s__Metschnikowia_pulcherrima(100); | Metschnikowia | 13 | Saprotroph | \|Undefined Saprotroph\| | Yeast | NULL | Probable | As nectar-tap_saprotroph (Põlme et al. 2020); Decay sugar-rich substrates (Põlme et al. 2020) | Põlme S, et al. 2020. Fungal Diversity 105:1-16 ((https://doi.org/10.1007/s13225-020-00466-2)) |
| Otu51820 | 1 | k__Fungi(100);p__Ascomycota(100);c__Saccharomycetes(100);o__Saccharomycetales(100);f__Metschnikowiaceae(100);g__Metschnikowia(100);s__Metschnikowia_pulcherrima(100); | Metschnikowia | 13 | Saprotroph | \|Undefined Saprotroph\| | Yeast | NULL | Probable | As nectar-tap_saprotroph (Põlme et al. 2020); Decay sugar-rich substrates (Põlme et al. 2020) | Põlme S, et al. 2020. Fungal Diversity 105:1-16 ((https://doi.org/10.1007/s13225-020-00466-2)) |
| Otu51821 | 1 | k__Fungi(100);p__Ascomycota(100);c__Saccharomycetes(100);o__Saccharomycetales(100);f__Metschnikowiaceae(100);g__Metschnikowia(100);g__Metschnikowia_unclassified(100); | Metschnikowia | 13 | Saprotroph | \|Undefined Saprotroph\| | Yeast | NULL | Probable | As nectar-tap_saprotroph (Põlme et al. 2020); Decay sugar-rich substrates (Põlme et al. 2020) | Põlme S, et al. 2020. Fungal Diversity 105:1-16 ((https://doi.org/10.1007/s13225-020-00466-2)) |
| Otu51822 | 1 | k__Fungi(100);p__Ascomycota(100);c__Saccharomycetes(100);o__Saccharomycetales(100);f__Metschnikowiaceae(100);g__Metschnikowia(100);s__Metschnikowia_pulcherrima(100); | Metschnikowia | 13 | Saprotroph | \|Undefined Saprotroph\| | Yeast | NULL | Probable | As nectar-tap_saprotroph (Põlme et al. 2020); Decay sugar-rich substrates (Põlme et al. 2020) | Põlme S, et al. 2020. Fungal Diversity 105:1-16 ((https://doi.org/10.1007/s13225-020-00466-2)) |
| Otu51823 | 1 | k__Fungi(100);p__Ascomycota(100);c__Saccharomycetes(100);o__Saccharomycetales(100);f__Metschnikowiaceae(100);g__Metschnikowia(100);s__Metschnikowia_pulcherrima(100); | Metschnikowia | 13 | Saprotroph | \|Undefined Saprotroph\| | Yeast | NULL | Probable | As nectar-tap_saprotroph (Põlme et al. 2020); Decay sugar-rich substrates (Põlme et al. 2020) | Põlme S, et al. 2020. Fungal Diversity 105:1-16 ((https://doi.org/10.1007/s13225-020-00466-2)) |
| Otu51824 | 1 | k__Fungi(100);p__Ascomycota(100);c__Saccharomycetes(100);o__Saccharomycetales(100);f__Metschnikowiaceae(100);g__Metschnikowia(100);s__Metschnikowia_pulcherrima(100); | Metschnikowia | 13 | Saprotroph | \|Undefined Saprotroph\| | Yeast | NULL | Probable | As nectar-tap_saprotroph (Põlme et al. 2020); Decay sugar-rich substrates (Põlme et al. 2020) | Põlme S, et al. 2020. Fungal Diversity 105:1-16 ((https://doi.org/10.1007/s13225-020-00466-2)) |
| Otu51826 | 1 | k__Fungi(100);p__Ascomycota(100);c__Saccharomycetes(100);o__Saccharomycetales(100);f__Metschnikowiaceae(100);g__Metschnikowia(100);g__Metschnikowia_unclassified(100); | Metschnikowia | 13 | Saprotroph | \|Undefined Saprotroph\| | Yeast | NULL | Probable | As nectar-tap_saprotroph (Põlme et al. 2020); Decay sugar-rich substrates (Põlme et al. 2020) | Põlme S, et al. 2020. Fungal Diversity 105:1-16 ((https://doi.org/10.1007/s13225-020-00466-2)) |
| Otu51827 | 1 | k__Fungi(100);p__Ascomycota(100);c__Saccharomycetes(100);o__Saccharomycetales(100);f__Metschnikowiaceae(100);g__Metschnikowia(100);g__Metschnikowia_unclassified(100); | Metschnikowia | 13 | Saprotroph | \|Undefined Saprotroph\| | Yeast | NULL | Probable | As nectar-tap_saprotroph (Põlme et al. 2020); Decay sugar-rich substrates (Põlme et al. 2020) | Põlme S, et al. 2020. Fungal Diversity 105:1-16 ((https://doi.org/10.1007/s13225-020-00466-2)) |
| Otu51830 | 1 | k__Fungi(100);p__Ascomycota(100);c__Saccharomycetes(100);o__Saccharomycetales(100);f__Metschnikowiaceae(100);g__Metschnikowia(100);s__Metschnikowia_pulcherrima(100); | Metschnikowia | 13 | Saprotroph | \|Undefined Saprotroph\| | Yeast | NULL | Probable | As nectar-tap_saprotroph (Põlme et al. 2020); Decay sugar-rich substrates (Põlme et al. 2020) | Põlme S, et al. 2020. Fungal Diversity 105:1-16 ((https://doi.org/10.1007/s13225-020-00466-2)) |
| Otu51832 | 1 | k__Fungi(100);p__Ascomycota(100);c__Saccharomycetes(100);o__Saccharomycetales(100);f__Metschnikowiaceae(100);g__Metschnikowia(100);g__Metschnikowia_unclassified(100); | Metschnikowia | 13 | Saprotroph | \|Undefined Saprotroph\| | Yeast | NULL | Probable | As nectar-tap_saprotroph (Põlme et al. 2020); Decay sugar-rich substrates (Põlme et al. 2020) | Põlme S, et al. 2020. Fungal Diversity 105:1-16 ((https://doi.org/10.1007/s13225-020-00466-2)) |
| Otu51833 | 1 | k__Fungi(100);p__Ascomycota(100);c__Saccharomycetes(100);o__Saccharomycetales(100);f__Metschnikowiaceae(100);g__Metschnikowia(100);g__Metschnikowia_unclassified(100); | Metschnikowia | 13 | Saprotroph | \|Undefined Saprotroph\| | Yeast | NULL | Probable | As nectar-tap_saprotroph (Põlme et al. 2020); Decay sugar-rich substrates (Põlme et al. 2020) | Põlme S, et al. 2020. Fungal Diversity 105:1-16 ((https://doi.org/10.1007/s13225-020-00466-2)) |
| Otu51834 | 1 | k__Fungi(100);p__Ascomycota(100);c__Saccharomycetes(100);o__Saccharomycetales(100);f__Metschnikowiaceae(100);g__Metschnikowia(100);g__Metschnikowia_unclassified(100); | Metschnikowia | 13 | Saprotroph | \|Undefined Saprotroph\| | Yeast | NULL | Probable | As nectar-tap_saprotroph (Põlme et al. 2020); Decay sugar-rich substrates (Põlme et al. 2020) | Põlme S, et al. 2020. Fungal Diversity 105:1-16 ((https://doi.org/10.1007/s13225-020-00466-2)) |
| Otu51835 | 1 | k__Fungi(100);p__Ascomycota(100);c__Saccharomycetes(100);o__Saccharomycetales(100);f__Metschnikowiaceae(100);g__Metschnikowia(100);s__Metschnikowia_pulcherrima(100); | Metschnikowia | 13 | Saprotroph | \|Undefined Saprotroph\| | Yeast | NULL | Probable | As nectar-tap_saprotroph (Põlme et al. 2020); Decay sugar-rich substrates (Põlme et al. 2020) | Põlme S, et al. 2020. Fungal Diversity 105:1-16 ((https://doi.org/10.1007/s13225-020-00466-2)) |
| Otu51836 | 1 | k__Fungi(100);p__Ascomycota(100);c__Saccharomycetes(100);o__Saccharomycetales(100);f__Metschnikowiaceae(100);g__Metschnikowia(100);g__Metschnikowia_unclassified(100); | Metschnikowia | 13 | Saprotroph | \|Undefined Saprotroph\| | Yeast | NULL | Probable | As nectar-tap_saprotroph (Põlme et al. 2020); Decay sugar-rich substrates (Põlme et al. 2020) | Põlme S, et al. 2020. Fungal Diversity 105:1-16 ((https://doi.org/10.1007/s13225-020-00466-2)) |
| Otu51837 | 1 | k__Fungi(100);p__Ascomycota(100);c__Saccharomycetes(100);o__Saccharomycetales(100);f__Metschnikowiaceae(100);g__Metschnikowia(100);g__Metschnikowia_unclassified(100); | Metschnikowia | 13 | Saprotroph | \|Undefined Saprotroph\| | Yeast | NULL | Probable | As nectar-tap_saprotroph (Põlme et al. 2020); Decay sugar-rich substrates (Põlme et al. 2020) | Põlme S, et al. 2020. Fungal Diversity 105:1-16 ((https://doi.org/10.1007/s13225-020-00466-2)) |
| Otu51838 | 1 | k__Fungi(100);p__Ascomycota(100);c__Saccharomycetes(100);o__Saccharomycetales(100);f__Metschnikowiaceae(100);g__Metschnikowia(100);s__Metschnikowia_pulcherrima(100); | Metschnikowia | 13 | Saprotroph | \|Undefined Saprotroph\| | Yeast | NULL | Probable | As nectar-tap_saprotroph (Põlme et al. 2020); Decay sugar-rich substrates (Põlme et al. 2020) | Põlme S, et al. 2020. Fungal Diversity 105:1-16 ((https://doi.org/10.1007/s13225-020-00466-2)) |
| Otu51839 | 1 | k__Fungi(100);p__Ascomycota(100);c__Saccharomycetes(100);o__Saccharomycetales(100);f__Metschnikowiaceae(100);g__Metschnikowia(100);s__Metschnikowia_pulcherrima(100); | Metschnikowia | 13 | Saprotroph | \|Undefined Saprotroph\| | Yeast | NULL | Probable | As nectar-tap_saprotroph (Põlme et al. 2020); Decay sugar-rich substrates (Põlme et al. 2020) | Põlme S, et al. 2020. Fungal Diversity 105:1-16 ((https://doi.org/10.1007/s13225-020-00466-2)) |
| Otu51843 | 1 | k__Fungi(100);p__Ascomycota(100);c__Saccharomycetes(100);o__Saccharomycetales(100);f__Metschnikowiaceae(100);g__Metschnikowia(100);g__Metschnikowia_unclassified(100); | Metschnikowia | 13 | Saprotroph | \|Undefined Saprotroph\| | Yeast | NULL | Probable | As nectar-tap_saprotroph (Põlme et al. 2020); Decay sugar-rich substrates (Põlme et al. 2020) | Põlme S, et al. 2020. Fungal Diversity 105:1-16 ((https://doi.org/10.1007/s13225-020-00466-2)) |
| Otu51844 | 1 | k__Fungi(100);p__Ascomycota(100);c__Saccharomycetes(100);o__Saccharomycetales(100);f__Metschnikowiaceae(100);g__Metschnikowia(100);s__Metschnikowia_pulcherrima(100); | Metschnikowia | 13 | Saprotroph | \|Undefined Saprotroph\| | Yeast | NULL | Probable | As nectar-tap_saprotroph (Põlme et al. 2020); Decay sugar-rich substrates (Põlme et al. 2020) | Põlme S, et al. 2020. Fungal Diversity 105:1-16 ((https://doi.org/10.1007/s13225-020-00466-2)) |
| Otu51845 | 1 | k__Fungi(100);p__Ascomycota(100);c__Saccharomycetes(100);o__Saccharomycetales(100);f__Metschnikowiaceae(100);g__Metschnikowia(100);s__Metschnikowia_pulcherrima(100); | Metschnikowia | 13 | Saprotroph | \|Undefined Saprotroph\| | Yeast | NULL | Probable | As nectar-tap_saprotroph (Põlme et al. 2020); Decay sugar-rich substrates (Põlme et al. 2020) | Põlme S, et al. 2020. Fungal Diversity 105:1-16 ((https://doi.org/10.1007/s13225-020-00466-2)) |
| Otu51847 | 1 | k__Fungi(100);p__Ascomycota(100);c__Saccharomycetes(100);o__Saccharomycetales(100);f__Metschnikowiaceae(100);g__Metschnikowia(100);s__Metschnikowia_pulcherrima(100); | Metschnikowia | 13 | Saprotroph | \|Undefined Saprotroph\| | Yeast | NULL | Probable | As nectar-tap_saprotroph (Põlme et al. 2020); Decay sugar-rich substrates (Põlme et al. 2020) | Põlme S, et al. 2020. Fungal Diversity 105:1-16 ((https://doi.org/10.1007/s13225-020-00466-2)) |
| Otu51848 | 1 | k__Fungi(100);p__Ascomycota(100);c__Saccharomycetes(100);o__Saccharomycetales(100);f__Metschnikowiaceae(100);g__Metschnikowia(100);s__Metschnikowia_pulcherrima(100); | Metschnikowia | 13 | Saprotroph | \|Undefined Saprotroph\| | Yeast | NULL | Probable | As nectar-tap_saprotroph (Põlme et al. 2020); Decay sugar-rich substrates (Põlme et al. 2020) | Põlme S, et al. 2020. Fungal Diversity 105:1-16 ((https://doi.org/10.1007/s13225-020-00466-2)) |
| Otu51849 | 1 | k__Fungi(100);p__Ascomycota(100);c__Saccharomycetes(100);o__Saccharomycetales(100);f__Metschnikowiaceae(100);g__Metschnikowia(100);s__Metschnikowia_pulcherrima(100); | Metschnikowia | 13 | Saprotroph | \|Undefined Saprotroph\| | Yeast | NULL | Probable | As nectar-tap_saprotroph (Põlme et al. 2020); Decay sugar-rich substrates (Põlme et al. 2020) | Põlme S, et al. 2020. Fungal Diversity 105:1-16 ((https://doi.org/10.1007/s13225-020-00466-2)) |
| Otu51850 | 1 | k__Fungi(100);p__Ascomycota(100);c__Saccharomycetes(100);o__Saccharomycetales(100);f__Saccharomycetales_family_Incertae_sedis(100);g__Candida(100);g__Candida_unclassified(100); | Candida | 13 | Pathotroph-Saprotroph-Symbiotroph | Animal Pathogen-Endophyte-Endosymbiont-Epiphyte-\|Undefined Saprotroph\| | Dimorphic Yeast-Yeast | NULL | Probable | As nectar-tap_saprotroph (Põlme et al. 2020); Decay sugar-rich substrates (Põlme et al. 2020); This asexual genus is polyphyletic and contain species that function very differently depend on what they are. Caution should be taken when interpreting guilds for Candida species. | Manolakaki D, et al. 2010. Virulence. 1: 367-375 ((https://doi.org/10.4161/viru.1.5.12796)); Mekbib SB, et al. 2011. Tropical Plant Pathology 36:233-240 ((https://doi.org/10.1590/S1982-56762011000400004)); Põlme S, et al. 2020. Fungal Diversity 105:1-16 ((https://doi.org/10.1007/s13225-020-00466-2)) |
| Otu51851 | 1 | k__Fungi(100);p__Ascomycota(100);c__Saccharomycetes(100);o__Saccharomycetales(100);f__Saccharomycetales_family_Incertae_sedis(100);g__Candida(100);g__Candida_unclassified(100); | Candida | 13 | Pathotroph-Saprotroph-Symbiotroph | Animal Pathogen-Endophyte-Endosymbiont-Epiphyte-\|Undefined Saprotroph\| | Dimorphic Yeast-Yeast | NULL | Probable | As nectar-tap_saprotroph (Põlme et al. 2020); Decay sugar-rich substrates (Põlme et al. 2020); This asexual genus is polyphyletic and contain species that function very differently depend on what they are. Caution should be taken when interpreting guilds for Candida species. | Manolakaki D, et al. 2010. Virulence. 1: 367-375 ((https://doi.org/10.4161/viru.1.5.12796)); Mekbib SB, et al. 2011. Tropical Plant Pathology 36:233-240 ((https://doi.org/10.1590/S1982-56762011000400004)); Põlme S, et al. 2020. Fungal Diversity 105:1-16 ((https://doi.org/10.1007/s13225-020-00466-2)) |
| Otu51852 | 1 | k__Fungi(100);p__Ascomycota(100);c__Saccharomycetes(100);o__Saccharomycetales(100);f__Metschnikowiaceae(100);g__Metschnikowia(100);s__Metschnikowia_pulcherrima(100); | Metschnikowia | 13 | Saprotroph | \|Undefined Saprotroph\| | Yeast | NULL | Probable | As nectar-tap_saprotroph (Põlme et al. 2020); Decay sugar-rich substrates (Põlme et al. 2020) | Põlme S, et al. 2020. Fungal Diversity 105:1-16 ((https://doi.org/10.1007/s13225-020-00466-2)) |
| Otu51853 | 1 | k__Fungi(100);p__Ascomycota(100);c__Saccharomycetes(100);o__Saccharomycetales(100);f__Metschnikowiaceae(100);g__Metschnikowia(100);s__Metschnikowia_pulcherrima(100); | Metschnikowia | 13 | Saprotroph | \|Undefined Saprotroph\| | Yeast | NULL | Probable | As nectar-tap_saprotroph (Põlme et al. 2020); Decay sugar-rich substrates (Põlme et al. 2020) | Põlme S, et al. 2020. Fungal Diversity 105:1-16 ((https://doi.org/10.1007/s13225-020-00466-2)) |
| Otu51857 | 1 | k__Fungi(100);p__Ascomycota(100);c__Saccharomycetes(100);o__Saccharomycetales(100);f__Metschnikowiaceae(100);g__Metschnikowia(100);s__Metschnikowia_pulcherrima(100); | Metschnikowia | 13 | Saprotroph | \|Undefined Saprotroph\| | Yeast | NULL | Probable | As nectar-tap_saprotroph (Põlme et al. 2020); Decay sugar-rich substrates (Põlme et al. 2020) | Põlme S, et al. 2020. Fungal Diversity 105:1-16 ((https://doi.org/10.1007/s13225-020-00466-2)) |
| Otu51858 | 1 | k__Fungi(100);p__Ascomycota(100);c__Saccharomycetes(100);o__Saccharomycetales(100);f__Metschnikowiaceae(100);g__Metschnikowia(100);s__Metschnikowia_pulcherrima(100); | Metschnikowia | 13 | Saprotroph | \|Undefined Saprotroph\| | Yeast | NULL | Probable | As nectar-tap_saprotroph (Põlme et al. 2020); Decay sugar-rich substrates (Põlme et al. 2020) | Põlme S, et al. 2020. Fungal Diversity 105:1-16 ((https://doi.org/10.1007/s13225-020-00466-2)) |
| Otu51859 | 1 | k__Fungi(100);p__Ascomycota(100);c__Saccharomycetes(100);o__Saccharomycetales(100);f__Metschnikowiaceae(100);g__Metschnikowia(100);g__Metschnikowia_unclassified(100); | Metschnikowia | 13 | Saprotroph | \|Undefined Saprotroph\| | Yeast | NULL | Probable | As nectar-tap_saprotroph (Põlme et al. 2020); Decay sugar-rich substrates (Põlme et al. 2020) | Põlme S, et al. 2020. Fungal Diversity 105:1-16 ((https://doi.org/10.1007/s13225-020-00466-2)) |
| Otu51860 | 1 | k__Fungi(100);p__Ascomycota(100);c__Saccharomycetes(100);o__Saccharomycetales(100);f__Metschnikowiaceae(100);g__Metschnikowia(100);s__Metschnikowia_pulcherrima(100); | Metschnikowia | 13 | Saprotroph | \|Undefined Saprotroph\| | Yeast | NULL | Probable | As nectar-tap_saprotroph (Põlme et al. 2020); Decay sugar-rich substrates (Põlme et al. 2020) | Põlme S, et al. 2020. Fungal Diversity 105:1-16 ((https://doi.org/10.1007/s13225-020-00466-2)) |
| Otu51861 | 1 | k__Fungi(100);p__Ascomycota(100);c__Saccharomycetes(100);o__Saccharomycetales(100);f__Metschnikowiaceae(100);g__Metschnikowia(100);s__Metschnikowia_pulcherrima(100); | Metschnikowia | 13 | Saprotroph | \|Undefined Saprotroph\| | Yeast | NULL | Probable | As nectar-tap_saprotroph (Põlme et al. 2020); Decay sugar-rich substrates (Põlme et al. 2020) | Põlme S, et al. 2020. Fungal Diversity 105:1-16 ((https://doi.org/10.1007/s13225-020-00466-2)) |
| Otu51863 | 1 | k__Fungi(100);p__Ascomycota(100);c__Saccharomycetes(100);o__Saccharomycetales(100);f__Metschnikowiaceae(100);g__Metschnikowia(100);g__Metschnikowia_unclassified(100); | Metschnikowia | 13 | Saprotroph | \|Undefined Saprotroph\| | Yeast | NULL | Probable | As nectar-tap_saprotroph (Põlme et al. 2020); Decay sugar-rich substrates (Põlme et al. 2020) | Põlme S, et al. 2020. Fungal Diversity 105:1-16 ((https://doi.org/10.1007/s13225-020-00466-2)) |
| Otu51864 | 1 | k__Fungi(100);p__Ascomycota(100);c__Saccharomycetes(100);o__Saccharomycetales(100);f__Metschnikowiaceae(100);g__Metschnikowia(100);s__Metschnikowia_pulcherrima(100); | Metschnikowia | 13 | Saprotroph | \|Undefined Saprotroph\| | Yeast | NULL | Probable | As nectar-tap_saprotroph (Põlme et al. 2020); Decay sugar-rich substrates (Põlme et al. 2020) | Põlme S, et al. 2020. Fungal Diversity 105:1-16 ((https://doi.org/10.1007/s13225-020-00466-2)) |
| Otu51865 | 1 | k__Fungi(100);p__Ascomycota(100);c__Saccharomycetes(100);o__Saccharomycetales(100);f__Metschnikowiaceae(100);g__Metschnikowia(100);s__Metschnikowia_pulcherrima(100); | Metschnikowia | 13 | Saprotroph | \|Undefined Saprotroph\| | Yeast | NULL | Probable | As nectar-tap_saprotroph (Põlme et al. 2020); Decay sugar-rich substrates (Põlme et al. 2020) | Põlme S, et al. 2020. Fungal Diversity 105:1-16 ((https://doi.org/10.1007/s13225-020-00466-2)) |
| Otu51866 | 1 | k__Fungi(100);p__Ascomycota(100);c__Saccharomycetes(100);o__Saccharomycetales(100);f__Metschnikowiaceae(100);g__Metschnikowia(100);g__Metschnikowia_unclassified(100); | Metschnikowia | 13 | Saprotroph | \|Undefined Saprotroph\| | Yeast | NULL | Probable | As nectar-tap_saprotroph (Põlme et al. 2020); Decay sugar-rich substrates (Põlme et al. 2020) | Põlme S, et al. 2020. Fungal Diversity 105:1-16 ((https://doi.org/10.1007/s13225-020-00466-2)) |
| Otu51867 | 1 | k__Fungi(100);p__Ascomycota(100);c__Saccharomycetes(100);o__Saccharomycetales(100);f__Metschnikowiaceae(100);g__Metschnikowia(100);s__Metschnikowia_pulcherrima(100); | Metschnikowia | 13 | Saprotroph | \|Undefined Saprotroph\| | Yeast | NULL | Probable | As nectar-tap_saprotroph (Põlme et al. 2020); Decay sugar-rich substrates (Põlme et al. 2020) | Põlme S, et al. 2020. Fungal Diversity 105:1-16 ((https://doi.org/10.1007/s13225-020-00466-2)) |
| Otu51868 | 1 | k__Fungi(100);p__Ascomycota(100);c__Saccharomycetes(100);o__Saccharomycetales(100);f__Saccharomycetales_family_Incertae_sedis(100);g__Candida(100);g__Candida_unclassified(100); | Candida | 13 | Pathotroph-Saprotroph-Symbiotroph | Animal Pathogen-Endophyte-Endosymbiont-Epiphyte-\|Undefined Saprotroph\| | Dimorphic Yeast-Yeast | NULL | Probable | As nectar-tap_saprotroph (Põlme et al. 2020); Decay sugar-rich substrates (Põlme et al. 2020); This asexual genus is polyphyletic and contain species that function very differently depend on what they are. Caution should be taken when interpreting guilds for Candida species. | Manolakaki D, et al. 2010. Virulence. 1: 367-375 ((https://doi.org/10.4161/viru.1.5.12796)); Mekbib SB, et al. 2011. Tropical Plant Pathology 36:233-240 ((https://doi.org/10.1590/S1982-56762011000400004)); Põlme S, et al. 2020. Fungal Diversity 105:1-16 ((https://doi.org/10.1007/s13225-020-00466-2)) |
| Otu51870 | 1 | k__Fungi(100);p__Ascomycota(100);c__Saccharomycetes(100);o__Saccharomycetales(100);f__Metschnikowiaceae(100);g__Metschnikowia(100);g__Metschnikowia_unclassified(100); | Metschnikowia | 13 | Saprotroph | \|Undefined Saprotroph\| | Yeast | NULL | Probable | As nectar-tap_saprotroph (Põlme et al. 2020); Decay sugar-rich substrates (Põlme et al. 2020) | Põlme S, et al. 2020. Fungal Diversity 105:1-16 ((https://doi.org/10.1007/s13225-020-00466-2)) |
| Otu51871 | 1 | k__Fungi(100);p__Ascomycota(100);c__Saccharomycetes(100);o__Saccharomycetales(100);f__Metschnikowiaceae(100);g__Metschnikowia(100);g__Metschnikowia_unclassified(100); | Metschnikowia | 13 | Saprotroph | \|Undefined Saprotroph\| | Yeast | NULL | Probable | As nectar-tap_saprotroph (Põlme et al. 2020); Decay sugar-rich substrates (Põlme et al. 2020) | Põlme S, et al. 2020. Fungal Diversity 105:1-16 ((https://doi.org/10.1007/s13225-020-00466-2)) |
| Otu51872 | 1 | k__Fungi(100);p__Ascomycota(100);c__Saccharomycetes(100);o__Saccharomycetales(100);f__Metschnikowiaceae(100);g__Metschnikowia(100);s__Metschnikowia_pulcherrima(100); | Metschnikowia | 13 | Saprotroph | \|Undefined Saprotroph\| | Yeast | NULL | Probable | As nectar-tap_saprotroph (Põlme et al. 2020); Decay sugar-rich substrates (Põlme et al. 2020) | Põlme S, et al. 2020. Fungal Diversity 105:1-16 ((https://doi.org/10.1007/s13225-020-00466-2)) |
| Otu51873 | 1 | k__Fungi(100);p__Ascomycota(100);c__Saccharomycetes(100);o__Saccharomycetales(100);f__Metschnikowiaceae(100);g__Metschnikowia(100);s__Metschnikowia_pulcherrima(100); | Metschnikowia | 13 | Saprotroph | \|Undefined Saprotroph\| | Yeast | NULL | Probable | As nectar-tap_saprotroph (Põlme et al. 2020); Decay sugar-rich substrates (Põlme et al. 2020) | Põlme S, et al. 2020. Fungal Diversity 105:1-16 ((https://doi.org/10.1007/s13225-020-00466-2)) |
| Otu51874 | 1 | k__Fungi(100);p__Ascomycota(100);c__Saccharomycetes(100);o__Saccharomycetales(100);f__Metschnikowiaceae(100);g__Metschnikowia(100);s__Metschnikowia_pulcherrima(100); | Metschnikowia | 13 | Saprotroph | \|Undefined Saprotroph\| | Yeast | NULL | Probable | As nectar-tap_saprotroph (Põlme et al. 2020); Decay sugar-rich substrates (Põlme et al. 2020) | Põlme S, et al. 2020. Fungal Diversity 105:1-16 ((https://doi.org/10.1007/s13225-020-00466-2)) |
| Otu51876 | 1 | k__Fungi(100);p__Ascomycota(100);c__Saccharomycetes(100);o__Saccharomycetales(100);f__Saccharomycetales_family_Incertae_sedis(100);g__Candida(100);g__Candida_unclassified(100); | Candida | 13 | Pathotroph-Saprotroph-Symbiotroph | Animal Pathogen-Endophyte-Endosymbiont-Epiphyte-\|Undefined Saprotroph\| | Dimorphic Yeast-Yeast | NULL | Probable | As nectar-tap_saprotroph (Põlme et al. 2020); Decay sugar-rich substrates (Põlme et al. 2020); This asexual genus is polyphyletic and contain species that function very differently depend on what they are. Caution should be taken when interpreting guilds for Candida species. | Manolakaki D, et al. 2010. Virulence. 1: 367-375 ((https://doi.org/10.4161/viru.1.5.12796)); Mekbib SB, et al. 2011. Tropical Plant Pathology 36:233-240 ((https://doi.org/10.1590/S1982-56762011000400004)); Põlme S, et al. 2020. Fungal Diversity 105:1-16 ((https://doi.org/10.1007/s13225-020-00466-2)) |
| Otu51877 | 1 | k__Fungi(100);p__Ascomycota(100);c__Saccharomycetes(100);o__Saccharomycetales(100);f__Metschnikowiaceae(100);g__Metschnikowia(100);g__Metschnikowia_unclassified(100); | Metschnikowia | 13 | Saprotroph | \|Undefined Saprotroph\| | Yeast | NULL | Probable | As nectar-tap_saprotroph (Põlme et al. 2020); Decay sugar-rich substrates (Põlme et al. 2020) | Põlme S, et al. 2020. Fungal Diversity 105:1-16 ((https://doi.org/10.1007/s13225-020-00466-2)) |
| Otu51878 | 1 | k__Fungi(100);p__Ascomycota(100);c__Saccharomycetes(100);o__Saccharomycetales(100);f__Metschnikowiaceae(100);g__Metschnikowia(100);g__Metschnikowia_unclassified(100); | Metschnikowia | 13 | Saprotroph | \|Undefined Saprotroph\| | Yeast | NULL | Probable | As nectar-tap_saprotroph (Põlme et al. 2020); Decay sugar-rich substrates (Põlme et al. 2020) | Põlme S, et al. 2020. Fungal Diversity 105:1-16 ((https://doi.org/10.1007/s13225-020-00466-2)) |
| Otu51879 | 1 | k__Fungi(100);p__Ascomycota(100);c__Saccharomycetes(100);o__Saccharomycetales(100);f__Metschnikowiaceae(100);g__Metschnikowia(100);s__Metschnikowia_pulcherrima(100); | Metschnikowia | 13 | Saprotroph | \|Undefined Saprotroph\| | Yeast | NULL | Probable | As nectar-tap_saprotroph (Põlme et al. 2020); Decay sugar-rich substrates (Põlme et al. 2020) | Põlme S, et al. 2020. Fungal Diversity 105:1-16 ((https://doi.org/10.1007/s13225-020-00466-2)) |
| Otu51880 | 1 | k__Fungi(100);p__Ascomycota(100);c__Saccharomycetes(100);o__Saccharomycetales(100);f__Metschnikowiaceae(100);g__Metschnikowia(100);s__Metschnikowia_pulcherrima(100); | Metschnikowia | 13 | Saprotroph | \|Undefined Saprotroph\| | Yeast | NULL | Probable | As nectar-tap_saprotroph (Põlme et al. 2020); Decay sugar-rich substrates (Põlme et al. 2020) | Põlme S, et al. 2020. Fungal Diversity 105:1-16 ((https://doi.org/10.1007/s13225-020-00466-2)) |
| Otu51881 | 1 | k__Fungi(100);p__Ascomycota(100);c__Saccharomycetes(100);o__Saccharomycetales(100);f__Metschnikowiaceae(100);g__Metschnikowia(100);s__Metschnikowia_pulcherrima(100); | Metschnikowia | 13 | Saprotroph | \|Undefined Saprotroph\| | Yeast | NULL | Probable | As nectar-tap_saprotroph (Põlme et al. 2020); Decay sugar-rich substrates (Põlme et al. 2020) | Põlme S, et al. 2020. Fungal Diversity 105:1-16 ((https://doi.org/10.1007/s13225-020-00466-2)) |
| Otu51882 | 1 | k__Fungi(100);p__Ascomycota(100);c__Saccharomycetes(100);o__Saccharomycetales(100);f__Metschnikowiaceae(100);g__Metschnikowia(100);s__Metschnikowia_pulcherrima(100); | Metschnikowia | 13 | Saprotroph | \|Undefined Saprotroph\| | Yeast | NULL | Probable | As nectar-tap_saprotroph (Põlme et al. 2020); Decay sugar-rich substrates (Põlme et al. 2020) | Põlme S, et al. 2020. Fungal Diversity 105:1-16 ((https://doi.org/10.1007/s13225-020-00466-2)) |
| Otu51884 | 1 | k__Fungi(100);p__Ascomycota(100);c__Saccharomycetes(100);o__Saccharomycetales(100);f__Metschnikowiaceae(100);g__Metschnikowia(100);s__Metschnikowia_pulcherrima(100); | Metschnikowia | 13 | Saprotroph | \|Undefined Saprotroph\| | Yeast | NULL | Probable | As nectar-tap_saprotroph (Põlme et al. 2020); Decay sugar-rich substrates (Põlme et al. 2020) | Põlme S, et al. 2020. Fungal Diversity 105:1-16 ((https://doi.org/10.1007/s13225-020-00466-2)) |
| Otu51885 | 1 | k__Fungi(100);p__Ascomycota(100);c__Saccharomycetes(100);o__Saccharomycetales(100);f__Metschnikowiaceae(100);g__Metschnikowia(100);g__Metschnikowia_unclassified(100); | Metschnikowia | 13 | Saprotroph | \|Undefined Saprotroph\| | Yeast | NULL | Probable | As nectar-tap_saprotroph (Põlme et al. 2020); Decay sugar-rich substrates (Põlme et al. 2020) | Põlme S, et al. 2020. Fungal Diversity 105:1-16 ((https://doi.org/10.1007/s13225-020-00466-2)) |
| Otu51886 | 1 | k__Fungi(100);p__Ascomycota(100);c__Saccharomycetes(100);o__Saccharomycetales(100);f__Metschnikowiaceae(100);g__Metschnikowia(100);s__Metschnikowia_pulcherrima(100); | Metschnikowia | 13 | Saprotroph | \|Undefined Saprotroph\| | Yeast | NULL | Probable | As nectar-tap_saprotroph (Põlme et al. 2020); Decay sugar-rich substrates (Põlme et al. 2020) | Põlme S, et al. 2020. Fungal Diversity 105:1-16 ((https://doi.org/10.1007/s13225-020-00466-2)) |
| Otu51887 | 1 | k__Fungi(100);p__Ascomycota(100);c__Saccharomycetes(100);o__Saccharomycetales(100);f__Metschnikowiaceae(100);g__Metschnikowia(100);s__Metschnikowia_pulcherrima(100); | Metschnikowia | 13 | Saprotroph | \|Undefined Saprotroph\| | Yeast | NULL | Probable | As nectar-tap_saprotroph (Põlme et al. 2020); Decay sugar-rich substrates (Põlme et al. 2020) | Põlme S, et al. 2020. Fungal Diversity 105:1-16 ((https://doi.org/10.1007/s13225-020-00466-2)) |
| Otu51891 | 1 | k__Fungi(100);p__Ascomycota(100);c__Saccharomycetes(100);o__Saccharomycetales(100);f__Metschnikowiaceae(100);g__Metschnikowia(100);s__Metschnikowia_pulcherrima(100); | Metschnikowia | 13 | Saprotroph | \|Undefined Saprotroph\| | Yeast | NULL | Probable | As nectar-tap_saprotroph (Põlme et al. 2020); Decay sugar-rich substrates (Põlme et al. 2020) | Põlme S, et al. 2020. Fungal Diversity 105:1-16 ((https://doi.org/10.1007/s13225-020-00466-2)) |
| Otu51892 | 1 | k__Fungi(100);p__Ascomycota(100);c__Saccharomycetes(100);o__Saccharomycetales(100);f__Metschnikowiaceae(100);g__Metschnikowia(100);s__Metschnikowia_pulcherrima(100); | Metschnikowia | 13 | Saprotroph | \|Undefined Saprotroph\| | Yeast | NULL | Probable | As nectar-tap_saprotroph (Põlme et al. 2020); Decay sugar-rich substrates (Põlme et al. 2020) | Põlme S, et al. 2020. Fungal Diversity 105:1-16 ((https://doi.org/10.1007/s13225-020-00466-2)) |
| Otu51893 | 1 | k__Fungi(100);p__Ascomycota(100);c__Saccharomycetes(100);o__Saccharomycetales(100);f__Metschnikowiaceae(100);g__Metschnikowia(100);s__Metschnikowia_pulcherrima(100); | Metschnikowia | 13 | Saprotroph | \|Undefined Saprotroph\| | Yeast | NULL | Probable | As nectar-tap_saprotroph (Põlme et al. 2020); Decay sugar-rich substrates (Põlme et al. 2020) | Põlme S, et al. 2020. Fungal Diversity 105:1-16 ((https://doi.org/10.1007/s13225-020-00466-2)) |
| Otu51894 | 1 | k__Fungi(100);p__Ascomycota(100);c__Saccharomycetes(100);o__Saccharomycetales(100);f__Metschnikowiaceae(100);g__Metschnikowia(100);g__Metschnikowia_unclassified(100); | Metschnikowia | 13 | Saprotroph | \|Undefined Saprotroph\| | Yeast | NULL | Probable | As nectar-tap_saprotroph (Põlme et al. 2020); Decay sugar-rich substrates (Põlme et al. 2020) | Põlme S, et al. 2020. Fungal Diversity 105:1-16 ((https://doi.org/10.1007/s13225-020-00466-2)) |
| Otu51896 | 1 | k__Fungi(100);p__Ascomycota(100);c__Saccharomycetes(100);o__Saccharomycetales(100);f__Metschnikowiaceae(100);g__Metschnikowia(100);s__Metschnikowia_pulcherrima(100); | Metschnikowia | 13 | Saprotroph | \|Undefined Saprotroph\| | Yeast | NULL | Probable | As nectar-tap_saprotroph (Põlme et al. 2020); Decay sugar-rich substrates (Põlme et al. 2020) | Põlme S, et al. 2020. Fungal Diversity 105:1-16 ((https://doi.org/10.1007/s13225-020-00466-2)) |
| Otu51898 | 1 | k__Fungi(100);p__Ascomycota(100);c__Saccharomycetes(100);o__Saccharomycetales(100);f__Metschnikowiaceae(100);g__Metschnikowia(100);s__Metschnikowia_pulcherrima(100); | Metschnikowia | 13 | Saprotroph | \|Undefined Saprotroph\| | Yeast | NULL | Probable | As nectar-tap_saprotroph (Põlme et al. 2020); Decay sugar-rich substrates (Põlme et al. 2020) | Põlme S, et al. 2020. Fungal Diversity 105:1-16 ((https://doi.org/10.1007/s13225-020-00466-2)) |
| Otu51899 | 1 | k__Fungi(100);p__Ascomycota(100);c__Saccharomycetes(100);o__Saccharomycetales(100);f__Metschnikowiaceae(100);g__Metschnikowia(100);s__Metschnikowia_pulcherrima(100); | Metschnikowia | 13 | Saprotroph | \|Undefined Saprotroph\| | Yeast | NULL | Probable | As nectar-tap_saprotroph (Põlme et al. 2020); Decay sugar-rich substrates (Põlme et al. 2020) | Põlme S, et al. 2020. Fungal Diversity 105:1-16 ((https://doi.org/10.1007/s13225-020-00466-2)) |
| Otu51901 | 1 | k__Fungi(100);p__Ascomycota(100);c__Saccharomycetes(100);o__Saccharomycetales(100);f__Saccharomycetales_family_Incertae_sedis(100);g__Candida(100);g__Candida_unclassified(100); | Candida | 13 | Pathotroph-Saprotroph-Symbiotroph | Animal Pathogen-Endophyte-Endosymbiont-Epiphyte-\|Undefined Saprotroph\| | Dimorphic Yeast-Yeast | NULL | Probable | As nectar-tap_saprotroph (Põlme et al. 2020); Decay sugar-rich substrates (Põlme et al. 2020); This asexual genus is polyphyletic and contain species that function very differently depend on what they are. Caution should be taken when interpreting guilds for Candida species. | Manolakaki D, et al. 2010. Virulence. 1: 367-375 ((https://doi.org/10.4161/viru.1.5.12796)); Mekbib SB, et al. 2011. Tropical Plant Pathology 36:233-240 ((https://doi.org/10.1590/S1982-56762011000400004)); Põlme S, et al. 2020. Fungal Diversity 105:1-16 ((https://doi.org/10.1007/s13225-020-00466-2)) |
| Otu51902 | 1 | k__Fungi(100);p__Ascomycota(100);c__Saccharomycetes(100);o__Saccharomycetales(100);f__Metschnikowiaceae(100);g__Metschnikowia(100);s__Metschnikowia_pulcherrima(100); | Metschnikowia | 13 | Saprotroph | \|Undefined Saprotroph\| | Yeast | NULL | Probable | As nectar-tap_saprotroph (Põlme et al. 2020); Decay sugar-rich substrates (Põlme et al. 2020) | Põlme S, et al. 2020. Fungal Diversity 105:1-16 ((https://doi.org/10.1007/s13225-020-00466-2)) |
| Otu51903 | 1 | k__Fungi(100);p__Ascomycota(100);c__Saccharomycetes(100);o__Saccharomycetales(100);f__Metschnikowiaceae(100);g__Metschnikowia(100);s__Metschnikowia_pulcherrima(100); | Metschnikowia | 13 | Saprotroph | \|Undefined Saprotroph\| | Yeast | NULL | Probable | As nectar-tap_saprotroph (Põlme et al. 2020); Decay sugar-rich substrates (Põlme et al. 2020) | Põlme S, et al. 2020. Fungal Diversity 105:1-16 ((https://doi.org/10.1007/s13225-020-00466-2)) |
| Otu51904 | 1 | k__Fungi(100);p__Ascomycota(100);c__Saccharomycetes(100);o__Saccharomycetales(100);f__Metschnikowiaceae(100);g__Metschnikowia(100);s__Metschnikowia_pulcherrima(100); | Metschnikowia | 13 | Saprotroph | \|Undefined Saprotroph\| | Yeast | NULL | Probable | As nectar-tap_saprotroph (Põlme et al. 2020); Decay sugar-rich substrates (Põlme et al. 2020) | Põlme S, et al. 2020. Fungal Diversity 105:1-16 ((https://doi.org/10.1007/s13225-020-00466-2)) |
| Otu51906 | 1 | k__Fungi(100);p__Ascomycota(100);c__Saccharomycetes(100);o__Saccharomycetales(100);f__Saccharomycetales_family_Incertae_sedis(100);g__Candida(100);g__Candida_unclassified(100); | Candida | 13 | Pathotroph-Saprotroph-Symbiotroph | Animal Pathogen-Endophyte-Endosymbiont-Epiphyte-\|Undefined Saprotroph\| | Dimorphic Yeast-Yeast | NULL | Probable | As nectar-tap_saprotroph (Põlme et al. 2020); Decay sugar-rich substrates (Põlme et al. 2020); This asexual genus is polyphyletic and contain species that function very differently depend on what they are. Caution should be taken when interpreting guilds for Candida species. | Manolakaki D, et al. 2010. Virulence. 1: 367-375 ((https://doi.org/10.4161/viru.1.5.12796)); Mekbib SB, et al. 2011. Tropical Plant Pathology 36:233-240 ((https://doi.org/10.1590/S1982-56762011000400004)); Põlme S, et al. 2020. Fungal Diversity 105:1-16 ((https://doi.org/10.1007/s13225-020-00466-2)) |
| Otu51907 | 1 | k__Fungi(100);p__Ascomycota(100);c__Saccharomycetes(100);o__Saccharomycetales(100);f__Metschnikowiaceae(100);g__Metschnikowia(100);s__Metschnikowia_pulcherrima(100); | Metschnikowia | 13 | Saprotroph | \|Undefined Saprotroph\| | Yeast | NULL | Probable | As nectar-tap_saprotroph (Põlme et al. 2020); Decay sugar-rich substrates (Põlme et al. 2020) | Põlme S, et al. 2020. Fungal Diversity 105:1-16 ((https://doi.org/10.1007/s13225-020-00466-2)) |
| Otu51908 | 1 | k__Fungi(100);p__Ascomycota(100);c__Saccharomycetes(100);o__Saccharomycetales(100);f__Metschnikowiaceae(100);g__Metschnikowia(100);s__Metschnikowia_pulcherrima(100); | Metschnikowia | 13 | Saprotroph | \|Undefined Saprotroph\| | Yeast | NULL | Probable | As nectar-tap_saprotroph (Põlme et al. 2020); Decay sugar-rich substrates (Põlme et al. 2020) | Põlme S, et al. 2020. Fungal Diversity 105:1-16 ((https://doi.org/10.1007/s13225-020-00466-2)) |
| Otu51909 | 1 | k__Fungi(100);p__Ascomycota(100);c__Saccharomycetes(100);o__Saccharomycetales(100);f__Metschnikowiaceae(100);g__Metschnikowia(100);g__Metschnikowia_unclassified(100); | Metschnikowia | 13 | Saprotroph | \|Undefined Saprotroph\| | Yeast | NULL | Probable | As nectar-tap_saprotroph (Põlme et al. 2020); Decay sugar-rich substrates (Põlme et al. 2020) | Põlme S, et al. 2020. Fungal Diversity 105:1-16 ((https://doi.org/10.1007/s13225-020-00466-2)) |
| Otu51911 | 1 | k__Fungi(100);p__Ascomycota(100);c__Saccharomycetes(100);o__Saccharomycetales(100);f__Metschnikowiaceae(100);g__Metschnikowia(100);g__Metschnikowia_unclassified(100); | Metschnikowia | 13 | Saprotroph | \|Undefined Saprotroph\| | Yeast | NULL | Probable | As nectar-tap_saprotroph (Põlme et al. 2020); Decay sugar-rich substrates (Põlme et al. 2020) | Põlme S, et al. 2020. Fungal Diversity 105:1-16 ((https://doi.org/10.1007/s13225-020-00466-2)) |
| Otu51912 | 1 | k__Fungi(100);p__Ascomycota(100);c__Saccharomycetes(100);o__Saccharomycetales(100);f__Metschnikowiaceae(100);g__Metschnikowia(100);g__Metschnikowia_unclassified(100); | Metschnikowia | 13 | Saprotroph | \|Undefined Saprotroph\| | Yeast | NULL | Probable | As nectar-tap_saprotroph (Põlme et al. 2020); Decay sugar-rich substrates (Põlme et al. 2020) | Põlme S, et al. 2020. Fungal Diversity 105:1-16 ((https://doi.org/10.1007/s13225-020-00466-2)) |
| Otu51913 | 1 | k__Fungi(100);p__Ascomycota(100);c__Saccharomycetes(100);o__Saccharomycetales(100);f__Metschnikowiaceae(100);g__Metschnikowia(100);s__Metschnikowia_pulcherrima(100); | Metschnikowia | 13 | Saprotroph | \|Undefined Saprotroph\| | Yeast | NULL | Probable | As nectar-tap_saprotroph (Põlme et al. 2020); Decay sugar-rich substrates (Põlme et al. 2020) | Põlme S, et al. 2020. Fungal Diversity 105:1-16 ((https://doi.org/10.1007/s13225-020-00466-2)) |
| Otu51914 | 1 | k__Fungi(100);p__Ascomycota(100);c__Saccharomycetes(100);o__Saccharomycetales(100);f__Metschnikowiaceae(100);g__Metschnikowia(100);s__Metschnikowia_pulcherrima(100); | Metschnikowia | 13 | Saprotroph | \|Undefined Saprotroph\| | Yeast | NULL | Probable | As nectar-tap_saprotroph (Põlme et al. 2020); Decay sugar-rich substrates (Põlme et al. 2020) | Põlme S, et al. 2020. Fungal Diversity 105:1-16 ((https://doi.org/10.1007/s13225-020-00466-2)) |
| Otu51915 | 1 | k__Fungi(100);p__Ascomycota(100);c__Saccharomycetes(100);o__Saccharomycetales(100);f__Metschnikowiaceae(100);g__Metschnikowia(100);s__Metschnikowia_pulcherrima(100); | Metschnikowia | 13 | Saprotroph | \|Undefined Saprotroph\| | Yeast | NULL | Probable | As nectar-tap_saprotroph (Põlme et al. 2020); Decay sugar-rich substrates (Põlme et al. 2020) | Põlme S, et al. 2020. Fungal Diversity 105:1-16 ((https://doi.org/10.1007/s13225-020-00466-2)) |
| Otu51916 | 1 | k__Fungi(100);p__Ascomycota(100);c__Saccharomycetes(100);o__Saccharomycetales(100);f__Metschnikowiaceae(100);g__Metschnikowia(100);s__Metschnikowia_pulcherrima(100); | Metschnikowia | 13 | Saprotroph | \|Undefined Saprotroph\| | Yeast | NULL | Probable | As nectar-tap_saprotroph (Põlme et al. 2020); Decay sugar-rich substrates (Põlme et al. 2020) | Põlme S, et al. 2020. Fungal Diversity 105:1-16 ((https://doi.org/10.1007/s13225-020-00466-2)) |
| Otu51917 | 1 | k__Fungi(100);p__Ascomycota(100);c__Saccharomycetes(100);o__Saccharomycetales(100);f__Metschnikowiaceae(100);g__Metschnikowia(100);s__Metschnikowia_pulcherrima(100); | Metschnikowia | 13 | Saprotroph | \|Undefined Saprotroph\| | Yeast | NULL | Probable | As nectar-tap_saprotroph (Põlme et al. 2020); Decay sugar-rich substrates (Põlme et al. 2020) | Põlme S, et al. 2020. Fungal Diversity 105:1-16 ((https://doi.org/10.1007/s13225-020-00466-2)) |
| Otu51918 | 1 | k__Fungi(100);p__Ascomycota(100);c__Saccharomycetes(100);o__Saccharomycetales(100);f__Metschnikowiaceae(100);g__Metschnikowia(100);s__Metschnikowia_pulcherrima(100); | Metschnikowia | 13 | Saprotroph | \|Undefined Saprotroph\| | Yeast | NULL | Probable | As nectar-tap_saprotroph (Põlme et al. 2020); Decay sugar-rich substrates (Põlme et al. 2020) | Põlme S, et al. 2020. Fungal Diversity 105:1-16 ((https://doi.org/10.1007/s13225-020-00466-2)) |
| Otu51919 | 1 | k__Fungi(100);p__Ascomycota(100);c__Saccharomycetes(100);o__Saccharomycetales(100);f__Metschnikowiaceae(100);g__Metschnikowia(100);g__Metschnikowia_unclassified(100); | Metschnikowia | 13 | Saprotroph | \|Undefined Saprotroph\| | Yeast | NULL | Probable | As nectar-tap_saprotroph (Põlme et al. 2020); Decay sugar-rich substrates (Põlme et al. 2020) | Põlme S, et al. 2020. Fungal Diversity 105:1-16 ((https://doi.org/10.1007/s13225-020-00466-2)) |
| Otu51920 | 1 | k__Fungi(100);p__Ascomycota(100);c__Saccharomycetes(100);o__Saccharomycetales(100);f__Metschnikowiaceae(100);g__Metschnikowia(100);s__Metschnikowia_pulcherrima(100); | Metschnikowia | 13 | Saprotroph | \|Undefined Saprotroph\| | Yeast | NULL | Probable | As nectar-tap_saprotroph (Põlme et al. 2020); Decay sugar-rich substrates (Põlme et al. 2020) | Põlme S, et al. 2020. Fungal Diversity 105:1-16 ((https://doi.org/10.1007/s13225-020-00466-2)) |
| Otu51921 | 1 | k__Fungi(100);p__Ascomycota(100);c__Saccharomycetes(100);o__Saccharomycetales(100);f__Metschnikowiaceae(100);g__Metschnikowia(100);s__Metschnikowia_pulcherrima(100); | Metschnikowia | 13 | Saprotroph | \|Undefined Saprotroph\| | Yeast | NULL | Probable | As nectar-tap_saprotroph (Põlme et al. 2020); Decay sugar-rich substrates (Põlme et al. 2020) | Põlme S, et al. 2020. Fungal Diversity 105:1-16 ((https://doi.org/10.1007/s13225-020-00466-2)) |
| Otu51923 | 1 | k__Fungi(100);p__Ascomycota(100);c__Saccharomycetes(100);o__Saccharomycetales(100);f__Metschnikowiaceae(100);g__Metschnikowia(100);g__Metschnikowia_unclassified(100); | Metschnikowia | 13 | Saprotroph | \|Undefined Saprotroph\| | Yeast | NULL | Probable | As nectar-tap_saprotroph (Põlme et al. 2020); Decay sugar-rich substrates (Põlme et al. 2020) | Põlme S, et al. 2020. Fungal Diversity 105:1-16 ((https://doi.org/10.1007/s13225-020-00466-2)) |
| Otu51924 | 1 | k__Fungi(100);p__Ascomycota(100);c__Saccharomycetes(100);o__Saccharomycetales(100);f__Metschnikowiaceae(100);g__Metschnikowia(100);s__Metschnikowia_pulcherrima(100); | Metschnikowia | 13 | Saprotroph | \|Undefined Saprotroph\| | Yeast | NULL | Probable | As nectar-tap_saprotroph (Põlme et al. 2020); Decay sugar-rich substrates (Põlme et al. 2020) | Põlme S, et al. 2020. Fungal Diversity 105:1-16 ((https://doi.org/10.1007/s13225-020-00466-2)) |
| Otu51925 | 1 | k__Fungi(100);p__Ascomycota(100);c__Saccharomycetes(100);o__Saccharomycetales(100);f__Metschnikowiaceae(100);g__Metschnikowia(100);s__Metschnikowia_pulcherrima(100); | Metschnikowia | 13 | Saprotroph | \|Undefined Saprotroph\| | Yeast | NULL | Probable | As nectar-tap_saprotroph (Põlme et al. 2020); Decay sugar-rich substrates (Põlme et al. 2020) | Põlme S, et al. 2020. Fungal Diversity 105:1-16 ((https://doi.org/10.1007/s13225-020-00466-2)) |
| Otu51926 | 1 | k__Fungi(100);p__Ascomycota(100);c__Saccharomycetes(100);o__Saccharomycetales(100);f__Metschnikowiaceae(100);g__Metschnikowia(100);s__Metschnikowia_pulcherrima(100); | Metschnikowia | 13 | Saprotroph | \|Undefined Saprotroph\| | Yeast | NULL | Probable | As nectar-tap_saprotroph (Põlme et al. 2020); Decay sugar-rich substrates (Põlme et al. 2020) | Põlme S, et al. 2020. Fungal Diversity 105:1-16 ((https://doi.org/10.1007/s13225-020-00466-2)) |
| Otu51927 | 1 | k__Fungi(100);p__Ascomycota(100);c__Saccharomycetes(100);o__Saccharomycetales(100);f__Metschnikowiaceae(100);g__Metschnikowia(100);s__Metschnikowia_pulcherrima(100); | Metschnikowia | 13 | Saprotroph | \|Undefined Saprotroph\| | Yeast | NULL | Probable | As nectar-tap_saprotroph (Põlme et al. 2020); Decay sugar-rich substrates (Põlme et al. 2020) | Põlme S, et al. 2020. Fungal Diversity 105:1-16 ((https://doi.org/10.1007/s13225-020-00466-2)) |
| Otu51928 | 1 | k__Fungi(100);p__Ascomycota(100);c__Saccharomycetes(100);o__Saccharomycetales(100);f__Metschnikowiaceae(100);g__Metschnikowia(100);s__Metschnikowia_pulcherrima(100); | Metschnikowia | 13 | Saprotroph | \|Undefined Saprotroph\| | Yeast | NULL | Probable | As nectar-tap_saprotroph (Põlme et al. 2020); Decay sugar-rich substrates (Põlme et al. 2020) | Põlme S, et al. 2020. Fungal Diversity 105:1-16 ((https://doi.org/10.1007/s13225-020-00466-2)) |
| Otu51929 | 1 | k__Fungi(100);p__Ascomycota(100);c__Saccharomycetes(100);o__Saccharomycetales(100);f__Metschnikowiaceae(100);g__Metschnikowia(100);s__Metschnikowia_pulcherrima(100); | Metschnikowia | 13 | Saprotroph | \|Undefined Saprotroph\| | Yeast | NULL | Probable | As nectar-tap_saprotroph (Põlme et al. 2020); Decay sugar-rich substrates (Põlme et al. 2020) | Põlme S, et al. 2020. Fungal Diversity 105:1-16 ((https://doi.org/10.1007/s13225-020-00466-2)) |
| Otu51930 | 1 | k__Fungi(100);p__Ascomycota(100);c__Saccharomycetes(100);o__Saccharomycetales(100);f__Metschnikowiaceae(100);g__Metschnikowia(100);g__Metschnikowia_unclassified(100); | Metschnikowia | 13 | Saprotroph | \|Undefined Saprotroph\| | Yeast | NULL | Probable | As nectar-tap_saprotroph (Põlme et al. 2020); Decay sugar-rich substrates (Põlme et al. 2020) | Põlme S, et al. 2020. Fungal Diversity 105:1-16 ((https://doi.org/10.1007/s13225-020-00466-2)) |
| Otu51931 | 1 | k__Fungi(100);p__Ascomycota(100);c__Saccharomycetes(100);o__Saccharomycetales(100);f__Metschnikowiaceae(100);g__Metschnikowia(100);s__Metschnikowia_pulcherrima(100); | Metschnikowia | 13 | Saprotroph | \|Undefined Saprotroph\| | Yeast | NULL | Probable | As nectar-tap_saprotroph (Põlme et al. 2020); Decay sugar-rich substrates (Põlme et al. 2020) | Põlme S, et al. 2020. Fungal Diversity 105:1-16 ((https://doi.org/10.1007/s13225-020-00466-2)) |
| Otu51932 | 1 | k__Fungi(100);p__Ascomycota(100);c__Saccharomycetes(100);o__Saccharomycetales(100);f__Metschnikowiaceae(100);g__Metschnikowia(100);s__Metschnikowia_pulcherrima(100); | Metschnikowia | 13 | Saprotroph | \|Undefined Saprotroph\| | Yeast | NULL | Probable | As nectar-tap_saprotroph (Põlme et al. 2020); Decay sugar-rich substrates (Põlme et al. 2020) | Põlme S, et al. 2020. Fungal Diversity 105:1-16 ((https://doi.org/10.1007/s13225-020-00466-2)) |
| Otu51933 | 1 | k__Fungi(100);p__Ascomycota(100);c__Saccharomycetes(100);o__Saccharomycetales(100);f__Metschnikowiaceae(100);g__Metschnikowia(100);s__Metschnikowia_pulcherrima(100); | Metschnikowia | 13 | Saprotroph | \|Undefined Saprotroph\| | Yeast | NULL | Probable | As nectar-tap_saprotroph (Põlme et al. 2020); Decay sugar-rich substrates (Põlme et al. 2020) | Põlme S, et al. 2020. Fungal Diversity 105:1-16 ((https://doi.org/10.1007/s13225-020-00466-2)) |

| | | | | | | | | | | | |
|---|---|---|---|---|---|---|---|---|---|---|---|
| Otu51934 | 1 | k__Fungi(100);p__Ascomycota(100);c__Saccharomycetes(100);o__Saccharomycetales(100);f__Metschnikowiaceae(100);g__Metschnikowia(100);s__Metschnikowia_pulcherrima(100); | Metschnikowia | 13 | Saprotroph | \|Undefined Saprotroph\| | Yeast | NULL | Probable | As nectar-tap_saprotroph (Põlme et al. 2020); Decay sugar-rich substrates (Põlme et al. 2020) | Põlme S, et al. 2020. Fungal Diversity 105:1-16 ((https://doi.org/10.1007/s13225-020-00466-2)) |
| Otu51935 | 1 | k__Fungi(100);p__Ascomycota(100);c__Saccharomycetes(100);o__Saccharomycetales(100);f__Metschnikowiaceae(100);g__Metschnikowia(100);s__Metschnikowia_pulcherrima(100); | Metschnikowia | 13 | Saprotroph | \|Undefined Saprotroph\| | Yeast | NULL | Probable | As nectar-tap_saprotroph (Põlme et al. 2020); Decay sugar-rich substrates (Põlme et al. 2020) | Põlme S, et al. 2020. Fungal Diversity 105:1-16 ((https://doi.org/10.1007/s13225-020-00466-2)) |
| Otu51936 | 1 | k__Fungi(100);p__Ascomycota(100);c__Saccharomycetes(100);o__Saccharomycetales(100);f__Metschnikowiaceae(100);g__Metschnikowia(100);s__Metschnikowia_pulcherrima(100); | Metschnikowia | 13 | Saprotroph | \|Undefined Saprotroph\| | Yeast | NULL | Probable | As nectar-tap_saprotroph (Põlme et al. 2020); Decay sugar-rich substrates (Põlme et al. 2020) | Põlme S, et al. 2020. Fungal Diversity 105:1-16 ((https://doi.org/10.1007/s13225-020-00466-2)) |
| Otu51937 | 1 | k__Fungi(100);p__Ascomycota(100);c__Saccharomycetes(100);o__Saccharomycetales(100);f__Metschnikowiaceae(100);g__Metschnikowia(100);g__Metschnikowia_unclassified(100); | Metschnikowia | 13 | Saprotroph | \|Undefined Saprotroph\| | Yeast | NULL | Probable | As nectar-tap_saprotroph (Põlme et al. 2020); Decay sugar-rich substrates (Põlme et al. 2020) | Põlme S, et al. 2020. Fungal Diversity 105:1-16 ((https://doi.org/10.1007/s13225-020-00466-2)) |
| Otu51938 | 1 | k__Fungi(100);p__Ascomycota(100);c__Saccharomycetes(100);o__Saccharomycetales(100);f__Metschnikowiaceae(100);g__Metschnikowia(100);g__Metschnikowia_unclassified(100); | Metschnikowia | 13 | Saprotroph | \|Undefined Saprotroph\| | Yeast | NULL | Probable | As nectar-tap_saprotroph (Põlme et al. 2020); Decay sugar-rich substrates (Põlme et al. 2020) | Põlme S, et al. 2020. Fungal Diversity 105:1-16 ((https://doi.org/10.1007/s13225-020-00466-2)) |
| Otu51939 | 1 | k__Fungi(100);p__Ascomycota(100);c__Saccharomycetes(100);o__Saccharomycetales(100);f__Metschnikowiaceae(100);g__Metschnikowia(100);s__Metschnikowia_pulcherrima(100); | Metschnikowia | 13 | Saprotroph | \|Undefined Saprotroph\| | Yeast | NULL | Probable | As nectar-tap_saprotroph (Põlme et al. 2020); Decay sugar-rich substrates (Põlme et al. 2020) | Põlme S, et al. 2020. Fungal Diversity 105:1-16 ((https://doi.org/10.1007/s13225-020-00466-2)) |
| Otu51940 | 1 | k__Fungi(100);p__Ascomycota(100);c__Saccharomycetes(100);o__Saccharomycetales(100);f__Metschnikowiaceae(100);g__Metschnikowia(100);s__Metschnikowia_pulcherrima(100); | Metschnikowia | 13 | Saprotroph | \|Undefined Saprotroph\| | Yeast | NULL | Probable | As nectar-tap_saprotroph (Põlme et al. 2020); Decay sugar-rich substrates (Põlme et al. 2020) | Põlme S, et al. 2020. Fungal Diversity 105:1-16 ((https://doi.org/10.1007/s13225-020-00466-2)) |
| Otu51941 | 1 | k__Fungi(100);p__Ascomycota(100);c__Saccharomycetes(100);o__Saccharomycetales(100);f__Metschnikowiaceae(100);g__Metschnikowia(100);s__Metschnikowia_pulcherrima(100); | Metschnikowia | 13 | Saprotroph | \|Undefined Saprotroph\| | Yeast | NULL | Probable | As nectar-tap_saprotroph (Põlme et al. 2020); Decay sugar-rich substrates (Põlme et al. 2020) | Põlme S, et al. 2020. Fungal Diversity 105:1-16 ((https://doi.org/10.1007/s13225-020-00466-2)) |
| Otu51942 | 1 | k__Fungi(100);p__Ascomycota(100);c__Saccharomycetes(100);o__Saccharomycetales(100);f__Metschnikowiaceae(100);g__Metschnikowia(100);g__Metschnikowia_unclassified(100); | Metschnikowia | 13 | Saprotroph | \|Undefined Saprotroph\| | Yeast | NULL | Probable | As nectar-tap_saprotroph (Põlme et al. 2020); Decay sugar-rich substrates (Põlme et al. 2020) | Põlme S, et al. 2020. Fungal Diversity 105:1-16 ((https://doi.org/10.1007/s13225-020-00466-2)) |
| Otu51943 | 1 | k__Fungi(100);p__Ascomycota(100);c__Saccharomycetes(100);o__Saccharomycetales(100);f__Metschnikowiaceae(100);g__Metschnikowia(100);g__Metschnikowia_unclassified(100); | Metschnikowia | 13 | Saprotroph | \|Undefined Saprotroph\| | Yeast | NULL | Probable | As nectar-tap_saprotroph (Põlme et al. 2020); Decay sugar-rich substrates (Põlme et al. 2020) | Põlme S, et al. 2020. Fungal Diversity 105:1-16 ((https://doi.org/10.1007/s13225-020-00466-2)) |
| Otu51944 | 1 | k__Fungi(100);p__Ascomycota(100);c__Saccharomycetes(100);o__Saccharomycetales(100);f__Metschnikowiaceae(100);g__Metschnikowia(100);g__Metschnikowia_unclassified(100); | Metschnikowia | 13 | Saprotroph | \|Undefined Saprotroph\| | Yeast | NULL | Probable | As nectar-tap_saprotroph (Põlme et al. 2020); Decay sugar-rich substrates (Põlme et al. 2020) | Põlme S, et al. 2020. Fungal Diversity 105:1-16 ((https://doi.org/10.1007/s13225-020-00466-2)) |
| Otu51945 | 1 | k__Fungi(100);p__Ascomycota(100);c__Saccharomycetes(100);o__Saccharomycetales(100);f__Metschnikowiaceae(100);g__Metschnikowia(100);s__Metschnikowia_pulcherrima(100); | Metschnikowia | 13 | Saprotroph | \|Undefined Saprotroph\| | Yeast | NULL | Probable | As nectar-tap_saprotroph (Põlme et al. 2020); Decay sugar-rich substrates (Põlme et al. 2020) | Põlme S, et al. 2020. Fungal Diversity 105:1-16 ((https://doi.org/10.1007/s13225-020-00466-2)) |
| Otu51946 | 1 | k__Fungi(100);p__Ascomycota(100);c__Saccharomycetes(100);o__Saccharomycetales(100);f__Metschnikowiaceae(100);g__Metschnikowia(100);s__Metschnikowia_pulcherrima(100); | Metschnikowia | 13 | Saprotroph | \|Undefined Saprotroph\| | Yeast | NULL | Probable | As nectar-tap_saprotroph (Põlme et al. 2020); Decay sugar-rich substrates (Põlme et al. 2020) | Põlme S, et al. 2020. Fungal Diversity 105:1-16 ((https://doi.org/10.1007/s13225-020-00466-2)) |
| Otu51948 | 1 | k__Fungi(100);p__Ascomycota(100);c__Saccharomycetes(100);o__Saccharomycetales(100);f__Metschnikowiaceae(100);g__Metschnikowia(100);s__Metschnikowia_pulcherrima(100); | Metschnikowia | 13 | Saprotroph | \|Undefined Saprotroph\| | Yeast | NULL | Probable | As nectar-tap_saprotroph (Põlme et al. 2020); Decay sugar-rich substrates (Põlme et al. 2020) | Põlme S, et al. 2020. Fungal Diversity 105:1-16 ((https://doi.org/10.1007/s13225-020-00466-2)) |
| Otu51949 | 1 | k__Fungi(100);p__Ascomycota(100);c__Saccharomycetes(100);o__Saccharomycetales(100);f__Metschnikowiaceae(100);g__Metschnikowia(100);g__Metschnikowia_unclassified(100); | Metschnikowia | 13 | Saprotroph | \|Undefined Saprotroph\| | Yeast | NULL | Probable | As nectar-tap_saprotroph (Põlme et al. 2020); Decay sugar-rich substrates (Põlme et al. 2020) | Põlme S, et al. 2020. Fungal Diversity 105:1-16 ((https://doi.org/10.1007/s13225-020-00466-2)) |
| Otu51951 | 1 | k__Fungi(100);p__Ascomycota(100);c__Saccharomycetes(100);o__Saccharomycetales(100);f__Metschnikowiaceae(100);g__Metschnikowia(100);g__Metschnikowia_unclassified(100); | Metschnikowia | 13 | Saprotroph | \|Undefined Saprotroph\| | Yeast | NULL | Probable | As nectar-tap_saprotroph (Põlme et al. 2020); Decay sugar-rich substrates (Põlme et al. 2020) | Põlme S, et al. 2020. Fungal Diversity 105:1-16 ((https://doi.org/10.1007/s13225-020-00466-2)) |
| Otu51952 | 1 | k__Fungi(100);p__Ascomycota(100);c__Saccharomycetes(100);o__Saccharomycetales(100);f__Metschnikowiaceae(100);g__Metschnikowia(100);s__Metschnikowia_pulcherrima(100); | Metschnikowia | 13 | Saprotroph | \|Undefined Saprotroph\| | Yeast | NULL | Probable | As nectar-tap_saprotroph (Põlme et al. 2020); Decay sugar-rich substrates (Põlme et al. 2020) | Põlme S, et al. 2020. Fungal Diversity 105:1-16 ((https://doi.org/10.1007/s13225-020-00466-2)) |
| Otu51953 | 1 | k__Fungi(100);p__Ascomycota(100);c__Saccharomycetes(100);o__Saccharomycetales(100);f__Metschnikowiaceae(100);g__Metschnikowia(100);g__Metschnikowia_unclassified(100); | Metschnikowia | 13 | Saprotroph | \|Undefined Saprotroph\| | Yeast | NULL | Probable | As nectar-tap_saprotroph (Põlme et al. 2020); Decay sugar-rich substrates (Põlme et al. 2020) | Põlme S, et al. 2020. Fungal Diversity 105:1-16 ((https://doi.org/10.1007/s13225-020-00466-2)) |
| Otu51954 | 1 | k__Fungi(100);p__Ascomycota(100);c__Saccharomycetes(100);o__Saccharomycetales(100);o__Saccharomycetales_unclassified(100);o__Saccharomycetales_unclassified(100);o__Saccharomycetales_unclassified(100); | Saccharomycetales | 7 | Saprotroph | Undefined Saprotroph | Yeast | NULL | Possible | NULL | Sterkenburg E, et al. 2015. New Phytologist 207:1145-1158 ((https://doi.org/10.1111/nph.13426)) |
| Otu51955 | 1 | k__Fungi(100);p__Ascomycota(100);c__Sordariomycetes(100);o__Sordariales(100);f__Chaetomiaceae(100);g__Humicola(100);g__Humicola_unclassified(100); | Humicola | 13 | Pathotroph-Saprotroph | Plant Pathogen-Undefined Saprotroph-\|Wood Saprotroph\| | Microfungus | Soft Rot | Probable | Decay wood (Põlme et al. 2020) | Seehann G, et al. 1975. List of Fungi in Soft-Rot Tissue Tests. Institute fur Holzbiologie und Holzschutz, Bundesforschungsanstalt fur Forst - und Holzwirtshaft, Hamburg - Lohbrugge, Germany; Costa IPMW, et al. 2012. Mycotaxon 119:e26 ((http://mycotaxon.com/resources/checklists/costa_v119_checklist.pdf)); Tedersoo L, et al. 2014. Science 346:e1256688 ((https://doi.org/10.1126/science.1256688)); Põlme S, et al. 2020. Fungal Diversity 105:1-16 ((https://doi.org/10.1007/s13225-020-00466-2)) |
| Otu51956 | 1 | k__Fungi(100);p__Ascomycota(100);c__Saccharomycetes(100);o__Saccharomycetales(100);f__Metschnikowiaceae(100);g__Metschnikowia(100);s__Metschnikowia_pulcherrima(100); | Metschnikowia | 13 | Saprotroph | \|Undefined Saprotroph\| | Yeast | NULL | Probable | As nectar-tap_saprotroph (Põlme et al. 2020); Decay sugar-rich substrates (Põlme et al. 2020) | Põlme S, et al. 2020. Fungal Diversity 105:1-16 ((https://doi.org/10.1007/s13225-020-00466-2)) |
| Otu51957 | 1 | k__Fungi(100);p__Ascomycota(100);c__Saccharomycetes(100);o__Saccharomycetales(100);f__Metschnikowiaceae(100);g__Metschnikowia(100);g__Metschnikowia_unclassified(100); | Metschnikowia | 13 | Saprotroph | \|Undefined Saprotroph\| | Yeast | NULL | Probable | As nectar-tap_saprotroph (Põlme et al. 2020); Decay sugar-rich substrates (Põlme et al. 2020) | Põlme S, et al. 2020. Fungal Diversity 105:1-16 ((https://doi.org/10.1007/s13225-020-00466-2)) |
| Otu51958 | 1 | k__Fungi(100);p__Ascomycota(100);c__Saccharomycetes(100);o__Saccharomycetales(100);f__Metschnikowiaceae(100);g__Metschnikowia(100);g__Metschnikowia_unclassified(100); | Metschnikowia | 13 | Saprotroph | \|Undefined Saprotroph\| | Yeast | NULL | Probable | As nectar-tap_saprotroph (Põlme et al. 2020); Decay sugar-rich substrates (Põlme et al. 2020) | Põlme S, et al. 2020. Fungal Diversity 105:1-16 ((https://doi.org/10.1007/s13225-020-00466-2)) |
| Otu51959 | 1 | k__Fungi(100);p__Ascomycota(100);c__Saccharomycetes(100);o__Saccharomycetales(100);f__Metschnikowiaceae(100);g__Metschnikowia(100);s__Metschnikowia_pulcherrima(100); | Metschnikowia | 13 | Saprotroph | \|Undefined Saprotroph\| | Yeast | NULL | Probable | As nectar-tap_saprotroph (Põlme et al. 2020); Decay sugar-rich substrates (Põlme et al. 2020) | Põlme S, et al. 2020. Fungal Diversity 105:1-16 ((https://doi.org/10.1007/s13225-020-00466-2)) |
| Otu51960 | 1 | k__Fungi(100);p__Ascomycota(100);c__Saccharomycetes(100);o__Saccharomycetales(100);f__Metschnikowiaceae(100);g__Metschnikowia(100);s__Metschnikowia_pulcherrima(100); | Metschnikowia | 13 | Saprotroph | \|Undefined Saprotroph\| | Yeast | NULL | Probable | As nectar-tap_saprotroph (Põlme et al. 2020); Decay sugar-rich substrates (Põlme et al. 2020) | Põlme S, et al. 2020. Fungal Diversity 105:1-16 ((https://doi.org/10.1007/s13225-020-00466-2)) |
| Otu51961 | 1 | k__Fungi(100);p__Ascomycota(100);c__Saccharomycetes(100);o__Saccharomycetales(100);f__Metschnikowiaceae(100);g__Metschnikowia(100);s__Metschnikowia_pulcherrima(100); | Metschnikowia | 13 | Saprotroph | \|Undefined Saprotroph\| | Yeast | NULL | Probable | As nectar-tap_saprotroph (Põlme et al. 2020); Decay sugar-rich substrates (Põlme et al. 2020) | Põlme S, et al. 2020. Fungal Diversity 105:1-16 ((https://doi.org/10.1007/s13225-020-00466-2)) |
| Otu51962 | 1 | k__Fungi(100);p__Ascomycota(100);c__Saccharomycetes(100);o__Saccharomycetales(100);f__Metschnikowiaceae(100);g__Metschnikowia(100);s__Metschnikowia_pulcherrima(100); | Metschnikowia | 13 | Saprotroph | \|Undefined Saprotroph\| | Yeast | NULL | Probable | As nectar-tap_saprotroph (Põlme et al. 2020); Decay sugar-rich substrates (Põlme et al. 2020) | Põlme S, et al. 2020. Fungal Diversity 105:1-16 ((https://doi.org/10.1007/s13225-020-00466-2)) |
| Otu51963 | 1 | k__Fungi(100);p__Ascomycota(100);c__Saccharomycetes(100);o__Saccharomycetales(100);f__Metschnikowiaceae(100);g__Metschnikowia(100);g__Metschnikowia_unclassified(100); | Metschnikowia | 13 | Saprotroph | \|Undefined Saprotroph\| | Yeast | NULL | Probable | As nectar-tap_saprotroph (Põlme et al. 2020); Decay sugar-rich substrates (Põlme et al. 2020) | Põlme S, et al. 2020. Fungal Diversity 105:1-16 ((https://doi.org/10.1007/s13225-020-00466-2)) |
| Otu51964 | 1 | k__Fungi(100);p__Ascomycota(100);c__Saccharomycetes(100);o__Saccharomycetales(100);f__Metschnikowiaceae(100);g__Metschnikowia(100);g__Metschnikowia_unclassified(100); | Metschnikowia | 13 | Saprotroph | \|Undefined Saprotroph\| | Yeast | NULL | Probable | As nectar-tap_saprotroph (Põlme et al. 2020); Decay sugar-rich substrates (Põlme et al. 2020) | Põlme S, et al. 2020. Fungal Diversity 105:1-16 ((https://doi.org/10.1007/s13225-020-00466-2)) |
| Otu51965 | 1 | k__Fungi(100);p__Ascomycota(100);c__Saccharomycetes(100);o__Saccharomycetales(100);f__Metschnikowiaceae(100);g__Metschnikowia(100);s__Metschnikowia_pulcherrima(100); | Metschnikowia | 13 | Saprotroph | \|Undefined Saprotroph\| | Yeast | NULL | Probable | As nectar-tap_saprotroph (Põlme et al. 2020); Decay sugar-rich substrates (Põlme et al. 2020) | Põlme S, et al. 2020. Fungal Diversity 105:1-16 ((https://doi.org/10.1007/s13225-020-00466-2)) |
| Otu51966 | 1 | k__Fungi(100);p__Ascomycota(100);c__Saccharomycetes(100);o__Saccharomycetales(100);f__Metschnikowiaceae(100);g__Metschnikowia(100);g__Metschnikowia_unclassified(100); | Metschnikowia | 13 | Saprotroph | \|Undefined Saprotroph\| | Yeast | NULL | Probable | As nectar-tap_saprotroph (Põlme et al. 2020); Decay sugar-rich substrates (Põlme et al. 2020) | Põlme S, et al. 2020. Fungal Diversity 105:1-16 ((https://doi.org/10.1007/s13225-020-00466-2)) |
| Otu51967 | 1 | k__Fungi(100);p__Ascomycota(100);c__Saccharomycetes(100);o__Saccharomycetales(100);f__Metschnikowiaceae(100);g__Metschnikowia(100);s__Metschnikowia_pulcherrima(100); | Metschnikowia | 13 | Saprotroph | \|Undefined Saprotroph\| | Yeast | NULL | Probable | As nectar-tap_saprotroph (Põlme et al. 2020); Decay sugar-rich substrates (Põlme et al. 2020) | Põlme S, et al. 2020. Fungal Diversity 105:1-16 ((https://doi.org/10.1007/s13225-020-00466-2)) |
| Otu51968 | 1 | k__Fungi(100);p__Ascomycota(100);c__Saccharomycetes(100);o__Saccharomycetales(100);f__Metschnikowiaceae(100);g__Metschnikowia(100);s__Metschnikowia_pulcherrima(100); | Metschnikowia | 13 | Saprotroph | \|Undefined Saprotroph\| | Yeast | NULL | Probable | As nectar-tap_saprotroph (Põlme et al. 2020); Decay sugar-rich substrates (Põlme et al. 2020) | Põlme S, et al. 2020. Fungal Diversity 105:1-16 ((https://doi.org/10.1007/s13225-020-00466-2)) |
| Otu51969 | 1 | k__Fungi(100);p__Ascomycota(100);c__Saccharomycetes(100);o__Saccharomycetales(100);f__Metschnikowiaceae(100);g__Metschnikowia(100);s__Metschnikowia_pulcherrima(100); | Metschnikowia | 13 | Saprotroph | \|Undefined Saprotroph\| | Yeast | NULL | Probable | As nectar-tap_saprotroph (Põlme et al. 2020); Decay sugar-rich substrates (Põlme et al. 2020) | Põlme S, et al. 2020. Fungal Diversity 105:1-16 ((https://doi.org/10.1007/s13225-020-00466-2)) |
| Otu51972 | 1 | k__Fungi(100);p__Ascomycota(100);c__Saccharomycetes(100);o__Saccharomycetales(100);f__Metschnikowiaceae(100);g__Metschnikowia(100);s__Metschnikowia_pulcherrima(100); | Metschnikowia | 13 | Saprotroph | \|Undefined Saprotroph\| | Yeast | NULL | Probable | As nectar-tap_saprotroph (Põlme et al. 2020); Decay sugar-rich substrates (Põlme et al. 2020) | Põlme S, et al. 2020. Fungal Diversity 105:1-16 ((https://doi.org/10.1007/s13225-020-00466-2)) |
| Otu51973 | 1 | k__Fungi(100);p__Ascomycota(100);c__Saccharomycetes(100);o__Saccharomycetales(100);f__Metschnikowiaceae(100);g__Metschnikowia(100);s__Metschnikowia_pulcherrima(100); | Metschnikowia | 13 | Saprotroph | \|Undefined Saprotroph\| | Yeast | NULL | Probable | As nectar-tap_saprotroph (Põlme et al. 2020); Decay sugar-rich substrates (Põlme et al. 2020) | Põlme S, et al. 2020. Fungal Diversity 105:1-16 ((https://doi.org/10.1007/s13225-020-00466-2)) |
| Otu51974 | 1 | k__Fungi(100);p__Ascomycota(100);c__Saccharomycetes(100);o__Saccharomycetales(100);f__Metschnikowiaceae(100);g__Metschnikowia(100);s__Metschnikowia_pulcherrima(100); | Metschnikowia | 13 | Saprotroph | \|Undefined Saprotroph\| | Yeast | NULL | Probable | As nectar-tap_saprotroph (Põlme et al. 2020); Decay sugar-rich substrates (Põlme et al. 2020) | Põlme S, et al. 2020. Fungal Diversity 105:1-16 ((https://doi.org/10.1007/s13225-020-00466-2)) |
| Otu51975 | 1 | k__Fungi(100);p__Ascomycota(100);c__Saccharomycetes(100);o__Saccharomycetales(100);f__Metschnikowiaceae(100);g__Metschnikowia(100);s__Metschnikowia_pulcherrima(100); | Metschnikowia | 13 | Saprotroph | \|Undefined Saprotroph\| | Yeast | NULL | Probable | As nectar-tap_saprotroph (Põlme et al. 2020); Decay sugar-rich substrates (Põlme et al. 2020) | Põlme S, et al. 2020. Fungal Diversity 105:1-16 ((https://doi.org/10.1007/s13225-020-00466-2)) |
| Otu51976 | 1 | k__Fungi(100);p__Ascomycota(100);c__Saccharomycetes(100);o__Saccharomycetales(100);f__Metschnikowiaceae(100);g__Metschnikowia(100);s__Metschnikowia_pulcherrima(100); | Metschnikowia | 13 | Saprotroph | \|Undefined Saprotroph\| | Yeast | NULL | Probable | As nectar-tap_saprotroph (Põlme et al. 2020); Decay sugar-rich substrates (Põlme et al. 2020) | Põlme S, et al. 2020. Fungal Diversity 105:1-16 ((https://doi.org/10.1007/s13225-020-00466-2)) |
| Otu51977 | 1 | k__Fungi(100);p__Ascomycota(100);c__Saccharomycetes(100);o__Saccharomycetales(100);f__Metschnikowiaceae(100);g__Metschnikowia(100);s__Metschnikowia_pulcherrima(100); | Metschnikowia | 13 | Saprotroph | \|Undefined Saprotroph\| | Yeast | NULL | Probable | As nectar-tap_saprotroph (Põlme et al. 2020); Decay sugar-rich substrates (Põlme et al. 2020) | Põlme S, et al. 2020. Fungal Diversity 105:1-16 ((https://doi.org/10.1007/s13225-020-00466-2)) |
| Otu51980 | 1 | k__Fungi(100);p__Ascomycota(100);c__Saccharomycetes(100);o__Saccharomycetales(100);f__Metschnikowiaceae(100);g__Metschnikowia(100);g__Metschnikowia_unclassified(100); | Metschnikowia | 13 | Saprotroph | \|Undefined Saprotroph\| | Yeast | NULL | Probable | As nectar-tap_saprotroph (Põlme et al. 2020); Decay sugar-rich substrates (Põlme et al. 2020) | Põlme S, et al. 2020. Fungal Diversity 105:1-16 ((https://doi.org/10.1007/s13225-020-00466-2)) |
| Otu51982 | 1 | k__Fungi(100);p__Ascomycota(100);c__Saccharomycetes(100);o__Saccharomycetales(100);f__Metschnikowiaceae(100);g__Metschnikowia(100);s__Metschnikowia_pulcherrima(100); | Metschnikowia | 13 | Saprotroph | \|Undefined Saprotroph\| | Yeast | NULL | Probable | As nectar-tap_saprotroph (Põlme et al. 2020); Decay sugar-rich substrates (Põlme et al. 2020) | Põlme S, et al. 2020. Fungal Diversity 105:1-16 ((https://doi.org/10.1007/s13225-020-00466-2)) |
| Otu51983 | 1 | k__Fungi(100);p__Ascomycota(100);c__Saccharomycetes(100);o__Saccharomycetales(100);f__Metschnikowiaceae(100);g__Metschnikowia(100);g__Metschnikowia_unclassified(100); | Metschnikowia | 13 | Saprotroph | \|Undefined Saprotroph\| | Yeast | NULL | Probable | As nectar-tap_saprotroph (Põlme et al. 2020); Decay sugar-rich substrates (Põlme et al. 2020) | Põlme S, et al. 2020. Fungal Diversity 105:1-16 ((https://doi.org/10.1007/s13225-020-00466-2)) |
| Otu51984 | 1 | k__Fungi(100);p__Ascomycota(100);c__Saccharomycetes(100);o__Saccharomycetales(100);f__Metschnikowiaceae(100);g__Metschnikowia(100);g__Metschnikowia_unclassified(100); | Metschnikowia | 13 | Saprotroph | \|Undefined Saprotroph\| | Yeast | NULL | Probable | As nectar-tap_saprotroph (Põlme et al. 2020); Decay sugar-rich substrates (Põlme et al. 2020) | Põlme S, et al. 2020. Fungal Diversity 105:1-16 ((https://doi.org/10.1007/s13225-020-00466-2)) |
| Otu51985 | 1 | k__Fungi(100);p__Ascomycota(100);c__Saccharomycetes(100);o__Saccharomycetales(100);f__Metschnikowiaceae(100);g__Metschnikowia(100);g__Metschnikowia_unclassified(100); | Metschnikowia | 13 | Saprotroph | \|Undefined Saprotroph\| | Yeast | NULL | Probable | As nectar-tap_saprotroph (Põlme et al. 2020); Decay sugar-rich substrates (Põlme et al. 2020) | Põlme S, et al. 2020. Fungal Diversity 105:1-16 ((https://doi.org/10.1007/s13225-020-00466-2)) |
| Otu51986 | 1 | k__Fungi(100);p__Ascomycota(100);c__Saccharomycetes(100);o__Saccharomycetales(100);f__Metschnikowiaceae(100);g__Metschnikowia(100);s__Metschnikowia_pulcherrima(100); | Metschnikowia | 13 | Saprotroph | \|Undefined Saprotroph\| | Yeast | NULL | Probable | As nectar-tap_saprotroph (Põlme et al. 2020); Decay sugar-rich substrates (Põlme et al. 2020) | Põlme S, et al. 2020. Fungal Diversity 105:1-16 ((https://doi.org/10.1007/s13225-020-00466-2)) |
| Otu51987 | 1 | k__Fungi(100);p__Ascomycota(100);c__Saccharomycetes(100);o__Saccharomycetales(100);f__Metschnikowiaceae(100);g__Metschnikowia(100);s__Metschnikowia_pulcherrima(100); | Metschnikowia | 13 | Saprotroph | \|Undefined Saprotroph\| | Yeast | NULL | Probable | As nectar-tap_saprotroph (Põlme et al. 2020); Decay sugar-rich substrates (Põlme et al. 2020) | Põlme S, et al. 2020. Fungal Diversity 105:1-16 ((https://doi.org/10.1007/s13225-020-00466-2)) |
| Otu51988 | 1 | k__Fungi(100);p__Ascomycota(100);c__Saccharomycetes(100);o__Saccharomycetales(100);f__Metschnikowiaceae(100);g__Metschnikowia(100);s__Metschnikowia_pulcherrima(100); | Metschnikowia | 13 | Saprotroph | \|Undefined Saprotroph\| | Yeast | NULL | Probable | As nectar-tap_saprotroph (Põlme et al. 2020); Decay sugar-rich substrates (Põlme et al. 2020) | Põlme S, et al. 2020. Fungal Diversity 105:1-16 ((https://doi.org/10.1007/s13225-020-00466-2)) |
| Otu51989 | 1 | k__Fungi(100);p__Ascomycota(100);c__Saccharomycetes(100);o__Saccharomycetales(100);f__Metschnikowiaceae(100);g__Metschnikowia(100);g__Metschnikowia_unclassified(100); | Metschnikowia | 13 | Saprotroph | \|Undefined Saprotroph\| | Yeast | NULL | Probable | As nectar-tap_saprotroph (Põlme et al. 2020); Decay sugar-rich substrates (Põlme et al. 2020) | Põlme S, et al. 2020. Fungal Diversity 105:1-16 ((https://doi.org/10.1007/s13225-020-00466-2)) |
| Otu51990 | 1 | k__Fungi(100);p__Ascomycota(100);c__Saccharomycetes(100);o__Saccharomycetales(100);f__Metschnikowiaceae(100);g__Metschnikowia(100);s__Metschnikowia_pulcherrima(100); | Metschnikowia | 13 | Saprotroph | \|Undefined Saprotroph\| | Yeast | NULL | Probable | As nectar-tap_saprotroph (Põlme et al. 2020); Decay sugar-rich substrates (Põlme et al. 2020) | Põlme S, et al. 2020. Fungal Diversity 105:1-16 ((https://doi.org/10.1007/s13225-020-00466-2)) |
| Otu51991 | 1 | k__Fungi(100);p__Ascomycota(100);c__Saccharomycetes(100);o__Saccharomycetales(100);f__Metschnikowiaceae(100);g__Metschnikowia(100);g__Metschnikowia_unclassified(100); | Metschnikowia | 13 | Saprotroph | \|Undefined Saprotroph\| | Yeast | NULL | Probable | As nectar-tap_saprotroph (Põlme et al. 2020); Decay sugar-rich substrates (Põlme et al. 2020) | Põlme S, et al. 2020. Fungal Diversity 105:1-16 ((https://doi.org/10.1007/s13225-020-00466-2)) |
| Otu51992 | 1 | k__Fungi(100);p__Ascomycota(100);c__Saccharomycetes(100);o__Saccharomycetales(100);f__Metschnikowiaceae(100);g__Metschnikowia(100);s__Metschnikowia_pulcherrima(100); | Metschnikowia | 13 | Saprotroph | \|Undefined Saprotroph\| | Yeast | NULL | Probable | As nectar-tap_saprotroph (Põlme et al. 2020); Decay sugar-rich substrates (Põlme et al. 2020) | Põlme S, et al. 2020. Fungal Diversity 105:1-16 ((https://doi.org/10.1007/s13225-020-00466-2)) |
| Otu51993 | 1 | k__Fungi(100);p__Ascomycota(100);c__Saccharomycetes(100);o__Saccharomycetales(100);f__Metschnikowiaceae(100);g__Metschnikowia(100);s__Metschnikowia_pulcherrima(100); | Metschnikowia | 13 | Saprotroph | \|Undefined Saprotroph\| | Yeast | NULL | Probable | As nectar-tap_saprotroph (Põlme et al. 2020); Decay sugar-rich substrates (Põlme et al. 2020) | Põlme S, et al. 2020. Fungal Diversity 105:1-16 ((https://doi.org/10.1007/s13225-020-00466-2)) |
| Otu51994 | 1 | k__Fungi(100);p__Ascomycota(100);c__Saccharomycetes(100);o__Saccharomycetales(100);f__Metschnikowiaceae(100);g__Metschnikowia(100);s__Metschnikowia_pulcherrima(100); | Metschnikowia | 13 | Saprotroph | \|Undefined Saprotroph\| | Yeast | NULL | Probable | As nectar-tap_saprotroph (Põlme et al. 2020); Decay sugar-rich substrates (Põlme et al. 2020) | Põlme S, et al. 2020. Fungal Diversity 105:1-16 ((https://doi.org/10.1007/s13225-020-00466-2)) |
| Otu51995 | 1 | k__Fungi(100);p__Ascomycota(100);c__Saccharomycetes(100);o__Saccharomycetales(100);f__Metschnikowiaceae(100);g__Metschnikowia(100);g__Metschnikowia_unclassified(100); | Metschnikowia | 13 | Saprotroph | \|Undefined Saprotroph\| | Yeast | NULL | Probable | As nectar-tap_saprotroph (Põlme et al. 2020); Decay sugar-rich substrates (Põlme et al. 2020) | Põlme S, et al. 2020. Fungal Diversity 105:1-16 ((https://doi.org/10.1007/s13225-020-00466-2)) |
| Otu51996 | 1 | k__Fungi(100);p__Ascomycota(100);c__Saccharomycetes(100);o__Saccharomycetales(100);f__Metschnikowiaceae(100);g__Metschnikowia(100);g__Metschnikowia_unclassified(100); | Metschnikowia | 13 | Saprotroph | \|Undefined Saprotroph\| | Yeast | NULL | Probable | As nectar-tap_saprotroph (Põlme et al. 2020); Decay sugar-rich substrates (Põlme et al. 2020) | Põlme S, et al. 2020. Fungal Diversity 105:1-16 ((https://doi.org/10.1007/s13225-020-00466-2)) |
| Otu51997 | 1 | k__Fungi(100);p__Ascomycota(100);c__Saccharomycetes(100);o__Saccharomycetales(100);f__Metschnikowiaceae(100);g__Metschnikowia(100);s__Metschnikowia_pulcherrima(100); | Metschnikowia | 13 | Saprotroph | \|Undefined Saprotroph\| | Yeast | NULL | Probable | As nectar-tap_saprotroph (Põlme et al. 2020); Decay sugar-rich substrates (Põlme et al. 2020) | Põlme S, et al. 2020. Fungal Diversity 105:1-16 ((https://doi.org/10.1007/s13225-020-00466-2)) |
| Otu51998 | 1 | k__Fungi(100);p__Ascomycota(100);c__Saccharomycetes(100);o__Saccharomycetales(100);f__Metschnikowiaceae(100);g__Metschnikowia(100);g__Metschnikowia_unclassified(100); | Metschnikowia | 13 | Saprotroph | \|Undefined Saprotroph\| | Yeast | NULL | Probable | As nectar-tap_saprotroph (Põlme et al. 2020); Decay sugar-rich substrates (Põlme et al. 2020) | Põlme S, et al. 2020. Fungal Diversity 105:1-16 ((https://doi.org/10.1007/s13225-020-00466-2)) |
| Otu51999 | 1 | k__Fungi(100);p__Ascomycota(100);c__Saccharomycetes(100);o__Saccharomycetales(100);f__Metschnikowiaceae(100);g__Metschnikowia(100);s__Metschnikowia_pulcherrima(100); | Metschnikowia | 13 | Saprotroph | \|Undefined Saprotroph\| | Yeast | NULL | Probable | As nectar-tap_saprotroph (Põlme et al. 2020); Decay sugar-rich substrates (Põlme et al. 2020) | Põlme S, et al. 2020. Fungal Diversity 105:1-16 ((https://doi.org/10.1007/s13225-020-00466-2)) |
| Otu52000 | 1 | k__Fungi(100);p__Ascomycota(100);c__Saccharomycetes(100);o__Saccharomycetales(100);f__Metschnikowiaceae(100);g__Metschnikowia(100);s__Metschnikowia_pulcherrima(100); | Metschnikowia | 13 | Saprotroph | \|Undefined Saprotroph\| | Yeast | NULL | Probable | As nectar-tap_saprotroph (Põlme et al. 2020); Decay sugar-rich substrates (Põlme et al. 2020) | Põlme S, et al. 2020. Fungal Diversity 105:1-16 ((https://doi.org/10.1007/s13225-020-00466-2)) |
| Otu52001 | 1 | k__Fungi(100);p__Ascomycota(100);c__Saccharomycetes(100);o__Saccharomycetales(100);f__Metschnikowiaceae(100);g__Metschnikowia(100);s__Metschnikowia_pulcherrima(100); | Metschnikowia | 13 | Saprotroph | \|Undefined Saprotroph\| | Yeast | NULL | Probable | As nectar-tap_saprotroph (Põlme et al. 2020); Decay sugar-rich substrates (Põlme et al. 2020) | Põlme S, et al. 2020. Fungal Diversity 105:1-16 ((https://doi.org/10.1007/s13225-020-00466-2)) |
| Otu52002 | 1 | k__Fungi(100);p__Ascomycota(100);c__Saccharomycetes(100);o__Saccharomycetales(100);f__Metschnikowiaceae(100);g__Metschnikowia(100);s__Metschnikowia_pulcherrima(100); | Metschnikowia | 13 | Saprotroph | \|Undefined Saprotroph\| | Yeast | NULL | Probable | As nectar-tap_saprotroph (Põlme et al. 2020); Decay sugar-rich substrates (Põlme et al. 2020) | Põlme S, et al. 2020. Fungal Diversity 105:1-16 ((https://doi.org/10.1007/s13225-020-00466-2)) |
| Otu52003 | 1 | k__Fungi(100);p__Ascomycota(100);c__Saccharomycetes(100);o__Saccharomycetales(100);f__Metschnikowiaceae(100);g__Metschnikowia(100);s__Metschnikowia_pulcherrima(100); | Metschnikowia | 13 | Saprotroph | \|Undefined Saprotroph\| | Yeast | NULL | Probable | As nectar-tap_saprotroph (Põlme et al. 2020); Decay sugar-rich substrates (Põlme et al. 2020) | Põlme S, et al. 2020. Fungal Diversity 105:1-16 ((https://doi.org/10.1007/s13225-020-00466-2)) |
| Otu52004 | 1 | k__Fungi(100);p__Ascomycota(100);c__Saccharomycetes(100);o__Saccharomycetales(100);f__Metschnikowiaceae(100);g__Metschnikowia(100);g__Metschnikowia_unclassified(100); | Metschnikowia | 13 | Saprotroph | \|Undefined Saprotroph\| | Yeast | NULL | Probable | As nectar-tap_saprotroph (Põlme et al. 2020); Decay sugar-rich substrates (Põlme et al. 2020) | Põlme S, et al. 2020. Fungal Diversity 105:1-16 ((https://doi.org/10.1007/s13225-020-00466-2)) |
| Otu52005 | 1 | k__Fungi(100);p__Ascomycota(100);c__Saccharomycetes(100);o__Saccharomycetales(100);f__Metschnikowiaceae(100);g__Metschnikowia(100);s__Metschnikowia_pulcherrima(100); | Metschnikowia | 13 | Saprotroph | \|Undefined Saprotroph\| | Yeast | NULL | Probable | As nectar-tap_saprotroph (Põlme et al. 2020); Decay sugar-rich substrates (Põlme et al. 2020) | Põlme S, et al. 2020. Fungal Diversity 105:1-16 ((https://doi.org/10.1007/s13225-020-00466-2)) |
| Otu52006 | 1 | k__Fungi(100);p__Ascomycota(100);c__Saccharomycetes(100);o__Saccharomycetales(100);f__Metschnikowiaceae(100);g__Metschnikowia(100);s__Metschnikowia_pulcherrima(100); | Metschnikowia | 13 | Saprotroph | \|Undefined Saprotroph\| | Yeast | NULL | Probable | As nectar-tap_saprotroph (Põlme et al. 2020); Decay sugar-rich substrates (Põlme et al. 2020) | Põlme S, et al. 2020. Fungal Diversity 105:1-16 ((https://doi.org/10.1007/s13225-020-00466-2)) |
| Otu52007 | 1 | k__Fungi(100);p__Ascomycota(100);c__Saccharomycetes(100);o__Saccharomycetales(100);f__Metschnikowiaceae(100);g__Metschnikowia(100);s__Metschnikowia_pulcherrima(100); | Metschnikowia | 13 | Saprotroph | \|Undefined Saprotroph\| | Yeast | NULL | Probable | As nectar-tap_saprotroph (Põlme et al. 2020); Decay sugar-rich substrates (Põlme et al. 2020) | Põlme S, et al. 2020. Fungal Diversity 105:1-16 ((https://doi.org/10.1007/s13225-020-00466-2)) |
| Otu52008 | 1 | k__Fungi(100);p__Ascomycota(100);c__Saccharomycetes(100);o__Saccharomycetales(100);f__Metschnikowiaceae(100);g__Metschnikowia(100);s__Metschnikowia_pulcherrima(100); | Metschnikowia | 13 | Saprotroph | \|Undefined Saprotroph\| | Yeast | NULL | Probable | As nectar-tap_saprotroph (Põlme et al. 2020); Decay sugar-rich substrates (Põlme et al. 2020) | Põlme S, et al. 2020. Fungal Diversity 105:1-16 ((https://doi.org/10.1007/s13225-020-00466-2)) |
| Otu52009 | 1 | k__Fungi(100);p__Ascomycota(100);c__Saccharomycetes(100);o__Saccharomycetales(100);f__Metschnikowiaceae(100);g__Metschnikowia(100);s__Metschnikowia_pulcherrima(100); | Metschnikowia | 13 | Saprotroph | \|Undefined Saprotroph\| | Yeast | NULL | Probable | As nectar-tap_saprotroph (Põlme et al. 2020); Decay sugar-rich substrates (Põlme et al. 2020) | Põlme S, et al. 2020. Fungal Diversity 105:1-16 ((https://doi.org/10.1007/s13225-020-00466-2)) |
| Otu52010 | 1 | k__Fungi(100);p__Ascomycota(100);c__Saccharomycetes(100);o__Saccharomycetales(100);f__Metschnikowiaceae(100);g__Metschnikowia(100);s__Metschnikowia_pulcherrima(100); | Metschnikowia | 13 | Saprotroph | \|Undefined Saprotroph\| | Yeast | NULL | Probable | As nectar-tap_saprotroph (Põlme et al. 2020); Decay sugar-rich substrates (Põlme et al. 2020) | Põlme S, et al. 2020. Fungal Diversity 105:1-16 ((https://doi.org/10.1007/s13225-020-00466-2)) |
| Otu52011 | 1 | k__Fungi(100);p__Ascomycota(100);c__Saccharomycetes(100);o__Saccharomycetales(100);f__Metschnikowiaceae(100);g__Metschnikowia(100);s__Metschnikowia_pulcherrima(100); | Metschnikowia | 13 | Saprotroph | \|Undefined Saprotroph\| | Yeast | NULL | Probable | As nectar-tap_saprotroph (Põlme et al. 2020); Decay sugar-rich substrates (Põlme et al. 2020) | Põlme S, et al. 2020. Fungal Diversity 105:1-16 ((https://doi.org/10.1007/s13225-020-00466-2)) |
| Otu52012 | 1 | k__Fungi(100);p__Ascomycota(100);c__Saccharomycetes(100);o__Saccharomycetales(100);f__Metschnikowiaceae(100);g__Metschnikowia(100);s__Metschnikowia_pulcherrima(100); | Metschnikowia | 13 | Saprotroph | \|Undefined Saprotroph\| | Yeast | NULL | Probable | As nectar-tap_saprotroph (Põlme et al. 2020); Decay sugar-rich substrates (Põlme et al. 2020) | Põlme S, et al. 2020. Fungal Diversity 105:1-16 ((https://doi.org/10.1007/s13225-020-00466-2)) |
| Otu52013 | 1 | k__Fungi(100);p__Ascomycota(100);c__Saccharomycetes(100);o__Saccharomycetales(100);f__Metschnikowiaceae(100);g__Metschnikowia(100);s__Metschnikowia_pulcherrima(100); | Metschnikowia | 13 | Saprotroph | \|Undefined Saprotroph\| | Yeast | NULL | Probable | As nectar-tap_saprotroph (Põlme et al. 2020); Decay sugar-rich substrates (Põlme et al. 2020) | Põlme S, et al. 2020. Fungal Diversity 105:1-16 ((https://doi.org/10.1007/s13225-020-00466-2)) |
| Otu52015 | 1 | k__Fungi(100);p__Ascomycota(100);c__Saccharomycetes(100);o__Saccharomycetales(100);f__Metschnikowiaceae(100);g__Metschnikowia(100);s__Metschnikowia_pulcherrima(100); | Metschnikowia | 13 | Saprotroph | \|Undefined Saprotroph\| | Yeast | NULL | Probable | As nectar-tap_saprotroph (Põlme et al. 2020); Decay sugar-rich substrates (Põlme et al. 2020) | Põlme S, et al. 2020. Fungal Diversity 105:1-16 ((https://doi.org/10.1007/s13225-020-00466-2)) |
| Otu52016 | 1 | k__Fungi(100);p__Ascomycota(100);c__Saccharomycetes(100);o__Saccharomycetales(100);f__Metschnikowiaceae(100);g__Metschnikowia(100);g__Metschnikowia_unclassified(100); | Metschnikowia | 13 | Saprotroph | \|Undefined Saprotroph\| | Yeast | NULL | Probable | As nectar-tap_saprotroph (Põlme et al. 2020); Decay sugar-rich substrates (Põlme et al. 2020) | Põlme S, et al. 2020. Fungal Diversity 105:1-16 ((https://doi.org/10.1007/s13225-020-00466-2)) |
| Otu52017 | 1 | k__Fungi(100);p__Ascomycota(100);c__Saccharomycetes(100);o__Saccharomycetales(100);f__Metschnikowiaceae(100);g__Metschnikowia(100);s__Metschnikowia_pulcherrima(100); | Metschnikowia | 13 | Saprotroph | \|Undefined Saprotroph\| | Yeast | NULL | Probable | As nectar-tap_saprotroph (Põlme et al. 2020); Decay sugar-rich substrates (Põlme et al. 2020) | Põlme S, et al. 2020. Fungal Diversity 105:1-16 ((https://doi.org/10.1007/s13225-020-00466-2)) |
| Otu52019 | 1 | k__Fungi(100);p__Ascomycota(100);c__Saccharomycetes(100);o__Saccharomycetales(100);f__Metschnikowiaceae(100);g__Metschnikowia(100);g__Metschnikowia_unclassified(100); | Metschnikowia | 13 | Saprotroph | \|Undefined Saprotroph\| | Yeast | NULL | Probable | As nectar-tap_saprotroph (Põlme et al. 2020); Decay sugar-rich substrates (Põlme et al. 2020) | Põlme S, et al. 2020. Fungal Diversity 105:1-16 ((https://doi.org/10.1007/s13225-020-00466-2)) |
| Otu52022 | 1 | k__Fungi(100);p__Ascomycota(100);c__Saccharomycetes(100);o__Saccharomycetales(100);f__Metschnikowiaceae(100);g__Metschnikowia(100);s__Metschnikowia_pulcherrima(100); | Metschnikowia | 13 | Saprotroph | \|Undefined Saprotroph\| | Yeast | NULL | Probable | As nectar-tap_saprotroph (Põlme et al. 2020); Decay sugar-rich substrates (Põlme et al. 2020) | Põlme S, et al. 2020. Fungal Diversity 105:1-16 ((https://doi.org/10.1007/s13225-020-00466-2)) |
| Otu52028 | 1 | k__Fungi(100);p__Ascomycota(100);c__Saccharomycetes(100);o__Saccharomycetales(100);f__Metschnikowiaceae(100);g__Metschnikowia(100);s__Metschnikowia_pulcherrima(100); | Metschnikowia | 13 | Saprotroph | \|Undefined Saprotroph\| | Yeast | NULL | Probable | As nectar-tap_saprotroph (Põlme et al. 2020); Decay sugar-rich substrates (Põlme et al. 2020) | Põlme S, et al. 2020. Fungal Diversity 105:1-16 ((https://doi.org/10.1007/s13225-020-00466-2)) |
| Otu52029 | 1 | k__Fungi(100);p__Ascomycota(100);c__Saccharomycetes(100);o__Saccharomycetales(100);f__Metschnikowiaceae(100);g__Metschnikowia(100);s__Metschnikowia_pulcherrima(100); | Metschnikowia | 13 | Saprotroph | \|Undefined Saprotroph\| | Yeast | NULL | Probable | As nectar-tap_saprotroph (Põlme et al. 2020); Decay sugar-rich substrates (Põlme et al. 2020) | Põlme S, et al. 2020. Fungal Diversity 105:1-16 ((https://doi.org/10.1007/s13225-020-00466-2)) |
| Otu52030 | 1 | k__Fungi(100);p__Ascomycota(100);c__Saccharomycetes(100);o__Saccharomycetales(100);f__Metschnikowiaceae(100);g__Metschnikowia(100);s__Metschnikowia_pulcherrima(100); | Metschnikowia | 13 | Saprotroph | \|Undefined Saprotroph\| | Yeast | NULL | Probable | As nectar-tap_saprotroph (Põlme et al. 2020); Decay sugar-rich substrates (Põlme et al. 2020) | Põlme S, et al. 2020. Fungal Diversity 105:1-16 ((https://doi.org/10.1007/s13225-020-00466-2)) |
| Otu52031 | 1 | k__Fungi(100);p__Ascomycota(100);c__Saccharomycetes(100);o__Saccharomycetales(100);f__Metschnikowiaceae(100);g__Metschnikowia(100);s__Metschnikowia_pulcherrima(100); | Metschnikowia | 13 | Saprotroph | \|Undefined Saprotroph\| | Yeast | NULL | Probable | As nectar-tap_saprotroph (Põlme et al. 2020); Decay sugar-rich substrates (Põlme et al. 2020) | Põlme S, et al. 2020. Fungal Diversity 105:1-16 ((https://doi.org/10.1007/s13225-020-00466-2)) |
| Otu52032 | 1 | k__Fungi(100);p__Ascomycota(100);c__Saccharomycetes(100);o__Saccharomycetales(100);f__Metschnikowiaceae(100);g__Metschnikowia(100);g__Metschnikowia_unclassified(100); | Metschnikowia | 13 | Saprotroph | \|Undefined Saprotroph\| | Yeast | NULL | Probable | As nectar-tap_saprotroph (Põlme et al. 2020); Decay sugar-rich substrates (Põlme et al. 2020) | Põlme S, et al. 2020. Fungal Diversity 105:1-16 ((https://doi.org/10.1007/s13225-020-00466-2)) |
| Otu52033 | 1 | k__Fungi(100);p__Ascomycota(100);c__Saccharomycetes(100);o__Saccharomycetales(100);f__Metschnikowiaceae(100);g__Metschnikowia(100);s__Metschnikowia_pulcherrima(100); | Metschnikowia | 13 | Saprotroph | \|Undefined Saprotroph\| | Yeast | NULL | Probable | As nectar-tap_saprotroph (Põlme et al. 2020); Decay sugar-rich substrates (Põlme et al. 2020) | Põlme S, et al. 2020. Fungal Diversity 105:1-16 ((https://doi.org/10.1007/s13225-020-00466-2)) |
| Otu52034 | 1 | k__Fungi(100);p__Ascomycota(100);c__Saccharomycetes(100);o__Saccharomycetales(100);f__Metschnikowiaceae(100);g__Metschnikowia(100);s__Metschnikowia_pulcherrima(100); | Metschnikowia | 13 | Saprotroph | \|Undefined Saprotroph\| | Yeast | NULL | Probable | As nectar-tap_saprotroph (Põlme et al. 2020); Decay sugar-rich substrates (Põlme et al. 2020) | Põlme S, et al. 2020. Fungal Diversity 105:1-16 ((https://doi.org/10.1007/s13225-020-00466-2)) |
| Otu52036 | 1 | k__Fungi(100);p__Ascomycota(100);c__Saccharomycetes(100);o__Saccharomycetales(100);f__Metschnikowiaceae(100);g__Metschnikowia(100);s__Metschnikowia_pulcherrima(100); | Metschnikowia | 13 | Saprotroph | \|Undefined Saprotroph\| | Yeast | NULL | Probable | As nectar-tap_saprotroph (Põlme et al. 2020); Decay sugar-rich substrates (Põlme et al. 2020) | Põlme S, et al. 2020. Fungal Diversity 105:1-16 ((https://doi.org/10.1007/s13225-020-00466-2)) |
| Otu52037 | 1 | k__Fungi(100);p__Ascomycota(100);c__Saccharomycetes(100);o__Saccharomycetales(100);f__Metschnikowiaceae(100);g__Metschnikowia(100);s__Metschnikowia_pulcherrima(100); | Metschnikowia | 13 | Saprotroph | \|Undefined Saprotroph\| | Yeast | NULL | Probable | As nectar-tap_saprotroph (Põlme et al. 2020); Decay sugar-rich substrates (Põlme et al. 2020) | Põlme S, et al. 2020. Fungal Diversity 105:1-16 ((https://doi.org/10.1007/s13225-020-00466-2)) |
| Otu52038 | 1 | k__Fungi(100);p__Ascomycota(100);c__Saccharomycetes(100);o__Saccharomycetales(100);f__Metschnikowiaceae(100);g__Metschnikowia(100);s__Metschnikowia_pulcherrima(100); | Metschnikowia | 13 | Saprotroph | \|Undefined Saprotroph\| | Yeast | NULL | Probable | As nectar-tap_saprotroph (Põlme et al. 2020); Decay sugar-rich substrates (Põlme et al. 2020) | Põlme S, et al. 2020. Fungal Diversity 105:1-16 ((https://doi.org/10.1007/s13225-020-00466-2)) |
| Otu52039 | 1 | k__Fungi(100);p__Ascomycota(100);c__Saccharomycetes(100);o__Saccharomycetales(100);f__Metschnikowiaceae(100);g__Metschnikowia(100);s__Metschnikowia_pulcherrima(100); | Metschnikowia | 13 | Saprotroph | \|Undefined Saprotroph\| | Yeast | NULL | Probable | As nectar-tap_saprotroph (Põlme et al. 2020); Decay sugar-rich substrates (Põlme et al. 2020) | Põlme S, et al. 2020. Fungal Diversity 105:1-16 ((https://doi.org/10.1007/s13225-020-00466-2)) |
| Otu52042 | 1 | k__Fungi(100);p__Ascomycota(100);c__Saccharomycetes(100);o__Saccharomycetales(100);o__Saccharomycetales_unclassified(100);o__Saccharomycetales_unclassified(100);o__Saccharomycetales_unclassified(100); | Saccharomycetales | 7 | Saprotroph | Undefined Saprotroph | Yeast | NULL | Possible | NULL | Sterkenburg E, et al. 2015. New Phytologist 207:1145-1158 ((https://doi.org/10.1111/nph.13426)) |
| Otu52043 | 1 | k__Fungi(100);p__Ascomycota(100);c__Saccharomycetes(100);o__Saccharomycetales(100);f__Metschnikowiaceae(100);g__Metschnikowia(100);g__Metschnikowia_unclassified(100); | Metschnikowia | 13 | Saprotroph | \|Undefined Saprotroph\| | Yeast | NULL | Probable | As nectar-tap_saprotroph (Põlme et al. 2020); Decay sugar-rich substrates (Põlme et al. 2020) | Põlme S, et al. 2020. Fungal Diversity 105:1-16 ((https://doi.org/10.1007/s13225-020-00466-2)) |
| Otu52044 | 1 | k__Fungi(100);p__Ascomycota(100);c__Saccharomycetes(100);o__Saccharomycetales(100);f__Metschnikowiaceae(100);g__Metschnikowia(100);s__Metschnikowia_pulcherrima(100); | Metschnikowia | 13 | Saprotroph | \|Undefined Saprotroph\| | Yeast | NULL | Probable | As nectar-tap_saprotroph (Põlme et al. 2020); Decay sugar-rich substrates (Põlme et al. 2020) | Põlme S, et al. 2020. Fungal Diversity 105:1-16 ((https://doi.org/10.1007/s13225-020-00466-2)) |
| Otu52045 | 1 | k__Fungi(100);p__Ascomycota(100);c__Saccharomycetes(100);o__Saccharomycetales(100);f__Metschnikowiaceae(100);g__Metschnikowia(100);s__Metschnikowia_pulcherrima(100); | Metschnikowia | 13 | Saprotroph | \|Undefined Saprotroph\| | Yeast | NULL | Probable | As nectar-tap_saprotroph (Põlme et al. 2020); Decay sugar-rich substrates (Põlme et al. 2020) | Põlme S, et al. 2020. Fungal Diversity 105:1-16 ((https://doi.org/10.1007/s13225-020-00466-2)) |
| Otu52046 | 1 | k__Fungi(100);p__Ascomycota(100);c__Saccharomycetes(100);o__Saccharomycetales(100);f__Metschnikowiaceae(100);g__Metschnikowia(100);s__Metschnikowia_pulcherrima(100); | Metschnikowia | 13 | Saprotroph | \|Undefined Saprotroph\| | Yeast | NULL | Probable | As nectar-tap_saprotroph (Põlme et al. 2020); Decay sugar-rich substrates (Põlme et al. 2020) | Põlme S, et al. 2020. Fungal Diversity 105:1-16 ((https://doi.org/10.1007/s13225-020-00466-2)) |
| Otu52047 | 1 | k__Fungi(100);p__Ascomycota(100);c__Saccharomycetes(100);o__Saccharomycetales(100);f__Metschnikowiaceae(100);g__Metschnikowia(100);s__Metschnikowia_pulcherrima(100); | Metschnikowia | 13 | Saprotroph | \|Undefined Saprotroph\| | Yeast | NULL | Probable | As nectar-tap_saprotroph (Põlme et al. 2020); Decay sugar-rich substrates (Põlme et al. 2020) | Põlme S, et al. 2020. Fungal Diversity 105:1-16 ((https://doi.org/10.1007/s13225-020-00466-2)) |
| Otu52048 | 1 | k__Fungi(100);p__Ascomycota(100);c__Saccharomycetes(100);o__Saccharomycetales(100);f__Metschnikowiaceae(100);g__Metschnikowia(100);s__Metschnikowia_pulcherrima(100); | Metschnikowia | 13 | Saprotroph | \|Undefined Saprotroph\| | Yeast | NULL | Probable | As nectar-tap_saprotroph (Põlme et al. 2020); Decay sugar-rich substrates (Põlme et al. 2020) | Põlme S, et al. 2020. Fungal Diversity 105:1-16 ((https://doi.org/10.1007/s13225-020-00466-2)) |
| Otu52049 | 1 | k__Fungi(100);p__Ascomycota(100);c__Saccharomycetes(100);o__Saccharomycetales(100);f__Metschnikowiaceae(100);g__Metschnikowia(100);g__Metschnikowia_unclassified(100); | Metschnikowia | 13 | Saprotroph | \|Undefined Saprotroph\| | Yeast | NULL | Probable | As nectar-tap_saprotroph (Põlme et al. 2020); Decay sugar-rich substrates (Põlme et al. 2020) | Põlme S, et al. 2020. Fungal Diversity 105:1-16 ((https://doi.org/10.1007/s13225-020-00466-2)) |
| Otu52050 | 1 | k__Fungi(100);p__Ascomycota(100);c__Saccharomycetes(100);o__Saccharomycetales(100);f__Metschnikowiaceae(100);g__Metschnikowia(100);s__Metschnikowia_pulcherrima(100); | Metschnikowia | 13 | Saprotroph | \|Undefined Saprotroph\| | Yeast | NULL | Probable | As nectar-tap_saprotroph (Põlme et al. 2020); Decay sugar-rich substrates (Põlme et al. 2020) | Põlme S, et al. 2020. Fungal Diversity 105:1-16 ((https://doi.org/10.1007/s13225-020-00466-2)) |
| Otu52051 | 1 | k__Fungi(100);p__Ascomycota(100);c__Saccharomycetes(100);o__Saccharomycetales(100);f__Metschnikowiaceae(100);g__Metschnikowia(100);g__Metschnikowia_unclassified(100); | Metschnikowia | 13 | Saprotroph | \|Undefined Saprotroph\| | Yeast | NULL | Probable | As nectar-tap_saprotroph (Põlme et al. 2020); Decay sugar-rich substrates (Põlme et al. 2020) | Põlme S, et al. 2020. Fungal Diversity 105:1-16 ((https://doi.org/10.1007/s13225-020-00466-2)) |
| Otu52053 | 1 | k__Fungi(100);p__Ascomycota(100);c__Saccharomycetes(100);o__Saccharomycetales(100);f__Metschnikowiaceae(100);g__Metschnikowia(100);g__Metschnikowia_unclassified(100); | Metschnikowia | 13 | Saprotroph | \|Undefined Saprotroph\| | Yeast | NULL | Probable | As nectar-tap_saprotroph (Põlme et al. 2020); Decay sugar-rich substrates (Põlme et al. 2020) | Põlme S, et al. 2020. Fungal Diversity 105:1-16 ((https://doi.org/10.1007/s13225-020-00466-2)) |
| Otu52054 | 1 | k__Fungi(100);p__Ascomycota(100);c__Saccharomycetes(100);o__Saccharomycetales(100);f__Metschnikowiaceae(100);g__Metschnikowia(100);s__Metschnikowia_pulcherrima(100); | Metschnikowia | 13 | Saprotroph | \|Undefined Saprotroph\| | Yeast | NULL | Probable | As nectar-tap_saprotroph (Põlme et al. 2020); Decay sugar-rich substrates (Põlme et al. 2020) | Põlme S, et al. 2020. Fungal Diversity 105:1-16 ((https://doi.org/10.1007/s13225-020-00466-2)) |
| Otu52055 | 1 | k__Fungi(100);p__Ascomycota(100);c__Saccharomycetes(100);o__Saccharomycetales(100);f__Metschnikowiaceae(100);g__Metschnikowia(100);s__Metschnikowia_pulcherrima(100); | Metschnikowia | 13 | Saprotroph | \|Undefined Saprotroph\| | Yeast | NULL | Probable | As nectar-tap_saprotroph (Põlme et al. 2020); Decay sugar-rich substrates (Põlme et al. 2020) | Põlme S, et al. 2020. Fungal Diversity 105:1-16 ((https://doi.org/10.1007/s13225-020-00466-2)) |
| Otu52056 | 1 | k__Fungi(100);p__Ascomycota(100);c__Saccharomycetes(100);o__Saccharomycetales(100);f__Metschnikowiaceae(100);g__Metschnikowia(100);s__Metschnikowia_pulcherrima(100); | Metschnikowia | 13 | Saprotroph | \|Undefined Saprotroph\| | Yeast | NULL | Probable | As nectar-tap_saprotroph (Põlme et al. 2020); Decay sugar-rich substrates (Põlme et al. 2020) | Põlme S, et al. 2020. Fungal Diversity 105:1-16 ((https://doi.org/10.1007/s13225-020-00466-2)) |
| Otu52057 | 1 | k__Fungi(100);p__Ascomycota(100);c__Saccharomycetes(100);o__Saccharomycetales(100);f__Metschnikowiaceae(100);g__Metschnikowia(100);g__Metschnikowia_unclassified(100); | Metschnikowia | 13 | Saprotroph | \|Undefined Saprotroph\| | Yeast | NULL | Probable | As nectar-tap_saprotroph (Põlme et al. 2020); Decay sugar-rich substrates (Põlme et al. 2020) | Põlme S, et al. 2020. Fungal Diversity 105:1-16 ((https://doi.org/10.1007/s13225-020-00466-2)) |
| Otu52058 | 1 | k__Fungi(100);p__Ascomycota(100);c__Saccharomycetes(100);o__Saccharomycetales(100);f__Metschnikowiaceae(100);g__Metschnikowia(100);s__Metschnikowia_pulcherrima(100); | Metschnikowia | 13 | Saprotroph | \|Undefined Saprotroph\| | Yeast | NULL | Probable | As nectar-tap_saprotroph (Põlme et al. 2020); Decay sugar-rich substrates (Põlme et al. 2020) | Põlme S, et al. 2020. Fungal Diversity 105:1-16 ((https://doi.org/10.1007/s13225-020-00466-2)) |
| Otu52059 | 1 | k__Fungi(100);p__Ascomycota(100);c__Saccharomycetes(100);o__Saccharomycetales(100);f__Metschnikowiaceae(100);g__Metschnikowia(100);g__Metschnikowia_unclassified(100); | Metschnikowia | 13 | Saprotroph | \|Undefined Saprotroph\| | Yeast | NULL | Probable | As nectar-tap_saprotroph (Põlme et al. 2020); Decay sugar-rich substrates (Põlme et al. 2020) | Põlme S, et al. 2020. Fungal Diversity 105:1-16 ((https://doi.org/10.1007/s13225-020-00466-2)) |
| Otu52060 | 1 | k__Fungi(100);p__Ascomycota(100);c__Saccharomycetes(100);o__Saccharomycetales(100);f__Metschnikowiaceae(100);g__Metschnikowia(100);s__Metschnikowia_pulcherrima(100); | Metschnikowia | 13 | Saprotroph | \|Undefined Saprotroph\| | Yeast | NULL | Probable | As nectar-tap_saprotroph (Põlme et al. 2020); Decay sugar-rich substrates (Põlme et al. 2020) | Põlme S, et al. 2020. Fungal Diversity 105:1-16 ((https://doi.org/10.1007/s13225-020-00466-2)) |
| Otu52061 | 1 | k__Fungi(100);p__Ascomycota(100);c__Saccharomycetes(100);o__Saccharomycetales(100);f__Metschnikowiaceae(100);g__Metschnikowia(100);g__Metschnikowia_unclassified(100); | Metschnikowia | 13 | Saprotroph | \|Undefined Saprotroph\| | Yeast | NULL | Probable | As nectar-tap_saprotroph (Põlme et al. 2020); Decay sugar-rich substrates (Põlme et al. 2020) | Põlme S, et al. 2020. Fungal Diversity 105:1-16 ((https://doi.org/10.1007/s13225-020-00466-2)) |
| Otu52062 | 1 | k__Fungi(100);p__Ascomycota(100);c__Saccharomycetes(100);o__Saccharomycetales(100);f__Metschnikowiaceae(100);g__Metschnikowia(100);s__Metschnikowia_pulcherrima(100); | Metschnikowia | 13 | Saprotroph | \|Undefined Saprotroph\| | Yeast | NULL | Probable | As nectar-tap_saprotroph (Põlme et al. 2020); Decay sugar-rich substrates (Põlme et al. 2020) | Põlme S, et al. 2020. Fungal Diversity 105:1-16 ((https://doi.org/10.1007/s13225-020-00466-2)) |
| Otu52063 | 1 | k__Fungi(100);p__Ascomycota(100);c__Saccharomycetes(100);o__Saccharomycetales(100);f__Metschnikowiaceae(100);g__Metschnikowia(100);s__Metschnikowia_pulcherrima(100); | Metschnikowia | 13 | Saprotroph | \|Undefined Saprotroph\| | Yeast | NULL | Probable | As nectar-tap_saprotroph (Põlme et al. 2020); Decay sugar-rich substrates (Põlme et al. 2020) | Põlme S, et al. 2020. Fungal Diversity 105:1-16 ((https://doi.org/10.1007/s13225-020-00466-2)) |
| Otu52064 | 1 | k__Fungi(100);p__Ascomycota(100);c__Saccharomycetes(100);o__Saccharomycetales(100);f__Metschnikowiaceae(100);g__Metschnikowia(100);s__Metschnikowia_pulcherrima(100); | Metschnikowia | 13 | Saprotroph | \|Undefined Saprotroph\| | Yeast | NULL | Probable | As nectar-tap_saprotroph (Põlme et al. 2020); Decay sugar-rich substrates (Põlme et al. 2020) | Põlme S, et al. 2020. Fungal Diversity 105:1-16 ((https://doi.org/10.1007/s13225-020-00466-2)) |
| Otu52065 | 1 | k__Fungi(100);p__Ascomycota(100);c__Saccharomycetes(100);o__Saccharomycetales(100);f__Metschnikowiaceae(100);g__Metschnikowia(100);s__Metschnikowia_viticola(100); | Metschnikowia | 13 | Saprotroph | \|Undefined Saprotroph\| | Yeast | NULL | Probable | As nectar-tap_saprotroph (Põlme et al. 2020); Decay sugar-rich substrates (Põlme et al. 2020) | Põlme S, et al. 2020. Fungal Diversity 105:1-16 ((https://doi.org/10.1007/s13225-020-00466-2)) |
| Otu52066 | 1 | k__Fungi(100);p__Ascomycota(100);c__Saccharomycetes(100);o__Saccharomycetales(100);f__Metschnikowiaceae(100);g__Metschnikowia(100);s__Metschnikowia_pulcherrima(100); | Metschnikowia | 13 | Saprotroph | \|Undefined Saprotroph\| | Yeast | NULL | Probable | As nectar-tap_saprotroph (Põlme et al. 2020); Decay sugar-rich substrates (Põlme et al. 2020) | Põlme S, et al. 2020. Fungal Diversity 105:1-16 ((https://doi.org/10.1007/s13225-020-00466-2)) |
| Otu52067 | 1 | k__Fungi(100);p__Ascomycota(100);c__Saccharomycetes(100);o__Saccharomycetales(100);f__Metschnikowiaceae(100);g__Metschnikowia(100);s__Metschnikowia_pulcherrima(100); | Metschnikowia | 13 | Saprotroph | \|Undefined Saprotroph\| | Yeast | NULL | Probable | As nectar-tap_saprotroph (Põlme et al. 2020); Decay sugar-rich substrates (Põlme et al. 2020) | Põlme S, et al. 2020. Fungal Diversity 105:1-16 ((https://doi.org/10.1007/s13225-020-00466-2)) |
| Otu52068 | 1 | k__Fungi(100);p__Ascomycota(100);c__Saccharomycetes(100);o__Saccharomycetales(100);f__Metschnikowiaceae(100);g__Metschnikowia(100);s__Metschnikowia_pulcherrima(100); | Metschnikowia | 13 | Saprotroph | \|Undefined Saprotroph\| | Yeast | NULL | Probable | As nectar-tap_saprotroph (Põlme et al. 2020); Decay sugar-rich substrates (Põlme et al. 2020) | Põlme S, et al. 2020. Fungal Diversity 105:1-16 ((https://doi.org/10.1007/s13225-020-00466-2)) |
| Otu52069 | 1 | k__Fungi(100);p__Ascomycota(100);c__Saccharomycetes(100);o__Saccharomycetales(100);f__Metschnikowiaceae(100);g__Metschnikowia(100);s__Metschnikowia_pulcherrima(100); | Metschnikowia | 13 | Saprotroph | \|Undefined Saprotroph\| | Yeast | NULL | Probable | As nectar-tap_saprotroph (Põlme et al. 2020); Decay sugar-rich substrates (Põlme et al. 2020) | Põlme S, et al. 2020. Fungal Diversity 105:1-16 ((https://doi.org/10.1007/s13225-020-00466-2)) |
| Otu52070 | 1 | k__Fungi(100);p__Ascomycota(100);c__Saccharomycetes(100);o__Saccharomycetales(100);f__Metschnikowiaceae(100);g__Metschnikowia(100);s__Metschnikowia_pulcherrima(100); | Metschnikowia | 13 | Saprotroph | \|Undefined Saprotroph\| | Yeast | NULL | Probable | As nectar-tap_saprotroph (Põlme et al. 2020); Decay sugar-rich substrates (Põlme et al. 2020) | Põlme S, et al. 2020. Fungal Diversity 105:1-16 ((https://doi.org/10.1007/s13225-020-00466-2)) |
| Otu52071 | 1 | k__Fungi(100);p__Ascomycota(100);c__Saccharomycetes(100);o__Saccharomycetales(100);f__Metschnikowiaceae(100);g__Metschnikowia(100);g__Metschnikowia_unclassified(100); | Metschnikowia | 13 | Saprotroph | \|Undefined Saprotroph\| | Yeast | NULL | Probable | As nectar-tap_saprotroph (Põlme et al. 2020); Decay sugar-rich substrates (Põlme et al. 2020) | Põlme S, et al. 2020. Fungal Diversity 105:1-16 ((https://doi.org/10.1007/s13225-020-00466-2)) |
| Otu52072 | 1 | k__Fungi(100);p__Ascomycota(100);c__Saccharomycetes(100);o__Saccharomycetales(100);f__Metschnikowiaceae(100);g__Metschnikowia(100);s__Metschnikowia_pulcherrima(100); | Metschnikowia | 13 | Saprotroph | \|Undefined Saprotroph\| | Yeast | NULL | Probable | As nectar-tap_saprotroph (Põlme et al. 2020); Decay sugar-rich substrates (Põlme et al. 2020) | Põlme S, et al. 2020. Fungal Diversity 105:1-16 ((https://doi.org/10.1007/s13225-020-00466-2)) |
| Otu52074 | 1 | k__Fungi(100);p__Ascomycota(100);c__Saccharomycetes(100);o__Saccharomycetales(100);f__Metschnikowiaceae(100);g__Metschnikowia(100);s__Metschnikowia_pulcherrima(100); | Metschnikowia | 13 | Saprotroph | \|Undefined Saprotroph\| | Yeast | NULL | Probable | As nectar-tap_saprotroph (Põlme et al. 2020); Decay sugar-rich substrates (Põlme et al. 2020) | Põlme S, et al. 2020. Fungal Diversity 105:1-16 ((https://doi.org/10.1007/s13225-020-00466-2)) |
| Otu52075 | 1 | k__Fungi(100);p__Ascomycota(100);c__Saccharomycetes(100);o__Saccharomycetales(100);f__Metschnikowiaceae(100);g__Metschnikowia(100);s__Metschnikowia_pulcherrima(100); | Metschnikowia | 13 | Saprotroph | \|Undefined Saprotroph\| | Yeast | NULL | Probable | As nectar-tap_saprotroph (Põlme et al. 2020); Decay sugar-rich substrates (Põlme et al. 2020) | Põlme S, et al. 2020. Fungal Diversity 105:1-16 ((https://doi.org/10.1007/s13225-020-00466-2)) |
| Otu52076 | 1 | k__Fungi(100);p__Ascomycota(100);c__Saccharomycetes(100);o__Saccharomycetales(100);f__Metschnikowiaceae(100);g__Metschnikowia(100);s__Metschnikowia_pulcherrima(100); | Metschnikowia | 13 | Saprotroph | \|Undefined Saprotroph\| | Yeast | NULL | Probable | As nectar-tap_saprotroph (Põlme et al. 2020); Decay sugar-rich substrates (Põlme et al. 2020) | Põlme S, et al. 2020. Fungal Diversity 105:1-16 ((https://doi.org/10.1007/s13225-020-00466-2)) |
| Otu52077 | 1 | k__Fungi(100);p__Ascomycota(100);c__Saccharomycetes(100);o__Saccharomycetales(100);f__Metschnikowiaceae(100);g__Metschnikowia(100);s__Metschnikowia_pulcherrima(100); | Metschnikowia | 13 | Saprotroph | \|Undefined Saprotroph\| | Yeast | NULL | Probable | As nectar-tap_saprotroph (Põlme et al. 2020); Decay sugar-rich substrates (Põlme et al. 2020) | Põlme S, et al. 2020. Fungal Diversity 105:1-16 ((https://doi.org/10.1007/s13225-020-00466-2)) |
| Otu52078 | 1 | k__Fungi(100);p__Ascomycota(100);c__Saccharomycetes(100);o__Saccharomycetales(100);f__Metschnikowiaceae(100);g__Metschnikowia(100);s__Metschnikowia_pulcherrima(100); | Metschnikowia | 13 | Saprotroph | \|Undefined Saprotroph\| | Yeast | NULL | Probable | As nectar-tap_saprotroph (Põlme et al. 2020); Decay sugar-rich substrates (Põlme et al. 2020) | Põlme S, et al. 2020. Fungal Diversity 105:1-16 ((https://doi.org/10.1007/s13225-020-00466-2)) |
| Otu52079 | 1 | k__Fungi(100);p__Ascomycota(100);c__Saccharomycetes(100);o__Saccharomycetales(100);f__Metschnikowiaceae(100);g__Metschnikowia(100);s__Metschnikowia_pulcherrima(100); | Metschnikowia | 13 | Saprotroph | \|Undefined Saprotroph\| | Yeast | NULL | Probable | As nectar-tap_saprotroph (Põlme et al. 2020); Decay sugar-rich substrates (Põlme et al. 2020) | Põlme S, et al. 2020. Fungal Diversity 105:1-16 ((https://doi.org/10.1007/s13225-020-00466-2)) |
| Otu52080 | 1 | k__Fungi(100);p__Ascomycota(100);c__Saccharomycetes(100);o__Saccharomycetales(100);f__Metschnikowiaceae(100);g__Metschnikowia(100);s__Metschnikowia_pulcherrima(100); | Metschnikowia | 13 | Saprotroph | \|Undefined Saprotroph\| | Yeast | NULL | Probable | As nectar-tap_saprotroph (Põlme et al. 2020); Decay sugar-rich substrates (Põlme et al. 2020) | Põlme S, et al. 2020. Fungal Diversity 105:1-16 ((https://doi.org/10.1007/s13225-020-00466-2)) |
| Otu52081 | 1 | k__Fungi(100);p__Ascomycota(100);c__Saccharomycetes(100);o__Saccharomycetales(100);f__Metschnikowiaceae(100);g__Metschnikowia(100);s__Metschnikowia_pulcherrima(100); | Metschnikowia | 13 | Saprotroph | \|Undefined Saprotroph\| | Yeast | NULL | Probable | As nectar-tap_saprotroph (Põlme et al. 2020); Decay sugar-rich substrates (Põlme et al. 2020) | Põlme S, et al. 2020. Fungal Diversity 105:1-16 ((https://doi.org/10.1007/s13225-020-00466-2)) |
| Otu52082 | 1 | k__Fungi(100);p__Ascomycota(100);c__Saccharomycetes(100);o__Saccharomycetales(100);f__Metschnikowiaceae(100);g__Metschnikowia(100);g__Metschnikowia_unclassified(100); | Metschnikowia | 13 | Saprotroph | \|Undefined Saprotroph\| | Yeast | NULL | Probable | As nectar-tap_saprotroph (Põlme et al. 2020); Decay sugar-rich substrates (Põlme et al. 2020) | Põlme S, et al. 2020. Fungal Diversity 105:1-16 ((https://doi.org/10.1007/s13225-020-00466-2)) |
| Otu52083 | 1 | k__Fungi(100);p__Ascomycota(100);c__Saccharomycetes(100);o__Saccharomycetales(100);f__Metschnikowiaceae(100);g__Metschnikowia(100);g__Metschnikowia_unclassified(100); | Metschnikowia | 13 | Saprotroph | \|Undefined Saprotroph\| | Yeast | NULL | Probable | As nectar-tap_saprotroph (Põlme et al. 2020); Decay sugar-rich substrates (Põlme et al. 2020) | Põlme S, et al. 2020. Fungal Diversity 105:1-16 ((https://doi.org/10.1007/s13225-020-00466-2)) |
| Otu52084 | 1 | k__Fungi(100);p__Ascomycota(100);c__Saccharomycetes(100);o__Saccharomycetales(100);f__Metschnikowiaceae(100);g__Metschnikowia(100);s__Metschnikowia_pulcherrima(100); | Metschnikowia | 13 | Saprotroph | \|Undefined Saprotroph\| | Yeast | NULL | Probable | As nectar-tap_saprotroph (Põlme et al. 2020); Decay sugar-rich substrates (Põlme et al. 2020) | Põlme S, et al. 2020. Fungal Diversity 105:1-16 ((https://doi.org/10.1007/s13225-020-00466-2)) |
| Otu52085 | 1 | k__Fungi(100);p__Ascomycota(100);c__Saccharomycetes(100);o__Saccharomycetales(100);f__Metschnikowiaceae(100);g__Metschnikowia(100);g__Metschnikowia_unclassified(100); | Metschnikowia | 13 | Saprotroph | \|Undefined Saprotroph\| | Yeast | NULL | Probable | As nectar-tap_saprotroph (Põlme et al. 2020); Decay sugar-rich substrates (Põlme et al. 2020) | Põlme S, et al. 2020. Fungal Diversity 105:1-16 ((https://doi.org/10.1007/s13225-020-00466-2)) |
| Otu52086 | 1 | k__Fungi(100);p__Ascomycota(100);c__Saccharomycetes(100);o__Saccharomycetales(100);f__Metschnikowiaceae(100);g__Metschnikowia(100);g__Metschnikowia_unclassified(100); | Metschnikowia | 13 | Saprotroph | \|Undefined Saprotroph\| | Yeast | NULL | Probable | As nectar-tap_saprotroph (Põlme et al. 2020); Decay sugar-rich substrates (Põlme et al. 2020) | Põlme S, et al. 2020. Fungal Diversity 105:1-16 ((https://doi.org/10.1007/s13225-020-00466-2)) |
| Otu52087 | 1 | k__Fungi(100);p__Ascomycota(100);c__Saccharomycetes(100);o__Saccharomycetales(100);f__Metschnikowiaceae(100);g__Metschnikowia(100);s__Metschnikowia_pulcherrima(100); | Metschnikowia | 13 | Saprotroph | \|Undefined Saprotroph\| | Yeast | NULL | Probable | As nectar-tap_saprotroph (Põlme et al. 2020); Decay sugar-rich substrates (Põlme et al. 2020) | Põlme S, et al. 2020. Fungal Diversity 105:1-16 ((https://doi.org/10.1007/s13225-020-00466-2)) |
| Otu52088 | 1 | k__Fungi(100);p__Ascomycota(100);c__Saccharomycetes(100);o__Saccharomycetales(100);f__Metschnikowiaceae(100);g__Metschnikowia(100);s__Metschnikowia_pulcherrima(100); | Metschnikowia | 13 | Saprotroph | \|Undefined Saprotroph\| | Yeast | NULL | Probable | As nectar-tap_saprotroph (Põlme et al. 2020); Decay sugar-rich substrates (Põlme et al. 2020) | Põlme S, et al. 2020. Fungal Diversity 105:1-16 ((https://doi.org/10.1007/s13225-020-00466-2)) |
| Otu52089 | 1 | k__Fungi(100);p__Ascomycota(100);c__Saccharomycetes(100);o__Saccharomycetales(100);f__Metschnikowiaceae(100);g__Metschnikowia(100);s__Metschnikowia_pulcherrima(100); | Metschnikowia | 13 | Saprotroph | \|Undefined Saprotroph\| | Yeast | NULL | Probable | As nectar-tap_saprotroph (Põlme et al. 2020); Decay sugar-rich substrates (Põlme et al. 2020) | Põlme S, et al. 2020. Fungal Diversity 105:1-16 ((https://doi.org/10.1007/s13225-020-00466-2)) |
| Otu52090 | 1 | k__Fungi(100);p__Ascomycota(100);c__Saccharomycetes(100);o__Saccharomycetales(100);f__Metschnikowiaceae(100);g__Metschnikowia(100);s__Metschnikowia_pulcherrima(100); | Metschnikowia | 13 | Saprotroph | \|Undefined Saprotroph\| | Yeast | NULL | Probable | As nectar-tap_saprotroph (Põlme et al. 2020); Decay sugar-rich substrates (Põlme et al. 2020) | Põlme S, et al. 2020. Fungal Diversity 105:1-16 ((https://doi.org/10.1007/s13225-020-00466-2)) |
| Otu52091 | 1 | k__Fungi(100);p__Ascomycota(100);c__Saccharomycetes(100);o__Saccharomycetales(100);f__Metschnikowiaceae(100);g__Metschnikowia(100);s__Metschnikowia_pulcherrima(100); | Metschnikowia | 13 | Saprotroph | \|Undefined Saprotroph\| | Yeast | NULL | Probable | As nectar-tap_saprotroph (Põlme et al. 2020); Decay sugar-rich substrates (Põlme et al. 2020) | Põlme S, et al. 2020. Fungal Diversity 105:1-16 ((https://doi.org/10.1007/s13225-020-00466-2)) |
| Otu52092 | 1 | k__Fungi(100);p__Ascomycota(100);c__Saccharomycetes(100);o__Saccharomycetales(100);f__Metschnikowiaceae(100);g__Metschnikowia(100);s__Metschnikowia_pulcherrima(100); | Metschnikowia | 13 | Saprotroph | \|Undefined Saprotroph\| | Yeast | NULL | Probable | As nectar-tap_saprotroph (Põlme et al. 2020); Decay sugar-rich substrates (Põlme et al. 2020) | Põlme S, et al. 2020. Fungal Diversity 105:1-16 ((https://doi.org/10.1007/s13225-020-00466-2)) |
| Otu52093 | 1 | k__Fungi(100);p__Ascomycota(100);c__Saccharomycetes(100);o__Saccharomycetales(100);f__Metschnikowiaceae(100);g__Metschnikowia(100);s__Metschnikowia_pulcherrima(100); | Metschnikowia | 13 | Saprotroph | \|Undefined Saprotroph\| | Yeast | NULL | Probable | As nectar-tap_saprotroph (Põlme et al. 2020); Decay sugar-rich substrates (Põlme et al. 2020) | Põlme S, et al. 2020. Fungal Diversity 105:1-16 ((https://doi.org/10.1007/s13225-020-00466-2)) |
| Otu52094 | 1 | k__Fungi(100);p__Ascomycota(100);c__Saccharomycetes(100);o__Saccharomycetales(100);f__Metschnikowiaceae(100);g__Metschnikowia(100);s__Metschnikowia_pulcherrima(100); | Metschnikowia | 13 | Saprotroph | \|Undefined Saprotroph\| | Yeast | NULL | Probable | As nectar-tap_saprotroph (Põlme et al. 2020); Decay sugar-rich substrates (Põlme et al. 2020) | Põlme S, et al. 2020. Fungal Diversity 105:1-16 ((https://doi.org/10.1007/s13225-020-00466-2)) |
| Otu52095 | 1 | k__Fungi(100);p__Ascomycota(100);c__Saccharomycetes(100);o__Saccharomycetales(100);f__Metschnikowiaceae(100);g__Metschnikowia(100);s__Metschnikowia_pulcherrima(100); | Metschnikowia | 13 | Saprotroph | \|Undefined Saprotroph\| | Yeast | NULL | Probable | As nectar-tap_saprotroph (Põlme et al. 2020); Decay sugar-rich substrates (Põlme et al. 2020) | Põlme S, et al. 2020. Fungal Diversity 105:1-16 ((https://doi.org/10.1007/s13225-020-00466-2)) |
| Otu52097 | 1 | k__Fungi(100);p__Ascomycota(100);c__Saccharomycetes(100);o__Saccharomycetales(100);f__Metschnikowiaceae(100);g__Metschnikowia(100);g__Metschnikowia_unclassified(100); | Metschnikowia | 13 | Saprotroph | \|Undefined Saprotroph\| | Yeast | NULL | Probable | As nectar-tap_saprotroph (Põlme et al. 2020); Decay sugar-rich substrates (Põlme et al. 2020) | Põlme S, et al. 2020. Fungal Diversity 105:1-16 ((https://doi.org/10.1007/s13225-020-00466-2)) |
| Otu52098 | 1 | k__Fungi(100);p__Ascomycota(100);c__Saccharomycetes(100);o__Saccharomycetales(100);f__Metschnikowiaceae(100);g__Metschnikowia(100);s__Metschnikowia_pulcherrima(100); | Metschnikowia | 13 | Saprotroph | \|Undefined Saprotroph\| | Yeast | NULL | Probable | As nectar-tap_saprotroph (Põlme et al. 2020); Decay sugar-rich substrates (Põlme et al. 2020) | Põlme S, et al. 2020. Fungal Diversity 105:1-16 ((https://doi.org/10.1007/s13225-020-00466-2)) |
| Otu52100 | 1 | k__Fungi(100);p__Ascomycota(100);c__Saccharomycetes(100);o__Saccharomycetales(100);f__Metschnikowiaceae(100);g__Metschnikowia(100);g__Metschnikowia_unclassified(100); | Metschnikowia | 13 | Saprotroph | \|Undefined Saprotroph\| | Yeast | NULL | Probable | As nectar-tap_saprotroph (Põlme et al. 2020); Decay sugar-rich substrates (Põlme et al. 2020) | Põlme S, et al. 2020. Fungal Diversity 105:1-16 ((https://doi.org/10.1007/s13225-020-00466-2)) |
| Otu52101 | 1 | k__Fungi(100);p__Ascomycota(100);c__Saccharomycetes(100);o__Saccharomycetales(100);f__Metschnikowiaceae(100);g__Metschnikowia(100);s__Metschnikowia_pulcherrima(100); | Metschnikowia | 13 | Saprotroph | \|Undefined Saprotroph\| | Yeast | NULL | Probable | As nectar-tap_saprotroph (Põlme et al. 2020); Decay sugar-rich substrates (Põlme et al. 2020) | Põlme S, et al. 2020. Fungal Diversity 105:1-16 ((https://doi.org/10.1007/s13225-020-00466-2)) |
| Otu52102 | 1 | k__Fungi(100);p__Ascomycota(100);c__Saccharomycetes(100);o__Saccharomycetales(100);f__Metschnikowiaceae(100);g__Metschnikowia(100);s__Metschnikowia_pulcherrima(100); | Metschnikowia | 13 | Saprotroph | \|Undefined Saprotroph\| | Yeast | NULL | Probable | As nectar-tap_saprotroph (Põlme et al. 2020); Decay sugar-rich substrates (Põlme et al. 2020) | Põlme S, et al. 2020. Fungal Diversity 105:1-16 ((https://doi.org/10.1007/s13225-020-00466-2)) |
| Otu52103 | 1 | k__Fungi(100);p__Ascomycota(100);c__Saccharomycetes(100);o__Saccharomycetales(100);f__Metschnikowiaceae(100);g__Metschnikowia(100);s__Metschnikowia_pulcherrima(100); | Metschnikowia | 13 | Saprotroph | \|Undefined Saprotroph\| | Yeast | NULL | Probable | As nectar-tap_saprotroph (Põlme et al. 2020); Decay sugar-rich substrates (Põlme et al. 2020) | Põlme S, et al. 2020. Fungal Diversity 105:1-16 ((https://doi.org/10.1007/s13225-020-00466-2)) |
| Otu52104 | 1 | k__Fungi(100);p__Ascomycota(100);c__Saccharomycetes(100);o__Saccharomycetales(100);f__Metschnikowiaceae(100);g__Metschnikowia(100);s__Metschnikowia_pulcherrima(100); | Metschnikowia | 13 | Saprotroph | \|Undefined Saprotroph\| | Yeast | NULL | Probable | As nectar-tap_saprotroph (Põlme et al. 2020); Decay sugar-rich substrates (Põlme et al. 2020) | Põlme S, et al. 2020. Fungal Diversity 105:1-16 ((https://doi.org/10.1007/s13225-020-00466-2)) |
| Otu52105 | 1 | k__Fungi(100);p__Ascomycota(100);c__Saccharomycetes(100);o__Saccharomycetales(100);f__Metschnikowiaceae(100);g__Metschnikowia(100);s__Metschnikowia_pulcherrima(100); | Metschnikowia | 13 | Saprotroph | \|Undefined Saprotroph\| | Yeast | NULL | Probable | As nectar-tap_saprotroph (Põlme et al. 2020); Decay sugar-rich substrates (Põlme et al. 2020) | Põlme S, et al. 2020. Fungal Diversity 105:1-16 ((https://doi.org/10.1007/s13225-020-00466-2)) |
| Otu52106 | 1 | k__Fungi(100);p__Ascomycota(100);c__Saccharomycetes(100);o__Saccharomycetales(100);f__Metschnikowiaceae(100);g__Metschnikowia(100);s__Metschnikowia_pulcherrima(100); | Metschnikowia | 13 | Saprotroph | \|Undefined Saprotroph\| | Yeast | NULL | Probable | As nectar-tap_saprotroph (Põlme et al. 2020); Decay sugar-rich substrates (Põlme et al. 2020) | Põlme S, et al. 2020. Fungal Diversity 105:1-16 ((https://doi.org/10.1007/s13225-020-00466-2)) |
| Otu52107 | 1 | k__Fungi(100);p__Ascomycota(100);c__Saccharomycetes(100);o__Saccharomycetales(100);f__Metschnikowiaceae(100);g__Metschnikowia(100);s__Metschnikowia_pulcherrima(100); | Metschnikowia | 13 | Saprotroph | \|Undefined Saprotroph\| | Yeast | NULL | Probable | As nectar-tap_saprotroph (Põlme et al. 2020); Decay sugar-rich substrates (Põlme et al. 2020) | Põlme S, et al. 2020. Fungal Diversity 105:1-16 ((https://doi.org/10.1007/s13225-020-00466-2)) |
| Otu52108 | 1 | k__Fungi(100);p__Ascomycota(100);c__Saccharomycetes(100);o__Saccharomycetales(100);f__Metschnikowiaceae(100);g__Metschnikowia(100);s__Metschnikowia_pulcherrima(100); | Metschnikowia | 13 | Saprotroph | \|Undefined Saprotroph\| | Yeast | NULL | Probable | As nectar-tap_saprotroph (Põlme et al. 2020); Decay sugar-rich substrates (Põlme et al. 2020) | Põlme S, et al. 2020. Fungal Diversity 105:1-16 ((https://doi.org/10.1007/s13225-020-00466-2)) |
| Otu52109 | 1 | k__Fungi(100);p__Ascomycota(100);c__Saccharomycetes(100);o__Saccharomycetales(100);f__Metschnikowiaceae(100);g__Metschnikowia(100);s__Metschnikowia_pulcherrima(100); | Metschnikowia | 13 | Saprotroph | \|Undefined Saprotroph\| | Yeast | NULL | Probable | As nectar-tap_saprotroph (Põlme et al. 2020); Decay sugar-rich substrates (Põlme et al. 2020) | Põlme S, et al. 2020. Fungal Diversity 105:1-16 ((https://doi.org/10.1007/s13225-020-00466-2)) |
| Otu52110 | 1 | k__Fungi(100);p__Ascomycota(100);c__Saccharomycetes(100);o__Saccharomycetales(100);f__Metschnikowiaceae(100);g__Metschnikowia(100);s__Metschnikowia_pulcherrima(100); | Metschnikowia | 13 | Saprotroph | \|Undefined Saprotroph\| | Yeast | NULL | Probable | As nectar-tap_saprotroph (Põlme et al. 2020); Decay sugar-rich substrates (Põlme et al. 2020) | Põlme S, et al. 2020. Fungal Diversity 105:1-16 ((https://doi.org/10.1007/s13225-020-00466-2)) |
| Otu52111 | 1 | k__Fungi(100);p__Ascomycota(100);c__Saccharomycetes(100);o__Saccharomycetales(100);f__Metschnikowiaceae(100);g__Metschnikowia(100);g__Metschnikowia_unclassified(100); | Metschnikowia | 13 | Saprotroph | \|Undefined Saprotroph\| | Yeast | NULL | Probable | As nectar-tap_saprotroph (Põlme et al. 2020); Decay sugar-rich substrates (Põlme et al. 2020) | Põlme S, et al. 2020. Fungal Diversity 105:1-16 ((https://doi.org/10.1007/s13225-020-00466-2)) |
| Otu52114 | 1 | k__Fungi(100);p__Ascomycota(100);c__Saccharomycetes(100);o__Saccharomycetales(100);f__Metschnikowiaceae(100);g__Metschnikowia(100);s__Metschnikowia_pulcherrima(100); | Metschnikowia | 13 | Saprotroph | \|Undefined Saprotroph\| | Yeast | NULL | Probable | As nectar-tap_saprotroph (Põlme et al. 2020); Decay sugar-rich substrates (Põlme et al. 2020) | Põlme S, et al. 2020. Fungal Diversity 105:1-16 ((https://doi.org/10.1007/s13225-020-00466-2)) |
| Otu52117 | 1 | k__Fungi(100);p__Ascomycota(100);c__Saccharomycetes(100);o__Saccharomycetales(100);f__Metschnikowiaceae(100);g__Metschnikowia(100);s__Metschnikowia_pulcherrima(100); | Metschnikowia | 13 | Saprotroph | \|Undefined Saprotroph\| | Yeast | NULL | Probable | As nectar-tap_saprotroph (Põlme et al. 2020); Decay sugar-rich substrates (Põlme et al. 2020) | Põlme S, et al. 2020. Fungal Diversity 105:1-16 ((https://doi.org/10.1007/s13225-020-00466-2)) |
| Otu52118 | 1 | k__Fungi(100);p__Ascomycota(100);c__Saccharomycetes(100);o__Saccharomycetales(100);f__Metschnikowiaceae(100);g__Metschnikowia(100);s__Metschnikowia_pulcherrima(100); | Metschnikowia | 13 | Saprotroph | \|Undefined Saprotroph\| | Yeast | NULL | Probable | As nectar-tap_saprotroph (Põlme et al. 2020); Decay sugar-rich substrates (Põlme et al. 2020) | Põlme S, et al. 2020. Fungal Diversity 105:1-16 ((https://doi.org/10.1007/s13225-020-00466-2)) |
| Otu52119 | 1 | k__Fungi(100);p__Ascomycota(100);c__Saccharomycetes(100);o__Saccharomycetales(100);f__Metschnikowiaceae(100);g__Metschnikowia(100);g__Metschnikowia_unclassified(100); | Metschnikowia | 13 | Saprotroph | \|Undefined Saprotroph\| | Yeast | NULL | Probable | As nectar-tap_saprotroph (Põlme et al. 2020); Decay sugar-rich substrates (Põlme et al. 2020) | Põlme S, et al. 2020. Fungal Diversity 105:1-16 ((https://doi.org/10.1007/s13225-020-00466-2)) |
| Otu52120 | 1 | k__Fungi(100);p__Ascomycota(100);c__Saccharomycetes(100);o__Saccharomycetales(100);f__Metschnikowiaceae(100);g__Metschnikowia(100);s__Metschnikowia_pulcherrima(100); | Metschnikowia | 13 | Saprotroph | \|Undefined Saprotroph\| | Yeast | NULL | Probable | As nectar-tap_saprotroph (Põlme et al. 2020); Decay sugar-rich substrates (Põlme et al. 2020) | Põlme S, et al. 2020. Fungal Diversity 105:1-16 ((https://doi.org/10.1007/s13225-020-00466-2)) |
| Otu52121 | 1 | k__Fungi(100);p__Ascomycota(100);c__Saccharomycetes(100);o__Saccharomycetales(100);f__Metschnikowiaceae(100);g__Metschnikowia(100);g__Metschnikowia_unclassified(100); | Metschnikowia | 13 | Saprotroph | \|Undefined Saprotroph\| | Yeast | NULL | Probable | As nectar-tap_saprotroph (Põlme et al. 2020); Decay sugar-rich substrates (Põlme et al. 2020) | Põlme S, et al. 2020. Fungal Diversity 105:1-16 ((https://doi.org/10.1007/s13225-020-00466-2)) |
| Otu52122 | 1 | k__Fungi(100);p__Ascomycota(100);c__Saccharomycetes(100);o__Saccharomycetales(100);f__Metschnikowiaceae(100);g__Metschnikowia(100);s__Metschnikowia_pulcherrima(100); | Metschnikowia | 13 | Saprotroph | \|Undefined Saprotroph\| | Yeast | NULL | Probable | As nectar-tap_saprotroph (Põlme et al. 2020); Decay sugar-rich substrates (Põlme et al. 2020) | Põlme S, et al. 2020. Fungal Diversity 105:1-16 ((https://doi.org/10.1007/s13225-020-00466-2)) |
| Otu52123 | 1 | k__Fungi(100);p__Ascomycota(100);c__Saccharomycetes(100);o__Saccharomycetales(100);f__Metschnikowiaceae(100);g__Metschnikowia(100);s__Metschnikowia_pulcherrima(100); | Metschnikowia | 13 | Saprotroph | \|Undefined Saprotroph\| | Yeast | NULL | Probable | As nectar-tap_saprotroph (Põlme et al. 2020); Decay sugar-rich substrates (Põlme et al. 2020) | Põlme S, et al. 2020. Fungal Diversity 105:1-16 ((https://doi.org/10.1007/s13225-020-00466-2)) |
| Otu52124 | 1 | k__Fungi(100);p__Ascomycota(100);c__Saccharomycetes(100);o__Saccharomycetales(100);f__Metschnikowiaceae(100);g__Metschnikowia(100);g__Metschnikowia_unclassified(100); | Metschnikowia | 13 | Saprotroph | \|Undefined Saprotroph\| | Yeast | NULL | Probable | As nectar-tap_saprotroph (Põlme et al. 2020); Decay sugar-rich substrates (Põlme et al. 2020) | Põlme S, et al. 2020. Fungal Diversity 105:1-16 ((https://doi.org/10.1007/s13225-020-00466-2)) |
| Otu52125 | 1 | k__Fungi(100);p__Ascomycota(100);c__Saccharomycetes(100);o__Saccharomycetales(100);f__Metschnikowiaceae(100);g__Metschnikowia(100);g__Metschnikowia_unclassified(100); | Metschnikowia | 13 | Saprotroph | \|Undefined Saprotroph\| | Yeast | NULL | Probable | As nectar-tap_saprotroph (Põlme et al. 2020); Decay sugar-rich substrates (Põlme et al. 2020) | Põlme S, et al. 2020. Fungal Diversity 105:1-16 ((https://doi.org/10.1007/s13225-020-00466-2)) |
| Otu52126 | 1 | k__Fungi(100);p__Ascomycota(100);c__Saccharomycetes(100);o__Saccharomycetales(100);f__Metschnikowiaceae(100);g__Metschnikowia(100);g__Metschnikowia_unclassified(100); | Metschnikowia | 13 | Saprotroph | \|Undefined Saprotroph\| | Yeast | NULL | Probable | As nectar-tap_saprotroph (Põlme et al. 2020); Decay sugar-rich substrates (Põlme et al. 2020) | Põlme S, et al. 2020. Fungal Diversity 105:1-16 ((https://doi.org/10.1007/s13225-020-00466-2)) |
| Otu52127 | 1 | k__Fungi(100);p__Ascomycota(100);c__Saccharomycetes(100);o__Saccharomycetales(100);f__Metschnikowiaceae(100);g__Metschnikowia(100);s__Metschnikowia_pulcherrima(100); | Metschnikowia | 13 | Saprotroph | \|Undefined Saprotroph\| | Yeast | NULL | Probable | As nectar-tap_saprotroph (Põlme et al. 2020); Decay sugar-rich substrates (Põlme et al. 2020) | Põlme S, et al. 2020. Fungal Diversity 105:1-16 ((https://doi.org/10.1007/s13225-020-00466-2)) |
| Otu52128 | 1 | k__Fungi(100);p__Ascomycota(100);c__Saccharomycetes(100);o__Saccharomycetales(100);f__Metschnikowiaceae(100);g__Metschnikowia(100);s__Metschnikowia_pulcherrima(100); | Metschnikowia | 13 | Saprotroph | \|Undefined Saprotroph\| | Yeast | NULL | Probable | As nectar-tap_saprotroph (Põlme et al. 2020); Decay sugar-rich substrates (Põlme et al. 2020) | Põlme S, et al. 2020. Fungal Diversity 105:1-16 ((https://doi.org/10.1007/s13225-020-00466-2)) |
| Otu52129 | 1 | k__Fungi(100);p__Ascomycota(100);c__Saccharomycetes(100);o__Saccharomycetales(100);f__Metschnikowiaceae(100);g__Metschnikowia(100);s__Metschnikowia_pulcherrima(100); | Metschnikowia | 13 | Saprotroph | \|Undefined Saprotroph\| | Yeast | NULL | Probable | As nectar-tap_saprotroph (Põlme et al. 2020); Decay sugar-rich substrates (Põlme et al. 2020) | Põlme S, et al. 2020. Fungal Diversity 105:1-16 ((https://doi.org/10.1007/s13225-020-00466-2)) |
| Otu52130 | 1 | k__Fungi(100);p__Ascomycota(100);c__Saccharomycetes(100);o__Saccharomycetales(100);f__Metschnikowiaceae(100);g__Metschnikowia(100);s__Metschnikowia_pulcherrima(100); | Metschnikowia | 13 | Saprotroph | \|Undefined Saprotroph\| | Yeast | NULL | Probable | As nectar-tap_saprotroph (Põlme et al. 2020); Decay sugar-rich substrates (Põlme et al. 2020) | Põlme S, et al. 2020. Fungal Diversity 105:1-16 ((https://doi.org/10.1007/s13225-020-00466-2)) |
| Otu52131 | 1 | k__Fungi(100);p__Ascomycota(100);c__Saccharomycetes(100);o__Saccharomycetales(100);f__Metschnikowiaceae(100);g__Metschnikowia(100);g__Metschnikowia_unclassified(100); | Metschnikowia | 13 | Saprotroph | \|Undefined Saprotroph\| | Yeast | NULL | Probable | As nectar-tap_saprotroph (Põlme et al. 2020); Decay sugar-rich substrates (Põlme et al. 2020) | Põlme S, et al. 2020. Fungal Diversity 105:1-16 ((https://doi.org/10.1007/s13225-020-00466-2)) |
| Otu52132 | 1 | k__Fungi(100);p__Ascomycota(100);c__Saccharomycetes(100);o__Saccharomycetales(100);f__Metschnikowiaceae(100);g__Metschnikowia(100);s__Metschnikowia_pulcherrima(100); | Metschnikowia | 13 | Saprotroph | \|Undefined Saprotroph\| | Yeast | NULL | Probable | As nectar-tap_saprotroph (Põlme et al. 2020); Decay sugar-rich substrates (Põlme et al. 2020) | Põlme S, et al. 2020. Fungal Diversity 105:1-16 ((https://doi.org/10.1007/s13225-020-00466-2)) |
| Otu52133 | 1 | k__Fungi(100);p__Ascomycota(100);c__Saccharomycetes(100);o__Saccharomycetales(100);f__Metschnikowiaceae(100);g__Metschnikowia(100);s__Metschnikowia_pulcherrima(100); | Metschnikowia | 13 | Saprotroph | \|Undefined Saprotroph\| | Yeast | NULL | Probable | As nectar-tap_saprotroph (Põlme et al. 2020); Decay sugar-rich substrates (Põlme et al. 2020) | Põlme S, et al. 2020. Fungal Diversity 105:1-16 ((https://doi.org/10.1007/s13225-020-00466-2)) |
| Otu52134 | 1 | k__Fungi(100);p__Ascomycota(100);c__Saccharomycetes(100);o__Saccharomycetales(100);f__Metschnikowiaceae(100);g__Metschnikowia(100);s__Metschnikowia_pulcherrima(100); | Metschnikowia | 13 | Saprotroph | \|Undefined Saprotroph\| | Yeast | NULL | Probable | As nectar-tap_saprotroph (Põlme et al. 2020); Decay sugar-rich substrates (Põlme et al. 2020) | Põlme S, et al. 2020. Fungal Diversity 105:1-16 ((https://doi.org/10.1007/s13225-020-00466-2)) |
| Otu52135 | 1 | k__Fungi(100);p__Ascomycota(100);c__Saccharomycetes(100);o__Saccharomycetales(100);f__Metschnikowiaceae(100);g__Metschnikowia(100);s__Metschnikowia_pulcherrima(100); | Metschnikowia | 13 | Saprotroph | \|Undefined Saprotroph\| | Yeast | NULL | Probable | As nectar-tap_saprotroph (Põlme et al. 2020); Decay sugar-rich substrates (Põlme et al. 2020) | Põlme S, et al. 2020. Fungal Diversity 105:1-16 ((https://doi.org/10.1007/s13225-020-00466-2)) |
| Otu52136 | 1 | k__Fungi(100);p__Ascomycota(100);c__Saccharomycetes(100);o__Saccharomycetales(100);f__Metschnikowiaceae(100);g__Metschnikowia(100);s__Metschnikowia_pulcherrima(100); | Metschnikowia | 13 | Saprotroph | \|Undefined Saprotroph\| | Yeast | NULL | Probable | As nectar-tap_saprotroph (Põlme et al. 2020); Decay sugar-rich substrates (Põlme et al. 2020) | Põlme S, et al. 2020. Fungal Diversity 105:1-16 ((https://doi.org/10.1007/s13225-020-00466-2)) |
| Otu52137 | 1 | k__Fungi(100);p__Ascomycota(100);c__Saccharomycetes(100);o__Saccharomycetales(100);f__Metschnikowiaceae(100);g__Metschnikowia(100);s__Metschnikowia_pulcherrima(100); | Metschnikowia | 13 | Saprotroph | \|Undefined Saprotroph\| | Yeast | NULL | Probable | As nectar-tap_saprotroph (Põlme et al. 2020); Decay sugar-rich substrates (Põlme et al. 2020) | Põlme S, et al. 2020. Fungal Diversity 105:1-16 ((https://doi.org/10.1007/s13225-020-00466-2)) |
| Otu52138 | 1 | k__Fungi(100);p__Ascomycota(100);c__Saccharomycetes(100);o__Saccharomycetales(100);f__Metschnikowiaceae(100);g__Metschnikowia(100);s__Metschnikowia_pulcherrima(100); | Metschnikowia | 13 | Saprotroph | \|Undefined Saprotroph\| | Yeast | NULL | Probable | As nectar-tap_saprotroph (Põlme et al. 2020); Decay sugar-rich substrates (Põlme et al. 2020) | Põlme S, et al. 2020. Fungal Diversity 105:1-16 ((https://doi.org/10.1007/s13225-020-00466-2)) |
| Otu52139 | 1 | k__Fungi(100);p__Ascomycota(100);c__Saccharomycetes(100);o__Saccharomycetales(100);f__Metschnikowiaceae(100);g__Metschnikowia(100);g__Metschnikowia_unclassified(100); | Metschnikowia | 13 | Saprotroph | \|Undefined Saprotroph\| | Yeast | NULL | Probable | As nectar-tap_saprotroph (Põlme et al. 2020); Decay sugar-rich substrates (Põlme et al. 2020) | Põlme S, et al. 2020. Fungal Diversity 105:1-16 ((https://doi.org/10.1007/s13225-020-00466-2)) |
| Otu52140 | 1 | k__Fungi(100);p__Ascomycota(100);c__Saccharomycetes(100);o__Saccharomycetales(100);f__Metschnikowiaceae(100);g__Metschnikowia(100);s__Metschnikowia_pulcherrima(100); | Metschnikowia | 13 | Saprotroph | \|Undefined Saprotroph\| | Yeast | NULL | Probable | As nectar-tap_saprotroph (Põlme et al. 2020); Decay sugar-rich substrates (Põlme et al. 2020) | Põlme S, et al. 2020. Fungal Diversity 105:1-16 ((https://doi.org/10.1007/s13225-020-00466-2)) |
| Otu52141 | 1 | k__Fungi(100);p__Ascomycota(100);c__Saccharomycetes(100);o__Saccharomycetales(100);f__Metschnikowiaceae(100);g__Metschnikowia(100);s__Metschnikowia_pulcherrima(100); | Metschnikowia | 13 | Saprotroph | \|Undefined Saprotroph\| | Yeast | NULL | Probable | As nectar-tap_saprotroph (Põlme et al. 2020); Decay sugar-rich substrates (Põlme et al. 2020) | Põlme S, et al. 2020. Fungal Diversity 105:1-16 ((https://doi.org/10.1007/s13225-020-00466-2)) |
| Otu52142 | 1 | k__Fungi(100);p__Ascomycota(100);c__Saccharomycetes(100);o__Saccharomycetales(100);f__Metschnikowiaceae(100);g__Metschnikowia(100);s__Metschnikowia_pulcherrima(100); | Metschnikowia | 13 | Saprotroph | \|Undefined Saprotroph\| | Yeast | NULL | Probable | As nectar-tap_saprotroph (Põlme et al. 2020); Decay sugar-rich substrates (Põlme et al. 2020) | Põlme S, et al. 2020. Fungal Diversity 105:1-16 ((https://doi.org/10.1007/s13225-020-00466-2)) |
| Otu52143 | 1 | k__Fungi(100);p__Ascomycota(100);c__Saccharomycetes(100);o__Saccharomycetales(100);f__Metschnikowiaceae(100);g__Metschnikowia(100);g__Metschnikowia_unclassified(100); | Metschnikowia | 13 | Saprotroph | \|Undefined Saprotroph\| | Yeast | NULL | Probable | As nectar-tap_saprotroph (Põlme et al. 2020); Decay sugar-rich substrates (Põlme et al. 2020) | Põlme S, et al. 2020. Fungal Diversity 105:1-16 ((https://doi.org/10.1007/s13225-020-00466-2)) |
| Otu52144 | 1 | k__Fungi(100);p__Ascomycota(100);c__Saccharomycetes(100);o__Saccharomycetales(100);f__Metschnikowiaceae(100);g__Metschnikowia(100);s__Metschnikowia_pulcherrima(100); | Metschnikowia | 13 | Saprotroph | \|Undefined Saprotroph\| | Yeast | NULL | Probable | As nectar-tap_saprotroph (Põlme et al. 2020); Decay sugar-rich substrates (Põlme et al. 2020) | Põlme S, et al. 2020. Fungal Diversity 105:1-16 ((https://doi.org/10.1007/s13225-020-00466-2)) |
| Otu52145 | 1 | k__Fungi(100);p__Ascomycota(100);c__Saccharomycetes(100);o__Saccharomycetales(100);f__Metschnikowiaceae(100);g__Metschnikowia(100);s__Metschnikowia_pulcherrima(100); | Metschnikowia | 13 | Saprotroph | \|Undefined Saprotroph\| | Yeast | NULL | Probable | As nectar-tap_saprotroph (Põlme et al. 2020); Decay sugar-rich substrates (Põlme et al. 2020) | Põlme S, et al. 2020. Fungal Diversity 105:1-16 ((https://doi.org/10.1007/s13225-020-00466-2)) |
| Otu52146 | 1 | k__Fungi(100);p__Ascomycota(100);c__Saccharomycetes(100);o__Saccharomycetales(100);f__Metschnikowiaceae(100);g__Metschnikowia(100);s__Metschnikowia_pulcherrima(100); | Metschnikowia | 13 | Saprotroph | \|Undefined Saprotroph\| | Yeast | NULL | Probable | As nectar-tap_saprotroph (Põlme et al. 2020); Decay sugar-rich substrates (Põlme et al. 2020) | Põlme S, et al. 2020. Fungal Diversity 105:1-16 ((https://doi.org/10.1007/s13225-020-00466-2)) |
| Otu52148 | 1 | k__Fungi(100);p__Ascomycota(100);c__Saccharomycetes(100);o__Saccharomycetales(100);f__Metschnikowiaceae(100);g__Metschnikowia(100);s__Metschnikowia_pulcherrima(100); | Metschnikowia | 13 | Saprotroph | \|Undefined Saprotroph\| | Yeast | NULL | Probable | As nectar-tap_saprotroph (Põlme et al. 2020); Decay sugar-rich substrates (Põlme et al. 2020) | Põlme S, et al. 2020. Fungal Diversity 105:1-16 ((https://doi.org/10.1007/s13225-020-00466-2)) |
| Otu52149 | 1 | k__Fungi(100);p__Ascomycota(100);c__Saccharomycetes(100);o__Saccharomycetales(100);f__Metschnikowiaceae(100);g__Metschnikowia(100);g__Metschnikowia_unclassified(100); | Metschnikowia | 13 | Saprotroph | \|Undefined Saprotroph\| | Yeast | NULL | Probable | As nectar-tap_saprotroph (Põlme et al. 2020); Decay sugar-rich substrates (Põlme et al. 2020) | Põlme S, et al. 2020. Fungal Diversity 105:1-16 ((https://doi.org/10.1007/s13225-020-00466-2)) |
| Otu52150 | 1 | k__Fungi(100);p__Ascomycota(100);c__Saccharomycetes(100);o__Saccharomycetales(100);f__Metschnikowiaceae(100);g__Metschnikowia(100);s__Metschnikowia_pulcherrima(100); | Metschnikowia | 13 | Saprotroph | \|Undefined Saprotroph\| | Yeast | NULL | Probable | As nectar-tap_saprotroph (Põlme et al. 2020); Decay sugar-rich substrates (Põlme et al. 2020) | Põlme S, et al. 2020. Fungal Diversity 105:1-16 ((https://doi.org/10.1007/s13225-020-00466-2)) |
| Otu52151 | 1 | k__Fungi(100);p__Ascomycota(100);c__Saccharomycetes(100);o__Saccharomycetales(100);f__Metschnikowiaceae(100);g__Metschnikowia(100);s__Metschnikowia_pulcherrima(100); | Metschnikowia | 13 | Saprotroph | \|Undefined Saprotroph\| | Yeast | NULL | Probable | As nectar-tap_saprotroph (Põlme et al. 2020); Decay sugar-rich substrates (Põlme et al. 2020) | Põlme S, et al. 2020. Fungal Diversity 105:1-16 ((https://doi.org/10.1007/s13225-020-00466-2)) |
| Otu52152 | 1 | k__Fungi(100);p__Ascomycota(100);c__Saccharomycetes(100);o__Saccharomycetales(100);f__Metschnikowiaceae(100);g__Metschnikowia(100);g__Metschnikowia_unclassified(100); | Metschnikowia | 13 | Saprotroph | \|Undefined Saprotroph\| | Yeast | NULL | Probable | As nectar-tap_saprotroph (Põlme et al. 2020); Decay sugar-rich substrates (Põlme et al. 2020) | Põlme S, et al. 2020. Fungal Diversity 105:1-16 ((https://doi.org/10.1007/s13225-020-00466-2)) |
| Otu52153 | 1 | k__Fungi(100);p__Ascomycota(100);c__Saccharomycetes(100);o__Saccharomycetales(100);f__Metschnikowiaceae(100);g__Metschnikowia(100);s__Metschnikowia_pulcherrima(100); | Metschnikowia | 13 | Saprotroph | \|Undefined Saprotroph\| | Yeast | NULL | Probable | As nectar-tap_saprotroph (Põlme et al. 2020); Decay sugar-rich substrates (Põlme et al. 2020) | Põlme S, et al. 2020. Fungal Diversity 105:1-16 ((https://doi.org/10.1007/s13225-020-00466-2)) |
| Otu52154 | 1 | k__Fungi(100);p__Ascomycota(100);c__Saccharomycetes(100);o__Saccharomycetales(100);f__Metschnikowiaceae(100);g__Metschnikowia(100);s__Metschnikowia_pulcherrima(100); | Metschnikowia | 13 | Saprotroph | \|Undefined Saprotroph\| | Yeast | NULL | Probable | As nectar-tap_saprotroph (Põlme et al. 2020); Decay sugar-rich substrates (Põlme et al. 2020) | Põlme S, et al. 2020. Fungal Diversity 105:1-16 ((https://doi.org/10.1007/s13225-020-00466-2)) |
| Otu52155 | 1 | k__Fungi(100);p__Ascomycota(100);c__Saccharomycetes(100);o__Saccharomycetales(100);f__Metschnikowiaceae(100);g__Metschnikowia(100);s__Metschnikowia_pulcherrima(100); | Metschnikowia | 13 | Saprotroph | \|Undefined Saprotroph\| | Yeast | NULL | Probable | As nectar-tap_saprotroph (Põlme et al. 2020); Decay sugar-rich substrates (Põlme et al. 2020) | Põlme S, et al. 2020. Fungal Diversity 105:1-16 ((https://doi.org/10.1007/s13225-020-00466-2)) |
| Otu52156 | 1 | k__Fungi(100);p__Ascomycota(100);c__Saccharomycetes(100);o__Saccharomycetales(100);f__Metschnikowiaceae(100);g__Metschnikowia(100);s__Metschnikowia_pulcherrima(100); | Metschnikowia | 13 | Saprotroph | \|Undefined Saprotroph\| | Yeast | NULL | Probable | As nectar-tap_saprotroph (Põlme et al. 2020); Decay sugar-rich substrates (Põlme et al. 2020) | Põlme S, et al. 2020. Fungal Diversity 105:1-16 ((https://doi.org/10.1007/s13225-020-00466-2)) |
| Otu52157 | 1 | k__Fungi(100);p__Ascomycota(100);c__Saccharomycetes(100);o__Saccharomycetales(100);f__Metschnikowiaceae(100);g__Metschnikowia(100);s__Metschnikowia_pulcherrima(100); | Metschnikowia | 13 | Saprotroph | \|Undefined Saprotroph\| | Yeast | NULL | Probable | As nectar-tap_saprotroph (Põlme et al. 2020); Decay sugar-rich substrates (Põlme et al. 2020) | Põlme S, et al. 2020. Fungal Diversity 105:1-16 ((https://doi.org/10.1007/s13225-020-00466-2)) |
| Otu52158 | 1 | k__Fungi(100);p__Ascomycota(100);c__Saccharomycetes(100);o__Saccharomycetales(100);f__Metschnikowiaceae(100);g__Metschnikowia(100);s__Metschnikowia_pulcherrima(100); | Metschnikowia | 13 | Saprotroph | \|Undefined Saprotroph\| | Yeast | NULL | Probable | As nectar-tap_saprotroph (Põlme et al. 2020); Decay sugar-rich substrates (Põlme et al. 2020) | Põlme S, et al. 2020. Fungal Diversity 105:1-16 ((https://doi.org/10.1007/s13225-020-00466-2)) |
| Otu52159 | 1 | k__Fungi(100);p__Ascomycota(100);c__Saccharomycetes(100);o__Saccharomycetales(100);f__Metschnikowiaceae(100);g__Metschnikowia(100);s__Metschnikowia_pulcherrima(100); | Metschnikowia | 13 | Saprotroph | \|Undefined Saprotroph\| | Yeast | NULL | Probable | As nectar-tap_saprotroph (Põlme et al. 2020); Decay sugar-rich substrates (Põlme et al. 2020) | Põlme S, et al. 2020. Fungal Diversity 105:1-16 ((https://doi.org/10.1007/s13225-020-00466-2)) |
| Otu52160 | 1 | k__Fungi(100);p__Ascomycota(100);c__Saccharomycetes(100);o__Saccharomycetales(100);f__Metschnikowiaceae(100);g__Metschnikowia(100);s__Metschnikowia_pulcherrima(100); | Metschnikowia | 13 | Saprotroph | \|Undefined Saprotroph\| | Yeast | NULL | Probable | As nectar-tap_saprotroph (Põlme et al. 2020); Decay sugar-rich substrates (Põlme et al. 2020) | Põlme S, et al. 2020. Fungal Diversity 105:1-16 ((https://doi.org/10.1007/s13225-020-00466-2)) |
| Otu52161 | 1 | k__Fungi(100);p__Ascomycota(100);c__Saccharomycetes(100);o__Saccharomycetales(100);f__Metschnikowiaceae(100);g__Metschnikowia(100);g__Metschnikowia_unclassified(100); | Metschnikowia | 13 | Saprotroph | \|Undefined Saprotroph\| | Yeast | NULL | Probable | As nectar-tap_saprotroph (Põlme et al. 2020); Decay sugar-rich substrates (Põlme et al. 2020) | Põlme S, et al. 2020. Fungal Diversity 105:1-16 ((https://doi.org/10.1007/s13225-020-00466-2)) |
| Otu52162 | 1 | k__Fungi(100);p__Ascomycota(100);c__Saccharomycetes(100);o__Saccharomycetales(100);f__Metschnikowiaceae(100);g__Metschnikowia(100);s__Metschnikowia_pulcherrima(100); | Metschnikowia | 13 | Saprotroph | \|Undefined Saprotroph\| | Yeast | NULL | Probable | As nectar-tap_saprotroph (Põlme et al. 2020); Decay sugar-rich substrates (Põlme et al. 2020) | Põlme S, et al. 2020. Fungal Diversity 105:1-16 ((https://doi.org/10.1007/s13225-020-00466-2)) |
| Otu52164 | 1 | k__Fungi(100);p__Ascomycota(100);c__Saccharomycetes(100);o__Saccharomycetales(100);f__Metschnikowiaceae(100);g__Metschnikowia(100);g__Metschnikowia_unclassified(100); | Metschnikowia | 13 | Saprotroph | \|Undefined Saprotroph\| | Yeast | NULL | Probable | As nectar-tap_saprotroph (Põlme et al. 2020); Decay sugar-rich substrates (Põlme et al. 2020) | Põlme S, et al. 2020. Fungal Diversity 105:1-16 ((https://doi.org/10.1007/s13225-020-00466-2)) |
| Otu52165 | 1 | k__Fungi(100);p__Ascomycota(100);c__Saccharomycetes(100);o__Saccharomycetales(100);f__Metschnikowiaceae(100);g__Metschnikowia(100);s__Metschnikowia_pulcherrima(100); | Metschnikowia | 13 | Saprotroph | \|Undefined Saprotroph\| | Yeast | NULL | Probable | As nectar-tap_saprotroph (Põlme et al. 2020); Decay sugar-rich substrates (Põlme et al. 2020) | Põlme S, et al. 2020. Fungal Diversity 105:1-16 ((https://doi.org/10.1007/s13225-020-00466-2)) |
| Otu52166 | 1 | k__Fungi(100);p__Ascomycota(100);c__Saccharomycetes(100);o__Saccharomycetales(100);f__Metschnikowiaceae(100);g__Metschnikowia(100);s__Metschnikowia_pulcherrima(100); | Metschnikowia | 13 | Saprotroph | \|Undefined Saprotroph\| | Yeast | NULL | Probable | As nectar-tap_saprotroph (Põlme et al. 2020); Decay sugar-rich substrates (Põlme et al. 2020) | Põlme S, et al. 2020. Fungal Diversity 105:1-16 ((https://doi.org/10.1007/s13225-020-00466-2)) |
| Otu52167 | 1 | k__Fungi(100);p__Ascomycota(100);c__Saccharomycetes(100);o__Saccharomycetales(100);f__Metschnikowiaceae(100);g__Metschnikowia(100);s__Metschnikowia_pulcherrima(100); | Metschnikowia | 13 | Saprotroph | \|Undefined Saprotroph\| | Yeast | NULL | Probable | As nectar-tap_saprotroph (Põlme et al. 2020); Decay sugar-rich substrates (Põlme et al. 2020) | Põlme S, et al. 2020. Fungal Diversity 105:1-16 ((https://doi.org/10.1007/s13225-020-00466-2)) |
| Otu52168 | 1 | k__Fungi(100);p__Ascomycota(100);c__Saccharomycetes(100);o__Saccharomycetales(100);f__Metschnikowiaceae(100);g__Metschnikowia(100);g__Metschnikowia_unclassified(100); | Metschnikowia | 13 | Saprotroph | \|Undefined Saprotroph\| | Yeast | NULL | Probable | As nectar-tap_saprotroph (Põlme et al. 2020); Decay sugar-rich substrates (Põlme et al. 2020) | Põlme S, et al. 2020. Fungal Diversity 105:1-16 ((https://doi.org/10.1007/s13225-020-00466-2)) |
| Otu52170 | 1 | k__Fungi(100);p__Ascomycota(100);c__Saccharomycetes(100);o__Saccharomycetales(100);f__Metschnikowiaceae(100);g__Metschnikowia(100);s__Metschnikowia_pulcherrima(100); | Metschnikowia | 13 | Saprotroph | \|Undefined Saprotroph\| | Yeast | NULL | Probable | As nectar-tap_saprotroph (Põlme et al. 2020); Decay sugar-rich substrates (Põlme et al. 2020) | Põlme S, et al. 2020. Fungal Diversity 105:1-16 ((https://doi.org/10.1007/s13225-020-00466-2)) |
| Otu52171 | 1 | k__Fungi(100);p__Ascomycota(100);c__Saccharomycetes(100);o__Saccharomycetales(100);f__Metschnikowiaceae(100);g__Metschnikowia(100);s__Metschnikowia_pulcherrima(100); | Metschnikowia | 13 | Saprotroph | \|Undefined Saprotroph\| | Yeast | NULL | Probable | As nectar-tap_saprotroph (Põlme et al. 2020); Decay sugar-rich substrates (Põlme et al. 2020) | Põlme S, et al. 2020. Fungal Diversity 105:1-16 ((https://doi.org/10.1007/s13225-020-00466-2)) |
| Otu52172 | 1 | k__Fungi(100);p__Ascomycota(100);c__Saccharomycetes(100);o__Saccharomycetales(100);f__Metschnikowiaceae(100);g__Metschnikowia(100);s__Metschnikowia_pulcherrima(100); | Metschnikowia | 13 | Saprotroph | \|Undefined Saprotroph\| | Yeast | NULL | Probable | As nectar-tap_saprotroph (Põlme et al. 2020); Decay sugar-rich substrates (Põlme et al. 2020) | Põlme S, et al. 2020. Fungal Diversity 105:1-16 ((https://doi.org/10.1007/s13225-020-00466-2)) |
| Otu52173 | 1 | k__Fungi(100);p__Ascomycota(100);c__Saccharomycetes(100);o__Saccharomycetales(100);f__Metschnikowiaceae(100);g__Metschnikowia(100);g__Metschnikowia_unclassified(100); | Metschnikowia | 13 | Saprotroph | \|Undefined Saprotroph\| | Yeast | NULL | Probable | As nectar-tap_saprotroph (Põlme et al. 2020); Decay sugar-rich substrates (Põlme et al. 2020) | Põlme S, et al. 2020. Fungal Diversity 105:1-16 ((https://doi.org/10.1007/s13225-020-00466-2)) |
| Otu52174 | 1 | k__Fungi(100);p__Ascomycota(100);c__Saccharomycetes(100);o__Saccharomycetales(100);f__Metschnikowiaceae(100);g__Metschnikowia(100);s__Metschnikowia_pulcherrima(100); | Metschnikowia | 13 | Saprotroph | \|Undefined Saprotroph\| | Yeast | NULL | Probable | As nectar-tap_saprotroph (Põlme et al. 2020); Decay sugar-rich substrates (Põlme et al. 2020) | Põlme S, et al. 2020. Fungal Diversity 105:1-16 ((https://doi.org/10.1007/s13225-020-00466-2)) |
| Otu52175 | 1 | k__Fungi(100);p__Ascomycota(100);c__Saccharomycetes(100);o__Saccharomycetales(100);f__Metschnikowiaceae(100);g__Metschnikowia(100);s__Metschnikowia_pulcherrima(100); | Metschnikowia | 13 | Saprotroph | \|Undefined Saprotroph\| | Yeast | NULL | Probable | As nectar-tap_saprotroph (Põlme et al. 2020); Decay sugar-rich substrates (Põlme et al. 2020) | Põlme S, et al. 2020. Fungal Diversity 105:1-16 ((https://doi.org/10.1007/s13225-020-00466-2)) |
| Otu52176 | 1 | k__Fungi(100);p__Ascomycota(100);c__Saccharomycetes(100);o__Saccharomycetales(100);f__Metschnikowiaceae(100);g__Metschnikowia(100);g__Metschnikowia_unclassified(100); | Metschnikowia | 13 | Saprotroph | \|Undefined Saprotroph\| | Yeast | NULL | Probable | As nectar-tap_saprotroph (Põlme et al. 2020); Decay sugar-rich substrates (Põlme et al. 2020) | Põlme S, et al. 2020. Fungal Diversity 105:1-16 ((https://doi.org/10.1007/s13225-020-00466-2)) |
| Otu52177 | 1 | k__Fungi(100);p__Ascomycota(100);c__Saccharomycetes(100);o__Saccharomycetales(100);f__Metschnikowiaceae(100);g__Metschnikowia(100);s__Metschnikowia_pulcherrima(100); | Metschnikowia | 13 | Saprotroph | \|Undefined Saprotroph\| | Yeast | NULL | Probable | As nectar-tap_saprotroph (Põlme et al. 2020); Decay sugar-rich substrates (Põlme et al. 2020) | Põlme S, et al. 2020. Fungal Diversity 105:1-16 ((https://doi.org/10.1007/s13225-020-00466-2)) |

| | | | | | | | | | | | |
|---|---|---|---|---|---|---|---|---|---|---|---|
| Otu52178 | 1 | k__Fungi(100);p__Ascomycota(100);c__Saccharomycetes(100);o__Saccharomycetales(100);f__Metschnikowiaceae(100);g__Metschnikowia(100);g__Metschnikowia_unclassified(100); | Metschnikowia | 13 | Saprotroph | \|Undefined Saprotroph\| | Yeast | NULL | Probable | As nectar-tap_saprotroph (Põlme et al. 2020); Decay sugar-rich substrates (Põlme et al. 2020) | Põlme S, et al. 2020. Fungal Diversity 105:1-16 ((https://doi.org/10.1007/s13225-020-00466-2)) |
| Otu52179 | 1 | k__Fungi(100);p__Ascomycota(100);c__Saccharomycetes(100);o__Saccharomycetales(100);f__Metschnikowiaceae(100);g__Metschnikowia(100);g__Metschnikowia_unclassified(100); | Metschnikowia | 13 | Saprotroph | \|Undefined Saprotroph\| | Yeast | NULL | Probable | As nectar-tap_saprotroph (Põlme et al. 2020); Decay sugar-rich substrates (Põlme et al. 2020) | Põlme S, et al. 2020. Fungal Diversity 105:1-16 ((https://doi.org/10.1007/s13225-020-00466-2)) |
| Otu52180 | 1 | k__Fungi(100);p__Ascomycota(100);c__Saccharomycetes(100);o__Saccharomycetales(100);f__Metschnikowiaceae(100);g__Metschnikowia(100);g__Metschnikowia_unclassified(100); | Metschnikowia | 13 | Saprotroph | \|Undefined Saprotroph\| | Yeast | NULL | Probable | As nectar-tap_saprotroph (Põlme et al. 2020); Decay sugar-rich substrates (Põlme et al. 2020) | Põlme S, et al. 2020. Fungal Diversity 105:1-16 ((https://doi.org/10.1007/s13225-020-00466-2)) |
| Otu52181 | 1 | k__Fungi(100);p__Ascomycota(100);c__Saccharomycetes(100);o__Saccharomycetales(100);f__Metschnikowiaceae(100);g__Metschnikowia(100);s__Metschnikowia_pulcherrima(100); | Metschnikowia | 13 | Saprotroph | \|Undefined Saprotroph\| | Yeast | NULL | Probable | As nectar-tap_saprotroph (Põlme et al. 2020); Decay sugar-rich substrates (Põlme et al. 2020) | Põlme S, et al. 2020. Fungal Diversity 105:1-16 ((https://doi.org/10.1007/s13225-020-00466-2)) |
| Otu52182 | 1 | k__Fungi(100);p__Ascomycota(100);c__Saccharomycetes(100);o__Saccharomycetales(100);f__Metschnikowiaceae(100);g__Metschnikowia(100);s__Metschnikowia_pulcherrima(100); | Metschnikowia | 13 | Saprotroph | \|Undefined Saprotroph\| | Yeast | NULL | Probable | As nectar-tap_saprotroph (Põlme et al. 2020); Decay sugar-rich substrates (Põlme et al. 2020) | Põlme S, et al. 2020. Fungal Diversity 105:1-16 ((https://doi.org/10.1007/s13225-020-00466-2)) |
| Otu52183 | 1 | k__Fungi(100);p__Ascomycota(100);c__Saccharomycetes(100);o__Saccharomycetales(100);f__Metschnikowiaceae(100);g__Metschnikowia(100);s__Metschnikowia_pulcherrima(100); | Metschnikowia | 13 | Saprotroph | \|Undefined Saprotroph\| | Yeast | NULL | Probable | As nectar-tap_saprotroph (Põlme et al. 2020); Decay sugar-rich substrates (Põlme et al. 2020) | Põlme S, et al. 2020. Fungal Diversity 105:1-16 ((https://doi.org/10.1007/s13225-020-00466-2)) |
| Otu52184 | 1 | k__Fungi(100);p__Ascomycota(100);c__Saccharomycetes(100);o__Saccharomycetales(100);f__Metschnikowiaceae(100);g__Metschnikowia(100);s__Metschnikowia_pulcherrima(100); | Metschnikowia | 13 | Saprotroph | \|Undefined Saprotroph\| | Yeast | NULL | Probable | As nectar-tap_saprotroph (Põlme et al. 2020); Decay sugar-rich substrates (Põlme et al. 2020) | Põlme S, et al. 2020. Fungal Diversity 105:1-16 ((https://doi.org/10.1007/s13225-020-00466-2)) |
| Otu52185 | 1 | k__Fungi(100);p__Ascomycota(100);c__Saccharomycetes(100);o__Saccharomycetales(100);f__Metschnikowiaceae(100);g__Metschnikowia(100);s__Metschnikowia_pulcherrima(100); | Metschnikowia | 13 | Saprotroph | \|Undefined Saprotroph\| | Yeast | NULL | Probable | As nectar-tap_saprotroph (Põlme et al. 2020); Decay sugar-rich substrates (Põlme et al. 2020) | Põlme S, et al. 2020. Fungal Diversity 105:1-16 ((https://doi.org/10.1007/s13225-020-00466-2)) |
| Otu52186 | 1 | k__Fungi(100);p__Ascomycota(100);c__Saccharomycetes(100);o__Saccharomycetales(100);f__Metschnikowiaceae(100);g__Metschnikowia(100);g__Metschnikowia_unclassified(100); | Metschnikowia | 13 | Saprotroph | \|Undefined Saprotroph\| | Yeast | NULL | Probable | As nectar-tap_saprotroph (Põlme et al. 2020); Decay sugar-rich substrates (Põlme et al. 2020) | Põlme S, et al. 2020. Fungal Diversity 105:1-16 ((https://doi.org/10.1007/s13225-020-00466-2)) |
| Otu52187 | 1 | k__Fungi(100);p__Ascomycota(100);c__Saccharomycetes(100);o__Saccharomycetales(100);f__Metschnikowiaceae(100);g__Metschnikowia(100);s__Metschnikowia_pulcherrima(100); | Metschnikowia | 13 | Saprotroph | \|Undefined Saprotroph\| | Yeast | NULL | Probable | As nectar-tap_saprotroph (Põlme et al. 2020); Decay sugar-rich substrates (Põlme et al. 2020) | Põlme S, et al. 2020. Fungal Diversity 105:1-16 ((https://doi.org/10.1007/s13225-020-00466-2)) |
| Otu52188 | 1 | k__Fungi(100);p__Ascomycota(100);c__Saccharomycetes(100);o__Saccharomycetales(100);f__Metschnikowiaceae(100);g__Metschnikowia(100);s__Metschnikowia_pulcherrima(100); | Metschnikowia | 13 | Saprotroph | \|Undefined Saprotroph\| | Yeast | NULL | Probable | As nectar-tap_saprotroph (Põlme et al. 2020); Decay sugar-rich substrates (Põlme et al. 2020) | Põlme S, et al. 2020. Fungal Diversity 105:1-16 ((https://doi.org/10.1007/s13225-020-00466-2)) |
| Otu52189 | 1 | k__Fungi(100);p__Ascomycota(100);c__Saccharomycetes(100);o__Saccharomycetales(100);f__Metschnikowiaceae(100);g__Metschnikowia(100);s__Metschnikowia_pulcherrima(100); | Metschnikowia | 13 | Saprotroph | \|Undefined Saprotroph\| | Yeast | NULL | Probable | As nectar-tap_saprotroph (Põlme et al. 2020); Decay sugar-rich substrates (Põlme et al. 2020) | Põlme S, et al. 2020. Fungal Diversity 105:1-16 ((https://doi.org/10.1007/s13225-020-00466-2)) |
| Otu52191 | 1 | k__Fungi(100);p__Ascomycota(100);c__Saccharomycetes(100);o__Saccharomycetales(100);f__Metschnikowiaceae(100);g__Metschnikowia(100);g__Metschnikowia_unclassified(100); | Metschnikowia | 13 | Saprotroph | \|Undefined Saprotroph\| | Yeast | NULL | Probable | As nectar-tap_saprotroph (Põlme et al. 2020); Decay sugar-rich substrates (Põlme et al. 2020) | Põlme S, et al. 2020. Fungal Diversity 105:1-16 ((https://doi.org/10.1007/s13225-020-00466-2)) |
| Otu52192 | 1 | k__Fungi(100);p__Ascomycota(100);c__Saccharomycetes(100);o__Saccharomycetales(100);f__Metschnikowiaceae(100);g__Metschnikowia(100);g__Metschnikowia_unclassified(100); | Metschnikowia | 13 | Saprotroph | \|Undefined Saprotroph\| | Yeast | NULL | Probable | As nectar-tap_saprotroph (Põlme et al. 2020); Decay sugar-rich substrates (Põlme et al. 2020) | Põlme S, et al. 2020. Fungal Diversity 105:1-16 ((https://doi.org/10.1007/s13225-020-00466-2)) |
| Otu52193 | 1 | k__Fungi(100);p__Ascomycota(100);c__Saccharomycetes(100);o__Saccharomycetales(100);f__Metschnikowiaceae(100);g__Metschnikowia(100);s__Metschnikowia_pulcherrima(100); | Metschnikowia | 13 | Saprotroph | \|Undefined Saprotroph\| | Yeast | NULL | Probable | As nectar-tap_saprotroph (Põlme et al. 2020); Decay sugar-rich substrates (Põlme et al. 2020) | Põlme S, et al. 2020. Fungal Diversity 105:1-16 ((https://doi.org/10.1007/s13225-020-00466-2)) |
| Otu52194 | 1 | k__Fungi(100);p__Ascomycota(100);c__Saccharomycetes(100);o__Saccharomycetales(100);f__Metschnikowiaceae(100);g__Metschnikowia(100);s__Metschnikowia_pulcherrima(100); | Metschnikowia | 13 | Saprotroph | \|Undefined Saprotroph\| | Yeast | NULL | Probable | As nectar-tap_saprotroph (Põlme et al. 2020); Decay sugar-rich substrates (Põlme et al. 2020) | Põlme S, et al. 2020. Fungal Diversity 105:1-16 ((https://doi.org/10.1007/s13225-020-00466-2)) |
| Otu52195 | 1 | k__Fungi(100);p__Ascomycota(100);c__Saccharomycetes(100);o__Saccharomycetales(100);f__Metschnikowiaceae(100);g__Metschnikowia(100);s__Metschnikowia_pulcherrima(100); | Metschnikowia | 13 | Saprotroph | \|Undefined Saprotroph\| | Yeast | NULL | Probable | As nectar-tap_saprotroph (Põlme et al. 2020); Decay sugar-rich substrates (Põlme et al. 2020) | Põlme S, et al. 2020. Fungal Diversity 105:1-16 ((https://doi.org/10.1007/s13225-020-00466-2)) |
| Otu52196 | 1 | k__Fungi(100);p__Ascomycota(100);c__Saccharomycetes(100);o__Saccharomycetales(100);f__Metschnikowiaceae(100);g__Metschnikowia(100);s__Metschnikowia_pulcherrima(100); | Metschnikowia | 13 | Saprotroph | \|Undefined Saprotroph\| | Yeast | NULL | Probable | As nectar-tap_saprotroph (Põlme et al. 2020); Decay sugar-rich substrates (Põlme et al. 2020) | Põlme S, et al. 2020. Fungal Diversity 105:1-16 ((https://doi.org/10.1007/s13225-020-00466-2)) |
| Otu52197 | 1 | k__Fungi(100);p__Ascomycota(100);c__Saccharomycetes(100);o__Saccharomycetales(100);f__Metschnikowiaceae(100);g__Metschnikowia(100);s__Metschnikowia_pulcherrima(100); | Metschnikowia | 13 | Saprotroph | \|Undefined Saprotroph\| | Yeast | NULL | Probable | As nectar-tap_saprotroph (Põlme et al. 2020); Decay sugar-rich substrates (Põlme et al. 2020) | Põlme S, et al. 2020. Fungal Diversity 105:1-16 ((https://doi.org/10.1007/s13225-020-00466-2)) |
| Otu52198 | 1 | k__Fungi(100);p__Ascomycota(100);c__Saccharomycetes(100);o__Saccharomycetales(100);f__Metschnikowiaceae(100);g__Metschnikowia(100);g__Metschnikowia_unclassified(100); | Metschnikowia | 13 | Saprotroph | \|Undefined Saprotroph\| | Yeast | NULL | Probable | As nectar-tap_saprotroph (Põlme et al. 2020); Decay sugar-rich substrates (Põlme et al. 2020) | Põlme S, et al. 2020. Fungal Diversity 105:1-16 ((https://doi.org/10.1007/s13225-020-00466-2)) |
| Otu52199 | 1 | k__Fungi(100);p__Ascomycota(100);c__Saccharomycetes(100);o__Saccharomycetales(100);f__Metschnikowiaceae(100);g__Metschnikowia(100);s__Metschnikowia_pulcherrima(100); | Metschnikowia | 13 | Saprotroph | \|Undefined Saprotroph\| | Yeast | NULL | Probable | As nectar-tap_saprotroph (Põlme et al. 2020); Decay sugar-rich substrates (Põlme et al. 2020) | Põlme S, et al. 2020. Fungal Diversity 105:1-16 ((https://doi.org/10.1007/s13225-020-00466-2)) |
| Otu52200 | 1 | k__Fungi(100);p__Ascomycota(100);c__Saccharomycetes(100);o__Saccharomycetales(100);f__Metschnikowiaceae(100);g__Metschnikowia(100);s__Metschnikowia_pulcherrima(100); | Metschnikowia | 13 | Saprotroph | \|Undefined Saprotroph\| | Yeast | NULL | Probable | As nectar-tap_saprotroph (Põlme et al. 2020); Decay sugar-rich substrates (Põlme et al. 2020) | Põlme S, et al. 2020. Fungal Diversity 105:1-16 ((https://doi.org/10.1007/s13225-020-00466-2)) |
| Otu52201 | 1 | k__Fungi(100);p__Ascomycota(100);c__Saccharomycetes(100);o__Saccharomycetales(100);f__Metschnikowiaceae(100);g__Metschnikowia(100);s__Metschnikowia_pulcherrima(100); | Metschnikowia | 13 | Saprotroph | \|Undefined Saprotroph\| | Yeast | NULL | Probable | As nectar-tap_saprotroph (Põlme et al. 2020); Decay sugar-rich substrates (Põlme et al. 2020) | Põlme S, et al. 2020. Fungal Diversity 105:1-16 ((https://doi.org/10.1007/s13225-020-00466-2)) |
| Otu52202 | 1 | k__Fungi(100);p__Ascomycota(100);c__Saccharomycetes(100);o__Saccharomycetales(100);f__Metschnikowiaceae(100);g__Metschnikowia(100);s__Metschnikowia_pulcherrima(100); | Metschnikowia | 13 | Saprotroph | \|Undefined Saprotroph\| | Yeast | NULL | Probable | As nectar-tap_saprotroph (Põlme et al. 2020); Decay sugar-rich substrates (Põlme et al. 2020) | Põlme S, et al. 2020. Fungal Diversity 105:1-16 ((https://doi.org/10.1007/s13225-020-00466-2)) |
| Otu52204 | 1 | k__Fungi(100);p__Ascomycota(100);c__Saccharomycetes(100);o__Saccharomycetales(100);f__Metschnikowiaceae(100);g__Metschnikowia(100);s__Metschnikowia_pulcherrima(100); | Metschnikowia | 13 | Saprotroph | \|Undefined Saprotroph\| | Yeast | NULL | Probable | As nectar-tap_saprotroph (Põlme et al. 2020); Decay sugar-rich substrates (Põlme et al. 2020) | Põlme S, et al. 2020. Fungal Diversity 105:1-16 ((https://doi.org/10.1007/s13225-020-00466-2)) |
| Otu52205 | 1 | k__Fungi(100);p__Ascomycota(100);c__Saccharomycetes(100);o__Saccharomycetales(100);f__Metschnikowiaceae(100);g__Metschnikowia(100);s__Metschnikowia_pulcherrima(100); | Metschnikowia | 13 | Saprotroph | \|Undefined Saprotroph\| | Yeast | NULL | Probable | As nectar-tap_saprotroph (Põlme et al. 2020); Decay sugar-rich substrates (Põlme et al. 2020) | Põlme S, et al. 2020. Fungal Diversity 105:1-16 ((https://doi.org/10.1007/s13225-020-00466-2)) |
| Otu52208 | 1 | k__Fungi(100);p__Ascomycota(100);c__Saccharomycetes(100);o__Saccharomycetales(100);f__Metschnikowiaceae(100);g__Metschnikowia(100);s__Metschnikowia_pulcherrima(100); | Metschnikowia | 13 | Saprotroph | \|Undefined Saprotroph\| | Yeast | NULL | Probable | As nectar-tap_saprotroph (Põlme et al. 2020); Decay sugar-rich substrates (Põlme et al. 2020) | Põlme S, et al. 2020. Fungal Diversity 105:1-16 ((https://doi.org/10.1007/s13225-020-00466-2)) |
| Otu52210 | 1 | k__Fungi(100);p__Ascomycota(100);c__Saccharomycetes(100);o__Saccharomycetales(100);f__Metschnikowiaceae(100);g__Metschnikowia(100);s__Metschnikowia_pulcherrima(100); | Metschnikowia | 13 | Saprotroph | \|Undefined Saprotroph\| | Yeast | NULL | Probable | As nectar-tap_saprotroph (Põlme et al. 2020); Decay sugar-rich substrates (Põlme et al. 2020) | Põlme S, et al. 2020. Fungal Diversity 105:1-16 ((https://doi.org/10.1007/s13225-020-00466-2)) |
| Otu52211 | 1 | k__Fungi(100);p__Ascomycota(100);c__Saccharomycetes(100);o__Saccharomycetales(100);f__Metschnikowiaceae(100);g__Metschnikowia(100);s__Metschnikowia_pulcherrima(100); | Metschnikowia | 13 | Saprotroph | \|Undefined Saprotroph\| | Yeast | NULL | Probable | As nectar-tap_saprotroph (Põlme et al. 2020); Decay sugar-rich substrates (Põlme et al. 2020) | Põlme S, et al. 2020. Fungal Diversity 105:1-16 ((https://doi.org/10.1007/s13225-020-00466-2)) |
| Otu52212 | 1 | k__Fungi(100);p__Ascomycota(100);c__Saccharomycetes(100);o__Saccharomycetales(100);f__Metschnikowiaceae(100);g__Metschnikowia(100);s__Metschnikowia_pulcherrima(100); | Metschnikowia | 13 | Saprotroph | \|Undefined Saprotroph\| | Yeast | NULL | Probable | As nectar-tap_saprotroph (Põlme et al. 2020); Decay sugar-rich substrates (Põlme et al. 2020) | Põlme S, et al. 2020. Fungal Diversity 105:1-16 ((https://doi.org/10.1007/s13225-020-00466-2)) |
| Otu52213 | 1 | k__Fungi(100);p__Ascomycota(100);c__Saccharomycetes(100);o__Saccharomycetales(100);f__Metschnikowiaceae(100);g__Metschnikowia(100);s__Metschnikowia_pulcherrima(100); | Metschnikowia | 13 | Saprotroph | \|Undefined Saprotroph\| | Yeast | NULL | Probable | As nectar-tap_saprotroph (Põlme et al. 2020); Decay sugar-rich substrates (Põlme et al. 2020) | Põlme S, et al. 2020. Fungal Diversity 105:1-16 ((https://doi.org/10.1007/s13225-020-00466-2)) |
| Otu52214 | 1 | k__Fungi(100);p__Ascomycota(100);c__Saccharomycetes(100);o__Saccharomycetales(100);f__Metschnikowiaceae(100);g__Metschnikowia(100);s__Metschnikowia_pulcherrima(100); | Metschnikowia | 13 | Saprotroph | \|Undefined Saprotroph\| | Yeast | NULL | Probable | As nectar-tap_saprotroph (Põlme et al. 2020); Decay sugar-rich substrates (Põlme et al. 2020) | Põlme S, et al. 2020. Fungal Diversity 105:1-16 ((https://doi.org/10.1007/s13225-020-00466-2)) |
| Otu52215 | 1 | k__Fungi(100);p__Ascomycota(100);c__Saccharomycetes(100);o__Saccharomycetales(100);f__Metschnikowiaceae(100);g__Metschnikowia(100);g__Metschnikowia_unclassified(100); | Metschnikowia | 13 | Saprotroph | \|Undefined Saprotroph\| | Yeast | NULL | Probable | As nectar-tap_saprotroph (Põlme et al. 2020); Decay sugar-rich substrates (Põlme et al. 2020) | Põlme S, et al. 2020. Fungal Diversity 105:1-16 ((https://doi.org/10.1007/s13225-020-00466-2)) |
| Otu52216 | 1 | k__Fungi(100);p__Ascomycota(100);c__Saccharomycetes(100);o__Saccharomycetales(100);f__Metschnikowiaceae(100);g__Metschnikowia(100);g__Metschnikowia_unclassified(100); | Metschnikowia | 13 | Saprotroph | \|Undefined Saprotroph\| | Yeast | NULL | Probable | As nectar-tap_saprotroph (Põlme et al. 2020); Decay sugar-rich substrates (Põlme et al. 2020) | Põlme S, et al. 2020. Fungal Diversity 105:1-16 ((https://doi.org/10.1007/s13225-020-00466-2)) |
| Otu52217 | 1 | k__Fungi(100);p__Ascomycota(100);c__Saccharomycetes(100);o__Saccharomycetales(100);f__Metschnikowiaceae(100);g__Metschnikowia(100);g__Metschnikowia_unclassified(100); | Metschnikowia | 13 | Saprotroph | \|Undefined Saprotroph\| | Yeast | NULL | Probable | As nectar-tap_saprotroph (Põlme et al. 2020); Decay sugar-rich substrates (Põlme et al. 2020) | Põlme S, et al. 2020. Fungal Diversity 105:1-16 ((https://doi.org/10.1007/s13225-020-00466-2)) |
| Otu52218 | 1 | k__Fungi(100);p__Ascomycota(100);c__Saccharomycetes(100);o__Saccharomycetales(100);f__Metschnikowiaceae(100);g__Metschnikowia(100);s__Metschnikowia_pulcherrima(100); | Metschnikowia | 13 | Saprotroph | \|Undefined Saprotroph\| | Yeast | NULL | Probable | As nectar-tap_saprotroph (Põlme et al. 2020); Decay sugar-rich substrates (Põlme et al. 2020) | Põlme S, et al. 2020. Fungal Diversity 105:1-16 ((https://doi.org/10.1007/s13225-020-00466-2)) |
| Otu52220 | 1 | k__Fungi(100);p__Ascomycota(100);c__Saccharomycetes(100);o__Saccharomycetales(100);f__Metschnikowiaceae(100);g__Metschnikowia(100);g__Metschnikowia_unclassified(100); | Metschnikowia | 13 | Saprotroph | \|Undefined Saprotroph\| | Yeast | NULL | Probable | As nectar-tap_saprotroph (Põlme et al. 2020); Decay sugar-rich substrates (Põlme et al. 2020) | Põlme S, et al. 2020. Fungal Diversity 105:1-16 ((https://doi.org/10.1007/s13225-020-00466-2)) |
| Otu52221 | 1 | k__Fungi(100);p__Ascomycota(100);c__Saccharomycetes(100);o__Saccharomycetales(100);f__Metschnikowiaceae(100);g__Metschnikowia(100);s__Metschnikowia_pulcherrima(100); | Metschnikowia | 13 | Saprotroph | \|Undefined Saprotroph\| | Yeast | NULL | Probable | As nectar-tap_saprotroph (Põlme et al. 2020); Decay sugar-rich substrates (Põlme et al. 2020) | Põlme S, et al. 2020. Fungal Diversity 105:1-16 ((https://doi.org/10.1007/s13225-020-00466-2)) |
| Otu52222 | 1 | k__Fungi(100);p__Ascomycota(100);c__Saccharomycetes(100);o__Saccharomycetales(100);f__Metschnikowiaceae(100);g__Metschnikowia(100);s__Metschnikowia_pulcherrima(100); | Metschnikowia | 13 | Saprotroph | \|Undefined Saprotroph\| | Yeast | NULL | Probable | As nectar-tap_saprotroph (Põlme et al. 2020); Decay sugar-rich substrates (Põlme et al. 2020) | Põlme S, et al. 2020. Fungal Diversity 105:1-16 ((https://doi.org/10.1007/s13225-020-00466-2)) |
| Otu52223 | 1 | k__Fungi(100);p__Ascomycota(100);c__Saccharomycetes(100);o__Saccharomycetales(100);f__Metschnikowiaceae(100);g__Metschnikowia(100);s__Metschnikowia_pulcherrima(100); | Metschnikowia | 13 | Saprotroph | \|Undefined Saprotroph\| | Yeast | NULL | Probable | As nectar-tap_saprotroph (Põlme et al. 2020); Decay sugar-rich substrates (Põlme et al. 2020) | Põlme S, et al. 2020. Fungal Diversity 105:1-16 ((https://doi.org/10.1007/s13225-020-00466-2)) |
| Otu52224 | 1 | k__Fungi(100);p__Ascomycota(100);c__Saccharomycetes(100);o__Saccharomycetales(100);f__Metschnikowiaceae(100);g__Metschnikowia(100);s__Metschnikowia_pulcherrima(100); | Metschnikowia | 13 | Saprotroph | \|Undefined Saprotroph\| | Yeast | NULL | Probable | As nectar-tap_saprotroph (Põlme et al. 2020); Decay sugar-rich substrates (Põlme et al. 2020) | Põlme S, et al. 2020. Fungal Diversity 105:1-16 ((https://doi.org/10.1007/s13225-020-00466-2)) |
| Otu52225 | 1 | k__Fungi(100);p__Ascomycota(100);c__Saccharomycetes(100);o__Saccharomycetales(100);f__Metschnikowiaceae(100);g__Metschnikowia(100);s__Metschnikowia_pulcherrima(100); | Metschnikowia | 13 | Saprotroph | \|Undefined Saprotroph\| | Yeast | NULL | Probable | As nectar-tap_saprotroph (Põlme et al. 2020); Decay sugar-rich substrates (Põlme et al. 2020) | Põlme S, et al. 2020. Fungal Diversity 105:1-16 ((https://doi.org/10.1007/s13225-020-00466-2)) |
| Otu52226 | 1 | k__Fungi(100);p__Ascomycota(100);c__Saccharomycetes(100);o__Saccharomycetales(100);f__Metschnikowiaceae(100);g__Metschnikowia(100);g__Metschnikowia_unclassified(100); | Metschnikowia | 13 | Saprotroph | \|Undefined Saprotroph\| | Yeast | NULL | Probable | As nectar-tap_saprotroph (Põlme et al. 2020); Decay sugar-rich substrates (Põlme et al. 2020) | Põlme S, et al. 2020. Fungal Diversity 105:1-16 ((https://doi.org/10.1007/s13225-020-00466-2)) |
| Otu52227 | 1 | k__Fungi(100);p__Ascomycota(100);c__Saccharomycetes(100);o__Saccharomycetales(100);f__Metschnikowiaceae(100);g__Metschnikowia(100);g__Metschnikowia_unclassified(100); | Metschnikowia | 13 | Saprotroph | \|Undefined Saprotroph\| | Yeast | NULL | Probable | As nectar-tap_saprotroph (Põlme et al. 2020); Decay sugar-rich substrates (Põlme et al. 2020) | Põlme S, et al. 2020. Fungal Diversity 105:1-16 ((https://doi.org/10.1007/s13225-020-00466-2)) |
| Otu52232 | 1 | k__Fungi(100);p__Ascomycota(100);c__Saccharomycetes(100);o__Saccharomycetales(100);f__Metschnikowiaceae(100);g__Metschnikowia(100);s__Metschnikowia_pulcherrima(100); | Metschnikowia | 13 | Saprotroph | \|Undefined Saprotroph\| | Yeast | NULL | Probable | As nectar-tap_saprotroph (Põlme et al. 2020); Decay sugar-rich substrates (Põlme et al. 2020) | Põlme S, et al. 2020. Fungal Diversity 105:1-16 ((https://doi.org/10.1007/s13225-020-00466-2)) |
| Otu52233 | 1 | k__Fungi(100);p__Ascomycota(100);c__Saccharomycetes(100);o__Saccharomycetales(100);f__Metschnikowiaceae(100);g__Metschnikowia(100);g__Metschnikowia_unclassified(100); | Metschnikowia | 13 | Saprotroph | \|Undefined Saprotroph\| | Yeast | NULL | Probable | As nectar-tap_saprotroph (Põlme et al. 2020); Decay sugar-rich substrates (Põlme et al. 2020) | Põlme S, et al. 2020. Fungal Diversity 105:1-16 ((https://doi.org/10.1007/s13225-020-00466-2)) |
| Otu52234 | 1 | k__Fungi(100);p__Ascomycota(100);c__Saccharomycetes(100);o__Saccharomycetales(100);f__Metschnikowiaceae(100);g__Metschnikowia(100);s__Metschnikowia_pulcherrima(100); | Metschnikowia | 13 | Saprotroph | \|Undefined Saprotroph\| | Yeast | NULL | Probable | As nectar-tap_saprotroph (Põlme et al. 2020); Decay sugar-rich substrates (Põlme et al. 2020) | Põlme S, et al. 2020. Fungal Diversity 105:1-16 ((https://doi.org/10.1007/s13225-020-00466-2)) |
| Otu52235 | 1 | k__Fungi(100);p__Ascomycota(100);c__Saccharomycetes(100);o__Saccharomycetales(100);f__Metschnikowiaceae(100);g__Metschnikowia(100);g__Metschnikowia_unclassified(100); | Metschnikowia | 13 | Saprotroph | \|Undefined Saprotroph\| | Yeast | NULL | Probable | As nectar-tap_saprotroph (Põlme et al. 2020); Decay sugar-rich substrates (Põlme et al. 2020) | Põlme S, et al. 2020. Fungal Diversity 105:1-16 ((https://doi.org/10.1007/s13225-020-00466-2)) |
| Otu52236 | 1 | k__Fungi(100);p__Ascomycota(100);c__Saccharomycetes(100);o__Saccharomycetales(100);f__Metschnikowiaceae(100);g__Metschnikowia(100);s__Metschnikowia_pulcherrima(100); | Metschnikowia | 13 | Saprotroph | \|Undefined Saprotroph\| | Yeast | NULL | Probable | As nectar-tap_saprotroph (Põlme et al. 2020); Decay sugar-rich substrates (Põlme et al. 2020) | Põlme S, et al. 2020. Fungal Diversity 105:1-16 ((https://doi.org/10.1007/s13225-020-00466-2)) |
| Otu52237 | 1 | k__Fungi(100);p__Ascomycota(100);c__Saccharomycetes(100);o__Saccharomycetales(100);f__Metschnikowiaceae(100);g__Metschnikowia(100);s__Metschnikowia_pulcherrima(100); | Metschnikowia | 13 | Saprotroph | \|Undefined Saprotroph\| | Yeast | NULL | Probable | As nectar-tap_saprotroph (Põlme et al. 2020); Decay sugar-rich substrates (Põlme et al. 2020) | Põlme S, et al. 2020. Fungal Diversity 105:1-16 ((https://doi.org/10.1007/s13225-020-00466-2)) |
| Otu52238 | 1 | k__Fungi(100);p__Ascomycota(100);c__Saccharomycetes(100);o__Saccharomycetales(100);f__Metschnikowiaceae(100);g__Metschnikowia(100);s__Metschnikowia_pulcherrima(100); | Metschnikowia | 13 | Saprotroph | \|Undefined Saprotroph\| | Yeast | NULL | Probable | As nectar-tap_saprotroph (Põlme et al. 2020); Decay sugar-rich substrates (Põlme et al. 2020) | Põlme S, et al. 2020. Fungal Diversity 105:1-16 ((https://doi.org/10.1007/s13225-020-00466-2)) |
| Otu52239 | 1 | k__Fungi(100);p__Ascomycota(100);c__Saccharomycetes(100);o__Saccharomycetales(100);f__Metschnikowiaceae(100);g__Metschnikowia(100);s__Metschnikowia_pulcherrima(100); | Metschnikowia | 13 | Saprotroph | \|Undefined Saprotroph\| | Yeast | NULL | Probable | As nectar-tap_saprotroph (Põlme et al. 2020); Decay sugar-rich substrates (Põlme et al. 2020) | Põlme S, et al. 2020. Fungal Diversity 105:1-16 ((https://doi.org/10.1007/s13225-020-00466-2)) |
| Otu52243 | 1 | k__Fungi(100);p__Ascomycota(100);c__Saccharomycetes(100);o__Saccharomycetales(100);f__Metschnikowiaceae(100);g__Metschnikowia(100);s__Metschnikowia_pulcherrima(100); | Metschnikowia | 13 | Saprotroph | \|Undefined Saprotroph\| | Yeast | NULL | Probable | As nectar-tap_saprotroph (Põlme et al. 2020); Decay sugar-rich substrates (Põlme et al. 2020) | Põlme S, et al. 2020. Fungal Diversity 105:1-16 ((https://doi.org/10.1007/s13225-020-00466-2)) |
| Otu52244 | 1 | k__Fungi(100);p__Ascomycota(100);c__Saccharomycetes(100);o__Saccharomycetales(100);f__Metschnikowiaceae(100);g__Metschnikowia(100);g__Metschnikowia_unclassified(100); | Metschnikowia | 13 | Saprotroph | \|Undefined Saprotroph\| | Yeast | NULL | Probable | As nectar-tap_saprotroph (Põlme et al. 2020); Decay sugar-rich substrates (Põlme et al. 2020) | Põlme S, et al. 2020. Fungal Diversity 105:1-16 ((https://doi.org/10.1007/s13225-020-00466-2)) |
| Otu52246 | 1 | k__Fungi(100);p__Ascomycota(100);c__Saccharomycetes(100);o__Saccharomycetales(100);f__Metschnikowiaceae(100);g__Metschnikowia(100);s__Metschnikowia_pulcherrima(100); | Metschnikowia | 13 | Saprotroph | \|Undefined Saprotroph\| | Yeast | NULL | Probable | As nectar-tap_saprotroph (Põlme et al. 2020); Decay sugar-rich substrates (Põlme et al. 2020) | Põlme S, et al. 2020. Fungal Diversity 105:1-16 ((https://doi.org/10.1007/s13225-020-00466-2)) |
| Otu52247 | 1 | k__Fungi(100);p__Ascomycota(100);c__Saccharomycetes(100);o__Saccharomycetales(100);f__Metschnikowiaceae(100);g__Metschnikowia(100);s__Metschnikowia_pulcherrima(100); | Metschnikowia | 13 | Saprotroph | \|Undefined Saprotroph\| | Yeast | NULL | Probable | As nectar-tap_saprotroph (Põlme et al. 2020); Decay sugar-rich substrates (Põlme et al. 2020) | Põlme S, et al. 2020. Fungal Diversity 105:1-16 ((https://doi.org/10.1007/s13225-020-00466-2)) |
| Otu52250 | 1 | k__Fungi(100);p__Ascomycota(100);c__Saccharomycetes(100);o__Saccharomycetales(100);f__Metschnikowiaceae(100);g__Metschnikowia(100);s__Metschnikowia_pulcherrima(100); | Metschnikowia | 13 | Saprotroph | \|Undefined Saprotroph\| | Yeast | NULL | Probable | As nectar-tap_saprotroph (Põlme et al. 2020); Decay sugar-rich substrates (Põlme et al. 2020) | Põlme S, et al. 2020. Fungal Diversity 105:1-16 ((https://doi.org/10.1007/s13225-020-00466-2)) |
| Otu52251 | 1 | k__Fungi(100);p__Ascomycota(100);c__Saccharomycetes(100);o__Saccharomycetales(100);f__Metschnikowiaceae(100);g__Metschnikowia(100);s__Metschnikowia_pulcherrima(100); | Metschnikowia | 13 | Saprotroph | \|Undefined Saprotroph\| | Yeast | NULL | Probable | As nectar-tap_saprotroph (Põlme et al. 2020); Decay sugar-rich substrates (Põlme et al. 2020) | Põlme S, et al. 2020. Fungal Diversity 105:1-16 ((https://doi.org/10.1007/s13225-020-00466-2)) |
| Otu52252 | 1 | k__Fungi(100);p__Ascomycota(100);c__Saccharomycetes(100);o__Saccharomycetales(100);f__Metschnikowiaceae(100);g__Metschnikowia(100);s__Metschnikowia_pulcherrima(100); | Metschnikowia | 13 | Saprotroph | \|Undefined Saprotroph\| | Yeast | NULL | Probable | As nectar-tap_saprotroph (Põlme et al. 2020); Decay sugar-rich substrates (Põlme et al. 2020) | Põlme S, et al. 2020. Fungal Diversity 105:1-16 ((https://doi.org/10.1007/s13225-020-00466-2)) |
| Otu52253 | 1 | k__Fungi(100);p__Ascomycota(100);c__Saccharomycetes(100);o__Saccharomycetales(100);f__Metschnikowiaceae(100);g__Metschnikowia(100);s__Metschnikowia_pulcherrima(100); | Metschnikowia | 13 | Saprotroph | \|Undefined Saprotroph\| | Yeast | NULL | Probable | As nectar-tap_saprotroph (Põlme et al. 2020); Decay sugar-rich substrates (Põlme et al. 2020) | Põlme S, et al. 2020. Fungal Diversity 105:1-16 ((https://doi.org/10.1007/s13225-020-00466-2)) |
| Otu52254 | 1 | k__Fungi(100);p__Ascomycota(100);c__Saccharomycetes(100);o__Saccharomycetales(100);f__Metschnikowiaceae(100);g__Metschnikowia(100);s__Metschnikowia_pulcherrima(100); | Metschnikowia | 13 | Saprotroph | \|Undefined Saprotroph\| | Yeast | NULL | Probable | As nectar-tap_saprotroph (Põlme et al. 2020); Decay sugar-rich substrates (Põlme et al. 2020) | Põlme S, et al. 2020. Fungal Diversity 105:1-16 ((https://doi.org/10.1007/s13225-020-00466-2)) |
| Otu52255 | 1 | k__Fungi(100);p__Ascomycota(100);c__Saccharomycetes(100);o__Saccharomycetales(100);f__Metschnikowiaceae(100);g__Metschnikowia(100);g__Metschnikowia_unclassified(100); | Metschnikowia | 13 | Saprotroph | \|Undefined Saprotroph\| | Yeast | NULL | Probable | As nectar-tap_saprotroph (Põlme et al. 2020); Decay sugar-rich substrates (Põlme et al. 2020) | Põlme S, et al. 2020. Fungal Diversity 105:1-16 ((https://doi.org/10.1007/s13225-020-00466-2)) |
| Otu52256 | 1 | k__Fungi(100);p__Ascomycota(100);c__Saccharomycetes(100);o__Saccharomycetales(100);f__Metschnikowiaceae(100);g__Metschnikowia(100);s__Metschnikowia_pulcherrima(100); | Metschnikowia | 13 | Saprotroph | \|Undefined Saprotroph\| | Yeast | NULL | Probable | As nectar-tap_saprotroph (Põlme et al. 2020); Decay sugar-rich substrates (Põlme et al. 2020) | Põlme S, et al. 2020. Fungal Diversity 105:1-16 ((https://doi.org/10.1007/s13225-020-00466-2)) |
| Otu52257 | 1 | k__Fungi(100);p__Ascomycota(100);c__Saccharomycetes(100);o__Saccharomycetales(100);f__Metschnikowiaceae(100);g__Metschnikowia(100);g__Metschnikowia_unclassified(100); | Metschnikowia | 13 | Saprotroph | \|Undefined Saprotroph\| | Yeast | NULL | Probable | As nectar-tap_saprotroph (Põlme et al. 2020); Decay sugar-rich substrates (Põlme et al. 2020) | Põlme S, et al. 2020. Fungal Diversity 105:1-16 ((https://doi.org/10.1007/s13225-020-00466-2)) |
| Otu52258 | 1 | k__Fungi(100);p__Ascomycota(100);c__Saccharomycetes(100);o__Saccharomycetales(100);f__Metschnikowiaceae(100);g__Metschnikowia(100);s__Metschnikowia_pulcherrima(100); | Metschnikowia | 13 | Saprotroph | \|Undefined Saprotroph\| | Yeast | NULL | Probable | As nectar-tap_saprotroph (Põlme et al. 2020); Decay sugar-rich substrates (Põlme et al. 2020) | Põlme S, et al. 2020. Fungal Diversity 105:1-16 ((https://doi.org/10.1007/s13225-020-00466-2)) |
| Otu52259 | 1 | k__Fungi(100);p__Ascomycota(100);c__Saccharomycetes(100);o__Saccharomycetales(100);f__Metschnikowiaceae(100);g__Metschnikowia(100);s__Metschnikowia_pulcherrima(100); | Metschnikowia | 13 | Saprotroph | \|Undefined Saprotroph\| | Yeast | NULL | Probable | As nectar-tap_saprotroph (Põlme et al. 2020); Decay sugar-rich substrates (Põlme et al. 2020) | Põlme S, et al. 2020. Fungal Diversity 105:1-16 ((https://doi.org/10.1007/s13225-020-00466-2)) |
| Otu52260 | 1 | k__Fungi(100);p__Ascomycota(100);c__Saccharomycetes(100);o__Saccharomycetales(100);f__Metschnikowiaceae(100);g__Metschnikowia(100);s__Metschnikowia_pulcherrima(100); | Metschnikowia | 13 | Saprotroph | \|Undefined Saprotroph\| | Yeast | NULL | Probable | As nectar-tap_saprotroph (Põlme et al. 2020); Decay sugar-rich substrates (Põlme et al. 2020) | Põlme S, et al. 2020. Fungal Diversity 105:1-16 ((https://doi.org/10.1007/s13225-020-00466-2)) |
| Otu52261 | 1 | k__Fungi(100);p__Ascomycota(100);c__Saccharomycetes(100);o__Saccharomycetales(100);f__Metschnikowiaceae(100);g__Metschnikowia(100);s__Metschnikowia_pulcherrima(100); | Metschnikowia | 13 | Saprotroph | \|Undefined Saprotroph\| | Yeast | NULL | Probable | As nectar-tap_saprotroph (Põlme et al. 2020); Decay sugar-rich substrates (Põlme et al. 2020) | Põlme S, et al. 2020. Fungal Diversity 105:1-16 ((https://doi.org/10.1007/s13225-020-00466-2)) |
| Otu52262 | 1 | k__Fungi(100);p__Ascomycota(100);c__Saccharomycetes(100);o__Saccharomycetales(100);f__Metschnikowiaceae(100);g__Metschnikowia(100);s__Metschnikowia_pulcherrima(100); | Metschnikowia | 13 | Saprotroph | \|Undefined Saprotroph\| | Yeast | NULL | Probable | As nectar-tap_saprotroph (Põlme et al. 2020); Decay sugar-rich substrates (Põlme et al. 2020) | Põlme S, et al. 2020. Fungal Diversity 105:1-16 ((https://doi.org/10.1007/s13225-020-00466-2)) |
| Otu52263 | 1 | k__Fungi(100);p__Ascomycota(100);c__Saccharomycetes(100);o__Saccharomycetales(100);f__Metschnikowiaceae(100);g__Metschnikowia(100);s__Metschnikowia_pulcherrima(100); | Metschnikowia | 13 | Saprotroph | \|Undefined Saprotroph\| | Yeast | NULL | Probable | As nectar-tap_saprotroph (Põlme et al. 2020); Decay sugar-rich substrates (Põlme et al. 2020) | Põlme S, et al. 2020. Fungal Diversity 105:1-16 ((https://doi.org/10.1007/s13225-020-00466-2)) |
| Otu52264 | 1 | k__Fungi(100);p__Ascomycota(100);c__Saccharomycetes(100);o__Saccharomycetales(100);f__Metschnikowiaceae(100);g__Metschnikowia(100);s__Metschnikowia_pulcherrima(100); | Metschnikowia | 13 | Saprotroph | \|Undefined Saprotroph\| | Yeast | NULL | Probable | As nectar-tap_saprotroph (Põlme et al. 2020); Decay sugar-rich substrates (Põlme et al. 2020) | Põlme S, et al. 2020. Fungal Diversity 105:1-16 ((https://doi.org/10.1007/s13225-020-00466-2)) |
| Otu52265 | 1 | k__Fungi(100);p__Ascomycota(100);c__Saccharomycetes(100);o__Saccharomycetales(100);f__Metschnikowiaceae(100);g__Metschnikowia(100);s__Metschnikowia_pulcherrima(100); | Metschnikowia | 13 | Saprotroph | \|Undefined Saprotroph\| | Yeast | NULL | Probable | As nectar-tap_saprotroph (Põlme et al. 2020); Decay sugar-rich substrates (Põlme et al. 2020) | Põlme S, et al. 2020. Fungal Diversity 105:1-16 ((https://doi.org/10.1007/s13225-020-00466-2)) |
| Otu52266 | 1 | k__Fungi(100);p__Ascomycota(100);c__Saccharomycetes(100);o__Saccharomycetales(100);f__Metschnikowiaceae(100);g__Metschnikowia(100);s__Metschnikowia_pulcherrima(100); | Metschnikowia | 13 | Saprotroph | \|Undefined Saprotroph\| | Yeast | NULL | Probable | As nectar-tap_saprotroph (Põlme et al. 2020); Decay sugar-rich substrates (Põlme et al. 2020) | Põlme S, et al. 2020. Fungal Diversity 105:1-16 ((https://doi.org/10.1007/s13225-020-00466-2)) |
| Otu52267 | 1 | k__Fungi(100);p__Ascomycota(100);c__Saccharomycetes(100);o__Saccharomycetales(100);f__Metschnikowiaceae(100);g__Metschnikowia(100);s__Metschnikowia_pulcherrima(100); | Metschnikowia | 13 | Saprotroph | \|Undefined Saprotroph\| | Yeast | NULL | Probable | As nectar-tap_saprotroph (Põlme et al. 2020); Decay sugar-rich substrates (Põlme et al. 2020) | Põlme S, et al. 2020. Fungal Diversity 105:1-16 ((https://doi.org/10.1007/s13225-020-00466-2)) |
| Otu52269 | 1 | k__Fungi(100);p__Ascomycota(100);c__Saccharomycetes(100);o__Saccharomycetales(100);f__Metschnikowiaceae(100);g__Metschnikowia(100);s__Metschnikowia_pulcherrima(100); | Metschnikowia | 13 | Saprotroph | \|Undefined Saprotroph\| | Yeast | NULL | Probable | As nectar-tap_saprotroph (Põlme et al. 2020); Decay sugar-rich substrates (Põlme et al. 2020) | Põlme S, et al. 2020. Fungal Diversity 105:1-16 ((https://doi.org/10.1007/s13225-020-00466-2)) |
| Otu52280 | 1 | k__Fungi(100);p__Ascomycota(100);c__Saccharomycetes(100);o__Saccharomycetales(100);f__Metschnikowiaceae(100);g__Metschnikowia(100);s__Metschnikowia_pulcherrima(100); | Metschnikowia | 13 | Saprotroph | \|Undefined Saprotroph\| | Yeast | NULL | Probable | As nectar-tap_saprotroph (Põlme et al. 2020); Decay sugar-rich substrates (Põlme et al. 2020) | Põlme S, et al. 2020. Fungal Diversity 105:1-16 ((https://doi.org/10.1007/s13225-020-00466-2)) |
| Otu52281 | 1 | k__Fungi(100);p__Ascomycota(100);c__Saccharomycetes(100);o__Saccharomycetales(100);f__Metschnikowiaceae(100);g__Metschnikowia(100);s__Metschnikowia_pulcherrima(100); | Metschnikowia | 13 | Saprotroph | \|Undefined Saprotroph\| | Yeast | NULL | Probable | As nectar-tap_saprotroph (Põlme et al. 2020); Decay sugar-rich substrates (Põlme et al. 2020) | Põlme S, et al. 2020. Fungal Diversity 105:1-16 ((https://doi.org/10.1007/s13225-020-00466-2)) |
| Otu52282 | 1 | k__Fungi(100);p__Ascomycota(100);c__Saccharomycetes(100);o__Saccharomycetales(100);f__Metschnikowiaceae(100);g__Metschnikowia(100);g__Metschnikowia_unclassified(100); | Metschnikowia | 13 | Saprotroph | \|Undefined Saprotroph\| | Yeast | NULL | Probable | As nectar-tap_saprotroph (Põlme et al. 2020); Decay sugar-rich substrates (Põlme et al. 2020) | Põlme S, et al. 2020. Fungal Diversity 105:1-16 ((https://doi.org/10.1007/s13225-020-00466-2)) |
| Otu52283 | 1 | k__Fungi(100);p__Ascomycota(100);c__Saccharomycetes(100);o__Saccharomycetales(100);f__Metschnikowiaceae(100);g__Metschnikowia(100);g__Metschnikowia_unclassified(100); | Metschnikowia | 13 | Saprotroph | \|Undefined Saprotroph\| | Yeast | NULL | Probable | As nectar-tap_saprotroph (Põlme et al. 2020); Decay sugar-rich substrates (Põlme et al. 2020) | Põlme S, et al. 2020. Fungal Diversity 105:1-16 ((https://doi.org/10.1007/s13225-020-00466-2)) |
| Otu52284 | 1 | k__Fungi(100);p__Ascomycota(100);c__Saccharomycetes(100);o__Saccharomycetales(100);f__Metschnikowiaceae(100);g__Metschnikowia(100);s__Metschnikowia_pulcherrima(100); | Metschnikowia | 13 | Saprotroph | \|Undefined Saprotroph\| | Yeast | NULL | Probable | As nectar-tap_saprotroph (Põlme et al. 2020); Decay sugar-rich substrates (Põlme et al. 2020) | Põlme S, et al. 2020. Fungal Diversity 105:1-16 ((https://doi.org/10.1007/s13225-020-00466-2)) |
| Otu52285 | 1 | k__Fungi(100);p__Ascomycota(100);c__Saccharomycetes(100);o__Saccharomycetales(100);f__Metschnikowiaceae(100);g__Metschnikowia(100);s__Metschnikowia_pulcherrima(100); | Metschnikowia | 13 | Saprotroph | \|Undefined Saprotroph\| | Yeast | NULL | Probable | As nectar-tap_saprotroph (Põlme et al. 2020); Decay sugar-rich substrates (Põlme et al. 2020) | Põlme S, et al. 2020. Fungal Diversity 105:1-16 ((https://doi.org/10.1007/s13225-020-00466-2)) |
| Otu52286 | 1 | k__Fungi(100);p__Ascomycota(100);c__Saccharomycetes(100);o__Saccharomycetales(100);f__Metschnikowiaceae(100);g__Metschnikowia(100);g__Metschnikowia_unclassified(100); | Metschnikowia | 13 | Saprotroph | \|Undefined Saprotroph\| | Yeast | NULL | Probable | As nectar-tap_saprotroph (Põlme et al. 2020); Decay sugar-rich substrates (Põlme et al. 2020) | Põlme S, et al. 2020. Fungal Diversity 105:1-16 ((https://doi.org/10.1007/s13225-020-00466-2)) |
| Otu52287 | 1 | k__Fungi(100);p__Ascomycota(100);c__Saccharomycetes(100);o__Saccharomycetales(100);f__Metschnikowiaceae(100);g__Metschnikowia(100);s__Metschnikowia_pulcherrima(100); | Metschnikowia | 13 | Saprotroph | \|Undefined Saprotroph\| | Yeast | NULL | Probable | As nectar-tap_saprotroph (Põlme et al. 2020); Decay sugar-rich substrates (Põlme et al. 2020) | Põlme S, et al. 2020. Fungal Diversity 105:1-16 ((https://doi.org/10.1007/s13225-020-00466-2)) |
| Otu52288 | 1 | k__Fungi(100);p__Ascomycota(100);c__Saccharomycetes(100);o__Saccharomycetales(100);f__Metschnikowiaceae(100);g__Metschnikowia(100);g__Metschnikowia_unclassified(100); | Metschnikowia | 13 | Saprotroph | \|Undefined Saprotroph\| | Yeast | NULL | Probable | As nectar-tap_saprotroph (Põlme et al. 2020); Decay sugar-rich substrates (Põlme et al. 2020) | Põlme S, et al. 2020. Fungal Diversity 105:1-16 ((https://doi.org/10.1007/s13225-020-00466-2)) |
| Otu52289 | 1 | k__Fungi(100);p__Ascomycota(100);c__Saccharomycetes(100);o__Saccharomycetales(100);f__Metschnikowiaceae(100);g__Metschnikowia(100);s__Metschnikowia_pulcherrima(100); | Metschnikowia | 13 | Saprotroph | \|Undefined Saprotroph\| | Yeast | NULL | Probable | As nectar-tap_saprotroph (Põlme et al. 2020); Decay sugar-rich substrates (Põlme et al. 2020) | Põlme S, et al. 2020. Fungal Diversity 105:1-16 ((https://doi.org/10.1007/s13225-020-00466-2)) |
| Otu52290 | 1 | k__Fungi(100);p__Ascomycota(100);c__Saccharomycetes(100);o__Saccharomycetales(100);f__Metschnikowiaceae(100);g__Metschnikowia(100);s__Metschnikowia_pulcherrima(100); | Metschnikowia | 13 | Saprotroph | \|Undefined Saprotroph\| | Yeast | NULL | Probable | As nectar-tap_saprotroph (Põlme et al. 2020); Decay sugar-rich substrates (Põlme et al. 2020) | Põlme S, et al. 2020. Fungal Diversity 105:1-16 ((https://doi.org/10.1007/s13225-020-00466-2)) |
| Otu52291 | 1 | k__Fungi(100);p__Ascomycota(100);c__Saccharomycetes(100);o__Saccharomycetales(100);f__Metschnikowiaceae(100);g__Metschnikowia(100);s__Metschnikowia_pulcherrima(100); | Metschnikowia | 13 | Saprotroph | \|Undefined Saprotroph\| | Yeast | NULL | Probable | As nectar-tap_saprotroph (Põlme et al. 2020); Decay sugar-rich substrates (Põlme et al. 2020) | Põlme S, et al. 2020. Fungal Diversity 105:1-16 ((https://doi.org/10.1007/s13225-020-00466-2)) |
| Otu52292 | 1 | k__Fungi(100);p__Ascomycota(100);c__Saccharomycetes(100);o__Saccharomycetales(100);f__Metschnikowiaceae(100);g__Metschnikowia(100);s__Metschnikowia_pulcherrima(100); | Metschnikowia | 13 | Saprotroph | \|Undefined Saprotroph\| | Yeast | NULL | Probable | As nectar-tap_saprotroph (Põlme et al. 2020); Decay sugar-rich substrates (Põlme et al. 2020) | Põlme S, et al. 2020. Fungal Diversity 105:1-16 ((https://doi.org/10.1007/s13225-020-00466-2)) |
| Otu52293 | 1 | k__Fungi(100);p__Ascomycota(100);c__Saccharomycetes(100);o__Saccharomycetales(100);f__Metschnikowiaceae(100);g__Metschnikowia(100);g__Metschnikowia_unclassified(100); | Metschnikowia | 13 | Saprotroph | \|Undefined Saprotroph\| | Yeast | NULL | Probable | As nectar-tap_saprotroph (Põlme et al. 2020); Decay sugar-rich substrates (Põlme et al. 2020) | Põlme S, et al. 2020. Fungal Diversity 105:1-16 ((https://doi.org/10.1007/s13225-020-00466-2)) |
| Otu52297 | 1 | k__Fungi(100);p__Ascomycota(100);c__Saccharomycetes(100);o__Saccharomycetales(100);f__Metschnikowiaceae(100);g__Metschnikowia(100);g__Metschnikowia_unclassified(100); | Metschnikowia | 13 | Saprotroph | \|Undefined Saprotroph\| | Yeast | NULL | Probable | As nectar-tap_saprotroph (Põlme et al. 2020); Decay sugar-rich substrates (Põlme et al. 2020) | Põlme S, et al. 2020. Fungal Diversity 105:1-16 ((https://doi.org/10.1007/s13225-020-00466-2)) |
| Otu52298 | 1 | k__Fungi(100);p__Ascomycota(100);c__Saccharomycetes(100);o__Saccharomycetales(100);f__Metschnikowiaceae(100);g__Metschnikowia(100);s__Metschnikowia_pulcherrima(100); | Metschnikowia | 13 | Saprotroph | \|Undefined Saprotroph\| | Yeast | NULL | Probable | As nectar-tap_saprotroph (Põlme et al. 2020); Decay sugar-rich substrates (Põlme et al. 2020) | Põlme S, et al. 2020. Fungal Diversity 105:1-16 ((https://doi.org/10.1007/s13225-020-00466-2)) |
| Otu52300 | 1 | k__Fungi(100);p__Ascomycota(100);c__Saccharomycetes(100);o__Saccharomycetales(100);f__Metschnikowiaceae(100);g__Metschnikowia(100);s__Metschnikowia_pulcherrima(100); | Metschnikowia | 13 | Saprotroph | \|Undefined Saprotroph\| | Yeast | NULL | Probable | As nectar-tap_saprotroph (Põlme et al. 2020); Decay sugar-rich substrates (Põlme et al. 2020) | Põlme S, et al. 2020. Fungal Diversity 105:1-16 ((https://doi.org/10.1007/s13225-020-00466-2)) |
| Otu52302 | 1 | k__Fungi(100);p__Ascomycota(100);c__Saccharomycetes(100);o__Saccharomycetales(100);f__Metschnikowiaceae(100);g__Metschnikowia(100);s__Metschnikowia_pulcherrima(100); | Metschnikowia | 13 | Saprotroph | \|Undefined Saprotroph\| | Yeast | NULL | Probable | As nectar-tap_saprotroph (Põlme et al. 2020); Decay sugar-rich substrates (Põlme et al. 2020) | Põlme S, et al. 2020. Fungal Diversity 105:1-16 ((https://doi.org/10.1007/s13225-020-00466-2)) |
| Otu52303 | 1 | k__Fungi(100);p__Ascomycota(100);c__Saccharomycetes(100);o__Saccharomycetales(100);f__Metschnikowiaceae(100);g__Metschnikowia(100);s__Metschnikowia_pulcherrima(100); | Metschnikowia | 13 | Saprotroph | \|Undefined Saprotroph\| | Yeast | NULL | Probable | As nectar-tap_saprotroph (Põlme et al. 2020); Decay sugar-rich substrates (Põlme et al. 2020) | Põlme S, et al. 2020. Fungal Diversity 105:1-16 ((https://doi.org/10.1007/s13225-020-00466-2)) |
| Otu52304 | 1 | k__Fungi(100);p__Ascomycota(100);c__Saccharomycetes(100);o__Saccharomycetales(100);f__Metschnikowiaceae(100);g__Metschnikowia(100);s__Metschnikowia_pulcherrima(100); | Metschnikowia | 13 | Saprotroph | \|Undefined Saprotroph\| | Yeast | NULL | Probable | As nectar-tap_saprotroph (Põlme et al. 2020); Decay sugar-rich substrates (Põlme et al. 2020) | Põlme S, et al. 2020. Fungal Diversity 105:1-16 ((https://doi.org/10.1007/s13225-020-00466-2)) |
| Otu52305 | 1 | k__Fungi(100);p__Ascomycota(100);c__Saccharomycetes(100);o__Saccharomycetales(100);f__Metschnikowiaceae(100);g__Metschnikowia(100);s__Metschnikowia_pulcherrima(100); | Metschnikowia | 13 | Saprotroph | \|Undefined Saprotroph\| | Yeast | NULL | Probable | As nectar-tap_saprotroph (Põlme et al. 2020); Decay sugar-rich substrates (Põlme et al. 2020) | Põlme S, et al. 2020. Fungal Diversity 105:1-16 ((https://doi.org/10.1007/s13225-020-00466-2)) |
| Otu52306 | 1 | k__Fungi(100);p__Ascomycota(100);c__Saccharomycetes(100);o__Saccharomycetales(100);f__Metschnikowiaceae(100);g__Metschnikowia(100);g__Metschnikowia_unclassified(100); | Metschnikowia | 13 | Saprotroph | \|Undefined Saprotroph\| | Yeast | NULL | Probable | As nectar-tap_saprotroph (Põlme et al. 2020); Decay sugar-rich substrates (Põlme et al. 2020) | Põlme S, et al. 2020. Fungal Diversity 105:1-16 ((https://doi.org/10.1007/s13225-020-00466-2)) |
| Otu52307 | 1 | k__Fungi(100);p__Ascomycota(100);c__Saccharomycetes(100);o__Saccharomycetales(100);f__Metschnikowiaceae(100);g__Metschnikowia(100);g__Metschnikowia_unclassified(100); | Metschnikowia | 13 | Saprotroph | \|Undefined Saprotroph\| | Yeast | NULL | Probable | As nectar-tap_saprotroph (Põlme et al. 2020); Decay sugar-rich substrates (Põlme et al. 2020) | Põlme S, et al. 2020. Fungal Diversity 105:1-16 ((https://doi.org/10.1007/s13225-020-00466-2)) |
| Otu52308 | 1 | k__Fungi(100);p__Ascomycota(100);c__Saccharomycetes(100);o__Saccharomycetales(100);f__Metschnikowiaceae(100);g__Metschnikowia(100);s__Metschnikowia_pulcherrima(100); | Metschnikowia | 13 | Saprotroph | \|Undefined Saprotroph\| | Yeast | NULL | Probable | As nectar-tap_saprotroph (Põlme et al. 2020); Decay sugar-rich substrates (Põlme et al. 2020) | Põlme S, et al. 2020. Fungal Diversity 105:1-16 ((https://doi.org/10.1007/s13225-020-00466-2)) |
| Otu52309 | 1 | k__Fungi(100);p__Ascomycota(100);c__Saccharomycetes(100);o__Saccharomycetales(100);f__Metschnikowiaceae(100);g__Metschnikowia(100);s__Metschnikowia_pulcherrima(100); | Metschnikowia | 13 | Saprotroph | \|Undefined Saprotroph\| | Yeast | NULL | Probable | As nectar-tap_saprotroph (Põlme et al. 2020); Decay sugar-rich substrates (Põlme et al. 2020) | Põlme S, et al. 2020. Fungal Diversity 105:1-16 ((https://doi.org/10.1007/s13225-020-00466-2)) |
| Otu52310 | 1 | k__Fungi(100);p__Ascomycota(100);c__Saccharomycetes(100);o__Saccharomycetales(100);f__Metschnikowiaceae(100);g__Metschnikowia(100);s__Metschnikowia_pulcherrima(100); | Metschnikowia | 13 | Saprotroph | \|Undefined Saprotroph\| | Yeast | NULL | Probable | As nectar-tap_saprotroph (Põlme et al. 2020); Decay sugar-rich substrates (Põlme et al. 2020) | Põlme S, et al. 2020. Fungal Diversity 105:1-16 ((https://doi.org/10.1007/s13225-020-00466-2)) |
| Otu52311 | 1 | k__Fungi(100);p__Ascomycota(100);c__Saccharomycetes(100);o__Saccharomycetales(100);f__Metschnikowiaceae(100);g__Metschnikowia(100);s__Metschnikowia_pulcherrima(100); | Metschnikowia | 13 | Saprotroph | \|Undefined Saprotroph\| | Yeast | NULL | Probable | As nectar-tap_saprotroph (Põlme et al. 2020); Decay sugar-rich substrates (Põlme et al. 2020) | Põlme S, et al. 2020. Fungal Diversity 105:1-16 ((https://doi.org/10.1007/s13225-020-00466-2)) |
| Otu52312 | 1 | k__Fungi(100);p__Ascomycota(100);c__Saccharomycetes(100);o__Saccharomycetales(100);f__Metschnikowiaceae(100);g__Metschnikowia(100);s__Metschnikowia_pulcherrima(100); | Metschnikowia | 13 | Saprotroph | \|Undefined Saprotroph\| | Yeast | NULL | Probable | As nectar-tap_saprotroph (Põlme et al. 2020); Decay sugar-rich substrates (Põlme et al. 2020) | Põlme S, et al. 2020. Fungal Diversity 105:1-16 ((https://doi.org/10.1007/s13225-020-00466-2)) |
| Otu52313 | 1 | k__Fungi(100);p__Ascomycota(100);c__Saccharomycetes(100);o__Saccharomycetales(100);f__Metschnikowiaceae(100);g__Metschnikowia(100);s__Metschnikowia_pulcherrima(100); | Metschnikowia | 13 | Saprotroph | \|Undefined Saprotroph\| | Yeast | NULL | Probable | As nectar-tap_saprotroph (Põlme et al. 2020); Decay sugar-rich substrates (Põlme et al. 2020) | Põlme S, et al. 2020. Fungal Diversity 105:1-16 ((https://doi.org/10.1007/s13225-020-00466-2)) |
| Otu52314 | 1 | k__Fungi(100);p__Ascomycota(100);c__Saccharomycetes(100);o__Saccharomycetales(100);f__Metschnikowiaceae(100);g__Metschnikowia(100);s__Metschnikowia_pulcherrima(100); | Metschnikowia | 13 | Saprotroph | \|Undefined Saprotroph\| | Yeast | NULL | Probable | As nectar-tap_saprotroph (Põlme et al. 2020); Decay sugar-rich substrates (Põlme et al. 2020) | Põlme S, et al. 2020. Fungal Diversity 105:1-16 ((https://doi.org/10.1007/s13225-020-00466-2)) |
| Otu52315 | 1 | k__Fungi(100);p__Ascomycota(100);c__Saccharomycetes(100);o__Saccharomycetales(100);f__Metschnikowiaceae(100);g__Metschnikowia(100);g__Metschnikowia_unclassified(100); | Metschnikowia | 13 | Saprotroph | \|Undefined Saprotroph\| | Yeast | NULL | Probable | As nectar-tap_saprotroph (Põlme et al. 2020); Decay sugar-rich substrates (Põlme et al. 2020) | Põlme S, et al. 2020. Fungal Diversity 105:1-16 ((https://doi.org/10.1007/s13225-020-00466-2)) |
| Otu52316 | 1 | k__Fungi(100);p__Ascomycota(100);c__Saccharomycetes(100);o__Saccharomycetales(100);f__Metschnikowiaceae(100);g__Metschnikowia(100);g__Metschnikowia_unclassified(100); | Metschnikowia | 13 | Saprotroph | \|Undefined Saprotroph\| | Yeast | NULL | Probable | As nectar-tap_saprotroph (Põlme et al. 2020); Decay sugar-rich substrates (Põlme et al. 2020) | Põlme S, et al. 2020. Fungal Diversity 105:1-16 ((https://doi.org/10.1007/s13225-020-00466-2)) |
| Otu52317 | 1 | k__Fungi(100);p__Ascomycota(100);c__Saccharomycetes(100);o__Saccharomycetales(100);f__Metschnikowiaceae(100);g__Metschnikowia(100);g__Metschnikowia_unclassified(100); | Metschnikowia | 13 | Saprotroph | \|Undefined Saprotroph\| | Yeast | NULL | Probable | As nectar-tap_saprotroph (Põlme et al. 2020); Decay sugar-rich substrates (Põlme et al. 2020) | Põlme S, et al. 2020. Fungal Diversity 105:1-16 ((https://doi.org/10.1007/s13225-020-00466-2)) |
| Otu52318 | 1 | k__Fungi(100);p__Ascomycota(100);c__Saccharomycetes(100);o__Saccharomycetales(100);f__Metschnikowiaceae(100);g__Metschnikowia(100);s__Metschnikowia_pulcherrima(100); | Metschnikowia | 13 | Saprotroph | \|Undefined Saprotroph\| | Yeast | NULL | Probable | As nectar-tap_saprotroph (Põlme et al. 2020); Decay sugar-rich substrates (Põlme et al. 2020) | Põlme S, et al. 2020. Fungal Diversity 105:1-16 ((https://doi.org/10.1007/s13225-020-00466-2)) |
| Otu52320 | 1 | k__Fungi(100);p__Ascomycota(100);c__Saccharomycetes(100);o__Saccharomycetales(100);f__Metschnikowiaceae(100);g__Metschnikowia(100);g__Metschnikowia_unclassified(100); | Metschnikowia | 13 | Saprotroph | \|Undefined Saprotroph\| | Yeast | NULL | Probable | As nectar-tap_saprotroph (Põlme et al. 2020); Decay sugar-rich substrates (Põlme et al. 2020) | Põlme S, et al. 2020. Fungal Diversity 105:1-16 ((https://doi.org/10.1007/s13225-020-00466-2)) |
| Otu52321 | 1 | k__Fungi(100);p__Ascomycota(100);c__Saccharomycetes(100);o__Saccharomycetales(100);f__Saccharomycetaceae(100);g__Kazachstania(100);s__Kazachstania_intestinalis(100); | Kazachstania | 13 | Saprotroph | \|Undefined Saprotroph\| | Yeast | NULL | Probable | As nectar-tap_saprotroph (Põlme et al. 2020); Decay sugar-rich substrates (Põlme et al. 2020) | Põlme S, et al. 2020. Fungal Diversity 105:1-16 ((https://doi.org/10.1007/s13225-020-00466-2)) |
| Otu52322 | 1 | k__Fungi(100);p__Ascomycota(100);c__Saccharomycetes(100);o__Saccharomycetales(100);f__Metschnikowiaceae(100);g__Metschnikowia(100);s__Metschnikowia_pulcherrima(100); | Metschnikowia | 13 | Saprotroph | \|Undefined Saprotroph\| | Yeast | NULL | Probable | As nectar-tap_saprotroph (Põlme et al. 2020); Decay sugar-rich substrates (Põlme et al. 2020) | Põlme S, et al. 2020. Fungal Diversity 105:1-16 ((https://doi.org/10.1007/s13225-020-00466-2)) |
| Otu52323 | 1 | k__Fungi(100);p__Ascomycota(100);c__Saccharomycetes(100);o__Saccharomycetales(100);f__Metschnikowiaceae(100);g__Metschnikowia(100);s__Metschnikowia_pulcherrima(100); | Metschnikowia | 13 | Saprotroph | \|Undefined Saprotroph\| | Yeast | NULL | Probable | As nectar-tap_saprotroph (Põlme et al. 2020); Decay sugar-rich substrates (Põlme et al. 2020) | Põlme S, et al. 2020. Fungal Diversity 105:1-16 ((https://doi.org/10.1007/s13225-020-00466-2)) |
| Otu52324 | 1 | k__Fungi(100);p__Ascomycota(100);c__Saccharomycetes(100);o__Saccharomycetales(100);f__Metschnikowiaceae(100);g__Metschnikowia(100);s__Metschnikowia_pulcherrima(100); | Metschnikowia | 13 | Saprotroph | \|Undefined Saprotroph\| | Yeast | NULL | Probable | As nectar-tap_saprotroph (Põlme et al. 2020); Decay sugar-rich substrates (Põlme et al. 2020) | Põlme S, et al. 2020. Fungal Diversity 105:1-16 ((https://doi.org/10.1007/s13225-020-00466-2)) |
| Otu52325 | 1 | k__Fungi(100);p__Ascomycota(100);c__Saccharomycetes(100);o__Saccharomycetales(100);f__Metschnikowiaceae(100);g__Metschnikowia(100);s__Metschnikowia_pulcherrima(100); | Metschnikowia | 13 | Saprotroph | \|Undefined Saprotroph\| | Yeast | NULL | Probable | As nectar-tap_saprotroph (Põlme et al. 2020); Decay sugar-rich substrates (Põlme et al. 2020) | Põlme S, et al. 2020. Fungal Diversity 105:1-16 ((https://doi.org/10.1007/s13225-020-00466-2)) |
| Otu52327 | 1 | k__Fungi(100);p__Ascomycota(100);c__Saccharomycetes(100);o__Saccharomycetales(100);f__Metschnikowiaceae(100);g__Metschnikowia(100);s__Metschnikowia_pulcherrima(100); | Metschnikowia | 13 | Saprotroph | \|Undefined Saprotroph\| | Yeast | NULL | Probable | As nectar-tap_saprotroph (Põlme et al. 2020); Decay sugar-rich substrates (Põlme et al. 2020) | Põlme S, et al. 2020. Fungal Diversity 105:1-16 ((https://doi.org/10.1007/s13225-020-00466-2)) |
| Otu52328 | 1 | k__Fungi(100);p__Ascomycota(100);c__Saccharomycetes(100);o__Saccharomycetales(100);f__Metschnikowiaceae(100);g__Metschnikowia(100);s__Metschnikowia_pulcherrima(100); | Metschnikowia | 13 | Saprotroph | \|Undefined Saprotroph\| | Yeast | NULL | Probable | As nectar-tap_saprotroph (Põlme et al. 2020); Decay sugar-rich substrates (Põlme et al. 2020) | Põlme S, et al. 2020. Fungal Diversity 105:1-16 ((https://doi.org/10.1007/s13225-020-00466-2)) |
| Otu52329 | 1 | k__Fungi(100);p__Ascomycota(100);c__Saccharomycetes(100);o__Saccharomycetales(100);f__Metschnikowiaceae(100);g__Metschnikowia(100);g__Metschnikowia_unclassified(100); | Metschnikowia | 13 | Saprotroph | \|Undefined Saprotroph\| | Yeast | NULL | Probable | As nectar-tap_saprotroph (Põlme et al. 2020); Decay sugar-rich substrates (Põlme et al. 2020) | Põlme S, et al. 2020. Fungal Diversity 105:1-16 ((https://doi.org/10.1007/s13225-020-00466-2)) |
| Otu52330 | 1 | k__Fungi(100);p__Ascomycota(100);c__Saccharomycetes(100);o__Saccharomycetales(100);f__Metschnikowiaceae(100);g__Metschnikowia(100);g__Metschnikowia_unclassified(100); | Metschnikowia | 13 | Saprotroph | \|Undefined Saprotroph\| | Yeast | NULL | Probable | As nectar-tap_saprotroph (Põlme et al. 2020); Decay sugar-rich substrates (Põlme et al. 2020) | Põlme S, et al. 2020. Fungal Diversity 105:1-16 ((https://doi.org/10.1007/s13225-020-00466-2)) |
| Otu52331 | 1 | k__Fungi(100);p__Ascomycota(100);c__Saccharomycetes(100);o__Saccharomycetales(100);f__Metschnikowiaceae(100);g__Metschnikowia(100);s__Metschnikowia_pulcherrima(100); | Metschnikowia | 13 | Saprotroph | \|Undefined Saprotroph\| | Yeast | NULL | Probable | As nectar-tap_saprotroph (Põlme et al. 2020); Decay sugar-rich substrates (Põlme et al. 2020) | Põlme S, et al. 2020. Fungal Diversity 105:1-16 ((https://doi.org/10.1007/s13225-020-00466-2)) |
| Otu52332 | 1 | k__Fungi(100);p__Ascomycota(100);c__Saccharomycetes(100);o__Saccharomycetales(100);f__Metschnikowiaceae(100);g__Metschnikowia(100);g__Metschnikowia_unclassified(100); | Metschnikowia | 13 | Saprotroph | \|Undefined Saprotroph\| | Yeast | NULL | Probable | As nectar-tap_saprotroph (Põlme et al. 2020); Decay sugar-rich substrates (Põlme et al. 2020) | Põlme S, et al. 2020. Fungal Diversity 105:1-16 ((https://doi.org/10.1007/s13225-020-00466-2)) |
| Otu52333 | 1 | k__Fungi(100);p__Ascomycota(100);c__Saccharomycetes(100);o__Saccharomycetales(100);f__Metschnikowiaceae(100);g__Metschnikowia(100);g__Metschnikowia_unclassified(100); | Metschnikowia | 13 | Saprotroph | \|Undefined Saprotroph\| | Yeast | NULL | Probable | As nectar-tap_saprotroph (Põlme et al. 2020); Decay sugar-rich substrates (Põlme et al. 2020) | Põlme S, et al. 2020. Fungal Diversity 105:1-16 ((https://doi.org/10.1007/s13225-020-00466-2)) |
| Otu52334 | 1 | k__Fungi(100);p__Ascomycota(100);c__Saccharomycetes(100);o__Saccharomycetales(100);f__Metschnikowiaceae(100);g__Metschnikowia(100);s__Metschnikowia_pulcherrima(100); | Metschnikowia | 13 | Saprotroph | \|Undefined Saprotroph\| | Yeast | NULL | Probable | As nectar-tap_saprotroph (Põlme et al. 2020); Decay sugar-rich substrates (Põlme et al. 2020) | Põlme S, et al. 2020. Fungal Diversity 105:1-16 ((https://doi.org/10.1007/s13225-020-00466-2)) |
| Otu52335 | 1 | k__Fungi(100);p__Ascomycota(100);c__Saccharomycetes(100);o__Saccharomycetales(100);f__Metschnikowiaceae(100);g__Metschnikowia(100);s__Metschnikowia_pulcherrima(100); | Metschnikowia | 13 | Saprotroph | \|Undefined Saprotroph\| | Yeast | NULL | Probable | As nectar-tap_saprotroph (Põlme et al. 2020); Decay sugar-rich substrates (Põlme et al. 2020) | Põlme S, et al. 2020. Fungal Diversity 105:1-16 ((https://doi.org/10.1007/s13225-020-00466-2)) |
| Otu52336 | 1 | k__Fungi(100);p__Ascomycota(100);c__Saccharomycetes(100);o__Saccharomycetales(100);f__Metschnikowiaceae(100);g__Metschnikowia(100);s__Metschnikowia_pulcherrima(100); | Metschnikowia | 13 | Saprotroph | \|Undefined Saprotroph\| | Yeast | NULL | Probable | As nectar-tap_saprotroph (Põlme et al. 2020); Decay sugar-rich substrates (Põlme et al. 2020) | Põlme S, et al. 2020. Fungal Diversity 105:1-16 ((https://doi.org/10.1007/s13225-020-00466-2)) |
| Otu52337 | 1 | k__Fungi(100);p__Ascomycota(100);c__Saccharomycetes(100);o__Saccharomycetales(100);f__Metschnikowiaceae(100);g__Metschnikowia(100);g__Metschnikowia_unclassified(100); | Metschnikowia | 13 | Saprotroph | \|Undefined Saprotroph\| | Yeast | NULL | Probable | As nectar-tap_saprotroph (Põlme et al. 2020); Decay sugar-rich substrates (Põlme et al. 2020) | Põlme S, et al. 2020. Fungal Diversity 105:1-16 ((https://doi.org/10.1007/s13225-020-00466-2)) |
| Otu52338 | 1 | k__Fungi(100);p__Ascomycota(100);c__Saccharomycetes(100);o__Saccharomycetales(100);f__Metschnikowiaceae(100);g__Metschnikowia(100);s__Metschnikowia_pulcherrima(100); | Metschnikowia | 13 | Saprotroph | \|Undefined Saprotroph\| | Yeast | NULL | Probable | As nectar-tap_saprotroph (Põlme et al. 2020); Decay sugar-rich substrates (Põlme et al. 2020) | Põlme S, et al. 2020. Fungal Diversity 105:1-16 ((https://doi.org/10.1007/s13225-020-00466-2)) |
| Otu52339 | 1 | k__Fungi(100);p__Ascomycota(100);c__Saccharomycetes(100);o__Saccharomycetales(100);f__Metschnikowiaceae(100);g__Metschnikowia(100);g__Metschnikowia_unclassified(100); | Metschnikowia | 13 | Saprotroph | \|Undefined Saprotroph\| | Yeast | NULL | Probable | As nectar-tap_saprotroph (Põlme et al. 2020); Decay sugar-rich substrates (Põlme et al. 2020) | Põlme S, et al. 2020. Fungal Diversity 105:1-16 ((https://doi.org/10.1007/s13225-020-00466-2)) |
| Otu52341 | 1 | k__Fungi(100);p__Ascomycota(100);c__Saccharomycetes(100);o__Saccharomycetales(100);f__Metschnikowiaceae(100);g__Metschnikowia(100);g__Metschnikowia_unclassified(100); | Metschnikowia | 13 | Saprotroph | \|Undefined Saprotroph\| | Yeast | NULL | Probable | As nectar-tap_saprotroph (Põlme et al. 2020); Decay sugar-rich substrates (Põlme et al. 2020) | Põlme S, et al. 2020. Fungal Diversity 105:1-16 ((https://doi.org/10.1007/s13225-020-00466-2)) |
| Otu52342 | 1 | k__Fungi(100);p__Ascomycota(100);c__Saccharomycetes(100);o__Saccharomycetales(100);f__Metschnikowiaceae(100);g__Metschnikowia(100);g__Metschnikowia_unclassified(100); | Metschnikowia | 13 | Saprotroph | \|Undefined Saprotroph\| | Yeast | NULL | Probable | As nectar-tap_saprotroph (Põlme et al. 2020); Decay sugar-rich substrates (Põlme et al. 2020) | Põlme S, et al. 2020. Fungal Diversity 105:1-16 ((https://doi.org/10.1007/s13225-020-00466-2)) |
| Otu52343 | 1 | k__Fungi(100);p__Ascomycota(100);c__Saccharomycetes(100);o__Saccharomycetales(100);f__Metschnikowiaceae(100);g__Metschnikowia(100);g__Metschnikowia_unclassified(100); | Metschnikowia | 13 | Saprotroph | \|Undefined Saprotroph\| | Yeast | NULL | Probable | As nectar-tap_saprotroph (Põlme et al. 2020); Decay sugar-rich substrates (Põlme et al. 2020) | Põlme S, et al. 2020. Fungal Diversity 105:1-16 ((https://doi.org/10.1007/s13225-020-00466-2)) |
| Otu52344 | 1 | k__Fungi(100);p__Ascomycota(100);c__Saccharomycetes(100);o__Saccharomycetales(100);f__Metschnikowiaceae(100);g__Metschnikowia(100);s__Metschnikowia_pulcherrima(100); | Metschnikowia | 13 | Saprotroph | \|Undefined Saprotroph\| | Yeast | NULL | Probable | As nectar-tap_saprotroph (Põlme et al. 2020); Decay sugar-rich substrates (Põlme et al. 2020) | Põlme S, et al. 2020. Fungal Diversity 105:1-16 ((https://doi.org/10.1007/s13225-020-00466-2)) |
| Otu52345 | 1 | k__Fungi(100);p__Ascomycota(100);c__Saccharomycetes(100);o__Saccharomycetales(100);f__Metschnikowiaceae(100);g__Metschnikowia(100);s__Metschnikowia_pulcherrima(100); | Metschnikowia | 13 | Saprotroph | \|Undefined Saprotroph\| | Yeast | NULL | Probable | As nectar-tap_saprotroph (Põlme et al. 2020); Decay sugar-rich substrates (Põlme et al. 2020) | Põlme S, et al. 2020. Fungal Diversity 105:1-16 ((https://doi.org/10.1007/s13225-020-00466-2)) |
| Otu52346 | 1 | k__Fungi(100);p__Ascomycota(100);c__Saccharomycetes(100);o__Saccharomycetales(100);f__Metschnikowiaceae(100);g__Metschnikowia(100);s__Metschnikowia_pulcherrima(100); | Metschnikowia | 13 | Saprotroph | \|Undefined Saprotroph\| | Yeast | NULL | Probable | As nectar-tap_saprotroph (Põlme et al. 2020); Decay sugar-rich substrates (Põlme et al. 2020) | Põlme S, et al. 2020. Fungal Diversity 105:1-16 ((https://doi.org/10.1007/s13225-020-00466-2)) |
| Otu52347 | 1 | k__Fungi(100);p__Ascomycota(100);c__Saccharomycetes(100);o__Saccharomycetales(100);f__Metschnikowiaceae(100);g__Metschnikowia(100);s__Metschnikowia_pulcherrima(100); | Metschnikowia | 13 | Saprotroph | \|Undefined Saprotroph\| | Yeast | NULL | Probable | As nectar-tap_saprotroph (Põlme et al. 2020); Decay sugar-rich substrates (Põlme et al. 2020) | Põlme S, et al. 2020. Fungal Diversity 105:1-16 ((https://doi.org/10.1007/s13225-020-00466-2)) |
| Otu52348 | 1 | k__Fungi(100);p__Ascomycota(100);c__Saccharomycetes(100);o__Saccharomycetales(100);f__Metschnikowiaceae(100);g__Metschnikowia(100);s__Metschnikowia_pulcherrima(100); | Metschnikowia | 13 | Saprotroph | \|Undefined Saprotroph\| | Yeast | NULL | Probable | As nectar-tap_saprotroph (Põlme et al. 2020); Decay sugar-rich substrates (Põlme et al. 2020) | Põlme S, et al. 2020. Fungal Diversity 105:1-16 ((https://doi.org/10.1007/s13225-020-00466-2)) |
| Otu52349 | 1 | k__Fungi(100);p__Ascomycota(100);c__Saccharomycetes(100);o__Saccharomycetales(100);f__Metschnikowiaceae(100);g__Metschnikowia(100);g__Metschnikowia_unclassified(100); | Metschnikowia | 13 | Saprotroph | \|Undefined Saprotroph\| | Yeast | NULL | Probable | As nectar-tap_saprotroph (Põlme et al. 2020); Decay sugar-rich substrates (Põlme et al. 2020) | Põlme S, et al. 2020. Fungal Diversity 105:1-16 ((https://doi.org/10.1007/s13225-020-00466-2)) |
| Otu52350 | 1 | k__Fungi(100);p__Ascomycota(100);c__Saccharomycetes(100);o__Saccharomycetales(100);f__Metschnikowiaceae(100);g__Metschnikowia(100);s__Metschnikowia_pulcherrima(100); | Metschnikowia | 13 | Saprotroph | \|Undefined Saprotroph\| | Yeast | NULL | Probable | As nectar-tap_saprotroph (Põlme et al. 2020); Decay sugar-rich substrates (Põlme et al. 2020) | Põlme S, et al. 2020. Fungal Diversity 105:1-16 ((https://doi.org/10.1007/s13225-020-00466-2)) |
| Otu52351 | 1 | k__Fungi(100);p__Ascomycota(100);c__Saccharomycetes(100);o__Saccharomycetales(100);f__Metschnikowiaceae(100);g__Metschnikowia(100);s__Metschnikowia_pulcherrima(100); | Metschnikowia | 13 | Saprotroph | \|Undefined Saprotroph\| | Yeast | NULL | Probable | As nectar-tap_saprotroph (Põlme et al. 2020); Decay sugar-rich substrates (Põlme et al. 2020) | Põlme S, et al. 2020. Fungal Diversity 105:1-16 ((https://doi.org/10.1007/s13225-020-00466-2)) |
| Otu52352 | 1 | k__Fungi(100);p__Ascomycota(100);c__Saccharomycetes(100);o__Saccharomycetales(100);f__Metschnikowiaceae(100);g__Metschnikowia(100);s__Metschnikowia_pulcherrima(100); | Metschnikowia | 13 | Saprotroph | \|Undefined Saprotroph\| | Yeast | NULL | Probable | As nectar-tap_saprotroph (Põlme et al. 2020); Decay sugar-rich substrates (Põlme et al. 2020) | Põlme S, et al. 2020. Fungal Diversity 105:1-16 ((https://doi.org/10.1007/s13225-020-00466-2)) |
| Otu52353 | 1 | k__Fungi(100);p__Ascomycota(100);c__Saccharomycetes(100);o__Saccharomycetales(100);f__Metschnikowiaceae(100);g__Metschnikowia(100);s__Metschnikowia_pulcherrima(100); | Metschnikowia | 13 | Saprotroph | \|Undefined Saprotroph\| | Yeast | NULL | Probable | As nectar-tap_saprotroph (Põlme et al. 2020); Decay sugar-rich substrates (Põlme et al. 2020) | Põlme S, et al. 2020. Fungal Diversity 105:1-16 ((https://doi.org/10.1007/s13225-020-00466-2)) |
| Otu52354 | 1 | k__Fungi(100);p__Ascomycota(100);c__Saccharomycetes(100);o__Saccharomycetales(100);f__Metschnikowiaceae(100);g__Metschnikowia(100);g__Metschnikowia_unclassified(100); | Metschnikowia | 13 | Saprotroph | \|Undefined Saprotroph\| | Yeast | NULL | Probable | As nectar-tap_saprotroph (Põlme et al. 2020); Decay sugar-rich substrates (Põlme et al. 2020) | Põlme S, et al. 2020. Fungal Diversity 105:1-16 ((https://doi.org/10.1007/s13225-020-00466-2)) |
| Otu52355 | 1 | k__Fungi(100);p__Basidiomycota(100);c__Microbotryomycetes(100);o__Sporidiobolales(100);f__Sporidiobolales_family_Incertae_sedis(100);g__Sporobolomyces(100);s__Sporobolomyces_lactosus(100); | Sporobolomyces | 13 | Pathotroph-Saprotroph | \|Fungal Parasite\|-Plant Saprotroph | Yeast | NULL | Probable | As litter_saprotroph (Põlme et al. 2020); As mycoparasite (Põlme et al. 2020); Decay fungal material (Põlme et al. 2020) | Tedersoo L, et al. 2014. Science 346:e1256688 ((https://doi.org/10.1126/science.1256688)); Purahong W, et al. 2016. Molecular Ecology 25:4059-4074 ((https://doi.org/10.1111/mec.13739)); Põlme S, et al. 2020. Fungal Diversity 105:1-16 ((https://doi.org/10.1007/s13225-020-00466-2)) |
| Otu52356 | 1 | k__Fungi(100);p__Ascomycota(100);c__Saccharomycetes(100);o__Saccharomycetales(100);f__Metschnikowiaceae(100);g__Metschnikowia(100);s__Metschnikowia_pulcherrima(100); | Metschnikowia | 13 | Saprotroph | \|Undefined Saprotroph\| | Yeast | NULL | Probable | As nectar-tap_saprotroph (Põlme et al. 2020); Decay sugar-rich substrates (Põlme et al. 2020) | Põlme S, et al. 2020. Fungal Diversity 105:1-16 ((https://doi.org/10.1007/s13225-020-00466-2)) |
| Otu52357 | 1 | k__Fungi(100);p__Ascomycota(100);c__Saccharomycetes(100);o__Saccharomycetales(100);f__Metschnikowiaceae(100);g__Metschnikowia(100);s__Metschnikowia_pulcherrima(100); | Metschnikowia | 13 | Saprotroph | \|Undefined Saprotroph\| | Yeast | NULL | Probable | As nectar-tap_saprotroph (Põlme et al. 2020); Decay sugar-rich substrates (Põlme et al. 2020) | Põlme S, et al. 2020. Fungal Diversity 105:1-16 ((https://doi.org/10.1007/s13225-020-00466-2)) |
| Otu52358 | 1 | k__Fungi(100);p__Ascomycota(100);c__Saccharomycetes(100);o__Saccharomycetales(100);f__Metschnikowiaceae(100);g__Metschnikowia(100);g__Metschnikowia_unclassified(100); | Metschnikowia | 13 | Saprotroph | \|Undefined Saprotroph\| | Yeast | NULL | Probable | As nectar-tap_saprotroph (Põlme et al. 2020); Decay sugar-rich substrates (Põlme et al. 2020) | Põlme S, et al. 2020. Fungal Diversity 105:1-16 ((https://doi.org/10.1007/s13225-020-00466-2)) |
| Otu52359 | 1 | k__Fungi(100);p__Ascomycota(100);c__Saccharomycetes(100);o__Saccharomycetales(100);f__Metschnikowiaceae(100);g__Metschnikowia(100);s__Metschnikowia_pulcherrima(100); | Metschnikowia | 13 | Saprotroph | \|Undefined Saprotroph\| | Yeast | NULL | Probable | As nectar-tap_saprotroph (Põlme et al. 2020); Decay sugar-rich substrates (Põlme et al. 2020) | Põlme S, et al. 2020. Fungal Diversity 105:1-16 ((https://doi.org/10.1007/s13225-020-00466-2)) |
| Otu52360 | 1 | k__Fungi(100);p__Ascomycota(100);c__Saccharomycetes(100);o__Saccharomycetales(100);f__Metschnikowiaceae(100);g__Metschnikowia(100);s__Metschnikowia_pulcherrima(100); | Metschnikowia | 13 | Saprotroph | \|Undefined Saprotroph\| | Yeast | NULL | Probable | As nectar-tap_saprotroph (Põlme et al. 2020); Decay sugar-rich substrates (Põlme et al. 2020) | Põlme S, et al. 2020. Fungal Diversity 105:1-16 ((https://doi.org/10.1007/s13225-020-00466-2)) |
| Otu52361 | 1 | k__Fungi(100);p__Ascomycota(100);c__Saccharomycetes(100);o__Saccharomycetales(100);f__Saccharomycetales_family_Incertae_sedis(100);g__Candida(100);g__Candida_unclassified(100); | Candida | 13 | Pathotroph-Saprotroph-Symbiotroph | Animal Pathogen-Endophyte-Endosymbiont-Epiphyte-\|Undefined Saprotroph\| | Dimorphic Yeast-Yeast | NULL | Probable | As nectar-tap_saprotroph (Põlme et al. 2020); Decay sugar-rich substrates (Põlme et al. 2020); This asexual genus is polyphyletic and contain species that function very differently depend on what they are. Caution should be taken when interpreting guilds for Candida species. | Manolakaki D, et al. 2010. Virulence. 1: 367-375 ((https://doi.org/10.4161/viru.1.5.12796)); Mekbib SB, et al. 2011. Tropical Plant Pathology 36:233-240 ((https://doi.org/10.1590/S1982-56762011000400004)); Põlme S, et al. 2020. Fungal Diversity 105:1-16 ((https://doi.org/10.1007/s13225-020-00466-2)) |
| Otu52363 | 1 | k__Fungi(100);p__Ascomycota(100);c__Saccharomycetes(100);o__Saccharomycetales(100);f__Metschnikowiaceae(100);g__Metschnikowia(100);s__Metschnikowia_pulcherrima(100); | Metschnikowia | 13 | Saprotroph | \|Undefined Saprotroph\| | Yeast | NULL | Probable | As nectar-tap_saprotroph (Põlme et al. 2020); Decay sugar-rich substrates (Põlme et al. 2020) | Põlme S, et al. 2020. Fungal Diversity 105:1-16 ((https://doi.org/10.1007/s13225-020-00466-2)) |
| Otu52364 | 1 | k__Fungi(100);p__Ascomycota(100);c__Saccharomycetes(100);o__Saccharomycetales(100);f__Metschnikowiaceae(100);g__Metschnikowia(100);s__Metschnikowia_pulcherrima(100); | Metschnikowia | 13 | Saprotroph | \|Undefined Saprotroph\| | Yeast | NULL | Probable | As nectar-tap_saprotroph (Põlme et al. 2020); Decay sugar-rich substrates (Põlme et al. 2020) | Põlme S, et al. 2020. Fungal Diversity 105:1-16 ((https://doi.org/10.1007/s13225-020-00466-2)) |
| Otu52365 | 1 | k__Fungi(100);p__Ascomycota(100);c__Saccharomycetes(100);o__Saccharomycetales(100);f__Metschnikowiaceae(100);g__Metschnikowia(100);g__Metschnikowia_unclassified(100); | Metschnikowia | 13 | Saprotroph | \|Undefined Saprotroph\| | Yeast | NULL | Probable | As nectar-tap_saprotroph (Põlme et al. 2020); Decay sugar-rich substrates (Põlme et al. 2020) | Põlme S, et al. 2020. Fungal Diversity 105:1-16 ((https://doi.org/10.1007/s13225-020-00466-2)) |
| Otu52366 | 1 | k__Fungi(100);p__Ascomycota(100);c__Saccharomycetes(100);o__Saccharomycetales(100);f__Metschnikowiaceae(100);g__Metschnikowia(100);g__Metschnikowia_unclassified(100); | Metschnikowia | 13 | Saprotroph | \|Undefined Saprotroph\| | Yeast | NULL | Probable | As nectar-tap_saprotroph (Põlme et al. 2020); Decay sugar-rich substrates (Põlme et al. 2020) | Põlme S, et al. 2020. Fungal Diversity 105:1-16 ((https://doi.org/10.1007/s13225-020-00466-2)) |
| Otu52367 | 1 | k__Fungi(100);p__Ascomycota(100);c__Saccharomycetes(100);o__Saccharomycetales(100);f__Metschnikowiaceae(100);g__Metschnikowia(100);g__Metschnikowia_unclassified(100); | Metschnikowia | 13 | Saprotroph | \|Undefined Saprotroph\| | Yeast | NULL | Probable | As nectar-tap_saprotroph (Põlme et al. 2020); Decay sugar-rich substrates (Põlme et al. 2020) | Põlme S, et al. 2020. Fungal Diversity 105:1-16 ((https://doi.org/10.1007/s13225-020-00466-2)) |
| Otu52368 | 1 | k__Fungi(100);p__Ascomycota(100);c__Saccharomycetes(100);o__Saccharomycetales(100);f__Metschnikowiaceae(100);g__Metschnikowia(100);s__Metschnikowia_pulcherrima(100); | Metschnikowia | 13 | Saprotroph | \|Undefined Saprotroph\| | Yeast | NULL | Probable | As nectar-tap_saprotroph (Põlme et al. 2020); Decay sugar-rich substrates (Põlme et al. 2020) | Põlme S, et al. 2020. Fungal Diversity 105:1-16 ((https://doi.org/10.1007/s13225-020-00466-2)) |
| Otu52369 | 1 | k__Fungi(100);p__Ascomycota(100);c__Saccharomycetes(100);o__Saccharomycetales(100);f__Metschnikowiaceae(100);g__Metschnikowia(100);g__Metschnikowia_unclassified(100); | Metschnikowia | 13 | Saprotroph | \|Undefined Saprotroph\| | Yeast | NULL | Probable | As nectar-tap_saprotroph (Põlme et al. 2020); Decay sugar-rich substrates (Põlme et al. 2020) | Põlme S, et al. 2020. Fungal Diversity 105:1-16 ((https://doi.org/10.1007/s13225-020-00466-2)) |
| Otu52370 | 1 | k__Fungi(100);p__Ascomycota(100);c__Saccharomycetes(100);o__Saccharomycetales(100);f__Metschnikowiaceae(100);g__Metschnikowia(100);s__Metschnikowia_pulcherrima(100); | Metschnikowia | 13 | Saprotroph | \|Undefined Saprotroph\| | Yeast | NULL | Probable | As nectar-tap_saprotroph (Põlme et al. 2020); Decay sugar-rich substrates (Põlme et al. 2020) | Põlme S, et al. 2020. Fungal Diversity 105:1-16 ((https://doi.org/10.1007/s13225-020-00466-2)) |
| Otu52371 | 1 | k__Fungi(100);p__Ascomycota(100);c__Saccharomycetes(100);o__Saccharomycetales(100);f__Metschnikowiaceae(100);g__Metschnikowia(100);s__Metschnikowia_pulcherrima(100); | Metschnikowia | 13 | Saprotroph | \|Undefined Saprotroph\| | Yeast | NULL | Probable | As nectar-tap_saprotroph (Põlme et al. 2020); Decay sugar-rich substrates (Põlme et al. 2020) | Põlme S, et al. 2020. Fungal Diversity 105:1-16 ((https://doi.org/10.1007/s13225-020-00466-2)) |
| Otu52372 | 1 | k__Fungi(100);p__Ascomycota(100);c__Saccharomycetes(100);o__Saccharomycetales(100);f__Metschnikowiaceae(100);g__Metschnikowia(100);s__Metschnikowia_pulcherrima(100); | Metschnikowia | 13 | Saprotroph | \|Undefined Saprotroph\| | Yeast | NULL | Probable | As nectar-tap_saprotroph (Põlme et al. 2020); Decay sugar-rich substrates (Põlme et al. 2020) | Põlme S, et al. 2020. Fungal Diversity 105:1-16 ((https://doi.org/10.1007/s13225-020-00466-2)) |
| Otu52373 | 1 | k__Fungi(100);p__Ascomycota(100);c__Saccharomycetes(100);o__Saccharomycetales(100);f__Metschnikowiaceae(100);g__Metschnikowia(100);s__Metschnikowia_pulcherrima(100); | Metschnikowia | 13 | Saprotroph | \|Undefined Saprotroph\| | Yeast | NULL | Probable | As nectar-tap_saprotroph (Põlme et al. 2020); Decay sugar-rich substrates (Põlme et al. 2020) | Põlme S, et al. 2020. Fungal Diversity 105:1-16 ((https://doi.org/10.1007/s13225-020-00466-2)) |
| Otu52374 | 1 | k__Fungi(100);p__Ascomycota(100);c__Saccharomycetes(100);o__Saccharomycetales(100);f__Metschnikowiaceae(100);g__Metschnikowia(100);g__Metschnikowia_unclassified(100); | Metschnikowia | 13 | Saprotroph | \|Undefined Saprotroph\| | Yeast | NULL | Probable | As nectar-tap_saprotroph (Põlme et al. 2020); Decay sugar-rich substrates (Põlme et al. 2020) | Põlme S, et al. 2020. Fungal Diversity 105:1-16 ((https://doi.org/10.1007/s13225-020-00466-2)) |
| Otu52375 | 1 | k__Fungi(100);p__Ascomycota(100);c__Saccharomycetes(100);o__Saccharomycetales(100);f__Metschnikowiaceae(100);g__Metschnikowia(100);g__Metschnikowia_unclassified(100); | Metschnikowia | 13 | Saprotroph | \|Undefined Saprotroph\| | Yeast | NULL | Probable | As nectar-tap_saprotroph (Põlme et al. 2020); Decay sugar-rich substrates (Põlme et al. 2020) | Põlme S, et al. 2020. Fungal Diversity 105:1-16 ((https://doi.org/10.1007/s13225-020-00466-2)) |
| Otu52376 | 1 | k__Fungi(100);p__Ascomycota(100);c__Saccharomycetes(100);o__Saccharomycetales(100);f__Metschnikowiaceae(100);g__Metschnikowia(100);s__Metschnikowia_pulcherrima(100); | Metschnikowia | 13 | Saprotroph | \|Undefined Saprotroph\| | Yeast | NULL | Probable | As nectar-tap_saprotroph (Põlme et al. 2020); Decay sugar-rich substrates (Põlme et al. 2020) | Põlme S, et al. 2020. Fungal Diversity 105:1-16 ((https://doi.org/10.1007/s13225-020-00466-2)) |
| Otu52377 | 1 | k__Fungi(100);p__Ascomycota(100);c__Saccharomycetes(100);o__Saccharomycetales(100);f__Metschnikowiaceae(100);g__Metschnikowia(100);g__Metschnikowia_unclassified(100); | Metschnikowia | 13 | Saprotroph | \|Undefined Saprotroph\| | Yeast | NULL | Probable | As nectar-tap_saprotroph (Põlme et al. 2020); Decay sugar-rich substrates (Põlme et al. 2020) | Põlme S, et al. 2020. Fungal Diversity 105:1-16 ((https://doi.org/10.1007/s13225-020-00466-2)) |
| Otu52379 | 1 | k__Fungi(100);p__Ascomycota(100);c__Saccharomycetes(100);o__Saccharomycetales(100);f__Metschnikowiaceae(100);g__Metschnikowia(100);s__Metschnikowia_pulcherrima(100); | Metschnikowia | 13 | Saprotroph | \|Undefined Saprotroph\| | Yeast | NULL | Probable | As nectar-tap_saprotroph (Põlme et al. 2020); Decay sugar-rich substrates (Põlme et al. 2020) | Põlme S, et al. 2020. Fungal Diversity 105:1-16 ((https://doi.org/10.1007/s13225-020-00466-2)) |
| Otu52380 | 1 | k__Fungi(100);p__Ascomycota(100);c__Saccharomycetes(100);o__Saccharomycetales(100);f__Metschnikowiaceae(100);g__Metschnikowia(100);s__Metschnikowia_pulcherrima(100); | Metschnikowia | 13 | Saprotroph | \|Undefined Saprotroph\| | Yeast | NULL | Probable | As nectar-tap_saprotroph (Põlme et al. 2020); Decay sugar-rich substrates (Põlme et al. 2020) | Põlme S, et al. 2020. Fungal Diversity 105:1-16 ((https://doi.org/10.1007/s13225-020-00466-2)) |
| Otu52381 | 1 | k__Fungi(100);p__Ascomycota(100);c__Saccharomycetes(100);o__Saccharomycetales(100);f__Metschnikowiaceae(100);g__Metschnikowia(100);s__Metschnikowia_pulcherrima(100); | Metschnikowia | 13 | Saprotroph | \|Undefined Saprotroph\| | Yeast | NULL | Probable | As nectar-tap_saprotroph (Põlme et al. 2020); Decay sugar-rich substrates (Põlme et al. 2020) | Põlme S, et al. 2020. Fungal Diversity 105:1-16 ((https://doi.org/10.1007/s13225-020-00466-2)) |
| Otu52382 | 1 | k__Fungi(100);p__Ascomycota(100);c__Saccharomycetes(100);o__Saccharomycetales(100);f__Metschnikowiaceae(100);g__Metschnikowia(100);s__Metschnikowia_pulcherrima(100); | Metschnikowia | 13 | Saprotroph | \|Undefined Saprotroph\| | Yeast | NULL | Probable | As nectar-tap_saprotroph (Põlme et al. 2020); Decay sugar-rich substrates (Põlme et al. 2020) | Põlme S, et al. 2020. Fungal Diversity 105:1-16 ((https://doi.org/10.1007/s13225-020-00466-2)) |
| Otu52383 | 1 | k__Fungi(100);p__Ascomycota(100);c__Saccharomycetes(100);o__Saccharomycetales(100);f__Metschnikowiaceae(100);g__Metschnikowia(100);s__Metschnikowia_pulcherrima(100); | Metschnikowia | 13 | Saprotroph | \|Undefined Saprotroph\| | Yeast | NULL | Probable | As nectar-tap_saprotroph (Põlme et al. 2020); Decay sugar-rich substrates (Põlme et al. 2020) | Põlme S, et al. 2020. Fungal Diversity 105:1-16 ((https://doi.org/10.1007/s13225-020-00466-2)) |
| Otu52384 | 1 | k__Fungi(100);p__Ascomycota(100);c__Saccharomycetes(100);o__Saccharomycetales(100);f__Metschnikowiaceae(100);g__Metschnikowia(100);s__Metschnikowia_pulcherrima(100); | Metschnikowia | 13 | Saprotroph | \|Undefined Saprotroph\| | Yeast | NULL | Probable | As nectar-tap_saprotroph (Põlme et al. 2020); Decay sugar-rich substrates (Põlme et al. 2020) | Põlme S, et al. 2020. Fungal Diversity 105:1-16 ((https://doi.org/10.1007/s13225-020-00466-2)) |
| Otu52385 | 1 | k__Fungi(100);p__Ascomycota(100);c__Saccharomycetes(100);o__Saccharomycetales(100);f__Metschnikowiaceae(100);g__Metschnikowia(100);s__Metschnikowia_pulcherrima(100); | Metschnikowia | 13 | Saprotroph | \|Undefined Saprotroph\| | Yeast | NULL | Probable | As nectar-tap_saprotroph (Põlme et al. 2020); Decay sugar-rich substrates (Põlme et al. 2020) | Põlme S, et al. 2020. Fungal Diversity 105:1-16 ((https://doi.org/10.1007/s13225-020-00466-2)) |
| Otu52386 | 1 | k__Fungi(100);p__Ascomycota(100);c__Saccharomycetes(100);o__Saccharomycetales(100);f__Metschnikowiaceae(100);g__Metschnikowia(100);s__Metschnikowia_pulcherrima(100); | Metschnikowia | 13 | Saprotroph | \|Undefined Saprotroph\| | Yeast | NULL | Probable | As nectar-tap_saprotroph (Põlme et al. 2020); Decay sugar-rich substrates (Põlme et al. 2020) | Põlme S, et al. 2020. Fungal Diversity 105:1-16 ((https://doi.org/10.1007/s13225-020-00466-2)) |
| Otu52388 | 1 | k__Fungi(100);p__Ascomycota(100);c__Saccharomycetes(100);o__Saccharomycetales(100);f__Metschnikowiaceae(100);g__Metschnikowia(100);g__Metschnikowia_unclassified(100); | Metschnikowia | 13 | Saprotroph | \|Undefined Saprotroph\| | Yeast | NULL | Probable | As nectar-tap_saprotroph (Põlme et al. 2020); Decay sugar-rich substrates (Põlme et al. 2020) | Põlme S, et al. 2020. Fungal Diversity 105:1-16 ((https://doi.org/10.1007/s13225-020-00466-2)) |
| Otu52389 | 1 | k__Fungi(100);p__Ascomycota(100);c__Saccharomycetes(100);o__Saccharomycetales(100);f__Metschnikowiaceae(100);g__Metschnikowia(100);g__Metschnikowia_unclassified(100); | Metschnikowia | 13 | Saprotroph | \|Undefined Saprotroph\| | Yeast | NULL | Probable | As nectar-tap_saprotroph (Põlme et al. 2020); Decay sugar-rich substrates (Põlme et al. 2020) | Põlme S, et al. 2020. Fungal Diversity 105:1-16 ((https://doi.org/10.1007/s13225-020-00466-2)) |
| Otu52390 | 1 | k__Fungi(100);p__Ascomycota(100);c__Saccharomycetes(100);o__Saccharomycetales(100);f__Metschnikowiaceae(100);g__Metschnikowia(100);s__Metschnikowia_pulcherrima(100); | Metschnikowia | 13 | Saprotroph | \|Undefined Saprotroph\| | Yeast | NULL | Probable | As nectar-tap_saprotroph (Põlme et al. 2020); Decay sugar-rich substrates (Põlme et al. 2020) | Põlme S, et al. 2020. Fungal Diversity 105:1-16 ((https://doi.org/10.1007/s13225-020-00466-2)) |
| Otu52391 | 1 | k__Fungi(100);p__Ascomycota(100);c__Saccharomycetes(100);o__Saccharomycetales(100);f__Metschnikowiaceae(100);g__Metschnikowia(100);s__Metschnikowia_pulcherrima(100); | Metschnikowia | 13 | Saprotroph | \|Undefined Saprotroph\| | Yeast | NULL | Probable | As nectar-tap_saprotroph (Põlme et al. 2020); Decay sugar-rich substrates (Põlme et al. 2020) | Põlme S, et al. 2020. Fungal Diversity 105:1-16 ((https://doi.org/10.1007/s13225-020-00466-2)) |
| Otu52392 | 1 | k__Fungi(100);p__Ascomycota(100);c__Saccharomycetes(100);o__Saccharomycetales(100);f__Metschnikowiaceae(100);g__Metschnikowia(100);g__Metschnikowia_unclassified(100); | Metschnikowia | 13 | Saprotroph | \|Undefined Saprotroph\| | Yeast | NULL | Probable | As nectar-tap_saprotroph (Põlme et al. 2020); Decay sugar-rich substrates (Põlme et al. 2020) | Põlme S, et al. 2020. Fungal Diversity 105:1-16 ((https://doi.org/10.1007/s13225-020-00466-2)) |
| Otu52393 | 1 | k__Fungi(100);p__Ascomycota(100);c__Saccharomycetes(100);o__Saccharomycetales(100);f__Metschnikowiaceae(100);g__Metschnikowia(100);g__Metschnikowia_unclassified(100); | Metschnikowia | 13 | Saprotroph | \|Undefined Saprotroph\| | Yeast | NULL | Probable | As nectar-tap_saprotroph (Põlme et al. 2020); Decay sugar-rich substrates (Põlme et al. 2020) | Põlme S, et al. 2020. Fungal Diversity 105:1-16 ((https://doi.org/10.1007/s13225-020-00466-2)) |
| Otu52394 | 1 | k__Fungi(100);p__Ascomycota(100);c__Saccharomycetes(100);o__Saccharomycetales(100);f__Metschnikowiaceae(100);g__Metschnikowia(100);s__Metschnikowia_pulcherrima(100); | Metschnikowia | 13 | Saprotroph | \|Undefined Saprotroph\| | Yeast | NULL | Probable | As nectar-tap_saprotroph (Põlme et al. 2020); Decay sugar-rich substrates (Põlme et al. 2020) | Põlme S, et al. 2020. Fungal Diversity 105:1-16 ((https://doi.org/10.1007/s13225-020-00466-2)) |
| Otu52395 | 1 | k__Fungi(100);p__Ascomycota(100);c__Saccharomycetes(100);o__Saccharomycetales(100);f__Metschnikowiaceae(100);g__Metschnikowia(100);s__Metschnikowia_pulcherrima(100); | Metschnikowia | 13 | Saprotroph | \|Undefined Saprotroph\| | Yeast | NULL | Probable | As nectar-tap_saprotroph (Põlme et al. 2020); Decay sugar-rich substrates (Põlme et al. 2020) | Põlme S, et al. 2020. Fungal Diversity 105:1-16 ((https://doi.org/10.1007/s13225-020-00466-2)) |
| Otu52396 | 1 | k__Fungi(100);p__Ascomycota(100);c__Saccharomycetes(100);o__Saccharomycetales(100);f__Metschnikowiaceae(100);g__Metschnikowia(100);g__Metschnikowia_unclassified(100); | Metschnikowia | 13 | Saprotroph | \|Undefined Saprotroph\| | Yeast | NULL | Probable | As nectar-tap_saprotroph (Põlme et al. 2020); Decay sugar-rich substrates (Põlme et al. 2020) | Põlme S, et al. 2020. Fungal Diversity 105:1-16 ((https://doi.org/10.1007/s13225-020-00466-2)) |
| Otu52397 | 1 | k__Fungi(100);p__Ascomycota(100);c__Saccharomycetes(100);o__Saccharomycetales(100);f__Metschnikowiaceae(100);g__Metschnikowia(100);s__Metschnikowia_pulcherrima(100); | Metschnikowia | 13 | Saprotroph | \|Undefined Saprotroph\| | Yeast | NULL | Probable | As nectar-tap_saprotroph (Põlme et al. 2020); Decay sugar-rich substrates (Põlme et al. 2020) | Põlme S, et al. 2020. Fungal Diversity 105:1-16 ((https://doi.org/10.1007/s13225-020-00466-2)) |
| Otu52398 | 1 | k__Fungi(100);p__Ascomycota(100);c__Saccharomycetes(100);o__Saccharomycetales(100);f__Metschnikowiaceae(100);g__Metschnikowia(100);g__Metschnikowia_unclassified(100); | Metschnikowia | 13 | Saprotroph | \|Undefined Saprotroph\| | Yeast | NULL | Probable | As nectar-tap_saprotroph (Põlme et al. 2020); Decay sugar-rich substrates (Põlme et al. 2020) | Põlme S, et al. 2020. Fungal Diversity 105:1-16 ((https://doi.org/10.1007/s13225-020-00466-2)) |
| Otu52399 | 1 | k__Fungi(100);p__Ascomycota(100);c__Saccharomycetes(100);o__Saccharomycetales(100);f__Metschnikowiaceae(100);g__Metschnikowia(100);s__Metschnikowia_pulcherrima(100); | Metschnikowia | 13 | Saprotroph | \|Undefined Saprotroph\| | Yeast | NULL | Probable | As nectar-tap_saprotroph (Põlme et al. 2020); Decay sugar-rich substrates (Põlme et al. 2020) | Põlme S, et al. 2020. Fungal Diversity 105:1-16 ((https://doi.org/10.1007/s13225-020-00466-2)) |
| Otu52400 | 1 | k__Fungi(100);p__Ascomycota(100);c__Saccharomycetes(100);o__Saccharomycetales(100);f__Metschnikowiaceae(100);g__Metschnikowia(100);s__Metschnikowia_pulcherrima(100); | Metschnikowia | 13 | Saprotroph | \|Undefined Saprotroph\| | Yeast | NULL | Probable | As nectar-tap_saprotroph (Põlme et al. 2020); Decay sugar-rich substrates (Põlme et al. 2020) | Põlme S, et al. 2020. Fungal Diversity 105:1-16 ((https://doi.org/10.1007/s13225-020-00466-2)) |
| Otu52401 | 1 | k__Fungi(100);p__Ascomycota(100);c__Saccharomycetes(100);o__Saccharomycetales(100);f__Metschnikowiaceae(100);g__Metschnikowia(100);s__Metschnikowia_pulcherrima(100); | Metschnikowia | 13 | Saprotroph | \|Undefined Saprotroph\| | Yeast | NULL | Probable | As nectar-tap_saprotroph (Põlme et al. 2020); Decay sugar-rich substrates (Põlme et al. 2020) | Põlme S, et al. 2020. Fungal Diversity 105:1-16 ((https://doi.org/10.1007/s13225-020-00466-2)) |
| Otu52402 | 1 | k__Fungi(100);p__Ascomycota(100);c__Saccharomycetes(100);o__Saccharomycetales(100);f__Metschnikowiaceae(100);g__Metschnikowia(100);s__Metschnikowia_pulcherrima(100); | Metschnikowia | 13 | Saprotroph | \|Undefined Saprotroph\| | Yeast | NULL | Probable | As nectar-tap_saprotroph (Põlme et al. 2020); Decay sugar-rich substrates (Põlme et al. 2020) | Põlme S, et al. 2020. Fungal Diversity 105:1-16 ((https://doi.org/10.1007/s13225-020-00466-2)) |
| Otu52403 | 1 | k__Fungi(100);p__Ascomycota(100);c__Saccharomycetes(100);o__Saccharomycetales(100);f__Metschnikowiaceae(100);g__Metschnikowia(100);s__Metschnikowia_pulcherrima(100); | Metschnikowia | 13 | Saprotroph | \|Undefined Saprotroph\| | Yeast | NULL | Probable | As nectar-tap_saprotroph (Põlme et al. 2020); Decay sugar-rich substrates (Põlme et al. 2020) | Põlme S, et al. 2020. Fungal Diversity 105:1-16 ((https://doi.org/10.1007/s13225-020-00466-2)) |
| Otu52404 | 1 | k__Fungi(100);p__Ascomycota(100);c__Saccharomycetes(100);o__Saccharomycetales(100);f__Metschnikowiaceae(100);g__Metschnikowia(100);s__Metschnikowia_pulcherrima(100); | Metschnikowia | 13 | Saprotroph | \|Undefined Saprotroph\| | Yeast | NULL | Probable | As nectar-tap_saprotroph (Põlme et al. 2020); Decay sugar-rich substrates (Põlme et al. 2020) | Põlme S, et al. 2020. Fungal Diversity 105:1-16 ((https://doi.org/10.1007/s13225-020-00466-2)) |
| Otu52405 | 1 | k__Fungi(100);p__Ascomycota(100);c__Saccharomycetes(100);o__Saccharomycetales(100);f__Metschnikowiaceae(100);g__Metschnikowia(100);g__Metschnikowia_unclassified(100); | Metschnikowia | 13 | Saprotroph | \|Undefined Saprotroph\| | Yeast | NULL | Probable | As nectar-tap_saprotroph (Põlme et al. 2020); Decay sugar-rich substrates (Põlme et al. 2020) | Põlme S, et al. 2020. Fungal Diversity 105:1-16 ((https://doi.org/10.1007/s13225-020-00466-2)) |
| Otu52406 | 1 | k__Fungi(100);p__Ascomycota(100);c__Saccharomycetes(100);o__Saccharomycetales(100);f__Metschnikowiaceae(100);g__Metschnikowia(100);s__Metschnikowia_pulcherrima(100); | Metschnikowia | 13 | Saprotroph | \|Undefined Saprotroph\| | Yeast | NULL | Probable | As nectar-tap_saprotroph (Põlme et al. 2020); Decay sugar-rich substrates (Põlme et al. 2020) | Põlme S, et al. 2020. Fungal Diversity 105:1-16 ((https://doi.org/10.1007/s13225-020-00466-2)) |
| Otu52407 | 1 | k__Fungi(100);p__Ascomycota(100);c__Saccharomycetes(100);o__Saccharomycetales(100);f__Metschnikowiaceae(100);g__Metschnikowia(100);s__Metschnikowia_pulcherrima(100); | Metschnikowia | 13 | Saprotroph | \|Undefined Saprotroph\| | Yeast | NULL | Probable | As nectar-tap_saprotroph (Põlme et al. 2020); Decay sugar-rich substrates (Põlme et al. 2020) | Põlme S, et al. 2020. Fungal Diversity 105:1-16 ((https://doi.org/10.1007/s13225-020-00466-2)) |
| Otu52408 | 1 | k__Fungi(100);p__Ascomycota(100);c__Saccharomycetes(100);o__Saccharomycetales(100);f__Metschnikowiaceae(100);g__Metschnikowia(100);s__Metschnikowia_pulcherrima(100); | Metschnikowia | 13 | Saprotroph | \|Undefined Saprotroph\| | Yeast | NULL | Probable | As nectar-tap_saprotroph (Põlme et al. 2020); Decay sugar-rich substrates (Põlme et al. 2020) | Põlme S, et al. 2020. Fungal Diversity 105:1-16 ((https://doi.org/10.1007/s13225-020-00466-2)) |
| Otu52409 | 1 | k__Fungi(100);p__Ascomycota(100);c__Saccharomycetes(100);o__Saccharomycetales(100);f__Metschnikowiaceae(100);g__Metschnikowia(100);s__Metschnikowia_pulcherrima(100); | Metschnikowia | 13 | Saprotroph | \|Undefined Saprotroph\| | Yeast | NULL | Probable | As nectar-tap_saprotroph (Põlme et al. 2020); Decay sugar-rich substrates (Põlme et al. 2020) | Põlme S, et al. 2020. Fungal Diversity 105:1-16 ((https://doi.org/10.1007/s13225-020-00466-2)) |
| Otu52410 | 1 | k__Fungi(100);p__Ascomycota(100);c__Saccharomycetes(100);o__Saccharomycetales(100);f__Metschnikowiaceae(100);g__Metschnikowia(100);s__Metschnikowia_pulcherrima(100); | Metschnikowia | 13 | Saprotroph | \|Undefined Saprotroph\| | Yeast | NULL | Probable | As nectar-tap_saprotroph (Põlme et al. 2020); Decay sugar-rich substrates (Põlme et al. 2020) | Põlme S, et al. 2020. Fungal Diversity 105:1-16 ((https://doi.org/10.1007/s13225-020-00466-2)) |
| Otu52411 | 1 | k__Fungi(100);p__Ascomycota(100);c__Saccharomycetes(100);o__Saccharomycetales(100);f__Metschnikowiaceae(100);g__Metschnikowia(100);g__Metschnikowia_unclassified(100); | Metschnikowia | 13 | Saprotroph | \|Undefined Saprotroph\| | Yeast | NULL | Probable | As nectar-tap_saprotroph (Põlme et al. 2020); Decay sugar-rich substrates (Põlme et al. 2020) | Põlme S, et al. 2020. Fungal Diversity 105:1-16 ((https://doi.org/10.1007/s13225-020-00466-2)) |
| Otu52412 | 1 | k__Fungi(100);p__Ascomycota(100);c__Saccharomycetes(100);o__Saccharomycetales(100);f__Metschnikowiaceae(100);g__Metschnikowia(100);g__Metschnikowia_unclassified(100); | Metschnikowia | 13 | Saprotroph | \|Undefined Saprotroph\| | Yeast | NULL | Probable | As nectar-tap_saprotroph (Põlme et al. 2020); Decay sugar-rich substrates (Põlme et al. 2020) | Põlme S, et al. 2020. Fungal Diversity 105:1-16 ((https://doi.org/10.1007/s13225-020-00466-2)) |
| Otu52413 | 1 | k__Fungi(100);p__Ascomycota(100);c__Saccharomycetes(100);o__Saccharomycetales(100);f__Metschnikowiaceae(100);g__Metschnikowia(100);s__Metschnikowia_pulcherrima(100); | Metschnikowia | 13 | Saprotroph | \|Undefined Saprotroph\| | Yeast | NULL | Probable | As nectar-tap_saprotroph (Põlme et al. 2020); Decay sugar-rich substrates (Põlme et al. 2020) | Põlme S, et al. 2020. Fungal Diversity 105:1-16 ((https://doi.org/10.1007/s13225-020-00466-2)) |
| Otu52414 | 1 | k__Fungi(100);p__Ascomycota(100);c__Saccharomycetes(100);o__Saccharomycetales(100);f__Metschnikowiaceae(100);g__Metschnikowia(100);s__Metschnikowia_pulcherrima(100); | Metschnikowia | 13 | Saprotroph | \|Undefined Saprotroph\| | Yeast | NULL | Probable | As nectar-tap_saprotroph (Põlme et al. 2020); Decay sugar-rich substrates (Põlme et al. 2020) | Põlme S, et al. 2020. Fungal Diversity 105:1-16 ((https://doi.org/10.1007/s13225-020-00466-2)) |
| Otu52415 | 1 | k__Fungi(100);p__Ascomycota(100);c__Saccharomycetes(100);o__Saccharomycetales(100);f__Metschnikowiaceae(100);g__Metschnikowia(100);s__Metschnikowia_pulcherrima(100); | Metschnikowia | 13 | Saprotroph | \|Undefined Saprotroph\| | Yeast | NULL | Probable | As nectar-tap_saprotroph (Põlme et al. 2020); Decay sugar-rich substrates (Põlme et al. 2020) | Põlme S, et al. 2020. Fungal Diversity 105:1-16 ((https://doi.org/10.1007/s13225-020-00466-2)) |
| Otu52416 | 1 | k__Fungi(100);p__Ascomycota(100);c__Saccharomycetes(100);o__Saccharomycetales(100);f__Metschnikowiaceae(100);g__Metschnikowia(100);s__Metschnikowia_pulcherrima(100); | Metschnikowia | 13 | Saprotroph | \|Undefined Saprotroph\| | Yeast | NULL | Probable | As nectar-tap_saprotroph (Põlme et al. 2020); Decay sugar-rich substrates (Põlme et al. 2020) | Põlme S, et al. 2020. Fungal Diversity 105:1-16 ((https://doi.org/10.1007/s13225-020-00466-2)) |
| Otu52417 | 1 | k__Fungi(100);p__Ascomycota(100);c__Saccharomycetes(100);o__Saccharomycetales(100);f__Metschnikowiaceae(100);g__Metschnikowia(100);s__Metschnikowia_pulcherrima(100); | Metschnikowia | 13 | Saprotroph | \|Undefined Saprotroph\| | Yeast | NULL | Probable | As nectar-tap_saprotroph (Põlme et al. 2020); Decay sugar-rich substrates (Põlme et al. 2020) | Põlme S, et al. 2020. Fungal Diversity 105:1-16 ((https://doi.org/10.1007/s13225-020-00466-2)) |
| Otu52418 | 1 | k__Fungi(100);p__Ascomycota(100);c__Saccharomycetes(100);o__Saccharomycetales(100);f__Metschnikowiaceae(100);g__Metschnikowia(100);g__Metschnikowia_unclassified(100); | Metschnikowia | 13 | Saprotroph | \|Undefined Saprotroph\| | Yeast | NULL | Probable | As nectar-tap_saprotroph (Põlme et al. 2020); Decay sugar-rich substrates (Põlme et al. 2020) | Põlme S, et al. 2020. Fungal Diversity 105:1-16 ((https://doi.org/10.1007/s13225-020-00466-2)) |
| Otu52419 | 1 | k__Fungi(100);p__Ascomycota(100);c__Saccharomycetes(100);o__Saccharomycetales(100);f__Metschnikowiaceae(100);g__Metschnikowia(100);s__Metschnikowia_pulcherrima(100); | Metschnikowia | 13 | Saprotroph | \|Undefined Saprotroph\| | Yeast | NULL | Probable | As nectar-tap_saprotroph (Põlme et al. 2020); Decay sugar-rich substrates (Põlme et al. 2020) | Põlme S, et al. 2020. Fungal Diversity 105:1-16 ((https://doi.org/10.1007/s13225-020-00466-2)) |
| Otu52420 | 1 | k__Fungi(100);p__Ascomycota(100);c__Saccharomycetes(100);o__Saccharomycetales(100);f__Metschnikowiaceae(100);g__Metschnikowia(100);s__Metschnikowia_pulcherrima(100); | Metschnikowia | 13 | Saprotroph | \|Undefined Saprotroph\| | Yeast | NULL | Probable | As nectar-tap_saprotroph (Põlme et al. 2020); Decay sugar-rich substrates (Põlme et al. 2020) | Põlme S, et al. 2020. Fungal Diversity 105:1-16 ((https://doi.org/10.1007/s13225-020-00466-2)) |
| Otu52421 | 1 | k__Fungi(100);p__Ascomycota(100);c__Saccharomycetes(100);o__Saccharomycetales(100);f__Metschnikowiaceae(100);g__Metschnikowia(100);s__Metschnikowia_pulcherrima(100); | Metschnikowia | 13 | Saprotroph | \|Undefined Saprotroph\| | Yeast | NULL | Probable | As nectar-tap_saprotroph (Põlme et al. 2020); Decay sugar-rich substrates (Põlme et al. 2020) | Põlme S, et al. 2020. Fungal Diversity 105:1-16 ((https://doi.org/10.1007/s13225-020-00466-2)) |
| Otu52422 | 1 | k__Fungi(100);p__Ascomycota(100);c__Saccharomycetes(100);o__Saccharomycetales(100);f__Metschnikowiaceae(100);g__Metschnikowia(100);s__Metschnikowia_pulcherrima(100); | Metschnikowia | 13 | Saprotroph | \|Undefined Saprotroph\| | Yeast | NULL | Probable | As nectar-tap_saprotroph (Põlme et al. 2020); Decay sugar-rich substrates (Põlme et al. 2020) | Põlme S, et al. 2020. Fungal Diversity 105:1-16 ((https://doi.org/10.1007/s13225-020-00466-2)) |
| Otu52423 | 1 | k__Fungi(100);p__Ascomycota(100);c__Saccharomycetes(100);o__Saccharomycetales(100);f__Metschnikowiaceae(100);g__Metschnikowia(100);s__Metschnikowia_pulcherrima(100); | Metschnikowia | 13 | Saprotroph | \|Undefined Saprotroph\| | Yeast | NULL | Probable | As nectar-tap_saprotroph (Põlme et al. 2020); Decay sugar-rich substrates (Põlme et al. 2020) | Põlme S, et al. 2020. Fungal Diversity 105:1-16 ((https://doi.org/10.1007/s13225-020-00466-2)) |
| Otu52425 | 1 | k__Fungi(100);p__Ascomycota(100);c__Saccharomycetes(100);o__Saccharomycetales(100);f__Metschnikowiaceae(100);g__Metschnikowia(100);g__Metschnikowia_unclassified(100); | Metschnikowia | 13 | Saprotroph | \|Undefined Saprotroph\| | Yeast | NULL | Probable | As nectar-tap_saprotroph (Põlme et al. 2020); Decay sugar-rich substrates (Põlme et al. 2020) | Põlme S, et al. 2020. Fungal Diversity 105:1-16 ((https://doi.org/10.1007/s13225-020-00466-2)) |
| Otu52426 | 1 | k__Fungi(100);p__Ascomycota(100);c__Saccharomycetes(100);o__Saccharomycetales(100);f__Metschnikowiaceae(100);g__Metschnikowia(100);s__Metschnikowia_pulcherrima(100); | Metschnikowia | 13 | Saprotroph | \|Undefined Saprotroph\| | Yeast | NULL | Probable | As nectar-tap_saprotroph (Põlme et al. 2020); Decay sugar-rich substrates (Põlme et al. 2020) | Põlme S, et al. 2020. Fungal Diversity 105:1-16 ((https://doi.org/10.1007/s13225-020-00466-2)) |
| Otu52427 | 1 | k__Fungi(100);p__Ascomycota(100);c__Saccharomycetes(100);o__Saccharomycetales(100);f__Metschnikowiaceae(100);g__Metschnikowia(100);g__Metschnikowia_unclassified(100); | Metschnikowia | 13 | Saprotroph | \|Undefined Saprotroph\| | Yeast | NULL | Probable | As nectar-tap_saprotroph (Põlme et al. 2020); Decay sugar-rich substrates (Põlme et al. 2020) | Põlme S, et al. 2020. Fungal Diversity 105:1-16 ((https://doi.org/10.1007/s13225-020-00466-2)) |
| Otu52428 | 1 | k__Fungi(100);p__Ascomycota(100);c__Saccharomycetes(100);o__Saccharomycetales(100);f__Metschnikowiaceae(100);g__Metschnikowia(100);s__Metschnikowia_pulcherrima(100); | Metschnikowia | 13 | Saprotroph | \|Undefined Saprotroph\| | Yeast | NULL | Probable | As nectar-tap_saprotroph (Põlme et al. 2020); Decay sugar-rich substrates (Põlme et al. 2020) | Põlme S, et al. 2020. Fungal Diversity 105:1-16 ((https://doi.org/10.1007/s13225-020-00466-2)) |
| Otu52429 | 1 | k__Fungi(100);p__Ascomycota(100);c__Saccharomycetes(100);o__Saccharomycetales(100);f__Metschnikowiaceae(100);g__Metschnikowia(100);s__Metschnikowia_pulcherrima(100); | Metschnikowia | 13 | Saprotroph | \|Undefined Saprotroph\| | Yeast | NULL | Probable | As nectar-tap_saprotroph (Põlme et al. 2020); Decay sugar-rich substrates (Põlme et al. 2020) | Põlme S, et al. 2020. Fungal Diversity 105:1-16 ((https://doi.org/10.1007/s13225-020-00466-2)) |
| Otu52430 | 1 | k__Fungi(100);p__Ascomycota(100);c__Saccharomycetes(100);o__Saccharomycetales(100);f__Metschnikowiaceae(100);g__Metschnikowia(100);s__Metschnikowia_pulcherrima(100); | Metschnikowia | 13 | Saprotroph | \|Undefined Saprotroph\| | Yeast | NULL | Probable | As nectar-tap_saprotroph (Põlme et al. 2020); Decay sugar-rich substrates (Põlme et al. 2020) | Põlme S, et al. 2020. Fungal Diversity 105:1-16 ((https://doi.org/10.1007/s13225-020-00466-2)) |

| | | | | | | | | | | | |
|---|---|---|---|---|---|---|---|---|---|---|---|
| Otu52431 | 1 | k__Fungi(100);p__Ascomycota(100);c__Saccharomycetes(100);o__Saccharomycetales(100);f__Metschnikowiaceae(100);g__Metschnikowia(100);g__Metschnikowia_unclassified(100); | Metschnikowia | 13 | Saprotroph | \|Undefined Saprotroph\| | Yeast | NULL | Probable | As nectar-tap_saprotroph (Põlme et al. 2020); Decay sugar-rich substrates (Põlme et al. 2020) | Põlme S, et al. 2020. Fungal Diversity 105:1-16 ((https://doi.org/10.1007/s13225-020-00466-2)) |
| Otu52432 | 1 | k__Fungi(100);p__Ascomycota(100);c__Saccharomycetes(100);o__Saccharomycetales(100);f__Metschnikowiaceae(100);g__Metschnikowia(100);s__Metschnikowia_pulcherrima(100); | Metschnikowia | 13 | Saprotroph | \|Undefined Saprotroph\| | Yeast | NULL | Probable | As nectar-tap_saprotroph (Põlme et al. 2020); Decay sugar-rich substrates (Põlme et al. 2020) | Põlme S, et al. 2020. Fungal Diversity 105:1-16 ((https://doi.org/10.1007/s13225-020-00466-2)) |
| Otu52433 | 1 | k__Fungi(100);p__Ascomycota(100);c__Saccharomycetes(100);o__Saccharomycetales(100);f__Metschnikowiaceae(100);g__Metschnikowia(100);s__Metschnikowia_pulcherrima(100); | Metschnikowia | 13 | Saprotroph | \|Undefined Saprotroph\| | Yeast | NULL | Probable | As nectar-tap_saprotroph (Põlme et al. 2020); Decay sugar-rich substrates (Põlme et al. 2020) | Põlme S, et al. 2020. Fungal Diversity 105:1-16 ((https://doi.org/10.1007/s13225-020-00466-2)) |
| Otu52434 | 1 | k__Fungi(100);p__Ascomycota(100);c__Saccharomycetes(100);o__Saccharomycetales(100);f__Metschnikowiaceae(100);g__Metschnikowia(100);g__Metschnikowia_unclassified(100); | Metschnikowia | 13 | Saprotroph | \|Undefined Saprotroph\| | Yeast | NULL | Probable | As nectar-tap_saprotroph (Põlme et al. 2020); Decay sugar-rich substrates (Põlme et al. 2020) | Põlme S, et al. 2020. Fungal Diversity 105:1-16 ((https://doi.org/10.1007/s13225-020-00466-2)) |
| Otu52435 | 1 | k__Fungi(100);p__Ascomycota(100);c__Saccharomycetes(100);o__Saccharomycetales(100);f__Metschnikowiaceae(100);g__Metschnikowia(100);s__Metschnikowia_pulcherrima(100); | Metschnikowia | 13 | Saprotroph | \|Undefined Saprotroph\| | Yeast | NULL | Probable | As nectar-tap_saprotroph (Põlme et al. 2020); Decay sugar-rich substrates (Põlme et al. 2020) | Põlme S, et al. 2020. Fungal Diversity 105:1-16 ((https://doi.org/10.1007/s13225-020-00466-2)) |
| Otu52436 | 1 | k__Fungi(100);p__Ascomycota(100);c__Saccharomycetes(100);o__Saccharomycetales(100);f__Metschnikowiaceae(100);g__Metschnikowia(100);s__Metschnikowia_pulcherrima(100); | Metschnikowia | 13 | Saprotroph | \|Undefined Saprotroph\| | Yeast | NULL | Probable | As nectar-tap_saprotroph (Põlme et al. 2020); Decay sugar-rich substrates (Põlme et al. 2020) | Põlme S, et al. 2020. Fungal Diversity 105:1-16 ((https://doi.org/10.1007/s13225-020-00466-2)) |
| Otu52437 | 1 | k__Fungi(100);p__Ascomycota(100);c__Saccharomycetes(100);o__Saccharomycetales(100);f__Metschnikowiaceae(100);g__Metschnikowia(100);s__Metschnikowia_pulcherrima(100); | Metschnikowia | 13 | Saprotroph | \|Undefined Saprotroph\| | Yeast | NULL | Probable | As nectar-tap_saprotroph (Põlme et al. 2020); Decay sugar-rich substrates (Põlme et al. 2020) | Põlme S, et al. 2020. Fungal Diversity 105:1-16 ((https://doi.org/10.1007/s13225-020-00466-2)) |
| Otu52438 | 1 | k__Fungi(100);p__Ascomycota(100);c__Saccharomycetes(100);o__Saccharomycetales(100);f__Metschnikowiaceae(100);g__Metschnikowia(100);s__Metschnikowia_pulcherrima(100); | Metschnikowia | 13 | Saprotroph | \|Undefined Saprotroph\| | Yeast | NULL | Probable | As nectar-tap_saprotroph (Põlme et al. 2020); Decay sugar-rich substrates (Põlme et al. 2020) | Põlme S, et al. 2020. Fungal Diversity 105:1-16 ((https://doi.org/10.1007/s13225-020-00466-2)) |
| Otu52439 | 1 | k__Fungi(100);p__Ascomycota(100);c__Saccharomycetes(100);o__Saccharomycetales(100);f__Metschnikowiaceae(100);g__Metschnikowia(100);g__Metschnikowia_unclassified(100); | Metschnikowia | 13 | Saprotroph | \|Undefined Saprotroph\| | Yeast | NULL | Probable | As nectar-tap_saprotroph (Põlme et al. 2020); Decay sugar-rich substrates (Põlme et al. 2020) | Põlme S, et al. 2020. Fungal Diversity 105:1-16 ((https://doi.org/10.1007/s13225-020-00466-2)) |
| Otu52440 | 1 | k__Fungi(100);p__Ascomycota(100);c__Saccharomycetes(100);o__Saccharomycetales(100);f__Metschnikowiaceae(100);g__Metschnikowia(100);s__Metschnikowia_pulcherrima(100); | Metschnikowia | 13 | Saprotroph | \|Undefined Saprotroph\| | Yeast | NULL | Probable | As nectar-tap_saprotroph (Põlme et al. 2020); Decay sugar-rich substrates (Põlme et al. 2020) | Põlme S, et al. 2020. Fungal Diversity 105:1-16 ((https://doi.org/10.1007/s13225-020-00466-2)) |
| Otu52441 | 1 | k__Fungi(100);p__Ascomycota(100);c__Saccharomycetes(100);o__Saccharomycetales(100);f__Metschnikowiaceae(100);g__Metschnikowia(100);s__Metschnikowia_pulcherrima(100); | Metschnikowia | 13 | Saprotroph | \|Undefined Saprotroph\| | Yeast | NULL | Probable | As nectar-tap_saprotroph (Põlme et al. 2020); Decay sugar-rich substrates (Põlme et al. 2020) | Põlme S, et al. 2020. Fungal Diversity 105:1-16 ((https://doi.org/10.1007/s13225-020-00466-2)) |
| Otu52442 | 1 | k__Fungi(100);p__Ascomycota(100);c__Saccharomycetes(100);o__Saccharomycetales(100);f__Metschnikowiaceae(100);g__Metschnikowia(100);g__Metschnikowia_unclassified(100); | Metschnikowia | 13 | Saprotroph | \|Undefined Saprotroph\| | Yeast | NULL | Probable | As nectar-tap_saprotroph (Põlme et al. 2020); Decay sugar-rich substrates (Põlme et al. 2020) | Põlme S, et al. 2020. Fungal Diversity 105:1-16 ((https://doi.org/10.1007/s13225-020-00466-2)) |
| Otu52443 | 1 | k__Fungi(100);p__Ascomycota(100);c__Saccharomycetes(100);o__Saccharomycetales(100);f__Metschnikowiaceae(100);g__Metschnikowia(100);s__Metschnikowia_pulcherrima(100); | Metschnikowia | 13 | Saprotroph | \|Undefined Saprotroph\| | Yeast | NULL | Probable | As nectar-tap_saprotroph (Põlme et al. 2020); Decay sugar-rich substrates (Põlme et al. 2020) | Põlme S, et al. 2020. Fungal Diversity 105:1-16 ((https://doi.org/10.1007/s13225-020-00466-2)) |
| Otu52444 | 1 | k__Fungi(100);p__Ascomycota(100);c__Saccharomycetes(100);o__Saccharomycetales(100);f__Metschnikowiaceae(100);g__Metschnikowia(100);s__Metschnikowia_pulcherrima(100); | Metschnikowia | 13 | Saprotroph | \|Undefined Saprotroph\| | Yeast | NULL | Probable | As nectar-tap_saprotroph (Põlme et al. 2020); Decay sugar-rich substrates (Põlme et al. 2020) | Põlme S, et al. 2020. Fungal Diversity 105:1-16 ((https://doi.org/10.1007/s13225-020-00466-2)) |
| Otu52445 | 1 | k__Fungi(100);p__Ascomycota(100);c__Saccharomycetes(100);o__Saccharomycetales(100);f__Metschnikowiaceae(100);g__Metschnikowia(100);g__Metschnikowia_unclassified(100); | Metschnikowia | 13 | Saprotroph | \|Undefined Saprotroph\| | Yeast | NULL | Probable | As nectar-tap_saprotroph (Põlme et al. 2020); Decay sugar-rich substrates (Põlme et al. 2020) | Põlme S, et al. 2020. Fungal Diversity 105:1-16 ((https://doi.org/10.1007/s13225-020-00466-2)) |
| Otu52446 | 1 | k__Fungi(100);p__Ascomycota(100);c__Saccharomycetes(100);o__Saccharomycetales(100);f__Metschnikowiaceae(100);g__Metschnikowia(100);s__Metschnikowia_pulcherrima(100); | Metschnikowia | 13 | Saprotroph | \|Undefined Saprotroph\| | Yeast | NULL | Probable | As nectar-tap_saprotroph (Põlme et al. 2020); Decay sugar-rich substrates (Põlme et al. 2020) | Põlme S, et al. 2020. Fungal Diversity 105:1-16 ((https://doi.org/10.1007/s13225-020-00466-2)) |
| Otu52447 | 1 | k__Fungi(100);p__Ascomycota(100);c__Saccharomycetes(100);o__Saccharomycetales(100);f__Metschnikowiaceae(100);g__Metschnikowia(100);g__Metschnikowia_unclassified(100); | Metschnikowia | 13 | Saprotroph | \|Undefined Saprotroph\| | Yeast | NULL | Probable | As nectar-tap_saprotroph (Põlme et al. 2020); Decay sugar-rich substrates (Põlme et al. 2020) | Põlme S, et al. 2020. Fungal Diversity 105:1-16 ((https://doi.org/10.1007/s13225-020-00466-2)) |
| Otu52448 | 1 | k__Fungi(100);p__Ascomycota(100);c__Saccharomycetes(100);o__Saccharomycetales(100);f__Metschnikowiaceae(100);g__Metschnikowia(100);s__Metschnikowia_pulcherrima(100); | Metschnikowia | 13 | Saprotroph | \|Undefined Saprotroph\| | Yeast | NULL | Probable | As nectar-tap_saprotroph (Põlme et al. 2020); Decay sugar-rich substrates (Põlme et al. 2020) | Põlme S, et al. 2020. Fungal Diversity 105:1-16 ((https://doi.org/10.1007/s13225-020-00466-2)) |
| Otu52449 | 1 | k__Fungi(100);p__Ascomycota(100);c__Saccharomycetes(100);o__Saccharomycetales(100);f__Metschnikowiaceae(100);g__Metschnikowia(100);s__Metschnikowia_pulcherrima(100); | Metschnikowia | 13 | Saprotroph | \|Undefined Saprotroph\| | Yeast | NULL | Probable | As nectar-tap_saprotroph (Põlme et al. 2020); Decay sugar-rich substrates (Põlme et al. 2020) | Põlme S, et al. 2020. Fungal Diversity 105:1-16 ((https://doi.org/10.1007/s13225-020-00466-2)) |
| Otu52451 | 1 | k__Fungi(100);p__Ascomycota(100);c__Saccharomycetes(100);o__Saccharomycetales(100);f__Metschnikowiaceae(100);g__Metschnikowia(100);s__Metschnikowia_pulcherrima(100); | Metschnikowia | 13 | Saprotroph | \|Undefined Saprotroph\| | Yeast | NULL | Probable | As nectar-tap_saprotroph (Põlme et al. 2020); Decay sugar-rich substrates (Põlme et al. 2020) | Põlme S, et al. 2020. Fungal Diversity 105:1-16 ((https://doi.org/10.1007/s13225-020-00466-2)) |
| Otu52452 | 1 | k__Fungi(100);p__Ascomycota(100);c__Saccharomycetes(100);o__Saccharomycetales(100);f__Metschnikowiaceae(100);g__Metschnikowia(100);g__Metschnikowia_unclassified(100); | Metschnikowia | 13 | Saprotroph | \|Undefined Saprotroph\| | Yeast | NULL | Probable | As nectar-tap_saprotroph (Põlme et al. 2020); Decay sugar-rich substrates (Põlme et al. 2020) | Põlme S, et al. 2020. Fungal Diversity 105:1-16 ((https://doi.org/10.1007/s13225-020-00466-2)) |
| Otu52453 | 1 | k__Fungi(100);p__Ascomycota(100);c__Saccharomycetes(100);o__Saccharomycetales(100);f__Metschnikowiaceae(100);g__Metschnikowia(100);s__Metschnikowia_pulcherrima(100); | Metschnikowia | 13 | Saprotroph | \|Undefined Saprotroph\| | Yeast | NULL | Probable | As nectar-tap_saprotroph (Põlme et al. 2020); Decay sugar-rich substrates (Põlme et al. 2020) | Põlme S, et al. 2020. Fungal Diversity 105:1-16 ((https://doi.org/10.1007/s13225-020-00466-2)) |
| Otu52454 | 1 | k__Fungi(100);p__Ascomycota(100);c__Saccharomycetes(100);o__Saccharomycetales(100);f__Metschnikowiaceae(100);g__Metschnikowia(100);s__Metschnikowia_pulcherrima(100); | Metschnikowia | 13 | Saprotroph | \|Undefined Saprotroph\| | Yeast | NULL | Probable | As nectar-tap_saprotroph (Põlme et al. 2020); Decay sugar-rich substrates (Põlme et al. 2020) | Põlme S, et al. 2020. Fungal Diversity 105:1-16 ((https://doi.org/10.1007/s13225-020-00466-2)) |
| Otu52455 | 1 | k__Fungi(100);p__Ascomycota(100);c__Saccharomycetes(100);o__Saccharomycetales(100);f__Metschnikowiaceae(100);g__Metschnikowia(100);s__Metschnikowia_pulcherrima(100); | Metschnikowia | 13 | Saprotroph | \|Undefined Saprotroph\| | Yeast | NULL | Probable | As nectar-tap_saprotroph (Põlme et al. 2020); Decay sugar-rich substrates (Põlme et al. 2020) | Põlme S, et al. 2020. Fungal Diversity 105:1-16 ((https://doi.org/10.1007/s13225-020-00466-2)) |
| Otu52456 | 1 | k__Fungi(100);p__Ascomycota(100);c__Saccharomycetes(100);o__Saccharomycetales(100);f__Metschnikowiaceae(100);g__Metschnikowia(100);g__Metschnikowia_unclassified(100); | Metschnikowia | 13 | Saprotroph | \|Undefined Saprotroph\| | Yeast | NULL | Probable | As nectar-tap_saprotroph (Põlme et al. 2020); Decay sugar-rich substrates (Põlme et al. 2020) | Põlme S, et al. 2020. Fungal Diversity 105:1-16 ((https://doi.org/10.1007/s13225-020-00466-2)) |
| Otu52457 | 1 | k__Fungi(100);p__Ascomycota(100);c__Saccharomycetes(100);o__Saccharomycetales(100);f__Metschnikowiaceae(100);g__Metschnikowia(100);s__Metschnikowia_pulcherrima(100); | Metschnikowia | 13 | Saprotroph | \|Undefined Saprotroph\| | Yeast | NULL | Probable | As nectar-tap_saprotroph (Põlme et al. 2020); Decay sugar-rich substrates (Põlme et al. 2020) | Põlme S, et al. 2020. Fungal Diversity 105:1-16 ((https://doi.org/10.1007/s13225-020-00466-2)) |
| Otu52458 | 1 | k__Fungi(100);p__Ascomycota(100);c__Saccharomycetes(100);o__Saccharomycetales(100);f__Metschnikowiaceae(100);g__Metschnikowia(100);s__Metschnikowia_pulcherrima(100); | Metschnikowia | 13 | Saprotroph | \|Undefined Saprotroph\| | Yeast | NULL | Probable | As nectar-tap_saprotroph (Põlme et al. 2020); Decay sugar-rich substrates (Põlme et al. 2020) | Põlme S, et al. 2020. Fungal Diversity 105:1-16 ((https://doi.org/10.1007/s13225-020-00466-2)) |
| Otu52459 | 1 | k__Fungi(100);p__Ascomycota(100);c__Saccharomycetes(100);o__Saccharomycetales(100);f__Metschnikowiaceae(100);g__Metschnikowia(100);s__Metschnikowia_pulcherrima(100); | Metschnikowia | 13 | Saprotroph | \|Undefined Saprotroph\| | Yeast | NULL | Probable | As nectar-tap_saprotroph (Põlme et al. 2020); Decay sugar-rich substrates (Põlme et al. 2020) | Põlme S, et al. 2020. Fungal Diversity 105:1-16 ((https://doi.org/10.1007/s13225-020-00466-2)) |
| Otu52460 | 1 | k__Fungi(100);p__Ascomycota(100);c__Saccharomycetes(100);o__Saccharomycetales(100);f__Metschnikowiaceae(100);g__Metschnikowia(100);s__Metschnikowia_pulcherrima(100); | Metschnikowia | 13 | Saprotroph | \|Undefined Saprotroph\| | Yeast | NULL | Probable | As nectar-tap_saprotroph (Põlme et al. 2020); Decay sugar-rich substrates (Põlme et al. 2020) | Põlme S, et al. 2020. Fungal Diversity 105:1-16 ((https://doi.org/10.1007/s13225-020-00466-2)) |
| Otu52461 | 1 | k__Fungi(100);p__Ascomycota(100);c__Saccharomycetes(100);o__Saccharomycetales(100);f__Metschnikowiaceae(100);g__Metschnikowia(100);s__Metschnikowia_pulcherrima(100); | Metschnikowia | 13 | Saprotroph | \|Undefined Saprotroph\| | Yeast | NULL | Probable | As nectar-tap_saprotroph (Põlme et al. 2020); Decay sugar-rich substrates (Põlme et al. 2020) | Põlme S, et al. 2020. Fungal Diversity 105:1-16 ((https://doi.org/10.1007/s13225-020-00466-2)) |
| Otu52462 | 1 | k__Fungi(100);p__Ascomycota(100);c__Saccharomycetes(100);o__Saccharomycetales(100);f__Metschnikowiaceae(100);g__Metschnikowia(100);s__Metschnikowia_pulcherrima(100); | Metschnikowia | 13 | Saprotroph | \|Undefined Saprotroph\| | Yeast | NULL | Probable | As nectar-tap_saprotroph (Põlme et al. 2020); Decay sugar-rich substrates (Põlme et al. 2020) | Põlme S, et al. 2020. Fungal Diversity 105:1-16 ((https://doi.org/10.1007/s13225-020-00466-2)) |
| Otu52463 | 1 | k__Fungi(100);p__Ascomycota(100);c__Saccharomycetes(100);o__Saccharomycetales(100);f__Metschnikowiaceae(100);g__Metschnikowia(100);g__Metschnikowia_unclassified(100); | Metschnikowia | 13 | Saprotroph | \|Undefined Saprotroph\| | Yeast | NULL | Probable | As nectar-tap_saprotroph (Põlme et al. 2020); Decay sugar-rich substrates (Põlme et al. 2020) | Põlme S, et al. 2020. Fungal Diversity 105:1-16 ((https://doi.org/10.1007/s13225-020-00466-2)) |
| Otu52464 | 1 | k__Fungi(100);p__Ascomycota(100);c__Saccharomycetes(100);o__Saccharomycetales(100);f__Metschnikowiaceae(100);g__Metschnikowia(100);s__Metschnikowia_pulcherrima(100); | Metschnikowia | 13 | Saprotroph | \|Undefined Saprotroph\| | Yeast | NULL | Probable | As nectar-tap_saprotroph (Põlme et al. 2020); Decay sugar-rich substrates (Põlme et al. 2020) | Põlme S, et al. 2020. Fungal Diversity 105:1-16 ((https://doi.org/10.1007/s13225-020-00466-2)) |
| Otu52465 | 1 | k__Fungi(100);p__Ascomycota(100);c__Saccharomycetes(100);o__Saccharomycetales(100);f__Metschnikowiaceae(100);g__Metschnikowia(100);s__Metschnikowia_pulcherrima(100); | Metschnikowia | 13 | Saprotroph | \|Undefined Saprotroph\| | Yeast | NULL | Probable | As nectar-tap_saprotroph (Põlme et al. 2020); Decay sugar-rich substrates (Põlme et al. 2020) | Põlme S, et al. 2020. Fungal Diversity 105:1-16 ((https://doi.org/10.1007/s13225-020-00466-2)) |
| Otu52466 | 1 | k__Fungi(100);p__Ascomycota(100);c__Saccharomycetes(100);o__Saccharomycetales(100);f__Metschnikowiaceae(100);g__Metschnikowia(100);s__Metschnikowia_pulcherrima(100); | Metschnikowia | 13 | Saprotroph | \|Undefined Saprotroph\| | Yeast | NULL | Probable | As nectar-tap_saprotroph (Põlme et al. 2020); Decay sugar-rich substrates (Põlme et al. 2020) | Põlme S, et al. 2020. Fungal Diversity 105:1-16 ((https://doi.org/10.1007/s13225-020-00466-2)) |
| Otu52467 | 1 | k__Fungi(100);p__Ascomycota(100);c__Saccharomycetes(100);o__Saccharomycetales(100);f__Metschnikowiaceae(100);g__Metschnikowia(100);s__Metschnikowia_pulcherrima(100); | Metschnikowia | 13 | Saprotroph | \|Undefined Saprotroph\| | Yeast | NULL | Probable | As nectar-tap_saprotroph (Põlme et al. 2020); Decay sugar-rich substrates (Põlme et al. 2020) | Põlme S, et al. 2020. Fungal Diversity 105:1-16 ((https://doi.org/10.1007/s13225-020-00466-2)) |
| Otu52468 | 1 | k__Fungi(100);p__Ascomycota(100);c__Saccharomycetes(100);o__Saccharomycetales(100);f__Metschnikowiaceae(100);g__Metschnikowia(100);s__Metschnikowia_pulcherrima(100); | Metschnikowia | 13 | Saprotroph | \|Undefined Saprotroph\| | Yeast | NULL | Probable | As nectar-tap_saprotroph (Põlme et al. 2020); Decay sugar-rich substrates (Põlme et al. 2020) | Põlme S, et al. 2020. Fungal Diversity 105:1-16 ((https://doi.org/10.1007/s13225-020-00466-2)) |
| Otu52470 | 1 | k__Fungi(100);p__Ascomycota(100);c__Saccharomycetes(100);o__Saccharomycetales(100);f__Metschnikowiaceae(100);g__Metschnikowia(100);s__Metschnikowia_pulcherrima(100); | Metschnikowia | 13 | Saprotroph | \|Undefined Saprotroph\| | Yeast | NULL | Probable | As nectar-tap_saprotroph (Põlme et al. 2020); Decay sugar-rich substrates (Põlme et al. 2020) | Põlme S, et al. 2020. Fungal Diversity 105:1-16 ((https://doi.org/10.1007/s13225-020-00466-2)) |
| Otu52471 | 1 | k__Fungi(100);p__Ascomycota(100);c__Saccharomycetes(100);o__Saccharomycetales(100);f__Metschnikowiaceae(100);g__Metschnikowia(100);s__Metschnikowia_pulcherrima(100); | Metschnikowia | 13 | Saprotroph | \|Undefined Saprotroph\| | Yeast | NULL | Probable | As nectar-tap_saprotroph (Põlme et al. 2020); Decay sugar-rich substrates (Põlme et al. 2020) | Põlme S, et al. 2020. Fungal Diversity 105:1-16 ((https://doi.org/10.1007/s13225-020-00466-2)) |
| Otu52472 | 1 | k__Fungi(100);p__Ascomycota(100);c__Saccharomycetes(100);o__Saccharomycetales(100);f__Metschnikowiaceae(100);g__Metschnikowia(100);g__Metschnikowia_unclassified(100); | Metschnikowia | 13 | Saprotroph | \|Undefined Saprotroph\| | Yeast | NULL | Probable | As nectar-tap_saprotroph (Põlme et al. 2020); Decay sugar-rich substrates (Põlme et al. 2020) | Põlme S, et al. 2020. Fungal Diversity 105:1-16 ((https://doi.org/10.1007/s13225-020-00466-2)) |
| Otu52473 | 1 | k__Fungi(100);p__Ascomycota(100);c__Saccharomycetes(100);o__Saccharomycetales(100);f__Metschnikowiaceae(100);g__Metschnikowia(100);s__Metschnikowia_pulcherrima(100); | Metschnikowia | 13 | Saprotroph | \|Undefined Saprotroph\| | Yeast | NULL | Probable | As nectar-tap_saprotroph (Põlme et al. 2020); Decay sugar-rich substrates (Põlme et al. 2020) | Põlme S, et al. 2020. Fungal Diversity 105:1-16 ((https://doi.org/10.1007/s13225-020-00466-2)) |
| Otu52474 | 1 | k__Fungi(100);p__Ascomycota(100);c__Saccharomycetes(100);o__Saccharomycetales(100);f__Metschnikowiaceae(100);g__Metschnikowia(100);s__Metschnikowia_pulcherrima(100); | Metschnikowia | 13 | Saprotroph | \|Undefined Saprotroph\| | Yeast | NULL | Probable | As nectar-tap_saprotroph (Põlme et al. 2020); Decay sugar-rich substrates (Põlme et al. 2020) | Põlme S, et al. 2020. Fungal Diversity 105:1-16 ((https://doi.org/10.1007/s13225-020-00466-2)) |
| Otu52475 | 1 | k__Fungi(100);p__Ascomycota(100);c__Saccharomycetes(100);o__Saccharomycetales(100);f__Metschnikowiaceae(100);g__Metschnikowia(100);s__Metschnikowia_pulcherrima(100); | Metschnikowia | 13 | Saprotroph | \|Undefined Saprotroph\| | Yeast | NULL | Probable | As nectar-tap_saprotroph (Põlme et al. 2020); Decay sugar-rich substrates (Põlme et al. 2020) | Põlme S, et al. 2020. Fungal Diversity 105:1-16 ((https://doi.org/10.1007/s13225-020-00466-2)) |
| Otu52476 | 1 | k__Fungi(100);p__Ascomycota(100);c__Saccharomycetes(100);o__Saccharomycetales(100);f__Metschnikowiaceae(100);g__Metschnikowia(100);s__Metschnikowia_pulcherrima(100); | Metschnikowia | 13 | Saprotroph | \|Undefined Saprotroph\| | Yeast | NULL | Probable | As nectar-tap_saprotroph (Põlme et al. 2020); Decay sugar-rich substrates (Põlme et al. 2020) | Põlme S, et al. 2020. Fungal Diversity 105:1-16 ((https://doi.org/10.1007/s13225-020-00466-2)) |
| Otu52477 | 1 | k__Fungi(100);p__Ascomycota(100);c__Saccharomycetes(100);o__Saccharomycetales(100);f__Metschnikowiaceae(100);g__Metschnikowia(100);g__Metschnikowia_unclassified(100); | Metschnikowia | 13 | Saprotroph | \|Undefined Saprotroph\| | Yeast | NULL | Probable | As nectar-tap_saprotroph (Põlme et al. 2020); Decay sugar-rich substrates (Põlme et al. 2020) | Põlme S, et al. 2020. Fungal Diversity 105:1-16 ((https://doi.org/10.1007/s13225-020-00466-2)) |
| Otu52478 | 1 | k__Fungi(100);p__Ascomycota(100);c__Saccharomycetes(100);o__Saccharomycetales(100);f__Metschnikowiaceae(100);g__Metschnikowia(100);s__Metschnikowia_pulcherrima(100); | Metschnikowia | 13 | Saprotroph | \|Undefined Saprotroph\| | Yeast | NULL | Probable | As nectar-tap_saprotroph (Põlme et al. 2020); Decay sugar-rich substrates (Põlme et al. 2020) | Põlme S, et al. 2020. Fungal Diversity 105:1-16 ((https://doi.org/10.1007/s13225-020-00466-2)) |
| Otu52479 | 1 | k__Fungi(100);p__Ascomycota(100);c__Saccharomycetes(100);o__Saccharomycetales(100);f__Metschnikowiaceae(100);g__Metschnikowia(100);s__Metschnikowia_pulcherrima(100); | Metschnikowia | 13 | Saprotroph | \|Undefined Saprotroph\| | Yeast | NULL | Probable | As nectar-tap_saprotroph (Põlme et al. 2020); Decay sugar-rich substrates (Põlme et al. 2020) | Põlme S, et al. 2020. Fungal Diversity 105:1-16 ((https://doi.org/10.1007/s13225-020-00466-2)) |
| Otu52480 | 1 | k__Fungi(100);p__Ascomycota(100);c__Saccharomycetes(100);o__Saccharomycetales(100);f__Metschnikowiaceae(100);g__Metschnikowia(100);g__Metschnikowia_unclassified(100); | Metschnikowia | 13 | Saprotroph | \|Undefined Saprotroph\| | Yeast | NULL | Probable | As nectar-tap_saprotroph (Põlme et al. 2020); Decay sugar-rich substrates (Põlme et al. 2020) | Põlme S, et al. 2020. Fungal Diversity 105:1-16 ((https://doi.org/10.1007/s13225-020-00466-2)) |
| Otu52481 | 1 | k__Fungi(100);p__Ascomycota(100);c__Saccharomycetes(100);o__Saccharomycetales(100);f__Metschnikowiaceae(100);g__Metschnikowia(100);s__Metschnikowia_pulcherrima(100); | Metschnikowia | 13 | Saprotroph | \|Undefined Saprotroph\| | Yeast | NULL | Probable | As nectar-tap_saprotroph (Põlme et al. 2020); Decay sugar-rich substrates (Põlme et al. 2020) | Põlme S, et al. 2020. Fungal Diversity 105:1-16 ((https://doi.org/10.1007/s13225-020-00466-2)) |
| Otu52482 | 1 | k__Fungi(100);p__Ascomycota(100);c__Saccharomycetes(100);o__Saccharomycetales(100);f__Metschnikowiaceae(100);g__Metschnikowia(100);s__Metschnikowia_pulcherrima(100); | Metschnikowia | 13 | Saprotroph | \|Undefined Saprotroph\| | Yeast | NULL | Probable | As nectar-tap_saprotroph (Põlme et al. 2020); Decay sugar-rich substrates (Põlme et al. 2020) | Põlme S, et al. 2020. Fungal Diversity 105:1-16 ((https://doi.org/10.1007/s13225-020-00466-2)) |
| Otu52483 | 1 | k__Fungi(100);p__Ascomycota(100);c__Saccharomycetes(100);o__Saccharomycetales(100);f__Metschnikowiaceae(100);g__Metschnikowia(100);g__Metschnikowia_unclassified(100); | Metschnikowia | 13 | Saprotroph | \|Undefined Saprotroph\| | Yeast | NULL | Probable | As nectar-tap_saprotroph (Põlme et al. 2020); Decay sugar-rich substrates (Põlme et al. 2020) | Põlme S, et al. 2020. Fungal Diversity 105:1-16 ((https://doi.org/10.1007/s13225-020-00466-2)) |
| Otu52484 | 1 | k__Fungi(100);p__Ascomycota(100);c__Saccharomycetes(100);o__Saccharomycetales(100);f__Metschnikowiaceae(100);g__Metschnikowia(100);s__Metschnikowia_pulcherrima(100); | Metschnikowia | 13 | Saprotroph | \|Undefined Saprotroph\| | Yeast | NULL | Probable | As nectar-tap_saprotroph (Põlme et al. 2020); Decay sugar-rich substrates (Põlme et al. 2020) | Põlme S, et al. 2020. Fungal Diversity 105:1-16 ((https://doi.org/10.1007/s13225-020-00466-2)) |
| Otu52487 | 1 | k__Fungi(100);p__Ascomycota(100);c__Saccharomycetes(100);o__Saccharomycetales(100);f__Metschnikowiaceae(100);g__Metschnikowia(100);s__Metschnikowia_pulcherrima(100); | Metschnikowia | 13 | Saprotroph | \|Undefined Saprotroph\| | Yeast | NULL | Probable | As nectar-tap_saprotroph (Põlme et al. 2020); Decay sugar-rich substrates (Põlme et al. 2020) | Põlme S, et al. 2020. Fungal Diversity 105:1-16 ((https://doi.org/10.1007/s13225-020-00466-2)) |
| Otu52488 | 1 | k__Fungi(100);p__Ascomycota(100);c__Saccharomycetes(100);o__Saccharomycetales(100);f__Metschnikowiaceae(100);g__Metschnikowia(100);s__Metschnikowia_pulcherrima(100); | Metschnikowia | 13 | Saprotroph | \|Undefined Saprotroph\| | Yeast | NULL | Probable | As nectar-tap_saprotroph (Põlme et al. 2020); Decay sugar-rich substrates (Põlme et al. 2020) | Põlme S, et al. 2020. Fungal Diversity 105:1-16 ((https://doi.org/10.1007/s13225-020-00466-2)) |
| Otu52489 | 1 | k__Fungi(100);p__Ascomycota(100);c__Saccharomycetes(100);o__Saccharomycetales(100);f__Metschnikowiaceae(100);g__Metschnikowia(100);s__Metschnikowia_pulcherrima(100); | Metschnikowia | 13 | Saprotroph | \|Undefined Saprotroph\| | Yeast | NULL | Probable | As nectar-tap_saprotroph (Põlme et al. 2020); Decay sugar-rich substrates (Põlme et al. 2020) | Põlme S, et al. 2020. Fungal Diversity 105:1-16 ((https://doi.org/10.1007/s13225-020-00466-2)) |
| Otu52490 | 1 | k__Fungi(100);p__Ascomycota(100);c__Saccharomycetes(100);o__Saccharomycetales(100);f__Metschnikowiaceae(100);g__Metschnikowia(100);s__Metschnikowia_pulcherrima(100); | Metschnikowia | 13 | Saprotroph | \|Undefined Saprotroph\| | Yeast | NULL | Probable | As nectar-tap_saprotroph (Põlme et al. 2020); Decay sugar-rich substrates (Põlme et al. 2020) | Põlme S, et al. 2020. Fungal Diversity 105:1-16 ((https://doi.org/10.1007/s13225-020-00466-2)) |
| Otu52491 | 1 | k__Fungi(100);p__Ascomycota(100);c__Saccharomycetes(100);o__Saccharomycetales(100);f__Metschnikowiaceae(100);g__Metschnikowia(100);g__Metschnikowia_unclassified(100); | Metschnikowia | 13 | Saprotroph | \|Undefined Saprotroph\| | Yeast | NULL | Probable | As nectar-tap_saprotroph (Põlme et al. 2020); Decay sugar-rich substrates (Põlme et al. 2020) | Põlme S, et al. 2020. Fungal Diversity 105:1-16 ((https://doi.org/10.1007/s13225-020-00466-2)) |
| Otu52492 | 1 | k__Fungi(100);p__Ascomycota(100);c__Saccharomycetes(100);o__Saccharomycetales(100);f__Metschnikowiaceae(100);g__Metschnikowia(100);g__Metschnikowia_unclassified(100); | Metschnikowia | 13 | Saprotroph | \|Undefined Saprotroph\| | Yeast | NULL | Probable | As nectar-tap_saprotroph (Põlme et al. 2020); Decay sugar-rich substrates (Põlme et al. 2020) | Põlme S, et al. 2020. Fungal Diversity 105:1-16 ((https://doi.org/10.1007/s13225-020-00466-2)) |
| Otu52493 | 1 | k__Fungi(100);p__Ascomycota(100);c__Saccharomycetes(100);o__Saccharomycetales(100);f__Metschnikowiaceae(100);g__Metschnikowia(100);g__Metschnikowia_unclassified(100); | Metschnikowia | 13 | Saprotroph | \|Undefined Saprotroph\| | Yeast | NULL | Probable | As nectar-tap_saprotroph (Põlme et al. 2020); Decay sugar-rich substrates (Põlme et al. 2020) | Põlme S, et al. 2020. Fungal Diversity 105:1-16 ((https://doi.org/10.1007/s13225-020-00466-2)) |
| Otu52494 | 1 | k__Fungi(100);p__Ascomycota(100);c__Saccharomycetes(100);o__Saccharomycetales(100);f__Metschnikowiaceae(100);g__Metschnikowia(100);s__Metschnikowia_pulcherrima(100); | Metschnikowia | 13 | Saprotroph | \|Undefined Saprotroph\| | Yeast | NULL | Probable | As nectar-tap_saprotroph (Põlme et al. 2020); Decay sugar-rich substrates (Põlme et al. 2020) | Põlme S, et al. 2020. Fungal Diversity 105:1-16 ((https://doi.org/10.1007/s13225-020-00466-2)) |
| Otu52495 | 1 | k__Fungi(100);p__Ascomycota(100);c__Saccharomycetes(100);o__Saccharomycetales(100);f__Metschnikowiaceae(100);g__Metschnikowia(100);s__Metschnikowia_pulcherrima(100); | Metschnikowia | 13 | Saprotroph | \|Undefined Saprotroph\| | Yeast | NULL | Probable | As nectar-tap_saprotroph (Põlme et al. 2020); Decay sugar-rich substrates (Põlme et al. 2020) | Põlme S, et al. 2020. Fungal Diversity 105:1-16 ((https://doi.org/10.1007/s13225-020-00466-2)) |
| Otu52496 | 1 | k__Fungi(100);p__Ascomycota(100);c__Saccharomycetes(100);o__Saccharomycetales(100);f__Metschnikowiaceae(100);g__Metschnikowia(100);s__Metschnikowia_pulcherrima(100); | Metschnikowia | 13 | Saprotroph | \|Undefined Saprotroph\| | Yeast | NULL | Probable | As nectar-tap_saprotroph (Põlme et al. 2020); Decay sugar-rich substrates (Põlme et al. 2020) | Põlme S, et al. 2020. Fungal Diversity 105:1-16 ((https://doi.org/10.1007/s13225-020-00466-2)) |
| Otu52497 | 1 | k__Fungi(100);p__Ascomycota(100);c__Saccharomycetes(100);o__Saccharomycetales(100);f__Metschnikowiaceae(100);g__Metschnikowia(100);g__Metschnikowia_unclassified(100); | Metschnikowia | 13 | Saprotroph | \|Undefined Saprotroph\| | Yeast | NULL | Probable | As nectar-tap_saprotroph (Põlme et al. 2020); Decay sugar-rich substrates (Põlme et al. 2020) | Põlme S, et al. 2020. Fungal Diversity 105:1-16 ((https://doi.org/10.1007/s13225-020-00466-2)) |
| Otu52498 | 1 | k__Fungi(100);p__Ascomycota(100);c__Saccharomycetes(100);o__Saccharomycetales(100);f__Metschnikowiaceae(100);g__Metschnikowia(100);g__Metschnikowia_unclassified(100); | Metschnikowia | 13 | Saprotroph | \|Undefined Saprotroph\| | Yeast | NULL | Probable | As nectar-tap_saprotroph (Põlme et al. 2020); Decay sugar-rich substrates (Põlme et al. 2020) | Põlme S, et al. 2020. Fungal Diversity 105:1-16 ((https://doi.org/10.1007/s13225-020-00466-2)) |
| Otu52499 | 1 | k__Fungi(100);p__Ascomycota(100);c__Saccharomycetes(100);o__Saccharomycetales(100);f__Metschnikowiaceae(100);g__Metschnikowia(100);s__Metschnikowia_pulcherrima(100); | Metschnikowia | 13 | Saprotroph | \|Undefined Saprotroph\| | Yeast | NULL | Probable | As nectar-tap_saprotroph (Põlme et al. 2020); Decay sugar-rich substrates (Põlme et al. 2020) | Põlme S, et al. 2020. Fungal Diversity 105:1-16 ((https://doi.org/10.1007/s13225-020-00466-2)) |
| Otu52500 | 1 | k__Fungi(100);p__Ascomycota(100);c__Saccharomycetes(100);o__Saccharomycetales(100);f__Metschnikowiaceae(100);g__Metschnikowia(100);g__Metschnikowia_unclassified(100); | Metschnikowia | 13 | Saprotroph | \|Undefined Saprotroph\| | Yeast | NULL | Probable | As nectar-tap_saprotroph (Põlme et al. 2020); Decay sugar-rich substrates (Põlme et al. 2020) | Põlme S, et al. 2020. Fungal Diversity 105:1-16 ((https://doi.org/10.1007/s13225-020-00466-2)) |
| Otu52501 | 1 | k__Fungi(100);p__Ascomycota(100);c__Saccharomycetes(100);o__Saccharomycetales(100);f__Metschnikowiaceae(100);g__Metschnikowia(100);s__Metschnikowia_pulcherrima(100); | Metschnikowia | 13 | Saprotroph | \|Undefined Saprotroph\| | Yeast | NULL | Probable | As nectar-tap_saprotroph (Põlme et al. 2020); Decay sugar-rich substrates (Põlme et al. 2020) | Põlme S, et al. 2020. Fungal Diversity 105:1-16 ((https://doi.org/10.1007/s13225-020-00466-2)) |
| Otu52502 | 1 | k__Fungi(100);p__Ascomycota(100);c__Saccharomycetes(100);o__Saccharomycetales(100);f__Metschnikowiaceae(100);g__Metschnikowia(100);s__Metschnikowia_pulcherrima(100); | Metschnikowia | 13 | Saprotroph | \|Undefined Saprotroph\| | Yeast | NULL | Probable | As nectar-tap_saprotroph (Põlme et al. 2020); Decay sugar-rich substrates (Põlme et al. 2020) | Põlme S, et al. 2020. Fungal Diversity 105:1-16 ((https://doi.org/10.1007/s13225-020-00466-2)) |
| Otu52503 | 1 | k__Fungi(100);p__Ascomycota(100);c__Saccharomycetes(100);o__Saccharomycetales(100);f__Metschnikowiaceae(100);g__Metschnikowia(100);s__Metschnikowia_pulcherrima(100); | Metschnikowia | 13 | Saprotroph | \|Undefined Saprotroph\| | Yeast | NULL | Probable | As nectar-tap_saprotroph (Põlme et al. 2020); Decay sugar-rich substrates (Põlme et al. 2020) | Põlme S, et al. 2020. Fungal Diversity 105:1-16 ((https://doi.org/10.1007/s13225-020-00466-2)) |
| Otu52504 | 1 | k__Fungi(100);p__Ascomycota(100);c__Saccharomycetes(100);o__Saccharomycetales(100);f__Metschnikowiaceae(100);g__Metschnikowia(100);s__Metschnikowia_pulcherrima(100); | Metschnikowia | 13 | Saprotroph | \|Undefined Saprotroph\| | Yeast | NULL | Probable | As nectar-tap_saprotroph (Põlme et al. 2020); Decay sugar-rich substrates (Põlme et al. 2020) | Põlme S, et al. 2020. Fungal Diversity 105:1-16 ((https://doi.org/10.1007/s13225-020-00466-2)) |
| Otu52505 | 1 | k__Fungi(100);p__Ascomycota(100);c__Saccharomycetes(100);o__Saccharomycetales(100);f__Metschnikowiaceae(100);g__Metschnikowia(100);s__Metschnikowia_pulcherrima(100); | Metschnikowia | 13 | Saprotroph | \|Undefined Saprotroph\| | Yeast | NULL | Probable | As nectar-tap_saprotroph (Põlme et al. 2020); Decay sugar-rich substrates (Põlme et al. 2020) | Põlme S, et al. 2020. Fungal Diversity 105:1-16 ((https://doi.org/10.1007/s13225-020-00466-2)) |
| Otu52506 | 1 | k__Fungi(100);p__Ascomycota(100);c__Saccharomycetes(100);o__Saccharomycetales(100);f__Metschnikowiaceae(100);g__Metschnikowia(100);s__Metschnikowia_pulcherrima(100); | Metschnikowia | 13 | Saprotroph | \|Undefined Saprotroph\| | Yeast | NULL | Probable | As nectar-tap_saprotroph (Põlme et al. 2020); Decay sugar-rich substrates (Põlme et al. 2020) | Põlme S, et al. 2020. Fungal Diversity 105:1-16 ((https://doi.org/10.1007/s13225-020-00466-2)) |
| Otu52507 | 1 | k__Fungi(100);p__Ascomycota(100);c__Saccharomycetes(100);o__Saccharomycetales(100);f__Metschnikowiaceae(100);g__Metschnikowia(100);s__Metschnikowia_pulcherrima(100); | Metschnikowia | 13 | Saprotroph | \|Undefined Saprotroph\| | Yeast | NULL | Probable | As nectar-tap_saprotroph (Põlme et al. 2020); Decay sugar-rich substrates (Põlme et al. 2020) | Põlme S, et al. 2020. Fungal Diversity 105:1-16 ((https://doi.org/10.1007/s13225-020-00466-2)) |
| Otu52508 | 1 | k__Fungi(100);p__Ascomycota(100);c__Saccharomycetes(100);o__Saccharomycetales(100);f__Metschnikowiaceae(100);g__Metschnikowia(100);s__Metschnikowia_pulcherrima(100); | Metschnikowia | 13 | Saprotroph | \|Undefined Saprotroph\| | Yeast | NULL | Probable | As nectar-tap_saprotroph (Põlme et al. 2020); Decay sugar-rich substrates (Põlme et al. 2020) | Põlme S, et al. 2020. Fungal Diversity 105:1-16 ((https://doi.org/10.1007/s13225-020-00466-2)) |
| Otu52509 | 1 | k__Fungi(100);p__Ascomycota(100);c__Saccharomycetes(100);o__Saccharomycetales(100);f__Metschnikowiaceae(100);g__Metschnikowia(100);s__Metschnikowia_pulcherrima(100); | Metschnikowia | 13 | Saprotroph | \|Undefined Saprotroph\| | Yeast | NULL | Probable | As nectar-tap_saprotroph (Põlme et al. 2020); Decay sugar-rich substrates (Põlme et al. 2020) | Põlme S, et al. 2020. Fungal Diversity 105:1-16 ((https://doi.org/10.1007/s13225-020-00466-2)) |
| Otu52510 | 1 | k__Fungi(100);p__Ascomycota(100);c__Saccharomycetes(100);o__Saccharomycetales(100);f__Metschnikowiaceae(100);g__Metschnikowia(100);s__Metschnikowia_pulcherrima(100); | Metschnikowia | 13 | Saprotroph | \|Undefined Saprotroph\| | Yeast | NULL | Probable | As nectar-tap_saprotroph (Põlme et al. 2020); Decay sugar-rich substrates (Põlme et al. 2020) | Põlme S, et al. 2020. Fungal Diversity 105:1-16 ((https://doi.org/10.1007/s13225-020-00466-2)) |
| Otu52511 | 1 | k__Fungi(100);p__Ascomycota(100);c__Saccharomycetes(100);o__Saccharomycetales(100);f__Metschnikowiaceae(100);g__Metschnikowia(100);s__Metschnikowia_pulcherrima(100); | Metschnikowia | 13 | Saprotroph | \|Undefined Saprotroph\| | Yeast | NULL | Probable | As nectar-tap_saprotroph (Põlme et al. 2020); Decay sugar-rich substrates (Põlme et al. 2020) | Põlme S, et al. 2020. Fungal Diversity 105:1-16 ((https://doi.org/10.1007/s13225-020-00466-2)) |
| Otu52512 | 1 | k__Fungi(100);p__Ascomycota(100);c__Saccharomycetes(100);o__Saccharomycetales(100);f__Metschnikowiaceae(100);g__Metschnikowia(100);s__Metschnikowia_pulcherrima(100); | Metschnikowia | 13 | Saprotroph | \|Undefined Saprotroph\| | Yeast | NULL | Probable | As nectar-tap_saprotroph (Põlme et al. 2020); Decay sugar-rich substrates (Põlme et al. 2020) | Põlme S, et al. 2020. Fungal Diversity 105:1-16 ((https://doi.org/10.1007/s13225-020-00466-2)) |
| Otu52513 | 1 | k__Fungi(100);p__Ascomycota(100);c__Saccharomycetes(100);o__Saccharomycetales(100);f__Metschnikowiaceae(100);g__Metschnikowia(100);g__Metschnikowia_unclassified(100); | Metschnikowia | 13 | Saprotroph | \|Undefined Saprotroph\| | Yeast | NULL | Probable | As nectar-tap_saprotroph (Põlme et al. 2020); Decay sugar-rich substrates (Põlme et al. 2020) | Põlme S, et al. 2020. Fungal Diversity 105:1-16 ((https://doi.org/10.1007/s13225-020-00466-2)) |
| Otu52514 | 1 | k__Fungi(100);p__Ascomycota(100);c__Saccharomycetes(100);o__Saccharomycetales(100);f__Metschnikowiaceae(100);g__Metschnikowia(100);s__Metschnikowia_pulcherrima(100); | Metschnikowia | 13 | Saprotroph | \|Undefined Saprotroph\| | Yeast | NULL | Probable | As nectar-tap_saprotroph (Põlme et al. 2020); Decay sugar-rich substrates (Põlme et al. 2020) | Põlme S, et al. 2020. Fungal Diversity 105:1-16 ((https://doi.org/10.1007/s13225-020-00466-2)) |
| Otu52515 | 1 | k__Fungi(100);p__Ascomycota(100);c__Saccharomycetes(100);o__Saccharomycetales(100);f__Metschnikowiaceae(100);g__Metschnikowia(100);s__Metschnikowia_pulcherrima(100); | Metschnikowia | 13 | Saprotroph | \|Undefined Saprotroph\| | Yeast | NULL | Probable | As nectar-tap_saprotroph (Põlme et al. 2020); Decay sugar-rich substrates (Põlme et al. 2020) | Põlme S, et al. 2020. Fungal Diversity 105:1-16 ((https://doi.org/10.1007/s13225-020-00466-2)) |
| Otu52516 | 1 | k__Fungi(100);p__Ascomycota(100);c__Saccharomycetes(100);o__Saccharomycetales(100);f__Metschnikowiaceae(100);g__Metschnikowia(100);s__Metschnikowia_pulcherrima(100); | Metschnikowia | 13 | Saprotroph | \|Undefined Saprotroph\| | Yeast | NULL | Probable | As nectar-tap_saprotroph (Põlme et al. 2020); Decay sugar-rich substrates (Põlme et al. 2020) | Põlme S, et al. 2020. Fungal Diversity 105:1-16 ((https://doi.org/10.1007/s13225-020-00466-2)) |
| Otu52517 | 1 | k__Fungi(100);p__Ascomycota(100);c__Saccharomycetes(100);o__Saccharomycetales(100);f__Metschnikowiaceae(100);g__Metschnikowia(100);s__Metschnikowia_pulcherrima(100); | Metschnikowia | 13 | Saprotroph | \|Undefined Saprotroph\| | Yeast | NULL | Probable | As nectar-tap_saprotroph (Põlme et al. 2020); Decay sugar-rich substrates (Põlme et al. 2020) | Põlme S, et al. 2020. Fungal Diversity 105:1-16 ((https://doi.org/10.1007/s13225-020-00466-2)) |
| Otu52518 | 1 | k__Fungi(100);p__Ascomycota(100);c__Saccharomycetes(100);o__Saccharomycetales(100);f__Metschnikowiaceae(100);g__Metschnikowia(100);s__Metschnikowia_pulcherrima(100); | Metschnikowia | 13 | Saprotroph | \|Undefined Saprotroph\| | Yeast | NULL | Probable | As nectar-tap_saprotroph (Põlme et al. 2020); Decay sugar-rich substrates (Põlme et al. 2020) | Põlme S, et al. 2020. Fungal Diversity 105:1-16 ((https://doi.org/10.1007/s13225-020-00466-2)) |
| Otu52519 | 1 | k__Fungi(100);p__Ascomycota(100);c__Saccharomycetes(100);o__Saccharomycetales(100);f__Metschnikowiaceae(100);g__Metschnikowia(100);s__Metschnikowia_pulcherrima(100); | Metschnikowia | 13 | Saprotroph | \|Undefined Saprotroph\| | Yeast | NULL | Probable | As nectar-tap_saprotroph (Põlme et al. 2020); Decay sugar-rich substrates (Põlme et al. 2020) | Põlme S, et al. 2020. Fungal Diversity 105:1-16 ((https://doi.org/10.1007/s13225-020-00466-2)) |
| Otu52520 | 1 | k__Fungi(100);p__Ascomycota(100);c__Saccharomycetes(100);o__Saccharomycetales(100);f__Metschnikowiaceae(100);g__Metschnikowia(100);s__Metschnikowia_pulcherrima(100); | Metschnikowia | 13 | Saprotroph | \|Undefined Saprotroph\| | Yeast | NULL | Probable | As nectar-tap_saprotroph (Põlme et al. 2020); Decay sugar-rich substrates (Põlme et al. 2020) | Põlme S, et al. 2020. Fungal Diversity 105:1-16 ((https://doi.org/10.1007/s13225-020-00466-2)) |
| Otu52521 | 1 | k__Fungi(100);p__Ascomycota(100);c__Saccharomycetes(100);o__Saccharomycetales(100);f__Metschnikowiaceae(100);g__Metschnikowia(100);s__Metschnikowia_pulcherrima(100); | Metschnikowia | 13 | Saprotroph | \|Undefined Saprotroph\| | Yeast | NULL | Probable | As nectar-tap_saprotroph (Põlme et al. 2020); Decay sugar-rich substrates (Põlme et al. 2020) | Põlme S, et al. 2020. Fungal Diversity 105:1-16 ((https://doi.org/10.1007/s13225-020-00466-2)) |
| Otu52522 | 1 | k__Fungi(100);p__Ascomycota(100);c__Saccharomycetes(100);o__Saccharomycetales(100);f__Metschnikowiaceae(100);g__Metschnikowia(100);g__Metschnikowia_unclassified(100); | Metschnikowia | 13 | Saprotroph | \|Undefined Saprotroph\| | Yeast | NULL | Probable | As nectar-tap_saprotroph (Põlme et al. 2020); Decay sugar-rich substrates (Põlme et al. 2020) | Põlme S, et al. 2020. Fungal Diversity 105:1-16 ((https://doi.org/10.1007/s13225-020-00466-2)) |
| Otu52523 | 1 | k__Fungi(100);p__Ascomycota(100);c__Saccharomycetes(100);o__Saccharomycetales(100);f__Metschnikowiaceae(100);g__Metschnikowia(100);g__Metschnikowia_unclassified(100); | Metschnikowia | 13 | Saprotroph | \|Undefined Saprotroph\| | Yeast | NULL | Probable | As nectar-tap_saprotroph (Põlme et al. 2020); Decay sugar-rich substrates (Põlme et al. 2020) | Põlme S, et al. 2020. Fungal Diversity 105:1-16 ((https://doi.org/10.1007/s13225-020-00466-2)) |
| Otu52524 | 1 | k__Fungi(100);p__Ascomycota(100);c__Saccharomycetes(100);o__Saccharomycetales(100);f__Metschnikowiaceae(100);g__Metschnikowia(100);g__Metschnikowia_unclassified(100); | Metschnikowia | 13 | Saprotroph | \|Undefined Saprotroph\| | Yeast | NULL | Probable | As nectar-tap_saprotroph (Põlme et al. 2020); Decay sugar-rich substrates (Põlme et al. 2020) | Põlme S, et al. 2020. Fungal Diversity 105:1-16 ((https://doi.org/10.1007/s13225-020-00466-2)) |
| Otu52525 | 1 | k__Fungi(100);p__Ascomycota(100);c__Saccharomycetes(100);o__Saccharomycetales(100);f__Metschnikowiaceae(100);g__Metschnikowia(100);s__Metschnikowia_pulcherrima(100); | Metschnikowia | 13 | Saprotroph | \|Undefined Saprotroph\| | Yeast | NULL | Probable | As nectar-tap_saprotroph (Põlme et al. 2020); Decay sugar-rich substrates (Põlme et al. 2020) | Põlme S, et al. 2020. Fungal Diversity 105:1-16 ((https://doi.org/10.1007/s13225-020-00466-2)) |
| Otu52526 | 1 | k__Fungi(100);p__Ascomycota(100);c__Saccharomycetes(100);o__Saccharomycetales(100);f__Metschnikowiaceae(100);g__Metschnikowia(100);s__Metschnikowia_pulcherrima(100); | Metschnikowia | 13 | Saprotroph | \|Undefined Saprotroph\| | Yeast | NULL | Probable | As nectar-tap_saprotroph (Põlme et al. 2020); Decay sugar-rich substrates (Põlme et al. 2020) | Põlme S, et al. 2020. Fungal Diversity 105:1-16 ((https://doi.org/10.1007/s13225-020-00466-2)) |
| Otu52527 | 1 | k__Fungi(100);p__Ascomycota(100);c__Saccharomycetes(100);o__Saccharomycetales(100);f__Metschnikowiaceae(100);g__Metschnikowia(100);s__Metschnikowia_pulcherrima(100); | Metschnikowia | 13 | Saprotroph | \|Undefined Saprotroph\| | Yeast | NULL | Probable | As nectar-tap_saprotroph (Põlme et al. 2020); Decay sugar-rich substrates (Põlme et al. 2020) | Põlme S, et al. 2020. Fungal Diversity 105:1-16 ((https://doi.org/10.1007/s13225-020-00466-2)) |
| Otu52528 | 1 | k__Fungi(100);p__Ascomycota(100);c__Saccharomycetes(100);o__Saccharomycetales(100);f__Metschnikowiaceae(100);g__Metschnikowia(100);g__Metschnikowia_unclassified(100); | Metschnikowia | 13 | Saprotroph | \|Undefined Saprotroph\| | Yeast | NULL | Probable | As nectar-tap_saprotroph (Põlme et al. 2020); Decay sugar-rich substrates (Põlme et al. 2020) | Põlme S, et al. 2020. Fungal Diversity 105:1-16 ((https://doi.org/10.1007/s13225-020-00466-2)) |
| Otu52529 | 1 | k__Fungi(100);p__Ascomycota(100);c__Saccharomycetes(100);o__Saccharomycetales(100);f__Metschnikowiaceae(100);g__Metschnikowia(100);s__Metschnikowia_pulcherrima(100); | Metschnikowia | 13 | Saprotroph | \|Undefined Saprotroph\| | Yeast | NULL | Probable | As nectar-tap_saprotroph (Põlme et al. 2020); Decay sugar-rich substrates (Põlme et al. 2020) | Põlme S, et al. 2020. Fungal Diversity 105:1-16 ((https://doi.org/10.1007/s13225-020-00466-2)) |
| Otu52530 | 1 | k__Fungi(100);p__Ascomycota(100);c__Saccharomycetes(100);o__Saccharomycetales(100);f__Metschnikowiaceae(100);g__Metschnikowia(100);s__Metschnikowia_pulcherrima(100); | Metschnikowia | 13 | Saprotroph | \|Undefined Saprotroph\| | Yeast | NULL | Probable | As nectar-tap_saprotroph (Põlme et al. 2020); Decay sugar-rich substrates (Põlme et al. 2020) | Põlme S, et al. 2020. Fungal Diversity 105:1-16 ((https://doi.org/10.1007/s13225-020-00466-2)) |
| Otu52533 | 1 | k__Fungi(100);p__Ascomycota(100);c__Saccharomycetes(100);o__Saccharomycetales(100);f__Metschnikowiaceae(100);g__Metschnikowia(100);s__Metschnikowia_pulcherrima(100); | Metschnikowia | 13 | Saprotroph | \|Undefined Saprotroph\| | Yeast | NULL | Probable | As nectar-tap_saprotroph (Põlme et al. 2020); Decay sugar-rich substrates (Põlme et al. 2020) | Põlme S, et al. 2020. Fungal Diversity 105:1-16 ((https://doi.org/10.1007/s13225-020-00466-2)) |
| Otu52534 | 1 | k__Fungi(100);p__Ascomycota(100);c__Saccharomycetes(100);o__Saccharomycetales(100);f__Metschnikowiaceae(100);g__Metschnikowia(100);s__Metschnikowia_pulcherrima(100); | Metschnikowia | 13 | Saprotroph | \|Undefined Saprotroph\| | Yeast | NULL | Probable | As nectar-tap_saprotroph (Põlme et al. 2020); Decay sugar-rich substrates (Põlme et al. 2020) | Põlme S, et al. 2020. Fungal Diversity 105:1-16 ((https://doi.org/10.1007/s13225-020-00466-2)) |
| Otu52535 | 1 | k__Fungi(100);p__Ascomycota(100);c__Saccharomycetes(100);o__Saccharomycetales(100);f__Metschnikowiaceae(100);g__Metschnikowia(100);g__Metschnikowia_unclassified(100); | Metschnikowia | 13 | Saprotroph | \|Undefined Saprotroph\| | Yeast | NULL | Probable | As nectar-tap_saprotroph (Põlme et al. 2020); Decay sugar-rich substrates (Põlme et al. 2020) | Põlme S, et al. 2020. Fungal Diversity 105:1-16 ((https://doi.org/10.1007/s13225-020-00466-2)) |
| Otu52536 | 1 | k__Fungi(100);p__Ascomycota(100);c__Saccharomycetes(100);o__Saccharomycetales(100);f__Metschnikowiaceae(100);g__Metschnikowia(100);s__Metschnikowia_pulcherrima(100); | Metschnikowia | 13 | Saprotroph | \|Undefined Saprotroph\| | Yeast | NULL | Probable | As nectar-tap_saprotroph (Põlme et al. 2020); Decay sugar-rich substrates (Põlme et al. 2020) | Põlme S, et al. 2020. Fungal Diversity 105:1-16 ((https://doi.org/10.1007/s13225-020-00466-2)) |
| Otu52537 | 1 | k__Fungi(100);p__Ascomycota(100);c__Saccharomycetes(100);o__Saccharomycetales(100);f__Metschnikowiaceae(100);g__Metschnikowia(100);g__Metschnikowia_unclassified(100); | Metschnikowia | 13 | Saprotroph | \|Undefined Saprotroph\| | Yeast | NULL | Probable | As nectar-tap_saprotroph (Põlme et al. 2020); Decay sugar-rich substrates (Põlme et al. 2020) | Põlme S, et al. 2020. Fungal Diversity 105:1-16 ((https://doi.org/10.1007/s13225-020-00466-2)) |
| Otu52538 | 1 | k__Fungi(100);p__Ascomycota(100);c__Saccharomycetes(100);o__Saccharomycetales(100);f__Metschnikowiaceae(100);g__Metschnikowia(100);s__Metschnikowia_pulcherrima(100); | Metschnikowia | 13 | Saprotroph | \|Undefined Saprotroph\| | Yeast | NULL | Probable | As nectar-tap_saprotroph (Põlme et al. 2020); Decay sugar-rich substrates (Põlme et al. 2020) | Põlme S, et al. 2020. Fungal Diversity 105:1-16 ((https://doi.org/10.1007/s13225-020-00466-2)) |
| Otu52540 | 1 | k__Fungi(100);p__Ascomycota(100);c__Saccharomycetes(100);o__Saccharomycetales(100);f__Metschnikowiaceae(100);g__Metschnikowia(100);s__Metschnikowia_pulcherrima(100); | Metschnikowia | 13 | Saprotroph | \|Undefined Saprotroph\| | Yeast | NULL | Probable | As nectar-tap_saprotroph (Põlme et al. 2020); Decay sugar-rich substrates (Põlme et al. 2020) | Põlme S, et al. 2020. Fungal Diversity 105:1-16 ((https://doi.org/10.1007/s13225-020-00466-2)) |
| Otu52541 | 1 | k__Fungi(100);p__Ascomycota(100);c__Saccharomycetes(100);o__Saccharomycetales(100);f__Metschnikowiaceae(100);g__Metschnikowia(100);s__Metschnikowia_pulcherrima(100); | Metschnikowia | 13 | Saprotroph | \|Undefined Saprotroph\| | Yeast | NULL | Probable | As nectar-tap_saprotroph (Põlme et al. 2020); Decay sugar-rich substrates (Põlme et al. 2020) | Põlme S, et al. 2020. Fungal Diversity 105:1-16 ((https://doi.org/10.1007/s13225-020-00466-2)) |
| Otu52542 | 1 | k__Fungi(100);p__Ascomycota(100);c__Saccharomycetes(100);o__Saccharomycetales(100);f__Metschnikowiaceae(100);g__Metschnikowia(100);s__Metschnikowia_pulcherrima(100); | Metschnikowia | 13 | Saprotroph | \|Undefined Saprotroph\| | Yeast | NULL | Probable | As nectar-tap_saprotroph (Põlme et al. 2020); Decay sugar-rich substrates (Põlme et al. 2020) | Põlme S, et al. 2020. Fungal Diversity 105:1-16 ((https://doi.org/10.1007/s13225-020-00466-2)) |
| Otu52543 | 1 | k__Fungi(100);p__Ascomycota(100);c__Saccharomycetes(100);o__Saccharomycetales(100);f__Metschnikowiaceae(100);g__Metschnikowia(100);s__Metschnikowia_pulcherrima(100); | Metschnikowia | 13 | Saprotroph | \|Undefined Saprotroph\| | Yeast | NULL | Probable | As nectar-tap_saprotroph (Põlme et al. 2020); Decay sugar-rich substrates (Põlme et al. 2020) | Põlme S, et al. 2020. Fungal Diversity 105:1-16 ((https://doi.org/10.1007/s13225-020-00466-2)) |
| Otu52544 | 1 | k__Fungi(100);p__Ascomycota(100);c__Saccharomycetes(100);o__Saccharomycetales(100);f__Metschnikowiaceae(100);g__Metschnikowia(100);s__Metschnikowia_pulcherrima(100); | Metschnikowia | 13 | Saprotroph | \|Undefined Saprotroph\| | Yeast | NULL | Probable | As nectar-tap_saprotroph (Põlme et al. 2020); Decay sugar-rich substrates (Põlme et al. 2020) | Põlme S, et al. 2020. Fungal Diversity 105:1-16 ((https://doi.org/10.1007/s13225-020-00466-2)) |
| Otu52545 | 1 | k__Fungi(100);p__Ascomycota(100);c__Saccharomycetes(100);o__Saccharomycetales(100);f__Metschnikowiaceae(100);g__Metschnikowia(100);s__Metschnikowia_pulcherrima(100); | Metschnikowia | 13 | Saprotroph | \|Undefined Saprotroph\| | Yeast | NULL | Probable | As nectar-tap_saprotroph (Põlme et al. 2020); Decay sugar-rich substrates (Põlme et al. 2020) | Põlme S, et al. 2020. Fungal Diversity 105:1-16 ((https://doi.org/10.1007/s13225-020-00466-2)) |
| Otu52546 | 1 | k__Fungi(100);p__Ascomycota(100);c__Saccharomycetes(100);o__Saccharomycetales(100);f__Metschnikowiaceae(100);g__Metschnikowia(100);s__Metschnikowia_pulcherrima(100); | Metschnikowia | 13 | Saprotroph | \|Undefined Saprotroph\| | Yeast | NULL | Probable | As nectar-tap_saprotroph (Põlme et al. 2020); Decay sugar-rich substrates (Põlme et al. 2020) | Põlme S, et al. 2020. Fungal Diversity 105:1-16 ((https://doi.org/10.1007/s13225-020-00466-2)) |
| Otu52547 | 1 | k__Fungi(100);p__Ascomycota(100);c__Saccharomycetes(100);o__Saccharomycetales(100);f__Metschnikowiaceae(100);g__Metschnikowia(100);g__Metschnikowia_unclassified(100); | Metschnikowia | 13 | Saprotroph | \|Undefined Saprotroph\| | Yeast | NULL | Probable | As nectar-tap_saprotroph (Põlme et al. 2020); Decay sugar-rich substrates (Põlme et al. 2020) | Põlme S, et al. 2020. Fungal Diversity 105:1-16 ((https://doi.org/10.1007/s13225-020-00466-2)) |
| Otu52548 | 1 | k__Fungi(100);p__Ascomycota(100);c__Saccharomycetes(100);o__Saccharomycetales(100);f__Metschnikowiaceae(100);g__Metschnikowia(100);s__Metschnikowia_pulcherrima(100); | Metschnikowia | 13 | Saprotroph | \|Undefined Saprotroph\| | Yeast | NULL | Probable | As nectar-tap_saprotroph (Põlme et al. 2020); Decay sugar-rich substrates (Põlme et al. 2020) | Põlme S, et al. 2020. Fungal Diversity 105:1-16 ((https://doi.org/10.1007/s13225-020-00466-2)) |
| Otu52550 | 1 | k__Fungi(100);p__Ascomycota(100);c__Saccharomycetes(100);o__Saccharomycetales(100);f__Metschnikowiaceae(100);g__Metschnikowia(100);s__Metschnikowia_pulcherrima(100); | Metschnikowia | 13 | Saprotroph | \|Undefined Saprotroph\| | Yeast | NULL | Probable | As nectar-tap_saprotroph (Põlme et al. 2020); Decay sugar-rich substrates (Põlme et al. 2020) | Põlme S, et al. 2020. Fungal Diversity 105:1-16 ((https://doi.org/10.1007/s13225-020-00466-2)) |
| Otu52551 | 1 | k__Fungi(100);p__Ascomycota(100);c__Saccharomycetes(100);o__Saccharomycetales(100);f__Metschnikowiaceae(100);g__Metschnikowia(100);s__Metschnikowia_pulcherrima(100); | Metschnikowia | 13 | Saprotroph | \|Undefined Saprotroph\| | Yeast | NULL | Probable | As nectar-tap_saprotroph (Põlme et al. 2020); Decay sugar-rich substrates (Põlme et al. 2020) | Põlme S, et al. 2020. Fungal Diversity 105:1-16 ((https://doi.org/10.1007/s13225-020-00466-2)) |
| Otu52552 | 1 | k__Fungi(100);p__Ascomycota(100);c__Saccharomycetes(100);o__Saccharomycetales(100);f__Metschnikowiaceae(100);g__Metschnikowia(100);s__Metschnikowia_pulcherrima(100); | Metschnikowia | 13 | Saprotroph | \|Undefined Saprotroph\| | Yeast | NULL | Probable | As nectar-tap_saprotroph (Põlme et al. 2020); Decay sugar-rich substrates (Põlme et al. 2020) | Põlme S, et al. 2020. Fungal Diversity 105:1-16 ((https://doi.org/10.1007/s13225-020-00466-2)) |
| Otu52554 | 1 | k__Fungi(100);p__Ascomycota(100);c__Saccharomycetes(100);o__Saccharomycetales(100);f__Metschnikowiaceae(100);g__Metschnikowia(100);s__Metschnikowia_pulcherrima(100); | Metschnikowia | 13 | Saprotroph | \|Undefined Saprotroph\| | Yeast | NULL | Probable | As nectar-tap_saprotroph (Põlme et al. 2020); Decay sugar-rich substrates (Põlme et al. 2020) | Põlme S, et al. 2020. Fungal Diversity 105:1-16 ((https://doi.org/10.1007/s13225-020-00466-2)) |
| Otu52555 | 1 | k__Fungi(100);p__Ascomycota(100);c__Saccharomycetes(100);o__Saccharomycetales(100);f__Metschnikowiaceae(100);g__Metschnikowia(100);g__Metschnikowia_unclassified(100); | Metschnikowia | 13 | Saprotroph | \|Undefined Saprotroph\| | Yeast | NULL | Probable | As nectar-tap_saprotroph (Põlme et al. 2020); Decay sugar-rich substrates (Põlme et al. 2020) | Põlme S, et al. 2020. Fungal Diversity 105:1-16 ((https://doi.org/10.1007/s13225-020-00466-2)) |
| Otu52556 | 1 | k__Fungi(100);p__Ascomycota(100);c__Saccharomycetes(100);o__Saccharomycetales(100);f__Metschnikowiaceae(100);g__Metschnikowia(100);s__Metschnikowia_pulcherrima(100); | Metschnikowia | 13 | Saprotroph | \|Undefined Saprotroph\| | Yeast | NULL | Probable | As nectar-tap_saprotroph (Põlme et al. 2020); Decay sugar-rich substrates (Põlme et al. 2020) | Põlme S, et al. 2020. Fungal Diversity 105:1-16 ((https://doi.org/10.1007/s13225-020-00466-2)) |
| Otu52567 | 1 | k__Fungi(100);p__Ascomycota(100);c__Saccharomycetes(100);o__Saccharomycetales(100);f__Metschnikowiaceae(100);g__Metschnikowia(100);s__Metschnikowia_pulcherrima(100); | Metschnikowia | 13 | Saprotroph | \|Undefined Saprotroph\| | Yeast | NULL | Probable | As nectar-tap_saprotroph (Põlme et al. 2020); Decay sugar-rich substrates (Põlme et al. 2020) | Põlme S, et al. 2020. Fungal Diversity 105:1-16 ((https://doi.org/10.1007/s13225-020-00466-2)) |
| Otu52568 | 1 | k__Fungi(100);p__Ascomycota(100);c__Saccharomycetes(100);o__Saccharomycetales(100);f__Metschnikowiaceae(100);g__Metschnikowia(100);s__Metschnikowia_pulcherrima(100); | Metschnikowia | 13 | Saprotroph | \|Undefined Saprotroph\| | Yeast | NULL | Probable | As nectar-tap_saprotroph (Põlme et al. 2020); Decay sugar-rich substrates (Põlme et al. 2020) | Põlme S, et al. 2020. Fungal Diversity 105:1-16 ((https://doi.org/10.1007/s13225-020-00466-2)) |
| Otu52569 | 1 | k__Fungi(100);p__Ascomycota(100);c__Saccharomycetes(100);o__Saccharomycetales(100);f__Metschnikowiaceae(100);g__Metschnikowia(100);g__Metschnikowia_unclassified(100); | Metschnikowia | 13 | Saprotroph | \|Undefined Saprotroph\| | Yeast | NULL | Probable | As nectar-tap_saprotroph (Põlme et al. 2020); Decay sugar-rich substrates (Põlme et al. 2020) | Põlme S, et al. 2020. Fungal Diversity 105:1-16 ((https://doi.org/10.1007/s13225-020-00466-2)) |
| Otu52570 | 1 | k__Fungi(100);p__Ascomycota(100);c__Saccharomycetes(100);o__Saccharomycetales(100);f__Metschnikowiaceae(100);g__Metschnikowia(100);s__Metschnikowia_pulcherrima(100); | Metschnikowia | 13 | Saprotroph | \|Undefined Saprotroph\| | Yeast | NULL | Probable | As nectar-tap_saprotroph (Põlme et al. 2020); Decay sugar-rich substrates (Põlme et al. 2020) | Põlme S, et al. 2020. Fungal Diversity 105:1-16 ((https://doi.org/10.1007/s13225-020-00466-2)) |
| Otu52571 | 1 | k__Fungi(100);p__Ascomycota(100);c__Saccharomycetes(100);o__Saccharomycetales(100);f__Metschnikowiaceae(100);g__Metschnikowia(100);s__Metschnikowia_pulcherrima(100); | Metschnikowia | 13 | Saprotroph | \|Undefined Saprotroph\| | Yeast | NULL | Probable | As nectar-tap_saprotroph (Põlme et al. 2020); Decay sugar-rich substrates (Põlme et al. 2020) | Põlme S, et al. 2020. Fungal Diversity 105:1-16 ((https://doi.org/10.1007/s13225-020-00466-2)) |
| Otu52572 | 1 | k__Fungi(100);p__Ascomycota(100);c__Saccharomycetes(100);o__Saccharomycetales(100);f__Metschnikowiaceae(100);g__Metschnikowia(100);s__Metschnikowia_pulcherrima(100); | Metschnikowia | 13 | Saprotroph | \|Undefined Saprotroph\| | Yeast | NULL | Probable | As nectar-tap_saprotroph (Põlme et al. 2020); Decay sugar-rich substrates (Põlme et al. 2020) | Põlme S, et al. 2020. Fungal Diversity 105:1-16 ((https://doi.org/10.1007/s13225-020-00466-2)) |
| Otu52573 | 1 | k__Fungi(100);p__Ascomycota(100);c__Saccharomycetes(100);o__Saccharomycetales(100);f__Metschnikowiaceae(100);g__Metschnikowia(100);g__Metschnikowia_unclassified(100); | Metschnikowia | 13 | Saprotroph | \|Undefined Saprotroph\| | Yeast | NULL | Probable | As nectar-tap_saprotroph (Põlme et al. 2020); Decay sugar-rich substrates (Põlme et al. 2020) | Põlme S, et al. 2020. Fungal Diversity 105:1-16 ((https://doi.org/10.1007/s13225-020-00466-2)) |
| Otu52574 | 1 | k__Fungi(100);p__Ascomycota(100);c__Saccharomycetes(100);o__Saccharomycetales(100);f__Metschnikowiaceae(100);g__Metschnikowia(100);g__Metschnikowia_unclassified(100); | Metschnikowia | 13 | Saprotroph | \|Undefined Saprotroph\| | Yeast | NULL | Probable | As nectar-tap_saprotroph (Põlme et al. 2020); Decay sugar-rich substrates (Põlme et al. 2020) | Põlme S, et al. 2020. Fungal Diversity 105:1-16 ((https://doi.org/10.1007/s13225-020-00466-2)) |
| Otu52575 | 1 | k__Fungi(100);p__Ascomycota(100);c__Saccharomycetes(100);o__Saccharomycetales(100);f__Metschnikowiaceae(100);g__Metschnikowia(100);s__Metschnikowia_pulcherrima(100); | Metschnikowia | 13 | Saprotroph | \|Undefined Saprotroph\| | Yeast | NULL | Probable | As nectar-tap_saprotroph (Põlme et al. 2020); Decay sugar-rich substrates (Põlme et al. 2020) | Põlme S, et al. 2020. Fungal Diversity 105:1-16 ((https://doi.org/10.1007/s13225-020-00466-2)) |
| Otu52576 | 1 | k__Fungi(100);p__Ascomycota(100);c__Saccharomycetes(100);o__Saccharomycetales(100);f__Metschnikowiaceae(100);g__Metschnikowia(100);s__Metschnikowia_pulcherrima(100); | Metschnikowia | 13 | Saprotroph | \|Undefined Saprotroph\| | Yeast | NULL | Probable | As nectar-tap_saprotroph (Põlme et al. 2020); Decay sugar-rich substrates (Põlme et al. 2020) | Põlme S, et al. 2020. Fungal Diversity 105:1-16 ((https://doi.org/10.1007/s13225-020-00466-2)) |
| Otu52577 | 1 | k__Fungi(100);p__Ascomycota(100);c__Saccharomycetes(100);o__Saccharomycetales(100);f__Metschnikowiaceae(100);g__Metschnikowia(100);g__Metschnikowia_unclassified(100); | Metschnikowia | 13 | Saprotroph | \|Undefined Saprotroph\| | Yeast | NULL | Probable | As nectar-tap_saprotroph (Põlme et al. 2020); Decay sugar-rich substrates (Põlme et al. 2020) | Põlme S, et al. 2020. Fungal Diversity 105:1-16 ((https://doi.org/10.1007/s13225-020-00466-2)) |
| Otu52578 | 1 | k__Fungi(100);p__Ascomycota(100);c__Saccharomycetes(100);o__Saccharomycetales(100);f__Metschnikowiaceae(100);g__Metschnikowia(100);s__Metschnikowia_pulcherrima(100); | Metschnikowia | 13 | Saprotroph | \|Undefined Saprotroph\| | Yeast | NULL | Probable | As nectar-tap_saprotroph (Põlme et al. 2020); Decay sugar-rich substrates (Põlme et al. 2020) | Põlme S, et al. 2020. Fungal Diversity 105:1-16 ((https://doi.org/10.1007/s13225-020-00466-2)) |
| Otu52579 | 1 | k__Fungi(100);p__Ascomycota(100);c__Saccharomycetes(100);o__Saccharomycetales(100);f__Metschnikowiaceae(100);g__Metschnikowia(100);g__Metschnikowia_unclassified(100); | Metschnikowia | 13 | Saprotroph | \|Undefined Saprotroph\| | Yeast | NULL | Probable | As nectar-tap_saprotroph (Põlme et al. 2020); Decay sugar-rich substrates (Põlme et al. 2020) | Põlme S, et al. 2020. Fungal Diversity 105:1-16 ((https://doi.org/10.1007/s13225-020-00466-2)) |
| Otu52580 | 1 | k__Fungi(100);p__Ascomycota(100);c__Saccharomycetes(100);o__Saccharomycetales(100);f__Metschnikowiaceae(100);g__Metschnikowia(100);g__Metschnikowia_unclassified(100); | Metschnikowia | 13 | Saprotroph | \|Undefined Saprotroph\| | Yeast | NULL | Probable | As nectar-tap_saprotroph (Põlme et al. 2020); Decay sugar-rich substrates (Põlme et al. 2020) | Põlme S, et al. 2020. Fungal Diversity 105:1-16 ((https://doi.org/10.1007/s13225-020-00466-2)) |
| Otu52581 | 1 | k__Fungi(100);p__Ascomycota(100);c__Saccharomycetes(100);o__Saccharomycetales(100);f__Metschnikowiaceae(100);g__Metschnikowia(100);g__Metschnikowia_unclassified(100); | Metschnikowia | 13 | Saprotroph | \|Undefined Saprotroph\| | Yeast | NULL | Probable | As nectar-tap_saprotroph (Põlme et al. 2020); Decay sugar-rich substrates (Põlme et al. 2020) | Põlme S, et al. 2020. Fungal Diversity 105:1-16 ((https://doi.org/10.1007/s13225-020-00466-2)) |
| Otu52582 | 1 | k__Fungi(100);p__Ascomycota(100);c__Saccharomycetes(100);o__Saccharomycetales(100);f__Metschnikowiaceae(100);g__Metschnikowia(100);s__Metschnikowia_pulcherrima(100); | Metschnikowia | 13 | Saprotroph | \|Undefined Saprotroph\| | Yeast | NULL | Probable | As nectar-tap_saprotroph (Põlme et al. 2020); Decay sugar-rich substrates (Põlme et al. 2020) | Põlme S, et al. 2020. Fungal Diversity 105:1-16 ((https://doi.org/10.1007/s13225-020-00466-2)) |
| Otu52583 | 1 | k__Fungi(100);p__Ascomycota(100);c__Saccharomycetes(100);o__Saccharomycetales(100);f__Metschnikowiaceae(100);g__Metschnikowia(100);g__Metschnikowia_unclassified(100); | Metschnikowia | 13 | Saprotroph | \|Undefined Saprotroph\| | Yeast | NULL | Probable | As nectar-tap_saprotroph (Põlme et al. 2020); Decay sugar-rich substrates (Põlme et al. 2020) | Põlme S, et al. 2020. Fungal Diversity 105:1-16 ((https://doi.org/10.1007/s13225-020-00466-2)) |
| Otu52584 | 1 | k__Fungi(100);p__Ascomycota(100);c__Saccharomycetes(100);o__Saccharomycetales(100);f__Metschnikowiaceae(100);g__Metschnikowia(100);s__Metschnikowia_pulcherrima(100); | Metschnikowia | 13 | Saprotroph | \|Undefined Saprotroph\| | Yeast | NULL | Probable | As nectar-tap_saprotroph (Põlme et al. 2020); Decay sugar-rich substrates (Põlme et al. 2020) | Põlme S, et al. 2020. Fungal Diversity 105:1-16 ((https://doi.org/10.1007/s13225-020-00466-2)) |
| Otu52585 | 1 | k__Fungi(100);p__Ascomycota(100);c__Saccharomycetes(100);o__Saccharomycetales(100);f__Metschnikowiaceae(100);g__Metschnikowia(100);s__Metschnikowia_pulcherrima(100); | Metschnikowia | 13 | Saprotroph | \|Undefined Saprotroph\| | Yeast | NULL | Probable | As nectar-tap_saprotroph (Põlme et al. 2020); Decay sugar-rich substrates (Põlme et al. 2020) | Põlme S, et al. 2020. Fungal Diversity 105:1-16 ((https://doi.org/10.1007/s13225-020-00466-2)) |
| Otu52586 | 1 | k__Fungi(100);p__Ascomycota(100);c__Saccharomycetes(100);o__Saccharomycetales(100);f__Metschnikowiaceae(100);g__Metschnikowia(100);s__Metschnikowia_pulcherrima(100); | Metschnikowia | 13 | Saprotroph | \|Undefined Saprotroph\| | Yeast | NULL | Probable | As nectar-tap_saprotroph (Põlme et al. 2020); Decay sugar-rich substrates (Põlme et al. 2020) | Põlme S, et al. 2020. Fungal Diversity 105:1-16 ((https://doi.org/10.1007/s13225-020-00466-2)) |
| Otu52587 | 1 | k__Fungi(100);p__Ascomycota(100);c__Saccharomycetes(100);o__Saccharomycetales(100);f__Metschnikowiaceae(100);g__Metschnikowia(100);s__Metschnikowia_pulcherrima(100); | Metschnikowia | 13 | Saprotroph | \|Undefined Saprotroph\| | Yeast | NULL | Probable | As nectar-tap_saprotroph (Põlme et al. 2020); Decay sugar-rich substrates (Põlme et al. 2020) | Põlme S, et al. 2020. Fungal Diversity 105:1-16 ((https://doi.org/10.1007/s13225-020-00466-2)) |
| Otu52588 | 1 | k__Fungi(100);p__Ascomycota(100);c__Saccharomycetes(100);o__Saccharomycetales(100);f__Metschnikowiaceae(100);g__Metschnikowia(100);s__Metschnikowia_pulcherrima(100); | Metschnikowia | 13 | Saprotroph | \|Undefined Saprotroph\| | Yeast | NULL | Probable | As nectar-tap_saprotroph (Põlme et al. 2020); Decay sugar-rich substrates (Põlme et al. 2020) | Põlme S, et al. 2020. Fungal Diversity 105:1-16 ((https://doi.org/10.1007/s13225-020-00466-2)) |
| Otu52589 | 1 | k__Fungi(100);p__Ascomycota(100);c__Saccharomycetes(100);o__Saccharomycetales(100);f__Metschnikowiaceae(100);g__Metschnikowia(100);s__Metschnikowia_pulcherrima(100); | Metschnikowia | 13 | Saprotroph | \|Undefined Saprotroph\| | Yeast | NULL | Probable | As nectar-tap_saprotroph (Põlme et al. 2020); Decay sugar-rich substrates (Põlme et al. 2020) | Põlme S, et al. 2020. Fungal Diversity 105:1-16 ((https://doi.org/10.1007/s13225-020-00466-2)) |
| Otu52590 | 1 | k__Fungi(100);p__Ascomycota(100);c__Saccharomycetes(100);o__Saccharomycetales(100);f__Metschnikowiaceae(100);g__Metschnikowia(100);g__Metschnikowia_unclassified(100); | Metschnikowia | 13 | Saprotroph | \|Undefined Saprotroph\| | Yeast | NULL | Probable | As nectar-tap_saprotroph (Põlme et al. 2020); Decay sugar-rich substrates (Põlme et al. 2020) | Põlme S, et al. 2020. Fungal Diversity 105:1-16 ((https://doi.org/10.1007/s13225-020-00466-2)) |
| Otu52591 | 1 | k__Fungi(100);p__Ascomycota(100);c__Saccharomycetes(100);o__Saccharomycetales(100);f__Metschnikowiaceae(100);g__Metschnikowia(100);s__Metschnikowia_pulcherrima(100); | Metschnikowia | 13 | Saprotroph | \|Undefined Saprotroph\| | Yeast | NULL | Probable | As nectar-tap_saprotroph (Põlme et al. 2020); Decay sugar-rich substrates (Põlme et al. 2020) | Põlme S, et al. 2020. Fungal Diversity 105:1-16 ((https://doi.org/10.1007/s13225-020-00466-2)) |
| Otu52592 | 1 | k__Fungi(100);p__Ascomycota(100);c__Saccharomycetes(100);o__Saccharomycetales(100);f__Saccharomycetales_family_Incertae_sedis(100);f__Saccharomycetales_family_Incertae_sedis_unclassified(100);f__Saccharomycetales_family_Incertae_sedis_unclassified(100); | Saccharomycetales | 7 | Saprotroph | Undefined Saprotroph | Yeast | NULL | Possible | NULL | Sterkenburg E, et al. 2015. New Phytologist 207:1145-1158 ((https://doi.org/10.1111/nph.13426)) |
| Otu52597 | 1 | k__Fungi(100);p__Ascomycota(100);c__Saccharomycetes(100);o__Saccharomycetales(100);f__Metschnikowiaceae(100);g__Metschnikowia(100);g__Metschnikowia_unclassified(100); | Metschnikowia | 13 | Saprotroph | \|Undefined Saprotroph\| | Yeast | NULL | Probable | As nectar-tap_saprotroph (Põlme et al. 2020); Decay sugar-rich substrates (Põlme et al. 2020) | Põlme S, et al. 2020. Fungal Diversity 105:1-16 ((https://doi.org/10.1007/s13225-020-00466-2)) |
| Otu52598 | 1 | k__Fungi(100);p__Ascomycota(100);c__Saccharomycetes(100);o__Saccharomycetales(100);f__Metschnikowiaceae(100);g__Metschnikowia(100);s__Metschnikowia_pulcherrima(100); | Metschnikowia | 13 | Saprotroph | \|Undefined Saprotroph\| | Yeast | NULL | Probable | As nectar-tap_saprotroph (Põlme et al. 2020); Decay sugar-rich substrates (Põlme et al. 2020) | Põlme S, et al. 2020. Fungal Diversity 105:1-16 ((https://doi.org/10.1007/s13225-020-00466-2)) |
| Otu52599 | 1 | k__Fungi(100);p__Ascomycota(100);c__Saccharomycetes(100);o__Saccharomycetales(100);f__Metschnikowiaceae(100);g__Metschnikowia(100);s__Metschnikowia_pulcherrima(100); | Metschnikowia | 13 | Saprotroph | \|Undefined Saprotroph\| | Yeast | NULL | Probable | As nectar-tap_saprotroph (Põlme et al. 2020); Decay sugar-rich substrates (Põlme et al. 2020) | Põlme S, et al. 2020. Fungal Diversity 105:1-16 ((https://doi.org/10.1007/s13225-020-00466-2)) |
| Otu52600 | 1 | k__Fungi(100);p__Ascomycota(100);c__Saccharomycetes(100);o__Saccharomycetales(100);f__Saccharomycetales_family_Incertae_sedis(100);g__Candida(100);g__Candida_unclassified(100); | Candida | 13 | Pathotroph-Saprotroph-Symbiotroph | Animal Pathogen-Endophyte-Endosymbiont-Epiphyte-\|Undefined Saprotroph\| | Dimorphic Yeast-Yeast | NULL | Probable | As nectar-tap_saprotroph (Põlme et al. 2020); Decay sugar-rich substrates (Põlme et al. 2020); This asexual genus is polyphyletic and contain species that function very differently depend on what they are. Caution should be taken when interpreting guilds for Candida species. | Manolakaki D, et al. 2010. Virulence. 1: 367-375 ((https://doi.org/10.4161/viru.1.5.12796)); Mekbib SB, et al. 2011. Tropical Plant Pathology 36:233-240 ((https://doi.org/10.1590/S1982-56762011000400004)); Põlme S, et al. 2020. Fungal Diversity 105:1-16 ((https://doi.org/10.1007/s13225-020-00466-2)) |
| Otu52601 | 1 | k__Fungi(100);p__Ascomycota(100);c__Saccharomycetes(100);o__Saccharomycetales(100);f__Metschnikowiaceae(100);g__Metschnikowia(100);s__Metschnikowia_pulcherrima(100); | Metschnikowia | 13 | Saprotroph | \|Undefined Saprotroph\| | Yeast | NULL | Probable | As nectar-tap_saprotroph (Põlme et al. 2020); Decay sugar-rich substrates (Põlme et al. 2020) | Põlme S, et al. 2020. Fungal Diversity 105:1-16 ((https://doi.org/10.1007/s13225-020-00466-2)) |
| Otu52602 | 1 | k__Fungi(100);p__Ascomycota(100);c__Saccharomycetes(100);o__Saccharomycetales(100);f__Metschnikowiaceae(100);g__Metschnikowia(100);g__Metschnikowia_unclassified(100); | Metschnikowia | 13 | Saprotroph | \|Undefined Saprotroph\| | Yeast | NULL | Probable | As nectar-tap_saprotroph (Põlme et al. 2020); Decay sugar-rich substrates (Põlme et al. 2020) | Põlme S, et al. 2020. Fungal Diversity 105:1-16 ((https://doi.org/10.1007/s13225-020-00466-2)) |
| Otu52604 | 1 | k__Fungi(100);p__Ascomycota(100);c__Saccharomycetes(100);o__Saccharomycetales(100);f__Metschnikowiaceae(100);g__Metschnikowia(100);s__Metschnikowia_pulcherrima(100); | Metschnikowia | 13 | Saprotroph | \|Undefined Saprotroph\| | Yeast | NULL | Probable | As nectar-tap_saprotroph (Põlme et al. 2020); Decay sugar-rich substrates (Põlme et al. 2020) | Põlme S, et al. 2020. Fungal Diversity 105:1-16 ((https://doi.org/10.1007/s13225-020-00466-2)) |
| Otu52605 | 1 | k__Fungi(100);p__Ascomycota(100);c__Saccharomycetes(100);o__Saccharomycetales(100);f__Metschnikowiaceae(100);g__Metschnikowia(100);s__Metschnikowia_pulcherrima(100); | Metschnikowia | 13 | Saprotroph | \|Undefined Saprotroph\| | Yeast | NULL | Probable | As nectar-tap_saprotroph (Põlme et al. 2020); Decay sugar-rich substrates (Põlme et al. 2020) | Põlme S, et al. 2020. Fungal Diversity 105:1-16 ((https://doi.org/10.1007/s13225-020-00466-2)) |
| Otu52606 | 1 | k__Fungi(100);p__Ascomycota(100);c__Saccharomycetes(100);o__Saccharomycetales(100);f__Metschnikowiaceae(100);g__Metschnikowia(100);s__Metschnikowia_pulcherrima(100); | Metschnikowia | 13 | Saprotroph | \|Undefined Saprotroph\| | Yeast | NULL | Probable | As nectar-tap_saprotroph (Põlme et al. 2020); Decay sugar-rich substrates (Põlme et al. 2020) | Põlme S, et al. 2020. Fungal Diversity 105:1-16 ((https://doi.org/10.1007/s13225-020-00466-2)) |
| Otu52607 | 1 | k__Fungi(100);p__Ascomycota(100);c__Saccharomycetes(100);o__Saccharomycetales(100);f__Metschnikowiaceae(100);g__Metschnikowia(100);g__Metschnikowia_unclassified(100); | Metschnikowia | 13 | Saprotroph | \|Undefined Saprotroph\| | Yeast | NULL | Probable | As nectar-tap_saprotroph (Põlme et al. 2020); Decay sugar-rich substrates (Põlme et al. 2020) | Põlme S, et al. 2020. Fungal Diversity 105:1-16 ((https://doi.org/10.1007/s13225-020-00466-2)) |
| Otu52608 | 1 | k__Fungi(100);p__Ascomycota(100);c__Saccharomycetes(100);o__Saccharomycetales(100);f__Metschnikowiaceae(100);g__Metschnikowia(100);s__Metschnikowia_pulcherrima(100); | Metschnikowia | 13 | Saprotroph | \|Undefined Saprotroph\| | Yeast | NULL | Probable | As nectar-tap_saprotroph (Põlme et al. 2020); Decay sugar-rich substrates (Põlme et al. 2020) | Põlme S, et al. 2020. Fungal Diversity 105:1-16 ((https://doi.org/10.1007/s13225-020-00466-2)) |
| Otu52609 | 1 | k__Fungi(100);p__Ascomycota(100);c__Saccharomycetes(100);o__Saccharomycetales(100);f__Metschnikowiaceae(100);g__Metschnikowia(100);s__Metschnikowia_pulcherrima(100); | Metschnikowia | 13 | Saprotroph | \|Undefined Saprotroph\| | Yeast | NULL | Probable | As nectar-tap_saprotroph (Põlme et al. 2020); Decay sugar-rich substrates (Põlme et al. 2020) | Põlme S, et al. 2020. Fungal Diversity 105:1-16 ((https://doi.org/10.1007/s13225-020-00466-2)) |
| Otu52610 | 1 | k__Fungi(100);p__Ascomycota(100);c__Saccharomycetes(100);o__Saccharomycetales(100);f__Metschnikowiaceae(100);g__Metschnikowia(100);s__Metschnikowia_pulcherrima(100); | Metschnikowia | 13 | Saprotroph | \|Undefined Saprotroph\| | Yeast | NULL | Probable | As nectar-tap_saprotroph (Põlme et al. 2020); Decay sugar-rich substrates (Põlme et al. 2020) | Põlme S, et al. 2020. Fungal Diversity 105:1-16 ((https://doi.org/10.1007/s13225-020-00466-2)) |
| Otu52612 | 1 | k__Fungi(100);p__Ascomycota(100);c__Saccharomycetes(100);o__Saccharomycetales(100);f__Metschnikowiaceae(100);g__Metschnikowia(100);s__Metschnikowia_pulcherrima(100); | Metschnikowia | 13 | Saprotroph | \|Undefined Saprotroph\| | Yeast | NULL | Probable | As nectar-tap_saprotroph (Põlme et al. 2020); Decay sugar-rich substrates (Põlme et al. 2020) | Põlme S, et al. 2020. Fungal Diversity 105:1-16 ((https://doi.org/10.1007/s13225-020-00466-2)) |
| Otu52613 | 1 | k__Fungi(100);p__Ascomycota(100);c__Saccharomycetes(100);o__Saccharomycetales(100);f__Metschnikowiaceae(100);g__Metschnikowia(100);s__Metschnikowia_pulcherrima(100); | Metschnikowia | 13 | Saprotroph | \|Undefined Saprotroph\| | Yeast | NULL | Probable | As nectar-tap_saprotroph (Põlme et al. 2020); Decay sugar-rich substrates (Põlme et al. 2020) | Põlme S, et al. 2020. Fungal Diversity 105:1-16 ((https://doi.org/10.1007/s13225-020-00466-2)) |
| Otu52614 | 1 | k__Fungi(100);p__Ascomycota(100);c__Sordariomycetes(100);o__Chaetosphaeriales(100);f__Chaetosphaeriaceae(100);g__Chaetosphaeria(100);s__Chaetosphaeria_sp_KRCF710(100); | Chaetosphaeria | 13 | Saprotroph-Symbiotroph | Endophyte-\|Plant Saprotroph\|-Wood Saprotroph | Microfungus | NULL | Probable | As foliar_endophyte (Põlme et al. 2020); As litter_saprotroph (Põlme et al. 2020); Decay wood, leaf, fruit, seed (Põlme et al. 2020) | Réblová M, et al. 2006. Mycological Research 110:104-109 ((https://doi.org/10.1016/j.mycres.2005.09.003)); Tedersoo L, et al. 2014. Science 346:e1256688 ((https://doi.org/10.1126/science.1256688)); Põlme S, et al. 2020. Fungal Diversity 105:1-16 ((https://doi.org/10.1007/s13225-020-00466-2)) |
| Otu52615 | 1 | k__Fungi(100);p__Ascomycota(100);c__Saccharomycetes(100);o__Saccharomycetales(100);f__Metschnikowiaceae(100);g__Metschnikowia(100);s__Metschnikowia_pulcherrima(100); | Metschnikowia | 13 | Saprotroph | \|Undefined Saprotroph\| | Yeast | NULL | Probable | As nectar-tap_saprotroph (Põlme et al. 2020); Decay sugar-rich substrates (Põlme et al. 2020) | Põlme S, et al. 2020. Fungal Diversity 105:1-16 ((https://doi.org/10.1007/s13225-020-00466-2)) |
| Otu52616 | 1 | k__Fungi(100);p__Ascomycota(100);c__Saccharomycetes(100);o__Saccharomycetales(100);f__Metschnikowiaceae(100);g__Metschnikowia(100);s__Metschnikowia_pulcherrima(100); | Metschnikowia | 13 | Saprotroph | \|Undefined Saprotroph\| | Yeast | NULL | Probable | As nectar-tap_saprotroph (Põlme et al. 2020); Decay sugar-rich substrates (Põlme et al. 2020) | Põlme S, et al. 2020. Fungal Diversity 105:1-16 ((https://doi.org/10.1007/s13225-020-00466-2)) |
| Otu52618 | 1 | k__Fungi(100);p__Ascomycota(100);c__Saccharomycetes(100);o__Saccharomycetales(100);f__Metschnikowiaceae(100);g__Metschnikowia(100);s__Metschnikowia_pulcherrima(100); | Metschnikowia | 13 | Saprotroph | \|Undefined Saprotroph\| | Yeast | NULL | Probable | As nectar-tap_saprotroph (Põlme et al. 2020); Decay sugar-rich substrates (Põlme et al. 2020) | Põlme S, et al. 2020. Fungal Diversity 105:1-16 ((https://doi.org/10.1007/s13225-020-00466-2)) |
| Otu52619 | 1 | k__Fungi(100);p__Ascomycota(100);c__Saccharomycetes(100);o__Saccharomycetales(100);f__Metschnikowiaceae(100);g__Metschnikowia(100);s__Metschnikowia_pulcherrima(100); | Metschnikowia | 13 | Saprotroph | \|Undefined Saprotroph\| | Yeast | NULL | Probable | As nectar-tap_saprotroph (Põlme et al. 2020); Decay sugar-rich substrates (Põlme et al. 2020) | Põlme S, et al. 2020. Fungal Diversity 105:1-16 ((https://doi.org/10.1007/s13225-020-00466-2)) |
| Otu52620 | 1 | k__Fungi(100);p__Ascomycota(100);c__Saccharomycetes(100);o__Saccharomycetales(100);f__Metschnikowiaceae(100);g__Metschnikowia(100);s__Metschnikowia_pulcherrima(100); | Metschnikowia | 13 | Saprotroph | \|Undefined Saprotroph\| | Yeast | NULL | Probable | As nectar-tap_saprotroph (Põlme et al. 2020); Decay sugar-rich substrates (Põlme et al. 2020) | Põlme S, et al. 2020. Fungal Diversity 105:1-16 ((https://doi.org/10.1007/s13225-020-00466-2)) |
| Otu52621 | 1 | k__Fungi(100);p__Ascomycota(100);c__Saccharomycetes(100);o__Saccharomycetales(100);f__Metschnikowiaceae(100);g__Metschnikowia(100);s__Metschnikowia_pulcherrima(100); | Metschnikowia | 13 | Saprotroph | \|Undefined Saprotroph\| | Yeast | NULL | Probable | As nectar-tap_saprotroph (Põlme et al. 2020); Decay sugar-rich substrates (Põlme et al. 2020) | Põlme S, et al. 2020. Fungal Diversity 105:1-16 ((https://doi.org/10.1007/s13225-020-00466-2)) |
| Otu52622 | 1 | k__Fungi(100);p__Ascomycota(100);c__Saccharomycetes(100);o__Saccharomycetales(100);f__Metschnikowiaceae(100);g__Metschnikowia(100);g__Metschnikowia_unclassified(100); | Metschnikowia | 13 | Saprotroph | \|Undefined Saprotroph\| | Yeast | NULL | Probable | As nectar-tap_saprotroph (Põlme et al. 2020); Decay sugar-rich substrates (Põlme et al. 2020) | Põlme S, et al. 2020. Fungal Diversity 105:1-16 ((https://doi.org/10.1007/s13225-020-00466-2)) |
| Otu52623 | 1 | k__Fungi(100);p__Ascomycota(100);c__Saccharomycetes(100);o__Saccharomycetales(100);f__Metschnikowiaceae(100);g__Metschnikowia(100);g__Metschnikowia_unclassified(100); | Metschnikowia | 13 | Saprotroph | \|Undefined Saprotroph\| | Yeast | NULL | Probable | As nectar-tap_saprotroph (Põlme et al. 2020); Decay sugar-rich substrates (Põlme et al. 2020) | Põlme S, et al. 2020. Fungal Diversity 105:1-16 ((https://doi.org/10.1007/s13225-020-00466-2)) |
| Otu52624 | 1 | k__Fungi(100);p__Ascomycota(100);c__Saccharomycetes(100);o__Saccharomycetales(100);f__Metschnikowiaceae(100);g__Metschnikowia(100);s__Metschnikowia_pulcherrima(100); | Metschnikowia | 13 | Saprotroph | \|Undefined Saprotroph\| | Yeast | NULL | Probable | As nectar-tap_saprotroph (Põlme et al. 2020); Decay sugar-rich substrates (Põlme et al. 2020) | Põlme S, et al. 2020. Fungal Diversity 105:1-16 ((https://doi.org/10.1007/s13225-020-00466-2)) |
| Otu52625 | 1 | k__Fungi(100);p__Ascomycota(100);c__Saccharomycetes(100);o__Saccharomycetales(100);f__Metschnikowiaceae(100);g__Metschnikowia(100);g__Metschnikowia_unclassified(100); | Metschnikowia | 13 | Saprotroph | \|Undefined Saprotroph\| | Yeast | NULL | Probable | As nectar-tap_saprotroph (Põlme et al. 2020); Decay sugar-rich substrates (Põlme et al. 2020) | Põlme S, et al. 2020. Fungal Diversity 105:1-16 ((https://doi.org/10.1007/s13225-020-00466-2)) |
| Otu52626 | 1 | k__Fungi(100);p__Ascomycota(100);c__Saccharomycetes(100);o__Saccharomycetales(100);f__Metschnikowiaceae(100);g__Metschnikowia(100);g__Metschnikowia_unclassified(100); | Metschnikowia | 13 | Saprotroph | \|Undefined Saprotroph\| | Yeast | NULL | Probable | As nectar-tap_saprotroph (Põlme et al. 2020); Decay sugar-rich substrates (Põlme et al. 2020) | Põlme S, et al. 2020. Fungal Diversity 105:1-16 ((https://doi.org/10.1007/s13225-020-00466-2)) |
| Otu52628 | 1 | k__Fungi(100);p__Ascomycota(100);c__Saccharomycetes(100);o__Saccharomycetales(100);f__Metschnikowiaceae(100);g__Metschnikowia(100);s__Metschnikowia_pulcherrima(100); | Metschnikowia | 13 | Saprotroph | \|Undefined Saprotroph\| | Yeast | NULL | Probable | As nectar-tap_saprotroph (Põlme et al. 2020); Decay sugar-rich substrates (Põlme et al. 2020) | Põlme S, et al. 2020. Fungal Diversity 105:1-16 ((https://doi.org/10.1007/s13225-020-00466-2)) |
| Otu52630 | 1 | k__Fungi(100);p__Ascomycota(100);c__Saccharomycetes(100);o__Saccharomycetales(100);f__Metschnikowiaceae(100);g__Metschnikowia(100);g__Metschnikowia_unclassified(100); | Metschnikowia | 13 | Saprotroph | \|Undefined Saprotroph\| | Yeast | NULL | Probable | As nectar-tap_saprotroph (Põlme et al. 2020); Decay sugar-rich substrates (Põlme et al. 2020) | Põlme S, et al. 2020. Fungal Diversity 105:1-16 ((https://doi.org/10.1007/s13225-020-00466-2)) |
| Otu52631 | 1 | k__Fungi(100);p__Ascomycota(100);c__Saccharomycetes(100);o__Saccharomycetales(100);f__Metschnikowiaceae(100);g__Metschnikowia(100);s__Metschnikowia_pulcherrima(100); | Metschnikowia | 13 | Saprotroph | \|Undefined Saprotroph\| | Yeast | NULL | Probable | As nectar-tap_saprotroph (Põlme et al. 2020); Decay sugar-rich substrates (Põlme et al. 2020) | Põlme S, et al. 2020. Fungal Diversity 105:1-16 ((https://doi.org/10.1007/s13225-020-00466-2)) |
| Otu52632 | 1 | k__Fungi(100);p__Ascomycota(100);c__Saccharomycetes(100);o__Saccharomycetales(100);f__Metschnikowiaceae(100);g__Metschnikowia(100);s__Metschnikowia_pulcherrima(100); | Metschnikowia | 13 | Saprotroph | \|Undefined Saprotroph\| | Yeast | NULL | Probable | As nectar-tap_saprotroph (Põlme et al. 2020); Decay sugar-rich substrates (Põlme et al. 2020) | Põlme S, et al. 2020. Fungal Diversity 105:1-16 ((https://doi.org/10.1007/s13225-020-00466-2)) |
| Otu52633 | 1 | k__Fungi(100);p__Ascomycota(100);c__Saccharomycetes(100);o__Saccharomycetales(100);f__Metschnikowiaceae(100);g__Metschnikowia(100);s__Metschnikowia_pulcherrima(100); | Metschnikowia | 13 | Saprotroph | \|Undefined Saprotroph\| | Yeast | NULL | Probable | As nectar-tap_saprotroph (Põlme et al. 2020); Decay sugar-rich substrates (Põlme et al. 2020) | Põlme S, et al. 2020. Fungal Diversity 105:1-16 ((https://doi.org/10.1007/s13225-020-00466-2)) |
| Otu52635 | 1 | k__Fungi(100);p__Ascomycota(100);c__Saccharomycetes(100);o__Saccharomycetales(100);f__Metschnikowiaceae(100);g__Metschnikowia(100);s__Metschnikowia_pulcherrima(100); | Metschnikowia | 13 | Saprotroph | \|Undefined Saprotroph\| | Yeast | NULL | Probable | As nectar-tap_saprotroph (Põlme et al. 2020); Decay sugar-rich substrates (Põlme et al. 2020) | Põlme S, et al. 2020. Fungal Diversity 105:1-16 ((https://doi.org/10.1007/s13225-020-00466-2)) |
| Otu52636 | 1 | k__Fungi(100);p__Ascomycota(100);c__Saccharomycetes(100);o__Saccharomycetales(100);f__Metschnikowiaceae(100);g__Metschnikowia(100);s__Metschnikowia_pulcherrima(100); | Metschnikowia | 13 | Saprotroph | \|Undefined Saprotroph\| | Yeast | NULL | Probable | As nectar-tap_saprotroph (Põlme et al. 2020); Decay sugar-rich substrates (Põlme et al. 2020) | Põlme S, et al. 2020. Fungal Diversity 105:1-16 ((https://doi.org/10.1007/s13225-020-00466-2)) |
| Otu52637 | 1 | k__Fungi(100);p__Ascomycota(100);c__Saccharomycetes(100);o__Saccharomycetales(100);f__Metschnikowiaceae(100);g__Metschnikowia(100);g__Metschnikowia_unclassified(100); | Metschnikowia | 13 | Saprotroph | \|Undefined Saprotroph\| | Yeast | NULL | Probable | As nectar-tap_saprotroph (Põlme et al. 2020); Decay sugar-rich substrates (Põlme et al. 2020) | Põlme S, et al. 2020. Fungal Diversity 105:1-16 ((https://doi.org/10.1007/s13225-020-00466-2)) |
| Otu52640 | 1 | k__Fungi(100);p__Ascomycota(100);c__Saccharomycetes(100);o__Saccharomycetales(100);f__Metschnikowiaceae(100);g__Metschnikowia(100);s__Metschnikowia_pulcherrima(100); | Metschnikowia | 13 | Saprotroph | \|Undefined Saprotroph\| | Yeast | NULL | Probable | As nectar-tap_saprotroph (Põlme et al. 2020); Decay sugar-rich substrates (Põlme et al. 2020) | Põlme S, et al. 2020. Fungal Diversity 105:1-16 ((https://doi.org/10.1007/s13225-020-00466-2)) |
| Otu52641 | 1 | k__Fungi(100);p__Ascomycota(100);c__Saccharomycetes(100);o__Saccharomycetales(100);f__Metschnikowiaceae(100);g__Metschnikowia(100);s__Metschnikowia_pulcherrima(100); | Metschnikowia | 13 | Saprotroph | \|Undefined Saprotroph\| | Yeast | NULL | Probable | As nectar-tap_saprotroph (Põlme et al. 2020); Decay sugar-rich substrates (Põlme et al. 2020) | Põlme S, et al. 2020. Fungal Diversity 105:1-16 ((https://doi.org/10.1007/s13225-020-00466-2)) |
| Otu52642 | 1 | k__Fungi(100);p__Ascomycota(100);c__Saccharomycetes(100);o__Saccharomycetales(100);f__Metschnikowiaceae(100);g__Metschnikowia(100);s__Metschnikowia_pulcherrima(100); | Metschnikowia | 13 | Saprotroph | \|Undefined Saprotroph\| | Yeast | NULL | Probable | As nectar-tap_saprotroph (Põlme et al. 2020); Decay sugar-rich substrates (Põlme et al. 2020) | Põlme S, et al. 2020. Fungal Diversity 105:1-16 ((https://doi.org/10.1007/s13225-020-00466-2)) |
| Otu52643 | 1 | k__Fungi(100);p__Ascomycota(100);c__Saccharomycetes(100);o__Saccharomycetales(100);f__Metschnikowiaceae(100);g__Metschnikowia(100);s__Metschnikowia_pulcherrima(100); | Metschnikowia | 13 | Saprotroph | \|Undefined Saprotroph\| | Yeast | NULL | Probable | As nectar-tap_saprotroph (Põlme et al. 2020); Decay sugar-rich substrates (Põlme et al. 2020) | Põlme S, et al. 2020. Fungal Diversity 105:1-16 ((https://doi.org/10.1007/s13225-020-00466-2)) |
| Otu52644 | 1 | k__Fungi(100);p__Ascomycota(100);c__Saccharomycetes(100);o__Saccharomycetales(100);f__Metschnikowiaceae(100);g__Metschnikowia(100);s__Metschnikowia_pulcherrima(100); | Metschnikowia | 13 | Saprotroph | \|Undefined Saprotroph\| | Yeast | NULL | Probable | As nectar-tap_saprotroph (Põlme et al. 2020); Decay sugar-rich substrates (Põlme et al. 2020) | Põlme S, et al. 2020. Fungal Diversity 105:1-16 ((https://doi.org/10.1007/s13225-020-00466-2)) |
| Otu52645 | 1 | k__Fungi(100);p__Ascomycota(100);c__Leotiomycetes(100);o__Helotiales(100);f__Sclerotiniaceae(100);g__Sclerotinia(100);s__Sclerotinia_sclerotiorum(100); | Sclerotinia | 13 | Pathotroph-Saprotroph | \|Plant Pathogen\|-Plant Saprotroph | Microfungus | NULL | Probable | As litter_saprotroph (Põlme et al. 2020); Decay leaf, fruit, seed (Põlme et al. 2020) | Tedersoo L, et al. 2014. Science 346:e1256688 ((https://doi.org/10.1126/science.1256688)); Põlme S, et al. 2020. Fungal Diversity 105:1-16 ((https://doi.org/10.1007/s13225-020-00466-2)) |
| Otu52646 | 1 | k__Fungi(100);p__Ascomycota(100);c__Saccharomycetes(100);o__Saccharomycetales(100);f__Metschnikowiaceae(100);g__Metschnikowia(100);s__Metschnikowia_pulcherrima(100); | Metschnikowia | 13 | Saprotroph | \|Undefined Saprotroph\| | Yeast | NULL | Probable | As nectar-tap_saprotroph (Põlme et al. 2020); Decay sugar-rich substrates (Põlme et al. 2020) | Põlme S, et al. 2020. Fungal Diversity 105:1-16 ((https://doi.org/10.1007/s13225-020-00466-2)) |
| Otu52647 | 1 | k__Fungi(100);p__Ascomycota(100);c__Saccharomycetes(100);o__Saccharomycetales(100);f__Metschnikowiaceae(100);g__Metschnikowia(100);s__Metschnikowia_pulcherrima(100); | Metschnikowia | 13 | Saprotroph | \|Undefined Saprotroph\| | Yeast | NULL | Probable | As nectar-tap_saprotroph (Põlme et al. 2020); Decay sugar-rich substrates (Põlme et al. 2020) | Põlme S, et al. 2020. Fungal Diversity 105:1-16 ((https://doi.org/10.1007/s13225-020-00466-2)) |
| Otu52648 | 1 | k__Fungi(100);p__Ascomycota(100);c__Saccharomycetes(100);o__Saccharomycetales(100);f__Metschnikowiaceae(100);g__Metschnikowia(100);g__Metschnikowia_unclassified(100); | Metschnikowia | 13 | Saprotroph | \|Undefined Saprotroph\| | Yeast | NULL | Probable | As nectar-tap_saprotroph (Põlme et al. 2020); Decay sugar-rich substrates (Põlme et al. 2020) | Põlme S, et al. 2020. Fungal Diversity 105:1-16 ((https://doi.org/10.1007/s13225-020-00466-2)) |
| Otu52649 | 1 | k__Fungi(100);p__Ascomycota(100);c__Saccharomycetes(100);o__Saccharomycetales(100);f__Metschnikowiaceae(100);g__Metschnikowia(100);g__Metschnikowia_unclassified(100); | Metschnikowia | 13 | Saprotroph | \|Undefined Saprotroph\| | Yeast | NULL | Probable | As nectar-tap_saprotroph (Põlme et al. 2020); Decay sugar-rich substrates (Põlme et al. 2020) | Põlme S, et al. 2020. Fungal Diversity 105:1-16 ((https://doi.org/10.1007/s13225-020-00466-2)) |
| Otu52650 | 1 | k__Fungi(100);p__Ascomycota(100);c__Saccharomycetes(100);o__Saccharomycetales(100);f__Metschnikowiaceae(100);g__Metschnikowia(100);g__Metschnikowia_unclassified(100); | Metschnikowia | 13 | Saprotroph | \|Undefined Saprotroph\| | Yeast | NULL | Probable | As nectar-tap_saprotroph (Põlme et al. 2020); Decay sugar-rich substrates (Põlme et al. 2020) | Põlme S, et al. 2020. Fungal Diversity 105:1-16 ((https://doi.org/10.1007/s13225-020-00466-2)) |
| Otu52651 | 1 | k__Fungi(100);p__Ascomycota(100);c__Saccharomycetes(100);o__Saccharomycetales(100);f__Metschnikowiaceae(100);g__Metschnikowia(100);s__Metschnikowia_pulcherrima(100); | Metschnikowia | 13 | Saprotroph | \|Undefined Saprotroph\| | Yeast | NULL | Probable | As nectar-tap_saprotroph (Põlme et al. 2020); Decay sugar-rich substrates (Põlme et al. 2020) | Põlme S, et al. 2020. Fungal Diversity 105:1-16 ((https://doi.org/10.1007/s13225-020-00466-2)) |
| Otu52652 | 1 | k__Fungi(100);p__Ascomycota(100);c__Saccharomycetes(100);o__Saccharomycetales(100);f__Metschnikowiaceae(100);g__Metschnikowia(100);s__Metschnikowia_pulcherrima(100); | Metschnikowia | 13 | Saprotroph | \|Undefined Saprotroph\| | Yeast | NULL | Probable | As nectar-tap_saprotroph (Põlme et al. 2020); Decay sugar-rich substrates (Põlme et al. 2020) | Põlme S, et al. 2020. Fungal Diversity 105:1-16 ((https://doi.org/10.1007/s13225-020-00466-2)) |
| Otu52653 | 1 | k__Fungi(100);p__Ascomycota(100);c__Saccharomycetes(100);o__Saccharomycetales(100);f__Metschnikowiaceae(100);g__Metschnikowia(100);s__Metschnikowia_pulcherrima(100); | Metschnikowia | 13 | Saprotroph | \|Undefined Saprotroph\| | Yeast | NULL | Probable | As nectar-tap_saprotroph (Põlme et al. 2020); Decay sugar-rich substrates (Põlme et al. 2020) | Põlme S, et al. 2020. Fungal Diversity 105:1-16 ((https://doi.org/10.1007/s13225-020-00466-2)) |
| Otu52654 | 1 | k__Fungi(100);p__Ascomycota(100);c__Saccharomycetes(100);o__Saccharomycetales(100);f__Metschnikowiaceae(100);g__Metschnikowia(100);s__Metschnikowia_pulcherrima(100); | Metschnikowia | 13 | Saprotroph | \|Undefined Saprotroph\| | Yeast | NULL | Probable | As nectar-tap_saprotroph (Põlme et al. 2020); Decay sugar-rich substrates (Põlme et al. 2020) | Põlme S, et al. 2020. Fungal Diversity 105:1-16 ((https://doi.org/10.1007/s13225-020-00466-2)) |
| Otu52655 | 1 | k__Fungi(100);p__Ascomycota(100);c__Saccharomycetes(100);o__Saccharomycetales(100);f__Metschnikowiaceae(100);g__Metschnikowia(100);s__Metschnikowia_pulcherrima(100); | Metschnikowia | 13 | Saprotroph | \|Undefined Saprotroph\| | Yeast | NULL | Probable | As nectar-tap_saprotroph (Põlme et al. 2020); Decay sugar-rich substrates (Põlme et al. 2020) | Põlme S, et al. 2020. Fungal Diversity 105:1-16 ((https://doi.org/10.1007/s13225-020-00466-2)) |
| Otu52656 | 1 | k__Fungi(100);p__Ascomycota(100);c__Saccharomycetes(100);o__Saccharomycetales(100);f__Metschnikowiaceae(100);g__Metschnikowia(100);s__Metschnikowia_pulcherrima(100); | Metschnikowia | 13 | Saprotroph | \|Undefined Saprotroph\| | Yeast | NULL | Probable | As nectar-tap_saprotroph (Põlme et al. 2020); Decay sugar-rich substrates (Põlme et al. 2020) | Põlme S, et al. 2020. Fungal Diversity 105:1-16 ((https://doi.org/10.1007/s13225-020-00466-2)) |
| Otu52657 | 1 | k__Fungi(100);p__Ascomycota(100);c__Saccharomycetes(100);o__Saccharomycetales(100);f__Metschnikowiaceae(100);g__Metschnikowia(100);g__Metschnikowia_unclassified(100); | Metschnikowia | 13 | Saprotroph | \|Undefined Saprotroph\| | Yeast | NULL | Probable | As nectar-tap_saprotroph (Põlme et al. 2020); Decay sugar-rich substrates (Põlme et al. 2020) | Põlme S, et al. 2020. Fungal Diversity 105:1-16 ((https://doi.org/10.1007/s13225-020-00466-2)) |
| Otu52658 | 1 | k__Fungi(100);p__Ascomycota(100);c__Saccharomycetes(100);o__Saccharomycetales(100);f__Metschnikowiaceae(100);g__Metschnikowia(100);s__Metschnikowia_pulcherrima(100); | Metschnikowia | 13 | Saprotroph | \|Undefined Saprotroph\| | Yeast | NULL | Probable | As nectar-tap_saprotroph (Põlme et al. 2020); Decay sugar-rich substrates (Põlme et al. 2020) | Põlme S, et al. 2020. Fungal Diversity 105:1-16 ((https://doi.org/10.1007/s13225-020-00466-2)) |
| Otu52659 | 1 | k__Fungi(100);p__Ascomycota(100);c__Saccharomycetes(100);o__Saccharomycetales(100);f__Metschnikowiaceae(100);g__Metschnikowia(100);g__Metschnikowia_unclassified(100); | Metschnikowia | 13 | Saprotroph | \|Undefined Saprotroph\| | Yeast | NULL | Probable | As nectar-tap_saprotroph (Põlme et al. 2020); Decay sugar-rich substrates (Põlme et al. 2020) | Põlme S, et al. 2020. Fungal Diversity 105:1-16 ((https://doi.org/10.1007/s13225-020-00466-2)) |
| Otu52660 | 1 | k__Fungi(100);p__Ascomycota(100);c__Saccharomycetes(100);o__Saccharomycetales(100);f__Metschnikowiaceae(100);g__Metschnikowia(100);s__Metschnikowia_pulcherrima(100); | Metschnikowia | 13 | Saprotroph | \|Undefined Saprotroph\| | Yeast | NULL | Probable | As nectar-tap_saprotroph (Põlme et al. 2020); Decay sugar-rich substrates (Põlme et al. 2020) | Põlme S, et al. 2020. Fungal Diversity 105:1-16 ((https://doi.org/10.1007/s13225-020-00466-2)) |
| Otu52661 | 1 | k__Fungi(100);p__Ascomycota(100);c__Saccharomycetes(100);o__Saccharomycetales(100);f__Metschnikowiaceae(100);g__Metschnikowia(100);g__Metschnikowia_unclassified(100); | Metschnikowia | 13 | Saprotroph | \|Undefined Saprotroph\| | Yeast | NULL | Probable | As nectar-tap_saprotroph (Põlme et al. 2020); Decay sugar-rich substrates (Põlme et al. 2020) | Põlme S, et al. 2020. Fungal Diversity 105:1-16 ((https://doi.org/10.1007/s13225-020-00466-2)) |
| Otu52662 | 1 | k__Fungi(100);p__Ascomycota(100);c__Saccharomycetes(100);o__Saccharomycetales(100);f__Metschnikowiaceae(100);g__Metschnikowia(100);s__Metschnikowia_pulcherrima(100); | Metschnikowia | 13 | Saprotroph | \|Undefined Saprotroph\| | Yeast | NULL | Probable | As nectar-tap_saprotroph (Põlme et al. 2020); Decay sugar-rich substrates (Põlme et al. 2020) | Põlme S, et al. 2020. Fungal Diversity 105:1-16 ((https://doi.org/10.1007/s13225-020-00466-2)) |
| Otu52663 | 1 | k__Fungi(100);p__Ascomycota(100);c__Saccharomycetes(100);o__Saccharomycetales(100);f__Metschnikowiaceae(100);g__Metschnikowia(100);s__Metschnikowia_pulcherrima(100); | Metschnikowia | 13 | Saprotroph | \|Undefined Saprotroph\| | Yeast | NULL | Probable | As nectar-tap_saprotroph (Põlme et al. 2020); Decay sugar-rich substrates (Põlme et al. 2020) | Põlme S, et al. 2020. Fungal Diversity 105:1-16 ((https://doi.org/10.1007/s13225-020-00466-2)) |
| Otu52664 | 1 | k__Fungi(100);p__Ascomycota(100);c__Saccharomycetes(100);o__Saccharomycetales(100);f__Metschnikowiaceae(100);g__Metschnikowia(100);s__Metschnikowia_pulcherrima(100); | Metschnikowia | 13 | Saprotroph | \|Undefined Saprotroph\| | Yeast | NULL | Probable | As nectar-tap_saprotroph (Põlme et al. 2020); Decay sugar-rich substrates (Põlme et al. 2020) | Põlme S, et al. 2020. Fungal Diversity 105:1-16 ((https://doi.org/10.1007/s13225-020-00466-2)) |
| Otu52665 | 1 | k__Fungi(100);p__Ascomycota(100);c__Saccharomycetes(100);o__Saccharomycetales(100);f__Metschnikowiaceae(100);g__Metschnikowia(100);s__Metschnikowia_pulcherrima(100); | Metschnikowia | 13 | Saprotroph | \|Undefined Saprotroph\| | Yeast | NULL | Probable | As nectar-tap_saprotroph (Põlme et al. 2020); Decay sugar-rich substrates (Põlme et al. 2020) | Põlme S, et al. 2020. Fungal Diversity 105:1-16 ((https://doi.org/10.1007/s13225-020-00466-2)) |
| Otu52666 | 1 | k__Fungi(100);p__Ascomycota(100);c__Saccharomycetes(100);o__Saccharomycetales(100);f__Metschnikowiaceae(100);g__Metschnikowia(100);s__Metschnikowia_pulcherrima(100); | Metschnikowia | 13 | Saprotroph | \|Undefined Saprotroph\| | Yeast | NULL | Probable | As nectar-tap_saprotroph (Põlme et al. 2020); Decay sugar-rich substrates (Põlme et al. 2020) | Põlme S, et al. 2020. Fungal Diversity 105:1-16 ((https://doi.org/10.1007/s13225-020-00466-2)) |
| Otu52667 | 1 | k__Fungi(100);p__Ascomycota(100);c__Saccharomycetes(100);o__Saccharomycetales(100);f__Metschnikowiaceae(100);g__Metschnikowia(100);g__Metschnikowia_unclassified(100); | Metschnikowia | 13 | Saprotroph | \|Undefined Saprotroph\| | Yeast | NULL | Probable | As nectar-tap_saprotroph (Põlme et al. 2020); Decay sugar-rich substrates (Põlme et al. 2020) | Põlme S, et al. 2020. Fungal Diversity 105:1-16 ((https://doi.org/10.1007/s13225-020-00466-2)) |
| Otu52668 | 1 | k__Fungi(100);p__Ascomycota(100);c__Saccharomycetes(100);o__Saccharomycetales(100);f__Metschnikowiaceae(100);g__Metschnikowia(100);s__Metschnikowia_pulcherrima(100); | Metschnikowia | 13 | Saprotroph | \|Undefined Saprotroph\| | Yeast | NULL | Probable | As nectar-tap_saprotroph (Põlme et al. 2020); Decay sugar-rich substrates (Põlme et al. 2020) | Põlme S, et al. 2020. Fungal Diversity 105:1-16 ((https://doi.org/10.1007/s13225-020-00466-2)) |
| Otu52669 | 1 | k__Fungi(100);p__Ascomycota(100);c__Saccharomycetes(100);o__Saccharomycetales(100);f__Metschnikowiaceae(100);g__Metschnikowia(100);s__Metschnikowia_pulcherrima(100); | Metschnikowia | 13 | Saprotroph | \|Undefined Saprotroph\| | Yeast | NULL | Probable | As nectar-tap_saprotroph (Põlme et al. 2020); Decay sugar-rich substrates (Põlme et al. 2020) | Põlme S, et al. 2020. Fungal Diversity 105:1-16 ((https://doi.org/10.1007/s13225-020-00466-2)) |
| Otu52670 | 1 | k__Fungi(100);p__Ascomycota(100);c__Saccharomycetes(100);o__Saccharomycetales(100);f__Metschnikowiaceae(100);g__Metschnikowia(100);s__Metschnikowia_pulcherrima(100); | Metschnikowia | 13 | Saprotroph | \|Undefined Saprotroph\| | Yeast | NULL | Probable | As nectar-tap_saprotroph (Põlme et al. 2020); Decay sugar-rich substrates (Põlme et al. 2020) | Põlme S, et al. 2020. Fungal Diversity 105:1-16 ((https://doi.org/10.1007/s13225-020-00466-2)) |
| Otu52671 | 1 | k__Fungi(100);p__Ascomycota(100);c__Saccharomycetes(100);o__Saccharomycetales(100);f__Metschnikowiaceae(100);g__Metschnikowia(100);s__Metschnikowia_pulcherrima(100); | Metschnikowia | 13 | Saprotroph | \|Undefined Saprotroph\| | Yeast | NULL | Probable | As nectar-tap_saprotroph (Põlme et al. 2020); Decay sugar-rich substrates (Põlme et al. 2020) | Põlme S, et al. 2020. Fungal Diversity 105:1-16 ((https://doi.org/10.1007/s13225-020-00466-2)) |
| Otu52672 | 1 | k__Fungi(100);p__Ascomycota(100);c__Saccharomycetes(100);o__Saccharomycetales(100);f__Metschnikowiaceae(100);g__Metschnikowia(100);g__Metschnikowia_unclassified(100); | Metschnikowia | 13 | Saprotroph | \|Undefined Saprotroph\| | Yeast | NULL | Probable | As nectar-tap_saprotroph (Põlme et al. 2020); Decay sugar-rich substrates (Põlme et al. 2020) | Põlme S, et al. 2020. Fungal Diversity 105:1-16 ((https://doi.org/10.1007/s13225-020-00466-2)) |
| Otu52673 | 1 | k__Fungi(100);p__Ascomycota(100);c__Saccharomycetes(100);o__Saccharomycetales(100);f__Metschnikowiaceae(100);g__Metschnikowia(100);s__Metschnikowia_pulcherrima(100); | Metschnikowia | 13 | Saprotroph | \|Undefined Saprotroph\| | Yeast | NULL | Probable | As nectar-tap_saprotroph (Põlme et al. 2020); Decay sugar-rich substrates (Põlme et al. 2020) | Põlme S, et al. 2020. Fungal Diversity 105:1-16 ((https://doi.org/10.1007/s13225-020-00466-2)) |
| Otu52674 | 1 | k__Fungi(100);p__Ascomycota(100);c__Saccharomycetes(100);o__Saccharomycetales(100);f__Metschnikowiaceae(100);g__Metschnikowia(100);s__Metschnikowia_pulcherrima(100); | Metschnikowia | 13 | Saprotroph | \|Undefined Saprotroph\| | Yeast | NULL | Probable | As nectar-tap_saprotroph (Põlme et al. 2020); Decay sugar-rich substrates (Põlme et al. 2020) | Põlme S, et al. 2020. Fungal Diversity 105:1-16 ((https://doi.org/10.1007/s13225-020-00466-2)) |

| | | | | | | | | | | | |
|---|---|---|---|---|---|---|---|---|---|---|---|
| Otu52675 | 1 | k__Fungi(100);p__Ascomycota(100);c__Saccharomycetes(100);o__Saccharomycetales(100);f__Metschnikowiaceae(100);g__Metschnikowia(100);s__Metschnikowia_pulcherrima(100); | Metschnikowia | 13 | Saprotroph | \|Undefined Saprotroph\| | Yeast | NULL | Probable | As nectar-tap_saprotroph (Põlme et al. 2020); Decay sugar-rich substrates (Põlme et al. 2020) | Põlme S, et al. 2020. Fungal Diversity 105:1-16 ((https://doi.org/10.1007/s13225-020-00466-2)) |
| Otu52676 | 1 | k__Fungi(100);p__Ascomycota(100);c__Saccharomycetes(100);o__Saccharomycetales(100);f__Metschnikowiaceae(100);g__Metschnikowia(100);s__Metschnikowia_pulcherrima(100); | Metschnikowia | 13 | Saprotroph | \|Undefined Saprotroph\| | Yeast | NULL | Probable | As nectar-tap_saprotroph (Põlme et al. 2020); Decay sugar-rich substrates (Põlme et al. 2020) | Põlme S, et al. 2020. Fungal Diversity 105:1-16 ((https://doi.org/10.1007/s13225-020-00466-2)) |
| Otu52677 | 1 | k__Fungi(100);p__Ascomycota(100);c__Saccharomycetes(100);o__Saccharomycetales(100);f__Metschnikowiaceae(100);g__Metschnikowia(100);s__Metschnikowia_pulcherrima(100); | Metschnikowia | 13 | Saprotroph | \|Undefined Saprotroph\| | Yeast | NULL | Probable | As nectar-tap_saprotroph (Põlme et al. 2020); Decay sugar-rich substrates (Põlme et al. 2020) | Põlme S, et al. 2020. Fungal Diversity 105:1-16 ((https://doi.org/10.1007/s13225-020-00466-2)) |
| Otu52678 | 1 | k__Fungi(100);p__Ascomycota(100);c__Saccharomycetes(100);o__Saccharomycetales(100);f__Metschnikowiaceae(100);g__Metschnikowia(100);g__Metschnikowia_unclassified(100); | Metschnikowia | 13 | Saprotroph | \|Undefined Saprotroph\| | Yeast | NULL | Probable | As nectar-tap_saprotroph (Põlme et al. 2020); Decay sugar-rich substrates (Põlme et al. 2020) | Põlme S, et al. 2020. Fungal Diversity 105:1-16 ((https://doi.org/10.1007/s13225-020-00466-2)) |
| Otu52679 | 1 | k__Fungi(100);p__Ascomycota(100);c__Saccharomycetes(100);o__Saccharomycetales(100);f__Metschnikowiaceae(100);g__Metschnikowia(100);s__Metschnikowia_pulcherrima(100); | Metschnikowia | 13 | Saprotroph | \|Undefined Saprotroph\| | Yeast | NULL | Probable | As nectar-tap_saprotroph (Põlme et al. 2020); Decay sugar-rich substrates (Põlme et al. 2020) | Põlme S, et al. 2020. Fungal Diversity 105:1-16 ((https://doi.org/10.1007/s13225-020-00466-2)) |
| Otu52680 | 1 | k__Fungi(100);p__Ascomycota(100);c__Saccharomycetes(100);o__Saccharomycetales(100);f__Metschnikowiaceae(100);g__Metschnikowia(100);g__Metschnikowia_unclassified(100); | Metschnikowia | 13 | Saprotroph | \|Undefined Saprotroph\| | Yeast | NULL | Probable | As nectar-tap_saprotroph (Põlme et al. 2020); Decay sugar-rich substrates (Põlme et al. 2020) | Põlme S, et al. 2020. Fungal Diversity 105:1-16 ((https://doi.org/10.1007/s13225-020-00466-2)) |
| Otu52681 | 1 | k__Fungi(100);p__Ascomycota(100);c__Saccharomycetes(100);o__Saccharomycetales(100);f__Metschnikowiaceae(100);g__Metschnikowia(100);s__Metschnikowia_pulcherrima(100); | Metschnikowia | 13 | Saprotroph | \|Undefined Saprotroph\| | Yeast | NULL | Probable | As nectar-tap_saprotroph (Põlme et al. 2020); Decay sugar-rich substrates (Põlme et al. 2020) | Põlme S, et al. 2020. Fungal Diversity 105:1-16 ((https://doi.org/10.1007/s13225-020-00466-2)) |
| Otu52682 | 1 | k__Fungi(100);p__Ascomycota(100);c__Saccharomycetes(100);o__Saccharomycetales(100);f__Metschnikowiaceae(100);g__Metschnikowia(100);s__Metschnikowia_pulcherrima(100); | Metschnikowia | 13 | Saprotroph | \|Undefined Saprotroph\| | Yeast | NULL | Probable | As nectar-tap_saprotroph (Põlme et al. 2020); Decay sugar-rich substrates (Põlme et al. 2020) | Põlme S, et al. 2020. Fungal Diversity 105:1-16 ((https://doi.org/10.1007/s13225-020-00466-2)) |
| Otu52685 | 1 | k__Fungi(100);p__Ascomycota(100);c__Saccharomycetes(100);o__Saccharomycetales(100);f__Metschnikowiaceae(100);g__Metschnikowia(100);s__Metschnikowia_pulcherrima(100); | Metschnikowia | 13 | Saprotroph | \|Undefined Saprotroph\| | Yeast | NULL | Probable | As nectar-tap_saprotroph (Põlme et al. 2020); Decay sugar-rich substrates (Põlme et al. 2020) | Põlme S, et al. 2020. Fungal Diversity 105:1-16 ((https://doi.org/10.1007/s13225-020-00466-2)) |
| Otu52686 | 1 | k__Fungi(100);p__Ascomycota(100);c__Saccharomycetes(100);o__Saccharomycetales(100);f__Metschnikowiaceae(100);g__Metschnikowia(100);g__Metschnikowia_unclassified(100); | Metschnikowia | 13 | Saprotroph | \|Undefined Saprotroph\| | Yeast | NULL | Probable | As nectar-tap_saprotroph (Põlme et al. 2020); Decay sugar-rich substrates (Põlme et al. 2020) | Põlme S, et al. 2020. Fungal Diversity 105:1-16 ((https://doi.org/10.1007/s13225-020-00466-2)) |
| Otu52687 | 1 | k__Fungi(100);p__Ascomycota(100);c__Saccharomycetes(100);o__Saccharomycetales(100);f__Metschnikowiaceae(100);g__Metschnikowia(100);s__Metschnikowia_pulcherrima(100); | Metschnikowia | 13 | Saprotroph | \|Undefined Saprotroph\| | Yeast | NULL | Probable | As nectar-tap_saprotroph (Põlme et al. 2020); Decay sugar-rich substrates (Põlme et al. 2020) | Põlme S, et al. 2020. Fungal Diversity 105:1-16 ((https://doi.org/10.1007/s13225-020-00466-2)) |
| Otu52688 | 1 | k__Fungi(100);p__Ascomycota(100);c__Saccharomycetes(100);o__Saccharomycetales(100);f__Metschnikowiaceae(100);g__Metschnikowia(100);g__Metschnikowia_unclassified(100); | Metschnikowia | 13 | Saprotroph | \|Undefined Saprotroph\| | Yeast | NULL | Probable | As nectar-tap_saprotroph (Põlme et al. 2020); Decay sugar-rich substrates (Põlme et al. 2020) | Põlme S, et al. 2020. Fungal Diversity 105:1-16 ((https://doi.org/10.1007/s13225-020-00466-2)) |
| Otu52689 | 1 | k__Fungi(100);p__Ascomycota(100);c__Saccharomycetes(100);o__Saccharomycetales(100);f__Metschnikowiaceae(100);g__Metschnikowia(100);s__Metschnikowia_pulcherrima(100); | Metschnikowia | 13 | Saprotroph | \|Undefined Saprotroph\| | Yeast | NULL | Probable | As nectar-tap_saprotroph (Põlme et al. 2020); Decay sugar-rich substrates (Põlme et al. 2020) | Põlme S, et al. 2020. Fungal Diversity 105:1-16 ((https://doi.org/10.1007/s13225-020-00466-2)) |
| Otu52690 | 1 | k__Fungi(100);p__Ascomycota(100);c__Saccharomycetes(100);o__Saccharomycetales(100);f__Metschnikowiaceae(100);g__Metschnikowia(100);g__Metschnikowia_unclassified(100); | Metschnikowia | 13 | Saprotroph | \|Undefined Saprotroph\| | Yeast | NULL | Probable | As nectar-tap_saprotroph (Põlme et al. 2020); Decay sugar-rich substrates (Põlme et al. 2020) | Põlme S, et al. 2020. Fungal Diversity 105:1-16 ((https://doi.org/10.1007/s13225-020-00466-2)) |
| Otu52691 | 1 | k__Fungi(100);p__Ascomycota(100);c__Saccharomycetes(100);o__Saccharomycetales(100);f__Metschnikowiaceae(100);g__Metschnikowia(100);g__Metschnikowia_unclassified(100); | Metschnikowia | 13 | Saprotroph | \|Undefined Saprotroph\| | Yeast | NULL | Probable | As nectar-tap_saprotroph (Põlme et al. 2020); Decay sugar-rich substrates (Põlme et al. 2020) | Põlme S, et al. 2020. Fungal Diversity 105:1-16 ((https://doi.org/10.1007/s13225-020-00466-2)) |
| Otu52692 | 1 | k__Fungi(100);p__Ascomycota(100);c__Saccharomycetes(100);o__Saccharomycetales(100);f__Metschnikowiaceae(100);g__Metschnikowia(100);s__Metschnikowia_pulcherrima(100); | Metschnikowia | 13 | Saprotroph | \|Undefined Saprotroph\| | Yeast | NULL | Probable | As nectar-tap_saprotroph (Põlme et al. 2020); Decay sugar-rich substrates (Põlme et al. 2020) | Põlme S, et al. 2020. Fungal Diversity 105:1-16 ((https://doi.org/10.1007/s13225-020-00466-2)) |
| Otu52693 | 1 | k__Fungi(100);p__Ascomycota(100);c__Saccharomycetes(100);o__Saccharomycetales(100);f__Metschnikowiaceae(100);g__Metschnikowia(100);g__Metschnikowia_unclassified(100); | Metschnikowia | 13 | Saprotroph | \|Undefined Saprotroph\| | Yeast | NULL | Probable | As nectar-tap_saprotroph (Põlme et al. 2020); Decay sugar-rich substrates (Põlme et al. 2020) | Põlme S, et al. 2020. Fungal Diversity 105:1-16 ((https://doi.org/10.1007/s13225-020-00466-2)) |
| Otu52694 | 1 | k__Fungi(100);p__Ascomycota(100);c__Saccharomycetes(100);o__Saccharomycetales(100);f__Metschnikowiaceae(100);g__Metschnikowia(100);g__Metschnikowia_unclassified(100); | Metschnikowia | 13 | Saprotroph | \|Undefined Saprotroph\| | Yeast | NULL | Probable | As nectar-tap_saprotroph (Põlme et al. 2020); Decay sugar-rich substrates (Põlme et al. 2020) | Põlme S, et al. 2020. Fungal Diversity 105:1-16 ((https://doi.org/10.1007/s13225-020-00466-2)) |
| Otu52695 | 1 | k__Fungi(100);p__Ascomycota(100);c__Saccharomycetes(100);o__Saccharomycetales(100);f__Metschnikowiaceae(100);g__Metschnikowia(100);s__Metschnikowia_pulcherrima(100); | Metschnikowia | 13 | Saprotroph | \|Undefined Saprotroph\| | Yeast | NULL | Probable | As nectar-tap_saprotroph (Põlme et al. 2020); Decay sugar-rich substrates (Põlme et al. 2020) | Põlme S, et al. 2020. Fungal Diversity 105:1-16 ((https://doi.org/10.1007/s13225-020-00466-2)) |
| Otu52696 | 1 | k__Fungi(100);p__Ascomycota(100);c__Saccharomycetes(100);o__Saccharomycetales(100);f__Metschnikowiaceae(100);g__Metschnikowia(100);s__Metschnikowia_pulcherrima(100); | Metschnikowia | 13 | Saprotroph | \|Undefined Saprotroph\| | Yeast | NULL | Probable | As nectar-tap_saprotroph (Põlme et al. 2020); Decay sugar-rich substrates (Põlme et al. 2020) | Põlme S, et al. 2020. Fungal Diversity 105:1-16 ((https://doi.org/10.1007/s13225-020-00466-2)) |
| Otu52697 | 1 | k__Fungi(100);p__Ascomycota(100);c__Saccharomycetes(100);o__Saccharomycetales(100);f__Metschnikowiaceae(100);g__Metschnikowia(100);s__Metschnikowia_pulcherrima(100); | Metschnikowia | 13 | Saprotroph | \|Undefined Saprotroph\| | Yeast | NULL | Probable | As nectar-tap_saprotroph (Põlme et al. 2020); Decay sugar-rich substrates (Põlme et al. 2020) | Põlme S, et al. 2020. Fungal Diversity 105:1-16 ((https://doi.org/10.1007/s13225-020-00466-2)) |
| Otu52698 | 1 | k__Fungi(100);p__Ascomycota(100);c__Saccharomycetes(100);o__Saccharomycetales(100);f__Metschnikowiaceae(100);g__Metschnikowia(100);g__Metschnikowia_unclassified(100); | Metschnikowia | 13 | Saprotroph | \|Undefined Saprotroph\| | Yeast | NULL | Probable | As nectar-tap_saprotroph (Põlme et al. 2020); Decay sugar-rich substrates (Põlme et al. 2020) | Põlme S, et al. 2020. Fungal Diversity 105:1-16 ((https://doi.org/10.1007/s13225-020-00466-2)) |
| Otu52699 | 1 | k__Fungi(100);p__Ascomycota(100);c__Saccharomycetes(100);o__Saccharomycetales(100);f__Metschnikowiaceae(100);g__Metschnikowia(100);s__Metschnikowia_pulcherrima(100); | Metschnikowia | 13 | Saprotroph | \|Undefined Saprotroph\| | Yeast | NULL | Probable | As nectar-tap_saprotroph (Põlme et al. 2020); Decay sugar-rich substrates (Põlme et al. 2020) | Põlme S, et al. 2020. Fungal Diversity 105:1-16 ((https://doi.org/10.1007/s13225-020-00466-2)) |
| Otu52700 | 1 | k__Fungi(100);p__Ascomycota(100);c__Saccharomycetes(100);o__Saccharomycetales(100);f__Metschnikowiaceae(100);g__Metschnikowia(100);g__Metschnikowia_unclassified(100); | Metschnikowia | 13 | Saprotroph | \|Undefined Saprotroph\| | Yeast | NULL | Probable | As nectar-tap_saprotroph (Põlme et al. 2020); Decay sugar-rich substrates (Põlme et al. 2020) | Põlme S, et al. 2020. Fungal Diversity 105:1-16 ((https://doi.org/10.1007/s13225-020-00466-2)) |
| Otu52701 | 1 | k__Fungi(100);p__Ascomycota(100);c__Saccharomycetes(100);o__Saccharomycetales(100);f__Metschnikowiaceae(100);g__Metschnikowia(100);s__Metschnikowia_pulcherrima(100); | Metschnikowia | 13 | Saprotroph | \|Undefined Saprotroph\| | Yeast | NULL | Probable | As nectar-tap_saprotroph (Põlme et al. 2020); Decay sugar-rich substrates (Põlme et al. 2020) | Põlme S, et al. 2020. Fungal Diversity 105:1-16 ((https://doi.org/10.1007/s13225-020-00466-2)) |
| Otu52702 | 1 | k__Fungi(100);p__Ascomycota(100);c__Saccharomycetes(100);o__Saccharomycetales(100);f__Metschnikowiaceae(100);g__Metschnikowia(100);s__Metschnikowia_pulcherrima(100); | Metschnikowia | 13 | Saprotroph | \|Undefined Saprotroph\| | Yeast | NULL | Probable | As nectar-tap_saprotroph (Põlme et al. 2020); Decay sugar-rich substrates (Põlme et al. 2020) | Põlme S, et al. 2020. Fungal Diversity 105:1-16 ((https://doi.org/10.1007/s13225-020-00466-2)) |
| Otu52703 | 1 | k__Fungi(100);p__Ascomycota(100);c__Saccharomycetes(100);o__Saccharomycetales(100);f__Metschnikowiaceae(100);g__Metschnikowia(100);g__Metschnikowia_unclassified(100); | Metschnikowia | 13 | Saprotroph | \|Undefined Saprotroph\| | Yeast | NULL | Probable | As nectar-tap_saprotroph (Põlme et al. 2020); Decay sugar-rich substrates (Põlme et al. 2020) | Põlme S, et al. 2020. Fungal Diversity 105:1-16 ((https://doi.org/10.1007/s13225-020-00466-2)) |
| Otu52704 | 1 | k__Fungi(100);p__Ascomycota(100);c__Saccharomycetes(100);o__Saccharomycetales(100);f__Metschnikowiaceae(100);g__Metschnikowia(100);s__Metschnikowia_pulcherrima(100); | Metschnikowia | 13 | Saprotroph | \|Undefined Saprotroph\| | Yeast | NULL | Probable | As nectar-tap_saprotroph (Põlme et al. 2020); Decay sugar-rich substrates (Põlme et al. 2020) | Põlme S, et al. 2020. Fungal Diversity 105:1-16 ((https://doi.org/10.1007/s13225-020-00466-2)) |
| Otu52705 | 1 | k__Fungi(100);p__Ascomycota(100);c__Saccharomycetes(100);o__Saccharomycetales(100);f__Metschnikowiaceae(100);g__Metschnikowia(100);s__Metschnikowia_pulcherrima(100); | Metschnikowia | 13 | Saprotroph | \|Undefined Saprotroph\| | Yeast | NULL | Probable | As nectar-tap_saprotroph (Põlme et al. 2020); Decay sugar-rich substrates (Põlme et al. 2020) | Põlme S, et al. 2020. Fungal Diversity 105:1-16 ((https://doi.org/10.1007/s13225-020-00466-2)) |
| Otu52706 | 1 | k__Fungi(100);p__Ascomycota(100);c__Saccharomycetes(100);o__Saccharomycetales(100);f__Metschnikowiaceae(100);g__Metschnikowia(100);s__Metschnikowia_pulcherrima(100); | Metschnikowia | 13 | Saprotroph | \|Undefined Saprotroph\| | Yeast | NULL | Probable | As nectar-tap_saprotroph (Põlme et al. 2020); Decay sugar-rich substrates (Põlme et al. 2020) | Põlme S, et al. 2020. Fungal Diversity 105:1-16 ((https://doi.org/10.1007/s13225-020-00466-2)) |
| Otu52707 | 1 | k__Fungi(100);p__Ascomycota(100);c__Saccharomycetes(100);o__Saccharomycetales(100);f__Metschnikowiaceae(100);g__Metschnikowia(100);s__Metschnikowia_pulcherrima(100); | Metschnikowia | 13 | Saprotroph | \|Undefined Saprotroph\| | Yeast | NULL | Probable | As nectar-tap_saprotroph (Põlme et al. 2020); Decay sugar-rich substrates (Põlme et al. 2020) | Põlme S, et al. 2020. Fungal Diversity 105:1-16 ((https://doi.org/10.1007/s13225-020-00466-2)) |
| Otu52708 | 1 | k__Fungi(100);p__Ascomycota(100);c__Saccharomycetes(100);o__Saccharomycetales(100);f__Metschnikowiaceae(100);g__Metschnikowia(100);g__Metschnikowia_unclassified(100); | Metschnikowia | 13 | Saprotroph | \|Undefined Saprotroph\| | Yeast | NULL | Probable | As nectar-tap_saprotroph (Põlme et al. 2020); Decay sugar-rich substrates (Põlme et al. 2020) | Põlme S, et al. 2020. Fungal Diversity 105:1-16 ((https://doi.org/10.1007/s13225-020-00466-2)) |
| Otu52709 | 1 | k__Fungi(100);p__Ascomycota(100);c__Saccharomycetes(100);o__Saccharomycetales(100);f__Metschnikowiaceae(100);g__Metschnikowia(100);s__Metschnikowia_pulcherrima(100); | Metschnikowia | 13 | Saprotroph | \|Undefined Saprotroph\| | Yeast | NULL | Probable | As nectar-tap_saprotroph (Põlme et al. 2020); Decay sugar-rich substrates (Põlme et al. 2020) | Põlme S, et al. 2020. Fungal Diversity 105:1-16 ((https://doi.org/10.1007/s13225-020-00466-2)) |
| Otu52710 | 1 | k__Fungi(100);p__Ascomycota(100);c__Saccharomycetes(100);o__Saccharomycetales(100);f__Metschnikowiaceae(100);g__Metschnikowia(100);g__Metschnikowia_unclassified(100); | Metschnikowia | 13 | Saprotroph | \|Undefined Saprotroph\| | Yeast | NULL | Probable | As nectar-tap_saprotroph (Põlme et al. 2020); Decay sugar-rich substrates (Põlme et al. 2020) | Põlme S, et al. 2020. Fungal Diversity 105:1-16 ((https://doi.org/10.1007/s13225-020-00466-2)) |
| Otu52711 | 1 | k__Fungi(100);p__Ascomycota(100);c__Saccharomycetes(100);o__Saccharomycetales(100);f__Metschnikowiaceae(100);g__Metschnikowia(100);s__Metschnikowia_pulcherrima(100); | Metschnikowia | 13 | Saprotroph | \|Undefined Saprotroph\| | Yeast | NULL | Probable | As nectar-tap_saprotroph (Põlme et al. 2020); Decay sugar-rich substrates (Põlme et al. 2020) | Põlme S, et al. 2020. Fungal Diversity 105:1-16 ((https://doi.org/10.1007/s13225-020-00466-2)) |
| Otu52712 | 1 | k__Fungi(100);p__Ascomycota(100);c__Saccharomycetes(100);o__Saccharomycetales(100);f__Metschnikowiaceae(100);g__Metschnikowia(100);s__Metschnikowia_pulcherrima(100); | Metschnikowia | 13 | Saprotroph | \|Undefined Saprotroph\| | Yeast | NULL | Probable | As nectar-tap_saprotroph (Põlme et al. 2020); Decay sugar-rich substrates (Põlme et al. 2020) | Põlme S, et al. 2020. Fungal Diversity 105:1-16 ((https://doi.org/10.1007/s13225-020-00466-2)) |
| Otu52713 | 1 | k__Fungi(100);p__Ascomycota(100);c__Saccharomycetes(100);o__Saccharomycetales(100);f__Metschnikowiaceae(100);g__Metschnikowia(100);g__Metschnikowia_unclassified(100); | Metschnikowia | 13 | Saprotroph | \|Undefined Saprotroph\| | Yeast | NULL | Probable | As nectar-tap_saprotroph (Põlme et al. 2020); Decay sugar-rich substrates (Põlme et al. 2020) | Põlme S, et al. 2020. Fungal Diversity 105:1-16 ((https://doi.org/10.1007/s13225-020-00466-2)) |
| Otu52714 | 1 | k__Fungi(100);p__Ascomycota(100);c__Saccharomycetes(100);o__Saccharomycetales(100);f__Metschnikowiaceae(100);g__Metschnikowia(100);s__Metschnikowia_pulcherrima(100); | Metschnikowia | 13 | Saprotroph | \|Undefined Saprotroph\| | Yeast | NULL | Probable | As nectar-tap_saprotroph (Põlme et al. 2020); Decay sugar-rich substrates (Põlme et al. 2020) | Põlme S, et al. 2020. Fungal Diversity 105:1-16 ((https://doi.org/10.1007/s13225-020-00466-2)) |
| Otu52715 | 1 | k__Fungi(100);p__Ascomycota(100);c__Saccharomycetes(100);o__Saccharomycetales(100);f__Metschnikowiaceae(100);g__Metschnikowia(100);s__Metschnikowia_pulcherrima(100); | Metschnikowia | 13 | Saprotroph | \|Undefined Saprotroph\| | Yeast | NULL | Probable | As nectar-tap_saprotroph (Põlme et al. 2020); Decay sugar-rich substrates (Põlme et al. 2020) | Põlme S, et al. 2020. Fungal Diversity 105:1-16 ((https://doi.org/10.1007/s13225-020-00466-2)) |
| Otu52716 | 1 | k__Fungi(100);p__Ascomycota(100);c__Saccharomycetes(100);o__Saccharomycetales(100);f__Metschnikowiaceae(100);g__Metschnikowia(100);s__Metschnikowia_pulcherrima(100); | Metschnikowia | 13 | Saprotroph | \|Undefined Saprotroph\| | Yeast | NULL | Probable | As nectar-tap_saprotroph (Põlme et al. 2020); Decay sugar-rich substrates (Põlme et al. 2020) | Põlme S, et al. 2020. Fungal Diversity 105:1-16 ((https://doi.org/10.1007/s13225-020-00466-2)) |
| Otu52717 | 1 | k__Fungi(100);p__Ascomycota(100);c__Saccharomycetes(100);o__Saccharomycetales(100);f__Metschnikowiaceae(100);g__Metschnikowia(100);s__Metschnikowia_pulcherrima(100); | Metschnikowia | 13 | Saprotroph | \|Undefined Saprotroph\| | Yeast | NULL | Probable | As nectar-tap_saprotroph (Põlme et al. 2020); Decay sugar-rich substrates (Põlme et al. 2020) | Põlme S, et al. 2020. Fungal Diversity 105:1-16 ((https://doi.org/10.1007/s13225-020-00466-2)) |
| Otu52718 | 1 | k__Fungi(100);p__Ascomycota(100);c__Saccharomycetes(100);o__Saccharomycetales(100);f__Metschnikowiaceae(100);g__Metschnikowia(100);s__Metschnikowia_pulcherrima(100); | Metschnikowia | 13 | Saprotroph | \|Undefined Saprotroph\| | Yeast | NULL | Probable | As nectar-tap_saprotroph (Põlme et al. 2020); Decay sugar-rich substrates (Põlme et al. 2020) | Põlme S, et al. 2020. Fungal Diversity 105:1-16 ((https://doi.org/10.1007/s13225-020-00466-2)) |
| Otu52721 | 1 | k__Fungi(100);p__Ascomycota(100);c__Saccharomycetes(100);o__Saccharomycetales(100);f__Metschnikowiaceae(100);g__Metschnikowia(100);s__Metschnikowia_pulcherrima(100); | Metschnikowia | 13 | Saprotroph | \|Undefined Saprotroph\| | Yeast | NULL | Probable | As nectar-tap_saprotroph (Põlme et al. 2020); Decay sugar-rich substrates (Põlme et al. 2020) | Põlme S, et al. 2020. Fungal Diversity 105:1-16 ((https://doi.org/10.1007/s13225-020-00466-2)) |
| Otu52723 | 1 | k__Fungi(100);p__Ascomycota(100);c__Saccharomycetes(100);o__Saccharomycetales(100);f__Metschnikowiaceae(100);g__Metschnikowia(100);s__Metschnikowia_pulcherrima(100); | Metschnikowia | 13 | Saprotroph | \|Undefined Saprotroph\| | Yeast | NULL | Probable | As nectar-tap_saprotroph (Põlme et al. 2020); Decay sugar-rich substrates (Põlme et al. 2020) | Põlme S, et al. 2020. Fungal Diversity 105:1-16 ((https://doi.org/10.1007/s13225-020-00466-2)) |
| Otu52724 | 1 | k__Fungi(100);p__Ascomycota(100);c__Saccharomycetes(100);o__Saccharomycetales(100);f__Saccharomycetales_family_Incertae_sedis(100);g__Candida(100);g__Candida_unclassified(100); | Candida | 13 | Pathotroph-Saprotroph-Symbiotroph | Animal Pathogen-Endophyte-Endosymbiont-Epiphyte-\|Undefined Saprotroph\| | Dimorphic Yeast-Yeast | NULL | Probable | As nectar-tap_saprotroph (Põlme et al. 2020); Decay sugar-rich substrates (Põlme et al. 2020); This asexual genus is polyphyletic and contain species that function very differently depend on what they are. Caution should be taken when interpreting guilds for Candida species. | Manolakaki D, et al. 2010. Virulence. 1: 367-375 ((https://doi.org/10.4161/viru.1.5.12796)); Mekbib SB, et al. 2011. Tropical Plant Pathology 36:233-240 ((https://doi.org/10.1590/S1982-56762011000400004)); Põlme S, et al. 2020. Fungal Diversity 105:1-16 ((https://doi.org/10.1007/s13225-020-00466-2)) |
| Otu52725 | 1 | k__Fungi(100);p__Ascomycota(100);c__Saccharomycetes(100);o__Saccharomycetales(100);o__Saccharomycetales_unclassified(100);o__Saccharomycetales_unclassified(100);o__Saccharomycetales_unclassified(100); | Saccharomycetales | 7 | Saprotroph | Undefined Saprotroph | Yeast | NULL | Possible | NULL | Sterkenburg E, et al. 2015. New Phytologist 207:1145-1158 ((https://doi.org/10.1111/nph.13426)) |
| Otu52726 | 1 | k__Fungi(100);p__Ascomycota(100);c__Saccharomycetes(100);o__Saccharomycetales(100);f__Metschnikowiaceae(100);g__Metschnikowia(100);g__Metschnikowia_unclassified(100); | Metschnikowia | 13 | Saprotroph | \|Undefined Saprotroph\| | Yeast | NULL | Probable | As nectar-tap_saprotroph (Põlme et al. 2020); Decay sugar-rich substrates (Põlme et al. 2020) | Põlme S, et al. 2020. Fungal Diversity 105:1-16 ((https://doi.org/10.1007/s13225-020-00466-2)) |
| Otu52727 | 1 | k__Fungi(100);p__Ascomycota(100);c__Saccharomycetes(100);o__Saccharomycetales(100);f__Metschnikowiaceae(100);g__Metschnikowia(100);g__Metschnikowia_unclassified(100); | Metschnikowia | 13 | Saprotroph | \|Undefined Saprotroph\| | Yeast | NULL | Probable | As nectar-tap_saprotroph (Põlme et al. 2020); Decay sugar-rich substrates (Põlme et al. 2020) | Põlme S, et al. 2020. Fungal Diversity 105:1-16 ((https://doi.org/10.1007/s13225-020-00466-2)) |
| Otu52728 | 1 | k__Fungi(100);p__Ascomycota(100);c__Saccharomycetes(100);o__Saccharomycetales(100);f__Metschnikowiaceae(100);g__Metschnikowia(100);g__Metschnikowia_unclassified(100); | Metschnikowia | 13 | Saprotroph | \|Undefined Saprotroph\| | Yeast | NULL | Probable | As nectar-tap_saprotroph (Põlme et al. 2020); Decay sugar-rich substrates (Põlme et al. 2020) | Põlme S, et al. 2020. Fungal Diversity 105:1-16 ((https://doi.org/10.1007/s13225-020-00466-2)) |
| Otu52729 | 1 | k__Fungi(100);p__Ascomycota(100);c__Saccharomycetes(100);o__Saccharomycetales(100);f__Metschnikowiaceae(100);g__Metschnikowia(100);s__Metschnikowia_pulcherrima(100); | Metschnikowia | 13 | Saprotroph | \|Undefined Saprotroph\| | Yeast | NULL | Probable | As nectar-tap_saprotroph (Põlme et al. 2020); Decay sugar-rich substrates (Põlme et al. 2020) | Põlme S, et al. 2020. Fungal Diversity 105:1-16 ((https://doi.org/10.1007/s13225-020-00466-2)) |
| Otu52730 | 1 | k__Fungi(100);p__Ascomycota(100);c__Saccharomycetes(100);o__Saccharomycetales(100);f__Metschnikowiaceae(100);g__Metschnikowia(100);g__Metschnikowia_unclassified(100); | Metschnikowia | 13 | Saprotroph | \|Undefined Saprotroph\| | Yeast | NULL | Probable | As nectar-tap_saprotroph (Põlme et al. 2020); Decay sugar-rich substrates (Põlme et al. 2020) | Põlme S, et al. 2020. Fungal Diversity 105:1-16 ((https://doi.org/10.1007/s13225-020-00466-2)) |
| Otu52731 | 1 | k__Fungi(100);p__Ascomycota(100);c__Saccharomycetes(100);o__Saccharomycetales(100);f__Metschnikowiaceae(100);g__Metschnikowia(100);s__Metschnikowia_pulcherrima(100); | Metschnikowia | 13 | Saprotroph | \|Undefined Saprotroph\| | Yeast | NULL | Probable | As nectar-tap_saprotroph (Põlme et al. 2020); Decay sugar-rich substrates (Põlme et al. 2020) | Põlme S, et al. 2020. Fungal Diversity 105:1-16 ((https://doi.org/10.1007/s13225-020-00466-2)) |
| Otu52732 | 1 | k__Fungi(100);p__Ascomycota(100);c__Saccharomycetes(100);o__Saccharomycetales(100);f__Metschnikowiaceae(100);g__Metschnikowia(100);s__Metschnikowia_pulcherrima(100); | Metschnikowia | 13 | Saprotroph | \|Undefined Saprotroph\| | Yeast | NULL | Probable | As nectar-tap_saprotroph (Põlme et al. 2020); Decay sugar-rich substrates (Põlme et al. 2020) | Põlme S, et al. 2020. Fungal Diversity 105:1-16 ((https://doi.org/10.1007/s13225-020-00466-2)) |
| Otu52733 | 1 | k__Fungi(100);p__Ascomycota(100);c__Saccharomycetes(100);o__Saccharomycetales(100);f__Metschnikowiaceae(100);g__Metschnikowia(100);s__Metschnikowia_pulcherrima(100); | Metschnikowia | 13 | Saprotroph | \|Undefined Saprotroph\| | Yeast | NULL | Probable | As nectar-tap_saprotroph (Põlme et al. 2020); Decay sugar-rich substrates (Põlme et al. 2020) | Põlme S, et al. 2020. Fungal Diversity 105:1-16 ((https://doi.org/10.1007/s13225-020-00466-2)) |
| Otu52734 | 1 | k__Fungi(100);p__Ascomycota(100);c__Saccharomycetes(100);o__Saccharomycetales(100);f__Metschnikowiaceae(100);g__Metschnikowia(100);s__Metschnikowia_pulcherrima(100); | Metschnikowia | 13 | Saprotroph | \|Undefined Saprotroph\| | Yeast | NULL | Probable | As nectar-tap_saprotroph (Põlme et al. 2020); Decay sugar-rich substrates (Põlme et al. 2020) | Põlme S, et al. 2020. Fungal Diversity 105:1-16 ((https://doi.org/10.1007/s13225-020-00466-2)) |
| Otu52735 | 1 | k__Fungi(100);p__Ascomycota(100);c__Saccharomycetes(100);o__Saccharomycetales(100);f__Metschnikowiaceae(100);g__Metschnikowia(100);s__Metschnikowia_pulcherrima(100); | Metschnikowia | 13 | Saprotroph | \|Undefined Saprotroph\| | Yeast | NULL | Probable | As nectar-tap_saprotroph (Põlme et al. 2020); Decay sugar-rich substrates (Põlme et al. 2020) | Põlme S, et al. 2020. Fungal Diversity 105:1-16 ((https://doi.org/10.1007/s13225-020-00466-2)) |
| Otu52736 | 1 | k__Fungi(100);p__Ascomycota(100);c__Saccharomycetes(100);o__Saccharomycetales(100);f__Metschnikowiaceae(100);g__Metschnikowia(100);s__Metschnikowia_pulcherrima(100); | Metschnikowia | 13 | Saprotroph | \|Undefined Saprotroph\| | Yeast | NULL | Probable | As nectar-tap_saprotroph (Põlme et al. 2020); Decay sugar-rich substrates (Põlme et al. 2020) | Põlme S, et al. 2020. Fungal Diversity 105:1-16 ((https://doi.org/10.1007/s13225-020-00466-2)) |
| Otu52737 | 1 | k__Fungi(100);p__Ascomycota(100);c__Saccharomycetes(100);o__Saccharomycetales(100);f__Metschnikowiaceae(100);g__Metschnikowia(100);s__Metschnikowia_pulcherrima(100); | Metschnikowia | 13 | Saprotroph | \|Undefined Saprotroph\| | Yeast | NULL | Probable | As nectar-tap_saprotroph (Põlme et al. 2020); Decay sugar-rich substrates (Põlme et al. 2020) | Põlme S, et al. 2020. Fungal Diversity 105:1-16 ((https://doi.org/10.1007/s13225-020-00466-2)) |
| Otu52738 | 1 | k__Fungi(100);p__Ascomycota(100);c__Saccharomycetes(100);o__Saccharomycetales(100);f__Metschnikowiaceae(100);g__Metschnikowia(100);s__Metschnikowia_pulcherrima(100); | Metschnikowia | 13 | Saprotroph | \|Undefined Saprotroph\| | Yeast | NULL | Probable | As nectar-tap_saprotroph (Põlme et al. 2020); Decay sugar-rich substrates (Põlme et al. 2020) | Põlme S, et al. 2020. Fungal Diversity 105:1-16 ((https://doi.org/10.1007/s13225-020-00466-2)) |
| Otu52739 | 1 | k__Fungi(100);p__Ascomycota(100);c__Saccharomycetes(100);o__Saccharomycetales(100);f__Metschnikowiaceae(100);g__Metschnikowia(100);g__Metschnikowia_unclassified(100); | Metschnikowia | 13 | Saprotroph | \|Undefined Saprotroph\| | Yeast | NULL | Probable | As nectar-tap_saprotroph (Põlme et al. 2020); Decay sugar-rich substrates (Põlme et al. 2020) | Põlme S, et al. 2020. Fungal Diversity 105:1-16 ((https://doi.org/10.1007/s13225-020-00466-2)) |
| Otu52740 | 1 | k__Fungi(100);p__Ascomycota(100);c__Saccharomycetes(100);o__Saccharomycetales(100);f__Metschnikowiaceae(100);g__Metschnikowia(100);g__Metschnikowia_unclassified(100); | Metschnikowia | 13 | Saprotroph | \|Undefined Saprotroph\| | Yeast | NULL | Probable | As nectar-tap_saprotroph (Põlme et al. 2020); Decay sugar-rich substrates (Põlme et al. 2020) | Põlme S, et al. 2020. Fungal Diversity 105:1-16 ((https://doi.org/10.1007/s13225-020-00466-2)) |
| Otu52741 | 1 | k__Fungi(100);p__Ascomycota(100);c__Saccharomycetes(100);o__Saccharomycetales(100);f__Metschnikowiaceae(100);g__Metschnikowia(100);s__Metschnikowia_pulcherrima(100); | Metschnikowia | 13 | Saprotroph | \|Undefined Saprotroph\| | Yeast | NULL | Probable | As nectar-tap_saprotroph (Põlme et al. 2020); Decay sugar-rich substrates (Põlme et al. 2020) | Põlme S, et al. 2020. Fungal Diversity 105:1-16 ((https://doi.org/10.1007/s13225-020-00466-2)) |
| Otu52742 | 1 | k__Fungi(100);p__Ascomycota(100);c__Saccharomycetes(100);o__Saccharomycetales(100);f__Metschnikowiaceae(100);g__Metschnikowia(100);g__Metschnikowia_unclassified(100); | Metschnikowia | 13 | Saprotroph | \|Undefined Saprotroph\| | Yeast | NULL | Probable | As nectar-tap_saprotroph (Põlme et al. 2020); Decay sugar-rich substrates (Põlme et al. 2020) | Põlme S, et al. 2020. Fungal Diversity 105:1-16 ((https://doi.org/10.1007/s13225-020-00466-2)) |
| Otu52743 | 1 | k__Fungi(100);p__Ascomycota(100);c__Saccharomycetes(100);o__Saccharomycetales(100);f__Metschnikowiaceae(100);g__Metschnikowia(100);s__Metschnikowia_pulcherrima(100); | Metschnikowia | 13 | Saprotroph | \|Undefined Saprotroph\| | Yeast | NULL | Probable | As nectar-tap_saprotroph (Põlme et al. 2020); Decay sugar-rich substrates (Põlme et al. 2020) | Põlme S, et al. 2020. Fungal Diversity 105:1-16 ((https://doi.org/10.1007/s13225-020-00466-2)) |
| Otu52745 | 1 | k__Fungi(100);p__Ascomycota(100);c__Saccharomycetes(100);o__Saccharomycetales(100);f__Metschnikowiaceae(100);g__Metschnikowia(100);g__Metschnikowia_unclassified(100); | Metschnikowia | 13 | Saprotroph | \|Undefined Saprotroph\| | Yeast | NULL | Probable | As nectar-tap_saprotroph (Põlme et al. 2020); Decay sugar-rich substrates (Põlme et al. 2020) | Põlme S, et al. 2020. Fungal Diversity 105:1-16 ((https://doi.org/10.1007/s13225-020-00466-2)) |
| Otu52746 | 1 | k__Fungi(100);p__Ascomycota(100);c__Saccharomycetes(100);o__Saccharomycetales(100);f__Metschnikowiaceae(100);g__Metschnikowia(100);s__Metschnikowia_pulcherrima(100); | Metschnikowia | 13 | Saprotroph | \|Undefined Saprotroph\| | Yeast | NULL | Probable | As nectar-tap_saprotroph (Põlme et al. 2020); Decay sugar-rich substrates (Põlme et al. 2020) | Põlme S, et al. 2020. Fungal Diversity 105:1-16 ((https://doi.org/10.1007/s13225-020-00466-2)) |
| Otu52748 | 1 | k__Fungi(100);p__Ascomycota(100);c__Saccharomycetes(100);o__Saccharomycetales(100);f__Metschnikowiaceae(100);g__Metschnikowia(100);g__Metschnikowia_unclassified(100); | Metschnikowia | 13 | Saprotroph | \|Undefined Saprotroph\| | Yeast | NULL | Probable | As nectar-tap_saprotroph (Põlme et al. 2020); Decay sugar-rich substrates (Põlme et al. 2020) | Põlme S, et al. 2020. Fungal Diversity 105:1-16 ((https://doi.org/10.1007/s13225-020-00466-2)) |
| Otu52749 | 1 | k__Fungi(100);p__Ascomycota(100);c__Saccharomycetes(100);o__Saccharomycetales(100);f__Metschnikowiaceae(100);g__Metschnikowia(100);g__Metschnikowia_unclassified(100); | Metschnikowia | 13 | Saprotroph | \|Undefined Saprotroph\| | Yeast | NULL | Probable | As nectar-tap_saprotroph (Põlme et al. 2020); Decay sugar-rich substrates (Põlme et al. 2020) | Põlme S, et al. 2020. Fungal Diversity 105:1-16 ((https://doi.org/10.1007/s13225-020-00466-2)) |
| Otu52750 | 1 | k__Fungi(100);p__Ascomycota(100);c__Saccharomycetes(100);o__Saccharomycetales(100);f__Metschnikowiaceae(100);g__Metschnikowia(100);s__Metschnikowia_pulcherrima(100); | Metschnikowia | 13 | Saprotroph | \|Undefined Saprotroph\| | Yeast | NULL | Probable | As nectar-tap_saprotroph (Põlme et al. 2020); Decay sugar-rich substrates (Põlme et al. 2020) | Põlme S, et al. 2020. Fungal Diversity 105:1-16 ((https://doi.org/10.1007/s13225-020-00466-2)) |
| Otu52751 | 1 | k__Fungi(100);p__Ascomycota(100);c__Saccharomycetes(100);o__Saccharomycetales(100);f__Metschnikowiaceae(100);g__Metschnikowia(100);s__Metschnikowia_pulcherrima(100); | Metschnikowia | 13 | Saprotroph | \|Undefined Saprotroph\| | Yeast | NULL | Probable | As nectar-tap_saprotroph (Põlme et al. 2020); Decay sugar-rich substrates (Põlme et al. 2020) | Põlme S, et al. 2020. Fungal Diversity 105:1-16 ((https://doi.org/10.1007/s13225-020-00466-2)) |
| Otu52752 | 1 | k__Fungi(100);p__Ascomycota(100);c__Saccharomycetes(100);o__Saccharomycetales(100);f__Metschnikowiaceae(100);g__Metschnikowia(100);s__Metschnikowia_pulcherrima(100); | Metschnikowia | 13 | Saprotroph | \|Undefined Saprotroph\| | Yeast | NULL | Probable | As nectar-tap_saprotroph (Põlme et al. 2020); Decay sugar-rich substrates (Põlme et al. 2020) | Põlme S, et al. 2020. Fungal Diversity 105:1-16 ((https://doi.org/10.1007/s13225-020-00466-2)) |
| Otu52753 | 1 | k__Fungi(100);p__Ascomycota(100);c__Saccharomycetes(100);o__Saccharomycetales(100);f__Metschnikowiaceae(100);g__Metschnikowia(100);s__Metschnikowia_pulcherrima(100); | Metschnikowia | 13 | Saprotroph | \|Undefined Saprotroph\| | Yeast | NULL | Probable | As nectar-tap_saprotroph (Põlme et al. 2020); Decay sugar-rich substrates (Põlme et al. 2020) | Põlme S, et al. 2020. Fungal Diversity 105:1-16 ((https://doi.org/10.1007/s13225-020-00466-2)) |
| Otu52754 | 1 | k__Fungi(100);p__Ascomycota(100);c__Saccharomycetes(100);o__Saccharomycetales(100);f__Metschnikowiaceae(100);g__Metschnikowia(100);g__Metschnikowia_unclassified(100); | Metschnikowia | 13 | Saprotroph | \|Undefined Saprotroph\| | Yeast | NULL | Probable | As nectar-tap_saprotroph (Põlme et al. 2020); Decay sugar-rich substrates (Põlme et al. 2020) | Põlme S, et al. 2020. Fungal Diversity 105:1-16 ((https://doi.org/10.1007/s13225-020-00466-2)) |
| Otu52755 | 1 | k__Fungi(100);p__Ascomycota(100);c__Saccharomycetes(100);o__Saccharomycetales(100);f__Metschnikowiaceae(100);g__Metschnikowia(100);s__Metschnikowia_pulcherrima(100); | Metschnikowia | 13 | Saprotroph | \|Undefined Saprotroph\| | Yeast | NULL | Probable | As nectar-tap_saprotroph (Põlme et al. 2020); Decay sugar-rich substrates (Põlme et al. 2020) | Põlme S, et al. 2020. Fungal Diversity 105:1-16 ((https://doi.org/10.1007/s13225-020-00466-2)) |
| Otu52756 | 1 | k__Fungi(100);p__Ascomycota(100);c__Saccharomycetes(100);o__Saccharomycetales(100);f__Metschnikowiaceae(100);g__Metschnikowia(100);g__Metschnikowia_unclassified(100); | Metschnikowia | 13 | Saprotroph | \|Undefined Saprotroph\| | Yeast | NULL | Probable | As nectar-tap_saprotroph (Põlme et al. 2020); Decay sugar-rich substrates (Põlme et al. 2020) | Põlme S, et al. 2020. Fungal Diversity 105:1-16 ((https://doi.org/10.1007/s13225-020-00466-2)) |
| Otu52757 | 1 | k__Fungi(100);p__Ascomycota(100);c__Saccharomycetes(100);o__Saccharomycetales(100);f__Metschnikowiaceae(100);g__Metschnikowia(100);g__Metschnikowia_unclassified(100); | Metschnikowia | 13 | Saprotroph | \|Undefined Saprotroph\| | Yeast | NULL | Probable | As nectar-tap_saprotroph (Põlme et al. 2020); Decay sugar-rich substrates (Põlme et al. 2020) | Põlme S, et al. 2020. Fungal Diversity 105:1-16 ((https://doi.org/10.1007/s13225-020-00466-2)) |
| Otu52758 | 1 | k__Fungi(100);p__Ascomycota(100);c__Saccharomycetes(100);o__Saccharomycetales(100);f__Metschnikowiaceae(100);g__Metschnikowia(100);s__Metschnikowia_pulcherrima(100); | Metschnikowia | 13 | Saprotroph | \|Undefined Saprotroph\| | Yeast | NULL | Probable | As nectar-tap_saprotroph (Põlme et al. 2020); Decay sugar-rich substrates (Põlme et al. 2020) | Põlme S, et al. 2020. Fungal Diversity 105:1-16 ((https://doi.org/10.1007/s13225-020-00466-2)) |
| Otu52759 | 1 | k__Fungi(100);p__Ascomycota(100);c__Saccharomycetes(100);o__Saccharomycetales(100);f__Metschnikowiaceae(100);g__Metschnikowia(100);s__Metschnikowia_pulcherrima(100); | Metschnikowia | 13 | Saprotroph | \|Undefined Saprotroph\| | Yeast | NULL | Probable | As nectar-tap_saprotroph (Põlme et al. 2020); Decay sugar-rich substrates (Põlme et al. 2020) | Põlme S, et al. 2020. Fungal Diversity 105:1-16 ((https://doi.org/10.1007/s13225-020-00466-2)) |
| Otu52761 | 1 | k__Fungi(100);p__Ascomycota(100);c__Saccharomycetes(100);o__Saccharomycetales(100);f__Metschnikowiaceae(100);g__Metschnikowia(100);s__Metschnikowia_pulcherrima(100); | Metschnikowia | 13 | Saprotroph | \|Undefined Saprotroph\| | Yeast | NULL | Probable | As nectar-tap_saprotroph (Põlme et al. 2020); Decay sugar-rich substrates (Põlme et al. 2020) | Põlme S, et al. 2020. Fungal Diversity 105:1-16 ((https://doi.org/10.1007/s13225-020-00466-2)) |
| Otu52762 | 1 | k__Fungi(100);p__Ascomycota(100);c__Saccharomycetes(100);o__Saccharomycetales(100);f__Metschnikowiaceae(100);g__Metschnikowia(100);s__Metschnikowia_pulcherrima(100); | Metschnikowia | 13 | Saprotroph | \|Undefined Saprotroph\| | Yeast | NULL | Probable | As nectar-tap_saprotroph (Põlme et al. 2020); Decay sugar-rich substrates (Põlme et al. 2020) | Põlme S, et al. 2020. Fungal Diversity 105:1-16 ((https://doi.org/10.1007/s13225-020-00466-2)) |
| Otu52763 | 1 | k__Fungi(100);p__Ascomycota(100);c__Saccharomycetes(100);o__Saccharomycetales(100);f__Metschnikowiaceae(100);g__Metschnikowia(100);s__Metschnikowia_pulcherrima(100); | Metschnikowia | 13 | Saprotroph | \|Undefined Saprotroph\| | Yeast | NULL | Probable | As nectar-tap_saprotroph (Põlme et al. 2020); Decay sugar-rich substrates (Põlme et al. 2020) | Põlme S, et al. 2020. Fungal Diversity 105:1-16 ((https://doi.org/10.1007/s13225-020-00466-2)) |
| Otu52764 | 1 | k__Fungi(100);p__Ascomycota(100);c__Saccharomycetes(100);o__Saccharomycetales(100);f__Metschnikowiaceae(100);g__Metschnikowia(100);s__Metschnikowia_pulcherrima(100); | Metschnikowia | 13 | Saprotroph | \|Undefined Saprotroph\| | Yeast | NULL | Probable | As nectar-tap_saprotroph (Põlme et al. 2020); Decay sugar-rich substrates (Põlme et al. 2020) | Põlme S, et al. 2020. Fungal Diversity 105:1-16 ((https://doi.org/10.1007/s13225-020-00466-2)) |
| Otu52765 | 1 | k__Fungi(100);p__Ascomycota(100);c__Saccharomycetes(100);o__Saccharomycetales(100);f__Metschnikowiaceae(100);g__Metschnikowia(100);g__Metschnikowia_unclassified(100); | Metschnikowia | 13 | Saprotroph | \|Undefined Saprotroph\| | Yeast | NULL | Probable | As nectar-tap_saprotroph (Põlme et al. 2020); Decay sugar-rich substrates (Põlme et al. 2020) | Põlme S, et al. 2020. Fungal Diversity 105:1-16 ((https://doi.org/10.1007/s13225-020-00466-2)) |
| Otu52766 | 1 | k__Fungi(100);p__Ascomycota(100);c__Saccharomycetes(100);o__Saccharomycetales(100);f__Metschnikowiaceae(100);g__Metschnikowia(100);g__Metschnikowia_unclassified(100); | Metschnikowia | 13 | Saprotroph | \|Undefined Saprotroph\| | Yeast | NULL | Probable | As nectar-tap_saprotroph (Põlme et al. 2020); Decay sugar-rich substrates (Põlme et al. 2020) | Põlme S, et al. 2020. Fungal Diversity 105:1-16 ((https://doi.org/10.1007/s13225-020-00466-2)) |
| Otu52767 | 1 | k__Fungi(100);p__Ascomycota(100);c__Saccharomycetes(100);o__Saccharomycetales(100);f__Metschnikowiaceae(100);g__Metschnikowia(100);g__Metschnikowia_unclassified(100); | Metschnikowia | 13 | Saprotroph | \|Undefined Saprotroph\| | Yeast | NULL | Probable | As nectar-tap_saprotroph (Põlme et al. 2020); Decay sugar-rich substrates (Põlme et al. 2020) | Põlme S, et al. 2020. Fungal Diversity 105:1-16 ((https://doi.org/10.1007/s13225-020-00466-2)) |
| Otu52768 | 1 | k__Fungi(100);p__Ascomycota(100);c__Saccharomycetes(100);o__Saccharomycetales(100);f__Metschnikowiaceae(100);g__Metschnikowia(100);s__Metschnikowia_pulcherrima(100); | Metschnikowia | 13 | Saprotroph | \|Undefined Saprotroph\| | Yeast | NULL | Probable | As nectar-tap_saprotroph (Põlme et al. 2020); Decay sugar-rich substrates (Põlme et al. 2020) | Põlme S, et al. 2020. Fungal Diversity 105:1-16 ((https://doi.org/10.1007/s13225-020-00466-2)) |
| Otu52769 | 1 | k__Fungi(100);p__Ascomycota(100);c__Saccharomycetes(100);o__Saccharomycetales(100);f__Metschnikowiaceae(100);g__Metschnikowia(100);s__Metschnikowia_pulcherrima(100); | Metschnikowia | 13 | Saprotroph | \|Undefined Saprotroph\| | Yeast | NULL | Probable | As nectar-tap_saprotroph (Põlme et al. 2020); Decay sugar-rich substrates (Põlme et al. 2020) | Põlme S, et al. 2020. Fungal Diversity 105:1-16 ((https://doi.org/10.1007/s13225-020-00466-2)) |
| Otu52770 | 1 | k__Fungi(100);p__Ascomycota(100);c__Saccharomycetes(100);o__Saccharomycetales(100);f__Metschnikowiaceae(100);g__Metschnikowia(100);s__Metschnikowia_pulcherrima(100); | Metschnikowia | 13 | Saprotroph | \|Undefined Saprotroph\| | Yeast | NULL | Probable | As nectar-tap_saprotroph (Põlme et al. 2020); Decay sugar-rich substrates (Põlme et al. 2020) | Põlme S, et al. 2020. Fungal Diversity 105:1-16 ((https://doi.org/10.1007/s13225-020-00466-2)) |
| Otu52771 | 1 | k__Fungi(100);p__Ascomycota(100);c__Saccharomycetes(100);o__Saccharomycetales(100);f__Metschnikowiaceae(100);g__Metschnikowia(100);g__Metschnikowia_unclassified(100); | Metschnikowia | 13 | Saprotroph | \|Undefined Saprotroph\| | Yeast | NULL | Probable | As nectar-tap_saprotroph (Põlme et al. 2020); Decay sugar-rich substrates (Põlme et al. 2020) | Põlme S, et al. 2020. Fungal Diversity 105:1-16 ((https://doi.org/10.1007/s13225-020-00466-2)) |
| Otu52772 | 1 | k__Fungi(100);p__Ascomycota(100);c__Saccharomycetes(100);o__Saccharomycetales(100);f__Metschnikowiaceae(100);g__Metschnikowia(100);g__Metschnikowia_unclassified(100); | Metschnikowia | 13 | Saprotroph | \|Undefined Saprotroph\| | Yeast | NULL | Probable | As nectar-tap_saprotroph (Põlme et al. 2020); Decay sugar-rich substrates (Põlme et al. 2020) | Põlme S, et al. 2020. Fungal Diversity 105:1-16 ((https://doi.org/10.1007/s13225-020-00466-2)) |
| Otu52773 | 1 | k__Fungi(100);p__Ascomycota(100);c__Saccharomycetes(100);o__Saccharomycetales(100);f__Metschnikowiaceae(100);g__Metschnikowia(100);s__Metschnikowia_pulcherrima(100); | Metschnikowia | 13 | Saprotroph | \|Undefined Saprotroph\| | Yeast | NULL | Probable | As nectar-tap_saprotroph (Põlme et al. 2020); Decay sugar-rich substrates (Põlme et al. 2020) | Põlme S, et al. 2020. Fungal Diversity 105:1-16 ((https://doi.org/10.1007/s13225-020-00466-2)) |
| Otu52775 | 1 | k__Fungi(100);p__Ascomycota(100);c__Saccharomycetes(100);o__Saccharomycetales(100);f__Metschnikowiaceae(100);g__Metschnikowia(100);s__Metschnikowia_pulcherrima(100); | Metschnikowia | 13 | Saprotroph | \|Undefined Saprotroph\| | Yeast | NULL | Probable | As nectar-tap_saprotroph (Põlme et al. 2020); Decay sugar-rich substrates (Põlme et al. 2020) | Põlme S, et al. 2020. Fungal Diversity 105:1-16 ((https://doi.org/10.1007/s13225-020-00466-2)) |
| Otu52776 | 1 | k__Fungi(100);p__Ascomycota(100);c__Saccharomycetes(100);o__Saccharomycetales(100);f__Metschnikowiaceae(100);g__Metschnikowia(100);s__Metschnikowia_pulcherrima(100); | Metschnikowia | 13 | Saprotroph | \|Undefined Saprotroph\| | Yeast | NULL | Probable | As nectar-tap_saprotroph (Põlme et al. 2020); Decay sugar-rich substrates (Põlme et al. 2020) | Põlme S, et al. 2020. Fungal Diversity 105:1-16 ((https://doi.org/10.1007/s13225-020-00466-2)) |
| Otu52777 | 1 | k__Fungi(100);p__Ascomycota(100);c__Saccharomycetes(100);o__Saccharomycetales(100);f__Metschnikowiaceae(100);g__Metschnikowia(100);s__Metschnikowia_pulcherrima(100); | Metschnikowia | 13 | Saprotroph | \|Undefined Saprotroph\| | Yeast | NULL | Probable | As nectar-tap_saprotroph (Põlme et al. 2020); Decay sugar-rich substrates (Põlme et al. 2020) | Põlme S, et al. 2020. Fungal Diversity 105:1-16 ((https://doi.org/10.1007/s13225-020-00466-2)) |
| Otu52778 | 1 | k__Fungi(100);p__Ascomycota(100);c__Saccharomycetes(100);o__Saccharomycetales(100);f__Metschnikowiaceae(100);g__Metschnikowia(100);s__Metschnikowia_pulcherrima(100); | Metschnikowia | 13 | Saprotroph | \|Undefined Saprotroph\| | Yeast | NULL | Probable | As nectar-tap_saprotroph (Põlme et al. 2020); Decay sugar-rich substrates (Põlme et al. 2020) | Põlme S, et al. 2020. Fungal Diversity 105:1-16 ((https://doi.org/10.1007/s13225-020-00466-2)) |
| Otu52779 | 1 | k__Fungi(100);p__Ascomycota(100);c__Saccharomycetes(100);o__Saccharomycetales(100);f__Metschnikowiaceae(100);g__Metschnikowia(100);s__Metschnikowia_pulcherrima(100); | Metschnikowia | 13 | Saprotroph | \|Undefined Saprotroph\| | Yeast | NULL | Probable | As nectar-tap_saprotroph (Põlme et al. 2020); Decay sugar-rich substrates (Põlme et al. 2020) | Põlme S, et al. 2020. Fungal Diversity 105:1-16 ((https://doi.org/10.1007/s13225-020-00466-2)) |
| Otu52780 | 1 | k__Fungi(100);p__Ascomycota(100);c__Saccharomycetes(100);o__Saccharomycetales(100);f__Metschnikowiaceae(100);g__Metschnikowia(100);s__Metschnikowia_pulcherrima(100); | Metschnikowia | 13 | Saprotroph | \|Undefined Saprotroph\| | Yeast | NULL | Probable | As nectar-tap_saprotroph (Põlme et al. 2020); Decay sugar-rich substrates (Põlme et al. 2020) | Põlme S, et al. 2020. Fungal Diversity 105:1-16 ((https://doi.org/10.1007/s13225-020-00466-2)) |
| Otu52781 | 1 | k__Fungi(100);p__Ascomycota(100);c__Saccharomycetes(100);o__Saccharomycetales(100);f__Metschnikowiaceae(100);g__Metschnikowia(100);s__Metschnikowia_pulcherrima(100); | Metschnikowia | 13 | Saprotroph | \|Undefined Saprotroph\| | Yeast | NULL | Probable | As nectar-tap_saprotroph (Põlme et al. 2020); Decay sugar-rich substrates (Põlme et al. 2020) | Põlme S, et al. 2020. Fungal Diversity 105:1-16 ((https://doi.org/10.1007/s13225-020-00466-2)) |
| Otu52783 | 1 | k__Fungi(100);p__Ascomycota(100);c__Saccharomycetes(100);o__Saccharomycetales(100);f__Metschnikowiaceae(100);g__Metschnikowia(100);g__Metschnikowia_unclassified(100); | Metschnikowia | 13 | Saprotroph | \|Undefined Saprotroph\| | Yeast | NULL | Probable | As nectar-tap_saprotroph (Põlme et al. 2020); Decay sugar-rich substrates (Põlme et al. 2020) | Põlme S, et al. 2020. Fungal Diversity 105:1-16 ((https://doi.org/10.1007/s13225-020-00466-2)) |
| Otu52784 | 1 | k__Fungi(100);p__Ascomycota(100);c__Saccharomycetes(100);o__Saccharomycetales(100);f__Metschnikowiaceae(100);g__Metschnikowia(100);g__Metschnikowia_unclassified(100); | Metschnikowia | 13 | Saprotroph | \|Undefined Saprotroph\| | Yeast | NULL | Probable | As nectar-tap_saprotroph (Põlme et al. 2020); Decay sugar-rich substrates (Põlme et al. 2020) | Põlme S, et al. 2020. Fungal Diversity 105:1-16 ((https://doi.org/10.1007/s13225-020-00466-2)) |
| Otu52785 | 1 | k__Fungi(100);p__Ascomycota(100);c__Saccharomycetes(100);o__Saccharomycetales(100);f__Metschnikowiaceae(100);g__Metschnikowia(100);g__Metschnikowia_unclassified(100); | Metschnikowia | 13 | Saprotroph | \|Undefined Saprotroph\| | Yeast | NULL | Probable | As nectar-tap_saprotroph (Põlme et al. 2020); Decay sugar-rich substrates (Põlme et al. 2020) | Põlme S, et al. 2020. Fungal Diversity 105:1-16 ((https://doi.org/10.1007/s13225-020-00466-2)) |
| Otu52787 | 1 | k__Fungi(100);p__Ascomycota(100);c__Saccharomycetes(100);o__Saccharomycetales(100);f__Metschnikowiaceae(100);g__Metschnikowia(100);s__Metschnikowia_pulcherrima(100); | Metschnikowia | 13 | Saprotroph | \|Undefined Saprotroph\| | Yeast | NULL | Probable | As nectar-tap_saprotroph (Põlme et al. 2020); Decay sugar-rich substrates (Põlme et al. 2020) | Põlme S, et al. 2020. Fungal Diversity 105:1-16 ((https://doi.org/10.1007/s13225-020-00466-2)) |
| Otu52789 | 1 | k__Fungi(100);p__Ascomycota(100);c__Saccharomycetes(100);o__Saccharomycetales(100);f__Metschnikowiaceae(100);g__Metschnikowia(100);g__Metschnikowia_unclassified(100); | Metschnikowia | 13 | Saprotroph | \|Undefined Saprotroph\| | Yeast | NULL | Probable | As nectar-tap_saprotroph (Põlme et al. 2020); Decay sugar-rich substrates (Põlme et al. 2020) | Põlme S, et al. 2020. Fungal Diversity 105:1-16 ((https://doi.org/10.1007/s13225-020-00466-2)) |
| Otu52790 | 1 | k__Fungi(100);p__Ascomycota(100);c__Saccharomycetes(100);o__Saccharomycetales(100);f__Metschnikowiaceae(100);g__Metschnikowia(100);g__Metschnikowia_unclassified(100); | Metschnikowia | 13 | Saprotroph | \|Undefined Saprotroph\| | Yeast | NULL | Probable | As nectar-tap_saprotroph (Põlme et al. 2020); Decay sugar-rich substrates (Põlme et al. 2020) | Põlme S, et al. 2020. Fungal Diversity 105:1-16 ((https://doi.org/10.1007/s13225-020-00466-2)) |
| Otu52791 | 1 | k__Fungi(100);p__Ascomycota(100);c__Saccharomycetes(100);o__Saccharomycetales(100);f__Metschnikowiaceae(100);g__Metschnikowia(100);s__Metschnikowia_pulcherrima(100); | Metschnikowia | 13 | Saprotroph | \|Undefined Saprotroph\| | Yeast | NULL | Probable | As nectar-tap_saprotroph (Põlme et al. 2020); Decay sugar-rich substrates (Põlme et al. 2020) | Põlme S, et al. 2020. Fungal Diversity 105:1-16 ((https://doi.org/10.1007/s13225-020-00466-2)) |
| Otu52792 | 1 | k__Fungi(100);p__Ascomycota(100);c__Saccharomycetes(100);o__Saccharomycetales(100);f__Metschnikowiaceae(100);g__Metschnikowia(100);g__Metschnikowia_unclassified(100); | Metschnikowia | 13 | Saprotroph | \|Undefined Saprotroph\| | Yeast | NULL | Probable | As nectar-tap_saprotroph (Põlme et al. 2020); Decay sugar-rich substrates (Põlme et al. 2020) | Põlme S, et al. 2020. Fungal Diversity 105:1-16 ((https://doi.org/10.1007/s13225-020-00466-2)) |
| Otu52793 | 1 | k__Fungi(100);p__Ascomycota(100);c__Saccharomycetes(100);o__Saccharomycetales(100);f__Metschnikowiaceae(100);g__Metschnikowia(100);g__Metschnikowia_unclassified(100); | Metschnikowia | 13 | Saprotroph | \|Undefined Saprotroph\| | Yeast | NULL | Probable | As nectar-tap_saprotroph (Põlme et al. 2020); Decay sugar-rich substrates (Põlme et al. 2020) | Põlme S, et al. 2020. Fungal Diversity 105:1-16 ((https://doi.org/10.1007/s13225-020-00466-2)) |
| Otu52794 | 1 | k__Fungi(100);p__Ascomycota(100);c__Saccharomycetes(100);o__Saccharomycetales(100);f__Metschnikowiaceae(100);g__Metschnikowia(100);s__Metschnikowia_pulcherrima(100); | Metschnikowia | 13 | Saprotroph | \|Undefined Saprotroph\| | Yeast | NULL | Probable | As nectar-tap_saprotroph (Põlme et al. 2020); Decay sugar-rich substrates (Põlme et al. 2020) | Põlme S, et al. 2020. Fungal Diversity 105:1-16 ((https://doi.org/10.1007/s13225-020-00466-2)) |
| Otu52795 | 1 | k__Fungi(100);p__Ascomycota(100);c__Saccharomycetes(100);o__Saccharomycetales(100);f__Metschnikowiaceae(100);g__Metschnikowia(100);s__Metschnikowia_pulcherrima(100); | Metschnikowia | 13 | Saprotroph | \|Undefined Saprotroph\| | Yeast | NULL | Probable | As nectar-tap_saprotroph (Põlme et al. 2020); Decay sugar-rich substrates (Põlme et al. 2020) | Põlme S, et al. 2020. Fungal Diversity 105:1-16 ((https://doi.org/10.1007/s13225-020-00466-2)) |
| Otu52797 | 1 | k__Fungi(100);p__Ascomycota(100);c__Saccharomycetes(100);o__Saccharomycetales(100);f__Metschnikowiaceae(100);g__Metschnikowia(100);s__Metschnikowia_pulcherrima(100); | Metschnikowia | 13 | Saprotroph | \|Undefined Saprotroph\| | Yeast | NULL | Probable | As nectar-tap_saprotroph (Põlme et al. 2020); Decay sugar-rich substrates (Põlme et al. 2020) | Põlme S, et al. 2020. Fungal Diversity 105:1-16 ((https://doi.org/10.1007/s13225-020-00466-2)) |
| Otu52798 | 1 | k__Fungi(100);p__Ascomycota(100);c__Saccharomycetes(100);o__Saccharomycetales(100);f__Metschnikowiaceae(100);g__Metschnikowia(100);s__Metschnikowia_pulcherrima(100); | Metschnikowia | 13 | Saprotroph | \|Undefined Saprotroph\| | Yeast | NULL | Probable | As nectar-tap_saprotroph (Põlme et al. 2020); Decay sugar-rich substrates (Põlme et al. 2020) | Põlme S, et al. 2020. Fungal Diversity 105:1-16 ((https://doi.org/10.1007/s13225-020-00466-2)) |
| Otu52799 | 1 | k__Fungi(100);p__Ascomycota(100);c__Saccharomycetes(100);o__Saccharomycetales(100);f__Metschnikowiaceae(100);g__Metschnikowia(100);s__Metschnikowia_pulcherrima(100); | Metschnikowia | 13 | Saprotroph | \|Undefined Saprotroph\| | Yeast | NULL | Probable | As nectar-tap_saprotroph (Põlme et al. 2020); Decay sugar-rich substrates (Põlme et al. 2020) | Põlme S, et al. 2020. Fungal Diversity 105:1-16 ((https://doi.org/10.1007/s13225-020-00466-2)) |
| Otu52800 | 1 | k__Fungi(100);p__Ascomycota(100);c__Saccharomycetes(100);o__Saccharomycetales(100);f__Metschnikowiaceae(100);g__Metschnikowia(100);s__Metschnikowia_pulcherrima(100); | Metschnikowia | 13 | Saprotroph | \|Undefined Saprotroph\| | Yeast | NULL | Probable | As nectar-tap_saprotroph (Põlme et al. 2020); Decay sugar-rich substrates (Põlme et al. 2020) | Põlme S, et al. 2020. Fungal Diversity 105:1-16 ((https://doi.org/10.1007/s13225-020-00466-2)) |
| Otu52801 | 1 | k__Fungi(100);p__Ascomycota(100);c__Saccharomycetes(100);o__Saccharomycetales(100);f__Metschnikowiaceae(100);g__Metschnikowia(100);s__Metschnikowia_pulcherrima(100); | Metschnikowia | 13 | Saprotroph | \|Undefined Saprotroph\| | Yeast | NULL | Probable | As nectar-tap_saprotroph (Põlme et al. 2020); Decay sugar-rich substrates (Põlme et al. 2020) | Põlme S, et al. 2020. Fungal Diversity 105:1-16 ((https://doi.org/10.1007/s13225-020-00466-2)) |
| Otu52802 | 1 | k__Fungi(100);p__Ascomycota(100);c__Saccharomycetes(100);o__Saccharomycetales(100);f__Metschnikowiaceae(100);g__Metschnikowia(100);s__Metschnikowia_pulcherrima(100); | Metschnikowia | 13 | Saprotroph | \|Undefined Saprotroph\| | Yeast | NULL | Probable | As nectar-tap_saprotroph (Põlme et al. 2020); Decay sugar-rich substrates (Põlme et al. 2020) | Põlme S, et al. 2020. Fungal Diversity 105:1-16 ((https://doi.org/10.1007/s13225-020-00466-2)) |
| Otu52803 | 1 | k__Fungi(100);p__Ascomycota(100);c__Saccharomycetes(100);o__Saccharomycetales(100);f__Metschnikowiaceae(100);g__Metschnikowia(100);g__Metschnikowia_unclassified(100); | Metschnikowia | 13 | Saprotroph | \|Undefined Saprotroph\| | Yeast | NULL | Probable | As nectar-tap_saprotroph (Põlme et al. 2020); Decay sugar-rich substrates (Põlme et al. 2020) | Põlme S, et al. 2020. Fungal Diversity 105:1-16 ((https://doi.org/10.1007/s13225-020-00466-2)) |
| Otu52804 | 1 | k__Fungi(100);p__Ascomycota(100);c__Saccharomycetes(100);o__Saccharomycetales(100);f__Metschnikowiaceae(100);g__Metschnikowia(100);s__Metschnikowia_pulcherrima(100); | Metschnikowia | 13 | Saprotroph | \|Undefined Saprotroph\| | Yeast | NULL | Probable | As nectar-tap_saprotroph (Põlme et al. 2020); Decay sugar-rich substrates (Põlme et al. 2020) | Põlme S, et al. 2020. Fungal Diversity 105:1-16 ((https://doi.org/10.1007/s13225-020-00466-2)) |
| Otu52805 | 1 | k__Fungi(100);p__Ascomycota(100);c__Saccharomycetes(100);o__Saccharomycetales(100);f__Metschnikowiaceae(100);g__Metschnikowia(100);g__Metschnikowia_unclassified(100); | Metschnikowia | 13 | Saprotroph | \|Undefined Saprotroph\| | Yeast | NULL | Probable | As nectar-tap_saprotroph (Põlme et al. 2020); Decay sugar-rich substrates (Põlme et al. 2020) | Põlme S, et al. 2020. Fungal Diversity 105:1-16 ((https://doi.org/10.1007/s13225-020-00466-2)) |
| Otu52806 | 1 | k__Fungi(100);p__Ascomycota(100);c__Saccharomycetes(100);o__Saccharomycetales(100);f__Metschnikowiaceae(100);g__Metschnikowia(100);s__Metschnikowia_pulcherrima(100); | Metschnikowia | 13 | Saprotroph | \|Undefined Saprotroph\| | Yeast | NULL | Probable | As nectar-tap_saprotroph (Põlme et al. 2020); Decay sugar-rich substrates (Põlme et al. 2020) | Põlme S, et al. 2020. Fungal Diversity 105:1-16 ((https://doi.org/10.1007/s13225-020-00466-2)) |
| Otu52807 | 1 | k__Fungi(100);p__Ascomycota(100);c__Saccharomycetes(100);o__Saccharomycetales(100);f__Metschnikowiaceae(100);g__Metschnikowia(100);g__Metschnikowia_unclassified(100); | Metschnikowia | 13 | Saprotroph | \|Undefined Saprotroph\| | Yeast | NULL | Probable | As nectar-tap_saprotroph (Põlme et al. 2020); Decay sugar-rich substrates (Põlme et al. 2020) | Põlme S, et al. 2020. Fungal Diversity 105:1-16 ((https://doi.org/10.1007/s13225-020-00466-2)) |
| Otu52808 | 1 | k__Fungi(100);p__Ascomycota(100);c__Saccharomycetes(100);o__Saccharomycetales(100);f__Metschnikowiaceae(100);g__Metschnikowia(100);g__Metschnikowia_unclassified(100); | Metschnikowia | 13 | Saprotroph | \|Undefined Saprotroph\| | Yeast | NULL | Probable | As nectar-tap_saprotroph (Põlme et al. 2020); Decay sugar-rich substrates (Põlme et al. 2020) | Põlme S, et al. 2020. Fungal Diversity 105:1-16 ((https://doi.org/10.1007/s13225-020-00466-2)) |
| Otu52809 | 1 | k__Fungi(100);p__Ascomycota(100);c__Saccharomycetes(100);o__Saccharomycetales(100);f__Metschnikowiaceae(100);g__Metschnikowia(100);s__Metschnikowia_pulcherrima(100); | Metschnikowia | 13 | Saprotroph | \|Undefined Saprotroph\| | Yeast | NULL | Probable | As nectar-tap_saprotroph (Põlme et al. 2020); Decay sugar-rich substrates (Põlme et al. 2020) | Põlme S, et al. 2020. Fungal Diversity 105:1-16 ((https://doi.org/10.1007/s13225-020-00466-2)) |
| Otu52810 | 1 | k__Fungi(100);p__Ascomycota(100);c__Saccharomycetes(100);o__Saccharomycetales(100);f__Metschnikowiaceae(100);g__Metschnikowia(100);s__Metschnikowia_pulcherrima(100); | Metschnikowia | 13 | Saprotroph | \|Undefined Saprotroph\| | Yeast | NULL | Probable | As nectar-tap_saprotroph (Põlme et al. 2020); Decay sugar-rich substrates (Põlme et al. 2020) | Põlme S, et al. 2020. Fungal Diversity 105:1-16 ((https://doi.org/10.1007/s13225-020-00466-2)) |
| Otu52811 | 1 | k__Fungi(100);p__Ascomycota(100);c__Saccharomycetes(100);o__Saccharomycetales(100);f__Metschnikowiaceae(100);g__Metschnikowia(100);s__Metschnikowia_pulcherrima(100); | Metschnikowia | 13 | Saprotroph | \|Undefined Saprotroph\| | Yeast | NULL | Probable | As nectar-tap_saprotroph (Põlme et al. 2020); Decay sugar-rich substrates (Põlme et al. 2020) | Põlme S, et al. 2020. Fungal Diversity 105:1-16 ((https://doi.org/10.1007/s13225-020-00466-2)) |
| Otu52812 | 1 | k__Fungi(100);p__Ascomycota(100);c__Saccharomycetes(100);o__Saccharomycetales(100);f__Metschnikowiaceae(100);g__Metschnikowia(100);s__Metschnikowia_pulcherrima(100); | Metschnikowia | 13 | Saprotroph | \|Undefined Saprotroph\| | Yeast | NULL | Probable | As nectar-tap_saprotroph (Põlme et al. 2020); Decay sugar-rich substrates (Põlme et al. 2020) | Põlme S, et al. 2020. Fungal Diversity 105:1-16 ((https://doi.org/10.1007/s13225-020-00466-2)) |
| Otu52813 | 1 | k__Fungi(100);p__Ascomycota(100);c__Saccharomycetes(100);o__Saccharomycetales(100);f__Metschnikowiaceae(100);g__Metschnikowia(100);s__Metschnikowia_pulcherrima(100); | Metschnikowia | 13 | Saprotroph | \|Undefined Saprotroph\| | Yeast | NULL | Probable | As nectar-tap_saprotroph (Põlme et al. 2020); Decay sugar-rich substrates (Põlme et al. 2020) | Põlme S, et al. 2020. Fungal Diversity 105:1-16 ((https://doi.org/10.1007/s13225-020-00466-2)) |
| Otu52814 | 1 | k__Fungi(100);p__Ascomycota(100);c__Saccharomycetes(100);o__Saccharomycetales(100);f__Metschnikowiaceae(100);g__Metschnikowia(100);g__Metschnikowia_unclassified(100); | Metschnikowia | 13 | Saprotroph | \|Undefined Saprotroph\| | Yeast | NULL | Probable | As nectar-tap_saprotroph (Põlme et al. 2020); Decay sugar-rich substrates (Põlme et al. 2020) | Põlme S, et al. 2020. Fungal Diversity 105:1-16 ((https://doi.org/10.1007/s13225-020-00466-2)) |
| Otu52816 | 1 | k__Fungi(100);p__Ascomycota(100);c__Saccharomycetes(100);o__Saccharomycetales(100);f__Metschnikowiaceae(100);g__Metschnikowia(100);s__Metschnikowia_pulcherrima(100); | Metschnikowia | 13 | Saprotroph | \|Undefined Saprotroph\| | Yeast | NULL | Probable | As nectar-tap_saprotroph (Põlme et al. 2020); Decay sugar-rich substrates (Põlme et al. 2020) | Põlme S, et al. 2020. Fungal Diversity 105:1-16 ((https://doi.org/10.1007/s13225-020-00466-2)) |
| Otu52820 | 1 | k__Fungi(100);p__Ascomycota(100);c__Saccharomycetes(100);o__Saccharomycetales(100);f__Metschnikowiaceae(100);g__Metschnikowia(100);g__Metschnikowia_unclassified(100); | Metschnikowia | 13 | Saprotroph | \|Undefined Saprotroph\| | Yeast | NULL | Probable | As nectar-tap_saprotroph (Põlme et al. 2020); Decay sugar-rich substrates (Põlme et al. 2020) | Põlme S, et al. 2020. Fungal Diversity 105:1-16 ((https://doi.org/10.1007/s13225-020-00466-2)) |
| Otu52821 | 1 | k__Fungi(100);p__Ascomycota(100);c__Saccharomycetes(100);o__Saccharomycetales(100);f__Metschnikowiaceae(100);g__Metschnikowia(100);g__Metschnikowia_unclassified(100); | Metschnikowia | 13 | Saprotroph | \|Undefined Saprotroph\| | Yeast | NULL | Probable | As nectar-tap_saprotroph (Põlme et al. 2020); Decay sugar-rich substrates (Põlme et al. 2020) | Põlme S, et al. 2020. Fungal Diversity 105:1-16 ((https://doi.org/10.1007/s13225-020-00466-2)) |
| Otu52822 | 1 | k__Fungi(100);p__Ascomycota(100);c__Saccharomycetes(100);o__Saccharomycetales(100);f__Metschnikowiaceae(100);g__Metschnikowia(100);s__Metschnikowia_pulcherrima(100); | Metschnikowia | 13 | Saprotroph | \|Undefined Saprotroph\| | Yeast | NULL | Probable | As nectar-tap_saprotroph (Põlme et al. 2020); Decay sugar-rich substrates (Põlme et al. 2020) | Põlme S, et al. 2020. Fungal Diversity 105:1-16 ((https://doi.org/10.1007/s13225-020-00466-2)) |
| Otu52823 | 1 | k__Fungi(100);p__Ascomycota(100);c__Saccharomycetes(100);o__Saccharomycetales(100);f__Metschnikowiaceae(100);g__Metschnikowia(100);g__Metschnikowia_unclassified(100); | Metschnikowia | 13 | Saprotroph | \|Undefined Saprotroph\| | Yeast | NULL | Probable | As nectar-tap_saprotroph (Põlme et al. 2020); Decay sugar-rich substrates (Põlme et al. 2020) | Põlme S, et al. 2020. Fungal Diversity 105:1-16 ((https://doi.org/10.1007/s13225-020-00466-2)) |
| Otu52824 | 1 | k__Fungi(100);p__Ascomycota(100);c__Saccharomycetes(100);o__Saccharomycetales(100);f__Metschnikowiaceae(100);g__Metschnikowia(100);s__Metschnikowia_pulcherrima(100); | Metschnikowia | 13 | Saprotroph | \|Undefined Saprotroph\| | Yeast | NULL | Probable | As nectar-tap_saprotroph (Põlme et al. 2020); Decay sugar-rich substrates (Põlme et al. 2020) | Põlme S, et al. 2020. Fungal Diversity 105:1-16 ((https://doi.org/10.1007/s13225-020-00466-2)) |
| Otu52826 | 1 | k__Fungi(100);p__Ascomycota(100);c__Saccharomycetes(100);o__Saccharomycetales(100);f__Metschnikowiaceae(100);g__Metschnikowia(100);g__Metschnikowia_unclassified(100); | Metschnikowia | 13 | Saprotroph | \|Undefined Saprotroph\| | Yeast | NULL | Probable | As nectar-tap_saprotroph (Põlme et al. 2020); Decay sugar-rich substrates (Põlme et al. 2020) | Põlme S, et al. 2020. Fungal Diversity 105:1-16 ((https://doi.org/10.1007/s13225-020-00466-2)) |
| Otu52833 | 1 | k__Fungi(100);p__Ascomycota(100);c__Saccharomycetes(100);o__Saccharomycetales(100);f__Metschnikowiaceae(100);g__Metschnikowia(100);s__Metschnikowia_pulcherrima(100); | Metschnikowia | 13 | Saprotroph | \|Undefined Saprotroph\| | Yeast | NULL | Probable | As nectar-tap_saprotroph (Põlme et al. 2020); Decay sugar-rich substrates (Põlme et al. 2020) | Põlme S, et al. 2020. Fungal Diversity 105:1-16 ((https://doi.org/10.1007/s13225-020-00466-2)) |
| Otu52844 | 1 | k__Fungi(100);p__Ascomycota(100);c__Saccharomycetes(100);o__Saccharomycetales(100);f__Metschnikowiaceae(100);g__Metschnikowia(100);s__Metschnikowia_pulcherrima(100); | Metschnikowia | 13 | Saprotroph | \|Undefined Saprotroph\| | Yeast | NULL | Probable | As nectar-tap_saprotroph (Põlme et al. 2020); Decay sugar-rich substrates (Põlme et al. 2020) | Põlme S, et al. 2020. Fungal Diversity 105:1-16 ((https://doi.org/10.1007/s13225-020-00466-2)) |
| Otu52845 | 1 | k__Fungi(100);p__Ascomycota(100);c__Saccharomycetes(100);o__Saccharomycetales(100);f__Metschnikowiaceae(100);g__Metschnikowia(100);g__Metschnikowia_unclassified(100); | Metschnikowia | 13 | Saprotroph | \|Undefined Saprotroph\| | Yeast | NULL | Probable | As nectar-tap_saprotroph (Põlme et al. 2020); Decay sugar-rich substrates (Põlme et al. 2020) | Põlme S, et al. 2020. Fungal Diversity 105:1-16 ((https://doi.org/10.1007/s13225-020-00466-2)) |
| Otu52846 | 1 | k__Fungi(100);p__Ascomycota(100);c__Saccharomycetes(100);o__Saccharomycetales(100);f__Metschnikowiaceae(100);g__Metschnikowia(100);s__Metschnikowia_pulcherrima(100); | Metschnikowia | 13 | Saprotroph | \|Undefined Saprotroph\| | Yeast | NULL | Probable | As nectar-tap_saprotroph (Põlme et al. 2020); Decay sugar-rich substrates (Põlme et al. 2020) | Põlme S, et al. 2020. Fungal Diversity 105:1-16 ((https://doi.org/10.1007/s13225-020-00466-2)) |
| Otu52847 | 1 | k__Fungi(100);p__Ascomycota(100);c__Saccharomycetes(100);o__Saccharomycetales(100);f__Metschnikowiaceae(100);g__Metschnikowia(100);g__Metschnikowia_unclassified(100); | Metschnikowia | 13 | Saprotroph | \|Undefined Saprotroph\| | Yeast | NULL | Probable | As nectar-tap_saprotroph (Põlme et al. 2020); Decay sugar-rich substrates (Põlme et al. 2020) | Põlme S, et al. 2020. Fungal Diversity 105:1-16 ((https://doi.org/10.1007/s13225-020-00466-2)) |
| Otu52848 | 1 | k__Fungi(100);p__Ascomycota(100);c__Saccharomycetes(100);o__Saccharomycetales(100);f__Metschnikowiaceae(100);g__Metschnikowia(100);s__Metschnikowia_pulcherrima(100); | Metschnikowia | 13 | Saprotroph | \|Undefined Saprotroph\| | Yeast | NULL | Probable | As nectar-tap_saprotroph (Põlme et al. 2020); Decay sugar-rich substrates (Põlme et al. 2020) | Põlme S, et al. 2020. Fungal Diversity 105:1-16 ((https://doi.org/10.1007/s13225-020-00466-2)) |
| Otu52849 | 1 | k__Fungi(100);p__Ascomycota(100);c__Saccharomycetes(100);o__Saccharomycetales(100);f__Metschnikowiaceae(100);g__Metschnikowia(100);s__Metschnikowia_pulcherrima(100); | Metschnikowia | 13 | Saprotroph | \|Undefined Saprotroph\| | Yeast | NULL | Probable | As nectar-tap_saprotroph (Põlme et al. 2020); Decay sugar-rich substrates (Põlme et al. 2020) | Põlme S, et al. 2020. Fungal Diversity 105:1-16 ((https://doi.org/10.1007/s13225-020-00466-2)) |
| Otu52850 | 1 | k__Fungi(100);p__Ascomycota(100);c__Saccharomycetes(100);o__Saccharomycetales(100);f__Metschnikowiaceae(100);g__Metschnikowia(100);s__Metschnikowia_pulcherrima(100); | Metschnikowia | 13 | Saprotroph | \|Undefined Saprotroph\| | Yeast | NULL | Probable | As nectar-tap_saprotroph (Põlme et al. 2020); Decay sugar-rich substrates (Põlme et al. 2020) | Põlme S, et al. 2020. Fungal Diversity 105:1-16 ((https://doi.org/10.1007/s13225-020-00466-2)) |
| Otu52851 | 1 | k__Fungi(100);p__Ascomycota(100);c__Saccharomycetes(100);o__Saccharomycetales(100);f__Metschnikowiaceae(100);g__Metschnikowia(100);g__Metschnikowia_unclassified(100); | Metschnikowia | 13 | Saprotroph | \|Undefined Saprotroph\| | Yeast | NULL | Probable | As nectar-tap_saprotroph (Põlme et al. 2020); Decay sugar-rich substrates (Põlme et al. 2020) | Põlme S, et al. 2020. Fungal Diversity 105:1-16 ((https://doi.org/10.1007/s13225-020-00466-2)) |
| Otu52852 | 1 | k__Fungi(100);p__Ascomycota(100);c__Saccharomycetes(100);o__Saccharomycetales(100);f__Metschnikowiaceae(100);g__Metschnikowia(100);g__Metschnikowia_unclassified(100); | Metschnikowia | 13 | Saprotroph | \|Undefined Saprotroph\| | Yeast | NULL | Probable | As nectar-tap_saprotroph (Põlme et al. 2020); Decay sugar-rich substrates (Põlme et al. 2020) | Põlme S, et al. 2020. Fungal Diversity 105:1-16 ((https://doi.org/10.1007/s13225-020-00466-2)) |
| Otu52853 | 1 | k__Fungi(100);p__Ascomycota(100);c__Saccharomycetes(100);o__Saccharomycetales(100);f__Metschnikowiaceae(100);g__Metschnikowia(100);g__Metschnikowia_unclassified(100); | Metschnikowia | 13 | Saprotroph | \|Undefined Saprotroph\| | Yeast | NULL | Probable | As nectar-tap_saprotroph (Põlme et al. 2020); Decay sugar-rich substrates (Põlme et al. 2020) | Põlme S, et al. 2020. Fungal Diversity 105:1-16 ((https://doi.org/10.1007/s13225-020-00466-2)) |
| Otu52854 | 1 | k__Fungi(100);p__Ascomycota(100);c__Saccharomycetes(100);o__Saccharomycetales(100);f__Metschnikowiaceae(100);g__Metschnikowia(100);s__Metschnikowia_pulcherrima(100); | Metschnikowia | 13 | Saprotroph | \|Undefined Saprotroph\| | Yeast | NULL | Probable | As nectar-tap_saprotroph (Põlme et al. 2020); Decay sugar-rich substrates (Põlme et al. 2020) | Põlme S, et al. 2020. Fungal Diversity 105:1-16 ((https://doi.org/10.1007/s13225-020-00466-2)) |
| Otu52855 | 1 | k__Fungi(100);p__Ascomycota(100);c__Saccharomycetes(100);o__Saccharomycetales(100);f__Saccharomycetales_family_Incertae_sedis(100);g__Candida(100);s__Candida_cretensis(100); | Candida | 13 | Pathotroph-Saprotroph-Symbiotroph | Animal Pathogen-Endophyte-Endosymbiont-Epiphyte-\|Undefined Saprotroph\| | Dimorphic Yeast-Yeast | NULL | Probable | As nectar-tap_saprotroph (Põlme et al. 2020); Decay sugar-rich substrates (Põlme et al. 2020); This asexual genus is polyphyletic and contain species that function very differently depend on what they are. Caution should be taken when interpreting guilds for Candida species. | Manolakaki D, et al. 2010. Virulence. 1: 367-375 ((https://doi.org/10.4161/viru.1.5.12796)); Mekbib SB, et al. 2011. Tropical Plant Pathology 36:233-240 ((https://doi.org/10.1590/S1982-56762011000400004)); Põlme S, et al. 2020. Fungal Diversity 105:1-16 ((https://doi.org/10.1007/s13225-020-00466-2)) |
| Otu52860 | 1 | k__Fungi(100);p__Ascomycota(100);c__Saccharomycetes(100);o__Saccharomycetales(100);f__Metschnikowiaceae(100);g__Metschnikowia(100);g__Metschnikowia_unclassified(100); | Metschnikowia | 13 | Saprotroph | \|Undefined Saprotroph\| | Yeast | NULL | Probable | As nectar-tap_saprotroph (Põlme et al. 2020); Decay sugar-rich substrates (Põlme et al. 2020) | Põlme S, et al. 2020. Fungal Diversity 105:1-16 ((https://doi.org/10.1007/s13225-020-00466-2)) |
| Otu52861 | 1 | k__Fungi(100);p__Ascomycota(100);c__Saccharomycetes(100);o__Saccharomycetales(100);f__Metschnikowiaceae(100);g__Metschnikowia(100);s__Metschnikowia_pulcherrima(100); | Metschnikowia | 13 | Saprotroph | \|Undefined Saprotroph\| | Yeast | NULL | Probable | As nectar-tap_saprotroph (Põlme et al. 2020); Decay sugar-rich substrates (Põlme et al. 2020) | Põlme S, et al. 2020. Fungal Diversity 105:1-16 ((https://doi.org/10.1007/s13225-020-00466-2)) |
| Otu52862 | 1 | k__Fungi(100);p__Ascomycota(100);c__Saccharomycetes(100);o__Saccharomycetales(100);f__Metschnikowiaceae(100);g__Metschnikowia(100);s__Metschnikowia_pulcherrima(100); | Metschnikowia | 13 | Saprotroph | \|Undefined Saprotroph\| | Yeast | NULL | Probable | As nectar-tap_saprotroph (Põlme et al. 2020); Decay sugar-rich substrates (Põlme et al. 2020) | Põlme S, et al. 2020. Fungal Diversity 105:1-16 ((https://doi.org/10.1007/s13225-020-00466-2)) |
| Otu52863 | 1 | k__Fungi(100);p__Ascomycota(100);c__Saccharomycetes(100);o__Saccharomycetales(100);f__Metschnikowiaceae(100);g__Metschnikowia(100);s__Metschnikowia_pulcherrima(100); | Metschnikowia | 13 | Saprotroph | \|Undefined Saprotroph\| | Yeast | NULL | Probable | As nectar-tap_saprotroph (Põlme et al. 2020); Decay sugar-rich substrates (Põlme et al. 2020) | Põlme S, et al. 2020. Fungal Diversity 105:1-16 ((https://doi.org/10.1007/s13225-020-00466-2)) |
| Otu52865 | 1 | k__Fungi(100);p__Ascomycota(100);c__Saccharomycetes(100);o__Saccharomycetales(100);f__Metschnikowiaceae(100);g__Metschnikowia(100);g__Metschnikowia_unclassified(100); | Metschnikowia | 13 | Saprotroph | \|Undefined Saprotroph\| | Yeast | NULL | Probable | As nectar-tap_saprotroph (Põlme et al. 2020); Decay sugar-rich substrates (Põlme et al. 2020) | Põlme S, et al. 2020. Fungal Diversity 105:1-16 ((https://doi.org/10.1007/s13225-020-00466-2)) |
| Otu52866 | 1 | k__Fungi(100);p__Ascomycota(100);c__Saccharomycetes(100);o__Saccharomycetales(100);f__Metschnikowiaceae(100);g__Metschnikowia(100);g__Metschnikowia_unclassified(100); | Metschnikowia | 13 | Saprotroph | \|Undefined Saprotroph\| | Yeast | NULL | Probable | As nectar-tap_saprotroph (Põlme et al. 2020); Decay sugar-rich substrates (Põlme et al. 2020) | Põlme S, et al. 2020. Fungal Diversity 105:1-16 ((https://doi.org/10.1007/s13225-020-00466-2)) |
| Otu52867 | 1 | k__Fungi(100);p__Ascomycota(100);c__Saccharomycetes(100);o__Saccharomycetales(100);f__Metschnikowiaceae(100);g__Metschnikowia(100);s__Metschnikowia_pulcherrima(100); | Metschnikowia | 13 | Saprotroph | \|Undefined Saprotroph\| | Yeast | NULL | Probable | As nectar-tap_saprotroph (Põlme et al. 2020); Decay sugar-rich substrates (Põlme et al. 2020) | Põlme S, et al. 2020. Fungal Diversity 105:1-16 ((https://doi.org/10.1007/s13225-020-00466-2)) |
| Otu52868 | 1 | k__Fungi(100);p__Ascomycota(100);c__Saccharomycetes(100);o__Saccharomycetales(100);f__Metschnikowiaceae(100);g__Metschnikowia(100);s__Metschnikowia_pulcherrima(100); | Metschnikowia | 13 | Saprotroph | \|Undefined Saprotroph\| | Yeast | NULL | Probable | As nectar-tap_saprotroph (Põlme et al. 2020); Decay sugar-rich substrates (Põlme et al. 2020) | Põlme S, et al. 2020. Fungal Diversity 105:1-16 ((https://doi.org/10.1007/s13225-020-00466-2)) |
| Otu52869 | 1 | k__Fungi(100);p__Ascomycota(100);c__Saccharomycetes(100);o__Saccharomycetales(100);f__Metschnikowiaceae(100);g__Metschnikowia(100);s__Metschnikowia_pulcherrima(100); | Metschnikowia | 13 | Saprotroph | \|Undefined Saprotroph\| | Yeast | NULL | Probable | As nectar-tap_saprotroph (Põlme et al. 2020); Decay sugar-rich substrates (Põlme et al. 2020) | Põlme S, et al. 2020. Fungal Diversity 105:1-16 ((https://doi.org/10.1007/s13225-020-00466-2)) |
| Otu52871 | 1 | k__Fungi(100);p__Ascomycota(100);c__Saccharomycetes(100);o__Saccharomycetales(100);f__Metschnikowiaceae(100);g__Metschnikowia(100);s__Metschnikowia_pulcherrima(100); | Metschnikowia | 13 | Saprotroph | \|Undefined Saprotroph\| | Yeast | NULL | Probable | As nectar-tap_saprotroph (Põlme et al. 2020); Decay sugar-rich substrates (Põlme et al. 2020) | Põlme S, et al. 2020. Fungal Diversity 105:1-16 ((https://doi.org/10.1007/s13225-020-00466-2)) |
| Otu52873 | 1 | k__Fungi(100);p__Ascomycota(100);c__Saccharomycetes(100);o__Saccharomycetales(100);f__Saccharomycetales_family_Incertae_sedis(100);g__Candida(100);g__Candida_unclassified(100); | Candida | 13 | Pathotroph-Saprotroph-Symbiotroph | Animal Pathogen-Endophyte-Endosymbiont-Epiphyte-\|Undefined Saprotroph\| | Dimorphic Yeast-Yeast | NULL | Probable | As nectar-tap_saprotroph (Põlme et al. 2020); Decay sugar-rich substrates (Põlme et al. 2020); This asexual genus is polyphyletic and contain species that function very differently depend on what they are. Caution should be taken when interpreting guilds for Candida species. | Manolakaki D, et al. 2010. Virulence. 1: 367-375 ((https://doi.org/10.4161/viru.1.5.12796)); Mekbib SB, et al. 2011. Tropical Plant Pathology 36:233-240 ((https://doi.org/10.1590/S1982-56762011000400004)); Põlme S, et al. 2020. Fungal Diversity 105:1-16 ((https://doi.org/10.1007/s13225-020-00466-2)) |
| Otu52875 | 1 | k__Fungi(100);p__Ascomycota(100);c__Saccharomycetes(100);o__Saccharomycetales(100);f__Saccharomycetales_family_Incertae_sedis(100);g__Candida(100);s__Candida_cretensis(100); | Candida | 13 | Pathotroph-Saprotroph-Symbiotroph | Animal Pathogen-Endophyte-Endosymbiont-Epiphyte-\|Undefined Saprotroph\| | Dimorphic Yeast-Yeast | NULL | Probable | As nectar-tap_saprotroph (Põlme et al. 2020); Decay sugar-rich substrates (Põlme et al. 2020); This asexual genus is polyphyletic and contain species that function very differently depend on what they are. Caution should be taken when interpreting guilds for Candida species. | Manolakaki D, et al. 2010. Virulence. 1: 367-375 ((https://doi.org/10.4161/viru.1.5.12796)); Mekbib SB, et al. 2011. Tropical Plant Pathology 36:233-240 ((https://doi.org/10.1590/S1982-56762011000400004)); Põlme S, et al. 2020. Fungal Diversity 105:1-16 ((https://doi.org/10.1007/s13225-020-00466-2)) |
| Otu52876 | 1 | k__Fungi(100);p__Ascomycota(100);c__Saccharomycetes(100);o__Saccharomycetales(100);f__Saccharomycetales_family_Incertae_sedis(100);g__Candida(100);g__Candida_unclassified(100); | Candida | 13 | Pathotroph-Saprotroph-Symbiotroph | Animal Pathogen-Endophyte-Endosymbiont-Epiphyte-\|Undefined Saprotroph\| | Dimorphic Yeast-Yeast | NULL | Probable | As nectar-tap_saprotroph (Põlme et al. 2020); Decay sugar-rich substrates (Põlme et al. 2020); This asexual genus is polyphyletic and contain species that function very differently depend on what they are. Caution should be taken when interpreting guilds for Candida species. | Manolakaki D, et al. 2010. Virulence. 1: 367-375 ((https://doi.org/10.4161/viru.1.5.12796)); Mekbib SB, et al. 2011. Tropical Plant Pathology 36:233-240 ((https://doi.org/10.1590/S1982-56762011000400004)); Põlme S, et al. 2020. Fungal Diversity 105:1-16 ((https://doi.org/10.1007/s13225-020-00466-2)) |
| Otu52878 | 1 | k__Fungi(100);p__Ascomycota(100);c__Saccharomycetes(100);o__Saccharomycetales(100);f__Metschnikowiaceae(100);g__Metschnikowia(100);s__Metschnikowia_pulcherrima(100); | Metschnikowia | 13 | Saprotroph | \|Undefined Saprotroph\| | Yeast | NULL | Probable | As nectar-tap_saprotroph (Põlme et al. 2020); Decay sugar-rich substrates (Põlme et al. 2020) | Põlme S, et al. 2020. Fungal Diversity 105:1-16 ((https://doi.org/10.1007/s13225-020-00466-2)) |
| Otu52881 | 1 | k__Fungi(100);p__Ascomycota(100);c__Saccharomycetes(100);o__Saccharomycetales(100);f__Metschnikowiaceae(100);g__Metschnikowia(100);g__Metschnikowia_unclassified(100); | Metschnikowia | 13 | Saprotroph | \|Undefined Saprotroph\| | Yeast | NULL | Probable | As nectar-tap_saprotroph (Põlme et al. 2020); Decay sugar-rich substrates (Põlme et al. 2020) | Põlme S, et al. 2020. Fungal Diversity 105:1-16 ((https://doi.org/10.1007/s13225-020-00466-2)) |
| Otu52882 | 1 | k__Fungi(100);p__Ascomycota(100);c__Saccharomycetes(100);o__Saccharomycetales(100);f__Metschnikowiaceae(100);g__Metschnikowia(100);g__Metschnikowia_unclassified(100); | Metschnikowia | 13 | Saprotroph | \|Undefined Saprotroph\| | Yeast | NULL | Probable | As nectar-tap_saprotroph (Põlme et al. 2020); Decay sugar-rich substrates (Põlme et al. 2020) | Põlme S, et al. 2020. Fungal Diversity 105:1-16 ((https://doi.org/10.1007/s13225-020-00466-2)) |
| Otu52883 | 1 | k__Fungi(100);p__Ascomycota(100);c__Saccharomycetes(100);o__Saccharomycetales(100);f__Metschnikowiaceae(100);g__Metschnikowia(100);g__Metschnikowia_unclassified(100); | Metschnikowia | 13 | Saprotroph | \|Undefined Saprotroph\| | Yeast | NULL | Probable | As nectar-tap_saprotroph (Põlme et al. 2020); Decay sugar-rich substrates (Põlme et al. 2020) | Põlme S, et al. 2020. Fungal Diversity 105:1-16 ((https://doi.org/10.1007/s13225-020-00466-2)) |
| Otu52884 | 1 | k__Fungi(100);p__Ascomycota(100);c__Saccharomycetes(100);o__Saccharomycetales(100);f__Metschnikowiaceae(100);g__Metschnikowia(100);g__Metschnikowia_unclassified(100); | Metschnikowia | 13 | Saprotroph | \|Undefined Saprotroph\| | Yeast | NULL | Probable | As nectar-tap_saprotroph (Põlme et al. 2020); Decay sugar-rich substrates (Põlme et al. 2020) | Põlme S, et al. 2020. Fungal Diversity 105:1-16 ((https://doi.org/10.1007/s13225-020-00466-2)) |
| Otu52885 | 1 | k__Fungi(100);p__Ascomycota(100);c__Saccharomycetes(100);o__Saccharomycetales(100);f__Metschnikowiaceae(100);g__Metschnikowia(100);s__Metschnikowia_pulcherrima(100); | Metschnikowia | 13 | Saprotroph | \|Undefined Saprotroph\| | Yeast | NULL | Probable | As nectar-tap_saprotroph (Põlme et al. 2020); Decay sugar-rich substrates (Põlme et al. 2020) | Põlme S, et al. 2020. Fungal Diversity 105:1-16 ((https://doi.org/10.1007/s13225-020-00466-2)) |
| Otu52886 | 1 | k__Fungi(100);p__Ascomycota(100);c__Saccharomycetes(100);o__Saccharomycetales(100);f__Metschnikowiaceae(100);g__Metschnikowia(100);g__Metschnikowia_unclassified(100); | Metschnikowia | 13 | Saprotroph | \|Undefined Saprotroph\| | Yeast | NULL | Probable | As nectar-tap_saprotroph (Põlme et al. 2020); Decay sugar-rich substrates (Põlme et al. 2020) | Põlme S, et al. 2020. Fungal Diversity 105:1-16 ((https://doi.org/10.1007/s13225-020-00466-2)) |
| Otu52887 | 1 | k__Fungi(100);p__Ascomycota(100);c__Saccharomycetes(100);o__Saccharomycetales(100);f__Metschnikowiaceae(100);g__Metschnikowia(100);s__Metschnikowia_pulcherrima(100); | Metschnikowia | 13 | Saprotroph | \|Undefined Saprotroph\| | Yeast | NULL | Probable | As nectar-tap_saprotroph (Põlme et al. 2020); Decay sugar-rich substrates (Põlme et al. 2020) | Põlme S, et al. 2020. Fungal Diversity 105:1-16 ((https://doi.org/10.1007/s13225-020-00466-2)) |
| Otu52888 | 1 | k__Fungi(100);p__Ascomycota(100);c__Saccharomycetes(100);o__Saccharomycetales(100);f__Metschnikowiaceae(100);g__Metschnikowia(100);s__Metschnikowia_pulcherrima(100); | Metschnikowia | 13 | Saprotroph | \|Undefined Saprotroph\| | Yeast | NULL | Probable | As nectar-tap_saprotroph (Põlme et al. 2020); Decay sugar-rich substrates (Põlme et al. 2020) | Põlme S, et al. 2020. Fungal Diversity 105:1-16 ((https://doi.org/10.1007/s13225-020-00466-2)) |
| Otu52889 | 1 | k__Fungi(100);p__Ascomycota(100);c__Saccharomycetes(100);o__Saccharomycetales(100);f__Saccharomycetales_family_Incertae_sedis(100);g__Candida(100);g__Candida_unclassified(100); | Candida | 13 | Pathotroph-Saprotroph-Symbiotroph | Animal Pathogen-Endophyte-Endosymbiont-Epiphyte-\|Undefined Saprotroph\| | Dimorphic Yeast-Yeast | NULL | Probable | As nectar-tap_saprotroph (Põlme et al. 2020); Decay sugar-rich substrates (Põlme et al. 2020); This asexual genus is polyphyletic and contain species that function very differently depend on what they are. Caution should be taken when interpreting guilds for Candida species. | Manolakaki D, et al. 2010. Virulence. 1: 367-375 ((https://doi.org/10.4161/viru.1.5.12796)); Mekbib SB, et al. 2011. Tropical Plant Pathology 36:233-240 ((https://doi.org/10.1590/S1982-56762011000400004)); Põlme S, et al. 2020. Fungal Diversity 105:1-16 ((https://doi.org/10.1007/s13225-020-00466-2)) |
| Otu52890 | 1 | k__Fungi(100);p__Ascomycota(100);c__Saccharomycetes(100);o__Saccharomycetales(100);f__Metschnikowiaceae(100);g__Metschnikowia(100);s__Metschnikowia_pulcherrima(100); | Metschnikowia | 13 | Saprotroph | \|Undefined Saprotroph\| | Yeast | NULL | Probable | As nectar-tap_saprotroph (Põlme et al. 2020); Decay sugar-rich substrates (Põlme et al. 2020) | Põlme S, et al. 2020. Fungal Diversity 105:1-16 ((https://doi.org/10.1007/s13225-020-00466-2)) |
| Otu52891 | 1 | k__Fungi(100);p__Ascomycota(100);c__Saccharomycetes(100);o__Saccharomycetales(100);f__Metschnikowiaceae(100);g__Metschnikowia(100);s__Metschnikowia_pulcherrima(100); | Metschnikowia | 13 | Saprotroph | \|Undefined Saprotroph\| | Yeast | NULL | Probable | As nectar-tap_saprotroph (Põlme et al. 2020); Decay sugar-rich substrates (Põlme et al. 2020) | Põlme S, et al. 2020. Fungal Diversity 105:1-16 ((https://doi.org/10.1007/s13225-020-00466-2)) |
| Otu52892 | 1 | k__Fungi(100);p__Ascomycota(100);c__Saccharomycetes(100);o__Saccharomycetales(100);f__Metschnikowiaceae(100);g__Metschnikowia(100);s__Metschnikowia_pulcherrima(100); | Metschnikowia | 13 | Saprotroph | \|Undefined Saprotroph\| | Yeast | NULL | Probable | As nectar-tap_saprotroph (Põlme et al. 2020); Decay sugar-rich substrates (Põlme et al. 2020) | Põlme S, et al. 2020. Fungal Diversity 105:1-16 ((https://doi.org/10.1007/s13225-020-00466-2)) |
| Otu52893 | 1 | k__Fungi(100);p__Ascomycota(100);c__Saccharomycetes(100);o__Saccharomycetales(100);f__Metschnikowiaceae(100);g__Metschnikowia(100);s__Metschnikowia_pulcherrima(100); | Metschnikowia | 13 | Saprotroph | \|Undefined Saprotroph\| | Yeast | NULL | Probable | As nectar-tap_saprotroph (Põlme et al. 2020); Decay sugar-rich substrates (Põlme et al. 2020) | Põlme S, et al. 2020. Fungal Diversity 105:1-16 ((https://doi.org/10.1007/s13225-020-00466-2)) |
| Otu52894 | 1 | k__Fungi(100);p__Ascomycota(100);c__Saccharomycetes(100);o__Saccharomycetales(100);f__Metschnikowiaceae(100);g__Metschnikowia(100);g__Metschnikowia_unclassified(100); | Metschnikowia | 13 | Saprotroph | \|Undefined Saprotroph\| | Yeast | NULL | Probable | As nectar-tap_saprotroph (Põlme et al. 2020); Decay sugar-rich substrates (Põlme et al. 2020) | Põlme S, et al. 2020. Fungal Diversity 105:1-16 ((https://doi.org/10.1007/s13225-020-00466-2)) |
| Otu52896 | 1 | k__Fungi(100);p__Ascomycota(100);c__Saccharomycetes(100);o__Saccharomycetales(100);o__Saccharomycetales_unclassified(100);o__Saccharomycetales_unclassified(100);o__Saccharomycetales_unclassified(100); | Saccharomycetales | 7 | Saprotroph | Undefined Saprotroph | Yeast | NULL | Possible | NULL | Sterkenburg E, et al. 2015. New Phytologist 207:1145-1158 ((https://doi.org/10.1111/nph.13426)) |
| Otu52898 | 1 | k__Fungi(100);p__Ascomycota(100);c__Saccharomycetes(100);o__Saccharomycetales(100);f__Metschnikowiaceae(100);g__Metschnikowia(100);s__Metschnikowia_pulcherrima(100); | Metschnikowia | 13 | Saprotroph | \|Undefined Saprotroph\| | Yeast | NULL | Probable | As nectar-tap_saprotroph (Põlme et al. 2020); Decay sugar-rich substrates (Põlme et al. 2020) | Põlme S, et al. 2020. Fungal Diversity 105:1-16 ((https://doi.org/10.1007/s13225-020-00466-2)) |
| Otu52899 | 1 | k__Fungi(100);p__Ascomycota(100);c__Saccharomycetes(100);o__Saccharomycetales(100);f__Metschnikowiaceae(100);g__Metschnikowia(100);s__Metschnikowia_pulcherrima(100); | Metschnikowia | 13 | Saprotroph | \|Undefined Saprotroph\| | Yeast | NULL | Probable | As nectar-tap_saprotroph (Põlme et al. 2020); Decay sugar-rich substrates (Põlme et al. 2020) | Põlme S, et al. 2020. Fungal Diversity 105:1-16 ((https://doi.org/10.1007/s13225-020-00466-2)) |
| Otu52900 | 1 | k__Fungi(100);p__Ascomycota(100);c__Saccharomycetes(100);o__Saccharomycetales(100);f__Metschnikowiaceae(100);g__Metschnikowia(100);s__Metschnikowia_pulcherrima(100); | Metschnikowia | 13 | Saprotroph | \|Undefined Saprotroph\| | Yeast | NULL | Probable | As nectar-tap_saprotroph (Põlme et al. 2020); Decay sugar-rich substrates (Põlme et al. 2020) | Põlme S, et al. 2020. Fungal Diversity 105:1-16 ((https://doi.org/10.1007/s13225-020-00466-2)) |
| Otu52901 | 1 | k__Fungi(100);p__Ascomycota(100);c__Saccharomycetes(100);o__Saccharomycetales(100);f__Metschnikowiaceae(100);g__Metschnikowia(100);g__Metschnikowia_unclassified(100); | Metschnikowia | 13 | Saprotroph | \|Undefined Saprotroph\| | Yeast | NULL | Probable | As nectar-tap_saprotroph (Põlme et al. 2020); Decay sugar-rich substrates (Põlme et al. 2020) | Põlme S, et al. 2020. Fungal Diversity 105:1-16 ((https://doi.org/10.1007/s13225-020-00466-2)) |
| Otu52902 | 1 | k__Fungi(100);p__Ascomycota(100);c__Saccharomycetes(100);o__Saccharomycetales(100);f__Metschnikowiaceae(100);g__Metschnikowia(100);s__Metschnikowia_pulcherrima(100); | Metschnikowia | 13 | Saprotroph | \|Undefined Saprotroph\| | Yeast | NULL | Probable | As nectar-tap_saprotroph (Põlme et al. 2020); Decay sugar-rich substrates (Põlme et al. 2020) | Põlme S, et al. 2020. Fungal Diversity 105:1-16 ((https://doi.org/10.1007/s13225-020-00466-2)) |
| Otu52903 | 1 | k__Fungi(100);p__Ascomycota(100);c__Saccharomycetes(100);o__Saccharomycetales(100);f__Metschnikowiaceae(100);g__Metschnikowia(100);s__Metschnikowia_pulcherrima(100); | Metschnikowia | 13 | Saprotroph | \|Undefined Saprotroph\| | Yeast | NULL | Probable | As nectar-tap_saprotroph (Põlme et al. 2020); Decay sugar-rich substrates (Põlme et al. 2020) | Põlme S, et al. 2020. Fungal Diversity 105:1-16 ((https://doi.org/10.1007/s13225-020-00466-2)) |
| Otu52904 | 1 | k__Fungi(100);p__Ascomycota(100);c__Saccharomycetes(100);o__Saccharomycetales(100);f__Metschnikowiaceae(100);g__Metschnikowia(100);s__Metschnikowia_pulcherrima(100); | Metschnikowia | 13 | Saprotroph | \|Undefined Saprotroph\| | Yeast | NULL | Probable | As nectar-tap_saprotroph (Põlme et al. 2020); Decay sugar-rich substrates (Põlme et al. 2020) | Põlme S, et al. 2020. Fungal Diversity 105:1-16 ((https://doi.org/10.1007/s13225-020-00466-2)) |
| Otu52905 | 1 | k__Fungi(100);p__Ascomycota(100);c__Saccharomycetes(100);o__Saccharomycetales(100);f__Metschnikowiaceae(100);g__Metschnikowia(100);s__Metschnikowia_pulcherrima(100); | Metschnikowia | 13 | Saprotroph | \|Undefined Saprotroph\| | Yeast | NULL | Probable | As nectar-tap_saprotroph (Põlme et al. 2020); Decay sugar-rich substrates (Põlme et al. 2020) | Põlme S, et al. 2020. Fungal Diversity 105:1-16 ((https://doi.org/10.1007/s13225-020-00466-2)) |
| Otu52906 | 1 | k__Fungi(100);p__Ascomycota(100);c__Saccharomycetes(100);o__Saccharomycetales(100);f__Metschnikowiaceae(100);g__Metschnikowia(100);g__Metschnikowia_unclassified(100); | Metschnikowia | 13 | Saprotroph | \|Undefined Saprotroph\| | Yeast | NULL | Probable | As nectar-tap_saprotroph (Põlme et al. 2020); Decay sugar-rich substrates (Põlme et al. 2020) | Põlme S, et al. 2020. Fungal Diversity 105:1-16 ((https://doi.org/10.1007/s13225-020-00466-2)) |
| Otu52907 | 1 | k__Fungi(100);p__Ascomycota(100);c__Saccharomycetes(100);o__Saccharomycetales(100);f__Metschnikowiaceae(100);g__Metschnikowia(100);s__Metschnikowia_pulcherrima(100); | Metschnikowia | 13 | Saprotroph | \|Undefined Saprotroph\| | Yeast | NULL | Probable | As nectar-tap_saprotroph (Põlme et al. 2020); Decay sugar-rich substrates (Põlme et al. 2020) | Põlme S, et al. 2020. Fungal Diversity 105:1-16 ((https://doi.org/10.1007/s13225-020-00466-2)) |
| Otu52908 | 1 | k__Fungi(100);p__Ascomycota(100);c__Saccharomycetes(100);o__Saccharomycetales(100);f__Metschnikowiaceae(100);g__Metschnikowia(100);g__Metschnikowia_unclassified(100); | Metschnikowia | 13 | Saprotroph | \|Undefined Saprotroph\| | Yeast | NULL | Probable | As nectar-tap_saprotroph (Põlme et al. 2020); Decay sugar-rich substrates (Põlme et al. 2020) | Põlme S, et al. 2020. Fungal Diversity 105:1-16 ((https://doi.org/10.1007/s13225-020-00466-2)) |
| Otu52909 | 1 | k__Fungi(100);p__Ascomycota(100);c__Saccharomycetes(100);o__Saccharomycetales(100);f__Metschnikowiaceae(100);g__Metschnikowia(100);s__Metschnikowia_pulcherrima(100); | Metschnikowia | 13 | Saprotroph | \|Undefined Saprotroph\| | Yeast | NULL | Probable | As nectar-tap_saprotroph (Põlme et al. 2020); Decay sugar-rich substrates (Põlme et al. 2020) | Põlme S, et al. 2020. Fungal Diversity 105:1-16 ((https://doi.org/10.1007/s13225-020-00466-2)) |
| Otu52910 | 1 | k__Fungi(100);p__Ascomycota(100);c__Saccharomycetes(100);o__Saccharomycetales(100);f__Metschnikowiaceae(100);g__Metschnikowia(100);g__Metschnikowia_unclassified(100); | Metschnikowia | 13 | Saprotroph | \|Undefined Saprotroph\| | Yeast | NULL | Probable | As nectar-tap_saprotroph (Põlme et al. 2020); Decay sugar-rich substrates (Põlme et al. 2020) | Põlme S, et al. 2020. Fungal Diversity 105:1-16 ((https://doi.org/10.1007/s13225-020-00466-2)) |
| Otu52911 | 1 | k__Fungi(100);p__Ascomycota(100);c__Saccharomycetes(100);o__Saccharomycetales(100);f__Metschnikowiaceae(100);g__Metschnikowia(100);s__Metschnikowia_pulcherrima(100); | Metschnikowia | 13 | Saprotroph | \|Undefined Saprotroph\| | Yeast | NULL | Probable | As nectar-tap_saprotroph (Põlme et al. 2020); Decay sugar-rich substrates (Põlme et al. 2020) | Põlme S, et al. 2020. Fungal Diversity 105:1-16 ((https://doi.org/10.1007/s13225-020-00466-2)) |
| Otu52912 | 1 | k__Fungi(100);p__Ascomycota(100);c__Saccharomycetes(100);o__Saccharomycetales(100);f__Metschnikowiaceae(100);g__Metschnikowia(100);s__Metschnikowia_pulcherrima(100); | Metschnikowia | 13 | Saprotroph | \|Undefined Saprotroph\| | Yeast | NULL | Probable | As nectar-tap_saprotroph (Põlme et al. 2020); Decay sugar-rich substrates (Põlme et al. 2020) | Põlme S, et al. 2020. Fungal Diversity 105:1-16 ((https://doi.org/10.1007/s13225-020-00466-2)) |
| Otu52913 | 1 | k__Fungi(100);p__Ascomycota(100);c__Saccharomycetes(100);o__Saccharomycetales(100);f__Metschnikowiaceae(100);g__Metschnikowia(100);s__Metschnikowia_pulcherrima(100); | Metschnikowia | 13 | Saprotroph | \|Undefined Saprotroph\| | Yeast | NULL | Probable | As nectar-tap_saprotroph (Põlme et al. 2020); Decay sugar-rich substrates (Põlme et al. 2020) | Põlme S, et al. 2020. Fungal Diversity 105:1-16 ((https://doi.org/10.1007/s13225-020-00466-2)) |
| Otu52914 | 1 | k__Fungi(100);p__Ascomycota(100);c__Saccharomycetes(100);o__Saccharomycetales(100);f__Metschnikowiaceae(100);g__Metschnikowia(100);g__Metschnikowia_unclassified(100); | Metschnikowia | 13 | Saprotroph | \|Undefined Saprotroph\| | Yeast | NULL | Probable | As nectar-tap_saprotroph (Põlme et al. 2020); Decay sugar-rich substrates (Põlme et al. 2020) | Põlme S, et al. 2020. Fungal Diversity 105:1-16 ((https://doi.org/10.1007/s13225-020-00466-2)) |
| Otu52915 | 1 | k__Fungi(100);p__Ascomycota(100);c__Saccharomycetes(100);o__Saccharomycetales(100);f__Metschnikowiaceae(100);g__Metschnikowia(100);s__Metschnikowia_pulcherrima(100); | Metschnikowia | 13 | Saprotroph | \|Undefined Saprotroph\| | Yeast | NULL | Probable | As nectar-tap_saprotroph (Põlme et al. 2020); Decay sugar-rich substrates (Põlme et al. 2020) | Põlme S, et al. 2020. Fungal Diversity 105:1-16 ((https://doi.org/10.1007/s13225-020-00466-2)) |
| Otu52916 | 1 | k__Fungi(100);p__Ascomycota(100);c__Saccharomycetes(100);o__Saccharomycetales(100);f__Metschnikowiaceae(100);g__Metschnikowia(100);s__Metschnikowia_pulcherrima(100); | Metschnikowia | 13 | Saprotroph | \|Undefined Saprotroph\| | Yeast | NULL | Probable | As nectar-tap_saprotroph (Põlme et al. 2020); Decay sugar-rich substrates (Põlme et al. 2020) | Põlme S, et al. 2020. Fungal Diversity 105:1-16 ((https://doi.org/10.1007/s13225-020-00466-2)) |
| Otu52917 | 1 | k__Fungi(100);p__Ascomycota(100);c__Saccharomycetes(100);o__Saccharomycetales(100);f__Metschnikowiaceae(100);g__Metschnikowia(100);s__Metschnikowia_pulcherrima(100); | Metschnikowia | 13 | Saprotroph | \|Undefined Saprotroph\| | Yeast | NULL | Probable | As nectar-tap_saprotroph (Põlme et al. 2020); Decay sugar-rich substrates (Põlme et al. 2020) | Põlme S, et al. 2020. Fungal Diversity 105:1-16 ((https://doi.org/10.1007/s13225-020-00466-2)) |
| Otu52918 | 1 | k__Fungi(100);p__Ascomycota(100);c__Saccharomycetes(100);o__Saccharomycetales(100);f__Metschnikowiaceae(100);g__Metschnikowia(100);g__Metschnikowia_unclassified(100); | Metschnikowia | 13 | Saprotroph | \|Undefined Saprotroph\| | Yeast | NULL | Probable | As nectar-tap_saprotroph (Põlme et al. 2020); Decay sugar-rich substrates (Põlme et al. 2020) | Põlme S, et al. 2020. Fungal Diversity 105:1-16 ((https://doi.org/10.1007/s13225-020-00466-2)) |
| Otu52919 | 1 | k__Fungi(100);p__Ascomycota(100);c__Saccharomycetes(100);o__Saccharomycetales(100);f__Metschnikowiaceae(100);g__Metschnikowia(100);s__Metschnikowia_pulcherrima(100); | Metschnikowia | 13 | Saprotroph | \|Undefined Saprotroph\| | Yeast | NULL | Probable | As nectar-tap_saprotroph (Põlme et al. 2020); Decay sugar-rich substrates (Põlme et al. 2020) | Põlme S, et al. 2020. Fungal Diversity 105:1-16 ((https://doi.org/10.1007/s13225-020-00466-2)) |
| Otu52920 | 1 | k__Fungi(100);p__Ascomycota(100);c__Saccharomycetes(100);o__Saccharomycetales(100);f__Metschnikowiaceae(100);g__Metschnikowia(100);s__Metschnikowia_pulcherrima(100); | Metschnikowia | 13 | Saprotroph | \|Undefined Saprotroph\| | Yeast | NULL | Probable | As nectar-tap_saprotroph (Põlme et al. 2020); Decay sugar-rich substrates (Põlme et al. 2020) | Põlme S, et al. 2020. Fungal Diversity 105:1-16 ((https://doi.org/10.1007/s13225-020-00466-2)) |
| Otu52921 | 1 | k__Fungi(100);p__Ascomycota(100);c__Saccharomycetes(100);o__Saccharomycetales(100);f__Metschnikowiaceae(100);g__Metschnikowia(100);g__Metschnikowia_unclassified(100); | Metschnikowia | 13 | Saprotroph | \|Undefined Saprotroph\| | Yeast | NULL | Probable | As nectar-tap_saprotroph (Põlme et al. 2020); Decay sugar-rich substrates (Põlme et al. 2020) | Põlme S, et al. 2020. Fungal Diversity 105:1-16 ((https://doi.org/10.1007/s13225-020-00466-2)) |
| Otu52922 | 1 | k__Fungi(100);p__Ascomycota(100);c__Saccharomycetes(100);o__Saccharomycetales(100);f__Metschnikowiaceae(100);g__Metschnikowia(100);g__Metschnikowia_unclassified(100); | Metschnikowia | 13 | Saprotroph | \|Undefined Saprotroph\| | Yeast | NULL | Probable | As nectar-tap_saprotroph (Põlme et al. 2020); Decay sugar-rich substrates (Põlme et al. 2020) | Põlme S, et al. 2020. Fungal Diversity 105:1-16 ((https://doi.org/10.1007/s13225-020-00466-2)) |
| Otu52923 | 1 | k__Fungi(100);p__Ascomycota(100);c__Saccharomycetes(100);o__Saccharomycetales(100);f__Metschnikowiaceae(100);g__Metschnikowia(100);g__Metschnikowia_unclassified(100); | Metschnikowia | 13 | Saprotroph | \|Undefined Saprotroph\| | Yeast | NULL | Probable | As nectar-tap_saprotroph (Põlme et al. 2020); Decay sugar-rich substrates (Põlme et al. 2020) | Põlme S, et al. 2020. Fungal Diversity 105:1-16 ((https://doi.org/10.1007/s13225-020-00466-2)) |
| Otu52924 | 1 | k__Fungi(100);p__Ascomycota(100);c__Saccharomycetes(100);o__Saccharomycetales(100);f__Metschnikowiaceae(100);g__Metschnikowia(100);s__Metschnikowia_pulcherrima(100); | Metschnikowia | 13 | Saprotroph | \|Undefined Saprotroph\| | Yeast | NULL | Probable | As nectar-tap_saprotroph (Põlme et al. 2020); Decay sugar-rich substrates (Põlme et al. 2020) | Põlme S, et al. 2020. Fungal Diversity 105:1-16 ((https://doi.org/10.1007/s13225-020-00466-2)) |
| Otu52925 | 1 | k__Fungi(100);p__Ascomycota(100);c__Saccharomycetes(100);o__Saccharomycetales(100);f__Metschnikowiaceae(100);g__Metschnikowia(100);s__Metschnikowia_pulcherrima(100); | Metschnikowia | 13 | Saprotroph | \|Undefined Saprotroph\| | Yeast | NULL | Probable | As nectar-tap_saprotroph (Põlme et al. 2020); Decay sugar-rich substrates (Põlme et al. 2020) | Põlme S, et al. 2020. Fungal Diversity 105:1-16 ((https://doi.org/10.1007/s13225-020-00466-2)) |
| Otu52926 | 1 | k__Fungi(100);p__Ascomycota(100);c__Saccharomycetes(100);o__Saccharomycetales(100);f__Metschnikowiaceae(100);g__Metschnikowia(100);s__Metschnikowia_pulcherrima(100); | Metschnikowia | 13 | Saprotroph | \|Undefined Saprotroph\| | Yeast | NULL | Probable | As nectar-tap_saprotroph (Põlme et al. 2020); Decay sugar-rich substrates (Põlme et al. 2020) | Põlme S, et al. 2020. Fungal Diversity 105:1-16 ((https://doi.org/10.1007/s13225-020-00466-2)) |
| Otu52927 | 1 | k__Fungi(100);p__Ascomycota(100);c__Saccharomycetes(100);o__Saccharomycetales(100);f__Metschnikowiaceae(100);g__Metschnikowia(100);s__Metschnikowia_pulcherrima(100); | Metschnikowia | 13 | Saprotroph | \|Undefined Saprotroph\| | Yeast | NULL | Probable | As nectar-tap_saprotroph (Põlme et al. 2020); Decay sugar-rich substrates (Põlme et al. 2020) | Põlme S, et al. 2020. Fungal Diversity 105:1-16 ((https://doi.org/10.1007/s13225-020-00466-2)) |
| Otu52928 | 1 | k__Fungi(100);p__Ascomycota(100);c__Saccharomycetes(100);o__Saccharomycetales(100);f__Metschnikowiaceae(100);g__Metschnikowia(100);s__Metschnikowia_pulcherrima(100); | Metschnikowia | 13 | Saprotroph | \|Undefined Saprotroph\| | Yeast | NULL | Probable | As nectar-tap_saprotroph (Põlme et al. 2020); Decay sugar-rich substrates (Põlme et al. 2020) | Põlme S, et al. 2020. Fungal Diversity 105:1-16 ((https://doi.org/10.1007/s13225-020-00466-2)) |
| Otu52929 | 1 | k__Fungi(100);p__Ascomycota(100);c__Saccharomycetes(100);o__Saccharomycetales(100);f__Metschnikowiaceae(100);g__Metschnikowia(100);s__Metschnikowia_pulcherrima(100); | Metschnikowia | 13 | Saprotroph | \|Undefined Saprotroph\| | Yeast | NULL | Probable | As nectar-tap_saprotroph (Põlme et al. 2020); Decay sugar-rich substrates (Põlme et al. 2020) | Põlme S, et al. 2020. Fungal Diversity 105:1-16 ((https://doi.org/10.1007/s13225-020-00466-2)) |

| | | | | | | | | | | | |
|---|---|---|---|---|---|---|---|---|---|---|---|
| Otu52930 | 1 | k__Fungi(100);p__Ascomycota(100);c__Saccharomycetes(100);o__Saccharomycetales(100);f__Metschnikowiaceae(100);g__Metschnikowia(100);s__Metschnikowia_pulcherrima(100); | Metschnikowia | 13 | Saprotroph | \|Undefined Saprotroph\| | Yeast | NULL | Probable | As nectar-tap_saprotroph (Põlme et al. 2020); Decay sugar-rich substrates (Põlme et al. 2020) | Põlme S, et al. 2020. Fungal Diversity 105:1-16 ((https://doi.org/10.1007/s13225-020-00466-2)) |
| Otu52931 | 1 | k__Fungi(100);p__Ascomycota(100);c__Saccharomycetes(100);o__Saccharomycetales(100);f__Metschnikowiaceae(100);g__Metschnikowia(100);s__Metschnikowia_pulcherrima(100); | Metschnikowia | 13 | Saprotroph | \|Undefined Saprotroph\| | Yeast | NULL | Probable | As nectar-tap_saprotroph (Põlme et al. 2020); Decay sugar-rich substrates (Põlme et al. 2020) | Põlme S, et al. 2020. Fungal Diversity 105:1-16 ((https://doi.org/10.1007/s13225-020-00466-2)) |
| Otu52932 | 1 | k__Fungi(100);p__Ascomycota(100);c__Saccharomycetes(100);o__Saccharomycetales(100);f__Metschnikowiaceae(100);g__Metschnikowia(100);s__Metschnikowia_pulcherrima(100); | Metschnikowia | 13 | Saprotroph | \|Undefined Saprotroph\| | Yeast | NULL | Probable | As nectar-tap_saprotroph (Põlme et al. 2020); Decay sugar-rich substrates (Põlme et al. 2020) | Põlme S, et al. 2020. Fungal Diversity 105:1-16 ((https://doi.org/10.1007/s13225-020-00466-2)) |
| Otu52933 | 1 | k__Fungi(100);p__Ascomycota(100);c__Saccharomycetes(100);o__Saccharomycetales(100);f__Metschnikowiaceae(100);g__Metschnikowia(100);s__Metschnikowia_pulcherrima(100); | Metschnikowia | 13 | Saprotroph | \|Undefined Saprotroph\| | Yeast | NULL | Probable | As nectar-tap_saprotroph (Põlme et al. 2020); Decay sugar-rich substrates (Põlme et al. 2020) | Põlme S, et al. 2020. Fungal Diversity 105:1-16 ((https://doi.org/10.1007/s13225-020-00466-2)) |
| Otu52934 | 1 | k__Fungi(100);p__Ascomycota(100);c__Saccharomycetes(100);o__Saccharomycetales(100);f__Metschnikowiaceae(100);g__Metschnikowia(100);g__Metschnikowia_unclassified(100); | Metschnikowia | 13 | Saprotroph | \|Undefined Saprotroph\| | Yeast | NULL | Probable | As nectar-tap_saprotroph (Põlme et al. 2020); Decay sugar-rich substrates (Põlme et al. 2020) | Põlme S, et al. 2020. Fungal Diversity 105:1-16 ((https://doi.org/10.1007/s13225-020-00466-2)) |
| Otu52935 | 1 | k__Fungi(100);p__Ascomycota(100);c__Saccharomycetes(100);o__Saccharomycetales(100);f__Metschnikowiaceae(100);g__Metschnikowia(100);s__Metschnikowia_pulcherrima(100); | Metschnikowia | 13 | Saprotroph | \|Undefined Saprotroph\| | Yeast | NULL | Probable | As nectar-tap_saprotroph (Põlme et al. 2020); Decay sugar-rich substrates (Põlme et al. 2020) | Põlme S, et al. 2020. Fungal Diversity 105:1-16 ((https://doi.org/10.1007/s13225-020-00466-2)) |
| Otu52936 | 1 | k__Fungi(100);p__Ascomycota(100);c__Saccharomycetes(100);o__Saccharomycetales(100);f__Metschnikowiaceae(100);g__Metschnikowia(100);s__Metschnikowia_pulcherrima(100); | Metschnikowia | 13 | Saprotroph | \|Undefined Saprotroph\| | Yeast | NULL | Probable | As nectar-tap_saprotroph (Põlme et al. 2020); Decay sugar-rich substrates (Põlme et al. 2020) | Põlme S, et al. 2020. Fungal Diversity 105:1-16 ((https://doi.org/10.1007/s13225-020-00466-2)) |
| Otu52937 | 1 | k__Fungi(100);p__Ascomycota(100);c__Saccharomycetes(100);o__Saccharomycetales(100);f__Metschnikowiaceae(100);g__Metschnikowia(100);s__Metschnikowia_pulcherrima(100); | Metschnikowia | 13 | Saprotroph | \|Undefined Saprotroph\| | Yeast | NULL | Probable | As nectar-tap_saprotroph (Põlme et al. 2020); Decay sugar-rich substrates (Põlme et al. 2020) | Põlme S, et al. 2020. Fungal Diversity 105:1-16 ((https://doi.org/10.1007/s13225-020-00466-2)) |
| Otu52938 | 1 | k__Fungi(100);p__Ascomycota(100);c__Saccharomycetes(100);o__Saccharomycetales(100);f__Metschnikowiaceae(100);g__Metschnikowia(100);s__Metschnikowia_pulcherrima(100); | Metschnikowia | 13 | Saprotroph | \|Undefined Saprotroph\| | Yeast | NULL | Probable | As nectar-tap_saprotroph (Põlme et al. 2020); Decay sugar-rich substrates (Põlme et al. 2020) | Põlme S, et al. 2020. Fungal Diversity 105:1-16 ((https://doi.org/10.1007/s13225-020-00466-2)) |
| Otu52939 | 1 | k__Fungi(100);p__Ascomycota(100);c__Saccharomycetes(100);o__Saccharomycetales(100);f__Metschnikowiaceae(100);g__Metschnikowia(100);s__Metschnikowia_pulcherrima(100); | Metschnikowia | 13 | Saprotroph | \|Undefined Saprotroph\| | Yeast | NULL | Probable | As nectar-tap_saprotroph (Põlme et al. 2020); Decay sugar-rich substrates (Põlme et al. 2020) | Põlme S, et al. 2020. Fungal Diversity 105:1-16 ((https://doi.org/10.1007/s13225-020-00466-2)) |
| Otu52940 | 1 | k__Fungi(100);p__Ascomycota(100);c__Saccharomycetes(100);o__Saccharomycetales(100);f__Metschnikowiaceae(100);g__Metschnikowia(100);g__Metschnikowia_unclassified(100); | Metschnikowia | 13 | Saprotroph | \|Undefined Saprotroph\| | Yeast | NULL | Probable | As nectar-tap_saprotroph (Põlme et al. 2020); Decay sugar-rich substrates (Põlme et al. 2020) | Põlme S, et al. 2020. Fungal Diversity 105:1-16 ((https://doi.org/10.1007/s13225-020-00466-2)) |
| Otu52941 | 1 | k__Fungi(100);p__Ascomycota(100);c__Saccharomycetes(100);o__Saccharomycetales(100);f__Metschnikowiaceae(100);g__Metschnikowia(100);s__Metschnikowia_pulcherrima(100); | Metschnikowia | 13 | Saprotroph | \|Undefined Saprotroph\| | Yeast | NULL | Probable | As nectar-tap_saprotroph (Põlme et al. 2020); Decay sugar-rich substrates (Põlme et al. 2020) | Põlme S, et al. 2020. Fungal Diversity 105:1-16 ((https://doi.org/10.1007/s13225-020-00466-2)) |
| Otu52942 | 1 | k__Fungi(100);p__Ascomycota(100);c__Saccharomycetes(100);o__Saccharomycetales(100);f__Metschnikowiaceae(100);g__Metschnikowia(100);g__Metschnikowia_unclassified(100); | Metschnikowia | 13 | Saprotroph | \|Undefined Saprotroph\| | Yeast | NULL | Probable | As nectar-tap_saprotroph (Põlme et al. 2020); Decay sugar-rich substrates (Põlme et al. 2020) | Põlme S, et al. 2020. Fungal Diversity 105:1-16 ((https://doi.org/10.1007/s13225-020-00466-2)) |
| Otu52943 | 1 | k__Fungi(100);p__Ascomycota(100);c__Saccharomycetes(100);o__Saccharomycetales(100);f__Metschnikowiaceae(100);g__Metschnikowia(100);s__Metschnikowia_pulcherrima(100); | Metschnikowia | 13 | Saprotroph | \|Undefined Saprotroph\| | Yeast | NULL | Probable | As nectar-tap_saprotroph (Põlme et al. 2020); Decay sugar-rich substrates (Põlme et al. 2020) | Põlme S, et al. 2020. Fungal Diversity 105:1-16 ((https://doi.org/10.1007/s13225-020-00466-2)) |
| Otu52944 | 1 | k__Fungi(100);p__Ascomycota(100);c__Saccharomycetes(100);o__Saccharomycetales(100);f__Metschnikowiaceae(100);g__Metschnikowia(100);s__Metschnikowia_pulcherrima(100); | Metschnikowia | 13 | Saprotroph | \|Undefined Saprotroph\| | Yeast | NULL | Probable | As nectar-tap_saprotroph (Põlme et al. 2020); Decay sugar-rich substrates (Põlme et al. 2020) | Põlme S, et al. 2020. Fungal Diversity 105:1-16 ((https://doi.org/10.1007/s13225-020-00466-2)) |
| Otu52945 | 1 | k__Fungi(100);p__Ascomycota(100);c__Saccharomycetes(100);o__Saccharomycetales(100);f__Metschnikowiaceae(100);g__Metschnikowia(100);g__Metschnikowia_unclassified(100); | Metschnikowia | 13 | Saprotroph | \|Undefined Saprotroph\| | Yeast | NULL | Probable | As nectar-tap_saprotroph (Põlme et al. 2020); Decay sugar-rich substrates (Põlme et al. 2020) | Põlme S, et al. 2020. Fungal Diversity 105:1-16 ((https://doi.org/10.1007/s13225-020-00466-2)) |
| Otu52946 | 1 | k__Fungi(100);p__Ascomycota(100);c__Saccharomycetes(100);o__Saccharomycetales(100);f__Metschnikowiaceae(100);g__Metschnikowia(100);g__Metschnikowia_unclassified(100); | Metschnikowia | 13 | Saprotroph | \|Undefined Saprotroph\| | Yeast | NULL | Probable | As nectar-tap_saprotroph (Põlme et al. 2020); Decay sugar-rich substrates (Põlme et al. 2020) | Põlme S, et al. 2020. Fungal Diversity 105:1-16 ((https://doi.org/10.1007/s13225-020-00466-2)) |
| Otu52947 | 1 | k__Fungi(100);p__Ascomycota(100);c__Saccharomycetes(100);o__Saccharomycetales(100);f__Metschnikowiaceae(100);g__Metschnikowia(100);s__Metschnikowia_pulcherrima(100); | Metschnikowia | 13 | Saprotroph | \|Undefined Saprotroph\| | Yeast | NULL | Probable | As nectar-tap_saprotroph (Põlme et al. 2020); Decay sugar-rich substrates (Põlme et al. 2020) | Põlme S, et al. 2020. Fungal Diversity 105:1-16 ((https://doi.org/10.1007/s13225-020-00466-2)) |
| Otu52948 | 1 | k__Fungi(100);p__Ascomycota(100);c__Saccharomycetes(100);o__Saccharomycetales(100);f__Metschnikowiaceae(100);g__Metschnikowia(100);s__Metschnikowia_pulcherrima(100); | Metschnikowia | 13 | Saprotroph | \|Undefined Saprotroph\| | Yeast | NULL | Probable | As nectar-tap_saprotroph (Põlme et al. 2020); Decay sugar-rich substrates (Põlme et al. 2020) | Põlme S, et al. 2020. Fungal Diversity 105:1-16 ((https://doi.org/10.1007/s13225-020-00466-2)) |
| Otu52949 | 1 | k__Fungi(100);p__Ascomycota(100);c__Saccharomycetes(100);o__Saccharomycetales(100);f__Metschnikowiaceae(100);g__Metschnikowia(100);g__Metschnikowia_unclassified(100); | Metschnikowia | 13 | Saprotroph | \|Undefined Saprotroph\| | Yeast | NULL | Probable | As nectar-tap_saprotroph (Põlme et al. 2020); Decay sugar-rich substrates (Põlme et al. 2020) | Põlme S, et al. 2020. Fungal Diversity 105:1-16 ((https://doi.org/10.1007/s13225-020-00466-2)) |
| Otu52950 | 1 | k__Fungi(100);p__Ascomycota(100);c__Saccharomycetes(100);o__Saccharomycetales(100);f__Metschnikowiaceae(100);g__Metschnikowia(100);s__Metschnikowia_pulcherrima(100); | Metschnikowia | 13 | Saprotroph | \|Undefined Saprotroph\| | Yeast | NULL | Probable | As nectar-tap_saprotroph (Põlme et al. 2020); Decay sugar-rich substrates (Põlme et al. 2020) | Põlme S, et al. 2020. Fungal Diversity 105:1-16 ((https://doi.org/10.1007/s13225-020-00466-2)) |
| Otu52951 | 1 | k__Fungi(100);p__Ascomycota(100);c__Saccharomycetes(100);o__Saccharomycetales(100);f__Metschnikowiaceae(100);g__Metschnikowia(100);g__Metschnikowia_unclassified(100); | Metschnikowia | 13 | Saprotroph | \|Undefined Saprotroph\| | Yeast | NULL | Probable | As nectar-tap_saprotroph (Põlme et al. 2020); Decay sugar-rich substrates (Põlme et al. 2020) | Põlme S, et al. 2020. Fungal Diversity 105:1-16 ((https://doi.org/10.1007/s13225-020-00466-2)) |
| Otu52952 | 1 | k__Fungi(100);p__Ascomycota(100);c__Saccharomycetes(100);o__Saccharomycetales(100);f__Metschnikowiaceae(100);g__Metschnikowia(100);s__Metschnikowia_pulcherrima(100); | Metschnikowia | 13 | Saprotroph | \|Undefined Saprotroph\| | Yeast | NULL | Probable | As nectar-tap_saprotroph (Põlme et al. 2020); Decay sugar-rich substrates (Põlme et al. 2020) | Põlme S, et al. 2020. Fungal Diversity 105:1-16 ((https://doi.org/10.1007/s13225-020-00466-2)) |
| Otu52953 | 1 | k__Fungi(100);p__Ascomycota(100);c__Saccharomycetes(100);o__Saccharomycetales(100);f__Metschnikowiaceae(100);g__Metschnikowia(100);g__Metschnikowia_unclassified(100); | Metschnikowia | 13 | Saprotroph | \|Undefined Saprotroph\| | Yeast | NULL | Probable | As nectar-tap_saprotroph (Põlme et al. 2020); Decay sugar-rich substrates (Põlme et al. 2020) | Põlme S, et al. 2020. Fungal Diversity 105:1-16 ((https://doi.org/10.1007/s13225-020-00466-2)) |
| Otu52955 | 1 | k__Fungi(100);p__Ascomycota(100);c__Saccharomycetes(100);o__Saccharomycetales(100);f__Metschnikowiaceae(100);g__Metschnikowia(100);s__Metschnikowia_pulcherrima(100); | Metschnikowia | 13 | Saprotroph | \|Undefined Saprotroph\| | Yeast | NULL | Probable | As nectar-tap_saprotroph (Põlme et al. 2020); Decay sugar-rich substrates (Põlme et al. 2020) | Põlme S, et al. 2020. Fungal Diversity 105:1-16 ((https://doi.org/10.1007/s13225-020-00466-2)) |
| Otu52956 | 1 | k__Fungi(100);p__Ascomycota(100);c__Saccharomycetes(100);o__Saccharomycetales(100);f__Metschnikowiaceae(100);g__Metschnikowia(100);s__Metschnikowia_pulcherrima(100); | Metschnikowia | 13 | Saprotroph | \|Undefined Saprotroph\| | Yeast | NULL | Probable | As nectar-tap_saprotroph (Põlme et al. 2020); Decay sugar-rich substrates (Põlme et al. 2020) | Põlme S, et al. 2020. Fungal Diversity 105:1-16 ((https://doi.org/10.1007/s13225-020-00466-2)) |
| Otu52957 | 1 | k__Fungi(100);p__Ascomycota(100);c__Saccharomycetes(100);o__Saccharomycetales(100);f__Metschnikowiaceae(100);g__Metschnikowia(100);g__Metschnikowia_unclassified(100); | Metschnikowia | 13 | Saprotroph | \|Undefined Saprotroph\| | Yeast | NULL | Probable | As nectar-tap_saprotroph (Põlme et al. 2020); Decay sugar-rich substrates (Põlme et al. 2020) | Põlme S, et al. 2020. Fungal Diversity 105:1-16 ((https://doi.org/10.1007/s13225-020-00466-2)) |
| Otu52958 | 1 | k__Fungi(100);p__Ascomycota(100);c__Saccharomycetes(100);o__Saccharomycetales(100);f__Metschnikowiaceae(100);g__Metschnikowia(100);s__Metschnikowia_pulcherrima(100); | Metschnikowia | 13 | Saprotroph | \|Undefined Saprotroph\| | Yeast | NULL | Probable | As nectar-tap_saprotroph (Põlme et al. 2020); Decay sugar-rich substrates (Põlme et al. 2020) | Põlme S, et al. 2020. Fungal Diversity 105:1-16 ((https://doi.org/10.1007/s13225-020-00466-2)) |
| Otu52959 | 1 | k__Fungi(100);p__Ascomycota(100);c__Saccharomycetes(100);o__Saccharomycetales(100);f__Metschnikowiaceae(100);g__Metschnikowia(100);s__Metschnikowia_pulcherrima(100); | Metschnikowia | 13 | Saprotroph | \|Undefined Saprotroph\| | Yeast | NULL | Probable | As nectar-tap_saprotroph (Põlme et al. 2020); Decay sugar-rich substrates (Põlme et al. 2020) | Põlme S, et al. 2020. Fungal Diversity 105:1-16 ((https://doi.org/10.1007/s13225-020-00466-2)) |
| Otu52960 | 1 | k__Fungi(100);p__Ascomycota(100);c__Saccharomycetes(100);o__Saccharomycetales(100);f__Metschnikowiaceae(100);g__Metschnikowia(100);s__Metschnikowia_pulcherrima(100); | Metschnikowia | 13 | Saprotroph | \|Undefined Saprotroph\| | Yeast | NULL | Probable | As nectar-tap_saprotroph (Põlme et al. 2020); Decay sugar-rich substrates (Põlme et al. 2020) | Põlme S, et al. 2020. Fungal Diversity 105:1-16 ((https://doi.org/10.1007/s13225-020-00466-2)) |
| Otu52961 | 1 | k__Fungi(100);p__Ascomycota(100);c__Saccharomycetes(100);o__Saccharomycetales(100);f__Metschnikowiaceae(100);g__Metschnikowia(100);g__Metschnikowia_unclassified(100); | Metschnikowia | 13 | Saprotroph | \|Undefined Saprotroph\| | Yeast | NULL | Probable | As nectar-tap_saprotroph (Põlme et al. 2020); Decay sugar-rich substrates (Põlme et al. 2020) | Põlme S, et al. 2020. Fungal Diversity 105:1-16 ((https://doi.org/10.1007/s13225-020-00466-2)) |
| Otu52962 | 1 | k__Fungi(100);p__Ascomycota(100);c__Saccharomycetes(100);o__Saccharomycetales(100);f__Metschnikowiaceae(100);g__Metschnikowia(100);g__Metschnikowia_unclassified(100); | Metschnikowia | 13 | Saprotroph | \|Undefined Saprotroph\| | Yeast | NULL | Probable | As nectar-tap_saprotroph (Põlme et al. 2020); Decay sugar-rich substrates (Põlme et al. 2020) | Põlme S, et al. 2020. Fungal Diversity 105:1-16 ((https://doi.org/10.1007/s13225-020-00466-2)) |
| Otu52963 | 1 | k__Fungi(100);p__Ascomycota(100);c__Saccharomycetes(100);o__Saccharomycetales(100);f__Metschnikowiaceae(100);g__Metschnikowia(100);g__Metschnikowia_unclassified(100); | Metschnikowia | 13 | Saprotroph | \|Undefined Saprotroph\| | Yeast | NULL | Probable | As nectar-tap_saprotroph (Põlme et al. 2020); Decay sugar-rich substrates (Põlme et al. 2020) | Põlme S, et al. 2020. Fungal Diversity 105:1-16 ((https://doi.org/10.1007/s13225-020-00466-2)) |
| Otu52964 | 1 | k__Fungi(100);p__Ascomycota(100);c__Saccharomycetes(100);o__Saccharomycetales(100);f__Metschnikowiaceae(100);g__Metschnikowia(100);g__Metschnikowia_unclassified(100); | Metschnikowia | 13 | Saprotroph | \|Undefined Saprotroph\| | Yeast | NULL | Probable | As nectar-tap_saprotroph (Põlme et al. 2020); Decay sugar-rich substrates (Põlme et al. 2020) | Põlme S, et al. 2020. Fungal Diversity 105:1-16 ((https://doi.org/10.1007/s13225-020-00466-2)) |
| Otu52965 | 1 | k__Fungi(100);p__Ascomycota(100);c__Saccharomycetes(100);o__Saccharomycetales(100);f__Metschnikowiaceae(100);g__Metschnikowia(100);s__Metschnikowia_pulcherrima(100); | Metschnikowia | 13 | Saprotroph | \|Undefined Saprotroph\| | Yeast | NULL | Probable | As nectar-tap_saprotroph (Põlme et al. 2020); Decay sugar-rich substrates (Põlme et al. 2020) | Põlme S, et al. 2020. Fungal Diversity 105:1-16 ((https://doi.org/10.1007/s13225-020-00466-2)) |
| Otu52967 | 1 | k__Fungi(100);p__Ascomycota(100);c__Saccharomycetes(100);o__Saccharomycetales(100);f__Metschnikowiaceae(100);g__Metschnikowia(100);s__Metschnikowia_pulcherrima(100); | Metschnikowia | 13 | Saprotroph | \|Undefined Saprotroph\| | Yeast | NULL | Probable | As nectar-tap_saprotroph (Põlme et al. 2020); Decay sugar-rich substrates (Põlme et al. 2020) | Põlme S, et al. 2020. Fungal Diversity 105:1-16 ((https://doi.org/10.1007/s13225-020-00466-2)) |
| Otu52968 | 1 | k__Fungi(100);p__Ascomycota(100);c__Saccharomycetes(100);o__Saccharomycetales(100);f__Metschnikowiaceae(100);g__Metschnikowia(100);s__Metschnikowia_pulcherrima(100); | Metschnikowia | 13 | Saprotroph | \|Undefined Saprotroph\| | Yeast | NULL | Probable | As nectar-tap_saprotroph (Põlme et al. 2020); Decay sugar-rich substrates (Põlme et al. 2020) | Põlme S, et al. 2020. Fungal Diversity 105:1-16 ((https://doi.org/10.1007/s13225-020-00466-2)) |
| Otu52970 | 1 | k__Fungi(100);p__Ascomycota(100);c__Saccharomycetes(100);o__Saccharomycetales(100);f__Metschnikowiaceae(100);g__Metschnikowia(100);g__Metschnikowia_unclassified(100); | Metschnikowia | 13 | Saprotroph | \|Undefined Saprotroph\| | Yeast | NULL | Probable | As nectar-tap_saprotroph (Põlme et al. 2020); Decay sugar-rich substrates (Põlme et al. 2020) | Põlme S, et al. 2020. Fungal Diversity 105:1-16 ((https://doi.org/10.1007/s13225-020-00466-2)) |
| Otu52971 | 1 | k__Fungi(100);p__Ascomycota(100);c__Saccharomycetes(100);o__Saccharomycetales(100);f__Metschnikowiaceae(100);g__Metschnikowia(100);s__Metschnikowia_pulcherrima(100); | Metschnikowia | 13 | Saprotroph | \|Undefined Saprotroph\| | Yeast | NULL | Probable | As nectar-tap_saprotroph (Põlme et al. 2020); Decay sugar-rich substrates (Põlme et al. 2020) | Põlme S, et al. 2020. Fungal Diversity 105:1-16 ((https://doi.org/10.1007/s13225-020-00466-2)) |
| Otu52972 | 1 | k__Fungi(100);p__Ascomycota(100);c__Saccharomycetes(100);o__Saccharomycetales(100);f__Metschnikowiaceae(100);g__Metschnikowia(100);s__Metschnikowia_pulcherrima(100); | Metschnikowia | 13 | Saprotroph | \|Undefined Saprotroph\| | Yeast | NULL | Probable | As nectar-tap_saprotroph (Põlme et al. 2020); Decay sugar-rich substrates (Põlme et al. 2020) | Põlme S, et al. 2020. Fungal Diversity 105:1-16 ((https://doi.org/10.1007/s13225-020-00466-2)) |
| Otu52973 | 1 | k__Fungi(100);p__Ascomycota(100);c__Saccharomycetes(100);o__Saccharomycetales(100);f__Metschnikowiaceae(100);g__Metschnikowia(100);s__Metschnikowia_pulcherrima(100); | Metschnikowia | 13 | Saprotroph | \|Undefined Saprotroph\| | Yeast | NULL | Probable | As nectar-tap_saprotroph (Põlme et al. 2020); Decay sugar-rich substrates (Põlme et al. 2020) | Põlme S, et al. 2020. Fungal Diversity 105:1-16 ((https://doi.org/10.1007/s13225-020-00466-2)) |
| Otu52974 | 1 | k__Fungi(100);p__Ascomycota(100);c__Saccharomycetes(100);o__Saccharomycetales(100);f__Metschnikowiaceae(100);g__Metschnikowia(100);s__Metschnikowia_pulcherrima(100); | Metschnikowia | 13 | Saprotroph | \|Undefined Saprotroph\| | Yeast | NULL | Probable | As nectar-tap_saprotroph (Põlme et al. 2020); Decay sugar-rich substrates (Põlme et al. 2020) | Põlme S, et al. 2020. Fungal Diversity 105:1-16 ((https://doi.org/10.1007/s13225-020-00466-2)) |
| Otu52975 | 1 | k__Fungi(100);p__Ascomycota(100);c__Saccharomycetes(100);o__Saccharomycetales(100);f__Saccharomycetales_family_Incertae_sedis(100);g__Candida(100);g__Candida_unclassified(100); | Candida | 13 | Pathotroph-Saprotroph-Symbiotroph | Animal Pathogen-Endophyte-Endosymbiont-Epiphyte-\|Undefined Saprotroph\| | Dimorphic Yeast-Yeast | NULL | Probable | As nectar-tap_saprotroph (Põlme et al. 2020); Decay sugar-rich substrates (Põlme et al. 2020); This asexual genus is polyphyletic and contain species that function very differently depend on what they are. Caution should be taken when interpreting guilds for Candida species. | Manolakaki D, et al. 2010. Virulence. 1: 367-375 ((https://doi.org/10.4161/viru.1.5.12796)); Mekbib SB, et al. 2011. Tropical Plant Pathology 36:233-240 ((https://doi.org/10.1590/S1982-56762011000400004)); Põlme S, et al. 2020. Fungal Diversity 105:1-16 ((https://doi.org/10.1007/s13225-020-00466-2)) |
| Otu52976 | 1 | k__Fungi(100);p__Ascomycota(100);c__Saccharomycetes(100);o__Saccharomycetales(100);f__Metschnikowiaceae(100);g__Metschnikowia(100);g__Metschnikowia_unclassified(100); | Metschnikowia | 13 | Saprotroph | \|Undefined Saprotroph\| | Yeast | NULL | Probable | As nectar-tap_saprotroph (Põlme et al. 2020); Decay sugar-rich substrates (Põlme et al. 2020) | Põlme S, et al. 2020. Fungal Diversity 105:1-16 ((https://doi.org/10.1007/s13225-020-00466-2)) |
| Otu52977 | 1 | k__Fungi(100);p__Ascomycota(100);c__Saccharomycetes(100);o__Saccharomycetales(100);f__Metschnikowiaceae(100);g__Metschnikowia(100);s__Metschnikowia_pulcherrima(100); | Metschnikowia | 13 | Saprotroph | \|Undefined Saprotroph\| | Yeast | NULL | Probable | As nectar-tap_saprotroph (Põlme et al. 2020); Decay sugar-rich substrates (Põlme et al. 2020) | Põlme S, et al. 2020. Fungal Diversity 105:1-16 ((https://doi.org/10.1007/s13225-020-00466-2)) |
| Otu52978 | 1 | k__Fungi(100);p__Ascomycota(100);c__Saccharomycetes(100);o__Saccharomycetales(100);f__Metschnikowiaceae(100);g__Metschnikowia(100);s__Metschnikowia_pulcherrima(100); | Metschnikowia | 13 | Saprotroph | \|Undefined Saprotroph\| | Yeast | NULL | Probable | As nectar-tap_saprotroph (Põlme et al. 2020); Decay sugar-rich substrates (Põlme et al. 2020) | Põlme S, et al. 2020. Fungal Diversity 105:1-16 ((https://doi.org/10.1007/s13225-020-00466-2)) |
| Otu52979 | 1 | k__Fungi(100);p__Ascomycota(100);c__Saccharomycetes(100);o__Saccharomycetales(100);f__Metschnikowiaceae(100);g__Metschnikowia(100);g__Metschnikowia_unclassified(100); | Metschnikowia | 13 | Saprotroph | \|Undefined Saprotroph\| | Yeast | NULL | Probable | As nectar-tap_saprotroph (Põlme et al. 2020); Decay sugar-rich substrates (Põlme et al. 2020) | Põlme S, et al. 2020. Fungal Diversity 105:1-16 ((https://doi.org/10.1007/s13225-020-00466-2)) |
| Otu52980 | 1 | k__Fungi(100);p__Ascomycota(100);c__Saccharomycetes(100);o__Saccharomycetales(100);f__Metschnikowiaceae(100);g__Metschnikowia(100);g__Metschnikowia_unclassified(100); | Metschnikowia | 13 | Saprotroph | \|Undefined Saprotroph\| | Yeast | NULL | Probable | As nectar-tap_saprotroph (Põlme et al. 2020); Decay sugar-rich substrates (Põlme et al. 2020) | Põlme S, et al. 2020. Fungal Diversity 105:1-16 ((https://doi.org/10.1007/s13225-020-00466-2)) |
| Otu52981 | 1 | k__Fungi(100);p__Ascomycota(100);c__Saccharomycetes(100);o__Saccharomycetales(100);f__Metschnikowiaceae(100);g__Metschnikowia(100);s__Metschnikowia_pulcherrima(100); | Metschnikowia | 13 | Saprotroph | \|Undefined Saprotroph\| | Yeast | NULL | Probable | As nectar-tap_saprotroph (Põlme et al. 2020); Decay sugar-rich substrates (Põlme et al. 2020) | Põlme S, et al. 2020. Fungal Diversity 105:1-16 ((https://doi.org/10.1007/s13225-020-00466-2)) |
| Otu52982 | 1 | k__Fungi(100);p__Ascomycota(100);c__Saccharomycetes(100);o__Saccharomycetales(100);f__Metschnikowiaceae(100);g__Metschnikowia(100);s__Metschnikowia_pulcherrima(100); | Metschnikowia | 13 | Saprotroph | \|Undefined Saprotroph\| | Yeast | NULL | Probable | As nectar-tap_saprotroph (Põlme et al. 2020); Decay sugar-rich substrates (Põlme et al. 2020) | Põlme S, et al. 2020. Fungal Diversity 105:1-16 ((https://doi.org/10.1007/s13225-020-00466-2)) |
| Otu52983 | 1 | k__Fungi(100);p__Ascomycota(100);c__Saccharomycetes(100);o__Saccharomycetales(100);f__Metschnikowiaceae(100);g__Metschnikowia(100);s__Metschnikowia_pulcherrima(100); | Metschnikowia | 13 | Saprotroph | \|Undefined Saprotroph\| | Yeast | NULL | Probable | As nectar-tap_saprotroph (Põlme et al. 2020); Decay sugar-rich substrates (Põlme et al. 2020) | Põlme S, et al. 2020. Fungal Diversity 105:1-16 ((https://doi.org/10.1007/s13225-020-00466-2)) |
| Otu52984 | 1 | k__Fungi(100);p__Ascomycota(100);c__Saccharomycetes(100);o__Saccharomycetales(100);f__Metschnikowiaceae(100);g__Metschnikowia(100);s__Metschnikowia_pulcherrima(100); | Metschnikowia | 13 | Saprotroph | \|Undefined Saprotroph\| | Yeast | NULL | Probable | As nectar-tap_saprotroph (Põlme et al. 2020); Decay sugar-rich substrates (Põlme et al. 2020) | Põlme S, et al. 2020. Fungal Diversity 105:1-16 ((https://doi.org/10.1007/s13225-020-00466-2)) |
| Otu52985 | 1 | k__Fungi(100);p__Ascomycota(100);c__Saccharomycetes(100);o__Saccharomycetales(100);f__Metschnikowiaceae(100);g__Metschnikowia(100);s__Metschnikowia_pulcherrima(100); | Metschnikowia | 13 | Saprotroph | \|Undefined Saprotroph\| | Yeast | NULL | Probable | As nectar-tap_saprotroph (Põlme et al. 2020); Decay sugar-rich substrates (Põlme et al. 2020) | Põlme S, et al. 2020. Fungal Diversity 105:1-16 ((https://doi.org/10.1007/s13225-020-00466-2)) |
| Otu52986 | 1 | k__Fungi(100);p__Ascomycota(100);c__Saccharomycetes(100);o__Saccharomycetales(100);f__Metschnikowiaceae(100);g__Metschnikowia(100);g__Metschnikowia_unclassified(100); | Metschnikowia | 13 | Saprotroph | \|Undefined Saprotroph\| | Yeast | NULL | Probable | As nectar-tap_saprotroph (Põlme et al. 2020); Decay sugar-rich substrates (Põlme et al. 2020) | Põlme S, et al. 2020. Fungal Diversity 105:1-16 ((https://doi.org/10.1007/s13225-020-00466-2)) |
| Otu52987 | 1 | k__Fungi(100);p__Ascomycota(100);c__Saccharomycetes(100);o__Saccharomycetales(100);f__Metschnikowiaceae(100);g__Metschnikowia(100);g__Metschnikowia_unclassified(100); | Metschnikowia | 13 | Saprotroph | \|Undefined Saprotroph\| | Yeast | NULL | Probable | As nectar-tap_saprotroph (Põlme et al. 2020); Decay sugar-rich substrates (Põlme et al. 2020) | Põlme S, et al. 2020. Fungal Diversity 105:1-16 ((https://doi.org/10.1007/s13225-020-00466-2)) |
| Otu52988 | 1 | k__Fungi(100);p__Ascomycota(100);c__Saccharomycetes(100);o__Saccharomycetales(100);f__Metschnikowiaceae(100);g__Metschnikowia(100);s__Metschnikowia_pulcherrima(100); | Metschnikowia | 13 | Saprotroph | \|Undefined Saprotroph\| | Yeast | NULL | Probable | As nectar-tap_saprotroph (Põlme et al. 2020); Decay sugar-rich substrates (Põlme et al. 2020) | Põlme S, et al. 2020. Fungal Diversity 105:1-16 ((https://doi.org/10.1007/s13225-020-00466-2)) |
| Otu52990 | 1 | k__Fungi(100);p__Ascomycota(100);c__Saccharomycetes(100);o__Saccharomycetales(100);f__Metschnikowiaceae(100);g__Metschnikowia(100);s__Metschnikowia_pulcherrima(100); | Metschnikowia | 13 | Saprotroph | \|Undefined Saprotroph\| | Yeast | NULL | Probable | As nectar-tap_saprotroph (Põlme et al. 2020); Decay sugar-rich substrates (Põlme et al. 2020) | Põlme S, et al. 2020. Fungal Diversity 105:1-16 ((https://doi.org/10.1007/s13225-020-00466-2)) |
| Otu52992 | 1 | k__Fungi(100);p__Ascomycota(100);c__Saccharomycetes(100);o__Saccharomycetales(100);f__Metschnikowiaceae(100);g__Metschnikowia(100);s__Metschnikowia_pulcherrima(100); | Metschnikowia | 13 | Saprotroph | \|Undefined Saprotroph\| | Yeast | NULL | Probable | As nectar-tap_saprotroph (Põlme et al. 2020); Decay sugar-rich substrates (Põlme et al. 2020) | Põlme S, et al. 2020. Fungal Diversity 105:1-16 ((https://doi.org/10.1007/s13225-020-00466-2)) |
| Otu52993 | 1 | k__Fungi(100);p__Ascomycota(100);c__Saccharomycetes(100);o__Saccharomycetales(100);f__Metschnikowiaceae(100);g__Metschnikowia(100);s__Metschnikowia_pulcherrima(100); | Metschnikowia | 13 | Saprotroph | \|Undefined Saprotroph\| | Yeast | NULL | Probable | As nectar-tap_saprotroph (Põlme et al. 2020); Decay sugar-rich substrates (Põlme et al. 2020) | Põlme S, et al. 2020. Fungal Diversity 105:1-16 ((https://doi.org/10.1007/s13225-020-00466-2)) |
| Otu52995 | 1 | k__Fungi(100);p__Ascomycota(100);c__Saccharomycetes(100);o__Saccharomycetales(100);f__Metschnikowiaceae(100);g__Metschnikowia(100);g__Metschnikowia_unclassified(100); | Metschnikowia | 13 | Saprotroph | \|Undefined Saprotroph\| | Yeast | NULL | Probable | As nectar-tap_saprotroph (Põlme et al. 2020); Decay sugar-rich substrates (Põlme et al. 2020) | Põlme S, et al. 2020. Fungal Diversity 105:1-16 ((https://doi.org/10.1007/s13225-020-00466-2)) |
| Otu52996 | 1 | k__Fungi(100);p__Ascomycota(100);c__Saccharomycetes(100);o__Saccharomycetales(100);f__Metschnikowiaceae(100);g__Metschnikowia(100);g__Metschnikowia_unclassified(100); | Metschnikowia | 13 | Saprotroph | \|Undefined Saprotroph\| | Yeast | NULL | Probable | As nectar-tap_saprotroph (Põlme et al. 2020); Decay sugar-rich substrates (Põlme et al. 2020) | Põlme S, et al. 2020. Fungal Diversity 105:1-16 ((https://doi.org/10.1007/s13225-020-00466-2)) |
| Otu52997 | 1 | k__Fungi(100);p__Ascomycota(100);c__Saccharomycetes(100);o__Saccharomycetales(100);f__Metschnikowiaceae(100);g__Metschnikowia(100);g__Metschnikowia_unclassified(100); | Metschnikowia | 13 | Saprotroph | \|Undefined Saprotroph\| | Yeast | NULL | Probable | As nectar-tap_saprotroph (Põlme et al. 2020); Decay sugar-rich substrates (Põlme et al. 2020) | Põlme S, et al. 2020. Fungal Diversity 105:1-16 ((https://doi.org/10.1007/s13225-020-00466-2)) |
| Otu52998 | 1 | k__Fungi(100);p__Ascomycota(100);c__Saccharomycetes(100);o__Saccharomycetales(100);f__Metschnikowiaceae(100);g__Metschnikowia(100);g__Metschnikowia_unclassified(100); | Metschnikowia | 13 | Saprotroph | \|Undefined Saprotroph\| | Yeast | NULL | Probable | As nectar-tap_saprotroph (Põlme et al. 2020); Decay sugar-rich substrates (Põlme et al. 2020) | Põlme S, et al. 2020. Fungal Diversity 105:1-16 ((https://doi.org/10.1007/s13225-020-00466-2)) |
| Otu52999 | 1 | k__Fungi(100);p__Ascomycota(100);c__Saccharomycetes(100);o__Saccharomycetales(100);f__Metschnikowiaceae(100);g__Metschnikowia(100);s__Metschnikowia_pulcherrima(100); | Metschnikowia | 13 | Saprotroph | \|Undefined Saprotroph\| | Yeast | NULL | Probable | As nectar-tap_saprotroph (Põlme et al. 2020); Decay sugar-rich substrates (Põlme et al. 2020) | Põlme S, et al. 2020. Fungal Diversity 105:1-16 ((https://doi.org/10.1007/s13225-020-00466-2)) |
| Otu53000 | 1 | k__Fungi(100);p__Ascomycota(100);c__Saccharomycetes(100);o__Saccharomycetales(100);f__Metschnikowiaceae(100);g__Metschnikowia(100);s__Metschnikowia_pulcherrima(100); | Metschnikowia | 13 | Saprotroph | \|Undefined Saprotroph\| | Yeast | NULL | Probable | As nectar-tap_saprotroph (Põlme et al. 2020); Decay sugar-rich substrates (Põlme et al. 2020) | Põlme S, et al. 2020. Fungal Diversity 105:1-16 ((https://doi.org/10.1007/s13225-020-00466-2)) |
| Otu53001 | 1 | k__Fungi(100);p__Ascomycota(100);c__Saccharomycetes(100);o__Saccharomycetales(100);f__Metschnikowiaceae(100);g__Metschnikowia(100);s__Metschnikowia_pulcherrima(100); | Metschnikowia | 13 | Saprotroph | \|Undefined Saprotroph\| | Yeast | NULL | Probable | As nectar-tap_saprotroph (Põlme et al. 2020); Decay sugar-rich substrates (Põlme et al. 2020) | Põlme S, et al. 2020. Fungal Diversity 105:1-16 ((https://doi.org/10.1007/s13225-020-00466-2)) |
| Otu53002 | 1 | k__Fungi(100);p__Ascomycota(100);c__Saccharomycetes(100);o__Saccharomycetales(100);f__Metschnikowiaceae(100);g__Metschnikowia(100);g__Metschnikowia_unclassified(100); | Metschnikowia | 13 | Saprotroph | \|Undefined Saprotroph\| | Yeast | NULL | Probable | As nectar-tap_saprotroph (Põlme et al. 2020); Decay sugar-rich substrates (Põlme et al. 2020) | Põlme S, et al. 2020. Fungal Diversity 105:1-16 ((https://doi.org/10.1007/s13225-020-00466-2)) |
| Otu53003 | 1 | k__Fungi(100);p__Ascomycota(100);c__Saccharomycetes(100);o__Saccharomycetales(100);f__Metschnikowiaceae(100);g__Metschnikowia(100);g__Metschnikowia_unclassified(100); | Metschnikowia | 13 | Saprotroph | \|Undefined Saprotroph\| | Yeast | NULL | Probable | As nectar-tap_saprotroph (Põlme et al. 2020); Decay sugar-rich substrates (Põlme et al. 2020) | Põlme S, et al. 2020. Fungal Diversity 105:1-16 ((https://doi.org/10.1007/s13225-020-00466-2)) |
| Otu53004 | 1 | k__Fungi(100);p__Ascomycota(100);c__Saccharomycetes(100);o__Saccharomycetales(100);f__Metschnikowiaceae(100);g__Metschnikowia(100);s__Metschnikowia_pulcherrima(100); | Metschnikowia | 13 | Saprotroph | \|Undefined Saprotroph\| | Yeast | NULL | Probable | As nectar-tap_saprotroph (Põlme et al. 2020); Decay sugar-rich substrates (Põlme et al. 2020) | Põlme S, et al. 2020. Fungal Diversity 105:1-16 ((https://doi.org/10.1007/s13225-020-00466-2)) |
| Otu53005 | 1 | k__Fungi(100);p__Ascomycota(100);c__Saccharomycetes(100);o__Saccharomycetales(100);f__Metschnikowiaceae(100);g__Metschnikowia(100);g__Metschnikowia_unclassified(100); | Metschnikowia | 13 | Saprotroph | \|Undefined Saprotroph\| | Yeast | NULL | Probable | As nectar-tap_saprotroph (Põlme et al. 2020); Decay sugar-rich substrates (Põlme et al. 2020) | Põlme S, et al. 2020. Fungal Diversity 105:1-16 ((https://doi.org/10.1007/s13225-020-00466-2)) |
| Otu53006 | 1 | k__Fungi(100);p__Ascomycota(100);c__Saccharomycetes(100);o__Saccharomycetales(100);f__Metschnikowiaceae(100);g__Metschnikowia(100);g__Metschnikowia_unclassified(100); | Metschnikowia | 13 | Saprotroph | \|Undefined Saprotroph\| | Yeast | NULL | Probable | As nectar-tap_saprotroph (Põlme et al. 2020); Decay sugar-rich substrates (Põlme et al. 2020) | Põlme S, et al. 2020. Fungal Diversity 105:1-16 ((https://doi.org/10.1007/s13225-020-00466-2)) |
| Otu53007 | 1 | k__Fungi(100);p__Ascomycota(100);c__Saccharomycetes(100);o__Saccharomycetales(100);f__Metschnikowiaceae(100);g__Metschnikowia(100);s__Metschnikowia_pulcherrima(100); | Metschnikowia | 13 | Saprotroph | \|Undefined Saprotroph\| | Yeast | NULL | Probable | As nectar-tap_saprotroph (Põlme et al. 2020); Decay sugar-rich substrates (Põlme et al. 2020) | Põlme S, et al. 2020. Fungal Diversity 105:1-16 ((https://doi.org/10.1007/s13225-020-00466-2)) |
| Otu53008 | 1 | k__Fungi(100);p__Ascomycota(100);c__Saccharomycetes(100);o__Saccharomycetales(100);f__Metschnikowiaceae(100);g__Metschnikowia(100);g__Metschnikowia_unclassified(100); | Metschnikowia | 13 | Saprotroph | \|Undefined Saprotroph\| | Yeast | NULL | Probable | As nectar-tap_saprotroph (Põlme et al. 2020); Decay sugar-rich substrates (Põlme et al. 2020) | Põlme S, et al. 2020. Fungal Diversity 105:1-16 ((https://doi.org/10.1007/s13225-020-00466-2)) |
| Otu53009 | 1 | k__Fungi(100);p__Ascomycota(100);c__Saccharomycetes(100);o__Saccharomycetales(100);f__Metschnikowiaceae(100);g__Metschnikowia(100);s__Metschnikowia_pulcherrima(100); | Metschnikowia | 13 | Saprotroph | \|Undefined Saprotroph\| | Yeast | NULL | Probable | As nectar-tap_saprotroph (Põlme et al. 2020); Decay sugar-rich substrates (Põlme et al. 2020) | Põlme S, et al. 2020. Fungal Diversity 105:1-16 ((https://doi.org/10.1007/s13225-020-00466-2)) |
| Otu53010 | 1 | k__Fungi(100);p__Ascomycota(100);c__Saccharomycetes(100);o__Saccharomycetales(100);f__Metschnikowiaceae(100);g__Metschnikowia(100);s__Metschnikowia_pulcherrima(100); | Metschnikowia | 13 | Saprotroph | \|Undefined Saprotroph\| | Yeast | NULL | Probable | As nectar-tap_saprotroph (Põlme et al. 2020); Decay sugar-rich substrates (Põlme et al. 2020) | Põlme S, et al. 2020. Fungal Diversity 105:1-16 ((https://doi.org/10.1007/s13225-020-00466-2)) |
| Otu53011 | 1 | k__Fungi(100);p__Ascomycota(100);c__Saccharomycetes(100);o__Saccharomycetales(100);f__Metschnikowiaceae(100);g__Metschnikowia(100);s__Metschnikowia_pulcherrima(100); | Metschnikowia | 13 | Saprotroph | \|Undefined Saprotroph\| | Yeast | NULL | Probable | As nectar-tap_saprotroph (Põlme et al. 2020); Decay sugar-rich substrates (Põlme et al. 2020) | Põlme S, et al. 2020. Fungal Diversity 105:1-16 ((https://doi.org/10.1007/s13225-020-00466-2)) |
| Otu53012 | 1 | k__Fungi(100);p__Ascomycota(100);c__Saccharomycetes(100);o__Saccharomycetales(100);f__Metschnikowiaceae(100);g__Metschnikowia(100);s__Metschnikowia_pulcherrima(100); | Metschnikowia | 13 | Saprotroph | \|Undefined Saprotroph\| | Yeast | NULL | Probable | As nectar-tap_saprotroph (Põlme et al. 2020); Decay sugar-rich substrates (Põlme et al. 2020) | Põlme S, et al. 2020. Fungal Diversity 105:1-16 ((https://doi.org/10.1007/s13225-020-00466-2)) |
| Otu53013 | 1 | k__Fungi(100);p__Ascomycota(100);c__Saccharomycetes(100);o__Saccharomycetales(100);f__Metschnikowiaceae(100);g__Metschnikowia(100);g__Metschnikowia_unclassified(100); | Metschnikowia | 13 | Saprotroph | \|Undefined Saprotroph\| | Yeast | NULL | Probable | As nectar-tap_saprotroph (Põlme et al. 2020); Decay sugar-rich substrates (Põlme et al. 2020) | Põlme S, et al. 2020. Fungal Diversity 105:1-16 ((https://doi.org/10.1007/s13225-020-00466-2)) |
| Otu53014 | 1 | k__Fungi(100);p__Ascomycota(100);c__Saccharomycetes(100);o__Saccharomycetales(100);f__Metschnikowiaceae(100);g__Metschnikowia(100);g__Metschnikowia_unclassified(100); | Metschnikowia | 13 | Saprotroph | \|Undefined Saprotroph\| | Yeast | NULL | Probable | As nectar-tap_saprotroph (Põlme et al. 2020); Decay sugar-rich substrates (Põlme et al. 2020) | Põlme S, et al. 2020. Fungal Diversity 105:1-16 ((https://doi.org/10.1007/s13225-020-00466-2)) |
| Otu53015 | 1 | k__Fungi(100);p__Ascomycota(100);c__Saccharomycetes(100);o__Saccharomycetales(100);f__Metschnikowiaceae(100);g__Metschnikowia(100);s__Metschnikowia_pulcherrima(100); | Metschnikowia | 13 | Saprotroph | \|Undefined Saprotroph\| | Yeast | NULL | Probable | As nectar-tap_saprotroph (Põlme et al. 2020); Decay sugar-rich substrates (Põlme et al. 2020) | Põlme S, et al. 2020. Fungal Diversity 105:1-16 ((https://doi.org/10.1007/s13225-020-00466-2)) |
| Otu53016 | 1 | k__Fungi(100);p__Ascomycota(100);c__Saccharomycetes(100);o__Saccharomycetales(100);f__Metschnikowiaceae(100);g__Metschnikowia(100);s__Metschnikowia_pulcherrima(100); | Metschnikowia | 13 | Saprotroph | \|Undefined Saprotroph\| | Yeast | NULL | Probable | As nectar-tap_saprotroph (Põlme et al. 2020); Decay sugar-rich substrates (Põlme et al. 2020) | Põlme S, et al. 2020. Fungal Diversity 105:1-16 ((https://doi.org/10.1007/s13225-020-00466-2)) |
| Otu53017 | 1 | k__Fungi(100);p__Ascomycota(100);c__Saccharomycetes(100);o__Saccharomycetales(100);f__Metschnikowiaceae(100);g__Metschnikowia(100);s__Metschnikowia_pulcherrima(100); | Metschnikowia | 13 | Saprotroph | \|Undefined Saprotroph\| | Yeast | NULL | Probable | As nectar-tap_saprotroph (Põlme et al. 2020); Decay sugar-rich substrates (Põlme et al. 2020) | Põlme S, et al. 2020. Fungal Diversity 105:1-16 ((https://doi.org/10.1007/s13225-020-00466-2)) |
| Otu53018 | 1 | k__Fungi(100);p__Ascomycota(100);c__Saccharomycetes(100);o__Saccharomycetales(100);f__Metschnikowiaceae(100);g__Metschnikowia(100);s__Metschnikowia_pulcherrima(100); | Metschnikowia | 13 | Saprotroph | \|Undefined Saprotroph\| | Yeast | NULL | Probable | As nectar-tap_saprotroph (Põlme et al. 2020); Decay sugar-rich substrates (Põlme et al. 2020) | Põlme S, et al. 2020. Fungal Diversity 105:1-16 ((https://doi.org/10.1007/s13225-020-00466-2)) |
| Otu53019 | 1 | k__Fungi(100);p__Ascomycota(100);c__Saccharomycetes(100);o__Saccharomycetales(100);f__Metschnikowiaceae(100);g__Metschnikowia(100);s__Metschnikowia_pulcherrima(100); | Metschnikowia | 13 | Saprotroph | \|Undefined Saprotroph\| | Yeast | NULL | Probable | As nectar-tap_saprotroph (Põlme et al. 2020); Decay sugar-rich substrates (Põlme et al. 2020) | Põlme S, et al. 2020. Fungal Diversity 105:1-16 ((https://doi.org/10.1007/s13225-020-00466-2)) |
| Otu53020 | 1 | k__Fungi(100);p__Ascomycota(100);c__Saccharomycetes(100);o__Saccharomycetales(100);f__Metschnikowiaceae(100);g__Metschnikowia(100);s__Metschnikowia_pulcherrima(100); | Metschnikowia | 13 | Saprotroph | \|Undefined Saprotroph\| | Yeast | NULL | Probable | As nectar-tap_saprotroph (Põlme et al. 2020); Decay sugar-rich substrates (Põlme et al. 2020) | Põlme S, et al. 2020. Fungal Diversity 105:1-16 ((https://doi.org/10.1007/s13225-020-00466-2)) |
| Otu53021 | 1 | k__Fungi(100);p__Ascomycota(100);c__Saccharomycetes(100);o__Saccharomycetales(100);f__Metschnikowiaceae(100);g__Metschnikowia(100);s__Metschnikowia_pulcherrima(100); | Metschnikowia | 13 | Saprotroph | \|Undefined Saprotroph\| | Yeast | NULL | Probable | As nectar-tap_saprotroph (Põlme et al. 2020); Decay sugar-rich substrates (Põlme et al. 2020) | Põlme S, et al. 2020. Fungal Diversity 105:1-16 ((https://doi.org/10.1007/s13225-020-00466-2)) |
| Otu53022 | 1 | k__Fungi(100);p__Ascomycota(100);c__Saccharomycetes(100);o__Saccharomycetales(100);f__Metschnikowiaceae(100);g__Metschnikowia(100);s__Metschnikowia_pulcherrima(100); | Metschnikowia | 13 | Saprotroph | \|Undefined Saprotroph\| | Yeast | NULL | Probable | As nectar-tap_saprotroph (Põlme et al. 2020); Decay sugar-rich substrates (Põlme et al. 2020) | Põlme S, et al. 2020. Fungal Diversity 105:1-16 ((https://doi.org/10.1007/s13225-020-00466-2)) |
| Otu53023 | 1 | k__Fungi(100);p__Ascomycota(100);c__Saccharomycetes(100);o__Saccharomycetales(100);f__Metschnikowiaceae(100);g__Metschnikowia(100);g__Metschnikowia_unclassified(100); | Metschnikowia | 13 | Saprotroph | \|Undefined Saprotroph\| | Yeast | NULL | Probable | As nectar-tap_saprotroph (Põlme et al. 2020); Decay sugar-rich substrates (Põlme et al. 2020) | Põlme S, et al. 2020. Fungal Diversity 105:1-16 ((https://doi.org/10.1007/s13225-020-00466-2)) |
| Otu53024 | 1 | k__Fungi(100);p__Ascomycota(100);c__Saccharomycetes(100);o__Saccharomycetales(100);f__Metschnikowiaceae(100);g__Metschnikowia(100);g__Metschnikowia_unclassified(100); | Metschnikowia | 13 | Saprotroph | \|Undefined Saprotroph\| | Yeast | NULL | Probable | As nectar-tap_saprotroph (Põlme et al. 2020); Decay sugar-rich substrates (Põlme et al. 2020) | Põlme S, et al. 2020. Fungal Diversity 105:1-16 ((https://doi.org/10.1007/s13225-020-00466-2)) |
| Otu53025 | 1 | k__Fungi(100);p__Ascomycota(100);c__Saccharomycetes(100);o__Saccharomycetales(100);f__Metschnikowiaceae(100);g__Metschnikowia(100);s__Metschnikowia_pulcherrima(100); | Metschnikowia | 13 | Saprotroph | \|Undefined Saprotroph\| | Yeast | NULL | Probable | As nectar-tap_saprotroph (Põlme et al. 2020); Decay sugar-rich substrates (Põlme et al. 2020) | Põlme S, et al. 2020. Fungal Diversity 105:1-16 ((https://doi.org/10.1007/s13225-020-00466-2)) |
| Otu53026 | 1 | k__Fungi(100);p__Ascomycota(100);c__Saccharomycetes(100);o__Saccharomycetales(100);f__Metschnikowiaceae(100);g__Metschnikowia(100);s__Metschnikowia_pulcherrima(100); | Metschnikowia | 13 | Saprotroph | \|Undefined Saprotroph\| | Yeast | NULL | Probable | As nectar-tap_saprotroph (Põlme et al. 2020); Decay sugar-rich substrates (Põlme et al. 2020) | Põlme S, et al. 2020. Fungal Diversity 105:1-16 ((https://doi.org/10.1007/s13225-020-00466-2)) |
| Otu53027 | 1 | k__Fungi(100);p__Ascomycota(100);c__Saccharomycetes(100);o__Saccharomycetales(100);f__Metschnikowiaceae(100);g__Metschnikowia(100);s__Metschnikowia_pulcherrima(100); | Metschnikowia | 13 | Saprotroph | \|Undefined Saprotroph\| | Yeast | NULL | Probable | As nectar-tap_saprotroph (Põlme et al. 2020); Decay sugar-rich substrates (Põlme et al. 2020) | Põlme S, et al. 2020. Fungal Diversity 105:1-16 ((https://doi.org/10.1007/s13225-020-00466-2)) |
| Otu53028 | 1 | k__Fungi(100);p__Ascomycota(100);c__Saccharomycetes(100);o__Saccharomycetales(100);f__Metschnikowiaceae(100);g__Metschnikowia(100);s__Metschnikowia_pulcherrima(100); | Metschnikowia | 13 | Saprotroph | \|Undefined Saprotroph\| | Yeast | NULL | Probable | As nectar-tap_saprotroph (Põlme et al. 2020); Decay sugar-rich substrates (Põlme et al. 2020) | Põlme S, et al. 2020. Fungal Diversity 105:1-16 ((https://doi.org/10.1007/s13225-020-00466-2)) |
| Otu53029 | 1 | k__Fungi(100);p__Ascomycota(100);c__Saccharomycetes(100);o__Saccharomycetales(100);f__Metschnikowiaceae(100);g__Metschnikowia(100);g__Metschnikowia_unclassified(100); | Metschnikowia | 13 | Saprotroph | \|Undefined Saprotroph\| | Yeast | NULL | Probable | As nectar-tap_saprotroph (Põlme et al. 2020); Decay sugar-rich substrates (Põlme et al. 2020) | Põlme S, et al. 2020. Fungal Diversity 105:1-16 ((https://doi.org/10.1007/s13225-020-00466-2)) |
| Otu53030 | 1 | k__Fungi(100);p__Ascomycota(100);c__Saccharomycetes(100);o__Saccharomycetales(100);f__Metschnikowiaceae(100);g__Metschnikowia(100);s__Metschnikowia_pulcherrima(100); | Metschnikowia | 13 | Saprotroph | \|Undefined Saprotroph\| | Yeast | NULL | Probable | As nectar-tap_saprotroph (Põlme et al. 2020); Decay sugar-rich substrates (Põlme et al. 2020) | Põlme S, et al. 2020. Fungal Diversity 105:1-16 ((https://doi.org/10.1007/s13225-020-00466-2)) |
| Otu53031 | 1 | k__Fungi(100);p__Ascomycota(100);c__Saccharomycetes(100);o__Saccharomycetales(100);f__Metschnikowiaceae(100);g__Metschnikowia(100);s__Metschnikowia_pulcherrima(100); | Metschnikowia | 13 | Saprotroph | \|Undefined Saprotroph\| | Yeast | NULL | Probable | As nectar-tap_saprotroph (Põlme et al. 2020); Decay sugar-rich substrates (Põlme et al. 2020) | Põlme S, et al. 2020. Fungal Diversity 105:1-16 ((https://doi.org/10.1007/s13225-020-00466-2)) |
| Otu53032 | 1 | k__Fungi(100);p__Ascomycota(100);c__Saccharomycetes(100);o__Saccharomycetales(100);f__Metschnikowiaceae(100);g__Metschnikowia(100);s__Metschnikowia_pulcherrima(100); | Metschnikowia | 13 | Saprotroph | \|Undefined Saprotroph\| | Yeast | NULL | Probable | As nectar-tap_saprotroph (Põlme et al. 2020); Decay sugar-rich substrates (Põlme et al. 2020) | Põlme S, et al. 2020. Fungal Diversity 105:1-16 ((https://doi.org/10.1007/s13225-020-00466-2)) |
| Otu53034 | 1 | k__Fungi(100);p__Ascomycota(100);c__Saccharomycetes(100);o__Saccharomycetales(100);f__Metschnikowiaceae(100);g__Metschnikowia(100);g__Metschnikowia_unclassified(100); | Metschnikowia | 13 | Saprotroph | \|Undefined Saprotroph\| | Yeast | NULL | Probable | As nectar-tap_saprotroph (Põlme et al. 2020); Decay sugar-rich substrates (Põlme et al. 2020) | Põlme S, et al. 2020. Fungal Diversity 105:1-16 ((https://doi.org/10.1007/s13225-020-00466-2)) |
| Otu53035 | 1 | k__Fungi(100);p__Ascomycota(100);c__Saccharomycetes(100);o__Saccharomycetales(100);f__Metschnikowiaceae(100);g__Metschnikowia(100);s__Metschnikowia_pulcherrima(100); | Metschnikowia | 13 | Saprotroph | \|Undefined Saprotroph\| | Yeast | NULL | Probable | As nectar-tap_saprotroph (Põlme et al. 2020); Decay sugar-rich substrates (Põlme et al. 2020) | Põlme S, et al. 2020. Fungal Diversity 105:1-16 ((https://doi.org/10.1007/s13225-020-00466-2)) |
| Otu53036 | 1 | k__Fungi(100);p__Ascomycota(100);c__Saccharomycetes(100);o__Saccharomycetales(100);f__Metschnikowiaceae(100);g__Metschnikowia(100);s__Metschnikowia_pulcherrima(100); | Metschnikowia | 13 | Saprotroph | \|Undefined Saprotroph\| | Yeast | NULL | Probable | As nectar-tap_saprotroph (Põlme et al. 2020); Decay sugar-rich substrates (Põlme et al. 2020) | Põlme S, et al. 2020. Fungal Diversity 105:1-16 ((https://doi.org/10.1007/s13225-020-00466-2)) |
| Otu53037 | 1 | k__Fungi(100);p__Ascomycota(100);c__Saccharomycetes(100);o__Saccharomycetales(100);f__Metschnikowiaceae(100);g__Metschnikowia(100);s__Metschnikowia_pulcherrima(100); | Metschnikowia | 13 | Saprotroph | \|Undefined Saprotroph\| | Yeast | NULL | Probable | As nectar-tap_saprotroph (Põlme et al. 2020); Decay sugar-rich substrates (Põlme et al. 2020) | Põlme S, et al. 2020. Fungal Diversity 105:1-16 ((https://doi.org/10.1007/s13225-020-00466-2)) |
| Otu53038 | 1 | k__Fungi(100);p__Ascomycota(100);c__Saccharomycetes(100);o__Saccharomycetales(100);f__Metschnikowiaceae(100);g__Metschnikowia(100);s__Metschnikowia_pulcherrima(100); | Metschnikowia | 13 | Saprotroph | \|Undefined Saprotroph\| | Yeast | NULL | Probable | As nectar-tap_saprotroph (Põlme et al. 2020); Decay sugar-rich substrates (Põlme et al. 2020) | Põlme S, et al. 2020. Fungal Diversity 105:1-16 ((https://doi.org/10.1007/s13225-020-00466-2)) |
| Otu53039 | 1 | k__Fungi(100);p__Ascomycota(100);c__Saccharomycetes(100);o__Saccharomycetales(100);f__Metschnikowiaceae(100);g__Metschnikowia(100);g__Metschnikowia_unclassified(100); | Metschnikowia | 13 | Saprotroph | \|Undefined Saprotroph\| | Yeast | NULL | Probable | As nectar-tap_saprotroph (Põlme et al. 2020); Decay sugar-rich substrates (Põlme et al. 2020) | Põlme S, et al. 2020. Fungal Diversity 105:1-16 ((https://doi.org/10.1007/s13225-020-00466-2)) |
| Otu53040 | 1 | k__Fungi(100);p__Ascomycota(100);c__Saccharomycetes(100);o__Saccharomycetales(100);f__Metschnikowiaceae(100);g__Metschnikowia(100);s__Metschnikowia_pulcherrima(100); | Metschnikowia | 13 | Saprotroph | \|Undefined Saprotroph\| | Yeast | NULL | Probable | As nectar-tap_saprotroph (Põlme et al. 2020); Decay sugar-rich substrates (Põlme et al. 2020) | Põlme S, et al. 2020. Fungal Diversity 105:1-16 ((https://doi.org/10.1007/s13225-020-00466-2)) |
| Otu53041 | 1 | k__Fungi(100);p__Ascomycota(100);c__Saccharomycetes(100);o__Saccharomycetales(100);f__Metschnikowiaceae(100);g__Metschnikowia(100);s__Metschnikowia_pulcherrima(100); | Metschnikowia | 13 | Saprotroph | \|Undefined Saprotroph\| | Yeast | NULL | Probable | As nectar-tap_saprotroph (Põlme et al. 2020); Decay sugar-rich substrates (Põlme et al. 2020) | Põlme S, et al. 2020. Fungal Diversity 105:1-16 ((https://doi.org/10.1007/s13225-020-00466-2)) |
| Otu53042 | 1 | k__Fungi(100);p__Ascomycota(100);c__Saccharomycetes(100);o__Saccharomycetales(100);f__Metschnikowiaceae(100);g__Metschnikowia(100);s__Metschnikowia_pulcherrima(100); | Metschnikowia | 13 | Saprotroph | \|Undefined Saprotroph\| | Yeast | NULL | Probable | As nectar-tap_saprotroph (Põlme et al. 2020); Decay sugar-rich substrates (Põlme et al. 2020) | Põlme S, et al. 2020. Fungal Diversity 105:1-16 ((https://doi.org/10.1007/s13225-020-00466-2)) |
| Otu53043 | 1 | k__Fungi(100);p__Ascomycota(100);c__Saccharomycetes(100);o__Saccharomycetales(100);f__Metschnikowiaceae(100);g__Metschnikowia(100);g__Metschnikowia_unclassified(100); | Metschnikowia | 13 | Saprotroph | \|Undefined Saprotroph\| | Yeast | NULL | Probable | As nectar-tap_saprotroph (Põlme et al. 2020); Decay sugar-rich substrates (Põlme et al. 2020) | Põlme S, et al. 2020. Fungal Diversity 105:1-16 ((https://doi.org/10.1007/s13225-020-00466-2)) |
| Otu53044 | 1 | k__Fungi(100);p__Ascomycota(100);c__Saccharomycetes(100);o__Saccharomycetales(100);f__Metschnikowiaceae(100);g__Metschnikowia(100);s__Metschnikowia_pulcherrima(100); | Metschnikowia | 13 | Saprotroph | \|Undefined Saprotroph\| | Yeast | NULL | Probable | As nectar-tap_saprotroph (Põlme et al. 2020); Decay sugar-rich substrates (Põlme et al. 2020) | Põlme S, et al. 2020. Fungal Diversity 105:1-16 ((https://doi.org/10.1007/s13225-020-00466-2)) |
| Otu53045 | 1 | k__Fungi(100);p__Ascomycota(100);c__Saccharomycetes(100);o__Saccharomycetales(100);f__Metschnikowiaceae(100);g__Metschnikowia(100);s__Metschnikowia_pulcherrima(100); | Metschnikowia | 13 | Saprotroph | \|Undefined Saprotroph\| | Yeast | NULL | Probable | As nectar-tap_saprotroph (Põlme et al. 2020); Decay sugar-rich substrates (Põlme et al. 2020) | Põlme S, et al. 2020. Fungal Diversity 105:1-16 ((https://doi.org/10.1007/s13225-020-00466-2)) |
| Otu53046 | 1 | k__Fungi(100);p__Ascomycota(100);c__Saccharomycetes(100);o__Saccharomycetales(100);f__Metschnikowiaceae(100);g__Metschnikowia(100);g__Metschnikowia_unclassified(100); | Metschnikowia | 13 | Saprotroph | \|Undefined Saprotroph\| | Yeast | NULL | Probable | As nectar-tap_saprotroph (Põlme et al. 2020); Decay sugar-rich substrates (Põlme et al. 2020) | Põlme S, et al. 2020. Fungal Diversity 105:1-16 ((https://doi.org/10.1007/s13225-020-00466-2)) |
| Otu53047 | 1 | k__Fungi(100);p__Ascomycota(100);c__Saccharomycetes(100);o__Saccharomycetales(100);f__Metschnikowiaceae(100);g__Metschnikowia(100);s__Metschnikowia_pulcherrima(100); | Metschnikowia | 13 | Saprotroph | \|Undefined Saprotroph\| | Yeast | NULL | Probable | As nectar-tap_saprotroph (Põlme et al. 2020); Decay sugar-rich substrates (Põlme et al. 2020) | Põlme S, et al. 2020. Fungal Diversity 105:1-16 ((https://doi.org/10.1007/s13225-020-00466-2)) |
| Otu53048 | 1 | k__Fungi(100);p__Ascomycota(100);c__Saccharomycetes(100);o__Saccharomycetales(100);f__Metschnikowiaceae(100);g__Metschnikowia(100);s__Metschnikowia_pulcherrima(100); | Metschnikowia | 13 | Saprotroph | \|Undefined Saprotroph\| | Yeast | NULL | Probable | As nectar-tap_saprotroph (Põlme et al. 2020); Decay sugar-rich substrates (Põlme et al. 2020) | Põlme S, et al. 2020. Fungal Diversity 105:1-16 ((https://doi.org/10.1007/s13225-020-00466-2)) |
| Otu53049 | 1 | k__Fungi(100);p__Ascomycota(100);c__Saccharomycetes(100);o__Saccharomycetales(100);f__Metschnikowiaceae(100);g__Metschnikowia(100);s__Metschnikowia_pulcherrima(100); | Metschnikowia | 13 | Saprotroph | \|Undefined Saprotroph\| | Yeast | NULL | Probable | As nectar-tap_saprotroph (Põlme et al. 2020); Decay sugar-rich substrates (Põlme et al. 2020) | Põlme S, et al. 2020. Fungal Diversity 105:1-16 ((https://doi.org/10.1007/s13225-020-00466-2)) |
| Otu53050 | 1 | k__Fungi(100);p__Ascomycota(100);c__Saccharomycetes(100);o__Saccharomycetales(100);f__Metschnikowiaceae(100);g__Metschnikowia(100);s__Metschnikowia_pulcherrima(100); | Metschnikowia | 13 | Saprotroph | \|Undefined Saprotroph\| | Yeast | NULL | Probable | As nectar-tap_saprotroph (Põlme et al. 2020); Decay sugar-rich substrates (Põlme et al. 2020) | Põlme S, et al. 2020. Fungal Diversity 105:1-16 ((https://doi.org/10.1007/s13225-020-00466-2)) |
| Otu53051 | 1 | k__Fungi(100);p__Ascomycota(100);c__Saccharomycetes(100);o__Saccharomycetales(100);f__Metschnikowiaceae(100);g__Metschnikowia(100);s__Metschnikowia_pulcherrima(100); | Metschnikowia | 13 | Saprotroph | \|Undefined Saprotroph\| | Yeast | NULL | Probable | As nectar-tap_saprotroph (Põlme et al. 2020); Decay sugar-rich substrates (Põlme et al. 2020) | Põlme S, et al. 2020. Fungal Diversity 105:1-16 ((https://doi.org/10.1007/s13225-020-00466-2)) |
| Otu53052 | 1 | k__Fungi(100);p__Ascomycota(100);c__Saccharomycetes(100);o__Saccharomycetales(100);f__Metschnikowiaceae(100);g__Metschnikowia(100);s__Metschnikowia_pulcherrima(100); | Metschnikowia | 13 | Saprotroph | \|Undefined Saprotroph\| | Yeast | NULL | Probable | As nectar-tap_saprotroph (Põlme et al. 2020); Decay sugar-rich substrates (Põlme et al. 2020) | Põlme S, et al. 2020. Fungal Diversity 105:1-16 ((https://doi.org/10.1007/s13225-020-00466-2)) |
| Otu53053 | 1 | k__Fungi(100);p__Ascomycota(100);c__Saccharomycetes(100);o__Saccharomycetales(100);f__Metschnikowiaceae(100);g__Metschnikowia(100);s__Metschnikowia_pulcherrima(100); | Metschnikowia | 13 | Saprotroph | \|Undefined Saprotroph\| | Yeast | NULL | Probable | As nectar-tap_saprotroph (Põlme et al. 2020); Decay sugar-rich substrates (Põlme et al. 2020) | Põlme S, et al. 2020. Fungal Diversity 105:1-16 ((https://doi.org/10.1007/s13225-020-00466-2)) |
| Otu53054 | 1 | k__Fungi(100);p__Ascomycota(100);c__Saccharomycetes(100);o__Saccharomycetales(100);f__Metschnikowiaceae(100);g__Metschnikowia(100);s__Metschnikowia_pulcherrima(100); | Metschnikowia | 13 | Saprotroph | \|Undefined Saprotroph\| | Yeast | NULL | Probable | As nectar-tap_saprotroph (Põlme et al. 2020); Decay sugar-rich substrates (Põlme et al. 2020) | Põlme S, et al. 2020. Fungal Diversity 105:1-16 ((https://doi.org/10.1007/s13225-020-00466-2)) |
| Otu53055 | 1 | k__Fungi(100);p__Ascomycota(100);c__Saccharomycetes(100);o__Saccharomycetales(100);f__Metschnikowiaceae(100);g__Metschnikowia(100);s__Metschnikowia_pulcherrima(100); | Metschnikowia | 13 | Saprotroph | \|Undefined Saprotroph\| | Yeast | NULL | Probable | As nectar-tap_saprotroph (Põlme et al. 2020); Decay sugar-rich substrates (Põlme et al. 2020) | Põlme S, et al. 2020. Fungal Diversity 105:1-16 ((https://doi.org/10.1007/s13225-020-00466-2)) |
| Otu53056 | 1 | k__Fungi(100);p__Ascomycota(100);c__Saccharomycetes(100);o__Saccharomycetales(100);f__Metschnikowiaceae(100);g__Metschnikowia(100);g__Metschnikowia_unclassified(100); | Metschnikowia | 13 | Saprotroph | \|Undefined Saprotroph\| | Yeast | NULL | Probable | As nectar-tap_saprotroph (Põlme et al. 2020); Decay sugar-rich substrates (Põlme et al. 2020) | Põlme S, et al. 2020. Fungal Diversity 105:1-16 ((https://doi.org/10.1007/s13225-020-00466-2)) |
| Otu53057 | 1 | k__Fungi(100);p__Ascomycota(100);c__Saccharomycetes(100);o__Saccharomycetales(100);f__Metschnikowiaceae(100);g__Metschnikowia(100);g__Metschnikowia_unclassified(100); | Metschnikowia | 13 | Saprotroph | \|Undefined Saprotroph\| | Yeast | NULL | Probable | As nectar-tap_saprotroph (Põlme et al. 2020); Decay sugar-rich substrates (Põlme et al. 2020) | Põlme S, et al. 2020. Fungal Diversity 105:1-16 ((https://doi.org/10.1007/s13225-020-00466-2)) |
| Otu53058 | 1 | k__Fungi(100);p__Ascomycota(100);c__Saccharomycetes(100);o__Saccharomycetales(100);f__Metschnikowiaceae(100);g__Metschnikowia(100);g__Metschnikowia_unclassified(100); | Metschnikowia | 13 | Saprotroph | \|Undefined Saprotroph\| | Yeast | NULL | Probable | As nectar-tap_saprotroph (Põlme et al. 2020); Decay sugar-rich substrates (Põlme et al. 2020) | Põlme S, et al. 2020. Fungal Diversity 105:1-16 ((https://doi.org/10.1007/s13225-020-00466-2)) |
| Otu53059 | 1 | k__Fungi(100);p__Ascomycota(100);c__Saccharomycetes(100);o__Saccharomycetales(100);f__Metschnikowiaceae(100);g__Metschnikowia(100);s__Metschnikowia_pulcherrima(100); | Metschnikowia | 13 | Saprotroph | \|Undefined Saprotroph\| | Yeast | NULL | Probable | As nectar-tap_saprotroph (Põlme et al. 2020); Decay sugar-rich substrates (Põlme et al. 2020) | Põlme S, et al. 2020. Fungal Diversity 105:1-16 ((https://doi.org/10.1007/s13225-020-00466-2)) |
| Otu53060 | 1 | k__Fungi(100);p__Ascomycota(100);c__Saccharomycetes(100);o__Saccharomycetales(100);f__Metschnikowiaceae(100);g__Metschnikowia(100);s__Metschnikowia_pulcherrima(100); | Metschnikowia | 13 | Saprotroph | \|Undefined Saprotroph\| | Yeast | NULL | Probable | As nectar-tap_saprotroph (Põlme et al. 2020); Decay sugar-rich substrates (Põlme et al. 2020) | Põlme S, et al. 2020. Fungal Diversity 105:1-16 ((https://doi.org/10.1007/s13225-020-00466-2)) |
| Otu53061 | 1 | k__Fungi(100);p__Ascomycota(100);c__Saccharomycetes(100);o__Saccharomycetales(100);f__Metschnikowiaceae(100);g__Metschnikowia(100);s__Metschnikowia_pulcherrima(100); | Metschnikowia | 13 | Saprotroph | \|Undefined Saprotroph\| | Yeast | NULL | Probable | As nectar-tap_saprotroph (Põlme et al. 2020); Decay sugar-rich substrates (Põlme et al. 2020) | Põlme S, et al. 2020. Fungal Diversity 105:1-16 ((https://doi.org/10.1007/s13225-020-00466-2)) |
| Otu53062 | 1 | k__Fungi(100);p__Ascomycota(100);c__Saccharomycetes(100);o__Saccharomycetales(100);f__Metschnikowiaceae(100);g__Metschnikowia(100);s__Metschnikowia_pulcherrima(100); | Metschnikowia | 13 | Saprotroph | \|Undefined Saprotroph\| | Yeast | NULL | Probable | As nectar-tap_saprotroph (Põlme et al. 2020); Decay sugar-rich substrates (Põlme et al. 2020) | Põlme S, et al. 2020. Fungal Diversity 105:1-16 ((https://doi.org/10.1007/s13225-020-00466-2)) |
| Otu53063 | 1 | k__Fungi(100);p__Ascomycota(100);c__Saccharomycetes(100);o__Saccharomycetales(100);f__Metschnikowiaceae(100);g__Metschnikowia(100);s__Metschnikowia_pulcherrima(100); | Metschnikowia | 13 | Saprotroph | \|Undefined Saprotroph\| | Yeast | NULL | Probable | As nectar-tap_saprotroph (Põlme et al. 2020); Decay sugar-rich substrates (Põlme et al. 2020) | Põlme S, et al. 2020. Fungal Diversity 105:1-16 ((https://doi.org/10.1007/s13225-020-00466-2)) |
| Otu53064 | 1 | k__Fungi(100);p__Ascomycota(100);c__Saccharomycetes(100);o__Saccharomycetales(100);f__Metschnikowiaceae(100);g__Metschnikowia(100);g__Metschnikowia_unclassified(100); | Metschnikowia | 13 | Saprotroph | \|Undefined Saprotroph\| | Yeast | NULL | Probable | As nectar-tap_saprotroph (Põlme et al. 2020); Decay sugar-rich substrates (Põlme et al. 2020) | Põlme S, et al. 2020. Fungal Diversity 105:1-16 ((https://doi.org/10.1007/s13225-020-00466-2)) |
| Otu53065 | 1 | k__Fungi(100);p__Ascomycota(100);c__Saccharomycetes(100);o__Saccharomycetales(100);f__Metschnikowiaceae(100);g__Metschnikowia(100);s__Metschnikowia_pulcherrima(100); | Metschnikowia | 13 | Saprotroph | \|Undefined Saprotroph\| | Yeast | NULL | Probable | As nectar-tap_saprotroph (Põlme et al. 2020); Decay sugar-rich substrates (Põlme et al. 2020) | Põlme S, et al. 2020. Fungal Diversity 105:1-16 ((https://doi.org/10.1007/s13225-020-00466-2)) |
| Otu53066 | 1 | k__Fungi(100);p__Ascomycota(100);c__Saccharomycetes(100);o__Saccharomycetales(100);f__Metschnikowiaceae(100);g__Metschnikowia(100);s__Metschnikowia_pulcherrima(100); | Metschnikowia | 13 | Saprotroph | \|Undefined Saprotroph\| | Yeast | NULL | Probable | As nectar-tap_saprotroph (Põlme et al. 2020); Decay sugar-rich substrates (Põlme et al. 2020) | Põlme S, et al. 2020. Fungal Diversity 105:1-16 ((https://doi.org/10.1007/s13225-020-00466-2)) |
| Otu53067 | 1 | k__Fungi(100);p__Ascomycota(100);c__Saccharomycetes(100);o__Saccharomycetales(100);f__Metschnikowiaceae(100);g__Metschnikowia(100);g__Metschnikowia_unclassified(100); | Metschnikowia | 13 | Saprotroph | \|Undefined Saprotroph\| | Yeast | NULL | Probable | As nectar-tap_saprotroph (Põlme et al. 2020); Decay sugar-rich substrates (Põlme et al. 2020) | Põlme S, et al. 2020. Fungal Diversity 105:1-16 ((https://doi.org/10.1007/s13225-020-00466-2)) |
| Otu53068 | 1 | k__Fungi(100);p__Ascomycota(100);c__Saccharomycetes(100);o__Saccharomycetales(100);f__Metschnikowiaceae(100);g__Metschnikowia(100);g__Metschnikowia_unclassified(100); | Metschnikowia | 13 | Saprotroph | \|Undefined Saprotroph\| | Yeast | NULL | Probable | As nectar-tap_saprotroph (Põlme et al. 2020); Decay sugar-rich substrates (Põlme et al. 2020) | Põlme S, et al. 2020. Fungal Diversity 105:1-16 ((https://doi.org/10.1007/s13225-020-00466-2)) |
| Otu53069 | 1 | k__Fungi(100);p__Ascomycota(100);c__Saccharomycetes(100);o__Saccharomycetales(100);f__Metschnikowiaceae(100);g__Metschnikowia(100);g__Metschnikowia_unclassified(100); | Metschnikowia | 13 | Saprotroph | \|Undefined Saprotroph\| | Yeast | NULL | Probable | As nectar-tap_saprotroph (Põlme et al. 2020); Decay sugar-rich substrates (Põlme et al. 2020) | Põlme S, et al. 2020. Fungal Diversity 105:1-16 ((https://doi.org/10.1007/s13225-020-00466-2)) |
| Otu53070 | 1 | k__Fungi(100);p__Ascomycota(100);c__Saccharomycetes(100);o__Saccharomycetales(100);f__Metschnikowiaceae(100);g__Metschnikowia(100);g__Metschnikowia_unclassified(100); | Metschnikowia | 13 | Saprotroph | \|Undefined Saprotroph\| | Yeast | NULL | Probable | As nectar-tap_saprotroph (Põlme et al. 2020); Decay sugar-rich substrates (Põlme et al. 2020) | Põlme S, et al. 2020. Fungal Diversity 105:1-16 ((https://doi.org/10.1007/s13225-020-00466-2)) |
| Otu53071 | 1 | k__Fungi(100);p__Ascomycota(100);c__Saccharomycetes(100);o__Saccharomycetales(100);f__Metschnikowiaceae(100);g__Metschnikowia(100);s__Metschnikowia_pulcherrima(100); | Metschnikowia | 13 | Saprotroph | \|Undefined Saprotroph\| | Yeast | NULL | Probable | As nectar-tap_saprotroph (Põlme et al. 2020); Decay sugar-rich substrates (Põlme et al. 2020) | Põlme S, et al. 2020. Fungal Diversity 105:1-16 ((https://doi.org/10.1007/s13225-020-00466-2)) |
| Otu53072 | 1 | k__Fungi(100);p__Ascomycota(100);c__Saccharomycetes(100);o__Saccharomycetales(100);f__Metschnikowiaceae(100);g__Metschnikowia(100);g__Metschnikowia_unclassified(100); | Metschnikowia | 13 | Saprotroph | \|Undefined Saprotroph\| | Yeast | NULL | Probable | As nectar-tap_saprotroph (Põlme et al. 2020); Decay sugar-rich substrates (Põlme et al. 2020) | Põlme S, et al. 2020. Fungal Diversity 105:1-16 ((https://doi.org/10.1007/s13225-020-00466-2)) |
| Otu53073 | 1 | k__Fungi(100);p__Ascomycota(100);c__Saccharomycetes(100);o__Saccharomycetales(100);f__Metschnikowiaceae(100);g__Metschnikowia(100);g__Metschnikowia_unclassified(100); | Metschnikowia | 13 | Saprotroph | \|Undefined Saprotroph\| | Yeast | NULL | Probable | As nectar-tap_saprotroph (Põlme et al. 2020); Decay sugar-rich substrates (Põlme et al. 2020) | Põlme S, et al. 2020. Fungal Diversity 105:1-16 ((https://doi.org/10.1007/s13225-020-00466-2)) |
| Otu53074 | 1 | k__Fungi(100);p__Ascomycota(100);c__Saccharomycetes(100);o__Saccharomycetales(100);f__Metschnikowiaceae(100);g__Metschnikowia(100);s__Metschnikowia_pulcherrima(100); | Metschnikowia | 13 | Saprotroph | \|Undefined Saprotroph\| | Yeast | NULL | Probable | As nectar-tap_saprotroph (Põlme et al. 2020); Decay sugar-rich substrates (Põlme et al. 2020) | Põlme S, et al. 2020. Fungal Diversity 105:1-16 ((https://doi.org/10.1007/s13225-020-00466-2)) |
| Otu53075 | 1 | k__Fungi(100);p__Ascomycota(100);c__Saccharomycetes(100);o__Saccharomycetales(100);f__Metschnikowiaceae(100);g__Metschnikowia(100);s__Metschnikowia_pulcherrima(100); | Metschnikowia | 13 | Saprotroph | \|Undefined Saprotroph\| | Yeast | NULL | Probable | As nectar-tap_saprotroph (Põlme et al. 2020); Decay sugar-rich substrates (Põlme et al. 2020) | Põlme S, et al. 2020. Fungal Diversity 105:1-16 ((https://doi.org/10.1007/s13225-020-00466-2)) |
| Otu53076 | 1 | k__Fungi(100);p__Ascomycota(100);c__Saccharomycetes(100);o__Saccharomycetales(100);f__Metschnikowiaceae(100);g__Metschnikowia(100);s__Metschnikowia_pulcherrima(100); | Metschnikowia | 13 | Saprotroph | \|Undefined Saprotroph\| | Yeast | NULL | Probable | As nectar-tap_saprotroph (Põlme et al. 2020); Decay sugar-rich substrates (Põlme et al. 2020) | Põlme S, et al. 2020. Fungal Diversity 105:1-16 ((https://doi.org/10.1007/s13225-020-00466-2)) |
| Otu53077 | 1 | k__Fungi(100);p__Ascomycota(100);c__Saccharomycetes(100);o__Saccharomycetales(100);f__Metschnikowiaceae(100);g__Metschnikowia(100);s__Metschnikowia_pulcherrima(100); | Metschnikowia | 13 | Saprotroph | \|Undefined Saprotroph\| | Yeast | NULL | Probable | As nectar-tap_saprotroph (Põlme et al. 2020); Decay sugar-rich substrates (Põlme et al. 2020) | Põlme S, et al. 2020. Fungal Diversity 105:1-16 ((https://doi.org/10.1007/s13225-020-00466-2)) |
| Otu53078 | 1 | k__Fungi(100);p__Ascomycota(100);c__Saccharomycetes(100);o__Saccharomycetales(100);f__Metschnikowiaceae(100);g__Metschnikowia(100);s__Metschnikowia_pulcherrima(100); | Metschnikowia | 13 | Saprotroph | \|Undefined Saprotroph\| | Yeast | NULL | Probable | As nectar-tap_saprotroph (Põlme et al. 2020); Decay sugar-rich substrates (Põlme et al. 2020) | Põlme S, et al. 2020. Fungal Diversity 105:1-16 ((https://doi.org/10.1007/s13225-020-00466-2)) |
| Otu53079 | 1 | k__Fungi(100);p__Ascomycota(100);c__Saccharomycetes(100);o__Saccharomycetales(100);f__Metschnikowiaceae(100);g__Metschnikowia(100);s__Metschnikowia_pulcherrima(100); | Metschnikowia | 13 | Saprotroph | \|Undefined Saprotroph\| | Yeast | NULL | Probable | As nectar-tap_saprotroph (Põlme et al. 2020); Decay sugar-rich substrates (Põlme et al. 2020) | Põlme S, et al. 2020. Fungal Diversity 105:1-16 ((https://doi.org/10.1007/s13225-020-00466-2)) |
| Otu53080 | 1 | k__Fungi(100);p__Ascomycota(100);c__Saccharomycetes(100);o__Saccharomycetales(100);f__Metschnikowiaceae(100);g__Metschnikowia(100);s__Metschnikowia_pulcherrima(100); | Metschnikowia | 13 | Saprotroph | \|Undefined Saprotroph\| | Yeast | NULL | Probable | As nectar-tap_saprotroph (Põlme et al. 2020); Decay sugar-rich substrates (Põlme et al. 2020) | Põlme S, et al. 2020. Fungal Diversity 105:1-16 ((https://doi.org/10.1007/s13225-020-00466-2)) |
| Otu53081 | 1 | k__Fungi(100);p__Ascomycota(100);c__Saccharomycetes(100);o__Saccharomycetales(100);f__Metschnikowiaceae(100);g__Metschnikowia(100);s__Metschnikowia_pulcherrima(100); | Metschnikowia | 13 | Saprotroph | \|Undefined Saprotroph\| | Yeast | NULL | Probable | As nectar-tap_saprotroph (Põlme et al. 2020); Decay sugar-rich substrates (Põlme et al. 2020) | Põlme S, et al. 2020. Fungal Diversity 105:1-16 ((https://doi.org/10.1007/s13225-020-00466-2)) |
| Otu53082 | 1 | k__Fungi(100);p__Ascomycota(100);c__Saccharomycetes(100);o__Saccharomycetales(100);f__Metschnikowiaceae(100);g__Metschnikowia(100);s__Metschnikowia_pulcherrima(100); | Metschnikowia | 13 | Saprotroph | \|Undefined Saprotroph\| | Yeast | NULL | Probable | As nectar-tap_saprotroph (Põlme et al. 2020); Decay sugar-rich substrates (Põlme et al. 2020) | Põlme S, et al. 2020. Fungal Diversity 105:1-16 ((https://doi.org/10.1007/s13225-020-00466-2)) |
| Otu53083 | 1 | k__Fungi(100);p__Ascomycota(100);c__Saccharomycetes(100);o__Saccharomycetales(100);f__Metschnikowiaceae(100);g__Metschnikowia(100);s__Metschnikowia_pulcherrima(100); | Metschnikowia | 13 | Saprotroph | \|Undefined Saprotroph\| | Yeast | NULL | Probable | As nectar-tap_saprotroph (Põlme et al. 2020); Decay sugar-rich substrates (Põlme et al. 2020) | Põlme S, et al. 2020. Fungal Diversity 105:1-16 ((https://doi.org/10.1007/s13225-020-00466-2)) |
| Otu53084 | 1 | k__Fungi(100);p__Ascomycota(100);c__Saccharomycetes(100);o__Saccharomycetales(100);f__Metschnikowiaceae(100);g__Metschnikowia(100);s__Metschnikowia_pulcherrima(100); | Metschnikowia | 13 | Saprotroph | \|Undefined Saprotroph\| | Yeast | NULL | Probable | As nectar-tap_saprotroph (Põlme et al. 2020); Decay sugar-rich substrates (Põlme et al. 2020) | Põlme S, et al. 2020. Fungal Diversity 105:1-16 ((https://doi.org/10.1007/s13225-020-00466-2)) |
| Otu53085 | 1 | k__Fungi(100);p__Ascomycota(100);c__Saccharomycetes(100);o__Saccharomycetales(100);f__Metschnikowiaceae(100);g__Metschnikowia(100);g__Metschnikowia_unclassified(100); | Metschnikowia | 13 | Saprotroph | \|Undefined Saprotroph\| | Yeast | NULL | Probable | As nectar-tap_saprotroph (Põlme et al. 2020); Decay sugar-rich substrates (Põlme et al. 2020) | Põlme S, et al. 2020. Fungal Diversity 105:1-16 ((https://doi.org/10.1007/s13225-020-00466-2)) |
| Otu53086 | 1 | k__Fungi(100);p__Ascomycota(100);c__Saccharomycetes(100);o__Saccharomycetales(100);f__Metschnikowiaceae(100);g__Metschnikowia(100);s__Metschnikowia_pulcherrima(100); | Metschnikowia | 13 | Saprotroph | \|Undefined Saprotroph\| | Yeast | NULL | Probable | As nectar-tap_saprotroph (Põlme et al. 2020); Decay sugar-rich substrates (Põlme et al. 2020) | Põlme S, et al. 2020. Fungal Diversity 105:1-16 ((https://doi.org/10.1007/s13225-020-00466-2)) |
| Otu53087 | 1 | k__Fungi(100);p__Ascomycota(100);c__Saccharomycetes(100);o__Saccharomycetales(100);f__Metschnikowiaceae(100);g__Metschnikowia(100);g__Metschnikowia_unclassified(100); | Metschnikowia | 13 | Saprotroph | \|Undefined Saprotroph\| | Yeast | NULL | Probable | As nectar-tap_saprotroph (Põlme et al. 2020); Decay sugar-rich substrates (Põlme et al. 2020) | Põlme S, et al. 2020. Fungal Diversity 105:1-16 ((https://doi.org/10.1007/s13225-020-00466-2)) |
| Otu53088 | 1 | k__Fungi(100);p__Ascomycota(100);c__Saccharomycetes(100);o__Saccharomycetales(100);f__Metschnikowiaceae(100);g__Metschnikowia(100);s__Metschnikowia_pulcherrima(100); | Metschnikowia | 13 | Saprotroph | \|Undefined Saprotroph\| | Yeast | NULL | Probable | As nectar-tap_saprotroph (Põlme et al. 2020); Decay sugar-rich substrates (Põlme et al. 2020) | Põlme S, et al. 2020. Fungal Diversity 105:1-16 ((https://doi.org/10.1007/s13225-020-00466-2)) |
| Otu53089 | 1 | k__Fungi(100);p__Ascomycota(100);c__Saccharomycetes(100);o__Saccharomycetales(100);f__Metschnikowiaceae(100);g__Metschnikowia(100);s__Metschnikowia_pulcherrima(100); | Metschnikowia | 13 | Saprotroph | \|Undefined Saprotroph\| | Yeast | NULL | Probable | As nectar-tap_saprotroph (Põlme et al. 2020); Decay sugar-rich substrates (Põlme et al. 2020) | Põlme S, et al. 2020. Fungal Diversity 105:1-16 ((https://doi.org/10.1007/s13225-020-00466-2)) |
| Otu53090 | 1 | k__Fungi(100);p__Ascomycota(100);c__Saccharomycetes(100);o__Saccharomycetales(100);f__Metschnikowiaceae(100);g__Metschnikowia(100);s__Metschnikowia_pulcherrima(100); | Metschnikowia | 13 | Saprotroph | \|Undefined Saprotroph\| | Yeast | NULL | Probable | As nectar-tap_saprotroph (Põlme et al. 2020); Decay sugar-rich substrates (Põlme et al. 2020) | Põlme S, et al. 2020. Fungal Diversity 105:1-16 ((https://doi.org/10.1007/s13225-020-00466-2)) |
| Otu53091 | 1 | k__Fungi(100);p__Ascomycota(100);c__Saccharomycetes(100);o__Saccharomycetales(100);f__Metschnikowiaceae(100);g__Metschnikowia(100);s__Metschnikowia_pulcherrima(100); | Metschnikowia | 13 | Saprotroph | \|Undefined Saprotroph\| | Yeast | NULL | Probable | As nectar-tap_saprotroph (Põlme et al. 2020); Decay sugar-rich substrates (Põlme et al. 2020) | Põlme S, et al. 2020. Fungal Diversity 105:1-16 ((https://doi.org/10.1007/s13225-020-00466-2)) |
| Otu53092 | 1 | k__Fungi(100);p__Ascomycota(100);c__Sordariomycetes(100);o__Hypocreales(100);f__Nectriaceae(100);f__Nectriaceae_unclassified(100);f__Nectriaceae_unclassified(100); | Nectriaceae | 9 | Pathotroph-Saprotroph-Symbiotroph | Animal Pathogen-Endophyte-Fungal Parasite-Lichen Parasite-Plant Pathogen-Wood Saprotroph | Microfungus | NULL | Possible | Fusarium species are amongst the economically significant plant pathogens (Cannon & Kirk 2007) | Cannon PF, Kirk PM. 2007. Fungal Families of the World. CAB International, Cambridge (ISBN: 978-0851998275); Hirooka Y, et al. 2012. Studies in Mycology 711-210 ((https://doi.org/10.3114/sim0001)); Tedersoo L, et al. 2014. Science 346:e1256688 ((https://doi.org/10.1126/science.1256688)); The American Phytopathological Society Common Names of Plant Diseases ((http://www.apsnet.org/publications/commonnames/Pages/default.aspx)); https://www.cdc.gov ((https://www.cdc.gov)) |
| Otu53093 | 1 | k__Fungi(100);p__Ascomycota(100);c__Saccharomycetes(100);o__Saccharomycetales(100);f__Metschnikowiaceae(100);g__Metschnikowia(100);s__Metschnikowia_pulcherrima(100); | Metschnikowia | 13 | Saprotroph | \|Undefined Saprotroph\| | Yeast | NULL | Probable | As nectar-tap_saprotroph (Põlme et al. 2020); Decay sugar-rich substrates (Põlme et al. 2020) | Põlme S, et al. 2020. Fungal Diversity 105:1-16 ((https://doi.org/10.1007/s13225-020-00466-2)) |
| Otu53094 | 1 | k__Fungi(100);p__Ascomycota(100);c__Saccharomycetes(100);o__Saccharomycetales(100);f__Metschnikowiaceae(100);g__Metschnikowia(100);g__Metschnikowia_unclassified(100); | Metschnikowia | 13 | Saprotroph | \|Undefined Saprotroph\| | Yeast | NULL | Probable | As nectar-tap_saprotroph (Põlme et al. 2020); Decay sugar-rich substrates (Põlme et al. 2020) | Põlme S, et al. 2020. Fungal Diversity 105:1-16 ((https://doi.org/10.1007/s13225-020-00466-2)) |
| Otu53095 | 1 | k__Fungi(100);p__Ascomycota(100);c__Saccharomycetes(100);o__Saccharomycetales(100);f__Metschnikowiaceae(100);g__Metschnikowia(100);s__Metschnikowia_pulcherrima(100); | Metschnikowia | 13 | Saprotroph | \|Undefined Saprotroph\| | Yeast | NULL | Probable | As nectar-tap_saprotroph (Põlme et al. 2020); Decay sugar-rich substrates (Põlme et al. 2020) | Põlme S, et al. 2020. Fungal Diversity 105:1-16 ((https://doi.org/10.1007/s13225-020-00466-2)) |
| Otu53096 | 1 | k__Fungi(100);p__Ascomycota(100);c__Saccharomycetes(100);o__Saccharomycetales(100);f__Metschnikowiaceae(100);g__Metschnikowia(100);s__Metschnikowia_pulcherrima(100); | Metschnikowia | 13 | Saprotroph | \|Undefined Saprotroph\| | Yeast | NULL | Probable | As nectar-tap_saprotroph (Põlme et al. 2020); Decay sugar-rich substrates (Põlme et al. 2020) | Põlme S, et al. 2020. Fungal Diversity 105:1-16 ((https://doi.org/10.1007/s13225-020-00466-2)) |
| Otu53097 | 1 | k__Fungi(100);p__Ascomycota(100);c__Saccharomycetes(100);o__Saccharomycetales(100);f__Metschnikowiaceae(100);g__Metschnikowia(100);s__Metschnikowia_pulcherrima(100); | Metschnikowia | 13 | Saprotroph | \|Undefined Saprotroph\| | Yeast | NULL | Probable | As nectar-tap_saprotroph (Põlme et al. 2020); Decay sugar-rich substrates (Põlme et al. 2020) | Põlme S, et al. 2020. Fungal Diversity 105:1-16 ((https://doi.org/10.1007/s13225-020-00466-2)) |
| Otu53100 | 1 | k__Fungi(100);p__Ascomycota(100);c__Saccharomycetes(100);o__Saccharomycetales(100);f__Metschnikowiaceae(100);g__Metschnikowia(100);s__Metschnikowia_pulcherrima(100); | Metschnikowia | 13 | Saprotroph | \|Undefined Saprotroph\| | Yeast | NULL | Probable | As nectar-tap_saprotroph (Põlme et al. 2020); Decay sugar-rich substrates (Põlme et al. 2020) | Põlme S, et al. 2020. Fungal Diversity 105:1-16 ((https://doi.org/10.1007/s13225-020-00466-2)) |
| Otu53101 | 1 | k__Fungi(100);p__Ascomycota(100);c__Saccharomycetes(100);o__Saccharomycetales(100);f__Metschnikowiaceae(100);g__Metschnikowia(100);s__Metschnikowia_pulcherrima(100); | Metschnikowia | 13 | Saprotroph | \|Undefined Saprotroph\| | Yeast | NULL | Probable | As nectar-tap_saprotroph (Põlme et al. 2020); Decay sugar-rich substrates (Põlme et al. 2020) | Põlme S, et al. 2020. Fungal Diversity 105:1-16 ((https://doi.org/10.1007/s13225-020-00466-2)) |
| Otu53102 | 1 | k__Fungi(100);p__Ascomycota(100);c__Saccharomycetes(100);o__Saccharomycetales(100);f__Metschnikowiaceae(100);g__Metschnikowia(100);s__Metschnikowia_pulcherrima(100); | Metschnikowia | 13 | Saprotroph | \|Undefined Saprotroph\| | Yeast | NULL | Probable | As nectar-tap_saprotroph (Põlme et al. 2020); Decay sugar-rich substrates (Põlme et al. 2020) | Põlme S, et al. 2020. Fungal Diversity 105:1-16 ((https://doi.org/10.1007/s13225-020-00466-2)) |
| Otu53103 | 1 | k__Fungi(100);p__Ascomycota(100);c__Saccharomycetes(100);o__Saccharomycetales(100);f__Metschnikowiaceae(100);g__Metschnikowia(100);s__Metschnikowia_pulcherrima(100); | Metschnikowia | 13 | Saprotroph | \|Undefined Saprotroph\| | Yeast | NULL | Probable | As nectar-tap_saprotroph (Põlme et al. 2020); Decay sugar-rich substrates (Põlme et al. 2020) | Põlme S, et al. 2020. Fungal Diversity 105:1-16 ((https://doi.org/10.1007/s13225-020-00466-2)) |
| Otu53105 | 1 | k__Fungi(100);p__Ascomycota(100);c__Saccharomycetes(100);o__Saccharomycetales(100);f__Metschnikowiaceae(100);g__Metschnikowia(100);s__Metschnikowia_pulcherrima(100); | Metschnikowia | 13 | Saprotroph | \|Undefined Saprotroph\| | Yeast | NULL | Probable | As nectar-tap_saprotroph (Põlme et al. 2020); Decay sugar-rich substrates (Põlme et al. 2020) | Põlme S, et al. 2020. Fungal Diversity 105:1-16 ((https://doi.org/10.1007/s13225-020-00466-2)) |
| Otu53106 | 1 | k__Fungi(100);p__Ascomycota(100);c__Saccharomycetes(100);o__Saccharomycetales(100);f__Metschnikowiaceae(100);g__Metschnikowia(100);s__Metschnikowia_pulcherrima(100); | Metschnikowia | 13 | Saprotroph | \|Undefined Saprotroph\| | Yeast | NULL | Probable | As nectar-tap_saprotroph (Põlme et al. 2020); Decay sugar-rich substrates (Põlme et al. 2020) | Põlme S, et al. 2020. Fungal Diversity 105:1-16 ((https://doi.org/10.1007/s13225-020-00466-2)) |
| Otu53107 | 1 | k__Fungi(100);p__Ascomycota(100);c__Saccharomycetes(100);o__Saccharomycetales(100);f__Metschnikowiaceae(100);g__Metschnikowia(100);s__Metschnikowia_pulcherrima(100); | Metschnikowia | 13 | Saprotroph | \|Undefined Saprotroph\| | Yeast | NULL | Probable | As nectar-tap_saprotroph (Põlme et al. 2020); Decay sugar-rich substrates (Põlme et al. 2020) | Põlme S, et al. 2020. Fungal Diversity 105:1-16 ((https://doi.org/10.1007/s13225-020-00466-2)) |
| Otu53108 | 1 | k__Fungi(100);p__Ascomycota(100);c__Saccharomycetes(100);o__Saccharomycetales(100);f__Metschnikowiaceae(100);g__Metschnikowia(100);g__Metschnikowia_unclassified(100); | Metschnikowia | 13 | Saprotroph | \|Undefined Saprotroph\| | Yeast | NULL | Probable | As nectar-tap_saprotroph (Põlme et al. 2020); Decay sugar-rich substrates (Põlme et al. 2020) | Põlme S, et al. 2020. Fungal Diversity 105:1-16 ((https://doi.org/10.1007/s13225-020-00466-2)) |
| Otu53109 | 1 | k__Fungi(100);p__Ascomycota(100);c__Saccharomycetes(100);o__Saccharomycetales(100);f__Metschnikowiaceae(100);g__Metschnikowia(100);s__Metschnikowia_pulcherrima(100); | Metschnikowia | 13 | Saprotroph | \|Undefined Saprotroph\| | Yeast | NULL | Probable | As nectar-tap_saprotroph (Põlme et al. 2020); Decay sugar-rich substrates (Põlme et al. 2020) | Põlme S, et al. 2020. Fungal Diversity 105:1-16 ((https://doi.org/10.1007/s13225-020-00466-2)) |
| Otu53110 | 1 | k__Fungi(100);p__Ascomycota(100);c__Saccharomycetes(100);o__Saccharomycetales(100);f__Metschnikowiaceae(100);g__Metschnikowia(100);s__Metschnikowia_pulcherrima(100); | Metschnikowia | 13 | Saprotroph | \|Undefined Saprotroph\| | Yeast | NULL | Probable | As nectar-tap_saprotroph (Põlme et al. 2020); Decay sugar-rich substrates (Põlme et al. 2020) | Põlme S, et al. 2020. Fungal Diversity 105:1-16 ((https://doi.org/10.1007/s13225-020-00466-2)) |
| Otu53111 | 1 | k__Fungi(100);p__Ascomycota(100);c__Saccharomycetes(100);o__Saccharomycetales(100);f__Metschnikowiaceae(100);g__Metschnikowia(100);s__Metschnikowia_pulcherrima(100); | Metschnikowia | 13 | Saprotroph | \|Undefined Saprotroph\| | Yeast | NULL | Probable | As nectar-tap_saprotroph (Põlme et al. 2020); Decay sugar-rich substrates (Põlme et al. 2020) | Põlme S, et al. 2020. Fungal Diversity 105:1-16 ((https://doi.org/10.1007/s13225-020-00466-2)) |
| Otu53112 | 1 | k__Fungi(100);p__Ascomycota(100);c__Saccharomycetes(100);o__Saccharomycetales(100);f__Metschnikowiaceae(100);g__Metschnikowia(100);s__Metschnikowia_pulcherrima(100); | Metschnikowia | 13 | Saprotroph | \|Undefined Saprotroph\| | Yeast | NULL | Probable | As nectar-tap_saprotroph (Põlme et al. 2020); Decay sugar-rich substrates (Põlme et al. 2020) | Põlme S, et al. 2020. Fungal Diversity 105:1-16 ((https://doi.org/10.1007/s13225-020-00466-2)) |
| Otu53113 | 1 | k__Fungi(100);p__Ascomycota(100);c__Saccharomycetes(100);o__Saccharomycetales(100);f__Metschnikowiaceae(100);g__Metschnikowia(100);s__Metschnikowia_pulcherrima(100); | Metschnikowia | 13 | Saprotroph | \|Undefined Saprotroph\| | Yeast | NULL | Probable | As nectar-tap_saprotroph (Põlme et al. 2020); Decay sugar-rich substrates (Põlme et al. 2020) | Põlme S, et al. 2020. Fungal Diversity 105:1-16 ((https://doi.org/10.1007/s13225-020-00466-2)) |
| Otu53114 | 1 | k__Fungi(100);p__Ascomycota(100);c__Saccharomycetes(100);o__Saccharomycetales(100);f__Metschnikowiaceae(100);g__Metschnikowia(100);g__Metschnikowia_unclassified(100); | Metschnikowia | 13 | Saprotroph | \|Undefined Saprotroph\| | Yeast | NULL | Probable | As nectar-tap_saprotroph (Põlme et al. 2020); Decay sugar-rich substrates (Põlme et al. 2020) | Põlme S, et al. 2020. Fungal Diversity 105:1-16 ((https://doi.org/10.1007/s13225-020-00466-2)) |
| Otu53115 | 1 | k__Fungi(100);p__Ascomycota(100);c__Saccharomycetes(100);o__Saccharomycetales(100);f__Metschnikowiaceae(100);g__Metschnikowia(100);g__Metschnikowia_unclassified(100); | Metschnikowia | 13 | Saprotroph | \|Undefined Saprotroph\| | Yeast | NULL | Probable | As nectar-tap_saprotroph (Põlme et al. 2020); Decay sugar-rich substrates (Põlme et al. 2020) | Põlme S, et al. 2020. Fungal Diversity 105:1-16 ((https://doi.org/10.1007/s13225-020-00466-2)) |
| Otu53116 | 1 | k__Fungi(100);p__Ascomycota(100);c__Saccharomycetes(100);o__Saccharomycetales(100);f__Metschnikowiaceae(100);g__Metschnikowia(100);g__Metschnikowia_unclassified(100); | Metschnikowia | 13 | Saprotroph | \|Undefined Saprotroph\| | Yeast | NULL | Probable | As nectar-tap_saprotroph (Põlme et al. 2020); Decay sugar-rich substrates (Põlme et al. 2020) | Põlme S, et al. 2020. Fungal Diversity 105:1-16 ((https://doi.org/10.1007/s13225-020-00466-2)) |
| Otu53117 | 1 | k__Fungi(100);p__Ascomycota(100);c__Saccharomycetes(100);o__Saccharomycetales(100);f__Metschnikowiaceae(100);g__Metschnikowia(100);g__Metschnikowia_unclassified(100); | Metschnikowia | 13 | Saprotroph | \|Undefined Saprotroph\| | Yeast | NULL | Probable | As nectar-tap_saprotroph (Põlme et al. 2020); Decay sugar-rich substrates (Põlme et al. 2020) | Põlme S, et al. 2020. Fungal Diversity 105:1-16 ((https://doi.org/10.1007/s13225-020-00466-2)) |
| Otu53118 | 1 | k__Fungi(100);p__Ascomycota(100);c__Saccharomycetes(100);o__Saccharomycetales(100);f__Metschnikowiaceae(100);g__Metschnikowia(100);s__Metschnikowia_pulcherrima(100); | Metschnikowia | 13 | Saprotroph | \|Undefined Saprotroph\| | Yeast | NULL | Probable | As nectar-tap_saprotroph (Põlme et al. 2020); Decay sugar-rich substrates (Põlme et al. 2020) | Põlme S, et al. 2020. Fungal Diversity 105:1-16 ((https://doi.org/10.1007/s13225-020-00466-2)) |
| Otu53119 | 1 | k__Fungi(100);p__Ascomycota(100);c__Saccharomycetes(100);o__Saccharomycetales(100);f__Metschnikowiaceae(100);g__Metschnikowia(100);s__Metschnikowia_pulcherrima(100); | Metschnikowia | 13 | Saprotroph | \|Undefined Saprotroph\| | Yeast | NULL | Probable | As nectar-tap_saprotroph (Põlme et al. 2020); Decay sugar-rich substrates (Põlme et al. 2020) | Põlme S, et al. 2020. Fungal Diversity 105:1-16 ((https://doi.org/10.1007/s13225-020-00466-2)) |
| Otu53120 | 1 | k__Fungi(100);p__Ascomycota(100);c__Saccharomycetes(100);o__Saccharomycetales(100);f__Metschnikowiaceae(100);g__Metschnikowia(100);s__Metschnikowia_pulcherrima(100); | Metschnikowia | 13 | Saprotroph | \|Undefined Saprotroph\| | Yeast | NULL | Probable | As nectar-tap_saprotroph (Põlme et al. 2020); Decay sugar-rich substrates (Põlme et al. 2020) | Põlme S, et al. 2020. Fungal Diversity 105:1-16 ((https://doi.org/10.1007/s13225-020-00466-2)) |
| Otu53121 | 1 | k__Fungi(100);p__Ascomycota(100);c__Saccharomycetes(100);o__Saccharomycetales(100);f__Metschnikowiaceae(100);g__Metschnikowia(100);s__Metschnikowia_pulcherrima(100); | Metschnikowia | 13 | Saprotroph | \|Undefined Saprotroph\| | Yeast | NULL | Probable | As nectar-tap_saprotroph (Põlme et al. 2020); Decay sugar-rich substrates (Põlme et al. 2020) | Põlme S, et al. 2020. Fungal Diversity 105:1-16 ((https://doi.org/10.1007/s13225-020-00466-2)) |
| Otu53122 | 1 | k__Fungi(100);p__Ascomycota(100);c__Saccharomycetes(100);o__Saccharomycetales(100);f__Metschnikowiaceae(100);g__Metschnikowia(100);s__Metschnikowia_pulcherrima(100); | Metschnikowia | 13 | Saprotroph | \|Undefined Saprotroph\| | Yeast | NULL | Probable | As nectar-tap_saprotroph (Põlme et al. 2020); Decay sugar-rich substrates (Põlme et al. 2020) | Põlme S, et al. 2020. Fungal Diversity 105:1-16 ((https://doi.org/10.1007/s13225-020-00466-2)) |
| Otu53123 | 1 | k__Fungi(100);p__Ascomycota(100);c__Saccharomycetes(100);o__Saccharomycetales(100);f__Metschnikowiaceae(100);g__Metschnikowia(100);s__Metschnikowia_pulcherrima(100); | Metschnikowia | 13 | Saprotroph | \|Undefined Saprotroph\| | Yeast | NULL | Probable | As nectar-tap_saprotroph (Põlme et al. 2020); Decay sugar-rich substrates (Põlme et al. 2020) | Põlme S, et al. 2020. Fungal Diversity 105:1-16 ((https://doi.org/10.1007/s13225-020-00466-2)) |
| Otu53124 | 1 | k__Fungi(100);p__Ascomycota(100);c__Saccharomycetes(100);o__Saccharomycetales(100);f__Metschnikowiaceae(100);g__Metschnikowia(100);g__Metschnikowia_unclassified(100); | Metschnikowia | 13 | Saprotroph | \|Undefined Saprotroph\| | Yeast | NULL | Probable | As nectar-tap_saprotroph (Põlme et al. 2020); Decay sugar-rich substrates (Põlme et al. 2020) | Põlme S, et al. 2020. Fungal Diversity 105:1-16 ((https://doi.org/10.1007/s13225-020-00466-2)) |
| Otu53125 | 1 | k__Fungi(100);p__Ascomycota(100);c__Saccharomycetes(100);o__Saccharomycetales(100);f__Metschnikowiaceae(100);g__Metschnikowia(100);s__Metschnikowia_pulcherrima(100); | Metschnikowia | 13 | Saprotroph | \|Undefined Saprotroph\| | Yeast | NULL | Probable | As nectar-tap_saprotroph (Põlme et al. 2020); Decay sugar-rich substrates (Põlme et al. 2020) | Põlme S, et al. 2020. Fungal Diversity 105:1-16 ((https://doi.org/10.1007/s13225-020-00466-2)) |
| Otu53126 | 1 | k__Fungi(100);p__Ascomycota(100);c__Saccharomycetes(100);o__Saccharomycetales(100);f__Metschnikowiaceae(100);g__Metschnikowia(100);g__Metschnikowia_unclassified(100); | Metschnikowia | 13 | Saprotroph | \|Undefined Saprotroph\| | Yeast | NULL | Probable | As nectar-tap_saprotroph (Põlme et al. 2020); Decay sugar-rich substrates (Põlme et al. 2020) | Põlme S, et al. 2020. Fungal Diversity 105:1-16 ((https://doi.org/10.1007/s13225-020-00466-2)) |
| Otu53127 | 1 | k__Fungi(100);p__Ascomycota(100);c__Saccharomycetes(100);o__Saccharomycetales(100);f__Metschnikowiaceae(100);g__Metschnikowia(100);g__Metschnikowia_unclassified(100); | Metschnikowia | 13 | Saprotroph | \|Undefined Saprotroph\| | Yeast | NULL | Probable | As nectar-tap_saprotroph (Põlme et al. 2020); Decay sugar-rich substrates (Põlme et al. 2020) | Põlme S, et al. 2020. Fungal Diversity 105:1-16 ((https://doi.org/10.1007/s13225-020-00466-2)) |
| Otu53128 | 1 | k__Fungi(100);p__Ascomycota(100);c__Saccharomycetes(100);o__Saccharomycetales(100);f__Metschnikowiaceae(100);g__Metschnikowia(100);g__Metschnikowia_unclassified(100); | Metschnikowia | 13 | Saprotroph | \|Undefined Saprotroph\| | Yeast | NULL | Probable | As nectar-tap_saprotroph (Põlme et al. 2020); Decay sugar-rich substrates (Põlme et al. 2020) | Põlme S, et al. 2020. Fungal Diversity 105:1-16 ((https://doi.org/10.1007/s13225-020-00466-2)) |
| Otu53129 | 1 | k__Fungi(100);p__Ascomycota(100);c__Saccharomycetes(100);o__Saccharomycetales(100);f__Metschnikowiaceae(100);g__Metschnikowia(100);s__Metschnikowia_pulcherrima(100); | Metschnikowia | 13 | Saprotroph | \|Undefined Saprotroph\| | Yeast | NULL | Probable | As nectar-tap_saprotroph (Põlme et al. 2020); Decay sugar-rich substrates (Põlme et al. 2020) | Põlme S, et al. 2020. Fungal Diversity 105:1-16 ((https://doi.org/10.1007/s13225-020-00466-2)) |
| Otu53130 | 1 | k__Fungi(100);p__Ascomycota(100);c__Saccharomycetes(100);o__Saccharomycetales(100);f__Metschnikowiaceae(100);g__Metschnikowia(100);s__Metschnikowia_pulcherrima(100); | Metschnikowia | 13 | Saprotroph | \|Undefined Saprotroph\| | Yeast | NULL | Probable | As nectar-tap_saprotroph (Põlme et al. 2020); Decay sugar-rich substrates (Põlme et al. 2020) | Põlme S, et al. 2020. Fungal Diversity 105:1-16 ((https://doi.org/10.1007/s13225-020-00466-2)) |
| Otu53131 | 1 | k__Fungi(100);p__Ascomycota(100);c__Saccharomycetes(100);o__Saccharomycetales(100);f__Metschnikowiaceae(100);g__Metschnikowia(100);s__Metschnikowia_pulcherrima(100); | Metschnikowia | 13 | Saprotroph | \|Undefined Saprotroph\| | Yeast | NULL | Probable | As nectar-tap_saprotroph (Põlme et al. 2020); Decay sugar-rich substrates (Põlme et al. 2020) | Põlme S, et al. 2020. Fungal Diversity 105:1-16 ((https://doi.org/10.1007/s13225-020-00466-2)) |
| Otu53132 | 1 | k__Fungi(100);p__Ascomycota(100);c__Saccharomycetes(100);o__Saccharomycetales(100);f__Metschnikowiaceae(100);g__Metschnikowia(100);g__Metschnikowia_unclassified(100); | Metschnikowia | 13 | Saprotroph | \|Undefined Saprotroph\| | Yeast | NULL | Probable | As nectar-tap_saprotroph (Põlme et al. 2020); Decay sugar-rich substrates (Põlme et al. 2020) | Põlme S, et al. 2020. Fungal Diversity 105:1-16 ((https://doi.org/10.1007/s13225-020-00466-2)) |
| Otu53133 | 1 | k__Fungi(100);p__Ascomycota(100);c__Saccharomycetes(100);o__Saccharomycetales(100);f__Metschnikowiaceae(100);g__Metschnikowia(100);g__Metschnikowia_unclassified(100); | Metschnikowia | 13 | Saprotroph | \|Undefined Saprotroph\| | Yeast | NULL | Probable | As nectar-tap_saprotroph (Põlme et al. 2020); Decay sugar-rich substrates (Põlme et al. 2020) | Põlme S, et al. 2020. Fungal Diversity 105:1-16 ((https://doi.org/10.1007/s13225-020-00466-2)) |
| Otu53134 | 1 | k__Fungi(100);p__Ascomycota(100);c__Saccharomycetes(100);o__Saccharomycetales(100);f__Metschnikowiaceae(100);g__Metschnikowia(100);s__Metschnikowia_pulcherrima(100); | Metschnikowia | 13 | Saprotroph | \|Undefined Saprotroph\| | Yeast | NULL | Probable | As nectar-tap_saprotroph (Põlme et al. 2020); Decay sugar-rich substrates (Põlme et al. 2020) | Põlme S, et al. 2020. Fungal Diversity 105:1-16 ((https://doi.org/10.1007/s13225-020-00466-2)) |
| Otu53135 | 1 | k__Fungi(100);p__Ascomycota(100);c__Saccharomycetes(100);o__Saccharomycetales(100);f__Metschnikowiaceae(100);g__Metschnikowia(100);g__Metschnikowia_unclassified(100); | Metschnikowia | 13 | Saprotroph | \|Undefined Saprotroph\| | Yeast | NULL | Probable | As nectar-tap_saprotroph (Põlme et al. 2020); Decay sugar-rich substrates (Põlme et al. 2020) | Põlme S, et al. 2020. Fungal Diversity 105:1-16 ((https://doi.org/10.1007/s13225-020-00466-2)) |
| Otu53137 | 1 | k__Fungi(100);p__Ascomycota(100);c__Saccharomycetes(100);o__Saccharomycetales(100);f__Metschnikowiaceae(100);g__Metschnikowia(100);s__Metschnikowia_pulcherrima(100); | Metschnikowia | 13 | Saprotroph | \|Undefined Saprotroph\| | Yeast | NULL | Probable | As nectar-tap_saprotroph (Põlme et al. 2020); Decay sugar-rich substrates (Põlme et al. 2020) | Põlme S, et al. 2020. Fungal Diversity 105:1-16 ((https://doi.org/10.1007/s13225-020-00466-2)) |
| Otu53138 | 1 | k__Fungi(100);p__Ascomycota(100);c__Saccharomycetes(100);o__Saccharomycetales(100);f__Metschnikowiaceae(100);g__Metschnikowia(100);g__Metschnikowia_unclassified(100); | Metschnikowia | 13 | Saprotroph | \|Undefined Saprotroph\| | Yeast | NULL | Probable | As nectar-tap_saprotroph (Põlme et al. 2020); Decay sugar-rich substrates (Põlme et al. 2020) | Põlme S, et al. 2020. Fungal Diversity 105:1-16 ((https://doi.org/10.1007/s13225-020-00466-2)) |
| Otu53139 | 1 | k__Fungi(100);p__Ascomycota(100);c__Saccharomycetes(100);o__Saccharomycetales(100);f__Metschnikowiaceae(100);g__Metschnikowia(100);g__Metschnikowia_unclassified(100); | Metschnikowia | 13 | Saprotroph | \|Undefined Saprotroph\| | Yeast | NULL | Probable | As nectar-tap_saprotroph (Põlme et al. 2020); Decay sugar-rich substrates (Põlme et al. 2020) | Põlme S, et al. 2020. Fungal Diversity 105:1-16 ((https://doi.org/10.1007/s13225-020-00466-2)) |
| Otu53140 | 1 | k__Fungi(100);p__Ascomycota(100);c__Saccharomycetes(100);o__Saccharomycetales(100);f__Metschnikowiaceae(100);g__Metschnikowia(100);s__Metschnikowia_pulcherrima(100); | Metschnikowia | 13 | Saprotroph | \|Undefined Saprotroph\| | Yeast | NULL | Probable | As nectar-tap_saprotroph (Põlme et al. 2020); Decay sugar-rich substrates (Põlme et al. 2020) | Põlme S, et al. 2020. Fungal Diversity 105:1-16 ((https://doi.org/10.1007/s13225-020-00466-2)) |
| Otu53141 | 1 | k__Fungi(100);p__Ascomycota(100);c__Saccharomycetes(100);o__Saccharomycetales(100);f__Metschnikowiaceae(100);g__Metschnikowia(100);s__Metschnikowia_pulcherrima(100); | Metschnikowia | 13 | Saprotroph | \|Undefined Saprotroph\| | Yeast | NULL | Probable | As nectar-tap_saprotroph (Põlme et al. 2020); Decay sugar-rich substrates (Põlme et al. 2020) | Põlme S, et al. 2020. Fungal Diversity 105:1-16 ((https://doi.org/10.1007/s13225-020-00466-2)) |
| Otu53143 | 1 | k__Fungi(100);p__Ascomycota(100);c__Saccharomycetes(100);o__Saccharomycetales(100);f__Metschnikowiaceae(100);g__Metschnikowia(100);g__Metschnikowia_unclassified(100); | Metschnikowia | 13 | Saprotroph | \|Undefined Saprotroph\| | Yeast | NULL | Probable | As nectar-tap_saprotroph (Põlme et al. 2020); Decay sugar-rich substrates (Põlme et al. 2020) | Põlme S, et al. 2020. Fungal Diversity 105:1-16 ((https://doi.org/10.1007/s13225-020-00466-2)) |
| Otu53144 | 1 | k__Fungi(100);p__Ascomycota(100);c__Saccharomycetes(100);o__Saccharomycetales(100);f__Metschnikowiaceae(100);g__Metschnikowia(100);g__Metschnikowia_unclassified(100); | Metschnikowia | 13 | Saprotroph | \|Undefined Saprotroph\| | Yeast | NULL | Probable | As nectar-tap_saprotroph (Põlme et al. 2020); Decay sugar-rich substrates (Põlme et al. 2020) | Põlme S, et al. 2020. Fungal Diversity 105:1-16 ((https://doi.org/10.1007/s13225-020-00466-2)) |
| Otu53145 | 1 | k__Fungi(100);p__Ascomycota(100);c__Saccharomycetes(100);o__Saccharomycetales(100);f__Metschnikowiaceae(100);g__Metschnikowia(100);s__Metschnikowia_pulcherrima(100); | Metschnikowia | 13 | Saprotroph | \|Undefined Saprotroph\| | Yeast | NULL | Probable | As nectar-tap_saprotroph (Põlme et al. 2020); Decay sugar-rich substrates (Põlme et al. 2020) | Põlme S, et al. 2020. Fungal Diversity 105:1-16 ((https://doi.org/10.1007/s13225-020-00466-2)) |
| Otu53146 | 1 | k__Fungi(100);p__Ascomycota(100);c__Saccharomycetes(100);o__Saccharomycetales(100);f__Metschnikowiaceae(100);g__Metschnikowia(100);g__Metschnikowia_unclassified(100); | Metschnikowia | 13 | Saprotroph | \|Undefined Saprotroph\| | Yeast | NULL | Probable | As nectar-tap_saprotroph (Põlme et al. 2020); Decay sugar-rich substrates (Põlme et al. 2020) | Põlme S, et al. 2020. Fungal Diversity 105:1-16 ((https://doi.org/10.1007/s13225-020-00466-2)) |
| Otu53147 | 1 | k__Fungi(100);p__Ascomycota(100);c__Saccharomycetes(100);o__Saccharomycetales(100);f__Metschnikowiaceae(100);g__Metschnikowia(100);s__Metschnikowia_pulcherrima(100); | Metschnikowia | 13 | Saprotroph | \|Undefined Saprotroph\| | Yeast | NULL | Probable | As nectar-tap_saprotroph (Põlme et al. 2020); Decay sugar-rich substrates (Põlme et al. 2020) | Põlme S, et al. 2020. Fungal Diversity 105:1-16 ((https://doi.org/10.1007/s13225-020-00466-2)) |
| Otu53148 | 1 | k__Fungi(100);p__Ascomycota(100);c__Saccharomycetes(100);o__Saccharomycetales(100);f__Metschnikowiaceae(100);g__Metschnikowia(100);s__Metschnikowia_pulcherrima(100); | Metschnikowia | 13 | Saprotroph | \|Undefined Saprotroph\| | Yeast | NULL | Probable | As nectar-tap_saprotroph (Põlme et al. 2020); Decay sugar-rich substrates (Põlme et al. 2020) | Põlme S, et al. 2020. Fungal Diversity 105:1-16 ((https://doi.org/10.1007/s13225-020-00466-2)) |
| Otu53149 | 1 | k__Fungi(100);p__Ascomycota(100);c__Saccharomycetes(100);o__Saccharomycetales(100);f__Metschnikowiaceae(100);g__Metschnikowia(100);g__Metschnikowia_unclassified(100); | Metschnikowia | 13 | Saprotroph | \|Undefined Saprotroph\| | Yeast | NULL | Probable | As nectar-tap_saprotroph (Põlme et al. 2020); Decay sugar-rich substrates (Põlme et al. 2020) | Põlme S, et al. 2020. Fungal Diversity 105:1-16 ((https://doi.org/10.1007/s13225-020-00466-2)) |
| Otu53150 | 1 | k__Fungi(100);p__Ascomycota(100);c__Saccharomycetes(100);o__Saccharomycetales(100);f__Metschnikowiaceae(100);g__Metschnikowia(100);s__Metschnikowia_pulcherrima(100); | Metschnikowia | 13 | Saprotroph | \|Undefined Saprotroph\| | Yeast | NULL | Probable | As nectar-tap_saprotroph (Põlme et al. 2020); Decay sugar-rich substrates (Põlme et al. 2020) | Põlme S, et al. 2020. Fungal Diversity 105:1-16 ((https://doi.org/10.1007/s13225-020-00466-2)) |
| Otu53151 | 1 | k__Fungi(100);p__Ascomycota(100);c__Saccharomycetes(100);o__Saccharomycetales(100);f__Metschnikowiaceae(100);g__Metschnikowia(100);s__Metschnikowia_pulcherrima(100); | Metschnikowia | 13 | Saprotroph | \|Undefined Saprotroph\| | Yeast | NULL | Probable | As nectar-tap_saprotroph (Põlme et al. 2020); Decay sugar-rich substrates (Põlme et al. 2020) | Põlme S, et al. 2020. Fungal Diversity 105:1-16 ((https://doi.org/10.1007/s13225-020-00466-2)) |
| Otu53152 | 1 | k__Fungi(100);p__Ascomycota(100);c__Saccharomycetes(100);o__Saccharomycetales(100);f__Metschnikowiaceae(100);g__Metschnikowia(100);g__Metschnikowia_unclassified(100); | Metschnikowia | 13 | Saprotroph | \|Undefined Saprotroph\| | Yeast | NULL | Probable | As nectar-tap_saprotroph (Põlme et al. 2020); Decay sugar-rich substrates (Põlme et al. 2020) | Põlme S, et al. 2020. Fungal Diversity 105:1-16 ((https://doi.org/10.1007/s13225-020-00466-2)) |
| Otu53153 | 1 | k__Fungi(100);p__Ascomycota(100);c__Saccharomycetes(100);o__Saccharomycetales(100);f__Metschnikowiaceae(100);g__Metschnikowia(100);s__Metschnikowia_pulcherrima(100); | Metschnikowia | 13 | Saprotroph | \|Undefined Saprotroph\| | Yeast | NULL | Probable | As nectar-tap_saprotroph (Põlme et al. 2020); Decay sugar-rich substrates (Põlme et al. 2020) | Põlme S, et al. 2020. Fungal Diversity 105:1-16 ((https://doi.org/10.1007/s13225-020-00466-2)) |
| Otu53154 | 1 | k__Fungi(100);p__Ascomycota(100);c__Saccharomycetes(100);o__Saccharomycetales(100);f__Metschnikowiaceae(100);g__Metschnikowia(100);s__Metschnikowia_pulcherrima(100); | Metschnikowia | 13 | Saprotroph | \|Undefined Saprotroph\| | Yeast | NULL | Probable | As nectar-tap_saprotroph (Põlme et al. 2020); Decay sugar-rich substrates (Põlme et al. 2020) | Põlme S, et al. 2020. Fungal Diversity 105:1-16 ((https://doi.org/10.1007/s13225-020-00466-2)) |

| | | | | | | | | | | | |
|---|---|---|---|---|---|---|---|---|---|---|---|
| Otu53155 | 1 | k__Fungi(100);p__Ascomycota(100);c__Saccharomycetes(100);o__Saccharomycetales(100);f__Metschnikowiaceae(100);g__Metschnikowia(100);s__Metschnikowia_pulcherrima(100); | Metschnikowia | 13 | Saprotroph | \|Undefined Saprotroph\| | Yeast | NULL | Probable | As nectar-tap_saprotroph (Põlme et al. 2020); Decay sugar-rich substrates (Põlme et al. 2020) | Põlme S, et al. 2020. Fungal Diversity 105:1-16 ((https://doi.org/10.1007/s13225-020-00466-2)) |
| Otu53156 | 1 | k__Fungi(100);p__Ascomycota(100);c__Saccharomycetes(100);o__Saccharomycetales(100);f__Metschnikowiaceae(100);g__Metschnikowia(100);g__Metschnikowia_unclassified(100); | Metschnikowia | 13 | Saprotroph | \|Undefined Saprotroph\| | Yeast | NULL | Probable | As nectar-tap_saprotroph (Põlme et al. 2020); Decay sugar-rich substrates (Põlme et al. 2020) | Põlme S, et al. 2020. Fungal Diversity 105:1-16 ((https://doi.org/10.1007/s13225-020-00466-2)) |
| Otu53157 | 1 | k__Fungi(100);p__Ascomycota(100);c__Saccharomycetes(100);o__Saccharomycetales(100);f__Metschnikowiaceae(100);g__Metschnikowia(100);s__Metschnikowia_pulcherrima(100); | Metschnikowia | 13 | Saprotroph | \|Undefined Saprotroph\| | Yeast | NULL | Probable | As nectar-tap_saprotroph (Põlme et al. 2020); Decay sugar-rich substrates (Põlme et al. 2020) | Põlme S, et al. 2020. Fungal Diversity 105:1-16 ((https://doi.org/10.1007/s13225-020-00466-2)) |
| Otu53158 | 1 | k__Fungi(100);p__Ascomycota(100);c__Saccharomycetes(100);o__Saccharomycetales(100);f__Metschnikowiaceae(100);g__Metschnikowia(100);s__Metschnikowia_pulcherrima(100); | Metschnikowia | 13 | Saprotroph | \|Undefined Saprotroph\| | Yeast | NULL | Probable | As nectar-tap_saprotroph (Põlme et al. 2020); Decay sugar-rich substrates (Põlme et al. 2020) | Põlme S, et al. 2020. Fungal Diversity 105:1-16 ((https://doi.org/10.1007/s13225-020-00466-2)) |
| Otu53159 | 1 | k__Fungi(100);p__Ascomycota(100);c__Saccharomycetes(100);o__Saccharomycetales(100);f__Metschnikowiaceae(100);g__Metschnikowia(100);s__Metschnikowia_pulcherrima(100); | Metschnikowia | 13 | Saprotroph | \|Undefined Saprotroph\| | Yeast | NULL | Probable | As nectar-tap_saprotroph (Põlme et al. 2020); Decay sugar-rich substrates (Põlme et al. 2020) | Põlme S, et al. 2020. Fungal Diversity 105:1-16 ((https://doi.org/10.1007/s13225-020-00466-2)) |
| Otu53160 | 1 | k__Fungi(100);p__Ascomycota(100);c__Saccharomycetes(100);o__Saccharomycetales(100);f__Metschnikowiaceae(100);g__Metschnikowia(100);s__Metschnikowia_pulcherrima(100); | Metschnikowia | 13 | Saprotroph | \|Undefined Saprotroph\| | Yeast | NULL | Probable | As nectar-tap_saprotroph (Põlme et al. 2020); Decay sugar-rich substrates (Põlme et al. 2020) | Põlme S, et al. 2020. Fungal Diversity 105:1-16 ((https://doi.org/10.1007/s13225-020-00466-2)) |
| Otu53161 | 1 | k__Fungi(100);p__Ascomycota(100);c__Saccharomycetes(100);o__Saccharomycetales(100);f__Metschnikowiaceae(100);g__Metschnikowia(100);g__Metschnikowia_unclassified(100); | Metschnikowia | 13 | Saprotroph | \|Undefined Saprotroph\| | Yeast | NULL | Probable | As nectar-tap_saprotroph (Põlme et al. 2020); Decay sugar-rich substrates (Põlme et al. 2020) | Põlme S, et al. 2020. Fungal Diversity 105:1-16 ((https://doi.org/10.1007/s13225-020-00466-2)) |
| Otu53163 | 1 | k__Fungi(100);p__Ascomycota(100);c__Saccharomycetes(100);o__Saccharomycetales(100);f__Metschnikowiaceae(100);g__Metschnikowia(100);g__Metschnikowia_unclassified(100); | Metschnikowia | 13 | Saprotroph | \|Undefined Saprotroph\| | Yeast | NULL | Probable | As nectar-tap_saprotroph (Põlme et al. 2020); Decay sugar-rich substrates (Põlme et al. 2020) | Põlme S, et al. 2020. Fungal Diversity 105:1-16 ((https://doi.org/10.1007/s13225-020-00466-2)) |
| Otu53164 | 1 | k__Fungi(100);p__Ascomycota(100);c__Saccharomycetes(100);o__Saccharomycetales(100);f__Metschnikowiaceae(100);g__Metschnikowia(100);s__Metschnikowia_pulcherrima(100); | Metschnikowia | 13 | Saprotroph | \|Undefined Saprotroph\| | Yeast | NULL | Probable | As nectar-tap_saprotroph (Põlme et al. 2020); Decay sugar-rich substrates (Põlme et al. 2020) | Põlme S, et al. 2020. Fungal Diversity 105:1-16 ((https://doi.org/10.1007/s13225-020-00466-2)) |
| Otu53165 | 1 | k__Fungi(100);p__Ascomycota(100);c__Saccharomycetes(100);o__Saccharomycetales(100);f__Metschnikowiaceae(100);g__Metschnikowia(100);s__Metschnikowia_pulcherrima(100); | Metschnikowia | 13 | Saprotroph | \|Undefined Saprotroph\| | Yeast | NULL | Probable | As nectar-tap_saprotroph (Põlme et al. 2020); Decay sugar-rich substrates (Põlme et al. 2020) | Põlme S, et al. 2020. Fungal Diversity 105:1-16 ((https://doi.org/10.1007/s13225-020-00466-2)) |
| Otu53166 | 1 | k__Fungi(100);p__Ascomycota(100);c__Saccharomycetes(100);o__Saccharomycetales(100);f__Metschnikowiaceae(100);g__Metschnikowia(100);s__Metschnikowia_pulcherrima(100); | Metschnikowia | 13 | Saprotroph | \|Undefined Saprotroph\| | Yeast | NULL | Probable | As nectar-tap_saprotroph (Põlme et al. 2020); Decay sugar-rich substrates (Põlme et al. 2020) | Põlme S, et al. 2020. Fungal Diversity 105:1-16 ((https://doi.org/10.1007/s13225-020-00466-2)) |
| Otu53167 | 1 | k__Fungi(100);p__Ascomycota(100);c__Saccharomycetes(100);o__Saccharomycetales(100);f__Metschnikowiaceae(100);g__Metschnikowia(100);g__Metschnikowia_unclassified(100); | Metschnikowia | 13 | Saprotroph | \|Undefined Saprotroph\| | Yeast | NULL | Probable | As nectar-tap_saprotroph (Põlme et al. 2020); Decay sugar-rich substrates (Põlme et al. 2020) | Põlme S, et al. 2020. Fungal Diversity 105:1-16 ((https://doi.org/10.1007/s13225-020-00466-2)) |
| Otu53168 | 1 | k__Fungi(100);p__Ascomycota(100);c__Saccharomycetes(100);o__Saccharomycetales(100);f__Metschnikowiaceae(100);g__Metschnikowia(100);g__Metschnikowia_unclassified(100); | Metschnikowia | 13 | Saprotroph | \|Undefined Saprotroph\| | Yeast | NULL | Probable | As nectar-tap_saprotroph (Põlme et al. 2020); Decay sugar-rich substrates (Põlme et al. 2020) | Põlme S, et al. 2020. Fungal Diversity 105:1-16 ((https://doi.org/10.1007/s13225-020-00466-2)) |
| Otu53169 | 1 | k__Fungi(100);p__Ascomycota(100);c__Saccharomycetes(100);o__Saccharomycetales(100);f__Metschnikowiaceae(100);g__Metschnikowia(100);s__Metschnikowia_pulcherrima(100); | Metschnikowia | 13 | Saprotroph | \|Undefined Saprotroph\| | Yeast | NULL | Probable | As nectar-tap_saprotroph (Põlme et al. 2020); Decay sugar-rich substrates (Põlme et al. 2020) | Põlme S, et al. 2020. Fungal Diversity 105:1-16 ((https://doi.org/10.1007/s13225-020-00466-2)) |
| Otu53170 | 1 | k__Fungi(100);p__Ascomycota(100);c__Saccharomycetes(100);o__Saccharomycetales(100);f__Metschnikowiaceae(100);g__Metschnikowia(100);s__Metschnikowia_pulcherrima(100); | Metschnikowia | 13 | Saprotroph | \|Undefined Saprotroph\| | Yeast | NULL | Probable | As nectar-tap_saprotroph (Põlme et al. 2020); Decay sugar-rich substrates (Põlme et al. 2020) | Põlme S, et al. 2020. Fungal Diversity 105:1-16 ((https://doi.org/10.1007/s13225-020-00466-2)) |
| Otu53171 | 1 | k__Fungi(100);p__Ascomycota(100);c__Saccharomycetes(100);o__Saccharomycetales(100);f__Metschnikowiaceae(100);g__Metschnikowia(100);s__Metschnikowia_pulcherrima(100); | Metschnikowia | 13 | Saprotroph | \|Undefined Saprotroph\| | Yeast | NULL | Probable | As nectar-tap_saprotroph (Põlme et al. 2020); Decay sugar-rich substrates (Põlme et al. 2020) | Põlme S, et al. 2020. Fungal Diversity 105:1-16 ((https://doi.org/10.1007/s13225-020-00466-2)) |
| Otu53172 | 1 | k__Fungi(100);p__Ascomycota(100);c__Saccharomycetes(100);o__Saccharomycetales(100);f__Metschnikowiaceae(100);g__Metschnikowia(100);s__Metschnikowia_pulcherrima(100); | Metschnikowia | 13 | Saprotroph | \|Undefined Saprotroph\| | Yeast | NULL | Probable | As nectar-tap_saprotroph (Põlme et al. 2020); Decay sugar-rich substrates (Põlme et al. 2020) | Põlme S, et al. 2020. Fungal Diversity 105:1-16 ((https://doi.org/10.1007/s13225-020-00466-2)) |
| Otu53173 | 1 | k__Fungi(100);p__Ascomycota(100);c__Saccharomycetes(100);o__Saccharomycetales(100);f__Metschnikowiaceae(100);g__Metschnikowia(100);g__Metschnikowia_unclassified(100); | Metschnikowia | 13 | Saprotroph | \|Undefined Saprotroph\| | Yeast | NULL | Probable | As nectar-tap_saprotroph (Põlme et al. 2020); Decay sugar-rich substrates (Põlme et al. 2020) | Põlme S, et al. 2020. Fungal Diversity 105:1-16 ((https://doi.org/10.1007/s13225-020-00466-2)) |
| Otu53175 | 1 | k__Fungi(100);p__Ascomycota(100);c__Saccharomycetes(100);o__Saccharomycetales(100);f__Metschnikowiaceae(100);g__Metschnikowia(100);g__Metschnikowia_unclassified(100); | Metschnikowia | 13 | Saprotroph | \|Undefined Saprotroph\| | Yeast | NULL | Probable | As nectar-tap_saprotroph (Põlme et al. 2020); Decay sugar-rich substrates (Põlme et al. 2020) | Põlme S, et al. 2020. Fungal Diversity 105:1-16 ((https://doi.org/10.1007/s13225-020-00466-2)) |
| Otu53180 | 1 | k__Fungi(100);p__Ascomycota(100);c__Saccharomycetes(100);o__Saccharomycetales(100);f__Metschnikowiaceae(100);g__Metschnikowia(100);s__Metschnikowia_pulcherrima(100); | Metschnikowia | 13 | Saprotroph | \|Undefined Saprotroph\| | Yeast | NULL | Probable | As nectar-tap_saprotroph (Põlme et al. 2020); Decay sugar-rich substrates (Põlme et al. 2020) | Põlme S, et al. 2020. Fungal Diversity 105:1-16 ((https://doi.org/10.1007/s13225-020-00466-2)) |
| Otu53182 | 1 | k__Fungi(100);p__Ascomycota(100);c__Saccharomycetes(100);o__Saccharomycetales(100);f__Metschnikowiaceae(100);g__Metschnikowia(100);s__Metschnikowia_pulcherrima(100); | Metschnikowia | 13 | Saprotroph | \|Undefined Saprotroph\| | Yeast | NULL | Probable | As nectar-tap_saprotroph (Põlme et al. 2020); Decay sugar-rich substrates (Põlme et al. 2020) | Põlme S, et al. 2020. Fungal Diversity 105:1-16 ((https://doi.org/10.1007/s13225-020-00466-2)) |
| Otu53183 | 1 | k__Fungi(100);p__Ascomycota(100);c__Saccharomycetes(100);o__Saccharomycetales(100);f__Metschnikowiaceae(100);g__Metschnikowia(100);s__Metschnikowia_pulcherrima(100); | Metschnikowia | 13 | Saprotroph | \|Undefined Saprotroph\| | Yeast | NULL | Probable | As nectar-tap_saprotroph (Põlme et al. 2020); Decay sugar-rich substrates (Põlme et al. 2020) | Põlme S, et al. 2020. Fungal Diversity 105:1-16 ((https://doi.org/10.1007/s13225-020-00466-2)) |
| Otu53184 | 1 | k__Fungi(100);p__Ascomycota(100);c__Saccharomycetes(100);o__Saccharomycetales(100);f__Metschnikowiaceae(100);g__Metschnikowia(100);s__Metschnikowia_pulcherrima(100); | Metschnikowia | 13 | Saprotroph | \|Undefined Saprotroph\| | Yeast | NULL | Probable | As nectar-tap_saprotroph (Põlme et al. 2020); Decay sugar-rich substrates (Põlme et al. 2020) | Põlme S, et al. 2020. Fungal Diversity 105:1-16 ((https://doi.org/10.1007/s13225-020-00466-2)) |
| Otu53185 | 1 | k__Fungi(100);p__Ascomycota(100);c__Saccharomycetes(100);o__Saccharomycetales(100);f__Metschnikowiaceae(100);g__Metschnikowia(100);s__Metschnikowia_pulcherrima(100); | Metschnikowia | 13 | Saprotroph | \|Undefined Saprotroph\| | Yeast | NULL | Probable | As nectar-tap_saprotroph (Põlme et al. 2020); Decay sugar-rich substrates (Põlme et al. 2020) | Põlme S, et al. 2020. Fungal Diversity 105:1-16 ((https://doi.org/10.1007/s13225-020-00466-2)) |
| Otu53186 | 1 | k__Fungi(100);p__Ascomycota(100);c__Saccharomycetes(100);o__Saccharomycetales(100);f__Metschnikowiaceae(100);g__Metschnikowia(100);s__Metschnikowia_pulcherrima(100); | Metschnikowia | 13 | Saprotroph | \|Undefined Saprotroph\| | Yeast | NULL | Probable | As nectar-tap_saprotroph (Põlme et al. 2020); Decay sugar-rich substrates (Põlme et al. 2020) | Põlme S, et al. 2020. Fungal Diversity 105:1-16 ((https://doi.org/10.1007/s13225-020-00466-2)) |
| Otu53187 | 1 | k__Fungi(100);p__Ascomycota(100);c__Saccharomycetes(100);o__Saccharomycetales(100);f__Metschnikowiaceae(100);g__Metschnikowia(100);s__Metschnikowia_pulcherrima(100); | Metschnikowia | 13 | Saprotroph | \|Undefined Saprotroph\| | Yeast | NULL | Probable | As nectar-tap_saprotroph (Põlme et al. 2020); Decay sugar-rich substrates (Põlme et al. 2020) | Põlme S, et al. 2020. Fungal Diversity 105:1-16 ((https://doi.org/10.1007/s13225-020-00466-2)) |
| Otu53188 | 1 | k__Fungi(100);p__Ascomycota(100);c__Saccharomycetes(100);o__Saccharomycetales(100);f__Metschnikowiaceae(100);g__Metschnikowia(100);s__Metschnikowia_pulcherrima(100); | Metschnikowia | 13 | Saprotroph | \|Undefined Saprotroph\| | Yeast | NULL | Probable | As nectar-tap_saprotroph (Põlme et al. 2020); Decay sugar-rich substrates (Põlme et al. 2020) | Põlme S, et al. 2020. Fungal Diversity 105:1-16 ((https://doi.org/10.1007/s13225-020-00466-2)) |
| Otu53189 | 1 | k__Fungi(100);p__Ascomycota(100);c__Saccharomycetes(100);o__Saccharomycetales(100);f__Metschnikowiaceae(100);g__Metschnikowia(100);s__Metschnikowia_pulcherrima(100); | Metschnikowia | 13 | Saprotroph | \|Undefined Saprotroph\| | Yeast | NULL | Probable | As nectar-tap_saprotroph (Põlme et al. 2020); Decay sugar-rich substrates (Põlme et al. 2020) | Põlme S, et al. 2020. Fungal Diversity 105:1-16 ((https://doi.org/10.1007/s13225-020-00466-2)) |
| Otu53190 | 1 | k__Fungi(100);p__Ascomycota(100);c__Saccharomycetes(100);o__Saccharomycetales(100);f__Metschnikowiaceae(100);g__Metschnikowia(100);s__Metschnikowia_pulcherrima(100); | Metschnikowia | 13 | Saprotroph | \|Undefined Saprotroph\| | Yeast | NULL | Probable | As nectar-tap_saprotroph (Põlme et al. 2020); Decay sugar-rich substrates (Põlme et al. 2020) | Põlme S, et al. 2020. Fungal Diversity 105:1-16 ((https://doi.org/10.1007/s13225-020-00466-2)) |
| Otu53191 | 1 | k__Fungi(100);p__Ascomycota(100);c__Saccharomycetes(100);o__Saccharomycetales(100);f__Metschnikowiaceae(100);g__Metschnikowia(100);s__Metschnikowia_pulcherrima(100); | Metschnikowia | 13 | Saprotroph | \|Undefined Saprotroph\| | Yeast | NULL | Probable | As nectar-tap_saprotroph (Põlme et al. 2020); Decay sugar-rich substrates (Põlme et al. 2020) | Põlme S, et al. 2020. Fungal Diversity 105:1-16 ((https://doi.org/10.1007/s13225-020-00466-2)) |
| Otu53192 | 1 | k__Fungi(100);p__Ascomycota(100);c__Saccharomycetes(100);o__Saccharomycetales(100);f__Metschnikowiaceae(100);g__Metschnikowia(100);g__Metschnikowia_unclassified(100); | Metschnikowia | 13 | Saprotroph | \|Undefined Saprotroph\| | Yeast | NULL | Probable | As nectar-tap_saprotroph (Põlme et al. 2020); Decay sugar-rich substrates (Põlme et al. 2020) | Põlme S, et al. 2020. Fungal Diversity 105:1-16 ((https://doi.org/10.1007/s13225-020-00466-2)) |
| Otu53193 | 1 | k__Fungi(100);p__Ascomycota(100);c__Saccharomycetes(100);o__Saccharomycetales(100);f__Metschnikowiaceae(100);g__Metschnikowia(100);g__Metschnikowia_unclassified(100); | Metschnikowia | 13 | Saprotroph | \|Undefined Saprotroph\| | Yeast | NULL | Probable | As nectar-tap_saprotroph (Põlme et al. 2020); Decay sugar-rich substrates (Põlme et al. 2020) | Põlme S, et al. 2020. Fungal Diversity 105:1-16 ((https://doi.org/10.1007/s13225-020-00466-2)) |
| Otu53194 | 1 | k__Fungi(100);p__Ascomycota(100);c__Saccharomycetes(100);o__Saccharomycetales(100);f__Metschnikowiaceae(100);g__Metschnikowia(100);s__Metschnikowia_pulcherrima(100); | Metschnikowia | 13 | Saprotroph | \|Undefined Saprotroph\| | Yeast | NULL | Probable | As nectar-tap_saprotroph (Põlme et al. 2020); Decay sugar-rich substrates (Põlme et al. 2020) | Põlme S, et al. 2020. Fungal Diversity 105:1-16 ((https://doi.org/10.1007/s13225-020-00466-2)) |
| Otu53195 | 1 | k__Fungi(100);p__Ascomycota(100);c__Saccharomycetes(100);o__Saccharomycetales(100);f__Metschnikowiaceae(100);g__Metschnikowia(100);g__Metschnikowia_unclassified(100); | Metschnikowia | 13 | Saprotroph | \|Undefined Saprotroph\| | Yeast | NULL | Probable | As nectar-tap_saprotroph (Põlme et al. 2020); Decay sugar-rich substrates (Põlme et al. 2020) | Põlme S, et al. 2020. Fungal Diversity 105:1-16 ((https://doi.org/10.1007/s13225-020-00466-2)) |
| Otu53197 | 1 | k__Fungi(100);p__Ascomycota(100);c__Saccharomycetes(100);o__Saccharomycetales(100);f__Metschnikowiaceae(100);g__Metschnikowia(100);s__Metschnikowia_pulcherrima(100); | Metschnikowia | 13 | Saprotroph | \|Undefined Saprotroph\| | Yeast | NULL | Probable | As nectar-tap_saprotroph (Põlme et al. 2020); Decay sugar-rich substrates (Põlme et al. 2020) | Põlme S, et al. 2020. Fungal Diversity 105:1-16 ((https://doi.org/10.1007/s13225-020-00466-2)) |
| Otu53198 | 1 | k__Fungi(100);p__Ascomycota(100);c__Saccharomycetes(100);o__Saccharomycetales(100);f__Metschnikowiaceae(100);g__Metschnikowia(100);s__Metschnikowia_pulcherrima(100); | Metschnikowia | 13 | Saprotroph | \|Undefined Saprotroph\| | Yeast | NULL | Probable | As nectar-tap_saprotroph (Põlme et al. 2020); Decay sugar-rich substrates (Põlme et al. 2020) | Põlme S, et al. 2020. Fungal Diversity 105:1-16 ((https://doi.org/10.1007/s13225-020-00466-2)) |
| Otu53199 | 1 | k__Fungi(100);p__Ascomycota(100);c__Saccharomycetes(100);o__Saccharomycetales(100);f__Metschnikowiaceae(100);g__Metschnikowia(100);s__Metschnikowia_pulcherrima(100); | Metschnikowia | 13 | Saprotroph | \|Undefined Saprotroph\| | Yeast | NULL | Probable | As nectar-tap_saprotroph (Põlme et al. 2020); Decay sugar-rich substrates (Põlme et al. 2020) | Põlme S, et al. 2020. Fungal Diversity 105:1-16 ((https://doi.org/10.1007/s13225-020-00466-2)) |
| Otu53200 | 1 | k__Fungi(100);p__Ascomycota(100);c__Saccharomycetes(100);o__Saccharomycetales(100);f__Metschnikowiaceae(100);g__Metschnikowia(100);s__Metschnikowia_pulcherrima(100); | Metschnikowia | 13 | Saprotroph | \|Undefined Saprotroph\| | Yeast | NULL | Probable | As nectar-tap_saprotroph (Põlme et al. 2020); Decay sugar-rich substrates (Põlme et al. 2020) | Põlme S, et al. 2020. Fungal Diversity 105:1-16 ((https://doi.org/10.1007/s13225-020-00466-2)) |
| Otu53201 | 1 | k__Fungi(100);p__Ascomycota(100);c__Saccharomycetes(100);o__Saccharomycetales(100);f__Metschnikowiaceae(100);g__Metschnikowia(100);g__Metschnikowia_unclassified(100); | Metschnikowia | 13 | Saprotroph | \|Undefined Saprotroph\| | Yeast | NULL | Probable | As nectar-tap_saprotroph (Põlme et al. 2020); Decay sugar-rich substrates (Põlme et al. 2020) | Põlme S, et al. 2020. Fungal Diversity 105:1-16 ((https://doi.org/10.1007/s13225-020-00466-2)) |
| Otu53202 | 1 | k__Fungi(100);p__Ascomycota(100);c__Saccharomycetes(100);o__Saccharomycetales(100);f__Metschnikowiaceae(100);g__Metschnikowia(100);s__Metschnikowia_pulcherrima(100); | Metschnikowia | 13 | Saprotroph | \|Undefined Saprotroph\| | Yeast | NULL | Probable | As nectar-tap_saprotroph (Põlme et al. 2020); Decay sugar-rich substrates (Põlme et al. 2020) | Põlme S, et al. 2020. Fungal Diversity 105:1-16 ((https://doi.org/10.1007/s13225-020-00466-2)) |
| Otu53203 | 1 | k__Fungi(100);p__Ascomycota(100);c__Saccharomycetes(100);o__Saccharomycetales(100);f__Metschnikowiaceae(100);g__Metschnikowia(100);s__Metschnikowia_pulcherrima(100); | Metschnikowia | 13 | Saprotroph | \|Undefined Saprotroph\| | Yeast | NULL | Probable | As nectar-tap_saprotroph (Põlme et al. 2020); Decay sugar-rich substrates (Põlme et al. 2020) | Põlme S, et al. 2020. Fungal Diversity 105:1-16 ((https://doi.org/10.1007/s13225-020-00466-2)) |
| Otu53204 | 1 | k__Fungi(100);p__Ascomycota(100);c__Saccharomycetes(100);o__Saccharomycetales(100);f__Metschnikowiaceae(100);g__Metschnikowia(100);s__Metschnikowia_pulcherrima(100); | Metschnikowia | 13 | Saprotroph | \|Undefined Saprotroph\| | Yeast | NULL | Probable | As nectar-tap_saprotroph (Põlme et al. 2020); Decay sugar-rich substrates (Põlme et al. 2020) | Põlme S, et al. 2020. Fungal Diversity 105:1-16 ((https://doi.org/10.1007/s13225-020-00466-2)) |
| Otu53205 | 1 | k__Fungi(100);p__Ascomycota(100);c__Saccharomycetes(100);o__Saccharomycetales(100);f__Metschnikowiaceae(100);g__Metschnikowia(100);s__Metschnikowia_pulcherrima(100); | Metschnikowia | 13 | Saprotroph | \|Undefined Saprotroph\| | Yeast | NULL | Probable | As nectar-tap_saprotroph (Põlme et al. 2020); Decay sugar-rich substrates (Põlme et al. 2020) | Põlme S, et al. 2020. Fungal Diversity 105:1-16 ((https://doi.org/10.1007/s13225-020-00466-2)) |
| Otu53206 | 1 | k__Fungi(100);p__Ascomycota(100);c__Saccharomycetes(100);o__Saccharomycetales(100);f__Metschnikowiaceae(100);g__Metschnikowia(100);g__Metschnikowia_unclassified(100); | Metschnikowia | 13 | Saprotroph | \|Undefined Saprotroph\| | Yeast | NULL | Probable | As nectar-tap_saprotroph (Põlme et al. 2020); Decay sugar-rich substrates (Põlme et al. 2020) | Põlme S, et al. 2020. Fungal Diversity 105:1-16 ((https://doi.org/10.1007/s13225-020-00466-2)) |
| Otu53207 | 1 | k__Fungi(100);p__Ascomycota(100);c__Saccharomycetes(100);o__Saccharomycetales(100);f__Metschnikowiaceae(100);g__Metschnikowia(100);s__Metschnikowia_pulcherrima(100); | Metschnikowia | 13 | Saprotroph | \|Undefined Saprotroph\| | Yeast | NULL | Probable | As nectar-tap_saprotroph (Põlme et al. 2020); Decay sugar-rich substrates (Põlme et al. 2020) | Põlme S, et al. 2020. Fungal Diversity 105:1-16 ((https://doi.org/10.1007/s13225-020-00466-2)) |
| Otu53208 | 1 | k__Fungi(100);p__Ascomycota(100);c__Saccharomycetes(100);o__Saccharomycetales(100);f__Metschnikowiaceae(100);g__Metschnikowia(100);s__Metschnikowia_pulcherrima(100); | Metschnikowia | 13 | Saprotroph | \|Undefined Saprotroph\| | Yeast | NULL | Probable | As nectar-tap_saprotroph (Põlme et al. 2020); Decay sugar-rich substrates (Põlme et al. 2020) | Põlme S, et al. 2020. Fungal Diversity 105:1-16 ((https://doi.org/10.1007/s13225-020-00466-2)) |
| Otu53210 | 1 | k__Fungi(100);p__Ascomycota(100);c__Saccharomycetes(100);o__Saccharomycetales(100);f__Metschnikowiaceae(100);g__Metschnikowia(100);s__Metschnikowia_pulcherrima(100); | Metschnikowia | 13 | Saprotroph | \|Undefined Saprotroph\| | Yeast | NULL | Probable | As nectar-tap_saprotroph (Põlme et al. 2020); Decay sugar-rich substrates (Põlme et al. 2020) | Põlme S, et al. 2020. Fungal Diversity 105:1-16 ((https://doi.org/10.1007/s13225-020-00466-2)) |
| Otu53212 | 1 | k__Fungi(100);p__Ascomycota(100);c__Saccharomycetes(100);o__Saccharomycetales(100);f__Metschnikowiaceae(100);g__Metschnikowia(100);s__Metschnikowia_pulcherrima(100); | Metschnikowia | 13 | Saprotroph | \|Undefined Saprotroph\| | Yeast | NULL | Probable | As nectar-tap_saprotroph (Põlme et al. 2020); Decay sugar-rich substrates (Põlme et al. 2020) | Põlme S, et al. 2020. Fungal Diversity 105:1-16 ((https://doi.org/10.1007/s13225-020-00466-2)) |
| Otu53213 | 1 | k__Fungi(100);p__Ascomycota(100);c__Saccharomycetes(100);o__Saccharomycetales(100);f__Metschnikowiaceae(100);g__Metschnikowia(100);s__Metschnikowia_pulcherrima(100); | Metschnikowia | 13 | Saprotroph | \|Undefined Saprotroph\| | Yeast | NULL | Probable | As nectar-tap_saprotroph (Põlme et al. 2020); Decay sugar-rich substrates (Põlme et al. 2020) | Põlme S, et al. 2020. Fungal Diversity 105:1-16 ((https://doi.org/10.1007/s13225-020-00466-2)) |
| Otu53214 | 1 | k__Fungi(100);p__Ascomycota(100);c__Saccharomycetes(100);o__Saccharomycetales(100);f__Metschnikowiaceae(100);g__Metschnikowia(100);s__Metschnikowia_pulcherrima(100); | Metschnikowia | 13 | Saprotroph | \|Undefined Saprotroph\| | Yeast | NULL | Probable | As nectar-tap_saprotroph (Põlme et al. 2020); Decay sugar-rich substrates (Põlme et al. 2020) | Põlme S, et al. 2020. Fungal Diversity 105:1-16 ((https://doi.org/10.1007/s13225-020-00466-2)) |
| Otu53215 | 1 | k__Fungi(100);p__Ascomycota(100);c__Saccharomycetes(100);o__Saccharomycetales(100);f__Metschnikowiaceae(100);g__Metschnikowia(100);g__Metschnikowia_unclassified(100); | Metschnikowia | 13 | Saprotroph | \|Undefined Saprotroph\| | Yeast | NULL | Probable | As nectar-tap_saprotroph (Põlme et al. 2020); Decay sugar-rich substrates (Põlme et al. 2020) | Põlme S, et al. 2020. Fungal Diversity 105:1-16 ((https://doi.org/10.1007/s13225-020-00466-2)) |
| Otu53216 | 1 | k__Fungi(100);p__Ascomycota(100);c__Saccharomycetes(100);o__Saccharomycetales(100);f__Metschnikowiaceae(100);g__Metschnikowia(100);s__Metschnikowia_pulcherrima(100); | Metschnikowia | 13 | Saprotroph | \|Undefined Saprotroph\| | Yeast | NULL | Probable | As nectar-tap_saprotroph (Põlme et al. 2020); Decay sugar-rich substrates (Põlme et al. 2020) | Põlme S, et al. 2020. Fungal Diversity 105:1-16 ((https://doi.org/10.1007/s13225-020-00466-2)) |
| Otu53217 | 1 | k__Fungi(100);p__Ascomycota(100);c__Saccharomycetes(100);o__Saccharomycetales(100);f__Metschnikowiaceae(100);g__Metschnikowia(100);s__Metschnikowia_pulcherrima(100); | Metschnikowia | 13 | Saprotroph | \|Undefined Saprotroph\| | Yeast | NULL | Probable | As nectar-tap_saprotroph (Põlme et al. 2020); Decay sugar-rich substrates (Põlme et al. 2020) | Põlme S, et al. 2020. Fungal Diversity 105:1-16 ((https://doi.org/10.1007/s13225-020-00466-2)) |
| Otu53218 | 1 | k__Fungi(100);p__Ascomycota(100);c__Saccharomycetes(100);o__Saccharomycetales(100);f__Metschnikowiaceae(100);g__Metschnikowia(100);s__Metschnikowia_pulcherrima(100); | Metschnikowia | 13 | Saprotroph | \|Undefined Saprotroph\| | Yeast | NULL | Probable | As nectar-tap_saprotroph (Põlme et al. 2020); Decay sugar-rich substrates (Põlme et al. 2020) | Põlme S, et al. 2020. Fungal Diversity 105:1-16 ((https://doi.org/10.1007/s13225-020-00466-2)) |
| Otu53219 | 1 | k__Fungi(100);p__Ascomycota(100);c__Saccharomycetes(100);o__Saccharomycetales(100);f__Metschnikowiaceae(100);g__Metschnikowia(100);s__Metschnikowia_pulcherrima(100); | Metschnikowia | 13 | Saprotroph | \|Undefined Saprotroph\| | Yeast | NULL | Probable | As nectar-tap_saprotroph (Põlme et al. 2020); Decay sugar-rich substrates (Põlme et al. 2020) | Põlme S, et al. 2020. Fungal Diversity 105:1-16 ((https://doi.org/10.1007/s13225-020-00466-2)) |
| Otu53220 | 1 | k__Fungi(100);p__Ascomycota(100);c__Saccharomycetes(100);o__Saccharomycetales(100);f__Metschnikowiaceae(100);g__Metschnikowia(100);s__Metschnikowia_pulcherrima(100); | Metschnikowia | 13 | Saprotroph | \|Undefined Saprotroph\| | Yeast | NULL | Probable | As nectar-tap_saprotroph (Põlme et al. 2020); Decay sugar-rich substrates (Põlme et al. 2020) | Põlme S, et al. 2020. Fungal Diversity 105:1-16 ((https://doi.org/10.1007/s13225-020-00466-2)) |
| Otu53221 | 1 | k__Fungi(100);p__Ascomycota(100);c__Saccharomycetes(100);o__Saccharomycetales(100);f__Metschnikowiaceae(100);g__Metschnikowia(100);s__Metschnikowia_pulcherrima(100); | Metschnikowia | 13 | Saprotroph | \|Undefined Saprotroph\| | Yeast | NULL | Probable | As nectar-tap_saprotroph (Põlme et al. 2020); Decay sugar-rich substrates (Põlme et al. 2020) | Põlme S, et al. 2020. Fungal Diversity 105:1-16 ((https://doi.org/10.1007/s13225-020-00466-2)) |
| Otu53222 | 1 | k__Fungi(100);p__Ascomycota(100);c__Saccharomycetes(100);o__Saccharomycetales(100);f__Metschnikowiaceae(100);g__Metschnikowia(100);s__Metschnikowia_pulcherrima(100); | Metschnikowia | 13 | Saprotroph | \|Undefined Saprotroph\| | Yeast | NULL | Probable | As nectar-tap_saprotroph (Põlme et al. 2020); Decay sugar-rich substrates (Põlme et al. 2020) | Põlme S, et al. 2020. Fungal Diversity 105:1-16 ((https://doi.org/10.1007/s13225-020-00466-2)) |
| Otu53223 | 1 | k__Fungi(100);p__Ascomycota(100);c__Saccharomycetes(100);o__Saccharomycetales(100);f__Metschnikowiaceae(100);g__Metschnikowia(100);s__Metschnikowia_pulcherrima(100); | Metschnikowia | 13 | Saprotroph | \|Undefined Saprotroph\| | Yeast | NULL | Probable | As nectar-tap_saprotroph (Põlme et al. 2020); Decay sugar-rich substrates (Põlme et al. 2020) | Põlme S, et al. 2020. Fungal Diversity 105:1-16 ((https://doi.org/10.1007/s13225-020-00466-2)) |
| Otu53224 | 1 | k__Fungi(100);p__Ascomycota(100);c__Saccharomycetes(100);o__Saccharomycetales(100);f__Metschnikowiaceae(100);g__Metschnikowia(100);s__Metschnikowia_pulcherrima(100); | Metschnikowia | 13 | Saprotroph | \|Undefined Saprotroph\| | Yeast | NULL | Probable | As nectar-tap_saprotroph (Põlme et al. 2020); Decay sugar-rich substrates (Põlme et al. 2020) | Põlme S, et al. 2020. Fungal Diversity 105:1-16 ((https://doi.org/10.1007/s13225-020-00466-2)) |
| Otu53225 | 1 | k__Fungi(100);p__Ascomycota(100);c__Saccharomycetes(100);o__Saccharomycetales(100);f__Metschnikowiaceae(100);g__Metschnikowia(100);s__Metschnikowia_pulcherrima(100); | Metschnikowia | 13 | Saprotroph | \|Undefined Saprotroph\| | Yeast | NULL | Probable | As nectar-tap_saprotroph (Põlme et al. 2020); Decay sugar-rich substrates (Põlme et al. 2020) | Põlme S, et al. 2020. Fungal Diversity 105:1-16 ((https://doi.org/10.1007/s13225-020-00466-2)) |
| Otu53226 | 1 | k__Fungi(100);p__Ascomycota(100);c__Saccharomycetes(100);o__Saccharomycetales(100);f__Metschnikowiaceae(100);g__Metschnikowia(100);g__Metschnikowia_unclassified(100); | Metschnikowia | 13 | Saprotroph | \|Undefined Saprotroph\| | Yeast | NULL | Probable | As nectar-tap_saprotroph (Põlme et al. 2020); Decay sugar-rich substrates (Põlme et al. 2020) | Põlme S, et al. 2020. Fungal Diversity 105:1-16 ((https://doi.org/10.1007/s13225-020-00466-2)) |
| Otu53227 | 1 | k__Fungi(100);p__Ascomycota(100);c__Saccharomycetes(100);o__Saccharomycetales(100);f__Metschnikowiaceae(100);g__Metschnikowia(100);s__Metschnikowia_pulcherrima(100); | Metschnikowia | 13 | Saprotroph | \|Undefined Saprotroph\| | Yeast | NULL | Probable | As nectar-tap_saprotroph (Põlme et al. 2020); Decay sugar-rich substrates (Põlme et al. 2020) | Põlme S, et al. 2020. Fungal Diversity 105:1-16 ((https://doi.org/10.1007/s13225-020-00466-2)) |
| Otu53229 | 1 | k__Fungi(100);p__Ascomycota(100);c__Saccharomycetes(100);o__Saccharomycetales(100);f__Metschnikowiaceae(100);g__Metschnikowia(100);s__Metschnikowia_pulcherrima(100); | Metschnikowia | 13 | Saprotroph | \|Undefined Saprotroph\| | Yeast | NULL | Probable | As nectar-tap_saprotroph (Põlme et al. 2020); Decay sugar-rich substrates (Põlme et al. 2020) | Põlme S, et al. 2020. Fungal Diversity 105:1-16 ((https://doi.org/10.1007/s13225-020-00466-2)) |
| Otu53230 | 1 | k__Fungi(100);p__Ascomycota(100);c__Saccharomycetes(100);o__Saccharomycetales(100);f__Metschnikowiaceae(100);g__Metschnikowia(100);s__Metschnikowia_pulcherrima(100); | Metschnikowia | 13 | Saprotroph | \|Undefined Saprotroph\| | Yeast | NULL | Probable | As nectar-tap_saprotroph (Põlme et al. 2020); Decay sugar-rich substrates (Põlme et al. 2020) | Põlme S, et al. 2020. Fungal Diversity 105:1-16 ((https://doi.org/10.1007/s13225-020-00466-2)) |
| Otu53231 | 1 | k__Fungi(100);p__Ascomycota(100);c__Saccharomycetes(100);o__Saccharomycetales(100);f__Metschnikowiaceae(100);g__Metschnikowia(100);s__Metschnikowia_pulcherrima(100); | Metschnikowia | 13 | Saprotroph | \|Undefined Saprotroph\| | Yeast | NULL | Probable | As nectar-tap_saprotroph (Põlme et al. 2020); Decay sugar-rich substrates (Põlme et al. 2020) | Põlme S, et al. 2020. Fungal Diversity 105:1-16 ((https://doi.org/10.1007/s13225-020-00466-2)) |
| Otu53232 | 1 | k__Fungi(100);p__Ascomycota(100);c__Saccharomycetes(100);o__Saccharomycetales(100);f__Metschnikowiaceae(100);g__Metschnikowia(100);s__Metschnikowia_pulcherrima(100); | Metschnikowia | 13 | Saprotroph | \|Undefined Saprotroph\| | Yeast | NULL | Probable | As nectar-tap_saprotroph (Põlme et al. 2020); Decay sugar-rich substrates (Põlme et al. 2020) | Põlme S, et al. 2020. Fungal Diversity 105:1-16 ((https://doi.org/10.1007/s13225-020-00466-2)) |
| Otu53233 | 1 | k__Fungi(100);p__Ascomycota(100);c__Saccharomycetes(100);o__Saccharomycetales(100);f__Metschnikowiaceae(100);g__Metschnikowia(100);s__Metschnikowia_pulcherrima(100); | Metschnikowia | 13 | Saprotroph | \|Undefined Saprotroph\| | Yeast | NULL | Probable | As nectar-tap_saprotroph (Põlme et al. 2020); Decay sugar-rich substrates (Põlme et al. 2020) | Põlme S, et al. 2020. Fungal Diversity 105:1-16 ((https://doi.org/10.1007/s13225-020-00466-2)) |
| Otu53234 | 1 | k__Fungi(100);p__Ascomycota(100);c__Saccharomycetes(100);o__Saccharomycetales(100);f__Metschnikowiaceae(100);g__Metschnikowia(100);s__Metschnikowia_pulcherrima(100); | Metschnikowia | 13 | Saprotroph | \|Undefined Saprotroph\| | Yeast | NULL | Probable | As nectar-tap_saprotroph (Põlme et al. 2020); Decay sugar-rich substrates (Põlme et al. 2020) | Põlme S, et al. 2020. Fungal Diversity 105:1-16 ((https://doi.org/10.1007/s13225-020-00466-2)) |
| Otu53235 | 1 | k__Fungi(100);p__Ascomycota(100);c__Sordariomycetes(100);o__Hypocreales(100);f__Nectriaceae(100);f__Nectriaceae_unclassified(100);f__Nectriaceae_unclassified(100); | Nectriaceae | 9 | Pathotroph-Saprotroph-Symbiotroph | Animal Pathogen-Endophyte-Fungal Parasite-Lichen Parasite-Plant Pathogen-Wood Saprotroph | Microfungus | NULL | Possible | Fusarium species are amongst the economically significant plant pathogens (Cannon & Kirk 2007) | Cannon PF, Kirk PM. 2007. Fungal Families of the World. CAB International, Cambridge ((ISBN: 978-0851998275)); Hirooka Y, et al. 2012. Studies in Mycology 71:1-210 ((https://doi.org/10.3114/sim0001)); Tedersoo L, et al. 2014. Science 346:e1256688 ((https://doi.org/10.1126/science.1256688)); The American Phytopathological Society Common Names of Plant Diseases ((http://www.apsnet.org/publications/commonnames/Pages/default.aspx)); https://www.cdc.gov ((https://www.cdc.gov)) |
| Otu53236 | 1 | k__Fungi(100);p__Ascomycota(100);c__Saccharomycetes(100);o__Saccharomycetales(100);f__Metschnikowiaceae(100);g__Metschnikowia(100);s__Metschnikowia_pulcherrima(100); | Metschnikowia | 13 | Saprotroph | \|Undefined Saprotroph\| | Yeast | NULL | Probable | As nectar-tap_saprotroph (Põlme et al. 2020); Decay sugar-rich substrates (Põlme et al. 2020) | Põlme S, et al. 2020. Fungal Diversity 105:1-16 ((https://doi.org/10.1007/s13225-020-00466-2)) |
| Otu53237 | 1 | k__Fungi(100);p__Ascomycota(100);c__Saccharomycetes(100);o__Saccharomycetales(100);f__Metschnikowiaceae(100);g__Metschnikowia(100);g__Metschnikowia_unclassified(100); | Metschnikowia | 13 | Saprotroph | \|Undefined Saprotroph\| | Yeast | NULL | Probable | As nectar-tap_saprotroph (Põlme et al. 2020); Decay sugar-rich substrates (Põlme et al. 2020) | Põlme S, et al. 2020. Fungal Diversity 105:1-16 ((https://doi.org/10.1007/s13225-020-00466-2)) |
| Otu53238 | 1 | k__Fungi(100);p__Ascomycota(100);c__Saccharomycetes(100);o__Saccharomycetales(100);f__Metschnikowiaceae(100);g__Metschnikowia(100);g__Metschnikowia_unclassified(100); | Metschnikowia | 13 | Saprotroph | \|Undefined Saprotroph\| | Yeast | NULL | Probable | As nectar-tap_saprotroph (Põlme et al. 2020); Decay sugar-rich substrates (Põlme et al. 2020) | Põlme S, et al. 2020. Fungal Diversity 105:1-16 ((https://doi.org/10.1007/s13225-020-00466-2)) |
| Otu53239 | 1 | k__Fungi(100);p__Ascomycota(100);c__Saccharomycetes(100);o__Saccharomycetales(100);f__Metschnikowiaceae(100);g__Metschnikowia(100);s__Metschnikowia_pulcherrima(100); | Metschnikowia | 13 | Saprotroph | \|Undefined Saprotroph\| | Yeast | NULL | Probable | As nectar-tap_saprotroph (Põlme et al. 2020); Decay sugar-rich substrates (Põlme et al. 2020) | Põlme S, et al. 2020. Fungal Diversity 105:1-16 ((https://doi.org/10.1007/s13225-020-00466-2)) |
| Otu53240 | 1 | k__Fungi(100);p__Ascomycota(100);c__Saccharomycetes(100);o__Saccharomycetales(100);f__Metschnikowiaceae(100);g__Metschnikowia(100);s__Metschnikowia_pulcherrima(100); | Metschnikowia | 13 | Saprotroph | \|Undefined Saprotroph\| | Yeast | NULL | Probable | As nectar-tap_saprotroph (Põlme et al. 2020); Decay sugar-rich substrates (Põlme et al. 2020) | Põlme S, et al. 2020. Fungal Diversity 105:1-16 ((https://doi.org/10.1007/s13225-020-00466-2)) |
| Otu53241 | 1 | k__Fungi(100);p__Ascomycota(100);c__Saccharomycetes(100);o__Saccharomycetales(100);f__Metschnikowiaceae(100);g__Metschnikowia(100);g__Metschnikowia_unclassified(100); | Metschnikowia | 13 | Saprotroph | \|Undefined Saprotroph\| | Yeast | NULL | Probable | As nectar-tap_saprotroph (Põlme et al. 2020); Decay sugar-rich substrates (Põlme et al. 2020) | Põlme S, et al. 2020. Fungal Diversity 105:1-16 ((https://doi.org/10.1007/s13225-020-00466-2)) |
| Otu53242 | 1 | k__Fungi(100);p__Ascomycota(100);c__Saccharomycetes(100);o__Saccharomycetales(100);f__Metschnikowiaceae(100);g__Metschnikowia(100);s__Metschnikowia_pulcherrima(100); | Metschnikowia | 13 | Saprotroph | \|Undefined Saprotroph\| | Yeast | NULL | Probable | As nectar-tap_saprotroph (Põlme et al. 2020); Decay sugar-rich substrates (Põlme et al. 2020) | Põlme S, et al. 2020. Fungal Diversity 105:1-16 ((https://doi.org/10.1007/s13225-020-00466-2)) |
| Otu53243 | 1 | k__Fungi(100);p__Ascomycota(100);c__Saccharomycetes(100);o__Saccharomycetales(100);f__Metschnikowiaceae(100);g__Metschnikowia(100);s__Metschnikowia_pulcherrima(100); | Metschnikowia | 13 | Saprotroph | \|Undefined Saprotroph\| | Yeast | NULL | Probable | As nectar-tap_saprotroph (Põlme et al. 2020); Decay sugar-rich substrates (Põlme et al. 2020) | Põlme S, et al. 2020. Fungal Diversity 105:1-16 ((https://doi.org/10.1007/s13225-020-00466-2)) |
| Otu53245 | 1 | k__Fungi(100);p__Ascomycota(100);c__Saccharomycetes(100);o__Saccharomycetales(100);f__Metschnikowiaceae(100);g__Metschnikowia(100);s__Metschnikowia_pulcherrima(100); | Metschnikowia | 13 | Saprotroph | \|Undefined Saprotroph\| | Yeast | NULL | Probable | As nectar-tap_saprotroph (Põlme et al. 2020); Decay sugar-rich substrates (Põlme et al. 2020) | Põlme S, et al. 2020. Fungal Diversity 105:1-16 ((https://doi.org/10.1007/s13225-020-00466-2)) |
| Otu53246 | 1 | k__Fungi(100);p__Ascomycota(100);c__Saccharomycetes(100);o__Saccharomycetales(100);f__Metschnikowiaceae(100);g__Metschnikowia(100);s__Metschnikowia_pulcherrima(100); | Metschnikowia | 13 | Saprotroph | \|Undefined Saprotroph\| | Yeast | NULL | Probable | As nectar-tap_saprotroph (Põlme et al. 2020); Decay sugar-rich substrates (Põlme et al. 2020) | Põlme S, et al. 2020. Fungal Diversity 105:1-16 ((https://doi.org/10.1007/s13225-020-00466-2)) |
| Otu53247 | 1 | k__Fungi(100);p__Ascomycota(100);c__Saccharomycetes(100);o__Saccharomycetales(100);f__Metschnikowiaceae(100);g__Metschnikowia(100);g__Metschnikowia_unclassified(100); | Metschnikowia | 13 | Saprotroph | \|Undefined Saprotroph\| | Yeast | NULL | Probable | As nectar-tap_saprotroph (Põlme et al. 2020); Decay sugar-rich substrates (Põlme et al. 2020) | Põlme S, et al. 2020. Fungal Diversity 105:1-16 ((https://doi.org/10.1007/s13225-020-00466-2)) |
| Otu53248 | 1 | k__Fungi(100);p__Ascomycota(100);c__Saccharomycetes(100);o__Saccharomycetales(100);f__Metschnikowiaceae(100);g__Metschnikowia(100);s__Metschnikowia_pulcherrima(100); | Metschnikowia | 13 | Saprotroph | \|Undefined Saprotroph\| | Yeast | NULL | Probable | As nectar-tap_saprotroph (Põlme et al. 2020); Decay sugar-rich substrates (Põlme et al. 2020) | Põlme S, et al. 2020. Fungal Diversity 105:1-16 ((https://doi.org/10.1007/s13225-020-00466-2)) |
| Otu53249 | 1 | k__Fungi(100);p__Ascomycota(100);c__Saccharomycetes(100);o__Saccharomycetales(100);f__Metschnikowiaceae(100);g__Metschnikowia(100);s__Metschnikowia_pulcherrima(100); | Metschnikowia | 13 | Saprotroph | \|Undefined Saprotroph\| | Yeast | NULL | Probable | As nectar-tap_saprotroph (Põlme et al. 2020); Decay sugar-rich substrates (Põlme et al. 2020) | Põlme S, et al. 2020. Fungal Diversity 105:1-16 ((https://doi.org/10.1007/s13225-020-00466-2)) |
| Otu53250 | 1 | k__Fungi(100);p__Ascomycota(100);c__Saccharomycetes(100);o__Saccharomycetales(100);f__Metschnikowiaceae(100);g__Metschnikowia(100);s__Metschnikowia_pulcherrima(100); | Metschnikowia | 13 | Saprotroph | \|Undefined Saprotroph\| | Yeast | NULL | Probable | As nectar-tap_saprotroph (Põlme et al. 2020); Decay sugar-rich substrates (Põlme et al. 2020) | Põlme S, et al. 2020. Fungal Diversity 105:1-16 ((https://doi.org/10.1007/s13225-020-00466-2)) |
| Otu53251 | 1 | k__Fungi(100);p__Ascomycota(100);c__Saccharomycetes(100);o__Saccharomycetales(100);f__Metschnikowiaceae(100);g__Metschnikowia(100);s__Metschnikowia_pulcherrima(100); | Metschnikowia | 13 | Saprotroph | \|Undefined Saprotroph\| | Yeast | NULL | Probable | As nectar-tap_saprotroph (Põlme et al. 2020); Decay sugar-rich substrates (Põlme et al. 2020) | Põlme S, et al. 2020. Fungal Diversity 105:1-16 ((https://doi.org/10.1007/s13225-020-00466-2)) |
| Otu53252 | 1 | k__Fungi(100);p__Ascomycota(100);c__Saccharomycetes(100);o__Saccharomycetales(100);f__Metschnikowiaceae(100);g__Metschnikowia(100);g__Metschnikowia_unclassified(100); | Metschnikowia | 13 | Saprotroph | \|Undefined Saprotroph\| | Yeast | NULL | Probable | As nectar-tap_saprotroph (Põlme et al. 2020); Decay sugar-rich substrates (Põlme et al. 2020) | Põlme S, et al. 2020. Fungal Diversity 105:1-16 ((https://doi.org/10.1007/s13225-020-00466-2)) |
| Otu53253 | 1 | k__Fungi(100);p__Ascomycota(100);c__Saccharomycetes(100);o__Saccharomycetales(100);f__Metschnikowiaceae(100);g__Metschnikowia(100);g__Metschnikowia_unclassified(100); | Metschnikowia | 13 | Saprotroph | \|Undefined Saprotroph\| | Yeast | NULL | Probable | As nectar-tap_saprotroph (Põlme et al. 2020); Decay sugar-rich substrates (Põlme et al. 2020) | Põlme S, et al. 2020. Fungal Diversity 105:1-16 ((https://doi.org/10.1007/s13225-020-00466-2)) |
| Otu53254 | 1 | k__Fungi(100);p__Ascomycota(100);c__Saccharomycetes(100);o__Saccharomycetales(100);f__Metschnikowiaceae(100);g__Metschnikowia(100);g__Metschnikowia_unclassified(100); | Metschnikowia | 13 | Saprotroph | \|Undefined Saprotroph\| | Yeast | NULL | Probable | As nectar-tap_saprotroph (Põlme et al. 2020); Decay sugar-rich substrates (Põlme et al. 2020) | Põlme S, et al. 2020. Fungal Diversity 105:1-16 ((https://doi.org/10.1007/s13225-020-00466-2)) |
| Otu53255 | 1 | k__Fungi(100);p__Ascomycota(100);c__Saccharomycetes(100);o__Saccharomycetales(100);f__Metschnikowiaceae(100);g__Metschnikowia(100);g__Metschnikowia_unclassified(100); | Metschnikowia | 13 | Saprotroph | \|Undefined Saprotroph\| | Yeast | NULL | Probable | As nectar-tap_saprotroph (Põlme et al. 2020); Decay sugar-rich substrates (Põlme et al. 2020) | Põlme S, et al. 2020. Fungal Diversity 105:1-16 ((https://doi.org/10.1007/s13225-020-00466-2)) |
| Otu53256 | 1 | k__Fungi(100);p__Ascomycota(100);c__Saccharomycetes(100);o__Saccharomycetales(100);f__Metschnikowiaceae(100);g__Metschnikowia(100);s__Metschnikowia_pulcherrima(100); | Metschnikowia | 13 | Saprotroph | \|Undefined Saprotroph\| | Yeast | NULL | Probable | As nectar-tap_saprotroph (Põlme et al. 2020); Decay sugar-rich substrates (Põlme et al. 2020) | Põlme S, et al. 2020. Fungal Diversity 105:1-16 ((https://doi.org/10.1007/s13225-020-00466-2)) |
| Otu53257 | 1 | k__Fungi(100);p__Ascomycota(100);c__Saccharomycetes(100);o__Saccharomycetales(100);f__Metschnikowiaceae(100);g__Metschnikowia(100);s__Metschnikowia_pulcherrima(100); | Metschnikowia | 13 | Saprotroph | \|Undefined Saprotroph\| | Yeast | NULL | Probable | As nectar-tap_saprotroph (Põlme et al. 2020); Decay sugar-rich substrates (Põlme et al. 2020) | Põlme S, et al. 2020. Fungal Diversity 105:1-16 ((https://doi.org/10.1007/s13225-020-00466-2)) |
| Otu53258 | 1 | k__Fungi(100);p__Ascomycota(100);c__Saccharomycetes(100);o__Saccharomycetales(100);f__Metschnikowiaceae(100);g__Metschnikowia(100);s__Metschnikowia_pulcherrima(100); | Metschnikowia | 13 | Saprotroph | \|Undefined Saprotroph\| | Yeast | NULL | Probable | As nectar-tap_saprotroph (Põlme et al. 2020); Decay sugar-rich substrates (Põlme et al. 2020) | Põlme S, et al. 2020. Fungal Diversity 105:1-16 ((https://doi.org/10.1007/s13225-020-00466-2)) |
| Otu53259 | 1 | k__Fungi(100);p__Ascomycota(100);c__Saccharomycetes(100);o__Saccharomycetales(100);f__Metschnikowiaceae(100);g__Metschnikowia(100);s__Metschnikowia_pulcherrima(100); | Metschnikowia | 13 | Saprotroph | \|Undefined Saprotroph\| | Yeast | NULL | Probable | As nectar-tap_saprotroph (Põlme et al. 2020); Decay sugar-rich substrates (Põlme et al. 2020) | Põlme S, et al. 2020. Fungal Diversity 105:1-16 ((https://doi.org/10.1007/s13225-020-00466-2)) |
| Otu53260 | 1 | k__Fungi(100);p__Ascomycota(100);c__Saccharomycetes(100);o__Saccharomycetales(100);f__Metschnikowiaceae(100);g__Metschnikowia(100);s__Metschnikowia_pulcherrima(100); | Metschnikowia | 13 | Saprotroph | \|Undefined Saprotroph\| | Yeast | NULL | Probable | As nectar-tap_saprotroph (Põlme et al. 2020); Decay sugar-rich substrates (Põlme et al. 2020) | Põlme S, et al. 2020. Fungal Diversity 105:1-16 ((https://doi.org/10.1007/s13225-020-00466-2)) |
| Otu53261 | 1 | k__Fungi(100);p__Ascomycota(100);c__Saccharomycetes(100);o__Saccharomycetales(100);f__Metschnikowiaceae(100);g__Metschnikowia(100);s__Metschnikowia_pulcherrima(100); | Metschnikowia | 13 | Saprotroph | \|Undefined Saprotroph\| | Yeast | NULL | Probable | As nectar-tap_saprotroph (Põlme et al. 2020); Decay sugar-rich substrates (Põlme et al. 2020) | Põlme S, et al. 2020. Fungal Diversity 105:1-16 ((https://doi.org/10.1007/s13225-020-00466-2)) |
| Otu53262 | 1 | k__Fungi(100);p__Ascomycota(100);c__Saccharomycetes(100);o__Saccharomycetales(100);f__Metschnikowiaceae(100);g__Metschnikowia(100);s__Metschnikowia_pulcherrima(100); | Metschnikowia | 13 | Saprotroph | \|Undefined Saprotroph\| | Yeast | NULL | Probable | As nectar-tap_saprotroph (Põlme et al. 2020); Decay sugar-rich substrates (Põlme et al. 2020) | Põlme S, et al. 2020. Fungal Diversity 105:1-16 ((https://doi.org/10.1007/s13225-020-00466-2)) |
| Otu53263 | 1 | k__Fungi(100);p__Ascomycota(100);c__Saccharomycetes(100);o__Saccharomycetales(100);f__Metschnikowiaceae(100);g__Metschnikowia(100);g__Metschnikowia_unclassified(100); | Metschnikowia | 13 | Saprotroph | \|Undefined Saprotroph\| | Yeast | NULL | Probable | As nectar-tap_saprotroph (Põlme et al. 2020); Decay sugar-rich substrates (Põlme et al. 2020) | Põlme S, et al. 2020. Fungal Diversity 105:1-16 ((https://doi.org/10.1007/s13225-020-00466-2)) |
| Otu53264 | 1 | k__Fungi(100);p__Ascomycota(100);c__Saccharomycetes(100);o__Saccharomycetales(100);f__Metschnikowiaceae(100);g__Metschnikowia(100);s__Metschnikowia_pulcherrima(100); | Metschnikowia | 13 | Saprotroph | \|Undefined Saprotroph\| | Yeast | NULL | Probable | As nectar-tap_saprotroph (Põlme et al. 2020); Decay sugar-rich substrates (Põlme et al. 2020) | Põlme S, et al. 2020. Fungal Diversity 105:1-16 ((https://doi.org/10.1007/s13225-020-00466-2)) |
| Otu53265 | 1 | k__Fungi(100);p__Ascomycota(100);c__Saccharomycetes(100);o__Saccharomycetales(100);f__Metschnikowiaceae(100);g__Metschnikowia(100);s__Metschnikowia_pulcherrima(100); | Metschnikowia | 13 | Saprotroph | \|Undefined Saprotroph\| | Yeast | NULL | Probable | As nectar-tap_saprotroph (Põlme et al. 2020); Decay sugar-rich substrates (Põlme et al. 2020) | Põlme S, et al. 2020. Fungal Diversity 105:1-16 ((https://doi.org/10.1007/s13225-020-00466-2)) |
| Otu53266 | 1 | k__Fungi(100);p__Ascomycota(100);c__Saccharomycetes(100);o__Saccharomycetales(100);f__Metschnikowiaceae(100);g__Metschnikowia(100);g__Metschnikowia_unclassified(100); | Metschnikowia | 13 | Saprotroph | \|Undefined Saprotroph\| | Yeast | NULL | Probable | As nectar-tap_saprotroph (Põlme et al. 2020); Decay sugar-rich substrates (Põlme et al. 2020) | Põlme S, et al. 2020. Fungal Diversity 105:1-16 ((https://doi.org/10.1007/s13225-020-00466-2)) |
| Otu53267 | 1 | k__Fungi(100);p__Ascomycota(100);c__Saccharomycetes(100);o__Saccharomycetales(100);f__Metschnikowiaceae(100);g__Metschnikowia(100);s__Metschnikowia_pulcherrima(100); | Metschnikowia | 13 | Saprotroph | \|Undefined Saprotroph\| | Yeast | NULL | Probable | As nectar-tap_saprotroph (Põlme et al. 2020); Decay sugar-rich substrates (Põlme et al. 2020) | Põlme S, et al. 2020. Fungal Diversity 105:1-16 ((https://doi.org/10.1007/s13225-020-00466-2)) |
| Otu53268 | 1 | k__Fungi(100);p__Ascomycota(100);c__Saccharomycetes(100);o__Saccharomycetales(100);f__Metschnikowiaceae(100);g__Metschnikowia(100);s__Metschnikowia_pulcherrima(100); | Metschnikowia | 13 | Saprotroph | \|Undefined Saprotroph\| | Yeast | NULL | Probable | As nectar-tap_saprotroph (Põlme et al. 2020); Decay sugar-rich substrates (Põlme et al. 2020) | Põlme S, et al. 2020. Fungal Diversity 105:1-16 ((https://doi.org/10.1007/s13225-020-00466-2)) |
| Otu53269 | 1 | k__Fungi(100);p__Ascomycota(100);c__Saccharomycetes(100);o__Saccharomycetales(100);f__Metschnikowiaceae(100);g__Metschnikowia(100);s__Metschnikowia_pulcherrima(100); | Metschnikowia | 13 | Saprotroph | \|Undefined Saprotroph\| | Yeast | NULL | Probable | As nectar-tap_saprotroph (Põlme et al. 2020); Decay sugar-rich substrates (Põlme et al. 2020) | Põlme S, et al. 2020. Fungal Diversity 105:1-16 ((https://doi.org/10.1007/s13225-020-00466-2)) |
| Otu53270 | 1 | k__Fungi(100);p__Ascomycota(100);c__Saccharomycetes(100);o__Saccharomycetales(100);f__Metschnikowiaceae(100);g__Metschnikowia(100);s__Metschnikowia_pulcherrima(100); | Metschnikowia | 13 | Saprotroph | \|Undefined Saprotroph\| | Yeast | NULL | Probable | As nectar-tap_saprotroph (Põlme et al. 2020); Decay sugar-rich substrates (Põlme et al. 2020) | Põlme S, et al. 2020. Fungal Diversity 105:1-16 ((https://doi.org/10.1007/s13225-020-00466-2)) |
| Otu53271 | 1 | k__Fungi(100);p__Ascomycota(100);c__Saccharomycetes(100);o__Saccharomycetales(100);f__Metschnikowiaceae(100);g__Metschnikowia(100);s__Metschnikowia_pulcherrima(100); | Metschnikowia | 13 | Saprotroph | \|Undefined Saprotroph\| | Yeast | NULL | Probable | As nectar-tap_saprotroph (Põlme et al. 2020); Decay sugar-rich substrates (Põlme et al. 2020) | Põlme S, et al. 2020. Fungal Diversity 105:1-16 ((https://doi.org/10.1007/s13225-020-00466-2)) |
| Otu53272 | 1 | k__Fungi(100);p__Ascomycota(100);c__Saccharomycetes(100);o__Saccharomycetales(100);f__Metschnikowiaceae(100);g__Metschnikowia(100);s__Metschnikowia_pulcherrima(100); | Metschnikowia | 13 | Saprotroph | \|Undefined Saprotroph\| | Yeast | NULL | Probable | As nectar-tap_saprotroph (Põlme et al. 2020); Decay sugar-rich substrates (Põlme et al. 2020) | Põlme S, et al. 2020. Fungal Diversity 105:1-16 ((https://doi.org/10.1007/s13225-020-00466-2)) |
| Otu53273 | 1 | k__Fungi(100);p__Ascomycota(100);c__Saccharomycetes(100);o__Saccharomycetales(100);f__Metschnikowiaceae(100);g__Metschnikowia(100);g__Metschnikowia_unclassified(100); | Metschnikowia | 13 | Saprotroph | \|Undefined Saprotroph\| | Yeast | NULL | Probable | As nectar-tap_saprotroph (Põlme et al. 2020); Decay sugar-rich substrates (Põlme et al. 2020) | Põlme S, et al. 2020. Fungal Diversity 105:1-16 ((https://doi.org/10.1007/s13225-020-00466-2)) |
| Otu53274 | 1 | k__Fungi(100);p__Ascomycota(100);c__Saccharomycetes(100);o__Saccharomycetales(100);f__Metschnikowiaceae(100);g__Metschnikowia(100);s__Metschnikowia_pulcherrima(100); | Metschnikowia | 13 | Saprotroph | \|Undefined Saprotroph\| | Yeast | NULL | Probable | As nectar-tap_saprotroph (Põlme et al. 2020); Decay sugar-rich substrates (Põlme et al. 2020) | Põlme S, et al. 2020. Fungal Diversity 105:1-16 ((https://doi.org/10.1007/s13225-020-00466-2)) |
| Otu53275 | 1 | k__Fungi(100);p__Ascomycota(100);c__Saccharomycetes(100);o__Saccharomycetales(100);f__Metschnikowiaceae(100);g__Metschnikowia(100);g__Metschnikowia_unclassified(100); | Metschnikowia | 13 | Saprotroph | \|Undefined Saprotroph\| | Yeast | NULL | Probable | As nectar-tap_saprotroph (Põlme et al. 2020); Decay sugar-rich substrates (Põlme et al. 2020) | Põlme S, et al. 2020. Fungal Diversity 105:1-16 ((https://doi.org/10.1007/s13225-020-00466-2)) |
| Otu53276 | 1 | k__Fungi(100);p__Ascomycota(100);c__Saccharomycetes(100);o__Saccharomycetales(100);f__Metschnikowiaceae(100);g__Metschnikowia(100);g__Metschnikowia_unclassified(100); | Metschnikowia | 13 | Saprotroph | \|Undefined Saprotroph\| | Yeast | NULL | Probable | As nectar-tap_saprotroph (Põlme et al. 2020); Decay sugar-rich substrates (Põlme et al. 2020) | Põlme S, et al. 2020. Fungal Diversity 105:1-16 ((https://doi.org/10.1007/s13225-020-00466-2)) |
| Otu53277 | 1 | k__Fungi(100);p__Ascomycota(100);c__Saccharomycetes(100);o__Saccharomycetales(100);f__Metschnikowiaceae(100);g__Metschnikowia(100);s__Metschnikowia_pulcherrima(100); | Metschnikowia | 13 | Saprotroph | \|Undefined Saprotroph\| | Yeast | NULL | Probable | As nectar-tap_saprotroph (Põlme et al. 2020); Decay sugar-rich substrates (Põlme et al. 2020) | Põlme S, et al. 2020. Fungal Diversity 105:1-16 ((https://doi.org/10.1007/s13225-020-00466-2)) |
| Otu53278 | 1 | k__Fungi(100);p__Ascomycota(100);c__Saccharomycetes(100);o__Saccharomycetales(100);f__Metschnikowiaceae(100);g__Metschnikowia(100);s__Metschnikowia_pulcherrima(100); | Metschnikowia | 13 | Saprotroph | \|Undefined Saprotroph\| | Yeast | NULL | Probable | As nectar-tap_saprotroph (Põlme et al. 2020); Decay sugar-rich substrates (Põlme et al. 2020) | Põlme S, et al. 2020. Fungal Diversity 105:1-16 ((https://doi.org/10.1007/s13225-020-00466-2)) |
| Otu53279 | 1 | k__Fungi(100);p__Ascomycota(100);c__Saccharomycetes(100);o__Saccharomycetales(100);f__Metschnikowiaceae(100);g__Metschnikowia(100);g__Metschnikowia_unclassified(100); | Metschnikowia | 13 | Saprotroph | \|Undefined Saprotroph\| | Yeast | NULL | Probable | As nectar-tap_saprotroph (Põlme et al. 2020); Decay sugar-rich substrates (Põlme et al. 2020) | Põlme S, et al. 2020. Fungal Diversity 105:1-16 ((https://doi.org/10.1007/s13225-020-00466-2)) |
| Otu53280 | 1 | k__Fungi(100);p__Ascomycota(100);c__Saccharomycetes(100);o__Saccharomycetales(100);f__Metschnikowiaceae(100);g__Metschnikowia(100);s__Metschnikowia_pulcherrima(100); | Metschnikowia | 13 | Saprotroph | \|Undefined Saprotroph\| | Yeast | NULL | Probable | As nectar-tap_saprotroph (Põlme et al. 2020); Decay sugar-rich substrates (Põlme et al. 2020) | Põlme S, et al. 2020. Fungal Diversity 105:1-16 ((https://doi.org/10.1007/s13225-020-00466-2)) |
| Otu53281 | 1 | k__Fungi(100);p__Ascomycota(100);c__Saccharomycetes(100);o__Saccharomycetales(100);f__Metschnikowiaceae(100);g__Metschnikowia(100);s__Metschnikowia_pulcherrima(100); | Metschnikowia | 13 | Saprotroph | \|Undefined Saprotroph\| | Yeast | NULL | Probable | As nectar-tap_saprotroph (Põlme et al. 2020); Decay sugar-rich substrates (Põlme et al. 2020) | Põlme S, et al. 2020. Fungal Diversity 105:1-16 ((https://doi.org/10.1007/s13225-020-00466-2)) |
| Otu53282 | 1 | k__Fungi(100);p__Ascomycota(100);c__Saccharomycetes(100);o__Saccharomycetales(100);f__Metschnikowiaceae(100);g__Metschnikowia(100);s__Metschnikowia_pulcherrima(100); | Metschnikowia | 13 | Saprotroph | \|Undefined Saprotroph\| | Yeast | NULL | Probable | As nectar-tap_saprotroph (Põlme et al. 2020); Decay sugar-rich substrates (Põlme et al. 2020) | Põlme S, et al. 2020. Fungal Diversity 105:1-16 ((https://doi.org/10.1007/s13225-020-00466-2)) |
| Otu53283 | 1 | k__Fungi(100);p__Ascomycota(100);c__Saccharomycetes(100);o__Saccharomycetales(100);f__Metschnikowiaceae(100);g__Metschnikowia(100);s__Metschnikowia_pulcherrima(100); | Metschnikowia | 13 | Saprotroph | \|Undefined Saprotroph\| | Yeast | NULL | Probable | As nectar-tap_saprotroph (Põlme et al. 2020); Decay sugar-rich substrates (Põlme et al. 2020) | Põlme S, et al. 2020. Fungal Diversity 105:1-16 ((https://doi.org/10.1007/s13225-020-00466-2)) |
| Otu53284 | 1 | k__Fungi(100);p__Ascomycota(100);c__Saccharomycetes(100);o__Saccharomycetales(100);f__Metschnikowiaceae(100);g__Metschnikowia(100);s__Metschnikowia_pulcherrima(100); | Metschnikowia | 13 | Saprotroph | \|Undefined Saprotroph\| | Yeast | NULL | Probable | As nectar-tap_saprotroph (Põlme et al. 2020); Decay sugar-rich substrates (Põlme et al. 2020) | Põlme S, et al. 2020. Fungal Diversity 105:1-16 ((https://doi.org/10.1007/s13225-020-00466-2)) |
| Otu53285 | 1 | k__Fungi(100);p__Ascomycota(100);c__Saccharomycetes(100);o__Saccharomycetales(100);f__Metschnikowiaceae(100);g__Metschnikowia(100);s__Metschnikowia_pulcherrima(100); | Metschnikowia | 13 | Saprotroph | \|Undefined Saprotroph\| | Yeast | NULL | Probable | As nectar-tap_saprotroph (Põlme et al. 2020); Decay sugar-rich substrates (Põlme et al. 2020) | Põlme S, et al. 2020. Fungal Diversity 105:1-16 ((https://doi.org/10.1007/s13225-020-00466-2)) |
| Otu53286 | 1 | k__Fungi(100);p__Ascomycota(100);c__Saccharomycetes(100);o__Saccharomycetales(100);f__Metschnikowiaceae(100);g__Metschnikowia(100);s__Metschnikowia_pulcherrima(100); | Metschnikowia | 13 | Saprotroph | \|Undefined Saprotroph\| | Yeast | NULL | Probable | As nectar-tap_saprotroph (Põlme et al. 2020); Decay sugar-rich substrates (Põlme et al. 2020) | Põlme S, et al. 2020. Fungal Diversity 105:1-16 ((https://doi.org/10.1007/s13225-020-00466-2)) |
| Otu53287 | 1 | k__Fungi(100);p__Ascomycota(100);c__Saccharomycetes(100);o__Saccharomycetales(100);f__Metschnikowiaceae(100);g__Metschnikowia(100);s__Metschnikowia_pulcherrima(100); | Metschnikowia | 13 | Saprotroph | \|Undefined Saprotroph\| | Yeast | NULL | Probable | As nectar-tap_saprotroph (Põlme et al. 2020); Decay sugar-rich substrates (Põlme et al. 2020) | Põlme S, et al. 2020. Fungal Diversity 105:1-16 ((https://doi.org/10.1007/s13225-020-00466-2)) |
| Otu53288 | 1 | k__Fungi(100);p__Ascomycota(100);c__Saccharomycetes(100);o__Saccharomycetales(100);f__Metschnikowiaceae(100);g__Metschnikowia(100);s__Metschnikowia_pulcherrima(100); | Metschnikowia | 13 | Saprotroph | \|Undefined Saprotroph\| | Yeast | NULL | Probable | As nectar-tap_saprotroph (Põlme et al. 2020); Decay sugar-rich substrates (Põlme et al. 2020) | Põlme S, et al. 2020. Fungal Diversity 105:1-16 ((https://doi.org/10.1007/s13225-020-00466-2)) |
| Otu53289 | 1 | k__Fungi(100);p__Ascomycota(100);c__Saccharomycetes(100);o__Saccharomycetales(100);f__Metschnikowiaceae(100);g__Metschnikowia(100);s__Metschnikowia_pulcherrima(100); | Metschnikowia | 13 | Saprotroph | \|Undefined Saprotroph\| | Yeast | NULL | Probable | As nectar-tap_saprotroph (Põlme et al. 2020); Decay sugar-rich substrates (Põlme et al. 2020) | Põlme S, et al. 2020. Fungal Diversity 105:1-16 ((https://doi.org/10.1007/s13225-020-00466-2)) |
| Otu53290 | 1 | k__Fungi(100);p__Ascomycota(100);c__Saccharomycetes(100);o__Saccharomycetales(100);f__Metschnikowiaceae(100);g__Metschnikowia(100);g__Metschnikowia_unclassified(100); | Metschnikowia | 13 | Saprotroph | \|Undefined Saprotroph\| | Yeast | NULL | Probable | As nectar-tap_saprotroph (Põlme et al. 2020); Decay sugar-rich substrates (Põlme et al. 2020) | Põlme S, et al. 2020. Fungal Diversity 105:1-16 ((https://doi.org/10.1007/s13225-020-00466-2)) |
| Otu53291 | 1 | k__Fungi(100);p__Ascomycota(100);c__Saccharomycetes(100);o__Saccharomycetales(100);f__Metschnikowiaceae(100);g__Metschnikowia(100);g__Metschnikowia_unclassified(100); | Metschnikowia | 13 | Saprotroph | \|Undefined Saprotroph\| | Yeast | NULL | Probable | As nectar-tap_saprotroph (Põlme et al. 2020); Decay sugar-rich substrates (Põlme et al. 2020) | Põlme S, et al. 2020. Fungal Diversity 105:1-16 ((https://doi.org/10.1007/s13225-020-00466-2)) |
| Otu53292 | 1 | k__Fungi(100);p__Ascomycota(100);c__Saccharomycetes(100);o__Saccharomycetales(100);f__Metschnikowiaceae(100);g__Metschnikowia(100);s__Metschnikowia_pulcherrima(100); | Metschnikowia | 13 | Saprotroph | \|Undefined Saprotroph\| | Yeast | NULL | Probable | As nectar-tap_saprotroph (Põlme et al. 2020); Decay sugar-rich substrates (Põlme et al. 2020) | Põlme S, et al. 2020. Fungal Diversity 105:1-16 ((https://doi.org/10.1007/s13225-020-00466-2)) |
| Otu53293 | 1 | k__Fungi(100);p__Ascomycota(100);c__Saccharomycetes(100);o__Saccharomycetales(100);f__Metschnikowiaceae(100);g__Metschnikowia(100);s__Metschnikowia_pulcherrima(100); | Metschnikowia | 13 | Saprotroph | \|Undefined Saprotroph\| | Yeast | NULL | Probable | As nectar-tap_saprotroph (Põlme et al. 2020); Decay sugar-rich substrates (Põlme et al. 2020) | Põlme S, et al. 2020. Fungal Diversity 105:1-16 ((https://doi.org/10.1007/s13225-020-00466-2)) |
| Otu53294 | 1 | k__Fungi(100);p__Ascomycota(100);c__Saccharomycetes(100);o__Saccharomycetales(100);f__Metschnikowiaceae(100);g__Metschnikowia(100);s__Metschnikowia_pulcherrima(100); | Metschnikowia | 13 | Saprotroph | \|Undefined Saprotroph\| | Yeast | NULL | Probable | As nectar-tap_saprotroph (Põlme et al. 2020); Decay sugar-rich substrates (Põlme et al. 2020) | Põlme S, et al. 2020. Fungal Diversity 105:1-16 ((https://doi.org/10.1007/s13225-020-00466-2)) |
| Otu53295 | 1 | k__Fungi(100);p__Ascomycota(100);c__Saccharomycetes(100);o__Saccharomycetales(100);f__Metschnikowiaceae(100);g__Metschnikowia(100);s__Metschnikowia_pulcherrima(100); | Metschnikowia | 13 | Saprotroph | \|Undefined Saprotroph\| | Yeast | NULL | Probable | As nectar-tap_saprotroph (Põlme et al. 2020); Decay sugar-rich substrates (Põlme et al. 2020) | Põlme S, et al. 2020. Fungal Diversity 105:1-16 ((https://doi.org/10.1007/s13225-020-00466-2)) |
| Otu53296 | 1 | k__Fungi(100);p__Ascomycota(100);c__Saccharomycetes(100);o__Saccharomycetales(100);f__Metschnikowiaceae(100);g__Metschnikowia(100);s__Metschnikowia_pulcherrima(100); | Metschnikowia | 13 | Saprotroph | \|Undefined Saprotroph\| | Yeast | NULL | Probable | As nectar-tap_saprotroph (Põlme et al. 2020); Decay sugar-rich substrates (Põlme et al. 2020) | Põlme S, et al. 2020. Fungal Diversity 105:1-16 ((https://doi.org/10.1007/s13225-020-00466-2)) |
| Otu53297 | 1 | k__Fungi(100);p__Ascomycota(100);c__Saccharomycetes(100);o__Saccharomycetales(100);f__Metschnikowiaceae(100);g__Metschnikowia(100);s__Metschnikowia_pulcherrima(100); | Metschnikowia | 13 | Saprotroph | \|Undefined Saprotroph\| | Yeast | NULL | Probable | As nectar-tap_saprotroph (Põlme et al. 2020); Decay sugar-rich substrates (Põlme et al. 2020) | Põlme S, et al. 2020. Fungal Diversity 105:1-16 ((https://doi.org/10.1007/s13225-020-00466-2)) |
| Otu53298 | 1 | k__Fungi(100);p__Ascomycota(100);c__Saccharomycetes(100);o__Saccharomycetales(100);f__Metschnikowiaceae(100);g__Metschnikowia(100);s__Metschnikowia_pulcherrima(100); | Metschnikowia | 13 | Saprotroph | \|Undefined Saprotroph\| | Yeast | NULL | Probable | As nectar-tap_saprotroph (Põlme et al. 2020); Decay sugar-rich substrates (Põlme et al. 2020) | Põlme S, et al. 2020. Fungal Diversity 105:1-16 ((https://doi.org/10.1007/s13225-020-00466-2)) |
| Otu53299 | 1 | k__Fungi(100);p__Ascomycota(100);c__Saccharomycetes(100);o__Saccharomycetales(100);f__Metschnikowiaceae(100);g__Metschnikowia(100);s__Metschnikowia_pulcherrima(100); | Metschnikowia | 13 | Saprotroph | \|Undefined Saprotroph\| | Yeast | NULL | Probable | As nectar-tap_saprotroph (Põlme et al. 2020); Decay sugar-rich substrates (Põlme et al. 2020) | Põlme S, et al. 2020. Fungal Diversity 105:1-16 ((https://doi.org/10.1007/s13225-020-00466-2)) |
| Otu53300 | 1 | k__Fungi(100);p__Ascomycota(100);c__Saccharomycetes(100);o__Saccharomycetales(100);f__Metschnikowiaceae(100);g__Metschnikowia(100);g__Metschnikowia_unclassified(100); | Metschnikowia | 13 | Saprotroph | \|Undefined Saprotroph\| | Yeast | NULL | Probable | As nectar-tap_saprotroph (Põlme et al. 2020); Decay sugar-rich substrates (Põlme et al. 2020) | Põlme S, et al. 2020. Fungal Diversity 105:1-16 ((https://doi.org/10.1007/s13225-020-00466-2)) |
| Otu53301 | 1 | k__Fungi(100);p__Ascomycota(100);c__Saccharomycetes(100);o__Saccharomycetales(100);f__Metschnikowiaceae(100);g__Metschnikowia(100);s__Metschnikowia_pulcherrima(100); | Metschnikowia | 13 | Saprotroph | \|Undefined Saprotroph\| | Yeast | NULL | Probable | As nectar-tap_saprotroph (Põlme et al. 2020); Decay sugar-rich substrates (Põlme et al. 2020) | Põlme S, et al. 2020. Fungal Diversity 105:1-16 ((https://doi.org/10.1007/s13225-020-00466-2)) |
| Otu53302 | 1 | k__Fungi(100);p__Ascomycota(100);c__Saccharomycetes(100);o__Saccharomycetales(100);f__Metschnikowiaceae(100);g__Metschnikowia(100);s__Metschnikowia_pulcherrima(100); | Metschnikowia | 13 | Saprotroph | \|Undefined Saprotroph\| | Yeast | NULL | Probable | As nectar-tap_saprotroph (Põlme et al. 2020); Decay sugar-rich substrates (Põlme et al. 2020) | Põlme S, et al. 2020. Fungal Diversity 105:1-16 ((https://doi.org/10.1007/s13225-020-00466-2)) |
| Otu53303 | 1 | k__Fungi(100);p__Ascomycota(100);c__Saccharomycetes(100);o__Saccharomycetales(100);f__Metschnikowiaceae(100);g__Metschnikowia(100);g__Metschnikowia_unclassified(100); | Metschnikowia | 13 | Saprotroph | \|Undefined Saprotroph\| | Yeast | NULL | Probable | As nectar-tap_saprotroph (Põlme et al. 2020); Decay sugar-rich substrates (Põlme et al. 2020) | Põlme S, et al. 2020. Fungal Diversity 105:1-16 ((https://doi.org/10.1007/s13225-020-00466-2)) |
| Otu53304 | 1 | k__Fungi(100);p__Ascomycota(100);c__Saccharomycetes(100);o__Saccharomycetales(100);f__Metschnikowiaceae(100);g__Metschnikowia(100);g__Metschnikowia_unclassified(100); | Metschnikowia | 13 | Saprotroph | \|Undefined Saprotroph\| | Yeast | NULL | Probable | As nectar-tap_saprotroph (Põlme et al. 2020); Decay sugar-rich substrates (Põlme et al. 2020) | Põlme S, et al. 2020. Fungal Diversity 105:1-16 ((https://doi.org/10.1007/s13225-020-00466-2)) |
| Otu53305 | 1 | k__Fungi(100);p__Ascomycota(100);c__Saccharomycetes(100);o__Saccharomycetales(100);f__Metschnikowiaceae(100);g__Metschnikowia(100);s__Metschnikowia_pulcherrima(100); | Metschnikowia | 13 | Saprotroph | \|Undefined Saprotroph\| | Yeast | NULL | Probable | As nectar-tap_saprotroph (Põlme et al. 2020); Decay sugar-rich substrates (Põlme et al. 2020) | Põlme S, et al. 2020. Fungal Diversity 105:1-16 ((https://doi.org/10.1007/s13225-020-00466-2)) |
| Otu53306 | 1 | k__Fungi(100);p__Ascomycota(100);c__Saccharomycetes(100);o__Saccharomycetales(100);f__Metschnikowiaceae(100);g__Metschnikowia(100);s__Metschnikowia_pulcherrima(100); | Metschnikowia | 13 | Saprotroph | \|Undefined Saprotroph\| | Yeast | NULL | Probable | As nectar-tap_saprotroph (Põlme et al. 2020); Decay sugar-rich substrates (Põlme et al. 2020) | Põlme S, et al. 2020. Fungal Diversity 105:1-16 ((https://doi.org/10.1007/s13225-020-00466-2)) |
| Otu53307 | 1 | k__Fungi(100);p__Ascomycota(100);c__Saccharomycetes(100);o__Saccharomycetales(100);f__Metschnikowiaceae(100);g__Metschnikowia(100);g__Metschnikowia_unclassified(100); | Metschnikowia | 13 | Saprotroph | \|Undefined Saprotroph\| | Yeast | NULL | Probable | As nectar-tap_saprotroph (Põlme et al. 2020); Decay sugar-rich substrates (Põlme et al. 2020) | Põlme S, et al. 2020. Fungal Diversity 105:1-16 ((https://doi.org/10.1007/s13225-020-00466-2)) |
| Otu53309 | 1 | k__Fungi(100);p__Ascomycota(100);c__Saccharomycetes(100);o__Saccharomycetales(100);f__Metschnikowiaceae(100);g__Metschnikowia(100);g__Metschnikowia_unclassified(100); | Metschnikowia | 13 | Saprotroph | \|Undefined Saprotroph\| | Yeast | NULL | Probable | As nectar-tap_saprotroph (Põlme et al. 2020); Decay sugar-rich substrates (Põlme et al. 2020) | Põlme S, et al. 2020. Fungal Diversity 105:1-16 ((https://doi.org/10.1007/s13225-020-00466-2)) |
| Otu53310 | 1 | k__Fungi(100);p__Ascomycota(100);c__Saccharomycetes(100);o__Saccharomycetales(100);f__Metschnikowiaceae(100);g__Metschnikowia(100);s__Metschnikowia_pulcherrima(100); | Metschnikowia | 13 | Saprotroph | \|Undefined Saprotroph\| | Yeast | NULL | Probable | As nectar-tap_saprotroph (Põlme et al. 2020); Decay sugar-rich substrates (Põlme et al. 2020) | Põlme S, et al. 2020. Fungal Diversity 105:1-16 ((https://doi.org/10.1007/s13225-020-00466-2)) |
| Otu53311 | 1 | k__Fungi(100);p__Ascomycota(100);c__Saccharomycetes(100);o__Saccharomycetales(100);f__Metschnikowiaceae(100);g__Metschnikowia(100);s__Metschnikowia_pulcherrima(100); | Metschnikowia | 13 | Saprotroph | \|Undefined Saprotroph\| | Yeast | NULL | Probable | As nectar-tap_saprotroph (Põlme et al. 2020); Decay sugar-rich substrates (Põlme et al. 2020) | Põlme S, et al. 2020. Fungal Diversity 105:1-16 ((https://doi.org/10.1007/s13225-020-00466-2)) |
| Otu53312 | 1 | k__Fungi(100);p__Ascomycota(100);c__Saccharomycetes(100);o__Saccharomycetales(100);f__Metschnikowiaceae(100);g__Metschnikowia(100);s__Metschnikowia_pulcherrima(100); | Metschnikowia | 13 | Saprotroph | \|Undefined Saprotroph\| | Yeast | NULL | Probable | As nectar-tap_saprotroph (Põlme et al. 2020); Decay sugar-rich substrates (Põlme et al. 2020) | Põlme S, et al. 2020. Fungal Diversity 105:1-16 ((https://doi.org/10.1007/s13225-020-00466-2)) |
| Otu53313 | 1 | k__Fungi(100);p__Ascomycota(100);c__Saccharomycetes(100);o__Saccharomycetales(100);f__Metschnikowiaceae(100);g__Metschnikowia(100);s__Metschnikowia_pulcherrima(100); | Metschnikowia | 13 | Saprotroph | \|Undefined Saprotroph\| | Yeast | NULL | Probable | As nectar-tap_saprotroph (Põlme et al. 2020); Decay sugar-rich substrates (Põlme et al. 2020) | Põlme S, et al. 2020. Fungal Diversity 105:1-16 ((https://doi.org/10.1007/s13225-020-00466-2)) |
| Otu53314 | 1 | k__Fungi(100);p__Ascomycota(100);c__Saccharomycetes(100);o__Saccharomycetales(100);f__Metschnikowiaceae(100);g__Metschnikowia(100);g__Metschnikowia_unclassified(100); | Metschnikowia | 13 | Saprotroph | \|Undefined Saprotroph\| | Yeast | NULL | Probable | As nectar-tap_saprotroph (Põlme et al. 2020); Decay sugar-rich substrates (Põlme et al. 2020) | Põlme S, et al. 2020. Fungal Diversity 105:1-16 ((https://doi.org/10.1007/s13225-020-00466-2)) |
| Otu53315 | 1 | k__Fungi(100);p__Ascomycota(100);c__Saccharomycetes(100);o__Saccharomycetales(100);f__Metschnikowiaceae(100);g__Metschnikowia(100);g__Metschnikowia_unclassified(100); | Metschnikowia | 13 | Saprotroph | \|Undefined Saprotroph\| | Yeast | NULL | Probable | As nectar-tap_saprotroph (Põlme et al. 2020); Decay sugar-rich substrates (Põlme et al. 2020) | Põlme S, et al. 2020. Fungal Diversity 105:1-16 ((https://doi.org/10.1007/s13225-020-00466-2)) |
| Otu53316 | 1 | k__Fungi(100);p__Ascomycota(100);c__Saccharomycetes(100);o__Saccharomycetales(100);f__Metschnikowiaceae(100);g__Metschnikowia(100);g__Metschnikowia_unclassified(100); | Metschnikowia | 13 | Saprotroph | \|Undefined Saprotroph\| | Yeast | NULL | Probable | As nectar-tap_saprotroph (Põlme et al. 2020); Decay sugar-rich substrates (Põlme et al. 2020) | Põlme S, et al. 2020. Fungal Diversity 105:1-16 ((https://doi.org/10.1007/s13225-020-00466-2)) |
| Otu53317 | 1 | k__Fungi(100);p__Ascomycota(100);c__Saccharomycetes(100);o__Saccharomycetales(100);f__Metschnikowiaceae(100);g__Metschnikowia(100);g__Metschnikowia_unclassified(100); | Metschnikowia | 13 | Saprotroph | \|Undefined Saprotroph\| | Yeast | NULL | Probable | As nectar-tap_saprotroph (Põlme et al. 2020); Decay sugar-rich substrates (Põlme et al. 2020) | Põlme S, et al. 2020. Fungal Diversity 105:1-16 ((https://doi.org/10.1007/s13225-020-00466-2)) |
| Otu53319 | 1 | k__Fungi(100);p__Ascomycota(100);c__Saccharomycetes(100);o__Saccharomycetales(100);f__Metschnikowiaceae(100);g__Metschnikowia(100);s__Metschnikowia_pulcherrima(100); | Metschnikowia | 13 | Saprotroph | \|Undefined Saprotroph\| | Yeast | NULL | Probable | As nectar-tap_saprotroph (Põlme et al. 2020); Decay sugar-rich substrates (Põlme et al. 2020) | Põlme S, et al. 2020. Fungal Diversity 105:1-16 ((https://doi.org/10.1007/s13225-020-00466-2)) |
| Otu53320 | 1 | k__Fungi(100);p__Ascomycota(100);c__Saccharomycetes(100);o__Saccharomycetales(100);f__Metschnikowiaceae(100);g__Metschnikowia(100);s__Metschnikowia_pulcherrima(100); | Metschnikowia | 13 | Saprotroph | \|Undefined Saprotroph\| | Yeast | NULL | Probable | As nectar-tap_saprotroph (Põlme et al. 2020); Decay sugar-rich substrates (Põlme et al. 2020) | Põlme S, et al. 2020. Fungal Diversity 105:1-16 ((https://doi.org/10.1007/s13225-020-00466-2)) |
| Otu53321 | 1 | k__Fungi(100);p__Ascomycota(100);c__Saccharomycetes(100);o__Saccharomycetales(100);f__Metschnikowiaceae(100);g__Metschnikowia(100);s__Metschnikowia_pulcherrima(100); | Metschnikowia | 13 | Saprotroph | \|Undefined Saprotroph\| | Yeast | NULL | Probable | As nectar-tap_saprotroph (Põlme et al. 2020); Decay sugar-rich substrates (Põlme et al. 2020) | Põlme S, et al. 2020. Fungal Diversity 105:1-16 ((https://doi.org/10.1007/s13225-020-00466-2)) |
| Otu53322 | 1 | k__Fungi(100);p__Ascomycota(100);c__Saccharomycetes(100);o__Saccharomycetales(100);f__Metschnikowiaceae(100);g__Metschnikowia(100);s__Metschnikowia_pulcherrima(100); | Metschnikowia | 13 | Saprotroph | \|Undefined Saprotroph\| | Yeast | NULL | Probable | As nectar-tap_saprotroph (Põlme et al. 2020); Decay sugar-rich substrates (Põlme et al. 2020) | Põlme S, et al. 2020. Fungal Diversity 105:1-16 ((https://doi.org/10.1007/s13225-020-00466-2)) |
| Otu53323 | 1 | k__Fungi(100);p__Ascomycota(100);c__Saccharomycetes(100);o__Saccharomycetales(100);f__Metschnikowiaceae(100);g__Metschnikowia(100);s__Metschnikowia_pulcherrima(100); | Metschnikowia | 13 | Saprotroph | \|Undefined Saprotroph\| | Yeast | NULL | Probable | As nectar-tap_saprotroph (Põlme et al. 2020); Decay sugar-rich substrates (Põlme et al. 2020) | Põlme S, et al. 2020. Fungal Diversity 105:1-16 ((https://doi.org/10.1007/s13225-020-00466-2)) |
| Otu53324 | 1 | k__Fungi(100);p__Ascomycota(100);c__Saccharomycetes(100);o__Saccharomycetales(100);f__Metschnikowiaceae(100);g__Metschnikowia(100);s__Metschnikowia_pulcherrima(100); | Metschnikowia | 13 | Saprotroph | \|Undefined Saprotroph\| | Yeast | NULL | Probable | As nectar-tap_saprotroph (Põlme et al. 2020); Decay sugar-rich substrates (Põlme et al. 2020) | Põlme S, et al. 2020. Fungal Diversity 105:1-16 ((https://doi.org/10.1007/s13225-020-00466-2)) |
| Otu53325 | 1 | k__Fungi(100);p__Ascomycota(100);c__Saccharomycetes(100);o__Saccharomycetales(100);f__Metschnikowiaceae(100);g__Metschnikowia(100);s__Metschnikowia_pulcherrima(100); | Metschnikowia | 13 | Saprotroph | \|Undefined Saprotroph\| | Yeast | NULL | Probable | As nectar-tap_saprotroph (Põlme et al. 2020); Decay sugar-rich substrates (Põlme et al. 2020) | Põlme S, et al. 2020. Fungal Diversity 105:1-16 ((https://doi.org/10.1007/s13225-020-00466-2)) |
| Otu53326 | 1 | k__Fungi(100);p__Ascomycota(100);c__Saccharomycetes(100);o__Saccharomycetales(100);f__Metschnikowiaceae(100);g__Metschnikowia(100);s__Metschnikowia_pulcherrima(100); | Metschnikowia | 13 | Saprotroph | \|Undefined Saprotroph\| | Yeast | NULL | Probable | As nectar-tap_saprotroph (Põlme et al. 2020); Decay sugar-rich substrates (Põlme et al. 2020) | Põlme S, et al. 2020. Fungal Diversity 105:1-16 ((https://doi.org/10.1007/s13225-020-00466-2)) |
| Otu53328 | 1 | k__Fungi(100);p__Ascomycota(100);c__Saccharomycetes(100);o__Saccharomycetales(100);f__Metschnikowiaceae(100);g__Metschnikowia(100);s__Metschnikowia_pulcherrima(100); | Metschnikowia | 13 | Saprotroph | \|Undefined Saprotroph\| | Yeast | NULL | Probable | As nectar-tap_saprotroph (Põlme et al. 2020); Decay sugar-rich substrates (Põlme et al. 2020) | Põlme S, et al. 2020. Fungal Diversity 105:1-16 ((https://doi.org/10.1007/s13225-020-00466-2)) |
| Otu53329 | 1 | k__Fungi(100);p__Ascomycota(100);c__Saccharomycetes(100);o__Saccharomycetales(100);f__Metschnikowiaceae(100);g__Metschnikowia(100);g__Metschnikowia_unclassified(100); | Metschnikowia | 13 | Saprotroph | \|Undefined Saprotroph\| | Yeast | NULL | Probable | As nectar-tap_saprotroph (Põlme et al. 2020); Decay sugar-rich substrates (Põlme et al. 2020) | Põlme S, et al. 2020. Fungal Diversity 105:1-16 ((https://doi.org/10.1007/s13225-020-00466-2)) |
| Otu53330 | 1 | k__Fungi(100);p__Ascomycota(100);c__Saccharomycetes(100);o__Saccharomycetales(100);f__Metschnikowiaceae(100);g__Metschnikowia(100);s__Metschnikowia_pulcherrima(100); | Metschnikowia | 13 | Saprotroph | \|Undefined Saprotroph\| | Yeast | NULL | Probable | As nectar-tap_saprotroph (Põlme et al. 2020); Decay sugar-rich substrates (Põlme et al. 2020) | Põlme S, et al. 2020. Fungal Diversity 105:1-16 ((https://doi.org/10.1007/s13225-020-00466-2)) |
| Otu53331 | 1 | k__Fungi(100);p__Ascomycota(100);c__Saccharomycetes(100);o__Saccharomycetales(100);f__Metschnikowiaceae(100);g__Metschnikowia(100);s__Metschnikowia_pulcherrima(100); | Metschnikowia | 13 | Saprotroph | \|Undefined Saprotroph\| | Yeast | NULL | Probable | As nectar-tap_saprotroph (Põlme et al. 2020); Decay sugar-rich substrates (Põlme et al. 2020) | Põlme S, et al. 2020. Fungal Diversity 105:1-16 ((https://doi.org/10.1007/s13225-020-00466-2)) |
| Otu53332 | 1 | k__Fungi(100);p__Ascomycota(100);c__Saccharomycetes(100);o__Saccharomycetales(100);f__Metschnikowiaceae(100);g__Metschnikowia(100);s__Metschnikowia_pulcherrima(100); | Metschnikowia | 13 | Saprotroph | \|Undefined Saprotroph\| | Yeast | NULL | Probable | As nectar-tap_saprotroph (Põlme et al. 2020); Decay sugar-rich substrates (Põlme et al. 2020) | Põlme S, et al. 2020. Fungal Diversity 105:1-16 ((https://doi.org/10.1007/s13225-020-00466-2)) |
| Otu53333 | 1 | k__Fungi(100);p__Ascomycota(100);c__Saccharomycetes(100);o__Saccharomycetales(100);f__Metschnikowiaceae(100);g__Metschnikowia(100);g__Metschnikowia_unclassified(100); | Metschnikowia | 13 | Saprotroph | \|Undefined Saprotroph\| | Yeast | NULL | Probable | As nectar-tap_saprotroph (Põlme et al. 2020); Decay sugar-rich substrates (Põlme et al. 2020) | Põlme S, et al. 2020. Fungal Diversity 105:1-16 ((https://doi.org/10.1007/s13225-020-00466-2)) |
| Otu53334 | 1 | k__Fungi(100);p__Ascomycota(100);c__Saccharomycetes(100);o__Saccharomycetales(100);f__Metschnikowiaceae(100);g__Metschnikowia(100);g__Metschnikowia_unclassified(100); | Metschnikowia | 13 | Saprotroph | \|Undefined Saprotroph\| | Yeast | NULL | Probable | As nectar-tap_saprotroph (Põlme et al. 2020); Decay sugar-rich substrates (Põlme et al. 2020) | Põlme S, et al. 2020. Fungal Diversity 105:1-16 ((https://doi.org/10.1007/s13225-020-00466-2)) |
| Otu53335 | 1 | k__Fungi(100);p__Ascomycota(100);c__Saccharomycetes(100);o__Saccharomycetales(100);f__Metschnikowiaceae(100);g__Metschnikowia(100);s__Metschnikowia_pulcherrima(100); | Metschnikowia | 13 | Saprotroph | \|Undefined Saprotroph\| | Yeast | NULL | Probable | As nectar-tap_saprotroph (Põlme et al. 2020); Decay sugar-rich substrates (Põlme et al. 2020) | Põlme S, et al. 2020. Fungal Diversity 105:1-16 ((https://doi.org/10.1007/s13225-020-00466-2)) |
| Otu53336 | 1 | k__Fungi(100);p__Ascomycota(100);c__Saccharomycetes(100);o__Saccharomycetales(100);f__Metschnikowiaceae(100);g__Metschnikowia(100);s__Metschnikowia_pulcherrima(100); | Metschnikowia | 13 | Saprotroph | \|Undefined Saprotroph\| | Yeast | NULL | Probable | As nectar-tap_saprotroph (Põlme et al. 2020); Decay sugar-rich substrates (Põlme et al. 2020) | Põlme S, et al. 2020. Fungal Diversity 105:1-16 ((https://doi.org/10.1007/s13225-020-00466-2)) |
| Otu53337 | 1 | k__Fungi(100);p__Ascomycota(100);c__Saccharomycetes(100);o__Saccharomycetales(100);f__Metschnikowiaceae(100);g__Metschnikowia(100);g__Metschnikowia_unclassified(100); | Metschnikowia | 13 | Saprotroph | \|Undefined Saprotroph\| | Yeast | NULL | Probable | As nectar-tap_saprotroph (Põlme et al. 2020); Decay sugar-rich substrates (Põlme et al. 2020) | Põlme S, et al. 2020. Fungal Diversity 105:1-16 ((https://doi.org/10.1007/s13225-020-00466-2)) |
| Otu53338 | 1 | k__Fungi(100);p__Ascomycota(100);c__Saccharomycetes(100);o__Saccharomycetales(100);f__Metschnikowiaceae(100);g__Metschnikowia(100);s__Metschnikowia_pulcherrima(100); | Metschnikowia | 13 | Saprotroph | \|Undefined Saprotroph\| | Yeast | NULL | Probable | As nectar-tap_saprotroph (Põlme et al. 2020); Decay sugar-rich substrates (Põlme et al. 2020) | Põlme S, et al. 2020. Fungal Diversity 105:1-16 ((https://doi.org/10.1007/s13225-020-00466-2)) |
| Otu53340 | 1 | k__Fungi(100);p__Ascomycota(100);c__Saccharomycetes(100);o__Saccharomycetales(100);f__Metschnikowiaceae(100);g__Metschnikowia(100);s__Metschnikowia_pulcherrima(100); | Metschnikowia | 13 | Saprotroph | \|Undefined Saprotroph\| | Yeast | NULL | Probable | As nectar-tap_saprotroph (Põlme et al. 2020); Decay sugar-rich substrates (Põlme et al. 2020) | Põlme S, et al. 2020. Fungal Diversity 105:1-16 ((https://doi.org/10.1007/s13225-020-00466-2)) |
| Otu53341 | 1 | k__Fungi(100);p__Ascomycota(100);c__Saccharomycetes(100);o__Saccharomycetales(100);f__Metschnikowiaceae(100);g__Metschnikowia(100);g__Metschnikowia_unclassified(100); | Metschnikowia | 13 | Saprotroph | \|Undefined Saprotroph\| | Yeast | NULL | Probable | As nectar-tap_saprotroph (Põlme et al. 2020); Decay sugar-rich substrates (Põlme et al. 2020) | Põlme S, et al. 2020. Fungal Diversity 105:1-16 ((https://doi.org/10.1007/s13225-020-00466-2)) |
| Otu53342 | 1 | k__Fungi(100);p__Ascomycota(100);c__Saccharomycetes(100);o__Saccharomycetales(100);f__Saccharomycetales_family_Incertae_sedis(100);g__Candida(100);g__Candida_unclassified(100); | Candida | 13 | Pathotroph-Saprotroph-Symbiotroph | Animal Pathogen-Endophyte-Endosymbiont-Epiphyte-\|Undefined Saprotroph\| | Dimorphic Yeast-Yeast | NULL | Probable | As nectar-tap_saprotroph (Põlme et al. 2020); Decay sugar-rich substrates (Põlme et al. 2020); This asexual genus is polyphyletic and contain species that function very differently depend on what they are. Caution should be taken when interpreting guilds for Candida species. | Manolakaki D, et al. 2010. Virulence. 1: 367-375 ((https://doi.org/10.4161/viru.1.5.12796)); Mekbib SB, et al. 2011. Tropical Plant Pathology 36:233-240 ((https://doi.org/10.1590/S1982-56762011000400004)); Põlme S, et al. 2020. Fungal Diversity 105:1-16 ((https://doi.org/10.1007/s13225-020-00466-2)) |
| Otu53344 | 1 | k__Fungi(100);p__Ascomycota(100);c__Saccharomycetes(100);o__Saccharomycetales(100);f__Metschnikowiaceae(100);g__Metschnikowia(100);s__Metschnikowia_pulcherrima(100); | Metschnikowia | 13 | Saprotroph | \|Undefined Saprotroph\| | Yeast | NULL | Probable | As nectar-tap_saprotroph (Põlme et al. 2020); Decay sugar-rich substrates (Põlme et al. 2020) | Põlme S, et al. 2020. Fungal Diversity 105:1-16 ((https://doi.org/10.1007/s13225-020-00466-2)) |
| Otu53345 | 1 | k__Fungi(100);p__Ascomycota(100);c__Saccharomycetes(100);o__Saccharomycetales(100);f__Metschnikowiaceae(100);g__Metschnikowia(100);s__Metschnikowia_pulcherrima(100); | Metschnikowia | 13 | Saprotroph | \|Undefined Saprotroph\| | Yeast | NULL | Probable | As nectar-tap_saprotroph (Põlme et al. 2020); Decay sugar-rich substrates (Põlme et al. 2020) | Põlme S, et al. 2020. Fungal Diversity 105:1-16 ((https://doi.org/10.1007/s13225-020-00466-2)) |
| Otu53346 | 1 | k__Fungi(100);p__Ascomycota(100);c__Saccharomycetes(100);o__Saccharomycetales(100);f__Metschnikowiaceae(100);g__Metschnikowia(100);s__Metschnikowia_pulcherrima(100); | Metschnikowia | 13 | Saprotroph | \|Undefined Saprotroph\| | Yeast | NULL | Probable | As nectar-tap_saprotroph (Põlme et al. 2020); Decay sugar-rich substrates (Põlme et al. 2020) | Põlme S, et al. 2020. Fungal Diversity 105:1-16 ((https://doi.org/10.1007/s13225-020-00466-2)) |
| Otu53348 | 1 | k__Fungi(100);p__Ascomycota(100);c__Saccharomycetes(100);o__Saccharomycetales(100);f__Metschnikowiaceae(100);g__Metschnikowia(100);g__Metschnikowia_unclassified(100); | Metschnikowia | 13 | Saprotroph | \|Undefined Saprotroph\| | Yeast | NULL | Probable | As nectar-tap_saprotroph (Põlme et al. 2020); Decay sugar-rich substrates (Põlme et al. 2020) | Põlme S, et al. 2020. Fungal Diversity 105:1-16 ((https://doi.org/10.1007/s13225-020-00466-2)) |
| Otu53349 | 1 | k__Fungi(100);p__Ascomycota(100);c__Saccharomycetes(100);o__Saccharomycetales(100);f__Metschnikowiaceae(100);g__Metschnikowia(100);s__Metschnikowia_pulcherrima(100); | Metschnikowia | 13 | Saprotroph | \|Undefined Saprotroph\| | Yeast | NULL | Probable | As nectar-tap_saprotroph (Põlme et al. 2020); Decay sugar-rich substrates (Põlme et al. 2020) | Põlme S, et al. 2020. Fungal Diversity 105:1-16 ((https://doi.org/10.1007/s13225-020-00466-2)) |
| Otu53350 | 1 | k__Fungi(100);p__Ascomycota(100);c__Saccharomycetes(100);o__Saccharomycetales(100);f__Metschnikowiaceae(100);g__Metschnikowia(100);s__Metschnikowia_pulcherrima(100); | Metschnikowia | 13 | Saprotroph | \|Undefined Saprotroph\| | Yeast | NULL | Probable | As nectar-tap_saprotroph (Põlme et al. 2020); Decay sugar-rich substrates (Põlme et al. 2020) | Põlme S, et al. 2020. Fungal Diversity 105:1-16 ((https://doi.org/10.1007/s13225-020-00466-2)) |
| Otu53352 | 1 | k__Fungi(100);p__Ascomycota(100);c__Saccharomycetes(100);o__Saccharomycetales(100);f__Metschnikowiaceae(100);g__Metschnikowia(100);s__Metschnikowia_pulcherrima(100); | Metschnikowia | 13 | Saprotroph | \|Undefined Saprotroph\| | Yeast | NULL | Probable | As nectar-tap_saprotroph (Põlme et al. 2020); Decay sugar-rich substrates (Põlme et al. 2020) | Põlme S, et al. 2020. Fungal Diversity 105:1-16 ((https://doi.org/10.1007/s13225-020-00466-2)) |
| Otu53353 | 1 | k__Fungi(100);p__Ascomycota(100);c__Saccharomycetes(100);o__Saccharomycetales(100);f__Metschnikowiaceae(100);g__Metschnikowia(100);g__Metschnikowia_unclassified(100); | Metschnikowia | 13 | Saprotroph | \|Undefined Saprotroph\| | Yeast | NULL | Probable | As nectar-tap_saprotroph (Põlme et al. 2020); Decay sugar-rich substrates (Põlme et al. 2020) | Põlme S, et al. 2020. Fungal Diversity 105:1-16 ((https://doi.org/10.1007/s13225-020-00466-2)) |
| Otu53354 | 1 | k__Fungi(100);p__Ascomycota(100);c__Saccharomycetes(100);o__Saccharomycetales(100);f__Metschnikowiaceae(100);g__Metschnikowia(100);s__Metschnikowia_pulcherrima(100); | Metschnikowia | 13 | Saprotroph | \|Undefined Saprotroph\| | Yeast | NULL | Probable | As nectar-tap_saprotroph (Põlme et al. 2020); Decay sugar-rich substrates (Põlme et al. 2020) | Põlme S, et al. 2020. Fungal Diversity 105:1-16 ((https://doi.org/10.1007/s13225-020-00466-2)) |
| Otu53355 | 1 | k__Fungi(100);p__Basidiomycota(100);c__Microbotryomycetes(100);o__Sporidiobolales(100);f__Sporidiobolales_family_Incertae_sedis(100);g__Sporobolomyces(100);s__Sporobolomyces_lactosus(100); | Sporobolomyces | 13 | Pathotroph-Saprotroph | \|Fungal Parasite\|-Plant Saprotroph | Yeast | NULL | Probable | As litter_saprotroph (Põlme et al. 2020); As mycoparasite (Põlme et al. 2020); Decay fungal material (Põlme et al. 2020) | Tedersoo L, et al. 2014. Science 346:e1256688 ((https://doi.org/10.1126/science.1256688)); Purahong W, et al. 2016. Molecular Ecology 25:4059-4074 ((https://doi.org/10.1111/mec.13739)); Põlme S, et al. 2020. Fungal Diversity 105:1-16 ((https://doi.org/10.1007/s13225-020-00466-2)) |
| Otu53357 | 1 | k__Fungi(100);p__Ascomycota(100);c__Saccharomycetes(100);o__Saccharomycetales(100);f__Metschnikowiaceae(100);g__Metschnikowia(100);s__Metschnikowia_pulcherrima(100); | Metschnikowia | 13 | Saprotroph | \|Undefined Saprotroph\| | Yeast | NULL | Probable | As nectar-tap_saprotroph (Põlme et al. 2020); Decay sugar-rich substrates (Põlme et al. 2020) | Põlme S, et al. 2020. Fungal Diversity 105:1-16 ((https://doi.org/10.1007/s13225-020-00466-2)) |
| Otu53358 | 1 | k__Fungi(100);p__Ascomycota(100);c__Saccharomycetes(100);o__Saccharomycetales(100);f__Metschnikowiaceae(100);g__Metschnikowia(100);s__Metschnikowia_pulcherrima(100); | Metschnikowia | 13 | Saprotroph | \|Undefined Saprotroph\| | Yeast | NULL | Probable | As nectar-tap_saprotroph (Põlme et al. 2020); Decay sugar-rich substrates (Põlme et al. 2020) | Põlme S, et al. 2020. Fungal Diversity 105:1-16 ((https://doi.org/10.1007/s13225-020-00466-2)) |
| Otu53359 | 1 | k__Fungi(100);p__Basidiomycota(100);c__Microbotryomycetes(100);o__Sporidiobolales(100);f__Sporidiobolales_family_Incertae_sedis(100);g__Sporobolomyces(100);s__Sporobolomyces_lactosus(100); | Sporobolomyces | 13 | Pathotroph-Saprotroph | \|Fungal Parasite\|-Plant Saprotroph | Yeast | NULL | Probable | As litter_saprotroph (Põlme et al. 2020); As mycoparasite (Põlme et al. 2020); Decay fungal material (Põlme et al. 2020) | Tedersoo L, et al. 2014. Science 346:e1256688 ((https://doi.org/10.1126/science.1256688)); Purahong W, et al. 2016. Molecular Ecology 25:4059-4074 ((https://doi.org/10.1111/mec.13739)); Põlme S, et al. 2020. Fungal Diversity 105:1-16 ((https://doi.org/10.1007/s13225-020-00466-2)) |
| Otu53361 | 1 | k__Fungi(100);p__Ascomycota(100);c__Saccharomycetes(100);o__Saccharomycetales(100);f__Saccharomycetales_family_Incertae_sedis(100);g__Candida(100);s__Candida_cretensis(100); | Candida | 13 | Pathotroph-Saprotroph-Symbiotroph | Animal Pathogen-Endophyte-Endosymbiont-Epiphyte-\|Undefined Saprotroph\| | Dimorphic Yeast-Yeast | NULL | Probable | As nectar-tap_saprotroph (Põlme et al. 2020); Decay sugar-rich substrates (Põlme et al. 2020); This asexual genus is polyphyletic and contain species that function very differently depend on what they are. Caution should be taken when interpreting guilds for Candida species. | Manolakaki D, et al. 2010. Virulence. 1: 367-375 ((https://doi.org/10.4161/viru.1.5.12796)); Mekbib SB, et al. 2011. Tropical Plant Pathology 36:233-240 ((https://doi.org/10.1590/S1982-56762011000400004)); Põlme S, et al. 2020. Fungal Diversity 105:1-16 ((https://doi.org/10.1007/s13225-020-00466-2)) |
| Otu53362 | 1 | k__Fungi(100);p__Ascomycota(100);c__Saccharomycetes(100);o__Saccharomycetales(100);f__Metschnikowiaceae(100);g__Metschnikowia(100);s__Metschnikowia_pulcherrima(100); | Metschnikowia | 13 | Saprotroph | \|Undefined Saprotroph\| | Yeast | NULL | Probable | As nectar-tap_saprotroph (Põlme et al. 2020); Decay sugar-rich substrates (Põlme et al. 2020) | Põlme S, et al. 2020. Fungal Diversity 105:1-16 ((https://doi.org/10.1007/s13225-020-00466-2)) |
| Otu53363 | 1 | k__Fungi(100);p__Ascomycota(100);c__Saccharomycetes(100);o__Saccharomycetales(100);f__Metschnikowiaceae(100);g__Metschnikowia(100);s__Metschnikowia_pulcherrima(100); | Metschnikowia | 13 | Saprotroph | \|Undefined Saprotroph\| | Yeast | NULL | Probable | As nectar-tap_saprotroph (Põlme et al. 2020); Decay sugar-rich substrates (Põlme et al. 2020) | Põlme S, et al. 2020. Fungal Diversity 105:1-16 ((https://doi.org/10.1007/s13225-020-00466-2)) |
| Otu53365 | 1 | k__Fungi(100);p__Ascomycota(100);c__Saccharomycetes(100);o__Saccharomycetales(100);o__Saccharomycetales_unclassified(100);o__Saccharomycetales_unclassified(100);o__Saccharomycetales_unclassified(100); | Saccharomycetales | 7 | Saprotroph | Undefined Saprotroph | Yeast | NULL | Possible | NULL | Sterkenburg E, et al. 2015. New Phytologist 207:1145-1158 ((https://doi.org/10.1111/nph.13426)) |
| Otu53366 | 1 | k__Fungi(100);p__Ascomycota(100);c__Saccharomycetes(100);o__Saccharomycetales(100);f__Metschnikowiaceae(100);g__Metschnikowia(100);g__Metschnikowia_unclassified(100); | Metschnikowia | 13 | Saprotroph | \|Undefined Saprotroph\| | Yeast | NULL | Probable | As nectar-tap_saprotroph (Põlme et al. 2020); Decay sugar-rich substrates (Põlme et al. 2020) | Põlme S, et al. 2020. Fungal Diversity 105:1-16 ((https://doi.org/10.1007/s13225-020-00466-2)) |
| Otu53367 | 1 | k__Fungi(100);p__Ascomycota(100);c__Saccharomycetes(100);o__Saccharomycetales(100);f__Metschnikowiaceae(100);g__Metschnikowia(100);s__Metschnikowia_pulcherrima(100); | Metschnikowia | 13 | Saprotroph | \|Undefined Saprotroph\| | Yeast | NULL | Probable | As nectar-tap_saprotroph (Põlme et al. 2020); Decay sugar-rich substrates (Põlme et al. 2020) | Põlme S, et al. 2020. Fungal Diversity 105:1-16 ((https://doi.org/10.1007/s13225-020-00466-2)) |
| Otu53368 | 1 | k__Fungi(100);p__Ascomycota(100);c__Saccharomycetes(100);o__Saccharomycetales(100);f__Metschnikowiaceae(100);g__Metschnikowia(100);s__Metschnikowia_pulcherrima(100); | Metschnikowia | 13 | Saprotroph | \|Undefined Saprotroph\| | Yeast | NULL | Probable | As nectar-tap_saprotroph (Põlme et al. 2020); Decay sugar-rich substrates (Põlme et al. 2020) | Põlme S, et al. 2020. Fungal Diversity 105:1-16 ((https://doi.org/10.1007/s13225-020-00466-2)) |
| Otu53370 | 1 | k__Fungi(100);p__Ascomycota(100);c__Saccharomycetes(100);o__Saccharomycetales(100);f__Saccharomycetales_family_Incertae_sedis(100);f__Saccharomycetales_family_Incertae_sedis_unclassified(100);f__Saccharomycetales_family_Incertae_sedis_unclassified(100); | Saccharomycetales | 7 | Saprotroph | Undefined Saprotroph | Yeast | NULL | Possible | NULL | Sterkenburg E, et al. 2015. New Phytologist 207:1145-1158 ((https://doi.org/10.1111/nph.13426)) |
| Otu53371 | 1 | k__Fungi(100);p__Ascomycota(100);c__Saccharomycetes(100);o__Saccharomycetales(100);f__Metschnikowiaceae(100);g__Metschnikowia(100);s__Metschnikowia_pulcherrima(100); | Metschnikowia | 13 | Saprotroph | \|Undefined Saprotroph\| | Yeast | NULL | Probable | As nectar-tap_saprotroph (Põlme et al. 2020); Decay sugar-rich substrates (Põlme et al. 2020) | Põlme S, et al. 2020. Fungal Diversity 105:1-16 ((https://doi.org/10.1007/s13225-020-00466-2)) |
| Otu53374 | 1 | k__Fungi(100);p__Ascomycota(100);c__Saccharomycetes(100);o__Saccharomycetales(100);f__Metschnikowiaceae(100);g__Metschnikowia(100);g__Metschnikowia_unclassified(100); | Metschnikowia | 13 | Saprotroph | \|Undefined Saprotroph\| | Yeast | NULL | Probable | As nectar-tap_saprotroph (Põlme et al. 2020); Decay sugar-rich substrates (Põlme et al. 2020) | Põlme S, et al. 2020. Fungal Diversity 105:1-16 ((https://doi.org/10.1007/s13225-020-00466-2)) |
| Otu53378 | 1 | k__Fungi(100);p__Ascomycota(100);c__Saccharomycetes(100);o__Saccharomycetales(100);f__Metschnikowiaceae(100);g__Metschnikowia(100);s__Metschnikowia_pulcherrima(100); | Metschnikowia | 13 | Saprotroph | \|Undefined Saprotroph\| | Yeast | NULL | Probable | As nectar-tap_saprotroph (Põlme et al. 2020); Decay sugar-rich substrates (Põlme et al. 2020) | Põlme S, et al. 2020. Fungal Diversity 105:1-16 ((https://doi.org/10.1007/s13225-020-00466-2)) |
| Otu53380 | 1 | k__Fungi(100);p__Ascomycota(100);c__Saccharomycetes(100);o__Saccharomycetales(100);f__Metschnikowiaceae(100);g__Metschnikowia(100);s__Metschnikowia_pulcherrima(100); | Metschnikowia | 13 | Saprotroph | \|Undefined Saprotroph\| | Yeast | NULL | Probable | As nectar-tap_saprotroph (Põlme et al. 2020); Decay sugar-rich substrates (Põlme et al. 2020) | Põlme S, et al. 2020. Fungal Diversity 105:1-16 ((https://doi.org/10.1007/s13225-020-00466-2)) |
| Otu53387 | 1 | k__Fungi(100);p__Ascomycota(100);c__Saccharomycetes(100);o__Saccharomycetales(100);f__Metschnikowiaceae(100);g__Metschnikowia(100);s__Metschnikowia_pulcherrima(100); | Metschnikowia | 13 | Saprotroph | \|Undefined Saprotroph\| | Yeast | NULL | Probable | As nectar-tap_saprotroph (Põlme et al. 2020); Decay sugar-rich substrates (Põlme et al. 2020) | Põlme S, et al. 2020. Fungal Diversity 105:1-16 ((https://doi.org/10.1007/s13225-020-00466-2)) |
| Otu53388 | 1 | k__Fungi(100);p__Ascomycota(100);c__Saccharomycetes(100);o__Saccharomycetales(100);f__Metschnikowiaceae(100);g__Metschnikowia(100);s__Metschnikowia_pulcherrima(100); | Metschnikowia | 13 | Saprotroph | \|Undefined Saprotroph\| | Yeast | NULL | Probable | As nectar-tap_saprotroph (Põlme et al. 2020); Decay sugar-rich substrates (Põlme et al. 2020) | Põlme S, et al. 2020. Fungal Diversity 105:1-16 ((https://doi.org/10.1007/s13225-020-00466-2)) |
| Otu53389 | 1 | k__Fungi(100);p__Ascomycota(100);c__Saccharomycetes(100);o__Saccharomycetales(100);f__Metschnikowiaceae(100);g__Metschnikowia(100);s__Metschnikowia_pulcherrima(100); | Metschnikowia | 13 | Saprotroph | \|Undefined Saprotroph\| | Yeast | NULL | Probable | As nectar-tap_saprotroph (Põlme et al. 2020); Decay sugar-rich substrates (Põlme et al. 2020) | Põlme S, et al. 2020. Fungal Diversity 105:1-16 ((https://doi.org/10.1007/s13225-020-00466-2)) |
| Otu53390 | 1 | k__Fungi(100);p__Ascomycota(100);c__Saccharomycetes(100);o__Saccharomycetales(100);f__Metschnikowiaceae(100);g__Metschnikowia(100);s__Metschnikowia_pulcherrima(100); | Metschnikowia | 13 | Saprotroph | \|Undefined Saprotroph\| | Yeast | NULL | Probable | As nectar-tap_saprotroph (Põlme et al. 2020); Decay sugar-rich substrates (Põlme et al. 2020) | Põlme S, et al. 2020. Fungal Diversity 105:1-16 ((https://doi.org/10.1007/s13225-020-00466-2)) |
| Otu53393 | 1 | k__Fungi(100);p__Ascomycota(100);c__Saccharomycetes(100);o__Saccharomycetales(100);o__Saccharomycetales_unclassified(100);o__Saccharomycetales_unclassified(100);o__Saccharomycetales_unclassified(100); | Saccharomycetales | 7 | Saprotroph | Undefined Saprotroph | Yeast | NULL | Possible | NULL | Sterkenburg E, et al. 2015. New Phytologist 207:1145-1158 ((https://doi.org/10.1111/nph.13426)) |
| Otu53395 | 1 | k__Fungi(100);p__Ascomycota(100);c__Saccharomycetes(100);o__Saccharomycetales(100);f__Metschnikowiaceae(100);g__Metschnikowia(100);s__Metschnikowia_pulcherrima(100); | Metschnikowia | 13 | Saprotroph | \|Undefined Saprotroph\| | Yeast | NULL | Probable | As nectar-tap_saprotroph (Põlme et al. 2020); Decay sugar-rich substrates (Põlme et al. 2020) | Põlme S, et al. 2020. Fungal Diversity 105:1-16 ((https://doi.org/10.1007/s13225-020-00466-2)) |
| Otu53396 | 1 | k__Fungi(100);p__Ascomycota(100);c__Saccharomycetes(100);o__Saccharomycetales(100);f__Metschnikowiaceae(100);g__Metschnikowia(100);g__Metschnikowia_unclassified(100); | Metschnikowia | 13 | Saprotroph | \|Undefined Saprotroph\| | Yeast | NULL | Probable | As nectar-tap_saprotroph (Põlme et al. 2020); Decay sugar-rich substrates (Põlme et al. 2020) | Põlme S, et al. 2020. Fungal Diversity 105:1-16 ((https://doi.org/10.1007/s13225-020-00466-2)) |
| Otu53397 | 1 | k__Fungi(100);p__Ascomycota(100);c__Saccharomycetes(100);o__Saccharomycetales(100);f__Metschnikowiaceae(100);g__Metschnikowia(100);s__Metschnikowia_pulcherrima(100); | Metschnikowia | 13 | Saprotroph | \|Undefined Saprotroph\| | Yeast | NULL | Probable | As nectar-tap_saprotroph (Põlme et al. 2020); Decay sugar-rich substrates (Põlme et al. 2020) | Põlme S, et al. 2020. Fungal Diversity 105:1-16 ((https://doi.org/10.1007/s13225-020-00466-2)) |
| Otu53398 | 1 | k__Fungi(100);p__Ascomycota(100);c__Saccharomycetes(100);o__Saccharomycetales(100);f__Metschnikowiaceae(100);g__Metschnikowia(100);s__Metschnikowia_pulcherrima(100); | Metschnikowia | 13 | Saprotroph | \|Undefined Saprotroph\| | Yeast | NULL | Probable | As nectar-tap_saprotroph (Põlme et al. 2020); Decay sugar-rich substrates (Põlme et al. 2020) | Põlme S, et al. 2020. Fungal Diversity 105:1-16 ((https://doi.org/10.1007/s13225-020-00466-2)) |
| Otu53399 | 1 | k__Fungi(100);p__Ascomycota(100);c__Saccharomycetes(100);o__Saccharomycetales(100);f__Metschnikowiaceae(100);g__Metschnikowia(100);s__Metschnikowia_pulcherrima(100); | Metschnikowia | 13 | Saprotroph | \|Undefined Saprotroph\| | Yeast | NULL | Probable | As nectar-tap_saprotroph (Põlme et al. 2020); Decay sugar-rich substrates (Põlme et al. 2020) | Põlme S, et al. 2020. Fungal Diversity 105:1-16 ((https://doi.org/10.1007/s13225-020-00466-2)) |
| Otu53400 | 1 | k__Fungi(100);p__Ascomycota(100);c__Saccharomycetes(100);o__Saccharomycetales(100);f__Metschnikowiaceae(100);g__Metschnikowia(100);s__Metschnikowia_pulcherrima(100); | Metschnikowia | 13 | Saprotroph | \|Undefined Saprotroph\| | Yeast | NULL | Probable | As nectar-tap_saprotroph (Põlme et al. 2020); Decay sugar-rich substrates (Põlme et al. 2020) | Põlme S, et al. 2020. Fungal Diversity 105:1-16 ((https://doi.org/10.1007/s13225-020-00466-2)) |
| Otu53401 | 1 | k__Fungi(100);p__Ascomycota(100);c__Saccharomycetes(100);o__Saccharomycetales(100);f__Metschnikowiaceae(100);g__Metschnikowia(100);s__Metschnikowia_pulcherrima(100); | Metschnikowia | 13 | Saprotroph | \|Undefined Saprotroph\| | Yeast | NULL | Probable | As nectar-tap_saprotroph (Põlme et al. 2020); Decay sugar-rich substrates (Põlme et al. 2020) | Põlme S, et al. 2020. Fungal Diversity 105:1-16 ((https://doi.org/10.1007/s13225-020-00466-2)) |

| | | | | | | | | | | | |
|---|---|---|---|---|---|---|---|---|---|---|---|
| Otu53402 | 1 | k__Fungi(100);p__Ascomycota(100);c__Saccharomycetes(100);o__Saccharomycetales(100);f__Metschnikowiaceae(100);g__Metschnikowia(100);s__Metschnikowia_pulcherrima(100); | Metschnikowia | 13 | Saprotroph | \|Undefined Saprotroph\| | Yeast | NULL | Probable | As nectar-tap_saprotroph (Põlme et al. 2020); Decay sugar-rich substrates (Põlme et al. 2020) | Põlme S, et al. 2020. Fungal Diversity 105:1-16 ((https://doi.org/10.1007/s13225-020-00466-2)) |
| Otu53403 | 1 | k__Fungi(100);p__Ascomycota(100);c__Saccharomycetes(100);o__Saccharomycetales(100);f__Metschnikowiaceae(100);g__Metschnikowia(100);s__Metschnikowia_pulcherrima(100); | Metschnikowia | 13 | Saprotroph | \|Undefined Saprotroph\| | Yeast | NULL | Probable | As nectar-tap_saprotroph (Põlme et al. 2020); Decay sugar-rich substrates (Põlme et al. 2020) | Põlme S, et al. 2020. Fungal Diversity 105:1-16 ((https://doi.org/10.1007/s13225-020-00466-2)) |
| Otu53404 | 1 | k__Fungi(100);p__Ascomycota(100);c__Saccharomycetes(100);o__Saccharomycetales(100);f__Metschnikowiaceae(100);g__Metschnikowia(100);s__Metschnikowia_pulcherrima(100); | Metschnikowia | 13 | Saprotroph | \|Undefined Saprotroph\| | Yeast | NULL | Probable | As nectar-tap_saprotroph (Põlme et al. 2020); Decay sugar-rich substrates (Põlme et al. 2020) | Põlme S, et al. 2020. Fungal Diversity 105:1-16 ((https://doi.org/10.1007/s13225-020-00466-2)) |
| Otu53405 | 1 | k__Fungi(100);p__Ascomycota(100);c__Saccharomycetes(100);o__Saccharomycetales(100);f__Metschnikowiaceae(100);g__Metschnikowia(100);s__Metschnikowia_pulcherrima(100); | Metschnikowia | 13 | Saprotroph | \|Undefined Saprotroph\| | Yeast | NULL | Probable | As nectar-tap_saprotroph (Põlme et al. 2020); Decay sugar-rich substrates (Põlme et al. 2020) | Põlme S, et al. 2020. Fungal Diversity 105:1-16 ((https://doi.org/10.1007/s13225-020-00466-2)) |
| Otu53406 | 1 | k__Fungi(100);p__Ascomycota(100);c__Saccharomycetes(100);o__Saccharomycetales(100);f__Metschnikowiaceae(100);g__Metschnikowia(100);s__Metschnikowia_pulcherrima(100); | Metschnikowia | 13 | Saprotroph | \|Undefined Saprotroph\| | Yeast | NULL | Probable | As nectar-tap_saprotroph (Põlme et al. 2020); Decay sugar-rich substrates (Põlme et al. 2020) | Põlme S, et al. 2020. Fungal Diversity 105:1-16 ((https://doi.org/10.1007/s13225-020-00466-2)) |
| Otu53408 | 1 | k__Fungi(100);p__Ascomycota(100);c__Saccharomycetes(100);o__Saccharomycetales(100);f__Metschnikowiaceae(100);g__Metschnikowia(100);g__Metschnikowia_unclassified(100); | Metschnikowia | 13 | Saprotroph | \|Undefined Saprotroph\| | Yeast | NULL | Probable | As nectar-tap_saprotroph (Põlme et al. 2020); Decay sugar-rich substrates (Põlme et al. 2020) | Põlme S, et al. 2020. Fungal Diversity 105:1-16 ((https://doi.org/10.1007/s13225-020-00466-2)) |
| Otu53409 | 1 | k__Fungi(100);p__Ascomycota(100);c__Saccharomycetes(100);o__Saccharomycetales(100);f__Metschnikowiaceae(100);g__Metschnikowia(100);g__Metschnikowia_unclassified(100); | Metschnikowia | 13 | Saprotroph | \|Undefined Saprotroph\| | Yeast | NULL | Probable | As nectar-tap_saprotroph (Põlme et al. 2020); Decay sugar-rich substrates (Põlme et al. 2020) | Põlme S, et al. 2020. Fungal Diversity 105:1-16 ((https://doi.org/10.1007/s13225-020-00466-2)) |
| Otu53410 | 1 | k__Fungi(100);p__Ascomycota(100);c__Saccharomycetes(100);o__Saccharomycetales(100);f__Metschnikowiaceae(100);g__Metschnikowia(100);s__Metschnikowia_pulcherrima(100); | Metschnikowia | 13 | Saprotroph | \|Undefined Saprotroph\| | Yeast | NULL | Probable | As nectar-tap_saprotroph (Põlme et al. 2020); Decay sugar-rich substrates (Põlme et al. 2020) | Põlme S, et al. 2020. Fungal Diversity 105:1-16 ((https://doi.org/10.1007/s13225-020-00466-2)) |
| Otu53411 | 1 | k__Fungi(100);p__Ascomycota(100);c__Saccharomycetes(100);o__Saccharomycetales(100);f__Metschnikowiaceae(100);g__Metschnikowia(100);s__Metschnikowia_pulcherrima(100); | Metschnikowia | 13 | Saprotroph | \|Undefined Saprotroph\| | Yeast | NULL | Probable | As nectar-tap_saprotroph (Põlme et al. 2020); Decay sugar-rich substrates (Põlme et al. 2020) | Põlme S, et al. 2020. Fungal Diversity 105:1-16 ((https://doi.org/10.1007/s13225-020-00466-2)) |
| Otu53412 | 1 | k__Fungi(100);p__Ascomycota(100);c__Saccharomycetes(100);o__Saccharomycetales(100);f__Metschnikowiaceae(100);g__Metschnikowia(100);s__Metschnikowia_pulcherrima(100); | Metschnikowia | 13 | Saprotroph | \|Undefined Saprotroph\| | Yeast | NULL | Probable | As nectar-tap_saprotroph (Põlme et al. 2020); Decay sugar-rich substrates (Põlme et al. 2020) | Põlme S, et al. 2020. Fungal Diversity 105:1-16 ((https://doi.org/10.1007/s13225-020-00466-2)) |
| Otu53413 | 1 | k__Fungi(100);p__Ascomycota(100);c__Saccharomycetes(100);o__Saccharomycetales(100);f__Metschnikowiaceae(100);g__Metschnikowia(100);s__Metschnikowia_pulcherrima(100); | Metschnikowia | 13 | Saprotroph | \|Undefined Saprotroph\| | Yeast | NULL | Probable | As nectar-tap_saprotroph (Põlme et al. 2020); Decay sugar-rich substrates (Põlme et al. 2020) | Põlme S, et al. 2020. Fungal Diversity 105:1-16 ((https://doi.org/10.1007/s13225-020-00466-2)) |
| Otu53414 | 1 | k__Fungi(100);p__Ascomycota(100);c__Saccharomycetes(100);o__Saccharomycetales(100);f__Metschnikowiaceae(100);g__Metschnikowia(100);s__Metschnikowia_pulcherrima(100); | Metschnikowia | 13 | Saprotroph | \|Undefined Saprotroph\| | Yeast | NULL | Probable | As nectar-tap_saprotroph (Põlme et al. 2020); Decay sugar-rich substrates (Põlme et al. 2020) | Põlme S, et al. 2020. Fungal Diversity 105:1-16 ((https://doi.org/10.1007/s13225-020-00466-2)) |
| Otu53415 | 1 | k__Fungi(100);p__Ascomycota(100);c__Saccharomycetes(100);o__Saccharomycetales(100);f__Saccharomycetales_family_Incertae_sedis(100);g__Candida(100);g__Candida_unclassified(100); | Candida | 13 | Pathotroph-Saprotroph-Symbiotroph | Animal Pathogen-Endophyte-Endosymbiont-Epiphyte-\|Undefined Saprotroph\| | Dimorphic Yeast-Yeast | NULL | Probable | As nectar-tap_saprotroph (Põlme et al. 2020); Decay sugar-rich substrates (Põlme et al. 2020); This asexual genus is polyphyletic and contain species that function very differently depend on what they are. Caution should be taken when interpreting guilds for Candida species. | Manolakaki D, et al. 2010. Virulence. 1: 367-375 ((https://doi.org/10.4161/viru.1.5.12796)); Mekbib SB, et al. 2011. Tropical Plant Pathology 36:233-240 ((https://doi.org/10.1590/S1982-56762011000400004)); Põlme S, et al. 2020. Fungal Diversity 105:1-16 ((https://doi.org/10.1007/s13225-020-00466-2)) |
| Otu53416 | 1 | k__Fungi(100);p__Ascomycota(100);c__Saccharomycetes(100);o__Saccharomycetales(100);f__Metschnikowiaceae(100);g__Metschnikowia(100);s__Metschnikowia_pulcherrima(100); | Metschnikowia | 13 | Saprotroph | \|Undefined Saprotroph\| | Yeast | NULL | Probable | As nectar-tap_saprotroph (Põlme et al. 2020); Decay sugar-rich substrates (Põlme et al. 2020) | Põlme S, et al. 2020. Fungal Diversity 105:1-16 ((https://doi.org/10.1007/s13225-020-00466-2)) |
| Otu53417 | 1 | k__Fungi(100);p__Ascomycota(100);c__Saccharomycetes(100);o__Saccharomycetales(100);f__Metschnikowiaceae(100);g__Metschnikowia(100);s__Metschnikowia_pulcherrima(100); | Metschnikowia | 13 | Saprotroph | \|Undefined Saprotroph\| | Yeast | NULL | Probable | As nectar-tap_saprotroph (Põlme et al. 2020); Decay sugar-rich substrates (Põlme et al. 2020) | Põlme S, et al. 2020. Fungal Diversity 105:1-16 ((https://doi.org/10.1007/s13225-020-00466-2)) |
| Otu53425 | 1 | k__Fungi(100);p__Ascomycota(100);c__Saccharomycetes(100);o__Saccharomycetales(100);f__Metschnikowiaceae(100);g__Metschnikowia(100);g__Metschnikowia_unclassified(100); | Metschnikowia | 13 | Saprotroph | \|Undefined Saprotroph\| | Yeast | NULL | Probable | As nectar-tap_saprotroph (Põlme et al. 2020); Decay sugar-rich substrates (Põlme et al. 2020) | Põlme S, et al. 2020. Fungal Diversity 105:1-16 ((https://doi.org/10.1007/s13225-020-00466-2)) |
| Otu53427 | 1 | k__Fungi(100);p__Ascomycota(100);c__Saccharomycetes(100);o__Saccharomycetales(100);f__Saccharomycetales_family_Incertae_sedis(100);g__Candida(100);g__Candida_unclassified(100); | Candida | 13 | Pathotroph-Saprotroph-Symbiotroph | Animal Pathogen-Endophyte-Endosymbiont-Epiphyte-\|Undefined Saprotroph\| | Dimorphic Yeast-Yeast | NULL | Probable | As nectar-tap_saprotroph (Põlme et al. 2020); Decay sugar-rich substrates (Põlme et al. 2020); This asexual genus is polyphyletic and contain species that function very differently depend on what they are. Caution should be taken when interpreting guilds for Candida species. | Manolakaki D, et al. 2010. Virulence. 1: 367-375 ((https://doi.org/10.4161/viru.1.5.12796)); Mekbib SB, et al. 2011. Tropical Plant Pathology 36:233-240 ((https://doi.org/10.1590/S1982-56762011000400004)); Põlme S, et al. 2020. Fungal Diversity 105:1-16 ((https://doi.org/10.1007/s13225-020-00466-2)) |
| Otu53428 | 1 | k__Fungi(100);p__Ascomycota(100);c__Saccharomycetes(100);o__Saccharomycetales(100);f__Metschnikowiaceae(100);g__Metschnikowia(100);s__Metschnikowia_pulcherrima(100); | Metschnikowia | 13 | Saprotroph | \|Undefined Saprotroph\| | Yeast | NULL | Probable | As nectar-tap_saprotroph (Põlme et al. 2020); Decay sugar-rich substrates (Põlme et al. 2020) | Põlme S, et al. 2020. Fungal Diversity 105:1-16 ((https://doi.org/10.1007/s13225-020-00466-2)) |
| Otu53434 | 1 | k__Fungi(100);p__Ascomycota(100);c__Saccharomycetes(100);o__Saccharomycetales(100);f__Saccharomycetales_family_Incertae_sedis(100);g__Candida(100);g__Candida_unclassified(100); | Candida | 13 | Pathotroph-Saprotroph-Symbiotroph | Animal Pathogen-Endophyte-Endosymbiont-Epiphyte-\|Undefined Saprotroph\| | Dimorphic Yeast-Yeast | NULL | Probable | As nectar-tap_saprotroph (Põlme et al. 2020); Decay sugar-rich substrates (Põlme et al. 2020); This asexual genus is polyphyletic and contain species that function very differently depend on what they are. Caution should be taken when interpreting guilds for Candida species. | Manolakaki D, et al. 2010. Virulence. 1: 367-375 ((https://doi.org/10.4161/viru.1.5.12796)); Mekbib SB, et al. 2011. Tropical Plant Pathology 36:233-240 ((https://doi.org/10.1590/S1982-56762011000400004)); Põlme S, et al. 2020. Fungal Diversity 105:1-16 ((https://doi.org/10.1007/s13225-020-00466-2)) |
| Otu53440 | 1 | k__Fungi(100);p__Ascomycota(100);c__Saccharomycetes(100);o__Saccharomycetales(100);f__Saccharomycetales_family_Incertae_sedis(100);g__Candida(100);g__Candida_unclassified(100); | Candida | 13 | Pathotroph-Saprotroph-Symbiotroph | Animal Pathogen-Endophyte-Endosymbiont-Epiphyte-\|Undefined Saprotroph\| | Dimorphic Yeast-Yeast | NULL | Probable | As nectar-tap_saprotroph (Põlme et al. 2020); Decay sugar-rich substrates (Põlme et al. 2020); This asexual genus is polyphyletic and contain species that function very differently depend on what they are. Caution should be taken when interpreting guilds for Candida species. | Manolakaki D, et al. 2010. Virulence. 1: 367-375 ((https://doi.org/10.4161/viru.1.5.12796)); Mekbib SB, et al. 2011. Tropical Plant Pathology 36:233-240 ((https://doi.org/10.1590/S1982-56762011000400004)); Põlme S, et al. 2020. Fungal Diversity 105:1-16 ((https://doi.org/10.1007/s13225-020-00466-2)) |
| Otu53443 | 1 | k__Fungi(100);p__Ascomycota(100);c__Saccharomycetes(100);o__Saccharomycetales(100);f__Metschnikowiaceae(100);g__Metschnikowia(100);s__Metschnikowia_pulcherrima(100); | Metschnikowia | 13 | Saprotroph | \|Undefined Saprotroph\| | Yeast | NULL | Probable | As nectar-tap_saprotroph (Põlme et al. 2020); Decay sugar-rich substrates (Põlme et al. 2020) | Põlme S, et al. 2020. Fungal Diversity 105:1-16 ((https://doi.org/10.1007/s13225-020-00466-2)) |
| Otu53444 | 1 | k__Fungi(100);p__Ascomycota(100);c__Saccharomycetes(100);o__Saccharomycetales(100);f__Saccharomycetales_family_Incertae_sedis(100);f__Saccharomycetales_family_Incertae_sedis_unclassified(100);f__Saccharomycetales_family_Incertae_sedis_unclassified(100); | Saccharomycetales | 7 | Saprotroph | Undefined Saprotroph | Yeast | NULL | Possible | NULL | Sterkenburg E, et al. 2015. New Phytologist 207:1145-1158 ((https://doi.org/10.1111/nph.13426)) |
| Otu53445 | 1 | k__Fungi(100);p__Ascomycota(100);c__Saccharomycetes(100);o__Saccharomycetales(100);f__Metschnikowiaceae(100);g__Metschnikowia(100);s__Metschnikowia_pulcherrima(100); | Metschnikowia | 13 | Saprotroph | \|Undefined Saprotroph\| | Yeast | NULL | Probable | As nectar-tap_saprotroph (Põlme et al. 2020); Decay sugar-rich substrates (Põlme et al. 2020) | Põlme S, et al. 2020. Fungal Diversity 105:1-16 ((https://doi.org/10.1007/s13225-020-00466-2)) |
| Otu53446 | 1 | k__Fungi(100);p__Ascomycota(100);c__Saccharomycetes(100);o__Saccharomycetales(100);f__Metschnikowiaceae(100);g__Metschnikowia(100);g__Metschnikowia_unclassified(100); | Metschnikowia | 13 | Saprotroph | \|Undefined Saprotroph\| | Yeast | NULL | Probable | As nectar-tap_saprotroph (Põlme et al. 2020); Decay sugar-rich substrates (Põlme et al. 2020) | Põlme S, et al. 2020. Fungal Diversity 105:1-16 ((https://doi.org/10.1007/s13225-020-00466-2)) |
| Otu53447 | 1 | k__Fungi(100);p__Ascomycota(100);c__Saccharomycetes(100);o__Saccharomycetales(100);f__Metschnikowiaceae(100);g__Metschnikowia(100);s__Metschnikowia_pulcherrima(100); | Metschnikowia | 13 | Saprotroph | \|Undefined Saprotroph\| | Yeast | NULL | Probable | As nectar-tap_saprotroph (Põlme et al. 2020); Decay sugar-rich substrates (Põlme et al. 2020) | Põlme S, et al. 2020. Fungal Diversity 105:1-16 ((https://doi.org/10.1007/s13225-020-00466-2)) |
| Otu53448 | 1 | k__Fungi(100);p__Ascomycota(100);c__Saccharomycetes(100);o__Saccharomycetales(100);f__Metschnikowiaceae(100);g__Metschnikowia(100);s__Metschnikowia_pulcherrima(100); | Metschnikowia | 13 | Saprotroph | \|Undefined Saprotroph\| | Yeast | NULL | Probable | As nectar-tap_saprotroph (Põlme et al. 2020); Decay sugar-rich substrates (Põlme et al. 2020) | Põlme S, et al. 2020. Fungal Diversity 105:1-16 ((https://doi.org/10.1007/s13225-020-00466-2)) |
| Otu53449 | 1 | k__Fungi(100);p__Ascomycota(100);c__Saccharomycetes(100);o__Saccharomycetales(100);f__Metschnikowiaceae(100);g__Metschnikowia(100);g__Metschnikowia_unclassified(100); | Metschnikowia | 13 | Saprotroph | \|Undefined Saprotroph\| | Yeast | NULL | Probable | As nectar-tap_saprotroph (Põlme et al. 2020); Decay sugar-rich substrates (Põlme et al. 2020) | Põlme S, et al. 2020. Fungal Diversity 105:1-16 ((https://doi.org/10.1007/s13225-020-00466-2)) |
| Otu53451 | 1 | k__Fungi(100);p__Ascomycota(100);c__Saccharomycetes(100);o__Saccharomycetales(100);f__Metschnikowiaceae(100);g__Metschnikowia(100);s__Metschnikowia_pulcherrima(100); | Metschnikowia | 13 | Saprotroph | \|Undefined Saprotroph\| | Yeast | NULL | Probable | As nectar-tap_saprotroph (Põlme et al. 2020); Decay sugar-rich substrates (Põlme et al. 2020) | Põlme S, et al. 2020. Fungal Diversity 105:1-16 ((https://doi.org/10.1007/s13225-020-00466-2)) |
| Otu53452 | 1 | k__Fungi(100);p__Ascomycota(100);c__Saccharomycetes(100);o__Saccharomycetales(100);f__Metschnikowiaceae(100);g__Metschnikowia(100);s__Metschnikowia_pulcherrima(100); | Metschnikowia | 13 | Saprotroph | \|Undefined Saprotroph\| | Yeast | NULL | Probable | As nectar-tap_saprotroph (Põlme et al. 2020); Decay sugar-rich substrates (Põlme et al. 2020) | Põlme S, et al. 2020. Fungal Diversity 105:1-16 ((https://doi.org/10.1007/s13225-020-00466-2)) |
| Otu53454 | 1 | k__Fungi(100);p__Ascomycota(100);c__Saccharomycetes(100);o__Saccharomycetales(100);f__Metschnikowiaceae(100);g__Metschnikowia(100);s__Metschnikowia_pulcherrima(100); | Metschnikowia | 13 | Saprotroph | \|Undefined Saprotroph\| | Yeast | NULL | Probable | As nectar-tap_saprotroph (Põlme et al. 2020); Decay sugar-rich substrates (Põlme et al. 2020) | Põlme S, et al. 2020. Fungal Diversity 105:1-16 ((https://doi.org/10.1007/s13225-020-00466-2)) |
| Otu53455 | 1 | k__Fungi(100);p__Ascomycota(100);c__Saccharomycetes(100);o__Saccharomycetales(100);f__Saccharomycetales_family_Incertae_sedis(100);g__Candida(100);g__Candida_unclassified(100); | Candida | 13 | Pathotroph-Saprotroph-Symbiotroph | Animal Pathogen-Endophyte-Endosymbiont-Epiphyte-\|Undefined Saprotroph\| | Dimorphic Yeast-Yeast | NULL | Probable | As nectar-tap_saprotroph (Põlme et al. 2020); Decay sugar-rich substrates (Põlme et al. 2020); This asexual genus is polyphyletic and contain species that function very differently depend on what they are. Caution should be taken when interpreting guilds for Candida species. | Manolakaki D, et al. 2010. Virulence. 1: 367-375 ((https://doi.org/10.4161/viru.1.5.12796)); Mekbib SB, et al. 2011. Tropical Plant Pathology 36:233-240 ((https://doi.org/10.1590/S1982-56762011000400004)); Põlme S, et al. 2020. Fungal Diversity 105:1-16 ((https://doi.org/10.1007/s13225-020-00466-2)) |
| Otu53460 | 1 | k__Fungi(100);p__Ascomycota(100);c__Saccharomycetes(100);o__Saccharomycetales(100);f__Metschnikowiaceae(100);g__Metschnikowia(100);g__Metschnikowia_unclassified(100); | Metschnikowia | 13 | Saprotroph | \|Undefined Saprotroph\| | Yeast | NULL | Probable | As nectar-tap_saprotroph (Põlme et al. 2020); Decay sugar-rich substrates (Põlme et al. 2020) | Põlme S, et al. 2020. Fungal Diversity 105:1-16 ((https://doi.org/10.1007/s13225-020-00466-2)) |
| Otu53469 | 1 | k__Fungi(100);p__Ascomycota(100);c__Saccharomycetes(100);o__Saccharomycetales(100);f__Metschnikowiaceae(100);g__Metschnikowia(100);s__Metschnikowia_pulcherrima(100); | Metschnikowia | 13 | Saprotroph | \|Undefined Saprotroph\| | Yeast | NULL | Probable | As nectar-tap_saprotroph (Põlme et al. 2020); Decay sugar-rich substrates (Põlme et al. 2020) | Põlme S, et al. 2020. Fungal Diversity 105:1-16 ((https://doi.org/10.1007/s13225-020-00466-2)) |
| Otu53470 | 1 | k__Fungi(100);p__Ascomycota(100);c__Saccharomycetes(100);o__Saccharomycetales(100);f__Metschnikowiaceae(100);g__Metschnikowia(100);s__Metschnikowia_pulcherrima(100); | Metschnikowia | 13 | Saprotroph | \|Undefined Saprotroph\| | Yeast | NULL | Probable | As nectar-tap_saprotroph (Põlme et al. 2020); Decay sugar-rich substrates (Põlme et al. 2020) | Põlme S, et al. 2020. Fungal Diversity 105:1-16 ((https://doi.org/10.1007/s13225-020-00466-2)) |
| Otu53471 | 1 | k__Fungi(100);p__Ascomycota(100);c__Saccharomycetes(100);o__Saccharomycetales(100);f__Saccharomycetales_family_Incertae_sedis(100);f__Saccharomycetales_family_Incertae_sedis_unclassified(100);f__Saccharomycetales_family_Incertae_sedis_unclassified(100); | Saccharomycetales | 7 | Saprotroph | Undefined Saprotroph | Yeast | NULL | Possible | NULL | Sterkenburg E, et al. 2015. New Phytologist 207:1145-1158 ((https://doi.org/10.1111/nph.13426)) |
| Otu53472 | 1 | k__Fungi(100);p__Ascomycota(100);c__Saccharomycetes(100);o__Saccharomycetales(100);f__Metschnikowiaceae(100);g__Metschnikowia(100);s__Metschnikowia_pulcherrima(100); | Metschnikowia | 13 | Saprotroph | \|Undefined Saprotroph\| | Yeast | NULL | Probable | As nectar-tap_saprotroph (Põlme et al. 2020); Decay sugar-rich substrates (Põlme et al. 2020) | Põlme S, et al. 2020. Fungal Diversity 105:1-16 ((https://doi.org/10.1007/s13225-020-00466-2)) |
| Otu53474 | 1 | k__Fungi(100);p__Ascomycota(100);c__Saccharomycetes(100);o__Saccharomycetales(100);f__Metschnikowiaceae(100);g__Metschnikowia(100);s__Metschnikowia_pulcherrima(100); | Metschnikowia | 13 | Saprotroph | \|Undefined Saprotroph\| | Yeast | NULL | Probable | As nectar-tap_saprotroph (Põlme et al. 2020); Decay sugar-rich substrates (Põlme et al. 2020) | Põlme S, et al. 2020. Fungal Diversity 105:1-16 ((https://doi.org/10.1007/s13225-020-00466-2)) |
| Otu53475 | 1 | k__Fungi(100);p__Ascomycota(100);c__Saccharomycetes(100);o__Saccharomycetales(100);f__Metschnikowiaceae(100);g__Metschnikowia(100);g__Metschnikowia_unclassified(100); | Metschnikowia | 13 | Saprotroph | \|Undefined Saprotroph\| | Yeast | NULL | Probable | As nectar-tap_saprotroph (Põlme et al. 2020); Decay sugar-rich substrates (Põlme et al. 2020) | Põlme S, et al. 2020. Fungal Diversity 105:1-16 ((https://doi.org/10.1007/s13225-020-00466-2)) |
| Otu53476 | 1 | k__Fungi(100);p__Ascomycota(100);c__Saccharomycetes(100);o__Saccharomycetales(100);f__Metschnikowiaceae(100);g__Metschnikowia(100);s__Metschnikowia_pulcherrima(100); | Metschnikowia | 13 | Saprotroph | \|Undefined Saprotroph\| | Yeast | NULL | Probable | As nectar-tap_saprotroph (Põlme et al. 2020); Decay sugar-rich substrates (Põlme et al. 2020) | Põlme S, et al. 2020. Fungal Diversity 105:1-16 ((https://doi.org/10.1007/s13225-020-00466-2)) |
| Otu53478 | 1 | k__Fungi(100);p__Ascomycota(100);c__Saccharomycetes(100);o__Saccharomycetales(100);f__Metschnikowiaceae(100);g__Metschnikowia(100);g__Metschnikowia_unclassified(100); | Metschnikowia | 13 | Saprotroph | \|Undefined Saprotroph\| | Yeast | NULL | Probable | As nectar-tap_saprotroph (Põlme et al. 2020); Decay sugar-rich substrates (Põlme et al. 2020) | Põlme S, et al. 2020. Fungal Diversity 105:1-16 ((https://doi.org/10.1007/s13225-020-00466-2)) |
| Otu53479 | 1 | k__Fungi(100);p__Ascomycota(100);c__Saccharomycetes(100);o__Saccharomycetales(100);f__Metschnikowiaceae(100);g__Metschnikowia(100);s__Metschnikowia_pulcherrima(100); | Metschnikowia | 13 | Saprotroph | \|Undefined Saprotroph\| | Yeast | NULL | Probable | As nectar-tap_saprotroph (Põlme et al. 2020); Decay sugar-rich substrates (Põlme et al. 2020) | Põlme S, et al. 2020. Fungal Diversity 105:1-16 ((https://doi.org/10.1007/s13225-020-00466-2)) |
| Otu53480 | 1 | k__Fungi(100);p__Ascomycota(100);c__Saccharomycetes(100);o__Saccharomycetales(100);f__Metschnikowiaceae(100);g__Metschnikowia(100);s__Metschnikowia_pulcherrima(100); | Metschnikowia | 13 | Saprotroph | \|Undefined Saprotroph\| | Yeast | NULL | Probable | As nectar-tap_saprotroph (Põlme et al. 2020); Decay sugar-rich substrates (Põlme et al. 2020) | Põlme S, et al. 2020. Fungal Diversity 105:1-16 ((https://doi.org/10.1007/s13225-020-00466-2)) |
| Otu53482 | 1 | k__Fungi(100);p__Ascomycota(100);c__Saccharomycetes(100);o__Saccharomycetales(100);f__Saccharomycetales_family_Incertae_sedis(100);g__Candida(100);g__Candida_unclassified(100); | Candida | 13 | Pathotroph-Saprotroph-Symbiotroph | Animal Pathogen-Endophyte-Endosymbiont-Epiphyte-\|Undefined Saprotroph\| | Dimorphic Yeast-Yeast | NULL | Probable | As nectar-tap_saprotroph (Põlme et al. 2020); Decay sugar-rich substrates (Põlme et al. 2020); This asexual genus is polyphyletic and contain species that function very differently depend on what they are. Caution should be taken when interpreting guilds for Candida species. | Manolakaki D, et al. 2010. Virulence. 1: 367-375 ((https://doi.org/10.4161/viru.1.5.12796)); Mekbib SB, et al. 2011. Tropical Plant Pathology 36:233-240 ((https://doi.org/10.1590/S1982-56762011000400004)); Põlme S, et al. 2020. Fungal Diversity 105:1-16 ((https://doi.org/10.1007/s13225-020-00466-2)) |
| Otu53484 | 1 | k__Fungi(100);p__Ascomycota(100);c__Saccharomycetes(100);o__Saccharomycetales(100);f__Saccharomycetales_family_Incertae_sedis(100);g__Candida(100);g__Candida_unclassified(100); | Candida | 13 | Pathotroph-Saprotroph-Symbiotroph | Animal Pathogen-Endophyte-Endosymbiont-Epiphyte-\|Undefined Saprotroph\| | Dimorphic Yeast-Yeast | NULL | Probable | As nectar-tap_saprotroph (Põlme et al. 2020); Decay sugar-rich substrates (Põlme et al. 2020); This asexual genus is polyphyletic and contain species that function very differently depend on what they are. Caution should be taken when interpreting guilds for Candida species. | Manolakaki D, et al. 2010. Virulence. 1: 367-375 ((https://doi.org/10.4161/viru.1.5.12796)); Mekbib SB, et al. 2011. Tropical Plant Pathology 36:233-240 ((https://doi.org/10.1590/S1982-56762011000400004)); Põlme S, et al. 2020. Fungal Diversity 105:1-16 ((https://doi.org/10.1007/s13225-020-00466-2)) |
| Otu53485 | 1 | k__Fungi(100);p__Ascomycota(100);c__Saccharomycetes(100);o__Saccharomycetales(100);f__Saccharomycetales_family_Incertae_sedis(100);g__Candida(100);g__Candida_unclassified(100); | Candida | 13 | Pathotroph-Saprotroph-Symbiotroph | Animal Pathogen-Endophyte-Endosymbiont-Epiphyte-\|Undefined Saprotroph\| | Dimorphic Yeast-Yeast | NULL | Probable | As nectar-tap_saprotroph (Põlme et al. 2020); Decay sugar-rich substrates (Põlme et al. 2020); This asexual genus is polyphyletic and contain species that function very differently depend on what they are. Caution should be taken when interpreting guilds for Candida species. | Manolakaki D, et al. 2010. Virulence. 1: 367-375 ((https://doi.org/10.4161/viru.1.5.12796)); Mekbib SB, et al. 2011. Tropical Plant Pathology 36:233-240 ((https://doi.org/10.1590/S1982-56762011000400004)); Põlme S, et al. 2020. Fungal Diversity 105:1-16 ((https://doi.org/10.1007/s13225-020-00466-2)) |
| Otu53487 | 1 | k__Fungi(100);p__Ascomycota(100);c__Saccharomycetes(100);o__Saccharomycetales(100);f__Saccharomycetales_family_Incertae_sedis(100);g__Candida(100);g__Candida_unclassified(100); | Candida | 13 | Pathotroph-Saprotroph-Symbiotroph | Animal Pathogen-Endophyte-Endosymbiont-Epiphyte-\|Undefined Saprotroph\| | Dimorphic Yeast-Yeast | NULL | Probable | As nectar-tap_saprotroph (Põlme et al. 2020); Decay sugar-rich substrates (Põlme et al. 2020); This asexual genus is polyphyletic and contain species that function very differently depend on what they are. Caution should be taken when interpreting guilds for Candida species. | Manolakaki D, et al. 2010. Virulence. 1: 367-375 ((https://doi.org/10.4161/viru.1.5.12796)); Mekbib SB, et al. 2011. Tropical Plant Pathology 36:233-240 ((https://doi.org/10.1590/S1982-56762011000400004)); Põlme S, et al. 2020. Fungal Diversity 105:1-16 ((https://doi.org/10.1007/s13225-020-00466-2)) |
| Otu53488 | 1 | k__Fungi(100);p__Ascomycota(100);c__Saccharomycetes(100);o__Saccharomycetales(100);f__Metschnikowiaceae(100);g__Metschnikowia(100);s__Metschnikowia_pulcherrima(100); | Metschnikowia | 13 | Saprotroph | \|Undefined Saprotroph\| | Yeast | NULL | Probable | As nectar-tap_saprotroph (Põlme et al. 2020); Decay sugar-rich substrates (Põlme et al. 2020) | Põlme S, et al. 2020. Fungal Diversity 105:1-16 ((https://doi.org/10.1007/s13225-020-00466-2)) |
| Otu53489 | 1 | k__Fungi(100);p__Ascomycota(100);c__Saccharomycetes(100);o__Saccharomycetales(100);f__Metschnikowiaceae(100);g__Metschnikowia(100);s__Metschnikowia_pulcherrima(100); | Metschnikowia | 13 | Saprotroph | \|Undefined Saprotroph\| | Yeast | NULL | Probable | As nectar-tap_saprotroph (Põlme et al. 2020); Decay sugar-rich substrates (Põlme et al. 2020) | Põlme S, et al. 2020. Fungal Diversity 105:1-16 ((https://doi.org/10.1007/s13225-020-00466-2)) |
| Otu53491 | 1 | k__Fungi(100);p__Ascomycota(100);c__Saccharomycetes(100);o__Saccharomycetales(100);f__Metschnikowiaceae(100);g__Metschnikowia(100);s__Metschnikowia_pulcherrima(100); | Metschnikowia | 13 | Saprotroph | \|Undefined Saprotroph\| | Yeast | NULL | Probable | As nectar-tap_saprotroph (Põlme et al. 2020); Decay sugar-rich substrates (Põlme et al. 2020) | Põlme S, et al. 2020. Fungal Diversity 105:1-16 ((https://doi.org/10.1007/s13225-020-00466-2)) |
| Otu53492 | 1 | k__Fungi(100);p__Ascomycota(100);c__Saccharomycetes(100);o__Saccharomycetales(100);f__Metschnikowiaceae(100);g__Metschnikowia(100);s__Metschnikowia_pulcherrima(100); | Metschnikowia | 13 | Saprotroph | \|Undefined Saprotroph\| | Yeast | NULL | Probable | As nectar-tap_saprotroph (Põlme et al. 2020); Decay sugar-rich substrates (Põlme et al. 2020) | Põlme S, et al. 2020. Fungal Diversity 105:1-16 ((https://doi.org/10.1007/s13225-020-00466-2)) |
| Otu53494 | 1 | k__Fungi(100);p__Ascomycota(100);c__Saccharomycetes(100);o__Saccharomycetales(100);f__Metschnikowiaceae(100);g__Metschnikowia(100);s__Metschnikowia_pulcherrima(100); | Metschnikowia | 13 | Saprotroph | \|Undefined Saprotroph\| | Yeast | NULL | Probable | As nectar-tap_saprotroph (Põlme et al. 2020); Decay sugar-rich substrates (Põlme et al. 2020) | Põlme S, et al. 2020. Fungal Diversity 105:1-16 ((https://doi.org/10.1007/s13225-020-00466-2)) |
| Otu53495 | 1 | k__Fungi(100);p__Ascomycota(100);c__Saccharomycetes(100);o__Saccharomycetales(100);f__Metschnikowiaceae(100);g__Metschnikowia(100);s__Metschnikowia_pulcherrima(100); | Metschnikowia | 13 | Saprotroph | \|Undefined Saprotroph\| | Yeast | NULL | Probable | As nectar-tap_saprotroph (Põlme et al. 2020); Decay sugar-rich substrates (Põlme et al. 2020) | Põlme S, et al. 2020. Fungal Diversity 105:1-16 ((https://doi.org/10.1007/s13225-020-00466-2)) |
| Otu53496 | 1 | k__Fungi(100);p__Ascomycota(100);c__Saccharomycetes(100);o__Saccharomycetales(100);f__Metschnikowiaceae(100);g__Metschnikowia(100);s__Metschnikowia_pulcherrima(100); | Metschnikowia | 13 | Saprotroph | \|Undefined Saprotroph\| | Yeast | NULL | Probable | As nectar-tap_saprotroph (Põlme et al. 2020); Decay sugar-rich substrates (Põlme et al. 2020) | Põlme S, et al. 2020. Fungal Diversity 105:1-16 ((https://doi.org/10.1007/s13225-020-00466-2)) |
| Otu53497 | 1 | k__Fungi(100);p__Ascomycota(100);c__Saccharomycetes(100);o__Saccharomycetales(100);f__Metschnikowiaceae(100);g__Metschnikowia(100);s__Metschnikowia_pulcherrima(100); | Metschnikowia | 13 | Saprotroph | \|Undefined Saprotroph\| | Yeast | NULL | Probable | As nectar-tap_saprotroph (Põlme et al. 2020); Decay sugar-rich substrates (Põlme et al. 2020) | Põlme S, et al. 2020. Fungal Diversity 105:1-16 ((https://doi.org/10.1007/s13225-020-00466-2)) |
| Otu53498 | 1 | k__Fungi(100);p__Ascomycota(100);c__Saccharomycetes(100);o__Saccharomycetales(100);f__Metschnikowiaceae(100);g__Metschnikowia(100);s__Metschnikowia_pulcherrima(100); | Metschnikowia | 13 | Saprotroph | \|Undefined Saprotroph\| | Yeast | NULL | Probable | As nectar-tap_saprotroph (Põlme et al. 2020); Decay sugar-rich substrates (Põlme et al. 2020) | Põlme S, et al. 2020. Fungal Diversity 105:1-16 ((https://doi.org/10.1007/s13225-020-00466-2)) |
| Otu53499 | 1 | k__Fungi(100);p__Ascomycota(100);c__Saccharomycetes(100);o__Saccharomycetales(100);f__Metschnikowiaceae(100);g__Metschnikowia(100);s__Metschnikowia_pulcherrima(100); | Metschnikowia | 13 | Saprotroph | \|Undefined Saprotroph\| | Yeast | NULL | Probable | As nectar-tap_saprotroph (Põlme et al. 2020); Decay sugar-rich substrates (Põlme et al. 2020) | Põlme S, et al. 2020. Fungal Diversity 105:1-16 ((https://doi.org/10.1007/s13225-020-00466-2)) |
| Otu53500 | 1 | k__Fungi(100);p__Ascomycota(100);c__Saccharomycetes(100);o__Saccharomycetales(100);f__Saccharomycetales_family_Incertae_sedis(100);g__Candida(100);g__Candida_unclassified(100); | Candida | 13 | Pathotroph-Saprotroph-Symbiotroph | Animal Pathogen-Endophyte-Endosymbiont-Epiphyte-\|Undefined Saprotroph\| | Dimorphic Yeast-Yeast | NULL | Probable | As nectar-tap_saprotroph (Põlme et al. 2020); Decay sugar-rich substrates (Põlme et al. 2020); This asexual genus is polyphyletic and contain species that function very differently depend on what they are. Caution should be taken when interpreting guilds for Candida species. | Manolakaki D, et al. 2010. Virulence. 1: 367-375 ((https://doi.org/10.4161/viru.1.5.12796)); Mekbib SB, et al. 2011. Tropical Plant Pathology 36:233-240 ((https://doi.org/10.1590/S1982-56762011000400004)); Põlme S, et al. 2020. Fungal Diversity 105:1-16 ((https://doi.org/10.1007/s13225-020-00466-2)) |
| Otu53501 | 1 | k__Fungi(100);p__Ascomycota(100);c__Saccharomycetes(100);o__Saccharomycetales(100);f__Metschnikowiaceae(100);g__Metschnikowia(100);s__Metschnikowia_pulcherrima(100); | Metschnikowia | 13 | Saprotroph | \|Undefined Saprotroph\| | Yeast | NULL | Probable | As nectar-tap_saprotroph (Põlme et al. 2020); Decay sugar-rich substrates (Põlme et al. 2020) | Põlme S, et al. 2020. Fungal Diversity 105:1-16 ((https://doi.org/10.1007/s13225-020-00466-2)) |
| Otu53502 | 1 | k__Fungi(100);p__Ascomycota(100);c__Saccharomycetes(100);o__Saccharomycetales(100);f__Metschnikowiaceae(100);g__Metschnikowia(100);s__Metschnikowia_pulcherrima(100); | Metschnikowia | 13 | Saprotroph | \|Undefined Saprotroph\| | Yeast | NULL | Probable | As nectar-tap_saprotroph (Põlme et al. 2020); Decay sugar-rich substrates (Põlme et al. 2020) | Põlme S, et al. 2020. Fungal Diversity 105:1-16 ((https://doi.org/10.1007/s13225-020-00466-2)) |
| Otu53503 | 1 | k__Fungi(100);p__Ascomycota(100);c__Saccharomycetes(100);o__Saccharomycetales(100);f__Metschnikowiaceae(100);g__Metschnikowia(100);g__Metschnikowia_unclassified(100); | Metschnikowia | 13 | Saprotroph | \|Undefined Saprotroph\| | Yeast | NULL | Probable | As nectar-tap_saprotroph (Põlme et al. 2020); Decay sugar-rich substrates (Põlme et al. 2020) | Põlme S, et al. 2020. Fungal Diversity 105:1-16 ((https://doi.org/10.1007/s13225-020-00466-2)) |
| Otu53505 | 1 | k__Fungi(100);p__Ascomycota(100);c__Saccharomycetes(100);o__Saccharomycetales(100);f__Saccharomycetales_family_Incertae_sedis(100);g__Candida(100);g__Candida_unclassified(100); | Candida | 13 | Pathotroph-Saprotroph-Symbiotroph | Animal Pathogen-Endophyte-Endosymbiont-Epiphyte-\|Undefined Saprotroph\| | Dimorphic Yeast-Yeast | NULL | Probable | As nectar-tap_saprotroph (Põlme et al. 2020); Decay sugar-rich substrates (Põlme et al. 2020); This asexual genus is polyphyletic and contain species that function very differently depend on what they are. Caution should be taken when interpreting guilds for Candida species. | Manolakaki D, et al. 2010. Virulence. 1: 367-375 ((https://doi.org/10.4161/viru.1.5.12796)); Mekbib SB, et al. 2011. Tropical Plant Pathology 36:233-240 ((https://doi.org/10.1590/S1982-56762011000400004)); Põlme S, et al. 2020. Fungal Diversity 105:1-16 ((https://doi.org/10.1007/s13225-020-00466-2)) |
| Otu53509 | 1 | k__Fungi(100);p__Ascomycota(100);c__Saccharomycetes(100);o__Saccharomycetales(100);f__Saccharomycetales_family_Incertae_sedis(100);f__Saccharomycetales_family_Incertae_sedis_unclassified(100);f__Saccharomycetales_family_Incertae_sedis_unclassified(100); | Saccharomycetales | 7 | Saprotroph | Undefined Saprotroph | Yeast | NULL | Possible | NULL | Sterkenburg E, et al. 2015. New Phytologist 207:1145-1158 ((https://doi.org/10.1111/nph.13426)) |
| Otu53511 | 1 | k__Fungi(100);p__Ascomycota(100);c__Saccharomycetes(100);o__Saccharomycetales(100);f__Saccharomycetales_family_Incertae_sedis(100);g__Candida(100);s__Candida_cretensis(100); | Candida | 13 | Pathotroph-Saprotroph-Symbiotroph | Animal Pathogen-Endophyte-Endosymbiont-Epiphyte-\|Undefined Saprotroph\| | Dimorphic Yeast-Yeast | NULL | Probable | As nectar-tap_saprotroph (Põlme et al. 2020); Decay sugar-rich substrates (Põlme et al. 2020); This asexual genus is polyphyletic and contain species that function very differently depend on what they are. Caution should be taken when interpreting guilds for Candida species. | Manolakaki D, et al. 2010. Virulence. 1: 367-375 ((https://doi.org/10.4161/viru.1.5.12796)); Mekbib SB, et al. 2011. Tropical Plant Pathology 36:233-240 ((https://doi.org/10.1590/S1982-56762011000400004)); Põlme S, et al. 2020. Fungal Diversity 105:1-16 ((https://doi.org/10.1007/s13225-020-00466-2)) |
| Otu53514 | 1 | k__Fungi(100);p__Ascomycota(100);c__Saccharomycetes(100);o__Saccharomycetales(100);f__Saccharomycetales_family_Incertae_sedis(100);g__Candida(100);g__Candida_unclassified(100); | Candida | 13 | Pathotroph-Saprotroph-Symbiotroph | Animal Pathogen-Endophyte-Endosymbiont-Epiphyte-\|Undefined Saprotroph\| | Dimorphic Yeast-Yeast | NULL | Probable | As nectar-tap_saprotroph (Põlme et al. 2020); Decay sugar-rich substrates (Põlme et al. 2020); This asexual genus is polyphyletic and contain species that function very differently depend on what they are. Caution should be taken when interpreting guilds for Candida species. | Manolakaki D, et al. 2010. Virulence. 1: 367-375 ((https://doi.org/10.4161/viru.1.5.12796)); Mekbib SB, et al. 2011. Tropical Plant Pathology 36:233-240 ((https://doi.org/10.1590/S1982-56762011000400004)); Põlme S, et al. 2020. Fungal Diversity 105:1-16 ((https://doi.org/10.1007/s13225-020-00466-2)) |
| Otu53516 | 1 | k__Fungi(100);p__Ascomycota(100);c__Saccharomycetes(100);o__Saccharomycetales(100);f__Saccharomycetales_family_Incertae_sedis(100);g__Candida(100);g__Candida_unclassified(100); | Candida | 13 | Pathotroph-Saprotroph-Symbiotroph | Animal Pathogen-Endophyte-Endosymbiont-Epiphyte-\|Undefined Saprotroph\| | Dimorphic Yeast-Yeast | NULL | Probable | As nectar-tap_saprotroph (Põlme et al. 2020); Decay sugar-rich substrates (Põlme et al. 2020); This asexual genus is polyphyletic and contain species that function very differently depend on what they are. Caution should be taken when interpreting guilds for Candida species. | Manolakaki D, et al. 2010. Virulence. 1: 367-375 ((https://doi.org/10.4161/viru.1.5.12796)); Mekbib SB, et al. 2011. Tropical Plant Pathology 36:233-240 ((https://doi.org/10.1590/S1982-56762011000400004)); Põlme S, et al. 2020. Fungal Diversity 105:1-16 ((https://doi.org/10.1007/s13225-020-00466-2)) |
| Otu53517 | 1 | k__Fungi(100);p__Ascomycota(100);c__Saccharomycetes(100);o__Saccharomycetales(100);f__Metschnikowiaceae(100);g__Metschnikowia(100);s__Metschnikowia_pulcherrima(100); | Metschnikowia | 13 | Saprotroph | \|Undefined Saprotroph\| | Yeast | NULL | Probable | As nectar-tap_saprotroph (Põlme et al. 2020); Decay sugar-rich substrates (Põlme et al. 2020) | Põlme S, et al. 2020. Fungal Diversity 105:1-16 ((https://doi.org/10.1007/s13225-020-00466-2)) |
| Otu53518 | 1 | k__Fungi(100);p__Ascomycota(100);c__Saccharomycetes(100);o__Saccharomycetales(100);f__Metschnikowiaceae(100);g__Metschnikowia(100);s__Metschnikowia_pulcherrima(100); | Metschnikowia | 13 | Saprotroph | \|Undefined Saprotroph\| | Yeast | NULL | Probable | As nectar-tap_saprotroph (Põlme et al. 2020); Decay sugar-rich substrates (Põlme et al. 2020) | Põlme S, et al. 2020. Fungal Diversity 105:1-16 ((https://doi.org/10.1007/s13225-020-00466-2)) |
| Otu53519 | 1 | k__Fungi(100);p__Ascomycota(100);c__Saccharomycetes(100);o__Saccharomycetales(100);f__Metschnikowiaceae(100);g__Metschnikowia(100);g__Metschnikowia_unclassified(100); | Metschnikowia | 13 | Saprotroph | \|Undefined Saprotroph\| | Yeast | NULL | Probable | As nectar-tap_saprotroph (Põlme et al. 2020); Decay sugar-rich substrates (Põlme et al. 2020) | Põlme S, et al. 2020. Fungal Diversity 105:1-16 ((https://doi.org/10.1007/s13225-020-00466-2)) |
| Otu53520 | 1 | k__Fungi(100);p__Ascomycota(100);c__Saccharomycetes(100);o__Saccharomycetales(100);f__Metschnikowiaceae(100);g__Metschnikowia(100);s__Metschnikowia_pulcherrima(100); | Metschnikowia | 13 | Saprotroph | \|Undefined Saprotroph\| | Yeast | NULL | Probable | As nectar-tap_saprotroph (Põlme et al. 2020); Decay sugar-rich substrates (Põlme et al. 2020) | Põlme S, et al. 2020. Fungal Diversity 105:1-16 ((https://doi.org/10.1007/s13225-020-00466-2)) |
| Otu53521 | 1 | k__Fungi(100);p__Ascomycota(100);c__Saccharomycetes(100);o__Saccharomycetales(100);f__Metschnikowiaceae(100);g__Metschnikowia(100);s__Metschnikowia_pulcherrima(100); | Metschnikowia | 13 | Saprotroph | \|Undefined Saprotroph\| | Yeast | NULL | Probable | As nectar-tap_saprotroph (Põlme et al. 2020); Decay sugar-rich substrates (Põlme et al. 2020) | Põlme S, et al. 2020. Fungal Diversity 105:1-16 ((https://doi.org/10.1007/s13225-020-00466-2)) |
| Otu53522 | 1 | k__Fungi(100);p__Ascomycota(100);c__Saccharomycetes(100);o__Saccharomycetales(100);f__Metschnikowiaceae(100);g__Metschnikowia(100);s__Metschnikowia_pulcherrima(100); | Metschnikowia | 13 | Saprotroph | \|Undefined Saprotroph\| | Yeast | NULL | Probable | As nectar-tap_saprotroph (Põlme et al. 2020); Decay sugar-rich substrates (Põlme et al. 2020) | Põlme S, et al. 2020. Fungal Diversity 105:1-16 ((https://doi.org/10.1007/s13225-020-00466-2)) |
| Otu53523 | 1 | k__Fungi(100);p__Ascomycota(100);c__Saccharomycetes(100);o__Saccharomycetales(100);f__Metschnikowiaceae(100);g__Metschnikowia(100);s__Metschnikowia_pulcherrima(100); | Metschnikowia | 13 | Saprotroph | \|Undefined Saprotroph\| | Yeast | NULL | Probable | As nectar-tap_saprotroph (Põlme et al. 2020); Decay sugar-rich substrates (Põlme et al. 2020) | Põlme S, et al. 2020. Fungal Diversity 105:1-16 ((https://doi.org/10.1007/s13225-020-00466-2)) |
| Otu53527 | 1 | k__Fungi(100);p__Ascomycota(100);c__Saccharomycetes(100);o__Saccharomycetales(100);f__Saccharomycetales_family_Incertae_sedis(100);g__Candida(100);g__Candida_unclassified(100); | Candida | 13 | Pathotroph-Saprotroph-Symbiotroph | Animal Pathogen-Endophyte-Endosymbiont-Epiphyte-\|Undefined Saprotroph\| | Dimorphic Yeast-Yeast | NULL | Probable | As nectar-tap_saprotroph (Põlme et al. 2020); Decay sugar-rich substrates (Põlme et al. 2020); This asexual genus is polyphyletic and contain species that function very differently depend on what they are. Caution should be taken when interpreting guilds for Candida species. | Manolakaki D, et al. 2010. Virulence. 1: 367-375 ((https://doi.org/10.4161/viru.1.5.12796)); Mekbib SB, et al. 2011. Tropical Plant Pathology 36:233-240 ((https://doi.org/10.1590/S1982-56762011000400004)); Põlme S, et al. 2020. Fungal Diversity 105:1-16 ((https://doi.org/10.1007/s13225-020-00466-2)) |
| Otu53528 | 1 | k__Fungi(100);p__Ascomycota(100);c__Saccharomycetes(100);o__Saccharomycetales(100);f__Saccharomycetales_family_Incertae_sedis(100);g__Candida(100);g__Candida_unclassified(100); | Candida | 13 | Pathotroph-Saprotroph-Symbiotroph | Animal Pathogen-Endophyte-Endosymbiont-Epiphyte-\|Undefined Saprotroph\| | Dimorphic Yeast-Yeast | NULL | Probable | As nectar-tap_saprotroph (Põlme et al. 2020); Decay sugar-rich substrates (Põlme et al. 2020); This asexual genus is polyphyletic and contain species that function very differently depend on what they are. Caution should be taken when interpreting guilds for Candida species. | Manolakaki D, et al. 2010. Virulence. 1: 367-375 ((https://doi.org/10.4161/viru.1.5.12796)); Mekbib SB, et al. 2011. Tropical Plant Pathology 36:233-240 ((https://doi.org/10.1590/S1982-56762011000400004)); Põlme S, et al. 2020. Fungal Diversity 105:1-16 ((https://doi.org/10.1007/s13225-020-00466-2)) |
| Otu53537 | 1 | k__Fungi(100);p__Ascomycota(100);c__Saccharomycetes(100);o__Saccharomycetales(100);f__Metschnikowiaceae(100);g__Metschnikowia(100);s__Metschnikowia_pulcherrima(100); | Metschnikowia | 13 | Saprotroph | \|Undefined Saprotroph\| | Yeast | NULL | Probable | As nectar-tap_saprotroph (Põlme et al. 2020); Decay sugar-rich substrates (Põlme et al. 2020) | Põlme S, et al. 2020. Fungal Diversity 105:1-16 ((https://doi.org/10.1007/s13225-020-00466-2)) |
| Otu53539 | 1 | k__Fungi(100);p__Ascomycota(100);c__Saccharomycetes(100);o__Saccharomycetales(100);f__Saccharomycetales_family_Incertae_sedis(100);f__Saccharomycetales_family_Incertae_sedis_unclassified(100);f__Saccharomycetales_family_Incertae_sedis_unclassified(100); | Saccharomycetales | 7 | Saprotroph | Undefined Saprotroph | Yeast | NULL | Possible | NULL | Sterkenburg E, et al. 2015. New Phytologist 207:1145-1158 ((https://doi.org/10.1111/nph.13426)) |
| Otu53545 | 1 | k__Fungi(100);p__Ascomycota(100);c__Saccharomycetes(100);o__Saccharomycetales(100);f__Saccharomycetales_family_Incertae_sedis(100);g__Candida(100);g__Candida_unclassified(100); | Candida | 13 | Pathotroph-Saprotroph-Symbiotroph | Animal Pathogen-Endophyte-Endosymbiont-Epiphyte-\|Undefined Saprotroph\| | Dimorphic Yeast-Yeast | NULL | Probable | As nectar-tap_saprotroph (Põlme et al. 2020); Decay sugar-rich substrates (Põlme et al. 2020); This asexual genus is polyphyletic and contain species that function very differently depend on what they are. Caution should be taken when interpreting guilds for Candida species. | Manolakaki D, et al. 2010. Virulence. 1: 367-375 ((https://doi.org/10.4161/viru.1.5.12796)); Mekbib SB, et al. 2011. Tropical Plant Pathology 36:233-240 ((https://doi.org/10.1590/S1982-56762011000400004)); Põlme S, et al. 2020. Fungal Diversity 105:1-16 ((https://doi.org/10.1007/s13225-020-00466-2)) |
| Otu53547 | 1 | k__Fungi(100);p__Ascomycota(100);c__Saccharomycetes(100);o__Saccharomycetales(100);f__Metschnikowiaceae(100);g__Metschnikowia(100);s__Metschnikowia_pulcherrima(100); | Metschnikowia | 13 | Saprotroph | \|Undefined Saprotroph\| | Yeast | NULL | Probable | As nectar-tap_saprotroph (Põlme et al. 2020); Decay sugar-rich substrates (Põlme et al. 2020) | Põlme S, et al. 2020. Fungal Diversity 105:1-16 ((https://doi.org/10.1007/s13225-020-00466-2)) |
| Otu53549 | 1 | k__Fungi(100);p__Ascomycota(100);c__Saccharomycetes(100);o__Saccharomycetales(100);f__Saccharomycetales_family_Incertae_sedis(100);g__Candida(100);g__Candida_unclassified(100); | Candida | 13 | Pathotroph-Saprotroph-Symbiotroph | Animal Pathogen-Endophyte-Endosymbiont-Epiphyte-\|Undefined Saprotroph\| | Dimorphic Yeast-Yeast | NULL | Probable | As nectar-tap_saprotroph (Põlme et al. 2020); Decay sugar-rich substrates (Põlme et al. 2020); This asexual genus is polyphyletic and contain species that function very differently depend on what they are. Caution should be taken when interpreting guilds for Candida species. | Manolakaki D, et al. 2010. Virulence. 1: 367-375 ((https://doi.org/10.4161/viru.1.5.12796)); Mekbib SB, et al. 2011. Tropical Plant Pathology 36:233-240 ((https://doi.org/10.1590/S1982-56762011000400004)); Põlme S, et al. 2020. Fungal Diversity 105:1-16 ((https://doi.org/10.1007/s13225-020-00466-2)) |
| Otu53550 | 1 | k__Fungi(100);p__Ascomycota(100);c__Saccharomycetes(100);o__Saccharomycetales(100);f__Saccharomycetales_family_Incertae_sedis(100);g__Candida(100);g__Candida_unclassified(100); | Candida | 13 | Pathotroph-Saprotroph-Symbiotroph | Animal Pathogen-Endophyte-Endosymbiont-Epiphyte-\|Undefined Saprotroph\| | Dimorphic Yeast-Yeast | NULL | Probable | As nectar-tap_saprotroph (Põlme et al. 2020); Decay sugar-rich substrates (Põlme et al. 2020); This asexual genus is polyphyletic and contain species that function very differently depend on what they are. Caution should be taken when interpreting guilds for Candida species. | Manolakaki D, et al. 2010. Virulence. 1: 367-375 ((https://doi.org/10.4161/viru.1.5.12796)); Mekbib SB, et al. 2011. Tropical Plant Pathology 36:233-240 ((https://doi.org/10.1590/S1982-56762011000400004)); Põlme S, et al. 2020. Fungal Diversity 105:1-16 ((https://doi.org/10.1007/s13225-020-00466-2)) |
| Otu53553 | 1 | k__Fungi(100);p__Ascomycota(100);c__Saccharomycetes(100);o__Saccharomycetales(100);f__Saccharomycetales_family_Incertae_sedis(100);f__Saccharomycetales_family_Incertae_sedis_unclassified(100);f__Saccharomycetales_family_Incertae_sedis_unclassified(100); | Saccharomycetales | 7 | Saprotroph | Undefined Saprotroph | Yeast | NULL | Possible | NULL | Sterkenburg E, et al. 2015. New Phytologist 207:1145-1158 ((https://doi.org/10.1111/nph.13426)) |
| Otu53554 | 1 | k__Fungi(100);p__Ascomycota(100);c__Saccharomycetes(100);o__Saccharomycetales(100);f__Saccharomycetales_family_Incertae_sedis(100);g__Candida(100);g__Candida_unclassified(100); | Candida | 13 | Pathotroph-Saprotroph-Symbiotroph | Animal Pathogen-Endophyte-Endosymbiont-Epiphyte-\|Undefined Saprotroph\| | Dimorphic Yeast-Yeast | NULL | Probable | As nectar-tap_saprotroph (Põlme et al. 2020); Decay sugar-rich substrates (Põlme et al. 2020); This asexual genus is polyphyletic and contain species that function very differently depend on what they are. Caution should be taken when interpreting guilds for Candida species. | Manolakaki D, et al. 2010. Virulence. 1: 367-375 ((https://doi.org/10.4161/viru.1.5.12796)); Mekbib SB, et al. 2011. Tropical Plant Pathology 36:233-240 ((https://doi.org/10.1590/S1982-56762011000400004)); Põlme S, et al. 2020. Fungal Diversity 105:1-16 ((https://doi.org/10.1007/s13225-020-00466-2)) |
| Otu53555 | 1 | k__Fungi(100);p__Ascomycota(100);c__Saccharomycetes(100);o__Saccharomycetales(100);f__Saccharomycetales_family_Incertae_sedis(100);f__Saccharomycetales_family_Incertae_sedis_unclassified(100);f__Saccharomycetales_family_Incertae_sedis_unclassified(100); | Saccharomycetales | 7 | Saprotroph | Undefined Saprotroph | Yeast | NULL | Possible | NULL | Sterkenburg E, et al. 2015. New Phytologist 207:1145-1158 ((https://doi.org/10.1111/nph.13426)) |
| Otu53562 | 1 | k__Fungi(100);p__Ascomycota(100);c__Saccharomycetes(100);o__Saccharomycetales(100);f__Saccharomycetales_family_Incertae_sedis(100);g__Candida(100);g__Candida_unclassified(100); | Candida | 13 | Pathotroph-Saprotroph-Symbiotroph | Animal Pathogen-Endophyte-Endosymbiont-Epiphyte-\|Undefined Saprotroph\| | Dimorphic Yeast-Yeast | NULL | Probable | As nectar-tap_saprotroph (Põlme et al. 2020); Decay sugar-rich substrates (Põlme et al. 2020); This asexual genus is polyphyletic and contain species that function very differently depend on what they are. Caution should be taken when interpreting guilds for Candida species. | Manolakaki D, et al. 2010. Virulence. 1: 367-375 ((https://doi.org/10.4161/viru.1.5.12796)); Mekbib SB, et al. 2011. Tropical Plant Pathology 36:233-240 ((https://doi.org/10.1590/S1982-56762011000400004)); Põlme S, et al. 2020. Fungal Diversity 105:1-16 ((https://doi.org/10.1007/s13225-020-00466-2)) |
| Otu53564 | 1 | k__Fungi(100);p__Ascomycota(100);c__Saccharomycetes(100);o__Saccharomycetales(100);f__Saccharomycetales_family_Incertae_sedis(100);g__Candida(100);g__Candida_unclassified(100); | Candida | 13 | Pathotroph-Saprotroph-Symbiotroph | Animal Pathogen-Endophyte-Endosymbiont-Epiphyte-\|Undefined Saprotroph\| | Dimorphic Yeast-Yeast | NULL | Probable | As nectar-tap_saprotroph (Põlme et al. 2020); Decay sugar-rich substrates (Põlme et al. 2020); This asexual genus is polyphyletic and contain species that function very differently depend on what they are. Caution should be taken when interpreting guilds for Candida species. | Manolakaki D, et al. 2010. Virulence. 1: 367-375 ((https://doi.org/10.4161/viru.1.5.12796)); Mekbib SB, et al. 2011. Tropical Plant Pathology 36:233-240 ((https://doi.org/10.1590/S1982-56762011000400004)); Põlme S, et al. 2020. Fungal Diversity 105:1-16 ((https://doi.org/10.1007/s13225-020-00466-2)) |
| Otu53569 | 1 | k__Fungi(100);p__Ascomycota(100);c__Saccharomycetes(100);o__Saccharomycetales(100);f__Saccharomycetales_family_Incertae_sedis(100);f__Saccharomycetales_family_Incertae_sedis_unclassified(100);f__Saccharomycetales_family_Incertae_sedis_unclassified(100); | Saccharomycetales | 7 | Saprotroph | Undefined Saprotroph | Yeast | NULL | Possible | NULL | Sterkenburg E, et al. 2015. New Phytologist 207:1145-1158 ((https://doi.org/10.1111/nph.13426)) |
| Otu53576 | 1 | k__Fungi(100);p__Ascomycota(100);c__Saccharomycetes(100);o__Saccharomycetales(100);f__Saccharomycetales_family_Incertae_sedis(100);f__Saccharomycetales_family_Incertae_sedis_unclassified(100);f__Saccharomycetales_family_Incertae_sedis_unclassified(100); | Saccharomycetales | 7 | Saprotroph | Undefined Saprotroph | Yeast | NULL | Possible | NULL | Sterkenburg E, et al. 2015. New Phytologist 207:1145-1158 ((https://doi.org/10.1111/nph.13426)) |
| Otu53578 | 1 | k__Fungi(100);p__Ascomycota(100);c__Saccharomycetes(100);o__Saccharomycetales(100);f__Saccharomycetales_family_Incertae_sedis(100);g__Candida(100);g__Candida_unclassified(100); | Candida | 13 | Pathotroph-Saprotroph-Symbiotroph | Animal Pathogen-Endophyte-Endosymbiont-Epiphyte-\|Undefined Saprotroph\| | Dimorphic Yeast-Yeast | NULL | Probable | As nectar-tap_saprotroph (Põlme et al. 2020); Decay sugar-rich substrates (Põlme et al. 2020); This asexual genus is polyphyletic and contain species that function very differently depend on what they are. Caution should be taken when interpreting guilds for Candida species. | Manolakaki D, et al. 2010. Virulence. 1: 367-375 ((https://doi.org/10.4161/viru.1.5.12796)); Mekbib SB, et al. 2011. Tropical Plant Pathology 36:233-240 ((https://doi.org/10.1590/S1982-56762011000400004)); Põlme S, et al. 2020. Fungal Diversity 105:1-16 ((https://doi.org/10.1007/s13225-020-00466-2)) |
| Otu53580 | 1 | k__Fungi(100);p__Ascomycota(100);c__Saccharomycetes(100);o__Saccharomycetales(100);f__Saccharomycetales_family_Incertae_sedis(100);g__Candida(100);s__Candida_cretensis(100); | Candida | 13 | Pathotroph-Saprotroph-Symbiotroph | Animal Pathogen-Endophyte-Endosymbiont-Epiphyte-\|Undefined Saprotroph\| | Dimorphic Yeast-Yeast | NULL | Probable | As nectar-tap_saprotroph (Põlme et al. 2020); Decay sugar-rich substrates (Põlme et al. 2020); This asexual genus is polyphyletic and contain species that function very differently depend on what they are. Caution should be taken when interpreting guilds for Candida species. | Manolakaki D, et al. 2010. Virulence. 1: 367-375 ((https://doi.org/10.4161/viru.1.5.12796)); Mekbib SB, et al. 2011. Tropical Plant Pathology 36:233-240 ((https://doi.org/10.1590/S1982-56762011000400004)); Põlme S, et al. 2020. Fungal Diversity 105:1-16 ((https://doi.org/10.1007/s13225-020-00466-2)) |
| Otu53582 | 1 | k__Fungi(100);p__Ascomycota(100);c__Saccharomycetes(100);o__Saccharomycetales(100);f__Saccharomycetales_family_Incertae_sedis(100);g__Candida(100);g__Candida_unclassified(100); | Candida | 13 | Pathotroph-Saprotroph-Symbiotroph | Animal Pathogen-Endophyte-Endosymbiont-Epiphyte-\|Undefined Saprotroph\| | Dimorphic Yeast-Yeast | NULL | Probable | As nectar-tap_saprotroph (Põlme et al. 2020); Decay sugar-rich substrates (Põlme et al. 2020); This asexual genus is polyphyletic and contain species that function very differently depend on what they are. Caution should be taken when interpreting guilds for Candida species. | Manolakaki D, et al. 2010. Virulence. 1: 367-375 ((https://doi.org/10.4161/viru.1.5.12796)); Mekbib SB, et al. 2011. Tropical Plant Pathology 36:233-240 ((https://doi.org/10.1590/S1982-56762011000400004)); Põlme S, et al. 2020. Fungal Diversity 105:1-16 ((https://doi.org/10.1007/s13225-020-00466-2)) |
| Otu53583 | 1 | k__Fungi(100);p__Ascomycota(100);c__Saccharomycetes(100);o__Saccharomycetales(100);f__Saccharomycetales_family_Incertae_sedis(100);g__Candida(100);s__Candida_cretensis(100); | Candida | 13 | Pathotroph-Saprotroph-Symbiotroph | Animal Pathogen-Endophyte-Endosymbiont-Epiphyte-\|Undefined Saprotroph\| | Dimorphic Yeast-Yeast | NULL | Probable | As nectar-tap_saprotroph (Põlme et al. 2020); Decay sugar-rich substrates (Põlme et al. 2020); This asexual genus is polyphyletic and contain species that function very differently depend on what they are. Caution should be taken when interpreting guilds for Candida species. | Manolakaki D, et al. 2010. Virulence. 1: 367-375 ((https://doi.org/10.4161/viru.1.5.12796)); Mekbib SB, et al. 2011. Tropical Plant Pathology 36:233-240 ((https://doi.org/10.1590/S1982-56762011000400004)); Põlme S, et al. 2020. Fungal Diversity 105:1-16 ((https://doi.org/10.1007/s13225-020-00466-2)) |
| Otu53586 | 1 | k__Fungi(100);p__Ascomycota(100);c__Saccharomycetes(100);o__Saccharomycetales(100);o__Saccharomycetales_unclassified(100);o__Saccharomycetales_unclassified(100);o__Saccharomycetales_unclassified(100); | Saccharomycetales | 7 | Saprotroph | Undefined Saprotroph | Yeast | NULL | Possible | NULL | Sterkenburg E, et al. 2015. New Phytologist 207:1145-1158 ((https://doi.org/10.1111/nph.13426)) |
| Otu53590 | 1 | k__Fungi(100);p__Ascomycota(100);c__Saccharomycetes(100);o__Saccharomycetales(100);f__Saccharomycetales_family_Incertae_sedis(100);f__Saccharomycetales_family_Incertae_sedis_unclassified(100);f__Saccharomycetales_family_Incertae_sedis_unclassified(100); | Saccharomycetales | 7 | Saprotroph | Undefined Saprotroph | Yeast | NULL | Possible | NULL | Sterkenburg E, et al. 2015. New Phytologist 207:1145-1158 ((https://doi.org/10.1111/nph.13426)) |
| Otu53592 | 1 | k__Fungi(100);p__Ascomycota(100);c__Saccharomycetes(100);o__Saccharomycetales(100);f__Saccharomycetales_family_Incertae_sedis(100);g__Candida(100);g__Candida_unclassified(100); | Candida | 13 | Pathotroph-Saprotroph-Symbiotroph | Animal Pathogen-Endophyte-Endosymbiont-Epiphyte-\|Undefined Saprotroph\| | Dimorphic Yeast-Yeast | NULL | Probable | As nectar-tap_saprotroph (Põlme et al. 2020); Decay sugar-rich substrates (Põlme et al. 2020); This asexual genus is polyphyletic and contain species that function very differently depend on what they are. Caution should be taken when interpreting guilds for Candida species. | Manolakaki D, et al. 2010. Virulence. 1: 367-375 ((https://doi.org/10.4161/viru.1.5.12796)); Mekbib SB, et al. 2011. Tropical Plant Pathology 36:233-240 ((https://doi.org/10.1590/S1982-56762011000400004)); Põlme S, et al. 2020. Fungal Diversity 105:1-16 ((https://doi.org/10.1007/s13225-020-00466-2)) |
| Otu53594 | 1 | k__Fungi(100);p__Ascomycota(100);c__Saccharomycetes(100);o__Saccharomycetales(100);f__Saccharomycetales_family_Incertae_sedis(100);g__Candida(100);g__Candida_unclassified(100); | Candida | 13 | Pathotroph-Saprotroph-Symbiotroph | Animal Pathogen-Endophyte-Endosymbiont-Epiphyte-\|Undefined Saprotroph\| | Dimorphic Yeast-Yeast | NULL | Probable | As nectar-tap_saprotroph (Põlme et al. 2020); Decay sugar-rich substrates (Põlme et al. 2020); This asexual genus is polyphyletic and contain species that function very differently depend on what they are. Caution should be taken when interpreting guilds for Candida species. | Manolakaki D, et al. 2010. Virulence. 1: 367-375 ((https://doi.org/10.4161/viru.1.5.12796)); Mekbib SB, et al. 2011. Tropical Plant Pathology 36:233-240 ((https://doi.org/10.1590/S1982-56762011000400004)); Põlme S, et al. 2020. Fungal Diversity 105:1-16 ((https://doi.org/10.1007/s13225-020-00466-2)) |
| Otu53596 | 1 | k__Fungi(100);p__Ascomycota(100);c__Saccharomycetes(100);o__Saccharomycetales(100);f__Saccharomycetales_family_Incertae_sedis(100);g__Candida(100);g__Candida_unclassified(100); | Candida | 13 | Pathotroph-Saprotroph-Symbiotroph | Animal Pathogen-Endophyte-Endosymbiont-Epiphyte-\|Undefined Saprotroph\| | Dimorphic Yeast-Yeast | NULL | Probable | As nectar-tap_saprotroph (Põlme et al. 2020); Decay sugar-rich substrates (Põlme et al. 2020); This asexual genus is polyphyletic and contain species that function very differently depend on what they are. Caution should be taken when interpreting guilds for Candida species. | Manolakaki D, et al. 2010. Virulence. 1: 367-375 ((https://doi.org/10.4161/viru.1.5.12796)); Mekbib SB, et al. 2011. Tropical Plant Pathology 36:233-240 ((https://doi.org/10.1590/S1982-56762011000400004)); Põlme S, et al. 2020. Fungal Diversity 105:1-16 ((https://doi.org/10.1007/s13225-020-00466-2)) |
| Otu53598 | 1 | k__Fungi(100);p__Ascomycota(100);c__Saccharomycetes(100);o__Saccharomycetales(100);f__Saccharomycetales_family_Incertae_sedis(100);f__Saccharomycetales_family_Incertae_sedis_unclassified(100);f__Saccharomycetales_family_Incertae_sedis_unclassified(100); | Saccharomycetales | 7 | Saprotroph | Undefined Saprotroph | Yeast | NULL | Possible | NULL | Sterkenburg E, et al. 2015. New Phytologist 207:1145-1158 ((https://doi.org/10.1111/nph.13426)) |
| Otu53600 | 1 | k__Fungi(100);p__Ascomycota(100);c__Saccharomycetes(100);o__Saccharomycetales(100);f__Saccharomycetales_family_Incertae_sedis(100);g__Candida(100);g__Candida_unclassified(100); | Candida | 13 | Pathotroph-Saprotroph-Symbiotroph | Animal Pathogen-Endophyte-Endosymbiont-Epiphyte-\|Undefined Saprotroph\| | Dimorphic Yeast-Yeast | NULL | Probable | As nectar-tap_saprotroph (Põlme et al. 2020); Decay sugar-rich substrates (Põlme et al. 2020); This asexual genus is polyphyletic and contain species that function very differently depend on what they are. Caution should be taken when interpreting guilds for Candida species. | Manolakaki D, et al. 2010. Virulence. 1: 367-375 ((https://doi.org/10.4161/viru.1.5.12796)); Mekbib SB, et al. 2011. Tropical Plant Pathology 36:233-240 ((https://doi.org/10.1590/S1982-56762011000400004)); Põlme S, et al. 2020. Fungal Diversity 105:1-16 ((https://doi.org/10.1007/s13225-020-00466-2)) |
| Otu53602 | 1 | k__Fungi(100);p__Ascomycota(100);c__Saccharomycetes(100);o__Saccharomycetales(100);f__Saccharomycetales_family_Incertae_sedis(100);g__Candida(100);g__Candida_unclassified(100); | Candida | 13 | Pathotroph-Saprotroph-Symbiotroph | Animal Pathogen-Endophyte-Endosymbiont-Epiphyte-\|Undefined Saprotroph\| | Dimorphic Yeast-Yeast | NULL | Probable | As nectar-tap_saprotroph (Põlme et al. 2020); Decay sugar-rich substrates (Põlme et al. 2020); This asexual genus is polyphyletic and contain species that function very differently depend on what they are. Caution should be taken when interpreting guilds for Candida species. | Manolakaki D, et al. 2010. Virulence. 1: 367-375 ((https://doi.org/10.4161/viru.1.5.12796)); Mekbib SB, et al. 2011. Tropical Plant Pathology 36:233-240 ((https://doi.org/10.1590/S1982-56762011000400004)); Põlme S, et al. 2020. Fungal Diversity 105:1-16 ((https://doi.org/10.1007/s13225-020-00466-2)) |
| Otu53608 | 1 | k__Fungi(100);p__Ascomycota(100);c__Saccharomycetes(100);o__Saccharomycetales(100);f__Saccharomycetales_family_Incertae_sedis(100);g__Candida(100);g__Candida_unclassified(100); | Candida | 13 | Pathotroph-Saprotroph-Symbiotroph | Animal Pathogen-Endophyte-Endosymbiont-Epiphyte-\|Undefined Saprotroph\| | Dimorphic Yeast-Yeast | NULL | Probable | As nectar-tap_saprotroph (Põlme et al. 2020); Decay sugar-rich substrates (Põlme et al. 2020); This asexual genus is polyphyletic and contain species that function very differently depend on what they are. Caution should be taken when interpreting guilds for Candida species. | Manolakaki D, et al. 2010. Virulence. 1: 367-375 ((https://doi.org/10.4161/viru.1.5.12796)); Mekbib SB, et al. 2011. Tropical Plant Pathology 36:233-240 ((https://doi.org/10.1590/S1982-56762011000400004)); Põlme S, et al. 2020. Fungal Diversity 105:1-16 ((https://doi.org/10.1007/s13225-020-00466-2)) |
| Otu53609 | 1 | k__Fungi(100);p__Ascomycota(100);c__Saccharomycetes(100);o__Saccharomycetales(100);f__Metschnikowiaceae(100);g__Metschnikowia(100);s__Metschnikowia_pulcherrima(100); | Metschnikowia | 13 | Saprotroph | \|Undefined Saprotroph\| | Yeast | NULL | Probable | As nectar-tap_saprotroph (Põlme et al. 2020); Decay sugar-rich substrates (Põlme et al. 2020) | Põlme S, et al. 2020. Fungal Diversity 105:1-16 ((https://doi.org/10.1007/s13225-020-00466-2)) |
| Otu53610 | 1 | k__Fungi(100);p__Ascomycota(100);c__Saccharomycetes(100);o__Saccharomycetales(100);f__Metschnikowiaceae(100);g__Metschnikowia(100);s__Metschnikowia_pulcherrima(100); | Metschnikowia | 13 | Saprotroph | \|Undefined Saprotroph\| | Yeast | NULL | Probable | As nectar-tap_saprotroph (Põlme et al. 2020); Decay sugar-rich substrates (Põlme et al. 2020) | Põlme S, et al. 2020. Fungal Diversity 105:1-16 ((https://doi.org/10.1007/s13225-020-00466-2)) |
| Otu53611 | 1 | k__Fungi(100);p__Ascomycota(100);c__Saccharomycetes(100);o__Saccharomycetales(100);f__Saccharomycetales_family_Incertae_sedis(100);g__Candida(100);g__Candida_unclassified(100); | Candida | 13 | Pathotroph-Saprotroph-Symbiotroph | Animal Pathogen-Endophyte-Endosymbiont-Epiphyte-\|Undefined Saprotroph\| | Dimorphic Yeast-Yeast | NULL | Probable | As nectar-tap_saprotroph (Põlme et al. 2020); Decay sugar-rich substrates (Põlme et al. 2020); This asexual genus is polyphyletic and contain species that function very differently depend on what they are. Caution should be taken when interpreting guilds for Candida species. | Manolakaki D, et al. 2010. Virulence. 1: 367-375 ((https://doi.org/10.4161/viru.1.5.12796)); Mekbib SB, et al. 2011. Tropical Plant Pathology 36:233-240 ((https://doi.org/10.1590/S1982-56762011000400004)); Põlme S, et al. 2020. Fungal Diversity 105:1-16 ((https://doi.org/10.1007/s13225-020-00466-2)) |
| Otu53613 | 1 | k__Fungi(100);p__Ascomycota(100);c__Saccharomycetes(100);o__Saccharomycetales(100);f__Saccharomycetales_family_Incertae_sedis(100);f__Saccharomycetales_family_Incertae_sedis_unclassified(100);f__Saccharomycetales_family_Incertae_sedis_unclassified(100); | Saccharomycetales | 7 | Saprotroph | Undefined Saprotroph | Yeast | NULL | Possible | NULL | Sterkenburg E, et al. 2015. New Phytologist 207:1145-1158 ((https://doi.org/10.1111/nph.13426)) |
| Otu53615 | 1 | k__Fungi(100);p__Ascomycota(100);c__Saccharomycetes(100);o__Saccharomycetales(100);f__Saccharomycetales_family_Incertae_sedis(100);g__Candida(100);g__Candida_unclassified(100); | Candida | 13 | Pathotroph-Saprotroph-Symbiotroph | Animal Pathogen-Endophyte-Endosymbiont-Epiphyte-\|Undefined Saprotroph\| | Dimorphic Yeast-Yeast | NULL | Probable | As nectar-tap_saprotroph (Põlme et al. 2020); Decay sugar-rich substrates (Põlme et al. 2020); This asexual genus is polyphyletic and contain species that function very differently depend on what they are. Caution should be taken when interpreting guilds for Candida species. | Manolakaki D, et al. 2010. Virulence. 1: 367-375 ((https://doi.org/10.4161/viru.1.5.12796)); Mekbib SB, et al. 2011. Tropical Plant Pathology 36:233-240 ((https://doi.org/10.1590/S1982-56762011000400004)); Põlme S, et al. 2020. Fungal Diversity 105:1-16 ((https://doi.org/10.1007/s13225-020-00466-2)) |
| Otu53616 | 1 | k__Fungi(100);p__Ascomycota(100);c__Saccharomycetes(100);o__Saccharomycetales(100);f__Saccharomycetales_family_Incertae_sedis(100);g__Candida(100);g__Candida_unclassified(100); | Candida | 13 | Pathotroph-Saprotroph-Symbiotroph | Animal Pathogen-Endophyte-Endosymbiont-Epiphyte-\|Undefined Saprotroph\| | Dimorphic Yeast-Yeast | NULL | Probable | As nectar-tap_saprotroph (Põlme et al. 2020); Decay sugar-rich substrates (Põlme et al. 2020); This asexual genus is polyphyletic and contain species that function very differently depend on what they are. Caution should be taken when interpreting guilds for Candida species. | Manolakaki D, et al. 2010. Virulence. 1: 367-375 ((https://doi.org/10.4161/viru.1.5.12796)); Mekbib SB, et al. 2011. Tropical Plant Pathology 36:233-240 ((https://doi.org/10.1590/S1982-56762011000400004)); Põlme S, et al. 2020. Fungal Diversity 105:1-16 ((https://doi.org/10.1007/s13225-020-00466-2)) |
| Otu53618 | 1 | k__Fungi(100);p__Ascomycota(100);c__Saccharomycetes(100);o__Saccharomycetales(100);f__Saccharomycetales_family_Incertae_sedis(100);f__Saccharomycetales_family_Incertae_sedis_unclassified(100);f__Saccharomycetales_family_Incertae_sedis_unclassified(100); | Saccharomycetales | 7 | Saprotroph | Undefined Saprotroph | Yeast | NULL | Possible | NULL | Sterkenburg E, et al. 2015. New Phytologist 207:1145-1158 ((https://doi.org/10.1111/nph.13426)) |
| Otu53620 | 1 | k__Fungi(100);p__Ascomycota(100);c__Saccharomycetes(100);o__Saccharomycetales(100);f__Saccharomycetales_family_Incertae_sedis(100);g__Candida(100);g__Candida_unclassified(100); | Candida | 13 | Pathotroph-Saprotroph-Symbiotroph | Animal Pathogen-Endophyte-Endosymbiont-Epiphyte-\|Undefined Saprotroph\| | Dimorphic Yeast-Yeast | NULL | Probable | As nectar-tap_saprotroph (Põlme et al. 2020); Decay sugar-rich substrates (Põlme et al. 2020); This asexual genus is polyphyletic and contain species that function very differently depend on what they are. Caution should be taken when interpreting guilds for Candida species. | Manolakaki D, et al. 2010. Virulence. 1: 367-375 ((https://doi.org/10.4161/viru.1.5.12796)); Mekbib SB, et al. 2011. Tropical Plant Pathology 36:233-240 ((https://doi.org/10.1590/S1982-56762011000400004)); Põlme S, et al. 2020. Fungal Diversity 105:1-16 ((https://doi.org/10.1007/s13225-020-00466-2)) |
| Otu53621 | 1 | k__Fungi(100);p__Ascomycota(100);c__Saccharomycetes(100);o__Saccharomycetales(100);f__Saccharomycetales_family_Incertae_sedis(100);g__Candida(100);g__Candida_unclassified(100); | Candida | 13 | Pathotroph-Saprotroph-Symbiotroph | Animal Pathogen-Endophyte-Endosymbiont-Epiphyte-\|Undefined Saprotroph\| | Dimorphic Yeast-Yeast | NULL | Probable | As nectar-tap_saprotroph (Põlme et al. 2020); Decay sugar-rich substrates (Põlme et al. 2020); This asexual genus is polyphyletic and contain species that function very differently depend on what they are. Caution should be taken when interpreting guilds for Candida species. | Manolakaki D, et al. 2010. Virulence. 1: 367-375 ((https://doi.org/10.4161/viru.1.5.12796)); Mekbib SB, et al. 2011. Tropical Plant Pathology 36:233-240 ((https://doi.org/10.1590/S1982-56762011000400004)); Põlme S, et al. 2020. Fungal Diversity 105:1-16 ((https://doi.org/10.1007/s13225-020-00466-2)) |
| Otu53622 | 1 | k__Fungi(100);p__Ascomycota(100);c__Saccharomycetes(100);o__Saccharomycetales(100);f__Saccharomycetales_family_Incertae_sedis(100);g__Candida(100);g__Candida_unclassified(100); | Candida | 13 | Pathotroph-Saprotroph-Symbiotroph | Animal Pathogen-Endophyte-Endosymbiont-Epiphyte-\|Undefined Saprotroph\| | Dimorphic Yeast-Yeast | NULL | Probable | As nectar-tap_saprotroph (Põlme et al. 2020); Decay sugar-rich substrates (Põlme et al. 2020); This asexual genus is polyphyletic and contain species that function very differently depend on what they are. Caution should be taken when interpreting guilds for Candida species. | Manolakaki D, et al. 2010. Virulence. 1: 367-375 ((https://doi.org/10.4161/viru.1.5.12796)); Mekbib SB, et al. 2011. Tropical Plant Pathology 36:233-240 ((https://doi.org/10.1590/S1982-56762011000400004)); Põlme S, et al. 2020. Fungal Diversity 105:1-16 ((https://doi.org/10.1007/s13225-020-00466-2)) |
| Otu53624 | 1 | k__Fungi(100);p__Ascomycota(100);c__Saccharomycetes(100);o__Saccharomycetales(100);f__Metschnikowiaceae(100);g__Metschnikowia(100);s__Metschnikowia_pulcherrima(100); | Metschnikowia | 13 | Saprotroph | \|Undefined Saprotroph\| | Yeast | NULL | Probable | As nectar-tap_saprotroph (Põlme et al. 2020); Decay sugar-rich substrates (Põlme et al. 2020) | Põlme S, et al. 2020. Fungal Diversity 105:1-16 ((https://doi.org/10.1007/s13225-020-00466-2)) |
| Otu53625 | 1 | k__Fungi(100);p__Ascomycota(100);c__Saccharomycetes(100);o__Saccharomycetales(100);f__Saccharomycetales_family_Incertae_sedis(100);g__Candida(100);g__Candida_unclassified(100); | Candida | 13 | Pathotroph-Saprotroph-Symbiotroph | Animal Pathogen-Endophyte-Endosymbiont-Epiphyte-\|Undefined Saprotroph\| | Dimorphic Yeast-Yeast | NULL | Probable | As nectar-tap_saprotroph (Põlme et al. 2020); Decay sugar-rich substrates (Põlme et al. 2020); This asexual genus is polyphyletic and contain species that function very differently depend on what they are. Caution should be taken when interpreting guilds for Candida species. | Manolakaki D, et al. 2010. Virulence. 1: 367-375 ((https://doi.org/10.4161/viru.1.5.12796)); Mekbib SB, et al. 2011. Tropical Plant Pathology 36:233-240 ((https://doi.org/10.1590/S1982-56762011000400004)); Põlme S, et al. 2020. Fungal Diversity 105:1-16 ((https://doi.org/10.1007/s13225-020-00466-2)) |
| Otu53628 | 1 | k__Fungi(100);p__Ascomycota(100);c__Saccharomycetes(100);o__Saccharomycetales(100);f__Saccharomycetales_family_Incertae_sedis(100);f__Saccharomycetales_family_Incertae_sedis_unclassified(100);f__Saccharomycetales_family_Incertae_sedis_unclassified(100); | Saccharomycetales | 7 | Saprotroph | Undefined Saprotroph | Yeast | NULL | Possible | NULL | Sterkenburg E, et al. 2015. New Phytologist 207:1145-1158 ((https://doi.org/10.1111/nph.13426)) |
| Otu53629 | 1 | k__Fungi(100);p__Ascomycota(100);c__Saccharomycetes(100);o__Saccharomycetales(100);f__Metschnikowiaceae(100);g__Metschnikowia(100);g__Metschnikowia_unclassified(100); | Metschnikowia | 13 | Saprotroph | \|Undefined Saprotroph\| | Yeast | NULL | Probable | As nectar-tap_saprotroph (Põlme et al. 2020); Decay sugar-rich substrates (Põlme et al. 2020) | Põlme S, et al. 2020. Fungal Diversity 105:1-16 ((https://doi.org/10.1007/s13225-020-00466-2)) |
| Otu53631 | 1 | k__Fungi(100);p__Ascomycota(100);c__Saccharomycetes(100);o__Saccharomycetales(100);f__Metschnikowiaceae(100);g__Metschnikowia(100);s__Metschnikowia_pulcherrima(100); | Metschnikowia | 13 | Saprotroph | \|Undefined Saprotroph\| | Yeast | NULL | Probable | As nectar-tap_saprotroph (Põlme et al. 2020); Decay sugar-rich substrates (Põlme et al. 2020) | Põlme S, et al. 2020. Fungal Diversity 105:1-16 ((https://doi.org/10.1007/s13225-020-00466-2)) |
| Otu53635 | 1 | k__Fungi(100);p__Ascomycota(100);c__Saccharomycetes(100);o__Saccharomycetales(100);f__Saccharomycetaceae(100);g__Kazachstania(100);s__Kazachstania_intestinalis(100); | Kazachstania | 13 | Saprotroph | \|Undefined Saprotroph\| | Yeast | NULL | Probable | As nectar-tap_saprotroph (Põlme et al. 2020); Decay sugar-rich substrates (Põlme et al. 2020) | Põlme S, et al. 2020. Fungal Diversity 105:1-16 ((https://doi.org/10.1007/s13225-020-00466-2)) |
| Otu53636 | 1 | k__Fungi(100);p__Ascomycota(100);c__Saccharomycetes(100);o__Saccharomycetales(100);f__Metschnikowiaceae(100);g__Metschnikowia(100);s__Metschnikowia_pulcherrima(100); | Metschnikowia | 13 | Saprotroph | \|Undefined Saprotroph\| | Yeast | NULL | Probable | As nectar-tap_saprotroph (Põlme et al. 2020); Decay sugar-rich substrates (Põlme et al. 2020) | Põlme S, et al. 2020. Fungal Diversity 105:1-16 ((https://doi.org/10.1007/s13225-020-00466-2)) |
| Otu53638 | 1 | k__Fungi(100);p__Ascomycota(100);c__Saccharomycetes(100);o__Saccharomycetales(100);f__Metschnikowiaceae(100);g__Metschnikowia(100);g__Metschnikowia_unclassified(100); | Metschnikowia | 13 | Saprotroph | \|Undefined Saprotroph\| | Yeast | NULL | Probable | As nectar-tap_saprotroph (Põlme et al. 2020); Decay sugar-rich substrates (Põlme et al. 2020) | Põlme S, et al. 2020. Fungal Diversity 105:1-16 ((https://doi.org/10.1007/s13225-020-00466-2)) |
| Otu53641 | 1 | k__Fungi(100);p__Ascomycota(100);c__Saccharomycetes(100);o__Saccharomycetales(100);f__Metschnikowiaceae(100);g__Metschnikowia(100);g__Metschnikowia_unclassified(100); | Metschnikowia | 13 | Saprotroph | \|Undefined Saprotroph\| | Yeast | NULL | Probable | As nectar-tap_saprotroph (Põlme et al. 2020); Decay sugar-rich substrates (Põlme et al. 2020) | Põlme S, et al. 2020. Fungal Diversity 105:1-16 ((https://doi.org/10.1007/s13225-020-00466-2)) |
| Otu53642 | 1 | k__Fungi(100);p__Ascomycota(100);c__Saccharomycetes(100);o__Saccharomycetales(100);f__Metschnikowiaceae(100);g__Metschnikowia(100);s__Metschnikowia_pulcherrima(100); | Metschnikowia | 13 | Saprotroph | \|Undefined Saprotroph\| | Yeast | NULL | Probable | As nectar-tap_saprotroph (Põlme et al. 2020); Decay sugar-rich substrates (Põlme et al. 2020) | Põlme S, et al. 2020. Fungal Diversity 105:1-16 ((https://doi.org/10.1007/s13225-020-00466-2)) |
| Otu53644 | 1 | k__Fungi(100);p__Ascomycota(100);c__Saccharomycetes(100);o__Saccharomycetales(100);f__Metschnikowiaceae(100);g__Metschnikowia(100);s__Metschnikowia_pulcherrima(100); | Metschnikowia | 13 | Saprotroph | \|Undefined Saprotroph\| | Yeast | NULL | Probable | As nectar-tap_saprotroph (Põlme et al. 2020); Decay sugar-rich substrates (Põlme et al. 2020) | Põlme S, et al. 2020. Fungal Diversity 105:1-16 ((https://doi.org/10.1007/s13225-020-00466-2)) |
| Otu53645 | 1 | k__Fungi(100);p__Ascomycota(100);c__Saccharomycetes(100);o__Saccharomycetales(100);f__Saccharomycetales_family_Incertae_sedis(100);g__Candida(100);g__Candida_unclassified(100); | Candida | 13 | Pathotroph-Saprotroph-Symbiotroph | Animal Pathogen-Endophyte-Endosymbiont-Epiphyte-\|Undefined Saprotroph\| | Dimorphic Yeast-Yeast | NULL | Probable | As nectar-tap_saprotroph (Põlme et al. 2020); Decay sugar-rich substrates (Põlme et al. 2020); This asexual genus is polyphyletic and contain species that function very differently depend on what they are. Caution should be taken when interpreting guilds for Candida species. | Manolakaki D, et al. 2010. Virulence. 1: 367-375 ((https://doi.org/10.4161/viru.1.5.12796)); Mekbib SB, et al. 2011. Tropical Plant Pathology 36:233-240 ((https://doi.org/10.1590/S1982-56762011000400004)); Põlme S, et al. 2020. Fungal Diversity 105:1-16 ((https://doi.org/10.1007/s13225-020-00466-2)) |
| Otu53646 | 1 | k__Fungi(100);p__Ascomycota(100);c__Saccharomycetes(100);o__Saccharomycetales(100);f__Saccharomycetales_family_Incertae_sedis(100);f__Saccharomycetales_family_Incertae_sedis_unclassified(100);f__Saccharomycetales_family_Incertae_sedis_unclassified(100); | Saccharomycetales | 7 | Saprotroph | Undefined Saprotroph | Yeast | NULL | Possible | NULL | Sterkenburg E, et al. 2015. New Phytologist 207:1145-1158 ((https://doi.org/10.1111/nph.13426)) |
| Otu53648 | 1 | k__Fungi(100);p__Ascomycota(100);c__Saccharomycetes(100);o__Saccharomycetales(100);f__Saccharomycetales_family_Incertae_sedis(100);g__Candida(100);g__Candida_unclassified(100); | Candida | 13 | Pathotroph-Saprotroph-Symbiotroph | Animal Pathogen-Endophyte-Endosymbiont-Epiphyte-\|Undefined Saprotroph\| | Dimorphic Yeast-Yeast | NULL | Probable | As nectar-tap_saprotroph (Põlme et al. 2020); Decay sugar-rich substrates (Põlme et al. 2020); This asexual genus is polyphyletic and contain species that function very differently depend on what they are. Caution should be taken when interpreting guilds for Candida species. | Manolakaki D, et al. 2010. Virulence. 1: 367-375 ((https://doi.org/10.4161/viru.1.5.12796)); Mekbib SB, et al. 2011. Tropical Plant Pathology 36:233-240 ((https://doi.org/10.1590/S1982-56762011000400004)); Põlme S, et al. 2020. Fungal Diversity 105:1-16 ((https://doi.org/10.1007/s13225-020-00466-2)) |
| Otu53649 | 1 | k__Fungi(100);p__Ascomycota(100);c__Saccharomycetes(100);o__Saccharomycetales(100);f__Metschnikowiaceae(100);g__Metschnikowia(100);s__Metschnikowia_pulcherrima(100); | Metschnikowia | 13 | Saprotroph | \|Undefined Saprotroph\| | Yeast | NULL | Probable | As nectar-tap_saprotroph (Põlme et al. 2020); Decay sugar-rich substrates (Põlme et al. 2020) | Põlme S, et al. 2020. Fungal Diversity 105:1-16 ((https://doi.org/10.1007/s13225-020-00466-2)) |
| Otu53650 | 1 | k__Fungi(100);p__Ascomycota(100);c__Saccharomycetes(100);o__Saccharomycetales(100);f__Metschnikowiaceae(100);g__Metschnikowia(100);s__Metschnikowia_pulcherrima(100); | Metschnikowia | 13 | Saprotroph | \|Undefined Saprotroph\| | Yeast | NULL | Probable | As nectar-tap_saprotroph (Põlme et al. 2020); Decay sugar-rich substrates (Põlme et al. 2020) | Põlme S, et al. 2020. Fungal Diversity 105:1-16 ((https://doi.org/10.1007/s13225-020-00466-2)) |
| Otu53652 | 1 | k__Fungi(100);p__Ascomycota(100);c__Saccharomycetes(100);o__Saccharomycetales(100);f__Metschnikowiaceae(100);g__Metschnikowia(100);s__Metschnikowia_pulcherrima(100); | Metschnikowia | 13 | Saprotroph | \|Undefined Saprotroph\| | Yeast | NULL | Probable | As nectar-tap_saprotroph (Põlme et al. 2020); Decay sugar-rich substrates (Põlme et al. 2020) | Põlme S, et al. 2020. Fungal Diversity 105:1-16 ((https://doi.org/10.1007/s13225-020-00466-2)) |
| Otu53653 | 1 | k__Fungi(100);p__Ascomycota(100);c__Saccharomycetes(100);o__Saccharomycetales(100);f__Saccharomycetales_family_Incertae_sedis(100);g__Candida(100);s__Candida_cretensis(100); | Candida | 13 | Pathotroph-Saprotroph-Symbiotroph | Animal Pathogen-Endophyte-Endosymbiont-Epiphyte-\|Undefined Saprotroph\| | Dimorphic Yeast-Yeast | NULL | Probable | As nectar-tap_saprotroph (Põlme et al. 2020); Decay sugar-rich substrates (Põlme et al. 2020); This asexual genus is polyphyletic and contain species that function very differently depend on what they are. Caution should be taken when interpreting guilds for Candida species. | Manolakaki D, et al. 2010. Virulence. 1: 367-375 ((https://doi.org/10.4161/viru.1.5.12796)); Mekbib SB, et al. 2011. Tropical Plant Pathology 36:233-240 ((https://doi.org/10.1590/S1982-56762011000400004)); Põlme S, et al. 2020. Fungal Diversity 105:1-16 ((https://doi.org/10.1007/s13225-020-00466-2)) |
| Otu53655 | 1 | k__Fungi(100);p__Ascomycota(100);c__Saccharomycetes(100);o__Saccharomycetales(100);f__Metschnikowiaceae(100);g__Metschnikowia(100);s__Metschnikowia_pulcherrima(100); | Metschnikowia | 13 | Saprotroph | \|Undefined Saprotroph\| | Yeast | NULL | Probable | As nectar-tap_saprotroph (Põlme et al. 2020); Decay sugar-rich substrates (Põlme et al. 2020) | Põlme S, et al. 2020. Fungal Diversity 105:1-16 ((https://doi.org/10.1007/s13225-020-00466-2)) |
| Otu53656 | 1 | k__Fungi(100);p__Ascomycota(100);c__Saccharomycetes(100);o__Saccharomycetales(100);f__Saccharomycetales_family_Incertae_sedis(100);g__Candida(100);g__Candida_unclassified(100); | Candida | 13 | Pathotroph-Saprotroph-Symbiotroph | Animal Pathogen-Endophyte-Endosymbiont-Epiphyte-\|Undefined Saprotroph\| | Dimorphic Yeast-Yeast | NULL | Probable | As nectar-tap_saprotroph (Põlme et al. 2020); Decay sugar-rich substrates (Põlme et al. 2020); This asexual genus is polyphyletic and contain species that function very differently depend on what they are. Caution should be taken when interpreting guilds for Candida species. | Manolakaki D, et al. 2010. Virulence. 1: 367-375 ((https://doi.org/10.4161/viru.1.5.12796)); Mekbib SB, et al. 2011. Tropical Plant Pathology 36:233-240 ((https://doi.org/10.1590/S1982-56762011000400004)); Põlme S, et al. 2020. Fungal Diversity 105:1-16 ((https://doi.org/10.1007/s13225-020-00466-2)) |
| Otu53657 | 1 | k__Fungi(100);p__Ascomycota(100);c__Saccharomycetes(100);o__Saccharomycetales(100);f__Metschnikowiaceae(100);g__Metschnikowia(100);s__Metschnikowia_pulcherrima(100); | Metschnikowia | 13 | Saprotroph | \|Undefined Saprotroph\| | Yeast | NULL | Probable | As nectar-tap_saprotroph (Põlme et al. 2020); Decay sugar-rich substrates (Põlme et al. 2020) | Põlme S, et al. 2020. Fungal Diversity 105:1-16 ((https://doi.org/10.1007/s13225-020-00466-2)) |
| Otu53661 | 1 | k__Fungi(100);p__Ascomycota(100);c__Saccharomycetes(100);o__Saccharomycetales(100);f__Saccharomycetales_family_Incertae_sedis(100);g__Candida(100);g__Candida_unclassified(100); | Candida | 13 | Pathotroph-Saprotroph-Symbiotroph | Animal Pathogen-Endophyte-Endosymbiont-Epiphyte-\|Undefined Saprotroph\| | Dimorphic Yeast-Yeast | NULL | Probable | As nectar-tap_saprotroph (Põlme et al. 2020); Decay sugar-rich substrates (Põlme et al. 2020); This asexual genus is polyphyletic and contain species that function very differently depend on what they are. Caution should be taken when interpreting guilds for Candida species. | Manolakaki D, et al. 2010. Virulence. 1: 367-375 ((https://doi.org/10.4161/viru.1.5.12796)); Mekbib SB, et al. 2011. Tropical Plant Pathology 36:233-240 ((https://doi.org/10.1590/S1982-56762011000400004)); Põlme S, et al. 2020. Fungal Diversity 105:1-16 ((https://doi.org/10.1007/s13225-020-00466-2)) |
| Otu53663 | 1 | k__Fungi(100);p__Ascomycota(100);c__Saccharomycetes(100);o__Saccharomycetales(100);f__Metschnikowiaceae(100);g__Metschnikowia(100);s__Metschnikowia_pulcherrima(100); | Metschnikowia | 13 | Saprotroph | \|Undefined Saprotroph\| | Yeast | NULL | Probable | As nectar-tap_saprotroph (Põlme et al. 2020); Decay sugar-rich substrates (Põlme et al. 2020) | Põlme S, et al. 2020. Fungal Diversity 105:1-16 ((https://doi.org/10.1007/s13225-020-00466-2)) |
| Otu53665 | 1 | k__Fungi(100);p__Ascomycota(100);c__Saccharomycetes(100);o__Saccharomycetales(100);f__Metschnikowiaceae(100);g__Metschnikowia(100);s__Metschnikowia_pulcherrima(100); | Metschnikowia | 13 | Saprotroph | \|Undefined Saprotroph\| | Yeast | NULL | Probable | As nectar-tap_saprotroph (Põlme et al. 2020); Decay sugar-rich substrates (Põlme et al. 2020) | Põlme S, et al. 2020. Fungal Diversity 105:1-16 ((https://doi.org/10.1007/s13225-020-00466-2)) |
| Otu53667 | 1 | k__Fungi(100);p__Ascomycota(100);c__Saccharomycetes(100);o__Saccharomycetales(100);f__Metschnikowiaceae(100);g__Metschnikowia(100);g__Metschnikowia_unclassified(100); | Metschnikowia | 13 | Saprotroph | \|Undefined Saprotroph\| | Yeast | NULL | Probable | As nectar-tap_saprotroph (Põlme et al. 2020); Decay sugar-rich substrates (Põlme et al. 2020) | Põlme S, et al. 2020. Fungal Diversity 105:1-16 ((https://doi.org/10.1007/s13225-020-00466-2)) |
| Otu53668 | 1 | k__Fungi(100);p__Ascomycota(100);c__Saccharomycetes(100);o__Saccharomycetales(100);f__Metschnikowiaceae(100);g__Metschnikowia(100);s__Metschnikowia_pulcherrima(100); | Metschnikowia | 13 | Saprotroph | \|Undefined Saprotroph\| | Yeast | NULL | Probable | As nectar-tap_saprotroph (Põlme et al. 2020); Decay sugar-rich substrates (Põlme et al. 2020) | Põlme S, et al. 2020. Fungal Diversity 105:1-16 ((https://doi.org/10.1007/s13225-020-00466-2)) |
| Otu53669 | 1 | k__Fungi(100);p__Ascomycota(100);c__Saccharomycetes(100);o__Saccharomycetales(100);f__Metschnikowiaceae(100);g__Metschnikowia(100);s__Metschnikowia_pulcherrima(100); | Metschnikowia | 13 | Saprotroph | \|Undefined Saprotroph\| | Yeast | NULL | Probable | As nectar-tap_saprotroph (Põlme et al. 2020); Decay sugar-rich substrates (Põlme et al. 2020) | Põlme S, et al. 2020. Fungal Diversity 105:1-16 ((https://doi.org/10.1007/s13225-020-00466-2)) |
| Otu53671 | 1 | k__Fungi(100);p__Ascomycota(100);c__Saccharomycetes(100);o__Saccharomycetales(100);f__Metschnikowiaceae(100);g__Metschnikowia(100);s__Metschnikowia_pulcherrima(100); | Metschnikowia | 13 | Saprotroph | \|Undefined Saprotroph\| | Yeast | NULL | Probable | As nectar-tap_saprotroph (Põlme et al. 2020); Decay sugar-rich substrates (Põlme et al. 2020) | Põlme S, et al. 2020. Fungal Diversity 105:1-16 ((https://doi.org/10.1007/s13225-020-00466-2)) |
| Otu53672 | 1 | k__Fungi(100);p__Ascomycota(100);c__Saccharomycetes(100);o__Saccharomycetales(100);f__Metschnikowiaceae(100);g__Metschnikowia(100);g__Metschnikowia_unclassified(100); | Metschnikowia | 13 | Saprotroph | \|Undefined Saprotroph\| | Yeast | NULL | Probable | As nectar-tap_saprotroph (Põlme et al. 2020); Decay sugar-rich substrates (Põlme et al. 2020) | Põlme S, et al. 2020. Fungal Diversity 105:1-16 ((https://doi.org/10.1007/s13225-020-00466-2)) |
| Otu53673 | 1 | k__Fungi(100);p__Ascomycota(100);c__Saccharomycetes(100);o__Saccharomycetales(100);f__Saccharomycetales_family_Incertae_sedis(100);f__Saccharomycetales_family_Incertae_sedis_unclassified(100);f__Saccharomycetales_family_Incertae_sedis_unclassified(100); | Saccharomycetales | 7 | Saprotroph | Undefined Saprotroph | Yeast | NULL | Possible | NULL | Sterkenburg E, et al. 2015. New Phytologist 207:1145-1158 ((https://doi.org/10.1111/nph.13426)) |
| Otu53676 | 1 | k__Fungi(100);p__Ascomycota(100);c__Saccharomycetes(100);o__Saccharomycetales(100);f__Saccharomycetales_family_Incertae_sedis(100);g__Candida(100);g__Candida_unclassified(100); | Candida | 13 | Pathotroph-Saprotroph-Symbiotroph | Animal Pathogen-Endophyte-Endosymbiont-Epiphyte-\|Undefined Saprotroph\| | Dimorphic Yeast-Yeast | NULL | Probable | As nectar-tap_saprotroph (Põlme et al. 2020); Decay sugar-rich substrates (Põlme et al. 2020); This asexual genus is polyphyletic and contain species that function very differently depend on what they are. Caution should be taken when interpreting guilds for Candida species. | Manolakaki D, et al. 2010. Virulence. 1: 367-375 ((https://doi.org/10.4161/viru.1.5.12796)); Mekbib SB, et al. 2011. Tropical Plant Pathology 36:233-240 ((https://doi.org/10.1590/S1982-56762011000400004)); Põlme S, et al. 2020. Fungal Diversity 105:1-16 ((https://doi.org/10.1007/s13225-020-00466-2)) |
| Otu53680 | 1 | k__Fungi(100);p__Ascomycota(100);c__Saccharomycetes(100);o__Saccharomycetales(100);f__Saccharomycetales_family_Incertae_sedis(100);g__Candida(100);g__Candida_unclassified(100); | Candida | 13 | Pathotroph-Saprotroph-Symbiotroph | Animal Pathogen-Endophyte-Endosymbiont-Epiphyte-\|Undefined Saprotroph\| | Dimorphic Yeast-Yeast | NULL | Probable | As nectar-tap_saprotroph (Põlme et al. 2020); Decay sugar-rich substrates (Põlme et al. 2020); This asexual genus is polyphyletic and contain species that function very differently depend on what they are. Caution should be taken when interpreting guilds for Candida species. | Manolakaki D, et al. 2010. Virulence. 1: 367-375 ((https://doi.org/10.4161/viru.1.5.12796)); Mekbib SB, et al. 2011. Tropical Plant Pathology 36:233-240 ((https://doi.org/10.1590/S1982-56762011000400004)); Põlme S, et al. 2020. Fungal Diversity 105:1-16 ((https://doi.org/10.1007/s13225-020-00466-2)) |
| Otu53681 | 1 | k__Fungi(100);p__Ascomycota(100);c__Saccharomycetes(100);o__Saccharomycetales(100);f__Saccharomycetales_family_Incertae_sedis(100);g__Candida(100);g__Candida_unclassified(100); | Candida | 13 | Pathotroph-Saprotroph-Symbiotroph | Animal Pathogen-Endophyte-Endosymbiont-Epiphyte-\|Undefined Saprotroph\| | Dimorphic Yeast-Yeast | NULL | Probable | As nectar-tap_saprotroph (Põlme et al. 2020); Decay sugar-rich substrates (Põlme et al. 2020); This asexual genus is polyphyletic and contain species that function very differently depend on what they are. Caution should be taken when interpreting guilds for Candida species. | Manolakaki D, et al. 2010. Virulence. 1: 367-375 ((https://doi.org/10.4161/viru.1.5.12796)); Mekbib SB, et al. 2011. Tropical Plant Pathology 36:233-240 ((https://doi.org/10.1590/S1982-56762011000400004)); Põlme S, et al. 2020. Fungal Diversity 105:1-16 ((https://doi.org/10.1007/s13225-020-00466-2)) |
| Otu53686 | 1 | k__Fungi(100);p__Ascomycota(100);c__Saccharomycetes(100);o__Saccharomycetales(100);o__Saccharomycetales_unclassified(100);o__Saccharomycetales_unclassified(100);o__Saccharomycetales_unclassified(100); | Saccharomycetales | 7 | Saprotroph | Undefined Saprotroph | Yeast | NULL | Possible | NULL | Sterkenburg E, et al. 2015. New Phytologist 207:1145-1158 ((https://doi.org/10.1111/nph.13426)) |
| Otu53687 | 1 | k__Fungi(100);p__Ascomycota(100);c__Saccharomycetes(100);o__Saccharomycetales(100);f__Metschnikowiaceae(100);g__Metschnikowia(100);s__Metschnikowia_pulcherrima(100); | Metschnikowia | 13 | Saprotroph | \|Undefined Saprotroph\| | Yeast | NULL | Probable | As nectar-tap_saprotroph (Põlme et al. 2020); Decay sugar-rich substrates (Põlme et al. 2020) | Põlme S, et al. 2020. Fungal Diversity 105:1-16 ((https://doi.org/10.1007/s13225-020-00466-2)) |
| Otu53688 | 1 | k__Fungi(100);p__Ascomycota(100);c__Saccharomycetes(100);o__Saccharomycetales(100);f__Saccharomycetales_family_Incertae_sedis(100);g__Candida(100);s__Candida_cretensis(100); | Candida | 13 | Pathotroph-Saprotroph-Symbiotroph | Animal Pathogen-Endophyte-Endosymbiont-Epiphyte-\|Undefined Saprotroph\| | Dimorphic Yeast-Yeast | NULL | Probable | As nectar-tap_saprotroph (Põlme et al. 2020); Decay sugar-rich substrates (Põlme et al. 2020); This asexual genus is polyphyletic and contain species that function very differently depend on what they are. Caution should be taken when interpreting guilds for Candida species. | Manolakaki D, et al. 2010. Virulence. 1: 367-375 ((https://doi.org/10.4161/viru.1.5.12796)); Mekbib SB, et al. 2011. Tropical Plant Pathology 36:233-240 ((https://doi.org/10.1590/S1982-56762011000400004)); Põlme S, et al. 2020. Fungal Diversity 105:1-16 ((https://doi.org/10.1007/s13225-020-00466-2)) |
| Otu53689 | 1 | k__Fungi(100);p__Ascomycota(100);c__Saccharomycetes(100);o__Saccharomycetales(100);f__Metschnikowiaceae(100);g__Metschnikowia(100);s__Metschnikowia_pulcherrima(100); | Metschnikowia | 13 | Saprotroph | \|Undefined Saprotroph\| | Yeast | NULL | Probable | As nectar-tap_saprotroph (Põlme et al. 2020); Decay sugar-rich substrates (Põlme et al. 2020) | Põlme S, et al. 2020. Fungal Diversity 105:1-16 ((https://doi.org/10.1007/s13225-020-00466-2)) |
| Otu53696 | 1 | k__Fungi(100);p__Ascomycota(100);c__Saccharomycetes(100);o__Saccharomycetales(100);f__Saccharomycetales_family_Incertae_sedis(100);g__Candida(100);g__Candida_unclassified(100); | Candida | 13 | Pathotroph-Saprotroph-Symbiotroph | Animal Pathogen-Endophyte-Endosymbiont-Epiphyte-\|Undefined Saprotroph\| | Dimorphic Yeast-Yeast | NULL | Probable | As nectar-tap_saprotroph (Põlme et al. 2020); Decay sugar-rich substrates (Põlme et al. 2020); This asexual genus is polyphyletic and contain species that function very differently depend on what they are. Caution should be taken when interpreting guilds for Candida species. | Manolakaki D, et al. 2010. Virulence. 1: 367-375 ((https://doi.org/10.4161/viru.1.5.12796)); Mekbib SB, et al. 2011. Tropical Plant Pathology 36:233-240 ((https://doi.org/10.1590/S1982-56762011000400004)); Põlme S, et al. 2020. Fungal Diversity 105:1-16 ((https://doi.org/10.1007/s13225-020-00466-2)) |
| Otu53697 | 1 | k__Fungi(100);p__Ascomycota(100);c__Saccharomycetes(100);o__Saccharomycetales(100);o__Saccharomycetales_unclassified(100);o__Saccharomycetales_unclassified(100);o__Saccharomycetales_unclassified(100); | Saccharomycetales | 7 | Saprotroph | Undefined Saprotroph | Yeast | NULL | Possible | NULL | Sterkenburg E, et al. 2015. New Phytologist 207:1145-1158 ((https://doi.org/10.1111/nph.13426)) |

| | | | | | | | | | | | |
|---|---|---|---|---|---|---|---|---|---|---|---|
| Otu53698 | 1 | k__Fungi(100);p__Ascomycota(100);c__Saccharomycetes(100);o__Saccharomycetales(100);f__Saccharomycetales_family_Incertae_sedis(100);g__Candida(100);g__Candida_unclassified(100); | Candida | 13 | Pathotroph-Saprotroph-Symbiotroph | Animal Pathogen-Endophyte-Endosymbiont-Epiphyte-\|Undefined Saprotroph\| | Dimorphic Yeast-Yeast | NULL | Probable | As nectar-tap_saprotroph (Põlme et al. 2020); Decay sugar-rich substrates (Põlme et al. 2020); This asexual genus is polyphyletic and contain species that function very differently depend on what they are. Caution should be taken when interpreting guilds for Candida species. | Manolakaki D, et al. 2010. Virulence. 1: 367-375 ((https://doi.org/10.4161/viru.1.5.12796)); Mekbib SB, et al. 2011. Tropical Plant Pathology 36:233-240 ((https://doi.org/10.1590/S1982-56762011000400004)); Põlme S, et al. 2020. Fungal Diversity 105:1-16 ((https://doi.org/10.1007/s13225-020-00466-2)) |
| Otu53699 | 1 | k__Fungi(100);p__Ascomycota(100);c__Saccharomycetes(100);o__Saccharomycetales(100);f__Saccharomycetales_family_Incertae_sedis(100);g__Candida(100);g__Candida_unclassified(100); | Candida | 13 | Pathotroph-Saprotroph-Symbiotroph | Animal Pathogen-Endophyte-Endosymbiont-Epiphyte-\|Undefined Saprotroph\| | Dimorphic Yeast-Yeast | NULL | Probable | As nectar-tap_saprotroph (Põlme et al. 2020); Decay sugar-rich substrates (Põlme et al. 2020); This asexual genus is polyphyletic and contain species that function very differently depend on what they are. Caution should be taken when interpreting guilds for Candida species. | Manolakaki D, et al. 2010. Virulence. 1: 367-375 ((https://doi.org/10.4161/viru.1.5.12796)); Mekbib SB, et al. 2011. Tropical Plant Pathology 36:233-240 ((https://doi.org/10.1590/S1982-56762011000400004)); Põlme S, et al. 2020. Fungal Diversity 105:1-16 ((https://doi.org/10.1007/s13225-020-00466-2)) |
| Otu53700 | 1 | k__Fungi(100);p__Ascomycota(100);c__Saccharomycetes(100);o__Saccharomycetales(100);f__Saccharomycetales_family_Incertae_sedis(100);g__Candida(100);g__Candida_unclassified(100); | Candida | 13 | Pathotroph-Saprotroph-Symbiotroph | Animal Pathogen-Endophyte-Endosymbiont-Epiphyte-\|Undefined Saprotroph\| | Dimorphic Yeast-Yeast | NULL | Probable | As nectar-tap_saprotroph (Põlme et al. 2020); Decay sugar-rich substrates (Põlme et al. 2020); This asexual genus is polyphyletic and contain species that function very differently depend on what they are. Caution should be taken when interpreting guilds for Candida species. | Manolakaki D, et al. 2010. Virulence. 1: 367-375 ((https://doi.org/10.4161/viru.1.5.12796)); Mekbib SB, et al. 2011. Tropical Plant Pathology 36:233-240 ((https://doi.org/10.1590/S1982-56762011000400004)); Põlme S, et al. 2020. Fungal Diversity 105:1-16 ((https://doi.org/10.1007/s13225-020-00466-2)) |
| Otu53701 | 1 | k__Fungi(100);p__Ascomycota(100);c__Saccharomycetes(100);o__Saccharomycetales(100);f__Saccharomycetales_family_Incertae_sedis(100);f__Saccharomycetales_family_Incertae_sedis_unclassified(100);f__Saccharomycetales_family_Incertae_sedis_unclassified(100); | Saccharomycetales | 7 | Saprotroph | Undefined Saprotroph | Yeast | NULL | Possible | NULL | Sterkenburg E, et al. 2015. New Phytologist 207:1145-1158 ((https://doi.org/10.1111/nph.13426)) |
| Otu53702 | 1 | k__Fungi(100);p__Ascomycota(100);c__Saccharomycetes(100);o__Saccharomycetales(100);f__Saccharomycetales_family_Incertae_sedis(100);g__Candida(100);s__Candida_cretensis(100); | Candida | 13 | Pathotroph-Saprotroph-Symbiotroph | Animal Pathogen-Endophyte-Endosymbiont-Epiphyte-\|Undefined Saprotroph\| | Dimorphic Yeast-Yeast | NULL | Probable | As nectar-tap_saprotroph (Põlme et al. 2020); Decay sugar-rich substrates (Põlme et al. 2020); This asexual genus is polyphyletic and contain species that function very differently depend on what they are. Caution should be taken when interpreting guilds for Candida species. | Manolakaki D, et al. 2010. Virulence. 1: 367-375 ((https://doi.org/10.4161/viru.1.5.12796)); Mekbib SB, et al. 2011. Tropical Plant Pathology 36:233-240 ((https://doi.org/10.1590/S1982-56762011000400004)); Põlme S, et al. 2020. Fungal Diversity 105:1-16 ((https://doi.org/10.1007/s13225-020-00466-2)) |
| Otu53705 | 1 | k__Fungi(100);p__Ascomycota(100);c__Saccharomycetes(100);o__Saccharomycetales(100);f__Saccharomycetales_family_Incertae_sedis(100);g__Candida(100);g__Candida_unclassified(100); | Candida | 13 | Pathotroph-Saprotroph-Symbiotroph | Animal Pathogen-Endophyte-Endosymbiont-Epiphyte-\|Undefined Saprotroph\| | Dimorphic Yeast-Yeast | NULL | Probable | As nectar-tap_saprotroph (Põlme et al. 2020); Decay sugar-rich substrates (Põlme et al. 2020); This asexual genus is polyphyletic and contain species that function very differently depend on what they are. Caution should be taken when interpreting guilds for Candida species. | Manolakaki D, et al. 2010. Virulence. 1: 367-375 ((https://doi.org/10.4161/viru.1.5.12796)); Mekbib SB, et al. 2011. Tropical Plant Pathology 36:233-240 ((https://doi.org/10.1590/S1982-56762011000400004)); Põlme S, et al. 2020. Fungal Diversity 105:1-16 ((https://doi.org/10.1007/s13225-020-00466-2)) |
| Otu53706 | 1 | k__Fungi(100);p__Ascomycota(100);c__Saccharomycetes(100);o__Saccharomycetales(100);f__Saccharomycetales_family_Incertae_sedis(100);g__Candida(100);s__Candida_cretensis(100); | Candida | 13 | Pathotroph-Saprotroph-Symbiotroph | Animal Pathogen-Endophyte-Endosymbiont-Epiphyte-\|Undefined Saprotroph\| | Dimorphic Yeast-Yeast | NULL | Probable | As nectar-tap_saprotroph (Põlme et al. 2020); Decay sugar-rich substrates (Põlme et al. 2020); This asexual genus is polyphyletic and contain species that function very differently depend on what they are. Caution should be taken when interpreting guilds for Candida species. | Manolakaki D, et al. 2010. Virulence. 1: 367-375 ((https://doi.org/10.4161/viru.1.5.12796)); Mekbib SB, et al. 2011. Tropical Plant Pathology 36:233-240 ((https://doi.org/10.1590/S1982-56762011000400004)); Põlme S, et al. 2020. Fungal Diversity 105:1-16 ((https://doi.org/10.1007/s13225-020-00466-2)) |
| Otu53707 | 1 | k__Fungi(100);p__Ascomycota(100);c__Saccharomycetes(100);o__Saccharomycetales(100);o__Saccharomycetales_unclassified(100);o__Saccharomycetales_unclassified(100);o__Saccharomycetales_unclassified(100); | Saccharomycetales | 7 | Saprotroph | Undefined Saprotroph | Yeast | NULL | Possible | NULL | Sterkenburg E, et al. 2015. New Phytologist 207:1145-1158 ((https://doi.org/10.1111/nph.13426)) |
| Otu53708 | 1 | k__Fungi(100);p__Ascomycota(100);c__Saccharomycetes(100);o__Saccharomycetales(100);f__Metschnikowiaceae(100);g__Metschnikowia(100);s__Metschnikowia_pulcherrima(100); | Metschnikowia | 13 | Saprotroph | \|Undefined Saprotroph\| | Yeast | NULL | Probable | As nectar-tap_saprotroph (Põlme et al. 2020); Decay sugar-rich substrates (Põlme et al. 2020) | Põlme S, et al. 2020. Fungal Diversity 105:1-16 ((https://doi.org/10.1007/s13225-020-00466-2)) |
| Otu53709 | 1 | k__Fungi(100);p__Ascomycota(100);c__Saccharomycetes(100);o__Saccharomycetales(100);f__Metschnikowiaceae(100);g__Metschnikowia(100);s__Metschnikowia_pulcherrima(100); | Metschnikowia | 13 | Saprotroph | \|Undefined Saprotroph\| | Yeast | NULL | Probable | As nectar-tap_saprotroph (Põlme et al. 2020); Decay sugar-rich substrates (Põlme et al. 2020) | Põlme S, et al. 2020. Fungal Diversity 105:1-16 ((https://doi.org/10.1007/s13225-020-00466-2)) |
| Otu53710 | 1 | k__Fungi(100);p__Ascomycota(100);c__Saccharomycetes(100);o__Saccharomycetales(100);f__Metschnikowiaceae(100);g__Metschnikowia(100);s__Metschnikowia_pulcherrima(100); | Metschnikowia | 13 | Saprotroph | \|Undefined Saprotroph\| | Yeast | NULL | Probable | As nectar-tap_saprotroph (Põlme et al. 2020); Decay sugar-rich substrates (Põlme et al. 2020) | Põlme S, et al. 2020. Fungal Diversity 105:1-16 ((https://doi.org/10.1007/s13225-020-00466-2)) |
| Otu53712 | 1 | k__Fungi(100);p__Ascomycota(100);c__Saccharomycetes(100);o__Saccharomycetales(100);f__Saccharomycetales_family_Incertae_sedis(100);g__Candida(100);g__Candida_unclassified(100); | Candida | 13 | Pathotroph-Saprotroph-Symbiotroph | Animal Pathogen-Endophyte-Endosymbiont-Epiphyte-\|Undefined Saprotroph\| | Dimorphic Yeast-Yeast | NULL | Probable | As nectar-tap_saprotroph (Põlme et al. 2020); Decay sugar-rich substrates (Põlme et al. 2020); This asexual genus is polyphyletic and contain species that function very differently depend on what they are. Caution should be taken when interpreting guilds for Candida species. | Manolakaki D, et al. 2010. Virulence. 1: 367-375 ((https://doi.org/10.4161/viru.1.5.12796)); Mekbib SB, et al. 2011. Tropical Plant Pathology 36:233-240 ((https://doi.org/10.1590/S1982-56762011000400004)); Põlme S, et al. 2020. Fungal Diversity 105:1-16 ((https://doi.org/10.1007/s13225-020-00466-2)) |
| Otu53714 | 1 | k__Fungi(100);p__Ascomycota(100);c__Saccharomycetes(100);o__Saccharomycetales(100);f__Saccharomycetales_family_Incertae_sedis(100);g__Candida(100);g__Candida_unclassified(100); | Candida | 13 | Pathotroph-Saprotroph-Symbiotroph | Animal Pathogen-Endophyte-Endosymbiont-Epiphyte-\|Undefined Saprotroph\| | Dimorphic Yeast-Yeast | NULL | Probable | As nectar-tap_saprotroph (Põlme et al. 2020); Decay sugar-rich substrates (Põlme et al. 2020); This asexual genus is polyphyletic and contain species that function very differently depend on what they are. Caution should be taken when interpreting guilds for Candida species. | Manolakaki D, et al. 2010. Virulence. 1: 367-375 ((https://doi.org/10.4161/viru.1.5.12796)); Mekbib SB, et al. 2011. Tropical Plant Pathology 36:233-240 ((https://doi.org/10.1590/S1982-56762011000400004)); Põlme S, et al. 2020. Fungal Diversity 105:1-16 ((https://doi.org/10.1007/s13225-020-00466-2)) |
| Otu53719 | 1 | k__Fungi(100);p__Ascomycota(100);c__Saccharomycetes(100);o__Saccharomycetales(100);o__Saccharomycetales_unclassified(100);o__Saccharomycetales_unclassified(100);o__Saccharomycetales_unclassified(100); | Saccharomycetales | 7 | Saprotroph | Undefined Saprotroph | Yeast | NULL | Possible | NULL | Sterkenburg E, et al. 2015. New Phytologist 207:1145-1158 ((https://doi.org/10.1111/nph.13426)) |
| Otu53720 | 1 | k__Fungi(100);p__Ascomycota(100);c__Saccharomycetes(100);o__Saccharomycetales(100);f__Metschnikowiaceae(100);g__Metschnikowia(100);s__Metschnikowia_pulcherrima(100); | Metschnikowia | 13 | Saprotroph | \|Undefined Saprotroph\| | Yeast | NULL | Probable | As nectar-tap_saprotroph (Põlme et al. 2020); Decay sugar-rich substrates (Põlme et al. 2020) | Põlme S, et al. 2020. Fungal Diversity 105:1-16 ((https://doi.org/10.1007/s13225-020-00466-2)) |
| Otu53724 | 1 | k__Fungi(100);p__Ascomycota(100);c__Saccharomycetes(100);o__Saccharomycetales(100);f__Saccharomycetales_family_Incertae_sedis(100);g__Candida(100);g__Candida_unclassified(100); | Candida | 13 | Pathotroph-Saprotroph-Symbiotroph | Animal Pathogen-Endophyte-Endosymbiont-Epiphyte-\|Undefined Saprotroph\| | Dimorphic Yeast-Yeast | NULL | Probable | As nectar-tap_saprotroph (Põlme et al. 2020); Decay sugar-rich substrates (Põlme et al. 2020); This asexual genus is polyphyletic and contain species that function very differently depend on what they are. Caution should be taken when interpreting guilds for Candida species. | Manolakaki D, et al. 2010. Virulence. 1: 367-375 ((https://doi.org/10.4161/viru.1.5.12796)); Mekbib SB, et al. 2011. Tropical Plant Pathology 36:233-240 ((https://doi.org/10.1590/S1982-56762011000400004)); Põlme S, et al. 2020. Fungal Diversity 105:1-16 ((https://doi.org/10.1007/s13225-020-00466-2)) |
| Otu53731 | 1 | k__Fungi(100);p__Ascomycota(100);c__Saccharomycetes(100);o__Saccharomycetales(100);f__Saccharomycetales_family_Incertae_sedis(100);g__Candida(100);g__Candida_unclassified(100); | Candida | 13 | Pathotroph-Saprotroph-Symbiotroph | Animal Pathogen-Endophyte-Endosymbiont-Epiphyte-\|Undefined Saprotroph\| | Dimorphic Yeast-Yeast | NULL | Probable | As nectar-tap_saprotroph (Põlme et al. 2020); Decay sugar-rich substrates (Põlme et al. 2020); This asexual genus is polyphyletic and contain species that function very differently depend on what they are. Caution should be taken when interpreting guilds for Candida species. | Manolakaki D, et al. 2010. Virulence. 1: 367-375 ((https://doi.org/10.4161/viru.1.5.12796)); Mekbib SB, et al. 2011. Tropical Plant Pathology 36:233-240 ((https://doi.org/10.1590/S1982-56762011000400004)); Põlme S, et al. 2020. Fungal Diversity 105:1-16 ((https://doi.org/10.1007/s13225-020-00466-2)) |
| Otu53733 | 1 | k__Fungi(100);p__Ascomycota(100);c__Saccharomycetes(100);o__Saccharomycetales(100);f__Metschnikowiaceae(100);g__Metschnikowia(100);g__Metschnikowia_unclassified(100); | Metschnikowia | 13 | Saprotroph | \|Undefined Saprotroph\| | Yeast | NULL | Probable | As nectar-tap_saprotroph (Põlme et al. 2020); Decay sugar-rich substrates (Põlme et al. 2020) | Põlme S, et al. 2020. Fungal Diversity 105:1-16 ((https://doi.org/10.1007/s13225-020-00466-2)) |
| Otu53735 | 1 | k__Fungi(100);p__Ascomycota(100);c__Saccharomycetes(100);o__Saccharomycetales(100);f__Metschnikowiaceae(100);g__Metschnikowia(100);s__Metschnikowia_pulcherrima(100); | Metschnikowia | 13 | Saprotroph | \|Undefined Saprotroph\| | Yeast | NULL | Probable | As nectar-tap_saprotroph (Põlme et al. 2020); Decay sugar-rich substrates (Põlme et al. 2020) | Põlme S, et al. 2020. Fungal Diversity 105:1-16 ((https://doi.org/10.1007/s13225-020-00466-2)) |
| Otu53736 | 1 | k__Fungi(100);p__Ascomycota(100);c__Saccharomycetes(100);o__Saccharomycetales(100);f__Metschnikowiaceae(100);g__Metschnikowia(100);s__Metschnikowia_pulcherrima(100); | Metschnikowia | 13 | Saprotroph | \|Undefined Saprotroph\| | Yeast | NULL | Probable | As nectar-tap_saprotroph (Põlme et al. 2020); Decay sugar-rich substrates (Põlme et al. 2020) | Põlme S, et al. 2020. Fungal Diversity 105:1-16 ((https://doi.org/10.1007/s13225-020-00466-2)) |
| Otu53740 | 1 | k__Fungi(100);p__Ascomycota(100);c__Saccharomycetes(100);o__Saccharomycetales(100);f__Saccharomycetales_family_Incertae_sedis(100);g__Candida(100);g__Candida_unclassified(100); | Candida | 13 | Pathotroph-Saprotroph-Symbiotroph | Animal Pathogen-Endophyte-Endosymbiont-Epiphyte-\|Undefined Saprotroph\| | Dimorphic Yeast-Yeast | NULL | Probable | As nectar-tap_saprotroph (Põlme et al. 2020); Decay sugar-rich substrates (Põlme et al. 2020); This asexual genus is polyphyletic and contain species that function very differently depend on what they are. Caution should be taken when interpreting guilds for Candida species. | Manolakaki D, et al. 2010. Virulence. 1: 367-375 ((https://doi.org/10.4161/viru.1.5.12796)); Mekbib SB, et al. 2011. Tropical Plant Pathology 36:233-240 ((https://doi.org/10.1590/S1982-56762011000400004)); Põlme S, et al. 2020. Fungal Diversity 105:1-16 ((https://doi.org/10.1007/s13225-020-00466-2)) |
| Otu53741 | 1 | k__Fungi(100);p__Ascomycota(100);c__Saccharomycetes(100);o__Saccharomycetales(100);f__Metschnikowiaceae(100);g__Metschnikowia(100);s__Metschnikowia_pulcherrima(100); | Metschnikowia | 13 | Saprotroph | \|Undefined Saprotroph\| | Yeast | NULL | Probable | As nectar-tap_saprotroph (Põlme et al. 2020); Decay sugar-rich substrates (Põlme et al. 2020) | Põlme S, et al. 2020. Fungal Diversity 105:1-16 ((https://doi.org/10.1007/s13225-020-00466-2)) |
| Otu53745 | 1 | k__Fungi(100);p__Ascomycota(100);c__Saccharomycetes(100);o__Saccharomycetales(100);f__Metschnikowiaceae(100);g__Metschnikowia(100);s__Metschnikowia_pulcherrima(100); | Metschnikowia | 13 | Saprotroph | \|Undefined Saprotroph\| | Yeast | NULL | Probable | As nectar-tap_saprotroph (Põlme et al. 2020); Decay sugar-rich substrates (Põlme et al. 2020) | Põlme S, et al. 2020. Fungal Diversity 105:1-16 ((https://doi.org/10.1007/s13225-020-00466-2)) |
| Otu53746 | 1 | k__Fungi(100);p__Ascomycota(100);c__Saccharomycetes(100);o__Saccharomycetales(100);f__Metschnikowiaceae(100);g__Metschnikowia(100);s__Metschnikowia_pulcherrima(100); | Metschnikowia | 13 | Saprotroph | \|Undefined Saprotroph\| | Yeast | NULL | Probable | As nectar-tap_saprotroph (Põlme et al. 2020); Decay sugar-rich substrates (Põlme et al. 2020) | Põlme S, et al. 2020. Fungal Diversity 105:1-16 ((https://doi.org/10.1007/s13225-020-00466-2)) |
| Otu53747 | 1 | k__Fungi(100);p__Ascomycota(100);c__Saccharomycetes(100);o__Saccharomycetales(100);f__Metschnikowiaceae(100);g__Metschnikowia(100);s__Metschnikowia_pulcherrima(100); | Metschnikowia | 13 | Saprotroph | \|Undefined Saprotroph\| | Yeast | NULL | Probable | As nectar-tap_saprotroph (Põlme et al. 2020); Decay sugar-rich substrates (Põlme et al. 2020) | Põlme S, et al. 2020. Fungal Diversity 105:1-16 ((https://doi.org/10.1007/s13225-020-00466-2)) |
| Otu53748 | 1 | k__Fungi(100);p__Ascomycota(100);c__Saccharomycetes(100);o__Saccharomycetales(100);f__Metschnikowiaceae(100);g__Metschnikowia(100);g__Metschnikowia_unclassified(100); | Metschnikowia | 13 | Saprotroph | \|Undefined Saprotroph\| | Yeast | NULL | Probable | As nectar-tap_saprotroph (Põlme et al. 2020); Decay sugar-rich substrates (Põlme et al. 2020) | Põlme S, et al. 2020. Fungal Diversity 105:1-16 ((https://doi.org/10.1007/s13225-020-00466-2)) |
| Otu53749 | 1 | k__Fungi(100);p__Ascomycota(100);c__Saccharomycetes(100);o__Saccharomycetales(100);f__Metschnikowiaceae(100);g__Metschnikowia(100);s__Metschnikowia_pulcherrima(100); | Metschnikowia | 13 | Saprotroph | \|Undefined Saprotroph\| | Yeast | NULL | Probable | As nectar-tap_saprotroph (Põlme et al. 2020); Decay sugar-rich substrates (Põlme et al. 2020) | Põlme S, et al. 2020. Fungal Diversity 105:1-16 ((https://doi.org/10.1007/s13225-020-00466-2)) |
| Otu53753 | 1 | k__Fungi(100);p__Ascomycota(100);c__Saccharomycetes(100);o__Saccharomycetales(100);f__Saccharomycetales_family_Incertae_sedis(100);g__Candida(100);g__Candida_unclassified(100); | Candida | 13 | Pathotroph-Saprotroph-Symbiotroph | Animal Pathogen-Endophyte-Endosymbiont-Epiphyte-\|Undefined Saprotroph\| | Dimorphic Yeast-Yeast | NULL | Probable | As nectar-tap_saprotroph (Põlme et al. 2020); Decay sugar-rich substrates (Põlme et al. 2020); This asexual genus is polyphyletic and contain species that function very differently depend on what they are. Caution should be taken when interpreting guilds for Candida species. | Manolakaki D, et al. 2010. Virulence. 1: 367-375 ((https://doi.org/10.4161/viru.1.5.12796)); Mekbib SB, et al. 2011. Tropical Plant Pathology 36:233-240 ((https://doi.org/10.1590/S1982-56762011000400004)); Põlme S, et al. 2020. Fungal Diversity 105:1-16 ((https://doi.org/10.1007/s13225-020-00466-2)) |
| Otu53755 | 1 | k__Fungi(100);p__Ascomycota(100);c__Saccharomycetes(100);o__Saccharomycetales(100);f__Metschnikowiaceae(100);g__Metschnikowia(100);g__Metschnikowia_unclassified(100); | Metschnikowia | 13 | Saprotroph | \|Undefined Saprotroph\| | Yeast | NULL | Probable | As nectar-tap_saprotroph (Põlme et al. 2020); Decay sugar-rich substrates (Põlme et al. 2020) | Põlme S, et al. 2020. Fungal Diversity 105:1-16 ((https://doi.org/10.1007/s13225-020-00466-2)) |
| Otu53756 | 1 | k__Fungi(100);p__Ascomycota(100);c__Saccharomycetes(100);o__Saccharomycetales(100);f__Saccharomycetales_family_Incertae_sedis(100);g__Candida(100);g__Candida_unclassified(100); | Candida | 13 | Pathotroph-Saprotroph-Symbiotroph | Animal Pathogen-Endophyte-Endosymbiont-Epiphyte-\|Undefined Saprotroph\| | Dimorphic Yeast-Yeast | NULL | Probable | As nectar-tap_saprotroph (Põlme et al. 2020); Decay sugar-rich substrates (Põlme et al. 2020); This asexual genus is polyphyletic and contain species that function very differently depend on what they are. Caution should be taken when interpreting guilds for Candida species. | Manolakaki D, et al. 2010. Virulence. 1: 367-375 ((https://doi.org/10.4161/viru.1.5.12796)); Mekbib SB, et al. 2011. Tropical Plant Pathology 36:233-240 ((https://doi.org/10.1590/S1982-56762011000400004)); Põlme S, et al. 2020. Fungal Diversity 105:1-16 ((https://doi.org/10.1007/s13225-020-00466-2)) |
| Otu53757 | 1 | k__Fungi(100);p__Ascomycota(100);c__Saccharomycetes(100);o__Saccharomycetales(100);f__Metschnikowiaceae(100);g__Metschnikowia(100);s__Metschnikowia_pulcherrima(100); | Metschnikowia | 13 | Saprotroph | \|Undefined Saprotroph\| | Yeast | NULL | Probable | As nectar-tap_saprotroph (Põlme et al. 2020); Decay sugar-rich substrates (Põlme et al. 2020) | Põlme S, et al. 2020. Fungal Diversity 105:1-16 ((https://doi.org/10.1007/s13225-020-00466-2)) |
| Otu53758 | 1 | k__Fungi(100);p__Ascomycota(100);c__Saccharomycetes(100);o__Saccharomycetales(100);f__Metschnikowiaceae(100);g__Metschnikowia(100);s__Metschnikowia_pulcherrima(100); | Metschnikowia | 13 | Saprotroph | \|Undefined Saprotroph\| | Yeast | NULL | Probable | As nectar-tap_saprotroph (Põlme et al. 2020); Decay sugar-rich substrates (Põlme et al. 2020) | Põlme S, et al. 2020. Fungal Diversity 105:1-16 ((https://doi.org/10.1007/s13225-020-00466-2)) |
| Otu53759 | 1 | k__Fungi(100);p__Ascomycota(100);c__Saccharomycetes(100);o__Saccharomycetales(100);f__Metschnikowiaceae(100);g__Metschnikowia(100);s__Metschnikowia_pulcherrima(100); | Metschnikowia | 13 | Saprotroph | \|Undefined Saprotroph\| | Yeast | NULL | Probable | As nectar-tap_saprotroph (Põlme et al. 2020); Decay sugar-rich substrates (Põlme et al. 2020) | Põlme S, et al. 2020. Fungal Diversity 105:1-16 ((https://doi.org/10.1007/s13225-020-00466-2)) |
| Otu53760 | 1 | k__Fungi(100);p__Ascomycota(100);c__Saccharomycetes(100);o__Saccharomycetales(100);f__Metschnikowiaceae(100);g__Metschnikowia(100);s__Metschnikowia_pulcherrima(100); | Metschnikowia | 13 | Saprotroph | \|Undefined Saprotroph\| | Yeast | NULL | Probable | As nectar-tap_saprotroph (Põlme et al. 2020); Decay sugar-rich substrates (Põlme et al. 2020) | Põlme S, et al. 2020. Fungal Diversity 105:1-16 ((https://doi.org/10.1007/s13225-020-00466-2)) |
| Otu53761 | 1 | k__Fungi(100);p__Ascomycota(100);c__Saccharomycetes(100);o__Saccharomycetales(100);f__Metschnikowiaceae(100);g__Metschnikowia(100);s__Metschnikowia_pulcherrima(100); | Metschnikowia | 13 | Saprotroph | \|Undefined Saprotroph\| | Yeast | NULL | Probable | As nectar-tap_saprotroph (Põlme et al. 2020); Decay sugar-rich substrates (Põlme et al. 2020) | Põlme S, et al. 2020. Fungal Diversity 105:1-16 ((https://doi.org/10.1007/s13225-020-00466-2)) |
| Otu53762 | 1 | k__Fungi(100);p__Ascomycota(100);c__Saccharomycetes(100);o__Saccharomycetales(100);f__Metschnikowiaceae(100);g__Metschnikowia(100);s__Metschnikowia_pulcherrima(100); | Metschnikowia | 13 | Saprotroph | \|Undefined Saprotroph\| | Yeast | NULL | Probable | As nectar-tap_saprotroph (Põlme et al. 2020); Decay sugar-rich substrates (Põlme et al. 2020) | Põlme S, et al. 2020. Fungal Diversity 105:1-16 ((https://doi.org/10.1007/s13225-020-00466-2)) |
| Otu53763 | 1 | k__Fungi(100);p__Ascomycota(100);c__Saccharomycetes(100);o__Saccharomycetales(100);f__Metschnikowiaceae(100);g__Metschnikowia(100);s__Metschnikowia_pulcherrima(100); | Metschnikowia | 13 | Saprotroph | \|Undefined Saprotroph\| | Yeast | NULL | Probable | As nectar-tap_saprotroph (Põlme et al. 2020); Decay sugar-rich substrates (Põlme et al. 2020) | Põlme S, et al. 2020. Fungal Diversity 105:1-16 ((https://doi.org/10.1007/s13225-020-00466-2)) |
| Otu53764 | 1 | k__Fungi(100);p__Ascomycota(100);c__Saccharomycetes(100);o__Saccharomycetales(100);f__Saccharomycetales_family_Incertae_sedis(100);g__Candida(100);g__Candida_unclassified(100); | Candida | 13 | Pathotroph-Saprotroph-Symbiotroph | Animal Pathogen-Endophyte-Endosymbiont-Epiphyte-\|Undefined Saprotroph\| | Dimorphic Yeast-Yeast | NULL | Probable | As nectar-tap_saprotroph (Põlme et al. 2020); Decay sugar-rich substrates (Põlme et al. 2020); This asexual genus is polyphyletic and contain species that function very differently depend on what they are. Caution should be taken when interpreting guilds for Candida species. | Manolakaki D, et al. 2010. Virulence. 1: 367-375 ((https://doi.org/10.4161/viru.1.5.12796)); Mekbib SB, et al. 2011. Tropical Plant Pathology 36:233-240 ((https://doi.org/10.1590/S1982-56762011000400004)); Põlme S, et al. 2020. Fungal Diversity 105:1-16 ((https://doi.org/10.1007/s13225-020-00466-2)) |
| Otu53765 | 1 | k__Fungi(100);p__Ascomycota(100);c__Saccharomycetes(100);o__Saccharomycetales(100);f__Metschnikowiaceae(100);g__Metschnikowia(100);s__Metschnikowia_pulcherrima(100); | Metschnikowia | 13 | Saprotroph | \|Undefined Saprotroph\| | Yeast | NULL | Probable | As nectar-tap_saprotroph (Põlme et al. 2020); Decay sugar-rich substrates (Põlme et al. 2020) | Põlme S, et al. 2020. Fungal Diversity 105:1-16 ((https://doi.org/10.1007/s13225-020-00466-2)) |
| Otu53766 | 1 | k__Fungi(100);p__Ascomycota(100);c__Saccharomycetes(100);o__Saccharomycetales(100);f__Metschnikowiaceae(100);g__Metschnikowia(100);s__Metschnikowia_pulcherrima(100); | Metschnikowia | 13 | Saprotroph | \|Undefined Saprotroph\| | Yeast | NULL | Probable | As nectar-tap_saprotroph (Põlme et al. 2020); Decay sugar-rich substrates (Põlme et al. 2020) | Põlme S, et al. 2020. Fungal Diversity 105:1-16 ((https://doi.org/10.1007/s13225-020-00466-2)) |
| Otu53767 | 1 | k__Fungi(100);p__Ascomycota(100);c__Saccharomycetes(100);o__Saccharomycetales(100);f__Metschnikowiaceae(100);g__Metschnikowia(100);g__Metschnikowia_unclassified(100); | Metschnikowia | 13 | Saprotroph | \|Undefined Saprotroph\| | Yeast | NULL | Probable | As nectar-tap_saprotroph (Põlme et al. 2020); Decay sugar-rich substrates (Põlme et al. 2020) | Põlme S, et al. 2020. Fungal Diversity 105:1-16 ((https://doi.org/10.1007/s13225-020-00466-2)) |
| Otu53768 | 1 | k__Fungi(100);p__Ascomycota(100);c__Saccharomycetes(100);o__Saccharomycetales(100);f__Metschnikowiaceae(100);g__Metschnikowia(100);s__Metschnikowia_pulcherrima(100); | Metschnikowia | 13 | Saprotroph | \|Undefined Saprotroph\| | Yeast | NULL | Probable | As nectar-tap_saprotroph (Põlme et al. 2020); Decay sugar-rich substrates (Põlme et al. 2020) | Põlme S, et al. 2020. Fungal Diversity 105:1-16 ((https://doi.org/10.1007/s13225-020-00466-2)) |
| Otu53769 | 1 | k__Fungi(100);p__Ascomycota(100);c__Saccharomycetes(100);o__Saccharomycetales(100);f__Metschnikowiaceae(100);g__Metschnikowia(100);s__Metschnikowia_pulcherrima(100); | Metschnikowia | 13 | Saprotroph | \|Undefined Saprotroph\| | Yeast | NULL | Probable | As nectar-tap_saprotroph (Põlme et al. 2020); Decay sugar-rich substrates (Põlme et al. 2020) | Põlme S, et al. 2020. Fungal Diversity 105:1-16 ((https://doi.org/10.1007/s13225-020-00466-2)) |
| Otu53770 | 1 | k__Fungi(100);p__Ascomycota(100);c__Saccharomycetes(100);o__Saccharomycetales(100);f__Metschnikowiaceae(100);g__Metschnikowia(100);g__Metschnikowia_unclassified(100); | Metschnikowia | 13 | Saprotroph | \|Undefined Saprotroph\| | Yeast | NULL | Probable | As nectar-tap_saprotroph (Põlme et al. 2020); Decay sugar-rich substrates (Põlme et al. 2020) | Põlme S, et al. 2020. Fungal Diversity 105:1-16 ((https://doi.org/10.1007/s13225-020-00466-2)) |
| Otu53771 | 1 | k__Fungi(100);p__Ascomycota(100);c__Saccharomycetes(100);o__Saccharomycetales(100);f__Metschnikowiaceae(100);g__Metschnikowia(100);s__Metschnikowia_pulcherrima(100); | Metschnikowia | 13 | Saprotroph | \|Undefined Saprotroph\| | Yeast | NULL | Probable | As nectar-tap_saprotroph (Põlme et al. 2020); Decay sugar-rich substrates (Põlme et al. 2020) | Põlme S, et al. 2020. Fungal Diversity 105:1-16 ((https://doi.org/10.1007/s13225-020-00466-2)) |
| Otu53772 | 1 | k__Fungi(100);p__Ascomycota(100);c__Saccharomycetes(100);o__Saccharomycetales(100);f__Saccharomycetales_family_Incertae_sedis(100);g__Candida(100);g__Candida_unclassified(100); | Candida | 13 | Pathotroph-Saprotroph-Symbiotroph | Animal Pathogen-Endophyte-Endosymbiont-Epiphyte-\|Undefined Saprotroph\| | Dimorphic Yeast-Yeast | NULL | Probable | As nectar-tap_saprotroph (Põlme et al. 2020); Decay sugar-rich substrates (Põlme et al. 2020); This asexual genus is polyphyletic and contain species that function very differently depend on what they are. Caution should be taken when interpreting guilds for Candida species. | Manolakaki D, et al. 2010. Virulence. 1: 367-375 ((https://doi.org/10.4161/viru.1.5.12796)); Mekbib SB, et al. 2011. Tropical Plant Pathology 36:233-240 ((https://doi.org/10.1590/S1982-56762011000400004)); Põlme S, et al. 2020. Fungal Diversity 105:1-16 ((https://doi.org/10.1007/s13225-020-00466-2)) |
| Otu53774 | 1 | k__Fungi(100);p__Ascomycota(100);c__Saccharomycetes(100);o__Saccharomycetales(100);f__Saccharomycetales_family_Incertae_sedis(100);g__Candida(100);g__Candida_unclassified(100); | Candida | 13 | Pathotroph-Saprotroph-Symbiotroph | Animal Pathogen-Endophyte-Endosymbiont-Epiphyte-\|Undefined Saprotroph\| | Dimorphic Yeast-Yeast | NULL | Probable | As nectar-tap_saprotroph (Põlme et al. 2020); Decay sugar-rich substrates (Põlme et al. 2020); This asexual genus is polyphyletic and contain species that function very differently depend on what they are. Caution should be taken when interpreting guilds for Candida species. | Manolakaki D, et al. 2010. Virulence. 1: 367-375 ((https://doi.org/10.4161/viru.1.5.12796)); Mekbib SB, et al. 2011. Tropical Plant Pathology 36:233-240 ((https://doi.org/10.1590/S1982-56762011000400004)); Põlme S, et al. 2020. Fungal Diversity 105:1-16 ((https://doi.org/10.1007/s13225-020-00466-2)) |
| Otu53776 | 1 | k__Fungi(100);p__Basidiomycota(100);c__Microbotryomycetes(100);o__Sporidiobolales(100);f__Sporidiobolales_family_Incertae_sedis(100);g__Sporobolomyces(100);s__Sporobolomyces_lactosus(100); | Sporobolomyces | 13 | Pathotroph-Saprotroph | \|Fungal Parasite\|-Plant Saprotroph | Yeast | NULL | Probable | As litter_saprotroph (Põlme et al. 2020); As mycoparasite (Põlme et al. 2020); Decay fungal material (Põlme et al. 2020) | Tedersoo L, et al. 2014. Science 346:e1256688 ((https://doi.org/10.1126/science.1256688)); Purahong W, et al. 2016. Molecular Ecology 25:4059-4074 ((https://doi.org/10.1111/mec.13739)); Põlme S, et al. 2020. Fungal Diversity 105:1-16 ((https://doi.org/10.1007/s13225-020-00466-2)) |
| Otu53777 | 1 | k__Fungi(100);p__Ascomycota(100);c__Saccharomycetes(100);o__Saccharomycetales(100);f__Metschnikowiaceae(100);g__Metschnikowia(100);s__Metschnikowia_pulcherrima(100); | Metschnikowia | 13 | Saprotroph | \|Undefined Saprotroph\| | Yeast | NULL | Probable | As nectar-tap_saprotroph (Põlme et al. 2020); Decay sugar-rich substrates (Põlme et al. 2020) | Põlme S, et al. 2020. Fungal Diversity 105:1-16 ((https://doi.org/10.1007/s13225-020-00466-2)) |
| Otu53778 | 1 | k__Fungi(100);p__Ascomycota(100);c__Saccharomycetes(100);o__Saccharomycetales(100);f__Metschnikowiaceae(100);g__Metschnikowia(100);g__Metschnikowia_unclassified(100); | Metschnikowia | 13 | Saprotroph | \|Undefined Saprotroph\| | Yeast | NULL | Probable | As nectar-tap_saprotroph (Põlme et al. 2020); Decay sugar-rich substrates (Põlme et al. 2020) | Põlme S, et al. 2020. Fungal Diversity 105:1-16 ((https://doi.org/10.1007/s13225-020-00466-2)) |
| Otu53779 | 1 | k__Fungi(100);p__Ascomycota(100);c__Saccharomycetes(100);o__Saccharomycetales(100);f__Metschnikowiaceae(100);g__Metschnikowia(100);s__Metschnikowia_pulcherrima(100); | Metschnikowia | 13 | Saprotroph | \|Undefined Saprotroph\| | Yeast | NULL | Probable | As nectar-tap_saprotroph (Põlme et al. 2020); Decay sugar-rich substrates (Põlme et al. 2020) | Põlme S, et al. 2020. Fungal Diversity 105:1-16 ((https://doi.org/10.1007/s13225-020-00466-2)) |
| Otu53780 | 1 | k__Fungi(100);p__Ascomycota(100);c__Saccharomycetes(100);o__Saccharomycetales(100);f__Metschnikowiaceae(100);g__Metschnikowia(100);s__Metschnikowia_pulcherrima(100); | Metschnikowia | 13 | Saprotroph | \|Undefined Saprotroph\| | Yeast | NULL | Probable | As nectar-tap_saprotroph (Põlme et al. 2020); Decay sugar-rich substrates (Põlme et al. 2020) | Põlme S, et al. 2020. Fungal Diversity 105:1-16 ((https://doi.org/10.1007/s13225-020-00466-2)) |
| Otu53781 | 1 | k__Fungi(100);p__Ascomycota(100);c__Saccharomycetes(100);o__Saccharomycetales(100);f__Metschnikowiaceae(100);g__Metschnikowia(100);s__Metschnikowia_pulcherrima(100); | Metschnikowia | 13 | Saprotroph | \|Undefined Saprotroph\| | Yeast | NULL | Probable | As nectar-tap_saprotroph (Põlme et al. 2020); Decay sugar-rich substrates (Põlme et al. 2020) | Põlme S, et al. 2020. Fungal Diversity 105:1-16 ((https://doi.org/10.1007/s13225-020-00466-2)) |
| Otu53783 | 1 | k__Fungi(100);p__Ascomycota(100);c__Saccharomycetes(100);o__Saccharomycetales(100);f__Metschnikowiaceae(100);g__Metschnikowia(100);s__Metschnikowia_pulcherrima(100); | Metschnikowia | 13 | Saprotroph | \|Undefined Saprotroph\| | Yeast | NULL | Probable | As nectar-tap_saprotroph (Põlme et al. 2020); Decay sugar-rich substrates (Põlme et al. 2020) | Põlme S, et al. 2020. Fungal Diversity 105:1-16 ((https://doi.org/10.1007/s13225-020-00466-2)) |
| Otu53785 | 1 | k__Fungi(100);p__Ascomycota(100);c__Saccharomycetes(100);o__Saccharomycetales(100);f__Saccharomycetales_family_Incertae_sedis(100);g__Candida(100);g__Candida_unclassified(100); | Candida | 13 | Pathotroph-Saprotroph-Symbiotroph | Animal Pathogen-Endophyte-Endosymbiont-Epiphyte-\|Undefined Saprotroph\| | Dimorphic Yeast-Yeast | NULL | Probable | As nectar-tap_saprotroph (Põlme et al. 2020); Decay sugar-rich substrates (Põlme et al. 2020); This asexual genus is polyphyletic and contain species that function very differently depend on what they are. Caution should be taken when interpreting guilds for Candida species. | Manolakaki D, et al. 2010. Virulence. 1: 367-375 ((https://doi.org/10.4161/viru.1.5.12796)); Mekbib SB, et al. 2011. Tropical Plant Pathology 36:233-240 ((https://doi.org/10.1590/S1982-56762011000400004)); Põlme S, et al. 2020. Fungal Diversity 105:1-16 ((https://doi.org/10.1007/s13225-020-00466-2)) |
| Otu53788 | 1 | k__Fungi(100);p__Ascomycota(100);c__Saccharomycetes(100);o__Saccharomycetales(100);f__Saccharomycetales_family_Incertae_sedis(100);g__Candida(100);g__Candida_unclassified(100); | Candida | 13 | Pathotroph-Saprotroph-Symbiotroph | Animal Pathogen-Endophyte-Endosymbiont-Epiphyte-\|Undefined Saprotroph\| | Dimorphic Yeast-Yeast | NULL | Probable | As nectar-tap_saprotroph (Põlme et al. 2020); Decay sugar-rich substrates (Põlme et al. 2020); This asexual genus is polyphyletic and contain species that function very differently depend on what they are. Caution should be taken when interpreting guilds for Candida species. | Manolakaki D, et al. 2010. Virulence. 1: 367-375 ((https://doi.org/10.4161/viru.1.5.12796)); Mekbib SB, et al. 2011. Tropical Plant Pathology 36:233-240 ((https://doi.org/10.1590/S1982-56762011000400004)); Põlme S, et al. 2020. Fungal Diversity 105:1-16 ((https://doi.org/10.1007/s13225-020-00466-2)) |
| Otu53790 | 1 | k__Fungi(100);p__Ascomycota(100);c__Saccharomycetes(100);o__Saccharomycetales(100);f__Saccharomycetales_family_Incertae_sedis(100);g__Candida(100);g__Candida_unclassified(100); | Candida | 13 | Pathotroph-Saprotroph-Symbiotroph | Animal Pathogen-Endophyte-Endosymbiont-Epiphyte-\|Undefined Saprotroph\| | Dimorphic Yeast-Yeast | NULL | Probable | As nectar-tap_saprotroph (Põlme et al. 2020); Decay sugar-rich substrates (Põlme et al. 2020); This asexual genus is polyphyletic and contain species that function very differently depend on what they are. Caution should be taken when interpreting guilds for Candida species. | Manolakaki D, et al. 2010. Virulence. 1: 367-375 ((https://doi.org/10.4161/viru.1.5.12796)); Mekbib SB, et al. 2011. Tropical Plant Pathology 36:233-240 ((https://doi.org/10.1590/S1982-56762011000400004)); Põlme S, et al. 2020. Fungal Diversity 105:1-16 ((https://doi.org/10.1007/s13225-020-00466-2)) |
| Otu53791 | 1 | k__Fungi(100);p__Ascomycota(100);c__Saccharomycetes(100);o__Saccharomycetales(100);f__Metschnikowiaceae(100);g__Metschnikowia(100);g__Metschnikowia_unclassified(100); | Metschnikowia | 13 | Saprotroph | \|Undefined Saprotroph\| | Yeast | NULL | Probable | As nectar-tap_saprotroph (Põlme et al. 2020); Decay sugar-rich substrates (Põlme et al. 2020) | Põlme S, et al. 2020. Fungal Diversity 105:1-16 ((https://doi.org/10.1007/s13225-020-00466-2)) |
| Otu53792 | 1 | k__Fungi(100);p__Ascomycota(100);c__Saccharomycetes(100);o__Saccharomycetales(100);f__Metschnikowiaceae(100);g__Metschnikowia(100);s__Metschnikowia_pulcherrima(100); | Metschnikowia | 13 | Saprotroph | \|Undefined Saprotroph\| | Yeast | NULL | Probable | As nectar-tap_saprotroph (Põlme et al. 2020); Decay sugar-rich substrates (Põlme et al. 2020) | Põlme S, et al. 2020. Fungal Diversity 105:1-16 ((https://doi.org/10.1007/s13225-020-00466-2)) |
| Otu53793 | 1 | k__Fungi(100);p__Ascomycota(100);c__Saccharomycetes(100);o__Saccharomycetales(100);f__Metschnikowiaceae(100);g__Metschnikowia(100);s__Metschnikowia_pulcherrima(100); | Metschnikowia | 13 | Saprotroph | \|Undefined Saprotroph\| | Yeast | NULL | Probable | As nectar-tap_saprotroph (Põlme et al. 2020); Decay sugar-rich substrates (Põlme et al. 2020) | Põlme S, et al. 2020. Fungal Diversity 105:1-16 ((https://doi.org/10.1007/s13225-020-00466-2)) |
| Otu53796 | 1 | k__Fungi(100);p__Ascomycota(100);c__Saccharomycetes(100);o__Saccharomycetales(100);f__Metschnikowiaceae(100);g__Metschnikowia(100);s__Metschnikowia_pulcherrima(100); | Metschnikowia | 13 | Saprotroph | \|Undefined Saprotroph\| | Yeast | NULL | Probable | As nectar-tap_saprotroph (Põlme et al. 2020); Decay sugar-rich substrates (Põlme et al. 2020) | Põlme S, et al. 2020. Fungal Diversity 105:1-16 ((https://doi.org/10.1007/s13225-020-00466-2)) |
| Otu53799 | 1 | k__Fungi(100);p__Ascomycota(100);c__Saccharomycetes(100);o__Saccharomycetales(100);f__Metschnikowiaceae(100);g__Metschnikowia(100);s__Metschnikowia_pulcherrima(100); | Metschnikowia | 13 | Saprotroph | \|Undefined Saprotroph\| | Yeast | NULL | Probable | As nectar-tap_saprotroph (Põlme et al. 2020); Decay sugar-rich substrates (Põlme et al. 2020) | Põlme S, et al. 2020. Fungal Diversity 105:1-16 ((https://doi.org/10.1007/s13225-020-00466-2)) |
| Otu53800 | 1 | k__Fungi(100);p__Ascomycota(100);c__Saccharomycetes(100);o__Saccharomycetales(100);f__Saccharomycetales_family_Incertae_sedis(100);g__Candida(100);g__Candida_unclassified(100); | Candida | 13 | Pathotroph-Saprotroph-Symbiotroph | Animal Pathogen-Endophyte-Endosymbiont-Epiphyte-\|Undefined Saprotroph\| | Dimorphic Yeast-Yeast | NULL | Probable | As nectar-tap_saprotroph (Põlme et al. 2020); Decay sugar-rich substrates (Põlme et al. 2020); This asexual genus is polyphyletic and contain species that function very differently depend on what they are. Caution should be taken when interpreting guilds for Candida species. | Manolakaki D, et al. 2010. Virulence. 1: 367-375 ((https://doi.org/10.4161/viru.1.5.12796)); Mekbib SB, et al. 2011. Tropical Plant Pathology 36:233-240 ((https://doi.org/10.1590/S1982-56762011000400004)); Põlme S, et al. 2020. Fungal Diversity 105:1-16 ((https://doi.org/10.1007/s13225-020-00466-2)) |
| Otu53801 | 1 | k__Fungi(100);p__Ascomycota(100);c__Saccharomycetes(100);o__Saccharomycetales(100);f__Metschnikowiaceae(100);g__Metschnikowia(100);g__Metschnikowia_unclassified(100); | Metschnikowia | 13 | Saprotroph | \|Undefined Saprotroph\| | Yeast | NULL | Probable | As nectar-tap_saprotroph (Põlme et al. 2020); Decay sugar-rich substrates (Põlme et al. 2020) | Põlme S, et al. 2020. Fungal Diversity 105:1-16 ((https://doi.org/10.1007/s13225-020-00466-2)) |
| Otu53802 | 1 | k__Fungi(100);p__Ascomycota(100);c__Saccharomycetes(100);o__Saccharomycetales(100);f__Metschnikowiaceae(100);g__Metschnikowia(100);s__Metschnikowia_pulcherrima(100); | Metschnikowia | 13 | Saprotroph | \|Undefined Saprotroph\| | Yeast | NULL | Probable | As nectar-tap_saprotroph (Põlme et al. 2020); Decay sugar-rich substrates (Põlme et al. 2020) | Põlme S, et al. 2020. Fungal Diversity 105:1-16 ((https://doi.org/10.1007/s13225-020-00466-2)) |
| Otu53803 | 1 | k__Fungi(100);p__Ascomycota(100);c__Saccharomycetes(100);o__Saccharomycetales(100);f__Saccharomycetales_family_Incertae_sedis(100);g__Candida(100);g__Candida_unclassified(100); | Candida | 13 | Pathotroph-Saprotroph-Symbiotroph | Animal Pathogen-Endophyte-Endosymbiont-Epiphyte-\|Undefined Saprotroph\| | Dimorphic Yeast-Yeast | NULL | Probable | As nectar-tap_saprotroph (Põlme et al. 2020); Decay sugar-rich substrates (Põlme et al. 2020); This asexual genus is polyphyletic and contain species that function very differently depend on what they are. Caution should be taken when interpreting guilds for Candida species. | Manolakaki D, et al. 2010. Virulence. 1: 367-375 ((https://doi.org/10.4161/viru.1.5.12796)); Mekbib SB, et al. 2011. Tropical Plant Pathology 36:233-240 ((https://doi.org/10.1590/S1982-56762011000400004)); Põlme S, et al. 2020. Fungal Diversity 105:1-16 ((https://doi.org/10.1007/s13225-020-00466-2)) |
| Otu53804 | 1 | k__Fungi(100);p__Ascomycota(100);c__Saccharomycetes(100);o__Saccharomycetales(100);f__Metschnikowiaceae(100);g__Metschnikowia(100);g__Metschnikowia_unclassified(100); | Metschnikowia | 13 | Saprotroph | \|Undefined Saprotroph\| | Yeast | NULL | Probable | As nectar-tap_saprotroph (Põlme et al. 2020); Decay sugar-rich substrates (Põlme et al. 2020) | Põlme S, et al. 2020. Fungal Diversity 105:1-16 ((https://doi.org/10.1007/s13225-020-00466-2)) |
| Otu53805 | 1 | k__Fungi(100);p__Ascomycota(100);c__Saccharomycetes(100);o__Saccharomycetales(100);f__Metschnikowiaceae(100);g__Metschnikowia(100);s__Metschnikowia_pulcherrima(100); | Metschnikowia | 13 | Saprotroph | \|Undefined Saprotroph\| | Yeast | NULL | Probable | As nectar-tap_saprotroph (Põlme et al. 2020); Decay sugar-rich substrates (Põlme et al. 2020) | Põlme S, et al. 2020. Fungal Diversity 105:1-16 ((https://doi.org/10.1007/s13225-020-00466-2)) |
| Otu53806 | 1 | k__Fungi(100);p__Ascomycota(100);c__Saccharomycetes(100);o__Saccharomycetales(100);f__Metschnikowiaceae(100);g__Metschnikowia(100);s__Metschnikowia_pulcherrima(100); | Metschnikowia | 13 | Saprotroph | \|Undefined Saprotroph\| | Yeast | NULL | Probable | As nectar-tap_saprotroph (Põlme et al. 2020); Decay sugar-rich substrates (Põlme et al. 2020) | Põlme S, et al. 2020. Fungal Diversity 105:1-16 ((https://doi.org/10.1007/s13225-020-00466-2)) |
| Otu53808 | 1 | k__Fungi(100);p__Ascomycota(100);c__Saccharomycetes(100);o__Saccharomycetales(100);f__Metschnikowiaceae(100);g__Metschnikowia(100);s__Metschnikowia_pulcherrima(100); | Metschnikowia | 13 | Saprotroph | \|Undefined Saprotroph\| | Yeast | NULL | Probable | As nectar-tap_saprotroph (Põlme et al. 2020); Decay sugar-rich substrates (Põlme et al. 2020) | Põlme S, et al. 2020. Fungal Diversity 105:1-16 ((https://doi.org/10.1007/s13225-020-00466-2)) |
| Otu53812 | 1 | k__Fungi(100);p__Ascomycota(100);c__Saccharomycetes(100);o__Saccharomycetales(100);f__Metschnikowiaceae(100);g__Metschnikowia(100);g__Metschnikowia_unclassified(100); | Metschnikowia | 13 | Saprotroph | \|Undefined Saprotroph\| | Yeast | NULL | Probable | As nectar-tap_saprotroph (Põlme et al. 2020); Decay sugar-rich substrates (Põlme et al. 2020) | Põlme S, et al. 2020. Fungal Diversity 105:1-16 ((https://doi.org/10.1007/s13225-020-00466-2)) |
| Otu53813 | 1 | k__Fungi(100);p__Ascomycota(100);c__Saccharomycetes(100);o__Saccharomycetales(100);f__Saccharomycetales_family_Incertae_sedis(100);g__Candida(100);g__Candida_unclassified(100); | Candida | 13 | Pathotroph-Saprotroph-Symbiotroph | Animal Pathogen-Endophyte-Endosymbiont-Epiphyte-\|Undefined Saprotroph\| | Dimorphic Yeast-Yeast | NULL | Probable | As nectar-tap_saprotroph (Põlme et al. 2020); Decay sugar-rich substrates (Põlme et al. 2020); This asexual genus is polyphyletic and contain species that function very differently depend on what they are. Caution should be taken when interpreting guilds for Candida species. | Manolakaki D, et al. 2010. Virulence. 1: 367-375 ((https://doi.org/10.4161/viru.1.5.12796)); Mekbib SB, et al. 2011. Tropical Plant Pathology 36:233-240 ((https://doi.org/10.1590/S1982-56762011000400004)); Põlme S, et al. 2020. Fungal Diversity 105:1-16 ((https://doi.org/10.1007/s13225-020-00466-2)) |
| Otu53814 | 1 | k__Fungi(100);p__Ascomycota(100);c__Saccharomycetes(100);o__Saccharomycetales(100);f__Metschnikowiaceae(100);g__Metschnikowia(100);g__Metschnikowia_unclassified(100); | Metschnikowia | 13 | Saprotroph | \|Undefined Saprotroph\| | Yeast | NULL | Probable | As nectar-tap_saprotroph (Põlme et al. 2020); Decay sugar-rich substrates (Põlme et al. 2020) | Põlme S, et al. 2020. Fungal Diversity 105:1-16 ((https://doi.org/10.1007/s13225-020-00466-2)) |
| Otu53815 | 1 | k__Fungi(100);p__Ascomycota(100);c__Saccharomycetes(100);o__Saccharomycetales(100);f__Metschnikowiaceae(100);g__Metschnikowia(100);s__Metschnikowia_pulcherrima(100); | Metschnikowia | 13 | Saprotroph | \|Undefined Saprotroph\| | Yeast | NULL | Probable | As nectar-tap_saprotroph (Põlme et al. 2020); Decay sugar-rich substrates (Põlme et al. 2020) | Põlme S, et al. 2020. Fungal Diversity 105:1-16 ((https://doi.org/10.1007/s13225-020-00466-2)) |
| Otu53816 | 1 | k__Fungi(100);p__Ascomycota(100);c__Saccharomycetes(100);o__Saccharomycetales(100);f__Metschnikowiaceae(100);g__Metschnikowia(100);g__Metschnikowia_unclassified(100); | Metschnikowia | 13 | Saprotroph | \|Undefined Saprotroph\| | Yeast | NULL | Probable | As nectar-tap_saprotroph (Põlme et al. 2020); Decay sugar-rich substrates (Põlme et al. 2020) | Põlme S, et al. 2020. Fungal Diversity 105:1-16 ((https://doi.org/10.1007/s13225-020-00466-2)) |
| Otu53819 | 1 | k__Fungi(100);p__Ascomycota(100);c__Saccharomycetes(100);o__Saccharomycetales(100);f__Metschnikowiaceae(100);g__Metschnikowia(100);s__Metschnikowia_pulcherrima(100); | Metschnikowia | 13 | Saprotroph | \|Undefined Saprotroph\| | Yeast | NULL | Probable | As nectar-tap_saprotroph (Põlme et al. 2020); Decay sugar-rich substrates (Põlme et al. 2020) | Põlme S, et al. 2020. Fungal Diversity 105:1-16 ((https://doi.org/10.1007/s13225-020-00466-2)) |
| Otu53820 | 1 | k__Fungi(100);p__Ascomycota(100);c__Saccharomycetes(100);o__Saccharomycetales(100);o__Saccharomycetales_unclassified(100);o__Saccharomycetales_unclassified(100);o__Saccharomycetales_unclassified(100); | Saccharomycetales | 7 | Saprotroph | Undefined Saprotroph | Yeast | NULL | Possible | NULL | Sterkenburg E, et al. 2015. New Phytologist 207:1145-1158 ((https://doi.org/10.1111/nph.13426)) |
| Otu53821 | 1 | k__Fungi(100);p__Ascomycota(100);c__Saccharomycetes(100);o__Saccharomycetales(100);f__Metschnikowiaceae(100);g__Metschnikowia(100);g__Metschnikowia_unclassified(100); | Metschnikowia | 13 | Saprotroph | \|Undefined Saprotroph\| | Yeast | NULL | Probable | As nectar-tap_saprotroph (Põlme et al. 2020); Decay sugar-rich substrates (Põlme et al. 2020) | Põlme S, et al. 2020. Fungal Diversity 105:1-16 ((https://doi.org/10.1007/s13225-020-00466-2)) |
| Otu53822 | 1 | k__Fungi(100);p__Ascomycota(100);c__Saccharomycetes(100);o__Saccharomycetales(100);o__Saccharomycetales_unclassified(100);o__Saccharomycetales_unclassified(100);o__Saccharomycetales_unclassified(100); | Saccharomycetales | 7 | Saprotroph | Undefined Saprotroph | Yeast | NULL | Possible | NULL | Sterkenburg E, et al. 2015. New Phytologist 207:1145-1158 ((https://doi.org/10.1111/nph.13426)) |
| Otu53823 | 1 | k__Fungi(100);p__Ascomycota(100);c__Saccharomycetes(100);o__Saccharomycetales(100);f__Metschnikowiaceae(100);g__Metschnikowia(100);s__Metschnikowia_pulcherrima(100); | Metschnikowia | 13 | Saprotroph | \|Undefined Saprotroph\| | Yeast | NULL | Probable | As nectar-tap_saprotroph (Põlme et al. 2020); Decay sugar-rich substrates (Põlme et al. 2020) | Põlme S, et al. 2020. Fungal Diversity 105:1-16 ((https://doi.org/10.1007/s13225-020-00466-2)) |
| Otu53826 | 1 | k__Fungi(100);p__Ascomycota(100);c__Saccharomycetes(100);o__Saccharomycetales(100);f__Saccharomycetales_family_Incertae_sedis(100);g__Candida(100);g__Candida_unclassified(100); | Candida | 13 | Pathotroph-Saprotroph-Symbiotroph | Animal Pathogen-Endophyte-Endosymbiont-Epiphyte-\|Undefined Saprotroph\| | Dimorphic Yeast-Yeast | NULL | Probable | As nectar-tap_saprotroph (Põlme et al. 2020); Decay sugar-rich substrates (Põlme et al. 2020); This asexual genus is polyphyletic and contain species that function very differently depend on what they are. Caution should be taken when interpreting guilds for Candida species. | Manolakaki D, et al. 2010. Virulence. 1: 367-375 ((https://doi.org/10.4161/viru.1.5.12796)); Mekbib SB, et al. 2011. Tropical Plant Pathology 36:233-240 ((https://doi.org/10.1590/S1982-56762011000400004)); Põlme S, et al. 2020. Fungal Diversity 105:1-16 ((https://doi.org/10.1007/s13225-020-00466-2)) |
| Otu53829 | 1 | k__Fungi(100);p__Ascomycota(100);c__Saccharomycetes(100);o__Saccharomycetales(100);f__Metschnikowiaceae(100);g__Metschnikowia(100);s__Metschnikowia_pulcherrima(100); | Metschnikowia | 13 | Saprotroph | \|Undefined Saprotroph\| | Yeast | NULL | Probable | As nectar-tap_saprotroph (Põlme et al. 2020); Decay sugar-rich substrates (Põlme et al. 2020) | Põlme S, et al. 2020. Fungal Diversity 105:1-16 ((https://doi.org/10.1007/s13225-020-00466-2)) |
| Otu53830 | 1 | k__Fungi(100);p__Ascomycota(100);c__Saccharomycetes(100);o__Saccharomycetales(100);f__Metschnikowiaceae(100);g__Metschnikowia(100);s__Metschnikowia_pulcherrima(100); | Metschnikowia | 13 | Saprotroph | \|Undefined Saprotroph\| | Yeast | NULL | Probable | As nectar-tap_saprotroph (Põlme et al. 2020); Decay sugar-rich substrates (Põlme et al. 2020) | Põlme S, et al. 2020. Fungal Diversity 105:1-16 ((https://doi.org/10.1007/s13225-020-00466-2)) |
| Otu53834 | 1 | k__Fungi(100);p__Ascomycota(100);c__Saccharomycetes(100);o__Saccharomycetales(100);f__Metschnikowiaceae(100);g__Metschnikowia(100);s__Metschnikowia_pulcherrima(100); | Metschnikowia | 13 | Saprotroph | \|Undefined Saprotroph\| | Yeast | NULL | Probable | As nectar-tap_saprotroph (Põlme et al. 2020); Decay sugar-rich substrates (Põlme et al. 2020) | Põlme S, et al. 2020. Fungal Diversity 105:1-16 ((https://doi.org/10.1007/s13225-020-00466-2)) |
| Otu53837 | 1 | k__Fungi(100);p__Ascomycota(100);c__Saccharomycetes(100);o__Saccharomycetales(100);f__Metschnikowiaceae(100);g__Metschnikowia(100);s__Metschnikowia_pulcherrima(100); | Metschnikowia | 13 | Saprotroph | \|Undefined Saprotroph\| | Yeast | NULL | Probable | As nectar-tap_saprotroph (Põlme et al. 2020); Decay sugar-rich substrates (Põlme et al. 2020) | Põlme S, et al. 2020. Fungal Diversity 105:1-16 ((https://doi.org/10.1007/s13225-020-00466-2)) |
| Otu53838 | 1 | k__Fungi(100);p__Ascomycota(100);c__Saccharomycetes(100);o__Saccharomycetales(100);f__Metschnikowiaceae(100);g__Metschnikowia(100);s__Metschnikowia_pulcherrima(100); | Metschnikowia | 13 | Saprotroph | \|Undefined Saprotroph\| | Yeast | NULL | Probable | As nectar-tap_saprotroph (Põlme et al. 2020); Decay sugar-rich substrates (Põlme et al. 2020) | Põlme S, et al. 2020. Fungal Diversity 105:1-16 ((https://doi.org/10.1007/s13225-020-00466-2)) |
| Otu53839 | 1 | k__Fungi(100);p__Ascomycota(100);c__Saccharomycetes(100);o__Saccharomycetales(100);f__Metschnikowiaceae(100);g__Metschnikowia(100);s__Metschnikowia_pulcherrima(100); | Metschnikowia | 13 | Saprotroph | \|Undefined Saprotroph\| | Yeast | NULL | Probable | As nectar-tap_saprotroph (Põlme et al. 2020); Decay sugar-rich substrates (Põlme et al. 2020) | Põlme S, et al. 2020. Fungal Diversity 105:1-16 ((https://doi.org/10.1007/s13225-020-00466-2)) |
| Otu53840 | 1 | k__Fungi(100);p__Ascomycota(100);c__Saccharomycetes(100);o__Saccharomycetales(100);f__Metschnikowiaceae(100);g__Metschnikowia(100);s__Metschnikowia_pulcherrima(100); | Metschnikowia | 13 | Saprotroph | \|Undefined Saprotroph\| | Yeast | NULL | Probable | As nectar-tap_saprotroph (Põlme et al. 2020); Decay sugar-rich substrates (Põlme et al. 2020) | Põlme S, et al. 2020. Fungal Diversity 105:1-16 ((https://doi.org/10.1007/s13225-020-00466-2)) |
| Otu53841 | 1 | k__Fungi(100);p__Ascomycota(100);c__Saccharomycetes(100);o__Saccharomycetales(100);f__Metschnikowiaceae(100);g__Metschnikowia(100);g__Metschnikowia_unclassified(100); | Metschnikowia | 13 | Saprotroph | \|Undefined Saprotroph\| | Yeast | NULL | Probable | As nectar-tap_saprotroph (Põlme et al. 2020); Decay sugar-rich substrates (Põlme et al. 2020) | Põlme S, et al. 2020. Fungal Diversity 105:1-16 ((https://doi.org/10.1007/s13225-020-00466-2)) |
| Otu53842 | 1 | k__Fungi(100);p__Ascomycota(100);c__Saccharomycetes(100);o__Saccharomycetales(100);f__Metschnikowiaceae(100);g__Metschnikowia(100);s__Metschnikowia_pulcherrima(100); | Metschnikowia | 13 | Saprotroph | \|Undefined Saprotroph\| | Yeast | NULL | Probable | As nectar-tap_saprotroph (Põlme et al. 2020); Decay sugar-rich substrates (Põlme et al. 2020) | Põlme S, et al. 2020. Fungal Diversity 105:1-16 ((https://doi.org/10.1007/s13225-020-00466-2)) |
| Otu53843 | 1 | k__Fungi(100);p__Ascomycota(100);c__Saccharomycetes(100);o__Saccharomycetales(100);f__Metschnikowiaceae(100);g__Metschnikowia(100);s__Metschnikowia_pulcherrima(100); | Metschnikowia | 13 | Saprotroph | \|Undefined Saprotroph\| | Yeast | NULL | Probable | As nectar-tap_saprotroph (Põlme et al. 2020); Decay sugar-rich substrates (Põlme et al. 2020) | Põlme S, et al. 2020. Fungal Diversity 105:1-16 ((https://doi.org/10.1007/s13225-020-00466-2)) |
| Otu53844 | 1 | k__Fungi(100);p__Ascomycota(100);c__Saccharomycetes(100);o__Saccharomycetales(100);f__Metschnikowiaceae(100);g__Metschnikowia(100);s__Metschnikowia_pulcherrima(100); | Metschnikowia | 13 | Saprotroph | \|Undefined Saprotroph\| | Yeast | NULL | Probable | As nectar-tap_saprotroph (Põlme et al. 2020); Decay sugar-rich substrates (Põlme et al. 2020) | Põlme S, et al. 2020. Fungal Diversity 105:1-16 ((https://doi.org/10.1007/s13225-020-00466-2)) |
| Otu53845 | 1 | k__Fungi(100);p__Ascomycota(100);c__Saccharomycetes(100);o__Saccharomycetales(100);f__Metschnikowiaceae(100);g__Metschnikowia(100);s__Metschnikowia_pulcherrima(100); | Metschnikowia | 13 | Saprotroph | \|Undefined Saprotroph\| | Yeast | NULL | Probable | As nectar-tap_saprotroph (Põlme et al. 2020); Decay sugar-rich substrates (Põlme et al. 2020) | Põlme S, et al. 2020. Fungal Diversity 105:1-16 ((https://doi.org/10.1007/s13225-020-00466-2)) |
| Otu53846 | 1 | k__Fungi(100);p__Ascomycota(100);c__Saccharomycetes(100);o__Saccharomycetales(100);f__Metschnikowiaceae(100);g__Metschnikowia(100);s__Metschnikowia_pulcherrima(100); | Metschnikowia | 13 | Saprotroph | \|Undefined Saprotroph\| | Yeast | NULL | Probable | As nectar-tap_saprotroph (Põlme et al. 2020); Decay sugar-rich substrates (Põlme et al. 2020) | Põlme S, et al. 2020. Fungal Diversity 105:1-16 ((https://doi.org/10.1007/s13225-020-00466-2)) |
| Otu53847 | 1 | k__Fungi(100);p__Ascomycota(100);c__Saccharomycetes(100);o__Saccharomycetales(100);f__Metschnikowiaceae(100);g__Metschnikowia(100);s__Metschnikowia_pulcherrima(100); | Metschnikowia | 13 | Saprotroph | \|Undefined Saprotroph\| | Yeast | NULL | Probable | As nectar-tap_saprotroph (Põlme et al. 2020); Decay sugar-rich substrates (Põlme et al. 2020) | Põlme S, et al. 2020. Fungal Diversity 105:1-16 ((https://doi.org/10.1007/s13225-020-00466-2)) |
| Otu53848 | 1 | k__Fungi(100);p__Ascomycota(100);c__Saccharomycetes(100);o__Saccharomycetales(100);f__Metschnikowiaceae(100);g__Metschnikowia(100);s__Metschnikowia_pulcherrima(100); | Metschnikowia | 13 | Saprotroph | \|Undefined Saprotroph\| | Yeast | NULL | Probable | As nectar-tap_saprotroph (Põlme et al. 2020); Decay sugar-rich substrates (Põlme et al. 2020) | Põlme S, et al. 2020. Fungal Diversity 105:1-16 ((https://doi.org/10.1007/s13225-020-00466-2)) |
| Otu53849 | 1 | k__Fungi(100);p__Ascomycota(100);c__Saccharomycetes(100);o__Saccharomycetales(100);f__Metschnikowiaceae(100);g__Metschnikowia(100);g__Metschnikowia_unclassified(100); | Metschnikowia | 13 | Saprotroph | \|Undefined Saprotroph\| | Yeast | NULL | Probable | As nectar-tap_saprotroph (Põlme et al. 2020); Decay sugar-rich substrates (Põlme et al. 2020) | Põlme S, et al. 2020. Fungal Diversity 105:1-16 ((https://doi.org/10.1007/s13225-020-00466-2)) |
| Otu53852 | 1 | k__Fungi(100);p__Ascomycota(100);c__Saccharomycetes(100);o__Saccharomycetales(100);f__Metschnikowiaceae(100);g__Metschnikowia(100);s__Metschnikowia_pulcherrima(100); | Metschnikowia | 13 | Saprotroph | \|Undefined Saprotroph\| | Yeast | NULL | Probable | As nectar-tap_saprotroph (Põlme et al. 2020); Decay sugar-rich substrates (Põlme et al. 2020) | Põlme S, et al. 2020. Fungal Diversity 105:1-16 ((https://doi.org/10.1007/s13225-020-00466-2)) |
| Otu53853 | 1 | k__Fungi(100);p__Ascomycota(100);c__Saccharomycetes(100);o__Saccharomycetales(100);f__Metschnikowiaceae(100);g__Metschnikowia(100);g__Metschnikowia_unclassified(100); | Metschnikowia | 13 | Saprotroph | \|Undefined Saprotroph\| | Yeast | NULL | Probable | As nectar-tap_saprotroph (Põlme et al. 2020); Decay sugar-rich substrates (Põlme et al. 2020) | Põlme S, et al. 2020. Fungal Diversity 105:1-16 ((https://doi.org/10.1007/s13225-020-00466-2)) |
| Otu53854 | 1 | k__Fungi(100);p__Ascomycota(100);c__Saccharomycetes(100);o__Saccharomycetales(100);f__Metschnikowiaceae(100);g__Metschnikowia(100);s__Metschnikowia_pulcherrima(100); | Metschnikowia | 13 | Saprotroph | \|Undefined Saprotroph\| | Yeast | NULL | Probable | As nectar-tap_saprotroph (Põlme et al. 2020); Decay sugar-rich substrates (Põlme et al. 2020) | Põlme S, et al. 2020. Fungal Diversity 105:1-16 ((https://doi.org/10.1007/s13225-020-00466-2)) |
| Otu53857 | 1 | k__Fungi(100);p__Ascomycota(100);c__Saccharomycetes(100);o__Saccharomycetales(100);f__Metschnikowiaceae(100);g__Metschnikowia(100);s__Metschnikowia_pulcherrima(100); | Metschnikowia | 13 | Saprotroph | \|Undefined Saprotroph\| | Yeast | NULL | Probable | As nectar-tap_saprotroph (Põlme et al. 2020); Decay sugar-rich substrates (Põlme et al. 2020) | Põlme S, et al. 2020. Fungal Diversity 105:1-16 ((https://doi.org/10.1007/s13225-020-00466-2)) |
| Otu53859 | 1 | k__Fungi(100);p__Ascomycota(100);c__Saccharomycetes(100);o__Saccharomycetales(100);f__Metschnikowiaceae(100);g__Metschnikowia(100);s__Metschnikowia_pulcherrima(100); | Metschnikowia | 13 | Saprotroph | \|Undefined Saprotroph\| | Yeast | NULL | Probable | As nectar-tap_saprotroph (Põlme et al. 2020); Decay sugar-rich substrates (Põlme et al. 2020) | Põlme S, et al. 2020. Fungal Diversity 105:1-16 ((https://doi.org/10.1007/s13225-020-00466-2)) |
| Otu53863 | 1 | k__Fungi(100);p__Ascomycota(100);c__Saccharomycetes(100);o__Saccharomycetales(100);f__Metschnikowiaceae(100);g__Metschnikowia(100);s__Metschnikowia_pulcherrima(100); | Metschnikowia | 13 | Saprotroph | \|Undefined Saprotroph\| | Yeast | NULL | Probable | As nectar-tap_saprotroph (Põlme et al. 2020); Decay sugar-rich substrates (Põlme et al. 2020) | Põlme S, et al. 2020. Fungal Diversity 105:1-16 ((https://doi.org/10.1007/s13225-020-00466-2)) |
| Otu53864 | 1 | k__Fungi(100);p__Ascomycota(100);c__Saccharomycetes(100);o__Saccharomycetales(100);f__Metschnikowiaceae(100);g__Metschnikowia(100);s__Metschnikowia_pulcherrima(100); | Metschnikowia | 13 | Saprotroph | \|Undefined Saprotroph\| | Yeast | NULL | Probable | As nectar-tap_saprotroph (Põlme et al. 2020); Decay sugar-rich substrates (Põlme et al. 2020) | Põlme S, et al. 2020. Fungal Diversity 105:1-16 ((https://doi.org/10.1007/s13225-020-00466-2)) |
| Otu53865 | 1 | k__Fungi(100);p__Ascomycota(100);c__Saccharomycetes(100);o__Saccharomycetales(100);f__Metschnikowiaceae(100);g__Metschnikowia(100);s__Metschnikowia_pulcherrima(100); | Metschnikowia | 13 | Saprotroph | \|Undefined Saprotroph\| | Yeast | NULL | Probable | As nectar-tap_saprotroph (Põlme et al. 2020); Decay sugar-rich substrates (Põlme et al. 2020) | Põlme S, et al. 2020. Fungal Diversity 105:1-16 ((https://doi.org/10.1007/s13225-020-00466-2)) |
| Otu53866 | 1 | k__Fungi(100);p__Ascomycota(100);c__Saccharomycetes(100);o__Saccharomycetales(100);f__Metschnikowiaceae(100);g__Metschnikowia(100);s__Metschnikowia_pulcherrima(100); | Metschnikowia | 13 | Saprotroph | \|Undefined Saprotroph\| | Yeast | NULL | Probable | As nectar-tap_saprotroph (Põlme et al. 2020); Decay sugar-rich substrates (Põlme et al. 2020) | Põlme S, et al. 2020. Fungal Diversity 105:1-16 ((https://doi.org/10.1007/s13225-020-00466-2)) |
| Otu53867 | 1 | k__Fungi(100);p__Ascomycota(100);c__Saccharomycetes(100);o__Saccharomycetales(100);f__Metschnikowiaceae(100);g__Metschnikowia(100);g__Metschnikowia_unclassified(100); | Metschnikowia | 13 | Saprotroph | \|Undefined Saprotroph\| | Yeast | NULL | Probable | As nectar-tap_saprotroph (Põlme et al. 2020); Decay sugar-rich substrates (Põlme et al. 2020) | Põlme S, et al. 2020. Fungal Diversity 105:1-16 ((https://doi.org/10.1007/s13225-020-00466-2)) |
| Otu53868 | 1 | k__Fungi(100);p__Ascomycota(100);c__Saccharomycetes(100);o__Saccharomycetales(100);f__Metschnikowiaceae(100);g__Metschnikowia(100);s__Metschnikowia_pulcherrima(100); | Metschnikowia | 13 | Saprotroph | \|Undefined Saprotroph\| | Yeast | NULL | Probable | As nectar-tap_saprotroph (Põlme et al. 2020); Decay sugar-rich substrates (Põlme et al. 2020) | Põlme S, et al. 2020. Fungal Diversity 105:1-16 ((https://doi.org/10.1007/s13225-020-00466-2)) |
| Otu53869 | 1 | k__Fungi(100);p__Ascomycota(100);c__Saccharomycetes(100);o__Saccharomycetales(100);f__Metschnikowiaceae(100);g__Metschnikowia(100);s__Metschnikowia_pulcherrima(100); | Metschnikowia | 13 | Saprotroph | \|Undefined Saprotroph\| | Yeast | NULL | Probable | As nectar-tap_saprotroph (Põlme et al. 2020); Decay sugar-rich substrates (Põlme et al. 2020) | Põlme S, et al. 2020. Fungal Diversity 105:1-16 ((https://doi.org/10.1007/s13225-020-00466-2)) |
| Otu53870 | 1 | k__Fungi(100);p__Ascomycota(100);c__Saccharomycetes(100);o__Saccharomycetales(100);f__Metschnikowiaceae(100);g__Metschnikowia(100);g__Metschnikowia_unclassified(100); | Metschnikowia | 13 | Saprotroph | \|Undefined Saprotroph\| | Yeast | NULL | Probable | As nectar-tap_saprotroph (Põlme et al. 2020); Decay sugar-rich substrates (Põlme et al. 2020) | Põlme S, et al. 2020. Fungal Diversity 105:1-16 ((https://doi.org/10.1007/s13225-020-00466-2)) |
| Otu53871 | 1 | k__Fungi(100);p__Ascomycota(100);c__Saccharomycetes(100);o__Saccharomycetales(100);f__Metschnikowiaceae(100);g__Metschnikowia(100);s__Metschnikowia_pulcherrima(100); | Metschnikowia | 13 | Saprotroph | \|Undefined Saprotroph\| | Yeast | NULL | Probable | As nectar-tap_saprotroph (Põlme et al. 2020); Decay sugar-rich substrates (Põlme et al. 2020) | Põlme S, et al. 2020. Fungal Diversity 105:1-16 ((https://doi.org/10.1007/s13225-020-00466-2)) |
| Otu53872 | 1 | k__Fungi(100);p__Ascomycota(100);c__Saccharomycetes(100);o__Saccharomycetales(100);f__Metschnikowiaceae(100);g__Metschnikowia(100);s__Metschnikowia_pulcherrima(100); | Metschnikowia | 13 | Saprotroph | \|Undefined Saprotroph\| | Yeast | NULL | Probable | As nectar-tap_saprotroph (Põlme et al. 2020); Decay sugar-rich substrates (Põlme et al. 2020) | Põlme S, et al. 2020. Fungal Diversity 105:1-16 ((https://doi.org/10.1007/s13225-020-00466-2)) |
| Otu53873 | 1 | k__Fungi(100);p__Ascomycota(100);c__Saccharomycetes(100);o__Saccharomycetales(100);f__Metschnikowiaceae(100);g__Metschnikowia(100);s__Metschnikowia_pulcherrima(100); | Metschnikowia | 13 | Saprotroph | \|Undefined Saprotroph\| | Yeast | NULL | Probable | As nectar-tap_saprotroph (Põlme et al. 2020); Decay sugar-rich substrates (Põlme et al. 2020) | Põlme S, et al. 2020. Fungal Diversity 105:1-16 ((https://doi.org/10.1007/s13225-020-00466-2)) |
| Otu53876 | 1 | k__Fungi(100);p__Ascomycota(100);c__Saccharomycetes(100);o__Saccharomycetales(100);f__Metschnikowiaceae(100);g__Metschnikowia(100);s__Metschnikowia_pulcherrima(100); | Metschnikowia | 13 | Saprotroph | \|Undefined Saprotroph\| | Yeast | NULL | Probable | As nectar-tap_saprotroph (Põlme et al. 2020); Decay sugar-rich substrates (Põlme et al. 2020) | Põlme S, et al. 2020. Fungal Diversity 105:1-16 ((https://doi.org/10.1007/s13225-020-00466-2)) |
| Otu53877 | 1 | k__Fungi(100);p__Ascomycota(100);c__Saccharomycetes(100);o__Saccharomycetales(100);f__Metschnikowiaceae(100);g__Metschnikowia(100);g__Metschnikowia_unclassified(100); | Metschnikowia | 13 | Saprotroph | \|Undefined Saprotroph\| | Yeast | NULL | Probable | As nectar-tap_saprotroph (Põlme et al. 2020); Decay sugar-rich substrates (Põlme et al. 2020) | Põlme S, et al. 2020. Fungal Diversity 105:1-16 ((https://doi.org/10.1007/s13225-020-00466-2)) |
| Otu53879 | 1 | k__Fungi(100);p__Ascomycota(100);c__Saccharomycetes(100);o__Saccharomycetales(100);f__Saccharomycetales_family_Incertae_sedis(100);f__Saccharomycetales_family_Incertae_sedis_unclassified(100);f__Saccharomycetales_family_Incertae_sedis_unclassified(100); | Saccharomycetales | 7 | Saprotroph | Undefined Saprotroph | Yeast | NULL | Possible | NULL | Sterkenburg E, et al. 2015. New Phytologist 207:1145-1158 ((https://doi.org/10.1111/nph.13426)) |
| Otu53880 | 1 | k__Fungi(100);p__Ascomycota(100);c__Saccharomycetes(100);o__Saccharomycetales(100);f__Metschnikowiaceae(100);g__Metschnikowia(100);s__Metschnikowia_pulcherrima(100); | Metschnikowia | 13 | Saprotroph | \|Undefined Saprotroph\| | Yeast | NULL | Probable | As nectar-tap_saprotroph (Põlme et al. 2020); Decay sugar-rich substrates (Põlme et al. 2020) | Põlme S, et al. 2020. Fungal Diversity 105:1-16 ((https://doi.org/10.1007/s13225-020-00466-2)) |
| Otu53881 | 1 | k__Fungi(100);p__Basidiomycota(100);c__Microbotryomycetes(100);o__Sporidiobolales(100);f__Sporidiobolales_family_Incertae_sedis(100);g__Sporobolomyces(100);s__Sporobolomyces_lactosus(100); | Sporobolomyces | 13 | Pathotroph-Saprotroph | \|Fungal Parasite\|-Plant Saprotroph | Yeast | NULL | Probable | As litter_saprotroph (Põlme et al. 2020); As mycoparasite (Põlme et al. 2020); Decay fungal material (Põlme et al. 2020) | Tedersoo L, et al. 2014. Science 346:e1256688 ((https://doi.org/10.1126/science.1256688)); Purahong W, et al. 2016. Molecular Ecology 25:4059-4074 ((https://doi.org/10.1111/mec.13739)); Põlme S, et al. 2020. Fungal Diversity 105:1-16 ((https://doi.org/10.1007/s13225-020-00466-2)) |
| Otu53882 | 1 | k__Fungi(100);p__Ascomycota(100);c__Saccharomycetes(100);o__Saccharomycetales(100);f__Metschnikowiaceae(100);g__Metschnikowia(100);g__Metschnikowia_unclassified(100); | Metschnikowia | 13 | Saprotroph | \|Undefined Saprotroph\| | Yeast | NULL | Probable | As nectar-tap_saprotroph (Põlme et al. 2020); Decay sugar-rich substrates (Põlme et al. 2020) | Põlme S, et al. 2020. Fungal Diversity 105:1-16 ((https://doi.org/10.1007/s13225-020-00466-2)) |
| Otu53883 | 1 | k__Fungi(100);p__Ascomycota(100);c__Saccharomycetes(100);o__Saccharomycetales(100);f__Metschnikowiaceae(100);g__Metschnikowia(100);s__Metschnikowia_pulcherrima(100); | Metschnikowia | 13 | Saprotroph | \|Undefined Saprotroph\| | Yeast | NULL | Probable | As nectar-tap_saprotroph (Põlme et al. 2020); Decay sugar-rich substrates (Põlme et al. 2020) | Põlme S, et al. 2020. Fungal Diversity 105:1-16 ((https://doi.org/10.1007/s13225-020-00466-2)) |
| Otu53884 | 1 | k__Fungi(100);p__Ascomycota(100);c__Saccharomycetes(100);o__Saccharomycetales(100);f__Metschnikowiaceae(100);g__Metschnikowia(100);g__Metschnikowia_unclassified(100); | Metschnikowia | 13 | Saprotroph | \|Undefined Saprotroph\| | Yeast | NULL | Probable | As nectar-tap_saprotroph (Põlme et al. 2020); Decay sugar-rich substrates (Põlme et al. 2020) | Põlme S, et al. 2020. Fungal Diversity 105:1-16 ((https://doi.org/10.1007/s13225-020-00466-2)) |
| Otu53885 | 1 | k__Fungi(100);p__Ascomycota(100);c__Saccharomycetes(100);o__Saccharomycetales(100);f__Metschnikowiaceae(100);g__Metschnikowia(100);s__Metschnikowia_pulcherrima(100); | Metschnikowia | 13 | Saprotroph | \|Undefined Saprotroph\| | Yeast | NULL | Probable | As nectar-tap_saprotroph (Põlme et al. 2020); Decay sugar-rich substrates (Põlme et al. 2020) | Põlme S, et al. 2020. Fungal Diversity 105:1-16 ((https://doi.org/10.1007/s13225-020-00466-2)) |
| Otu53886 | 1 | k__Fungi(100);p__Ascomycota(100);c__Saccharomycetes(100);o__Saccharomycetales(100);f__Metschnikowiaceae(100);g__Metschnikowia(100);g__Metschnikowia_unclassified(100); | Metschnikowia | 13 | Saprotroph | \|Undefined Saprotroph\| | Yeast | NULL | Probable | As nectar-tap_saprotroph (Põlme et al. 2020); Decay sugar-rich substrates (Põlme et al. 2020) | Põlme S, et al. 2020. Fungal Diversity 105:1-16 ((https://doi.org/10.1007/s13225-020-00466-2)) |
| Otu53887 | 1 | k__Fungi(100);p__Ascomycota(100);c__Saccharomycetes(100);o__Saccharomycetales(100);f__Metschnikowiaceae(100);g__Metschnikowia(100);s__Metschnikowia_pulcherrima(100); | Metschnikowia | 13 | Saprotroph | \|Undefined Saprotroph\| | Yeast | NULL | Probable | As nectar-tap_saprotroph (Põlme et al. 2020); Decay sugar-rich substrates (Põlme et al. 2020) | Põlme S, et al. 2020. Fungal Diversity 105:1-16 ((https://doi.org/10.1007/s13225-020-00466-2)) |
| Otu53888 | 1 | k__Fungi(100);p__Ascomycota(100);c__Saccharomycetes(100);o__Saccharomycetales(100);f__Metschnikowiaceae(100);g__Metschnikowia(100);s__Metschnikowia_pulcherrima(100); | Metschnikowia | 13 | Saprotroph | \|Undefined Saprotroph\| | Yeast | NULL | Probable | As nectar-tap_saprotroph (Põlme et al. 2020); Decay sugar-rich substrates (Põlme et al. 2020) | Põlme S, et al. 2020. Fungal Diversity 105:1-16 ((https://doi.org/10.1007/s13225-020-00466-2)) |
| Otu53896 | 1 | k__Fungi(100);p__Ascomycota(100);c__Saccharomycetes(100);o__Saccharomycetales(100);f__Metschnikowiaceae(100);g__Metschnikowia(100);g__Metschnikowia_unclassified(100); | Metschnikowia | 13 | Saprotroph | \|Undefined Saprotroph\| | Yeast | NULL | Probable | As nectar-tap_saprotroph (Põlme et al. 2020); Decay sugar-rich substrates (Põlme et al. 2020) | Põlme S, et al. 2020. Fungal Diversity 105:1-16 ((https://doi.org/10.1007/s13225-020-00466-2)) |
| Otu53898 | 1 | k__Fungi(100);p__Ascomycota(100);c__Saccharomycetes(100);o__Saccharomycetales(100);f__Metschnikowiaceae(100);g__Metschnikowia(100);g__Metschnikowia_unclassified(100); | Metschnikowia | 13 | Saprotroph | \|Undefined Saprotroph\| | Yeast | NULL | Probable | As nectar-tap_saprotroph (Põlme et al. 2020); Decay sugar-rich substrates (Põlme et al. 2020) | Põlme S, et al. 2020. Fungal Diversity 105:1-16 ((https://doi.org/10.1007/s13225-020-00466-2)) |
| Otu53899 | 1 | k__Fungi(100);p__Ascomycota(100);c__Saccharomycetes(100);o__Saccharomycetales(100);f__Metschnikowiaceae(100);g__Metschnikowia(100);g__Metschnikowia_unclassified(100); | Metschnikowia | 13 | Saprotroph | \|Undefined Saprotroph\| | Yeast | NULL | Probable | As nectar-tap_saprotroph (Põlme et al. 2020); Decay sugar-rich substrates (Põlme et al. 2020) | Põlme S, et al. 2020. Fungal Diversity 105:1-16 ((https://doi.org/10.1007/s13225-020-00466-2)) |
| Otu53900 | 1 | k__Fungi(100);p__Ascomycota(100);c__Saccharomycetes(100);o__Saccharomycetales(100);f__Metschnikowiaceae(100);g__Metschnikowia(100);s__Metschnikowia_pulcherrima(100); | Metschnikowia | 13 | Saprotroph | \|Undefined Saprotroph\| | Yeast | NULL | Probable | As nectar-tap_saprotroph (Põlme et al. 2020); Decay sugar-rich substrates (Põlme et al. 2020) | Põlme S, et al. 2020. Fungal Diversity 105:1-16 ((https://doi.org/10.1007/s13225-020-00466-2)) |
| Otu53908 | 1 | k__Fungi(100);p__Ascomycota(100);c__Saccharomycetes(100);o__Saccharomycetales(100);f__Metschnikowiaceae(100);g__Metschnikowia(100);g__Metschnikowia_unclassified(100); | Metschnikowia | 13 | Saprotroph | \|Undefined Saprotroph\| | Yeast | NULL | Probable | As nectar-tap_saprotroph (Põlme et al. 2020); Decay sugar-rich substrates (Põlme et al. 2020) | Põlme S, et al. 2020. Fungal Diversity 105:1-16 ((https://doi.org/10.1007/s13225-020-00466-2)) |
| Otu53909 | 1 | k__Fungi(100);p__Ascomycota(100);c__Saccharomycetes(100);o__Saccharomycetales(100);f__Metschnikowiaceae(100);g__Metschnikowia(100);g__Metschnikowia_unclassified(100); | Metschnikowia | 13 | Saprotroph | \|Undefined Saprotroph\| | Yeast | NULL | Probable | As nectar-tap_saprotroph (Põlme et al. 2020); Decay sugar-rich substrates (Põlme et al. 2020) | Põlme S, et al. 2020. Fungal Diversity 105:1-16 ((https://doi.org/10.1007/s13225-020-00466-2)) |
| Otu53911 | 1 | k__Fungi(100);p__Ascomycota(100);c__Saccharomycetes(100);o__Saccharomycetales(100);f__Metschnikowiaceae(100);g__Metschnikowia(100);s__Metschnikowia_pulcherrima(100); | Metschnikowia | 13 | Saprotroph | \|Undefined Saprotroph\| | Yeast | NULL | Probable | As nectar-tap_saprotroph (Põlme et al. 2020); Decay sugar-rich substrates (Põlme et al. 2020) | Põlme S, et al. 2020. Fungal Diversity 105:1-16 ((https://doi.org/10.1007/s13225-020-00466-2)) |
| Otu53912 | 1 | k__Fungi(100);p__Ascomycota(100);c__Saccharomycetes(100);o__Saccharomycetales(100);f__Metschnikowiaceae(100);g__Metschnikowia(100);s__Metschnikowia_pulcherrima(100); | Metschnikowia | 13 | Saprotroph | \|Undefined Saprotroph\| | Yeast | NULL | Probable | As nectar-tap_saprotroph (Põlme et al. 2020); Decay sugar-rich substrates (Põlme et al. 2020) | Põlme S, et al. 2020. Fungal Diversity 105:1-16 ((https://doi.org/10.1007/s13225-020-00466-2)) |
| Otu53913 | 1 | k__Fungi(100);p__Ascomycota(100);c__Saccharomycetes(100);o__Saccharomycetales(100);f__Metschnikowiaceae(100);g__Metschnikowia(100);s__Metschnikowia_pulcherrima(100); | Metschnikowia | 13 | Saprotroph | \|Undefined Saprotroph\| | Yeast | NULL | Probable | As nectar-tap_saprotroph (Põlme et al. 2020); Decay sugar-rich substrates (Põlme et al. 2020) | Põlme S, et al. 2020. Fungal Diversity 105:1-16 ((https://doi.org/10.1007/s13225-020-00466-2)) |
| Otu53920 | 1 | k__Fungi(100);p__Ascomycota(100);c__Saccharomycetes(100);o__Saccharomycetales(100);f__Metschnikowiaceae(100);g__Metschnikowia(100);s__Metschnikowia_pulcherrima(100); | Metschnikowia | 13 | Saprotroph | \|Undefined Saprotroph\| | Yeast | NULL | Probable | As nectar-tap_saprotroph (Põlme et al. 2020); Decay sugar-rich substrates (Põlme et al. 2020) | Põlme S, et al. 2020. Fungal Diversity 105:1-16 ((https://doi.org/10.1007/s13225-020-00466-2)) |
| Otu53923 | 1 | k__Fungi(100);p__Ascomycota(100);c__Saccharomycetes(100);o__Saccharomycetales(100);f__Saccharomycetales_family_Incertae_sedis(100);g__Candida(100);g__Candida_unclassified(100); | Candida | 13 | Pathotroph-Saprotroph-Symbiotroph | Animal Pathogen-Endophyte-Endosymbiont-Epiphyte-\|Undefined Saprotroph\| | Dimorphic Yeast-Yeast | NULL | Probable | As nectar-tap_saprotroph (Põlme et al. 2020); Decay sugar-rich substrates (Põlme et al. 2020); This asexual genus is polyphyletic and contain species that function very differently depend on what they are. Caution should be taken when interpreting guilds for Candida species. | Manolakaki D, et al. 2010. Virulence. 1: 367-375 ((https://doi.org/10.4161/viru.1.5.12796)); Mekbib SB, et al. 2011. Tropical Plant Pathology 36:233-240 ((https://doi.org/10.1590/S1982-56762011000400004)); Põlme S, et al. 2020. Fungal Diversity 105:1-16 ((https://doi.org/10.1007/s13225-020-00466-2)) |
| Otu53928 | 1 | k__Fungi(100);p__Ascomycota(100);c__Saccharomycetes(100);o__Saccharomycetales(100);f__Metschnikowiaceae(100);g__Metschnikowia(100);s__Metschnikowia_pulcherrima(100); | Metschnikowia | 13 | Saprotroph | \|Undefined Saprotroph\| | Yeast | NULL | Probable | As nectar-tap_saprotroph (Põlme et al. 2020); Decay sugar-rich substrates (Põlme et al. 2020) | Põlme S, et al. 2020. Fungal Diversity 105:1-16 ((https://doi.org/10.1007/s13225-020-00466-2)) |
| Otu53929 | 1 | k__Fungi(100);p__Ascomycota(100);c__Saccharomycetes(100);o__Saccharomycetales(100);f__Metschnikowiaceae(100);g__Metschnikowia(100);s__Metschnikowia_pulcherrima(100); | Metschnikowia | 13 | Saprotroph | \|Undefined Saprotroph\| | Yeast | NULL | Probable | As nectar-tap_saprotroph (Põlme et al. 2020); Decay sugar-rich substrates (Põlme et al. 2020) | Põlme S, et al. 2020. Fungal Diversity 105:1-16 ((https://doi.org/10.1007/s13225-020-00466-2)) |
| Otu53933 | 1 | k__Fungi(100);p__Ascomycota(100);c__Saccharomycetes(100);o__Saccharomycetales(100);f__Metschnikowiaceae(100);g__Metschnikowia(100);g__Metschnikowia_unclassified(100); | Metschnikowia | 13 | Saprotroph | \|Undefined Saprotroph\| | Yeast | NULL | Probable | As nectar-tap_saprotroph (Põlme et al. 2020); Decay sugar-rich substrates (Põlme et al. 2020) | Põlme S, et al. 2020. Fungal Diversity 105:1-16 ((https://doi.org/10.1007/s13225-020-00466-2)) |
| Otu53935 | 1 | k__Fungi(100);p__Ascomycota(100);c__Saccharomycetes(100);o__Saccharomycetales(100);f__Metschnikowiaceae(100);g__Metschnikowia(100);s__Metschnikowia_pulcherrima(100); | Metschnikowia | 13 | Saprotroph | \|Undefined Saprotroph\| | Yeast | NULL | Probable | As nectar-tap_saprotroph (Põlme et al. 2020); Decay sugar-rich substrates (Põlme et al. 2020) | Põlme S, et al. 2020. Fungal Diversity 105:1-16 ((https://doi.org/10.1007/s13225-020-00466-2)) |
| Otu53937 | 1 | k__Fungi(100);p__Ascomycota(100);c__Saccharomycetes(100);o__Saccharomycetales(100);f__Saccharomycetales_family_Incertae_sedis(100);g__Candida(100);g__Candida_unclassified(100); | Candida | 13 | Pathotroph-Saprotroph-Symbiotroph | Animal Pathogen-Endophyte-Endosymbiont-Epiphyte-\|Undefined Saprotroph\| | Dimorphic Yeast-Yeast | NULL | Probable | As nectar-tap_saprotroph (Põlme et al. 2020); Decay sugar-rich substrates (Põlme et al. 2020); This asexual genus is polyphyletic and contain species that function very differently depend on what they are. Caution should be taken when interpreting guilds for Candida species. | Manolakaki D, et al. 2010. Virulence. 1: 367-375 ((https://doi.org/10.4161/viru.1.5.12796)); Mekbib SB, et al. 2011. Tropical Plant Pathology 36:233-240 ((https://doi.org/10.1590/S1982-56762011000400004)); Põlme S, et al. 2020. Fungal Diversity 105:1-16 ((https://doi.org/10.1007/s13225-020-00466-2)) |
| Otu53939 | 1 | k__Fungi(100);p__Ascomycota(100);c__Saccharomycetes(100);o__Saccharomycetales(100);f__Metschnikowiaceae(100);g__Metschnikowia(100);s__Metschnikowia_pulcherrima(100); | Metschnikowia | 13 | Saprotroph | \|Undefined Saprotroph\| | Yeast | NULL | Probable | As nectar-tap_saprotroph (Põlme et al. 2020); Decay sugar-rich substrates (Põlme et al. 2020) | Põlme S, et al. 2020. Fungal Diversity 105:1-16 ((https://doi.org/10.1007/s13225-020-00466-2)) |
| Otu53941 | 1 | k__Fungi(100);p__Ascomycota(100);c__Saccharomycetes(100);o__Saccharomycetales(100);f__Metschnikowiaceae(100);g__Metschnikowia(100);s__Metschnikowia_pulcherrima(100); | Metschnikowia | 13 | Saprotroph | \|Undefined Saprotroph\| | Yeast | NULL | Probable | As nectar-tap_saprotroph (Põlme et al. 2020); Decay sugar-rich substrates (Põlme et al. 2020) | Põlme S, et al. 2020. Fungal Diversity 105:1-16 ((https://doi.org/10.1007/s13225-020-00466-2)) |
| Otu53942 | 1 | k__Fungi(100);p__Ascomycota(100);c__Saccharomycetes(100);o__Saccharomycetales(100);f__Metschnikowiaceae(100);g__Metschnikowia(100);s__Metschnikowia_pulcherrima(100); | Metschnikowia | 13 | Saprotroph | \|Undefined Saprotroph\| | Yeast | NULL | Probable | As nectar-tap_saprotroph (Põlme et al. 2020); Decay sugar-rich substrates (Põlme et al. 2020) | Põlme S, et al. 2020. Fungal Diversity 105:1-16 ((https://doi.org/10.1007/s13225-020-00466-2)) |
| Otu53943 | 1 | k__Fungi(100);p__Ascomycota(100);c__Saccharomycetes(100);o__Saccharomycetales(100);f__Metschnikowiaceae(100);g__Metschnikowia(100);s__Metschnikowia_pulcherrima(100); | Metschnikowia | 13 | Saprotroph | \|Undefined Saprotroph\| | Yeast | NULL | Probable | As nectar-tap_saprotroph (Põlme et al. 2020); Decay sugar-rich substrates (Põlme et al. 2020) | Põlme S, et al. 2020. Fungal Diversity 105:1-16 ((https://doi.org/10.1007/s13225-020-00466-2)) |
| Otu53944 | 1 | k__Fungi(100);p__Ascomycota(100);c__Saccharomycetes(100);o__Saccharomycetales(100);f__Metschnikowiaceae(100);g__Metschnikowia(100);s__Metschnikowia_pulcherrima(100); | Metschnikowia | 13 | Saprotroph | \|Undefined Saprotroph\| | Yeast | NULL | Probable | As nectar-tap_saprotroph (Põlme et al. 2020); Decay sugar-rich substrates (Põlme et al. 2020) | Põlme S, et al. 2020. Fungal Diversity 105:1-16 ((https://doi.org/10.1007/s13225-020-00466-2)) |
| Otu53945 | 1 | k__Fungi(100);p__Ascomycota(100);c__Saccharomycetes(100);o__Saccharomycetales(100);f__Metschnikowiaceae(100);g__Metschnikowia(100);s__Metschnikowia_pulcherrima(100); | Metschnikowia | 13 | Saprotroph | \|Undefined Saprotroph\| | Yeast | NULL | Probable | As nectar-tap_saprotroph (Põlme et al. 2020); Decay sugar-rich substrates (Põlme et al. 2020) | Põlme S, et al. 2020. Fungal Diversity 105:1-16 ((https://doi.org/10.1007/s13225-020-00466-2)) |
| Otu53946 | 1 | k__Fungi(100);p__Ascomycota(100);c__Saccharomycetes(100);o__Saccharomycetales(100);f__Metschnikowiaceae(100);g__Metschnikowia(100);s__Metschnikowia_pulcherrima(100); | Metschnikowia | 13 | Saprotroph | \|Undefined Saprotroph\| | Yeast | NULL | Probable | As nectar-tap_saprotroph (Põlme et al. 2020); Decay sugar-rich substrates (Põlme et al. 2020) | Põlme S, et al. 2020. Fungal Diversity 105:1-16 ((https://doi.org/10.1007/s13225-020-00466-2)) |
| Otu53948 | 1 | k__Fungi(100);p__Ascomycota(100);c__Saccharomycetes(100);o__Saccharomycetales(100);f__Metschnikowiaceae(100);g__Metschnikowia(100);g__Metschnikowia_unclassified(100); | Metschnikowia | 13 | Saprotroph | \|Undefined Saprotroph\| | Yeast | NULL | Probable | As nectar-tap_saprotroph (Põlme et al. 2020); Decay sugar-rich substrates (Põlme et al. 2020) | Põlme S, et al. 2020. Fungal Diversity 105:1-16 ((https://doi.org/10.1007/s13225-020-00466-2)) |
| Otu53949 | 1 | k__Fungi(100);p__Ascomycota(100);c__Saccharomycetes(100);o__Saccharomycetales(100);f__Metschnikowiaceae(100);g__Metschnikowia(100);g__Metschnikowia_unclassified(100); | Metschnikowia | 13 | Saprotroph | \|Undefined Saprotroph\| | Yeast | NULL | Probable | As nectar-tap_saprotroph (Põlme et al. 2020); Decay sugar-rich substrates (Põlme et al. 2020) | Põlme S, et al. 2020. Fungal Diversity 105:1-16 ((https://doi.org/10.1007/s13225-020-00466-2)) |
| Otu53950 | 1 | k__Fungi(100);p__Ascomycota(100);c__Saccharomycetes(100);o__Saccharomycetales(100);f__Metschnikowiaceae(100);g__Metschnikowia(100);s__Metschnikowia_pulcherrima(100); | Metschnikowia | 13 | Saprotroph | \|Undefined Saprotroph\| | Yeast | NULL | Probable | As nectar-tap_saprotroph (Põlme et al. 2020); Decay sugar-rich substrates (Põlme et al. 2020) | Põlme S, et al. 2020. Fungal Diversity 105:1-16 ((https://doi.org/10.1007/s13225-020-00466-2)) |
| Otu53952 | 1 | k__Fungi(100);p__Ascomycota(100);c__Saccharomycetes(100);o__Saccharomycetales(100);f__Saccharomycetales_family_Incertae_sedis(100);g__Candida(100);g__Candida_unclassified(100); | Candida | 13 | Pathotroph-Saprotroph-Symbiotroph | Animal Pathogen-Endophyte-Endosymbiont-Epiphyte-\|Undefined Saprotroph\| | Dimorphic Yeast-Yeast | NULL | Probable | As nectar-tap_saprotroph (Põlme et al. 2020); Decay sugar-rich substrates (Põlme et al. 2020); This asexual genus is polyphyletic and contain species that function very differently depend on what they are. Caution should be taken when interpreting guilds for Candida species. | Manolakaki D, et al. 2010. Virulence. 1: 367-375 ((https://doi.org/10.4161/viru.1.5.12796)); Mekbib SB, et al. 2011. Tropical Plant Pathology 36:233-240 ((https://doi.org/10.1590/S1982-56762011000400004)); Põlme S, et al. 2020. Fungal Diversity 105:1-16 ((https://doi.org/10.1007/s13225-020-00466-2)) |
| Otu53953 | 1 | k__Fungi(100);p__Ascomycota(100);c__Saccharomycetes(100);o__Saccharomycetales(100);f__Metschnikowiaceae(100);g__Metschnikowia(100);s__Metschnikowia_pulcherrima(100); | Metschnikowia | 13 | Saprotroph | \|Undefined Saprotroph\| | Yeast | NULL | Probable | As nectar-tap_saprotroph (Põlme et al. 2020); Decay sugar-rich substrates (Põlme et al. 2020) | Põlme S, et al. 2020. Fungal Diversity 105:1-16 ((https://doi.org/10.1007/s13225-020-00466-2)) |
| Otu53954 | 1 | k__Fungi(100);p__Ascomycota(100);c__Saccharomycetes(100);o__Saccharomycetales(100);f__Metschnikowiaceae(100);g__Metschnikowia(100);g__Metschnikowia_unclassified(100); | Metschnikowia | 13 | Saprotroph | \|Undefined Saprotroph\| | Yeast | NULL | Probable | As nectar-tap_saprotroph (Põlme et al. 2020); Decay sugar-rich substrates (Põlme et al. 2020) | Põlme S, et al. 2020. Fungal Diversity 105:1-16 ((https://doi.org/10.1007/s13225-020-00466-2)) |
| Otu53956 | 1 | k__Fungi(100);p__Ascomycota(100);c__Saccharomycetes(100);o__Saccharomycetales(100);f__Saccharomycetales_family_Incertae_sedis(100);g__Candida(100);g__Candida_unclassified(100); | Candida | 13 | Pathotroph-Saprotroph-Symbiotroph | Animal Pathogen-Endophyte-Endosymbiont-Epiphyte-\|Undefined Saprotroph\| | Dimorphic Yeast-Yeast | NULL | Probable | As nectar-tap_saprotroph (Põlme et al. 2020); Decay sugar-rich substrates (Põlme et al. 2020); This asexual genus is polyphyletic and contain species that function very differently depend on what they are. Caution should be taken when interpreting guilds for Candida species. | Manolakaki D, et al. 2010. Virulence. 1: 367-375 ((https://doi.org/10.4161/viru.1.5.12796)); Mekbib SB, et al. 2011. Tropical Plant Pathology 36:233-240 ((https://doi.org/10.1590/S1982-56762011000400004)); Põlme S, et al. 2020. Fungal Diversity 105:1-16 ((https://doi.org/10.1007/s13225-020-00466-2)) |
| Otu53958 | 1 | k__Fungi(100);p__Ascomycota(100);c__Saccharomycetes(100);o__Saccharomycetales(100);f__Metschnikowiaceae(100);g__Metschnikowia(100);s__Metschnikowia_pulcherrima(100); | Metschnikowia | 13 | Saprotroph | \|Undefined Saprotroph\| | Yeast | NULL | Probable | As nectar-tap_saprotroph (Põlme et al. 2020); Decay sugar-rich substrates (Põlme et al. 2020) | Põlme S, et al. 2020. Fungal Diversity 105:1-16 ((https://doi.org/10.1007/s13225-020-00466-2)) |
| Otu53960 | 1 | k__Fungi(100);p__Ascomycota(100);c__Saccharomycetes(100);o__Saccharomycetales(100);f__Metschnikowiaceae(100);g__Metschnikowia(100);s__Metschnikowia_pulcherrima(100); | Metschnikowia | 13 | Saprotroph | \|Undefined Saprotroph\| | Yeast | NULL | Probable | As nectar-tap_saprotroph (Põlme et al. 2020); Decay sugar-rich substrates (Põlme et al. 2020) | Põlme S, et al. 2020. Fungal Diversity 105:1-16 ((https://doi.org/10.1007/s13225-020-00466-2)) |
| Otu53961 | 1 | k__Fungi(100);p__Ascomycota(100);c__Saccharomycetes(100);o__Saccharomycetales(100);f__Metschnikowiaceae(100);g__Metschnikowia(100);s__Metschnikowia_pulcherrima(100); | Metschnikowia | 13 | Saprotroph | \|Undefined Saprotroph\| | Yeast | NULL | Probable | As nectar-tap_saprotroph (Põlme et al. 2020); Decay sugar-rich substrates (Põlme et al. 2020) | Põlme S, et al. 2020. Fungal Diversity 105:1-16 ((https://doi.org/10.1007/s13225-020-00466-2)) |
| Otu53962 | 1 | k__Fungi(100);p__Ascomycota(100);c__Saccharomycetes(100);o__Saccharomycetales(100);f__Metschnikowiaceae(100);g__Metschnikowia(100);s__Metschnikowia_pulcherrima(100); | Metschnikowia | 13 | Saprotroph | \|Undefined Saprotroph\| | Yeast | NULL | Probable | As nectar-tap_saprotroph (Põlme et al. 2020); Decay sugar-rich substrates (Põlme et al. 2020) | Põlme S, et al. 2020. Fungal Diversity 105:1-16 ((https://doi.org/10.1007/s13225-020-00466-2)) |
| Otu53963 | 1 | k__Fungi(100);p__Ascomycota(100);c__Saccharomycetes(100);o__Saccharomycetales(100);f__Metschnikowiaceae(100);g__Metschnikowia(100);s__Metschnikowia_pulcherrima(100); | Metschnikowia | 13 | Saprotroph | \|Undefined Saprotroph\| | Yeast | NULL | Probable | As nectar-tap_saprotroph (Põlme et al. 2020); Decay sugar-rich substrates (Põlme et al. 2020) | Põlme S, et al. 2020. Fungal Diversity 105:1-16 ((https://doi.org/10.1007/s13225-020-00466-2)) |
| Otu53964 | 1 | k__Fungi(100);p__Ascomycota(100);c__Saccharomycetes(100);o__Saccharomycetales(100);f__Metschnikowiaceae(100);g__Metschnikowia(100);g__Metschnikowia_unclassified(100); | Metschnikowia | 13 | Saprotroph | \|Undefined Saprotroph\| | Yeast | NULL | Probable | As nectar-tap_saprotroph (Põlme et al. 2020); Decay sugar-rich substrates (Põlme et al. 2020) | Põlme S, et al. 2020. Fungal Diversity 105:1-16 ((https://doi.org/10.1007/s13225-020-00466-2)) |
| Otu53965 | 1 | k__Fungi(100);p__Ascomycota(100);c__Saccharomycetes(100);o__Saccharomycetales(100);f__Metschnikowiaceae(100);g__Metschnikowia(100);s__Metschnikowia_pulcherrima(100); | Metschnikowia | 13 | Saprotroph | \|Undefined Saprotroph\| | Yeast | NULL | Probable | As nectar-tap_saprotroph (Põlme et al. 2020); Decay sugar-rich substrates (Põlme et al. 2020) | Põlme S, et al. 2020. Fungal Diversity 105:1-16 ((https://doi.org/10.1007/s13225-020-00466-2)) |
| Otu53966 | 1 | k__Fungi(100);p__Ascomycota(100);c__Saccharomycetes(100);o__Saccharomycetales(100);f__Metschnikowiaceae(100);g__Metschnikowia(100);s__Metschnikowia_pulcherrima(100); | Metschnikowia | 13 | Saprotroph | \|Undefined Saprotroph\| | Yeast | NULL | Probable | As nectar-tap_saprotroph (Põlme et al. 2020); Decay sugar-rich substrates (Põlme et al. 2020) | Põlme S, et al. 2020. Fungal Diversity 105:1-16 ((https://doi.org/10.1007/s13225-020-00466-2)) |
| Otu53967 | 1 | k__Fungi(100);p__Ascomycota(100);c__Saccharomycetes(100);o__Saccharomycetales(100);f__Metschnikowiaceae(100);g__Metschnikowia(100);s__Metschnikowia_pulcherrima(100); | Metschnikowia | 13 | Saprotroph | \|Undefined Saprotroph\| | Yeast | NULL | Probable | As nectar-tap_saprotroph (Põlme et al. 2020); Decay sugar-rich substrates (Põlme et al. 2020) | Põlme S, et al. 2020. Fungal Diversity 105:1-16 ((https://doi.org/10.1007/s13225-020-00466-2)) |
| Otu53968 | 1 | k__Fungi(100);p__Ascomycota(100);c__Saccharomycetes(100);o__Saccharomycetales(100);f__Metschnikowiaceae(100);g__Metschnikowia(100);g__Metschnikowia_unclassified(100); | Metschnikowia | 13 | Saprotroph | \|Undefined Saprotroph\| | Yeast | NULL | Probable | As nectar-tap_saprotroph (Põlme et al. 2020); Decay sugar-rich substrates (Põlme et al. 2020) | Põlme S, et al. 2020. Fungal Diversity 105:1-16 ((https://doi.org/10.1007/s13225-020-00466-2)) |
| Otu53969 | 1 | k__Fungi(100);p__Ascomycota(100);c__Saccharomycetes(100);o__Saccharomycetales(100);f__Metschnikowiaceae(100);g__Metschnikowia(100);s__Metschnikowia_pulcherrima(100); | Metschnikowia | 13 | Saprotroph | \|Undefined Saprotroph\| | Yeast | NULL | Probable | As nectar-tap_saprotroph (Põlme et al. 2020); Decay sugar-rich substrates (Põlme et al. 2020) | Põlme S, et al. 2020. Fungal Diversity 105:1-16 ((https://doi.org/10.1007/s13225-020-00466-2)) |
| Otu53972 | 1 | k__Fungi(100);p__Ascomycota(100);c__Saccharomycetes(100);o__Saccharomycetales(100);f__Metschnikowiaceae(100);g__Metschnikowia(100);s__Metschnikowia_pulcherrima(100); | Metschnikowia | 13 | Saprotroph | \|Undefined Saprotroph\| | Yeast | NULL | Probable | As nectar-tap_saprotroph (Põlme et al. 2020); Decay sugar-rich substrates (Põlme et al. 2020) | Põlme S, et al. 2020. Fungal Diversity 105:1-16 ((https://doi.org/10.1007/s13225-020-00466-2)) |
| Otu53973 | 1 | k__Fungi(100);p__Ascomycota(100);c__Saccharomycetes(100);o__Saccharomycetales(100);f__Metschnikowiaceae(100);g__Metschnikowia(100);s__Metschnikowia_pulcherrima(100); | Metschnikowia | 13 | Saprotroph | \|Undefined Saprotroph\| | Yeast | NULL | Probable | As nectar-tap_saprotroph (Põlme et al. 2020); Decay sugar-rich substrates (Põlme et al. 2020) | Põlme S, et al. 2020. Fungal Diversity 105:1-16 ((https://doi.org/10.1007/s13225-020-00466-2)) |
| Otu53979 | 1 | k__Fungi(100);p__Ascomycota(100);c__Saccharomycetes(100);o__Saccharomycetales(100);f__Metschnikowiaceae(100);g__Metschnikowia(100);g__Metschnikowia_unclassified(100); | Metschnikowia | 13 | Saprotroph | \|Undefined Saprotroph\| | Yeast | NULL | Probable | As nectar-tap_saprotroph (Põlme et al. 2020); Decay sugar-rich substrates (Põlme et al. 2020) | Põlme S, et al. 2020. Fungal Diversity 105:1-16 ((https://doi.org/10.1007/s13225-020-00466-2)) |
| Otu53980 | 1 | k__Fungi(100);p__Ascomycota(100);c__Saccharomycetes(100);o__Saccharomycetales(100);f__Saccharomycetales_family_Incertae_sedis(100);g__Candida(100);s__Candida_cretensis(100); | Candida | 13 | Pathotroph-Saprotroph-Symbiotroph | Animal Pathogen-Endophyte-Endosymbiont-Epiphyte-\|Undefined Saprotroph\| | Dimorphic Yeast-Yeast | NULL | Probable | As nectar-tap_saprotroph (Põlme et al. 2020); Decay sugar-rich substrates (Põlme et al. 2020); This asexual genus is polyphyletic and contain species that function very differently depend on what they are. Caution should be taken when interpreting guilds for Candida species. | Manolakaki D, et al. 2010. Virulence. 1: 367-375 ((https://doi.org/10.4161/viru.1.5.12796)); Mekbib SB, et al. 2011. Tropical Plant Pathology 36:233-240 ((https://doi.org/10.1590/S1982-56762011000400004)); Põlme S, et al. 2020. Fungal Diversity 105:1-16 ((https://doi.org/10.1007/s13225-020-00466-2)) |
| Otu53981 | 1 | k__Fungi(100);p__Ascomycota(100);c__Saccharomycetes(100);o__Saccharomycetales(100);f__Metschnikowiaceae(100);g__Metschnikowia(100);s__Metschnikowia_pulcherrima(100); | Metschnikowia | 13 | Saprotroph | \|Undefined Saprotroph\| | Yeast | NULL | Probable | As nectar-tap_saprotroph (Põlme et al. 2020); Decay sugar-rich substrates (Põlme et al. 2020) | Põlme S, et al. 2020. Fungal Diversity 105:1-16 ((https://doi.org/10.1007/s13225-020-00466-2)) |
| Otu53982 | 1 | k__Fungi(100);p__Ascomycota(100);c__Saccharomycetes(100);o__Saccharomycetales(100);f__Saccharomycetales_family_Incertae_sedis(100);g__Candida(100);g__Candida_unclassified(100); | Candida | 13 | Pathotroph-Saprotroph-Symbiotroph | Animal Pathogen-Endophyte-Endosymbiont-Epiphyte-\|Undefined Saprotroph\| | Dimorphic Yeast-Yeast | NULL | Probable | As nectar-tap_saprotroph (Põlme et al. 2020); Decay sugar-rich substrates (Põlme et al. 2020); This asexual genus is polyphyletic and contain species that function very differently depend on what they are. Caution should be taken when interpreting guilds for Candida species. | Manolakaki D, et al. 2010. Virulence. 1: 367-375 ((https://doi.org/10.4161/viru.1.5.12796)); Mekbib SB, et al. 2011. Tropical Plant Pathology 36:233-240 ((https://doi.org/10.1590/S1982-56762011000400004)); Põlme S, et al. 2020. Fungal Diversity 105:1-16 ((https://doi.org/10.1007/s13225-020-00466-2)) |
| Otu53989 | 1 | k__Fungi(100);p__Ascomycota(100);c__Saccharomycetes(100);o__Saccharomycetales(100);f__Metschnikowiaceae(100);g__Metschnikowia(100);s__Metschnikowia_pulcherrima(100); | Metschnikowia | 13 | Saprotroph | \|Undefined Saprotroph\| | Yeast | NULL | Probable | As nectar-tap_saprotroph (Põlme et al. 2020); Decay sugar-rich substrates (Põlme et al. 2020) | Põlme S, et al. 2020. Fungal Diversity 105:1-16 ((https://doi.org/10.1007/s13225-020-00466-2)) |
| Otu53990 | 1 | k__Fungi(100);p__Ascomycota(100);c__Saccharomycetes(100);o__Saccharomycetales(100);f__Metschnikowiaceae(100);g__Metschnikowia(100);s__Metschnikowia_pulcherrima(100); | Metschnikowia | 13 | Saprotroph | \|Undefined Saprotroph\| | Yeast | NULL | Probable | As nectar-tap_saprotroph (Põlme et al. 2020); Decay sugar-rich substrates (Põlme et al. 2020) | Põlme S, et al. 2020. Fungal Diversity 105:1-16 ((https://doi.org/10.1007/s13225-020-00466-2)) |

| | | | | | | | | | | | |
|---|---|---|---|---|---|---|---|---|---|---|---|
| Otu53991 | 1 | k__Fungi(100);p__Ascomycota(100);c__Saccharomycetes(100);o__Saccharomycetales(100);f__Saccharomycetales_family_Incertae_sedis(100);g__Candida(100);g__Candida_unclassified(100); | Candida | 13 | Pathotroph-Saprotroph-Symbiotroph | Animal Pathogen-Endophyte-Endosymbiont-Epiphyte-\|Undefined Saprotroph\| | Dimorphic Yeast-Yeast | NULL | Probable | As nectar-tap_saprotroph (Põlme et al. 2020); Decay sugar-rich substrates (Põlme et al. 2020); This asexual genus is polyphyletic and contain species that function very differently depend on what they are. Caution should be taken when interpreting guilds for Candida species. | Manolakaki D, et al. 2010. Virulence. 1: 367-375 ((https://doi.org/10.4161/viru.1.5.12796)); Mekbib SB, et al. 2011. Tropical Plant Pathology 36:233-240 ((https://doi.org/10.1590/S1982-56762011000400004)); Põlme S, et al. 2020. Fungal Diversity 105:1-16 ((https://doi.org/10.1007/s13225-020-00466-2)) |
| Otu53992 | 1 | k__Fungi(100);p__Ascomycota(100);c__Saccharomycetes(100);o__Saccharomycetales(100);f__Metschnikowiaceae(100);g__Metschnikowia(100);g__Metschnikowia_unclassified(100); | Metschnikowia | 13 | Saprotroph | \|Undefined Saprotroph\| | Yeast | NULL | Probable | As nectar-tap_saprotroph (Põlme et al. 2020); Decay sugar-rich substrates (Põlme et al. 2020) | Põlme S, et al. 2020. Fungal Diversity 105:1-16 ((https://doi.org/10.1007/s13225-020-00466-2)) |
| Otu53993 | 1 | k__Fungi(100);p__Ascomycota(100);c__Saccharomycetes(100);o__Saccharomycetales(100);f__Metschnikowiaceae(100);g__Metschnikowia(100);s__Metschnikowia_pulcherrima(100); | Metschnikowia | 13 | Saprotroph | \|Undefined Saprotroph\| | Yeast | NULL | Probable | As nectar-tap_saprotroph (Põlme et al. 2020); Decay sugar-rich substrates (Põlme et al. 2020) | Põlme S, et al. 2020. Fungal Diversity 105:1-16 ((https://doi.org/10.1007/s13225-020-00466-2)) |
| Otu53994 | 1 | k__Fungi(100);p__Ascomycota(100);c__Saccharomycetes(100);o__Saccharomycetales(100);f__Metschnikowiaceae(100);g__Metschnikowia(100);g__Metschnikowia_unclassified(100); | Metschnikowia | 13 | Saprotroph | \|Undefined Saprotroph\| | Yeast | NULL | Probable | As nectar-tap_saprotroph (Põlme et al. 2020); Decay sugar-rich substrates (Põlme et al. 2020) | Põlme S, et al. 2020. Fungal Diversity 105:1-16 ((https://doi.org/10.1007/s13225-020-00466-2)) |
| Otu53995 | 1 | k__Fungi(100);p__Ascomycota(100);c__Saccharomycetes(100);o__Saccharomycetales(100);f__Metschnikowiaceae(100);g__Metschnikowia(100);s__Metschnikowia_pulcherrima(100); | Metschnikowia | 13 | Saprotroph | \|Undefined Saprotroph\| | Yeast | NULL | Probable | As nectar-tap_saprotroph (Põlme et al. 2020); Decay sugar-rich substrates (Põlme et al. 2020) | Põlme S, et al. 2020. Fungal Diversity 105:1-16 ((https://doi.org/10.1007/s13225-020-00466-2)) |
| Otu53998 | 1 | k__Fungi(100);p__Ascomycota(100);c__Saccharomycetes(100);o__Saccharomycetales(100);f__Metschnikowiaceae(100);g__Metschnikowia(100);g__Metschnikowia_unclassified(100); | Metschnikowia | 13 | Saprotroph | \|Undefined Saprotroph\| | Yeast | NULL | Probable | As nectar-tap_saprotroph (Põlme et al. 2020); Decay sugar-rich substrates (Põlme et al. 2020) | Põlme S, et al. 2020. Fungal Diversity 105:1-16 ((https://doi.org/10.1007/s13225-020-00466-2)) |
| Otu53999 | 1 | k__Fungi(100);p__Ascomycota(100);c__Saccharomycetes(100);o__Saccharomycetales(100);f__Metschnikowiaceae(100);g__Metschnikowia(100);s__Metschnikowia_pulcherrima(100); | Metschnikowia | 13 | Saprotroph | \|Undefined Saprotroph\| | Yeast | NULL | Probable | As nectar-tap_saprotroph (Põlme et al. 2020); Decay sugar-rich substrates (Põlme et al. 2020) | Põlme S, et al. 2020. Fungal Diversity 105:1-16 ((https://doi.org/10.1007/s13225-020-00466-2)) |
| Otu54000 | 1 | k__Fungi(100);p__Ascomycota(100);c__Saccharomycetes(100);o__Saccharomycetales(100);f__Metschnikowiaceae(100);g__Metschnikowia(100);s__Metschnikowia_pulcherrima(100); | Metschnikowia | 13 | Saprotroph | \|Undefined Saprotroph\| | Yeast | NULL | Probable | As nectar-tap_saprotroph (Põlme et al. 2020); Decay sugar-rich substrates (Põlme et al. 2020) | Põlme S, et al. 2020. Fungal Diversity 105:1-16 ((https://doi.org/10.1007/s13225-020-00466-2)) |
| Otu54001 | 1 | k__Fungi(100);p__Ascomycota(100);c__Saccharomycetes(100);o__Saccharomycetales(100);f__Metschnikowiaceae(100);g__Metschnikowia(100);s__Metschnikowia_pulcherrima(100); | Metschnikowia | 13 | Saprotroph | \|Undefined Saprotroph\| | Yeast | NULL | Probable | As nectar-tap_saprotroph (Põlme et al. 2020); Decay sugar-rich substrates (Põlme et al. 2020) | Põlme S, et al. 2020. Fungal Diversity 105:1-16 ((https://doi.org/10.1007/s13225-020-00466-2)) |
| Otu54002 | 1 | k__Fungi(100);p__Ascomycota(100);c__Saccharomycetes(100);o__Saccharomycetales(100);f__Metschnikowiaceae(100);g__Metschnikowia(100);s__Metschnikowia_pulcherrima(100); | Metschnikowia | 13 | Saprotroph | \|Undefined Saprotroph\| | Yeast | NULL | Probable | As nectar-tap_saprotroph (Põlme et al. 2020); Decay sugar-rich substrates (Põlme et al. 2020) | Põlme S, et al. 2020. Fungal Diversity 105:1-16 ((https://doi.org/10.1007/s13225-020-00466-2)) |
| Otu54003 | 1 | k__Fungi(100);p__Ascomycota(100);c__Saccharomycetes(100);o__Saccharomycetales(100);f__Metschnikowiaceae(100);g__Metschnikowia(100);g__Metschnikowia_unclassified(100); | Metschnikowia | 13 | Saprotroph | \|Undefined Saprotroph\| | Yeast | NULL | Probable | As nectar-tap_saprotroph (Põlme et al. 2020); Decay sugar-rich substrates (Põlme et al. 2020) | Põlme S, et al. 2020. Fungal Diversity 105:1-16 ((https://doi.org/10.1007/s13225-020-00466-2)) |
| Otu54004 | 1 | k__Fungi(100);p__Ascomycota(100);c__Saccharomycetes(100);o__Saccharomycetales(100);f__Metschnikowiaceae(100);g__Metschnikowia(100);s__Metschnikowia_pulcherrima(100); | Metschnikowia | 13 | Saprotroph | \|Undefined Saprotroph\| | Yeast | NULL | Probable | As nectar-tap_saprotroph (Põlme et al. 2020); Decay sugar-rich substrates (Põlme et al. 2020) | Põlme S, et al. 2020. Fungal Diversity 105:1-16 ((https://doi.org/10.1007/s13225-020-00466-2)) |
| Otu54005 | 1 | k__Fungi(100);p__Ascomycota(100);c__Saccharomycetes(100);o__Saccharomycetales(100);f__Metschnikowiaceae(100);g__Metschnikowia(100);g__Metschnikowia_unclassified(100); | Metschnikowia | 13 | Saprotroph | \|Undefined Saprotroph\| | Yeast | NULL | Probable | As nectar-tap_saprotroph (Põlme et al. 2020); Decay sugar-rich substrates (Põlme et al. 2020) | Põlme S, et al. 2020. Fungal Diversity 105:1-16 ((https://doi.org/10.1007/s13225-020-00466-2)) |
| Otu54006 | 1 | k__Fungi(100);p__Ascomycota(100);c__Saccharomycetes(100);o__Saccharomycetales(100);f__Metschnikowiaceae(100);g__Metschnikowia(100);s__Metschnikowia_pulcherrima(100); | Metschnikowia | 13 | Saprotroph | \|Undefined Saprotroph\| | Yeast | NULL | Probable | As nectar-tap_saprotroph (Põlme et al. 2020); Decay sugar-rich substrates (Põlme et al. 2020) | Põlme S, et al. 2020. Fungal Diversity 105:1-16 ((https://doi.org/10.1007/s13225-020-00466-2)) |
| Otu54007 | 1 | k__Fungi(100);p__Ascomycota(100);c__Saccharomycetes(100);o__Saccharomycetales(100);f__Metschnikowiaceae(100);g__Metschnikowia(100);s__Metschnikowia_pulcherrima(100); | Metschnikowia | 13 | Saprotroph | \|Undefined Saprotroph\| | Yeast | NULL | Probable | As nectar-tap_saprotroph (Põlme et al. 2020); Decay sugar-rich substrates (Põlme et al. 2020) | Põlme S, et al. 2020. Fungal Diversity 105:1-16 ((https://doi.org/10.1007/s13225-020-00466-2)) |
| Otu54008 | 1 | k__Fungi(100);p__Ascomycota(100);c__Saccharomycetes(100);o__Saccharomycetales(100);f__Metschnikowiaceae(100);g__Metschnikowia(100);g__Metschnikowia_unclassified(100); | Metschnikowia | 13 | Saprotroph | \|Undefined Saprotroph\| | Yeast | NULL | Probable | As nectar-tap_saprotroph (Põlme et al. 2020); Decay sugar-rich substrates (Põlme et al. 2020) | Põlme S, et al. 2020. Fungal Diversity 105:1-16 ((https://doi.org/10.1007/s13225-020-00466-2)) |
| Otu54010 | 1 | k__Fungi(100);p__Ascomycota(100);c__Saccharomycetes(100);o__Saccharomycetales(100);f__Metschnikowiaceae(100);g__Metschnikowia(100);g__Metschnikowia_unclassified(100); | Metschnikowia | 13 | Saprotroph | \|Undefined Saprotroph\| | Yeast | NULL | Probable | As nectar-tap_saprotroph (Põlme et al. 2020); Decay sugar-rich substrates (Põlme et al. 2020) | Põlme S, et al. 2020. Fungal Diversity 105:1-16 ((https://doi.org/10.1007/s13225-020-00466-2)) |
| Otu54011 | 1 | k__Fungi(100);p__Ascomycota(100);c__Leotiomycetes(100);o__Helotiales(100);f__Vibrisseaceae(100);g__Vibrissea(100);g__Vibrissea_unclassified(100); | Vibrissea | 13 | Saprotroph | \|Plant Saprotroph\|-Undefined Saprotroph | Microfungus | NULL | Probable | As litter_saprotroph (Põlme et al. 2020); Decay leaf, fruit, seed (Põlme et al. 2020) | Tedersoo L, et al. 2014. Science 346:e1256688 ((https://doi.org/10.1126/science.1256688)); Põlme S, et al. 2020. Fungal Diversity 105:1-16 ((https://doi.org/10.1007/s13225-020-00466-2)) |
| Otu54012 | 1 | k__Fungi(100);p__Ascomycota(100);c__Saccharomycetes(100);o__Saccharomycetales(100);f__Metschnikowiaceae(100);g__Metschnikowia(100);s__Metschnikowia_pulcherrima(100); | Metschnikowia | 13 | Saprotroph | \|Undefined Saprotroph\| | Yeast | NULL | Probable | As nectar-tap_saprotroph (Põlme et al. 2020); Decay sugar-rich substrates (Põlme et al. 2020) | Põlme S, et al. 2020. Fungal Diversity 105:1-16 ((https://doi.org/10.1007/s13225-020-00466-2)) |
| Otu54013 | 1 | k__Fungi(100);p__Ascomycota(100);c__Saccharomycetes(100);o__Saccharomycetales(100);f__Metschnikowiaceae(100);g__Metschnikowia(100);s__Metschnikowia_pulcherrima(100); | Metschnikowia | 13 | Saprotroph | \|Undefined Saprotroph\| | Yeast | NULL | Probable | As nectar-tap_saprotroph (Põlme et al. 2020); Decay sugar-rich substrates (Põlme et al. 2020) | Põlme S, et al. 2020. Fungal Diversity 105:1-16 ((https://doi.org/10.1007/s13225-020-00466-2)) |
| Otu54015 | 1 | k__Fungi(100);p__Ascomycota(100);c__Saccharomycetes(100);o__Saccharomycetales(100);f__Metschnikowiaceae(100);g__Metschnikowia(100);g__Metschnikowia_unclassified(100); | Metschnikowia | 13 | Saprotroph | \|Undefined Saprotroph\| | Yeast | NULL | Probable | As nectar-tap_saprotroph (Põlme et al. 2020); Decay sugar-rich substrates (Põlme et al. 2020) | Põlme S, et al. 2020. Fungal Diversity 105:1-16 ((https://doi.org/10.1007/s13225-020-00466-2)) |
| Otu54016 | 1 | k__Fungi(100);p__Ascomycota(100);c__Saccharomycetes(100);o__Saccharomycetales(100);f__Metschnikowiaceae(100);g__Metschnikowia(100);s__Metschnikowia_pulcherrima(100); | Metschnikowia | 13 | Saprotroph | \|Undefined Saprotroph\| | Yeast | NULL | Probable | As nectar-tap_saprotroph (Põlme et al. 2020); Decay sugar-rich substrates (Põlme et al. 2020) | Põlme S, et al. 2020. Fungal Diversity 105:1-16 ((https://doi.org/10.1007/s13225-020-00466-2)) |
| Otu54017 | 1 | k__Fungi(100);p__Ascomycota(100);c__Saccharomycetes(100);o__Saccharomycetales(100);f__Metschnikowiaceae(100);g__Metschnikowia(100);g__Metschnikowia_unclassified(100); | Metschnikowia | 13 | Saprotroph | \|Undefined Saprotroph\| | Yeast | NULL | Probable | As nectar-tap_saprotroph (Põlme et al. 2020); Decay sugar-rich substrates (Põlme et al. 2020) | Põlme S, et al. 2020. Fungal Diversity 105:1-16 ((https://doi.org/10.1007/s13225-020-00466-2)) |
| Otu54018 | 1 | k__Fungi(100);p__Ascomycota(100);c__Saccharomycetes(100);o__Saccharomycetales(100);f__Metschnikowiaceae(100);g__Metschnikowia(100);g__Metschnikowia_unclassified(100); | Metschnikowia | 13 | Saprotroph | \|Undefined Saprotroph\| | Yeast | NULL | Probable | As nectar-tap_saprotroph (Põlme et al. 2020); Decay sugar-rich substrates (Põlme et al. 2020) | Põlme S, et al. 2020. Fungal Diversity 105:1-16 ((https://doi.org/10.1007/s13225-020-00466-2)) |
| Otu54022 | 1 | k__Fungi(100);p__Ascomycota(100);c__Saccharomycetes(100);o__Saccharomycetales(100);f__Metschnikowiaceae(100);g__Metschnikowia(100);s__Metschnikowia_pulcherrima(100); | Metschnikowia | 13 | Saprotroph | \|Undefined Saprotroph\| | Yeast | NULL | Probable | As nectar-tap_saprotroph (Põlme et al. 2020); Decay sugar-rich substrates (Põlme et al. 2020) | Põlme S, et al. 2020. Fungal Diversity 105:1-16 ((https://doi.org/10.1007/s13225-020-00466-2)) |
| Otu54023 | 1 | k__Fungi(100);p__Ascomycota(100);c__Saccharomycetes(100);o__Saccharomycetales(100);f__Metschnikowiaceae(100);g__Metschnikowia(100);s__Metschnikowia_pulcherrima(100); | Metschnikowia | 13 | Saprotroph | \|Undefined Saprotroph\| | Yeast | NULL | Probable | As nectar-tap_saprotroph (Põlme et al. 2020); Decay sugar-rich substrates (Põlme et al. 2020) | Põlme S, et al. 2020. Fungal Diversity 105:1-16 ((https://doi.org/10.1007/s13225-020-00466-2)) |
| Otu54024 | 1 | k__Fungi(100);p__Ascomycota(100);c__Saccharomycetes(100);o__Saccharomycetales(100);f__Metschnikowiaceae(100);g__Metschnikowia(100);g__Metschnikowia_unclassified(100); | Metschnikowia | 13 | Saprotroph | \|Undefined Saprotroph\| | Yeast | NULL | Probable | As nectar-tap_saprotroph (Põlme et al. 2020); Decay sugar-rich substrates (Põlme et al. 2020) | Põlme S, et al. 2020. Fungal Diversity 105:1-16 ((https://doi.org/10.1007/s13225-020-00466-2)) |
| Otu54025 | 1 | k__Fungi(100);p__Ascomycota(100);c__Saccharomycetes(100);o__Saccharomycetales(100);f__Metschnikowiaceae(100);g__Metschnikowia(100);g__Metschnikowia_unclassified(100); | Metschnikowia | 13 | Saprotroph | \|Undefined Saprotroph\| | Yeast | NULL | Probable | As nectar-tap_saprotroph (Põlme et al. 2020); Decay sugar-rich substrates (Põlme et al. 2020) | Põlme S, et al. 2020. Fungal Diversity 105:1-16 ((https://doi.org/10.1007/s13225-020-00466-2)) |
| Otu54026 | 1 | k__Fungi(100);p__Ascomycota(100);c__Saccharomycetes(100);o__Saccharomycetales(100);f__Metschnikowiaceae(100);g__Metschnikowia(100);s__Metschnikowia_pulcherrima(100); | Metschnikowia | 13 | Saprotroph | \|Undefined Saprotroph\| | Yeast | NULL | Probable | As nectar-tap_saprotroph (Põlme et al. 2020); Decay sugar-rich substrates (Põlme et al. 2020) | Põlme S, et al. 2020. Fungal Diversity 105:1-16 ((https://doi.org/10.1007/s13225-020-00466-2)) |
| Otu54027 | 1 | k__Fungi(100);p__Ascomycota(100);c__Saccharomycetes(100);o__Saccharomycetales(100);f__Metschnikowiaceae(100);g__Metschnikowia(100);s__Metschnikowia_pulcherrima(100); | Metschnikowia | 13 | Saprotroph | \|Undefined Saprotroph\| | Yeast | NULL | Probable | As nectar-tap_saprotroph (Põlme et al. 2020); Decay sugar-rich substrates (Põlme et al. 2020) | Põlme S, et al. 2020. Fungal Diversity 105:1-16 ((https://doi.org/10.1007/s13225-020-00466-2)) |
| Otu54028 | 1 | k__Fungi(100);p__Ascomycota(100);c__Saccharomycetes(100);o__Saccharomycetales(100);f__Metschnikowiaceae(100);g__Metschnikowia(100);s__Metschnikowia_pulcherrima(100); | Metschnikowia | 13 | Saprotroph | \|Undefined Saprotroph\| | Yeast | NULL | Probable | As nectar-tap_saprotroph (Põlme et al. 2020); Decay sugar-rich substrates (Põlme et al. 2020) | Põlme S, et al. 2020. Fungal Diversity 105:1-16 ((https://doi.org/10.1007/s13225-020-00466-2)) |
| Otu54029 | 1 | k__Fungi(100);p__Ascomycota(100);c__Saccharomycetes(100);o__Saccharomycetales(100);f__Metschnikowiaceae(100);g__Metschnikowia(100);s__Metschnikowia_pulcherrima(100); | Metschnikowia | 13 | Saprotroph | \|Undefined Saprotroph\| | Yeast | NULL | Probable | As nectar-tap_saprotroph (Põlme et al. 2020); Decay sugar-rich substrates (Põlme et al. 2020) | Põlme S, et al. 2020. Fungal Diversity 105:1-16 ((https://doi.org/10.1007/s13225-020-00466-2)) |
| Otu54030 | 1 | k__Fungi(100);p__Ascomycota(100);c__Saccharomycetes(100);o__Saccharomycetales(100);f__Metschnikowiaceae(100);g__Metschnikowia(100);g__Metschnikowia_unclassified(100); | Metschnikowia | 13 | Saprotroph | \|Undefined Saprotroph\| | Yeast | NULL | Probable | As nectar-tap_saprotroph (Põlme et al. 2020); Decay sugar-rich substrates (Põlme et al. 2020) | Põlme S, et al. 2020. Fungal Diversity 105:1-16 ((https://doi.org/10.1007/s13225-020-00466-2)) |
| Otu54031 | 1 | k__Fungi(100);p__Ascomycota(100);c__Saccharomycetes(100);o__Saccharomycetales(100);f__Metschnikowiaceae(100);g__Metschnikowia(100);s__Metschnikowia_pulcherrima(100); | Metschnikowia | 13 | Saprotroph | \|Undefined Saprotroph\| | Yeast | NULL | Probable | As nectar-tap_saprotroph (Põlme et al. 2020); Decay sugar-rich substrates (Põlme et al. 2020) | Põlme S, et al. 2020. Fungal Diversity 105:1-16 ((https://doi.org/10.1007/s13225-020-00466-2)) |
| Otu54032 | 1 | k__Fungi(100);p__Ascomycota(100);c__Saccharomycetes(100);o__Saccharomycetales(100);f__Metschnikowiaceae(100);g__Metschnikowia(100);s__Metschnikowia_pulcherrima(100); | Metschnikowia | 13 | Saprotroph | \|Undefined Saprotroph\| | Yeast | NULL | Probable | As nectar-tap_saprotroph (Põlme et al. 2020); Decay sugar-rich substrates (Põlme et al. 2020) | Põlme S, et al. 2020. Fungal Diversity 105:1-16 ((https://doi.org/10.1007/s13225-020-00466-2)) |
| Otu54034 | 1 | k__Fungi(100);p__Ascomycota(100);c__Saccharomycetes(100);o__Saccharomycetales(100);f__Metschnikowiaceae(100);g__Metschnikowia(100);s__Metschnikowia_pulcherrima(100); | Metschnikowia | 13 | Saprotroph | \|Undefined Saprotroph\| | Yeast | NULL | Probable | As nectar-tap_saprotroph (Põlme et al. 2020); Decay sugar-rich substrates (Põlme et al. 2020) | Põlme S, et al. 2020. Fungal Diversity 105:1-16 ((https://doi.org/10.1007/s13225-020-00466-2)) |
| Otu54035 | 1 | k__Fungi(100);p__Ascomycota(100);c__Saccharomycetes(100);o__Saccharomycetales(100);f__Metschnikowiaceae(100);g__Metschnikowia(100);g__Metschnikowia_unclassified(100); | Metschnikowia | 13 | Saprotroph | \|Undefined Saprotroph\| | Yeast | NULL | Probable | As nectar-tap_saprotroph (Põlme et al. 2020); Decay sugar-rich substrates (Põlme et al. 2020) | Põlme S, et al. 2020. Fungal Diversity 105:1-16 ((https://doi.org/10.1007/s13225-020-00466-2)) |
| Otu54036 | 1 | k__Fungi(100);p__Ascomycota(100);c__Saccharomycetes(100);o__Saccharomycetales(100);f__Metschnikowiaceae(100);g__Metschnikowia(100);s__Metschnikowia_pulcherrima(100); | Metschnikowia | 13 | Saprotroph | \|Undefined Saprotroph\| | Yeast | NULL | Probable | As nectar-tap_saprotroph (Põlme et al. 2020); Decay sugar-rich substrates (Põlme et al. 2020) | Põlme S, et al. 2020. Fungal Diversity 105:1-16 ((https://doi.org/10.1007/s13225-020-00466-2)) |
| Otu54037 | 1 | k__Fungi(100);p__Ascomycota(100);c__Saccharomycetes(100);o__Saccharomycetales(100);f__Metschnikowiaceae(100);g__Metschnikowia(100);g__Metschnikowia_unclassified(100); | Metschnikowia | 13 | Saprotroph | \|Undefined Saprotroph\| | Yeast | NULL | Probable | As nectar-tap_saprotroph (Põlme et al. 2020); Decay sugar-rich substrates (Põlme et al. 2020) | Põlme S, et al. 2020. Fungal Diversity 105:1-16 ((https://doi.org/10.1007/s13225-020-00466-2)) |
| Otu54038 | 1 | k__Fungi(100);p__Ascomycota(100);c__Saccharomycetes(100);o__Saccharomycetales(100);f__Metschnikowiaceae(100);g__Metschnikowia(100);s__Metschnikowia_pulcherrima(100); | Metschnikowia | 13 | Saprotroph | \|Undefined Saprotroph\| | Yeast | NULL | Probable | As nectar-tap_saprotroph (Põlme et al. 2020); Decay sugar-rich substrates (Põlme et al. 2020) | Põlme S, et al. 2020. Fungal Diversity 105:1-16 ((https://doi.org/10.1007/s13225-020-00466-2)) |
| Otu54040 | 1 | k__Fungi(100);p__Ascomycota(100);c__Saccharomycetes(100);o__Saccharomycetales(100);f__Metschnikowiaceae(100);g__Metschnikowia(100);g__Metschnikowia_unclassified(100); | Metschnikowia | 13 | Saprotroph | \|Undefined Saprotroph\| | Yeast | NULL | Probable | As nectar-tap_saprotroph (Põlme et al. 2020); Decay sugar-rich substrates (Põlme et al. 2020) | Põlme S, et al. 2020. Fungal Diversity 105:1-16 ((https://doi.org/10.1007/s13225-020-00466-2)) |
| Otu54041 | 1 | k__Fungi(100);p__Ascomycota(100);c__Saccharomycetes(100);o__Saccharomycetales(100);f__Metschnikowiaceae(100);g__Metschnikowia(100);s__Metschnikowia_pulcherrima(100); | Metschnikowia | 13 | Saprotroph | \|Undefined Saprotroph\| | Yeast | NULL | Probable | As nectar-tap_saprotroph (Põlme et al. 2020); Decay sugar-rich substrates (Põlme et al. 2020) | Põlme S, et al. 2020. Fungal Diversity 105:1-16 ((https://doi.org/10.1007/s13225-020-00466-2)) |
| Otu54042 | 1 | k__Fungi(100);p__Ascomycota(100);c__Saccharomycetes(100);o__Saccharomycetales(100);f__Metschnikowiaceae(100);g__Metschnikowia(100);g__Metschnikowia_unclassified(100); | Metschnikowia | 13 | Saprotroph | \|Undefined Saprotroph\| | Yeast | NULL | Probable | As nectar-tap_saprotroph (Põlme et al. 2020); Decay sugar-rich substrates (Põlme et al. 2020) | Põlme S, et al. 2020. Fungal Diversity 105:1-16 ((https://doi.org/10.1007/s13225-020-00466-2)) |
| Otu54044 | 1 | k__Fungi(100);p__Ascomycota(100);c__Saccharomycetes(100);o__Saccharomycetales(100);f__Metschnikowiaceae(100);g__Metschnikowia(100);s__Metschnikowia_pulcherrima(100); | Metschnikowia | 13 | Saprotroph | \|Undefined Saprotroph\| | Yeast | NULL | Probable | As nectar-tap_saprotroph (Põlme et al. 2020); Decay sugar-rich substrates (Põlme et al. 2020) | Põlme S, et al. 2020. Fungal Diversity 105:1-16 ((https://doi.org/10.1007/s13225-020-00466-2)) |
| Otu54045 | 1 | k__Fungi(100);p__Ascomycota(100);c__Saccharomycetes(100);o__Saccharomycetales(100);f__Metschnikowiaceae(100);g__Metschnikowia(100);g__Metschnikowia_unclassified(100); | Metschnikowia | 13 | Saprotroph | \|Undefined Saprotroph\| | Yeast | NULL | Probable | As nectar-tap_saprotroph (Põlme et al. 2020); Decay sugar-rich substrates (Põlme et al. 2020) | Põlme S, et al. 2020. Fungal Diversity 105:1-16 ((https://doi.org/10.1007/s13225-020-00466-2)) |
| Otu54046 | 1 | k__Fungi(100);p__Ascomycota(100);c__Saccharomycetes(100);o__Saccharomycetales(100);f__Metschnikowiaceae(100);g__Metschnikowia(100);g__Metschnikowia_unclassified(100); | Metschnikowia | 13 | Saprotroph | \|Undefined Saprotroph\| | Yeast | NULL | Probable | As nectar-tap_saprotroph (Põlme et al. 2020); Decay sugar-rich substrates (Põlme et al. 2020) | Põlme S, et al. 2020. Fungal Diversity 105:1-16 ((https://doi.org/10.1007/s13225-020-00466-2)) |
| Otu54047 | 1 | k__Fungi(100);p__Ascomycota(100);c__Saccharomycetes(100);o__Saccharomycetales(100);f__Metschnikowiaceae(100);g__Metschnikowia(100);s__Metschnikowia_pulcherrima(100); | Metschnikowia | 13 | Saprotroph | \|Undefined Saprotroph\| | Yeast | NULL | Probable | As nectar-tap_saprotroph (Põlme et al. 2020); Decay sugar-rich substrates (Põlme et al. 2020) | Põlme S, et al. 2020. Fungal Diversity 105:1-16 ((https://doi.org/10.1007/s13225-020-00466-2)) |
| Otu54048 | 1 | k__Fungi(100);p__Ascomycota(100);c__Saccharomycetes(100);o__Saccharomycetales(100);f__Metschnikowiaceae(100);g__Metschnikowia(100);s__Metschnikowia_pulcherrima(100); | Metschnikowia | 13 | Saprotroph | \|Undefined Saprotroph\| | Yeast | NULL | Probable | As nectar-tap_saprotroph (Põlme et al. 2020); Decay sugar-rich substrates (Põlme et al. 2020) | Põlme S, et al. 2020. Fungal Diversity 105:1-16 ((https://doi.org/10.1007/s13225-020-00466-2)) |
| Otu54049 | 1 | k__Fungi(100);p__Ascomycota(100);c__Saccharomycetes(100);o__Saccharomycetales(100);f__Metschnikowiaceae(100);g__Metschnikowia(100);s__Metschnikowia_pulcherrima(100); | Metschnikowia | 13 | Saprotroph | \|Undefined Saprotroph\| | Yeast | NULL | Probable | As nectar-tap_saprotroph (Põlme et al. 2020); Decay sugar-rich substrates (Põlme et al. 2020) | Põlme S, et al. 2020. Fungal Diversity 105:1-16 ((https://doi.org/10.1007/s13225-020-00466-2)) |
| Otu54052 | 1 | k__Fungi(100);p__Ascomycota(100);c__Saccharomycetes(100);o__Saccharomycetales(100);f__Metschnikowiaceae(100);g__Metschnikowia(100);s__Metschnikowia_pulcherrima(100); | Metschnikowia | 13 | Saprotroph | \|Undefined Saprotroph\| | Yeast | NULL | Probable | As nectar-tap_saprotroph (Põlme et al. 2020); Decay sugar-rich substrates (Põlme et al. 2020) | Põlme S, et al. 2020. Fungal Diversity 105:1-16 ((https://doi.org/10.1007/s13225-020-00466-2)) |
| Otu54053 | 1 | k__Fungi(100);p__Ascomycota(100);c__Saccharomycetes(100);o__Saccharomycetales(100);f__Saccharomycetaceae(100);g__Kazachstania(100);s__Kazachstania_intestinalis(100); | Kazachstania | 13 | Saprotroph | \|Undefined Saprotroph\| | Yeast | NULL | Probable | As nectar-tap_saprotroph (Põlme et al. 2020); Decay sugar-rich substrates (Põlme et al. 2020) | Põlme S, et al. 2020. Fungal Diversity 105:1-16 ((https://doi.org/10.1007/s13225-020-00466-2)) |
| Otu54055 | 1 | k__Fungi(100);p__Ascomycota(100);c__Saccharomycetes(100);o__Saccharomycetales(100);f__Saccharomycetales_family_Incertae_sedis(100);g__Candida(100);g__Candida_unclassified(100); | Candida | 13 | Pathotroph-Saprotroph-Symbiotroph | Animal Pathogen-Endophyte-Endosymbiont-Epiphyte-\|Undefined Saprotroph\| | Dimorphic Yeast-Yeast | NULL | Probable | As nectar-tap_saprotroph (Põlme et al. 2020); Decay sugar-rich substrates (Põlme et al. 2020); This asexual genus is polyphyletic and contain species that function very differently depend on what they are. Caution should be taken when interpreting guilds for Candida species. | Manolakaki D, et al. 2010. Virulence. 1: 367-375 ((https://doi.org/10.4161/viru.1.5.12796)); Mekbib SB, et al. 2011. Tropical Plant Pathology 36:233-240 ((https://doi.org/10.1590/S1982-56762011000400004)); Põlme S, et al. 2020. Fungal Diversity 105:1-16 ((https://doi.org/10.1007/s13225-020-00466-2)) |
| Otu54057 | 1 | k__Fungi(100);p__Ascomycota(100);c__Saccharomycetes(100);o__Saccharomycetales(100);f__Metschnikowiaceae(100);g__Metschnikowia(100);g__Metschnikowia_unclassified(100); | Metschnikowia | 13 | Saprotroph | \|Undefined Saprotroph\| | Yeast | NULL | Probable | As nectar-tap_saprotroph (Põlme et al. 2020); Decay sugar-rich substrates (Põlme et al. 2020) | Põlme S, et al. 2020. Fungal Diversity 105:1-16 ((https://doi.org/10.1007/s13225-020-00466-2)) |
| Otu54059 | 1 | k__Fungi(100);p__Ascomycota(100);c__Saccharomycetes(100);o__Saccharomycetales(100);f__Metschnikowiaceae(100);g__Metschnikowia(100);s__Metschnikowia_pulcherrima(100); | Metschnikowia | 13 | Saprotroph | \|Undefined Saprotroph\| | Yeast | NULL | Probable | As nectar-tap_saprotroph (Põlme et al. 2020); Decay sugar-rich substrates (Põlme et al. 2020) | Põlme S, et al. 2020. Fungal Diversity 105:1-16 ((https://doi.org/10.1007/s13225-020-00466-2)) |
| Otu54061 | 1 | k__Fungi(100);p__Ascomycota(100);c__Saccharomycetes(100);o__Saccharomycetales(100);f__Saccharomycetales_family_Incertae_sedis(100);g__Candida(100);g__Candida_unclassified(100); | Candida | 13 | Pathotroph-Saprotroph-Symbiotroph | Animal Pathogen-Endophyte-Endosymbiont-Epiphyte-\|Undefined Saprotroph\| | Dimorphic Yeast-Yeast | NULL | Probable | As nectar-tap_saprotroph (Põlme et al. 2020); Decay sugar-rich substrates (Põlme et al. 2020); This asexual genus is polyphyletic and contain species that function very differently depend on what they are. Caution should be taken when interpreting guilds for Candida species. | Manolakaki D, et al. 2010. Virulence. 1: 367-375 ((https://doi.org/10.4161/viru.1.5.12796)); Mekbib SB, et al. 2011. Tropical Plant Pathology 36:233-240 ((https://doi.org/10.1590/S1982-56762011000400004)); Põlme S, et al. 2020. Fungal Diversity 105:1-16 ((https://doi.org/10.1007/s13225-020-00466-2)) |
| Otu54063 | 1 | k__Fungi(100);p__Ascomycota(100);c__Saccharomycetes(100);o__Saccharomycetales(100);f__Metschnikowiaceae(100);g__Metschnikowia(100);s__Metschnikowia_pulcherrima(100); | Metschnikowia | 13 | Saprotroph | \|Undefined Saprotroph\| | Yeast | NULL | Probable | As nectar-tap_saprotroph (Põlme et al. 2020); Decay sugar-rich substrates (Põlme et al. 2020) | Põlme S, et al. 2020. Fungal Diversity 105:1-16 ((https://doi.org/10.1007/s13225-020-00466-2)) |
| Otu54064 | 1 | k__Fungi(100);p__Ascomycota(100);c__Saccharomycetes(100);o__Saccharomycetales(100);f__Metschnikowiaceae(100);g__Metschnikowia(100);g__Metschnikowia_unclassified(100); | Metschnikowia | 13 | Saprotroph | \|Undefined Saprotroph\| | Yeast | NULL | Probable | As nectar-tap_saprotroph (Põlme et al. 2020); Decay sugar-rich substrates (Põlme et al. 2020) | Põlme S, et al. 2020. Fungal Diversity 105:1-16 ((https://doi.org/10.1007/s13225-020-00466-2)) |
| Otu54065 | 1 | k__Fungi(100);p__Ascomycota(100);c__Saccharomycetes(100);o__Saccharomycetales(100);f__Metschnikowiaceae(100);g__Metschnikowia(100);s__Metschnikowia_pulcherrima(100); | Metschnikowia | 13 | Saprotroph | \|Undefined Saprotroph\| | Yeast | NULL | Probable | As nectar-tap_saprotroph (Põlme et al. 2020); Decay sugar-rich substrates (Põlme et al. 2020) | Põlme S, et al. 2020. Fungal Diversity 105:1-16 ((https://doi.org/10.1007/s13225-020-00466-2)) |
| Otu54066 | 1 | k__Fungi(100);p__Ascomycota(100);c__Saccharomycetes(100);o__Saccharomycetales(100);f__Metschnikowiaceae(100);g__Metschnikowia(100);s__Metschnikowia_pulcherrima(100); | Metschnikowia | 13 | Saprotroph | \|Undefined Saprotroph\| | Yeast | NULL | Probable | As nectar-tap_saprotroph (Põlme et al. 2020); Decay sugar-rich substrates (Põlme et al. 2020) | Põlme S, et al. 2020. Fungal Diversity 105:1-16 ((https://doi.org/10.1007/s13225-020-00466-2)) |
| Otu54067 | 1 | k__Fungi(100);p__Basidiomycota(100);c__Microbotryomycetes(100);o__Sporidiobolales(100);f__Sporidiobolales_family_Incertae_sedis(100);g__Sporobolomyces(100);s__Sporobolomyces_lactosus(100); | Sporobolomyces | 13 | Pathotroph-Saprotroph | \|Fungal Parasite\|-Plant Saprotroph | Yeast | NULL | Probable | As litter_saprotroph (Põlme et al. 2020); As mycoparasite (Põlme et al. 2020); Decay fungal material (Põlme et al. 2020) | Tedersoo L, et al. 2014. Science 346:e1256688 ((https://doi.org/10.1126/science.1256688)); Purahong W, et al. 2016. Molecular Ecology 25:4059-4074 ((https://doi.org/10.1111/mec.13739)); Põlme S, et al. 2020. Fungal Diversity 105:1-16 ((https://doi.org/10.1007/s13225-020-00466-2)) |
| Otu54068 | 1 | k__Fungi(100);p__Ascomycota(100);c__Saccharomycetes(100);o__Saccharomycetales(100);f__Metschnikowiaceae(100);g__Metschnikowia(100);s__Metschnikowia_pulcherrima(100); | Metschnikowia | 13 | Saprotroph | \|Undefined Saprotroph\| | Yeast | NULL | Probable | As nectar-tap_saprotroph (Põlme et al. 2020); Decay sugar-rich substrates (Põlme et al. 2020) | Põlme S, et al. 2020. Fungal Diversity 105:1-16 ((https://doi.org/10.1007/s13225-020-00466-2)) |
| Otu54069 | 1 | k__Fungi(100);p__Ascomycota(100);c__Saccharomycetes(100);o__Saccharomycetales(100);f__Metschnikowiaceae(100);g__Metschnikowia(100);s__Metschnikowia_pulcherrima(100); | Metschnikowia | 13 | Saprotroph | \|Undefined Saprotroph\| | Yeast | NULL | Probable | As nectar-tap_saprotroph (Põlme et al. 2020); Decay sugar-rich substrates (Põlme et al. 2020) | Põlme S, et al. 2020. Fungal Diversity 105:1-16 ((https://doi.org/10.1007/s13225-020-00466-2)) |
| Otu54072 | 1 | k__Fungi(100);p__Ascomycota(100);c__Saccharomycetes(100);o__Saccharomycetales(100);f__Metschnikowiaceae(100);g__Metschnikowia(100);g__Metschnikowia_unclassified(100); | Metschnikowia | 13 | Saprotroph | \|Undefined Saprotroph\| | Yeast | NULL | Probable | As nectar-tap_saprotroph (Põlme et al. 2020); Decay sugar-rich substrates (Põlme et al. 2020) | Põlme S, et al. 2020. Fungal Diversity 105:1-16 ((https://doi.org/10.1007/s13225-020-00466-2)) |
| Otu54074 | 1 | k__Fungi(100);p__Ascomycota(100);c__Saccharomycetes(100);o__Saccharomycetales(100);f__Saccharomycetales_family_Incertae_sedis(100);g__Candida(100);g__Candida_unclassified(100); | Candida | 13 | Pathotroph-Saprotroph-Symbiotroph | Animal Pathogen-Endophyte-Endosymbiont-Epiphyte-\|Undefined Saprotroph\| | Dimorphic Yeast-Yeast | NULL | Probable | As nectar-tap_saprotroph (Põlme et al. 2020); Decay sugar-rich substrates (Põlme et al. 2020); This asexual genus is polyphyletic and contain species that function very differently depend on what they are. Caution should be taken when interpreting guilds for Candida species. | Manolakaki D, et al. 2010. Virulence. 1: 367-375 ((https://doi.org/10.4161/viru.1.5.12796)); Mekbib SB, et al. 2011. Tropical Plant Pathology 36:233-240 ((https://doi.org/10.1590/S1982-56762011000400004)); Põlme S, et al. 2020. Fungal Diversity 105:1-16 ((https://doi.org/10.1007/s13225-020-00466-2)) |
| Otu54075 | 1 | k__Fungi(100);p__Ascomycota(100);c__Saccharomycetes(100);o__Saccharomycetales(100);f__Metschnikowiaceae(100);g__Metschnikowia(100);s__Metschnikowia_pulcherrima(100); | Metschnikowia | 13 | Saprotroph | \|Undefined Saprotroph\| | Yeast | NULL | Probable | As nectar-tap_saprotroph (Põlme et al. 2020); Decay sugar-rich substrates (Põlme et al. 2020) | Põlme S, et al. 2020. Fungal Diversity 105:1-16 ((https://doi.org/10.1007/s13225-020-00466-2)) |
| Otu54076 | 1 | k__Fungi(100);p__Ascomycota(100);c__Saccharomycetes(100);o__Saccharomycetales(100);f__Metschnikowiaceae(100);g__Metschnikowia(100);s__Metschnikowia_pulcherrima(100); | Metschnikowia | 13 | Saprotroph | \|Undefined Saprotroph\| | Yeast | NULL | Probable | As nectar-tap_saprotroph (Põlme et al. 2020); Decay sugar-rich substrates (Põlme et al. 2020) | Põlme S, et al. 2020. Fungal Diversity 105:1-16 ((https://doi.org/10.1007/s13225-020-00466-2)) |
| Otu54077 | 1 | k__Fungi(100);p__Ascomycota(100);c__Saccharomycetes(100);o__Saccharomycetales(100);f__Saccharomycetales_family_Incertae_sedis(100);f__Saccharomycetales_family_Incertae_sedis_unclassified(100);f__Saccharomycetales_family_Incertae_sedis_unclassified(100); | Saccharomycetales | 7 | Saprotroph | Undefined Saprotroph | Yeast | NULL | Possible | NULL | Sterkenburg E, et al. 2015. New Phytologist 207:1145-1158 ((https://doi.org/10.1111/nph.13426)) |
| Otu54080 | 1 | k__Fungi(100);p__Ascomycota(100);c__Saccharomycetes(100);o__Saccharomycetales(100);f__Saccharomycetales_family_Incertae_sedis(100);g__Candida(100);g__Candida_unclassified(100); | Candida | 13 | Pathotroph-Saprotroph-Symbiotroph | Animal Pathogen-Endophyte-Endosymbiont-Epiphyte-\|Undefined Saprotroph\| | Dimorphic Yeast-Yeast | NULL | Probable | As nectar-tap_saprotroph (Põlme et al. 2020); Decay sugar-rich substrates (Põlme et al. 2020); This asexual genus is polyphyletic and contain species that function very differently depend on what they are. Caution should be taken when interpreting guilds for Candida species. | Manolakaki D, et al. 2010. Virulence. 1: 367-375 ((https://doi.org/10.4161/viru.1.5.12796)); Mekbib SB, et al. 2011. Tropical Plant Pathology 36:233-240 ((https://doi.org/10.1590/S1982-56762011000400004)); Põlme S, et al. 2020. Fungal Diversity 105:1-16 ((https://doi.org/10.1007/s13225-020-00466-2)) |
| Otu54083 | 1 | k__Fungi(100);p__Ascomycota(100);c__Saccharomycetes(100);o__Saccharomycetales(100);f__Metschnikowiaceae(100);g__Metschnikowia(100);s__Metschnikowia_pulcherrima(100); | Metschnikowia | 13 | Saprotroph | \|Undefined Saprotroph\| | Yeast | NULL | Probable | As nectar-tap_saprotroph (Põlme et al. 2020); Decay sugar-rich substrates (Põlme et al. 2020) | Põlme S, et al. 2020. Fungal Diversity 105:1-16 ((https://doi.org/10.1007/s13225-020-00466-2)) |
| Otu54084 | 1 | k__Fungi(100);p__Ascomycota(100);c__Saccharomycetes(100);o__Saccharomycetales(100);f__Saccharomycetales_family_Incertae_sedis(100);g__Candida(100);g__Candida_unclassified(100); | Candida | 13 | Pathotroph-Saprotroph-Symbiotroph | Animal Pathogen-Endophyte-Endosymbiont-Epiphyte-\|Undefined Saprotroph\| | Dimorphic Yeast-Yeast | NULL | Probable | As nectar-tap_saprotroph (Põlme et al. 2020); Decay sugar-rich substrates (Põlme et al. 2020); This asexual genus is polyphyletic and contain species that function very differently depend on what they are. Caution should be taken when interpreting guilds for Candida species. | Manolakaki D, et al. 2010. Virulence. 1: 367-375 ((https://doi.org/10.4161/viru.1.5.12796)); Mekbib SB, et al. 2011. Tropical Plant Pathology 36:233-240 ((https://doi.org/10.1590/S1982-56762011000400004)); Põlme S, et al. 2020. Fungal Diversity 105:1-16 ((https://doi.org/10.1007/s13225-020-00466-2)) |
| Otu54086 | 1 | k__Fungi(100);p__Ascomycota(100);c__Saccharomycetes(100);o__Saccharomycetales(100);f__Saccharomycetales_family_Incertae_sedis(100);g__Candida(100);s__Candida_cretensis(100); | Candida | 13 | Pathotroph-Saprotroph-Symbiotroph | Animal Pathogen-Endophyte-Endosymbiont-Epiphyte-\|Undefined Saprotroph\| | Dimorphic Yeast-Yeast | NULL | Probable | As nectar-tap_saprotroph (Põlme et al. 2020); Decay sugar-rich substrates (Põlme et al. 2020); This asexual genus is polyphyletic and contain species that function very differently depend on what they are. Caution should be taken when interpreting guilds for Candida species. | Manolakaki D, et al. 2010. Virulence. 1: 367-375 ((https://doi.org/10.4161/viru.1.5.12796)); Mekbib SB, et al. 2011. Tropical Plant Pathology 36:233-240 ((https://doi.org/10.1590/S1982-56762011000400004)); Põlme S, et al. 2020. Fungal Diversity 105:1-16 ((https://doi.org/10.1007/s13225-020-00466-2)) |
| Otu54087 | 1 | k__Fungi(100);p__Ascomycota(100);c__Saccharomycetes(100);o__Saccharomycetales(100);f__Metschnikowiaceae(100);g__Metschnikowia(100);s__Metschnikowia_pulcherrima(100); | Metschnikowia | 13 | Saprotroph | \|Undefined Saprotroph\| | Yeast | NULL | Probable | As nectar-tap_saprotroph (Põlme et al. 2020); Decay sugar-rich substrates (Põlme et al. 2020) | Põlme S, et al. 2020. Fungal Diversity 105:1-16 ((https://doi.org/10.1007/s13225-020-00466-2)) |
| Otu54090 | 1 | k__Fungi(100);p__Ascomycota(100);c__Saccharomycetes(100);o__Saccharomycetales(100);f__Saccharomycetales_family_Incertae_sedis(100);g__Candida(100);g__Candida_unclassified(100); | Candida | 13 | Pathotroph-Saprotroph-Symbiotroph | Animal Pathogen-Endophyte-Endosymbiont-Epiphyte-\|Undefined Saprotroph\| | Dimorphic Yeast-Yeast | NULL | Probable | As nectar-tap_saprotroph (Põlme et al. 2020); Decay sugar-rich substrates (Põlme et al. 2020); This asexual genus is polyphyletic and contain species that function very differently depend on what they are. Caution should be taken when interpreting guilds for Candida species. | Manolakaki D, et al. 2010. Virulence. 1: 367-375 ((https://doi.org/10.4161/viru.1.5.12796)); Mekbib SB, et al. 2011. Tropical Plant Pathology 36:233-240 ((https://doi.org/10.1590/S1982-56762011000400004)); Põlme S, et al. 2020. Fungal Diversity 105:1-16 ((https://doi.org/10.1007/s13225-020-00466-2)) |
| Otu54091 | 1 | k__Fungi(100);p__Ascomycota(100);c__Saccharomycetes(100);o__Saccharomycetales(100);f__Metschnikowiaceae(100);g__Metschnikowia(100);s__Metschnikowia_pulcherrima(100); | Metschnikowia | 13 | Saprotroph | \|Undefined Saprotroph\| | Yeast | NULL | Probable | As nectar-tap_saprotroph (Põlme et al. 2020); Decay sugar-rich substrates (Põlme et al. 2020) | Põlme S, et al. 2020. Fungal Diversity 105:1-16 ((https://doi.org/10.1007/s13225-020-00466-2)) |
| Otu54093 | 1 | k__Fungi(100);p__Ascomycota(100);c__Saccharomycetes(100);o__Saccharomycetales(100);f__Metschnikowiaceae(100);g__Metschnikowia(100);s__Metschnikowia_pulcherrima(100); | Metschnikowia | 13 | Saprotroph | \|Undefined Saprotroph\| | Yeast | NULL | Probable | As nectar-tap_saprotroph (Põlme et al. 2020); Decay sugar-rich substrates (Põlme et al. 2020) | Põlme S, et al. 2020. Fungal Diversity 105:1-16 ((https://doi.org/10.1007/s13225-020-00466-2)) |
| Otu54094 | 1 | k__Fungi(100);p__Ascomycota(100);c__Saccharomycetes(100);o__Saccharomycetales(100);f__Saccharomycetales_family_Incertae_sedis(100);g__Candida(100);s__Candida_cretensis(100); | Candida | 13 | Pathotroph-Saprotroph-Symbiotroph | Animal Pathogen-Endophyte-Endosymbiont-Epiphyte-\|Undefined Saprotroph\| | Dimorphic Yeast-Yeast | NULL | Probable | As nectar-tap_saprotroph (Põlme et al. 2020); Decay sugar-rich substrates (Põlme et al. 2020); This asexual genus is polyphyletic and contain species that function very differently depend on what they are. Caution should be taken when interpreting guilds for Candida species. | Manolakaki D, et al. 2010. Virulence. 1: 367-375 ((https://doi.org/10.4161/viru.1.5.12796)); Mekbib SB, et al. 2011. Tropical Plant Pathology 36:233-240 ((https://doi.org/10.1590/S1982-56762011000400004)); Põlme S, et al. 2020. Fungal Diversity 105:1-16 ((https://doi.org/10.1007/s13225-020-00466-2)) |
| Otu54096 | 1 | k__Fungi(100);p__Ascomycota(100);c__Saccharomycetes(100);o__Saccharomycetales(100);f__Metschnikowiaceae(100);g__Metschnikowia(100);s__Metschnikowia_pulcherrima(100); | Metschnikowia | 13 | Saprotroph | \|Undefined Saprotroph\| | Yeast | NULL | Probable | As nectar-tap_saprotroph (Põlme et al. 2020); Decay sugar-rich substrates (Põlme et al. 2020) | Põlme S, et al. 2020. Fungal Diversity 105:1-16 ((https://doi.org/10.1007/s13225-020-00466-2)) |
| Otu54097 | 1 | k__Fungi(100);p__Ascomycota(100);c__Saccharomycetes(100);o__Saccharomycetales(100);f__Metschnikowiaceae(100);g__Metschnikowia(100);g__Metschnikowia_unclassified(100); | Metschnikowia | 13 | Saprotroph | \|Undefined Saprotroph\| | Yeast | NULL | Probable | As nectar-tap_saprotroph (Põlme et al. 2020); Decay sugar-rich substrates (Põlme et al. 2020) | Põlme S, et al. 2020. Fungal Diversity 105:1-16 ((https://doi.org/10.1007/s13225-020-00466-2)) |
| Otu54098 | 1 | k__Fungi(100);p__Ascomycota(100);c__Saccharomycetes(100);o__Saccharomycetales(100);f__Metschnikowiaceae(100);g__Metschnikowia(100);s__Metschnikowia_pulcherrima(100); | Metschnikowia | 13 | Saprotroph | \|Undefined Saprotroph\| | Yeast | NULL | Probable | As nectar-tap_saprotroph (Põlme et al. 2020); Decay sugar-rich substrates (Põlme et al. 2020) | Põlme S, et al. 2020. Fungal Diversity 105:1-16 ((https://doi.org/10.1007/s13225-020-00466-2)) |
| Otu54100 | 1 | k__Fungi(100);p__Ascomycota(100);c__Saccharomycetes(100);o__Saccharomycetales(100);f__Metschnikowiaceae(100);g__Metschnikowia(100);s__Metschnikowia_pulcherrima(100); | Metschnikowia | 13 | Saprotroph | \|Undefined Saprotroph\| | Yeast | NULL | Probable | As nectar-tap_saprotroph (Põlme et al. 2020); Decay sugar-rich substrates (Põlme et al. 2020) | Põlme S, et al. 2020. Fungal Diversity 105:1-16 ((https://doi.org/10.1007/s13225-020-00466-2)) |
| Otu54101 | 1 | k__Fungi(100);p__Ascomycota(100);c__Saccharomycetes(100);o__Saccharomycetales(100);f__Metschnikowiaceae(100);g__Metschnikowia(100);s__Metschnikowia_pulcherrima(100); | Metschnikowia | 13 | Saprotroph | \|Undefined Saprotroph\| | Yeast | NULL | Probable | As nectar-tap_saprotroph (Põlme et al. 2020); Decay sugar-rich substrates (Põlme et al. 2020) | Põlme S, et al. 2020. Fungal Diversity 105:1-16 ((https://doi.org/10.1007/s13225-020-00466-2)) |
| Otu54103 | 1 | k__Fungi(100);p__Ascomycota(100);c__Saccharomycetes(100);o__Saccharomycetales(100);f__Metschnikowiaceae(100);g__Metschnikowia(100);s__Metschnikowia_pulcherrima(100); | Metschnikowia | 13 | Saprotroph | \|Undefined Saprotroph\| | Yeast | NULL | Probable | As nectar-tap_saprotroph (Põlme et al. 2020); Decay sugar-rich substrates (Põlme et al. 2020) | Põlme S, et al. 2020. Fungal Diversity 105:1-16 ((https://doi.org/10.1007/s13225-020-00466-2)) |
| Otu54104 | 1 | k__Fungi(100);p__Ascomycota(100);c__Saccharomycetes(100);o__Saccharomycetales(100);f__Metschnikowiaceae(100);g__Metschnikowia(100);g__Metschnikowia_unclassified(100); | Metschnikowia | 13 | Saprotroph | \|Undefined Saprotroph\| | Yeast | NULL | Probable | As nectar-tap_saprotroph (Põlme et al. 2020); Decay sugar-rich substrates (Põlme et al. 2020) | Põlme S, et al. 2020. Fungal Diversity 105:1-16 ((https://doi.org/10.1007/s13225-020-00466-2)) |
| Otu54106 | 1 | k__Fungi(100);p__Ascomycota(100);c__Saccharomycetes(100);o__Saccharomycetales(100);f__Metschnikowiaceae(100);g__Metschnikowia(100);s__Metschnikowia_pulcherrima(100); | Metschnikowia | 13 | Saprotroph | \|Undefined Saprotroph\| | Yeast | NULL | Probable | As nectar-tap_saprotroph (Põlme et al. 2020); Decay sugar-rich substrates (Põlme et al. 2020) | Põlme S, et al. 2020. Fungal Diversity 105:1-16 ((https://doi.org/10.1007/s13225-020-00466-2)) |
| Otu54110 | 1 | k__Fungi(100);p__Ascomycota(100);c__Saccharomycetes(100);o__Saccharomycetales(100);f__Metschnikowiaceae(100);g__Metschnikowia(100);s__Metschnikowia_pulcherrima(100); | Metschnikowia | 13 | Saprotroph | \|Undefined Saprotroph\| | Yeast | NULL | Probable | As nectar-tap_saprotroph (Põlme et al. 2020); Decay sugar-rich substrates (Põlme et al. 2020) | Põlme S, et al. 2020. Fungal Diversity 105:1-16 ((https://doi.org/10.1007/s13225-020-00466-2)) |
| Otu54111 | 1 | k__Fungi(100);p__Ascomycota(100);c__Saccharomycetes(100);o__Saccharomycetales(100);f__Metschnikowiaceae(100);g__Metschnikowia(100);s__Metschnikowia_pulcherrima(100); | Metschnikowia | 13 | Saprotroph | \|Undefined Saprotroph\| | Yeast | NULL | Probable | As nectar-tap_saprotroph (Põlme et al. 2020); Decay sugar-rich substrates (Põlme et al. 2020) | Põlme S, et al. 2020. Fungal Diversity 105:1-16 ((https://doi.org/10.1007/s13225-020-00466-2)) |
| Otu54112 | 1 | k__Fungi(100);p__Ascomycota(100);c__Saccharomycetes(100);o__Saccharomycetales(100);f__Metschnikowiaceae(100);g__Metschnikowia(100);g__Metschnikowia_unclassified(100); | Metschnikowia | 13 | Saprotroph | \|Undefined Saprotroph\| | Yeast | NULL | Probable | As nectar-tap_saprotroph (Põlme et al. 2020); Decay sugar-rich substrates (Põlme et al. 2020) | Põlme S, et al. 2020. Fungal Diversity 105:1-16 ((https://doi.org/10.1007/s13225-020-00466-2)) |
| Otu54114 | 1 | k__Fungi(100);p__Ascomycota(100);c__Saccharomycetes(100);o__Saccharomycetales(100);f__Metschnikowiaceae(100);g__Metschnikowia(100);s__Metschnikowia_pulcherrima(100); | Metschnikowia | 13 | Saprotroph | \|Undefined Saprotroph\| | Yeast | NULL | Probable | As nectar-tap_saprotroph (Põlme et al. 2020); Decay sugar-rich substrates (Põlme et al. 2020) | Põlme S, et al. 2020. Fungal Diversity 105:1-16 ((https://doi.org/10.1007/s13225-020-00466-2)) |
| Otu54115 | 1 | k__Fungi(100);p__Ascomycota(100);c__Saccharomycetes(100);o__Saccharomycetales(100);f__Saccharomycetales_family_Incertae_sedis(100);g__Candida(100);s__Candida_cretensis(100); | Candida | 13 | Pathotroph-Saprotroph-Symbiotroph | Animal Pathogen-Endophyte-Endosymbiont-Epiphyte-\|Undefined Saprotroph\| | Dimorphic Yeast-Yeast | NULL | Probable | As nectar-tap_saprotroph (Põlme et al. 2020); Decay sugar-rich substrates (Põlme et al. 2020); This asexual genus is polyphyletic and contain species that function very differently depend on what they are. Caution should be taken when interpreting guilds for Candida species. | Manolakaki D, et al. 2010. Virulence. 1: 367-375 ((https://doi.org/10.4161/viru.1.5.12796)); Mekbib SB, et al. 2011. Tropical Plant Pathology 36:233-240 ((https://doi.org/10.1590/S1982-56762011000400004)); Põlme S, et al. 2020. Fungal Diversity 105:1-16 ((https://doi.org/10.1007/s13225-020-00466-2)) |
| Otu54116 | 1 | k__Fungi(100);p__Ascomycota(100);c__Saccharomycetes(100);o__Saccharomycetales(100);f__Metschnikowiaceae(100);g__Metschnikowia(100);s__Metschnikowia_pulcherrima(100); | Metschnikowia | 13 | Saprotroph | \|Undefined Saprotroph\| | Yeast | NULL | Probable | As nectar-tap_saprotroph (Põlme et al. 2020); Decay sugar-rich substrates (Põlme et al. 2020) | Põlme S, et al. 2020. Fungal Diversity 105:1-16 ((https://doi.org/10.1007/s13225-020-00466-2)) |
| Otu54117 | 1 | k__Fungi(100);p__Ascomycota(100);c__Saccharomycetes(100);o__Saccharomycetales(100);f__Metschnikowiaceae(100);g__Metschnikowia(100);s__Metschnikowia_pulcherrima(100); | Metschnikowia | 13 | Saprotroph | \|Undefined Saprotroph\| | Yeast | NULL | Probable | As nectar-tap_saprotroph (Põlme et al. 2020); Decay sugar-rich substrates (Põlme et al. 2020) | Põlme S, et al. 2020. Fungal Diversity 105:1-16 ((https://doi.org/10.1007/s13225-020-00466-2)) |
| Otu54118 | 1 | k__Fungi(100);p__Ascomycota(100);c__Saccharomycetes(100);o__Saccharomycetales(100);f__Metschnikowiaceae(100);g__Metschnikowia(100);g__Metschnikowia_unclassified(100); | Metschnikowia | 13 | Saprotroph | \|Undefined Saprotroph\| | Yeast | NULL | Probable | As nectar-tap_saprotroph (Põlme et al. 2020); Decay sugar-rich substrates (Põlme et al. 2020) | Põlme S, et al. 2020. Fungal Diversity 105:1-16 ((https://doi.org/10.1007/s13225-020-00466-2)) |
| Otu54119 | 1 | k__Fungi(100);p__Ascomycota(100);c__Saccharomycetes(100);o__Saccharomycetales(100);f__Metschnikowiaceae(100);g__Metschnikowia(100);s__Metschnikowia_pulcherrima(100); | Metschnikowia | 13 | Saprotroph | \|Undefined Saprotroph\| | Yeast | NULL | Probable | As nectar-tap_saprotroph (Põlme et al. 2020); Decay sugar-rich substrates (Põlme et al. 2020) | Põlme S, et al. 2020. Fungal Diversity 105:1-16 ((https://doi.org/10.1007/s13225-020-00466-2)) |
| Otu54120 | 1 | k__Fungi(100);p__Ascomycota(100);c__Saccharomycetes(100);o__Saccharomycetales(100);f__Metschnikowiaceae(100);g__Metschnikowia(100);s__Metschnikowia_pulcherrima(100); | Metschnikowia | 13 | Saprotroph | \|Undefined Saprotroph\| | Yeast | NULL | Probable | As nectar-tap_saprotroph (Põlme et al. 2020); Decay sugar-rich substrates (Põlme et al. 2020) | Põlme S, et al. 2020. Fungal Diversity 105:1-16 ((https://doi.org/10.1007/s13225-020-00466-2)) |
| Otu54121 | 1 | k__Fungi(100);p__Ascomycota(100);c__Saccharomycetes(100);o__Saccharomycetales(100);f__Metschnikowiaceae(100);g__Metschnikowia(100);s__Metschnikowia_pulcherrima(100); | Metschnikowia | 13 | Saprotroph | \|Undefined Saprotroph\| | Yeast | NULL | Probable | As nectar-tap_saprotroph (Põlme et al. 2020); Decay sugar-rich substrates (Põlme et al. 2020) | Põlme S, et al. 2020. Fungal Diversity 105:1-16 ((https://doi.org/10.1007/s13225-020-00466-2)) |
| Otu54122 | 1 | k__Fungi(100);p__Ascomycota(100);c__Saccharomycetes(100);o__Saccharomycetales(100);f__Metschnikowiaceae(100);g__Metschnikowia(100);g__Metschnikowia_unclassified(100); | Metschnikowia | 13 | Saprotroph | \|Undefined Saprotroph\| | Yeast | NULL | Probable | As nectar-tap_saprotroph (Põlme et al. 2020); Decay sugar-rich substrates (Põlme et al. 2020) | Põlme S, et al. 2020. Fungal Diversity 105:1-16 ((https://doi.org/10.1007/s13225-020-00466-2)) |
| Otu54123 | 1 | k__Fungi(100);p__Ascomycota(100);c__Saccharomycetes(100);o__Saccharomycetales(100);f__Metschnikowiaceae(100);g__Metschnikowia(100);s__Metschnikowia_pulcherrima(100); | Metschnikowia | 13 | Saprotroph | \|Undefined Saprotroph\| | Yeast | NULL | Probable | As nectar-tap_saprotroph (Põlme et al. 2020); Decay sugar-rich substrates (Põlme et al. 2020) | Põlme S, et al. 2020. Fungal Diversity 105:1-16 ((https://doi.org/10.1007/s13225-020-00466-2)) |
| Otu54124 | 1 | k__Fungi(100);p__Ascomycota(100);c__Saccharomycetes(100);o__Saccharomycetales(100);f__Metschnikowiaceae(100);g__Metschnikowia(100);s__Metschnikowia_pulcherrima(100); | Metschnikowia | 13 | Saprotroph | \|Undefined Saprotroph\| | Yeast | NULL | Probable | As nectar-tap_saprotroph (Põlme et al. 2020); Decay sugar-rich substrates (Põlme et al. 2020) | Põlme S, et al. 2020. Fungal Diversity 105:1-16 ((https://doi.org/10.1007/s13225-020-00466-2)) |
| Otu54125 | 1 | k__Fungi(100);p__Ascomycota(100);c__Saccharomycetes(100);o__Saccharomycetales(100);f__Metschnikowiaceae(100);g__Metschnikowia(100);g__Metschnikowia_unclassified(100); | Metschnikowia | 13 | Saprotroph | \|Undefined Saprotroph\| | Yeast | NULL | Probable | As nectar-tap_saprotroph (Põlme et al. 2020); Decay sugar-rich substrates (Põlme et al. 2020) | Põlme S, et al. 2020. Fungal Diversity 105:1-16 ((https://doi.org/10.1007/s13225-020-00466-2)) |
| Otu54126 | 1 | k__Fungi(100);p__Ascomycota(100);c__Saccharomycetes(100);o__Saccharomycetales(100);f__Metschnikowiaceae(100);g__Metschnikowia(100);s__Metschnikowia_pulcherrima(100); | Metschnikowia | 13 | Saprotroph | \|Undefined Saprotroph\| | Yeast | NULL | Probable | As nectar-tap_saprotroph (Põlme et al. 2020); Decay sugar-rich substrates (Põlme et al. 2020) | Põlme S, et al. 2020. Fungal Diversity 105:1-16 ((https://doi.org/10.1007/s13225-020-00466-2)) |
| Otu54127 | 1 | k__Fungi(100);p__Ascomycota(100);c__Saccharomycetes(100);o__Saccharomycetales(100);f__Metschnikowiaceae(100);g__Metschnikowia(100);s__Metschnikowia_pulcherrima(100); | Metschnikowia | 13 | Saprotroph | \|Undefined Saprotroph\| | Yeast | NULL | Probable | As nectar-tap_saprotroph (Põlme et al. 2020); Decay sugar-rich substrates (Põlme et al. 2020) | Põlme S, et al. 2020. Fungal Diversity 105:1-16 ((https://doi.org/10.1007/s13225-020-00466-2)) |
| Otu54128 | 1 | k__Fungi(100);p__Ascomycota(100);c__Saccharomycetes(100);o__Saccharomycetales(100);f__Metschnikowiaceae(100);g__Metschnikowia(100);s__Metschnikowia_pulcherrima(100); | Metschnikowia | 13 | Saprotroph | \|Undefined Saprotroph\| | Yeast | NULL | Probable | As nectar-tap_saprotroph (Põlme et al. 2020); Decay sugar-rich substrates (Põlme et al. 2020) | Põlme S, et al. 2020. Fungal Diversity 105:1-16 ((https://doi.org/10.1007/s13225-020-00466-2)) |
| Otu54129 | 1 | k__Fungi(100);p__Ascomycota(100);c__Saccharomycetes(100);o__Saccharomycetales(100);f__Metschnikowiaceae(100);g__Metschnikowia(100);s__Metschnikowia_pulcherrima(100); | Metschnikowia | 13 | Saprotroph | \|Undefined Saprotroph\| | Yeast | NULL | Probable | As nectar-tap_saprotroph (Põlme et al. 2020); Decay sugar-rich substrates (Põlme et al. 2020) | Põlme S, et al. 2020. Fungal Diversity 105:1-16 ((https://doi.org/10.1007/s13225-020-00466-2)) |
| Otu54130 | 1 | k__Fungi(100);p__Ascomycota(100);c__Saccharomycetes(100);o__Saccharomycetales(100);f__Metschnikowiaceae(100);g__Metschnikowia(100);s__Metschnikowia_pulcherrima(100); | Metschnikowia | 13 | Saprotroph | \|Undefined Saprotroph\| | Yeast | NULL | Probable | As nectar-tap_saprotroph (Põlme et al. 2020); Decay sugar-rich substrates (Põlme et al. 2020) | Põlme S, et al. 2020. Fungal Diversity 105:1-16 ((https://doi.org/10.1007/s13225-020-00466-2)) |
| Otu54131 | 1 | k__Fungi(100);p__Ascomycota(100);c__Saccharomycetes(100);o__Saccharomycetales(100);f__Metschnikowiaceae(100);g__Metschnikowia(100);s__Metschnikowia_pulcherrima(100); | Metschnikowia | 13 | Saprotroph | \|Undefined Saprotroph\| | Yeast | NULL | Probable | As nectar-tap_saprotroph (Põlme et al. 2020); Decay sugar-rich substrates (Põlme et al. 2020) | Põlme S, et al. 2020. Fungal Diversity 105:1-16 ((https://doi.org/10.1007/s13225-020-00466-2)) |
| Otu54132 | 1 | k__Fungi(100);p__Ascomycota(100);c__Saccharomycetes(100);o__Saccharomycetales(100);f__Metschnikowiaceae(100);g__Metschnikowia(100);s__Metschnikowia_pulcherrima(100); | Metschnikowia | 13 | Saprotroph | \|Undefined Saprotroph\| | Yeast | NULL | Probable | As nectar-tap_saprotroph (Põlme et al. 2020); Decay sugar-rich substrates (Põlme et al. 2020) | Põlme S, et al. 2020. Fungal Diversity 105:1-16 ((https://doi.org/10.1007/s13225-020-00466-2)) |
| Otu54133 | 1 | k__Fungi(100);p__Ascomycota(100);c__Saccharomycetes(100);o__Saccharomycetales(100);f__Metschnikowiaceae(100);g__Metschnikowia(100);s__Metschnikowia_pulcherrima(100); | Metschnikowia | 13 | Saprotroph | \|Undefined Saprotroph\| | Yeast | NULL | Probable | As nectar-tap_saprotroph (Põlme et al. 2020); Decay sugar-rich substrates (Põlme et al. 2020) | Põlme S, et al. 2020. Fungal Diversity 105:1-16 ((https://doi.org/10.1007/s13225-020-00466-2)) |
| Otu54134 | 1 | k__Fungi(100);p__Ascomycota(100);c__Saccharomycetes(100);o__Saccharomycetales(100);f__Metschnikowiaceae(100);g__Metschnikowia(100);s__Metschnikowia_pulcherrima(100); | Metschnikowia | 13 | Saprotroph | \|Undefined Saprotroph\| | Yeast | NULL | Probable | As nectar-tap_saprotroph (Põlme et al. 2020); Decay sugar-rich substrates (Põlme et al. 2020) | Põlme S, et al. 2020. Fungal Diversity 105:1-16 ((https://doi.org/10.1007/s13225-020-00466-2)) |
| Otu54136 | 1 | k__Fungi(100);p__Ascomycota(100);c__Saccharomycetes(100);o__Saccharomycetales(100);f__Metschnikowiaceae(100);g__Metschnikowia(100);s__Metschnikowia_pulcherrima(100); | Metschnikowia | 13 | Saprotroph | \|Undefined Saprotroph\| | Yeast | NULL | Probable | As nectar-tap_saprotroph (Põlme et al. 2020); Decay sugar-rich substrates (Põlme et al. 2020) | Põlme S, et al. 2020. Fungal Diversity 105:1-16 ((https://doi.org/10.1007/s13225-020-00466-2)) |
| Otu54137 | 1 | k__Fungi(100);p__Ascomycota(100);c__Saccharomycetes(100);o__Saccharomycetales(100);f__Metschnikowiaceae(100);g__Metschnikowia(100);s__Metschnikowia_pulcherrima(100); | Metschnikowia | 13 | Saprotroph | \|Undefined Saprotroph\| | Yeast | NULL | Probable | As nectar-tap_saprotroph (Põlme et al. 2020); Decay sugar-rich substrates (Põlme et al. 2020) | Põlme S, et al. 2020. Fungal Diversity 105:1-16 ((https://doi.org/10.1007/s13225-020-00466-2)) |
| Otu54138 | 1 | k__Fungi(100);p__Ascomycota(100);c__Saccharomycetes(100);o__Saccharomycetales(100);f__Metschnikowiaceae(100);g__Metschnikowia(100);s__Metschnikowia_pulcherrima(100); | Metschnikowia | 13 | Saprotroph | \|Undefined Saprotroph\| | Yeast | NULL | Probable | As nectar-tap_saprotroph (Põlme et al. 2020); Decay sugar-rich substrates (Põlme et al. 2020) | Põlme S, et al. 2020. Fungal Diversity 105:1-16 ((https://doi.org/10.1007/s13225-020-00466-2)) |
| Otu54139 | 1 | k__Fungi(100);p__Ascomycota(100);c__Saccharomycetes(100);o__Saccharomycetales(100);f__Metschnikowiaceae(100);g__Metschnikowia(100);s__Metschnikowia_pulcherrima(100); | Metschnikowia | 13 | Saprotroph | \|Undefined Saprotroph\| | Yeast | NULL | Probable | As nectar-tap_saprotroph (Põlme et al. 2020); Decay sugar-rich substrates (Põlme et al. 2020) | Põlme S, et al. 2020. Fungal Diversity 105:1-16 ((https://doi.org/10.1007/s13225-020-00466-2)) |
| Otu54140 | 1 | k__Fungi(100);p__Ascomycota(100);c__Saccharomycetes(100);o__Saccharomycetales(100);f__Metschnikowiaceae(100);g__Metschnikowia(100);g__Metschnikowia_unclassified(100); | Metschnikowia | 13 | Saprotroph | \|Undefined Saprotroph\| | Yeast | NULL | Probable | As nectar-tap_saprotroph (Põlme et al. 2020); Decay sugar-rich substrates (Põlme et al. 2020) | Põlme S, et al. 2020. Fungal Diversity 105:1-16 ((https://doi.org/10.1007/s13225-020-00466-2)) |
| Otu54141 | 1 | k__Fungi(100);p__Ascomycota(100);c__Saccharomycetes(100);o__Saccharomycetales(100);f__Metschnikowiaceae(100);g__Metschnikowia(100);s__Metschnikowia_pulcherrima(100); | Metschnikowia | 13 | Saprotroph | \|Undefined Saprotroph\| | Yeast | NULL | Probable | As nectar-tap_saprotroph (Põlme et al. 2020); Decay sugar-rich substrates (Põlme et al. 2020) | Põlme S, et al. 2020. Fungal Diversity 105:1-16 ((https://doi.org/10.1007/s13225-020-00466-2)) |
| Otu54142 | 1 | k__Fungi(100);p__Ascomycota(100);c__Saccharomycetes(100);o__Saccharomycetales(100);f__Metschnikowiaceae(100);g__Metschnikowia(100);s__Metschnikowia_pulcherrima(100); | Metschnikowia | 13 | Saprotroph | \|Undefined Saprotroph\| | Yeast | NULL | Probable | As nectar-tap_saprotroph (Põlme et al. 2020); Decay sugar-rich substrates (Põlme et al. 2020) | Põlme S, et al. 2020. Fungal Diversity 105:1-16 ((https://doi.org/10.1007/s13225-020-00466-2)) |
| Otu54143 | 1 | k__Fungi(100);p__Ascomycota(100);c__Saccharomycetes(100);o__Saccharomycetales(100);f__Metschnikowiaceae(100);g__Metschnikowia(100);s__Metschnikowia_pulcherrima(100); | Metschnikowia | 13 | Saprotroph | \|Undefined Saprotroph\| | Yeast | NULL | Probable | As nectar-tap_saprotroph (Põlme et al. 2020); Decay sugar-rich substrates (Põlme et al. 2020) | Põlme S, et al. 2020. Fungal Diversity 105:1-16 ((https://doi.org/10.1007/s13225-020-00466-2)) |
| Otu54144 | 1 | k__Fungi(100);p__Ascomycota(100);c__Saccharomycetes(100);o__Saccharomycetales(100);f__Metschnikowiaceae(100);g__Metschnikowia(100);s__Metschnikowia_pulcherrima(100); | Metschnikowia | 13 | Saprotroph | \|Undefined Saprotroph\| | Yeast | NULL | Probable | As nectar-tap_saprotroph (Põlme et al. 2020); Decay sugar-rich substrates (Põlme et al. 2020) | Põlme S, et al. 2020. Fungal Diversity 105:1-16 ((https://doi.org/10.1007/s13225-020-00466-2)) |
| Otu54145 | 1 | k__Fungi(100);p__Ascomycota(100);c__Saccharomycetes(100);o__Saccharomycetales(100);f__Metschnikowiaceae(100);g__Metschnikowia(100);s__Metschnikowia_pulcherrima(100); | Metschnikowia | 13 | Saprotroph | \|Undefined Saprotroph\| | Yeast | NULL | Probable | As nectar-tap_saprotroph (Põlme et al. 2020); Decay sugar-rich substrates (Põlme et al. 2020) | Põlme S, et al. 2020. Fungal Diversity 105:1-16 ((https://doi.org/10.1007/s13225-020-00466-2)) |
| Otu54146 | 1 | k__Fungi(100);p__Ascomycota(100);c__Saccharomycetes(100);o__Saccharomycetales(100);f__Metschnikowiaceae(100);g__Metschnikowia(100);s__Metschnikowia_pulcherrima(100); | Metschnikowia | 13 | Saprotroph | \|Undefined Saprotroph\| | Yeast | NULL | Probable | As nectar-tap_saprotroph (Põlme et al. 2020); Decay sugar-rich substrates (Põlme et al. 2020) | Põlme S, et al. 2020. Fungal Diversity 105:1-16 ((https://doi.org/10.1007/s13225-020-00466-2)) |
| Otu54148 | 1 | k__Fungi(100);p__Ascomycota(100);c__Saccharomycetes(100);o__Saccharomycetales(100);f__Metschnikowiaceae(100);g__Metschnikowia(100);s__Metschnikowia_pulcherrima(100); | Metschnikowia | 13 | Saprotroph | \|Undefined Saprotroph\| | Yeast | NULL | Probable | As nectar-tap_saprotroph (Põlme et al. 2020); Decay sugar-rich substrates (Põlme et al. 2020) | Põlme S, et al. 2020. Fungal Diversity 105:1-16 ((https://doi.org/10.1007/s13225-020-00466-2)) |
| Otu54149 | 1 | k__Fungi(100);p__Ascomycota(100);c__Saccharomycetes(100);o__Saccharomycetales(100);f__Metschnikowiaceae(100);g__Metschnikowia(100);s__Metschnikowia_pulcherrima(100); | Metschnikowia | 13 | Saprotroph | \|Undefined Saprotroph\| | Yeast | NULL | Probable | As nectar-tap_saprotroph (Põlme et al. 2020); Decay sugar-rich substrates (Põlme et al. 2020) | Põlme S, et al. 2020. Fungal Diversity 105:1-16 ((https://doi.org/10.1007/s13225-020-00466-2)) |
| Otu54150 | 1 | k__Fungi(100);p__Ascomycota(100);c__Saccharomycetes(100);o__Saccharomycetales(100);f__Metschnikowiaceae(100);g__Metschnikowia(100);s__Metschnikowia_pulcherrima(100); | Metschnikowia | 13 | Saprotroph | \|Undefined Saprotroph\| | Yeast | NULL | Probable | As nectar-tap_saprotroph (Põlme et al. 2020); Decay sugar-rich substrates (Põlme et al. 2020) | Põlme S, et al. 2020. Fungal Diversity 105:1-16 ((https://doi.org/10.1007/s13225-020-00466-2)) |
| Otu54151 | 1 | k__Fungi(100);p__Ascomycota(100);c__Saccharomycetes(100);o__Saccharomycetales(100);f__Metschnikowiaceae(100);g__Metschnikowia(100);s__Metschnikowia_pulcherrima(100); | Metschnikowia | 13 | Saprotroph | \|Undefined Saprotroph\| | Yeast | NULL | Probable | As nectar-tap_saprotroph (Põlme et al. 2020); Decay sugar-rich substrates (Põlme et al. 2020) | Põlme S, et al. 2020. Fungal Diversity 105:1-16 ((https://doi.org/10.1007/s13225-020-00466-2)) |
| Otu54152 | 1 | k__Fungi(100);p__Ascomycota(100);c__Saccharomycetes(100);o__Saccharomycetales(100);f__Metschnikowiaceae(100);g__Metschnikowia(100);s__Metschnikowia_pulcherrima(100); | Metschnikowia | 13 | Saprotroph | \|Undefined Saprotroph\| | Yeast | NULL | Probable | As nectar-tap_saprotroph (Põlme et al. 2020); Decay sugar-rich substrates (Põlme et al. 2020) | Põlme S, et al. 2020. Fungal Diversity 105:1-16 ((https://doi.org/10.1007/s13225-020-00466-2)) |
| Otu54153 | 1 | k__Fungi(100);p__Ascomycota(100);c__Saccharomycetes(100);o__Saccharomycetales(100);f__Metschnikowiaceae(100);g__Metschnikowia(100);s__Metschnikowia_pulcherrima(100); | Metschnikowia | 13 | Saprotroph | \|Undefined Saprotroph\| | Yeast | NULL | Probable | As nectar-tap_saprotroph (Põlme et al. 2020); Decay sugar-rich substrates (Põlme et al. 2020) | Põlme S, et al. 2020. Fungal Diversity 105:1-16 ((https://doi.org/10.1007/s13225-020-00466-2)) |
| Otu54154 | 1 | k__Fungi(100);p__Ascomycota(100);c__Saccharomycetes(100);o__Saccharomycetales(100);f__Metschnikowiaceae(100);g__Metschnikowia(100);s__Metschnikowia_pulcherrima(100); | Metschnikowia | 13 | Saprotroph | \|Undefined Saprotroph\| | Yeast | NULL | Probable | As nectar-tap_saprotroph (Põlme et al. 2020); Decay sugar-rich substrates (Põlme et al. 2020) | Põlme S, et al. 2020. Fungal Diversity 105:1-16 ((https://doi.org/10.1007/s13225-020-00466-2)) |
| Otu54155 | 1 | k__Fungi(100);p__Ascomycota(100);c__Saccharomycetes(100);o__Saccharomycetales(100);f__Metschnikowiaceae(100);g__Metschnikowia(100);s__Metschnikowia_pulcherrima(100); | Metschnikowia | 13 | Saprotroph | \|Undefined Saprotroph\| | Yeast | NULL | Probable | As nectar-tap_saprotroph (Põlme et al. 2020); Decay sugar-rich substrates (Põlme et al. 2020) | Põlme S, et al. 2020. Fungal Diversity 105:1-16 ((https://doi.org/10.1007/s13225-020-00466-2)) |
| Otu54156 | 1 | k__Fungi(100);p__Ascomycota(100);c__Saccharomycetes(100);o__Saccharomycetales(100);f__Metschnikowiaceae(100);g__Metschnikowia(100);s__Metschnikowia_pulcherrima(100); | Metschnikowia | 13 | Saprotroph | \|Undefined Saprotroph\| | Yeast | NULL | Probable | As nectar-tap_saprotroph (Põlme et al. 2020); Decay sugar-rich substrates (Põlme et al. 2020) | Põlme S, et al. 2020. Fungal Diversity 105:1-16 ((https://doi.org/10.1007/s13225-020-00466-2)) |
| Otu54157 | 1 | k__Fungi(100);p__Ascomycota(100);c__Saccharomycetes(100);o__Saccharomycetales(100);f__Metschnikowiaceae(100);g__Metschnikowia(100);s__Metschnikowia_pulcherrima(100); | Metschnikowia | 13 | Saprotroph | \|Undefined Saprotroph\| | Yeast | NULL | Probable | As nectar-tap_saprotroph (Põlme et al. 2020); Decay sugar-rich substrates (Põlme et al. 2020) | Põlme S, et al. 2020. Fungal Diversity 105:1-16 ((https://doi.org/10.1007/s13225-020-00466-2)) |
| Otu54158 | 1 | k__Fungi(100);p__Ascomycota(100);c__Saccharomycetes(100);o__Saccharomycetales(100);f__Metschnikowiaceae(100);g__Metschnikowia(100);g__Metschnikowia_unclassified(100); | Metschnikowia | 13 | Saprotroph | \|Undefined Saprotroph\| | Yeast | NULL | Probable | As nectar-tap_saprotroph (Põlme et al. 2020); Decay sugar-rich substrates (Põlme et al. 2020) | Põlme S, et al. 2020. Fungal Diversity 105:1-16 ((https://doi.org/10.1007/s13225-020-00466-2)) |
| Otu54159 | 1 | k__Fungi(100);p__Ascomycota(100);c__Saccharomycetes(100);o__Saccharomycetales(100);f__Metschnikowiaceae(100);g__Metschnikowia(100);g__Metschnikowia_unclassified(100); | Metschnikowia | 13 | Saprotroph | \|Undefined Saprotroph\| | Yeast | NULL | Probable | As nectar-tap_saprotroph (Põlme et al. 2020); Decay sugar-rich substrates (Põlme et al. 2020) | Põlme S, et al. 2020. Fungal Diversity 105:1-16 ((https://doi.org/10.1007/s13225-020-00466-2)) |
| Otu54161 | 1 | k__Fungi(100);p__Ascomycota(100);c__Saccharomycetes(100);o__Saccharomycetales(100);f__Metschnikowiaceae(100);g__Metschnikowia(100);s__Metschnikowia_pulcherrima(100); | Metschnikowia | 13 | Saprotroph | \|Undefined Saprotroph\| | Yeast | NULL | Probable | As nectar-tap_saprotroph (Põlme et al. 2020); Decay sugar-rich substrates (Põlme et al. 2020) | Põlme S, et al. 2020. Fungal Diversity 105:1-16 ((https://doi.org/10.1007/s13225-020-00466-2)) |
| Otu54162 | 1 | k__Fungi(100);p__Ascomycota(100);c__Saccharomycetes(100);o__Saccharomycetales(100);f__Metschnikowiaceae(100);g__Metschnikowia(100);g__Metschnikowia_unclassified(100); | Metschnikowia | 13 | Saprotroph | \|Undefined Saprotroph\| | Yeast | NULL | Probable | As nectar-tap_saprotroph (Põlme et al. 2020); Decay sugar-rich substrates (Põlme et al. 2020) | Põlme S, et al. 2020. Fungal Diversity 105:1-16 ((https://doi.org/10.1007/s13225-020-00466-2)) |
| Otu54163 | 1 | k__Fungi(100);p__Ascomycota(100);c__Saccharomycetes(100);o__Saccharomycetales(100);f__Metschnikowiaceae(100);g__Metschnikowia(100);s__Metschnikowia_pulcherrima(100); | Metschnikowia | 13 | Saprotroph | \|Undefined Saprotroph\| | Yeast | NULL | Probable | As nectar-tap_saprotroph (Põlme et al. 2020); Decay sugar-rich substrates (Põlme et al. 2020) | Põlme S, et al. 2020. Fungal Diversity 105:1-16 ((https://doi.org/10.1007/s13225-020-00466-2)) |
| Otu54164 | 1 | k__Fungi(100);p__Ascomycota(100);c__Saccharomycetes(100);o__Saccharomycetales(100);f__Metschnikowiaceae(100);g__Metschnikowia(100);s__Metschnikowia_pulcherrima(100); | Metschnikowia | 13 | Saprotroph | \|Undefined Saprotroph\| | Yeast | NULL | Probable | As nectar-tap_saprotroph (Põlme et al. 2020); Decay sugar-rich substrates (Põlme et al. 2020) | Põlme S, et al. 2020. Fungal Diversity 105:1-16 ((https://doi.org/10.1007/s13225-020-00466-2)) |
| Otu54166 | 1 | k__Fungi(100);p__Ascomycota(100);c__Saccharomycetes(100);o__Saccharomycetales(100);f__Metschnikowiaceae(100);g__Metschnikowia(100);g__Metschnikowia_unclassified(100); | Metschnikowia | 13 | Saprotroph | \|Undefined Saprotroph\| | Yeast | NULL | Probable | As nectar-tap_saprotroph (Põlme et al. 2020); Decay sugar-rich substrates (Põlme et al. 2020) | Põlme S, et al. 2020. Fungal Diversity 105:1-16 ((https://doi.org/10.1007/s13225-020-00466-2)) |
| Otu54167 | 1 | k__Fungi(100);p__Ascomycota(100);c__Saccharomycetes(100);o__Saccharomycetales(100);f__Metschnikowiaceae(100);g__Metschnikowia(100);s__Metschnikowia_pulcherrima(100); | Metschnikowia | 13 | Saprotroph | \|Undefined Saprotroph\| | Yeast | NULL | Probable | As nectar-tap_saprotroph (Põlme et al. 2020); Decay sugar-rich substrates (Põlme et al. 2020) | Põlme S, et al. 2020. Fungal Diversity 105:1-16 ((https://doi.org/10.1007/s13225-020-00466-2)) |
| Otu54168 | 1 | k__Fungi(100);p__Ascomycota(100);c__Saccharomycetes(100);o__Saccharomycetales(100);f__Metschnikowiaceae(100);g__Metschnikowia(100);s__Metschnikowia_pulcherrima(100); | Metschnikowia | 13 | Saprotroph | \|Undefined Saprotroph\| | Yeast | NULL | Probable | As nectar-tap_saprotroph (Põlme et al. 2020); Decay sugar-rich substrates (Põlme et al. 2020) | Põlme S, et al. 2020. Fungal Diversity 105:1-16 ((https://doi.org/10.1007/s13225-020-00466-2)) |
| Otu54169 | 1 | k__Fungi(100);p__Ascomycota(100);c__Saccharomycetes(100);o__Saccharomycetales(100);f__Metschnikowiaceae(100);g__Metschnikowia(100);g__Metschnikowia_unclassified(100); | Metschnikowia | 13 | Saprotroph | \|Undefined Saprotroph\| | Yeast | NULL | Probable | As nectar-tap_saprotroph (Põlme et al. 2020); Decay sugar-rich substrates (Põlme et al. 2020) | Põlme S, et al. 2020. Fungal Diversity 105:1-16 ((https://doi.org/10.1007/s13225-020-00466-2)) |
| Otu54170 | 1 | k__Fungi(100);p__Ascomycota(100);c__Saccharomycetes(100);o__Saccharomycetales(100);f__Metschnikowiaceae(100);g__Metschnikowia(100);g__Metschnikowia_unclassified(100); | Metschnikowia | 13 | Saprotroph | \|Undefined Saprotroph\| | Yeast | NULL | Probable | As nectar-tap_saprotroph (Põlme et al. 2020); Decay sugar-rich substrates (Põlme et al. 2020) | Põlme S, et al. 2020. Fungal Diversity 105:1-16 ((https://doi.org/10.1007/s13225-020-00466-2)) |
| Otu54171 | 1 | k__Fungi(100);p__Ascomycota(100);c__Saccharomycetes(100);o__Saccharomycetales(100);f__Metschnikowiaceae(100);g__Metschnikowia(100);g__Metschnikowia_unclassified(100); | Metschnikowia | 13 | Saprotroph | \|Undefined Saprotroph\| | Yeast | NULL | Probable | As nectar-tap_saprotroph (Põlme et al. 2020); Decay sugar-rich substrates (Põlme et al. 2020) | Põlme S, et al. 2020. Fungal Diversity 105:1-16 ((https://doi.org/10.1007/s13225-020-00466-2)) |
| Otu54172 | 1 | k__Fungi(100);p__Ascomycota(100);c__Saccharomycetes(100);o__Saccharomycetales(100);f__Metschnikowiaceae(100);g__Metschnikowia(100);g__Metschnikowia_unclassified(100); | Metschnikowia | 13 | Saprotroph | \|Undefined Saprotroph\| | Yeast | NULL | Probable | As nectar-tap_saprotroph (Põlme et al. 2020); Decay sugar-rich substrates (Põlme et al. 2020) | Põlme S, et al. 2020. Fungal Diversity 105:1-16 ((https://doi.org/10.1007/s13225-020-00466-2)) |
| Otu54173 | 1 | k__Fungi(100);p__Ascomycota(100);c__Saccharomycetes(100);o__Saccharomycetales(100);f__Metschnikowiaceae(100);g__Metschnikowia(100);s__Metschnikowia_pulcherrima(100); | Metschnikowia | 13 | Saprotroph | \|Undefined Saprotroph\| | Yeast | NULL | Probable | As nectar-tap_saprotroph (Põlme et al. 2020); Decay sugar-rich substrates (Põlme et al. 2020) | Põlme S, et al. 2020. Fungal Diversity 105:1-16 ((https://doi.org/10.1007/s13225-020-00466-2)) |
| Otu54174 | 1 | k__Fungi(100);p__Ascomycota(100);c__Saccharomycetes(100);o__Saccharomycetales(100);f__Metschnikowiaceae(100);g__Metschnikowia(100);s__Metschnikowia_pulcherrima(100); | Metschnikowia | 13 | Saprotroph | \|Undefined Saprotroph\| | Yeast | NULL | Probable | As nectar-tap_saprotroph (Põlme et al. 2020); Decay sugar-rich substrates (Põlme et al. 2020) | Põlme S, et al. 2020. Fungal Diversity 105:1-16 ((https://doi.org/10.1007/s13225-020-00466-2)) |
| Otu54175 | 1 | k__Fungi(100);p__Ascomycota(100);c__Saccharomycetes(100);o__Saccharomycetales(100);f__Metschnikowiaceae(100);g__Metschnikowia(100);s__Metschnikowia_pulcherrima(100); | Metschnikowia | 13 | Saprotroph | \|Undefined Saprotroph\| | Yeast | NULL | Probable | As nectar-tap_saprotroph (Põlme et al. 2020); Decay sugar-rich substrates (Põlme et al. 2020) | Põlme S, et al. 2020. Fungal Diversity 105:1-16 ((https://doi.org/10.1007/s13225-020-00466-2)) |
| Otu54176 | 1 | k__Fungi(100);p__Ascomycota(100);c__Saccharomycetes(100);o__Saccharomycetales(100);f__Metschnikowiaceae(100);g__Metschnikowia(100);s__Metschnikowia_pulcherrima(100); | Metschnikowia | 13 | Saprotroph | \|Undefined Saprotroph\| | Yeast | NULL | Probable | As nectar-tap_saprotroph (Põlme et al. 2020); Decay sugar-rich substrates (Põlme et al. 2020) | Põlme S, et al. 2020. Fungal Diversity 105:1-16 ((https://doi.org/10.1007/s13225-020-00466-2)) |
| Otu54177 | 1 | k__Fungi(100);p__Ascomycota(100);c__Saccharomycetes(100);o__Saccharomycetales(100);f__Metschnikowiaceae(100);g__Metschnikowia(100);s__Metschnikowia_pulcherrima(100); | Metschnikowia | 13 | Saprotroph | \|Undefined Saprotroph\| | Yeast | NULL | Probable | As nectar-tap_saprotroph (Põlme et al. 2020); Decay sugar-rich substrates (Põlme et al. 2020) | Põlme S, et al. 2020. Fungal Diversity 105:1-16 ((https://doi.org/10.1007/s13225-020-00466-2)) |
| Otu54178 | 1 | k__Fungi(100);p__Ascomycota(100);c__Saccharomycetes(100);o__Saccharomycetales(100);f__Metschnikowiaceae(100);g__Metschnikowia(100);s__Metschnikowia_pulcherrima(100); | Metschnikowia | 13 | Saprotroph | \|Undefined Saprotroph\| | Yeast | NULL | Probable | As nectar-tap_saprotroph (Põlme et al. 2020); Decay sugar-rich substrates (Põlme et al. 2020) | Põlme S, et al. 2020. Fungal Diversity 105:1-16 ((https://doi.org/10.1007/s13225-020-00466-2)) |
| Otu54179 | 1 | k__Fungi(100);p__Ascomycota(100);c__Saccharomycetes(100);o__Saccharomycetales(100);f__Metschnikowiaceae(100);g__Metschnikowia(100);s__Metschnikowia_pulcherrima(100); | Metschnikowia | 13 | Saprotroph | \|Undefined Saprotroph\| | Yeast | NULL | Probable | As nectar-tap_saprotroph (Põlme et al. 2020); Decay sugar-rich substrates (Põlme et al. 2020) | Põlme S, et al. 2020. Fungal Diversity 105:1-16 ((https://doi.org/10.1007/s13225-020-00466-2)) |
| Otu54180 | 1 | k__Fungi(100);p__Ascomycota(100);c__Saccharomycetes(100);o__Saccharomycetales(100);f__Metschnikowiaceae(100);g__Metschnikowia(100);s__Metschnikowia_pulcherrima(100); | Metschnikowia | 13 | Saprotroph | \|Undefined Saprotroph\| | Yeast | NULL | Probable | As nectar-tap_saprotroph (Põlme et al. 2020); Decay sugar-rich substrates (Põlme et al. 2020) | Põlme S, et al. 2020. Fungal Diversity 105:1-16 ((https://doi.org/10.1007/s13225-020-00466-2)) |
| Otu54181 | 1 | k__Fungi(100);p__Ascomycota(100);c__Saccharomycetes(100);o__Saccharomycetales(100);f__Metschnikowiaceae(100);g__Metschnikowia(100);s__Metschnikowia_pulcherrima(100); | Metschnikowia | 13 | Saprotroph | \|Undefined Saprotroph\| | Yeast | NULL | Probable | As nectar-tap_saprotroph (Põlme et al. 2020); Decay sugar-rich substrates (Põlme et al. 2020) | Põlme S, et al. 2020. Fungal Diversity 105:1-16 ((https://doi.org/10.1007/s13225-020-00466-2)) |
| Otu54182 | 1 | k__Fungi(100);p__Ascomycota(100);c__Saccharomycetes(100);o__Saccharomycetales(100);f__Metschnikowiaceae(100);g__Metschnikowia(100);g__Metschnikowia_unclassified(100); | Metschnikowia | 13 | Saprotroph | \|Undefined Saprotroph\| | Yeast | NULL | Probable | As nectar-tap_saprotroph (Põlme et al. 2020); Decay sugar-rich substrates (Põlme et al. 2020) | Põlme S, et al. 2020. Fungal Diversity 105:1-16 ((https://doi.org/10.1007/s13225-020-00466-2)) |
| Otu54183 | 1 | k__Fungi(100);p__Ascomycota(100);c__Saccharomycetes(100);o__Saccharomycetales(100);f__Metschnikowiaceae(100);g__Metschnikowia(100);s__Metschnikowia_pulcherrima(100); | Metschnikowia | 13 | Saprotroph | \|Undefined Saprotroph\| | Yeast | NULL | Probable | As nectar-tap_saprotroph (Põlme et al. 2020); Decay sugar-rich substrates (Põlme et al. 2020) | Põlme S, et al. 2020. Fungal Diversity 105:1-16 ((https://doi.org/10.1007/s13225-020-00466-2)) |
| Otu54184 | 1 | k__Fungi(100);p__Ascomycota(100);c__Saccharomycetes(100);o__Saccharomycetales(100);f__Metschnikowiaceae(100);g__Metschnikowia(100);s__Metschnikowia_pulcherrima(100); | Metschnikowia | 13 | Saprotroph | \|Undefined Saprotroph\| | Yeast | NULL | Probable | As nectar-tap_saprotroph (Põlme et al. 2020); Decay sugar-rich substrates (Põlme et al. 2020) | Põlme S, et al. 2020. Fungal Diversity 105:1-16 ((https://doi.org/10.1007/s13225-020-00466-2)) |
| Otu54185 | 1 | k__Fungi(100);p__Ascomycota(100);c__Saccharomycetes(100);o__Saccharomycetales(100);f__Metschnikowiaceae(100);g__Metschnikowia(100);s__Metschnikowia_pulcherrima(100); | Metschnikowia | 13 | Saprotroph | \|Undefined Saprotroph\| | Yeast | NULL | Probable | As nectar-tap_saprotroph (Põlme et al. 2020); Decay sugar-rich substrates (Põlme et al. 2020) | Põlme S, et al. 2020. Fungal Diversity 105:1-16 ((https://doi.org/10.1007/s13225-020-00466-2)) |
| Otu54186 | 1 | k__Fungi(100);p__Ascomycota(100);c__Saccharomycetes(100);o__Saccharomycetales(100);f__Metschnikowiaceae(100);g__Metschnikowia(100);s__Metschnikowia_pulcherrima(100); | Metschnikowia | 13 | Saprotroph | \|Undefined Saprotroph\| | Yeast | NULL | Probable | As nectar-tap_saprotroph (Põlme et al. 2020); Decay sugar-rich substrates (Põlme et al. 2020) | Põlme S, et al. 2020. Fungal Diversity 105:1-16 ((https://doi.org/10.1007/s13225-020-00466-2)) |
| Otu54187 | 1 | k__Fungi(100);p__Ascomycota(100);c__Saccharomycetes(100);o__Saccharomycetales(100);f__Metschnikowiaceae(100);g__Metschnikowia(100);g__Metschnikowia_unclassified(100); | Metschnikowia | 13 | Saprotroph | \|Undefined Saprotroph\| | Yeast | NULL | Probable | As nectar-tap_saprotroph (Põlme et al. 2020); Decay sugar-rich substrates (Põlme et al. 2020) | Põlme S, et al. 2020. Fungal Diversity 105:1-16 ((https://doi.org/10.1007/s13225-020-00466-2)) |
| Otu54188 | 1 | k__Fungi(100);p__Ascomycota(100);c__Saccharomycetes(100);o__Saccharomycetales(100);f__Metschnikowiaceae(100);g__Metschnikowia(100);s__Metschnikowia_pulcherrima(100); | Metschnikowia | 13 | Saprotroph | \|Undefined Saprotroph\| | Yeast | NULL | Probable | As nectar-tap_saprotroph (Põlme et al. 2020); Decay sugar-rich substrates (Põlme et al. 2020) | Põlme S, et al. 2020. Fungal Diversity 105:1-16 ((https://doi.org/10.1007/s13225-020-00466-2)) |
| Otu54189 | 1 | k__Fungi(100);p__Ascomycota(100);c__Saccharomycetes(100);o__Saccharomycetales(100);f__Metschnikowiaceae(100);g__Metschnikowia(100);s__Metschnikowia_pulcherrima(100); | Metschnikowia | 13 | Saprotroph | \|Undefined Saprotroph\| | Yeast | NULL | Probable | As nectar-tap_saprotroph (Põlme et al. 2020); Decay sugar-rich substrates (Põlme et al. 2020) | Põlme S, et al. 2020. Fungal Diversity 105:1-16 ((https://doi.org/10.1007/s13225-020-00466-2)) |
| Otu54190 | 1 | k__Fungi(100);p__Ascomycota(100);c__Saccharomycetes(100);o__Saccharomycetales(100);f__Metschnikowiaceae(100);g__Metschnikowia(100);s__Metschnikowia_pulcherrima(100); | Metschnikowia | 13 | Saprotroph | \|Undefined Saprotroph\| | Yeast | NULL | Probable | As nectar-tap_saprotroph (Põlme et al. 2020); Decay sugar-rich substrates (Põlme et al. 2020) | Põlme S, et al. 2020. Fungal Diversity 105:1-16 ((https://doi.org/10.1007/s13225-020-00466-2)) |
| Otu54191 | 1 | k__Fungi(100);p__Ascomycota(100);c__Saccharomycetes(100);o__Saccharomycetales(100);f__Metschnikowiaceae(100);g__Metschnikowia(100);s__Metschnikowia_pulcherrima(100); | Metschnikowia | 13 | Saprotroph | \|Undefined Saprotroph\| | Yeast | NULL | Probable | As nectar-tap_saprotroph (Põlme et al. 2020); Decay sugar-rich substrates (Põlme et al. 2020) | Põlme S, et al. 2020. Fungal Diversity 105:1-16 ((https://doi.org/10.1007/s13225-020-00466-2)) |
| Otu54192 | 1 | k__Fungi(100);p__Ascomycota(100);c__Saccharomycetes(100);o__Saccharomycetales(100);f__Metschnikowiaceae(100);g__Metschnikowia(100);s__Metschnikowia_pulcherrima(100); | Metschnikowia | 13 | Saprotroph | \|Undefined Saprotroph\| | Yeast | NULL | Probable | As nectar-tap_saprotroph (Põlme et al. 2020); Decay sugar-rich substrates (Põlme et al. 2020) | Põlme S, et al. 2020. Fungal Diversity 105:1-16 ((https://doi.org/10.1007/s13225-020-00466-2)) |
| Otu54193 | 1 | k__Fungi(100);p__Ascomycota(100);c__Saccharomycetes(100);o__Saccharomycetales(100);f__Metschnikowiaceae(100);g__Metschnikowia(100);g__Metschnikowia_unclassified(100); | Metschnikowia | 13 | Saprotroph | \|Undefined Saprotroph\| | Yeast | NULL | Probable | As nectar-tap_saprotroph (Põlme et al. 2020); Decay sugar-rich substrates (Põlme et al. 2020) | Põlme S, et al. 2020. Fungal Diversity 105:1-16 ((https://doi.org/10.1007/s13225-020-00466-2)) |
| Otu54194 | 1 | k__Fungi(100);p__Ascomycota(100);c__Saccharomycetes(100);o__Saccharomycetales(100);f__Metschnikowiaceae(100);g__Metschnikowia(100);s__Metschnikowia_pulcherrima(100); | Metschnikowia | 13 | Saprotroph | \|Undefined Saprotroph\| | Yeast | NULL | Probable | As nectar-tap_saprotroph (Põlme et al. 2020); Decay sugar-rich substrates (Põlme et al. 2020) | Põlme S, et al. 2020. Fungal Diversity 105:1-16 ((https://doi.org/10.1007/s13225-020-00466-2)) |
| Otu54195 | 1 | k__Fungi(100);p__Ascomycota(100);c__Saccharomycetes(100);o__Saccharomycetales(100);f__Metschnikowiaceae(100);g__Metschnikowia(100);s__Metschnikowia_pulcherrima(100); | Metschnikowia | 13 | Saprotroph | \|Undefined Saprotroph\| | Yeast | NULL | Probable | As nectar-tap_saprotroph (Põlme et al. 2020); Decay sugar-rich substrates (Põlme et al. 2020) | Põlme S, et al. 2020. Fungal Diversity 105:1-16 ((https://doi.org/10.1007/s13225-020-00466-2)) |
| Otu54196 | 1 | k__Fungi(100);p__Ascomycota(100);c__Saccharomycetes(100);o__Saccharomycetales(100);f__Metschnikowiaceae(100);g__Metschnikowia(100);g__Metschnikowia_unclassified(100); | Metschnikowia | 13 | Saprotroph | \|Undefined Saprotroph\| | Yeast | NULL | Probable | As nectar-tap_saprotroph (Põlme et al. 2020); Decay sugar-rich substrates (Põlme et al. 2020) | Põlme S, et al. 2020. Fungal Diversity 105:1-16 ((https://doi.org/10.1007/s13225-020-00466-2)) |
| Otu54197 | 1 | k__Fungi(100);p__Ascomycota(100);c__Saccharomycetes(100);o__Saccharomycetales(100);f__Metschnikowiaceae(100);g__Metschnikowia(100);s__Metschnikowia_pulcherrima(100); | Metschnikowia | 13 | Saprotroph | \|Undefined Saprotroph\| | Yeast | NULL | Probable | As nectar-tap_saprotroph (Põlme et al. 2020); Decay sugar-rich substrates (Põlme et al. 2020) | Põlme S, et al. 2020. Fungal Diversity 105:1-16 ((https://doi.org/10.1007/s13225-020-00466-2)) |
| Otu54198 | 1 | k__Fungi(100);p__Ascomycota(100);c__Saccharomycetes(100);o__Saccharomycetales(100);f__Metschnikowiaceae(100);g__Metschnikowia(100);s__Metschnikowia_pulcherrima(100); | Metschnikowia | 13 | Saprotroph | \|Undefined Saprotroph\| | Yeast | NULL | Probable | As nectar-tap_saprotroph (Põlme et al. 2020); Decay sugar-rich substrates (Põlme et al. 2020) | Põlme S, et al. 2020. Fungal Diversity 105:1-16 ((https://doi.org/10.1007/s13225-020-00466-2)) |
| Otu54199 | 1 | k__Fungi(100);p__Ascomycota(100);c__Saccharomycetes(100);o__Saccharomycetales(100);f__Metschnikowiaceae(100);g__Metschnikowia(100);s__Metschnikowia_pulcherrima(100); | Metschnikowia | 13 | Saprotroph | \|Undefined Saprotroph\| | Yeast | NULL | Probable | As nectar-tap_saprotroph (Põlme et al. 2020); Decay sugar-rich substrates (Põlme et al. 2020) | Põlme S, et al. 2020. Fungal Diversity 105:1-16 ((https://doi.org/10.1007/s13225-020-00466-2)) |
| Otu54200 | 1 | k__Fungi(100);p__Ascomycota(100);c__Saccharomycetes(100);o__Saccharomycetales(100);f__Metschnikowiaceae(100);g__Metschnikowia(100);g__Metschnikowia_unclassified(100); | Metschnikowia | 13 | Saprotroph | \|Undefined Saprotroph\| | Yeast | NULL | Probable | As nectar-tap_saprotroph (Põlme et al. 2020); Decay sugar-rich substrates (Põlme et al. 2020) | Põlme S, et al. 2020. Fungal Diversity 105:1-16 ((https://doi.org/10.1007/s13225-020-00466-2)) |
| Otu54201 | 1 | k__Fungi(100);p__Ascomycota(100);c__Saccharomycetes(100);o__Saccharomycetales(100);f__Metschnikowiaceae(100);g__Metschnikowia(100);g__Metschnikowia_unclassified(100); | Metschnikowia | 13 | Saprotroph | \|Undefined Saprotroph\| | Yeast | NULL | Probable | As nectar-tap_saprotroph (Põlme et al. 2020); Decay sugar-rich substrates (Põlme et al. 2020) | Põlme S, et al. 2020. Fungal Diversity 105:1-16 ((https://doi.org/10.1007/s13225-020-00466-2)) |
| Otu54202 | 1 | k__Fungi(100);p__Ascomycota(100);c__Saccharomycetes(100);o__Saccharomycetales(100);f__Metschnikowiaceae(100);g__Metschnikowia(100);g__Metschnikowia_unclassified(100); | Metschnikowia | 13 | Saprotroph | \|Undefined Saprotroph\| | Yeast | NULL | Probable | As nectar-tap_saprotroph (Põlme et al. 2020); Decay sugar-rich substrates (Põlme et al. 2020) | Põlme S, et al. 2020. Fungal Diversity 105:1-16 ((https://doi.org/10.1007/s13225-020-00466-2)) |
| Otu54203 | 1 | k__Fungi(100);p__Ascomycota(100);c__Saccharomycetes(100);o__Saccharomycetales(100);f__Metschnikowiaceae(100);g__Metschnikowia(100);s__Metschnikowia_pulcherrima(100); | Metschnikowia | 13 | Saprotroph | \|Undefined Saprotroph\| | Yeast | NULL | Probable | As nectar-tap_saprotroph (Põlme et al. 2020); Decay sugar-rich substrates (Põlme et al. 2020) | Põlme S, et al. 2020. Fungal Diversity 105:1-16 ((https://doi.org/10.1007/s13225-020-00466-2)) |
| Otu54204 | 1 | k__Fungi(100);p__Ascomycota(100);c__Saccharomycetes(100);o__Saccharomycetales(100);f__Metschnikowiaceae(100);g__Metschnikowia(100);g__Metschnikowia_unclassified(100); | Metschnikowia | 13 | Saprotroph | \|Undefined Saprotroph\| | Yeast | NULL | Probable | As nectar-tap_saprotroph (Põlme et al. 2020); Decay sugar-rich substrates (Põlme et al. 2020) | Põlme S, et al. 2020. Fungal Diversity 105:1-16 ((https://doi.org/10.1007/s13225-020-00466-2)) |
| Otu54205 | 1 | k__Fungi(100);p__Ascomycota(100);c__Saccharomycetes(100);o__Saccharomycetales(100);f__Metschnikowiaceae(100);g__Metschnikowia(100);s__Metschnikowia_pulcherrima(100); | Metschnikowia | 13 | Saprotroph | \|Undefined Saprotroph\| | Yeast | NULL | Probable | As nectar-tap_saprotroph (Põlme et al. 2020); Decay sugar-rich substrates (Põlme et al. 2020) | Põlme S, et al. 2020. Fungal Diversity 105:1-16 ((https://doi.org/10.1007/s13225-020-00466-2)) |
| Otu54206 | 1 | k__Fungi(100);p__Ascomycota(100);c__Saccharomycetes(100);o__Saccharomycetales(100);f__Metschnikowiaceae(100);g__Metschnikowia(100);s__Metschnikowia_pulcherrima(100); | Metschnikowia | 13 | Saprotroph | \|Undefined Saprotroph\| | Yeast | NULL | Probable | As nectar-tap_saprotroph (Põlme et al. 2020); Decay sugar-rich substrates (Põlme et al. 2020) | Põlme S, et al. 2020. Fungal Diversity 105:1-16 ((https://doi.org/10.1007/s13225-020-00466-2)) |
| Otu54207 | 1 | k__Fungi(100);p__Ascomycota(100);c__Saccharomycetes(100);o__Saccharomycetales(100);f__Metschnikowiaceae(100);g__Metschnikowia(100);s__Metschnikowia_pulcherrima(100); | Metschnikowia | 13 | Saprotroph | \|Undefined Saprotroph\| | Yeast | NULL | Probable | As nectar-tap_saprotroph (Põlme et al. 2020); Decay sugar-rich substrates (Põlme et al. 2020) | Põlme S, et al. 2020. Fungal Diversity 105:1-16 ((https://doi.org/10.1007/s13225-020-00466-2)) |
| Otu54208 | 1 | k__Fungi(100);p__Ascomycota(100);c__Saccharomycetes(100);o__Saccharomycetales(100);f__Metschnikowiaceae(100);g__Metschnikowia(100);s__Metschnikowia_pulcherrima(100); | Metschnikowia | 13 | Saprotroph | \|Undefined Saprotroph\| | Yeast | NULL | Probable | As nectar-tap_saprotroph (Põlme et al. 2020); Decay sugar-rich substrates (Põlme et al. 2020) | Põlme S, et al. 2020. Fungal Diversity 105:1-16 ((https://doi.org/10.1007/s13225-020-00466-2)) |
| Otu54209 | 1 | k__Fungi(100);p__Ascomycota(100);c__Saccharomycetes(100);o__Saccharomycetales(100);f__Metschnikowiaceae(100);g__Metschnikowia(100);s__Metschnikowia_pulcherrima(100); | Metschnikowia | 13 | Saprotroph | \|Undefined Saprotroph\| | Yeast | NULL | Probable | As nectar-tap_saprotroph (Põlme et al. 2020); Decay sugar-rich substrates (Põlme et al. 2020) | Põlme S, et al. 2020. Fungal Diversity 105:1-16 ((https://doi.org/10.1007/s13225-020-00466-2)) |
| Otu54210 | 1 | k__Fungi(100);p__Ascomycota(100);c__Saccharomycetes(100);o__Saccharomycetales(100);f__Metschnikowiaceae(100);g__Metschnikowia(100);s__Metschnikowia_pulcherrima(100); | Metschnikowia | 13 | Saprotroph | \|Undefined Saprotroph\| | Yeast | NULL | Probable | As nectar-tap_saprotroph (Põlme et al. 2020); Decay sugar-rich substrates (Põlme et al. 2020) | Põlme S, et al. 2020. Fungal Diversity 105:1-16 ((https://doi.org/10.1007/s13225-020-00466-2)) |
| Otu54211 | 1 | k__Fungi(100);p__Ascomycota(100);c__Saccharomycetes(100);o__Saccharomycetales(100);f__Metschnikowiaceae(100);g__Metschnikowia(100);s__Metschnikowia_pulcherrima(100); | Metschnikowia | 13 | Saprotroph | \|Undefined Saprotroph\| | Yeast | NULL | Probable | As nectar-tap_saprotroph (Põlme et al. 2020); Decay sugar-rich substrates (Põlme et al. 2020) | Põlme S, et al. 2020. Fungal Diversity 105:1-16 ((https://doi.org/10.1007/s13225-020-00466-2)) |
| Otu54212 | 1 | k__Fungi(100);p__Ascomycota(100);c__Saccharomycetes(100);o__Saccharomycetales(100);f__Metschnikowiaceae(100);g__Metschnikowia(100);s__Metschnikowia_pulcherrima(100); | Metschnikowia | 13 | Saprotroph | \|Undefined Saprotroph\| | Yeast | NULL | Probable | As nectar-tap_saprotroph (Põlme et al. 2020); Decay sugar-rich substrates (Põlme et al. 2020) | Põlme S, et al. 2020. Fungal Diversity 105:1-16 ((https://doi.org/10.1007/s13225-020-00466-2)) |
| Otu54213 | 1 | k__Fungi(100);p__Ascomycota(100);c__Saccharomycetes(100);o__Saccharomycetales(100);f__Metschnikowiaceae(100);g__Metschnikowia(100);g__Metschnikowia_unclassified(100); | Metschnikowia | 13 | Saprotroph | \|Undefined Saprotroph\| | Yeast | NULL | Probable | As nectar-tap_saprotroph (Põlme et al. 2020); Decay sugar-rich substrates (Põlme et al. 2020) | Põlme S, et al. 2020. Fungal Diversity 105:1-16 ((https://doi.org/10.1007/s13225-020-00466-2)) |
| Otu54214 | 1 | k__Fungi(100);p__Ascomycota(100);c__Saccharomycetes(100);o__Saccharomycetales(100);f__Metschnikowiaceae(100);g__Metschnikowia(100);s__Metschnikowia_pulcherrima(100); | Metschnikowia | 13 | Saprotroph | \|Undefined Saprotroph\| | Yeast | NULL | Probable | As nectar-tap_saprotroph (Põlme et al. 2020); Decay sugar-rich substrates (Põlme et al. 2020) | Põlme S, et al. 2020. Fungal Diversity 105:1-16 ((https://doi.org/10.1007/s13225-020-00466-2)) |
| Otu54215 | 1 | k__Fungi(100);p__Ascomycota(100);c__Saccharomycetes(100);o__Saccharomycetales(100);f__Metschnikowiaceae(100);g__Metschnikowia(100);s__Metschnikowia_pulcherrima(100); | Metschnikowia | 13 | Saprotroph | \|Undefined Saprotroph\| | Yeast | NULL | Probable | As nectar-tap_saprotroph (Põlme et al. 2020); Decay sugar-rich substrates (Põlme et al. 2020) | Põlme S, et al. 2020. Fungal Diversity 105:1-16 ((https://doi.org/10.1007/s13225-020-00466-2)) |
| Otu54216 | 1 | k__Fungi(100);p__Ascomycota(100);c__Saccharomycetes(100);o__Saccharomycetales(100);f__Metschnikowiaceae(100);g__Metschnikowia(100);s__Metschnikowia_pulcherrima(100); | Metschnikowia | 13 | Saprotroph | \|Undefined Saprotroph\| | Yeast | NULL | Probable | As nectar-tap_saprotroph (Põlme et al. 2020); Decay sugar-rich substrates (Põlme et al. 2020) | Põlme S, et al. 2020. Fungal Diversity 105:1-16 ((https://doi.org/10.1007/s13225-020-00466-2)) |
| Otu54217 | 1 | k__Fungi(100);p__Ascomycota(100);c__Saccharomycetes(100);o__Saccharomycetales(100);f__Metschnikowiaceae(100);g__Metschnikowia(100);s__Metschnikowia_pulcherrima(100); | Metschnikowia | 13 | Saprotroph | \|Undefined Saprotroph\| | Yeast | NULL | Probable | As nectar-tap_saprotroph (Põlme et al. 2020); Decay sugar-rich substrates (Põlme et al. 2020) | Põlme S, et al. 2020. Fungal Diversity 105:1-16 ((https://doi.org/10.1007/s13225-020-00466-2)) |
| Otu54218 | 1 | k__Fungi(100);p__Ascomycota(100);c__Saccharomycetes(100);o__Saccharomycetales(100);f__Metschnikowiaceae(100);g__Metschnikowia(100);s__Metschnikowia_pulcherrima(100); | Metschnikowia | 13 | Saprotroph | \|Undefined Saprotroph\| | Yeast | NULL | Probable | As nectar-tap_saprotroph (Põlme et al. 2020); Decay sugar-rich substrates (Põlme et al. 2020) | Põlme S, et al. 2020. Fungal Diversity 105:1-16 ((https://doi.org/10.1007/s13225-020-00466-2)) |
| Otu54219 | 1 | k__Fungi(100);p__Ascomycota(100);c__Saccharomycetes(100);o__Saccharomycetales(100);f__Metschnikowiaceae(100);g__Metschnikowia(100);s__Metschnikowia_pulcherrima(100); | Metschnikowia | 13 | Saprotroph | \|Undefined Saprotroph\| | Yeast | NULL | Probable | As nectar-tap_saprotroph (Põlme et al. 2020); Decay sugar-rich substrates (Põlme et al. 2020) | Põlme S, et al. 2020. Fungal Diversity 105:1-16 ((https://doi.org/10.1007/s13225-020-00466-2)) |
| Otu54220 | 1 | k__Fungi(100);p__Ascomycota(100);c__Saccharomycetes(100);o__Saccharomycetales(100);f__Metschnikowiaceae(100);g__Metschnikowia(100);g__Metschnikowia_unclassified(100); | Metschnikowia | 13 | Saprotroph | \|Undefined Saprotroph\| | Yeast | NULL | Probable | As nectar-tap_saprotroph (Põlme et al. 2020); Decay sugar-rich substrates (Põlme et al. 2020) | Põlme S, et al. 2020. Fungal Diversity 105:1-16 ((https://doi.org/10.1007/s13225-020-00466-2)) |
| Otu54221 | 1 | k__Fungi(100);p__Ascomycota(100);c__Saccharomycetes(100);o__Saccharomycetales(100);f__Metschnikowiaceae(100);g__Metschnikowia(100);s__Metschnikowia_pulcherrima(100); | Metschnikowia | 13 | Saprotroph | \|Undefined Saprotroph\| | Yeast | NULL | Probable | As nectar-tap_saprotroph (Põlme et al. 2020); Decay sugar-rich substrates (Põlme et al. 2020) | Põlme S, et al. 2020. Fungal Diversity 105:1-16 ((https://doi.org/10.1007/s13225-020-00466-2)) |
| Otu54223 | 1 | k__Fungi(100);p__Ascomycota(100);c__Saccharomycetes(100);o__Saccharomycetales(100);f__Metschnikowiaceae(100);g__Metschnikowia(100);g__Metschnikowia_unclassified(100); | Metschnikowia | 13 | Saprotroph | \|Undefined Saprotroph\| | Yeast | NULL | Probable | As nectar-tap_saprotroph (Põlme et al. 2020); Decay sugar-rich substrates (Põlme et al. 2020) | Põlme S, et al. 2020. Fungal Diversity 105:1-16 ((https://doi.org/10.1007/s13225-020-00466-2)) |
| Otu54224 | 1 | k__Fungi(100);p__Ascomycota(100);c__Saccharomycetes(100);o__Saccharomycetales(100);f__Metschnikowiaceae(100);g__Metschnikowia(100);g__Metschnikowia_unclassified(100); | Metschnikowia | 13 | Saprotroph | \|Undefined Saprotroph\| | Yeast | NULL | Probable | As nectar-tap_saprotroph (Põlme et al. 2020); Decay sugar-rich substrates (Põlme et al. 2020) | Põlme S, et al. 2020. Fungal Diversity 105:1-16 ((https://doi.org/10.1007/s13225-020-00466-2)) |
| Otu54225 | 1 | k__Fungi(100);p__Ascomycota(100);c__Saccharomycetes(100);o__Saccharomycetales(100);f__Metschnikowiaceae(100);g__Metschnikowia(100);s__Metschnikowia_pulcherrima(100); | Metschnikowia | 13 | Saprotroph | \|Undefined Saprotroph\| | Yeast | NULL | Probable | As nectar-tap_saprotroph (Põlme et al. 2020); Decay sugar-rich substrates (Põlme et al. 2020) | Põlme S, et al. 2020. Fungal Diversity 105:1-16 ((https://doi.org/10.1007/s13225-020-00466-2)) |
| Otu54226 | 1 | k__Fungi(100);p__Ascomycota(100);c__Saccharomycetes(100);o__Saccharomycetales(100);f__Metschnikowiaceae(100);g__Metschnikowia(100);s__Metschnikowia_pulcherrima(100); | Metschnikowia | 13 | Saprotroph | \|Undefined Saprotroph\| | Yeast | NULL | Probable | As nectar-tap_saprotroph (Põlme et al. 2020); Decay sugar-rich substrates (Põlme et al. 2020) | Põlme S, et al. 2020. Fungal Diversity 105:1-16 ((https://doi.org/10.1007/s13225-020-00466-2)) |
| Otu54227 | 1 | k__Fungi(100);p__Ascomycota(100);c__Saccharomycetes(100);o__Saccharomycetales(100);f__Metschnikowiaceae(100);g__Metschnikowia(100);s__Metschnikowia_pulcherrima(100); | Metschnikowia | 13 | Saprotroph | \|Undefined Saprotroph\| | Yeast | NULL | Probable | As nectar-tap_saprotroph (Põlme et al. 2020); Decay sugar-rich substrates (Põlme et al. 2020) | Põlme S, et al. 2020. Fungal Diversity 105:1-16 ((https://doi.org/10.1007/s13225-020-00466-2)) |
| Otu54228 | 1 | k__Fungi(100);p__Ascomycota(100);c__Saccharomycetes(100);o__Saccharomycetales(100);f__Metschnikowiaceae(100);g__Metschnikowia(100);s__Metschnikowia_pulcherrima(100); | Metschnikowia | 13 | Saprotroph | \|Undefined Saprotroph\| | Yeast | NULL | Probable | As nectar-tap_saprotroph (Põlme et al. 2020); Decay sugar-rich substrates (Põlme et al. 2020) | Põlme S, et al. 2020. Fungal Diversity 105:1-16 ((https://doi.org/10.1007/s13225-020-00466-2)) |
| Otu54229 | 1 | k__Fungi(100);p__Ascomycota(100);c__Saccharomycetes(100);o__Saccharomycetales(100);f__Metschnikowiaceae(100);g__Metschnikowia(100);g__Metschnikowia_unclassified(100); | Metschnikowia | 13 | Saprotroph | \|Undefined Saprotroph\| | Yeast | NULL | Probable | As nectar-tap_saprotroph (Põlme et al. 2020); Decay sugar-rich substrates (Põlme et al. 2020) | Põlme S, et al. 2020. Fungal Diversity 105:1-16 ((https://doi.org/10.1007/s13225-020-00466-2)) |
| Otu54230 | 1 | k__Fungi(100);p__Ascomycota(100);c__Saccharomycetes(100);o__Saccharomycetales(100);f__Metschnikowiaceae(100);g__Metschnikowia(100);s__Metschnikowia_pulcherrima(100); | Metschnikowia | 13 | Saprotroph | \|Undefined Saprotroph\| | Yeast | NULL | Probable | As nectar-tap_saprotroph (Põlme et al. 2020); Decay sugar-rich substrates (Põlme et al. 2020) | Põlme S, et al. 2020. Fungal Diversity 105:1-16 ((https://doi.org/10.1007/s13225-020-00466-2)) |
| Otu54231 | 1 | k__Fungi(100);p__Ascomycota(100);c__Saccharomycetes(100);o__Saccharomycetales(100);f__Metschnikowiaceae(100);g__Metschnikowia(100);s__Metschnikowia_pulcherrima(100); | Metschnikowia | 13 | Saprotroph | \|Undefined Saprotroph\| | Yeast | NULL | Probable | As nectar-tap_saprotroph (Põlme et al. 2020); Decay sugar-rich substrates (Põlme et al. 2020) | Põlme S, et al. 2020. Fungal Diversity 105:1-16 ((https://doi.org/10.1007/s13225-020-00466-2)) |
| Otu54232 | 1 | k__Fungi(100);p__Ascomycota(100);c__Saccharomycetes(100);o__Saccharomycetales(100);f__Metschnikowiaceae(100);g__Metschnikowia(100);s__Metschnikowia_pulcherrima(100); | Metschnikowia | 13 | Saprotroph | \|Undefined Saprotroph\| | Yeast | NULL | Probable | As nectar-tap_saprotroph (Põlme et al. 2020); Decay sugar-rich substrates (Põlme et al. 2020) | Põlme S, et al. 2020. Fungal Diversity 105:1-16 ((https://doi.org/10.1007/s13225-020-00466-2)) |

| | | | | | | | | | | | |
|---|---|---|---|---|---|---|---|---|---|---|---|
| Otu54234 | 1 | k__Fungi(100);p__Ascomycota(100);c__Saccharomycetes(100);o__Saccharomycetales(100);f__Metschnikowiaceae(100);g__Metschnikowia(100);s__Metschnikowia_pulcherrima(100); | Metschnikowia | 13 | Saprotroph | (Undefined Saprotroph) | Yeast | NULL | Probable | As nectar-tap_saprotroph (Põlme et al. 2020); Decay sugar-rich substrates (Põlme et al. 2020) | Põlme S, et al. 2020. Fungal Diversity 105:1-16 ((https://doi.org/10.1007/s13225-020-00466-2)) |
| Otu54235 | 1 | k__Fungi(100);p__Ascomycota(100);c__Saccharomycetes(100);o__Saccharomycetales(100);f__Metschnikowiaceae(100);g__Metschnikowia(100);s__Metschnikowia_pulcherrima(100); | Metschnikowia | 13 | Saprotroph | (Undefined Saprotroph) | Yeast | NULL | Probable | As nectar-tap_saprotroph (Põlme et al. 2020); Decay sugar-rich substrates (Põlme et al. 2020) | Põlme S, et al. 2020. Fungal Diversity 105:1-16 ((https://doi.org/10.1007/s13225-020-00466-2)) |
| Otu54236 | 1 | k__Fungi(100);p__Ascomycota(100);c__Saccharomycetes(100);o__Saccharomycetales(100);f__Metschnikowiaceae(100);g__Metschnikowia(100);s__Metschnikowia_pulcherrima(100); | Metschnikowia | 13 | Saprotroph | (Undefined Saprotroph) | Yeast | NULL | Probable | As nectar-tap_saprotroph (Põlme et al. 2020); Decay sugar-rich substrates (Põlme et al. 2020) | Põlme S, et al. 2020. Fungal Diversity 105:1-16 ((https://doi.org/10.1007/s13225-020-00466-2)) |
| Otu54237 | 1 | k__Fungi(100);p__Ascomycota(100);c__Saccharomycetes(100);o__Saccharomycetales(100);f__Metschnikowiaceae(100);g__Metschnikowia(100);s__Metschnikowia_pulcherrima(100); | Metschnikowia | 13 | Saprotroph | (Undefined Saprotroph) | Yeast | NULL | Probable | As nectar-tap_saprotroph (Põlme et al. 2020); Decay sugar-rich substrates (Põlme et al. 2020) | Põlme S, et al. 2020. Fungal Diversity 105:1-16 ((https://doi.org/10.1007/s13225-020-00466-2)) |
| Otu54238 | 1 | k__Fungi(100);p__Ascomycota(100);c__Saccharomycetes(100);o__Saccharomycetales(100);f__Metschnikowiaceae(100);g__Metschnikowia(100);g__Metschnikowia_unclassified(100); | Metschnikowia | 13 | Saprotroph | (Undefined Saprotroph) | Yeast | NULL | Probable | As nectar-tap_saprotroph (Põlme et al. 2020); Decay sugar-rich substrates (Põlme et al. 2020) | Põlme S, et al. 2020. Fungal Diversity 105:1-16 ((https://doi.org/10.1007/s13225-020-00466-2)) |
| Otu54239 | 1 | k__Fungi(100);p__Ascomycota(100);c__Saccharomycetes(100);o__Saccharomycetales(100);f__Metschnikowiaceae(100);g__Metschnikowia(100);s__Metschnikowia_pulcherrima(100); | Metschnikowia | 13 | Saprotroph | (Undefined Saprotroph) | Yeast | NULL | Probable | As nectar-tap_saprotroph (Põlme et al. 2020); Decay sugar-rich substrates (Põlme et al. 2020) | Põlme S, et al. 2020. Fungal Diversity 105:1-16 ((https://doi.org/10.1007/s13225-020-00466-2)) |
| Otu54240 | 1 | k__Fungi(100);p__Ascomycota(100);c__Saccharomycetes(100);o__Saccharomycetales(100);f__Metschnikowiaceae(100);g__Metschnikowia(100);s__Metschnikowia_pulcherrima(100); | Metschnikowia | 13 | Saprotroph | (Undefined Saprotroph) | Yeast | NULL | Probable | As nectar-tap_saprotroph (Põlme et al. 2020); Decay sugar-rich substrates (Põlme et al. 2020) | Põlme S, et al. 2020. Fungal Diversity 105:1-16 ((https://doi.org/10.1007/s13225-020-00466-2)) |
| Otu54241 | 1 | k__Fungi(100);p__Ascomycota(100);c__Saccharomycetes(100);o__Saccharomycetales(100);f__Metschnikowiaceae(100);g__Metschnikowia(100);s__Metschnikowia_pulcherrima(100); | Metschnikowia | 13 | Saprotroph | (Undefined Saprotroph) | Yeast | NULL | Probable | As nectar-tap_saprotroph (Põlme et al. 2020); Decay sugar-rich substrates (Põlme et al. 2020) | Põlme S, et al. 2020. Fungal Diversity 105:1-16 ((https://doi.org/10.1007/s13225-020-00466-2)) |
| Otu54245 | 1 | k__Fungi(100);p__Ascomycota(100);c__Saccharomycetes(100);o__Saccharomycetales(100);f__Metschnikowiaceae(100);g__Metschnikowia(100);s__Metschnikowia_pulcherrima(100); | Metschnikowia | 13 | Saprotroph | (Undefined Saprotroph) | Yeast | NULL | Probable | As nectar-tap_saprotroph (Põlme et al. 2020); Decay sugar-rich substrates (Põlme et al. 2020) | Põlme S, et al. 2020. Fungal Diversity 105:1-16 ((https://doi.org/10.1007/s13225-020-00466-2)) |
| Otu54246 | 1 | k__Fungi(100);p__Ascomycota(100);c__Saccharomycetes(100);o__Saccharomycetales(100);f__Metschnikowiaceae(100);g__Metschnikowia(100);g__Metschnikowia_unclassified(100); | Metschnikowia | 13 | Saprotroph | (Undefined Saprotroph) | Yeast | NULL | Probable | As nectar-tap_saprotroph (Põlme et al. 2020); Decay sugar-rich substrates (Põlme et al. 2020) | Põlme S, et al. 2020. Fungal Diversity 105:1-16 ((https://doi.org/10.1007/s13225-020-00466-2)) |
| Otu54247 | 1 | k__Fungi(100);p__Ascomycota(100);c__Saccharomycetes(100);o__Saccharomycetales(100);f__Metschnikowiaceae(100);g__Metschnikowia(100);g__Metschnikowia_unclassified(100); | Metschnikowia | 13 | Saprotroph | (Undefined Saprotroph) | Yeast | NULL | Probable | As nectar-tap_saprotroph (Põlme et al. 2020); Decay sugar-rich substrates (Põlme et al. 2020) | Põlme S, et al. 2020. Fungal Diversity 105:1-16 ((https://doi.org/10.1007/s13225-020-00466-2)) |
| Otu54248 | 1 | k__Fungi(100);p__Ascomycota(100);c__Saccharomycetes(100);o__Saccharomycetales(100);f__Metschnikowiaceae(100);g__Metschnikowia(100);s__Metschnikowia_pulcherrima(100); | Metschnikowia | 13 | Saprotroph | (Undefined Saprotroph) | Yeast | NULL | Probable | As nectar-tap_saprotroph (Põlme et al. 2020); Decay sugar-rich substrates (Põlme et al. 2020) | Põlme S, et al. 2020. Fungal Diversity 105:1-16 ((https://doi.org/10.1007/s13225-020-00466-2)) |
| Otu54249 | 1 | k__Fungi(100);p__Ascomycota(100);c__Saccharomycetes(100);o__Saccharomycetales(100);f__Metschnikowiaceae(100);g__Metschnikowia(100);s__Metschnikowia_pulcherrima(100); | Metschnikowia | 13 | Saprotroph | (Undefined Saprotroph) | Yeast | NULL | Probable | As nectar-tap_saprotroph (Põlme et al. 2020); Decay sugar-rich substrates (Põlme et al. 2020) | Põlme S, et al. 2020. Fungal Diversity 105:1-16 ((https://doi.org/10.1007/s13225-020-00466-2)) |
| Otu54250 | 1 | k__Fungi(100);p__Ascomycota(100);c__Saccharomycetes(100);o__Saccharomycetales(100);f__Metschnikowiaceae(100);g__Metschnikowia(100);s__Metschnikowia_pulcherrima(100); | Metschnikowia | 13 | Saprotroph | (Undefined Saprotroph) | Yeast | NULL | Probable | As nectar-tap_saprotroph (Põlme et al. 2020); Decay sugar-rich substrates (Põlme et al. 2020) | Põlme S, et al. 2020. Fungal Diversity 105:1-16 ((https://doi.org/10.1007/s13225-020-00466-2)) |
| Otu54251 | 1 | k__Fungi(100);p__Ascomycota(100);c__Saccharomycetes(100);o__Saccharomycetales(100);f__Metschnikowiaceae(100);g__Metschnikowia(100);g__Metschnikowia_unclassified(100); | Metschnikowia | 13 | Saprotroph | (Undefined Saprotroph) | Yeast | NULL | Probable | As nectar-tap_saprotroph (Põlme et al. 2020); Decay sugar-rich substrates (Põlme et al. 2020) | Põlme S, et al. 2020. Fungal Diversity 105:1-16 ((https://doi.org/10.1007/s13225-020-00466-2)) |
| Otu54252 | 1 | k__Fungi(100);p__Ascomycota(100);c__Saccharomycetes(100);o__Saccharomycetales(100);f__Metschnikowiaceae(100);g__Metschnikowia(100);g__Metschnikowia_unclassified(100); | Metschnikowia | 13 | Saprotroph | (Undefined Saprotroph) | Yeast | NULL | Probable | As nectar-tap_saprotroph (Põlme et al. 2020); Decay sugar-rich substrates (Põlme et al. 2020) | Põlme S, et al. 2020. Fungal Diversity 105:1-16 ((https://doi.org/10.1007/s13225-020-00466-2)) |
| Otu54253 | 1 | k__Fungi(100);p__Ascomycota(100);c__Saccharomycetes(100);o__Saccharomycetales(100);f__Metschnikowiaceae(100);g__Metschnikowia(100);g__Metschnikowia_unclassified(100); | Metschnikowia | 13 | Saprotroph | (Undefined Saprotroph) | Yeast | NULL | Probable | As nectar-tap_saprotroph (Põlme et al. 2020); Decay sugar-rich substrates (Põlme et al. 2020) | Põlme S, et al. 2020. Fungal Diversity 105:1-16 ((https://doi.org/10.1007/s13225-020-00466-2)) |
| Otu54254 | 1 | k__Fungi(100);p__Ascomycota(100);c__Saccharomycetes(100);o__Saccharomycetales(100);f__Metschnikowiaceae(100);g__Metschnikowia(100);s__Metschnikowia_pulcherrima(100); | Metschnikowia | 13 | Saprotroph | (Undefined Saprotroph) | Yeast | NULL | Probable | As nectar-tap_saprotroph (Põlme et al. 2020); Decay sugar-rich substrates (Põlme et al. 2020) | Põlme S, et al. 2020. Fungal Diversity 105:1-16 ((https://doi.org/10.1007/s13225-020-00466-2)) |
| Otu54255 | 1 | k__Fungi(100);p__Ascomycota(100);c__Saccharomycetes(100);o__Saccharomycetales(100);f__Metschnikowiaceae(100);g__Metschnikowia(100);g__Metschnikowia_unclassified(100); | Metschnikowia | 13 | Saprotroph | (Undefined Saprotroph) | Yeast | NULL | Probable | As nectar-tap_saprotroph (Põlme et al. 2020); Decay sugar-rich substrates (Põlme et al. 2020) | Põlme S, et al. 2020. Fungal Diversity 105:1-16 ((https://doi.org/10.1007/s13225-020-00466-2)) |
| Otu54256 | 1 | k__Fungi(100);p__Ascomycota(100);c__Saccharomycetes(100);o__Saccharomycetales(100);f__Metschnikowiaceae(100);g__Metschnikowia(100);s__Metschnikowia_pulcherrima(100); | Metschnikowia | 13 | Saprotroph | (Undefined Saprotroph) | Yeast | NULL | Probable | As nectar-tap_saprotroph (Põlme et al. 2020); Decay sugar-rich substrates (Põlme et al. 2020) | Põlme S, et al. 2020. Fungal Diversity 105:1-16 ((https://doi.org/10.1007/s13225-020-00466-2)) |
| Otu54257 | 1 | k__Fungi(100);p__Ascomycota(100);c__Saccharomycetes(100);o__Saccharomycetales(100);f__Metschnikowiaceae(100);g__Metschnikowia(100);s__Metschnikowia_pulcherrima(100); | Metschnikowia | 13 | Saprotroph | (Undefined Saprotroph) | Yeast | NULL | Probable | As nectar-tap_saprotroph (Põlme et al. 2020); Decay sugar-rich substrates (Põlme et al. 2020) | Põlme S, et al. 2020. Fungal Diversity 105:1-16 ((https://doi.org/10.1007/s13225-020-00466-2)) |
| Otu54258 | 1 | k__Fungi(100);p__Ascomycota(100);c__Saccharomycetes(100);o__Saccharomycetales(100);f__Metschnikowiaceae(100);g__Metschnikowia(100);g__Metschnikowia_unclassified(100); | Metschnikowia | 13 | Saprotroph | (Undefined Saprotroph) | Yeast | NULL | Probable | As nectar-tap_saprotroph (Põlme et al. 2020); Decay sugar-rich substrates (Põlme et al. 2020) | Põlme S, et al. 2020. Fungal Diversity 105:1-16 ((https://doi.org/10.1007/s13225-020-00466-2)) |
| Otu54259 | 1 | k__Fungi(100);p__Ascomycota(100);c__Saccharomycetes(100);o__Saccharomycetales(100);f__Metschnikowiaceae(100);g__Metschnikowia(100);g__Metschnikowia_unclassified(100); | Metschnikowia | 13 | Saprotroph | (Undefined Saprotroph) | Yeast | NULL | Probable | As nectar-tap_saprotroph (Põlme et al. 2020); Decay sugar-rich substrates (Põlme et al. 2020) | Põlme S, et al. 2020. Fungal Diversity 105:1-16 ((https://doi.org/10.1007/s13225-020-00466-2)) |
| Otu54260 | 1 | k__Fungi(100);p__Ascomycota(100);c__Saccharomycetes(100);o__Saccharomycetales(100);f__Metschnikowiaceae(100);g__Metschnikowia(100);s__Metschnikowia_pulcherrima(100); | Metschnikowia | 13 | Saprotroph | (Undefined Saprotroph) | Yeast | NULL | Probable | As nectar-tap_saprotroph (Põlme et al. 2020); Decay sugar-rich substrates (Põlme et al. 2020) | Põlme S, et al. 2020. Fungal Diversity 105:1-16 ((https://doi.org/10.1007/s13225-020-00466-2)) |
| Otu54261 | 1 | k__Fungi(100);p__Ascomycota(100);c__Saccharomycetes(100);o__Saccharomycetales(100);f__Metschnikowiaceae(100);g__Metschnikowia(100);s__Metschnikowia_pulcherrima(100); | Metschnikowia | 13 | Saprotroph | (Undefined Saprotroph) | Yeast | NULL | Probable | As nectar-tap_saprotroph (Põlme et al. 2020); Decay sugar-rich substrates (Põlme et al. 2020) | Põlme S, et al. 2020. Fungal Diversity 105:1-16 ((https://doi.org/10.1007/s13225-020-00466-2)) |
| Otu54262 | 1 | k__Fungi(100);p__Ascomycota(100);c__Saccharomycetes(100);o__Saccharomycetales(100);f__Metschnikowiaceae(100);g__Metschnikowia(100);s__Metschnikowia_pulcherrima(100); | Metschnikowia | 13 | Saprotroph | (Undefined Saprotroph) | Yeast | NULL | Probable | As nectar-tap_saprotroph (Põlme et al. 2020); Decay sugar-rich substrates (Põlme et al. 2020) | Põlme S, et al. 2020. Fungal Diversity 105:1-16 ((https://doi.org/10.1007/s13225-020-00466-2)) |
| Otu54263 | 1 | k__Fungi(100);p__Ascomycota(100);c__Saccharomycetes(100);o__Saccharomycetales(100);f__Metschnikowiaceae(100);g__Metschnikowia(100);g__Metschnikowia_unclassified(100); | Metschnikowia | 13 | Saprotroph | (Undefined Saprotroph) | Yeast | NULL | Probable | As nectar-tap_saprotroph (Põlme et al. 2020); Decay sugar-rich substrates (Põlme et al. 2020) | Põlme S, et al. 2020. Fungal Diversity 105:1-16 ((https://doi.org/10.1007/s13225-020-00466-2)) |
| Otu54264 | 1 | k__Fungi(100);p__Ascomycota(100);c__Saccharomycetes(100);o__Saccharomycetales(100);f__Metschnikowiaceae(100);g__Metschnikowia(100);s__Metschnikowia_pulcherrima(100); | Metschnikowia | 13 | Saprotroph | (Undefined Saprotroph) | Yeast | NULL | Probable | As nectar-tap_saprotroph (Põlme et al. 2020); Decay sugar-rich substrates (Põlme et al. 2020) | Põlme S, et al. 2020. Fungal Diversity 105:1-16 ((https://doi.org/10.1007/s13225-020-00466-2)) |
| Otu54265 | 1 | k__Fungi(100);p__Ascomycota(100);c__Saccharomycetes(100);o__Saccharomycetales(100);f__Metschnikowiaceae(100);g__Metschnikowia(100);g__Metschnikowia_unclassified(100); | Metschnikowia | 13 | Saprotroph | (Undefined Saprotroph) | Yeast | NULL | Probable | As nectar-tap_saprotroph (Põlme et al. 2020); Decay sugar-rich substrates (Põlme et al. 2020) | Põlme S, et al. 2020. Fungal Diversity 105:1-16 ((https://doi.org/10.1007/s13225-020-00466-2)) |
| Otu54266 | 1 | k__Fungi(100);p__Ascomycota(100);c__Saccharomycetes(100);o__Saccharomycetales(100);f__Metschnikowiaceae(100);g__Metschnikowia(100);s__Metschnikowia_pulcherrima(100); | Metschnikowia | 13 | Saprotroph | (Undefined Saprotroph) | Yeast | NULL | Probable | As nectar-tap_saprotroph (Põlme et al. 2020); Decay sugar-rich substrates (Põlme et al. 2020) | Põlme S, et al. 2020. Fungal Diversity 105:1-16 ((https://doi.org/10.1007/s13225-020-00466-2)) |
| Otu54267 | 1 | k__Fungi(100);p__Ascomycota(100);c__Saccharomycetes(100);o__Saccharomycetales(100);f__Metschnikowiaceae(100);g__Metschnikowia(100);s__Metschnikowia_pulcherrima(100); | Metschnikowia | 13 | Saprotroph | (Undefined Saprotroph) | Yeast | NULL | Probable | As nectar-tap_saprotroph (Põlme et al. 2020); Decay sugar-rich substrates (Põlme et al. 2020) | Põlme S, et al. 2020. Fungal Diversity 105:1-16 ((https://doi.org/10.1007/s13225-020-00466-2)) |
| Otu54268 | 1 | k__Fungi(100);p__Ascomycota(100);c__Saccharomycetes(100);o__Saccharomycetales(100);f__Metschnikowiaceae(100);g__Metschnikowia(100);s__Metschnikowia_pulcherrima(100); | Metschnikowia | 13 | Saprotroph | (Undefined Saprotroph) | Yeast | NULL | Probable | As nectar-tap_saprotroph (Põlme et al. 2020); Decay sugar-rich substrates (Põlme et al. 2020) | Põlme S, et al. 2020. Fungal Diversity 105:1-16 ((https://doi.org/10.1007/s13225-020-00466-2)) |
| Otu54269 | 1 | k__Fungi(100);p__Ascomycota(100);c__Saccharomycetes(100);o__Saccharomycetales(100);f__Metschnikowiaceae(100);g__Metschnikowia(100);g__Metschnikowia_unclassified(100); | Metschnikowia | 13 | Saprotroph | (Undefined Saprotroph) | Yeast | NULL | Probable | As nectar-tap_saprotroph (Põlme et al. 2020); Decay sugar-rich substrates (Põlme et al. 2020) | Põlme S, et al. 2020. Fungal Diversity 105:1-16 ((https://doi.org/10.1007/s13225-020-00466-2)) |
| Otu54270 | 1 | k__Fungi(100);p__Ascomycota(100);c__Saccharomycetes(100);o__Saccharomycetales(100);f__Metschnikowiaceae(100);g__Metschnikowia(100);g__Metschnikowia_unclassified(100); | Metschnikowia | 13 | Saprotroph | (Undefined Saprotroph) | Yeast | NULL | Probable | As nectar-tap_saprotroph (Põlme et al. 2020); Decay sugar-rich substrates (Põlme et al. 2020) | Põlme S, et al. 2020. Fungal Diversity 105:1-16 ((https://doi.org/10.1007/s13225-020-00466-2)) |
| Otu54271 | 1 | k__Fungi(100);p__Ascomycota(100);c__Saccharomycetes(100);o__Saccharomycetales(100);f__Metschnikowiaceae(100);g__Metschnikowia(100);s__Metschnikowia_pulcherrima(100); | Metschnikowia | 13 | Saprotroph | (Undefined Saprotroph) | Yeast | NULL | Probable | As nectar-tap_saprotroph (Põlme et al. 2020); Decay sugar-rich substrates (Põlme et al. 2020) | Põlme S, et al. 2020. Fungal Diversity 105:1-16 ((https://doi.org/10.1007/s13225-020-00466-2)) |
| Otu54272 | 1 | k__Fungi(100);p__Ascomycota(100);c__Saccharomycetes(100);o__Saccharomycetales(100);f__Metschnikowiaceae(100);g__Metschnikowia(100);s__Metschnikowia_pulcherrima(100); | Metschnikowia | 13 | Saprotroph | (Undefined Saprotroph) | Yeast | NULL | Probable | As nectar-tap_saprotroph (Põlme et al. 2020); Decay sugar-rich substrates (Põlme et al. 2020) | Põlme S, et al. 2020. Fungal Diversity 105:1-16 ((https://doi.org/10.1007/s13225-020-00466-2)) |
| Otu54273 | 1 | k__Fungi(100);p__Ascomycota(100);c__Saccharomycetes(100);o__Saccharomycetales(100);f__Metschnikowiaceae(100);g__Metschnikowia(100);s__Metschnikowia_pulcherrima(100); | Metschnikowia | 13 | Saprotroph | (Undefined Saprotroph) | Yeast | NULL | Probable | As nectar-tap_saprotroph (Põlme et al. 2020); Decay sugar-rich substrates (Põlme et al. 2020) | Põlme S, et al. 2020. Fungal Diversity 105:1-16 ((https://doi.org/10.1007/s13225-020-00466-2)) |
| Otu54274 | 1 | k__Fungi(100);p__Ascomycota(100);c__Saccharomycetes(100);o__Saccharomycetales(100);f__Metschnikowiaceae(100);g__Metschnikowia(100);s__Metschnikowia_pulcherrima(100); | Metschnikowia | 13 | Saprotroph | (Undefined Saprotroph) | Yeast | NULL | Probable | As nectar-tap_saprotroph (Põlme et al. 2020); Decay sugar-rich substrates (Põlme et al. 2020) | Põlme S, et al. 2020. Fungal Diversity 105:1-16 ((https://doi.org/10.1007/s13225-020-00466-2)) |
| Otu54275 | 1 | k__Fungi(100);p__Ascomycota(100);c__Saccharomycetes(100);o__Saccharomycetales(100);f__Metschnikowiaceae(100);g__Metschnikowia(100);g__Metschnikowia_unclassified(100); | Metschnikowia | 13 | Saprotroph | (Undefined Saprotroph) | Yeast | NULL | Probable | As nectar-tap_saprotroph (Põlme et al. 2020); Decay sugar-rich substrates (Põlme et al. 2020) | Põlme S, et al. 2020. Fungal Diversity 105:1-16 ((https://doi.org/10.1007/s13225-020-00466-2)) |
| Otu54276 | 1 | k__Fungi(100);p__Ascomycota(100);c__Saccharomycetes(100);o__Saccharomycetales(100);f__Metschnikowiaceae(100);g__Metschnikowia(100);s__Metschnikowia_pulcherrima(100); | Metschnikowia | 13 | Saprotroph | (Undefined Saprotroph) | Yeast | NULL | Probable | As nectar-tap_saprotroph (Põlme et al. 2020); Decay sugar-rich substrates (Põlme et al. 2020) | Põlme S, et al. 2020. Fungal Diversity 105:1-16 ((https://doi.org/10.1007/s13225-020-00466-2)) |
| Otu54278 | 1 | k__Fungi(100);p__Ascomycota(100);c__Saccharomycetes(100);o__Saccharomycetales(100);f__Metschnikowiaceae(100);g__Metschnikowia(100);s__Metschnikowia_pulcherrima(100); | Metschnikowia | 13 | Saprotroph | (Undefined Saprotroph) | Yeast | NULL | Probable | As nectar-tap_saprotroph (Põlme et al. 2020); Decay sugar-rich substrates (Põlme et al. 2020) | Põlme S, et al. 2020. Fungal Diversity 105:1-16 ((https://doi.org/10.1007/s13225-020-00466-2)) |
| Otu54279 | 1 | k__Fungi(100);p__Ascomycota(100);c__Saccharomycetes(100);o__Saccharomycetales(100);f__Metschnikowiaceae(100);g__Metschnikowia(100);s__Metschnikowia_pulcherrima(100); | Metschnikowia | 13 | Saprotroph | (Undefined Saprotroph) | Yeast | NULL | Probable | As nectar-tap_saprotroph (Põlme et al. 2020); Decay sugar-rich substrates (Põlme et al. 2020) | Põlme S, et al. 2020. Fungal Diversity 105:1-16 ((https://doi.org/10.1007/s13225-020-00466-2)) |
| Otu54280 | 1 | k__Fungi(100);p__Ascomycota(100);c__Saccharomycetes(100);o__Saccharomycetales(100);f__Metschnikowiaceae(100);g__Metschnikowia(100);s__Metschnikowia_pulcherrima(100); | Metschnikowia | 13 | Saprotroph | (Undefined Saprotroph) | Yeast | NULL | Probable | As nectar-tap_saprotroph (Põlme et al. 2020); Decay sugar-rich substrates (Põlme et al. 2020) | Põlme S, et al. 2020. Fungal Diversity 105:1-16 ((https://doi.org/10.1007/s13225-020-00466-2)) |
| Otu54281 | 1 | k__Fungi(100);p__Ascomycota(100);c__Saccharomycetes(100);o__Saccharomycetales(100);f__Metschnikowiaceae(100);g__Metschnikowia(100);s__Metschnikowia_pulcherrima(100); | Metschnikowia | 13 | Saprotroph | (Undefined Saprotroph) | Yeast | NULL | Probable | As nectar-tap_saprotroph (Põlme et al. 2020); Decay sugar-rich substrates (Põlme et al. 2020) | Põlme S, et al. 2020. Fungal Diversity 105:1-16 ((https://doi.org/10.1007/s13225-020-00466-2)) |
| Otu54282 | 1 | k__Fungi(100);p__Ascomycota(100);c__Saccharomycetes(100);o__Saccharomycetales(100);f__Metschnikowiaceae(100);g__Metschnikowia(100);s__Metschnikowia_pulcherrima(100); | Metschnikowia | 13 | Saprotroph | (Undefined Saprotroph) | Yeast | NULL | Probable | As nectar-tap_saprotroph (Põlme et al. 2020); Decay sugar-rich substrates (Põlme et al. 2020) | Põlme S, et al. 2020. Fungal Diversity 105:1-16 ((https://doi.org/10.1007/s13225-020-00466-2)) |
| Otu54283 | 1 | k__Fungi(100);p__Ascomycota(100);c__Saccharomycetes(100);o__Saccharomycetales(100);f__Metschnikowiaceae(100);g__Metschnikowia(100);g__Metschnikowia_unclassified(100); | Metschnikowia | 13 | Saprotroph | (Undefined Saprotroph) | Yeast | NULL | Probable | As nectar-tap_saprotroph (Põlme et al. 2020); Decay sugar-rich substrates (Põlme et al. 2020) | Põlme S, et al. 2020. Fungal Diversity 105:1-16 ((https://doi.org/10.1007/s13225-020-00466-2)) |
| Otu54284 | 1 | k__Fungi(100);p__Ascomycota(100);c__Saccharomycetes(100);o__Saccharomycetales(100);f__Metschnikowiaceae(100);g__Metschnikowia(100);s__Metschnikowia_pulcherrima(100); | Metschnikowia | 13 | Saprotroph | (Undefined Saprotroph) | Yeast | NULL | Probable | As nectar-tap_saprotroph (Põlme et al. 2020); Decay sugar-rich substrates (Põlme et al. 2020) | Põlme S, et al. 2020. Fungal Diversity 105:1-16 ((https://doi.org/10.1007/s13225-020-00466-2)) |
| Otu54286 | 1 | k__Fungi(100);p__Ascomycota(100);c__Saccharomycetes(100);o__Saccharomycetales(100);f__Metschnikowiaceae(100);g__Metschnikowia(100);s__Metschnikowia_pulcherrima(100); | Metschnikowia | 13 | Saprotroph | (Undefined Saprotroph) | Yeast | NULL | Probable | As nectar-tap_saprotroph (Põlme et al. 2020); Decay sugar-rich substrates (Põlme et al. 2020) | Põlme S, et al. 2020. Fungal Diversity 105:1-16 ((https://doi.org/10.1007/s13225-020-00466-2)) |
| Otu54287 | 1 | k__Fungi(100);p__Ascomycota(100);c__Saccharomycetes(100);o__Saccharomycetales(100);f__Metschnikowiaceae(100);g__Metschnikowia(100);s__Metschnikowia_pulcherrima(100); | Metschnikowia | 13 | Saprotroph | (Undefined Saprotroph) | Yeast | NULL | Probable | As nectar-tap_saprotroph (Põlme et al. 2020); Decay sugar-rich substrates (Põlme et al. 2020) | Põlme S, et al. 2020. Fungal Diversity 105:1-16 ((https://doi.org/10.1007/s13225-020-00466-2)) |
| Otu54288 | 1 | k__Fungi(100);p__Ascomycota(100);c__Saccharomycetes(100);o__Saccharomycetales(100);f__Metschnikowiaceae(100);g__Metschnikowia(100);s__Metschnikowia_pulcherrima(100); | Metschnikowia | 13 | Saprotroph | (Undefined Saprotroph) | Yeast | NULL | Probable | As nectar-tap_saprotroph (Põlme et al. 2020); Decay sugar-rich substrates (Põlme et al. 2020) | Põlme S, et al. 2020. Fungal Diversity 105:1-16 ((https://doi.org/10.1007/s13225-020-00466-2)) |
| Otu54289 | 1 | k__Fungi(100);p__Ascomycota(100);c__Saccharomycetes(100);o__Saccharomycetales(100);f__Metschnikowiaceae(100);g__Metschnikowia(100);s__Metschnikowia_pulcherrima(100); | Metschnikowia | 13 | Saprotroph | (Undefined Saprotroph) | Yeast | NULL | Probable | As nectar-tap_saprotroph (Põlme et al. 2020); Decay sugar-rich substrates (Põlme et al. 2020) | Põlme S, et al. 2020. Fungal Diversity 105:1-16 ((https://doi.org/10.1007/s13225-020-00466-2)) |
| Otu54290 | 1 | k__Fungi(100);p__Ascomycota(100);c__Saccharomycetes(100);o__Saccharomycetales(100);f__Metschnikowiaceae(100);g__Metschnikowia(100);g__Metschnikowia_unclassified(100); | Metschnikowia | 13 | Saprotroph | (Undefined Saprotroph) | Yeast | NULL | Probable | As nectar-tap_saprotroph (Põlme et al. 2020); Decay sugar-rich substrates (Põlme et al. 2020) | Põlme S, et al. 2020. Fungal Diversity 105:1-16 ((https://doi.org/10.1007/s13225-020-00466-2)) |
| Otu54291 | 1 | k__Fungi(100);p__Ascomycota(100);c__Saccharomycetes(100);o__Saccharomycetales(100);f__Metschnikowiaceae(100);g__Metschnikowia(100);s__Metschnikowia_pulcherrima(100); | Metschnikowia | 13 | Saprotroph | (Undefined Saprotroph) | Yeast | NULL | Probable | As nectar-tap_saprotroph (Põlme et al. 2020); Decay sugar-rich substrates (Põlme et al. 2020) | Põlme S, et al. 2020. Fungal Diversity 105:1-16 ((https://doi.org/10.1007/s13225-020-00466-2)) |
| Otu54292 | 1 | k__Fungi(100);p__Ascomycota(100);c__Saccharomycetes(100);o__Saccharomycetales(100);f__Metschnikowiaceae(100);g__Metschnikowia(100);s__Metschnikowia_pulcherrima(100); | Metschnikowia | 13 | Saprotroph | (Undefined Saprotroph) | Yeast | NULL | Probable | As nectar-tap_saprotroph (Põlme et al. 2020); Decay sugar-rich substrates (Põlme et al. 2020) | Põlme S, et al. 2020. Fungal Diversity 105:1-16 ((https://doi.org/10.1007/s13225-020-00466-2)) |
| Otu54293 | 1 | k__Fungi(100);p__Ascomycota(100);c__Saccharomycetes(100);o__Saccharomycetales(100);f__Metschnikowiaceae(100);g__Metschnikowia(100);s__Metschnikowia_pulcherrima(100); | Metschnikowia | 13 | Saprotroph | (Undefined Saprotroph) | Yeast | NULL | Probable | As nectar-tap_saprotroph (Põlme et al. 2020); Decay sugar-rich substrates (Põlme et al. 2020) | Põlme S, et al. 2020. Fungal Diversity 105:1-16 ((https://doi.org/10.1007/s13225-020-00466-2)) |
| Otu54294 | 1 | k__Fungi(100);p__Ascomycota(100);c__Saccharomycetes(100);o__Saccharomycetales(100);f__Metschnikowiaceae(100);g__Metschnikowia(100);g__Metschnikowia_unclassified(100); | Metschnikowia | 13 | Saprotroph | (Undefined Saprotroph) | Yeast | NULL | Probable | As nectar-tap_saprotroph (Põlme et al. 2020); Decay sugar-rich substrates (Põlme et al. 2020) | Põlme S, et al. 2020. Fungal Diversity 105:1-16 ((https://doi.org/10.1007/s13225-020-00466-2)) |
| Otu54295 | 1 | k__Fungi(100);p__Ascomycota(100);c__Saccharomycetes(100);o__Saccharomycetales(100);f__Metschnikowiaceae(100);g__Metschnikowia(100);g__Metschnikowia_unclassified(100); | Metschnikowia | 13 | Saprotroph | (Undefined Saprotroph) | Yeast | NULL | Probable | As nectar-tap_saprotroph (Põlme et al. 2020); Decay sugar-rich substrates (Põlme et al. 2020) | Põlme S, et al. 2020. Fungal Diversity 105:1-16 ((https://doi.org/10.1007/s13225-020-00466-2)) |
| Otu54296 | 1 | k__Fungi(100);p__Ascomycota(100);c__Saccharomycetes(100);o__Saccharomycetales(100);f__Metschnikowiaceae(100);g__Metschnikowia(100);g__Metschnikowia_unclassified(100); | Metschnikowia | 13 | Saprotroph | (Undefined Saprotroph) | Yeast | NULL | Probable | As nectar-tap_saprotroph (Põlme et al. 2020); Decay sugar-rich substrates (Põlme et al. 2020) | Põlme S, et al. 2020. Fungal Diversity 105:1-16 ((https://doi.org/10.1007/s13225-020-00466-2)) |
| Otu54297 | 1 | k__Fungi(100);p__Ascomycota(100);c__Saccharomycetes(100);o__Saccharomycetales(100);f__Metschnikowiaceae(100);g__Metschnikowia(100);s__Metschnikowia_pulcherrima(100); | Metschnikowia | 13 | Saprotroph | (Undefined Saprotroph) | Yeast | NULL | Probable | As nectar-tap_saprotroph (Põlme et al. 2020); Decay sugar-rich substrates (Põlme et al. 2020) | Põlme S, et al. 2020. Fungal Diversity 105:1-16 ((https://doi.org/10.1007/s13225-020-00466-2)) |
| Otu54298 | 1 | k__Fungi(100);p__Ascomycota(100);c__Saccharomycetes(100);o__Saccharomycetales(100);f__Metschnikowiaceae(100);g__Metschnikowia(100);s__Metschnikowia_pulcherrima(100); | Metschnikowia | 13 | Saprotroph | (Undefined Saprotroph) | Yeast | NULL | Probable | As nectar-tap_saprotroph (Põlme et al. 2020); Decay sugar-rich substrates (Põlme et al. 2020) | Põlme S, et al. 2020. Fungal Diversity 105:1-16 ((https://doi.org/10.1007/s13225-020-00466-2)) |
| Otu54299 | 1 | k__Fungi(100);p__Ascomycota(100);c__Saccharomycetes(100);o__Saccharomycetales(100);f__Metschnikowiaceae(100);g__Metschnikowia(100);s__Metschnikowia_pulcherrima(100); | Metschnikowia | 13 | Saprotroph | (Undefined Saprotroph) | Yeast | NULL | Probable | As nectar-tap_saprotroph (Põlme et al. 2020); Decay sugar-rich substrates (Põlme et al. 2020) | Põlme S, et al. 2020. Fungal Diversity 105:1-16 ((https://doi.org/10.1007/s13225-020-00466-2)) |
| Otu54300 | 1 | k__Fungi(100);p__Ascomycota(100);c__Saccharomycetes(100);o__Saccharomycetales(100);f__Metschnikowiaceae(100);g__Metschnikowia(100);s__Metschnikowia_pulcherrima(100); | Metschnikowia | 13 | Saprotroph | (Undefined Saprotroph) | Yeast | NULL | Probable | As nectar-tap_saprotroph (Põlme et al. 2020); Decay sugar-rich substrates (Põlme et al. 2020) | Põlme S, et al. 2020. Fungal Diversity 105:1-16 ((https://doi.org/10.1007/s13225-020-00466-2)) |
| Otu54301 | 1 | k__Fungi(100);p__Ascomycota(100);c__Saccharomycetes(100);o__Saccharomycetales(100);f__Metschnikowiaceae(100);g__Metschnikowia(100);s__Metschnikowia_pulcherrima(100); | Metschnikowia | 13 | Saprotroph | (Undefined Saprotroph) | Yeast | NULL | Probable | As nectar-tap_saprotroph (Põlme et al. 2020); Decay sugar-rich substrates (Põlme et al. 2020) | Põlme S, et al. 2020. Fungal Diversity 105:1-16 ((https://doi.org/10.1007/s13225-020-00466-2)) |
| Otu54302 | 1 | k__Fungi(100);p__Ascomycota(100);c__Saccharomycetes(100);o__Saccharomycetales(100);f__Metschnikowiaceae(100);g__Metschnikowia(100);s__Metschnikowia_pulcherrima(100); | Metschnikowia | 13 | Saprotroph | (Undefined Saprotroph) | Yeast | NULL | Probable | As nectar-tap_saprotroph (Põlme et al. 2020); Decay sugar-rich substrates (Põlme et al. 2020) | Põlme S, et al. 2020. Fungal Diversity 105:1-16 ((https://doi.org/10.1007/s13225-020-00466-2)) |
| Otu54303 | 1 | k__Fungi(100);p__Ascomycota(100);c__Saccharomycetes(100);o__Saccharomycetales(100);f__Metschnikowiaceae(100);g__Metschnikowia(100);s__Metschnikowia_pulcherrima(100); | Metschnikowia | 13 | Saprotroph | (Undefined Saprotroph) | Yeast | NULL | Probable | As nectar-tap_saprotroph (Põlme et al. 2020); Decay sugar-rich substrates (Põlme et al. 2020) | Põlme S, et al. 2020. Fungal Diversity 105:1-16 ((https://doi.org/10.1007/s13225-020-00466-2)) |
| Otu54304 | 1 | k__Fungi(100);p__Ascomycota(100);c__Saccharomycetes(100);o__Saccharomycetales(100);f__Metschnikowiaceae(100);g__Metschnikowia(100);s__Metschnikowia_pulcherrima(100); | Metschnikowia | 13 | Saprotroph | (Undefined Saprotroph) | Yeast | NULL | Probable | As nectar-tap_saprotroph (Põlme et al. 2020); Decay sugar-rich substrates (Põlme et al. 2020) | Põlme S, et al. 2020. Fungal Diversity 105:1-16 ((https://doi.org/10.1007/s13225-020-00466-2)) |
| Otu54305 | 1 | k__Fungi(100);p__Ascomycota(100);c__Saccharomycetes(100);o__Saccharomycetales(100);f__Metschnikowiaceae(100);g__Metschnikowia(100);s__Metschnikowia_pulcherrima(100); | Metschnikowia | 13 | Saprotroph | (Undefined Saprotroph) | Yeast | NULL | Probable | As nectar-tap_saprotroph (Põlme et al. 2020); Decay sugar-rich substrates (Põlme et al. 2020) | Põlme S, et al. 2020. Fungal Diversity 105:1-16 ((https://doi.org/10.1007/s13225-020-00466-2)) |
| Otu54306 | 1 | k__Fungi(100);p__Ascomycota(100);c__Saccharomycetes(100);o__Saccharomycetales(100);f__Metschnikowiaceae(100);g__Metschnikowia(100);s__Metschnikowia_pulcherrima(100); | Metschnikowia | 13 | Saprotroph | (Undefined Saprotroph) | Yeast | NULL | Probable | As nectar-tap_saprotroph (Põlme et al. 2020); Decay sugar-rich substrates (Põlme et al. 2020) | Põlme S, et al. 2020. Fungal Diversity 105:1-16 ((https://doi.org/10.1007/s13225-020-00466-2)) |
| Otu54307 | 1 | k__Fungi(100);p__Ascomycota(100);c__Saccharomycetes(100);o__Saccharomycetales(100);f__Metschnikowiaceae(100);g__Metschnikowia(100);s__Metschnikowia_pulcherrima(100); | Metschnikowia | 13 | Saprotroph | (Undefined Saprotroph) | Yeast | NULL | Probable | As nectar-tap_saprotroph (Põlme et al. 2020); Decay sugar-rich substrates (Põlme et al. 2020) | Põlme S, et al. 2020. Fungal Diversity 105:1-16 ((https://doi.org/10.1007/s13225-020-00466-2)) |
| Otu54308 | 1 | k__Fungi(100);p__Ascomycota(100);c__Saccharomycetes(100);o__Saccharomycetales(100);f__Metschnikowiaceae(100);g__Metschnikowia(100);g__Metschnikowia_unclassified(100); | Metschnikowia | 13 | Saprotroph | (Undefined Saprotroph) | Yeast | NULL | Probable | As nectar-tap_saprotroph (Põlme et al. 2020); Decay sugar-rich substrates (Põlme et al. 2020) | Põlme S, et al. 2020. Fungal Diversity 105:1-16 ((https://doi.org/10.1007/s13225-020-00466-2)) |
| Otu54309 | 1 | k__Fungi(100);p__Ascomycota(100);c__Saccharomycetes(100);o__Saccharomycetales(100);f__Metschnikowiaceae(100);g__Metschnikowia(100);s__Metschnikowia_pulcherrima(100); | Metschnikowia | 13 | Saprotroph | (Undefined Saprotroph) | Yeast | NULL | Probable | As nectar-tap_saprotroph (Põlme et al. 2020); Decay sugar-rich substrates (Põlme et al. 2020) | Põlme S, et al. 2020. Fungal Diversity 105:1-16 ((https://doi.org/10.1007/s13225-020-00466-2)) |
| Otu54310 | 1 | k__Fungi(100);p__Ascomycota(100);c__Saccharomycetes(100);o__Saccharomycetales(100);f__Metschnikowiaceae(100);g__Metschnikowia(100);s__Metschnikowia_pulcherrima(100); | Metschnikowia | 13 | Saprotroph | (Undefined Saprotroph) | Yeast | NULL | Probable | As nectar-tap_saprotroph (Põlme et al. 2020); Decay sugar-rich substrates (Põlme et al. 2020) | Põlme S, et al. 2020. Fungal Diversity 105:1-16 ((https://doi.org/10.1007/s13225-020-00466-2)) |
| Otu54311 | 1 | k__Fungi(100);p__Ascomycota(100);c__Saccharomycetes(100);o__Saccharomycetales(100);f__Metschnikowiaceae(100);g__Metschnikowia(100);g__Metschnikowia_unclassified(100); | Metschnikowia | 13 | Saprotroph | (Undefined Saprotroph) | Yeast | NULL | Probable | As nectar-tap_saprotroph (Põlme et al. 2020); Decay sugar-rich substrates (Põlme et al. 2020) | Põlme S, et al. 2020. Fungal Diversity 105:1-16 ((https://doi.org/10.1007/s13225-020-00466-2)) |
| Otu54312 | 1 | k__Fungi(100);p__Ascomycota(100);c__Saccharomycetes(100);o__Saccharomycetales(100);f__Metschnikowiaceae(100);g__Metschnikowia(100);s__Metschnikowia_pulcherrima(100); | Metschnikowia | 13 | Saprotroph | (Undefined Saprotroph) | Yeast | NULL | Probable | As nectar-tap_saprotroph (Põlme et al. 2020); Decay sugar-rich substrates (Põlme et al. 2020) | Põlme S, et al. 2020. Fungal Diversity 105:1-16 ((https://doi.org/10.1007/s13225-020-00466-2)) |
| Otu54313 | 1 | k__Fungi(100);p__Ascomycota(100);c__Saccharomycetes(100);o__Saccharomycetales(100);f__Metschnikowiaceae(100);g__Metschnikowia(100);s__Metschnikowia_pulcherrima(100); | Metschnikowia | 13 | Saprotroph | (Undefined Saprotroph) | Yeast | NULL | Probable | As nectar-tap_saprotroph (Põlme et al. 2020); Decay sugar-rich substrates (Põlme et al. 2020) | Põlme S, et al. 2020. Fungal Diversity 105:1-16 ((https://doi.org/10.1007/s13225-020-00466-2)) |
| Otu54314 | 1 | k__Fungi(100);p__Ascomycota(100);c__Saccharomycetes(100);o__Saccharomycetales(100);f__Metschnikowiaceae(100);g__Metschnikowia(100);s__Metschnikowia_pulcherrima(100); | Metschnikowia | 13 | Saprotroph | (Undefined Saprotroph) | Yeast | NULL | Probable | As nectar-tap_saprotroph (Põlme et al. 2020); Decay sugar-rich substrates (Põlme et al. 2020) | Põlme S, et al. 2020. Fungal Diversity 105:1-16 ((https://doi.org/10.1007/s13225-020-00466-2)) |
| Otu54315 | 1 | k__Fungi(100);p__Ascomycota(100);c__Saccharomycetes(100);o__Saccharomycetales(100);f__Metschnikowiaceae(100);g__Metschnikowia(100);s__Metschnikowia_pulcherrima(100); | Metschnikowia | 13 | Saprotroph | (Undefined Saprotroph) | Yeast | NULL | Probable | As nectar-tap_saprotroph (Põlme et al. 2020); Decay sugar-rich substrates (Põlme et al. 2020) | Põlme S, et al. 2020. Fungal Diversity 105:1-16 ((https://doi.org/10.1007/s13225-020-00466-2)) |
| Otu54316 | 1 | k__Fungi(100);p__Ascomycota(100);c__Saccharomycetes(100);o__Saccharomycetales(100);f__Metschnikowiaceae(100);g__Metschnikowia(100);g__Metschnikowia_unclassified(100); | Metschnikowia | 13 | Saprotroph | (Undefined Saprotroph) | Yeast | NULL | Probable | As nectar-tap_saprotroph (Põlme et al. 2020); Decay sugar-rich substrates (Põlme et al. 2020) | Põlme S, et al. 2020. Fungal Diversity 105:1-16 ((https://doi.org/10.1007/s13225-020-00466-2)) |
| Otu54317 | 1 | k__Fungi(100);p__Ascomycota(100);c__Saccharomycetes(100);o__Saccharomycetales(100);f__Metschnikowiaceae(100);g__Metschnikowia(100);g__Metschnikowia_unclassified(100); | Metschnikowia | 13 | Saprotroph | (Undefined Saprotroph) | Yeast | NULL | Probable | As nectar-tap_saprotroph (Põlme et al. 2020); Decay sugar-rich substrates (Põlme et al. 2020) | Põlme S, et al. 2020. Fungal Diversity 105:1-16 ((https://doi.org/10.1007/s13225-020-00466-2)) |
| Otu54318 | 1 | k__Fungi(100);p__Ascomycota(100);c__Saccharomycetes(100);o__Saccharomycetales(100);f__Metschnikowiaceae(100);g__Metschnikowia(100);s__Metschnikowia_pulcherrima(100); | Metschnikowia | 13 | Saprotroph | (Undefined Saprotroph) | Yeast | NULL | Probable | As nectar-tap_saprotroph (Põlme et al. 2020); Decay sugar-rich substrates (Põlme et al. 2020) | Põlme S, et al. 2020. Fungal Diversity 105:1-16 ((https://doi.org/10.1007/s13225-020-00466-2)) |
| Otu54319 | 1 | k__Fungi(100);p__Ascomycota(100);c__Saccharomycetes(100);o__Saccharomycetales(100);f__Metschnikowiaceae(100);g__Metschnikowia(100);g__Metschnikowia_unclassified(100); | Metschnikowia | 13 | Saprotroph | (Undefined Saprotroph) | Yeast | NULL | Probable | As nectar-tap_saprotroph (Põlme et al. 2020); Decay sugar-rich substrates (Põlme et al. 2020) | Põlme S, et al. 2020. Fungal Diversity 105:1-16 ((https://doi.org/10.1007/s13225-020-00466-2)) |
| Otu54320 | 1 | k__Fungi(100);p__Ascomycota(100);c__Saccharomycetes(100);o__Saccharomycetales(100);f__Metschnikowiaceae(100);g__Metschnikowia(100);g__Metschnikowia_unclassified(100); | Metschnikowia | 13 | Saprotroph | (Undefined Saprotroph) | Yeast | NULL | Probable | As nectar-tap_saprotroph (Põlme et al. 2020); Decay sugar-rich substrates (Põlme et al. 2020) | Põlme S, et al. 2020. Fungal Diversity 105:1-16 ((https://doi.org/10.1007/s13225-020-00466-2)) |
| Otu54321 | 1 | k__Fungi(100);p__Ascomycota(100);c__Saccharomycetes(100);o__Saccharomycetales(100);f__Metschnikowiaceae(100);g__Metschnikowia(100);g__Metschnikowia_unclassified(100); | Metschnikowia | 13 | Saprotroph | (Undefined Saprotroph) | Yeast | NULL | Probable | As nectar-tap_saprotroph (Põlme et al. 2020); Decay sugar-rich substrates (Põlme et al. 2020) | Põlme S, et al. 2020. Fungal Diversity 105:1-16 ((https://doi.org/10.1007/s13225-020-00466-2)) |
| Otu54322 | 1 | k__Fungi(100);p__Ascomycota(100);c__Saccharomycetes(100);o__Saccharomycetales(100);f__Metschnikowiaceae(100);g__Metschnikowia(100);s__Metschnikowia_pulcherrima(100); | Metschnikowia | 13 | Saprotroph | (Undefined Saprotroph) | Yeast | NULL | Probable | As nectar-tap_saprotroph (Põlme et al. 2020); Decay sugar-rich substrates (Põlme et al. 2020) | Põlme S, et al. 2020. Fungal Diversity 105:1-16 ((https://doi.org/10.1007/s13225-020-00466-2)) |
| Otu54323 | 1 | k__Fungi(100);p__Ascomycota(100);c__Saccharomycetes(100);o__Saccharomycetales(100);f__Metschnikowiaceae(100);g__Metschnikowia(100);g__Metschnikowia_unclassified(100); | Metschnikowia | 13 | Saprotroph | (Undefined Saprotroph) | Yeast | NULL | Probable | As nectar-tap_saprotroph (Põlme et al. 2020); Decay sugar-rich substrates (Põlme et al. 2020) | Põlme S, et al. 2020. Fungal Diversity 105:1-16 ((https://doi.org/10.1007/s13225-020-00466-2)) |
| Otu54324 | 1 | k__Fungi(100);p__Ascomycota(100);c__Saccharomycetes(100);o__Saccharomycetales(100);f__Metschnikowiaceae(100);g__Metschnikowia(100);g__Metschnikowia_unclassified(100); | Metschnikowia | 13 | Saprotroph | (Undefined Saprotroph) | Yeast | NULL | Probable | As nectar-tap_saprotroph (Põlme et al. 2020); Decay sugar-rich substrates (Põlme et al. 2020) | Põlme S, et al. 2020. Fungal Diversity 105:1-16 ((https://doi.org/10.1007/s13225-020-00466-2)) |
| Otu54325 | 1 | k__Fungi(100);p__Ascomycota(100);c__Saccharomycetes(100);o__Saccharomycetales(100);f__Metschnikowiaceae(100);g__Metschnikowia(100);s__Metschnikowia_pulcherrima(100); | Metschnikowia | 13 | Saprotroph | (Undefined Saprotroph) | Yeast | NULL | Probable | As nectar-tap_saprotroph (Põlme et al. 2020); Decay sugar-rich substrates (Põlme et al. 2020) | Põlme S, et al. 2020. Fungal Diversity 105:1-16 ((https://doi.org/10.1007/s13225-020-00466-2)) |
| Otu54326 | 1 | k__Fungi(100);p__Ascomycota(100);c__Saccharomycetes(100);o__Saccharomycetales(100);f__Metschnikowiaceae(100);g__Metschnikowia(100);s__Metschnikowia_pulcherrima(100); | Metschnikowia | 13 | Saprotroph | (Undefined Saprotroph) | Yeast | NULL | Probable | As nectar-tap_saprotroph (Põlme et al. 2020); Decay sugar-rich substrates (Põlme et al. 2020) | Põlme S, et al. 2020. Fungal Diversity 105:1-16 ((https://doi.org/10.1007/s13225-020-00466-2)) |
| Otu54327 | 1 | k__Fungi(100);p__Ascomycota(100);c__Saccharomycetes(100);o__Saccharomycetales(100);f__Metschnikowiaceae(100);g__Metschnikowia(100);g__Metschnikowia_unclassified(100); | Metschnikowia | 13 | Saprotroph | (Undefined Saprotroph) | Yeast | NULL | Probable | As nectar-tap_saprotroph (Põlme et al. 2020); Decay sugar-rich substrates (Põlme et al. 2020) | Põlme S, et al. 2020. Fungal Diversity 105:1-16 ((https://doi.org/10.1007/s13225-020-00466-2)) |
| Otu54328 | 1 | k__Fungi(100);p__Ascomycota(100);c__Saccharomycetes(100);o__Saccharomycetales(100);f__Metschnikowiaceae(100);g__Metschnikowia(100);s__Metschnikowia_pulcherrima(100); | Metschnikowia | 13 | Saprotroph | (Undefined Saprotroph) | Yeast | NULL | Probable | As nectar-tap_saprotroph (Põlme et al. 2020); Decay sugar-rich substrates (Põlme et al. 2020) | Põlme S, et al. 2020. Fungal Diversity 105:1-16 ((https://doi.org/10.1007/s13225-020-00466-2)) |
| Otu54329 | 1 | k__Fungi(100);p__Ascomycota(100);c__Saccharomycetes(100);o__Saccharomycetales(100);f__Metschnikowiaceae(100);g__Metschnikowia(100);s__Metschnikowia_pulcherrima(100); | Metschnikowia | 13 | Saprotroph | (Undefined Saprotroph) | Yeast | NULL | Probable | As nectar-tap_saprotroph (Põlme et al. 2020); Decay sugar-rich substrates (Põlme et al. 2020) | Põlme S, et al. 2020. Fungal Diversity 105:1-16 ((https://doi.org/10.1007/s13225-020-00466-2)) |
| Otu54330 | 1 | k__Fungi(100);p__Ascomycota(100);c__Saccharomycetes(100);o__Saccharomycetales(100);f__Metschnikowiaceae(100);g__Metschnikowia(100);s__Metschnikowia_pulcherrima(100); | Metschnikowia | 13 | Saprotroph | (Undefined Saprotroph) | Yeast | NULL | Probable | As nectar-tap_saprotroph (Põlme et al. 2020); Decay sugar-rich substrates (Põlme et al. 2020) | Põlme S, et al. 2020. Fungal Diversity 105:1-16 ((https://doi.org/10.1007/s13225-020-00466-2)) |
| Otu54331 | 1 | k__Fungi(100);p__Ascomycota(100);c__Saccharomycetes(100);o__Saccharomycetales(100);f__Metschnikowiaceae(100);g__Metschnikowia(100);s__Metschnikowia_pulcherrima(100); | Metschnikowia | 13 | Saprotroph | (Undefined Saprotroph) | Yeast | NULL | Probable | As nectar-tap_saprotroph (Põlme et al. 2020); Decay sugar-rich substrates (Põlme et al. 2020) | Põlme S, et al. 2020. Fungal Diversity 105:1-16 ((https://doi.org/10.1007/s13225-020-00466-2)) |
| Otu54332 | 1 | k__Fungi(100);p__Ascomycota(100);c__Saccharomycetes(100);o__Saccharomycetales(100);f__Metschnikowiaceae(100);g__Metschnikowia(100);s__Metschnikowia_pulcherrima(100); | Metschnikowia | 13 | Saprotroph | (Undefined Saprotroph) | Yeast | NULL | Probable | As nectar-tap_saprotroph (Põlme et al. 2020); Decay sugar-rich substrates (Põlme et al. 2020) | Põlme S, et al. 2020. Fungal Diversity 105:1-16 ((https://doi.org/10.1007/s13225-020-00466-2)) |
| Otu54333 | 1 | k__Fungi(100);p__Ascomycota(100);c__Saccharomycetes(100);o__Saccharomycetales(100);f__Metschnikowiaceae(100);g__Metschnikowia(100);s__Metschnikowia_pulcherrima(100); | Metschnikowia | 13 | Saprotroph | (Undefined Saprotroph) | Yeast | NULL | Probable | As nectar-tap_saprotroph (Põlme et al. 2020); Decay sugar-rich substrates (Põlme et al. 2020) | Põlme S, et al. 2020. Fungal Diversity 105:1-16 ((https://doi.org/10.1007/s13225-020-00466-2)) |
| Otu54334 | 1 | k__Fungi(100);p__Ascomycota(100);c__Saccharomycetes(100);o__Saccharomycetales(100);f__Metschnikowiaceae(100);g__Metschnikowia(100);s__Metschnikowia_pulcherrima(100); | Metschnikowia | 13 | Saprotroph | (Undefined Saprotroph) | Yeast | NULL | Probable | As nectar-tap_saprotroph (Põlme et al. 2020); Decay sugar-rich substrates (Põlme et al. 2020) | Põlme S, et al. 2020. Fungal Diversity 105:1-16 ((https://doi.org/10.1007/s13225-020-00466-2)) |
| Otu54335 | 1 | k__Fungi(100);p__Ascomycota(100);c__Saccharomycetes(100);o__Saccharomycetales(100);f__Metschnikowiaceae(100);g__Metschnikowia(100);s__Metschnikowia_pulcherrima(100); | Metschnikowia | 13 | Saprotroph | (Undefined Saprotroph) | Yeast | NULL | Probable | As nectar-tap_saprotroph (Põlme et al. 2020); Decay sugar-rich substrates (Põlme et al. 2020) | Põlme S, et al. 2020. Fungal Diversity 105:1-16 ((https://doi.org/10.1007/s13225-020-00466-2)) |
| Otu54336 | 1 | k__Fungi(100);p__Ascomycota(100);c__Saccharomycetes(100);o__Saccharomycetales(100);f__Metschnikowiaceae(100);g__Metschnikowia(100);s__Metschnikowia_pulcherrima(100); | Metschnikowia | 13 | Saprotroph | (Undefined Saprotroph) | Yeast | NULL | Probable | As nectar-tap_saprotroph (Põlme et al. 2020); Decay sugar-rich substrates (Põlme et al. 2020) | Põlme S, et al. 2020. Fungal Diversity 105:1-16 ((https://doi.org/10.1007/s13225-020-00466-2)) |
| Otu54337 | 1 | k__Fungi(100);p__Ascomycota(100);c__Saccharomycetes(100);o__Saccharomycetales(100);f__Metschnikowiaceae(100);g__Metschnikowia(100);g__Metschnikowia_unclassified(100); | Metschnikowia | 13 | Saprotroph | (Undefined Saprotroph) | Yeast | NULL | Probable | As nectar-tap_saprotroph (Põlme et al. 2020); Decay sugar-rich substrates (Põlme et al. 2020) | Põlme S, et al. 2020. Fungal Diversity 105:1-16 ((https://doi.org/10.1007/s13225-020-00466-2)) |
| Otu54338 | 1 | k__Fungi(100);p__Ascomycota(100);c__Saccharomycetes(100);o__Saccharomycetales(100);f__Metschnikowiaceae(100);g__Metschnikowia(100);g__Metschnikowia_unclassified(100); | Metschnikowia | 13 | Saprotroph | (Undefined Saprotroph) | Yeast | NULL | Probable | As nectar-tap_saprotroph (Põlme et al. 2020); Decay sugar-rich substrates (Põlme et al. 2020) | Põlme S, et al. 2020. Fungal Diversity 105:1-16 ((https://doi.org/10.1007/s13225-020-00466-2)) |
| Otu54339 | 1 | k__Fungi(100);p__Ascomycota(100);c__Saccharomycetes(100);o__Saccharomycetales(100);f__Metschnikowiaceae(100);g__Metschnikowia(100);s__Metschnikowia_pulcherrima(100); | Metschnikowia | 13 | Saprotroph | (Undefined Saprotroph) | Yeast | NULL | Probable | As nectar-tap_saprotroph (Põlme et al. 2020); Decay sugar-rich substrates (Põlme et al. 2020) | Põlme S, et al. 2020. Fungal Diversity 105:1-16 ((https://doi.org/10.1007/s13225-020-00466-2)) |
| Otu54340 | 1 | k__Fungi(100);p__Ascomycota(100);c__Saccharomycetes(100);o__Saccharomycetales(100);f__Metschnikowiaceae(100);g__Metschnikowia(100);s__Metschnikowia_pulcherrima(100); | Metschnikowia | 13 | Saprotroph | (Undefined Saprotroph) | Yeast | NULL | Probable | As nectar-tap_saprotroph (Põlme et al. 2020); Decay sugar-rich substrates (Põlme et al. 2020) | Põlme S, et al. 2020. Fungal Diversity 105:1-16 ((https://doi.org/10.1007/s13225-020-00466-2)) |
| Otu54341 | 1 | k__Fungi(100);p__Ascomycota(100);c__Saccharomycetes(100);o__Saccharomycetales(100);f__Metschnikowiaceae(100);g__Metschnikowia(100);s__Metschnikowia_pulcherrima(100); | Metschnikowia | 13 | Saprotroph | (Undefined Saprotroph) | Yeast | NULL | Probable | As nectar-tap_saprotroph (Põlme et al. 2020); Decay sugar-rich substrates (Põlme et al. 2020) | Põlme S, et al. 2020. Fungal Diversity 105:1-16 ((https://doi.org/10.1007/s13225-020-00466-2)) |
| Otu54342 | 1 | k__Fungi(100);p__Ascomycota(100);c__Saccharomycetes(100);o__Saccharomycetales(100);f__Metschnikowiaceae(100);g__Metschnikowia(100);s__Metschnikowia_pulcherrima(100); | Metschnikowia | 13 | Saprotroph | (Undefined Saprotroph) | Yeast | NULL | Probable | As nectar-tap_saprotroph (Põlme et al. 2020); Decay sugar-rich substrates (Põlme et al. 2020) | Põlme S, et al. 2020. Fungal Diversity 105:1-16 ((https://doi.org/10.1007/s13225-020-00466-2)) |
| Otu54343 | 1 | k__Fungi(100);p__Ascomycota(100);c__Saccharomycetes(100);o__Saccharomycetales(100);f__Metschnikowiaceae(100);g__Metschnikowia(100);s__Metschnikowia_pulcherrima(100); | Metschnikowia | 13 | Saprotroph | (Undefined Saprotroph) | Yeast | NULL | Probable | As nectar-tap_saprotroph (Põlme et al. 2020); Decay sugar-rich substrates (Põlme et al. 2020) | Põlme S, et al. 2020. Fungal Diversity 105:1-16 ((https://doi.org/10.1007/s13225-020-00466-2)) |
| Otu54344 | 1 | k__Fungi(100);p__Ascomycota(100);c__Saccharomycetes(100);o__Saccharomycetales(100);f__Metschnikowiaceae(100);g__Metschnikowia(100);s__Metschnikowia_pulcherrima(100); | Metschnikowia | 13 | Saprotroph | (Undefined Saprotroph) | Yeast | NULL | Probable | As nectar-tap_saprotroph (Põlme et al. 2020); Decay sugar-rich substrates (Põlme et al. 2020) | Põlme S, et al. 2020. Fungal Diversity 105:1-16 ((https://doi.org/10.1007/s13225-020-00466-2)) |
| Otu54345 | 1 | k__Fungi(100);p__Ascomycota(100);c__Saccharomycetes(100);o__Saccharomycetales(100);f__Metschnikowiaceae(100);g__Metschnikowia(100);g__Metschnikowia_unclassified(100); | Metschnikowia | 13 | Saprotroph | (Undefined Saprotroph) | Yeast | NULL | Probable | As nectar-tap_saprotroph (Põlme et al. 2020); Decay sugar-rich substrates (Põlme et al. 2020) | Põlme S, et al. 2020. Fungal Diversity 105:1-16 ((https://doi.org/10.1007/s13225-020-00466-2)) |
| Otu54346 | 1 | k__Fungi(100);p__Ascomycota(100);c__Saccharomycetes(100);o__Saccharomycetales(100);f__Metschnikowiaceae(100);g__Metschnikowia(100);g__Metschnikowia_unclassified(100); | Metschnikowia | 13 | Saprotroph | (Undefined Saprotroph) | Yeast | NULL | Probable | As nectar-tap_saprotroph (Põlme et al. 2020); Decay sugar-rich substrates (Põlme et al. 2020) | Põlme S, et al. 2020. Fungal Diversity 105:1-16 ((https://doi.org/10.1007/s13225-020-00466-2)) |
| Otu54347 | 1 | k__Fungi(100);p__Ascomycota(100);c__Saccharomycetes(100);o__Saccharomycetales(100);f__Metschnikowiaceae(100);g__Metschnikowia(100);s__Metschnikowia_pulcherrima(100); | Metschnikowia | 13 | Saprotroph | (Undefined Saprotroph) | Yeast | NULL | Probable | As nectar-tap_saprotroph (Põlme et al. 2020); Decay sugar-rich substrates (Põlme et al. 2020) | Põlme S, et al. 2020. Fungal Diversity 105:1-16 ((https://doi.org/10.1007/s13225-020-00466-2)) |
| Otu54348 | 1 | k__Fungi(100);p__Ascomycota(100);c__Saccharomycetes(100);o__Saccharomycetales(100);f__Metschnikowiaceae(100);g__Metschnikowia(100);s__Metschnikowia_pulcherrima(100); | Metschnikowia | 13 | Saprotroph | (Undefined Saprotroph) | Yeast | NULL | Probable | As nectar-tap_saprotroph (Põlme et al. 2020); Decay sugar-rich substrates (Põlme et al. 2020) | Põlme S, et al. 2020. Fungal Diversity 105:1-16 ((https://doi.org/10.1007/s13225-020-00466-2)) |
| Otu54350 | 1 | k__Fungi(100);p__Ascomycota(100);c__Saccharomycetes(100);o__Saccharomycetales(100);f__Metschnikowiaceae(100);g__Metschnikowia(100);g__Metschnikowia_unclassified(100); | Metschnikowia | 13 | Saprotroph | (Undefined Saprotroph) | Yeast | NULL | Probable | As nectar-tap_saprotroph (Põlme et al. 2020); Decay sugar-rich substrates (Põlme et al. 2020) | Põlme S, et al. 2020. Fungal Diversity 105:1-16 ((https://doi.org/10.1007/s13225-020-00466-2)) |
| Otu54351 | 1 | k__Fungi(100);p__Ascomycota(100);c__Saccharomycetes(100);o__Saccharomycetales(100);f__Metschnikowiaceae(100);g__Metschnikowia(100);g__Metschnikowia_unclassified(100); | Metschnikowia | 13 | Saprotroph | (Undefined Saprotroph) | Yeast | NULL | Probable | As nectar-tap_saprotroph (Põlme et al. 2020); Decay sugar-rich substrates (Põlme et al. 2020) | Põlme S, et al. 2020. Fungal Diversity 105:1-16 ((https://doi.org/10.1007/s13225-020-00466-2)) |
| Otu54352 | 1 | k__Fungi(100);p__Ascomycota(100);c__Saccharomycetes(100);o__Saccharomycetales(100);f__Metschnikowiaceae(100);g__Metschnikowia(100);s__Metschnikowia_pulcherrima(100); | Metschnikowia | 13 | Saprotroph | (Undefined Saprotroph) | Yeast | NULL | Probable | As nectar-tap_saprotroph (Põlme et al. 2020); Decay sugar-rich substrates (Põlme et al. 2020) | Põlme S, et al. 2020. Fungal Diversity 105:1-16 ((https://doi.org/10.1007/s13225-020-00466-2)) |
| Otu54353 | 1 | k__Fungi(100);p__Ascomycota(100);c__Saccharomycetes(100);o__Saccharomycetales(100);f__Metschnikowiaceae(100);g__Metschnikowia(100);g__Metschnikowia_unclassified(100); | Metschnikowia | 13 | Saprotroph | (Undefined Saprotroph) | Yeast | NULL | Probable | As nectar-tap_saprotroph (Põlme et al. 2020); Decay sugar-rich substrates (Põlme et al. 2020) | Põlme S, et al. 2020. Fungal Diversity 105:1-16 ((https://doi.org/10.1007/s13225-020-00466-2)) |
| Otu54354 | 1 | k__Fungi(100);p__Ascomycota(100);c__Saccharomycetes(100);o__Saccharomycetales(100);f__Metschnikowiaceae(100);g__Metschnikowia(100);s__Metschnikowia_pulcherrima(100); | Metschnikowia | 13 | Saprotroph | (Undefined Saprotroph) | Yeast | NULL | Probable | As nectar-tap_saprotroph (Põlme et al. 2020); Decay sugar-rich substrates (Põlme et al. 2020) | Põlme S, et al. 2020. Fungal Diversity 105:1-16 ((https://doi.org/10.1007/s13225-020-00466-2)) |
| Otu54355 | 1 | k__Fungi(100);p__Ascomycota(100);c__Saccharomycetes(100);o__Saccharomycetales(100);f__Metschnikowiaceae(100);g__Metschnikowia(100);g__Metschnikowia_unclassified(100); | Metschnikowia | 13 | Saprotroph | (Undefined Saprotroph) | Yeast | NULL | Probable | As nectar-tap_saprotroph (Põlme et al. 2020); Decay sugar-rich substrates (Põlme et al. 2020) | Põlme S, et al. 2020. Fungal Diversity 105:1-16 ((https://doi.org/10.1007/s13225-020-00466-2)) |
| Otu54356 | 1 | k__Fungi(100);p__Ascomycota(100);c__Saccharomycetes(100);o__Saccharomycetales(100);f__Metschnikowiaceae(100);g__Metschnikowia(100);g__Metschnikowia_unclassified(100); | Metschnikowia | 13 | Saprotroph | (Undefined Saprotroph) | Yeast | NULL | Probable | As nectar-tap_saprotroph (Põlme et al. 2020); Decay sugar-rich substrates (Põlme et al. 2020) | Põlme S, et al. 2020. Fungal Diversity 105:1-16 ((https://doi.org/10.1007/s13225-020-00466-2)) |
| Otu54357 | 1 | k__Fungi(100);p__Ascomycota(100);c__Saccharomycetes(100);o__Saccharomycetales(100);f__Metschnikowiaceae(100);g__Metschnikowia(100);s__Metschnikowia_pulcherrima(100); | Metschnikowia | 13 | Saprotroph | (Undefined Saprotroph) | Yeast | NULL | Probable | As nectar-tap_saprotroph (Põlme et al. 2020); Decay sugar-rich substrates (Põlme et al. 2020) | Põlme S, et al. 2020. Fungal Diversity 105:1-16 ((https://doi.org/10.1007/s13225-020-00466-2)) |
| Otu54358 | 1 | k__Fungi(100);p__Ascomycota(100);c__Saccharomycetes(100);o__Saccharomycetales(100);f__Metschnikowiaceae(100);g__Metschnikowia(100);s__Metschnikowia_pulcherrima(100); | Metschnikowia | 13 | Saprotroph | (Undefined Saprotroph) | Yeast | NULL | Probable | As nectar-tap_saprotroph (Põlme et al. 2020); Decay sugar-rich substrates (Põlme et al. 2020) | Põlme S, et al. 2020. Fungal Diversity 105:1-16 ((https://doi.org/10.1007/s13225-020-00466-2)) |
| Otu54359 | 1 | k__Fungi(100);p__Ascomycota(100);c__Saccharomycetes(100);o__Saccharomycetales(100);f__Metschnikowiaceae(100);g__Metschnikowia(100);s__Metschnikowia_pulcherrima(100); | Metschnikowia | 13 | Saprotroph | (Undefined Saprotroph) | Yeast | NULL | Probable | As nectar-tap_saprotroph (Põlme et al. 2020); Decay sugar-rich substrates (Põlme et al. 2020) | Põlme S, et al. 2020. Fungal Diversity 105:1-16 ((https://doi.org/10.1007/s13225-020-00466-2)) |
| Otu54360 | 1 | k__Fungi(100);p__Ascomycota(100);c__Saccharomycetes(100);o__Saccharomycetales(100);f__Metschnikowiaceae(100);g__Metschnikowia(100);g__Metschnikowia_unclassified(100); | Metschnikowia | 13 | Saprotroph | (Undefined Saprotroph) | Yeast | NULL | Probable | As nectar-tap_saprotroph (Põlme et al. 2020); Decay sugar-rich substrates (Põlme et al. 2020) | Põlme S, et al. 2020. Fungal Diversity 105:1-16 ((https://doi.org/10.1007/s13225-020-00466-2)) |
| Otu54361 | 1 | k__Fungi(100);p__Ascomycota(100);c__Saccharomycetes(100);o__Saccharomycetales(100);f__Metschnikowiaceae(100);g__Metschnikowia(100);g__Metschnikowia_unclassified(100); | Metschnikowia | 13 | Saprotroph | (Undefined Saprotroph) | Yeast | NULL | Probable | As nectar-tap_saprotroph (Põlme et al. 2020); Decay sugar-rich substrates (Põlme et al. 2020) | Põlme S, et al. 2020. Fungal Diversity 105:1-16 ((https://doi.org/10.1007/s13225-020-00466-2)) |
| Otu54362 | 1 | k__Fungi(100);p__Ascomycota(100);c__Saccharomycetes(100);o__Saccharomycetales(100);f__Metschnikowiaceae(100);g__Metschnikowia(100);s__Metschnikowia_pulcherrima(100); | Metschnikowia | 13 | Saprotroph | (Undefined Saprotroph) | Yeast | NULL | Probable | As nectar-tap_saprotroph (Põlme et al. 2020); Decay sugar-rich substrates (Põlme et al. 2020) | Põlme S, et al. 2020. Fungal Diversity 105:1-16 ((https://doi.org/10.1007/s13225-020-00466-2)) |
| Otu54363 | 1 | k__Fungi(100);p__Ascomycota(100);c__Saccharomycetes(100);o__Saccharomycetales(100);f__Metschnikowiaceae(100);g__Metschnikowia(100);s__Metschnikowia_pulcherrima(100); | Metschnikowia | 13 | Saprotroph | (Undefined Saprotroph) | Yeast | NULL | Probable | As nectar-tap_saprotroph (Põlme et al. 2020); Decay sugar-rich substrates (Põlme et al. 2020) | Põlme S, et al. 2020. Fungal Diversity 105:1-16 ((https://doi.org/10.1007/s13225-020-00466-2)) |
| Otu54364 | 1 | k__Fungi(100);p__Ascomycota(100);c__Saccharomycetes(100);o__Saccharomycetales(100);f__Metschnikowiaceae(100);g__Metschnikowia(100);s__Metschnikowia_pulcherrima(100); | Metschnikowia | 13 | Saprotroph | (Undefined Saprotroph) | Yeast | NULL | Probable | As nectar-tap_saprotroph (Põlme et al. 2020); Decay sugar-rich substrates (Põlme et al. 2020) | Põlme S, et al. 2020. Fungal Diversity 105:1-16 ((https://doi.org/10.1007/s13225-020-00466-2)) |
| Otu54365 | 1 | k__Fungi(100);p__Ascomycota(100);c__Saccharomycetes(100);o__Saccharomycetales(100);f__Metschnikowiaceae(100);g__Metschnikowia(100);s__Metschnikowia_pulcherrima(100); | Metschnikowia | 13 | Saprotroph | (Undefined Saprotroph) | Yeast | NULL | Probable | As nectar-tap_saprotroph (Põlme et al. 2020); Decay sugar-rich substrates (Põlme et al. 2020) | Põlme S, et al. 2020. Fungal Diversity 105:1-16 ((https://doi.org/10.1007/s13225-020-00466-2)) |
| Otu54366 | 1 | k__Fungi(100);p__Ascomycota(100);c__Saccharomycetes(100);o__Saccharomycetales(100);f__Metschnikowiaceae(100);g__Metschnikowia(100);s__Metschnikowia_pulcherrima(100); | Metschnikowia | 13 | Saprotroph | (Undefined Saprotroph) | Yeast | NULL | Probable | As nectar-tap_saprotroph (Põlme et al. 2020); Decay sugar-rich substrates (Põlme et al. 2020) | Põlme S, et al. 2020. Fungal Diversity 105:1-16 ((https://doi.org/10.1007/s13225-020-00466-2)) |
| Otu54367 | 1 | k__Fungi(100);p__Ascomycota(100);c__Saccharomycetes(100);o__Saccharomycetales(100);f__Metschnikowiaceae(100);g__Metschnikowia(100);s__Metschnikowia_pulcherrima(100); | Metschnikowia | 13 | Saprotroph | (Undefined Saprotroph) | Yeast | NULL | Probable | As nectar-tap_saprotroph (Põlme et al. 2020); Decay sugar-rich substrates (Põlme et al. 2020) | Põlme S, et al. 2020. Fungal Diversity 105:1-16 ((https://doi.org/10.1007/s13225-020-00466-2)) |
| Otu54368 | 1 | k__Fungi(100);p__Ascomycota(100);c__Saccharomycetes(100);o__Saccharomycetales(100);f__Metschnikowiaceae(100);g__Metschnikowia(100);g__Metschnikowia_unclassified(100); | Metschnikowia | 13 | Saprotroph | (Undefined Saprotroph) | Yeast | NULL | Probable | As nectar-tap_saprotroph (Põlme et al. 2020); Decay sugar-rich substrates (Põlme et al. 2020) | Põlme S, et al. 2020. Fungal Diversity 105:1-16 ((https://doi.org/10.1007/s13225-020-00466-2)) |
| Otu54370 | 1 | k__Fungi(100);p__Ascomycota(100);c__Saccharomycetes(100);o__Saccharomycetales(100);f__Metschnikowiaceae(100);g__Metschnikowia(100);s__Metschnikowia_pulcherrima(100); | Metschnikowia | 13 | Saprotroph | (Undefined Saprotroph) | Yeast | NULL | Probable | As nectar-tap_saprotroph (Põlme et al. 2020); Decay sugar-rich substrates (Põlme et al. 2020) | Põlme S, et al. 2020. Fungal Diversity 105:1-16 ((https://doi.org/10.1007/s13225-020-00466-2)) |
| Otu54371 | 1 | k__Fungi(100);p__Ascomycota(100);c__Saccharomycetes(100);o__Saccharomycetales(100);f__Metschnikowiaceae(100);g__Metschnikowia(100);s__Metschnikowia_pulcherrima(100); | Metschnikowia | 13 | Saprotroph | (Undefined Saprotroph) | Yeast | NULL | Probable | As nectar-tap_saprotroph (Põlme et al. 2020); Decay sugar-rich substrates (Põlme et al. 2020) | Põlme S, et al. 2020. Fungal Diversity 105:1-16 ((https://doi.org/10.1007/s13225-020-00466-2)) |
| Otu54372 | 1 | k__Fungi(100);p__Ascomycota(100);c__Saccharomycetes(100);o__Saccharomycetales(100);f__Metschnikowiaceae(100);g__Metschnikowia(100);s__Metschnikowia_pulcherrima(100); | Metschnikowia | 13 | Saprotroph | (Undefined Saprotroph) | Yeast | NULL | Probable | As nectar-tap_saprotroph (Põlme et al. 2020); Decay sugar-rich substrates (Põlme et al. 2020) | Põlme S, et al. 2020. Fungal Diversity 105:1-16 ((https://doi.org/10.1007/s13225-020-00466-2)) |
| Otu54373 | 1 | k__Fungi(100);p__Ascomycota(100);c__Saccharomycetes(100);o__Saccharomycetales(100);f__Metschnikowiaceae(100);g__Metschnikowia(100);s__Metschnikowia_pulcherrima(100); | Metschnikowia | 13 | Saprotroph | (Undefined Saprotroph) | Yeast | NULL | Probable | As nectar-tap_saprotroph (Põlme et al. 2020); Decay sugar-rich substrates (Põlme et al. 2020) | Põlme S, et al. 2020. Fungal Diversity 105:1-16 ((https://doi.org/10.1007/s13225-020-00466-2)) |
| Otu54374 | 1 | k__Fungi(100);p__Ascomycota(100);c__Saccharomycetes(100);o__Saccharomycetales(100);f__Metschnikowiaceae(100);g__Metschnikowia(100);g__Metschnikowia_unclassified(100); | Metschnikowia | 13 | Saprotroph | (Undefined Saprotroph) | Yeast | NULL | Probable | As nectar-tap_saprotroph (Põlme et al. 2020); Decay sugar-rich substrates (Põlme et al. 2020) | Põlme S, et al. 2020. Fungal Diversity 105:1-16 ((https://doi.org/10.1007/s13225-020-00466-2)) |
| Otu54377 | 1 | k__Fungi(100);p__Ascomycota(100);c__Saccharomycetes(100);o__Saccharomycetales(100);f__Metschnikowiaceae(100);g__Metschnikowia(100);s__Metschnikowia_pulcherrima(100); | Metschnikowia | 13 | Saprotroph | (Undefined Saprotroph) | Yeast | NULL | Probable | As nectar-tap_saprotroph (Põlme et al. 2020); Decay sugar-rich substrates (Põlme et al. 2020) | Põlme S, et al. 2020. Fungal Diversity 105:1-16 ((https://doi.org/10.1007/s13225-020-00466-2)) |
| Otu54378 | 1 | k__Fungi(100);p__Ascomycota(100);c__Saccharomycetes(100);o__Saccharomycetales(100);f__Metschnikowiaceae(100);g__Metschnikowia(100);g__Metschnikowia_unclassified(100); | Metschnikowia | 13 | Saprotroph | (Undefined Saprotroph) | Yeast | NULL | Probable | As nectar-tap_saprotroph (Põlme et al. 2020); Decay sugar-rich substrates (Põlme et al. 2020) | Põlme S, et al. 2020. Fungal Diversity 105:1-16 ((https://doi.org/10.1007/s13225-020-00466-2)) |
| Otu54379 | 1 | k__Fungi(100);p__Ascomycota(100);c__Saccharomycetes(100);o__Saccharomycetales(100);f__Metschnikowiaceae(100);g__Metschnikowia(100);s__Metschnikowia_pulcherrima(100); | Metschnikowia | 13 | Saprotroph | (Undefined Saprotroph) | Yeast | NULL | Probable | As nectar-tap_saprotroph (Põlme et al. 2020); Decay sugar-rich substrates (Põlme et al. 2020) | Põlme S, et al. 2020. Fungal Diversity 105:1-16 ((https://doi.org/10.1007/s13225-020-00466-2)) |
| Otu54380 | 1 | k__Fungi(100);p__Ascomycota(100);c__Saccharomycetes(100);o__Saccharomycetales(100);f__Metschnikowiaceae(100);g__Metschnikowia(100);s__Metschnikowia_pulcherrima(100); | Metschnikowia | 13 | Saprotroph | (Undefined Saprotroph) | Yeast | NULL | Probable | As nectar-tap_saprotroph (Põlme et al. 2020); Decay sugar-rich substrates (Põlme et al. 2020) | Põlme S, et al. 2020. Fungal Diversity 105:1-16 ((https://doi.org/10.1007/s13225-020-00466-2)) |
| Otu54381 | 1 | k__Fungi(100);p__Ascomycota(100);c__Lecanoromycetes(100);o__Lecanorales(100);o__Lecanorales_unclassified(100);o__Lecanorales_unclassified(100);o__Lecanorales_unclassified(100); | Lecanorales | 7 | Pathotroph-Symbiotroph | Lichen Parasite-Lichenized | Microfungus-Thallus | NULL | Probable | Lichenized with green algae, occasionally lichenicolous, on rocks, mosses and wood (Cannon & Kirk 2007); ...an order of mostly lichen-forming fungi (Kirk et al. 2008) | Cannon PF, Kirk PM. 2007. Fungal Families of the World. CAB International, Cambridge (ISBN: 978-0851998275); Kirk PM et al. 2008. Dictionary of the Fungi. Tenth Edition. CAB International, Wallingford (ISBN: 978-0851998268); Esslinger TL. 2014. North Dakota State University ((http://www.ndsu.edu/pubweb/~esslinge/chcklst/chcklst7.htm)) |
| Otu54382 | 1 | k__Fungi(100);p__Ascomycota(100);c__Saccharomycetes(100);o__Saccharomycetales(100);f__Metschnikowiaceae(100);g__Metschnikowia(100);s__Metschnikowia_pulcherrima(100); | Metschnikowia | 13 | Saprotroph | (Undefined Saprotroph) | Yeast | NULL | Probable | As nectar-tap_saprotroph (Põlme et al. 2020); Decay sugar-rich substrates (Põlme et al. 2020) | Põlme S, et al. 2020. Fungal Diversity 105:1-16 ((https://doi.org/10.1007/s13225-020-00466-2)) |
| Otu54383 | 1 | k__Fungi(100);p__Ascomycota(100);c__Saccharomycetes(100);o__Saccharomycetales(100);f__Metschnikowiaceae(100);g__Metschnikowia(100);s__Metschnikowia_pulcherrima(100); | Metschnikowia | 13 | Saprotroph | (Undefined Saprotroph) | Yeast | NULL | Probable | As nectar-tap_saprotroph (Põlme et al. 2020); Decay sugar-rich substrates (Põlme et al. 2020) | Põlme S, et al. 2020. Fungal Diversity 105:1-16 ((https://doi.org/10.1007/s13225-020-00466-2)) |
| Otu54384 | 1 | k__Fungi(100);p__Ascomycota(100);c__Saccharomycetes(100);o__Saccharomycetales(100);f__Metschnikowiaceae(100);g__Metschnikowia(100);s__Metschnikowia_pulcherrima(100); | Metschnikowia | 13 | Saprotroph | (Undefined Saprotroph) | Yeast | NULL | Probable | As nectar-tap_saprotroph (Põlme et al. 2020); Decay sugar-rich substrates (Põlme et al. 2020) | Põlme S, et al. 2020. Fungal Diversity 105:1-16 ((https://doi.org/10.1007/s13225-020-00466-2)) |
| Otu54385 | 1 | k__Fungi(100);p__Ascomycota(100);c__Saccharomycetes(100);o__Saccharomycetales(100);f__Metschnikowiaceae(100);g__Metschnikowia(100);g__Metschnikowia_unclassified(100); | Metschnikowia | 13 | Saprotroph | (Undefined Saprotroph) | Yeast | NULL | Probable | As nectar-tap_saprotroph (Põlme et al. 2020); Decay sugar-rich substrates (Põlme et al. 2020) | Põlme S, et al. 2020. Fungal Diversity 105:1-16 ((https://doi.org/10.1007/s13225-020-00466-2)) |
| Otu54386 | 1 | k__Fungi(100);p__Ascomycota(100);c__Saccharomycetes(100);o__Saccharomycetales(100);f__Metschnikowiaceae(100);g__Metschnikowia(100);s__Metschnikowia_pulcherrima(100); | Metschnikowia | 13 | Saprotroph | (Undefined Saprotroph) | Yeast | NULL | Probable | As nectar-tap_saprotroph (Põlme et al. 2020); Decay sugar-rich substrates (Põlme et al. 2020) | Põlme S, et al. 2020. Fungal Diversity 105:1-16 ((https://doi.org/10.1007/s13225-020-00466-2)) |
| Otu54387 | 1 | k__Fungi(100);p__Ascomycota(100);c__Saccharomycetes(100);o__Saccharomycetales(100);f__Metschnikowiaceae(100);g__Metschnikowia(100);s__Metschnikowia_pulcherrima(100); | Metschnikowia | 13 | Saprotroph | (Undefined Saprotroph) | Yeast | NULL | Probable | As nectar-tap_saprotroph (Põlme et al. 2020); Decay sugar-rich substrates (Põlme et al. 2020) | Põlme S, et al. 2020. Fungal Diversity 105:1-16 ((https://doi.org/10.1007/s13225-020-00466-2)) |
| Otu54388 | 1 | k__Fungi(100);p__Ascomycota(100);c__Saccharomycetes(100);o__Saccharomycetales(100);f__Metschnikowiaceae(100);g__Metschnikowia(100);s__Metschnikowia_pulcherrima(100); | Metschnikowia | 13 | Saprotroph | (Undefined Saprotroph) | Yeast | NULL | Probable | As nectar-tap_saprotroph (Põlme et al. 2020); Decay sugar-rich substrates (Põlme et al. 2020) | Põlme S, et al. 2020. Fungal Diversity 105:1-16 ((https://doi.org/10.1007/s13225-020-00466-2)) |
| Otu54389 | 1 | k__Fungi(100);p__Ascomycota(100);c__Saccharomycetes(100);o__Saccharomycetales(100);f__Metschnikowiaceae(100);g__Metschnikowia(100);s__Metschnikowia_pulcherrima(100); | Metschnikowia | 13 | Saprotroph | (Undefined Saprotroph) | Yeast | NULL | Probable | As nectar-tap_saprotroph (Põlme et al. 2020); Decay sugar-rich substrates (Põlme et al. 2020) | Põlme S, et al. 2020. Fungal Diversity 105:1-16 ((https://doi.org/10.1007/s13225-020-00466-2)) |
| Otu54391 | 1 | k__Fungi(100);p__Ascomycota(100);c__Saccharomycetes(100);o__Saccharomycetales(100);f__Metschnikowiaceae(100);g__Metschnikowia(100);g__Metschnikowia_unclassified(100); | Metschnikowia | 13 | Saprotroph | (Undefined Saprotroph) | Yeast | NULL | Probable | As nectar-tap_saprotroph (Põlme et al. 2020); Decay sugar-rich substrates (Põlme et al. 2020) | Põlme S, et al. 2020. Fungal Diversity 105:1-16 ((https://doi.org/10.1007/s13225-020-00466-2)) |
| Otu54392 | 1 | k__Fungi(100);p__Ascomycota(100);c__Saccharomycetes(100);o__Saccharomycetales(100);f__Metschnikowiaceae(100);g__Metschnikowia(100);s__Metschnikowia_pulcherrima(100); | Metschnikowia | 13 | Saprotroph | (Undefined Saprotroph) | Yeast | NULL | Probable | As nectar-tap_saprotroph (Põlme et al. 2020); Decay sugar-rich substrates (Põlme et al. 2020) | Põlme S, et al. 2020. Fungal Diversity 105:1-16 ((https://doi.org/10.1007/s13225-020-00466-2)) |
| Otu54393 | 1 | k__Fungi(100);p__Ascomycota(100);c__Saccharomycetes(100);o__Saccharomycetales(100);f__Metschnikowiaceae(100);g__Metschnikowia(100);s__Metschnikowia_pulcherrima(100); | Metschnikowia | 13 | Saprotroph | (Undefined Saprotroph) | Yeast | NULL | Probable | As nectar-tap_saprotroph (Põlme et al. 2020); Decay sugar-rich substrates (Põlme et al. 2020) | Põlme S, et al. 2020. Fungal Diversity 105:1-16 ((https://doi.org/10.1007/s13225-020-00466-2)) |
| Otu54394 | 1 | k__Fungi(100);p__Ascomycota(100);c__Saccharomycetes(100);o__Saccharomycetales(100);f__Metschnikowiaceae(100);g__Metschnikowia(100);s__Metschnikowia_pulcherrima(100); | Metschnikowia | 13 | Saprotroph | (Undefined Saprotroph) | Yeast | NULL | Probable | As nectar-tap_saprotroph (Põlme et al. 2020); Decay sugar-rich substrates (Põlme et al. 2020) | Põlme S, et al. 2020. Fungal Diversity 105:1-16 ((https://doi.org/10.1007/s13225-020-00466-2)) |
| Otu54395 | 1 | k__Fungi(100);p__Ascomycota(100);c__Saccharomycetes(100);o__Saccharomycetales(100);f__Metschnikowiaceae(100);g__Metschnikowia(100);g__Metschnikowia_unclassified(100); | Metschnikowia | 13 | Saprotroph | (Undefined Saprotroph) | Yeast | NULL | Probable | As nectar-tap_saprotroph (Põlme et al. 2020); Decay sugar-rich substrates (Põlme et al. 2020) | Põlme S, et al. 2020. Fungal Diversity 105:1-16 ((https://doi.org/10.1007/s13225-020-00466-2)) |
| Otu54396 | 1 | k__Fungi(100);p__Ascomycota(100);c__Saccharomycetes(100);o__Saccharomycetales(100);f__Metschnikowiaceae(100);g__Metschnikowia(100);s__Metschnikowia_sp(100); | Metschnikowia | 13 | Saprotroph | (Undefined Saprotroph) | Yeast | NULL | Probable | As nectar-tap_saprotroph (Põlme et al. 2020); Decay sugar-rich substrates (Põlme et al. 2020) | Põlme S, et al. 2020. Fungal Diversity 105:1-16 ((https://doi.org/10.1007/s13225-020-00466-2)) |
| Otu54397 | 1 | k__Fungi(100);p__Ascomycota(100);c__Saccharomycetes(100);o__Saccharomycetales(100);f__Metschnikowiaceae(100);g__Metschnikowia(100);s__Metschnikowia_pulcherrima(100); | Metschnikowia | 13 | Saprotroph | (Undefined Saprotroph) | Yeast | NULL | Probable | As nectar-tap_saprotroph (Põlme et al. 2020); Decay sugar-rich substrates (Põlme et al. 2020) | Põlme S, et al. 2020. Fungal Diversity 105:1-16 ((https://doi.org/10.1007/s13225-020-00466-2)) |
| Otu54398 | 1 | k__Fungi(100);p__Ascomycota(100);c__Saccharomycetes(100);o__Saccharomycetales(100);f__Metschnikowiaceae(100);g__Metschnikowia(100);s__Metschnikowia_pulcherrima(100); | Metschnikowia | 13 | Saprotroph | (Undefined Saprotroph) | Yeast | NULL | Probable | As nectar-tap_saprotroph (Põlme et al. 2020); Decay sugar-rich substrates (Põlme et al. 2020) | Põlme S, et al. 2020. Fungal Diversity 105:1-16 ((https://doi.org/10.1007/s13225-020-00466-2)) |
| Otu54399 | 1 | k__Fungi(100);p__Ascomycota(100);c__Saccharomycetes(100);o__Saccharomycetales(100);f__Metschnikowiaceae(100);g__Metschnikowia(100);s__Metschnikowia_pulcherrima(100); | Metschnikowia | 13 | Saprotroph | (Undefined Saprotroph) | Yeast | NULL | Probable | As nectar-tap_saprotroph (Põlme et al. 2020); Decay sugar-rich substrates (Põlme et al. 2020) | Põlme S, et al. 2020. Fungal Diversity 105:1-16 ((https://doi.org/10.1007/s13225-020-00466-2)) |
| Otu54400 | 1 | k__Fungi(100);p__Ascomycota(100);c__Saccharomycetes(100);o__Saccharomycetales(100);f__Metschnikowiaceae(100);g__Metschnikowia(100);s__Metschnikowia_pulcherrima(100); | Metschnikowia | 13 | Saprotroph | (Undefined Saprotroph) | Yeast | NULL | Probable | As nectar-tap_saprotroph (Põlme et al. 2020); Decay sugar-rich substrates (Põlme et al. 2020) | Põlme S, et al. 2020. Fungal Diversity 105:1-16 ((https://doi.org/10.1007/s13225-020-00466-2)) |
| Otu54401 | 1 | k__Fungi(100);p__Ascomycota(100);c__Saccharomycetes(100);o__Saccharomycetales(100);f__Metschnikowiaceae(100);g__Metschnikowia(100);s__Metschnikowia_pulcherrima(100); | Metschnikowia | 13 | Saprotroph | (Undefined Saprotroph) | Yeast | NULL | Probable | As nectar-tap_saprotroph (Põlme et al. 2020); Decay sugar-rich substrates (Põlme et al. 2020) | Põlme S, et al. 2020. Fungal Diversity 105:1-16 ((https://doi.org/10.1007/s13225-020-00466-2)) |
| Otu54402 | 1 | k__Fungi(100);p__Ascomycota(100);c__Saccharomycetes(100);o__Saccharomycetales(100);f__Metschnikowiaceae(100);g__Metschnikowia(100);s__Metschnikowia_pulcherrima(100); | Metschnikowia | 13 | Saprotroph | (Undefined Saprotroph) | Yeast | NULL | Probable | As nectar-tap_saprotroph (Põlme et al. 2020); Decay sugar-rich substrates (Põlme et al. 2020) | Põlme S, et al. 2020. Fungal Diversity 105:1-16 ((https://doi.org/10.1007/s13225-020-00466-2)) |
| Otu54403 | 1 | k__Fungi(100);p__Ascomycota(100);c__Saccharomycetes(100);o__Saccharomycetales(100);f__Metschnikowiaceae(100);g__Metschnikowia(100);s__Metschnikowia_pulcherrima(100); | Metschnikowia | 13 | Saprotroph | (Undefined Saprotroph) | Yeast | NULL | Probable | As nectar-tap_saprotroph (Põlme et al. 2020); Decay sugar-rich substrates (Põlme et al. 2020) | Põlme S, et al. 2020. Fungal Diversity 105:1-16 ((https://doi.org/10.1007/s13225-020-00466-2)) |
| Otu54404 | 1 | k__Fungi(100);p__Ascomycota(100);c__Saccharomycetes(100);o__Saccharomycetales(100);f__Metschnikowiaceae(100);g__Metschnikowia(100);g__Metschnikowia_unclassified(100); | Metschnikowia | 13 | Saprotroph | (Undefined Saprotroph) | Yeast | NULL | Probable | As nectar-tap_saprotroph (Põlme et al. 2020); Decay sugar-rich substrates (Põlme et al. 2020) | Põlme S, et al. 2020. Fungal Diversity 105:1-16 ((https://doi.org/10.1007/s13225-020-00466-2)) |
| Otu54405 | 1 | k__Fungi(100);p__Ascomycota(100);c__Saccharomycetes(100);o__Saccharomycetales(100);f__Metschnikowiaceae(100);g__Metschnikowia(100);s__Metschnikowia_pulcherrima(100); | Metschnikowia | 13 | Saprotroph | (Undefined Saprotroph) | Yeast | NULL | Probable | As nectar-tap_saprotroph (Põlme et al. 2020); Decay sugar-rich substrates (Põlme et al. 2020) | Põlme S, et al. 2020. Fungal Diversity 105:1-16 ((https://doi.org/10.1007/s13225-020-00466-2)) |
| Otu54406 | 1 | k__Fungi(100);p__Ascomycota(100);c__Saccharomycetes(100);o__Saccharomycetales(100);f__Metschnikowiaceae(100);g__Metschnikowia(100);g__Metschnikowia_unclassified(100); | Metschnikowia | 13 | Saprotroph | (Undefined Saprotroph) | Yeast | NULL | Probable | As nectar-tap_saprotroph (Põlme et al. 2020); Decay sugar-rich substrates (Põlme et al. 2020) | Põlme S, et al. 2020. Fungal Diversity 105:1-16 ((https://doi.org/10.1007/s13225-020-00466-2)) |
| Otu54407 | 1 | k__Fungi(100);p__Ascomycota(100);c__Saccharomycetes(100);o__Saccharomycetales(100);f__Metschnikowiaceae(100);g__Metschnikowia(100);s__Metschnikowia_pulcherrima(100); | Metschnikowia | 13 | Saprotroph | (Undefined Saprotroph) | Yeast | NULL | Probable | As nectar-tap_saprotroph (Põlme et al. 2020); Decay sugar-rich substrates (Põlme et al. 2020) | Põlme S, et al. 2020. Fungal Diversity 105:1-16 ((https://doi.org/10.1007/s13225-020-00466-2)) |
| Otu54408 | 1 | k__Fungi(100);p__Ascomycota(100);c__Saccharomycetes(100);o__Saccharomycetales(100);f__Metschnikowiaceae(100);g__Metschnikowia(100);g__Metschnikowia_unclassified(100); | Metschnikowia | 13 | Saprotroph | (Undefined Saprotroph) | Yeast | NULL | Probable | As nectar-tap_saprotroph (Põlme et al. 2020); Decay sugar-rich substrates (Põlme et al. 2020) | Põlme S, et al. 2020. Fungal Diversity 105:1-16 ((https://doi.org/10.1007/s13225-020-00466-2)) |
| Otu54414 | 1 | k__Fungi(100);p__Ascomycota(100);c__Saccharomycetes(100);o__Saccharomycetales(100);f__Metschnikowiaceae(100);g__Metschnikowia(100);g__Metschnikowia_unclassified(100); | Metschnikowia | 13 | Saprotroph | (Undefined Saprotroph) | Yeast | NULL | Probable | As nectar-tap_saprotroph (Põlme et al. 2020); Decay sugar-rich substrates (Põlme et al. 2020) | Põlme S, et al. 2020. Fungal Diversity 105:1-16 ((https://doi.org/10.1007/s13225-020-00466-2)) |
| Otu54415 | 1 | k__Fungi(100);p__Ascomycota(100);c__Saccharomycetes(100);o__Saccharomycetales(100);f__Metschnikowiaceae(100);g__Metschnikowia(100);s__Metschnikowia_pulcherrima(100); | Metschnikowia | 13 | Saprotroph | (Undefined Saprotroph) | Yeast | NULL | Probable | As nectar-tap_saprotroph (Põlme et al. 2020); Decay sugar-rich substrates (Põlme et al. 2020) | Põlme S, et al. 2020. Fungal Diversity 105:1-16 ((https://doi.org/10.1007/s13225-020-00466-2)) |
| Otu54416 | 1 | k__Fungi(100);p__Ascomycota(100);c__Saccharomycetes(100);o__Saccharomycetales(100);f__Metschnikowiaceae(100);g__Metschnikowia(100);s__Metschnikowia_pulcherrima(100); | Metschnikowia | 13 | Saprotroph | (Undefined Saprotroph) | Yeast | NULL | Probable | As nectar-tap_saprotroph (Põlme et al. 2020); Decay sugar-rich substrates (Põlme et al. 2020) | Põlme S, et al. 2020. Fungal Diversity 105:1-16 ((https://doi.org/10.1007/s13225-020-00466-2)) |
| Otu54417 | 1 | k__Fungi(100);p__Ascomycota(100);c__Saccharomycetes(100);o__Saccharomycetales(100);f__Metschnikowiaceae(100);g__Metschnikowia(100);g__Metschnikowia_unclassified(100); | Metschnikowia | 13 | Saprotroph | (Undefined Saprotroph) | Yeast | NULL | Probable | As nectar-tap_saprotroph (Põlme et al. 2020); Decay sugar-rich substrates (Põlme et al. 2020) | Põlme S, et al. 2020. Fungal Diversity 105:1-16 ((https://doi.org/10.1007/s13225-020-00466-2)) |
| Otu54418 | 1 | k__Fungi(100);p__Ascomycota(100);c__Saccharomycetes(100);o__Saccharomycetales(100);f__Metschnikowiaceae(100);g__Metschnikowia(100);s__Metschnikowia_pulcherrima(100); | Metschnikowia | 13 | Saprotroph | (Undefined Saprotroph) | Yeast | NULL | Probable | As nectar-tap_saprotroph (Põlme et al. 2020); Decay sugar-rich substrates (Põlme et al. 2020) | Põlme S, et al. 2020. Fungal Diversity 105:1-16 ((https://doi.org/10.1007/s13225-020-00466-2)) |
| Otu54419 | 1 | k__Fungi(100);p__Ascomycota(100);c__Saccharomycetes(100);o__Saccharomycetales(100);f__Metschnikowiaceae(100);g__Metschnikowia(100);s__Metschnikowia_pulcherrima(100); | Metschnikowia | 13 | Saprotroph | (Undefined Saprotroph) | Yeast | NULL | Probable | As nectar-tap_saprotroph (Põlme et al. 2020); Decay sugar-rich substrates (Põlme et al. 2020) | Põlme S, et al. 2020. Fungal Diversity 105:1-16 ((https://doi.org/10.1007/s13225-020-00466-2)) |
| Otu54420 | 1 | k__Fungi(100);p__Ascomycota(100);c__Saccharomycetes(100);o__Saccharomycetales(100);f__Metschnikowiaceae(100);g__Metschnikowia(100);s__Metschnikowia_pulcherrima(100); | Metschnikowia | 13 | Saprotroph | (Undefined Saprotroph) | Yeast | NULL | Probable | As nectar-tap_saprotroph (Põlme et al. 2020); Decay sugar-rich substrates (Põlme et al. 2020) | Põlme S, et al. 2020. Fungal Diversity 105:1-16 ((https://doi.org/10.1007/s13225-020-00466-2)) |
| Otu54421 | 1 | k__Fungi(100);p__Ascomycota(100);c__Saccharomycetes(100);o__Saccharomycetales(100);f__Metschnikowiaceae(100);g__Metschnikowia(100);g__Metschnikowia_unclassified(100); | Metschnikowia | 13 | Saprotroph | (Undefined Saprotroph) | Yeast | NULL | Probable | As nectar-tap_saprotroph (Põlme et al. 2020); Decay sugar-rich substrates (Põlme et al. 2020) | Põlme S, et al. 2020. Fungal Diversity 105:1-16 ((https://doi.org/10.1007/s13225-020-00466-2)) |
| Otu54422 | 1 | k__Fungi(100);p__Ascomycota(100);c__Saccharomycetes(100);o__Saccharomycetales(100);f__Metschnikowiaceae(100);g__Metschnikowia(100);g__Metschnikowia_unclassified(100); | Metschnikowia | 13 | Saprotroph | (Undefined Saprotroph) | Yeast | NULL | Probable | As nectar-tap_saprotroph (Põlme et al. 2020); Decay sugar-rich substrates (Põlme et al. 2020) | Põlme S, et al. 2020. Fungal Diversity 105:1-16 ((https://doi.org/10.1007/s13225-020-00466-2)) |
| Otu54423 | 1 | k__Fungi(100);p__Ascomycota(100);c__Saccharomycetes(100);o__Saccharomycetales(100);f__Metschnikowiaceae(100);g__Metschnikowia(100);s__Metschnikowia_pulcherrima(100); | Metschnikowia | 13 | Saprotroph | (Undefined Saprotroph) | Yeast | NULL | Probable | As nectar-tap_saprotroph (Põlme et al. 2020); Decay sugar-rich substrates (Põlme et al. 2020) | Põlme S, et al. 2020. Fungal Diversity 105:1-16 ((https://doi.org/10.1007/s13225-020-00466-2)) |
| Otu54424 | 1 | k__Fungi(100);p__Ascomycota(100);c__Saccharomycetes(100);o__Saccharomycetales(100);f__Metschnikowiaceae(100);g__Metschnikowia(100);s__Metschnikowia_pulcherrima(100); | Metschnikowia | 13 | Saprotroph | (Undefined Saprotroph) | Yeast | NULL | Probable | As nectar-tap_saprotroph (Põlme et al. 2020); Decay sugar-rich substrates (Põlme et al. 2020) | Põlme S, et al. 2020. Fungal Diversity 105:1-16 ((https://doi.org/10.1007/s13225-020-00466-2)) |
| Otu54425 | 1 | k__Fungi(100);p__Ascomycota(100);c__Saccharomycetes(100);o__Saccharomycetales(100);f__Metschnikowiaceae(100);g__Metschnikowia(100);s__Metschnikowia_pulcherrima(100); | Metschnikowia | 13 | Saprotroph | (Undefined Saprotroph) | Yeast | NULL | Probable | As nectar-tap_saprotroph (Põlme et al. 2020); Decay sugar-rich substrates (Põlme et al. 2020) | Põlme S, et al. 2020. Fungal Diversity 105:1-16 ((https://doi.org/10.1007/s13225-020-00466-2)) |
| Otu54426 | 1 | k__Fungi(100);p__Ascomycota(100);c__Saccharomycetes(100);o__Saccharomycetales(100);f__Metschnikowiaceae(100);g__Metschnikowia(100);s__Metschnikowia_pulcherrima(100); | Metschnikowia | 13 | Saprotroph | (Undefined Saprotroph) | Yeast | NULL | Probable | As nectar-tap_saprotroph (Põlme et al. 2020); Decay sugar-rich substrates (Põlme et al. 2020) | Põlme S, et al. 2020. Fungal Diversity 105:1-16 ((https://doi.org/10.1007/s13225-020-00466-2)) |
| Otu54427 | 1 | k__Fungi(100);p__Ascomycota(100);c__Saccharomycetes(100);o__Saccharomycetales(100);f__Metschnikowiaceae(100);g__Metschnikowia(100);s__Metschnikowia_pulcherrima(100); | Metschnikowia | 13 | Saprotroph | (Undefined Saprotroph) | Yeast | NULL | Probable | As nectar-tap_saprotroph (Põlme et al. 2020); Decay sugar-rich substrates (Põlme et al. 2020) | Põlme S, et al. 2020. Fungal Diversity 105:1-16 ((https://doi.org/10.1007/s13225-020-00466-2)) |
| Otu54428 | 1 | k__Fungi(100);p__Ascomycota(100);c__Saccharomycetes(100);o__Saccharomycetales(100);f__Metschnikowiaceae(100);g__Metschnikowia(100);s__Metschnikowia_pulcherrima(100); | Metschnikowia | 13 | Saprotroph | (Undefined Saprotroph) | Yeast | NULL | Probable | As nectar-tap_saprotroph (Põlme et al. 2020); Decay sugar-rich substrates (Põlme et al. 2020) | Põlme S, et al. 2020. Fungal Diversity 105:1-16 ((https://doi.org/10.1007/s13225-020-00466-2)) |
| Otu54430 | 1 | k__Fungi(100);p__Ascomycota(100);c__Saccharomycetes(100);o__Saccharomycetales(100);f__Metschnikowiaceae(100);g__Metschnikowia(100);g__Metschnikowia_unclassified(100); | Metschnikowia | 13 | Saprotroph | (Undefined Saprotroph) | Yeast | NULL | Probable | As nectar-tap_saprotroph (Põlme et al. 2020); Decay sugar-rich substrates (Põlme et al. 2020) | Põlme S, et al. 2020. Fungal Diversity 105:1-16 ((https://doi.org/10.1007/s13225-020-00466-2)) |
| Otu54431 | 1 | k__Fungi(100);p__Ascomycota(100);c__Saccharomycetes(100);o__Saccharomycetales(100);f__Metschnikowiaceae(100);g__Metschnikowia(100);s__Metschnikowia_pulcherrima(100); | Metschnikowia | 13 | Saprotroph | (Undefined Saprotroph) | Yeast | NULL | Probable | As nectar-tap_saprotroph (Põlme et al. 2020); Decay sugar-rich substrates (Põlme et al. 2020) | Põlme S, et al. 2020. Fungal Diversity 105:1-16 ((https://doi.org/10.1007/s13225-020-00466-2)) |
| Otu54432 | 1 | k__Fungi(100);p__Ascomycota(100);c__Saccharomycetes(100);o__Saccharomycetales(100);f__Metschnikowiaceae(100);g__Metschnikowia(100);s__Metschnikowia_pulcherrima(100); | Metschnikowia | 13 | Saprotroph | (Undefined Saprotroph) | Yeast | NULL | Probable | As nectar-tap_saprotroph (Põlme et al. 2020); Decay sugar-rich substrates (Põlme et al. 2020) | Põlme S, et al. 2020. Fungal Diversity 105:1-16 ((https://doi.org/10.1007/s13225-020-00466-2)) |
| Otu54433 | 1 | k__Fungi(100);p__Ascomycota(100);c__Saccharomycetes(100);o__Saccharomycetales(100);f__Metschnikowiaceae(100);g__Metschnikowia(100);g__Metschnikowia_unclassified(100); | Metschnikowia | 13 | Saprotroph | (Undefined Saprotroph) | Yeast | NULL | Probable | As nectar-tap_saprotroph (Põlme et al. 2020); Decay sugar-rich substrates (Põlme et al. 2020) | Põlme S, et al. 2020. Fungal Diversity 105:1-16 ((https://doi.org/10.1007/s13225-020-00466-2)) |
| Otu54434 | 1 | k__Fungi(100);p__Ascomycota(100);c__Saccharomycetes(100);o__Saccharomycetales(100);f__Metschnikowiaceae(100);g__Metschnikowia(100);s__Metschnikowia_pulcherrima(100); | Metschnikowia | 13 | Saprotroph | (Undefined Saprotroph) | Yeast | NULL | Probable | As nectar-tap_saprotroph (Põlme et al. 2020); Decay sugar-rich substrates (Põlme et al. 2020) | Põlme S, et al. 2020. Fungal Diversity 105:1-16 ((https://doi.org/10.1007/s13225-020-00466-2)) |
| Otu54435 | 1 | k__Fungi(100);p__Ascomycota(100);c__Saccharomycetes(100);o__Saccharomycetales(100);f__Metschnikowiaceae(100);g__Metschnikowia(100);s__Metschnikowia_pulcherrima(100); | Metschnikowia | 13 | Saprotroph | (Undefined Saprotroph) | Yeast | NULL | Probable | As nectar-tap_saprotroph (Põlme et al. 2020); Decay sugar-rich substrates (Põlme et al. 2020) | Põlme S, et al. 2020. Fungal Diversity 105:1-16 ((https://doi.org/10.1007/s13225-020-00466-2)) |
| Otu54436 | 1 | k__Fungi(100);p__Ascomycota(100);c__Saccharomycetes(100);o__Saccharomycetales(100);f__Metschnikowiaceae(100);g__Metschnikowia(100);g__Metschnikowia_unclassified(100); | Metschnikowia | 13 | Saprotroph | (Undefined Saprotroph) | Yeast | NULL | Probable | As nectar-tap_saprotroph (Põlme et al. 2020); Decay sugar-rich substrates (Põlme et al. 2020) | Põlme S, et al. 2020. Fungal Diversity 105:1-16 ((https://doi.org/10.1007/s13225-020-00466-2)) |
| Otu54437 | 1 | k__Fungi(100);p__Ascomycota(100);c__Saccharomycetes(100);o__Saccharomycetales(100);f__Metschnikowiaceae(100);g__Metschnikowia(100);g__Metschnikowia_unclassified(100); | Metschnikowia | 13 | Saprotroph | (Undefined Saprotroph) | Yeast | NULL | Probable | As nectar-tap_saprotroph (Põlme et al. 2020); Decay sugar-rich substrates (Põlme et al. 2020) | Põlme S, et al. 2020. Fungal Diversity 105:1-16 ((https://doi.org/10.1007/s13225-020-00466-2)) |
| Otu54438 | 1 | k__Fungi(100);p__Ascomycota(100);c__Saccharomycetes(100);o__Saccharomycetales(100);f__Metschnikowiaceae(100);g__Metschnikowia(100);g__Metschnikowia_unclassified(100); | Metschnikowia | 13 | Saprotroph | (Undefined Saprotroph) | Yeast | NULL | Probable | As nectar-tap_saprotroph (Põlme et al. 2020); Decay sugar-rich substrates (Põlme et al. 2020) | Põlme S, et al. 2020. Fungal Diversity 105:1-16 ((https://doi.org/10.1007/s13225-020-00466-2)) |
| Otu54439 | 1 | k__Fungi(100);p__Ascomycota(100);c__Saccharomycetes(100);o__Saccharomycetales(100);f__Metschnikowiaceae(100);g__Metschnikowia(100);g__Metschnikowia_unclassified(100); | Metschnikowia | 13 | Saprotroph | (Undefined Saprotroph) | Yeast | NULL | Probable | As nectar-tap_saprotroph (Põlme et al. 2020); Decay sugar-rich substrates (Põlme et al. 2020) | Põlme S, et al. 2020. Fungal Diversity 105:1-16 ((https://doi.org/10.1007/s13225-020-00466-2)) |
| Otu54440 | 1 | k__Fungi(100);p__Ascomycota(100);c__Saccharomycetes(100);o__Saccharomycetales(100);f__Metschnikowiaceae(100);g__Metschnikowia(100);g__Metschnikowia_unclassified(100); | Metschnikowia | 13 | Saprotroph | (Undefined Saprotroph) | Yeast | NULL | Probable | As nectar-tap_saprotroph (Põlme et al. 2020); Decay sugar-rich substrates (Põlme et al. 2020) | Põlme S, et al. 2020. Fungal Diversity 105:1-16 ((https://doi.org/10.1007/s13225-020-00466-2)) |
| Otu54441 | 1 | k__Fungi(100);p__Ascomycota(100);c__Saccharomycetes(100);o__Saccharomycetales(100);f__Metschnikowiaceae(100);g__Metschnikowia(100);s__Metschnikowia_pulcherrima(100); | Metschnikowia | 13 | Saprotroph | (Undefined Saprotroph) | Yeast | NULL | Probable | As nectar-tap_saprotroph (Põlme et al. 2020); Decay sugar-rich substrates (Põlme et al. 2020) | Põlme S, et al. 2020. Fungal Diversity 105:1-16 ((https://doi.org/10.1007/s13225-020-00466-2)) |
| Otu54442 | 1 | k__Fungi(100);p__Ascomycota(100);c__Saccharomycetes(100);o__Saccharomycetales(100);f__Metschnikowiaceae(100);g__Metschnikowia(100);g__Metschnikowia_unclassified(100); | Metschnikowia | 13 | Saprotroph | (Undefined Saprotroph) | Yeast | NULL | Probable | As nectar-tap_saprotroph (Põlme et al. 2020); Decay sugar-rich substrates (Põlme et al. 2020) | Põlme S, et al. 2020. Fungal Diversity 105:1-16 ((https://doi.org/10.1007/s13225-020-00466-2)) |
| Otu54443 | 1 | k__Fungi(100);p__Ascomycota(100);c__Saccharomycetes(100);o__Saccharomycetales(100);f__Metschnikowiaceae(100);g__Metschnikowia(100);s__Metschnikowia_pulcherrima(100); | Metschnikowia | 13 | Saprotroph | (Undefined Saprotroph) | Yeast | NULL | Probable | As nectar-tap_saprotroph (Põlme et al. 2020); Decay sugar-rich substrates (Põlme et al. 2020) | Põlme S, et al. 2020. Fungal Diversity 105:1-16 ((https://doi.org/10.1007/s13225-020-00466-2)) |
| Otu54444 | 1 | k__Fungi(100);p__Ascomycota(100);c__Saccharomycetes(100);o__Saccharomycetales(100);f__Metschnikowiaceae(100);g__Metschnikowia(100);s__Metschnikowia_pulcherrima(100); | Metschnikowia | 13 | Saprotroph | (Undefined Saprotroph) | Yeast | NULL | Probable | As nectar-tap_saprotroph (Põlme et al. 2020); Decay sugar-rich substrates (Põlme et al. 2020) | Põlme S, et al. 2020. Fungal Diversity 105:1-16 ((https://doi.org/10.1007/s13225-020-00466-2)) |
| Otu54445 | 1 | k__Fungi(100);p__Ascomycota(100);c__Saccharomycetes(100);o__Saccharomycetales(100);f__Metschnikowiaceae(100);g__Metschnikowia(100);s__Metschnikowia_pulcherrima(100); | Metschnikowia | 13 | Saprotroph | (Undefined Saprotroph) | Yeast | NULL | Probable | As nectar-tap_saprotroph (Põlme et al. 2020); Decay sugar-rich substrates (Põlme et al. 2020) | Põlme S, et al. 2020. Fungal Diversity 105:1-16 ((https://doi.org/10.1007/s13225-020-00466-2)) |
| Otu54446 | 1 | k__Fungi(100);p__Ascomycota(100);c__Saccharomycetes(100);o__Saccharomycetales(100);f__Metschnikowiaceae(100);g__Metschnikowia(100);s__Metschnikowia_pulcherrima(100); | Metschnikowia | 13 | Saprotroph | (Undefined Saprotroph) | Yeast | NULL | Probable | As nectar-tap_saprotroph (Põlme et al. 2020); Decay sugar-rich substrates (Põlme et al. 2020) | Põlme S, et al. 2020. Fungal Diversity 105:1-16 ((https://doi.org/10.1007/s13225-020-00466-2)) |
| Otu54447 | 1 | k__Fungi(100);p__Ascomycota(100);c__Saccharomycetes(100);o__Saccharomycetales(100);f__Metschnikowiaceae(100);g__Metschnikowia(100);s__Metschnikowia_pulcherrima(100); | Metschnikowia | 13 | Saprotroph | (Undefined Saprotroph) | Yeast | NULL | Probable | As nectar-tap_saprotroph (Põlme et al. 2020); Decay sugar-rich substrates (Põlme et al. 2020) | Põlme S, et al. 2020. Fungal Diversity 105:1-16 ((https://doi.org/10.1007/s13225-020-00466-2)) |
| Otu54448 | 1 | k__Fungi(100);p__Ascomycota(100);c__Saccharomycetes(100);o__Saccharomycetales(100);f__Metschnikowiaceae(100);g__Metschnikowia(100);s__Metschnikowia_pulcherrima(100); | Metschnikowia | 13 | Saprotroph | (Undefined Saprotroph) | Yeast | NULL | Probable | As nectar-tap_saprotroph (Põlme et al. 2020); Decay sugar-rich substrates (Põlme et al. 2020) | Põlme S, et al. 2020. Fungal Diversity 105:1-16 ((https://doi.org/10.1007/s13225-020-00466-2)) |
| Otu54449 | 1 | k__Fungi(100);p__Ascomycota(100);c__Saccharomycetes(100);o__Saccharomycetales(100);f__Metschnikowiaceae(100);g__Metschnikowia(100);s__Metschnikowia_pulcherrima(100); | Metschnikowia | 13 | Saprotroph | (Undefined Saprotroph) | Yeast | NULL | Probable | As nectar-tap_saprotroph (Põlme et al. 2020); Decay sugar-rich substrates (Põlme et al. 2020) | Põlme S, et al. 2020. Fungal Diversity 105:1-16 ((https://doi.org/10.1007/s13225-020-00466-2)) |
| Otu54450 | 1 | k__Fungi(100);p__Ascomycota(100);c__Saccharomycetes(100);o__Saccharomycetales(100);f__Metschnikowiaceae(100);g__Metschnikowia(100);s__Metschnikowia_pulcherrima(100); | Metschnikowia | 13 | Saprotroph | (Undefined Saprotroph) | Yeast | NULL | Probable | As nectar-tap_saprotroph (Põlme et al. 2020); Decay sugar-rich substrates (Põlme et al. 2020) | Põlme S, et al. 2020. Fungal Diversity 105:1-16 ((https://doi.org/10.1007/s13225-020-00466-2)) |
| Otu54451 | 1 | k__Fungi(100);p__Ascomycota(100);c__Saccharomycetes(100);o__Saccharomycetales(100);f__Metschnikowiaceae(100);g__Metschnikowia(100);s__Metschnikowia_pulcherrima(100); | Metschnikowia | 13 | Saprotroph | (Undefined Saprotroph) | Yeast | NULL | Probable | As nectar-tap_saprotroph (Põlme et al. 2020); Decay sugar-rich substrates (Põlme et al. 2020) | Põlme S, et al. 2020. Fungal Diversity 105:1-16 ((https://doi.org/10.1007/s13225-020-00466-2)) |
| Otu54452 | 1 | k__Fungi(100);p__Ascomycota(100);c__Saccharomycetes(100);o__Saccharomycetales(100);f__Metschnikowiaceae(100);g__Metschnikowia(100);s__Metschnikowia_pulcherrima(100); | Metschnikowia | 13 | Saprotroph | (Undefined Saprotroph) | Yeast | NULL | Probable | As nectar-tap_saprotroph (Põlme et al. 2020); Decay sugar-rich substrates (Põlme et al. 2020) | Põlme S, et al. 2020. Fungal Diversity 105:1-16 ((https://doi.org/10.1007/s13225-020-00466-2)) |
| Otu54453 | 1 | k__Fungi(100);p__Ascomycota(100);c__Saccharomycetes(100);o__Saccharomycetales(100);f__Metschnikowiaceae(100);g__Metschnikowia(100);s__Metschnikowia_pulcherrima(100); | Metschnikowia | 13 | Saprotroph | (Undefined Saprotroph) | Yeast | NULL | Probable | As nectar-tap_saprotroph (Põlme et al. 2020); Decay sugar-rich substrates (Põlme et al. 2020) | Põlme S, et al. 2020. Fungal Diversity 105:1-16 ((https://doi.org/10.1007/s13225-020-00466-2)) |
| Otu54454 | 1 | k__Fungi(100);p__Ascomycota(100);c__Saccharomycetes(100);o__Saccharomycetales(100);f__Metschnikowiaceae(100);g__Metschnikowia(100);s__Metschnikowia_pulcherrima(100); | Metschnikowia | 13 | Saprotroph | (Undefined Saprotroph) | Yeast | NULL | Probable | As nectar-tap_saprotroph (Põlme et al. 2020); Decay sugar-rich substrates (Põlme et al. 2020) | Põlme S, et al. 2020. Fungal Diversity 105:1-16 ((https://doi.org/10.1007/s13225-020-00466-2)) |
| Otu54455 | 1 | k__Fungi(100);p__Ascomycota(100);c__Saccharomycetes(100);o__Saccharomycetales(100);f__Metschnikowiaceae(100);g__Metschnikowia(100);g__Metschnikowia_unclassified(100); | Metschnikowia | 13 | Saprotroph | (Undefined Saprotroph) | Yeast | NULL | Probable | As nectar-tap_saprotroph (Põlme et al. 2020); Decay sugar-rich substrates (Põlme et al. 2020) | Põlme S, et al. 2020. Fungal Diversity 105:1-16 ((https://doi.org/10.1007/s13225-020-00466-2)) |
| Otu54456 | 1 | k__Fungi(100);p__Ascomycota(100);c__Saccharomycetes(100);o__Saccharomycetales(100);f__Metschnikowiaceae(100);g__Metschnikowia(100);s__Metschnikowia_pulcherrima(100); | Metschnikowia | 13 | Saprotroph | (Undefined Saprotroph) | Yeast | NULL | Probable | As nectar-tap_saprotroph (Põlme et al. 2020); Decay sugar-rich substrates (Põlme et al. 2020) | Põlme S, et al. 2020. Fungal Diversity 105:1-16 ((https://doi.org/10.1007/s13225-020-00466-2)) |
| Otu54457 | 1 | k__Fungi(100);p__Ascomycota(100);c__Saccharomycetes(100);o__Saccharomycetales(100);f__Metschnikowiaceae(100);g__Metschnikowia(100);s__Metschnikowia_pulcherrima(100); | Metschnikowia | 13 | Saprotroph | (Undefined Saprotroph) | Yeast | NULL | Probable | As nectar-tap_saprotroph (Põlme et al. 2020); Decay sugar-rich substrates (Põlme et al. 2020) | Põlme S, et al. 2020. Fungal Diversity 105:1-16 ((https://doi.org/10.1007/s13225-020-00466-2)) |
| Otu54458 | 1 | k__Fungi(100);p__Ascomycota(100);c__Saccharomycetes(100);o__Saccharomycetales(100);f__Metschnikowiaceae(100);g__Metschnikowia(100);s__Metschnikowia_pulcherrima(100); | Metschnikowia | 13 | Saprotroph | (Undefined Saprotroph) | Yeast | NULL | Probable | As nectar-tap_saprotroph (Põlme et al. 2020); Decay sugar-rich substrates (Põlme et al. 2020) | Põlme S, et al. 2020. Fungal Diversity 105:1-16 ((https://doi.org/10.1007/s13225-020-00466-2)) |
| Otu54459 | 1 | k__Fungi(100);p__Ascomycota(100);c__Saccharomycetes(100);o__Saccharomycetales(100);f__Metschnikowiaceae(100);g__Metschnikowia(100);g__Metschnikowia_unclassified(100); | Metschnikowia | 13 | Saprotroph | (Undefined Saprotroph) | Yeast | NULL | Probable | As nectar-tap_saprotroph (Põlme et al. 2020); Decay sugar-rich substrates (Põlme et al. 2020) | Põlme S, et al. 2020. Fungal Diversity 105:1-16 ((https://doi.org/10.1007/s13225-020-00466-2)) |
| Otu54460 | 1 | k__Fungi(100);p__Ascomycota(100);c__Saccharomycetes(100);o__Saccharomycetales(100);f__Metschnikowiaceae(100);g__Metschnikowia(100);s__Metschnikowia_pulcherrima(100); | Metschnikowia | 13 | Saprotroph | (Undefined Saprotroph) | Yeast | NULL | Probable | As nectar-tap_saprotroph (Põlme et al. 2020); Decay sugar-rich substrates (Põlme et al. 2020) | Põlme S, et al. 2020. Fungal Diversity 105:1-16 ((https://doi.org/10.1007/s13225-020-00466-2)) |
| Otu54461 | 1 | k__Fungi(100);p__Ascomycota(100);c__Saccharomycetes(100);o__Saccharomycetales(100);f__Metschnikowiaceae(100);g__Metschnikowia(100);s__Metschnikowia_pulcherrima(100); | Metschnikowia | 13 | Saprotroph | (Undefined Saprotroph) | Yeast | NULL | Probable | As nectar-tap_saprotroph (Põlme et al. 2020); Decay sugar-rich substrates (Põlme et al. 2020) | Põlme S, et al. 2020. Fungal Diversity 105:1-16 ((https://doi.org/10.1007/s13225-020-00466-2)) |
| Otu54462 | 1 | k__Fungi(100);p__Ascomycota(100);c__Saccharomycetes(100);o__Saccharomycetales(100);f__Metschnikowiaceae(100);g__Metschnikowia(100);s__Metschnikowia_pulcherrima(100); | Metschnikowia | 13 | Saprotroph | (Undefined Saprotroph) | Yeast | NULL | Probable | As nectar-tap_saprotroph (Põlme et al. 2020); Decay sugar-rich substrates (Põlme et al. 2020) | Põlme S, et al. 2020. Fungal Diversity 105:1-16 ((https://doi.org/10.1007/s13225-020-00466-2)) |
| Otu54463 | 1 | k__Fungi(100);p__Ascomycota(100);c__Saccharomycetes(100);o__Saccharomycetales(100);f__Metschnikowiaceae(100);g__Metschnikowia(100);s__Metschnikowia_pulcherrima(100); | Metschnikowia | 13 | Saprotroph | (Undefined Saprotroph) | Yeast | NULL | Probable | As nectar-tap_saprotroph (Põlme et al. 2020); Decay sugar-rich substrates (Põlme et al. 2020) | Põlme S, et al. 2020. Fungal Diversity 105:1-16 ((https://doi.org/10.1007/s13225-020-00466-2)) |
| Otu54464 | 1 | k__Fungi(100);p__Ascomycota(100);c__Saccharomycetes(100);o__Saccharomycetales(100);f__Metschnikowiaceae(100);g__Metschnikowia(100);s__Metschnikowia_pulcherrima(100); | Metschnikowia | 13 | Saprotroph | (Undefined Saprotroph) | Yeast | NULL | Probable | As nectar-tap_saprotroph (Põlme et al. 2020); Decay sugar-rich substrates (Põlme et al. 2020) | Põlme S, et al. 2020. Fungal Diversity 105:1-16 ((https://doi.org/10.1007/s13225-020-00466-2)) |

| | | | | | | | | | | | |
|---|---|---|---|---|---|---|---|---|---|---|---|
| Otu54465 | 1 | k__Fungi(100);p__Ascomycota(100);c__Saccharomycetes(100);o__Saccharomycetales(100);f__Metschnikowiaceae(100);g__Metschnikowia(100);s__Metschnikowia_pulcherrima(100); | Metschnikowia | 13 | Saprotroph | \|Undefined Saprotroph\| | Yeast | NULL | Probable | As nectar-tap_saprotroph (Põlme et al. 2020); Decay sugar-rich substrates (Põlme et al. 2020) | Põlme S, et al. 2020. Fungal Diversity 105:1-16 ((https://doi.org/10.1007/s13225-020-00466-2)) |
| Otu54466 | 1 | k__Fungi(100);p__Ascomycota(100);c__Saccharomycetes(100);o__Saccharomycetales(100);f__Metschnikowiaceae(100);g__Metschnikowia(100);s__Metschnikowia_pulcherrima(100); | Metschnikowia | 13 | Saprotroph | \|Undefined Saprotroph\| | Yeast | NULL | Probable | As nectar-tap_saprotroph (Põlme et al. 2020); Decay sugar-rich substrates (Põlme et al. 2020) | Põlme S, et al. 2020. Fungal Diversity 105:1-16 ((https://doi.org/10.1007/s13225-020-00466-2)) |
| Otu54467 | 1 | k__Fungi(100);p__Ascomycota(100);c__Saccharomycetes(100);o__Saccharomycetales(100);f__Metschnikowiaceae(100);g__Metschnikowia(100);s__Metschnikowia_pulcherrima(100); | Metschnikowia | 13 | Saprotroph | \|Undefined Saprotroph\| | Yeast | NULL | Probable | As nectar-tap_saprotroph (Põlme et al. 2020); Decay sugar-rich substrates (Põlme et al. 2020) | Põlme S, et al. 2020. Fungal Diversity 105:1-16 ((https://doi.org/10.1007/s13225-020-00466-2)) |
| Otu54468 | 1 | k__Fungi(100);p__Ascomycota(100);c__Saccharomycetes(100);o__Saccharomycetales(100);f__Metschnikowiaceae(100);g__Metschnikowia(100);g__Metschnikowia_unclassified(100); | Metschnikowia | 13 | Saprotroph | \|Undefined Saprotroph\| | Yeast | NULL | Probable | As nectar-tap_saprotroph (Põlme et al. 2020); Decay sugar-rich substrates (Põlme et al. 2020) | Põlme S, et al. 2020. Fungal Diversity 105:1-16 ((https://doi.org/10.1007/s13225-020-00466-2)) |
| Otu54469 | 1 | k__Fungi(100);p__Ascomycota(100);c__Saccharomycetes(100);o__Saccharomycetales(100);f__Metschnikowiaceae(100);g__Metschnikowia(100);s__Metschnikowia_pulcherrima(100); | Metschnikowia | 13 | Saprotroph | \|Undefined Saprotroph\| | Yeast | NULL | Probable | As nectar-tap_saprotroph (Põlme et al. 2020); Decay sugar-rich substrates (Põlme et al. 2020) | Põlme S, et al. 2020. Fungal Diversity 105:1-16 ((https://doi.org/10.1007/s13225-020-00466-2)) |
| Otu54470 | 1 | k__Fungi(100);p__Ascomycota(100);c__Saccharomycetes(100);o__Saccharomycetales(100);f__Metschnikowiaceae(100);g__Metschnikowia(100);s__Metschnikowia_pulcherrima(100); | Metschnikowia | 13 | Saprotroph | \|Undefined Saprotroph\| | Yeast | NULL | Probable | As nectar-tap_saprotroph (Põlme et al. 2020); Decay sugar-rich substrates (Põlme et al. 2020) | Põlme S, et al. 2020. Fungal Diversity 105:1-16 ((https://doi.org/10.1007/s13225-020-00466-2)) |
| Otu54471 | 1 | k__Fungi(100);p__Ascomycota(100);c__Saccharomycetes(100);o__Saccharomycetales(100);f__Metschnikowiaceae(100);g__Metschnikowia(100);s__Metschnikowia_pulcherrima(100); | Metschnikowia | 13 | Saprotroph | \|Undefined Saprotroph\| | Yeast | NULL | Probable | As nectar-tap_saprotroph (Põlme et al. 2020); Decay sugar-rich substrates (Põlme et al. 2020) | Põlme S, et al. 2020. Fungal Diversity 105:1-16 ((https://doi.org/10.1007/s13225-020-00466-2)) |
| Otu54472 | 1 | k__Fungi(100);p__Ascomycota(100);c__Saccharomycetes(100);o__Saccharomycetales(100);f__Metschnikowiaceae(100);g__Metschnikowia(100);s__Metschnikowia_pulcherrima(100); | Metschnikowia | 13 | Saprotroph | \|Undefined Saprotroph\| | Yeast | NULL | Probable | As nectar-tap_saprotroph (Põlme et al. 2020); Decay sugar-rich substrates (Põlme et al. 2020) | Põlme S, et al. 2020. Fungal Diversity 105:1-16 ((https://doi.org/10.1007/s13225-020-00466-2)) |
| Otu54473 | 1 | k__Fungi(100);p__Ascomycota(100);c__Saccharomycetes(100);o__Saccharomycetales(100);f__Metschnikowiaceae(100);g__Metschnikowia(100);g__Metschnikowia_unclassified(100); | Metschnikowia | 13 | Saprotroph | \|Undefined Saprotroph\| | Yeast | NULL | Probable | As nectar-tap_saprotroph (Põlme et al. 2020); Decay sugar-rich substrates (Põlme et al. 2020) | Põlme S, et al. 2020. Fungal Diversity 105:1-16 ((https://doi.org/10.1007/s13225-020-00466-2)) |
| Otu54474 | 1 | k__Fungi(100);p__Ascomycota(100);c__Saccharomycetes(100);o__Saccharomycetales(100);f__Metschnikowiaceae(100);g__Metschnikowia(100);s__Metschnikowia_pulcherrima(100); | Metschnikowia | 13 | Saprotroph | \|Undefined Saprotroph\| | Yeast | NULL | Probable | As nectar-tap_saprotroph (Põlme et al. 2020); Decay sugar-rich substrates (Põlme et al. 2020) | Põlme S, et al. 2020. Fungal Diversity 105:1-16 ((https://doi.org/10.1007/s13225-020-00466-2)) |
| Otu54476 | 1 | k__Fungi(100);p__Ascomycota(100);c__Saccharomycetes(100);o__Saccharomycetales(100);f__Metschnikowiaceae(100);g__Metschnikowia(100);s__Metschnikowia_pulcherrima(100); | Metschnikowia | 13 | Saprotroph | \|Undefined Saprotroph\| | Yeast | NULL | Probable | As nectar-tap_saprotroph (Põlme et al. 2020); Decay sugar-rich substrates (Põlme et al. 2020) | Põlme S, et al. 2020. Fungal Diversity 105:1-16 ((https://doi.org/10.1007/s13225-020-00466-2)) |
| Otu54477 | 1 | k__Fungi(100);p__Ascomycota(100);c__Saccharomycetes(100);o__Saccharomycetales(100);f__Metschnikowiaceae(100);g__Metschnikowia(100);s__Metschnikowia_pulcherrima(100); | Metschnikowia | 13 | Saprotroph | \|Undefined Saprotroph\| | Yeast | NULL | Probable | As nectar-tap_saprotroph (Põlme et al. 2020); Decay sugar-rich substrates (Põlme et al. 2020) | Põlme S, et al. 2020. Fungal Diversity 105:1-16 ((https://doi.org/10.1007/s13225-020-00466-2)) |
| Otu54478 | 1 | k__Fungi(100);p__Ascomycota(100);c__Saccharomycetes(100);o__Saccharomycetales(100);f__Metschnikowiaceae(100);g__Metschnikowia(100);s__Metschnikowia_pulcherrima(100); | Metschnikowia | 13 | Saprotroph | \|Undefined Saprotroph\| | Yeast | NULL | Probable | As nectar-tap_saprotroph (Põlme et al. 2020); Decay sugar-rich substrates (Põlme et al. 2020) | Põlme S, et al. 2020. Fungal Diversity 105:1-16 ((https://doi.org/10.1007/s13225-020-00466-2)) |
| Otu54479 | 1 | k__Fungi(100);p__Ascomycota(100);c__Saccharomycetes(100);o__Saccharomycetales(100);f__Metschnikowiaceae(100);g__Metschnikowia(100);s__Metschnikowia_pulcherrima(100); | Metschnikowia | 13 | Saprotroph | \|Undefined Saprotroph\| | Yeast | NULL | Probable | As nectar-tap_saprotroph (Põlme et al. 2020); Decay sugar-rich substrates (Põlme et al. 2020) | Põlme S, et al. 2020. Fungal Diversity 105:1-16 ((https://doi.org/10.1007/s13225-020-00466-2)) |
| Otu54480 | 1 | k__Fungi(100);p__Ascomycota(100);c__Saccharomycetes(100);o__Saccharomycetales(100);f__Metschnikowiaceae(100);g__Metschnikowia(100);s__Metschnikowia_pulcherrima(100); | Metschnikowia | 13 | Saprotroph | \|Undefined Saprotroph\| | Yeast | NULL | Probable | As nectar-tap_saprotroph (Põlme et al. 2020); Decay sugar-rich substrates (Põlme et al. 2020) | Põlme S, et al. 2020. Fungal Diversity 105:1-16 ((https://doi.org/10.1007/s13225-020-00466-2)) |
| Otu54481 | 1 | k__Fungi(100);p__Ascomycota(100);c__Saccharomycetes(100);o__Saccharomycetales(100);f__Metschnikowiaceae(100);g__Metschnikowia(100);s__Metschnikowia_pulcherrima(100); | Metschnikowia | 13 | Saprotroph | \|Undefined Saprotroph\| | Yeast | NULL | Probable | As nectar-tap_saprotroph (Põlme et al. 2020); Decay sugar-rich substrates (Põlme et al. 2020) | Põlme S, et al. 2020. Fungal Diversity 105:1-16 ((https://doi.org/10.1007/s13225-020-00466-2)) |
| Otu54482 | 1 | k__Fungi(100);p__Ascomycota(100);c__Saccharomycetes(100);o__Saccharomycetales(100);f__Metschnikowiaceae(100);g__Metschnikowia(100);s__Metschnikowia_pulcherrima(100); | Metschnikowia | 13 | Saprotroph | \|Undefined Saprotroph\| | Yeast | NULL | Probable | As nectar-tap_saprotroph (Põlme et al. 2020); Decay sugar-rich substrates (Põlme et al. 2020) | Põlme S, et al. 2020. Fungal Diversity 105:1-16 ((https://doi.org/10.1007/s13225-020-00466-2)) |
| Otu54483 | 1 | k__Fungi(100);p__Ascomycota(100);c__Saccharomycetes(100);o__Saccharomycetales(100);f__Metschnikowiaceae(100);g__Metschnikowia(100);s__Metschnikowia_pulcherrima(100); | Metschnikowia | 13 | Saprotroph | \|Undefined Saprotroph\| | Yeast | NULL | Probable | As nectar-tap_saprotroph (Põlme et al. 2020); Decay sugar-rich substrates (Põlme et al. 2020) | Põlme S, et al. 2020. Fungal Diversity 105:1-16 ((https://doi.org/10.1007/s13225-020-00466-2)) |
| Otu54484 | 1 | k__Fungi(100);p__Ascomycota(100);c__Saccharomycetes(100);o__Saccharomycetales(100);f__Metschnikowiaceae(100);g__Metschnikowia(100);s__Metschnikowia_pulcherrima(100); | Metschnikowia | 13 | Saprotroph | \|Undefined Saprotroph\| | Yeast | NULL | Probable | As nectar-tap_saprotroph (Põlme et al. 2020); Decay sugar-rich substrates (Põlme et al. 2020) | Põlme S, et al. 2020. Fungal Diversity 105:1-16 ((https://doi.org/10.1007/s13225-020-00466-2)) |
| Otu54485 | 1 | k__Fungi(100);p__Ascomycota(100);c__Saccharomycetes(100);o__Saccharomycetales(100);f__Metschnikowiaceae(100);g__Metschnikowia(100);s__Metschnikowia_pulcherrima(100); | Metschnikowia | 13 | Saprotroph | \|Undefined Saprotroph\| | Yeast | NULL | Probable | As nectar-tap_saprotroph (Põlme et al. 2020); Decay sugar-rich substrates (Põlme et al. 2020) | Põlme S, et al. 2020. Fungal Diversity 105:1-16 ((https://doi.org/10.1007/s13225-020-00466-2)) |
| Otu54486 | 1 | k__Fungi(100);p__Ascomycota(100);c__Saccharomycetes(100);o__Saccharomycetales(100);f__Metschnikowiaceae(100);g__Metschnikowia(100);g__Metschnikowia_unclassified(100); | Metschnikowia | 13 | Saprotroph | \|Undefined Saprotroph\| | Yeast | NULL | Probable | As nectar-tap_saprotroph (Põlme et al. 2020); Decay sugar-rich substrates (Põlme et al. 2020) | Põlme S, et al. 2020. Fungal Diversity 105:1-16 ((https://doi.org/10.1007/s13225-020-00466-2)) |
| Otu54487 | 1 | k__Fungi(100);p__Ascomycota(100);c__Saccharomycetes(100);o__Saccharomycetales(100);f__Metschnikowiaceae(100);g__Metschnikowia(100);g__Metschnikowia_unclassified(100); | Metschnikowia | 13 | Saprotroph | \|Undefined Saprotroph\| | Yeast | NULL | Probable | As nectar-tap_saprotroph (Põlme et al. 2020); Decay sugar-rich substrates (Põlme et al. 2020) | Põlme S, et al. 2020. Fungal Diversity 105:1-16 ((https://doi.org/10.1007/s13225-020-00466-2)) |
| Otu54488 | 1 | k__Fungi(100);p__Ascomycota(100);c__Saccharomycetes(100);o__Saccharomycetales(100);f__Metschnikowiaceae(100);g__Metschnikowia(100);s__Metschnikowia_pulcherrima(100); | Metschnikowia | 13 | Saprotroph | \|Undefined Saprotroph\| | Yeast | NULL | Probable | As nectar-tap_saprotroph (Põlme et al. 2020); Decay sugar-rich substrates (Põlme et al. 2020) | Põlme S, et al. 2020. Fungal Diversity 105:1-16 ((https://doi.org/10.1007/s13225-020-00466-2)) |
| Otu54489 | 1 | k__Fungi(100);p__Ascomycota(100);c__Saccharomycetes(100);o__Saccharomycetales(100);f__Metschnikowiaceae(100);g__Metschnikowia(100);g__Metschnikowia_unclassified(100); | Metschnikowia | 13 | Saprotroph | \|Undefined Saprotroph\| | Yeast | NULL | Probable | As nectar-tap_saprotroph (Põlme et al. 2020); Decay sugar-rich substrates (Põlme et al. 2020) | Põlme S, et al. 2020. Fungal Diversity 105:1-16 ((https://doi.org/10.1007/s13225-020-00466-2)) |
| Otu54490 | 1 | k__Fungi(100);p__Ascomycota(100);c__Saccharomycetes(100);o__Saccharomycetales(100);f__Metschnikowiaceae(100);g__Metschnikowia(100);s__Metschnikowia_pulcherrima(100); | Metschnikowia | 13 | Saprotroph | \|Undefined Saprotroph\| | Yeast | NULL | Probable | As nectar-tap_saprotroph (Põlme et al. 2020); Decay sugar-rich substrates (Põlme et al. 2020) | Põlme S, et al. 2020. Fungal Diversity 105:1-16 ((https://doi.org/10.1007/s13225-020-00466-2)) |
| Otu54491 | 1 | k__Fungi(100);p__Ascomycota(100);c__Saccharomycetes(100);o__Saccharomycetales(100);f__Metschnikowiaceae(100);g__Metschnikowia(100);s__Metschnikowia_pulcherrima(100); | Metschnikowia | 13 | Saprotroph | \|Undefined Saprotroph\| | Yeast | NULL | Probable | As nectar-tap_saprotroph (Põlme et al. 2020); Decay sugar-rich substrates (Põlme et al. 2020) | Põlme S, et al. 2020. Fungal Diversity 105:1-16 ((https://doi.org/10.1007/s13225-020-00466-2)) |
| Otu54492 | 1 | k__Fungi(100);p__Ascomycota(100);c__Saccharomycetes(100);o__Saccharomycetales(100);f__Metschnikowiaceae(100);g__Metschnikowia(100);s__Metschnikowia_pulcherrima(100); | Metschnikowia | 13 | Saprotroph | \|Undefined Saprotroph\| | Yeast | NULL | Probable | As nectar-tap_saprotroph (Põlme et al. 2020); Decay sugar-rich substrates (Põlme et al. 2020) | Põlme S, et al. 2020. Fungal Diversity 105:1-16 ((https://doi.org/10.1007/s13225-020-00466-2)) |
| Otu54493 | 1 | k__Fungi(100);p__Ascomycota(100);c__Saccharomycetes(100);o__Saccharomycetales(100);f__Metschnikowiaceae(100);g__Metschnikowia(100);s__Metschnikowia_pulcherrima(100); | Metschnikowia | 13 | Saprotroph | \|Undefined Saprotroph\| | Yeast | NULL | Probable | As nectar-tap_saprotroph (Põlme et al. 2020); Decay sugar-rich substrates (Põlme et al. 2020) | Põlme S, et al. 2020. Fungal Diversity 105:1-16 ((https://doi.org/10.1007/s13225-020-00466-2)) |
| Otu54494 | 1 | k__Fungi(100);p__Ascomycota(100);c__Saccharomycetes(100);o__Saccharomycetales(100);f__Metschnikowiaceae(100);g__Metschnikowia(100);s__Metschnikowia_pulcherrima(100); | Metschnikowia | 13 | Saprotroph | \|Undefined Saprotroph\| | Yeast | NULL | Probable | As nectar-tap_saprotroph (Põlme et al. 2020); Decay sugar-rich substrates (Põlme et al. 2020) | Põlme S, et al. 2020. Fungal Diversity 105:1-16 ((https://doi.org/10.1007/s13225-020-00466-2)) |
| Otu54495 | 1 | k__Fungi(100);p__Ascomycota(100);c__Saccharomycetes(100);o__Saccharomycetales(100);f__Metschnikowiaceae(100);g__Metschnikowia(100);g__Metschnikowia_unclassified(100); | Metschnikowia | 13 | Saprotroph | \|Undefined Saprotroph\| | Yeast | NULL | Probable | As nectar-tap_saprotroph (Põlme et al. 2020); Decay sugar-rich substrates (Põlme et al. 2020) | Põlme S, et al. 2020. Fungal Diversity 105:1-16 ((https://doi.org/10.1007/s13225-020-00466-2)) |
| Otu54496 | 1 | k__Fungi(100);p__Ascomycota(100);c__Saccharomycetes(100);o__Saccharomycetales(100);f__Metschnikowiaceae(100);g__Metschnikowia(100);s__Metschnikowia_pulcherrima(100); | Metschnikowia | 13 | Saprotroph | \|Undefined Saprotroph\| | Yeast | NULL | Probable | As nectar-tap_saprotroph (Põlme et al. 2020); Decay sugar-rich substrates (Põlme et al. 2020) | Põlme S, et al. 2020. Fungal Diversity 105:1-16 ((https://doi.org/10.1007/s13225-020-00466-2)) |
| Otu54497 | 1 | k__Fungi(100);p__Ascomycota(100);c__Saccharomycetes(100);o__Saccharomycetales(100);f__Metschnikowiaceae(100);g__Metschnikowia(100);s__Metschnikowia_pulcherrima(100); | Metschnikowia | 13 | Saprotroph | \|Undefined Saprotroph\| | Yeast | NULL | Probable | As nectar-tap_saprotroph (Põlme et al. 2020); Decay sugar-rich substrates (Põlme et al. 2020) | Põlme S, et al. 2020. Fungal Diversity 105:1-16 ((https://doi.org/10.1007/s13225-020-00466-2)) |
| Otu54498 | 1 | k__Fungi(100);p__Ascomycota(100);c__Saccharomycetes(100);o__Saccharomycetales(100);f__Metschnikowiaceae(100);g__Metschnikowia(100);s__Metschnikowia_pulcherrima(100); | Metschnikowia | 13 | Saprotroph | \|Undefined Saprotroph\| | Yeast | NULL | Probable | As nectar-tap_saprotroph (Põlme et al. 2020); Decay sugar-rich substrates (Põlme et al. 2020) | Põlme S, et al. 2020. Fungal Diversity 105:1-16 ((https://doi.org/10.1007/s13225-020-00466-2)) |
| Otu54499 | 1 | k__Fungi(100);p__Ascomycota(100);c__Saccharomycetes(100);o__Saccharomycetales(100);f__Metschnikowiaceae(100);g__Metschnikowia(100);s__Metschnikowia_pulcherrima(100); | Metschnikowia | 13 | Saprotroph | \|Undefined Saprotroph\| | Yeast | NULL | Probable | As nectar-tap_saprotroph (Põlme et al. 2020); Decay sugar-rich substrates (Põlme et al. 2020) | Põlme S, et al. 2020. Fungal Diversity 105:1-16 ((https://doi.org/10.1007/s13225-020-00466-2)) |
| Otu54500 | 1 | k__Fungi(100);p__Ascomycota(100);c__Saccharomycetes(100);o__Saccharomycetales(100);f__Metschnikowiaceae(100);g__Metschnikowia(100);s__Metschnikowia_pulcherrima(100); | Metschnikowia | 13 | Saprotroph | \|Undefined Saprotroph\| | Yeast | NULL | Probable | As nectar-tap_saprotroph (Põlme et al. 2020); Decay sugar-rich substrates (Põlme et al. 2020) | Põlme S, et al. 2020. Fungal Diversity 105:1-16 ((https://doi.org/10.1007/s13225-020-00466-2)) |
| Otu54501 | 1 | k__Fungi(100);p__Ascomycota(100);c__Saccharomycetes(100);o__Saccharomycetales(100);f__Metschnikowiaceae(100);g__Metschnikowia(100);s__Metschnikowia_pulcherrima(100); | Metschnikowia | 13 | Saprotroph | \|Undefined Saprotroph\| | Yeast | NULL | Probable | As nectar-tap_saprotroph (Põlme et al. 2020); Decay sugar-rich substrates (Põlme et al. 2020) | Põlme S, et al. 2020. Fungal Diversity 105:1-16 ((https://doi.org/10.1007/s13225-020-00466-2)) |
| Otu54502 | 1 | k__Fungi(100);p__Ascomycota(100);c__Saccharomycetes(100);o__Saccharomycetales(100);f__Metschnikowiaceae(100);g__Metschnikowia(100);g__Metschnikowia_unclassified(100); | Metschnikowia | 13 | Saprotroph | \|Undefined Saprotroph\| | Yeast | NULL | Probable | As nectar-tap_saprotroph (Põlme et al. 2020); Decay sugar-rich substrates (Põlme et al. 2020) | Põlme S, et al. 2020. Fungal Diversity 105:1-16 ((https://doi.org/10.1007/s13225-020-00466-2)) |
| Otu54503 | 1 | k__Fungi(100);p__Ascomycota(100);c__Saccharomycetes(100);o__Saccharomycetales(100);f__Metschnikowiaceae(100);g__Metschnikowia(100);s__Metschnikowia_pulcherrima(100); | Metschnikowia | 13 | Saprotroph | \|Undefined Saprotroph\| | Yeast | NULL | Probable | As nectar-tap_saprotroph (Põlme et al. 2020); Decay sugar-rich substrates (Põlme et al. 2020) | Põlme S, et al. 2020. Fungal Diversity 105:1-16 ((https://doi.org/10.1007/s13225-020-00466-2)) |
| Otu54504 | 1 | k__Fungi(100);p__Ascomycota(100);c__Saccharomycetes(100);o__Saccharomycetales(100);f__Metschnikowiaceae(100);g__Metschnikowia(100);s__Metschnikowia_pulcherrima(100); | Metschnikowia | 13 | Saprotroph | \|Undefined Saprotroph\| | Yeast | NULL | Probable | As nectar-tap_saprotroph (Põlme et al. 2020); Decay sugar-rich substrates (Põlme et al. 2020) | Põlme S, et al. 2020. Fungal Diversity 105:1-16 ((https://doi.org/10.1007/s13225-020-00466-2)) |
| Otu54505 | 1 | k__Fungi(100);p__Ascomycota(100);c__Saccharomycetes(100);o__Saccharomycetales(100);f__Metschnikowiaceae(100);g__Metschnikowia(100);s__Metschnikowia_pulcherrima(100); | Metschnikowia | 13 | Saprotroph | \|Undefined Saprotroph\| | Yeast | NULL | Probable | As nectar-tap_saprotroph (Põlme et al. 2020); Decay sugar-rich substrates (Põlme et al. 2020) | Põlme S, et al. 2020. Fungal Diversity 105:1-16 ((https://doi.org/10.1007/s13225-020-00466-2)) |
| Otu54506 | 1 | k__Fungi(100);p__Ascomycota(100);c__Saccharomycetes(100);o__Saccharomycetales(100);f__Metschnikowiaceae(100);g__Metschnikowia(100);s__Metschnikowia_pulcherrima(100); | Metschnikowia | 13 | Saprotroph | \|Undefined Saprotroph\| | Yeast | NULL | Probable | As nectar-tap_saprotroph (Põlme et al. 2020); Decay sugar-rich substrates (Põlme et al. 2020) | Põlme S, et al. 2020. Fungal Diversity 105:1-16 ((https://doi.org/10.1007/s13225-020-00466-2)) |
| Otu54507 | 1 | k__Fungi(100);p__Ascomycota(100);c__Saccharomycetes(100);o__Saccharomycetales(100);f__Metschnikowiaceae(100);g__Metschnikowia(100);g__Metschnikowia_unclassified(100); | Metschnikowia | 13 | Saprotroph | \|Undefined Saprotroph\| | Yeast | NULL | Probable | As nectar-tap_saprotroph (Põlme et al. 2020); Decay sugar-rich substrates (Põlme et al. 2020) | Põlme S, et al. 2020. Fungal Diversity 105:1-16 ((https://doi.org/10.1007/s13225-020-00466-2)) |
| Otu54508 | 1 | k__Fungi(100);p__Ascomycota(100);c__Saccharomycetes(100);o__Saccharomycetales(100);f__Metschnikowiaceae(100);g__Metschnikowia(100);s__Metschnikowia_pulcherrima(100); | Metschnikowia | 13 | Saprotroph | \|Undefined Saprotroph\| | Yeast | NULL | Probable | As nectar-tap_saprotroph (Põlme et al. 2020); Decay sugar-rich substrates (Põlme et al. 2020) | Põlme S, et al. 2020. Fungal Diversity 105:1-16 ((https://doi.org/10.1007/s13225-020-00466-2)) |
| Otu54509 | 1 | k__Fungi(100);p__Ascomycota(100);c__Saccharomycetes(100);o__Saccharomycetales(100);f__Metschnikowiaceae(100);g__Metschnikowia(100);g__Metschnikowia_unclassified(100); | Metschnikowia | 13 | Saprotroph | \|Undefined Saprotroph\| | Yeast | NULL | Probable | As nectar-tap_saprotroph (Põlme et al. 2020); Decay sugar-rich substrates (Põlme et al. 2020) | Põlme S, et al. 2020. Fungal Diversity 105:1-16 ((https://doi.org/10.1007/s13225-020-00466-2)) |
| Otu54510 | 1 | k__Fungi(100);p__Ascomycota(100);c__Saccharomycetes(100);o__Saccharomycetales(100);f__Metschnikowiaceae(100);g__Metschnikowia(100);s__Metschnikowia_pulcherrima(100); | Metschnikowia | 13 | Saprotroph | \|Undefined Saprotroph\| | Yeast | NULL | Probable | As nectar-tap_saprotroph (Põlme et al. 2020); Decay sugar-rich substrates (Põlme et al. 2020) | Põlme S, et al. 2020. Fungal Diversity 105:1-16 ((https://doi.org/10.1007/s13225-020-00466-2)) |
| Otu54511 | 1 | k__Fungi(100);p__Ascomycota(100);c__Saccharomycetes(100);o__Saccharomycetales(100);f__Metschnikowiaceae(100);g__Metschnikowia(100);g__Metschnikowia_unclassified(100); | Metschnikowia | 13 | Saprotroph | \|Undefined Saprotroph\| | Yeast | NULL | Probable | As nectar-tap_saprotroph (Põlme et al. 2020); Decay sugar-rich substrates (Põlme et al. 2020) | Põlme S, et al. 2020. Fungal Diversity 105:1-16 ((https://doi.org/10.1007/s13225-020-00466-2)) |
| Otu54512 | 1 | k__Fungi(100);p__Ascomycota(100);c__Saccharomycetes(100);o__Saccharomycetales(100);f__Metschnikowiaceae(100);g__Metschnikowia(100);s__Metschnikowia_pulcherrima(100); | Metschnikowia | 13 | Saprotroph | \|Undefined Saprotroph\| | Yeast | NULL | Probable | As nectar-tap_saprotroph (Põlme et al. 2020); Decay sugar-rich substrates (Põlme et al. 2020) | Põlme S, et al. 2020. Fungal Diversity 105:1-16 ((https://doi.org/10.1007/s13225-020-00466-2)) |
| Otu54513 | 1 | k__Fungi(100);p__Ascomycota(100);c__Saccharomycetes(100);o__Saccharomycetales(100);f__Metschnikowiaceae(100);g__Metschnikowia(100);g__Metschnikowia_unclassified(100); | Metschnikowia | 13 | Saprotroph | \|Undefined Saprotroph\| | Yeast | NULL | Probable | As nectar-tap_saprotroph (Põlme et al. 2020); Decay sugar-rich substrates (Põlme et al. 2020) | Põlme S, et al. 2020. Fungal Diversity 105:1-16 ((https://doi.org/10.1007/s13225-020-00466-2)) |
| Otu54514 | 1 | k__Fungi(100);p__Ascomycota(100);c__Saccharomycetes(100);o__Saccharomycetales(100);f__Metschnikowiaceae(100);g__Metschnikowia(100);s__Metschnikowia_pulcherrima(100); | Metschnikowia | 13 | Saprotroph | \|Undefined Saprotroph\| | Yeast | NULL | Probable | As nectar-tap_saprotroph (Põlme et al. 2020); Decay sugar-rich substrates (Põlme et al. 2020) | Põlme S, et al. 2020. Fungal Diversity 105:1-16 ((https://doi.org/10.1007/s13225-020-00466-2)) |
| Otu54515 | 1 | k__Fungi(100);p__Ascomycota(100);c__Saccharomycetes(100);o__Saccharomycetales(100);f__Metschnikowiaceae(100);g__Metschnikowia(100);g__Metschnikowia_unclassified(100); | Metschnikowia | 13 | Saprotroph | \|Undefined Saprotroph\| | Yeast | NULL | Probable | As nectar-tap_saprotroph (Põlme et al. 2020); Decay sugar-rich substrates (Põlme et al. 2020) | Põlme S, et al. 2020. Fungal Diversity 105:1-16 ((https://doi.org/10.1007/s13225-020-00466-2)) |
| Otu54516 | 1 | k__Fungi(100);p__Ascomycota(100);c__Saccharomycetes(100);o__Saccharomycetales(100);f__Metschnikowiaceae(100);g__Metschnikowia(100);s__Metschnikowia_pulcherrima(100); | Metschnikowia | 13 | Saprotroph | \|Undefined Saprotroph\| | Yeast | NULL | Probable | As nectar-tap_saprotroph (Põlme et al. 2020); Decay sugar-rich substrates (Põlme et al. 2020) | Põlme S, et al. 2020. Fungal Diversity 105:1-16 ((https://doi.org/10.1007/s13225-020-00466-2)) |
| Otu54517 | 1 | k__Fungi(100);p__Ascomycota(100);c__Saccharomycetes(100);o__Saccharomycetales(100);f__Metschnikowiaceae(100);g__Metschnikowia(100);s__Metschnikowia_pulcherrima(100); | Metschnikowia | 13 | Saprotroph | \|Undefined Saprotroph\| | Yeast | NULL | Probable | As nectar-tap_saprotroph (Põlme et al. 2020); Decay sugar-rich substrates (Põlme et al. 2020) | Põlme S, et al. 2020. Fungal Diversity 105:1-16 ((https://doi.org/10.1007/s13225-020-00466-2)) |
| Otu54518 | 1 | k__Fungi(100);p__Ascomycota(100);c__Saccharomycetes(100);o__Saccharomycetales(100);f__Metschnikowiaceae(100);g__Metschnikowia(100);s__Metschnikowia_pulcherrima(100); | Metschnikowia | 13 | Saprotroph | \|Undefined Saprotroph\| | Yeast | NULL | Probable | As nectar-tap_saprotroph (Põlme et al. 2020); Decay sugar-rich substrates (Põlme et al. 2020) | Põlme S, et al. 2020. Fungal Diversity 105:1-16 ((https://doi.org/10.1007/s13225-020-00466-2)) |
| Otu54519 | 1 | k__Fungi(100);p__Ascomycota(100);c__Saccharomycetes(100);o__Saccharomycetales(100);f__Metschnikowiaceae(100);g__Metschnikowia(100);g__Metschnikowia_unclassified(100); | Metschnikowia | 13 | Saprotroph | \|Undefined Saprotroph\| | Yeast | NULL | Probable | As nectar-tap_saprotroph (Põlme et al. 2020); Decay sugar-rich substrates (Põlme et al. 2020) | Põlme S, et al. 2020. Fungal Diversity 105:1-16 ((https://doi.org/10.1007/s13225-020-00466-2)) |
| Otu54520 | 1 | k__Fungi(100);p__Ascomycota(100);c__Saccharomycetes(100);o__Saccharomycetales(100);f__Metschnikowiaceae(100);g__Metschnikowia(100);s__Metschnikowia_pulcherrima(100); | Metschnikowia | 13 | Saprotroph | \|Undefined Saprotroph\| | Yeast | NULL | Probable | As nectar-tap_saprotroph (Põlme et al. 2020); Decay sugar-rich substrates (Põlme et al. 2020) | Põlme S, et al. 2020. Fungal Diversity 105:1-16 ((https://doi.org/10.1007/s13225-020-00466-2)) |
| Otu54521 | 1 | k__Fungi(100);p__Ascomycota(100);c__Saccharomycetes(100);o__Saccharomycetales(100);f__Metschnikowiaceae(100);g__Metschnikowia(100);s__Metschnikowia_pulcherrima(100); | Metschnikowia | 13 | Saprotroph | \|Undefined Saprotroph\| | Yeast | NULL | Probable | As nectar-tap_saprotroph (Põlme et al. 2020); Decay sugar-rich substrates (Põlme et al. 2020) | Põlme S, et al. 2020. Fungal Diversity 105:1-16 ((https://doi.org/10.1007/s13225-020-00466-2)) |
| Otu54522 | 1 | k__Fungi(100);p__Ascomycota(100);c__Saccharomycetes(100);o__Saccharomycetales(100);f__Metschnikowiaceae(100);g__Metschnikowia(100);g__Metschnikowia_unclassified(100); | Metschnikowia | 13 | Saprotroph | \|Undefined Saprotroph\| | Yeast | NULL | Probable | As nectar-tap_saprotroph (Põlme et al. 2020); Decay sugar-rich substrates (Põlme et al. 2020) | Põlme S, et al. 2020. Fungal Diversity 105:1-16 ((https://doi.org/10.1007/s13225-020-00466-2)) |
| Otu54523 | 1 | k__Fungi(100);p__Ascomycota(100);c__Saccharomycetes(100);o__Saccharomycetales(100);f__Metschnikowiaceae(100);g__Metschnikowia(100);s__Metschnikowia_pulcherrima(100); | Metschnikowia | 13 | Saprotroph | \|Undefined Saprotroph\| | Yeast | NULL | Probable | As nectar-tap_saprotroph (Põlme et al. 2020); Decay sugar-rich substrates (Põlme et al. 2020) | Põlme S, et al. 2020. Fungal Diversity 105:1-16 ((https://doi.org/10.1007/s13225-020-00466-2)) |
| Otu54524 | 1 | k__Fungi(100);p__Ascomycota(100);c__Saccharomycetes(100);o__Saccharomycetales(100);f__Metschnikowiaceae(100);g__Metschnikowia(100);s__Metschnikowia_pulcherrima(100); | Metschnikowia | 13 | Saprotroph | \|Undefined Saprotroph\| | Yeast | NULL | Probable | As nectar-tap_saprotroph (Põlme et al. 2020); Decay sugar-rich substrates (Põlme et al. 2020) | Põlme S, et al. 2020. Fungal Diversity 105:1-16 ((https://doi.org/10.1007/s13225-020-00466-2)) |
| Otu54525 | 1 | k__Fungi(100);p__Ascomycota(100);c__Saccharomycetes(100);o__Saccharomycetales(100);f__Metschnikowiaceae(100);g__Metschnikowia(100);s__Metschnikowia_pulcherrima(100); | Metschnikowia | 13 | Saprotroph | \|Undefined Saprotroph\| | Yeast | NULL | Probable | As nectar-tap_saprotroph (Põlme et al. 2020); Decay sugar-rich substrates (Põlme et al. 2020) | Põlme S, et al. 2020. Fungal Diversity 105:1-16 ((https://doi.org/10.1007/s13225-020-00466-2)) |
| Otu54526 | 1 | k__Fungi(100);p__Ascomycota(100);c__Saccharomycetes(100);o__Saccharomycetales(100);f__Metschnikowiaceae(100);g__Metschnikowia(100);s__Metschnikowia_pulcherrima(100); | Metschnikowia | 13 | Saprotroph | \|Undefined Saprotroph\| | Yeast | NULL | Probable | As nectar-tap_saprotroph (Põlme et al. 2020); Decay sugar-rich substrates (Põlme et al. 2020) | Põlme S, et al. 2020. Fungal Diversity 105:1-16 ((https://doi.org/10.1007/s13225-020-00466-2)) |
| Otu54527 | 1 | k__Fungi(100);p__Ascomycota(100);c__Saccharomycetes(100);o__Saccharomycetales(100);f__Metschnikowiaceae(100);g__Metschnikowia(100);s__Metschnikowia_pulcherrima(100); | Metschnikowia | 13 | Saprotroph | \|Undefined Saprotroph\| | Yeast | NULL | Probable | As nectar-tap_saprotroph (Põlme et al. 2020); Decay sugar-rich substrates (Põlme et al. 2020) | Põlme S, et al. 2020. Fungal Diversity 105:1-16 ((https://doi.org/10.1007/s13225-020-00466-2)) |
| Otu54528 | 1 | k__Fungi(100);p__Ascomycota(100);c__Saccharomycetes(100);o__Saccharomycetales(100);f__Metschnikowiaceae(100);g__Metschnikowia(100);s__Metschnikowia_pulcherrima(100); | Metschnikowia | 13 | Saprotroph | \|Undefined Saprotroph\| | Yeast | NULL | Probable | As nectar-tap_saprotroph (Põlme et al. 2020); Decay sugar-rich substrates (Põlme et al. 2020) | Põlme S, et al. 2020. Fungal Diversity 105:1-16 ((https://doi.org/10.1007/s13225-020-00466-2)) |
| Otu54529 | 1 | k__Fungi(100);p__Ascomycota(100);c__Saccharomycetes(100);o__Saccharomycetales(100);f__Metschnikowiaceae(100);g__Metschnikowia(100);s__Metschnikowia_pulcherrima(100); | Metschnikowia | 13 | Saprotroph | \|Undefined Saprotroph\| | Yeast | NULL | Probable | As nectar-tap_saprotroph (Põlme et al. 2020); Decay sugar-rich substrates (Põlme et al. 2020) | Põlme S, et al. 2020. Fungal Diversity 105:1-16 ((https://doi.org/10.1007/s13225-020-00466-2)) |
| Otu54530 | 1 | k__Fungi(100);p__Ascomycota(100);c__Saccharomycetes(100);o__Saccharomycetales(100);f__Metschnikowiaceae(100);g__Metschnikowia(100);s__Metschnikowia_pulcherrima(100); | Metschnikowia | 13 | Saprotroph | \|Undefined Saprotroph\| | Yeast | NULL | Probable | As nectar-tap_saprotroph (Põlme et al. 2020); Decay sugar-rich substrates (Põlme et al. 2020) | Põlme S, et al. 2020. Fungal Diversity 105:1-16 ((https://doi.org/10.1007/s13225-020-00466-2)) |
| Otu54531 | 1 | k__Fungi(100);p__Ascomycota(100);c__Saccharomycetes(100);o__Saccharomycetales(100);f__Metschnikowiaceae(100);g__Metschnikowia(100);s__Metschnikowia_pulcherrima(100); | Metschnikowia | 13 | Saprotroph | \|Undefined Saprotroph\| | Yeast | NULL | Probable | As nectar-tap_saprotroph (Põlme et al. 2020); Decay sugar-rich substrates (Põlme et al. 2020) | Põlme S, et al. 2020. Fungal Diversity 105:1-16 ((https://doi.org/10.1007/s13225-020-00466-2)) |
| Otu54532 | 1 | k__Fungi(100);p__Ascomycota(100);c__Saccharomycetes(100);o__Saccharomycetales(100);f__Metschnikowiaceae(100);g__Metschnikowia(100);s__Metschnikowia_pulcherrima(100); | Metschnikowia | 13 | Saprotroph | \|Undefined Saprotroph\| | Yeast | NULL | Probable | As nectar-tap_saprotroph (Põlme et al. 2020); Decay sugar-rich substrates (Põlme et al. 2020) | Põlme S, et al. 2020. Fungal Diversity 105:1-16 ((https://doi.org/10.1007/s13225-020-00466-2)) |
| Otu54533 | 1 | k__Fungi(100);p__Ascomycota(100);c__Saccharomycetes(100);o__Saccharomycetales(100);f__Metschnikowiaceae(100);g__Metschnikowia(100);s__Metschnikowia_pulcherrima(100); | Metschnikowia | 13 | Saprotroph | \|Undefined Saprotroph\| | Yeast | NULL | Probable | As nectar-tap_saprotroph (Põlme et al. 2020); Decay sugar-rich substrates (Põlme et al. 2020) | Põlme S, et al. 2020. Fungal Diversity 105:1-16 ((https://doi.org/10.1007/s13225-020-00466-2)) |
| Otu54534 | 1 | k__Fungi(100);p__Ascomycota(100);c__Saccharomycetes(100);o__Saccharomycetales(100);f__Metschnikowiaceae(100);g__Metschnikowia(100);s__Metschnikowia_pulcherrima(100); | Metschnikowia | 13 | Saprotroph | \|Undefined Saprotroph\| | Yeast | NULL | Probable | As nectar-tap_saprotroph (Põlme et al. 2020); Decay sugar-rich substrates (Põlme et al. 2020) | Põlme S, et al. 2020. Fungal Diversity 105:1-16 ((https://doi.org/10.1007/s13225-020-00466-2)) |
| Otu54535 | 1 | k__Fungi(100);p__Ascomycota(100);c__Saccharomycetes(100);o__Saccharomycetales(100);f__Metschnikowiaceae(100);g__Metschnikowia(100);s__Metschnikowia_pulcherrima(100); | Metschnikowia | 13 | Saprotroph | \|Undefined Saprotroph\| | Yeast | NULL | Probable | As nectar-tap_saprotroph (Põlme et al. 2020); Decay sugar-rich substrates (Põlme et al. 2020) | Põlme S, et al. 2020. Fungal Diversity 105:1-16 ((https://doi.org/10.1007/s13225-020-00466-2)) |
| Otu54536 | 1 | k__Fungi(100);p__Ascomycota(100);c__Saccharomycetes(100);o__Saccharomycetales(100);f__Metschnikowiaceae(100);g__Metschnikowia(100);s__Metschnikowia_pulcherrima(100); | Metschnikowia | 13 | Saprotroph | \|Undefined Saprotroph\| | Yeast | NULL | Probable | As nectar-tap_saprotroph (Põlme et al. 2020); Decay sugar-rich substrates (Põlme et al. 2020) | Põlme S, et al. 2020. Fungal Diversity 105:1-16 ((https://doi.org/10.1007/s13225-020-00466-2)) |
| Otu54537 | 1 | k__Fungi(100);p__Ascomycota(100);c__Saccharomycetes(100);o__Saccharomycetales(100);f__Metschnikowiaceae(100);g__Metschnikowia(100);s__Metschnikowia_pulcherrima(100); | Metschnikowia | 13 | Saprotroph | \|Undefined Saprotroph\| | Yeast | NULL | Probable | As nectar-tap_saprotroph (Põlme et al. 2020); Decay sugar-rich substrates (Põlme et al. 2020) | Põlme S, et al. 2020. Fungal Diversity 105:1-16 ((https://doi.org/10.1007/s13225-020-00466-2)) |
| Otu54538 | 1 | k__Fungi(100);p__Ascomycota(100);c__Saccharomycetes(100);o__Saccharomycetales(100);f__Metschnikowiaceae(100);g__Metschnikowia(100);g__Metschnikowia_unclassified(100); | Metschnikowia | 13 | Saprotroph | \|Undefined Saprotroph\| | Yeast | NULL | Probable | As nectar-tap_saprotroph (Põlme et al. 2020); Decay sugar-rich substrates (Põlme et al. 2020) | Põlme S, et al. 2020. Fungal Diversity 105:1-16 ((https://doi.org/10.1007/s13225-020-00466-2)) |
| Otu54539 | 1 | k__Fungi(100);p__Ascomycota(100);c__Saccharomycetes(100);o__Saccharomycetales(100);f__Metschnikowiaceae(100);g__Metschnikowia(100);s__Metschnikowia_pulcherrima(100); | Metschnikowia | 13 | Saprotroph | \|Undefined Saprotroph\| | Yeast | NULL | Probable | As nectar-tap_saprotroph (Põlme et al. 2020); Decay sugar-rich substrates (Põlme et al. 2020) | Põlme S, et al. 2020. Fungal Diversity 105:1-16 ((https://doi.org/10.1007/s13225-020-00466-2)) |
| Otu54540 | 1 | k__Fungi(100);p__Ascomycota(100);c__Saccharomycetes(100);o__Saccharomycetales(100);f__Metschnikowiaceae(100);g__Metschnikowia(100);s__Metschnikowia_pulcherrima(100); | Metschnikowia | 13 | Saprotroph | \|Undefined Saprotroph\| | Yeast | NULL | Probable | As nectar-tap_saprotroph (Põlme et al. 2020); Decay sugar-rich substrates (Põlme et al. 2020) | Põlme S, et al. 2020. Fungal Diversity 105:1-16 ((https://doi.org/10.1007/s13225-020-00466-2)) |
| Otu54541 | 1 | k__Fungi(100);p__Ascomycota(100);c__Saccharomycetes(100);o__Saccharomycetales(100);f__Metschnikowiaceae(100);g__Metschnikowia(100);g__Metschnikowia_unclassified(100); | Metschnikowia | 13 | Saprotroph | \|Undefined Saprotroph\| | Yeast | NULL | Probable | As nectar-tap_saprotroph (Põlme et al. 2020); Decay sugar-rich substrates (Põlme et al. 2020) | Põlme S, et al. 2020. Fungal Diversity 105:1-16 ((https://doi.org/10.1007/s13225-020-00466-2)) |
| Otu54542 | 1 | k__Fungi(100);p__Ascomycota(100);c__Saccharomycetes(100);o__Saccharomycetales(100);f__Metschnikowiaceae(100);g__Metschnikowia(100);s__Metschnikowia_pulcherrima(100); | Metschnikowia | 13 | Saprotroph | \|Undefined Saprotroph\| | Yeast | NULL | Probable | As nectar-tap_saprotroph (Põlme et al. 2020); Decay sugar-rich substrates (Põlme et al. 2020) | Põlme S, et al. 2020. Fungal Diversity 105:1-16 ((https://doi.org/10.1007/s13225-020-00466-2)) |
| Otu54543 | 1 | k__Fungi(100);p__Ascomycota(100);c__Saccharomycetes(100);o__Saccharomycetales(100);f__Metschnikowiaceae(100);g__Metschnikowia(100);s__Metschnikowia_pulcherrima(100); | Metschnikowia | 13 | Saprotroph | \|Undefined Saprotroph\| | Yeast | NULL | Probable | As nectar-tap_saprotroph (Põlme et al. 2020); Decay sugar-rich substrates (Põlme et al. 2020) | Põlme S, et al. 2020. Fungal Diversity 105:1-16 ((https://doi.org/10.1007/s13225-020-00466-2)) |
| Otu54544 | 1 | k__Fungi(100);p__Ascomycota(100);c__Saccharomycetes(100);o__Saccharomycetales(100);f__Metschnikowiaceae(100);g__Metschnikowia(100);s__Metschnikowia_pulcherrima(100); | Metschnikowia | 13 | Saprotroph | \|Undefined Saprotroph\| | Yeast | NULL | Probable | As nectar-tap_saprotroph (Põlme et al. 2020); Decay sugar-rich substrates (Põlme et al. 2020) | Põlme S, et al. 2020. Fungal Diversity 105:1-16 ((https://doi.org/10.1007/s13225-020-00466-2)) |
| Otu54545 | 1 | k__Fungi(100);p__Ascomycota(100);c__Saccharomycetes(100);o__Saccharomycetales(100);f__Metschnikowiaceae(100);g__Metschnikowia(100);s__Metschnikowia_pulcherrima(100); | Metschnikowia | 13 | Saprotroph | \|Undefined Saprotroph\| | Yeast | NULL | Probable | As nectar-tap_saprotroph (Põlme et al. 2020); Decay sugar-rich substrates (Põlme et al. 2020) | Põlme S, et al. 2020. Fungal Diversity 105:1-16 ((https://doi.org/10.1007/s13225-020-00466-2)) |
| Otu54546 | 1 | k__Fungi(100);p__Ascomycota(100);c__Saccharomycetes(100);o__Saccharomycetales(100);f__Metschnikowiaceae(100);g__Metschnikowia(100);s__Metschnikowia_pulcherrima(100); | Metschnikowia | 13 | Saprotroph | \|Undefined Saprotroph\| | Yeast | NULL | Probable | As nectar-tap_saprotroph (Põlme et al. 2020); Decay sugar-rich substrates (Põlme et al. 2020) | Põlme S, et al. 2020. Fungal Diversity 105:1-16 ((https://doi.org/10.1007/s13225-020-00466-2)) |
| Otu54547 | 1 | k__Fungi(100);p__Ascomycota(100);c__Saccharomycetes(100);o__Saccharomycetales(100);f__Metschnikowiaceae(100);g__Metschnikowia(100);s__Metschnikowia_pulcherrima(100); | Metschnikowia | 13 | Saprotroph | \|Undefined Saprotroph\| | Yeast | NULL | Probable | As nectar-tap_saprotroph (Põlme et al. 2020); Decay sugar-rich substrates (Põlme et al. 2020) | Põlme S, et al. 2020. Fungal Diversity 105:1-16 ((https://doi.org/10.1007/s13225-020-00466-2)) |
| Otu54548 | 1 | k__Fungi(100);p__Ascomycota(100);c__Saccharomycetes(100);o__Saccharomycetales(100);f__Metschnikowiaceae(100);g__Metschnikowia(100);s__Metschnikowia_pulcherrima(100); | Metschnikowia | 13 | Saprotroph | \|Undefined Saprotroph\| | Yeast | NULL | Probable | As nectar-tap_saprotroph (Põlme et al. 2020); Decay sugar-rich substrates (Põlme et al. 2020) | Põlme S, et al. 2020. Fungal Diversity 105:1-16 ((https://doi.org/10.1007/s13225-020-00466-2)) |
| Otu54549 | 1 | k__Fungi(100);p__Ascomycota(100);c__Saccharomycetes(100);o__Saccharomycetales(100);f__Metschnikowiaceae(100);g__Metschnikowia(100);s__Metschnikowia_pulcherrima(100); | Metschnikowia | 13 | Saprotroph | \|Undefined Saprotroph\| | Yeast | NULL | Probable | As nectar-tap_saprotroph (Põlme et al. 2020); Decay sugar-rich substrates (Põlme et al. 2020) | Põlme S, et al. 2020. Fungal Diversity 105:1-16 ((https://doi.org/10.1007/s13225-020-00466-2)) |
| Otu54550 | 1 | k__Fungi(100);p__Ascomycota(100);c__Saccharomycetes(100);o__Saccharomycetales(100);f__Metschnikowiaceae(100);g__Metschnikowia(100);s__Metschnikowia_pulcherrima(100); | Metschnikowia | 13 | Saprotroph | \|Undefined Saprotroph\| | Yeast | NULL | Probable | As nectar-tap_saprotroph (Põlme et al. 2020); Decay sugar-rich substrates (Põlme et al. 2020) | Põlme S, et al. 2020. Fungal Diversity 105:1-16 ((https://doi.org/10.1007/s13225-020-00466-2)) |
| Otu54551 | 1 | k__Fungi(100);p__Ascomycota(100);c__Saccharomycetes(100);o__Saccharomycetales(100);f__Metschnikowiaceae(100);g__Metschnikowia(100);g__Metschnikowia_unclassified(100); | Metschnikowia | 13 | Saprotroph | \|Undefined Saprotroph\| | Yeast | NULL | Probable | As nectar-tap_saprotroph (Põlme et al. 2020); Decay sugar-rich substrates (Põlme et al. 2020) | Põlme S, et al. 2020. Fungal Diversity 105:1-16 ((https://doi.org/10.1007/s13225-020-00466-2)) |
| Otu54552 | 1 | k__Fungi(100);p__Ascomycota(100);c__Saccharomycetes(100);o__Saccharomycetales(100);f__Metschnikowiaceae(100);g__Metschnikowia(100);g__Metschnikowia_unclassified(100); | Metschnikowia | 13 | Saprotroph | \|Undefined Saprotroph\| | Yeast | NULL | Probable | As nectar-tap_saprotroph (Põlme et al. 2020); Decay sugar-rich substrates (Põlme et al. 2020) | Põlme S, et al. 2020. Fungal Diversity 105:1-16 ((https://doi.org/10.1007/s13225-020-00466-2)) |
| Otu54553 | 1 | k__Fungi(100);p__Ascomycota(100);c__Saccharomycetes(100);o__Saccharomycetales(100);f__Metschnikowiaceae(100);g__Metschnikowia(100);g__Metschnikowia_unclassified(100); | Metschnikowia | 13 | Saprotroph | \|Undefined Saprotroph\| | Yeast | NULL | Probable | As nectar-tap_saprotroph (Põlme et al. 2020); Decay sugar-rich substrates (Põlme et al. 2020) | Põlme S, et al. 2020. Fungal Diversity 105:1-16 ((https://doi.org/10.1007/s13225-020-00466-2)) |
| Otu54554 | 1 | k__Fungi(100);p__Ascomycota(100);c__Saccharomycetes(100);o__Saccharomycetales(100);f__Metschnikowiaceae(100);g__Metschnikowia(100);s__Metschnikowia_pulcherrima(100); | Metschnikowia | 13 | Saprotroph | \|Undefined Saprotroph\| | Yeast | NULL | Probable | As nectar-tap_saprotroph (Põlme et al. 2020); Decay sugar-rich substrates (Põlme et al. 2020) | Põlme S, et al. 2020. Fungal Diversity 105:1-16 ((https://doi.org/10.1007/s13225-020-00466-2)) |
| Otu54555 | 1 | k__Fungi(100);p__Ascomycota(100);c__Saccharomycetes(100);o__Saccharomycetales(100);f__Metschnikowiaceae(100);g__Metschnikowia(100);g__Metschnikowia_unclassified(100); | Metschnikowia | 13 | Saprotroph | \|Undefined Saprotroph\| | Yeast | NULL | Probable | As nectar-tap_saprotroph (Põlme et al. 2020); Decay sugar-rich substrates (Põlme et al. 2020) | Põlme S, et al. 2020. Fungal Diversity 105:1-16 ((https://doi.org/10.1007/s13225-020-00466-2)) |
| Otu54556 | 1 | k__Fungi(100);p__Ascomycota(100);c__Saccharomycetes(100);o__Saccharomycetales(100);f__Metschnikowiaceae(100);g__Metschnikowia(100);s__Metschnikowia_pulcherrima(100); | Metschnikowia | 13 | Saprotroph | \|Undefined Saprotroph\| | Yeast | NULL | Probable | As nectar-tap_saprotroph (Põlme et al. 2020); Decay sugar-rich substrates (Põlme et al. 2020) | Põlme S, et al. 2020. Fungal Diversity 105:1-16 ((https://doi.org/10.1007/s13225-020-00466-2)) |
| Otu54557 | 1 | k__Fungi(100);p__Ascomycota(100);c__Saccharomycetes(100);o__Saccharomycetales(100);f__Metschnikowiaceae(100);g__Metschnikowia(100);g__Metschnikowia_unclassified(100); | Metschnikowia | 13 | Saprotroph | \|Undefined Saprotroph\| | Yeast | NULL | Probable | As nectar-tap_saprotroph (Põlme et al. 2020); Decay sugar-rich substrates (Põlme et al. 2020) | Põlme S, et al. 2020. Fungal Diversity 105:1-16 ((https://doi.org/10.1007/s13225-020-00466-2)) |
| Otu54558 | 1 | k__Fungi(100);p__Ascomycota(100);c__Saccharomycetes(100);o__Saccharomycetales(100);f__Metschnikowiaceae(100);g__Metschnikowia(100);s__Metschnikowia_pulcherrima(100); | Metschnikowia | 13 | Saprotroph | \|Undefined Saprotroph\| | Yeast | NULL | Probable | As nectar-tap_saprotroph (Põlme et al. 2020); Decay sugar-rich substrates (Põlme et al. 2020) | Põlme S, et al. 2020. Fungal Diversity 105:1-16 ((https://doi.org/10.1007/s13225-020-00466-2)) |
| Otu54559 | 1 | k__Fungi(100);p__Ascomycota(100);c__Saccharomycetes(100);o__Saccharomycetales(100);f__Metschnikowiaceae(100);g__Metschnikowia(100);s__Metschnikowia_pulcherrima(100); | Metschnikowia | 13 | Saprotroph | \|Undefined Saprotroph\| | Yeast | NULL | Probable | As nectar-tap_saprotroph (Põlme et al. 2020); Decay sugar-rich substrates (Põlme et al. 2020) | Põlme S, et al. 2020. Fungal Diversity 105:1-16 ((https://doi.org/10.1007/s13225-020-00466-2)) |
| Otu54560 | 1 | k__Fungi(100);p__Ascomycota(100);c__Saccharomycetes(100);o__Saccharomycetales(100);f__Metschnikowiaceae(100);g__Metschnikowia(100);s__Metschnikowia_pulcherrima(100); | Metschnikowia | 13 | Saprotroph | \|Undefined Saprotroph\| | Yeast | NULL | Probable | As nectar-tap_saprotroph (Põlme et al. 2020); Decay sugar-rich substrates (Põlme et al. 2020) | Põlme S, et al. 2020. Fungal Diversity 105:1-16 ((https://doi.org/10.1007/s13225-020-00466-2)) |
| Otu54561 | 1 | k__Fungi(100);p__Ascomycota(100);c__Saccharomycetes(100);o__Saccharomycetales(100);f__Metschnikowiaceae(100);g__Metschnikowia(100);s__Metschnikowia_pulcherrima(100); | Metschnikowia | 13 | Saprotroph | \|Undefined Saprotroph\| | Yeast | NULL | Probable | As nectar-tap_saprotroph (Põlme et al. 2020); Decay sugar-rich substrates (Põlme et al. 2020) | Põlme S, et al. 2020. Fungal Diversity 105:1-16 ((https://doi.org/10.1007/s13225-020-00466-2)) |
| Otu54562 | 1 | k__Fungi(100);p__Basidiomycota(100);c__Microbotryomycetes(100);o__Sporidiobolales(100);f__Sporidiobolales_family_Incertae_sedis(100);g__Sporobolomyces(100);s__Sporobolomyces_lactosus(100); | Sporobolomyces | 13 | Pathotroph-Saprotroph | \|Fungal Parasite\|-Plant Saprotroph | Yeast | NULL | Probable | As litter_saprotroph (Põlme et al. 2020); As mycoparasite (Põlme et al. 2020); Decay fungal material (Põlme et al. 2020) | Tedersoo L, et al. 2014. Science 346:e1256688 ((https://doi.org/10.1126/science.1256688)); Purahong W, et al. 2016. Molecular Ecology 25:4059-4074 ((https://doi.org/10.1111/mec.13739)); Põlme S, et al. 2020. Fungal Diversity 105:1-16 ((https://doi.org/10.1007/s13225-020-00466-2)) |
| Otu54566 | 1 | k__Fungi(100);p__Ascomycota(100);c__Saccharomycetes(100);o__Saccharomycetales(100);f__Metschnikowiaceae(100);g__Metschnikowia(100);s__Metschnikowia_pulcherrima(100); | Metschnikowia | 13 | Saprotroph | \|Undefined Saprotroph\| | Yeast | NULL | Probable | As nectar-tap_saprotroph (Põlme et al. 2020); Decay sugar-rich substrates (Põlme et al. 2020) | Põlme S, et al. 2020. Fungal Diversity 105:1-16 ((https://doi.org/10.1007/s13225-020-00466-2)) |
| Otu54567 | 1 | k__Fungi(100);p__Ascomycota(100);c__Saccharomycetes(100);o__Saccharomycetales(100);f__Saccharomycetales_family_Incertae_sedis(100);g__Candida(100);g__Candida_unclassified(100); | Candida | 13 | Pathotroph-Saprotroph-Symbiotroph | Animal Pathogen-Endophyte-Endosymbiont-Epiphyte-\|Undefined Saprotroph\| | Dimorphic Yeast-Yeast | NULL | Probable | As nectar-tap_saprotroph (Põlme et al. 2020); Decay sugar-rich substrates (Põlme et al. 2020); This asexual genus is polyphyletic and contain species that function very differently depend on what they are. Caution should be taken when interpreting guilds for Candida species. | Manolakaki D, et al. 2010. Virulence. 1: 367-375 ((https://doi.org/10.4161/viru.1.5.12796)); Mekbib SB, et al. 2011. Tropical Plant Pathology 36:233-240 ((https://doi.org/10.1590/S1982-56762011000400004)); Põlme S, et al. 2020. Fungal Diversity 105:1-16 ((https://doi.org/10.1007/s13225-020-00466-2)) |
| Otu54568 | 1 | k__Fungi(100);p__Ascomycota(100);c__Saccharomycetes(100);o__Saccharomycetales(100);f__Saccharomycetales_family_Incertae_sedis(100);g__Candida(100);g__Candida_unclassified(100); | Candida | 13 | Pathotroph-Saprotroph-Symbiotroph | Animal Pathogen-Endophyte-Endosymbiont-Epiphyte-\|Undefined Saprotroph\| | Dimorphic Yeast-Yeast | NULL | Probable | As nectar-tap_saprotroph (Põlme et al. 2020); Decay sugar-rich substrates (Põlme et al. 2020); This asexual genus is polyphyletic and contain species that function very differently depend on what they are. Caution should be taken when interpreting guilds for Candida species. | Manolakaki D, et al. 2010. Virulence. 1: 367-375 ((https://doi.org/10.4161/viru.1.5.12796)); Mekbib SB, et al. 2011. Tropical Plant Pathology 36:233-240 ((https://doi.org/10.1590/S1982-56762011000400004)); Põlme S, et al. 2020. Fungal Diversity 105:1-16 ((https://doi.org/10.1007/s13225-020-00466-2)) |
| Otu54570 | 1 | k__Fungi(100);p__Ascomycota(100);c__Saccharomycetes(100);o__Saccharomycetales(100);f__Saccharomycetales_family_Incertae_sedis(100);g__Candida(100);g__Candida_unclassified(100); | Candida | 13 | Pathotroph-Saprotroph-Symbiotroph | Animal Pathogen-Endophyte-Endosymbiont-Epiphyte-\|Undefined Saprotroph\| | Dimorphic Yeast-Yeast | NULL | Probable | As nectar-tap_saprotroph (Põlme et al. 2020); Decay sugar-rich substrates (Põlme et al. 2020); This asexual genus is polyphyletic and contain species that function very differently depend on what they are. Caution should be taken when interpreting guilds for Candida species. | Manolakaki D, et al. 2010. Virulence. 1: 367-375 ((https://doi.org/10.4161/viru.1.5.12796)); Mekbib SB, et al. 2011. Tropical Plant Pathology 36:233-240 ((https://doi.org/10.1590/S1982-56762011000400004)); Põlme S, et al. 2020. Fungal Diversity 105:1-16 ((https://doi.org/10.1007/s13225-020-00466-2)) |
| Otu54573 | 1 | k__Fungi(100);p__Ascomycota(100);c__Saccharomycetes(100);o__Saccharomycetales(100);f__Metschnikowiaceae(100);g__Metschnikowia(100);g__Metschnikowia_unclassified(100); | Metschnikowia | 13 | Saprotroph | \|Undefined Saprotroph\| | Yeast | NULL | Probable | As nectar-tap_saprotroph (Põlme et al. 2020); Decay sugar-rich substrates (Põlme et al. 2020) | Põlme S, et al. 2020. Fungal Diversity 105:1-16 ((https://doi.org/10.1007/s13225-020-00466-2)) |
| Otu54574 | 1 | k__Fungi(100);p__Ascomycota(100);c__Saccharomycetes(100);o__Saccharomycetales(100);f__Metschnikowiaceae(100);g__Metschnikowia(100);s__Metschnikowia_pulcherrima(100); | Metschnikowia | 13 | Saprotroph | \|Undefined Saprotroph\| | Yeast | NULL | Probable | As nectar-tap_saprotroph (Põlme et al. 2020); Decay sugar-rich substrates (Põlme et al. 2020) | Põlme S, et al. 2020. Fungal Diversity 105:1-16 ((https://doi.org/10.1007/s13225-020-00466-2)) |
| Otu54577 | 1 | k__Fungi(100);p__Ascomycota(100);c__Leotiomycetes(100);o__Helotiales(100);f__Helotiaceae(100);g__Hymenoscyphus(100);s__Hymenoscyphus_sp_1558(100); | Hymenoscyphus | 13 | Pathotroph-Saprotroph-Symbiotroph | Bryophyte Parasite-Ectomycorrhizal-Ericoid Mycorrhizal-Plant Pathogen-\|Plant Saprotroph\|-Undefined Saprotroph | Microfungus | NULL | Possible | As litter_saprotroph (Põlme et al. 2020); Dark Septate Microfungus; Syn. Pezoloma among other (see e.g., Pezoloma ericae); Decay leaf, fruit, seed (Põlme et al. 2020) | Baral HO, Krieglsteiner L. 2006. Acta Mycologica 41:11-20 ((https://doi.org/10.5586/am.2006.003)) (as a bryophyte parasite); Huhtinen S, et al. 2010. Nova Hedwigia 90:413-431 ((https://doi.org/10.1127/0029-5035/2010/0090-0413)); Bent E, et al. 2011. Applied and Environmental Microbiology 77:3351-3359 ((https://doi.org/10.1128/AEM.02575-10)) (as ectomycorrhizae); Tedersoo L, et al. 2014. Science 346:1256688 ((https://doi.org/10.1126/science.1256688)) (as saprotroph); Põlme S, et al. 2020. Fungal Diversity 105:1-16 ((https://doi.org/10.1007/s13225-020-00466-2)) |
| Otu54578 | 1 | k__Fungi(100);p__Ascomycota(100);c__Saccharomycetes(100);o__Saccharomycetales(100);f__Metschnikowiaceae(100);g__Metschnikowia(100);s__Metschnikowia_pulcherrima(100); | Metschnikowia | 13 | Saprotroph | \|Undefined Saprotroph\| | Yeast | NULL | Probable | As nectar-tap_saprotroph (Põlme et al. 2020); Decay sugar-rich substrates (Põlme et al. 2020) | Põlme S, et al. 2020. Fungal Diversity 105:1-16 ((https://doi.org/10.1007/s13225-020-00466-2)) |
| Otu54579 | 1 | k__Fungi(100);p__Ascomycota(100);c__Saccharomycetes(100);o__Saccharomycetales(100);f__Metschnikowiaceae(100);g__Metschnikowia(100);g__Metschnikowia_unclassified(100); | Metschnikowia | 13 | Saprotroph | \|Undefined Saprotroph\| | Yeast | NULL | Probable | As nectar-tap_saprotroph (Põlme et al. 2020); Decay sugar-rich substrates (Põlme et al. 2020) | Põlme S, et al. 2020. Fungal Diversity 105:1-16 ((https://doi.org/10.1007/s13225-020-00466-2)) |
| Otu54580 | 1 | k__Fungi(100);p__Ascomycota(100);c__Saccharomycetes(100);o__Saccharomycetales(100);f__Metschnikowiaceae(100);g__Metschnikowia(100);s__Metschnikowia_pulcherrima(100); | Metschnikowia | 13 | Saprotroph | \|Undefined Saprotroph\| | Yeast | NULL | Probable | As nectar-tap_saprotroph (Põlme et al. 2020); Decay sugar-rich substrates (Põlme et al. 2020) | Põlme S, et al. 2020. Fungal Diversity 105:1-16 ((https://doi.org/10.1007/s13225-020-00466-2)) |
| Otu54581 | 1 | k__Fungi(100);p__Ascomycota(100);c__Saccharomycetes(100);o__Saccharomycetales(100);f__Metschnikowiaceae(100);g__Metschnikowia(100);s__Metschnikowia_pulcherrima(100); | Metschnikowia | 13 | Saprotroph | \|Undefined Saprotroph\| | Yeast | NULL | Probable | As nectar-tap_saprotroph (Põlme et al. 2020); Decay sugar-rich substrates (Põlme et al. 2020) | Põlme S, et al. 2020. Fungal Diversity 105:1-16 ((https://doi.org/10.1007/s13225-020-00466-2)) |
| Otu54582 | 1 | k__Fungi(100);p__Ascomycota(100);c__Saccharomycetes(100);o__Saccharomycetales(100);f__Metschnikowiaceae(100);g__Metschnikowia(100);s__Metschnikowia_pulcherrima(100); | Metschnikowia | 13 | Saprotroph | \|Undefined Saprotroph\| | Yeast | NULL | Probable | As nectar-tap_saprotroph (Põlme et al. 2020); Decay sugar-rich substrates (Põlme et al. 2020) | Põlme S, et al. 2020. Fungal Diversity 105:1-16 ((https://doi.org/10.1007/s13225-020-00466-2)) |
| Otu54583 | 1 | k__Fungi(100);p__Ascomycota(100);c__Saccharomycetes(100);o__Saccharomycetales(100);f__Metschnikowiaceae(100);g__Metschnikowia(100);g__Metschnikowia_unclassified(100); | Metschnikowia | 13 | Saprotroph | \|Undefined Saprotroph\| | Yeast | NULL | Probable | As nectar-tap_saprotroph (Põlme et al. 2020); Decay sugar-rich substrates (Põlme et al. 2020) | Põlme S, et al. 2020. Fungal Diversity 105:1-16 ((https://doi.org/10.1007/s13225-020-00466-2)) |
| Otu54584 | 1 | k__Fungi(100);p__Ascomycota(100);c__Saccharomycetes(100);o__Saccharomycetales(100);f__Metschnikowiaceae(100);g__Metschnikowia(100);g__Metschnikowia_unclassified(100); | Metschnikowia | 13 | Saprotroph | \|Undefined Saprotroph\| | Yeast | NULL | Probable | As nectar-tap_saprotroph (Põlme et al. 2020); Decay sugar-rich substrates (Põlme et al. 2020) | Põlme S, et al. 2020. Fungal Diversity 105:1-16 ((https://doi.org/10.1007/s13225-020-00466-2)) |
| Otu54585 | 1 | k__Fungi(100);p__Ascomycota(100);c__Saccharomycetes(100);o__Saccharomycetales(100);f__Metschnikowiaceae(100);g__Metschnikowia(100);s__Metschnikowia_pulcherrima(100); | Metschnikowia | 13 | Saprotroph | \|Undefined Saprotroph\| | Yeast | NULL | Probable | As nectar-tap_saprotroph (Põlme et al. 2020); Decay sugar-rich substrates (Põlme et al. 2020) | Põlme S, et al. 2020. Fungal Diversity 105:1-16 ((https://doi.org/10.1007/s13225-020-00466-2)) |
| Otu54586 | 1 | k__Fungi(100);p__Ascomycota(100);c__Saccharomycetes(100);o__Saccharomycetales(100);f__Metschnikowiaceae(100);g__Metschnikowia(100);s__Metschnikowia_pulcherrima(100); | Metschnikowia | 13 | Saprotroph | \|Undefined Saprotroph\| | Yeast | NULL | Probable | As nectar-tap_saprotroph (Põlme et al. 2020); Decay sugar-rich substrates (Põlme et al. 2020) | Põlme S, et al. 2020. Fungal Diversity 105:1-16 ((https://doi.org/10.1007/s13225-020-00466-2)) |
| Otu54587 | 1 | k__Fungi(100);p__Basidiomycota(100);c__Microbotryomycetes(100);o__Sporidiobolales(100);f__Sporidiobolales_family_Incertae_sedis(100);g__Sporobolomyces(100);s__Sporobolomyces_lactosus(100); | Sporobolomyces | 13 | Pathotroph-Saprotroph | \|Fungal Parasite\|-Plant Saprotroph | Yeast | NULL | Probable | As litter_saprotroph (Põlme et al. 2020); As mycoparasite (Põlme et al. 2020); Decay fungal material (Põlme et al. 2020) | Tedersoo L, et al. 2014. Science 346:e1256688 ((https://doi.org/10.1126/science.1256688)); Purahong W, et al. 2016. Molecular Ecology 25:4059-4074 ((https://doi.org/10.1111/mec.13739)); Põlme S, et al. 2020. Fungal Diversity 105:1-16 ((https://doi.org/10.1007/s13225-020-00466-2)) |
| Otu54588 | 1 | k__Fungi(100);p__Ascomycota(100);c__Saccharomycetes(100);o__Saccharomycetales(100);o__Saccharomycetales_unclassified(100);o__Saccharomycetales_unclassified(100);o__Saccharomycetales_unclassified(100); | Saccharomycetales | 7 | Saprotroph | Undefined Saprotroph | Yeast | NULL | Possible | NULL | Sterkenburg E, et al. 2015. New Phytologist 207:1145-1158 ((https://doi.org/10.1111/nph.13426)) |
| Otu54590 | 1 | k__Fungi(100);p__Ascomycota(100);c__Saccharomycetes(100);o__Saccharomycetales(100);f__Saccharomycetales_family_Incertae_sedis(100);g__Candida(100);s__Candida_cretensis(100); | Candida | 13 | Pathotroph-Saprotroph-Symbiotroph | Animal Pathogen-Endophyte-Endosymbiont-Epiphyte-\|Undefined Saprotroph\| | Dimorphic Yeast-Yeast | NULL | Probable | As nectar-tap_saprotroph (Põlme et al. 2020); Decay sugar-rich substrates (Põlme et al. 2020); This asexual genus is polyphyletic and contain species that function very differently depend on what they are. Caution should be taken when interpreting guilds for Candida species. | Manolakaki D, et al. 2010. Virulence. 1: 367-375 ((https://doi.org/10.4161/viru.1.5.12796)); Mekbib SB, et al. 2011. Tropical Plant Pathology 36:233-240 ((https://doi.org/10.1590/S1982-56762011000400004)); Põlme S, et al. 2020. Fungal Diversity 105:1-16 ((https://doi.org/10.1007/s13225-020-00466-2)) |
| Otu54591 | 1 | k__Fungi(100);p__Ascomycota(100);c__Saccharomycetes(100);o__Saccharomycetales(100);f__Saccharomycetales_family_Incertae_sedis(100);g__Candida(100);g__Candida_unclassified(100); | Candida | 13 | Pathotroph-Saprotroph-Symbiotroph | Animal Pathogen-Endophyte-Endosymbiont-Epiphyte-\|Undefined Saprotroph\| | Dimorphic Yeast-Yeast | NULL | Probable | As nectar-tap_saprotroph (Põlme et al. 2020); Decay sugar-rich substrates (Põlme et al. 2020); This asexual genus is polyphyletic and contain species that function very differently depend on what they are. Caution should be taken when interpreting guilds for Candida species. | Manolakaki D, et al. 2010. Virulence. 1: 367-375 ((https://doi.org/10.4161/viru.1.5.12796)); Mekbib SB, et al. 2011. Tropical Plant Pathology 36:233-240 ((https://doi.org/10.1590/S1982-56762011000400004)); Põlme S, et al. 2020. Fungal Diversity 105:1-16 ((https://doi.org/10.1007/s13225-020-00466-2)) |
| Otu54593 | 1 | k__Fungi(100);p__Ascomycota(100);c__Saccharomycetes(100);o__Saccharomycetales(100);f__Metschnikowiaceae(100);g__Metschnikowia(100);g__Metschnikowia_unclassified(100); | Metschnikowia | 13 | Saprotroph | \|Undefined Saprotroph\| | Yeast | NULL | Probable | As nectar-tap_saprotroph (Põlme et al. 2020); Decay sugar-rich substrates (Põlme et al. 2020) | Põlme S, et al. 2020. Fungal Diversity 105:1-16 ((https://doi.org/10.1007/s13225-020-00466-2)) |
| Otu54594 | 1 | k__Fungi(100);p__Ascomycota(100);c__Saccharomycetes(100);o__Saccharomycetales(100);f__Metschnikowiaceae(100);g__Metschnikowia(100);s__Metschnikowia_pulcherrima(100); | Metschnikowia | 13 | Saprotroph | \|Undefined Saprotroph\| | Yeast | NULL | Probable | As nectar-tap_saprotroph (Põlme et al. 2020); Decay sugar-rich substrates (Põlme et al. 2020) | Põlme S, et al. 2020. Fungal Diversity 105:1-16 ((https://doi.org/10.1007/s13225-020-00466-2)) |
| Otu54595 | 1 | k__Fungi(100);p__Ascomycota(100);c__Saccharomycetes(100);o__Saccharomycetales(100);f__Metschnikowiaceae(100);g__Metschnikowia(100);s__Metschnikowia_pulcherrima(100); | Metschnikowia | 13 | Saprotroph | \|Undefined Saprotroph\| | Yeast | NULL | Probable | As nectar-tap_saprotroph (Põlme et al. 2020); Decay sugar-rich substrates (Põlme et al. 2020) | Põlme S, et al. 2020. Fungal Diversity 105:1-16 ((https://doi.org/10.1007/s13225-020-00466-2)) |
| Otu54596 | 1 | k__Fungi(100);p__Ascomycota(100);c__Saccharomycetes(100);o__Saccharomycetales(100);f__Metschnikowiaceae(100);g__Metschnikowia(100);s__Metschnikowia_pulcherrima(100); | Metschnikowia | 13 | Saprotroph | \|Undefined Saprotroph\| | Yeast | NULL | Probable | As nectar-tap_saprotroph (Põlme et al. 2020); Decay sugar-rich substrates (Põlme et al. 2020) | Põlme S, et al. 2020. Fungal Diversity 105:1-16 ((https://doi.org/10.1007/s13225-020-00466-2)) |
| Otu54597 | 1 | k__Fungi(100);p__Ascomycota(100);c__Saccharomycetes(100);o__Saccharomycetales(100);f__Saccharomycetales_family_Incertae_sedis(100);g__Candida(100);g__Candida_unclassified(100); | Candida | 13 | Pathotroph-Saprotroph-Symbiotroph | Animal Pathogen-Endophyte-Endosymbiont-Epiphyte-\|Undefined Saprotroph\| | Dimorphic Yeast-Yeast | NULL | Probable | As nectar-tap_saprotroph (Põlme et al. 2020); Decay sugar-rich substrates (Põlme et al. 2020); This asexual genus is polyphyletic and contain species that function very differently depend on what they are. Caution should be taken when interpreting guilds for Candida species. | Manolakaki D, et al. 2010. Virulence. 1: 367-375 ((https://doi.org/10.4161/viru.1.5.12796)); Mekbib SB, et al. 2011. Tropical Plant Pathology 36:233-240 ((https://doi.org/10.1590/S1982-56762011000400004)); Põlme S, et al. 2020. Fungal Diversity 105:1-16 ((https://doi.org/10.1007/s13225-020-00466-2)) |
| Otu54598 | 1 | k__Fungi(100);p__Ascomycota(100);c__Saccharomycetes(100);o__Saccharomycetales(100);f__Metschnikowiaceae(100);g__Metschnikowia(100);g__Metschnikowia_unclassified(100); | Metschnikowia | 13 | Saprotroph | \|Undefined Saprotroph\| | Yeast | NULL | Probable | As nectar-tap_saprotroph (Põlme et al. 2020); Decay sugar-rich substrates (Põlme et al. 2020) | Põlme S, et al. 2020. Fungal Diversity 105:1-16 ((https://doi.org/10.1007/s13225-020-00466-2)) |
| Otu54600 | 1 | k__Fungi(100);p__Ascomycota(100);c__Saccharomycetes(100);o__Saccharomycetales(100);f__Metschnikowiaceae(100);g__Metschnikowia(100);g__Metschnikowia_unclassified(100); | Metschnikowia | 13 | Saprotroph | \|Undefined Saprotroph\| | Yeast | NULL | Probable | As nectar-tap_saprotroph (Põlme et al. 2020); Decay sugar-rich substrates (Põlme et al. 2020) | Põlme S, et al. 2020. Fungal Diversity 105:1-16 ((https://doi.org/10.1007/s13225-020-00466-2)) |
| Otu54602 | 1 | k__Fungi(100);p__Ascomycota(100);c__Saccharomycetes(100);o__Saccharomycetales(100);f__Saccharomycetales_family_Incertae_sedis(100);g__Candida(100);g__Candida_unclassified(100); | Candida | 13 | Pathotroph-Saprotroph-Symbiotroph | Animal Pathogen-Endophyte-Endosymbiont-Epiphyte-\|Undefined Saprotroph\| | Dimorphic Yeast-Yeast | NULL | Probable | As nectar-tap_saprotroph (Põlme et al. 2020); Decay sugar-rich substrates (Põlme et al. 2020); This asexual genus is polyphyletic and contain species that function very differently depend on what they are. Caution should be taken when interpreting guilds for Candida species. | Manolakaki D, et al. 2010. Virulence. 1: 367-375 ((https://doi.org/10.4161/viru.1.5.12796)); Mekbib SB, et al. 2011. Tropical Plant Pathology 36:233-240 ((https://doi.org/10.1590/S1982-56762011000400004)); Põlme S, et al. 2020. Fungal Diversity 105:1-16 ((https://doi.org/10.1007/s13225-020-00466-2)) |
| Otu54604 | 1 | k__Fungi(100);p__Ascomycota(100);c__Saccharomycetes(100);o__Saccharomycetales(100);f__Saccharomycetales_family_Incertae_sedis(100);g__Candida(100);g__Candida_unclassified(100); | Candida | 13 | Pathotroph-Saprotroph-Symbiotroph | Animal Pathogen-Endophyte-Endosymbiont-Epiphyte-\|Undefined Saprotroph\| | Dimorphic Yeast-Yeast | NULL | Probable | As nectar-tap_saprotroph (Põlme et al. 2020); Decay sugar-rich substrates (Põlme et al. 2020); This asexual genus is polyphyletic and contain species that function very differently depend on what they are. Caution should be taken when interpreting guilds for Candida species. | Manolakaki D, et al. 2010. Virulence. 1: 367-375 ((https://doi.org/10.4161/viru.1.5.12796)); Mekbib SB, et al. 2011. Tropical Plant Pathology 36:233-240 ((https://doi.org/10.1590/S1982-56762011000400004)); Põlme S, et al. 2020. Fungal Diversity 105:1-16 ((https://doi.org/10.1007/s13225-020-00466-2)) |
| Otu54609 | 1 | k__Fungi(100);p__Ascomycota(100);c__Saccharomycetes(100);o__Saccharomycetales(100);f__Saccharomycetales_family_Incertae_sedis(100);f__Saccharomycetales_family_Incertae_sedis_unclassified(100);f__Saccharomycetales_family_Incertae_sedis_unclassified(100); | Saccharomycetales | 7 | Saprotroph | Undefined Saprotroph | Yeast | NULL | Possible | NULL | Sterkenburg E, et al. 2015. New Phytologist 207:1145-1158 ((https://doi.org/10.1111/nph.13426)) |
| Otu54614 | 1 | k__Fungi(100);p__Basidiomycota(100);c__Microbotryomycetes(100);o__Sporidiobolales(100);f__Sporidiobolales_family_Incertae_sedis(100);g__Sporobolomyces(100);s__Sporobolomyces_lactosus(100); | Sporobolomyces | 13 | Pathotroph-Saprotroph | \|Fungal Parasite\|-Plant Saprotroph | Yeast | NULL | Probable | As litter_saprotroph (Põlme et al. 2020); As mycoparasite (Põlme et al. 2020); Decay fungal material (Põlme et al. 2020) | Tedersoo L, et al. 2014. Science 346:e1256688 ((https://doi.org/10.1126/science.1256688)); Purahong W, et al. 2016. Molecular Ecology 25:4059-4074 ((https://doi.org/10.1111/mec.13739)); Põlme S, et al. 2020. Fungal Diversity 105:1-16 ((https://doi.org/10.1007/s13225-020-00466-2)) |
| Otu54615 | 1 | k__Fungi(100);p__Basidiomycota(100);c__Microbotryomycetes(100);o__Sporidiobolales(100);f__Sporidiobolales_family_Incertae_sedis(100);g__Sporobolomyces(100);s__Sporobolomyces_lactosus(100); | Sporobolomyces | 13 | Pathotroph-Saprotroph | \|Fungal Parasite\|-Plant Saprotroph | Yeast | NULL | Probable | As litter_saprotroph (Põlme et al. 2020); As mycoparasite (Põlme et al. 2020); Decay fungal material (Põlme et al. 2020) | Tedersoo L, et al. 2014. Science 346:e1256688 ((https://doi.org/10.1126/science.1256688)); Purahong W, et al. 2016. Molecular Ecology 25:4059-4074 ((https://doi.org/10.1111/mec.13739)); Põlme S, et al. 2020. Fungal Diversity 105:1-16 ((https://doi.org/10.1007/s13225-020-00466-2)) |
| Otu54616 | 1 | k__Fungi(100);p__Ascomycota(100);c__Dothideomycetes(100);o__Dothideomycetes_order_Incertae_sedis(100);f__Pseudeurotiaceae(100);g__Pseudeurotium(100);g__Pseudeurotium_unclassified(100); | Pseudeurotium | 13 | Saprotroph | \|Dung Saprotroph\|-Undefined Saprotroph | Microfungus | NULL | Probable | As soil_saprotroph (Põlme et al. 2020); Decay soil, dung (Põlme et al. 2020) | Tedersoo L, et al. 2014. Science 346:e1256688 ((https://doi.org/10.1126/science.1256688)); Põlme S, et al. 2020. Fungal Diversity 105:1-16 ((https://doi.org/10.1007/s13225-020-00466-2)) |
| Otu54617 | 1 | k__Fungi(100);p__Basidiomycota(100);c__Microbotryomycetes(100);o__Sporidiobolales(100);f__Sporidiobolales_family_Incertae_sedis(100);g__Sporobolomyces(100);s__Sporobolomyces_lactosus(100); | Sporobolomyces | 13 | Pathotroph-Saprotroph | \|Fungal Parasite\|-Plant Saprotroph | Yeast | NULL | Probable | As litter_saprotroph (Põlme et al. 2020); As mycoparasite (Põlme et al. 2020); Decay fungal material (Põlme et al. 2020) | Tedersoo L, et al. 2014. Science 346:e1256688 ((https://doi.org/10.1126/science.1256688)); Purahong W, et al. 2016. Molecular Ecology 25:4059-4074 ((https://doi.org/10.1111/mec.13739)); Põlme S, et al. 2020. Fungal Diversity 105:1-16 ((https://doi.org/10.1007/s13225-020-00466-2)) |
| Otu54618 | 1 | k__Fungi(100);p__Basidiomycota(100);c__Microbotryomycetes(100);o__Sporidiobolales(100);f__Sporidiobolales_family_Incertae_sedis(100);g__Sporobolomyces(100);s__Sporobolomyces_lactosus(100); | Sporobolomyces | 13 | Pathotroph-Saprotroph | \|Fungal Parasite\|-Plant Saprotroph | Yeast | NULL | Probable | As litter_saprotroph (Põlme et al. 2020); As mycoparasite (Põlme et al. 2020); Decay fungal material (Põlme et al. 2020) | Tedersoo L, et al. 2014. Science 346:e1256688 ((https://doi.org/10.1126/science.1256688)); Purahong W, et al. 2016. Molecular Ecology 25:4059-4074 ((https://doi.org/10.1111/mec.13739)); Põlme S, et al. 2020. Fungal Diversity 105:1-16 ((https://doi.org/10.1007/s13225-020-00466-2)) |
| Otu54619 | 1 | k__Fungi(100);p__Basidiomycota(100);c__Microbotryomycetes(100);o__Sporidiobolales(100);f__Sporidiobolales_family_Incertae_sedis(100);g__Sporobolomyces(100);s__Sporobolomyces_lactosus(100); | Sporobolomyces | 13 | Pathotroph-Saprotroph | \|Fungal Parasite\|-Plant Saprotroph | Yeast | NULL | Probable | As litter_saprotroph (Põlme et al. 2020); As mycoparasite (Põlme et al. 2020); Decay fungal material (Põlme et al. 2020) | Tedersoo L, et al. 2014. Science 346:e1256688 ((https://doi.org/10.1126/science.1256688)); Purahong W, et al. 2016. Molecular Ecology 25:4059-4074 ((https://doi.org/10.1111/mec.13739)); Põlme S, et al. 2020. Fungal Diversity 105:1-16 ((https://doi.org/10.1007/s13225-020-00466-2)) |
| Otu54620 | 1 | k__Fungi(100);p__Basidiomycota(100);c__Microbotryomycetes(100);o__Sporidiobolales(100);f__Sporidiobolales_family_Incertae_sedis(100);g__Sporobolomyces(100);s__Sporobolomyces_lactosus(100); | Sporobolomyces | 13 | Pathotroph-Saprotroph | \|Fungal Parasite\|-Plant Saprotroph | Yeast | NULL | Probable | As litter_saprotroph (Põlme et al. 2020); As mycoparasite (Põlme et al. 2020); Decay fungal material (Põlme et al. 2020) | Tedersoo L, et al. 2014. Science 346:e1256688 ((https://doi.org/10.1126/science.1256688)); Purahong W, et al. 2016. Molecular Ecology 25:4059-4074 ((https://doi.org/10.1111/mec.13739)); Põlme S, et al. 2020. Fungal Diversity 105:1-16 ((https://doi.org/10.1007/s13225-020-00466-2)) |
| Otu54621 | 1 | k__Fungi(100);p__Basidiomycota(100);c__Microbotryomycetes(100);o__Sporidiobolales(100);f__Sporidiobolales_family_Incertae_sedis(100);g__Sporobolomyces(100);s__Sporobolomyces_lactosus(100); | Sporobolomyces | 13 | Pathotroph-Saprotroph | \|Fungal Parasite\|-Plant Saprotroph | Yeast | NULL | Probable | As litter_saprotroph (Põlme et al. 2020); As mycoparasite (Põlme et al. 2020); Decay fungal material (Põlme et al. 2020) | Tedersoo L, et al. 2014. Science 346:e1256688 ((https://doi.org/10.1126/science.1256688)); Purahong W, et al. 2016. Molecular Ecology 25:4059-4074 ((https://doi.org/10.1111/mec.13739)); Põlme S, et al. 2020. Fungal Diversity 105:1-16 ((https://doi.org/10.1007/s13225-020-00466-2)) |
| Otu54622 | 1 | k__Fungi(100);p__Basidiomycota(100);c__Microbotryomycetes(100);o__Sporidiobolales(100);f__Sporidiobolales_family_Incertae_sedis(100);g__Sporobolomyces(100);s__Sporobolomyces_lactosus(100); | Sporobolomyces | 13 | Pathotroph-Saprotroph | \|Fungal Parasite\|-Plant Saprotroph | Yeast | NULL | Probable | As litter_saprotroph (Põlme et al. 2020); As mycoparasite (Põlme et al. 2020); Decay fungal material (Põlme et al. 2020) | Tedersoo L, et al. 2014. Science 346:e1256688 ((https://doi.org/10.1126/science.1256688)); Purahong W, et al. 2016. Molecular Ecology 25:4059-4074 ((https://doi.org/10.1111/mec.13739)); Põlme S, et al. 2020. Fungal Diversity 105:1-16 ((https://doi.org/10.1007/s13225-020-00466-2)) |
| Otu54623 | 1 | k__Fungi(100);p__Basidiomycota(100);c__Microbotryomycetes(100);o__Sporidiobolales(100);f__Sporidiobolales_family_Incertae_sedis(100);g__Sporobolomyces(100);s__Sporobolomyces_lactosus(100); | Sporobolomyces | 13 | Pathotroph-Saprotroph | \|Fungal Parasite\|-Plant Saprotroph | Yeast | NULL | Probable | As litter_saprotroph (Põlme et al. 2020); As mycoparasite (Põlme et al. 2020); Decay fungal material (Põlme et al. 2020) | Tedersoo L, et al. 2014. Science 346:e1256688 ((https://doi.org/10.1126/science.1256688)); Purahong W, et al. 2016. Molecular Ecology 25:4059-4074 ((https://doi.org/10.1111/mec.13739)); Põlme S, et al. 2020. Fungal Diversity 105:1-16 ((https://doi.org/10.1007/s13225-020-00466-2)) |
| Otu54624 | 1 | k__Fungi(100);p__Basidiomycota(100);c__Microbotryomycetes(100);o__Sporidiobolales(100);f__Sporidiobolales_family_Incertae_sedis(100);g__Sporobolomyces(100);s__Sporobolomyces_lactosus(100); | Sporobolomyces | 13 | Pathotroph-Saprotroph | \|Fungal Parasite\|-Plant Saprotroph | Yeast | NULL | Probable | As litter_saprotroph (Põlme et al. 2020); As mycoparasite (Põlme et al. 2020); Decay fungal material (Põlme et al. 2020) | Tedersoo L, et al. 2014. Science 346:e1256688 ((https://doi.org/10.1126/science.1256688)); Purahong W, et al. 2016. Molecular Ecology 25:4059-4074 ((https://doi.org/10.1111/mec.13739)); Põlme S, et al. 2020. Fungal Diversity 105:1-16 ((https://doi.org/10.1007/s13225-020-00466-2)) |
| Otu54625 | 1 | k__Fungi(100);p__Basidiomycota(100);c__Microbotryomycetes(100);o__Sporidiobolales(100);f__Sporidiobolales_family_Incertae_sedis(100);g__Sporobolomyces(100);s__Sporobolomyces_lactosus(100); | Sporobolomyces | 13 | Pathotroph-Saprotroph | \|Fungal Parasite\|-Plant Saprotroph | Yeast | NULL | Probable | As litter_saprotroph (Põlme et al. 2020); As mycoparasite (Põlme et al. 2020); Decay fungal material (Põlme et al. 2020) | Tedersoo L, et al. 2014. Science 346:e1256688 ((https://doi.org/10.1126/science.1256688)); Purahong W, et al. 2016. Molecular Ecology 25:4059-4074 ((https://doi.org/10.1111/mec.13739)); Põlme S, et al. 2020. Fungal Diversity 105:1-16 ((https://doi.org/10.1007/s13225-020-00466-2)) |
| Otu54627 | 1 | k__Fungi(100);p__Basidiomycota(100);c__Microbotryomycetes(100);o__Sporidiobolales(100);f__Sporidiobolales_family_Incertae_sedis(100);g__Sporobolomyces(100);s__Sporobolomyces_lactosus(100); | Sporobolomyces | 13 | Pathotroph-Saprotroph | \|Fungal Parasite\|-Plant Saprotroph | Yeast | NULL | Probable | As litter_saprotroph (Põlme et al. 2020); As mycoparasite (Põlme et al. 2020); Decay fungal material (Põlme et al. 2020) | Tedersoo L, et al. 2014. Science 346:e1256688 ((https://doi.org/10.1126/science.1256688)); Purahong W, et al. 2016. Molecular Ecology 25:4059-4074 ((https://doi.org/10.1111/mec.13739)); Põlme S, et al. 2020. Fungal Diversity 105:1-16 ((https://doi.org/10.1007/s13225-020-00466-2)) |
| Otu54628 | 1 | k__Fungi(100);p__Basidiomycota(100);c__Microbotryomycetes(100);o__Sporidiobolales(100);f__Sporidiobolales_family_Incertae_sedis(100);g__Sporobolomyces(100);s__Sporobolomyces_lactosus(100); | Sporobolomyces | 13 | Pathotroph-Saprotroph | \|Fungal Parasite\|-Plant Saprotroph | Yeast | NULL | Probable | As litter_saprotroph (Põlme et al. 2020); As mycoparasite (Põlme et al. 2020); Decay fungal material (Põlme et al. 2020) | Tedersoo L, et al. 2014. Science 346:e1256688 ((https://doi.org/10.1126/science.1256688)); Purahong W, et al. 2016. Molecular Ecology 25:4059-4074 ((https://doi.org/10.1111/mec.13739)); Põlme S, et al. 2020. Fungal Diversity 105:1-16 ((https://doi.org/10.1007/s13225-020-00466-2)) |
| Otu54629 | 1 | k__Fungi(100);p__Basidiomycota(100);c__Microbotryomycetes(100);o__Sporidiobolales(100);f__Sporidiobolales_family_Incertae_sedis(100);g__Sporobolomyces(100);s__Sporobolomyces_lactosus(100); | Sporobolomyces | 13 | Pathotroph-Saprotroph | \|Fungal Parasite\|-Plant Saprotroph | Yeast | NULL | Probable | As litter_saprotroph (Põlme et al. 2020); As mycoparasite (Põlme et al. 2020); Decay fungal material (Põlme et al. 2020) | Tedersoo L, et al. 2014. Science 346:e1256688 ((https://doi.org/10.1126/science.1256688)); Purahong W, et al. 2016. Molecular Ecology 25:4059-4074 ((https://doi.org/10.1111/mec.13739)); Põlme S, et al. 2020. Fungal Diversity 105:1-16 ((https://doi.org/10.1007/s13225-020-00466-2)) |
| Otu54630 | 1 | k__Fungi(100);p__Basidiomycota(100);c__Microbotryomycetes(100);o__Sporidiobolales(100);f__Sporidiobolales_family_Incertae_sedis(100);g__Sporobolomyces(100);s__Sporobolomyces_lactosus(100); | Sporobolomyces | 13 | Pathotroph-Saprotroph | \|Fungal Parasite\|-Plant Saprotroph | Yeast | NULL | Probable | As litter_saprotroph (Põlme et al. 2020); As mycoparasite (Põlme et al. 2020); Decay fungal material (Põlme et al. 2020) | Tedersoo L, et al. 2014. Science 346:e1256688 ((https://doi.org/10.1126/science.1256688)); Purahong W, et al. 2016. Molecular Ecology 25:4059-4074 ((https://doi.org/10.1111/mec.13739)); Põlme S, et al. 2020. Fungal Diversity 105:1-16 ((https://doi.org/10.1007/s13225-020-00466-2)) |
| Otu54632 | 1 | k__Fungi(100);p__Ascomycota(100);c__Saccharomycetes(100);o__Saccharomycetales(100);f__Metschnikowiaceae(100);g__Metschnikowia(100);g__Metschnikowia_unclassified(100); | Metschnikowia | 13 | Saprotroph | \|Undefined Saprotroph\| | Yeast | NULL | Probable | As nectar-tap_saprotroph (Põlme et al. 2020); Decay sugar-rich substrates (Põlme et al. 2020) | Põlme S, et al. 2020. Fungal Diversity 105:1-16 ((https://doi.org/10.1007/s13225-020-00466-2)) |
| Otu54633 | 1 | k__Fungi(100);p__Ascomycota(100);c__Dothideomycetes(100);o__Dothideomycetes_order_Incertae_sedis(100);f__Pseudeurotiaceae(100);g__Pseudeurotium(100);g__Pseudeurotium_unclassified(100); | Pseudeurotium | 13 | Saprotroph | \|Dung Saprotroph\|-Undefined Saprotroph | Microfungus | NULL | Probable | As soil_saprotroph (Põlme et al. 2020); Decay soil, dung (Põlme et al. 2020) | Tedersoo L, et al. 2014. Science 346:e1256688 ((https://doi.org/10.1126/science.1256688)); Põlme S, et al. 2020. Fungal Diversity 105:1-16 ((https://doi.org/10.1007/s13225-020-00466-2)) |
| Otu54634 | 1 | k__Fungi(100);p__Basidiomycota(100);c__Microbotryomycetes(100);o__Sporidiobolales(100);f__Sporidiobolales_family_Incertae_sedis(100);g__Sporobolomyces(100);s__Sporobolomyces_lactosus(100); | Sporobolomyces | 13 | Pathotroph-Saprotroph | \|Fungal Parasite\|-Plant Saprotroph | Yeast | NULL | Probable | As litter_saprotroph (Põlme et al. 2020); As mycoparasite (Põlme et al. 2020); Decay fungal material (Põlme et al. 2020) | Tedersoo L, et al. 2014. Science 346:e1256688 ((https://doi.org/10.1126/science.1256688)); Purahong W, et al. 2016. Molecular Ecology 25:4059-4074 ((https://doi.org/10.1111/mec.13739)); Põlme S, et al. 2020. Fungal Diversity 105:1-16 ((https://doi.org/10.1007/s13225-020-00466-2)) |
| Otu54635 | 1 | k__Fungi(100);p__Basidiomycota(100);c__Microbotryomycetes(100);o__Sporidiobolales(100);f__Sporidiobolales_family_Incertae_sedis(100);g__Sporobolomyces(100);s__Sporobolomyces_lactosus(100); | Sporobolomyces | 13 | Pathotroph-Saprotroph | \|Fungal Parasite\|-Plant Saprotroph | Yeast | NULL | Probable | As litter_saprotroph (Põlme et al. 2020); As mycoparasite (Põlme et al. 2020); Decay fungal material (Põlme et al. 2020) | Tedersoo L, et al. 2014. Science 346:e1256688 ((https://doi.org/10.1126/science.1256688)); Purahong W, et al. 2016. Molecular Ecology 25:4059-4074 ((https://doi.org/10.1111/mec.13739)); Põlme S, et al. 2020. Fungal Diversity 105:1-16 ((https://doi.org/10.1007/s13225-020-00466-2)) |
| Otu54636 | 1 | k__Fungi(100);p__Basidiomycota(100);c__Microbotryomycetes(100);o__Sporidiobolales(100);f__Sporidiobolales_family_Incertae_sedis(100);g__Sporobolomyces(100);s__Sporobolomyces_lactosus(100); | Sporobolomyces | 13 | Pathotroph-Saprotroph | \|Fungal Parasite\|-Plant Saprotroph | Yeast | NULL | Probable | As litter_saprotroph (Põlme et al. 2020); As mycoparasite (Põlme et al. 2020); Decay fungal material (Põlme et al. 2020) | Tedersoo L, et al. 2014. Science 346:e1256688 ((https://doi.org/10.1126/science.1256688)); Purahong W, et al. 2016. Molecular Ecology 25:4059-4074 ((https://doi.org/10.1111/mec.13739)); Põlme S, et al. 2020. Fungal Diversity 105:1-16 ((https://doi.org/10.1007/s13225-020-00466-2)) |
| Otu54637 | 1 | k__Fungi(100);p__Ascomycota(100);c__Saccharomycetes(100);o__Saccharomycetales(100);f__Metschnikowiaceae(100);g__Metschnikowia(100);s__Metschnikowia_pulcherrima(100); | Metschnikowia | 13 | Saprotroph | \|Undefined Saprotroph\| | Yeast | NULL | Probable | As nectar-tap_saprotroph (Põlme et al. 2020); Decay sugar-rich substrates (Põlme et al. 2020) | Põlme S, et al. 2020. Fungal Diversity 105:1-16 ((https://doi.org/10.1007/s13225-020-00466-2)) |
| Otu54638 | 1 | k__Fungi(100);p__Basidiomycota(100);c__Microbotryomycetes(100);o__Sporidiobolales(100);f__Sporidiobolales_family_Incertae_sedis(100);g__Sporobolomyces(100);s__Sporobolomyces_lactosus(100); | Sporobolomyces | 13 | Pathotroph-Saprotroph | \|Fungal Parasite\|-Plant Saprotroph | Yeast | NULL | Probable | As litter_saprotroph (Põlme et al. 2020); As mycoparasite (Põlme et al. 2020); Decay fungal material (Põlme et al. 2020) | Tedersoo L, et al. 2014. Science 346:e1256688 ((https://doi.org/10.1126/science.1256688)); Purahong W, et al. 2016. Molecular Ecology 25:4059-4074 ((https://doi.org/10.1111/mec.13739)); Põlme S, et al. 2020. Fungal Diversity 105:1-16 ((https://doi.org/10.1007/s13225-020-00466-2)) |
| Otu54639 | 1 | k__Fungi(100);p__Ascomycota(100);c__Saccharomycetes(100);o__Saccharomycetales(100);f__Metschnikowiaceae(100);g__Metschnikowia(100);g__Metschnikowia_unclassified(100); | Metschnikowia | 13 | Saprotroph | \|Undefined Saprotroph\| | Yeast | NULL | Probable | As nectar-tap_saprotroph (Põlme et al. 2020); Decay sugar-rich substrates (Põlme et al. 2020) | Põlme S, et al. 2020. Fungal Diversity 105:1-16 ((https://doi.org/10.1007/s13225-020-00466-2)) |
| Otu54640 | 1 | k__Fungi(100);p__Basidiomycota(100);c__Microbotryomycetes(100);o__Sporidiobolales(100);f__Sporidiobolales_family_Incertae_sedis(100);g__Sporobolomyces(100);s__Sporobolomyces_lactosus(100); | Sporobolomyces | 13 | Pathotroph-Saprotroph | \|Fungal Parasite\|-Plant Saprotroph | Yeast | NULL | Probable | As litter_saprotroph (Põlme et al. 2020); As mycoparasite (Põlme et al. 2020); Decay fungal material (Põlme et al. 2020) | Tedersoo L, et al. 2014. Science 346:e1256688 ((https://doi.org/10.1126/science.1256688)); Purahong W, et al. 2016. Molecular Ecology 25:4059-4074 ((https://doi.org/10.1111/mec.13739)); Põlme S, et al. 2020. Fungal Diversity 105:1-16 ((https://doi.org/10.1007/s13225-020-00466-2)) |
| Otu54641 | 1 | k__Fungi(100);p__Ascomycota(100);c__Saccharomycetes(100);o__Saccharomycetales(100);f__Metschnikowiaceae(100);g__Metschnikowia(100);s__Metschnikowia_pulcherrima(100); | Metschnikowia | 13 | Saprotroph | \|Undefined Saprotroph\| | Yeast | NULL | Probable | As nectar-tap_saprotroph (Põlme et al. 2020); Decay sugar-rich substrates (Põlme et al. 2020) | Põlme S, et al. 2020. Fungal Diversity 105:1-16 ((https://doi.org/10.1007/s13225-020-00466-2)) |
| Otu54643 | 1 | k__Fungi(100);p__Ascomycota(100);c__Saccharomycetes(100);o__Saccharomycetales(100);f__Metschnikowiaceae(100);g__Metschnikowia(100);s__Metschnikowia_pulcherrima(100); | Metschnikowia | 13 | Saprotroph | \|Undefined Saprotroph\| | Yeast | NULL | Probable | As nectar-tap_saprotroph (Põlme et al. 2020); Decay sugar-rich substrates (Põlme et al. 2020) | Põlme S, et al. 2020. Fungal Diversity 105:1-16 ((https://doi.org/10.1007/s13225-020-00466-2)) |
| Otu54644 | 1 | k__Fungi(100);p__Ascomycota(100);c__Saccharomycetes(100);o__Saccharomycetales(100);f__Metschnikowiaceae(100);g__Metschnikowia(100);s__Metschnikowia_pulcherrima(100); | Metschnikowia | 13 | Saprotroph | \|Undefined Saprotroph\| | Yeast | NULL | Probable | As nectar-tap_saprotroph (Põlme et al. 2020); Decay sugar-rich substrates (Põlme et al. 2020) | Põlme S, et al. 2020. Fungal Diversity 105:1-16 ((https://doi.org/10.1007/s13225-020-00466-2)) |
| Otu54645 | 1 | k__Fungi(100);p__Basidiomycota(100);c__Microbotryomycetes(100);o__Sporidiobolales(100);f__Sporidiobolales_family_Incertae_sedis(100);g__Sporobolomyces(100);s__Sporobolomyces_lactosus(100); | Sporobolomyces | 13 | Pathotroph-Saprotroph | \|Fungal Parasite\|-Plant Saprotroph | Yeast | NULL | Probable | As litter_saprotroph (Põlme et al. 2020); As mycoparasite (Põlme et al. 2020); Decay fungal material (Põlme et al. 2020) | Tedersoo L, et al. 2014. Science 346:e1256688 ((https://doi.org/10.1126/science.1256688)); Purahong W, et al. 2016. Molecular Ecology 25:4059-4074 ((https://doi.org/10.1111/mec.13739)); Põlme S, et al. 2020. Fungal Diversity 105:1-16 ((https://doi.org/10.1007/s13225-020-00466-2)) |
| Otu54646 | 1 | k__Fungi(100);p__Ascomycota(100);c__Dothideomycetes(100);o__Capnodiales(100);f__Mycosphaerellaceae(100);g__Passalora(100);s__Passalora_zambiae(100); | Passalora | 13 | Pathotroph | \|Plant Pathogen\| | Microfungus | NULL | Probable | NULL | Tedersoo L, et al. 2014. Science 346:e1256688 ((https://doi.org/10.1126/science.1256688)); Põlme S, et al. 2020. Fungal Diversity 105:1-16 ((https://doi.org/10.1007/s13225-020-00466-2)) |
| Otu54648 | 1 | k__Fungi(100);p__Basidiomycota(100);c__Microbotryomycetes(100);o__Sporidiobolales(100);f__Sporidiobolales_family_Incertae_sedis(100);g__Sporobolomyces(100);s__Sporobolomyces_lactosus(100); | Sporobolomyces | 13 | Pathotroph-Saprotroph | \|Fungal Parasite\|-Plant Saprotroph | Yeast | NULL | Probable | As litter_saprotroph (Põlme et al. 2020); As mycoparasite (Põlme et al. 2020); Decay fungal material (Põlme et al. 2020) | Tedersoo L, et al. 2014. Science 346:e1256688 ((https://doi.org/10.1126/science.1256688)); Purahong W, et al. 2016. Molecular Ecology 25:4059-4074 ((https://doi.org/10.1111/mec.13739)); Põlme S, et al. 2020. Fungal Diversity 105:1-16 ((https://doi.org/10.1007/s13225-020-00466-2)) |
| Otu54649 | 1 | k__Fungi(100);p__Basidiomycota(100);c__Microbotryomycetes(100);o__Sporidiobolales(100);f__Sporidiobolales_family_Incertae_sedis(100);g__Sporobolomyces(100);s__Sporobolomyces_lactosus(100); | Sporobolomyces | 13 | Pathotroph-Saprotroph | \|Fungal Parasite\|-Plant Saprotroph | Yeast | NULL | Probable | As litter_saprotroph (Põlme et al. 2020); As mycoparasite (Põlme et al. 2020); Decay fungal material (Põlme et al. 2020) | Tedersoo L, et al. 2014. Science 346:e1256688 ((https://doi.org/10.1126/science.1256688)); Purahong W, et al. 2016. Molecular Ecology 25:4059-4074 ((https://doi.org/10.1111/mec.13739)); Põlme S, et al. 2020. Fungal Diversity 105:1-16 ((https://doi.org/10.1007/s13225-020-00466-2)) |
| Otu54650 | 1 | k__Fungi(100);p__Basidiomycota(100);c__Microbotryomycetes(100);o__Sporidiobolales(100);f__Sporidiobolales_family_Incertae_sedis(100);g__Sporobolomyces(100);s__Sporobolomyces_lactosus(100); | Sporobolomyces | 13 | Pathotroph-Saprotroph | \|Fungal Parasite\|-Plant Saprotroph | Yeast | NULL | Probable | As litter_saprotroph (Põlme et al. 2020); As mycoparasite (Põlme et al. 2020); Decay fungal material (Põlme et al. 2020) | Tedersoo L, et al. 2014. Science 346:e1256688 ((https://doi.org/10.1126/science.1256688)); Purahong W, et al. 2016. Molecular Ecology 25:4059-4074 ((https://doi.org/10.1111/mec.13739)); Põlme S, et al. 2020. Fungal Diversity 105:1-16 ((https://doi.org/10.1007/s13225-020-00466-2)) |
| Otu54652 | 1 | k__Fungi(100);p__Basidiomycota(100);c__Microbotryomycetes(100);o__Sporidiobolales(100);f__Sporidiobolales_family_Incertae_sedis(100);g__Sporobolomyces(100);s__Sporobolomyces_lactosus(100); | Sporobolomyces | 13 | Pathotroph-Saprotroph | \|Fungal Parasite\|-Plant Saprotroph | Yeast | NULL | Probable | As litter_saprotroph (Põlme et al. 2020); As mycoparasite (Põlme et al. 2020); Decay fungal material (Põlme et al. 2020) | Tedersoo L, et al. 2014. Science 346:e1256688 ((https://doi.org/10.1126/science.1256688)); Purahong W, et al. 2016. Molecular Ecology 25:4059-4074 ((https://doi.org/10.1111/mec.13739)); Põlme S, et al. 2020. Fungal Diversity 105:1-16 ((https://doi.org/10.1007/s13225-020-00466-2)) |
| Otu54655 | 1 | k__Fungi(100);p__Ascomycota(100);c__Leotiomycetes(100);o__Thelebolales(100);f__Thelebolaceae(100);g__Antarctomyces(100);s__Antarctomyces_psychrotrophicus(100); | Antarctomyces | 13 | Saprotroph | \|Undefined Saprotroph\| | Microfungus | NULL | Probable | As soil_saprotroph (Põlme et al. 2020); Decay soil (Põlme et al. 2020) | Tedersoo L, et al. 2014. Science 346:e1256688 ((https://doi.org/10.1126/science.1256688)); Põlme S, et al. 2020. Fungal Diversity 105:1-16 ((https://doi.org/10.1007/s13225-020-00466-2)) |

| | | | | | | | | | | | |
|---|---|---|---|---|---|---|---|---|---|---|---|
| Otu54656 | 1 | k__Fungi(100);p__Basidiomycota(100);c__Microbotryomycetes(100);o__Sporidiobolales(100);f__Sporidiobolales_family_Incertae_sedis(100);g__Sporobolomyces(100);s__Sporobolomyces_lactosus(100); | Sporobolomyces | 13 | Pathotroph-Saprotroph | \|Fungal Parasite\|-Plant Saprotroph | Yeast | NULL | Probable | As litter_saprotroph (Põlme et al. 2020); As mycoparasite (Põlme et al. 2020); Decay fungal material (Põlme et al. 2020) | Tedersoo L, et al. 2014. Science 346:e1256688 ((https://doi.org/10.1126/science.1256688)); Purahong W, et al. 2016. Molecular Ecology 25:4059-4074 ((https://doi.org/10.1111/mec.13739)); Põlme S, et al. 2020. Fungal Diversity 105:1-16 ((https://doi.org/10.1007/s13225-020-00466-2)) |
| Otu54660 | 1 | k__Fungi(100);p__Ascomycota(100);c__Leotiomycetes(100);o__Helotiales(100);f__Hyaloscyphaceae(100);g__Hyaloscypha(100);s__Hyaloscypha_albohyalina_var_albohyalina(100); | Hyaloscypha | 13 | Pathotroph-Saprotroph-Symbiotroph | Ectomycorrhizal-Ericoid Mycorrhizal-Plant Pathogen-\|Plant Saprotroph\|-Undefined Saprotroph-Wood Saprotroph | Microfungus | NULL | Probable | As litter_saprotroph (Põlme et al. 2020); Decay wood, leaf, fruit, seed (Põlme et al. 2020); Host - mosses (Põlme et al. 2020) | Tedersoo L, et al. 2014. Science 346:e1256688 ((https://doi.org/10.1126/science.1256688)); Põlme S, et al. 2020. Fungal Diversity 105:1-16 ((https://doi.org/10.1007/s13225-020-00466-2)) |
| Otu54661 | 1 | k__Fungi(100);p__Basidiomycota(100);c__Microbotryomycetes(100);o__Sporidiobolales(100);f__Sporidiobolales_family_Incertae_sedis(100);g__Sporobolomyces(100);s__Sporobolomyces_lactosus(100); | Sporobolomyces | 13 | Pathotroph-Saprotroph | \|Fungal Parasite\|-Plant Saprotroph | Yeast | NULL | Probable | As litter_saprotroph (Põlme et al. 2020); As mycoparasite (Põlme et al. 2020); Decay fungal material (Põlme et al. 2020) | Tedersoo L, et al. 2014. Science 346:e1256688 ((https://doi.org/10.1126/science.1256688)); Purahong W, et al. 2016. Molecular Ecology 25:4059-4074 ((https://doi.org/10.1111/mec.13739)); Põlme S, et al. 2020. Fungal Diversity 105:1-16 ((https://doi.org/10.1007/s13225-020-00466-2)) |
| Otu54663 | 1 | k__Fungi(100);p__Ascomycota(100);c__Dothideomycetes(100);o__Capnodiales(100);f__Mycosphaerellaceae(100);f__Mycosphaerellaceae_unclassified(100);f__Mycosphaerellaceae_unclassified(100); | Mycosphaerellaceae | 9 | Pathotroph-Saprotroph | Plant Pathogen-Undefined Saprotroph | Microfungus | NULL | Probable | Biotrophic, necrotrophic or saprobic on various plant tissues (Cannon & Kirk 2007) | Cannon PF, Kirk PM. 2007. Fungal Families of the World. CAB International, Cambridge (ISBN: 978-0851998275) |
| Otu54664 | 1 | k__Fungi(100);p__Ascomycota(100);c__Dothideomycetes(100);o__Pleosporales(100);f__Venturiaceae(100);g__Fusicladium(100);g__Fusicladium_unclassified(100); | Fusicladium | 13 | Pathotroph-Symbiotroph | Endophyte-Lichen Parasite-\|Plant Pathogen\| | Microfungus | NULL | Probable | NULL | Tedersoo L, et al. 2014. Science 346:e1256688 ((https://doi.org/10.1126/science.1256688)); Põlme S, et al. 2020. Fungal Diversity 105:1-16 ((https://doi.org/10.1007/s13225-020-00466-2)) |
| Otu54666 | 1 | k__Fungi(100);p__Ascomycota(100);c__Saccharomycetes(100);o__Saccharomycetales(100);f__Metschnikowiaceae(100);g__Metschnikowia(100);s__Metschnikowia_pulcherrima(100); | Metschnikowia | 13 | Saprotroph | \|Undefined Saprotroph\| | Yeast | NULL | Probable | As nectar-tap_saprotroph (Põlme et al. 2020); Decay sugar-rich substrates (Põlme et al. 2020) | Põlme S, et al. 2020. Fungal Diversity 105:1-16 ((https://doi.org/10.1007/s13225-020-00466-2)) |
| Otu54667 | 1 | k__Fungi(100);p__Ascomycota(100);c__Saccharomycetes(100);o__Saccharomycetales(100);f__Saccharomycetales_family_Incertae_sedis(100);g__Candida(100);g__Candida_unclassified(100); | Candida | 13 | Pathotroph-Saprotroph-Symbiotroph | Animal Pathogen-Endophyte-Endosymbiont-Epiphyte-\|Undefined Saprotroph\| | Dimorphic Yeast-Yeast | NULL | Probable | As nectar-tap_saprotroph (Põlme et al. 2020); Decay sugar-rich substrates (Põlme et al. 2020); This asexual genus is polyphyletic and contain species that function very differently depend on what they are. Caution should be taken when interpreting guilds for Candida species. | Manolakaki D, et al. 2010. Virulence. 1: 367-375 ((https://doi.org/10.4161/viru.1.5.12796)); Mekbib SB, et al. 2011. Tropical Plant Pathology 36:233-240 ((https://doi.org/10.1590/S1982-56762011000400004)); Põlme S, et al. 2020. Fungal Diversity 105:1-16 ((https://doi.org/10.1007/s13225-020-00466-2)) |
| Otu54668 | 1 | k__Fungi(100);p__Ascomycota(100);c__Saccharomycetes(100);o__Saccharomycetales(100);f__Metschnikowiaceae(100);g__Metschnikowia(100);g__Metschnikowia_unclassified(100); | Metschnikowia | 13 | Saprotroph | \|Undefined Saprotroph\| | Yeast | NULL | Probable | As nectar-tap_saprotroph (Põlme et al. 2020); Decay sugar-rich substrates (Põlme et al. 2020) | Põlme S, et al. 2020. Fungal Diversity 105:1-16 ((https://doi.org/10.1007/s13225-020-00466-2)) |
| Otu54669 | 1 | k__Fungi(100);p__Ascomycota(100);c__Saccharomycetes(100);o__Saccharomycetales(100);f__Metschnikowiaceae(100);g__Metschnikowia(100);s__Metschnikowia_pulcherrima(100); | Metschnikowia | 13 | Saprotroph | \|Undefined Saprotroph\| | Yeast | NULL | Probable | As nectar-tap_saprotroph (Põlme et al. 2020); Decay sugar-rich substrates (Põlme et al. 2020) | Põlme S, et al. 2020. Fungal Diversity 105:1-16 ((https://doi.org/10.1007/s13225-020-00466-2)) |
| Otu54670 | 1 | k__Fungi(100);p__Ascomycota(100);c__Saccharomycetes(100);o__Saccharomycetales(100);f__Metschnikowiaceae(100);g__Metschnikowia(100);s__Metschnikowia_pulcherrima(100); | Metschnikowia | 13 | Saprotroph | \|Undefined Saprotroph\| | Yeast | NULL | Probable | As nectar-tap_saprotroph (Põlme et al. 2020); Decay sugar-rich substrates (Põlme et al. 2020) | Põlme S, et al. 2020. Fungal Diversity 105:1-16 ((https://doi.org/10.1007/s13225-020-00466-2)) |
| Otu54672 | 1 | k__Fungi(100);p__Ascomycota(100);c__Saccharomycetes(100);o__Saccharomycetales(100);f__Metschnikowiaceae(100);g__Metschnikowia(100);s__Metschnikowia_pulcherrima(100); | Metschnikowia | 13 | Saprotroph | \|Undefined Saprotroph\| | Yeast | NULL | Probable | As nectar-tap_saprotroph (Põlme et al. 2020); Decay sugar-rich substrates (Põlme et al. 2020) | Põlme S, et al. 2020. Fungal Diversity 105:1-16 ((https://doi.org/10.1007/s13225-020-00466-2)) |
| Otu54673 | 1 | k__Fungi(100);p__Ascomycota(100);c__Saccharomycetes(100);o__Saccharomycetales(100);f__Metschnikowiaceae(100);g__Metschnikowia(100);s__Metschnikowia_pulcherrima(100); | Metschnikowia | 13 | Saprotroph | \|Undefined Saprotroph\| | Yeast | NULL | Probable | As nectar-tap_saprotroph (Põlme et al. 2020); Decay sugar-rich substrates (Põlme et al. 2020) | Põlme S, et al. 2020. Fungal Diversity 105:1-16 ((https://doi.org/10.1007/s13225-020-00466-2)) |
| Otu54674 | 1 | k__Fungi(100);p__Ascomycota(100);c__Saccharomycetes(100);o__Saccharomycetales(100);f__Metschnikowiaceae(100);g__Metschnikowia(100);s__Metschnikowia_pulcherrima(100); | Metschnikowia | 13 | Saprotroph | \|Undefined Saprotroph\| | Yeast | NULL | Probable | As nectar-tap_saprotroph (Põlme et al. 2020); Decay sugar-rich substrates (Põlme et al. 2020) | Põlme S, et al. 2020. Fungal Diversity 105:1-16 ((https://doi.org/10.1007/s13225-020-00466-2)) |
| Otu54675 | 1 | k__Fungi(100);p__Ascomycota(100);c__Saccharomycetes(100);o__Saccharomycetales(100);f__Metschnikowiaceae(100);g__Metschnikowia(100);g__Metschnikowia_unclassified(100); | Metschnikowia | 13 | Saprotroph | \|Undefined Saprotroph\| | Yeast | NULL | Probable | As nectar-tap_saprotroph (Põlme et al. 2020); Decay sugar-rich substrates (Põlme et al. 2020) | Põlme S, et al. 2020. Fungal Diversity 105:1-16 ((https://doi.org/10.1007/s13225-020-00466-2)) |
| Otu54676 | 1 | k__Fungi(100);p__Ascomycota(100);c__Saccharomycetes(100);o__Saccharomycetales(100);f__Metschnikowiaceae(100);g__Metschnikowia(100);g__Metschnikowia_unclassified(100); | Metschnikowia | 13 | Saprotroph | \|Undefined Saprotroph\| | Yeast | NULL | Probable | As nectar-tap_saprotroph (Põlme et al. 2020); Decay sugar-rich substrates (Põlme et al. 2020) | Põlme S, et al. 2020. Fungal Diversity 105:1-16 ((https://doi.org/10.1007/s13225-020-00466-2)) |
| Otu54677 | 1 | k__Fungi(100);p__Ascomycota(100);c__Saccharomycetes(100);o__Saccharomycetales(100);f__Metschnikowiaceae(100);g__Metschnikowia(100);s__Metschnikowia_pulcherrima(100); | Metschnikowia | 13 | Saprotroph | \|Undefined Saprotroph\| | Yeast | NULL | Probable | As nectar-tap_saprotroph (Põlme et al. 2020); Decay sugar-rich substrates (Põlme et al. 2020) | Põlme S, et al. 2020. Fungal Diversity 105:1-16 ((https://doi.org/10.1007/s13225-020-00466-2)) |
| Otu54679 | 1 | k__Fungi(100);p__Ascomycota(100);c__Saccharomycetes(100);o__Saccharomycetales(100);f__Saccharomycetales_family_Incertae_sedis(100);g__Candida(100);g__Candida_unclassified(100); | Candida | 13 | Pathotroph-Saprotroph-Symbiotroph | Animal Pathogen-Endophyte-Endosymbiont-Epiphyte-\|Undefined Saprotroph\| | Dimorphic Yeast-Yeast | NULL | Probable | As nectar-tap_saprotroph (Põlme et al. 2020); Decay sugar-rich substrates (Põlme et al. 2020); This asexual genus is polyphyletic and contain species that function very differently depend on what they are. Caution should be taken when interpreting guilds for Candida species. | Manolakaki D, et al. 2010. Virulence. 1: 367-375 ((https://doi.org/10.4161/viru.1.5.12796)); Mekbib SB, et al. 2011. Tropical Plant Pathology 36:233-240 ((https://doi.org/10.1590/S1982-56762011000400004)); Põlme S, et al. 2020. Fungal Diversity 105:1-16 ((https://doi.org/10.1007/s13225-020-00466-2)) |
| Otu54680 | 1 | k__Fungi(100);p__Ascomycota(100);c__Saccharomycetes(100);o__Saccharomycetales(100);f__Saccharomycetales_family_Incertae_sedis(100);f__Saccharomycetales_family_Incertae_sedis_unclassified(100);f__Saccharomycetales_family_Incertae_sedis_unclassified(100); | Saccharomycetales | 7 | Saprotroph | Undefined Saprotroph | Yeast | NULL | Possible | NULL | Sterkenburg E, et al. 2015. New Phytologist 207:1145-1158 ((https://doi.org/10.1111/nph.13426)) |
| Otu54681 | 1 | k__Fungi(100);p__Ascomycota(100);c__Saccharomycetes(100);o__Saccharomycetales(100);f__Metschnikowiaceae(100);g__Metschnikowia(100);g__Metschnikowia_unclassified(100); | Metschnikowia | 13 | Saprotroph | \|Undefined Saprotroph\| | Yeast | NULL | Probable | As nectar-tap_saprotroph (Põlme et al. 2020); Decay sugar-rich substrates (Põlme et al. 2020) | Põlme S, et al. 2020. Fungal Diversity 105:1-16 ((https://doi.org/10.1007/s13225-020-00466-2)) |
| Otu54683 | 1 | k__Fungi(100);p__Ascomycota(100);c__Saccharomycetes(100);o__Saccharomycetales(100);f__Metschnikowiaceae(100);g__Metschnikowia(100);s__Metschnikowia_pulcherrima(100); | Metschnikowia | 13 | Saprotroph | \|Undefined Saprotroph\| | Yeast | NULL | Probable | As nectar-tap_saprotroph (Põlme et al. 2020); Decay sugar-rich substrates (Põlme et al. 2020) | Põlme S, et al. 2020. Fungal Diversity 105:1-16 ((https://doi.org/10.1007/s13225-020-00466-2)) |
| Otu54686 | 1 | k__Fungi(100);p__Ascomycota(100);c__Saccharomycetes(100);o__Saccharomycetales(100);f__Saccharomycetales_family_Incertae_sedis(100);f__Saccharomycetales_family_Incertae_sedis_unclassified(100);f__Saccharomycetales_family_Incertae_sedis_unclassified(100); | Saccharomycetales | 7 | Saprotroph | Undefined Saprotroph | Yeast | NULL | Possible | NULL | Sterkenburg E, et al. 2015. New Phytologist 207:1145-1158 ((https://doi.org/10.1111/nph.13426)) |
| Otu54688 | 1 | k__Fungi(100);p__Ascomycota(100);c__Saccharomycetes(100);o__Saccharomycetales(100);f__Metschnikowiaceae(100);g__Metschnikowia(100);s__Metschnikowia_pulcherrima(100); | Metschnikowia | 13 | Saprotroph | \|Undefined Saprotroph\| | Yeast | NULL | Probable | As nectar-tap_saprotroph (Põlme et al. 2020); Decay sugar-rich substrates (Põlme et al. 2020) | Põlme S, et al. 2020. Fungal Diversity 105:1-16 ((https://doi.org/10.1007/s13225-020-00466-2)) |
| Otu54689 | 1 | k__Fungi(100);p__Ascomycota(100);c__Saccharomycetes(100);o__Saccharomycetales(100);f__Saccharomycetales_family_Incertae_sedis(100);f__Saccharomycetales_family_Incertae_sedis_unclassified(100);f__Saccharomycetales_family_Incertae_sedis_unclassified(100); | Saccharomycetales | 7 | Saprotroph | Undefined Saprotroph | Yeast | NULL | Possible | NULL | Sterkenburg E, et al. 2015. New Phytologist 207:1145-1158 ((https://doi.org/10.1111/nph.13426)) |
| Otu54690 | 1 | k__Fungi(100);p__Ascomycota(100);c__Saccharomycetes(100);o__Saccharomycetales(100);f__Metschnikowiaceae(100);g__Metschnikowia(100);s__Metschnikowia_pulcherrima(100); | Metschnikowia | 13 | Saprotroph | \|Undefined Saprotroph\| | Yeast | NULL | Probable | As nectar-tap_saprotroph (Põlme et al. 2020); Decay sugar-rich substrates (Põlme et al. 2020) | Põlme S, et al. 2020. Fungal Diversity 105:1-16 ((https://doi.org/10.1007/s13225-020-00466-2)) |
| Otu54692 | 1 | k__Fungi(100);p__Ascomycota(100);c__Saccharomycetes(100);o__Saccharomycetales(100);f__Metschnikowiaceae(100);g__Metschnikowia(100);g__Metschnikowia_unclassified(100); | Metschnikowia | 13 | Saprotroph | \|Undefined Saprotroph\| | Yeast | NULL | Probable | As nectar-tap_saprotroph (Põlme et al. 2020); Decay sugar-rich substrates (Põlme et al. 2020) | Põlme S, et al. 2020. Fungal Diversity 105:1-16 ((https://doi.org/10.1007/s13225-020-00466-2)) |
| Otu54701 | 1 | k__Fungi(100);p__Ascomycota(100);c__Saccharomycetes(100);o__Saccharomycetales(100);f__Saccharomycetales_family_Incertae_sedis(100);g__Candida(100);g__Candida_unclassified(100); | Candida | 13 | Pathotroph-Saprotroph-Symbiotroph | Animal Pathogen-Endophyte-Endosymbiont-Epiphyte-\|Undefined Saprotroph\| | Dimorphic Yeast-Yeast | NULL | Probable | As nectar-tap_saprotroph (Põlme et al. 2020); Decay sugar-rich substrates (Põlme et al. 2020); This asexual genus is polyphyletic and contain species that function very differently depend on what they are. Caution should be taken when interpreting guilds for Candida species. | Manolakaki D, et al. 2010. Virulence. 1: 367-375 ((https://doi.org/10.4161/viru.1.5.12796)); Mekbib SB, et al. 2011. Tropical Plant Pathology 36:233-240 ((https://doi.org/10.1590/S1982-56762011000400004)); Põlme S, et al. 2020. Fungal Diversity 105:1-16 ((https://doi.org/10.1007/s13225-020-00466-2)) |
| Otu54703 | 1 | k__Fungi(100);p__Ascomycota(100);c__Saccharomycetes(100);o__Saccharomycetales(100);f__Metschnikowiaceae(100);g__Metschnikowia(100);g__Metschnikowia_unclassified(100); | Metschnikowia | 13 | Saprotroph | \|Undefined Saprotroph\| | Yeast | NULL | Probable | As nectar-tap_saprotroph (Põlme et al. 2020); Decay sugar-rich substrates (Põlme et al. 2020) | Põlme S, et al. 2020. Fungal Diversity 105:1-16 ((https://doi.org/10.1007/s13225-020-00466-2)) |
| Otu54704 | 1 | k__Fungi(100);p__Ascomycota(100);c__Saccharomycetes(100);o__Saccharomycetales(100);f__Metschnikowiaceae(100);g__Metschnikowia(100);s__Metschnikowia_pulcherrima(100); | Metschnikowia | 13 | Saprotroph | \|Undefined Saprotroph\| | Yeast | NULL | Probable | As nectar-tap_saprotroph (Põlme et al. 2020); Decay sugar-rich substrates (Põlme et al. 2020) | Põlme S, et al. 2020. Fungal Diversity 105:1-16 ((https://doi.org/10.1007/s13225-020-00466-2)) |
| Otu54706 | 1 | k__Fungi(100);p__Ascomycota(100);c__Saccharomycetes(100);o__Saccharomycetales(100);f__Metschnikowiaceae(100);g__Metschnikowia(100);s__Metschnikowia_pulcherrima(100); | Metschnikowia | 13 | Saprotroph | \|Undefined Saprotroph\| | Yeast | NULL | Probable | As nectar-tap_saprotroph (Põlme et al. 2020); Decay sugar-rich substrates (Põlme et al. 2020) | Põlme S, et al. 2020. Fungal Diversity 105:1-16 ((https://doi.org/10.1007/s13225-020-00466-2)) |
| Otu54707 | 1 | k__Fungi(100);p__Ascomycota(100);c__Saccharomycetes(100);o__Saccharomycetales(100);f__Metschnikowiaceae(100);g__Metschnikowia(100);s__Metschnikowia_pulcherrima(100); | Metschnikowia | 13 | Saprotroph | \|Undefined Saprotroph\| | Yeast | NULL | Probable | As nectar-tap_saprotroph (Põlme et al. 2020); Decay sugar-rich substrates (Põlme et al. 2020) | Põlme S, et al. 2020. Fungal Diversity 105:1-16 ((https://doi.org/10.1007/s13225-020-00466-2)) |
| Otu54708 | 1 | k__Fungi(100);p__Ascomycota(100);c__Saccharomycetes(100);o__Saccharomycetales(100);f__Metschnikowiaceae(100);g__Metschnikowia(100);s__Metschnikowia_pulcherrima(100); | Metschnikowia | 13 | Saprotroph | \|Undefined Saprotroph\| | Yeast | NULL | Probable | As nectar-tap_saprotroph (Põlme et al. 2020); Decay sugar-rich substrates (Põlme et al. 2020) | Põlme S, et al. 2020. Fungal Diversity 105:1-16 ((https://doi.org/10.1007/s13225-020-00466-2)) |
| Otu54709 | 1 | k__Fungi(100);p__Ascomycota(100);c__Saccharomycetes(100);o__Saccharomycetales(100);f__Metschnikowiaceae(100);g__Metschnikowia(100);g__Metschnikowia_unclassified(100); | Metschnikowia | 13 | Saprotroph | \|Undefined Saprotroph\| | Yeast | NULL | Probable | As nectar-tap_saprotroph (Põlme et al. 2020); Decay sugar-rich substrates (Põlme et al. 2020) | Põlme S, et al. 2020. Fungal Diversity 105:1-16 ((https://doi.org/10.1007/s13225-020-00466-2)) |
| Otu54710 | 1 | k__Fungi(100);p__Ascomycota(100);c__Saccharomycetes(100);o__Saccharomycetales(100);f__Metschnikowiaceae(100);g__Metschnikowia(100);g__Metschnikowia_unclassified(100); | Metschnikowia | 13 | Saprotroph | \|Undefined Saprotroph\| | Yeast | NULL | Probable | As nectar-tap_saprotroph (Põlme et al. 2020); Decay sugar-rich substrates (Põlme et al. 2020) | Põlme S, et al. 2020. Fungal Diversity 105:1-16 ((https://doi.org/10.1007/s13225-020-00466-2)) |
| Otu54711 | 1 | k__Fungi(100);p__Ascomycota(100);c__Saccharomycetes(100);o__Saccharomycetales(100);f__Metschnikowiaceae(100);g__Metschnikowia(100);s__Metschnikowia_pulcherrima(100); | Metschnikowia | 13 | Saprotroph | \|Undefined Saprotroph\| | Yeast | NULL | Probable | As nectar-tap_saprotroph (Põlme et al. 2020); Decay sugar-rich substrates (Põlme et al. 2020) | Põlme S, et al. 2020. Fungal Diversity 105:1-16 ((https://doi.org/10.1007/s13225-020-00466-2)) |
| Otu54712 | 1 | k__Fungi(100);p__Ascomycota(100);c__Saccharomycetes(100);o__Saccharomycetales(100);f__Metschnikowiaceae(100);g__Metschnikowia(100);g__Metschnikowia_unclassified(100); | Metschnikowia | 13 | Saprotroph | \|Undefined Saprotroph\| | Yeast | NULL | Probable | As nectar-tap_saprotroph (Põlme et al. 2020); Decay sugar-rich substrates (Põlme et al. 2020) | Põlme S, et al. 2020. Fungal Diversity 105:1-16 ((https://doi.org/10.1007/s13225-020-00466-2)) |
| Otu54713 | 1 | k__Fungi(100);p__Ascomycota(100);c__Saccharomycetes(100);o__Saccharomycetales(100);f__Metschnikowiaceae(100);g__Metschnikowia(100);s__Metschnikowia_pulcherrima(100); | Metschnikowia | 13 | Saprotroph | \|Undefined Saprotroph\| | Yeast | NULL | Probable | As nectar-tap_saprotroph (Põlme et al. 2020); Decay sugar-rich substrates (Põlme et al. 2020) | Põlme S, et al. 2020. Fungal Diversity 105:1-16 ((https://doi.org/10.1007/s13225-020-00466-2)) |
| Otu54714 | 1 | k__Fungi(100);p__Ascomycota(100);c__Saccharomycetes(100);o__Saccharomycetales(100);f__Metschnikowiaceae(100);g__Metschnikowia(100);s__Metschnikowia_pulcherrima(100); | Metschnikowia | 13 | Saprotroph | \|Undefined Saprotroph\| | Yeast | NULL | Probable | As nectar-tap_saprotroph (Põlme et al. 2020); Decay sugar-rich substrates (Põlme et al. 2020) | Põlme S, et al. 2020. Fungal Diversity 105:1-16 ((https://doi.org/10.1007/s13225-020-00466-2)) |
| Otu54715 | 1 | k__Fungi(100);p__Ascomycota(100);c__Saccharomycetes(100);o__Saccharomycetales(100);f__Metschnikowiaceae(100);g__Metschnikowia(100);s__Metschnikowia_pulcherrima(100); | Metschnikowia | 13 | Saprotroph | \|Undefined Saprotroph\| | Yeast | NULL | Probable | As nectar-tap_saprotroph (Põlme et al. 2020); Decay sugar-rich substrates (Põlme et al. 2020) | Põlme S, et al. 2020. Fungal Diversity 105:1-16 ((https://doi.org/10.1007/s13225-020-00466-2)) |
| Otu54716 | 1 | k__Fungi(100);p__Ascomycota(100);c__Saccharomycetes(100);o__Saccharomycetales(100);f__Metschnikowiaceae(100);g__Metschnikowia(100);g__Metschnikowia_unclassified(100); | Metschnikowia | 13 | Saprotroph | \|Undefined Saprotroph\| | Yeast | NULL | Probable | As nectar-tap_saprotroph (Põlme et al. 2020); Decay sugar-rich substrates (Põlme et al. 2020) | Põlme S, et al. 2020. Fungal Diversity 105:1-16 ((https://doi.org/10.1007/s13225-020-00466-2)) |
| Otu54717 | 1 | k__Fungi(100);p__Ascomycota(100);c__Saccharomycetes(100);o__Saccharomycetales(100);f__Metschnikowiaceae(100);g__Metschnikowia(100);s__Metschnikowia_pulcherrima(100); | Metschnikowia | 13 | Saprotroph | \|Undefined Saprotroph\| | Yeast | NULL | Probable | As nectar-tap_saprotroph (Põlme et al. 2020); Decay sugar-rich substrates (Põlme et al. 2020) | Põlme S, et al. 2020. Fungal Diversity 105:1-16 ((https://doi.org/10.1007/s13225-020-00466-2)) |
| Otu54718 | 1 | k__Fungi(100);p__Ascomycota(100);c__Saccharomycetes(100);o__Saccharomycetales(100);f__Metschnikowiaceae(100);g__Metschnikowia(100);g__Metschnikowia_unclassified(100); | Metschnikowia | 13 | Saprotroph | \|Undefined Saprotroph\| | Yeast | NULL | Probable | As nectar-tap_saprotroph (Põlme et al. 2020); Decay sugar-rich substrates (Põlme et al. 2020) | Põlme S, et al. 2020. Fungal Diversity 105:1-16 ((https://doi.org/10.1007/s13225-020-00466-2)) |
| Otu54719 | 1 | k__Fungi(100);p__Ascomycota(100);c__Saccharomycetes(100);o__Saccharomycetales(100);f__Saccharomycetales_family_Incertae_sedis(100);g__Candida(100);g__Candida_unclassified(100); | Candida | 13 | Pathotroph-Saprotroph-Symbiotroph | Animal Pathogen-Endophyte-Endosymbiont-Epiphyte-\|Undefined Saprotroph\| | Dimorphic Yeast-Yeast | NULL | Probable | As nectar-tap_saprotroph (Põlme et al. 2020); Decay sugar-rich substrates (Põlme et al. 2020); This asexual genus is polyphyletic and contain species that function very differently depend on what they are. Caution should be taken when interpreting guilds for Candida species. | Manolakaki D, et al. 2010. Virulence. 1: 367-375 ((https://doi.org/10.4161/viru.1.5.12796)); Mekbib SB, et al. 2011. Tropical Plant Pathology 36:233-240 ((https://doi.org/10.1590/S1982-56762011000400004)); Põlme S, et al. 2020. Fungal Diversity 105:1-16 ((https://doi.org/10.1007/s13225-020-00466-2)) |
| Otu54725 | 1 | k__Fungi(100);p__Ascomycota(100);c__Saccharomycetes(100);o__Saccharomycetales(100);f__Saccharomycetales_family_Incertae_sedis(100);g__Candida(100);g__Candida_unclassified(100); | Candida | 13 | Pathotroph-Saprotroph-Symbiotroph | Animal Pathogen-Endophyte-Endosymbiont-Epiphyte-\|Undefined Saprotroph\| | Dimorphic Yeast-Yeast | NULL | Probable | As nectar-tap_saprotroph (Põlme et al. 2020); Decay sugar-rich substrates (Põlme et al. 2020); This asexual genus is polyphyletic and contain species that function very differently depend on what they are. Caution should be taken when interpreting guilds for Candida species. | Manolakaki D, et al. 2010. Virulence. 1: 367-375 ((https://doi.org/10.4161/viru.1.5.12796)); Mekbib SB, et al. 2011. Tropical Plant Pathology 36:233-240 ((https://doi.org/10.1590/S1982-56762011000400004)); Põlme S, et al. 2020. Fungal Diversity 105:1-16 ((https://doi.org/10.1007/s13225-020-00466-2)) |
| Otu54727 | 1 | k__Fungi(100);p__Ascomycota(100);c__Saccharomycetes(100);o__Saccharomycetales(100);f__Metschnikowiaceae(100);g__Metschnikowia(100);s__Metschnikowia_pulcherrima(100); | Metschnikowia | 13 | Saprotroph | \|Undefined Saprotroph\| | Yeast | NULL | Probable | As nectar-tap_saprotroph (Põlme et al. 2020); Decay sugar-rich substrates (Põlme et al. 2020) | Põlme S, et al. 2020. Fungal Diversity 105:1-16 ((https://doi.org/10.1007/s13225-020-00466-2)) |
| Otu54729 | 1 | k__Fungi(100);p__Ascomycota(100);c__Saccharomycetes(100);o__Saccharomycetales(100);o__Saccharomycetales_unclassified(100);o__Saccharomycetales_unclassified(100);o__Saccharomycetales_unclassified(100); | Saccharomycetales | 7 | Saprotroph | Undefined Saprotroph | Yeast | NULL | Possible | NULL | Sterkenburg E, et al. 2015. New Phytologist 207:1145-1158 ((https://doi.org/10.1111/nph.13426)) |
| Otu54731 | 1 | k__Fungi(100);p__Ascomycota(100);c__Saccharomycetes(100);o__Saccharomycetales(100);f__Saccharomycetales_family_Incertae_sedis(100);g__Candida(100);g__Candida_unclassified(100); | Candida | 13 | Pathotroph-Saprotroph-Symbiotroph | Animal Pathogen-Endophyte-Endosymbiont-Epiphyte-\|Undefined Saprotroph\| | Dimorphic Yeast-Yeast | NULL | Probable | As nectar-tap_saprotroph (Põlme et al. 2020); Decay sugar-rich substrates (Põlme et al. 2020); This asexual genus is polyphyletic and contain species that function very differently depend on what they are. Caution should be taken when interpreting guilds for Candida species. | Manolakaki D, et al. 2010. Virulence. 1: 367-375 ((https://doi.org/10.4161/viru.1.5.12796)); Mekbib SB, et al. 2011. Tropical Plant Pathology 36:233-240 ((https://doi.org/10.1590/S1982-56762011000400004)); Põlme S, et al. 2020. Fungal Diversity 105:1-16 ((https://doi.org/10.1007/s13225-020-00466-2)) |
| Otu54734 | 1 | k__Fungi(100);p__Ascomycota(100);c__Saccharomycetes(100);o__Saccharomycetales(100);f__Metschnikowiaceae(100);g__Metschnikowia(100);s__Metschnikowia_pulcherrima(100); | Metschnikowia | 13 | Saprotroph | \|Undefined Saprotroph\| | Yeast | NULL | Probable | As nectar-tap_saprotroph (Põlme et al. 2020); Decay sugar-rich substrates (Põlme et al. 2020) | Põlme S, et al. 2020. Fungal Diversity 105:1-16 ((https://doi.org/10.1007/s13225-020-00466-2)) |
| Otu54735 | 1 | k__Fungi(100);p__Ascomycota(100);c__Saccharomycetes(100);o__Saccharomycetales(100);f__Metschnikowiaceae(100);g__Metschnikowia(100);s__Metschnikowia_pulcherrima(100); | Metschnikowia | 13 | Saprotroph | \|Undefined Saprotroph\| | Yeast | NULL | Probable | As nectar-tap_saprotroph (Põlme et al. 2020); Decay sugar-rich substrates (Põlme et al. 2020) | Põlme S, et al. 2020. Fungal Diversity 105:1-16 ((https://doi.org/10.1007/s13225-020-00466-2)) |
| Otu54736 | 1 | k__Fungi(100);p__Ascomycota(100);c__Saccharomycetes(100);o__Saccharomycetales(100);f__Metschnikowiaceae(100);g__Metschnikowia(100);g__Metschnikowia_unclassified(100); | Metschnikowia | 13 | Saprotroph | \|Undefined Saprotroph\| | Yeast | NULL | Probable | As nectar-tap_saprotroph (Põlme et al. 2020); Decay sugar-rich substrates (Põlme et al. 2020) | Põlme S, et al. 2020. Fungal Diversity 105:1-16 ((https://doi.org/10.1007/s13225-020-00466-2)) |
| Otu54742 | 1 | k__Fungi(100);p__Ascomycota(100);c__Saccharomycetes(100);o__Saccharomycetales(100);f__Saccharomycetales_family_Incertae_sedis(100);g__Candida(100);g__Candida_unclassified(100); | Candida | 13 | Pathotroph-Saprotroph-Symbiotroph | Animal Pathogen-Endophyte-Endosymbiont-Epiphyte-\|Undefined Saprotroph\| | Dimorphic Yeast-Yeast | NULL | Probable | As nectar-tap_saprotroph (Põlme et al. 2020); Decay sugar-rich substrates (Põlme et al. 2020); This asexual genus is polyphyletic and contain species that function very differently depend on what they are. Caution should be taken when interpreting guilds for Candida species. | Manolakaki D, et al. 2010. Virulence. 1: 367-375 ((https://doi.org/10.4161/viru.1.5.12796)); Mekbib SB, et al. 2011. Tropical Plant Pathology 36:233-240 ((https://doi.org/10.1590/S1982-56762011000400004)); Põlme S, et al. 2020. Fungal Diversity 105:1-16 ((https://doi.org/10.1007/s13225-020-00466-2)) |
| Otu54744 | 1 | k__Fungi(100);p__Ascomycota(100);c__Saccharomycetes(100);o__Saccharomycetales(100);f__Metschnikowiaceae(100);g__Metschnikowia(100);s__Metschnikowia_pulcherrima(100); | Metschnikowia | 13 | Saprotroph | \|Undefined Saprotroph\| | Yeast | NULL | Probable | As nectar-tap_saprotroph (Põlme et al. 2020); Decay sugar-rich substrates (Põlme et al. 2020) | Põlme S, et al. 2020. Fungal Diversity 105:1-16 ((https://doi.org/10.1007/s13225-020-00466-2)) |
| Otu54747 | 1 | k__Fungi(100);p__Ascomycota(100);c__Saccharomycetes(100);o__Saccharomycetales(100);f__Metschnikowiaceae(100);g__Metschnikowia(100);g__Metschnikowia_unclassified(100); | Metschnikowia | 13 | Saprotroph | \|Undefined Saprotroph\| | Yeast | NULL | Probable | As nectar-tap_saprotroph (Põlme et al. 2020); Decay sugar-rich substrates (Põlme et al. 2020) | Põlme S, et al. 2020. Fungal Diversity 105:1-16 ((https://doi.org/10.1007/s13225-020-00466-2)) |
| Otu54748 | 1 | k__Fungi(100);p__Ascomycota(100);c__Dothideomycetes(100);o__Dothideomycetes_order_Incertae_sedis(100);f__Pseudeurotiaceae(100);g__Pseudeurotium(100);g__Pseudeurotium_unclassified(100); | Pseudeurotium | 13 | Saprotroph | \|Dung Saprotroph\|-Undefined Saprotroph | Microfungus | NULL | Probable | As soil_saprotroph (Põlme et al. 2020); Decay soil, dung (Põlme et al. 2020) | Tedersoo L, et al. 2014. Science 346:e1256688 ((https://doi.org/10.1126/science.1256688)); Põlme S, et al. 2020. Fungal Diversity 105:1-16 ((https://doi.org/10.1007/s13225-020-00466-2)) |
| Otu54749 | 1 | k__Fungi(100);p__Ascomycota(100);c__Saccharomycetes(100);o__Saccharomycetales(100);f__Metschnikowiaceae(100);g__Metschnikowia(100);s__Metschnikowia_pulcherrima(100); | Metschnikowia | 13 | Saprotroph | \|Undefined Saprotroph\| | Yeast | NULL | Probable | As nectar-tap_saprotroph (Põlme et al. 2020); Decay sugar-rich substrates (Põlme et al. 2020) | Põlme S, et al. 2020. Fungal Diversity 105:1-16 ((https://doi.org/10.1007/s13225-020-00466-2)) |
| Otu54750 | 1 | k__Fungi(100);p__Ascomycota(100);c__Saccharomycetes(100);o__Saccharomycetales(100);f__Metschnikowiaceae(100);g__Metschnikowia(100);s__Metschnikowia_pulcherrima(100); | Metschnikowia | 13 | Saprotroph | \|Undefined Saprotroph\| | Yeast | NULL | Probable | As nectar-tap_saprotroph (Põlme et al. 2020); Decay sugar-rich substrates (Põlme et al. 2020) | Põlme S, et al. 2020. Fungal Diversity 105:1-16 ((https://doi.org/10.1007/s13225-020-00466-2)) |
| Otu54751 | 1 | k__Fungi(100);p__Ascomycota(100);c__Saccharomycetes(100);o__Saccharomycetales(100);f__Saccharomycetales_family_Incertae_sedis(100);g__Candida(100);s__Candida_cretensis(100); | Candida | 13 | Pathotroph-Saprotroph-Symbiotroph | Animal Pathogen-Endophyte-Endosymbiont-Epiphyte-\|Undefined Saprotroph\| | Dimorphic Yeast-Yeast | NULL | Probable | As nectar-tap_saprotroph (Põlme et al. 2020); Decay sugar-rich substrates (Põlme et al. 2020); This asexual genus is polyphyletic and contain species that function very differently depend on what they are. Caution should be taken when interpreting guilds for Candida species. | Manolakaki D, et al. 2010. Virulence. 1: 367-375 ((https://doi.org/10.4161/viru.1.5.12796)); Mekbib SB, et al. 2011. Tropical Plant Pathology 36:233-240 ((https://doi.org/10.1590/S1982-56762011000400004)); Põlme S, et al. 2020. Fungal Diversity 105:1-16 ((https://doi.org/10.1007/s13225-020-00466-2)) |
| Otu54752 | 1 | k__Fungi(100);p__Ascomycota(100);c__Saccharomycetes(100);o__Saccharomycetales(100);f__Metschnikowiaceae(100);g__Metschnikowia(100);s__Metschnikowia_pulcherrima(100); | Metschnikowia | 13 | Saprotroph | \|Undefined Saprotroph\| | Yeast | NULL | Probable | As nectar-tap_saprotroph (Põlme et al. 2020); Decay sugar-rich substrates (Põlme et al. 2020) | Põlme S, et al. 2020. Fungal Diversity 105:1-16 ((https://doi.org/10.1007/s13225-020-00466-2)) |
| Otu54754 | 1 | k__Fungi(100);p__Ascomycota(100);c__Saccharomycetes(100);o__Saccharomycetales(100);f__Metschnikowiaceae(100);g__Metschnikowia(100);s__Metschnikowia_pulcherrima(100); | Metschnikowia | 13 | Saprotroph | \|Undefined Saprotroph\| | Yeast | NULL | Probable | As nectar-tap_saprotroph (Põlme et al. 2020); Decay sugar-rich substrates (Põlme et al. 2020) | Põlme S, et al. 2020. Fungal Diversity 105:1-16 ((https://doi.org/10.1007/s13225-020-00466-2)) |
| Otu54755 | 1 | k__Fungi(100);p__Ascomycota(100);c__Saccharomycetes(100);o__Saccharomycetales(100);f__Metschnikowiaceae(100);g__Metschnikowia(100);s__Metschnikowia_pulcherrima(100); | Metschnikowia | 13 | Saprotroph | \|Undefined Saprotroph\| | Yeast | NULL | Probable | As nectar-tap_saprotroph (Põlme et al. 2020); Decay sugar-rich substrates (Põlme et al. 2020) | Põlme S, et al. 2020. Fungal Diversity 105:1-16 ((https://doi.org/10.1007/s13225-020-00466-2)) |
| Otu54756 | 1 | k__Fungi(100);p__Ascomycota(100);c__Saccharomycetes(100);o__Saccharomycetales(100);f__Metschnikowiaceae(100);g__Metschnikowia(100);g__Metschnikowia_unclassified(100); | Metschnikowia | 13 | Saprotroph | \|Undefined Saprotroph\| | Yeast | NULL | Probable | As nectar-tap_saprotroph (Põlme et al. 2020); Decay sugar-rich substrates (Põlme et al. 2020) | Põlme S, et al. 2020. Fungal Diversity 105:1-16 ((https://doi.org/10.1007/s13225-020-00466-2)) |
| Otu54762 | 1 | k__Fungi(100);p__Basidiomycota(100);c__Microbotryomycetes(100);o__Sporidiobolales(100);f__Sporidiobolales_family_Incertae_sedis(100);g__Sporobolomyces(100);s__Sporobolomyces_lactosus(100); | Sporobolomyces | 13 | Pathotroph-Saprotroph | \|Fungal Parasite\|-Plant Saprotroph | Yeast | NULL | Probable | As litter_saprotroph (Põlme et al. 2020); As mycoparasite (Põlme et al. 2020); Decay fungal material (Põlme et al. 2020) | Tedersoo L, et al. 2014. Science 346:e1256688 ((https://doi.org/10.1126/science.1256688)); Purahong W, et al. 2016. Molecular Ecology 25:4059-4074 ((https://doi.org/10.1111/mec.13739)); Põlme S, et al. 2020. Fungal Diversity 105:1-16 ((https://doi.org/10.1007/s13225-020-00466-2)) |
| Otu54763 | 1 | k__Fungi(100);p__Basidiomycota(100);c__Microbotryomycetes(100);o__Sporidiobolales(100);f__Sporidiobolales_family_Incertae_sedis(100);g__Sporobolomyces(100);s__Sporobolomyces_lactosus(100); | Sporobolomyces | 13 | Pathotroph-Saprotroph | \|Fungal Parasite\|-Plant Saprotroph | Yeast | NULL | Probable | As litter_saprotroph (Põlme et al. 2020); As mycoparasite (Põlme et al. 2020); Decay fungal material (Põlme et al. 2020) | Tedersoo L, et al. 2014. Science 346:e1256688 ((https://doi.org/10.1126/science.1256688)); Purahong W, et al. 2016. Molecular Ecology 25:4059-4074 ((https://doi.org/10.1111/mec.13739)); Põlme S, et al. 2020. Fungal Diversity 105:1-16 ((https://doi.org/10.1007/s13225-020-00466-2)) |
| Otu54764 | 1 | k__Fungi(100);p__Ascomycota(100);c__Saccharomycetes(100);o__Saccharomycetales(100);f__Saccharomycetales_family_Incertae_sedis(100);g__Candida(100);g__Candida_unclassified(100); | Candida | 13 | Pathotroph-Saprotroph-Symbiotroph | Animal Pathogen-Endophyte-Endosymbiont-Epiphyte-\|Undefined Saprotroph\| | Dimorphic Yeast-Yeast | NULL | Probable | As nectar-tap_saprotroph (Põlme et al. 2020); Decay sugar-rich substrates (Põlme et al. 2020); This asexual genus is polyphyletic and contain species that function very differently depend on what they are. Caution should be taken when interpreting guilds for Candida species. | Manolakaki D, et al. 2010. Virulence. 1: 367-375 ((https://doi.org/10.4161/viru.1.5.12796)); Mekbib SB, et al. 2011. Tropical Plant Pathology 36:233-240 ((https://doi.org/10.1590/S1982-56762011000400004)); Põlme S, et al. 2020. Fungal Diversity 105:1-16 ((https://doi.org/10.1007/s13225-020-00466-2)) |
| Otu54765 | 1 | k__Fungi(100);p__Ascomycota(100);c__Saccharomycetes(100);o__Saccharomycetales(100);f__Saccharomycetales_family_Incertae_sedis(100);g__Candida(100);g__Candida_unclassified(100); | Candida | 13 | Pathotroph-Saprotroph-Symbiotroph | Animal Pathogen-Endophyte-Endosymbiont-Epiphyte-\|Undefined Saprotroph\| | Dimorphic Yeast-Yeast | NULL | Probable | As nectar-tap_saprotroph (Põlme et al. 2020); Decay sugar-rich substrates (Põlme et al. 2020); This asexual genus is polyphyletic and contain species that function very differently depend on what they are. Caution should be taken when interpreting guilds for Candida species. | Manolakaki D, et al. 2010. Virulence. 1: 367-375 ((https://doi.org/10.4161/viru.1.5.12796)); Mekbib SB, et al. 2011. Tropical Plant Pathology 36:233-240 ((https://doi.org/10.1590/S1982-56762011000400004)); Põlme S, et al. 2020. Fungal Diversity 105:1-16 ((https://doi.org/10.1007/s13225-020-00466-2)) |
| Otu54766 | 1 | k__Fungi(100);p__Ascomycota(100);c__Saccharomycetes(100);o__Saccharomycetales(100);f__Saccharomycetales_family_Incertae_sedis(100);g__Candida(100);g__Candida_unclassified(100); | Candida | 13 | Pathotroph-Saprotroph-Symbiotroph | Animal Pathogen-Endophyte-Endosymbiont-Epiphyte-\|Undefined Saprotroph\| | Dimorphic Yeast-Yeast | NULL | Probable | As nectar-tap_saprotroph (Põlme et al. 2020); Decay sugar-rich substrates (Põlme et al. 2020); This asexual genus is polyphyletic and contain species that function very differently depend on what they are. Caution should be taken when interpreting guilds for Candida species. | Manolakaki D, et al. 2010. Virulence. 1: 367-375 ((https://doi.org/10.4161/viru.1.5.12796)); Mekbib SB, et al. 2011. Tropical Plant Pathology 36:233-240 ((https://doi.org/10.1590/S1982-56762011000400004)); Põlme S, et al. 2020. Fungal Diversity 105:1-16 ((https://doi.org/10.1007/s13225-020-00466-2)) |
| Otu54773 | 1 | k__Fungi(100);p__Ascomycota(100);c__Saccharomycetes(100);o__Saccharomycetales(100);f__Saccharomycetales_family_Incertae_sedis(100);g__Candida(100);s__Candida_cretensis(100); | Candida | 13 | Pathotroph-Saprotroph-Symbiotroph | Animal Pathogen-Endophyte-Endosymbiont-Epiphyte-\|Undefined Saprotroph\| | Dimorphic Yeast-Yeast | NULL | Probable | As nectar-tap_saprotroph (Põlme et al. 2020); Decay sugar-rich substrates (Põlme et al. 2020); This asexual genus is polyphyletic and contain species that function very differently depend on what they are. Caution should be taken when interpreting guilds for Candida species. | Manolakaki D, et al. 2010. Virulence. 1: 367-375 ((https://doi.org/10.4161/viru.1.5.12796)); Mekbib SB, et al. 2011. Tropical Plant Pathology 36:233-240 ((https://doi.org/10.1590/S1982-56762011000400004)); Põlme S, et al. 2020. Fungal Diversity 105:1-16 ((https://doi.org/10.1007/s13225-020-00466-2)) |
| Otu54775 | 1 | k__Fungi(100);p__Ascomycota(100);c__Saccharomycetes(100);o__Saccharomycetales(100);f__Saccharomycetales_family_Incertae_sedis(100);g__Candida(100);g__Candida_unclassified(100); | Candida | 13 | Pathotroph-Saprotroph-Symbiotroph | Animal Pathogen-Endophyte-Endosymbiont-Epiphyte-\|Undefined Saprotroph\| | Dimorphic Yeast-Yeast | NULL | Probable | As nectar-tap_saprotroph (Põlme et al. 2020); Decay sugar-rich substrates (Põlme et al. 2020); This asexual genus is polyphyletic and contain species that function very differently depend on what they are. Caution should be taken when interpreting guilds for Candida species. | Manolakaki D, et al. 2010. Virulence. 1: 367-375 ((https://doi.org/10.4161/viru.1.5.12796)); Mekbib SB, et al. 2011. Tropical Plant Pathology 36:233-240 ((https://doi.org/10.1590/S1982-56762011000400004)); Põlme S, et al. 2020. Fungal Diversity 105:1-16 ((https://doi.org/10.1007/s13225-020-00466-2)) |
| Otu54779 | 1 | k__Fungi(100);p__Ascomycota(100);c__Saccharomycetes(100);o__Saccharomycetales(100);f__Saccharomycetales_family_Incertae_sedis(100);g__Candida(100);g__Candida_unclassified(100); | Candida | 13 | Pathotroph-Saprotroph-Symbiotroph | Animal Pathogen-Endophyte-Endosymbiont-Epiphyte-\|Undefined Saprotroph\| | Dimorphic Yeast-Yeast | NULL | Probable | As nectar-tap_saprotroph (Põlme et al. 2020); Decay sugar-rich substrates (Põlme et al. 2020); This asexual genus is polyphyletic and contain species that function very differently depend on what they are. Caution should be taken when interpreting guilds for Candida species. | Manolakaki D, et al. 2010. Virulence. 1: 367-375 ((https://doi.org/10.4161/viru.1.5.12796)); Mekbib SB, et al. 2011. Tropical Plant Pathology 36:233-240 ((https://doi.org/10.1590/S1982-56762011000400004)); Põlme S, et al. 2020. Fungal Diversity 105:1-16 ((https://doi.org/10.1007/s13225-020-00466-2)) |
| Otu54780 | 1 | k__Fungi(100);p__Ascomycota(100);c__Saccharomycetes(100);o__Saccharomycetales(100);o__Saccharomycetales_unclassified(100);o__Saccharomycetales_unclassified(100);o__Saccharomycetales_unclassified(100); | Saccharomycetales | 7 | Saprotroph | Undefined Saprotroph | Yeast | NULL | Possible | NULL | Sterkenburg E, et al. 2015. New Phytologist 207:1145-1158 ((https://doi.org/10.1111/nph.13426)) |
| Otu54782 | 1 | k__Fungi(100);p__Ascomycota(100);c__Saccharomycetes(100);o__Saccharomycetales(100);f__Saccharomycetales_family_Incertae_sedis(100);g__Candida(100);s__Candida_cretensis(100); | Candida | 13 | Pathotroph-Saprotroph-Symbiotroph | Animal Pathogen-Endophyte-Endosymbiont-Epiphyte-\|Undefined Saprotroph\| | Dimorphic Yeast-Yeast | NULL | Probable | As nectar-tap_saprotroph (Põlme et al. 2020); Decay sugar-rich substrates (Põlme et al. 2020); This asexual genus is polyphyletic and contain species that function very differently depend on what they are. Caution should be taken when interpreting guilds for Candida species. | Manolakaki D, et al. 2010. Virulence. 1: 367-375 ((https://doi.org/10.4161/viru.1.5.12796)); Mekbib SB, et al. 2011. Tropical Plant Pathology 36:233-240 ((https://doi.org/10.1590/S1982-56762011000400004)); Põlme S, et al. 2020. Fungal Diversity 105:1-16 ((https://doi.org/10.1007/s13225-020-00466-2)) |
| Otu54785 | 1 | k__Fungi(100);p__Ascomycota(100);c__Saccharomycetes(100);o__Saccharomycetales(100);f__Metschnikowiaceae(100);g__Metschnikowia(100);s__Metschnikowia_pulcherrima(100); | Metschnikowia | 13 | Saprotroph | \|Undefined Saprotroph\| | Yeast | NULL | Probable | As nectar-tap_saprotroph (Põlme et al. 2020); Decay sugar-rich substrates (Põlme et al. 2020) | Põlme S, et al. 2020. Fungal Diversity 105:1-16 ((https://doi.org/10.1007/s13225-020-00466-2)) |
| Otu54786 | 1 | k__Fungi(100);p__Ascomycota(100);c__Saccharomycetes(100);o__Saccharomycetales(100);f__Metschnikowiaceae(100);g__Metschnikowia(100);s__Metschnikowia_pulcherrima(100); | Metschnikowia | 13 | Saprotroph | \|Undefined Saprotroph\| | Yeast | NULL | Probable | As nectar-tap_saprotroph (Põlme et al. 2020); Decay sugar-rich substrates (Põlme et al. 2020) | Põlme S, et al. 2020. Fungal Diversity 105:1-16 ((https://doi.org/10.1007/s13225-020-00466-2)) |
| Otu54792 | 1 | k__Fungi(100);p__Ascomycota(100);c__Saccharomycetes(100);o__Saccharomycetales(100);f__Saccharomycetales_family_Incertae_sedis(100);f__Saccharomycetales_family_Incertae_sedis_unclassified(100);f__Saccharomycetales_family_Incertae_sedis_unclassified(100); | Saccharomycetales | 7 | Saprotroph | Undefined Saprotroph | Yeast | NULL | Possible | NULL | Sterkenburg E, et al. 2015. New Phytologist 207:1145-1158 ((https://doi.org/10.1111/nph.13426)) |
| Otu54793 | 1 | k__Fungi(100);p__Ascomycota(100);c__Saccharomycetes(100);o__Saccharomycetales(100);f__Saccharomycetales_family_Incertae_sedis(100);g__Candida(100);g__Candida_unclassified(100); | Candida | 13 | Pathotroph-Saprotroph-Symbiotroph | Animal Pathogen-Endophyte-Endosymbiont-Epiphyte-\|Undefined Saprotroph\| | Dimorphic Yeast-Yeast | NULL | Probable | As nectar-tap_saprotroph (Põlme et al. 2020); Decay sugar-rich substrates (Põlme et al. 2020); This asexual genus is polyphyletic and contain species that function very differently depend on what they are. Caution should be taken when interpreting guilds for Candida species. | Manolakaki D, et al. 2010. Virulence. 1: 367-375 ((https://doi.org/10.4161/viru.1.5.12796)); Mekbib SB, et al. 2011. Tropical Plant Pathology 36:233-240 ((https://doi.org/10.1590/S1982-56762011000400004)); Põlme S, et al. 2020. Fungal Diversity 105:1-16 ((https://doi.org/10.1007/s13225-020-00466-2)) |
| Otu54796 | 1 | k__Fungi(100);p__Basidiomycota(100);c__Microbotryomycetes(100);o__Sporidiobolales(100);f__Sporidiobolales_family_Incertae_sedis(100);g__Sporobolomyces(100);s__Sporobolomyces_lactosus(100); | Sporobolomyces | 13 | Pathotroph-Saprotroph | \|Fungal Parasite\|-Plant Saprotroph | Yeast | NULL | Probable | As litter_saprotroph (Põlme et al. 2020); As mycoparasite (Põlme et al. 2020); Decay fungal material (Põlme et al. 2020) | Tedersoo L, et al. 2014. Science 346:e1256688 ((https://doi.org/10.1126/science.1256688)); Purahong W, et al. 2016. Molecular Ecology 25:4059-4074 ((https://doi.org/10.1111/mec.13739)); Põlme S, et al. 2020. Fungal Diversity 105:1-16 ((https://doi.org/10.1007/s13225-020-00466-2)) |
| Otu54798 | 1 | k__Fungi(100);p__Ascomycota(100);c__Saccharomycetes(100);o__Saccharomycetales(100);f__Metschnikowiaceae(100);g__Metschnikowia(100);s__Metschnikowia_pulcherrima(100); | Metschnikowia | 13 | Saprotroph | \|Undefined Saprotroph\| | Yeast | NULL | Probable | As nectar-tap_saprotroph (Põlme et al. 2020); Decay sugar-rich substrates (Põlme et al. 2020) | Põlme S, et al. 2020. Fungal Diversity 105:1-16 ((https://doi.org/10.1007/s13225-020-00466-2)) |
| Otu54803 | 1 | k__Fungi(100);p__Basidiomycota(100);c__Microbotryomycetes(100);o__Sporidiobolales(100);f__Sporidiobolales_family_Incertae_sedis(100);g__Sporobolomyces(100);s__Sporobolomyces_lactosus(100); | Sporobolomyces | 13 | Pathotroph-Saprotroph | \|Fungal Parasite\|-Plant Saprotroph | Yeast | NULL | Probable | As litter_saprotroph (Põlme et al. 2020); As mycoparasite (Põlme et al. 2020); Decay fungal material (Põlme et al. 2020) | Tedersoo L, et al. 2014. Science 346:e1256688 ((https://doi.org/10.1126/science.1256688)); Purahong W, et al. 2016. Molecular Ecology 25:4059-4074 ((https://doi.org/10.1111/mec.13739)); Põlme S, et al. 2020. Fungal Diversity 105:1-16 ((https://doi.org/10.1007/s13225-020-00466-2)) |
| Otu54804 | 1 | k__Fungi(100);p__Basidiomycota(100);c__Microbotryomycetes(100);o__Sporidiobolales(100);f__Sporidiobolales_family_Incertae_sedis(100);g__Sporobolomyces(100);s__Sporobolomyces_lactosus(100); | Sporobolomyces | 13 | Pathotroph-Saprotroph | \|Fungal Parasite\|-Plant Saprotroph | Yeast | NULL | Probable | As litter_saprotroph (Põlme et al. 2020); As mycoparasite (Põlme et al. 2020); Decay fungal material (Põlme et al. 2020) | Tedersoo L, et al. 2014. Science 346:e1256688 ((https://doi.org/10.1126/science.1256688)); Purahong W, et al. 2016. Molecular Ecology 25:4059-4074 ((https://doi.org/10.1111/mec.13739)); Põlme S, et al. 2020. Fungal Diversity 105:1-16 ((https://doi.org/10.1007/s13225-020-00466-2)) |
| Otu54805 | 1 | k__Fungi(100);p__Basidiomycota(100);c__Microbotryomycetes(100);o__Sporidiobolales(100);f__Sporidiobolales_family_Incertae_sedis(100);g__Sporobolomyces(100);s__Sporobolomyces_lactosus(100); | Sporobolomyces | 13 | Pathotroph-Saprotroph | \|Fungal Parasite\|-Plant Saprotroph | Yeast | NULL | Probable | As litter_saprotroph (Põlme et al. 2020); As mycoparasite (Põlme et al. 2020); Decay fungal material (Põlme et al. 2020) | Tedersoo L, et al. 2014. Science 346:e1256688 ((https://doi.org/10.1126/science.1256688)); Purahong W, et al. 2016. Molecular Ecology 25:4059-4074 ((https://doi.org/10.1111/mec.13739)); Põlme S, et al. 2020. Fungal Diversity 105:1-16 ((https://doi.org/10.1007/s13225-020-00466-2)) |
| Otu54807 | 1 | k__Fungi(100);p__Basidiomycota(100);c__Microbotryomycetes(100);o__Sporidiobolales(100);f__Sporidiobolales_family_Incertae_sedis(100);g__Sporobolomyces(100);s__Sporobolomyces_lactosus(100); | Sporobolomyces | 13 | Pathotroph-Saprotroph | \|Fungal Parasite\|-Plant Saprotroph | Yeast | NULL | Probable | As litter_saprotroph (Põlme et al. 2020); As mycoparasite (Põlme et al. 2020); Decay fungal material (Põlme et al. 2020) | Tedersoo L, et al. 2014. Science 346:e1256688 ((https://doi.org/10.1126/science.1256688)); Purahong W, et al. 2016. Molecular Ecology 25:4059-4074 ((https://doi.org/10.1111/mec.13739)); Põlme S, et al. 2020. Fungal Diversity 105:1-16 ((https://doi.org/10.1007/s13225-020-00466-2)) |
| Otu54808 | 1 | k__Fungi(100);p__Ascomycota(100);c__Saccharomycetes(100);o__Saccharomycetales(100);f__Metschnikowiaceae(100);g__Metschnikowia(100);s__Metschnikowia_pulcherrima(100); | Metschnikowia | 13 | Saprotroph | \|Undefined Saprotroph\| | Yeast | NULL | Probable | As nectar-tap_saprotroph (Põlme et al. 2020); Decay sugar-rich substrates (Põlme et al. 2020) | Põlme S, et al. 2020. Fungal Diversity 105:1-16 ((https://doi.org/10.1007/s13225-020-00466-2)) |
| Otu54809 | 1 | k__Fungi(100);p__Ascomycota(100);c__Saccharomycetes(100);o__Saccharomycetales(100);f__Metschnikowiaceae(100);g__Metschnikowia(100);s__Metschnikowia_pulcherrima(100); | Metschnikowia | 13 | Saprotroph | \|Undefined Saprotroph\| | Yeast | NULL | Probable | As nectar-tap_saprotroph (Põlme et al. 2020); Decay sugar-rich substrates (Põlme et al. 2020) | Põlme S, et al. 2020. Fungal Diversity 105:1-16 ((https://doi.org/10.1007/s13225-020-00466-2)) |
| Otu54810 | 1 | k__Fungi(100);p__Ascomycota(100);c__Saccharomycetes(100);o__Saccharomycetales(100);f__Metschnikowiaceae(100);g__Metschnikowia(100);g__Metschnikowia_unclassified(100); | Metschnikowia | 13 | Saprotroph | \|Undefined Saprotroph\| | Yeast | NULL | Probable | As nectar-tap_saprotroph (Põlme et al. 2020); Decay sugar-rich substrates (Põlme et al. 2020) | Põlme S, et al. 2020. Fungal Diversity 105:1-16 ((https://doi.org/10.1007/s13225-020-00466-2)) |
| Otu54811 | 1 | k__Fungi(100);p__Ascomycota(100);c__Saccharomycetes(100);o__Saccharomycetales(100);f__Metschnikowiaceae(100);g__Metschnikowia(100);s__Metschnikowia_pulcherrima(100); | Metschnikowia | 13 | Saprotroph | \|Undefined Saprotroph\| | Yeast | NULL | Probable | As nectar-tap_saprotroph (Põlme et al. 2020); Decay sugar-rich substrates (Põlme et al. 2020) | Põlme S, et al. 2020. Fungal Diversity 105:1-16 ((https://doi.org/10.1007/s13225-020-00466-2)) |
| Otu54813 | 1 | k__Fungi(100);p__Ascomycota(100);c__Saccharomycetes(100);o__Saccharomycetales(100);f__Metschnikowiaceae(100);g__Metschnikowia(100);g__Metschnikowia_unclassified(100); | Metschnikowia | 13 | Saprotroph | \|Undefined Saprotroph\| | Yeast | NULL | Probable | As nectar-tap_saprotroph (Põlme et al. 2020); Decay sugar-rich substrates (Põlme et al. 2020) | Põlme S, et al. 2020. Fungal Diversity 105:1-16 ((https://doi.org/10.1007/s13225-020-00466-2)) |
| Otu54817 | 1 | k__Fungi(100);p__Ascomycota(100);c__Saccharomycetes(100);o__Saccharomycetales(100);o__Saccharomycetales_unclassified(100);o__Saccharomycetales_unclassified(100);o__Saccharomycetales_unclassified(100); | Saccharomycetales | 7 | Saprotroph | Undefined Saprotroph | Yeast | NULL | Possible | NULL | Sterkenburg E, et al. 2015. New Phytologist 207:1145-1158 ((https://doi.org/10.1111/nph.13426)) |
| Otu54819 | 1 | k__Fungi(100);p__Basidiomycota(100);c__Microbotryomycetes(100);o__Sporidiobolales(100);f__Sporidiobolales_family_Incertae_sedis(100);g__Sporobolomyces(100);s__Sporobolomyces_lactosus(100); | Sporobolomyces | 13 | Pathotroph-Saprotroph | \|Fungal Parasite\|-Plant Saprotroph | Yeast | NULL | Probable | As litter_saprotroph (Põlme et al. 2020); As mycoparasite (Põlme et al. 2020); Decay fungal material (Põlme et al. 2020) | Tedersoo L, et al. 2014. Science 346:e1256688 ((https://doi.org/10.1126/science.1256688)); Purahong W, et al. 2016. Molecular Ecology 25:4059-4074 ((https://doi.org/10.1111/mec.13739)); Põlme S, et al. 2020. Fungal Diversity 105:1-16 ((https://doi.org/10.1007/s13225-020-00466-2)) |
| Otu54820 | 1 | k__Fungi(100);p__Ascomycota(100);c__Saccharomycetes(100);o__Saccharomycetales(100);f__Saccharomycetales_family_Incertae_sedis(100);g__Candida(100);g__Candida_unclassified(100); | Candida | 13 | Pathotroph-Saprotroph-Symbiotroph | Animal Pathogen-Endophyte-Endosymbiont-Epiphyte-\|Undefined Saprotroph\| | Dimorphic Yeast-Yeast | NULL | Probable | As nectar-tap_saprotroph (Põlme et al. 2020); Decay sugar-rich substrates (Põlme et al. 2020); This asexual genus is polyphyletic and contain species that function very differently depend on what they are. Caution should be taken when interpreting guilds for Candida species. | Manolakaki D, et al. 2010. Virulence. 1: 367-375 ((https://doi.org/10.4161/viru.1.5.12796)); Mekbib SB, et al. 2011. Tropical Plant Pathology 36:233-240 ((https://doi.org/10.1590/S1982-56762011000400004)); Põlme S, et al. 2020. Fungal Diversity 105:1-16 ((https://doi.org/10.1007/s13225-020-00466-2)) |
| Otu54821 | 1 | k__Fungi(100);p__Ascomycota(100);c__Saccharomycetes(100);o__Saccharomycetales(100);f__Saccharomycetales_family_Incertae_sedis(100);g__Candida(100);g__Candida_unclassified(100); | Candida | 13 | Pathotroph-Saprotroph-Symbiotroph | Animal Pathogen-Endophyte-Endosymbiont-Epiphyte-\|Undefined Saprotroph\| | Dimorphic Yeast-Yeast | NULL | Probable | As nectar-tap_saprotroph (Põlme et al. 2020); Decay sugar-rich substrates (Põlme et al. 2020); This asexual genus is polyphyletic and contain species that function very differently depend on what they are. Caution should be taken when interpreting guilds for Candida species. | Manolakaki D, et al. 2010. Virulence. 1: 367-375 ((https://doi.org/10.4161/viru.1.5.12796)); Mekbib SB, et al. 2011. Tropical Plant Pathology 36:233-240 ((https://doi.org/10.1590/S1982-56762011000400004)); Põlme S, et al. 2020. Fungal Diversity 105:1-16 ((https://doi.org/10.1007/s13225-020-00466-2)) |
| Otu54823 | 1 | k__Fungi(100);p__Ascomycota(100);c__Saccharomycetes(100);o__Saccharomycetales(100);f__Metschnikowiaceae(100);g__Metschnikowia(100);s__Metschnikowia_pulcherrima(100); | Metschnikowia | 13 | Saprotroph | \|Undefined Saprotroph\| | Yeast | NULL | Probable | As nectar-tap_saprotroph (Põlme et al. 2020); Decay sugar-rich substrates (Põlme et al. 2020) | Põlme S, et al. 2020. Fungal Diversity 105:1-16 ((https://doi.org/10.1007/s13225-020-00466-2)) |
| Otu54824 | 1 | k__Fungi(100);p__Ascomycota(100);c__Saccharomycetes(100);o__Saccharomycetales(100);f__Metschnikowiaceae(100);g__Metschnikowia(100);s__Metschnikowia_pulcherrima(100); | Metschnikowia | 13 | Saprotroph | \|Undefined Saprotroph\| | Yeast | NULL | Probable | As nectar-tap_saprotroph (Põlme et al. 2020); Decay sugar-rich substrates (Põlme et al. 2020) | Põlme S, et al. 2020. Fungal Diversity 105:1-16 ((https://doi.org/10.1007/s13225-020-00466-2)) |
| Otu54825 | 1 | k__Fungi(100);p__Ascomycota(100);c__Saccharomycetes(100);o__Saccharomycetales(100);f__Metschnikowiaceae(100);g__Metschnikowia(100);s__Metschnikowia_pulcherrima(100); | Metschnikowia | 13 | Saprotroph | \|Undefined Saprotroph\| | Yeast | NULL | Probable | As nectar-tap_saprotroph (Põlme et al. 2020); Decay sugar-rich substrates (Põlme et al. 2020) | Põlme S, et al. 2020. Fungal Diversity 105:1-16 ((https://doi.org/10.1007/s13225-020-00466-2)) |
| Otu54826 | 1 | k__Fungi(100);p__Ascomycota(100);c__Saccharomycetes(100);o__Saccharomycetales(100);f__Metschnikowiaceae(100);g__Metschnikowia(100);s__Metschnikowia_pulcherrima(100); | Metschnikowia | 13 | Saprotroph | \|Undefined Saprotroph\| | Yeast | NULL | Probable | As nectar-tap_saprotroph (Põlme et al. 2020); Decay sugar-rich substrates (Põlme et al. 2020) | Põlme S, et al. 2020. Fungal Diversity 105:1-16 ((https://doi.org/10.1007/s13225-020-00466-2)) |
| Otu54828 | 1 | k__Fungi(100);p__Basidiomycota(100);c__Microbotryomycetes(100);o__Sporidiobolales(100);f__Sporidiobolales_family_Incertae_sedis(100);g__Sporobolomyces(100);s__Sporobolomyces_lactosus(100); | Sporobolomyces | 13 | Pathotroph-Saprotroph | \|Fungal Parasite\|-Plant Saprotroph | Yeast | NULL | Probable | As litter_saprotroph (Põlme et al. 2020); As mycoparasite (Põlme et al. 2020); Decay fungal material (Põlme et al. 2020) | Tedersoo L, et al. 2014. Science 346:e1256688 ((https://doi.org/10.1126/science.1256688)); Purahong W, et al. 2016. Molecular Ecology 25:4059-4074 ((https://doi.org/10.1111/mec.13739)); Põlme S, et al. 2020. Fungal Diversity 105:1-16 ((https://doi.org/10.1007/s13225-020-00466-2)) |
| Otu54829 | 1 | k__Fungi(100);p__Basidiomycota(100);c__Microbotryomycetes(100);o__Sporidiobolales(100);f__Sporidiobolales_family_Incertae_sedis(100);g__Sporobolomyces(100);s__Sporobolomyces_lactosus(100); | Sporobolomyces | 13 | Pathotroph-Saprotroph | \|Fungal Parasite\|-Plant Saprotroph | Yeast | NULL | Probable | As litter_saprotroph (Põlme et al. 2020); As mycoparasite (Põlme et al. 2020); Decay fungal material (Põlme et al. 2020) | Tedersoo L, et al. 2014. Science 346:e1256688 ((https://doi.org/10.1126/science.1256688)); Purahong W, et al. 2016. Molecular Ecology 25:4059-4074 ((https://doi.org/10.1111/mec.13739)); Põlme S, et al. 2020. Fungal Diversity 105:1-16 ((https://doi.org/10.1007/s13225-020-00466-2)) |
| Otu54830 | 1 | k__Fungi(100);p__Basidiomycota(100);c__Microbotryomycetes(100);o__Sporidiobolales(100);f__Sporidiobolales_family_Incertae_sedis(100);g__Sporobolomyces(100);s__Sporobolomyces_lactosus(100); | Sporobolomyces | 13 | Pathotroph-Saprotroph | \|Fungal Parasite\|-Plant Saprotroph | Yeast | NULL | Probable | As litter_saprotroph (Põlme et al. 2020); As mycoparasite (Põlme et al. 2020); Decay fungal material (Põlme et al. 2020) | Tedersoo L, et al. 2014. Science 346:e1256688 ((https://doi.org/10.1126/science.1256688)); Purahong W, et al. 2016. Molecular Ecology 25:4059-4074 ((https://doi.org/10.1111/mec.13739)); Põlme S, et al. 2020. Fungal Diversity 105:1-16 ((https://doi.org/10.1007/s13225-020-00466-2)) |
| Otu54832 | 1 | k__Fungi(100);p__Basidiomycota(100);c__Microbotryomycetes(100);o__Sporidiobolales(100);f__Sporidiobolales_family_Incertae_sedis(100);g__Sporobolomyces(100);s__Sporobolomyces_lactosus(100); | Sporobolomyces | 13 | Pathotroph-Saprotroph | \|Fungal Parasite\|-Plant Saprotroph | Yeast | NULL | Probable | As litter_saprotroph (Põlme et al. 2020); As mycoparasite (Põlme et al. 2020); Decay fungal material (Põlme et al. 2020) | Tedersoo L, et al. 2014. Science 346:e1256688 ((https://doi.org/10.1126/science.1256688)); Purahong W, et al. 2016. Molecular Ecology 25:4059-4074 ((https://doi.org/10.1111/mec.13739)); Põlme S, et al. 2020. Fungal Diversity 105:1-16 ((https://doi.org/10.1007/s13225-020-00466-2)) |
| Otu54833 | 1 | k__Fungi(100);p__Ascomycota(100);c__Saccharomycetes(100);o__Saccharomycetales(100);f__Saccharomycetales_family_Incertae_sedis(100);f__Saccharomycetales_family_Incertae_sedis_unclassified(100);f__Saccharomycetales_family_Incertae_sedis_unclassified(100); | Saccharomycetales | 7 | Saprotroph | Undefined Saprotroph | Yeast | NULL | Possible | NULL | Sterkenburg E, et al. 2015. New Phytologist 207:1145-1158 ((https://doi.org/10.1111/nph.13426)) |
| Otu54834 | 1 | k__Fungi(100);p__Ascomycota(100);c__Saccharomycetes(100);o__Saccharomycetales(100);f__Metschnikowiaceae(100);g__Metschnikowia(100);s__Metschnikowia_pulcherrima(100); | Metschnikowia | 13 | Saprotroph | \|Undefined Saprotroph\| | Yeast | NULL | Probable | As nectar-tap_saprotroph (Põlme et al. 2020); Decay sugar-rich substrates (Põlme et al. 2020) | Põlme S, et al. 2020. Fungal Diversity 105:1-16 ((https://doi.org/10.1007/s13225-020-00466-2)) |
| Otu54835 | 1 | k__Fungi(100);p__Ascomycota(100);c__Saccharomycetes(100);o__Saccharomycetales(100);f__Metschnikowiaceae(100);g__Metschnikowia(100);s__Metschnikowia_pulcherrima(100); | Metschnikowia | 13 | Saprotroph | \|Undefined Saprotroph\| | Yeast | NULL | Probable | As nectar-tap_saprotroph (Põlme et al. 2020); Decay sugar-rich substrates (Põlme et al. 2020) | Põlme S, et al. 2020. Fungal Diversity 105:1-16 ((https://doi.org/10.1007/s13225-020-00466-2)) |
| Otu54839 | 1 | k__Fungi(100);p__Ascomycota(100);c__Saccharomycetes(100);o__Saccharomycetales(100);f__Metschnikowiaceae(100);g__Metschnikowia(100);s__Metschnikowia_pulcherrima(100); | Metschnikowia | 13 | Saprotroph | \|Undefined Saprotroph\| | Yeast | NULL | Probable | As nectar-tap_saprotroph (Põlme et al. 2020); Decay sugar-rich substrates (Põlme et al. 2020) | Põlme S, et al. 2020. Fungal Diversity 105:1-16 ((https://doi.org/10.1007/s13225-020-00466-2)) |
| Otu54840 | 1 | k__Fungi(100);p__Ascomycota(100);c__Saccharomycetes(100);o__Saccharomycetales(100);f__Metschnikowiaceae(100);g__Metschnikowia(100);g__Metschnikowia_unclassified(100); | Metschnikowia | 13 | Saprotroph | \|Undefined Saprotroph\| | Yeast | NULL | Probable | As nectar-tap_saprotroph (Põlme et al. 2020); Decay sugar-rich substrates (Põlme et al. 2020) | Põlme S, et al. 2020. Fungal Diversity 105:1-16 ((https://doi.org/10.1007/s13225-020-00466-2)) |
| Otu54842 | 1 | k__Fungi(100);p__Ascomycota(100);c__Saccharomycetes(100);o__Saccharomycetales(100);f__Metschnikowiaceae(100);g__Metschnikowia(100);g__Metschnikowia_unclassified(100); | Metschnikowia | 13 | Saprotroph | \|Undefined Saprotroph\| | Yeast | NULL | Probable | As nectar-tap_saprotroph (Põlme et al. 2020); Decay sugar-rich substrates (Põlme et al. 2020) | Põlme S, et al. 2020. Fungal Diversity 105:1-16 ((https://doi.org/10.1007/s13225-020-00466-2)) |
| Otu54843 | 1 | k__Fungi(100);p__Ascomycota(100);c__Saccharomycetes(100);o__Saccharomycetales(100);f__Metschnikowiaceae(100);g__Metschnikowia(100);g__Metschnikowia_unclassified(100); | Metschnikowia | 13 | Saprotroph | \|Undefined Saprotroph\| | Yeast | NULL | Probable | As nectar-tap_saprotroph (Põlme et al. 2020); Decay sugar-rich substrates (Põlme et al. 2020) | Põlme S, et al. 2020. Fungal Diversity 105:1-16 ((https://doi.org/10.1007/s13225-020-00466-2)) |
| Otu54844 | 1 | k__Fungi(100);p__Ascomycota(100);c__Saccharomycetes(100);o__Saccharomycetales(100);f__Metschnikowiaceae(100);g__Metschnikowia(100);g__Metschnikowia_unclassified(100); | Metschnikowia | 13 | Saprotroph | \|Undefined Saprotroph\| | Yeast | NULL | Probable | As nectar-tap_saprotroph (Põlme et al. 2020); Decay sugar-rich substrates (Põlme et al. 2020) | Põlme S, et al. 2020. Fungal Diversity 105:1-16 ((https://doi.org/10.1007/s13225-020-00466-2)) |
| Otu54845 | 1 | k__Fungi(100);p__Ascomycota(100);c__Saccharomycetes(100);o__Saccharomycetales(100);f__Metschnikowiaceae(100);g__Metschnikowia(100);g__Metschnikowia_unclassified(100); | Metschnikowia | 13 | Saprotroph | \|Undefined Saprotroph\| | Yeast | NULL | Probable | As nectar-tap_saprotroph (Põlme et al. 2020); Decay sugar-rich substrates (Põlme et al. 2020) | Põlme S, et al. 2020. Fungal Diversity 105:1-16 ((https://doi.org/10.1007/s13225-020-00466-2)) |
| Otu54846 | 1 | k__Fungi(100);p__Ascomycota(100);c__Saccharomycetes(100);o__Saccharomycetales(100);f__Metschnikowiaceae(100);g__Metschnikowia(100);g__Metschnikowia_unclassified(100); | Metschnikowia | 13 | Saprotroph | \|Undefined Saprotroph\| | Yeast | NULL | Probable | As nectar-tap_saprotroph (Põlme et al. 2020); Decay sugar-rich substrates (Põlme et al. 2020) | Põlme S, et al. 2020. Fungal Diversity 105:1-16 ((https://doi.org/10.1007/s13225-020-00466-2)) |
| Otu54847 | 1 | k__Fungi(100);p__Ascomycota(100);c__Saccharomycetes(100);o__Saccharomycetales(100);f__Metschnikowiaceae(100);g__Metschnikowia(100);g__Metschnikowia_unclassified(100); | Metschnikowia | 13 | Saprotroph | \|Undefined Saprotroph\| | Yeast | NULL | Probable | As nectar-tap_saprotroph (Põlme et al. 2020); Decay sugar-rich substrates (Põlme et al. 2020) | Põlme S, et al. 2020. Fungal Diversity 105:1-16 ((https://doi.org/10.1007/s13225-020-00466-2)) |
| Otu54850 | 1 | k__Fungi(100);p__Ascomycota(100);c__Saccharomycetes(100);o__Saccharomycetales(100);f__Metschnikowiaceae(100);g__Metschnikowia(100);s__Metschnikowia_pulcherrima(100); | Metschnikowia | 13 | Saprotroph | \|Undefined Saprotroph\| | Yeast | NULL | Probable | As nectar-tap_saprotroph (Põlme et al. 2020); Decay sugar-rich substrates (Põlme et al. 2020) | Põlme S, et al. 2020. Fungal Diversity 105:1-16 ((https://doi.org/10.1007/s13225-020-00466-2)) |
| Otu54852 | 1 | k__Fungi(100);p__Ascomycota(100);c__Saccharomycetes(100);o__Saccharomycetales(100);f__Metschnikowiaceae(100);g__Metschnikowia(100);g__Metschnikowia_unclassified(100); | Metschnikowia | 13 | Saprotroph | \|Undefined Saprotroph\| | Yeast | NULL | Probable | As nectar-tap_saprotroph (Põlme et al. 2020); Decay sugar-rich substrates (Põlme et al. 2020) | Põlme S, et al. 2020. Fungal Diversity 105:1-16 ((https://doi.org/10.1007/s13225-020-00466-2)) |
| Otu54858 | 1 | k__Fungi(100);p__Ascomycota(100);c__Saccharomycetes(100);o__Saccharomycetales(100);f__Saccharomycetales_family_Incertae_sedis(100);g__Candida(100);g__Candida_unclassified(100); | Candida | 13 | Pathotroph-Saprotroph-Symbiotroph | Animal Pathogen-Endophyte-Endosymbiont-Epiphyte-\|Undefined Saprotroph\| | Dimorphic Yeast-Yeast | NULL | Probable | As nectar-tap_saprotroph (Põlme et al. 2020); Decay sugar-rich substrates (Põlme et al. 2020); This asexual genus is polyphyletic and contain species that function very differently depend on what they are. Caution should be taken when interpreting guilds for Candida species. | Manolakaki D, et al. 2010. Virulence. 1: 367-375 ((https://doi.org/10.4161/viru.1.5.12796)); Mekbib SB, et al. 2011. Tropical Plant Pathology 36:233-240 ((https://doi.org/10.1590/S1982-56762011000400004)); Põlme S, et al. 2020. Fungal Diversity 105:1-16 ((https://doi.org/10.1007/s13225-020-00466-2)) |
| Otu54867 | 1 | k__Fungi(100);p__Ascomycota(100);c__Saccharomycetes(100);o__Saccharomycetales(100);f__Metschnikowiaceae(100);g__Metschnikowia(100);s__Metschnikowia_pulcherrima(100); | Metschnikowia | 13 | Saprotroph | \|Undefined Saprotroph\| | Yeast | NULL | Probable | As nectar-tap_saprotroph (Põlme et al. 2020); Decay sugar-rich substrates (Põlme et al. 2020) | Põlme S, et al. 2020. Fungal Diversity 105:1-16 ((https://doi.org/10.1007/s13225-020-00466-2)) |
| Otu54870 | 1 | k__Fungi(100);p__Ascomycota(100);c__Saccharomycetes(100);o__Saccharomycetales(100);f__Saccharomycetales_family_Incertae_sedis(100);g__Candida(100);s__Candida_cretensis(100); | Candida | 13 | Pathotroph-Saprotroph-Symbiotroph | Animal Pathogen-Endophyte-Endosymbiont-Epiphyte-\|Undefined Saprotroph\| | Dimorphic Yeast-Yeast | NULL | Probable | As nectar-tap_saprotroph (Põlme et al. 2020); Decay sugar-rich substrates (Põlme et al. 2020); This asexual genus is polyphyletic and contain species that function very differently depend on what they are. Caution should be taken when interpreting guilds for Candida species. | Manolakaki D, et al. 2010. Virulence. 1: 367-375 ((https://doi.org/10.4161/viru.1.5.12796)); Mekbib SB, et al. 2011. Tropical Plant Pathology 36:233-240 ((https://doi.org/10.1590/S1982-56762011000400004)); Põlme S, et al. 2020. Fungal Diversity 105:1-16 ((https://doi.org/10.1007/s13225-020-00466-2)) |
| Otu54874 | 1 | k__Fungi(100);p__Basidiomycota(100);c__Microbotryomycetes(100);o__Sporidiobolales(100);f__Sporidiobolales_family_Incertae_sedis(100);g__Sporobolomyces(100);s__Sporobolomyces_lactosus(100); | Sporobolomyces | 13 | Pathotroph-Saprotroph | \|Fungal Parasite\|-Plant Saprotroph | Yeast | NULL | Probable | As litter_saprotroph (Põlme et al. 2020); As mycoparasite (Põlme et al. 2020); Decay fungal material (Põlme et al. 2020) | Tedersoo L, et al. 2014. Science 346:e1256688 ((https://doi.org/10.1126/science.1256688)); Purahong W, et al. 2016. Molecular Ecology 25:4059-4074 ((https://doi.org/10.1111/mec.13739)); Põlme S, et al. 2020. Fungal Diversity 105:1-16 ((https://doi.org/10.1007/s13225-020-00466-2)) |
| Otu54875 | 1 | k__Fungi(100);p__Ascomycota(100);c__Saccharomycetes(100);o__Saccharomycetales(100);f__Metschnikowiaceae(100);g__Metschnikowia(100);s__Metschnikowia_pulcherrima(100); | Metschnikowia | 13 | Saprotroph | \|Undefined Saprotroph\| | Yeast | NULL | Probable | As nectar-tap_saprotroph (Põlme et al. 2020); Decay sugar-rich substrates (Põlme et al. 2020) | Põlme S, et al. 2020. Fungal Diversity 105:1-16 ((https://doi.org/10.1007/s13225-020-00466-2)) |
| Otu54879 | 1 | k__Fungi(100);p__Ascomycota(100);c__Saccharomycetes(100);o__Saccharomycetales(100);f__Metschnikowiaceae(100);g__Metschnikowia(100);s__Metschnikowia_pulcherrima(100); | Metschnikowia | 13 | Saprotroph | \|Undefined Saprotroph\| | Yeast | NULL | Probable | As nectar-tap_saprotroph (Põlme et al. 2020); Decay sugar-rich substrates (Põlme et al. 2020) | Põlme S, et al. 2020. Fungal Diversity 105:1-16 ((https://doi.org/10.1007/s13225-020-00466-2)) |
| Otu54882 | 1 | k__Fungi(100);p__Ascomycota(100);c__Saccharomycetes(100);o__Saccharomycetales(100);f__Metschnikowiaceae(100);g__Metschnikowia(100);s__Metschnikowia_pulcherrima(100); | Metschnikowia | 13 | Saprotroph | \|Undefined Saprotroph\| | Yeast | NULL | Probable | As nectar-tap_saprotroph (Põlme et al. 2020); Decay sugar-rich substrates (Põlme et al. 2020) | Põlme S, et al. 2020. Fungal Diversity 105:1-16 ((https://doi.org/10.1007/s13225-020-00466-2)) |
| Otu54883 | 1 | k__Fungi(100);p__Ascomycota(100);c__Saccharomycetes(100);o__Saccharomycetales(100);f__Metschnikowiaceae(100);g__Metschnikowia(100);s__Metschnikowia_pulcherrima(100); | Metschnikowia | 13 | Saprotroph | \|Undefined Saprotroph\| | Yeast | NULL | Probable | As nectar-tap_saprotroph (Põlme et al. 2020); Decay sugar-rich substrates (Põlme et al. 2020) | Põlme S, et al. 2020. Fungal Diversity 105:1-16 ((https://doi.org/10.1007/s13225-020-00466-2)) |
| Otu54884 | 1 | k__Fungi(100);p__Ascomycota(100);c__Saccharomycetes(100);o__Saccharomycetales(100);f__Metschnikowiaceae(100);g__Metschnikowia(100);s__Metschnikowia_pulcherrima(100); | Metschnikowia | 13 | Saprotroph | \|Undefined Saprotroph\| | Yeast | NULL | Probable | As nectar-tap_saprotroph (Põlme et al. 2020); Decay sugar-rich substrates (Põlme et al. 2020) | Põlme S, et al. 2020. Fungal Diversity 105:1-16 ((https://doi.org/10.1007/s13225-020-00466-2)) |
| Otu54885 | 1 | k__Fungi(100);p__Ascomycota(100);c__Saccharomycetes(100);o__Saccharomycetales(100);f__Metschnikowiaceae(100);g__Metschnikowia(100);g__Metschnikowia_unclassified(100); | Metschnikowia | 13 | Saprotroph | \|Undefined Saprotroph\| | Yeast | NULL | Probable | As nectar-tap_saprotroph (Põlme et al. 2020); Decay sugar-rich substrates (Põlme et al. 2020) | Põlme S, et al. 2020. Fungal Diversity 105:1-16 ((https://doi.org/10.1007/s13225-020-00466-2)) |
| Otu54886 | 1 | k__Fungi(100);p__Ascomycota(100);c__Saccharomycetes(100);o__Saccharomycetales(100);f__Metschnikowiaceae(100);g__Metschnikowia(100);s__Metschnikowia_pulcherrima(100); | Metschnikowia | 13 | Saprotroph | \|Undefined Saprotroph\| | Yeast | NULL | Probable | As nectar-tap_saprotroph (Põlme et al. 2020); Decay sugar-rich substrates (Põlme et al. 2020) | Põlme S, et al. 2020. Fungal Diversity 105:1-16 ((https://doi.org/10.1007/s13225-020-00466-2)) |
| Otu54890 | 1 | k__Fungi(100);p__Ascomycota(100);c__Saccharomycetes(100);o__Saccharomycetales(100);f__Metschnikowiaceae(100);g__Metschnikowia(100);s__Metschnikowia_pulcherrima(100); | Metschnikowia | 13 | Saprotroph | \|Undefined Saprotroph\| | Yeast | NULL | Probable | As nectar-tap_saprotroph (Põlme et al. 2020); Decay sugar-rich substrates (Põlme et al. 2020) | Põlme S, et al. 2020. Fungal Diversity 105:1-16 ((https://doi.org/10.1007/s13225-020-00466-2)) |
| Otu54891 | 1 | k__Fungi(100);p__Basidiomycota(100);c__Microbotryomycetes(100);o__Sporidiobolales(100);f__Sporidiobolales_family_Incertae_sedis(100);g__Sporobolomyces(100);s__Sporobolomyces_lactosus(100); | Sporobolomyces | 13 | Pathotroph-Saprotroph | \|Fungal Parasite\|-Plant Saprotroph | Yeast | NULL | Probable | As litter_saprotroph (Põlme et al. 2020); As mycoparasite (Põlme et al. 2020); Decay fungal material (Põlme et al. 2020) | Tedersoo L, et al. 2014. Science 346:e1256688 ((https://doi.org/10.1126/science.1256688)); Purahong W, et al. 2016. Molecular Ecology 25:4059-4074 ((https://doi.org/10.1111/mec.13739)); Põlme S, et al. 2020. Fungal Diversity 105:1-16 ((https://doi.org/10.1007/s13225-020-00466-2)) |
| Otu54892 | 1 | k__Fungi(100);p__Basidiomycota(100);c__Microbotryomycetes(100);o__Sporidiobolales(100);f__Sporidiobolales_family_Incertae_sedis(100);g__Sporobolomyces(100);s__Sporobolomyces_lactosus(100); | Sporobolomyces | 13 | Pathotroph-Saprotroph | \|Fungal Parasite\|-Plant Saprotroph | Yeast | NULL | Probable | As litter_saprotroph (Põlme et al. 2020); As mycoparasite (Põlme et al. 2020); Decay fungal material (Põlme et al. 2020) | Tedersoo L, et al. 2014. Science 346:e1256688 ((https://doi.org/10.1126/science.1256688)); Purahong W, et al. 2016. Molecular Ecology 25:4059-4074 ((https://doi.org/10.1111/mec.13739)); Põlme S, et al. 2020. Fungal Diversity 105:1-16 ((https://doi.org/10.1007/s13225-020-00466-2)) |
| Otu54893 | 1 | k__Fungi(100);p__Basidiomycota(100);c__Microbotryomycetes(100);o__Sporidiobolales(100);f__Sporidiobolales_family_Incertae_sedis(100);g__Sporobolomyces(100);s__Sporobolomyces_lactosus(100); | Sporobolomyces | 13 | Pathotroph-Saprotroph | \|Fungal Parasite\|-Plant Saprotroph | Yeast | NULL | Probable | As litter_saprotroph (Põlme et al. 2020); As mycoparasite (Põlme et al. 2020); Decay fungal material (Põlme et al. 2020) | Tedersoo L, et al. 2014. Science 346:e1256688 ((https://doi.org/10.1126/science.1256688)); Purahong W, et al. 2016. Molecular Ecology 25:4059-4074 ((https://doi.org/10.1111/mec.13739)); Põlme S, et al. 2020. Fungal Diversity 105:1-16 ((https://doi.org/10.1007/s13225-020-00466-2)) |
| Otu54895 | 1 | k__Fungi(100);p__Basidiomycota(100);c__Microbotryomycetes(100);o__Sporidiobolales(100);f__Sporidiobolales_family_Incertae_sedis(100);g__Sporobolomyces(100);s__Sporobolomyces_lactosus(100); | Sporobolomyces | 13 | Pathotroph-Saprotroph | \|Fungal Parasite\|-Plant Saprotroph | Yeast | NULL | Probable | As litter_saprotroph (Põlme et al. 2020); As mycoparasite (Põlme et al. 2020); Decay fungal material (Põlme et al. 2020) | Tedersoo L, et al. 2014. Science 346:e1256688 ((https://doi.org/10.1126/science.1256688)); Purahong W, et al. 2016. Molecular Ecology 25:4059-4074 ((https://doi.org/10.1111/mec.13739)); Põlme S, et al. 2020. Fungal Diversity 105:1-16 ((https://doi.org/10.1007/s13225-020-00466-2)) |
| Otu54897 | 1 | k__Fungi(100);p__Basidiomycota(100);c__Microbotryomycetes(100);o__Sporidiobolales(100);f__Sporidiobolales_family_Incertae_sedis(100);g__Sporobolomyces(100);s__Sporobolomyces_lactosus(100); | Sporobolomyces | 13 | Pathotroph-Saprotroph | \|Fungal Parasite\|-Plant Saprotroph | Yeast | NULL | Probable | As litter_saprotroph (Põlme et al. 2020); As mycoparasite (Põlme et al. 2020); Decay fungal material (Põlme et al. 2020) | Tedersoo L, et al. 2014. Science 346:e1256688 ((https://doi.org/10.1126/science.1256688)); Purahong W, et al. 2016. Molecular Ecology 25:4059-4074 ((https://doi.org/10.1111/mec.13739)); Põlme S, et al. 2020. Fungal Diversity 105:1-16 ((https://doi.org/10.1007/s13225-020-00466-2)) |
| Otu54898 | 1 | k__Fungi(100);p__Ascomycota(100);c__Saccharomycetes(100);o__Saccharomycetales(100);f__Metschnikowiaceae(100);g__Metschnikowia(100);g__Metschnikowia_unclassified(100); | Metschnikowia | 13 | Saprotroph | \|Undefined Saprotroph\| | Yeast | NULL | Probable | As nectar-tap_saprotroph (Põlme et al. 2020); Decay sugar-rich substrates (Põlme et al. 2020) | Põlme S, et al. 2020. Fungal Diversity 105:1-16 ((https://doi.org/10.1007/s13225-020-00466-2)) |
| Otu54900 | 1 | k__Fungi(100);p__Ascomycota(100);c__Saccharomycetes(100);o__Saccharomycetales(100);f__Metschnikowiaceae(100);g__Metschnikowia(100);s__Metschnikowia_pulcherrima(100); | Metschnikowia | 13 | Saprotroph | \|Undefined Saprotroph\| | Yeast | NULL | Probable | As nectar-tap_saprotroph (Põlme et al. 2020); Decay sugar-rich substrates (Põlme et al. 2020) | Põlme S, et al. 2020. Fungal Diversity 105:1-16 ((https://doi.org/10.1007/s13225-020-00466-2)) |
| Otu54901 | 1 | k__Fungi(100);p__Ascomycota(100);c__Saccharomycetes(100);o__Saccharomycetales(100);f__Metschnikowiaceae(100);g__Metschnikowia(100);s__Metschnikowia_pulcherrima(100); | Metschnikowia | 13 | Saprotroph | \|Undefined Saprotroph\| | Yeast | NULL | Probable | As nectar-tap_saprotroph (Põlme et al. 2020); Decay sugar-rich substrates (Põlme et al. 2020) | Põlme S, et al. 2020. Fungal Diversity 105:1-16 ((https://doi.org/10.1007/s13225-020-00466-2)) |
| Otu54902 | 1 | k__Fungi(100);p__Ascomycota(100);c__Saccharomycetes(100);o__Saccharomycetales(100);f__Metschnikowiaceae(100);g__Metschnikowia(100);s__Metschnikowia_pulcherrima(100); | Metschnikowia | 13 | Saprotroph | \|Undefined Saprotroph\| | Yeast | NULL | Probable | As nectar-tap_saprotroph (Põlme et al. 2020); Decay sugar-rich substrates (Põlme et al. 2020) | Põlme S, et al. 2020. Fungal Diversity 105:1-16 ((https://doi.org/10.1007/s13225-020-00466-2)) |
| Otu54904 | 1 | k__Fungi(100);p__Ascomycota(100);c__Saccharomycetes(100);o__Saccharomycetales(100);f__Metschnikowiaceae(100);g__Metschnikowia(100);s__Metschnikowia_pulcherrima(100); | Metschnikowia | 13 | Saprotroph | \|Undefined Saprotroph\| | Yeast | NULL | Probable | As nectar-tap_saprotroph (Põlme et al. 2020); Decay sugar-rich substrates (Põlme et al. 2020) | Põlme S, et al. 2020. Fungal Diversity 105:1-16 ((https://doi.org/10.1007/s13225-020-00466-2)) |
| Otu54905 | 1 | k__Fungi(100);p__Ascomycota(100);c__Saccharomycetes(100);o__Saccharomycetales(100);f__Metschnikowiaceae(100);g__Metschnikowia(100);s__Metschnikowia_pulcherrima(100); | Metschnikowia | 13 | Saprotroph | \|Undefined Saprotroph\| | Yeast | NULL | Probable | As nectar-tap_saprotroph (Põlme et al. 2020); Decay sugar-rich substrates (Põlme et al. 2020) | Põlme S, et al. 2020. Fungal Diversity 105:1-16 ((https://doi.org/10.1007/s13225-020-00466-2)) |
| Otu54910 | 1 | k__Fungi(100);p__Ascomycota(100);c__Saccharomycetes(100);o__Saccharomycetales(100);f__Metschnikowiaceae(100);g__Metschnikowia(100);s__Metschnikowia_pulcherrima(100); | Metschnikowia | 13 | Saprotroph | \|Undefined Saprotroph\| | Yeast | NULL | Probable | As nectar-tap_saprotroph (Põlme et al. 2020); Decay sugar-rich substrates (Põlme et al. 2020) | Põlme S, et al. 2020. Fungal Diversity 105:1-16 ((https://doi.org/10.1007/s13225-020-00466-2)) |
| Otu54914 | 1 | k__Fungi(100);p__Ascomycota(100);c__Saccharomycetes(100);o__Saccharomycetales(100);f__Metschnikowiaceae(100);g__Metschnikowia(100);s__Metschnikowia_pulcherrima(100); | Metschnikowia | 13 | Saprotroph | \|Undefined Saprotroph\| | Yeast | NULL | Probable | As nectar-tap_saprotroph (Põlme et al. 2020); Decay sugar-rich substrates (Põlme et al. 2020) | Põlme S, et al. 2020. Fungal Diversity 105:1-16 ((https://doi.org/10.1007/s13225-020-00466-2)) |
| Otu54919 | 1 | k__Fungi(100);p__Ascomycota(100);c__Saccharomycetes(100);o__Saccharomycetales(100);f__Saccharomycetales_family_Incertae_sedis(100);g__Candida(100);s__Candida_cretensis(100); | Candida | 13 | Pathotroph-Saprotroph-Symbiotroph | Animal Pathogen-Endophyte-Endosymbiont-Epiphyte-\|Undefined Saprotroph\| | Dimorphic Yeast-Yeast | NULL | Probable | As nectar-tap_saprotroph (Põlme et al. 2020); Decay sugar-rich substrates (Põlme et al. 2020); This asexual genus is polyphyletic and contain species that function very differently depend on what they are. Caution should be taken when interpreting guilds for Candida species. | Manolakaki D, et al. 2010. Virulence. 1: 367-375 ((https://doi.org/10.4161/viru.1.5.12796)); Mekbib SB, et al. 2011. Tropical Plant Pathology 36:233-240 ((https://doi.org/10.1590/S1982-56762011000400004)); Põlme S, et al. 2020. Fungal Diversity 105:1-16 ((https://doi.org/10.1007/s13225-020-00466-2)) |
| Otu54922 | 1 | k__Fungi(100);p__Ascomycota(100);c__Saccharomycetes(100);o__Saccharomycetales(100);f__Metschnikowiaceae(100);g__Metschnikowia(100);s__Metschnikowia_pulcherrima(100); | Metschnikowia | 13 | Saprotroph | \|Undefined Saprotroph\| | Yeast | NULL | Probable | As nectar-tap_saprotroph (Põlme et al. 2020); Decay sugar-rich substrates (Põlme et al. 2020) | Põlme S, et al. 2020. Fungal Diversity 105:1-16 ((https://doi.org/10.1007/s13225-020-00466-2)) |
| Otu54923 | 1 | k__Fungi(100);p__Ascomycota(100);c__Saccharomycetes(100);o__Saccharomycetales(100);f__Metschnikowiaceae(100);g__Metschnikowia(100);s__Metschnikowia_pulcherrima(100); | Metschnikowia | 13 | Saprotroph | \|Undefined Saprotroph\| | Yeast | NULL | Probable | As nectar-tap_saprotroph (Põlme et al. 2020); Decay sugar-rich substrates (Põlme et al. 2020) | Põlme S, et al. 2020. Fungal Diversity 105:1-16 ((https://doi.org/10.1007/s13225-020-00466-2)) |
| Otu54925 | 1 | k__Fungi(100);p__Ascomycota(100);c__Saccharomycetes(100);o__Saccharomycetales(100);f__Metschnikowiaceae(100);g__Metschnikowia(100);s__Metschnikowia_pulcherrima(100); | Metschnikowia | 13 | Saprotroph | \|Undefined Saprotroph\| | Yeast | NULL | Probable | As nectar-tap_saprotroph (Põlme et al. 2020); Decay sugar-rich substrates (Põlme et al. 2020) | Põlme S, et al. 2020. Fungal Diversity 105:1-16 ((https://doi.org/10.1007/s13225-020-00466-2)) |
| Otu54929 | 1 | k__Fungi(100);p__Ascomycota(100);c__Saccharomycetes(100);o__Saccharomycetales(100);f__Metschnikowiaceae(100);g__Metschnikowia(100);g__Metschnikowia_unclassified(100); | Metschnikowia | 13 | Saprotroph | \|Undefined Saprotroph\| | Yeast | NULL | Probable | As nectar-tap_saprotroph (Põlme et al. 2020); Decay sugar-rich substrates (Põlme et al. 2020) | Põlme S, et al. 2020. Fungal Diversity 105:1-16 ((https://doi.org/10.1007/s13225-020-00466-2)) |
| Otu54934 | 1 | k__Fungi(100);p__Ascomycota(100);c__Saccharomycetes(100);o__Saccharomycetales(100);f__Metschnikowiaceae(100);g__Metschnikowia(100);s__Metschnikowia_pulcherrima(100); | Metschnikowia | 13 | Saprotroph | \|Undefined Saprotroph\| | Yeast | NULL | Probable | As nectar-tap_saprotroph (Põlme et al. 2020); Decay sugar-rich substrates (Põlme et al. 2020) | Põlme S, et al. 2020. Fungal Diversity 105:1-16 ((https://doi.org/10.1007/s13225-020-00466-2)) |
| Otu54936 | 1 | k__Fungi(100);p__Ascomycota(100);c__Saccharomycetes(100);o__Saccharomycetales(100);f__Saccharomycetales_family_Incertae_sedis(100);g__Candida(100);g__Candida_unclassified(100); | Candida | 13 | Pathotroph-Saprotroph-Symbiotroph | Animal Pathogen-Endophyte-Endosymbiont-Epiphyte-\|Undefined Saprotroph\| | Dimorphic Yeast-Yeast | NULL | Probable | As nectar-tap_saprotroph (Põlme et al. 2020); Decay sugar-rich substrates (Põlme et al. 2020); This asexual genus is polyphyletic and contain species that function very differently depend on what they are. Caution should be taken when interpreting guilds for Candida species. | Manolakaki D, et al. 2010. Virulence. 1: 367-375 ((https://doi.org/10.4161/viru.1.5.12796)); Mekbib SB, et al. 2011. Tropical Plant Pathology 36:233-240 ((https://doi.org/10.1590/S1982-56762011000400004)); Põlme S, et al. 2020. Fungal Diversity 105:1-16 ((https://doi.org/10.1007/s13225-020-00466-2)) |
| Otu54938 | 1 | k__Fungi(100);p__Ascomycota(100);c__Saccharomycetes(100);o__Saccharomycetales(100);f__Metschnikowiaceae(100);g__Metschnikowia(100);s__Metschnikowia_pulcherrima(100); | Metschnikowia | 13 | Saprotroph | \|Undefined Saprotroph\| | Yeast | NULL | Probable | As nectar-tap_saprotroph (Põlme et al. 2020); Decay sugar-rich substrates (Põlme et al. 2020) | Põlme S, et al. 2020. Fungal Diversity 105:1-16 ((https://doi.org/10.1007/s13225-020-00466-2)) |
| Otu54939 | 1 | k__Fungi(100);p__Ascomycota(100);c__Saccharomycetes(100);o__Saccharomycetales(100);f__Saccharomycetales_family_Incertae_sedis(100);g__Candida(100);g__Candida_unclassified(100); | Candida | 13 | Pathotroph-Saprotroph-Symbiotroph | Animal Pathogen-Endophyte-Endosymbiont-Epiphyte-\|Undefined Saprotroph\| | Dimorphic Yeast-Yeast | NULL | Probable | As nectar-tap_saprotroph (Põlme et al. 2020); Decay sugar-rich substrates (Põlme et al. 2020); This asexual genus is polyphyletic and contain species that function very differently depend on what they are. Caution should be taken when interpreting guilds for Candida species. | Manolakaki D, et al. 2010. Virulence. 1: 367-375 ((https://doi.org/10.4161/viru.1.5.12796)); Mekbib SB, et al. 2011. Tropical Plant Pathology 36:233-240 ((https://doi.org/10.1590/S1982-56762011000400004)); Põlme S, et al. 2020. Fungal Diversity 105:1-16 ((https://doi.org/10.1007/s13225-020-00466-2)) |
| Otu54940 | 1 | k__Fungi(100);p__Ascomycota(100);c__Saccharomycetes(100);o__Saccharomycetales(100);o__Saccharomycetales_unclassified(100);o__Saccharomycetales_unclassified(100);o__Saccharomycetales_unclassified(100); | Saccharomycetales | 7 | Saprotroph | Undefined Saprotroph | Yeast | NULL | Possible | NULL | Sterkenburg E, et al. 2015. New Phytologist 207:1145-1158 ((https://doi.org/10.1111/nph.13426)) |
| Otu54942 | 1 | k__Fungi(100);p__Ascomycota(100);c__Saccharomycetes(100);o__Saccharomycetales(100);f__Saccharomycetales_family_Incertae_sedis(100);f__Saccharomycetales_family_Incertae_sedis_unclassified(100);f__Saccharomycetales_family_Incertae_sedis_unclassified(100); | Saccharomycetales | 7 | Saprotroph | Undefined Saprotroph | Yeast | NULL | Possible | NULL | Sterkenburg E, et al. 2015. New Phytologist 207:1145-1158 ((https://doi.org/10.1111/nph.13426)) |
| Otu54947 | 1 | k__Fungi(100);p__Ascomycota(100);c__Saccharomycetes(100);o__Saccharomycetales(100);f__Saccharomycetales_family_Incertae_sedis(100);g__Candida(100);g__Candida_unclassified(100); | Candida | 13 | Pathotroph-Saprotroph-Symbiotroph | Animal Pathogen-Endophyte-Endosymbiont-Epiphyte-\|Undefined Saprotroph\| | Dimorphic Yeast-Yeast | NULL | Probable | As nectar-tap_saprotroph (Põlme et al. 2020); Decay sugar-rich substrates (Põlme et al. 2020); This asexual genus is polyphyletic and contain species that function very differently depend on what they are. Caution should be taken when interpreting guilds for Candida species. | Manolakaki D, et al. 2010. Virulence. 1: 367-375 ((https://doi.org/10.4161/viru.1.5.12796)); Mekbib SB, et al. 2011. Tropical Plant Pathology 36:233-240 ((https://doi.org/10.1590/S1982-56762011000400004)); Põlme S, et al. 2020. Fungal Diversity 105:1-16 ((https://doi.org/10.1007/s13225-020-00466-2)) |
| Otu54949 | 1 | k__Fungi(100);p__Ascomycota(100);c__Dothideomycetes(100);o__Pleosporales(100);f__Montagnulaceae(100);g__Paraphaeosphaeria(100);s__Paraphaeosphaeria_michotii(100); | Paraphaeosphaeria | 13 | Saprotroph | Undefined Saprotroph-\|Wood Saprotroph\| | Microfungus | NULL | Probable | Decay wood (Põlme et al. 2020) | Tedersoo L, et al. 2014. Science 346:e1256688 ((https://doi.org/10.1126/science.1256688)); Põlme S, et al. 2020. Fungal Diversity 105:1-16 ((https://doi.org/10.1007/s13225-020-00466-2)) |
| Otu54956 | 1 | k__Fungi(100);p__Ascomycota(100);c__Saccharomycetes(100);o__Saccharomycetales(100);f__Metschnikowiaceae(100);g__Metschnikowia(100);s__Metschnikowia_pulcherrima(100); | Metschnikowia | 13 | Saprotroph | \|Undefined Saprotroph\| | Yeast | NULL | Probable | As nectar-tap_saprotroph (Põlme et al. 2020); Decay sugar-rich substrates (Põlme et al. 2020) | Põlme S, et al. 2020. Fungal Diversity 105:1-16 ((https://doi.org/10.1007/s13225-020-00466-2)) |
| Otu54958 | 1 | k__Fungi(100);p__Ascomycota(100);c__Saccharomycetes(100);o__Saccharomycetales(100);f__Metschnikowiaceae(100);g__Metschnikowia(100);s__Metschnikowia_pulcherrima(100); | Metschnikowia | 13 | Saprotroph | \|Undefined Saprotroph\| | Yeast | NULL | Probable | As nectar-tap_saprotroph (Põlme et al. 2020); Decay sugar-rich substrates (Põlme et al. 2020) | Põlme S, et al. 2020. Fungal Diversity 105:1-16 ((https://doi.org/10.1007/s13225-020-00466-2)) |
| Otu54959 | 1 | k__Fungi(100);p__Ascomycota(100);c__Saccharomycetes(100);o__Saccharomycetales(100);f__Metschnikowiaceae(100);g__Metschnikowia(100);g__Metschnikowia_unclassified(100); | Metschnikowia | 13 | Saprotroph | \|Undefined Saprotroph\| | Yeast | NULL | Probable | As nectar-tap_saprotroph (Põlme et al. 2020); Decay sugar-rich substrates (Põlme et al. 2020) | Põlme S, et al. 2020. Fungal Diversity 105:1-16 ((https://doi.org/10.1007/s13225-020-00466-2)) |
| Otu54960 | 1 | k__Fungi(100);p__Ascomycota(100);c__Saccharomycetes(100);o__Saccharomycetales(100);f__Metschnikowiaceae(100);g__Metschnikowia(100);s__Metschnikowia_pulcherrima(100); | Metschnikowia | 13 | Saprotroph | \|Undefined Saprotroph\| | Yeast | NULL | Probable | As nectar-tap_saprotroph (Põlme et al. 2020); Decay sugar-rich substrates (Põlme et al. 2020) | Põlme S, et al. 2020. Fungal Diversity 105:1-16 ((https://doi.org/10.1007/s13225-020-00466-2)) |
| Otu54961 | 1 | k__Fungi(100);p__Ascomycota(100);c__Saccharomycetes(100);o__Saccharomycetales(100);f__Metschnikowiaceae(100);g__Metschnikowia(100);g__Metschnikowia_unclassified(100); | Metschnikowia | 13 | Saprotroph | \|Undefined Saprotroph\| | Yeast | NULL | Probable | As nectar-tap_saprotroph (Põlme et al. 2020); Decay sugar-rich substrates (Põlme et al. 2020) | Põlme S, et al. 2020. Fungal Diversity 105:1-16 ((https://doi.org/10.1007/s13225-020-00466-2)) |

| | | | | | | | | | | | |
|---|---|---|---|---|---|---|---|---|---|---|---|
| Otu54962 | 1 | k__Fungi(100);p__Ascomycota(100);c__Saccharomycetes(100);o__Saccharomycetales(100);f__Metschnikowiaceae(100);g__Metschnikowia(100);g__Metschnikowia_unclassified(100); | Metschnikowia | 13 | Saprotroph | \|Undefined Saprotroph\| | Yeast | NULL | Probable | As nectar-tap_saprotroph (Põlme et al. 2020); Decay sugar-rich substrates (Põlme et al. 2020) | Põlme S, et al. 2020. Fungal Diversity 105:1-16 ((https://doi.org/10.1007/s13225-020-00466-2)) |
| Otu54963 | 1 | k__Fungi(100);p__Ascomycota(100);c__Saccharomycetes(100);o__Saccharomycetales(100);f__Metschnikowiaceae(100);g__Metschnikowia(100);s__Metschnikowia_pulcherrima(100); | Metschnikowia | 13 | Saprotroph | \|Undefined Saprotroph\| | Yeast | NULL | Probable | As nectar-tap_saprotroph (Põlme et al. 2020); Decay sugar-rich substrates (Põlme et al. 2020) | Põlme S, et al. 2020. Fungal Diversity 105:1-16 ((https://doi.org/10.1007/s13225-020-00466-2)) |
| Otu54964 | 1 | k__Fungi(100);p__Basidiomycota(100);c__Microbotryomycetes(100);o__Sporidiobolales(100);f__Sporidiobolales_family_Incertae_sedis(100);g__Sporobolomyces(100);s__Sporobolomyces_lactosus(100); | Sporobolomyces | 13 | Pathotroph-Saprotroph | \|Fungal Parasite\|-Plant Saprotroph | Yeast | NULL | Probable | As litter_saprotroph (Põlme et al. 2020); As mycoparasite (Põlme et al. 2020); Decay fungal material (Põlme et al. 2020) | Tedersoo L, et al. 2014. Science 346:e1256688 ((https://doi.org/10.1126/science.1256688)); Purahong W, et al. 2016. Molecular Ecology 25:4059-4074 ((https://doi.org/10.1111/mec.13739)); Põlme S, et al. 2020. Fungal Diversity 105:1-16 ((https://doi.org/10.1007/s13225-020-00466-2)) |
| Otu54965 | 1 | k__Fungi(100);p__Ascomycota(100);c__Saccharomycetes(100);o__Saccharomycetales(100);f__Metschnikowiaceae(100);g__Metschnikowia(100);s__Metschnikowia_pulcherrima(100); | Metschnikowia | 13 | Saprotroph | \|Undefined Saprotroph\| | Yeast | NULL | Probable | As nectar-tap_saprotroph (Põlme et al. 2020); Decay sugar-rich substrates (Põlme et al. 2020) | Põlme S, et al. 2020. Fungal Diversity 105:1-16 ((https://doi.org/10.1007/s13225-020-00466-2)) |
| Otu54968 | 1 | k__Fungi(100);p__Ascomycota(100);c__Saccharomycetes(100);o__Saccharomycetales(100);f__Metschnikowiaceae(100);g__Metschnikowia(100);s__Metschnikowia_pulcherrima(100); | Metschnikowia | 13 | Saprotroph | \|Undefined Saprotroph\| | Yeast | NULL | Probable | As nectar-tap_saprotroph (Põlme et al. 2020); Decay sugar-rich substrates (Põlme et al. 2020) | Põlme S, et al. 2020. Fungal Diversity 105:1-16 ((https://doi.org/10.1007/s13225-020-00466-2)) |
| Otu54969 | 1 | k__Fungi(100);p__Ascomycota(100);c__Saccharomycetes(100);o__Saccharomycetales(100);f__Metschnikowiaceae(100);g__Metschnikowia(100);s__Metschnikowia_pulcherrima(100); | Metschnikowia | 13 | Saprotroph | \|Undefined Saprotroph\| | Yeast | NULL | Probable | As nectar-tap_saprotroph (Põlme et al. 2020); Decay sugar-rich substrates (Põlme et al. 2020) | Põlme S, et al. 2020. Fungal Diversity 105:1-16 ((https://doi.org/10.1007/s13225-020-00466-2)) |
| Otu54970 | 1 | k__Fungi(100);p__Ascomycota(100);c__Saccharomycetes(100);o__Saccharomycetales(100);f__Metschnikowiaceae(100);g__Metschnikowia(100);s__Metschnikowia_pulcherrima(100); | Metschnikowia | 13 | Saprotroph | \|Undefined Saprotroph\| | Yeast | NULL | Probable | As nectar-tap_saprotroph (Põlme et al. 2020); Decay sugar-rich substrates (Põlme et al. 2020) | Põlme S, et al. 2020. Fungal Diversity 105:1-16 ((https://doi.org/10.1007/s13225-020-00466-2)) |
| Otu54971 | 1 | k__Fungi(100);p__Ascomycota(100);c__Saccharomycetes(100);o__Saccharomycetales(100);f__Metschnikowiaceae(100);g__Metschnikowia(100);s__Metschnikowia_pulcherrima(100); | Metschnikowia | 13 | Saprotroph | \|Undefined Saprotroph\| | Yeast | NULL | Probable | As nectar-tap_saprotroph (Põlme et al. 2020); Decay sugar-rich substrates (Põlme et al. 2020) | Põlme S, et al. 2020. Fungal Diversity 105:1-16 ((https://doi.org/10.1007/s13225-020-00466-2)) |
| Otu54972 | 1 | k__Fungi(100);p__Ascomycota(100);c__Saccharomycetes(100);o__Saccharomycetales(100);f__Metschnikowiaceae(100);g__Metschnikowia(100);s__Metschnikowia_pulcherrima(100); | Metschnikowia | 13 | Saprotroph | \|Undefined Saprotroph\| | Yeast | NULL | Probable | As nectar-tap_saprotroph (Põlme et al. 2020); Decay sugar-rich substrates (Põlme et al. 2020) | Põlme S, et al. 2020. Fungal Diversity 105:1-16 ((https://doi.org/10.1007/s13225-020-00466-2)) |
| Otu54973 | 1 | k__Fungi(100);p__Ascomycota(100);c__Saccharomycetes(100);o__Saccharomycetales(100);f__Metschnikowiaceae(100);g__Metschnikowia(100);g__Metschnikowia_unclassified(100); | Metschnikowia | 13 | Saprotroph | \|Undefined Saprotroph\| | Yeast | NULL | Probable | As nectar-tap_saprotroph (Põlme et al. 2020); Decay sugar-rich substrates (Põlme et al. 2020) | Põlme S, et al. 2020. Fungal Diversity 105:1-16 ((https://doi.org/10.1007/s13225-020-00466-2)) |
| Otu54977 | 1 | k__Fungi(100);p__Ascomycota(100);c__Saccharomycetes(100);o__Saccharomycetales(100);f__Metschnikowiaceae(100);g__Metschnikowia(100);s__Metschnikowia_pulcherrima(100); | Metschnikowia | 13 | Saprotroph | \|Undefined Saprotroph\| | Yeast | NULL | Probable | As nectar-tap_saprotroph (Põlme et al. 2020); Decay sugar-rich substrates (Põlme et al. 2020) | Põlme S, et al. 2020. Fungal Diversity 105:1-16 ((https://doi.org/10.1007/s13225-020-00466-2)) |
| Otu54978 | 1 | k__Fungi(100);p__Ascomycota(100);c__Saccharomycetes(100);o__Saccharomycetales(100);f__Metschnikowiaceae(100);g__Metschnikowia(100);g__Metschnikowia_unclassified(100); | Metschnikowia | 13 | Saprotroph | \|Undefined Saprotroph\| | Yeast | NULL | Probable | As nectar-tap_saprotroph (Põlme et al. 2020); Decay sugar-rich substrates (Põlme et al. 2020) | Põlme S, et al. 2020. Fungal Diversity 105:1-16 ((https://doi.org/10.1007/s13225-020-00466-2)) |
| Otu54979 | 1 | k__Fungi(100);p__Ascomycota(100);c__Saccharomycetes(100);o__Saccharomycetales(100);f__Metschnikowiaceae(100);g__Metschnikowia(100);s__Metschnikowia_pulcherrima(100); | Metschnikowia | 13 | Saprotroph | \|Undefined Saprotroph\| | Yeast | NULL | Probable | As nectar-tap_saprotroph (Põlme et al. 2020); Decay sugar-rich substrates (Põlme et al. 2020) | Põlme S, et al. 2020. Fungal Diversity 105:1-16 ((https://doi.org/10.1007/s13225-020-00466-2)) |
| Otu54980 | 1 | k__Fungi(100);p__Ascomycota(100);c__Saccharomycetes(100);o__Saccharomycetales(100);f__Metschnikowiaceae(100);g__Metschnikowia(100);s__Metschnikowia_pulcherrima(100); | Metschnikowia | 13 | Saprotroph | \|Undefined Saprotroph\| | Yeast | NULL | Probable | As nectar-tap_saprotroph (Põlme et al. 2020); Decay sugar-rich substrates (Põlme et al. 2020) | Põlme S, et al. 2020. Fungal Diversity 105:1-16 ((https://doi.org/10.1007/s13225-020-00466-2)) |
| Otu54981 | 1 | k__Fungi(100);p__Ascomycota(100);c__Saccharomycetes(100);o__Saccharomycetales(100);f__Metschnikowiaceae(100);g__Metschnikowia(100);s__Metschnikowia_pulcherrima(100); | Metschnikowia | 13 | Saprotroph | \|Undefined Saprotroph\| | Yeast | NULL | Probable | As nectar-tap_saprotroph (Põlme et al. 2020); Decay sugar-rich substrates (Põlme et al. 2020) | Põlme S, et al. 2020. Fungal Diversity 105:1-16 ((https://doi.org/10.1007/s13225-020-00466-2)) |
| Otu54982 | 1 | k__Fungi(100);p__Ascomycota(100);c__Saccharomycetes(100);o__Saccharomycetales(100);f__Metschnikowiaceae(100);g__Metschnikowia(100);s__Metschnikowia_pulcherrima(100); | Metschnikowia | 13 | Saprotroph | \|Undefined Saprotroph\| | Yeast | NULL | Probable | As nectar-tap_saprotroph (Põlme et al. 2020); Decay sugar-rich substrates (Põlme et al. 2020) | Põlme S, et al. 2020. Fungal Diversity 105:1-16 ((https://doi.org/10.1007/s13225-020-00466-2)) |
| Otu54983 | 1 | k__Fungi(100);p__Ascomycota(100);c__Saccharomycetes(100);o__Saccharomycetales(100);f__Metschnikowiaceae(100);g__Metschnikowia(100);s__Metschnikowia_pulcherrima(100); | Metschnikowia | 13 | Saprotroph | \|Undefined Saprotroph\| | Yeast | NULL | Probable | As nectar-tap_saprotroph (Põlme et al. 2020); Decay sugar-rich substrates (Põlme et al. 2020) | Põlme S, et al. 2020. Fungal Diversity 105:1-16 ((https://doi.org/10.1007/s13225-020-00466-2)) |
| Otu54984 | 1 | k__Fungi(100);p__Ascomycota(100);c__Saccharomycetes(100);o__Saccharomycetales(100);f__Metschnikowiaceae(100);g__Metschnikowia(100);s__Metschnikowia_pulcherrima(100); | Metschnikowia | 13 | Saprotroph | \|Undefined Saprotroph\| | Yeast | NULL | Probable | As nectar-tap_saprotroph (Põlme et al. 2020); Decay sugar-rich substrates (Põlme et al. 2020) | Põlme S, et al. 2020. Fungal Diversity 105:1-16 ((https://doi.org/10.1007/s13225-020-00466-2)) |
| Otu54985 | 1 | k__Fungi(100);p__Ascomycota(100);c__Saccharomycetes(100);o__Saccharomycetales(100);f__Metschnikowiaceae(100);g__Metschnikowia(100);g__Metschnikowia_unclassified(100); | Metschnikowia | 13 | Saprotroph | \|Undefined Saprotroph\| | Yeast | NULL | Probable | As nectar-tap_saprotroph (Põlme et al. 2020); Decay sugar-rich substrates (Põlme et al. 2020) | Põlme S, et al. 2020. Fungal Diversity 105:1-16 ((https://doi.org/10.1007/s13225-020-00466-2)) |
| Otu54987 | 1 | k__Fungi(100);p__Ascomycota(100);c__Saccharomycetes(100);o__Saccharomycetales(100);f__Metschnikowiaceae(100);g__Metschnikowia(100);s__Metschnikowia_pulcherrima(100); | Metschnikowia | 13 | Saprotroph | \|Undefined Saprotroph\| | Yeast | NULL | Probable | As nectar-tap_saprotroph (Põlme et al. 2020); Decay sugar-rich substrates (Põlme et al. 2020) | Põlme S, et al. 2020. Fungal Diversity 105:1-16 ((https://doi.org/10.1007/s13225-020-00466-2)) |
| Otu54988 | 1 | k__Fungi(100);p__Ascomycota(100);c__Saccharomycetes(100);o__Saccharomycetales(100);f__Metschnikowiaceae(100);g__Metschnikowia(100);g__Metschnikowia_unclassified(100); | Metschnikowia | 13 | Saprotroph | \|Undefined Saprotroph\| | Yeast | NULL | Probable | As nectar-tap_saprotroph (Põlme et al. 2020); Decay sugar-rich substrates (Põlme et al. 2020) | Põlme S, et al. 2020. Fungal Diversity 105:1-16 ((https://doi.org/10.1007/s13225-020-00466-2)) |
| Otu54989 | 1 | k__Fungi(100);p__Ascomycota(100);c__Saccharomycetes(100);o__Saccharomycetales(100);f__Metschnikowiaceae(100);g__Metschnikowia(100);s__Metschnikowia_pulcherrima(100); | Metschnikowia | 13 | Saprotroph | \|Undefined Saprotroph\| | Yeast | NULL | Probable | As nectar-tap_saprotroph (Põlme et al. 2020); Decay sugar-rich substrates (Põlme et al. 2020) | Põlme S, et al. 2020. Fungal Diversity 105:1-16 ((https://doi.org/10.1007/s13225-020-00466-2)) |
| Otu54990 | 1 | k__Fungi(100);p__Ascomycota(100);c__Saccharomycetes(100);o__Saccharomycetales(100);f__Metschnikowiaceae(100);g__Metschnikowia(100);s__Metschnikowia_pulcherrima(100); | Metschnikowia | 13 | Saprotroph | \|Undefined Saprotroph\| | Yeast | NULL | Probable | As nectar-tap_saprotroph (Põlme et al. 2020); Decay sugar-rich substrates (Põlme et al. 2020) | Põlme S, et al. 2020. Fungal Diversity 105:1-16 ((https://doi.org/10.1007/s13225-020-00466-2)) |
| Otu54991 | 1 | k__Fungi(100);p__Ascomycota(100);c__Saccharomycetes(100);o__Saccharomycetales(100);f__Metschnikowiaceae(100);g__Metschnikowia(100);s__Metschnikowia_pulcherrima(100); | Metschnikowia | 13 | Saprotroph | \|Undefined Saprotroph\| | Yeast | NULL | Probable | As nectar-tap_saprotroph (Põlme et al. 2020); Decay sugar-rich substrates (Põlme et al. 2020) | Põlme S, et al. 2020. Fungal Diversity 105:1-16 ((https://doi.org/10.1007/s13225-020-00466-2)) |
| Otu54992 | 1 | k__Fungi(100);p__Ascomycota(100);c__Saccharomycetes(100);o__Saccharomycetales(100);f__Metschnikowiaceae(100);g__Metschnikowia(100);s__Metschnikowia_pulcherrima(100); | Metschnikowia | 13 | Saprotroph | \|Undefined Saprotroph\| | Yeast | NULL | Probable | As nectar-tap_saprotroph (Põlme et al. 2020); Decay sugar-rich substrates (Põlme et al. 2020) | Põlme S, et al. 2020. Fungal Diversity 105:1-16 ((https://doi.org/10.1007/s13225-020-00466-2)) |
| Otu54993 | 1 | k__Fungi(100);p__Ascomycota(100);c__Saccharomycetes(100);o__Saccharomycetales(100);f__Metschnikowiaceae(100);g__Metschnikowia(100);g__Metschnikowia_unclassified(100); | Metschnikowia | 13 | Saprotroph | \|Undefined Saprotroph\| | Yeast | NULL | Probable | As nectar-tap_saprotroph (Põlme et al. 2020); Decay sugar-rich substrates (Põlme et al. 2020) | Põlme S, et al. 2020. Fungal Diversity 105:1-16 ((https://doi.org/10.1007/s13225-020-00466-2)) |
| Otu54994 | 1 | k__Fungi(100);p__Ascomycota(100);c__Saccharomycetes(100);o__Saccharomycetales(100);f__Metschnikowiaceae(100);g__Metschnikowia(100);s__Metschnikowia_pulcherrima(100); | Metschnikowia | 13 | Saprotroph | \|Undefined Saprotroph\| | Yeast | NULL | Probable | As nectar-tap_saprotroph (Põlme et al. 2020); Decay sugar-rich substrates (Põlme et al. 2020) | Põlme S, et al. 2020. Fungal Diversity 105:1-16 ((https://doi.org/10.1007/s13225-020-00466-2)) |
| Otu54995 | 1 | k__Fungi(100);p__Ascomycota(100);c__Saccharomycetes(100);o__Saccharomycetales(100);f__Metschnikowiaceae(100);g__Metschnikowia(100);g__Metschnikowia_unclassified(100); | Metschnikowia | 13 | Saprotroph | \|Undefined Saprotroph\| | Yeast | NULL | Probable | As nectar-tap_saprotroph (Põlme et al. 2020); Decay sugar-rich substrates (Põlme et al. 2020) | Põlme S, et al. 2020. Fungal Diversity 105:1-16 ((https://doi.org/10.1007/s13225-020-00466-2)) |
| Otu54996 | 1 | k__Fungi(100);p__Ascomycota(100);c__Saccharomycetes(100);o__Saccharomycetales(100);f__Metschnikowiaceae(100);g__Metschnikowia(100);s__Metschnikowia_pulcherrima(100); | Metschnikowia | 13 | Saprotroph | \|Undefined Saprotroph\| | Yeast | NULL | Probable | As nectar-tap_saprotroph (Põlme et al. 2020); Decay sugar-rich substrates (Põlme et al. 2020) | Põlme S, et al. 2020. Fungal Diversity 105:1-16 ((https://doi.org/10.1007/s13225-020-00466-2)) |
| Otu54997 | 1 | k__Fungi(100);p__Ascomycota(100);c__Saccharomycetes(100);o__Saccharomycetales(100);f__Metschnikowiaceae(100);g__Metschnikowia(100);s__Metschnikowia_pulcherrima(100); | Metschnikowia | 13 | Saprotroph | \|Undefined Saprotroph\| | Yeast | NULL | Probable | As nectar-tap_saprotroph (Põlme et al. 2020); Decay sugar-rich substrates (Põlme et al. 2020) | Põlme S, et al. 2020. Fungal Diversity 105:1-16 ((https://doi.org/10.1007/s13225-020-00466-2)) |
| Otu54998 | 1 | k__Fungi(100);p__Ascomycota(100);c__Saccharomycetes(100);o__Saccharomycetales(100);f__Metschnikowiaceae(100);g__Metschnikowia(100);s__Metschnikowia_pulcherrima(100); | Metschnikowia | 13 | Saprotroph | \|Undefined Saprotroph\| | Yeast | NULL | Probable | As nectar-tap_saprotroph (Põlme et al. 2020); Decay sugar-rich substrates (Põlme et al. 2020) | Põlme S, et al. 2020. Fungal Diversity 105:1-16 ((https://doi.org/10.1007/s13225-020-00466-2)) |
| Otu54999 | 1 | k__Fungi(100);p__Ascomycota(100);c__Saccharomycetes(100);o__Saccharomycetales(100);f__Metschnikowiaceae(100);g__Metschnikowia(100);s__Metschnikowia_pulcherrima(100); | Metschnikowia | 13 | Saprotroph | \|Undefined Saprotroph\| | Yeast | NULL | Probable | As nectar-tap_saprotroph (Põlme et al. 2020); Decay sugar-rich substrates (Põlme et al. 2020) | Põlme S, et al. 2020. Fungal Diversity 105:1-16 ((https://doi.org/10.1007/s13225-020-00466-2)) |
| Otu55000 | 1 | k__Fungi(100);p__Ascomycota(100);c__Saccharomycetes(100);o__Saccharomycetales(100);f__Metschnikowiaceae(100);g__Metschnikowia(100);s__Metschnikowia_pulcherrima(100); | Metschnikowia | 13 | Saprotroph | \|Undefined Saprotroph\| | Yeast | NULL | Probable | As nectar-tap_saprotroph (Põlme et al. 2020); Decay sugar-rich substrates (Põlme et al. 2020) | Põlme S, et al. 2020. Fungal Diversity 105:1-16 ((https://doi.org/10.1007/s13225-020-00466-2)) |
| Otu55001 | 1 | k__Fungi(100);p__Ascomycota(100);c__Saccharomycetes(100);o__Saccharomycetales(100);f__Metschnikowiaceae(100);g__Metschnikowia(100);s__Metschnikowia_pulcherrima(100); | Metschnikowia | 13 | Saprotroph | \|Undefined Saprotroph\| | Yeast | NULL | Probable | As nectar-tap_saprotroph (Põlme et al. 2020); Decay sugar-rich substrates (Põlme et al. 2020) | Põlme S, et al. 2020. Fungal Diversity 105:1-16 ((https://doi.org/10.1007/s13225-020-00466-2)) |
| Otu55002 | 1 | k__Fungi(100);p__Ascomycota(100);c__Saccharomycetes(100);o__Saccharomycetales(100);f__Metschnikowiaceae(100);g__Metschnikowia(100);s__Metschnikowia_pulcherrima(100); | Metschnikowia | 13 | Saprotroph | \|Undefined Saprotroph\| | Yeast | NULL | Probable | As nectar-tap_saprotroph (Põlme et al. 2020); Decay sugar-rich substrates (Põlme et al. 2020) | Põlme S, et al. 2020. Fungal Diversity 105:1-16 ((https://doi.org/10.1007/s13225-020-00466-2)) |
| Otu55004 | 1 | k__Fungi(100);p__Ascomycota(100);c__Saccharomycetes(100);o__Saccharomycetales(100);f__Metschnikowiaceae(100);g__Metschnikowia(100);s__Metschnikowia_pulcherrima(100); | Metschnikowia | 13 | Saprotroph | \|Undefined Saprotroph\| | Yeast | NULL | Probable | As nectar-tap_saprotroph (Põlme et al. 2020); Decay sugar-rich substrates (Põlme et al. 2020) | Põlme S, et al. 2020. Fungal Diversity 105:1-16 ((https://doi.org/10.1007/s13225-020-00466-2)) |
| Otu55005 | 1 | k__Fungi(100);p__Ascomycota(100);c__Saccharomycetes(100);o__Saccharomycetales(100);f__Metschnikowiaceae(100);g__Metschnikowia(100);g__Metschnikowia_unclassified(100); | Metschnikowia | 13 | Saprotroph | \|Undefined Saprotroph\| | Yeast | NULL | Probable | As nectar-tap_saprotroph (Põlme et al. 2020); Decay sugar-rich substrates (Põlme et al. 2020) | Põlme S, et al. 2020. Fungal Diversity 105:1-16 ((https://doi.org/10.1007/s13225-020-00466-2)) |
| Otu55006 | 1 | k__Fungi(100);p__Ascomycota(100);c__Saccharomycetes(100);o__Saccharomycetales(100);f__Metschnikowiaceae(100);g__Metschnikowia(100);g__Metschnikowia_unclassified(100); | Metschnikowia | 13 | Saprotroph | \|Undefined Saprotroph\| | Yeast | NULL | Probable | As nectar-tap_saprotroph (Põlme et al. 2020); Decay sugar-rich substrates (Põlme et al. 2020) | Põlme S, et al. 2020. Fungal Diversity 105:1-16 ((https://doi.org/10.1007/s13225-020-00466-2)) |
| Otu55007 | 1 | k__Fungi(100);p__Ascomycota(100);c__Saccharomycetes(100);o__Saccharomycetales(100);f__Metschnikowiaceae(100);g__Metschnikowia(100);s__Metschnikowia_pulcherrima(100); | Metschnikowia | 13 | Saprotroph | \|Undefined Saprotroph\| | Yeast | NULL | Probable | As nectar-tap_saprotroph (Põlme et al. 2020); Decay sugar-rich substrates (Põlme et al. 2020) | Põlme S, et al. 2020. Fungal Diversity 105:1-16 ((https://doi.org/10.1007/s13225-020-00466-2)) |
| Otu55008 | 1 | k__Fungi(100);p__Ascomycota(100);c__Saccharomycetes(100);o__Saccharomycetales(100);f__Metschnikowiaceae(100);g__Metschnikowia(100);g__Metschnikowia_unclassified(100); | Metschnikowia | 13 | Saprotroph | \|Undefined Saprotroph\| | Yeast | NULL | Probable | As nectar-tap_saprotroph (Põlme et al. 2020); Decay sugar-rich substrates (Põlme et al. 2020) | Põlme S, et al. 2020. Fungal Diversity 105:1-16 ((https://doi.org/10.1007/s13225-020-00466-2)) |
| Otu55009 | 1 | k__Fungi(100);p__Ascomycota(100);c__Saccharomycetes(100);o__Saccharomycetales(100);f__Metschnikowiaceae(100);g__Metschnikowia(100);g__Metschnikowia_unclassified(100); | Metschnikowia | 13 | Saprotroph | \|Undefined Saprotroph\| | Yeast | NULL | Probable | As nectar-tap_saprotroph (Põlme et al. 2020); Decay sugar-rich substrates (Põlme et al. 2020) | Põlme S, et al. 2020. Fungal Diversity 105:1-16 ((https://doi.org/10.1007/s13225-020-00466-2)) |
| Otu55010 | 1 | k__Fungi(100);p__Ascomycota(100);c__Saccharomycetes(100);o__Saccharomycetales(100);f__Saccharomycetales_family_Incertae_sedis(100);g__Candida(100);g__Candida_unclassified(100); | Candida | 13 | Pathotroph-Saprotroph-Symbiotroph | Animal Pathogen-Endophyte-Endosymbiont-Epiphyte-\|Undefined Saprotroph\| | Dimorphic Yeast-Yeast | NULL | Probable | As nectar-tap_saprotroph (Põlme et al. 2020); Decay sugar-rich substrates (Põlme et al. 2020); This asexual genus is polyphyletic and contain species that function very differently depend on what they are. Caution should be taken when interpreting guilds for Candida species. | Manolakaki D, et al. 2010. Virulence. 1: 367-375 ((https://doi.org/10.4161/viru.1.5.12796)); Mekbib SB, et al. 2011. Tropical Plant Pathology 36:233-240 ((https://doi.org/10.1590/S1982-56762011000400004)); Põlme S, et al. 2020. Fungal Diversity 105:1-16 ((https://doi.org/10.1007/s13225-020-00466-2)) |
| Otu55011 | 1 | k__Fungi(100);p__Ascomycota(100);c__Saccharomycetes(100);o__Saccharomycetales(100);f__Metschnikowiaceae(100);g__Metschnikowia(100);s__Metschnikowia_pulcherrima(100); | Metschnikowia | 13 | Saprotroph | \|Undefined Saprotroph\| | Yeast | NULL | Probable | As nectar-tap_saprotroph (Põlme et al. 2020); Decay sugar-rich substrates (Põlme et al. 2020) | Põlme S, et al. 2020. Fungal Diversity 105:1-16 ((https://doi.org/10.1007/s13225-020-00466-2)) |
| Otu55012 | 1 | k__Fungi(100);p__Ascomycota(100);c__Saccharomycetes(100);o__Saccharomycetales(100);f__Metschnikowiaceae(100);g__Metschnikowia(100);g__Metschnikowia_unclassified(100); | Metschnikowia | 13 | Saprotroph | \|Undefined Saprotroph\| | Yeast | NULL | Probable | As nectar-tap_saprotroph (Põlme et al. 2020); Decay sugar-rich substrates (Põlme et al. 2020) | Põlme S, et al. 2020. Fungal Diversity 105:1-16 ((https://doi.org/10.1007/s13225-020-00466-2)) |
| Otu55013 | 1 | k__Fungi(100);p__Ascomycota(100);c__Saccharomycetes(100);o__Saccharomycetales(100);o__Saccharomycetales_unclassified(100);o__Saccharomycetales_unclassified(100);o__Saccharomycetales_unclassified(100); | Saccharomycetales | 7 | Saprotroph | Undefined Saprotroph | Yeast | NULL | Possible | NULL | Sterkenburg E, et al. 2015. New Phytologist 207:1145-1158 ((https://doi.org/10.1111/nph.13426)) |
| Otu55014 | 1 | k__Fungi(100);p__Ascomycota(100);c__Saccharomycetes(100);o__Saccharomycetales(100);f__Metschnikowiaceae(100);g__Metschnikowia(100);s__Metschnikowia_pulcherrima(100); | Metschnikowia | 13 | Saprotroph | \|Undefined Saprotroph\| | Yeast | NULL | Probable | As nectar-tap_saprotroph (Põlme et al. 2020); Decay sugar-rich substrates (Põlme et al. 2020) | Põlme S, et al. 2020. Fungal Diversity 105:1-16 ((https://doi.org/10.1007/s13225-020-00466-2)) |
| Otu55015 | 1 | k__Fungi(100);p__Ascomycota(100);c__Saccharomycetes(100);o__Saccharomycetales(100);f__Metschnikowiaceae(100);g__Metschnikowia(100);s__Metschnikowia_pulcherrima(100); | Metschnikowia | 13 | Saprotroph | \|Undefined Saprotroph\| | Yeast | NULL | Probable | As nectar-tap_saprotroph (Põlme et al. 2020); Decay sugar-rich substrates (Põlme et al. 2020) | Põlme S, et al. 2020. Fungal Diversity 105:1-16 ((https://doi.org/10.1007/s13225-020-00466-2)) |
| Otu55016 | 1 | k__Fungi(100);p__Ascomycota(100);c__Saccharomycetes(100);o__Saccharomycetales(100);f__Metschnikowiaceae(100);g__Metschnikowia(100);s__Metschnikowia_pulcherrima(100); | Metschnikowia | 13 | Saprotroph | \|Undefined Saprotroph\| | Yeast | NULL | Probable | As nectar-tap_saprotroph (Põlme et al. 2020); Decay sugar-rich substrates (Põlme et al. 2020) | Põlme S, et al. 2020. Fungal Diversity 105:1-16 ((https://doi.org/10.1007/s13225-020-00466-2)) |
| Otu55017 | 1 | k__Fungi(100);p__Ascomycota(100);c__Saccharomycetes(100);o__Saccharomycetales(100);f__Metschnikowiaceae(100);g__Metschnikowia(100);g__Metschnikowia_unclassified(100); | Metschnikowia | 13 | Saprotroph | \|Undefined Saprotroph\| | Yeast | NULL | Probable | As nectar-tap_saprotroph (Põlme et al. 2020); Decay sugar-rich substrates (Põlme et al. 2020) | Põlme S, et al. 2020. Fungal Diversity 105:1-16 ((https://doi.org/10.1007/s13225-020-00466-2)) |
| Otu55018 | 1 | k__Fungi(100);p__Ascomycota(100);c__Saccharomycetes(100);o__Saccharomycetales(100);f__Metschnikowiaceae(100);g__Metschnikowia(100);s__Metschnikowia_pulcherrima(100); | Metschnikowia | 13 | Saprotroph | \|Undefined Saprotroph\| | Yeast | NULL | Probable | As nectar-tap_saprotroph (Põlme et al. 2020); Decay sugar-rich substrates (Põlme et al. 2020) | Põlme S, et al. 2020. Fungal Diversity 105:1-16 ((https://doi.org/10.1007/s13225-020-00466-2)) |
| Otu55019 | 1 | k__Fungi(100);p__Ascomycota(100);c__Saccharomycetes(100);o__Saccharomycetales(100);f__Saccharomycetales_family_Incertae_sedis(100);g__Candida(100);g__Candida_unclassified(100); | Candida | 13 | Pathotroph-Saprotroph-Symbiotroph | Animal Pathogen-Endophyte-Endosymbiont-Epiphyte-\|Undefined Saprotroph\| | Dimorphic Yeast-Yeast | NULL | Probable | As nectar-tap_saprotroph (Põlme et al. 2020); Decay sugar-rich substrates (Põlme et al. 2020); This asexual genus is polyphyletic and contain species that function very differently depend on what they are. Caution should be taken when interpreting guilds for Candida species. | Manolakaki D, et al. 2010. Virulence. 1: 367-375 ((https://doi.org/10.4161/viru.1.5.12796)); Mekbib SB, et al. 2011. Tropical Plant Pathology 36:233-240 ((https://doi.org/10.1590/S1982-56762011000400004)); Põlme S, et al. 2020. Fungal Diversity 105:1-16 ((https://doi.org/10.1007/s13225-020-00466-2)) |
| Otu55022 | 1 | k__Fungi(100);p__Ascomycota(100);c__Saccharomycetes(100);o__Saccharomycetales(100);o__Saccharomycetales_unclassified(100);o__Saccharomycetales_unclassified(100);o__Saccharomycetales_unclassified(100); | Saccharomycetales | 7 | Saprotroph | Undefined Saprotroph | Yeast | NULL | Possible | NULL | Sterkenburg E, et al. 2015. New Phytologist 207:1145-1158 ((https://doi.org/10.1111/nph.13426)) |
| Otu55024 | 1 | k__Fungi(100);p__Ascomycota(100);c__Saccharomycetes(100);o__Saccharomycetales(100);f__Metschnikowiaceae(100);g__Metschnikowia(100);s__Metschnikowia_pulcherrima(100); | Metschnikowia | 13 | Saprotroph | \|Undefined Saprotroph\| | Yeast | NULL | Probable | As nectar-tap_saprotroph (Põlme et al. 2020); Decay sugar-rich substrates (Põlme et al. 2020) | Põlme S, et al. 2020. Fungal Diversity 105:1-16 ((https://doi.org/10.1007/s13225-020-00466-2)) |
| Otu55025 | 1 | k__Fungi(100);p__Ascomycota(100);c__Saccharomycetes(100);o__Saccharomycetales(100);f__Saccharomycetales_family_Incertae_sedis(100);g__Candida(100);g__Candida_unclassified(100); | Candida | 13 | Pathotroph-Saprotroph-Symbiotroph | Animal Pathogen-Endophyte-Endosymbiont-Epiphyte-\|Undefined Saprotroph\| | Dimorphic Yeast-Yeast | NULL | Probable | As nectar-tap_saprotroph (Põlme et al. 2020); Decay sugar-rich substrates (Põlme et al. 2020); This asexual genus is polyphyletic and contain species that function very differently depend on what they are. Caution should be taken when interpreting guilds for Candida species. | Manolakaki D, et al. 2010. Virulence. 1: 367-375 ((https://doi.org/10.4161/viru.1.5.12796)); Mekbib SB, et al. 2011. Tropical Plant Pathology 36:233-240 ((https://doi.org/10.1590/S1982-56762011000400004)); Põlme S, et al. 2020. Fungal Diversity 105:1-16 ((https://doi.org/10.1007/s13225-020-00466-2)) |
| Otu55026 | 1 | k__Fungi(100);p__Ascomycota(100);c__Saccharomycetes(100);o__Saccharomycetales(100);f__Metschnikowiaceae(100);g__Metschnikowia(100);s__Metschnikowia_pulcherrima(100); | Metschnikowia | 13 | Saprotroph | \|Undefined Saprotroph\| | Yeast | NULL | Probable | As nectar-tap_saprotroph (Põlme et al. 2020); Decay sugar-rich substrates (Põlme et al. 2020) | Põlme S, et al. 2020. Fungal Diversity 105:1-16 ((https://doi.org/10.1007/s13225-020-00466-2)) |
| Otu55027 | 1 | k__Fungi(100);p__Ascomycota(100);c__Saccharomycetes(100);o__Saccharomycetales(100);o__Saccharomycetales_unclassified(100);o__Saccharomycetales_unclassified(100);o__Saccharomycetales_unclassified(100); | Saccharomycetales | 7 | Saprotroph | Undefined Saprotroph | Yeast | NULL | Possible | NULL | Sterkenburg E, et al. 2015. New Phytologist 207:1145-1158 ((https://doi.org/10.1111/nph.13426)) |
| Otu55029 | 1 | k__Fungi(100);p__Ascomycota(100);c__Saccharomycetes(100);o__Saccharomycetales(100);f__Metschnikowiaceae(100);g__Metschnikowia(100);s__Metschnikowia_pulcherrima(100); | Metschnikowia | 13 | Saprotroph | \|Undefined Saprotroph\| | Yeast | NULL | Probable | As nectar-tap_saprotroph (Põlme et al. 2020); Decay sugar-rich substrates (Põlme et al. 2020) | Põlme S, et al. 2020. Fungal Diversity 105:1-16 ((https://doi.org/10.1007/s13225-020-00466-2)) |
| Otu55030 | 1 | k__Fungi(100);p__Ascomycota(100);c__Saccharomycetes(100);o__Saccharomycetales(100);f__Metschnikowiaceae(100);g__Metschnikowia(100);s__Metschnikowia_pulcherrima(100); | Metschnikowia | 13 | Saprotroph | \|Undefined Saprotroph\| | Yeast | NULL | Probable | As nectar-tap_saprotroph (Põlme et al. 2020); Decay sugar-rich substrates (Põlme et al. 2020) | Põlme S, et al. 2020. Fungal Diversity 105:1-16 ((https://doi.org/10.1007/s13225-020-00466-2)) |
| Otu55031 | 1 | k__Fungi(100);p__Ascomycota(100);c__Saccharomycetes(100);o__Saccharomycetales(100);f__Metschnikowiaceae(100);g__Metschnikowia(100);s__Metschnikowia_pulcherrima(100); | Metschnikowia | 13 | Saprotroph | \|Undefined Saprotroph\| | Yeast | NULL | Probable | As nectar-tap_saprotroph (Põlme et al. 2020); Decay sugar-rich substrates (Põlme et al. 2020) | Põlme S, et al. 2020. Fungal Diversity 105:1-16 ((https://doi.org/10.1007/s13225-020-00466-2)) |
| Otu55032 | 1 | k__Fungi(100);p__Ascomycota(100);c__Saccharomycetes(100);o__Saccharomycetales(100);f__Metschnikowiaceae(100);g__Metschnikowia(100);s__Metschnikowia_pulcherrima(100); | Metschnikowia | 13 | Saprotroph | \|Undefined Saprotroph\| | Yeast | NULL | Probable | As nectar-tap_saprotroph (Põlme et al. 2020); Decay sugar-rich substrates (Põlme et al. 2020) | Põlme S, et al. 2020. Fungal Diversity 105:1-16 ((https://doi.org/10.1007/s13225-020-00466-2)) |
| Otu55033 | 1 | k__Fungi(100);p__Ascomycota(100);c__Saccharomycetes(100);o__Saccharomycetales(100);f__Metschnikowiaceae(100);g__Metschnikowia(100);s__Metschnikowia_pulcherrima(100); | Metschnikowia | 13 | Saprotroph | \|Undefined Saprotroph\| | Yeast | NULL | Probable | As nectar-tap_saprotroph (Põlme et al. 2020); Decay sugar-rich substrates (Põlme et al. 2020) | Põlme S, et al. 2020. Fungal Diversity 105:1-16 ((https://doi.org/10.1007/s13225-020-00466-2)) |
| Otu55034 | 1 | k__Fungi(100);p__Ascomycota(100);c__Saccharomycetes(100);o__Saccharomycetales(100);f__Metschnikowiaceae(100);g__Metschnikowia(100);s__Metschnikowia_pulcherrima(100); | Metschnikowia | 13 | Saprotroph | \|Undefined Saprotroph\| | Yeast | NULL | Probable | As nectar-tap_saprotroph (Põlme et al. 2020); Decay sugar-rich substrates (Põlme et al. 2020) | Põlme S, et al. 2020. Fungal Diversity 105:1-16 ((https://doi.org/10.1007/s13225-020-00466-2)) |
| Otu55035 | 1 | k__Fungi(100);p__Ascomycota(100);c__Saccharomycetes(100);o__Saccharomycetales(100);f__Metschnikowiaceae(100);g__Metschnikowia(100);s__Metschnikowia_pulcherrima(100); | Metschnikowia | 13 | Saprotroph | \|Undefined Saprotroph\| | Yeast | NULL | Probable | As nectar-tap_saprotroph (Põlme et al. 2020); Decay sugar-rich substrates (Põlme et al. 2020) | Põlme S, et al. 2020. Fungal Diversity 105:1-16 ((https://doi.org/10.1007/s13225-020-00466-2)) |
| Otu55036 | 1 | k__Fungi(100);p__Ascomycota(100);c__Saccharomycetes(100);o__Saccharomycetales(100);f__Metschnikowiaceae(100);g__Metschnikowia(100);s__Metschnikowia_pulcherrima(100); | Metschnikowia | 13 | Saprotroph | \|Undefined Saprotroph\| | Yeast | NULL | Probable | As nectar-tap_saprotroph (Põlme et al. 2020); Decay sugar-rich substrates (Põlme et al. 2020) | Põlme S, et al. 2020. Fungal Diversity 105:1-16 ((https://doi.org/10.1007/s13225-020-00466-2)) |
| Otu55037 | 1 | k__Fungi(100);p__Ascomycota(100);c__Saccharomycetes(100);o__Saccharomycetales(100);f__Metschnikowiaceae(100);g__Metschnikowia(100);s__Metschnikowia_pulcherrima(100); | Metschnikowia | 13 | Saprotroph | \|Undefined Saprotroph\| | Yeast | NULL | Probable | As nectar-tap_saprotroph (Põlme et al. 2020); Decay sugar-rich substrates (Põlme et al. 2020) | Põlme S, et al. 2020. Fungal Diversity 105:1-16 ((https://doi.org/10.1007/s13225-020-00466-2)) |
| Otu55038 | 1 | k__Fungi(100);p__Ascomycota(100);c__Saccharomycetes(100);o__Saccharomycetales(100);f__Metschnikowiaceae(100);g__Metschnikowia(100);g__Metschnikowia_unclassified(100); | Metschnikowia | 13 | Saprotroph | \|Undefined Saprotroph\| | Yeast | NULL | Probable | As nectar-tap_saprotroph (Põlme et al. 2020); Decay sugar-rich substrates (Põlme et al. 2020) | Põlme S, et al. 2020. Fungal Diversity 105:1-16 ((https://doi.org/10.1007/s13225-020-00466-2)) |
| Otu55039 | 1 | k__Fungi(100);p__Ascomycota(100);c__Saccharomycetes(100);o__Saccharomycetales(100);f__Metschnikowiaceae(100);g__Metschnikowia(100);g__Metschnikowia_unclassified(100); | Metschnikowia | 13 | Saprotroph | \|Undefined Saprotroph\| | Yeast | NULL | Probable | As nectar-tap_saprotroph (Põlme et al. 2020); Decay sugar-rich substrates (Põlme et al. 2020) | Põlme S, et al. 2020. Fungal Diversity 105:1-16 ((https://doi.org/10.1007/s13225-020-00466-2)) |
| Otu55040 | 1 | k__Fungi(100);p__Ascomycota(100);c__Saccharomycetes(100);o__Saccharomycetales(100);f__Metschnikowiaceae(100);g__Metschnikowia(100);s__Metschnikowia_pulcherrima(100); | Metschnikowia | 13 | Saprotroph | \|Undefined Saprotroph\| | Yeast | NULL | Probable | As nectar-tap_saprotroph (Põlme et al. 2020); Decay sugar-rich substrates (Põlme et al. 2020) | Põlme S, et al. 2020. Fungal Diversity 105:1-16 ((https://doi.org/10.1007/s13225-020-00466-2)) |
| Otu55041 | 1 | k__Fungi(100);p__Ascomycota(100);c__Saccharomycetes(100);o__Saccharomycetales(100);f__Metschnikowiaceae(100);g__Metschnikowia(100);s__Metschnikowia_pulcherrima(100); | Metschnikowia | 13 | Saprotroph | \|Undefined Saprotroph\| | Yeast | NULL | Probable | As nectar-tap_saprotroph (Põlme et al. 2020); Decay sugar-rich substrates (Põlme et al. 2020) | Põlme S, et al. 2020. Fungal Diversity 105:1-16 ((https://doi.org/10.1007/s13225-020-00466-2)) |
| Otu55042 | 1 | k__Fungi(100);p__Ascomycota(100);c__Saccharomycetes(100);o__Saccharomycetales(100);f__Metschnikowiaceae(100);g__Metschnikowia(100);s__Metschnikowia_pulcherrima(100); | Metschnikowia | 13 | Saprotroph | \|Undefined Saprotroph\| | Yeast | NULL | Probable | As nectar-tap_saprotroph (Põlme et al. 2020); Decay sugar-rich substrates (Põlme et al. 2020) | Põlme S, et al. 2020. Fungal Diversity 105:1-16 ((https://doi.org/10.1007/s13225-020-00466-2)) |
| Otu55045 | 1 | k__Fungi(100);p__Ascomycota(100);c__Saccharomycetes(100);o__Saccharomycetales(100);f__Metschnikowiaceae(100);g__Metschnikowia(100);s__Metschnikowia_pulcherrima(100); | Metschnikowia | 13 | Saprotroph | \|Undefined Saprotroph\| | Yeast | NULL | Probable | As nectar-tap_saprotroph (Põlme et al. 2020); Decay sugar-rich substrates (Põlme et al. 2020) | Põlme S, et al. 2020. Fungal Diversity 105:1-16 ((https://doi.org/10.1007/s13225-020-00466-2)) |
| Otu55046 | 1 | k__Fungi(100);p__Ascomycota(100);c__Saccharomycetes(100);o__Saccharomycetales(100);f__Metschnikowiaceae(100);g__Metschnikowia(100);g__Metschnikowia_unclassified(100); | Metschnikowia | 13 | Saprotroph | \|Undefined Saprotroph\| | Yeast | NULL | Probable | As nectar-tap_saprotroph (Põlme et al. 2020); Decay sugar-rich substrates (Põlme et al. 2020) | Põlme S, et al. 2020. Fungal Diversity 105:1-16 ((https://doi.org/10.1007/s13225-020-00466-2)) |
| Otu55047 | 1 | k__Fungi(100);p__Ascomycota(100);c__Saccharomycetes(100);o__Saccharomycetales(100);f__Metschnikowiaceae(100);g__Metschnikowia(100);g__Metschnikowia_unclassified(100); | Metschnikowia | 13 | Saprotroph | \|Undefined Saprotroph\| | Yeast | NULL | Probable | As nectar-tap_saprotroph (Põlme et al. 2020); Decay sugar-rich substrates (Põlme et al. 2020) | Põlme S, et al. 2020. Fungal Diversity 105:1-16 ((https://doi.org/10.1007/s13225-020-00466-2)) |
| Otu55048 | 1 | k__Fungi(100);p__Ascomycota(100);c__Saccharomycetes(100);o__Saccharomycetales(100);f__Metschnikowiaceae(100);g__Metschnikowia(100);s__Metschnikowia_pulcherrima(100); | Metschnikowia | 13 | Saprotroph | \|Undefined Saprotroph\| | Yeast | NULL | Probable | As nectar-tap_saprotroph (Põlme et al. 2020); Decay sugar-rich substrates (Põlme et al. 2020) | Põlme S, et al. 2020. Fungal Diversity 105:1-16 ((https://doi.org/10.1007/s13225-020-00466-2)) |
| Otu55049 | 1 | k__Fungi(100);p__Ascomycota(100);c__Saccharomycetes(100);o__Saccharomycetales(100);f__Metschnikowiaceae(100);g__Metschnikowia(100);s__Metschnikowia_pulcherrima(100); | Metschnikowia | 13 | Saprotroph | \|Undefined Saprotroph\| | Yeast | NULL | Probable | As nectar-tap_saprotroph (Põlme et al. 2020); Decay sugar-rich substrates (Põlme et al. 2020) | Põlme S, et al. 2020. Fungal Diversity 105:1-16 ((https://doi.org/10.1007/s13225-020-00466-2)) |
| Otu55050 | 1 | k__Fungi(100);p__Ascomycota(100);c__Saccharomycetes(100);o__Saccharomycetales(100);f__Metschnikowiaceae(100);g__Metschnikowia(100);s__Metschnikowia_pulcherrima(100); | Metschnikowia | 13 | Saprotroph | \|Undefined Saprotroph\| | Yeast | NULL | Probable | As nectar-tap_saprotroph (Põlme et al. 2020); Decay sugar-rich substrates (Põlme et al. 2020) | Põlme S, et al. 2020. Fungal Diversity 105:1-16 ((https://doi.org/10.1007/s13225-020-00466-2)) |
| Otu55051 | 1 | k__Fungi(100);p__Ascomycota(100);c__Saccharomycetes(100);o__Saccharomycetales(100);f__Metschnikowiaceae(100);g__Metschnikowia(100);g__Metschnikowia_unclassified(100); | Metschnikowia | 13 | Saprotroph | \|Undefined Saprotroph\| | Yeast | NULL | Probable | As nectar-tap_saprotroph (Põlme et al. 2020); Decay sugar-rich substrates (Põlme et al. 2020) | Põlme S, et al. 2020. Fungal Diversity 105:1-16 ((https://doi.org/10.1007/s13225-020-00466-2)) |
| Otu55052 | 1 | k__Fungi(100);p__Ascomycota(100);c__Saccharomycetes(100);o__Saccharomycetales(100);f__Metschnikowiaceae(100);g__Metschnikowia(100);s__Metschnikowia_pulcherrima(100); | Metschnikowia | 13 | Saprotroph | \|Undefined Saprotroph\| | Yeast | NULL | Probable | As nectar-tap_saprotroph (Põlme et al. 2020); Decay sugar-rich substrates (Põlme et al. 2020) | Põlme S, et al. 2020. Fungal Diversity 105:1-16 ((https://doi.org/10.1007/s13225-020-00466-2)) |
| Otu55053 | 1 | k__Fungi(100);p__Ascomycota(100);c__Saccharomycetes(100);o__Saccharomycetales(100);f__Metschnikowiaceae(100);g__Metschnikowia(100);s__Metschnikowia_pulcherrima(100); | Metschnikowia | 13 | Saprotroph | \|Undefined Saprotroph\| | Yeast | NULL | Probable | As nectar-tap_saprotroph (Põlme et al. 2020); Decay sugar-rich substrates (Põlme et al. 2020) | Põlme S, et al. 2020. Fungal Diversity 105:1-16 ((https://doi.org/10.1007/s13225-020-00466-2)) |
| Otu55055 | 1 | k__Fungi(100);p__Ascomycota(100);c__Saccharomycetes(100);o__Saccharomycetales(100);f__Metschnikowiaceae(100);g__Metschnikowia(100);s__Metschnikowia_pulcherrima(100); | Metschnikowia | 13 | Saprotroph | \|Undefined Saprotroph\| | Yeast | NULL | Probable | As nectar-tap_saprotroph (Põlme et al. 2020); Decay sugar-rich substrates (Põlme et al. 2020) | Põlme S, et al. 2020. Fungal Diversity 105:1-16 ((https://doi.org/10.1007/s13225-020-00466-2)) |
| Otu55056 | 1 | k__Fungi(100);p__Ascomycota(100);c__Saccharomycetes(100);o__Saccharomycetales(100);f__Metschnikowiaceae(100);g__Metschnikowia(100);s__Metschnikowia_pulcherrima(100); | Metschnikowia | 13 | Saprotroph | \|Undefined Saprotroph\| | Yeast | NULL | Probable | As nectar-tap_saprotroph (Põlme et al. 2020); Decay sugar-rich substrates (Põlme et al. 2020) | Põlme S, et al. 2020. Fungal Diversity 105:1-16 ((https://doi.org/10.1007/s13225-020-00466-2)) |
| Otu55057 | 1 | k__Fungi(100);p__Ascomycota(100);c__Saccharomycetes(100);o__Saccharomycetales(100);f__Metschnikowiaceae(100);g__Metschnikowia(100);s__Metschnikowia_pulcherrima(100); | Metschnikowia | 13 | Saprotroph | \|Undefined Saprotroph\| | Yeast | NULL | Probable | As nectar-tap_saprotroph (Põlme et al. 2020); Decay sugar-rich substrates (Põlme et al. 2020) | Põlme S, et al. 2020. Fungal Diversity 105:1-16 ((https://doi.org/10.1007/s13225-020-00466-2)) |
| Otu55058 | 1 | k__Fungi(100);p__Ascomycota(100);c__Saccharomycetes(100);o__Saccharomycetales(100);f__Metschnikowiaceae(100);g__Metschnikowia(100);s__Metschnikowia_pulcherrima(100); | Metschnikowia | 13 | Saprotroph | \|Undefined Saprotroph\| | Yeast | NULL | Probable | As nectar-tap_saprotroph (Põlme et al. 2020); Decay sugar-rich substrates (Põlme et al. 2020) | Põlme S, et al. 2020. Fungal Diversity 105:1-16 ((https://doi.org/10.1007/s13225-020-00466-2)) |
| Otu55059 | 1 | k__Fungi(100);p__Ascomycota(100);c__Saccharomycetes(100);o__Saccharomycetales(100);f__Metschnikowiaceae(100);g__Metschnikowia(100);s__Metschnikowia_pulcherrima(100); | Metschnikowia | 13 | Saprotroph | \|Undefined Saprotroph\| | Yeast | NULL | Probable | As nectar-tap_saprotroph (Põlme et al. 2020); Decay sugar-rich substrates (Põlme et al. 2020) | Põlme S, et al. 2020. Fungal Diversity 105:1-16 ((https://doi.org/10.1007/s13225-020-00466-2)) |
| Otu55060 | 1 | k__Fungi(100);p__Ascomycota(100);c__Saccharomycetes(100);o__Saccharomycetales(100);f__Metschnikowiaceae(100);g__Metschnikowia(100);s__Metschnikowia_pulcherrima(100); | Metschnikowia | 13 | Saprotroph | \|Undefined Saprotroph\| | Yeast | NULL | Probable | As nectar-tap_saprotroph (Põlme et al. 2020); Decay sugar-rich substrates (Põlme et al. 2020) | Põlme S, et al. 2020. Fungal Diversity 105:1-16 ((https://doi.org/10.1007/s13225-020-00466-2)) |
| Otu55061 | 1 | k__Fungi(100);p__Ascomycota(100);c__Saccharomycetes(100);o__Saccharomycetales(100);f__Metschnikowiaceae(100);g__Metschnikowia(100);s__Metschnikowia_pulcherrima(100); | Metschnikowia | 13 | Saprotroph | \|Undefined Saprotroph\| | Yeast | NULL | Probable | As nectar-tap_saprotroph (Põlme et al. 2020); Decay sugar-rich substrates (Põlme et al. 2020) | Põlme S, et al. 2020. Fungal Diversity 105:1-16 ((https://doi.org/10.1007/s13225-020-00466-2)) |
| Otu55062 | 1 | k__Fungi(100);p__Ascomycota(100);c__Saccharomycetes(100);o__Saccharomycetales(100);f__Metschnikowiaceae(100);g__Metschnikowia(100);g__Metschnikowia_unclassified(100); | Metschnikowia | 13 | Saprotroph | \|Undefined Saprotroph\| | Yeast | NULL | Probable | As nectar-tap_saprotroph (Põlme et al. 2020); Decay sugar-rich substrates (Põlme et al. 2020) | Põlme S, et al. 2020. Fungal Diversity 105:1-16 ((https://doi.org/10.1007/s13225-020-00466-2)) |
| Otu55063 | 1 | k__Fungi(100);p__Ascomycota(100);c__Saccharomycetes(100);o__Saccharomycetales(100);f__Metschnikowiaceae(100);g__Metschnikowia(100);s__Metschnikowia_pulcherrima(100); | Metschnikowia | 13 | Saprotroph | \|Undefined Saprotroph\| | Yeast | NULL | Probable | As nectar-tap_saprotroph (Põlme et al. 2020); Decay sugar-rich substrates (Põlme et al. 2020) | Põlme S, et al. 2020. Fungal Diversity 105:1-16 ((https://doi.org/10.1007/s13225-020-00466-2)) |
| Otu55064 | 1 | k__Fungi(100);p__Ascomycota(100);c__Saccharomycetes(100);o__Saccharomycetales(100);f__Metschnikowiaceae(100);g__Metschnikowia(100);s__Metschnikowia_pulcherrima(100); | Metschnikowia | 13 | Saprotroph | \|Undefined Saprotroph\| | Yeast | NULL | Probable | As nectar-tap_saprotroph (Põlme et al. 2020); Decay sugar-rich substrates (Põlme et al. 2020) | Põlme S, et al. 2020. Fungal Diversity 105:1-16 ((https://doi.org/10.1007/s13225-020-00466-2)) |
| Otu55065 | 1 | k__Fungi(100);p__Ascomycota(100);c__Saccharomycetes(100);o__Saccharomycetales(100);f__Metschnikowiaceae(100);g__Metschnikowia(100);s__Metschnikowia_pulcherrima(100); | Metschnikowia | 13 | Saprotroph | \|Undefined Saprotroph\| | Yeast | NULL | Probable | As nectar-tap_saprotroph (Põlme et al. 2020); Decay sugar-rich substrates (Põlme et al. 2020) | Põlme S, et al. 2020. Fungal Diversity 105:1-16 ((https://doi.org/10.1007/s13225-020-00466-2)) |
| Otu55066 | 1 | k__Fungi(100);p__Ascomycota(100);c__Saccharomycetes(100);o__Saccharomycetales(100);f__Metschnikowiaceae(100);g__Metschnikowia(100);g__Metschnikowia_unclassified(100); | Metschnikowia | 13 | Saprotroph | \|Undefined Saprotroph\| | Yeast | NULL | Probable | As nectar-tap_saprotroph (Põlme et al. 2020); Decay sugar-rich substrates (Põlme et al. 2020) | Põlme S, et al. 2020. Fungal Diversity 105:1-16 ((https://doi.org/10.1007/s13225-020-00466-2)) |
| Otu55067 | 1 | k__Fungi(100);p__Ascomycota(100);c__Saccharomycetes(100);o__Saccharomycetales(100);f__Metschnikowiaceae(100);g__Metschnikowia(100);s__Metschnikowia_pulcherrima(100); | Metschnikowia | 13 | Saprotroph | \|Undefined Saprotroph\| | Yeast | NULL | Probable | As nectar-tap_saprotroph (Põlme et al. 2020); Decay sugar-rich substrates (Põlme et al. 2020) | Põlme S, et al. 2020. Fungal Diversity 105:1-16 ((https://doi.org/10.1007/s13225-020-00466-2)) |
| Otu55068 | 1 | k__Fungi(100);p__Ascomycota(100);c__Saccharomycetes(100);o__Saccharomycetales(100);f__Metschnikowiaceae(100);g__Metschnikowia(100);g__Metschnikowia_unclassified(100); | Metschnikowia | 13 | Saprotroph | \|Undefined Saprotroph\| | Yeast | NULL | Probable | As nectar-tap_saprotroph (Põlme et al. 2020); Decay sugar-rich substrates (Põlme et al. 2020) | Põlme S, et al. 2020. Fungal Diversity 105:1-16 ((https://doi.org/10.1007/s13225-020-00466-2)) |
| Otu55069 | 1 | k__Fungi(100);p__Ascomycota(100);c__Saccharomycetes(100);o__Saccharomycetales(100);f__Metschnikowiaceae(100);g__Metschnikowia(100);g__Metschnikowia_unclassified(100); | Metschnikowia | 13 | Saprotroph | \|Undefined Saprotroph\| | Yeast | NULL | Probable | As nectar-tap_saprotroph (Põlme et al. 2020); Decay sugar-rich substrates (Põlme et al. 2020) | Põlme S, et al. 2020. Fungal Diversity 105:1-16 ((https://doi.org/10.1007/s13225-020-00466-2)) |
| Otu55070 | 1 | k__Fungi(100);p__Ascomycota(100);c__Saccharomycetes(100);o__Saccharomycetales(100);f__Metschnikowiaceae(100);g__Metschnikowia(100);s__Metschnikowia_pulcherrima(100); | Metschnikowia | 13 | Saprotroph | \|Undefined Saprotroph\| | Yeast | NULL | Probable | As nectar-tap_saprotroph (Põlme et al. 2020); Decay sugar-rich substrates (Põlme et al. 2020) | Põlme S, et al. 2020. Fungal Diversity 105:1-16 ((https://doi.org/10.1007/s13225-020-00466-2)) |
| Otu55071 | 1 | k__Fungi(100);p__Ascomycota(100);c__Saccharomycetes(100);o__Saccharomycetales(100);f__Metschnikowiaceae(100);g__Metschnikowia(100);s__Metschnikowia_pulcherrima(100); | Metschnikowia | 13 | Saprotroph | \|Undefined Saprotroph\| | Yeast | NULL | Probable | As nectar-tap_saprotroph (Põlme et al. 2020); Decay sugar-rich substrates (Põlme et al. 2020) | Põlme S, et al. 2020. Fungal Diversity 105:1-16 ((https://doi.org/10.1007/s13225-020-00466-2)) |
| Otu55072 | 1 | k__Fungi(100);p__Ascomycota(100);c__Saccharomycetes(100);o__Saccharomycetales(100);f__Metschnikowiaceae(100);g__Metschnikowia(100);s__Metschnikowia_pulcherrima(100); | Metschnikowia | 13 | Saprotroph | \|Undefined Saprotroph\| | Yeast | NULL | Probable | As nectar-tap_saprotroph (Põlme et al. 2020); Decay sugar-rich substrates (Põlme et al. 2020) | Põlme S, et al. 2020. Fungal Diversity 105:1-16 ((https://doi.org/10.1007/s13225-020-00466-2)) |
| Otu55073 | 1 | k__Fungi(100);p__Ascomycota(100);c__Saccharomycetes(100);o__Saccharomycetales(100);f__Metschnikowiaceae(100);g__Metschnikowia(100);g__Metschnikowia_unclassified(100); | Metschnikowia | 13 | Saprotroph | \|Undefined Saprotroph\| | Yeast | NULL | Probable | As nectar-tap_saprotroph (Põlme et al. 2020); Decay sugar-rich substrates (Põlme et al. 2020) | Põlme S, et al. 2020. Fungal Diversity 105:1-16 ((https://doi.org/10.1007/s13225-020-00466-2)) |
| Otu55074 | 1 | k__Fungi(100);p__Ascomycota(100);c__Saccharomycetes(100);o__Saccharomycetales(100);f__Metschnikowiaceae(100);g__Metschnikowia(100);s__Metschnikowia_pulcherrima(100); | Metschnikowia | 13 | Saprotroph | \|Undefined Saprotroph\| | Yeast | NULL | Probable | As nectar-tap_saprotroph (Põlme et al. 2020); Decay sugar-rich substrates (Põlme et al. 2020) | Põlme S, et al. 2020. Fungal Diversity 105:1-16 ((https://doi.org/10.1007/s13225-020-00466-2)) |
| Otu55075 | 1 | k__Fungi(100);p__Ascomycota(100);c__Saccharomycetes(100);o__Saccharomycetales(100);f__Metschnikowiaceae(100);g__Metschnikowia(100);s__Metschnikowia_pulcherrima(100); | Metschnikowia | 13 | Saprotroph | \|Undefined Saprotroph\| | Yeast | NULL | Probable | As nectar-tap_saprotroph (Põlme et al. 2020); Decay sugar-rich substrates (Põlme et al. 2020) | Põlme S, et al. 2020. Fungal Diversity 105:1-16 ((https://doi.org/10.1007/s13225-020-00466-2)) |
| Otu55076 | 1 | k__Fungi(100);p__Ascomycota(100);c__Saccharomycetes(100);o__Saccharomycetales(100);f__Metschnikowiaceae(100);g__Metschnikowia(100);s__Metschnikowia_pulcherrima(100); | Metschnikowia | 13 | Saprotroph | \|Undefined Saprotroph\| | Yeast | NULL | Probable | As nectar-tap_saprotroph (Põlme et al. 2020); Decay sugar-rich substrates (Põlme et al. 2020) | Põlme S, et al. 2020. Fungal Diversity 105:1-16 ((https://doi.org/10.1007/s13225-020-00466-2)) |
| Otu55077 | 1 | k__Fungi(100);p__Ascomycota(100);c__Saccharomycetes(100);o__Saccharomycetales(100);f__Metschnikowiaceae(100);g__Metschnikowia(100);s__Metschnikowia_pulcherrima(100); | Metschnikowia | 13 | Saprotroph | \|Undefined Saprotroph\| | Yeast | NULL | Probable | As nectar-tap_saprotroph (Põlme et al. 2020); Decay sugar-rich substrates (Põlme et al. 2020) | Põlme S, et al. 2020. Fungal Diversity 105:1-16 ((https://doi.org/10.1007/s13225-020-00466-2)) |
| Otu55078 | 1 | k__Fungi(100);p__Ascomycota(100);c__Saccharomycetes(100);o__Saccharomycetales(100);f__Metschnikowiaceae(100);g__Metschnikowia(100);g__Metschnikowia_unclassified(100); | Metschnikowia | 13 | Saprotroph | \|Undefined Saprotroph\| | Yeast | NULL | Probable | As nectar-tap_saprotroph (Põlme et al. 2020); Decay sugar-rich substrates (Põlme et al. 2020) | Põlme S, et al. 2020. Fungal Diversity 105:1-16 ((https://doi.org/10.1007/s13225-020-00466-2)) |
| Otu55079 | 1 | k__Fungi(100);p__Ascomycota(100);c__Saccharomycetes(100);o__Saccharomycetales(100);f__Metschnikowiaceae(100);g__Metschnikowia(100);g__Metschnikowia_unclassified(100); | Metschnikowia | 13 | Saprotroph | \|Undefined Saprotroph\| | Yeast | NULL | Probable | As nectar-tap_saprotroph (Põlme et al. 2020); Decay sugar-rich substrates (Põlme et al. 2020) | Põlme S, et al. 2020. Fungal Diversity 105:1-16 ((https://doi.org/10.1007/s13225-020-00466-2)) |
| Otu55080 | 1 | k__Fungi(100);p__Ascomycota(100);c__Saccharomycetes(100);o__Saccharomycetales(100);f__Metschnikowiaceae(100);g__Metschnikowia(100);g__Metschnikowia_unclassified(100); | Metschnikowia | 13 | Saprotroph | \|Undefined Saprotroph\| | Yeast | NULL | Probable | As nectar-tap_saprotroph (Põlme et al. 2020); Decay sugar-rich substrates (Põlme et al. 2020) | Põlme S, et al. 2020. Fungal Diversity 105:1-16 ((https://doi.org/10.1007/s13225-020-00466-2)) |
| Otu55082 | 1 | k__Fungi(100);p__Ascomycota(100);c__Saccharomycetes(100);o__Saccharomycetales(100);f__Metschnikowiaceae(100);g__Metschnikowia(100);g__Metschnikowia_unclassified(100); | Metschnikowia | 13 | Saprotroph | \|Undefined Saprotroph\| | Yeast | NULL | Probable | As nectar-tap_saprotroph (Põlme et al. 2020); Decay sugar-rich substrates (Põlme et al. 2020) | Põlme S, et al. 2020. Fungal Diversity 105:1-16 ((https://doi.org/10.1007/s13225-020-00466-2)) |
| Otu55083 | 1 | k__Fungi(100);p__Ascomycota(100);c__Saccharomycetes(100);o__Saccharomycetales(100);f__Metschnikowiaceae(100);g__Metschnikowia(100);s__Metschnikowia_pulcherrima(100); | Metschnikowia | 13 | Saprotroph | \|Undefined Saprotroph\| | Yeast | NULL | Probable | As nectar-tap_saprotroph (Põlme et al. 2020); Decay sugar-rich substrates (Põlme et al. 2020) | Põlme S, et al. 2020. Fungal Diversity 105:1-16 ((https://doi.org/10.1007/s13225-020-00466-2)) |
| Otu55085 | 1 | k__Fungi(100);p__Ascomycota(100);c__Saccharomycetes(100);o__Saccharomycetales(100);f__Metschnikowiaceae(100);g__Metschnikowia(100);s__Metschnikowia_pulcherrima(100); | Metschnikowia | 13 | Saprotroph | \|Undefined Saprotroph\| | Yeast | NULL | Probable | As nectar-tap_saprotroph (Põlme et al. 2020); Decay sugar-rich substrates (Põlme et al. 2020) | Põlme S, et al. 2020. Fungal Diversity 105:1-16 ((https://doi.org/10.1007/s13225-020-00466-2)) |
| Otu55087 | 1 | k__Fungi(100);p__Ascomycota(100);c__Saccharomycetes(100);o__Saccharomycetales(100);f__Metschnikowiaceae(100);g__Metschnikowia(100);s__Metschnikowia_pulcherrima(100); | Metschnikowia | 13 | Saprotroph | \|Undefined Saprotroph\| | Yeast | NULL | Probable | As nectar-tap_saprotroph (Põlme et al. 2020); Decay sugar-rich substrates (Põlme et al. 2020) | Põlme S, et al. 2020. Fungal Diversity 105:1-16 ((https://doi.org/10.1007/s13225-020-00466-2)) |
| Otu55093 | 1 | k__Fungi(100);p__Ascomycota(100);c__Saccharomycetes(100);o__Saccharomycetales(100);f__Metschnikowiaceae(100);g__Metschnikowia(100);s__Metschnikowia_pulcherrima(100); | Metschnikowia | 13 | Saprotroph | \|Undefined Saprotroph\| | Yeast | NULL | Probable | As nectar-tap_saprotroph (Põlme et al. 2020); Decay sugar-rich substrates (Põlme et al. 2020) | Põlme S, et al. 2020. Fungal Diversity 105:1-16 ((https://doi.org/10.1007/s13225-020-00466-2)) |
| Otu55096 | 1 | k__Fungi(100);p__Ascomycota(100);c__Saccharomycetes(100);o__Saccharomycetales(100);f__Saccharomycetales_family_Incertae_sedis(100);g__Candida(100);g__Candida_unclassified(100); | Candida | 13 | Pathotroph-Saprotroph-Symbiotroph | Animal Pathogen-Endophyte-Endosymbiont-Epiphyte-\|Undefined Saprotroph\| | Dimorphic Yeast-Yeast | NULL | Probable | As nectar-tap_saprotroph (Põlme et al. 2020); Decay sugar-rich substrates (Põlme et al. 2020); This asexual genus is polyphyletic and contain species that function very differently depend on what they are. Caution should be taken when interpreting guilds for Candida species. | Manolakaki D, et al. 2010. Virulence. 1: 367-375 ((https://doi.org/10.4161/viru.1.5.12796)); Mekbib SB, et al. 2011. Tropical Plant Pathology 36:233-240 ((https://doi.org/10.1590/S1982-56762011000400004)); Põlme S, et al. 2020. Fungal Diversity 105:1-16 ((https://doi.org/10.1007/s13225-020-00466-2)) |
| Otu55097 | 1 | k__Fungi(100);p__Ascomycota(100);c__Saccharomycetes(100);o__Saccharomycetales(100);f__Metschnikowiaceae(100);g__Metschnikowia(100);s__Metschnikowia_pulcherrima(100); | Metschnikowia | 13 | Saprotroph | \|Undefined Saprotroph\| | Yeast | NULL | Probable | As nectar-tap_saprotroph (Põlme et al. 2020); Decay sugar-rich substrates (Põlme et al. 2020) | Põlme S, et al. 2020. Fungal Diversity 105:1-16 ((https://doi.org/10.1007/s13225-020-00466-2)) |
| Otu55098 | 1 | k__Fungi(100);p__Ascomycota(100);c__Saccharomycetes(100);o__Saccharomycetales(100);f__Metschnikowiaceae(100);g__Metschnikowia(100);s__Metschnikowia_pulcherrima(100); | Metschnikowia | 13 | Saprotroph | \|Undefined Saprotroph\| | Yeast | NULL | Probable | As nectar-tap_saprotroph (Põlme et al. 2020); Decay sugar-rich substrates (Põlme et al. 2020) | Põlme S, et al. 2020. Fungal Diversity 105:1-16 ((https://doi.org/10.1007/s13225-020-00466-2)) |
| Otu55102 | 1 | k__Fungi(100);p__Ascomycota(100);c__Saccharomycetes(100);o__Saccharomycetales(100);f__Metschnikowiaceae(100);g__Metschnikowia(100);s__Metschnikowia_pulcherrima(100); | Metschnikowia | 13 | Saprotroph | \|Undefined Saprotroph\| | Yeast | NULL | Probable | As nectar-tap_saprotroph (Põlme et al. 2020); Decay sugar-rich substrates (Põlme et al. 2020) | Põlme S, et al. 2020. Fungal Diversity 105:1-16 ((https://doi.org/10.1007/s13225-020-00466-2)) |
| Otu55103 | 1 | k__Fungi(100);p__Ascomycota(100);c__Saccharomycetes(100);o__Saccharomycetales(100);f__Saccharomycetales_family_Incertae_sedis(100);g__Candida(100);g__Candida_unclassified(100); | Candida | 13 | Pathotroph-Saprotroph-Symbiotroph | Animal Pathogen-Endophyte-Endosymbiont-Epiphyte-\|Undefined Saprotroph\| | Dimorphic Yeast-Yeast | NULL | Probable | As nectar-tap_saprotroph (Põlme et al. 2020); Decay sugar-rich substrates (Põlme et al. 2020); This asexual genus is polyphyletic and contain species that function very differently depend on what they are. Caution should be taken when interpreting guilds for Candida species. | Manolakaki D, et al. 2010. Virulence. 1: 367-375 ((https://doi.org/10.4161/viru.1.5.12796)); Mekbib SB, et al. 2011. Tropical Plant Pathology 36:233-240 ((https://doi.org/10.1590/S1982-56762011000400004)); Põlme S, et al. 2020. Fungal Diversity 105:1-16 ((https://doi.org/10.1007/s13225-020-00466-2)) |
| Otu55105 | 1 | k__Fungi(100);p__Ascomycota(100);c__Saccharomycetes(100);o__Saccharomycetales(100);f__Saccharomycetales_family_Incertae_sedis(100);g__Candida(100);g__Candida_unclassified(100); | Candida | 13 | Pathotroph-Saprotroph-Symbiotroph | Animal Pathogen-Endophyte-Endosymbiont-Epiphyte-\|Undefined Saprotroph\| | Dimorphic Yeast-Yeast | NULL | Probable | As nectar-tap_saprotroph (Põlme et al. 2020); Decay sugar-rich substrates (Põlme et al. 2020); This asexual genus is polyphyletic and contain species that function very differently depend on what they are. Caution should be taken when interpreting guilds for Candida species. | Manolakaki D, et al. 2010. Virulence. 1: 367-375 ((https://doi.org/10.4161/viru.1.5.12796)); Mekbib SB, et al. 2011. Tropical Plant Pathology 36:233-240 ((https://doi.org/10.1590/S1982-56762011000400004)); Põlme S, et al. 2020. Fungal Diversity 105:1-16 ((https://doi.org/10.1007/s13225-020-00466-2)) |
| Otu55107 | 1 | k__Fungi(100);p__Ascomycota(100);c__Saccharomycetes(100);o__Saccharomycetales(100);f__Metschnikowiaceae(100);g__Metschnikowia(100);s__Metschnikowia_pulcherrima(100); | Metschnikowia | 13 | Saprotroph | \|Undefined Saprotroph\| | Yeast | NULL | Probable | As nectar-tap_saprotroph (Põlme et al. 2020); Decay sugar-rich substrates (Põlme et al. 2020) | Põlme S, et al. 2020. Fungal Diversity 105:1-16 ((https://doi.org/10.1007/s13225-020-00466-2)) |
| Otu55108 | 1 | k__Fungi(100);p__Ascomycota(100);c__Saccharomycetes(100);o__Saccharomycetales(100);f__Metschnikowiaceae(100);g__Metschnikowia(100);s__Metschnikowia_pulcherrima(100); | Metschnikowia | 13 | Saprotroph | \|Undefined Saprotroph\| | Yeast | NULL | Probable | As nectar-tap_saprotroph (Põlme et al. 2020); Decay sugar-rich substrates (Põlme et al. 2020) | Põlme S, et al. 2020. Fungal Diversity 105:1-16 ((https://doi.org/10.1007/s13225-020-00466-2)) |
| Otu55109 | 1 | k__Fungi(100);p__Ascomycota(100);c__Saccharomycetes(100);o__Saccharomycetales(100);f__Metschnikowiaceae(100);g__Metschnikowia(100);s__Metschnikowia_pulcherrima(100); | Metschnikowia | 13 | Saprotroph | \|Undefined Saprotroph\| | Yeast | NULL | Probable | As nectar-tap_saprotroph (Põlme et al. 2020); Decay sugar-rich substrates (Põlme et al. 2020) | Põlme S, et al. 2020. Fungal Diversity 105:1-16 ((https://doi.org/10.1007/s13225-020-00466-2)) |
| Otu55110 | 1 | k__Fungi(100);p__Ascomycota(100);c__Saccharomycetes(100);o__Saccharomycetales(100);f__Metschnikowiaceae(100);g__Metschnikowia(100);s__Metschnikowia_pulcherrima(100); | Metschnikowia | 13 | Saprotroph | \|Undefined Saprotroph\| | Yeast | NULL | Probable | As nectar-tap_saprotroph (Põlme et al. 2020); Decay sugar-rich substrates (Põlme et al. 2020) | Põlme S, et al. 2020. Fungal Diversity 105:1-16 ((https://doi.org/10.1007/s13225-020-00466-2)) |
| Otu55111 | 1 | k__Fungi(100);p__Ascomycota(100);c__Saccharomycetes(100);o__Saccharomycetales(100);f__Metschnikowiaceae(100);g__Metschnikowia(100);g__Metschnikowia_unclassified(100); | Metschnikowia | 13 | Saprotroph | \|Undefined Saprotroph\| | Yeast | NULL | Probable | As nectar-tap_saprotroph (Põlme et al. 2020); Decay sugar-rich substrates (Põlme et al. 2020) | Põlme S, et al. 2020. Fungal Diversity 105:1-16 ((https://doi.org/10.1007/s13225-020-00466-2)) |
| Otu55112 | 1 | k__Fungi(100);p__Ascomycota(100);c__Saccharomycetes(100);o__Saccharomycetales(100);f__Metschnikowiaceae(100);g__Metschnikowia(100);s__Metschnikowia_pulcherrima(100); | Metschnikowia | 13 | Saprotroph | \|Undefined Saprotroph\| | Yeast | NULL | Probable | As nectar-tap_saprotroph (Põlme et al. 2020); Decay sugar-rich substrates (Põlme et al. 2020) | Põlme S, et al. 2020. Fungal Diversity 105:1-16 ((https://doi.org/10.1007/s13225-020-00466-2)) |
| Otu55115 | 1 | k__Fungi(100);p__Ascomycota(100);c__Saccharomycetes(100);o__Saccharomycetales(100);f__Metschnikowiaceae(100);g__Metschnikowia(100);g__Metschnikowia_unclassified(100); | Metschnikowia | 13 | Saprotroph | \|Undefined Saprotroph\| | Yeast | NULL | Probable | As nectar-tap_saprotroph (Põlme et al. 2020); Decay sugar-rich substrates (Põlme et al. 2020) | Põlme S, et al. 2020. Fungal Diversity 105:1-16 ((https://doi.org/10.1007/s13225-020-00466-2)) |
| Otu55116 | 1 | k__Fungi(100);p__Ascomycota(100);c__Saccharomycetes(100);o__Saccharomycetales(100);f__Metschnikowiaceae(100);g__Metschnikowia(100);s__Metschnikowia_pulcherrima(100); | Metschnikowia | 13 | Saprotroph | \|Undefined Saprotroph\| | Yeast | NULL | Probable | As nectar-tap_saprotroph (Põlme et al. 2020); Decay sugar-rich substrates (Põlme et al. 2020) | Põlme S, et al. 2020. Fungal Diversity 105:1-16 ((https://doi.org/10.1007/s13225-020-00466-2)) |
| Otu55117 | 1 | k__Fungi(100);p__Ascomycota(100);c__Saccharomycetes(100);o__Saccharomycetales(100);f__Metschnikowiaceae(100);g__Metschnikowia(100);s__Metschnikowia_pulcherrima(100); | Metschnikowia | 13 | Saprotroph | \|Undefined Saprotroph\| | Yeast | NULL | Probable | As nectar-tap_saprotroph (Põlme et al. 2020); Decay sugar-rich substrates (Põlme et al. 2020) | Põlme S, et al. 2020. Fungal Diversity 105:1-16 ((https://doi.org/10.1007/s13225-020-00466-2)) |
| Otu55118 | 1 | k__Fungi(100);p__Ascomycota(100);c__Saccharomycetes(100);o__Saccharomycetales(100);f__Metschnikowiaceae(100);g__Metschnikowia(100);s__Metschnikowia_pulcherrima(100); | Metschnikowia | 13 | Saprotroph | \|Undefined Saprotroph\| | Yeast | NULL | Probable | As nectar-tap_saprotroph (Põlme et al. 2020); Decay sugar-rich substrates (Põlme et al. 2020) | Põlme S, et al. 2020. Fungal Diversity 105:1-16 ((https://doi.org/10.1007/s13225-020-00466-2)) |
| Otu55119 | 1 | k__Fungi(100);p__Ascomycota(100);c__Saccharomycetes(100);o__Saccharomycetales(100);f__Metschnikowiaceae(100);g__Metschnikowia(100);g__Metschnikowia_unclassified(100); | Metschnikowia | 13 | Saprotroph | \|Undefined Saprotroph\| | Yeast | NULL | Probable | As nectar-tap_saprotroph (Põlme et al. 2020); Decay sugar-rich substrates (Põlme et al. 2020) | Põlme S, et al. 2020. Fungal Diversity 105:1-16 ((https://doi.org/10.1007/s13225-020-00466-2)) |
| Otu55120 | 1 | k__Fungi(100);p__Ascomycota(100);c__Saccharomycetes(100);o__Saccharomycetales(100);f__Saccharomycetales_family_Incertae_sedis(100);g__Candida(100);g__Candida_unclassified(100); | Candida | 13 | Pathotroph-Saprotroph-Symbiotroph | Animal Pathogen-Endophyte-Endosymbiont-Epiphyte-\|Undefined Saprotroph\| | Dimorphic Yeast-Yeast | NULL | Probable | As nectar-tap_saprotroph (Põlme et al. 2020); Decay sugar-rich substrates (Põlme et al. 2020); This asexual genus is polyphyletic and contain species that function very differently depend on what they are. Caution should be taken when interpreting guilds for Candida species. | Manolakaki D, et al. 2010. Virulence. 1: 367-375 ((https://doi.org/10.4161/viru.1.5.12796)); Mekbib SB, et al. 2011. Tropical Plant Pathology 36:233-240 ((https://doi.org/10.1590/S1982-56762011000400004)); Põlme S, et al. 2020. Fungal Diversity 105:1-16 ((https://doi.org/10.1007/s13225-020-00466-2)) |
| Otu55121 | 1 | k__Fungi(100);p__Ascomycota(100);c__Saccharomycetes(100);o__Saccharomycetales(100);f__Metschnikowiaceae(100);g__Metschnikowia(100);g__Metschnikowia_unclassified(100); | Metschnikowia | 13 | Saprotroph | \|Undefined Saprotroph\| | Yeast | NULL | Probable | As nectar-tap_saprotroph (Põlme et al. 2020); Decay sugar-rich substrates (Põlme et al. 2020) | Põlme S, et al. 2020. Fungal Diversity 105:1-16 ((https://doi.org/10.1007/s13225-020-00466-2)) |
| Otu55122 | 1 | k__Fungi(100);p__Ascomycota(100);c__Saccharomycetes(100);o__Saccharomycetales(100);f__Metschnikowiaceae(100);g__Metschnikowia(100);g__Metschnikowia_unclassified(100); | Metschnikowia | 13 | Saprotroph | \|Undefined Saprotroph\| | Yeast | NULL | Probable | As nectar-tap_saprotroph (Põlme et al. 2020); Decay sugar-rich substrates (Põlme et al. 2020) | Põlme S, et al. 2020. Fungal Diversity 105:1-16 ((https://doi.org/10.1007/s13225-020-00466-2)) |
| Otu55123 | 1 | k__Fungi(100);p__Ascomycota(100);c__Saccharomycetes(100);o__Saccharomycetales(100);f__Metschnikowiaceae(100);g__Metschnikowia(100);s__Metschnikowia_pulcherrima(100); | Metschnikowia | 13 | Saprotroph | \|Undefined Saprotroph\| | Yeast | NULL | Probable | As nectar-tap_saprotroph (Põlme et al. 2020); Decay sugar-rich substrates (Põlme et al. 2020) | Põlme S, et al. 2020. Fungal Diversity 105:1-16 ((https://doi.org/10.1007/s13225-020-00466-2)) |
| Otu55124 | 1 | k__Fungi(100);p__Ascomycota(100);c__Saccharomycetes(100);o__Saccharomycetales(100);f__Metschnikowiaceae(100);g__Metschnikowia(100);s__Metschnikowia_pulcherrima(100); | Metschnikowia | 13 | Saprotroph | \|Undefined Saprotroph\| | Yeast | NULL | Probable | As nectar-tap_saprotroph (Põlme et al. 2020); Decay sugar-rich substrates (Põlme et al. 2020) | Põlme S, et al. 2020. Fungal Diversity 105:1-16 ((https://doi.org/10.1007/s13225-020-00466-2)) |
| Otu55125 | 1 | k__Fungi(100);p__Ascomycota(100);c__Saccharomycetes(100);o__Saccharomycetales(100);f__Metschnikowiaceae(100);g__Metschnikowia(100);s__Metschnikowia_pulcherrima(100); | Metschnikowia | 13 | Saprotroph | \|Undefined Saprotroph\| | Yeast | NULL | Probable | As nectar-tap_saprotroph (Põlme et al. 2020); Decay sugar-rich substrates (Põlme et al. 2020) | Põlme S, et al. 2020. Fungal Diversity 105:1-16 ((https://doi.org/10.1007/s13225-020-00466-2)) |
| Otu55126 | 1 | k__Fungi(100);p__Ascomycota(100);c__Saccharomycetes(100);o__Saccharomycetales(100);f__Metschnikowiaceae(100);g__Metschnikowia(100);s__Metschnikowia_pulcherrima(100); | Metschnikowia | 13 | Saprotroph | \|Undefined Saprotroph\| | Yeast | NULL | Probable | As nectar-tap_saprotroph (Põlme et al. 2020); Decay sugar-rich substrates (Põlme et al. 2020) | Põlme S, et al. 2020. Fungal Diversity 105:1-16 ((https://doi.org/10.1007/s13225-020-00466-2)) |
| Otu55127 | 1 | k__Fungi(100);p__Ascomycota(100);c__Saccharomycetes(100);o__Saccharomycetales(100);f__Metschnikowiaceae(100);g__Metschnikowia(100);s__Metschnikowia_pulcherrima(100); | Metschnikowia | 13 | Saprotroph | \|Undefined Saprotroph\| | Yeast | NULL | Probable | As nectar-tap_saprotroph (Põlme et al. 2020); Decay sugar-rich substrates (Põlme et al. 2020) | Põlme S, et al. 2020. Fungal Diversity 105:1-16 ((https://doi.org/10.1007/s13225-020-00466-2)) |
| Otu55128 | 1 | k__Fungi(100);p__Ascomycota(100);c__Saccharomycetes(100);o__Saccharomycetales(100);f__Metschnikowiaceae(100);g__Metschnikowia(100);s__Metschnikowia_pulcherrima(100); | Metschnikowia | 13 | Saprotroph | \|Undefined Saprotroph\| | Yeast | NULL | Probable | As nectar-tap_saprotroph (Põlme et al. 2020); Decay sugar-rich substrates (Põlme et al. 2020) | Põlme S, et al. 2020. Fungal Diversity 105:1-16 ((https://doi.org/10.1007/s13225-020-00466-2)) |
| Otu55129 | 1 | k__Fungi(100);p__Ascomycota(100);c__Saccharomycetes(100);o__Saccharomycetales(100);f__Metschnikowiaceae(100);g__Metschnikowia(100);s__Metschnikowia_pulcherrima(100); | Metschnikowia | 13 | Saprotroph | \|Undefined Saprotroph\| | Yeast | NULL | Probable | As nectar-tap_saprotroph (Põlme et al. 2020); Decay sugar-rich substrates (Põlme et al. 2020) | Põlme S, et al. 2020. Fungal Diversity 105:1-16 ((https://doi.org/10.1007/s13225-020-00466-2)) |
| Otu55130 | 1 | k__Fungi(100);p__Ascomycota(100);c__Saccharomycetes(100);o__Saccharomycetales(100);f__Metschnikowiaceae(100);g__Metschnikowia(100);s__Metschnikowia_pulcherrima(100); | Metschnikowia | 13 | Saprotroph | \|Undefined Saprotroph\| | Yeast | NULL | Probable | As nectar-tap_saprotroph (Põlme et al. 2020); Decay sugar-rich substrates (Põlme et al. 2020) | Põlme S, et al. 2020. Fungal Diversity 105:1-16 ((https://doi.org/10.1007/s13225-020-00466-2)) |
| Otu55131 | 1 | k__Fungi(100);p__Ascomycota(100);c__Saccharomycetes(100);o__Saccharomycetales(100);f__Metschnikowiaceae(100);g__Metschnikowia(100);s__Metschnikowia_pulcherrima(100); | Metschnikowia | 13 | Saprotroph | \|Undefined Saprotroph\| | Yeast | NULL | Probable | As nectar-tap_saprotroph (Põlme et al. 2020); Decay sugar-rich substrates (Põlme et al. 2020) | Põlme S, et al. 2020. Fungal Diversity 105:1-16 ((https://doi.org/10.1007/s13225-020-00466-2)) |
| Otu55132 | 1 | k__Fungi(100);p__Ascomycota(100);c__Saccharomycetes(100);o__Saccharomycetales(100);f__Metschnikowiaceae(100);g__Metschnikowia(100);s__Metschnikowia_pulcherrima(100); | Metschnikowia | 13 | Saprotroph | \|Undefined Saprotroph\| | Yeast | NULL | Probable | As nectar-tap_saprotroph (Põlme et al. 2020); Decay sugar-rich substrates (Põlme et al. 2020) | Põlme S, et al. 2020. Fungal Diversity 105:1-16 ((https://doi.org/10.1007/s13225-020-00466-2)) |
| Otu55133 | 1 | k__Fungi(100);p__Ascomycota(100);c__Saccharomycetes(100);o__Saccharomycetales(100);f__Metschnikowiaceae(100);g__Metschnikowia(100);s__Metschnikowia_pulcherrima(100); | Metschnikowia | 13 | Saprotroph | \|Undefined Saprotroph\| | Yeast | NULL | Probable | As nectar-tap_saprotroph (Põlme et al. 2020); Decay sugar-rich substrates (Põlme et al. 2020) | Põlme S, et al. 2020. Fungal Diversity 105:1-16 ((https://doi.org/10.1007/s13225-020-00466-2)) |
| Otu55134 | 1 | k__Fungi(100);p__Ascomycota(100);c__Saccharomycetes(100);o__Saccharomycetales(100);f__Metschnikowiaceae(100);g__Metschnikowia(100);s__Metschnikowia_pulcherrima(100); | Metschnikowia | 13 | Saprotroph | \|Undefined Saprotroph\| | Yeast | NULL | Probable | As nectar-tap_saprotroph (Põlme et al. 2020); Decay sugar-rich substrates (Põlme et al. 2020) | Põlme S, et al. 2020. Fungal Diversity 105:1-16 ((https://doi.org/10.1007/s13225-020-00466-2)) |
| Otu55135 | 1 | k__Fungi(100);p__Ascomycota(100);c__Saccharomycetes(100);o__Saccharomycetales(100);f__Metschnikowiaceae(100);g__Metschnikowia(100);s__Metschnikowia_pulcherrima(100); | Metschnikowia | 13 | Saprotroph | \|Undefined Saprotroph\| | Yeast | NULL | Probable | As nectar-tap_saprotroph (Põlme et al. 2020); Decay sugar-rich substrates (Põlme et al. 2020) | Põlme S, et al. 2020. Fungal Diversity 105:1-16 ((https://doi.org/10.1007/s13225-020-00466-2)) |
| Otu55136 | 1 | k__Fungi(100);p__Ascomycota(100);c__Saccharomycetes(100);o__Saccharomycetales(100);f__Metschnikowiaceae(100);g__Metschnikowia(100);s__Metschnikowia_pulcherrima(100); | Metschnikowia | 13 | Saprotroph | \|Undefined Saprotroph\| | Yeast | NULL | Probable | As nectar-tap_saprotroph (Põlme et al. 2020); Decay sugar-rich substrates (Põlme et al. 2020) | Põlme S, et al. 2020. Fungal Diversity 105:1-16 ((https://doi.org/10.1007/s13225-020-00466-2)) |
| Otu55137 | 1 | k__Fungi(100);p__Ascomycota(100);c__Saccharomycetes(100);o__Saccharomycetales(100);f__Metschnikowiaceae(100);g__Metschnikowia(100);s__Metschnikowia_pulcherrima(100); | Metschnikowia | 13 | Saprotroph | \|Undefined Saprotroph\| | Yeast | NULL | Probable | As nectar-tap_saprotroph (Põlme et al. 2020); Decay sugar-rich substrates (Põlme et al. 2020) | Põlme S, et al. 2020. Fungal Diversity 105:1-16 ((https://doi.org/10.1007/s13225-020-00466-2)) |
| Otu55139 | 1 | k__Fungi(100);p__Ascomycota(100);c__Saccharomycetes(100);o__Saccharomycetales(100);f__Metschnikowiaceae(100);g__Metschnikowia(100);s__Metschnikowia_pulcherrima(100); | Metschnikowia | 13 | Saprotroph | \|Undefined Saprotroph\| | Yeast | NULL | Probable | As nectar-tap_saprotroph (Põlme et al. 2020); Decay sugar-rich substrates (Põlme et al. 2020) | Põlme S, et al. 2020. Fungal Diversity 105:1-16 ((https://doi.org/10.1007/s13225-020-00466-2)) |
| Otu55140 | 1 | k__Fungi(100);p__Ascomycota(100);c__Saccharomycetes(100);o__Saccharomycetales(100);f__Metschnikowiaceae(100);g__Metschnikowia(100);g__Metschnikowia_unclassified(100); | Metschnikowia | 13 | Saprotroph | \|Undefined Saprotroph\| | Yeast | NULL | Probable | As nectar-tap_saprotroph (Põlme et al. 2020); Decay sugar-rich substrates (Põlme et al. 2020) | Põlme S, et al. 2020. Fungal Diversity 105:1-16 ((https://doi.org/10.1007/s13225-020-00466-2)) |
| Otu55141 | 1 | k__Fungi(100);p__Ascomycota(100);c__Saccharomycetes(100);o__Saccharomycetales(100);f__Metschnikowiaceae(100);g__Metschnikowia(100);s__Metschnikowia_pulcherrima(100); | Metschnikowia | 13 | Saprotroph | \|Undefined Saprotroph\| | Yeast | NULL | Probable | As nectar-tap_saprotroph (Põlme et al. 2020); Decay sugar-rich substrates (Põlme et al. 2020) | Põlme S, et al. 2020. Fungal Diversity 105:1-16 ((https://doi.org/10.1007/s13225-020-00466-2)) |
| Otu55142 | 1 | k__Fungi(100);p__Ascomycota(100);c__Saccharomycetes(100);o__Saccharomycetales(100);f__Metschnikowiaceae(100);g__Metschnikowia(100);s__Metschnikowia_pulcherrima(100); | Metschnikowia | 13 | Saprotroph | \|Undefined Saprotroph\| | Yeast | NULL | Probable | As nectar-tap_saprotroph (Põlme et al. 2020); Decay sugar-rich substrates (Põlme et al. 2020) | Põlme S, et al. 2020. Fungal Diversity 105:1-16 ((https://doi.org/10.1007/s13225-020-00466-2)) |
| Otu55143 | 1 | k__Fungi(100);p__Ascomycota(100);c__Saccharomycetes(100);o__Saccharomycetales(100);f__Metschnikowiaceae(100);g__Metschnikowia(100);s__Metschnikowia_pulcherrima(100); | Metschnikowia | 13 | Saprotroph | \|Undefined Saprotroph\| | Yeast | NULL | Probable | As nectar-tap_saprotroph (Põlme et al. 2020); Decay sugar-rich substrates (Põlme et al. 2020) | Põlme S, et al. 2020. Fungal Diversity 105:1-16 ((https://doi.org/10.1007/s13225-020-00466-2)) |
| Otu55144 | 1 | k__Fungi(100);p__Ascomycota(100);c__Saccharomycetes(100);o__Saccharomycetales(100);f__Metschnikowiaceae(100);g__Metschnikowia(100);s__Metschnikowia_pulcherrima(100); | Metschnikowia | 13 | Saprotroph | \|Undefined Saprotroph\| | Yeast | NULL | Probable | As nectar-tap_saprotroph (Põlme et al. 2020); Decay sugar-rich substrates (Põlme et al. 2020) | Põlme S, et al. 2020. Fungal Diversity 105:1-16 ((https://doi.org/10.1007/s13225-020-00466-2)) |
| Otu55145 | 1 | k__Fungi(100);p__Ascomycota(100);c__Saccharomycetes(100);o__Saccharomycetales(100);f__Metschnikowiaceae(100);g__Metschnikowia(100);s__Metschnikowia_pulcherrima(100); | Metschnikowia | 13 | Saprotroph | \|Undefined Saprotroph\| | Yeast | NULL | Probable | As nectar-tap_saprotroph (Põlme et al. 2020); Decay sugar-rich substrates (Põlme et al. 2020) | Põlme S, et al. 2020. Fungal Diversity 105:1-16 ((https://doi.org/10.1007/s13225-020-00466-2)) |
| Otu55146 | 1 | k__Fungi(100);p__Ascomycota(100);c__Saccharomycetes(100);o__Saccharomycetales(100);f__Metschnikowiaceae(100);g__Metschnikowia(100);s__Metschnikowia_pulcherrima(100); | Metschnikowia | 13 | Saprotroph | \|Undefined Saprotroph\| | Yeast | NULL | Probable | As nectar-tap_saprotroph (Põlme et al. 2020); Decay sugar-rich substrates (Põlme et al. 2020) | Põlme S, et al. 2020. Fungal Diversity 105:1-16 ((https://doi.org/10.1007/s13225-020-00466-2)) |
| Otu55147 | 1 | k__Fungi(100);p__Ascomycota(100);c__Saccharomycetes(100);o__Saccharomycetales(100);f__Metschnikowiaceae(100);g__Metschnikowia(100);s__Metschnikowia_pulcherrima(100); | Metschnikowia | 13 | Saprotroph | \|Undefined Saprotroph\| | Yeast | NULL | Probable | As nectar-tap_saprotroph (Põlme et al. 2020); Decay sugar-rich substrates (Põlme et al. 2020) | Põlme S, et al. 2020. Fungal Diversity 105:1-16 ((https://doi.org/10.1007/s13225-020-00466-2)) |
| Otu55151 | 1 | k__Fungi(100);p__Ascomycota(100);c__Saccharomycetes(100);o__Saccharomycetales(100);f__Metschnikowiaceae(100);g__Metschnikowia(100);s__Metschnikowia_pulcherrima(100); | Metschnikowia | 13 | Saprotroph | \|Undefined Saprotroph\| | Yeast | NULL | Probable | As nectar-tap_saprotroph (Põlme et al. 2020); Decay sugar-rich substrates (Põlme et al. 2020) | Põlme S, et al. 2020. Fungal Diversity 105:1-16 ((https://doi.org/10.1007/s13225-020-00466-2)) |
| Otu55152 | 1 | k__Fungi(100);p__Ascomycota(100);c__Saccharomycetes(100);o__Saccharomycetales(100);f__Saccharomycetales_family_Incertae_sedis(100);g__Candida(100);s__Candida_cretensis(100); | Candida | 13 | Pathotroph-Saprotroph-Symbiotroph | Animal Pathogen-Endophyte-Endosymbiont-Epiphyte-\|Undefined Saprotroph\| | Dimorphic Yeast-Yeast | NULL | Probable | As nectar-tap_saprotroph (Põlme et al. 2020); Decay sugar-rich substrates (Põlme et al. 2020); This asexual genus is polyphyletic and contain species that function very differently depend on what they are. Caution should be taken when interpreting guilds for Candida species. | Manolakaki D, et al. 2010. Virulence. 1: 367-375 ((https://doi.org/10.4161/viru.1.5.12796)); Mekbib SB, et al. 2011. Tropical Plant Pathology 36:233-240 ((https://doi.org/10.1590/S1982-56762011000400004)); Põlme S, et al. 2020. Fungal Diversity 105:1-16 ((https://doi.org/10.1007/s13225-020-00466-2)) |
| Otu55154 | 1 | k__Fungi(100);p__Ascomycota(100);c__Saccharomycetes(100);o__Saccharomycetales(100);f__Metschnikowiaceae(100);g__Metschnikowia(100);s__Metschnikowia_pulcherrima(100); | Metschnikowia | 13 | Saprotroph | \|Undefined Saprotroph\| | Yeast | NULL | Probable | As nectar-tap_saprotroph (Põlme et al. 2020); Decay sugar-rich substrates (Põlme et al. 2020) | Põlme S, et al. 2020. Fungal Diversity 105:1-16 ((https://doi.org/10.1007/s13225-020-00466-2)) |
| Otu55155 | 1 | k__Fungi(100);p__Ascomycota(100);c__Saccharomycetes(100);o__Saccharomycetales(100);f__Metschnikowiaceae(100);g__Metschnikowia(100);g__Metschnikowia_unclassified(100); | Metschnikowia | 13 | Saprotroph | \|Undefined Saprotroph\| | Yeast | NULL | Probable | As nectar-tap_saprotroph (Põlme et al. 2020); Decay sugar-rich substrates (Põlme et al. 2020) | Põlme S, et al. 2020. Fungal Diversity 105:1-16 ((https://doi.org/10.1007/s13225-020-00466-2)) |
| Otu55156 | 1 | k__Fungi(100);p__Ascomycota(100);c__Saccharomycetes(100);o__Saccharomycetales(100);f__Metschnikowiaceae(100);g__Metschnikowia(100);s__Metschnikowia_pulcherrima(100); | Metschnikowia | 13 | Saprotroph | \|Undefined Saprotroph\| | Yeast | NULL | Probable | As nectar-tap_saprotroph (Põlme et al. 2020); Decay sugar-rich substrates (Põlme et al. 2020) | Põlme S, et al. 2020. Fungal Diversity 105:1-16 ((https://doi.org/10.1007/s13225-020-00466-2)) |
| Otu55157 | 1 | k__Fungi(100);p__Ascomycota(100);c__Saccharomycetes(100);o__Saccharomycetales(100);f__Metschnikowiaceae(100);g__Metschnikowia(100);s__Metschnikowia_pulcherrima(100); | Metschnikowia | 13 | Saprotroph | \|Undefined Saprotroph\| | Yeast | NULL | Probable | As nectar-tap_saprotroph (Põlme et al. 2020); Decay sugar-rich substrates (Põlme et al. 2020) | Põlme S, et al. 2020. Fungal Diversity 105:1-16 ((https://doi.org/10.1007/s13225-020-00466-2)) |
| Otu55158 | 1 | k__Fungi(100);p__Ascomycota(100);c__Saccharomycetes(100);o__Saccharomycetales(100);f__Saccharomycetales_family_Incertae_sedis(100);g__Candida(100);g__Candida_unclassified(100); | Candida | 13 | Pathotroph-Saprotroph-Symbiotroph | Animal Pathogen-Endophyte-Endosymbiont-Epiphyte-\|Undefined Saprotroph\| | Dimorphic Yeast-Yeast | NULL | Probable | As nectar-tap_saprotroph (Põlme et al. 2020); Decay sugar-rich substrates (Põlme et al. 2020); This asexual genus is polyphyletic and contain species that function very differently depend on what they are. Caution should be taken when interpreting guilds for Candida species. | Manolakaki D, et al. 2010. Virulence. 1: 367-375 ((https://doi.org/10.4161/viru.1.5.12796)); Mekbib SB, et al. 2011. Tropical Plant Pathology 36:233-240 ((https://doi.org/10.1590/S1982-56762011000400004)); Põlme S, et al. 2020. Fungal Diversity 105:1-16 ((https://doi.org/10.1007/s13225-020-00466-2)) |
| Otu55161 | 1 | k__Fungi(100);p__Ascomycota(100);c__Saccharomycetes(100);o__Saccharomycetales(100);f__Saccharomycetales_family_Incertae_sedis(100);g__Candida(100);g__Candida_unclassified(100); | Candida | 13 | Pathotroph-Saprotroph-Symbiotroph | Animal Pathogen-Endophyte-Endosymbiont-Epiphyte-\|Undefined Saprotroph\| | Dimorphic Yeast-Yeast | NULL | Probable | As nectar-tap_saprotroph (Põlme et al. 2020); Decay sugar-rich substrates (Põlme et al. 2020); This asexual genus is polyphyletic and contain species that function very differently depend on what they are. Caution should be taken when interpreting guilds for Candida species. | Manolakaki D, et al. 2010. Virulence. 1: 367-375 ((https://doi.org/10.4161/viru.1.5.12796)); Mekbib SB, et al. 2011. Tropical Plant Pathology 36:233-240 ((https://doi.org/10.1590/S1982-56762011000400004)); Põlme S, et al. 2020. Fungal Diversity 105:1-16 ((https://doi.org/10.1007/s13225-020-00466-2)) |
| Otu55162 | 1 | k__Fungi(100);p__Ascomycota(100);c__Saccharomycetes(100);o__Saccharomycetales(100);f__Saccharomycetales_family_Incertae_sedis(100);g__Candida(100);g__Candida_unclassified(100); | Candida | 13 | Pathotroph-Saprotroph-Symbiotroph | Animal Pathogen-Endophyte-Endosymbiont-Epiphyte-\|Undefined Saprotroph\| | Dimorphic Yeast-Yeast | NULL | Probable | As nectar-tap_saprotroph (Põlme et al. 2020); Decay sugar-rich substrates (Põlme et al. 2020); This asexual genus is polyphyletic and contain species that function very differently depend on what they are. Caution should be taken when interpreting guilds for Candida species. | Manolakaki D, et al. 2010. Virulence. 1: 367-375 ((https://doi.org/10.4161/viru.1.5.12796)); Mekbib SB, et al. 2011. Tropical Plant Pathology 36:233-240 ((https://doi.org/10.1590/S1982-56762011000400004)); Põlme S, et al. 2020. Fungal Diversity 105:1-16 ((https://doi.org/10.1007/s13225-020-00466-2)) |
| Otu55163 | 1 | k__Fungi(100);p__Ascomycota(100);c__Saccharomycetes(100);o__Saccharomycetales(100);f__Metschnikowiaceae(100);g__Metschnikowia(100);g__Metschnikowia_unclassified(100); | Metschnikowia | 13 | Saprotroph | \|Undefined Saprotroph\| | Yeast | NULL | Probable | As nectar-tap_saprotroph (Põlme et al. 2020); Decay sugar-rich substrates (Põlme et al. 2020) | Põlme S, et al. 2020. Fungal Diversity 105:1-16 ((https://doi.org/10.1007/s13225-020-00466-2)) |
| Otu55165 | 1 | k__Fungi(100);p__Ascomycota(100);c__Saccharomycetes(100);o__Saccharomycetales(100);f__Saccharomycetales_family_Incertae_sedis(100);g__Candida(100);g__Candida_unclassified(100); | Candida | 13 | Pathotroph-Saprotroph-Symbiotroph | Animal Pathogen-Endophyte-Endosymbiont-Epiphyte-\|Undefined Saprotroph\| | Dimorphic Yeast-Yeast | NULL | Probable | As nectar-tap_saprotroph (Põlme et al. 2020); Decay sugar-rich substrates (Põlme et al. 2020); This asexual genus is polyphyletic and contain species that function very differently depend on what they are. Caution should be taken when interpreting guilds for Candida species. | Manolakaki D, et al. 2010. Virulence. 1: 367-375 ((https://doi.org/10.4161/viru.1.5.12796)); Mekbib SB, et al. 2011. Tropical Plant Pathology 36:233-240 ((https://doi.org/10.1590/S1982-56762011000400004)); Põlme S, et al. 2020. Fungal Diversity 105:1-16 ((https://doi.org/10.1007/s13225-020-00466-2)) |
| Otu55168 | 1 | k__Fungi(100);p__Ascomycota(100);c__Saccharomycetes(100);o__Saccharomycetales(100);f__Metschnikowiaceae(100);g__Metschnikowia(100);g__Metschnikowia_unclassified(100); | Metschnikowia | 13 | Saprotroph | \|Undefined Saprotroph\| | Yeast | NULL | Probable | As nectar-tap_saprotroph (Põlme et al. 2020); Decay sugar-rich substrates (Põlme et al. 2020) | Põlme S, et al. 2020. Fungal Diversity 105:1-16 ((https://doi.org/10.1007/s13225-020-00466-2)) |
| Otu55170 | 1 | k__Fungi(100);p__Ascomycota(100);c__Saccharomycetes(100);o__Saccharomycetales(100);f__Metschnikowiaceae(100);g__Metschnikowia(100);g__Metschnikowia_unclassified(100); | Metschnikowia | 13 | Saprotroph | \|Undefined Saprotroph\| | Yeast | NULL | Probable | As nectar-tap_saprotroph (Põlme et al. 2020); Decay sugar-rich substrates (Põlme et al. 2020) | Põlme S, et al. 2020. Fungal Diversity 105:1-16 ((https://doi.org/10.1007/s13225-020-00466-2)) |
| Otu55171 | 1 | k__Fungi(100);p__Ascomycota(100);c__Saccharomycetes(100);o__Saccharomycetales(100);f__Metschnikowiaceae(100);g__Metschnikowia(100);s__Metschnikowia_pulcherrima(100); | Metschnikowia | 13 | Saprotroph | \|Undefined Saprotroph\| | Yeast | NULL | Probable | As nectar-tap_saprotroph (Põlme et al. 2020); Decay sugar-rich substrates (Põlme et al. 2020) | Põlme S, et al. 2020. Fungal Diversity 105:1-16 ((https://doi.org/10.1007/s13225-020-00466-2)) |
| Otu55172 | 1 | k__Fungi(100);p__Ascomycota(100);c__Saccharomycetes(100);o__Saccharomycetales(100);f__Metschnikowiaceae(100);g__Metschnikowia(100);s__Metschnikowia_pulcherrima(100); | Metschnikowia | 13 | Saprotroph | \|Undefined Saprotroph\| | Yeast | NULL | Probable | As nectar-tap_saprotroph (Põlme et al. 2020); Decay sugar-rich substrates (Põlme et al. 2020) | Põlme S, et al. 2020. Fungal Diversity 105:1-16 ((https://doi.org/10.1007/s13225-020-00466-2)) |
| Otu55173 | 1 | k__Fungi(100);p__Ascomycota(100);c__Saccharomycetes(100);o__Saccharomycetales(100);f__Metschnikowiaceae(100);g__Metschnikowia(100);s__Metschnikowia_pulcherrima(100); | Metschnikowia | 13 | Saprotroph | \|Undefined Saprotroph\| | Yeast | NULL | Probable | As nectar-tap_saprotroph (Põlme et al. 2020); Decay sugar-rich substrates (Põlme et al. 2020) | Põlme S, et al. 2020. Fungal Diversity 105:1-16 ((https://doi.org/10.1007/s13225-020-00466-2)) |
| Otu55174 | 1 | k__Fungi(100);p__Ascomycota(100);c__Saccharomycetes(100);o__Saccharomycetales(100);f__Metschnikowiaceae(100);g__Metschnikowia(100);g__Metschnikowia_unclassified(100); | Metschnikowia | 13 | Saprotroph | \|Undefined Saprotroph\| | Yeast | NULL | Probable | As nectar-tap_saprotroph (Põlme et al. 2020); Decay sugar-rich substrates (Põlme et al. 2020) | Põlme S, et al. 2020. Fungal Diversity 105:1-16 ((https://doi.org/10.1007/s13225-020-00466-2)) |
| Otu55176 | 1 | k__Fungi(100);p__Ascomycota(100);c__Saccharomycetes(100);o__Saccharomycetales(100);f__Saccharomycetales_family_Incertae_sedis(100);g__Candida(100);g__Candida_unclassified(100); | Candida | 13 | Pathotroph-Saprotroph-Symbiotroph | Animal Pathogen-Endophyte-Endosymbiont-Epiphyte-\|Undefined Saprotroph\| | Dimorphic Yeast-Yeast | NULL | Probable | As nectar-tap_saprotroph (Põlme et al. 2020); Decay sugar-rich substrates (Põlme et al. 2020); This asexual genus is polyphyletic and contain species that function very differently depend on what they are. Caution should be taken when interpreting guilds for Candida species. | Manolakaki D, et al. 2010. Virulence. 1: 367-375 ((https://doi.org/10.4161/viru.1.5.12796)); Mekbib SB, et al. 2011. Tropical Plant Pathology 36:233-240 ((https://doi.org/10.1590/S1982-56762011000400004)); Põlme S, et al. 2020. Fungal Diversity 105:1-16 ((https://doi.org/10.1007/s13225-020-00466-2)) |
| Otu55178 | 1 | k__Fungi(100);p__Ascomycota(100);c__Saccharomycetes(100);o__Saccharomycetales(100);f__Saccharomycetales_family_Incertae_sedis(100);g__Candida(100);g__Candida_unclassified(100); | Candida | 13 | Pathotroph-Saprotroph-Symbiotroph | Animal Pathogen-Endophyte-Endosymbiont-Epiphyte-\|Undefined Saprotroph\| | Dimorphic Yeast-Yeast | NULL | Probable | As nectar-tap_saprotroph (Põlme et al. 2020); Decay sugar-rich substrates (Põlme et al. 2020); This asexual genus is polyphyletic and contain species that function very differently depend on what they are. Caution should be taken when interpreting guilds for Candida species. | Manolakaki D, et al. 2010. Virulence. 1: 367-375 ((https://doi.org/10.4161/viru.1.5.12796)); Mekbib SB, et al. 2011. Tropical Plant Pathology 36:233-240 ((https://doi.org/10.1590/S1982-56762011000400004)); Põlme S, et al. 2020. Fungal Diversity 105:1-16 ((https://doi.org/10.1007/s13225-020-00466-2)) |
| Otu55181 | 1 | k__Fungi(100);p__Ascomycota(100);c__Saccharomycetes(100);o__Saccharomycetales(100);f__Metschnikowiaceae(100);g__Metschnikowia(100);s__Metschnikowia_pulcherrima(100); | Metschnikowia | 13 | Saprotroph | \|Undefined Saprotroph\| | Yeast | NULL | Probable | As nectar-tap_saprotroph (Põlme et al. 2020); Decay sugar-rich substrates (Põlme et al. 2020) | Põlme S, et al. 2020. Fungal Diversity 105:1-16 ((https://doi.org/10.1007/s13225-020-00466-2)) |
| Otu55182 | 1 | k__Fungi(100);p__Ascomycota(100);c__Saccharomycetes(100);o__Saccharomycetales(100);f__Metschnikowiaceae(100);g__Metschnikowia(100);s__Metschnikowia_pulcherrima(100); | Metschnikowia | 13 | Saprotroph | \|Undefined Saprotroph\| | Yeast | NULL | Probable | As nectar-tap_saprotroph (Põlme et al. 2020); Decay sugar-rich substrates (Põlme et al. 2020) | Põlme S, et al. 2020. Fungal Diversity 105:1-16 ((https://doi.org/10.1007/s13225-020-00466-2)) |
| Otu55183 | 1 | k__Fungi(100);p__Ascomycota(100);c__Saccharomycetes(100);o__Saccharomycetales(100);f__Metschnikowiaceae(100);g__Metschnikowia(100);s__Metschnikowia_pulcherrima(100); | Metschnikowia | 13 | Saprotroph | \|Undefined Saprotroph\| | Yeast | NULL | Probable | As nectar-tap_saprotroph (Põlme et al. 2020); Decay sugar-rich substrates (Põlme et al. 2020) | Põlme S, et al. 2020. Fungal Diversity 105:1-16 ((https://doi.org/10.1007/s13225-020-00466-2)) |
| Otu55184 | 1 | k__Fungi(100);p__Ascomycota(100);c__Saccharomycetes(100);o__Saccharomycetales(100);f__Metschnikowiaceae(100);g__Metschnikowia(100);g__Metschnikowia_unclassified(100); | Metschnikowia | 13 | Saprotroph | \|Undefined Saprotroph\| | Yeast | NULL | Probable | As nectar-tap_saprotroph (Põlme et al. 2020); Decay sugar-rich substrates (Põlme et al. 2020) | Põlme S, et al. 2020. Fungal Diversity 105:1-16 ((https://doi.org/10.1007/s13225-020-00466-2)) |
| Otu55185 | 1 | k__Fungi(100);p__Ascomycota(100);c__Saccharomycetes(100);o__Saccharomycetales(100);f__Metschnikowiaceae(100);g__Metschnikowia(100);s__Metschnikowia_pulcherrima(100); | Metschnikowia | 13 | Saprotroph | \|Undefined Saprotroph\| | Yeast | NULL | Probable | As nectar-tap_saprotroph (Põlme et al. 2020); Decay sugar-rich substrates (Põlme et al. 2020) | Põlme S, et al. 2020. Fungal Diversity 105:1-16 ((https://doi.org/10.1007/s13225-020-00466-2)) |
| Otu55186 | 1 | k__Fungi(100);p__Ascomycota(100);c__Saccharomycetes(100);o__Saccharomycetales(100);f__Metschnikowiaceae(100);g__Metschnikowia(100);s__Metschnikowia_pulcherrima(100); | Metschnikowia | 13 | Saprotroph | \|Undefined Saprotroph\| | Yeast | NULL | Probable | As nectar-tap_saprotroph (Põlme et al. 2020); Decay sugar-rich substrates (Põlme et al. 2020) | Põlme S, et al. 2020. Fungal Diversity 105:1-16 ((https://doi.org/10.1007/s13225-020-00466-2)) |
| Otu55187 | 1 | k__Fungi(100);p__Ascomycota(100);c__Saccharomycetes(100);o__Saccharomycetales(100);f__Metschnikowiaceae(100);g__Metschnikowia(100);s__Metschnikowia_pulcherrima(100); | Metschnikowia | 13 | Saprotroph | \|Undefined Saprotroph\| | Yeast | NULL | Probable | As nectar-tap_saprotroph (Põlme et al. 2020); Decay sugar-rich substrates (Põlme et al. 2020) | Põlme S, et al. 2020. Fungal Diversity 105:1-16 ((https://doi.org/10.1007/s13225-020-00466-2)) |
| Otu55188 | 1 | k__Fungi(100);p__Ascomycota(100);c__Saccharomycetes(100);o__Saccharomycetales(100);f__Metschnikowiaceae(100);g__Metschnikowia(100);s__Metschnikowia_pulcherrima(100); | Metschnikowia | 13 | Saprotroph | \|Undefined Saprotroph\| | Yeast | NULL | Probable | As nectar-tap_saprotroph (Põlme et al. 2020); Decay sugar-rich substrates (Põlme et al. 2020) | Põlme S, et al. 2020. Fungal Diversity 105:1-16 ((https://doi.org/10.1007/s13225-020-00466-2)) |
| Otu55189 | 1 | k__Fungi(100);p__Ascomycota(100);c__Saccharomycetes(100);o__Saccharomycetales(100);f__Metschnikowiaceae(100);g__Metschnikowia(100);s__Metschnikowia_pulcherrima(100); | Metschnikowia | 13 | Saprotroph | \|Undefined Saprotroph\| | Yeast | NULL | Probable | As nectar-tap_saprotroph (Põlme et al. 2020); Decay sugar-rich substrates (Põlme et al. 2020) | Põlme S, et al. 2020. Fungal Diversity 105:1-16 ((https://doi.org/10.1007/s13225-020-00466-2)) |
| Otu55190 | 1 | k__Fungi(100);p__Ascomycota(100);c__Saccharomycetes(100);o__Saccharomycetales(100);f__Metschnikowiaceae(100);g__Metschnikowia(100);g__Metschnikowia_unclassified(100); | Metschnikowia | 13 | Saprotroph | \|Undefined Saprotroph\| | Yeast | NULL | Probable | As nectar-tap_saprotroph (Põlme et al. 2020); Decay sugar-rich substrates (Põlme et al. 2020) | Põlme S, et al. 2020. Fungal Diversity 105:1-16 ((https://doi.org/10.1007/s13225-020-00466-2)) |
| Otu55191 | 1 | k__Fungi(100);p__Ascomycota(100);c__Saccharomycetes(100);o__Saccharomycetales(100);f__Metschnikowiaceae(100);g__Metschnikowia(100);s__Metschnikowia_pulcherrima(100); | Metschnikowia | 13 | Saprotroph | \|Undefined Saprotroph\| | Yeast | NULL | Probable | As nectar-tap_saprotroph (Põlme et al. 2020); Decay sugar-rich substrates (Põlme et al. 2020) | Põlme S, et al. 2020. Fungal Diversity 105:1-16 ((https://doi.org/10.1007/s13225-020-00466-2)) |
| Otu55192 | 1 | k__Fungi(100);p__Ascomycota(100);c__Saccharomycetes(100);o__Saccharomycetales(100);f__Metschnikowiaceae(100);g__Metschnikowia(100);s__Metschnikowia_pulcherrima(100); | Metschnikowia | 13 | Saprotroph | \|Undefined Saprotroph\| | Yeast | NULL | Probable | As nectar-tap_saprotroph (Põlme et al. 2020); Decay sugar-rich substrates (Põlme et al. 2020) | Põlme S, et al. 2020. Fungal Diversity 105:1-16 ((https://doi.org/10.1007/s13225-020-00466-2)) |
| Otu55193 | 1 | k__Fungi(100);p__Ascomycota(100);c__Saccharomycetes(100);o__Saccharomycetales(100);f__Metschnikowiaceae(100);g__Metschnikowia(100);s__Metschnikowia_pulcherrima(100); | Metschnikowia | 13 | Saprotroph | \|Undefined Saprotroph\| | Yeast | NULL | Probable | As nectar-tap_saprotroph (Põlme et al. 2020); Decay sugar-rich substrates (Põlme et al. 2020) | Põlme S, et al. 2020. Fungal Diversity 105:1-16 ((https://doi.org/10.1007/s13225-020-00466-2)) |
| Otu55194 | 1 | k__Fungi(100);p__Ascomycota(100);c__Saccharomycetes(100);o__Saccharomycetales(100);f__Metschnikowiaceae(100);g__Metschnikowia(100);s__Metschnikowia_pulcherrima(100); | Metschnikowia | 13 | Saprotroph | \|Undefined Saprotroph\| | Yeast | NULL | Probable | As nectar-tap_saprotroph (Põlme et al. 2020); Decay sugar-rich substrates (Põlme et al. 2020) | Põlme S, et al. 2020. Fungal Diversity 105:1-16 ((https://doi.org/10.1007/s13225-020-00466-2)) |
| Otu55195 | 1 | k__Fungi(100);p__Ascomycota(100);c__Saccharomycetes(100);o__Saccharomycetales(100);f__Metschnikowiaceae(100);g__Metschnikowia(100);s__Metschnikowia_pulcherrima(100); | Metschnikowia | 13 | Saprotroph | \|Undefined Saprotroph\| | Yeast | NULL | Probable | As nectar-tap_saprotroph (Põlme et al. 2020); Decay sugar-rich substrates (Põlme et al. 2020) | Põlme S, et al. 2020. Fungal Diversity 105:1-16 ((https://doi.org/10.1007/s13225-020-00466-2)) |
| Otu55196 | 1 | k__Fungi(100);p__Ascomycota(100);c__Saccharomycetes(100);o__Saccharomycetales(100);f__Metschnikowiaceae(100);g__Metschnikowia(100);s__Metschnikowia_pulcherrima(100); | Metschnikowia | 13 | Saprotroph | \|Undefined Saprotroph\| | Yeast | NULL | Probable | As nectar-tap_saprotroph (Põlme et al. 2020); Decay sugar-rich substrates (Põlme et al. 2020) | Põlme S, et al. 2020. Fungal Diversity 105:1-16 ((https://doi.org/10.1007/s13225-020-00466-2)) |
| Otu55198 | 1 | k__Fungi(100);p__Ascomycota(100);c__Saccharomycetes(100);o__Saccharomycetales(100);f__Metschnikowiaceae(100);g__Metschnikowia(100);s__Metschnikowia_pulcherrima(100); | Metschnikowia | 13 | Saprotroph | \|Undefined Saprotroph\| | Yeast | NULL | Probable | As nectar-tap_saprotroph (Põlme et al. 2020); Decay sugar-rich substrates (Põlme et al. 2020) | Põlme S, et al. 2020. Fungal Diversity 105:1-16 ((https://doi.org/10.1007/s13225-020-00466-2)) |
| Otu55201 | 1 | k__Fungi(100);p__Ascomycota(100);c__Saccharomycetes(100);o__Saccharomycetales(100);f__Metschnikowiaceae(100);g__Metschnikowia(100);s__Metschnikowia_pulcherrima(100); | Metschnikowia | 13 | Saprotroph | \|Undefined Saprotroph\| | Yeast | NULL | Probable | As nectar-tap_saprotroph (Põlme et al. 2020); Decay sugar-rich substrates (Põlme et al. 2020) | Põlme S, et al. 2020. Fungal Diversity 105:1-16 ((https://doi.org/10.1007/s13225-020-00466-2)) |
| Otu55203 | 1 | k__Fungi(100);p__Ascomycota(100);c__Saccharomycetes(100);o__Saccharomycetales(100);f__Metschnikowiaceae(100);g__Metschnikowia(100);s__Metschnikowia_pulcherrima(100); | Metschnikowia | 13 | Saprotroph | \|Undefined Saprotroph\| | Yeast | NULL | Probable | As nectar-tap_saprotroph (Põlme et al. 2020); Decay sugar-rich substrates (Põlme et al. 2020) | Põlme S, et al. 2020. Fungal Diversity 105:1-16 ((https://doi.org/10.1007/s13225-020-00466-2)) |
| Otu55204 | 1 | k__Fungi(100);p__Ascomycota(100);c__Saccharomycetes(100);o__Saccharomycetales(100);f__Metschnikowiaceae(100);g__Metschnikowia(100);s__Metschnikowia_pulcherrima(100); | Metschnikowia | 13 | Saprotroph | \|Undefined Saprotroph\| | Yeast | NULL | Probable | As nectar-tap_saprotroph (Põlme et al. 2020); Decay sugar-rich substrates (Põlme et al. 2020) | Põlme S, et al. 2020. Fungal Diversity 105:1-16 ((https://doi.org/10.1007/s13225-020-00466-2)) |
| Otu55206 | 1 | k__Fungi(100);p__Ascomycota(100);c__Saccharomycetes(100);o__Saccharomycetales(100);f__Metschnikowiaceae(100);g__Metschnikowia(100);s__Metschnikowia_pulcherrima(100); | Metschnikowia | 13 | Saprotroph | \|Undefined Saprotroph\| | Yeast | NULL | Probable | As nectar-tap_saprotroph (Põlme et al. 2020); Decay sugar-rich substrates (Põlme et al. 2020) | Põlme S, et al. 2020. Fungal Diversity 105:1-16 ((https://doi.org/10.1007/s13225-020-00466-2)) |
| Otu55211 | 1 | k__Fungi(100);p__Ascomycota(100);c__Saccharomycetes(100);o__Saccharomycetales(100);f__Metschnikowiaceae(100);g__Metschnikowia(100);s__Metschnikowia_pulcherrima(100); | Metschnikowia | 13 | Saprotroph | \|Undefined Saprotroph\| | Yeast | NULL | Probable | As nectar-tap_saprotroph (Põlme et al. 2020); Decay sugar-rich substrates (Põlme et al. 2020) | Põlme S, et al. 2020. Fungal Diversity 105:1-16 ((https://doi.org/10.1007/s13225-020-00466-2)) |
| Otu55212 | 1 | k__Fungi(100);p__Ascomycota(100);c__Saccharomycetes(100);o__Saccharomycetales(100);f__Metschnikowiaceae(100);g__Metschnikowia(100);g__Metschnikowia_unclassified(100); | Metschnikowia | 13 | Saprotroph | \|Undefined Saprotroph\| | Yeast | NULL | Probable | As nectar-tap_saprotroph (Põlme et al. 2020); Decay sugar-rich substrates (Põlme et al. 2020) | Põlme S, et al. 2020. Fungal Diversity 105:1-16 ((https://doi.org/10.1007/s13225-020-00466-2)) |

| | | | | | | | | | | | |
|---|---|---|---|---|---|---|---|---|---|---|---|
| Otu55213 | 1 | k__Fungi(100);p__Ascomycota(100);c__Saccharomycetes(100);o__Saccharomycetales(100);f__Metschnikowiaceae(100);g__Metschnikowia(100);s__Metschnikowia_pulcherrima(100); | Metschnikowia | 13 | Saprotroph | \|Undefined Saprotroph\| | Yeast | NULL | Probable | As nectar-tap_saprotroph (Põlme et al. 2020); Decay sugar-rich substrates (Põlme et al. 2020) | Põlme S, et al. 2020. Fungal Diversity 105:1-16 ((https://doi.org/10.1007/s13225-020-00466-2)) |
| Otu55214 | 1 | k__Fungi(100);p__Ascomycota(100);c__Saccharomycetes(100);o__Saccharomycetales(100);f__Metschnikowiaceae(100);g__Metschnikowia(100);s__Metschnikowia_pulcherrima(100); | Metschnikowia | 13 | Saprotroph | \|Undefined Saprotroph\| | Yeast | NULL | Probable | As nectar-tap_saprotroph (Põlme et al. 2020); Decay sugar-rich substrates (Põlme et al. 2020) | Põlme S, et al. 2020. Fungal Diversity 105:1-16 ((https://doi.org/10.1007/s13225-020-00466-2)) |
| Otu55215 | 1 | k__Fungi(100);p__Ascomycota(100);c__Saccharomycetes(100);o__Saccharomycetales(100);f__Metschnikowiaceae(100);g__Metschnikowia(100);s__Metschnikowia_pulcherrima(100); | Metschnikowia | 13 | Saprotroph | \|Undefined Saprotroph\| | Yeast | NULL | Probable | As nectar-tap_saprotroph (Põlme et al. 2020); Decay sugar-rich substrates (Põlme et al. 2020) | Põlme S, et al. 2020. Fungal Diversity 105:1-16 ((https://doi.org/10.1007/s13225-020-00466-2)) |
| Otu55216 | 1 | k__Fungi(100);p__Ascomycota(100);c__Saccharomycetes(100);o__Saccharomycetales(100);f__Metschnikowiaceae(100);g__Metschnikowia(100);g__Metschnikowia_unclassified(100); | Metschnikowia | 13 | Saprotroph | \|Undefined Saprotroph\| | Yeast | NULL | Probable | As nectar-tap_saprotroph (Põlme et al. 2020); Decay sugar-rich substrates (Põlme et al. 2020) | Põlme S, et al. 2020. Fungal Diversity 105:1-16 ((https://doi.org/10.1007/s13225-020-00466-2)) |
| Otu55217 | 1 | k__Fungi(100);p__Ascomycota(100);c__Saccharomycetes(100);o__Saccharomycetales(100);f__Metschnikowiaceae(100);g__Metschnikowia(100);s__Metschnikowia_pulcherrima(100); | Metschnikowia | 13 | Saprotroph | \|Undefined Saprotroph\| | Yeast | NULL | Probable | As nectar-tap_saprotroph (Põlme et al. 2020); Decay sugar-rich substrates (Põlme et al. 2020) | Põlme S, et al. 2020. Fungal Diversity 105:1-16 ((https://doi.org/10.1007/s13225-020-00466-2)) |
| Otu55218 | 1 | k__Fungi(100);p__Ascomycota(100);c__Saccharomycetes(100);o__Saccharomycetales(100);f__Metschnikowiaceae(100);g__Metschnikowia(100);s__Metschnikowia_pulcherrima(100); | Metschnikowia | 13 | Saprotroph | \|Undefined Saprotroph\| | Yeast | NULL | Probable | As nectar-tap_saprotroph (Põlme et al. 2020); Decay sugar-rich substrates (Põlme et al. 2020) | Põlme S, et al. 2020. Fungal Diversity 105:1-16 ((https://doi.org/10.1007/s13225-020-00466-2)) |
| Otu55219 | 1 | k__Fungi(100);p__Ascomycota(100);c__Saccharomycetes(100);o__Saccharomycetales(100);o__Saccharomycetales_unclassified(100);o__Saccharomycetales_unclassified(100);o__Saccharomycetales_unclassified(100); | Saccharomycetales | 7 | Saprotroph | Undefined Saprotroph | Yeast | NULL | Possible | NULL | Sterkenburg E, et al. 2015. New Phytologist 207:1145-1158 ((https://doi.org/10.1111/nph.13426)) |
| Otu55221 | 1 | k__Fungi(100);p__Ascomycota(100);c__Saccharomycetes(100);o__Saccharomycetales(100);f__Metschnikowiaceae(100);g__Metschnikowia(100);s__Metschnikowia_pulcherrima(100); | Metschnikowia | 13 | Saprotroph | \|Undefined Saprotroph\| | Yeast | NULL | Probable | As nectar-tap_saprotroph (Põlme et al. 2020); Decay sugar-rich substrates (Põlme et al. 2020) | Põlme S, et al. 2020. Fungal Diversity 105:1-16 ((https://doi.org/10.1007/s13225-020-00466-2)) |
| Otu55223 | 1 | k__Fungi(100);p__Ascomycota(100);c__Saccharomycetes(100);o__Saccharomycetales(100);f__Metschnikowiaceae(100);g__Metschnikowia(100);g__Metschnikowia_unclassified(100); | Metschnikowia | 13 | Saprotroph | \|Undefined Saprotroph\| | Yeast | NULL | Probable | As nectar-tap_saprotroph (Põlme et al. 2020); Decay sugar-rich substrates (Põlme et al. 2020) | Põlme S, et al. 2020. Fungal Diversity 105:1-16 ((https://doi.org/10.1007/s13225-020-00466-2)) |
| Otu55225 | 1 | k__Fungi(100);p__Ascomycota(100);c__Saccharomycetes(100);o__Saccharomycetales(100);f__Metschnikowiaceae(100);g__Metschnikowia(100);s__Metschnikowia_pulcherrima(100); | Metschnikowia | 13 | Saprotroph | \|Undefined Saprotroph\| | Yeast | NULL | Probable | As nectar-tap_saprotroph (Põlme et al. 2020); Decay sugar-rich substrates (Põlme et al. 2020) | Põlme S, et al. 2020. Fungal Diversity 105:1-16 ((https://doi.org/10.1007/s13225-020-00466-2)) |
| Otu55226 | 1 | k__Fungi(100);p__Ascomycota(100);c__Saccharomycetes(100);o__Saccharomycetales(100);f__Metschnikowiaceae(100);g__Metschnikowia(100);s__Metschnikowia_pulcherrima(100); | Metschnikowia | 13 | Saprotroph | \|Undefined Saprotroph\| | Yeast | NULL | Probable | As nectar-tap_saprotroph (Põlme et al. 2020); Decay sugar-rich substrates (Põlme et al. 2020) | Põlme S, et al. 2020. Fungal Diversity 105:1-16 ((https://doi.org/10.1007/s13225-020-00466-2)) |
| Otu55227 | 1 | k__Fungi(100);p__Ascomycota(100);c__Saccharomycetes(100);o__Saccharomycetales(100);f__Metschnikowiaceae(100);g__Metschnikowia(100);g__Metschnikowia_unclassified(100); | Metschnikowia | 13 | Saprotroph | \|Undefined Saprotroph\| | Yeast | NULL | Probable | As nectar-tap_saprotroph (Põlme et al. 2020); Decay sugar-rich substrates (Põlme et al. 2020) | Põlme S, et al. 2020. Fungal Diversity 105:1-16 ((https://doi.org/10.1007/s13225-020-00466-2)) |
| Otu55228 | 1 | k__Fungi(100);p__Ascomycota(100);c__Saccharomycetes(100);o__Saccharomycetales(100);f__Metschnikowiaceae(100);g__Metschnikowia(100);s__Metschnikowia_pulcherrima(100); | Metschnikowia | 13 | Saprotroph | \|Undefined Saprotroph\| | Yeast | NULL | Probable | As nectar-tap_saprotroph (Põlme et al. 2020); Decay sugar-rich substrates (Põlme et al. 2020) | Põlme S, et al. 2020. Fungal Diversity 105:1-16 ((https://doi.org/10.1007/s13225-020-00466-2)) |
| Otu55230 | 1 | k__Fungi(100);p__Ascomycota(100);c__Saccharomycetes(100);o__Saccharomycetales(100);f__Metschnikowiaceae(100);g__Metschnikowia(100);s__Metschnikowia_pulcherrima(100); | Metschnikowia | 13 | Saprotroph | \|Undefined Saprotroph\| | Yeast | NULL | Probable | As nectar-tap_saprotroph (Põlme et al. 2020); Decay sugar-rich substrates (Põlme et al. 2020) | Põlme S, et al. 2020. Fungal Diversity 105:1-16 ((https://doi.org/10.1007/s13225-020-00466-2)) |
| Otu55231 | 1 | k__Fungi(100);p__Ascomycota(100);c__Saccharomycetes(100);o__Saccharomycetales(100);f__Saccharomycetales_family_Incertae_sedis(100);f__Saccharomycetales_family_Incertae_sedis_unclassified(100);f__Saccharomycetales_family_Incertae_sedis_unclassified(100); | Saccharomycetales | 7 | Saprotroph | Undefined Saprotroph | Yeast | NULL | Possible | NULL | Sterkenburg E, et al. 2015. New Phytologist 207:1145-1158 ((https://doi.org/10.1111/nph.13426)) |
| Otu55232 | 1 | k__Fungi(100);p__Ascomycota(100);c__Saccharomycetes(100);o__Saccharomycetales(100);f__Metschnikowiaceae(100);g__Metschnikowia(100);g__Metschnikowia_unclassified(100); | Metschnikowia | 13 | Saprotroph | \|Undefined Saprotroph\| | Yeast | NULL | Probable | As nectar-tap_saprotroph (Põlme et al. 2020); Decay sugar-rich substrates (Põlme et al. 2020) | Põlme S, et al. 2020. Fungal Diversity 105:1-16 ((https://doi.org/10.1007/s13225-020-00466-2)) |
| Otu55233 | 1 | k__Fungi(100);p__Ascomycota(100);c__Saccharomycetes(100);o__Saccharomycetales(100);f__Metschnikowiaceae(100);g__Metschnikowia(100);s__Metschnikowia_pulcherrima(100); | Metschnikowia | 13 | Saprotroph | \|Undefined Saprotroph\| | Yeast | NULL | Probable | As nectar-tap_saprotroph (Põlme et al. 2020); Decay sugar-rich substrates (Põlme et al. 2020) | Põlme S, et al. 2020. Fungal Diversity 105:1-16 ((https://doi.org/10.1007/s13225-020-00466-2)) |
| Otu55235 | 1 | k__Fungi(100);p__Ascomycota(100);c__Saccharomycetes(100);o__Saccharomycetales(100);f__Metschnikowiaceae(100);g__Metschnikowia(100);g__Metschnikowia_unclassified(100); | Metschnikowia | 13 | Saprotroph | \|Undefined Saprotroph\| | Yeast | NULL | Probable | As nectar-tap_saprotroph (Põlme et al. 2020); Decay sugar-rich substrates (Põlme et al. 2020) | Põlme S, et al. 2020. Fungal Diversity 105:1-16 ((https://doi.org/10.1007/s13225-020-00466-2)) |
| Otu55236 | 1 | k__Fungi(100);p__Ascomycota(100);c__Saccharomycetes(100);o__Saccharomycetales(100);f__Metschnikowiaceae(100);g__Metschnikowia(100);s__Metschnikowia_pulcherrima(100); | Metschnikowia | 13 | Saprotroph | \|Undefined Saprotroph\| | Yeast | NULL | Probable | As nectar-tap_saprotroph (Põlme et al. 2020); Decay sugar-rich substrates (Põlme et al. 2020) | Põlme S, et al. 2020. Fungal Diversity 105:1-16 ((https://doi.org/10.1007/s13225-020-00466-2)) |
| Otu55237 | 1 | k__Fungi(100);p__Ascomycota(100);c__Saccharomycetes(100);o__Saccharomycetales(100);f__Metschnikowiaceae(100);g__Metschnikowia(100);s__Metschnikowia_pulcherrima(100); | Metschnikowia | 13 | Saprotroph | \|Undefined Saprotroph\| | Yeast | NULL | Probable | As nectar-tap_saprotroph (Põlme et al. 2020); Decay sugar-rich substrates (Põlme et al. 2020) | Põlme S, et al. 2020. Fungal Diversity 105:1-16 ((https://doi.org/10.1007/s13225-020-00466-2)) |
| Otu55238 | 1 | k__Fungi(100);p__Ascomycota(100);c__Saccharomycetes(100);o__Saccharomycetales(100);f__Metschnikowiaceae(100);g__Metschnikowia(100);s__Metschnikowia_pulcherrima(100); | Metschnikowia | 13 | Saprotroph | \|Undefined Saprotroph\| | Yeast | NULL | Probable | As nectar-tap_saprotroph (Põlme et al. 2020); Decay sugar-rich substrates (Põlme et al. 2020) | Põlme S, et al. 2020. Fungal Diversity 105:1-16 ((https://doi.org/10.1007/s13225-020-00466-2)) |
| Otu55239 | 1 | k__Fungi(100);p__Ascomycota(100);c__Saccharomycetes(100);o__Saccharomycetales(100);f__Metschnikowiaceae(100);g__Metschnikowia(100);s__Metschnikowia_pulcherrima(100); | Metschnikowia | 13 | Saprotroph | \|Undefined Saprotroph\| | Yeast | NULL | Probable | As nectar-tap_saprotroph (Põlme et al. 2020); Decay sugar-rich substrates (Põlme et al. 2020) | Põlme S, et al. 2020. Fungal Diversity 105:1-16 ((https://doi.org/10.1007/s13225-020-00466-2)) |
| Otu55241 | 1 | k__Fungi(100);p__Ascomycota(100);c__Saccharomycetes(100);o__Saccharomycetales(100);f__Metschnikowiaceae(100);g__Metschnikowia(100);s__Metschnikowia_pulcherrima(100); | Metschnikowia | 13 | Saprotroph | \|Undefined Saprotroph\| | Yeast | NULL | Probable | As nectar-tap_saprotroph (Põlme et al. 2020); Decay sugar-rich substrates (Põlme et al. 2020) | Põlme S, et al. 2020. Fungal Diversity 105:1-16 ((https://doi.org/10.1007/s13225-020-00466-2)) |
| Otu55242 | 1 | k__Fungi(100);p__Ascomycota(100);c__Saccharomycetes(100);o__Saccharomycetales(100);f__Metschnikowiaceae(100);g__Metschnikowia(100);s__Metschnikowia_pulcherrima(100); | Metschnikowia | 13 | Saprotroph | \|Undefined Saprotroph\| | Yeast | NULL | Probable | As nectar-tap_saprotroph (Põlme et al. 2020); Decay sugar-rich substrates (Põlme et al. 2020) | Põlme S, et al. 2020. Fungal Diversity 105:1-16 ((https://doi.org/10.1007/s13225-020-00466-2)) |
| Otu55244 | 1 | k__Fungi(100);p__Ascomycota(100);c__Saccharomycetes(100);o__Saccharomycetales(100);f__Metschnikowiaceae(100);g__Metschnikowia(100);s__Metschnikowia_pulcherrima(100); | Metschnikowia | 13 | Saprotroph | \|Undefined Saprotroph\| | Yeast | NULL | Probable | As nectar-tap_saprotroph (Põlme et al. 2020); Decay sugar-rich substrates (Põlme et al. 2020) | Põlme S, et al. 2020. Fungal Diversity 105:1-16 ((https://doi.org/10.1007/s13225-020-00466-2)) |
| Otu55245 | 1 | k__Fungi(100);p__Ascomycota(100);c__Saccharomycetes(100);o__Saccharomycetales(100);f__Metschnikowiaceae(100);g__Metschnikowia(100);s__Metschnikowia_pulcherrima(100); | Metschnikowia | 13 | Saprotroph | \|Undefined Saprotroph\| | Yeast | NULL | Probable | As nectar-tap_saprotroph (Põlme et al. 2020); Decay sugar-rich substrates (Põlme et al. 2020) | Põlme S, et al. 2020. Fungal Diversity 105:1-16 ((https://doi.org/10.1007/s13225-020-00466-2)) |
| Otu55246 | 1 | k__Fungi(100);p__Ascomycota(100);c__Saccharomycetes(100);o__Saccharomycetales(100);f__Metschnikowiaceae(100);g__Metschnikowia(100);s__Metschnikowia_pulcherrima(100); | Metschnikowia | 13 | Saprotroph | \|Undefined Saprotroph\| | Yeast | NULL | Probable | As nectar-tap_saprotroph (Põlme et al. 2020); Decay sugar-rich substrates (Põlme et al. 2020) | Põlme S, et al. 2020. Fungal Diversity 105:1-16 ((https://doi.org/10.1007/s13225-020-00466-2)) |
| Otu55247 | 1 | k__Fungi(100);p__Ascomycota(100);c__Saccharomycetes(100);o__Saccharomycetales(100);f__Metschnikowiaceae(100);g__Metschnikowia(100);s__Metschnikowia_pulcherrima(100); | Metschnikowia | 13 | Saprotroph | \|Undefined Saprotroph\| | Yeast | NULL | Probable | As nectar-tap_saprotroph (Põlme et al. 2020); Decay sugar-rich substrates (Põlme et al. 2020) | Põlme S, et al. 2020. Fungal Diversity 105:1-16 ((https://doi.org/10.1007/s13225-020-00466-2)) |
| Otu55248 | 1 | k__Fungi(100);p__Ascomycota(100);c__Saccharomycetes(100);o__Saccharomycetales(100);f__Metschnikowiaceae(100);g__Metschnikowia(100);g__Metschnikowia_unclassified(100); | Metschnikowia | 13 | Saprotroph | \|Undefined Saprotroph\| | Yeast | NULL | Probable | As nectar-tap_saprotroph (Põlme et al. 2020); Decay sugar-rich substrates (Põlme et al. 2020) | Põlme S, et al. 2020. Fungal Diversity 105:1-16 ((https://doi.org/10.1007/s13225-020-00466-2)) |
| Otu55249 | 1 | k__Fungi(100);p__Ascomycota(100);c__Saccharomycetes(100);o__Saccharomycetales(100);f__Metschnikowiaceae(100);g__Metschnikowia(100);s__Metschnikowia_pulcherrima(100); | Metschnikowia | 13 | Saprotroph | \|Undefined Saprotroph\| | Yeast | NULL | Probable | As nectar-tap_saprotroph (Põlme et al. 2020); Decay sugar-rich substrates (Põlme et al. 2020) | Põlme S, et al. 2020. Fungal Diversity 105:1-16 ((https://doi.org/10.1007/s13225-020-00466-2)) |
| Otu55250 | 1 | k__Fungi(100);p__Ascomycota(100);c__Saccharomycetes(100);o__Saccharomycetales(100);f__Metschnikowiaceae(100);g__Metschnikowia(100);s__Metschnikowia_pulcherrima(100); | Metschnikowia | 13 | Saprotroph | \|Undefined Saprotroph\| | Yeast | NULL | Probable | As nectar-tap_saprotroph (Põlme et al. 2020); Decay sugar-rich substrates (Põlme et al. 2020) | Põlme S, et al. 2020. Fungal Diversity 105:1-16 ((https://doi.org/10.1007/s13225-020-00466-2)) |
| Otu55252 | 1 | k__Fungi(100);p__Ascomycota(100);c__Saccharomycetes(100);o__Saccharomycetales(100);f__Metschnikowiaceae(100);g__Metschnikowia(100);s__Metschnikowia_pulcherrima(100); | Metschnikowia | 13 | Saprotroph | \|Undefined Saprotroph\| | Yeast | NULL | Probable | As nectar-tap_saprotroph (Põlme et al. 2020); Decay sugar-rich substrates (Põlme et al. 2020) | Põlme S, et al. 2020. Fungal Diversity 105:1-16 ((https://doi.org/10.1007/s13225-020-00466-2)) |
| Otu55253 | 1 | k__Fungi(100);p__Ascomycota(100);c__Saccharomycetes(100);o__Saccharomycetales(100);f__Metschnikowiaceae(100);g__Metschnikowia(100);s__Metschnikowia_pulcherrima(100); | Metschnikowia | 13 | Saprotroph | \|Undefined Saprotroph\| | Yeast | NULL | Probable | As nectar-tap_saprotroph (Põlme et al. 2020); Decay sugar-rich substrates (Põlme et al. 2020) | Põlme S, et al. 2020. Fungal Diversity 105:1-16 ((https://doi.org/10.1007/s13225-020-00466-2)) |
| Otu55254 | 1 | k__Fungi(100);p__Ascomycota(100);c__Saccharomycetes(100);o__Saccharomycetales(100);f__Metschnikowiaceae(100);g__Metschnikowia(100);s__Metschnikowia_pulcherrima(100); | Metschnikowia | 13 | Saprotroph | \|Undefined Saprotroph\| | Yeast | NULL | Probable | As nectar-tap_saprotroph (Põlme et al. 2020); Decay sugar-rich substrates (Põlme et al. 2020) | Põlme S, et al. 2020. Fungal Diversity 105:1-16 ((https://doi.org/10.1007/s13225-020-00466-2)) |
| Otu55255 | 1 | k__Fungi(100);p__Ascomycota(100);c__Saccharomycetes(100);o__Saccharomycetales(100);o__Saccharomycetales_unclassified(100);o__Saccharomycetales_unclassified(100);o__Saccharomycetales_unclassified(100); | Saccharomycetales | 7 | Saprotroph | Undefined Saprotroph | Yeast | NULL | Possible | NULL | Sterkenburg E, et al. 2015. New Phytologist 207:1145-1158 ((https://doi.org/10.1111/nph.13426)) |
| Otu55257 | 1 | k__Fungi(100);p__Ascomycota(100);c__Saccharomycetes(100);o__Saccharomycetales(100);f__Metschnikowiaceae(100);g__Metschnikowia(100);s__Metschnikowia_pulcherrima(100); | Metschnikowia | 13 | Saprotroph | \|Undefined Saprotroph\| | Yeast | NULL | Probable | As nectar-tap_saprotroph (Põlme et al. 2020); Decay sugar-rich substrates (Põlme et al. 2020) | Põlme S, et al. 2020. Fungal Diversity 105:1-16 ((https://doi.org/10.1007/s13225-020-00466-2)) |
| Otu55259 | 1 | k__Fungi(100);p__Ascomycota(100);c__Saccharomycetes(100);o__Saccharomycetales(100);f__Metschnikowiaceae(100);g__Metschnikowia(100);s__Metschnikowia_pulcherrima(100); | Metschnikowia | 13 | Saprotroph | \|Undefined Saprotroph\| | Yeast | NULL | Probable | As nectar-tap_saprotroph (Põlme et al. 2020); Decay sugar-rich substrates (Põlme et al. 2020) | Põlme S, et al. 2020. Fungal Diversity 105:1-16 ((https://doi.org/10.1007/s13225-020-00466-2)) |
| Otu55260 | 1 | k__Fungi(100);p__Ascomycota(100);c__Saccharomycetes(100);o__Saccharomycetales(100);f__Metschnikowiaceae(100);g__Metschnikowia(100);s__Metschnikowia_pulcherrima(100); | Metschnikowia | 13 | Saprotroph | \|Undefined Saprotroph\| | Yeast | NULL | Probable | As nectar-tap_saprotroph (Põlme et al. 2020); Decay sugar-rich substrates (Põlme et al. 2020) | Põlme S, et al. 2020. Fungal Diversity 105:1-16 ((https://doi.org/10.1007/s13225-020-00466-2)) |
| Otu55261 | 1 | k__Fungi(100);p__Ascomycota(100);c__Saccharomycetes(100);o__Saccharomycetales(100);f__Metschnikowiaceae(100);g__Metschnikowia(100);s__Metschnikowia_pulcherrima(100); | Metschnikowia | 13 | Saprotroph | \|Undefined Saprotroph\| | Yeast | NULL | Probable | As nectar-tap_saprotroph (Põlme et al. 2020); Decay sugar-rich substrates (Põlme et al. 2020) | Põlme S, et al. 2020. Fungal Diversity 105:1-16 ((https://doi.org/10.1007/s13225-020-00466-2)) |
| Otu55262 | 1 | k__Fungi(100);p__Ascomycota(100);c__Saccharomycetes(100);o__Saccharomycetales(100);f__Metschnikowiaceae(100);g__Metschnikowia(100);s__Metschnikowia_pulcherrima(100); | Metschnikowia | 13 | Saprotroph | \|Undefined Saprotroph\| | Yeast | NULL | Probable | As nectar-tap_saprotroph (Põlme et al. 2020); Decay sugar-rich substrates (Põlme et al. 2020) | Põlme S, et al. 2020. Fungal Diversity 105:1-16 ((https://doi.org/10.1007/s13225-020-00466-2)) |
| Otu55267 | 1 | k__Fungi(100);p__Ascomycota(100);c__Saccharomycetes(100);o__Saccharomycetales(100);f__Metschnikowiaceae(100);g__Metschnikowia(100);s__Metschnikowia_pulcherrima(100); | Metschnikowia | 13 | Saprotroph | \|Undefined Saprotroph\| | Yeast | NULL | Probable | As nectar-tap_saprotroph (Põlme et al. 2020); Decay sugar-rich substrates (Põlme et al. 2020) | Põlme S, et al. 2020. Fungal Diversity 105:1-16 ((https://doi.org/10.1007/s13225-020-00466-2)) |
| Otu55269 | 1 | k__Fungi(100);p__Ascomycota(100);c__Saccharomycetes(100);o__Saccharomycetales(100);f__Metschnikowiaceae(100);g__Metschnikowia(100);s__Metschnikowia_pulcherrima(100); | Metschnikowia | 13 | Saprotroph | \|Undefined Saprotroph\| | Yeast | NULL | Probable | As nectar-tap_saprotroph (Põlme et al. 2020); Decay sugar-rich substrates (Põlme et al. 2020) | Põlme S, et al. 2020. Fungal Diversity 105:1-16 ((https://doi.org/10.1007/s13225-020-00466-2)) |
| Otu55277 | 1 | k__Fungi(100);p__Ascomycota(100);c__Saccharomycetes(100);o__Saccharomycetales(100);f__Metschnikowiaceae(100);g__Metschnikowia(100);s__Metschnikowia_pulcherrima(100); | Metschnikowia | 13 | Saprotroph | \|Undefined Saprotroph\| | Yeast | NULL | Probable | As nectar-tap_saprotroph (Põlme et al. 2020); Decay sugar-rich substrates (Põlme et al. 2020) | Põlme S, et al. 2020. Fungal Diversity 105:1-16 ((https://doi.org/10.1007/s13225-020-00466-2)) |
| Otu55278 | 1 | k__Fungi(100);p__Ascomycota(100);c__Saccharomycetes(100);o__Saccharomycetales(100);f__Metschnikowiaceae(100);g__Metschnikowia(100);s__Metschnikowia_pulcherrima(100); | Metschnikowia | 13 | Saprotroph | \|Undefined Saprotroph\| | Yeast | NULL | Probable | As nectar-tap_saprotroph (Põlme et al. 2020); Decay sugar-rich substrates (Põlme et al. 2020) | Põlme S, et al. 2020. Fungal Diversity 105:1-16 ((https://doi.org/10.1007/s13225-020-00466-2)) |
| Otu55279 | 1 | k__Fungi(100);p__Ascomycota(100);c__Saccharomycetes(100);o__Saccharomycetales(100);f__Metschnikowiaceae(100);g__Metschnikowia(100);s__Metschnikowia_pulcherrima(100); | Metschnikowia | 13 | Saprotroph | \|Undefined Saprotroph\| | Yeast | NULL | Probable | As nectar-tap_saprotroph (Põlme et al. 2020); Decay sugar-rich substrates (Põlme et al. 2020) | Põlme S, et al. 2020. Fungal Diversity 105:1-16 ((https://doi.org/10.1007/s13225-020-00466-2)) |
| Otu55280 | 1 | k__Fungi(100);p__Ascomycota(100);c__Saccharomycetes(100);o__Saccharomycetales(100);f__Metschnikowiaceae(100);g__Metschnikowia(100);s__Metschnikowia_pulcherrima(100); | Metschnikowia | 13 | Saprotroph | \|Undefined Saprotroph\| | Yeast | NULL | Probable | As nectar-tap_saprotroph (Põlme et al. 2020); Decay sugar-rich substrates (Põlme et al. 2020) | Põlme S, et al. 2020. Fungal Diversity 105:1-16 ((https://doi.org/10.1007/s13225-020-00466-2)) |
| Otu55284 | 1 | k__Fungi(100);p__Ascomycota(100);c__Saccharomycetes(100);o__Saccharomycetales(100);f__Metschnikowiaceae(100);g__Metschnikowia(100);s__Metschnikowia_pulcherrima(100); | Metschnikowia | 13 | Saprotroph | \|Undefined Saprotroph\| | Yeast | NULL | Probable | As nectar-tap_saprotroph (Põlme et al. 2020); Decay sugar-rich substrates (Põlme et al. 2020) | Põlme S, et al. 2020. Fungal Diversity 105:1-16 ((https://doi.org/10.1007/s13225-020-00466-2)) |
| Otu55285 | 1 | k__Fungi(100);p__Ascomycota(100);c__Saccharomycetes(100);o__Saccharomycetales(100);f__Metschnikowiaceae(100);g__Metschnikowia(100);s__Metschnikowia_pulcherrima(100); | Metschnikowia | 13 | Saprotroph | \|Undefined Saprotroph\| | Yeast | NULL | Probable | As nectar-tap_saprotroph (Põlme et al. 2020); Decay sugar-rich substrates (Põlme et al. 2020) | Põlme S, et al. 2020. Fungal Diversity 105:1-16 ((https://doi.org/10.1007/s13225-020-00466-2)) |
| Otu55287 | 1 | k__Fungi(100);p__Ascomycota(100);c__Saccharomycetes(100);o__Saccharomycetales(100);f__Metschnikowiaceae(100);g__Metschnikowia(100);g__Metschnikowia_unclassified(100); | Metschnikowia | 13 | Saprotroph | \|Undefined Saprotroph\| | Yeast | NULL | Probable | As nectar-tap_saprotroph (Põlme et al. 2020); Decay sugar-rich substrates (Põlme et al. 2020) | Põlme S, et al. 2020. Fungal Diversity 105:1-16 ((https://doi.org/10.1007/s13225-020-00466-2)) |
| Otu55288 | 1 | k__Fungi(100);p__Ascomycota(100);c__Saccharomycetes(100);o__Saccharomycetales(100);f__Metschnikowiaceae(100);g__Metschnikowia(100);s__Metschnikowia_pulcherrima(100); | Metschnikowia | 13 | Saprotroph | \|Undefined Saprotroph\| | Yeast | NULL | Probable | As nectar-tap_saprotroph (Põlme et al. 2020); Decay sugar-rich substrates (Põlme et al. 2020) | Põlme S, et al. 2020. Fungal Diversity 105:1-16 ((https://doi.org/10.1007/s13225-020-00466-2)) |
| Otu55289 | 1 | k__Fungi(100);p__Ascomycota(100);c__Saccharomycetes(100);o__Saccharomycetales(100);f__Metschnikowiaceae(100);g__Metschnikowia(100);s__Metschnikowia_pulcherrima(100); | Metschnikowia | 13 | Saprotroph | \|Undefined Saprotroph\| | Yeast | NULL | Probable | As nectar-tap_saprotroph (Põlme et al. 2020); Decay sugar-rich substrates (Põlme et al. 2020) | Põlme S, et al. 2020. Fungal Diversity 105:1-16 ((https://doi.org/10.1007/s13225-020-00466-2)) |
| Otu55290 | 1 | k__Fungi(100);p__Ascomycota(100);c__Saccharomycetes(100);o__Saccharomycetales(100);f__Metschnikowiaceae(100);g__Metschnikowia(100);g__Metschnikowia_unclassified(100); | Metschnikowia | 13 | Saprotroph | \|Undefined Saprotroph\| | Yeast | NULL | Probable | As nectar-tap_saprotroph (Põlme et al. 2020); Decay sugar-rich substrates (Põlme et al. 2020) | Põlme S, et al. 2020. Fungal Diversity 105:1-16 ((https://doi.org/10.1007/s13225-020-00466-2)) |
| Otu55291 | 1 | k__Fungi(100);p__Ascomycota(100);c__Saccharomycetes(100);o__Saccharomycetales(100);f__Metschnikowiaceae(100);g__Metschnikowia(100);s__Metschnikowia_pulcherrima(100); | Metschnikowia | 13 | Saprotroph | \|Undefined Saprotroph\| | Yeast | NULL | Probable | As nectar-tap_saprotroph (Põlme et al. 2020); Decay sugar-rich substrates (Põlme et al. 2020) | Põlme S, et al. 2020. Fungal Diversity 105:1-16 ((https://doi.org/10.1007/s13225-020-00466-2)) |
| Otu55292 | 1 | k__Fungi(100);p__Ascomycota(100);c__Saccharomycetes(100);o__Saccharomycetales(100);f__Metschnikowiaceae(100);g__Metschnikowia(100);g__Metschnikowia_unclassified(100); | Metschnikowia | 13 | Saprotroph | \|Undefined Saprotroph\| | Yeast | NULL | Probable | As nectar-tap_saprotroph (Põlme et al. 2020); Decay sugar-rich substrates (Põlme et al. 2020) | Põlme S, et al. 2020. Fungal Diversity 105:1-16 ((https://doi.org/10.1007/s13225-020-00466-2)) |
| Otu55293 | 1 | k__Fungi(100);p__Ascomycota(100);c__Saccharomycetes(100);o__Saccharomycetales(100);f__Metschnikowiaceae(100);g__Metschnikowia(100);s__Metschnikowia_pulcherrima(100); | Metschnikowia | 13 | Saprotroph | \|Undefined Saprotroph\| | Yeast | NULL | Probable | As nectar-tap_saprotroph (Põlme et al. 2020); Decay sugar-rich substrates (Põlme et al. 2020) | Põlme S, et al. 2020. Fungal Diversity 105:1-16 ((https://doi.org/10.1007/s13225-020-00466-2)) |
| Otu55294 | 1 | k__Fungi(100);p__Ascomycota(100);c__Saccharomycetes(100);o__Saccharomycetales(100);f__Metschnikowiaceae(100);g__Metschnikowia(100);s__Metschnikowia_pulcherrima(100); | Metschnikowia | 13 | Saprotroph | \|Undefined Saprotroph\| | Yeast | NULL | Probable | As nectar-tap_saprotroph (Põlme et al. 2020); Decay sugar-rich substrates (Põlme et al. 2020) | Põlme S, et al. 2020. Fungal Diversity 105:1-16 ((https://doi.org/10.1007/s13225-020-00466-2)) |
| Otu55295 | 1 | k__Fungi(100);p__Ascomycota(100);c__Saccharomycetes(100);o__Saccharomycetales(100);f__Metschnikowiaceae(100);g__Metschnikowia(100);s__Metschnikowia_pulcherrima(100); | Metschnikowia | 13 | Saprotroph | \|Undefined Saprotroph\| | Yeast | NULL | Probable | As nectar-tap_saprotroph (Põlme et al. 2020); Decay sugar-rich substrates (Põlme et al. 2020) | Põlme S, et al. 2020. Fungal Diversity 105:1-16 ((https://doi.org/10.1007/s13225-020-00466-2)) |
| Otu55296 | 1 | k__Fungi(100);p__Ascomycota(100);c__Saccharomycetes(100);o__Saccharomycetales(100);f__Metschnikowiaceae(100);g__Metschnikowia(100);s__Metschnikowia_pulcherrima(100); | Metschnikowia | 13 | Saprotroph | \|Undefined Saprotroph\| | Yeast | NULL | Probable | As nectar-tap_saprotroph (Põlme et al. 2020); Decay sugar-rich substrates (Põlme et al. 2020) | Põlme S, et al. 2020. Fungal Diversity 105:1-16 ((https://doi.org/10.1007/s13225-020-00466-2)) |
| Otu55297 | 1 | k__Fungi(100);p__Ascomycota(100);c__Saccharomycetes(100);o__Saccharomycetales(100);f__Metschnikowiaceae(100);g__Metschnikowia(100);s__Metschnikowia_pulcherrima(100); | Metschnikowia | 13 | Saprotroph | \|Undefined Saprotroph\| | Yeast | NULL | Probable | As nectar-tap_saprotroph (Põlme et al. 2020); Decay sugar-rich substrates (Põlme et al. 2020) | Põlme S, et al. 2020. Fungal Diversity 105:1-16 ((https://doi.org/10.1007/s13225-020-00466-2)) |
| Otu55298 | 1 | k__Fungi(100);p__Ascomycota(100);c__Saccharomycetes(100);o__Saccharomycetales(100);f__Metschnikowiaceae(100);g__Metschnikowia(100);s__Metschnikowia_pulcherrima(100); | Metschnikowia | 13 | Saprotroph | \|Undefined Saprotroph\| | Yeast | NULL | Probable | As nectar-tap_saprotroph (Põlme et al. 2020); Decay sugar-rich substrates (Põlme et al. 2020) | Põlme S, et al. 2020. Fungal Diversity 105:1-16 ((https://doi.org/10.1007/s13225-020-00466-2)) |
| Otu55299 | 1 | k__Fungi(100);p__Ascomycota(100);c__Saccharomycetes(100);o__Saccharomycetales(100);f__Metschnikowiaceae(100);g__Metschnikowia(100);s__Metschnikowia_pulcherrima(100); | Metschnikowia | 13 | Saprotroph | \|Undefined Saprotroph\| | Yeast | NULL | Probable | As nectar-tap_saprotroph (Põlme et al. 2020); Decay sugar-rich substrates (Põlme et al. 2020) | Põlme S, et al. 2020. Fungal Diversity 105:1-16 ((https://doi.org/10.1007/s13225-020-00466-2)) |
| Otu55300 | 1 | k__Fungi(100);p__Ascomycota(100);c__Saccharomycetes(100);o__Saccharomycetales(100);f__Metschnikowiaceae(100);g__Metschnikowia(100);s__Metschnikowia_pulcherrima(100); | Metschnikowia | 13 | Saprotroph | \|Undefined Saprotroph\| | Yeast | NULL | Probable | As nectar-tap_saprotroph (Põlme et al. 2020); Decay sugar-rich substrates (Põlme et al. 2020) | Põlme S, et al. 2020. Fungal Diversity 105:1-16 ((https://doi.org/10.1007/s13225-020-00466-2)) |
| Otu55301 | 1 | k__Fungi(100);p__Ascomycota(100);c__Saccharomycetes(100);o__Saccharomycetales(100);f__Metschnikowiaceae(100);g__Metschnikowia(100);s__Metschnikowia_pulcherrima(100); | Metschnikowia | 13 | Saprotroph | \|Undefined Saprotroph\| | Yeast | NULL | Probable | As nectar-tap_saprotroph (Põlme et al. 2020); Decay sugar-rich substrates (Põlme et al. 2020) | Põlme S, et al. 2020. Fungal Diversity 105:1-16 ((https://doi.org/10.1007/s13225-020-00466-2)) |
| Otu55302 | 1 | k__Fungi(100);p__Ascomycota(100);c__Saccharomycetes(100);o__Saccharomycetales(100);f__Metschnikowiaceae(100);g__Metschnikowia(100);s__Metschnikowia_pulcherrima(100); | Metschnikowia | 13 | Saprotroph | \|Undefined Saprotroph\| | Yeast | NULL | Probable | As nectar-tap_saprotroph (Põlme et al. 2020); Decay sugar-rich substrates (Põlme et al. 2020) | Põlme S, et al. 2020. Fungal Diversity 105:1-16 ((https://doi.org/10.1007/s13225-020-00466-2)) |
| Otu55304 | 1 | k__Fungi(100);p__Ascomycota(100);c__Saccharomycetes(100);o__Saccharomycetales(100);f__Metschnikowiaceae(100);g__Metschnikowia(100);s__Metschnikowia_pulcherrima(100); | Metschnikowia | 13 | Saprotroph | \|Undefined Saprotroph\| | Yeast | NULL | Probable | As nectar-tap_saprotroph (Põlme et al. 2020); Decay sugar-rich substrates (Põlme et al. 2020) | Põlme S, et al. 2020. Fungal Diversity 105:1-16 ((https://doi.org/10.1007/s13225-020-00466-2)) |
| Otu55305 | 1 | k__Fungi(100);p__Ascomycota(100);c__Saccharomycetes(100);o__Saccharomycetales(100);f__Metschnikowiaceae(100);g__Metschnikowia(100);s__Metschnikowia_pulcherrima(100); | Metschnikowia | 13 | Saprotroph | \|Undefined Saprotroph\| | Yeast | NULL | Probable | As nectar-tap_saprotroph (Põlme et al. 2020); Decay sugar-rich substrates (Põlme et al. 2020) | Põlme S, et al. 2020. Fungal Diversity 105:1-16 ((https://doi.org/10.1007/s13225-020-00466-2)) |
| Otu55306 | 1 | k__Fungi(100);p__Ascomycota(100);c__Saccharomycetes(100);o__Saccharomycetales(100);f__Metschnikowiaceae(100);g__Metschnikowia(100);s__Metschnikowia_pulcherrima(100); | Metschnikowia | 13 | Saprotroph | \|Undefined Saprotroph\| | Yeast | NULL | Probable | As nectar-tap_saprotroph (Põlme et al. 2020); Decay sugar-rich substrates (Põlme et al. 2020) | Põlme S, et al. 2020. Fungal Diversity 105:1-16 ((https://doi.org/10.1007/s13225-020-00466-2)) |
| Otu55307 | 1 | k__Fungi(100);p__Ascomycota(100);c__Saccharomycetes(100);o__Saccharomycetales(100);f__Metschnikowiaceae(100);g__Metschnikowia(100);s__Metschnikowia_pulcherrima(100); | Metschnikowia | 13 | Saprotroph | \|Undefined Saprotroph\| | Yeast | NULL | Probable | As nectar-tap_saprotroph (Põlme et al. 2020); Decay sugar-rich substrates (Põlme et al. 2020) | Põlme S, et al. 2020. Fungal Diversity 105:1-16 ((https://doi.org/10.1007/s13225-020-00466-2)) |
| Otu55308 | 1 | k__Fungi(100);p__Ascomycota(100);c__Saccharomycetes(100);o__Saccharomycetales(100);f__Metschnikowiaceae(100);g__Metschnikowia(100);g__Metschnikowia_unclassified(100); | Metschnikowia | 13 | Saprotroph | \|Undefined Saprotroph\| | Yeast | NULL | Probable | As nectar-tap_saprotroph (Põlme et al. 2020); Decay sugar-rich substrates (Põlme et al. 2020) | Põlme S, et al. 2020. Fungal Diversity 105:1-16 ((https://doi.org/10.1007/s13225-020-00466-2)) |
| Otu55309 | 1 | k__Fungi(100);p__Ascomycota(100);c__Saccharomycetes(100);o__Saccharomycetales(100);f__Metschnikowiaceae(100);g__Metschnikowia(100);s__Metschnikowia_pulcherrima(100); | Metschnikowia | 13 | Saprotroph | \|Undefined Saprotroph\| | Yeast | NULL | Probable | As nectar-tap_saprotroph (Põlme et al. 2020); Decay sugar-rich substrates (Põlme et al. 2020) | Põlme S, et al. 2020. Fungal Diversity 105:1-16 ((https://doi.org/10.1007/s13225-020-00466-2)) |
| Otu55310 | 1 | k__Fungi(100);p__Ascomycota(100);c__Saccharomycetes(100);o__Saccharomycetales(100);f__Metschnikowiaceae(100);g__Metschnikowia(100);s__Metschnikowia_pulcherrima(100); | Metschnikowia | 13 | Saprotroph | \|Undefined Saprotroph\| | Yeast | NULL | Probable | As nectar-tap_saprotroph (Põlme et al. 2020); Decay sugar-rich substrates (Põlme et al. 2020) | Põlme S, et al. 2020. Fungal Diversity 105:1-16 ((https://doi.org/10.1007/s13225-020-00466-2)) |
| Otu55311 | 1 | k__Fungi(100);p__Ascomycota(100);c__Saccharomycetes(100);o__Saccharomycetales(100);f__Metschnikowiaceae(100);g__Metschnikowia(100);s__Metschnikowia_pulcherrima(100); | Metschnikowia | 13 | Saprotroph | \|Undefined Saprotroph\| | Yeast | NULL | Probable | As nectar-tap_saprotroph (Põlme et al. 2020); Decay sugar-rich substrates (Põlme et al. 2020) | Põlme S, et al. 2020. Fungal Diversity 105:1-16 ((https://doi.org/10.1007/s13225-020-00466-2)) |
| Otu55312 | 1 | k__Fungi(100);p__Ascomycota(100);c__Saccharomycetes(100);o__Saccharomycetales(100);f__Metschnikowiaceae(100);g__Metschnikowia(100);s__Metschnikowia_pulcherrima(100); | Metschnikowia | 13 | Saprotroph | \|Undefined Saprotroph\| | Yeast | NULL | Probable | As nectar-tap_saprotroph (Põlme et al. 2020); Decay sugar-rich substrates (Põlme et al. 2020) | Põlme S, et al. 2020. Fungal Diversity 105:1-16 ((https://doi.org/10.1007/s13225-020-00466-2)) |
| Otu55313 | 1 | k__Fungi(100);p__Ascomycota(100);c__Saccharomycetes(100);o__Saccharomycetales(100);f__Metschnikowiaceae(100);g__Metschnikowia(100);g__Metschnikowia_unclassified(100); | Metschnikowia | 13 | Saprotroph | \|Undefined Saprotroph\| | Yeast | NULL | Probable | As nectar-tap_saprotroph (Põlme et al. 2020); Decay sugar-rich substrates (Põlme et al. 2020) | Põlme S, et al. 2020. Fungal Diversity 105:1-16 ((https://doi.org/10.1007/s13225-020-00466-2)) |
| Otu55314 | 1 | k__Fungi(100);p__Ascomycota(100);c__Saccharomycetes(100);o__Saccharomycetales(100);f__Metschnikowiaceae(100);g__Metschnikowia(100);s__Metschnikowia_pulcherrima(100); | Metschnikowia | 13 | Saprotroph | \|Undefined Saprotroph\| | Yeast | NULL | Probable | As nectar-tap_saprotroph (Põlme et al. 2020); Decay sugar-rich substrates (Põlme et al. 2020) | Põlme S, et al. 2020. Fungal Diversity 105:1-16 ((https://doi.org/10.1007/s13225-020-00466-2)) |
| Otu55315 | 1 | k__Fungi(100);p__Ascomycota(100);c__Saccharomycetes(100);o__Saccharomycetales(100);f__Metschnikowiaceae(100);g__Metschnikowia(100);s__Metschnikowia_pulcherrima(100); | Metschnikowia | 13 | Saprotroph | \|Undefined Saprotroph\| | Yeast | NULL | Probable | As nectar-tap_saprotroph (Põlme et al. 2020); Decay sugar-rich substrates (Põlme et al. 2020) | Põlme S, et al. 2020. Fungal Diversity 105:1-16 ((https://doi.org/10.1007/s13225-020-00466-2)) |
| Otu55316 | 1 | k__Fungi(100);p__Ascomycota(100);c__Saccharomycetes(100);o__Saccharomycetales(100);f__Metschnikowiaceae(100);g__Metschnikowia(100);s__Metschnikowia_pulcherrima(100); | Metschnikowia | 13 | Saprotroph | \|Undefined Saprotroph\| | Yeast | NULL | Probable | As nectar-tap_saprotroph (Põlme et al. 2020); Decay sugar-rich substrates (Põlme et al. 2020) | Põlme S, et al. 2020. Fungal Diversity 105:1-16 ((https://doi.org/10.1007/s13225-020-00466-2)) |
| Otu55317 | 1 | k__Fungi(100);p__Ascomycota(100);c__Saccharomycetes(100);o__Saccharomycetales(100);f__Metschnikowiaceae(100);g__Metschnikowia(100);g__Metschnikowia_unclassified(100); | Metschnikowia | 13 | Saprotroph | \|Undefined Saprotroph\| | Yeast | NULL | Probable | As nectar-tap_saprotroph (Põlme et al. 2020); Decay sugar-rich substrates (Põlme et al. 2020) | Põlme S, et al. 2020. Fungal Diversity 105:1-16 ((https://doi.org/10.1007/s13225-020-00466-2)) |
| Otu55318 | 1 | k__Fungi(100);p__Ascomycota(100);c__Saccharomycetes(100);o__Saccharomycetales(100);f__Metschnikowiaceae(100);g__Metschnikowia(100);s__Metschnikowia_pulcherrima(100); | Metschnikowia | 13 | Saprotroph | \|Undefined Saprotroph\| | Yeast | NULL | Probable | As nectar-tap_saprotroph (Põlme et al. 2020); Decay sugar-rich substrates (Põlme et al. 2020) | Põlme S, et al. 2020. Fungal Diversity 105:1-16 ((https://doi.org/10.1007/s13225-020-00466-2)) |
| Otu55319 | 1 | k__Fungi(100);p__Ascomycota(100);c__Saccharomycetes(100);o__Saccharomycetales(100);f__Metschnikowiaceae(100);g__Metschnikowia(100);s__Metschnikowia_pulcherrima(100); | Metschnikowia | 13 | Saprotroph | \|Undefined Saprotroph\| | Yeast | NULL | Probable | As nectar-tap_saprotroph (Põlme et al. 2020); Decay sugar-rich substrates (Põlme et al. 2020) | Põlme S, et al. 2020. Fungal Diversity 105:1-16 ((https://doi.org/10.1007/s13225-020-00466-2)) |
| Otu55320 | 1 | k__Fungi(100);p__Ascomycota(100);c__Saccharomycetes(100);o__Saccharomycetales(100);f__Metschnikowiaceae(100);g__Metschnikowia(100);g__Metschnikowia_unclassified(100); | Metschnikowia | 13 | Saprotroph | \|Undefined Saprotroph\| | Yeast | NULL | Probable | As nectar-tap_saprotroph (Põlme et al. 2020); Decay sugar-rich substrates (Põlme et al. 2020) | Põlme S, et al. 2020. Fungal Diversity 105:1-16 ((https://doi.org/10.1007/s13225-020-00466-2)) |
| Otu55322 | 1 | k__Fungi(100);p__Ascomycota(100);c__Saccharomycetes(100);o__Saccharomycetales(100);f__Metschnikowiaceae(100);g__Metschnikowia(100);s__Metschnikowia_pulcherrima(100); | Metschnikowia | 13 | Saprotroph | \|Undefined Saprotroph\| | Yeast | NULL | Probable | As nectar-tap_saprotroph (Põlme et al. 2020); Decay sugar-rich substrates (Põlme et al. 2020) | Põlme S, et al. 2020. Fungal Diversity 105:1-16 ((https://doi.org/10.1007/s13225-020-00466-2)) |
| Otu55323 | 1 | k__Fungi(100);p__Ascomycota(100);c__Saccharomycetes(100);o__Saccharomycetales(100);f__Metschnikowiaceae(100);g__Metschnikowia(100);g__Metschnikowia_unclassified(100); | Metschnikowia | 13 | Saprotroph | \|Undefined Saprotroph\| | Yeast | NULL | Probable | As nectar-tap_saprotroph (Põlme et al. 2020); Decay sugar-rich substrates (Põlme et al. 2020) | Põlme S, et al. 2020. Fungal Diversity 105:1-16 ((https://doi.org/10.1007/s13225-020-00466-2)) |
| Otu55324 | 1 | k__Fungi(100);p__Ascomycota(100);c__Saccharomycetes(100);o__Saccharomycetales(100);f__Metschnikowiaceae(100);g__Metschnikowia(100);s__Metschnikowia_pulcherrima(100); | Metschnikowia | 13 | Saprotroph | \|Undefined Saprotroph\| | Yeast | NULL | Probable | As nectar-tap_saprotroph (Põlme et al. 2020); Decay sugar-rich substrates (Põlme et al. 2020) | Põlme S, et al. 2020. Fungal Diversity 105:1-16 ((https://doi.org/10.1007/s13225-020-00466-2)) |
| Otu55325 | 1 | k__Fungi(100);p__Ascomycota(100);c__Saccharomycetes(100);o__Saccharomycetales(100);f__Metschnikowiaceae(100);g__Metschnikowia(100);s__Metschnikowia_pulcherrima(100); | Metschnikowia | 13 | Saprotroph | \|Undefined Saprotroph\| | Yeast | NULL | Probable | As nectar-tap_saprotroph (Põlme et al. 2020); Decay sugar-rich substrates (Põlme et al. 2020) | Põlme S, et al. 2020. Fungal Diversity 105:1-16 ((https://doi.org/10.1007/s13225-020-00466-2)) |
| Otu55326 | 1 | k__Fungi(100);p__Ascomycota(100);c__Saccharomycetes(100);o__Saccharomycetales(100);f__Metschnikowiaceae(100);g__Metschnikowia(100);s__Metschnikowia_pulcherrima(100); | Metschnikowia | 13 | Saprotroph | \|Undefined Saprotroph\| | Yeast | NULL | Probable | As nectar-tap_saprotroph (Põlme et al. 2020); Decay sugar-rich substrates (Põlme et al. 2020) | Põlme S, et al. 2020. Fungal Diversity 105:1-16 ((https://doi.org/10.1007/s13225-020-00466-2)) |
| Otu55327 | 1 | k__Fungi(100);p__Ascomycota(100);c__Saccharomycetes(100);o__Saccharomycetales(100);f__Metschnikowiaceae(100);g__Metschnikowia(100);g__Metschnikowia_unclassified(100); | Metschnikowia | 13 | Saprotroph | \|Undefined Saprotroph\| | Yeast | NULL | Probable | As nectar-tap_saprotroph (Põlme et al. 2020); Decay sugar-rich substrates (Põlme et al. 2020) | Põlme S, et al. 2020. Fungal Diversity 105:1-16 ((https://doi.org/10.1007/s13225-020-00466-2)) |
| Otu55328 | 1 | k__Fungi(100);p__Ascomycota(100);c__Saccharomycetes(100);o__Saccharomycetales(100);f__Metschnikowiaceae(100);g__Metschnikowia(100);s__Metschnikowia_pulcherrima(100); | Metschnikowia | 13 | Saprotroph | \|Undefined Saprotroph\| | Yeast | NULL | Probable | As nectar-tap_saprotroph (Põlme et al. 2020); Decay sugar-rich substrates (Põlme et al. 2020) | Põlme S, et al. 2020. Fungal Diversity 105:1-16 ((https://doi.org/10.1007/s13225-020-00466-2)) |
| Otu55329 | 1 | k__Fungi(100);p__Ascomycota(100);c__Saccharomycetes(100);o__Saccharomycetales(100);f__Metschnikowiaceae(100);g__Metschnikowia(100);g__Metschnikowia_unclassified(100); | Metschnikowia | 13 | Saprotroph | \|Undefined Saprotroph\| | Yeast | NULL | Probable | As nectar-tap_saprotroph (Põlme et al. 2020); Decay sugar-rich substrates (Põlme et al. 2020) | Põlme S, et al. 2020. Fungal Diversity 105:1-16 ((https://doi.org/10.1007/s13225-020-00466-2)) |
| Otu55330 | 1 | k__Fungi(100);p__Ascomycota(100);c__Saccharomycetes(100);o__Saccharomycetales(100);f__Metschnikowiaceae(100);g__Metschnikowia(100);s__Metschnikowia_pulcherrima(100); | Metschnikowia | 13 | Saprotroph | \|Undefined Saprotroph\| | Yeast | NULL | Probable | As nectar-tap_saprotroph (Põlme et al. 2020); Decay sugar-rich substrates (Põlme et al. 2020) | Põlme S, et al. 2020. Fungal Diversity 105:1-16 ((https://doi.org/10.1007/s13225-020-00466-2)) |
| Otu55331 | 1 | k__Fungi(100);p__Ascomycota(100);c__Saccharomycetes(100);o__Saccharomycetales(100);f__Metschnikowiaceae(100);g__Metschnikowia(100);s__Metschnikowia_pulcherrima(100); | Metschnikowia | 13 | Saprotroph | \|Undefined Saprotroph\| | Yeast | NULL | Probable | As nectar-tap_saprotroph (Põlme et al. 2020); Decay sugar-rich substrates (Põlme et al. 2020) | Põlme S, et al. 2020. Fungal Diversity 105:1-16 ((https://doi.org/10.1007/s13225-020-00466-2)) |
| Otu55332 | 1 | k__Fungi(100);p__Ascomycota(100);c__Saccharomycetes(100);o__Saccharomycetales(100);f__Metschnikowiaceae(100);g__Metschnikowia(100);g__Metschnikowia_unclassified(100); | Metschnikowia | 13 | Saprotroph | \|Undefined Saprotroph\| | Yeast | NULL | Probable | As nectar-tap_saprotroph (Põlme et al. 2020); Decay sugar-rich substrates (Põlme et al. 2020) | Põlme S, et al. 2020. Fungal Diversity 105:1-16 ((https://doi.org/10.1007/s13225-020-00466-2)) |
| Otu55333 | 1 | k__Fungi(100);p__Ascomycota(100);c__Saccharomycetes(100);o__Saccharomycetales(100);f__Metschnikowiaceae(100);g__Metschnikowia(100);s__Metschnikowia_pulcherrima(100); | Metschnikowia | 13 | Saprotroph | \|Undefined Saprotroph\| | Yeast | NULL | Probable | As nectar-tap_saprotroph (Põlme et al. 2020); Decay sugar-rich substrates (Põlme et al. 2020) | Põlme S, et al. 2020. Fungal Diversity 105:1-16 ((https://doi.org/10.1007/s13225-020-00466-2)) |
| Otu55334 | 1 | k__Fungi(100);p__Ascomycota(100);c__Saccharomycetes(100);o__Saccharomycetales(100);f__Metschnikowiaceae(100);g__Metschnikowia(100);g__Metschnikowia_unclassified(100); | Metschnikowia | 13 | Saprotroph | \|Undefined Saprotroph\| | Yeast | NULL | Probable | As nectar-tap_saprotroph (Põlme et al. 2020); Decay sugar-rich substrates (Põlme et al. 2020) | Põlme S, et al. 2020. Fungal Diversity 105:1-16 ((https://doi.org/10.1007/s13225-020-00466-2)) |
| Otu55335 | 1 | k__Fungi(100);p__Ascomycota(100);c__Saccharomycetes(100);o__Saccharomycetales(100);f__Metschnikowiaceae(100);g__Metschnikowia(100);s__Metschnikowia_pulcherrima(100); | Metschnikowia | 13 | Saprotroph | \|Undefined Saprotroph\| | Yeast | NULL | Probable | As nectar-tap_saprotroph (Põlme et al. 2020); Decay sugar-rich substrates (Põlme et al. 2020) | Põlme S, et al. 2020. Fungal Diversity 105:1-16 ((https://doi.org/10.1007/s13225-020-00466-2)) |
| Otu55336 | 1 | k__Fungi(100);p__Ascomycota(100);c__Saccharomycetes(100);o__Saccharomycetales(100);f__Metschnikowiaceae(100);g__Metschnikowia(100);s__Metschnikowia_pulcherrima(100); | Metschnikowia | 13 | Saprotroph | \|Undefined Saprotroph\| | Yeast | NULL | Probable | As nectar-tap_saprotroph (Põlme et al. 2020); Decay sugar-rich substrates (Põlme et al. 2020) | Põlme S, et al. 2020. Fungal Diversity 105:1-16 ((https://doi.org/10.1007/s13225-020-00466-2)) |
| Otu55337 | 1 | k__Fungi(100);p__Ascomycota(100);c__Saccharomycetes(100);o__Saccharomycetales(100);f__Metschnikowiaceae(100);g__Metschnikowia(100);g__Metschnikowia_unclassified(100); | Metschnikowia | 13 | Saprotroph | \|Undefined Saprotroph\| | Yeast | NULL | Probable | As nectar-tap_saprotroph (Põlme et al. 2020); Decay sugar-rich substrates (Põlme et al. 2020) | Põlme S, et al. 2020. Fungal Diversity 105:1-16 ((https://doi.org/10.1007/s13225-020-00466-2)) |
| Otu55338 | 1 | k__Fungi(100);p__Ascomycota(100);c__Saccharomycetes(100);o__Saccharomycetales(100);f__Metschnikowiaceae(100);g__Metschnikowia(100);s__Metschnikowia_pulcherrima(100); | Metschnikowia | 13 | Saprotroph | \|Undefined Saprotroph\| | Yeast | NULL | Probable | As nectar-tap_saprotroph (Põlme et al. 2020); Decay sugar-rich substrates (Põlme et al. 2020) | Põlme S, et al. 2020. Fungal Diversity 105:1-16 ((https://doi.org/10.1007/s13225-020-00466-2)) |
| Otu55339 | 1 | k__Fungi(100);p__Ascomycota(100);c__Saccharomycetes(100);o__Saccharomycetales(100);f__Metschnikowiaceae(100);g__Metschnikowia(100);s__Metschnikowia_pulcherrima(100); | Metschnikowia | 13 | Saprotroph | \|Undefined Saprotroph\| | Yeast | NULL | Probable | As nectar-tap_saprotroph (Põlme et al. 2020); Decay sugar-rich substrates (Põlme et al. 2020) | Põlme S, et al. 2020. Fungal Diversity 105:1-16 ((https://doi.org/10.1007/s13225-020-00466-2)) |
| Otu55340 | 1 | k__Fungi(100);p__Ascomycota(100);c__Saccharomycetes(100);o__Saccharomycetales(100);f__Metschnikowiaceae(100);g__Metschnikowia(100);g__Metschnikowia_unclassified(100); | Metschnikowia | 13 | Saprotroph | \|Undefined Saprotroph\| | Yeast | NULL | Probable | As nectar-tap_saprotroph (Põlme et al. 2020); Decay sugar-rich substrates (Põlme et al. 2020) | Põlme S, et al. 2020. Fungal Diversity 105:1-16 ((https://doi.org/10.1007/s13225-020-00466-2)) |
| Otu55342 | 1 | k__Fungi(100);p__Ascomycota(100);c__Saccharomycetes(100);o__Saccharomycetales(100);f__Metschnikowiaceae(100);g__Metschnikowia(100);s__Metschnikowia_pulcherrima(100); | Metschnikowia | 13 | Saprotroph | \|Undefined Saprotroph\| | Yeast | NULL | Probable | As nectar-tap_saprotroph (Põlme et al. 2020); Decay sugar-rich substrates (Põlme et al. 2020) | Põlme S, et al. 2020. Fungal Diversity 105:1-16 ((https://doi.org/10.1007/s13225-020-00466-2)) |
| Otu55343 | 1 | k__Fungi(100);p__Ascomycota(100);c__Saccharomycetes(100);o__Saccharomycetales(100);f__Metschnikowiaceae(100);g__Metschnikowia(100);s__Metschnikowia_pulcherrima(100); | Metschnikowia | 13 | Saprotroph | \|Undefined Saprotroph\| | Yeast | NULL | Probable | As nectar-tap_saprotroph (Põlme et al. 2020); Decay sugar-rich substrates (Põlme et al. 2020) | Põlme S, et al. 2020. Fungal Diversity 105:1-16 ((https://doi.org/10.1007/s13225-020-00466-2)) |
| Otu55344 | 1 | k__Fungi(100);p__Ascomycota(100);c__Saccharomycetes(100);o__Saccharomycetales(100);f__Metschnikowiaceae(100);g__Metschnikowia(100);s__Metschnikowia_pulcherrima(100); | Metschnikowia | 13 | Saprotroph | \|Undefined Saprotroph\| | Yeast | NULL | Probable | As nectar-tap_saprotroph (Põlme et al. 2020); Decay sugar-rich substrates (Põlme et al. 2020) | Põlme S, et al. 2020. Fungal Diversity 105:1-16 ((https://doi.org/10.1007/s13225-020-00466-2)) |
| Otu55345 | 1 | k__Fungi(100);p__Ascomycota(100);c__Saccharomycetes(100);o__Saccharomycetales(100);f__Metschnikowiaceae(100);g__Metschnikowia(100);s__Metschnikowia_pulcherrima(100); | Metschnikowia | 13 | Saprotroph | \|Undefined Saprotroph\| | Yeast | NULL | Probable | As nectar-tap_saprotroph (Põlme et al. 2020); Decay sugar-rich substrates (Põlme et al. 2020) | Põlme S, et al. 2020. Fungal Diversity 105:1-16 ((https://doi.org/10.1007/s13225-020-00466-2)) |
| Otu55346 | 1 | k__Fungi(100);p__Ascomycota(100);c__Saccharomycetes(100);o__Saccharomycetales(100);f__Metschnikowiaceae(100);g__Metschnikowia(100);s__Metschnikowia_pulcherrima(100); | Metschnikowia | 13 | Saprotroph | \|Undefined Saprotroph\| | Yeast | NULL | Probable | As nectar-tap_saprotroph (Põlme et al. 2020); Decay sugar-rich substrates (Põlme et al. 2020) | Põlme S, et al. 2020. Fungal Diversity 105:1-16 ((https://doi.org/10.1007/s13225-020-00466-2)) |
| Otu55347 | 1 | k__Fungi(100);p__Ascomycota(100);c__Saccharomycetes(100);o__Saccharomycetales(100);f__Metschnikowiaceae(100);g__Metschnikowia(100);s__Metschnikowia_pulcherrima(100); | Metschnikowia | 13 | Saprotroph | \|Undefined Saprotroph\| | Yeast | NULL | Probable | As nectar-tap_saprotroph (Põlme et al. 2020); Decay sugar-rich substrates (Põlme et al. 2020) | Põlme S, et al. 2020. Fungal Diversity 105:1-16 ((https://doi.org/10.1007/s13225-020-00466-2)) |
| Otu55348 | 1 | k__Fungi(100);p__Ascomycota(100);c__Saccharomycetes(100);o__Saccharomycetales(100);f__Metschnikowiaceae(100);g__Metschnikowia(100);s__Metschnikowia_pulcherrima(100); | Metschnikowia | 13 | Saprotroph | \|Undefined Saprotroph\| | Yeast | NULL | Probable | As nectar-tap_saprotroph (Põlme et al. 2020); Decay sugar-rich substrates (Põlme et al. 2020) | Põlme S, et al. 2020. Fungal Diversity 105:1-16 ((https://doi.org/10.1007/s13225-020-00466-2)) |
| Otu55349 | 1 | k__Fungi(100);p__Ascomycota(100);c__Saccharomycetes(100);o__Saccharomycetales(100);f__Metschnikowiaceae(100);g__Metschnikowia(100);s__Metschnikowia_pulcherrima(100); | Metschnikowia | 13 | Saprotroph | \|Undefined Saprotroph\| | Yeast | NULL | Probable | As nectar-tap_saprotroph (Põlme et al. 2020); Decay sugar-rich substrates (Põlme et al. 2020) | Põlme S, et al. 2020. Fungal Diversity 105:1-16 ((https://doi.org/10.1007/s13225-020-00466-2)) |
| Otu55350 | 1 | k__Fungi(100);p__Ascomycota(100);c__Saccharomycetes(100);o__Saccharomycetales(100);f__Metschnikowiaceae(100);g__Metschnikowia(100);s__Metschnikowia_pulcherrima(100); | Metschnikowia | 13 | Saprotroph | \|Undefined Saprotroph\| | Yeast | NULL | Probable | As nectar-tap_saprotroph (Põlme et al. 2020); Decay sugar-rich substrates (Põlme et al. 2020) | Põlme S, et al. 2020. Fungal Diversity 105:1-16 ((https://doi.org/10.1007/s13225-020-00466-2)) |
| Otu55354 | 1 | k__Fungi(100);p__Ascomycota(100);c__Saccharomycetes(100);o__Saccharomycetales(100);f__Metschnikowiaceae(100);g__Metschnikowia(100);s__Metschnikowia_pulcherrima(100); | Metschnikowia | 13 | Saprotroph | \|Undefined Saprotroph\| | Yeast | NULL | Probable | As nectar-tap_saprotroph (Põlme et al. 2020); Decay sugar-rich substrates (Põlme et al. 2020) | Põlme S, et al. 2020. Fungal Diversity 105:1-16 ((https://doi.org/10.1007/s13225-020-00466-2)) |
| Otu55355 | 1 | k__Fungi(100);p__Ascomycota(100);c__Saccharomycetes(100);o__Saccharomycetales(100);f__Metschnikowiaceae(100);g__Metschnikowia(100);s__Metschnikowia_pulcherrima(100); | Metschnikowia | 13 | Saprotroph | \|Undefined Saprotroph\| | Yeast | NULL | Probable | As nectar-tap_saprotroph (Põlme et al. 2020); Decay sugar-rich substrates (Põlme et al. 2020) | Põlme S, et al. 2020. Fungal Diversity 105:1-16 ((https://doi.org/10.1007/s13225-020-00466-2)) |
| Otu55356 | 1 | k__Fungi(100);p__Ascomycota(100);c__Saccharomycetes(100);o__Saccharomycetales(100);f__Metschnikowiaceae(100);g__Metschnikowia(100);s__Metschnikowia_pulcherrima(100); | Metschnikowia | 13 | Saprotroph | \|Undefined Saprotroph\| | Yeast | NULL | Probable | As nectar-tap_saprotroph (Põlme et al. 2020); Decay sugar-rich substrates (Põlme et al. 2020) | Põlme S, et al. 2020. Fungal Diversity 105:1-16 ((https://doi.org/10.1007/s13225-020-00466-2)) |
| Otu55357 | 1 | k__Fungi(100);p__Ascomycota(100);c__Saccharomycetes(100);o__Saccharomycetales(100);f__Metschnikowiaceae(100);g__Metschnikowia(100);s__Metschnikowia_pulcherrima(100); | Metschnikowia | 13 | Saprotroph | \|Undefined Saprotroph\| | Yeast | NULL | Probable | As nectar-tap_saprotroph (Põlme et al. 2020); Decay sugar-rich substrates (Põlme et al. 2020) | Põlme S, et al. 2020. Fungal Diversity 105:1-16 ((https://doi.org/10.1007/s13225-020-00466-2)) |
| Otu55358 | 1 | k__Fungi(100);p__Ascomycota(100);c__Saccharomycetes(100);o__Saccharomycetales(100);f__Metschnikowiaceae(100);g__Metschnikowia(100);s__Metschnikowia_pulcherrima(100); | Metschnikowia | 13 | Saprotroph | \|Undefined Saprotroph\| | Yeast | NULL | Probable | As nectar-tap_saprotroph (Põlme et al. 2020); Decay sugar-rich substrates (Põlme et al. 2020) | Põlme S, et al. 2020. Fungal Diversity 105:1-16 ((https://doi.org/10.1007/s13225-020-00466-2)) |
| Otu55360 | 1 | k__Fungi(100);p__Ascomycota(100);c__Saccharomycetes(100);o__Saccharomycetales(100);f__Metschnikowiaceae(100);g__Metschnikowia(100);s__Metschnikowia_pulcherrima(100); | Metschnikowia | 13 | Saprotroph | \|Undefined Saprotroph\| | Yeast | NULL | Probable | As nectar-tap_saprotroph (Põlme et al. 2020); Decay sugar-rich substrates (Põlme et al. 2020) | Põlme S, et al. 2020. Fungal Diversity 105:1-16 ((https://doi.org/10.1007/s13225-020-00466-2)) |
| Otu55361 | 1 | k__Fungi(100);p__Ascomycota(100);c__Saccharomycetes(100);o__Saccharomycetales(100);f__Metschnikowiaceae(100);g__Metschnikowia(100);s__Metschnikowia_pulcherrima(100); | Metschnikowia | 13 | Saprotroph | \|Undefined Saprotroph\| | Yeast | NULL | Probable | As nectar-tap_saprotroph (Põlme et al. 2020); Decay sugar-rich substrates (Põlme et al. 2020) | Põlme S, et al. 2020. Fungal Diversity 105:1-16 ((https://doi.org/10.1007/s13225-020-00466-2)) |
| Otu55362 | 1 | k__Fungi(100);p__Ascomycota(100);c__Saccharomycetes(100);o__Saccharomycetales(100);f__Metschnikowiaceae(100);g__Metschnikowia(100);s__Metschnikowia_pulcherrima(100); | Metschnikowia | 13 | Saprotroph | \|Undefined Saprotroph\| | Yeast | NULL | Probable | As nectar-tap_saprotroph (Põlme et al. 2020); Decay sugar-rich substrates (Põlme et al. 2020) | Põlme S, et al. 2020. Fungal Diversity 105:1-16 ((https://doi.org/10.1007/s13225-020-00466-2)) |
| Otu55364 | 1 | k__Fungi(100);p__Ascomycota(100);c__Saccharomycetes(100);o__Saccharomycetales(100);f__Saccharomycetales_family_Incertae_sedis(100);g__Candida(100);g__Candida_unclassified(100); | Candida | 13 | Pathotroph-Saprotroph-Symbiotroph | Animal Pathogen-Endophyte-Endosymbiont-Epiphyte-\|Undefined Saprotroph\| | Dimorphic Yeast-Yeast | NULL | Probable | As nectar-tap_saprotroph (Põlme et al. 2020); Decay sugar-rich substrates (Põlme et al. 2020); This asexual genus is polyphyletic and contain species that function very differently depend on what they are. Caution should be taken when interpreting guilds for Candida species. | Manolakaki D, et al. 2010. Virulence. 1: 367-375 ((https://doi.org/10.4161/viru.1.5.12796)); Mekbib SB, et al. 2011. Tropical Plant Pathology 36:233-240 ((https://doi.org/10.1590/S1982-56762011000400004)); Põlme S, et al. 2020. Fungal Diversity 105:1-16 ((https://doi.org/10.1007/s13225-020-00466-2)) |
| Otu55365 | 1 | k__Fungi(100);p__Ascomycota(100);c__Saccharomycetes(100);o__Saccharomycetales(100);f__Metschnikowiaceae(100);g__Metschnikowia(100);s__Metschnikowia_pulcherrima(100); | Metschnikowia | 13 | Saprotroph | \|Undefined Saprotroph\| | Yeast | NULL | Probable | As nectar-tap_saprotroph (Põlme et al. 2020); Decay sugar-rich substrates (Põlme et al. 2020) | Põlme S, et al. 2020. Fungal Diversity 105:1-16 ((https://doi.org/10.1007/s13225-020-00466-2)) |
| Otu55366 | 1 | k__Fungi(100);p__Ascomycota(100);c__Saccharomycetes(100);o__Saccharomycetales(100);f__Metschnikowiaceae(100);g__Metschnikowia(100);g__Metschnikowia_unclassified(100); | Metschnikowia | 13 | Saprotroph | \|Undefined Saprotroph\| | Yeast | NULL | Probable | As nectar-tap_saprotroph (Põlme et al. 2020); Decay sugar-rich substrates (Põlme et al. 2020) | Põlme S, et al. 2020. Fungal Diversity 105:1-16 ((https://doi.org/10.1007/s13225-020-00466-2)) |
| Otu55368 | 1 | k__Fungi(100);p__Ascomycota(100);c__Saccharomycetes(100);o__Saccharomycetales(100);f__Metschnikowiaceae(100);g__Metschnikowia(100);g__Metschnikowia_unclassified(100); | Metschnikowia | 13 | Saprotroph | \|Undefined Saprotroph\| | Yeast | NULL | Probable | As nectar-tap_saprotroph (Põlme et al. 2020); Decay sugar-rich substrates (Põlme et al. 2020) | Põlme S, et al. 2020. Fungal Diversity 105:1-16 ((https://doi.org/10.1007/s13225-020-00466-2)) |
| Otu55369 | 1 | k__Fungi(100);p__Ascomycota(100);c__Saccharomycetes(100);o__Saccharomycetales(100);f__Metschnikowiaceae(100);g__Metschnikowia(100);g__Metschnikowia_unclassified(100); | Metschnikowia | 13 | Saprotroph | \|Undefined Saprotroph\| | Yeast | NULL | Probable | As nectar-tap_saprotroph (Põlme et al. 2020); Decay sugar-rich substrates (Põlme et al. 2020) | Põlme S, et al. 2020. Fungal Diversity 105:1-16 ((https://doi.org/10.1007/s13225-020-00466-2)) |
| Otu55370 | 1 | k__Fungi(100);p__Ascomycota(100);c__Saccharomycetes(100);o__Saccharomycetales(100);f__Metschnikowiaceae(100);g__Metschnikowia(100);s__Metschnikowia_pulcherrima(100); | Metschnikowia | 13 | Saprotroph | \|Undefined Saprotroph\| | Yeast | NULL | Probable | As nectar-tap_saprotroph (Põlme et al. 2020); Decay sugar-rich substrates (Põlme et al. 2020) | Põlme S, et al. 2020. Fungal Diversity 105:1-16 ((https://doi.org/10.1007/s13225-020-00466-2)) |
| Otu55371 | 1 | k__Fungi(100);p__Ascomycota(100);c__Saccharomycetes(100);o__Saccharomycetales(100);f__Metschnikowiaceae(100);g__Metschnikowia(100);s__Metschnikowia_pulcherrima(100); | Metschnikowia | 13 | Saprotroph | \|Undefined Saprotroph\| | Yeast | NULL | Probable | As nectar-tap_saprotroph (Põlme et al. 2020); Decay sugar-rich substrates (Põlme et al. 2020) | Põlme S, et al. 2020. Fungal Diversity 105:1-16 ((https://doi.org/10.1007/s13225-020-00466-2)) |
| Otu55372 | 1 | k__Fungi(100);p__Ascomycota(100);c__Saccharomycetes(100);o__Saccharomycetales(100);f__Metschnikowiaceae(100);g__Metschnikowia(100);s__Metschnikowia_pulcherrima(100); | Metschnikowia | 13 | Saprotroph | \|Undefined Saprotroph\| | Yeast | NULL | Probable | As nectar-tap_saprotroph (Põlme et al. 2020); Decay sugar-rich substrates (Põlme et al. 2020) | Põlme S, et al. 2020. Fungal Diversity 105:1-16 ((https://doi.org/10.1007/s13225-020-00466-2)) |
| Otu55373 | 1 | k__Fungi(100);p__Ascomycota(100);c__Saccharomycetes(100);o__Saccharomycetales(100);f__Metschnikowiaceae(100);g__Metschnikowia(100);s__Metschnikowia_pulcherrima(100); | Metschnikowia | 13 | Saprotroph | \|Undefined Saprotroph\| | Yeast | NULL | Probable | As nectar-tap_saprotroph (Põlme et al. 2020); Decay sugar-rich substrates (Põlme et al. 2020) | Põlme S, et al. 2020. Fungal Diversity 105:1-16 ((https://doi.org/10.1007/s13225-020-00466-2)) |
| Otu55374 | 1 | k__Fungi(100);p__Ascomycota(100);c__Saccharomycetes(100);o__Saccharomycetales(100);f__Metschnikowiaceae(100);g__Metschnikowia(100);s__Metschnikowia_pulcherrima(100); | Metschnikowia | 13 | Saprotroph | \|Undefined Saprotroph\| | Yeast | NULL | Probable | As nectar-tap_saprotroph (Põlme et al. 2020); Decay sugar-rich substrates (Põlme et al. 2020) | Põlme S, et al. 2020. Fungal Diversity 105:1-16 ((https://doi.org/10.1007/s13225-020-00466-2)) |
| Otu55375 | 1 | k__Fungi(100);p__Ascomycota(100);c__Saccharomycetes(100);o__Saccharomycetales(100);f__Metschnikowiaceae(100);g__Metschnikowia(100);s__Metschnikowia_pulcherrima(100); | Metschnikowia | 13 | Saprotroph | \|Undefined Saprotroph\| | Yeast | NULL | Probable | As nectar-tap_saprotroph (Põlme et al. 2020); Decay sugar-rich substrates (Põlme et al. 2020) | Põlme S, et al. 2020. Fungal Diversity 105:1-16 ((https://doi.org/10.1007/s13225-020-00466-2)) |
| Otu55376 | 1 | k__Fungi(100);p__Ascomycota(100);c__Saccharomycetes(100);o__Saccharomycetales(100);f__Metschnikowiaceae(100);g__Metschnikowia(100);s__Metschnikowia_pulcherrima(100); | Metschnikowia | 13 | Saprotroph | \|Undefined Saprotroph\| | Yeast | NULL | Probable | As nectar-tap_saprotroph (Põlme et al. 2020); Decay sugar-rich substrates (Põlme et al. 2020) | Põlme S, et al. 2020. Fungal Diversity 105:1-16 ((https://doi.org/10.1007/s13225-020-00466-2)) |
| Otu55377 | 1 | k__Fungi(100);p__Ascomycota(100);c__Saccharomycetes(100);o__Saccharomycetales(100);f__Metschnikowiaceae(100);g__Metschnikowia(100);s__Metschnikowia_pulcherrima(100); | Metschnikowia | 13 | Saprotroph | \|Undefined Saprotroph\| | Yeast | NULL | Probable | As nectar-tap_saprotroph (Põlme et al. 2020); Decay sugar-rich substrates (Põlme et al. 2020) | Põlme S, et al. 2020. Fungal Diversity 105:1-16 ((https://doi.org/10.1007/s13225-020-00466-2)) |
| Otu55378 | 1 | k__Fungi(100);p__Ascomycota(100);c__Saccharomycetes(100);o__Saccharomycetales(100);f__Metschnikowiaceae(100);g__Metschnikowia(100);s__Metschnikowia_pulcherrima(100); | Metschnikowia | 13 | Saprotroph | \|Undefined Saprotroph\| | Yeast | NULL | Probable | As nectar-tap_saprotroph (Põlme et al. 2020); Decay sugar-rich substrates (Põlme et al. 2020) | Põlme S, et al. 2020. Fungal Diversity 105:1-16 ((https://doi.org/10.1007/s13225-020-00466-2)) |
| Otu55379 | 1 | k__Fungi(100);p__Ascomycota(100);c__Saccharomycetes(100);o__Saccharomycetales(100);f__Metschnikowiaceae(100);g__Metschnikowia(100);g__Metschnikowia_unclassified(100); | Metschnikowia | 13 | Saprotroph | \|Undefined Saprotroph\| | Yeast | NULL | Probable | As nectar-tap_saprotroph (Põlme et al. 2020); Decay sugar-rich substrates (Põlme et al. 2020) | Põlme S, et al. 2020. Fungal Diversity 105:1-16 ((https://doi.org/10.1007/s13225-020-00466-2)) |
| Otu55381 | 1 | k__Fungi(100);p__Ascomycota(100);c__Saccharomycetes(100);o__Saccharomycetales(100);f__Metschnikowiaceae(100);g__Metschnikowia(100);s__Metschnikowia_pulcherrima(100); | Metschnikowia | 13 | Saprotroph | \|Undefined Saprotroph\| | Yeast | NULL | Probable | As nectar-tap_saprotroph (Põlme et al. 2020); Decay sugar-rich substrates (Põlme et al. 2020) | Põlme S, et al. 2020. Fungal Diversity 105:1-16 ((https://doi.org/10.1007/s13225-020-00466-2)) |
| Otu55382 | 1 | k__Fungi(100);p__Ascomycota(100);c__Saccharomycetes(100);o__Saccharomycetales(100);f__Metschnikowiaceae(100);g__Metschnikowia(100);s__Metschnikowia_pulcherrima(100); | Metschnikowia | 13 | Saprotroph | \|Undefined Saprotroph\| | Yeast | NULL | Probable | As nectar-tap_saprotroph (Põlme et al. 2020); Decay sugar-rich substrates (Põlme et al. 2020) | Põlme S, et al. 2020. Fungal Diversity 105:1-16 ((https://doi.org/10.1007/s13225-020-00466-2)) |
| Otu55383 | 1 | k__Fungi(100);p__Ascomycota(100);c__Saccharomycetes(100);o__Saccharomycetales(100);f__Metschnikowiaceae(100);g__Metschnikowia(100);s__Metschnikowia_pulcherrima(100); | Metschnikowia | 13 | Saprotroph | \|Undefined Saprotroph\| | Yeast | NULL | Probable | As nectar-tap_saprotroph (Põlme et al. 2020); Decay sugar-rich substrates (Põlme et al. 2020) | Põlme S, et al. 2020. Fungal Diversity 105:1-16 ((https://doi.org/10.1007/s13225-020-00466-2)) |
| Otu55384 | 1 | k__Fungi(100);p__Ascomycota(100);c__Saccharomycetes(100);o__Saccharomycetales(100);f__Metschnikowiaceae(100);g__Metschnikowia(100);s__Metschnikowia_pulcherrima(100); | Metschnikowia | 13 | Saprotroph | \|Undefined Saprotroph\| | Yeast | NULL | Probable | As nectar-tap_saprotroph (Põlme et al. 2020); Decay sugar-rich substrates (Põlme et al. 2020) | Põlme S, et al. 2020. Fungal Diversity 105:1-16 ((https://doi.org/10.1007/s13225-020-00466-2)) |
| Otu55385 | 1 | k__Fungi(100);p__Ascomycota(100);c__Saccharomycetes(100);o__Saccharomycetales(100);f__Metschnikowiaceae(100);g__Metschnikowia(100);s__Metschnikowia_pulcherrima(100); | Metschnikowia | 13 | Saprotroph | \|Undefined Saprotroph\| | Yeast | NULL | Probable | As nectar-tap_saprotroph (Põlme et al. 2020); Decay sugar-rich substrates (Põlme et al. 2020) | Põlme S, et al. 2020. Fungal Diversity 105:1-16 ((https://doi.org/10.1007/s13225-020-00466-2)) |
| Otu55386 | 1 | k__Fungi(100);p__Ascomycota(100);c__Saccharomycetes(100);o__Saccharomycetales(100);f__Metschnikowiaceae(100);g__Metschnikowia(100);g__Metschnikowia_unclassified(100); | Metschnikowia | 13 | Saprotroph | \|Undefined Saprotroph\| | Yeast | NULL | Probable | As nectar-tap_saprotroph (Põlme et al. 2020); Decay sugar-rich substrates (Põlme et al. 2020) | Põlme S, et al. 2020. Fungal Diversity 105:1-16 ((https://doi.org/10.1007/s13225-020-00466-2)) |
| Otu55387 | 1 | k__Fungi(100);p__Ascomycota(100);c__Saccharomycetes(100);o__Saccharomycetales(100);f__Metschnikowiaceae(100);g__Metschnikowia(100);s__Metschnikowia_pulcherrima(100); | Metschnikowia | 13 | Saprotroph | \|Undefined Saprotroph\| | Yeast | NULL | Probable | As nectar-tap_saprotroph (Põlme et al. 2020); Decay sugar-rich substrates (Põlme et al. 2020) | Põlme S, et al. 2020. Fungal Diversity 105:1-16 ((https://doi.org/10.1007/s13225-020-00466-2)) |
| Otu55388 | 1 | k__Fungi(100);p__Ascomycota(100);c__Saccharomycetes(100);o__Saccharomycetales(100);f__Metschnikowiaceae(100);g__Metschnikowia(100);s__Metschnikowia_pulcherrima(100); | Metschnikowia | 13 | Saprotroph | \|Undefined Saprotroph\| | Yeast | NULL | Probable | As nectar-tap_saprotroph (Põlme et al. 2020); Decay sugar-rich substrates (Põlme et al. 2020) | Põlme S, et al. 2020. Fungal Diversity 105:1-16 ((https://doi.org/10.1007/s13225-020-00466-2)) |
| Otu55389 | 1 | k__Fungi(100);p__Ascomycota(100);c__Saccharomycetes(100);o__Saccharomycetales(100);f__Metschnikowiaceae(100);g__Metschnikowia(100);g__Metschnikowia_unclassified(100); | Metschnikowia | 13 | Saprotroph | \|Undefined Saprotroph\| | Yeast | NULL | Probable | As nectar-tap_saprotroph (Põlme et al. 2020); Decay sugar-rich substrates (Põlme et al. 2020) | Põlme S, et al. 2020. Fungal Diversity 105:1-16 ((https://doi.org/10.1007/s13225-020-00466-2)) |
| Otu55390 | 1 | k__Fungi(100);p__Ascomycota(100);c__Saccharomycetes(100);o__Saccharomycetales(100);f__Metschnikowiaceae(100);g__Metschnikowia(100);s__Metschnikowia_pulcherrima(100); | Metschnikowia | 13 | Saprotroph | \|Undefined Saprotroph\| | Yeast | NULL | Probable | As nectar-tap_saprotroph (Põlme et al. 2020); Decay sugar-rich substrates (Põlme et al. 2020) | Põlme S, et al. 2020. Fungal Diversity 105:1-16 ((https://doi.org/10.1007/s13225-020-00466-2)) |
| Otu55391 | 1 | k__Fungi(100);p__Ascomycota(100);c__Saccharomycetes(100);o__Saccharomycetales(100);f__Metschnikowiaceae(100);g__Metschnikowia(100);s__Metschnikowia_pulcherrima(100); | Metschnikowia | 13 | Saprotroph | \|Undefined Saprotroph\| | Yeast | NULL | Probable | As nectar-tap_saprotroph (Põlme et al. 2020); Decay sugar-rich substrates (Põlme et al. 2020) | Põlme S, et al. 2020. Fungal Diversity 105:1-16 ((https://doi.org/10.1007/s13225-020-00466-2)) |
| Otu55392 | 1 | k__Fungi(100);p__Ascomycota(100);c__Saccharomycetes(100);o__Saccharomycetales(100);f__Metschnikowiaceae(100);g__Metschnikowia(100);s__Metschnikowia_pulcherrima(100); | Metschnikowia | 13 | Saprotroph | \|Undefined Saprotroph\| | Yeast | NULL | Probable | As nectar-tap_saprotroph (Põlme et al. 2020); Decay sugar-rich substrates (Põlme et al. 2020) | Põlme S, et al. 2020. Fungal Diversity 105:1-16 ((https://doi.org/10.1007/s13225-020-00466-2)) |
| Otu55393 | 1 | k__Fungi(100);p__Ascomycota(100);c__Saccharomycetes(100);o__Saccharomycetales(100);f__Metschnikowiaceae(100);g__Metschnikowia(100);s__Metschnikowia_pulcherrima(100); | Metschnikowia | 13 | Saprotroph | \|Undefined Saprotroph\| | Yeast | NULL | Probable | As nectar-tap_saprotroph (Põlme et al. 2020); Decay sugar-rich substrates (Põlme et al. 2020) | Põlme S, et al. 2020. Fungal Diversity 105:1-16 ((https://doi.org/10.1007/s13225-020-00466-2)) |
| Otu55394 | 1 | k__Fungi(100);p__Ascomycota(100);c__Saccharomycetes(100);o__Saccharomycetales(100);f__Metschnikowiaceae(100);g__Metschnikowia(100);s__Metschnikowia_pulcherrima(100); | Metschnikowia | 13 | Saprotroph | \|Undefined Saprotroph\| | Yeast | NULL | Probable | As nectar-tap_saprotroph (Põlme et al. 2020); Decay sugar-rich substrates (Põlme et al. 2020) | Põlme S, et al. 2020. Fungal Diversity 105:1-16 ((https://doi.org/10.1007/s13225-020-00466-2)) |
| Otu55395 | 1 | k__Fungi(100);p__Ascomycota(100);c__Saccharomycetes(100);o__Saccharomycetales(100);f__Metschnikowiaceae(100);g__Metschnikowia(100);s__Metschnikowia_pulcherrima(100); | Metschnikowia | 13 | Saprotroph | \|Undefined Saprotroph\| | Yeast | NULL | Probable | As nectar-tap_saprotroph (Põlme et al. 2020); Decay sugar-rich substrates (Põlme et al. 2020) | Põlme S, et al. 2020. Fungal Diversity 105:1-16 ((https://doi.org/10.1007/s13225-020-00466-2)) |
| Otu55396 | 1 | k__Fungi(100);p__Ascomycota(100);c__Saccharomycetes(100);o__Saccharomycetales(100);f__Metschnikowiaceae(100);g__Metschnikowia(100);g__Metschnikowia_unclassified(100); | Metschnikowia | 13 | Saprotroph | \|Undefined Saprotroph\| | Yeast | NULL | Probable | As nectar-tap_saprotroph (Põlme et al. 2020); Decay sugar-rich substrates (Põlme et al. 2020) | Põlme S, et al. 2020. Fungal Diversity 105:1-16 ((https://doi.org/10.1007/s13225-020-00466-2)) |
| Otu55397 | 1 | k__Fungi(100);p__Ascomycota(100);c__Saccharomycetes(100);o__Saccharomycetales(100);f__Metschnikowiaceae(100);g__Metschnikowia(100);s__Metschnikowia_pulcherrima(100); | Metschnikowia | 13 | Saprotroph | \|Undefined Saprotroph\| | Yeast | NULL | Probable | As nectar-tap_saprotroph (Põlme et al. 2020); Decay sugar-rich substrates (Põlme et al. 2020) | Põlme S, et al. 2020. Fungal Diversity 105:1-16 ((https://doi.org/10.1007/s13225-020-00466-2)) |
| Otu55398 | 1 | k__Fungi(100);p__Ascomycota(100);c__Saccharomycetes(100);o__Saccharomycetales(100);f__Metschnikowiaceae(100);g__Metschnikowia(100);s__Metschnikowia_pulcherrima(100); | Metschnikowia | 13 | Saprotroph | \|Undefined Saprotroph\| | Yeast | NULL | Probable | As nectar-tap_saprotroph (Põlme et al. 2020); Decay sugar-rich substrates (Põlme et al. 2020) | Põlme S, et al. 2020. Fungal Diversity 105:1-16 ((https://doi.org/10.1007/s13225-020-00466-2)) |
| Otu55399 | 1 | k__Fungi(100);p__Ascomycota(100);c__Saccharomycetes(100);o__Saccharomycetales(100);f__Metschnikowiaceae(100);g__Metschnikowia(100);s__Metschnikowia_pulcherrima(100); | Metschnikowia | 13 | Saprotroph | \|Undefined Saprotroph\| | Yeast | NULL | Probable | As nectar-tap_saprotroph (Põlme et al. 2020); Decay sugar-rich substrates (Põlme et al. 2020) | Põlme S, et al. 2020. Fungal Diversity 105:1-16 ((https://doi.org/10.1007/s13225-020-00466-2)) |
| Otu55400 | 1 | k__Fungi(100);p__Ascomycota(100);c__Saccharomycetes(100);o__Saccharomycetales(100);f__Metschnikowiaceae(100);g__Metschnikowia(100);s__Metschnikowia_pulcherrima(100); | Metschnikowia | 13 | Saprotroph | \|Undefined Saprotroph\| | Yeast | NULL | Probable | As nectar-tap_saprotroph (Põlme et al. 2020); Decay sugar-rich substrates (Põlme et al. 2020) | Põlme S, et al. 2020. Fungal Diversity 105:1-16 ((https://doi.org/10.1007/s13225-020-00466-2)) |
| Otu55401 | 1 | k__Fungi(100);p__Ascomycota(100);c__Saccharomycetes(100);o__Saccharomycetales(100);f__Metschnikowiaceae(100);g__Metschnikowia(100);s__Metschnikowia_pulcherrima(100); | Metschnikowia | 13 | Saprotroph | \|Undefined Saprotroph\| | Yeast | NULL | Probable | As nectar-tap_saprotroph (Põlme et al. 2020); Decay sugar-rich substrates (Põlme et al. 2020) | Põlme S, et al. 2020. Fungal Diversity 105:1-16 ((https://doi.org/10.1007/s13225-020-00466-2)) |
| Otu55405 | 1 | k__Fungi(100);p__Ascomycota(100);c__Saccharomycetes(100);o__Saccharomycetales(100);f__Metschnikowiaceae(100);g__Metschnikowia(100);s__Metschnikowia_pulcherrima(100); | Metschnikowia | 13 | Saprotroph | \|Undefined Saprotroph\| | Yeast | NULL | Probable | As nectar-tap_saprotroph (Põlme et al. 2020); Decay sugar-rich substrates (Põlme et al. 2020) | Põlme S, et al. 2020. Fungal Diversity 105:1-16 ((https://doi.org/10.1007/s13225-020-00466-2)) |
| Otu55406 | 1 | k__Fungi(100);p__Ascomycota(100);c__Saccharomycetes(100);o__Saccharomycetales(100);f__Metschnikowiaceae(100);g__Metschnikowia(100);g__Metschnikowia_unclassified(100); | Metschnikowia | 13 | Saprotroph | \|Undefined Saprotroph\| | Yeast | NULL | Probable | As nectar-tap_saprotroph (Põlme et al. 2020); Decay sugar-rich substrates (Põlme et al. 2020) | Põlme S, et al. 2020. Fungal Diversity 105:1-16 ((https://doi.org/10.1007/s13225-020-00466-2)) |
| Otu55407 | 1 | k__Fungi(100);p__Ascomycota(100);c__Saccharomycetes(100);o__Saccharomycetales(100);f__Metschnikowiaceae(100);g__Metschnikowia(100);s__Metschnikowia_pulcherrima(100); | Metschnikowia | 13 | Saprotroph | \|Undefined Saprotroph\| | Yeast | NULL | Probable | As nectar-tap_saprotroph (Põlme et al. 2020); Decay sugar-rich substrates (Põlme et al. 2020) | Põlme S, et al. 2020. Fungal Diversity 105:1-16 ((https://doi.org/10.1007/s13225-020-00466-2)) |
| Otu55409 | 1 | k__Fungi(100);p__Ascomycota(100);c__Saccharomycetes(100);o__Saccharomycetales(100);f__Metschnikowiaceae(100);g__Metschnikowia(100);s__Metschnikowia_pulcherrima(100); | Metschnikowia | 13 | Saprotroph | \|Undefined Saprotroph\| | Yeast | NULL | Probable | As nectar-tap_saprotroph (Põlme et al. 2020); Decay sugar-rich substrates (Põlme et al. 2020) | Põlme S, et al. 2020. Fungal Diversity 105:1-16 ((https://doi.org/10.1007/s13225-020-00466-2)) |
| Otu55410 | 1 | k__Fungi(100);p__Ascomycota(100);c__Saccharomycetes(100);o__Saccharomycetales(100);f__Metschnikowiaceae(100);g__Metschnikowia(100);s__Metschnikowia_pulcherrima(100); | Metschnikowia | 13 | Saprotroph | \|Undefined Saprotroph\| | Yeast | NULL | Probable | As nectar-tap_saprotroph (Põlme et al. 2020); Decay sugar-rich substrates (Põlme et al. 2020) | Põlme S, et al. 2020. Fungal Diversity 105:1-16 ((https://doi.org/10.1007/s13225-020-00466-2)) |
| Otu55411 | 1 | k__Fungi(100);p__Ascomycota(100);c__Saccharomycetes(100);o__Saccharomycetales(100);f__Metschnikowiaceae(100);g__Metschnikowia(100);g__Metschnikowia_unclassified(100); | Metschnikowia | 13 | Saprotroph | \|Undefined Saprotroph\| | Yeast | NULL | Probable | As nectar-tap_saprotroph (Põlme et al. 2020); Decay sugar-rich substrates (Põlme et al. 2020) | Põlme S, et al. 2020. Fungal Diversity 105:1-16 ((https://doi.org/10.1007/s13225-020-00466-2)) |
| Otu55412 | 1 | k__Fungi(100);p__Ascomycota(100);c__Saccharomycetes(100);o__Saccharomycetales(100);f__Metschnikowiaceae(100);g__Metschnikowia(100);s__Metschnikowia_pulcherrima(100); | Metschnikowia | 13 | Saprotroph | \|Undefined Saprotroph\| | Yeast | NULL | Probable | As nectar-tap_saprotroph (Põlme et al. 2020); Decay sugar-rich substrates (Põlme et al. 2020) | Põlme S, et al. 2020. Fungal Diversity 105:1-16 ((https://doi.org/10.1007/s13225-020-00466-2)) |
| Otu55413 | 1 | k__Fungi(100);p__Ascomycota(100);c__Saccharomycetes(100);o__Saccharomycetales(100);f__Metschnikowiaceae(100);g__Metschnikowia(100);s__Metschnikowia_pulcherrima(100); | Metschnikowia | 13 | Saprotroph | \|Undefined Saprotroph\| | Yeast | NULL | Probable | As nectar-tap_saprotroph (Põlme et al. 2020); Decay sugar-rich substrates (Põlme et al. 2020) | Põlme S, et al. 2020. Fungal Diversity 105:1-16 ((https://doi.org/10.1007/s13225-020-00466-2)) |
| Otu55414 | 1 | k__Fungi(100);p__Ascomycota(100);c__Saccharomycetes(100);o__Saccharomycetales(100);f__Metschnikowiaceae(100);g__Metschnikowia(100);s__Metschnikowia_pulcherrima(100); | Metschnikowia | 13 | Saprotroph | \|Undefined Saprotroph\| | Yeast | NULL | Probable | As nectar-tap_saprotroph (Põlme et al. 2020); Decay sugar-rich substrates (Põlme et al. 2020) | Põlme S, et al. 2020. Fungal Diversity 105:1-16 ((https://doi.org/10.1007/s13225-020-00466-2)) |
| Otu55415 | 1 | k__Fungi(100);p__Ascomycota(100);c__Saccharomycetes(100);o__Saccharomycetales(100);f__Metschnikowiaceae(100);g__Metschnikowia(100);s__Metschnikowia_pulcherrima(100); | Metschnikowia | 13 | Saprotroph | \|Undefined Saprotroph\| | Yeast | NULL | Probable | As nectar-tap_saprotroph (Põlme et al. 2020); Decay sugar-rich substrates (Põlme et al. 2020) | Põlme S, et al. 2020. Fungal Diversity 105:1-16 ((https://doi.org/10.1007/s13225-020-00466-2)) |
| Otu55416 | 1 | k__Fungi(100);p__Ascomycota(100);c__Saccharomycetes(100);o__Saccharomycetales(100);f__Metschnikowiaceae(100);g__Metschnikowia(100);s__Metschnikowia_pulcherrima(100); | Metschnikowia | 13 | Saprotroph | \|Undefined Saprotroph\| | Yeast | NULL | Probable | As nectar-tap_saprotroph (Põlme et al. 2020); Decay sugar-rich substrates (Põlme et al. 2020) | Põlme S, et al. 2020. Fungal Diversity 105:1-16 ((https://doi.org/10.1007/s13225-020-00466-2)) |
| Otu55417 | 1 | k__Fungi(100);p__Ascomycota(100);c__Saccharomycetes(100);o__Saccharomycetales(100);f__Metschnikowiaceae(100);g__Metschnikowia(100);s__Metschnikowia_pulcherrima(100); | Metschnikowia | 13 | Saprotroph | \|Undefined Saprotroph\| | Yeast | NULL | Probable | As nectar-tap_saprotroph (Põlme et al. 2020); Decay sugar-rich substrates (Põlme et al. 2020) | Põlme S, et al. 2020. Fungal Diversity 105:1-16 ((https://doi.org/10.1007/s13225-020-00466-2)) |
| Otu55418 | 1 | k__Fungi(100);p__Ascomycota(100);c__Saccharomycetes(100);o__Saccharomycetales(100);f__Metschnikowiaceae(100);g__Metschnikowia(100);s__Metschnikowia_pulcherrima(100); | Metschnikowia | 13 | Saprotroph | \|Undefined Saprotroph\| | Yeast | NULL | Probable | As nectar-tap_saprotroph (Põlme et al. 2020); Decay sugar-rich substrates (Põlme et al. 2020) | Põlme S, et al. 2020. Fungal Diversity 105:1-16 ((https://doi.org/10.1007/s13225-020-00466-2)) |
| Otu55419 | 1 | k__Fungi(100);p__Ascomycota(100);c__Saccharomycetes(100);o__Saccharomycetales(100);f__Metschnikowiaceae(100);g__Metschnikowia(100);s__Metschnikowia_pulcherrima(100); | Metschnikowia | 13 | Saprotroph | \|Undefined Saprotroph\| | Yeast | NULL | Probable | As nectar-tap_saprotroph (Põlme et al. 2020); Decay sugar-rich substrates (Põlme et al. 2020) | Põlme S, et al. 2020. Fungal Diversity 105:1-16 ((https://doi.org/10.1007/s13225-020-00466-2)) |
| Otu55421 | 1 | k__Fungi(100);p__Ascomycota(100);c__Saccharomycetes(100);o__Saccharomycetales(100);f__Metschnikowiaceae(100);g__Metschnikowia(100);g__Metschnikowia_unclassified(100); | Metschnikowia | 13 | Saprotroph | \|Undefined Saprotroph\| | Yeast | NULL | Probable | As nectar-tap_saprotroph (Põlme et al. 2020); Decay sugar-rich substrates (Põlme et al. 2020) | Põlme S, et al. 2020. Fungal Diversity 105:1-16 ((https://doi.org/10.1007/s13225-020-00466-2)) |
| Otu55423 | 1 | k__Fungi(100);p__Ascomycota(100);c__Saccharomycetes(100);o__Saccharomycetales(100);f__Metschnikowiaceae(100);g__Metschnikowia(100);s__Metschnikowia_pulcherrima(100); | Metschnikowia | 13 | Saprotroph | \|Undefined Saprotroph\| | Yeast | NULL | Probable | As nectar-tap_saprotroph (Põlme et al. 2020); Decay sugar-rich substrates (Põlme et al. 2020) | Põlme S, et al. 2020. Fungal Diversity 105:1-16 ((https://doi.org/10.1007/s13225-020-00466-2)) |
| Otu55424 | 1 | k__Fungi(100);p__Ascomycota(100);c__Saccharomycetes(100);o__Saccharomycetales(100);f__Metschnikowiaceae(100);g__Metschnikowia(100);g__Metschnikowia_unclassified(100); | Metschnikowia | 13 | Saprotroph | \|Undefined Saprotroph\| | Yeast | NULL | Probable | As nectar-tap_saprotroph (Põlme et al. 2020); Decay sugar-rich substrates (Põlme et al. 2020) | Põlme S, et al. 2020. Fungal Diversity 105:1-16 ((https://doi.org/10.1007/s13225-020-00466-2)) |
| Otu55425 | 1 | k__Fungi(100);p__Ascomycota(100);c__Saccharomycetes(100);o__Saccharomycetales(100);f__Metschnikowiaceae(100);g__Metschnikowia(100);s__Metschnikowia_pulcherrima(100); | Metschnikowia | 13 | Saprotroph | \|Undefined Saprotroph\| | Yeast | NULL | Probable | As nectar-tap_saprotroph (Põlme et al. 2020); Decay sugar-rich substrates (Põlme et al. 2020) | Põlme S, et al. 2020. Fungal Diversity 105:1-16 ((https://doi.org/10.1007/s13225-020-00466-2)) |
| Otu55426 | 1 | k__Fungi(100);p__Ascomycota(100);c__Saccharomycetes(100);o__Saccharomycetales(100);f__Metschnikowiaceae(100);g__Metschnikowia(100);s__Metschnikowia_pulcherrima(100); | Metschnikowia | 13 | Saprotroph | \|Undefined Saprotroph\| | Yeast | NULL | Probable | As nectar-tap_saprotroph (Põlme et al. 2020); Decay sugar-rich substrates (Põlme et al. 2020) | Põlme S, et al. 2020. Fungal Diversity 105:1-16 ((https://doi.org/10.1007/s13225-020-00466-2)) |
| Otu55427 | 1 | k__Fungi(100);p__Ascomycota(100);c__Saccharomycetes(100);o__Saccharomycetales(100);f__Metschnikowiaceae(100);g__Metschnikowia(100);s__Metschnikowia_pulcherrima(100); | Metschnikowia | 13 | Saprotroph | \|Undefined Saprotroph\| | Yeast | NULL | Probable | As nectar-tap_saprotroph (Põlme et al. 2020); Decay sugar-rich substrates (Põlme et al. 2020) | Põlme S, et al. 2020. Fungal Diversity 105:1-16 ((https://doi.org/10.1007/s13225-020-00466-2)) |
| Otu55428 | 1 | k__Fungi(100);p__Ascomycota(100);c__Saccharomycetes(100);o__Saccharomycetales(100);f__Metschnikowiaceae(100);g__Metschnikowia(100);s__Metschnikowia_pulcherrima(100); | Metschnikowia | 13 | Saprotroph | \|Undefined Saprotroph\| | Yeast | NULL | Probable | As nectar-tap_saprotroph (Põlme et al. 2020); Decay sugar-rich substrates (Põlme et al. 2020) | Põlme S, et al. 2020. Fungal Diversity 105:1-16 ((https://doi.org/10.1007/s13225-020-00466-2)) |
| Otu55429 | 1 | k__Fungi(100);p__Ascomycota(100);c__Saccharomycetes(100);o__Saccharomycetales(100);f__Metschnikowiaceae(100);g__Metschnikowia(100);s__Metschnikowia_pulcherrima(100); | Metschnikowia | 13 | Saprotroph | \|Undefined Saprotroph\| | Yeast | NULL | Probable | As nectar-tap_saprotroph (Põlme et al. 2020); Decay sugar-rich substrates (Põlme et al. 2020) | Põlme S, et al. 2020. Fungal Diversity 105:1-16 ((https://doi.org/10.1007/s13225-020-00466-2)) |
| Otu55430 | 1 | k__Fungi(100);p__Ascomycota(100);c__Saccharomycetes(100);o__Saccharomycetales(100);f__Metschnikowiaceae(100);g__Metschnikowia(100);s__Metschnikowia_pulcherrima(100); | Metschnikowia | 13 | Saprotroph | \|Undefined Saprotroph\| | Yeast | NULL | Probable | As nectar-tap_saprotroph (Põlme et al. 2020); Decay sugar-rich substrates (Põlme et al. 2020) | Põlme S, et al. 2020. Fungal Diversity 105:1-16 ((https://doi.org/10.1007/s13225-020-00466-2)) |
| Otu55431 | 1 | k__Fungi(100);p__Ascomycota(100);c__Saccharomycetes(100);o__Saccharomycetales(100);f__Metschnikowiaceae(100);g__Metschnikowia(100);s__Metschnikowia_pulcherrima(100); | Metschnikowia | 13 | Saprotroph | \|Undefined Saprotroph\| | Yeast | NULL | Probable | As nectar-tap_saprotroph (Põlme et al. 2020); Decay sugar-rich substrates (Põlme et al. 2020) | Põlme S, et al. 2020. Fungal Diversity 105:1-16 ((https://doi.org/10.1007/s13225-020-00466-2)) |
| Otu55432 | 1 | k__Fungi(100);p__Ascomycota(100);c__Saccharomycetes(100);o__Saccharomycetales(100);f__Metschnikowiaceae(100);g__Metschnikowia(100);g__Metschnikowia_unclassified(100); | Metschnikowia | 13 | Saprotroph | \|Undefined Saprotroph\| | Yeast | NULL | Probable | As nectar-tap_saprotroph (Põlme et al. 2020); Decay sugar-rich substrates (Põlme et al. 2020) | Põlme S, et al. 2020. Fungal Diversity 105:1-16 ((https://doi.org/10.1007/s13225-020-00466-2)) |
| Otu55433 | 1 | k__Fungi(100);p__Ascomycota(100);c__Saccharomycetes(100);o__Saccharomycetales(100);f__Metschnikowiaceae(100);g__Metschnikowia(100);s__Metschnikowia_pulcherrima(100); | Metschnikowia | 13 | Saprotroph | \|Undefined Saprotroph\| | Yeast | NULL | Probable | As nectar-tap_saprotroph (Põlme et al. 2020); Decay sugar-rich substrates (Põlme et al. 2020) | Põlme S, et al. 2020. Fungal Diversity 105:1-16 ((https://doi.org/10.1007/s13225-020-00466-2)) |
| Otu55434 | 1 | k__Fungi(100);p__Ascomycota(100);c__Saccharomycetes(100);o__Saccharomycetales(100);f__Metschnikowiaceae(100);g__Metschnikowia(100);g__Metschnikowia_unclassified(100); | Metschnikowia | 13 | Saprotroph | \|Undefined Saprotroph\| | Yeast | NULL | Probable | As nectar-tap_saprotroph (Põlme et al. 2020); Decay sugar-rich substrates (Põlme et al. 2020) | Põlme S, et al. 2020. Fungal Diversity 105:1-16 ((https://doi.org/10.1007/s13225-020-00466-2)) |
| Otu55435 | 1 | k__Fungi(100);p__Ascomycota(100);c__Saccharomycetes(100);o__Saccharomycetales(100);f__Metschnikowiaceae(100);g__Metschnikowia(100);s__Metschnikowia_pulcherrima(100); | Metschnikowia | 13 | Saprotroph | \|Undefined Saprotroph\| | Yeast | NULL | Probable | As nectar-tap_saprotroph (Põlme et al. 2020); Decay sugar-rich substrates (Põlme et al. 2020) | Põlme S, et al. 2020. Fungal Diversity 105:1-16 ((https://doi.org/10.1007/s13225-020-00466-2)) |
| Otu55436 | 1 | k__Fungi(100);p__Ascomycota(100);c__Saccharomycetes(100);o__Saccharomycetales(100);f__Metschnikowiaceae(100);g__Metschnikowia(100);g__Metschnikowia_unclassified(100); | Metschnikowia | 13 | Saprotroph | \|Undefined Saprotroph\| | Yeast | NULL | Probable | As nectar-tap_saprotroph (Põlme et al. 2020); Decay sugar-rich substrates (Põlme et al. 2020) | Põlme S, et al. 2020. Fungal Diversity 105:1-16 ((https://doi.org/10.1007/s13225-020-00466-2)) |
| Otu55437 | 1 | k__Fungi(100);p__Ascomycota(100);c__Saccharomycetes(100);o__Saccharomycetales(100);f__Metschnikowiaceae(100);g__Metschnikowia(100);s__Metschnikowia_pulcherrima(100); | Metschnikowia | 13 | Saprotroph | \|Undefined Saprotroph\| | Yeast | NULL | Probable | As nectar-tap_saprotroph (Põlme et al. 2020); Decay sugar-rich substrates (Põlme et al. 2020) | Põlme S, et al. 2020. Fungal Diversity 105:1-16 ((https://doi.org/10.1007/s13225-020-00466-2)) |
| Otu55438 | 1 | k__Fungi(100);p__Ascomycota(100);c__Saccharomycetes(100);o__Saccharomycetales(100);f__Metschnikowiaceae(100);g__Metschnikowia(100);s__Metschnikowia_pulcherrima(100); | Metschnikowia | 13 | Saprotroph | \|Undefined Saprotroph\| | Yeast | NULL | Probable | As nectar-tap_saprotroph (Põlme et al. 2020); Decay sugar-rich substrates (Põlme et al. 2020) | Põlme S, et al. 2020. Fungal Diversity 105:1-16 ((https://doi.org/10.1007/s13225-020-00466-2)) |
| Otu55439 | 1 | k__Fungi(100);p__Ascomycota(100);c__Saccharomycetes(100);o__Saccharomycetales(100);f__Metschnikowiaceae(100);g__Metschnikowia(100);s__Metschnikowia_pulcherrima(100); | Metschnikowia | 13 | Saprotroph | \|Undefined Saprotroph\| | Yeast | NULL | Probable | As nectar-tap_saprotroph (Põlme et al. 2020); Decay sugar-rich substrates (Põlme et al. 2020) | Põlme S, et al. 2020. Fungal Diversity 105:1-16 ((https://doi.org/10.1007/s13225-020-00466-2)) |
| Otu55440 | 1 | k__Fungi(100);p__Ascomycota(100);c__Saccharomycetes(100);o__Saccharomycetales(100);f__Metschnikowiaceae(100);g__Metschnikowia(100);s__Metschnikowia_pulcherrima(100); | Metschnikowia | 13 | Saprotroph | \|Undefined Saprotroph\| | Yeast | NULL | Probable | As nectar-tap_saprotroph (Põlme et al. 2020); Decay sugar-rich substrates (Põlme et al. 2020) | Põlme S, et al. 2020. Fungal Diversity 105:1-16 ((https://doi.org/10.1007/s13225-020-00466-2)) |
| Otu55441 | 1 | k__Fungi(100);p__Ascomycota(100);c__Saccharomycetes(100);o__Saccharomycetales(100);f__Metschnikowiaceae(100);g__Metschnikowia(100);g__Metschnikowia_unclassified(100); | Metschnikowia | 13 | Saprotroph | \|Undefined Saprotroph\| | Yeast | NULL | Probable | As nectar-tap_saprotroph (Põlme et al. 2020); Decay sugar-rich substrates (Põlme et al. 2020) | Põlme S, et al. 2020. Fungal Diversity 105:1-16 ((https://doi.org/10.1007/s13225-020-00466-2)) |
| Otu55442 | 1 | k__Fungi(100);p__Ascomycota(100);c__Saccharomycetes(100);o__Saccharomycetales(100);f__Metschnikowiaceae(100);g__Metschnikowia(100);s__Metschnikowia_pulcherrima(100); | Metschnikowia | 13 | Saprotroph | \|Undefined Saprotroph\| | Yeast | NULL | Probable | As nectar-tap_saprotroph (Põlme et al. 2020); Decay sugar-rich substrates (Põlme et al. 2020) | Põlme S, et al. 2020. Fungal Diversity 105:1-16 ((https://doi.org/10.1007/s13225-020-00466-2)) |
| Otu55443 | 1 | k__Fungi(100);p__Ascomycota(100);c__Saccharomycetes(100);o__Saccharomycetales(100);f__Metschnikowiaceae(100);g__Metschnikowia(100);s__Metschnikowia_pulcherrima(100); | Metschnikowia | 13 | Saprotroph | \|Undefined Saprotroph\| | Yeast | NULL | Probable | As nectar-tap_saprotroph (Põlme et al. 2020); Decay sugar-rich substrates (Põlme et al. 2020) | Põlme S, et al. 2020. Fungal Diversity 105:1-16 ((https://doi.org/10.1007/s13225-020-00466-2)) |
| Otu55444 | 1 | k__Fungi(100);p__Ascomycota(100);c__Saccharomycetes(100);o__Saccharomycetales(100);f__Metschnikowiaceae(100);g__Metschnikowia(100);s__Metschnikowia_pulcherrima(100); | Metschnikowia | 13 | Saprotroph | \|Undefined Saprotroph\| | Yeast | NULL | Probable | As nectar-tap_saprotroph (Põlme et al. 2020); Decay sugar-rich substrates (Põlme et al. 2020) | Põlme S, et al. 2020. Fungal Diversity 105:1-16 ((https://doi.org/10.1007/s13225-020-00466-2)) |
| Otu55445 | 1 | k__Fungi(100);p__Ascomycota(100);c__Saccharomycetes(100);o__Saccharomycetales(100);f__Metschnikowiaceae(100);g__Metschnikowia(100);s__Metschnikowia_pulcherrima(100); | Metschnikowia | 13 | Saprotroph | \|Undefined Saprotroph\| | Yeast | NULL | Probable | As nectar-tap_saprotroph (Põlme et al. 2020); Decay sugar-rich substrates (Põlme et al. 2020) | Põlme S, et al. 2020. Fungal Diversity 105:1-16 ((https://doi.org/10.1007/s13225-020-00466-2)) |
| Otu55446 | 1 | k__Fungi(100);p__Ascomycota(100);c__Saccharomycetes(100);o__Saccharomycetales(100);f__Metschnikowiaceae(100);g__Metschnikowia(100);s__Metschnikowia_pulcherrima(100); | Metschnikowia | 13 | Saprotroph | \|Undefined Saprotroph\| | Yeast | NULL | Probable | As nectar-tap_saprotroph (Põlme et al. 2020); Decay sugar-rich substrates (Põlme et al. 2020) | Põlme S, et al. 2020. Fungal Diversity 105:1-16 ((https://doi.org/10.1007/s13225-020-00466-2)) |
| Otu55447 | 1 | k__Fungi(100);p__Ascomycota(100);c__Saccharomycetes(100);o__Saccharomycetales(100);f__Metschnikowiaceae(100);g__Metschnikowia(100);s__Metschnikowia_pulcherrima(100); | Metschnikowia | 13 | Saprotroph | \|Undefined Saprotroph\| | Yeast | NULL | Probable | As nectar-tap_saprotroph (Põlme et al. 2020); Decay sugar-rich substrates (Põlme et al. 2020) | Põlme S, et al. 2020. Fungal Diversity 105:1-16 ((https://doi.org/10.1007/s13225-020-00466-2)) |
| Otu55448 | 1 | k__Fungi(100);p__Ascomycota(100);c__Saccharomycetes(100);o__Saccharomycetales(100);f__Metschnikowiaceae(100);g__Metschnikowia(100);s__Metschnikowia_pulcherrima(100); | Metschnikowia | 13 | Saprotroph | \|Undefined Saprotroph\| | Yeast | NULL | Probable | As nectar-tap_saprotroph (Põlme et al. 2020); Decay sugar-rich substrates (Põlme et al. 2020) | Põlme S, et al. 2020. Fungal Diversity 105:1-16 ((https://doi.org/10.1007/s13225-020-00466-2)) |
| Otu55449 | 1 | k__Fungi(100);p__Ascomycota(100);c__Saccharomycetes(100);o__Saccharomycetales(100);f__Metschnikowiaceae(100);g__Metschnikowia(100);s__Metschnikowia_pulcherrima(100); | Metschnikowia | 13 | Saprotroph | \|Undefined Saprotroph\| | Yeast | NULL | Probable | As nectar-tap_saprotroph (Põlme et al. 2020); Decay sugar-rich substrates (Põlme et al. 2020) | Põlme S, et al. 2020. Fungal Diversity 105:1-16 ((https://doi.org/10.1007/s13225-020-00466-2)) |
| Otu55451 | 1 | k__Fungi(100);p__Ascomycota(100);c__Saccharomycetes(100);o__Saccharomycetales(100);f__Metschnikowiaceae(100);g__Metschnikowia(100);g__Metschnikowia_unclassified(100); | Metschnikowia | 13 | Saprotroph | \|Undefined Saprotroph\| | Yeast | NULL | Probable | As nectar-tap_saprotroph (Põlme et al. 2020); Decay sugar-rich substrates (Põlme et al. 2020) | Põlme S, et al. 2020. Fungal Diversity 105:1-16 ((https://doi.org/10.1007/s13225-020-00466-2)) |
| Otu55452 | 1 | k__Fungi(100);p__Ascomycota(100);c__Saccharomycetes(100);o__Saccharomycetales(100);f__Metschnikowiaceae(100);g__Metschnikowia(100);s__Metschnikowia_pulcherrima(100); | Metschnikowia | 13 | Saprotroph | \|Undefined Saprotroph\| | Yeast | NULL | Probable | As nectar-tap_saprotroph (Põlme et al. 2020); Decay sugar-rich substrates (Põlme et al. 2020) | Põlme S, et al. 2020. Fungal Diversity 105:1-16 ((https://doi.org/10.1007/s13225-020-00466-2)) |
| Otu55453 | 1 | k__Fungi(100);p__Ascomycota(100);c__Saccharomycetes(100);o__Saccharomycetales(100);f__Metschnikowiaceae(100);g__Metschnikowia(100);g__Metschnikowia_unclassified(100); | Metschnikowia | 13 | Saprotroph | \|Undefined Saprotroph\| | Yeast | NULL | Probable | As nectar-tap_saprotroph (Põlme et al. 2020); Decay sugar-rich substrates (Põlme et al. 2020) | Põlme S, et al. 2020. Fungal Diversity 105:1-16 ((https://doi.org/10.1007/s13225-020-00466-2)) |
| Otu55454 | 1 | k__Fungi(100);p__Ascomycota(100);c__Saccharomycetes(100);o__Saccharomycetales(100);f__Metschnikowiaceae(100);g__Metschnikowia(100);s__Metschnikowia_pulcherrima(100); | Metschnikowia | 13 | Saprotroph | \|Undefined Saprotroph\| | Yeast | NULL | Probable | As nectar-tap_saprotroph (Põlme et al. 2020); Decay sugar-rich substrates (Põlme et al. 2020) | Põlme S, et al. 2020. Fungal Diversity 105:1-16 ((https://doi.org/10.1007/s13225-020-00466-2)) |
| Otu55455 | 1 | k__Fungi(100);p__Ascomycota(100);c__Saccharomycetes(100);o__Saccharomycetales(100);f__Metschnikowiaceae(100);g__Metschnikowia(100);s__Metschnikowia_pulcherrima(100); | Metschnikowia | 13 | Saprotroph | \|Undefined Saprotroph\| | Yeast | NULL | Probable | As nectar-tap_saprotroph (Põlme et al. 2020); Decay sugar-rich substrates (Põlme et al. 2020) | Põlme S, et al. 2020. Fungal Diversity 105:1-16 ((https://doi.org/10.1007/s13225-020-00466-2)) |
| Otu55456 | 1 | k__Fungi(100);p__Ascomycota(100);c__Saccharomycetes(100);o__Saccharomycetales(100);f__Metschnikowiaceae(100);g__Metschnikowia(100);g__Metschnikowia_unclassified(100); | Metschnikowia | 13 | Saprotroph | \|Undefined Saprotroph\| | Yeast | NULL | Probable | As nectar-tap_saprotroph (Põlme et al. 2020); Decay sugar-rich substrates (Põlme et al. 2020) | Põlme S, et al. 2020. Fungal Diversity 105:1-16 ((https://doi.org/10.1007/s13225-020-00466-2)) |
| Otu55457 | 1 | k__Fungi(100);p__Ascomycota(100);c__Saccharomycetes(100);o__Saccharomycetales(100);f__Metschnikowiaceae(100);g__Metschnikowia(100);g__Metschnikowia_unclassified(100); | Metschnikowia | 13 | Saprotroph | \|Undefined Saprotroph\| | Yeast | NULL | Probable | As nectar-tap_saprotroph (Põlme et al. 2020); Decay sugar-rich substrates (Põlme et al. 2020) | Põlme S, et al. 2020. Fungal Diversity 105:1-16 ((https://doi.org/10.1007/s13225-020-00466-2)) |
| Otu55458 | 1 | k__Fungi(100);p__Ascomycota(100);c__Saccharomycetes(100);o__Saccharomycetales(100);f__Metschnikowiaceae(100);g__Metschnikowia(100);s__Metschnikowia_pulcherrima(100); | Metschnikowia | 13 | Saprotroph | \|Undefined Saprotroph\| | Yeast | NULL | Probable | As nectar-tap_saprotroph (Põlme et al. 2020); Decay sugar-rich substrates (Põlme et al. 2020) | Põlme S, et al. 2020. Fungal Diversity 105:1-16 ((https://doi.org/10.1007/s13225-020-00466-2)) |
| Otu55459 | 1 | k__Fungi(100);p__Ascomycota(100);c__Saccharomycetes(100);o__Saccharomycetales(100);f__Metschnikowiaceae(100);g__Metschnikowia(100);s__Metschnikowia_pulcherrima(100); | Metschnikowia | 13 | Saprotroph | \|Undefined Saprotroph\| | Yeast | NULL | Probable | As nectar-tap_saprotroph (Põlme et al. 2020); Decay sugar-rich substrates (Põlme et al. 2020) | Põlme S, et al. 2020. Fungal Diversity 105:1-16 ((https://doi.org/10.1007/s13225-020-00466-2)) |
| Otu55460 | 1 | k__Fungi(100);p__Ascomycota(100);c__Saccharomycetes(100);o__Saccharomycetales(100);f__Metschnikowiaceae(100);g__Metschnikowia(100);s__Metschnikowia_pulcherrima(100); | Metschnikowia | 13 | Saprotroph | \|Undefined Saprotroph\| | Yeast | NULL | Probable | As nectar-tap_saprotroph (Põlme et al. 2020); Decay sugar-rich substrates (Põlme et al. 2020) | Põlme S, et al. 2020. Fungal Diversity 105:1-16 ((https://doi.org/10.1007/s13225-020-00466-2)) |
| Otu55461 | 1 | k__Fungi(100);p__Ascomycota(100);c__Saccharomycetes(100);o__Saccharomycetales(100);f__Metschnikowiaceae(100);g__Metschnikowia(100);s__Metschnikowia_pulcherrima(100); | Metschnikowia | 13 | Saprotroph | \|Undefined Saprotroph\| | Yeast | NULL | Probable | As nectar-tap_saprotroph (Põlme et al. 2020); Decay sugar-rich substrates (Põlme et al. 2020) | Põlme S, et al. 2020. Fungal Diversity 105:1-16 ((https://doi.org/10.1007/s13225-020-00466-2)) |
| Otu55462 | 1 | k__Fungi(100);p__Ascomycota(100);c__Saccharomycetes(100);o__Saccharomycetales(100);f__Metschnikowiaceae(100);g__Metschnikowia(100);s__Metschnikowia_pulcherrima(100); | Metschnikowia | 13 | Saprotroph | \|Undefined Saprotroph\| | Yeast | NULL | Probable | As nectar-tap_saprotroph (Põlme et al. 2020); Decay sugar-rich substrates (Põlme et al. 2020) | Põlme S, et al. 2020. Fungal Diversity 105:1-16 ((https://doi.org/10.1007/s13225-020-00466-2)) |
| Otu55463 | 1 | k__Fungi(100);p__Ascomycota(100);c__Saccharomycetes(100);o__Saccharomycetales(100);f__Metschnikowiaceae(100);g__Metschnikowia(100);s__Metschnikowia_pulcherrima(100); | Metschnikowia | 13 | Saprotroph | \|Undefined Saprotroph\| | Yeast | NULL | Probable | As nectar-tap_saprotroph (Põlme et al. 2020); Decay sugar-rich substrates (Põlme et al. 2020) | Põlme S, et al. 2020. Fungal Diversity 105:1-16 ((https://doi.org/10.1007/s13225-020-00466-2)) |
| Otu55464 | 1 | k__Fungi(100);p__Ascomycota(100);c__Saccharomycetes(100);o__Saccharomycetales(100);f__Metschnikowiaceae(100);g__Metschnikowia(100);s__Metschnikowia_pulcherrima(100); | Metschnikowia | 13 | Saprotroph | \|Undefined Saprotroph\| | Yeast | NULL | Probable | As nectar-tap_saprotroph (Põlme et al. 2020); Decay sugar-rich substrates (Põlme et al. 2020) | Põlme S, et al. 2020. Fungal Diversity 105:1-16 ((https://doi.org/10.1007/s13225-020-00466-2)) |
| Otu55465 | 1 | k__Fungi(100);p__Ascomycota(100);c__Saccharomycetes(100);o__Saccharomycetales(100);f__Metschnikowiaceae(100);g__Metschnikowia(100);g__Metschnikowia_unclassified(100); | Metschnikowia | 13 | Saprotroph | \|Undefined Saprotroph\| | Yeast | NULL | Probable | As nectar-tap_saprotroph (Põlme et al. 2020); Decay sugar-rich substrates (Põlme et al. 2020) | Põlme S, et al. 2020. Fungal Diversity 105:1-16 ((https://doi.org/10.1007/s13225-020-00466-2)) |
| Otu55466 | 1 | k__Fungi(100);p__Ascomycota(100);c__Saccharomycetes(100);o__Saccharomycetales(100);f__Metschnikowiaceae(100);g__Metschnikowia(100);s__Metschnikowia_pulcherrima(100); | Metschnikowia | 13 | Saprotroph | \|Undefined Saprotroph\| | Yeast | NULL | Probable | As nectar-tap_saprotroph (Põlme et al. 2020); Decay sugar-rich substrates (Põlme et al. 2020) | Põlme S, et al. 2020. Fungal Diversity 105:1-16 ((https://doi.org/10.1007/s13225-020-00466-2)) |
| Otu55467 | 1 | k__Fungi(100);p__Ascomycota(100);c__Saccharomycetes(100);o__Saccharomycetales(100);f__Metschnikowiaceae(100);g__Metschnikowia(100);s__Metschnikowia_pulcherrima(100); | Metschnikowia | 13 | Saprotroph | \|Undefined Saprotroph\| | Yeast | NULL | Probable | As nectar-tap_saprotroph (Põlme et al. 2020); Decay sugar-rich substrates (Põlme et al. 2020) | Põlme S, et al. 2020. Fungal Diversity 105:1-16 ((https://doi.org/10.1007/s13225-020-00466-2)) |
| Otu55468 | 1 | k__Fungi(100);p__Ascomycota(100);c__Saccharomycetes(100);o__Saccharomycetales(100);f__Metschnikowiaceae(100);g__Metschnikowia(100);s__Metschnikowia_pulcherrima(100); | Metschnikowia | 13 | Saprotroph | \|Undefined Saprotroph\| | Yeast | NULL | Probable | As nectar-tap_saprotroph (Põlme et al. 2020); Decay sugar-rich substrates (Põlme et al. 2020) | Põlme S, et al. 2020. Fungal Diversity 105:1-16 ((https://doi.org/10.1007/s13225-020-00466-2)) |
| Otu55469 | 1 | k__Fungi(100);p__Ascomycota(100);c__Saccharomycetes(100);o__Saccharomycetales(100);f__Metschnikowiaceae(100);g__Metschnikowia(100);s__Metschnikowia_pulcherrima(100); | Metschnikowia | 13 | Saprotroph | \|Undefined Saprotroph\| | Yeast | NULL | Probable | As nectar-tap_saprotroph (Põlme et al. 2020); Decay sugar-rich substrates (Põlme et al. 2020) | Põlme S, et al. 2020. Fungal Diversity 105:1-16 ((https://doi.org/10.1007/s13225-020-00466-2)) |
| Otu55470 | 1 | k__Fungi(100);p__Ascomycota(100);c__Saccharomycetes(100);o__Saccharomycetales(100);f__Metschnikowiaceae(100);g__Metschnikowia(100);s__Metschnikowia_pulcherrima(100); | Metschnikowia | 13 | Saprotroph | \|Undefined Saprotroph\| | Yeast | NULL | Probable | As nectar-tap_saprotroph (Põlme et al. 2020); Decay sugar-rich substrates (Põlme et al. 2020) | Põlme S, et al. 2020. Fungal Diversity 105:1-16 ((https://doi.org/10.1007/s13225-020-00466-2)) |

| | | | | | | | | | | | |
|---|---|---|---|---|---|---|---|---|---|---|---|
| Otu55471 | 1 | k__Fungi(100);p__Ascomycota(100);c__Saccharomycetes(100);o__Saccharomycetales(100);f__Metschnikowiaceae(100);g__Metschnikowia(100);g__Metschnikowia_unclassified(100); | Metschnikowia | 13 | Saprotroph | \|Undefined Saprotroph\| | Yeast | NULL | Probable | As nectar-tap_saprotroph (Põlme et al. 2020); Decay sugar-rich substrates (Põlme et al. 2020) | Põlme S, et al. 2020. Fungal Diversity 105:1-16 ((https://doi.org/10.1007/s13225-020-00466-2)) |
| Otu55472 | 1 | k__Fungi(100);p__Ascomycota(100);c__Saccharomycetes(100);o__Saccharomycetales(100);f__Metschnikowiaceae(100);g__Metschnikowia(100);g__Metschnikowia_unclassified(100); | Metschnikowia | 13 | Saprotroph | \|Undefined Saprotroph\| | Yeast | NULL | Probable | As nectar-tap_saprotroph (Põlme et al. 2020); Decay sugar-rich substrates (Põlme et al. 2020) | Põlme S, et al. 2020. Fungal Diversity 105:1-16 ((https://doi.org/10.1007/s13225-020-00466-2)) |
| Otu55473 | 1 | k__Fungi(100);p__Ascomycota(100);c__Saccharomycetes(100);o__Saccharomycetales(100);f__Metschnikowiaceae(100);g__Metschnikowia(100);s__Metschnikowia_pulcherrima(100); | Metschnikowia | 13 | Saprotroph | \|Undefined Saprotroph\| | Yeast | NULL | Probable | As nectar-tap_saprotroph (Põlme et al. 2020); Decay sugar-rich substrates (Põlme et al. 2020) | Põlme S, et al. 2020. Fungal Diversity 105:1-16 ((https://doi.org/10.1007/s13225-020-00466-2)) |
| Otu55474 | 1 | k__Fungi(100);p__Ascomycota(100);c__Saccharomycetes(100);o__Saccharomycetales(100);f__Metschnikowiaceae(100);g__Metschnikowia(100);s__Metschnikowia_pulcherrima(100); | Metschnikowia | 13 | Saprotroph | \|Undefined Saprotroph\| | Yeast | NULL | Probable | As nectar-tap_saprotroph (Põlme et al. 2020); Decay sugar-rich substrates (Põlme et al. 2020) | Põlme S, et al. 2020. Fungal Diversity 105:1-16 ((https://doi.org/10.1007/s13225-020-00466-2)) |
| Otu55475 | 1 | k__Fungi(100);p__Ascomycota(100);c__Saccharomycetes(100);o__Saccharomycetales(100);f__Metschnikowiaceae(100);g__Metschnikowia(100);s__Metschnikowia_pulcherrima(100); | Metschnikowia | 13 | Saprotroph | \|Undefined Saprotroph\| | Yeast | NULL | Probable | As nectar-tap_saprotroph (Põlme et al. 2020); Decay sugar-rich substrates (Põlme et al. 2020) | Põlme S, et al. 2020. Fungal Diversity 105:1-16 ((https://doi.org/10.1007/s13225-020-00466-2)) |
| Otu55476 | 1 | k__Fungi(100);p__Ascomycota(100);c__Saccharomycetes(100);o__Saccharomycetales(100);f__Metschnikowiaceae(100);g__Metschnikowia(100);s__Metschnikowia_pulcherrima(100); | Metschnikowia | 13 | Saprotroph | \|Undefined Saprotroph\| | Yeast | NULL | Probable | As nectar-tap_saprotroph (Põlme et al. 2020); Decay sugar-rich substrates (Põlme et al. 2020) | Põlme S, et al. 2020. Fungal Diversity 105:1-16 ((https://doi.org/10.1007/s13225-020-00466-2)) |
| Otu55477 | 1 | k__Fungi(100);p__Ascomycota(100);c__Saccharomycetes(100);o__Saccharomycetales(100);f__Metschnikowiaceae(100);g__Metschnikowia(100);s__Metschnikowia_pulcherrima(100); | Metschnikowia | 13 | Saprotroph | \|Undefined Saprotroph\| | Yeast | NULL | Probable | As nectar-tap_saprotroph (Põlme et al. 2020); Decay sugar-rich substrates (Põlme et al. 2020) | Põlme S, et al. 2020. Fungal Diversity 105:1-16 ((https://doi.org/10.1007/s13225-020-00466-2)) |
| Otu55478 | 1 | k__Fungi(100);p__Ascomycota(100);c__Saccharomycetes(100);o__Saccharomycetales(100);f__Metschnikowiaceae(100);g__Metschnikowia(100);g__Metschnikowia_unclassified(100); | Metschnikowia | 13 | Saprotroph | \|Undefined Saprotroph\| | Yeast | NULL | Probable | As nectar-tap_saprotroph (Põlme et al. 2020); Decay sugar-rich substrates (Põlme et al. 2020) | Põlme S, et al. 2020. Fungal Diversity 105:1-16 ((https://doi.org/10.1007/s13225-020-00466-2)) |
| Otu55479 | 1 | k__Fungi(100);p__Ascomycota(100);c__Saccharomycetes(100);o__Saccharomycetales(100);f__Metschnikowiaceae(100);g__Metschnikowia(100);s__Metschnikowia_pulcherrima(100); | Metschnikowia | 13 | Saprotroph | \|Undefined Saprotroph\| | Yeast | NULL | Probable | As nectar-tap_saprotroph (Põlme et al. 2020); Decay sugar-rich substrates (Põlme et al. 2020) | Põlme S, et al. 2020. Fungal Diversity 105:1-16 ((https://doi.org/10.1007/s13225-020-00466-2)) |
| Otu55481 | 1 | k__Fungi(100);p__Ascomycota(100);c__Saccharomycetes(100);o__Saccharomycetales(100);f__Metschnikowiaceae(100);g__Metschnikowia(100);s__Metschnikowia_pulcherrima(100); | Metschnikowia | 13 | Saprotroph | \|Undefined Saprotroph\| | Yeast | NULL | Probable | As nectar-tap_saprotroph (Põlme et al. 2020); Decay sugar-rich substrates (Põlme et al. 2020) | Põlme S, et al. 2020. Fungal Diversity 105:1-16 ((https://doi.org/10.1007/s13225-020-00466-2)) |
| Otu55482 | 1 | k__Fungi(100);p__Ascomycota(100);c__Saccharomycetes(100);o__Saccharomycetales(100);f__Metschnikowiaceae(100);g__Metschnikowia(100);s__Metschnikowia_pulcherrima(100); | Metschnikowia | 13 | Saprotroph | \|Undefined Saprotroph\| | Yeast | NULL | Probable | As nectar-tap_saprotroph (Põlme et al. 2020); Decay sugar-rich substrates (Põlme et al. 2020) | Põlme S, et al. 2020. Fungal Diversity 105:1-16 ((https://doi.org/10.1007/s13225-020-00466-2)) |
| Otu55483 | 1 | k__Fungi(100);p__Ascomycota(100);c__Saccharomycetes(100);o__Saccharomycetales(100);f__Metschnikowiaceae(100);g__Metschnikowia(100);s__Metschnikowia_pulcherrima(100); | Metschnikowia | 13 | Saprotroph | \|Undefined Saprotroph\| | Yeast | NULL | Probable | As nectar-tap_saprotroph (Põlme et al. 2020); Decay sugar-rich substrates (Põlme et al. 2020) | Põlme S, et al. 2020. Fungal Diversity 105:1-16 ((https://doi.org/10.1007/s13225-020-00466-2)) |
| Otu55484 | 1 | k__Fungi(100);p__Ascomycota(100);c__Saccharomycetes(100);o__Saccharomycetales(100);f__Metschnikowiaceae(100);g__Metschnikowia(100);s__Metschnikowia_pulcherrima(100); | Metschnikowia | 13 | Saprotroph | \|Undefined Saprotroph\| | Yeast | NULL | Probable | As nectar-tap_saprotroph (Põlme et al. 2020); Decay sugar-rich substrates (Põlme et al. 2020) | Põlme S, et al. 2020. Fungal Diversity 105:1-16 ((https://doi.org/10.1007/s13225-020-00466-2)) |
| Otu55485 | 1 | k__Fungi(100);p__Ascomycota(100);c__Saccharomycetes(100);o__Saccharomycetales(100);f__Metschnikowiaceae(100);g__Metschnikowia(100);s__Metschnikowia_pulcherrima(100); | Metschnikowia | 13 | Saprotroph | \|Undefined Saprotroph\| | Yeast | NULL | Probable | As nectar-tap_saprotroph (Põlme et al. 2020); Decay sugar-rich substrates (Põlme et al. 2020) | Põlme S, et al. 2020. Fungal Diversity 105:1-16 ((https://doi.org/10.1007/s13225-020-00466-2)) |
| Otu55487 | 1 | k__Fungi(100);p__Ascomycota(100);c__Saccharomycetes(100);o__Saccharomycetales(100);f__Metschnikowiaceae(100);g__Metschnikowia(100);s__Metschnikowia_pulcherrima(100); | Metschnikowia | 13 | Saprotroph | \|Undefined Saprotroph\| | Yeast | NULL | Probable | As nectar-tap_saprotroph (Põlme et al. 2020); Decay sugar-rich substrates (Põlme et al. 2020) | Põlme S, et al. 2020. Fungal Diversity 105:1-16 ((https://doi.org/10.1007/s13225-020-00466-2)) |
| Otu55488 | 1 | k__Fungi(100);p__Ascomycota(100);c__Saccharomycetes(100);o__Saccharomycetales(100);f__Metschnikowiaceae(100);g__Metschnikowia(100);g__Metschnikowia_unclassified(100); | Metschnikowia | 13 | Saprotroph | \|Undefined Saprotroph\| | Yeast | NULL | Probable | As nectar-tap_saprotroph (Põlme et al. 2020); Decay sugar-rich substrates (Põlme et al. 2020) | Põlme S, et al. 2020. Fungal Diversity 105:1-16 ((https://doi.org/10.1007/s13225-020-00466-2)) |
| Otu55489 | 1 | k__Fungi(100);p__Ascomycota(100);c__Saccharomycetes(100);o__Saccharomycetales(100);f__Metschnikowiaceae(100);g__Metschnikowia(100);s__Metschnikowia_pulcherrima(100); | Metschnikowia | 13 | Saprotroph | \|Undefined Saprotroph\| | Yeast | NULL | Probable | As nectar-tap_saprotroph (Põlme et al. 2020); Decay sugar-rich substrates (Põlme et al. 2020) | Põlme S, et al. 2020. Fungal Diversity 105:1-16 ((https://doi.org/10.1007/s13225-020-00466-2)) |
| Otu55490 | 1 | k__Fungi(100);p__Ascomycota(100);c__Saccharomycetes(100);o__Saccharomycetales(100);f__Metschnikowiaceae(100);g__Metschnikowia(100);s__Metschnikowia_pulcherrima(100); | Metschnikowia | 13 | Saprotroph | \|Undefined Saprotroph\| | Yeast | NULL | Probable | As nectar-tap_saprotroph (Põlme et al. 2020); Decay sugar-rich substrates (Põlme et al. 2020) | Põlme S, et al. 2020. Fungal Diversity 105:1-16 ((https://doi.org/10.1007/s13225-020-00466-2)) |
| Otu55491 | 1 | k__Fungi(100);p__Ascomycota(100);c__Saccharomycetes(100);o__Saccharomycetales(100);f__Metschnikowiaceae(100);g__Metschnikowia(100);s__Metschnikowia_pulcherrima(100); | Metschnikowia | 13 | Saprotroph | \|Undefined Saprotroph\| | Yeast | NULL | Probable | As nectar-tap_saprotroph (Põlme et al. 2020); Decay sugar-rich substrates (Põlme et al. 2020) | Põlme S, et al. 2020. Fungal Diversity 105:1-16 ((https://doi.org/10.1007/s13225-020-00466-2)) |
| Otu55492 | 1 | k__Fungi(100);p__Ascomycota(100);c__Saccharomycetes(100);o__Saccharomycetales(100);f__Metschnikowiaceae(100);g__Metschnikowia(100);s__Metschnikowia_pulcherrima(100); | Metschnikowia | 13 | Saprotroph | \|Undefined Saprotroph\| | Yeast | NULL | Probable | As nectar-tap_saprotroph (Põlme et al. 2020); Decay sugar-rich substrates (Põlme et al. 2020) | Põlme S, et al. 2020. Fungal Diversity 105:1-16 ((https://doi.org/10.1007/s13225-020-00466-2)) |
| Otu55493 | 1 | k__Fungi(100);p__Ascomycota(100);c__Saccharomycetes(100);o__Saccharomycetales(100);f__Metschnikowiaceae(100);g__Metschnikowia(100);s__Metschnikowia_pulcherrima(100); | Metschnikowia | 13 | Saprotroph | \|Undefined Saprotroph\| | Yeast | NULL | Probable | As nectar-tap_saprotroph (Põlme et al. 2020); Decay sugar-rich substrates (Põlme et al. 2020) | Põlme S, et al. 2020. Fungal Diversity 105:1-16 ((https://doi.org/10.1007/s13225-020-00466-2)) |
| Otu55494 | 1 | k__Fungi(100);p__Ascomycota(100);c__Saccharomycetes(100);o__Saccharomycetales(100);f__Metschnikowiaceae(100);g__Metschnikowia(100);s__Metschnikowia_pulcherrima(100); | Metschnikowia | 13 | Saprotroph | \|Undefined Saprotroph\| | Yeast | NULL | Probable | As nectar-tap_saprotroph (Põlme et al. 2020); Decay sugar-rich substrates (Põlme et al. 2020) | Põlme S, et al. 2020. Fungal Diversity 105:1-16 ((https://doi.org/10.1007/s13225-020-00466-2)) |
| Otu55495 | 1 | k__Fungi(100);p__Ascomycota(100);c__Saccharomycetes(100);o__Saccharomycetales(100);f__Metschnikowiaceae(100);g__Metschnikowia(100);s__Metschnikowia_pulcherrima(100); | Metschnikowia | 13 | Saprotroph | \|Undefined Saprotroph\| | Yeast | NULL | Probable | As nectar-tap_saprotroph (Põlme et al. 2020); Decay sugar-rich substrates (Põlme et al. 2020) | Põlme S, et al. 2020. Fungal Diversity 105:1-16 ((https://doi.org/10.1007/s13225-020-00466-2)) |
| Otu55497 | 1 | k__Fungi(100);p__Ascomycota(100);c__Saccharomycetes(100);o__Saccharomycetales(100);f__Metschnikowiaceae(100);g__Metschnikowia(100);s__Metschnikowia_pulcherrima(100); | Metschnikowia | 13 | Saprotroph | \|Undefined Saprotroph\| | Yeast | NULL | Probable | As nectar-tap_saprotroph (Põlme et al. 2020); Decay sugar-rich substrates (Põlme et al. 2020) | Põlme S, et al. 2020. Fungal Diversity 105:1-16 ((https://doi.org/10.1007/s13225-020-00466-2)) |
| Otu55498 | 1 | k__Fungi(100);p__Ascomycota(100);c__Saccharomycetes(100);o__Saccharomycetales(100);f__Metschnikowiaceae(100);g__Metschnikowia(100);s__Metschnikowia_pulcherrima(100); | Metschnikowia | 13 | Saprotroph | \|Undefined Saprotroph\| | Yeast | NULL | Probable | As nectar-tap_saprotroph (Põlme et al. 2020); Decay sugar-rich substrates (Põlme et al. 2020) | Põlme S, et al. 2020. Fungal Diversity 105:1-16 ((https://doi.org/10.1007/s13225-020-00466-2)) |
| Otu55507 | 1 | k__Fungi(100);p__Ascomycota(100);c__Saccharomycetes(100);o__Saccharomycetales(100);f__Metschnikowiaceae(100);g__Metschnikowia(100);s__Metschnikowia_pulcherrima(100); | Metschnikowia | 13 | Saprotroph | \|Undefined Saprotroph\| | Yeast | NULL | Probable | As nectar-tap_saprotroph (Põlme et al. 2020); Decay sugar-rich substrates (Põlme et al. 2020) | Põlme S, et al. 2020. Fungal Diversity 105:1-16 ((https://doi.org/10.1007/s13225-020-00466-2)) |
| Otu55509 | 1 | k__Fungi(100);p__Ascomycota(100);c__Saccharomycetes(100);o__Saccharomycetales(100);f__Metschnikowiaceae(100);g__Metschnikowia(100);s__Metschnikowia_pulcherrima(100); | Metschnikowia | 13 | Saprotroph | \|Undefined Saprotroph\| | Yeast | NULL | Probable | As nectar-tap_saprotroph (Põlme et al. 2020); Decay sugar-rich substrates (Põlme et al. 2020) | Põlme S, et al. 2020. Fungal Diversity 105:1-16 ((https://doi.org/10.1007/s13225-020-00466-2)) |
| Otu55510 | 1 | k__Fungi(100);p__Ascomycota(100);c__Saccharomycetes(100);o__Saccharomycetales(100);f__Metschnikowiaceae(100);g__Metschnikowia(100);g__Metschnikowia_unclassified(100); | Metschnikowia | 13 | Saprotroph | \|Undefined Saprotroph\| | Yeast | NULL | Probable | As nectar-tap_saprotroph (Põlme et al. 2020); Decay sugar-rich substrates (Põlme et al. 2020) | Põlme S, et al. 2020. Fungal Diversity 105:1-16 ((https://doi.org/10.1007/s13225-020-00466-2)) |
| Otu55511 | 1 | k__Fungi(100);p__Ascomycota(100);c__Saccharomycetes(100);o__Saccharomycetales(100);f__Metschnikowiaceae(100);g__Metschnikowia(100);g__Metschnikowia_unclassified(100); | Metschnikowia | 13 | Saprotroph | \|Undefined Saprotroph\| | Yeast | NULL | Probable | As nectar-tap_saprotroph (Põlme et al. 2020); Decay sugar-rich substrates (Põlme et al. 2020) | Põlme S, et al. 2020. Fungal Diversity 105:1-16 ((https://doi.org/10.1007/s13225-020-00466-2)) |
| Otu55512 | 1 | k__Fungi(100);p__Ascomycota(100);c__Saccharomycetes(100);o__Saccharomycetales(100);f__Metschnikowiaceae(100);g__Metschnikowia(100);g__Metschnikowia_unclassified(100); | Metschnikowia | 13 | Saprotroph | \|Undefined Saprotroph\| | Yeast | NULL | Probable | As nectar-tap_saprotroph (Põlme et al. 2020); Decay sugar-rich substrates (Põlme et al. 2020) | Põlme S, et al. 2020. Fungal Diversity 105:1-16 ((https://doi.org/10.1007/s13225-020-00466-2)) |
| Otu55513 | 1 | k__Fungi(100);p__Ascomycota(100);c__Saccharomycetes(100);o__Saccharomycetales(100);f__Metschnikowiaceae(100);g__Metschnikowia(100);g__Metschnikowia_unclassified(100); | Metschnikowia | 13 | Saprotroph | \|Undefined Saprotroph\| | Yeast | NULL | Probable | As nectar-tap_saprotroph (Põlme et al. 2020); Decay sugar-rich substrates (Põlme et al. 2020) | Põlme S, et al. 2020. Fungal Diversity 105:1-16 ((https://doi.org/10.1007/s13225-020-00466-2)) |
| Otu55514 | 1 | k__Fungi(100);p__Ascomycota(100);c__Saccharomycetes(100);o__Saccharomycetales(100);f__Metschnikowiaceae(100);g__Metschnikowia(100);s__Metschnikowia_pulcherrima(100); | Metschnikowia | 13 | Saprotroph | \|Undefined Saprotroph\| | Yeast | NULL | Probable | As nectar-tap_saprotroph (Põlme et al. 2020); Decay sugar-rich substrates (Põlme et al. 2020) | Põlme S, et al. 2020. Fungal Diversity 105:1-16 ((https://doi.org/10.1007/s13225-020-00466-2)) |
| Otu55515 | 1 | k__Fungi(100);p__Ascomycota(100);c__Saccharomycetes(100);o__Saccharomycetales(100);f__Metschnikowiaceae(100);g__Metschnikowia(100);s__Metschnikowia_pulcherrima(100); | Metschnikowia | 13 | Saprotroph | \|Undefined Saprotroph\| | Yeast | NULL | Probable | As nectar-tap_saprotroph (Põlme et al. 2020); Decay sugar-rich substrates (Põlme et al. 2020) | Põlme S, et al. 2020. Fungal Diversity 105:1-16 ((https://doi.org/10.1007/s13225-020-00466-2)) |
| Otu55517 | 1 | k__Fungi(100);p__Ascomycota(100);c__Saccharomycetes(100);o__Saccharomycetales(100);f__Metschnikowiaceae(100);g__Metschnikowia(100);s__Metschnikowia_pulcherrima(100); | Metschnikowia | 13 | Saprotroph | \|Undefined Saprotroph\| | Yeast | NULL | Probable | As nectar-tap_saprotroph (Põlme et al. 2020); Decay sugar-rich substrates (Põlme et al. 2020) | Põlme S, et al. 2020. Fungal Diversity 105:1-16 ((https://doi.org/10.1007/s13225-020-00466-2)) |
| Otu55518 | 1 | k__Fungi(100);p__Ascomycota(100);c__Saccharomycetes(100);o__Saccharomycetales(100);f__Metschnikowiaceae(100);g__Metschnikowia(100);g__Metschnikowia_unclassified(100); | Metschnikowia | 13 | Saprotroph | \|Undefined Saprotroph\| | Yeast | NULL | Probable | As nectar-tap_saprotroph (Põlme et al. 2020); Decay sugar-rich substrates (Põlme et al. 2020) | Põlme S, et al. 2020. Fungal Diversity 105:1-16 ((https://doi.org/10.1007/s13225-020-00466-2)) |
| Otu55519 | 1 | k__Fungi(100);p__Ascomycota(100);c__Saccharomycetes(100);o__Saccharomycetales(100);f__Metschnikowiaceae(100);g__Metschnikowia(100);s__Metschnikowia_pulcherrima(100); | Metschnikowia | 13 | Saprotroph | \|Undefined Saprotroph\| | Yeast | NULL | Probable | As nectar-tap_saprotroph (Põlme et al. 2020); Decay sugar-rich substrates (Põlme et al. 2020) | Põlme S, et al. 2020. Fungal Diversity 105:1-16 ((https://doi.org/10.1007/s13225-020-00466-2)) |
| Otu55520 | 1 | k__Fungi(100);p__Ascomycota(100);c__Saccharomycetes(100);o__Saccharomycetales(100);f__Metschnikowiaceae(100);g__Metschnikowia(100);g__Metschnikowia_unclassified(100); | Metschnikowia | 13 | Saprotroph | \|Undefined Saprotroph\| | Yeast | NULL | Probable | As nectar-tap_saprotroph (Põlme et al. 2020); Decay sugar-rich substrates (Põlme et al. 2020) | Põlme S, et al. 2020. Fungal Diversity 105:1-16 ((https://doi.org/10.1007/s13225-020-00466-2)) |
| Otu55522 | 1 | k__Fungi(100);p__Ascomycota(100);c__Saccharomycetes(100);o__Saccharomycetales(100);f__Metschnikowiaceae(100);g__Metschnikowia(100);g__Metschnikowia_unclassified(100); | Metschnikowia | 13 | Saprotroph | \|Undefined Saprotroph\| | Yeast | NULL | Probable | As nectar-tap_saprotroph (Põlme et al. 2020); Decay sugar-rich substrates (Põlme et al. 2020) | Põlme S, et al. 2020. Fungal Diversity 105:1-16 ((https://doi.org/10.1007/s13225-020-00466-2)) |
| Otu55523 | 1 | k__Fungi(100);p__Ascomycota(100);c__Saccharomycetes(100);o__Saccharomycetales(100);f__Metschnikowiaceae(100);g__Metschnikowia(100);s__Metschnikowia_pulcherrima(100); | Metschnikowia | 13 | Saprotroph | \|Undefined Saprotroph\| | Yeast | NULL | Probable | As nectar-tap_saprotroph (Põlme et al. 2020); Decay sugar-rich substrates (Põlme et al. 2020) | Põlme S, et al. 2020. Fungal Diversity 105:1-16 ((https://doi.org/10.1007/s13225-020-00466-2)) |
| Otu55524 | 1 | k__Fungi(100);p__Ascomycota(100);c__Saccharomycetes(100);o__Saccharomycetales(100);f__Metschnikowiaceae(100);g__Metschnikowia(100);s__Metschnikowia_pulcherrima(100); | Metschnikowia | 13 | Saprotroph | \|Undefined Saprotroph\| | Yeast | NULL | Probable | As nectar-tap_saprotroph (Põlme et al. 2020); Decay sugar-rich substrates (Põlme et al. 2020) | Põlme S, et al. 2020. Fungal Diversity 105:1-16 ((https://doi.org/10.1007/s13225-020-00466-2)) |
| Otu55525 | 1 | k__Fungi(100);p__Ascomycota(100);c__Saccharomycetes(100);o__Saccharomycetales(100);f__Metschnikowiaceae(100);g__Metschnikowia(100);s__Metschnikowia_pulcherrima(100); | Metschnikowia | 13 | Saprotroph | \|Undefined Saprotroph\| | Yeast | NULL | Probable | As nectar-tap_saprotroph (Põlme et al. 2020); Decay sugar-rich substrates (Põlme et al. 2020) | Põlme S, et al. 2020. Fungal Diversity 105:1-16 ((https://doi.org/10.1007/s13225-020-00466-2)) |
| Otu55527 | 1 | k__Fungi(100);p__Ascomycota(100);c__Saccharomycetes(100);o__Saccharomycetales(100);f__Metschnikowiaceae(100);g__Metschnikowia(100);s__Metschnikowia_pulcherrima(100); | Metschnikowia | 13 | Saprotroph | \|Undefined Saprotroph\| | Yeast | NULL | Probable | As nectar-tap_saprotroph (Põlme et al. 2020); Decay sugar-rich substrates (Põlme et al. 2020) | Põlme S, et al. 2020. Fungal Diversity 105:1-16 ((https://doi.org/10.1007/s13225-020-00466-2)) |
| Otu55528 | 1 | k__Fungi(100);p__Ascomycota(100);c__Saccharomycetes(100);o__Saccharomycetales(100);f__Metschnikowiaceae(100);g__Metschnikowia(100);g__Metschnikowia_unclassified(100); | Metschnikowia | 13 | Saprotroph | \|Undefined Saprotroph\| | Yeast | NULL | Probable | As nectar-tap_saprotroph (Põlme et al. 2020); Decay sugar-rich substrates (Põlme et al. 2020) | Põlme S, et al. 2020. Fungal Diversity 105:1-16 ((https://doi.org/10.1007/s13225-020-00466-2)) |
| Otu55529 | 1 | k__Fungi(100);p__Ascomycota(100);c__Saccharomycetes(100);o__Saccharomycetales(100);f__Metschnikowiaceae(100);g__Metschnikowia(100);g__Metschnikowia_unclassified(100); | Metschnikowia | 13 | Saprotroph | \|Undefined Saprotroph\| | Yeast | NULL | Probable | As nectar-tap_saprotroph (Põlme et al. 2020); Decay sugar-rich substrates (Põlme et al. 2020) | Põlme S, et al. 2020. Fungal Diversity 105:1-16 ((https://doi.org/10.1007/s13225-020-00466-2)) |
| Otu55530 | 1 | k__Fungi(100);p__Ascomycota(100);c__Saccharomycetes(100);o__Saccharomycetales(100);f__Metschnikowiaceae(100);g__Metschnikowia(100);g__Metschnikowia_unclassified(100); | Metschnikowia | 13 | Saprotroph | \|Undefined Saprotroph\| | Yeast | NULL | Probable | As nectar-tap_saprotroph (Põlme et al. 2020); Decay sugar-rich substrates (Põlme et al. 2020) | Põlme S, et al. 2020. Fungal Diversity 105:1-16 ((https://doi.org/10.1007/s13225-020-00466-2)) |
| Otu55531 | 1 | k__Fungi(100);p__Ascomycota(100);c__Saccharomycetes(100);o__Saccharomycetales(100);f__Metschnikowiaceae(100);g__Metschnikowia(100);s__Metschnikowia_pulcherrima(100); | Metschnikowia | 13 | Saprotroph | \|Undefined Saprotroph\| | Yeast | NULL | Probable | As nectar-tap_saprotroph (Põlme et al. 2020); Decay sugar-rich substrates (Põlme et al. 2020) | Põlme S, et al. 2020. Fungal Diversity 105:1-16 ((https://doi.org/10.1007/s13225-020-00466-2)) |
| Otu55532 | 1 | k__Fungi(100);p__Ascomycota(100);c__Saccharomycetes(100);o__Saccharomycetales(100);f__Metschnikowiaceae(100);g__Metschnikowia(100);s__Metschnikowia_pulcherrima(100); | Metschnikowia | 13 | Saprotroph | \|Undefined Saprotroph\| | Yeast | NULL | Probable | As nectar-tap_saprotroph (Põlme et al. 2020); Decay sugar-rich substrates (Põlme et al. 2020) | Põlme S, et al. 2020. Fungal Diversity 105:1-16 ((https://doi.org/10.1007/s13225-020-00466-2)) |
| Otu55533 | 1 | k__Fungi(100);p__Ascomycota(100);c__Saccharomycetes(100);o__Saccharomycetales(100);f__Metschnikowiaceae(100);g__Metschnikowia(100);s__Metschnikowia_pulcherrima(100); | Metschnikowia | 13 | Saprotroph | \|Undefined Saprotroph\| | Yeast | NULL | Probable | As nectar-tap_saprotroph (Põlme et al. 2020); Decay sugar-rich substrates (Põlme et al. 2020) | Põlme S, et al. 2020. Fungal Diversity 105:1-16 ((https://doi.org/10.1007/s13225-020-00466-2)) |
| Otu55534 | 1 | k__Fungi(100);p__Ascomycota(100);c__Saccharomycetes(100);o__Saccharomycetales(100);f__Metschnikowiaceae(100);g__Metschnikowia(100);s__Metschnikowia_pulcherrima(100); | Metschnikowia | 13 | Saprotroph | \|Undefined Saprotroph\| | Yeast | NULL | Probable | As nectar-tap_saprotroph (Põlme et al. 2020); Decay sugar-rich substrates (Põlme et al. 2020) | Põlme S, et al. 2020. Fungal Diversity 105:1-16 ((https://doi.org/10.1007/s13225-020-00466-2)) |
| Otu55536 | 1 | k__Fungi(100);p__Ascomycota(100);c__Saccharomycetes(100);o__Saccharomycetales(100);f__Metschnikowiaceae(100);g__Metschnikowia(100);s__Metschnikowia_pulcherrima(100); | Metschnikowia | 13 | Saprotroph | \|Undefined Saprotroph\| | Yeast | NULL | Probable | As nectar-tap_saprotroph (Põlme et al. 2020); Decay sugar-rich substrates (Põlme et al. 2020) | Põlme S, et al. 2020. Fungal Diversity 105:1-16 ((https://doi.org/10.1007/s13225-020-00466-2)) |
| Otu55537 | 1 | k__Fungi(100);p__Ascomycota(100);c__Saccharomycetes(100);o__Saccharomycetales(100);f__Metschnikowiaceae(100);g__Metschnikowia(100);s__Metschnikowia_pulcherrima(100); | Metschnikowia | 13 | Saprotroph | \|Undefined Saprotroph\| | Yeast | NULL | Probable | As nectar-tap_saprotroph (Põlme et al. 2020); Decay sugar-rich substrates (Põlme et al. 2020) | Põlme S, et al. 2020. Fungal Diversity 105:1-16 ((https://doi.org/10.1007/s13225-020-00466-2)) |
| Otu55538 | 1 | k__Fungi(100);p__Ascomycota(100);c__Saccharomycetes(100);o__Saccharomycetales(100);f__Metschnikowiaceae(100);g__Metschnikowia(100);g__Metschnikowia_unclassified(100); | Metschnikowia | 13 | Saprotroph | \|Undefined Saprotroph\| | Yeast | NULL | Probable | As nectar-tap_saprotroph (Põlme et al. 2020); Decay sugar-rich substrates (Põlme et al. 2020) | Põlme S, et al. 2020. Fungal Diversity 105:1-16 ((https://doi.org/10.1007/s13225-020-00466-2)) |
| Otu55539 | 1 | k__Fungi(100);p__Ascomycota(100);c__Saccharomycetes(100);o__Saccharomycetales(100);f__Metschnikowiaceae(100);g__Metschnikowia(100);s__Metschnikowia_pulcherrima(100); | Metschnikowia | 13 | Saprotroph | \|Undefined Saprotroph\| | Yeast | NULL | Probable | As nectar-tap_saprotroph (Põlme et al. 2020); Decay sugar-rich substrates (Põlme et al. 2020) | Põlme S, et al. 2020. Fungal Diversity 105:1-16 ((https://doi.org/10.1007/s13225-020-00466-2)) |
| Otu55540 | 1 | k__Fungi(100);p__Ascomycota(100);c__Saccharomycetes(100);o__Saccharomycetales(100);f__Metschnikowiaceae(100);g__Metschnikowia(100);s__Metschnikowia_pulcherrima(100); | Metschnikowia | 13 | Saprotroph | \|Undefined Saprotroph\| | Yeast | NULL | Probable | As nectar-tap_saprotroph (Põlme et al. 2020); Decay sugar-rich substrates (Põlme et al. 2020) | Põlme S, et al. 2020. Fungal Diversity 105:1-16 ((https://doi.org/10.1007/s13225-020-00466-2)) |
| Otu55541 | 1 | k__Fungi(100);p__Ascomycota(100);c__Saccharomycetes(100);o__Saccharomycetales(100);f__Metschnikowiaceae(100);g__Metschnikowia(100);g__Metschnikowia_unclassified(100); | Metschnikowia | 13 | Saprotroph | \|Undefined Saprotroph\| | Yeast | NULL | Probable | As nectar-tap_saprotroph (Põlme et al. 2020); Decay sugar-rich substrates (Põlme et al. 2020) | Põlme S, et al. 2020. Fungal Diversity 105:1-16 ((https://doi.org/10.1007/s13225-020-00466-2)) |
| Otu55542 | 1 | k__Fungi(100);p__Ascomycota(100);c__Saccharomycetes(100);o__Saccharomycetales(100);f__Metschnikowiaceae(100);g__Metschnikowia(100);g__Metschnikowia_unclassified(100); | Metschnikowia | 13 | Saprotroph | \|Undefined Saprotroph\| | Yeast | NULL | Probable | As nectar-tap_saprotroph (Põlme et al. 2020); Decay sugar-rich substrates (Põlme et al. 2020) | Põlme S, et al. 2020. Fungal Diversity 105:1-16 ((https://doi.org/10.1007/s13225-020-00466-2)) |
| Otu55545 | 1 | k__Fungi(100);p__Ascomycota(100);c__Saccharomycetes(100);o__Saccharomycetales(100);f__Metschnikowiaceae(100);g__Metschnikowia(100);s__Metschnikowia_pulcherrima(100); | Metschnikowia | 13 | Saprotroph | \|Undefined Saprotroph\| | Yeast | NULL | Probable | As nectar-tap_saprotroph (Põlme et al. 2020); Decay sugar-rich substrates (Põlme et al. 2020) | Põlme S, et al. 2020. Fungal Diversity 105:1-16 ((https://doi.org/10.1007/s13225-020-00466-2)) |
| Otu55546 | 1 | k__Fungi(100);p__Ascomycota(100);c__Saccharomycetes(100);o__Saccharomycetales(100);f__Metschnikowiaceae(100);g__Metschnikowia(100);s__Metschnikowia_pulcherrima(100); | Metschnikowia | 13 | Saprotroph | \|Undefined Saprotroph\| | Yeast | NULL | Probable | As nectar-tap_saprotroph (Põlme et al. 2020); Decay sugar-rich substrates (Põlme et al. 2020) | Põlme S, et al. 2020. Fungal Diversity 105:1-16 ((https://doi.org/10.1007/s13225-020-00466-2)) |
| Otu55547 | 1 | k__Fungi(100);p__Ascomycota(100);c__Saccharomycetes(100);o__Saccharomycetales(100);f__Metschnikowiaceae(100);g__Metschnikowia(100);s__Metschnikowia_pulcherrima(100); | Metschnikowia | 13 | Saprotroph | \|Undefined Saprotroph\| | Yeast | NULL | Probable | As nectar-tap_saprotroph (Põlme et al. 2020); Decay sugar-rich substrates (Põlme et al. 2020) | Põlme S, et al. 2020. Fungal Diversity 105:1-16 ((https://doi.org/10.1007/s13225-020-00466-2)) |
| Otu55548 | 1 | k__Fungi(100);p__Ascomycota(100);c__Saccharomycetes(100);o__Saccharomycetales(100);f__Metschnikowiaceae(100);g__Metschnikowia(100);g__Metschnikowia_unclassified(100); | Metschnikowia | 13 | Saprotroph | \|Undefined Saprotroph\| | Yeast | NULL | Probable | As nectar-tap_saprotroph (Põlme et al. 2020); Decay sugar-rich substrates (Põlme et al. 2020) | Põlme S, et al. 2020. Fungal Diversity 105:1-16 ((https://doi.org/10.1007/s13225-020-00466-2)) |
| Otu55549 | 1 | k__Fungi(100);p__Ascomycota(100);c__Saccharomycetes(100);o__Saccharomycetales(100);f__Metschnikowiaceae(100);g__Metschnikowia(100);s__Metschnikowia_pulcherrima(100); | Metschnikowia | 13 | Saprotroph | \|Undefined Saprotroph\| | Yeast | NULL | Probable | As nectar-tap_saprotroph (Põlme et al. 2020); Decay sugar-rich substrates (Põlme et al. 2020) | Põlme S, et al. 2020. Fungal Diversity 105:1-16 ((https://doi.org/10.1007/s13225-020-00466-2)) |
| Otu55550 | 1 | k__Fungi(100);p__Ascomycota(100);c__Saccharomycetes(100);o__Saccharomycetales(100);f__Metschnikowiaceae(100);g__Metschnikowia(100);s__Metschnikowia_pulcherrima(100); | Metschnikowia | 13 | Saprotroph | \|Undefined Saprotroph\| | Yeast | NULL | Probable | As nectar-tap_saprotroph (Põlme et al. 2020); Decay sugar-rich substrates (Põlme et al. 2020) | Põlme S, et al. 2020. Fungal Diversity 105:1-16 ((https://doi.org/10.1007/s13225-020-00466-2)) |
| Otu55551 | 1 | k__Fungi(100);p__Ascomycota(100);c__Saccharomycetes(100);o__Saccharomycetales(100);f__Metschnikowiaceae(100);g__Metschnikowia(100);s__Metschnikowia_pulcherrima(100); | Metschnikowia | 13 | Saprotroph | \|Undefined Saprotroph\| | Yeast | NULL | Probable | As nectar-tap_saprotroph (Põlme et al. 2020); Decay sugar-rich substrates (Põlme et al. 2020) | Põlme S, et al. 2020. Fungal Diversity 105:1-16 ((https://doi.org/10.1007/s13225-020-00466-2)) |
| Otu55554 | 1 | k__Fungi(100);p__Ascomycota(100);c__Saccharomycetes(100);o__Saccharomycetales(100);f__Metschnikowiaceae(100);g__Metschnikowia(100);g__Metschnikowia_unclassified(100); | Metschnikowia | 13 | Saprotroph | \|Undefined Saprotroph\| | Yeast | NULL | Probable | As nectar-tap_saprotroph (Põlme et al. 2020); Decay sugar-rich substrates (Põlme et al. 2020) | Põlme S, et al. 2020. Fungal Diversity 105:1-16 ((https://doi.org/10.1007/s13225-020-00466-2)) |
| Otu55556 | 1 | k__Fungi(100);p__Ascomycota(100);c__Saccharomycetes(100);o__Saccharomycetales(100);f__Metschnikowiaceae(100);g__Metschnikowia(100);s__Metschnikowia_pulcherrima(100); | Metschnikowia | 13 | Saprotroph | \|Undefined Saprotroph\| | Yeast | NULL | Probable | As nectar-tap_saprotroph (Põlme et al. 2020); Decay sugar-rich substrates (Põlme et al. 2020) | Põlme S, et al. 2020. Fungal Diversity 105:1-16 ((https://doi.org/10.1007/s13225-020-00466-2)) |
| Otu55557 | 1 | k__Fungi(100);p__Ascomycota(100);c__Saccharomycetes(100);o__Saccharomycetales(100);f__Metschnikowiaceae(100);g__Metschnikowia(100);s__Metschnikowia_pulcherrima(100); | Metschnikowia | 13 | Saprotroph | \|Undefined Saprotroph\| | Yeast | NULL | Probable | As nectar-tap_saprotroph (Põlme et al. 2020); Decay sugar-rich substrates (Põlme et al. 2020) | Põlme S, et al. 2020. Fungal Diversity 105:1-16 ((https://doi.org/10.1007/s13225-020-00466-2)) |
| Otu55558 | 1 | k__Fungi(100);p__Ascomycota(100);c__Saccharomycetes(100);o__Saccharomycetales(100);f__Metschnikowiaceae(100);g__Metschnikowia(100);g__Metschnikowia_unclassified(100); | Metschnikowia | 13 | Saprotroph | \|Undefined Saprotroph\| | Yeast | NULL | Probable | As nectar-tap_saprotroph (Põlme et al. 2020); Decay sugar-rich substrates (Põlme et al. 2020) | Põlme S, et al. 2020. Fungal Diversity 105:1-16 ((https://doi.org/10.1007/s13225-020-00466-2)) |
| Otu55559 | 1 | k__Fungi(100);p__Ascomycota(100);c__Saccharomycetes(100);o__Saccharomycetales(100);f__Metschnikowiaceae(100);g__Metschnikowia(100);s__Metschnikowia_pulcherrima(100); | Metschnikowia | 13 | Saprotroph | \|Undefined Saprotroph\| | Yeast | NULL | Probable | As nectar-tap_saprotroph (Põlme et al. 2020); Decay sugar-rich substrates (Põlme et al. 2020) | Põlme S, et al. 2020. Fungal Diversity 105:1-16 ((https://doi.org/10.1007/s13225-020-00466-2)) |
| Otu55560 | 1 | k__Fungi(100);p__Ascomycota(100);c__Saccharomycetes(100);o__Saccharomycetales(100);f__Metschnikowiaceae(100);g__Metschnikowia(100);s__Metschnikowia_pulcherrima(100); | Metschnikowia | 13 | Saprotroph | \|Undefined Saprotroph\| | Yeast | NULL | Probable | As nectar-tap_saprotroph (Põlme et al. 2020); Decay sugar-rich substrates (Põlme et al. 2020) | Põlme S, et al. 2020. Fungal Diversity 105:1-16 ((https://doi.org/10.1007/s13225-020-00466-2)) |
| Otu55562 | 1 | k__Fungi(100);p__Ascomycota(100);c__Saccharomycetes(100);o__Saccharomycetales(100);f__Metschnikowiaceae(100);g__Metschnikowia(100);s__Metschnikowia_pulcherrima(100); | Metschnikowia | 13 | Saprotroph | \|Undefined Saprotroph\| | Yeast | NULL | Probable | As nectar-tap_saprotroph (Põlme et al. 2020); Decay sugar-rich substrates (Põlme et al. 2020) | Põlme S, et al. 2020. Fungal Diversity 105:1-16 ((https://doi.org/10.1007/s13225-020-00466-2)) |
| Otu55564 | 1 | k__Fungi(100);p__Ascomycota(100);c__Saccharomycetes(100);o__Saccharomycetales(100);f__Metschnikowiaceae(100);g__Metschnikowia(100);s__Metschnikowia_pulcherrima(100); | Metschnikowia | 13 | Saprotroph | \|Undefined Saprotroph\| | Yeast | NULL | Probable | As nectar-tap_saprotroph (Põlme et al. 2020); Decay sugar-rich substrates (Põlme et al. 2020) | Põlme S, et al. 2020. Fungal Diversity 105:1-16 ((https://doi.org/10.1007/s13225-020-00466-2)) |
| Otu55565 | 1 | k__Fungi(100);p__Ascomycota(100);c__Saccharomycetes(100);o__Saccharomycetales(100);f__Metschnikowiaceae(100);g__Metschnikowia(100);s__Metschnikowia_pulcherrima(100); | Metschnikowia | 13 | Saprotroph | \|Undefined Saprotroph\| | Yeast | NULL | Probable | As nectar-tap_saprotroph (Põlme et al. 2020); Decay sugar-rich substrates (Põlme et al. 2020) | Põlme S, et al. 2020. Fungal Diversity 105:1-16 ((https://doi.org/10.1007/s13225-020-00466-2)) |
| Otu55566 | 1 | k__Fungi(100);p__Ascomycota(100);c__Saccharomycetes(100);o__Saccharomycetales(100);f__Metschnikowiaceae(100);g__Metschnikowia(100);s__Metschnikowia_pulcherrima(100); | Metschnikowia | 13 | Saprotroph | \|Undefined Saprotroph\| | Yeast | NULL | Probable | As nectar-tap_saprotroph (Põlme et al. 2020); Decay sugar-rich substrates (Põlme et al. 2020) | Põlme S, et al. 2020. Fungal Diversity 105:1-16 ((https://doi.org/10.1007/s13225-020-00466-2)) |
| Otu55567 | 1 | k__Fungi(100);p__Ascomycota(100);c__Saccharomycetes(100);o__Saccharomycetales(100);f__Metschnikowiaceae(100);g__Metschnikowia(100);s__Metschnikowia_pulcherrima(100); | Metschnikowia | 13 | Saprotroph | \|Undefined Saprotroph\| | Yeast | NULL | Probable | As nectar-tap_saprotroph (Põlme et al. 2020); Decay sugar-rich substrates (Põlme et al. 2020) | Põlme S, et al. 2020. Fungal Diversity 105:1-16 ((https://doi.org/10.1007/s13225-020-00466-2)) |
| Otu55568 | 1 | k__Fungi(100);p__Ascomycota(100);c__Saccharomycetes(100);o__Saccharomycetales(100);f__Metschnikowiaceae(100);g__Metschnikowia(100);s__Metschnikowia_pulcherrima(100); | Metschnikowia | 13 | Saprotroph | \|Undefined Saprotroph\| | Yeast | NULL | Probable | As nectar-tap_saprotroph (Põlme et al. 2020); Decay sugar-rich substrates (Põlme et al. 2020) | Põlme S, et al. 2020. Fungal Diversity 105:1-16 ((https://doi.org/10.1007/s13225-020-00466-2)) |
| Otu55569 | 1 | k__Fungi(100);p__Ascomycota(100);c__Saccharomycetes(100);o__Saccharomycetales(100);f__Metschnikowiaceae(100);g__Metschnikowia(100);s__Metschnikowia_pulcherrima(100); | Metschnikowia | 13 | Saprotroph | \|Undefined Saprotroph\| | Yeast | NULL | Probable | As nectar-tap_saprotroph (Põlme et al. 2020); Decay sugar-rich substrates (Põlme et al. 2020) | Põlme S, et al. 2020. Fungal Diversity 105:1-16 ((https://doi.org/10.1007/s13225-020-00466-2)) |
| Otu55570 | 1 | k__Fungi(100);p__Ascomycota(100);c__Saccharomycetes(100);o__Saccharomycetales(100);f__Metschnikowiaceae(100);g__Metschnikowia(100);s__Metschnikowia_pulcherrima(100); | Metschnikowia | 13 | Saprotroph | \|Undefined Saprotroph\| | Yeast | NULL | Probable | As nectar-tap_saprotroph (Põlme et al. 2020); Decay sugar-rich substrates (Põlme et al. 2020) | Põlme S, et al. 2020. Fungal Diversity 105:1-16 ((https://doi.org/10.1007/s13225-020-00466-2)) |
| Otu55571 | 1 | k__Fungi(100);p__Ascomycota(100);c__Saccharomycetes(100);o__Saccharomycetales(100);f__Metschnikowiaceae(100);g__Metschnikowia(100);s__Metschnikowia_pulcherrima(100); | Metschnikowia | 13 | Saprotroph | \|Undefined Saprotroph\| | Yeast | NULL | Probable | As nectar-tap_saprotroph (Põlme et al. 2020); Decay sugar-rich substrates (Põlme et al. 2020) | Põlme S, et al. 2020. Fungal Diversity 105:1-16 ((https://doi.org/10.1007/s13225-020-00466-2)) |
| Otu55572 | 1 | k__Fungi(100);p__Ascomycota(100);c__Saccharomycetes(100);o__Saccharomycetales(100);f__Metschnikowiaceae(100);g__Metschnikowia(100);s__Metschnikowia_pulcherrima(100); | Metschnikowia | 13 | Saprotroph | \|Undefined Saprotroph\| | Yeast | NULL | Probable | As nectar-tap_saprotroph (Põlme et al. 2020); Decay sugar-rich substrates (Põlme et al. 2020) | Põlme S, et al. 2020. Fungal Diversity 105:1-16 ((https://doi.org/10.1007/s13225-020-00466-2)) |
| Otu55573 | 1 | k__Fungi(100);p__Ascomycota(100);c__Saccharomycetes(100);o__Saccharomycetales(100);f__Metschnikowiaceae(100);g__Metschnikowia(100);g__Metschnikowia_unclassified(100); | Metschnikowia | 13 | Saprotroph | \|Undefined Saprotroph\| | Yeast | NULL | Probable | As nectar-tap_saprotroph (Põlme et al. 2020); Decay sugar-rich substrates (Põlme et al. 2020) | Põlme S, et al. 2020. Fungal Diversity 105:1-16 ((https://doi.org/10.1007/s13225-020-00466-2)) |
| Otu55577 | 1 | k__Fungi(100);p__Ascomycota(100);c__Saccharomycetes(100);o__Saccharomycetales(100);f__Metschnikowiaceae(100);g__Metschnikowia(100);s__Metschnikowia_pulcherrima(100); | Metschnikowia | 13 | Saprotroph | \|Undefined Saprotroph\| | Yeast | NULL | Probable | As nectar-tap_saprotroph (Põlme et al. 2020); Decay sugar-rich substrates (Põlme et al. 2020) | Põlme S, et al. 2020. Fungal Diversity 105:1-16 ((https://doi.org/10.1007/s13225-020-00466-2)) |
| Otu55578 | 1 | k__Fungi(100);p__Ascomycota(100);c__Saccharomycetes(100);o__Saccharomycetales(100);f__Saccharomycetales_family_Incertae_sedis(100);g__Candida(100);g__Candida_unclassified(100); | Candida | 13 | Pathotroph-Saprotroph-Symbiotroph | Animal Pathogen-Endophyte-Endosymbiont-Epiphyte-\|Undefined Saprotroph\| | Dimorphic Yeast-Yeast | NULL | Probable | As nectar-tap_saprotroph (Põlme et al. 2020); Decay sugar-rich substrates (Põlme et al. 2020); This asexual genus is polyphyletic and contain species that function very differently depend on what they are. Caution should be taken when interpreting guilds for Candida species. | Manolakaki D, et al. 2010. Virulence. 1: 367-375 ((https://doi.org/10.4161/viru.1.5.12796)); Mekbib SB, et al. 2011. Tropical Plant Pathology 36:233-240 ((https://doi.org/10.1590/S1982-56762011000400004)); Põlme S, et al. 2020. Fungal Diversity 105:1-16 ((https://doi.org/10.1007/s13225-020-00466-2)) |
| Otu55579 | 1 | k__Fungi(100);p__Ascomycota(100);c__Saccharomycetes(100);o__Saccharomycetales(100);f__Metschnikowiaceae(100);g__Metschnikowia(100);s__Metschnikowia_pulcherrima(100); | Metschnikowia | 13 | Saprotroph | \|Undefined Saprotroph\| | Yeast | NULL | Probable | As nectar-tap_saprotroph (Põlme et al. 2020); Decay sugar-rich substrates (Põlme et al. 2020) | Põlme S, et al. 2020. Fungal Diversity 105:1-16 ((https://doi.org/10.1007/s13225-020-00466-2)) |
| Otu55580 | 1 | k__Fungi(100);p__Ascomycota(100);c__Saccharomycetes(100);o__Saccharomycetales(100);f__Metschnikowiaceae(100);g__Metschnikowia(100);g__Metschnikowia_unclassified(100); | Metschnikowia | 13 | Saprotroph | \|Undefined Saprotroph\| | Yeast | NULL | Probable | As nectar-tap_saprotroph (Põlme et al. 2020); Decay sugar-rich substrates (Põlme et al. 2020) | Põlme S, et al. 2020. Fungal Diversity 105:1-16 ((https://doi.org/10.1007/s13225-020-00466-2)) |
| Otu55581 | 1 | k__Fungi(100);p__Ascomycota(100);c__Saccharomycetes(100);o__Saccharomycetales(100);f__Metschnikowiaceae(100);g__Metschnikowia(100);g__Metschnikowia_unclassified(100); | Metschnikowia | 13 | Saprotroph | \|Undefined Saprotroph\| | Yeast | NULL | Probable | As nectar-tap_saprotroph (Põlme et al. 2020); Decay sugar-rich substrates (Põlme et al. 2020) | Põlme S, et al. 2020. Fungal Diversity 105:1-16 ((https://doi.org/10.1007/s13225-020-00466-2)) |
| Otu55583 | 1 | k__Fungi(100);p__Ascomycota(100);c__Saccharomycetes(100);o__Saccharomycetales(100);f__Metschnikowiaceae(100);g__Metschnikowia(100);g__Metschnikowia_unclassified(100); | Metschnikowia | 13 | Saprotroph | \|Undefined Saprotroph\| | Yeast | NULL | Probable | As nectar-tap_saprotroph (Põlme et al. 2020); Decay sugar-rich substrates (Põlme et al. 2020) | Põlme S, et al. 2020. Fungal Diversity 105:1-16 ((https://doi.org/10.1007/s13225-020-00466-2)) |
| Otu55584 | 1 | k__Fungi(100);p__Ascomycota(100);c__Saccharomycetes(100);o__Saccharomycetales(100);f__Metschnikowiaceae(100);g__Metschnikowia(100);g__Metschnikowia_unclassified(100); | Metschnikowia | 13 | Saprotroph | \|Undefined Saprotroph\| | Yeast | NULL | Probable | As nectar-tap_saprotroph (Põlme et al. 2020); Decay sugar-rich substrates (Põlme et al. 2020) | Põlme S, et al. 2020. Fungal Diversity 105:1-16 ((https://doi.org/10.1007/s13225-020-00466-2)) |
| Otu55586 | 1 | k__Fungi(100);p__Ascomycota(100);c__Saccharomycetes(100);o__Saccharomycetales(100);f__Metschnikowiaceae(100);g__Metschnikowia(100);s__Metschnikowia_pulcherrima(100); | Metschnikowia | 13 | Saprotroph | \|Undefined Saprotroph\| | Yeast | NULL | Probable | As nectar-tap_saprotroph (Põlme et al. 2020); Decay sugar-rich substrates (Põlme et al. 2020) | Põlme S, et al. 2020. Fungal Diversity 105:1-16 ((https://doi.org/10.1007/s13225-020-00466-2)) |
| Otu55587 | 1 | k__Fungi(100);p__Ascomycota(100);c__Saccharomycetes(100);o__Saccharomycetales(100);f__Metschnikowiaceae(100);g__Metschnikowia(100);s__Metschnikowia_pulcherrima(100); | Metschnikowia | 13 | Saprotroph | \|Undefined Saprotroph\| | Yeast | NULL | Probable | As nectar-tap_saprotroph (Põlme et al. 2020); Decay sugar-rich substrates (Põlme et al. 2020) | Põlme S, et al. 2020. Fungal Diversity 105:1-16 ((https://doi.org/10.1007/s13225-020-00466-2)) |
| Otu55588 | 1 | k__Fungi(100);p__Ascomycota(100);c__Saccharomycetes(100);o__Saccharomycetales(100);f__Metschnikowiaceae(100);g__Metschnikowia(100);s__Metschnikowia_pulcherrima(100); | Metschnikowia | 13 | Saprotroph | \|Undefined Saprotroph\| | Yeast | NULL | Probable | As nectar-tap_saprotroph (Põlme et al. 2020); Decay sugar-rich substrates (Põlme et al. 2020) | Põlme S, et al. 2020. Fungal Diversity 105:1-16 ((https://doi.org/10.1007/s13225-020-00466-2)) |
| Otu55589 | 1 | k__Fungi(100);p__Ascomycota(100);c__Saccharomycetes(100);o__Saccharomycetales(100);f__Metschnikowiaceae(100);g__Metschnikowia(100);g__Metschnikowia_unclassified(100); | Metschnikowia | 13 | Saprotroph | \|Undefined Saprotroph\| | Yeast | NULL | Probable | As nectar-tap_saprotroph (Põlme et al. 2020); Decay sugar-rich substrates (Põlme et al. 2020) | Põlme S, et al. 2020. Fungal Diversity 105:1-16 ((https://doi.org/10.1007/s13225-020-00466-2)) |
| Otu55590 | 1 | k__Fungi(100);p__Ascomycota(100);c__Saccharomycetes(100);o__Saccharomycetales(100);f__Metschnikowiaceae(100);g__Metschnikowia(100);s__Metschnikowia_pulcherrima(100); | Metschnikowia | 13 | Saprotroph | \|Undefined Saprotroph\| | Yeast | NULL | Probable | As nectar-tap_saprotroph (Põlme et al. 2020); Decay sugar-rich substrates (Põlme et al. 2020) | Põlme S, et al. 2020. Fungal Diversity 105:1-16 ((https://doi.org/10.1007/s13225-020-00466-2)) |
| Otu55591 | 1 | k__Fungi(100);p__Ascomycota(100);c__Saccharomycetes(100);o__Saccharomycetales(100);f__Metschnikowiaceae(100);g__Metschnikowia(100);g__Metschnikowia_unclassified(100); | Metschnikowia | 13 | Saprotroph | \|Undefined Saprotroph\| | Yeast | NULL | Probable | As nectar-tap_saprotroph (Põlme et al. 2020); Decay sugar-rich substrates (Põlme et al. 2020) | Põlme S, et al. 2020. Fungal Diversity 105:1-16 ((https://doi.org/10.1007/s13225-020-00466-2)) |
| Otu55592 | 1 | k__Fungi(100);p__Ascomycota(100);c__Saccharomycetes(100);o__Saccharomycetales(100);f__Saccharomycetales_family_Incertae_sedis(100);g__Candida(100);s__Candida_cretensis(100); | Candida | 13 | Pathotroph-Saprotroph-Symbiotroph | Animal Pathogen-Endophyte-Endosymbiont-Epiphyte-\|Undefined Saprotroph\| | Dimorphic Yeast-Yeast | NULL | Probable | As nectar-tap_saprotroph (Põlme et al. 2020); Decay sugar-rich substrates (Põlme et al. 2020); This asexual genus is polyphyletic and contain species that function very differently depend on what they are. Caution should be taken when interpreting guilds for Candida species. | Manolakaki D, et al. 2010. Virulence. 1: 367-375 ((https://doi.org/10.4161/viru.1.5.12796)); Mekbib SB, et al. 2011. Tropical Plant Pathology 36:233-240 ((https://doi.org/10.1590/S1982-56762011000400004)); Põlme S, et al. 2020. Fungal Diversity 105:1-16 ((https://doi.org/10.1007/s13225-020-00466-2)) |
| Otu55593 | 1 | k__Fungi(100);p__Ascomycota(100);c__Saccharomycetes(100);o__Saccharomycetales(100);f__Metschnikowiaceae(100);g__Metschnikowia(100);s__Metschnikowia_pulcherrima(100); | Metschnikowia | 13 | Saprotroph | \|Undefined Saprotroph\| | Yeast | NULL | Probable | As nectar-tap_saprotroph (Põlme et al. 2020); Decay sugar-rich substrates (Põlme et al. 2020) | Põlme S, et al. 2020. Fungal Diversity 105:1-16 ((https://doi.org/10.1007/s13225-020-00466-2)) |
| Otu55594 | 1 | k__Fungi(100);p__Ascomycota(100);c__Saccharomycetes(100);o__Saccharomycetales(100);f__Metschnikowiaceae(100);g__Metschnikowia(100);s__Metschnikowia_pulcherrima(100); | Metschnikowia | 13 | Saprotroph | \|Undefined Saprotroph\| | Yeast | NULL | Probable | As nectar-tap_saprotroph (Põlme et al. 2020); Decay sugar-rich substrates (Põlme et al. 2020) | Põlme S, et al. 2020. Fungal Diversity 105:1-16 ((https://doi.org/10.1007/s13225-020-00466-2)) |
| Otu55596 | 1 | k__Fungi(100);p__Ascomycota(100);c__Saccharomycetes(100);o__Saccharomycetales(100);f__Metschnikowiaceae(100);g__Metschnikowia(100);s__Metschnikowia_pulcherrima(100); | Metschnikowia | 13 | Saprotroph | \|Undefined Saprotroph\| | Yeast | NULL | Probable | As nectar-tap_saprotroph (Põlme et al. 2020); Decay sugar-rich substrates (Põlme et al. 2020) | Põlme S, et al. 2020. Fungal Diversity 105:1-16 ((https://doi.org/10.1007/s13225-020-00466-2)) |
| Otu55597 | 1 | k__Fungi(100);p__Ascomycota(100);c__Saccharomycetes(100);o__Saccharomycetales(100);f__Metschnikowiaceae(100);g__Metschnikowia(100);s__Metschnikowia_pulcherrima(100); | Metschnikowia | 13 | Saprotroph | \|Undefined Saprotroph\| | Yeast | NULL | Probable | As nectar-tap_saprotroph (Põlme et al. 2020); Decay sugar-rich substrates (Põlme et al. 2020) | Põlme S, et al. 2020. Fungal Diversity 105:1-16 ((https://doi.org/10.1007/s13225-020-00466-2)) |
| Otu55598 | 1 | k__Fungi(100);p__Ascomycota(100);c__Saccharomycetes(100);o__Saccharomycetales(100);f__Metschnikowiaceae(100);g__Metschnikowia(100);s__Metschnikowia_pulcherrima(100); | Metschnikowia | 13 | Saprotroph | \|Undefined Saprotroph\| | Yeast | NULL | Probable | As nectar-tap_saprotroph (Põlme et al. 2020); Decay sugar-rich substrates (Põlme et al. 2020) | Põlme S, et al. 2020. Fungal Diversity 105:1-16 ((https://doi.org/10.1007/s13225-020-00466-2)) |
| Otu55600 | 1 | k__Fungi(100);p__Ascomycota(100);c__Saccharomycetes(100);o__Saccharomycetales(100);f__Metschnikowiaceae(100);g__Metschnikowia(100);s__Metschnikowia_pulcherrima(100); | Metschnikowia | 13 | Saprotroph | \|Undefined Saprotroph\| | Yeast | NULL | Probable | As nectar-tap_saprotroph (Põlme et al. 2020); Decay sugar-rich substrates (Põlme et al. 2020) | Põlme S, et al. 2020. Fungal Diversity 105:1-16 ((https://doi.org/10.1007/s13225-020-00466-2)) |
| Otu55601 | 1 | k__Fungi(100);p__Ascomycota(100);c__Saccharomycetes(100);o__Saccharomycetales(100);f__Metschnikowiaceae(100);g__Metschnikowia(100);g__Metschnikowia_unclassified(100); | Metschnikowia | 13 | Saprotroph | \|Undefined Saprotroph\| | Yeast | NULL | Probable | As nectar-tap_saprotroph (Põlme et al. 2020); Decay sugar-rich substrates (Põlme et al. 2020) | Põlme S, et al. 2020. Fungal Diversity 105:1-16 ((https://doi.org/10.1007/s13225-020-00466-2)) |
| Otu55602 | 1 | k__Fungi(100);p__Ascomycota(100);c__Saccharomycetes(100);o__Saccharomycetales(100);f__Metschnikowiaceae(100);g__Metschnikowia(100);g__Metschnikowia_unclassified(100); | Metschnikowia | 13 | Saprotroph | \|Undefined Saprotroph\| | Yeast | NULL | Probable | As nectar-tap_saprotroph (Põlme et al. 2020); Decay sugar-rich substrates (Põlme et al. 2020) | Põlme S, et al. 2020. Fungal Diversity 105:1-16 ((https://doi.org/10.1007/s13225-020-00466-2)) |
| Otu55603 | 1 | k__Fungi(100);p__Ascomycota(100);c__Saccharomycetes(100);o__Saccharomycetales(100);f__Metschnikowiaceae(100);g__Metschnikowia(100);s__Metschnikowia_pulcherrima(100); | Metschnikowia | 13 | Saprotroph | \|Undefined Saprotroph\| | Yeast | NULL | Probable | As nectar-tap_saprotroph (Põlme et al. 2020); Decay sugar-rich substrates (Põlme et al. 2020) | Põlme S, et al. 2020. Fungal Diversity 105:1-16 ((https://doi.org/10.1007/s13225-020-00466-2)) |
| Otu55604 | 1 | k__Fungi(100);p__Ascomycota(100);c__Saccharomycetes(100);o__Saccharomycetales(100);f__Metschnikowiaceae(100);g__Metschnikowia(100);g__Metschnikowia_unclassified(100); | Metschnikowia | 13 | Saprotroph | \|Undefined Saprotroph\| | Yeast | NULL | Probable | As nectar-tap_saprotroph (Põlme et al. 2020); Decay sugar-rich substrates (Põlme et al. 2020) | Põlme S, et al. 2020. Fungal Diversity 105:1-16 ((https://doi.org/10.1007/s13225-020-00466-2)) |
| Otu55605 | 1 | k__Fungi(100);p__Ascomycota(100);c__Saccharomycetes(100);o__Saccharomycetales(100);f__Metschnikowiaceae(100);g__Metschnikowia(100);s__Metschnikowia_pulcherrima(100); | Metschnikowia | 13 | Saprotroph | \|Undefined Saprotroph\| | Yeast | NULL | Probable | As nectar-tap_saprotroph (Põlme et al. 2020); Decay sugar-rich substrates (Põlme et al. 2020) | Põlme S, et al. 2020. Fungal Diversity 105:1-16 ((https://doi.org/10.1007/s13225-020-00466-2)) |
| Otu55606 | 1 | k__Fungi(100);p__Ascomycota(100);c__Saccharomycetes(100);o__Saccharomycetales(100);f__Metschnikowiaceae(100);g__Metschnikowia(100);s__Metschnikowia_pulcherrima(100); | Metschnikowia | 13 | Saprotroph | \|Undefined Saprotroph\| | Yeast | NULL | Probable | As nectar-tap_saprotroph (Põlme et al. 2020); Decay sugar-rich substrates (Põlme et al. 2020) | Põlme S, et al. 2020. Fungal Diversity 105:1-16 ((https://doi.org/10.1007/s13225-020-00466-2)) |
| Otu55607 | 1 | k__Fungi(100);p__Ascomycota(100);c__Saccharomycetes(100);o__Saccharomycetales(100);f__Metschnikowiaceae(100);g__Metschnikowia(100);s__Metschnikowia_pulcherrima(100); | Metschnikowia | 13 | Saprotroph | \|Undefined Saprotroph\| | Yeast | NULL | Probable | As nectar-tap_saprotroph (Põlme et al. 2020); Decay sugar-rich substrates (Põlme et al. 2020) | Põlme S, et al. 2020. Fungal Diversity 105:1-16 ((https://doi.org/10.1007/s13225-020-00466-2)) |
| Otu55609 | 1 | k__Fungi(100);p__Ascomycota(100);c__Saccharomycetes(100);o__Saccharomycetales(100);f__Saccharomycetales_family_Incertae_sedis(100);g__Candida(100);g__Candida_unclassified(100); | Candida | 13 | Pathotroph-Saprotroph-Symbiotroph | Animal Pathogen-Endophyte-Endosymbiont-Epiphyte-\|Undefined Saprotroph\| | Dimorphic Yeast-Yeast | NULL | Probable | As nectar-tap_saprotroph (Põlme et al. 2020); Decay sugar-rich substrates (Põlme et al. 2020); This asexual genus is polyphyletic and contain species that function very differently depend on what they are. Caution should be taken when interpreting guilds for Candida species. | Manolakaki D, et al. 2010. Virulence. 1: 367-375 ((https://doi.org/10.4161/viru.1.5.12796)); Mekbib SB, et al. 2011. Tropical Plant Pathology 36:233-240 ((https://doi.org/10.1590/S1982-56762011000400004)); Põlme S, et al. 2020. Fungal Diversity 105:1-16 ((https://doi.org/10.1007/s13225-020-00466-2)) |
| Otu55610 | 1 | k__Fungi(100);p__Ascomycota(100);c__Saccharomycetes(100);o__Saccharomycetales(100);f__Metschnikowiaceae(100);g__Metschnikowia(100);s__Metschnikowia_pulcherrima(100); | Metschnikowia | 13 | Saprotroph | \|Undefined Saprotroph\| | Yeast | NULL | Probable | As nectar-tap_saprotroph (Põlme et al. 2020); Decay sugar-rich substrates (Põlme et al. 2020) | Põlme S, et al. 2020. Fungal Diversity 105:1-16 ((https://doi.org/10.1007/s13225-020-00466-2)) |
| Otu55611 | 1 | k__Fungi(100);p__Ascomycota(100);c__Saccharomycetes(100);o__Saccharomycetales(100);f__Metschnikowiaceae(100);g__Metschnikowia(100);s__Metschnikowia_pulcherrima(100); | Metschnikowia | 13 | Saprotroph | \|Undefined Saprotroph\| | Yeast | NULL | Probable | As nectar-tap_saprotroph (Põlme et al. 2020); Decay sugar-rich substrates (Põlme et al. 2020) | Põlme S, et al. 2020. Fungal Diversity 105:1-16 ((https://doi.org/10.1007/s13225-020-00466-2)) |
| Otu55612 | 1 | k__Fungi(100);p__Ascomycota(100);c__Saccharomycetes(100);o__Saccharomycetales(100);f__Metschnikowiaceae(100);g__Metschnikowia(100);s__Metschnikowia_pulcherrima(100); | Metschnikowia | 13 | Saprotroph | \|Undefined Saprotroph\| | Yeast | NULL | Probable | As nectar-tap_saprotroph (Põlme et al. 2020); Decay sugar-rich substrates (Põlme et al. 2020) | Põlme S, et al. 2020. Fungal Diversity 105:1-16 ((https://doi.org/10.1007/s13225-020-00466-2)) |
| Otu55613 | 1 | k__Fungi(100);p__Ascomycota(100);c__Saccharomycetes(100);o__Saccharomycetales(100);f__Metschnikowiaceae(100);g__Metschnikowia(100);s__Metschnikowia_pulcherrima(100); | Metschnikowia | 13 | Saprotroph | \|Undefined Saprotroph\| | Yeast | NULL | Probable | As nectar-tap_saprotroph (Põlme et al. 2020); Decay sugar-rich substrates (Põlme et al. 2020) | Põlme S, et al. 2020. Fungal Diversity 105:1-16 ((https://doi.org/10.1007/s13225-020-00466-2)) |
| Otu55614 | 1 | k__Fungi(100);p__Ascomycota(100);c__Saccharomycetes(100);o__Saccharomycetales(100);f__Metschnikowiaceae(100);g__Metschnikowia(100);g__Metschnikowia_unclassified(100); | Metschnikowia | 13 | Saprotroph | \|Undefined Saprotroph\| | Yeast | NULL | Probable | As nectar-tap_saprotroph (Põlme et al. 2020); Decay sugar-rich substrates (Põlme et al. 2020) | Põlme S, et al. 2020. Fungal Diversity 105:1-16 ((https://doi.org/10.1007/s13225-020-00466-2)) |
| Otu55616 | 1 | k__Fungi(100);p__Ascomycota(100);c__Saccharomycetes(100);o__Saccharomycetales(100);f__Metschnikowiaceae(100);g__Metschnikowia(100);s__Metschnikowia_pulcherrima(100); | Metschnikowia | 13 | Saprotroph | \|Undefined Saprotroph\| | Yeast | NULL | Probable | As nectar-tap_saprotroph (Põlme et al. 2020); Decay sugar-rich substrates (Põlme et al. 2020) | Põlme S, et al. 2020. Fungal Diversity 105:1-16 ((https://doi.org/10.1007/s13225-020-00466-2)) |
| Otu55617 | 1 | k__Fungi(100);p__Ascomycota(100);c__Saccharomycetes(100);o__Saccharomycetales(100);f__Saccharomycetales_family_Incertae_sedis(100);f__Saccharomycetales_family_Incertae_sedis_unclassified(100);f__Saccharomycetales_family_Incertae_sedis_unclassified(100); | Saccharomycetales | 7 | Saprotroph | Undefined Saprotroph | Yeast | NULL | Possible | NULL | Sterkenburg E, et al. 2015. New Phytologist 207:1145-1158 ((https://doi.org/10.1111/nph.13426)) |
| Otu55618 | 1 | k__Fungi(100);p__Ascomycota(100);c__Saccharomycetes(100);o__Saccharomycetales(100);f__Metschnikowiaceae(100);g__Metschnikowia(100);s__Metschnikowia_pulcherrima(100); | Metschnikowia | 13 | Saprotroph | \|Undefined Saprotroph\| | Yeast | NULL | Probable | As nectar-tap_saprotroph (Põlme et al. 2020); Decay sugar-rich substrates (Põlme et al. 2020) | Põlme S, et al. 2020. Fungal Diversity 105:1-16 ((https://doi.org/10.1007/s13225-020-00466-2)) |
| Otu55619 | 1 | k__Fungi(100);p__Ascomycota(100);c__Saccharomycetes(100);o__Saccharomycetales(100);f__Metschnikowiaceae(100);g__Metschnikowia(100);s__Metschnikowia_pulcherrima(100); | Metschnikowia | 13 | Saprotroph | \|Undefined Saprotroph\| | Yeast | NULL | Probable | As nectar-tap_saprotroph (Põlme et al. 2020); Decay sugar-rich substrates (Põlme et al. 2020) | Põlme S, et al. 2020. Fungal Diversity 105:1-16 ((https://doi.org/10.1007/s13225-020-00466-2)) |
| Otu55620 | 1 | k__Fungi(100);p__Ascomycota(100);c__Saccharomycetes(100);o__Saccharomycetales(100);f__Metschnikowiaceae(100);g__Metschnikowia(100);s__Metschnikowia_pulcherrima(100); | Metschnikowia | 13 | Saprotroph | \|Undefined Saprotroph\| | Yeast | NULL | Probable | As nectar-tap_saprotroph (Põlme et al. 2020); Decay sugar-rich substrates (Põlme et al. 2020) | Põlme S, et al. 2020. Fungal Diversity 105:1-16 ((https://doi.org/10.1007/s13225-020-00466-2)) |
| Otu55621 | 1 | k__Fungi(100);p__Ascomycota(100);c__Saccharomycetes(100);o__Saccharomycetales(100);f__Metschnikowiaceae(100);g__Metschnikowia(100);s__Metschnikowia_pulcherrima(100); | Metschnikowia | 13 | Saprotroph | \|Undefined Saprotroph\| | Yeast | NULL | Probable | As nectar-tap_saprotroph (Põlme et al. 2020); Decay sugar-rich substrates (Põlme et al. 2020) | Põlme S, et al. 2020. Fungal Diversity 105:1-16 ((https://doi.org/10.1007/s13225-020-00466-2)) |
| Otu55622 | 1 | k__Fungi(100);p__Ascomycota(100);c__Saccharomycetes(100);o__Saccharomycetales(100);f__Metschnikowiaceae(100);g__Metschnikowia(100);s__Metschnikowia_pulcherrima(100); | Metschnikowia | 13 | Saprotroph | \|Undefined Saprotroph\| | Yeast | NULL | Probable | As nectar-tap_saprotroph (Põlme et al. 2020); Decay sugar-rich substrates (Põlme et al. 2020) | Põlme S, et al. 2020. Fungal Diversity 105:1-16 ((https://doi.org/10.1007/s13225-020-00466-2)) |
| Otu55623 | 1 | k__Fungi(100);p__Ascomycota(100);c__Saccharomycetes(100);o__Saccharomycetales(100);f__Metschnikowiaceae(100);g__Metschnikowia(100);s__Metschnikowia_pulcherrima(100); | Metschnikowia | 13 | Saprotroph | \|Undefined Saprotroph\| | Yeast | NULL | Probable | As nectar-tap_saprotroph (Põlme et al. 2020); Decay sugar-rich substrates (Põlme et al. 2020) | Põlme S, et al. 2020. Fungal Diversity 105:1-16 ((https://doi.org/10.1007/s13225-020-00466-2)) |
| Otu55624 | 1 | k__Fungi(100);p__Ascomycota(100);c__Saccharomycetes(100);o__Saccharomycetales(100);f__Metschnikowiaceae(100);g__Metschnikowia(100);s__Metschnikowia_pulcherrima(100); | Metschnikowia | 13 | Saprotroph | \|Undefined Saprotroph\| | Yeast | NULL | Probable | As nectar-tap_saprotroph (Põlme et al. 2020); Decay sugar-rich substrates (Põlme et al. 2020) | Põlme S, et al. 2020. Fungal Diversity 105:1-16 ((https://doi.org/10.1007/s13225-020-00466-2)) |
| Otu55625 | 1 | k__Fungi(100);p__Ascomycota(100);c__Saccharomycetes(100);o__Saccharomycetales(100);f__Metschnikowiaceae(100);g__Metschnikowia(100);s__Metschnikowia_pulcherrima(100); | Metschnikowia | 13 | Saprotroph | \|Undefined Saprotroph\| | Yeast | NULL | Probable | As nectar-tap_saprotroph (Põlme et al. 2020); Decay sugar-rich substrates (Põlme et al. 2020) | Põlme S, et al. 2020. Fungal Diversity 105:1-16 ((https://doi.org/10.1007/s13225-020-00466-2)) |
| Otu55626 | 1 | k__Fungi(100);p__Ascomycota(100);c__Saccharomycetes(100);o__Saccharomycetales(100);f__Metschnikowiaceae(100);g__Metschnikowia(100);s__Metschnikowia_pulcherrima(100); | Metschnikowia | 13 | Saprotroph | \|Undefined Saprotroph\| | Yeast | NULL | Probable | As nectar-tap_saprotroph (Põlme et al. 2020); Decay sugar-rich substrates (Põlme et al. 2020) | Põlme S, et al. 2020. Fungal Diversity 105:1-16 ((https://doi.org/10.1007/s13225-020-00466-2)) |
| Otu55627 | 1 | k__Fungi(100);p__Ascomycota(100);c__Saccharomycetes(100);o__Saccharomycetales(100);f__Metschnikowiaceae(100);g__Metschnikowia(100);s__Metschnikowia_pulcherrima(100); | Metschnikowia | 13 | Saprotroph | \|Undefined Saprotroph\| | Yeast | NULL | Probable | As nectar-tap_saprotroph (Põlme et al. 2020); Decay sugar-rich substrates (Põlme et al. 2020) | Põlme S, et al. 2020. Fungal Diversity 105:1-16 ((https://doi.org/10.1007/s13225-020-00466-2)) |
| Otu55628 | 1 | k__Fungi(100);p__Ascomycota(100);c__Saccharomycetes(100);o__Saccharomycetales(100);f__Metschnikowiaceae(100);g__Metschnikowia(100);s__Metschnikowia_pulcherrima(100); | Metschnikowia | 13 | Saprotroph | \|Undefined Saprotroph\| | Yeast | NULL | Probable | As nectar-tap_saprotroph (Põlme et al. 2020); Decay sugar-rich substrates (Põlme et al. 2020) | Põlme S, et al. 2020. Fungal Diversity 105:1-16 ((https://doi.org/10.1007/s13225-020-00466-2)) |
| Otu55629 | 1 | k__Fungi(100);p__Ascomycota(100);c__Saccharomycetes(100);o__Saccharomycetales(100);f__Metschnikowiaceae(100);g__Metschnikowia(100);s__Metschnikowia_pulcherrima(100); | Metschnikowia | 13 | Saprotroph | \|Undefined Saprotroph\| | Yeast | NULL | Probable | As nectar-tap_saprotroph (Põlme et al. 2020); Decay sugar-rich substrates (Põlme et al. 2020) | Põlme S, et al. 2020. Fungal Diversity 105:1-16 ((https://doi.org/10.1007/s13225-020-00466-2)) |
| Otu55631 | 1 | k__Fungi(100);p__Ascomycota(100);c__Saccharomycetes(100);o__Saccharomycetales(100);f__Metschnikowiaceae(100);g__Metschnikowia(100);s__Metschnikowia_pulcherrima(100); | Metschnikowia | 13 | Saprotroph | \|Undefined Saprotroph\| | Yeast | NULL | Probable | As nectar-tap_saprotroph (Põlme et al. 2020); Decay sugar-rich substrates (Põlme et al. 2020) | Põlme S, et al. 2020. Fungal Diversity 105:1-16 ((https://doi.org/10.1007/s13225-020-00466-2)) |
| Otu55632 | 1 | k__Fungi(100);p__Ascomycota(100);c__Saccharomycetes(100);o__Saccharomycetales(100);f__Metschnikowiaceae(100);g__Metschnikowia(100);g__Metschnikowia_unclassified(100); | Metschnikowia | 13 | Saprotroph | \|Undefined Saprotroph\| | Yeast | NULL | Probable | As nectar-tap_saprotroph (Põlme et al. 2020); Decay sugar-rich substrates (Põlme et al. 2020) | Põlme S, et al. 2020. Fungal Diversity 105:1-16 ((https://doi.org/10.1007/s13225-020-00466-2)) |
| Otu55633 | 1 | k__Fungi(100);p__Ascomycota(100);c__Saccharomycetes(100);o__Saccharomycetales(100);f__Metschnikowiaceae(100);g__Metschnikowia(100);g__Metschnikowia_unclassified(100); | Metschnikowia | 13 | Saprotroph | \|Undefined Saprotroph\| | Yeast | NULL | Probable | As nectar-tap_saprotroph (Põlme et al. 2020); Decay sugar-rich substrates (Põlme et al. 2020) | Põlme S, et al. 2020. Fungal Diversity 105:1-16 ((https://doi.org/10.1007/s13225-020-00466-2)) |
| Otu55634 | 1 | k__Fungi(100);p__Ascomycota(100);c__Saccharomycetes(100);o__Saccharomycetales(100);f__Metschnikowiaceae(100);g__Metschnikowia(100);g__Metschnikowia_unclassified(100); | Metschnikowia | 13 | Saprotroph | \|Undefined Saprotroph\| | Yeast | NULL | Probable | As nectar-tap_saprotroph (Põlme et al. 2020); Decay sugar-rich substrates (Põlme et al. 2020) | Põlme S, et al. 2020. Fungal Diversity 105:1-16 ((https://doi.org/10.1007/s13225-020-00466-2)) |
| Otu55635 | 1 | k__Fungi(100);p__Ascomycota(100);c__Saccharomycetes(100);o__Saccharomycetales(100);f__Metschnikowiaceae(100);g__Metschnikowia(100);g__Metschnikowia_unclassified(100); | Metschnikowia | 13 | Saprotroph | \|Undefined Saprotroph\| | Yeast | NULL | Probable | As nectar-tap_saprotroph (Põlme et al. 2020); Decay sugar-rich substrates (Põlme et al. 2020) | Põlme S, et al. 2020. Fungal Diversity 105:1-16 ((https://doi.org/10.1007/s13225-020-00466-2)) |
| Otu55636 | 1 | k__Fungi(100);p__Ascomycota(100);c__Saccharomycetes(100);o__Saccharomycetales(100);f__Metschnikowiaceae(100);g__Metschnikowia(100);s__Metschnikowia_pulcherrima(100); | Metschnikowia | 13 | Saprotroph | \|Undefined Saprotroph\| | Yeast | NULL | Probable | As nectar-tap_saprotroph (Põlme et al. 2020); Decay sugar-rich substrates (Põlme et al. 2020) | Põlme S, et al. 2020. Fungal Diversity 105:1-16 ((https://doi.org/10.1007/s13225-020-00466-2)) |
| Otu55637 | 1 | k__Fungi(100);p__Ascomycota(100);c__Saccharomycetes(100);o__Saccharomycetales(100);f__Metschnikowiaceae(100);g__Metschnikowia(100);s__Metschnikowia_pulcherrima(100); | Metschnikowia | 13 | Saprotroph | \|Undefined Saprotroph\| | Yeast | NULL | Probable | As nectar-tap_saprotroph (Põlme et al. 2020); Decay sugar-rich substrates (Põlme et al. 2020) | Põlme S, et al. 2020. Fungal Diversity 105:1-16 ((https://doi.org/10.1007/s13225-020-00466-2)) |
| Otu55638 | 1 | k__Fungi(100);p__Ascomycota(100);c__Saccharomycetes(100);o__Saccharomycetales(100);f__Metschnikowiaceae(100);g__Metschnikowia(100);s__Metschnikowia_pulcherrima(100); | Metschnikowia | 13 | Saprotroph | \|Undefined Saprotroph\| | Yeast | NULL | Probable | As nectar-tap_saprotroph (Põlme et al. 2020); Decay sugar-rich substrates (Põlme et al. 2020) | Põlme S, et al. 2020. Fungal Diversity 105:1-16 ((https://doi.org/10.1007/s13225-020-00466-2)) |
| Otu55639 | 1 | k__Fungi(100);p__Ascomycota(100);c__Saccharomycetes(100);o__Saccharomycetales(100);f__Metschnikowiaceae(100);g__Metschnikowia(100);s__Metschnikowia_pulcherrima(100); | Metschnikowia | 13 | Saprotroph | \|Undefined Saprotroph\| | Yeast | NULL | Probable | As nectar-tap_saprotroph (Põlme et al. 2020); Decay sugar-rich substrates (Põlme et al. 2020) | Põlme S, et al. 2020. Fungal Diversity 105:1-16 ((https://doi.org/10.1007/s13225-020-00466-2)) |
| Otu55640 | 1 | k__Fungi(100);p__Ascomycota(100);c__Saccharomycetes(100);o__Saccharomycetales(100);f__Metschnikowiaceae(100);g__Metschnikowia(100);s__Metschnikowia_pulcherrima(100); | Metschnikowia | 13 | Saprotroph | \|Undefined Saprotroph\| | Yeast | NULL | Probable | As nectar-tap_saprotroph (Põlme et al. 2020); Decay sugar-rich substrates (Põlme et al. 2020) | Põlme S, et al. 2020. Fungal Diversity 105:1-16 ((https://doi.org/10.1007/s13225-020-00466-2)) |
| Otu55641 | 1 | k__Fungi(100);p__Ascomycota(100);c__Saccharomycetes(100);o__Saccharomycetales(100);f__Metschnikowiaceae(100);g__Metschnikowia(100);s__Metschnikowia_pulcherrima(100); | Metschnikowia | 13 | Saprotroph | \|Undefined Saprotroph\| | Yeast | NULL | Probable | As nectar-tap_saprotroph (Põlme et al. 2020); Decay sugar-rich substrates (Põlme et al. 2020) | Põlme S, et al. 2020. Fungal Diversity 105:1-16 ((https://doi.org/10.1007/s13225-020-00466-2)) |
| Otu55642 | 1 | k__Fungi(100);p__Ascomycota(100);c__Saccharomycetes(100);o__Saccharomycetales(100);f__Metschnikowiaceae(100);g__Metschnikowia(100);s__Metschnikowia_pulcherrima(100); | Metschnikowia | 13 | Saprotroph | \|Undefined Saprotroph\| | Yeast | NULL | Probable | As nectar-tap_saprotroph (Põlme et al. 2020); Decay sugar-rich substrates (Põlme et al. 2020) | Põlme S, et al. 2020. Fungal Diversity 105:1-16 ((https://doi.org/10.1007/s13225-020-00466-2)) |
| Otu55643 | 1 | k__Fungi(100);p__Ascomycota(100);c__Saccharomycetes(100);o__Saccharomycetales(100);f__Metschnikowiaceae(100);g__Metschnikowia(100);s__Metschnikowia_pulcherrima(100); | Metschnikowia | 13 | Saprotroph | \|Undefined Saprotroph\| | Yeast | NULL | Probable | As nectar-tap_saprotroph (Põlme et al. 2020); Decay sugar-rich substrates (Põlme et al. 2020) | Põlme S, et al. 2020. Fungal Diversity 105:1-16 ((https://doi.org/10.1007/s13225-020-00466-2)) |
| Otu55644 | 1 | k__Fungi(100);p__Ascomycota(100);c__Saccharomycetes(100);o__Saccharomycetales(100);f__Metschnikowiaceae(100);g__Metschnikowia(100);s__Metschnikowia_pulcherrima(100); | Metschnikowia | 13 | Saprotroph | \|Undefined Saprotroph\| | Yeast | NULL | Probable | As nectar-tap_saprotroph (Põlme et al. 2020); Decay sugar-rich substrates (Põlme et al. 2020) | Põlme S, et al. 2020. Fungal Diversity 105:1-16 ((https://doi.org/10.1007/s13225-020-00466-2)) |
| Otu55645 | 1 | k__Fungi(100);p__Ascomycota(100);c__Saccharomycetes(100);o__Saccharomycetales(100);f__Metschnikowiaceae(100);g__Metschnikowia(100);s__Metschnikowia_pulcherrima(100); | Metschnikowia | 13 | Saprotroph | \|Undefined Saprotroph\| | Yeast | NULL | Probable | As nectar-tap_saprotroph (Põlme et al. 2020); Decay sugar-rich substrates (Põlme et al. 2020) | Põlme S, et al. 2020. Fungal Diversity 105:1-16 ((https://doi.org/10.1007/s13225-020-00466-2)) |
| Otu55646 | 1 | k__Fungi(100);p__Ascomycota(100);c__Saccharomycetes(100);o__Saccharomycetales(100);f__Metschnikowiaceae(100);g__Metschnikowia(100);s__Metschnikowia_pulcherrima(100); | Metschnikowia | 13 | Saprotroph | \|Undefined Saprotroph\| | Yeast | NULL | Probable | As nectar-tap_saprotroph (Põlme et al. 2020); Decay sugar-rich substrates (Põlme et al. 2020) | Põlme S, et al. 2020. Fungal Diversity 105:1-16 ((https://doi.org/10.1007/s13225-020-00466-2)) |
| Otu55647 | 1 | k__Fungi(100);p__Ascomycota(100);c__Saccharomycetes(100);o__Saccharomycetales(100);f__Metschnikowiaceae(100);g__Metschnikowia(100);s__Metschnikowia_pulcherrima(100); | Metschnikowia | 13 | Saprotroph | \|Undefined Saprotroph\| | Yeast | NULL | Probable | As nectar-tap_saprotroph (Põlme et al. 2020); Decay sugar-rich substrates (Põlme et al. 2020) | Põlme S, et al. 2020. Fungal Diversity 105:1-16 ((https://doi.org/10.1007/s13225-020-00466-2)) |
| Otu55648 | 1 | k__Fungi(100);p__Ascomycota(100);c__Saccharomycetes(100);o__Saccharomycetales(100);f__Metschnikowiaceae(100);g__Metschnikowia(100);s__Metschnikowia_pulcherrima(100); | Metschnikowia | 13 | Saprotroph | \|Undefined Saprotroph\| | Yeast | NULL | Probable | As nectar-tap_saprotroph (Põlme et al. 2020); Decay sugar-rich substrates (Põlme et al. 2020) | Põlme S, et al. 2020. Fungal Diversity 105:1-16 ((https://doi.org/10.1007/s13225-020-00466-2)) |
| Otu55649 | 1 | k__Fungi(100);p__Ascomycota(100);c__Saccharomycetes(100);o__Saccharomycetales(100);f__Metschnikowiaceae(100);g__Metschnikowia(100);s__Metschnikowia_pulcherrima(100); | Metschnikowia | 13 | Saprotroph | \|Undefined Saprotroph\| | Yeast | NULL | Probable | As nectar-tap_saprotroph (Põlme et al. 2020); Decay sugar-rich substrates (Põlme et al. 2020) | Põlme S, et al. 2020. Fungal Diversity 105:1-16 ((https://doi.org/10.1007/s13225-020-00466-2)) |
| Otu55650 | 1 | k__Fungi(100);p__Ascomycota(100);c__Saccharomycetes(100);o__Saccharomycetales(100);f__Metschnikowiaceae(100);g__Metschnikowia(100);s__Metschnikowia_pulcherrima(100); | Metschnikowia | 13 | Saprotroph | \|Undefined Saprotroph\| | Yeast | NULL | Probable | As nectar-tap_saprotroph (Põlme et al. 2020); Decay sugar-rich substrates (Põlme et al. 2020) | Põlme S, et al. 2020. Fungal Diversity 105:1-16 ((https://doi.org/10.1007/s13225-020-00466-2)) |
| Otu55651 | 1 | k__Fungi(100);p__Ascomycota(100);c__Saccharomycetes(100);o__Saccharomycetales(100);f__Metschnikowiaceae(100);g__Metschnikowia(100);s__Metschnikowia_pulcherrima(100); | Metschnikowia | 13 | Saprotroph | \|Undefined Saprotroph\| | Yeast | NULL | Probable | As nectar-tap_saprotroph (Põlme et al. 2020); Decay sugar-rich substrates (Põlme et al. 2020) | Põlme S, et al. 2020. Fungal Diversity 105:1-16 ((https://doi.org/10.1007/s13225-020-00466-2)) |
| Otu55653 | 1 | k__Fungi(100);p__Ascomycota(100);c__Saccharomycetes(100);o__Saccharomycetales(100);f__Metschnikowiaceae(100);g__Metschnikowia(100);s__Metschnikowia_pulcherrima(100); | Metschnikowia | 13 | Saprotroph | \|Undefined Saprotroph\| | Yeast | NULL | Probable | As nectar-tap_saprotroph (Põlme et al. 2020); Decay sugar-rich substrates (Põlme et al. 2020) | Põlme S, et al. 2020. Fungal Diversity 105:1-16 ((https://doi.org/10.1007/s13225-020-00466-2)) |
| Otu55654 | 1 | k__Fungi(100);p__Ascomycota(100);c__Saccharomycetes(100);o__Saccharomycetales(100);f__Metschnikowiaceae(100);g__Metschnikowia(100);s__Metschnikowia_pulcherrima(100); | Metschnikowia | 13 | Saprotroph | \|Undefined Saprotroph\| | Yeast | NULL | Probable | As nectar-tap_saprotroph (Põlme et al. 2020); Decay sugar-rich substrates (Põlme et al. 2020) | Põlme S, et al. 2020. Fungal Diversity 105:1-16 ((https://doi.org/10.1007/s13225-020-00466-2)) |
| Otu55656 | 1 | k__Fungi(100);p__Ascomycota(100);c__Saccharomycetes(100);o__Saccharomycetales(100);f__Metschnikowiaceae(100);g__Metschnikowia(100);s__Metschnikowia_pulcherrima(100); | Metschnikowia | 13 | Saprotroph | \|Undefined Saprotroph\| | Yeast | NULL | Probable | As nectar-tap_saprotroph (Põlme et al. 2020); Decay sugar-rich substrates (Põlme et al. 2020) | Põlme S, et al. 2020. Fungal Diversity 105:1-16 ((https://doi.org/10.1007/s13225-020-00466-2)) |
| Otu55658 | 1 | k__Fungi(100);p__Ascomycota(100);c__Saccharomycetes(100);o__Saccharomycetales(100);f__Metschnikowiaceae(100);g__Metschnikowia(100);s__Metschnikowia_pulcherrima(100); | Metschnikowia | 13 | Saprotroph | \|Undefined Saprotroph\| | Yeast | NULL | Probable | As nectar-tap_saprotroph (Põlme et al. 2020); Decay sugar-rich substrates (Põlme et al. 2020) | Põlme S, et al. 2020. Fungal Diversity 105:1-16 ((https://doi.org/10.1007/s13225-020-00466-2)) |
| Otu55659 | 1 | k__Fungi(100);p__Ascomycota(100);c__Saccharomycetes(100);o__Saccharomycetales(100);f__Metschnikowiaceae(100);g__Metschnikowia(100);s__Metschnikowia_pulcherrima(100); | Metschnikowia | 13 | Saprotroph | \|Undefined Saprotroph\| | Yeast | NULL | Probable | As nectar-tap_saprotroph (Põlme et al. 2020); Decay sugar-rich substrates (Põlme et al. 2020) | Põlme S, et al. 2020. Fungal Diversity 105:1-16 ((https://doi.org/10.1007/s13225-020-00466-2)) |
| Otu55660 | 1 | k__Fungi(100);p__Ascomycota(100);c__Saccharomycetes(100);o__Saccharomycetales(100);f__Metschnikowiaceae(100);g__Metschnikowia(100);g__Metschnikowia_unclassified(100); | Metschnikowia | 13 | Saprotroph | \|Undefined Saprotroph\| | Yeast | NULL | Probable | As nectar-tap_saprotroph (Põlme et al. 2020); Decay sugar-rich substrates (Põlme et al. 2020) | Põlme S, et al. 2020. Fungal Diversity 105:1-16 ((https://doi.org/10.1007/s13225-020-00466-2)) |
| Otu55661 | 1 | k__Fungi(100);p__Ascomycota(100);c__Saccharomycetes(100);o__Saccharomycetales(100);f__Metschnikowiaceae(100);g__Metschnikowia(100);s__Metschnikowia_pulcherrima(100); | Metschnikowia | 13 | Saprotroph | \|Undefined Saprotroph\| | Yeast | NULL | Probable | As nectar-tap_saprotroph (Põlme et al. 2020); Decay sugar-rich substrates (Põlme et al. 2020) | Põlme S, et al. 2020. Fungal Diversity 105:1-16 ((https://doi.org/10.1007/s13225-020-00466-2)) |
| Otu55664 | 1 | k__Fungi(100);p__Ascomycota(100);c__Saccharomycetes(100);o__Saccharomycetales(100);f__Metschnikowiaceae(100);g__Metschnikowia(100);s__Metschnikowia_pulcherrima(100); | Metschnikowia | 13 | Saprotroph | \|Undefined Saprotroph\| | Yeast | NULL | Probable | As nectar-tap_saprotroph (Põlme et al. 2020); Decay sugar-rich substrates (Põlme et al. 2020) | Põlme S, et al. 2020. Fungal Diversity 105:1-16 ((https://doi.org/10.1007/s13225-020-00466-2)) |
| Otu55668 | 1 | k__Fungi(100);p__Ascomycota(100);c__Saccharomycetes(100);o__Saccharomycetales(100);f__Metschnikowiaceae(100);g__Metschnikowia(100);s__Metschnikowia_pulcherrima(100); | Metschnikowia | 13 | Saprotroph | \|Undefined Saprotroph\| | Yeast | NULL | Probable | As nectar-tap_saprotroph (Põlme et al. 2020); Decay sugar-rich substrates (Põlme et al. 2020) | Põlme S, et al. 2020. Fungal Diversity 105:1-16 ((https://doi.org/10.1007/s13225-020-00466-2)) |
| Otu55669 | 1 | k__Fungi(100);p__Ascomycota(100);c__Saccharomycetes(100);o__Saccharomycetales(100);f__Saccharomycetales_family_Incertae_sedis(100);g__Candida(100);g__Candida_unclassified(100); | Candida | 13 | Pathotroph-Saprotroph-Symbiotroph | Animal Pathogen-Endophyte-Endosymbiont-Epiphyte-\|Undefined Saprotroph\| | Dimorphic Yeast-Yeast | NULL | Probable | As nectar-tap_saprotroph (Põlme et al. 2020); Decay sugar-rich substrates (Põlme et al. 2020); This asexual genus is polyphyletic and contain species that function very differently depend on what they are. Caution should be taken when interpreting guilds for Candida species. | Manolakaki D, et al. 2010. Virulence. 1: 367-375 ((https://doi.org/10.4161/viru.1.5.12796)); Mekbib SB, et al. 2011. Tropical Plant Pathology 36:233-240 ((https://doi.org/10.1590/S1982-56762011000400004)); Põlme S, et al. 2020. Fungal Diversity 105:1-16 ((https://doi.org/10.1007/s13225-020-00466-2)) |
| Otu55672 | 1 | k__Fungi(100);p__Ascomycota(100);c__Saccharomycetes(100);o__Saccharomycetales(100);f__Metschnikowiaceae(100);g__Metschnikowia(100);s__Metschnikowia_pulcherrima(100); | Metschnikowia | 13 | Saprotroph | \|Undefined Saprotroph\| | Yeast | NULL | Probable | As nectar-tap_saprotroph (Põlme et al. 2020); Decay sugar-rich substrates (Põlme et al. 2020) | Põlme S, et al. 2020. Fungal Diversity 105:1-16 ((https://doi.org/10.1007/s13225-020-00466-2)) |
| Otu55674 | 1 | k__Fungi(100);p__Ascomycota(100);c__Saccharomycetes(100);o__Saccharomycetales(100);f__Metschnikowiaceae(100);g__Metschnikowia(100);g__Metschnikowia_unclassified(100); | Metschnikowia | 13 | Saprotroph | \|Undefined Saprotroph\| | Yeast | NULL | Probable | As nectar-tap_saprotroph (Põlme et al. 2020); Decay sugar-rich substrates (Põlme et al. 2020) | Põlme S, et al. 2020. Fungal Diversity 105:1-16 ((https://doi.org/10.1007/s13225-020-00466-2)) |
| Otu55675 | 1 | k__Fungi(100);p__Ascomycota(100);c__Saccharomycetes(100);o__Saccharomycetales(100);f__Metschnikowiaceae(100);g__Metschnikowia(100);s__Metschnikowia_pulcherrima(100); | Metschnikowia | 13 | Saprotroph | \|Undefined Saprotroph\| | Yeast | NULL | Probable | As nectar-tap_saprotroph (Põlme et al. 2020); Decay sugar-rich substrates (Põlme et al. 2020) | Põlme S, et al. 2020. Fungal Diversity 105:1-16 ((https://doi.org/10.1007/s13225-020-00466-2)) |
| Otu55678 | 1 | k__Fungi(100);p__Ascomycota(100);c__Saccharomycetes(100);o__Saccharomycetales(100);f__Metschnikowiaceae(100);g__Metschnikowia(100);s__Metschnikowia_pulcherrima(100); | Metschnikowia | 13 | Saprotroph | \|Undefined Saprotroph\| | Yeast | NULL | Probable | As nectar-tap_saprotroph (Põlme et al. 2020); Decay sugar-rich substrates (Põlme et al. 2020) | Põlme S, et al. 2020. Fungal Diversity 105:1-16 ((https://doi.org/10.1007/s13225-020-00466-2)) |
| Otu55679 | 1 | k__Fungi(100);p__Ascomycota(100);c__Saccharomycetes(100);o__Saccharomycetales(100);f__Metschnikowiaceae(100);g__Metschnikowia(100);s__Metschnikowia_pulcherrima(100); | Metschnikowia | 13 | Saprotroph | \|Undefined Saprotroph\| | Yeast | NULL | Probable | As nectar-tap_saprotroph (Põlme et al. 2020); Decay sugar-rich substrates (Põlme et al. 2020) | Põlme S, et al. 2020. Fungal Diversity 105:1-16 ((https://doi.org/10.1007/s13225-020-00466-2)) |
| Otu55680 | 1 | k__Fungi(100);p__Ascomycota(100);c__Saccharomycetes(100);o__Saccharomycetales(100);f__Metschnikowiaceae(100);g__Metschnikowia(100);s__Metschnikowia_pulcherrima(100); | Metschnikowia | 13 | Saprotroph | \|Undefined Saprotroph\| | Yeast | NULL | Probable | As nectar-tap_saprotroph (Põlme et al. 2020); Decay sugar-rich substrates (Põlme et al. 2020) | Põlme S, et al. 2020. Fungal Diversity 105:1-16 ((https://doi.org/10.1007/s13225-020-00466-2)) |
| Otu55681 | 1 | k__Fungi(100);p__Ascomycota(100);c__Saccharomycetes(100);o__Saccharomycetales(100);f__Metschnikowiaceae(100);g__Metschnikowia(100);s__Metschnikowia_pulcherrima(100); | Metschnikowia | 13 | Saprotroph | \|Undefined Saprotroph\| | Yeast | NULL | Probable | As nectar-tap_saprotroph (Põlme et al. 2020); Decay sugar-rich substrates (Põlme et al. 2020) | Põlme S, et al. 2020. Fungal Diversity 105:1-16 ((https://doi.org/10.1007/s13225-020-00466-2)) |
| Otu55683 | 1 | k__Fungi(100);p__Ascomycota(100);c__Saccharomycetes(100);o__Saccharomycetales(100);f__Metschnikowiaceae(100);g__Metschnikowia(100);g__Metschnikowia_unclassified(100); | Metschnikowia | 13 | Saprotroph | \|Undefined Saprotroph\| | Yeast | NULL | Probable | As nectar-tap_saprotroph (Põlme et al. 2020); Decay sugar-rich substrates (Põlme et al. 2020) | Põlme S, et al. 2020. Fungal Diversity 105:1-16 ((https://doi.org/10.1007/s13225-020-00466-2)) |
| Otu55684 | 1 | k__Fungi(100);p__Ascomycota(100);c__Saccharomycetes(100);o__Saccharomycetales(100);f__Metschnikowiaceae(100);g__Metschnikowia(100);g__Metschnikowia_unclassified(100); | Metschnikowia | 13 | Saprotroph | \|Undefined Saprotroph\| | Yeast | NULL | Probable | As nectar-tap_saprotroph (Põlme et al. 2020); Decay sugar-rich substrates (Põlme et al. 2020) | Põlme S, et al. 2020. Fungal Diversity 105:1-16 ((https://doi.org/10.1007/s13225-020-00466-2)) |
| Otu55685 | 1 | k__Fungi(100);p__Ascomycota(100);c__Saccharomycetes(100);o__Saccharomycetales(100);f__Metschnikowiaceae(100);g__Metschnikowia(100);s__Metschnikowia_pulcherrima(100); | Metschnikowia | 13 | Saprotroph | \|Undefined Saprotroph\| | Yeast | NULL | Probable | As nectar-tap_saprotroph (Põlme et al. 2020); Decay sugar-rich substrates (Põlme et al. 2020) | Põlme S, et al. 2020. Fungal Diversity 105:1-16 ((https://doi.org/10.1007/s13225-020-00466-2)) |
| Otu55686 | 1 | k__Fungi(100);p__Ascomycota(100);c__Saccharomycetes(100);o__Saccharomycetales(100);f__Metschnikowiaceae(100);g__Metschnikowia(100);s__Metschnikowia_pulcherrima(100); | Metschnikowia | 13 | Saprotroph | \|Undefined Saprotroph\| | Yeast | NULL | Probable | As nectar-tap_saprotroph (Põlme et al. 2020); Decay sugar-rich substrates (Põlme et al. 2020) | Põlme S, et al. 2020. Fungal Diversity 105:1-16 ((https://doi.org/10.1007/s13225-020-00466-2)) |
| Otu55687 | 1 | k__Fungi(100);p__Ascomycota(100);c__Saccharomycetes(100);o__Saccharomycetales(100);f__Metschnikowiaceae(100);g__Metschnikowia(100);s__Metschnikowia_pulcherrima(100); | Metschnikowia | 13 | Saprotroph | \|Undefined Saprotroph\| | Yeast | NULL | Probable | As nectar-tap_saprotroph (Põlme et al. 2020); Decay sugar-rich substrates (Põlme et al. 2020) | Põlme S, et al. 2020. Fungal Diversity 105:1-16 ((https://doi.org/10.1007/s13225-020-00466-2)) |
| Otu55688 | 1 | k__Fungi(100);p__Ascomycota(100);c__Saccharomycetes(100);o__Saccharomycetales(100);f__Metschnikowiaceae(100);g__Metschnikowia(100);s__Metschnikowia_pulcherrima(100); | Metschnikowia | 13 | Saprotroph | \|Undefined Saprotroph\| | Yeast | NULL | Probable | As nectar-tap_saprotroph (Põlme et al. 2020); Decay sugar-rich substrates (Põlme et al. 2020) | Põlme S, et al. 2020. Fungal Diversity 105:1-16 ((https://doi.org/10.1007/s13225-020-00466-2)) |
| Otu55689 | 1 | k__Fungi(100);p__Ascomycota(100);c__Saccharomycetes(100);o__Saccharomycetales(100);f__Metschnikowiaceae(100);g__Metschnikowia(100);s__Metschnikowia_pulcherrima(100); | Metschnikowia | 13 | Saprotroph | \|Undefined Saprotroph\| | Yeast | NULL | Probable | As nectar-tap_saprotroph (Põlme et al. 2020); Decay sugar-rich substrates (Põlme et al. 2020) | Põlme S, et al. 2020. Fungal Diversity 105:1-16 ((https://doi.org/10.1007/s13225-020-00466-2)) |
| Otu55690 | 1 | k__Fungi(100);p__Ascomycota(100);c__Saccharomycetes(100);o__Saccharomycetales(100);f__Metschnikowiaceae(100);g__Metschnikowia(100);s__Metschnikowia_pulcherrima(100); | Metschnikowia | 13 | Saprotroph | \|Undefined Saprotroph\| | Yeast | NULL | Probable | As nectar-tap_saprotroph (Põlme et al. 2020); Decay sugar-rich substrates (Põlme et al. 2020) | Põlme S, et al. 2020. Fungal Diversity 105:1-16 ((https://doi.org/10.1007/s13225-020-00466-2)) |
| Otu55692 | 1 | k__Fungi(100);p__Ascomycota(100);c__Saccharomycetes(100);o__Saccharomycetales(100);f__Metschnikowiaceae(100);g__Metschnikowia(100);s__Metschnikowia_pulcherrima(100); | Metschnikowia | 13 | Saprotroph | \|Undefined Saprotroph\| | Yeast | NULL | Probable | As nectar-tap_saprotroph (Põlme et al. 2020); Decay sugar-rich substrates (Põlme et al. 2020) | Põlme S, et al. 2020. Fungal Diversity 105:1-16 ((https://doi.org/10.1007/s13225-020-00466-2)) |
| Otu55693 | 1 | k__Fungi(100);p__Ascomycota(100);c__Saccharomycetes(100);o__Saccharomycetales(100);f__Metschnikowiaceae(100);g__Metschnikowia(100);s__Metschnikowia_pulcherrima(100); | Metschnikowia | 13 | Saprotroph | \|Undefined Saprotroph\| | Yeast | NULL | Probable | As nectar-tap_saprotroph (Põlme et al. 2020); Decay sugar-rich substrates (Põlme et al. 2020) | Põlme S, et al. 2020. Fungal Diversity 105:1-16 ((https://doi.org/10.1007/s13225-020-00466-2)) |
| Otu55694 | 1 | k__Fungi(100);p__Ascomycota(100);c__Saccharomycetes(100);o__Saccharomycetales(100);f__Metschnikowiaceae(100);g__Metschnikowia(100);s__Metschnikowia_pulcherrima(100); | Metschnikowia | 13 | Saprotroph | \|Undefined Saprotroph\| | Yeast | NULL | Probable | As nectar-tap_saprotroph (Põlme et al. 2020); Decay sugar-rich substrates (Põlme et al. 2020) | Põlme S, et al. 2020. Fungal Diversity 105:1-16 ((https://doi.org/10.1007/s13225-020-00466-2)) |
| Otu55695 | 1 | k__Fungi(100);p__Ascomycota(100);c__Saccharomycetes(100);o__Saccharomycetales(100);f__Metschnikowiaceae(100);g__Metschnikowia(100);s__Metschnikowia_pulcherrima(100); | Metschnikowia | 13 | Saprotroph | \|Undefined Saprotroph\| | Yeast | NULL | Probable | As nectar-tap_saprotroph (Põlme et al. 2020); Decay sugar-rich substrates (Põlme et al. 2020) | Põlme S, et al. 2020. Fungal Diversity 105:1-16 ((https://doi.org/10.1007/s13225-020-00466-2)) |
| Otu55696 | 1 | k__Fungi(100);p__Ascomycota(100);c__Saccharomycetes(100);o__Saccharomycetales(100);f__Metschnikowiaceae(100);g__Metschnikowia(100);s__Metschnikowia_pulcherrima(100); | Metschnikowia | 13 | Saprotroph | \|Undefined Saprotroph\| | Yeast | NULL | Probable | As nectar-tap_saprotroph (Põlme et al. 2020); Decay sugar-rich substrates (Põlme et al. 2020) | Põlme S, et al. 2020. Fungal Diversity 105:1-16 ((https://doi.org/10.1007/s13225-020-00466-2)) |
| Otu55697 | 1 | k__Fungi(100);p__Ascomycota(100);c__Saccharomycetes(100);o__Saccharomycetales(100);f__Metschnikowiaceae(100);g__Metschnikowia(100);s__Metschnikowia_pulcherrima(100); | Metschnikowia | 13 | Saprotroph | \|Undefined Saprotroph\| | Yeast | NULL | Probable | As nectar-tap_saprotroph (Põlme et al. 2020); Decay sugar-rich substrates (Põlme et al. 2020) | Põlme S, et al. 2020. Fungal Diversity 105:1-16 ((https://doi.org/10.1007/s13225-020-00466-2)) |
| Otu55698 | 1 | k__Fungi(100);p__Ascomycota(100);c__Saccharomycetes(100);o__Saccharomycetales(100);f__Metschnikowiaceae(100);g__Metschnikowia(100);s__Metschnikowia_pulcherrima(100); | Metschnikowia | 13 | Saprotroph | \|Undefined Saprotroph\| | Yeast | NULL | Probable | As nectar-tap_saprotroph (Põlme et al. 2020); Decay sugar-rich substrates (Põlme et al. 2020) | Põlme S, et al. 2020. Fungal Diversity 105:1-16 ((https://doi.org/10.1007/s13225-020-00466-2)) |
| Otu55699 | 1 | k__Fungi(100);p__Ascomycota(100);c__Saccharomycetes(100);o__Saccharomycetales(100);f__Metschnikowiaceae(100);g__Metschnikowia(100);s__Metschnikowia_pulcherrima(100); | Metschnikowia | 13 | Saprotroph | \|Undefined Saprotroph\| | Yeast | NULL | Probable | As nectar-tap_saprotroph (Põlme et al. 2020); Decay sugar-rich substrates (Põlme et al. 2020) | Põlme S, et al. 2020. Fungal Diversity 105:1-16 ((https://doi.org/10.1007/s13225-020-00466-2)) |
| Otu55703 | 1 | k__Fungi(100);p__Ascomycota(100);c__Saccharomycetes(100);o__Saccharomycetales(100);f__Metschnikowiaceae(100);g__Metschnikowia(100);g__Metschnikowia_unclassified(100); | Metschnikowia | 13 | Saprotroph | \|Undefined Saprotroph\| | Yeast | NULL | Probable | As nectar-tap_saprotroph (Põlme et al. 2020); Decay sugar-rich substrates (Põlme et al. 2020) | Põlme S, et al. 2020. Fungal Diversity 105:1-16 ((https://doi.org/10.1007/s13225-020-00466-2)) |
| Otu55707 | 1 | k__Fungi(100);p__Ascomycota(100);c__Saccharomycetes(100);o__Saccharomycetales(100);f__Metschnikowiaceae(100);g__Metschnikowia(100);s__Metschnikowia_pulcherrima(100); | Metschnikowia | 13 | Saprotroph | \|Undefined Saprotroph\| | Yeast | NULL | Probable | As nectar-tap_saprotroph (Põlme et al. 2020); Decay sugar-rich substrates (Põlme et al. 2020) | Põlme S, et al. 2020. Fungal Diversity 105:1-16 ((https://doi.org/10.1007/s13225-020-00466-2)) |
| Otu55709 | 1 | k__Fungi(100);p__Ascomycota(100);c__Saccharomycetes(100);o__Saccharomycetales(100);f__Metschnikowiaceae(100);g__Metschnikowia(100);s__Metschnikowia_pulcherrima(100); | Metschnikowia | 13 | Saprotroph | \|Undefined Saprotroph\| | Yeast | NULL | Probable | As nectar-tap_saprotroph (Põlme et al. 2020); Decay sugar-rich substrates (Põlme et al. 2020) | Põlme S, et al. 2020. Fungal Diversity 105:1-16 ((https://doi.org/10.1007/s13225-020-00466-2)) |
| Otu55711 | 1 | k__Fungi(100);p__Ascomycota(100);c__Saccharomycetes(100);o__Saccharomycetales(100);f__Metschnikowiaceae(100);g__Metschnikowia(100);s__Metschnikowia_pulcherrima(100); | Metschnikowia | 13 | Saprotroph | \|Undefined Saprotroph\| | Yeast | NULL | Probable | As nectar-tap_saprotroph (Põlme et al. 2020); Decay sugar-rich substrates (Põlme et al. 2020) | Põlme S, et al. 2020. Fungal Diversity 105:1-16 ((https://doi.org/10.1007/s13225-020-00466-2)) |
| Otu55717 | 1 | k__Fungi(100);p__Ascomycota(100);c__Saccharomycetes(100);o__Saccharomycetales(100);f__Metschnikowiaceae(100);g__Metschnikowia(100);s__Metschnikowia_pulcherrima(100); | Metschnikowia | 13 | Saprotroph | \|Undefined Saprotroph\| | Yeast | NULL | Probable | As nectar-tap_saprotroph (Põlme et al. 2020); Decay sugar-rich substrates (Põlme et al. 2020) | Põlme S, et al. 2020. Fungal Diversity 105:1-16 ((https://doi.org/10.1007/s13225-020-00466-2)) |
| Otu55718 | 1 | k__Fungi(100);p__Ascomycota(100);c__Saccharomycetes(100);o__Saccharomycetales(100);f__Metschnikowiaceae(100);g__Metschnikowia(100);s__Metschnikowia_pulcherrima(100); | Metschnikowia | 13 | Saprotroph | \|Undefined Saprotroph\| | Yeast | NULL | Probable | As nectar-tap_saprotroph (Põlme et al. 2020); Decay sugar-rich substrates (Põlme et al. 2020) | Põlme S, et al. 2020. Fungal Diversity 105:1-16 ((https://doi.org/10.1007/s13225-020-00466-2)) |
| Otu55719 | 1 | k__Fungi(100);p__Ascomycota(100);c__Saccharomycetes(100);o__Saccharomycetales(100);f__Metschnikowiaceae(100);g__Metschnikowia(100);s__Metschnikowia_pulcherrima(100); | Metschnikowia | 13 | Saprotroph | \|Undefined Saprotroph\| | Yeast | NULL | Probable | As nectar-tap_saprotroph (Põlme et al. 2020); Decay sugar-rich substrates (Põlme et al. 2020) | Põlme S, et al. 2020. Fungal Diversity 105:1-16 ((https://doi.org/10.1007/s13225-020-00466-2)) |
| Otu55720 | 1 | k__Fungi(100);p__Ascomycota(100);c__Saccharomycetes(100);o__Saccharomycetales(100);f__Metschnikowiaceae(100);g__Metschnikowia(100);g__Metschnikowia_unclassified(100); | Metschnikowia | 13 | Saprotroph | \|Undefined Saprotroph\| | Yeast | NULL | Probable | As nectar-tap_saprotroph (Põlme et al. 2020); Decay sugar-rich substrates (Põlme et al. 2020) | Põlme S, et al. 2020. Fungal Diversity 105:1-16 ((https://doi.org/10.1007/s13225-020-00466-2)) |
| Otu55721 | 1 | k__Fungi(100);p__Ascomycota(100);c__Saccharomycetes(100);o__Saccharomycetales(100);f__Metschnikowiaceae(100);g__Metschnikowia(100);g__Metschnikowia_unclassified(100); | Metschnikowia | 13 | Saprotroph | \|Undefined Saprotroph\| | Yeast | NULL | Probable | As nectar-tap_saprotroph (Põlme et al. 2020); Decay sugar-rich substrates (Põlme et al. 2020) | Põlme S, et al. 2020. Fungal Diversity 105:1-16 ((https://doi.org/10.1007/s13225-020-00466-2)) |
| Otu55722 | 1 | k__Fungi(100);p__Ascomycota(100);c__Saccharomycetes(100);o__Saccharomycetales(100);f__Metschnikowiaceae(100);g__Metschnikowia(100);s__Metschnikowia_pulcherrima(100); | Metschnikowia | 13 | Saprotroph | \|Undefined Saprotroph\| | Yeast | NULL | Probable | As nectar-tap_saprotroph (Põlme et al. 2020); Decay sugar-rich substrates (Põlme et al. 2020) | Põlme S, et al. 2020. Fungal Diversity 105:1-16 ((https://doi.org/10.1007/s13225-020-00466-2)) |
| Otu55723 | 1 | k__Fungi(100);p__Ascomycota(100);c__Saccharomycetes(100);o__Saccharomycetales(100);f__Metschnikowiaceae(100);g__Metschnikowia(100);s__Metschnikowia_pulcherrima(100); | Metschnikowia | 13 | Saprotroph | \|Undefined Saprotroph\| | Yeast | NULL | Probable | As nectar-tap_saprotroph (Põlme et al. 2020); Decay sugar-rich substrates (Põlme et al. 2020) | Põlme S, et al. 2020. Fungal Diversity 105:1-16 ((https://doi.org/10.1007/s13225-020-00466-2)) |
| Otu55726 | 1 | k__Fungi(100);p__Ascomycota(100);c__Saccharomycetes(100);o__Saccharomycetales(100);f__Metschnikowiaceae(100);g__Metschnikowia(100);s__Metschnikowia_pulcherrima(100); | Metschnikowia | 13 | Saprotroph | \|Undefined Saprotroph\| | Yeast | NULL | Probable | As nectar-tap_saprotroph (Põlme et al. 2020); Decay sugar-rich substrates (Põlme et al. 2020) | Põlme S, et al. 2020. Fungal Diversity 105:1-16 ((https://doi.org/10.1007/s13225-020-00466-2)) |
| Otu55727 | 1 | k__Fungi(100);p__Ascomycota(100);c__Saccharomycetes(100);o__Saccharomycetales(100);f__Metschnikowiaceae(100);g__Metschnikowia(100);s__Metschnikowia_pulcherrima(100); | Metschnikowia | 13 | Saprotroph | \|Undefined Saprotroph\| | Yeast | NULL | Probable | As nectar-tap_saprotroph (Põlme et al. 2020); Decay sugar-rich substrates (Põlme et al. 2020) | Põlme S, et al. 2020. Fungal Diversity 105:1-16 ((https://doi.org/10.1007/s13225-020-00466-2)) |
| Otu55728 | 1 | k__Fungi(100);p__Ascomycota(100);c__Saccharomycetes(100);o__Saccharomycetales(100);f__Metschnikowiaceae(100);g__Metschnikowia(100);g__Metschnikowia_unclassified(100); | Metschnikowia | 13 | Saprotroph | \|Undefined Saprotroph\| | Yeast | NULL | Probable | As nectar-tap_saprotroph (Põlme et al. 2020); Decay sugar-rich substrates (Põlme et al. 2020) | Põlme S, et al. 2020. Fungal Diversity 105:1-16 ((https://doi.org/10.1007/s13225-020-00466-2)) |
| Otu55729 | 1 | k__Fungi(100);p__Ascomycota(100);c__Saccharomycetes(100);o__Saccharomycetales(100);f__Metschnikowiaceae(100);g__Metschnikowia(100);s__Metschnikowia_pulcherrima(100); | Metschnikowia | 13 | Saprotroph | \|Undefined Saprotroph\| | Yeast | NULL | Probable | As nectar-tap_saprotroph (Põlme et al. 2020); Decay sugar-rich substrates (Põlme et al. 2020) | Põlme S, et al. 2020. Fungal Diversity 105:1-16 ((https://doi.org/10.1007/s13225-020-00466-2)) |
| Otu55730 | 1 | k__Fungi(100);p__Ascomycota(100);c__Saccharomycetes(100);o__Saccharomycetales(100);f__Metschnikowiaceae(100);g__Metschnikowia(100);s__Metschnikowia_pulcherrima(100); | Metschnikowia | 13 | Saprotroph | \|Undefined Saprotroph\| | Yeast | NULL | Probable | As nectar-tap_saprotroph (Põlme et al. 2020); Decay sugar-rich substrates (Põlme et al. 2020) | Põlme S, et al. 2020. Fungal Diversity 105:1-16 ((https://doi.org/10.1007/s13225-020-00466-2)) |
| Otu55731 | 1 | k__Fungi(100);p__Ascomycota(100);c__Saccharomycetes(100);o__Saccharomycetales(100);f__Metschnikowiaceae(100);g__Metschnikowia(100);s__Metschnikowia_pulcherrima(100); | Metschnikowia | 13 | Saprotroph | \|Undefined Saprotroph\| | Yeast | NULL | Probable | As nectar-tap_saprotroph (Põlme et al. 2020); Decay sugar-rich substrates (Põlme et al. 2020) | Põlme S, et al. 2020. Fungal Diversity 105:1-16 ((https://doi.org/10.1007/s13225-020-00466-2)) |
| Otu55732 | 1 | k__Fungi(100);p__Ascomycota(100);c__Saccharomycetes(100);o__Saccharomycetales(100);f__Metschnikowiaceae(100);g__Metschnikowia(100);s__Metschnikowia_pulcherrima(100); | Metschnikowia | 13 | Saprotroph | \|Undefined Saprotroph\| | Yeast | NULL | Probable | As nectar-tap_saprotroph (Põlme et al. 2020); Decay sugar-rich substrates (Põlme et al. 2020) | Põlme S, et al. 2020. Fungal Diversity 105:1-16 ((https://doi.org/10.1007/s13225-020-00466-2)) |
| Otu55733 | 1 | k__Fungi(100);p__Ascomycota(100);c__Saccharomycetes(100);o__Saccharomycetales(100);f__Metschnikowiaceae(100);g__Metschnikowia(100);g__Metschnikowia_unclassified(100); | Metschnikowia | 13 | Saprotroph | \|Undefined Saprotroph\| | Yeast | NULL | Probable | As nectar-tap_saprotroph (Põlme et al. 2020); Decay sugar-rich substrates (Põlme et al. 2020) | Põlme S, et al. 2020. Fungal Diversity 105:1-16 ((https://doi.org/10.1007/s13225-020-00466-2)) |
| Otu55734 | 1 | k__Fungi(100);p__Ascomycota(100);c__Saccharomycetes(100);o__Saccharomycetales(100);f__Metschnikowiaceae(100);g__Metschnikowia(100);s__Metschnikowia_pulcherrima(100); | Metschnikowia | 13 | Saprotroph | \|Undefined Saprotroph\| | Yeast | NULL | Probable | As nectar-tap_saprotroph (Põlme et al. 2020); Decay sugar-rich substrates (Põlme et al. 2020) | Põlme S, et al. 2020. Fungal Diversity 105:1-16 ((https://doi.org/10.1007/s13225-020-00466-2)) |
| Otu55735 | 1 | k__Fungi(100);p__Ascomycota(100);c__Saccharomycetes(100);o__Saccharomycetales(100);f__Metschnikowiaceae(100);g__Metschnikowia(100);s__Metschnikowia_pulcherrima(100); | Metschnikowia | 13 | Saprotroph | \|Undefined Saprotroph\| | Yeast | NULL | Probable | As nectar-tap_saprotroph (Põlme et al. 2020); Decay sugar-rich substrates (Põlme et al. 2020) | Põlme S, et al. 2020. Fungal Diversity 105:1-16 ((https://doi.org/10.1007/s13225-020-00466-2)) |
| Otu55736 | 1 | k__Fungi(100);p__Ascomycota(100);c__Saccharomycetes(100);o__Saccharomycetales(100);f__Metschnikowiaceae(100);g__Metschnikowia(100);s__Metschnikowia_pulcherrima(100); | Metschnikowia | 13 | Saprotroph | \|Undefined Saprotroph\| | Yeast | NULL | Probable | As nectar-tap_saprotroph (Põlme et al. 2020); Decay sugar-rich substrates (Põlme et al. 2020) | Põlme S, et al. 2020. Fungal Diversity 105:1-16 ((https://doi.org/10.1007/s13225-020-00466-2)) |
| Otu55737 | 1 | k__Fungi(100);p__Ascomycota(100);c__Saccharomycetes(100);o__Saccharomycetales(100);f__Metschnikowiaceae(100);g__Metschnikowia(100);s__Metschnikowia_pulcherrima(100); | Metschnikowia | 13 | Saprotroph | \|Undefined Saprotroph\| | Yeast | NULL | Probable | As nectar-tap_saprotroph (Põlme et al. 2020); Decay sugar-rich substrates (Põlme et al. 2020) | Põlme S, et al. 2020. Fungal Diversity 105:1-16 ((https://doi.org/10.1007/s13225-020-00466-2)) |
| Otu55738 | 1 | k__Fungi(100);p__Ascomycota(100);c__Saccharomycetes(100);o__Saccharomycetales(100);f__Metschnikowiaceae(100);g__Metschnikowia(100);s__Metschnikowia_pulcherrima(100); | Metschnikowia | 13 | Saprotroph | \|Undefined Saprotroph\| | Yeast | NULL | Probable | As nectar-tap_saprotroph (Põlme et al. 2020); Decay sugar-rich substrates (Põlme et al. 2020) | Põlme S, et al. 2020. Fungal Diversity 105:1-16 ((https://doi.org/10.1007/s13225-020-00466-2)) |
| Otu55739 | 1 | k__Fungi(100);p__Ascomycota(100);c__Saccharomycetes(100);o__Saccharomycetales(100);f__Metschnikowiaceae(100);g__Metschnikowia(100);s__Metschnikowia_pulcherrima(100); | Metschnikowia | 13 | Saprotroph | \|Undefined Saprotroph\| | Yeast | NULL | Probable | As nectar-tap_saprotroph (Põlme et al. 2020); Decay sugar-rich substrates (Põlme et al. 2020) | Põlme S, et al. 2020. Fungal Diversity 105:1-16 ((https://doi.org/10.1007/s13225-020-00466-2)) |
| Otu55740 | 1 | k__Fungi(100);p__Ascomycota(100);c__Saccharomycetes(100);o__Saccharomycetales(100);f__Metschnikowiaceae(100);g__Metschnikowia(100);g__Metschnikowia_unclassified(100); | Metschnikowia | 13 | Saprotroph | \|Undefined Saprotroph\| | Yeast | NULL | Probable | As nectar-tap_saprotroph (Põlme et al. 2020); Decay sugar-rich substrates (Põlme et al. 2020) | Põlme S, et al. 2020. Fungal Diversity 105:1-16 ((https://doi.org/10.1007/s13225-020-00466-2)) |
| Otu55741 | 1 | k__Fungi(100);p__Ascomycota(100);c__Saccharomycetes(100);o__Saccharomycetales(100);f__Metschnikowiaceae(100);g__Metschnikowia(100);s__Metschnikowia_pulcherrima(100); | Metschnikowia | 13 | Saprotroph | \|Undefined Saprotroph\| | Yeast | NULL | Probable | As nectar-tap_saprotroph (Põlme et al. 2020); Decay sugar-rich substrates (Põlme et al. 2020) | Põlme S, et al. 2020. Fungal Diversity 105:1-16 ((https://doi.org/10.1007/s13225-020-00466-2)) |
| Otu55742 | 1 | k__Fungi(100);p__Ascomycota(100);c__Saccharomycetes(100);o__Saccharomycetales(100);f__Metschnikowiaceae(100);g__Metschnikowia(100);s__Metschnikowia_pulcherrima(100); | Metschnikowia | 13 | Saprotroph | \|Undefined Saprotroph\| | Yeast | NULL | Probable | As nectar-tap_saprotroph (Põlme et al. 2020); Decay sugar-rich substrates (Põlme et al. 2020) | Põlme S, et al. 2020. Fungal Diversity 105:1-16 ((https://doi.org/10.1007/s13225-020-00466-2)) |
| Otu55745 | 1 | k__Fungi(100);p__Ascomycota(100);c__Saccharomycetes(100);o__Saccharomycetales(100);f__Metschnikowiaceae(100);g__Metschnikowia(100);g__Metschnikowia_unclassified(100); | Metschnikowia | 13 | Saprotroph | \|Undefined Saprotroph\| | Yeast | NULL | Probable | As nectar-tap_saprotroph (Põlme et al. 2020); Decay sugar-rich substrates (Põlme et al. 2020) | Põlme S, et al. 2020. Fungal Diversity 105:1-16 ((https://doi.org/10.1007/s13225-020-00466-2)) |
| Otu55746 | 1 | k__Fungi(100);p__Ascomycota(100);c__Saccharomycetes(100);o__Saccharomycetales(100);f__Metschnikowiaceae(100);g__Metschnikowia(100);g__Metschnikowia_unclassified(100); | Metschnikowia | 13 | Saprotroph | \|Undefined Saprotroph\| | Yeast | NULL | Probable | As nectar-tap_saprotroph (Põlme et al. 2020); Decay sugar-rich substrates (Põlme et al. 2020) | Põlme S, et al. 2020. Fungal Diversity 105:1-16 ((https://doi.org/10.1007/s13225-020-00466-2)) |

| | | | | | | | | | | | |
|---|---|---|---|---|---|---|---|---|---|---|---|
| Otu55747 | 1 | k__Fungi(100);p__Ascomycota(100);c__Saccharomycetes(100);o__Saccharomycetales(100);f__Metschnikowiaceae(100);g__Metschnikowia(100);s__Metschnikowia_pulcherrima(100); | Metschnikowia | 13 | Saprotroph | \|Undefined Saprotroph\| | Yeast | NULL | Probable | As nectar-tap_saprotroph (Põlme et al. 2020); Decay sugar-rich substrates (Põlme et al. 2020) | Põlme S, et al. 2020. Fungal Diversity 105:1-16 ((https://doi.org/10.1007/s13225-020-00466-2)) |
| Otu55748 | 1 | k__Fungi(100);p__Ascomycota(100);c__Saccharomycetes(100);o__Saccharomycetales(100);f__Metschnikowiaceae(100);g__Metschnikowia(100);s__Metschnikowia_pulcherrima(100); | Metschnikowia | 13 | Saprotroph | \|Undefined Saprotroph\| | Yeast | NULL | Probable | As nectar-tap_saprotroph (Põlme et al. 2020); Decay sugar-rich substrates (Põlme et al. 2020) | Põlme S, et al. 2020. Fungal Diversity 105:1-16 ((https://doi.org/10.1007/s13225-020-00466-2)) |
| Otu55749 | 1 | k__Fungi(100);p__Ascomycota(100);c__Saccharomycetes(100);o__Saccharomycetales(100);f__Metschnikowiaceae(100);g__Metschnikowia(100);g__Metschnikowia_unclassified(100); | Metschnikowia | 13 | Saprotroph | \|Undefined Saprotroph\| | Yeast | NULL | Probable | As nectar-tap_saprotroph (Põlme et al. 2020); Decay sugar-rich substrates (Põlme et al. 2020) | Põlme S, et al. 2020. Fungal Diversity 105:1-16 ((https://doi.org/10.1007/s13225-020-00466-2)) |
| Otu55750 | 1 | k__Fungi(100);p__Ascomycota(100);c__Saccharomycetes(100);o__Saccharomycetales(100);f__Metschnikowiaceae(100);g__Metschnikowia(100);s__Metschnikowia_pulcherrima(100); | Metschnikowia | 13 | Saprotroph | \|Undefined Saprotroph\| | Yeast | NULL | Probable | As nectar-tap_saprotroph (Põlme et al. 2020); Decay sugar-rich substrates (Põlme et al. 2020) | Põlme S, et al. 2020. Fungal Diversity 105:1-16 ((https://doi.org/10.1007/s13225-020-00466-2)) |
| Otu55751 | 1 | k__Fungi(100);p__Ascomycota(100);c__Saccharomycetes(100);o__Saccharomycetales(100);f__Metschnikowiaceae(100);g__Metschnikowia(100);g__Metschnikowia_unclassified(100); | Metschnikowia | 13 | Saprotroph | \|Undefined Saprotroph\| | Yeast | NULL | Probable | As nectar-tap_saprotroph (Põlme et al. 2020); Decay sugar-rich substrates (Põlme et al. 2020) | Põlme S, et al. 2020. Fungal Diversity 105:1-16 ((https://doi.org/10.1007/s13225-020-00466-2)) |
| Otu55752 | 1 | k__Fungi(100);p__Ascomycota(100);c__Saccharomycetes(100);o__Saccharomycetales(100);f__Metschnikowiaceae(100);g__Metschnikowia(100);s__Metschnikowia_pulcherrima(100); | Metschnikowia | 13 | Saprotroph | \|Undefined Saprotroph\| | Yeast | NULL | Probable | As nectar-tap_saprotroph (Põlme et al. 2020); Decay sugar-rich substrates (Põlme et al. 2020) | Põlme S, et al. 2020. Fungal Diversity 105:1-16 ((https://doi.org/10.1007/s13225-020-00466-2)) |
| Otu55753 | 1 | k__Fungi(100);p__Ascomycota(100);c__Saccharomycetes(100);o__Saccharomycetales(100);f__Metschnikowiaceae(100);g__Metschnikowia(100);s__Metschnikowia_pulcherrima(100); | Metschnikowia | 13 | Saprotroph | \|Undefined Saprotroph\| | Yeast | NULL | Probable | As nectar-tap_saprotroph (Põlme et al. 2020); Decay sugar-rich substrates (Põlme et al. 2020) | Põlme S, et al. 2020. Fungal Diversity 105:1-16 ((https://doi.org/10.1007/s13225-020-00466-2)) |
| Otu55754 | 1 | k__Fungi(100);p__Ascomycota(100);c__Saccharomycetes(100);o__Saccharomycetales(100);f__Metschnikowiaceae(100);g__Metschnikowia(100);s__Metschnikowia_pulcherrima(100); | Metschnikowia | 13 | Saprotroph | \|Undefined Saprotroph\| | Yeast | NULL | Probable | As nectar-tap_saprotroph (Põlme et al. 2020); Decay sugar-rich substrates (Põlme et al. 2020) | Põlme S, et al. 2020. Fungal Diversity 105:1-16 ((https://doi.org/10.1007/s13225-020-00466-2)) |
| Otu55755 | 1 | k__Fungi(100);p__Ascomycota(100);c__Saccharomycetes(100);o__Saccharomycetales(100);f__Metschnikowiaceae(100);g__Metschnikowia(100);s__Metschnikowia_pulcherrima(100); | Metschnikowia | 13 | Saprotroph | \|Undefined Saprotroph\| | Yeast | NULL | Probable | As nectar-tap_saprotroph (Põlme et al. 2020); Decay sugar-rich substrates (Põlme et al. 2020) | Põlme S, et al. 2020. Fungal Diversity 105:1-16 ((https://doi.org/10.1007/s13225-020-00466-2)) |
| Otu55756 | 1 | k__Fungi(100);p__Ascomycota(100);c__Saccharomycetes(100);o__Saccharomycetales(100);f__Metschnikowiaceae(100);g__Metschnikowia(100);g__Metschnikowia_unclassified(100); | Metschnikowia | 13 | Saprotroph | \|Undefined Saprotroph\| | Yeast | NULL | Probable | As nectar-tap_saprotroph (Põlme et al. 2020); Decay sugar-rich substrates (Põlme et al. 2020) | Põlme S, et al. 2020. Fungal Diversity 105:1-16 ((https://doi.org/10.1007/s13225-020-00466-2)) |
| Otu55759 | 1 | k__Fungi(100);p__Ascomycota(100);c__Saccharomycetes(100);o__Saccharomycetales(100);o__Saccharomycetales_unclassified(100);o__Saccharomycetales_unclassified(100);o__Saccharomycetales_unclassified(100); | Saccharomycetales | 7 | Saprotroph | Undefined Saprotroph | Yeast | NULL | Possible | NULL | Sterkenburg E, et al. 2015. New Phytologist 207:1145-1158 ((https://doi.org/10.1111/nph.13426)) |
| Otu55760 | 1 | k__Fungi(100);p__Ascomycota(100);c__Saccharomycetes(100);o__Saccharomycetales(100);f__Metschnikowiaceae(100);g__Metschnikowia(100);s__Metschnikowia_pulcherrima(100); | Metschnikowia | 13 | Saprotroph | \|Undefined Saprotroph\| | Yeast | NULL | Probable | As nectar-tap_saprotroph (Põlme et al. 2020); Decay sugar-rich substrates (Põlme et al. 2020) | Põlme S, et al. 2020. Fungal Diversity 105:1-16 ((https://doi.org/10.1007/s13225-020-00466-2)) |
| Otu55761 | 1 | k__Fungi(100);p__Ascomycota(100);c__Saccharomycetes(100);o__Saccharomycetales(100);f__Metschnikowiaceae(100);g__Metschnikowia(100);s__Metschnikowia_pulcherrima(100); | Metschnikowia | 13 | Saprotroph | \|Undefined Saprotroph\| | Yeast | NULL | Probable | As nectar-tap_saprotroph (Põlme et al. 2020); Decay sugar-rich substrates (Põlme et al. 2020) | Põlme S, et al. 2020. Fungal Diversity 105:1-16 ((https://doi.org/10.1007/s13225-020-00466-2)) |
| Otu55762 | 1 | k__Fungi(100);p__Ascomycota(100);c__Saccharomycetes(100);o__Saccharomycetales(100);f__Metschnikowiaceae(100);g__Metschnikowia(100);s__Metschnikowia_pulcherrima(100); | Metschnikowia | 13 | Saprotroph | \|Undefined Saprotroph\| | Yeast | NULL | Probable | As nectar-tap_saprotroph (Põlme et al. 2020); Decay sugar-rich substrates (Põlme et al. 2020) | Põlme S, et al. 2020. Fungal Diversity 105:1-16 ((https://doi.org/10.1007/s13225-020-00466-2)) |
| Otu55763 | 1 | k__Fungi(100);p__Ascomycota(100);c__Saccharomycetes(100);o__Saccharomycetales(100);f__Metschnikowiaceae(100);g__Metschnikowia(100);s__Metschnikowia_pulcherrima(100); | Metschnikowia | 13 | Saprotroph | \|Undefined Saprotroph\| | Yeast | NULL | Probable | As nectar-tap_saprotroph (Põlme et al. 2020); Decay sugar-rich substrates (Põlme et al. 2020) | Põlme S, et al. 2020. Fungal Diversity 105:1-16 ((https://doi.org/10.1007/s13225-020-00466-2)) |
| Otu55764 | 1 | k__Fungi(100);p__Ascomycota(100);c__Saccharomycetes(100);o__Saccharomycetales(100);f__Metschnikowiaceae(100);g__Metschnikowia(100);s__Metschnikowia_pulcherrima(100); | Metschnikowia | 13 | Saprotroph | \|Undefined Saprotroph\| | Yeast | NULL | Probable | As nectar-tap_saprotroph (Põlme et al. 2020); Decay sugar-rich substrates (Põlme et al. 2020) | Põlme S, et al. 2020. Fungal Diversity 105:1-16 ((https://doi.org/10.1007/s13225-020-00466-2)) |
| Otu55765 | 1 | k__Fungi(100);p__Ascomycota(100);c__Saccharomycetes(100);o__Saccharomycetales(100);f__Metschnikowiaceae(100);g__Metschnikowia(100);s__Metschnikowia_pulcherrima(100); | Metschnikowia | 13 | Saprotroph | \|Undefined Saprotroph\| | Yeast | NULL | Probable | As nectar-tap_saprotroph (Põlme et al. 2020); Decay sugar-rich substrates (Põlme et al. 2020) | Põlme S, et al. 2020. Fungal Diversity 105:1-16 ((https://doi.org/10.1007/s13225-020-00466-2)) |
| Otu55766 | 1 | k__Fungi(100);p__Ascomycota(100);c__Saccharomycetes(100);o__Saccharomycetales(100);f__Metschnikowiaceae(100);g__Metschnikowia(100);s__Metschnikowia_pulcherrima(100); | Metschnikowia | 13 | Saprotroph | \|Undefined Saprotroph\| | Yeast | NULL | Probable | As nectar-tap_saprotroph (Põlme et al. 2020); Decay sugar-rich substrates (Põlme et al. 2020) | Põlme S, et al. 2020. Fungal Diversity 105:1-16 ((https://doi.org/10.1007/s13225-020-00466-2)) |
| Otu55768 | 1 | k__Fungi(100);p__Ascomycota(100);c__Saccharomycetes(100);o__Saccharomycetales(100);f__Metschnikowiaceae(100);g__Metschnikowia(100);s__Metschnikowia_pulcherrima(100); | Metschnikowia | 13 | Saprotroph | \|Undefined Saprotroph\| | Yeast | NULL | Probable | As nectar-tap_saprotroph (Põlme et al. 2020); Decay sugar-rich substrates (Põlme et al. 2020) | Põlme S, et al. 2020. Fungal Diversity 105:1-16 ((https://doi.org/10.1007/s13225-020-00466-2)) |
| Otu55769 | 1 | k__Fungi(100);p__Ascomycota(100);c__Saccharomycetes(100);o__Saccharomycetales(100);f__Metschnikowiaceae(100);g__Metschnikowia(100);s__Metschnikowia_pulcherrima(100); | Metschnikowia | 13 | Saprotroph | \|Undefined Saprotroph\| | Yeast | NULL | Probable | As nectar-tap_saprotroph (Põlme et al. 2020); Decay sugar-rich substrates (Põlme et al. 2020) | Põlme S, et al. 2020. Fungal Diversity 105:1-16 ((https://doi.org/10.1007/s13225-020-00466-2)) |
| Otu55770 | 1 | k__Fungi(100);p__Ascomycota(100);c__Saccharomycetes(100);o__Saccharomycetales(100);f__Metschnikowiaceae(100);g__Metschnikowia(100);s__Metschnikowia_pulcherrima(100); | Metschnikowia | 13 | Saprotroph | \|Undefined Saprotroph\| | Yeast | NULL | Probable | As nectar-tap_saprotroph (Põlme et al. 2020); Decay sugar-rich substrates (Põlme et al. 2020) | Põlme S, et al. 2020. Fungal Diversity 105:1-16 ((https://doi.org/10.1007/s13225-020-00466-2)) |
| Otu55771 | 1 | k__Fungi(100);p__Ascomycota(100);c__Saccharomycetes(100);o__Saccharomycetales(100);f__Metschnikowiaceae(100);g__Metschnikowia(100);s__Metschnikowia_pulcherrima(100); | Metschnikowia | 13 | Saprotroph | \|Undefined Saprotroph\| | Yeast | NULL | Probable | As nectar-tap_saprotroph (Põlme et al. 2020); Decay sugar-rich substrates (Põlme et al. 2020) | Põlme S, et al. 2020. Fungal Diversity 105:1-16 ((https://doi.org/10.1007/s13225-020-00466-2)) |
| Otu55772 | 1 | k__Fungi(100);p__Ascomycota(100);c__Saccharomycetes(100);o__Saccharomycetales(100);f__Metschnikowiaceae(100);g__Metschnikowia(100);g__Metschnikowia_unclassified(100); | Metschnikowia | 13 | Saprotroph | \|Undefined Saprotroph\| | Yeast | NULL | Probable | As nectar-tap_saprotroph (Põlme et al. 2020); Decay sugar-rich substrates (Põlme et al. 2020) | Põlme S, et al. 2020. Fungal Diversity 105:1-16 ((https://doi.org/10.1007/s13225-020-00466-2)) |
| Otu55773 | 1 | k__Fungi(100);p__Ascomycota(100);c__Saccharomycetes(100);o__Saccharomycetales(100);f__Metschnikowiaceae(100);g__Metschnikowia(100);s__Metschnikowia_pulcherrima(100); | Metschnikowia | 13 | Saprotroph | \|Undefined Saprotroph\| | Yeast | NULL | Probable | As nectar-tap_saprotroph (Põlme et al. 2020); Decay sugar-rich substrates (Põlme et al. 2020) | Põlme S, et al. 2020. Fungal Diversity 105:1-16 ((https://doi.org/10.1007/s13225-020-00466-2)) |
| Otu55774 | 1 | k__Fungi(100);p__Ascomycota(100);c__Saccharomycetes(100);o__Saccharomycetales(100);f__Metschnikowiaceae(100);g__Metschnikowia(100);s__Metschnikowia_pulcherrima(100); | Metschnikowia | 13 | Saprotroph | \|Undefined Saprotroph\| | Yeast | NULL | Probable | As nectar-tap_saprotroph (Põlme et al. 2020); Decay sugar-rich substrates (Põlme et al. 2020) | Põlme S, et al. 2020. Fungal Diversity 105:1-16 ((https://doi.org/10.1007/s13225-020-00466-2)) |
| Otu55775 | 1 | k__Fungi(100);p__Ascomycota(100);c__Saccharomycetes(100);o__Saccharomycetales(100);f__Metschnikowiaceae(100);g__Metschnikowia(100);s__Metschnikowia_pulcherrima(100); | Metschnikowia | 13 | Saprotroph | \|Undefined Saprotroph\| | Yeast | NULL | Probable | As nectar-tap_saprotroph (Põlme et al. 2020); Decay sugar-rich substrates (Põlme et al. 2020) | Põlme S, et al. 2020. Fungal Diversity 105:1-16 ((https://doi.org/10.1007/s13225-020-00466-2)) |
| Otu55776 | 1 | k__Fungi(100);p__Ascomycota(100);c__Saccharomycetes(100);o__Saccharomycetales(100);f__Metschnikowiaceae(100);g__Metschnikowia(100);s__Metschnikowia_pulcherrima(100); | Metschnikowia | 13 | Saprotroph | \|Undefined Saprotroph\| | Yeast | NULL | Probable | As nectar-tap_saprotroph (Põlme et al. 2020); Decay sugar-rich substrates (Põlme et al. 2020) | Põlme S, et al. 2020. Fungal Diversity 105:1-16 ((https://doi.org/10.1007/s13225-020-00466-2)) |
| Otu55777 | 1 | k__Fungi(100);p__Ascomycota(100);c__Saccharomycetes(100);o__Saccharomycetales(100);f__Metschnikowiaceae(100);g__Metschnikowia(100);g__Metschnikowia_unclassified(100); | Metschnikowia | 13 | Saprotroph | \|Undefined Saprotroph\| | Yeast | NULL | Probable | As nectar-tap_saprotroph (Põlme et al. 2020); Decay sugar-rich substrates (Põlme et al. 2020) | Põlme S, et al. 2020. Fungal Diversity 105:1-16 ((https://doi.org/10.1007/s13225-020-00466-2)) |
| Otu55779 | 1 | k__Fungi(100);p__Ascomycota(100);c__Saccharomycetes(100);o__Saccharomycetales(100);f__Metschnikowiaceae(100);g__Metschnikowia(100);s__Metschnikowia_pulcherrima(100); | Metschnikowia | 13 | Saprotroph | \|Undefined Saprotroph\| | Yeast | NULL | Probable | As nectar-tap_saprotroph (Põlme et al. 2020); Decay sugar-rich substrates (Põlme et al. 2020) | Põlme S, et al. 2020. Fungal Diversity 105:1-16 ((https://doi.org/10.1007/s13225-020-00466-2)) |
| Otu55780 | 1 | k__Fungi(100);p__Ascomycota(100);c__Saccharomycetes(100);o__Saccharomycetales(100);f__Metschnikowiaceae(100);g__Metschnikowia(100);s__Metschnikowia_pulcherrima(100); | Metschnikowia | 13 | Saprotroph | \|Undefined Saprotroph\| | Yeast | NULL | Probable | As nectar-tap_saprotroph (Põlme et al. 2020); Decay sugar-rich substrates (Põlme et al. 2020) | Põlme S, et al. 2020. Fungal Diversity 105:1-16 ((https://doi.org/10.1007/s13225-020-00466-2)) |
| Otu55781 | 1 | k__Fungi(100);p__Ascomycota(100);c__Saccharomycetes(100);o__Saccharomycetales(100);f__Metschnikowiaceae(100);g__Metschnikowia(100);s__Metschnikowia_pulcherrima(100); | Metschnikowia | 13 | Saprotroph | \|Undefined Saprotroph\| | Yeast | NULL | Probable | As nectar-tap_saprotroph (Põlme et al. 2020); Decay sugar-rich substrates (Põlme et al. 2020) | Põlme S, et al. 2020. Fungal Diversity 105:1-16 ((https://doi.org/10.1007/s13225-020-00466-2)) |
| Otu55791 | 1 | k__Fungi(100);p__Ascomycota(100);c__Saccharomycetes(100);o__Saccharomycetales(100);f__Metschnikowiaceae(100);g__Metschnikowia(100);s__Metschnikowia_pulcherrima(100); | Metschnikowia | 13 | Saprotroph | \|Undefined Saprotroph\| | Yeast | NULL | Probable | As nectar-tap_saprotroph (Põlme et al. 2020); Decay sugar-rich substrates (Põlme et al. 2020) | Põlme S, et al. 2020. Fungal Diversity 105:1-16 ((https://doi.org/10.1007/s13225-020-00466-2)) |
| Otu55794 | 1 | k__Fungi(100);p__Ascomycota(100);c__Saccharomycetes(100);o__Saccharomycetales(100);f__Metschnikowiaceae(100);g__Metschnikowia(100);s__Metschnikowia_pulcherrima(100); | Metschnikowia | 13 | Saprotroph | \|Undefined Saprotroph\| | Yeast | NULL | Probable | As nectar-tap_saprotroph (Põlme et al. 2020); Decay sugar-rich substrates (Põlme et al. 2020) | Põlme S, et al. 2020. Fungal Diversity 105:1-16 ((https://doi.org/10.1007/s13225-020-00466-2)) |
| Otu55795 | 1 | k__Fungi(100);p__Ascomycota(100);c__Saccharomycetes(100);o__Saccharomycetales(100);f__Metschnikowiaceae(100);g__Metschnikowia(100);s__Metschnikowia_pulcherrima(100); | Metschnikowia | 13 | Saprotroph | \|Undefined Saprotroph\| | Yeast | NULL | Probable | As nectar-tap_saprotroph (Põlme et al. 2020); Decay sugar-rich substrates (Põlme et al. 2020) | Põlme S, et al. 2020. Fungal Diversity 105:1-16 ((https://doi.org/10.1007/s13225-020-00466-2)) |
| Otu55796 | 1 | k__Fungi(100);p__Ascomycota(100);c__Saccharomycetes(100);o__Saccharomycetales(100);f__Metschnikowiaceae(100);g__Metschnikowia(100);s__Metschnikowia_pulcherrima(100); | Metschnikowia | 13 | Saprotroph | \|Undefined Saprotroph\| | Yeast | NULL | Probable | As nectar-tap_saprotroph (Põlme et al. 2020); Decay sugar-rich substrates (Põlme et al. 2020) | Põlme S, et al. 2020. Fungal Diversity 105:1-16 ((https://doi.org/10.1007/s13225-020-00466-2)) |
| Otu55797 | 1 | k__Fungi(100);p__Ascomycota(100);c__Saccharomycetes(100);o__Saccharomycetales(100);f__Metschnikowiaceae(100);g__Metschnikowia(100);s__Metschnikowia_pulcherrima(100); | Metschnikowia | 13 | Saprotroph | \|Undefined Saprotroph\| | Yeast | NULL | Probable | As nectar-tap_saprotroph (Põlme et al. 2020); Decay sugar-rich substrates (Põlme et al. 2020) | Põlme S, et al. 2020. Fungal Diversity 105:1-16 ((https://doi.org/10.1007/s13225-020-00466-2)) |
| Otu55798 | 1 | k__Fungi(100);p__Ascomycota(100);c__Saccharomycetes(100);o__Saccharomycetales(100);f__Metschnikowiaceae(100);g__Metschnikowia(100);s__Metschnikowia_pulcherrima(100); | Metschnikowia | 13 | Saprotroph | \|Undefined Saprotroph\| | Yeast | NULL | Probable | As nectar-tap_saprotroph (Põlme et al. 2020); Decay sugar-rich substrates (Põlme et al. 2020) | Põlme S, et al. 2020. Fungal Diversity 105:1-16 ((https://doi.org/10.1007/s13225-020-00466-2)) |
| Otu55799 | 1 | k__Fungi(100);p__Ascomycota(100);c__Saccharomycetes(100);o__Saccharomycetales(100);f__Metschnikowiaceae(100);g__Metschnikowia(100);s__Metschnikowia_pulcherrima(100); | Metschnikowia | 13 | Saprotroph | \|Undefined Saprotroph\| | Yeast | NULL | Probable | As nectar-tap_saprotroph (Põlme et al. 2020); Decay sugar-rich substrates (Põlme et al. 2020) | Põlme S, et al. 2020. Fungal Diversity 105:1-16 ((https://doi.org/10.1007/s13225-020-00466-2)) |
| Otu55800 | 1 | k__Fungi(100);p__Ascomycota(100);c__Saccharomycetes(100);o__Saccharomycetales(100);f__Metschnikowiaceae(100);g__Metschnikowia(100);s__Metschnikowia_pulcherrima(100); | Metschnikowia | 13 | Saprotroph | \|Undefined Saprotroph\| | Yeast | NULL | Probable | As nectar-tap_saprotroph (Põlme et al. 2020); Decay sugar-rich substrates (Põlme et al. 2020) | Põlme S, et al. 2020. Fungal Diversity 105:1-16 ((https://doi.org/10.1007/s13225-020-00466-2)) |
| Otu55802 | 1 | k__Fungi(100);p__Ascomycota(100);c__Saccharomycetes(100);o__Saccharomycetales(100);f__Metschnikowiaceae(100);g__Metschnikowia(100);s__Metschnikowia_pulcherrima(100); | Metschnikowia | 13 | Saprotroph | \|Undefined Saprotroph\| | Yeast | NULL | Probable | As nectar-tap_saprotroph (Põlme et al. 2020); Decay sugar-rich substrates (Põlme et al. 2020) | Põlme S, et al. 2020. Fungal Diversity 105:1-16 ((https://doi.org/10.1007/s13225-020-00466-2)) |
| Otu55803 | 1 | k__Fungi(100);p__Ascomycota(100);c__Saccharomycetes(100);o__Saccharomycetales(100);f__Metschnikowiaceae(100);g__Metschnikowia(100);s__Metschnikowia_pulcherrima(100); | Metschnikowia | 13 | Saprotroph | \|Undefined Saprotroph\| | Yeast | NULL | Probable | As nectar-tap_saprotroph (Põlme et al. 2020); Decay sugar-rich substrates (Põlme et al. 2020) | Põlme S, et al. 2020. Fungal Diversity 105:1-16 ((https://doi.org/10.1007/s13225-020-00466-2)) |
| Otu55804 | 1 | k__Fungi(100);p__Ascomycota(100);c__Saccharomycetes(100);o__Saccharomycetales(100);f__Metschnikowiaceae(100);g__Metschnikowia(100);s__Metschnikowia_pulcherrima(100); | Metschnikowia | 13 | Saprotroph | \|Undefined Saprotroph\| | Yeast | NULL | Probable | As nectar-tap_saprotroph (Põlme et al. 2020); Decay sugar-rich substrates (Põlme et al. 2020) | Põlme S, et al. 2020. Fungal Diversity 105:1-16 ((https://doi.org/10.1007/s13225-020-00466-2)) |
| Otu55805 | 1 | k__Fungi(100);p__Ascomycota(100);c__Saccharomycetes(100);o__Saccharomycetales(100);f__Metschnikowiaceae(100);g__Metschnikowia(100);s__Metschnikowia_pulcherrima(100); | Metschnikowia | 13 | Saprotroph | \|Undefined Saprotroph\| | Yeast | NULL | Probable | As nectar-tap_saprotroph (Põlme et al. 2020); Decay sugar-rich substrates (Põlme et al. 2020) | Põlme S, et al. 2020. Fungal Diversity 105:1-16 ((https://doi.org/10.1007/s13225-020-00466-2)) |
| Otu55807 | 1 | k__Fungi(100);p__Ascomycota(100);c__Saccharomycetes(100);o__Saccharomycetales(100);f__Metschnikowiaceae(100);g__Metschnikowia(100);s__Metschnikowia_pulcherrima(100); | Metschnikowia | 13 | Saprotroph | \|Undefined Saprotroph\| | Yeast | NULL | Probable | As nectar-tap_saprotroph (Põlme et al. 2020); Decay sugar-rich substrates (Põlme et al. 2020) | Põlme S, et al. 2020. Fungal Diversity 105:1-16 ((https://doi.org/10.1007/s13225-020-00466-2)) |
| Otu55808 | 1 | k__Fungi(100);p__Ascomycota(100);c__Saccharomycetes(100);o__Saccharomycetales(100);f__Metschnikowiaceae(100);g__Metschnikowia(100);g__Metschnikowia_unclassified(100); | Metschnikowia | 13 | Saprotroph | \|Undefined Saprotroph\| | Yeast | NULL | Probable | As nectar-tap_saprotroph (Põlme et al. 2020); Decay sugar-rich substrates (Põlme et al. 2020) | Põlme S, et al. 2020. Fungal Diversity 105:1-16 ((https://doi.org/10.1007/s13225-020-00466-2)) |
| Otu55809 | 1 | k__Fungi(100);p__Ascomycota(100);c__Saccharomycetes(100);o__Saccharomycetales(100);f__Metschnikowiaceae(100);g__Metschnikowia(100);s__Metschnikowia_pulcherrima(100); | Metschnikowia | 13 | Saprotroph | \|Undefined Saprotroph\| | Yeast | NULL | Probable | As nectar-tap_saprotroph (Põlme et al. 2020); Decay sugar-rich substrates (Põlme et al. 2020) | Põlme S, et al. 2020. Fungal Diversity 105:1-16 ((https://doi.org/10.1007/s13225-020-00466-2)) |
| Otu55810 | 1 | k__Fungi(100);p__Ascomycota(100);c__Saccharomycetes(100);o__Saccharomycetales(100);f__Metschnikowiaceae(100);g__Metschnikowia(100);s__Metschnikowia_pulcherrima(100); | Metschnikowia | 13 | Saprotroph | \|Undefined Saprotroph\| | Yeast | NULL | Probable | As nectar-tap_saprotroph (Põlme et al. 2020); Decay sugar-rich substrates (Põlme et al. 2020) | Põlme S, et al. 2020. Fungal Diversity 105:1-16 ((https://doi.org/10.1007/s13225-020-00466-2)) |
| Otu55811 | 1 | k__Fungi(100);p__Ascomycota(100);c__Saccharomycetes(100);o__Saccharomycetales(100);f__Metschnikowiaceae(100);g__Metschnikowia(100);s__Metschnikowia_pulcherrima(100); | Metschnikowia | 13 | Saprotroph | \|Undefined Saprotroph\| | Yeast | NULL | Probable | As nectar-tap_saprotroph (Põlme et al. 2020); Decay sugar-rich substrates (Põlme et al. 2020) | Põlme S, et al. 2020. Fungal Diversity 105:1-16 ((https://doi.org/10.1007/s13225-020-00466-2)) |
| Otu55812 | 1 | k__Fungi(100);p__Ascomycota(100);c__Saccharomycetes(100);o__Saccharomycetales(100);f__Metschnikowiaceae(100);g__Metschnikowia(100);g__Metschnikowia_unclassified(100); | Metschnikowia | 13 | Saprotroph | \|Undefined Saprotroph\| | Yeast | NULL | Probable | As nectar-tap_saprotroph (Põlme et al. 2020); Decay sugar-rich substrates (Põlme et al. 2020) | Põlme S, et al. 2020. Fungal Diversity 105:1-16 ((https://doi.org/10.1007/s13225-020-00466-2)) |
| Otu55813 | 1 | k__Fungi(100);p__Ascomycota(100);c__Saccharomycetes(100);o__Saccharomycetales(100);f__Metschnikowiaceae(100);g__Metschnikowia(100);s__Metschnikowia_pulcherrima(100); | Metschnikowia | 13 | Saprotroph | \|Undefined Saprotroph\| | Yeast | NULL | Probable | As nectar-tap_saprotroph (Põlme et al. 2020); Decay sugar-rich substrates (Põlme et al. 2020) | Põlme S, et al. 2020. Fungal Diversity 105:1-16 ((https://doi.org/10.1007/s13225-020-00466-2)) |
| Otu55814 | 1 | k__Fungi(100);p__Ascomycota(100);c__Saccharomycetes(100);o__Saccharomycetales(100);f__Metschnikowiaceae(100);g__Metschnikowia(100);s__Metschnikowia_pulcherrima(100); | Metschnikowia | 13 | Saprotroph | \|Undefined Saprotroph\| | Yeast | NULL | Probable | As nectar-tap_saprotroph (Põlme et al. 2020); Decay sugar-rich substrates (Põlme et al. 2020) | Põlme S, et al. 2020. Fungal Diversity 105:1-16 ((https://doi.org/10.1007/s13225-020-00466-2)) |
| Otu55815 | 1 | k__Fungi(100);p__Ascomycota(100);c__Saccharomycetes(100);o__Saccharomycetales(100);f__Metschnikowiaceae(100);g__Metschnikowia(100);g__Metschnikowia_unclassified(100); | Metschnikowia | 13 | Saprotroph | \|Undefined Saprotroph\| | Yeast | NULL | Probable | As nectar-tap_saprotroph (Põlme et al. 2020); Decay sugar-rich substrates (Põlme et al. 2020) | Põlme S, et al. 2020. Fungal Diversity 105:1-16 ((https://doi.org/10.1007/s13225-020-00466-2)) |
| Otu55816 | 1 | k__Fungi(100);p__Ascomycota(100);c__Saccharomycetes(100);o__Saccharomycetales(100);f__Metschnikowiaceae(100);g__Metschnikowia(100);s__Metschnikowia_pulcherrima(100); | Metschnikowia | 13 | Saprotroph | \|Undefined Saprotroph\| | Yeast | NULL | Probable | As nectar-tap_saprotroph (Põlme et al. 2020); Decay sugar-rich substrates (Põlme et al. 2020) | Põlme S, et al. 2020. Fungal Diversity 105:1-16 ((https://doi.org/10.1007/s13225-020-00466-2)) |
| Otu55817 | 1 | k__Fungi(100);p__Ascomycota(100);c__Saccharomycetes(100);o__Saccharomycetales(100);f__Metschnikowiaceae(100);g__Metschnikowia(100);s__Metschnikowia_pulcherrima(100); | Metschnikowia | 13 | Saprotroph | \|Undefined Saprotroph\| | Yeast | NULL | Probable | As nectar-tap_saprotroph (Põlme et al. 2020); Decay sugar-rich substrates (Põlme et al. 2020) | Põlme S, et al. 2020. Fungal Diversity 105:1-16 ((https://doi.org/10.1007/s13225-020-00466-2)) |
| Otu55818 | 1 | k__Fungi(100);p__Ascomycota(100);c__Saccharomycetes(100);o__Saccharomycetales(100);f__Metschnikowiaceae(100);g__Metschnikowia(100);s__Metschnikowia_pulcherrima(100); | Metschnikowia | 13 | Saprotroph | \|Undefined Saprotroph\| | Yeast | NULL | Probable | As nectar-tap_saprotroph (Põlme et al. 2020); Decay sugar-rich substrates (Põlme et al. 2020) | Põlme S, et al. 2020. Fungal Diversity 105:1-16 ((https://doi.org/10.1007/s13225-020-00466-2)) |
| Otu55820 | 1 | k__Fungi(100);p__Ascomycota(100);c__Saccharomycetes(100);o__Saccharomycetales(100);f__Metschnikowiaceae(100);g__Metschnikowia(100);g__Metschnikowia_unclassified(100); | Metschnikowia | 13 | Saprotroph | \|Undefined Saprotroph\| | Yeast | NULL | Probable | As nectar-tap_saprotroph (Põlme et al. 2020); Decay sugar-rich substrates (Põlme et al. 2020) | Põlme S, et al. 2020. Fungal Diversity 105:1-16 ((https://doi.org/10.1007/s13225-020-00466-2)) |
| Otu55821 | 1 | k__Fungi(100);p__Ascomycota(100);c__Saccharomycetes(100);o__Saccharomycetales(100);f__Metschnikowiaceae(100);g__Metschnikowia(100);s__Metschnikowia_pulcherrima(100); | Metschnikowia | 13 | Saprotroph | \|Undefined Saprotroph\| | Yeast | NULL | Probable | As nectar-tap_saprotroph (Põlme et al. 2020); Decay sugar-rich substrates (Põlme et al. 2020) | Põlme S, et al. 2020. Fungal Diversity 105:1-16 ((https://doi.org/10.1007/s13225-020-00466-2)) |
| Otu55822 | 1 | k__Fungi(100);p__Ascomycota(100);c__Saccharomycetes(100);o__Saccharomycetales(100);f__Metschnikowiaceae(100);g__Metschnikowia(100);s__Metschnikowia_pulcherrima(100); | Metschnikowia | 13 | Saprotroph | \|Undefined Saprotroph\| | Yeast | NULL | Probable | As nectar-tap_saprotroph (Põlme et al. 2020); Decay sugar-rich substrates (Põlme et al. 2020) | Põlme S, et al. 2020. Fungal Diversity 105:1-16 ((https://doi.org/10.1007/s13225-020-00466-2)) |
| Otu55823 | 1 | k__Fungi(100);p__Ascomycota(100);c__Saccharomycetes(100);o__Saccharomycetales(100);f__Metschnikowiaceae(100);g__Metschnikowia(100);s__Metschnikowia_pulcherrima(100); | Metschnikowia | 13 | Saprotroph | \|Undefined Saprotroph\| | Yeast | NULL | Probable | As nectar-tap_saprotroph (Põlme et al. 2020); Decay sugar-rich substrates (Põlme et al. 2020) | Põlme S, et al. 2020. Fungal Diversity 105:1-16 ((https://doi.org/10.1007/s13225-020-00466-2)) |
| Otu55824 | 1 | k__Fungi(100);p__Ascomycota(100);c__Saccharomycetes(100);o__Saccharomycetales(100);f__Metschnikowiaceae(100);g__Metschnikowia(100);s__Metschnikowia_pulcherrima(100); | Metschnikowia | 13 | Saprotroph | \|Undefined Saprotroph\| | Yeast | NULL | Probable | As nectar-tap_saprotroph (Põlme et al. 2020); Decay sugar-rich substrates (Põlme et al. 2020) | Põlme S, et al. 2020. Fungal Diversity 105:1-16 ((https://doi.org/10.1007/s13225-020-00466-2)) |
| Otu55827 | 1 | k__Fungi(100);p__Ascomycota(100);c__Saccharomycetes(100);o__Saccharomycetales(100);f__Metschnikowiaceae(100);g__Metschnikowia(100);s__Metschnikowia_pulcherrima(100); | Metschnikowia | 13 | Saprotroph | \|Undefined Saprotroph\| | Yeast | NULL | Probable | As nectar-tap_saprotroph (Põlme et al. 2020); Decay sugar-rich substrates (Põlme et al. 2020) | Põlme S, et al. 2020. Fungal Diversity 105:1-16 ((https://doi.org/10.1007/s13225-020-00466-2)) |
| Otu55828 | 1 | k__Fungi(100);p__Ascomycota(100);c__Saccharomycetes(100);o__Saccharomycetales(100);f__Metschnikowiaceae(100);g__Metschnikowia(100);s__Metschnikowia_pulcherrima(100); | Metschnikowia | 13 | Saprotroph | \|Undefined Saprotroph\| | Yeast | NULL | Probable | As nectar-tap_saprotroph (Põlme et al. 2020); Decay sugar-rich substrates (Põlme et al. 2020) | Põlme S, et al. 2020. Fungal Diversity 105:1-16 ((https://doi.org/10.1007/s13225-020-00466-2)) |
| Otu55830 | 1 | k__Fungi(100);p__Ascomycota(100);c__Saccharomycetes(100);o__Saccharomycetales(100);f__Saccharomycetaceae(100);g__Kazachstania(100);s__Kazachstania_intestinalis(100); | Kazachstania | 13 | Saprotroph | \|Undefined Saprotroph\| | Yeast | NULL | Probable | As nectar-tap_saprotroph (Põlme et al. 2020); Decay sugar-rich substrates (Põlme et al. 2020) | Põlme S, et al. 2020. Fungal Diversity 105:1-16 ((https://doi.org/10.1007/s13225-020-00466-2)) |
| Otu55833 | 1 | k__Fungi(100);p__Ascomycota(100);c__Saccharomycetes(100);o__Saccharomycetales(100);f__Metschnikowiaceae(100);g__Metschnikowia(100);g__Metschnikowia_unclassified(100); | Metschnikowia | 13 | Saprotroph | \|Undefined Saprotroph\| | Yeast | NULL | Probable | As nectar-tap_saprotroph (Põlme et al. 2020); Decay sugar-rich substrates (Põlme et al. 2020) | Põlme S, et al. 2020. Fungal Diversity 105:1-16 ((https://doi.org/10.1007/s13225-020-00466-2)) |
| Otu55834 | 1 | k__Fungi(100);p__Ascomycota(100);c__Saccharomycetes(100);o__Saccharomycetales(100);f__Metschnikowiaceae(100);g__Metschnikowia(100);s__Metschnikowia_pulcherrima(100); | Metschnikowia | 13 | Saprotroph | \|Undefined Saprotroph\| | Yeast | NULL | Probable | As nectar-tap_saprotroph (Põlme et al. 2020); Decay sugar-rich substrates (Põlme et al. 2020) | Põlme S, et al. 2020. Fungal Diversity 105:1-16 ((https://doi.org/10.1007/s13225-020-00466-2)) |
| Otu55835 | 1 | k__Fungi(100);p__Ascomycota(100);c__Saccharomycetes(100);o__Saccharomycetales(100);f__Metschnikowiaceae(100);g__Metschnikowia(100);s__Metschnikowia_pulcherrima(100); | Metschnikowia | 13 | Saprotroph | \|Undefined Saprotroph\| | Yeast | NULL | Probable | As nectar-tap_saprotroph (Põlme et al. 2020); Decay sugar-rich substrates (Põlme et al. 2020) | Põlme S, et al. 2020. Fungal Diversity 105:1-16 ((https://doi.org/10.1007/s13225-020-00466-2)) |
| Otu55836 | 1 | k__Fungi(100);p__Ascomycota(100);c__Saccharomycetes(100);o__Saccharomycetales(100);f__Metschnikowiaceae(100);g__Metschnikowia(100);s__Metschnikowia_pulcherrima(100); | Metschnikowia | 13 | Saprotroph | \|Undefined Saprotroph\| | Yeast | NULL | Probable | As nectar-tap_saprotroph (Põlme et al. 2020); Decay sugar-rich substrates (Põlme et al. 2020) | Põlme S, et al. 2020. Fungal Diversity 105:1-16 ((https://doi.org/10.1007/s13225-020-00466-2)) |
| Otu55837 | 1 | k__Fungi(100);p__Ascomycota(100);c__Saccharomycetes(100);o__Saccharomycetales(100);f__Metschnikowiaceae(100);g__Metschnikowia(100);s__Metschnikowia_pulcherrima(100); | Metschnikowia | 13 | Saprotroph | \|Undefined Saprotroph\| | Yeast | NULL | Probable | As nectar-tap_saprotroph (Põlme et al. 2020); Decay sugar-rich substrates (Põlme et al. 2020) | Põlme S, et al. 2020. Fungal Diversity 105:1-16 ((https://doi.org/10.1007/s13225-020-00466-2)) |
| Otu55838 | 1 | k__Fungi(100);p__Ascomycota(100);c__Saccharomycetes(100);o__Saccharomycetales(100);f__Metschnikowiaceae(100);g__Metschnikowia(100);s__Metschnikowia_pulcherrima(100); | Metschnikowia | 13 | Saprotroph | \|Undefined Saprotroph\| | Yeast | NULL | Probable | As nectar-tap_saprotroph (Põlme et al. 2020); Decay sugar-rich substrates (Põlme et al. 2020) | Põlme S, et al. 2020. Fungal Diversity 105:1-16 ((https://doi.org/10.1007/s13225-020-00466-2)) |
| Otu55839 | 1 | k__Fungi(100);p__Ascomycota(100);c__Saccharomycetes(100);o__Saccharomycetales(100);f__Metschnikowiaceae(100);g__Metschnikowia(100);s__Metschnikowia_pulcherrima(100); | Metschnikowia | 13 | Saprotroph | \|Undefined Saprotroph\| | Yeast | NULL | Probable | As nectar-tap_saprotroph (Põlme et al. 2020); Decay sugar-rich substrates (Põlme et al. 2020) | Põlme S, et al. 2020. Fungal Diversity 105:1-16 ((https://doi.org/10.1007/s13225-020-00466-2)) |
| Otu55840 | 1 | k__Fungi(100);p__Ascomycota(100);c__Saccharomycetes(100);o__Saccharomycetales(100);f__Metschnikowiaceae(100);g__Metschnikowia(100);s__Metschnikowia_pulcherrima(100); | Metschnikowia | 13 | Saprotroph | \|Undefined Saprotroph\| | Yeast | NULL | Probable | As nectar-tap_saprotroph (Põlme et al. 2020); Decay sugar-rich substrates (Põlme et al. 2020) | Põlme S, et al. 2020. Fungal Diversity 105:1-16 ((https://doi.org/10.1007/s13225-020-00466-2)) |
| Otu55841 | 1 | k__Fungi(100);p__Ascomycota(100);c__Saccharomycetes(100);o__Saccharomycetales(100);f__Metschnikowiaceae(100);g__Metschnikowia(100);s__Metschnikowia_pulcherrima(100); | Metschnikowia | 13 | Saprotroph | \|Undefined Saprotroph\| | Yeast | NULL | Probable | As nectar-tap_saprotroph (Põlme et al. 2020); Decay sugar-rich substrates (Põlme et al. 2020) | Põlme S, et al. 2020. Fungal Diversity 105:1-16 ((https://doi.org/10.1007/s13225-020-00466-2)) |
| Otu55842 | 1 | k__Fungi(100);p__Ascomycota(100);c__Saccharomycetes(100);o__Saccharomycetales(100);f__Metschnikowiaceae(100);g__Metschnikowia(100);g__Metschnikowia_unclassified(100); | Metschnikowia | 13 | Saprotroph | \|Undefined Saprotroph\| | Yeast | NULL | Probable | As nectar-tap_saprotroph (Põlme et al. 2020); Decay sugar-rich substrates (Põlme et al. 2020) | Põlme S, et al. 2020. Fungal Diversity 105:1-16 ((https://doi.org/10.1007/s13225-020-00466-2)) |
| Otu55843 | 1 | k__Fungi(100);p__Ascomycota(100);c__Saccharomycetes(100);o__Saccharomycetales(100);f__Metschnikowiaceae(100);g__Metschnikowia(100);g__Metschnikowia_unclassified(100); | Metschnikowia | 13 | Saprotroph | \|Undefined Saprotroph\| | Yeast | NULL | Probable | As nectar-tap_saprotroph (Põlme et al. 2020); Decay sugar-rich substrates (Põlme et al. 2020) | Põlme S, et al. 2020. Fungal Diversity 105:1-16 ((https://doi.org/10.1007/s13225-020-00466-2)) |
| Otu55844 | 1 | k__Fungi(100);p__Ascomycota(100);c__Saccharomycetes(100);o__Saccharomycetales(100);f__Metschnikowiaceae(100);g__Metschnikowia(100);s__Metschnikowia_pulcherrima(100); | Metschnikowia | 13 | Saprotroph | \|Undefined Saprotroph\| | Yeast | NULL | Probable | As nectar-tap_saprotroph (Põlme et al. 2020); Decay sugar-rich substrates (Põlme et al. 2020) | Põlme S, et al. 2020. Fungal Diversity 105:1-16 ((https://doi.org/10.1007/s13225-020-00466-2)) |
| Otu55845 | 1 | k__Fungi(100);p__Ascomycota(100);c__Saccharomycetes(100);o__Saccharomycetales(100);f__Metschnikowiaceae(100);g__Metschnikowia(100);s__Metschnikowia_pulcherrima(100); | Metschnikowia | 13 | Saprotroph | \|Undefined Saprotroph\| | Yeast | NULL | Probable | As nectar-tap_saprotroph (Põlme et al. 2020); Decay sugar-rich substrates (Põlme et al. 2020) | Põlme S, et al. 2020. Fungal Diversity 105:1-16 ((https://doi.org/10.1007/s13225-020-00466-2)) |
| Otu55847 | 1 | k__Fungi(100);p__Ascomycota(100);c__Saccharomycetes(100);o__Saccharomycetales(100);f__Metschnikowiaceae(100);g__Metschnikowia(100);g__Metschnikowia_unclassified(100); | Metschnikowia | 13 | Saprotroph | \|Undefined Saprotroph\| | Yeast | NULL | Probable | As nectar-tap_saprotroph (Põlme et al. 2020); Decay sugar-rich substrates (Põlme et al. 2020) | Põlme S, et al. 2020. Fungal Diversity 105:1-16 ((https://doi.org/10.1007/s13225-020-00466-2)) |
| Otu55848 | 1 | k__Fungi(100);p__Ascomycota(100);c__Saccharomycetes(100);o__Saccharomycetales(100);f__Metschnikowiaceae(100);g__Metschnikowia(100);s__Metschnikowia_pulcherrima(100); | Metschnikowia | 13 | Saprotroph | \|Undefined Saprotroph\| | Yeast | NULL | Probable | As nectar-tap_saprotroph (Põlme et al. 2020); Decay sugar-rich substrates (Põlme et al. 2020) | Põlme S, et al. 2020. Fungal Diversity 105:1-16 ((https://doi.org/10.1007/s13225-020-00466-2)) |
| Otu55849 | 1 | k__Fungi(100);p__Ascomycota(100);c__Saccharomycetes(100);o__Saccharomycetales(100);f__Metschnikowiaceae(100);g__Metschnikowia(100);s__Metschnikowia_pulcherrima(100); | Metschnikowia | 13 | Saprotroph | \|Undefined Saprotroph\| | Yeast | NULL | Probable | As nectar-tap_saprotroph (Põlme et al. 2020); Decay sugar-rich substrates (Põlme et al. 2020) | Põlme S, et al. 2020. Fungal Diversity 105:1-16 ((https://doi.org/10.1007/s13225-020-00466-2)) |
| Otu55850 | 1 | k__Fungi(100);p__Ascomycota(100);c__Saccharomycetes(100);o__Saccharomycetales(100);f__Metschnikowiaceae(100);g__Metschnikowia(100);s__Metschnikowia_pulcherrima(100); | Metschnikowia | 13 | Saprotroph | \|Undefined Saprotroph\| | Yeast | NULL | Probable | As nectar-tap_saprotroph (Põlme et al. 2020); Decay sugar-rich substrates (Põlme et al. 2020) | Põlme S, et al. 2020. Fungal Diversity 105:1-16 ((https://doi.org/10.1007/s13225-020-00466-2)) |
| Otu55852 | 1 | k__Fungi(100);p__Ascomycota(100);c__Saccharomycetes(100);o__Saccharomycetales(100);f__Metschnikowiaceae(100);g__Metschnikowia(100);s__Metschnikowia_pulcherrima(100); | Metschnikowia | 13 | Saprotroph | \|Undefined Saprotroph\| | Yeast | NULL | Probable | As nectar-tap_saprotroph (Põlme et al. 2020); Decay sugar-rich substrates (Põlme et al. 2020) | Põlme S, et al. 2020. Fungal Diversity 105:1-16 ((https://doi.org/10.1007/s13225-020-00466-2)) |
| Otu55853 | 1 | k__Fungi(100);p__Ascomycota(100);c__Saccharomycetes(100);o__Saccharomycetales(100);f__Metschnikowiaceae(100);g__Metschnikowia(100);s__Metschnikowia_pulcherrima(100); | Metschnikowia | 13 | Saprotroph | \|Undefined Saprotroph\| | Yeast | NULL | Probable | As nectar-tap_saprotroph (Põlme et al. 2020); Decay sugar-rich substrates (Põlme et al. 2020) | Põlme S, et al. 2020. Fungal Diversity 105:1-16 ((https://doi.org/10.1007/s13225-020-00466-2)) |
| Otu55858 | 1 | k__Fungi(100);p__Ascomycota(100);c__Saccharomycetes(100);o__Saccharomycetales(100);f__Metschnikowiaceae(100);g__Metschnikowia(100);s__Metschnikowia_pulcherrima(100); | Metschnikowia | 13 | Saprotroph | \|Undefined Saprotroph\| | Yeast | NULL | Probable | As nectar-tap_saprotroph (Põlme et al. 2020); Decay sugar-rich substrates (Põlme et al. 2020) | Põlme S, et al. 2020. Fungal Diversity 105:1-16 ((https://doi.org/10.1007/s13225-020-00466-2)) |
| Otu55859 | 1 | k__Fungi(100);p__Ascomycota(100);c__Saccharomycetes(100);o__Saccharomycetales(100);f__Metschnikowiaceae(100);g__Metschnikowia(100);s__Metschnikowia_pulcherrima(100); | Metschnikowia | 13 | Saprotroph | \|Undefined Saprotroph\| | Yeast | NULL | Probable | As nectar-tap_saprotroph (Põlme et al. 2020); Decay sugar-rich substrates (Põlme et al. 2020) | Põlme S, et al. 2020. Fungal Diversity 105:1-16 ((https://doi.org/10.1007/s13225-020-00466-2)) |
| Otu55868 | 1 | k__Fungi(100);p__Ascomycota(100);c__Saccharomycetes(100);o__Saccharomycetales(100);f__Metschnikowiaceae(100);g__Metschnikowia(100);g__Metschnikowia_unclassified(100); | Metschnikowia | 13 | Saprotroph | \|Undefined Saprotroph\| | Yeast | NULL | Probable | As nectar-tap_saprotroph (Põlme et al. 2020); Decay sugar-rich substrates (Põlme et al. 2020) | Põlme S, et al. 2020. Fungal Diversity 105:1-16 ((https://doi.org/10.1007/s13225-020-00466-2)) |
| Otu55869 | 1 | k__Fungi(100);p__Ascomycota(100);c__Saccharomycetes(100);o__Saccharomycetales(100);f__Metschnikowiaceae(100);g__Metschnikowia(100);s__Metschnikowia_pulcherrima(100); | Metschnikowia | 13 | Saprotroph | \|Undefined Saprotroph\| | Yeast | NULL | Probable | As nectar-tap_saprotroph (Põlme et al. 2020); Decay sugar-rich substrates (Põlme et al. 2020) | Põlme S, et al. 2020. Fungal Diversity 105:1-16 ((https://doi.org/10.1007/s13225-020-00466-2)) |
| Otu55870 | 1 | k__Fungi(100);p__Ascomycota(100);c__Saccharomycetes(100);o__Saccharomycetales(100);f__Metschnikowiaceae(100);g__Metschnikowia(100);s__Metschnikowia_pulcherrima(100); | Metschnikowia | 13 | Saprotroph | \|Undefined Saprotroph\| | Yeast | NULL | Probable | As nectar-tap_saprotroph (Põlme et al. 2020); Decay sugar-rich substrates (Põlme et al. 2020) | Põlme S, et al. 2020. Fungal Diversity 105:1-16 ((https://doi.org/10.1007/s13225-020-00466-2)) |
| Otu55871 | 1 | k__Fungi(100);p__Ascomycota(100);c__Saccharomycetes(100);o__Saccharomycetales(100);f__Metschnikowiaceae(100);g__Metschnikowia(100);s__Metschnikowia_pulcherrima(100); | Metschnikowia | 13 | Saprotroph | \|Undefined Saprotroph\| | Yeast | NULL | Probable | As nectar-tap_saprotroph (Põlme et al. 2020); Decay sugar-rich substrates (Põlme et al. 2020) | Põlme S, et al. 2020. Fungal Diversity 105:1-16 ((https://doi.org/10.1007/s13225-020-00466-2)) |
| Otu55874 | 1 | k__Fungi(100);p__Ascomycota(100);c__Saccharomycetes(100);o__Saccharomycetales(100);f__Saccharomycetales_family_Incertae_sedis(100);g__Candida(100);g__Candida_unclassified(100); | Candida | 13 | Pathotroph-Saprotroph-Symbiotroph | Animal Pathogen-Endophyte-Endosymbiont-Epiphyte-\|Undefined Saprotroph\| | Dimorphic Yeast-Yeast | NULL | Probable | As nectar-tap_saprotroph (Põlme et al. 2020); Decay sugar-rich substrates (Põlme et al. 2020); This asexual genus is polyphyletic and contain species that function very differently depend on what they are. Caution should be taken when interpreting guilds for Candida species. | Manolakaki D, et al. 2010. Virulence. 1: 367-375 ((https://doi.org/10.4161/viru.1.5.12796)); Mekbib SB, et al. 2011. Tropical Plant Pathology 36:233-240 ((https://doi.org/10.1590/S1982-56762011000400004)); Põlme S, et al. 2020. Fungal Diversity 105:1-16 ((https://doi.org/10.1007/s13225-020-00466-2)) |
| Otu55875 | 1 | k__Fungi(100);p__Ascomycota(100);c__Saccharomycetes(100);o__Saccharomycetales(100);f__Metschnikowiaceae(100);g__Metschnikowia(100);s__Metschnikowia_pulcherrima(100); | Metschnikowia | 13 | Saprotroph | \|Undefined Saprotroph\| | Yeast | NULL | Probable | As nectar-tap_saprotroph (Põlme et al. 2020); Decay sugar-rich substrates (Põlme et al. 2020) | Põlme S, et al. 2020. Fungal Diversity 105:1-16 ((https://doi.org/10.1007/s13225-020-00466-2)) |
| Otu55877 | 1 | k__Fungi(100);p__Ascomycota(100);c__Saccharomycetes(100);o__Saccharomycetales(100);f__Metschnikowiaceae(100);g__Metschnikowia(100);g__Metschnikowia_unclassified(100); | Metschnikowia | 13 | Saprotroph | \|Undefined Saprotroph\| | Yeast | NULL | Probable | As nectar-tap_saprotroph (Põlme et al. 2020); Decay sugar-rich substrates (Põlme et al. 2020) | Põlme S, et al. 2020. Fungal Diversity 105:1-16 ((https://doi.org/10.1007/s13225-020-00466-2)) |
| Otu55878 | 1 | k__Fungi(100);p__Ascomycota(100);c__Saccharomycetes(100);o__Saccharomycetales(100);f__Metschnikowiaceae(100);g__Metschnikowia(100);g__Metschnikowia_unclassified(100); | Metschnikowia | 13 | Saprotroph | \|Undefined Saprotroph\| | Yeast | NULL | Probable | As nectar-tap_saprotroph (Põlme et al. 2020); Decay sugar-rich substrates (Põlme et al. 2020) | Põlme S, et al. 2020. Fungal Diversity 105:1-16 ((https://doi.org/10.1007/s13225-020-00466-2)) |
| Otu55881 | 1 | k__Fungi(100);p__Ascomycota(100);c__Saccharomycetes(100);o__Saccharomycetales(100);f__Saccharomycetales_family_Incertae_sedis(100);g__Candida(100);g__Candida_unclassified(100); | Candida | 13 | Pathotroph-Saprotroph-Symbiotroph | Animal Pathogen-Endophyte-Endosymbiont-Epiphyte-\|Undefined Saprotroph\| | Dimorphic Yeast-Yeast | NULL | Probable | As nectar-tap_saprotroph (Põlme et al. 2020); Decay sugar-rich substrates (Põlme et al. 2020); This asexual genus is polyphyletic and contain species that function very differently depend on what they are. Caution should be taken when interpreting guilds for Candida species. | Manolakaki D, et al. 2010. Virulence. 1: 367-375 ((https://doi.org/10.4161/viru.1.5.12796)); Mekbib SB, et al. 2011. Tropical Plant Pathology 36:233-240 ((https://doi.org/10.1590/S1982-56762011000400004)); Põlme S, et al. 2020. Fungal Diversity 105:1-16 ((https://doi.org/10.1007/s13225-020-00466-2)) |
| Otu55886 | 1 | k__Fungi(100);p__Ascomycota(100);c__Saccharomycetes(100);o__Saccharomycetales(100);f__Metschnikowiaceae(100);g__Metschnikowia(100);s__Metschnikowia_pulcherrima(100); | Metschnikowia | 13 | Saprotroph | \|Undefined Saprotroph\| | Yeast | NULL | Probable | As nectar-tap_saprotroph (Põlme et al. 2020); Decay sugar-rich substrates (Põlme et al. 2020) | Põlme S, et al. 2020. Fungal Diversity 105:1-16 ((https://doi.org/10.1007/s13225-020-00466-2)) |
| Otu55887 | 1 | k__Fungi(100);p__Ascomycota(100);c__Saccharomycetes(100);o__Saccharomycetales(100);f__Metschnikowiaceae(100);g__Metschnikowia(100);s__Metschnikowia_pulcherrima(100); | Metschnikowia | 13 | Saprotroph | \|Undefined Saprotroph\| | Yeast | NULL | Probable | As nectar-tap_saprotroph (Põlme et al. 2020); Decay sugar-rich substrates (Põlme et al. 2020) | Põlme S, et al. 2020. Fungal Diversity 105:1-16 ((https://doi.org/10.1007/s13225-020-00466-2)) |
| Otu55888 | 1 | k__Fungi(100);p__Ascomycota(100);c__Saccharomycetes(100);o__Saccharomycetales(100);f__Metschnikowiaceae(100);g__Metschnikowia(100);s__Metschnikowia_pulcherrima(100); | Metschnikowia | 13 | Saprotroph | \|Undefined Saprotroph\| | Yeast | NULL | Probable | As nectar-tap_saprotroph (Põlme et al. 2020); Decay sugar-rich substrates (Põlme et al. 2020) | Põlme S, et al. 2020. Fungal Diversity 105:1-16 ((https://doi.org/10.1007/s13225-020-00466-2)) |
| Otu55889 | 1 | k__Fungi(100);p__Ascomycota(100);c__Saccharomycetes(100);o__Saccharomycetales(100);f__Metschnikowiaceae(100);g__Metschnikowia(100);s__Metschnikowia_pulcherrima(100); | Metschnikowia | 13 | Saprotroph | \|Undefined Saprotroph\| | Yeast | NULL | Probable | As nectar-tap_saprotroph (Põlme et al. 2020); Decay sugar-rich substrates (Põlme et al. 2020) | Põlme S, et al. 2020. Fungal Diversity 105:1-16 ((https://doi.org/10.1007/s13225-020-00466-2)) |
| Otu55890 | 1 | k__Fungi(100);p__Ascomycota(100);c__Saccharomycetes(100);o__Saccharomycetales(100);f__Metschnikowiaceae(100);g__Metschnikowia(100);s__Metschnikowia_pulcherrima(100); | Metschnikowia | 13 | Saprotroph | \|Undefined Saprotroph\| | Yeast | NULL | Probable | As nectar-tap_saprotroph (Põlme et al. 2020); Decay sugar-rich substrates (Põlme et al. 2020) | Põlme S, et al. 2020. Fungal Diversity 105:1-16 ((https://doi.org/10.1007/s13225-020-00466-2)) |
| Otu55891 | 1 | k__Fungi(100);p__Ascomycota(100);c__Saccharomycetes(100);o__Saccharomycetales(100);f__Metschnikowiaceae(100);g__Metschnikowia(100);s__Metschnikowia_pulcherrima(100); | Metschnikowia | 13 | Saprotroph | \|Undefined Saprotroph\| | Yeast | NULL | Probable | As nectar-tap_saprotroph (Põlme et al. 2020); Decay sugar-rich substrates (Põlme et al. 2020) | Põlme S, et al. 2020. Fungal Diversity 105:1-16 ((https://doi.org/10.1007/s13225-020-00466-2)) |
| Otu55898 | 1 | k__Fungi(100);p__Ascomycota(100);c__Saccharomycetes(100);o__Saccharomycetales(100);f__Metschnikowiaceae(100);g__Metschnikowia(100);s__Metschnikowia_pulcherrima(100); | Metschnikowia | 13 | Saprotroph | \|Undefined Saprotroph\| | Yeast | NULL | Probable | As nectar-tap_saprotroph (Põlme et al. 2020); Decay sugar-rich substrates (Põlme et al. 2020) | Põlme S, et al. 2020. Fungal Diversity 105:1-16 ((https://doi.org/10.1007/s13225-020-00466-2)) |
| Otu55905 | 1 | k__Fungi(100);p__Ascomycota(100);c__Saccharomycetes(100);o__Saccharomycetales(100);f__Metschnikowiaceae(100);g__Metschnikowia(100);s__Metschnikowia_pulcherrima(100); | Metschnikowia | 13 | Saprotroph | \|Undefined Saprotroph\| | Yeast | NULL | Probable | As nectar-tap_saprotroph (Põlme et al. 2020); Decay sugar-rich substrates (Põlme et al. 2020) | Põlme S, et al. 2020. Fungal Diversity 105:1-16 ((https://doi.org/10.1007/s13225-020-00466-2)) |
| Otu55911 | 1 | k__Fungi(100);p__Ascomycota(100);c__Saccharomycetes(100);o__Saccharomycetales(100);f__Saccharomycetales_family_Incertae_sedis(100);g__Candida(100);g__Candida_unclassified(100); | Candida | 13 | Pathotroph-Saprotroph-Symbiotroph | Animal Pathogen-Endophyte-Endosymbiont-Epiphyte-\|Undefined Saprotroph\| | Dimorphic Yeast-Yeast | NULL | Probable | As nectar-tap_saprotroph (Põlme et al. 2020); Decay sugar-rich substrates (Põlme et al. 2020); This asexual genus is polyphyletic and contain species that function very differently depend on what they are. Caution should be taken when interpreting guilds for Candida species. | Manolakaki D, et al. 2010. Virulence. 1: 367-375 ((https://doi.org/10.4161/viru.1.5.12796)); Mekbib SB, et al. 2011. Tropical Plant Pathology 36:233-240 ((https://doi.org/10.1590/S1982-56762011000400004)); Põlme S, et al. 2020. Fungal Diversity 105:1-16 ((https://doi.org/10.1007/s13225-020-00466-2)) |
| Otu55912 | 1 | k__Fungi(100);p__Ascomycota(100);c__Saccharomycetes(100);o__Saccharomycetales(100);f__Metschnikowiaceae(100);g__Metschnikowia(100);s__Metschnikowia_pulcherrima(100); | Metschnikowia | 13 | Saprotroph | \|Undefined Saprotroph\| | Yeast | NULL | Probable | As nectar-tap_saprotroph (Põlme et al. 2020); Decay sugar-rich substrates (Põlme et al. 2020) | Põlme S, et al. 2020. Fungal Diversity 105:1-16 ((https://doi.org/10.1007/s13225-020-00466-2)) |
| Otu55913 | 1 | k__Fungi(100);p__Ascomycota(100);c__Saccharomycetes(100);o__Saccharomycetales(100);f__Metschnikowiaceae(100);g__Metschnikowia(100);s__Metschnikowia_pulcherrima(100); | Metschnikowia | 13 | Saprotroph | \|Undefined Saprotroph\| | Yeast | NULL | Probable | As nectar-tap_saprotroph (Põlme et al. 2020); Decay sugar-rich substrates (Põlme et al. 2020) | Põlme S, et al. 2020. Fungal Diversity 105:1-16 ((https://doi.org/10.1007/s13225-020-00466-2)) |
| Otu55914 | 1 | k__Fungi(100);p__Basidiomycota(100);c__Microbotryomycetes(100);o__Sporidiobolales(100);f__Sporidiobolales_family_Incertae_sedis(100);g__Sporobolomyces(100);s__Sporobolomyces_lactosus(100); | Sporobolomyces | 13 | Pathotroph-Saprotroph | \|Fungal Parasite\|-Plant Saprotroph | Yeast | NULL | Probable | As litter_saprotroph (Põlme et al. 2020); As mycoparasite (Põlme et al. 2020); Decay fungal material (Põlme et al. 2020) | Tedersoo L, et al. 2014. Science 346:e1256688 ((https://doi.org/10.1126/science.1256688)); Purahong W, et al. 2016. Molecular Ecology 25:4059-4074 ((https://doi.org/10.1111/mec.13739)); Põlme S, et al. 2020. Fungal Diversity 105:1-16 ((https://doi.org/10.1007/s13225-020-00466-2)) |
| Otu55915 | 1 | k__Fungi(100);p__Ascomycota(100);c__Saccharomycetes(100);o__Saccharomycetales(100);f__Metschnikowiaceae(100);g__Metschnikowia(100);s__Metschnikowia_pulcherrima(100); | Metschnikowia | 13 | Saprotroph | \|Undefined Saprotroph\| | Yeast | NULL | Probable | As nectar-tap_saprotroph (Põlme et al. 2020); Decay sugar-rich substrates (Põlme et al. 2020) | Põlme S, et al. 2020. Fungal Diversity 105:1-16 ((https://doi.org/10.1007/s13225-020-00466-2)) |
| Otu55916 | 1 | k__Fungi(100);p__Ascomycota(100);c__Saccharomycetes(100);o__Saccharomycetales(100);f__Metschnikowiaceae(100);g__Metschnikowia(100);s__Metschnikowia_pulcherrima(100); | Metschnikowia | 13 | Saprotroph | \|Undefined Saprotroph\| | Yeast | NULL | Probable | As nectar-tap_saprotroph (Põlme et al. 2020); Decay sugar-rich substrates (Põlme et al. 2020) | Põlme S, et al. 2020. Fungal Diversity 105:1-16 ((https://doi.org/10.1007/s13225-020-00466-2)) |
| Otu55917 | 1 | k__Fungi(100);p__Ascomycota(100);c__Saccharomycetes(100);o__Saccharomycetales(100);f__Metschnikowiaceae(100);g__Metschnikowia(100);s__Metschnikowia_pulcherrima(100); | Metschnikowia | 13 | Saprotroph | \|Undefined Saprotroph\| | Yeast | NULL | Probable | As nectar-tap_saprotroph (Põlme et al. 2020); Decay sugar-rich substrates (Põlme et al. 2020) | Põlme S, et al. 2020. Fungal Diversity 105:1-16 ((https://doi.org/10.1007/s13225-020-00466-2)) |
| Otu55918 | 1 | k__Fungi(100);p__Ascomycota(100);c__Saccharomycetes(100);o__Saccharomycetales(100);f__Metschnikowiaceae(100);g__Metschnikowia(100);s__Metschnikowia_pulcherrima(100); | Metschnikowia | 13 | Saprotroph | \|Undefined Saprotroph\| | Yeast | NULL | Probable | As nectar-tap_saprotroph (Põlme et al. 2020); Decay sugar-rich substrates (Põlme et al. 2020) | Põlme S, et al. 2020. Fungal Diversity 105:1-16 ((https://doi.org/10.1007/s13225-020-00466-2)) |
| Otu55921 | 1 | k__Fungi(100);p__Ascomycota(100);c__Saccharomycetes(100);o__Saccharomycetales(100);f__Saccharomycetales_family_Incertae_sedis(100);g__Candida(100);g__Candida_unclassified(100); | Candida | 13 | Pathotroph-Saprotroph-Symbiotroph | Animal Pathogen-Endophyte-Endosymbiont-Epiphyte-\|Undefined Saprotroph\| | Dimorphic Yeast-Yeast | NULL | Probable | As nectar-tap_saprotroph (Põlme et al. 2020); Decay sugar-rich substrates (Põlme et al. 2020); This asexual genus is polyphyletic and contain species that function very differently depend on what they are. Caution should be taken when interpreting guilds for Candida species. | Manolakaki D, et al. 2010. Virulence. 1: 367-375 ((https://doi.org/10.4161/viru.1.5.12796)); Mekbib SB, et al. 2011. Tropical Plant Pathology 36:233-240 ((https://doi.org/10.1590/S1982-56762011000400004)); Põlme S, et al. 2020. Fungal Diversity 105:1-16 ((https://doi.org/10.1007/s13225-020-00466-2)) |
| Otu55922 | 1 | k__Fungi(100);p__Ascomycota(100);c__Saccharomycetes(100);o__Saccharomycetales(100);f__Metschnikowiaceae(100);g__Metschnikowia(100);s__Metschnikowia_pulcherrima(100); | Metschnikowia | 13 | Saprotroph | \|Undefined Saprotroph\| | Yeast | NULL | Probable | As nectar-tap_saprotroph (Põlme et al. 2020); Decay sugar-rich substrates (Põlme et al. 2020) | Põlme S, et al. 2020. Fungal Diversity 105:1-16 ((https://doi.org/10.1007/s13225-020-00466-2)) |
| Otu55923 | 1 | k__Fungi(100);p__Ascomycota(100);c__Saccharomycetes(100);o__Saccharomycetales(100);f__Saccharomycetales_family_Incertae_sedis(100);g__Candida(100);g__Candida_unclassified(100); | Candida | 13 | Pathotroph-Saprotroph-Symbiotroph | Animal Pathogen-Endophyte-Endosymbiont-Epiphyte-\|Undefined Saprotroph\| | Dimorphic Yeast-Yeast | NULL | Probable | As nectar-tap_saprotroph (Põlme et al. 2020); Decay sugar-rich substrates (Põlme et al. 2020); This asexual genus is polyphyletic and contain species that function very differently depend on what they are. Caution should be taken when interpreting guilds for Candida species. | Manolakaki D, et al. 2010. Virulence. 1: 367-375 ((https://doi.org/10.4161/viru.1.5.12796)); Mekbib SB, et al. 2011. Tropical Plant Pathology 36:233-240 ((https://doi.org/10.1590/S1982-56762011000400004)); Põlme S, et al. 2020. Fungal Diversity 105:1-16 ((https://doi.org/10.1007/s13225-020-00466-2)) |
| Otu55924 | 1 | k__Fungi(100);p__Ascomycota(100);c__Saccharomycetes(100);o__Saccharomycetales(100);f__Metschnikowiaceae(100);g__Metschnikowia(100);s__Metschnikowia_pulcherrima(100); | Metschnikowia | 13 | Saprotroph | \|Undefined Saprotroph\| | Yeast | NULL | Probable | As nectar-tap_saprotroph (Põlme et al. 2020); Decay sugar-rich substrates (Põlme et al. 2020) | Põlme S, et al. 2020. Fungal Diversity 105:1-16 ((https://doi.org/10.1007/s13225-020-00466-2)) |
| Otu55926 | 1 | k__Fungi(100);p__Ascomycota(100);c__Saccharomycetes(100);o__Saccharomycetales(100);f__Metschnikowiaceae(100);g__Metschnikowia(100);s__Metschnikowia_pulcherrima(100); | Metschnikowia | 13 | Saprotroph | \|Undefined Saprotroph\| | Yeast | NULL | Probable | As nectar-tap_saprotroph (Põlme et al. 2020); Decay sugar-rich substrates (Põlme et al. 2020) | Põlme S, et al. 2020. Fungal Diversity 105:1-16 ((https://doi.org/10.1007/s13225-020-00466-2)) |
| Otu55927 | 1 | k__Fungi(100);p__Ascomycota(100);c__Saccharomycetes(100);o__Saccharomycetales(100);f__Metschnikowiaceae(100);g__Metschnikowia(100);s__Metschnikowia_pulcherrima(100); | Metschnikowia | 13 | Saprotroph | \|Undefined Saprotroph\| | Yeast | NULL | Probable | As nectar-tap_saprotroph (Põlme et al. 2020); Decay sugar-rich substrates (Põlme et al. 2020) | Põlme S, et al. 2020. Fungal Diversity 105:1-16 ((https://doi.org/10.1007/s13225-020-00466-2)) |
| Otu55928 | 1 | k__Fungi(100);p__Ascomycota(100);c__Saccharomycetes(100);o__Saccharomycetales(100);f__Metschnikowiaceae(100);g__Metschnikowia(100);s__Metschnikowia_pulcherrima(100); | Metschnikowia | 13 | Saprotroph | \|Undefined Saprotroph\| | Yeast | NULL | Probable | As nectar-tap_saprotroph (Põlme et al. 2020); Decay sugar-rich substrates (Põlme et al. 2020) | Põlme S, et al. 2020. Fungal Diversity 105:1-16 ((https://doi.org/10.1007/s13225-020-00466-2)) |
| Otu55929 | 1 | k__Fungi(100);p__Ascomycota(100);c__Saccharomycetes(100);o__Saccharomycetales(100);f__Metschnikowiaceae(100);g__Metschnikowia(100);s__Metschnikowia_pulcherrima(100); | Metschnikowia | 13 | Saprotroph | \|Undefined Saprotroph\| | Yeast | NULL | Probable | As nectar-tap_saprotroph (Põlme et al. 2020); Decay sugar-rich substrates (Põlme et al. 2020) | Põlme S, et al. 2020. Fungal Diversity 105:1-16 ((https://doi.org/10.1007/s13225-020-00466-2)) |
| Otu55930 | 1 | k__Fungi(100);p__Ascomycota(100);c__Saccharomycetes(100);o__Saccharomycetales(100);f__Metschnikowiaceae(100);g__Metschnikowia(100);g__Metschnikowia_unclassified(100); | Metschnikowia | 13 | Saprotroph | \|Undefined Saprotroph\| | Yeast | NULL | Probable | As nectar-tap_saprotroph (Põlme et al. 2020); Decay sugar-rich substrates (Põlme et al. 2020) | Põlme S, et al. 2020. Fungal Diversity 105:1-16 ((https://doi.org/10.1007/s13225-020-00466-2)) |
| Otu55931 | 1 | k__Fungi(100);p__Ascomycota(100);c__Saccharomycetes(100);o__Saccharomycetales(100);f__Metschnikowiaceae(100);g__Metschnikowia(100);s__Metschnikowia_pulcherrima(100); | Metschnikowia | 13 | Saprotroph | \|Undefined Saprotroph\| | Yeast | NULL | Probable | As nectar-tap_saprotroph (Põlme et al. 2020); Decay sugar-rich substrates (Põlme et al. 2020) | Põlme S, et al. 2020. Fungal Diversity 105:1-16 ((https://doi.org/10.1007/s13225-020-00466-2)) |
| Otu55932 | 1 | k__Fungi(100);p__Ascomycota(100);c__Saccharomycetes(100);o__Saccharomycetales(100);f__Metschnikowiaceae(100);g__Metschnikowia(100);s__Metschnikowia_pulcherrima(100); | Metschnikowia | 13 | Saprotroph | \|Undefined Saprotroph\| | Yeast | NULL | Probable | As nectar-tap_saprotroph (Põlme et al. 2020); Decay sugar-rich substrates (Põlme et al. 2020) | Põlme S, et al. 2020. Fungal Diversity 105:1-16 ((https://doi.org/10.1007/s13225-020-00466-2)) |
| Otu55933 | 1 | k__Fungi(100);p__Ascomycota(100);c__Saccharomycetes(100);o__Saccharomycetales(100);f__Metschnikowiaceae(100);g__Metschnikowia(100);s__Metschnikowia_pulcherrima(100); | Metschnikowia | 13 | Saprotroph | \|Undefined Saprotroph\| | Yeast | NULL | Probable | As nectar-tap_saprotroph (Põlme et al. 2020); Decay sugar-rich substrates (Põlme et al. 2020) | Põlme S, et al. 2020. Fungal Diversity 105:1-16 ((https://doi.org/10.1007/s13225-020-00466-2)) |
| Otu55934 | 1 | k__Fungi(100);p__Ascomycota(100);c__Saccharomycetes(100);o__Saccharomycetales(100);f__Metschnikowiaceae(100);g__Metschnikowia(100);g__Metschnikowia_unclassified(100); | Metschnikowia | 13 | Saprotroph | \|Undefined Saprotroph\| | Yeast | NULL | Probable | As nectar-tap_saprotroph (Põlme et al. 2020); Decay sugar-rich substrates (Põlme et al. 2020) | Põlme S, et al. 2020. Fungal Diversity 105:1-16 ((https://doi.org/10.1007/s13225-020-00466-2)) |
| Otu55935 | 1 | k__Fungi(100);p__Ascomycota(100);c__Saccharomycetes(100);o__Saccharomycetales(100);f__Metschnikowiaceae(100);g__Metschnikowia(100);s__Metschnikowia_pulcherrima(100); | Metschnikowia | 13 | Saprotroph | \|Undefined Saprotroph\| | Yeast | NULL | Probable | As nectar-tap_saprotroph (Põlme et al. 2020); Decay sugar-rich substrates (Põlme et al. 2020) | Põlme S, et al. 2020. Fungal Diversity 105:1-16 ((https://doi.org/10.1007/s13225-020-00466-2)) |
| Otu55936 | 1 | k__Fungi(100);p__Ascomycota(100);c__Saccharomycetes(100);o__Saccharomycetales(100);f__Metschnikowiaceae(100);g__Metschnikowia(100);s__Metschnikowia_pulcherrima(100); | Metschnikowia | 13 | Saprotroph | \|Undefined Saprotroph\| | Yeast | NULL | Probable | As nectar-tap_saprotroph (Põlme et al. 2020); Decay sugar-rich substrates (Põlme et al. 2020) | Põlme S, et al. 2020. Fungal Diversity 105:1-16 ((https://doi.org/10.1007/s13225-020-00466-2)) |
| Otu55937 | 1 | k__Fungi(100);p__Ascomycota(100);c__Saccharomycetes(100);o__Saccharomycetales(100);o__Saccharomycetales_unclassified(100);o__Saccharomycetales_unclassified(100);o__Saccharomycetales_unclassified(100); | Saccharomycetales | 7 | Saprotroph | Undefined Saprotroph | Yeast | NULL | Possible | NULL | Sterkenburg E, et al. 2015. New Phytologist 207:1145-1158 ((https://doi.org/10.1111/nph.13426)) |
| Otu55938 | 1 | k__Fungi(100);p__Ascomycota(100);c__Saccharomycetes(100);o__Saccharomycetales(100);f__Saccharomycetales_family_Incertae_sedis(100);g__Candida(100);g__Candida_unclassified(100); | Candida | 13 | Pathotroph-Saprotroph-Symbiotroph | Animal Pathogen-Endophyte-Endosymbiont-Epiphyte-\|Undefined Saprotroph\| | Dimorphic Yeast-Yeast | NULL | Probable | As nectar-tap_saprotroph (Põlme et al. 2020); Decay sugar-rich substrates (Põlme et al. 2020); This asexual genus is polyphyletic and contain species that function very differently depend on what they are. Caution should be taken when interpreting guilds for Candida species. | Manolakaki D, et al. 2010. Virulence. 1: 367-375 ((https://doi.org/10.4161/viru.1.5.12796)); Mekbib SB, et al. 2011. Tropical Plant Pathology 36:233-240 ((https://doi.org/10.1590/S1982-56762011000400004)); Põlme S, et al. 2020. Fungal Diversity 105:1-16 ((https://doi.org/10.1007/s13225-020-00466-2)) |
| Otu55939 | 1 | k__Fungi(100);p__Ascomycota(100);c__Saccharomycetes(100);o__Saccharomycetales(100);f__Metschnikowiaceae(100);g__Metschnikowia(100);s__Metschnikowia_pulcherrima(100); | Metschnikowia | 13 | Saprotroph | \|Undefined Saprotroph\| | Yeast | NULL | Probable | As nectar-tap_saprotroph (Põlme et al. 2020); Decay sugar-rich substrates (Põlme et al. 2020) | Põlme S, et al. 2020. Fungal Diversity 105:1-16 ((https://doi.org/10.1007/s13225-020-00466-2)) |
| Otu55940 | 1 | k__Fungi(100);p__Ascomycota(100);c__Saccharomycetes(100);o__Saccharomycetales(100);f__Metschnikowiaceae(100);g__Metschnikowia(100);s__Metschnikowia_pulcherrima(100); | Metschnikowia | 13 | Saprotroph | \|Undefined Saprotroph\| | Yeast | NULL | Probable | As nectar-tap_saprotroph (Põlme et al. 2020); Decay sugar-rich substrates (Põlme et al. 2020) | Põlme S, et al. 2020. Fungal Diversity 105:1-16 ((https://doi.org/10.1007/s13225-020-00466-2)) |
| Otu55947 | 1 | k__Fungi(100);p__Ascomycota(100);c__Saccharomycetes(100);o__Saccharomycetales(100);f__Saccharomycetales_family_Incertae_sedis(100);f__Saccharomycetales_family_Incertae_sedis_unclassified(100);f__Saccharomycetales_family_Incertae_sedis_unclassified(100); | Saccharomycetales | 7 | Saprotroph | Undefined Saprotroph | Yeast | NULL | Possible | NULL | Sterkenburg E, et al. 2015. New Phytologist 207:1145-1158 ((https://doi.org/10.1111/nph.13426)) |
| Otu55948 | 1 | k__Fungi(100);p__Ascomycota(100);c__Saccharomycetes(100);o__Saccharomycetales(100);f__Metschnikowiaceae(100);g__Metschnikowia(100);s__Metschnikowia_pulcherrima(100); | Metschnikowia | 13 | Saprotroph | \|Undefined Saprotroph\| | Yeast | NULL | Probable | As nectar-tap_saprotroph (Põlme et al. 2020); Decay sugar-rich substrates (Põlme et al. 2020) | Põlme S, et al. 2020. Fungal Diversity 105:1-16 ((https://doi.org/10.1007/s13225-020-00466-2)) |
| Otu55950 | 1 | k__Fungi(100);p__Ascomycota(100);c__Saccharomycetes(100);o__Saccharomycetales(100);f__Metschnikowiaceae(100);g__Metschnikowia(100);s__Metschnikowia_pulcherrima(100); | Metschnikowia | 13 | Saprotroph | \|Undefined Saprotroph\| | Yeast | NULL | Probable | As nectar-tap_saprotroph (Põlme et al. 2020); Decay sugar-rich substrates (Põlme et al. 2020) | Põlme S, et al. 2020. Fungal Diversity 105:1-16 ((https://doi.org/10.1007/s13225-020-00466-2)) |
| Otu55951 | 1 | k__Fungi(100);p__Ascomycota(100);c__Saccharomycetes(100);o__Saccharomycetales(100);f__Saccharomycetales_family_Incertae_sedis(100);g__Candida(100);g__Candida_unclassified(100); | Candida | 13 | Pathotroph-Saprotroph-Symbiotroph | Animal Pathogen-Endophyte-Endosymbiont-Epiphyte-\|Undefined Saprotroph\| | Dimorphic Yeast-Yeast | NULL | Probable | As nectar-tap_saprotroph (Põlme et al. 2020); Decay sugar-rich substrates (Põlme et al. 2020); This asexual genus is polyphyletic and contain species that function very differently depend on what they are. Caution should be taken when interpreting guilds for Candida species. | Manolakaki D, et al. 2010. Virulence. 1: 367-375 ((https://doi.org/10.4161/viru.1.5.12796)); Mekbib SB, et al. 2011. Tropical Plant Pathology 36:233-240 ((https://doi.org/10.1590/S1982-56762011000400004)); Põlme S, et al. 2020. Fungal Diversity 105:1-16 ((https://doi.org/10.1007/s13225-020-00466-2)) |
| Otu55952 | 1 | k__Fungi(100);p__Ascomycota(100);c__Saccharomycetes(100);o__Saccharomycetales(100);f__Saccharomycetales_family_Incertae_sedis(100);g__Candida(100);g__Candida_unclassified(100); | Candida | 13 | Pathotroph-Saprotroph-Symbiotroph | Animal Pathogen-Endophyte-Endosymbiont-Epiphyte-\|Undefined Saprotroph\| | Dimorphic Yeast-Yeast | NULL | Probable | As nectar-tap_saprotroph (Põlme et al. 2020); Decay sugar-rich substrates (Põlme et al. 2020); This asexual genus is polyphyletic and contain species that function very differently depend on what they are. Caution should be taken when interpreting guilds for Candida species. | Manolakaki D, et al. 2010. Virulence. 1: 367-375 ((https://doi.org/10.4161/viru.1.5.12796)); Mekbib SB, et al. 2011. Tropical Plant Pathology 36:233-240 ((https://doi.org/10.1590/S1982-56762011000400004)); Põlme S, et al. 2020. Fungal Diversity 105:1-16 ((https://doi.org/10.1007/s13225-020-00466-2)) |
| Otu55954 | 1 | k__Fungi(100);p__Ascomycota(100);c__Saccharomycetes(100);o__Saccharomycetales(100);f__Metschnikowiaceae(100);g__Metschnikowia(100);s__Metschnikowia_pulcherrima(100); | Metschnikowia | 13 | Saprotroph | \|Undefined Saprotroph\| | Yeast | NULL | Probable | As nectar-tap_saprotroph (Põlme et al. 2020); Decay sugar-rich substrates (Põlme et al. 2020) | Põlme S, et al. 2020. Fungal Diversity 105:1-16 ((https://doi.org/10.1007/s13225-020-00466-2)) |
| Otu55956 | 1 | k__Fungi(100);p__Ascomycota(100);c__Saccharomycetes(100);o__Saccharomycetales(100);f__Metschnikowiaceae(100);g__Metschnikowia(100);s__Metschnikowia_pulcherrima(100); | Metschnikowia | 13 | Saprotroph | \|Undefined Saprotroph\| | Yeast | NULL | Probable | As nectar-tap_saprotroph (Põlme et al. 2020); Decay sugar-rich substrates (Põlme et al. 2020) | Põlme S, et al. 2020. Fungal Diversity 105:1-16 ((https://doi.org/10.1007/s13225-020-00466-2)) |
| Otu55957 | 1 | k__Fungi(100);p__Ascomycota(100);c__Saccharomycetes(100);o__Saccharomycetales(100);f__Saccharomycetales_family_Incertae_sedis(100);g__Candida(100);s__Candida_cretensis(100); | Candida | 13 | Pathotroph-Saprotroph-Symbiotroph | Animal Pathogen-Endophyte-Endosymbiont-Epiphyte-\|Undefined Saprotroph\| | Dimorphic Yeast-Yeast | NULL | Probable | As nectar-tap_saprotroph (Põlme et al. 2020); Decay sugar-rich substrates (Põlme et al. 2020); This asexual genus is polyphyletic and contain species that function very differently depend on what they are. Caution should be taken when interpreting guilds for Candida species. | Manolakaki D, et al. 2010. Virulence. 1: 367-375 ((https://doi.org/10.4161/viru.1.5.12796)); Mekbib SB, et al. 2011. Tropical Plant Pathology 36:233-240 ((https://doi.org/10.1590/S1982-56762011000400004)); Põlme S, et al. 2020. Fungal Diversity 105:1-16 ((https://doi.org/10.1007/s13225-020-00466-2)) |
| Otu55958 | 1 | k__Fungi(100);p__Ascomycota(100);c__Saccharomycetes(100);o__Saccharomycetales(100);f__Saccharomycetales_family_Incertae_sedis(100);g__Candida(100);g__Candida_unclassified(100); | Candida | 13 | Pathotroph-Saprotroph-Symbiotroph | Animal Pathogen-Endophyte-Endosymbiont-Epiphyte-\|Undefined Saprotroph\| | Dimorphic Yeast-Yeast | NULL | Probable | As nectar-tap_saprotroph (Põlme et al. 2020); Decay sugar-rich substrates (Põlme et al. 2020); This asexual genus is polyphyletic and contain species that function very differently depend on what they are. Caution should be taken when interpreting guilds for Candida species. | Manolakaki D, et al. 2010. Virulence. 1: 367-375 ((https://doi.org/10.4161/viru.1.5.12796)); Mekbib SB, et al. 2011. Tropical Plant Pathology 36:233-240 ((https://doi.org/10.1590/S1982-56762011000400004)); Põlme S, et al. 2020. Fungal Diversity 105:1-16 ((https://doi.org/10.1007/s13225-020-00466-2)) |
| Otu55961 | 1 | k__Fungi(100);p__Basidiomycota(100);c__Microbotryomycetes(100);o__Sporidiobolales(100);f__Sporidiobolales_family_Incertae_sedis(100);g__Sporobolomyces(100);s__Sporobolomyces_lactosus(100); | Sporobolomyces | 13 | Pathotroph-Saprotroph | \|Fungal Parasite\|-Plant Saprotroph | Yeast | NULL | Probable | As litter_saprotroph (Põlme et al. 2020); As mycoparasite (Põlme et al. 2020); Decay fungal material (Põlme et al. 2020) | Tedersoo L, et al. 2014. Science 346:e1256688 ((https://doi.org/10.1126/science.1256688)); Purahong W, et al. 2016. Molecular Ecology 25:4059-4074 ((https://doi.org/10.1111/mec.13739)); Põlme S, et al. 2020. Fungal Diversity 105:1-16 ((https://doi.org/10.1007/s13225-020-00466-2)) |
| Otu55962 | 1 | k__Fungi(100);p__Ascomycota(100);c__Saccharomycetes(100);o__Saccharomycetales(100);f__Metschnikowiaceae(100);g__Metschnikowia(100);s__Metschnikowia_pulcherrima(100); | Metschnikowia | 13 | Saprotroph | \|Undefined Saprotroph\| | Yeast | NULL | Probable | As nectar-tap_saprotroph (Põlme et al. 2020); Decay sugar-rich substrates (Põlme et al. 2020) | Põlme S, et al. 2020. Fungal Diversity 105:1-16 ((https://doi.org/10.1007/s13225-020-00466-2)) |
| Otu55964 | 1 | k__Fungi(100);p__Ascomycota(100);c__Saccharomycetes(100);o__Saccharomycetales(100);f__Saccharomycetales_family_Incertae_sedis(100);g__Candida(100);g__Candida_unclassified(100); | Candida | 13 | Pathotroph-Saprotroph-Symbiotroph | Animal Pathogen-Endophyte-Endosymbiont-Epiphyte-\|Undefined Saprotroph\| | Dimorphic Yeast-Yeast | NULL | Probable | As nectar-tap_saprotroph (Põlme et al. 2020); Decay sugar-rich substrates (Põlme et al. 2020); This asexual genus is polyphyletic and contain species that function very differently depend on what they are. Caution should be taken when interpreting guilds for Candida species. | Manolakaki D, et al. 2010. Virulence. 1: 367-375 ((https://doi.org/10.4161/viru.1.5.12796)); Mekbib SB, et al. 2011. Tropical Plant Pathology 36:233-240 ((https://doi.org/10.1590/S1982-56762011000400004)); Põlme S, et al. 2020. Fungal Diversity 105:1-16 ((https://doi.org/10.1007/s13225-020-00466-2)) |
| Otu55966 | 1 | k__Fungi(100);p__Ascomycota(100);c__Saccharomycetes(100);o__Saccharomycetales(100);f__Saccharomycetales_family_Incertae_sedis(100);g__Candida(100);g__Candida_unclassified(100); | Candida | 13 | Pathotroph-Saprotroph-Symbiotroph | Animal Pathogen-Endophyte-Endosymbiont-Epiphyte-\|Undefined Saprotroph\| | Dimorphic Yeast-Yeast | NULL | Probable | As nectar-tap_saprotroph (Põlme et al. 2020); Decay sugar-rich substrates (Põlme et al. 2020); This asexual genus is polyphyletic and contain species that function very differently depend on what they are. Caution should be taken when interpreting guilds for Candida species. | Manolakaki D, et al. 2010. Virulence. 1: 367-375 ((https://doi.org/10.4161/viru.1.5.12796)); Mekbib SB, et al. 2011. Tropical Plant Pathology 36:233-240 ((https://doi.org/10.1590/S1982-56762011000400004)); Põlme S, et al. 2020. Fungal Diversity 105:1-16 ((https://doi.org/10.1007/s13225-020-00466-2)) |
| Otu55967 | 1 | k__Fungi(100);p__Ascomycota(100);c__Saccharomycetes(100);o__Saccharomycetales(100);f__Metschnikowiaceae(100);g__Metschnikowia(100);s__Metschnikowia_pulcherrima(100); | Metschnikowia | 13 | Saprotroph | \|Undefined Saprotroph\| | Yeast | NULL | Probable | As nectar-tap_saprotroph (Põlme et al. 2020); Decay sugar-rich substrates (Põlme et al. 2020) | Põlme S, et al. 2020. Fungal Diversity 105:1-16 ((https://doi.org/10.1007/s13225-020-00466-2)) |
| Otu55968 | 1 | k__Fungi(100);p__Ascomycota(100);c__Saccharomycetes(100);o__Saccharomycetales(100);f__Metschnikowiaceae(100);g__Metschnikowia(100);s__Metschnikowia_pulcherrima(100); | Metschnikowia | 13 | Saprotroph | \|Undefined Saprotroph\| | Yeast | NULL | Probable | As nectar-tap_saprotroph (Põlme et al. 2020); Decay sugar-rich substrates (Põlme et al. 2020) | Põlme S, et al. 2020. Fungal Diversity 105:1-16 ((https://doi.org/10.1007/s13225-020-00466-2)) |
| Otu55969 | 1 | k__Fungi(100);p__Ascomycota(100);c__Saccharomycetes(100);o__Saccharomycetales(100);f__Saccharomycetales_family_Incertae_sedis(100);g__Candida(100);g__Candida_unclassified(100); | Candida | 13 | Pathotroph-Saprotroph-Symbiotroph | Animal Pathogen-Endophyte-Endosymbiont-Epiphyte-\|Undefined Saprotroph\| | Dimorphic Yeast-Yeast | NULL | Probable | As nectar-tap_saprotroph (Põlme et al. 2020); Decay sugar-rich substrates (Põlme et al. 2020); This asexual genus is polyphyletic and contain species that function very differently depend on what they are. Caution should be taken when interpreting guilds for Candida species. | Manolakaki D, et al. 2010. Virulence. 1: 367-375 ((https://doi.org/10.4161/viru.1.5.12796)); Mekbib SB, et al. 2011. Tropical Plant Pathology 36:233-240 ((https://doi.org/10.1590/S1982-56762011000400004)); Põlme S, et al. 2020. Fungal Diversity 105:1-16 ((https://doi.org/10.1007/s13225-020-00466-2)) |
| Otu55970 | 1 | k__Fungi(100);p__Ascomycota(100);c__Saccharomycetes(100);o__Saccharomycetales(100);f__Metschnikowiaceae(100);g__Metschnikowia(100);s__Metschnikowia_pulcherrima(100); | Metschnikowia | 13 | Saprotroph | \|Undefined Saprotroph\| | Yeast | NULL | Probable | As nectar-tap_saprotroph (Põlme et al. 2020); Decay sugar-rich substrates (Põlme et al. 2020) | Põlme S, et al. 2020. Fungal Diversity 105:1-16 ((https://doi.org/10.1007/s13225-020-00466-2)) |
| Otu55971 | 1 | k__Fungi(100);p__Ascomycota(100);c__Saccharomycetes(100);o__Saccharomycetales(100);f__Metschnikowiaceae(100);g__Metschnikowia(100);s__Metschnikowia_pulcherrima(100); | Metschnikowia | 13 | Saprotroph | \|Undefined Saprotroph\| | Yeast | NULL | Probable | As nectar-tap_saprotroph (Põlme et al. 2020); Decay sugar-rich substrates (Põlme et al. 2020) | Põlme S, et al. 2020. Fungal Diversity 105:1-16 ((https://doi.org/10.1007/s13225-020-00466-2)) |
| Otu55972 | 1 | k__Fungi(100);p__Ascomycota(100);c__Saccharomycetes(100);o__Saccharomycetales(100);f__Saccharomycetales_family_Incertae_sedis(100);g__Candida(100);s__Candida_cretensis(100); | Candida | 13 | Pathotroph-Saprotroph-Symbiotroph | Animal Pathogen-Endophyte-Endosymbiont-Epiphyte-\|Undefined Saprotroph\| | Dimorphic Yeast-Yeast | NULL | Probable | As nectar-tap_saprotroph (Põlme et al. 2020); Decay sugar-rich substrates (Põlme et al. 2020); This asexual genus is polyphyletic and contain species that function very differently depend on what they are. Caution should be taken when interpreting guilds for Candida species. | Manolakaki D, et al. 2010. Virulence. 1: 367-375 ((https://doi.org/10.4161/viru.1.5.12796)); Mekbib SB, et al. 2011. Tropical Plant Pathology 36:233-240 ((https://doi.org/10.1590/S1982-56762011000400004)); Põlme S, et al. 2020. Fungal Diversity 105:1-16 ((https://doi.org/10.1007/s13225-020-00466-2)) |
| Otu55973 | 1 | k__Fungi(100);p__Ascomycota(100);c__Saccharomycetes(100);o__Saccharomycetales(100);f__Saccharomycetales_family_Incertae_sedis(100);g__Candida(100);g__Candida_unclassified(100); | Candida | 13 | Pathotroph-Saprotroph-Symbiotroph | Animal Pathogen-Endophyte-Endosymbiont-Epiphyte-\|Undefined Saprotroph\| | Dimorphic Yeast-Yeast | NULL | Probable | As nectar-tap_saprotroph (Põlme et al. 2020); Decay sugar-rich substrates (Põlme et al. 2020); This asexual genus is polyphyletic and contain species that function very differently depend on what they are. Caution should be taken when interpreting guilds for Candida species. | Manolakaki D, et al. 2010. Virulence. 1: 367-375 ((https://doi.org/10.4161/viru.1.5.12796)); Mekbib SB, et al. 2011. Tropical Plant Pathology 36:233-240 ((https://doi.org/10.1590/S1982-56762011000400004)); Põlme S, et al. 2020. Fungal Diversity 105:1-16 ((https://doi.org/10.1007/s13225-020-00466-2)) |
| Otu55974 | 1 | k__Fungi(100);p__Ascomycota(100);c__Saccharomycetes(100);o__Saccharomycetales(100);f__Metschnikowiaceae(100);g__Metschnikowia(100);g__Metschnikowia_unclassified(100); | Metschnikowia | 13 | Saprotroph | \|Undefined Saprotroph\| | Yeast | NULL | Probable | As nectar-tap_saprotroph (Põlme et al. 2020); Decay sugar-rich substrates (Põlme et al. 2020) | Põlme S, et al. 2020. Fungal Diversity 105:1-16 ((https://doi.org/10.1007/s13225-020-00466-2)) |
| Otu55975 | 1 | k__Fungi(100);p__Ascomycota(100);c__Saccharomycetes(100);o__Saccharomycetales(100);f__Saccharomycetales_family_Incertae_sedis(100);g__Candida(100);g__Candida_unclassified(100); | Candida | 13 | Pathotroph-Saprotroph-Symbiotroph | Animal Pathogen-Endophyte-Endosymbiont-Epiphyte-\|Undefined Saprotroph\| | Dimorphic Yeast-Yeast | NULL | Probable | As nectar-tap_saprotroph (Põlme et al. 2020); Decay sugar-rich substrates (Põlme et al. 2020); This asexual genus is polyphyletic and contain species that function very differently depend on what they are. Caution should be taken when interpreting guilds for Candida species. | Manolakaki D, et al. 2010. Virulence. 1: 367-375 ((https://doi.org/10.4161/viru.1.5.12796)); Mekbib SB, et al. 2011. Tropical Plant Pathology 36:233-240 ((https://doi.org/10.1590/S1982-56762011000400004)); Põlme S, et al. 2020. Fungal Diversity 105:1-16 ((https://doi.org/10.1007/s13225-020-00466-2)) |
| Otu55977 | 1 | k__Fungi(100);p__Ascomycota(100);c__Saccharomycetes(100);o__Saccharomycetales(100);f__Metschnikowiaceae(100);g__Metschnikowia(100);g__Metschnikowia_unclassified(100); | Metschnikowia | 13 | Saprotroph | \|Undefined Saprotroph\| | Yeast | NULL | Probable | As nectar-tap_saprotroph (Põlme et al. 2020); Decay sugar-rich substrates (Põlme et al. 2020) | Põlme S, et al. 2020. Fungal Diversity 105:1-16 ((https://doi.org/10.1007/s13225-020-00466-2)) |
| Otu55978 | 1 | k__Fungi(100);p__Ascomycota(100);c__Saccharomycetes(100);o__Saccharomycetales(100);f__Metschnikowiaceae(100);g__Metschnikowia(100);s__Metschnikowia_pulcherrima(100); | Metschnikowia | 13 | Saprotroph | \|Undefined Saprotroph\| | Yeast | NULL | Probable | As nectar-tap_saprotroph (Põlme et al. 2020); Decay sugar-rich substrates (Põlme et al. 2020) | Põlme S, et al. 2020. Fungal Diversity 105:1-16 ((https://doi.org/10.1007/s13225-020-00466-2)) |
| Otu55979 | 1 | k__Fungi(100);p__Ascomycota(100);c__Saccharomycetes(100);o__Saccharomycetales(100);f__Metschnikowiaceae(100);g__Metschnikowia(100);g__Metschnikowia_unclassified(100); | Metschnikowia | 13 | Saprotroph | \|Undefined Saprotroph\| | Yeast | NULL | Probable | As nectar-tap_saprotroph (Põlme et al. 2020); Decay sugar-rich substrates (Põlme et al. 2020) | Põlme S, et al. 2020. Fungal Diversity 105:1-16 ((https://doi.org/10.1007/s13225-020-00466-2)) |
| Otu55981 | 1 | k__Fungi(100);p__Ascomycota(100);c__Saccharomycetes(100);o__Saccharomycetales(100);f__Metschnikowiaceae(100);g__Metschnikowia(100);s__Metschnikowia_pulcherrima(100); | Metschnikowia | 13 | Saprotroph | \|Undefined Saprotroph\| | Yeast | NULL | Probable | As nectar-tap_saprotroph (Põlme et al. 2020); Decay sugar-rich substrates (Põlme et al. 2020) | Põlme S, et al. 2020. Fungal Diversity 105:1-16 ((https://doi.org/10.1007/s13225-020-00466-2)) |
| Otu55982 | 1 | k__Fungi(100);p__Ascomycota(100);c__Saccharomycetes(100);o__Saccharomycetales(100);f__Metschnikowiaceae(100);g__Metschnikowia(100);s__Metschnikowia_pulcherrima(100); | Metschnikowia | 13 | Saprotroph | \|Undefined Saprotroph\| | Yeast | NULL | Probable | As nectar-tap_saprotroph (Põlme et al. 2020); Decay sugar-rich substrates (Põlme et al. 2020) | Põlme S, et al. 2020. Fungal Diversity 105:1-16 ((https://doi.org/10.1007/s13225-020-00466-2)) |
| Otu55983 | 1 | k__Fungi(100);p__Ascomycota(100);c__Saccharomycetes(100);o__Saccharomycetales(100);f__Metschnikowiaceae(100);g__Metschnikowia(100);s__Metschnikowia_pulcherrima(100); | Metschnikowia | 13 | Saprotroph | \|Undefined Saprotroph\| | Yeast | NULL | Probable | As nectar-tap_saprotroph (Põlme et al. 2020); Decay sugar-rich substrates (Põlme et al. 2020) | Põlme S, et al. 2020. Fungal Diversity 105:1-16 ((https://doi.org/10.1007/s13225-020-00466-2)) |
| Otu55985 | 1 | k__Fungi(100);p__Ascomycota(100);c__Saccharomycetes(100);o__Saccharomycetales(100);f__Metschnikowiaceae(100);g__Metschnikowia(100);s__Metschnikowia_pulcherrima(100); | Metschnikowia | 13 | Saprotroph | \|Undefined Saprotroph\| | Yeast | NULL | Probable | As nectar-tap_saprotroph (Põlme et al. 2020); Decay sugar-rich substrates (Põlme et al. 2020) | Põlme S, et al. 2020. Fungal Diversity 105:1-16 ((https://doi.org/10.1007/s13225-020-00466-2)) |
| Otu55986 | 1 | k__Fungi(100);p__Ascomycota(100);c__Saccharomycetes(100);o__Saccharomycetales(100);f__Saccharomycetales_family_Incertae_sedis(100);g__Candida(100);g__Candida_unclassified(100); | Candida | 13 | Pathotroph-Saprotroph-Symbiotroph | Animal Pathogen-Endophyte-Endosymbiont-Epiphyte-\|Undefined Saprotroph\| | Dimorphic Yeast-Yeast | NULL | Probable | As nectar-tap_saprotroph (Põlme et al. 2020); Decay sugar-rich substrates (Põlme et al. 2020); This asexual genus is polyphyletic and contain species that function very differently depend on what they are. Caution should be taken when interpreting guilds for Candida species. | Manolakaki D, et al. 2010. Virulence. 1: 367-375 ((https://doi.org/10.4161/viru.1.5.12796)); Mekbib SB, et al. 2011. Tropical Plant Pathology 36:233-240 ((https://doi.org/10.1590/S1982-56762011000400004)); Põlme S, et al. 2020. Fungal Diversity 105:1-16 ((https://doi.org/10.1007/s13225-020-00466-2)) |
| Otu55987 | 1 | k__Fungi(100);p__Ascomycota(100);c__Saccharomycetes(100);o__Saccharomycetales(100);f__Metschnikowiaceae(100);g__Metschnikowia(100);s__Metschnikowia_pulcherrima(100); | Metschnikowia | 13 | Saprotroph | \|Undefined Saprotroph\| | Yeast | NULL | Probable | As nectar-tap_saprotroph (Põlme et al. 2020); Decay sugar-rich substrates (Põlme et al. 2020) | Põlme S, et al. 2020. Fungal Diversity 105:1-16 ((https://doi.org/10.1007/s13225-020-00466-2)) |
| Otu55988 | 1 | k__Fungi(100);p__Ascomycota(100);c__Saccharomycetes(100);o__Saccharomycetales(100);f__Metschnikowiaceae(100);g__Metschnikowia(100);s__Metschnikowia_pulcherrima(100); | Metschnikowia | 13 | Saprotroph | \|Undefined Saprotroph\| | Yeast | NULL | Probable | As nectar-tap_saprotroph (Põlme et al. 2020); Decay sugar-rich substrates (Põlme et al. 2020) | Põlme S, et al. 2020. Fungal Diversity 105:1-16 ((https://doi.org/10.1007/s13225-020-00466-2)) |
| Otu55990 | 1 | k__Fungi(100);p__Ascomycota(100);c__Saccharomycetes(100);o__Saccharomycetales(100);f__Metschnikowiaceae(100);g__Metschnikowia(100);s__Metschnikowia_pulcherrima(100); | Metschnikowia | 13 | Saprotroph | \|Undefined Saprotroph\| | Yeast | NULL | Probable | As nectar-tap_saprotroph (Põlme et al. 2020); Decay sugar-rich substrates (Põlme et al. 2020) | Põlme S, et al. 2020. Fungal Diversity 105:1-16 ((https://doi.org/10.1007/s13225-020-00466-2)) |
| Otu55991 | 1 | k__Fungi(100);p__Ascomycota(100);c__Saccharomycetes(100);o__Saccharomycetales(100);f__Metschnikowiaceae(100);g__Metschnikowia(100);s__Metschnikowia_pulcherrima(100); | Metschnikowia | 13 | Saprotroph | \|Undefined Saprotroph\| | Yeast | NULL | Probable | As nectar-tap_saprotroph (Põlme et al. 2020); Decay sugar-rich substrates (Põlme et al. 2020) | Põlme S, et al. 2020. Fungal Diversity 105:1-16 ((https://doi.org/10.1007/s13225-020-00466-2)) |
| Otu55992 | 1 | k__Fungi(100);p__Ascomycota(100);c__Saccharomycetes(100);o__Saccharomycetales(100);f__Metschnikowiaceae(100);g__Metschnikowia(100);s__Metschnikowia_pulcherrima(100); | Metschnikowia | 13 | Saprotroph | \|Undefined Saprotroph\| | Yeast | NULL | Probable | As nectar-tap_saprotroph (Põlme et al. 2020); Decay sugar-rich substrates (Põlme et al. 2020) | Põlme S, et al. 2020. Fungal Diversity 105:1-16 ((https://doi.org/10.1007/s13225-020-00466-2)) |
| Otu55993 | 1 | k__Fungi(100);p__Ascomycota(100);c__Saccharomycetes(100);o__Saccharomycetales(100);f__Metschnikowiaceae(100);g__Metschnikowia(100);s__Metschnikowia_pulcherrima(100); | Metschnikowia | 13 | Saprotroph | \|Undefined Saprotroph\| | Yeast | NULL | Probable | As nectar-tap_saprotroph (Põlme et al. 2020); Decay sugar-rich substrates (Põlme et al. 2020) | Põlme S, et al. 2020. Fungal Diversity 105:1-16 ((https://doi.org/10.1007/s13225-020-00466-2)) |
| Otu55994 | 1 | k__Fungi(100);p__Ascomycota(100);c__Saccharomycetes(100);o__Saccharomycetales(100);f__Metschnikowiaceae(100);g__Metschnikowia(100);g__Metschnikowia_unclassified(100); | Metschnikowia | 13 | Saprotroph | \|Undefined Saprotroph\| | Yeast | NULL | Probable | As nectar-tap_saprotroph (Põlme et al. 2020); Decay sugar-rich substrates (Põlme et al. 2020) | Põlme S, et al. 2020. Fungal Diversity 105:1-16 ((https://doi.org/10.1007/s13225-020-00466-2)) |
| Otu55995 | 1 | k__Fungi(100);p__Ascomycota(100);c__Saccharomycetes(100);o__Saccharomycetales(100);f__Metschnikowiaceae(100);g__Metschnikowia(100);s__Metschnikowia_pulcherrima(100); | Metschnikowia | 13 | Saprotroph | \|Undefined Saprotroph\| | Yeast | NULL | Probable | As nectar-tap_saprotroph (Põlme et al. 2020); Decay sugar-rich substrates (Põlme et al. 2020) | Põlme S, et al. 2020. Fungal Diversity 105:1-16 ((https://doi.org/10.1007/s13225-020-00466-2)) |
| Otu55996 | 1 | k__Fungi(100);p__Ascomycota(100);c__Saccharomycetes(100);o__Saccharomycetales(100);f__Metschnikowiaceae(100);g__Metschnikowia(100);g__Metschnikowia_unclassified(100); | Metschnikowia | 13 | Saprotroph | \|Undefined Saprotroph\| | Yeast | NULL | Probable | As nectar-tap_saprotroph (Põlme et al. 2020); Decay sugar-rich substrates (Põlme et al. 2020) | Põlme S, et al. 2020. Fungal Diversity 105:1-16 ((https://doi.org/10.1007/s13225-020-00466-2)) |
| Otu55997 | 1 | k__Fungi(100);p__Ascomycota(100);c__Saccharomycetes(100);o__Saccharomycetales(100);f__Metschnikowiaceae(100);g__Metschnikowia(100);s__Metschnikowia_pulcherrima(100); | Metschnikowia | 13 | Saprotroph | \|Undefined Saprotroph\| | Yeast | NULL | Probable | As nectar-tap_saprotroph (Põlme et al. 2020); Decay sugar-rich substrates (Põlme et al. 2020) | Põlme S, et al. 2020. Fungal Diversity 105:1-16 ((https://doi.org/10.1007/s13225-020-00466-2)) |
| Otu55998 | 1 | k__Fungi(100);p__Ascomycota(100);c__Saccharomycetes(100);o__Saccharomycetales(100);f__Metschnikowiaceae(100);g__Metschnikowia(100);g__Metschnikowia_unclassified(100); | Metschnikowia | 13 | Saprotroph | \|Undefined Saprotroph\| | Yeast | NULL | Probable | As nectar-tap_saprotroph (Põlme et al. 2020); Decay sugar-rich substrates (Põlme et al. 2020) | Põlme S, et al. 2020. Fungal Diversity 105:1-16 ((https://doi.org/10.1007/s13225-020-00466-2)) |
| Otu55999 | 1 | k__Fungi(100);p__Ascomycota(100);c__Saccharomycetes(100);o__Saccharomycetales(100);f__Metschnikowiaceae(100);g__Metschnikowia(100);s__Metschnikowia_pulcherrima(100); | Metschnikowia | 13 | Saprotroph | \|Undefined Saprotroph\| | Yeast | NULL | Probable | As nectar-tap_saprotroph (Põlme et al. 2020); Decay sugar-rich substrates (Põlme et al. 2020) | Põlme S, et al. 2020. Fungal Diversity 105:1-16 ((https://doi.org/10.1007/s13225-020-00466-2)) |
| Otu56000 | 1 | k__Fungi(100);p__Ascomycota(100);c__Saccharomycetes(100);o__Saccharomycetales(100);f__Metschnikowiaceae(100);g__Metschnikowia(100);s__Metschnikowia_pulcherrima(100); | Metschnikowia | 13 | Saprotroph | \|Undefined Saprotroph\| | Yeast | NULL | Probable | As nectar-tap_saprotroph (Põlme et al. 2020); Decay sugar-rich substrates (Põlme et al. 2020) | Põlme S, et al. 2020. Fungal Diversity 105:1-16 ((https://doi.org/10.1007/s13225-020-00466-2)) |
| Otu56001 | 1 | k__Fungi(100);p__Ascomycota(100);c__Saccharomycetes(100);o__Saccharomycetales(100);f__Metschnikowiaceae(100);g__Metschnikowia(100);s__Metschnikowia_pulcherrima(100); | Metschnikowia | 13 | Saprotroph | \|Undefined Saprotroph\| | Yeast | NULL | Probable | As nectar-tap_saprotroph (Põlme et al. 2020); Decay sugar-rich substrates (Põlme et al. 2020) | Põlme S, et al. 2020. Fungal Diversity 105:1-16 ((https://doi.org/10.1007/s13225-020-00466-2)) |
| Otu56004 | 1 | k__Fungi(100);p__Ascomycota(100);c__Saccharomycetes(100);o__Saccharomycetales(100);f__Metschnikowiaceae(100);g__Metschnikowia(100);s__Metschnikowia_pulcherrima(100); | Metschnikowia | 13 | Saprotroph | \|Undefined Saprotroph\| | Yeast | NULL | Probable | As nectar-tap_saprotroph (Põlme et al. 2020); Decay sugar-rich substrates (Põlme et al. 2020) | Põlme S, et al. 2020. Fungal Diversity 105:1-16 ((https://doi.org/10.1007/s13225-020-00466-2)) |
| Otu56005 | 1 | k__Fungi(100);p__Ascomycota(100);c__Saccharomycetes(100);o__Saccharomycetales(100);f__Metschnikowiaceae(100);g__Metschnikowia(100);s__Metschnikowia_pulcherrima(100); | Metschnikowia | 13 | Saprotroph | \|Undefined Saprotroph\| | Yeast | NULL | Probable | As nectar-tap_saprotroph (Põlme et al. 2020); Decay sugar-rich substrates (Põlme et al. 2020) | Põlme S, et al. 2020. Fungal Diversity 105:1-16 ((https://doi.org/10.1007/s13225-020-00466-2)) |
| Otu56008 | 1 | k__Fungi(100);p__Ascomycota(100);c__Saccharomycetes(100);o__Saccharomycetales(100);f__Metschnikowiaceae(100);g__Metschnikowia(100);s__Metschnikowia_pulcherrima(100); | Metschnikowia | 13 | Saprotroph | \|Undefined Saprotroph\| | Yeast | NULL | Probable | As nectar-tap_saprotroph (Põlme et al. 2020); Decay sugar-rich substrates (Põlme et al. 2020) | Põlme S, et al. 2020. Fungal Diversity 105:1-16 ((https://doi.org/10.1007/s13225-020-00466-2)) |
| Otu56010 | 1 | k__Fungi(100);p__Ascomycota(100);c__Saccharomycetes(100);o__Saccharomycetales(100);f__Metschnikowiaceae(100);g__Metschnikowia(100);g__Metschnikowia_unclassified(100); | Metschnikowia | 13 | Saprotroph | \|Undefined Saprotroph\| | Yeast | NULL | Probable | As nectar-tap_saprotroph (Põlme et al. 2020); Decay sugar-rich substrates (Põlme et al. 2020) | Põlme S, et al. 2020. Fungal Diversity 105:1-16 ((https://doi.org/10.1007/s13225-020-00466-2)) |
| Otu56011 | 1 | k__Fungi(100);p__Ascomycota(100);c__Saccharomycetes(100);o__Saccharomycetales(100);f__Metschnikowiaceae(100);g__Metschnikowia(100);s__Metschnikowia_pulcherrima(100); | Metschnikowia | 13 | Saprotroph | \|Undefined Saprotroph\| | Yeast | NULL | Probable | As nectar-tap_saprotroph (Põlme et al. 2020); Decay sugar-rich substrates (Põlme et al. 2020) | Põlme S, et al. 2020. Fungal Diversity 105:1-16 ((https://doi.org/10.1007/s13225-020-00466-2)) |
| Otu56012 | 1 | k__Fungi(100);p__Ascomycota(100);c__Saccharomycetes(100);o__Saccharomycetales(100);f__Metschnikowiaceae(100);g__Metschnikowia(100);s__Metschnikowia_pulcherrima(100); | Metschnikowia | 13 | Saprotroph | \|Undefined Saprotroph\| | Yeast | NULL | Probable | As nectar-tap_saprotroph (Põlme et al. 2020); Decay sugar-rich substrates (Põlme et al. 2020) | Põlme S, et al. 2020. Fungal Diversity 105:1-16 ((https://doi.org/10.1007/s13225-020-00466-2)) |
| Otu56014 | 1 | k__Fungi(100);p__Ascomycota(100);c__Saccharomycetes(100);o__Saccharomycetales(100);f__Metschnikowiaceae(100);g__Metschnikowia(100);s__Metschnikowia_pulcherrima(100); | Metschnikowia | 13 | Saprotroph | \|Undefined Saprotroph\| | Yeast | NULL | Probable | As nectar-tap_saprotroph (Põlme et al. 2020); Decay sugar-rich substrates (Põlme et al. 2020) | Põlme S, et al. 2020. Fungal Diversity 105:1-16 ((https://doi.org/10.1007/s13225-020-00466-2)) |
| Otu56016 | 1 | k__Fungi(100);p__Ascomycota(100);c__Saccharomycetes(100);o__Saccharomycetales(100);f__Saccharomycetales_family_Incertae_sedis(100);f__Saccharomycetales_family_Incertae_sedis_unclassified(100);f__Saccharomycetales_family_Incertae_sedis_unclassified(100); | Saccharomycetales | 7 | Saprotroph | Undefined Saprotroph | Yeast | NULL | Possible | NULL | Sterkenburg E, et al. 2015. New Phytologist 207:1145-1158 ((https://doi.org/10.1111/nph.13426)) |
| Otu56017 | 1 | k__Fungi(100);p__Ascomycota(100);c__Saccharomycetes(100);o__Saccharomycetales(100);f__Saccharomycetales_family_Incertae_sedis(100);g__Candida(100);g__Candida_unclassified(100); | Candida | 13 | Pathotroph-Saprotroph-Symbiotroph | Animal Pathogen-Endophyte-Endosymbiont-Epiphyte-\|Undefined Saprotroph\| | Dimorphic Yeast-Yeast | NULL | Probable | As nectar-tap_saprotroph (Põlme et al. 2020); Decay sugar-rich substrates (Põlme et al. 2020); This asexual genus is polyphyletic and contain species that function very differently depend on what they are. Caution should be taken when interpreting guilds for Candida species. | Manolakaki D, et al. 2010. Virulence. 1: 367-375 ((https://doi.org/10.4161/viru.1.5.12796)); Mekbib SB, et al. 2011. Tropical Plant Pathology 36:233-240 ((https://doi.org/10.1590/S1982-56762011000400004)); Põlme S, et al. 2020. Fungal Diversity 105:1-16 ((https://doi.org/10.1007/s13225-020-00466-2)) |
| Otu56018 | 1 | k__Fungi(100);p__Ascomycota(100);c__Saccharomycetes(100);o__Saccharomycetales(100);f__Saccharomycetales_family_Incertae_sedis(100);g__Candida(100);g__Candida_unclassified(100); | Candida | 13 | Pathotroph-Saprotroph-Symbiotroph | Animal Pathogen-Endophyte-Endosymbiont-Epiphyte-\|Undefined Saprotroph\| | Dimorphic Yeast-Yeast | NULL | Probable | As nectar-tap_saprotroph (Põlme et al. 2020); Decay sugar-rich substrates (Põlme et al. 2020); This asexual genus is polyphyletic and contain species that function very differently depend on what they are. Caution should be taken when interpreting guilds for Candida species. | Manolakaki D, et al. 2010. Virulence. 1: 367-375 ((https://doi.org/10.4161/viru.1.5.12796)); Mekbib SB, et al. 2011. Tropical Plant Pathology 36:233-240 ((https://doi.org/10.1590/S1982-56762011000400004)); Põlme S, et al. 2020. Fungal Diversity 105:1-16 ((https://doi.org/10.1007/s13225-020-00466-2)) |
| Otu56021 | 1 | k__Fungi(100);p__Ascomycota(100);c__Saccharomycetes(100);o__Saccharomycetales(100);o__Saccharomycetales_unclassified(100);o__Saccharomycetales_unclassified(100);o__Saccharomycetales_unclassified(100); | Saccharomycetales | 7 | Saprotroph | Undefined Saprotroph | Yeast | NULL | Possible | NULL | Sterkenburg E, et al. 2015. New Phytologist 207:1145-1158 ((https://doi.org/10.1111/nph.13426)) |
| Otu56023 | 1 | k__Fungi(100);p__Ascomycota(100);c__Saccharomycetes(100);o__Saccharomycetales(100);f__Metschnikowiaceae(100);g__Metschnikowia(100);s__Metschnikowia_pulcherrima(100); | Metschnikowia | 13 | Saprotroph | \|Undefined Saprotroph\| | Yeast | NULL | Probable | As nectar-tap_saprotroph (Põlme et al. 2020); Decay sugar-rich substrates (Põlme et al. 2020) | Põlme S, et al. 2020. Fungal Diversity 105:1-16 ((https://doi.org/10.1007/s13225-020-00466-2)) |
| Otu56024 | 1 | k__Fungi(100);p__Ascomycota(100);c__Saccharomycetes(100);o__Saccharomycetales(100);f__Metschnikowiaceae(100);g__Metschnikowia(100);s__Metschnikowia_pulcherrima(100); | Metschnikowia | 13 | Saprotroph | \|Undefined Saprotroph\| | Yeast | NULL | Probable | As nectar-tap_saprotroph (Põlme et al. 2020); Decay sugar-rich substrates (Põlme et al. 2020) | Põlme S, et al. 2020. Fungal Diversity 105:1-16 ((https://doi.org/10.1007/s13225-020-00466-2)) |
| Otu56025 | 1 | k__Fungi(100);p__Ascomycota(100);c__Saccharomycetes(100);o__Saccharomycetales(100);f__Saccharomycetales_family_Incertae_sedis(100);g__Candida(100);g__Candida_unclassified(100); | Candida | 13 | Pathotroph-Saprotroph-Symbiotroph | Animal Pathogen-Endophyte-Endosymbiont-Epiphyte-\|Undefined Saprotroph\| | Dimorphic Yeast-Yeast | NULL | Probable | As nectar-tap_saprotroph (Põlme et al. 2020); Decay sugar-rich substrates (Põlme et al. 2020); This asexual genus is polyphyletic and contain species that function very differently depend on what they are. Caution should be taken when interpreting guilds for Candida species. | Manolakaki D, et al. 2010. Virulence. 1: 367-375 ((https://doi.org/10.4161/viru.1.5.12796)); Mekbib SB, et al. 2011. Tropical Plant Pathology 36:233-240 ((https://doi.org/10.1590/S1982-56762011000400004)); Põlme S, et al. 2020. Fungal Diversity 105:1-16 ((https://doi.org/10.1007/s13225-020-00466-2)) |

| | | | | | | | | | | | |
|---|---|---|---|---|---|---|---|---|---|---|---|
| Otu56026 | 1 | k__Fungi(100);p__Ascomycota(100);c__Saccharomycetes(100);o__Saccharomycetales(100);f__Metschnikowiaceae(100);g__Metschnikowia(100);s__Metschnikowia_pulcherrima(100); | Metschnikowia | 13 | Saprotroph | \|Undefined Saprotroph\| | Yeast | NULL | Probable | As nectar-tap_saprotroph (Põlme et al. 2020); Decay sugar-rich substrates (Põlme et al. 2020) | Põlme S, et al. 2020. Fungal Diversity 105:1-16 ((https://doi.org/10.1007/s13225-020-00466-2)) |
| Otu56027 | 1 | k__Fungi(100);p__Ascomycota(100);c__Saccharomycetes(100);o__Saccharomycetales(100);f__Metschnikowiaceae(100);g__Metschnikowia(100);s__Metschnikowia_pulcherrima(100); | Metschnikowia | 13 | Saprotroph | \|Undefined Saprotroph\| | Yeast | NULL | Probable | As nectar-tap_saprotroph (Põlme et al. 2020); Decay sugar-rich substrates (Põlme et al. 2020) | Põlme S, et al. 2020. Fungal Diversity 105:1-16 ((https://doi.org/10.1007/s13225-020-00466-2)) |
| Otu56028 | 1 | k__Fungi(100);p__Ascomycota(100);c__Saccharomycetes(100);o__Saccharomycetales(100);f__Saccharomycetales_family_Incertae_sedis(100);g__Candida(100);g__Candida_unclassified(100); | Candida | 13 | Pathotroph-Saprotroph-Symbiotroph | Animal Pathogen-Endophyte-Endosymbiont-Epiphyte-\|Undefined Saprotroph\| | Dimorphic Yeast-Yeast | NULL | Probable | As nectar-tap_saprotroph (Põlme et al. 2020); Decay sugar-rich substrates (Põlme et al. 2020); This asexual genus is polyphyletic and contain species that function very differently depend on what they are. Caution should be taken when interpreting guilds for Candida species. | Manolakaki D, et al. 2010. Virulence. 1: 367-375 ((https://doi.org/10.4161/viru.1.5.12796)); Mekbib SB, et al. 2011. Tropical Plant Pathology 36:233-240 ((https://doi.org/10.1590/S1982-56762011000400004)); Põlme S, et al. 2020. Fungal Diversity 105:1-16 ((https://doi.org/10.1007/s13225-020-00466-2)) |
| Otu56029 | 1 | k__Fungi(100);p__Ascomycota(100);c__Saccharomycetes(100);o__Saccharomycetales(100);f__Saccharomycetales_family_Incertae_sedis(100);g__Candida(100);s__Candida_cretensis(100); | Candida | 13 | Pathotroph-Saprotroph-Symbiotroph | Animal Pathogen-Endophyte-Endosymbiont-Epiphyte-\|Undefined Saprotroph\| | Dimorphic Yeast-Yeast | NULL | Probable | As nectar-tap_saprotroph (Põlme et al. 2020); Decay sugar-rich substrates (Põlme et al. 2020); This asexual genus is polyphyletic and contain species that function very differently depend on what they are. Caution should be taken when interpreting guilds for Candida species. | Manolakaki D, et al. 2010. Virulence. 1: 367-375 ((https://doi.org/10.4161/viru.1.5.12796)); Mekbib SB, et al. 2011. Tropical Plant Pathology 36:233-240 ((https://doi.org/10.1590/S1982-56762011000400004)); Põlme S, et al. 2020. Fungal Diversity 105:1-16 ((https://doi.org/10.1007/s13225-020-00466-2)) |
| Otu56030 | 1 | k__Fungi(100);p__Ascomycota(100);c__Saccharomycetes(100);o__Saccharomycetales(100);f__Saccharomycetales_family_Incertae_sedis(100);f__Saccharomycetales_family_Incertae_sedis_unclassified(100);f__Saccharomycetales_family_Incertae_sedis_unclassified(100); | Saccharomycetales | 7 | Saprotroph | Undefined Saprotroph | Yeast | NULL | Possible | NULL | Sterkenburg E, et al. 2015. New Phytologist 207:1145-1158 ((https://doi.org/10.1111/nph.13426)) |
| Otu56031 | 1 | k__Fungi(100);p__Ascomycota(100);c__Saccharomycetes(100);o__Saccharomycetales(100);f__Saccharomycetales_family_Incertae_sedis(100);f__Saccharomycetales_family_Incertae_sedis_unclassified(100);f__Saccharomycetales_family_Incertae_sedis_unclassified(100); | Saccharomycetales | 7 | Saprotroph | Undefined Saprotroph | Yeast | NULL | Possible | NULL | Sterkenburg E, et al. 2015. New Phytologist 207:1145-1158 ((https://doi.org/10.1111/nph.13426)) |
| Otu56033 | 1 | k__Fungi(100);p__Ascomycota(100);c__Saccharomycetes(100);o__Saccharomycetales(100);f__Saccharomycetales_family_Incertae_sedis(100);f__Saccharomycetales_family_Incertae_sedis_unclassified(100);f__Saccharomycetales_family_Incertae_sedis_unclassified(100); | Saccharomycetales | 7 | Saprotroph | Undefined Saprotroph | Yeast | NULL | Possible | NULL | Sterkenburg E, et al. 2015. New Phytologist 207:1145-1158 ((https://doi.org/10.1111/nph.13426)) |
| Otu56034 | 1 | k__Fungi(100);p__Ascomycota(100);c__Saccharomycetes(100);o__Saccharomycetales(100);f__Saccharomycetales_family_Incertae_sedis(100);f__Saccharomycetales_family_Incertae_sedis_unclassified(100);f__Saccharomycetales_family_Incertae_sedis_unclassified(100); | Saccharomycetales | 7 | Saprotroph | Undefined Saprotroph | Yeast | NULL | Possible | NULL | Sterkenburg E, et al. 2015. New Phytologist 207:1145-1158 ((https://doi.org/10.1111/nph.13426)) |
| Otu56035 | 1 | k__Fungi(100);p__Ascomycota(100);c__Saccharomycetes(100);o__Saccharomycetales(100);f__Saccharomycetales_family_Incertae_sedis(100);g__Candida(100);g__Candida_unclassified(100); | Candida | 13 | Pathotroph-Saprotroph-Symbiotroph | Animal Pathogen-Endophyte-Endosymbiont-Epiphyte-\|Undefined Saprotroph\| | Dimorphic Yeast-Yeast | NULL | Probable | As nectar-tap_saprotroph (Põlme et al. 2020); Decay sugar-rich substrates (Põlme et al. 2020); This asexual genus is polyphyletic and contain species that function very differently depend on what they are. Caution should be taken when interpreting guilds for Candida species. | Manolakaki D, et al. 2010. Virulence. 1: 367-375 ((https://doi.org/10.4161/viru.1.5.12796)); Mekbib SB, et al. 2011. Tropical Plant Pathology 36:233-240 ((https://doi.org/10.1590/S1982-56762011000400004)); Põlme S, et al. 2020. Fungal Diversity 105:1-16 ((https://doi.org/10.1007/s13225-020-00466-2)) |
| Otu56036 | 1 | k__Fungi(100);p__Ascomycota(100);c__Saccharomycetes(100);o__Saccharomycetales(100);f__Saccharomycetales_family_Incertae_sedis(100);g__Candida(100);g__Candida_unclassified(100); | Candida | 13 | Pathotroph-Saprotroph-Symbiotroph | Animal Pathogen-Endophyte-Endosymbiont-Epiphyte-\|Undefined Saprotroph\| | Dimorphic Yeast-Yeast | NULL | Probable | As nectar-tap_saprotroph (Põlme et al. 2020); Decay sugar-rich substrates (Põlme et al. 2020); This asexual genus is polyphyletic and contain species that function very differently depend on what they are. Caution should be taken when interpreting guilds for Candida species. | Manolakaki D, et al. 2010. Virulence. 1: 367-375 ((https://doi.org/10.4161/viru.1.5.12796)); Mekbib SB, et al. 2011. Tropical Plant Pathology 36:233-240 ((https://doi.org/10.1590/S1982-56762011000400004)); Põlme S, et al. 2020. Fungal Diversity 105:1-16 ((https://doi.org/10.1007/s13225-020-00466-2)) |
| Otu56039 | 1 | k__Fungi(100);p__Ascomycota(100);c__Saccharomycetes(100);o__Saccharomycetales(100);f__Metschnikowiaceae(100);g__Metschnikowia(100);s__Metschnikowia_pulcherrima(100); | Metschnikowia | 13 | Saprotroph | \|Undefined Saprotroph\| | Yeast | NULL | Probable | As nectar-tap_saprotroph (Põlme et al. 2020); Decay sugar-rich substrates (Põlme et al. 2020) | Põlme S, et al. 2020. Fungal Diversity 105:1-16 ((https://doi.org/10.1007/s13225-020-00466-2)) |
| Otu56040 | 1 | k__Fungi(100);p__Ascomycota(100);c__Saccharomycetes(100);o__Saccharomycetales(100);f__Saccharomycetales_family_Incertae_sedis(100);g__Candida(100);g__Candida_unclassified(100); | Candida | 13 | Pathotroph-Saprotroph-Symbiotroph | Animal Pathogen-Endophyte-Endosymbiont-Epiphyte-\|Undefined Saprotroph\| | Dimorphic Yeast-Yeast | NULL | Probable | As nectar-tap_saprotroph (Põlme et al. 2020); Decay sugar-rich substrates (Põlme et al. 2020); This asexual genus is polyphyletic and contain species that function very differently depend on what they are. Caution should be taken when interpreting guilds for Candida species. | Manolakaki D, et al. 2010. Virulence. 1: 367-375 ((https://doi.org/10.4161/viru.1.5.12796)); Mekbib SB, et al. 2011. Tropical Plant Pathology 36:233-240 ((https://doi.org/10.1590/S1982-56762011000400004)); Põlme S, et al. 2020. Fungal Diversity 105:1-16 ((https://doi.org/10.1007/s13225-020-00466-2)) |
| Otu56042 | 1 | k__Fungi(100);p__Ascomycota(100);c__Saccharomycetes(100);o__Saccharomycetales(100);o__Saccharomycetales_unclassified(100);o__Saccharomycetales_unclassified(100);o__Saccharomycetales_unclassified(100); | Saccharomycetales | 7 | Saprotroph | Undefined Saprotroph | Yeast | NULL | Possible | NULL | Sterkenburg E, et al. 2015. New Phytologist 207:1145-1158 ((https://doi.org/10.1111/nph.13426)) |
| Otu56043 | 1 | k__Fungi(100);p__Ascomycota(100);c__Saccharomycetes(100);o__Saccharomycetales(100);f__Saccharomycetales_family_Incertae_sedis(100);g__Candida(100);g__Candida_unclassified(100); | Candida | 13 | Pathotroph-Saprotroph-Symbiotroph | Animal Pathogen-Endophyte-Endosymbiont-Epiphyte-\|Undefined Saprotroph\| | Dimorphic Yeast-Yeast | NULL | Probable | As nectar-tap_saprotroph (Põlme et al. 2020); Decay sugar-rich substrates (Põlme et al. 2020); This asexual genus is polyphyletic and contain species that function very differently depend on what they are. Caution should be taken when interpreting guilds for Candida species. | Manolakaki D, et al. 2010. Virulence. 1: 367-375 ((https://doi.org/10.4161/viru.1.5.12796)); Mekbib SB, et al. 2011. Tropical Plant Pathology 36:233-240 ((https://doi.org/10.1590/S1982-56762011000400004)); Põlme S, et al. 2020. Fungal Diversity 105:1-16 ((https://doi.org/10.1007/s13225-020-00466-2)) |
| Otu56046 | 1 | k__Fungi(100);p__Ascomycota(100);c__Saccharomycetes(100);o__Saccharomycetales(100);f__Saccharomycetales_family_Incertae_sedis(100);g__Candida(100);g__Candida_unclassified(100); | Candida | 13 | Pathotroph-Saprotroph-Symbiotroph | Animal Pathogen-Endophyte-Endosymbiont-Epiphyte-\|Undefined Saprotroph\| | Dimorphic Yeast-Yeast | NULL | Probable | As nectar-tap_saprotroph (Põlme et al. 2020); Decay sugar-rich substrates (Põlme et al. 2020); This asexual genus is polyphyletic and contain species that function very differently depend on what they are. Caution should be taken when interpreting guilds for Candida species. | Manolakaki D, et al. 2010. Virulence. 1: 367-375 ((https://doi.org/10.4161/viru.1.5.12796)); Mekbib SB, et al. 2011. Tropical Plant Pathology 36:233-240 ((https://doi.org/10.1590/S1982-56762011000400004)); Põlme S, et al. 2020. Fungal Diversity 105:1-16 ((https://doi.org/10.1007/s13225-020-00466-2)) |
| Otu56047 | 1 | k__Fungi(100);p__Ascomycota(100);c__Saccharomycetes(100);o__Saccharomycetales(100);f__Metschnikowiaceae(100);g__Metschnikowia(100);g__Metschnikowia_unclassified(100); | Metschnikowia | 13 | Saprotroph | \|Undefined Saprotroph\| | Yeast | NULL | Probable | As nectar-tap_saprotroph (Põlme et al. 2020); Decay sugar-rich substrates (Põlme et al. 2020) | Põlme S, et al. 2020. Fungal Diversity 105:1-16 ((https://doi.org/10.1007/s13225-020-00466-2)) |
| Otu56048 | 1 | k__Fungi(100);p__Ascomycota(100);c__Saccharomycetes(100);o__Saccharomycetales(100);f__Saccharomycetales_family_Incertae_sedis(100);g__Candida(100);g__Candida_unclassified(100); | Candida | 13 | Pathotroph-Saprotroph-Symbiotroph | Animal Pathogen-Endophyte-Endosymbiont-Epiphyte-\|Undefined Saprotroph\| | Dimorphic Yeast-Yeast | NULL | Probable | As nectar-tap_saprotroph (Põlme et al. 2020); Decay sugar-rich substrates (Põlme et al. 2020); This asexual genus is polyphyletic and contain species that function very differently depend on what they are. Caution should be taken when interpreting guilds for Candida species. | Manolakaki D, et al. 2010. Virulence. 1: 367-375 ((https://doi.org/10.4161/viru.1.5.12796)); Mekbib SB, et al. 2011. Tropical Plant Pathology 36:233-240 ((https://doi.org/10.1590/S1982-56762011000400004)); Põlme S, et al. 2020. Fungal Diversity 105:1-16 ((https://doi.org/10.1007/s13225-020-00466-2)) |
| Otu56051 | 1 | k__Fungi(100);p__Ascomycota(100);c__Saccharomycetes(100);o__Saccharomycetales(100);f__Saccharomycetales_family_Incertae_sedis(100);g__Candida(100);s__Candida_cretensis(100); | Candida | 13 | Pathotroph-Saprotroph-Symbiotroph | Animal Pathogen-Endophyte-Endosymbiont-Epiphyte-\|Undefined Saprotroph\| | Dimorphic Yeast-Yeast | NULL | Probable | As nectar-tap_saprotroph (Põlme et al. 2020); Decay sugar-rich substrates (Põlme et al. 2020); This asexual genus is polyphyletic and contain species that function very differently depend on what they are. Caution should be taken when interpreting guilds for Candida species. | Manolakaki D, et al. 2010. Virulence. 1: 367-375 ((https://doi.org/10.4161/viru.1.5.12796)); Mekbib SB, et al. 2011. Tropical Plant Pathology 36:233-240 ((https://doi.org/10.1590/S1982-56762011000400004)); Põlme S, et al. 2020. Fungal Diversity 105:1-16 ((https://doi.org/10.1007/s13225-020-00466-2)) |
| Otu56052 | 1 | k__Fungi(100);p__Ascomycota(100);c__Saccharomycetes(100);o__Saccharomycetales(100);f__Saccharomycetales_family_Incertae_sedis(100);g__Candida(100);g__Candida_unclassified(100); | Candida | 13 | Pathotroph-Saprotroph-Symbiotroph | Animal Pathogen-Endophyte-Endosymbiont-Epiphyte-\|Undefined Saprotroph\| | Dimorphic Yeast-Yeast | NULL | Probable | As nectar-tap_saprotroph (Põlme et al. 2020); Decay sugar-rich substrates (Põlme et al. 2020); This asexual genus is polyphyletic and contain species that function very differently depend on what they are. Caution should be taken when interpreting guilds for Candida species. | Manolakaki D, et al. 2010. Virulence. 1: 367-375 ((https://doi.org/10.4161/viru.1.5.12796)); Mekbib SB, et al. 2011. Tropical Plant Pathology 36:233-240 ((https://doi.org/10.1590/S1982-56762011000400004)); Põlme S, et al. 2020. Fungal Diversity 105:1-16 ((https://doi.org/10.1007/s13225-020-00466-2)) |
| Otu56054 | 1 | k__Fungi(100);p__Ascomycota(100);c__Saccharomycetes(100);o__Saccharomycetales(100);f__Saccharomycetales_family_Incertae_sedis(100);g__Candida(100);g__Candida_unclassified(100); | Candida | 13 | Pathotroph-Saprotroph-Symbiotroph | Animal Pathogen-Endophyte-Endosymbiont-Epiphyte-\|Undefined Saprotroph\| | Dimorphic Yeast-Yeast | NULL | Probable | As nectar-tap_saprotroph (Põlme et al. 2020); Decay sugar-rich substrates (Põlme et al. 2020); This asexual genus is polyphyletic and contain species that function very differently depend on what they are. Caution should be taken when interpreting guilds for Candida species. | Manolakaki D, et al. 2010. Virulence. 1: 367-375 ((https://doi.org/10.4161/viru.1.5.12796)); Mekbib SB, et al. 2011. Tropical Plant Pathology 36:233-240 ((https://doi.org/10.1590/S1982-56762011000400004)); Põlme S, et al. 2020. Fungal Diversity 105:1-16 ((https://doi.org/10.1007/s13225-020-00466-2)) |
| Otu56056 | 1 | k__Fungi(100);p__Ascomycota(100);c__Saccharomycetes(100);o__Saccharomycetales(100);f__Saccharomycetales_family_Incertae_sedis(100);f__Saccharomycetales_family_Incertae_sedis_unclassified(100);f__Saccharomycetales_family_Incertae_sedis_unclassified(100); | Saccharomycetales | 7 | Saprotroph | Undefined Saprotroph | Yeast | NULL | Possible | NULL | Sterkenburg E, et al. 2015. New Phytologist 207:1145-1158 ((https://doi.org/10.1111/nph.13426)) |
| Otu56057 | 1 | k__Fungi(100);p__Ascomycota(100);c__Saccharomycetes(100);o__Saccharomycetales(100);f__Saccharomycetales_family_Incertae_sedis(100);g__Candida(100);g__Candida_unclassified(100); | Candida | 13 | Pathotroph-Saprotroph-Symbiotroph | Animal Pathogen-Endophyte-Endosymbiont-Epiphyte-\|Undefined Saprotroph\| | Dimorphic Yeast-Yeast | NULL | Probable | As nectar-tap_saprotroph (Põlme et al. 2020); Decay sugar-rich substrates (Põlme et al. 2020); This asexual genus is polyphyletic and contain species that function very differently depend on what they are. Caution should be taken when interpreting guilds for Candida species. | Manolakaki D, et al. 2010. Virulence. 1: 367-375 ((https://doi.org/10.4161/viru.1.5.12796)); Mekbib SB, et al. 2011. Tropical Plant Pathology 36:233-240 ((https://doi.org/10.1590/S1982-56762011000400004)); Põlme S, et al. 2020. Fungal Diversity 105:1-16 ((https://doi.org/10.1007/s13225-020-00466-2)) |
| Otu56060 | 1 | k__Fungi(100);p__Ascomycota(100);c__Saccharomycetes(100);o__Saccharomycetales(100);f__Saccharomycetales_family_Incertae_sedis(100);g__Candida(100);g__Candida_unclassified(100); | Candida | 13 | Pathotroph-Saprotroph-Symbiotroph | Animal Pathogen-Endophyte-Endosymbiont-Epiphyte-\|Undefined Saprotroph\| | Dimorphic Yeast-Yeast | NULL | Probable | As nectar-tap_saprotroph (Põlme et al. 2020); Decay sugar-rich substrates (Põlme et al. 2020); This asexual genus is polyphyletic and contain species that function very differently depend on what they are. Caution should be taken when interpreting guilds for Candida species. | Manolakaki D, et al. 2010. Virulence. 1: 367-375 ((https://doi.org/10.4161/viru.1.5.12796)); Mekbib SB, et al. 2011. Tropical Plant Pathology 36:233-240 ((https://doi.org/10.1590/S1982-56762011000400004)); Põlme S, et al. 2020. Fungal Diversity 105:1-16 ((https://doi.org/10.1007/s13225-020-00466-2)) |
| Otu56061 | 1 | k__Fungi(100);p__Ascomycota(100);c__Saccharomycetes(100);o__Saccharomycetales(100);f__Saccharomycetales_family_Incertae_sedis(100);g__Candida(100);g__Candida_unclassified(100); | Candida | 13 | Pathotroph-Saprotroph-Symbiotroph | Animal Pathogen-Endophyte-Endosymbiont-Epiphyte-\|Undefined Saprotroph\| | Dimorphic Yeast-Yeast | NULL | Probable | As nectar-tap_saprotroph (Põlme et al. 2020); Decay sugar-rich substrates (Põlme et al. 2020); This asexual genus is polyphyletic and contain species that function very differently depend on what they are. Caution should be taken when interpreting guilds for Candida species. | Manolakaki D, et al. 2010. Virulence. 1: 367-375 ((https://doi.org/10.4161/viru.1.5.12796)); Mekbib SB, et al. 2011. Tropical Plant Pathology 36:233-240 ((https://doi.org/10.1590/S1982-56762011000400004)); Põlme S, et al. 2020. Fungal Diversity 105:1-16 ((https://doi.org/10.1007/s13225-020-00466-2)) |
| Otu56062 | 1 | k__Fungi(100);p__Ascomycota(100);c__Saccharomycetes(100);o__Saccharomycetales(100);f__Saccharomycetales_family_Incertae_sedis(100);g__Candida(100);g__Candida_unclassified(100); | Candida | 13 | Pathotroph-Saprotroph-Symbiotroph | Animal Pathogen-Endophyte-Endosymbiont-Epiphyte-\|Undefined Saprotroph\| | Dimorphic Yeast-Yeast | NULL | Probable | As nectar-tap_saprotroph (Põlme et al. 2020); Decay sugar-rich substrates (Põlme et al. 2020); This asexual genus is polyphyletic and contain species that function very differently depend on what they are. Caution should be taken when interpreting guilds for Candida species. | Manolakaki D, et al. 2010. Virulence. 1: 367-375 ((https://doi.org/10.4161/viru.1.5.12796)); Mekbib SB, et al. 2011. Tropical Plant Pathology 36:233-240 ((https://doi.org/10.1590/S1982-56762011000400004)); Põlme S, et al. 2020. Fungal Diversity 105:1-16 ((https://doi.org/10.1007/s13225-020-00466-2)) |
| Otu56063 | 1 | k__Fungi(100);p__Ascomycota(100);c__Saccharomycetes(100);o__Saccharomycetales(100);f__Saccharomycetales_family_Incertae_sedis(100);f__Saccharomycetales_family_Incertae_sedis_unclassified(100);f__Saccharomycetales_family_Incertae_sedis_unclassified(100); | Saccharomycetales | 7 | Saprotroph | Undefined Saprotroph | Yeast | NULL | Possible | NULL | Sterkenburg E, et al. 2015. New Phytologist 207:1145-1158 ((https://doi.org/10.1111/nph.13426)) |
| Otu56064 | 1 | k__Fungi(100);p__Ascomycota(100);c__Saccharomycetes(100);o__Saccharomycetales(100);f__Saccharomycetales_family_Incertae_sedis(100);g__Candida(100);g__Candida_unclassified(100); | Candida | 13 | Pathotroph-Saprotroph-Symbiotroph | Animal Pathogen-Endophyte-Endosymbiont-Epiphyte-\|Undefined Saprotroph\| | Dimorphic Yeast-Yeast | NULL | Probable | As nectar-tap_saprotroph (Põlme et al. 2020); Decay sugar-rich substrates (Põlme et al. 2020); This asexual genus is polyphyletic and contain species that function very differently depend on what they are. Caution should be taken when interpreting guilds for Candida species. | Manolakaki D, et al. 2010. Virulence. 1: 367-375 ((https://doi.org/10.4161/viru.1.5.12796)); Mekbib SB, et al. 2011. Tropical Plant Pathology 36:233-240 ((https://doi.org/10.1590/S1982-56762011000400004)); Põlme S, et al. 2020. Fungal Diversity 105:1-16 ((https://doi.org/10.1007/s13225-020-00466-2)) |
| Otu56065 | 1 | k__Fungi(100);p__Ascomycota(100);c__Saccharomycetes(100);o__Saccharomycetales(100);f__Saccharomycetales_family_Incertae_sedis(100);g__Candida(100);g__Candida_unclassified(100); | Candida | 13 | Pathotroph-Saprotroph-Symbiotroph | Animal Pathogen-Endophyte-Endosymbiont-Epiphyte-\|Undefined Saprotroph\| | Dimorphic Yeast-Yeast | NULL | Probable | As nectar-tap_saprotroph (Põlme et al. 2020); Decay sugar-rich substrates (Põlme et al. 2020); This asexual genus is polyphyletic and contain species that function very differently depend on what they are. Caution should be taken when interpreting guilds for Candida species. | Manolakaki D, et al. 2010. Virulence. 1: 367-375 ((https://doi.org/10.4161/viru.1.5.12796)); Mekbib SB, et al. 2011. Tropical Plant Pathology 36:233-240 ((https://doi.org/10.1590/S1982-56762011000400004)); Põlme S, et al. 2020. Fungal Diversity 105:1-16 ((https://doi.org/10.1007/s13225-020-00466-2)) |
| Otu56066 | 1 | k__Fungi(100);p__Ascomycota(100);c__Saccharomycetes(100);o__Saccharomycetales(100);f__Saccharomycetales_family_Incertae_sedis(100);g__Candida(100);s__Candida_cretensis(100); | Candida | 13 | Pathotroph-Saprotroph-Symbiotroph | Animal Pathogen-Endophyte-Endosymbiont-Epiphyte-\|Undefined Saprotroph\| | Dimorphic Yeast-Yeast | NULL | Probable | As nectar-tap_saprotroph (Põlme et al. 2020); Decay sugar-rich substrates (Põlme et al. 2020); This asexual genus is polyphyletic and contain species that function very differently depend on what they are. Caution should be taken when interpreting guilds for Candida species. | Manolakaki D, et al. 2010. Virulence. 1: 367-375 ((https://doi.org/10.4161/viru.1.5.12796)); Mekbib SB, et al. 2011. Tropical Plant Pathology 36:233-240 ((https://doi.org/10.1590/S1982-56762011000400004)); Põlme S, et al. 2020. Fungal Diversity 105:1-16 ((https://doi.org/10.1007/s13225-020-00466-2)) |
| Otu56067 | 1 | k__Fungi(100);p__Ascomycota(100);c__Saccharomycetes(100);o__Saccharomycetales(100);f__Metschnikowiaceae(100);g__Metschnikowia(100);s__Metschnikowia_pulcherrima(100); | Metschnikowia | 13 | Saprotroph | \|Undefined Saprotroph\| | Yeast | NULL | Probable | As nectar-tap_saprotroph (Põlme et al. 2020); Decay sugar-rich substrates (Põlme et al. 2020) | Põlme S, et al. 2020. Fungal Diversity 105:1-16 ((https://doi.org/10.1007/s13225-020-00466-2)) |
| Otu56069 | 1 | k__Fungi(100);p__Ascomycota(100);c__Saccharomycetes(100);o__Saccharomycetales(100);f__Saccharomycetales_family_Incertae_sedis(100);g__Candida(100);g__Candida_unclassified(100); | Candida | 13 | Pathotroph-Saprotroph-Symbiotroph | Animal Pathogen-Endophyte-Endosymbiont-Epiphyte-\|Undefined Saprotroph\| | Dimorphic Yeast-Yeast | NULL | Probable | As nectar-tap_saprotroph (Põlme et al. 2020); Decay sugar-rich substrates (Põlme et al. 2020); This asexual genus is polyphyletic and contain species that function very differently depend on what they are. Caution should be taken when interpreting guilds for Candida species. | Manolakaki D, et al. 2010. Virulence. 1: 367-375 ((https://doi.org/10.4161/viru.1.5.12796)); Mekbib SB, et al. 2011. Tropical Plant Pathology 36:233-240 ((https://doi.org/10.1590/S1982-56762011000400004)); Põlme S, et al. 2020. Fungal Diversity 105:1-16 ((https://doi.org/10.1007/s13225-020-00466-2)) |
| Otu56071 | 1 | k__Fungi(100);p__Ascomycota(100);c__Saccharomycetes(100);o__Saccharomycetales(100);f__Metschnikowiaceae(100);g__Metschnikowia(100);s__Metschnikowia_pulcherrima(100); | Metschnikowia | 13 | Saprotroph | \|Undefined Saprotroph\| | Yeast | NULL | Probable | As nectar-tap_saprotroph (Põlme et al. 2020); Decay sugar-rich substrates (Põlme et al. 2020) | Põlme S, et al. 2020. Fungal Diversity 105:1-16 ((https://doi.org/10.1007/s13225-020-00466-2)) |
| Otu56072 | 1 | k__Fungi(100);p__Ascomycota(100);c__Saccharomycetes(100);o__Saccharomycetales(100);f__Metschnikowiaceae(100);g__Metschnikowia(100);s__Metschnikowia_pulcherrima(100); | Metschnikowia | 13 | Saprotroph | \|Undefined Saprotroph\| | Yeast | NULL | Probable | As nectar-tap_saprotroph (Põlme et al. 2020); Decay sugar-rich substrates (Põlme et al. 2020) | Põlme S, et al. 2020. Fungal Diversity 105:1-16 ((https://doi.org/10.1007/s13225-020-00466-2)) |
| Otu56075 | 1 | k__Fungi(100);p__Ascomycota(100);c__Saccharomycetes(100);o__Saccharomycetales(100);f__Saccharomycetales_family_Incertae_sedis(100);g__Candida(100);g__Candida_unclassified(100); | Candida | 13 | Pathotroph-Saprotroph-Symbiotroph | Animal Pathogen-Endophyte-Endosymbiont-Epiphyte-\|Undefined Saprotroph\| | Dimorphic Yeast-Yeast | NULL | Probable | As nectar-tap_saprotroph (Põlme et al. 2020); Decay sugar-rich substrates (Põlme et al. 2020); This asexual genus is polyphyletic and contain species that function very differently depend on what they are. Caution should be taken when interpreting guilds for Candida species. | Manolakaki D, et al. 2010. Virulence. 1: 367-375 ((https://doi.org/10.4161/viru.1.5.12796)); Mekbib SB, et al. 2011. Tropical Plant Pathology 36:233-240 ((https://doi.org/10.1590/S1982-56762011000400004)); Põlme S, et al. 2020. Fungal Diversity 105:1-16 ((https://doi.org/10.1007/s13225-020-00466-2)) |
| Otu56076 | 1 | k__Fungi(100);p__Ascomycota(100);c__Saccharomycetes(100);o__Saccharomycetales(100);f__Saccharomycetales_family_Incertae_sedis(100);g__Candida(100);g__Candida_unclassified(100); | Candida | 13 | Pathotroph-Saprotroph-Symbiotroph | Animal Pathogen-Endophyte-Endosymbiont-Epiphyte-\|Undefined Saprotroph\| | Dimorphic Yeast-Yeast | NULL | Probable | As nectar-tap_saprotroph (Põlme et al. 2020); Decay sugar-rich substrates (Põlme et al. 2020); This asexual genus is polyphyletic and contain species that function very differently depend on what they are. Caution should be taken when interpreting guilds for Candida species. | Manolakaki D, et al. 2010. Virulence. 1: 367-375 ((https://doi.org/10.4161/viru.1.5.12796)); Mekbib SB, et al. 2011. Tropical Plant Pathology 36:233-240 ((https://doi.org/10.1590/S1982-56762011000400004)); Põlme S, et al. 2020. Fungal Diversity 105:1-16 ((https://doi.org/10.1007/s13225-020-00466-2)) |
| Otu56077 | 1 | k__Fungi(100);p__Ascomycota(100);c__Leotiomycetes(100);o__Helotiales(100);f__Sclerotiniaceae(100);g__Botrytis(100);s__Botrytis_caroliniana(100); | Botrytis | 13 | Pathotroph-Saprotroph | \|Plant Pathogen\|-Plant Saprotroph | Microfungus | NULL | Probable | As litter_saprotroph (Põlme et al. 2020); Decay leaf, fruit, seed (Põlme et al. 2020) | Põlme S, et al. 2020. Fungal Diversity 105:1-16 ((https://doi.org/10.1007/s13225-020-00466-2)); The American Phytopathological Society Common Names of Plant Diseases ((https://www.apsnet.org/edcenter/resources/commonnames/Pages/default.aspx)) |
| Otu56078 | 1 | k__Fungi(100);p__Ascomycota(100);c__Saccharomycetes(100);o__Saccharomycetales(100);f__Metschnikowiaceae(100);g__Metschnikowia(100);s__Metschnikowia_pulcherrima(100); | Metschnikowia | 13 | Saprotroph | \|Undefined Saprotroph\| | Yeast | NULL | Probable | As nectar-tap_saprotroph (Põlme et al. 2020); Decay sugar-rich substrates (Põlme et al. 2020) | Põlme S, et al. 2020. Fungal Diversity 105:1-16 ((https://doi.org/10.1007/s13225-020-00466-2)) |
| Otu56079 | 1 | k__Fungi(100);p__Ascomycota(100);c__Saccharomycetes(100);o__Saccharomycetales(100);f__Metschnikowiaceae(100);g__Metschnikowia(100);g__Metschnikowia_unclassified(100); | Metschnikowia | 13 | Saprotroph | \|Undefined Saprotroph\| | Yeast | NULL | Probable | As nectar-tap_saprotroph (Põlme et al. 2020); Decay sugar-rich substrates (Põlme et al. 2020) | Põlme S, et al. 2020. Fungal Diversity 105:1-16 ((https://doi.org/10.1007/s13225-020-00466-2)) |
| Otu56081 | 1 | k__Fungi(100);p__Ascomycota(100);c__Saccharomycetes(100);o__Saccharomycetales(100);f__Metschnikowiaceae(100);g__Metschnikowia(100);s__Metschnikowia_pulcherrima(100); | Metschnikowia | 13 | Saprotroph | \|Undefined Saprotroph\| | Yeast | NULL | Probable | As nectar-tap_saprotroph (Põlme et al. 2020); Decay sugar-rich substrates (Põlme et al. 2020) | Põlme S, et al. 2020. Fungal Diversity 105:1-16 ((https://doi.org/10.1007/s13225-020-00466-2)) |
| Otu56082 | 1 | k__Fungi(100);p__Ascomycota(100);c__Saccharomycetes(100);o__Saccharomycetales(100);f__Metschnikowiaceae(100);g__Metschnikowia(100);s__Metschnikowia_pulcherrima(100); | Metschnikowia | 13 | Saprotroph | \|Undefined Saprotroph\| | Yeast | NULL | Probable | As nectar-tap_saprotroph (Põlme et al. 2020); Decay sugar-rich substrates (Põlme et al. 2020) | Põlme S, et al. 2020. Fungal Diversity 105:1-16 ((https://doi.org/10.1007/s13225-020-00466-2)) |
| Otu56083 | 1 | k__Fungi(100);p__Ascomycota(100);c__Saccharomycetes(100);o__Saccharomycetales(100);f__Metschnikowiaceae(100);g__Metschnikowia(100);g__Metschnikowia_unclassified(100); | Metschnikowia | 13 | Saprotroph | \|Undefined Saprotroph\| | Yeast | NULL | Probable | As nectar-tap_saprotroph (Põlme et al. 2020); Decay sugar-rich substrates (Põlme et al. 2020) | Põlme S, et al. 2020. Fungal Diversity 105:1-16 ((https://doi.org/10.1007/s13225-020-00466-2)) |
| Otu56084 | 1 | k__Fungi(100);p__Ascomycota(100);c__Saccharomycetes(100);o__Saccharomycetales(100);f__Metschnikowiaceae(100);g__Metschnikowia(100);s__Metschnikowia_pulcherrima(100); | Metschnikowia | 13 | Saprotroph | \|Undefined Saprotroph\| | Yeast | NULL | Probable | As nectar-tap_saprotroph (Põlme et al. 2020); Decay sugar-rich substrates (Põlme et al. 2020) | Põlme S, et al. 2020. Fungal Diversity 105:1-16 ((https://doi.org/10.1007/s13225-020-00466-2)) |
| Otu56085 | 1 | k__Fungi(100);p__Ascomycota(100);c__Saccharomycetes(100);o__Saccharomycetales(100);f__Metschnikowiaceae(100);g__Metschnikowia(100);g__Metschnikowia_unclassified(100); | Metschnikowia | 13 | Saprotroph | \|Undefined Saprotroph\| | Yeast | NULL | Probable | As nectar-tap_saprotroph (Põlme et al. 2020); Decay sugar-rich substrates (Põlme et al. 2020) | Põlme S, et al. 2020. Fungal Diversity 105:1-16 ((https://doi.org/10.1007/s13225-020-00466-2)) |
| Otu56087 | 1 | k__Fungi(100);p__Ascomycota(100);c__Saccharomycetes(100);o__Saccharomycetales(100);f__Metschnikowiaceae(100);g__Metschnikowia(100);s__Metschnikowia_pulcherrima(100); | Metschnikowia | 13 | Saprotroph | \|Undefined Saprotroph\| | Yeast | NULL | Probable | As nectar-tap_saprotroph (Põlme et al. 2020); Decay sugar-rich substrates (Põlme et al. 2020) | Põlme S, et al. 2020. Fungal Diversity 105:1-16 ((https://doi.org/10.1007/s13225-020-00466-2)) |
| Otu56088 | 1 | k__Fungi(100);p__Ascomycota(100);c__Saccharomycetes(100);o__Saccharomycetales(100);f__Metschnikowiaceae(100);g__Metschnikowia(100);s__Metschnikowia_pulcherrima(100); | Metschnikowia | 13 | Saprotroph | \|Undefined Saprotroph\| | Yeast | NULL | Probable | As nectar-tap_saprotroph (Põlme et al. 2020); Decay sugar-rich substrates (Põlme et al. 2020) | Põlme S, et al. 2020. Fungal Diversity 105:1-16 ((https://doi.org/10.1007/s13225-020-00466-2)) |
| Otu56089 | 1 | k__Fungi(100);p__Ascomycota(100);c__Saccharomycetes(100);o__Saccharomycetales(100);f__Metschnikowiaceae(100);g__Metschnikowia(100);s__Metschnikowia_pulcherrima(100); | Metschnikowia | 13 | Saprotroph | \|Undefined Saprotroph\| | Yeast | NULL | Probable | As nectar-tap_saprotroph (Põlme et al. 2020); Decay sugar-rich substrates (Põlme et al. 2020) | Põlme S, et al. 2020. Fungal Diversity 105:1-16 ((https://doi.org/10.1007/s13225-020-00466-2)) |
| Otu56090 | 1 | k__Fungi(100);p__Ascomycota(100);c__Saccharomycetes(100);o__Saccharomycetales(100);f__Metschnikowiaceae(100);g__Metschnikowia(100);s__Metschnikowia_pulcherrima(100); | Metschnikowia | 13 | Saprotroph | \|Undefined Saprotroph\| | Yeast | NULL | Probable | As nectar-tap_saprotroph (Põlme et al. 2020); Decay sugar-rich substrates (Põlme et al. 2020) | Põlme S, et al. 2020. Fungal Diversity 105:1-16 ((https://doi.org/10.1007/s13225-020-00466-2)) |
| Otu56091 | 1 | k__Fungi(100);p__Ascomycota(100);c__Saccharomycetes(100);o__Saccharomycetales(100);f__Metschnikowiaceae(100);g__Metschnikowia(100);s__Metschnikowia_pulcherrima(100); | Metschnikowia | 13 | Saprotroph | \|Undefined Saprotroph\| | Yeast | NULL | Probable | As nectar-tap_saprotroph (Põlme et al. 2020); Decay sugar-rich substrates (Põlme et al. 2020) | Põlme S, et al. 2020. Fungal Diversity 105:1-16 ((https://doi.org/10.1007/s13225-020-00466-2)) |
| Otu56092 | 1 | k__Fungi(100);p__Ascomycota(100);c__Saccharomycetes(100);o__Saccharomycetales(100);f__Metschnikowiaceae(100);g__Metschnikowia(100);s__Metschnikowia_pulcherrima(100); | Metschnikowia | 13 | Saprotroph | \|Undefined Saprotroph\| | Yeast | NULL | Probable | As nectar-tap_saprotroph (Põlme et al. 2020); Decay sugar-rich substrates (Põlme et al. 2020) | Põlme S, et al. 2020. Fungal Diversity 105:1-16 ((https://doi.org/10.1007/s13225-020-00466-2)) |
| Otu56093 | 1 | k__Fungi(100);p__Ascomycota(100);c__Saccharomycetes(100);o__Saccharomycetales(100);f__Metschnikowiaceae(100);g__Metschnikowia(100);s__Metschnikowia_pulcherrima(100); | Metschnikowia | 13 | Saprotroph | \|Undefined Saprotroph\| | Yeast | NULL | Probable | As nectar-tap_saprotroph (Põlme et al. 2020); Decay sugar-rich substrates (Põlme et al. 2020) | Põlme S, et al. 2020. Fungal Diversity 105:1-16 ((https://doi.org/10.1007/s13225-020-00466-2)) |
| Otu56095 | 1 | k__Fungi(100);p__Ascomycota(100);c__Saccharomycetes(100);o__Saccharomycetales(100);f__Metschnikowiaceae(100);g__Metschnikowia(100);g__Metschnikowia_unclassified(100); | Metschnikowia | 13 | Saprotroph | \|Undefined Saprotroph\| | Yeast | NULL | Probable | As nectar-tap_saprotroph (Põlme et al. 2020); Decay sugar-rich substrates (Põlme et al. 2020) | Põlme S, et al. 2020. Fungal Diversity 105:1-16 ((https://doi.org/10.1007/s13225-020-00466-2)) |
| Otu56096 | 1 | k__Fungi(100);p__Ascomycota(100);c__Saccharomycetes(100);o__Saccharomycetales(100);f__Metschnikowiaceae(100);g__Metschnikowia(100);s__Metschnikowia_pulcherrima(100); | Metschnikowia | 13 | Saprotroph | \|Undefined Saprotroph\| | Yeast | NULL | Probable | As nectar-tap_saprotroph (Põlme et al. 2020); Decay sugar-rich substrates (Põlme et al. 2020) | Põlme S, et al. 2020. Fungal Diversity 105:1-16 ((https://doi.org/10.1007/s13225-020-00466-2)) |
| Otu56098 | 1 | k__Fungi(100);p__Ascomycota(100);c__Saccharomycetes(100);o__Saccharomycetales(100);f__Metschnikowiaceae(100);g__Metschnikowia(100);s__Metschnikowia_pulcherrima(100); | Metschnikowia | 13 | Saprotroph | \|Undefined Saprotroph\| | Yeast | NULL | Probable | As nectar-tap_saprotroph (Põlme et al. 2020); Decay sugar-rich substrates (Põlme et al. 2020) | Põlme S, et al. 2020. Fungal Diversity 105:1-16 ((https://doi.org/10.1007/s13225-020-00466-2)) |
| Otu56099 | 1 | k__Fungi(100);p__Ascomycota(100);c__Saccharomycetes(100);o__Saccharomycetales(100);f__Metschnikowiaceae(100);g__Metschnikowia(100);s__Metschnikowia_pulcherrima(100); | Metschnikowia | 13 | Saprotroph | \|Undefined Saprotroph\| | Yeast | NULL | Probable | As nectar-tap_saprotroph (Põlme et al. 2020); Decay sugar-rich substrates (Põlme et al. 2020) | Põlme S, et al. 2020. Fungal Diversity 105:1-16 ((https://doi.org/10.1007/s13225-020-00466-2)) |
| Otu56100 | 1 | k__Fungi(100);p__Ascomycota(100);c__Saccharomycetes(100);o__Saccharomycetales(100);f__Metschnikowiaceae(100);g__Metschnikowia(100);s__Metschnikowia_pulcherrima(100); | Metschnikowia | 13 | Saprotroph | \|Undefined Saprotroph\| | Yeast | NULL | Probable | As nectar-tap_saprotroph (Põlme et al. 2020); Decay sugar-rich substrates (Põlme et al. 2020) | Põlme S, et al. 2020. Fungal Diversity 105:1-16 ((https://doi.org/10.1007/s13225-020-00466-2)) |
| Otu56101 | 1 | k__Fungi(100);p__Ascomycota(100);c__Saccharomycetes(100);o__Saccharomycetales(100);f__Metschnikowiaceae(100);g__Metschnikowia(100);s__Metschnikowia_pulcherrima(100); | Metschnikowia | 13 | Saprotroph | \|Undefined Saprotroph\| | Yeast | NULL | Probable | As nectar-tap_saprotroph (Põlme et al. 2020); Decay sugar-rich substrates (Põlme et al. 2020) | Põlme S, et al. 2020. Fungal Diversity 105:1-16 ((https://doi.org/10.1007/s13225-020-00466-2)) |
| Otu56102 | 1 | k__Fungi(100);p__Ascomycota(100);c__Saccharomycetes(100);o__Saccharomycetales(100);f__Metschnikowiaceae(100);g__Metschnikowia(100);s__Metschnikowia_pulcherrima(100); | Metschnikowia | 13 | Saprotroph | \|Undefined Saprotroph\| | Yeast | NULL | Probable | As nectar-tap_saprotroph (Põlme et al. 2020); Decay sugar-rich substrates (Põlme et al. 2020) | Põlme S, et al. 2020. Fungal Diversity 105:1-16 ((https://doi.org/10.1007/s13225-020-00466-2)) |
| Otu56103 | 1 | k__Fungi(100);p__Ascomycota(100);c__Saccharomycetes(100);o__Saccharomycetales(100);f__Saccharomycetaceae(100);g__Kazachstania(100);s__Kazachstania_intestinalis(100); | Kazachstania | 13 | Saprotroph | \|Undefined Saprotroph\| | Yeast | NULL | Probable | As nectar-tap_saprotroph (Põlme et al. 2020); Decay sugar-rich substrates (Põlme et al. 2020) | Põlme S, et al. 2020. Fungal Diversity 105:1-16 ((https://doi.org/10.1007/s13225-020-00466-2)) |
| Otu56104 | 1 | k__Fungi(100);p__Ascomycota(100);c__Saccharomycetes(100);o__Saccharomycetales(100);f__Metschnikowiaceae(100);g__Metschnikowia(100);g__Metschnikowia_unclassified(100); | Metschnikowia | 13 | Saprotroph | \|Undefined Saprotroph\| | Yeast | NULL | Probable | As nectar-tap_saprotroph (Põlme et al. 2020); Decay sugar-rich substrates (Põlme et al. 2020) | Põlme S, et al. 2020. Fungal Diversity 105:1-16 ((https://doi.org/10.1007/s13225-020-00466-2)) |
| Otu56105 | 1 | k__Fungi(100);p__Ascomycota(100);c__Saccharomycetes(100);o__Saccharomycetales(100);f__Metschnikowiaceae(100);g__Metschnikowia(100);s__Metschnikowia_pulcherrima(100); | Metschnikowia | 13 | Saprotroph | \|Undefined Saprotroph\| | Yeast | NULL | Probable | As nectar-tap_saprotroph (Põlme et al. 2020); Decay sugar-rich substrates (Põlme et al. 2020) | Põlme S, et al. 2020. Fungal Diversity 105:1-16 ((https://doi.org/10.1007/s13225-020-00466-2)) |
| Otu56106 | 1 | k__Fungi(100);p__Ascomycota(100);c__Saccharomycetes(100);o__Saccharomycetales(100);f__Saccharomycetales_family_Incertae_sedis(100);f__Saccharomycetales_family_Incertae_sedis_unclassified(100);f__Saccharomycetales_family_Incertae_sedis_unclassified(100); | Saccharomycetales | 7 | Saprotroph | Undefined Saprotroph | Yeast | NULL | Possible | NULL | Sterkenburg E, et al. 2015. New Phytologist 207:1145-1158 ((https://doi.org/10.1111/nph.13426)) |
| Otu56107 | 1 | k__Fungi(100);p__Ascomycota(100);c__Saccharomycetes(100);o__Saccharomycetales(100);f__Metschnikowiaceae(100);g__Metschnikowia(100);s__Metschnikowia_pulcherrima(100); | Metschnikowia | 13 | Saprotroph | \|Undefined Saprotroph\| | Yeast | NULL | Probable | As nectar-tap_saprotroph (Põlme et al. 2020); Decay sugar-rich substrates (Põlme et al. 2020) | Põlme S, et al. 2020. Fungal Diversity 105:1-16 ((https://doi.org/10.1007/s13225-020-00466-2)) |
| Otu56108 | 1 | k__Fungi(100);p__Ascomycota(100);c__Saccharomycetes(100);o__Saccharomycetales(100);f__Metschnikowiaceae(100);g__Metschnikowia(100);s__Metschnikowia_pulcherrima(100); | Metschnikowia | 13 | Saprotroph | \|Undefined Saprotroph\| | Yeast | NULL | Probable | As nectar-tap_saprotroph (Põlme et al. 2020); Decay sugar-rich substrates (Põlme et al. 2020) | Põlme S, et al. 2020. Fungal Diversity 105:1-16 ((https://doi.org/10.1007/s13225-020-00466-2)) |
| Otu56111 | 1 | k__Fungi(100);p__Ascomycota(100);c__Saccharomycetes(100);o__Saccharomycetales(100);f__Metschnikowiaceae(100);g__Metschnikowia(100);g__Metschnikowia_unclassified(100); | Metschnikowia | 13 | Saprotroph | \|Undefined Saprotroph\| | Yeast | NULL | Probable | As nectar-tap_saprotroph (Põlme et al. 2020); Decay sugar-rich substrates (Põlme et al. 2020) | Põlme S, et al. 2020. Fungal Diversity 105:1-16 ((https://doi.org/10.1007/s13225-020-00466-2)) |
| Otu56112 | 1 | k__Fungi(100);p__Ascomycota(100);c__Saccharomycetes(100);o__Saccharomycetales(100);f__Saccharomycetales_family_Incertae_sedis(100);g__Candida(100);g__Candida_unclassified(100); | Candida | 13 | Pathotroph-Saprotroph-Symbiotroph | Animal Pathogen-Endophyte-Endosymbiont-Epiphyte-\|Undefined Saprotroph\| | Dimorphic Yeast-Yeast | NULL | Probable | As nectar-tap_saprotroph (Põlme et al. 2020); Decay sugar-rich substrates (Põlme et al. 2020); This asexual genus is polyphyletic and contain species that function very differently depend on what they are. Caution should be taken when interpreting guilds for Candida species. | Manolakaki D, et al. 2010. Virulence. 1: 367-375 ((https://doi.org/10.4161/viru.1.5.12796)); Mekbib SB, et al. 2011. Tropical Plant Pathology 36:233-240 ((https://doi.org/10.1590/S1982-56762011000400004)); Põlme S, et al. 2020. Fungal Diversity 105:1-16 ((https://doi.org/10.1007/s13225-020-00466-2)) |
| Otu56116 | 1 | k__Fungi(100);p__Ascomycota(100);c__Saccharomycetes(100);o__Saccharomycetales(100);f__Metschnikowiaceae(100);g__Metschnikowia(100);s__Metschnikowia_pulcherrima(100); | Metschnikowia | 13 | Saprotroph | \|Undefined Saprotroph\| | Yeast | NULL | Probable | As nectar-tap_saprotroph (Põlme et al. 2020); Decay sugar-rich substrates (Põlme et al. 2020) | Põlme S, et al. 2020. Fungal Diversity 105:1-16 ((https://doi.org/10.1007/s13225-020-00466-2)) |
| Otu56118 | 1 | k__Fungi(100);p__Ascomycota(100);c__Saccharomycetes(100);o__Saccharomycetales(100);f__Saccharomycetales_family_Incertae_sedis(100);g__Candida(100);g__Candida_unclassified(100); | Candida | 13 | Pathotroph-Saprotroph-Symbiotroph | Animal Pathogen-Endophyte-Endosymbiont-Epiphyte-\|Undefined Saprotroph\| | Dimorphic Yeast-Yeast | NULL | Probable | As nectar-tap_saprotroph (Põlme et al. 2020); Decay sugar-rich substrates (Põlme et al. 2020); This asexual genus is polyphyletic and contain species that function very differently depend on what they are. Caution should be taken when interpreting guilds for Candida species. | Manolakaki D, et al. 2010. Virulence. 1: 367-375 ((https://doi.org/10.4161/viru.1.5.12796)); Mekbib SB, et al. 2011. Tropical Plant Pathology 36:233-240 ((https://doi.org/10.1590/S1982-56762011000400004)); Põlme S, et al. 2020. Fungal Diversity 105:1-16 ((https://doi.org/10.1007/s13225-020-00466-2)) |
| Otu56119 | 1 | k__Fungi(100);p__Ascomycota(100);c__Saccharomycetes(100);o__Saccharomycetales(100);f__Saccharomycetales_family_Incertae_sedis(100);g__Candida(100);s__Candida_sp_AS23486(100); | Candida | 13 | Pathotroph-Saprotroph-Symbiotroph | Animal Pathogen-Endophyte-Endosymbiont-Epiphyte-\|Undefined Saprotroph\| | Dimorphic Yeast-Yeast | NULL | Probable | As nectar-tap_saprotroph (Põlme et al. 2020); Decay sugar-rich substrates (Põlme et al. 2020); This asexual genus is polyphyletic and contain species that function very differently depend on what they are. Caution should be taken when interpreting guilds for Candida species. | Manolakaki D, et al. 2010. Virulence. 1: 367-375 ((https://doi.org/10.4161/viru.1.5.12796)); Mekbib SB, et al. 2011. Tropical Plant Pathology 36:233-240 ((https://doi.org/10.1590/S1982-56762011000400004)); Põlme S, et al. 2020. Fungal Diversity 105:1-16 ((https://doi.org/10.1007/s13225-020-00466-2)) |
| Otu56120 | 1 | k__Fungi(100);p__Ascomycota(100);c__Saccharomycetes(100);o__Saccharomycetales(100);f__Saccharomycetales_family_Incertae_sedis(100);g__Candida(100);g__Candida_unclassified(100); | Candida | 13 | Pathotroph-Saprotroph-Symbiotroph | Animal Pathogen-Endophyte-Endosymbiont-Epiphyte-\|Undefined Saprotroph\| | Dimorphic Yeast-Yeast | NULL | Probable | As nectar-tap_saprotroph (Põlme et al. 2020); Decay sugar-rich substrates (Põlme et al. 2020); This asexual genus is polyphyletic and contain species that function very differently depend on what they are. Caution should be taken when interpreting guilds for Candida species. | Manolakaki D, et al. 2010. Virulence. 1: 367-375 ((https://doi.org/10.4161/viru.1.5.12796)); Mekbib SB, et al. 2011. Tropical Plant Pathology 36:233-240 ((https://doi.org/10.1590/S1982-56762011000400004)); Põlme S, et al. 2020. Fungal Diversity 105:1-16 ((https://doi.org/10.1007/s13225-020-00466-2)) |
| Otu56124 | 1 | k__Fungi(100);p__Ascomycota(100);c__Saccharomycetes(100);o__Saccharomycetales(100);f__Metschnikowiaceae(100);g__Metschnikowia(100);s__Metschnikowia_pulcherrima(100); | Metschnikowia | 13 | Saprotroph | \|Undefined Saprotroph\| | Yeast | NULL | Probable | As nectar-tap_saprotroph (Põlme et al. 2020); Decay sugar-rich substrates (Põlme et al. 2020) | Põlme S, et al. 2020. Fungal Diversity 105:1-16 ((https://doi.org/10.1007/s13225-020-00466-2)) |
| Otu56125 | 1 | k__Fungi(100);p__Ascomycota(100);c__Saccharomycetes(100);o__Saccharomycetales(100);f__Metschnikowiaceae(100);g__Metschnikowia(100);s__Metschnikowia_pulcherrima(100); | Metschnikowia | 13 | Saprotroph | \|Undefined Saprotroph\| | Yeast | NULL | Probable | As nectar-tap_saprotroph (Põlme et al. 2020); Decay sugar-rich substrates (Põlme et al. 2020) | Põlme S, et al. 2020. Fungal Diversity 105:1-16 ((https://doi.org/10.1007/s13225-020-00466-2)) |
| Otu56127 | 1 | k__Fungi(100);p__Ascomycota(100);c__Saccharomycetes(100);o__Saccharomycetales(100);f__Saccharomycetales_family_Incertae_sedis(100);g__Candida(100);s__Candida_sp_AS23486(100); | Candida | 13 | Pathotroph-Saprotroph-Symbiotroph | Animal Pathogen-Endophyte-Endosymbiont-Epiphyte-\|Undefined Saprotroph\| | Dimorphic Yeast-Yeast | NULL | Probable | As nectar-tap_saprotroph (Põlme et al. 2020); Decay sugar-rich substrates (Põlme et al. 2020); This asexual genus is polyphyletic and contain species that function very differently depend on what they are. Caution should be taken when interpreting guilds for Candida species. | Manolakaki D, et al. 2010. Virulence. 1: 367-375 ((https://doi.org/10.4161/viru.1.5.12796)); Mekbib SB, et al. 2011. Tropical Plant Pathology 36:233-240 ((https://doi.org/10.1590/S1982-56762011000400004)); Põlme S, et al. 2020. Fungal Diversity 105:1-16 ((https://doi.org/10.1007/s13225-020-00466-2)) |
| Otu56130 | 1 | k__Fungi(100);p__Ascomycota(100);c__Saccharomycetes(100);o__Saccharomycetales(100);f__Metschnikowiaceae(100);g__Metschnikowia(100);s__Metschnikowia_pulcherrima(100); | Metschnikowia | 13 | Saprotroph | \|Undefined Saprotroph\| | Yeast | NULL | Probable | As nectar-tap_saprotroph (Põlme et al. 2020); Decay sugar-rich substrates (Põlme et al. 2020) | Põlme S, et al. 2020. Fungal Diversity 105:1-16 ((https://doi.org/10.1007/s13225-020-00466-2)) |
| Otu56133 | 1 | k__Fungi(100);p__Ascomycota(100);c__Saccharomycetes(100);o__Saccharomycetales(100);f__Metschnikowiaceae(100);g__Metschnikowia(100);s__Metschnikowia_pulcherrima(100); | Metschnikowia | 13 | Saprotroph | \|Undefined Saprotroph\| | Yeast | NULL | Probable | As nectar-tap_saprotroph (Põlme et al. 2020); Decay sugar-rich substrates (Põlme et al. 2020) | Põlme S, et al. 2020. Fungal Diversity 105:1-16 ((https://doi.org/10.1007/s13225-020-00466-2)) |
| Otu56134 | 1 | k__Fungi(100);p__Ascomycota(100);c__Saccharomycetes(100);o__Saccharomycetales(100);f__Metschnikowiaceae(100);g__Metschnikowia(100);g__Metschnikowia_unclassified(100); | Metschnikowia | 13 | Saprotroph | \|Undefined Saprotroph\| | Yeast | NULL | Probable | As nectar-tap_saprotroph (Põlme et al. 2020); Decay sugar-rich substrates (Põlme et al. 2020) | Põlme S, et al. 2020. Fungal Diversity 105:1-16 ((https://doi.org/10.1007/s13225-020-00466-2)) |
| Otu56135 | 1 | k__Fungi(100);p__Ascomycota(100);c__Saccharomycetes(100);o__Saccharomycetales(100);f__Metschnikowiaceae(100);g__Metschnikowia(100);s__Metschnikowia_pulcherrima(100); | Metschnikowia | 13 | Saprotroph | \|Undefined Saprotroph\| | Yeast | NULL | Probable | As nectar-tap_saprotroph (Põlme et al. 2020); Decay sugar-rich substrates (Põlme et al. 2020) | Põlme S, et al. 2020. Fungal Diversity 105:1-16 ((https://doi.org/10.1007/s13225-020-00466-2)) |
| Otu56136 | 1 | k__Fungi(100);p__Ascomycota(100);c__Saccharomycetes(100);o__Saccharomycetales(100);f__Metschnikowiaceae(100);g__Metschnikowia(100);s__Metschnikowia_pulcherrima(100); | Metschnikowia | 13 | Saprotroph | \|Undefined Saprotroph\| | Yeast | NULL | Probable | As nectar-tap_saprotroph (Põlme et al. 2020); Decay sugar-rich substrates (Põlme et al. 2020) | Põlme S, et al. 2020. Fungal Diversity 105:1-16 ((https://doi.org/10.1007/s13225-020-00466-2)) |
| Otu56137 | 1 | k__Fungi(100);p__Ascomycota(100);c__Saccharomycetes(100);o__Saccharomycetales(100);f__Metschnikowiaceae(100);g__Metschnikowia(100);g__Metschnikowia_unclassified(100); | Metschnikowia | 13 | Saprotroph | \|Undefined Saprotroph\| | Yeast | NULL | Probable | As nectar-tap_saprotroph (Põlme et al. 2020); Decay sugar-rich substrates (Põlme et al. 2020) | Põlme S, et al. 2020. Fungal Diversity 105:1-16 ((https://doi.org/10.1007/s13225-020-00466-2)) |
| Otu56138 | 1 | k__Fungi(100);p__Ascomycota(100);c__Saccharomycetes(100);o__Saccharomycetales(100);f__Metschnikowiaceae(100);g__Metschnikowia(100);s__Metschnikowia_pulcherrima(100); | Metschnikowia | 13 | Saprotroph | \|Undefined Saprotroph\| | Yeast | NULL | Probable | As nectar-tap_saprotroph (Põlme et al. 2020); Decay sugar-rich substrates (Põlme et al. 2020) | Põlme S, et al. 2020. Fungal Diversity 105:1-16 ((https://doi.org/10.1007/s13225-020-00466-2)) |
| Otu56139 | 1 | k__Fungi(100);p__Ascomycota(100);c__Saccharomycetes(100);o__Saccharomycetales(100);f__Metschnikowiaceae(100);g__Metschnikowia(100);s__Metschnikowia_pulcherrima(100); | Metschnikowia | 13 | Saprotroph | \|Undefined Saprotroph\| | Yeast | NULL | Probable | As nectar-tap_saprotroph (Põlme et al. 2020); Decay sugar-rich substrates (Põlme et al. 2020) | Põlme S, et al. 2020. Fungal Diversity 105:1-16 ((https://doi.org/10.1007/s13225-020-00466-2)) |
| Otu56140 | 1 | k__Fungi(100);p__Ascomycota(100);c__Saccharomycetes(100);o__Saccharomycetales(100);f__Metschnikowiaceae(100);g__Metschnikowia(100);s__Metschnikowia_pulcherrima(100); | Metschnikowia | 13 | Saprotroph | \|Undefined Saprotroph\| | Yeast | NULL | Probable | As nectar-tap_saprotroph (Põlme et al. 2020); Decay sugar-rich substrates (Põlme et al. 2020) | Põlme S, et al. 2020. Fungal Diversity 105:1-16 ((https://doi.org/10.1007/s13225-020-00466-2)) |
| Otu56141 | 1 | k__Fungi(100);p__Ascomycota(100);c__Saccharomycetes(100);o__Saccharomycetales(100);o__Saccharomycetales_unclassified(100);o__Saccharomycetales_unclassified(100);o__Saccharomycetales_unclassified(100); | Saccharomycetales | 7 | Saprotroph | Undefined Saprotroph | Yeast | NULL | Possible | NULL | Sterkenburg E, et al. 2015. New Phytologist 207:1145-1158 ((https://doi.org/10.1111/nph.13426)) |
| Otu56142 | 1 | k__Fungi(100);p__Ascomycota(100);c__Saccharomycetes(100);o__Saccharomycetales(100);f__Metschnikowiaceae(100);g__Metschnikowia(100);g__Metschnikowia_unclassified(100); | Metschnikowia | 13 | Saprotroph | \|Undefined Saprotroph\| | Yeast | NULL | Probable | As nectar-tap_saprotroph (Põlme et al. 2020); Decay sugar-rich substrates (Põlme et al. 2020) | Põlme S, et al. 2020. Fungal Diversity 105:1-16 ((https://doi.org/10.1007/s13225-020-00466-2)) |
| Otu56143 | 1 | k__Fungi(100);p__Ascomycota(100);c__Saccharomycetes(100);o__Saccharomycetales(100);f__Metschnikowiaceae(100);g__Metschnikowia(100);s__Metschnikowia_pulcherrima(100); | Metschnikowia | 13 | Saprotroph | \|Undefined Saprotroph\| | Yeast | NULL | Probable | As nectar-tap_saprotroph (Põlme et al. 2020); Decay sugar-rich substrates (Põlme et al. 2020) | Põlme S, et al. 2020. Fungal Diversity 105:1-16 ((https://doi.org/10.1007/s13225-020-00466-2)) |
| Otu56144 | 1 | k__Fungi(100);p__Ascomycota(100);c__Saccharomycetes(100);o__Saccharomycetales(100);f__Metschnikowiaceae(100);g__Metschnikowia(100);g__Metschnikowia_unclassified(100); | Metschnikowia | 13 | Saprotroph | \|Undefined Saprotroph\| | Yeast | NULL | Probable | As nectar-tap_saprotroph (Põlme et al. 2020); Decay sugar-rich substrates (Põlme et al. 2020) | Põlme S, et al. 2020. Fungal Diversity 105:1-16 ((https://doi.org/10.1007/s13225-020-00466-2)) |
| Otu56145 | 1 | k__Fungi(100);p__Ascomycota(100);c__Saccharomycetes(100);o__Saccharomycetales(100);f__Metschnikowiaceae(100);g__Metschnikowia(100);s__Metschnikowia_pulcherrima(100); | Metschnikowia | 13 | Saprotroph | \|Undefined Saprotroph\| | Yeast | NULL | Probable | As nectar-tap_saprotroph (Põlme et al. 2020); Decay sugar-rich substrates (Põlme et al. 2020) | Põlme S, et al. 2020. Fungal Diversity 105:1-16 ((https://doi.org/10.1007/s13225-020-00466-2)) |
| Otu56148 | 1 | k__Fungi(100);p__Ascomycota(100);c__Saccharomycetes(100);o__Saccharomycetales(100);f__Metschnikowiaceae(100);g__Metschnikowia(100);s__Metschnikowia_pulcherrima(100); | Metschnikowia | 13 | Saprotroph | \|Undefined Saprotroph\| | Yeast | NULL | Probable | As nectar-tap_saprotroph (Põlme et al. 2020); Decay sugar-rich substrates (Põlme et al. 2020) | Põlme S, et al. 2020. Fungal Diversity 105:1-16 ((https://doi.org/10.1007/s13225-020-00466-2)) |
| Otu56149 | 1 | k__Fungi(100);p__Ascomycota(100);c__Saccharomycetes(100);o__Saccharomycetales(100);f__Metschnikowiaceae(100);g__Metschnikowia(100);s__Metschnikowia_pulcherrima(100); | Metschnikowia | 13 | Saprotroph | \|Undefined Saprotroph\| | Yeast | NULL | Probable | As nectar-tap_saprotroph (Põlme et al. 2020); Decay sugar-rich substrates (Põlme et al. 2020) | Põlme S, et al. 2020. Fungal Diversity 105:1-16 ((https://doi.org/10.1007/s13225-020-00466-2)) |
| Otu56151 | 1 | k__Fungi(100);p__Ascomycota(100);c__Saccharomycetes(100);o__Saccharomycetales(100);f__Metschnikowiaceae(100);g__Metschnikowia(100);s__Metschnikowia_pulcherrima(100); | Metschnikowia | 13 | Saprotroph | \|Undefined Saprotroph\| | Yeast | NULL | Probable | As nectar-tap_saprotroph (Põlme et al. 2020); Decay sugar-rich substrates (Põlme et al. 2020) | Põlme S, et al. 2020. Fungal Diversity 105:1-16 ((https://doi.org/10.1007/s13225-020-00466-2)) |
| Otu56153 | 1 | k__Fungi(100);p__Ascomycota(100);c__Saccharomycetes(100);o__Saccharomycetales(100);f__Metschnikowiaceae(100);g__Metschnikowia(100);s__Metschnikowia_pulcherrima(100); | Metschnikowia | 13 | Saprotroph | \|Undefined Saprotroph\| | Yeast | NULL | Probable | As nectar-tap_saprotroph (Põlme et al. 2020); Decay sugar-rich substrates (Põlme et al. 2020) | Põlme S, et al. 2020. Fungal Diversity 105:1-16 ((https://doi.org/10.1007/s13225-020-00466-2)) |
| Otu56158 | 1 | k__Fungi(100);p__Ascomycota(100);c__Saccharomycetes(100);o__Saccharomycetales(100);f__Saccharomycetales_family_Incertae_sedis(100);g__Candida(100);s__Candida_cretensis(100); | Candida | 13 | Pathotroph-Saprotroph-Symbiotroph | Animal Pathogen-Endophyte-Endosymbiont-Epiphyte-\|Undefined Saprotroph\| | Dimorphic Yeast-Yeast | NULL | Probable | As nectar-tap_saprotroph (Põlme et al. 2020); Decay sugar-rich substrates (Põlme et al. 2020); This asexual genus is polyphyletic and contain species that function very differently depend on what they are. Caution should be taken when interpreting guilds for Candida species. | Manolakaki D, et al. 2010. Virulence. 1: 367-375 ((https://doi.org/10.4161/viru.1.5.12796)); Mekbib SB, et al. 2011. Tropical Plant Pathology 36:233-240 ((https://doi.org/10.1590/S1982-56762011000400004)); Põlme S, et al. 2020. Fungal Diversity 105:1-16 ((https://doi.org/10.1007/s13225-020-00466-2)) |
| Otu56159 | 1 | k__Fungi(100);p__Ascomycota(100);c__Saccharomycetes(100);o__Saccharomycetales(100);f__Metschnikowiaceae(100);g__Metschnikowia(100);s__Metschnikowia_pulcherrima(100); | Metschnikowia | 13 | Saprotroph | \|Undefined Saprotroph\| | Yeast | NULL | Probable | As nectar-tap_saprotroph (Põlme et al. 2020); Decay sugar-rich substrates (Põlme et al. 2020) | Põlme S, et al. 2020. Fungal Diversity 105:1-16 ((https://doi.org/10.1007/s13225-020-00466-2)) |
| Otu56161 | 1 | k__Fungi(100);p__Ascomycota(100);c__Saccharomycetes(100);o__Saccharomycetales(100);f__Metschnikowiaceae(100);g__Metschnikowia(100);s__Metschnikowia_pulcherrima(100); | Metschnikowia | 13 | Saprotroph | \|Undefined Saprotroph\| | Yeast | NULL | Probable | As nectar-tap_saprotroph (Põlme et al. 2020); Decay sugar-rich substrates (Põlme et al. 2020) | Põlme S, et al. 2020. Fungal Diversity 105:1-16 ((https://doi.org/10.1007/s13225-020-00466-2)) |
| Otu56162 | 1 | k__Fungi(100);p__Ascomycota(100);c__Saccharomycetes(100);o__Saccharomycetales(100);f__Metschnikowiaceae(100);g__Metschnikowia(100);s__Metschnikowia_pulcherrima(100); | Metschnikowia | 13 | Saprotroph | \|Undefined Saprotroph\| | Yeast | NULL | Probable | As nectar-tap_saprotroph (Põlme et al. 2020); Decay sugar-rich substrates (Põlme et al. 2020) | Põlme S, et al. 2020. Fungal Diversity 105:1-16 ((https://doi.org/10.1007/s13225-020-00466-2)) |
| Otu56163 | 1 | k__Fungi(100);p__Ascomycota(100);c__Saccharomycetes(100);o__Saccharomycetales(100);f__Metschnikowiaceae(100);g__Metschnikowia(100);s__Metschnikowia_pulcherrima(100); | Metschnikowia | 13 | Saprotroph | \|Undefined Saprotroph\| | Yeast | NULL | Probable | As nectar-tap_saprotroph (Põlme et al. 2020); Decay sugar-rich substrates (Põlme et al. 2020) | Põlme S, et al. 2020. Fungal Diversity 105:1-16 ((https://doi.org/10.1007/s13225-020-00466-2)) |
| Otu56164 | 1 | k__Fungi(100);p__Ascomycota(100);c__Saccharomycetes(100);o__Saccharomycetales(100);f__Metschnikowiaceae(100);g__Metschnikowia(100);s__Metschnikowia_pulcherrima(100); | Metschnikowia | 13 | Saprotroph | \|Undefined Saprotroph\| | Yeast | NULL | Probable | As nectar-tap_saprotroph (Põlme et al. 2020); Decay sugar-rich substrates (Põlme et al. 2020) | Põlme S, et al. 2020. Fungal Diversity 105:1-16 ((https://doi.org/10.1007/s13225-020-00466-2)) |
| Otu56165 | 1 | k__Fungi(100);p__Ascomycota(100);c__Saccharomycetes(100);o__Saccharomycetales(100);f__Metschnikowiaceae(100);g__Metschnikowia(100);s__Metschnikowia_pulcherrima(100); | Metschnikowia | 13 | Saprotroph | \|Undefined Saprotroph\| | Yeast | NULL | Probable | As nectar-tap_saprotroph (Põlme et al. 2020); Decay sugar-rich substrates (Põlme et al. 2020) | Põlme S, et al. 2020. Fungal Diversity 105:1-16 ((https://doi.org/10.1007/s13225-020-00466-2)) |
| Otu56168 | 1 | k__Fungi(100);p__Ascomycota(100);c__Saccharomycetes(100);o__Saccharomycetales(100);f__Metschnikowiaceae(100);g__Metschnikowia(100);s__Metschnikowia_pulcherrima(100); | Metschnikowia | 13 | Saprotroph | \|Undefined Saprotroph\| | Yeast | NULL | Probable | As nectar-tap_saprotroph (Põlme et al. 2020); Decay sugar-rich substrates (Põlme et al. 2020) | Põlme S, et al. 2020. Fungal Diversity 105:1-16 ((https://doi.org/10.1007/s13225-020-00466-2)) |
| Otu56170 | 1 | k__Fungi(100);p__Ascomycota(100);c__Saccharomycetes(100);o__Saccharomycetales(100);f__Metschnikowiaceae(100);g__Metschnikowia(100);s__Metschnikowia_pulcherrima(100); | Metschnikowia | 13 | Saprotroph | \|Undefined Saprotroph\| | Yeast | NULL | Probable | As nectar-tap_saprotroph (Põlme et al. 2020); Decay sugar-rich substrates (Põlme et al. 2020) | Põlme S, et al. 2020. Fungal Diversity 105:1-16 ((https://doi.org/10.1007/s13225-020-00466-2)) |
| Otu56177 | 1 | k__Fungi(100);p__Ascomycota(100);c__Saccharomycetes(100);o__Saccharomycetales(100);f__Saccharomycetales_family_Incertae_sedis(100);g__Candida(100);g__Candida_unclassified(100); | Candida | 13 | Pathotroph-Saprotroph-Symbiotroph | Animal Pathogen-Endophyte-Endosymbiont-Epiphyte-\|Undefined Saprotroph\| | Dimorphic Yeast-Yeast | NULL | Probable | As nectar-tap_saprotroph (Põlme et al. 2020); Decay sugar-rich substrates (Põlme et al. 2020); This asexual genus is polyphyletic and contain species that function very differently depend on what they are. Caution should be taken when interpreting guilds for Candida species. | Manolakaki D, et al. 2010. Virulence. 1: 367-375 ((https://doi.org/10.4161/viru.1.5.12796)); Mekbib SB, et al. 2011. Tropical Plant Pathology 36:233-240 ((https://doi.org/10.1590/S1982-56762011000400004)); Põlme S, et al. 2020. Fungal Diversity 105:1-16 ((https://doi.org/10.1007/s13225-020-00466-2)) |
| Otu56179 | 1 | k__Fungi(100);p__Ascomycota(100);c__Saccharomycetes(100);o__Saccharomycetales(100);f__Saccharomycetales_family_Incertae_sedis(100);f__Saccharomycetales_family_Incertae_sedis_unclassified(100);f__Saccharomycetales_family_Incertae_sedis_unclassified(100); | Saccharomycetales | 7 | Saprotroph | Undefined Saprotroph | Yeast | NULL | Possible | NULL | Sterkenburg E, et al. 2015. New Phytologist 207:1145-1158 ((https://doi.org/10.1111/nph.13426)) |
| Otu56180 | 1 | k__Fungi(100);p__Ascomycota(100);c__Saccharomycetes(100);o__Saccharomycetales(100);f__Metschnikowiaceae(100);g__Metschnikowia(100);s__Metschnikowia_pulcherrima(100); | Metschnikowia | 13 | Saprotroph | \|Undefined Saprotroph\| | Yeast | NULL | Probable | As nectar-tap_saprotroph (Põlme et al. 2020); Decay sugar-rich substrates (Põlme et al. 2020) | Põlme S, et al. 2020. Fungal Diversity 105:1-16 ((https://doi.org/10.1007/s13225-020-00466-2)) |
| Otu56186 | 1 | k__Fungi(100);p__Ascomycota(100);c__Saccharomycetes(100);o__Saccharomycetales(100);f__Metschnikowiaceae(100);g__Metschnikowia(100);s__Metschnikowia_pulcherrima(100); | Metschnikowia | 13 | Saprotroph | \|Undefined Saprotroph\| | Yeast | NULL | Probable | As nectar-tap_saprotroph (Põlme et al. 2020); Decay sugar-rich substrates (Põlme et al. 2020) | Põlme S, et al. 2020. Fungal Diversity 105:1-16 ((https://doi.org/10.1007/s13225-020-00466-2)) |
| Otu56188 | 1 | k__Fungi(100);p__Ascomycota(100);c__Saccharomycetes(100);o__Saccharomycetales(100);f__Metschnikowiaceae(100);g__Metschnikowia(100);s__Metschnikowia_pulcherrima(100); | Metschnikowia | 13 | Saprotroph | \|Undefined Saprotroph\| | Yeast | NULL | Probable | As nectar-tap_saprotroph (Põlme et al. 2020); Decay sugar-rich substrates (Põlme et al. 2020) | Põlme S, et al. 2020. Fungal Diversity 105:1-16 ((https://doi.org/10.1007/s13225-020-00466-2)) |
| Otu56189 | 1 | k__Fungi(100);p__Ascomycota(100);c__Saccharomycetes(100);o__Saccharomycetales(100);f__Metschnikowiaceae(100);g__Metschnikowia(100);g__Metschnikowia_unclassified(100); | Metschnikowia | 13 | Saprotroph | \|Undefined Saprotroph\| | Yeast | NULL | Probable | As nectar-tap_saprotroph (Põlme et al. 2020); Decay sugar-rich substrates (Põlme et al. 2020) | Põlme S, et al. 2020. Fungal Diversity 105:1-16 ((https://doi.org/10.1007/s13225-020-00466-2)) |
| Otu56190 | 1 | k__Fungi(100);p__Ascomycota(100);c__Saccharomycetes(100);o__Saccharomycetales(100);f__Metschnikowiaceae(100);g__Metschnikowia(100);s__Metschnikowia_pulcherrima(100); | Metschnikowia | 13 | Saprotroph | \|Undefined Saprotroph\| | Yeast | NULL | Probable | As nectar-tap_saprotroph (Põlme et al. 2020); Decay sugar-rich substrates (Põlme et al. 2020) | Põlme S, et al. 2020. Fungal Diversity 105:1-16 ((https://doi.org/10.1007/s13225-020-00466-2)) |
| Otu56191 | 1 | k__Fungi(100);p__Ascomycota(100);c__Saccharomycetes(100);o__Saccharomycetales(100);f__Metschnikowiaceae(100);g__Metschnikowia(100);s__Metschnikowia_pulcherrima(100); | Metschnikowia | 13 | Saprotroph | \|Undefined Saprotroph\| | Yeast | NULL | Probable | As nectar-tap_saprotroph (Põlme et al. 2020); Decay sugar-rich substrates (Põlme et al. 2020) | Põlme S, et al. 2020. Fungal Diversity 105:1-16 ((https://doi.org/10.1007/s13225-020-00466-2)) |
| Otu56192 | 1 | k__Fungi(100);p__Ascomycota(100);c__Saccharomycetes(100);o__Saccharomycetales(100);f__Metschnikowiaceae(100);g__Metschnikowia(100);g__Metschnikowia_unclassified(100); | Metschnikowia | 13 | Saprotroph | \|Undefined Saprotroph\| | Yeast | NULL | Probable | As nectar-tap_saprotroph (Põlme et al. 2020); Decay sugar-rich substrates (Põlme et al. 2020) | Põlme S, et al. 2020. Fungal Diversity 105:1-16 ((https://doi.org/10.1007/s13225-020-00466-2)) |
| Otu56193 | 1 | k__Fungi(100);p__Ascomycota(100);c__Saccharomycetes(100);o__Saccharomycetales(100);f__Metschnikowiaceae(100);g__Metschnikowia(100);s__Metschnikowia_pulcherrima(100); | Metschnikowia | 13 | Saprotroph | \|Undefined Saprotroph\| | Yeast | NULL | Probable | As nectar-tap_saprotroph (Põlme et al. 2020); Decay sugar-rich substrates (Põlme et al. 2020) | Põlme S, et al. 2020. Fungal Diversity 105:1-16 ((https://doi.org/10.1007/s13225-020-00466-2)) |
| Otu56194 | 1 | k__Fungi(100);p__Ascomycota(100);c__Saccharomycetes(100);o__Saccharomycetales(100);f__Metschnikowiaceae(100);g__Metschnikowia(100);s__Metschnikowia_pulcherrima(100); | Metschnikowia | 13 | Saprotroph | \|Undefined Saprotroph\| | Yeast | NULL | Probable | As nectar-tap_saprotroph (Põlme et al. 2020); Decay sugar-rich substrates (Põlme et al. 2020) | Põlme S, et al. 2020. Fungal Diversity 105:1-16 ((https://doi.org/10.1007/s13225-020-00466-2)) |
| Otu56195 | 1 | k__Fungi(100);p__Ascomycota(100);c__Saccharomycetes(100);o__Saccharomycetales(100);f__Metschnikowiaceae(100);g__Metschnikowia(100);s__Metschnikowia_pulcherrima(100); | Metschnikowia | 13 | Saprotroph | \|Undefined Saprotroph\| | Yeast | NULL | Probable | As nectar-tap_saprotroph (Põlme et al. 2020); Decay sugar-rich substrates (Põlme et al. 2020) | Põlme S, et al. 2020. Fungal Diversity 105:1-16 ((https://doi.org/10.1007/s13225-020-00466-2)) |
| Otu56198 | 1 | k__Fungi(100);p__Ascomycota(100);c__Saccharomycetes(100);o__Saccharomycetales(100);f__Metschnikowiaceae(100);g__Metschnikowia(100);s__Metschnikowia_pulcherrima(100); | Metschnikowia | 13 | Saprotroph | \|Undefined Saprotroph\| | Yeast | NULL | Probable | As nectar-tap_saprotroph (Põlme et al. 2020); Decay sugar-rich substrates (Põlme et al. 2020) | Põlme S, et al. 2020. Fungal Diversity 105:1-16 ((https://doi.org/10.1007/s13225-020-00466-2)) |
| Otu56199 | 1 | k__Fungi(100);p__Ascomycota(100);c__Saccharomycetes(100);o__Saccharomycetales(100);f__Metschnikowiaceae(100);g__Metschnikowia(100);s__Metschnikowia_pulcherrima(100); | Metschnikowia | 13 | Saprotroph | \|Undefined Saprotroph\| | Yeast | NULL | Probable | As nectar-tap_saprotroph (Põlme et al. 2020); Decay sugar-rich substrates (Põlme et al. 2020) | Põlme S, et al. 2020. Fungal Diversity 105:1-16 ((https://doi.org/10.1007/s13225-020-00466-2)) |
| Otu56200 | 1 | k__Fungi(100);p__Ascomycota(100);c__Saccharomycetes(100);o__Saccharomycetales(100);f__Metschnikowiaceae(100);g__Metschnikowia(100);s__Metschnikowia_pulcherrima(100); | Metschnikowia | 13 | Saprotroph | \|Undefined Saprotroph\| | Yeast | NULL | Probable | As nectar-tap_saprotroph (Põlme et al. 2020); Decay sugar-rich substrates (Põlme et al. 2020) | Põlme S, et al. 2020. Fungal Diversity 105:1-16 ((https://doi.org/10.1007/s13225-020-00466-2)) |
| Otu56201 | 1 | k__Fungi(100);p__Ascomycota(100);c__Saccharomycetes(100);o__Saccharomycetales(100);f__Metschnikowiaceae(100);g__Metschnikowia(100);s__Metschnikowia_pulcherrima(100); | Metschnikowia | 13 | Saprotroph | \|Undefined Saprotroph\| | Yeast | NULL | Probable | As nectar-tap_saprotroph (Põlme et al. 2020); Decay sugar-rich substrates (Põlme et al. 2020) | Põlme S, et al. 2020. Fungal Diversity 105:1-16 ((https://doi.org/10.1007/s13225-020-00466-2)) |
| Otu56202 | 1 | k__Fungi(100);p__Ascomycota(100);c__Saccharomycetes(100);o__Saccharomycetales(100);f__Metschnikowiaceae(100);g__Metschnikowia(100);s__Metschnikowia_pulcherrima(100); | Metschnikowia | 13 | Saprotroph | \|Undefined Saprotroph\| | Yeast | NULL | Probable | As nectar-tap_saprotroph (Põlme et al. 2020); Decay sugar-rich substrates (Põlme et al. 2020) | Põlme S, et al. 2020. Fungal Diversity 105:1-16 ((https://doi.org/10.1007/s13225-020-00466-2)) |
| Otu56203 | 1 | k__Fungi(100);p__Ascomycota(100);c__Saccharomycetes(100);o__Saccharomycetales(100);f__Metschnikowiaceae(100);g__Metschnikowia(100);s__Metschnikowia_pulcherrima(100); | Metschnikowia | 13 | Saprotroph | \|Undefined Saprotroph\| | Yeast | NULL | Probable | As nectar-tap_saprotroph (Põlme et al. 2020); Decay sugar-rich substrates (Põlme et al. 2020) | Põlme S, et al. 2020. Fungal Diversity 105:1-16 ((https://doi.org/10.1007/s13225-020-00466-2)) |
| Otu56207 | 1 | k__Fungi(100);p__Ascomycota(100);c__Saccharomycetes(100);o__Saccharomycetales(100);f__Metschnikowiaceae(100);g__Metschnikowia(100);s__Metschnikowia_pulcherrima(100); | Metschnikowia | 13 | Saprotroph | \|Undefined Saprotroph\| | Yeast | NULL | Probable | As nectar-tap_saprotroph (Põlme et al. 2020); Decay sugar-rich substrates (Põlme et al. 2020) | Põlme S, et al. 2020. Fungal Diversity 105:1-16 ((https://doi.org/10.1007/s13225-020-00466-2)) |
| Otu56208 | 1 | k__Fungi(100);p__Ascomycota(100);c__Saccharomycetes(100);o__Saccharomycetales(100);f__Metschnikowiaceae(100);g__Metschnikowia(100);s__Metschnikowia_pulcherrima(100); | Metschnikowia | 13 | Saprotroph | \|Undefined Saprotroph\| | Yeast | NULL | Probable | As nectar-tap_saprotroph (Põlme et al. 2020); Decay sugar-rich substrates (Põlme et al. 2020) | Põlme S, et al. 2020. Fungal Diversity 105:1-16 ((https://doi.org/10.1007/s13225-020-00466-2)) |
| Otu56211 | 1 | k__Fungi(100);p__Ascomycota(100);c__Saccharomycetes(100);o__Saccharomycetales(100);f__Metschnikowiaceae(100);g__Metschnikowia(100);s__Metschnikowia_pulcherrima(100); | Metschnikowia | 13 | Saprotroph | \|Undefined Saprotroph\| | Yeast | NULL | Probable | As nectar-tap_saprotroph (Põlme et al. 2020); Decay sugar-rich substrates (Põlme et al. 2020) | Põlme S, et al. 2020. Fungal Diversity 105:1-16 ((https://doi.org/10.1007/s13225-020-00466-2)) |
| Otu56214 | 1 | k__Fungi(100);p__Ascomycota(100);c__Saccharomycetes(100);o__Saccharomycetales(100);f__Metschnikowiaceae(100);g__Metschnikowia(100);s__Metschnikowia_pulcherrima(100); | Metschnikowia | 13 | Saprotroph | \|Undefined Saprotroph\| | Yeast | NULL | Probable | As nectar-tap_saprotroph (Põlme et al. 2020); Decay sugar-rich substrates (Põlme et al. 2020) | Põlme S, et al. 2020. Fungal Diversity 105:1-16 ((https://doi.org/10.1007/s13225-020-00466-2)) |
| Otu56219 | 1 | k__Fungi(100);p__Ascomycota(100);c__Saccharomycetes(100);o__Saccharomycetales(100);f__Metschnikowiaceae(100);g__Metschnikowia(100);s__Metschnikowia_pulcherrima(100); | Metschnikowia | 13 | Saprotroph | \|Undefined Saprotroph\| | Yeast | NULL | Probable | As nectar-tap_saprotroph (Põlme et al. 2020); Decay sugar-rich substrates (Põlme et al. 2020) | Põlme S, et al. 2020. Fungal Diversity 105:1-16 ((https://doi.org/10.1007/s13225-020-00466-2)) |
| Otu56222 | 1 | k__Fungi(100);p__Ascomycota(100);c__Saccharomycetes(100);o__Saccharomycetales(100);f__Metschnikowiaceae(100);g__Metschnikowia(100);s__Metschnikowia_pulcherrima(100); | Metschnikowia | 13 | Saprotroph | \|Undefined Saprotroph\| | Yeast | NULL | Probable | As nectar-tap_saprotroph (Põlme et al. 2020); Decay sugar-rich substrates (Põlme et al. 2020) | Põlme S, et al. 2020. Fungal Diversity 105:1-16 ((https://doi.org/10.1007/s13225-020-00466-2)) |
| Otu56223 | 1 | k__Fungi(100);p__Ascomycota(100);c__Saccharomycetes(100);o__Saccharomycetales(100);f__Metschnikowiaceae(100);g__Metschnikowia(100);g__Metschnikowia_unclassified(100); | Metschnikowia | 13 | Saprotroph | \|Undefined Saprotroph\| | Yeast | NULL | Probable | As nectar-tap_saprotroph (Põlme et al. 2020); Decay sugar-rich substrates (Põlme et al. 2020) | Põlme S, et al. 2020. Fungal Diversity 105:1-16 ((https://doi.org/10.1007/s13225-020-00466-2)) |
| Otu56224 | 1 | k__Fungi(100);p__Ascomycota(100);c__Saccharomycetes(100);o__Saccharomycetales(100);f__Metschnikowiaceae(100);g__Metschnikowia(100);s__Metschnikowia_pulcherrima(100); | Metschnikowia | 13 | Saprotroph | \|Undefined Saprotroph\| | Yeast | NULL | Probable | As nectar-tap_saprotroph (Põlme et al. 2020); Decay sugar-rich substrates (Põlme et al. 2020) | Põlme S, et al. 2020. Fungal Diversity 105:1-16 ((https://doi.org/10.1007/s13225-020-00466-2)) |
| Otu56225 | 1 | k__Fungi(100);p__Ascomycota(100);c__Saccharomycetes(100);o__Saccharomycetales(100);f__Metschnikowiaceae(100);g__Metschnikowia(100);s__Metschnikowia_pulcherrima(100); | Metschnikowia | 13 | Saprotroph | \|Undefined Saprotroph\| | Yeast | NULL | Probable | As nectar-tap_saprotroph (Põlme et al. 2020); Decay sugar-rich substrates (Põlme et al. 2020) | Põlme S, et al. 2020. Fungal Diversity 105:1-16 ((https://doi.org/10.1007/s13225-020-00466-2)) |
| Otu56226 | 1 | k__Fungi(100);p__Ascomycota(100);c__Saccharomycetes(100);o__Saccharomycetales(100);f__Metschnikowiaceae(100);g__Metschnikowia(100);s__Metschnikowia_pulcherrima(100); | Metschnikowia | 13 | Saprotroph | \|Undefined Saprotroph\| | Yeast | NULL | Probable | As nectar-tap_saprotroph (Põlme et al. 2020); Decay sugar-rich substrates (Põlme et al. 2020) | Põlme S, et al. 2020. Fungal Diversity 105:1-16 ((https://doi.org/10.1007/s13225-020-00466-2)) |
| Otu56227 | 1 | k__Fungi(100);p__Ascomycota(100);c__Saccharomycetes(100);o__Saccharomycetales(100);f__Metschnikowiaceae(100);g__Metschnikowia(100);s__Metschnikowia_pulcherrima(100); | Metschnikowia | 13 | Saprotroph | \|Undefined Saprotroph\| | Yeast | NULL | Probable | As nectar-tap_saprotroph (Põlme et al. 2020); Decay sugar-rich substrates (Põlme et al. 2020) | Põlme S, et al. 2020. Fungal Diversity 105:1-16 ((https://doi.org/10.1007/s13225-020-00466-2)) |
| Otu56228 | 1 | k__Fungi(100);p__Ascomycota(100);c__Saccharomycetes(100);o__Saccharomycetales(100);f__Metschnikowiaceae(100);g__Metschnikowia(100);s__Metschnikowia_pulcherrima(100); | Metschnikowia | 13 | Saprotroph | \|Undefined Saprotroph\| | Yeast | NULL | Probable | As nectar-tap_saprotroph (Põlme et al. 2020); Decay sugar-rich substrates (Põlme et al. 2020) | Põlme S, et al. 2020. Fungal Diversity 105:1-16 ((https://doi.org/10.1007/s13225-020-00466-2)) |
| Otu56229 | 1 | k__Fungi(100);p__Ascomycota(100);c__Saccharomycetes(100);o__Saccharomycetales(100);f__Metschnikowiaceae(100);g__Metschnikowia(100);s__Metschnikowia_pulcherrima(100); | Metschnikowia | 13 | Saprotroph | \|Undefined Saprotroph\| | Yeast | NULL | Probable | As nectar-tap_saprotroph (Põlme et al. 2020); Decay sugar-rich substrates (Põlme et al. 2020) | Põlme S, et al. 2020. Fungal Diversity 105:1-16 ((https://doi.org/10.1007/s13225-020-00466-2)) |
| Otu56230 | 1 | k__Fungi(100);p__Ascomycota(100);c__Saccharomycetes(100);o__Saccharomycetales(100);f__Metschnikowiaceae(100);g__Metschnikowia(100);s__Metschnikowia_pulcherrima(100); | Metschnikowia | 13 | Saprotroph | \|Undefined Saprotroph\| | Yeast | NULL | Probable | As nectar-tap_saprotroph (Põlme et al. 2020); Decay sugar-rich substrates (Põlme et al. 2020) | Põlme S, et al. 2020. Fungal Diversity 105:1-16 ((https://doi.org/10.1007/s13225-020-00466-2)) |
| Otu56231 | 1 | k__Fungi(100);p__Ascomycota(100);c__Saccharomycetes(100);o__Saccharomycetales(100);f__Metschnikowiaceae(100);g__Metschnikowia(100);g__Metschnikowia_unclassified(100); | Metschnikowia | 13 | Saprotroph | \|Undefined Saprotroph\| | Yeast | NULL | Probable | As nectar-tap_saprotroph (Põlme et al. 2020); Decay sugar-rich substrates (Põlme et al. 2020) | Põlme S, et al. 2020. Fungal Diversity 105:1-16 ((https://doi.org/10.1007/s13225-020-00466-2)) |
| Otu56232 | 1 | k__Fungi(100);p__Ascomycota(100);c__Saccharomycetes(100);o__Saccharomycetales(100);f__Metschnikowiaceae(100);g__Metschnikowia(100);s__Metschnikowia_pulcherrima(100); | Metschnikowia | 13 | Saprotroph | \|Undefined Saprotroph\| | Yeast | NULL | Probable | As nectar-tap_saprotroph (Põlme et al. 2020); Decay sugar-rich substrates (Põlme et al. 2020) | Põlme S, et al. 2020. Fungal Diversity 105:1-16 ((https://doi.org/10.1007/s13225-020-00466-2)) |
| Otu56233 | 1 | k__Fungi(100);p__Ascomycota(100);c__Saccharomycetes(100);o__Saccharomycetales(100);f__Metschnikowiaceae(100);g__Metschnikowia(100);s__Metschnikowia_pulcherrima(100); | Metschnikowia | 13 | Saprotroph | \|Undefined Saprotroph\| | Yeast | NULL | Probable | As nectar-tap_saprotroph (Põlme et al. 2020); Decay sugar-rich substrates (Põlme et al. 2020) | Põlme S, et al. 2020. Fungal Diversity 105:1-16 ((https://doi.org/10.1007/s13225-020-00466-2)) |
| Otu56234 | 1 | k__Fungi(100);p__Basidiomycota(100);c__Tremellomycetes(100);o__Tremellales(100);f__Tremellales_family_Incertae_sedis(100);g__Tremella(100);s__Tremella_mesenterica(100); | Tremella | 13 | Pathotroph | \|Fungal Parasite\|-Lichen Parasite | Dimorphic-Facultative Yeast-Microfungus-Tremelloid | NULL | Probable | As mycoparasite (Põlme et al. 2020); Decay fungal material (Põlme et al. 2020) | Tedersoo L, et al. 2014. Science 346:e1256688 ((https://doi.org/10.1126/science.1256688)); Põlme S, et al. 2020. Fungal Diversity 105:1-16 ((https://doi.org/10.1007/s13225-020-00466-2)) |
| Otu56235 | 1 | k__Fungi(100);p__Ascomycota(100);c__Saccharomycetes(100);o__Saccharomycetales(100);f__Metschnikowiaceae(100);g__Metschnikowia(100);s__Metschnikowia_pulcherrima(100); | Metschnikowia | 13 | Saprotroph | \|Undefined Saprotroph\| | Yeast | NULL | Probable | As nectar-tap_saprotroph (Põlme et al. 2020); Decay sugar-rich substrates (Põlme et al. 2020) | Põlme S, et al. 2020. Fungal Diversity 105:1-16 ((https://doi.org/10.1007/s13225-020-00466-2)) |
| Otu56236 | 1 | k__Fungi(100);p__Ascomycota(100);c__Saccharomycetes(100);o__Saccharomycetales(100);f__Metschnikowiaceae(100);g__Metschnikowia(100);s__Metschnikowia_pulcherrima(100); | Metschnikowia | 13 | Saprotroph | \|Undefined Saprotroph\| | Yeast | NULL | Probable | As nectar-tap_saprotroph (Põlme et al. 2020); Decay sugar-rich substrates (Põlme et al. 2020) | Põlme S, et al. 2020. Fungal Diversity 105:1-16 ((https://doi.org/10.1007/s13225-020-00466-2)) |
| Otu56238 | 1 | k__Fungi(100);p__Ascomycota(100);c__Saccharomycetes(100);o__Saccharomycetales(100);f__Metschnikowiaceae(100);g__Metschnikowia(100);s__Metschnikowia_pulcherrima(100); | Metschnikowia | 13 | Saprotroph | \|Undefined Saprotroph\| | Yeast | NULL | Probable | As nectar-tap_saprotroph (Põlme et al. 2020); Decay sugar-rich substrates (Põlme et al. 2020) | Põlme S, et al. 2020. Fungal Diversity 105:1-16 ((https://doi.org/10.1007/s13225-020-00466-2)) |
| Otu56239 | 1 | k__Fungi(100);p__Ascomycota(100);c__Saccharomycetes(100);o__Saccharomycetales(100);f__Metschnikowiaceae(100);g__Metschnikowia(100);s__Metschnikowia_pulcherrima(100); | Metschnikowia | 13 | Saprotroph | \|Undefined Saprotroph\| | Yeast | NULL | Probable | As nectar-tap_saprotroph (Põlme et al. 2020); Decay sugar-rich substrates (Põlme et al. 2020) | Põlme S, et al. 2020. Fungal Diversity 105:1-16 ((https://doi.org/10.1007/s13225-020-00466-2)) |
| Otu56241 | 1 | k__Fungi(100);p__Ascomycota(100);c__Saccharomycetes(100);o__Saccharomycetales(100);f__Metschnikowiaceae(100);g__Metschnikowia(100);s__Metschnikowia_pulcherrima(100); | Metschnikowia | 13 | Saprotroph | \|Undefined Saprotroph\| | Yeast | NULL | Probable | As nectar-tap_saprotroph (Põlme et al. 2020); Decay sugar-rich substrates (Põlme et al. 2020) | Põlme S, et al. 2020. Fungal Diversity 105:1-16 ((https://doi.org/10.1007/s13225-020-00466-2)) |
| Otu56243 | 1 | k__Fungi(100);p__Ascomycota(100);c__Saccharomycetes(100);o__Saccharomycetales(100);f__Metschnikowiaceae(100);g__Metschnikowia(100);s__Metschnikowia_pulcherrima(100); | Metschnikowia | 13 | Saprotroph | \|Undefined Saprotroph\| | Yeast | NULL | Probable | As nectar-tap_saprotroph (Põlme et al. 2020); Decay sugar-rich substrates (Põlme et al. 2020) | Põlme S, et al. 2020. Fungal Diversity 105:1-16 ((https://doi.org/10.1007/s13225-020-00466-2)) |
| Otu56244 | 1 | k__Fungi(100);p__Ascomycota(100);c__Saccharomycetes(100);o__Saccharomycetales(100);f__Metschnikowiaceae(100);g__Metschnikowia(100);s__Metschnikowia_pulcherrima(100); | Metschnikowia | 13 | Saprotroph | \|Undefined Saprotroph\| | Yeast | NULL | Probable | As nectar-tap_saprotroph (Põlme et al. 2020); Decay sugar-rich substrates (Põlme et al. 2020) | Põlme S, et al. 2020. Fungal Diversity 105:1-16 ((https://doi.org/10.1007/s13225-020-00466-2)) |
| Otu56245 | 1 | k__Fungi(100);p__Ascomycota(100);c__Saccharomycetes(100);o__Saccharomycetales(100);f__Metschnikowiaceae(100);g__Metschnikowia(100);g__Metschnikowia_unclassified(100); | Metschnikowia | 13 | Saprotroph | \|Undefined Saprotroph\| | Yeast | NULL | Probable | As nectar-tap_saprotroph (Põlme et al. 2020); Decay sugar-rich substrates (Põlme et al. 2020) | Põlme S, et al. 2020. Fungal Diversity 105:1-16 ((https://doi.org/10.1007/s13225-020-00466-2)) |
| Otu56246 | 1 | k__Fungi(100);p__Ascomycota(100);c__Saccharomycetes(100);o__Saccharomycetales(100);f__Metschnikowiaceae(100);g__Metschnikowia(100);s__Metschnikowia_pulcherrima(100); | Metschnikowia | 13 | Saprotroph | \|Undefined Saprotroph\| | Yeast | NULL | Probable | As nectar-tap_saprotroph (Põlme et al. 2020); Decay sugar-rich substrates (Põlme et al. 2020) | Põlme S, et al. 2020. Fungal Diversity 105:1-16 ((https://doi.org/10.1007/s13225-020-00466-2)) |
| Otu56249 | 1 | k__Fungi(100);p__Ascomycota(100);c__Saccharomycetes(100);o__Saccharomycetales(100);f__Saccharomycetales_family_Incertae_sedis(100);g__Candida(100);s__Candida_cretensis(100); | Candida | 13 | Pathotroph-Saprotroph-Symbiotroph | Animal Pathogen-Endophyte-Endosymbiont-Epiphyte-\|Undefined Saprotroph\| | Dimorphic Yeast-Yeast | NULL | Probable | As nectar-tap_saprotroph (Põlme et al. 2020); Decay sugar-rich substrates (Põlme et al. 2020); This asexual genus is polyphyletic and contain species that function very differently depend on what they are. Caution should be taken when interpreting guilds for Candida species. | Manolakaki D, et al. 2010. Virulence. 1: 367-375 ((https://doi.org/10.4161/viru.1.5.12796)); Mekbib SB, et al. 2011. Tropical Plant Pathology 36:233-240 ((https://doi.org/10.1590/S1982-56762011000400004)); Põlme S, et al. 2020. Fungal Diversity 105:1-16 ((https://doi.org/10.1007/s13225-020-00466-2)) |
| Otu56251 | 1 | k__Fungi(100);p__Ascomycota(100);c__Saccharomycetes(100);o__Saccharomycetales(100);f__Metschnikowiaceae(100);g__Metschnikowia(100);s__Metschnikowia_pulcherrima(100); | Metschnikowia | 13 | Saprotroph | \|Undefined Saprotroph\| | Yeast | NULL | Probable | As nectar-tap_saprotroph (Põlme et al. 2020); Decay sugar-rich substrates (Põlme et al. 2020) | Põlme S, et al. 2020. Fungal Diversity 105:1-16 ((https://doi.org/10.1007/s13225-020-00466-2)) |
| Otu56252 | 1 | k__Fungi(100);p__Ascomycota(100);c__Saccharomycetes(100);o__Saccharomycetales(100);f__Metschnikowiaceae(100);g__Metschnikowia(100);g__Metschnikowia_unclassified(100); | Metschnikowia | 13 | Saprotroph | \|Undefined Saprotroph\| | Yeast | NULL | Probable | As nectar-tap_saprotroph (Põlme et al. 2020); Decay sugar-rich substrates (Põlme et al. 2020) | Põlme S, et al. 2020. Fungal Diversity 105:1-16 ((https://doi.org/10.1007/s13225-020-00466-2)) |
| Otu56253 | 1 | k__Fungi(100);p__Ascomycota(100);c__Saccharomycetes(100);o__Saccharomycetales(100);f__Metschnikowiaceae(100);g__Metschnikowia(100);s__Metschnikowia_pulcherrima(100); | Metschnikowia | 13 | Saprotroph | \|Undefined Saprotroph\| | Yeast | NULL | Probable | As nectar-tap_saprotroph (Põlme et al. 2020); Decay sugar-rich substrates (Põlme et al. 2020) | Põlme S, et al. 2020. Fungal Diversity 105:1-16 ((https://doi.org/10.1007/s13225-020-00466-2)) |
| Otu56254 | 1 | k__Fungi(100);p__Ascomycota(100);c__Saccharomycetes(100);o__Saccharomycetales(100);f__Metschnikowiaceae(100);g__Metschnikowia(100);s__Metschnikowia_pulcherrima(100); | Metschnikowia | 13 | Saprotroph | \|Undefined Saprotroph\| | Yeast | NULL | Probable | As nectar-tap_saprotroph (Põlme et al. 2020); Decay sugar-rich substrates (Põlme et al. 2020) | Põlme S, et al. 2020. Fungal Diversity 105:1-16 ((https://doi.org/10.1007/s13225-020-00466-2)) |
| Otu56255 | 1 | k__Fungi(100);p__Ascomycota(100);c__Saccharomycetes(100);o__Saccharomycetales(100);f__Saccharomycetales_family_Incertae_sedis(100);g__Candida(100);g__Candida_unclassified(100); | Candida | 13 | Pathotroph-Saprotroph-Symbiotroph | Animal Pathogen-Endophyte-Endosymbiont-Epiphyte-\|Undefined Saprotroph\| | Dimorphic Yeast-Yeast | NULL | Probable | As nectar-tap_saprotroph (Põlme et al. 2020); Decay sugar-rich substrates (Põlme et al. 2020); This asexual genus is polyphyletic and contain species that function very differently depend on what they are. Caution should be taken when interpreting guilds for Candida species. | Manolakaki D, et al. 2010. Virulence. 1: 367-375 ((https://doi.org/10.4161/viru.1.5.12796)); Mekbib SB, et al. 2011. Tropical Plant Pathology 36:233-240 ((https://doi.org/10.1590/S1982-56762011000400004)); Põlme S, et al. 2020. Fungal Diversity 105:1-16 ((https://doi.org/10.1007/s13225-020-00466-2)) |
| Otu56256 | 1 | k__Fungi(100);p__Ascomycota(100);c__Saccharomycetes(100);o__Saccharomycetales(100);f__Metschnikowiaceae(100);g__Metschnikowia(100);s__Metschnikowia_pulcherrima(100); | Metschnikowia | 13 | Saprotroph | \|Undefined Saprotroph\| | Yeast | NULL | Probable | As nectar-tap_saprotroph (Põlme et al. 2020); Decay sugar-rich substrates (Põlme et al. 2020) | Põlme S, et al. 2020. Fungal Diversity 105:1-16 ((https://doi.org/10.1007/s13225-020-00466-2)) |
| Otu56257 | 1 | k__Fungi(100);p__Ascomycota(100);c__Saccharomycetes(100);o__Saccharomycetales(100);f__Saccharomycetales_family_Incertae_sedis(100);g__Candida(100);g__Candida_unclassified(100); | Candida | 13 | Pathotroph-Saprotroph-Symbiotroph | Animal Pathogen-Endophyte-Endosymbiont-Epiphyte-\|Undefined Saprotroph\| | Dimorphic Yeast-Yeast | NULL | Probable | As nectar-tap_saprotroph (Põlme et al. 2020); Decay sugar-rich substrates (Põlme et al. 2020); This asexual genus is polyphyletic and contain species that function very differently depend on what they are. Caution should be taken when interpreting guilds for Candida species. | Manolakaki D, et al. 2010. Virulence. 1: 367-375 ((https://doi.org/10.4161/viru.1.5.12796)); Mekbib SB, et al. 2011. Tropical Plant Pathology 36:233-240 ((https://doi.org/10.1590/S1982-56762011000400004)); Põlme S, et al. 2020. Fungal Diversity 105:1-16 ((https://doi.org/10.1007/s13225-020-00466-2)) |
| Otu56258 | 1 | k__Fungi(100);p__Ascomycota(100);c__Saccharomycetes(100);o__Saccharomycetales(100);f__Metschnikowiaceae(100);g__Metschnikowia(100);s__Metschnikowia_pulcherrima(100); | Metschnikowia | 13 | Saprotroph | \|Undefined Saprotroph\| | Yeast | NULL | Probable | As nectar-tap_saprotroph (Põlme et al. 2020); Decay sugar-rich substrates (Põlme et al. 2020) | Põlme S, et al. 2020. Fungal Diversity 105:1-16 ((https://doi.org/10.1007/s13225-020-00466-2)) |
| Otu56260 | 1 | k__Fungi(100);p__Ascomycota(100);c__Saccharomycetes(100);o__Saccharomycetales(100);f__Saccharomycetales_family_Incertae_sedis(100);g__Candida(100);s__Candida_cretensis(100); | Candida | 13 | Pathotroph-Saprotroph-Symbiotroph | Animal Pathogen-Endophyte-Endosymbiont-Epiphyte-\|Undefined Saprotroph\| | Dimorphic Yeast-Yeast | NULL | Probable | As nectar-tap_saprotroph (Põlme et al. 2020); Decay sugar-rich substrates (Põlme et al. 2020); This asexual genus is polyphyletic and contain species that function very differently depend on what they are. Caution should be taken when interpreting guilds for Candida species. | Manolakaki D, et al. 2010. Virulence. 1: 367-375 ((https://doi.org/10.4161/viru.1.5.12796)); Mekbib SB, et al. 2011. Tropical Plant Pathology 36:233-240 ((https://doi.org/10.1590/S1982-56762011000400004)); Põlme S, et al. 2020. Fungal Diversity 105:1-16 ((https://doi.org/10.1007/s13225-020-00466-2)) |
| Otu56261 | 1 | k__Fungi(100);p__Basidiomycota(100);c__Tremellomycetes(100);o__Tremellales(100);f__Tremellales_family_Incertae_sedis(100);g__Hannaella(100);s__Hannaella_luteola(100); | Hannaella | 13 | Pathotroph-Saprotroph | \|Animal Parasite\|-Undefined Saprotroph | Yeast | NULL | Probable | As animal_decomposer (Põlme et al. 2020); Decay animal material (Põlme et al. 2020) | Põlme S, et al. 2020. Fungal Diversity 105:1-16 ((https://doi.org/10.1007/s13225-020-00466-2)) |
| Otu56263 | 1 | k__Fungi(100);p__Ascomycota(100);c__Saccharomycetes(100);o__Saccharomycetales(100);f__Saccharomycetales_family_Incertae_sedis(100);g__Candida(100);g__Candida_unclassified(100); | Candida | 13 | Pathotroph-Saprotroph-Symbiotroph | Animal Pathogen-Endophyte-Endosymbiont-Epiphyte-\|Undefined Saprotroph\| | Dimorphic Yeast-Yeast | NULL | Probable | As nectar-tap_saprotroph (Põlme et al. 2020); Decay sugar-rich substrates (Põlme et al. 2020); This asexual genus is polyphyletic and contain species that function very differently depend on what they are. Caution should be taken when interpreting guilds for Candida species. | Manolakaki D, et al. 2010. Virulence. 1: 367-375 ((https://doi.org/10.4161/viru.1.5.12796)); Mekbib SB, et al. 2011. Tropical Plant Pathology 36:233-240 ((https://doi.org/10.1590/S1982-56762011000400004)); Põlme S, et al. 2020. Fungal Diversity 105:1-16 ((https://doi.org/10.1007/s13225-020-00466-2)) |
| Otu56264 | 1 | k__Fungi(100);p__Ascomycota(100);c__Saccharomycetes(100);o__Saccharomycetales(100);f__Metschnikowiaceae(100);g__Metschnikowia(100);g__Metschnikowia_unclassified(100); | Metschnikowia | 13 | Saprotroph | \|Undefined Saprotroph\| | Yeast | NULL | Probable | As nectar-tap_saprotroph (Põlme et al. 2020); Decay sugar-rich substrates (Põlme et al. 2020) | Põlme S, et al. 2020. Fungal Diversity 105:1-16 ((https://doi.org/10.1007/s13225-020-00466-2)) |
| Otu56265 | 1 | k__Fungi(100);p__Ascomycota(100);c__Saccharomycetes(100);o__Saccharomycetales(100);f__Metschnikowiaceae(100);g__Metschnikowia(100);s__Metschnikowia_pulcherrima(100); | Metschnikowia | 13 | Saprotroph | \|Undefined Saprotroph\| | Yeast | NULL | Probable | As nectar-tap_saprotroph (Põlme et al. 2020); Decay sugar-rich substrates (Põlme et al. 2020) | Põlme S, et al. 2020. Fungal Diversity 105:1-16 ((https://doi.org/10.1007/s13225-020-00466-2)) |
| Otu56267 | 1 | k__Fungi(100);p__Ascomycota(100);c__Saccharomycetes(100);o__Saccharomycetales(100);f__Metschnikowiaceae(100);g__Metschnikowia(100);s__Metschnikowia_pulcherrima(100); | Metschnikowia | 13 | Saprotroph | \|Undefined Saprotroph\| | Yeast | NULL | Probable | As nectar-tap_saprotroph (Põlme et al. 2020); Decay sugar-rich substrates (Põlme et al. 2020) | Põlme S, et al. 2020. Fungal Diversity 105:1-16 ((https://doi.org/10.1007/s13225-020-00466-2)) |
| Otu56268 | 1 | k__Fungi(100);p__Ascomycota(100);c__Saccharomycetes(100);o__Saccharomycetales(100);f__Metschnikowiaceae(100);g__Metschnikowia(100);s__Metschnikowia_pulcherrima(100); | Metschnikowia | 13 | Saprotroph | \|Undefined Saprotroph\| | Yeast | NULL | Probable | As nectar-tap_saprotroph (Põlme et al. 2020); Decay sugar-rich substrates (Põlme et al. 2020) | Põlme S, et al. 2020. Fungal Diversity 105:1-16 ((https://doi.org/10.1007/s13225-020-00466-2)) |
| Otu56269 | 1 | k__Fungi(100);p__Ascomycota(100);c__Saccharomycetes(100);o__Saccharomycetales(100);f__Metschnikowiaceae(100);g__Metschnikowia(100);s__Metschnikowia_pulcherrima(100); | Metschnikowia | 13 | Saprotroph | \|Undefined Saprotroph\| | Yeast | NULL | Probable | As nectar-tap_saprotroph (Põlme et al. 2020); Decay sugar-rich substrates (Põlme et al. 2020) | Põlme S, et al. 2020. Fungal Diversity 105:1-16 ((https://doi.org/10.1007/s13225-020-00466-2)) |
| Otu56270 | 1 | k__Fungi(100);p__Ascomycota(100);c__Saccharomycetes(100);o__Saccharomycetales(100);f__Metschnikowiaceae(100);g__Metschnikowia(100);g__Metschnikowia_unclassified(100); | Metschnikowia | 13 | Saprotroph | \|Undefined Saprotroph\| | Yeast | NULL | Probable | As nectar-tap_saprotroph (Põlme et al. 2020); Decay sugar-rich substrates (Põlme et al. 2020) | Põlme S, et al. 2020. Fungal Diversity 105:1-16 ((https://doi.org/10.1007/s13225-020-00466-2)) |
| Otu56271 | 1 | k__Fungi(100);p__Ascomycota(100);c__Saccharomycetes(100);o__Saccharomycetales(100);f__Metschnikowiaceae(100);g__Metschnikowia(100);s__Metschnikowia_pulcherrima(100); | Metschnikowia | 13 | Saprotroph | \|Undefined Saprotroph\| | Yeast | NULL | Probable | As nectar-tap_saprotroph (Põlme et al. 2020); Decay sugar-rich substrates (Põlme et al. 2020) | Põlme S, et al. 2020. Fungal Diversity 105:1-16 ((https://doi.org/10.1007/s13225-020-00466-2)) |
| Otu56272 | 1 | k__Fungi(100);p__Ascomycota(100);c__Saccharomycetes(100);o__Saccharomycetales(100);f__Metschnikowiaceae(100);g__Metschnikowia(100);s__Metschnikowia_pulcherrima(100); | Metschnikowia | 13 | Saprotroph | \|Undefined Saprotroph\| | Yeast | NULL | Probable | As nectar-tap_saprotroph (Põlme et al. 2020); Decay sugar-rich substrates (Põlme et al. 2020) | Põlme S, et al. 2020. Fungal Diversity 105:1-16 ((https://doi.org/10.1007/s13225-020-00466-2)) |
| Otu56273 | 1 | k__Fungi(100);p__Ascomycota(100);c__Saccharomycetes(100);o__Saccharomycetales(100);f__Metschnikowiaceae(100);g__Metschnikowia(100);g__Metschnikowia_unclassified(100); | Metschnikowia | 13 | Saprotroph | \|Undefined Saprotroph\| | Yeast | NULL | Probable | As nectar-tap_saprotroph (Põlme et al. 2020); Decay sugar-rich substrates (Põlme et al. 2020) | Põlme S, et al. 2020. Fungal Diversity 105:1-16 ((https://doi.org/10.1007/s13225-020-00466-2)) |
| Otu56274 | 1 | k__Fungi(100);p__Ascomycota(100);c__Saccharomycetes(100);o__Saccharomycetales(100);f__Metschnikowiaceae(100);g__Metschnikowia(100);g__Metschnikowia_unclassified(100); | Metschnikowia | 13 | Saprotroph | \|Undefined Saprotroph\| | Yeast | NULL | Probable | As nectar-tap_saprotroph (Põlme et al. 2020); Decay sugar-rich substrates (Põlme et al. 2020) | Põlme S, et al. 2020. Fungal Diversity 105:1-16 ((https://doi.org/10.1007/s13225-020-00466-2)) |
| Otu56275 | 1 | k__Fungi(100);p__Ascomycota(100);c__Saccharomycetes(100);o__Saccharomycetales(100);f__Metschnikowiaceae(100);g__Metschnikowia(100);g__Metschnikowia_unclassified(100); | Metschnikowia | 13 | Saprotroph | \|Undefined Saprotroph\| | Yeast | NULL | Probable | As nectar-tap_saprotroph (Põlme et al. 2020); Decay sugar-rich substrates (Põlme et al. 2020) | Põlme S, et al. 2020. Fungal Diversity 105:1-16 ((https://doi.org/10.1007/s13225-020-00466-2)) |
| Otu56277 | 1 | k__Fungi(100);p__Ascomycota(100);c__Saccharomycetes(100);o__Saccharomycetales(100);f__Metschnikowiaceae(100);g__Metschnikowia(100);s__Metschnikowia_pulcherrima(100); | Metschnikowia | 13 | Saprotroph | \|Undefined Saprotroph\| | Yeast | NULL | Probable | As nectar-tap_saprotroph (Põlme et al. 2020); Decay sugar-rich substrates (Põlme et al. 2020) | Põlme S, et al. 2020. Fungal Diversity 105:1-16 ((https://doi.org/10.1007/s13225-020-00466-2)) |
| Otu56279 | 1 | k__Fungi(100);p__Ascomycota(100);c__Saccharomycetes(100);o__Saccharomycetales(100);f__Metschnikowiaceae(100);g__Metschnikowia(100);s__Metschnikowia_pulcherrima(100); | Metschnikowia | 13 | Saprotroph | \|Undefined Saprotroph\| | Yeast | NULL | Probable | As nectar-tap_saprotroph (Põlme et al. 2020); Decay sugar-rich substrates (Põlme et al. 2020) | Põlme S, et al. 2020. Fungal Diversity 105:1-16 ((https://doi.org/10.1007/s13225-020-00466-2)) |

| | | | | | | | | | | | |
|---|---|---|---|---|---|---|---|---|---|---|---|
| Otu56281 | 1 | k__Fungi(100);p__Ascomycota(100);c__Saccharomycetes(100);o__Saccharomycetales(100);f__Metschnikowiaceae(100);g__Metschnikowia(100);s__Metschnikowia_pulcherrima(100); | Metschnikowia | 13 | Saprotroph | \|Undefined Saprotroph\| | Yeast | NULL | Probable | As nectar-tap_saprotroph (Põlme et al. 2020); Decay sugar-rich substrates (Põlme et al. 2020) | Põlme S, et al. 2020. Fungal Diversity 105:1-16 ((https://doi.org/10.1007/s13225-020-00466-2)) |
| Otu56282 | 1 | k__Fungi(100);p__Ascomycota(100);c__Saccharomycetes(100);o__Saccharomycetales(100);f__Metschnikowiaceae(100);g__Metschnikowia(100);s__Metschnikowia_pulcherrima(100); | Metschnikowia | 13 | Saprotroph | \|Undefined Saprotroph\| | Yeast | NULL | Probable | As nectar-tap_saprotroph (Põlme et al. 2020); Decay sugar-rich substrates (Põlme et al. 2020) | Põlme S, et al. 2020. Fungal Diversity 105:1-16 ((https://doi.org/10.1007/s13225-020-00466-2)) |
| Otu56283 | 1 | k__Fungi(100);p__Ascomycota(100);c__Saccharomycetes(100);o__Saccharomycetales(100);f__Metschnikowiaceae(100);g__Metschnikowia(100);s__Metschnikowia_pulcherrima(100); | Metschnikowia | 13 | Saprotroph | \|Undefined Saprotroph\| | Yeast | NULL | Probable | As nectar-tap_saprotroph (Põlme et al. 2020); Decay sugar-rich substrates (Põlme et al. 2020) | Põlme S, et al. 2020. Fungal Diversity 105:1-16 ((https://doi.org/10.1007/s13225-020-00466-2)) |
| Otu56284 | 1 | k__Fungi(100);p__Ascomycota(100);c__Saccharomycetes(100);o__Saccharomycetales(100);f__Metschnikowiaceae(100);g__Metschnikowia(100);s__Metschnikowia_pulcherrima(100); | Metschnikowia | 13 | Saprotroph | \|Undefined Saprotroph\| | Yeast | NULL | Probable | As nectar-tap_saprotroph (Põlme et al. 2020); Decay sugar-rich substrates (Põlme et al. 2020) | Põlme S, et al. 2020. Fungal Diversity 105:1-16 ((https://doi.org/10.1007/s13225-020-00466-2)) |
| Otu56285 | 1 | k__Fungi(100);p__Ascomycota(100);c__Saccharomycetes(100);o__Saccharomycetales(100);f__Metschnikowiaceae(100);g__Metschnikowia(100);s__Metschnikowia_pulcherrima(100); | Metschnikowia | 13 | Saprotroph | \|Undefined Saprotroph\| | Yeast | NULL | Probable | As nectar-tap_saprotroph (Põlme et al. 2020); Decay sugar-rich substrates (Põlme et al. 2020) | Põlme S, et al. 2020. Fungal Diversity 105:1-16 ((https://doi.org/10.1007/s13225-020-00466-2)) |
| Otu56286 | 1 | k__Fungi(100);p__Ascomycota(100);c__Saccharomycetes(100);o__Saccharomycetales(100);f__Metschnikowiaceae(100);g__Metschnikowia(100);s__Metschnikowia_pulcherrima(100); | Metschnikowia | 13 | Saprotroph | \|Undefined Saprotroph\| | Yeast | NULL | Probable | As nectar-tap_saprotroph (Põlme et al. 2020); Decay sugar-rich substrates (Põlme et al. 2020) | Põlme S, et al. 2020. Fungal Diversity 105:1-16 ((https://doi.org/10.1007/s13225-020-00466-2)) |
| Otu56288 | 1 | k__Fungi(100);p__Ascomycota(100);c__Saccharomycetes(100);o__Saccharomycetales(100);f__Metschnikowiaceae(100);g__Metschnikowia(100);s__Metschnikowia_pulcherrima(100); | Metschnikowia | 13 | Saprotroph | \|Undefined Saprotroph\| | Yeast | NULL | Probable | As nectar-tap_saprotroph (Põlme et al. 2020); Decay sugar-rich substrates (Põlme et al. 2020) | Põlme S, et al. 2020. Fungal Diversity 105:1-16 ((https://doi.org/10.1007/s13225-020-00466-2)) |
| Otu56289 | 1 | k__Fungi(100);p__Ascomycota(100);c__Saccharomycetes(100);o__Saccharomycetales(100);f__Saccharomycetales_family_Incertae_sedis(100);f__Saccharomycetales_family_Incertae_sedis_unclassified(100);f__Saccharomycetales_family_Incertae_sedis_unclassified(100); | Saccharomycetales | 7 | Saprotroph | Undefined Saprotroph | Yeast | NULL | Possible | NULL | Sterkenburg E, et al. 2015. New Phytologist 207:1145-1158 ((https://doi.org/10.1111/nph.13426)) |
| Otu56291 | 1 | k__Fungi(100);p__Ascomycota(100);c__Saccharomycetes(100);o__Saccharomycetales(100);f__Metschnikowiaceae(100);g__Metschnikowia(100);s__Metschnikowia_pulcherrima(100); | Metschnikowia | 13 | Saprotroph | \|Undefined Saprotroph\| | Yeast | NULL | Probable | As nectar-tap_saprotroph (Põlme et al. 2020); Decay sugar-rich substrates (Põlme et al. 2020) | Põlme S, et al. 2020. Fungal Diversity 105:1-16 ((https://doi.org/10.1007/s13225-020-00466-2)) |
| Otu56293 | 1 | k__Fungi(100);p__Ascomycota(100);c__Saccharomycetes(100);o__Saccharomycetales(100);f__Metschnikowiaceae(100);g__Metschnikowia(100);s__Metschnikowia_pulcherrima(100); | Metschnikowia | 13 | Saprotroph | \|Undefined Saprotroph\| | Yeast | NULL | Probable | As nectar-tap_saprotroph (Põlme et al. 2020); Decay sugar-rich substrates (Põlme et al. 2020) | Põlme S, et al. 2020. Fungal Diversity 105:1-16 ((https://doi.org/10.1007/s13225-020-00466-2)) |
| Otu56294 | 1 | k__Fungi(100);p__Ascomycota(100);c__Saccharomycetes(100);o__Saccharomycetales(100);f__Metschnikowiaceae(100);g__Metschnikowia(100);g__Metschnikowia_unclassified(100); | Metschnikowia | 13 | Saprotroph | \|Undefined Saprotroph\| | Yeast | NULL | Probable | As nectar-tap_saprotroph (Põlme et al. 2020); Decay sugar-rich substrates (Põlme et al. 2020) | Põlme S, et al. 2020. Fungal Diversity 105:1-16 ((https://doi.org/10.1007/s13225-020-00466-2)) |
| Otu56297 | 1 | k__Fungi(100);p__Ascomycota(100);c__Saccharomycetes(100);o__Saccharomycetales(100);f__Metschnikowiaceae(100);g__Metschnikowia(100);s__Metschnikowia_pulcherrima(100); | Metschnikowia | 13 | Saprotroph | \|Undefined Saprotroph\| | Yeast | NULL | Probable | As nectar-tap_saprotroph (Põlme et al. 2020); Decay sugar-rich substrates (Põlme et al. 2020) | Põlme S, et al. 2020. Fungal Diversity 105:1-16 ((https://doi.org/10.1007/s13225-020-00466-2)) |
| Otu56298 | 1 | k__Fungi(100);p__Ascomycota(100);c__Saccharomycetes(100);o__Saccharomycetales(100);f__Metschnikowiaceae(100);g__Metschnikowia(100);s__Metschnikowia_pulcherrima(100); | Metschnikowia | 13 | Saprotroph | \|Undefined Saprotroph\| | Yeast | NULL | Probable | As nectar-tap_saprotroph (Põlme et al. 2020); Decay sugar-rich substrates (Põlme et al. 2020) | Põlme S, et al. 2020. Fungal Diversity 105:1-16 ((https://doi.org/10.1007/s13225-020-00466-2)) |
| Otu56299 | 1 | k__Fungi(100);p__Ascomycota(100);c__Saccharomycetes(100);o__Saccharomycetales(100);f__Metschnikowiaceae(100);g__Metschnikowia(100);s__Metschnikowia_pulcherrima(100); | Metschnikowia | 13 | Saprotroph | \|Undefined Saprotroph\| | Yeast | NULL | Probable | As nectar-tap_saprotroph (Põlme et al. 2020); Decay sugar-rich substrates (Põlme et al. 2020) | Põlme S, et al. 2020. Fungal Diversity 105:1-16 ((https://doi.org/10.1007/s13225-020-00466-2)) |
| Otu56301 | 1 | k__Fungi(100);p__Ascomycota(100);c__Saccharomycetes(100);o__Saccharomycetales(100);f__Metschnikowiaceae(100);g__Metschnikowia(100);s__Metschnikowia_pulcherrima(100); | Metschnikowia | 13 | Saprotroph | \|Undefined Saprotroph\| | Yeast | NULL | Probable | As nectar-tap_saprotroph (Põlme et al. 2020); Decay sugar-rich substrates (Põlme et al. 2020) | Põlme S, et al. 2020. Fungal Diversity 105:1-16 ((https://doi.org/10.1007/s13225-020-00466-2)) |
| Otu56302 | 1 | k__Fungi(100);p__Ascomycota(100);c__Saccharomycetes(100);o__Saccharomycetales(100);f__Metschnikowiaceae(100);g__Metschnikowia(100);g__Metschnikowia_unclassified(100); | Metschnikowia | 13 | Saprotroph | \|Undefined Saprotroph\| | Yeast | NULL | Probable | As nectar-tap_saprotroph (Põlme et al. 2020); Decay sugar-rich substrates (Põlme et al. 2020) | Põlme S, et al. 2020. Fungal Diversity 105:1-16 ((https://doi.org/10.1007/s13225-020-00466-2)) |
| Otu56304 | 1 | k__Fungi(100);p__Ascomycota(100);c__Saccharomycetes(100);o__Saccharomycetales(100);f__Metschnikowiaceae(100);g__Metschnikowia(100);s__Metschnikowia_pulcherrima(100); | Metschnikowia | 13 | Saprotroph | \|Undefined Saprotroph\| | Yeast | NULL | Probable | As nectar-tap_saprotroph (Põlme et al. 2020); Decay sugar-rich substrates (Põlme et al. 2020) | Põlme S, et al. 2020. Fungal Diversity 105:1-16 ((https://doi.org/10.1007/s13225-020-00466-2)) |
| Otu56305 | 1 | k__Fungi(100);p__Ascomycota(100);c__Saccharomycetes(100);o__Saccharomycetales(100);f__Metschnikowiaceae(100);g__Metschnikowia(100);s__Metschnikowia_pulcherrima(100); | Metschnikowia | 13 | Saprotroph | \|Undefined Saprotroph\| | Yeast | NULL | Probable | As nectar-tap_saprotroph (Põlme et al. 2020); Decay sugar-rich substrates (Põlme et al. 2020) | Põlme S, et al. 2020. Fungal Diversity 105:1-16 ((https://doi.org/10.1007/s13225-020-00466-2)) |
| Otu56306 | 1 | k__Fungi(100);p__Ascomycota(100);c__Saccharomycetes(100);o__Saccharomycetales(100);f__Saccharomycetales_family_Incertae_sedis(100);g__Candida(100);g__Candida_unclassified(100); | Candida | 13 | Pathotroph-Saprotroph-Symbiotroph | Animal Pathogen-Endophyte-Endosymbiont-Epiphyte-\|Undefined Saprotroph\| | Dimorphic Yeast-Yeast | NULL | Probable | As nectar-tap_saprotroph (Põlme et al. 2020); Decay sugar-rich substrates (Põlme et al. 2020); This asexual genus is polyphyletic and contain species that function very differently depend on what they are. Caution should be taken when interpreting guilds for Candida species. | Manolakaki D, et al. 2010. Virulence. 1: 367-375 ((https://doi.org/10.4161/viru.1.5.12796)); Mekbib SB, et al. 2011. Tropical Plant Pathology 36:233-240 ((https://doi.org/10.1590/S1982-56762011000400004)); Põlme S, et al. 2020. Fungal Diversity 105:1-16 ((https://doi.org/10.1007/s13225-020-00466-2)) |
| Otu56307 | 1 | k__Fungi(100);p__Ascomycota(100);c__Saccharomycetes(100);o__Saccharomycetales(100);f__Metschnikowiaceae(100);g__Metschnikowia(100);s__Metschnikowia_pulcherrima(100); | Metschnikowia | 13 | Saprotroph | \|Undefined Saprotroph\| | Yeast | NULL | Probable | As nectar-tap_saprotroph (Põlme et al. 2020); Decay sugar-rich substrates (Põlme et al. 2020) | Põlme S, et al. 2020. Fungal Diversity 105:1-16 ((https://doi.org/10.1007/s13225-020-00466-2)) |
| Otu56308 | 1 | k__Fungi(100);p__Ascomycota(100);c__Saccharomycetes(100);o__Saccharomycetales(100);f__Metschnikowiaceae(100);g__Metschnikowia(100);g__Metschnikowia_unclassified(100); | Metschnikowia | 13 | Saprotroph | \|Undefined Saprotroph\| | Yeast | NULL | Probable | As nectar-tap_saprotroph (Põlme et al. 2020); Decay sugar-rich substrates (Põlme et al. 2020) | Põlme S, et al. 2020. Fungal Diversity 105:1-16 ((https://doi.org/10.1007/s13225-020-00466-2)) |
| Otu56309 | 1 | k__Fungi(100);p__Ascomycota(100);c__Saccharomycetes(100);o__Saccharomycetales(100);f__Metschnikowiaceae(100);g__Metschnikowia(100);s__Metschnikowia_pulcherrima(100); | Metschnikowia | 13 | Saprotroph | \|Undefined Saprotroph\| | Yeast | NULL | Probable | As nectar-tap_saprotroph (Põlme et al. 2020); Decay sugar-rich substrates (Põlme et al. 2020) | Põlme S, et al. 2020. Fungal Diversity 105:1-16 ((https://doi.org/10.1007/s13225-020-00466-2)) |
| Otu56311 | 1 | k__Fungi(100);p__Ascomycota(100);c__Saccharomycetes(100);o__Saccharomycetales(100);f__Metschnikowiaceae(100);g__Metschnikowia(100);g__Metschnikowia_unclassified(100); | Metschnikowia | 13 | Saprotroph | \|Undefined Saprotroph\| | Yeast | NULL | Probable | As nectar-tap_saprotroph (Põlme et al. 2020); Decay sugar-rich substrates (Põlme et al. 2020) | Põlme S, et al. 2020. Fungal Diversity 105:1-16 ((https://doi.org/10.1007/s13225-020-00466-2)) |
| Otu56312 | 1 | k__Fungi(100);p__Ascomycota(100);c__Saccharomycetes(100);o__Saccharomycetales(100);f__Metschnikowiaceae(100);g__Metschnikowia(100);s__Metschnikowia_pulcherrima(100); | Metschnikowia | 13 | Saprotroph | \|Undefined Saprotroph\| | Yeast | NULL | Probable | As nectar-tap_saprotroph (Põlme et al. 2020); Decay sugar-rich substrates (Põlme et al. 2020) | Põlme S, et al. 2020. Fungal Diversity 105:1-16 ((https://doi.org/10.1007/s13225-020-00466-2)) |
| Otu56314 | 1 | k__Fungi(100);p__Ascomycota(100);c__Saccharomycetes(100);o__Saccharomycetales(100);f__Metschnikowiaceae(100);g__Metschnikowia(100);s__Metschnikowia_pulcherrima(100); | Metschnikowia | 13 | Saprotroph | \|Undefined Saprotroph\| | Yeast | NULL | Probable | As nectar-tap_saprotroph (Põlme et al. 2020); Decay sugar-rich substrates (Põlme et al. 2020) | Põlme S, et al. 2020. Fungal Diversity 105:1-16 ((https://doi.org/10.1007/s13225-020-00466-2)) |
| Otu56315 | 1 | k__Fungi(100);p__Ascomycota(100);c__Saccharomycetes(100);o__Saccharomycetales(100);f__Metschnikowiaceae(100);g__Metschnikowia(100);s__Metschnikowia_pulcherrima(100); | Metschnikowia | 13 | Saprotroph | \|Undefined Saprotroph\| | Yeast | NULL | Probable | As nectar-tap_saprotroph (Põlme et al. 2020); Decay sugar-rich substrates (Põlme et al. 2020) | Põlme S, et al. 2020. Fungal Diversity 105:1-16 ((https://doi.org/10.1007/s13225-020-00466-2)) |
| Otu56316 | 1 | k__Fungi(100);p__Ascomycota(100);c__Saccharomycetes(100);o__Saccharomycetales(100);f__Metschnikowiaceae(100);g__Metschnikowia(100);s__Metschnikowia_pulcherrima(100); | Metschnikowia | 13 | Saprotroph | \|Undefined Saprotroph\| | Yeast | NULL | Probable | As nectar-tap_saprotroph (Põlme et al. 2020); Decay sugar-rich substrates (Põlme et al. 2020) | Põlme S, et al. 2020. Fungal Diversity 105:1-16 ((https://doi.org/10.1007/s13225-020-00466-2)) |
| Otu56317 | 1 | k__Fungi(100);p__Ascomycota(100);c__Saccharomycetes(100);o__Saccharomycetales(100);f__Metschnikowiaceae(100);g__Metschnikowia(100);g__Metschnikowia_unclassified(100); | Metschnikowia | 13 | Saprotroph | \|Undefined Saprotroph\| | Yeast | NULL | Probable | As nectar-tap_saprotroph (Põlme et al. 2020); Decay sugar-rich substrates (Põlme et al. 2020) | Põlme S, et al. 2020. Fungal Diversity 105:1-16 ((https://doi.org/10.1007/s13225-020-00466-2)) |
| Otu56318 | 1 | k__Fungi(100);p__Ascomycota(100);c__Saccharomycetes(100);o__Saccharomycetales(100);f__Metschnikowiaceae(100);g__Metschnikowia(100);s__Metschnikowia_pulcherrima(100); | Metschnikowia | 13 | Saprotroph | \|Undefined Saprotroph\| | Yeast | NULL | Probable | As nectar-tap_saprotroph (Põlme et al. 2020); Decay sugar-rich substrates (Põlme et al. 2020) | Põlme S, et al. 2020. Fungal Diversity 105:1-16 ((https://doi.org/10.1007/s13225-020-00466-2)) |
| Otu56319 | 1 | k__Fungi(100);p__Ascomycota(100);c__Saccharomycetes(100);o__Saccharomycetales(100);f__Metschnikowiaceae(100);g__Metschnikowia(100);s__Metschnikowia_pulcherrima(100); | Metschnikowia | 13 | Saprotroph | \|Undefined Saprotroph\| | Yeast | NULL | Probable | As nectar-tap_saprotroph (Põlme et al. 2020); Decay sugar-rich substrates (Põlme et al. 2020) | Põlme S, et al. 2020. Fungal Diversity 105:1-16 ((https://doi.org/10.1007/s13225-020-00466-2)) |
| Otu56320 | 1 | k__Fungi(100);p__Ascomycota(100);c__Saccharomycetes(100);o__Saccharomycetales(100);f__Metschnikowiaceae(100);g__Metschnikowia(100);s__Metschnikowia_pulcherrima(100); | Metschnikowia | 13 | Saprotroph | \|Undefined Saprotroph\| | Yeast | NULL | Probable | As nectar-tap_saprotroph (Põlme et al. 2020); Decay sugar-rich substrates (Põlme et al. 2020) | Põlme S, et al. 2020. Fungal Diversity 105:1-16 ((https://doi.org/10.1007/s13225-020-00466-2)) |
| Otu56321 | 1 | k__Fungi(100);p__Ascomycota(100);c__Saccharomycetes(100);o__Saccharomycetales(100);f__Metschnikowiaceae(100);g__Metschnikowia(100);g__Metschnikowia_unclassified(100); | Metschnikowia | 13 | Saprotroph | \|Undefined Saprotroph\| | Yeast | NULL | Probable | As nectar-tap_saprotroph (Põlme et al. 2020); Decay sugar-rich substrates (Põlme et al. 2020) | Põlme S, et al. 2020. Fungal Diversity 105:1-16 ((https://doi.org/10.1007/s13225-020-00466-2)) |
| Otu56322 | 1 | k__Fungi(100);p__Ascomycota(100);c__Saccharomycetes(100);o__Saccharomycetales(100);f__Metschnikowiaceae(100);g__Metschnikowia(100);s__Metschnikowia_pulcherrima(100); | Metschnikowia | 13 | Saprotroph | \|Undefined Saprotroph\| | Yeast | NULL | Probable | As nectar-tap_saprotroph (Põlme et al. 2020); Decay sugar-rich substrates (Põlme et al. 2020) | Põlme S, et al. 2020. Fungal Diversity 105:1-16 ((https://doi.org/10.1007/s13225-020-00466-2)) |
| Otu56323 | 1 | k__Fungi(100);p__Ascomycota(100);c__Saccharomycetes(100);o__Saccharomycetales(100);f__Metschnikowiaceae(100);g__Metschnikowia(100);s__Metschnikowia_pulcherrima(100); | Metschnikowia | 13 | Saprotroph | \|Undefined Saprotroph\| | Yeast | NULL | Probable | As nectar-tap_saprotroph (Põlme et al. 2020); Decay sugar-rich substrates (Põlme et al. 2020) | Põlme S, et al. 2020. Fungal Diversity 105:1-16 ((https://doi.org/10.1007/s13225-020-00466-2)) |
| Otu56324 | 1 | k__Fungi(100);p__Ascomycota(100);c__Saccharomycetes(100);o__Saccharomycetales(100);f__Metschnikowiaceae(100);g__Metschnikowia(100);s__Metschnikowia_pulcherrima(100); | Metschnikowia | 13 | Saprotroph | \|Undefined Saprotroph\| | Yeast | NULL | Probable | As nectar-tap_saprotroph (Põlme et al. 2020); Decay sugar-rich substrates (Põlme et al. 2020) | Põlme S, et al. 2020. Fungal Diversity 105:1-16 ((https://doi.org/10.1007/s13225-020-00466-2)) |
| Otu56325 | 1 | k__Fungi(100);p__Ascomycota(100);c__Saccharomycetes(100);o__Saccharomycetales(100);f__Metschnikowiaceae(100);g__Metschnikowia(100);s__Metschnikowia_pulcherrima(100); | Metschnikowia | 13 | Saprotroph | \|Undefined Saprotroph\| | Yeast | NULL | Probable | As nectar-tap_saprotroph (Põlme et al. 2020); Decay sugar-rich substrates (Põlme et al. 2020) | Põlme S, et al. 2020. Fungal Diversity 105:1-16 ((https://doi.org/10.1007/s13225-020-00466-2)) |
| Otu56326 | 1 | k__Fungi(100);p__Ascomycota(100);c__Saccharomycetes(100);o__Saccharomycetales(100);f__Metschnikowiaceae(100);g__Metschnikowia(100);s__Metschnikowia_pulcherrima(100); | Metschnikowia | 13 | Saprotroph | \|Undefined Saprotroph\| | Yeast | NULL | Probable | As nectar-tap_saprotroph (Põlme et al. 2020); Decay sugar-rich substrates (Põlme et al. 2020) | Põlme S, et al. 2020. Fungal Diversity 105:1-16 ((https://doi.org/10.1007/s13225-020-00466-2)) |
| Otu56328 | 1 | k__Fungi(100);p__Ascomycota(100);c__Saccharomycetes(100);o__Saccharomycetales(100);f__Metschnikowiaceae(100);g__Metschnikowia(100);s__Metschnikowia_pulcherrima(100); | Metschnikowia | 13 | Saprotroph | \|Undefined Saprotroph\| | Yeast | NULL | Probable | As nectar-tap_saprotroph (Põlme et al. 2020); Decay sugar-rich substrates (Põlme et al. 2020) | Põlme S, et al. 2020. Fungal Diversity 105:1-16 ((https://doi.org/10.1007/s13225-020-00466-2)) |
| Otu56329 | 1 | k__Fungi(100);p__Ascomycota(100);c__Saccharomycetes(100);o__Saccharomycetales(100);f__Metschnikowiaceae(100);g__Metschnikowia(100);s__Metschnikowia_pulcherrima(100); | Metschnikowia | 13 | Saprotroph | \|Undefined Saprotroph\| | Yeast | NULL | Probable | As nectar-tap_saprotroph (Põlme et al. 2020); Decay sugar-rich substrates (Põlme et al. 2020) | Põlme S, et al. 2020. Fungal Diversity 105:1-16 ((https://doi.org/10.1007/s13225-020-00466-2)) |
| Otu56330 | 1 | k__Fungi(100);p__Ascomycota(100);c__Saccharomycetes(100);o__Saccharomycetales(100);f__Metschnikowiaceae(100);g__Metschnikowia(100);s__Metschnikowia_pulcherrima(100); | Metschnikowia | 13 | Saprotroph | \|Undefined Saprotroph\| | Yeast | NULL | Probable | As nectar-tap_saprotroph (Põlme et al. 2020); Decay sugar-rich substrates (Põlme et al. 2020) | Põlme S, et al. 2020. Fungal Diversity 105:1-16 ((https://doi.org/10.1007/s13225-020-00466-2)) |
| Otu56331 | 1 | k__Fungi(100);p__Ascomycota(100);c__Saccharomycetes(100);o__Saccharomycetales(100);f__Saccharomycetales_family_Incertae_sedis(100);g__Candida(100);g__Candida_unclassified(100); | Candida | 13 | Pathotroph-Saprotroph-Symbiotroph | Animal Pathogen-Endophyte-Endosymbiont-Epiphyte-\|Undefined Saprotroph\| | Dimorphic Yeast-Yeast | NULL | Probable | As nectar-tap_saprotroph (Põlme et al. 2020); Decay sugar-rich substrates (Põlme et al. 2020); This asexual genus is polyphyletic and contain species that function very differently depend on what they are. Caution should be taken when interpreting guilds for Candida species. | Manolakaki D, et al. 2010. Virulence. 1: 367-375 ((https://doi.org/10.4161/viru.1.5.12796)); Mekbib SB, et al. 2011. Tropical Plant Pathology 36:233-240 ((https://doi.org/10.1590/S1982-56762011000400004)); Põlme S, et al. 2020. Fungal Diversity 105:1-16 ((https://doi.org/10.1007/s13225-020-00466-2)) |
| Otu56332 | 1 | k__Fungi(100);p__Basidiomycota(100);c__Microbotryomycetes(100);o__Sporidiobolales(100);f__Sporidiobolales_family_Incertae_sedis(100);g__Sporobolomyces(100);s__Sporobolomyces_lactosus(100); | Sporobolomyces | 13 | Pathotroph-Saprotroph | \|Fungal Parasite\|-Plant Saprotroph | Yeast | NULL | Probable | As litter_saprotroph (Põlme et al. 2020); As mycoparasite (Põlme et al. 2020); Decay fungal material (Põlme et al. 2020) | Tedersoo L, et al. 2014. Science 346:e1256688 ((https://doi.org/10.1126/science.1256688)); Purahong W, et al. 2016. Molecular Ecology 25:4059-4074 ((https://doi.org/10.1111/mec.13739)); Põlme S, et al. 2020. Fungal Diversity 105:1-16 ((https://doi.org/10.1007/s13225-020-00466-2)) |
| Otu56333 | 1 | k__Fungi(100);p__Ascomycota(100);c__Saccharomycetes(100);o__Saccharomycetales(100);f__Metschnikowiaceae(100);g__Metschnikowia(100);s__Metschnikowia_pulcherrima(100); | Metschnikowia | 13 | Saprotroph | \|Undefined Saprotroph\| | Yeast | NULL | Probable | As nectar-tap_saprotroph (Põlme et al. 2020); Decay sugar-rich substrates (Põlme et al. 2020) | Põlme S, et al. 2020. Fungal Diversity 105:1-16 ((https://doi.org/10.1007/s13225-020-00466-2)) |
| Otu56334 | 1 | k__Fungi(100);p__Ascomycota(100);c__Saccharomycetes(100);o__Saccharomycetales(100);f__Metschnikowiaceae(100);g__Metschnikowia(100);s__Metschnikowia_pulcherrima(100); | Metschnikowia | 13 | Saprotroph | \|Undefined Saprotroph\| | Yeast | NULL | Probable | As nectar-tap_saprotroph (Põlme et al. 2020); Decay sugar-rich substrates (Põlme et al. 2020) | Põlme S, et al. 2020. Fungal Diversity 105:1-16 ((https://doi.org/10.1007/s13225-020-00466-2)) |
| Otu56335 | 1 | k__Fungi(100);p__Ascomycota(100);c__Saccharomycetes(100);o__Saccharomycetales(100);f__Metschnikowiaceae(100);g__Metschnikowia(100);s__Metschnikowia_pulcherrima(100); | Metschnikowia | 13 | Saprotroph | \|Undefined Saprotroph\| | Yeast | NULL | Probable | As nectar-tap_saprotroph (Põlme et al. 2020); Decay sugar-rich substrates (Põlme et al. 2020) | Põlme S, et al. 2020. Fungal Diversity 105:1-16 ((https://doi.org/10.1007/s13225-020-00466-2)) |
| Otu56336 | 1 | k__Fungi(100);p__Ascomycota(100);c__Saccharomycetes(100);o__Saccharomycetales(100);f__Metschnikowiaceae(100);g__Metschnikowia(100);s__Metschnikowia_pulcherrima(100); | Metschnikowia | 13 | Saprotroph | \|Undefined Saprotroph\| | Yeast | NULL | Probable | As nectar-tap_saprotroph (Põlme et al. 2020); Decay sugar-rich substrates (Põlme et al. 2020) | Põlme S, et al. 2020. Fungal Diversity 105:1-16 ((https://doi.org/10.1007/s13225-020-00466-2)) |
| Otu56337 | 1 | k__Fungi(100);p__Ascomycota(100);c__Saccharomycetes(100);o__Saccharomycetales(100);f__Metschnikowiaceae(100);g__Metschnikowia(100);g__Metschnikowia_unclassified(100); | Metschnikowia | 13 | Saprotroph | \|Undefined Saprotroph\| | Yeast | NULL | Probable | As nectar-tap_saprotroph (Põlme et al. 2020); Decay sugar-rich substrates (Põlme et al. 2020) | Põlme S, et al. 2020. Fungal Diversity 105:1-16 ((https://doi.org/10.1007/s13225-020-00466-2)) |
| Otu56338 | 1 | k__Fungi(100);p__Ascomycota(100);c__Saccharomycetes(100);o__Saccharomycetales(100);f__Metschnikowiaceae(100);g__Metschnikowia(100);s__Metschnikowia_pulcherrima(100); | Metschnikowia | 13 | Saprotroph | \|Undefined Saprotroph\| | Yeast | NULL | Probable | As nectar-tap_saprotroph (Põlme et al. 2020); Decay sugar-rich substrates (Põlme et al. 2020) | Põlme S, et al. 2020. Fungal Diversity 105:1-16 ((https://doi.org/10.1007/s13225-020-00466-2)) |
| Otu56339 | 1 | k__Fungi(100);p__Ascomycota(100);c__Saccharomycetes(100);o__Saccharomycetales(100);f__Metschnikowiaceae(100);g__Metschnikowia(100);s__Metschnikowia_pulcherrima(100); | Metschnikowia | 13 | Saprotroph | \|Undefined Saprotroph\| | Yeast | NULL | Probable | As nectar-tap_saprotroph (Põlme et al. 2020); Decay sugar-rich substrates (Põlme et al. 2020) | Põlme S, et al. 2020. Fungal Diversity 105:1-16 ((https://doi.org/10.1007/s13225-020-00466-2)) |
| Otu56340 | 1 | k__Fungi(100);p__Ascomycota(100);c__Saccharomycetes(100);o__Saccharomycetales(100);f__Metschnikowiaceae(100);g__Metschnikowia(100);g__Metschnikowia_unclassified(100); | Metschnikowia | 13 | Saprotroph | \|Undefined Saprotroph\| | Yeast | NULL | Probable | As nectar-tap_saprotroph (Põlme et al. 2020); Decay sugar-rich substrates (Põlme et al. 2020) | Põlme S, et al. 2020. Fungal Diversity 105:1-16 ((https://doi.org/10.1007/s13225-020-00466-2)) |
| Otu56341 | 1 | k__Fungi(100);p__Ascomycota(100);c__Saccharomycetes(100);o__Saccharomycetales(100);f__Metschnikowiaceae(100);g__Metschnikowia(100);s__Metschnikowia_pulcherrima(100); | Metschnikowia | 13 | Saprotroph | \|Undefined Saprotroph\| | Yeast | NULL | Probable | As nectar-tap_saprotroph (Põlme et al. 2020); Decay sugar-rich substrates (Põlme et al. 2020) | Põlme S, et al. 2020. Fungal Diversity 105:1-16 ((https://doi.org/10.1007/s13225-020-00466-2)) |
| Otu56342 | 1 | k__Fungi(100);p__Ascomycota(100);c__Saccharomycetes(100);o__Saccharomycetales(100);f__Metschnikowiaceae(100);g__Metschnikowia(100);s__Metschnikowia_pulcherrima(100); | Metschnikowia | 13 | Saprotroph | \|Undefined Saprotroph\| | Yeast | NULL | Probable | As nectar-tap_saprotroph (Põlme et al. 2020); Decay sugar-rich substrates (Põlme et al. 2020) | Põlme S, et al. 2020. Fungal Diversity 105:1-16 ((https://doi.org/10.1007/s13225-020-00466-2)) |
| Otu56343 | 1 | k__Fungi(100);p__Ascomycota(100);c__Saccharomycetes(100);o__Saccharomycetales(100);f__Metschnikowiaceae(100);g__Metschnikowia(100);s__Metschnikowia_pulcherrima(100); | Metschnikowia | 13 | Saprotroph | \|Undefined Saprotroph\| | Yeast | NULL | Probable | As nectar-tap_saprotroph (Põlme et al. 2020); Decay sugar-rich substrates (Põlme et al. 2020) | Põlme S, et al. 2020. Fungal Diversity 105:1-16 ((https://doi.org/10.1007/s13225-020-00466-2)) |
| Otu56346 | 1 | k__Fungi(100);p__Ascomycota(100);c__Saccharomycetes(100);o__Saccharomycetales(100);f__Metschnikowiaceae(100);g__Metschnikowia(100);g__Metschnikowia_unclassified(100); | Metschnikowia | 13 | Saprotroph | \|Undefined Saprotroph\| | Yeast | NULL | Probable | As nectar-tap_saprotroph (Põlme et al. 2020); Decay sugar-rich substrates (Põlme et al. 2020) | Põlme S, et al. 2020. Fungal Diversity 105:1-16 ((https://doi.org/10.1007/s13225-020-00466-2)) |
| Otu56349 | 1 | k__Fungi(100);p__Ascomycota(100);c__Saccharomycetes(100);o__Saccharomycetales(100);f__Metschnikowiaceae(100);g__Metschnikowia(100);s__Metschnikowia_pulcherrima(100); | Metschnikowia | 13 | Saprotroph | \|Undefined Saprotroph\| | Yeast | NULL | Probable | As nectar-tap_saprotroph (Põlme et al. 2020); Decay sugar-rich substrates (Põlme et al. 2020) | Põlme S, et al. 2020. Fungal Diversity 105:1-16 ((https://doi.org/10.1007/s13225-020-00466-2)) |
| Otu56350 | 1 | k__Fungi(100);p__Ascomycota(100);c__Saccharomycetes(100);o__Saccharomycetales(100);f__Metschnikowiaceae(100);g__Metschnikowia(100);s__Metschnikowia_pulcherrima(100); | Metschnikowia | 13 | Saprotroph | \|Undefined Saprotroph\| | Yeast | NULL | Probable | As nectar-tap_saprotroph (Põlme et al. 2020); Decay sugar-rich substrates (Põlme et al. 2020) | Põlme S, et al. 2020. Fungal Diversity 105:1-16 ((https://doi.org/10.1007/s13225-020-00466-2)) |
| Otu56352 | 1 | k__Fungi(100);p__Ascomycota(100);c__Saccharomycetes(100);o__Saccharomycetales(100);o__Saccharomycetales_unclassified(100);o__Saccharomycetales_unclassified(100);o__Saccharomycetales_unclassified(100); | Saccharomycetales | 7 | Saprotroph | Undefined Saprotroph | Yeast | NULL | Possible | NULL | Sterkenburg E, et al. 2015. New Phytologist 207:1145-1158 ((https://doi.org/10.1111/nph.13426)) |
| Otu56353 | 1 | k__Fungi(100);p__Ascomycota(100);c__Saccharomycetes(100);o__Saccharomycetales(100);f__Metschnikowiaceae(100);g__Metschnikowia(100);g__Metschnikowia_unclassified(100); | Metschnikowia | 13 | Saprotroph | \|Undefined Saprotroph\| | Yeast | NULL | Probable | As nectar-tap_saprotroph (Põlme et al. 2020); Decay sugar-rich substrates (Põlme et al. 2020) | Põlme S, et al. 2020. Fungal Diversity 105:1-16 ((https://doi.org/10.1007/s13225-020-00466-2)) |
| Otu56354 | 1 | k__Fungi(100);p__Ascomycota(100);c__Saccharomycetes(100);o__Saccharomycetales(100);f__Metschnikowiaceae(100);g__Metschnikowia(100);s__Metschnikowia_pulcherrima(100); | Metschnikowia | 13 | Saprotroph | \|Undefined Saprotroph\| | Yeast | NULL | Probable | As nectar-tap_saprotroph (Põlme et al. 2020); Decay sugar-rich substrates (Põlme et al. 2020) | Põlme S, et al. 2020. Fungal Diversity 105:1-16 ((https://doi.org/10.1007/s13225-020-00466-2)) |
| Otu56355 | 1 | k__Fungi(100);p__Ascomycota(100);c__Saccharomycetes(100);o__Saccharomycetales(100);f__Metschnikowiaceae(100);g__Metschnikowia(100);s__Metschnikowia_pulcherrima(100); | Metschnikowia | 13 | Saprotroph | \|Undefined Saprotroph\| | Yeast | NULL | Probable | As nectar-tap_saprotroph (Põlme et al. 2020); Decay sugar-rich substrates (Põlme et al. 2020) | Põlme S, et al. 2020. Fungal Diversity 105:1-16 ((https://doi.org/10.1007/s13225-020-00466-2)) |
| Otu56356 | 1 | k__Fungi(100);p__Ascomycota(100);c__Saccharomycetes(100);o__Saccharomycetales(100);f__Metschnikowiaceae(100);g__Metschnikowia(100);g__Metschnikowia_unclassified(100); | Metschnikowia | 13 | Saprotroph | \|Undefined Saprotroph\| | Yeast | NULL | Probable | As nectar-tap_saprotroph (Põlme et al. 2020); Decay sugar-rich substrates (Põlme et al. 2020) | Põlme S, et al. 2020. Fungal Diversity 105:1-16 ((https://doi.org/10.1007/s13225-020-00466-2)) |
| Otu56357 | 1 | k__Fungi(100);p__Ascomycota(100);c__Saccharomycetes(100);o__Saccharomycetales(100);f__Metschnikowiaceae(100);g__Metschnikowia(100);g__Metschnikowia_unclassified(100); | Metschnikowia | 13 | Saprotroph | \|Undefined Saprotroph\| | Yeast | NULL | Probable | As nectar-tap_saprotroph (Põlme et al. 2020); Decay sugar-rich substrates (Põlme et al. 2020) | Põlme S, et al. 2020. Fungal Diversity 105:1-16 ((https://doi.org/10.1007/s13225-020-00466-2)) |
| Otu56358 | 1 | k__Fungi(100);p__Ascomycota(100);c__Saccharomycetes(100);o__Saccharomycetales(100);f__Metschnikowiaceae(100);g__Metschnikowia(100);s__Metschnikowia_pulcherrima(100); | Metschnikowia | 13 | Saprotroph | \|Undefined Saprotroph\| | Yeast | NULL | Probable | As nectar-tap_saprotroph (Põlme et al. 2020); Decay sugar-rich substrates (Põlme et al. 2020) | Põlme S, et al. 2020. Fungal Diversity 105:1-16 ((https://doi.org/10.1007/s13225-020-00466-2)) |
| Otu56359 | 1 | k__Fungi(100);p__Ascomycota(100);c__Saccharomycetes(100);o__Saccharomycetales(100);f__Metschnikowiaceae(100);g__Metschnikowia(100);s__Metschnikowia_pulcherrima(100); | Metschnikowia | 13 | Saprotroph | \|Undefined Saprotroph\| | Yeast | NULL | Probable | As nectar-tap_saprotroph (Põlme et al. 2020); Decay sugar-rich substrates (Põlme et al. 2020) | Põlme S, et al. 2020. Fungal Diversity 105:1-16 ((https://doi.org/10.1007/s13225-020-00466-2)) |
| Otu56360 | 1 | k__Fungi(100);p__Ascomycota(100);c__Saccharomycetes(100);o__Saccharomycetales(100);f__Metschnikowiaceae(100);g__Metschnikowia(100);s__Metschnikowia_pulcherrima(100); | Metschnikowia | 13 | Saprotroph | \|Undefined Saprotroph\| | Yeast | NULL | Probable | As nectar-tap_saprotroph (Põlme et al. 2020); Decay sugar-rich substrates (Põlme et al. 2020) | Põlme S, et al. 2020. Fungal Diversity 105:1-16 ((https://doi.org/10.1007/s13225-020-00466-2)) |
| Otu56361 | 1 | k__Fungi(100);p__Ascomycota(100);c__Saccharomycetes(100);o__Saccharomycetales(100);f__Metschnikowiaceae(100);g__Metschnikowia(100);s__Metschnikowia_pulcherrima(100); | Metschnikowia | 13 | Saprotroph | \|Undefined Saprotroph\| | Yeast | NULL | Probable | As nectar-tap_saprotroph (Põlme et al. 2020); Decay sugar-rich substrates (Põlme et al. 2020) | Põlme S, et al. 2020. Fungal Diversity 105:1-16 ((https://doi.org/10.1007/s13225-020-00466-2)) |
| Otu56362 | 1 | k__Fungi(100);p__Ascomycota(100);c__Saccharomycetes(100);o__Saccharomycetales(100);f__Metschnikowiaceae(100);g__Metschnikowia(100);s__Metschnikowia_pulcherrima(100); | Metschnikowia | 13 | Saprotroph | \|Undefined Saprotroph\| | Yeast | NULL | Probable | As nectar-tap_saprotroph (Põlme et al. 2020); Decay sugar-rich substrates (Põlme et al. 2020) | Põlme S, et al. 2020. Fungal Diversity 105:1-16 ((https://doi.org/10.1007/s13225-020-00466-2)) |
| Otu56363 | 1 | k__Fungi(100);p__Ascomycota(100);c__Saccharomycetes(100);o__Saccharomycetales(100);f__Metschnikowiaceae(100);g__Metschnikowia(100);s__Metschnikowia_pulcherrima(100); | Metschnikowia | 13 | Saprotroph | \|Undefined Saprotroph\| | Yeast | NULL | Probable | As nectar-tap_saprotroph (Põlme et al. 2020); Decay sugar-rich substrates (Põlme et al. 2020) | Põlme S, et al. 2020. Fungal Diversity 105:1-16 ((https://doi.org/10.1007/s13225-020-00466-2)) |
| Otu56364 | 1 | k__Fungi(100);p__Ascomycota(100);c__Saccharomycetes(100);o__Saccharomycetales(100);f__Metschnikowiaceae(100);g__Metschnikowia(100);s__Metschnikowia_pulcherrima(100); | Metschnikowia | 13 | Saprotroph | \|Undefined Saprotroph\| | Yeast | NULL | Probable | As nectar-tap_saprotroph (Põlme et al. 2020); Decay sugar-rich substrates (Põlme et al. 2020) | Põlme S, et al. 2020. Fungal Diversity 105:1-16 ((https://doi.org/10.1007/s13225-020-00466-2)) |
| Otu56365 | 1 | k__Fungi(100);p__Ascomycota(100);c__Saccharomycetes(100);o__Saccharomycetales(100);f__Metschnikowiaceae(100);g__Metschnikowia(100);s__Metschnikowia_pulcherrima(100); | Metschnikowia | 13 | Saprotroph | \|Undefined Saprotroph\| | Yeast | NULL | Probable | As nectar-tap_saprotroph (Põlme et al. 2020); Decay sugar-rich substrates (Põlme et al. 2020) | Põlme S, et al. 2020. Fungal Diversity 105:1-16 ((https://doi.org/10.1007/s13225-020-00466-2)) |
| Otu56366 | 1 | k__Fungi(100);p__Ascomycota(100);c__Sordariomycetes(100);o__Hypocreales(100);f__Nectriaceae(100);g__Gibberella(100);s__Gibberella_zeae(100); | Gibberella | 13 | Pathotroph-Saprotroph-Symbiotroph | Endophyte-\|Plant Pathogen\|-Plant Saprotroph-Undefined Saprotroph | Microfungus | Soft Rot | Probable | As foliar_endophyte (Põlme et al. 2020); As litter_saprotroph (Põlme et al. 2020); Causing damping-off disease (Põlme et al. 2020); Decay leaf, fruit, seed, roots, soil, animal material (Põlme et al. 2020) | Tedersoo L, et al. 2014. Science 346:e1256688 ((https://doi.org/10.1126/science.1256688)); Põlme S, et al. 2020. Fungal Diversity 105:1-16 ((https://doi.org/10.1007/s13225-020-00466-2)) |
| Otu56367 | 1 | k__Fungi(100);p__Ascomycota(100);c__Saccharomycetes(100);o__Saccharomycetales(100);f__Metschnikowiaceae(100);g__Metschnikowia(100);s__Metschnikowia_pulcherrima(100); | Metschnikowia | 13 | Saprotroph | \|Undefined Saprotroph\| | Yeast | NULL | Probable | As nectar-tap_saprotroph (Põlme et al. 2020); Decay sugar-rich substrates (Põlme et al. 2020) | Põlme S, et al. 2020. Fungal Diversity 105:1-16 ((https://doi.org/10.1007/s13225-020-00466-2)) |
| Otu56368 | 1 | k__Fungi(100);p__Ascomycota(100);c__Saccharomycetes(100);o__Saccharomycetales(100);f__Metschnikowiaceae(100);g__Metschnikowia(100);g__Metschnikowia_unclassified(100); | Metschnikowia | 13 | Saprotroph | \|Undefined Saprotroph\| | Yeast | NULL | Probable | As nectar-tap_saprotroph (Põlme et al. 2020); Decay sugar-rich substrates (Põlme et al. 2020) | Põlme S, et al. 2020. Fungal Diversity 105:1-16 ((https://doi.org/10.1007/s13225-020-00466-2)) |
| Otu56369 | 1 | k__Fungi(100);p__Ascomycota(100);c__Saccharomycetes(100);o__Saccharomycetales(100);f__Metschnikowiaceae(100);g__Metschnikowia(100);s__Metschnikowia_pulcherrima(100); | Metschnikowia | 13 | Saprotroph | \|Undefined Saprotroph\| | Yeast | NULL | Probable | As nectar-tap_saprotroph (Põlme et al. 2020); Decay sugar-rich substrates (Põlme et al. 2020) | Põlme S, et al. 2020. Fungal Diversity 105:1-16 ((https://doi.org/10.1007/s13225-020-00466-2)) |
| Otu56370 | 1 | k__Fungi(100);p__Ascomycota(100);c__Saccharomycetes(100);o__Saccharomycetales(100);f__Metschnikowiaceae(100);g__Metschnikowia(100);s__Metschnikowia_pulcherrima(100); | Metschnikowia | 13 | Saprotroph | \|Undefined Saprotroph\| | Yeast | NULL | Probable | As nectar-tap_saprotroph (Põlme et al. 2020); Decay sugar-rich substrates (Põlme et al. 2020) | Põlme S, et al. 2020. Fungal Diversity 105:1-16 ((https://doi.org/10.1007/s13225-020-00466-2)) |
| Otu56371 | 1 | k__Fungi(100);p__Ascomycota(100);c__Saccharomycetes(100);o__Saccharomycetales(100);f__Metschnikowiaceae(100);g__Metschnikowia(100);s__Metschnikowia_pulcherrima(100); | Metschnikowia | 13 | Saprotroph | \|Undefined Saprotroph\| | Yeast | NULL | Probable | As nectar-tap_saprotroph (Põlme et al. 2020); Decay sugar-rich substrates (Põlme et al. 2020) | Põlme S, et al. 2020. Fungal Diversity 105:1-16 ((https://doi.org/10.1007/s13225-020-00466-2)) |
| Otu56372 | 1 | k__Fungi(100);p__Ascomycota(100);c__Saccharomycetes(100);o__Saccharomycetales(100);f__Metschnikowiaceae(100);g__Metschnikowia(100);s__Metschnikowia_pulcherrima(100); | Metschnikowia | 13 | Saprotroph | \|Undefined Saprotroph\| | Yeast | NULL | Probable | As nectar-tap_saprotroph (Põlme et al. 2020); Decay sugar-rich substrates (Põlme et al. 2020) | Põlme S, et al. 2020. Fungal Diversity 105:1-16 ((https://doi.org/10.1007/s13225-020-00466-2)) |
| Otu56373 | 1 | k__Fungi(100);p__Ascomycota(100);c__Saccharomycetes(100);o__Saccharomycetales(100);f__Metschnikowiaceae(100);g__Metschnikowia(100);s__Metschnikowia_pulcherrima(100); | Metschnikowia | 13 | Saprotroph | \|Undefined Saprotroph\| | Yeast | NULL | Probable | As nectar-tap_saprotroph (Põlme et al. 2020); Decay sugar-rich substrates (Põlme et al. 2020) | Põlme S, et al. 2020. Fungal Diversity 105:1-16 ((https://doi.org/10.1007/s13225-020-00466-2)) |
| Otu56374 | 1 | k__Fungi(100);p__Ascomycota(100);c__Saccharomycetes(100);o__Saccharomycetales(100);f__Metschnikowiaceae(100);g__Metschnikowia(100);s__Metschnikowia_pulcherrima(100); | Metschnikowia | 13 | Saprotroph | \|Undefined Saprotroph\| | Yeast | NULL | Probable | As nectar-tap_saprotroph (Põlme et al. 2020); Decay sugar-rich substrates (Põlme et al. 2020) | Põlme S, et al. 2020. Fungal Diversity 105:1-16 ((https://doi.org/10.1007/s13225-020-00466-2)) |
| Otu56375 | 1 | k__Fungi(100);p__Ascomycota(100);c__Saccharomycetes(100);o__Saccharomycetales(100);f__Metschnikowiaceae(100);g__Metschnikowia(100);s__Metschnikowia_pulcherrima(100); | Metschnikowia | 13 | Saprotroph | \|Undefined Saprotroph\| | Yeast | NULL | Probable | As nectar-tap_saprotroph (Põlme et al. 2020); Decay sugar-rich substrates (Põlme et al. 2020) | Põlme S, et al. 2020. Fungal Diversity 105:1-16 ((https://doi.org/10.1007/s13225-020-00466-2)) |
| Otu56376 | 1 | k__Fungi(100);p__Ascomycota(100);c__Saccharomycetes(100);o__Saccharomycetales(100);f__Metschnikowiaceae(100);g__Metschnikowia(100);s__Metschnikowia_pulcherrima(100); | Metschnikowia | 13 | Saprotroph | \|Undefined Saprotroph\| | Yeast | NULL | Probable | As nectar-tap_saprotroph (Põlme et al. 2020); Decay sugar-rich substrates (Põlme et al. 2020) | Põlme S, et al. 2020. Fungal Diversity 105:1-16 ((https://doi.org/10.1007/s13225-020-00466-2)) |
| Otu56377 | 1 | k__Fungi(100);p__Ascomycota(100);c__Saccharomycetes(100);o__Saccharomycetales(100);f__Metschnikowiaceae(100);g__Metschnikowia(100);s__Metschnikowia_pulcherrima(100); | Metschnikowia | 13 | Saprotroph | \|Undefined Saprotroph\| | Yeast | NULL | Probable | As nectar-tap_saprotroph (Põlme et al. 2020); Decay sugar-rich substrates (Põlme et al. 2020) | Põlme S, et al. 2020. Fungal Diversity 105:1-16 ((https://doi.org/10.1007/s13225-020-00466-2)) |
| Otu56380 | 1 | k__Fungi(100);p__Ascomycota(100);c__Saccharomycetes(100);o__Saccharomycetales(100);f__Metschnikowiaceae(100);g__Metschnikowia(100);g__Metschnikowia_unclassified(100); | Metschnikowia | 13 | Saprotroph | \|Undefined Saprotroph\| | Yeast | NULL | Probable | As nectar-tap_saprotroph (Põlme et al. 2020); Decay sugar-rich substrates (Põlme et al. 2020) | Põlme S, et al. 2020. Fungal Diversity 105:1-16 ((https://doi.org/10.1007/s13225-020-00466-2)) |
| Otu56381 | 1 | k__Fungi(100);p__Ascomycota(100);c__Saccharomycetes(100);o__Saccharomycetales(100);f__Metschnikowiaceae(100);g__Metschnikowia(100);s__Metschnikowia_pulcherrima(100); | Metschnikowia | 13 | Saprotroph | \|Undefined Saprotroph\| | Yeast | NULL | Probable | As nectar-tap_saprotroph (Põlme et al. 2020); Decay sugar-rich substrates (Põlme et al. 2020) | Põlme S, et al. 2020. Fungal Diversity 105:1-16 ((https://doi.org/10.1007/s13225-020-00466-2)) |
| Otu56382 | 1 | k__Fungi(100);p__Ascomycota(100);c__Saccharomycetes(100);o__Saccharomycetales(100);f__Metschnikowiaceae(100);g__Metschnikowia(100);s__Metschnikowia_pulcherrima(100); | Metschnikowia | 13 | Saprotroph | \|Undefined Saprotroph\| | Yeast | NULL | Probable | As nectar-tap_saprotroph (Põlme et al. 2020); Decay sugar-rich substrates (Põlme et al. 2020) | Põlme S, et al. 2020. Fungal Diversity 105:1-16 ((https://doi.org/10.1007/s13225-020-00466-2)) |
| Otu56383 | 1 | k__Fungi(100);p__Ascomycota(100);c__Saccharomycetes(100);o__Saccharomycetales(100);f__Metschnikowiaceae(100);g__Metschnikowia(100);s__Metschnikowia_pulcherrima(100); | Metschnikowia | 13 | Saprotroph | \|Undefined Saprotroph\| | Yeast | NULL | Probable | As nectar-tap_saprotroph (Põlme et al. 2020); Decay sugar-rich substrates (Põlme et al. 2020) | Põlme S, et al. 2020. Fungal Diversity 105:1-16 ((https://doi.org/10.1007/s13225-020-00466-2)) |
| Otu56387 | 1 | k__Fungi(100);p__Ascomycota(100);c__Saccharomycetes(100);o__Saccharomycetales(100);f__Metschnikowiaceae(100);g__Metschnikowia(100);s__Metschnikowia_pulcherrima(100); | Metschnikowia | 13 | Saprotroph | \|Undefined Saprotroph\| | Yeast | NULL | Probable | As nectar-tap_saprotroph (Põlme et al. 2020); Decay sugar-rich substrates (Põlme et al. 2020) | Põlme S, et al. 2020. Fungal Diversity 105:1-16 ((https://doi.org/10.1007/s13225-020-00466-2)) |
| Otu56389 | 1 | k__Fungi(100);p__Ascomycota(100);c__Saccharomycetes(100);o__Saccharomycetales(100);f__Saccharomycetales_family_Incertae_sedis(100);g__Candida(100);g__Candida_unclassified(100); | Candida | 13 | Pathotroph-Saprotroph-Symbiotroph | Animal Pathogen-Endophyte-Endosymbiont-Epiphyte-\|Undefined Saprotroph\| | Dimorphic Yeast-Yeast | NULL | Probable | As nectar-tap_saprotroph (Põlme et al. 2020); Decay sugar-rich substrates (Põlme et al. 2020); This asexual genus is polyphyletic and contain species that function very differently depend on what they are. Caution should be taken when interpreting guilds for Candida species. | Manolakaki D, et al. 2010. Virulence. 1: 367-375 ((https://doi.org/10.4161/viru.1.5.12796)); Mekbib SB, et al. 2011. Tropical Plant Pathology 36:233-240 ((https://doi.org/10.1590/S1982-56762011000400004)); Põlme S, et al. 2020. Fungal Diversity 105:1-16 ((https://doi.org/10.1007/s13225-020-00466-2)) |
| Otu56390 | 1 | k__Fungi(100);p__Ascomycota(100);c__Saccharomycetes(100);o__Saccharomycetales(100);f__Saccharomycetales_family_Incertae_sedis(100);f__Saccharomycetales_family_Incertae_sedis_unclassified(100);f__Saccharomycetales_family_Incertae_sedis_unclassified(100); | Saccharomycetales | 7 | Saprotroph | Undefined Saprotroph | Yeast | NULL | Possible | NULL | Sterkenburg E, et al. 2015. New Phytologist 207:1145-1158 ((https://doi.org/10.1111/nph.13426)) |
| Otu56392 | 1 | k__Fungi(100);p__Ascomycota(100);c__Saccharomycetes(100);o__Saccharomycetales(100);f__Metschnikowiaceae(100);g__Metschnikowia(100);g__Metschnikowia_unclassified(100); | Metschnikowia | 13 | Saprotroph | \|Undefined Saprotroph\| | Yeast | NULL | Probable | As nectar-tap_saprotroph (Põlme et al. 2020); Decay sugar-rich substrates (Põlme et al. 2020) | Põlme S, et al. 2020. Fungal Diversity 105:1-16 ((https://doi.org/10.1007/s13225-020-00466-2)) |
| Otu56394 | 1 | k__Fungi(100);p__Ascomycota(100);c__Saccharomycetes(100);o__Saccharomycetales(100);f__Metschnikowiaceae(100);g__Metschnikowia(100);s__Metschnikowia_pulcherrima(100); | Metschnikowia | 13 | Saprotroph | \|Undefined Saprotroph\| | Yeast | NULL | Probable | As nectar-tap_saprotroph (Põlme et al. 2020); Decay sugar-rich substrates (Põlme et al. 2020) | Põlme S, et al. 2020. Fungal Diversity 105:1-16 ((https://doi.org/10.1007/s13225-020-00466-2)) |
| Otu56395 | 1 | k__Fungi(100);p__Ascomycota(100);c__Saccharomycetes(100);o__Saccharomycetales(100);f__Metschnikowiaceae(100);g__Metschnikowia(100);s__Metschnikowia_pulcherrima(100); | Metschnikowia | 13 | Saprotroph | \|Undefined Saprotroph\| | Yeast | NULL | Probable | As nectar-tap_saprotroph (Põlme et al. 2020); Decay sugar-rich substrates (Põlme et al. 2020) | Põlme S, et al. 2020. Fungal Diversity 105:1-16 ((https://doi.org/10.1007/s13225-020-00466-2)) |
| Otu56396 | 1 | k__Fungi(100);p__Ascomycota(100);c__Saccharomycetes(100);o__Saccharomycetales(100);f__Metschnikowiaceae(100);g__Metschnikowia(100);s__Metschnikowia_pulcherrima(100); | Metschnikowia | 13 | Saprotroph | \|Undefined Saprotroph\| | Yeast | NULL | Probable | As nectar-tap_saprotroph (Põlme et al. 2020); Decay sugar-rich substrates (Põlme et al. 2020) | Põlme S, et al. 2020. Fungal Diversity 105:1-16 ((https://doi.org/10.1007/s13225-020-00466-2)) |
| Otu56397 | 1 | k__Fungi(100);p__Ascomycota(100);c__Saccharomycetes(100);o__Saccharomycetales(100);f__Metschnikowiaceae(100);g__Metschnikowia(100);s__Metschnikowia_pulcherrima(100); | Metschnikowia | 13 | Saprotroph | \|Undefined Saprotroph\| | Yeast | NULL | Probable | As nectar-tap_saprotroph (Põlme et al. 2020); Decay sugar-rich substrates (Põlme et al. 2020) | Põlme S, et al. 2020. Fungal Diversity 105:1-16 ((https://doi.org/10.1007/s13225-020-00466-2)) |
| Otu56400 | 1 | k__Fungi(100);p__Ascomycota(100);c__Saccharomycetes(100);o__Saccharomycetales(100);f__Metschnikowiaceae(100);g__Metschnikowia(100);g__Metschnikowia_unclassified(100); | Metschnikowia | 13 | Saprotroph | \|Undefined Saprotroph\| | Yeast | NULL | Probable | As nectar-tap_saprotroph (Põlme et al. 2020); Decay sugar-rich substrates (Põlme et al. 2020) | Põlme S, et al. 2020. Fungal Diversity 105:1-16 ((https://doi.org/10.1007/s13225-020-00466-2)) |
| Otu56401 | 1 | k__Fungi(100);p__Ascomycota(100);c__Saccharomycetes(100);o__Saccharomycetales(100);f__Metschnikowiaceae(100);g__Metschnikowia(100);g__Metschnikowia_unclassified(100); | Metschnikowia | 13 | Saprotroph | \|Undefined Saprotroph\| | Yeast | NULL | Probable | As nectar-tap_saprotroph (Põlme et al. 2020); Decay sugar-rich substrates (Põlme et al. 2020) | Põlme S, et al. 2020. Fungal Diversity 105:1-16 ((https://doi.org/10.1007/s13225-020-00466-2)) |
| Otu56402 | 1 | k__Fungi(100);p__Ascomycota(100);c__Saccharomycetes(100);o__Saccharomycetales(100);f__Metschnikowiaceae(100);g__Metschnikowia(100);s__Metschnikowia_pulcherrima(100); | Metschnikowia | 13 | Saprotroph | \|Undefined Saprotroph\| | Yeast | NULL | Probable | As nectar-tap_saprotroph (Põlme et al. 2020); Decay sugar-rich substrates (Põlme et al. 2020) | Põlme S, et al. 2020. Fungal Diversity 105:1-16 ((https://doi.org/10.1007/s13225-020-00466-2)) |
| Otu56403 | 1 | k__Fungi(100);p__Ascomycota(100);c__Saccharomycetes(100);o__Saccharomycetales(100);f__Metschnikowiaceae(100);g__Metschnikowia(100);s__Metschnikowia_pulcherrima(100); | Metschnikowia | 13 | Saprotroph | \|Undefined Saprotroph\| | Yeast | NULL | Probable | As nectar-tap_saprotroph (Põlme et al. 2020); Decay sugar-rich substrates (Põlme et al. 2020) | Põlme S, et al. 2020. Fungal Diversity 105:1-16 ((https://doi.org/10.1007/s13225-020-00466-2)) |
| Otu56406 | 1 | k__Fungi(100);p__Ascomycota(100);c__Saccharomycetes(100);o__Saccharomycetales(100);f__Metschnikowiaceae(100);g__Metschnikowia(100);g__Metschnikowia_unclassified(100); | Metschnikowia | 13 | Saprotroph | \|Undefined Saprotroph\| | Yeast | NULL | Probable | As nectar-tap_saprotroph (Põlme et al. 2020); Decay sugar-rich substrates (Põlme et al. 2020) | Põlme S, et al. 2020. Fungal Diversity 105:1-16 ((https://doi.org/10.1007/s13225-020-00466-2)) |
| Otu56407 | 1 | k__Fungi(100);p__Ascomycota(100);c__Saccharomycetes(100);o__Saccharomycetales(100);f__Metschnikowiaceae(100);g__Metschnikowia(100);s__Metschnikowia_pulcherrima(100); | Metschnikowia | 13 | Saprotroph | \|Undefined Saprotroph\| | Yeast | NULL | Probable | As nectar-tap_saprotroph (Põlme et al. 2020); Decay sugar-rich substrates (Põlme et al. 2020) | Põlme S, et al. 2020. Fungal Diversity 105:1-16 ((https://doi.org/10.1007/s13225-020-00466-2)) |
| Otu56408 | 1 | k__Fungi(100);p__Ascomycota(100);c__Saccharomycetes(100);o__Saccharomycetales(100);f__Metschnikowiaceae(100);g__Metschnikowia(100);s__Metschnikowia_pulcherrima(100); | Metschnikowia | 13 | Saprotroph | \|Undefined Saprotroph\| | Yeast | NULL | Probable | As nectar-tap_saprotroph (Põlme et al. 2020); Decay sugar-rich substrates (Põlme et al. 2020) | Põlme S, et al. 2020. Fungal Diversity 105:1-16 ((https://doi.org/10.1007/s13225-020-00466-2)) |
| Otu56409 | 1 | k__Fungi(100);p__Ascomycota(100);c__Saccharomycetes(100);o__Saccharomycetales(100);f__Metschnikowiaceae(100);g__Metschnikowia(100);s__Metschnikowia_pulcherrima(100); | Metschnikowia | 13 | Saprotroph | \|Undefined Saprotroph\| | Yeast | NULL | Probable | As nectar-tap_saprotroph (Põlme et al. 2020); Decay sugar-rich substrates (Põlme et al. 2020) | Põlme S, et al. 2020. Fungal Diversity 105:1-16 ((https://doi.org/10.1007/s13225-020-00466-2)) |
| Otu56410 | 1 | k__Fungi(100);p__Ascomycota(100);c__Saccharomycetes(100);o__Saccharomycetales(100);f__Saccharomycetales_family_Incertae_sedis(100);f__Saccharomycetales_family_Incertae_sedis_unclassified(100);f__Saccharomycetales_family_Incertae_sedis_unclassified(100); | Saccharomycetales | 7 | Saprotroph | Undefined Saprotroph | Yeast | NULL | Possible | NULL | Sterkenburg E, et al. 2015. New Phytologist 207:1145-1158 ((https://doi.org/10.1111/nph.13426)) |
| Otu56411 | 1 | k__Fungi(100);p__Ascomycota(100);c__Saccharomycetes(100);o__Saccharomycetales(100);f__Metschnikowiaceae(100);g__Metschnikowia(100);g__Metschnikowia_unclassified(100); | Metschnikowia | 13 | Saprotroph | \|Undefined Saprotroph\| | Yeast | NULL | Probable | As nectar-tap_saprotroph (Põlme et al. 2020); Decay sugar-rich substrates (Põlme et al. 2020) | Põlme S, et al. 2020. Fungal Diversity 105:1-16 ((https://doi.org/10.1007/s13225-020-00466-2)) |
| Otu56417 | 1 | k__Fungi(100);p__Ascomycota(100);c__Saccharomycetes(100);o__Saccharomycetales(100);f__Metschnikowiaceae(100);g__Metschnikowia(100);s__Metschnikowia_pulcherrima(100); | Metschnikowia | 13 | Saprotroph | \|Undefined Saprotroph\| | Yeast | NULL | Probable | As nectar-tap_saprotroph (Põlme et al. 2020); Decay sugar-rich substrates (Põlme et al. 2020) | Põlme S, et al. 2020. Fungal Diversity 105:1-16 ((https://doi.org/10.1007/s13225-020-00466-2)) |
| Otu56421 | 1 | k__Fungi(100);p__Ascomycota(100);c__Saccharomycetes(100);o__Saccharomycetales(100);f__Metschnikowiaceae(100);g__Metschnikowia(100);s__Metschnikowia_pulcherrima(100); | Metschnikowia | 13 | Saprotroph | \|Undefined Saprotroph\| | Yeast | NULL | Probable | As nectar-tap_saprotroph (Põlme et al. 2020); Decay sugar-rich substrates (Põlme et al. 2020) | Põlme S, et al. 2020. Fungal Diversity 105:1-16 ((https://doi.org/10.1007/s13225-020-00466-2)) |
| Otu56422 | 1 | k__Fungi(100);p__Ascomycota(100);c__Saccharomycetes(100);o__Saccharomycetales(100);f__Metschnikowiaceae(100);g__Metschnikowia(100);s__Metschnikowia_pulcherrima(100); | Metschnikowia | 13 | Saprotroph | \|Undefined Saprotroph\| | Yeast | NULL | Probable | As nectar-tap_saprotroph (Põlme et al. 2020); Decay sugar-rich substrates (Põlme et al. 2020) | Põlme S, et al. 2020. Fungal Diversity 105:1-16 ((https://doi.org/10.1007/s13225-020-00466-2)) |
| Otu56426 | 1 | k__Fungi(100);p__Ascomycota(100);c__Saccharomycetes(100);o__Saccharomycetales(100);f__Metschnikowiaceae(100);g__Metschnikowia(100);s__Metschnikowia_pulcherrima(100); | Metschnikowia | 13 | Saprotroph | \|Undefined Saprotroph\| | Yeast | NULL | Probable | As nectar-tap_saprotroph (Põlme et al. 2020); Decay sugar-rich substrates (Põlme et al. 2020) | Põlme S, et al. 2020. Fungal Diversity 105:1-16 ((https://doi.org/10.1007/s13225-020-00466-2)) |
| Otu56428 | 1 | k__Fungi(100);p__Ascomycota(100);c__Saccharomycetes(100);o__Saccharomycetales(100);f__Metschnikowiaceae(100);g__Metschnikowia(100);s__Metschnikowia_pulcherrima(100); | Metschnikowia | 13 | Saprotroph | \|Undefined Saprotroph\| | Yeast | NULL | Probable | As nectar-tap_saprotroph (Põlme et al. 2020); Decay sugar-rich substrates (Põlme et al. 2020) | Põlme S, et al. 2020. Fungal Diversity 105:1-16 ((https://doi.org/10.1007/s13225-020-00466-2)) |
| Otu56430 | 1 | k__Fungi(100);p__Ascomycota(100);c__Saccharomycetes(100);o__Saccharomycetales(100);f__Metschnikowiaceae(100);g__Metschnikowia(100);s__Metschnikowia_pulcherrima(100); | Metschnikowia | 13 | Saprotroph | \|Undefined Saprotroph\| | Yeast | NULL | Probable | As nectar-tap_saprotroph (Põlme et al. 2020); Decay sugar-rich substrates (Põlme et al. 2020) | Põlme S, et al. 2020. Fungal Diversity 105:1-16 ((https://doi.org/10.1007/s13225-020-00466-2)) |
| Otu56431 | 1 | k__Fungi(100);p__Ascomycota(100);c__Saccharomycetes(100);o__Saccharomycetales(100);f__Metschnikowiaceae(100);g__Metschnikowia(100);s__Metschnikowia_pulcherrima(100); | Metschnikowia | 13 | Saprotroph | \|Undefined Saprotroph\| | Yeast | NULL | Probable | As nectar-tap_saprotroph (Põlme et al. 2020); Decay sugar-rich substrates (Põlme et al. 2020) | Põlme S, et al. 2020. Fungal Diversity 105:1-16 ((https://doi.org/10.1007/s13225-020-00466-2)) |
| Otu56432 | 1 | k__Fungi(100);p__Ascomycota(100);c__Saccharomycetes(100);o__Saccharomycetales(100);f__Metschnikowiaceae(100);g__Metschnikowia(100);s__Metschnikowia_pulcherrima(100); | Metschnikowia | 13 | Saprotroph | \|Undefined Saprotroph\| | Yeast | NULL | Probable | As nectar-tap_saprotroph (Põlme et al. 2020); Decay sugar-rich substrates (Põlme et al. 2020) | Põlme S, et al. 2020. Fungal Diversity 105:1-16 ((https://doi.org/10.1007/s13225-020-00466-2)) |
| Otu56433 | 1 | k__Fungi(100);p__Ascomycota(100);c__Saccharomycetes(100);o__Saccharomycetales(100);f__Metschnikowiaceae(100);g__Metschnikowia(100);g__Metschnikowia_unclassified(100); | Metschnikowia | 13 | Saprotroph | \|Undefined Saprotroph\| | Yeast | NULL | Probable | As nectar-tap_saprotroph (Põlme et al. 2020); Decay sugar-rich substrates (Põlme et al. 2020) | Põlme S, et al. 2020. Fungal Diversity 105:1-16 ((https://doi.org/10.1007/s13225-020-00466-2)) |
| Otu56434 | 1 | k__Fungi(100);p__Ascomycota(100);c__Saccharomycetes(100);o__Saccharomycetales(100);f__Metschnikowiaceae(100);g__Metschnikowia(100);s__Metschnikowia_pulcherrima(100); | Metschnikowia | 13 | Saprotroph | \|Undefined Saprotroph\| | Yeast | NULL | Probable | As nectar-tap_saprotroph (Põlme et al. 2020); Decay sugar-rich substrates (Põlme et al. 2020) | Põlme S, et al. 2020. Fungal Diversity 105:1-16 ((https://doi.org/10.1007/s13225-020-00466-2)) |
| Otu56439 | 1 | k__Fungi(100);p__Ascomycota(100);c__Saccharomycetes(100);o__Saccharomycetales(100);f__Metschnikowiaceae(100);g__Metschnikowia(100);s__Metschnikowia_pulcherrima(100); | Metschnikowia | 13 | Saprotroph | \|Undefined Saprotroph\| | Yeast | NULL | Probable | As nectar-tap_saprotroph (Põlme et al. 2020); Decay sugar-rich substrates (Põlme et al. 2020) | Põlme S, et al. 2020. Fungal Diversity 105:1-16 ((https://doi.org/10.1007/s13225-020-00466-2)) |
| Otu56440 | 1 | k__Fungi(100);p__Ascomycota(100);c__Saccharomycetes(100);o__Saccharomycetales(100);f__Metschnikowiaceae(100);g__Metschnikowia(100);s__Metschnikowia_pulcherrima(100); | Metschnikowia | 13 | Saprotroph | \|Undefined Saprotroph\| | Yeast | NULL | Probable | As nectar-tap_saprotroph (Põlme et al. 2020); Decay sugar-rich substrates (Põlme et al. 2020) | Põlme S, et al. 2020. Fungal Diversity 105:1-16 ((https://doi.org/10.1007/s13225-020-00466-2)) |
| Otu56453 | 1 | k__Fungi(100);p__Ascomycota(100);c__Saccharomycetes(100);o__Saccharomycetales(100);f__Metschnikowiaceae(100);g__Metschnikowia(100);s__Metschnikowia_pulcherrima(100); | Metschnikowia | 13 | Saprotroph | \|Undefined Saprotroph\| | Yeast | NULL | Probable | As nectar-tap_saprotroph (Põlme et al. 2020); Decay sugar-rich substrates (Põlme et al. 2020) | Põlme S, et al. 2020. Fungal Diversity 105:1-16 ((https://doi.org/10.1007/s13225-020-00466-2)) |
| Otu56458 | 1 | k__Fungi(100);p__Ascomycota(100);c__Saccharomycetes(100);o__Saccharomycetales(100);f__Metschnikowiaceae(100);g__Metschnikowia(100);s__Metschnikowia_pulcherrima(100); | Metschnikowia | 13 | Saprotroph | \|Undefined Saprotroph\| | Yeast | NULL | Probable | As nectar-tap_saprotroph (Põlme et al. 2020); Decay sugar-rich substrates (Põlme et al. 2020) | Põlme S, et al. 2020. Fungal Diversity 105:1-16 ((https://doi.org/10.1007/s13225-020-00466-2)) |
| Otu56459 | 1 | k__Fungi(100);p__Ascomycota(100);c__Saccharomycetes(100);o__Saccharomycetales(100);f__Metschnikowiaceae(100);g__Metschnikowia(100);s__Metschnikowia_pulcherrima(100); | Metschnikowia | 13 | Saprotroph | \|Undefined Saprotroph\| | Yeast | NULL | Probable | As nectar-tap_saprotroph (Põlme et al. 2020); Decay sugar-rich substrates (Põlme et al. 2020) | Põlme S, et al. 2020. Fungal Diversity 105:1-16 ((https://doi.org/10.1007/s13225-020-00466-2)) |
| Otu56460 | 1 | k__Fungi(100);p__Ascomycota(100);c__Saccharomycetes(100);o__Saccharomycetales(100);f__Metschnikowiaceae(100);g__Metschnikowia(100);s__Metschnikowia_pulcherrima(100); | Metschnikowia | 13 | Saprotroph | \|Undefined Saprotroph\| | Yeast | NULL | Probable | As nectar-tap_saprotroph (Põlme et al. 2020); Decay sugar-rich substrates (Põlme et al. 2020) | Põlme S, et al. 2020. Fungal Diversity 105:1-16 ((https://doi.org/10.1007/s13225-020-00466-2)) |
| Otu56461 | 1 | k__Fungi(100);p__Ascomycota(100);c__Saccharomycetes(100);o__Saccharomycetales(100);f__Metschnikowiaceae(100);g__Metschnikowia(100);s__Metschnikowia_pulcherrima(100); | Metschnikowia | 13 | Saprotroph | \|Undefined Saprotroph\| | Yeast | NULL | Probable | As nectar-tap_saprotroph (Põlme et al. 2020); Decay sugar-rich substrates (Põlme et al. 2020) | Põlme S, et al. 2020. Fungal Diversity 105:1-16 ((https://doi.org/10.1007/s13225-020-00466-2)) |
| Otu56462 | 1 | k__Fungi(100);p__Ascomycota(100);c__Saccharomycetes(100);o__Saccharomycetales(100);f__Metschnikowiaceae(100);g__Metschnikowia(100);s__Metschnikowia_pulcherrima(100); | Metschnikowia | 13 | Saprotroph | \|Undefined Saprotroph\| | Yeast | NULL | Probable | As nectar-tap_saprotroph (Põlme et al. 2020); Decay sugar-rich substrates (Põlme et al. 2020) | Põlme S, et al. 2020. Fungal Diversity 105:1-16 ((https://doi.org/10.1007/s13225-020-00466-2)) |
| Otu56463 | 1 | k__Fungi(100);p__Ascomycota(100);c__Saccharomycetes(100);o__Saccharomycetales(100);f__Metschnikowiaceae(100);g__Metschnikowia(100);s__Metschnikowia_pulcherrima(100); | Metschnikowia | 13 | Saprotroph | \|Undefined Saprotroph\| | Yeast | NULL | Probable | As nectar-tap_saprotroph (Põlme et al. 2020); Decay sugar-rich substrates (Põlme et al. 2020) | Põlme S, et al. 2020. Fungal Diversity 105:1-16 ((https://doi.org/10.1007/s13225-020-00466-2)) |
| Otu56464 | 1 | k__Fungi(100);p__Ascomycota(100);c__Saccharomycetes(100);o__Saccharomycetales(100);f__Metschnikowiaceae(100);g__Metschnikowia(100);s__Metschnikowia_pulcherrima(100); | Metschnikowia | 13 | Saprotroph | \|Undefined Saprotroph\| | Yeast | NULL | Probable | As nectar-tap_saprotroph (Põlme et al. 2020); Decay sugar-rich substrates (Põlme et al. 2020) | Põlme S, et al. 2020. Fungal Diversity 105:1-16 ((https://doi.org/10.1007/s13225-020-00466-2)) |
| Otu56465 | 1 | k__Fungi(100);p__Ascomycota(100);c__Saccharomycetes(100);o__Saccharomycetales(100);f__Metschnikowiaceae(100);g__Metschnikowia(100);s__Metschnikowia_pulcherrima(100); | Metschnikowia | 13 | Saprotroph | \|Undefined Saprotroph\| | Yeast | NULL | Probable | As nectar-tap_saprotroph (Põlme et al. 2020); Decay sugar-rich substrates (Põlme et al. 2020) | Põlme S, et al. 2020. Fungal Diversity 105:1-16 ((https://doi.org/10.1007/s13225-020-00466-2)) |
| Otu56466 | 1 | k__Fungi(100);p__Ascomycota(100);c__Saccharomycetes(100);o__Saccharomycetales(100);f__Metschnikowiaceae(100);g__Metschnikowia(100);s__Metschnikowia_pulcherrima(100); | Metschnikowia | 13 | Saprotroph | \|Undefined Saprotroph\| | Yeast | NULL | Probable | As nectar-tap_saprotroph (Põlme et al. 2020); Decay sugar-rich substrates (Põlme et al. 2020) | Põlme S, et al. 2020. Fungal Diversity 105:1-16 ((https://doi.org/10.1007/s13225-020-00466-2)) |
| Otu56467 | 1 | k__Fungi(100);p__Ascomycota(100);c__Saccharomycetes(100);o__Saccharomycetales(100);f__Metschnikowiaceae(100);g__Metschnikowia(100);s__Metschnikowia_pulcherrima(100); | Metschnikowia | 13 | Saprotroph | \|Undefined Saprotroph\| | Yeast | NULL | Probable | As nectar-tap_saprotroph (Põlme et al. 2020); Decay sugar-rich substrates (Põlme et al. 2020) | Põlme S, et al. 2020. Fungal Diversity 105:1-16 ((https://doi.org/10.1007/s13225-020-00466-2)) |
| Otu56470 | 1 | k__Fungi(100);p__Ascomycota(100);c__Saccharomycetes(100);o__Saccharomycetales(100);f__Metschnikowiaceae(100);g__Metschnikowia(100);s__Metschnikowia_pulcherrima(100); | Metschnikowia | 13 | Saprotroph | \|Undefined Saprotroph\| | Yeast | NULL | Probable | As nectar-tap_saprotroph (Põlme et al. 2020); Decay sugar-rich substrates (Põlme et al. 2020) | Põlme S, et al. 2020. Fungal Diversity 105:1-16 ((https://doi.org/10.1007/s13225-020-00466-2)) |
| Otu56471 | 1 | k__Fungi(100);p__Ascomycota(100);c__Saccharomycetes(100);o__Saccharomycetales(100);f__Metschnikowiaceae(100);g__Metschnikowia(100);g__Metschnikowia_unclassified(100); | Metschnikowia | 13 | Saprotroph | \|Undefined Saprotroph\| | Yeast | NULL | Probable | As nectar-tap_saprotroph (Põlme et al. 2020); Decay sugar-rich substrates (Põlme et al. 2020) | Põlme S, et al. 2020. Fungal Diversity 105:1-16 ((https://doi.org/10.1007/s13225-020-00466-2)) |
| Otu56473 | 1 | k__Fungi(100);p__Ascomycota(100);c__Saccharomycetes(100);o__Saccharomycetales(100);f__Metschnikowiaceae(100);g__Metschnikowia(100);g__Metschnikowia_unclassified(100); | Metschnikowia | 13 | Saprotroph | \|Undefined Saprotroph\| | Yeast | NULL | Probable | As nectar-tap_saprotroph (Põlme et al. 2020); Decay sugar-rich substrates (Põlme et al. 2020) | Põlme S, et al. 2020. Fungal Diversity 105:1-16 ((https://doi.org/10.1007/s13225-020-00466-2)) |
| Otu56474 | 1 | k__Fungi(100);p__Ascomycota(100);c__Saccharomycetes(100);o__Saccharomycetales(100);f__Metschnikowiaceae(100);g__Metschnikowia(100);s__Metschnikowia_pulcherrima(100); | Metschnikowia | 13 | Saprotroph | \|Undefined Saprotroph\| | Yeast | NULL | Probable | As nectar-tap_saprotroph (Põlme et al. 2020); Decay sugar-rich substrates (Põlme et al. 2020) | Põlme S, et al. 2020. Fungal Diversity 105:1-16 ((https://doi.org/10.1007/s13225-020-00466-2)) |
| Otu56475 | 1 | k__Fungi(100);p__Ascomycota(100);c__Saccharomycetes(100);o__Saccharomycetales(100);f__Metschnikowiaceae(100);g__Metschnikowia(100);g__Metschnikowia_unclassified(100); | Metschnikowia | 13 | Saprotroph | \|Undefined Saprotroph\| | Yeast | NULL | Probable | As nectar-tap_saprotroph (Põlme et al. 2020); Decay sugar-rich substrates (Põlme et al. 2020) | Põlme S, et al. 2020. Fungal Diversity 105:1-16 ((https://doi.org/10.1007/s13225-020-00466-2)) |
| Otu56476 | 1 | k__Fungi(100);p__Ascomycota(100);c__Saccharomycetes(100);o__Saccharomycetales(100);f__Metschnikowiaceae(100);g__Metschnikowia(100);s__Metschnikowia_pulcherrima(100); | Metschnikowia | 13 | Saprotroph | \|Undefined Saprotroph\| | Yeast | NULL | Probable | As nectar-tap_saprotroph (Põlme et al. 2020); Decay sugar-rich substrates (Põlme et al. 2020) | Põlme S, et al. 2020. Fungal Diversity 105:1-16 ((https://doi.org/10.1007/s13225-020-00466-2)) |
| Otu56477 | 1 | k__Fungi(100);p__Ascomycota(100);c__Saccharomycetes(100);o__Saccharomycetales(100);f__Metschnikowiaceae(100);g__Metschnikowia(100);s__Metschnikowia_pulcherrima(100); | Metschnikowia | 13 | Saprotroph | \|Undefined Saprotroph\| | Yeast | NULL | Probable | As nectar-tap_saprotroph (Põlme et al. 2020); Decay sugar-rich substrates (Põlme et al. 2020) | Põlme S, et al. 2020. Fungal Diversity 105:1-16 ((https://doi.org/10.1007/s13225-020-00466-2)) |
| Otu56484 | 1 | k__Fungi(100);p__Ascomycota(100);c__Saccharomycetes(100);o__Saccharomycetales(100);f__Metschnikowiaceae(100);g__Metschnikowia(100);s__Metschnikowia_pulcherrima(100); | Metschnikowia | 13 | Saprotroph | \|Undefined Saprotroph\| | Yeast | NULL | Probable | As nectar-tap_saprotroph (Põlme et al. 2020); Decay sugar-rich substrates (Põlme et al. 2020) | Põlme S, et al. 2020. Fungal Diversity 105:1-16 ((https://doi.org/10.1007/s13225-020-00466-2)) |
| Otu56486 | 1 | k__Fungi(100);p__Ascomycota(100);c__Saccharomycetes(100);o__Saccharomycetales(100);f__Metschnikowiaceae(100);g__Metschnikowia(100);s__Metschnikowia_pulcherrima(100); | Metschnikowia | 13 | Saprotroph | \|Undefined Saprotroph\| | Yeast | NULL | Probable | As nectar-tap_saprotroph (Põlme et al. 2020); Decay sugar-rich substrates (Põlme et al. 2020) | Põlme S, et al. 2020. Fungal Diversity 105:1-16 ((https://doi.org/10.1007/s13225-020-00466-2)) |
| Otu56489 | 1 | k__Fungi(100);p__Ascomycota(100);c__Saccharomycetes(100);o__Saccharomycetales(100);f__Saccharomycetales_family_Incertae_sedis(100);g__Candida(100);g__Candida_unclassified(100); | Candida | 13 | Pathotroph-Saprotroph-Symbiotroph | Animal Pathogen-Endophyte-Endosymbiont-Epiphyte-\|Undefined Saprotroph\| | Dimorphic Yeast-Yeast | NULL | Probable | As nectar-tap_saprotroph (Põlme et al. 2020); Decay sugar-rich substrates (Põlme et al. 2020); This asexual genus is polyphyletic and contain species that function very differently depend on what they are. Caution should be taken when interpreting guilds for Candida species. | Manolakaki D, et al. 2010. Virulence. 1: 367-375 ((https://doi.org/10.4161/viru.1.5.12796)); Mekbib SB, et al. 2011. Tropical Plant Pathology 36:233-240 ((https://doi.org/10.1590/S1982-56762011000400004)); Põlme S, et al. 2020. Fungal Diversity 105:1-16 ((https://doi.org/10.1007/s13225-020-00466-2)) |
| Otu56490 | 1 | k__Fungi(100);p__Ascomycota(100);c__Saccharomycetes(100);o__Saccharomycetales(100);f__Saccharomycetales_family_Incertae_sedis(100);g__Candida(100);s__Candida_sp_AS23486(100); | Candida | 13 | Pathotroph-Saprotroph-Symbiotroph | Animal Pathogen-Endophyte-Endosymbiont-Epiphyte-\|Undefined Saprotroph\| | Dimorphic Yeast-Yeast | NULL | Probable | As nectar-tap_saprotroph (Põlme et al. 2020); Decay sugar-rich substrates (Põlme et al. 2020); This asexual genus is polyphyletic and contain species that function very differently depend on what they are. Caution should be taken when interpreting guilds for Candida species. | Manolakaki D, et al. 2010. Virulence. 1: 367-375 ((https://doi.org/10.4161/viru.1.5.12796)); Mekbib SB, et al. 2011. Tropical Plant Pathology 36:233-240 ((https://doi.org/10.1590/S1982-56762011000400004)); Põlme S, et al. 2020. Fungal Diversity 105:1-16 ((https://doi.org/10.1007/s13225-020-00466-2)) |
| Otu56495 | 1 | k__Fungi(100);p__Ascomycota(100);c__Saccharomycetes(100);o__Saccharomycetales(100);f__Metschnikowiaceae(100);g__Metschnikowia(100);s__Metschnikowia_pulcherrima(100); | Metschnikowia | 13 | Saprotroph | \|Undefined Saprotroph\| | Yeast | NULL | Probable | As nectar-tap_saprotroph (Põlme et al. 2020); Decay sugar-rich substrates (Põlme et al. 2020) | Põlme S, et al. 2020. Fungal Diversity 105:1-16 ((https://doi.org/10.1007/s13225-020-00466-2)) |
| Otu56496 | 1 | k__Fungi(100);p__Ascomycota(100);c__Saccharomycetes(100);o__Saccharomycetales(100);f__Metschnikowiaceae(100);g__Metschnikowia(100);s__Metschnikowia_pulcherrima(100); | Metschnikowia | 13 | Saprotroph | \|Undefined Saprotroph\| | Yeast | NULL | Probable | As nectar-tap_saprotroph (Põlme et al. 2020); Decay sugar-rich substrates (Põlme et al. 2020) | Põlme S, et al. 2020. Fungal Diversity 105:1-16 ((https://doi.org/10.1007/s13225-020-00466-2)) |
| Otu56497 | 1 | k__Fungi(100);p__Ascomycota(100);c__Saccharomycetes(100);o__Saccharomycetales(100);f__Metschnikowiaceae(100);g__Metschnikowia(100);s__Metschnikowia_pulcherrima(100); | Metschnikowia | 13 | Saprotroph | \|Undefined Saprotroph\| | Yeast | NULL | Probable | As nectar-tap_saprotroph (Põlme et al. 2020); Decay sugar-rich substrates (Põlme et al. 2020) | Põlme S, et al. 2020. Fungal Diversity 105:1-16 ((https://doi.org/10.1007/s13225-020-00466-2)) |
| Otu56498 | 1 | k__Fungi(100);p__Ascomycota(100);c__Saccharomycetes(100);o__Saccharomycetales(100);f__Metschnikowiaceae(100);g__Metschnikowia(100);s__Metschnikowia_pulcherrima(100); | Metschnikowia | 13 | Saprotroph | \|Undefined Saprotroph\| | Yeast | NULL | Probable | As nectar-tap_saprotroph (Põlme et al. 2020); Decay sugar-rich substrates (Põlme et al. 2020) | Põlme S, et al. 2020. Fungal Diversity 105:1-16 ((https://doi.org/10.1007/s13225-020-00466-2)) |
| Otu56499 | 1 | k__Fungi(100);p__Ascomycota(100);c__Saccharomycetes(100);o__Saccharomycetales(100);f__Metschnikowiaceae(100);g__Metschnikowia(100);g__Metschnikowia_unclassified(100); | Metschnikowia | 13 | Saprotroph | \|Undefined Saprotroph\| | Yeast | NULL | Probable | As nectar-tap_saprotroph (Põlme et al. 2020); Decay sugar-rich substrates (Põlme et al. 2020) | Põlme S, et al. 2020. Fungal Diversity 105:1-16 ((https://doi.org/10.1007/s13225-020-00466-2)) |
| Otu56500 | 1 | k__Fungi(100);p__Ascomycota(100);c__Saccharomycetes(100);o__Saccharomycetales(100);f__Metschnikowiaceae(100);g__Metschnikowia(100);s__Metschnikowia_pulcherrima(100); | Metschnikowia | 13 | Saprotroph | \|Undefined Saprotroph\| | Yeast | NULL | Probable | As nectar-tap_saprotroph (Põlme et al. 2020); Decay sugar-rich substrates (Põlme et al. 2020) | Põlme S, et al. 2020. Fungal Diversity 105:1-16 ((https://doi.org/10.1007/s13225-020-00466-2)) |
| Otu56501 | 1 | k__Fungi(100);p__Ascomycota(100);c__Saccharomycetes(100);o__Saccharomycetales(100);f__Metschnikowiaceae(100);g__Metschnikowia(100);s__Metschnikowia_pulcherrima(100); | Metschnikowia | 13 | Saprotroph | \|Undefined Saprotroph\| | Yeast | NULL | Probable | As nectar-tap_saprotroph (Põlme et al. 2020); Decay sugar-rich substrates (Põlme et al. 2020) | Põlme S, et al. 2020. Fungal Diversity 105:1-16 ((https://doi.org/10.1007/s13225-020-00466-2)) |
| Otu56510 | 1 | k__Fungi(100);p__Ascomycota(100);c__Saccharomycetes(100);o__Saccharomycetales(100);f__Metschnikowiaceae(100);g__Metschnikowia(100);s__Metschnikowia_pulcherrima(100); | Metschnikowia | 13 | Saprotroph | \|Undefined Saprotroph\| | Yeast | NULL | Probable | As nectar-tap_saprotroph (Põlme et al. 2020); Decay sugar-rich substrates (Põlme et al. 2020) | Põlme S, et al. 2020. Fungal Diversity 105:1-16 ((https://doi.org/10.1007/s13225-020-00466-2)) |
| Otu56511 | 1 | k__Fungi(100);p__Ascomycota(100);c__Saccharomycetes(100);o__Saccharomycetales(100);f__Saccharomycetales_family_Incertae_sedis(100);g__Candida(100);g__Candida_unclassified(100); | Candida | 13 | Pathotroph-Saprotroph-Symbiotroph | Animal Pathogen-Endophyte-Endosymbiont-Epiphyte-\|Undefined Saprotroph\| | Dimorphic Yeast-Yeast | NULL | Probable | As nectar-tap_saprotroph (Põlme et al. 2020); Decay sugar-rich substrates (Põlme et al. 2020); This asexual genus is polyphyletic and contain species that function very differently depend on what they are. Caution should be taken when interpreting guilds for Candida species. | Manolakaki D, et al. 2010. Virulence. 1: 367-375 ((https://doi.org/10.4161/viru.1.5.12796)); Mekbib SB, et al. 2011. Tropical Plant Pathology 36:233-240 ((https://doi.org/10.1590/S1982-56762011000400004)); Põlme S, et al. 2020. Fungal Diversity 105:1-16 ((https://doi.org/10.1007/s13225-020-00466-2)) |
| Otu56512 | 1 | k__Fungi(100);p__Ascomycota(100);c__Saccharomycetes(100);o__Saccharomycetales(100);f__Saccharomycetales_family_Incertae_sedis(100);g__Candida(100);s__Candida_stellata(100); | Candida | 13 | Pathotroph-Saprotroph-Symbiotroph | Animal Pathogen-Endophyte-Endosymbiont-Epiphyte-\|Undefined Saprotroph\| | Dimorphic Yeast-Yeast | NULL | Probable | As nectar-tap_saprotroph (Põlme et al. 2020); Decay sugar-rich substrates (Põlme et al. 2020); This asexual genus is polyphyletic and contain species that function very differently depend on what they are. Caution should be taken when interpreting guilds for Candida species. | Manolakaki D, et al. 2010. Virulence. 1: 367-375 ((https://doi.org/10.4161/viru.1.5.12796)); Mekbib SB, et al. 2011. Tropical Plant Pathology 36:233-240 ((https://doi.org/10.1590/S1982-56762011000400004)); Põlme S, et al. 2020. Fungal Diversity 105:1-16 ((https://doi.org/10.1007/s13225-020-00466-2)) |
| Otu56513 | 1 | k__Fungi(100);p__Ascomycota(100);c__Saccharomycetes(100);o__Saccharomycetales(100);f__Metschnikowiaceae(100);g__Metschnikowia(100);s__Metschnikowia_pulcherrima(100); | Metschnikowia | 13 | Saprotroph | \|Undefined Saprotroph\| | Yeast | NULL | Probable | As nectar-tap_saprotroph (Põlme et al. 2020); Decay sugar-rich substrates (Põlme et al. 2020) | Põlme S, et al. 2020. Fungal Diversity 105:1-16 ((https://doi.org/10.1007/s13225-020-00466-2)) |
| Otu56516 | 1 | k__Fungi(100);p__Ascomycota(100);c__Saccharomycetes(100);o__Saccharomycetales(100);f__Saccharomycetales_family_Incertae_sedis(100);f__Saccharomycetales_family_Incertae_sedis_unclassified(100);f__Saccharomycetales_family_Incertae_sedis_unclassified(100); | Saccharomycetales | 7 | Saprotroph | Undefined Saprotroph | Yeast | NULL | Possible | NULL | Sterkenburg E, et al. 2015. New Phytologist 207:1145-1158 ((https://doi.org/10.1111/nph.13426)) |
| Otu56519 | 1 | k__Fungi(100);p__Ascomycota(100);c__Saccharomycetes(100);o__Saccharomycetales(100);f__Metschnikowiaceae(100);g__Metschnikowia(100);s__Metschnikowia_pulcherrima(100); | Metschnikowia | 13 | Saprotroph | \|Undefined Saprotroph\| | Yeast | NULL | Probable | As nectar-tap_saprotroph (Põlme et al. 2020); Decay sugar-rich substrates (Põlme et al. 2020) | Põlme S, et al. 2020. Fungal Diversity 105:1-16 ((https://doi.org/10.1007/s13225-020-00466-2)) |
| Otu56520 | 1 | k__Fungi(100);p__Ascomycota(100);c__Saccharomycetes(100);o__Saccharomycetales(100);f__Metschnikowiaceae(100);g__Metschnikowia(100);s__Metschnikowia_pulcherrima(100); | Metschnikowia | 13 | Saprotroph | \|Undefined Saprotroph\| | Yeast | NULL | Probable | As nectar-tap_saprotroph (Põlme et al. 2020); Decay sugar-rich substrates (Põlme et al. 2020) | Põlme S, et al. 2020. Fungal Diversity 105:1-16 ((https://doi.org/10.1007/s13225-020-00466-2)) |
| Otu56521 | 1 | k__Fungi(100);p__Ascomycota(100);c__Saccharomycetes(100);o__Saccharomycetales(100);f__Metschnikowiaceae(100);g__Metschnikowia(100);s__Metschnikowia_pulcherrima(100); | Metschnikowia | 13 | Saprotroph | \|Undefined Saprotroph\| | Yeast | NULL | Probable | As nectar-tap_saprotroph (Põlme et al. 2020); Decay sugar-rich substrates (Põlme et al. 2020) | Põlme S, et al. 2020. Fungal Diversity 105:1-16 ((https://doi.org/10.1007/s13225-020-00466-2)) |
| Otu56522 | 1 | k__Fungi(100);p__Ascomycota(100);c__Saccharomycetes(100);o__Saccharomycetales(100);f__Metschnikowiaceae(100);g__Metschnikowia(100);s__Metschnikowia_pulcherrima(100); | Metschnikowia | 13 | Saprotroph | \|Undefined Saprotroph\| | Yeast | NULL | Probable | As nectar-tap_saprotroph (Põlme et al. 2020); Decay sugar-rich substrates (Põlme et al. 2020) | Põlme S, et al. 2020. Fungal Diversity 105:1-16 ((https://doi.org/10.1007/s13225-020-00466-2)) |
| Otu56524 | 1 | k__Fungi(100);p__Ascomycota(100);c__Saccharomycetes(100);o__Saccharomycetales(100);f__Metschnikowiaceae(100);g__Metschnikowia(100);s__Metschnikowia_pulcherrima(100); | Metschnikowia | 13 | Saprotroph | \|Undefined Saprotroph\| | Yeast | NULL | Probable | As nectar-tap_saprotroph (Põlme et al. 2020); Decay sugar-rich substrates (Põlme et al. 2020) | Põlme S, et al. 2020. Fungal Diversity 105:1-16 ((https://doi.org/10.1007/s13225-020-00466-2)) |
| Otu56525 | 1 | k__Fungi(100);p__Ascomycota(100);c__Saccharomycetes(100);o__Saccharomycetales(100);f__Metschnikowiaceae(100);g__Metschnikowia(100);s__Metschnikowia_pulcherrima(100); | Metschnikowia | 13 | Saprotroph | \|Undefined Saprotroph\| | Yeast | NULL | Probable | As nectar-tap_saprotroph (Põlme et al. 2020); Decay sugar-rich substrates (Põlme et al. 2020) | Põlme S, et al. 2020. Fungal Diversity 105:1-16 ((https://doi.org/10.1007/s13225-020-00466-2)) |
| Otu56526 | 1 | k__Fungi(100);p__Ascomycota(100);c__Saccharomycetes(100);o__Saccharomycetales(100);f__Metschnikowiaceae(100);g__Metschnikowia(100);s__Metschnikowia_pulcherrima(100); | Metschnikowia | 13 | Saprotroph | \|Undefined Saprotroph\| | Yeast | NULL | Probable | As nectar-tap_saprotroph (Põlme et al. 2020); Decay sugar-rich substrates (Põlme et al. 2020) | Põlme S, et al. 2020. Fungal Diversity 105:1-16 ((https://doi.org/10.1007/s13225-020-00466-2)) |
| Otu56527 | 1 | k__Fungi(100);p__Ascomycota(100);c__Saccharomycetes(100);o__Saccharomycetales(100);f__Metschnikowiaceae(100);g__Metschnikowia(100);s__Metschnikowia_pulcherrima(100); | Metschnikowia | 13 | Saprotroph | \|Undefined Saprotroph\| | Yeast | NULL | Probable | As nectar-tap_saprotroph (Põlme et al. 2020); Decay sugar-rich substrates (Põlme et al. 2020) | Põlme S, et al. 2020. Fungal Diversity 105:1-16 ((https://doi.org/10.1007/s13225-020-00466-2)) |
| Otu56528 | 1 | k__Fungi(100);p__Ascomycota(100);c__Saccharomycetes(100);o__Saccharomycetales(100);f__Metschnikowiaceae(100);g__Metschnikowia(100);s__Metschnikowia_pulcherrima(100); | Metschnikowia | 13 | Saprotroph | \|Undefined Saprotroph\| | Yeast | NULL | Probable | As nectar-tap_saprotroph (Põlme et al. 2020); Decay sugar-rich substrates (Põlme et al. 2020) | Põlme S, et al. 2020. Fungal Diversity 105:1-16 ((https://doi.org/10.1007/s13225-020-00466-2)) |
| Otu56530 | 1 | k__Fungi(100);p__Basidiomycota(100);c__Microbotryomycetes(100);o__Sporidiobolales(100);f__Sporidiobolales_family_Incertae_sedis(100);g__Sporobolomyces(100);s__Sporobolomyces_lactosus(100); | Sporobolomyces | 13 | Pathotroph-Saprotroph | \|Fungal Parasite\|-Plant Saprotroph | Yeast | NULL | Probable | As litter_saprotroph (Põlme et al. 2020); As mycoparasite (Põlme et al. 2020); Decay fungal material (Põlme et al. 2020) | Tedersoo L, et al. 2014. Science 346:e1256688 ((https://doi.org/10.1126/science.1256688)); Purahong W, et al. 2016. Molecular Ecology 25:4059-4074 ((https://doi.org/10.1111/mec.13739)); Põlme S, et al. 2020. Fungal Diversity 105:1-16 ((https://doi.org/10.1007/s13225-020-00466-2)) |
| Otu56531 | 1 | k__Fungi(100);p__Basidiomycota(100);c__Microbotryomycetes(100);o__Sporidiobolales(100);f__Sporidiobolales_family_Incertae_sedis(100);g__Sporobolomyces(100);s__Sporobolomyces_lactosus(100); | Sporobolomyces | 13 | Pathotroph-Saprotroph | \|Fungal Parasite\|-Plant Saprotroph | Yeast | NULL | Probable | As litter_saprotroph (Põlme et al. 2020); As mycoparasite (Põlme et al. 2020); Decay fungal material (Põlme et al. 2020) | Tedersoo L, et al. 2014. Science 346:e1256688 ((https://doi.org/10.1126/science.1256688)); Purahong W, et al. 2016. Molecular Ecology 25:4059-4074 ((https://doi.org/10.1111/mec.13739)); Põlme S, et al. 2020. Fungal Diversity 105:1-16 ((https://doi.org/10.1007/s13225-020-00466-2)) |
| Otu56532 | 1 | k__Fungi(100);p__Ascomycota(100);c__Saccharomycetes(100);o__Saccharomycetales(100);f__Metschnikowiaceae(100);g__Metschnikowia(100);s__Metschnikowia_pulcherrima(100); | Metschnikowia | 13 | Saprotroph | \|Undefined Saprotroph\| | Yeast | NULL | Probable | As nectar-tap_saprotroph (Põlme et al. 2020); Decay sugar-rich substrates (Põlme et al. 2020) | Põlme S, et al. 2020. Fungal Diversity 105:1-16 ((https://doi.org/10.1007/s13225-020-00466-2)) |
| Otu56536 | 1 | k__Fungi(100);p__Ascomycota(100);c__Saccharomycetes(100);o__Saccharomycetales(100);f__Saccharomycetales_family_Incertae_sedis(100);g__Candida(100);g__Candida_unclassified(100); | Candida | 13 | Pathotroph-Saprotroph-Symbiotroph | Animal Pathogen-Endophyte-Endosymbiont-Epiphyte-\|Undefined Saprotroph\| | Dimorphic Yeast-Yeast | NULL | Probable | As nectar-tap_saprotroph (Põlme et al. 2020); Decay sugar-rich substrates (Põlme et al. 2020); This asexual genus is polyphyletic and contain species that function very differently depend on what they are. Caution should be taken when interpreting guilds for Candida species. | Manolakaki D, et al. 2010. Virulence. 1: 367-375 ((https://doi.org/10.4161/viru.1.5.12796)); Mekbib SB, et al. 2011. Tropical Plant Pathology 36:233-240 ((https://doi.org/10.1590/S1982-56762011000400004)); Põlme S, et al. 2020. Fungal Diversity 105:1-16 ((https://doi.org/10.1007/s13225-020-00466-2)) |
| Otu56537 | 1 | k__Fungi(100);p__Ascomycota(100);c__Saccharomycetes(100);o__Saccharomycetales(100);f__Metschnikowiaceae(100);g__Metschnikowia(100);s__Metschnikowia_pulcherrima(100); | Metschnikowia | 13 | Saprotroph | \|Undefined Saprotroph\| | Yeast | NULL | Probable | As nectar-tap_saprotroph (Põlme et al. 2020); Decay sugar-rich substrates (Põlme et al. 2020) | Põlme S, et al. 2020. Fungal Diversity 105:1-16 ((https://doi.org/10.1007/s13225-020-00466-2)) |
| Otu56542 | 1 | k__Fungi(100);p__Ascomycota(100);c__Saccharomycetes(100);o__Saccharomycetales(100);f__Metschnikowiaceae(100);g__Metschnikowia(100);s__Metschnikowia_pulcherrima(100); | Metschnikowia | 13 | Saprotroph | \|Undefined Saprotroph\| | Yeast | NULL | Probable | As nectar-tap_saprotroph (Põlme et al. 2020); Decay sugar-rich substrates (Põlme et al. 2020) | Põlme S, et al. 2020. Fungal Diversity 105:1-16 ((https://doi.org/10.1007/s13225-020-00466-2)) |
| Otu56543 | 1 | k__Fungi(100);p__Ascomycota(100);c__Saccharomycetes(100);o__Saccharomycetales(100);f__Saccharomycetales_family_Incertae_sedis(100);g__Candida(100);g__Candida_unclassified(100); | Candida | 13 | Pathotroph-Saprotroph-Symbiotroph | Animal Pathogen-Endophyte-Endosymbiont-Epiphyte-\|Undefined Saprotroph\| | Dimorphic Yeast-Yeast | NULL | Probable | As nectar-tap_saprotroph (Põlme et al. 2020); Decay sugar-rich substrates (Põlme et al. 2020); This asexual genus is polyphyletic and contain species that function very differently depend on what they are. Caution should be taken when interpreting guilds for Candida species. | Manolakaki D, et al. 2010. Virulence. 1: 367-375 ((https://doi.org/10.4161/viru.1.5.12796)); Mekbib SB, et al. 2011. Tropical Plant Pathology 36:233-240 ((https://doi.org/10.1590/S1982-56762011000400004)); Põlme S, et al. 2020. Fungal Diversity 105:1-16 ((https://doi.org/10.1007/s13225-020-00466-2)) |
| Otu56545 | 1 | k__Fungi(100);p__Ascomycota(100);c__Saccharomycetes(100);o__Saccharomycetales(100);f__Metschnikowiaceae(100);g__Metschnikowia(100);g__Metschnikowia_unclassified(100); | Metschnikowia | 13 | Saprotroph | \|Undefined Saprotroph\| | Yeast | NULL | Probable | As nectar-tap_saprotroph (Põlme et al. 2020); Decay sugar-rich substrates (Põlme et al. 2020) | Põlme S, et al. 2020. Fungal Diversity 105:1-16 ((https://doi.org/10.1007/s13225-020-00466-2)) |
| Otu56546 | 1 | k__Fungi(100);p__Ascomycota(100);c__Saccharomycetes(100);o__Saccharomycetales(100);f__Metschnikowiaceae(100);g__Metschnikowia(100);s__Metschnikowia_pulcherrima(100); | Metschnikowia | 13 | Saprotroph | \|Undefined Saprotroph\| | Yeast | NULL | Probable | As nectar-tap_saprotroph (Põlme et al. 2020); Decay sugar-rich substrates (Põlme et al. 2020) | Põlme S, et al. 2020. Fungal Diversity 105:1-16 ((https://doi.org/10.1007/s13225-020-00466-2)) |
| Otu56549 | 1 | k__Fungi(100);p__Ascomycota(100);c__Saccharomycetes(100);o__Saccharomycetales(100);f__Saccharomycetales_family_Incertae_sedis(100);f__Saccharomycetales_family_Incertae_sedis_unclassified(100);f__Saccharomycetales_family_Incertae_sedis_unclassified(100); | Saccharomycetales | 7 | Saprotroph | Undefined Saprotroph | Yeast | NULL | Possible | NULL | Sterkenburg E, et al. 2015. New Phytologist 207:1145-1158 ((https://doi.org/10.1111/nph.13426)) |
| Otu56550 | 1 | k__Fungi(100);p__Ascomycota(100);c__Saccharomycetes(100);o__Saccharomycetales(100);f__Saccharomycetales_family_Incertae_sedis(100);g__Candida(100);g__Candida_unclassified(100); | Candida | 13 | Pathotroph-Saprotroph-Symbiotroph | Animal Pathogen-Endophyte-Endosymbiont-Epiphyte-\|Undefined Saprotroph\| | Dimorphic Yeast-Yeast | NULL | Probable | As nectar-tap_saprotroph (Põlme et al. 2020); Decay sugar-rich substrates (Põlme et al. 2020); This asexual genus is polyphyletic and contain species that function very differently depend on what they are. Caution should be taken when interpreting guilds for Candida species. | Manolakaki D, et al. 2010. Virulence. 1: 367-375 ((https://doi.org/10.4161/viru.1.5.12796)); Mekbib SB, et al. 2011. Tropical Plant Pathology 36:233-240 ((https://doi.org/10.1590/S1982-56762011000400004)); Põlme S, et al. 2020. Fungal Diversity 105:1-16 ((https://doi.org/10.1007/s13225-020-00466-2)) |
| Otu56551 | 1 | k__Fungi(100);p__Ascomycota(100);c__Saccharomycetes(100);o__Saccharomycetales(100);f__Saccharomycetales_family_Incertae_sedis(100);g__Candida(100);g__Candida_unclassified(100); | Candida | 13 | Pathotroph-Saprotroph-Symbiotroph | Animal Pathogen-Endophyte-Endosymbiont-Epiphyte-\|Undefined Saprotroph\| | Dimorphic Yeast-Yeast | NULL | Probable | As nectar-tap_saprotroph (Põlme et al. 2020); Decay sugar-rich substrates (Põlme et al. 2020); This asexual genus is polyphyletic and contain species that function very differently depend on what they are. Caution should be taken when interpreting guilds for Candida species. | Manolakaki D, et al. 2010. Virulence. 1: 367-375 ((https://doi.org/10.4161/viru.1.5.12796)); Mekbib SB, et al. 2011. Tropical Plant Pathology 36:233-240 ((https://doi.org/10.1590/S1982-56762011000400004)); Põlme S, et al. 2020. Fungal Diversity 105:1-16 ((https://doi.org/10.1007/s13225-020-00466-2)) |
| Otu56553 | 1 | k__Fungi(100);p__Ascomycota(100);c__Saccharomycetes(100);o__Saccharomycetales(100);f__Saccharomycetales_family_Incertae_sedis(100);g__Candida(100);g__Candida_unclassified(100); | Candida | 13 | Pathotroph-Saprotroph-Symbiotroph | Animal Pathogen-Endophyte-Endosymbiont-Epiphyte-\|Undefined Saprotroph\| | Dimorphic Yeast-Yeast | NULL | Probable | As nectar-tap_saprotroph (Põlme et al. 2020); Decay sugar-rich substrates (Põlme et al. 2020); This asexual genus is polyphyletic and contain species that function very differently depend on what they are. Caution should be taken when interpreting guilds for Candida species. | Manolakaki D, et al. 2010. Virulence. 1: 367-375 ((https://doi.org/10.4161/viru.1.5.12796)); Mekbib SB, et al. 2011. Tropical Plant Pathology 36:233-240 ((https://doi.org/10.1590/S1982-56762011000400004)); Põlme S, et al. 2020. Fungal Diversity 105:1-16 ((https://doi.org/10.1007/s13225-020-00466-2)) |
| Otu56554 | 1 | k__Fungi(100);p__Ascomycota(100);c__Saccharomycetes(100);o__Saccharomycetales(100);f__Saccharomycetales_family_Incertae_sedis(100);g__Candida(100);g__Candida_unclassified(100); | Candida | 13 | Pathotroph-Saprotroph-Symbiotroph | Animal Pathogen-Endophyte-Endosymbiont-Epiphyte-\|Undefined Saprotroph\| | Dimorphic Yeast-Yeast | NULL | Probable | As nectar-tap_saprotroph (Põlme et al. 2020); Decay sugar-rich substrates (Põlme et al. 2020); This asexual genus is polyphyletic and contain species that function very differently depend on what they are. Caution should be taken when interpreting guilds for Candida species. | Manolakaki D, et al. 2010. Virulence. 1: 367-375 ((https://doi.org/10.4161/viru.1.5.12796)); Mekbib SB, et al. 2011. Tropical Plant Pathology 36:233-240 ((https://doi.org/10.1590/S1982-56762011000400004)); Põlme S, et al. 2020. Fungal Diversity 105:1-16 ((https://doi.org/10.1007/s13225-020-00466-2)) |
| Otu56558 | 1 | k__Fungi(100);p__Ascomycota(100);c__Saccharomycetes(100);o__Saccharomycetales(100);f__Saccharomycetales_family_Incertae_sedis(100);g__Candida(100);s__Candida_cretensis(100); | Candida | 13 | Pathotroph-Saprotroph-Symbiotroph | Animal Pathogen-Endophyte-Endosymbiont-Epiphyte-\|Undefined Saprotroph\| | Dimorphic Yeast-Yeast | NULL | Probable | As nectar-tap_saprotroph (Põlme et al. 2020); Decay sugar-rich substrates (Põlme et al. 2020); This asexual genus is polyphyletic and contain species that function very differently depend on what they are. Caution should be taken when interpreting guilds for Candida species. | Manolakaki D, et al. 2010. Virulence. 1: 367-375 ((https://doi.org/10.4161/viru.1.5.12796)); Mekbib SB, et al. 2011. Tropical Plant Pathology 36:233-240 ((https://doi.org/10.1590/S1982-56762011000400004)); Põlme S, et al. 2020. Fungal Diversity 105:1-16 ((https://doi.org/10.1007/s13225-020-00466-2)) |
| Otu56560 | 1 | k__Fungi(100);p__Ascomycota(100);c__Saccharomycetes(100);o__Saccharomycetales(100);f__Metschnikowiaceae(100);g__Metschnikowia(100);s__Metschnikowia_pulcherrima(100); | Metschnikowia | 13 | Saprotroph | \|Undefined Saprotroph\| | Yeast | NULL | Probable | As nectar-tap_saprotroph (Põlme et al. 2020); Decay sugar-rich substrates (Põlme et al. 2020) | Põlme S, et al. 2020. Fungal Diversity 105:1-16 ((https://doi.org/10.1007/s13225-020-00466-2)) |
| Otu56561 | 1 | k__Fungi(100);p__Ascomycota(100);c__Saccharomycetes(100);o__Saccharomycetales(100);f__Metschnikowiaceae(100);g__Metschnikowia(100);s__Metschnikowia_pulcherrima(100); | Metschnikowia | 13 | Saprotroph | \|Undefined Saprotroph\| | Yeast | NULL | Probable | As nectar-tap_saprotroph (Põlme et al. 2020); Decay sugar-rich substrates (Põlme et al. 2020) | Põlme S, et al. 2020. Fungal Diversity 105:1-16 ((https://doi.org/10.1007/s13225-020-00466-2)) |
| Otu56562 | 1 | k__Fungi(100);p__Ascomycota(100);c__Saccharomycetes(100);o__Saccharomycetales(100);f__Metschnikowiaceae(100);g__Metschnikowia(100);s__Metschnikowia_pulcherrima(100); | Metschnikowia | 13 | Saprotroph | \|Undefined Saprotroph\| | Yeast | NULL | Probable | As nectar-tap_saprotroph (Põlme et al. 2020); Decay sugar-rich substrates (Põlme et al. 2020) | Põlme S, et al. 2020. Fungal Diversity 105:1-16 ((https://doi.org/10.1007/s13225-020-00466-2)) |
| Otu56563 | 1 | k__Fungi(100);p__Ascomycota(100);c__Saccharomycetes(100);o__Saccharomycetales(100);f__Metschnikowiaceae(100);g__Metschnikowia(100);s__Metschnikowia_pulcherrima(100); | Metschnikowia | 13 | Saprotroph | \|Undefined Saprotroph\| | Yeast | NULL | Probable | As nectar-tap_saprotroph (Põlme et al. 2020); Decay sugar-rich substrates (Põlme et al. 2020) | Põlme S, et al. 2020. Fungal Diversity 105:1-16 ((https://doi.org/10.1007/s13225-020-00466-2)) |
| Otu56564 | 1 | k__Fungi(100);p__Ascomycota(100);c__Saccharomycetes(100);o__Saccharomycetales(100);f__Metschnikowiaceae(100);g__Metschnikowia(100);s__Metschnikowia_pulcherrima(100); | Metschnikowia | 13 | Saprotroph | \|Undefined Saprotroph\| | Yeast | NULL | Probable | As nectar-tap_saprotroph (Põlme et al. 2020); Decay sugar-rich substrates (Põlme et al. 2020) | Põlme S, et al. 2020. Fungal Diversity 105:1-16 ((https://doi.org/10.1007/s13225-020-00466-2)) |
| Otu56566 | 1 | k__Fungi(100);p__Ascomycota(100);c__Saccharomycetes(100);o__Saccharomycetales(100);f__Metschnikowiaceae(100);g__Metschnikowia(100);s__Metschnikowia_pulcherrima(100); | Metschnikowia | 13 | Saprotroph | \|Undefined Saprotroph\| | Yeast | NULL | Probable | As nectar-tap_saprotroph (Põlme et al. 2020); Decay sugar-rich substrates (Põlme et al. 2020) | Põlme S, et al. 2020. Fungal Diversity 105:1-16 ((https://doi.org/10.1007/s13225-020-00466-2)) |
| Otu56567 | 1 | k__Fungi(100);p__Ascomycota(100);c__Saccharomycetes(100);o__Saccharomycetales(100);f__Metschnikowiaceae(100);g__Metschnikowia(100);s__Metschnikowia_pulcherrima(100); | Metschnikowia | 13 | Saprotroph | \|Undefined Saprotroph\| | Yeast | NULL | Probable | As nectar-tap_saprotroph (Põlme et al. 2020); Decay sugar-rich substrates (Põlme et al. 2020) | Põlme S, et al. 2020. Fungal Diversity 105:1-16 ((https://doi.org/10.1007/s13225-020-00466-2)) |
| Otu56568 | 1 | k__Fungi(100);p__Ascomycota(100);c__Saccharomycetes(100);o__Saccharomycetales(100);f__Metschnikowiaceae(100);g__Metschnikowia(100);s__Metschnikowia_pulcherrima(100); | Metschnikowia | 13 | Saprotroph | \|Undefined Saprotroph\| | Yeast | NULL | Probable | As nectar-tap_saprotroph (Põlme et al. 2020); Decay sugar-rich substrates (Põlme et al. 2020) | Põlme S, et al. 2020. Fungal Diversity 105:1-16 ((https://doi.org/10.1007/s13225-020-00466-2)) |
| Otu56569 | 1 | k__Fungi(100);p__Ascomycota(100);c__Saccharomycetes(100);o__Saccharomycetales(100);f__Metschnikowiaceae(100);g__Metschnikowia(100);s__Metschnikowia_pulcherrima(100); | Metschnikowia | 13 | Saprotroph | \|Undefined Saprotroph\| | Yeast | NULL | Probable | As nectar-tap_saprotroph (Põlme et al. 2020); Decay sugar-rich substrates (Põlme et al. 2020) | Põlme S, et al. 2020. Fungal Diversity 105:1-16 ((https://doi.org/10.1007/s13225-020-00466-2)) |
| Otu56574 | 1 | k__Fungi(100);p__Ascomycota(100);c__Saccharomycetes(100);o__Saccharomycetales(100);f__Metschnikowiaceae(100);g__Metschnikowia(100);s__Metschnikowia_pulcherrima(100); | Metschnikowia | 13 | Saprotroph | \|Undefined Saprotroph\| | Yeast | NULL | Probable | As nectar-tap_saprotroph (Põlme et al. 2020); Decay sugar-rich substrates (Põlme et al. 2020) | Põlme S, et al. 2020. Fungal Diversity 105:1-16 ((https://doi.org/10.1007/s13225-020-00466-2)) |
| Otu56575 | 1 | k__Fungi(100);p__Ascomycota(100);c__Saccharomycetes(100);o__Saccharomycetales(100);f__Metschnikowiaceae(100);g__Metschnikowia(100);s__Metschnikowia_pulcherrima(100); | Metschnikowia | 13 | Saprotroph | \|Undefined Saprotroph\| | Yeast | NULL | Probable | As nectar-tap_saprotroph (Põlme et al. 2020); Decay sugar-rich substrates (Põlme et al. 2020) | Põlme S, et al. 2020. Fungal Diversity 105:1-16 ((https://doi.org/10.1007/s13225-020-00466-2)) |
| Otu56576 | 1 | k__Fungi(100);p__Ascomycota(100);c__Saccharomycetes(100);o__Saccharomycetales(100);f__Saccharomycetales_family_Incertae_sedis(100);g__Candida(100);g__Candida_unclassified(100); | Candida | 13 | Pathotroph-Saprotroph-Symbiotroph | Animal Pathogen-Endophyte-Endosymbiont-Epiphyte-\|Undefined Saprotroph\| | Dimorphic Yeast-Yeast | NULL | Probable | As nectar-tap_saprotroph (Põlme et al. 2020); Decay sugar-rich substrates (Põlme et al. 2020); This asexual genus is polyphyletic and contain species that function very differently depend on what they are. Caution should be taken when interpreting guilds for Candida species. | Manolakaki D, et al. 2010. Virulence. 1: 367-375 ((https://doi.org/10.4161/viru.1.5.12796)); Mekbib SB, et al. 2011. Tropical Plant Pathology 36:233-240 ((https://doi.org/10.1590/S1982-56762011000400004)); Põlme S, et al. 2020. Fungal Diversity 105:1-16 ((https://doi.org/10.1007/s13225-020-00466-2)) |
| Otu56579 | 1 | k__Fungi(100);p__Ascomycota(100);c__Saccharomycetes(100);o__Saccharomycetales(100);f__Metschnikowiaceae(100);g__Metschnikowia(100);s__Metschnikowia_pulcherrima(100); | Metschnikowia | 13 | Saprotroph | \|Undefined Saprotroph\| | Yeast | NULL | Probable | As nectar-tap_saprotroph (Põlme et al. 2020); Decay sugar-rich substrates (Põlme et al. 2020) | Põlme S, et al. 2020. Fungal Diversity 105:1-16 ((https://doi.org/10.1007/s13225-020-00466-2)) |
| Otu56580 | 1 | k__Fungi(100);p__Ascomycota(100);c__Saccharomycetes(100);o__Saccharomycetales(100);f__Metschnikowiaceae(100);g__Metschnikowia(100);s__Metschnikowia_pulcherrima(100); | Metschnikowia | 13 | Saprotroph | \|Undefined Saprotroph\| | Yeast | NULL | Probable | As nectar-tap_saprotroph (Põlme et al. 2020); Decay sugar-rich substrates (Põlme et al. 2020) | Põlme S, et al. 2020. Fungal Diversity 105:1-16 ((https://doi.org/10.1007/s13225-020-00466-2)) |
| Otu56581 | 1 | k__Fungi(100);p__Ascomycota(100);c__Saccharomycetes(100);o__Saccharomycetales(100);f__Metschnikowiaceae(100);g__Metschnikowia(100);s__Metschnikowia_pulcherrima(100); | Metschnikowia | 13 | Saprotroph | \|Undefined Saprotroph\| | Yeast | NULL | Probable | As nectar-tap_saprotroph (Põlme et al. 2020); Decay sugar-rich substrates (Põlme et al. 2020) | Põlme S, et al. 2020. Fungal Diversity 105:1-16 ((https://doi.org/10.1007/s13225-020-00466-2)) |
| Otu56582 | 1 | k__Fungi(100);p__Ascomycota(100);c__Saccharomycetes(100);o__Saccharomycetales(100);f__Metschnikowiaceae(100);g__Metschnikowia(100);g__Metschnikowia_unclassified(100); | Metschnikowia | 13 | Saprotroph | \|Undefined Saprotroph\| | Yeast | NULL | Probable | As nectar-tap_saprotroph (Põlme et al. 2020); Decay sugar-rich substrates (Põlme et al. 2020) | Põlme S, et al. 2020. Fungal Diversity 105:1-16 ((https://doi.org/10.1007/s13225-020-00466-2)) |
| Otu56583 | 1 | k__Fungi(100);p__Ascomycota(100);c__Saccharomycetes(100);o__Saccharomycetales(100);f__Metschnikowiaceae(100);g__Metschnikowia(100);g__Metschnikowia_unclassified(100); | Metschnikowia | 13 | Saprotroph | \|Undefined Saprotroph\| | Yeast | NULL | Probable | As nectar-tap_saprotroph (Põlme et al. 2020); Decay sugar-rich substrates (Põlme et al. 2020) | Põlme S, et al. 2020. Fungal Diversity 105:1-16 ((https://doi.org/10.1007/s13225-020-00466-2)) |

| | | | | | | | | | | | |
|---|---|---|---|---|---|---|---|---|---|---|---|
| Otu56584 | 1 | k__Fungi(100);p__Ascomycota(100);c__Saccharomycetes(100);o__Saccharomycetales(100);f__Metschnikowiaceae(100);g__Metschnikowia(100);s__Metschnikowia_pulcherrima(100); | Metschnikowia | 13 | Saprotroph | \|Undefined Saprotroph\| | Yeast | NULL | Probable | As nectar-tap_saprotroph (Põlme et al. 2020); Decay sugar-rich substrates (Põlme et al. 2020) | Põlme S, et al. 2020. Fungal Diversity 105:1-16 ((https://doi.org/10.1007/s13225-020-00466-2)) |
| Otu56585 | 1 | k__Fungi(100);p__Ascomycota(100);c__Saccharomycetes(100);o__Saccharomycetales(100);f__Metschnikowiaceae(100);g__Metschnikowia(100);s__Metschnikowia_pulcherrima(100); | Metschnikowia | 13 | Saprotroph | \|Undefined Saprotroph\| | Yeast | NULL | Probable | As nectar-tap_saprotroph (Põlme et al. 2020); Decay sugar-rich substrates (Põlme et al. 2020) | Põlme S, et al. 2020. Fungal Diversity 105:1-16 ((https://doi.org/10.1007/s13225-020-00466-2)) |
| Otu56586 | 1 | k__Fungi(100);p__Ascomycota(100);c__Saccharomycetes(100);o__Saccharomycetales(100);f__Metschnikowiaceae(100);g__Metschnikowia(100);s__Metschnikowia_pulcherrima(100); | Metschnikowia | 13 | Saprotroph | \|Undefined Saprotroph\| | Yeast | NULL | Probable | As nectar-tap_saprotroph (Põlme et al. 2020); Decay sugar-rich substrates (Põlme et al. 2020) | Põlme S, et al. 2020. Fungal Diversity 105:1-16 ((https://doi.org/10.1007/s13225-020-00466-2)) |
| Otu56587 | 1 | k__Fungi(100);p__Ascomycota(100);c__Saccharomycetes(100);o__Saccharomycetales(100);f__Metschnikowiaceae(100);g__Metschnikowia(100);s__Metschnikowia_pulcherrima(100); | Metschnikowia | 13 | Saprotroph | \|Undefined Saprotroph\| | Yeast | NULL | Probable | As nectar-tap_saprotroph (Põlme et al. 2020); Decay sugar-rich substrates (Põlme et al. 2020) | Põlme S, et al. 2020. Fungal Diversity 105:1-16 ((https://doi.org/10.1007/s13225-020-00466-2)) |
| Otu56588 | 1 | k__Fungi(100);p__Ascomycota(100);c__Saccharomycetes(100);o__Saccharomycetales(100);f__Metschnikowiaceae(100);g__Metschnikowia(100);s__Metschnikowia_pulcherrima(100); | Metschnikowia | 13 | Saprotroph | \|Undefined Saprotroph\| | Yeast | NULL | Probable | As nectar-tap_saprotroph (Põlme et al. 2020); Decay sugar-rich substrates (Põlme et al. 2020) | Põlme S, et al. 2020. Fungal Diversity 105:1-16 ((https://doi.org/10.1007/s13225-020-00466-2)) |
| Otu56589 | 1 | k__Fungi(100);p__Ascomycota(100);c__Saccharomycetes(100);o__Saccharomycetales(100);f__Metschnikowiaceae(100);g__Metschnikowia(100);s__Metschnikowia_pulcherrima(100); | Metschnikowia | 13 | Saprotroph | \|Undefined Saprotroph\| | Yeast | NULL | Probable | As nectar-tap_saprotroph (Põlme et al. 2020); Decay sugar-rich substrates (Põlme et al. 2020) | Põlme S, et al. 2020. Fungal Diversity 105:1-16 ((https://doi.org/10.1007/s13225-020-00466-2)) |
| Otu56590 | 1 | k__Fungi(100);p__Ascomycota(100);c__Saccharomycetes(100);o__Saccharomycetales(100);f__Metschnikowiaceae(100);g__Metschnikowia(100);s__Metschnikowia_pulcherrima(100); | Metschnikowia | 13 | Saprotroph | \|Undefined Saprotroph\| | Yeast | NULL | Probable | As nectar-tap_saprotroph (Põlme et al. 2020); Decay sugar-rich substrates (Põlme et al. 2020) | Põlme S, et al. 2020. Fungal Diversity 105:1-16 ((https://doi.org/10.1007/s13225-020-00466-2)) |
| Otu56591 | 1 | k__Fungi(100);p__Ascomycota(100);c__Saccharomycetes(100);o__Saccharomycetales(100);f__Metschnikowiaceae(100);g__Metschnikowia(100);s__Metschnikowia_pulcherrima(100); | Metschnikowia | 13 | Saprotroph | \|Undefined Saprotroph\| | Yeast | NULL | Probable | As nectar-tap_saprotroph (Põlme et al. 2020); Decay sugar-rich substrates (Põlme et al. 2020) | Põlme S, et al. 2020. Fungal Diversity 105:1-16 ((https://doi.org/10.1007/s13225-020-00466-2)) |
| Otu56592 | 1 | k__Fungi(100);p__Ascomycota(100);c__Saccharomycetes(100);o__Saccharomycetales(100);f__Metschnikowiaceae(100);g__Metschnikowia(100);s__Metschnikowia_pulcherrima(100); | Metschnikowia | 13 | Saprotroph | \|Undefined Saprotroph\| | Yeast | NULL | Probable | As nectar-tap_saprotroph (Põlme et al. 2020); Decay sugar-rich substrates (Põlme et al. 2020) | Põlme S, et al. 2020. Fungal Diversity 105:1-16 ((https://doi.org/10.1007/s13225-020-00466-2)) |
| Otu56593 | 1 | k__Fungi(100);p__Ascomycota(100);c__Saccharomycetes(100);o__Saccharomycetales(100);f__Metschnikowiaceae(100);g__Metschnikowia(100);s__Metschnikowia_pulcherrima(100); | Metschnikowia | 13 | Saprotroph | \|Undefined Saprotroph\| | Yeast | NULL | Probable | As nectar-tap_saprotroph (Põlme et al. 2020); Decay sugar-rich substrates (Põlme et al. 2020) | Põlme S, et al. 2020. Fungal Diversity 105:1-16 ((https://doi.org/10.1007/s13225-020-00466-2)) |
| Otu56594 | 1 | k__Fungi(100);p__Ascomycota(100);c__Saccharomycetes(100);o__Saccharomycetales(100);f__Saccharomycetales_family_Incertae_sedis(100);g__Candida(100);s__Candida_cretensis(100); | Candida | 13 | Pathotroph-Saprotroph-Symbiotroph | Animal Pathogen-Endophyte-Endosymbiont-Epiphyte-\|Undefined Saprotroph\| | Dimorphic Yeast-Yeast | NULL | Probable | As nectar-tap_saprotroph (Põlme et al. 2020); Decay sugar-rich substrates (Põlme et al. 2020); This asexual genus is polyphyletic and contain species that function very differently depend on what they are. Caution should be taken when interpreting guilds for Candida species. | Manolakaki D, et al. 2010. Virulence. 1: 367-375 ((https://doi.org/10.4161/viru.1.5.12796)); Mekbib SB, et al. 2011. Tropical Plant Pathology 36:233-240 ((https://doi.org/10.1590/S1982-56762011000400004)); Põlme S, et al. 2020. Fungal Diversity 105:1-16 ((https://doi.org/10.1007/s13225-020-00466-2)) |
| Otu56595 | 1 | k__Fungi(100);p__Ascomycota(100);c__Saccharomycetes(100);o__Saccharomycetales(100);f__Metschnikowiaceae(100);g__Metschnikowia(100);s__Metschnikowia_pulcherrima(100); | Metschnikowia | 13 | Saprotroph | \|Undefined Saprotroph\| | Yeast | NULL | Probable | As nectar-tap_saprotroph (Põlme et al. 2020); Decay sugar-rich substrates (Põlme et al. 2020) | Põlme S, et al. 2020. Fungal Diversity 105:1-16 ((https://doi.org/10.1007/s13225-020-00466-2)) |
| Otu56597 | 1 | k__Fungi(100);p__Ascomycota(100);c__Saccharomycetes(100);o__Saccharomycetales(100);f__Metschnikowiaceae(100);g__Metschnikowia(100);s__Metschnikowia_pulcherrima(100); | Metschnikowia | 13 | Saprotroph | \|Undefined Saprotroph\| | Yeast | NULL | Probable | As nectar-tap_saprotroph (Põlme et al. 2020); Decay sugar-rich substrates (Põlme et al. 2020) | Põlme S, et al. 2020. Fungal Diversity 105:1-16 ((https://doi.org/10.1007/s13225-020-00466-2)) |
| Otu56598 | 1 | k__Fungi(100);p__Ascomycota(100);c__Saccharomycetes(100);o__Saccharomycetales(100);f__Metschnikowiaceae(100);g__Metschnikowia(100);s__Metschnikowia_pulcherrima(100); | Metschnikowia | 13 | Saprotroph | \|Undefined Saprotroph\| | Yeast | NULL | Probable | As nectar-tap_saprotroph (Põlme et al. 2020); Decay sugar-rich substrates (Põlme et al. 2020) | Põlme S, et al. 2020. Fungal Diversity 105:1-16 ((https://doi.org/10.1007/s13225-020-00466-2)) |
| Otu56599 | 1 | k__Fungi(100);p__Ascomycota(100);c__Saccharomycetes(100);o__Saccharomycetales(100);f__Metschnikowiaceae(100);g__Metschnikowia(100);s__Metschnikowia_pulcherrima(100); | Metschnikowia | 13 | Saprotroph | \|Undefined Saprotroph\| | Yeast | NULL | Probable | As nectar-tap_saprotroph (Põlme et al. 2020); Decay sugar-rich substrates (Põlme et al. 2020) | Põlme S, et al. 2020. Fungal Diversity 105:1-16 ((https://doi.org/10.1007/s13225-020-00466-2)) |
| Otu56600 | 1 | k__Fungi(100);p__Ascomycota(100);c__Saccharomycetes(100);o__Saccharomycetales(100);f__Metschnikowiaceae(100);g__Metschnikowia(100);s__Metschnikowia_pulcherrima(100); | Metschnikowia | 13 | Saprotroph | \|Undefined Saprotroph\| | Yeast | NULL | Probable | As nectar-tap_saprotroph (Põlme et al. 2020); Decay sugar-rich substrates (Põlme et al. 2020) | Põlme S, et al. 2020. Fungal Diversity 105:1-16 ((https://doi.org/10.1007/s13225-020-00466-2)) |
| Otu56601 | 1 | k__Fungi(100);p__Basidiomycota(100);c__Microbotryomycetes(100);o__Sporidiobolales(100);f__Sporidiobolales_family_Incertae_sedis(100);g__Sporobolomyces(100);s__Sporobolomyces_lactosus(100); | Sporobolomyces | 13 | Pathotroph-Saprotroph | \|Fungal Parasite\|-Plant Saprotroph | Yeast | NULL | Probable | As litter_saprotroph (Põlme et al. 2020); As mycoparasite (Põlme et al. 2020); Decay fungal material (Põlme et al. 2020) | Tedersoo L, et al. 2014. Science 346:e1256688 ((https://doi.org/10.1126/science.1256688)); Purahong W, et al. 2016. Molecular Ecology 25:4059-4074 ((https://doi.org/10.1111/mec.13739)); Põlme S, et al. 2020. Fungal Diversity 105:1-16 ((https://doi.org/10.1007/s13225-020-00466-2)) |
| Otu56602 | 1 | k__Fungi(100);p__Basidiomycota(100);c__Microbotryomycetes(100);o__Sporidiobolales(100);f__Sporidiobolales_family_Incertae_sedis(100);g__Sporobolomyces(100);s__Sporobolomyces_lactosus(100); | Sporobolomyces | 13 | Pathotroph-Saprotroph | \|Fungal Parasite\|-Plant Saprotroph | Yeast | NULL | Probable | As litter_saprotroph (Põlme et al. 2020); As mycoparasite (Põlme et al. 2020); Decay fungal material (Põlme et al. 2020) | Tedersoo L, et al. 2014. Science 346:e1256688 ((https://doi.org/10.1126/science.1256688)); Purahong W, et al. 2016. Molecular Ecology 25:4059-4074 ((https://doi.org/10.1111/mec.13739)); Põlme S, et al. 2020. Fungal Diversity 105:1-16 ((https://doi.org/10.1007/s13225-020-00466-2)) |
| Otu56603 | 1 | k__Fungi(100);p__Ascomycota(100);c__Saccharomycetes(100);o__Saccharomycetales(100);f__Metschnikowiaceae(100);g__Metschnikowia(100);s__Metschnikowia_pulcherrima(100); | Metschnikowia | 13 | Saprotroph | \|Undefined Saprotroph\| | Yeast | NULL | Probable | As nectar-tap_saprotroph (Põlme et al. 2020); Decay sugar-rich substrates (Põlme et al. 2020) | Põlme S, et al. 2020. Fungal Diversity 105:1-16 ((https://doi.org/10.1007/s13225-020-00466-2)) |
| Otu56604 | 1 | k__Fungi(100);p__Ascomycota(100);c__Saccharomycetes(100);o__Saccharomycetales(100);f__Metschnikowiaceae(100);g__Metschnikowia(100);s__Metschnikowia_pulcherrima(100); | Metschnikowia | 13 | Saprotroph | \|Undefined Saprotroph\| | Yeast | NULL | Probable | As nectar-tap_saprotroph (Põlme et al. 2020); Decay sugar-rich substrates (Põlme et al. 2020) | Põlme S, et al. 2020. Fungal Diversity 105:1-16 ((https://doi.org/10.1007/s13225-020-00466-2)) |
| Otu56605 | 1 | k__Fungi(100);p__Ascomycota(100);c__Saccharomycetes(100);o__Saccharomycetales(100);f__Metschnikowiaceae(100);g__Metschnikowia(100);s__Metschnikowia_pulcherrima(100); | Metschnikowia | 13 | Saprotroph | \|Undefined Saprotroph\| | Yeast | NULL | Probable | As nectar-tap_saprotroph (Põlme et al. 2020); Decay sugar-rich substrates (Põlme et al. 2020) | Põlme S, et al. 2020. Fungal Diversity 105:1-16 ((https://doi.org/10.1007/s13225-020-00466-2)) |
| Otu56606 | 1 | k__Fungi(100);p__Basidiomycota(100);c__Microbotryomycetes(100);o__Sporidiobolales(100);f__Sporidiobolales_family_Incertae_sedis(100);g__Sporobolomyces(100);s__Sporobolomyces_lactosus(100); | Sporobolomyces | 13 | Pathotroph-Saprotroph | \|Fungal Parasite\|-Plant Saprotroph | Yeast | NULL | Probable | As litter_saprotroph (Põlme et al. 2020); As mycoparasite (Põlme et al. 2020); Decay fungal material (Põlme et al. 2020) | Tedersoo L, et al. 2014. Science 346:e1256688 ((https://doi.org/10.1126/science.1256688)); Purahong W, et al. 2016. Molecular Ecology 25:4059-4074 ((https://doi.org/10.1111/mec.13739)); Põlme S, et al. 2020. Fungal Diversity 105:1-16 ((https://doi.org/10.1007/s13225-020-00466-2)) |
| Otu56607 | 1 | k__Fungi(100);p__Ascomycota(100);c__Saccharomycetes(100);o__Saccharomycetales(100);f__Metschnikowiaceae(100);g__Metschnikowia(100);s__Metschnikowia_pulcherrima(100); | Metschnikowia | 13 | Saprotroph | \|Undefined Saprotroph\| | Yeast | NULL | Probable | As nectar-tap_saprotroph (Põlme et al. 2020); Decay sugar-rich substrates (Põlme et al. 2020) | Põlme S, et al. 2020. Fungal Diversity 105:1-16 ((https://doi.org/10.1007/s13225-020-00466-2)) |
| Otu56608 | 1 | k__Fungi(100);p__Basidiomycota(100);c__Microbotryomycetes(100);o__Sporidiobolales(100);f__Sporidiobolales_family_Incertae_sedis(100);g__Sporobolomyces(100);s__Sporobolomyces_lactosus(100); | Sporobolomyces | 13 | Pathotroph-Saprotroph | \|Fungal Parasite\|-Plant Saprotroph | Yeast | NULL | Probable | As litter_saprotroph (Põlme et al. 2020); As mycoparasite (Põlme et al. 2020); Decay fungal material (Põlme et al. 2020) | Tedersoo L, et al. 2014. Science 346:e1256688 ((https://doi.org/10.1126/science.1256688)); Purahong W, et al. 2016. Molecular Ecology 25:4059-4074 ((https://doi.org/10.1111/mec.13739)); Põlme S, et al. 2020. Fungal Diversity 105:1-16 ((https://doi.org/10.1007/s13225-020-00466-2)) |
| Otu56609 | 1 | k__Fungi(100);p__Ascomycota(100);c__Saccharomycetes(100);o__Saccharomycetales(100);f__Metschnikowiaceae(100);g__Metschnikowia(100);g__Metschnikowia_unclassified(100); | Metschnikowia | 13 | Saprotroph | \|Undefined Saprotroph\| | Yeast | NULL | Probable | As nectar-tap_saprotroph (Põlme et al. 2020); Decay sugar-rich substrates (Põlme et al. 2020) | Põlme S, et al. 2020. Fungal Diversity 105:1-16 ((https://doi.org/10.1007/s13225-020-00466-2)) |
| Otu56610 | 1 | k__Fungi(100);p__Ascomycota(100);c__Saccharomycetes(100);o__Saccharomycetales(100);f__Metschnikowiaceae(100);g__Metschnikowia(100);s__Metschnikowia_pulcherrima(100); | Metschnikowia | 13 | Saprotroph | \|Undefined Saprotroph\| | Yeast | NULL | Probable | As nectar-tap_saprotroph (Põlme et al. 2020); Decay sugar-rich substrates (Põlme et al. 2020) | Põlme S, et al. 2020. Fungal Diversity 105:1-16 ((https://doi.org/10.1007/s13225-020-00466-2)) |
| Otu56611 | 1 | k__Fungi(100);p__Ascomycota(100);c__Saccharomycetes(100);o__Saccharomycetales(100);f__Metschnikowiaceae(100);g__Metschnikowia(100);s__Metschnikowia_pulcherrima(100); | Metschnikowia | 13 | Saprotroph | \|Undefined Saprotroph\| | Yeast | NULL | Probable | As nectar-tap_saprotroph (Põlme et al. 2020); Decay sugar-rich substrates (Põlme et al. 2020) | Põlme S, et al. 2020. Fungal Diversity 105:1-16 ((https://doi.org/10.1007/s13225-020-00466-2)) |
| Otu56612 | 1 | k__Fungi(100);p__Ascomycota(100);c__Saccharomycetes(100);o__Saccharomycetales(100);f__Metschnikowiaceae(100);g__Metschnikowia(100);s__Metschnikowia_pulcherrima(100); | Metschnikowia | 13 | Saprotroph | \|Undefined Saprotroph\| | Yeast | NULL | Probable | As nectar-tap_saprotroph (Põlme et al. 2020); Decay sugar-rich substrates (Põlme et al. 2020) | Põlme S, et al. 2020. Fungal Diversity 105:1-16 ((https://doi.org/10.1007/s13225-020-00466-2)) |
| Otu56618 | 1 | k__Fungi(100);p__Ascomycota(100);c__Saccharomycetes(100);o__Saccharomycetales(100);f__Metschnikowiaceae(100);g__Metschnikowia(100);s__Metschnikowia_pulcherrima(100); | Metschnikowia | 13 | Saprotroph | \|Undefined Saprotroph\| | Yeast | NULL | Probable | As nectar-tap_saprotroph (Põlme et al. 2020); Decay sugar-rich substrates (Põlme et al. 2020) | Põlme S, et al. 2020. Fungal Diversity 105:1-16 ((https://doi.org/10.1007/s13225-020-00466-2)) |
| Otu56620 | 1 | k__Fungi(100);p__Ascomycota(100);c__Saccharomycetes(100);o__Saccharomycetales(100);f__Metschnikowiaceae(100);g__Metschnikowia(100);s__Metschnikowia_pulcherrima(100); | Metschnikowia | 13 | Saprotroph | \|Undefined Saprotroph\| | Yeast | NULL | Probable | As nectar-tap_saprotroph (Põlme et al. 2020); Decay sugar-rich substrates (Põlme et al. 2020) | Põlme S, et al. 2020. Fungal Diversity 105:1-16 ((https://doi.org/10.1007/s13225-020-00466-2)) |
| Otu56621 | 1 | k__Fungi(100);p__Ascomycota(100);c__Saccharomycetes(100);o__Saccharomycetales(100);f__Metschnikowiaceae(100);g__Metschnikowia(100);s__Metschnikowia_pulcherrima(100); | Metschnikowia | 13 | Saprotroph | \|Undefined Saprotroph\| | Yeast | NULL | Probable | As nectar-tap_saprotroph (Põlme et al. 2020); Decay sugar-rich substrates (Põlme et al. 2020) | Põlme S, et al. 2020. Fungal Diversity 105:1-16 ((https://doi.org/10.1007/s13225-020-00466-2)) |
| Otu56624 | 1 | k__Fungi(100);p__Ascomycota(100);c__Saccharomycetes(100);o__Saccharomycetales(100);f__Metschnikowiaceae(100);g__Metschnikowia(100);g__Metschnikowia_unclassified(100); | Metschnikowia | 13 | Saprotroph | \|Undefined Saprotroph\| | Yeast | NULL | Probable | As nectar-tap_saprotroph (Põlme et al. 2020); Decay sugar-rich substrates (Põlme et al. 2020) | Põlme S, et al. 2020. Fungal Diversity 105:1-16 ((https://doi.org/10.1007/s13225-020-00466-2)) |
| Otu56625 | 1 | k__Fungi(100);p__Ascomycota(100);c__Saccharomycetes(100);o__Saccharomycetales(100);f__Saccharomycetales_family_Incertae_sedis(100);g__Candida(100);g__Candida_unclassified(100); | Candida | 13 | Pathotroph-Saprotroph-Symbiotroph | Animal Pathogen-Endophyte-Endosymbiont-Epiphyte-\|Undefined Saprotroph\| | Dimorphic Yeast-Yeast | NULL | Probable | As nectar-tap_saprotroph (Põlme et al. 2020); Decay sugar-rich substrates (Põlme et al. 2020); This asexual genus is polyphyletic and contain species that function very differently depend on what they are. Caution should be taken when interpreting guilds for Candida species. | Manolakaki D, et al. 2010. Virulence. 1: 367-375 ((https://doi.org/10.4161/viru.1.5.12796)); Mekbib SB, et al. 2011. Tropical Plant Pathology 36:233-240 ((https://doi.org/10.1590/S1982-56762011000400004)); Põlme S, et al. 2020. Fungal Diversity 105:1-16 ((https://doi.org/10.1007/s13225-020-00466-2)) |
| Otu56626 | 1 | k__Fungi(100);p__Ascomycota(100);c__Saccharomycetes(100);o__Saccharomycetales(100);f__Metschnikowiaceae(100);g__Metschnikowia(100);s__Metschnikowia_pulcherrima(100); | Metschnikowia | 13 | Saprotroph | \|Undefined Saprotroph\| | Yeast | NULL | Probable | As nectar-tap_saprotroph (Põlme et al. 2020); Decay sugar-rich substrates (Põlme et al. 2020) | Põlme S, et al. 2020. Fungal Diversity 105:1-16 ((https://doi.org/10.1007/s13225-020-00466-2)) |
| Otu56629 | 1 | k__Fungi(100);p__Ascomycota(100);c__Saccharomycetes(100);o__Saccharomycetales(100);f__Metschnikowiaceae(100);g__Metschnikowia(100);s__Metschnikowia_pulcherrima(100); | Metschnikowia | 13 | Saprotroph | \|Undefined Saprotroph\| | Yeast | NULL | Probable | As nectar-tap_saprotroph (Põlme et al. 2020); Decay sugar-rich substrates (Põlme et al. 2020) | Põlme S, et al. 2020. Fungal Diversity 105:1-16 ((https://doi.org/10.1007/s13225-020-00466-2)) |
| Otu56630 | 1 | k__Fungi(100);p__Ascomycota(100);c__Saccharomycetes(100);o__Saccharomycetales(100);f__Metschnikowiaceae(100);g__Metschnikowia(100);s__Metschnikowia_pulcherrima(100); | Metschnikowia | 13 | Saprotroph | \|Undefined Saprotroph\| | Yeast | NULL | Probable | As nectar-tap_saprotroph (Põlme et al. 2020); Decay sugar-rich substrates (Põlme et al. 2020) | Põlme S, et al. 2020. Fungal Diversity 105:1-16 ((https://doi.org/10.1007/s13225-020-00466-2)) |
| Otu56631 | 1 | k__Fungi(100);p__Ascomycota(100);c__Saccharomycetes(100);o__Saccharomycetales(100);f__Metschnikowiaceae(100);g__Metschnikowia(100);g__Metschnikowia_unclassified(100); | Metschnikowia | 13 | Saprotroph | \|Undefined Saprotroph\| | Yeast | NULL | Probable | As nectar-tap_saprotroph (Põlme et al. 2020); Decay sugar-rich substrates (Põlme et al. 2020) | Põlme S, et al. 2020. Fungal Diversity 105:1-16 ((https://doi.org/10.1007/s13225-020-00466-2)) |
| Otu56632 | 1 | k__Fungi(100);p__Ascomycota(100);c__Saccharomycetes(100);o__Saccharomycetales(100);f__Metschnikowiaceae(100);g__Metschnikowia(100);s__Metschnikowia_pulcherrima(100); | Metschnikowia | 13 | Saprotroph | \|Undefined Saprotroph\| | Yeast | NULL | Probable | As nectar-tap_saprotroph (Põlme et al. 2020); Decay sugar-rich substrates (Põlme et al. 2020) | Põlme S, et al. 2020. Fungal Diversity 105:1-16 ((https://doi.org/10.1007/s13225-020-00466-2)) |
| Otu56633 | 1 | k__Fungi(100);p__Ascomycota(100);c__Saccharomycetes(100);o__Saccharomycetales(100);f__Saccharomycetales_family_Incertae_sedis(100);g__Candida(100);s__Candida_cretensis(100); | Candida | 13 | Pathotroph-Saprotroph-Symbiotroph | Animal Pathogen-Endophyte-Endosymbiont-Epiphyte-\|Undefined Saprotroph\| | Dimorphic Yeast-Yeast | NULL | Probable | As nectar-tap_saprotroph (Põlme et al. 2020); Decay sugar-rich substrates (Põlme et al. 2020); This asexual genus is polyphyletic and contain species that function very differently depend on what they are. Caution should be taken when interpreting guilds for Candida species. | Manolakaki D, et al. 2010. Virulence. 1: 367-375 ((https://doi.org/10.4161/viru.1.5.12796)); Mekbib SB, et al. 2011. Tropical Plant Pathology 36:233-240 ((https://doi.org/10.1590/S1982-56762011000400004)); Põlme S, et al. 2020. Fungal Diversity 105:1-16 ((https://doi.org/10.1007/s13225-020-00466-2)) |
| Otu56634 | 1 | k__Fungi(100);p__Ascomycota(100);c__Saccharomycetes(100);o__Saccharomycetales(100);f__Metschnikowiaceae(100);g__Metschnikowia(100);g__Metschnikowia_unclassified(100); | Metschnikowia | 13 | Saprotroph | \|Undefined Saprotroph\| | Yeast | NULL | Probable | As nectar-tap_saprotroph (Põlme et al. 2020); Decay sugar-rich substrates (Põlme et al. 2020) | Põlme S, et al. 2020. Fungal Diversity 105:1-16 ((https://doi.org/10.1007/s13225-020-00466-2)) |
| Otu56635 | 1 | k__Fungi(100);p__Basidiomycota(100);c__Microbotryomycetes(100);o__Sporidiobolales(100);f__Sporidiobolales_family_Incertae_sedis(100);g__Sporobolomyces(100);s__Sporobolomyces_lactosus(100); | Sporobolomyces | 13 | Pathotroph-Saprotroph | \|Fungal Parasite\|-Plant Saprotroph | Yeast | NULL | Probable | As litter_saprotroph (Põlme et al. 2020); As mycoparasite (Põlme et al. 2020); Decay fungal material (Põlme et al. 2020) | Tedersoo L, et al. 2014. Science 346:e1256688 ((https://doi.org/10.1126/science.1256688)); Purahong W, et al. 2016. Molecular Ecology 25:4059-4074 ((https://doi.org/10.1111/mec.13739)); Põlme S, et al. 2020. Fungal Diversity 105:1-16 ((https://doi.org/10.1007/s13225-020-00466-2)) |
| Otu56639 | 1 | k__Fungi(100);p__Ascomycota(100);c__Saccharomycetes(100);o__Saccharomycetales(100);f__Metschnikowiaceae(100);g__Metschnikowia(100);g__Metschnikowia_unclassified(100); | Metschnikowia | 13 | Saprotroph | \|Undefined Saprotroph\| | Yeast | NULL | Probable | As nectar-tap_saprotroph (Põlme et al. 2020); Decay sugar-rich substrates (Põlme et al. 2020) | Põlme S, et al. 2020. Fungal Diversity 105:1-16 ((https://doi.org/10.1007/s13225-020-00466-2)) |
| Otu56640 | 1 | k__Fungi(100);p__Ascomycota(100);c__Saccharomycetes(100);o__Saccharomycetales(100);f__Metschnikowiaceae(100);g__Metschnikowia(100);g__Metschnikowia_unclassified(100); | Metschnikowia | 13 | Saprotroph | \|Undefined Saprotroph\| | Yeast | NULL | Probable | As nectar-tap_saprotroph (Põlme et al. 2020); Decay sugar-rich substrates (Põlme et al. 2020) | Põlme S, et al. 2020. Fungal Diversity 105:1-16 ((https://doi.org/10.1007/s13225-020-00466-2)) |
| Otu56641 | 1 | k__Fungi(100);p__Ascomycota(100);c__Saccharomycetes(100);o__Saccharomycetales(100);f__Metschnikowiaceae(100);g__Metschnikowia(100);s__Metschnikowia_pulcherrima(100); | Metschnikowia | 13 | Saprotroph | \|Undefined Saprotroph\| | Yeast | NULL | Probable | As nectar-tap_saprotroph (Põlme et al. 2020); Decay sugar-rich substrates (Põlme et al. 2020) | Põlme S, et al. 2020. Fungal Diversity 105:1-16 ((https://doi.org/10.1007/s13225-020-00466-2)) |
| Otu56643 | 1 | k__Fungi(100);p__Ascomycota(100);c__Saccharomycetes(100);o__Saccharomycetales(100);f__Metschnikowiaceae(100);g__Metschnikowia(100);s__Metschnikowia_pulcherrima(100); | Metschnikowia | 13 | Saprotroph | \|Undefined Saprotroph\| | Yeast | NULL | Probable | As nectar-tap_saprotroph (Põlme et al. 2020); Decay sugar-rich substrates (Põlme et al. 2020) | Põlme S, et al. 2020. Fungal Diversity 105:1-16 ((https://doi.org/10.1007/s13225-020-00466-2)) |
| Otu56646 | 1 | k__Fungi(100);p__Ascomycota(100);c__Saccharomycetes(100);o__Saccharomycetales(100);f__Metschnikowiaceae(100);g__Metschnikowia(100);s__Metschnikowia_pulcherrima(100); | Metschnikowia | 13 | Saprotroph | \|Undefined Saprotroph\| | Yeast | NULL | Probable | As nectar-tap_saprotroph (Põlme et al. 2020); Decay sugar-rich substrates (Põlme et al. 2020) | Põlme S, et al. 2020. Fungal Diversity 105:1-16 ((https://doi.org/10.1007/s13225-020-00466-2)) |
| Otu56650 | 1 | k__Fungi(100);p__Ascomycota(100);c__Saccharomycetes(100);o__Saccharomycetales(100);f__Metschnikowiaceae(100);g__Metschnikowia(100);s__Metschnikowia_pulcherrima(100); | Metschnikowia | 13 | Saprotroph | \|Undefined Saprotroph\| | Yeast | NULL | Probable | As nectar-tap_saprotroph (Põlme et al. 2020); Decay sugar-rich substrates (Põlme et al. 2020) | Põlme S, et al. 2020. Fungal Diversity 105:1-16 ((https://doi.org/10.1007/s13225-020-00466-2)) |
| Otu56651 | 1 | k__Fungi(100);p__Ascomycota(100);c__Saccharomycetes(100);o__Saccharomycetales(100);f__Metschnikowiaceae(100);g__Metschnikowia(100);s__Metschnikowia_pulcherrima(100); | Metschnikowia | 13 | Saprotroph | \|Undefined Saprotroph\| | Yeast | NULL | Probable | As nectar-tap_saprotroph (Põlme et al. 2020); Decay sugar-rich substrates (Põlme et al. 2020) | Põlme S, et al. 2020. Fungal Diversity 105:1-16 ((https://doi.org/10.1007/s13225-020-00466-2)) |
| Otu56652 | 1 | k__Fungi(100);p__Ascomycota(100);c__Saccharomycetes(100);o__Saccharomycetales(100);f__Metschnikowiaceae(100);g__Metschnikowia(100);g__Metschnikowia_unclassified(100); | Metschnikowia | 13 | Saprotroph | \|Undefined Saprotroph\| | Yeast | NULL | Probable | As nectar-tap_saprotroph (Põlme et al. 2020); Decay sugar-rich substrates (Põlme et al. 2020) | Põlme S, et al. 2020. Fungal Diversity 105:1-16 ((https://doi.org/10.1007/s13225-020-00466-2)) |
| Otu56653 | 1 | k__Fungi(100);p__Ascomycota(100);c__Saccharomycetes(100);o__Saccharomycetales(100);f__Metschnikowiaceae(100);g__Metschnikowia(100);s__Metschnikowia_pulcherrima(100); | Metschnikowia | 13 | Saprotroph | \|Undefined Saprotroph\| | Yeast | NULL | Probable | As nectar-tap_saprotroph (Põlme et al. 2020); Decay sugar-rich substrates (Põlme et al. 2020) | Põlme S, et al. 2020. Fungal Diversity 105:1-16 ((https://doi.org/10.1007/s13225-020-00466-2)) |
| Otu56654 | 1 | k__Fungi(100);p__Ascomycota(100);c__Saccharomycetes(100);o__Saccharomycetales(100);f__Metschnikowiaceae(100);g__Metschnikowia(100);g__Metschnikowia_unclassified(100); | Metschnikowia | 13 | Saprotroph | \|Undefined Saprotroph\| | Yeast | NULL | Probable | As nectar-tap_saprotroph (Põlme et al. 2020); Decay sugar-rich substrates (Põlme et al. 2020) | Põlme S, et al. 2020. Fungal Diversity 105:1-16 ((https://doi.org/10.1007/s13225-020-00466-2)) |
| Otu56655 | 1 | k__Fungi(100);p__Ascomycota(100);c__Saccharomycetes(100);o__Saccharomycetales(100);f__Metschnikowiaceae(100);g__Metschnikowia(100);s__Metschnikowia_pulcherrima(100); | Metschnikowia | 13 | Saprotroph | \|Undefined Saprotroph\| | Yeast | NULL | Probable | As nectar-tap_saprotroph (Põlme et al. 2020); Decay sugar-rich substrates (Põlme et al. 2020) | Põlme S, et al. 2020. Fungal Diversity 105:1-16 ((https://doi.org/10.1007/s13225-020-00466-2)) |
| Otu56656 | 1 | k__Fungi(100);p__Ascomycota(100);c__Saccharomycetes(100);o__Saccharomycetales(100);f__Metschnikowiaceae(100);g__Metschnikowia(100);s__Metschnikowia_pulcherrima(100); | Metschnikowia | 13 | Saprotroph | \|Undefined Saprotroph\| | Yeast | NULL | Probable | As nectar-tap_saprotroph (Põlme et al. 2020); Decay sugar-rich substrates (Põlme et al. 2020) | Põlme S, et al. 2020. Fungal Diversity 105:1-16 ((https://doi.org/10.1007/s13225-020-00466-2)) |
| Otu56657 | 1 | k__Fungi(100);p__Ascomycota(100);c__Saccharomycetes(100);o__Saccharomycetales(100);f__Metschnikowiaceae(100);g__Metschnikowia(100);s__Metschnikowia_pulcherrima(100); | Metschnikowia | 13 | Saprotroph | \|Undefined Saprotroph\| | Yeast | NULL | Probable | As nectar-tap_saprotroph (Põlme et al. 2020); Decay sugar-rich substrates (Põlme et al. 2020) | Põlme S, et al. 2020. Fungal Diversity 105:1-16 ((https://doi.org/10.1007/s13225-020-00466-2)) |
| Otu56658 | 1 | k__Fungi(100);p__Ascomycota(100);c__Saccharomycetes(100);o__Saccharomycetales(100);f__Metschnikowiaceae(100);g__Metschnikowia(100);g__Metschnikowia_unclassified(100); | Metschnikowia | 13 | Saprotroph | \|Undefined Saprotroph\| | Yeast | NULL | Probable | As nectar-tap_saprotroph (Põlme et al. 2020); Decay sugar-rich substrates (Põlme et al. 2020) | Põlme S, et al. 2020. Fungal Diversity 105:1-16 ((https://doi.org/10.1007/s13225-020-00466-2)) |
| Otu56659 | 1 | k__Fungi(100);p__Ascomycota(100);c__Saccharomycetes(100);o__Saccharomycetales(100);f__Metschnikowiaceae(100);g__Metschnikowia(100);s__Metschnikowia_pulcherrima(100); | Metschnikowia | 13 | Saprotroph | \|Undefined Saprotroph\| | Yeast | NULL | Probable | As nectar-tap_saprotroph (Põlme et al. 2020); Decay sugar-rich substrates (Põlme et al. 2020) | Põlme S, et al. 2020. Fungal Diversity 105:1-16 ((https://doi.org/10.1007/s13225-020-00466-2)) |
| Otu56660 | 1 | k__Fungi(100);p__Ascomycota(100);c__Saccharomycetes(100);o__Saccharomycetales(100);f__Metschnikowiaceae(100);g__Metschnikowia(100);g__Metschnikowia_unclassified(100); | Metschnikowia | 13 | Saprotroph | \|Undefined Saprotroph\| | Yeast | NULL | Probable | As nectar-tap_saprotroph (Põlme et al. 2020); Decay sugar-rich substrates (Põlme et al. 2020) | Põlme S, et al. 2020. Fungal Diversity 105:1-16 ((https://doi.org/10.1007/s13225-020-00466-2)) |
| Otu56661 | 1 | k__Fungi(100);p__Ascomycota(100);c__Saccharomycetes(100);o__Saccharomycetales(100);f__Metschnikowiaceae(100);g__Metschnikowia(100);s__Metschnikowia_pulcherrima(100); | Metschnikowia | 13 | Saprotroph | \|Undefined Saprotroph\| | Yeast | NULL | Probable | As nectar-tap_saprotroph (Põlme et al. 2020); Decay sugar-rich substrates (Põlme et al. 2020) | Põlme S, et al. 2020. Fungal Diversity 105:1-16 ((https://doi.org/10.1007/s13225-020-00466-2)) |
| Otu56663 | 1 | k__Fungi(100);p__Ascomycota(100);c__Saccharomycetes(100);o__Saccharomycetales(100);f__Metschnikowiaceae(100);g__Metschnikowia(100);s__Metschnikowia_pulcherrima(100); | Metschnikowia | 13 | Saprotroph | \|Undefined Saprotroph\| | Yeast | NULL | Probable | As nectar-tap_saprotroph (Põlme et al. 2020); Decay sugar-rich substrates (Põlme et al. 2020) | Põlme S, et al. 2020. Fungal Diversity 105:1-16 ((https://doi.org/10.1007/s13225-020-00466-2)) |
| Otu56664 | 1 | k__Fungi(100);p__Ascomycota(100);c__Saccharomycetes(100);o__Saccharomycetales(100);f__Metschnikowiaceae(100);g__Metschnikowia(100);g__Metschnikowia_unclassified(100); | Metschnikowia | 13 | Saprotroph | \|Undefined Saprotroph\| | Yeast | NULL | Probable | As nectar-tap_saprotroph (Põlme et al. 2020); Decay sugar-rich substrates (Põlme et al. 2020) | Põlme S, et al. 2020. Fungal Diversity 105:1-16 ((https://doi.org/10.1007/s13225-020-00466-2)) |
| Otu56665 | 1 | k__Fungi(100);p__Ascomycota(100);c__Saccharomycetes(100);o__Saccharomycetales(100);f__Metschnikowiaceae(100);g__Metschnikowia(100);g__Metschnikowia_unclassified(100); | Metschnikowia | 13 | Saprotroph | \|Undefined Saprotroph\| | Yeast | NULL | Probable | As nectar-tap_saprotroph (Põlme et al. 2020); Decay sugar-rich substrates (Põlme et al. 2020) | Põlme S, et al. 2020. Fungal Diversity 105:1-16 ((https://doi.org/10.1007/s13225-020-00466-2)) |
| Otu56666 | 1 | k__Fungi(100);p__Ascomycota(100);c__Saccharomycetes(100);o__Saccharomycetales(100);f__Metschnikowiaceae(100);g__Metschnikowia(100);s__Metschnikowia_pulcherrima(100); | Metschnikowia | 13 | Saprotroph | \|Undefined Saprotroph\| | Yeast | NULL | Probable | As nectar-tap_saprotroph (Põlme et al. 2020); Decay sugar-rich substrates (Põlme et al. 2020) | Põlme S, et al. 2020. Fungal Diversity 105:1-16 ((https://doi.org/10.1007/s13225-020-00466-2)) |
| Otu56667 | 1 | k__Fungi(100);p__Ascomycota(100);c__Saccharomycetes(100);o__Saccharomycetales(100);f__Metschnikowiaceae(100);g__Metschnikowia(100);s__Metschnikowia_pulcherrima(100); | Metschnikowia | 13 | Saprotroph | \|Undefined Saprotroph\| | Yeast | NULL | Probable | As nectar-tap_saprotroph (Põlme et al. 2020); Decay sugar-rich substrates (Põlme et al. 2020) | Põlme S, et al. 2020. Fungal Diversity 105:1-16 ((https://doi.org/10.1007/s13225-020-00466-2)) |
| Otu56668 | 1 | k__Fungi(100);p__Ascomycota(100);c__Saccharomycetes(100);o__Saccharomycetales(100);f__Metschnikowiaceae(100);g__Metschnikowia(100);g__Metschnikowia_unclassified(100); | Metschnikowia | 13 | Saprotroph | \|Undefined Saprotroph\| | Yeast | NULL | Probable | As nectar-tap_saprotroph (Põlme et al. 2020); Decay sugar-rich substrates (Põlme et al. 2020) | Põlme S, et al. 2020. Fungal Diversity 105:1-16 ((https://doi.org/10.1007/s13225-020-00466-2)) |
| Otu56669 | 1 | k__Fungi(100);p__Ascomycota(100);c__Saccharomycetes(100);o__Saccharomycetales(100);f__Metschnikowiaceae(100);g__Metschnikowia(100);g__Metschnikowia_unclassified(100); | Metschnikowia | 13 | Saprotroph | \|Undefined Saprotroph\| | Yeast | NULL | Probable | As nectar-tap_saprotroph (Põlme et al. 2020); Decay sugar-rich substrates (Põlme et al. 2020) | Põlme S, et al. 2020. Fungal Diversity 105:1-16 ((https://doi.org/10.1007/s13225-020-00466-2)) |
| Otu56670 | 1 | k__Fungi(100);p__Ascomycota(100);c__Saccharomycetes(100);o__Saccharomycetales(100);f__Metschnikowiaceae(100);g__Metschnikowia(100);s__Metschnikowia_pulcherrima(100); | Metschnikowia | 13 | Saprotroph | \|Undefined Saprotroph\| | Yeast | NULL | Probable | As nectar-tap_saprotroph (Põlme et al. 2020); Decay sugar-rich substrates (Põlme et al. 2020) | Põlme S, et al. 2020. Fungal Diversity 105:1-16 ((https://doi.org/10.1007/s13225-020-00466-2)) |
| Otu56672 | 1 | k__Fungi(100);p__Ascomycota(100);c__Saccharomycetes(100);o__Saccharomycetales(100);f__Metschnikowiaceae(100);g__Metschnikowia(100);s__Metschnikowia_pulcherrima(100); | Metschnikowia | 13 | Saprotroph | \|Undefined Saprotroph\| | Yeast | NULL | Probable | As nectar-tap_saprotroph (Põlme et al. 2020); Decay sugar-rich substrates (Põlme et al. 2020) | Põlme S, et al. 2020. Fungal Diversity 105:1-16 ((https://doi.org/10.1007/s13225-020-00466-2)) |
| Otu56673 | 1 | k__Fungi(100);p__Ascomycota(100);c__Saccharomycetes(100);o__Saccharomycetales(100);f__Metschnikowiaceae(100);g__Metschnikowia(100);s__Metschnikowia_pulcherrima(100); | Metschnikowia | 13 | Saprotroph | \|Undefined Saprotroph\| | Yeast | NULL | Probable | As nectar-tap_saprotroph (Põlme et al. 2020); Decay sugar-rich substrates (Põlme et al. 2020) | Põlme S, et al. 2020. Fungal Diversity 105:1-16 ((https://doi.org/10.1007/s13225-020-00466-2)) |
| Otu56674 | 1 | k__Fungi(100);p__Ascomycota(100);c__Saccharomycetes(100);o__Saccharomycetales(100);f__Metschnikowiaceae(100);g__Metschnikowia(100);s__Metschnikowia_pulcherrima(100); | Metschnikowia | 13 | Saprotroph | \|Undefined Saprotroph\| | Yeast | NULL | Probable | As nectar-tap_saprotroph (Põlme et al. 2020); Decay sugar-rich substrates (Põlme et al. 2020) | Põlme S, et al. 2020. Fungal Diversity 105:1-16 ((https://doi.org/10.1007/s13225-020-00466-2)) |
| Otu56675 | 1 | k__Fungi(100);p__Ascomycota(100);c__Saccharomycetes(100);o__Saccharomycetales(100);f__Metschnikowiaceae(100);g__Metschnikowia(100);s__Metschnikowia_pulcherrima(100); | Metschnikowia | 13 | Saprotroph | \|Undefined Saprotroph\| | Yeast | NULL | Probable | As nectar-tap_saprotroph (Põlme et al. 2020); Decay sugar-rich substrates (Põlme et al. 2020) | Põlme S, et al. 2020. Fungal Diversity 105:1-16 ((https://doi.org/10.1007/s13225-020-00466-2)) |
| Otu56676 | 1 | k__Fungi(100);p__Ascomycota(100);c__Saccharomycetes(100);o__Saccharomycetales(100);f__Metschnikowiaceae(100);g__Metschnikowia(100);s__Metschnikowia_pulcherrima(100); | Metschnikowia | 13 | Saprotroph | \|Undefined Saprotroph\| | Yeast | NULL | Probable | As nectar-tap_saprotroph (Põlme et al. 2020); Decay sugar-rich substrates (Põlme et al. 2020) | Põlme S, et al. 2020. Fungal Diversity 105:1-16 ((https://doi.org/10.1007/s13225-020-00466-2)) |
| Otu56677 | 1 | k__Fungi(100);p__Ascomycota(100);c__Saccharomycetes(100);o__Saccharomycetales(100);f__Metschnikowiaceae(100);g__Metschnikowia(100);g__Metschnikowia_unclassified(100); | Metschnikowia | 13 | Saprotroph | \|Undefined Saprotroph\| | Yeast | NULL | Probable | As nectar-tap_saprotroph (Põlme et al. 2020); Decay sugar-rich substrates (Põlme et al. 2020) | Põlme S, et al. 2020. Fungal Diversity 105:1-16 ((https://doi.org/10.1007/s13225-020-00466-2)) |
| Otu56678 | 1 | k__Fungi(100);p__Ascomycota(100);c__Saccharomycetes(100);o__Saccharomycetales(100);f__Metschnikowiaceae(100);g__Metschnikowia(100);s__Metschnikowia_pulcherrima(100); | Metschnikowia | 13 | Saprotroph | \|Undefined Saprotroph\| | Yeast | NULL | Probable | As nectar-tap_saprotroph (Põlme et al. 2020); Decay sugar-rich substrates (Põlme et al. 2020) | Põlme S, et al. 2020. Fungal Diversity 105:1-16 ((https://doi.org/10.1007/s13225-020-00466-2)) |
| Otu56679 | 1 | k__Fungi(100);p__Ascomycota(100);c__Saccharomycetes(100);o__Saccharomycetales(100);f__Metschnikowiaceae(100);g__Metschnikowia(100);g__Metschnikowia_unclassified(100); | Metschnikowia | 13 | Saprotroph | \|Undefined Saprotroph\| | Yeast | NULL | Probable | As nectar-tap_saprotroph (Põlme et al. 2020); Decay sugar-rich substrates (Põlme et al. 2020) | Põlme S, et al. 2020. Fungal Diversity 105:1-16 ((https://doi.org/10.1007/s13225-020-00466-2)) |
| Otu56680 | 1 | k__Fungi(100);p__Ascomycota(100);c__Saccharomycetes(100);o__Saccharomycetales(100);f__Metschnikowiaceae(100);g__Metschnikowia(100);s__Metschnikowia_pulcherrima(100); | Metschnikowia | 13 | Saprotroph | \|Undefined Saprotroph\| | Yeast | NULL | Probable | As nectar-tap_saprotroph (Põlme et al. 2020); Decay sugar-rich substrates (Põlme et al. 2020) | Põlme S, et al. 2020. Fungal Diversity 105:1-16 ((https://doi.org/10.1007/s13225-020-00466-2)) |
| Otu56681 | 1 | k__Fungi(100);p__Ascomycota(100);c__Saccharomycetes(100);o__Saccharomycetales(100);f__Metschnikowiaceae(100);g__Metschnikowia(100);s__Metschnikowia_pulcherrima(100); | Metschnikowia | 13 | Saprotroph | \|Undefined Saprotroph\| | Yeast | NULL | Probable | As nectar-tap_saprotroph (Põlme et al. 2020); Decay sugar-rich substrates (Põlme et al. 2020) | Põlme S, et al. 2020. Fungal Diversity 105:1-16 ((https://doi.org/10.1007/s13225-020-00466-2)) |
| Otu56682 | 1 | k__Fungi(100);p__Ascomycota(100);c__Saccharomycetes(100);o__Saccharomycetales(100);f__Metschnikowiaceae(100);g__Metschnikowia(100);s__Metschnikowia_pulcherrima(100); | Metschnikowia | 13 | Saprotroph | \|Undefined Saprotroph\| | Yeast | NULL | Probable | As nectar-tap_saprotroph (Põlme et al. 2020); Decay sugar-rich substrates (Põlme et al. 2020) | Põlme S, et al. 2020. Fungal Diversity 105:1-16 ((https://doi.org/10.1007/s13225-020-00466-2)) |
| Otu56683 | 1 | k__Fungi(100);p__Ascomycota(100);c__Saccharomycetes(100);o__Saccharomycetales(100);f__Metschnikowiaceae(100);g__Metschnikowia(100);s__Metschnikowia_pulcherrima(100); | Metschnikowia | 13 | Saprotroph | \|Undefined Saprotroph\| | Yeast | NULL | Probable | As nectar-tap_saprotroph (Põlme et al. 2020); Decay sugar-rich substrates (Põlme et al. 2020) | Põlme S, et al. 2020. Fungal Diversity 105:1-16 ((https://doi.org/10.1007/s13225-020-00466-2)) |
| Otu56686 | 1 | k__Fungi(100);p__Ascomycota(100);c__Saccharomycetes(100);o__Saccharomycetales(100);f__Metschnikowiaceae(100);g__Metschnikowia(100);s__Metschnikowia_pulcherrima(100); | Metschnikowia | 13 | Saprotroph | \|Undefined Saprotroph\| | Yeast | NULL | Probable | As nectar-tap_saprotroph (Põlme et al. 2020); Decay sugar-rich substrates (Põlme et al. 2020) | Põlme S, et al. 2020. Fungal Diversity 105:1-16 ((https://doi.org/10.1007/s13225-020-00466-2)) |
| Otu56688 | 1 | k__Fungi(100);p__Ascomycota(100);c__Saccharomycetes(100);o__Saccharomycetales(100);f__Metschnikowiaceae(100);g__Metschnikowia(100);s__Metschnikowia_pulcherrima(100); | Metschnikowia | 13 | Saprotroph | \|Undefined Saprotroph\| | Yeast | NULL | Probable | As nectar-tap_saprotroph (Põlme et al. 2020); Decay sugar-rich substrates (Põlme et al. 2020) | Põlme S, et al. 2020. Fungal Diversity 105:1-16 ((https://doi.org/10.1007/s13225-020-00466-2)) |
| Otu56689 | 1 | k__Fungi(100);p__Ascomycota(100);c__Saccharomycetes(100);o__Saccharomycetales(100);f__Metschnikowiaceae(100);g__Metschnikowia(100);s__Metschnikowia_pulcherrima(100); | Metschnikowia | 13 | Saprotroph | \|Undefined Saprotroph\| | Yeast | NULL | Probable | As nectar-tap_saprotroph (Põlme et al. 2020); Decay sugar-rich substrates (Põlme et al. 2020) | Põlme S, et al. 2020. Fungal Diversity 105:1-16 ((https://doi.org/10.1007/s13225-020-00466-2)) |
| Otu56690 | 1 | k__Fungi(100);p__Ascomycota(100);c__Saccharomycetes(100);o__Saccharomycetales(100);f__Metschnikowiaceae(100);g__Metschnikowia(100);g__Metschnikowia_unclassified(100); | Metschnikowia | 13 | Saprotroph | \|Undefined Saprotroph\| | Yeast | NULL | Probable | As nectar-tap_saprotroph (Põlme et al. 2020); Decay sugar-rich substrates (Põlme et al. 2020) | Põlme S, et al. 2020. Fungal Diversity 105:1-16 ((https://doi.org/10.1007/s13225-020-00466-2)) |
| Otu56691 | 1 | k__Fungi(100);p__Ascomycota(100);c__Saccharomycetes(100);o__Saccharomycetales(100);f__Metschnikowiaceae(100);g__Metschnikowia(100);s__Metschnikowia_pulcherrima(100); | Metschnikowia | 13 | Saprotroph | \|Undefined Saprotroph\| | Yeast | NULL | Probable | As nectar-tap_saprotroph (Põlme et al. 2020); Decay sugar-rich substrates (Põlme et al. 2020) | Põlme S, et al. 2020. Fungal Diversity 105:1-16 ((https://doi.org/10.1007/s13225-020-00466-2)) |
| Otu56693 | 1 | k__Fungi(100);p__Ascomycota(100);c__Saccharomycetes(100);o__Saccharomycetales(100);f__Metschnikowiaceae(100);g__Metschnikowia(100);s__Metschnikowia_pulcherrima(100); | Metschnikowia | 13 | Saprotroph | \|Undefined Saprotroph\| | Yeast | NULL | Probable | As nectar-tap_saprotroph (Põlme et al. 2020); Decay sugar-rich substrates (Põlme et al. 2020) | Põlme S, et al. 2020. Fungal Diversity 105:1-16 ((https://doi.org/10.1007/s13225-020-00466-2)) |
| Otu56694 | 1 | k__Fungi(100);p__Ascomycota(100);c__Saccharomycetes(100);o__Saccharomycetales(100);f__Metschnikowiaceae(100);g__Metschnikowia(100);s__Metschnikowia_pulcherrima(100); | Metschnikowia | 13 | Saprotroph | \|Undefined Saprotroph\| | Yeast | NULL | Probable | As nectar-tap_saprotroph (Põlme et al. 2020); Decay sugar-rich substrates (Põlme et al. 2020) | Põlme S, et al. 2020. Fungal Diversity 105:1-16 ((https://doi.org/10.1007/s13225-020-00466-2)) |
| Otu56695 | 1 | k__Fungi(100);p__Ascomycota(100);c__Saccharomycetes(100);o__Saccharomycetales(100);f__Metschnikowiaceae(100);g__Metschnikowia(100);s__Metschnikowia_pulcherrima(100); | Metschnikowia | 13 | Saprotroph | \|Undefined Saprotroph\| | Yeast | NULL | Probable | As nectar-tap_saprotroph (Põlme et al. 2020); Decay sugar-rich substrates (Põlme et al. 2020) | Põlme S, et al. 2020. Fungal Diversity 105:1-16 ((https://doi.org/10.1007/s13225-020-00466-2)) |
| Otu56696 | 1 | k__Fungi(100);p__Ascomycota(100);c__Saccharomycetes(100);o__Saccharomycetales(100);f__Metschnikowiaceae(100);g__Metschnikowia(100);g__Metschnikowia_unclassified(100); | Metschnikowia | 13 | Saprotroph | \|Undefined Saprotroph\| | Yeast | NULL | Probable | As nectar-tap_saprotroph (Põlme et al. 2020); Decay sugar-rich substrates (Põlme et al. 2020) | Põlme S, et al. 2020. Fungal Diversity 105:1-16 ((https://doi.org/10.1007/s13225-020-00466-2)) |
| Otu56698 | 1 | k__Fungi(100);p__Ascomycota(100);c__Saccharomycetes(100);o__Saccharomycetales(100);f__Metschnikowiaceae(100);g__Metschnikowia(100);s__Metschnikowia_pulcherrima(100); | Metschnikowia | 13 | Saprotroph | \|Undefined Saprotroph\| | Yeast | NULL | Probable | As nectar-tap_saprotroph (Põlme et al. 2020); Decay sugar-rich substrates (Põlme et al. 2020) | Põlme S, et al. 2020. Fungal Diversity 105:1-16 ((https://doi.org/10.1007/s13225-020-00466-2)) |
| Otu56700 | 1 | k__Fungi(100);p__Ascomycota(100);c__Saccharomycetes(100);o__Saccharomycetales(100);f__Metschnikowiaceae(100);g__Metschnikowia(100);s__Metschnikowia_pulcherrima(100); | Metschnikowia | 13 | Saprotroph | \|Undefined Saprotroph\| | Yeast | NULL | Probable | As nectar-tap_saprotroph (Põlme et al. 2020); Decay sugar-rich substrates (Põlme et al. 2020) | Põlme S, et al. 2020. Fungal Diversity 105:1-16 ((https://doi.org/10.1007/s13225-020-00466-2)) |
| Otu56701 | 1 | k__Fungi(100);p__Ascomycota(100);c__Saccharomycetes(100);o__Saccharomycetales(100);f__Metschnikowiaceae(100);g__Metschnikowia(100);s__Metschnikowia_pulcherrima(100); | Metschnikowia | 13 | Saprotroph | \|Undefined Saprotroph\| | Yeast | NULL | Probable | As nectar-tap_saprotroph (Põlme et al. 2020); Decay sugar-rich substrates (Põlme et al. 2020) | Põlme S, et al. 2020. Fungal Diversity 105:1-16 ((https://doi.org/10.1007/s13225-020-00466-2)) |
| Otu56702 | 1 | k__Fungi(100);p__Ascomycota(100);c__Saccharomycetes(100);o__Saccharomycetales(100);f__Metschnikowiaceae(100);g__Metschnikowia(100);s__Metschnikowia_pulcherrima(100); | Metschnikowia | 13 | Saprotroph | \|Undefined Saprotroph\| | Yeast | NULL | Probable | As nectar-tap_saprotroph (Põlme et al. 2020); Decay sugar-rich substrates (Põlme et al. 2020) | Põlme S, et al. 2020. Fungal Diversity 105:1-16 ((https://doi.org/10.1007/s13225-020-00466-2)) |
| Otu56703 | 1 | k__Fungi(100);p__Ascomycota(100);c__Saccharomycetes(100);o__Saccharomycetales(100);f__Metschnikowiaceae(100);g__Metschnikowia(100);g__Metschnikowia_unclassified(100); | Metschnikowia | 13 | Saprotroph | \|Undefined Saprotroph\| | Yeast | NULL | Probable | As nectar-tap_saprotroph (Põlme et al. 2020); Decay sugar-rich substrates (Põlme et al. 2020) | Põlme S, et al. 2020. Fungal Diversity 105:1-16 ((https://doi.org/10.1007/s13225-020-00466-2)) |
| Otu56704 | 1 | k__Fungi(100);p__Ascomycota(100);c__Saccharomycetes(100);o__Saccharomycetales(100);f__Metschnikowiaceae(100);g__Metschnikowia(100);s__Metschnikowia_pulcherrima(100); | Metschnikowia | 13 | Saprotroph | \|Undefined Saprotroph\| | Yeast | NULL | Probable | As nectar-tap_saprotroph (Põlme et al. 2020); Decay sugar-rich substrates (Põlme et al. 2020) | Põlme S, et al. 2020. Fungal Diversity 105:1-16 ((https://doi.org/10.1007/s13225-020-00466-2)) |
| Otu56706 | 1 | k__Fungi(100);p__Ascomycota(100);c__Saccharomycetes(100);o__Saccharomycetales(100);f__Metschnikowiaceae(100);g__Metschnikowia(100);g__Metschnikowia_unclassified(100); | Metschnikowia | 13 | Saprotroph | \|Undefined Saprotroph\| | Yeast | NULL | Probable | As nectar-tap_saprotroph (Põlme et al. 2020); Decay sugar-rich substrates (Põlme et al. 2020) | Põlme S, et al. 2020. Fungal Diversity 105:1-16 ((https://doi.org/10.1007/s13225-020-00466-2)) |
| Otu56709 | 1 | k__Fungi(100);p__Ascomycota(100);c__Saccharomycetes(100);o__Saccharomycetales(100);f__Metschnikowiaceae(100);g__Metschnikowia(100);s__Metschnikowia_pulcherrima(100); | Metschnikowia | 13 | Saprotroph | \|Undefined Saprotroph\| | Yeast | NULL | Probable | As nectar-tap_saprotroph (Põlme et al. 2020); Decay sugar-rich substrates (Põlme et al. 2020) | Põlme S, et al. 2020. Fungal Diversity 105:1-16 ((https://doi.org/10.1007/s13225-020-00466-2)) |
| Otu56710 | 1 | k__Fungi(100);p__Ascomycota(100);c__Saccharomycetes(100);o__Saccharomycetales(100);f__Metschnikowiaceae(100);g__Metschnikowia(100);s__Metschnikowia_pulcherrima(100); | Metschnikowia | 13 | Saprotroph | \|Undefined Saprotroph\| | Yeast | NULL | Probable | As nectar-tap_saprotroph (Põlme et al. 2020); Decay sugar-rich substrates (Põlme et al. 2020) | Põlme S, et al. 2020. Fungal Diversity 105:1-16 ((https://doi.org/10.1007/s13225-020-00466-2)) |
| Otu56711 | 1 | k__Fungi(100);p__Ascomycota(100);c__Saccharomycetes(100);o__Saccharomycetales(100);f__Metschnikowiaceae(100);g__Metschnikowia(100);s__Metschnikowia_pulcherrima(100); | Metschnikowia | 13 | Saprotroph | \|Undefined Saprotroph\| | Yeast | NULL | Probable | As nectar-tap_saprotroph (Põlme et al. 2020); Decay sugar-rich substrates (Põlme et al. 2020) | Põlme S, et al. 2020. Fungal Diversity 105:1-16 ((https://doi.org/10.1007/s13225-020-00466-2)) |
| Otu56712 | 1 | k__Fungi(100);p__Ascomycota(100);c__Saccharomycetes(100);o__Saccharomycetales(100);f__Metschnikowiaceae(100);g__Metschnikowia(100);s__Metschnikowia_pulcherrima(100); | Metschnikowia | 13 | Saprotroph | \|Undefined Saprotroph\| | Yeast | NULL | Probable | As nectar-tap_saprotroph (Põlme et al. 2020); Decay sugar-rich substrates (Põlme et al. 2020) | Põlme S, et al. 2020. Fungal Diversity 105:1-16 ((https://doi.org/10.1007/s13225-020-00466-2)) |
| Otu56713 | 1 | k__Fungi(100);p__Ascomycota(100);c__Saccharomycetes(100);o__Saccharomycetales(100);f__Metschnikowiaceae(100);g__Metschnikowia(100);s__Metschnikowia_pulcherrima(100); | Metschnikowia | 13 | Saprotroph | \|Undefined Saprotroph\| | Yeast | NULL | Probable | As nectar-tap_saprotroph (Põlme et al. 2020); Decay sugar-rich substrates (Põlme et al. 2020) | Põlme S, et al. 2020. Fungal Diversity 105:1-16 ((https://doi.org/10.1007/s13225-020-00466-2)) |
| Otu56714 | 1 | k__Fungi(100);p__Ascomycota(100);c__Saccharomycetes(100);o__Saccharomycetales(100);f__Metschnikowiaceae(100);g__Metschnikowia(100);s__Metschnikowia_pulcherrima(100); | Metschnikowia | 13 | Saprotroph | \|Undefined Saprotroph\| | Yeast | NULL | Probable | As nectar-tap_saprotroph (Põlme et al. 2020); Decay sugar-rich substrates (Põlme et al. 2020) | Põlme S, et al. 2020. Fungal Diversity 105:1-16 ((https://doi.org/10.1007/s13225-020-00466-2)) |
| Otu56715 | 1 | k__Fungi(100);p__Ascomycota(100);c__Saccharomycetes(100);o__Saccharomycetales(100);f__Metschnikowiaceae(100);g__Metschnikowia(100);g__Metschnikowia_unclassified(100); | Metschnikowia | 13 | Saprotroph | \|Undefined Saprotroph\| | Yeast | NULL | Probable | As nectar-tap_saprotroph (Põlme et al. 2020); Decay sugar-rich substrates (Põlme et al. 2020) | Põlme S, et al. 2020. Fungal Diversity 105:1-16 ((https://doi.org/10.1007/s13225-020-00466-2)) |
| Otu56716 | 1 | k__Fungi(100);p__Ascomycota(100);c__Saccharomycetes(100);o__Saccharomycetales(100);f__Metschnikowiaceae(100);g__Metschnikowia(100);g__Metschnikowia_unclassified(100); | Metschnikowia | 13 | Saprotroph | \|Undefined Saprotroph\| | Yeast | NULL | Probable | As nectar-tap_saprotroph (Põlme et al. 2020); Decay sugar-rich substrates (Põlme et al. 2020) | Põlme S, et al. 2020. Fungal Diversity 105:1-16 ((https://doi.org/10.1007/s13225-020-00466-2)) |
| Otu56717 | 1 | k__Fungi(100);p__Ascomycota(100);c__Saccharomycetes(100);o__Saccharomycetales(100);f__Metschnikowiaceae(100);g__Metschnikowia(100);s__Metschnikowia_pulcherrima(100); | Metschnikowia | 13 | Saprotroph | \|Undefined Saprotroph\| | Yeast | NULL | Probable | As nectar-tap_saprotroph (Põlme et al. 2020); Decay sugar-rich substrates (Põlme et al. 2020) | Põlme S, et al. 2020. Fungal Diversity 105:1-16 ((https://doi.org/10.1007/s13225-020-00466-2)) |
| Otu56718 | 1 | k__Fungi(100);p__Ascomycota(100);c__Saccharomycetes(100);o__Saccharomycetales(100);f__Metschnikowiaceae(100);g__Metschnikowia(100);s__Metschnikowia_pulcherrima(100); | Metschnikowia | 13 | Saprotroph | \|Undefined Saprotroph\| | Yeast | NULL | Probable | As nectar-tap_saprotroph (Põlme et al. 2020); Decay sugar-rich substrates (Põlme et al. 2020) | Põlme S, et al. 2020. Fungal Diversity 105:1-16 ((https://doi.org/10.1007/s13225-020-00466-2)) |
| Otu56719 | 1 | k__Fungi(100);p__Ascomycota(100);c__Saccharomycetes(100);o__Saccharomycetales(100);f__Metschnikowiaceae(100);g__Metschnikowia(100);s__Metschnikowia_pulcherrima(100); | Metschnikowia | 13 | Saprotroph | \|Undefined Saprotroph\| | Yeast | NULL | Probable | As nectar-tap_saprotroph (Põlme et al. 2020); Decay sugar-rich substrates (Põlme et al. 2020) | Põlme S, et al. 2020. Fungal Diversity 105:1-16 ((https://doi.org/10.1007/s13225-020-00466-2)) |
| Otu56720 | 1 | k__Fungi(100);p__Ascomycota(100);c__Saccharomycetes(100);o__Saccharomycetales(100);f__Metschnikowiaceae(100);g__Metschnikowia(100);g__Metschnikowia_unclassified(100); | Metschnikowia | 13 | Saprotroph | \|Undefined Saprotroph\| | Yeast | NULL | Probable | As nectar-tap_saprotroph (Põlme et al. 2020); Decay sugar-rich substrates (Põlme et al. 2020) | Põlme S, et al. 2020. Fungal Diversity 105:1-16 ((https://doi.org/10.1007/s13225-020-00466-2)) |
| Otu56721 | 1 | k__Fungi(100);p__Ascomycota(100);c__Saccharomycetes(100);o__Saccharomycetales(100);f__Metschnikowiaceae(100);g__Metschnikowia(100);s__Metschnikowia_pulcherrima(100); | Metschnikowia | 13 | Saprotroph | \|Undefined Saprotroph\| | Yeast | NULL | Probable | As nectar-tap_saprotroph (Põlme et al. 2020); Decay sugar-rich substrates (Põlme et al. 2020) | Põlme S, et al. 2020. Fungal Diversity 105:1-16 ((https://doi.org/10.1007/s13225-020-00466-2)) |
| Otu56723 | 1 | k__Fungi(100);p__Ascomycota(100);c__Saccharomycetes(100);o__Saccharomycetales(100);f__Metschnikowiaceae(100);g__Metschnikowia(100);g__Metschnikowia_unclassified(100); | Metschnikowia | 13 | Saprotroph | \|Undefined Saprotroph\| | Yeast | NULL | Probable | As nectar-tap_saprotroph (Põlme et al. 2020); Decay sugar-rich substrates (Põlme et al. 2020) | Põlme S, et al. 2020. Fungal Diversity 105:1-16 ((https://doi.org/10.1007/s13225-020-00466-2)) |
| Otu56724 | 1 | k__Fungi(100);p__Ascomycota(100);c__Saccharomycetes(100);o__Saccharomycetales(100);f__Metschnikowiaceae(100);g__Metschnikowia(100);s__Metschnikowia_pulcherrima(100); | Metschnikowia | 13 | Saprotroph | \|Undefined Saprotroph\| | Yeast | NULL | Probable | As nectar-tap_saprotroph (Põlme et al. 2020); Decay sugar-rich substrates (Põlme et al. 2020) | Põlme S, et al. 2020. Fungal Diversity 105:1-16 ((https://doi.org/10.1007/s13225-020-00466-2)) |
| Otu56725 | 1 | k__Fungi(100);p__Ascomycota(100);c__Saccharomycetes(100);o__Saccharomycetales(100);f__Metschnikowiaceae(100);g__Metschnikowia(100);s__Metschnikowia_pulcherrima(100); | Metschnikowia | 13 | Saprotroph | \|Undefined Saprotroph\| | Yeast | NULL | Probable | As nectar-tap_saprotroph (Põlme et al. 2020); Decay sugar-rich substrates (Põlme et al. 2020) | Põlme S, et al. 2020. Fungal Diversity 105:1-16 ((https://doi.org/10.1007/s13225-020-00466-2)) |
| Otu56726 | 1 | k__Fungi(100);p__Ascomycota(100);c__Saccharomycetes(100);o__Saccharomycetales(100);f__Metschnikowiaceae(100);g__Metschnikowia(100);g__Metschnikowia_unclassified(100); | Metschnikowia | 13 | Saprotroph | \|Undefined Saprotroph\| | Yeast | NULL | Probable | As nectar-tap_saprotroph (Põlme et al. 2020); Decay sugar-rich substrates (Põlme et al. 2020) | Põlme S, et al. 2020. Fungal Diversity 105:1-16 ((https://doi.org/10.1007/s13225-020-00466-2)) |
| Otu56727 | 1 | k__Fungi(100);p__Ascomycota(100);c__Saccharomycetes(100);o__Saccharomycetales(100);f__Metschnikowiaceae(100);g__Metschnikowia(100);s__Metschnikowia_pulcherrima(100); | Metschnikowia | 13 | Saprotroph | \|Undefined Saprotroph\| | Yeast | NULL | Probable | As nectar-tap_saprotroph (Põlme et al. 2020); Decay sugar-rich substrates (Põlme et al. 2020) | Põlme S, et al. 2020. Fungal Diversity 105:1-16 ((https://doi.org/10.1007/s13225-020-00466-2)) |
| Otu56728 | 1 | k__Fungi(100);p__Ascomycota(100);c__Saccharomycetes(100);o__Saccharomycetales(100);f__Metschnikowiaceae(100);g__Metschnikowia(100);s__Metschnikowia_pulcherrima(100); | Metschnikowia | 13 | Saprotroph | \|Undefined Saprotroph\| | Yeast | NULL | Probable | As nectar-tap_saprotroph (Põlme et al. 2020); Decay sugar-rich substrates (Põlme et al. 2020) | Põlme S, et al. 2020. Fungal Diversity 105:1-16 ((https://doi.org/10.1007/s13225-020-00466-2)) |
| Otu56729 | 1 | k__Fungi(100);p__Ascomycota(100);c__Saccharomycetes(100);o__Saccharomycetales(100);f__Metschnikowiaceae(100);g__Metschnikowia(100);s__Metschnikowia_pulcherrima(100); | Metschnikowia | 13 | Saprotroph | \|Undefined Saprotroph\| | Yeast | NULL | Probable | As nectar-tap_saprotroph (Põlme et al. 2020); Decay sugar-rich substrates (Põlme et al. 2020) | Põlme S, et al. 2020. Fungal Diversity 105:1-16 ((https://doi.org/10.1007/s13225-020-00466-2)) |
| Otu56730 | 1 | k__Fungi(100);p__Ascomycota(100);c__Saccharomycetes(100);o__Saccharomycetales(100);f__Metschnikowiaceae(100);g__Metschnikowia(100);s__Metschnikowia_pulcherrima(100); | Metschnikowia | 13 | Saprotroph | \|Undefined Saprotroph\| | Yeast | NULL | Probable | As nectar-tap_saprotroph (Põlme et al. 2020); Decay sugar-rich substrates (Põlme et al. 2020) | Põlme S, et al. 2020. Fungal Diversity 105:1-16 ((https://doi.org/10.1007/s13225-020-00466-2)) |
| Otu56731 | 1 | k__Fungi(100);p__Ascomycota(100);c__Saccharomycetes(100);o__Saccharomycetales(100);f__Metschnikowiaceae(100);g__Metschnikowia(100);g__Metschnikowia_unclassified(100); | Metschnikowia | 13 | Saprotroph | \|Undefined Saprotroph\| | Yeast | NULL | Probable | As nectar-tap_saprotroph (Põlme et al. 2020); Decay sugar-rich substrates (Põlme et al. 2020) | Põlme S, et al. 2020. Fungal Diversity 105:1-16 ((https://doi.org/10.1007/s13225-020-00466-2)) |
| Otu56732 | 1 | k__Fungi(100);p__Ascomycota(100);c__Saccharomycetes(100);o__Saccharomycetales(100);f__Metschnikowiaceae(100);g__Metschnikowia(100);g__Metschnikowia_unclassified(100); | Metschnikowia | 13 | Saprotroph | \|Undefined Saprotroph\| | Yeast | NULL | Probable | As nectar-tap_saprotroph (Põlme et al. 2020); Decay sugar-rich substrates (Põlme et al. 2020) | Põlme S, et al. 2020. Fungal Diversity 105:1-16 ((https://doi.org/10.1007/s13225-020-00466-2)) |
| Otu56733 | 1 | k__Fungi(100);p__Ascomycota(100);c__Saccharomycetes(100);o__Saccharomycetales(100);f__Metschnikowiaceae(100);g__Metschnikowia(100);s__Metschnikowia_pulcherrima(100); | Metschnikowia | 13 | Saprotroph | \|Undefined Saprotroph\| | Yeast | NULL | Probable | As nectar-tap_saprotroph (Põlme et al. 2020); Decay sugar-rich substrates (Põlme et al. 2020) | Põlme S, et al. 2020. Fungal Diversity 105:1-16 ((https://doi.org/10.1007/s13225-020-00466-2)) |
| Otu56734 | 1 | k__Fungi(100);p__Ascomycota(100);c__Saccharomycetes(100);o__Saccharomycetales(100);f__Metschnikowiaceae(100);g__Metschnikowia(100);g__Metschnikowia_unclassified(100); | Metschnikowia | 13 | Saprotroph | \|Undefined Saprotroph\| | Yeast | NULL | Probable | As nectar-tap_saprotroph (Põlme et al. 2020); Decay sugar-rich substrates (Põlme et al. 2020) | Põlme S, et al. 2020. Fungal Diversity 105:1-16 ((https://doi.org/10.1007/s13225-020-00466-2)) |
| Otu56735 | 1 | k__Fungi(100);p__Ascomycota(100);c__Saccharomycetes(100);o__Saccharomycetales(100);f__Metschnikowiaceae(100);g__Metschnikowia(100);s__Metschnikowia_pulcherrima(100); | Metschnikowia | 13 | Saprotroph | \|Undefined Saprotroph\| | Yeast | NULL | Probable | As nectar-tap_saprotroph (Põlme et al. 2020); Decay sugar-rich substrates (Põlme et al. 2020) | Põlme S, et al. 2020. Fungal Diversity 105:1-16 ((https://doi.org/10.1007/s13225-020-00466-2)) |
| Otu56736 | 1 | k__Fungi(100);p__Ascomycota(100);c__Saccharomycetes(100);o__Saccharomycetales(100);f__Metschnikowiaceae(100);g__Metschnikowia(100);g__Metschnikowia_unclassified(100); | Metschnikowia | 13 | Saprotroph | \|Undefined Saprotroph\| | Yeast | NULL | Probable | As nectar-tap_saprotroph (Põlme et al. 2020); Decay sugar-rich substrates (Põlme et al. 2020) | Põlme S, et al. 2020. Fungal Diversity 105:1-16 ((https://doi.org/10.1007/s13225-020-00466-2)) |
| Otu56737 | 1 | k__Fungi(100);p__Ascomycota(100);c__Dothideomycetes(100);o__Dothideomycetes_order_Incertae_sedis(100);f__Myxotrichaceae(100);g__Oidiodendron(100);g__Oidiodendron_unclassified(100); | Oidiodendron | 13 | Pathotroph-Saprotroph-Symbiotroph | Endophyte-Ericoid Mycorrhizal-\|Undefined Saprotroph\| | Microfungus | NULL | Probable | As root_endophyte (Põlme et al. 2020); As soil_saprotroph (Põlme et al. 2020); Dark Septate Endophyte; Decay soil (Põlme et al. 2020) | Newsham KK. 2011. New Phytologist 190:783-793 ((https://doi.org/10.1111/j.1469-8137.2010.03611.x)) (as dark septate root endophytes); Põlme S, et al. 2020. Fungal Diversity 105:1-16 ((https://doi.org/10.1007/s13225-020-00466-2)) |
| Otu56738 | 1 | k__Fungi(100);p__Ascomycota(100);c__Saccharomycetes(100);o__Saccharomycetales(100);f__Metschnikowiaceae(100);g__Metschnikowia(100);s__Metschnikowia_pulcherrima(100); | Metschnikowia | 13 | Saprotroph | \|Undefined Saprotroph\| | Yeast | NULL | Probable | As nectar-tap_saprotroph (Põlme et al. 2020); Decay sugar-rich substrates (Põlme et al. 2020) | Põlme S, et al. 2020. Fungal Diversity 105:1-16 ((https://doi.org/10.1007/s13225-020-00466-2)) |
| Otu56739 | 1 | k__Fungi(100);p__Ascomycota(100);c__Saccharomycetes(100);o__Saccharomycetales(100);f__Metschnikowiaceae(100);g__Metschnikowia(100);s__Metschnikowia_pulcherrima(100); | Metschnikowia | 13 | Saprotroph | \|Undefined Saprotroph\| | Yeast | NULL | Probable | As nectar-tap_saprotroph (Põlme et al. 2020); Decay sugar-rich substrates (Põlme et al. 2020) | Põlme S, et al. 2020. Fungal Diversity 105:1-16 ((https://doi.org/10.1007/s13225-020-00466-2)) |
| Otu56748 | 1 | k__Fungi(100);p__Ascomycota(100);c__Saccharomycetes(100);o__Saccharomycetales(100);f__Metschnikowiaceae(100);g__Metschnikowia(100);s__Metschnikowia_pulcherrima(100); | Metschnikowia | 13 | Saprotroph | \|Undefined Saprotroph\| | Yeast | NULL | Probable | As nectar-tap_saprotroph (Põlme et al. 2020); Decay sugar-rich substrates (Põlme et al. 2020) | Põlme S, et al. 2020. Fungal Diversity 105:1-16 ((https://doi.org/10.1007/s13225-020-00466-2)) |
| Otu56751 | 1 | k__Fungi(100);p__Ascomycota(100);c__Saccharomycetes(100);o__Saccharomycetales(100);f__Metschnikowiaceae(100);g__Metschnikowia(100);s__Metschnikowia_pulcherrima(100); | Metschnikowia | 13 | Saprotroph | \|Undefined Saprotroph\| | Yeast | NULL | Probable | As nectar-tap_saprotroph (Põlme et al. 2020); Decay sugar-rich substrates (Põlme et al. 2020) | Põlme S, et al. 2020. Fungal Diversity 105:1-16 ((https://doi.org/10.1007/s13225-020-00466-2)) |
| Otu56754 | 1 | k__Fungi(100);p__Ascomycota(100);c__Saccharomycetes(100);o__Saccharomycetales(100);f__Metschnikowiaceae(100);g__Metschnikowia(100);s__Metschnikowia_pulcherrima(100); | Metschnikowia | 13 | Saprotroph | \|Undefined Saprotroph\| | Yeast | NULL | Probable | As nectar-tap_saprotroph (Põlme et al. 2020); Decay sugar-rich substrates (Põlme et al. 2020) | Põlme S, et al. 2020. Fungal Diversity 105:1-16 ((https://doi.org/10.1007/s13225-020-00466-2)) |
| Otu56755 | 1 | k__Fungi(100);p__Ascomycota(100);c__Saccharomycetes(100);o__Saccharomycetales(100);f__Metschnikowiaceae(100);g__Metschnikowia(100);s__Metschnikowia_pulcherrima(100); | Metschnikowia | 13 | Saprotroph | \|Undefined Saprotroph\| | Yeast | NULL | Probable | As nectar-tap_saprotroph (Põlme et al. 2020); Decay sugar-rich substrates (Põlme et al. 2020) | Põlme S, et al. 2020. Fungal Diversity 105:1-16 ((https://doi.org/10.1007/s13225-020-00466-2)) |
| Otu56758 | 1 | k__Fungi(100);p__Ascomycota(100);c__Saccharomycetes(100);o__Saccharomycetales(100);f__Metschnikowiaceae(100);g__Metschnikowia(100);s__Metschnikowia_pulcherrima(100); | Metschnikowia | 13 | Saprotroph | \|Undefined Saprotroph\| | Yeast | NULL | Probable | As nectar-tap_saprotroph (Põlme et al. 2020); Decay sugar-rich substrates (Põlme et al. 2020) | Põlme S, et al. 2020. Fungal Diversity 105:1-16 ((https://doi.org/10.1007/s13225-020-00466-2)) |
| Otu56760 | 1 | k__Fungi(100);p__Ascomycota(100);c__Saccharomycetes(100);o__Saccharomycetales(100);f__Metschnikowiaceae(100);g__Metschnikowia(100);s__Metschnikowia_pulcherrima(100); | Metschnikowia | 13 | Saprotroph | \|Undefined Saprotroph\| | Yeast | NULL | Probable | As nectar-tap_saprotroph (Põlme et al. 2020); Decay sugar-rich substrates (Põlme et al. 2020) | Põlme S, et al. 2020. Fungal Diversity 105:1-16 ((https://doi.org/10.1007/s13225-020-00466-2)) |
| Otu56762 | 1 | k__Fungi(100);p__Ascomycota(100);c__Saccharomycetes(100);o__Saccharomycetales(100);f__Metschnikowiaceae(100);g__Metschnikowia(100);g__Metschnikowia_unclassified(100); | Metschnikowia | 13 | Saprotroph | \|Undefined Saprotroph\| | Yeast | NULL | Probable | As nectar-tap_saprotroph (Põlme et al. 2020); Decay sugar-rich substrates (Põlme et al. 2020) | Põlme S, et al. 2020. Fungal Diversity 105:1-16 ((https://doi.org/10.1007/s13225-020-00466-2)) |
| Otu56763 | 1 | k__Fungi(100);p__Ascomycota(100);c__Saccharomycetes(100);o__Saccharomycetales(100);f__Metschnikowiaceae(100);g__Metschnikowia(100);s__Metschnikowia_pulcherrima(100); | Metschnikowia | 13 | Saprotroph | \|Undefined Saprotroph\| | Yeast | NULL | Probable | As nectar-tap_saprotroph (Põlme et al. 2020); Decay sugar-rich substrates (Põlme et al. 2020) | Põlme S, et al. 2020. Fungal Diversity 105:1-16 ((https://doi.org/10.1007/s13225-020-00466-2)) |
| Otu56765 | 1 | k__Fungi(100);p__Ascomycota(100);c__Saccharomycetes(100);o__Saccharomycetales(100);f__Metschnikowiaceae(100);g__Metschnikowia(100);s__Metschnikowia_pulcherrima(100); | Metschnikowia | 13 | Saprotroph | \|Undefined Saprotroph\| | Yeast | NULL | Probable | As nectar-tap_saprotroph (Põlme et al. 2020); Decay sugar-rich substrates (Põlme et al. 2020) | Põlme S, et al. 2020. Fungal Diversity 105:1-16 ((https://doi.org/10.1007/s13225-020-00466-2)) |
| Otu56766 | 1 | k__Fungi(100);p__Ascomycota(100);c__Saccharomycetes(100);o__Saccharomycetales(100);f__Metschnikowiaceae(100);g__Metschnikowia(100);s__Metschnikowia_pulcherrima(100); | Metschnikowia | 13 | Saprotroph | \|Undefined Saprotroph\| | Yeast | NULL | Probable | As nectar-tap_saprotroph (Põlme et al. 2020); Decay sugar-rich substrates (Põlme et al. 2020) | Põlme S, et al. 2020. Fungal Diversity 105:1-16 ((https://doi.org/10.1007/s13225-020-00466-2)) |
| Otu56769 | 1 | k__Fungi(100);p__Ascomycota(100);c__Saccharomycetes(100);o__Saccharomycetales(100);f__Metschnikowiaceae(100);g__Metschnikowia(100);s__Metschnikowia_pulcherrima(100); | Metschnikowia | 13 | Saprotroph | \|Undefined Saprotroph\| | Yeast | NULL | Probable | As nectar-tap_saprotroph (Põlme et al. 2020); Decay sugar-rich substrates (Põlme et al. 2020) | Põlme S, et al. 2020. Fungal Diversity 105:1-16 ((https://doi.org/10.1007/s13225-020-00466-2)) |
| Otu56770 | 1 | k__Fungi(100);p__Ascomycota(100);c__Saccharomycetes(100);o__Saccharomycetales(100);f__Metschnikowiaceae(100);g__Metschnikowia(100);g__Metschnikowia_unclassified(100); | Metschnikowia | 13 | Saprotroph | \|Undefined Saprotroph\| | Yeast | NULL | Probable | As nectar-tap_saprotroph (Põlme et al. 2020); Decay sugar-rich substrates (Põlme et al. 2020) | Põlme S, et al. 2020. Fungal Diversity 105:1-16 ((https://doi.org/10.1007/s13225-020-00466-2)) |
| Otu56771 | 1 | k__Fungi(100);p__Ascomycota(100);c__Saccharomycetes(100);o__Saccharomycetales(100);f__Metschnikowiaceae(100);g__Metschnikowia(100);g__Metschnikowia_unclassified(100); | Metschnikowia | 13 | Saprotroph | \|Undefined Saprotroph\| | Yeast | NULL | Probable | As nectar-tap_saprotroph (Põlme et al. 2020); Decay sugar-rich substrates (Põlme et al. 2020) | Põlme S, et al. 2020. Fungal Diversity 105:1-16 ((https://doi.org/10.1007/s13225-020-00466-2)) |
| Otu56772 | 1 | k__Fungi(100);p__Ascomycota(100);c__Saccharomycetes(100);o__Saccharomycetales(100);f__Metschnikowiaceae(100);g__Metschnikowia(100);s__Metschnikowia_pulcherrima(100); | Metschnikowia | 13 | Saprotroph | \|Undefined Saprotroph\| | Yeast | NULL | Probable | As nectar-tap_saprotroph (Põlme et al. 2020); Decay sugar-rich substrates (Põlme et al. 2020) | Põlme S, et al. 2020. Fungal Diversity 105:1-16 ((https://doi.org/10.1007/s13225-020-00466-2)) |
| Otu56773 | 1 | k__Fungi(100);p__Ascomycota(100);c__Saccharomycetes(100);o__Saccharomycetales(100);f__Metschnikowiaceae(100);g__Metschnikowia(100);s__Metschnikowia_pulcherrima(100); | Metschnikowia | 13 | Saprotroph | \|Undefined Saprotroph\| | Yeast | NULL | Probable | As nectar-tap_saprotroph (Põlme et al. 2020); Decay sugar-rich substrates (Põlme et al. 2020) | Põlme S, et al. 2020. Fungal Diversity 105:1-16 ((https://doi.org/10.1007/s13225-020-00466-2)) |
| Otu56774 | 1 | k__Fungi(100);p__Ascomycota(100);c__Saccharomycetes(100);o__Saccharomycetales(100);f__Metschnikowiaceae(100);g__Metschnikowia(100);s__Metschnikowia_pulcherrima(100); | Metschnikowia | 13 | Saprotroph | \|Undefined Saprotroph\| | Yeast | NULL | Probable | As nectar-tap_saprotroph (Põlme et al. 2020); Decay sugar-rich substrates (Põlme et al. 2020) | Põlme S, et al. 2020. Fungal Diversity 105:1-16 ((https://doi.org/10.1007/s13225-020-00466-2)) |
| Otu56775 | 1 | k__Fungi(100);p__Ascomycota(100);c__Saccharomycetes(100);o__Saccharomycetales(100);f__Metschnikowiaceae(100);g__Metschnikowia(100);s__Metschnikowia_pulcherrima(100); | Metschnikowia | 13 | Saprotroph | \|Undefined Saprotroph\| | Yeast | NULL | Probable | As nectar-tap_saprotroph (Põlme et al. 2020); Decay sugar-rich substrates (Põlme et al. 2020) | Põlme S, et al. 2020. Fungal Diversity 105:1-16 ((https://doi.org/10.1007/s13225-020-00466-2)) |
| Otu56776 | 1 | k__Fungi(100);p__Ascomycota(100);c__Saccharomycetes(100);o__Saccharomycetales(100);f__Metschnikowiaceae(100);g__Metschnikowia(100);s__Metschnikowia_pulcherrima(100); | Metschnikowia | 13 | Saprotroph | \|Undefined Saprotroph\| | Yeast | NULL | Probable | As nectar-tap_saprotroph (Põlme et al. 2020); Decay sugar-rich substrates (Põlme et al. 2020) | Põlme S, et al. 2020. Fungal Diversity 105:1-16 ((https://doi.org/10.1007/s13225-020-00466-2)) |
| Otu56778 | 1 | k__Fungi(100);p__Ascomycota(100);c__Saccharomycetes(100);o__Saccharomycetales(100);f__Metschnikowiaceae(100);g__Metschnikowia(100);s__Metschnikowia_pulcherrima(100); | Metschnikowia | 13 | Saprotroph | \|Undefined Saprotroph\| | Yeast | NULL | Probable | As nectar-tap_saprotroph (Põlme et al. 2020); Decay sugar-rich substrates (Põlme et al. 2020) | Põlme S, et al. 2020. Fungal Diversity 105:1-16 ((https://doi.org/10.1007/s13225-020-00466-2)) |
| Otu56779 | 1 | k__Fungi(100);p__Ascomycota(100);c__Saccharomycetes(100);o__Saccharomycetales(100);f__Metschnikowiaceae(100);g__Metschnikowia(100);g__Metschnikowia_unclassified(100); | Metschnikowia | 13 | Saprotroph | \|Undefined Saprotroph\| | Yeast | NULL | Probable | As nectar-tap_saprotroph (Põlme et al. 2020); Decay sugar-rich substrates (Põlme et al. 2020) | Põlme S, et al. 2020. Fungal Diversity 105:1-16 ((https://doi.org/10.1007/s13225-020-00466-2)) |
| Otu56781 | 1 | k__Fungi(100);p__Ascomycota(100);c__Saccharomycetes(100);o__Saccharomycetales(100);f__Metschnikowiaceae(100);g__Metschnikowia(100);s__Metschnikowia_pulcherrima(100); | Metschnikowia | 13 | Saprotroph | \|Undefined Saprotroph\| | Yeast | NULL | Probable | As nectar-tap_saprotroph (Põlme et al. 2020); Decay sugar-rich substrates (Põlme et al. 2020) | Põlme S, et al. 2020. Fungal Diversity 105:1-16 ((https://doi.org/10.1007/s13225-020-00466-2)) |
| Otu56782 | 1 | k__Fungi(100);p__Ascomycota(100);c__Saccharomycetes(100);o__Saccharomycetales(100);f__Metschnikowiaceae(100);g__Metschnikowia(100);s__Metschnikowia_pulcherrima(100); | Metschnikowia | 13 | Saprotroph | \|Undefined Saprotroph\| | Yeast | NULL | Probable | As nectar-tap_saprotroph (Põlme et al. 2020); Decay sugar-rich substrates (Põlme et al. 2020) | Põlme S, et al. 2020. Fungal Diversity 105:1-16 ((https://doi.org/10.1007/s13225-020-00466-2)) |
| Otu56783 | 1 | k__Fungi(100);p__Ascomycota(100);c__Saccharomycetes(100);o__Saccharomycetales(100);f__Metschnikowiaceae(100);g__Metschnikowia(100);g__Metschnikowia_unclassified(100); | Metschnikowia | 13 | Saprotroph | \|Undefined Saprotroph\| | Yeast | NULL | Probable | As nectar-tap_saprotroph (Põlme et al. 2020); Decay sugar-rich substrates (Põlme et al. 2020) | Põlme S, et al. 2020. Fungal Diversity 105:1-16 ((https://doi.org/10.1007/s13225-020-00466-2)) |
| Otu56792 | 1 | k__Fungi(100);p__Ascomycota(100);c__Saccharomycetes(100);o__Saccharomycetales(100);f__Metschnikowiaceae(100);g__Metschnikowia(100);g__Metschnikowia_unclassified(100); | Metschnikowia | 13 | Saprotroph | \|Undefined Saprotroph\| | Yeast | NULL | Probable | As nectar-tap_saprotroph (Põlme et al. 2020); Decay sugar-rich substrates (Põlme et al. 2020) | Põlme S, et al. 2020. Fungal Diversity 105:1-16 ((https://doi.org/10.1007/s13225-020-00466-2)) |
| Otu56794 | 1 | k__Fungi(100);p__Ascomycota(100);c__Saccharomycetes(100);o__Saccharomycetales(100);f__Metschnikowiaceae(100);g__Metschnikowia(100);g__Metschnikowia_unclassified(100); | Metschnikowia | 13 | Saprotroph | \|Undefined Saprotroph\| | Yeast | NULL | Probable | As nectar-tap_saprotroph (Põlme et al. 2020); Decay sugar-rich substrates (Põlme et al. 2020) | Põlme S, et al. 2020. Fungal Diversity 105:1-16 ((https://doi.org/10.1007/s13225-020-00466-2)) |
| Otu56795 | 1 | k__Fungi(100);p__Ascomycota(100);c__Saccharomycetes(100);o__Saccharomycetales(100);f__Metschnikowiaceae(100);g__Metschnikowia(100);g__Metschnikowia_unclassified(100); | Metschnikowia | 13 | Saprotroph | \|Undefined Saprotroph\| | Yeast | NULL | Probable | As nectar-tap_saprotroph (Põlme et al. 2020); Decay sugar-rich substrates (Põlme et al. 2020) | Põlme S, et al. 2020. Fungal Diversity 105:1-16 ((https://doi.org/10.1007/s13225-020-00466-2)) |
| Otu56796 | 1 | k__Fungi(100);p__Ascomycota(100);c__Saccharomycetes(100);o__Saccharomycetales(100);f__Metschnikowiaceae(100);g__Metschnikowia(100);s__Metschnikowia_pulcherrima(100); | Metschnikowia | 13 | Saprotroph | \|Undefined Saprotroph\| | Yeast | NULL | Probable | As nectar-tap_saprotroph (Põlme et al. 2020); Decay sugar-rich substrates (Põlme et al. 2020) | Põlme S, et al. 2020. Fungal Diversity 105:1-16 ((https://doi.org/10.1007/s13225-020-00466-2)) |
| Otu56799 | 1 | k__Fungi(100);p__Ascomycota(100);c__Saccharomycetes(100);o__Saccharomycetales(100);f__Metschnikowiaceae(100);g__Metschnikowia(100);g__Metschnikowia_unclassified(100); | Metschnikowia | 13 | Saprotroph | \|Undefined Saprotroph\| | Yeast | NULL | Probable | As nectar-tap_saprotroph (Põlme et al. 2020); Decay sugar-rich substrates (Põlme et al. 2020) | Põlme S, et al. 2020. Fungal Diversity 105:1-16 ((https://doi.org/10.1007/s13225-020-00466-2)) |
| Otu56800 | 1 | k__Fungi(100);p__Ascomycota(100);c__Saccharomycetes(100);o__Saccharomycetales(100);f__Metschnikowiaceae(100);g__Metschnikowia(100);s__Metschnikowia_pulcherrima(100); | Metschnikowia | 13 | Saprotroph | \|Undefined Saprotroph\| | Yeast | NULL | Probable | As nectar-tap_saprotroph (Põlme et al. 2020); Decay sugar-rich substrates (Põlme et al. 2020) | Põlme S, et al. 2020. Fungal Diversity 105:1-16 ((https://doi.org/10.1007/s13225-020-00466-2)) |
| Otu56801 | 1 | k__Fungi(100);p__Ascomycota(100);c__Saccharomycetes(100);o__Saccharomycetales(100);f__Metschnikowiaceae(100);g__Metschnikowia(100);s__Metschnikowia_pulcherrima(100); | Metschnikowia | 13 | Saprotroph | \|Undefined Saprotroph\| | Yeast | NULL | Probable | As nectar-tap_saprotroph (Põlme et al. 2020); Decay sugar-rich substrates (Põlme et al. 2020) | Põlme S, et al. 2020. Fungal Diversity 105:1-16 ((https://doi.org/10.1007/s13225-020-00466-2)) |
| Otu56802 | 1 | k__Fungi(100);p__Ascomycota(100);c__Saccharomycetes(100);o__Saccharomycetales(100);f__Metschnikowiaceae(100);g__Metschnikowia(100);s__Metschnikowia_pulcherrima(100); | Metschnikowia | 13 | Saprotroph | \|Undefined Saprotroph\| | Yeast | NULL | Probable | As nectar-tap_saprotroph (Põlme et al. 2020); Decay sugar-rich substrates (Põlme et al. 2020) | Põlme S, et al. 2020. Fungal Diversity 105:1-16 ((https://doi.org/10.1007/s13225-020-00466-2)) |
| Otu56803 | 1 | k__Fungi(100);p__Ascomycota(100);c__Saccharomycetes(100);o__Saccharomycetales(100);f__Metschnikowiaceae(100);g__Metschnikowia(100);s__Metschnikowia_pulcherrima(100); | Metschnikowia | 13 | Saprotroph | \|Undefined Saprotroph\| | Yeast | NULL | Probable | As nectar-tap_saprotroph (Põlme et al. 2020); Decay sugar-rich substrates (Põlme et al. 2020) | Põlme S, et al. 2020. Fungal Diversity 105:1-16 ((https://doi.org/10.1007/s13225-020-00466-2)) |
| Otu56804 | 1 | k__Fungi(100);p__Ascomycota(100);c__Saccharomycetes(100);o__Saccharomycetales(100);f__Metschnikowiaceae(100);g__Metschnikowia(100);g__Metschnikowia_unclassified(100); | Metschnikowia | 13 | Saprotroph | \|Undefined Saprotroph\| | Yeast | NULL | Probable | As nectar-tap_saprotroph (Põlme et al. 2020); Decay sugar-rich substrates (Põlme et al. 2020) | Põlme S, et al. 2020. Fungal Diversity 105:1-16 ((https://doi.org/10.1007/s13225-020-00466-2)) |
| Otu56805 | 1 | k__Fungi(100);p__Ascomycota(100);c__Saccharomycetes(100);o__Saccharomycetales(100);f__Metschnikowiaceae(100);g__Metschnikowia(100);s__Metschnikowia_pulcherrima(100); | Metschnikowia | 13 | Saprotroph | \|Undefined Saprotroph\| | Yeast | NULL | Probable | As nectar-tap_saprotroph (Põlme et al. 2020); Decay sugar-rich substrates (Põlme et al. 2020) | Põlme S, et al. 2020. Fungal Diversity 105:1-16 ((https://doi.org/10.1007/s13225-020-00466-2)) |
| Otu56806 | 1 | k__Fungi(100);p__Ascomycota(100);c__Saccharomycetes(100);o__Saccharomycetales(100);f__Metschnikowiaceae(100);g__Metschnikowia(100);s__Metschnikowia_pulcherrima(100); | Metschnikowia | 13 | Saprotroph | \|Undefined Saprotroph\| | Yeast | NULL | Probable | As nectar-tap_saprotroph (Põlme et al. 2020); Decay sugar-rich substrates (Põlme et al. 2020) | Põlme S, et al. 2020. Fungal Diversity 105:1-16 ((https://doi.org/10.1007/s13225-020-00466-2)) |
| Otu56808 | 1 | k__Fungi(100);p__Ascomycota(100);c__Saccharomycetes(100);o__Saccharomycetales(100);f__Metschnikowiaceae(100);g__Metschnikowia(100);s__Metschnikowia_pulcherrima(100); | Metschnikowia | 13 | Saprotroph | \|Undefined Saprotroph\| | Yeast | NULL | Probable | As nectar-tap_saprotroph (Põlme et al. 2020); Decay sugar-rich substrates (Põlme et al. 2020) | Põlme S, et al. 2020. Fungal Diversity 105:1-16 ((https://doi.org/10.1007/s13225-020-00466-2)) |
| Otu56810 | 1 | k__Fungi(100);p__Ascomycota(100);c__Saccharomycetes(100);o__Saccharomycetales(100);f__Metschnikowiaceae(100);g__Metschnikowia(100);s__Metschnikowia_pulcherrima(100); | Metschnikowia | 13 | Saprotroph | \|Undefined Saprotroph\| | Yeast | NULL | Probable | As nectar-tap_saprotroph (Põlme et al. 2020); Decay sugar-rich substrates (Põlme et al. 2020) | Põlme S, et al. 2020. Fungal Diversity 105:1-16 ((https://doi.org/10.1007/s13225-020-00466-2)) |
| Otu56811 | 1 | k__Fungi(100);p__Ascomycota(100);c__Saccharomycetes(100);o__Saccharomycetales(100);f__Metschnikowiaceae(100);g__Metschnikowia(100);s__Metschnikowia_pulcherrima(100); | Metschnikowia | 13 | Saprotroph | \|Undefined Saprotroph\| | Yeast | NULL | Probable | As nectar-tap_saprotroph (Põlme et al. 2020); Decay sugar-rich substrates (Põlme et al. 2020) | Põlme S, et al. 2020. Fungal Diversity 105:1-16 ((https://doi.org/10.1007/s13225-020-00466-2)) |
| Otu56813 | 1 | k__Fungi(100);p__Ascomycota(100);c__Saccharomycetes(100);o__Saccharomycetales(100);f__Metschnikowiaceae(100);g__Metschnikowia(100);s__Metschnikowia_pulcherrima(100); | Metschnikowia | 13 | Saprotroph | \|Undefined Saprotroph\| | Yeast | NULL | Probable | As nectar-tap_saprotroph (Põlme et al. 2020); Decay sugar-rich substrates (Põlme et al. 2020) | Põlme S, et al. 2020. Fungal Diversity 105:1-16 ((https://doi.org/10.1007/s13225-020-00466-2)) |
| Otu56814 | 1 | k__Fungi(100);p__Ascomycota(100);c__Saccharomycetes(100);o__Saccharomycetales(100);f__Metschnikowiaceae(100);g__Metschnikowia(100);g__Metschnikowia_unclassified(100); | Metschnikowia | 13 | Saprotroph | \|Undefined Saprotroph\| | Yeast | NULL | Probable | As nectar-tap_saprotroph (Põlme et al. 2020); Decay sugar-rich substrates (Põlme et al. 2020) | Põlme S, et al. 2020. Fungal Diversity 105:1-16 ((https://doi.org/10.1007/s13225-020-00466-2)) |
| Otu56815 | 1 | k__Fungi(100);p__Ascomycota(100);c__Saccharomycetes(100);o__Saccharomycetales(100);f__Metschnikowiaceae(100);g__Metschnikowia(100);g__Metschnikowia_unclassified(100); | Metschnikowia | 13 | Saprotroph | \|Undefined Saprotroph\| | Yeast | NULL | Probable | As nectar-tap_saprotroph (Põlme et al. 2020); Decay sugar-rich substrates (Põlme et al. 2020) | Põlme S, et al. 2020. Fungal Diversity 105:1-16 ((https://doi.org/10.1007/s13225-020-00466-2)) |
| Otu56816 | 1 | k__Fungi(100);p__Ascomycota(100);c__Saccharomycetes(100);o__Saccharomycetales(100);f__Metschnikowiaceae(100);g__Metschnikowia(100);s__Metschnikowia_pulcherrima(100); | Metschnikowia | 13 | Saprotroph | \|Undefined Saprotroph\| | Yeast | NULL | Probable | As nectar-tap_saprotroph (Põlme et al. 2020); Decay sugar-rich substrates (Põlme et al. 2020) | Põlme S, et al. 2020. Fungal Diversity 105:1-16 ((https://doi.org/10.1007/s13225-020-00466-2)) |
| Otu56817 | 1 | k__Fungi(100);p__Ascomycota(100);c__Saccharomycetes(100);o__Saccharomycetales(100);f__Metschnikowiaceae(100);g__Metschnikowia(100);s__Metschnikowia_pulcherrima(100); | Metschnikowia | 13 | Saprotroph | \|Undefined Saprotroph\| | Yeast | NULL | Probable | As nectar-tap_saprotroph (Põlme et al. 2020); Decay sugar-rich substrates (Põlme et al. 2020) | Põlme S, et al. 2020. Fungal Diversity 105:1-16 ((https://doi.org/10.1007/s13225-020-00466-2)) |
| Otu56818 | 1 | k__Fungi(100);p__Ascomycota(100);c__Saccharomycetes(100);o__Saccharomycetales(100);f__Metschnikowiaceae(100);g__Metschnikowia(100);g__Metschnikowia_unclassified(100); | Metschnikowia | 13 | Saprotroph | \|Undefined Saprotroph\| | Yeast | NULL | Probable | As nectar-tap_saprotroph (Põlme et al. 2020); Decay sugar-rich substrates (Põlme et al. 2020) | Põlme S, et al. 2020. Fungal Diversity 105:1-16 ((https://doi.org/10.1007/s13225-020-00466-2)) |
| Otu56819 | 1 | k__Fungi(100);p__Ascomycota(100);c__Saccharomycetes(100);o__Saccharomycetales(100);f__Metschnikowiaceae(100);g__Metschnikowia(100);s__Metschnikowia_pulcherrima(100); | Metschnikowia | 13 | Saprotroph | \|Undefined Saprotroph\| | Yeast | NULL | Probable | As nectar-tap_saprotroph (Põlme et al. 2020); Decay sugar-rich substrates (Põlme et al. 2020) | Põlme S, et al. 2020. Fungal Diversity 105:1-16 ((https://doi.org/10.1007/s13225-020-00466-2)) |
| Otu56820 | 1 | k__Fungi(100);p__Ascomycota(100);c__Saccharomycetes(100);o__Saccharomycetales(100);f__Metschnikowiaceae(100);g__Metschnikowia(100);s__Metschnikowia_pulcherrima(100); | Metschnikowia | 13 | Saprotroph | \|Undefined Saprotroph\| | Yeast | NULL | Probable | As nectar-tap_saprotroph (Põlme et al. 2020); Decay sugar-rich substrates (Põlme et al. 2020) | Põlme S, et al. 2020. Fungal Diversity 105:1-16 ((https://doi.org/10.1007/s13225-020-00466-2)) |
| Otu56822 | 1 | k__Fungi(100);p__Ascomycota(100);c__Saccharomycetes(100);o__Saccharomycetales(100);f__Metschnikowiaceae(100);g__Metschnikowia(100);s__Metschnikowia_pulcherrima(100); | Metschnikowia | 13 | Saprotroph | \|Undefined Saprotroph\| | Yeast | NULL | Probable | As nectar-tap_saprotroph (Põlme et al. 2020); Decay sugar-rich substrates (Põlme et al. 2020) | Põlme S, et al. 2020. Fungal Diversity 105:1-16 ((https://doi.org/10.1007/s13225-020-00466-2)) |
| Otu56824 | 1 | k__Fungi(100);p__Ascomycota(100);c__Saccharomycetes(100);o__Saccharomycetales(100);f__Metschnikowiaceae(100);g__Metschnikowia(100);s__Metschnikowia_pulcherrima(100); | Metschnikowia | 13 | Saprotroph | \|Undefined Saprotroph\| | Yeast | NULL | Probable | As nectar-tap_saprotroph (Põlme et al. 2020); Decay sugar-rich substrates (Põlme et al. 2020) | Põlme S, et al. 2020. Fungal Diversity 105:1-16 ((https://doi.org/10.1007/s13225-020-00466-2)) |
| Otu56825 | 1 | k__Fungi(100);p__Ascomycota(100);c__Saccharomycetes(100);o__Saccharomycetales(100);f__Metschnikowiaceae(100);g__Metschnikowia(100);g__Metschnikowia_unclassified(100); | Metschnikowia | 13 | Saprotroph | \|Undefined Saprotroph\| | Yeast | NULL | Probable | As nectar-tap_saprotroph (Põlme et al. 2020); Decay sugar-rich substrates (Põlme et al. 2020) | Põlme S, et al. 2020. Fungal Diversity 105:1-16 ((https://doi.org/10.1007/s13225-020-00466-2)) |
| Otu56826 | 1 | k__Fungi(100);p__Ascomycota(100);c__Saccharomycetes(100);o__Saccharomycetales(100);f__Metschnikowiaceae(100);g__Metschnikowia(100);g__Metschnikowia_unclassified(100); | Metschnikowia | 13 | Saprotroph | \|Undefined Saprotroph\| | Yeast | NULL | Probable | As nectar-tap_saprotroph (Põlme et al. 2020); Decay sugar-rich substrates (Põlme et al. 2020) | Põlme S, et al. 2020. Fungal Diversity 105:1-16 ((https://doi.org/10.1007/s13225-020-00466-2)) |
| Otu56827 | 1 | k__Fungi(100);p__Ascomycota(100);c__Saccharomycetes(100);o__Saccharomycetales(100);f__Metschnikowiaceae(100);g__Metschnikowia(100);s__Metschnikowia_pulcherrima(100); | Metschnikowia | 13 | Saprotroph | \|Undefined Saprotroph\| | Yeast | NULL | Probable | As nectar-tap_saprotroph (Põlme et al. 2020); Decay sugar-rich substrates (Põlme et al. 2020) | Põlme S, et al. 2020. Fungal Diversity 105:1-16 ((https://doi.org/10.1007/s13225-020-00466-2)) |
| Otu56828 | 1 | k__Fungi(100);p__Ascomycota(100);c__Saccharomycetes(100);o__Saccharomycetales(100);f__Metschnikowiaceae(100);g__Metschnikowia(100);s__Metschnikowia_pulcherrima(100); | Metschnikowia | 13 | Saprotroph | \|Undefined Saprotroph\| | Yeast | NULL | Probable | As nectar-tap_saprotroph (Põlme et al. 2020); Decay sugar-rich substrates (Põlme et al. 2020) | Põlme S, et al. 2020. Fungal Diversity 105:1-16 ((https://doi.org/10.1007/s13225-020-00466-2)) |
| Otu56830 | 1 | k__Fungi(100);p__Ascomycota(100);c__Saccharomycetes(100);o__Saccharomycetales(100);f__Metschnikowiaceae(100);g__Metschnikowia(100);s__Metschnikowia_pulcherrima(100); | Metschnikowia | 13 | Saprotroph | \|Undefined Saprotroph\| | Yeast | NULL | Probable | As nectar-tap_saprotroph (Põlme et al. 2020); Decay sugar-rich substrates (Põlme et al. 2020) | Põlme S, et al. 2020. Fungal Diversity 105:1-16 ((https://doi.org/10.1007/s13225-020-00466-2)) |
| Otu56831 | 1 | k__Fungi(100);p__Ascomycota(100);c__Saccharomycetes(100);o__Saccharomycetales(100);f__Metschnikowiaceae(100);g__Metschnikowia(100);s__Metschnikowia_pulcherrima(100); | Metschnikowia | 13 | Saprotroph | \|Undefined Saprotroph\| | Yeast | NULL | Probable | As nectar-tap_saprotroph (Põlme et al. 2020); Decay sugar-rich substrates (Põlme et al. 2020) | Põlme S, et al. 2020. Fungal Diversity 105:1-16 ((https://doi.org/10.1007/s13225-020-00466-2)) |
| Otu56832 | 1 | k__Fungi(100);p__Ascomycota(100);c__Saccharomycetes(100);o__Saccharomycetales(100);f__Metschnikowiaceae(100);g__Metschnikowia(100);s__Metschnikowia_pulcherrima(100); | Metschnikowia | 13 | Saprotroph | \|Undefined Saprotroph\| | Yeast | NULL | Probable | As nectar-tap_saprotroph (Põlme et al. 2020); Decay sugar-rich substrates (Põlme et al. 2020) | Põlme S, et al. 2020. Fungal Diversity 105:1-16 ((https://doi.org/10.1007/s13225-020-00466-2)) |
| Otu56833 | 1 | k__Fungi(100);p__Ascomycota(100);c__Saccharomycetes(100);o__Saccharomycetales(100);f__Metschnikowiaceae(100);g__Metschnikowia(100);g__Metschnikowia_unclassified(100); | Metschnikowia | 13 | Saprotroph | \|Undefined Saprotroph\| | Yeast | NULL | Probable | As nectar-tap_saprotroph (Põlme et al. 2020); Decay sugar-rich substrates (Põlme et al. 2020) | Põlme S, et al. 2020. Fungal Diversity 105:1-16 ((https://doi.org/10.1007/s13225-020-00466-2)) |
| Otu56835 | 1 | k__Fungi(100);p__Ascomycota(100);c__Saccharomycetes(100);o__Saccharomycetales(100);f__Metschnikowiaceae(100);g__Metschnikowia(100);s__Metschnikowia_pulcherrima(100); | Metschnikowia | 13 | Saprotroph | \|Undefined Saprotroph\| | Yeast | NULL | Probable | As nectar-tap_saprotroph (Põlme et al. 2020); Decay sugar-rich substrates (Põlme et al. 2020) | Põlme S, et al. 2020. Fungal Diversity 105:1-16 ((https://doi.org/10.1007/s13225-020-00466-2)) |
| Otu56836 | 1 | k__Fungi(100);p__Ascomycota(100);c__Saccharomycetes(100);o__Saccharomycetales(100);f__Metschnikowiaceae(100);g__Metschnikowia(100);s__Metschnikowia_pulcherrima(100); | Metschnikowia | 13 | Saprotroph | \|Undefined Saprotroph\| | Yeast | NULL | Probable | As nectar-tap_saprotroph (Põlme et al. 2020); Decay sugar-rich substrates (Põlme et al. 2020) | Põlme S, et al. 2020. Fungal Diversity 105:1-16 ((https://doi.org/10.1007/s13225-020-00466-2)) |
| Otu56837 | 1 | k__Fungi(100);p__Ascomycota(100);c__Saccharomycetes(100);o__Saccharomycetales(100);f__Metschnikowiaceae(100);g__Metschnikowia(100);s__Metschnikowia_pulcherrima(100); | Metschnikowia | 13 | Saprotroph | \|Undefined Saprotroph\| | Yeast | NULL | Probable | As nectar-tap_saprotroph (Põlme et al. 2020); Decay sugar-rich substrates (Põlme et al. 2020) | Põlme S, et al. 2020. Fungal Diversity 105:1-16 ((https://doi.org/10.1007/s13225-020-00466-2)) |
| Otu56838 | 1 | k__Fungi(100);p__Ascomycota(100);c__Saccharomycetes(100);o__Saccharomycetales(100);f__Metschnikowiaceae(100);g__Metschnikowia(100);s__Metschnikowia_pulcherrima(100); | Metschnikowia | 13 | Saprotroph | \|Undefined Saprotroph\| | Yeast | NULL | Probable | As nectar-tap_saprotroph (Põlme et al. 2020); Decay sugar-rich substrates (Põlme et al. 2020) | Põlme S, et al. 2020. Fungal Diversity 105:1-16 ((https://doi.org/10.1007/s13225-020-00466-2)) |
| Otu56839 | 1 | k__Fungi(100);p__Ascomycota(100);c__Saccharomycetes(100);o__Saccharomycetales(100);f__Metschnikowiaceae(100);g__Metschnikowia(100);s__Metschnikowia_pulcherrima(100); | Metschnikowia | 13 | Saprotroph | \|Undefined Saprotroph\| | Yeast | NULL | Probable | As nectar-tap_saprotroph (Põlme et al. 2020); Decay sugar-rich substrates (Põlme et al. 2020) | Põlme S, et al. 2020. Fungal Diversity 105:1-16 ((https://doi.org/10.1007/s13225-020-00466-2)) |
| Otu56840 | 1 | k__Fungi(100);p__Ascomycota(100);c__Saccharomycetes(100);o__Saccharomycetales(100);f__Metschnikowiaceae(100);g__Metschnikowia(100);s__Metschnikowia_pulcherrima(100); | Metschnikowia | 13 | Saprotroph | \|Undefined Saprotroph\| | Yeast | NULL | Probable | As nectar-tap_saprotroph (Põlme et al. 2020); Decay sugar-rich substrates (Põlme et al. 2020) | Põlme S, et al. 2020. Fungal Diversity 105:1-16 ((https://doi.org/10.1007/s13225-020-00466-2)) |
| Otu56841 | 1 | k__Fungi(100);p__Ascomycota(100);c__Saccharomycetes(100);o__Saccharomycetales(100);f__Metschnikowiaceae(100);g__Metschnikowia(100);s__Metschnikowia_pulcherrima(100); | Metschnikowia | 13 | Saprotroph | \|Undefined Saprotroph\| | Yeast | NULL | Probable | As nectar-tap_saprotroph (Põlme et al. 2020); Decay sugar-rich substrates (Põlme et al. 2020) | Põlme S, et al. 2020. Fungal Diversity 105:1-16 ((https://doi.org/10.1007/s13225-020-00466-2)) |
| Otu56842 | 1 | k__Fungi(100);p__Ascomycota(100);c__Saccharomycetes(100);o__Saccharomycetales(100);f__Metschnikowiaceae(100);g__Metschnikowia(100);s__Metschnikowia_pulcherrima(100); | Metschnikowia | 13 | Saprotroph | \|Undefined Saprotroph\| | Yeast | NULL | Probable | As nectar-tap_saprotroph (Põlme et al. 2020); Decay sugar-rich substrates (Põlme et al. 2020) | Põlme S, et al. 2020. Fungal Diversity 105:1-16 ((https://doi.org/10.1007/s13225-020-00466-2)) |
| Otu56844 | 1 | k__Fungi(100);p__Ascomycota(100);c__Saccharomycetes(100);o__Saccharomycetales(100);f__Metschnikowiaceae(100);g__Metschnikowia(100);s__Metschnikowia_pulcherrima(100); | Metschnikowia | 13 | Saprotroph | \|Undefined Saprotroph\| | Yeast | NULL | Probable | As nectar-tap_saprotroph (Põlme et al. 2020); Decay sugar-rich substrates (Põlme et al. 2020) | Põlme S, et al. 2020. Fungal Diversity 105:1-16 ((https://doi.org/10.1007/s13225-020-00466-2)) |
| Otu56845 | 1 | k__Fungi(100);p__Ascomycota(100);c__Saccharomycetes(100);o__Saccharomycetales(100);f__Metschnikowiaceae(100);g__Metschnikowia(100);s__Metschnikowia_pulcherrima(100); | Metschnikowia | 13 | Saprotroph | \|Undefined Saprotroph\| | Yeast | NULL | Probable | As nectar-tap_saprotroph (Põlme et al. 2020); Decay sugar-rich substrates (Põlme et al. 2020) | Põlme S, et al. 2020. Fungal Diversity 105:1-16 ((https://doi.org/10.1007/s13225-020-00466-2)) |
| Otu56846 | 1 | k__Fungi(100);p__Ascomycota(100);c__Saccharomycetes(100);o__Saccharomycetales(100);f__Metschnikowiaceae(100);g__Metschnikowia(100);s__Metschnikowia_pulcherrima(100); | Metschnikowia | 13 | Saprotroph | \|Undefined Saprotroph\| | Yeast | NULL | Probable | As nectar-tap_saprotroph (Põlme et al. 2020); Decay sugar-rich substrates (Põlme et al. 2020) | Põlme S, et al. 2020. Fungal Diversity 105:1-16 ((https://doi.org/10.1007/s13225-020-00466-2)) |
| Otu56847 | 1 | k__Fungi(100);p__Ascomycota(100);c__Saccharomycetes(100);o__Saccharomycetales(100);f__Metschnikowiaceae(100);g__Metschnikowia(100);s__Metschnikowia_pulcherrima(100); | Metschnikowia | 13 | Saprotroph | \|Undefined Saprotroph\| | Yeast | NULL | Probable | As nectar-tap_saprotroph (Põlme et al. 2020); Decay sugar-rich substrates (Põlme et al. 2020) | Põlme S, et al. 2020. Fungal Diversity 105:1-16 ((https://doi.org/10.1007/s13225-020-00466-2)) |
| Otu56849 | 1 | k__Fungi(100);p__Ascomycota(100);c__Saccharomycetes(100);o__Saccharomycetales(100);f__Metschnikowiaceae(100);g__Metschnikowia(100);g__Metschnikowia_unclassified(100); | Metschnikowia | 13 | Saprotroph | \|Undefined Saprotroph\| | Yeast | NULL | Probable | As nectar-tap_saprotroph (Põlme et al. 2020); Decay sugar-rich substrates (Põlme et al. 2020) | Põlme S, et al. 2020. Fungal Diversity 105:1-16 ((https://doi.org/10.1007/s13225-020-00466-2)) |
| Otu56850 | 1 | k__Fungi(100);p__Ascomycota(100);c__Saccharomycetes(100);o__Saccharomycetales(100);f__Metschnikowiaceae(100);g__Metschnikowia(100);g__Metschnikowia_unclassified(100); | Metschnikowia | 13 | Saprotroph | \|Undefined Saprotroph\| | Yeast | NULL | Probable | As nectar-tap_saprotroph (Põlme et al. 2020); Decay sugar-rich substrates (Põlme et al. 2020) | Põlme S, et al. 2020. Fungal Diversity 105:1-16 ((https://doi.org/10.1007/s13225-020-00466-2)) |
| Otu56851 | 1 | k__Fungi(100);p__Ascomycota(100);c__Saccharomycetes(100);o__Saccharomycetales(100);f__Metschnikowiaceae(100);g__Metschnikowia(100);s__Metschnikowia_pulcherrima(100); | Metschnikowia | 13 | Saprotroph | \|Undefined Saprotroph\| | Yeast | NULL | Probable | As nectar-tap_saprotroph (Põlme et al. 2020); Decay sugar-rich substrates (Põlme et al. 2020) | Põlme S, et al. 2020. Fungal Diversity 105:1-16 ((https://doi.org/10.1007/s13225-020-00466-2)) |
| Otu56852 | 1 | k__Fungi(100);p__Ascomycota(100);c__Saccharomycetes(100);o__Saccharomycetales(100);f__Metschnikowiaceae(100);g__Metschnikowia(100);s__Metschnikowia_pulcherrima(100); | Metschnikowia | 13 | Saprotroph | \|Undefined Saprotroph\| | Yeast | NULL | Probable | As nectar-tap_saprotroph (Põlme et al. 2020); Decay sugar-rich substrates (Põlme et al. 2020) | Põlme S, et al. 2020. Fungal Diversity 105:1-16 ((https://doi.org/10.1007/s13225-020-00466-2)) |
| Otu56853 | 1 | k__Fungi(100);p__Ascomycota(100);c__Saccharomycetes(100);o__Saccharomycetales(100);f__Metschnikowiaceae(100);g__Metschnikowia(100);s__Metschnikowia_pulcherrima(100); | Metschnikowia | 13 | Saprotroph | \|Undefined Saprotroph\| | Yeast | NULL | Probable | As nectar-tap_saprotroph (Põlme et al. 2020); Decay sugar-rich substrates (Põlme et al. 2020) | Põlme S, et al. 2020. Fungal Diversity 105:1-16 ((https://doi.org/10.1007/s13225-020-00466-2)) |
| Otu56854 | 1 | k__Fungi(100);p__Ascomycota(100);c__Saccharomycetes(100);o__Saccharomycetales(100);f__Metschnikowiaceae(100);g__Metschnikowia(100);s__Metschnikowia_pulcherrima(100); | Metschnikowia | 13 | Saprotroph | \|Undefined Saprotroph\| | Yeast | NULL | Probable | As nectar-tap_saprotroph (Põlme et al. 2020); Decay sugar-rich substrates (Põlme et al. 2020) | Põlme S, et al. 2020. Fungal Diversity 105:1-16 ((https://doi.org/10.1007/s13225-020-00466-2)) |
| Otu56855 | 1 | k__Fungi(100);p__Ascomycota(100);c__Saccharomycetes(100);o__Saccharomycetales(100);f__Metschnikowiaceae(100);g__Metschnikowia(100);s__Metschnikowia_pulcherrima(100); | Metschnikowia | 13 | Saprotroph | \|Undefined Saprotroph\| | Yeast | NULL | Probable | As nectar-tap_saprotroph (Põlme et al. 2020); Decay sugar-rich substrates (Põlme et al. 2020) | Põlme S, et al. 2020. Fungal Diversity 105:1-16 ((https://doi.org/10.1007/s13225-020-00466-2)) |
| Otu56856 | 1 | k__Fungi(100);p__Ascomycota(100);c__Saccharomycetes(100);o__Saccharomycetales(100);f__Metschnikowiaceae(100);g__Metschnikowia(100);g__Metschnikowia_unclassified(100); | Metschnikowia | 13 | Saprotroph | \|Undefined Saprotroph\| | Yeast | NULL | Probable | As nectar-tap_saprotroph (Põlme et al. 2020); Decay sugar-rich substrates (Põlme et al. 2020) | Põlme S, et al. 2020. Fungal Diversity 105:1-16 ((https://doi.org/10.1007/s13225-020-00466-2)) |
| Otu56858 | 1 | k__Fungi(100);p__Ascomycota(100);c__Saccharomycetes(100);o__Saccharomycetales(100);f__Metschnikowiaceae(100);g__Metschnikowia(100);g__Metschnikowia_unclassified(100); | Metschnikowia | 13 | Saprotroph | \|Undefined Saprotroph\| | Yeast | NULL | Probable | As nectar-tap_saprotroph (Põlme et al. 2020); Decay sugar-rich substrates (Põlme et al. 2020) | Põlme S, et al. 2020. Fungal Diversity 105:1-16 ((https://doi.org/10.1007/s13225-020-00466-2)) |
| Otu56859 | 1 | k__Fungi(100);p__Ascomycota(100);c__Saccharomycetes(100);o__Saccharomycetales(100);f__Metschnikowiaceae(100);g__Metschnikowia(100);s__Metschnikowia_pulcherrima(100); | Metschnikowia | 13 | Saprotroph | \|Undefined Saprotroph\| | Yeast | NULL | Probable | As nectar-tap_saprotroph (Põlme et al. 2020); Decay sugar-rich substrates (Põlme et al. 2020) | Põlme S, et al. 2020. Fungal Diversity 105:1-16 ((https://doi.org/10.1007/s13225-020-00466-2)) |
| Otu56860 | 1 | k__Fungi(100);p__Ascomycota(100);c__Saccharomycetes(100);o__Saccharomycetales(100);f__Metschnikowiaceae(100);g__Metschnikowia(100);s__Metschnikowia_pulcherrima(100); | Metschnikowia | 13 | Saprotroph | \|Undefined Saprotroph\| | Yeast | NULL | Probable | As nectar-tap_saprotroph (Põlme et al. 2020); Decay sugar-rich substrates (Põlme et al. 2020) | Põlme S, et al. 2020. Fungal Diversity 105:1-16 ((https://doi.org/10.1007/s13225-020-00466-2)) |
| Otu56861 | 1 | k__Fungi(100);p__Ascomycota(100);c__Saccharomycetes(100);o__Saccharomycetales(100);f__Metschnikowiaceae(100);g__Metschnikowia(100);s__Metschnikowia_pulcherrima(100); | Metschnikowia | 13 | Saprotroph | \|Undefined Saprotroph\| | Yeast | NULL | Probable | As nectar-tap_saprotroph (Põlme et al. 2020); Decay sugar-rich substrates (Põlme et al. 2020) | Põlme S, et al. 2020. Fungal Diversity 105:1-16 ((https://doi.org/10.1007/s13225-020-00466-2)) |
| Otu56862 | 1 | k__Fungi(100);p__Ascomycota(100);c__Saccharomycetes(100);o__Saccharomycetales(100);f__Metschnikowiaceae(100);g__Metschnikowia(100);s__Metschnikowia_pulcherrima(100); | Metschnikowia | 13 | Saprotroph | \|Undefined Saprotroph\| | Yeast | NULL | Probable | As nectar-tap_saprotroph (Põlme et al. 2020); Decay sugar-rich substrates (Põlme et al. 2020) | Põlme S, et al. 2020. Fungal Diversity 105:1-16 ((https://doi.org/10.1007/s13225-020-00466-2)) |

| | | | | | | | | | | | |
|---|---|---|---|---|---|---|---|---|---|---|---|
| Otu56863 | 1 | k__Fungi(100);p__Ascomycota(100);c__Saccharomycetes(100);o__Saccharomycetales(100);f__Metschnikowiaceae(100);g__Metschnikowia(100);g__Metschnikowia_unclassified(100); | Metschnikowia | 13 | Saprotroph | \|Undefined Saprotroph\| | Yeast | NULL | Probable | As nectar-tap_saprotroph (Põlme et al. 2020); Decay sugar-rich substrates (Põlme et al. 2020) | Põlme S, et al. 2020. Fungal Diversity 105:1-16 ((https://doi.org/10.1007/s13225-020-00466-2)) |
| Otu56864 | 1 | k__Fungi(100);p__Ascomycota(100);c__Saccharomycetes(100);o__Saccharomycetales(100);f__Metschnikowiaceae(100);g__Metschnikowia(100);g__Metschnikowia_unclassified(100); | Metschnikowia | 13 | Saprotroph | \|Undefined Saprotroph\| | Yeast | NULL | Probable | As nectar-tap_saprotroph (Põlme et al. 2020); Decay sugar-rich substrates (Põlme et al. 2020) | Põlme S, et al. 2020. Fungal Diversity 105:1-16 ((https://doi.org/10.1007/s13225-020-00466-2)) |
| Otu56865 | 1 | k__Fungi(100);p__Ascomycota(100);c__Saccharomycetes(100);o__Saccharomycetales(100);f__Metschnikowiaceae(100);g__Metschnikowia(100);s__Metschnikowia_pulcherrima(100); | Metschnikowia | 13 | Saprotroph | \|Undefined Saprotroph\| | Yeast | NULL | Probable | As nectar-tap_saprotroph (Põlme et al. 2020); Decay sugar-rich substrates (Põlme et al. 2020) | Põlme S, et al. 2020. Fungal Diversity 105:1-16 ((https://doi.org/10.1007/s13225-020-00466-2)) |
| Otu56866 | 1 | k__Fungi(100);p__Ascomycota(100);c__Saccharomycetes(100);o__Saccharomycetales(100);f__Metschnikowiaceae(100);g__Metschnikowia(100);s__Metschnikowia_pulcherrima(100); | Metschnikowia | 13 | Saprotroph | \|Undefined Saprotroph\| | Yeast | NULL | Probable | As nectar-tap_saprotroph (Põlme et al. 2020); Decay sugar-rich substrates (Põlme et al. 2020) | Põlme S, et al. 2020. Fungal Diversity 105:1-16 ((https://doi.org/10.1007/s13225-020-00466-2)) |
| Otu56867 | 1 | k__Fungi(100);p__Ascomycota(100);c__Saccharomycetes(100);o__Saccharomycetales(100);f__Metschnikowiaceae(100);g__Metschnikowia(100);s__Metschnikowia_pulcherrima(100); | Metschnikowia | 13 | Saprotroph | \|Undefined Saprotroph\| | Yeast | NULL | Probable | As nectar-tap_saprotroph (Põlme et al. 2020); Decay sugar-rich substrates (Põlme et al. 2020) | Põlme S, et al. 2020. Fungal Diversity 105:1-16 ((https://doi.org/10.1007/s13225-020-00466-2)) |
| Otu56871 | 1 | k__Fungi(100);p__Ascomycota(100);c__Saccharomycetes(100);o__Saccharomycetales(100);f__Metschnikowiaceae(100);g__Metschnikowia(100);s__Metschnikowia_pulcherrima(100); | Metschnikowia | 13 | Saprotroph | \|Undefined Saprotroph\| | Yeast | NULL | Probable | As nectar-tap_saprotroph (Põlme et al. 2020); Decay sugar-rich substrates (Põlme et al. 2020) | Põlme S, et al. 2020. Fungal Diversity 105:1-16 ((https://doi.org/10.1007/s13225-020-00466-2)) |
| Otu56872 | 1 | k__Fungi(100);p__Ascomycota(100);c__Saccharomycetes(100);o__Saccharomycetales(100);f__Metschnikowiaceae(100);g__Metschnikowia(100);s__Metschnikowia_pulcherrima(100); | Metschnikowia | 13 | Saprotroph | \|Undefined Saprotroph\| | Yeast | NULL | Probable | As nectar-tap_saprotroph (Põlme et al. 2020); Decay sugar-rich substrates (Põlme et al. 2020) | Põlme S, et al. 2020. Fungal Diversity 105:1-16 ((https://doi.org/10.1007/s13225-020-00466-2)) |
| Otu56873 | 1 | k__Fungi(100);p__Ascomycota(100);c__Saccharomycetes(100);o__Saccharomycetales(100);f__Metschnikowiaceae(100);g__Metschnikowia(100);s__Metschnikowia_pulcherrima(100); | Metschnikowia | 13 | Saprotroph | \|Undefined Saprotroph\| | Yeast | NULL | Probable | As nectar-tap_saprotroph (Põlme et al. 2020); Decay sugar-rich substrates (Põlme et al. 2020) | Põlme S, et al. 2020. Fungal Diversity 105:1-16 ((https://doi.org/10.1007/s13225-020-00466-2)) |
| Otu56874 | 1 | k__Fungi(100);p__Ascomycota(100);c__Saccharomycetes(100);o__Saccharomycetales(100);f__Metschnikowiaceae(100);g__Metschnikowia(100);s__Metschnikowia_pulcherrima(100); | Metschnikowia | 13 | Saprotroph | \|Undefined Saprotroph\| | Yeast | NULL | Probable | As nectar-tap_saprotroph (Põlme et al. 2020); Decay sugar-rich substrates (Põlme et al. 2020) | Põlme S, et al. 2020. Fungal Diversity 105:1-16 ((https://doi.org/10.1007/s13225-020-00466-2)) |
| Otu56875 | 1 | k__Fungi(100);p__Ascomycota(100);c__Saccharomycetes(100);o__Saccharomycetales(100);f__Metschnikowiaceae(100);g__Metschnikowia(100);s__Metschnikowia_pulcherrima(100); | Metschnikowia | 13 | Saprotroph | \|Undefined Saprotroph\| | Yeast | NULL | Probable | As nectar-tap_saprotroph (Põlme et al. 2020); Decay sugar-rich substrates (Põlme et al. 2020) | Põlme S, et al. 2020. Fungal Diversity 105:1-16 ((https://doi.org/10.1007/s13225-020-00466-2)) |
| Otu56876 | 1 | k__Fungi(100);p__Ascomycota(100);c__Saccharomycetes(100);o__Saccharomycetales(100);f__Metschnikowiaceae(100);g__Metschnikowia(100);s__Metschnikowia_pulcherrima(100); | Metschnikowia | 13 | Saprotroph | \|Undefined Saprotroph\| | Yeast | NULL | Probable | As nectar-tap_saprotroph (Põlme et al. 2020); Decay sugar-rich substrates (Põlme et al. 2020) | Põlme S, et al. 2020. Fungal Diversity 105:1-16 ((https://doi.org/10.1007/s13225-020-00466-2)) |
| Otu56877 | 1 | k__Fungi(100);p__Ascomycota(100);c__Saccharomycetes(100);o__Saccharomycetales(100);f__Metschnikowiaceae(100);g__Metschnikowia(100);s__Metschnikowia_pulcherrima(100); | Metschnikowia | 13 | Saprotroph | \|Undefined Saprotroph\| | Yeast | NULL | Probable | As nectar-tap_saprotroph (Põlme et al. 2020); Decay sugar-rich substrates (Põlme et al. 2020) | Põlme S, et al. 2020. Fungal Diversity 105:1-16 ((https://doi.org/10.1007/s13225-020-00466-2)) |
| Otu56878 | 1 | k__Fungi(100);p__Ascomycota(100);c__Saccharomycetes(100);o__Saccharomycetales(100);f__Metschnikowiaceae(100);g__Metschnikowia(100);s__Metschnikowia_pulcherrima(100); | Metschnikowia | 13 | Saprotroph | \|Undefined Saprotroph\| | Yeast | NULL | Probable | As nectar-tap_saprotroph (Põlme et al. 2020); Decay sugar-rich substrates (Põlme et al. 2020) | Põlme S, et al. 2020. Fungal Diversity 105:1-16 ((https://doi.org/10.1007/s13225-020-00466-2)) |
| Otu56879 | 1 | k__Fungi(100);p__Ascomycota(100);c__Saccharomycetes(100);o__Saccharomycetales(100);f__Metschnikowiaceae(100);g__Metschnikowia(100);s__Metschnikowia_pulcherrima(100); | Metschnikowia | 13 | Saprotroph | \|Undefined Saprotroph\| | Yeast | NULL | Probable | As nectar-tap_saprotroph (Põlme et al. 2020); Decay sugar-rich substrates (Põlme et al. 2020) | Põlme S, et al. 2020. Fungal Diversity 105:1-16 ((https://doi.org/10.1007/s13225-020-00466-2)) |
| Otu56880 | 1 | k__Fungi(100);p__Ascomycota(100);c__Saccharomycetes(100);o__Saccharomycetales(100);f__Metschnikowiaceae(100);g__Metschnikowia(100);s__Metschnikowia_pulcherrima(100); | Metschnikowia | 13 | Saprotroph | \|Undefined Saprotroph\| | Yeast | NULL | Probable | As nectar-tap_saprotroph (Põlme et al. 2020); Decay sugar-rich substrates (Põlme et al. 2020) | Põlme S, et al. 2020. Fungal Diversity 105:1-16 ((https://doi.org/10.1007/s13225-020-00466-2)) |
| Otu56881 | 1 | k__Fungi(100);p__Ascomycota(100);c__Saccharomycetes(100);o__Saccharomycetales(100);f__Metschnikowiaceae(100);g__Metschnikowia(100);s__Metschnikowia_pulcherrima(100); | Metschnikowia | 13 | Saprotroph | \|Undefined Saprotroph\| | Yeast | NULL | Probable | As nectar-tap_saprotroph (Põlme et al. 2020); Decay sugar-rich substrates (Põlme et al. 2020) | Põlme S, et al. 2020. Fungal Diversity 105:1-16 ((https://doi.org/10.1007/s13225-020-00466-2)) |
| Otu56882 | 1 | k__Fungi(100);p__Ascomycota(100);c__Saccharomycetes(100);o__Saccharomycetales(100);f__Metschnikowiaceae(100);g__Metschnikowia(100);g__Metschnikowia_unclassified(100); | Metschnikowia | 13 | Saprotroph | \|Undefined Saprotroph\| | Yeast | NULL | Probable | As nectar-tap_saprotroph (Põlme et al. 2020); Decay sugar-rich substrates (Põlme et al. 2020) | Põlme S, et al. 2020. Fungal Diversity 105:1-16 ((https://doi.org/10.1007/s13225-020-00466-2)) |
| Otu56884 | 1 | k__Fungi(100);p__Ascomycota(100);c__Saccharomycetes(100);o__Saccharomycetales(100);f__Metschnikowiaceae(100);g__Metschnikowia(100);s__Metschnikowia_pulcherrima(100); | Metschnikowia | 13 | Saprotroph | \|Undefined Saprotroph\| | Yeast | NULL | Probable | As nectar-tap_saprotroph (Põlme et al. 2020); Decay sugar-rich substrates (Põlme et al. 2020) | Põlme S, et al. 2020. Fungal Diversity 105:1-16 ((https://doi.org/10.1007/s13225-020-00466-2)) |
| Otu56885 | 1 | k__Fungi(100);p__Ascomycota(100);c__Saccharomycetes(100);o__Saccharomycetales(100);f__Metschnikowiaceae(100);g__Metschnikowia(100);s__Metschnikowia_pulcherrima(100); | Metschnikowia | 13 | Saprotroph | \|Undefined Saprotroph\| | Yeast | NULL | Probable | As nectar-tap_saprotroph (Põlme et al. 2020); Decay sugar-rich substrates (Põlme et al. 2020) | Põlme S, et al. 2020. Fungal Diversity 105:1-16 ((https://doi.org/10.1007/s13225-020-00466-2)) |
| Otu56886 | 1 | k__Fungi(100);p__Ascomycota(100);c__Saccharomycetes(100);o__Saccharomycetales(100);f__Metschnikowiaceae(100);g__Metschnikowia(100);s__Metschnikowia_pulcherrima(100); | Metschnikowia | 13 | Saprotroph | \|Undefined Saprotroph\| | Yeast | NULL | Probable | As nectar-tap_saprotroph (Põlme et al. 2020); Decay sugar-rich substrates (Põlme et al. 2020) | Põlme S, et al. 2020. Fungal Diversity 105:1-16 ((https://doi.org/10.1007/s13225-020-00466-2)) |
| Otu56887 | 1 | k__Fungi(100);p__Ascomycota(100);c__Saccharomycetes(100);o__Saccharomycetales(100);f__Metschnikowiaceae(100);g__Metschnikowia(100);s__Metschnikowia_pulcherrima(100); | Metschnikowia | 13 | Saprotroph | \|Undefined Saprotroph\| | Yeast | NULL | Probable | As nectar-tap_saprotroph (Põlme et al. 2020); Decay sugar-rich substrates (Põlme et al. 2020) | Põlme S, et al. 2020. Fungal Diversity 105:1-16 ((https://doi.org/10.1007/s13225-020-00466-2)) |
| Otu56888 | 1 | k__Fungi(100);p__Ascomycota(100);c__Saccharomycetes(100);o__Saccharomycetales(100);f__Metschnikowiaceae(100);g__Metschnikowia(100);s__Metschnikowia_pulcherrima(100); | Metschnikowia | 13 | Saprotroph | \|Undefined Saprotroph\| | Yeast | NULL | Probable | As nectar-tap_saprotroph (Põlme et al. 2020); Decay sugar-rich substrates (Põlme et al. 2020) | Põlme S, et al. 2020. Fungal Diversity 105:1-16 ((https://doi.org/10.1007/s13225-020-00466-2)) |
| Otu56889 | 1 | k__Fungi(100);p__Ascomycota(100);c__Saccharomycetes(100);o__Saccharomycetales(100);f__Metschnikowiaceae(100);g__Metschnikowia(100);s__Metschnikowia_pulcherrima(100); | Metschnikowia | 13 | Saprotroph | \|Undefined Saprotroph\| | Yeast | NULL | Probable | As nectar-tap_saprotroph (Põlme et al. 2020); Decay sugar-rich substrates (Põlme et al. 2020) | Põlme S, et al. 2020. Fungal Diversity 105:1-16 ((https://doi.org/10.1007/s13225-020-00466-2)) |
| Otu56890 | 1 | k__Fungi(100);p__Ascomycota(100);c__Saccharomycetes(100);o__Saccharomycetales(100);f__Metschnikowiaceae(100);g__Metschnikowia(100);g__Metschnikowia_unclassified(100); | Metschnikowia | 13 | Saprotroph | \|Undefined Saprotroph\| | Yeast | NULL | Probable | As nectar-tap_saprotroph (Põlme et al. 2020); Decay sugar-rich substrates (Põlme et al. 2020) | Põlme S, et al. 2020. Fungal Diversity 105:1-16 ((https://doi.org/10.1007/s13225-020-00466-2)) |
| Otu56891 | 1 | k__Fungi(100);p__Ascomycota(100);c__Saccharomycetes(100);o__Saccharomycetales(100);f__Metschnikowiaceae(100);g__Metschnikowia(100);s__Metschnikowia_pulcherrima(100); | Metschnikowia | 13 | Saprotroph | \|Undefined Saprotroph\| | Yeast | NULL | Probable | As nectar-tap_saprotroph (Põlme et al. 2020); Decay sugar-rich substrates (Põlme et al. 2020) | Põlme S, et al. 2020. Fungal Diversity 105:1-16 ((https://doi.org/10.1007/s13225-020-00466-2)) |
| Otu56892 | 1 | k__Fungi(100);p__Ascomycota(100);c__Saccharomycetes(100);o__Saccharomycetales(100);f__Metschnikowiaceae(100);g__Metschnikowia(100);s__Metschnikowia_pulcherrima(100); | Metschnikowia | 13 | Saprotroph | \|Undefined Saprotroph\| | Yeast | NULL | Probable | As nectar-tap_saprotroph (Põlme et al. 2020); Decay sugar-rich substrates (Põlme et al. 2020) | Põlme S, et al. 2020. Fungal Diversity 105:1-16 ((https://doi.org/10.1007/s13225-020-00466-2)) |
| Otu56893 | 1 | k__Fungi(100);p__Ascomycota(100);c__Saccharomycetes(100);o__Saccharomycetales(100);f__Metschnikowiaceae(100);g__Metschnikowia(100);s__Metschnikowia_pulcherrima(100); | Metschnikowia | 13 | Saprotroph | \|Undefined Saprotroph\| | Yeast | NULL | Probable | As nectar-tap_saprotroph (Põlme et al. 2020); Decay sugar-rich substrates (Põlme et al. 2020) | Põlme S, et al. 2020. Fungal Diversity 105:1-16 ((https://doi.org/10.1007/s13225-020-00466-2)) |
| Otu56894 | 1 | k__Fungi(100);p__Ascomycota(100);c__Saccharomycetes(100);o__Saccharomycetales(100);f__Metschnikowiaceae(100);g__Metschnikowia(100);s__Metschnikowia_pulcherrima(100); | Metschnikowia | 13 | Saprotroph | \|Undefined Saprotroph\| | Yeast | NULL | Probable | As nectar-tap_saprotroph (Põlme et al. 2020); Decay sugar-rich substrates (Põlme et al. 2020) | Põlme S, et al. 2020. Fungal Diversity 105:1-16 ((https://doi.org/10.1007/s13225-020-00466-2)) |
| Otu56895 | 1 | k__Fungi(100);p__Ascomycota(100);c__Saccharomycetes(100);o__Saccharomycetales(100);f__Metschnikowiaceae(100);g__Metschnikowia(100);s__Metschnikowia_pulcherrima(100); | Metschnikowia | 13 | Saprotroph | \|Undefined Saprotroph\| | Yeast | NULL | Probable | As nectar-tap_saprotroph (Põlme et al. 2020); Decay sugar-rich substrates (Põlme et al. 2020) | Põlme S, et al. 2020. Fungal Diversity 105:1-16 ((https://doi.org/10.1007/s13225-020-00466-2)) |
| Otu56896 | 1 | k__Fungi(100);p__Ascomycota(100);c__Saccharomycetes(100);o__Saccharomycetales(100);f__Metschnikowiaceae(100);g__Metschnikowia(100);s__Metschnikowia_pulcherrima(100); | Metschnikowia | 13 | Saprotroph | \|Undefined Saprotroph\| | Yeast | NULL | Probable | As nectar-tap_saprotroph (Põlme et al. 2020); Decay sugar-rich substrates (Põlme et al. 2020) | Põlme S, et al. 2020. Fungal Diversity 105:1-16 ((https://doi.org/10.1007/s13225-020-00466-2)) |
| Otu56898 | 1 | k__Fungi(100);p__Ascomycota(100);c__Saccharomycetes(100);o__Saccharomycetales(100);f__Metschnikowiaceae(100);g__Metschnikowia(100);s__Metschnikowia_pulcherrima(100); | Metschnikowia | 13 | Saprotroph | \|Undefined Saprotroph\| | Yeast | NULL | Probable | As nectar-tap_saprotroph (Põlme et al. 2020); Decay sugar-rich substrates (Põlme et al. 2020) | Põlme S, et al. 2020. Fungal Diversity 105:1-16 ((https://doi.org/10.1007/s13225-020-00466-2)) |
| Otu56900 | 1 | k__Fungi(100);p__Ascomycota(100);c__Saccharomycetes(100);o__Saccharomycetales(100);f__Metschnikowiaceae(100);g__Metschnikowia(100);s__Metschnikowia_pulcherrima(100); | Metschnikowia | 13 | Saprotroph | \|Undefined Saprotroph\| | Yeast | NULL | Probable | As nectar-tap_saprotroph (Põlme et al. 2020); Decay sugar-rich substrates (Põlme et al. 2020) | Põlme S, et al. 2020. Fungal Diversity 105:1-16 ((https://doi.org/10.1007/s13225-020-00466-2)) |
| Otu56901 | 1 | k__Fungi(100);p__Ascomycota(100);c__Saccharomycetes(100);o__Saccharomycetales(100);f__Metschnikowiaceae(100);g__Metschnikowia(100);s__Metschnikowia_pulcherrima(100); | Metschnikowia | 13 | Saprotroph | \|Undefined Saprotroph\| | Yeast | NULL | Probable | As nectar-tap_saprotroph (Põlme et al. 2020); Decay sugar-rich substrates (Põlme et al. 2020) | Põlme S, et al. 2020. Fungal Diversity 105:1-16 ((https://doi.org/10.1007/s13225-020-00466-2)) |
| Otu56902 | 1 | k__Fungi(100);p__Ascomycota(100);c__Saccharomycetes(100);o__Saccharomycetales(100);f__Metschnikowiaceae(100);g__Metschnikowia(100);s__Metschnikowia_pulcherrima(100); | Metschnikowia | 13 | Saprotroph | \|Undefined Saprotroph\| | Yeast | NULL | Probable | As nectar-tap_saprotroph (Põlme et al. 2020); Decay sugar-rich substrates (Põlme et al. 2020) | Põlme S, et al. 2020. Fungal Diversity 105:1-16 ((https://doi.org/10.1007/s13225-020-00466-2)) |
| Otu56903 | 1 | k__Fungi(100);p__Ascomycota(100);c__Saccharomycetes(100);o__Saccharomycetales(100);f__Metschnikowiaceae(100);g__Metschnikowia(100);s__Metschnikowia_pulcherrima(100); | Metschnikowia | 13 | Saprotroph | \|Undefined Saprotroph\| | Yeast | NULL | Probable | As nectar-tap_saprotroph (Põlme et al. 2020); Decay sugar-rich substrates (Põlme et al. 2020) | Põlme S, et al. 2020. Fungal Diversity 105:1-16 ((https://doi.org/10.1007/s13225-020-00466-2)) |
| Otu56904 | 1 | k__Fungi(100);p__Ascomycota(100);c__Saccharomycetes(100);o__Saccharomycetales(100);f__Metschnikowiaceae(100);g__Metschnikowia(100);g__Metschnikowia_unclassified(100); | Metschnikowia | 13 | Saprotroph | \|Undefined Saprotroph\| | Yeast | NULL | Probable | As nectar-tap_saprotroph (Põlme et al. 2020); Decay sugar-rich substrates (Põlme et al. 2020) | Põlme S, et al. 2020. Fungal Diversity 105:1-16 ((https://doi.org/10.1007/s13225-020-00466-2)) |
| Otu56905 | 1 | k__Fungi(100);p__Ascomycota(100);c__Saccharomycetes(100);o__Saccharomycetales(100);f__Metschnikowiaceae(100);g__Metschnikowia(100);s__Metschnikowia_pulcherrima(100); | Metschnikowia | 13 | Saprotroph | \|Undefined Saprotroph\| | Yeast | NULL | Probable | As nectar-tap_saprotroph (Põlme et al. 2020); Decay sugar-rich substrates (Põlme et al. 2020) | Põlme S, et al. 2020. Fungal Diversity 105:1-16 ((https://doi.org/10.1007/s13225-020-00466-2)) |
| Otu56907 | 1 | k__Fungi(100);p__Ascomycota(100);c__Saccharomycetes(100);o__Saccharomycetales(100);f__Metschnikowiaceae(100);g__Metschnikowia(100);s__Metschnikowia_pulcherrima(100); | Metschnikowia | 13 | Saprotroph | \|Undefined Saprotroph\| | Yeast | NULL | Probable | As nectar-tap_saprotroph (Põlme et al. 2020); Decay sugar-rich substrates (Põlme et al. 2020) | Põlme S, et al. 2020. Fungal Diversity 105:1-16 ((https://doi.org/10.1007/s13225-020-00466-2)) |
| Otu56908 | 1 | k__Fungi(100);p__Ascomycota(100);c__Saccharomycetes(100);o__Saccharomycetales(100);f__Metschnikowiaceae(100);g__Metschnikowia(100);s__Metschnikowia_pulcherrima(100); | Metschnikowia | 13 | Saprotroph | \|Undefined Saprotroph\| | Yeast | NULL | Probable | As nectar-tap_saprotroph (Põlme et al. 2020); Decay sugar-rich substrates (Põlme et al. 2020) | Põlme S, et al. 2020. Fungal Diversity 105:1-16 ((https://doi.org/10.1007/s13225-020-00466-2)) |
| Otu56909 | 1 | k__Fungi(100);p__Ascomycota(100);c__Saccharomycetes(100);o__Saccharomycetales(100);f__Metschnikowiaceae(100);g__Metschnikowia(100);s__Metschnikowia_pulcherrima(100); | Metschnikowia | 13 | Saprotroph | \|Undefined Saprotroph\| | Yeast | NULL | Probable | As nectar-tap_saprotroph (Põlme et al. 2020); Decay sugar-rich substrates (Põlme et al. 2020) | Põlme S, et al. 2020. Fungal Diversity 105:1-16 ((https://doi.org/10.1007/s13225-020-00466-2)) |
| Otu56910 | 1 | k__Fungi(100);p__Ascomycota(100);c__Saccharomycetes(100);o__Saccharomycetales(100);f__Metschnikowiaceae(100);g__Metschnikowia(100);s__Metschnikowia_pulcherrima(100); | Metschnikowia | 13 | Saprotroph | \|Undefined Saprotroph\| | Yeast | NULL | Probable | As nectar-tap_saprotroph (Põlme et al. 2020); Decay sugar-rich substrates (Põlme et al. 2020) | Põlme S, et al. 2020. Fungal Diversity 105:1-16 ((https://doi.org/10.1007/s13225-020-00466-2)) |
| Otu56911 | 1 | k__Fungi(100);p__Ascomycota(100);c__Saccharomycetes(100);o__Saccharomycetales(100);f__Metschnikowiaceae(100);g__Metschnikowia(100);s__Metschnikowia_pulcherrima(100); | Metschnikowia | 13 | Saprotroph | \|Undefined Saprotroph\| | Yeast | NULL | Probable | As nectar-tap_saprotroph (Põlme et al. 2020); Decay sugar-rich substrates (Põlme et al. 2020) | Põlme S, et al. 2020. Fungal Diversity 105:1-16 ((https://doi.org/10.1007/s13225-020-00466-2)) |
| Otu56912 | 1 | k__Fungi(100);p__Ascomycota(100);c__Saccharomycetes(100);o__Saccharomycetales(100);f__Metschnikowiaceae(100);g__Metschnikowia(100);s__Metschnikowia_pulcherrima(100); | Metschnikowia | 13 | Saprotroph | \|Undefined Saprotroph\| | Yeast | NULL | Probable | As nectar-tap_saprotroph (Põlme et al. 2020); Decay sugar-rich substrates (Põlme et al. 2020) | Põlme S, et al. 2020. Fungal Diversity 105:1-16 ((https://doi.org/10.1007/s13225-020-00466-2)) |
| Otu56913 | 1 | k__Fungi(100);p__Ascomycota(100);c__Saccharomycetes(100);o__Saccharomycetales(100);f__Metschnikowiaceae(100);g__Metschnikowia(100);g__Metschnikowia_unclassified(100); | Metschnikowia | 13 | Saprotroph | \|Undefined Saprotroph\| | Yeast | NULL | Probable | As nectar-tap_saprotroph (Põlme et al. 2020); Decay sugar-rich substrates (Põlme et al. 2020) | Põlme S, et al. 2020. Fungal Diversity 105:1-16 ((https://doi.org/10.1007/s13225-020-00466-2)) |
| Otu56914 | 1 | k__Fungi(100);p__Ascomycota(100);c__Saccharomycetes(100);o__Saccharomycetales(100);f__Metschnikowiaceae(100);g__Metschnikowia(100);s__Metschnikowia_pulcherrima(100); | Metschnikowia | 13 | Saprotroph | \|Undefined Saprotroph\| | Yeast | NULL | Probable | As nectar-tap_saprotroph (Põlme et al. 2020); Decay sugar-rich substrates (Põlme et al. 2020) | Põlme S, et al. 2020. Fungal Diversity 105:1-16 ((https://doi.org/10.1007/s13225-020-00466-2)) |
| Otu56918 | 1 | k__Fungi(100);p__Ascomycota(100);c__Saccharomycetes(100);o__Saccharomycetales(100);f__Metschnikowiaceae(100);g__Metschnikowia(100);s__Metschnikowia_pulcherrima(100); | Metschnikowia | 13 | Saprotroph | \|Undefined Saprotroph\| | Yeast | NULL | Probable | As nectar-tap_saprotroph (Põlme et al. 2020); Decay sugar-rich substrates (Põlme et al. 2020) | Põlme S, et al. 2020. Fungal Diversity 105:1-16 ((https://doi.org/10.1007/s13225-020-00466-2)) |
| Otu56919 | 1 | k__Fungi(100);p__Ascomycota(100);c__Saccharomycetes(100);o__Saccharomycetales(100);f__Metschnikowiaceae(100);g__Metschnikowia(100);s__Metschnikowia_pulcherrima(100); | Metschnikowia | 13 | Saprotroph | \|Undefined Saprotroph\| | Yeast | NULL | Probable | As nectar-tap_saprotroph (Põlme et al. 2020); Decay sugar-rich substrates (Põlme et al. 2020) | Põlme S, et al. 2020. Fungal Diversity 105:1-16 ((https://doi.org/10.1007/s13225-020-00466-2)) |
| Otu56920 | 1 | k__Fungi(100);p__Ascomycota(100);c__Saccharomycetes(100);o__Saccharomycetales(100);f__Metschnikowiaceae(100);g__Metschnikowia(100);s__Metschnikowia_pulcherrima(100); | Metschnikowia | 13 | Saprotroph | \|Undefined Saprotroph\| | Yeast | NULL | Probable | As nectar-tap_saprotroph (Põlme et al. 2020); Decay sugar-rich substrates (Põlme et al. 2020) | Põlme S, et al. 2020. Fungal Diversity 105:1-16 ((https://doi.org/10.1007/s13225-020-00466-2)) |
| Otu56921 | 1 | k__Fungi(100);p__Ascomycota(100);c__Saccharomycetes(100);o__Saccharomycetales(100);f__Metschnikowiaceae(100);g__Metschnikowia(100);s__Metschnikowia_pulcherrima(100); | Metschnikowia | 13 | Saprotroph | \|Undefined Saprotroph\| | Yeast | NULL | Probable | As nectar-tap_saprotroph (Põlme et al. 2020); Decay sugar-rich substrates (Põlme et al. 2020) | Põlme S, et al. 2020. Fungal Diversity 105:1-16 ((https://doi.org/10.1007/s13225-020-00466-2)) |
| Otu56924 | 1 | k__Fungi(100);p__Ascomycota(100);c__Saccharomycetes(100);o__Saccharomycetales(100);f__Metschnikowiaceae(100);g__Metschnikowia(100);s__Metschnikowia_pulcherrima(100); | Metschnikowia | 13 | Saprotroph | \|Undefined Saprotroph\| | Yeast | NULL | Probable | As nectar-tap_saprotroph (Põlme et al. 2020); Decay sugar-rich substrates (Põlme et al. 2020) | Põlme S, et al. 2020. Fungal Diversity 105:1-16 ((https://doi.org/10.1007/s13225-020-00466-2)) |
| Otu56927 | 1 | k__Fungi(100);p__Ascomycota(100);c__Saccharomycetes(100);o__Saccharomycetales(100);f__Metschnikowiaceae(100);g__Metschnikowia(100);g__Metschnikowia_unclassified(100); | Metschnikowia | 13 | Saprotroph | \|Undefined Saprotroph\| | Yeast | NULL | Probable | As nectar-tap_saprotroph (Põlme et al. 2020); Decay sugar-rich substrates (Põlme et al. 2020) | Põlme S, et al. 2020. Fungal Diversity 105:1-16 ((https://doi.org/10.1007/s13225-020-00466-2)) |
| Otu56928 | 1 | k__Fungi(100);p__Ascomycota(100);c__Saccharomycetes(100);o__Saccharomycetales(100);f__Metschnikowiaceae(100);g__Metschnikowia(100);s__Metschnikowia_pulcherrima(100); | Metschnikowia | 13 | Saprotroph | \|Undefined Saprotroph\| | Yeast | NULL | Probable | As nectar-tap_saprotroph (Põlme et al. 2020); Decay sugar-rich substrates (Põlme et al. 2020) | Põlme S, et al. 2020. Fungal Diversity 105:1-16 ((https://doi.org/10.1007/s13225-020-00466-2)) |
| Otu56929 | 1 | k__Fungi(100);p__Ascomycota(100);c__Saccharomycetes(100);o__Saccharomycetales(100);f__Metschnikowiaceae(100);g__Metschnikowia(100);s__Metschnikowia_pulcherrima(100); | Metschnikowia | 13 | Saprotroph | \|Undefined Saprotroph\| | Yeast | NULL | Probable | As nectar-tap_saprotroph (Põlme et al. 2020); Decay sugar-rich substrates (Põlme et al. 2020) | Põlme S, et al. 2020. Fungal Diversity 105:1-16 ((https://doi.org/10.1007/s13225-020-00466-2)) |
| Otu56931 | 1 | k__Fungi(100);p__Ascomycota(100);c__Saccharomycetes(100);o__Saccharomycetales(100);f__Metschnikowiaceae(100);g__Metschnikowia(100);s__Metschnikowia_pulcherrima(100); | Metschnikowia | 13 | Saprotroph | \|Undefined Saprotroph\| | Yeast | NULL | Probable | As nectar-tap_saprotroph (Põlme et al. 2020); Decay sugar-rich substrates (Põlme et al. 2020) | Põlme S, et al. 2020. Fungal Diversity 105:1-16 ((https://doi.org/10.1007/s13225-020-00466-2)) |
| Otu56932 | 1 | k__Fungi(100);p__Ascomycota(100);c__Saccharomycetes(100);o__Saccharomycetales(100);f__Metschnikowiaceae(100);g__Metschnikowia(100);s__Metschnikowia_pulcherrima(100); | Metschnikowia | 13 | Saprotroph | \|Undefined Saprotroph\| | Yeast | NULL | Probable | As nectar-tap_saprotroph (Põlme et al. 2020); Decay sugar-rich substrates (Põlme et al. 2020) | Põlme S, et al. 2020. Fungal Diversity 105:1-16 ((https://doi.org/10.1007/s13225-020-00466-2)) |
| Otu56935 | 1 | k__Fungi(100);p__Ascomycota(100);c__Saccharomycetes(100);o__Saccharomycetales(100);f__Metschnikowiaceae(100);g__Metschnikowia(100);s__Metschnikowia_pulcherrima(100); | Metschnikowia | 13 | Saprotroph | \|Undefined Saprotroph\| | Yeast | NULL | Probable | As nectar-tap_saprotroph (Põlme et al. 2020); Decay sugar-rich substrates (Põlme et al. 2020) | Põlme S, et al. 2020. Fungal Diversity 105:1-16 ((https://doi.org/10.1007/s13225-020-00466-2)) |
| Otu56940 | 1 | k__Fungi(100);p__Ascomycota(100);c__Saccharomycetes(100);o__Saccharomycetales(100);f__Metschnikowiaceae(100);g__Metschnikowia(100);s__Metschnikowia_pulcherrima(100); | Metschnikowia | 13 | Saprotroph | \|Undefined Saprotroph\| | Yeast | NULL | Probable | As nectar-tap_saprotroph (Põlme et al. 2020); Decay sugar-rich substrates (Põlme et al. 2020) | Põlme S, et al. 2020. Fungal Diversity 105:1-16 ((https://doi.org/10.1007/s13225-020-00466-2)) |
| Otu56942 | 1 | k__Fungi(100);p__Ascomycota(100);c__Saccharomycetes(100);o__Saccharomycetales(100);f__Metschnikowiaceae(100);g__Metschnikowia(100);s__Metschnikowia_pulcherrima(100); | Metschnikowia | 13 | Saprotroph | \|Undefined Saprotroph\| | Yeast | NULL | Probable | As nectar-tap_saprotroph (Põlme et al. 2020); Decay sugar-rich substrates (Põlme et al. 2020) | Põlme S, et al. 2020. Fungal Diversity 105:1-16 ((https://doi.org/10.1007/s13225-020-00466-2)) |
| Otu56945 | 1 | k__Fungi(100);p__Ascomycota(100);c__Saccharomycetes(100);o__Saccharomycetales(100);f__Metschnikowiaceae(100);g__Metschnikowia(100);s__Metschnikowia_pulcherrima(100); | Metschnikowia | 13 | Saprotroph | \|Undefined Saprotroph\| | Yeast | NULL | Probable | As nectar-tap_saprotroph (Põlme et al. 2020); Decay sugar-rich substrates (Põlme et al. 2020) | Põlme S, et al. 2020. Fungal Diversity 105:1-16 ((https://doi.org/10.1007/s13225-020-00466-2)) |
| Otu56946 | 1 | k__Fungi(100);p__Ascomycota(100);c__Saccharomycetes(100);o__Saccharomycetales(100);f__Metschnikowiaceae(100);g__Metschnikowia(100);s__Metschnikowia_pulcherrima(100); | Metschnikowia | 13 | Saprotroph | \|Undefined Saprotroph\| | Yeast | NULL | Probable | As nectar-tap_saprotroph (Põlme et al. 2020); Decay sugar-rich substrates (Põlme et al. 2020) | Põlme S, et al. 2020. Fungal Diversity 105:1-16 ((https://doi.org/10.1007/s13225-020-00466-2)) |
| Otu56948 | 1 | k__Fungi(100);p__Ascomycota(100);c__Saccharomycetes(100);o__Saccharomycetales(100);f__Metschnikowiaceae(100);g__Metschnikowia(100);s__Metschnikowia_pulcherrima(100); | Metschnikowia | 13 | Saprotroph | \|Undefined Saprotroph\| | Yeast | NULL | Probable | As nectar-tap_saprotroph (Põlme et al. 2020); Decay sugar-rich substrates (Põlme et al. 2020) | Põlme S, et al. 2020. Fungal Diversity 105:1-16 ((https://doi.org/10.1007/s13225-020-00466-2)) |
| Otu56949 | 1 | k__Fungi(100);p__Ascomycota(100);c__Saccharomycetes(100);o__Saccharomycetales(100);f__Metschnikowiaceae(100);g__Metschnikowia(100);s__Metschnikowia_pulcherrima(100); | Metschnikowia | 13 | Saprotroph | \|Undefined Saprotroph\| | Yeast | NULL | Probable | As nectar-tap_saprotroph (Põlme et al. 2020); Decay sugar-rich substrates (Põlme et al. 2020) | Põlme S, et al. 2020. Fungal Diversity 105:1-16 ((https://doi.org/10.1007/s13225-020-00466-2)) |
| Otu56950 | 1 | k__Fungi(100);p__Ascomycota(100);c__Saccharomycetes(100);o__Saccharomycetales(100);f__Metschnikowiaceae(100);g__Metschnikowia(100);s__Metschnikowia_pulcherrima(100); | Metschnikowia | 13 | Saprotroph | \|Undefined Saprotroph\| | Yeast | NULL | Probable | As nectar-tap_saprotroph (Põlme et al. 2020); Decay sugar-rich substrates (Põlme et al. 2020) | Põlme S, et al. 2020. Fungal Diversity 105:1-16 ((https://doi.org/10.1007/s13225-020-00466-2)) |
| Otu56951 | 1 | k__Fungi(100);p__Ascomycota(100);c__Saccharomycetes(100);o__Saccharomycetales(100);f__Metschnikowiaceae(100);g__Metschnikowia(100);s__Metschnikowia_pulcherrima(100); | Metschnikowia | 13 | Saprotroph | \|Undefined Saprotroph\| | Yeast | NULL | Probable | As nectar-tap_saprotroph (Põlme et al. 2020); Decay sugar-rich substrates (Põlme et al. 2020) | Põlme S, et al. 2020. Fungal Diversity 105:1-16 ((https://doi.org/10.1007/s13225-020-00466-2)) |
| Otu56952 | 1 | k__Fungi(100);p__Ascomycota(100);c__Saccharomycetes(100);o__Saccharomycetales(100);f__Metschnikowiaceae(100);g__Metschnikowia(100);s__Metschnikowia_pulcherrima(100); | Metschnikowia | 13 | Saprotroph | \|Undefined Saprotroph\| | Yeast | NULL | Probable | As nectar-tap_saprotroph (Põlme et al. 2020); Decay sugar-rich substrates (Põlme et al. 2020) | Põlme S, et al. 2020. Fungal Diversity 105:1-16 ((https://doi.org/10.1007/s13225-020-00466-2)) |
| Otu56954 | 1 | k__Fungi(100);p__Ascomycota(100);c__Saccharomycetes(100);o__Saccharomycetales(100);f__Metschnikowiaceae(100);g__Metschnikowia(100);s__Metschnikowia_pulcherrima(100); | Metschnikowia | 13 | Saprotroph | \|Undefined Saprotroph\| | Yeast | NULL | Probable | As nectar-tap_saprotroph (Põlme et al. 2020); Decay sugar-rich substrates (Põlme et al. 2020) | Põlme S, et al. 2020. Fungal Diversity 105:1-16 ((https://doi.org/10.1007/s13225-020-00466-2)) |
| Otu56958 | 1 | k__Fungi(100);p__Ascomycota(100);c__Saccharomycetes(100);o__Saccharomycetales(100);f__Metschnikowiaceae(100);g__Metschnikowia(100);s__Metschnikowia_pulcherrima(100); | Metschnikowia | 13 | Saprotroph | \|Undefined Saprotroph\| | Yeast | NULL | Probable | As nectar-tap_saprotroph (Põlme et al. 2020); Decay sugar-rich substrates (Põlme et al. 2020) | Põlme S, et al. 2020. Fungal Diversity 105:1-16 ((https://doi.org/10.1007/s13225-020-00466-2)) |
| Otu56959 | 1 | k__Fungi(100);p__Ascomycota(100);c__Saccharomycetes(100);o__Saccharomycetales(100);f__Metschnikowiaceae(100);g__Metschnikowia(100);s__Metschnikowia_pulcherrima(100); | Metschnikowia | 13 | Saprotroph | \|Undefined Saprotroph\| | Yeast | NULL | Probable | As nectar-tap_saprotroph (Põlme et al. 2020); Decay sugar-rich substrates (Põlme et al. 2020) | Põlme S, et al. 2020. Fungal Diversity 105:1-16 ((https://doi.org/10.1007/s13225-020-00466-2)) |
| Otu56960 | 1 | k__Fungi(100);p__Ascomycota(100);c__Saccharomycetes(100);o__Saccharomycetales(100);f__Metschnikowiaceae(100);g__Metschnikowia(100);s__Metschnikowia_pulcherrima(100); | Metschnikowia | 13 | Saprotroph | \|Undefined Saprotroph\| | Yeast | NULL | Probable | As nectar-tap_saprotroph (Põlme et al. 2020); Decay sugar-rich substrates (Põlme et al. 2020) | Põlme S, et al. 2020. Fungal Diversity 105:1-16 ((https://doi.org/10.1007/s13225-020-00466-2)) |
| Otu56962 | 1 | k__Fungi(100);p__Ascomycota(100);c__Saccharomycetes(100);o__Saccharomycetales(100);f__Metschnikowiaceae(100);g__Metschnikowia(100);s__Metschnikowia_pulcherrima(100); | Metschnikowia | 13 | Saprotroph | \|Undefined Saprotroph\| | Yeast | NULL | Probable | As nectar-tap_saprotroph (Põlme et al. 2020); Decay sugar-rich substrates (Põlme et al. 2020) | Põlme S, et al. 2020. Fungal Diversity 105:1-16 ((https://doi.org/10.1007/s13225-020-00466-2)) |
| Otu56963 | 1 | k__Fungi(100);p__Ascomycota(100);c__Saccharomycetes(100);o__Saccharomycetales(100);f__Metschnikowiaceae(100);g__Metschnikowia(100);s__Metschnikowia_pulcherrima(100); | Metschnikowia | 13 | Saprotroph | \|Undefined Saprotroph\| | Yeast | NULL | Probable | As nectar-tap_saprotroph (Põlme et al. 2020); Decay sugar-rich substrates (Põlme et al. 2020) | Põlme S, et al. 2020. Fungal Diversity 105:1-16 ((https://doi.org/10.1007/s13225-020-00466-2)) |
| Otu56964 | 1 | k__Fungi(100);p__Ascomycota(100);c__Saccharomycetes(100);o__Saccharomycetales(100);f__Metschnikowiaceae(100);g__Metschnikowia(100);s__Metschnikowia_pulcherrima(100); | Metschnikowia | 13 | Saprotroph | \|Undefined Saprotroph\| | Yeast | NULL | Probable | As nectar-tap_saprotroph (Põlme et al. 2020); Decay sugar-rich substrates (Põlme et al. 2020) | Põlme S, et al. 2020. Fungal Diversity 105:1-16 ((https://doi.org/10.1007/s13225-020-00466-2)) |
| Otu56965 | 1 | k__Fungi(100);p__Ascomycota(100);c__Saccharomycetes(100);o__Saccharomycetales(100);f__Metschnikowiaceae(100);g__Metschnikowia(100);s__Metschnikowia_pulcherrima(100); | Metschnikowia | 13 | Saprotroph | \|Undefined Saprotroph\| | Yeast | NULL | Probable | As nectar-tap_saprotroph (Põlme et al. 2020); Decay sugar-rich substrates (Põlme et al. 2020) | Põlme S, et al. 2020. Fungal Diversity 105:1-16 ((https://doi.org/10.1007/s13225-020-00466-2)) |
| Otu56966 | 1 | k__Fungi(100);p__Ascomycota(100);c__Saccharomycetes(100);o__Saccharomycetales(100);f__Metschnikowiaceae(100);g__Metschnikowia(100);g__Metschnikowia_unclassified(100); | Metschnikowia | 13 | Saprotroph | \|Undefined Saprotroph\| | Yeast | NULL | Probable | As nectar-tap_saprotroph (Põlme et al. 2020); Decay sugar-rich substrates (Põlme et al. 2020) | Põlme S, et al. 2020. Fungal Diversity 105:1-16 ((https://doi.org/10.1007/s13225-020-00466-2)) |
| Otu56967 | 1 | k__Fungi(100);p__Ascomycota(100);c__Saccharomycetes(100);o__Saccharomycetales(100);f__Metschnikowiaceae(100);g__Metschnikowia(100);s__Metschnikowia_pulcherrima(100); | Metschnikowia | 13 | Saprotroph | \|Undefined Saprotroph\| | Yeast | NULL | Probable | As nectar-tap_saprotroph (Põlme et al. 2020); Decay sugar-rich substrates (Põlme et al. 2020) | Põlme S, et al. 2020. Fungal Diversity 105:1-16 ((https://doi.org/10.1007/s13225-020-00466-2)) |
| Otu56968 | 1 | k__Fungi(100);p__Ascomycota(100);c__Saccharomycetes(100);o__Saccharomycetales(100);f__Metschnikowiaceae(100);g__Metschnikowia(100);s__Metschnikowia_pulcherrima(100); | Metschnikowia | 13 | Saprotroph | \|Undefined Saprotroph\| | Yeast | NULL | Probable | As nectar-tap_saprotroph (Põlme et al. 2020); Decay sugar-rich substrates (Põlme et al. 2020) | Põlme S, et al. 2020. Fungal Diversity 105:1-16 ((https://doi.org/10.1007/s13225-020-00466-2)) |
| Otu56969 | 1 | k__Fungi(100);p__Ascomycota(100);c__Saccharomycetes(100);o__Saccharomycetales(100);f__Metschnikowiaceae(100);g__Metschnikowia(100);s__Metschnikowia_pulcherrima(100); | Metschnikowia | 13 | Saprotroph | \|Undefined Saprotroph\| | Yeast | NULL | Probable | As nectar-tap_saprotroph (Põlme et al. 2020); Decay sugar-rich substrates (Põlme et al. 2020) | Põlme S, et al. 2020. Fungal Diversity 105:1-16 ((https://doi.org/10.1007/s13225-020-00466-2)) |
| Otu56970 | 1 | k__Fungi(100);p__Ascomycota(100);c__Saccharomycetes(100);o__Saccharomycetales(100);f__Metschnikowiaceae(100);g__Metschnikowia(100);s__Metschnikowia_pulcherrima(100); | Metschnikowia | 13 | Saprotroph | \|Undefined Saprotroph\| | Yeast | NULL | Probable | As nectar-tap_saprotroph (Põlme et al. 2020); Decay sugar-rich substrates (Põlme et al. 2020) | Põlme S, et al. 2020. Fungal Diversity 105:1-16 ((https://doi.org/10.1007/s13225-020-00466-2)) |
| Otu56971 | 1 | k__Fungi(100);p__Ascomycota(100);c__Saccharomycetes(100);o__Saccharomycetales(100);f__Metschnikowiaceae(100);g__Metschnikowia(100);s__Metschnikowia_pulcherrima(100); | Metschnikowia | 13 | Saprotroph | \|Undefined Saprotroph\| | Yeast | NULL | Probable | As nectar-tap_saprotroph (Põlme et al. 2020); Decay sugar-rich substrates (Põlme et al. 2020) | Põlme S, et al. 2020. Fungal Diversity 105:1-16 ((https://doi.org/10.1007/s13225-020-00466-2)) |
| Otu56972 | 1 | k__Fungi(100);p__Ascomycota(100);c__Saccharomycetes(100);o__Saccharomycetales(100);f__Metschnikowiaceae(100);g__Metschnikowia(100);s__Metschnikowia_pulcherrima(100); | Metschnikowia | 13 | Saprotroph | \|Undefined Saprotroph\| | Yeast | NULL | Probable | As nectar-tap_saprotroph (Põlme et al. 2020); Decay sugar-rich substrates (Põlme et al. 2020) | Põlme S, et al. 2020. Fungal Diversity 105:1-16 ((https://doi.org/10.1007/s13225-020-00466-2)) |
| Otu56973 | 1 | k__Fungi(100);p__Ascomycota(100);c__Saccharomycetes(100);o__Saccharomycetales(100);f__Metschnikowiaceae(100);g__Metschnikowia(100);s__Metschnikowia_pulcherrima(100); | Metschnikowia | 13 | Saprotroph | \|Undefined Saprotroph\| | Yeast | NULL | Probable | As nectar-tap_saprotroph (Põlme et al. 2020); Decay sugar-rich substrates (Põlme et al. 2020) | Põlme S, et al. 2020. Fungal Diversity 105:1-16 ((https://doi.org/10.1007/s13225-020-00466-2)) |
| Otu56974 | 1 | k__Fungi(100);p__Ascomycota(100);c__Saccharomycetes(100);o__Saccharomycetales(100);f__Metschnikowiaceae(100);g__Metschnikowia(100);s__Metschnikowia_pulcherrima(100); | Metschnikowia | 13 | Saprotroph | \|Undefined Saprotroph\| | Yeast | NULL | Probable | As nectar-tap_saprotroph (Põlme et al. 2020); Decay sugar-rich substrates (Põlme et al. 2020) | Põlme S, et al. 2020. Fungal Diversity 105:1-16 ((https://doi.org/10.1007/s13225-020-00466-2)) |
| Otu56975 | 1 | k__Fungi(100);p__Ascomycota(100);c__Saccharomycetes(100);o__Saccharomycetales(100);f__Metschnikowiaceae(100);g__Metschnikowia(100);s__Metschnikowia_pulcherrima(100); | Metschnikowia | 13 | Saprotroph | \|Undefined Saprotroph\| | Yeast | NULL | Probable | As nectar-tap_saprotroph (Põlme et al. 2020); Decay sugar-rich substrates (Põlme et al. 2020) | Põlme S, et al. 2020. Fungal Diversity 105:1-16 ((https://doi.org/10.1007/s13225-020-00466-2)) |
| Otu56976 | 1 | k__Fungi(100);p__Ascomycota(100);c__Saccharomycetes(100);o__Saccharomycetales(100);f__Metschnikowiaceae(100);g__Metschnikowia(100);s__Metschnikowia_pulcherrima(100); | Metschnikowia | 13 | Saprotroph | \|Undefined Saprotroph\| | Yeast | NULL | Probable | As nectar-tap_saprotroph (Põlme et al. 2020); Decay sugar-rich substrates (Põlme et al. 2020) | Põlme S, et al. 2020. Fungal Diversity 105:1-16 ((https://doi.org/10.1007/s13225-020-00466-2)) |
| Otu56977 | 1 | k__Fungi(100);p__Ascomycota(100);c__Saccharomycetes(100);o__Saccharomycetales(100);f__Metschnikowiaceae(100);g__Metschnikowia(100);s__Metschnikowia_pulcherrima(100); | Metschnikowia | 13 | Saprotroph | \|Undefined Saprotroph\| | Yeast | NULL | Probable | As nectar-tap_saprotroph (Põlme et al. 2020); Decay sugar-rich substrates (Põlme et al. 2020) | Põlme S, et al. 2020. Fungal Diversity 105:1-16 ((https://doi.org/10.1007/s13225-020-00466-2)) |
| Otu56978 | 1 | k__Fungi(100);p__Ascomycota(100);c__Saccharomycetes(100);o__Saccharomycetales(100);f__Metschnikowiaceae(100);g__Metschnikowia(100);g__Metschnikowia_unclassified(100); | Metschnikowia | 13 | Saprotroph | \|Undefined Saprotroph\| | Yeast | NULL | Probable | As nectar-tap_saprotroph (Põlme et al. 2020); Decay sugar-rich substrates (Põlme et al. 2020) | Põlme S, et al. 2020. Fungal Diversity 105:1-16 ((https://doi.org/10.1007/s13225-020-00466-2)) |
| Otu56979 | 1 | k__Fungi(100);p__Ascomycota(100);c__Saccharomycetes(100);o__Saccharomycetales(100);f__Metschnikowiaceae(100);g__Metschnikowia(100);s__Metschnikowia_pulcherrima(100); | Metschnikowia | 13 | Saprotroph | \|Undefined Saprotroph\| | Yeast | NULL | Probable | As nectar-tap_saprotroph (Põlme et al. 2020); Decay sugar-rich substrates (Põlme et al. 2020) | Põlme S, et al. 2020. Fungal Diversity 105:1-16 ((https://doi.org/10.1007/s13225-020-00466-2)) |
| Otu56980 | 1 | k__Fungi(100);p__Ascomycota(100);c__Saccharomycetes(100);o__Saccharomycetales(100);f__Metschnikowiaceae(100);g__Metschnikowia(100);s__Metschnikowia_pulcherrima(100); | Metschnikowia | 13 | Saprotroph | \|Undefined Saprotroph\| | Yeast | NULL | Probable | As nectar-tap_saprotroph (Põlme et al. 2020); Decay sugar-rich substrates (Põlme et al. 2020) | Põlme S, et al. 2020. Fungal Diversity 105:1-16 ((https://doi.org/10.1007/s13225-020-00466-2)) |
| Otu56981 | 1 | k__Fungi(100);p__Ascomycota(100);c__Saccharomycetes(100);o__Saccharomycetales(100);f__Metschnikowiaceae(100);g__Metschnikowia(100);s__Metschnikowia_pulcherrima(100); | Metschnikowia | 13 | Saprotroph | \|Undefined Saprotroph\| | Yeast | NULL | Probable | As nectar-tap_saprotroph (Põlme et al. 2020); Decay sugar-rich substrates (Põlme et al. 2020) | Põlme S, et al. 2020. Fungal Diversity 105:1-16 ((https://doi.org/10.1007/s13225-020-00466-2)) |
| Otu56982 | 1 | k__Fungi(100);p__Ascomycota(100);c__Saccharomycetes(100);o__Saccharomycetales(100);f__Metschnikowiaceae(100);g__Metschnikowia(100);s__Metschnikowia_pulcherrima(100); | Metschnikowia | 13 | Saprotroph | \|Undefined Saprotroph\| | Yeast | NULL | Probable | As nectar-tap_saprotroph (Põlme et al. 2020); Decay sugar-rich substrates (Põlme et al. 2020) | Põlme S, et al. 2020. Fungal Diversity 105:1-16 ((https://doi.org/10.1007/s13225-020-00466-2)) |
| Otu56983 | 1 | k__Fungi(100);p__Ascomycota(100);c__Saccharomycetes(100);o__Saccharomycetales(100);f__Metschnikowiaceae(100);g__Metschnikowia(100);s__Metschnikowia_pulcherrima(100); | Metschnikowia | 13 | Saprotroph | \|Undefined Saprotroph\| | Yeast | NULL | Probable | As nectar-tap_saprotroph (Põlme et al. 2020); Decay sugar-rich substrates (Põlme et al. 2020) | Põlme S, et al. 2020. Fungal Diversity 105:1-16 ((https://doi.org/10.1007/s13225-020-00466-2)) |
| Otu56984 | 1 | k__Fungi(100);p__Ascomycota(100);c__Saccharomycetes(100);o__Saccharomycetales(100);f__Metschnikowiaceae(100);g__Metschnikowia(100);s__Metschnikowia_pulcherrima(100); | Metschnikowia | 13 | Saprotroph | \|Undefined Saprotroph\| | Yeast | NULL | Probable | As nectar-tap_saprotroph (Põlme et al. 2020); Decay sugar-rich substrates (Põlme et al. 2020) | Põlme S, et al. 2020. Fungal Diversity 105:1-16 ((https://doi.org/10.1007/s13225-020-00466-2)) |
| Otu56985 | 1 | k__Fungi(100);p__Ascomycota(100);c__Saccharomycetes(100);o__Saccharomycetales(100);f__Metschnikowiaceae(100);g__Metschnikowia(100);s__Metschnikowia_pulcherrima(100); | Metschnikowia | 13 | Saprotroph | \|Undefined Saprotroph\| | Yeast | NULL | Probable | As nectar-tap_saprotroph (Põlme et al. 2020); Decay sugar-rich substrates (Põlme et al. 2020) | Põlme S, et al. 2020. Fungal Diversity 105:1-16 ((https://doi.org/10.1007/s13225-020-00466-2)) |
| Otu56986 | 1 | k__Fungi(100);p__Ascomycota(100);c__Saccharomycetes(100);o__Saccharomycetales(100);f__Metschnikowiaceae(100);g__Metschnikowia(100);s__Metschnikowia_pulcherrima(100); | Metschnikowia | 13 | Saprotroph | \|Undefined Saprotroph\| | Yeast | NULL | Probable | As nectar-tap_saprotroph (Põlme et al. 2020); Decay sugar-rich substrates (Põlme et al. 2020) | Põlme S, et al. 2020. Fungal Diversity 105:1-16 ((https://doi.org/10.1007/s13225-020-00466-2)) |
| Otu56987 | 1 | k__Fungi(100);p__Ascomycota(100);c__Saccharomycetes(100);o__Saccharomycetales(100);f__Metschnikowiaceae(100);g__Metschnikowia(100);s__Metschnikowia_pulcherrima(100); | Metschnikowia | 13 | Saprotroph | \|Undefined Saprotroph\| | Yeast | NULL | Probable | As nectar-tap_saprotroph (Põlme et al. 2020); Decay sugar-rich substrates (Põlme et al. 2020) | Põlme S, et al. 2020. Fungal Diversity 105:1-16 ((https://doi.org/10.1007/s13225-020-00466-2)) |
| Otu56990 | 1 | k__Fungi(100);p__Ascomycota(100);c__Saccharomycetes(100);o__Saccharomycetales(100);f__Metschnikowiaceae(100);g__Metschnikowia(100);s__Metschnikowia_pulcherrima(100); | Metschnikowia | 13 | Saprotroph | \|Undefined Saprotroph\| | Yeast | NULL | Probable | As nectar-tap_saprotroph (Põlme et al. 2020); Decay sugar-rich substrates (Põlme et al. 2020) | Põlme S, et al. 2020. Fungal Diversity 105:1-16 ((https://doi.org/10.1007/s13225-020-00466-2)) |
| Otu56991 | 1 | k__Fungi(100);p__Ascomycota(100);c__Saccharomycetes(100);o__Saccharomycetales(100);f__Metschnikowiaceae(100);g__Metschnikowia(100);s__Metschnikowia_pulcherrima(100); | Metschnikowia | 13 | Saprotroph | \|Undefined Saprotroph\| | Yeast | NULL | Probable | As nectar-tap_saprotroph (Põlme et al. 2020); Decay sugar-rich substrates (Põlme et al. 2020) | Põlme S, et al. 2020. Fungal Diversity 105:1-16 ((https://doi.org/10.1007/s13225-020-00466-2)) |
| Otu56992 | 1 | k__Fungi(100);p__Ascomycota(100);c__Saccharomycetes(100);o__Saccharomycetales(100);f__Metschnikowiaceae(100);g__Metschnikowia(100);s__Metschnikowia_pulcherrima(100); | Metschnikowia | 13 | Saprotroph | \|Undefined Saprotroph\| | Yeast | NULL | Probable | As nectar-tap_saprotroph (Põlme et al. 2020); Decay sugar-rich substrates (Põlme et al. 2020) | Põlme S, et al. 2020. Fungal Diversity 105:1-16 ((https://doi.org/10.1007/s13225-020-00466-2)) |
| Otu56993 | 1 | k__Fungi(100);p__Ascomycota(100);c__Saccharomycetes(100);o__Saccharomycetales(100);f__Metschnikowiaceae(100);g__Metschnikowia(100);g__Metschnikowia_unclassified(100); | Metschnikowia | 13 | Saprotroph | \|Undefined Saprotroph\| | Yeast | NULL | Probable | As nectar-tap_saprotroph (Põlme et al. 2020); Decay sugar-rich substrates (Põlme et al. 2020) | Põlme S, et al. 2020. Fungal Diversity 105:1-16 ((https://doi.org/10.1007/s13225-020-00466-2)) |
| Otu56994 | 1 | k__Fungi(100);p__Ascomycota(100);c__Saccharomycetes(100);o__Saccharomycetales(100);f__Metschnikowiaceae(100);g__Metschnikowia(100);s__Metschnikowia_pulcherrima(100); | Metschnikowia | 13 | Saprotroph | \|Undefined Saprotroph\| | Yeast | NULL | Probable | As nectar-tap_saprotroph (Põlme et al. 2020); Decay sugar-rich substrates (Põlme et al. 2020) | Põlme S, et al. 2020. Fungal Diversity 105:1-16 ((https://doi.org/10.1007/s13225-020-00466-2)) |
| Otu56995 | 1 | k__Fungi(100);p__Ascomycota(100);c__Saccharomycetes(100);o__Saccharomycetales(100);f__Metschnikowiaceae(100);g__Metschnikowia(100);s__Metschnikowia_pulcherrima(100); | Metschnikowia | 13 | Saprotroph | \|Undefined Saprotroph\| | Yeast | NULL | Probable | As nectar-tap_saprotroph (Põlme et al. 2020); Decay sugar-rich substrates (Põlme et al. 2020) | Põlme S, et al. 2020. Fungal Diversity 105:1-16 ((https://doi.org/10.1007/s13225-020-00466-2)) |
| Otu56996 | 1 | k__Fungi(100);p__Ascomycota(100);c__Saccharomycetes(100);o__Saccharomycetales(100);f__Metschnikowiaceae(100);g__Metschnikowia(100);s__Metschnikowia_pulcherrima(100); | Metschnikowia | 13 | Saprotroph | \|Undefined Saprotroph\| | Yeast | NULL | Probable | As nectar-tap_saprotroph (Põlme et al. 2020); Decay sugar-rich substrates (Põlme et al. 2020) | Põlme S, et al. 2020. Fungal Diversity 105:1-16 ((https://doi.org/10.1007/s13225-020-00466-2)) |
| Otu56997 | 1 | k__Fungi(100);p__Ascomycota(100);c__Saccharomycetes(100);o__Saccharomycetales(100);f__Metschnikowiaceae(100);g__Metschnikowia(100);s__Metschnikowia_pulcherrima(100); | Metschnikowia | 13 | Saprotroph | \|Undefined Saprotroph\| | Yeast | NULL | Probable | As nectar-tap_saprotroph (Põlme et al. 2020); Decay sugar-rich substrates (Põlme et al. 2020) | Põlme S, et al. 2020. Fungal Diversity 105:1-16 ((https://doi.org/10.1007/s13225-020-00466-2)) |
| Otu56998 | 1 | k__Fungi(100);p__Ascomycota(100);c__Saccharomycetes(100);o__Saccharomycetales(100);f__Metschnikowiaceae(100);g__Metschnikowia(100);s__Metschnikowia_pulcherrima(100); | Metschnikowia | 13 | Saprotroph | \|Undefined Saprotroph\| | Yeast | NULL | Probable | As nectar-tap_saprotroph (Põlme et al. 2020); Decay sugar-rich substrates (Põlme et al. 2020) | Põlme S, et al. 2020. Fungal Diversity 105:1-16 ((https://doi.org/10.1007/s13225-020-00466-2)) |
| Otu56999 | 1 | k__Fungi(100);p__Ascomycota(100);c__Saccharomycetes(100);o__Saccharomycetales(100);f__Metschnikowiaceae(100);g__Metschnikowia(100);s__Metschnikowia_pulcherrima(100); | Metschnikowia | 13 | Saprotroph | \|Undefined Saprotroph\| | Yeast | NULL | Probable | As nectar-tap_saprotroph (Põlme et al. 2020); Decay sugar-rich substrates (Põlme et al. 2020) | Põlme S, et al. 2020. Fungal Diversity 105:1-16 ((https://doi.org/10.1007/s13225-020-00466-2)) |
| Otu57001 | 1 | k__Fungi(100);p__Ascomycota(100);c__Saccharomycetes(100);o__Saccharomycetales(100);f__Metschnikowiaceae(100);g__Metschnikowia(100);s__Metschnikowia_pulcherrima(100); | Metschnikowia | 13 | Saprotroph | \|Undefined Saprotroph\| | Yeast | NULL | Probable | As nectar-tap_saprotroph (Põlme et al. 2020); Decay sugar-rich substrates (Põlme et al. 2020) | Põlme S, et al. 2020. Fungal Diversity 105:1-16 ((https://doi.org/10.1007/s13225-020-00466-2)) |
| Otu57002 | 1 | k__Fungi(100);p__Ascomycota(100);c__Saccharomycetes(100);o__Saccharomycetales(100);f__Metschnikowiaceae(100);g__Metschnikowia(100);s__Metschnikowia_pulcherrima(100); | Metschnikowia | 13 | Saprotroph | \|Undefined Saprotroph\| | Yeast | NULL | Probable | As nectar-tap_saprotroph (Põlme et al. 2020); Decay sugar-rich substrates (Põlme et al. 2020) | Põlme S, et al. 2020. Fungal Diversity 105:1-16 ((https://doi.org/10.1007/s13225-020-00466-2)) |
| Otu57003 | 1 | k__Fungi(100);p__Ascomycota(100);c__Saccharomycetes(100);o__Saccharomycetales(100);f__Metschnikowiaceae(100);g__Metschnikowia(100);g__Metschnikowia_unclassified(100); | Metschnikowia | 13 | Saprotroph | \|Undefined Saprotroph\| | Yeast | NULL | Probable | As nectar-tap_saprotroph (Põlme et al. 2020); Decay sugar-rich substrates (Põlme et al. 2020) | Põlme S, et al. 2020. Fungal Diversity 105:1-16 ((https://doi.org/10.1007/s13225-020-00466-2)) |
| Otu57004 | 1 | k__Fungi(100);p__Ascomycota(100);c__Saccharomycetes(100);o__Saccharomycetales(100);f__Metschnikowiaceae(100);g__Metschnikowia(100);g__Metschnikowia_unclassified(100); | Metschnikowia | 13 | Saprotroph | \|Undefined Saprotroph\| | Yeast | NULL | Probable | As nectar-tap_saprotroph (Põlme et al. 2020); Decay sugar-rich substrates (Põlme et al. 2020) | Põlme S, et al. 2020. Fungal Diversity 105:1-16 ((https://doi.org/10.1007/s13225-020-00466-2)) |
| Otu57005 | 1 | k__Fungi(100);p__Ascomycota(100);c__Saccharomycetes(100);o__Saccharomycetales(100);f__Metschnikowiaceae(100);g__Metschnikowia(100);g__Metschnikowia_unclassified(100); | Metschnikowia | 13 | Saprotroph | \|Undefined Saprotroph\| | Yeast | NULL | Probable | As nectar-tap_saprotroph (Põlme et al. 2020); Decay sugar-rich substrates (Põlme et al. 2020) | Põlme S, et al. 2020. Fungal Diversity 105:1-16 ((https://doi.org/10.1007/s13225-020-00466-2)) |
| Otu57006 | 1 | k__Fungi(100);p__Ascomycota(100);c__Saccharomycetes(100);o__Saccharomycetales(100);f__Metschnikowiaceae(100);g__Metschnikowia(100);s__Metschnikowia_pulcherrima(100); | Metschnikowia | 13 | Saprotroph | \|Undefined Saprotroph\| | Yeast | NULL | Probable | As nectar-tap_saprotroph (Põlme et al. 2020); Decay sugar-rich substrates (Põlme et al. 2020) | Põlme S, et al. 2020. Fungal Diversity 105:1-16 ((https://doi.org/10.1007/s13225-020-00466-2)) |
| Otu57007 | 1 | k__Fungi(100);p__Ascomycota(100);c__Saccharomycetes(100);o__Saccharomycetales(100);f__Metschnikowiaceae(100);g__Metschnikowia(100);s__Metschnikowia_pulcherrima(100); | Metschnikowia | 13 | Saprotroph | \|Undefined Saprotroph\| | Yeast | NULL | Probable | As nectar-tap_saprotroph (Põlme et al. 2020); Decay sugar-rich substrates (Põlme et al. 2020) | Põlme S, et al. 2020. Fungal Diversity 105:1-16 ((https://doi.org/10.1007/s13225-020-00466-2)) |
| Otu57008 | 1 | k__Fungi(100);p__Ascomycota(100);c__Saccharomycetes(100);o__Saccharomycetales(100);f__Metschnikowiaceae(100);g__Metschnikowia(100);g__Metschnikowia_unclassified(100); | Metschnikowia | 13 | Saprotroph | \|Undefined Saprotroph\| | Yeast | NULL | Probable | As nectar-tap_saprotroph (Põlme et al. 2020); Decay sugar-rich substrates (Põlme et al. 2020) | Põlme S, et al. 2020. Fungal Diversity 105:1-16 ((https://doi.org/10.1007/s13225-020-00466-2)) |
| Otu57009 | 1 | k__Fungi(100);p__Ascomycota(100);c__Saccharomycetes(100);o__Saccharomycetales(100);f__Metschnikowiaceae(100);g__Metschnikowia(100);s__Metschnikowia_pulcherrima(100); | Metschnikowia | 13 | Saprotroph | \|Undefined Saprotroph\| | Yeast | NULL | Probable | As nectar-tap_saprotroph (Põlme et al. 2020); Decay sugar-rich substrates (Põlme et al. 2020) | Põlme S, et al. 2020. Fungal Diversity 105:1-16 ((https://doi.org/10.1007/s13225-020-00466-2)) |
| Otu57010 | 1 | k__Fungi(100);p__Ascomycota(100);c__Saccharomycetes(100);o__Saccharomycetales(100);f__Metschnikowiaceae(100);g__Metschnikowia(100);s__Metschnikowia_pulcherrima(100); | Metschnikowia | 13 | Saprotroph | \|Undefined Saprotroph\| | Yeast | NULL | Probable | As nectar-tap_saprotroph (Põlme et al. 2020); Decay sugar-rich substrates (Põlme et al. 2020) | Põlme S, et al. 2020. Fungal Diversity 105:1-16 ((https://doi.org/10.1007/s13225-020-00466-2)) |
| Otu57011 | 1 | k__Fungi(100);p__Ascomycota(100);c__Saccharomycetes(100);o__Saccharomycetales(100);f__Metschnikowiaceae(100);g__Metschnikowia(100);s__Metschnikowia_pulcherrima(100); | Metschnikowia | 13 | Saprotroph | \|Undefined Saprotroph\| | Yeast | NULL | Probable | As nectar-tap_saprotroph (Põlme et al. 2020); Decay sugar-rich substrates (Põlme et al. 2020) | Põlme S, et al. 2020. Fungal Diversity 105:1-16 ((https://doi.org/10.1007/s13225-020-00466-2)) |
| Otu57012 | 1 | k__Fungi(100);p__Ascomycota(100);c__Saccharomycetes(100);o__Saccharomycetales(100);f__Metschnikowiaceae(100);g__Metschnikowia(100);s__Metschnikowia_pulcherrima(100); | Metschnikowia | 13 | Saprotroph | \|Undefined Saprotroph\| | Yeast | NULL | Probable | As nectar-tap_saprotroph (Põlme et al. 2020); Decay sugar-rich substrates (Põlme et al. 2020) | Põlme S, et al. 2020. Fungal Diversity 105:1-16 ((https://doi.org/10.1007/s13225-020-00466-2)) |
| Otu57014 | 1 | k__Fungi(100);p__Ascomycota(100);c__Saccharomycetes(100);o__Saccharomycetales(100);f__Metschnikowiaceae(100);g__Metschnikowia(100);s__Metschnikowia_pulcherrima(100); | Metschnikowia | 13 | Saprotroph | \|Undefined Saprotroph\| | Yeast | NULL | Probable | As nectar-tap_saprotroph (Põlme et al. 2020); Decay sugar-rich substrates (Põlme et al. 2020) | Põlme S, et al. 2020. Fungal Diversity 105:1-16 ((https://doi.org/10.1007/s13225-020-00466-2)) |
| Otu57015 | 1 | k__Fungi(100);p__Ascomycota(100);c__Saccharomycetes(100);o__Saccharomycetales(100);f__Metschnikowiaceae(100);g__Metschnikowia(100);s__Metschnikowia_pulcherrima(100); | Metschnikowia | 13 | Saprotroph | \|Undefined Saprotroph\| | Yeast | NULL | Probable | As nectar-tap_saprotroph (Põlme et al. 2020); Decay sugar-rich substrates (Põlme et al. 2020) | Põlme S, et al. 2020. Fungal Diversity 105:1-16 ((https://doi.org/10.1007/s13225-020-00466-2)) |
| Otu57016 | 1 | k__Fungi(100);p__Ascomycota(100);c__Saccharomycetes(100);o__Saccharomycetales(100);f__Metschnikowiaceae(100);g__Metschnikowia(100);g__Metschnikowia_unclassified(100); | Metschnikowia | 13 | Saprotroph | \|Undefined Saprotroph\| | Yeast | NULL | Probable | As nectar-tap_saprotroph (Põlme et al. 2020); Decay sugar-rich substrates (Põlme et al. 2020) | Põlme S, et al. 2020. Fungal Diversity 105:1-16 ((https://doi.org/10.1007/s13225-020-00466-2)) |
| Otu57017 | 1 | k__Fungi(100);p__Ascomycota(100);c__Saccharomycetes(100);o__Saccharomycetales(100);f__Metschnikowiaceae(100);g__Metschnikowia(100);s__Metschnikowia_pulcherrima(100); | Metschnikowia | 13 | Saprotroph | \|Undefined Saprotroph\| | Yeast | NULL | Probable | As nectar-tap_saprotroph (Põlme et al. 2020); Decay sugar-rich substrates (Põlme et al. 2020) | Põlme S, et al. 2020. Fungal Diversity 105:1-16 ((https://doi.org/10.1007/s13225-020-00466-2)) |
| Otu57019 | 1 | k__Fungi(100);p__Ascomycota(100);c__Saccharomycetes(100);o__Saccharomycetales(100);f__Metschnikowiaceae(100);g__Metschnikowia(100);s__Metschnikowia_pulcherrima(100); | Metschnikowia | 13 | Saprotroph | \|Undefined Saprotroph\| | Yeast | NULL | Probable | As nectar-tap_saprotroph (Põlme et al. 2020); Decay sugar-rich substrates (Põlme et al. 2020) | Põlme S, et al. 2020. Fungal Diversity 105:1-16 ((https://doi.org/10.1007/s13225-020-00466-2)) |
| Otu57020 | 1 | k__Fungi(100);p__Ascomycota(100);c__Saccharomycetes(100);o__Saccharomycetales(100);f__Metschnikowiaceae(100);g__Metschnikowia(100);s__Metschnikowia_pulcherrima(100); | Metschnikowia | 13 | Saprotroph | \|Undefined Saprotroph\| | Yeast | NULL | Probable | As nectar-tap_saprotroph (Põlme et al. 2020); Decay sugar-rich substrates (Põlme et al. 2020) | Põlme S, et al. 2020. Fungal Diversity 105:1-16 ((https://doi.org/10.1007/s13225-020-00466-2)) |
| Otu57021 | 1 | k__Fungi(100);p__Ascomycota(100);c__Saccharomycetes(100);o__Saccharomycetales(100);f__Metschnikowiaceae(100);g__Metschnikowia(100);s__Metschnikowia_pulcherrima(100); | Metschnikowia | 13 | Saprotroph | \|Undefined Saprotroph\| | Yeast | NULL | Probable | As nectar-tap_saprotroph (Põlme et al. 2020); Decay sugar-rich substrates (Põlme et al. 2020) | Põlme S, et al. 2020. Fungal Diversity 105:1-16 ((https://doi.org/10.1007/s13225-020-00466-2)) |
| Otu57022 | 1 | k__Fungi(100);p__Ascomycota(100);c__Saccharomycetes(100);o__Saccharomycetales(100);f__Metschnikowiaceae(100);g__Metschnikowia(100);s__Metschnikowia_pulcherrima(100); | Metschnikowia | 13 | Saprotroph | \|Undefined Saprotroph\| | Yeast | NULL | Probable | As nectar-tap_saprotroph (Põlme et al. 2020); Decay sugar-rich substrates (Põlme et al. 2020) | Põlme S, et al. 2020. Fungal Diversity 105:1-16 ((https://doi.org/10.1007/s13225-020-00466-2)) |
| Otu57023 | 1 | k__Fungi(100);p__Ascomycota(100);c__Saccharomycetes(100);o__Saccharomycetales(100);f__Metschnikowiaceae(100);g__Metschnikowia(100);g__Metschnikowia_unclassified(100); | Metschnikowia | 13 | Saprotroph | \|Undefined Saprotroph\| | Yeast | NULL | Probable | As nectar-tap_saprotroph (Põlme et al. 2020); Decay sugar-rich substrates (Põlme et al. 2020) | Põlme S, et al. 2020. Fungal Diversity 105:1-16 ((https://doi.org/10.1007/s13225-020-00466-2)) |
| Otu57025 | 1 | k__Fungi(100);p__Ascomycota(100);c__Saccharomycetes(100);o__Saccharomycetales(100);f__Metschnikowiaceae(100);g__Metschnikowia(100);s__Metschnikowia_pulcherrima(100); | Metschnikowia | 13 | Saprotroph | \|Undefined Saprotroph\| | Yeast | NULL | Probable | As nectar-tap_saprotroph (Põlme et al. 2020); Decay sugar-rich substrates (Põlme et al. 2020) | Põlme S, et al. 2020. Fungal Diversity 105:1-16 ((https://doi.org/10.1007/s13225-020-00466-2)) |
| Otu57026 | 1 | k__Fungi(100);p__Ascomycota(100);c__Saccharomycetes(100);o__Saccharomycetales(100);f__Metschnikowiaceae(100);g__Metschnikowia(100);s__Metschnikowia_pulcherrima(100); | Metschnikowia | 13 | Saprotroph | \|Undefined Saprotroph\| | Yeast | NULL | Probable | As nectar-tap_saprotroph (Põlme et al. 2020); Decay sugar-rich substrates (Põlme et al. 2020) | Põlme S, et al. 2020. Fungal Diversity 105:1-16 ((https://doi.org/10.1007/s13225-020-00466-2)) |
| Otu57027 | 1 | k__Fungi(100);p__Ascomycota(100);c__Saccharomycetes(100);o__Saccharomycetales(100);f__Metschnikowiaceae(100);g__Metschnikowia(100);s__Metschnikowia_pulcherrima(100); | Metschnikowia | 13 | Saprotroph | \|Undefined Saprotroph\| | Yeast | NULL | Probable | As nectar-tap_saprotroph (Põlme et al. 2020); Decay sugar-rich substrates (Põlme et al. 2020) | Põlme S, et al. 2020. Fungal Diversity 105:1-16 ((https://doi.org/10.1007/s13225-020-00466-2)) |
| Otu57028 | 1 | k__Fungi(100);p__Ascomycota(100);c__Saccharomycetes(100);o__Saccharomycetales(100);f__Metschnikowiaceae(100);g__Metschnikowia(100);g__Metschnikowia_unclassified(100); | Metschnikowia | 13 | Saprotroph | \|Undefined Saprotroph\| | Yeast | NULL | Probable | As nectar-tap_saprotroph (Põlme et al. 2020); Decay sugar-rich substrates (Põlme et al. 2020) | Põlme S, et al. 2020. Fungal Diversity 105:1-16 ((https://doi.org/10.1007/s13225-020-00466-2)) |
| Otu57029 | 1 | k__Fungi(100);p__Ascomycota(100);c__Saccharomycetes(100);o__Saccharomycetales(100);f__Metschnikowiaceae(100);g__Metschnikowia(100);g__Metschnikowia_unclassified(100); | Metschnikowia | 13 | Saprotroph | \|Undefined Saprotroph\| | Yeast | NULL | Probable | As nectar-tap_saprotroph (Põlme et al. 2020); Decay sugar-rich substrates (Põlme et al. 2020) | Põlme S, et al. 2020. Fungal Diversity 105:1-16 ((https://doi.org/10.1007/s13225-020-00466-2)) |
| Otu57030 | 1 | k__Fungi(100);p__Ascomycota(100);c__Saccharomycetes(100);o__Saccharomycetales(100);f__Metschnikowiaceae(100);g__Metschnikowia(100);s__Metschnikowia_pulcherrima(100); | Metschnikowia | 13 | Saprotroph | \|Undefined Saprotroph\| | Yeast | NULL | Probable | As nectar-tap_saprotroph (Põlme et al. 2020); Decay sugar-rich substrates (Põlme et al. 2020) | Põlme S, et al. 2020. Fungal Diversity 105:1-16 ((https://doi.org/10.1007/s13225-020-00466-2)) |
| Otu57031 | 1 | k__Fungi(100);p__Ascomycota(100);c__Saccharomycetes(100);o__Saccharomycetales(100);f__Metschnikowiaceae(100);g__Metschnikowia(100);s__Metschnikowia_pulcherrima(100); | Metschnikowia | 13 | Saprotroph | \|Undefined Saprotroph\| | Yeast | NULL | Probable | As nectar-tap_saprotroph (Põlme et al. 2020); Decay sugar-rich substrates (Põlme et al. 2020) | Põlme S, et al. 2020. Fungal Diversity 105:1-16 ((https://doi.org/10.1007/s13225-020-00466-2)) |
| Otu57032 | 1 | k__Fungi(100);p__Ascomycota(100);c__Saccharomycetes(100);o__Saccharomycetales(100);f__Metschnikowiaceae(100);g__Metschnikowia(100);s__Metschnikowia_pulcherrima(100); | Metschnikowia | 13 | Saprotroph | \|Undefined Saprotroph\| | Yeast | NULL | Probable | As nectar-tap_saprotroph (Põlme et al. 2020); Decay sugar-rich substrates (Põlme et al. 2020) | Põlme S, et al. 2020. Fungal Diversity 105:1-16 ((https://doi.org/10.1007/s13225-020-00466-2)) |
| Otu57033 | 1 | k__Fungi(100);p__Ascomycota(100);c__Saccharomycetes(100);o__Saccharomycetales(100);f__Metschnikowiaceae(100);g__Metschnikowia(100);s__Metschnikowia_pulcherrima(100); | Metschnikowia | 13 | Saprotroph | \|Undefined Saprotroph\| | Yeast | NULL | Probable | As nectar-tap_saprotroph (Põlme et al. 2020); Decay sugar-rich substrates (Põlme et al. 2020) | Põlme S, et al. 2020. Fungal Diversity 105:1-16 ((https://doi.org/10.1007/s13225-020-00466-2)) |
| Otu57034 | 1 | k__Fungi(100);p__Ascomycota(100);c__Saccharomycetes(100);o__Saccharomycetales(100);f__Metschnikowiaceae(100);g__Metschnikowia(100);s__Metschnikowia_pulcherrima(100); | Metschnikowia | 13 | Saprotroph | \|Undefined Saprotroph\| | Yeast | NULL | Probable | As nectar-tap_saprotroph (Põlme et al. 2020); Decay sugar-rich substrates (Põlme et al. 2020) | Põlme S, et al. 2020. Fungal Diversity 105:1-16 ((https://doi.org/10.1007/s13225-020-00466-2)) |
| Otu57035 | 1 | k__Fungi(100);p__Ascomycota(100);c__Saccharomycetes(100);o__Saccharomycetales(100);f__Metschnikowiaceae(100);g__Metschnikowia(100);s__Metschnikowia_pulcherrima(100); | Metschnikowia | 13 | Saprotroph | \|Undefined Saprotroph\| | Yeast | NULL | Probable | As nectar-tap_saprotroph (Põlme et al. 2020); Decay sugar-rich substrates (Põlme et al. 2020) | Põlme S, et al. 2020. Fungal Diversity 105:1-16 ((https://doi.org/10.1007/s13225-020-00466-2)) |
| Otu57037 | 1 | k__Fungi(100);p__Ascomycota(100);c__Saccharomycetes(100);o__Saccharomycetales(100);f__Metschnikowiaceae(100);g__Metschnikowia(100);s__Metschnikowia_pulcherrima(100); | Metschnikowia | 13 | Saprotroph | \|Undefined Saprotroph\| | Yeast | NULL | Probable | As nectar-tap_saprotroph (Põlme et al. 2020); Decay sugar-rich substrates (Põlme et al. 2020) | Põlme S, et al. 2020. Fungal Diversity 105:1-16 ((https://doi.org/10.1007/s13225-020-00466-2)) |
| Otu57038 | 1 | k__Fungi(100);p__Ascomycota(100);c__Saccharomycetes(100);o__Saccharomycetales(100);f__Metschnikowiaceae(100);g__Metschnikowia(100);g__Metschnikowia_unclassified(100); | Metschnikowia | 13 | Saprotroph | \|Undefined Saprotroph\| | Yeast | NULL | Probable | As nectar-tap_saprotroph (Põlme et al. 2020); Decay sugar-rich substrates (Põlme et al. 2020) | Põlme S, et al. 2020. Fungal Diversity 105:1-16 ((https://doi.org/10.1007/s13225-020-00466-2)) |
| Otu57040 | 1 | k__Fungi(100);p__Ascomycota(100);c__Saccharomycetes(100);o__Saccharomycetales(100);f__Metschnikowiaceae(100);g__Metschnikowia(100);s__Metschnikowia_pulcherrima(100); | Metschnikowia | 13 | Saprotroph | \|Undefined Saprotroph\| | Yeast | NULL | Probable | As nectar-tap_saprotroph (Põlme et al. 2020); Decay sugar-rich substrates (Põlme et al. 2020) | Põlme S, et al. 2020. Fungal Diversity 105:1-16 ((https://doi.org/10.1007/s13225-020-00466-2)) |
| Otu57041 | 1 | k__Fungi(100);p__Ascomycota(100);c__Saccharomycetes(100);o__Saccharomycetales(100);f__Metschnikowiaceae(100);g__Metschnikowia(100);s__Metschnikowia_pulcherrima(100); | Metschnikowia | 13 | Saprotroph | \|Undefined Saprotroph\| | Yeast | NULL | Probable | As nectar-tap_saprotroph (Põlme et al. 2020); Decay sugar-rich substrates (Põlme et al. 2020) | Põlme S, et al. 2020. Fungal Diversity 105:1-16 ((https://doi.org/10.1007/s13225-020-00466-2)) |
| Otu57042 | 1 | k__Fungi(100);p__Ascomycota(100);c__Saccharomycetes(100);o__Saccharomycetales(100);f__Metschnikowiaceae(100);g__Metschnikowia(100);s__Metschnikowia_pulcherrima(100); | Metschnikowia | 13 | Saprotroph | \|Undefined Saprotroph\| | Yeast | NULL | Probable | As nectar-tap_saprotroph (Põlme et al. 2020); Decay sugar-rich substrates (Põlme et al. 2020) | Põlme S, et al. 2020. Fungal Diversity 105:1-16 ((https://doi.org/10.1007/s13225-020-00466-2)) |
| Otu57043 | 1 | k__Fungi(100);p__Ascomycota(100);c__Saccharomycetes(100);o__Saccharomycetales(100);f__Metschnikowiaceae(100);g__Metschnikowia(100);s__Metschnikowia_pulcherrima(100); | Metschnikowia | 13 | Saprotroph | \|Undefined Saprotroph\| | Yeast | NULL | Probable | As nectar-tap_saprotroph (Põlme et al. 2020); Decay sugar-rich substrates (Põlme et al. 2020) | Põlme S, et al. 2020. Fungal Diversity 105:1-16 ((https://doi.org/10.1007/s13225-020-00466-2)) |
| Otu57044 | 1 | k__Fungi(100);p__Ascomycota(100);c__Saccharomycetes(100);o__Saccharomycetales(100);f__Metschnikowiaceae(100);g__Metschnikowia(100);s__Metschnikowia_pulcherrima(100); | Metschnikowia | 13 | Saprotroph | \|Undefined Saprotroph\| | Yeast | NULL | Probable | As nectar-tap_saprotroph (Põlme et al. 2020); Decay sugar-rich substrates (Põlme et al. 2020) | Põlme S, et al. 2020. Fungal Diversity 105:1-16 ((https://doi.org/10.1007/s13225-020-00466-2)) |
| Otu57045 | 1 | k__Fungi(100);p__Ascomycota(100);c__Saccharomycetes(100);o__Saccharomycetales(100);f__Metschnikowiaceae(100);g__Metschnikowia(100);s__Metschnikowia_pulcherrima(100); | Metschnikowia | 13 | Saprotroph | \|Undefined Saprotroph\| | Yeast | NULL | Probable | As nectar-tap_saprotroph (Põlme et al. 2020); Decay sugar-rich substrates (Põlme et al. 2020) | Põlme S, et al. 2020. Fungal Diversity 105:1-16 ((https://doi.org/10.1007/s13225-020-00466-2)) |
| Otu57046 | 1 | k__Fungi(100);p__Ascomycota(100);c__Saccharomycetes(100);o__Saccharomycetales(100);f__Metschnikowiaceae(100);g__Metschnikowia(100);s__Metschnikowia_pulcherrima(100); | Metschnikowia | 13 | Saprotroph | \|Undefined Saprotroph\| | Yeast | NULL | Probable | As nectar-tap_saprotroph (Põlme et al. 2020); Decay sugar-rich substrates (Põlme et al. 2020) | Põlme S, et al. 2020. Fungal Diversity 105:1-16 ((https://doi.org/10.1007/s13225-020-00466-2)) |
| Otu57047 | 1 | k__Fungi(100);p__Ascomycota(100);c__Saccharomycetes(100);o__Saccharomycetales(100);f__Metschnikowiaceae(100);g__Metschnikowia(100);s__Metschnikowia_pulcherrima(100); | Metschnikowia | 13 | Saprotroph | \|Undefined Saprotroph\| | Yeast | NULL | Probable | As nectar-tap_saprotroph (Põlme et al. 2020); Decay sugar-rich substrates (Põlme et al. 2020) | Põlme S, et al. 2020. Fungal Diversity 105:1-16 ((https://doi.org/10.1007/s13225-020-00466-2)) |
| Otu57048 | 1 | k__Fungi(100);p__Ascomycota(100);c__Saccharomycetes(100);o__Saccharomycetales(100);f__Metschnikowiaceae(100);g__Metschnikowia(100);s__Metschnikowia_pulcherrima(100); | Metschnikowia | 13 | Saprotroph | \|Undefined Saprotroph\| | Yeast | NULL | Probable | As nectar-tap_saprotroph (Põlme et al. 2020); Decay sugar-rich substrates (Põlme et al. 2020) | Põlme S, et al. 2020. Fungal Diversity 105:1-16 ((https://doi.org/10.1007/s13225-020-00466-2)) |
| Otu57049 | 1 | k__Fungi(100);p__Ascomycota(100);c__Saccharomycetes(100);o__Saccharomycetales(100);f__Metschnikowiaceae(100);g__Metschnikowia(100);s__Metschnikowia_pulcherrima(100); | Metschnikowia | 13 | Saprotroph | \|Undefined Saprotroph\| | Yeast | NULL | Probable | As nectar-tap_saprotroph (Põlme et al. 2020); Decay sugar-rich substrates (Põlme et al. 2020) | Põlme S, et al. 2020. Fungal Diversity 105:1-16 ((https://doi.org/10.1007/s13225-020-00466-2)) |
| Otu57050 | 1 | k__Fungi(100);p__Ascomycota(100);c__Saccharomycetes(100);o__Saccharomycetales(100);f__Metschnikowiaceae(100);g__Metschnikowia(100);s__Metschnikowia_pulcherrima(100); | Metschnikowia | 13 | Saprotroph | \|Undefined Saprotroph\| | Yeast | NULL | Probable | As nectar-tap_saprotroph (Põlme et al. 2020); Decay sugar-rich substrates (Põlme et al. 2020) | Põlme S, et al. 2020. Fungal Diversity 105:1-16 ((https://doi.org/10.1007/s13225-020-00466-2)) |
| Otu57051 | 1 | k__Fungi(100);p__Ascomycota(100);c__Saccharomycetes(100);o__Saccharomycetales(100);f__Metschnikowiaceae(100);g__Metschnikowia(100);s__Metschnikowia_pulcherrima(100); | Metschnikowia | 13 | Saprotroph | \|Undefined Saprotroph\| | Yeast | NULL | Probable | As nectar-tap_saprotroph (Põlme et al. 2020); Decay sugar-rich substrates (Põlme et al. 2020) | Põlme S, et al. 2020. Fungal Diversity 105:1-16 ((https://doi.org/10.1007/s13225-020-00466-2)) |
| Otu57052 | 1 | k__Fungi(100);p__Ascomycota(100);c__Saccharomycetes(100);o__Saccharomycetales(100);f__Metschnikowiaceae(100);g__Metschnikowia(100);s__Metschnikowia_pulcherrima(100); | Metschnikowia | 13 | Saprotroph | \|Undefined Saprotroph\| | Yeast | NULL | Probable | As nectar-tap_saprotroph (Põlme et al. 2020); Decay sugar-rich substrates (Põlme et al. 2020) | Põlme S, et al. 2020. Fungal Diversity 105:1-16 ((https://doi.org/10.1007/s13225-020-00466-2)) |
| Otu57053 | 1 | k__Fungi(100);p__Ascomycota(100);c__Saccharomycetes(100);o__Saccharomycetales(100);f__Metschnikowiaceae(100);g__Metschnikowia(100);g__Metschnikowia_unclassified(100); | Metschnikowia | 13 | Saprotroph | \|Undefined Saprotroph\| | Yeast | NULL | Probable | As nectar-tap_saprotroph (Põlme et al. 2020); Decay sugar-rich substrates (Põlme et al. 2020) | Põlme S, et al. 2020. Fungal Diversity 105:1-16 ((https://doi.org/10.1007/s13225-020-00466-2)) |
| Otu57054 | 1 | k__Fungi(100);p__Ascomycota(100);c__Saccharomycetes(100);o__Saccharomycetales(100);f__Metschnikowiaceae(100);g__Metschnikowia(100);s__Metschnikowia_pulcherrima(100); | Metschnikowia | 13 | Saprotroph | \|Undefined Saprotroph\| | Yeast | NULL | Probable | As nectar-tap_saprotroph (Põlme et al. 2020); Decay sugar-rich substrates (Põlme et al. 2020) | Põlme S, et al. 2020. Fungal Diversity 105:1-16 ((https://doi.org/10.1007/s13225-020-00466-2)) |
| Otu57055 | 1 | k__Fungi(100);p__Ascomycota(100);c__Saccharomycetes(100);o__Saccharomycetales(100);f__Metschnikowiaceae(100);g__Metschnikowia(100);s__Metschnikowia_pulcherrima(100); | Metschnikowia | 13 | Saprotroph | \|Undefined Saprotroph\| | Yeast | NULL | Probable | As nectar-tap_saprotroph (Põlme et al. 2020); Decay sugar-rich substrates (Põlme et al. 2020) | Põlme S, et al. 2020. Fungal Diversity 105:1-16 ((https://doi.org/10.1007/s13225-020-00466-2)) |
| Otu57056 | 1 | k__Fungi(100);p__Ascomycota(100);c__Saccharomycetes(100);o__Saccharomycetales(100);f__Metschnikowiaceae(100);g__Metschnikowia(100);s__Metschnikowia_pulcherrima(100); | Metschnikowia | 13 | Saprotroph | \|Undefined Saprotroph\| | Yeast | NULL | Probable | As nectar-tap_saprotroph (Põlme et al. 2020); Decay sugar-rich substrates (Põlme et al. 2020) | Põlme S, et al. 2020. Fungal Diversity 105:1-16 ((https://doi.org/10.1007/s13225-020-00466-2)) |
| Otu57057 | 1 | k__Fungi(100);p__Ascomycota(100);c__Saccharomycetes(100);o__Saccharomycetales(100);f__Metschnikowiaceae(100);g__Metschnikowia(100);s__Metschnikowia_pulcherrima(100); | Metschnikowia | 13 | Saprotroph | \|Undefined Saprotroph\| | Yeast | NULL | Probable | As nectar-tap_saprotroph (Põlme et al. 2020); Decay sugar-rich substrates (Põlme et al. 2020) | Põlme S, et al. 2020. Fungal Diversity 105:1-16 ((https://doi.org/10.1007/s13225-020-00466-2)) |
| Otu57058 | 1 | k__Fungi(100);p__Ascomycota(100);c__Saccharomycetes(100);o__Saccharomycetales(100);f__Metschnikowiaceae(100);g__Metschnikowia(100);s__Metschnikowia_pulcherrima(100); | Metschnikowia | 13 | Saprotroph | \|Undefined Saprotroph\| | Yeast | NULL | Probable | As nectar-tap_saprotroph (Põlme et al. 2020); Decay sugar-rich substrates (Põlme et al. 2020) | Põlme S, et al. 2020. Fungal Diversity 105:1-16 ((https://doi.org/10.1007/s13225-020-00466-2)) |
| Otu57059 | 1 | k__Fungi(100);p__Ascomycota(100);c__Saccharomycetes(100);o__Saccharomycetales(100);f__Metschnikowiaceae(100);g__Metschnikowia(100);g__Metschnikowia_unclassified(100); | Metschnikowia | 13 | Saprotroph | \|Undefined Saprotroph\| | Yeast | NULL | Probable | As nectar-tap_saprotroph (Põlme et al. 2020); Decay sugar-rich substrates (Põlme et al. 2020) | Põlme S, et al. 2020. Fungal Diversity 105:1-16 ((https://doi.org/10.1007/s13225-020-00466-2)) |
| Otu57060 | 1 | k__Fungi(100);p__Ascomycota(100);c__Saccharomycetes(100);o__Saccharomycetales(100);f__Metschnikowiaceae(100);g__Metschnikowia(100);s__Metschnikowia_pulcherrima(100); | Metschnikowia | 13 | Saprotroph | \|Undefined Saprotroph\| | Yeast | NULL | Probable | As nectar-tap_saprotroph (Põlme et al. 2020); Decay sugar-rich substrates (Põlme et al. 2020) | Põlme S, et al. 2020. Fungal Diversity 105:1-16 ((https://doi.org/10.1007/s13225-020-00466-2)) |
| Otu57061 | 1 | k__Fungi(100);p__Ascomycota(100);c__Saccharomycetes(100);o__Saccharomycetales(100);f__Metschnikowiaceae(100);g__Metschnikowia(100);g__Metschnikowia_unclassified(100); | Metschnikowia | 13 | Saprotroph | \|Undefined Saprotroph\| | Yeast | NULL | Probable | As nectar-tap_saprotroph (Põlme et al. 2020); Decay sugar-rich substrates (Põlme et al. 2020) | Põlme S, et al. 2020. Fungal Diversity 105:1-16 ((https://doi.org/10.1007/s13225-020-00466-2)) |
| Otu57062 | 1 | k__Fungi(100);p__Ascomycota(100);c__Saccharomycetes(100);o__Saccharomycetales(100);f__Metschnikowiaceae(100);g__Metschnikowia(100);s__Metschnikowia_pulcherrima(100); | Metschnikowia | 13 | Saprotroph | \|Undefined Saprotroph\| | Yeast | NULL | Probable | As nectar-tap_saprotroph (Põlme et al. 2020); Decay sugar-rich substrates (Põlme et al. 2020) | Põlme S, et al. 2020. Fungal Diversity 105:1-16 ((https://doi.org/10.1007/s13225-020-00466-2)) |
| Otu57064 | 1 | k__Fungi(100);p__Ascomycota(100);c__Saccharomycetes(100);o__Saccharomycetales(100);f__Metschnikowiaceae(100);g__Metschnikowia(100);s__Metschnikowia_pulcherrima(100); | Metschnikowia | 13 | Saprotroph | \|Undefined Saprotroph\| | Yeast | NULL | Probable | As nectar-tap_saprotroph (Põlme et al. 2020); Decay sugar-rich substrates (Põlme et al. 2020) | Põlme S, et al. 2020. Fungal Diversity 105:1-16 ((https://doi.org/10.1007/s13225-020-00466-2)) |
| Otu57065 | 1 | k__Fungi(100);p__Ascomycota(100);c__Saccharomycetes(100);o__Saccharomycetales(100);f__Metschnikowiaceae(100);g__Metschnikowia(100);s__Metschnikowia_pulcherrima(100); | Metschnikowia | 13 | Saprotroph | \|Undefined Saprotroph\| | Yeast | NULL | Probable | As nectar-tap_saprotroph (Põlme et al. 2020); Decay sugar-rich substrates (Põlme et al. 2020) | Põlme S, et al. 2020. Fungal Diversity 105:1-16 ((https://doi.org/10.1007/s13225-020-00466-2)) |
| Otu57066 | 1 | k__Fungi(100);p__Ascomycota(100);c__Saccharomycetes(100);o__Saccharomycetales(100);f__Metschnikowiaceae(100);g__Metschnikowia(100);s__Metschnikowia_pulcherrima(100); | Metschnikowia | 13 | Saprotroph | \|Undefined Saprotroph\| | Yeast | NULL | Probable | As nectar-tap_saprotroph (Põlme et al. 2020); Decay sugar-rich substrates (Põlme et al. 2020) | Põlme S, et al. 2020. Fungal Diversity 105:1-16 ((https://doi.org/10.1007/s13225-020-00466-2)) |
| Otu57067 | 1 | k__Fungi(100);p__Ascomycota(100);c__Saccharomycetes(100);o__Saccharomycetales(100);f__Metschnikowiaceae(100);g__Metschnikowia(100);s__Metschnikowia_pulcherrima(100); | Metschnikowia | 13 | Saprotroph | \|Undefined Saprotroph\| | Yeast | NULL | Probable | As nectar-tap_saprotroph (Põlme et al. 2020); Decay sugar-rich substrates (Põlme et al. 2020) | Põlme S, et al. 2020. Fungal Diversity 105:1-16 ((https://doi.org/10.1007/s13225-020-00466-2)) |
| Otu57068 | 1 | k__Fungi(100);p__Ascomycota(100);c__Saccharomycetes(100);o__Saccharomycetales(100);f__Metschnikowiaceae(100);g__Metschnikowia(100);s__Metschnikowia_pulcherrima(100); | Metschnikowia | 13 | Saprotroph | \|Undefined Saprotroph\| | Yeast | NULL | Probable | As nectar-tap_saprotroph (Põlme et al. 2020); Decay sugar-rich substrates (Põlme et al. 2020) | Põlme S, et al. 2020. Fungal Diversity 105:1-16 ((https://doi.org/10.1007/s13225-020-00466-2)) |
| Otu57069 | 1 | k__Fungi(100);p__Ascomycota(100);c__Saccharomycetes(100);o__Saccharomycetales(100);f__Metschnikowiaceae(100);g__Metschnikowia(100);g__Metschnikowia_unclassified(100); | Metschnikowia | 13 | Saprotroph | \|Undefined Saprotroph\| | Yeast | NULL | Probable | As nectar-tap_saprotroph (Põlme et al. 2020); Decay sugar-rich substrates (Põlme et al. 2020) | Põlme S, et al. 2020. Fungal Diversity 105:1-16 ((https://doi.org/10.1007/s13225-020-00466-2)) |
| Otu57070 | 1 | k__Fungi(100);p__Ascomycota(100);c__Saccharomycetes(100);o__Saccharomycetales(100);f__Metschnikowiaceae(100);g__Metschnikowia(100);g__Metschnikowia_unclassified(100); | Metschnikowia | 13 | Saprotroph | \|Undefined Saprotroph\| | Yeast | NULL | Probable | As nectar-tap_saprotroph (Põlme et al. 2020); Decay sugar-rich substrates (Põlme et al. 2020) | Põlme S, et al. 2020. Fungal Diversity 105:1-16 ((https://doi.org/10.1007/s13225-020-00466-2)) |
| Otu57071 | 1 | k__Fungi(100);p__Ascomycota(100);c__Saccharomycetes(100);o__Saccharomycetales(100);f__Metschnikowiaceae(100);g__Metschnikowia(100);s__Metschnikowia_pulcherrima(100); | Metschnikowia | 13 | Saprotroph | \|Undefined Saprotroph\| | Yeast | NULL | Probable | As nectar-tap_saprotroph (Põlme et al. 2020); Decay sugar-rich substrates (Põlme et al. 2020) | Põlme S, et al. 2020. Fungal Diversity 105:1-16 ((https://doi.org/10.1007/s13225-020-00466-2)) |
| Otu57072 | 1 | k__Fungi(100);p__Ascomycota(100);c__Saccharomycetes(100);o__Saccharomycetales(100);f__Metschnikowiaceae(100);g__Metschnikowia(100);s__Metschnikowia_pulcherrima(100); | Metschnikowia | 13 | Saprotroph | \|Undefined Saprotroph\| | Yeast | NULL | Probable | As nectar-tap_saprotroph (Põlme et al. 2020); Decay sugar-rich substrates (Põlme et al. 2020) | Põlme S, et al. 2020. Fungal Diversity 105:1-16 ((https://doi.org/10.1007/s13225-020-00466-2)) |
| Otu57073 | 1 | k__Fungi(100);p__Ascomycota(100);c__Saccharomycetes(100);o__Saccharomycetales(100);f__Metschnikowiaceae(100);g__Metschnikowia(100);s__Metschnikowia_pulcherrima(100); | Metschnikowia | 13 | Saprotroph | \|Undefined Saprotroph\| | Yeast | NULL | Probable | As nectar-tap_saprotroph (Põlme et al. 2020); Decay sugar-rich substrates (Põlme et al. 2020) | Põlme S, et al. 2020. Fungal Diversity 105:1-16 ((https://doi.org/10.1007/s13225-020-00466-2)) |
| Otu57074 | 1 | k__Fungi(100);p__Ascomycota(100);c__Saccharomycetes(100);o__Saccharomycetales(100);f__Metschnikowiaceae(100);g__Metschnikowia(100);s__Metschnikowia_pulcherrima(100); | Metschnikowia | 13 | Saprotroph | \|Undefined Saprotroph\| | Yeast | NULL | Probable | As nectar-tap_saprotroph (Põlme et al. 2020); Decay sugar-rich substrates (Põlme et al. 2020) | Põlme S, et al. 2020. Fungal Diversity 105:1-16 ((https://doi.org/10.1007/s13225-020-00466-2)) |
| Otu57075 | 1 | k__Fungi(100);p__Ascomycota(100);c__Saccharomycetes(100);o__Saccharomycetales(100);f__Metschnikowiaceae(100);g__Metschnikowia(100);s__Metschnikowia_pulcherrima(100); | Metschnikowia | 13 | Saprotroph | \|Undefined Saprotroph\| | Yeast | NULL | Probable | As nectar-tap_saprotroph (Põlme et al. 2020); Decay sugar-rich substrates (Põlme et al. 2020) | Põlme S, et al. 2020. Fungal Diversity 105:1-16 ((https://doi.org/10.1007/s13225-020-00466-2)) |
| Otu57076 | 1 | k__Fungi(100);p__Ascomycota(100);c__Saccharomycetes(100);o__Saccharomycetales(100);f__Metschnikowiaceae(100);g__Metschnikowia(100);g__Metschnikowia_unclassified(100); | Metschnikowia | 13 | Saprotroph | \|Undefined Saprotroph\| | Yeast | NULL | Probable | As nectar-tap_saprotroph (Põlme et al. 2020); Decay sugar-rich substrates (Põlme et al. 2020) | Põlme S, et al. 2020. Fungal Diversity 105:1-16 ((https://doi.org/10.1007/s13225-020-00466-2)) |
| Otu57077 | 1 | k__Fungi(100);p__Ascomycota(100);c__Saccharomycetes(100);o__Saccharomycetales(100);f__Metschnikowiaceae(100);g__Metschnikowia(100);s__Metschnikowia_pulcherrima(100); | Metschnikowia | 13 | Saprotroph | \|Undefined Saprotroph\| | Yeast | NULL | Probable | As nectar-tap_saprotroph (Põlme et al. 2020); Decay sugar-rich substrates (Põlme et al. 2020) | Põlme S, et al. 2020. Fungal Diversity 105:1-16 ((https://doi.org/10.1007/s13225-020-00466-2)) |
| Otu57078 | 1 | k__Fungi(100);p__Ascomycota(100);c__Saccharomycetes(100);o__Saccharomycetales(100);f__Metschnikowiaceae(100);g__Metschnikowia(100);g__Metschnikowia_unclassified(100); | Metschnikowia | 13 | Saprotroph | \|Undefined Saprotroph\| | Yeast | NULL | Probable | As nectar-tap_saprotroph (Põlme et al. 2020); Decay sugar-rich substrates (Põlme et al. 2020) | Põlme S, et al. 2020. Fungal Diversity 105:1-16 ((https://doi.org/10.1007/s13225-020-00466-2)) |
| Otu57079 | 1 | k__Fungi(100);p__Ascomycota(100);c__Saccharomycetes(100);o__Saccharomycetales(100);f__Metschnikowiaceae(100);g__Metschnikowia(100);g__Metschnikowia_unclassified(100); | Metschnikowia | 13 | Saprotroph | \|Undefined Saprotroph\| | Yeast | NULL | Probable | As nectar-tap_saprotroph (Põlme et al. 2020); Decay sugar-rich substrates (Põlme et al. 2020) | Põlme S, et al. 2020. Fungal Diversity 105:1-16 ((https://doi.org/10.1007/s13225-020-00466-2)) |
| Otu57080 | 1 | k__Fungi(100);p__Ascomycota(100);c__Saccharomycetes(100);o__Saccharomycetales(100);f__Metschnikowiaceae(100);g__Metschnikowia(100);s__Metschnikowia_pulcherrima(100); | Metschnikowia | 13 | Saprotroph | \|Undefined Saprotroph\| | Yeast | NULL | Probable | As nectar-tap_saprotroph (Põlme et al. 2020); Decay sugar-rich substrates (Põlme et al. 2020) | Põlme S, et al. 2020. Fungal Diversity 105:1-16 ((https://doi.org/10.1007/s13225-020-00466-2)) |
| Otu57081 | 1 | k__Fungi(100);p__Ascomycota(100);c__Saccharomycetes(100);o__Saccharomycetales(100);f__Metschnikowiaceae(100);g__Metschnikowia(100);g__Metschnikowia_unclassified(100); | Metschnikowia | 13 | Saprotroph | \|Undefined Saprotroph\| | Yeast | NULL | Probable | As nectar-tap_saprotroph (Põlme et al. 2020); Decay sugar-rich substrates (Põlme et al. 2020) | Põlme S, et al. 2020. Fungal Diversity 105:1-16 ((https://doi.org/10.1007/s13225-020-00466-2)) |
| Otu57082 | 1 | k__Fungi(100);p__Ascomycota(100);c__Saccharomycetes(100);o__Saccharomycetales(100);f__Metschnikowiaceae(100);g__Metschnikowia(100);s__Metschnikowia_pulcherrima(100); | Metschnikowia | 13 | Saprotroph | \|Undefined Saprotroph\| | Yeast | NULL | Probable | As nectar-tap_saprotroph (Põlme et al. 2020); Decay sugar-rich substrates (Põlme et al. 2020) | Põlme S, et al. 2020. Fungal Diversity 105:1-16 ((https://doi.org/10.1007/s13225-020-00466-2)) |
| Otu57083 | 1 | k__Fungi(100);p__Ascomycota(100);c__Saccharomycetes(100);o__Saccharomycetales(100);f__Metschnikowiaceae(100);g__Metschnikowia(100);s__Metschnikowia_pulcherrima(100); | Metschnikowia | 13 | Saprotroph | \|Undefined Saprotroph\| | Yeast | NULL | Probable | As nectar-tap_saprotroph (Põlme et al. 2020); Decay sugar-rich substrates (Põlme et al. 2020) | Põlme S, et al. 2020. Fungal Diversity 105:1-16 ((https://doi.org/10.1007/s13225-020-00466-2)) |
| Otu57084 | 1 | k__Fungi(100);p__Ascomycota(100);c__Saccharomycetes(100);o__Saccharomycetales(100);f__Metschnikowiaceae(100);g__Metschnikowia(100);s__Metschnikowia_pulcherrima(100); | Metschnikowia | 13 | Saprotroph | \|Undefined Saprotroph\| | Yeast | NULL | Probable | As nectar-tap_saprotroph (Põlme et al. 2020); Decay sugar-rich substrates (Põlme et al. 2020) | Põlme S, et al. 2020. Fungal Diversity 105:1-16 ((https://doi.org/10.1007/s13225-020-00466-2)) |
| Otu57085 | 1 | k__Fungi(100);p__Ascomycota(100);c__Saccharomycetes(100);o__Saccharomycetales(100);f__Metschnikowiaceae(100);g__Metschnikowia(100);g__Metschnikowia_unclassified(100); | Metschnikowia | 13 | Saprotroph | \|Undefined Saprotroph\| | Yeast | NULL | Probable | As nectar-tap_saprotroph (Põlme et al. 2020); Decay sugar-rich substrates (Põlme et al. 2020) | Põlme S, et al. 2020. Fungal Diversity 105:1-16 ((https://doi.org/10.1007/s13225-020-00466-2)) |
| Otu57086 | 1 | k__Fungi(100);p__Ascomycota(100);c__Saccharomycetes(100);o__Saccharomycetales(100);f__Metschnikowiaceae(100);g__Metschnikowia(100);s__Metschnikowia_pulcherrima(100); | Metschnikowia | 13 | Saprotroph | \|Undefined Saprotroph\| | Yeast | NULL | Probable | As nectar-tap_saprotroph (Põlme et al. 2020); Decay sugar-rich substrates (Põlme et al. 2020) | Põlme S, et al. 2020. Fungal Diversity 105:1-16 ((https://doi.org/10.1007/s13225-020-00466-2)) |
| Otu57087 | 1 | k__Fungi(100);p__Ascomycota(100);c__Saccharomycetes(100);o__Saccharomycetales(100);f__Metschnikowiaceae(100);g__Metschnikowia(100);s__Metschnikowia_pulcherrima(100); | Metschnikowia | 13 | Saprotroph | \|Undefined Saprotroph\| | Yeast | NULL | Probable | As nectar-tap_saprotroph (Põlme et al. 2020); Decay sugar-rich substrates (Põlme et al. 2020) | Põlme S, et al. 2020. Fungal Diversity 105:1-16 ((https://doi.org/10.1007/s13225-020-00466-2)) |
| Otu57088 | 1 | k__Fungi(100);p__Ascomycota(100);c__Saccharomycetes(100);o__Saccharomycetales(100);f__Metschnikowiaceae(100);g__Metschnikowia(100);s__Metschnikowia_pulcherrima(100); | Metschnikowia | 13 | Saprotroph | \|Undefined Saprotroph\| | Yeast | NULL | Probable | As nectar-tap_saprotroph (Põlme et al. 2020); Decay sugar-rich substrates (Põlme et al. 2020) | Põlme S, et al. 2020. Fungal Diversity 105:1-16 ((https://doi.org/10.1007/s13225-020-00466-2)) |
| Otu57089 | 1 | k__Fungi(100);p__Ascomycota(100);c__Saccharomycetes(100);o__Saccharomycetales(100);f__Metschnikowiaceae(100);g__Metschnikowia(100);s__Metschnikowia_pulcherrima(100); | Metschnikowia | 13 | Saprotroph | \|Undefined Saprotroph\| | Yeast | NULL | Probable | As nectar-tap_saprotroph (Põlme et al. 2020); Decay sugar-rich substrates (Põlme et al. 2020) | Põlme S, et al. 2020. Fungal Diversity 105:1-16 ((https://doi.org/10.1007/s13225-020-00466-2)) |
| Otu57090 | 1 | k__Fungi(100);p__Ascomycota(100);c__Saccharomycetes(100);o__Saccharomycetales(100);f__Metschnikowiaceae(100);g__Metschnikowia(100);s__Metschnikowia_pulcherrima(100); | Metschnikowia | 13 | Saprotroph | \|Undefined Saprotroph\| | Yeast | NULL | Probable | As nectar-tap_saprotroph (Põlme et al. 2020); Decay sugar-rich substrates (Põlme et al. 2020) | Põlme S, et al. 2020. Fungal Diversity 105:1-16 ((https://doi.org/10.1007/s13225-020-00466-2)) |
| Otu57091 | 1 | k__Fungi(100);p__Ascomycota(100);c__Saccharomycetes(100);o__Saccharomycetales(100);f__Metschnikowiaceae(100);g__Metschnikowia(100);s__Metschnikowia_pulcherrima(100); | Metschnikowia | 13 | Saprotroph | \|Undefined Saprotroph\| | Yeast | NULL | Probable | As nectar-tap_saprotroph (Põlme et al. 2020); Decay sugar-rich substrates (Põlme et al. 2020) | Põlme S, et al. 2020. Fungal Diversity 105:1-16 ((https://doi.org/10.1007/s13225-020-00466-2)) |
| Otu57092 | 1 | k__Fungi(100);p__Ascomycota(100);c__Saccharomycetes(100);o__Saccharomycetales(100);f__Metschnikowiaceae(100);g__Metschnikowia(100);s__Metschnikowia_pulcherrima(100); | Metschnikowia | 13 | Saprotroph | \|Undefined Saprotroph\| | Yeast | NULL | Probable | As nectar-tap_saprotroph (Põlme et al. 2020); Decay sugar-rich substrates (Põlme et al. 2020) | Põlme S, et al. 2020. Fungal Diversity 105:1-16 ((https://doi.org/10.1007/s13225-020-00466-2)) |
| Otu57093 | 1 | k__Fungi(100);p__Ascomycota(100);c__Saccharomycetes(100);o__Saccharomycetales(100);f__Metschnikowiaceae(100);g__Metschnikowia(100);s__Metschnikowia_pulcherrima(100); | Metschnikowia | 13 | Saprotroph | \|Undefined Saprotroph\| | Yeast | NULL | Probable | As nectar-tap_saprotroph (Põlme et al. 2020); Decay sugar-rich substrates (Põlme et al. 2020) | Põlme S, et al. 2020. Fungal Diversity 105:1-16 ((https://doi.org/10.1007/s13225-020-00466-2)) |
| Otu57094 | 1 | k__Fungi(100);p__Ascomycota(100);c__Saccharomycetes(100);o__Saccharomycetales(100);f__Metschnikowiaceae(100);g__Metschnikowia(100);s__Metschnikowia_pulcherrima(100); | Metschnikowia | 13 | Saprotroph | \|Undefined Saprotroph\| | Yeast | NULL | Probable | As nectar-tap_saprotroph (Põlme et al. 2020); Decay sugar-rich substrates (Põlme et al. 2020) | Põlme S, et al. 2020. Fungal Diversity 105:1-16 ((https://doi.org/10.1007/s13225-020-00466-2)) |
| Otu57095 | 1 | k__Fungi(100);p__Ascomycota(100);c__Saccharomycetes(100);o__Saccharomycetales(100);f__Metschnikowiaceae(100);g__Metschnikowia(100);s__Metschnikowia_pulcherrima(100); | Metschnikowia | 13 | Saprotroph | \|Undefined Saprotroph\| | Yeast | NULL | Probable | As nectar-tap_saprotroph (Põlme et al. 2020); Decay sugar-rich substrates (Põlme et al. 2020) | Põlme S, et al. 2020. Fungal Diversity 105:1-16 ((https://doi.org/10.1007/s13225-020-00466-2)) |
| Otu57096 | 1 | k__Fungi(100);p__Ascomycota(100);c__Saccharomycetes(100);o__Saccharomycetales(100);f__Metschnikowiaceae(100);g__Metschnikowia(100);s__Metschnikowia_pulcherrima(100); | Metschnikowia | 13 | Saprotroph | \|Undefined Saprotroph\| | Yeast | NULL | Probable | As nectar-tap_saprotroph (Põlme et al. 2020); Decay sugar-rich substrates (Põlme et al. 2020) | Põlme S, et al. 2020. Fungal Diversity 105:1-16 ((https://doi.org/10.1007/s13225-020-00466-2)) |
| Otu57097 | 1 | k__Fungi(100);p__Ascomycota(100);c__Saccharomycetes(100);o__Saccharomycetales(100);f__Metschnikowiaceae(100);g__Metschnikowia(100);s__Metschnikowia_pulcherrima(100); | Metschnikowia | 13 | Saprotroph | \|Undefined Saprotroph\| | Yeast | NULL | Probable | As nectar-tap_saprotroph (Põlme et al. 2020); Decay sugar-rich substrates (Põlme et al. 2020) | Põlme S, et al. 2020. Fungal Diversity 105:1-16 ((https://doi.org/10.1007/s13225-020-00466-2)) |
| Otu57098 | 1 | k__Fungi(100);p__Ascomycota(100);c__Saccharomycetes(100);o__Saccharomycetales(100);f__Metschnikowiaceae(100);g__Metschnikowia(100);s__Metschnikowia_pulcherrima(100); | Metschnikowia | 13 | Saprotroph | \|Undefined Saprotroph\| | Yeast | NULL | Probable | As nectar-tap_saprotroph (Põlme et al. 2020); Decay sugar-rich substrates (Põlme et al. 2020) | Põlme S, et al. 2020. Fungal Diversity 105:1-16 ((https://doi.org/10.1007/s13225-020-00466-2)) |
| Otu57099 | 1 | k__Fungi(100);p__Ascomycota(100);c__Saccharomycetes(100);o__Saccharomycetales(100);f__Metschnikowiaceae(100);g__Metschnikowia(100);s__Metschnikowia_pulcherrima(100); | Metschnikowia | 13 | Saprotroph | \|Undefined Saprotroph\| | Yeast | NULL | Probable | As nectar-tap_saprotroph (Põlme et al. 2020); Decay sugar-rich substrates (Põlme et al. 2020) | Põlme S, et al. 2020. Fungal Diversity 105:1-16 ((https://doi.org/10.1007/s13225-020-00466-2)) |
| Otu57100 | 1 | k__Fungi(100);p__Ascomycota(100);c__Saccharomycetes(100);o__Saccharomycetales(100);f__Metschnikowiaceae(100);g__Metschnikowia(100);s__Metschnikowia_pulcherrima(100); | Metschnikowia | 13 | Saprotroph | \|Undefined Saprotroph\| | Yeast | NULL | Probable | As nectar-tap_saprotroph (Põlme et al. 2020); Decay sugar-rich substrates (Põlme et al. 2020) | Põlme S, et al. 2020. Fungal Diversity 105:1-16 ((https://doi.org/10.1007/s13225-020-00466-2)) |
| Otu57101 | 1 | k__Fungi(100);p__Ascomycota(100);c__Saccharomycetes(100);o__Saccharomycetales(100);f__Metschnikowiaceae(100);g__Metschnikowia(100);g__Metschnikowia_unclassified(100); | Metschnikowia | 13 | Saprotroph | \|Undefined Saprotroph\| | Yeast | NULL | Probable | As nectar-tap_saprotroph (Põlme et al. 2020); Decay sugar-rich substrates (Põlme et al. 2020) | Põlme S, et al. 2020. Fungal Diversity 105:1-16 ((https://doi.org/10.1007/s13225-020-00466-2)) |
| Otu57102 | 1 | k__Fungi(100);p__Ascomycota(100);c__Saccharomycetes(100);o__Saccharomycetales(100);f__Metschnikowiaceae(100);g__Metschnikowia(100);g__Metschnikowia_unclassified(100); | Metschnikowia | 13 | Saprotroph | \|Undefined Saprotroph\| | Yeast | NULL | Probable | As nectar-tap_saprotroph (Põlme et al. 2020); Decay sugar-rich substrates (Põlme et al. 2020) | Põlme S, et al. 2020. Fungal Diversity 105:1-16 ((https://doi.org/10.1007/s13225-020-00466-2)) |
| Otu57103 | 1 | k__Fungi(100);p__Ascomycota(100);c__Saccharomycetes(100);o__Saccharomycetales(100);f__Metschnikowiaceae(100);g__Metschnikowia(100);s__Metschnikowia_pulcherrima(100); | Metschnikowia | 13 | Saprotroph | \|Undefined Saprotroph\| | Yeast | NULL | Probable | As nectar-tap_saprotroph (Põlme et al. 2020); Decay sugar-rich substrates (Põlme et al. 2020) | Põlme S, et al. 2020. Fungal Diversity 105:1-16 ((https://doi.org/10.1007/s13225-020-00466-2)) |
| Otu57104 | 1 | k__Fungi(100);p__Ascomycota(100);c__Saccharomycetes(100);o__Saccharomycetales(100);f__Metschnikowiaceae(100);g__Metschnikowia(100);s__Metschnikowia_pulcherrima(100); | Metschnikowia | 13 | Saprotroph | \|Undefined Saprotroph\| | Yeast | NULL | Probable | As nectar-tap_saprotroph (Põlme et al. 2020); Decay sugar-rich substrates (Põlme et al. 2020) | Põlme S, et al. 2020. Fungal Diversity 105:1-16 ((https://doi.org/10.1007/s13225-020-00466-2)) |
| Otu57105 | 1 | k__Fungi(100);p__Ascomycota(100);c__Saccharomycetes(100);o__Saccharomycetales(100);f__Metschnikowiaceae(100);g__Metschnikowia(100);s__Metschnikowia_pulcherrima(100); | Metschnikowia | 13 | Saprotroph | \|Undefined Saprotroph\| | Yeast | NULL | Probable | As nectar-tap_saprotroph (Põlme et al. 2020); Decay sugar-rich substrates (Põlme et al. 2020) | Põlme S, et al. 2020. Fungal Diversity 105:1-16 ((https://doi.org/10.1007/s13225-020-00466-2)) |
| Otu57106 | 1 | k__Fungi(100);p__Ascomycota(100);c__Saccharomycetes(100);o__Saccharomycetales(100);f__Metschnikowiaceae(100);g__Metschnikowia(100);s__Metschnikowia_pulcherrima(100); | Metschnikowia | 13 | Saprotroph | \|Undefined Saprotroph\| | Yeast | NULL | Probable | As nectar-tap_saprotroph (Põlme et al. 2020); Decay sugar-rich substrates (Põlme et al. 2020) | Põlme S, et al. 2020. Fungal Diversity 105:1-16 ((https://doi.org/10.1007/s13225-020-00466-2)) |
| Otu57107 | 1 | k__Fungi(100);p__Ascomycota(100);c__Saccharomycetes(100);o__Saccharomycetales(100);f__Metschnikowiaceae(100);g__Metschnikowia(100);s__Metschnikowia_pulcherrima(100); | Metschnikowia | 13 | Saprotroph | \|Undefined Saprotroph\| | Yeast | NULL | Probable | As nectar-tap_saprotroph (Põlme et al. 2020); Decay sugar-rich substrates (Põlme et al. 2020) | Põlme S, et al. 2020. Fungal Diversity 105:1-16 ((https://doi.org/10.1007/s13225-020-00466-2)) |
| Otu57108 | 1 | k__Fungi(100);p__Ascomycota(100);c__Saccharomycetes(100);o__Saccharomycetales(100);f__Metschnikowiaceae(100);g__Metschnikowia(100);g__Metschnikowia_unclassified(100); | Metschnikowia | 13 | Saprotroph | \|Undefined Saprotroph\| | Yeast | NULL | Probable | As nectar-tap_saprotroph (Põlme et al. 2020); Decay sugar-rich substrates (Põlme et al. 2020) | Põlme S, et al. 2020. Fungal Diversity 105:1-16 ((https://doi.org/10.1007/s13225-020-00466-2)) |
| Otu57109 | 1 | k__Fungi(100);p__Ascomycota(100);c__Saccharomycetes(100);o__Saccharomycetales(100);f__Metschnikowiaceae(100);g__Metschnikowia(100);g__Metschnikowia_unclassified(100); | Metschnikowia | 13 | Saprotroph | \|Undefined Saprotroph\| | Yeast | NULL | Probable | As nectar-tap_saprotroph (Põlme et al. 2020); Decay sugar-rich substrates (Põlme et al. 2020) | Põlme S, et al. 2020. Fungal Diversity 105:1-16 ((https://doi.org/10.1007/s13225-020-00466-2)) |
| Otu57110 | 1 | k__Fungi(100);p__Ascomycota(100);c__Saccharomycetes(100);o__Saccharomycetales(100);f__Metschnikowiaceae(100);g__Metschnikowia(100);s__Metschnikowia_pulcherrima(100); | Metschnikowia | 13 | Saprotroph | \|Undefined Saprotroph\| | Yeast | NULL | Probable | As nectar-tap_saprotroph (Põlme et al. 2020); Decay sugar-rich substrates (Põlme et al. 2020) | Põlme S, et al. 2020. Fungal Diversity 105:1-16 ((https://doi.org/10.1007/s13225-020-00466-2)) |
| Otu57111 | 1 | k__Fungi(100);p__Ascomycota(100);c__Saccharomycetes(100);o__Saccharomycetales(100);f__Metschnikowiaceae(100);g__Metschnikowia(100);s__Metschnikowia_pulcherrima(100); | Metschnikowia | 13 | Saprotroph | \|Undefined Saprotroph\| | Yeast | NULL | Probable | As nectar-tap_saprotroph (Põlme et al. 2020); Decay sugar-rich substrates (Põlme et al. 2020) | Põlme S, et al. 2020. Fungal Diversity 105:1-16 ((https://doi.org/10.1007/s13225-020-00466-2)) |
| Otu57112 | 1 | k__Fungi(100);p__Ascomycota(100);c__Saccharomycetes(100);o__Saccharomycetales(100);f__Metschnikowiaceae(100);g__Metschnikowia(100);g__Metschnikowia_unclassified(100); | Metschnikowia | 13 | Saprotroph | \|Undefined Saprotroph\| | Yeast | NULL | Probable | As nectar-tap_saprotroph (Põlme et al. 2020); Decay sugar-rich substrates (Põlme et al. 2020) | Põlme S, et al. 2020. Fungal Diversity 105:1-16 ((https://doi.org/10.1007/s13225-020-00466-2)) |
| Otu57113 | 1 | k__Fungi(100);p__Ascomycota(100);c__Saccharomycetes(100);o__Saccharomycetales(100);f__Metschnikowiaceae(100);g__Metschnikowia(100);s__Metschnikowia_pulcherrima(100); | Metschnikowia | 13 | Saprotroph | \|Undefined Saprotroph\| | Yeast | NULL | Probable | As nectar-tap_saprotroph (Põlme et al. 2020); Decay sugar-rich substrates (Põlme et al. 2020) | Põlme S, et al. 2020. Fungal Diversity 105:1-16 ((https://doi.org/10.1007/s13225-020-00466-2)) |
| Otu57114 | 1 | k__Fungi(100);p__Ascomycota(100);c__Saccharomycetes(100);o__Saccharomycetales(100);f__Metschnikowiaceae(100);g__Metschnikowia(100);s__Metschnikowia_pulcherrima(100); | Metschnikowia | 13 | Saprotroph | \|Undefined Saprotroph\| | Yeast | NULL | Probable | As nectar-tap_saprotroph (Põlme et al. 2020); Decay sugar-rich substrates (Põlme et al. 2020) | Põlme S, et al. 2020. Fungal Diversity 105:1-16 ((https://doi.org/10.1007/s13225-020-00466-2)) |
| Otu57115 | 1 | k__Fungi(100);p__Ascomycota(100);c__Saccharomycetes(100);o__Saccharomycetales(100);f__Metschnikowiaceae(100);g__Metschnikowia(100);g__Metschnikowia_unclassified(100); | Metschnikowia | 13 | Saprotroph | \|Undefined Saprotroph\| | Yeast | NULL | Probable | As nectar-tap_saprotroph (Põlme et al. 2020); Decay sugar-rich substrates (Põlme et al. 2020) | Põlme S, et al. 2020. Fungal Diversity 105:1-16 ((https://doi.org/10.1007/s13225-020-00466-2)) |
| Otu57116 | 1 | k__Fungi(100);p__Ascomycota(100);c__Saccharomycetes(100);o__Saccharomycetales(100);f__Metschnikowiaceae(100);g__Metschnikowia(100);s__Metschnikowia_pulcherrima(100); | Metschnikowia | 13 | Saprotroph | \|Undefined Saprotroph\| | Yeast | NULL | Probable | As nectar-tap_saprotroph (Põlme et al. 2020); Decay sugar-rich substrates (Põlme et al. 2020) | Põlme S, et al. 2020. Fungal Diversity 105:1-16 ((https://doi.org/10.1007/s13225-020-00466-2)) |
| Otu57118 | 1 | k__Fungi(100);p__Ascomycota(100);c__Saccharomycetes(100);o__Saccharomycetales(100);f__Metschnikowiaceae(100);g__Metschnikowia(100);s__Metschnikowia_pulcherrima(100); | Metschnikowia | 13 | Saprotroph | \|Undefined Saprotroph\| | Yeast | NULL | Probable | As nectar-tap_saprotroph (Põlme et al. 2020); Decay sugar-rich substrates (Põlme et al. 2020) | Põlme S, et al. 2020. Fungal Diversity 105:1-16 ((https://doi.org/10.1007/s13225-020-00466-2)) |

| | | | | | | | | | | | |
|---|---|---|---|---|---|---|---|---|---|---|---|
| Otu57119 | 1 | k__Fungi(100);p__Ascomycota(100);c__Saccharomycetes(100);o__Saccharomycetales(100);f__Metschnikowiaceae(100);g__Metschnikowia(100);s__Metschnikowia_pulcherrima(100); | Metschnikowia | 13 | Saprotroph | \|Undefined Saprotroph\| | Yeast | NULL | Probable | As nectar-tap_saprotroph (Põlme et al. 2020); Decay sugar-rich substrates (Põlme et al. 2020) | Põlme S, et al. 2020. Fungal Diversity 105:1-16 ((https://doi.org/10.1007/s13225-020-00466-2)) |
| Otu57120 | 1 | k__Fungi(100);p__Ascomycota(100);c__Saccharomycetes(100);o__Saccharomycetales(100);f__Metschnikowiaceae(100);g__Metschnikowia(100);g__Metschnikowia_unclassified(100); | Metschnikowia | 13 | Saprotroph | \|Undefined Saprotroph\| | Yeast | NULL | Probable | As nectar-tap_saprotroph (Põlme et al. 2020); Decay sugar-rich substrates (Põlme et al. 2020) | Põlme S, et al. 2020. Fungal Diversity 105:1-16 ((https://doi.org/10.1007/s13225-020-00466-2)) |
| Otu57121 | 1 | k__Fungi(100);p__Ascomycota(100);c__Saccharomycetes(100);o__Saccharomycetales(100);f__Metschnikowiaceae(100);g__Metschnikowia(100);s__Metschnikowia_pulcherrima(100); | Metschnikowia | 13 | Saprotroph | \|Undefined Saprotroph\| | Yeast | NULL | Probable | As nectar-tap_saprotroph (Põlme et al. 2020); Decay sugar-rich substrates (Põlme et al. 2020) | Põlme S, et al. 2020. Fungal Diversity 105:1-16 ((https://doi.org/10.1007/s13225-020-00466-2)) |
| Otu57122 | 1 | k__Fungi(100);p__Ascomycota(100);c__Saccharomycetes(100);o__Saccharomycetales(100);f__Metschnikowiaceae(100);g__Metschnikowia(100);s__Metschnikowia_pulcherrima(100); | Metschnikowia | 13 | Saprotroph | \|Undefined Saprotroph\| | Yeast | NULL | Probable | As nectar-tap_saprotroph (Põlme et al. 2020); Decay sugar-rich substrates (Põlme et al. 2020) | Põlme S, et al. 2020. Fungal Diversity 105:1-16 ((https://doi.org/10.1007/s13225-020-00466-2)) |
| Otu57123 | 1 | k__Fungi(100);p__Ascomycota(100);c__Saccharomycetes(100);o__Saccharomycetales(100);f__Metschnikowiaceae(100);g__Metschnikowia(100);s__Metschnikowia_pulcherrima(100); | Metschnikowia | 13 | Saprotroph | \|Undefined Saprotroph\| | Yeast | NULL | Probable | As nectar-tap_saprotroph (Põlme et al. 2020); Decay sugar-rich substrates (Põlme et al. 2020) | Põlme S, et al. 2020. Fungal Diversity 105:1-16 ((https://doi.org/10.1007/s13225-020-00466-2)) |
| Otu57124 | 1 | k__Fungi(100);p__Ascomycota(100);c__Saccharomycetes(100);o__Saccharomycetales(100);f__Metschnikowiaceae(100);g__Metschnikowia(100);s__Metschnikowia_pulcherrima(100); | Metschnikowia | 13 | Saprotroph | \|Undefined Saprotroph\| | Yeast | NULL | Probable | As nectar-tap_saprotroph (Põlme et al. 2020); Decay sugar-rich substrates (Põlme et al. 2020) | Põlme S, et al. 2020. Fungal Diversity 105:1-16 ((https://doi.org/10.1007/s13225-020-00466-2)) |
| Otu57126 | 1 | k__Fungi(100);p__Ascomycota(100);c__Saccharomycetes(100);o__Saccharomycetales(100);f__Metschnikowiaceae(100);g__Metschnikowia(100);s__Metschnikowia_pulcherrima(100); | Metschnikowia | 13 | Saprotroph | \|Undefined Saprotroph\| | Yeast | NULL | Probable | As nectar-tap_saprotroph (Põlme et al. 2020); Decay sugar-rich substrates (Põlme et al. 2020) | Põlme S, et al. 2020. Fungal Diversity 105:1-16 ((https://doi.org/10.1007/s13225-020-00466-2)) |
| Otu57127 | 1 | k__Fungi(100);p__Ascomycota(100);c__Saccharomycetes(100);o__Saccharomycetales(100);f__Metschnikowiaceae(100);g__Metschnikowia(100);s__Metschnikowia_pulcherrima(100); | Metschnikowia | 13 | Saprotroph | \|Undefined Saprotroph\| | Yeast | NULL | Probable | As nectar-tap_saprotroph (Põlme et al. 2020); Decay sugar-rich substrates (Põlme et al. 2020) | Põlme S, et al. 2020. Fungal Diversity 105:1-16 ((https://doi.org/10.1007/s13225-020-00466-2)) |
| Otu57128 | 1 | k__Fungi(100);p__Ascomycota(100);c__Saccharomycetes(100);o__Saccharomycetales(100);f__Metschnikowiaceae(100);g__Metschnikowia(100);s__Metschnikowia_pulcherrima(100); | Metschnikowia | 13 | Saprotroph | \|Undefined Saprotroph\| | Yeast | NULL | Probable | As nectar-tap_saprotroph (Põlme et al. 2020); Decay sugar-rich substrates (Põlme et al. 2020) | Põlme S, et al. 2020. Fungal Diversity 105:1-16 ((https://doi.org/10.1007/s13225-020-00466-2)) |
| Otu57129 | 1 | k__Fungi(100);p__Ascomycota(100);c__Saccharomycetes(100);o__Saccharomycetales(100);f__Metschnikowiaceae(100);g__Metschnikowia(100);s__Metschnikowia_pulcherrima(100); | Metschnikowia | 13 | Saprotroph | \|Undefined Saprotroph\| | Yeast | NULL | Probable | As nectar-tap_saprotroph (Põlme et al. 2020); Decay sugar-rich substrates (Põlme et al. 2020) | Põlme S, et al. 2020. Fungal Diversity 105:1-16 ((https://doi.org/10.1007/s13225-020-00466-2)) |
| Otu57130 | 1 | k__Fungi(100);p__Ascomycota(100);c__Saccharomycetes(100);o__Saccharomycetales(100);f__Metschnikowiaceae(100);g__Metschnikowia(100);s__Metschnikowia_pulcherrima(100); | Metschnikowia | 13 | Saprotroph | \|Undefined Saprotroph\| | Yeast | NULL | Probable | As nectar-tap_saprotroph (Põlme et al. 2020); Decay sugar-rich substrates (Põlme et al. 2020) | Põlme S, et al. 2020. Fungal Diversity 105:1-16 ((https://doi.org/10.1007/s13225-020-00466-2)) |
| Otu57131 | 1 | k__Fungi(100);p__Ascomycota(100);c__Saccharomycetes(100);o__Saccharomycetales(100);f__Metschnikowiaceae(100);g__Metschnikowia(100);g__Metschnikowia_unclassified(100); | Metschnikowia | 13 | Saprotroph | \|Undefined Saprotroph\| | Yeast | NULL | Probable | As nectar-tap_saprotroph (Põlme et al. 2020); Decay sugar-rich substrates (Põlme et al. 2020) | Põlme S, et al. 2020. Fungal Diversity 105:1-16 ((https://doi.org/10.1007/s13225-020-00466-2)) |
| Otu57132 | 1 | k__Fungi(100);p__Ascomycota(100);c__Saccharomycetes(100);o__Saccharomycetales(100);f__Metschnikowiaceae(100);g__Metschnikowia(100);s__Metschnikowia_pulcherrima(100); | Metschnikowia | 13 | Saprotroph | \|Undefined Saprotroph\| | Yeast | NULL | Probable | As nectar-tap_saprotroph (Põlme et al. 2020); Decay sugar-rich substrates (Põlme et al. 2020) | Põlme S, et al. 2020. Fungal Diversity 105:1-16 ((https://doi.org/10.1007/s13225-020-00466-2)) |
| Otu57133 | 1 | k__Fungi(100);p__Ascomycota(100);c__Saccharomycetes(100);o__Saccharomycetales(100);f__Metschnikowiaceae(100);g__Metschnikowia(100);s__Metschnikowia_pulcherrima(100); | Metschnikowia | 13 | Saprotroph | \|Undefined Saprotroph\| | Yeast | NULL | Probable | As nectar-tap_saprotroph (Põlme et al. 2020); Decay sugar-rich substrates (Põlme et al. 2020) | Põlme S, et al. 2020. Fungal Diversity 105:1-16 ((https://doi.org/10.1007/s13225-020-00466-2)) |
| Otu57134 | 1 | k__Fungi(100);p__Ascomycota(100);c__Saccharomycetes(100);o__Saccharomycetales(100);f__Metschnikowiaceae(100);g__Metschnikowia(100);g__Metschnikowia_unclassified(100); | Metschnikowia | 13 | Saprotroph | \|Undefined Saprotroph\| | Yeast | NULL | Probable | As nectar-tap_saprotroph (Põlme et al. 2020); Decay sugar-rich substrates (Põlme et al. 2020) | Põlme S, et al. 2020. Fungal Diversity 105:1-16 ((https://doi.org/10.1007/s13225-020-00466-2)) |
| Otu57135 | 1 | k__Fungi(100);p__Ascomycota(100);c__Saccharomycetes(100);o__Saccharomycetales(100);f__Metschnikowiaceae(100);g__Metschnikowia(100);g__Metschnikowia_unclassified(100); | Metschnikowia | 13 | Saprotroph | \|Undefined Saprotroph\| | Yeast | NULL | Probable | As nectar-tap_saprotroph (Põlme et al. 2020); Decay sugar-rich substrates (Põlme et al. 2020) | Põlme S, et al. 2020. Fungal Diversity 105:1-16 ((https://doi.org/10.1007/s13225-020-00466-2)) |
| Otu57136 | 1 | k__Fungi(100);p__Ascomycota(100);c__Saccharomycetes(100);o__Saccharomycetales(100);f__Metschnikowiaceae(100);g__Metschnikowia(100);g__Metschnikowia_unclassified(100); | Metschnikowia | 13 | Saprotroph | \|Undefined Saprotroph\| | Yeast | NULL | Probable | As nectar-tap_saprotroph (Põlme et al. 2020); Decay sugar-rich substrates (Põlme et al. 2020) | Põlme S, et al. 2020. Fungal Diversity 105:1-16 ((https://doi.org/10.1007/s13225-020-00466-2)) |
| Otu57137 | 1 | k__Fungi(100);p__Ascomycota(100);c__Saccharomycetes(100);o__Saccharomycetales(100);f__Metschnikowiaceae(100);g__Metschnikowia(100);s__Metschnikowia_pulcherrima(100); | Metschnikowia | 13 | Saprotroph | \|Undefined Saprotroph\| | Yeast | NULL | Probable | As nectar-tap_saprotroph (Põlme et al. 2020); Decay sugar-rich substrates (Põlme et al. 2020) | Põlme S, et al. 2020. Fungal Diversity 105:1-16 ((https://doi.org/10.1007/s13225-020-00466-2)) |
| Otu57139 | 1 | k__Fungi(100);p__Ascomycota(100);c__Saccharomycetes(100);o__Saccharomycetales(100);f__Metschnikowiaceae(100);g__Metschnikowia(100);g__Metschnikowia_unclassified(100); | Metschnikowia | 13 | Saprotroph | \|Undefined Saprotroph\| | Yeast | NULL | Probable | As nectar-tap_saprotroph (Põlme et al. 2020); Decay sugar-rich substrates (Põlme et al. 2020) | Põlme S, et al. 2020. Fungal Diversity 105:1-16 ((https://doi.org/10.1007/s13225-020-00466-2)) |
| Otu57140 | 1 | k__Fungi(100);p__Ascomycota(100);c__Saccharomycetes(100);o__Saccharomycetales(100);f__Metschnikowiaceae(100);g__Metschnikowia(100);s__Metschnikowia_pulcherrima(100); | Metschnikowia | 13 | Saprotroph | \|Undefined Saprotroph\| | Yeast | NULL | Probable | As nectar-tap_saprotroph (Põlme et al. 2020); Decay sugar-rich substrates (Põlme et al. 2020) | Põlme S, et al. 2020. Fungal Diversity 105:1-16 ((https://doi.org/10.1007/s13225-020-00466-2)) |
| Otu57141 | 1 | k__Fungi(100);p__Ascomycota(100);c__Saccharomycetes(100);o__Saccharomycetales(100);f__Metschnikowiaceae(100);g__Metschnikowia(100);g__Metschnikowia_unclassified(100); | Metschnikowia | 13 | Saprotroph | \|Undefined Saprotroph\| | Yeast | NULL | Probable | As nectar-tap_saprotroph (Põlme et al. 2020); Decay sugar-rich substrates (Põlme et al. 2020) | Põlme S, et al. 2020. Fungal Diversity 105:1-16 ((https://doi.org/10.1007/s13225-020-00466-2)) |
| Otu57142 | 1 | k__Fungi(100);p__Ascomycota(100);c__Saccharomycetes(100);o__Saccharomycetales(100);f__Metschnikowiaceae(100);g__Metschnikowia(100);g__Metschnikowia_unclassified(100); | Metschnikowia | 13 | Saprotroph | \|Undefined Saprotroph\| | Yeast | NULL | Probable | As nectar-tap_saprotroph (Põlme et al. 2020); Decay sugar-rich substrates (Põlme et al. 2020) | Põlme S, et al. 2020. Fungal Diversity 105:1-16 ((https://doi.org/10.1007/s13225-020-00466-2)) |
| Otu57143 | 1 | k__Fungi(100);p__Ascomycota(100);c__Saccharomycetes(100);o__Saccharomycetales(100);f__Metschnikowiaceae(100);g__Metschnikowia(100);s__Metschnikowia_pulcherrima(100); | Metschnikowia | 13 | Saprotroph | \|Undefined Saprotroph\| | Yeast | NULL | Probable | As nectar-tap_saprotroph (Põlme et al. 2020); Decay sugar-rich substrates (Põlme et al. 2020) | Põlme S, et al. 2020. Fungal Diversity 105:1-16 ((https://doi.org/10.1007/s13225-020-00466-2)) |
| Otu57144 | 1 | k__Fungi(100);p__Ascomycota(100);c__Saccharomycetes(100);o__Saccharomycetales(100);f__Metschnikowiaceae(100);g__Metschnikowia(100);s__Metschnikowia_pulcherrima(100); | Metschnikowia | 13 | Saprotroph | \|Undefined Saprotroph\| | Yeast | NULL | Probable | As nectar-tap_saprotroph (Põlme et al. 2020); Decay sugar-rich substrates (Põlme et al. 2020) | Põlme S, et al. 2020. Fungal Diversity 105:1-16 ((https://doi.org/10.1007/s13225-020-00466-2)) |
| Otu57145 | 1 | k__Fungi(100);p__Ascomycota(100);c__Saccharomycetes(100);o__Saccharomycetales(100);f__Metschnikowiaceae(100);g__Metschnikowia(100);s__Metschnikowia_pulcherrima(100); | Metschnikowia | 13 | Saprotroph | \|Undefined Saprotroph\| | Yeast | NULL | Probable | As nectar-tap_saprotroph (Põlme et al. 2020); Decay sugar-rich substrates (Põlme et al. 2020) | Põlme S, et al. 2020. Fungal Diversity 105:1-16 ((https://doi.org/10.1007/s13225-020-00466-2)) |
| Otu57146 | 1 | k__Fungi(100);p__Ascomycota(100);c__Saccharomycetes(100);o__Saccharomycetales(100);f__Metschnikowiaceae(100);g__Metschnikowia(100);s__Metschnikowia_pulcherrima(100); | Metschnikowia | 13 | Saprotroph | \|Undefined Saprotroph\| | Yeast | NULL | Probable | As nectar-tap_saprotroph (Põlme et al. 2020); Decay sugar-rich substrates (Põlme et al. 2020) | Põlme S, et al. 2020. Fungal Diversity 105:1-16 ((https://doi.org/10.1007/s13225-020-00466-2)) |
| Otu57147 | 1 | k__Fungi(100);p__Ascomycota(100);c__Saccharomycetes(100);o__Saccharomycetales(100);f__Metschnikowiaceae(100);g__Metschnikowia(100);g__Metschnikowia_unclassified(100); | Metschnikowia | 13 | Saprotroph | \|Undefined Saprotroph\| | Yeast | NULL | Probable | As nectar-tap_saprotroph (Põlme et al. 2020); Decay sugar-rich substrates (Põlme et al. 2020) | Põlme S, et al. 2020. Fungal Diversity 105:1-16 ((https://doi.org/10.1007/s13225-020-00466-2)) |
| Otu57148 | 1 | k__Fungi(100);p__Ascomycota(100);c__Saccharomycetes(100);o__Saccharomycetales(100);f__Metschnikowiaceae(100);g__Metschnikowia(100);s__Metschnikowia_pulcherrima(100); | Metschnikowia | 13 | Saprotroph | \|Undefined Saprotroph\| | Yeast | NULL | Probable | As nectar-tap_saprotroph (Põlme et al. 2020); Decay sugar-rich substrates (Põlme et al. 2020) | Põlme S, et al. 2020. Fungal Diversity 105:1-16 ((https://doi.org/10.1007/s13225-020-00466-2)) |
| Otu57149 | 1 | k__Fungi(100);p__Ascomycota(100);c__Saccharomycetes(100);o__Saccharomycetales(100);f__Metschnikowiaceae(100);g__Metschnikowia(100);s__Metschnikowia_pulcherrima(100); | Metschnikowia | 13 | Saprotroph | \|Undefined Saprotroph\| | Yeast | NULL | Probable | As nectar-tap_saprotroph (Põlme et al. 2020); Decay sugar-rich substrates (Põlme et al. 2020) | Põlme S, et al. 2020. Fungal Diversity 105:1-16 ((https://doi.org/10.1007/s13225-020-00466-2)) |
| Otu57150 | 1 | k__Fungi(100);p__Ascomycota(100);c__Saccharomycetes(100);o__Saccharomycetales(100);f__Metschnikowiaceae(100);g__Metschnikowia(100);s__Metschnikowia_pulcherrima(100); | Metschnikowia | 13 | Saprotroph | \|Undefined Saprotroph\| | Yeast | NULL | Probable | As nectar-tap_saprotroph (Põlme et al. 2020); Decay sugar-rich substrates (Põlme et al. 2020) | Põlme S, et al. 2020. Fungal Diversity 105:1-16 ((https://doi.org/10.1007/s13225-020-00466-2)) |
| Otu57151 | 1 | k__Fungi(100);p__Ascomycota(100);c__Saccharomycetes(100);o__Saccharomycetales(100);f__Metschnikowiaceae(100);g__Metschnikowia(100);g__Metschnikowia_unclassified(100); | Metschnikowia | 13 | Saprotroph | \|Undefined Saprotroph\| | Yeast | NULL | Probable | As nectar-tap_saprotroph (Põlme et al. 2020); Decay sugar-rich substrates (Põlme et al. 2020) | Põlme S, et al. 2020. Fungal Diversity 105:1-16 ((https://doi.org/10.1007/s13225-020-00466-2)) |
| Otu57152 | 1 | k__Fungi(100);p__Ascomycota(100);c__Saccharomycetes(100);o__Saccharomycetales(100);f__Metschnikowiaceae(100);g__Metschnikowia(100);s__Metschnikowia_pulcherrima(100); | Metschnikowia | 13 | Saprotroph | \|Undefined Saprotroph\| | Yeast | NULL | Probable | As nectar-tap_saprotroph (Põlme et al. 2020); Decay sugar-rich substrates (Põlme et al. 2020) | Põlme S, et al. 2020. Fungal Diversity 105:1-16 ((https://doi.org/10.1007/s13225-020-00466-2)) |
| Otu57153 | 1 | k__Fungi(100);p__Ascomycota(100);c__Saccharomycetes(100);o__Saccharomycetales(100);f__Metschnikowiaceae(100);g__Metschnikowia(100);s__Metschnikowia_pulcherrima(100); | Metschnikowia | 13 | Saprotroph | \|Undefined Saprotroph\| | Yeast | NULL | Probable | As nectar-tap_saprotroph (Põlme et al. 2020); Decay sugar-rich substrates (Põlme et al. 2020) | Põlme S, et al. 2020. Fungal Diversity 105:1-16 ((https://doi.org/10.1007/s13225-020-00466-2)) |
| Otu57154 | 1 | k__Fungi(100);p__Ascomycota(100);c__Saccharomycetes(100);o__Saccharomycetales(100);f__Metschnikowiaceae(100);g__Metschnikowia(100);s__Metschnikowia_pulcherrima(100); | Metschnikowia | 13 | Saprotroph | \|Undefined Saprotroph\| | Yeast | NULL | Probable | As nectar-tap_saprotroph (Põlme et al. 2020); Decay sugar-rich substrates (Põlme et al. 2020) | Põlme S, et al. 2020. Fungal Diversity 105:1-16 ((https://doi.org/10.1007/s13225-020-00466-2)) |
| Otu57155 | 1 | k__Fungi(100);p__Ascomycota(100);c__Saccharomycetes(100);o__Saccharomycetales(100);f__Metschnikowiaceae(100);g__Metschnikowia(100);s__Metschnikowia_pulcherrima(100); | Metschnikowia | 13 | Saprotroph | \|Undefined Saprotroph\| | Yeast | NULL | Probable | As nectar-tap_saprotroph (Põlme et al. 2020); Decay sugar-rich substrates (Põlme et al. 2020) | Põlme S, et al. 2020. Fungal Diversity 105:1-16 ((https://doi.org/10.1007/s13225-020-00466-2)) |
| Otu57158 | 1 | k__Fungi(100);p__Ascomycota(100);c__Saccharomycetes(100);o__Saccharomycetales(100);f__Metschnikowiaceae(100);g__Metschnikowia(100);s__Metschnikowia_pulcherrima(100); | Metschnikowia | 13 | Saprotroph | \|Undefined Saprotroph\| | Yeast | NULL | Probable | As nectar-tap_saprotroph (Põlme et al. 2020); Decay sugar-rich substrates (Põlme et al. 2020) | Põlme S, et al. 2020. Fungal Diversity 105:1-16 ((https://doi.org/10.1007/s13225-020-00466-2)) |
| Otu57161 | 1 | k__Fungi(100);p__Ascomycota(100);c__Saccharomycetes(100);o__Saccharomycetales(100);f__Metschnikowiaceae(100);g__Metschnikowia(100);s__Metschnikowia_pulcherrima(100); | Metschnikowia | 13 | Saprotroph | \|Undefined Saprotroph\| | Yeast | NULL | Probable | As nectar-tap_saprotroph (Põlme et al. 2020); Decay sugar-rich substrates (Põlme et al. 2020) | Põlme S, et al. 2020. Fungal Diversity 105:1-16 ((https://doi.org/10.1007/s13225-020-00466-2)) |
| Otu57165 | 1 | k__Fungi(100);p__Ascomycota(100);c__Saccharomycetes(100);o__Saccharomycetales(100);f__Metschnikowiaceae(100);g__Metschnikowia(100);g__Metschnikowia_unclassified(100); | Metschnikowia | 13 | Saprotroph | \|Undefined Saprotroph\| | Yeast | NULL | Probable | As nectar-tap_saprotroph (Põlme et al. 2020); Decay sugar-rich substrates (Põlme et al. 2020) | Põlme S, et al. 2020. Fungal Diversity 105:1-16 ((https://doi.org/10.1007/s13225-020-00466-2)) |
| Otu57166 | 1 | k__Fungi(100);p__Ascomycota(100);c__Saccharomycetes(100);o__Saccharomycetales(100);f__Metschnikowiaceae(100);g__Metschnikowia(100);s__Metschnikowia_pulcherrima(100); | Metschnikowia | 13 | Saprotroph | \|Undefined Saprotroph\| | Yeast | NULL | Probable | As nectar-tap_saprotroph (Põlme et al. 2020); Decay sugar-rich substrates (Põlme et al. 2020) | Põlme S, et al. 2020. Fungal Diversity 105:1-16 ((https://doi.org/10.1007/s13225-020-00466-2)) |
| Otu57167 | 1 | k__Fungi(100);p__Ascomycota(100);c__Saccharomycetes(100);o__Saccharomycetales(100);f__Metschnikowiaceae(100);g__Metschnikowia(100);s__Metschnikowia_pulcherrima(100); | Metschnikowia | 13 | Saprotroph | \|Undefined Saprotroph\| | Yeast | NULL | Probable | As nectar-tap_saprotroph (Põlme et al. 2020); Decay sugar-rich substrates (Põlme et al. 2020) | Põlme S, et al. 2020. Fungal Diversity 105:1-16 ((https://doi.org/10.1007/s13225-020-00466-2)) |
| Otu57168 | 1 | k__Fungi(100);p__Ascomycota(100);c__Saccharomycetes(100);o__Saccharomycetales(100);f__Metschnikowiaceae(100);g__Metschnikowia(100);s__Metschnikowia_pulcherrima(100); | Metschnikowia | 13 | Saprotroph | \|Undefined Saprotroph\| | Yeast | NULL | Probable | As nectar-tap_saprotroph (Põlme et al. 2020); Decay sugar-rich substrates (Põlme et al. 2020) | Põlme S, et al. 2020. Fungal Diversity 105:1-16 ((https://doi.org/10.1007/s13225-020-00466-2)) |
| Otu57169 | 1 | k__Fungi(100);p__Ascomycota(100);c__Saccharomycetes(100);o__Saccharomycetales(100);f__Metschnikowiaceae(100);g__Metschnikowia(100);g__Metschnikowia_unclassified(100); | Metschnikowia | 13 | Saprotroph | \|Undefined Saprotroph\| | Yeast | NULL | Probable | As nectar-tap_saprotroph (Põlme et al. 2020); Decay sugar-rich substrates (Põlme et al. 2020) | Põlme S, et al. 2020. Fungal Diversity 105:1-16 ((https://doi.org/10.1007/s13225-020-00466-2)) |
| Otu57172 | 1 | k__Fungi(100);p__Ascomycota(100);c__Saccharomycetes(100);o__Saccharomycetales(100);f__Metschnikowiaceae(100);g__Metschnikowia(100);g__Metschnikowia_unclassified(100); | Metschnikowia | 13 | Saprotroph | \|Undefined Saprotroph\| | Yeast | NULL | Probable | As nectar-tap_saprotroph (Põlme et al. 2020); Decay sugar-rich substrates (Põlme et al. 2020) | Põlme S, et al. 2020. Fungal Diversity 105:1-16 ((https://doi.org/10.1007/s13225-020-00466-2)) |
| Otu57173 | 1 | k__Fungi(100);p__Ascomycota(100);c__Saccharomycetes(100);o__Saccharomycetales(100);f__Metschnikowiaceae(100);g__Metschnikowia(100);g__Metschnikowia_unclassified(100); | Metschnikowia | 13 | Saprotroph | \|Undefined Saprotroph\| | Yeast | NULL | Probable | As nectar-tap_saprotroph (Põlme et al. 2020); Decay sugar-rich substrates (Põlme et al. 2020) | Põlme S, et al. 2020. Fungal Diversity 105:1-16 ((https://doi.org/10.1007/s13225-020-00466-2)) |
| Otu57174 | 1 | k__Fungi(100);p__Ascomycota(100);c__Saccharomycetes(100);o__Saccharomycetales(100);f__Metschnikowiaceae(100);g__Metschnikowia(100);g__Metschnikowia_unclassified(100); | Metschnikowia | 13 | Saprotroph | \|Undefined Saprotroph\| | Yeast | NULL | Probable | As nectar-tap_saprotroph (Põlme et al. 2020); Decay sugar-rich substrates (Põlme et al. 2020) | Põlme S, et al. 2020. Fungal Diversity 105:1-16 ((https://doi.org/10.1007/s13225-020-00466-2)) |
| Otu57175 | 1 | k__Fungi(100);p__Ascomycota(100);c__Saccharomycetes(100);o__Saccharomycetales(100);f__Metschnikowiaceae(100);g__Metschnikowia(100);s__Metschnikowia_pulcherrima(100); | Metschnikowia | 13 | Saprotroph | \|Undefined Saprotroph\| | Yeast | NULL | Probable | As nectar-tap_saprotroph (Põlme et al. 2020); Decay sugar-rich substrates (Põlme et al. 2020) | Põlme S, et al. 2020. Fungal Diversity 105:1-16 ((https://doi.org/10.1007/s13225-020-00466-2)) |
| Otu57176 | 1 | k__Fungi(100);p__Ascomycota(100);c__Saccharomycetes(100);o__Saccharomycetales(100);f__Metschnikowiaceae(100);g__Metschnikowia(100);s__Metschnikowia_pulcherrima(100); | Metschnikowia | 13 | Saprotroph | \|Undefined Saprotroph\| | Yeast | NULL | Probable | As nectar-tap_saprotroph (Põlme et al. 2020); Decay sugar-rich substrates (Põlme et al. 2020) | Põlme S, et al. 2020. Fungal Diversity 105:1-16 ((https://doi.org/10.1007/s13225-020-00466-2)) |
| Otu57177 | 1 | k__Fungi(100);p__Ascomycota(100);c__Saccharomycetes(100);o__Saccharomycetales(100);f__Metschnikowiaceae(100);g__Metschnikowia(100);s__Metschnikowia_pulcherrima(100); | Metschnikowia | 13 | Saprotroph | \|Undefined Saprotroph\| | Yeast | NULL | Probable | As nectar-tap_saprotroph (Põlme et al. 2020); Decay sugar-rich substrates (Põlme et al. 2020) | Põlme S, et al. 2020. Fungal Diversity 105:1-16 ((https://doi.org/10.1007/s13225-020-00466-2)) |
| Otu57178 | 1 | k__Fungi(100);p__Ascomycota(100);c__Saccharomycetes(100);o__Saccharomycetales(100);f__Metschnikowiaceae(100);g__Metschnikowia(100);s__Metschnikowia_pulcherrima(100); | Metschnikowia | 13 | Saprotroph | \|Undefined Saprotroph\| | Yeast | NULL | Probable | As nectar-tap_saprotroph (Põlme et al. 2020); Decay sugar-rich substrates (Põlme et al. 2020) | Põlme S, et al. 2020. Fungal Diversity 105:1-16 ((https://doi.org/10.1007/s13225-020-00466-2)) |
| Otu57179 | 1 | k__Fungi(100);p__Ascomycota(100);c__Saccharomycetes(100);o__Saccharomycetales(100);f__Metschnikowiaceae(100);g__Metschnikowia(100);s__Metschnikowia_pulcherrima(100); | Metschnikowia | 13 | Saprotroph | \|Undefined Saprotroph\| | Yeast | NULL | Probable | As nectar-tap_saprotroph (Põlme et al. 2020); Decay sugar-rich substrates (Põlme et al. 2020) | Põlme S, et al. 2020. Fungal Diversity 105:1-16 ((https://doi.org/10.1007/s13225-020-00466-2)) |
| Otu57180 | 1 | k__Fungi(100);p__Ascomycota(100);c__Saccharomycetes(100);o__Saccharomycetales(100);f__Metschnikowiaceae(100);g__Metschnikowia(100);g__Metschnikowia_unclassified(100); | Metschnikowia | 13 | Saprotroph | \|Undefined Saprotroph\| | Yeast | NULL | Probable | As nectar-tap_saprotroph (Põlme et al. 2020); Decay sugar-rich substrates (Põlme et al. 2020) | Põlme S, et al. 2020. Fungal Diversity 105:1-16 ((https://doi.org/10.1007/s13225-020-00466-2)) |
| Otu57181 | 1 | k__Fungi(100);p__Ascomycota(100);c__Saccharomycetes(100);o__Saccharomycetales(100);f__Metschnikowiaceae(100);g__Metschnikowia(100);s__Metschnikowia_pulcherrima(100); | Metschnikowia | 13 | Saprotroph | \|Undefined Saprotroph\| | Yeast | NULL | Probable | As nectar-tap_saprotroph (Põlme et al. 2020); Decay sugar-rich substrates (Põlme et al. 2020) | Põlme S, et al. 2020. Fungal Diversity 105:1-16 ((https://doi.org/10.1007/s13225-020-00466-2)) |
| Otu57184 | 1 | k__Fungi(100);p__Ascomycota(100);c__Saccharomycetes(100);o__Saccharomycetales(100);f__Metschnikowiaceae(100);g__Metschnikowia(100);g__Metschnikowia_unclassified(100); | Metschnikowia | 13 | Saprotroph | \|Undefined Saprotroph\| | Yeast | NULL | Probable | As nectar-tap_saprotroph (Põlme et al. 2020); Decay sugar-rich substrates (Põlme et al. 2020) | Põlme S, et al. 2020. Fungal Diversity 105:1-16 ((https://doi.org/10.1007/s13225-020-00466-2)) |
| Otu57186 | 1 | k__Fungi(100);p__Ascomycota(100);c__Saccharomycetes(100);o__Saccharomycetales(100);f__Metschnikowiaceae(100);g__Metschnikowia(100);s__Metschnikowia_pulcherrima(100); | Metschnikowia | 13 | Saprotroph | \|Undefined Saprotroph\| | Yeast | NULL | Probable | As nectar-tap_saprotroph (Põlme et al. 2020); Decay sugar-rich substrates (Põlme et al. 2020) | Põlme S, et al. 2020. Fungal Diversity 105:1-16 ((https://doi.org/10.1007/s13225-020-00466-2)) |
| Otu57191 | 1 | k__Fungi(100);p__Ascomycota(100);c__Saccharomycetes(100);o__Saccharomycetales(100);f__Metschnikowiaceae(100);g__Metschnikowia(100);s__Metschnikowia_pulcherrima(100); | Metschnikowia | 13 | Saprotroph | \|Undefined Saprotroph\| | Yeast | NULL | Probable | As nectar-tap_saprotroph (Põlme et al. 2020); Decay sugar-rich substrates (Põlme et al. 2020) | Põlme S, et al. 2020. Fungal Diversity 105:1-16 ((https://doi.org/10.1007/s13225-020-00466-2)) |
| Otu57193 | 1 | k__Fungi(100);p__Ascomycota(100);c__Saccharomycetes(100);o__Saccharomycetales(100);f__Metschnikowiaceae(100);g__Metschnikowia(100);g__Metschnikowia_unclassified(100); | Metschnikowia | 13 | Saprotroph | \|Undefined Saprotroph\| | Yeast | NULL | Probable | As nectar-tap_saprotroph (Põlme et al. 2020); Decay sugar-rich substrates (Põlme et al. 2020) | Põlme S, et al. 2020. Fungal Diversity 105:1-16 ((https://doi.org/10.1007/s13225-020-00466-2)) |
| Otu57194 | 1 | k__Fungi(100);p__Ascomycota(100);c__Saccharomycetes(100);o__Saccharomycetales(100);f__Metschnikowiaceae(100);g__Metschnikowia(100);s__Metschnikowia_pulcherrima(100); | Metschnikowia | 13 | Saprotroph | \|Undefined Saprotroph\| | Yeast | NULL | Probable | As nectar-tap_saprotroph (Põlme et al. 2020); Decay sugar-rich substrates (Põlme et al. 2020) | Põlme S, et al. 2020. Fungal Diversity 105:1-16 ((https://doi.org/10.1007/s13225-020-00466-2)) |
| Otu57195 | 1 | k__Fungi(100);p__Ascomycota(100);c__Saccharomycetes(100);o__Saccharomycetales(100);f__Metschnikowiaceae(100);g__Metschnikowia(100);g__Metschnikowia_unclassified(100); | Metschnikowia | 13 | Saprotroph | \|Undefined Saprotroph\| | Yeast | NULL | Probable | As nectar-tap_saprotroph (Põlme et al. 2020); Decay sugar-rich substrates (Põlme et al. 2020) | Põlme S, et al. 2020. Fungal Diversity 105:1-16 ((https://doi.org/10.1007/s13225-020-00466-2)) |
| Otu57196 | 1 | k__Fungi(100);p__Ascomycota(100);c__Saccharomycetes(100);o__Saccharomycetales(100);f__Metschnikowiaceae(100);g__Metschnikowia(100);s__Metschnikowia_pulcherrima(100); | Metschnikowia | 13 | Saprotroph | \|Undefined Saprotroph\| | Yeast | NULL | Probable | As nectar-tap_saprotroph (Põlme et al. 2020); Decay sugar-rich substrates (Põlme et al. 2020) | Põlme S, et al. 2020. Fungal Diversity 105:1-16 ((https://doi.org/10.1007/s13225-020-00466-2)) |
| Otu57197 | 1 | k__Fungi(100);p__Ascomycota(100);c__Saccharomycetes(100);o__Saccharomycetales(100);f__Metschnikowiaceae(100);g__Metschnikowia(100);g__Metschnikowia_unclassified(100); | Metschnikowia | 13 | Saprotroph | \|Undefined Saprotroph\| | Yeast | NULL | Probable | As nectar-tap_saprotroph (Põlme et al. 2020); Decay sugar-rich substrates (Põlme et al. 2020) | Põlme S, et al. 2020. Fungal Diversity 105:1-16 ((https://doi.org/10.1007/s13225-020-00466-2)) |
| Otu57198 | 1 | k__Fungi(100);p__Ascomycota(100);c__Saccharomycetes(100);o__Saccharomycetales(100);f__Metschnikowiaceae(100);g__Metschnikowia(100);s__Metschnikowia_pulcherrima(100); | Metschnikowia | 13 | Saprotroph | \|Undefined Saprotroph\| | Yeast | NULL | Probable | As nectar-tap_saprotroph (Põlme et al. 2020); Decay sugar-rich substrates (Põlme et al. 2020) | Põlme S, et al. 2020. Fungal Diversity 105:1-16 ((https://doi.org/10.1007/s13225-020-00466-2)) |
| Otu57199 | 1 | k__Fungi(100);p__Ascomycota(100);c__Saccharomycetes(100);o__Saccharomycetales(100);f__Metschnikowiaceae(100);g__Metschnikowia(100);g__Metschnikowia_unclassified(100); | Metschnikowia | 13 | Saprotroph | \|Undefined Saprotroph\| | Yeast | NULL | Probable | As nectar-tap_saprotroph (Põlme et al. 2020); Decay sugar-rich substrates (Põlme et al. 2020) | Põlme S, et al. 2020. Fungal Diversity 105:1-16 ((https://doi.org/10.1007/s13225-020-00466-2)) |
| Otu57204 | 1 | k__Fungi(100);p__Ascomycota(100);c__Saccharomycetes(100);o__Saccharomycetales(100);f__Metschnikowiaceae(100);g__Metschnikowia(100);s__Metschnikowia_pulcherrima(100); | Metschnikowia | 13 | Saprotroph | \|Undefined Saprotroph\| | Yeast | NULL | Probable | As nectar-tap_saprotroph (Põlme et al. 2020); Decay sugar-rich substrates (Põlme et al. 2020) | Põlme S, et al. 2020. Fungal Diversity 105:1-16 ((https://doi.org/10.1007/s13225-020-00466-2)) |
| Otu57205 | 1 | k__Fungi(100);p__Ascomycota(100);c__Saccharomycetes(100);o__Saccharomycetales(100);f__Metschnikowiaceae(100);g__Metschnikowia(100);g__Metschnikowia_unclassified(100); | Metschnikowia | 13 | Saprotroph | \|Undefined Saprotroph\| | Yeast | NULL | Probable | As nectar-tap_saprotroph (Põlme et al. 2020); Decay sugar-rich substrates (Põlme et al. 2020) | Põlme S, et al. 2020. Fungal Diversity 105:1-16 ((https://doi.org/10.1007/s13225-020-00466-2)) |
| Otu57206 | 1 | k__Fungi(100);p__Ascomycota(100);c__Saccharomycetes(100);o__Saccharomycetales(100);f__Metschnikowiaceae(100);g__Metschnikowia(100);s__Metschnikowia_pulcherrima(100); | Metschnikowia | 13 | Saprotroph | \|Undefined Saprotroph\| | Yeast | NULL | Probable | As nectar-tap_saprotroph (Põlme et al. 2020); Decay sugar-rich substrates (Põlme et al. 2020) | Põlme S, et al. 2020. Fungal Diversity 105:1-16 ((https://doi.org/10.1007/s13225-020-00466-2)) |
| Otu57207 | 1 | k__Fungi(100);p__Ascomycota(100);c__Saccharomycetes(100);o__Saccharomycetales(100);f__Metschnikowiaceae(100);g__Metschnikowia(100);s__Metschnikowia_pulcherrima(100); | Metschnikowia | 13 | Saprotroph | \|Undefined Saprotroph\| | Yeast | NULL | Probable | As nectar-tap_saprotroph (Põlme et al. 2020); Decay sugar-rich substrates (Põlme et al. 2020) | Põlme S, et al. 2020. Fungal Diversity 105:1-16 ((https://doi.org/10.1007/s13225-020-00466-2)) |
| Otu57208 | 1 | k__Fungi(100);p__Ascomycota(100);c__Saccharomycetes(100);o__Saccharomycetales(100);f__Metschnikowiaceae(100);g__Metschnikowia(100);s__Metschnikowia_pulcherrima(100); | Metschnikowia | 13 | Saprotroph | \|Undefined Saprotroph\| | Yeast | NULL | Probable | As nectar-tap_saprotroph (Põlme et al. 2020); Decay sugar-rich substrates (Põlme et al. 2020) | Põlme S, et al. 2020. Fungal Diversity 105:1-16 ((https://doi.org/10.1007/s13225-020-00466-2)) |
| Otu57209 | 1 | k__Fungi(100);p__Ascomycota(100);c__Saccharomycetes(100);o__Saccharomycetales(100);f__Metschnikowiaceae(100);g__Metschnikowia(100);g__Metschnikowia_unclassified(100); | Metschnikowia | 13 | Saprotroph | \|Undefined Saprotroph\| | Yeast | NULL | Probable | As nectar-tap_saprotroph (Põlme et al. 2020); Decay sugar-rich substrates (Põlme et al. 2020) | Põlme S, et al. 2020. Fungal Diversity 105:1-16 ((https://doi.org/10.1007/s13225-020-00466-2)) |
| Otu57210 | 1 | k__Fungi(100);p__Ascomycota(100);c__Saccharomycetes(100);o__Saccharomycetales(100);f__Metschnikowiaceae(100);g__Metschnikowia(100);s__Metschnikowia_pulcherrima(100); | Metschnikowia | 13 | Saprotroph | \|Undefined Saprotroph\| | Yeast | NULL | Probable | As nectar-tap_saprotroph (Põlme et al. 2020); Decay sugar-rich substrates (Põlme et al. 2020) | Põlme S, et al. 2020. Fungal Diversity 105:1-16 ((https://doi.org/10.1007/s13225-020-00466-2)) |
| Otu57211 | 1 | k__Fungi(100);p__Ascomycota(100);c__Saccharomycetes(100);o__Saccharomycetales(100);f__Metschnikowiaceae(100);g__Metschnikowia(100);g__Metschnikowia_unclassified(100); | Metschnikowia | 13 | Saprotroph | \|Undefined Saprotroph\| | Yeast | NULL | Probable | As nectar-tap_saprotroph (Põlme et al. 2020); Decay sugar-rich substrates (Põlme et al. 2020) | Põlme S, et al. 2020. Fungal Diversity 105:1-16 ((https://doi.org/10.1007/s13225-020-00466-2)) |
| Otu57212 | 1 | k__Fungi(100);p__Ascomycota(100);c__Saccharomycetes(100);o__Saccharomycetales(100);f__Metschnikowiaceae(100);g__Metschnikowia(100);s__Metschnikowia_pulcherrima(100); | Metschnikowia | 13 | Saprotroph | \|Undefined Saprotroph\| | Yeast | NULL | Probable | As nectar-tap_saprotroph (Põlme et al. 2020); Decay sugar-rich substrates (Põlme et al. 2020) | Põlme S, et al. 2020. Fungal Diversity 105:1-16 ((https://doi.org/10.1007/s13225-020-00466-2)) |
| Otu57216 | 1 | k__Fungi(100);p__Ascomycota(100);c__Saccharomycetes(100);o__Saccharomycetales(100);f__Metschnikowiaceae(100);g__Metschnikowia(100);s__Metschnikowia_pulcherrima(100); | Metschnikowia | 13 | Saprotroph | \|Undefined Saprotroph\| | Yeast | NULL | Probable | As nectar-tap_saprotroph (Põlme et al. 2020); Decay sugar-rich substrates (Põlme et al. 2020) | Põlme S, et al. 2020. Fungal Diversity 105:1-16 ((https://doi.org/10.1007/s13225-020-00466-2)) |
| Otu57217 | 1 | k__Fungi(100);p__Ascomycota(100);c__Saccharomycetes(100);o__Saccharomycetales(100);f__Metschnikowiaceae(100);g__Metschnikowia(100);s__Metschnikowia_pulcherrima(100); | Metschnikowia | 13 | Saprotroph | \|Undefined Saprotroph\| | Yeast | NULL | Probable | As nectar-tap_saprotroph (Põlme et al. 2020); Decay sugar-rich substrates (Põlme et al. 2020) | Põlme S, et al. 2020. Fungal Diversity 105:1-16 ((https://doi.org/10.1007/s13225-020-00466-2)) |
| Otu57219 | 1 | k__Fungi(100);p__Ascomycota(100);c__Saccharomycetes(100);o__Saccharomycetales(100);f__Metschnikowiaceae(100);g__Metschnikowia(100);s__Metschnikowia_pulcherrima(100); | Metschnikowia | 13 | Saprotroph | \|Undefined Saprotroph\| | Yeast | NULL | Probable | As nectar-tap_saprotroph (Põlme et al. 2020); Decay sugar-rich substrates (Põlme et al. 2020) | Põlme S, et al. 2020. Fungal Diversity 105:1-16 ((https://doi.org/10.1007/s13225-020-00466-2)) |
| Otu57223 | 1 | k__Fungi(100);p__Ascomycota(100);c__Saccharomycetes(100);o__Saccharomycetales(100);f__Metschnikowiaceae(100);g__Metschnikowia(100);s__Metschnikowia_pulcherrima(100); | Metschnikowia | 13 | Saprotroph | \|Undefined Saprotroph\| | Yeast | NULL | Probable | As nectar-tap_saprotroph (Põlme et al. 2020); Decay sugar-rich substrates (Põlme et al. 2020) | Põlme S, et al. 2020. Fungal Diversity 105:1-16 ((https://doi.org/10.1007/s13225-020-00466-2)) |
| Otu57224 | 1 | k__Fungi(100);p__Ascomycota(100);c__Saccharomycetes(100);o__Saccharomycetales(100);f__Metschnikowiaceae(100);g__Metschnikowia(100);s__Metschnikowia_pulcherrima(100); | Metschnikowia | 13 | Saprotroph | \|Undefined Saprotroph\| | Yeast | NULL | Probable | As nectar-tap_saprotroph (Põlme et al. 2020); Decay sugar-rich substrates (Põlme et al. 2020) | Põlme S, et al. 2020. Fungal Diversity 105:1-16 ((https://doi.org/10.1007/s13225-020-00466-2)) |
| Otu57225 | 1 | k__Fungi(100);p__Ascomycota(100);c__Saccharomycetes(100);o__Saccharomycetales(100);f__Metschnikowiaceae(100);g__Metschnikowia(100);s__Metschnikowia_pulcherrima(100); | Metschnikowia | 13 | Saprotroph | \|Undefined Saprotroph\| | Yeast | NULL | Probable | As nectar-tap_saprotroph (Põlme et al. 2020); Decay sugar-rich substrates (Põlme et al. 2020) | Põlme S, et al. 2020. Fungal Diversity 105:1-16 ((https://doi.org/10.1007/s13225-020-00466-2)) |
| Otu57227 | 1 | k__Fungi(100);p__Ascomycota(100);c__Saccharomycetes(100);o__Saccharomycetales(100);f__Metschnikowiaceae(100);g__Metschnikowia(100);g__Metschnikowia_unclassified(100); | Metschnikowia | 13 | Saprotroph | \|Undefined Saprotroph\| | Yeast | NULL | Probable | As nectar-tap_saprotroph (Põlme et al. 2020); Decay sugar-rich substrates (Põlme et al. 2020) | Põlme S, et al. 2020. Fungal Diversity 105:1-16 ((https://doi.org/10.1007/s13225-020-00466-2)) |
| Otu57228 | 1 | k__Fungi(100);p__Ascomycota(100);c__Saccharomycetes(100);o__Saccharomycetales(100);f__Metschnikowiaceae(100);g__Metschnikowia(100);s__Metschnikowia_pulcherrima(100); | Metschnikowia | 13 | Saprotroph | \|Undefined Saprotroph\| | Yeast | NULL | Probable | As nectar-tap_saprotroph (Põlme et al. 2020); Decay sugar-rich substrates (Põlme et al. 2020) | Põlme S, et al. 2020. Fungal Diversity 105:1-16 ((https://doi.org/10.1007/s13225-020-00466-2)) |
| Otu57229 | 1 | k__Fungi(100);p__Ascomycota(100);c__Saccharomycetes(100);o__Saccharomycetales(100);f__Metschnikowiaceae(100);g__Metschnikowia(100);s__Metschnikowia_pulcherrima(100); | Metschnikowia | 13 | Saprotroph | \|Undefined Saprotroph\| | Yeast | NULL | Probable | As nectar-tap_saprotroph (Põlme et al. 2020); Decay sugar-rich substrates (Põlme et al. 2020) | Põlme S, et al. 2020. Fungal Diversity 105:1-16 ((https://doi.org/10.1007/s13225-020-00466-2)) |
| Otu57231 | 1 | k__Fungi(100);p__Ascomycota(100);c__Saccharomycetes(100);o__Saccharomycetales(100);f__Metschnikowiaceae(100);g__Metschnikowia(100);s__Metschnikowia_pulcherrima(100); | Metschnikowia | 13 | Saprotroph | \|Undefined Saprotroph\| | Yeast | NULL | Probable | As nectar-tap_saprotroph (Põlme et al. 2020); Decay sugar-rich substrates (Põlme et al. 2020) | Põlme S, et al. 2020. Fungal Diversity 105:1-16 ((https://doi.org/10.1007/s13225-020-00466-2)) |
| Otu57232 | 1 | k__Fungi(100);p__Ascomycota(100);c__Saccharomycetes(100);o__Saccharomycetales(100);f__Metschnikowiaceae(100);g__Metschnikowia(100);s__Metschnikowia_pulcherrima(100); | Metschnikowia | 13 | Saprotroph | \|Undefined Saprotroph\| | Yeast | NULL | Probable | As nectar-tap_saprotroph (Põlme et al. 2020); Decay sugar-rich substrates (Põlme et al. 2020) | Põlme S, et al. 2020. Fungal Diversity 105:1-16 ((https://doi.org/10.1007/s13225-020-00466-2)) |
| Otu57233 | 1 | k__Fungi(100);p__Ascomycota(100);c__Saccharomycetes(100);o__Saccharomycetales(100);f__Metschnikowiaceae(100);g__Metschnikowia(100);s__Metschnikowia_pulcherrima(100); | Metschnikowia | 13 | Saprotroph | \|Undefined Saprotroph\| | Yeast | NULL | Probable | As nectar-tap_saprotroph (Põlme et al. 2020); Decay sugar-rich substrates (Põlme et al. 2020) | Põlme S, et al. 2020. Fungal Diversity 105:1-16 ((https://doi.org/10.1007/s13225-020-00466-2)) |
| Otu57234 | 1 | k__Fungi(100);p__Ascomycota(100);c__Saccharomycetes(100);o__Saccharomycetales(100);f__Metschnikowiaceae(100);g__Metschnikowia(100);g__Metschnikowia_unclassified(100); | Metschnikowia | 13 | Saprotroph | \|Undefined Saprotroph\| | Yeast | NULL | Probable | As nectar-tap_saprotroph (Põlme et al. 2020); Decay sugar-rich substrates (Põlme et al. 2020) | Põlme S, et al. 2020. Fungal Diversity 105:1-16 ((https://doi.org/10.1007/s13225-020-00466-2)) |
| Otu57235 | 1 | k__Fungi(100);p__Ascomycota(100);c__Saccharomycetes(100);o__Saccharomycetales(100);f__Metschnikowiaceae(100);g__Metschnikowia(100);g__Metschnikowia_unclassified(100); | Metschnikowia | 13 | Saprotroph | \|Undefined Saprotroph\| | Yeast | NULL | Probable | As nectar-tap_saprotroph (Põlme et al. 2020); Decay sugar-rich substrates (Põlme et al. 2020) | Põlme S, et al. 2020. Fungal Diversity 105:1-16 ((https://doi.org/10.1007/s13225-020-00466-2)) |
| Otu57236 | 1 | k__Fungi(100);p__Ascomycota(100);c__Saccharomycetes(100);o__Saccharomycetales(100);f__Metschnikowiaceae(100);g__Metschnikowia(100);s__Metschnikowia_pulcherrima(100); | Metschnikowia | 13 | Saprotroph | \|Undefined Saprotroph\| | Yeast | NULL | Probable | As nectar-tap_saprotroph (Põlme et al. 2020); Decay sugar-rich substrates (Põlme et al. 2020) | Põlme S, et al. 2020. Fungal Diversity 105:1-16 ((https://doi.org/10.1007/s13225-020-00466-2)) |
| Otu57237 | 1 | k__Fungi(100);p__Ascomycota(100);c__Saccharomycetes(100);o__Saccharomycetales(100);f__Metschnikowiaceae(100);g__Metschnikowia(100);s__Metschnikowia_pulcherrima(100); | Metschnikowia | 13 | Saprotroph | \|Undefined Saprotroph\| | Yeast | NULL | Probable | As nectar-tap_saprotroph (Põlme et al. 2020); Decay sugar-rich substrates (Põlme et al. 2020) | Põlme S, et al. 2020. Fungal Diversity 105:1-16 ((https://doi.org/10.1007/s13225-020-00466-2)) |
| Otu57238 | 1 | k__Fungi(100);p__Ascomycota(100);c__Saccharomycetes(100);o__Saccharomycetales(100);f__Metschnikowiaceae(100);g__Metschnikowia(100);g__Metschnikowia_unclassified(100); | Metschnikowia | 13 | Saprotroph | \|Undefined Saprotroph\| | Yeast | NULL | Probable | As nectar-tap_saprotroph (Põlme et al. 2020); Decay sugar-rich substrates (Põlme et al. 2020) | Põlme S, et al. 2020. Fungal Diversity 105:1-16 ((https://doi.org/10.1007/s13225-020-00466-2)) |
| Otu57239 | 1 | k__Fungi(100);p__Ascomycota(100);c__Saccharomycetes(100);o__Saccharomycetales(100);f__Metschnikowiaceae(100);g__Metschnikowia(100);s__Metschnikowia_pulcherrima(100); | Metschnikowia | 13 | Saprotroph | \|Undefined Saprotroph\| | Yeast | NULL | Probable | As nectar-tap_saprotroph (Põlme et al. 2020); Decay sugar-rich substrates (Põlme et al. 2020) | Põlme S, et al. 2020. Fungal Diversity 105:1-16 ((https://doi.org/10.1007/s13225-020-00466-2)) |
| Otu57240 | 1 | k__Fungi(100);p__Ascomycota(100);c__Saccharomycetes(100);o__Saccharomycetales(100);f__Metschnikowiaceae(100);g__Metschnikowia(100);s__Metschnikowia_pulcherrima(100); | Metschnikowia | 13 | Saprotroph | \|Undefined Saprotroph\| | Yeast | NULL | Probable | As nectar-tap_saprotroph (Põlme et al. 2020); Decay sugar-rich substrates (Põlme et al. 2020) | Põlme S, et al. 2020. Fungal Diversity 105:1-16 ((https://doi.org/10.1007/s13225-020-00466-2)) |
| Otu57241 | 1 | k__Fungi(100);p__Ascomycota(100);c__Saccharomycetes(100);o__Saccharomycetales(100);f__Metschnikowiaceae(100);g__Metschnikowia(100);s__Metschnikowia_pulcherrima(100); | Metschnikowia | 13 | Saprotroph | \|Undefined Saprotroph\| | Yeast | NULL | Probable | As nectar-tap_saprotroph (Põlme et al. 2020); Decay sugar-rich substrates (Põlme et al. 2020) | Põlme S, et al. 2020. Fungal Diversity 105:1-16 ((https://doi.org/10.1007/s13225-020-00466-2)) |
| Otu57242 | 1 | k__Fungi(100);p__Ascomycota(100);c__Saccharomycetes(100);o__Saccharomycetales(100);f__Metschnikowiaceae(100);g__Metschnikowia(100);s__Metschnikowia_pulcherrima(100); | Metschnikowia | 13 | Saprotroph | \|Undefined Saprotroph\| | Yeast | NULL | Probable | As nectar-tap_saprotroph (Põlme et al. 2020); Decay sugar-rich substrates (Põlme et al. 2020) | Põlme S, et al. 2020. Fungal Diversity 105:1-16 ((https://doi.org/10.1007/s13225-020-00466-2)) |
| Otu57243 | 1 | k__Fungi(100);p__Ascomycota(100);c__Saccharomycetes(100);o__Saccharomycetales(100);f__Metschnikowiaceae(100);g__Metschnikowia(100);s__Metschnikowia_pulcherrima(100); | Metschnikowia | 13 | Saprotroph | \|Undefined Saprotroph\| | Yeast | NULL | Probable | As nectar-tap_saprotroph (Põlme et al. 2020); Decay sugar-rich substrates (Põlme et al. 2020) | Põlme S, et al. 2020. Fungal Diversity 105:1-16 ((https://doi.org/10.1007/s13225-020-00466-2)) |
| Otu57244 | 1 | k__Fungi(100);p__Ascomycota(100);c__Saccharomycetes(100);o__Saccharomycetales(100);f__Metschnikowiaceae(100);g__Metschnikowia(100);g__Metschnikowia_unclassified(100); | Metschnikowia | 13 | Saprotroph | \|Undefined Saprotroph\| | Yeast | NULL | Probable | As nectar-tap_saprotroph (Põlme et al. 2020); Decay sugar-rich substrates (Põlme et al. 2020) | Põlme S, et al. 2020. Fungal Diversity 105:1-16 ((https://doi.org/10.1007/s13225-020-00466-2)) |
| Otu57245 | 1 | k__Fungi(100);p__Ascomycota(100);c__Saccharomycetes(100);o__Saccharomycetales(100);f__Metschnikowiaceae(100);g__Metschnikowia(100);g__Metschnikowia_unclassified(100); | Metschnikowia | 13 | Saprotroph | \|Undefined Saprotroph\| | Yeast | NULL | Probable | As nectar-tap_saprotroph (Põlme et al. 2020); Decay sugar-rich substrates (Põlme et al. 2020) | Põlme S, et al. 2020. Fungal Diversity 105:1-16 ((https://doi.org/10.1007/s13225-020-00466-2)) |
| Otu57246 | 1 | k__Fungi(100);p__Ascomycota(100);c__Saccharomycetes(100);o__Saccharomycetales(100);f__Metschnikowiaceae(100);g__Metschnikowia(100);g__Metschnikowia_unclassified(100); | Metschnikowia | 13 | Saprotroph | \|Undefined Saprotroph\| | Yeast | NULL | Probable | As nectar-tap_saprotroph (Põlme et al. 2020); Decay sugar-rich substrates (Põlme et al. 2020) | Põlme S, et al. 2020. Fungal Diversity 105:1-16 ((https://doi.org/10.1007/s13225-020-00466-2)) |
| Otu57247 | 1 | k__Fungi(100);p__Ascomycota(100);c__Saccharomycetes(100);o__Saccharomycetales(100);f__Metschnikowiaceae(100);g__Metschnikowia(100);s__Metschnikowia_pulcherrima(100); | Metschnikowia | 13 | Saprotroph | \|Undefined Saprotroph\| | Yeast | NULL | Probable | As nectar-tap_saprotroph (Põlme et al. 2020); Decay sugar-rich substrates (Põlme et al. 2020) | Põlme S, et al. 2020. Fungal Diversity 105:1-16 ((https://doi.org/10.1007/s13225-020-00466-2)) |
| Otu57248 | 1 | k__Fungi(100);p__Ascomycota(100);c__Saccharomycetes(100);o__Saccharomycetales(100);f__Metschnikowiaceae(100);g__Metschnikowia(100);s__Metschnikowia_pulcherrima(100); | Metschnikowia | 13 | Saprotroph | \|Undefined Saprotroph\| | Yeast | NULL | Probable | As nectar-tap_saprotroph (Põlme et al. 2020); Decay sugar-rich substrates (Põlme et al. 2020) | Põlme S, et al. 2020. Fungal Diversity 105:1-16 ((https://doi.org/10.1007/s13225-020-00466-2)) |
| Otu57249 | 1 | k__Fungi(100);p__Ascomycota(100);c__Saccharomycetes(100);o__Saccharomycetales(100);f__Metschnikowiaceae(100);g__Metschnikowia(100);s__Metschnikowia_pulcherrima(100); | Metschnikowia | 13 | Saprotroph | \|Undefined Saprotroph\| | Yeast | NULL | Probable | As nectar-tap_saprotroph (Põlme et al. 2020); Decay sugar-rich substrates (Põlme et al. 2020) | Põlme S, et al. 2020. Fungal Diversity 105:1-16 ((https://doi.org/10.1007/s13225-020-00466-2)) |
| Otu57250 | 1 | k__Fungi(100);p__Ascomycota(100);c__Saccharomycetes(100);o__Saccharomycetales(100);f__Metschnikowiaceae(100);g__Metschnikowia(100);s__Metschnikowia_pulcherrima(100); | Metschnikowia | 13 | Saprotroph | \|Undefined Saprotroph\| | Yeast | NULL | Probable | As nectar-tap_saprotroph (Põlme et al. 2020); Decay sugar-rich substrates (Põlme et al. 2020) | Põlme S, et al. 2020. Fungal Diversity 105:1-16 ((https://doi.org/10.1007/s13225-020-00466-2)) |
| Otu57251 | 1 | k__Fungi(100);p__Ascomycota(100);c__Saccharomycetes(100);o__Saccharomycetales(100);f__Metschnikowiaceae(100);g__Metschnikowia(100);s__Metschnikowia_pulcherrima(100); | Metschnikowia | 13 | Saprotroph | \|Undefined Saprotroph\| | Yeast | NULL | Probable | As nectar-tap_saprotroph (Põlme et al. 2020); Decay sugar-rich substrates (Põlme et al. 2020) | Põlme S, et al. 2020. Fungal Diversity 105:1-16 ((https://doi.org/10.1007/s13225-020-00466-2)) |
| Otu57259 | 1 | k__Fungi(100);p__Ascomycota(100);c__Saccharomycetes(100);o__Saccharomycetales(100);f__Metschnikowiaceae(100);g__Metschnikowia(100);g__Metschnikowia_unclassified(100); | Metschnikowia | 13 | Saprotroph | \|Undefined Saprotroph\| | Yeast | NULL | Probable | As nectar-tap_saprotroph (Põlme et al. 2020); Decay sugar-rich substrates (Põlme et al. 2020) | Põlme S, et al. 2020. Fungal Diversity 105:1-16 ((https://doi.org/10.1007/s13225-020-00466-2)) |
| Otu57260 | 1 | k__Fungi(100);p__Ascomycota(100);c__Saccharomycetes(100);o__Saccharomycetales(100);f__Metschnikowiaceae(100);g__Metschnikowia(100);s__Metschnikowia_pulcherrima(100); | Metschnikowia | 13 | Saprotroph | \|Undefined Saprotroph\| | Yeast | NULL | Probable | As nectar-tap_saprotroph (Põlme et al. 2020); Decay sugar-rich substrates (Põlme et al. 2020) | Põlme S, et al. 2020. Fungal Diversity 105:1-16 ((https://doi.org/10.1007/s13225-020-00466-2)) |
| Otu57261 | 1 | k__Fungi(100);p__Ascomycota(100);c__Saccharomycetes(100);o__Saccharomycetales(100);f__Metschnikowiaceae(100);g__Metschnikowia(100);s__Metschnikowia_pulcherrima(100); | Metschnikowia | 13 | Saprotroph | \|Undefined Saprotroph\| | Yeast | NULL | Probable | As nectar-tap_saprotroph (Põlme et al. 2020); Decay sugar-rich substrates (Põlme et al. 2020) | Põlme S, et al. 2020. Fungal Diversity 105:1-16 ((https://doi.org/10.1007/s13225-020-00466-2)) |
| Otu57262 | 1 | k__Fungi(100);p__Ascomycota(100);c__Saccharomycetes(100);o__Saccharomycetales(100);f__Metschnikowiaceae(100);g__Metschnikowia(100);g__Metschnikowia_unclassified(100); | Metschnikowia | 13 | Saprotroph | \|Undefined Saprotroph\| | Yeast | NULL | Probable | As nectar-tap_saprotroph (Põlme et al. 2020); Decay sugar-rich substrates (Põlme et al. 2020) | Põlme S, et al. 2020. Fungal Diversity 105:1-16 ((https://doi.org/10.1007/s13225-020-00466-2)) |
| Otu57263 | 1 | k__Fungi(100);p__Ascomycota(100);c__Saccharomycetes(100);o__Saccharomycetales(100);f__Metschnikowiaceae(100);g__Metschnikowia(100);g__Metschnikowia_unclassified(100); | Metschnikowia | 13 | Saprotroph | \|Undefined Saprotroph\| | Yeast | NULL | Probable | As nectar-tap_saprotroph (Põlme et al. 2020); Decay sugar-rich substrates (Põlme et al. 2020) | Põlme S, et al. 2020. Fungal Diversity 105:1-16 ((https://doi.org/10.1007/s13225-020-00466-2)) |
| Otu57265 | 1 | k__Fungi(100);p__Ascomycota(100);c__Saccharomycetes(100);o__Saccharomycetales(100);f__Metschnikowiaceae(100);g__Metschnikowia(100);g__Metschnikowia_unclassified(100); | Metschnikowia | 13 | Saprotroph | \|Undefined Saprotroph\| | Yeast | NULL | Probable | As nectar-tap_saprotroph (Põlme et al. 2020); Decay sugar-rich substrates (Põlme et al. 2020) | Põlme S, et al. 2020. Fungal Diversity 105:1-16 ((https://doi.org/10.1007/s13225-020-00466-2)) |
| Otu57266 | 1 | k__Fungi(100);p__Ascomycota(100);c__Saccharomycetes(100);o__Saccharomycetales(100);f__Metschnikowiaceae(100);g__Metschnikowia(100);s__Metschnikowia_pulcherrima(100); | Metschnikowia | 13 | Saprotroph | \|Undefined Saprotroph\| | Yeast | NULL | Probable | As nectar-tap_saprotroph (Põlme et al. 2020); Decay sugar-rich substrates (Põlme et al. 2020) | Põlme S, et al. 2020. Fungal Diversity 105:1-16 ((https://doi.org/10.1007/s13225-020-00466-2)) |
| Otu57267 | 1 | k__Fungi(100);p__Ascomycota(100);c__Saccharomycetes(100);o__Saccharomycetales(100);f__Metschnikowiaceae(100);g__Metschnikowia(100);s__Metschnikowia_pulcherrima(100); | Metschnikowia | 13 | Saprotroph | \|Undefined Saprotroph\| | Yeast | NULL | Probable | As nectar-tap_saprotroph (Põlme et al. 2020); Decay sugar-rich substrates (Põlme et al. 2020) | Põlme S, et al. 2020. Fungal Diversity 105:1-16 ((https://doi.org/10.1007/s13225-020-00466-2)) |
| Otu57268 | 1 | k__Fungi(100);p__Ascomycota(100);c__Saccharomycetes(100);o__Saccharomycetales(100);f__Metschnikowiaceae(100);g__Metschnikowia(100);s__Metschnikowia_pulcherrima(100); | Metschnikowia | 13 | Saprotroph | \|Undefined Saprotroph\| | Yeast | NULL | Probable | As nectar-tap_saprotroph (Põlme et al. 2020); Decay sugar-rich substrates (Põlme et al. 2020) | Põlme S, et al. 2020. Fungal Diversity 105:1-16 ((https://doi.org/10.1007/s13225-020-00466-2)) |
| Otu57269 | 1 | k__Fungi(100);p__Ascomycota(100);c__Saccharomycetes(100);o__Saccharomycetales(100);f__Metschnikowiaceae(100);g__Metschnikowia(100);g__Metschnikowia_unclassified(100); | Metschnikowia | 13 | Saprotroph | \|Undefined Saprotroph\| | Yeast | NULL | Probable | As nectar-tap_saprotroph (Põlme et al. 2020); Decay sugar-rich substrates (Põlme et al. 2020) | Põlme S, et al. 2020. Fungal Diversity 105:1-16 ((https://doi.org/10.1007/s13225-020-00466-2)) |
| Otu57270 | 1 | k__Fungi(100);p__Ascomycota(100);c__Saccharomycetes(100);o__Saccharomycetales(100);f__Metschnikowiaceae(100);g__Metschnikowia(100);s__Metschnikowia_pulcherrima(100); | Metschnikowia | 13 | Saprotroph | \|Undefined Saprotroph\| | Yeast | NULL | Probable | As nectar-tap_saprotroph (Põlme et al. 2020); Decay sugar-rich substrates (Põlme et al. 2020) | Põlme S, et al. 2020. Fungal Diversity 105:1-16 ((https://doi.org/10.1007/s13225-020-00466-2)) |
| Otu57271 | 1 | k__Fungi(100);p__Ascomycota(100);c__Saccharomycetes(100);o__Saccharomycetales(100);f__Metschnikowiaceae(100);g__Metschnikowia(100);s__Metschnikowia_pulcherrima(100); | Metschnikowia | 13 | Saprotroph | \|Undefined Saprotroph\| | Yeast | NULL | Probable | As nectar-tap_saprotroph (Põlme et al. 2020); Decay sugar-rich substrates (Põlme et al. 2020) | Põlme S, et al. 2020. Fungal Diversity 105:1-16 ((https://doi.org/10.1007/s13225-020-00466-2)) |
| Otu57272 | 1 | k__Fungi(100);p__Ascomycota(100);c__Saccharomycetes(100);o__Saccharomycetales(100);f__Metschnikowiaceae(100);g__Metschnikowia(100);s__Metschnikowia_pulcherrima(100); | Metschnikowia | 13 | Saprotroph | \|Undefined Saprotroph\| | Yeast | NULL | Probable | As nectar-tap_saprotroph (Põlme et al. 2020); Decay sugar-rich substrates (Põlme et al. 2020) | Põlme S, et al. 2020. Fungal Diversity 105:1-16 ((https://doi.org/10.1007/s13225-020-00466-2)) |
| Otu57273 | 1 | k__Fungi(100);p__Ascomycota(100);c__Saccharomycetes(100);o__Saccharomycetales(100);f__Metschnikowiaceae(100);g__Metschnikowia(100);s__Metschnikowia_pulcherrima(100); | Metschnikowia | 13 | Saprotroph | \|Undefined Saprotroph\| | Yeast | NULL | Probable | As nectar-tap_saprotroph (Põlme et al. 2020); Decay sugar-rich substrates (Põlme et al. 2020) | Põlme S, et al. 2020. Fungal Diversity 105:1-16 ((https://doi.org/10.1007/s13225-020-00466-2)) |
| Otu57274 | 1 | k__Fungi(100);p__Ascomycota(100);c__Saccharomycetes(100);o__Saccharomycetales(100);f__Metschnikowiaceae(100);g__Metschnikowia(100);g__Metschnikowia_unclassified(100); | Metschnikowia | 13 | Saprotroph | \|Undefined Saprotroph\| | Yeast | NULL | Probable | As nectar-tap_saprotroph (Põlme et al. 2020); Decay sugar-rich substrates (Põlme et al. 2020) | Põlme S, et al. 2020. Fungal Diversity 105:1-16 ((https://doi.org/10.1007/s13225-020-00466-2)) |
| Otu57275 | 1 | k__Fungi(100);p__Ascomycota(100);c__Saccharomycetes(100);o__Saccharomycetales(100);f__Metschnikowiaceae(100);g__Metschnikowia(100);g__Metschnikowia_unclassified(100); | Metschnikowia | 13 | Saprotroph | \|Undefined Saprotroph\| | Yeast | NULL | Probable | As nectar-tap_saprotroph (Põlme et al. 2020); Decay sugar-rich substrates (Põlme et al. 2020) | Põlme S, et al. 2020. Fungal Diversity 105:1-16 ((https://doi.org/10.1007/s13225-020-00466-2)) |
| Otu57276 | 1 | k__Fungi(100);p__Ascomycota(100);c__Saccharomycetes(100);o__Saccharomycetales(100);f__Metschnikowiaceae(100);g__Metschnikowia(100);s__Metschnikowia_pulcherrima(100); | Metschnikowia | 13 | Saprotroph | \|Undefined Saprotroph\| | Yeast | NULL | Probable | As nectar-tap_saprotroph (Põlme et al. 2020); Decay sugar-rich substrates (Põlme et al. 2020) | Põlme S, et al. 2020. Fungal Diversity 105:1-16 ((https://doi.org/10.1007/s13225-020-00466-2)) |
| Otu57277 | 1 | k__Fungi(100);p__Ascomycota(100);c__Saccharomycetes(100);o__Saccharomycetales(100);f__Metschnikowiaceae(100);g__Metschnikowia(100);g__Metschnikowia_unclassified(100); | Metschnikowia | 13 | Saprotroph | \|Undefined Saprotroph\| | Yeast | NULL | Probable | As nectar-tap_saprotroph (Põlme et al. 2020); Decay sugar-rich substrates (Põlme et al. 2020) | Põlme S, et al. 2020. Fungal Diversity 105:1-16 ((https://doi.org/10.1007/s13225-020-00466-2)) |
| Otu57278 | 1 | k__Fungi(100);p__Ascomycota(100);c__Saccharomycetes(100);o__Saccharomycetales(100);f__Metschnikowiaceae(100);g__Metschnikowia(100);s__Metschnikowia_pulcherrima(100); | Metschnikowia | 13 | Saprotroph | \|Undefined Saprotroph\| | Yeast | NULL | Probable | As nectar-tap_saprotroph (Põlme et al. 2020); Decay sugar-rich substrates (Põlme et al. 2020) | Põlme S, et al. 2020. Fungal Diversity 105:1-16 ((https://doi.org/10.1007/s13225-020-00466-2)) |
| Otu57279 | 1 | k__Fungi(100);p__Ascomycota(100);c__Saccharomycetes(100);o__Saccharomycetales(100);f__Metschnikowiaceae(100);g__Metschnikowia(100);g__Metschnikowia_unclassified(100); | Metschnikowia | 13 | Saprotroph | \|Undefined Saprotroph\| | Yeast | NULL | Probable | As nectar-tap_saprotroph (Põlme et al. 2020); Decay sugar-rich substrates (Põlme et al. 2020) | Põlme S, et al. 2020. Fungal Diversity 105:1-16 ((https://doi.org/10.1007/s13225-020-00466-2)) |
| Otu57280 | 1 | k__Fungi(100);p__Ascomycota(100);c__Saccharomycetes(100);o__Saccharomycetales(100);f__Metschnikowiaceae(100);g__Metschnikowia(100);s__Metschnikowia_pulcherrima(100); | Metschnikowia | 13 | Saprotroph | \|Undefined Saprotroph\| | Yeast | NULL | Probable | As nectar-tap_saprotroph (Põlme et al. 2020); Decay sugar-rich substrates (Põlme et al. 2020) | Põlme S, et al. 2020. Fungal Diversity 105:1-16 ((https://doi.org/10.1007/s13225-020-00466-2)) |
| Otu57281 | 1 | k__Fungi(100);p__Ascomycota(100);c__Saccharomycetes(100);o__Saccharomycetales(100);f__Metschnikowiaceae(100);g__Metschnikowia(100);g__Metschnikowia_unclassified(100); | Metschnikowia | 13 | Saprotroph | \|Undefined Saprotroph\| | Yeast | NULL | Probable | As nectar-tap_saprotroph (Põlme et al. 2020); Decay sugar-rich substrates (Põlme et al. 2020) | Põlme S, et al. 2020. Fungal Diversity 105:1-16 ((https://doi.org/10.1007/s13225-020-00466-2)) |
| Otu57282 | 1 | k__Fungi(100);p__Ascomycota(100);c__Saccharomycetes(100);o__Saccharomycetales(100);f__Metschnikowiaceae(100);g__Metschnikowia(100);g__Metschnikowia_unclassified(100); | Metschnikowia | 13 | Saprotroph | \|Undefined Saprotroph\| | Yeast | NULL | Probable | As nectar-tap_saprotroph (Põlme et al. 2020); Decay sugar-rich substrates (Põlme et al. 2020) | Põlme S, et al. 2020. Fungal Diversity 105:1-16 ((https://doi.org/10.1007/s13225-020-00466-2)) |
| Otu57283 | 1 | k__Fungi(100);p__Ascomycota(100);c__Saccharomycetes(100);o__Saccharomycetales(100);f__Metschnikowiaceae(100);g__Metschnikowia(100);s__Metschnikowia_pulcherrima(100); | Metschnikowia | 13 | Saprotroph | \|Undefined Saprotroph\| | Yeast | NULL | Probable | As nectar-tap_saprotroph (Põlme et al. 2020); Decay sugar-rich substrates (Põlme et al. 2020) | Põlme S, et al. 2020. Fungal Diversity 105:1-16 ((https://doi.org/10.1007/s13225-020-00466-2)) |
| Otu57284 | 1 | k__Fungi(100);p__Ascomycota(100);c__Saccharomycetes(100);o__Saccharomycetales(100);f__Metschnikowiaceae(100);g__Metschnikowia(100);s__Metschnikowia_pulcherrima(100); | Metschnikowia | 13 | Saprotroph | \|Undefined Saprotroph\| | Yeast | NULL | Probable | As nectar-tap_saprotroph (Põlme et al. 2020); Decay sugar-rich substrates (Põlme et al. 2020) | Põlme S, et al. 2020. Fungal Diversity 105:1-16 ((https://doi.org/10.1007/s13225-020-00466-2)) |
| Otu57285 | 1 | k__Fungi(100);p__Ascomycota(100);c__Saccharomycetes(100);o__Saccharomycetales(100);f__Metschnikowiaceae(100);g__Metschnikowia(100);s__Metschnikowia_pulcherrima(100); | Metschnikowia | 13 | Saprotroph | \|Undefined Saprotroph\| | Yeast | NULL | Probable | As nectar-tap_saprotroph (Põlme et al. 2020); Decay sugar-rich substrates (Põlme et al. 2020) | Põlme S, et al. 2020. Fungal Diversity 105:1-16 ((https://doi.org/10.1007/s13225-020-00466-2)) |
| Otu57287 | 1 | k__Fungi(100);p__Ascomycota(100);c__Saccharomycetes(100);o__Saccharomycetales(100);f__Metschnikowiaceae(100);g__Metschnikowia(100);g__Metschnikowia_unclassified(100); | Metschnikowia | 13 | Saprotroph | \|Undefined Saprotroph\| | Yeast | NULL | Probable | As nectar-tap_saprotroph (Põlme et al. 2020); Decay sugar-rich substrates (Põlme et al. 2020) | Põlme S, et al. 2020. Fungal Diversity 105:1-16 ((https://doi.org/10.1007/s13225-020-00466-2)) |
| Otu57288 | 1 | k__Fungi(100);p__Ascomycota(100);c__Saccharomycetes(100);o__Saccharomycetales(100);f__Metschnikowiaceae(100);g__Metschnikowia(100);g__Metschnikowia_unclassified(100); | Metschnikowia | 13 | Saprotroph | \|Undefined Saprotroph\| | Yeast | NULL | Probable | As nectar-tap_saprotroph (Põlme et al. 2020); Decay sugar-rich substrates (Põlme et al. 2020) | Põlme S, et al. 2020. Fungal Diversity 105:1-16 ((https://doi.org/10.1007/s13225-020-00466-2)) |
| Otu57289 | 1 | k__Fungi(100);p__Ascomycota(100);c__Saccharomycetes(100);o__Saccharomycetales(100);f__Metschnikowiaceae(100);g__Metschnikowia(100);g__Metschnikowia_unclassified(100); | Metschnikowia | 13 | Saprotroph | \|Undefined Saprotroph\| | Yeast | NULL | Probable | As nectar-tap_saprotroph (Põlme et al. 2020); Decay sugar-rich substrates (Põlme et al. 2020) | Põlme S, et al. 2020. Fungal Diversity 105:1-16 ((https://doi.org/10.1007/s13225-020-00466-2)) |
| Otu57290 | 1 | k__Fungi(100);p__Ascomycota(100);c__Saccharomycetes(100);o__Saccharomycetales(100);f__Metschnikowiaceae(100);g__Metschnikowia(100);g__Metschnikowia_unclassified(100); | Metschnikowia | 13 | Saprotroph | \|Undefined Saprotroph\| | Yeast | NULL | Probable | As nectar-tap_saprotroph (Põlme et al. 2020); Decay sugar-rich substrates (Põlme et al. 2020) | Põlme S, et al. 2020. Fungal Diversity 105:1-16 ((https://doi.org/10.1007/s13225-020-00466-2)) |
| Otu57291 | 1 | k__Fungi(100);p__Ascomycota(100);c__Saccharomycetes(100);o__Saccharomycetales(100);f__Metschnikowiaceae(100);g__Metschnikowia(100);g__Metschnikowia_unclassified(100); | Metschnikowia | 13 | Saprotroph | \|Undefined Saprotroph\| | Yeast | NULL | Probable | As nectar-tap_saprotroph (Põlme et al. 2020); Decay sugar-rich substrates (Põlme et al. 2020) | Põlme S, et al. 2020. Fungal Diversity 105:1-16 ((https://doi.org/10.1007/s13225-020-00466-2)) |
| Otu57292 | 1 | k__Fungi(100);p__Ascomycota(100);c__Saccharomycetes(100);o__Saccharomycetales(100);f__Metschnikowiaceae(100);g__Metschnikowia(100);g__Metschnikowia_unclassified(100); | Metschnikowia | 13 | Saprotroph | \|Undefined Saprotroph\| | Yeast | NULL | Probable | As nectar-tap_saprotroph (Põlme et al. 2020); Decay sugar-rich substrates (Põlme et al. 2020) | Põlme S, et al. 2020. Fungal Diversity 105:1-16 ((https://doi.org/10.1007/s13225-020-00466-2)) |
| Otu57293 | 1 | k__Fungi(100);p__Ascomycota(100);c__Saccharomycetes(100);o__Saccharomycetales(100);f__Metschnikowiaceae(100);g__Metschnikowia(100);g__Metschnikowia_unclassified(100); | Metschnikowia | 13 | Saprotroph | \|Undefined Saprotroph\| | Yeast | NULL | Probable | As nectar-tap_saprotroph (Põlme et al. 2020); Decay sugar-rich substrates (Põlme et al. 2020) | Põlme S, et al. 2020. Fungal Diversity 105:1-16 ((https://doi.org/10.1007/s13225-020-00466-2)) |
| Otu57294 | 1 | k__Fungi(100);p__Ascomycota(100);c__Saccharomycetes(100);o__Saccharomycetales(100);f__Metschnikowiaceae(100);g__Metschnikowia(100);s__Metschnikowia_pulcherrima(100); | Metschnikowia | 13 | Saprotroph | \|Undefined Saprotroph\| | Yeast | NULL | Probable | As nectar-tap_saprotroph (Põlme et al. 2020); Decay sugar-rich substrates (Põlme et al. 2020) | Põlme S, et al. 2020. Fungal Diversity 105:1-16 ((https://doi.org/10.1007/s13225-020-00466-2)) |
| Otu57295 | 1 | k__Fungi(100);p__Ascomycota(100);c__Saccharomycetes(100);o__Saccharomycetales(100);f__Metschnikowiaceae(100);g__Metschnikowia(100);s__Metschnikowia_pulcherrima(100); | Metschnikowia | 13 | Saprotroph | \|Undefined Saprotroph\| | Yeast | NULL | Probable | As nectar-tap_saprotroph (Põlme et al. 2020); Decay sugar-rich substrates (Põlme et al. 2020) | Põlme S, et al. 2020. Fungal Diversity 105:1-16 ((https://doi.org/10.1007/s13225-020-00466-2)) |
| Otu57296 | 1 | k__Fungi(100);p__Ascomycota(100);c__Saccharomycetes(100);o__Saccharomycetales(100);f__Metschnikowiaceae(100);g__Metschnikowia(100);s__Metschnikowia_pulcherrima(100); | Metschnikowia | 13 | Saprotroph | \|Undefined Saprotroph\| | Yeast | NULL | Probable | As nectar-tap_saprotroph (Põlme et al. 2020); Decay sugar-rich substrates (Põlme et al. 2020) | Põlme S, et al. 2020. Fungal Diversity 105:1-16 ((https://doi.org/10.1007/s13225-020-00466-2)) |
| Otu57297 | 1 | k__Fungi(100);p__Ascomycota(100);c__Saccharomycetes(100);o__Saccharomycetales(100);f__Metschnikowiaceae(100);g__Metschnikowia(100);s__Metschnikowia_pulcherrima(100); | Metschnikowia | 13 | Saprotroph | \|Undefined Saprotroph\| | Yeast | NULL | Probable | As nectar-tap_saprotroph (Põlme et al. 2020); Decay sugar-rich substrates (Põlme et al. 2020) | Põlme S, et al. 2020. Fungal Diversity 105:1-16 ((https://doi.org/10.1007/s13225-020-00466-2)) |
| Otu57298 | 1 | k__Fungi(100);p__Ascomycota(100);c__Saccharomycetes(100);o__Saccharomycetales(100);f__Metschnikowiaceae(100);g__Metschnikowia(100);s__Metschnikowia_pulcherrima(100); | Metschnikowia | 13 | Saprotroph | \|Undefined Saprotroph\| | Yeast | NULL | Probable | As nectar-tap_saprotroph (Põlme et al. 2020); Decay sugar-rich substrates (Põlme et al. 2020) | Põlme S, et al. 2020. Fungal Diversity 105:1-16 ((https://doi.org/10.1007/s13225-020-00466-2)) |
| Otu57299 | 1 | k__Fungi(100);p__Ascomycota(100);c__Saccharomycetes(100);o__Saccharomycetales(100);f__Metschnikowiaceae(100);g__Metschnikowia(100);g__Metschnikowia_unclassified(100); | Metschnikowia | 13 | Saprotroph | \|Undefined Saprotroph\| | Yeast | NULL | Probable | As nectar-tap_saprotroph (Põlme et al. 2020); Decay sugar-rich substrates (Põlme et al. 2020) | Põlme S, et al. 2020. Fungal Diversity 105:1-16 ((https://doi.org/10.1007/s13225-020-00466-2)) |
| Otu57300 | 1 | k__Fungi(100);p__Ascomycota(100);c__Saccharomycetes(100);o__Saccharomycetales(100);f__Metschnikowiaceae(100);g__Metschnikowia(100);g__Metschnikowia_unclassified(100); | Metschnikowia | 13 | Saprotroph | \|Undefined Saprotroph\| | Yeast | NULL | Probable | As nectar-tap_saprotroph (Põlme et al. 2020); Decay sugar-rich substrates (Põlme et al. 2020) | Põlme S, et al. 2020. Fungal Diversity 105:1-16 ((https://doi.org/10.1007/s13225-020-00466-2)) |
| Otu57301 | 1 | k__Fungi(100);p__Ascomycota(100);c__Saccharomycetes(100);o__Saccharomycetales(100);f__Metschnikowiaceae(100);g__Metschnikowia(100);s__Metschnikowia_pulcherrima(100); | Metschnikowia | 13 | Saprotroph | \|Undefined Saprotroph\| | Yeast | NULL | Probable | As nectar-tap_saprotroph (Põlme et al. 2020); Decay sugar-rich substrates (Põlme et al. 2020) | Põlme S, et al. 2020. Fungal Diversity 105:1-16 ((https://doi.org/10.1007/s13225-020-00466-2)) |
| Otu57302 | 1 | k__Fungi(100);p__Ascomycota(100);c__Saccharomycetes(100);o__Saccharomycetales(100);f__Metschnikowiaceae(100);g__Metschnikowia(100);s__Metschnikowia_pulcherrima(100); | Metschnikowia | 13 | Saprotroph | \|Undefined Saprotroph\| | Yeast | NULL | Probable | As nectar-tap_saprotroph (Põlme et al. 2020); Decay sugar-rich substrates (Põlme et al. 2020) | Põlme S, et al. 2020. Fungal Diversity 105:1-16 ((https://doi.org/10.1007/s13225-020-00466-2)) |
| Otu57303 | 1 | k__Fungi(100);p__Ascomycota(100);c__Saccharomycetes(100);o__Saccharomycetales(100);f__Metschnikowiaceae(100);g__Metschnikowia(100);s__Metschnikowia_pulcherrima(100); | Metschnikowia | 13 | Saprotroph | \|Undefined Saprotroph\| | Yeast | NULL | Probable | As nectar-tap_saprotroph (Põlme et al. 2020); Decay sugar-rich substrates (Põlme et al. 2020) | Põlme S, et al. 2020. Fungal Diversity 105:1-16 ((https://doi.org/10.1007/s13225-020-00466-2)) |
| Otu57304 | 1 | k__Fungi(100);p__Ascomycota(100);c__Saccharomycetes(100);o__Saccharomycetales(100);f__Metschnikowiaceae(100);g__Metschnikowia(100);s__Metschnikowia_pulcherrima(100); | Metschnikowia | 13 | Saprotroph | \|Undefined Saprotroph\| | Yeast | NULL | Probable | As nectar-tap_saprotroph (Põlme et al. 2020); Decay sugar-rich substrates (Põlme et al. 2020) | Põlme S, et al. 2020. Fungal Diversity 105:1-16 ((https://doi.org/10.1007/s13225-020-00466-2)) |
| Otu57305 | 1 | k__Fungi(100);p__Ascomycota(100);c__Saccharomycetes(100);o__Saccharomycetales(100);f__Metschnikowiaceae(100);g__Metschnikowia(100);s__Metschnikowia_pulcherrima(100); | Metschnikowia | 13 | Saprotroph | \|Undefined Saprotroph\| | Yeast | NULL | Probable | As nectar-tap_saprotroph (Põlme et al. 2020); Decay sugar-rich substrates (Põlme et al. 2020) | Põlme S, et al. 2020. Fungal Diversity 105:1-16 ((https://doi.org/10.1007/s13225-020-00466-2)) |
| Otu57306 | 1 | k__Fungi(100);p__Ascomycota(100);c__Saccharomycetes(100);o__Saccharomycetales(100);f__Metschnikowiaceae(100);g__Metschnikowia(100);g__Metschnikowia_unclassified(100); | Metschnikowia | 13 | Saprotroph | \|Undefined Saprotroph\| | Yeast | NULL | Probable | As nectar-tap_saprotroph (Põlme et al. 2020); Decay sugar-rich substrates (Põlme et al. 2020) | Põlme S, et al. 2020. Fungal Diversity 105:1-16 ((https://doi.org/10.1007/s13225-020-00466-2)) |
| Otu57307 | 1 | k__Fungi(100);p__Ascomycota(100);c__Saccharomycetes(100);o__Saccharomycetales(100);f__Metschnikowiaceae(100);g__Metschnikowia(100);s__Metschnikowia_pulcherrima(100); | Metschnikowia | 13 | Saprotroph | \|Undefined Saprotroph\| | Yeast | NULL | Probable | As nectar-tap_saprotroph (Põlme et al. 2020); Decay sugar-rich substrates (Põlme et al. 2020) | Põlme S, et al. 2020. Fungal Diversity 105:1-16 ((https://doi.org/10.1007/s13225-020-00466-2)) |
| Otu57308 | 1 | k__Fungi(100);p__Ascomycota(100);c__Saccharomycetes(100);o__Saccharomycetales(100);f__Metschnikowiaceae(100);g__Metschnikowia(100);s__Metschnikowia_pulcherrima(100); | Metschnikowia | 13 | Saprotroph | \|Undefined Saprotroph\| | Yeast | NULL | Probable | As nectar-tap_saprotroph (Põlme et al. 2020); Decay sugar-rich substrates (Põlme et al. 2020) | Põlme S, et al. 2020. Fungal Diversity 105:1-16 ((https://doi.org/10.1007/s13225-020-00466-2)) |
| Otu57309 | 1 | k__Fungi(100);p__Ascomycota(100);c__Saccharomycetes(100);o__Saccharomycetales(100);f__Metschnikowiaceae(100);g__Metschnikowia(100);s__Metschnikowia_pulcherrima(100); | Metschnikowia | 13 | Saprotroph | \|Undefined Saprotroph\| | Yeast | NULL | Probable | As nectar-tap_saprotroph (Põlme et al. 2020); Decay sugar-rich substrates (Põlme et al. 2020) | Põlme S, et al. 2020. Fungal Diversity 105:1-16 ((https://doi.org/10.1007/s13225-020-00466-2)) |
| Otu57310 | 1 | k__Fungi(100);p__Ascomycota(100);c__Saccharomycetes(100);o__Saccharomycetales(100);f__Metschnikowiaceae(100);g__Metschnikowia(100);s__Metschnikowia_pulcherrima(100); | Metschnikowia | 13 | Saprotroph | \|Undefined Saprotroph\| | Yeast | NULL | Probable | As nectar-tap_saprotroph (Põlme et al. 2020); Decay sugar-rich substrates (Põlme et al. 2020) | Põlme S, et al. 2020. Fungal Diversity 105:1-16 ((https://doi.org/10.1007/s13225-020-00466-2)) |
| Otu57311 | 1 | k__Fungi(100);p__Ascomycota(100);c__Saccharomycetes(100);o__Saccharomycetales(100);f__Metschnikowiaceae(100);g__Metschnikowia(100);g__Metschnikowia_unclassified(100); | Metschnikowia | 13 | Saprotroph | \|Undefined Saprotroph\| | Yeast | NULL | Probable | As nectar-tap_saprotroph (Põlme et al. 2020); Decay sugar-rich substrates (Põlme et al. 2020) | Põlme S, et al. 2020. Fungal Diversity 105:1-16 ((https://doi.org/10.1007/s13225-020-00466-2)) |
| Otu57313 | 1 | k__Fungi(100);p__Ascomycota(100);c__Saccharomycetes(100);o__Saccharomycetales(100);f__Metschnikowiaceae(100);g__Metschnikowia(100);s__Metschnikowia_pulcherrima(100); | Metschnikowia | 13 | Saprotroph | \|Undefined Saprotroph\| | Yeast | NULL | Probable | As nectar-tap_saprotroph (Põlme et al. 2020); Decay sugar-rich substrates (Põlme et al. 2020) | Põlme S, et al. 2020. Fungal Diversity 105:1-16 ((https://doi.org/10.1007/s13225-020-00466-2)) |
| Otu57321 | 1 | k__Fungi(100);p__Ascomycota(100);c__Saccharomycetes(100);o__Saccharomycetales(100);f__Metschnikowiaceae(100);g__Metschnikowia(100);s__Metschnikowia_pulcherrima(100); | Metschnikowia | 13 | Saprotroph | \|Undefined Saprotroph\| | Yeast | NULL | Probable | As nectar-tap_saprotroph (Põlme et al. 2020); Decay sugar-rich substrates (Põlme et al. 2020) | Põlme S, et al. 2020. Fungal Diversity 105:1-16 ((https://doi.org/10.1007/s13225-020-00466-2)) |
| Otu57322 | 1 | k__Fungi(100);p__Ascomycota(100);c__Saccharomycetes(100);o__Saccharomycetales(100);f__Metschnikowiaceae(100);g__Metschnikowia(100);s__Metschnikowia_pulcherrima(100); | Metschnikowia | 13 | Saprotroph | \|Undefined Saprotroph\| | Yeast | NULL | Probable | As nectar-tap_saprotroph (Põlme et al. 2020); Decay sugar-rich substrates (Põlme et al. 2020) | Põlme S, et al. 2020. Fungal Diversity 105:1-16 ((https://doi.org/10.1007/s13225-020-00466-2)) |
| Otu57324 | 1 | k__Fungi(100);p__Ascomycota(100);c__Saccharomycetes(100);o__Saccharomycetales(100);f__Metschnikowiaceae(100);g__Metschnikowia(100);s__Metschnikowia_pulcherrima(100); | Metschnikowia | 13 | Saprotroph | \|Undefined Saprotroph\| | Yeast | NULL | Probable | As nectar-tap_saprotroph (Põlme et al. 2020); Decay sugar-rich substrates (Põlme et al. 2020) | Põlme S, et al. 2020. Fungal Diversity 105:1-16 ((https://doi.org/10.1007/s13225-020-00466-2)) |
| Otu57325 | 1 | k__Fungi(100);p__Ascomycota(100);c__Saccharomycetes(100);o__Saccharomycetales(100);f__Metschnikowiaceae(100);g__Metschnikowia(100);s__Metschnikowia_pulcherrima(100); | Metschnikowia | 13 | Saprotroph | \|Undefined Saprotroph\| | Yeast | NULL | Probable | As nectar-tap_saprotroph (Põlme et al. 2020); Decay sugar-rich substrates (Põlme et al. 2020) | Põlme S, et al. 2020. Fungal Diversity 105:1-16 ((https://doi.org/10.1007/s13225-020-00466-2)) |
| Otu57331 | 1 | k__Fungi(100);p__Ascomycota(100);c__Saccharomycetes(100);o__Saccharomycetales(100);f__Metschnikowiaceae(100);g__Metschnikowia(100);s__Metschnikowia_pulcherrima(100); | Metschnikowia | 13 | Saprotroph | \|Undefined Saprotroph\| | Yeast | NULL | Probable | As nectar-tap_saprotroph (Põlme et al. 2020); Decay sugar-rich substrates (Põlme et al. 2020) | Põlme S, et al. 2020. Fungal Diversity 105:1-16 ((https://doi.org/10.1007/s13225-020-00466-2)) |
| Otu57332 | 1 | k__Fungi(100);p__Ascomycota(100);c__Saccharomycetes(100);o__Saccharomycetales(100);f__Metschnikowiaceae(100);g__Metschnikowia(100);s__Metschnikowia_pulcherrima(100); | Metschnikowia | 13 | Saprotroph | \|Undefined Saprotroph\| | Yeast | NULL | Probable | As nectar-tap_saprotroph (Põlme et al. 2020); Decay sugar-rich substrates (Põlme et al. 2020) | Põlme S, et al. 2020. Fungal Diversity 105:1-16 ((https://doi.org/10.1007/s13225-020-00466-2)) |
| Otu57333 | 1 | k__Fungi(100);p__Ascomycota(100);c__Saccharomycetes(100);o__Saccharomycetales(100);f__Metschnikowiaceae(100);g__Metschnikowia(100);s__Metschnikowia_pulcherrima(100); | Metschnikowia | 13 | Saprotroph | \|Undefined Saprotroph\| | Yeast | NULL | Probable | As nectar-tap_saprotroph (Põlme et al. 2020); Decay sugar-rich substrates (Põlme et al. 2020) | Põlme S, et al. 2020. Fungal Diversity 105:1-16 ((https://doi.org/10.1007/s13225-020-00466-2)) |
| Otu57334 | 1 | k__Fungi(100);p__Ascomycota(100);c__Saccharomycetes(100);o__Saccharomycetales(100);f__Metschnikowiaceae(100);g__Metschnikowia(100);s__Metschnikowia_pulcherrima(100); | Metschnikowia | 13 | Saprotroph | \|Undefined Saprotroph\| | Yeast | NULL | Probable | As nectar-tap_saprotroph (Põlme et al. 2020); Decay sugar-rich substrates (Põlme et al. 2020) | Põlme S, et al. 2020. Fungal Diversity 105:1-16 ((https://doi.org/10.1007/s13225-020-00466-2)) |
| Otu57337 | 1 | k__Fungi(100);p__Ascomycota(100);c__Saccharomycetes(100);o__Saccharomycetales(100);f__Metschnikowiaceae(100);g__Metschnikowia(100);s__Metschnikowia_pulcherrima(100); | Metschnikowia | 13 | Saprotroph | \|Undefined Saprotroph\| | Yeast | NULL | Probable | As nectar-tap_saprotroph (Põlme et al. 2020); Decay sugar-rich substrates (Põlme et al. 2020) | Põlme S, et al. 2020. Fungal Diversity 105:1-16 ((https://doi.org/10.1007/s13225-020-00466-2)) |
| Otu57338 | 1 | k__Fungi(100);p__Ascomycota(100);c__Saccharomycetes(100);o__Saccharomycetales(100);f__Metschnikowiaceae(100);g__Metschnikowia(100);g__Metschnikowia_unclassified(100); | Metschnikowia | 13 | Saprotroph | \|Undefined Saprotroph\| | Yeast | NULL | Probable | As nectar-tap_saprotroph (Põlme et al. 2020); Decay sugar-rich substrates (Põlme et al. 2020) | Põlme S, et al. 2020. Fungal Diversity 105:1-16 ((https://doi.org/10.1007/s13225-020-00466-2)) |
| Otu57339 | 1 | k__Fungi(100);p__Ascomycota(100);c__Saccharomycetes(100);o__Saccharomycetales(100);f__Metschnikowiaceae(100);g__Metschnikowia(100);s__Metschnikowia_pulcherrima(100); | Metschnikowia | 13 | Saprotroph | \|Undefined Saprotroph\| | Yeast | NULL | Probable | As nectar-tap_saprotroph (Põlme et al. 2020); Decay sugar-rich substrates (Põlme et al. 2020) | Põlme S, et al. 2020. Fungal Diversity 105:1-16 ((https://doi.org/10.1007/s13225-020-00466-2)) |
| Otu57340 | 1 | k__Fungi(100);p__Ascomycota(100);c__Saccharomycetes(100);o__Saccharomycetales(100);f__Metschnikowiaceae(100);g__Metschnikowia(100);s__Metschnikowia_pulcherrima(100); | Metschnikowia | 13 | Saprotroph | \|Undefined Saprotroph\| | Yeast | NULL | Probable | As nectar-tap_saprotroph (Põlme et al. 2020); Decay sugar-rich substrates (Põlme et al. 2020) | Põlme S, et al. 2020. Fungal Diversity 105:1-16 ((https://doi.org/10.1007/s13225-020-00466-2)) |
| Otu57341 | 1 | k__Fungi(100);p__Ascomycota(100);c__Saccharomycetes(100);o__Saccharomycetales(100);f__Metschnikowiaceae(100);g__Metschnikowia(100);s__Metschnikowia_pulcherrima(100); | Metschnikowia | 13 | Saprotroph | \|Undefined Saprotroph\| | Yeast | NULL | Probable | As nectar-tap_saprotroph (Põlme et al. 2020); Decay sugar-rich substrates (Põlme et al. 2020) | Põlme S, et al. 2020. Fungal Diversity 105:1-16 ((https://doi.org/10.1007/s13225-020-00466-2)) |
| Otu57342 | 1 | k__Fungi(100);p__Ascomycota(100);c__Saccharomycetes(100);o__Saccharomycetales(100);f__Metschnikowiaceae(100);g__Metschnikowia(100);s__Metschnikowia_pulcherrima(100); | Metschnikowia | 13 | Saprotroph | \|Undefined Saprotroph\| | Yeast | NULL | Probable | As nectar-tap_saprotroph (Põlme et al. 2020); Decay sugar-rich substrates (Põlme et al. 2020) | Põlme S, et al. 2020. Fungal Diversity 105:1-16 ((https://doi.org/10.1007/s13225-020-00466-2)) |
| Otu57343 | 1 | k__Fungi(100);p__Ascomycota(100);c__Saccharomycetes(100);o__Saccharomycetales(100);f__Metschnikowiaceae(100);g__Metschnikowia(100);g__Metschnikowia_unclassified(100); | Metschnikowia | 13 | Saprotroph | \|Undefined Saprotroph\| | Yeast | NULL | Probable | As nectar-tap_saprotroph (Põlme et al. 2020); Decay sugar-rich substrates (Põlme et al. 2020) | Põlme S, et al. 2020. Fungal Diversity 105:1-16 ((https://doi.org/10.1007/s13225-020-00466-2)) |
| Otu57344 | 1 | k__Fungi(100);p__Ascomycota(100);c__Saccharomycetes(100);o__Saccharomycetales(100);f__Metschnikowiaceae(100);g__Metschnikowia(100);s__Metschnikowia_pulcherrima(100); | Metschnikowia | 13 | Saprotroph | \|Undefined Saprotroph\| | Yeast | NULL | Probable | As nectar-tap_saprotroph (Põlme et al. 2020); Decay sugar-rich substrates (Põlme et al. 2020) | Põlme S, et al. 2020. Fungal Diversity 105:1-16 ((https://doi.org/10.1007/s13225-020-00466-2)) |
| Otu57345 | 1 | k__Fungi(100);p__Ascomycota(100);c__Saccharomycetes(100);o__Saccharomycetales(100);f__Metschnikowiaceae(100);g__Metschnikowia(100);s__Metschnikowia_pulcherrima(100); | Metschnikowia | 13 | Saprotroph | \|Undefined Saprotroph\| | Yeast | NULL | Probable | As nectar-tap_saprotroph (Põlme et al. 2020); Decay sugar-rich substrates (Põlme et al. 2020) | Põlme S, et al. 2020. Fungal Diversity 105:1-16 ((https://doi.org/10.1007/s13225-020-00466-2)) |
| Otu57346 | 1 | k__Fungi(100);p__Ascomycota(100);c__Saccharomycetes(100);o__Saccharomycetales(100);f__Metschnikowiaceae(100);g__Metschnikowia(100);s__Metschnikowia_pulcherrima(100); | Metschnikowia | 13 | Saprotroph | \|Undefined Saprotroph\| | Yeast | NULL | Probable | As nectar-tap_saprotroph (Põlme et al. 2020); Decay sugar-rich substrates (Põlme et al. 2020) | Põlme S, et al. 2020. Fungal Diversity 105:1-16 ((https://doi.org/10.1007/s13225-020-00466-2)) |
| Otu57347 | 1 | k__Fungi(100);p__Ascomycota(100);c__Saccharomycetes(100);o__Saccharomycetales(100);f__Metschnikowiaceae(100);g__Metschnikowia(100);s__Metschnikowia_pulcherrima(100); | Metschnikowia | 13 | Saprotroph | \|Undefined Saprotroph\| | Yeast | NULL | Probable | As nectar-tap_saprotroph (Põlme et al. 2020); Decay sugar-rich substrates (Põlme et al. 2020) | Põlme S, et al. 2020. Fungal Diversity 105:1-16 ((https://doi.org/10.1007/s13225-020-00466-2)) |
| Otu57348 | 1 | k__Fungi(100);p__Ascomycota(100);c__Saccharomycetes(100);o__Saccharomycetales(100);f__Metschnikowiaceae(100);g__Metschnikowia(100);s__Metschnikowia_pulcherrima(100); | Metschnikowia | 13 | Saprotroph | \|Undefined Saprotroph\| | Yeast | NULL | Probable | As nectar-tap_saprotroph (Põlme et al. 2020); Decay sugar-rich substrates (Põlme et al. 2020) | Põlme S, et al. 2020. Fungal Diversity 105:1-16 ((https://doi.org/10.1007/s13225-020-00466-2)) |
| Otu57349 | 1 | k__Fungi(100);p__Ascomycota(100);c__Saccharomycetes(100);o__Saccharomycetales(100);f__Metschnikowiaceae(100);g__Metschnikowia(100);s__Metschnikowia_pulcherrima(100); | Metschnikowia | 13 | Saprotroph | \|Undefined Saprotroph\| | Yeast | NULL | Probable | As nectar-tap_saprotroph (Põlme et al. 2020); Decay sugar-rich substrates (Põlme et al. 2020) | Põlme S, et al. 2020. Fungal Diversity 105:1-16 ((https://doi.org/10.1007/s13225-020-00466-2)) |
| Otu57350 | 1 | k__Fungi(100);p__Ascomycota(100);c__Saccharomycetes(100);o__Saccharomycetales(100);f__Metschnikowiaceae(100);g__Metschnikowia(100);s__Metschnikowia_pulcherrima(100); | Metschnikowia | 13 | Saprotroph | \|Undefined Saprotroph\| | Yeast | NULL | Probable | As nectar-tap_saprotroph (Põlme et al. 2020); Decay sugar-rich substrates (Põlme et al. 2020) | Põlme S, et al. 2020. Fungal Diversity 105:1-16 ((https://doi.org/10.1007/s13225-020-00466-2)) |
| Otu57351 | 1 | k__Fungi(100);p__Ascomycota(100);c__Saccharomycetes(100);o__Saccharomycetales(100);f__Metschnikowiaceae(100);g__Metschnikowia(100);s__Metschnikowia_pulcherrima(100); | Metschnikowia | 13 | Saprotroph | \|Undefined Saprotroph\| | Yeast | NULL | Probable | As nectar-tap_saprotroph (Põlme et al. 2020); Decay sugar-rich substrates (Põlme et al. 2020) | Põlme S, et al. 2020. Fungal Diversity 105:1-16 ((https://doi.org/10.1007/s13225-020-00466-2)) |
| Otu57352 | 1 | k__Fungi(100);p__Ascomycota(100);c__Saccharomycetes(100);o__Saccharomycetales(100);f__Metschnikowiaceae(100);g__Metschnikowia(100);s__Metschnikowia_pulcherrima(100); | Metschnikowia | 13 | Saprotroph | \|Undefined Saprotroph\| | Yeast | NULL | Probable | As nectar-tap_saprotroph (Põlme et al. 2020); Decay sugar-rich substrates (Põlme et al. 2020) | Põlme S, et al. 2020. Fungal Diversity 105:1-16 ((https://doi.org/10.1007/s13225-020-00466-2)) |
| Otu57353 | 1 | k__Fungi(100);p__Ascomycota(100);c__Saccharomycetes(100);o__Saccharomycetales(100);f__Metschnikowiaceae(100);g__Metschnikowia(100);g__Metschnikowia_unclassified(100); | Metschnikowia | 13 | Saprotroph | \|Undefined Saprotroph\| | Yeast | NULL | Probable | As nectar-tap_saprotroph (Põlme et al. 2020); Decay sugar-rich substrates (Põlme et al. 2020) | Põlme S, et al. 2020. Fungal Diversity 105:1-16 ((https://doi.org/10.1007/s13225-020-00466-2)) |
| Otu57354 | 1 | k__Fungi(100);p__Ascomycota(100);c__Saccharomycetes(100);o__Saccharomycetales(100);f__Metschnikowiaceae(100);g__Metschnikowia(100);g__Metschnikowia_unclassified(100); | Metschnikowia | 13 | Saprotroph | \|Undefined Saprotroph\| | Yeast | NULL | Probable | As nectar-tap_saprotroph (Põlme et al. 2020); Decay sugar-rich substrates (Põlme et al. 2020) | Põlme S, et al. 2020. Fungal Diversity 105:1-16 ((https://doi.org/10.1007/s13225-020-00466-2)) |
| Otu57362 | 1 | k__Fungi(100);p__Ascomycota(100);c__Saccharomycetes(100);o__Saccharomycetales(100);f__Metschnikowiaceae(100);g__Metschnikowia(100);s__Metschnikowia_pulcherrima(100); | Metschnikowia | 13 | Saprotroph | \|Undefined Saprotroph\| | Yeast | NULL | Probable | As nectar-tap_saprotroph (Põlme et al. 2020); Decay sugar-rich substrates (Põlme et al. 2020) | Põlme S, et al. 2020. Fungal Diversity 105:1-16 ((https://doi.org/10.1007/s13225-020-00466-2)) |
| Otu57363 | 1 | k__Fungi(100);p__Ascomycota(100);c__Saccharomycetes(100);o__Saccharomycetales(100);f__Metschnikowiaceae(100);g__Metschnikowia(100);s__Metschnikowia_pulcherrima(100); | Metschnikowia | 13 | Saprotroph | \|Undefined Saprotroph\| | Yeast | NULL | Probable | As nectar-tap_saprotroph (Põlme et al. 2020); Decay sugar-rich substrates (Põlme et al. 2020) | Põlme S, et al. 2020. Fungal Diversity 105:1-16 ((https://doi.org/10.1007/s13225-020-00466-2)) |
| Otu57364 | 1 | k__Fungi(100);p__Ascomycota(100);c__Saccharomycetes(100);o__Saccharomycetales(100);f__Metschnikowiaceae(100);g__Metschnikowia(100);s__Metschnikowia_pulcherrima(100); | Metschnikowia | 13 | Saprotroph | \|Undefined Saprotroph\| | Yeast | NULL | Probable | As nectar-tap_saprotroph (Põlme et al. 2020); Decay sugar-rich substrates (Põlme et al. 2020) | Põlme S, et al. 2020. Fungal Diversity 105:1-16 ((https://doi.org/10.1007/s13225-020-00466-2)) |
| Otu57365 | 1 | k__Fungi(100);p__Ascomycota(100);c__Saccharomycetes(100);o__Saccharomycetales(100);f__Metschnikowiaceae(100);g__Metschnikowia(100);s__Metschnikowia_pulcherrima(100); | Metschnikowia | 13 | Saprotroph | \|Undefined Saprotroph\| | Yeast | NULL | Probable | As nectar-tap_saprotroph (Põlme et al. 2020); Decay sugar-rich substrates (Põlme et al. 2020) | Põlme S, et al. 2020. Fungal Diversity 105:1-16 ((https://doi.org/10.1007/s13225-020-00466-2)) |
| Otu57366 | 1 | k__Fungi(100);p__Ascomycota(100);c__Saccharomycetes(100);o__Saccharomycetales(100);f__Metschnikowiaceae(100);g__Metschnikowia(100);s__Metschnikowia_pulcherrima(100); | Metschnikowia | 13 | Saprotroph | \|Undefined Saprotroph\| | Yeast | NULL | Probable | As nectar-tap_saprotroph (Põlme et al. 2020); Decay sugar-rich substrates (Põlme et al. 2020) | Põlme S, et al. 2020. Fungal Diversity 105:1-16 ((https://doi.org/10.1007/s13225-020-00466-2)) |
| Otu57367 | 1 | k__Fungi(100);p__Ascomycota(100);c__Saccharomycetes(100);o__Saccharomycetales(100);f__Metschnikowiaceae(100);g__Metschnikowia(100);s__Metschnikowia_pulcherrima(100); | Metschnikowia | 13 | Saprotroph | \|Undefined Saprotroph\| | Yeast | NULL | Probable | As nectar-tap_saprotroph (Põlme et al. 2020); Decay sugar-rich substrates (Põlme et al. 2020) | Põlme S, et al. 2020. Fungal Diversity 105:1-16 ((https://doi.org/10.1007/s13225-020-00466-2)) |
| Otu57368 | 1 | k__Fungi(100);p__Ascomycota(100);c__Saccharomycetes(100);o__Saccharomycetales(100);f__Metschnikowiaceae(100);g__Metschnikowia(100);s__Metschnikowia_pulcherrima(100); | Metschnikowia | 13 | Saprotroph | \|Undefined Saprotroph\| | Yeast | NULL | Probable | As nectar-tap_saprotroph (Põlme et al. 2020); Decay sugar-rich substrates (Põlme et al. 2020) | Põlme S, et al. 2020. Fungal Diversity 105:1-16 ((https://doi.org/10.1007/s13225-020-00466-2)) |
| Otu57369 | 1 | k__Fungi(100);p__Ascomycota(100);c__Saccharomycetes(100);o__Saccharomycetales(100);f__Metschnikowiaceae(100);g__Metschnikowia(100);g__Metschnikowia_unclassified(100); | Metschnikowia | 13 | Saprotroph | \|Undefined Saprotroph\| | Yeast | NULL | Probable | As nectar-tap_saprotroph (Põlme et al. 2020); Decay sugar-rich substrates (Põlme et al. 2020) | Põlme S, et al. 2020. Fungal Diversity 105:1-16 ((https://doi.org/10.1007/s13225-020-00466-2)) |
| Otu57370 | 1 | k__Fungi(100);p__Ascomycota(100);c__Saccharomycetes(100);o__Saccharomycetales(100);f__Metschnikowiaceae(100);g__Metschnikowia(100);s__Metschnikowia_pulcherrima(100); | Metschnikowia | 13 | Saprotroph | \|Undefined Saprotroph\| | Yeast | NULL | Probable | As nectar-tap_saprotroph (Põlme et al. 2020); Decay sugar-rich substrates (Põlme et al. 2020) | Põlme S, et al. 2020. Fungal Diversity 105:1-16 ((https://doi.org/10.1007/s13225-020-00466-2)) |
| Otu57371 | 1 | k__Fungi(100);p__Ascomycota(100);c__Saccharomycetes(100);o__Saccharomycetales(100);f__Metschnikowiaceae(100);g__Metschnikowia(100);s__Metschnikowia_pulcherrima(100); | Metschnikowia | 13 | Saprotroph | \|Undefined Saprotroph\| | Yeast | NULL | Probable | As nectar-tap_saprotroph (Põlme et al. 2020); Decay sugar-rich substrates (Põlme et al. 2020) | Põlme S, et al. 2020. Fungal Diversity 105:1-16 ((https://doi.org/10.1007/s13225-020-00466-2)) |
| Otu57372 | 1 | k__Fungi(100);p__Ascomycota(100);c__Saccharomycetes(100);o__Saccharomycetales(100);f__Metschnikowiaceae(100);g__Metschnikowia(100);s__Metschnikowia_pulcherrima(100); | Metschnikowia | 13 | Saprotroph | \|Undefined Saprotroph\| | Yeast | NULL | Probable | As nectar-tap_saprotroph (Põlme et al. 2020); Decay sugar-rich substrates (Põlme et al. 2020) | Põlme S, et al. 2020. Fungal Diversity 105:1-16 ((https://doi.org/10.1007/s13225-020-00466-2)) |
| Otu57373 | 1 | k__Fungi(100);p__Ascomycota(100);c__Saccharomycetes(100);o__Saccharomycetales(100);f__Metschnikowiaceae(100);g__Metschnikowia(100);s__Metschnikowia_pulcherrima(100); | Metschnikowia | 13 | Saprotroph | \|Undefined Saprotroph\| | Yeast | NULL | Probable | As nectar-tap_saprotroph (Põlme et al. 2020); Decay sugar-rich substrates (Põlme et al. 2020) | Põlme S, et al. 2020. Fungal Diversity 105:1-16 ((https://doi.org/10.1007/s13225-020-00466-2)) |
| Otu57374 | 1 | k__Fungi(100);p__Ascomycota(100);c__Saccharomycetes(100);o__Saccharomycetales(100);f__Metschnikowiaceae(100);g__Metschnikowia(100);s__Metschnikowia_pulcherrima(100); | Metschnikowia | 13 | Saprotroph | \|Undefined Saprotroph\| | Yeast | NULL | Probable | As nectar-tap_saprotroph (Põlme et al. 2020); Decay sugar-rich substrates (Põlme et al. 2020) | Põlme S, et al. 2020. Fungal Diversity 105:1-16 ((https://doi.org/10.1007/s13225-020-00466-2)) |
| Otu57375 | 1 | k__Fungi(100);p__Ascomycota(100);c__Saccharomycetes(100);o__Saccharomycetales(100);f__Metschnikowiaceae(100);g__Metschnikowia(100);s__Metschnikowia_pulcherrima(100); | Metschnikowia | 13 | Saprotroph | \|Undefined Saprotroph\| | Yeast | NULL | Probable | As nectar-tap_saprotroph (Põlme et al. 2020); Decay sugar-rich substrates (Põlme et al. 2020) | Põlme S, et al. 2020. Fungal Diversity 105:1-16 ((https://doi.org/10.1007/s13225-020-00466-2)) |
| Otu57376 | 1 | k__Fungi(100);p__Ascomycota(100);c__Saccharomycetes(100);o__Saccharomycetales(100);f__Metschnikowiaceae(100);g__Metschnikowia(100);s__Metschnikowia_pulcherrima(100); | Metschnikowia | 13 | Saprotroph | \|Undefined Saprotroph\| | Yeast | NULL | Probable | As nectar-tap_saprotroph (Põlme et al. 2020); Decay sugar-rich substrates (Põlme et al. 2020) | Põlme S, et al. 2020. Fungal Diversity 105:1-16 ((https://doi.org/10.1007/s13225-020-00466-2)) |
| Otu57377 | 1 | k__Fungi(100);p__Ascomycota(100);c__Saccharomycetes(100);o__Saccharomycetales(100);f__Metschnikowiaceae(100);g__Metschnikowia(100);s__Metschnikowia_pulcherrima(100); | Metschnikowia | 13 | Saprotroph | \|Undefined Saprotroph\| | Yeast | NULL | Probable | As nectar-tap_saprotroph (Põlme et al. 2020); Decay sugar-rich substrates (Põlme et al. 2020) | Põlme S, et al. 2020. Fungal Diversity 105:1-16 ((https://doi.org/10.1007/s13225-020-00466-2)) |
| Otu57378 | 1 | k__Fungi(100);p__Ascomycota(100);c__Saccharomycetes(100);o__Saccharomycetales(100);f__Metschnikowiaceae(100);g__Metschnikowia(100);g__Metschnikowia_unclassified(100); | Metschnikowia | 13 | Saprotroph | \|Undefined Saprotroph\| | Yeast | NULL | Probable | As nectar-tap_saprotroph (Põlme et al. 2020); Decay sugar-rich substrates (Põlme et al. 2020) | Põlme S, et al. 2020. Fungal Diversity 105:1-16 ((https://doi.org/10.1007/s13225-020-00466-2)) |
| Otu57379 | 1 | k__Fungi(100);p__Ascomycota(100);c__Saccharomycetes(100);o__Saccharomycetales(100);f__Metschnikowiaceae(100);g__Metschnikowia(100);s__Metschnikowia_pulcherrima(100); | Metschnikowia | 13 | Saprotroph | \|Undefined Saprotroph\| | Yeast | NULL | Probable | As nectar-tap_saprotroph (Põlme et al. 2020); Decay sugar-rich substrates (Põlme et al. 2020) | Põlme S, et al. 2020. Fungal Diversity 105:1-16 ((https://doi.org/10.1007/s13225-020-00466-2)) |
| Otu57380 | 1 | k__Fungi(100);p__Ascomycota(100);c__Saccharomycetes(100);o__Saccharomycetales(100);f__Metschnikowiaceae(100);g__Metschnikowia(100);s__Metschnikowia_pulcherrima(100); | Metschnikowia | 13 | Saprotroph | \|Undefined Saprotroph\| | Yeast | NULL | Probable | As nectar-tap_saprotroph (Põlme et al. 2020); Decay sugar-rich substrates (Põlme et al. 2020) | Põlme S, et al. 2020. Fungal Diversity 105:1-16 ((https://doi.org/10.1007/s13225-020-00466-2)) |
| Otu57381 | 1 | k__Fungi(100);p__Ascomycota(100);c__Saccharomycetes(100);o__Saccharomycetales(100);f__Metschnikowiaceae(100);g__Metschnikowia(100);s__Metschnikowia_pulcherrima(100); | Metschnikowia | 13 | Saprotroph | \|Undefined Saprotroph\| | Yeast | NULL | Probable | As nectar-tap_saprotroph (Põlme et al. 2020); Decay sugar-rich substrates (Põlme et al. 2020) | Põlme S, et al. 2020. Fungal Diversity 105:1-16 ((https://doi.org/10.1007/s13225-020-00466-2)) |
| Otu57382 | 1 | k__Fungi(100);p__Ascomycota(100);c__Saccharomycetes(100);o__Saccharomycetales(100);f__Metschnikowiaceae(100);g__Metschnikowia(100);s__Metschnikowia_pulcherrima(100); | Metschnikowia | 13 | Saprotroph | \|Undefined Saprotroph\| | Yeast | NULL | Probable | As nectar-tap_saprotroph (Põlme et al. 2020); Decay sugar-rich substrates (Põlme et al. 2020) | Põlme S, et al. 2020. Fungal Diversity 105:1-16 ((https://doi.org/10.1007/s13225-020-00466-2)) |
| Otu57383 | 1 | k__Fungi(100);p__Ascomycota(100);c__Saccharomycetes(100);o__Saccharomycetales(100);f__Metschnikowiaceae(100);g__Metschnikowia(100);s__Metschnikowia_pulcherrima(100); | Metschnikowia | 13 | Saprotroph | \|Undefined Saprotroph\| | Yeast | NULL | Probable | As nectar-tap_saprotroph (Põlme et al. 2020); Decay sugar-rich substrates (Põlme et al. 2020) | Põlme S, et al. 2020. Fungal Diversity 105:1-16 ((https://doi.org/10.1007/s13225-020-00466-2)) |
| Otu57384 | 1 | k__Fungi(100);p__Ascomycota(100);c__Saccharomycetes(100);o__Saccharomycetales(100);f__Metschnikowiaceae(100);g__Metschnikowia(100);s__Metschnikowia_pulcherrima(100); | Metschnikowia | 13 | Saprotroph | \|Undefined Saprotroph\| | Yeast | NULL | Probable | As nectar-tap_saprotroph (Põlme et al. 2020); Decay sugar-rich substrates (Põlme et al. 2020) | Põlme S, et al. 2020. Fungal Diversity 105:1-16 ((https://doi.org/10.1007/s13225-020-00466-2)) |
| Otu57385 | 1 | k__Fungi(100);p__Ascomycota(100);c__Saccharomycetes(100);o__Saccharomycetales(100);f__Metschnikowiaceae(100);g__Metschnikowia(100);g__Metschnikowia_unclassified(100); | Metschnikowia | 13 | Saprotroph | \|Undefined Saprotroph\| | Yeast | NULL | Probable | As nectar-tap_saprotroph (Põlme et al. 2020); Decay sugar-rich substrates (Põlme et al. 2020) | Põlme S, et al. 2020. Fungal Diversity 105:1-16 ((https://doi.org/10.1007/s13225-020-00466-2)) |
| Otu57386 | 1 | k__Fungi(100);p__Ascomycota(100);c__Saccharomycetes(100);o__Saccharomycetales(100);f__Metschnikowiaceae(100);g__Metschnikowia(100);s__Metschnikowia_pulcherrima(100); | Metschnikowia | 13 | Saprotroph | \|Undefined Saprotroph\| | Yeast | NULL | Probable | As nectar-tap_saprotroph (Põlme et al. 2020); Decay sugar-rich substrates (Põlme et al. 2020) | Põlme S, et al. 2020. Fungal Diversity 105:1-16 ((https://doi.org/10.1007/s13225-020-00466-2)) |
| Otu57388 | 1 | k__Fungi(100);p__Ascomycota(100);c__Saccharomycetes(100);o__Saccharomycetales(100);f__Metschnikowiaceae(100);g__Metschnikowia(100);g__Metschnikowia_unclassified(100); | Metschnikowia | 13 | Saprotroph | \|Undefined Saprotroph\| | Yeast | NULL | Probable | As nectar-tap_saprotroph (Põlme et al. 2020); Decay sugar-rich substrates (Põlme et al. 2020) | Põlme S, et al. 2020. Fungal Diversity 105:1-16 ((https://doi.org/10.1007/s13225-020-00466-2)) |
| Otu57390 | 1 | k__Fungi(100);p__Ascomycota(100);c__Saccharomycetes(100);o__Saccharomycetales(100);f__Metschnikowiaceae(100);g__Metschnikowia(100);s__Metschnikowia_pulcherrima(100); | Metschnikowia | 13 | Saprotroph | \|Undefined Saprotroph\| | Yeast | NULL | Probable | As nectar-tap_saprotroph (Põlme et al. 2020); Decay sugar-rich substrates (Põlme et al. 2020) | Põlme S, et al. 2020. Fungal Diversity 105:1-16 ((https://doi.org/10.1007/s13225-020-00466-2)) |
| Otu57391 | 1 | k__Fungi(100);p__Ascomycota(100);c__Saccharomycetes(100);o__Saccharomycetales(100);f__Metschnikowiaceae(100);g__Metschnikowia(100);g__Metschnikowia_unclassified(100); | Metschnikowia | 13 | Saprotroph | \|Undefined Saprotroph\| | Yeast | NULL | Probable | As nectar-tap_saprotroph (Põlme et al. 2020); Decay sugar-rich substrates (Põlme et al. 2020) | Põlme S, et al. 2020. Fungal Diversity 105:1-16 ((https://doi.org/10.1007/s13225-020-00466-2)) |
| Otu57392 | 1 | k__Fungi(100);p__Ascomycota(100);c__Saccharomycetes(100);o__Saccharomycetales(100);f__Metschnikowiaceae(100);g__Metschnikowia(100);g__Metschnikowia_unclassified(100); | Metschnikowia | 13 | Saprotroph | \|Undefined Saprotroph\| | Yeast | NULL | Probable | As nectar-tap_saprotroph (Põlme et al. 2020); Decay sugar-rich substrates (Põlme et al. 2020) | Põlme S, et al. 2020. Fungal Diversity 105:1-16 ((https://doi.org/10.1007/s13225-020-00466-2)) |
| Otu57393 | 1 | k__Fungi(100);p__Ascomycota(100);c__Saccharomycetes(100);o__Saccharomycetales(100);f__Metschnikowiaceae(100);g__Metschnikowia(100);g__Metschnikowia_unclassified(100); | Metschnikowia | 13 | Saprotroph | \|Undefined Saprotroph\| | Yeast | NULL | Probable | As nectar-tap_saprotroph (Põlme et al. 2020); Decay sugar-rich substrates (Põlme et al. 2020) | Põlme S, et al. 2020. Fungal Diversity 105:1-16 ((https://doi.org/10.1007/s13225-020-00466-2)) |
| Otu57394 | 1 | k__Fungi(100);p__Ascomycota(100);c__Saccharomycetes(100);o__Saccharomycetales(100);f__Metschnikowiaceae(100);g__Metschnikowia(100);g__Metschnikowia_unclassified(100); | Metschnikowia | 13 | Saprotroph | \|Undefined Saprotroph\| | Yeast | NULL | Probable | As nectar-tap_saprotroph (Põlme et al. 2020); Decay sugar-rich substrates (Põlme et al. 2020) | Põlme S, et al. 2020. Fungal Diversity 105:1-16 ((https://doi.org/10.1007/s13225-020-00466-2)) |
| Otu57395 | 1 | k__Fungi(100);p__Ascomycota(100);c__Saccharomycetes(100);o__Saccharomycetales(100);f__Metschnikowiaceae(100);g__Metschnikowia(100);s__Metschnikowia_pulcherrima(100); | Metschnikowia | 13 | Saprotroph | \|Undefined Saprotroph\| | Yeast | NULL | Probable | As nectar-tap_saprotroph (Põlme et al. 2020); Decay sugar-rich substrates (Põlme et al. 2020) | Põlme S, et al. 2020. Fungal Diversity 105:1-16 ((https://doi.org/10.1007/s13225-020-00466-2)) |
| Otu57396 | 1 | k__Fungi(100);p__Ascomycota(100);c__Saccharomycetes(100);o__Saccharomycetales(100);f__Metschnikowiaceae(100);g__Metschnikowia(100);s__Metschnikowia_pulcherrima(100); | Metschnikowia | 13 | Saprotroph | \|Undefined Saprotroph\| | Yeast | NULL | Probable | As nectar-tap_saprotroph (Põlme et al. 2020); Decay sugar-rich substrates (Põlme et al. 2020) | Põlme S, et al. 2020. Fungal Diversity 105:1-16 ((https://doi.org/10.1007/s13225-020-00466-2)) |
| Otu57397 | 1 | k__Fungi(100);p__Ascomycota(100);c__Saccharomycetes(100);o__Saccharomycetales(100);f__Metschnikowiaceae(100);g__Metschnikowia(100);g__Metschnikowia_unclassified(100); | Metschnikowia | 13 | Saprotroph | \|Undefined Saprotroph\| | Yeast | NULL | Probable | As nectar-tap_saprotroph (Põlme et al. 2020); Decay sugar-rich substrates (Põlme et al. 2020) | Põlme S, et al. 2020. Fungal Diversity 105:1-16 ((https://doi.org/10.1007/s13225-020-00466-2)) |
| Otu57398 | 1 | k__Fungi(100);p__Ascomycota(100);c__Saccharomycetes(100);o__Saccharomycetales(100);f__Metschnikowiaceae(100);g__Metschnikowia(100);s__Metschnikowia_pulcherrima(100); | Metschnikowia | 13 | Saprotroph | \|Undefined Saprotroph\| | Yeast | NULL | Probable | As nectar-tap_saprotroph (Põlme et al. 2020); Decay sugar-rich substrates (Põlme et al. 2020) | Põlme S, et al. 2020. Fungal Diversity 105:1-16 ((https://doi.org/10.1007/s13225-020-00466-2)) |
| Otu57399 | 1 | k__Fungi(100);p__Ascomycota(100);c__Saccharomycetes(100);o__Saccharomycetales(100);f__Metschnikowiaceae(100);g__Metschnikowia(100);s__Metschnikowia_pulcherrima(100); | Metschnikowia | 13 | Saprotroph | \|Undefined Saprotroph\| | Yeast | NULL | Probable | As nectar-tap_saprotroph (Põlme et al. 2020); Decay sugar-rich substrates (Põlme et al. 2020) | Põlme S, et al. 2020. Fungal Diversity 105:1-16 ((https://doi.org/10.1007/s13225-020-00466-2)) |
| Otu57400 | 1 | k__Fungi(100);p__Ascomycota(100);c__Saccharomycetes(100);o__Saccharomycetales(100);f__Metschnikowiaceae(100);g__Metschnikowia(100);s__Metschnikowia_pulcherrima(100); | Metschnikowia | 13 | Saprotroph | \|Undefined Saprotroph\| | Yeast | NULL | Probable | As nectar-tap_saprotroph (Põlme et al. 2020); Decay sugar-rich substrates (Põlme et al. 2020) | Põlme S, et al. 2020. Fungal Diversity 105:1-16 ((https://doi.org/10.1007/s13225-020-00466-2)) |

| | | | | | | | | | | | |
|---|---|---|---|---|---|---|---|---|---|---|---|
| Otu57401 | 1 | k__Fungi(100);p__Ascomycota(100);c__Saccharomycetes(100);o__Saccharomycetales(100);f__Metschnikowiaceae(100);g__Metschnikowia(100);s__Metschnikowia_pulcherrima(100); | Metschnikowia | 13 | Saprotroph | \|Undefined Saprotroph\| | Yeast | NULL | Probable | As nectar-tap_saprotroph (Põlme et al. 2020); Decay sugar-rich substrates (Põlme et al. 2020) | Põlme S, et al. 2020. Fungal Diversity 105:1-16 ((https://doi.org/10.1007/s13225-020-00466-2)) |
| Otu57402 | 1 | k__Fungi(100);p__Ascomycota(100);c__Saccharomycetes(100);o__Saccharomycetales(100);f__Metschnikowiaceae(100);g__Metschnikowia(100);g__Metschnikowia_unclassified(100); | Metschnikowia | 13 | Saprotroph | \|Undefined Saprotroph\| | Yeast | NULL | Probable | As nectar-tap_saprotroph (Põlme et al. 2020); Decay sugar-rich substrates (Põlme et al. 2020) | Põlme S, et al. 2020. Fungal Diversity 105:1-16 ((https://doi.org/10.1007/s13225-020-00466-2)) |
| Otu57403 | 1 | k__Fungi(100);p__Ascomycota(100);c__Saccharomycetes(100);o__Saccharomycetales(100);f__Metschnikowiaceae(100);g__Metschnikowia(100);g__Metschnikowia_unclassified(100); | Metschnikowia | 13 | Saprotroph | \|Undefined Saprotroph\| | Yeast | NULL | Probable | As nectar-tap_saprotroph (Põlme et al. 2020); Decay sugar-rich substrates (Põlme et al. 2020) | Põlme S, et al. 2020. Fungal Diversity 105:1-16 ((https://doi.org/10.1007/s13225-020-00466-2)) |
| Otu57404 | 1 | k__Fungi(100);p__Ascomycota(100);c__Saccharomycetes(100);o__Saccharomycetales(100);f__Metschnikowiaceae(100);g__Metschnikowia(100);s__Metschnikowia_pulcherrima(100); | Metschnikowia | 13 | Saprotroph | \|Undefined Saprotroph\| | Yeast | NULL | Probable | As nectar-tap_saprotroph (Põlme et al. 2020); Decay sugar-rich substrates (Põlme et al. 2020) | Põlme S, et al. 2020. Fungal Diversity 105:1-16 ((https://doi.org/10.1007/s13225-020-00466-2)) |
| Otu57405 | 1 | k__Fungi(100);p__Ascomycota(100);c__Saccharomycetes(100);o__Saccharomycetales(100);f__Metschnikowiaceae(100);g__Metschnikowia(100);g__Metschnikowia_unclassified(100); | Metschnikowia | 13 | Saprotroph | \|Undefined Saprotroph\| | Yeast | NULL | Probable | As nectar-tap_saprotroph (Põlme et al. 2020); Decay sugar-rich substrates (Põlme et al. 2020) | Põlme S, et al. 2020. Fungal Diversity 105:1-16 ((https://doi.org/10.1007/s13225-020-00466-2)) |
| Otu57406 | 1 | k__Fungi(100);p__Ascomycota(100);c__Saccharomycetes(100);o__Saccharomycetales(100);f__Metschnikowiaceae(100);g__Metschnikowia(100);s__Metschnikowia_pulcherrima(100); | Metschnikowia | 13 | Saprotroph | \|Undefined Saprotroph\| | Yeast | NULL | Probable | As nectar-tap_saprotroph (Põlme et al. 2020); Decay sugar-rich substrates (Põlme et al. 2020) | Põlme S, et al. 2020. Fungal Diversity 105:1-16 ((https://doi.org/10.1007/s13225-020-00466-2)) |
| Otu57407 | 1 | k__Fungi(100);p__Ascomycota(100);c__Saccharomycetes(100);o__Saccharomycetales(100);f__Metschnikowiaceae(100);g__Metschnikowia(100);g__Metschnikowia_unclassified(100); | Metschnikowia | 13 | Saprotroph | \|Undefined Saprotroph\| | Yeast | NULL | Probable | As nectar-tap_saprotroph (Põlme et al. 2020); Decay sugar-rich substrates (Põlme et al. 2020) | Põlme S, et al. 2020. Fungal Diversity 105:1-16 ((https://doi.org/10.1007/s13225-020-00466-2)) |
| Otu57408 | 1 | k__Fungi(100);p__Ascomycota(100);c__Saccharomycetes(100);o__Saccharomycetales(100);f__Metschnikowiaceae(100);g__Metschnikowia(100);s__Metschnikowia_pulcherrima(100); | Metschnikowia | 13 | Saprotroph | \|Undefined Saprotroph\| | Yeast | NULL | Probable | As nectar-tap_saprotroph (Põlme et al. 2020); Decay sugar-rich substrates (Põlme et al. 2020) | Põlme S, et al. 2020. Fungal Diversity 105:1-16 ((https://doi.org/10.1007/s13225-020-00466-2)) |
| Otu57409 | 1 | k__Fungi(100);p__Ascomycota(100);c__Saccharomycetes(100);o__Saccharomycetales(100);f__Metschnikowiaceae(100);g__Metschnikowia(100);s__Metschnikowia_pulcherrima(100); | Metschnikowia | 13 | Saprotroph | \|Undefined Saprotroph\| | Yeast | NULL | Probable | As nectar-tap_saprotroph (Põlme et al. 2020); Decay sugar-rich substrates (Põlme et al. 2020) | Põlme S, et al. 2020. Fungal Diversity 105:1-16 ((https://doi.org/10.1007/s13225-020-00466-2)) |
| Otu57410 | 1 | k__Fungi(100);p__Ascomycota(100);c__Saccharomycetes(100);o__Saccharomycetales(100);f__Metschnikowiaceae(100);g__Metschnikowia(100);s__Metschnikowia_pulcherrima(100); | Metschnikowia | 13 | Saprotroph | \|Undefined Saprotroph\| | Yeast | NULL | Probable | As nectar-tap_saprotroph (Põlme et al. 2020); Decay sugar-rich substrates (Põlme et al. 2020) | Põlme S, et al. 2020. Fungal Diversity 105:1-16 ((https://doi.org/10.1007/s13225-020-00466-2)) |
| Otu57412 | 1 | k__Fungi(100);p__Ascomycota(100);c__Saccharomycetes(100);o__Saccharomycetales(100);f__Metschnikowiaceae(100);g__Metschnikowia(100);s__Metschnikowia_pulcherrima(100); | Metschnikowia | 13 | Saprotroph | \|Undefined Saprotroph\| | Yeast | NULL | Probable | As nectar-tap_saprotroph (Põlme et al. 2020); Decay sugar-rich substrates (Põlme et al. 2020) | Põlme S, et al. 2020. Fungal Diversity 105:1-16 ((https://doi.org/10.1007/s13225-020-00466-2)) |
| Otu57413 | 1 | k__Fungi(100);p__Ascomycota(100);c__Saccharomycetes(100);o__Saccharomycetales(100);f__Metschnikowiaceae(100);g__Metschnikowia(100);s__Metschnikowia_pulcherrima(100); | Metschnikowia | 13 | Saprotroph | \|Undefined Saprotroph\| | Yeast | NULL | Probable | As nectar-tap_saprotroph (Põlme et al. 2020); Decay sugar-rich substrates (Põlme et al. 2020) | Põlme S, et al. 2020. Fungal Diversity 105:1-16 ((https://doi.org/10.1007/s13225-020-00466-2)) |
| Otu57414 | 1 | k__Fungi(100);p__Ascomycota(100);c__Saccharomycetes(100);o__Saccharomycetales(100);f__Metschnikowiaceae(100);g__Metschnikowia(100);s__Metschnikowia_pulcherrima(100); | Metschnikowia | 13 | Saprotroph | \|Undefined Saprotroph\| | Yeast | NULL | Probable | As nectar-tap_saprotroph (Põlme et al. 2020); Decay sugar-rich substrates (Põlme et al. 2020) | Põlme S, et al. 2020. Fungal Diversity 105:1-16 ((https://doi.org/10.1007/s13225-020-00466-2)) |
| Otu57418 | 1 | k__Fungi(100);p__Ascomycota(100);c__Saccharomycetes(100);o__Saccharomycetales(100);f__Metschnikowiaceae(100);g__Metschnikowia(100);s__Metschnikowia_pulcherrima(100); | Metschnikowia | 13 | Saprotroph | \|Undefined Saprotroph\| | Yeast | NULL | Probable | As nectar-tap_saprotroph (Põlme et al. 2020); Decay sugar-rich substrates (Põlme et al. 2020) | Põlme S, et al. 2020. Fungal Diversity 105:1-16 ((https://doi.org/10.1007/s13225-020-00466-2)) |
| Otu57419 | 1 | k__Fungi(100);p__Ascomycota(100);c__Saccharomycetes(100);o__Saccharomycetales(100);f__Metschnikowiaceae(100);g__Metschnikowia(100);s__Metschnikowia_pulcherrima(100); | Metschnikowia | 13 | Saprotroph | \|Undefined Saprotroph\| | Yeast | NULL | Probable | As nectar-tap_saprotroph (Põlme et al. 2020); Decay sugar-rich substrates (Põlme et al. 2020) | Põlme S, et al. 2020. Fungal Diversity 105:1-16 ((https://doi.org/10.1007/s13225-020-00466-2)) |
| Otu57420 | 1 | k__Fungi(100);p__Ascomycota(100);c__Saccharomycetes(100);o__Saccharomycetales(100);f__Metschnikowiaceae(100);g__Metschnikowia(100);g__Metschnikowia_unclassified(100); | Metschnikowia | 13 | Saprotroph | \|Undefined Saprotroph\| | Yeast | NULL | Probable | As nectar-tap_saprotroph (Põlme et al. 2020); Decay sugar-rich substrates (Põlme et al. 2020) | Põlme S, et al. 2020. Fungal Diversity 105:1-16 ((https://doi.org/10.1007/s13225-020-00466-2)) |
| Otu57421 | 1 | k__Fungi(100);p__Ascomycota(100);c__Saccharomycetes(100);o__Saccharomycetales(100);f__Metschnikowiaceae(100);g__Metschnikowia(100);s__Metschnikowia_pulcherrima(100); | Metschnikowia | 13 | Saprotroph | \|Undefined Saprotroph\| | Yeast | NULL | Probable | As nectar-tap_saprotroph (Põlme et al. 2020); Decay sugar-rich substrates (Põlme et al. 2020) | Põlme S, et al. 2020. Fungal Diversity 105:1-16 ((https://doi.org/10.1007/s13225-020-00466-2)) |
| Otu57422 | 1 | k__Fungi(100);p__Ascomycota(100);c__Saccharomycetes(100);o__Saccharomycetales(100);f__Metschnikowiaceae(100);g__Metschnikowia(100);s__Metschnikowia_pulcherrima(100); | Metschnikowia | 13 | Saprotroph | \|Undefined Saprotroph\| | Yeast | NULL | Probable | As nectar-tap_saprotroph (Põlme et al. 2020); Decay sugar-rich substrates (Põlme et al. 2020) | Põlme S, et al. 2020. Fungal Diversity 105:1-16 ((https://doi.org/10.1007/s13225-020-00466-2)) |
| Otu57423 | 1 | k__Fungi(100);p__Ascomycota(100);c__Saccharomycetes(100);o__Saccharomycetales(100);f__Metschnikowiaceae(100);g__Metschnikowia(100);s__Metschnikowia_pulcherrima(100); | Metschnikowia | 13 | Saprotroph | \|Undefined Saprotroph\| | Yeast | NULL | Probable | As nectar-tap_saprotroph (Põlme et al. 2020); Decay sugar-rich substrates (Põlme et al. 2020) | Põlme S, et al. 2020. Fungal Diversity 105:1-16 ((https://doi.org/10.1007/s13225-020-00466-2)) |
| Otu57426 | 1 | k__Fungi(100);p__Ascomycota(100);c__Saccharomycetes(100);o__Saccharomycetales(100);f__Metschnikowiaceae(100);g__Metschnikowia(100);s__Metschnikowia_pulcherrima(100); | Metschnikowia | 13 | Saprotroph | \|Undefined Saprotroph\| | Yeast | NULL | Probable | As nectar-tap_saprotroph (Põlme et al. 2020); Decay sugar-rich substrates (Põlme et al. 2020) | Põlme S, et al. 2020. Fungal Diversity 105:1-16 ((https://doi.org/10.1007/s13225-020-00466-2)) |
| Otu57427 | 1 | k__Fungi(100);p__Ascomycota(100);c__Saccharomycetes(100);o__Saccharomycetales(100);f__Metschnikowiaceae(100);g__Metschnikowia(100);s__Metschnikowia_pulcherrima(100); | Metschnikowia | 13 | Saprotroph | \|Undefined Saprotroph\| | Yeast | NULL | Probable | As nectar-tap_saprotroph (Põlme et al. 2020); Decay sugar-rich substrates (Põlme et al. 2020) | Põlme S, et al. 2020. Fungal Diversity 105:1-16 ((https://doi.org/10.1007/s13225-020-00466-2)) |
| Otu57436 | 1 | k__Fungi(100);p__Ascomycota(100);c__Saccharomycetes(100);o__Saccharomycetales(100);f__Metschnikowiaceae(100);g__Metschnikowia(100);s__Metschnikowia_pulcherrima(100); | Metschnikowia | 13 | Saprotroph | \|Undefined Saprotroph\| | Yeast | NULL | Probable | As nectar-tap_saprotroph (Põlme et al. 2020); Decay sugar-rich substrates (Põlme et al. 2020) | Põlme S, et al. 2020. Fungal Diversity 105:1-16 ((https://doi.org/10.1007/s13225-020-00466-2)) |
| Otu57438 | 1 | k__Fungi(100);p__Ascomycota(100);c__Saccharomycetes(100);o__Saccharomycetales(100);f__Metschnikowiaceae(100);g__Metschnikowia(100);s__Metschnikowia_pulcherrima(100); | Metschnikowia | 13 | Saprotroph | \|Undefined Saprotroph\| | Yeast | NULL | Probable | As nectar-tap_saprotroph (Põlme et al. 2020); Decay sugar-rich substrates (Põlme et al. 2020) | Põlme S, et al. 2020. Fungal Diversity 105:1-16 ((https://doi.org/10.1007/s13225-020-00466-2)) |
| Otu57439 | 1 | k__Fungi(100);p__Ascomycota(100);c__Saccharomycetes(100);o__Saccharomycetales(100);f__Metschnikowiaceae(100);g__Metschnikowia(100);s__Metschnikowia_pulcherrima(100); | Metschnikowia | 13 | Saprotroph | \|Undefined Saprotroph\| | Yeast | NULL | Probable | As nectar-tap_saprotroph (Põlme et al. 2020); Decay sugar-rich substrates (Põlme et al. 2020) | Põlme S, et al. 2020. Fungal Diversity 105:1-16 ((https://doi.org/10.1007/s13225-020-00466-2)) |
| Otu57440 | 1 | k__Fungi(100);p__Ascomycota(100);c__Saccharomycetes(100);o__Saccharomycetales(100);f__Metschnikowiaceae(100);g__Metschnikowia(100);g__Metschnikowia_unclassified(100); | Metschnikowia | 13 | Saprotroph | \|Undefined Saprotroph\| | Yeast | NULL | Probable | As nectar-tap_saprotroph (Põlme et al. 2020); Decay sugar-rich substrates (Põlme et al. 2020) | Põlme S, et al. 2020. Fungal Diversity 105:1-16 ((https://doi.org/10.1007/s13225-020-00466-2)) |
| Otu57441 | 1 | k__Fungi(100);p__Ascomycota(100);c__Saccharomycetes(100);o__Saccharomycetales(100);f__Metschnikowiaceae(100);g__Metschnikowia(100);s__Metschnikowia_pulcherrima(100); | Metschnikowia | 13 | Saprotroph | \|Undefined Saprotroph\| | Yeast | NULL | Probable | As nectar-tap_saprotroph (Põlme et al. 2020); Decay sugar-rich substrates (Põlme et al. 2020) | Põlme S, et al. 2020. Fungal Diversity 105:1-16 ((https://doi.org/10.1007/s13225-020-00466-2)) |
| Otu57444 | 1 | k__Fungi(100);p__Ascomycota(100);c__Saccharomycetes(100);o__Saccharomycetales(100);f__Metschnikowiaceae(100);g__Metschnikowia(100);s__Metschnikowia_pulcherrima(100); | Metschnikowia | 13 | Saprotroph | \|Undefined Saprotroph\| | Yeast | NULL | Probable | As nectar-tap_saprotroph (Põlme et al. 2020); Decay sugar-rich substrates (Põlme et al. 2020) | Põlme S, et al. 2020. Fungal Diversity 105:1-16 ((https://doi.org/10.1007/s13225-020-00466-2)) |
| Otu57445 | 1 | k__Fungi(100);p__Ascomycota(100);c__Saccharomycetes(100);o__Saccharomycetales(100);f__Metschnikowiaceae(100);g__Metschnikowia(100);s__Metschnikowia_pulcherrima(100); | Metschnikowia | 13 | Saprotroph | \|Undefined Saprotroph\| | Yeast | NULL | Probable | As nectar-tap_saprotroph (Põlme et al. 2020); Decay sugar-rich substrates (Põlme et al. 2020) | Põlme S, et al. 2020. Fungal Diversity 105:1-16 ((https://doi.org/10.1007/s13225-020-00466-2)) |
| Otu57446 | 1 | k__Fungi(100);p__Ascomycota(100);c__Saccharomycetes(100);o__Saccharomycetales(100);f__Metschnikowiaceae(100);g__Metschnikowia(100);s__Metschnikowia_pulcherrima(100); | Metschnikowia | 13 | Saprotroph | \|Undefined Saprotroph\| | Yeast | NULL | Probable | As nectar-tap_saprotroph (Põlme et al. 2020); Decay sugar-rich substrates (Põlme et al. 2020) | Põlme S, et al. 2020. Fungal Diversity 105:1-16 ((https://doi.org/10.1007/s13225-020-00466-2)) |
| Otu57447 | 1 | k__Fungi(100);p__Ascomycota(100);c__Saccharomycetes(100);o__Saccharomycetales(100);f__Metschnikowiaceae(100);g__Metschnikowia(100);s__Metschnikowia_pulcherrima(100); | Metschnikowia | 13 | Saprotroph | \|Undefined Saprotroph\| | Yeast | NULL | Probable | As nectar-tap_saprotroph (Põlme et al. 2020); Decay sugar-rich substrates (Põlme et al. 2020) | Põlme S, et al. 2020. Fungal Diversity 105:1-16 ((https://doi.org/10.1007/s13225-020-00466-2)) |
| Otu57449 | 1 | k__Fungi(100);p__Ascomycota(100);c__Saccharomycetes(100);o__Saccharomycetales(100);f__Metschnikowiaceae(100);g__Metschnikowia(100);g__Metschnikowia_unclassified(100); | Metschnikowia | 13 | Saprotroph | \|Undefined Saprotroph\| | Yeast | NULL | Probable | As nectar-tap_saprotroph (Põlme et al. 2020); Decay sugar-rich substrates (Põlme et al. 2020) | Põlme S, et al. 2020. Fungal Diversity 105:1-16 ((https://doi.org/10.1007/s13225-020-00466-2)) |
| Otu57451 | 1 | k__Fungi(100);p__Ascomycota(100);c__Saccharomycetes(100);o__Saccharomycetales(100);f__Metschnikowiaceae(100);g__Metschnikowia(100);s__Metschnikowia_pulcherrima(100); | Metschnikowia | 13 | Saprotroph | \|Undefined Saprotroph\| | Yeast | NULL | Probable | As nectar-tap_saprotroph (Põlme et al. 2020); Decay sugar-rich substrates (Põlme et al. 2020) | Põlme S, et al. 2020. Fungal Diversity 105:1-16 ((https://doi.org/10.1007/s13225-020-00466-2)) |
| Otu57452 | 1 | k__Fungi(100);p__Ascomycota(100);c__Saccharomycetes(100);o__Saccharomycetales(100);f__Metschnikowiaceae(100);g__Metschnikowia(100);g__Metschnikowia_unclassified(100); | Metschnikowia | 13 | Saprotroph | \|Undefined Saprotroph\| | Yeast | NULL | Probable | As nectar-tap_saprotroph (Põlme et al. 2020); Decay sugar-rich substrates (Põlme et al. 2020) | Põlme S, et al. 2020. Fungal Diversity 105:1-16 ((https://doi.org/10.1007/s13225-020-00466-2)) |
| Otu57453 | 1 | k__Fungi(100);p__Ascomycota(100);c__Saccharomycetes(100);o__Saccharomycetales(100);f__Metschnikowiaceae(100);g__Metschnikowia(100);s__Metschnikowia_pulcherrima(100); | Metschnikowia | 13 | Saprotroph | \|Undefined Saprotroph\| | Yeast | NULL | Probable | As nectar-tap_saprotroph (Põlme et al. 2020); Decay sugar-rich substrates (Põlme et al. 2020) | Põlme S, et al. 2020. Fungal Diversity 105:1-16 ((https://doi.org/10.1007/s13225-020-00466-2)) |
| Otu57454 | 1 | k__Fungi(100);p__Ascomycota(100);c__Saccharomycetes(100);o__Saccharomycetales(100);f__Metschnikowiaceae(100);g__Metschnikowia(100);s__Metschnikowia_pulcherrima(100); | Metschnikowia | 13 | Saprotroph | \|Undefined Saprotroph\| | Yeast | NULL | Probable | As nectar-tap_saprotroph (Põlme et al. 2020); Decay sugar-rich substrates (Põlme et al. 2020) | Põlme S, et al. 2020. Fungal Diversity 105:1-16 ((https://doi.org/10.1007/s13225-020-00466-2)) |
| Otu57464 | 1 | k__Fungi(100);p__Ascomycota(100);c__Saccharomycetes(100);o__Saccharomycetales(100);f__Metschnikowiaceae(100);g__Metschnikowia(100);g__Metschnikowia_unclassified(100); | Metschnikowia | 13 | Saprotroph | \|Undefined Saprotroph\| | Yeast | NULL | Probable | As nectar-tap_saprotroph (Põlme et al. 2020); Decay sugar-rich substrates (Põlme et al. 2020) | Põlme S, et al. 2020. Fungal Diversity 105:1-16 ((https://doi.org/10.1007/s13225-020-00466-2)) |
| Otu57466 | 1 | k__Fungi(100);p__Ascomycota(100);c__Saccharomycetes(100);o__Saccharomycetales(100);f__Metschnikowiaceae(100);g__Metschnikowia(100);s__Metschnikowia_pulcherrima(100); | Metschnikowia | 13 | Saprotroph | \|Undefined Saprotroph\| | Yeast | NULL | Probable | As nectar-tap_saprotroph (Põlme et al. 2020); Decay sugar-rich substrates (Põlme et al. 2020) | Põlme S, et al. 2020. Fungal Diversity 105:1-16 ((https://doi.org/10.1007/s13225-020-00466-2)) |
| Otu57467 | 1 | k__Fungi(100);p__Ascomycota(100);c__Saccharomycetes(100);o__Saccharomycetales(100);f__Metschnikowiaceae(100);g__Metschnikowia(100);s__Metschnikowia_pulcherrima(100); | Metschnikowia | 13 | Saprotroph | \|Undefined Saprotroph\| | Yeast | NULL | Probable | As nectar-tap_saprotroph (Põlme et al. 2020); Decay sugar-rich substrates (Põlme et al. 2020) | Põlme S, et al. 2020. Fungal Diversity 105:1-16 ((https://doi.org/10.1007/s13225-020-00466-2)) |
| Otu57469 | 1 | k__Fungi(100);p__Ascomycota(100);c__Saccharomycetes(100);o__Saccharomycetales(100);o__Saccharomycetales_unclassified(100);o__Saccharomycetales_unclassified(100);o__Saccharomycetales_unclassified(100); | Saccharomycetales | 7 | Saprotroph | Undefined Saprotroph | Yeast | NULL | Possible | NULL | Sterkenburg E, et al. 2015. New Phytologist 207:1145-1158 ((https://doi.org/10.1111/nph.13426)) |
| Otu57470 | 1 | k__Fungi(100);p__Ascomycota(100);c__Saccharomycetes(100);o__Saccharomycetales(100);o__Saccharomycetales_unclassified(100);o__Saccharomycetales_unclassified(100);o__Saccharomycetales_unclassified(100); | Saccharomycetales | 7 | Saprotroph | Undefined Saprotroph | Yeast | NULL | Possible | NULL | Sterkenburg E, et al. 2015. New Phytologist 207:1145-1158 ((https://doi.org/10.1111/nph.13426)) |
| Otu57472 | 1 | k__Fungi(100);p__Ascomycota(100);c__Saccharomycetes(100);o__Saccharomycetales(100);f__Metschnikowiaceae(100);g__Metschnikowia(100);s__Metschnikowia_pulcherrima(100); | Metschnikowia | 13 | Saprotroph | \|Undefined Saprotroph\| | Yeast | NULL | Probable | As nectar-tap_saprotroph (Põlme et al. 2020); Decay sugar-rich substrates (Põlme et al. 2020) | Põlme S, et al. 2020. Fungal Diversity 105:1-16 ((https://doi.org/10.1007/s13225-020-00466-2)) |
| Otu57473 | 1 | k__Fungi(100);p__Ascomycota(100);c__Saccharomycetes(100);o__Saccharomycetales(100);f__Metschnikowiaceae(100);g__Metschnikowia(100);s__Metschnikowia_pulcherrima(100); | Metschnikowia | 13 | Saprotroph | \|Undefined Saprotroph\| | Yeast | NULL | Probable | As nectar-tap_saprotroph (Põlme et al. 2020); Decay sugar-rich substrates (Põlme et al. 2020) | Põlme S, et al. 2020. Fungal Diversity 105:1-16 ((https://doi.org/10.1007/s13225-020-00466-2)) |
| Otu57474 | 1 | k__Fungi(100);p__Ascomycota(100);c__Saccharomycetes(100);o__Saccharomycetales(100);f__Metschnikowiaceae(100);g__Metschnikowia(100);g__Metschnikowia_unclassified(100); | Metschnikowia | 13 | Saprotroph | \|Undefined Saprotroph\| | Yeast | NULL | Probable | As nectar-tap_saprotroph (Põlme et al. 2020); Decay sugar-rich substrates (Põlme et al. 2020) | Põlme S, et al. 2020. Fungal Diversity 105:1-16 ((https://doi.org/10.1007/s13225-020-00466-2)) |
| Otu57476 | 1 | k__Fungi(100);p__Ascomycota(100);c__Saccharomycetes(100);o__Saccharomycetales(100);f__Metschnikowiaceae(100);g__Metschnikowia(100);s__Metschnikowia_pulcherrima(100); | Metschnikowia | 13 | Saprotroph | \|Undefined Saprotroph\| | Yeast | NULL | Probable | As nectar-tap_saprotroph (Põlme et al. 2020); Decay sugar-rich substrates (Põlme et al. 2020) | Põlme S, et al. 2020. Fungal Diversity 105:1-16 ((https://doi.org/10.1007/s13225-020-00466-2)) |
| Otu57477 | 1 | k__Fungi(100);p__Ascomycota(100);c__Saccharomycetes(100);o__Saccharomycetales(100);f__Metschnikowiaceae(100);g__Metschnikowia(100);g__Metschnikowia_unclassified(100); | Metschnikowia | 13 | Saprotroph | \|Undefined Saprotroph\| | Yeast | NULL | Probable | As nectar-tap_saprotroph (Põlme et al. 2020); Decay sugar-rich substrates (Põlme et al. 2020) | Põlme S, et al. 2020. Fungal Diversity 105:1-16 ((https://doi.org/10.1007/s13225-020-00466-2)) |
| Otu57478 | 1 | k__Fungi(100);p__Ascomycota(100);c__Saccharomycetes(100);o__Saccharomycetales(100);f__Metschnikowiaceae(100);g__Metschnikowia(100);s__Metschnikowia_pulcherrima(100); | Metschnikowia | 13 | Saprotroph | \|Undefined Saprotroph\| | Yeast | NULL | Probable | As nectar-tap_saprotroph (Põlme et al. 2020); Decay sugar-rich substrates (Põlme et al. 2020) | Põlme S, et al. 2020. Fungal Diversity 105:1-16 ((https://doi.org/10.1007/s13225-020-00466-2)) |
| Otu57479 | 1 | k__Fungi(100);p__Ascomycota(100);c__Saccharomycetes(100);o__Saccharomycetales(100);f__Metschnikowiaceae(100);g__Metschnikowia(100);g__Metschnikowia_unclassified(100); | Metschnikowia | 13 | Saprotroph | \|Undefined Saprotroph\| | Yeast | NULL | Probable | As nectar-tap_saprotroph (Põlme et al. 2020); Decay sugar-rich substrates (Põlme et al. 2020) | Põlme S, et al. 2020. Fungal Diversity 105:1-16 ((https://doi.org/10.1007/s13225-020-00466-2)) |
| Otu57480 | 1 | k__Fungi(100);p__Ascomycota(100);c__Saccharomycetes(100);o__Saccharomycetales(100);f__Metschnikowiaceae(100);g__Metschnikowia(100);g__Metschnikowia_unclassified(100); | Metschnikowia | 13 | Saprotroph | \|Undefined Saprotroph\| | Yeast | NULL | Probable | As nectar-tap_saprotroph (Põlme et al. 2020); Decay sugar-rich substrates (Põlme et al. 2020) | Põlme S, et al. 2020. Fungal Diversity 105:1-16 ((https://doi.org/10.1007/s13225-020-00466-2)) |
| Otu57482 | 1 | k__Fungi(100);p__Ascomycota(100);c__Saccharomycetes(100);o__Saccharomycetales(100);f__Metschnikowiaceae(100);g__Metschnikowia(100);s__Metschnikowia_pulcherrima(100); | Metschnikowia | 13 | Saprotroph | \|Undefined Saprotroph\| | Yeast | NULL | Probable | As nectar-tap_saprotroph (Põlme et al. 2020); Decay sugar-rich substrates (Põlme et al. 2020) | Põlme S, et al. 2020. Fungal Diversity 105:1-16 ((https://doi.org/10.1007/s13225-020-00466-2)) |
| Otu57483 | 1 | k__Fungi(100);p__Ascomycota(100);c__Saccharomycetes(100);o__Saccharomycetales(100);f__Metschnikowiaceae(100);g__Metschnikowia(100);s__Metschnikowia_pulcherrima(100); | Metschnikowia | 13 | Saprotroph | \|Undefined Saprotroph\| | Yeast | NULL | Probable | As nectar-tap_saprotroph (Põlme et al. 2020); Decay sugar-rich substrates (Põlme et al. 2020) | Põlme S, et al. 2020. Fungal Diversity 105:1-16 ((https://doi.org/10.1007/s13225-020-00466-2)) |
| Otu57484 | 1 | k__Fungi(100);p__Ascomycota(100);c__Saccharomycetes(100);o__Saccharomycetales(100);f__Metschnikowiaceae(100);g__Metschnikowia(100);s__Metschnikowia_pulcherrima(100); | Metschnikowia | 13 | Saprotroph | \|Undefined Saprotroph\| | Yeast | NULL | Probable | As nectar-tap_saprotroph (Põlme et al. 2020); Decay sugar-rich substrates (Põlme et al. 2020) | Põlme S, et al. 2020. Fungal Diversity 105:1-16 ((https://doi.org/10.1007/s13225-020-00466-2)) |
| Otu57485 | 1 | k__Fungi(100);p__Ascomycota(100);c__Saccharomycetes(100);o__Saccharomycetales(100);f__Metschnikowiaceae(100);g__Metschnikowia(100);s__Metschnikowia_pulcherrima(100); | Metschnikowia | 13 | Saprotroph | \|Undefined Saprotroph\| | Yeast | NULL | Probable | As nectar-tap_saprotroph (Põlme et al. 2020); Decay sugar-rich substrates (Põlme et al. 2020) | Põlme S, et al. 2020. Fungal Diversity 105:1-16 ((https://doi.org/10.1007/s13225-020-00466-2)) |
| Otu57486 | 1 | k__Fungi(100);p__Ascomycota(100);c__Saccharomycetes(100);o__Saccharomycetales(100);f__Metschnikowiaceae(100);g__Metschnikowia(100);s__Metschnikowia_pulcherrima(100); | Metschnikowia | 13 | Saprotroph | \|Undefined Saprotroph\| | Yeast | NULL | Probable | As nectar-tap_saprotroph (Põlme et al. 2020); Decay sugar-rich substrates (Põlme et al. 2020) | Põlme S, et al. 2020. Fungal Diversity 105:1-16 ((https://doi.org/10.1007/s13225-020-00466-2)) |
| Otu57487 | 1 | k__Fungi(100);p__Ascomycota(100);c__Saccharomycetes(100);o__Saccharomycetales(100);f__Saccharomycetales_family_Incertae_sedis(100);g__Candida(100);g__Candida_unclassified(100); | Candida | 13 | Pathotroph-Saprotroph-Symbiotroph | Animal Pathogen-Endophyte-Endosymbiont-Epiphyte-\|Undefined Saprotroph\| | Dimorphic Yeast-Yeast | NULL | Probable | As nectar-tap_saprotroph (Põlme et al. 2020); Decay sugar-rich substrates (Põlme et al. 2020); This asexual genus is polyphyletic and contain species that function very differently depend on what they are. Caution should be taken when interpreting guilds for Candida species. | Manolakaki D, et al. 2010. Virulence. 1: 367-375 ((https://doi.org/10.4161/viru.1.5.12796)); Mekbib SB, et al. 2011. Tropical Plant Pathology 36:233-240 ((https://doi.org/10.1590/S1982-56762011000400004)); Põlme S, et al. 2020. Fungal Diversity 105:1-16 ((https://doi.org/10.1007/s13225-020-00466-2)) |
| Otu57488 | 1 | k__Fungi(100);p__Ascomycota(100);c__Saccharomycetes(100);o__Saccharomycetales(100);f__Metschnikowiaceae(100);g__Metschnikowia(100);s__Metschnikowia_pulcherrima(100); | Metschnikowia | 13 | Saprotroph | \|Undefined Saprotroph\| | Yeast | NULL | Probable | As nectar-tap_saprotroph (Põlme et al. 2020); Decay sugar-rich substrates (Põlme et al. 2020) | Põlme S, et al. 2020. Fungal Diversity 105:1-16 ((https://doi.org/10.1007/s13225-020-00466-2)) |
| Otu57489 | 1 | k__Fungi(100);p__Ascomycota(100);c__Saccharomycetes(100);o__Saccharomycetales(100);f__Metschnikowiaceae(100);g__Metschnikowia(100);g__Metschnikowia_unclassified(100); | Metschnikowia | 13 | Saprotroph | \|Undefined Saprotroph\| | Yeast | NULL | Probable | As nectar-tap_saprotroph (Põlme et al. 2020); Decay sugar-rich substrates (Põlme et al. 2020) | Põlme S, et al. 2020. Fungal Diversity 105:1-16 ((https://doi.org/10.1007/s13225-020-00466-2)) |
| Otu57490 | 1 | k__Fungi(100);p__Ascomycota(100);c__Saccharomycetes(100);o__Saccharomycetales(100);f__Metschnikowiaceae(100);g__Metschnikowia(100);s__Metschnikowia_pulcherrima(100); | Metschnikowia | 13 | Saprotroph | \|Undefined Saprotroph\| | Yeast | NULL | Probable | As nectar-tap_saprotroph (Põlme et al. 2020); Decay sugar-rich substrates (Põlme et al. 2020) | Põlme S, et al. 2020. Fungal Diversity 105:1-16 ((https://doi.org/10.1007/s13225-020-00466-2)) |
| Otu57493 | 1 | k__Fungi(100);p__Ascomycota(100);c__Saccharomycetes(100);o__Saccharomycetales(100);f__Metschnikowiaceae(100);g__Metschnikowia(100);s__Metschnikowia_pulcherrima(100); | Metschnikowia | 13 | Saprotroph | \|Undefined Saprotroph\| | Yeast | NULL | Probable | As nectar-tap_saprotroph (Põlme et al. 2020); Decay sugar-rich substrates (Põlme et al. 2020) | Põlme S, et al. 2020. Fungal Diversity 105:1-16 ((https://doi.org/10.1007/s13225-020-00466-2)) |
| Otu57494 | 1 | k__Fungi(100);p__Ascomycota(100);c__Saccharomycetes(100);o__Saccharomycetales(100);f__Metschnikowiaceae(100);g__Metschnikowia(100);s__Metschnikowia_pulcherrima(100); | Metschnikowia | 13 | Saprotroph | \|Undefined Saprotroph\| | Yeast | NULL | Probable | As nectar-tap_saprotroph (Põlme et al. 2020); Decay sugar-rich substrates (Põlme et al. 2020) | Põlme S, et al. 2020. Fungal Diversity 105:1-16 ((https://doi.org/10.1007/s13225-020-00466-2)) |
| Otu57495 | 1 | k__Fungi(100);p__Ascomycota(100);c__Saccharomycetes(100);o__Saccharomycetales(100);f__Metschnikowiaceae(100);g__Metschnikowia(100);g__Metschnikowia_unclassified(100); | Metschnikowia | 13 | Saprotroph | \|Undefined Saprotroph\| | Yeast | NULL | Probable | As nectar-tap_saprotroph (Põlme et al. 2020); Decay sugar-rich substrates (Põlme et al. 2020) | Põlme S, et al. 2020. Fungal Diversity 105:1-16 ((https://doi.org/10.1007/s13225-020-00466-2)) |
| Otu57496 | 1 | k__Fungi(100);p__Ascomycota(100);c__Saccharomycetes(100);o__Saccharomycetales(100);f__Metschnikowiaceae(100);g__Metschnikowia(100);g__Metschnikowia_unclassified(100); | Metschnikowia | 13 | Saprotroph | \|Undefined Saprotroph\| | Yeast | NULL | Probable | As nectar-tap_saprotroph (Põlme et al. 2020); Decay sugar-rich substrates (Põlme et al. 2020) | Põlme S, et al. 2020. Fungal Diversity 105:1-16 ((https://doi.org/10.1007/s13225-020-00466-2)) |
| Otu57497 | 1 | k__Fungi(100);p__Ascomycota(100);c__Saccharomycetes(100);o__Saccharomycetales(100);f__Metschnikowiaceae(100);g__Metschnikowia(100);s__Metschnikowia_pulcherrima(100); | Metschnikowia | 13 | Saprotroph | \|Undefined Saprotroph\| | Yeast | NULL | Probable | As nectar-tap_saprotroph (Põlme et al. 2020); Decay sugar-rich substrates (Põlme et al. 2020) | Põlme S, et al. 2020. Fungal Diversity 105:1-16 ((https://doi.org/10.1007/s13225-020-00466-2)) |
| Otu57498 | 1 | k__Fungi(100);p__Ascomycota(100);c__Saccharomycetes(100);o__Saccharomycetales(100);f__Saccharomycetales_family_Incertae_sedis(100);g__Candida(100);s__Candida_cretensis(100); | Candida | 13 | Pathotroph-Saprotroph-Symbiotroph | Animal Pathogen-Endophyte-Endosymbiont-Epiphyte-\|Undefined Saprotroph\| | Dimorphic Yeast-Yeast | NULL | Probable | As nectar-tap_saprotroph (Põlme et al. 2020); Decay sugar-rich substrates (Põlme et al. 2020); This asexual genus is polyphyletic and contain species that function very differently depend on what they are. Caution should be taken when interpreting guilds for Candida species. | Manolakaki D, et al. 2010. Virulence. 1: 367-375 ((https://doi.org/10.4161/viru.1.5.12796)); Mekbib SB, et al. 2011. Tropical Plant Pathology 36:233-240 ((https://doi.org/10.1590/S1982-56762011000400004)); Põlme S, et al. 2020. Fungal Diversity 105:1-16 ((https://doi.org/10.1007/s13225-020-00466-2)) |
| Otu57499 | 1 | k__Fungi(100);p__Ascomycota(100);c__Saccharomycetes(100);o__Saccharomycetales(100);f__Metschnikowiaceae(100);g__Metschnikowia(100);s__Metschnikowia_pulcherrima(100); | Metschnikowia | 13 | Saprotroph | \|Undefined Saprotroph\| | Yeast | NULL | Probable | As nectar-tap_saprotroph (Põlme et al. 2020); Decay sugar-rich substrates (Põlme et al. 2020) | Põlme S, et al. 2020. Fungal Diversity 105:1-16 ((https://doi.org/10.1007/s13225-020-00466-2)) |
| Otu57500 | 1 | k__Fungi(100);p__Ascomycota(100);c__Saccharomycetes(100);o__Saccharomycetales(100);f__Metschnikowiaceae(100);g__Metschnikowia(100);s__Metschnikowia_pulcherrima(100); | Metschnikowia | 13 | Saprotroph | \|Undefined Saprotroph\| | Yeast | NULL | Probable | As nectar-tap_saprotroph (Põlme et al. 2020); Decay sugar-rich substrates (Põlme et al. 2020) | Põlme S, et al. 2020. Fungal Diversity 105:1-16 ((https://doi.org/10.1007/s13225-020-00466-2)) |
| Otu57501 | 1 | k__Fungi(100);p__Ascomycota(100);c__Saccharomycetes(100);o__Saccharomycetales(100);f__Metschnikowiaceae(100);g__Metschnikowia(100);s__Metschnikowia_pulcherrima(100); | Metschnikowia | 13 | Saprotroph | \|Undefined Saprotroph\| | Yeast | NULL | Probable | As nectar-tap_saprotroph (Põlme et al. 2020); Decay sugar-rich substrates (Põlme et al. 2020) | Põlme S, et al. 2020. Fungal Diversity 105:1-16 ((https://doi.org/10.1007/s13225-020-00466-2)) |
| Otu57502 | 1 | k__Fungi(100);p__Ascomycota(100);c__Saccharomycetes(100);o__Saccharomycetales(100);f__Metschnikowiaceae(100);g__Metschnikowia(100);s__Metschnikowia_pulcherrima(100); | Metschnikowia | 13 | Saprotroph | \|Undefined Saprotroph\| | Yeast | NULL | Probable | As nectar-tap_saprotroph (Põlme et al. 2020); Decay sugar-rich substrates (Põlme et al. 2020) | Põlme S, et al. 2020. Fungal Diversity 105:1-16 ((https://doi.org/10.1007/s13225-020-00466-2)) |
| Otu57503 | 1 | k__Fungi(100);p__Ascomycota(100);c__Saccharomycetes(100);o__Saccharomycetales(100);f__Metschnikowiaceae(100);g__Metschnikowia(100);s__Metschnikowia_pulcherrima(100); | Metschnikowia | 13 | Saprotroph | \|Undefined Saprotroph\| | Yeast | NULL | Probable | As nectar-tap_saprotroph (Põlme et al. 2020); Decay sugar-rich substrates (Põlme et al. 2020) | Põlme S, et al. 2020. Fungal Diversity 105:1-16 ((https://doi.org/10.1007/s13225-020-00466-2)) |
| Otu57504 | 1 | k__Fungi(100);p__Ascomycota(100);c__Saccharomycetes(100);o__Saccharomycetales(100);f__Metschnikowiaceae(100);g__Metschnikowia(100);g__Metschnikowia_unclassified(100); | Metschnikowia | 13 | Saprotroph | \|Undefined Saprotroph\| | Yeast | NULL | Probable | As nectar-tap_saprotroph (Põlme et al. 2020); Decay sugar-rich substrates (Põlme et al. 2020) | Põlme S, et al. 2020. Fungal Diversity 105:1-16 ((https://doi.org/10.1007/s13225-020-00466-2)) |
| Otu57506 | 1 | k__Fungi(100);p__Ascomycota(100);c__Saccharomycetes(100);o__Saccharomycetales(100);f__Metschnikowiaceae(100);g__Metschnikowia(100);s__Metschnikowia_pulcherrima(100); | Metschnikowia | 13 | Saprotroph | \|Undefined Saprotroph\| | Yeast | NULL | Probable | As nectar-tap_saprotroph (Põlme et al. 2020); Decay sugar-rich substrates (Põlme et al. 2020) | Põlme S, et al. 2020. Fungal Diversity 105:1-16 ((https://doi.org/10.1007/s13225-020-00466-2)) |
| Otu57507 | 1 | k__Fungi(100);p__Ascomycota(100);c__Saccharomycetes(100);o__Saccharomycetales(100);f__Metschnikowiaceae(100);g__Metschnikowia(100);s__Metschnikowia_pulcherrima(100); | Metschnikowia | 13 | Saprotroph | \|Undefined Saprotroph\| | Yeast | NULL | Probable | As nectar-tap_saprotroph (Põlme et al. 2020); Decay sugar-rich substrates (Põlme et al. 2020) | Põlme S, et al. 2020. Fungal Diversity 105:1-16 ((https://doi.org/10.1007/s13225-020-00466-2)) |
| Otu57508 | 1 | k__Fungi(100);p__Ascomycota(100);c__Saccharomycetes(100);o__Saccharomycetales(100);f__Metschnikowiaceae(100);g__Metschnikowia(100);s__Metschnikowia_pulcherrima(100); | Metschnikowia | 13 | Saprotroph | \|Undefined Saprotroph\| | Yeast | NULL | Probable | As nectar-tap_saprotroph (Põlme et al. 2020); Decay sugar-rich substrates (Põlme et al. 2020) | Põlme S, et al. 2020. Fungal Diversity 105:1-16 ((https://doi.org/10.1007/s13225-020-00466-2)) |
| Otu57509 | 1 | k__Fungi(100);p__Ascomycota(100);c__Saccharomycetes(100);o__Saccharomycetales(100);f__Metschnikowiaceae(100);g__Metschnikowia(100);s__Metschnikowia_pulcherrima(100); | Metschnikowia | 13 | Saprotroph | \|Undefined Saprotroph\| | Yeast | NULL | Probable | As nectar-tap_saprotroph (Põlme et al. 2020); Decay sugar-rich substrates (Põlme et al. 2020) | Põlme S, et al. 2020. Fungal Diversity 105:1-16 ((https://doi.org/10.1007/s13225-020-00466-2)) |
| Otu57512 | 1 | k__Fungi(100);p__Ascomycota(100);c__Saccharomycetes(100);o__Saccharomycetales(100);f__Metschnikowiaceae(100);g__Metschnikowia(100);s__Metschnikowia_pulcherrima(100); | Metschnikowia | 13 | Saprotroph | \|Undefined Saprotroph\| | Yeast | NULL | Probable | As nectar-tap_saprotroph (Põlme et al. 2020); Decay sugar-rich substrates (Põlme et al. 2020) | Põlme S, et al. 2020. Fungal Diversity 105:1-16 ((https://doi.org/10.1007/s13225-020-00466-2)) |
| Otu57516 | 1 | k__Fungi(100);p__Ascomycota(100);c__Saccharomycetes(100);o__Saccharomycetales(100);f__Metschnikowiaceae(100);g__Metschnikowia(100);s__Metschnikowia_pulcherrima(100); | Metschnikowia | 13 | Saprotroph | \|Undefined Saprotroph\| | Yeast | NULL | Probable | As nectar-tap_saprotroph (Põlme et al. 2020); Decay sugar-rich substrates (Põlme et al. 2020) | Põlme S, et al. 2020. Fungal Diversity 105:1-16 ((https://doi.org/10.1007/s13225-020-00466-2)) |
| Otu57518 | 1 | k__Fungi(100);p__Ascomycota(100);c__Saccharomycetes(100);o__Saccharomycetales(100);f__Saccharomycetales_family_Incertae_sedis(100);g__Candida(100);g__Candida_unclassified(100); | Candida | 13 | Pathotroph-Saprotroph-Symbiotroph | Animal Pathogen-Endophyte-Endosymbiont-Epiphyte-\|Undefined Saprotroph\| | Dimorphic Yeast-Yeast | NULL | Probable | As nectar-tap_saprotroph (Põlme et al. 2020); Decay sugar-rich substrates (Põlme et al. 2020); This asexual genus is polyphyletic and contain species that function very differently depend on what they are. Caution should be taken when interpreting guilds for Candida species. | Manolakaki D, et al. 2010. Virulence. 1: 367-375 ((https://doi.org/10.4161/viru.1.5.12796)); Mekbib SB, et al. 2011. Tropical Plant Pathology 36:233-240 ((https://doi.org/10.1590/S1982-56762011000400004)); Põlme S, et al. 2020. Fungal Diversity 105:1-16 ((https://doi.org/10.1007/s13225-020-00466-2)) |
| Otu57522 | 1 | k__Fungi(100);p__Ascomycota(100);c__Saccharomycetes(100);o__Saccharomycetales(100);f__Metschnikowiaceae(100);g__Metschnikowia(100);s__Metschnikowia_pulcherrima(100); | Metschnikowia | 13 | Saprotroph | \|Undefined Saprotroph\| | Yeast | NULL | Probable | As nectar-tap_saprotroph (Põlme et al. 2020); Decay sugar-rich substrates (Põlme et al. 2020) | Põlme S, et al. 2020. Fungal Diversity 105:1-16 ((https://doi.org/10.1007/s13225-020-00466-2)) |
| Otu57525 | 1 | k__Fungi(100);p__Ascomycota(100);c__Saccharomycetes(100);o__Saccharomycetales(100);f__Metschnikowiaceae(100);g__Metschnikowia(100);s__Metschnikowia_pulcherrima(100); | Metschnikowia | 13 | Saprotroph | \|Undefined Saprotroph\| | Yeast | NULL | Probable | As nectar-tap_saprotroph (Põlme et al. 2020); Decay sugar-rich substrates (Põlme et al. 2020) | Põlme S, et al. 2020. Fungal Diversity 105:1-16 ((https://doi.org/10.1007/s13225-020-00466-2)) |
| Otu57526 | 1 | k__Fungi(100);p__Ascomycota(100);c__Saccharomycetes(100);o__Saccharomycetales(100);f__Metschnikowiaceae(100);g__Metschnikowia(100);s__Metschnikowia_pulcherrima(100); | Metschnikowia | 13 | Saprotroph | \|Undefined Saprotroph\| | Yeast | NULL | Probable | As nectar-tap_saprotroph (Põlme et al. 2020); Decay sugar-rich substrates (Põlme et al. 2020) | Põlme S, et al. 2020. Fungal Diversity 105:1-16 ((https://doi.org/10.1007/s13225-020-00466-2)) |
| Otu57528 | 1 | k__Fungi(100);p__Ascomycota(100);c__Saccharomycetes(100);o__Saccharomycetales(100);f__Metschnikowiaceae(100);g__Metschnikowia(100);s__Metschnikowia_pulcherrima(100); | Metschnikowia | 13 | Saprotroph | \|Undefined Saprotroph\| | Yeast | NULL | Probable | As nectar-tap_saprotroph (Põlme et al. 2020); Decay sugar-rich substrates (Põlme et al. 2020) | Põlme S, et al. 2020. Fungal Diversity 105:1-16 ((https://doi.org/10.1007/s13225-020-00466-2)) |
| Otu57536 | 1 | k__Fungi(100);p__Ascomycota(100);c__Saccharomycetes(100);o__Saccharomycetales(100);f__Metschnikowiaceae(100);g__Metschnikowia(100);s__Metschnikowia_pulcherrima(100); | Metschnikowia | 13 | Saprotroph | \|Undefined Saprotroph\| | Yeast | NULL | Probable | As nectar-tap_saprotroph (Põlme et al. 2020); Decay sugar-rich substrates (Põlme et al. 2020) | Põlme S, et al. 2020. Fungal Diversity 105:1-16 ((https://doi.org/10.1007/s13225-020-00466-2)) |
| Otu57537 | 1 | k__Fungi(100);p__Ascomycota(100);c__Saccharomycetes(100);o__Saccharomycetales(100);f__Metschnikowiaceae(100);g__Metschnikowia(100);s__Metschnikowia_pulcherrima(100); | Metschnikowia | 13 | Saprotroph | \|Undefined Saprotroph\| | Yeast | NULL | Probable | As nectar-tap_saprotroph (Põlme et al. 2020); Decay sugar-rich substrates (Põlme et al. 2020) | Põlme S, et al. 2020. Fungal Diversity 105:1-16 ((https://doi.org/10.1007/s13225-020-00466-2)) |
| Otu57538 | 1 | k__Fungi(100);p__Ascomycota(100);c__Saccharomycetes(100);o__Saccharomycetales(100);f__Metschnikowiaceae(100);g__Metschnikowia(100);g__Metschnikowia_unclassified(100); | Metschnikowia | 13 | Saprotroph | \|Undefined Saprotroph\| | Yeast | NULL | Probable | As nectar-tap_saprotroph (Põlme et al. 2020); Decay sugar-rich substrates (Põlme et al. 2020) | Põlme S, et al. 2020. Fungal Diversity 105:1-16 ((https://doi.org/10.1007/s13225-020-00466-2)) |
| Otu57539 | 1 | k__Fungi(100);p__Ascomycota(100);c__Saccharomycetes(100);o__Saccharomycetales(100);f__Metschnikowiaceae(100);g__Metschnikowia(100);s__Metschnikowia_pulcherrima(100); | Metschnikowia | 13 | Saprotroph | \|Undefined Saprotroph\| | Yeast | NULL | Probable | As nectar-tap_saprotroph (Põlme et al. 2020); Decay sugar-rich substrates (Põlme et al. 2020) | Põlme S, et al. 2020. Fungal Diversity 105:1-16 ((https://doi.org/10.1007/s13225-020-00466-2)) |
| Otu57541 | 1 | k__Fungi(100);p__Ascomycota(100);c__Saccharomycetes(100);o__Saccharomycetales(100);f__Metschnikowiaceae(100);g__Metschnikowia(100);s__Metschnikowia_pulcherrima(100); | Metschnikowia | 13 | Saprotroph | \|Undefined Saprotroph\| | Yeast | NULL | Probable | As nectar-tap_saprotroph (Põlme et al. 2020); Decay sugar-rich substrates (Põlme et al. 2020) | Põlme S, et al. 2020. Fungal Diversity 105:1-16 ((https://doi.org/10.1007/s13225-020-00466-2)) |
| Otu57542 | 1 | k__Fungi(100);p__Ascomycota(100);c__Saccharomycetes(100);o__Saccharomycetales(100);f__Metschnikowiaceae(100);g__Metschnikowia(100);s__Metschnikowia_pulcherrima(100); | Metschnikowia | 13 | Saprotroph | \|Undefined Saprotroph\| | Yeast | NULL | Probable | As nectar-tap_saprotroph (Põlme et al. 2020); Decay sugar-rich substrates (Põlme et al. 2020) | Põlme S, et al. 2020. Fungal Diversity 105:1-16 ((https://doi.org/10.1007/s13225-020-00466-2)) |
| Otu57545 | 1 | k__Fungi(100);p__Ascomycota(100);c__Saccharomycetes(100);o__Saccharomycetales(100);f__Metschnikowiaceae(100);g__Metschnikowia(100);s__Metschnikowia_pulcherrima(100); | Metschnikowia | 13 | Saprotroph | \|Undefined Saprotroph\| | Yeast | NULL | Probable | As nectar-tap_saprotroph (Põlme et al. 2020); Decay sugar-rich substrates (Põlme et al. 2020) | Põlme S, et al. 2020. Fungal Diversity 105:1-16 ((https://doi.org/10.1007/s13225-020-00466-2)) |
| Otu57546 | 1 | k__Fungi(100);p__Ascomycota(100);c__Saccharomycetes(100);o__Saccharomycetales(100);f__Metschnikowiaceae(100);g__Metschnikowia(100);s__Metschnikowia_pulcherrima(100); | Metschnikowia | 13 | Saprotroph | \|Undefined Saprotroph\| | Yeast | NULL | Probable | As nectar-tap_saprotroph (Põlme et al. 2020); Decay sugar-rich substrates (Põlme et al. 2020) | Põlme S, et al. 2020. Fungal Diversity 105:1-16 ((https://doi.org/10.1007/s13225-020-00466-2)) |
| Otu57548 | 1 | k__Fungi(100);p__Ascomycota(100);c__Saccharomycetes(100);o__Saccharomycetales(100);f__Metschnikowiaceae(100);g__Metschnikowia(100);s__Metschnikowia_pulcherrima(100); | Metschnikowia | 13 | Saprotroph | \|Undefined Saprotroph\| | Yeast | NULL | Probable | As nectar-tap_saprotroph (Põlme et al. 2020); Decay sugar-rich substrates (Põlme et al. 2020) | Põlme S, et al. 2020. Fungal Diversity 105:1-16 ((https://doi.org/10.1007/s13225-020-00466-2)) |
| Otu57549 | 1 | k__Fungi(100);p__Ascomycota(100);c__Saccharomycetes(100);o__Saccharomycetales(100);f__Metschnikowiaceae(100);g__Metschnikowia(100);s__Metschnikowia_pulcherrima(100); | Metschnikowia | 13 | Saprotroph | \|Undefined Saprotroph\| | Yeast | NULL | Probable | As nectar-tap_saprotroph (Põlme et al. 2020); Decay sugar-rich substrates (Põlme et al. 2020) | Põlme S, et al. 2020. Fungal Diversity 105:1-16 ((https://doi.org/10.1007/s13225-020-00466-2)) |
| Otu57550 | 1 | k__Fungi(100);p__Ascomycota(100);c__Saccharomycetes(100);o__Saccharomycetales(100);f__Metschnikowiaceae(100);g__Metschnikowia(100);s__Metschnikowia_pulcherrima(100); | Metschnikowia | 13 | Saprotroph | \|Undefined Saprotroph\| | Yeast | NULL | Probable | As nectar-tap_saprotroph (Põlme et al. 2020); Decay sugar-rich substrates (Põlme et al. 2020) | Põlme S, et al. 2020. Fungal Diversity 105:1-16 ((https://doi.org/10.1007/s13225-020-00466-2)) |
| Otu57551 | 1 | k__Fungi(100);p__Ascomycota(100);c__Saccharomycetes(100);o__Saccharomycetales(100);f__Metschnikowiaceae(100);g__Metschnikowia(100);s__Metschnikowia_pulcherrima(100); | Metschnikowia | 13 | Saprotroph | \|Undefined Saprotroph\| | Yeast | NULL | Probable | As nectar-tap_saprotroph (Põlme et al. 2020); Decay sugar-rich substrates (Põlme et al. 2020) | Põlme S, et al. 2020. Fungal Diversity 105:1-16 ((https://doi.org/10.1007/s13225-020-00466-2)) |
| Otu57552 | 1 | k__Fungi(100);p__Ascomycota(100);c__Saccharomycetes(100);o__Saccharomycetales(100);f__Metschnikowiaceae(100);g__Metschnikowia(100);g__Metschnikowia_unclassified(100); | Metschnikowia | 13 | Saprotroph | \|Undefined Saprotroph\| | Yeast | NULL | Probable | As nectar-tap_saprotroph (Põlme et al. 2020); Decay sugar-rich substrates (Põlme et al. 2020) | Põlme S, et al. 2020. Fungal Diversity 105:1-16 ((https://doi.org/10.1007/s13225-020-00466-2)) |
| Otu57556 | 1 | k__Fungi(100);p__Ascomycota(100);c__Saccharomycetes(100);o__Saccharomycetales(100);f__Metschnikowiaceae(100);g__Metschnikowia(100);g__Metschnikowia_unclassified(100); | Metschnikowia | 13 | Saprotroph | \|Undefined Saprotroph\| | Yeast | NULL | Probable | As nectar-tap_saprotroph (Põlme et al. 2020); Decay sugar-rich substrates (Põlme et al. 2020) | Põlme S, et al. 2020. Fungal Diversity 105:1-16 ((https://doi.org/10.1007/s13225-020-00466-2)) |
| Otu57559 | 1 | k__Fungi(100);p__Ascomycota(100);c__Saccharomycetes(100);o__Saccharomycetales(100);f__Metschnikowiaceae(100);g__Metschnikowia(100);s__Metschnikowia_pulcherrima(100); | Metschnikowia | 13 | Saprotroph | \|Undefined Saprotroph\| | Yeast | NULL | Probable | As nectar-tap_saprotroph (Põlme et al. 2020); Decay sugar-rich substrates (Põlme et al. 2020) | Põlme S, et al. 2020. Fungal Diversity 105:1-16 ((https://doi.org/10.1007/s13225-020-00466-2)) |
| Otu57561 | 1 | k__Fungi(100);p__Ascomycota(100);c__Saccharomycetes(100);o__Saccharomycetales(100);f__Metschnikowiaceae(100);g__Metschnikowia(100);s__Metschnikowia_pulcherrima(100); | Metschnikowia | 13 | Saprotroph | \|Undefined Saprotroph\| | Yeast | NULL | Probable | As nectar-tap_saprotroph (Põlme et al. 2020); Decay sugar-rich substrates (Põlme et al. 2020) | Põlme S, et al. 2020. Fungal Diversity 105:1-16 ((https://doi.org/10.1007/s13225-020-00466-2)) |
| Otu57562 | 1 | k__Fungi(100);p__Ascomycota(100);c__Saccharomycetes(100);o__Saccharomycetales(100);f__Metschnikowiaceae(100);g__Metschnikowia(100);s__Metschnikowia_pulcherrima(100); | Metschnikowia | 13 | Saprotroph | \|Undefined Saprotroph\| | Yeast | NULL | Probable | As nectar-tap_saprotroph (Põlme et al. 2020); Decay sugar-rich substrates (Põlme et al. 2020) | Põlme S, et al. 2020. Fungal Diversity 105:1-16 ((https://doi.org/10.1007/s13225-020-00466-2)) |
| Otu57563 | 1 | k__Fungi(100);p__Ascomycota(100);c__Saccharomycetes(100);o__Saccharomycetales(100);f__Metschnikowiaceae(100);g__Metschnikowia(100);s__Metschnikowia_pulcherrima(100); | Metschnikowia | 13 | Saprotroph | \|Undefined Saprotroph\| | Yeast | NULL | Probable | As nectar-tap_saprotroph (Põlme et al. 2020); Decay sugar-rich substrates (Põlme et al. 2020) | Põlme S, et al. 2020. Fungal Diversity 105:1-16 ((https://doi.org/10.1007/s13225-020-00466-2)) |
| Otu57567 | 1 | k__Fungi(100);p__Ascomycota(100);c__Saccharomycetes(100);o__Saccharomycetales(100);f__Metschnikowiaceae(100);g__Metschnikowia(100);s__Metschnikowia_pulcherrima(100); | Metschnikowia | 13 | Saprotroph | \|Undefined Saprotroph\| | Yeast | NULL | Probable | As nectar-tap_saprotroph (Põlme et al. 2020); Decay sugar-rich substrates (Põlme et al. 2020) | Põlme S, et al. 2020. Fungal Diversity 105:1-16 ((https://doi.org/10.1007/s13225-020-00466-2)) |
| Otu57568 | 1 | k__Fungi(100);p__Ascomycota(100);c__Dothideomycetes(100);o__Pleosporales(100);f__Pleosporaceae(100);g__Alternaria(100);g__Alternaria_unclassified(100); | Alternaria | 13 | Pathotroph-Saprotroph-Symbiotroph | Animal Pathogen-Endophyte-\|Plant Pathogen\|-Plant Saprotroph-Wood Saprotroph | Microfungus | Soft Rot | Possible | As foliar_endophyte (Põlme et al. 2020); As litter_saprotroph (Põlme et al. 2020); Decay leaf, fruit, seed (Põlme et al. 2020) | Duncan CG, Eslyn WE. 1966. Mycologia 58:642-664 ((https://doi.org/10.2307/3757045)); Seehann G, et al. 1975. List of Fungi in Soft-Rot Tissue Tests. Institute fur Holzbiologie und Holzschutz, Bundesforschungsanstalt fur Forst - und Holzwirtschaft, Hamburg - Lohbrugge, Germany ((https://www.irg-wp.com/irgdocs/details.php?d6493033-c9ea-40f3-8e3e-5697f32d640d)); Tedersoo L, et al. 2014. Science 346: e1256688 ((https://doi.org/10.1126/science.1256688)); Põlme S, et al. 2020. Fungal Diversity 105:1-16 ((https://doi.org/10.1007/s13225-020-00466-2)) |
| Otu57569 | 1 | k__Fungi(100);p__Ascomycota(100);c__Saccharomycetes(100);o__Saccharomycetales(100);f__Metschnikowiaceae(100);g__Metschnikowia(100);s__Metschnikowia_pulcherrima(100); | Metschnikowia | 13 | Saprotroph | \|Undefined Saprotroph\| | Yeast | NULL | Probable | As nectar-tap_saprotroph (Põlme et al. 2020); Decay sugar-rich substrates (Põlme et al. 2020) | Põlme S, et al. 2020. Fungal Diversity 105:1-16 ((https://doi.org/10.1007/s13225-020-00466-2)) |
| Otu57570 | 1 | k__Fungi(100);p__Ascomycota(100);c__Saccharomycetes(100);o__Saccharomycetales(100);f__Metschnikowiaceae(100);g__Metschnikowia(100);s__Metschnikowia_pulcherrima(100); | Metschnikowia | 13 | Saprotroph | \|Undefined Saprotroph\| | Yeast | NULL | Probable | As nectar-tap_saprotroph (Põlme et al. 2020); Decay sugar-rich substrates (Põlme et al. 2020) | Põlme S, et al. 2020. Fungal Diversity 105:1-16 ((https://doi.org/10.1007/s13225-020-00466-2)) |
| Otu57572 | 1 | k__Fungi(100);p__Ascomycota(100);c__Saccharomycetes(100);o__Saccharomycetales(100);f__Metschnikowiaceae(100);g__Metschnikowia(100);s__Metschnikowia_pulcherrima(100); | Metschnikowia | 13 | Saprotroph | \|Undefined Saprotroph\| | Yeast | NULL | Probable | As nectar-tap_saprotroph (Põlme et al. 2020); Decay sugar-rich substrates (Põlme et al. 2020) | Põlme S, et al. 2020. Fungal Diversity 105:1-16 ((https://doi.org/10.1007/s13225-020-00466-2)) |
| Otu57573 | 1 | k__Fungi(100);p__Ascomycota(100);c__Saccharomycetes(100);o__Saccharomycetales(100);f__Metschnikowiaceae(100);g__Metschnikowia(100);g__Metschnikowia_unclassified(100); | Metschnikowia | 13 | Saprotroph | \|Undefined Saprotroph\| | Yeast | NULL | Probable | As nectar-tap_saprotroph (Põlme et al. 2020); Decay sugar-rich substrates (Põlme et al. 2020) | Põlme S, et al. 2020. Fungal Diversity 105:1-16 ((https://doi.org/10.1007/s13225-020-00466-2)) |
| Otu57575 | 1 | k__Fungi(100);p__Ascomycota(100);c__Saccharomycetes(100);o__Saccharomycetales(100);f__Metschnikowiaceae(100);g__Metschnikowia(100);s__Metschnikowia_pulcherrima(100); | Metschnikowia | 13 | Saprotroph | \|Undefined Saprotroph\| | Yeast | NULL | Probable | As nectar-tap_saprotroph (Põlme et al. 2020); Decay sugar-rich substrates (Põlme et al. 2020) | Põlme S, et al. 2020. Fungal Diversity 105:1-16 ((https://doi.org/10.1007/s13225-020-00466-2)) |
| Otu57576 | 1 | k__Fungi(100);p__Ascomycota(100);c__Saccharomycetes(100);o__Saccharomycetales(100);f__Metschnikowiaceae(100);g__Metschnikowia(100);s__Metschnikowia_pulcherrima(100); | Metschnikowia | 13 | Saprotroph | \|Undefined Saprotroph\| | Yeast | NULL | Probable | As nectar-tap_saprotroph (Põlme et al. 2020); Decay sugar-rich substrates (Põlme et al. 2020) | Põlme S, et al. 2020. Fungal Diversity 105:1-16 ((https://doi.org/10.1007/s13225-020-00466-2)) |
| Otu57577 | 1 | k__Fungi(100);p__Ascomycota(100);c__Saccharomycetes(100);o__Saccharomycetales(100);f__Metschnikowiaceae(100);g__Metschnikowia(100);s__Metschnikowia_pulcherrima(100); | Metschnikowia | 13 | Saprotroph | \|Undefined Saprotroph\| | Yeast | NULL | Probable | As nectar-tap_saprotroph (Põlme et al. 2020); Decay sugar-rich substrates (Põlme et al. 2020) | Põlme S, et al. 2020. Fungal Diversity 105:1-16 ((https://doi.org/10.1007/s13225-020-00466-2)) |
| Otu57578 | 1 | k__Fungi(100);p__Ascomycota(100);c__Saccharomycetes(100);o__Saccharomycetales(100);f__Metschnikowiaceae(100);g__Metschnikowia(100);s__Metschnikowia_pulcherrima(100); | Metschnikowia | 13 | Saprotroph | \|Undefined Saprotroph\| | Yeast | NULL | Probable | As nectar-tap_saprotroph (Põlme et al. 2020); Decay sugar-rich substrates (Põlme et al. 2020) | Põlme S, et al. 2020. Fungal Diversity 105:1-16 ((https://doi.org/10.1007/s13225-020-00466-2)) |
| Otu57579 | 1 | k__Fungi(100);p__Ascomycota(100);c__Saccharomycetes(100);o__Saccharomycetales(100);f__Metschnikowiaceae(100);g__Metschnikowia(100);s__Metschnikowia_pulcherrima(100); | Metschnikowia | 13 | Saprotroph | \|Undefined Saprotroph\| | Yeast | NULL | Probable | As nectar-tap_saprotroph (Põlme et al. 2020); Decay sugar-rich substrates (Põlme et al. 2020) | Põlme S, et al. 2020. Fungal Diversity 105:1-16 ((https://doi.org/10.1007/s13225-020-00466-2)) |
| Otu57580 | 1 | k__Fungi(100);p__Ascomycota(100);c__Sordariomycetes(100);o__Chaetosphaeriales(100);f__Chaetosphaeriaceae(100);f__Chaetosphaeriaceae_unclassified(100);f__Chaetosphaeriaceae_unclassified(100); | Chaetosphaeriaceae | 9 | Saprotroph | Plant Saprotroph-Wood Saprotroph | Microfungus | NULL | Probable | Saprobes associated with decaying woody and herbaceous plant material, including timber and timber products in damp buildings (Cannon & Kirk 2007) | Cannon PF, Kirk PM. 2007. Fungal Families of the World. CAB International, Cambridge (ISBN: 978-0851998275) |
| Otu57581 | 1 | k__Fungi(100);p__Ascomycota(100);c__Saccharomycetes(100);o__Saccharomycetales(100);f__Metschnikowiaceae(100);g__Metschnikowia(100);g__Metschnikowia_unclassified(100); | Metschnikowia | 13 | Saprotroph | \|Undefined Saprotroph\| | Yeast | NULL | Probable | As nectar-tap_saprotroph (Põlme et al. 2020); Decay sugar-rich substrates (Põlme et al. 2020) | Põlme S, et al. 2020. Fungal Diversity 105:1-16 ((https://doi.org/10.1007/s13225-020-00466-2)) |
| Otu57582 | 1 | k__Fungi(100);p__Ascomycota(100);c__Saccharomycetes(100);o__Saccharomycetales(100);f__Metschnikowiaceae(100);g__Metschnikowia(100);s__Metschnikowia_pulcherrima(100); | Metschnikowia | 13 | Saprotroph | \|Undefined Saprotroph\| | Yeast | NULL | Probable | As nectar-tap_saprotroph (Põlme et al. 2020); Decay sugar-rich substrates (Põlme et al. 2020) | Põlme S, et al. 2020. Fungal Diversity 105:1-16 ((https://doi.org/10.1007/s13225-020-00466-2)) |
| Otu57583 | 1 | k__Fungi(100);p__Ascomycota(100);c__Saccharomycetes(100);o__Saccharomycetales(100);f__Metschnikowiaceae(100);g__Metschnikowia(100);s__Metschnikowia_pulcherrima(100); | Metschnikowia | 13 | Saprotroph | \|Undefined Saprotroph\| | Yeast | NULL | Probable | As nectar-tap_saprotroph (Põlme et al. 2020); Decay sugar-rich substrates (Põlme et al. 2020) | Põlme S, et al. 2020. Fungal Diversity 105:1-16 ((https://doi.org/10.1007/s13225-020-00466-2)) |
| Otu57588 | 1 | k__Fungi(100);p__Ascomycota(100);c__Saccharomycetes(100);o__Saccharomycetales(100);f__Metschnikowiaceae(100);g__Metschnikowia(100);s__Metschnikowia_pulcherrima(100); | Metschnikowia | 13 | Saprotroph | \|Undefined Saprotroph\| | Yeast | NULL | Probable | As nectar-tap_saprotroph (Põlme et al. 2020); Decay sugar-rich substrates (Põlme et al. 2020) | Põlme S, et al. 2020. Fungal Diversity 105:1-16 ((https://doi.org/10.1007/s13225-020-00466-2)) |
| Otu57590 | 1 | k__Fungi(100);p__Ascomycota(100);c__Saccharomycetes(100);o__Saccharomycetales(100);f__Saccharomycetales_family_Incertae_sedis(100);g__Candida(100);g__Candida_unclassified(100); | Candida | 13 | Pathotroph-Saprotroph-Symbiotroph | Animal Pathogen-Endophyte-Endosymbiont-Epiphyte-\|Undefined Saprotroph\| | Dimorphic Yeast-Yeast | NULL | Probable | As nectar-tap_saprotroph (Põlme et al. 2020); Decay sugar-rich substrates (Põlme et al. 2020); This asexual genus is polyphyletic and contain species that function very differently depend on what they are. Caution should be taken when interpreting guilds for Candida species. | Manolakaki D, et al. 2010. Virulence. 1: 367-375 ((https://doi.org/10.4161/viru.1.5.12796)); Mekbib SB, et al. 2011. Tropical Plant Pathology 36:233-240 ((https://doi.org/10.1590/S1982-56762011000400004)); Põlme S, et al. 2020. Fungal Diversity 105:1-16 ((https://doi.org/10.1007/s13225-020-00466-2)) |
| Otu57591 | 1 | k__Fungi(100);p__Ascomycota(100);c__Saccharomycetes(100);o__Saccharomycetales(100);f__Metschnikowiaceae(100);g__Metschnikowia(100);s__Metschnikowia_pulcherrima(100); | Metschnikowia | 13 | Saprotroph | \|Undefined Saprotroph\| | Yeast | NULL | Probable | As nectar-tap_saprotroph (Põlme et al. 2020); Decay sugar-rich substrates (Põlme et al. 2020) | Põlme S, et al. 2020. Fungal Diversity 105:1-16 ((https://doi.org/10.1007/s13225-020-00466-2)) |
| Otu57592 | 1 | k__Fungi(100);p__Ascomycota(100);c__Saccharomycetes(100);o__Saccharomycetales(100);f__Metschnikowiaceae(100);g__Metschnikowia(100);g__Metschnikowia_unclassified(100); | Metschnikowia | 13 | Saprotroph | \|Undefined Saprotroph\| | Yeast | NULL | Probable | As nectar-tap_saprotroph (Põlme et al. 2020); Decay sugar-rich substrates (Põlme et al. 2020) | Põlme S, et al. 2020. Fungal Diversity 105:1-16 ((https://doi.org/10.1007/s13225-020-00466-2)) |
| Otu57593 | 1 | k__Fungi(100);p__Ascomycota(100);c__Saccharomycetes(100);o__Saccharomycetales(100);f__Metschnikowiaceae(100);g__Metschnikowia(100);s__Metschnikowia_pulcherrima(100); | Metschnikowia | 13 | Saprotroph | \|Undefined Saprotroph\| | Yeast | NULL | Probable | As nectar-tap_saprotroph (Põlme et al. 2020); Decay sugar-rich substrates (Põlme et al. 2020) | Põlme S, et al. 2020. Fungal Diversity 105:1-16 ((https://doi.org/10.1007/s13225-020-00466-2)) |
| Otu57596 | 1 | k__Fungi(100);p__Ascomycota(100);c__Saccharomycetes(100);o__Saccharomycetales(100);f__Metschnikowiaceae(100);g__Metschnikowia(100);s__Metschnikowia_pulcherrima(100); | Metschnikowia | 13 | Saprotroph | \|Undefined Saprotroph\| | Yeast | NULL | Probable | As nectar-tap_saprotroph (Põlme et al. 2020); Decay sugar-rich substrates (Põlme et al. 2020) | Põlme S, et al. 2020. Fungal Diversity 105:1-16 ((https://doi.org/10.1007/s13225-020-00466-2)) |
| Otu57597 | 1 | k__Fungi(100);p__Ascomycota(100);c__Saccharomycetes(100);o__Saccharomycetales(100);f__Metschnikowiaceae(100);g__Metschnikowia(100);g__Metschnikowia_unclassified(100); | Metschnikowia | 13 | Saprotroph | \|Undefined Saprotroph\| | Yeast | NULL | Probable | As nectar-tap_saprotroph (Põlme et al. 2020); Decay sugar-rich substrates (Põlme et al. 2020) | Põlme S, et al. 2020. Fungal Diversity 105:1-16 ((https://doi.org/10.1007/s13225-020-00466-2)) |
| Otu57598 | 1 | k__Fungi(100);p__Ascomycota(100);c__Saccharomycetes(100);o__Saccharomycetales(100);f__Metschnikowiaceae(100);g__Metschnikowia(100);s__Metschnikowia_pulcherrima(100); | Metschnikowia | 13 | Saprotroph | \|Undefined Saprotroph\| | Yeast | NULL | Probable | As nectar-tap_saprotroph (Põlme et al. 2020); Decay sugar-rich substrates (Põlme et al. 2020) | Põlme S, et al. 2020. Fungal Diversity 105:1-16 ((https://doi.org/10.1007/s13225-020-00466-2)) |
| Otu57601 | 1 | k__Fungi(100);p__Ascomycota(100);c__Saccharomycetes(100);o__Saccharomycetales(100);f__Metschnikowiaceae(100);g__Metschnikowia(100);s__Metschnikowia_pulcherrima(100); | Metschnikowia | 13 | Saprotroph | \|Undefined Saprotroph\| | Yeast | NULL | Probable | As nectar-tap_saprotroph (Põlme et al. 2020); Decay sugar-rich substrates (Põlme et al. 2020) | Põlme S, et al. 2020. Fungal Diversity 105:1-16 ((https://doi.org/10.1007/s13225-020-00466-2)) |
| Otu57602 | 1 | k__Fungi(100);p__Ascomycota(100);c__Saccharomycetes(100);o__Saccharomycetales(100);f__Metschnikowiaceae(100);g__Metschnikowia(100);s__Metschnikowia_pulcherrima(100); | Metschnikowia | 13 | Saprotroph | \|Undefined Saprotroph\| | Yeast | NULL | Probable | As nectar-tap_saprotroph (Põlme et al. 2020); Decay sugar-rich substrates (Põlme et al. 2020) | Põlme S, et al. 2020. Fungal Diversity 105:1-16 ((https://doi.org/10.1007/s13225-020-00466-2)) |
| Otu57604 | 1 | k__Fungi(100);p__Ascomycota(100);c__Saccharomycetes(100);o__Saccharomycetales(100);f__Metschnikowiaceae(100);g__Metschnikowia(100);s__Metschnikowia_pulcherrima(100); | Metschnikowia | 13 | Saprotroph | \|Undefined Saprotroph\| | Yeast | NULL | Probable | As nectar-tap_saprotroph (Põlme et al. 2020); Decay sugar-rich substrates (Põlme et al. 2020) | Põlme S, et al. 2020. Fungal Diversity 105:1-16 ((https://doi.org/10.1007/s13225-020-00466-2)) |
| Otu57605 | 1 | k__Fungi(100);p__Ascomycota(100);c__Saccharomycetes(100);o__Saccharomycetales(100);f__Metschnikowiaceae(100);g__Metschnikowia(100);s__Metschnikowia_pulcherrima(100); | Metschnikowia | 13 | Saprotroph | \|Undefined Saprotroph\| | Yeast | NULL | Probable | As nectar-tap_saprotroph (Põlme et al. 2020); Decay sugar-rich substrates (Põlme et al. 2020) | Põlme S, et al. 2020. Fungal Diversity 105:1-16 ((https://doi.org/10.1007/s13225-020-00466-2)) |
| Otu57606 | 1 | k__Fungi(100);p__Ascomycota(100);c__Saccharomycetes(100);o__Saccharomycetales(100);f__Metschnikowiaceae(100);g__Metschnikowia(100);s__Metschnikowia_pulcherrima(100); | Metschnikowia | 13 | Saprotroph | \|Undefined Saprotroph\| | Yeast | NULL | Probable | As nectar-tap_saprotroph (Põlme et al. 2020); Decay sugar-rich substrates (Põlme et al. 2020) | Põlme S, et al. 2020. Fungal Diversity 105:1-16 ((https://doi.org/10.1007/s13225-020-00466-2)) |
| Otu57607 | 1 | k__Fungi(100);p__Ascomycota(100);c__Saccharomycetes(100);o__Saccharomycetales(100);f__Metschnikowiaceae(100);g__Metschnikowia(100);s__Metschnikowia_pulcherrima(100); | Metschnikowia | 13 | Saprotroph | \|Undefined Saprotroph\| | Yeast | NULL | Probable | As nectar-tap_saprotroph (Põlme et al. 2020); Decay sugar-rich substrates (Põlme et al. 2020) | Põlme S, et al. 2020. Fungal Diversity 105:1-16 ((https://doi.org/10.1007/s13225-020-00466-2)) |
| Otu57608 | 1 | k__Fungi(100);p__Ascomycota(100);c__Saccharomycetes(100);o__Saccharomycetales(100);f__Metschnikowiaceae(100);g__Metschnikowia(100);g__Metschnikowia_unclassified(100); | Metschnikowia | 13 | Saprotroph | \|Undefined Saprotroph\| | Yeast | NULL | Probable | As nectar-tap_saprotroph (Põlme et al. 2020); Decay sugar-rich substrates (Põlme et al. 2020) | Põlme S, et al. 2020. Fungal Diversity 105:1-16 ((https://doi.org/10.1007/s13225-020-00466-2)) |
| Otu57609 | 1 | k__Fungi(100);p__Ascomycota(100);c__Saccharomycetes(100);o__Saccharomycetales(100);f__Metschnikowiaceae(100);g__Metschnikowia(100);s__Metschnikowia_pulcherrima(100); | Metschnikowia | 13 | Saprotroph | \|Undefined Saprotroph\| | Yeast | NULL | Probable | As nectar-tap_saprotroph (Põlme et al. 2020); Decay sugar-rich substrates (Põlme et al. 2020) | Põlme S, et al. 2020. Fungal Diversity 105:1-16 ((https://doi.org/10.1007/s13225-020-00466-2)) |
| Otu57610 | 1 | k__Fungi(100);p__Ascomycota(100);c__Saccharomycetes(100);o__Saccharomycetales(100);f__Metschnikowiaceae(100);g__Metschnikowia(100);s__Metschnikowia_pulcherrima(100); | Metschnikowia | 13 | Saprotroph | \|Undefined Saprotroph\| | Yeast | NULL | Probable | As nectar-tap_saprotroph (Põlme et al. 2020); Decay sugar-rich substrates (Põlme et al. 2020) | Põlme S, et al. 2020. Fungal Diversity 105:1-16 ((https://doi.org/10.1007/s13225-020-00466-2)) |
| Otu57612 | 1 | k__Fungi(100);p__Ascomycota(100);c__Saccharomycetes(100);o__Saccharomycetales(100);f__Saccharomycetales_family_Incertae_sedis(100);g__Candida(100);g__Candida_unclassified(100); | Candida | 13 | Pathotroph-Saprotroph-Symbiotroph | Animal Pathogen-Endophyte-Endosymbiont-Epiphyte-\|Undefined Saprotroph\| | Dimorphic Yeast-Yeast | NULL | Probable | As nectar-tap_saprotroph (Põlme et al. 2020); Decay sugar-rich substrates (Põlme et al. 2020); This asexual genus is polyphyletic and contain species that function very differently depend on what they are. Caution should be taken when interpreting guilds for Candida species. | Manolakaki D, et al. 2010. Virulence. 1: 367-375 ((https://doi.org/10.4161/viru.1.5.12796)); Mekbib SB, et al. 2011. Tropical Plant Pathology 36:233-240 ((https://doi.org/10.1590/S1982-56762011000400004)); Põlme S, et al. 2020. Fungal Diversity 105:1-16 ((https://doi.org/10.1007/s13225-020-00466-2)) |
| Otu57613 | 1 | k__Fungi(100);p__Ascomycota(100);c__Saccharomycetes(100);o__Saccharomycetales(100);f__Metschnikowiaceae(100);g__Metschnikowia(100);s__Metschnikowia_pulcherrima(100); | Metschnikowia | 13 | Saprotroph | \|Undefined Saprotroph\| | Yeast | NULL | Probable | As nectar-tap_saprotroph (Põlme et al. 2020); Decay sugar-rich substrates (Põlme et al. 2020) | Põlme S, et al. 2020. Fungal Diversity 105:1-16 ((https://doi.org/10.1007/s13225-020-00466-2)) |
| Otu57614 | 1 | k__Fungi(100);p__Ascomycota(100);c__Saccharomycetes(100);o__Saccharomycetales(100);f__Metschnikowiaceae(100);g__Metschnikowia(100);s__Metschnikowia_pulcherrima(100); | Metschnikowia | 13 | Saprotroph | \|Undefined Saprotroph\| | Yeast | NULL | Probable | As nectar-tap_saprotroph (Põlme et al. 2020); Decay sugar-rich substrates (Põlme et al. 2020) | Põlme S, et al. 2020. Fungal Diversity 105:1-16 ((https://doi.org/10.1007/s13225-020-00466-2)) |
| Otu57616 | 1 | k__Fungi(100);p__Ascomycota(100);c__Saccharomycetes(100);o__Saccharomycetales(100);f__Metschnikowiaceae(100);g__Metschnikowia(100);g__Metschnikowia_unclassified(100); | Metschnikowia | 13 | Saprotroph | \|Undefined Saprotroph\| | Yeast | NULL | Probable | As nectar-tap_saprotroph (Põlme et al. 2020); Decay sugar-rich substrates (Põlme et al. 2020) | Põlme S, et al. 2020. Fungal Diversity 105:1-16 ((https://doi.org/10.1007/s13225-020-00466-2)) |
| Otu57617 | 1 | k__Fungi(100);p__Ascomycota(100);c__Saccharomycetes(100);o__Saccharomycetales(100);f__Metschnikowiaceae(100);g__Metschnikowia(100);s__Metschnikowia_pulcherrima(100); | Metschnikowia | 13 | Saprotroph | \|Undefined Saprotroph\| | Yeast | NULL | Probable | As nectar-tap_saprotroph (Põlme et al. 2020); Decay sugar-rich substrates (Põlme et al. 2020) | Põlme S, et al. 2020. Fungal Diversity 105:1-16 ((https://doi.org/10.1007/s13225-020-00466-2)) |
| Otu57618 | 1 | k__Fungi(100);p__Ascomycota(100);c__Saccharomycetes(100);o__Saccharomycetales(100);f__Metschnikowiaceae(100);g__Metschnikowia(100);s__Metschnikowia_pulcherrima(100); | Metschnikowia | 13 | Saprotroph | \|Undefined Saprotroph\| | Yeast | NULL | Probable | As nectar-tap_saprotroph (Põlme et al. 2020); Decay sugar-rich substrates (Põlme et al. 2020) | Põlme S, et al. 2020. Fungal Diversity 105:1-16 ((https://doi.org/10.1007/s13225-020-00466-2)) |
| Otu57619 | 1 | k__Fungi(100);p__Ascomycota(100);c__Saccharomycetes(100);o__Saccharomycetales(100);f__Metschnikowiaceae(100);g__Metschnikowia(100);s__Metschnikowia_pulcherrima(100); | Metschnikowia | 13 | Saprotroph | \|Undefined Saprotroph\| | Yeast | NULL | Probable | As nectar-tap_saprotroph (Põlme et al. 2020); Decay sugar-rich substrates (Põlme et al. 2020) | Põlme S, et al. 2020. Fungal Diversity 105:1-16 ((https://doi.org/10.1007/s13225-020-00466-2)) |
| Otu57620 | 1 | k__Fungi(100);p__Ascomycota(100);c__Saccharomycetes(100);o__Saccharomycetales(100);f__Metschnikowiaceae(100);g__Metschnikowia(100);g__Metschnikowia_unclassified(100); | Metschnikowia | 13 | Saprotroph | \|Undefined Saprotroph\| | Yeast | NULL | Probable | As nectar-tap_saprotroph (Põlme et al. 2020); Decay sugar-rich substrates (Põlme et al. 2020) | Põlme S, et al. 2020. Fungal Diversity 105:1-16 ((https://doi.org/10.1007/s13225-020-00466-2)) |
| Otu57621 | 1 | k__Fungi(100);p__Ascomycota(100);c__Saccharomycetes(100);o__Saccharomycetales(100);f__Metschnikowiaceae(100);g__Metschnikowia(100);s__Metschnikowia_pulcherrima(100); | Metschnikowia | 13 | Saprotroph | \|Undefined Saprotroph\| | Yeast | NULL | Probable | As nectar-tap_saprotroph (Põlme et al. 2020); Decay sugar-rich substrates (Põlme et al. 2020) | Põlme S, et al. 2020. Fungal Diversity 105:1-16 ((https://doi.org/10.1007/s13225-020-00466-2)) |
| Otu57622 | 1 | k__Fungi(100);p__Ascomycota(100);c__Saccharomycetes(100);o__Saccharomycetales(100);f__Metschnikowiaceae(100);g__Metschnikowia(100);s__Metschnikowia_pulcherrima(100); | Metschnikowia | 13 | Saprotroph | \|Undefined Saprotroph\| | Yeast | NULL | Probable | As nectar-tap_saprotroph (Põlme et al. 2020); Decay sugar-rich substrates (Põlme et al. 2020) | Põlme S, et al. 2020. Fungal Diversity 105:1-16 ((https://doi.org/10.1007/s13225-020-00466-2)) |
| Otu57623 | 1 | k__Fungi(100);p__Ascomycota(100);c__Saccharomycetes(100);o__Saccharomycetales(100);f__Saccharomycetaceae(100);g__Kazachstania(100);s__Kazachstania_intestinalis(100); | Kazachstania | 13 | Saprotroph | \|Undefined Saprotroph\| | Yeast | NULL | Probable | As nectar-tap_saprotroph (Põlme et al. 2020); Decay sugar-rich substrates (Põlme et al. 2020) | Põlme S, et al. 2020. Fungal Diversity 105:1-16 ((https://doi.org/10.1007/s13225-020-00466-2)) |
| Otu57625 | 1 | k__Fungi(100);p__Ascomycota(100);c__Saccharomycetes(100);o__Saccharomycetales(100);f__Metschnikowiaceae(100);g__Metschnikowia(100);s__Metschnikowia_pulcherrima(100); | Metschnikowia | 13 | Saprotroph | \|Undefined Saprotroph\| | Yeast | NULL | Probable | As nectar-tap_saprotroph (Põlme et al. 2020); Decay sugar-rich substrates (Põlme et al. 2020) | Põlme S, et al. 2020. Fungal Diversity 105:1-16 ((https://doi.org/10.1007/s13225-020-00466-2)) |
| Otu57626 | 1 | k__Fungi(100);p__Ascomycota(100);c__Saccharomycetes(100);o__Saccharomycetales(100);f__Metschnikowiaceae(100);g__Metschnikowia(100);s__Metschnikowia_pulcherrima(100); | Metschnikowia | 13 | Saprotroph | \|Undefined Saprotroph\| | Yeast | NULL | Probable | As nectar-tap_saprotroph (Põlme et al. 2020); Decay sugar-rich substrates (Põlme et al. 2020) | Põlme S, et al. 2020. Fungal Diversity 105:1-16 ((https://doi.org/10.1007/s13225-020-00466-2)) |
| Otu57628 | 1 | k__Fungi(100);p__Ascomycota(100);c__Saccharomycetes(100);o__Saccharomycetales(100);f__Metschnikowiaceae(100);g__Metschnikowia(100);s__Metschnikowia_pulcherrima(100); | Metschnikowia | 13 | Saprotroph | \|Undefined Saprotroph\| | Yeast | NULL | Probable | As nectar-tap_saprotroph (Põlme et al. 2020); Decay sugar-rich substrates (Põlme et al. 2020) | Põlme S, et al. 2020. Fungal Diversity 105:1-16 ((https://doi.org/10.1007/s13225-020-00466-2)) |
| Otu57629 | 1 | k__Fungi(100);p__Ascomycota(100);c__Saccharomycetes(100);o__Saccharomycetales(100);f__Metschnikowiaceae(100);g__Metschnikowia(100);g__Metschnikowia_unclassified(100); | Metschnikowia | 13 | Saprotroph | \|Undefined Saprotroph\| | Yeast | NULL | Probable | As nectar-tap_saprotroph (Põlme et al. 2020); Decay sugar-rich substrates (Põlme et al. 2020) | Põlme S, et al. 2020. Fungal Diversity 105:1-16 ((https://doi.org/10.1007/s13225-020-00466-2)) |
| Otu57631 | 1 | k__Fungi(100);p__Ascomycota(100);c__Saccharomycetes(100);o__Saccharomycetales(100);f__Metschnikowiaceae(100);g__Metschnikowia(100);s__Metschnikowia_pulcherrima(100); | Metschnikowia | 13 | Saprotroph | \|Undefined Saprotroph\| | Yeast | NULL | Probable | As nectar-tap_saprotroph (Põlme et al. 2020); Decay sugar-rich substrates (Põlme et al. 2020) | Põlme S, et al. 2020. Fungal Diversity 105:1-16 ((https://doi.org/10.1007/s13225-020-00466-2)) |
| Otu57632 | 1 | k__Fungi(100);p__Ascomycota(100);c__Saccharomycetes(100);o__Saccharomycetales(100);f__Metschnikowiaceae(100);g__Metschnikowia(100);s__Metschnikowia_pulcherrima(100); | Metschnikowia | 13 | Saprotroph | \|Undefined Saprotroph\| | Yeast | NULL | Probable | As nectar-tap_saprotroph (Põlme et al. 2020); Decay sugar-rich substrates (Põlme et al. 2020) | Põlme S, et al. 2020. Fungal Diversity 105:1-16 ((https://doi.org/10.1007/s13225-020-00466-2)) |
| Otu57633 | 1 | k__Fungi(100);p__Ascomycota(100);c__Saccharomycetes(100);o__Saccharomycetales(100);f__Metschnikowiaceae(100);g__Metschnikowia(100);s__Metschnikowia_pulcherrima(100); | Metschnikowia | 13 | Saprotroph | \|Undefined Saprotroph\| | Yeast | NULL | Probable | As nectar-tap_saprotroph (Põlme et al. 2020); Decay sugar-rich substrates (Põlme et al. 2020) | Põlme S, et al. 2020. Fungal Diversity 105:1-16 ((https://doi.org/10.1007/s13225-020-00466-2)) |
| Otu57634 | 1 | k__Fungi(100);p__Ascomycota(100);c__Saccharomycetes(100);o__Saccharomycetales(100);f__Metschnikowiaceae(100);g__Metschnikowia(100);s__Metschnikowia_pulcherrima(100); | Metschnikowia | 13 | Saprotroph | \|Undefined Saprotroph\| | Yeast | NULL | Probable | As nectar-tap_saprotroph (Põlme et al. 2020); Decay sugar-rich substrates (Põlme et al. 2020) | Põlme S, et al. 2020. Fungal Diversity 105:1-16 ((https://doi.org/10.1007/s13225-020-00466-2)) |
| Otu57635 | 1 | k__Fungi(100);p__Ascomycota(100);c__Saccharomycetes(100);o__Saccharomycetales(100);f__Metschnikowiaceae(100);g__Metschnikowia(100);s__Metschnikowia_pulcherrima(100); | Metschnikowia | 13 | Saprotroph | \|Undefined Saprotroph\| | Yeast | NULL | Probable | As nectar-tap_saprotroph (Põlme et al. 2020); Decay sugar-rich substrates (Põlme et al. 2020) | Põlme S, et al. 2020. Fungal Diversity 105:1-16 ((https://doi.org/10.1007/s13225-020-00466-2)) |
| Otu57638 | 1 | k__Fungi(100);p__Ascomycota(100);c__Saccharomycetes(100);o__Saccharomycetales(100);f__Metschnikowiaceae(100);g__Metschnikowia(100);s__Metschnikowia_pulcherrima(100); | Metschnikowia | 13 | Saprotroph | \|Undefined Saprotroph\| | Yeast | NULL | Probable | As nectar-tap_saprotroph (Põlme et al. 2020); Decay sugar-rich substrates (Põlme et al. 2020) | Põlme S, et al. 2020. Fungal Diversity 105:1-16 ((https://doi.org/10.1007/s13225-020-00466-2)) |
| Otu57639 | 1 | k__Fungi(100);p__Ascomycota(100);c__Saccharomycetes(100);o__Saccharomycetales(100);f__Metschnikowiaceae(100);g__Metschnikowia(100);s__Metschnikowia_pulcherrima(100); | Metschnikowia | 13 | Saprotroph | \|Undefined Saprotroph\| | Yeast | NULL | Probable | As nectar-tap_saprotroph (Põlme et al. 2020); Decay sugar-rich substrates (Põlme et al. 2020) | Põlme S, et al. 2020. Fungal Diversity 105:1-16 ((https://doi.org/10.1007/s13225-020-00466-2)) |
| Otu57642 | 1 | k__Fungi(100);p__Ascomycota(100);c__Saccharomycetes(100);o__Saccharomycetales(100);f__Metschnikowiaceae(100);g__Metschnikowia(100);s__Metschnikowia_pulcherrima(100); | Metschnikowia | 13 | Saprotroph | \|Undefined Saprotroph\| | Yeast | NULL | Probable | As nectar-tap_saprotroph (Põlme et al. 2020); Decay sugar-rich substrates (Põlme et al. 2020) | Põlme S, et al. 2020. Fungal Diversity 105:1-16 ((https://doi.org/10.1007/s13225-020-00466-2)) |
| Otu57645 | 1 | k__Fungi(100);p__Ascomycota(100);c__Saccharomycetes(100);o__Saccharomycetales(100);f__Metschnikowiaceae(100);g__Metschnikowia(100);s__Metschnikowia_pulcherrima(100); | Metschnikowia | 13 | Saprotroph | \|Undefined Saprotroph\| | Yeast | NULL | Probable | As nectar-tap_saprotroph (Põlme et al. 2020); Decay sugar-rich substrates (Põlme et al. 2020) | Põlme S, et al. 2020. Fungal Diversity 105:1-16 ((https://doi.org/10.1007/s13225-020-00466-2)) |
| Otu57649 | 1 | k__Fungi(100);p__Ascomycota(100);c__Saccharomycetes(100);o__Saccharomycetales(100);f__Metschnikowiaceae(100);g__Metschnikowia(100);s__Metschnikowia_pulcherrima(100); | Metschnikowia | 13 | Saprotroph | \|Undefined Saprotroph\| | Yeast | NULL | Probable | As nectar-tap_saprotroph (Põlme et al. 2020); Decay sugar-rich substrates (Põlme et al. 2020) | Põlme S, et al. 2020. Fungal Diversity 105:1-16 ((https://doi.org/10.1007/s13225-020-00466-2)) |
| Otu57650 | 1 | k__Fungi(100);p__Ascomycota(100);c__Saccharomycetes(100);o__Saccharomycetales(100);f__Metschnikowiaceae(100);g__Metschnikowia(100);s__Metschnikowia_pulcherrima(100); | Metschnikowia | 13 | Saprotroph | \|Undefined Saprotroph\| | Yeast | NULL | Probable | As nectar-tap_saprotroph (Põlme et al. 2020); Decay sugar-rich substrates (Põlme et al. 2020) | Põlme S, et al. 2020. Fungal Diversity 105:1-16 ((https://doi.org/10.1007/s13225-020-00466-2)) |
| Otu57651 | 1 | k__Fungi(100);p__Ascomycota(100);c__Saccharomycetes(100);o__Saccharomycetales(100);f__Metschnikowiaceae(100);g__Metschnikowia(100);s__Metschnikowia_pulcherrima(100); | Metschnikowia | 13 | Saprotroph | \|Undefined Saprotroph\| | Yeast | NULL | Probable | As nectar-tap_saprotroph (Põlme et al. 2020); Decay sugar-rich substrates (Põlme et al. 2020) | Põlme S, et al. 2020. Fungal Diversity 105:1-16 ((https://doi.org/10.1007/s13225-020-00466-2)) |
| Otu57652 | 1 | k__Fungi(100);p__Ascomycota(100);c__Saccharomycetes(100);o__Saccharomycetales(100);f__Metschnikowiaceae(100);g__Metschnikowia(100);s__Metschnikowia_pulcherrima(100); | Metschnikowia | 13 | Saprotroph | \|Undefined Saprotroph\| | Yeast | NULL | Probable | As nectar-tap_saprotroph (Põlme et al. 2020); Decay sugar-rich substrates (Põlme et al. 2020) | Põlme S, et al. 2020. Fungal Diversity 105:1-16 ((https://doi.org/10.1007/s13225-020-00466-2)) |
| Otu57653 | 1 | k__Fungi(100);p__Ascomycota(100);c__Saccharomycetes(100);o__Saccharomycetales(100);f__Metschnikowiaceae(100);g__Metschnikowia(100);s__Metschnikowia_pulcherrima(100); | Metschnikowia | 13 | Saprotroph | \|Undefined Saprotroph\| | Yeast | NULL | Probable | As nectar-tap_saprotroph (Põlme et al. 2020); Decay sugar-rich substrates (Põlme et al. 2020) | Põlme S, et al. 2020. Fungal Diversity 105:1-16 ((https://doi.org/10.1007/s13225-020-00466-2)) |
| Otu57654 | 1 | k__Fungi(100);p__Ascomycota(100);c__Saccharomycetes(100);o__Saccharomycetales(100);f__Metschnikowiaceae(100);g__Metschnikowia(100);s__Metschnikowia_pulcherrima(100); | Metschnikowia | 13 | Saprotroph | \|Undefined Saprotroph\| | Yeast | NULL | Probable | As nectar-tap_saprotroph (Põlme et al. 2020); Decay sugar-rich substrates (Põlme et al. 2020) | Põlme S, et al. 2020. Fungal Diversity 105:1-16 ((https://doi.org/10.1007/s13225-020-00466-2)) |
| Otu57657 | 1 | k__Fungi(100);p__Ascomycota(100);c__Saccharomycetes(100);o__Saccharomycetales(100);f__Metschnikowiaceae(100);g__Metschnikowia(100);s__Metschnikowia_pulcherrima(100); | Metschnikowia | 13 | Saprotroph | \|Undefined Saprotroph\| | Yeast | NULL | Probable | As nectar-tap_saprotroph (Põlme et al. 2020); Decay sugar-rich substrates (Põlme et al. 2020) | Põlme S, et al. 2020. Fungal Diversity 105:1-16 ((https://doi.org/10.1007/s13225-020-00466-2)) |
| Otu57661 | 1 | k__Fungi(100);p__Ascomycota(100);c__Saccharomycetes(100);o__Saccharomycetales(100);f__Metschnikowiaceae(100);g__Metschnikowia(100);s__Metschnikowia_pulcherrima(100); | Metschnikowia | 13 | Saprotroph | \|Undefined Saprotroph\| | Yeast | NULL | Probable | As nectar-tap_saprotroph (Põlme et al. 2020); Decay sugar-rich substrates (Põlme et al. 2020) | Põlme S, et al. 2020. Fungal Diversity 105:1-16 ((https://doi.org/10.1007/s13225-020-00466-2)) |
| Otu57663 | 1 | k__Fungi(100);p__Ascomycota(100);c__Saccharomycetes(100);o__Saccharomycetales(100);f__Metschnikowiaceae(100);g__Metschnikowia(100);s__Metschnikowia_pulcherrima(100); | Metschnikowia | 13 | Saprotroph | \|Undefined Saprotroph\| | Yeast | NULL | Probable | As nectar-tap_saprotroph (Põlme et al. 2020); Decay sugar-rich substrates (Põlme et al. 2020) | Põlme S, et al. 2020. Fungal Diversity 105:1-16 ((https://doi.org/10.1007/s13225-020-00466-2)) |
| Otu57664 | 1 | k__Fungi(100);p__Ascomycota(100);c__Saccharomycetes(100);o__Saccharomycetales(100);f__Saccharomycetales_family_Incertae_sedis(100);f__Saccharomycetales_family_Incertae_sedis_unclassified(100);f__Saccharomycetales_family_Incertae_sedis_unclassified(100); | Saccharomycetales | 7 | Saprotroph | Undefined Saprotroph | Yeast | NULL | Possible | NULL | Sterkenburg E, et al. 2015. New Phytologist 207:1145-1158 ((https://doi.org/10.1111/nph.13426)) |
| Otu57665 | 1 | k__Fungi(100);p__Ascomycota(100);c__Saccharomycetes(100);o__Saccharomycetales(100);f__Metschnikowiaceae(100);g__Metschnikowia(100);g__Metschnikowia_unclassified(100); | Metschnikowia | 13 | Saprotroph | \|Undefined Saprotroph\| | Yeast | NULL | Probable | As nectar-tap_saprotroph (Põlme et al. 2020); Decay sugar-rich substrates (Põlme et al. 2020) | Põlme S, et al. 2020. Fungal Diversity 105:1-16 ((https://doi.org/10.1007/s13225-020-00466-2)) |
| Otu57666 | 1 | k__Fungi(100);p__Ascomycota(100);c__Saccharomycetes(100);o__Saccharomycetales(100);f__Metschnikowiaceae(100);g__Metschnikowia(100);s__Metschnikowia_pulcherrima(100); | Metschnikowia | 13 | Saprotroph | \|Undefined Saprotroph\| | Yeast | NULL | Probable | As nectar-tap_saprotroph (Põlme et al. 2020); Decay sugar-rich substrates (Põlme et al. 2020) | Põlme S, et al. 2020. Fungal Diversity 105:1-16 ((https://doi.org/10.1007/s13225-020-00466-2)) |
| Otu57669 | 1 | k__Fungi(100);p__Ascomycota(100);c__Saccharomycetes(100);o__Saccharomycetales(100);f__Saccharomycetales_family_Incertae_sedis(100);g__Candida(100);g__Candida_unclassified(100); | Candida | 13 | Pathotroph-Saprotroph-Symbiotroph | Animal Pathogen-Endophyte-Endosymbiont-Epiphyte-\|Undefined Saprotroph\| | Dimorphic Yeast-Yeast | NULL | Probable | As nectar-tap_saprotroph (Põlme et al. 2020); Decay sugar-rich substrates (Põlme et al. 2020); This asexual genus is polyphyletic and contain species that function very differently depend on what they are. Caution should be taken when interpreting guilds for Candida species. | Manolakaki D, et al. 2010. Virulence. 1: 367-375 ((https://doi.org/10.4161/viru.1.5.12796)); Mekbib SB, et al. 2011. Tropical Plant Pathology 36:233-240 ((https://doi.org/10.1590/S1982-56762011000400004)); Põlme S, et al. 2020. Fungal Diversity 105:1-16 ((https://doi.org/10.1007/s13225-020-00466-2)) |
| Otu57671 | 1 | k__Fungi(100);p__Ascomycota(100);c__Saccharomycetes(100);o__Saccharomycetales(100);f__Metschnikowiaceae(100);g__Metschnikowia(100);g__Metschnikowia_unclassified(100); | Metschnikowia | 13 | Saprotroph | \|Undefined Saprotroph\| | Yeast | NULL | Probable | As nectar-tap_saprotroph (Põlme et al. 2020); Decay sugar-rich substrates (Põlme et al. 2020) | Põlme S, et al. 2020. Fungal Diversity 105:1-16 ((https://doi.org/10.1007/s13225-020-00466-2)) |
| Otu57672 | 1 | k__Fungi(100);p__Ascomycota(100);c__Saccharomycetes(100);o__Saccharomycetales(100);f__Metschnikowiaceae(100);g__Metschnikowia(100);s__Metschnikowia_pulcherrima(100); | Metschnikowia | 13 | Saprotroph | \|Undefined Saprotroph\| | Yeast | NULL | Probable | As nectar-tap_saprotroph (Põlme et al. 2020); Decay sugar-rich substrates (Põlme et al. 2020) | Põlme S, et al. 2020. Fungal Diversity 105:1-16 ((https://doi.org/10.1007/s13225-020-00466-2)) |
| Otu57673 | 1 | k__Fungi(100);p__Ascomycota(100);c__Saccharomycetes(100);o__Saccharomycetales(100);f__Metschnikowiaceae(100);g__Metschnikowia(100);s__Metschnikowia_pulcherrima(100); | Metschnikowia | 13 | Saprotroph | \|Undefined Saprotroph\| | Yeast | NULL | Probable | As nectar-tap_saprotroph (Põlme et al. 2020); Decay sugar-rich substrates (Põlme et al. 2020) | Põlme S, et al. 2020. Fungal Diversity 105:1-16 ((https://doi.org/10.1007/s13225-020-00466-2)) |
| Otu57686 | 1 | k__Fungi(100);p__Ascomycota(100);c__Saccharomycetes(100);o__Saccharomycetales(100);f__Metschnikowiaceae(100);g__Metschnikowia(100);g__Metschnikowia_unclassified(100); | Metschnikowia | 13 | Saprotroph | \|Undefined Saprotroph\| | Yeast | NULL | Probable | As nectar-tap_saprotroph (Põlme et al. 2020); Decay sugar-rich substrates (Põlme et al. 2020) | Põlme S, et al. 2020. Fungal Diversity 105:1-16 ((https://doi.org/10.1007/s13225-020-00466-2)) |
| Otu57688 | 1 | k__Fungi(100);p__Ascomycota(100);c__Saccharomycetes(100);o__Saccharomycetales(100);f__Metschnikowiaceae(100);g__Metschnikowia(100);s__Metschnikowia_pulcherrima(100); | Metschnikowia | 13 | Saprotroph | \|Undefined Saprotroph\| | Yeast | NULL | Probable | As nectar-tap_saprotroph (Põlme et al. 2020); Decay sugar-rich substrates (Põlme et al. 2020) | Põlme S, et al. 2020. Fungal Diversity 105:1-16 ((https://doi.org/10.1007/s13225-020-00466-2)) |
| Otu57691 | 1 | k__Fungi(100);p__Ascomycota(100);c__Saccharomycetes(100);o__Saccharomycetales(100);f__Saccharomycetales_family_Incertae_sedis(100);g__Candida(100);g__Candida_unclassified(100); | Candida | 13 | Pathotroph-Saprotroph-Symbiotroph | Animal Pathogen-Endophyte-Endosymbiont-Epiphyte-\|Undefined Saprotroph\| | Dimorphic Yeast-Yeast | NULL | Probable | As nectar-tap_saprotroph (Põlme et al. 2020); Decay sugar-rich substrates (Põlme et al. 2020); This asexual genus is polyphyletic and contain species that function very differently depend on what they are. Caution should be taken when interpreting guilds for Candida species. | Manolakaki D, et al. 2010. Virulence. 1: 367-375 ((https://doi.org/10.4161/viru.1.5.12796)); Mekbib SB, et al. 2011. Tropical Plant Pathology 36:233-240 ((https://doi.org/10.1590/S1982-56762011000400004)); Põlme S, et al. 2020. Fungal Diversity 105:1-16 ((https://doi.org/10.1007/s13225-020-00466-2)) |
| Otu57693 | 1 | k__Fungi(100);p__Ascomycota(100);c__Saccharomycetes(100);o__Saccharomycetales(100);f__Metschnikowiaceae(100);g__Metschnikowia(100);s__Metschnikowia_pulcherrima(100); | Metschnikowia | 13 | Saprotroph | \|Undefined Saprotroph\| | Yeast | NULL | Probable | As nectar-tap_saprotroph (Põlme et al. 2020); Decay sugar-rich substrates (Põlme et al. 2020) | Põlme S, et al. 2020. Fungal Diversity 105:1-16 ((https://doi.org/10.1007/s13225-020-00466-2)) |
| Otu57695 | 1 | k__Fungi(100);p__Ascomycota(100);c__Saccharomycetes(100);o__Saccharomycetales(100);f__Metschnikowiaceae(100);g__Metschnikowia(100);s__Metschnikowia_pulcherrima(100); | Metschnikowia | 13 | Saprotroph | \|Undefined Saprotroph\| | Yeast | NULL | Probable | As nectar-tap_saprotroph (Põlme et al. 2020); Decay sugar-rich substrates (Põlme et al. 2020) | Põlme S, et al. 2020. Fungal Diversity 105:1-16 ((https://doi.org/10.1007/s13225-020-00466-2)) |
| Otu57696 | 1 | k__Fungi(100);p__Ascomycota(100);c__Saccharomycetes(100);o__Saccharomycetales(100);f__Metschnikowiaceae(100);g__Metschnikowia(100);s__Metschnikowia_pulcherrima(100); | Metschnikowia | 13 | Saprotroph | \|Undefined Saprotroph\| | Yeast | NULL | Probable | As nectar-tap_saprotroph (Põlme et al. 2020); Decay sugar-rich substrates (Põlme et al. 2020) | Põlme S, et al. 2020. Fungal Diversity 105:1-16 ((https://doi.org/10.1007/s13225-020-00466-2)) |
| Otu57697 | 1 | k__Fungi(100);p__Ascomycota(100);c__Saccharomycetes(100);o__Saccharomycetales(100);f__Metschnikowiaceae(100);g__Metschnikowia(100);s__Metschnikowia_pulcherrima(100); | Metschnikowia | 13 | Saprotroph | \|Undefined Saprotroph\| | Yeast | NULL | Probable | As nectar-tap_saprotroph (Põlme et al. 2020); Decay sugar-rich substrates (Põlme et al. 2020) | Põlme S, et al. 2020. Fungal Diversity 105:1-16 ((https://doi.org/10.1007/s13225-020-00466-2)) |
| Otu57699 | 1 | k__Fungi(100);p__Ascomycota(100);c__Saccharomycetes(100);o__Saccharomycetales(100);f__Metschnikowiaceae(100);g__Metschnikowia(100);s__Metschnikowia_pulcherrima(100); | Metschnikowia | 13 | Saprotroph | \|Undefined Saprotroph\| | Yeast | NULL | Probable | As nectar-tap_saprotroph (Põlme et al. 2020); Decay sugar-rich substrates (Põlme et al. 2020) | Põlme S, et al. 2020. Fungal Diversity 105:1-16 ((https://doi.org/10.1007/s13225-020-00466-2)) |
| Otu57700 | 1 | k__Fungi(100);p__Ascomycota(100);c__Saccharomycetes(100);o__Saccharomycetales(100);f__Metschnikowiaceae(100);g__Metschnikowia(100);s__Metschnikowia_pulcherrima(100); | Metschnikowia | 13 | Saprotroph | \|Undefined Saprotroph\| | Yeast | NULL | Probable | As nectar-tap_saprotroph (Põlme et al. 2020); Decay sugar-rich substrates (Põlme et al. 2020) | Põlme S, et al. 2020. Fungal Diversity 105:1-16 ((https://doi.org/10.1007/s13225-020-00466-2)) |
| Otu57701 | 1 | k__Fungi(100);p__Ascomycota(100);c__Saccharomycetes(100);o__Saccharomycetales(100);f__Metschnikowiaceae(100);g__Metschnikowia(100);g__Metschnikowia_unclassified(100); | Metschnikowia | 13 | Saprotroph | \|Undefined Saprotroph\| | Yeast | NULL | Probable | As nectar-tap_saprotroph (Põlme et al. 2020); Decay sugar-rich substrates (Põlme et al. 2020) | Põlme S, et al. 2020. Fungal Diversity 105:1-16 ((https://doi.org/10.1007/s13225-020-00466-2)) |
| Otu57702 | 1 | k__Fungi(100);p__Ascomycota(100);c__Saccharomycetes(100);o__Saccharomycetales(100);f__Metschnikowiaceae(100);g__Metschnikowia(100);s__Metschnikowia_pulcherrima(100); | Metschnikowia | 13 | Saprotroph | \|Undefined Saprotroph\| | Yeast | NULL | Probable | As nectar-tap_saprotroph (Põlme et al. 2020); Decay sugar-rich substrates (Põlme et al. 2020) | Põlme S, et al. 2020. Fungal Diversity 105:1-16 ((https://doi.org/10.1007/s13225-020-00466-2)) |
| Otu57707 | 1 | k__Fungi(100);p__Ascomycota(100);c__Saccharomycetes(100);o__Saccharomycetales(100);f__Metschnikowiaceae(100);g__Metschnikowia(100);s__Metschnikowia_pulcherrima(100); | Metschnikowia | 13 | Saprotroph | \|Undefined Saprotroph\| | Yeast | NULL | Probable | As nectar-tap_saprotroph (Põlme et al. 2020); Decay sugar-rich substrates (Põlme et al. 2020) | Põlme S, et al. 2020. Fungal Diversity 105:1-16 ((https://doi.org/10.1007/s13225-020-00466-2)) |
| Otu57708 | 1 | k__Fungi(100);p__Ascomycota(100);c__Saccharomycetes(100);o__Saccharomycetales(100);f__Metschnikowiaceae(100);g__Metschnikowia(100);s__Metschnikowia_pulcherrima(100); | Metschnikowia | 13 | Saprotroph | \|Undefined Saprotroph\| | Yeast | NULL | Probable | As nectar-tap_saprotroph (Põlme et al. 2020); Decay sugar-rich substrates (Põlme et al. 2020) | Põlme S, et al. 2020. Fungal Diversity 105:1-16 ((https://doi.org/10.1007/s13225-020-00466-2)) |
| Otu57709 | 1 | k__Fungi(100);p__Ascomycota(100);c__Saccharomycetes(100);o__Saccharomycetales(100);f__Metschnikowiaceae(100);g__Metschnikowia(100);s__Metschnikowia_pulcherrima(100); | Metschnikowia | 13 | Saprotroph | \|Undefined Saprotroph\| | Yeast | NULL | Probable | As nectar-tap_saprotroph (Põlme et al. 2020); Decay sugar-rich substrates (Põlme et al. 2020) | Põlme S, et al. 2020. Fungal Diversity 105:1-16 ((https://doi.org/10.1007/s13225-020-00466-2)) |
| Otu57710 | 1 | k__Fungi(100);p__Ascomycota(100);c__Saccharomycetes(100);o__Saccharomycetales(100);f__Metschnikowiaceae(100);g__Metschnikowia(100);s__Metschnikowia_pulcherrima(100); | Metschnikowia | 13 | Saprotroph | \|Undefined Saprotroph\| | Yeast | NULL | Probable | As nectar-tap_saprotroph (Põlme et al. 2020); Decay sugar-rich substrates (Põlme et al. 2020) | Põlme S, et al. 2020. Fungal Diversity 105:1-16 ((https://doi.org/10.1007/s13225-020-00466-2)) |
| Otu57711 | 1 | k__Fungi(100);p__Ascomycota(100);c__Saccharomycetes(100);o__Saccharomycetales(100);f__Metschnikowiaceae(100);g__Metschnikowia(100);s__Metschnikowia_pulcherrima(100); | Metschnikowia | 13 | Saprotroph | \|Undefined Saprotroph\| | Yeast | NULL | Probable | As nectar-tap_saprotroph (Põlme et al. 2020); Decay sugar-rich substrates (Põlme et al. 2020) | Põlme S, et al. 2020. Fungal Diversity 105:1-16 ((https://doi.org/10.1007/s13225-020-00466-2)) |
| Otu57712 | 1 | k__Fungi(100);p__Ascomycota(100);c__Saccharomycetes(100);o__Saccharomycetales(100);f__Metschnikowiaceae(100);g__Metschnikowia(100);g__Metschnikowia_unclassified(100); | Metschnikowia | 13 | Saprotroph | \|Undefined Saprotroph\| | Yeast | NULL | Probable | As nectar-tap_saprotroph (Põlme et al. 2020); Decay sugar-rich substrates (Põlme et al. 2020) | Põlme S, et al. 2020. Fungal Diversity 105:1-16 ((https://doi.org/10.1007/s13225-020-00466-2)) |
| Otu57713 | 1 | k__Fungi(100);p__Ascomycota(100);c__Saccharomycetes(100);o__Saccharomycetales(100);f__Metschnikowiaceae(100);g__Metschnikowia(100);s__Metschnikowia_pulcherrima(100); | Metschnikowia | 13 | Saprotroph | \|Undefined Saprotroph\| | Yeast | NULL | Probable | As nectar-tap_saprotroph (Põlme et al. 2020); Decay sugar-rich substrates (Põlme et al. 2020) | Põlme S, et al. 2020. Fungal Diversity 105:1-16 ((https://doi.org/10.1007/s13225-020-00466-2)) |
| Otu57714 | 1 | k__Fungi(100);p__Ascomycota(100);c__Saccharomycetes(100);o__Saccharomycetales(100);f__Metschnikowiaceae(100);g__Metschnikowia(100);s__Metschnikowia_pulcherrima(100); | Metschnikowia | 13 | Saprotroph | \|Undefined Saprotroph\| | Yeast | NULL | Probable | As nectar-tap_saprotroph (Põlme et al. 2020); Decay sugar-rich substrates (Põlme et al. 2020) | Põlme S, et al. 2020. Fungal Diversity 105:1-16 ((https://doi.org/10.1007/s13225-020-00466-2)) |
| Otu57716 | 1 | k__Fungi(100);p__Zygomycota(100);c__Zygomycota_class_Incertae_sedis(100);o__Mortierellales(100);f__Mortierellaceae(100);g__Mortierella(100);g__Mortierella_unclassified(100); | Mortierella | 13 | Saprotroph-Symbiotroph | Endophyte-Plant Saprotroph-\|Undefined Saprotroph\| | Microfungus | NULL | Probable | As chitinolytic (Põlme et al. 2020); As soil_saprotroph (Põlme et al. 2020); Decay roots, soil (Põlme et al. 2020); Saprobic in soil and thus frequenctly isolated from soil and material contaminated with soil such as hay or rotten silage and spoil tips (Cannon & Kirk 2007) | Cannon PF, Kirk PM. 2007. Fungal Families of the World. CAB International, Cambridge (ISBN: 978-0851998275); Tedersoo L, et al. 2014. Science 346:e1256688 ((https://doi.org/10.1126/science.1256688)); Purahong W, et al. 2016. Molecular Ecology 25:4059-4074 ((https://doi.org/10.1111/mec.13739)); Põlme S, et al. 2020. Fungal Diversity 105:1-16 ((https://doi.org/10.1007/s13225-020-00466-2)) |
| Otu57721 | 1 | k__Fungi(100);p__Ascomycota(100);c__Saccharomycetes(100);o__Saccharomycetales(100);f__Metschnikowiaceae(100);g__Metschnikowia(100);g__Metschnikowia_unclassified(100); | Metschnikowia | 13 | Saprotroph | \|Undefined Saprotroph\| | Yeast | NULL | Probable | As nectar-tap_saprotroph (Põlme et al. 2020); Decay sugar-rich substrates (Põlme et al. 2020) | Põlme S, et al. 2020. Fungal Diversity 105:1-16 ((https://doi.org/10.1007/s13225-020-00466-2)) |
| Otu57731 | 1 | k__Fungi(100);p__Ascomycota(100);c__Saccharomycetes(100);o__Saccharomycetales(100);f__Metschnikowiaceae(100);g__Metschnikowia(100);s__Metschnikowia_pulcherrima(100); | Metschnikowia | 13 | Saprotroph | \|Undefined Saprotroph\| | Yeast | NULL | Probable | As nectar-tap_saprotroph (Põlme et al. 2020); Decay sugar-rich substrates (Põlme et al. 2020) | Põlme S, et al. 2020. Fungal Diversity 105:1-16 ((https://doi.org/10.1007/s13225-020-00466-2)) |
| Otu57735 | 1 | k__Fungi(100);p__Ascomycota(100);c__Saccharomycetes(100);o__Saccharomycetales(100);f__Saccharomycetales_family_Incertae_sedis(100);g__Candida(100);s__Candida_cretensis(100); | Candida | 13 | Pathotroph-Saprotroph-Symbiotroph | Animal Pathogen-Endophyte-Endosymbiont-Epiphyte-\|Undefined Saprotroph\| | Dimorphic Yeast-Yeast | NULL | Probable | As nectar-tap_saprotroph (Põlme et al. 2020); Decay sugar-rich substrates (Põlme et al. 2020); This asexual genus is polyphyletic and contain species that function very differently depend on what they are. Caution should be taken when interpreting guilds for Candida species. | Manolakaki D, et al. 2010. Virulence. 1: 367-375 ((https://doi.org/10.4161/viru.1.5.12796)); Mekbib SB, et al. 2011. Tropical Plant Pathology 36:233-240 ((https://doi.org/10.1590/S1982-56762011000400004)); Põlme S, et al. 2020. Fungal Diversity 105:1-16 ((https://doi.org/10.1007/s13225-020-00466-2)) |
| Otu57737 | 1 | k__Fungi(100);p__Ascomycota(100);c__Saccharomycetes(100);o__Saccharomycetales(100);f__Saccharomycetales_family_Incertae_sedis(100);g__Candida(100);g__Candida_unclassified(100); | Candida | 13 | Pathotroph-Saprotroph-Symbiotroph | Animal Pathogen-Endophyte-Endosymbiont-Epiphyte-\|Undefined Saprotroph\| | Dimorphic Yeast-Yeast | NULL | Probable | As nectar-tap_saprotroph (Põlme et al. 2020); Decay sugar-rich substrates (Põlme et al. 2020); This asexual genus is polyphyletic and contain species that function very differently depend on what they are. Caution should be taken when interpreting guilds for Candida species. | Manolakaki D, et al. 2010. Virulence. 1: 367-375 ((https://doi.org/10.4161/viru.1.5.12796)); Mekbib SB, et al. 2011. Tropical Plant Pathology 36:233-240 ((https://doi.org/10.1590/S1982-56762011000400004)); Põlme S, et al. 2020. Fungal Diversity 105:1-16 ((https://doi.org/10.1007/s13225-020-00466-2)) |
| Otu57739 | 1 | k__Fungi(100);p__Ascomycota(100);c__Saccharomycetes(100);o__Saccharomycetales(100);f__Metschnikowiaceae(100);g__Metschnikowia(100);s__Metschnikowia_pulcherrima(100); | Metschnikowia | 13 | Saprotroph | \|Undefined Saprotroph\| | Yeast | NULL | Probable | As nectar-tap_saprotroph (Põlme et al. 2020); Decay sugar-rich substrates (Põlme et al. 2020) | Põlme S, et al. 2020. Fungal Diversity 105:1-16 ((https://doi.org/10.1007/s13225-020-00466-2)) |
| Otu57740 | 1 | k__Fungi(100);p__Ascomycota(100);c__Saccharomycetes(100);o__Saccharomycetales(100);f__Metschnikowiaceae(100);g__Metschnikowia(100);g__Metschnikowia_unclassified(100); | Metschnikowia | 13 | Saprotroph | \|Undefined Saprotroph\| | Yeast | NULL | Probable | As nectar-tap_saprotroph (Põlme et al. 2020); Decay sugar-rich substrates (Põlme et al. 2020) | Põlme S, et al. 2020. Fungal Diversity 105:1-16 ((https://doi.org/10.1007/s13225-020-00466-2)) |
| Otu57741 | 1 | k__Fungi(100);p__Ascomycota(100);c__Saccharomycetes(100);o__Saccharomycetales(100);f__Metschnikowiaceae(100);g__Metschnikowia(100);s__Metschnikowia_pulcherrima(100); | Metschnikowia | 13 | Saprotroph | \|Undefined Saprotroph\| | Yeast | NULL | Probable | As nectar-tap_saprotroph (Põlme et al. 2020); Decay sugar-rich substrates (Põlme et al. 2020) | Põlme S, et al. 2020. Fungal Diversity 105:1-16 ((https://doi.org/10.1007/s13225-020-00466-2)) |
| Otu57747 | 1 | k__Fungi(100);p__Ascomycota(100);c__Saccharomycetes(100);o__Saccharomycetales(100);f__Metschnikowiaceae(100);g__Metschnikowia(100);s__Metschnikowia_pulcherrima(100); | Metschnikowia | 13 | Saprotroph | \|Undefined Saprotroph\| | Yeast | NULL | Probable | As nectar-tap_saprotroph (Põlme et al. 2020); Decay sugar-rich substrates (Põlme et al. 2020) | Põlme S, et al. 2020. Fungal Diversity 105:1-16 ((https://doi.org/10.1007/s13225-020-00466-2)) |
| Otu57748 | 1 | k__Fungi(100);p__Ascomycota(100);c__Saccharomycetes(100);o__Saccharomycetales(100);f__Metschnikowiaceae(100);g__Metschnikowia(100);s__Metschnikowia_pulcherrima(100); | Metschnikowia | 13 | Saprotroph | \|Undefined Saprotroph\| | Yeast | NULL | Probable | As nectar-tap_saprotroph (Põlme et al. 2020); Decay sugar-rich substrates (Põlme et al. 2020) | Põlme S, et al. 2020. Fungal Diversity 105:1-16 ((https://doi.org/10.1007/s13225-020-00466-2)) |
| Otu57749 | 1 | k__Fungi(100);p__Ascomycota(100);c__Saccharomycetes(100);o__Saccharomycetales(100);f__Metschnikowiaceae(100);g__Metschnikowia(100);s__Metschnikowia_pulcherrima(100); | Metschnikowia | 13 | Saprotroph | \|Undefined Saprotroph\| | Yeast | NULL | Probable | As nectar-tap_saprotroph (Põlme et al. 2020); Decay sugar-rich substrates (Põlme et al. 2020) | Põlme S, et al. 2020. Fungal Diversity 105:1-16 ((https://doi.org/10.1007/s13225-020-00466-2)) |

| | | | | | | | | | | | |
|---|---|---|---|---|---|---|---|---|---|---|---|
| Otu57750 | 1 | k__Fungi(100);p__Ascomycota(100);c__Saccharomycetes(100);o__Saccharomycetales(100);f__Metschnikowiaceae(100);g__Metschnikowia(100);s__Metschnikowia_pulcherrima(100); | Metschnikowia | 13 | Saprotroph | \|Undefined Saprotroph\| | Yeast | NULL | Probable | As nectar-tap_saprotroph (Põlme et al. 2020); Decay sugar-rich substrates (Põlme et al. 2020) | Põlme S, et al. 2020. Fungal Diversity 105:1-16 ((https://doi.org/10.1007/s13225-020-00466-2)) |
| Otu57751 | 1 | k__Fungi(100);p__Basidiomycota(100);c__Tremellomycetes(100);o__Tremellales(100);f__Tremellales_family_Incertae_sedis(100);g__Cryptococcus(100);s__Cryptococcus_sp_JCM_11356(100); | Cryptococcus | 13 | Pathotroph-Saprotroph-Symbiotroph | Animal Pathogen-Endophyte-Epiphyte-\|Undefined Saprotroph\| | Dimorphic Yeast-Yeast | NULL | Possible | As foliar_endophyte (Põlme et al. 2020); Decay leaf, fruit, seed, soil, dung, animal material (Põlme et al. 2020) | Andrews JH, et al. 1983. Phytopathology 73:228-234 ((https://doi.org/10.1094/Phyto-73-228)); Kurtzman CP, et al. (eds.) 2011. The Yeasts, a Taxonomic Study. Fifth Edition. Vols 1-3. Elsevier, San Diego (ISBN: 9780444521491); Põlme S, et al. 2020. Fungal Diversity 105:1-16 ((https://doi.org/10.1007/s13225-020-00466-2)) |
| Otu57753 | 1 | k__Fungi(100);p__Ascomycota(100);c__Saccharomycetes(100);o__Saccharomycetales(100);f__Metschnikowiaceae(100);g__Metschnikowia(100);s__Metschnikowia_pulcherrima(100); | Metschnikowia | 13 | Saprotroph | \|Undefined Saprotroph\| | Yeast | NULL | Probable | As nectar-tap_saprotroph (Põlme et al. 2020); Decay sugar-rich substrates (Põlme et al. 2020) | Põlme S, et al. 2020. Fungal Diversity 105:1-16 ((https://doi.org/10.1007/s13225-020-00466-2)) |
| Otu57755 | 1 | k__Fungi(100);p__Ascomycota(100);c__Saccharomycetes(100);o__Saccharomycetales(100);f__Metschnikowiaceae(100);g__Metschnikowia(100);g__Metschnikowia_unclassified(100); | Metschnikowia | 13 | Saprotroph | \|Undefined Saprotroph\| | Yeast | NULL | Probable | As nectar-tap_saprotroph (Põlme et al. 2020); Decay sugar-rich substrates (Põlme et al. 2020) | Põlme S, et al. 2020. Fungal Diversity 105:1-16 ((https://doi.org/10.1007/s13225-020-00466-2)) |
| Otu57756 | 1 | k__Fungi(100);p__Ascomycota(100);c__Saccharomycetes(100);o__Saccharomycetales(100);f__Metschnikowiaceae(100);g__Metschnikowia(100);s__Metschnikowia_pulcherrima(100); | Metschnikowia | 13 | Saprotroph | \|Undefined Saprotroph\| | Yeast | NULL | Probable | As nectar-tap_saprotroph (Põlme et al. 2020); Decay sugar-rich substrates (Põlme et al. 2020) | Põlme S, et al. 2020. Fungal Diversity 105:1-16 ((https://doi.org/10.1007/s13225-020-00466-2)) |
| Otu57758 | 1 | k__Fungi(100);p__Ascomycota(100);c__Saccharomycetes(100);o__Saccharomycetales(100);f__Metschnikowiaceae(100);g__Metschnikowia(100);s__Metschnikowia_pulcherrima(100); | Metschnikowia | 13 | Saprotroph | \|Undefined Saprotroph\| | Yeast | NULL | Probable | As nectar-tap_saprotroph (Põlme et al. 2020); Decay sugar-rich substrates (Põlme et al. 2020) | Põlme S, et al. 2020. Fungal Diversity 105:1-16 ((https://doi.org/10.1007/s13225-020-00466-2)) |
| Otu57759 | 1 | k__Fungi(100);p__Ascomycota(100);c__Saccharomycetes(100);o__Saccharomycetales(100);f__Metschnikowiaceae(100);g__Metschnikowia(100);s__Metschnikowia_pulcherrima(100); | Metschnikowia | 13 | Saprotroph | \|Undefined Saprotroph\| | Yeast | NULL | Probable | As nectar-tap_saprotroph (Põlme et al. 2020); Decay sugar-rich substrates (Põlme et al. 2020) | Põlme S, et al. 2020. Fungal Diversity 105:1-16 ((https://doi.org/10.1007/s13225-020-00466-2)) |
| Otu57760 | 1 | k__Fungi(100);p__Ascomycota(100);c__Saccharomycetes(100);o__Saccharomycetales(100);f__Metschnikowiaceae(100);g__Metschnikowia(100);s__Metschnikowia_pulcherrima(100); | Metschnikowia | 13 | Saprotroph | \|Undefined Saprotroph\| | Yeast | NULL | Probable | As nectar-tap_saprotroph (Põlme et al. 2020); Decay sugar-rich substrates (Põlme et al. 2020) | Põlme S, et al. 2020. Fungal Diversity 105:1-16 ((https://doi.org/10.1007/s13225-020-00466-2)) |
| Otu57761 | 1 | k__Fungi(100);p__Ascomycota(100);c__Saccharomycetes(100);o__Saccharomycetales(100);f__Metschnikowiaceae(100);g__Metschnikowia(100);g__Metschnikowia_unclassified(100); | Metschnikowia | 13 | Saprotroph | \|Undefined Saprotroph\| | Yeast | NULL | Probable | As nectar-tap_saprotroph (Põlme et al. 2020); Decay sugar-rich substrates (Põlme et al. 2020) | Põlme S, et al. 2020. Fungal Diversity 105:1-16 ((https://doi.org/10.1007/s13225-020-00466-2)) |
| Otu57762 | 1 | k__Fungi(100);p__Ascomycota(100);c__Saccharomycetes(100);o__Saccharomycetales(100);f__Metschnikowiaceae(100);g__Metschnikowia(100);s__Metschnikowia_pulcherrima(100); | Metschnikowia | 13 | Saprotroph | \|Undefined Saprotroph\| | Yeast | NULL | Probable | As nectar-tap_saprotroph (Põlme et al. 2020); Decay sugar-rich substrates (Põlme et al. 2020) | Põlme S, et al. 2020. Fungal Diversity 105:1-16 ((https://doi.org/10.1007/s13225-020-00466-2)) |
| Otu57763 | 1 | k__Fungi(100);p__Ascomycota(100);c__Saccharomycetes(100);o__Saccharomycetales(100);f__Metschnikowiaceae(100);g__Metschnikowia(100);s__Metschnikowia_pulcherrima(100); | Metschnikowia | 13 | Saprotroph | \|Undefined Saprotroph\| | Yeast | NULL | Probable | As nectar-tap_saprotroph (Põlme et al. 2020); Decay sugar-rich substrates (Põlme et al. 2020) | Põlme S, et al. 2020. Fungal Diversity 105:1-16 ((https://doi.org/10.1007/s13225-020-00466-2)) |
| Otu57765 | 1 | k__Fungi(100);p__Ascomycota(100);c__Saccharomycetes(100);o__Saccharomycetales(100);f__Metschnikowiaceae(100);g__Metschnikowia(100);s__Metschnikowia_pulcherrima(100); | Metschnikowia | 13 | Saprotroph | \|Undefined Saprotroph\| | Yeast | NULL | Probable | As nectar-tap_saprotroph (Põlme et al. 2020); Decay sugar-rich substrates (Põlme et al. 2020) | Põlme S, et al. 2020. Fungal Diversity 105:1-16 ((https://doi.org/10.1007/s13225-020-00466-2)) |
| Otu57766 | 1 | k__Fungi(100);p__Ascomycota(100);c__Saccharomycetes(100);o__Saccharomycetales(100);f__Metschnikowiaceae(100);g__Metschnikowia(100);s__Metschnikowia_pulcherrima(100); | Metschnikowia | 13 | Saprotroph | \|Undefined Saprotroph\| | Yeast | NULL | Probable | As nectar-tap_saprotroph (Põlme et al. 2020); Decay sugar-rich substrates (Põlme et al. 2020) | Põlme S, et al. 2020. Fungal Diversity 105:1-16 ((https://doi.org/10.1007/s13225-020-00466-2)) |
| Otu57769 | 1 | k__Fungi(100);p__Ascomycota(100);c__Saccharomycetes(100);o__Saccharomycetales(100);f__Metschnikowiaceae(100);g__Metschnikowia(100);s__Metschnikowia_pulcherrima(100); | Metschnikowia | 13 | Saprotroph | \|Undefined Saprotroph\| | Yeast | NULL | Probable | As nectar-tap_saprotroph (Põlme et al. 2020); Decay sugar-rich substrates (Põlme et al. 2020) | Põlme S, et al. 2020. Fungal Diversity 105:1-16 ((https://doi.org/10.1007/s13225-020-00466-2)) |
| Otu57771 | 1 | k__Fungi(100);p__Ascomycota(100);c__Saccharomycetes(100);o__Saccharomycetales(100);f__Metschnikowiaceae(100);g__Metschnikowia(100);s__Metschnikowia_pulcherrima(100); | Metschnikowia | 13 | Saprotroph | \|Undefined Saprotroph\| | Yeast | NULL | Probable | As nectar-tap_saprotroph (Põlme et al. 2020); Decay sugar-rich substrates (Põlme et al. 2020) | Põlme S, et al. 2020. Fungal Diversity 105:1-16 ((https://doi.org/10.1007/s13225-020-00466-2)) |
| Otu57773 | 1 | k__Fungi(100);p__Ascomycota(100);c__Saccharomycetes(100);o__Saccharomycetales(100);f__Metschnikowiaceae(100);g__Metschnikowia(100);s__Metschnikowia_pulcherrima(100); | Metschnikowia | 13 | Saprotroph | \|Undefined Saprotroph\| | Yeast | NULL | Probable | As nectar-tap_saprotroph (Põlme et al. 2020); Decay sugar-rich substrates (Põlme et al. 2020) | Põlme S, et al. 2020. Fungal Diversity 105:1-16 ((https://doi.org/10.1007/s13225-020-00466-2)) |
| Otu57781 | 1 | k__Fungi(100);p__Ascomycota(100);c__Saccharomycetes(100);o__Saccharomycetales(100);f__Metschnikowiaceae(100);g__Metschnikowia(100);s__Metschnikowia_pulcherrima(100); | Metschnikowia | 13 | Saprotroph | \|Undefined Saprotroph\| | Yeast | NULL | Probable | As nectar-tap_saprotroph (Põlme et al. 2020); Decay sugar-rich substrates (Põlme et al. 2020) | Põlme S, et al. 2020. Fungal Diversity 105:1-16 ((https://doi.org/10.1007/s13225-020-00466-2)) |
| Otu57783 | 1 | k__Fungi(100);p__Ascomycota(100);c__Saccharomycetes(100);o__Saccharomycetales(100);f__Metschnikowiaceae(100);g__Metschnikowia(100);s__Metschnikowia_pulcherrima(100); | Metschnikowia | 13 | Saprotroph | \|Undefined Saprotroph\| | Yeast | NULL | Probable | As nectar-tap_saprotroph (Põlme et al. 2020); Decay sugar-rich substrates (Põlme et al. 2020) | Põlme S, et al. 2020. Fungal Diversity 105:1-16 ((https://doi.org/10.1007/s13225-020-00466-2)) |
| Otu57788 | 1 | k__Fungi(100);p__Ascomycota(100);c__Saccharomycetes(100);o__Saccharomycetales(100);f__Metschnikowiaceae(100);g__Metschnikowia(100);s__Metschnikowia_pulcherrima(100); | Metschnikowia | 13 | Saprotroph | \|Undefined Saprotroph\| | Yeast | NULL | Probable | As nectar-tap_saprotroph (Põlme et al. 2020); Decay sugar-rich substrates (Põlme et al. 2020) | Põlme S, et al. 2020. Fungal Diversity 105:1-16 ((https://doi.org/10.1007/s13225-020-00466-2)) |
| Otu57795 | 1 | k__Fungi(100);p__Ascomycota(100);c__Saccharomycetes(100);o__Saccharomycetales(100);f__Saccharomycetaceae(100);g__Kazachstania(100);s__Kazachstania_intestinalis(100); | Kazachstania | 13 | Saprotroph | \|Undefined Saprotroph\| | Yeast | NULL | Probable | As nectar-tap_saprotroph (Põlme et al. 2020); Decay sugar-rich substrates (Põlme et al. 2020) | Põlme S, et al. 2020. Fungal Diversity 105:1-16 ((https://doi.org/10.1007/s13225-020-00466-2)) |
| Otu57797 | 1 | k__Fungi(100);p__Ascomycota(100);c__Saccharomycetes(100);o__Saccharomycetales(100);f__Metschnikowiaceae(100);g__Metschnikowia(100);g__Metschnikowia_unclassified(100); | Metschnikowia | 13 | Saprotroph | \|Undefined Saprotroph\| | Yeast | NULL | Probable | As nectar-tap_saprotroph (Põlme et al. 2020); Decay sugar-rich substrates (Põlme et al. 2020) | Põlme S, et al. 2020. Fungal Diversity 105:1-16 ((https://doi.org/10.1007/s13225-020-00466-2)) |
| Otu57798 | 1 | k__Fungi(100);p__Ascomycota(100);c__Saccharomycetes(100);o__Saccharomycetales(100);f__Metschnikowiaceae(100);g__Metschnikowia(100);s__Metschnikowia_pulcherrima(100); | Metschnikowia | 13 | Saprotroph | \|Undefined Saprotroph\| | Yeast | NULL | Probable | As nectar-tap_saprotroph (Põlme et al. 2020); Decay sugar-rich substrates (Põlme et al. 2020) | Põlme S, et al. 2020. Fungal Diversity 105:1-16 ((https://doi.org/10.1007/s13225-020-00466-2)) |
| Otu57800 | 1 | k__Fungi(100);p__Ascomycota(100);c__Saccharomycetes(100);o__Saccharomycetales(100);f__Metschnikowiaceae(100);g__Metschnikowia(100);s__Metschnikowia_pulcherrima(100); | Metschnikowia | 13 | Saprotroph | \|Undefined Saprotroph\| | Yeast | NULL | Probable | As nectar-tap_saprotroph (Põlme et al. 2020); Decay sugar-rich substrates (Põlme et al. 2020) | Põlme S, et al. 2020. Fungal Diversity 105:1-16 ((https://doi.org/10.1007/s13225-020-00466-2)) |
| Otu57801 | 1 | k__Fungi(100);p__Ascomycota(100);c__Saccharomycetes(100);o__Saccharomycetales(100);f__Metschnikowiaceae(100);g__Metschnikowia(100);s__Metschnikowia_pulcherrima(100); | Metschnikowia | 13 | Saprotroph | \|Undefined Saprotroph\| | Yeast | NULL | Probable | As nectar-tap_saprotroph (Põlme et al. 2020); Decay sugar-rich substrates (Põlme et al. 2020) | Põlme S, et al. 2020. Fungal Diversity 105:1-16 ((https://doi.org/10.1007/s13225-020-00466-2)) |
| Otu57802 | 1 | k__Fungi(100);p__Ascomycota(100);c__Saccharomycetes(100);o__Saccharomycetales(100);f__Metschnikowiaceae(100);g__Metschnikowia(100);s__Metschnikowia_pulcherrima(100); | Metschnikowia | 13 | Saprotroph | \|Undefined Saprotroph\| | Yeast | NULL | Probable | As nectar-tap_saprotroph (Põlme et al. 2020); Decay sugar-rich substrates (Põlme et al. 2020) | Põlme S, et al. 2020. Fungal Diversity 105:1-16 ((https://doi.org/10.1007/s13225-020-00466-2)) |
| Otu57803 | 1 | k__Fungi(100);p__Ascomycota(100);c__Saccharomycetes(100);o__Saccharomycetales(100);f__Metschnikowiaceae(100);g__Metschnikowia(100);g__Metschnikowia_unclassified(100); | Metschnikowia | 13 | Saprotroph | \|Undefined Saprotroph\| | Yeast | NULL | Probable | As nectar-tap_saprotroph (Põlme et al. 2020); Decay sugar-rich substrates (Põlme et al. 2020) | Põlme S, et al. 2020. Fungal Diversity 105:1-16 ((https://doi.org/10.1007/s13225-020-00466-2)) |
| Otu57805 | 1 | k__Fungi(100);p__Ascomycota(100);c__Saccharomycetes(100);o__Saccharomycetales(100);f__Metschnikowiaceae(100);g__Metschnikowia(100);s__Metschnikowia_pulcherrima(100); | Metschnikowia | 13 | Saprotroph | \|Undefined Saprotroph\| | Yeast | NULL | Probable | As nectar-tap_saprotroph (Põlme et al. 2020); Decay sugar-rich substrates (Põlme et al. 2020) | Põlme S, et al. 2020. Fungal Diversity 105:1-16 ((https://doi.org/10.1007/s13225-020-00466-2)) |
| Otu57806 | 1 | k__Fungi(100);p__Ascomycota(100);c__Dothideomycetes(100);o__Capnodiales(100);f__Mycosphaerellaceae(100);f__Mycosphaerellaceae_unclassified(100);f__Mycosphaerellaceae_unclassified(100); | Mycosphaerellaceae | 9 | Pathotroph-Saprotroph | Plant Pathogen-Undefined Saprotroph | Microfungus | NULL | Probable | Biotrophic, necrotrophic or saprobic on various plant tissues (Cannon & Kirk 2007) | Cannon PF, Kirk PM. 2007. Fungal Families of the World. CAB International, Cambridge (ISBN: 978-0851998275) |
| Otu57807 | 1 | k__Fungi(100);p__Basidiomycota(100);c__Dacrymycetes(100);o__Dacrymycetales(100);f__Dacrymycetaceae(100);f__Dacrymycetaceae_unclassified(100);f__Dacrymycetaceae_unclassified(100); | Dacrymycetaceae | 9 | Saprotroph | Wood Saprotroph | Tremelloid | NULL | Probable | Saprobic on rotten coniderous or broadleaved wood and bark (Cannon & Kirk 2007) | Cannon PF, Kirk PM. 2007. Fungal Families of the World. CAB International, Cambridge (ISBN: 978-0851998275) |
| Otu57811 | 1 | k__Fungi(100);p__Ascomycota(100);c__Leotiomycetes(100);o__Helotiales(100);f__Hyaloscyphaceae(100);g__Cistella(100);s__Cistella_sp_KUS_F52527(100); | Cistella | 13 | Saprotroph-Symbiotroph | Endophyte-Plant Saprotroph-Undefined Saprotroph-\|Wood Saprotroph\| | Microfungus | NULL | Probable | As foliar_endophyte (Põlme et al. 2020); As litter_saprotroph (Põlme et al. 2020); Decay wood, leaf, fruit, seed (Põlme et al. 2020) | Tedersoo L, et al. 2014. Science 346:e1256688 ((https://doi.org/10.1126/science.1256688)); Põlme S, et al. 2020. Fungal Diversity 105:1-16 ((https://doi.org/10.1007/s13225-020-00466-2)) |
| Otu57813 | 1 | k__Fungi(100);p__Ascomycota(100);c__Saccharomycetes(100);o__Saccharomycetales(100);f__Saccharomycetaceae(100);g__Kazachstania(100);s__Kazachstania_intestinalis(100); | Kazachstania | 13 | Saprotroph | \|Undefined Saprotroph\| | Yeast | NULL | Probable | As nectar-tap_saprotroph (Põlme et al. 2020); Decay sugar-rich substrates (Põlme et al. 2020) | Põlme S, et al. 2020. Fungal Diversity 105:1-16 ((https://doi.org/10.1007/s13225-020-00466-2)) |
| Otu57814 | 1 | k__Fungi(100);p__Ascomycota(100);c__Saccharomycetes(100);o__Saccharomycetales(100);f__Metschnikowiaceae(100);g__Metschnikowia(100);g__Metschnikowia_unclassified(100); | Metschnikowia | 13 | Saprotroph | \|Undefined Saprotroph\| | Yeast | NULL | Probable | As nectar-tap_saprotroph (Põlme et al. 2020); Decay sugar-rich substrates (Põlme et al. 2020) | Põlme S, et al. 2020. Fungal Diversity 105:1-16 ((https://doi.org/10.1007/s13225-020-00466-2)) |
| Otu57817 | 1 | k__Fungi(100);p__Ascomycota(100);c__Saccharomycetes(100);o__Saccharomycetales(100);f__Metschnikowiaceae(100);g__Metschnikowia(100);s__Metschnikowia_pulcherrima(100); | Metschnikowia | 13 | Saprotroph | \|Undefined Saprotroph\| | Yeast | NULL | Probable | As nectar-tap_saprotroph (Põlme et al. 2020); Decay sugar-rich substrates (Põlme et al. 2020) | Põlme S, et al. 2020. Fungal Diversity 105:1-16 ((https://doi.org/10.1007/s13225-020-00466-2)) |
| Otu57821 | 1 | k__Fungi(100);p__Ascomycota(100);c__Saccharomycetes(100);o__Saccharomycetales(100);f__Metschnikowiaceae(100);g__Metschnikowia(100);s__Metschnikowia_pulcherrima(100); | Metschnikowia | 13 | Saprotroph | \|Undefined Saprotroph\| | Yeast | NULL | Probable | As nectar-tap_saprotroph (Põlme et al. 2020); Decay sugar-rich substrates (Põlme et al. 2020) | Põlme S, et al. 2020. Fungal Diversity 105:1-16 ((https://doi.org/10.1007/s13225-020-00466-2)) |
| Otu57824 | 1 | k__Fungi(100);p__Ascomycota(100);c__Dothideomycetes(100);o__Capnodiales(100);f__Mycosphaerellaceae(100);g__Pseudocercospora(100);s__Pseudocercospora_sp_CPC_11592(100); | Pseudocercospora | 13 | Pathotroph | \|Plant Pathogen\| | Microfungus | NULL | Probable | NULL | Tedersoo L, et al. 2014. Science 346:e1256688 ((https://doi.org/10.1126/science.1256688)); Põlme S, et al. 2020. Fungal Diversity 105:1-16 ((https://doi.org/10.1007/s13225-020-00466-2)) |
| Otu57826 | 1 | k__Fungi(100);p__Ascomycota(100);c__Saccharomycetes(100);o__Saccharomycetales(100);f__Metschnikowiaceae(100);g__Metschnikowia(100);g__Metschnikowia_unclassified(100); | Metschnikowia | 13 | Saprotroph | \|Undefined Saprotroph\| | Yeast | NULL | Probable | As nectar-tap_saprotroph (Põlme et al. 2020); Decay sugar-rich substrates (Põlme et al. 2020) | Põlme S, et al. 2020. Fungal Diversity 105:1-16 ((https://doi.org/10.1007/s13225-020-00466-2)) |
| Otu57827 | 1 | k__Fungi(100);p__Ascomycota(100);c__Saccharomycetes(100);o__Saccharomycetales(100);f__Metschnikowiaceae(100);g__Metschnikowia(100);s__Metschnikowia_pulcherrima(100); | Metschnikowia | 13 | Saprotroph | \|Undefined Saprotroph\| | Yeast | NULL | Probable | As nectar-tap_saprotroph (Põlme et al. 2020); Decay sugar-rich substrates (Põlme et al. 2020) | Põlme S, et al. 2020. Fungal Diversity 105:1-16 ((https://doi.org/10.1007/s13225-020-00466-2)) |
| Otu57828 | 1 | k__Fungi(100);p__Ascomycota(100);c__Saccharomycetes(100);o__Saccharomycetales(100);f__Metschnikowiaceae(100);g__Metschnikowia(100);g__Metschnikowia_unclassified(100); | Metschnikowia | 13 | Saprotroph | \|Undefined Saprotroph\| | Yeast | NULL | Probable | As nectar-tap_saprotroph (Põlme et al. 2020); Decay sugar-rich substrates (Põlme et al. 2020) | Põlme S, et al. 2020. Fungal Diversity 105:1-16 ((https://doi.org/10.1007/s13225-020-00466-2)) |
| Otu57829 | 1 | k__Fungi(100);p__Ascomycota(100);c__Saccharomycetes(100);o__Saccharomycetales(100);f__Metschnikowiaceae(100);g__Metschnikowia(100);s__Metschnikowia_pulcherrima(100); | Metschnikowia | 13 | Saprotroph | \|Undefined Saprotroph\| | Yeast | NULL | Probable | As nectar-tap_saprotroph (Põlme et al. 2020); Decay sugar-rich substrates (Põlme et al. 2020) | Põlme S, et al. 2020. Fungal Diversity 105:1-16 ((https://doi.org/10.1007/s13225-020-00466-2)) |
| Otu57830 | 1 | k__Fungi(100);p__Ascomycota(100);c__Saccharomycetes(100);o__Saccharomycetales(100);f__Metschnikowiaceae(100);g__Metschnikowia(100);s__Metschnikowia_pulcherrima(100); | Metschnikowia | 13 | Saprotroph | \|Undefined Saprotroph\| | Yeast | NULL | Probable | As nectar-tap_saprotroph (Põlme et al. 2020); Decay sugar-rich substrates (Põlme et al. 2020) | Põlme S, et al. 2020. Fungal Diversity 105:1-16 ((https://doi.org/10.1007/s13225-020-00466-2)) |
| Otu57831 | 1 | k__Fungi(100);p__Ascomycota(100);c__Saccharomycetes(100);o__Saccharomycetales(100);f__Metschnikowiaceae(100);g__Metschnikowia(100);s__Metschnikowia_pulcherrima(100); | Metschnikowia | 13 | Saprotroph | \|Undefined Saprotroph\| | Yeast | NULL | Probable | As nectar-tap_saprotroph (Põlme et al. 2020); Decay sugar-rich substrates (Põlme et al. 2020) | Põlme S, et al. 2020. Fungal Diversity 105:1-16 ((https://doi.org/10.1007/s13225-020-00466-2)) |
| Otu57832 | 1 | k__Fungi(100);p__Ascomycota(100);c__Saccharomycetes(100);o__Saccharomycetales(100);f__Metschnikowiaceae(100);g__Metschnikowia(100);s__Metschnikowia_pulcherrima(100); | Metschnikowia | 13 | Saprotroph | \|Undefined Saprotroph\| | Yeast | NULL | Probable | As nectar-tap_saprotroph (Põlme et al. 2020); Decay sugar-rich substrates (Põlme et al. 2020) | Põlme S, et al. 2020. Fungal Diversity 105:1-16 ((https://doi.org/10.1007/s13225-020-00466-2)) |
| Otu57834 | 1 | k__Fungi(100);p__Ascomycota(100);c__Saccharomycetes(100);o__Saccharomycetales(100);f__Metschnikowiaceae(100);g__Metschnikowia(100);s__Metschnikowia_pulcherrima(100); | Metschnikowia | 13 | Saprotroph | \|Undefined Saprotroph\| | Yeast | NULL | Probable | As nectar-tap_saprotroph (Põlme et al. 2020); Decay sugar-rich substrates (Põlme et al. 2020) | Põlme S, et al. 2020. Fungal Diversity 105:1-16 ((https://doi.org/10.1007/s13225-020-00466-2)) |
| Otu57837 | 1 | k__Fungi(100);p__Ascomycota(100);c__Saccharomycetes(100);o__Saccharomycetales(100);f__Metschnikowiaceae(100);g__Metschnikowia(100);s__Metschnikowia_pulcherrima(100); | Metschnikowia | 13 | Saprotroph | \|Undefined Saprotroph\| | Yeast | NULL | Probable | As nectar-tap_saprotroph (Põlme et al. 2020); Decay sugar-rich substrates (Põlme et al. 2020) | Põlme S, et al. 2020. Fungal Diversity 105:1-16 ((https://doi.org/10.1007/s13225-020-00466-2)) |
| Otu57839 | 1 | k__Fungi(100);p__Ascomycota(100);c__Saccharomycetes(100);o__Saccharomycetales(100);f__Metschnikowiaceae(100);g__Metschnikowia(100);s__Metschnikowia_pulcherrima(100); | Metschnikowia | 13 | Saprotroph | \|Undefined Saprotroph\| | Yeast | NULL | Probable | As nectar-tap_saprotroph (Põlme et al. 2020); Decay sugar-rich substrates (Põlme et al. 2020) | Põlme S, et al. 2020. Fungal Diversity 105:1-16 ((https://doi.org/10.1007/s13225-020-00466-2)) |
| Otu57840 | 1 | k__Fungi(100);p__Ascomycota(100);c__Saccharomycetes(100);o__Saccharomycetales(100);f__Metschnikowiaceae(100);g__Metschnikowia(100);g__Metschnikowia_unclassified(100); | Metschnikowia | 13 | Saprotroph | \|Undefined Saprotroph\| | Yeast | NULL | Probable | As nectar-tap_saprotroph (Põlme et al. 2020); Decay sugar-rich substrates (Põlme et al. 2020) | Põlme S, et al. 2020. Fungal Diversity 105:1-16 ((https://doi.org/10.1007/s13225-020-00466-2)) |
| Otu57841 | 1 | k__Fungi(100);p__Ascomycota(100);c__Saccharomycetes(100);o__Saccharomycetales(100);f__Metschnikowiaceae(100);g__Metschnikowia(100);s__Metschnikowia_pulcherrima(100); | Metschnikowia | 13 | Saprotroph | \|Undefined Saprotroph\| | Yeast | NULL | Probable | As nectar-tap_saprotroph (Põlme et al. 2020); Decay sugar-rich substrates (Põlme et al. 2020) | Põlme S, et al. 2020. Fungal Diversity 105:1-16 ((https://doi.org/10.1007/s13225-020-00466-2)) |
| Otu57842 | 1 | k__Fungi(100);p__Ascomycota(100);c__Saccharomycetes(100);o__Saccharomycetales(100);f__Metschnikowiaceae(100);g__Metschnikowia(100);s__Metschnikowia_pulcherrima(100); | Metschnikowia | 13 | Saprotroph | \|Undefined Saprotroph\| | Yeast | NULL | Probable | As nectar-tap_saprotroph (Põlme et al. 2020); Decay sugar-rich substrates (Põlme et al. 2020) | Põlme S, et al. 2020. Fungal Diversity 105:1-16 ((https://doi.org/10.1007/s13225-020-00466-2)) |
| Otu57843 | 1 | k__Fungi(100);p__Ascomycota(100);c__Saccharomycetes(100);o__Saccharomycetales(100);f__Metschnikowiaceae(100);g__Metschnikowia(100);s__Metschnikowia_pulcherrima(100); | Metschnikowia | 13 | Saprotroph | \|Undefined Saprotroph\| | Yeast | NULL | Probable | As nectar-tap_saprotroph (Põlme et al. 2020); Decay sugar-rich substrates (Põlme et al. 2020) | Põlme S, et al. 2020. Fungal Diversity 105:1-16 ((https://doi.org/10.1007/s13225-020-00466-2)) |
| Otu57844 | 1 | k__Fungi(100);p__Ascomycota(100);c__Saccharomycetes(100);o__Saccharomycetales(100);f__Metschnikowiaceae(100);g__Metschnikowia(100);s__Metschnikowia_pulcherrima(100); | Metschnikowia | 13 | Saprotroph | \|Undefined Saprotroph\| | Yeast | NULL | Probable | As nectar-tap_saprotroph (Põlme et al. 2020); Decay sugar-rich substrates (Põlme et al. 2020) | Põlme S, et al. 2020. Fungal Diversity 105:1-16 ((https://doi.org/10.1007/s13225-020-00466-2)) |
| Otu57845 | 1 | k__Fungi(100);p__Ascomycota(100);c__Saccharomycetes(100);o__Saccharomycetales(100);f__Metschnikowiaceae(100);g__Metschnikowia(100);s__Metschnikowia_pulcherrima(100); | Metschnikowia | 13 | Saprotroph | \|Undefined Saprotroph\| | Yeast | NULL | Probable | As nectar-tap_saprotroph (Põlme et al. 2020); Decay sugar-rich substrates (Põlme et al. 2020) | Põlme S, et al. 2020. Fungal Diversity 105:1-16 ((https://doi.org/10.1007/s13225-020-00466-2)) |
| Otu57846 | 1 | k__Fungi(100);p__Ascomycota(100);c__Saccharomycetes(100);o__Saccharomycetales(100);f__Metschnikowiaceae(100);g__Metschnikowia(100);s__Metschnikowia_pulcherrima(100); | Metschnikowia | 13 | Saprotroph | \|Undefined Saprotroph\| | Yeast | NULL | Probable | As nectar-tap_saprotroph (Põlme et al. 2020); Decay sugar-rich substrates (Põlme et al. 2020) | Põlme S, et al. 2020. Fungal Diversity 105:1-16 ((https://doi.org/10.1007/s13225-020-00466-2)) |
| Otu57847 | 1 | k__Fungi(100);p__Ascomycota(100);c__Saccharomycetes(100);o__Saccharomycetales(100);f__Metschnikowiaceae(100);g__Metschnikowia(100);s__Metschnikowia_pulcherrima(100); | Metschnikowia | 13 | Saprotroph | \|Undefined Saprotroph\| | Yeast | NULL | Probable | As nectar-tap_saprotroph (Põlme et al. 2020); Decay sugar-rich substrates (Põlme et al. 2020) | Põlme S, et al. 2020. Fungal Diversity 105:1-16 ((https://doi.org/10.1007/s13225-020-00466-2)) |
| Otu57848 | 1 | k__Fungi(100);p__Ascomycota(100);c__Saccharomycetes(100);o__Saccharomycetales(100);f__Metschnikowiaceae(100);g__Metschnikowia(100);s__Metschnikowia_pulcherrima(100); | Metschnikowia | 13 | Saprotroph | \|Undefined Saprotroph\| | Yeast | NULL | Probable | As nectar-tap_saprotroph (Põlme et al. 2020); Decay sugar-rich substrates (Põlme et al. 2020) | Põlme S, et al. 2020. Fungal Diversity 105:1-16 ((https://doi.org/10.1007/s13225-020-00466-2)) |
| Otu57849 | 1 | k__Fungi(100);p__Ascomycota(100);c__Saccharomycetes(100);o__Saccharomycetales(100);f__Metschnikowiaceae(100);g__Metschnikowia(100);g__Metschnikowia_unclassified(100); | Metschnikowia | 13 | Saprotroph | \|Undefined Saprotroph\| | Yeast | NULL | Probable | As nectar-tap_saprotroph (Põlme et al. 2020); Decay sugar-rich substrates (Põlme et al. 2020) | Põlme S, et al. 2020. Fungal Diversity 105:1-16 ((https://doi.org/10.1007/s13225-020-00466-2)) |
| Otu57850 | 1 | k__Fungi(100);p__Ascomycota(100);c__Saccharomycetes(100);o__Saccharomycetales(100);f__Metschnikowiaceae(100);g__Metschnikowia(100);s__Metschnikowia_pulcherrima(100); | Metschnikowia | 13 | Saprotroph | \|Undefined Saprotroph\| | Yeast | NULL | Probable | As nectar-tap_saprotroph (Põlme et al. 2020); Decay sugar-rich substrates (Põlme et al. 2020) | Põlme S, et al. 2020. Fungal Diversity 105:1-16 ((https://doi.org/10.1007/s13225-020-00466-2)) |
| Otu57853 | 1 | k__Fungi(100);p__Ascomycota(100);c__Saccharomycetes(100);o__Saccharomycetales(100);f__Metschnikowiaceae(100);g__Metschnikowia(100);s__Metschnikowia_pulcherrima(100); | Metschnikowia | 13 | Saprotroph | \|Undefined Saprotroph\| | Yeast | NULL | Probable | As nectar-tap_saprotroph (Põlme et al. 2020); Decay sugar-rich substrates (Põlme et al. 2020) | Põlme S, et al. 2020. Fungal Diversity 105:1-16 ((https://doi.org/10.1007/s13225-020-00466-2)) |
| Otu57854 | 1 | k__Fungi(100);p__Ascomycota(100);c__Saccharomycetes(100);o__Saccharomycetales(100);f__Metschnikowiaceae(100);g__Metschnikowia(100);s__Metschnikowia_pulcherrima(100); | Metschnikowia | 13 | Saprotroph | \|Undefined Saprotroph\| | Yeast | NULL | Probable | As nectar-tap_saprotroph (Põlme et al. 2020); Decay sugar-rich substrates (Põlme et al. 2020) | Põlme S, et al. 2020. Fungal Diversity 105:1-16 ((https://doi.org/10.1007/s13225-020-00466-2)) |
| Otu57856 | 1 | k__Fungi(100);p__Ascomycota(100);c__Saccharomycetes(100);o__Saccharomycetales(100);f__Metschnikowiaceae(100);g__Metschnikowia(100);s__Metschnikowia_pulcherrima(100); | Metschnikowia | 13 | Saprotroph | \|Undefined Saprotroph\| | Yeast | NULL | Probable | As nectar-tap_saprotroph (Põlme et al. 2020); Decay sugar-rich substrates (Põlme et al. 2020) | Põlme S, et al. 2020. Fungal Diversity 105:1-16 ((https://doi.org/10.1007/s13225-020-00466-2)) |
| Otu57857 | 1 | k__Fungi(100);p__Ascomycota(100);c__Saccharomycetes(100);o__Saccharomycetales(100);f__Metschnikowiaceae(100);g__Metschnikowia(100);s__Metschnikowia_pulcherrima(100); | Metschnikowia | 13 | Saprotroph | \|Undefined Saprotroph\| | Yeast | NULL | Probable | As nectar-tap_saprotroph (Põlme et al. 2020); Decay sugar-rich substrates (Põlme et al. 2020) | Põlme S, et al. 2020. Fungal Diversity 105:1-16 ((https://doi.org/10.1007/s13225-020-00466-2)) |
| Otu57858 | 1 | k__Fungi(100);p__Ascomycota(100);c__Saccharomycetes(100);o__Saccharomycetales(100);f__Metschnikowiaceae(100);g__Metschnikowia(100);s__Metschnikowia_pulcherrima(100); | Metschnikowia | 13 | Saprotroph | \|Undefined Saprotroph\| | Yeast | NULL | Probable | As nectar-tap_saprotroph (Põlme et al. 2020); Decay sugar-rich substrates (Põlme et al. 2020) | Põlme S, et al. 2020. Fungal Diversity 105:1-16 ((https://doi.org/10.1007/s13225-020-00466-2)) |
| Otu57859 | 1 | k__Fungi(100);p__Ascomycota(100);c__Saccharomycetes(100);o__Saccharomycetales(100);f__Metschnikowiaceae(100);g__Metschnikowia(100);g__Metschnikowia_unclassified(100); | Metschnikowia | 13 | Saprotroph | \|Undefined Saprotroph\| | Yeast | NULL | Probable | As nectar-tap_saprotroph (Põlme et al. 2020); Decay sugar-rich substrates (Põlme et al. 2020) | Põlme S, et al. 2020. Fungal Diversity 105:1-16 ((https://doi.org/10.1007/s13225-020-00466-2)) |
| Otu57860 | 1 | k__Fungi(100);p__Ascomycota(100);c__Saccharomycetes(100);o__Saccharomycetales(100);f__Metschnikowiaceae(100);g__Metschnikowia(100);g__Metschnikowia_unclassified(100); | Metschnikowia | 13 | Saprotroph | \|Undefined Saprotroph\| | Yeast | NULL | Probable | As nectar-tap_saprotroph (Põlme et al. 2020); Decay sugar-rich substrates (Põlme et al. 2020) | Põlme S, et al. 2020. Fungal Diversity 105:1-16 ((https://doi.org/10.1007/s13225-020-00466-2)) |
| Otu57861 | 1 | k__Fungi(100);p__Ascomycota(100);c__Saccharomycetes(100);o__Saccharomycetales(100);f__Metschnikowiaceae(100);g__Metschnikowia(100);s__Metschnikowia_pulcherrima(100); | Metschnikowia | 13 | Saprotroph | \|Undefined Saprotroph\| | Yeast | NULL | Probable | As nectar-tap_saprotroph (Põlme et al. 2020); Decay sugar-rich substrates (Põlme et al. 2020) | Põlme S, et al. 2020. Fungal Diversity 105:1-16 ((https://doi.org/10.1007/s13225-020-00466-2)) |
| Otu57862 | 1 | k__Fungi(100);p__Ascomycota(100);c__Saccharomycetes(100);o__Saccharomycetales(100);f__Metschnikowiaceae(100);g__Metschnikowia(100);s__Metschnikowia_pulcherrima(100); | Metschnikowia | 13 | Saprotroph | \|Undefined Saprotroph\| | Yeast | NULL | Probable | As nectar-tap_saprotroph (Põlme et al. 2020); Decay sugar-rich substrates (Põlme et al. 2020) | Põlme S, et al. 2020. Fungal Diversity 105:1-16 ((https://doi.org/10.1007/s13225-020-00466-2)) |
| Otu57863 | 1 | k__Fungi(100);p__Ascomycota(100);c__Saccharomycetes(100);o__Saccharomycetales(100);f__Metschnikowiaceae(100);g__Metschnikowia(100);s__Metschnikowia_pulcherrima(100); | Metschnikowia | 13 | Saprotroph | \|Undefined Saprotroph\| | Yeast | NULL | Probable | As nectar-tap_saprotroph (Põlme et al. 2020); Decay sugar-rich substrates (Põlme et al. 2020) | Põlme S, et al. 2020. Fungal Diversity 105:1-16 ((https://doi.org/10.1007/s13225-020-00466-2)) |
| Otu57864 | 1 | k__Fungi(100);p__Ascomycota(100);c__Saccharomycetes(100);o__Saccharomycetales(100);f__Metschnikowiaceae(100);g__Metschnikowia(100);g__Metschnikowia_unclassified(100); | Metschnikowia | 13 | Saprotroph | \|Undefined Saprotroph\| | Yeast | NULL | Probable | As nectar-tap_saprotroph (Põlme et al. 2020); Decay sugar-rich substrates (Põlme et al. 2020) | Põlme S, et al. 2020. Fungal Diversity 105:1-16 ((https://doi.org/10.1007/s13225-020-00466-2)) |
| Otu57866 | 1 | k__Fungi(100);p__Ascomycota(100);c__Saccharomycetes(100);o__Saccharomycetales(100);f__Metschnikowiaceae(100);g__Metschnikowia(100);s__Metschnikowia_pulcherrima(100); | Metschnikowia | 13 | Saprotroph | \|Undefined Saprotroph\| | Yeast | NULL | Probable | As nectar-tap_saprotroph (Põlme et al. 2020); Decay sugar-rich substrates (Põlme et al. 2020) | Põlme S, et al. 2020. Fungal Diversity 105:1-16 ((https://doi.org/10.1007/s13225-020-00466-2)) |
| Otu57868 | 1 | k__Fungi(100);p__Ascomycota(100);c__Saccharomycetes(100);o__Saccharomycetales(100);f__Metschnikowiaceae(100);g__Metschnikowia(100);s__Metschnikowia_pulcherrima(100); | Metschnikowia | 13 | Saprotroph | \|Undefined Saprotroph\| | Yeast | NULL | Probable | As nectar-tap_saprotroph (Põlme et al. 2020); Decay sugar-rich substrates (Põlme et al. 2020) | Põlme S, et al. 2020. Fungal Diversity 105:1-16 ((https://doi.org/10.1007/s13225-020-00466-2)) |
| Otu57869 | 1 | k__Fungi(100);p__Ascomycota(100);c__Saccharomycetes(100);o__Saccharomycetales(100);f__Metschnikowiaceae(100);g__Metschnikowia(100);s__Metschnikowia_pulcherrima(100); | Metschnikowia | 13 | Saprotroph | \|Undefined Saprotroph\| | Yeast | NULL | Probable | As nectar-tap_saprotroph (Põlme et al. 2020); Decay sugar-rich substrates (Põlme et al. 2020) | Põlme S, et al. 2020. Fungal Diversity 105:1-16 ((https://doi.org/10.1007/s13225-020-00466-2)) |
| Otu57870 | 1 | k__Fungi(100);p__Ascomycota(100);c__Saccharomycetes(100);o__Saccharomycetales(100);f__Metschnikowiaceae(100);g__Metschnikowia(100);s__Metschnikowia_pulcherrima(100); | Metschnikowia | 13 | Saprotroph | \|Undefined Saprotroph\| | Yeast | NULL | Probable | As nectar-tap_saprotroph (Põlme et al. 2020); Decay sugar-rich substrates (Põlme et al. 2020) | Põlme S, et al. 2020. Fungal Diversity 105:1-16 ((https://doi.org/10.1007/s13225-020-00466-2)) |
| Otu57871 | 1 | k__Fungi(100);p__Ascomycota(100);c__Saccharomycetes(100);o__Saccharomycetales(100);f__Metschnikowiaceae(100);g__Metschnikowia(100);s__Metschnikowia_pulcherrima(100); | Metschnikowia | 13 | Saprotroph | \|Undefined Saprotroph\| | Yeast | NULL | Probable | As nectar-tap_saprotroph (Põlme et al. 2020); Decay sugar-rich substrates (Põlme et al. 2020) | Põlme S, et al. 2020. Fungal Diversity 105:1-16 ((https://doi.org/10.1007/s13225-020-00466-2)) |
| Otu57872 | 1 | k__Fungi(100);p__Ascomycota(100);c__Saccharomycetes(100);o__Saccharomycetales(100);f__Metschnikowiaceae(100);g__Metschnikowia(100);s__Metschnikowia_pulcherrima(100); | Metschnikowia | 13 | Saprotroph | \|Undefined Saprotroph\| | Yeast | NULL | Probable | As nectar-tap_saprotroph (Põlme et al. 2020); Decay sugar-rich substrates (Põlme et al. 2020) | Põlme S, et al. 2020. Fungal Diversity 105:1-16 ((https://doi.org/10.1007/s13225-020-00466-2)) |
| Otu57874 | 1 | k__Fungi(100);p__Ascomycota(100);c__Saccharomycetes(100);o__Saccharomycetales(100);f__Metschnikowiaceae(100);g__Metschnikowia(100);s__Metschnikowia_pulcherrima(100); | Metschnikowia | 13 | Saprotroph | \|Undefined Saprotroph\| | Yeast | NULL | Probable | As nectar-tap_saprotroph (Põlme et al. 2020); Decay sugar-rich substrates (Põlme et al. 2020) | Põlme S, et al. 2020. Fungal Diversity 105:1-16 ((https://doi.org/10.1007/s13225-020-00466-2)) |
| Otu57879 | 1 | k__Fungi(100);p__Ascomycota(100);c__Saccharomycetes(100);o__Saccharomycetales(100);f__Metschnikowiaceae(100);g__Metschnikowia(100);s__Metschnikowia_pulcherrima(100); | Metschnikowia | 13 | Saprotroph | \|Undefined Saprotroph\| | Yeast | NULL | Probable | As nectar-tap_saprotroph (Põlme et al. 2020); Decay sugar-rich substrates (Põlme et al. 2020) | Põlme S, et al. 2020. Fungal Diversity 105:1-16 ((https://doi.org/10.1007/s13225-020-00466-2)) |
| Otu57880 | 1 | k__Fungi(100);p__Ascomycota(100);c__Saccharomycetes(100);o__Saccharomycetales(100);f__Metschnikowiaceae(100);g__Metschnikowia(100);s__Metschnikowia_pulcherrima(100); | Metschnikowia | 13 | Saprotroph | \|Undefined Saprotroph\| | Yeast | NULL | Probable | As nectar-tap_saprotroph (Põlme et al. 2020); Decay sugar-rich substrates (Põlme et al. 2020) | Põlme S, et al. 2020. Fungal Diversity 105:1-16 ((https://doi.org/10.1007/s13225-020-00466-2)) |
| Otu57881 | 1 | k__Fungi(100);p__Ascomycota(100);c__Saccharomycetes(100);o__Saccharomycetales(100);f__Metschnikowiaceae(100);g__Metschnikowia(100);g__Metschnikowia_unclassified(100); | Metschnikowia | 13 | Saprotroph | \|Undefined Saprotroph\| | Yeast | NULL | Probable | As nectar-tap_saprotroph (Põlme et al. 2020); Decay sugar-rich substrates (Põlme et al. 2020) | Põlme S, et al. 2020. Fungal Diversity 105:1-16 ((https://doi.org/10.1007/s13225-020-00466-2)) |
| Otu57883 | 1 | k__Fungi(100);p__Ascomycota(100);c__Saccharomycetes(100);o__Saccharomycetales(100);f__Metschnikowiaceae(100);g__Metschnikowia(100);g__Metschnikowia_unclassified(100); | Metschnikowia | 13 | Saprotroph | \|Undefined Saprotroph\| | Yeast | NULL | Probable | As nectar-tap_saprotroph (Põlme et al. 2020); Decay sugar-rich substrates (Põlme et al. 2020) | Põlme S, et al. 2020. Fungal Diversity 105:1-16 ((https://doi.org/10.1007/s13225-020-00466-2)) |
| Otu57885 | 1 | k__Fungi(100);p__Ascomycota(100);c__Saccharomycetes(100);o__Saccharomycetales(100);f__Metschnikowiaceae(100);g__Metschnikowia(100);s__Metschnikowia_pulcherrima(100); | Metschnikowia | 13 | Saprotroph | \|Undefined Saprotroph\| | Yeast | NULL | Probable | As nectar-tap_saprotroph (Põlme et al. 2020); Decay sugar-rich substrates (Põlme et al. 2020) | Põlme S, et al. 2020. Fungal Diversity 105:1-16 ((https://doi.org/10.1007/s13225-020-00466-2)) |
| Otu57886 | 1 | k__Fungi(100);p__Ascomycota(100);c__Saccharomycetes(100);o__Saccharomycetales(100);f__Metschnikowiaceae(100);g__Metschnikowia(100);s__Metschnikowia_pulcherrima(100); | Metschnikowia | 13 | Saprotroph | \|Undefined Saprotroph\| | Yeast | NULL | Probable | As nectar-tap_saprotroph (Põlme et al. 2020); Decay sugar-rich substrates (Põlme et al. 2020) | Põlme S, et al. 2020. Fungal Diversity 105:1-16 ((https://doi.org/10.1007/s13225-020-00466-2)) |
| Otu57887 | 1 | k__Fungi(100);p__Ascomycota(100);c__Saccharomycetes(100);o__Saccharomycetales(100);f__Metschnikowiaceae(100);g__Metschnikowia(100);s__Metschnikowia_pulcherrima(100); | Metschnikowia | 13 | Saprotroph | \|Undefined Saprotroph\| | Yeast | NULL | Probable | As nectar-tap_saprotroph (Põlme et al. 2020); Decay sugar-rich substrates (Põlme et al. 2020) | Põlme S, et al. 2020. Fungal Diversity 105:1-16 ((https://doi.org/10.1007/s13225-020-00466-2)) |
| Otu57890 | 1 | k__Fungi(100);p__Ascomycota(100);c__Saccharomycetes(100);o__Saccharomycetales(100);f__Metschnikowiaceae(100);g__Metschnikowia(100);g__Metschnikowia_unclassified(100); | Metschnikowia | 13 | Saprotroph | \|Undefined Saprotroph\| | Yeast | NULL | Probable | As nectar-tap_saprotroph (Põlme et al. 2020); Decay sugar-rich substrates (Põlme et al. 2020) | Põlme S, et al. 2020. Fungal Diversity 105:1-16 ((https://doi.org/10.1007/s13225-020-00466-2)) |
| Otu57891 | 1 | k__Fungi(100);p__Ascomycota(100);c__Saccharomycetes(100);o__Saccharomycetales(100);f__Metschnikowiaceae(100);g__Metschnikowia(100);s__Metschnikowia_pulcherrima(100); | Metschnikowia | 13 | Saprotroph | \|Undefined Saprotroph\| | Yeast | NULL | Probable | As nectar-tap_saprotroph (Põlme et al. 2020); Decay sugar-rich substrates (Põlme et al. 2020) | Põlme S, et al. 2020. Fungal Diversity 105:1-16 ((https://doi.org/10.1007/s13225-020-00466-2)) |
| Otu57892 | 1 | k__Fungi(100);p__Ascomycota(100);c__Saccharomycetes(100);o__Saccharomycetales(100);f__Metschnikowiaceae(100);g__Metschnikowia(100);s__Metschnikowia_pulcherrima(100); | Metschnikowia | 13 | Saprotroph | \|Undefined Saprotroph\| | Yeast | NULL | Probable | As nectar-tap_saprotroph (Põlme et al. 2020); Decay sugar-rich substrates (Põlme et al. 2020) | Põlme S, et al. 2020. Fungal Diversity 105:1-16 ((https://doi.org/10.1007/s13225-020-00466-2)) |
| Otu57893 | 1 | k__Fungi(100);p__Ascomycota(100);c__Saccharomycetes(100);o__Saccharomycetales(100);f__Metschnikowiaceae(100);g__Metschnikowia(100);g__Metschnikowia_unclassified(100); | Metschnikowia | 13 | Saprotroph | \|Undefined Saprotroph\| | Yeast | NULL | Probable | As nectar-tap_saprotroph (Põlme et al. 2020); Decay sugar-rich substrates (Põlme et al. 2020) | Põlme S, et al. 2020. Fungal Diversity 105:1-16 ((https://doi.org/10.1007/s13225-020-00466-2)) |
| Otu57894 | 1 | k__Fungi(100);p__Ascomycota(100);c__Saccharomycetes(100);o__Saccharomycetales(100);f__Metschnikowiaceae(100);g__Metschnikowia(100);s__Metschnikowia_pulcherrima(100); | Metschnikowia | 13 | Saprotroph | \|Undefined Saprotroph\| | Yeast | NULL | Probable | As nectar-tap_saprotroph (Põlme et al. 2020); Decay sugar-rich substrates (Põlme et al. 2020) | Põlme S, et al. 2020. Fungal Diversity 105:1-16 ((https://doi.org/10.1007/s13225-020-00466-2)) |
| Otu57895 | 1 | k__Fungi(100);p__Ascomycota(100);c__Saccharomycetes(100);o__Saccharomycetales(100);f__Metschnikowiaceae(100);g__Metschnikowia(100);g__Metschnikowia_unclassified(100); | Metschnikowia | 13 | Saprotroph | \|Undefined Saprotroph\| | Yeast | NULL | Probable | As nectar-tap_saprotroph (Põlme et al. 2020); Decay sugar-rich substrates (Põlme et al. 2020) | Põlme S, et al. 2020. Fungal Diversity 105:1-16 ((https://doi.org/10.1007/s13225-020-00466-2)) |
| Otu57896 | 1 | k__Fungi(100);p__Ascomycota(100);c__Saccharomycetes(100);o__Saccharomycetales(100);f__Metschnikowiaceae(100);g__Metschnikowia(100);g__Metschnikowia_unclassified(100); | Metschnikowia | 13 | Saprotroph | \|Undefined Saprotroph\| | Yeast | NULL | Probable | As nectar-tap_saprotroph (Põlme et al. 2020); Decay sugar-rich substrates (Põlme et al. 2020) | Põlme S, et al. 2020. Fungal Diversity 105:1-16 ((https://doi.org/10.1007/s13225-020-00466-2)) |
| Otu57897 | 1 | k__Fungi(100);p__Ascomycota(100);c__Saccharomycetes(100);o__Saccharomycetales(100);f__Metschnikowiaceae(100);g__Metschnikowia(100);s__Metschnikowia_pulcherrima(100); | Metschnikowia | 13 | Saprotroph | \|Undefined Saprotroph\| | Yeast | NULL | Probable | As nectar-tap_saprotroph (Põlme et al. 2020); Decay sugar-rich substrates (Põlme et al. 2020) | Põlme S, et al. 2020. Fungal Diversity 105:1-16 ((https://doi.org/10.1007/s13225-020-00466-2)) |
| Otu57898 | 1 | k__Fungi(100);p__Ascomycota(100);c__Saccharomycetes(100);o__Saccharomycetales(100);f__Metschnikowiaceae(100);g__Metschnikowia(100);s__Metschnikowia_pulcherrima(100); | Metschnikowia | 13 | Saprotroph | \|Undefined Saprotroph\| | Yeast | NULL | Probable | As nectar-tap_saprotroph (Põlme et al. 2020); Decay sugar-rich substrates (Põlme et al. 2020) | Põlme S, et al. 2020. Fungal Diversity 105:1-16 ((https://doi.org/10.1007/s13225-020-00466-2)) |
| Otu57899 | 1 | k__Fungi(100);p__Ascomycota(100);c__Saccharomycetes(100);o__Saccharomycetales(100);f__Metschnikowiaceae(100);g__Metschnikowia(100);s__Metschnikowia_pulcherrima(100); | Metschnikowia | 13 | Saprotroph | \|Undefined Saprotroph\| | Yeast | NULL | Probable | As nectar-tap_saprotroph (Põlme et al. 2020); Decay sugar-rich substrates (Põlme et al. 2020) | Põlme S, et al. 2020. Fungal Diversity 105:1-16 ((https://doi.org/10.1007/s13225-020-00466-2)) |
| Otu57900 | 1 | k__Fungi(100);p__Ascomycota(100);c__Saccharomycetes(100);o__Saccharomycetales(100);f__Metschnikowiaceae(100);g__Metschnikowia(100);s__Metschnikowia_pulcherrima(100); | Metschnikowia | 13 | Saprotroph | \|Undefined Saprotroph\| | Yeast | NULL | Probable | As nectar-tap_saprotroph (Põlme et al. 2020); Decay sugar-rich substrates (Põlme et al. 2020) | Põlme S, et al. 2020. Fungal Diversity 105:1-16 ((https://doi.org/10.1007/s13225-020-00466-2)) |
| Otu57901 | 1 | k__Fungi(100);p__Ascomycota(100);c__Saccharomycetes(100);o__Saccharomycetales(100);f__Metschnikowiaceae(100);g__Metschnikowia(100);s__Metschnikowia_pulcherrima(100); | Metschnikowia | 13 | Saprotroph | \|Undefined Saprotroph\| | Yeast | NULL | Probable | As nectar-tap_saprotroph (Põlme et al. 2020); Decay sugar-rich substrates (Põlme et al. 2020) | Põlme S, et al. 2020. Fungal Diversity 105:1-16 ((https://doi.org/10.1007/s13225-020-00466-2)) |
| Otu57902 | 1 | k__Fungi(100);p__Ascomycota(100);c__Saccharomycetes(100);o__Saccharomycetales(100);f__Metschnikowiaceae(100);g__Metschnikowia(100);s__Metschnikowia_pulcherrima(100); | Metschnikowia | 13 | Saprotroph | \|Undefined Saprotroph\| | Yeast | NULL | Probable | As nectar-tap_saprotroph (Põlme et al. 2020); Decay sugar-rich substrates (Põlme et al. 2020) | Põlme S, et al. 2020. Fungal Diversity 105:1-16 ((https://doi.org/10.1007/s13225-020-00466-2)) |
| Otu57904 | 1 | k__Fungi(100);p__Ascomycota(100);c__Saccharomycetes(100);o__Saccharomycetales(100);f__Metschnikowiaceae(100);g__Metschnikowia(100);g__Metschnikowia_unclassified(100); | Metschnikowia | 13 | Saprotroph | \|Undefined Saprotroph\| | Yeast | NULL | Probable | As nectar-tap_saprotroph (Põlme et al. 2020); Decay sugar-rich substrates (Põlme et al. 2020) | Põlme S, et al. 2020. Fungal Diversity 105:1-16 ((https://doi.org/10.1007/s13225-020-00466-2)) |
| Otu57905 | 1 | k__Fungi(100);p__Ascomycota(100);c__Saccharomycetes(100);o__Saccharomycetales(100);f__Metschnikowiaceae(100);g__Metschnikowia(100);g__Metschnikowia_unclassified(100); | Metschnikowia | 13 | Saprotroph | \|Undefined Saprotroph\| | Yeast | NULL | Probable | As nectar-tap_saprotroph (Põlme et al. 2020); Decay sugar-rich substrates (Põlme et al. 2020) | Põlme S, et al. 2020. Fungal Diversity 105:1-16 ((https://doi.org/10.1007/s13225-020-00466-2)) |
| Otu57906 | 1 | k__Fungi(100);p__Ascomycota(100);c__Saccharomycetes(100);o__Saccharomycetales(100);f__Metschnikowiaceae(100);g__Metschnikowia(100);s__Metschnikowia_pulcherrima(100); | Metschnikowia | 13 | Saprotroph | \|Undefined Saprotroph\| | Yeast | NULL | Probable | As nectar-tap_saprotroph (Põlme et al. 2020); Decay sugar-rich substrates (Põlme et al. 2020) | Põlme S, et al. 2020. Fungal Diversity 105:1-16 ((https://doi.org/10.1007/s13225-020-00466-2)) |
| Otu57907 | 1 | k__Fungi(100);p__Ascomycota(100);c__Saccharomycetes(100);o__Saccharomycetales(100);f__Metschnikowiaceae(100);g__Metschnikowia(100);s__Metschnikowia_pulcherrima(100); | Metschnikowia | 13 | Saprotroph | \|Undefined Saprotroph\| | Yeast | NULL | Probable | As nectar-tap_saprotroph (Põlme et al. 2020); Decay sugar-rich substrates (Põlme et al. 2020) | Põlme S, et al. 2020. Fungal Diversity 105:1-16 ((https://doi.org/10.1007/s13225-020-00466-2)) |
| Otu57908 | 1 | k__Fungi(100);p__Ascomycota(100);c__Saccharomycetes(100);o__Saccharomycetales(100);f__Metschnikowiaceae(100);g__Metschnikowia(100);g__Metschnikowia_unclassified(100); | Metschnikowia | 13 | Saprotroph | \|Undefined Saprotroph\| | Yeast | NULL | Probable | As nectar-tap_saprotroph (Põlme et al. 2020); Decay sugar-rich substrates (Põlme et al. 2020) | Põlme S, et al. 2020. Fungal Diversity 105:1-16 ((https://doi.org/10.1007/s13225-020-00466-2)) |
| Otu57909 | 1 | k__Fungi(100);p__Ascomycota(100);c__Saccharomycetes(100);o__Saccharomycetales(100);f__Metschnikowiaceae(100);g__Metschnikowia(100);g__Metschnikowia_unclassified(100); | Metschnikowia | 13 | Saprotroph | \|Undefined Saprotroph\| | Yeast | NULL | Probable | As nectar-tap_saprotroph (Põlme et al. 2020); Decay sugar-rich substrates (Põlme et al. 2020) | Põlme S, et al. 2020. Fungal Diversity 105:1-16 ((https://doi.org/10.1007/s13225-020-00466-2)) |
| Otu57910 | 1 | k__Fungi(100);p__Ascomycota(100);c__Saccharomycetes(100);o__Saccharomycetales(100);f__Metschnikowiaceae(100);g__Metschnikowia(100);s__Metschnikowia_pulcherrima(100); | Metschnikowia | 13 | Saprotroph | \|Undefined Saprotroph\| | Yeast | NULL | Probable | As nectar-tap_saprotroph (Põlme et al. 2020); Decay sugar-rich substrates (Põlme et al. 2020) | Põlme S, et al. 2020. Fungal Diversity 105:1-16 ((https://doi.org/10.1007/s13225-020-00466-2)) |
| Otu57911 | 1 | k__Fungi(100);p__Ascomycota(100);c__Saccharomycetes(100);o__Saccharomycetales(100);f__Metschnikowiaceae(100);g__Metschnikowia(100);s__Metschnikowia_pulcherrima(100); | Metschnikowia | 13 | Saprotroph | \|Undefined Saprotroph\| | Yeast | NULL | Probable | As nectar-tap_saprotroph (Põlme et al. 2020); Decay sugar-rich substrates (Põlme et al. 2020) | Põlme S, et al. 2020. Fungal Diversity 105:1-16 ((https://doi.org/10.1007/s13225-020-00466-2)) |
| Otu57912 | 1 | k__Fungi(100);p__Ascomycota(100);c__Saccharomycetes(100);o__Saccharomycetales(100);f__Metschnikowiaceae(100);g__Metschnikowia(100);s__Metschnikowia_pulcherrima(100); | Metschnikowia | 13 | Saprotroph | \|Undefined Saprotroph\| | Yeast | NULL | Probable | As nectar-tap_saprotroph (Põlme et al. 2020); Decay sugar-rich substrates (Põlme et al. 2020) | Põlme S, et al. 2020. Fungal Diversity 105:1-16 ((https://doi.org/10.1007/s13225-020-00466-2)) |
| Otu57919 | 1 | k__Fungi(100);p__Ascomycota(100);c__Saccharomycetes(100);o__Saccharomycetales(100);f__Metschnikowiaceae(100);g__Metschnikowia(100);s__Metschnikowia_pulcherrima(100); | Metschnikowia | 13 | Saprotroph | \|Undefined Saprotroph\| | Yeast | NULL | Probable | As nectar-tap_saprotroph (Põlme et al. 2020); Decay sugar-rich substrates (Põlme et al. 2020) | Põlme S, et al. 2020. Fungal Diversity 105:1-16 ((https://doi.org/10.1007/s13225-020-00466-2)) |
| Otu57920 | 1 | k__Fungi(100);p__Ascomycota(100);c__Saccharomycetes(100);o__Saccharomycetales(100);f__Metschnikowiaceae(100);g__Metschnikowia(100);s__Metschnikowia_pulcherrima(100); | Metschnikowia | 13 | Saprotroph | \|Undefined Saprotroph\| | Yeast | NULL | Probable | As nectar-tap_saprotroph (Põlme et al. 2020); Decay sugar-rich substrates (Põlme et al. 2020) | Põlme S, et al. 2020. Fungal Diversity 105:1-16 ((https://doi.org/10.1007/s13225-020-00466-2)) |
| Otu57925 | 1 | k__Fungi(100);p__Ascomycota(100);c__Saccharomycetes(100);o__Saccharomycetales(100);f__Metschnikowiaceae(100);g__Metschnikowia(100);s__Metschnikowia_pulcherrima(100); | Metschnikowia | 13 | Saprotroph | \|Undefined Saprotroph\| | Yeast | NULL | Probable | As nectar-tap_saprotroph (Põlme et al. 2020); Decay sugar-rich substrates (Põlme et al. 2020) | Põlme S, et al. 2020. Fungal Diversity 105:1-16 ((https://doi.org/10.1007/s13225-020-00466-2)) |
| Otu57926 | 1 | k__Fungi(100);p__Ascomycota(100);c__Saccharomycetes(100);o__Saccharomycetales(100);f__Metschnikowiaceae(100);g__Metschnikowia(100);s__Metschnikowia_pulcherrima(100); | Metschnikowia | 13 | Saprotroph | \|Undefined Saprotroph\| | Yeast | NULL | Probable | As nectar-tap_saprotroph (Põlme et al. 2020); Decay sugar-rich substrates (Põlme et al. 2020) | Põlme S, et al. 2020. Fungal Diversity 105:1-16 ((https://doi.org/10.1007/s13225-020-00466-2)) |
| Otu57927 | 1 | k__Fungi(100);p__Ascomycota(100);c__Saccharomycetes(100);o__Saccharomycetales(100);f__Metschnikowiaceae(100);g__Metschnikowia(100);g__Metschnikowia_unclassified(100); | Metschnikowia | 13 | Saprotroph | \|Undefined Saprotroph\| | Yeast | NULL | Probable | As nectar-tap_saprotroph (Põlme et al. 2020); Decay sugar-rich substrates (Põlme et al. 2020) | Põlme S, et al. 2020. Fungal Diversity 105:1-16 ((https://doi.org/10.1007/s13225-020-00466-2)) |
| Otu57928 | 1 | k__Fungi(100);p__Ascomycota(100);c__Saccharomycetes(100);o__Saccharomycetales(100);f__Metschnikowiaceae(100);g__Metschnikowia(100);g__Metschnikowia_unclassified(100); | Metschnikowia | 13 | Saprotroph | \|Undefined Saprotroph\| | Yeast | NULL | Probable | As nectar-tap_saprotroph (Põlme et al. 2020); Decay sugar-rich substrates (Põlme et al. 2020) | Põlme S, et al. 2020. Fungal Diversity 105:1-16 ((https://doi.org/10.1007/s13225-020-00466-2)) |
| Otu57929 | 1 | k__Fungi(100);p__Ascomycota(100);c__Saccharomycetes(100);o__Saccharomycetales(100);f__Metschnikowiaceae(100);g__Metschnikowia(100);s__Metschnikowia_pulcherrima(100); | Metschnikowia | 13 | Saprotroph | \|Undefined Saprotroph\| | Yeast | NULL | Probable | As nectar-tap_saprotroph (Põlme et al. 2020); Decay sugar-rich substrates (Põlme et al. 2020) | Põlme S, et al. 2020. Fungal Diversity 105:1-16 ((https://doi.org/10.1007/s13225-020-00466-2)) |
| Otu57930 | 1 | k__Fungi(100);p__Ascomycota(100);c__Saccharomycetes(100);o__Saccharomycetales(100);f__Metschnikowiaceae(100);g__Metschnikowia(100);g__Metschnikowia_unclassified(100); | Metschnikowia | 13 | Saprotroph | \|Undefined Saprotroph\| | Yeast | NULL | Probable | As nectar-tap_saprotroph (Põlme et al. 2020); Decay sugar-rich substrates (Põlme et al. 2020) | Põlme S, et al. 2020. Fungal Diversity 105:1-16 ((https://doi.org/10.1007/s13225-020-00466-2)) |
| Otu57931 | 1 | k__Fungi(100);p__Ascomycota(100);c__Saccharomycetes(100);o__Saccharomycetales(100);f__Metschnikowiaceae(100);g__Metschnikowia(100);g__Metschnikowia_unclassified(100); | Metschnikowia | 13 | Saprotroph | \|Undefined Saprotroph\| | Yeast | NULL | Probable | As nectar-tap_saprotroph (Põlme et al. 2020); Decay sugar-rich substrates (Põlme et al. 2020) | Põlme S, et al. 2020. Fungal Diversity 105:1-16 ((https://doi.org/10.1007/s13225-020-00466-2)) |
| Otu57933 | 1 | k__Fungi(100);p__Ascomycota(100);c__Saccharomycetes(100);o__Saccharomycetales(100);f__Metschnikowiaceae(100);g__Metschnikowia(100);s__Metschnikowia_pulcherrima(100); | Metschnikowia | 13 | Saprotroph | \|Undefined Saprotroph\| | Yeast | NULL | Probable | As nectar-tap_saprotroph (Põlme et al. 2020); Decay sugar-rich substrates (Põlme et al. 2020) | Põlme S, et al. 2020. Fungal Diversity 105:1-16 ((https://doi.org/10.1007/s13225-020-00466-2)) |
| Otu57934 | 1 | k__Fungi(100);p__Ascomycota(100);c__Saccharomycetes(100);o__Saccharomycetales(100);f__Metschnikowiaceae(100);g__Metschnikowia(100);s__Metschnikowia_pulcherrima(100); | Metschnikowia | 13 | Saprotroph | \|Undefined Saprotroph\| | Yeast | NULL | Probable | As nectar-tap_saprotroph (Põlme et al. 2020); Decay sugar-rich substrates (Põlme et al. 2020) | Põlme S, et al. 2020. Fungal Diversity 105:1-16 ((https://doi.org/10.1007/s13225-020-00466-2)) |
| Otu57937 | 1 | k__Fungi(100);p__Ascomycota(100);c__Saccharomycetes(100);o__Saccharomycetales(100);f__Metschnikowiaceae(100);g__Metschnikowia(100);g__Metschnikowia_unclassified(100); | Metschnikowia | 13 | Saprotroph | \|Undefined Saprotroph\| | Yeast | NULL | Probable | As nectar-tap_saprotroph (Põlme et al. 2020); Decay sugar-rich substrates (Põlme et al. 2020) | Põlme S, et al. 2020. Fungal Diversity 105:1-16 ((https://doi.org/10.1007/s13225-020-00466-2)) |
| Otu57938 | 1 | k__Fungi(100);p__Ascomycota(100);c__Saccharomycetes(100);o__Saccharomycetales(100);f__Metschnikowiaceae(100);g__Metschnikowia(100);s__Metschnikowia_pulcherrima(100); | Metschnikowia | 13 | Saprotroph | \|Undefined Saprotroph\| | Yeast | NULL | Probable | As nectar-tap_saprotroph (Põlme et al. 2020); Decay sugar-rich substrates (Põlme et al. 2020) | Põlme S, et al. 2020. Fungal Diversity 105:1-16 ((https://doi.org/10.1007/s13225-020-00466-2)) |
| Otu57940 | 1 | k__Fungi(100);p__Ascomycota(100);c__Saccharomycetes(100);o__Saccharomycetales(100);f__Metschnikowiaceae(100);g__Metschnikowia(100);s__Metschnikowia_pulcherrima(100); | Metschnikowia | 13 | Saprotroph | \|Undefined Saprotroph\| | Yeast | NULL | Probable | As nectar-tap_saprotroph (Põlme et al. 2020); Decay sugar-rich substrates (Põlme et al. 2020) | Põlme S, et al. 2020. Fungal Diversity 105:1-16 ((https://doi.org/10.1007/s13225-020-00466-2)) |
| Otu57941 | 1 | k__Fungi(100);p__Ascomycota(100);c__Saccharomycetes(100);o__Saccharomycetales(100);f__Metschnikowiaceae(100);g__Metschnikowia(100);s__Metschnikowia_pulcherrima(100); | Metschnikowia | 13 | Saprotroph | \|Undefined Saprotroph\| | Yeast | NULL | Probable | As nectar-tap_saprotroph (Põlme et al. 2020); Decay sugar-rich substrates (Põlme et al. 2020) | Põlme S, et al. 2020. Fungal Diversity 105:1-16 ((https://doi.org/10.1007/s13225-020-00466-2)) |
| Otu57943 | 1 | k__Fungi(100);p__Ascomycota(100);c__Saccharomycetes(100);o__Saccharomycetales(100);f__Metschnikowiaceae(100);g__Metschnikowia(100);s__Metschnikowia_pulcherrima(100); | Metschnikowia | 13 | Saprotroph | \|Undefined Saprotroph\| | Yeast | NULL | Probable | As nectar-tap_saprotroph (Põlme et al. 2020); Decay sugar-rich substrates (Põlme et al. 2020) | Põlme S, et al. 2020. Fungal Diversity 105:1-16 ((https://doi.org/10.1007/s13225-020-00466-2)) |
| Otu57944 | 1 | k__Fungi(100);p__Ascomycota(100);c__Saccharomycetes(100);o__Saccharomycetales(100);f__Metschnikowiaceae(100);g__Metschnikowia(100);s__Metschnikowia_pulcherrima(100); | Metschnikowia | 13 | Saprotroph | \|Undefined Saprotroph\| | Yeast | NULL | Probable | As nectar-tap_saprotroph (Põlme et al. 2020); Decay sugar-rich substrates (Põlme et al. 2020) | Põlme S, et al. 2020. Fungal Diversity 105:1-16 ((https://doi.org/10.1007/s13225-020-00466-2)) |
| Otu57945 | 1 | k__Fungi(100);p__Ascomycota(100);c__Saccharomycetes(100);o__Saccharomycetales(100);f__Metschnikowiaceae(100);g__Metschnikowia(100);g__Metschnikowia_unclassified(100); | Metschnikowia | 13 | Saprotroph | \|Undefined Saprotroph\| | Yeast | NULL | Probable | As nectar-tap_saprotroph (Põlme et al. 2020); Decay sugar-rich substrates (Põlme et al. 2020) | Põlme S, et al. 2020. Fungal Diversity 105:1-16 ((https://doi.org/10.1007/s13225-020-00466-2)) |
| Otu57946 | 1 | k__Fungi(100);p__Ascomycota(100);c__Saccharomycetes(100);o__Saccharomycetales(100);f__Metschnikowiaceae(100);g__Metschnikowia(100);s__Metschnikowia_pulcherrima(100); | Metschnikowia | 13 | Saprotroph | \|Undefined Saprotroph\| | Yeast | NULL | Probable | As nectar-tap_saprotroph (Põlme et al. 2020); Decay sugar-rich substrates (Põlme et al. 2020) | Põlme S, et al. 2020. Fungal Diversity 105:1-16 ((https://doi.org/10.1007/s13225-020-00466-2)) |
| Otu57947 | 1 | k__Fungi(100);p__Ascomycota(100);c__Saccharomycetes(100);o__Saccharomycetales(100);f__Metschnikowiaceae(100);g__Metschnikowia(100);s__Metschnikowia_pulcherrima(100); | Metschnikowia | 13 | Saprotroph | \|Undefined Saprotroph\| | Yeast | NULL | Probable | As nectar-tap_saprotroph (Põlme et al. 2020); Decay sugar-rich substrates (Põlme et al. 2020) | Põlme S, et al. 2020. Fungal Diversity 105:1-16 ((https://doi.org/10.1007/s13225-020-00466-2)) |
| Otu57948 | 1 | k__Fungi(100);p__Ascomycota(100);c__Saccharomycetes(100);o__Saccharomycetales(100);f__Metschnikowiaceae(100);g__Metschnikowia(100);s__Metschnikowia_pulcherrima(100); | Metschnikowia | 13 | Saprotroph | \|Undefined Saprotroph\| | Yeast | NULL | Probable | As nectar-tap_saprotroph (Põlme et al. 2020); Decay sugar-rich substrates (Põlme et al. 2020) | Põlme S, et al. 2020. Fungal Diversity 105:1-16 ((https://doi.org/10.1007/s13225-020-00466-2)) |
| Otu57949 | 1 | k__Fungi(100);p__Ascomycota(100);c__Saccharomycetes(100);o__Saccharomycetales(100);f__Metschnikowiaceae(100);g__Metschnikowia(100);s__Metschnikowia_pulcherrima(100); | Metschnikowia | 13 | Saprotroph | \|Undefined Saprotroph\| | Yeast | NULL | Probable | As nectar-tap_saprotroph (Põlme et al. 2020); Decay sugar-rich substrates (Põlme et al. 2020) | Põlme S, et al. 2020. Fungal Diversity 105:1-16 ((https://doi.org/10.1007/s13225-020-00466-2)) |
| Otu57950 | 1 | k__Fungi(100);p__Ascomycota(100);c__Saccharomycetes(100);o__Saccharomycetales(100);f__Metschnikowiaceae(100);g__Metschnikowia(100);s__Metschnikowia_pulcherrima(100); | Metschnikowia | 13 | Saprotroph | \|Undefined Saprotroph\| | Yeast | NULL | Probable | As nectar-tap_saprotroph (Põlme et al. 2020); Decay sugar-rich substrates (Põlme et al. 2020) | Põlme S, et al. 2020. Fungal Diversity 105:1-16 ((https://doi.org/10.1007/s13225-020-00466-2)) |
| Otu57951 | 1 | k__Fungi(100);p__Ascomycota(100);c__Saccharomycetes(100);o__Saccharomycetales(100);f__Metschnikowiaceae(100);g__Metschnikowia(100);s__Metschnikowia_pulcherrima(100); | Metschnikowia | 13 | Saprotroph | \|Undefined Saprotroph\| | Yeast | NULL | Probable | As nectar-tap_saprotroph (Põlme et al. 2020); Decay sugar-rich substrates (Põlme et al. 2020) | Põlme S, et al. 2020. Fungal Diversity 105:1-16 ((https://doi.org/10.1007/s13225-020-00466-2)) |
| Otu57952 | 1 | k__Fungi(100);p__Ascomycota(100);c__Saccharomycetes(100);o__Saccharomycetales(100);f__Metschnikowiaceae(100);g__Metschnikowia(100);s__Metschnikowia_pulcherrima(100); | Metschnikowia | 13 | Saprotroph | \|Undefined Saprotroph\| | Yeast | NULL | Probable | As nectar-tap_saprotroph (Põlme et al. 2020); Decay sugar-rich substrates (Põlme et al. 2020) | Põlme S, et al. 2020. Fungal Diversity 105:1-16 ((https://doi.org/10.1007/s13225-020-00466-2)) |
| Otu57953 | 1 | k__Fungi(100);p__Ascomycota(100);c__Saccharomycetes(100);o__Saccharomycetales(100);f__Metschnikowiaceae(100);g__Metschnikowia(100);s__Metschnikowia_pulcherrima(100); | Metschnikowia | 13 | Saprotroph | \|Undefined Saprotroph\| | Yeast | NULL | Probable | As nectar-tap_saprotroph (Põlme et al. 2020); Decay sugar-rich substrates (Põlme et al. 2020) | Põlme S, et al. 2020. Fungal Diversity 105:1-16 ((https://doi.org/10.1007/s13225-020-00466-2)) |
| Otu57954 | 1 | k__Fungi(100);p__Ascomycota(100);c__Saccharomycetes(100);o__Saccharomycetales(100);f__Metschnikowiaceae(100);g__Metschnikowia(100);s__Metschnikowia_pulcherrima(100); | Metschnikowia | 13 | Saprotroph | \|Undefined Saprotroph\| | Yeast | NULL | Probable | As nectar-tap_saprotroph (Põlme et al. 2020); Decay sugar-rich substrates (Põlme et al. 2020) | Põlme S, et al. 2020. Fungal Diversity 105:1-16 ((https://doi.org/10.1007/s13225-020-00466-2)) |
| Otu57955 | 1 | k__Fungi(100);p__Ascomycota(100);c__Saccharomycetes(100);o__Saccharomycetales(100);f__Metschnikowiaceae(100);g__Metschnikowia(100);s__Metschnikowia_pulcherrima(100); | Metschnikowia | 13 | Saprotroph | \|Undefined Saprotroph\| | Yeast | NULL | Probable | As nectar-tap_saprotroph (Põlme et al. 2020); Decay sugar-rich substrates (Põlme et al. 2020) | Põlme S, et al. 2020. Fungal Diversity 105:1-16 ((https://doi.org/10.1007/s13225-020-00466-2)) |
| Otu57957 | 1 | k__Fungi(100);p__Ascomycota(100);c__Saccharomycetes(100);o__Saccharomycetales(100);f__Metschnikowiaceae(100);g__Metschnikowia(100);s__Metschnikowia_pulcherrima(100); | Metschnikowia | 13 | Saprotroph | \|Undefined Saprotroph\| | Yeast | NULL | Probable | As nectar-tap_saprotroph (Põlme et al. 2020); Decay sugar-rich substrates (Põlme et al. 2020) | Põlme S, et al. 2020. Fungal Diversity 105:1-16 ((https://doi.org/10.1007/s13225-020-00466-2)) |
| Otu57958 | 1 | k__Fungi(100);p__Ascomycota(100);c__Saccharomycetes(100);o__Saccharomycetales(100);f__Metschnikowiaceae(100);g__Metschnikowia(100);s__Metschnikowia_pulcherrima(100); | Metschnikowia | 13 | Saprotroph | \|Undefined Saprotroph\| | Yeast | NULL | Probable | As nectar-tap_saprotroph (Põlme et al. 2020); Decay sugar-rich substrates (Põlme et al. 2020) | Põlme S, et al. 2020. Fungal Diversity 105:1-16 ((https://doi.org/10.1007/s13225-020-00466-2)) |
| Otu57959 | 1 | k__Fungi(100);p__Ascomycota(100);c__Saccharomycetes(100);o__Saccharomycetales(100);f__Metschnikowiaceae(100);g__Metschnikowia(100);g__Metschnikowia_unclassified(100); | Metschnikowia | 13 | Saprotroph | \|Undefined Saprotroph\| | Yeast | NULL | Probable | As nectar-tap_saprotroph (Põlme et al. 2020); Decay sugar-rich substrates (Põlme et al. 2020) | Põlme S, et al. 2020. Fungal Diversity 105:1-16 ((https://doi.org/10.1007/s13225-020-00466-2)) |
| Otu57960 | 1 | k__Fungi(100);p__Ascomycota(100);c__Saccharomycetes(100);o__Saccharomycetales(100);f__Metschnikowiaceae(100);g__Metschnikowia(100);s__Metschnikowia_pulcherrima(100); | Metschnikowia | 13 | Saprotroph | \|Undefined Saprotroph\| | Yeast | NULL | Probable | As nectar-tap_saprotroph (Põlme et al. 2020); Decay sugar-rich substrates (Põlme et al. 2020) | Põlme S, et al. 2020. Fungal Diversity 105:1-16 ((https://doi.org/10.1007/s13225-020-00466-2)) |
| Otu57961 | 1 | k__Fungi(100);p__Ascomycota(100);c__Saccharomycetes(100);o__Saccharomycetales(100);f__Metschnikowiaceae(100);g__Metschnikowia(100);g__Metschnikowia_unclassified(100); | Metschnikowia | 13 | Saprotroph | \|Undefined Saprotroph\| | Yeast | NULL | Probable | As nectar-tap_saprotroph (Põlme et al. 2020); Decay sugar-rich substrates (Põlme et al. 2020) | Põlme S, et al. 2020. Fungal Diversity 105:1-16 ((https://doi.org/10.1007/s13225-020-00466-2)) |
| Otu57962 | 1 | k__Fungi(100);p__Ascomycota(100);c__Saccharomycetes(100);o__Saccharomycetales(100);f__Metschnikowiaceae(100);g__Metschnikowia(100);s__Metschnikowia_pulcherrima(100); | Metschnikowia | 13 | Saprotroph | \|Undefined Saprotroph\| | Yeast | NULL | Probable | As nectar-tap_saprotroph (Põlme et al. 2020); Decay sugar-rich substrates (Põlme et al. 2020) | Põlme S, et al. 2020. Fungal Diversity 105:1-16 ((https://doi.org/10.1007/s13225-020-00466-2)) |
| Otu57963 | 1 | k__Fungi(100);p__Ascomycota(100);c__Saccharomycetes(100);o__Saccharomycetales(100);f__Metschnikowiaceae(100);g__Metschnikowia(100);s__Metschnikowia_pulcherrima(100); | Metschnikowia | 13 | Saprotroph | \|Undefined Saprotroph\| | Yeast | NULL | Probable | As nectar-tap_saprotroph (Põlme et al. 2020); Decay sugar-rich substrates (Põlme et al. 2020) | Põlme S, et al. 2020. Fungal Diversity 105:1-16 ((https://doi.org/10.1007/s13225-020-00466-2)) |
| Otu57964 | 1 | k__Fungi(100);p__Ascomycota(100);c__Saccharomycetes(100);o__Saccharomycetales(100);f__Metschnikowiaceae(100);g__Metschnikowia(100);s__Metschnikowia_pulcherrima(100); | Metschnikowia | 13 | Saprotroph | \|Undefined Saprotroph\| | Yeast | NULL | Probable | As nectar-tap_saprotroph (Põlme et al. 2020); Decay sugar-rich substrates (Põlme et al. 2020) | Põlme S, et al. 2020. Fungal Diversity 105:1-16 ((https://doi.org/10.1007/s13225-020-00466-2)) |
| Otu57965 | 1 | k__Fungi(100);p__Ascomycota(100);c__Saccharomycetes(100);o__Saccharomycetales(100);f__Metschnikowiaceae(100);g__Metschnikowia(100);s__Metschnikowia_pulcherrima(100); | Metschnikowia | 13 | Saprotroph | \|Undefined Saprotroph\| | Yeast | NULL | Probable | As nectar-tap_saprotroph (Põlme et al. 2020); Decay sugar-rich substrates (Põlme et al. 2020) | Põlme S, et al. 2020. Fungal Diversity 105:1-16 ((https://doi.org/10.1007/s13225-020-00466-2)) |
| Otu57966 | 1 | k__Fungi(100);p__Ascomycota(100);c__Saccharomycetes(100);o__Saccharomycetales(100);f__Metschnikowiaceae(100);g__Metschnikowia(100);g__Metschnikowia_unclassified(100); | Metschnikowia | 13 | Saprotroph | \|Undefined Saprotroph\| | Yeast | NULL | Probable | As nectar-tap_saprotroph (Põlme et al. 2020); Decay sugar-rich substrates (Põlme et al. 2020) | Põlme S, et al. 2020. Fungal Diversity 105:1-16 ((https://doi.org/10.1007/s13225-020-00466-2)) |
| Otu57967 | 1 | k__Fungi(100);p__Ascomycota(100);c__Saccharomycetes(100);o__Saccharomycetales(100);f__Metschnikowiaceae(100);g__Metschnikowia(100);s__Metschnikowia_pulcherrima(100); | Metschnikowia | 13 | Saprotroph | \|Undefined Saprotroph\| | Yeast | NULL | Probable | As nectar-tap_saprotroph (Põlme et al. 2020); Decay sugar-rich substrates (Põlme et al. 2020) | Põlme S, et al. 2020. Fungal Diversity 105:1-16 ((https://doi.org/10.1007/s13225-020-00466-2)) |
| Otu57968 | 1 | k__Fungi(100);p__Ascomycota(100);c__Saccharomycetes(100);o__Saccharomycetales(100);f__Metschnikowiaceae(100);g__Metschnikowia(100);s__Metschnikowia_pulcherrima(100); | Metschnikowia | 13 | Saprotroph | \|Undefined Saprotroph\| | Yeast | NULL | Probable | As nectar-tap_saprotroph (Põlme et al. 2020); Decay sugar-rich substrates (Põlme et al. 2020) | Põlme S, et al. 2020. Fungal Diversity 105:1-16 ((https://doi.org/10.1007/s13225-020-00466-2)) |
| Otu57969 | 1 | k__Fungi(100);p__Ascomycota(100);c__Saccharomycetes(100);o__Saccharomycetales(100);f__Metschnikowiaceae(100);g__Metschnikowia(100);s__Metschnikowia_pulcherrima(100); | Metschnikowia | 13 | Saprotroph | \|Undefined Saprotroph\| | Yeast | NULL | Probable | As nectar-tap_saprotroph (Põlme et al. 2020); Decay sugar-rich substrates (Põlme et al. 2020) | Põlme S, et al. 2020. Fungal Diversity 105:1-16 ((https://doi.org/10.1007/s13225-020-00466-2)) |
| Otu57970 | 1 | k__Fungi(100);p__Ascomycota(100);c__Saccharomycetes(100);o__Saccharomycetales(100);f__Metschnikowiaceae(100);g__Metschnikowia(100);s__Metschnikowia_pulcherrima(100); | Metschnikowia | 13 | Saprotroph | \|Undefined Saprotroph\| | Yeast | NULL | Probable | As nectar-tap_saprotroph (Põlme et al. 2020); Decay sugar-rich substrates (Põlme et al. 2020) | Põlme S, et al. 2020. Fungal Diversity 105:1-16 ((https://doi.org/10.1007/s13225-020-00466-2)) |
| Otu57971 | 1 | k__Fungi(100);p__Ascomycota(100);c__Saccharomycetes(100);o__Saccharomycetales(100);f__Metschnikowiaceae(100);g__Metschnikowia(100);s__Metschnikowia_pulcherrima(100); | Metschnikowia | 13 | Saprotroph | \|Undefined Saprotroph\| | Yeast | NULL | Probable | As nectar-tap_saprotroph (Põlme et al. 2020); Decay sugar-rich substrates (Põlme et al. 2020) | Põlme S, et al. 2020. Fungal Diversity 105:1-16 ((https://doi.org/10.1007/s13225-020-00466-2)) |
| Otu57972 | 1 | k__Fungi(100);p__Ascomycota(100);c__Saccharomycetes(100);o__Saccharomycetales(100);f__Metschnikowiaceae(100);g__Metschnikowia(100);g__Metschnikowia_unclassified(100); | Metschnikowia | 13 | Saprotroph | \|Undefined Saprotroph\| | Yeast | NULL | Probable | As nectar-tap_saprotroph (Põlme et al. 2020); Decay sugar-rich substrates (Põlme et al. 2020) | Põlme S, et al. 2020. Fungal Diversity 105:1-16 ((https://doi.org/10.1007/s13225-020-00466-2)) |
| Otu57973 | 1 | k__Fungi(100);p__Ascomycota(100);c__Saccharomycetes(100);o__Saccharomycetales(100);f__Metschnikowiaceae(100);g__Metschnikowia(100);s__Metschnikowia_pulcherrima(100); | Metschnikowia | 13 | Saprotroph | \|Undefined Saprotroph\| | Yeast | NULL | Probable | As nectar-tap_saprotroph (Põlme et al. 2020); Decay sugar-rich substrates (Põlme et al. 2020) | Põlme S, et al. 2020. Fungal Diversity 105:1-16 ((https://doi.org/10.1007/s13225-020-00466-2)) |
| Otu57974 | 1 | k__Fungi(100);p__Ascomycota(100);c__Saccharomycetes(100);o__Saccharomycetales(100);f__Metschnikowiaceae(100);g__Metschnikowia(100);s__Metschnikowia_pulcherrima(100); | Metschnikowia | 13 | Saprotroph | \|Undefined Saprotroph\| | Yeast | NULL | Probable | As nectar-tap_saprotroph (Põlme et al. 2020); Decay sugar-rich substrates (Põlme et al. 2020) | Põlme S, et al. 2020. Fungal Diversity 105:1-16 ((https://doi.org/10.1007/s13225-020-00466-2)) |
| Otu57975 | 1 | k__Fungi(100);p__Ascomycota(100);c__Saccharomycetes(100);o__Saccharomycetales(100);f__Metschnikowiaceae(100);g__Metschnikowia(100);s__Metschnikowia_pulcherrima(100); | Metschnikowia | 13 | Saprotroph | \|Undefined Saprotroph\| | Yeast | NULL | Probable | As nectar-tap_saprotroph (Põlme et al. 2020); Decay sugar-rich substrates (Põlme et al. 2020) | Põlme S, et al. 2020. Fungal Diversity 105:1-16 ((https://doi.org/10.1007/s13225-020-00466-2)) |
| Otu57976 | 1 | k__Fungi(100);p__Ascomycota(100);c__Saccharomycetes(100);o__Saccharomycetales(100);f__Metschnikowiaceae(100);g__Metschnikowia(100);g__Metschnikowia_unclassified(100); | Metschnikowia | 13 | Saprotroph | \|Undefined Saprotroph\| | Yeast | NULL | Probable | As nectar-tap_saprotroph (Põlme et al. 2020); Decay sugar-rich substrates (Põlme et al. 2020) | Põlme S, et al. 2020. Fungal Diversity 105:1-16 ((https://doi.org/10.1007/s13225-020-00466-2)) |
| Otu57977 | 1 | k__Fungi(100);p__Ascomycota(100);c__Saccharomycetes(100);o__Saccharomycetales(100);f__Metschnikowiaceae(100);g__Metschnikowia(100);s__Metschnikowia_pulcherrima(100); | Metschnikowia | 13 | Saprotroph | \|Undefined Saprotroph\| | Yeast | NULL | Probable | As nectar-tap_saprotroph (Põlme et al. 2020); Decay sugar-rich substrates (Põlme et al. 2020) | Põlme S, et al. 2020. Fungal Diversity 105:1-16 ((https://doi.org/10.1007/s13225-020-00466-2)) |
| Otu57979 | 1 | k__Fungi(100);p__Ascomycota(100);c__Saccharomycetes(100);o__Saccharomycetales(100);f__Metschnikowiaceae(100);g__Metschnikowia(100);s__Metschnikowia_pulcherrima(100); | Metschnikowia | 13 | Saprotroph | \|Undefined Saprotroph\| | Yeast | NULL | Probable | As nectar-tap_saprotroph (Põlme et al. 2020); Decay sugar-rich substrates (Põlme et al. 2020) | Põlme S, et al. 2020. Fungal Diversity 105:1-16 ((https://doi.org/10.1007/s13225-020-00466-2)) |
| Otu57980 | 1 | k__Fungi(100);p__Ascomycota(100);c__Saccharomycetes(100);o__Saccharomycetales(100);f__Metschnikowiaceae(100);g__Metschnikowia(100);s__Metschnikowia_pulcherrima(100); | Metschnikowia | 13 | Saprotroph | \|Undefined Saprotroph\| | Yeast | NULL | Probable | As nectar-tap_saprotroph (Põlme et al. 2020); Decay sugar-rich substrates (Põlme et al. 2020) | Põlme S, et al. 2020. Fungal Diversity 105:1-16 ((https://doi.org/10.1007/s13225-020-00466-2)) |
| Otu57981 | 1 | k__Fungi(100);p__Ascomycota(100);c__Saccharomycetes(100);o__Saccharomycetales(100);f__Metschnikowiaceae(100);g__Metschnikowia(100);s__Metschnikowia_pulcherrima(100); | Metschnikowia | 13 | Saprotroph | \|Undefined Saprotroph\| | Yeast | NULL | Probable | As nectar-tap_saprotroph (Põlme et al. 2020); Decay sugar-rich substrates (Põlme et al. 2020) | Põlme S, et al. 2020. Fungal Diversity 105:1-16 ((https://doi.org/10.1007/s13225-020-00466-2)) |
| Otu57982 | 1 | k__Fungi(100);p__Ascomycota(100);c__Saccharomycetes(100);o__Saccharomycetales(100);f__Metschnikowiaceae(100);g__Metschnikowia(100);s__Metschnikowia_pulcherrima(100); | Metschnikowia | 13 | Saprotroph | \|Undefined Saprotroph\| | Yeast | NULL | Probable | As nectar-tap_saprotroph (Põlme et al. 2020); Decay sugar-rich substrates (Põlme et al. 2020) | Põlme S, et al. 2020. Fungal Diversity 105:1-16 ((https://doi.org/10.1007/s13225-020-00466-2)) |
| Otu57983 | 1 | k__Fungi(100);p__Ascomycota(100);c__Saccharomycetes(100);o__Saccharomycetales(100);f__Metschnikowiaceae(100);g__Metschnikowia(100);s__Metschnikowia_pulcherrima(100); | Metschnikowia | 13 | Saprotroph | \|Undefined Saprotroph\| | Yeast | NULL | Probable | As nectar-tap_saprotroph (Põlme et al. 2020); Decay sugar-rich substrates (Põlme et al. 2020) | Põlme S, et al. 2020. Fungal Diversity 105:1-16 ((https://doi.org/10.1007/s13225-020-00466-2)) |
| Otu57984 | 1 | k__Fungi(100);p__Ascomycota(100);c__Saccharomycetes(100);o__Saccharomycetales(100);f__Metschnikowiaceae(100);g__Metschnikowia(100);s__Metschnikowia_pulcherrima(100); | Metschnikowia | 13 | Saprotroph | \|Undefined Saprotroph\| | Yeast | NULL | Probable | As nectar-tap_saprotroph (Põlme et al. 2020); Decay sugar-rich substrates (Põlme et al. 2020) | Põlme S, et al. 2020. Fungal Diversity 105:1-16 ((https://doi.org/10.1007/s13225-020-00466-2)) |
| Otu57985 | 1 | k__Fungi(100);p__Ascomycota(100);c__Saccharomycetes(100);o__Saccharomycetales(100);f__Metschnikowiaceae(100);g__Metschnikowia(100);g__Metschnikowia_unclassified(100); | Metschnikowia | 13 | Saprotroph | \|Undefined Saprotroph\| | Yeast | NULL | Probable | As nectar-tap_saprotroph (Põlme et al. 2020); Decay sugar-rich substrates (Põlme et al. 2020) | Põlme S, et al. 2020. Fungal Diversity 105:1-16 ((https://doi.org/10.1007/s13225-020-00466-2)) |
| Otu57986 | 1 | k__Fungi(100);p__Ascomycota(100);c__Saccharomycetes(100);o__Saccharomycetales(100);f__Metschnikowiaceae(100);g__Metschnikowia(100);s__Metschnikowia_pulcherrima(100); | Metschnikowia | 13 | Saprotroph | \|Undefined Saprotroph\| | Yeast | NULL | Probable | As nectar-tap_saprotroph (Põlme et al. 2020); Decay sugar-rich substrates (Põlme et al. 2020) | Põlme S, et al. 2020. Fungal Diversity 105:1-16 ((https://doi.org/10.1007/s13225-020-00466-2)) |
| Otu57987 | 1 | k__Fungi(100);p__Ascomycota(100);c__Saccharomycetes(100);o__Saccharomycetales(100);f__Metschnikowiaceae(100);g__Metschnikowia(100);s__Metschnikowia_pulcherrima(100); | Metschnikowia | 13 | Saprotroph | \|Undefined Saprotroph\| | Yeast | NULL | Probable | As nectar-tap_saprotroph (Põlme et al. 2020); Decay sugar-rich substrates (Põlme et al. 2020) | Põlme S, et al. 2020. Fungal Diversity 105:1-16 ((https://doi.org/10.1007/s13225-020-00466-2)) |
| Otu57988 | 1 | k__Fungi(100);p__Ascomycota(100);c__Saccharomycetes(100);o__Saccharomycetales(100);f__Metschnikowiaceae(100);g__Metschnikowia(100);s__Metschnikowia_pulcherrima(100); | Metschnikowia | 13 | Saprotroph | \|Undefined Saprotroph\| | Yeast | NULL | Probable | As nectar-tap_saprotroph (Põlme et al. 2020); Decay sugar-rich substrates (Põlme et al. 2020) | Põlme S, et al. 2020. Fungal Diversity 105:1-16 ((https://doi.org/10.1007/s13225-020-00466-2)) |
| Otu57989 | 1 | k__Fungi(100);p__Ascomycota(100);c__Saccharomycetes(100);o__Saccharomycetales(100);f__Metschnikowiaceae(100);g__Metschnikowia(100);s__Metschnikowia_pulcherrima(100); | Metschnikowia | 13 | Saprotroph | \|Undefined Saprotroph\| | Yeast | NULL | Probable | As nectar-tap_saprotroph (Põlme et al. 2020); Decay sugar-rich substrates (Põlme et al. 2020) | Põlme S, et al. 2020. Fungal Diversity 105:1-16 ((https://doi.org/10.1007/s13225-020-00466-2)) |
| Otu57991 | 1 | k__Fungi(100);p__Ascomycota(100);c__Saccharomycetes(100);o__Saccharomycetales(100);f__Metschnikowiaceae(100);g__Metschnikowia(100);s__Metschnikowia_pulcherrima(100); | Metschnikowia | 13 | Saprotroph | \|Undefined Saprotroph\| | Yeast | NULL | Probable | As nectar-tap_saprotroph (Põlme et al. 2020); Decay sugar-rich substrates (Põlme et al. 2020) | Põlme S, et al. 2020. Fungal Diversity 105:1-16 ((https://doi.org/10.1007/s13225-020-00466-2)) |
| Otu57992 | 1 | k__Fungi(100);p__Ascomycota(100);c__Saccharomycetes(100);o__Saccharomycetales(100);f__Metschnikowiaceae(100);g__Metschnikowia(100);g__Metschnikowia_unclassified(100); | Metschnikowia | 13 | Saprotroph | \|Undefined Saprotroph\| | Yeast | NULL | Probable | As nectar-tap_saprotroph (Põlme et al. 2020); Decay sugar-rich substrates (Põlme et al. 2020) | Põlme S, et al. 2020. Fungal Diversity 105:1-16 ((https://doi.org/10.1007/s13225-020-00466-2)) |
| Otu57994 | 1 | k__Fungi(100);p__Ascomycota(100);c__Saccharomycetes(100);o__Saccharomycetales(100);f__Metschnikowiaceae(100);g__Metschnikowia(100);s__Metschnikowia_pulcherrima(100); | Metschnikowia | 13 | Saprotroph | \|Undefined Saprotroph\| | Yeast | NULL | Probable | As nectar-tap_saprotroph (Põlme et al. 2020); Decay sugar-rich substrates (Põlme et al. 2020) | Põlme S, et al. 2020. Fungal Diversity 105:1-16 ((https://doi.org/10.1007/s13225-020-00466-2)) |
| Otu57995 | 1 | k__Fungi(100);p__Ascomycota(100);c__Saccharomycetes(100);o__Saccharomycetales(100);f__Metschnikowiaceae(100);g__Metschnikowia(100);g__Metschnikowia_unclassified(100); | Metschnikowia | 13 | Saprotroph | \|Undefined Saprotroph\| | Yeast | NULL | Probable | As nectar-tap_saprotroph (Põlme et al. 2020); Decay sugar-rich substrates (Põlme et al. 2020) | Põlme S, et al. 2020. Fungal Diversity 105:1-16 ((https://doi.org/10.1007/s13225-020-00466-2)) |
| Otu57996 | 1 | k__Fungi(100);p__Ascomycota(100);c__Saccharomycetes(100);o__Saccharomycetales(100);f__Metschnikowiaceae(100);g__Metschnikowia(100);s__Metschnikowia_pulcherrima(100); | Metschnikowia | 13 | Saprotroph | \|Undefined Saprotroph\| | Yeast | NULL | Probable | As nectar-tap_saprotroph (Põlme et al. 2020); Decay sugar-rich substrates (Põlme et al. 2020) | Põlme S, et al. 2020. Fungal Diversity 105:1-16 ((https://doi.org/10.1007/s13225-020-00466-2)) |
| Otu57997 | 1 | k__Fungi(100);p__Ascomycota(100);c__Saccharomycetes(100);o__Saccharomycetales(100);f__Metschnikowiaceae(100);g__Metschnikowia(100);s__Metschnikowia_pulcherrima(100); | Metschnikowia | 13 | Saprotroph | \|Undefined Saprotroph\| | Yeast | NULL | Probable | As nectar-tap_saprotroph (Põlme et al. 2020); Decay sugar-rich substrates (Põlme et al. 2020) | Põlme S, et al. 2020. Fungal Diversity 105:1-16 ((https://doi.org/10.1007/s13225-020-00466-2)) |
| Otu57998 | 1 | k__Fungi(100);p__Ascomycota(100);c__Saccharomycetes(100);o__Saccharomycetales(100);f__Metschnikowiaceae(100);g__Metschnikowia(100);s__Metschnikowia_pulcherrima(100); | Metschnikowia | 13 | Saprotroph | \|Undefined Saprotroph\| | Yeast | NULL | Probable | As nectar-tap_saprotroph (Põlme et al. 2020); Decay sugar-rich substrates (Põlme et al. 2020) | Põlme S, et al. 2020. Fungal Diversity 105:1-16 ((https://doi.org/10.1007/s13225-020-00466-2)) |
| Otu57999 | 1 | k__Fungi(100);p__Ascomycota(100);c__Saccharomycetes(100);o__Saccharomycetales(100);f__Metschnikowiaceae(100);g__Metschnikowia(100);s__Metschnikowia_pulcherrima(100); | Metschnikowia | 13 | Saprotroph | \|Undefined Saprotroph\| | Yeast | NULL | Probable | As nectar-tap_saprotroph (Põlme et al. 2020); Decay sugar-rich substrates (Põlme et al. 2020) | Põlme S, et al. 2020. Fungal Diversity 105:1-16 ((https://doi.org/10.1007/s13225-020-00466-2)) |
| Otu58000 | 1 | k__Fungi(100);p__Ascomycota(100);c__Saccharomycetes(100);o__Saccharomycetales(100);f__Metschnikowiaceae(100);g__Metschnikowia(100);s__Metschnikowia_pulcherrima(100); | Metschnikowia | 13 | Saprotroph | \|Undefined Saprotroph\| | Yeast | NULL | Probable | As nectar-tap_saprotroph (Põlme et al. 2020); Decay sugar-rich substrates (Põlme et al. 2020) | Põlme S, et al. 2020. Fungal Diversity 105:1-16 ((https://doi.org/10.1007/s13225-020-00466-2)) |
| Otu58001 | 1 | k__Fungi(100);p__Ascomycota(100);c__Saccharomycetes(100);o__Saccharomycetales(100);f__Metschnikowiaceae(100);g__Metschnikowia(100);s__Metschnikowia_pulcherrima(100); | Metschnikowia | 13 | Saprotroph | \|Undefined Saprotroph\| | Yeast | NULL | Probable | As nectar-tap_saprotroph (Põlme et al. 2020); Decay sugar-rich substrates (Põlme et al. 2020) | Põlme S, et al. 2020. Fungal Diversity 105:1-16 ((https://doi.org/10.1007/s13225-020-00466-2)) |
| Otu58002 | 1 | k__Fungi(100);p__Ascomycota(100);c__Saccharomycetes(100);o__Saccharomycetales(100);f__Metschnikowiaceae(100);g__Metschnikowia(100);s__Metschnikowia_pulcherrima(100); | Metschnikowia | 13 | Saprotroph | \|Undefined Saprotroph\| | Yeast | NULL | Probable | As nectar-tap_saprotroph (Põlme et al. 2020); Decay sugar-rich substrates (Põlme et al. 2020) | Põlme S, et al. 2020. Fungal Diversity 105:1-16 ((https://doi.org/10.1007/s13225-020-00466-2)) |
| Otu58003 | 1 | k__Fungi(100);p__Ascomycota(100);c__Saccharomycetes(100);o__Saccharomycetales(100);f__Metschnikowiaceae(100);g__Metschnikowia(100);g__Metschnikowia_unclassified(100); | Metschnikowia | 13 | Saprotroph | \|Undefined Saprotroph\| | Yeast | NULL | Probable | As nectar-tap_saprotroph (Põlme et al. 2020); Decay sugar-rich substrates (Põlme et al. 2020) | Põlme S, et al. 2020. Fungal Diversity 105:1-16 ((https://doi.org/10.1007/s13225-020-00466-2)) |
| Otu58004 | 1 | k__Fungi(100);p__Ascomycota(100);c__Saccharomycetes(100);o__Saccharomycetales(100);f__Metschnikowiaceae(100);g__Metschnikowia(100);s__Metschnikowia_pulcherrima(100); | Metschnikowia | 13 | Saprotroph | \|Undefined Saprotroph\| | Yeast | NULL | Probable | As nectar-tap_saprotroph (Põlme et al. 2020); Decay sugar-rich substrates (Põlme et al. 2020) | Põlme S, et al. 2020. Fungal Diversity 105:1-16 ((https://doi.org/10.1007/s13225-020-00466-2)) |
| Otu58005 | 1 | k__Fungi(100);p__Ascomycota(100);c__Saccharomycetes(100);o__Saccharomycetales(100);f__Metschnikowiaceae(100);g__Metschnikowia(100);g__Metschnikowia_unclassified(100); | Metschnikowia | 13 | Saprotroph | \|Undefined Saprotroph\| | Yeast | NULL | Probable | As nectar-tap_saprotroph (Põlme et al. 2020); Decay sugar-rich substrates (Põlme et al. 2020) | Põlme S, et al. 2020. Fungal Diversity 105:1-16 ((https://doi.org/10.1007/s13225-020-00466-2)) |
| Otu58006 | 1 | k__Fungi(100);p__Ascomycota(100);c__Saccharomycetes(100);o__Saccharomycetales(100);f__Metschnikowiaceae(100);g__Metschnikowia(100);s__Metschnikowia_pulcherrima(100); | Metschnikowia | 13 | Saprotroph | \|Undefined Saprotroph\| | Yeast | NULL | Probable | As nectar-tap_saprotroph (Põlme et al. 2020); Decay sugar-rich substrates (Põlme et al. 2020) | Põlme S, et al. 2020. Fungal Diversity 105:1-16 ((https://doi.org/10.1007/s13225-020-00466-2)) |
| Otu58007 | 1 | k__Fungi(100);p__Ascomycota(100);c__Saccharomycetes(100);o__Saccharomycetales(100);f__Metschnikowiaceae(100);g__Metschnikowia(100);g__Metschnikowia_unclassified(100); | Metschnikowia | 13 | Saprotroph | \|Undefined Saprotroph\| | Yeast | NULL | Probable | As nectar-tap_saprotroph (Põlme et al. 2020); Decay sugar-rich substrates (Põlme et al. 2020) | Põlme S, et al. 2020. Fungal Diversity 105:1-16 ((https://doi.org/10.1007/s13225-020-00466-2)) |
| Otu58008 | 1 | k__Fungi(100);p__Ascomycota(100);c__Saccharomycetes(100);o__Saccharomycetales(100);f__Metschnikowiaceae(100);g__Metschnikowia(100);s__Metschnikowia_pulcherrima(100); | Metschnikowia | 13 | Saprotroph | \|Undefined Saprotroph\| | Yeast | NULL | Probable | As nectar-tap_saprotroph (Põlme et al. 2020); Decay sugar-rich substrates (Põlme et al. 2020) | Põlme S, et al. 2020. Fungal Diversity 105:1-16 ((https://doi.org/10.1007/s13225-020-00466-2)) |
| Otu58009 | 1 | k__Fungi(100);p__Ascomycota(100);c__Saccharomycetes(100);o__Saccharomycetales(100);f__Metschnikowiaceae(100);g__Metschnikowia(100);s__Metschnikowia_pulcherrima(100); | Metschnikowia | 13 | Saprotroph | \|Undefined Saprotroph\| | Yeast | NULL | Probable | As nectar-tap_saprotroph (Põlme et al. 2020); Decay sugar-rich substrates (Põlme et al. 2020) | Põlme S, et al. 2020. Fungal Diversity 105:1-16 ((https://doi.org/10.1007/s13225-020-00466-2)) |
| Otu58010 | 1 | k__Fungi(100);p__Ascomycota(100);c__Saccharomycetes(100);o__Saccharomycetales(100);f__Metschnikowiaceae(100);g__Metschnikowia(100);s__Metschnikowia_pulcherrima(100); | Metschnikowia | 13 | Saprotroph | \|Undefined Saprotroph\| | Yeast | NULL | Probable | As nectar-tap_saprotroph (Põlme et al. 2020); Decay sugar-rich substrates (Põlme et al. 2020) | Põlme S, et al. 2020. Fungal Diversity 105:1-16 ((https://doi.org/10.1007/s13225-020-00466-2)) |
| Otu58011 | 1 | k__Fungi(100);p__Ascomycota(100);c__Saccharomycetes(100);o__Saccharomycetales(100);f__Metschnikowiaceae(100);g__Metschnikowia(100);g__Metschnikowia_unclassified(100); | Metschnikowia | 13 | Saprotroph | \|Undefined Saprotroph\| | Yeast | NULL | Probable | As nectar-tap_saprotroph (Põlme et al. 2020); Decay sugar-rich substrates (Põlme et al. 2020) | Põlme S, et al. 2020. Fungal Diversity 105:1-16 ((https://doi.org/10.1007/s13225-020-00466-2)) |
| Otu58013 | 1 | k__Fungi(100);p__Ascomycota(100);c__Saccharomycetes(100);o__Saccharomycetales(100);f__Metschnikowiaceae(100);g__Metschnikowia(100);s__Metschnikowia_pulcherrima(100); | Metschnikowia | 13 | Saprotroph | \|Undefined Saprotroph\| | Yeast | NULL | Probable | As nectar-tap_saprotroph (Põlme et al. 2020); Decay sugar-rich substrates (Põlme et al. 2020) | Põlme S, et al. 2020. Fungal Diversity 105:1-16 ((https://doi.org/10.1007/s13225-020-00466-2)) |
| Otu58015 | 1 | k__Fungi(100);p__Ascomycota(100);c__Saccharomycetes(100);o__Saccharomycetales(100);f__Metschnikowiaceae(100);g__Metschnikowia(100);s__Metschnikowia_pulcherrima(100); | Metschnikowia | 13 | Saprotroph | \|Undefined Saprotroph\| | Yeast | NULL | Probable | As nectar-tap_saprotroph (Põlme et al. 2020); Decay sugar-rich substrates (Põlme et al. 2020) | Põlme S, et al. 2020. Fungal Diversity 105:1-16 ((https://doi.org/10.1007/s13225-020-00466-2)) |
| Otu58016 | 1 | k__Fungi(100);p__Ascomycota(100);c__Saccharomycetes(100);o__Saccharomycetales(100);f__Metschnikowiaceae(100);g__Metschnikowia(100);g__Metschnikowia_unclassified(100); | Metschnikowia | 13 | Saprotroph | \|Undefined Saprotroph\| | Yeast | NULL | Probable | As nectar-tap_saprotroph (Põlme et al. 2020); Decay sugar-rich substrates (Põlme et al. 2020) | Põlme S, et al. 2020. Fungal Diversity 105:1-16 ((https://doi.org/10.1007/s13225-020-00466-2)) |
| Otu58017 | 1 | k__Fungi(100);p__Ascomycota(100);c__Saccharomycetes(100);o__Saccharomycetales(100);f__Metschnikowiaceae(100);g__Metschnikowia(100);g__Metschnikowia_unclassified(100); | Metschnikowia | 13 | Saprotroph | \|Undefined Saprotroph\| | Yeast | NULL | Probable | As nectar-tap_saprotroph (Põlme et al. 2020); Decay sugar-rich substrates (Põlme et al. 2020) | Põlme S, et al. 2020. Fungal Diversity 105:1-16 ((https://doi.org/10.1007/s13225-020-00466-2)) |
| Otu58018 | 1 | k__Fungi(100);p__Ascomycota(100);c__Saccharomycetes(100);o__Saccharomycetales(100);f__Metschnikowiaceae(100);g__Metschnikowia(100);g__Metschnikowia_unclassified(100); | Metschnikowia | 13 | Saprotroph | \|Undefined Saprotroph\| | Yeast | NULL | Probable | As nectar-tap_saprotroph (Põlme et al. 2020); Decay sugar-rich substrates (Põlme et al. 2020) | Põlme S, et al. 2020. Fungal Diversity 105:1-16 ((https://doi.org/10.1007/s13225-020-00466-2)) |
| Otu58019 | 1 | k__Fungi(100);p__Ascomycota(100);c__Saccharomycetes(100);o__Saccharomycetales(100);f__Metschnikowiaceae(100);g__Metschnikowia(100);g__Metschnikowia_unclassified(100); | Metschnikowia | 13 | Saprotroph | \|Undefined Saprotroph\| | Yeast | NULL | Probable | As nectar-tap_saprotroph (Põlme et al. 2020); Decay sugar-rich substrates (Põlme et al. 2020) | Põlme S, et al. 2020. Fungal Diversity 105:1-16 ((https://doi.org/10.1007/s13225-020-00466-2)) |
| Otu58020 | 1 | k__Fungi(100);p__Ascomycota(100);c__Saccharomycetes(100);o__Saccharomycetales(100);f__Metschnikowiaceae(100);g__Metschnikowia(100);s__Metschnikowia_pulcherrima(100); | Metschnikowia | 13 | Saprotroph | \|Undefined Saprotroph\| | Yeast | NULL | Probable | As nectar-tap_saprotroph (Põlme et al. 2020); Decay sugar-rich substrates (Põlme et al. 2020) | Põlme S, et al. 2020. Fungal Diversity 105:1-16 ((https://doi.org/10.1007/s13225-020-00466-2)) |
| Otu58021 | 1 | k__Fungi(100);p__Ascomycota(100);c__Saccharomycetes(100);o__Saccharomycetales(100);f__Metschnikowiaceae(100);g__Metschnikowia(100);s__Metschnikowia_pulcherrima(100); | Metschnikowia | 13 | Saprotroph | \|Undefined Saprotroph\| | Yeast | NULL | Probable | As nectar-tap_saprotroph (Põlme et al. 2020); Decay sugar-rich substrates (Põlme et al. 2020) | Põlme S, et al. 2020. Fungal Diversity 105:1-16 ((https://doi.org/10.1007/s13225-020-00466-2)) |
| Otu58022 | 1 | k__Fungi(100);p__Ascomycota(100);c__Saccharomycetes(100);o__Saccharomycetales(100);f__Metschnikowiaceae(100);g__Metschnikowia(100);s__Metschnikowia_pulcherrima(100); | Metschnikowia | 13 | Saprotroph | \|Undefined Saprotroph\| | Yeast | NULL | Probable | As nectar-tap_saprotroph (Põlme et al. 2020); Decay sugar-rich substrates (Põlme et al. 2020) | Põlme S, et al. 2020. Fungal Diversity 105:1-16 ((https://doi.org/10.1007/s13225-020-00466-2)) |
| Otu58023 | 1 | k__Fungi(100);p__Ascomycota(100);c__Saccharomycetes(100);o__Saccharomycetales(100);f__Metschnikowiaceae(100);g__Metschnikowia(100);s__Metschnikowia_pulcherrima(100); | Metschnikowia | 13 | Saprotroph | \|Undefined Saprotroph\| | Yeast | NULL | Probable | As nectar-tap_saprotroph (Põlme et al. 2020); Decay sugar-rich substrates (Põlme et al. 2020) | Põlme S, et al. 2020. Fungal Diversity 105:1-16 ((https://doi.org/10.1007/s13225-020-00466-2)) |
| Otu58024 | 1 | k__Fungi(100);p__Ascomycota(100);c__Saccharomycetes(100);o__Saccharomycetales(100);f__Metschnikowiaceae(100);g__Metschnikowia(100);s__Metschnikowia_pulcherrima(100); | Metschnikowia | 13 | Saprotroph | \|Undefined Saprotroph\| | Yeast | NULL | Probable | As nectar-tap_saprotroph (Põlme et al. 2020); Decay sugar-rich substrates (Põlme et al. 2020) | Põlme S, et al. 2020. Fungal Diversity 105:1-16 ((https://doi.org/10.1007/s13225-020-00466-2)) |
| Otu58025 | 1 | k__Fungi(100);p__Ascomycota(100);c__Saccharomycetes(100);o__Saccharomycetales(100);f__Metschnikowiaceae(100);g__Metschnikowia(100);s__Metschnikowia_pulcherrima(100); | Metschnikowia | 13 | Saprotroph | \|Undefined Saprotroph\| | Yeast | NULL | Probable | As nectar-tap_saprotroph (Põlme et al. 2020); Decay sugar-rich substrates (Põlme et al. 2020) | Põlme S, et al. 2020. Fungal Diversity 105:1-16 ((https://doi.org/10.1007/s13225-020-00466-2)) |
| Otu58026 | 1 | k__Fungi(100);p__Ascomycota(100);c__Saccharomycetes(100);o__Saccharomycetales(100);f__Metschnikowiaceae(100);g__Metschnikowia(100);s__Metschnikowia_pulcherrima(100); | Metschnikowia | 13 | Saprotroph | \|Undefined Saprotroph\| | Yeast | NULL | Probable | As nectar-tap_saprotroph (Põlme et al. 2020); Decay sugar-rich substrates (Põlme et al. 2020) | Põlme S, et al. 2020. Fungal Diversity 105:1-16 ((https://doi.org/10.1007/s13225-020-00466-2)) |
| Otu58027 | 1 | k__Fungi(100);p__Ascomycota(100);c__Saccharomycetes(100);o__Saccharomycetales(100);f__Metschnikowiaceae(100);g__Metschnikowia(100);g__Metschnikowia_unclassified(100); | Metschnikowia | 13 | Saprotroph | \|Undefined Saprotroph\| | Yeast | NULL | Probable | As nectar-tap_saprotroph (Põlme et al. 2020); Decay sugar-rich substrates (Põlme et al. 2020) | Põlme S, et al. 2020. Fungal Diversity 105:1-16 ((https://doi.org/10.1007/s13225-020-00466-2)) |
| Otu58028 | 1 | k__Fungi(100);p__Ascomycota(100);c__Saccharomycetes(100);o__Saccharomycetales(100);f__Metschnikowiaceae(100);g__Metschnikowia(100);s__Metschnikowia_pulcherrima(100); | Metschnikowia | 13 | Saprotroph | \|Undefined Saprotroph\| | Yeast | NULL | Probable | As nectar-tap_saprotroph (Põlme et al. 2020); Decay sugar-rich substrates (Põlme et al. 2020) | Põlme S, et al. 2020. Fungal Diversity 105:1-16 ((https://doi.org/10.1007/s13225-020-00466-2)) |
| Otu58029 | 1 | k__Fungi(100);p__Ascomycota(100);c__Saccharomycetes(100);o__Saccharomycetales(100);f__Metschnikowiaceae(100);g__Metschnikowia(100);s__Metschnikowia_pulcherrima(100); | Metschnikowia | 13 | Saprotroph | \|Undefined Saprotroph\| | Yeast | NULL | Probable | As nectar-tap_saprotroph (Põlme et al. 2020); Decay sugar-rich substrates (Põlme et al. 2020) | Põlme S, et al. 2020. Fungal Diversity 105:1-16 ((https://doi.org/10.1007/s13225-020-00466-2)) |
| Otu58030 | 1 | k__Fungi(100);p__Ascomycota(100);c__Saccharomycetes(100);o__Saccharomycetales(100);f__Metschnikowiaceae(100);g__Metschnikowia(100);s__Metschnikowia_pulcherrima(100); | Metschnikowia | 13 | Saprotroph | \|Undefined Saprotroph\| | Yeast | NULL | Probable | As nectar-tap_saprotroph (Põlme et al. 2020); Decay sugar-rich substrates (Põlme et al. 2020) | Põlme S, et al. 2020. Fungal Diversity 105:1-16 ((https://doi.org/10.1007/s13225-020-00466-2)) |
| Otu58031 | 1 | k__Fungi(100);p__Ascomycota(100);c__Saccharomycetes(100);o__Saccharomycetales(100);f__Metschnikowiaceae(100);g__Metschnikowia(100);s__Metschnikowia_pulcherrima(100); | Metschnikowia | 13 | Saprotroph | \|Undefined Saprotroph\| | Yeast | NULL | Probable | As nectar-tap_saprotroph (Põlme et al. 2020); Decay sugar-rich substrates (Põlme et al. 2020) | Põlme S, et al. 2020. Fungal Diversity 105:1-16 ((https://doi.org/10.1007/s13225-020-00466-2)) |
| Otu58032 | 1 | k__Fungi(100);p__Ascomycota(100);c__Saccharomycetes(100);o__Saccharomycetales(100);f__Metschnikowiaceae(100);g__Metschnikowia(100);s__Metschnikowia_pulcherrima(100); | Metschnikowia | 13 | Saprotroph | \|Undefined Saprotroph\| | Yeast | NULL | Probable | As nectar-tap_saprotroph (Põlme et al. 2020); Decay sugar-rich substrates (Põlme et al. 2020) | Põlme S, et al. 2020. Fungal Diversity 105:1-16 ((https://doi.org/10.1007/s13225-020-00466-2)) |
| Otu58033 | 1 | k__Fungi(100);p__Ascomycota(100);c__Saccharomycetes(100);o__Saccharomycetales(100);f__Metschnikowiaceae(100);g__Metschnikowia(100);s__Metschnikowia_pulcherrima(100); | Metschnikowia | 13 | Saprotroph | \|Undefined Saprotroph\| | Yeast | NULL | Probable | As nectar-tap_saprotroph (Põlme et al. 2020); Decay sugar-rich substrates (Põlme et al. 2020) | Põlme S, et al. 2020. Fungal Diversity 105:1-16 ((https://doi.org/10.1007/s13225-020-00466-2)) |
| Otu58034 | 1 | k__Fungi(100);p__Ascomycota(100);c__Saccharomycetes(100);o__Saccharomycetales(100);f__Metschnikowiaceae(100);g__Metschnikowia(100);s__Metschnikowia_pulcherrima(100); | Metschnikowia | 13 | Saprotroph | \|Undefined Saprotroph\| | Yeast | NULL | Probable | As nectar-tap_saprotroph (Põlme et al. 2020); Decay sugar-rich substrates (Põlme et al. 2020) | Põlme S, et al. 2020. Fungal Diversity 105:1-16 ((https://doi.org/10.1007/s13225-020-00466-2)) |
| Otu58035 | 1 | k__Fungi(100);p__Ascomycota(100);c__Saccharomycetes(100);o__Saccharomycetales(100);f__Metschnikowiaceae(100);g__Metschnikowia(100);s__Metschnikowia_pulcherrima(100); | Metschnikowia | 13 | Saprotroph | \|Undefined Saprotroph\| | Yeast | NULL | Probable | As nectar-tap_saprotroph (Põlme et al. 2020); Decay sugar-rich substrates (Põlme et al. 2020) | Põlme S, et al. 2020. Fungal Diversity 105:1-16 ((https://doi.org/10.1007/s13225-020-00466-2)) |
| Otu58036 | 1 | k__Fungi(100);p__Ascomycota(100);c__Saccharomycetes(100);o__Saccharomycetales(100);f__Metschnikowiaceae(100);g__Metschnikowia(100);s__Metschnikowia_pulcherrima(100); | Metschnikowia | 13 | Saprotroph | \|Undefined Saprotroph\| | Yeast | NULL | Probable | As nectar-tap_saprotroph (Põlme et al. 2020); Decay sugar-rich substrates (Põlme et al. 2020) | Põlme S, et al. 2020. Fungal Diversity 105:1-16 ((https://doi.org/10.1007/s13225-020-00466-2)) |
| Otu58038 | 1 | k__Fungi(100);p__Ascomycota(100);c__Saccharomycetes(100);o__Saccharomycetales(100);f__Metschnikowiaceae(100);g__Metschnikowia(100);s__Metschnikowia_pulcherrima(100); | Metschnikowia | 13 | Saprotroph | \|Undefined Saprotroph\| | Yeast | NULL | Probable | As nectar-tap_saprotroph (Põlme et al. 2020); Decay sugar-rich substrates (Põlme et al. 2020) | Põlme S, et al. 2020. Fungal Diversity 105:1-16 ((https://doi.org/10.1007/s13225-020-00466-2)) |
| Otu58041 | 1 | k__Fungi(100);p__Ascomycota(100);c__Saccharomycetes(100);o__Saccharomycetales(100);f__Metschnikowiaceae(100);g__Metschnikowia(100);s__Metschnikowia_pulcherrima(100); | Metschnikowia | 13 | Saprotroph | \|Undefined Saprotroph\| | Yeast | NULL | Probable | As nectar-tap_saprotroph (Põlme et al. 2020); Decay sugar-rich substrates (Põlme et al. 2020) | Põlme S, et al. 2020. Fungal Diversity 105:1-16 ((https://doi.org/10.1007/s13225-020-00466-2)) |
| Otu58044 | 1 | k__Fungi(100);p__Ascomycota(100);c__Saccharomycetes(100);o__Saccharomycetales(100);f__Metschnikowiaceae(100);g__Metschnikowia(100);s__Metschnikowia_pulcherrima(100); | Metschnikowia | 13 | Saprotroph | \|Undefined Saprotroph\| | Yeast | NULL | Probable | As nectar-tap_saprotroph (Põlme et al. 2020); Decay sugar-rich substrates (Põlme et al. 2020) | Põlme S, et al. 2020. Fungal Diversity 105:1-16 ((https://doi.org/10.1007/s13225-020-00466-2)) |
| Otu58048 | 1 | k__Fungi(100);p__Ascomycota(100);c__Sordariomycetes(100);o__Hypocreales(100);f__Nectriaceae(100);g__Gibberella(100);s__Gibberella_intricans(100); | Gibberella | 13 | Pathotroph-Saprotroph-Symbiotroph | Endophyte-\|Plant Pathogen\|-Plant Saprotroph-Undefined Saprotroph | Microfungus | Soft Rot | Probable | As foliar_endophyte (Põlme et al. 2020); As litter_saprotroph (Põlme et al. 2020); Causing damping-off disease (Põlme et al. 2020); Decay leaf, fruit, seed, roots, soil, animal material (Põlme et al. 2020) | Tedersoo L, et al. 2014. Science 346:e1256688 ((https://doi.org/10.1126/science.1256688)); Põlme S, et al. 2020. Fungal Diversity 105:1-16 ((https://doi.org/10.1007/s13225-020-00466-2)) |
| Otu58052 | 1 | k__Fungi(100);p__Ascomycota(100);c__Saccharomycetes(100);o__Saccharomycetales(100);f__Metschnikowiaceae(100);g__Metschnikowia(100);s__Metschnikowia_pulcherrima(100); | Metschnikowia | 13 | Saprotroph | \|Undefined Saprotroph\| | Yeast | NULL | Probable | As nectar-tap_saprotroph (Põlme et al. 2020); Decay sugar-rich substrates (Põlme et al. 2020) | Põlme S, et al. 2020. Fungal Diversity 105:1-16 ((https://doi.org/10.1007/s13225-020-00466-2)) |
| Otu58056 | 1 | k__Fungi(100);p__Ascomycota(100);c__Saccharomycetes(100);o__Saccharomycetales(100);f__Metschnikowiaceae(100);g__Metschnikowia(100);s__Metschnikowia_pulcherrima(100); | Metschnikowia | 13 | Saprotroph | \|Undefined Saprotroph\| | Yeast | NULL | Probable | As nectar-tap_saprotroph (Põlme et al. 2020); Decay sugar-rich substrates (Põlme et al. 2020) | Põlme S, et al. 2020. Fungal Diversity 105:1-16 ((https://doi.org/10.1007/s13225-020-00466-2)) |
| Otu58057 | 1 | k__Fungi(100);p__Ascomycota(100);c__Saccharomycetes(100);o__Saccharomycetales(100);f__Metschnikowiaceae(100);g__Metschnikowia(100);s__Metschnikowia_pulcherrima(100); | Metschnikowia | 13 | Saprotroph | \|Undefined Saprotroph\| | Yeast | NULL | Probable | As nectar-tap_saprotroph (Põlme et al. 2020); Decay sugar-rich substrates (Põlme et al. 2020) | Põlme S, et al. 2020. Fungal Diversity 105:1-16 ((https://doi.org/10.1007/s13225-020-00466-2)) |

| | | | | | | | | | | | |
|---|---|---|---|---|---|---|---|---|---|---|---|
| Otu58058 | 1 | k__Fungi(100);p__Ascomycota(100);c__Saccharomycetes(100);o__Saccharomycetales(100);f__Metschnikowiaceae(100);g__Metschnikowia(100);g__Metschnikowia_unclassified(100); | Metschnikowia | 13 | Saprotroph | \|Undefined Saprotroph\| | Yeast | NULL | Probable | As nectar-tap_saprotroph (Põlme et al. 2020); Decay sugar-rich substrates (Põlme et al. 2020) | Põlme S, et al. 2020. Fungal Diversity 105:1-16 ((https://doi.org/10.1007/s13225-020-00466-2)) |
| Otu58059 | 1 | k__Fungi(100);p__Ascomycota(100);c__Saccharomycetes(100);o__Saccharomycetales(100);f__Metschnikowiaceae(100);g__Metschnikowia(100);g__Metschnikowia_unclassified(100); | Metschnikowia | 13 | Saprotroph | \|Undefined Saprotroph\| | Yeast | NULL | Probable | As nectar-tap_saprotroph (Põlme et al. 2020); Decay sugar-rich substrates (Põlme et al. 2020) | Põlme S, et al. 2020. Fungal Diversity 105:1-16 ((https://doi.org/10.1007/s13225-020-00466-2)) |
| Otu58060 | 1 | k__Fungi(100);p__Ascomycota(100);c__Saccharomycetes(100);o__Saccharomycetales(100);f__Metschnikowiaceae(100);g__Metschnikowia(100);g__Metschnikowia_unclassified(100); | Metschnikowia | 13 | Saprotroph | \|Undefined Saprotroph\| | Yeast | NULL | Probable | As nectar-tap_saprotroph (Põlme et al. 2020); Decay sugar-rich substrates (Põlme et al. 2020) | Põlme S, et al. 2020. Fungal Diversity 105:1-16 ((https://doi.org/10.1007/s13225-020-00466-2)) |
| Otu58061 | 1 | k__Fungi(100);p__Ascomycota(100);c__Saccharomycetes(100);o__Saccharomycetales(100);f__Metschnikowiaceae(100);g__Metschnikowia(100);s__Metschnikowia_pulcherrima(100); | Metschnikowia | 13 | Saprotroph | \|Undefined Saprotroph\| | Yeast | NULL | Probable | As nectar-tap_saprotroph (Põlme et al. 2020); Decay sugar-rich substrates (Põlme et al. 2020) | Põlme S, et al. 2020. Fungal Diversity 105:1-16 ((https://doi.org/10.1007/s13225-020-00466-2)) |
| Otu58062 | 1 | k__Fungi(100);p__Ascomycota(100);c__Saccharomycetes(100);o__Saccharomycetales(100);f__Metschnikowiaceae(100);g__Metschnikowia(100);g__Metschnikowia_unclassified(100); | Metschnikowia | 13 | Saprotroph | \|Undefined Saprotroph\| | Yeast | NULL | Probable | As nectar-tap_saprotroph (Põlme et al. 2020); Decay sugar-rich substrates (Põlme et al. 2020) | Põlme S, et al. 2020. Fungal Diversity 105:1-16 ((https://doi.org/10.1007/s13225-020-00466-2)) |
| Otu58063 | 1 | k__Fungi(100);p__Ascomycota(100);c__Saccharomycetes(100);o__Saccharomycetales(100);f__Metschnikowiaceae(100);g__Metschnikowia(100);s__Metschnikowia_pulcherrima(100); | Metschnikowia | 13 | Saprotroph | \|Undefined Saprotroph\| | Yeast | NULL | Probable | As nectar-tap_saprotroph (Põlme et al. 2020); Decay sugar-rich substrates (Põlme et al. 2020) | Põlme S, et al. 2020. Fungal Diversity 105:1-16 ((https://doi.org/10.1007/s13225-020-00466-2)) |
| Otu58064 | 1 | k__Fungi(100);p__Ascomycota(100);c__Saccharomycetes(100);o__Saccharomycetales(100);f__Metschnikowiaceae(100);g__Metschnikowia(100);s__Metschnikowia_pulcherrima(100); | Metschnikowia | 13 | Saprotroph | \|Undefined Saprotroph\| | Yeast | NULL | Probable | As nectar-tap_saprotroph (Põlme et al. 2020); Decay sugar-rich substrates (Põlme et al. 2020) | Põlme S, et al. 2020. Fungal Diversity 105:1-16 ((https://doi.org/10.1007/s13225-020-00466-2)) |
| Otu58065 | 1 | k__Fungi(100);p__Ascomycota(100);c__Saccharomycetes(100);o__Saccharomycetales(100);f__Metschnikowiaceae(100);g__Metschnikowia(100);g__Metschnikowia_unclassified(100); | Metschnikowia | 13 | Saprotroph | \|Undefined Saprotroph\| | Yeast | NULL | Probable | As nectar-tap_saprotroph (Põlme et al. 2020); Decay sugar-rich substrates (Põlme et al. 2020) | Põlme S, et al. 2020. Fungal Diversity 105:1-16 ((https://doi.org/10.1007/s13225-020-00466-2)) |
| Otu58066 | 1 | k__Fungi(100);p__Ascomycota(100);c__Saccharomycetes(100);o__Saccharomycetales(100);f__Metschnikowiaceae(100);g__Metschnikowia(100);s__Metschnikowia_pulcherrima(100); | Metschnikowia | 13 | Saprotroph | \|Undefined Saprotroph\| | Yeast | NULL | Probable | As nectar-tap_saprotroph (Põlme et al. 2020); Decay sugar-rich substrates (Põlme et al. 2020) | Põlme S, et al. 2020. Fungal Diversity 105:1-16 ((https://doi.org/10.1007/s13225-020-00466-2)) |
| Otu58067 | 1 | k__Fungi(100);p__Ascomycota(100);c__Saccharomycetes(100);o__Saccharomycetales(100);f__Metschnikowiaceae(100);g__Metschnikowia(100);g__Metschnikowia_unclassified(100); | Metschnikowia | 13 | Saprotroph | \|Undefined Saprotroph\| | Yeast | NULL | Probable | As nectar-tap_saprotroph (Põlme et al. 2020); Decay sugar-rich substrates (Põlme et al. 2020) | Põlme S, et al. 2020. Fungal Diversity 105:1-16 ((https://doi.org/10.1007/s13225-020-00466-2)) |
| Otu58068 | 1 | k__Fungi(100);p__Ascomycota(100);c__Saccharomycetes(100);o__Saccharomycetales(100);f__Metschnikowiaceae(100);g__Metschnikowia(100);s__Metschnikowia_pulcherrima(100); | Metschnikowia | 13 | Saprotroph | \|Undefined Saprotroph\| | Yeast | NULL | Probable | As nectar-tap_saprotroph (Põlme et al. 2020); Decay sugar-rich substrates (Põlme et al. 2020) | Põlme S, et al. 2020. Fungal Diversity 105:1-16 ((https://doi.org/10.1007/s13225-020-00466-2)) |
| Otu58070 | 1 | k__Fungi(100);p__Ascomycota(100);c__Saccharomycetes(100);o__Saccharomycetales(100);f__Metschnikowiaceae(100);g__Metschnikowia(100);g__Metschnikowia_unclassified(100); | Metschnikowia | 13 | Saprotroph | \|Undefined Saprotroph\| | Yeast | NULL | Probable | As nectar-tap_saprotroph (Põlme et al. 2020); Decay sugar-rich substrates (Põlme et al. 2020) | Põlme S, et al. 2020. Fungal Diversity 105:1-16 ((https://doi.org/10.1007/s13225-020-00466-2)) |
| Otu58071 | 1 | k__Fungi(100);p__Ascomycota(100);c__Saccharomycetes(100);o__Saccharomycetales(100);f__Metschnikowiaceae(100);g__Metschnikowia(100);s__Metschnikowia_pulcherrima(100); | Metschnikowia | 13 | Saprotroph | \|Undefined Saprotroph\| | Yeast | NULL | Probable | As nectar-tap_saprotroph (Põlme et al. 2020); Decay sugar-rich substrates (Põlme et al. 2020) | Põlme S, et al. 2020. Fungal Diversity 105:1-16 ((https://doi.org/10.1007/s13225-020-00466-2)) |
| Otu58072 | 1 | k__Fungi(100);p__Ascomycota(100);c__Saccharomycetes(100);o__Saccharomycetales(100);f__Metschnikowiaceae(100);g__Metschnikowia(100);g__Metschnikowia_unclassified(100); | Metschnikowia | 13 | Saprotroph | \|Undefined Saprotroph\| | Yeast | NULL | Probable | As nectar-tap_saprotroph (Põlme et al. 2020); Decay sugar-rich substrates (Põlme et al. 2020) | Põlme S, et al. 2020. Fungal Diversity 105:1-16 ((https://doi.org/10.1007/s13225-020-00466-2)) |
| Otu58073 | 1 | k__Fungi(100);p__Ascomycota(100);c__Saccharomycetes(100);o__Saccharomycetales(100);f__Metschnikowiaceae(100);g__Metschnikowia(100);g__Metschnikowia_unclassified(100); | Metschnikowia | 13 | Saprotroph | \|Undefined Saprotroph\| | Yeast | NULL | Probable | As nectar-tap_saprotroph (Põlme et al. 2020); Decay sugar-rich substrates (Põlme et al. 2020) | Põlme S, et al. 2020. Fungal Diversity 105:1-16 ((https://doi.org/10.1007/s13225-020-00466-2)) |
| Otu58074 | 1 | k__Fungi(100);p__Ascomycota(100);c__Saccharomycetes(100);o__Saccharomycetales(100);f__Metschnikowiaceae(100);g__Metschnikowia(100);g__Metschnikowia_unclassified(100); | Metschnikowia | 13 | Saprotroph | \|Undefined Saprotroph\| | Yeast | NULL | Probable | As nectar-tap_saprotroph (Põlme et al. 2020); Decay sugar-rich substrates (Põlme et al. 2020) | Põlme S, et al. 2020. Fungal Diversity 105:1-16 ((https://doi.org/10.1007/s13225-020-00466-2)) |
| Otu58075 | 1 | k__Fungi(100);p__Ascomycota(100);c__Saccharomycetes(100);o__Saccharomycetales(100);f__Metschnikowiaceae(100);g__Metschnikowia(100);s__Metschnikowia_pulcherrima(100); | Metschnikowia | 13 | Saprotroph | \|Undefined Saprotroph\| | Yeast | NULL | Probable | As nectar-tap_saprotroph (Põlme et al. 2020); Decay sugar-rich substrates (Põlme et al. 2020) | Põlme S, et al. 2020. Fungal Diversity 105:1-16 ((https://doi.org/10.1007/s13225-020-00466-2)) |
| Otu58076 | 1 | k__Fungi(100);p__Ascomycota(100);c__Saccharomycetes(100);o__Saccharomycetales(100);f__Metschnikowiaceae(100);g__Metschnikowia(100);s__Metschnikowia_pulcherrima(100); | Metschnikowia | 13 | Saprotroph | \|Undefined Saprotroph\| | Yeast | NULL | Probable | As nectar-tap_saprotroph (Põlme et al. 2020); Decay sugar-rich substrates (Põlme et al. 2020) | Põlme S, et al. 2020. Fungal Diversity 105:1-16 ((https://doi.org/10.1007/s13225-020-00466-2)) |
| Otu58077 | 1 | k__Fungi(100);p__Ascomycota(100);c__Saccharomycetes(100);o__Saccharomycetales(100);f__Metschnikowiaceae(100);g__Metschnikowia(100);s__Metschnikowia_pulcherrima(100); | Metschnikowia | 13 | Saprotroph | \|Undefined Saprotroph\| | Yeast | NULL | Probable | As nectar-tap_saprotroph (Põlme et al. 2020); Decay sugar-rich substrates (Põlme et al. 2020) | Põlme S, et al. 2020. Fungal Diversity 105:1-16 ((https://doi.org/10.1007/s13225-020-00466-2)) |
| Otu58078 | 1 | k__Fungi(100);p__Ascomycota(100);c__Saccharomycetes(100);o__Saccharomycetales(100);f__Metschnikowiaceae(100);g__Metschnikowia(100);s__Metschnikowia_pulcherrima(100); | Metschnikowia | 13 | Saprotroph | \|Undefined Saprotroph\| | Yeast | NULL | Probable | As nectar-tap_saprotroph (Põlme et al. 2020); Decay sugar-rich substrates (Põlme et al. 2020) | Põlme S, et al. 2020. Fungal Diversity 105:1-16 ((https://doi.org/10.1007/s13225-020-00466-2)) |
| Otu58079 | 1 | k__Fungi(100);p__Ascomycota(100);c__Saccharomycetes(100);o__Saccharomycetales(100);f__Metschnikowiaceae(100);g__Metschnikowia(100);s__Metschnikowia_pulcherrima(100); | Metschnikowia | 13 | Saprotroph | \|Undefined Saprotroph\| | Yeast | NULL | Probable | As nectar-tap_saprotroph (Põlme et al. 2020); Decay sugar-rich substrates (Põlme et al. 2020) | Põlme S, et al. 2020. Fungal Diversity 105:1-16 ((https://doi.org/10.1007/s13225-020-00466-2)) |
| Otu58080 | 1 | k__Fungi(100);p__Ascomycota(100);c__Saccharomycetes(100);o__Saccharomycetales(100);f__Metschnikowiaceae(100);g__Metschnikowia(100);s__Metschnikowia_pulcherrima(100); | Metschnikowia | 13 | Saprotroph | \|Undefined Saprotroph\| | Yeast | NULL | Probable | As nectar-tap_saprotroph (Põlme et al. 2020); Decay sugar-rich substrates (Põlme et al. 2020) | Põlme S, et al. 2020. Fungal Diversity 105:1-16 ((https://doi.org/10.1007/s13225-020-00466-2)) |
| Otu58081 | 1 | k__Fungi(100);p__Ascomycota(100);c__Saccharomycetes(100);o__Saccharomycetales(100);f__Metschnikowiaceae(100);g__Metschnikowia(100);s__Metschnikowia_pulcherrima(100); | Metschnikowia | 13 | Saprotroph | \|Undefined Saprotroph\| | Yeast | NULL | Probable | As nectar-tap_saprotroph (Põlme et al. 2020); Decay sugar-rich substrates (Põlme et al. 2020) | Põlme S, et al. 2020. Fungal Diversity 105:1-16 ((https://doi.org/10.1007/s13225-020-00466-2)) |
| Otu58082 | 1 | k__Fungi(100);p__Ascomycota(100);c__Saccharomycetes(100);o__Saccharomycetales(100);f__Metschnikowiaceae(100);g__Metschnikowia(100);s__Metschnikowia_pulcherrima(100); | Metschnikowia | 13 | Saprotroph | \|Undefined Saprotroph\| | Yeast | NULL | Probable | As nectar-tap_saprotroph (Põlme et al. 2020); Decay sugar-rich substrates (Põlme et al. 2020) | Põlme S, et al. 2020. Fungal Diversity 105:1-16 ((https://doi.org/10.1007/s13225-020-00466-2)) |
| Otu58084 | 1 | k__Fungi(100);p__Ascomycota(100);c__Saccharomycetes(100);o__Saccharomycetales(100);f__Metschnikowiaceae(100);g__Metschnikowia(100);s__Metschnikowia_pulcherrima(100); | Metschnikowia | 13 | Saprotroph | \|Undefined Saprotroph\| | Yeast | NULL | Probable | As nectar-tap_saprotroph (Põlme et al. 2020); Decay sugar-rich substrates (Põlme et al. 2020) | Põlme S, et al. 2020. Fungal Diversity 105:1-16 ((https://doi.org/10.1007/s13225-020-00466-2)) |
| Otu58089 | 1 | k__Fungi(100);p__Ascomycota(100);c__Saccharomycetes(100);o__Saccharomycetales(100);f__Metschnikowiaceae(100);g__Metschnikowia(100);g__Metschnikowia_unclassified(100); | Metschnikowia | 13 | Saprotroph | \|Undefined Saprotroph\| | Yeast | NULL | Probable | As nectar-tap_saprotroph (Põlme et al. 2020); Decay sugar-rich substrates (Põlme et al. 2020) | Põlme S, et al. 2020. Fungal Diversity 105:1-16 ((https://doi.org/10.1007/s13225-020-00466-2)) |
| Otu58091 | 1 | k__Fungi(100);p__Ascomycota(100);c__Saccharomycetes(100);o__Saccharomycetales(100);f__Metschnikowiaceae(100);g__Metschnikowia(100);g__Metschnikowia_unclassified(100); | Metschnikowia | 13 | Saprotroph | \|Undefined Saprotroph\| | Yeast | NULL | Probable | As nectar-tap_saprotroph (Põlme et al. 2020); Decay sugar-rich substrates (Põlme et al. 2020) | Põlme S, et al. 2020. Fungal Diversity 105:1-16 ((https://doi.org/10.1007/s13225-020-00466-2)) |
| Otu58092 | 1 | k__Fungi(100);p__Ascomycota(100);c__Saccharomycetes(100);o__Saccharomycetales(100);f__Metschnikowiaceae(100);g__Metschnikowia(100);g__Metschnikowia_unclassified(100); | Metschnikowia | 13 | Saprotroph | \|Undefined Saprotroph\| | Yeast | NULL | Probable | As nectar-tap_saprotroph (Põlme et al. 2020); Decay sugar-rich substrates (Põlme et al. 2020) | Põlme S, et al. 2020. Fungal Diversity 105:1-16 ((https://doi.org/10.1007/s13225-020-00466-2)) |
| Otu58094 | 1 | k__Fungi(100);p__Ascomycota(100);c__Saccharomycetes(100);o__Saccharomycetales(100);f__Metschnikowiaceae(100);g__Metschnikowia(100);g__Metschnikowia_unclassified(100); | Metschnikowia | 13 | Saprotroph | \|Undefined Saprotroph\| | Yeast | NULL | Probable | As nectar-tap_saprotroph (Põlme et al. 2020); Decay sugar-rich substrates (Põlme et al. 2020) | Põlme S, et al. 2020. Fungal Diversity 105:1-16 ((https://doi.org/10.1007/s13225-020-00466-2)) |
| Otu58095 | 1 | k__Fungi(100);p__Ascomycota(100);c__Saccharomycetes(100);o__Saccharomycetales(100);f__Metschnikowiaceae(100);g__Metschnikowia(100);s__Metschnikowia_pulcherrima(100); | Metschnikowia | 13 | Saprotroph | \|Undefined Saprotroph\| | Yeast | NULL | Probable | As nectar-tap_saprotroph (Põlme et al. 2020); Decay sugar-rich substrates (Põlme et al. 2020) | Põlme S, et al. 2020. Fungal Diversity 105:1-16 ((https://doi.org/10.1007/s13225-020-00466-2)) |
| Otu58096 | 1 | k__Fungi(100);p__Ascomycota(100);c__Saccharomycetes(100);o__Saccharomycetales(100);f__Metschnikowiaceae(100);g__Metschnikowia(100);g__Metschnikowia_unclassified(100); | Metschnikowia | 13 | Saprotroph | \|Undefined Saprotroph\| | Yeast | NULL | Probable | As nectar-tap_saprotroph (Põlme et al. 2020); Decay sugar-rich substrates (Põlme et al. 2020) | Põlme S, et al. 2020. Fungal Diversity 105:1-16 ((https://doi.org/10.1007/s13225-020-00466-2)) |
| Otu58097 | 1 | k__Fungi(100);p__Ascomycota(100);c__Saccharomycetes(100);o__Saccharomycetales(100);f__Metschnikowiaceae(100);g__Metschnikowia(100);s__Metschnikowia_pulcherrima(100); | Metschnikowia | 13 | Saprotroph | \|Undefined Saprotroph\| | Yeast | NULL | Probable | As nectar-tap_saprotroph (Põlme et al. 2020); Decay sugar-rich substrates (Põlme et al. 2020) | Põlme S, et al. 2020. Fungal Diversity 105:1-16 ((https://doi.org/10.1007/s13225-020-00466-2)) |
| Otu58098 | 1 | k__Fungi(100);p__Ascomycota(100);c__Saccharomycetes(100);o__Saccharomycetales(100);f__Metschnikowiaceae(100);g__Metschnikowia(100);g__Metschnikowia_unclassified(100); | Metschnikowia | 13 | Saprotroph | \|Undefined Saprotroph\| | Yeast | NULL | Probable | As nectar-tap_saprotroph (Põlme et al. 2020); Decay sugar-rich substrates (Põlme et al. 2020) | Põlme S, et al. 2020. Fungal Diversity 105:1-16 ((https://doi.org/10.1007/s13225-020-00466-2)) |
| Otu58099 | 1 | k__Fungi(100);p__Ascomycota(100);c__Saccharomycetes(100);o__Saccharomycetales(100);f__Metschnikowiaceae(100);g__Metschnikowia(100);s__Metschnikowia_pulcherrima(100); | Metschnikowia | 13 | Saprotroph | \|Undefined Saprotroph\| | Yeast | NULL | Probable | As nectar-tap_saprotroph (Põlme et al. 2020); Decay sugar-rich substrates (Põlme et al. 2020) | Põlme S, et al. 2020. Fungal Diversity 105:1-16 ((https://doi.org/10.1007/s13225-020-00466-2)) |
| Otu58100 | 1 | k__Fungi(100);p__Ascomycota(100);c__Saccharomycetes(100);o__Saccharomycetales(100);f__Metschnikowiaceae(100);g__Metschnikowia(100);s__Metschnikowia_pulcherrima(100); | Metschnikowia | 13 | Saprotroph | \|Undefined Saprotroph\| | Yeast | NULL | Probable | As nectar-tap_saprotroph (Põlme et al. 2020); Decay sugar-rich substrates (Põlme et al. 2020) | Põlme S, et al. 2020. Fungal Diversity 105:1-16 ((https://doi.org/10.1007/s13225-020-00466-2)) |
| Otu58101 | 1 | k__Fungi(100);p__Ascomycota(100);c__Saccharomycetes(100);o__Saccharomycetales(100);f__Metschnikowiaceae(100);g__Metschnikowia(100);s__Metschnikowia_pulcherrima(100); | Metschnikowia | 13 | Saprotroph | \|Undefined Saprotroph\| | Yeast | NULL | Probable | As nectar-tap_saprotroph (Põlme et al. 2020); Decay sugar-rich substrates (Põlme et al. 2020) | Põlme S, et al. 2020. Fungal Diversity 105:1-16 ((https://doi.org/10.1007/s13225-020-00466-2)) |
| Otu58103 | 1 | k__Fungi(100);p__Ascomycota(100);c__Saccharomycetes(100);o__Saccharomycetales(100);f__Metschnikowiaceae(100);g__Metschnikowia(100);s__Metschnikowia_pulcherrima(100); | Metschnikowia | 13 | Saprotroph | \|Undefined Saprotroph\| | Yeast | NULL | Probable | As nectar-tap_saprotroph (Põlme et al. 2020); Decay sugar-rich substrates (Põlme et al. 2020) | Põlme S, et al. 2020. Fungal Diversity 105:1-16 ((https://doi.org/10.1007/s13225-020-00466-2)) |
| Otu58104 | 1 | k__Fungi(100);p__Ascomycota(100);c__Saccharomycetes(100);o__Saccharomycetales(100);f__Metschnikowiaceae(100);g__Metschnikowia(100);g__Metschnikowia_unclassified(100); | Metschnikowia | 13 | Saprotroph | \|Undefined Saprotroph\| | Yeast | NULL | Probable | As nectar-tap_saprotroph (Põlme et al. 2020); Decay sugar-rich substrates (Põlme et al. 2020) | Põlme S, et al. 2020. Fungal Diversity 105:1-16 ((https://doi.org/10.1007/s13225-020-00466-2)) |
| Otu58105 | 1 | k__Fungi(100);p__Ascomycota(100);c__Saccharomycetes(100);o__Saccharomycetales(100);f__Metschnikowiaceae(100);g__Metschnikowia(100);s__Metschnikowia_pulcherrima(100); | Metschnikowia | 13 | Saprotroph | \|Undefined Saprotroph\| | Yeast | NULL | Probable | As nectar-tap_saprotroph (Põlme et al. 2020); Decay sugar-rich substrates (Põlme et al. 2020) | Põlme S, et al. 2020. Fungal Diversity 105:1-16 ((https://doi.org/10.1007/s13225-020-00466-2)) |
| Otu58106 | 1 | k__Fungi(100);p__Ascomycota(100);c__Saccharomycetes(100);o__Saccharomycetales(100);f__Metschnikowiaceae(100);g__Metschnikowia(100);s__Metschnikowia_pulcherrima(100); | Metschnikowia | 13 | Saprotroph | \|Undefined Saprotroph\| | Yeast | NULL | Probable | As nectar-tap_saprotroph (Põlme et al. 2020); Decay sugar-rich substrates (Põlme et al. 2020) | Põlme S, et al. 2020. Fungal Diversity 105:1-16 ((https://doi.org/10.1007/s13225-020-00466-2)) |
| Otu58107 | 1 | k__Fungi(100);p__Ascomycota(100);c__Saccharomycetes(100);o__Saccharomycetales(100);f__Metschnikowiaceae(100);g__Metschnikowia(100);s__Metschnikowia_pulcherrima(100); | Metschnikowia | 13 | Saprotroph | \|Undefined Saprotroph\| | Yeast | NULL | Probable | As nectar-tap_saprotroph (Põlme et al. 2020); Decay sugar-rich substrates (Põlme et al. 2020) | Põlme S, et al. 2020. Fungal Diversity 105:1-16 ((https://doi.org/10.1007/s13225-020-00466-2)) |
| Otu58108 | 1 | k__Fungi(100);p__Ascomycota(100);c__Saccharomycetes(100);o__Saccharomycetales(100);f__Metschnikowiaceae(100);g__Metschnikowia(100);g__Metschnikowia_unclassified(100); | Metschnikowia | 13 | Saprotroph | \|Undefined Saprotroph\| | Yeast | NULL | Probable | As nectar-tap_saprotroph (Põlme et al. 2020); Decay sugar-rich substrates (Põlme et al. 2020) | Põlme S, et al. 2020. Fungal Diversity 105:1-16 ((https://doi.org/10.1007/s13225-020-00466-2)) |
| Otu58109 | 1 | k__Fungi(100);p__Ascomycota(100);c__Saccharomycetes(100);o__Saccharomycetales(100);f__Metschnikowiaceae(100);g__Metschnikowia(100);s__Metschnikowia_pulcherrima(100); | Metschnikowia | 13 | Saprotroph | \|Undefined Saprotroph\| | Yeast | NULL | Probable | As nectar-tap_saprotroph (Põlme et al. 2020); Decay sugar-rich substrates (Põlme et al. 2020) | Põlme S, et al. 2020. Fungal Diversity 105:1-16 ((https://doi.org/10.1007/s13225-020-00466-2)) |
| Otu58111 | 1 | k__Fungi(100);p__Ascomycota(100);c__Saccharomycetes(100);o__Saccharomycetales(100);f__Metschnikowiaceae(100);g__Metschnikowia(100);s__Metschnikowia_pulcherrima(100); | Metschnikowia | 13 | Saprotroph | \|Undefined Saprotroph\| | Yeast | NULL | Probable | As nectar-tap_saprotroph (Põlme et al. 2020); Decay sugar-rich substrates (Põlme et al. 2020) | Põlme S, et al. 2020. Fungal Diversity 105:1-16 ((https://doi.org/10.1007/s13225-020-00466-2)) |
| Otu58112 | 1 | k__Fungi(100);p__Ascomycota(100);c__Saccharomycetes(100);o__Saccharomycetales(100);f__Metschnikowiaceae(100);g__Metschnikowia(100);g__Metschnikowia_unclassified(100); | Metschnikowia | 13 | Saprotroph | \|Undefined Saprotroph\| | Yeast | NULL | Probable | As nectar-tap_saprotroph (Põlme et al. 2020); Decay sugar-rich substrates (Põlme et al. 2020) | Põlme S, et al. 2020. Fungal Diversity 105:1-16 ((https://doi.org/10.1007/s13225-020-00466-2)) |
| Otu58113 | 1 | k__Fungi(100);p__Ascomycota(100);c__Saccharomycetes(100);o__Saccharomycetales(100);f__Metschnikowiaceae(100);g__Metschnikowia(100);s__Metschnikowia_pulcherrima(100); | Metschnikowia | 13 | Saprotroph | \|Undefined Saprotroph\| | Yeast | NULL | Probable | As nectar-tap_saprotroph (Põlme et al. 2020); Decay sugar-rich substrates (Põlme et al. 2020) | Põlme S, et al. 2020. Fungal Diversity 105:1-16 ((https://doi.org/10.1007/s13225-020-00466-2)) |
| Otu58114 | 1 | k__Fungi(100);p__Ascomycota(100);c__Saccharomycetes(100);o__Saccharomycetales(100);f__Metschnikowiaceae(100);g__Metschnikowia(100);s__Metschnikowia_pulcherrima(100); | Metschnikowia | 13 | Saprotroph | \|Undefined Saprotroph\| | Yeast | NULL | Probable | As nectar-tap_saprotroph (Põlme et al. 2020); Decay sugar-rich substrates (Põlme et al. 2020) | Põlme S, et al. 2020. Fungal Diversity 105:1-16 ((https://doi.org/10.1007/s13225-020-00466-2)) |
| Otu58115 | 1 | k__Fungi(100);p__Ascomycota(100);c__Saccharomycetes(100);o__Saccharomycetales(100);f__Metschnikowiaceae(100);g__Metschnikowia(100);s__Metschnikowia_pulcherrima(100); | Metschnikowia | 13 | Saprotroph | \|Undefined Saprotroph\| | Yeast | NULL | Probable | As nectar-tap_saprotroph (Põlme et al. 2020); Decay sugar-rich substrates (Põlme et al. 2020) | Põlme S, et al. 2020. Fungal Diversity 105:1-16 ((https://doi.org/10.1007/s13225-020-00466-2)) |
| Otu58118 | 1 | k__Fungi(100);p__Ascomycota(100);c__Saccharomycetes(100);o__Saccharomycetales(100);f__Metschnikowiaceae(100);g__Metschnikowia(100);s__Metschnikowia_pulcherrima(100); | Metschnikowia | 13 | Saprotroph | \|Undefined Saprotroph\| | Yeast | NULL | Probable | As nectar-tap_saprotroph (Põlme et al. 2020); Decay sugar-rich substrates (Põlme et al. 2020) | Põlme S, et al. 2020. Fungal Diversity 105:1-16 ((https://doi.org/10.1007/s13225-020-00466-2)) |
| Otu58120 | 1 | k__Fungi(100);p__Ascomycota(100);c__Saccharomycetes(100);o__Saccharomycetales(100);f__Metschnikowiaceae(100);g__Metschnikowia(100);s__Metschnikowia_pulcherrima(100); | Metschnikowia | 13 | Saprotroph | \|Undefined Saprotroph\| | Yeast | NULL | Probable | As nectar-tap_saprotroph (Põlme et al. 2020); Decay sugar-rich substrates (Põlme et al. 2020) | Põlme S, et al. 2020. Fungal Diversity 105:1-16 ((https://doi.org/10.1007/s13225-020-00466-2)) |
| Otu58121 | 1 | k__Fungi(100);p__Ascomycota(100);c__Saccharomycetes(100);o__Saccharomycetales(100);f__Metschnikowiaceae(100);g__Metschnikowia(100);s__Metschnikowia_pulcherrima(100); | Metschnikowia | 13 | Saprotroph | \|Undefined Saprotroph\| | Yeast | NULL | Probable | As nectar-tap_saprotroph (Põlme et al. 2020); Decay sugar-rich substrates (Põlme et al. 2020) | Põlme S, et al. 2020. Fungal Diversity 105:1-16 ((https://doi.org/10.1007/s13225-020-00466-2)) |
| Otu58122 | 1 | k__Fungi(100);p__Ascomycota(100);c__Saccharomycetes(100);o__Saccharomycetales(100);f__Metschnikowiaceae(100);g__Metschnikowia(100);s__Metschnikowia_pulcherrima(100); | Metschnikowia | 13 | Saprotroph | \|Undefined Saprotroph\| | Yeast | NULL | Probable | As nectar-tap_saprotroph (Põlme et al. 2020); Decay sugar-rich substrates (Põlme et al. 2020) | Põlme S, et al. 2020. Fungal Diversity 105:1-16 ((https://doi.org/10.1007/s13225-020-00466-2)) |
| Otu58123 | 1 | k__Fungi(100);p__Ascomycota(100);c__Saccharomycetes(100);o__Saccharomycetales(100);f__Metschnikowiaceae(100);g__Metschnikowia(100);s__Metschnikowia_pulcherrima(100); | Metschnikowia | 13 | Saprotroph | \|Undefined Saprotroph\| | Yeast | NULL | Probable | As nectar-tap_saprotroph (Põlme et al. 2020); Decay sugar-rich substrates (Põlme et al. 2020) | Põlme S, et al. 2020. Fungal Diversity 105:1-16 ((https://doi.org/10.1007/s13225-020-00466-2)) |
| Otu58124 | 1 | k__Fungi(100);p__Ascomycota(100);c__Saccharomycetes(100);o__Saccharomycetales(100);f__Metschnikowiaceae(100);g__Metschnikowia(100);g__Metschnikowia_unclassified(100); | Metschnikowia | 13 | Saprotroph | \|Undefined Saprotroph\| | Yeast | NULL | Probable | As nectar-tap_saprotroph (Põlme et al. 2020); Decay sugar-rich substrates (Põlme et al. 2020) | Põlme S, et al. 2020. Fungal Diversity 105:1-16 ((https://doi.org/10.1007/s13225-020-00466-2)) |
| Otu58125 | 1 | k__Fungi(100);p__Ascomycota(100);c__Saccharomycetes(100);o__Saccharomycetales(100);f__Metschnikowiaceae(100);g__Metschnikowia(100);s__Metschnikowia_pulcherrima(100); | Metschnikowia | 13 | Saprotroph | \|Undefined Saprotroph\| | Yeast | NULL | Probable | As nectar-tap_saprotroph (Põlme et al. 2020); Decay sugar-rich substrates (Põlme et al. 2020) | Põlme S, et al. 2020. Fungal Diversity 105:1-16 ((https://doi.org/10.1007/s13225-020-00466-2)) |
| Otu58126 | 1 | k__Fungi(100);p__Ascomycota(100);c__Saccharomycetes(100);o__Saccharomycetales(100);f__Metschnikowiaceae(100);g__Metschnikowia(100);s__Metschnikowia_pulcherrima(100); | Metschnikowia | 13 | Saprotroph | \|Undefined Saprotroph\| | Yeast | NULL | Probable | As nectar-tap_saprotroph (Põlme et al. 2020); Decay sugar-rich substrates (Põlme et al. 2020) | Põlme S, et al. 2020. Fungal Diversity 105:1-16 ((https://doi.org/10.1007/s13225-020-00466-2)) |
| Otu58128 | 1 | k__Fungi(100);p__Ascomycota(100);c__Saccharomycetes(100);o__Saccharomycetales(100);f__Metschnikowiaceae(100);g__Metschnikowia(100);s__Metschnikowia_pulcherrima(100); | Metschnikowia | 13 | Saprotroph | \|Undefined Saprotroph\| | Yeast | NULL | Probable | As nectar-tap_saprotroph (Põlme et al. 2020); Decay sugar-rich substrates (Põlme et al. 2020) | Põlme S, et al. 2020. Fungal Diversity 105:1-16 ((https://doi.org/10.1007/s13225-020-00466-2)) |
| Otu58129 | 1 | k__Fungi(100);p__Ascomycota(100);c__Saccharomycetes(100);o__Saccharomycetales(100);f__Metschnikowiaceae(100);g__Metschnikowia(100);s__Metschnikowia_pulcherrima(100); | Metschnikowia | 13 | Saprotroph | \|Undefined Saprotroph\| | Yeast | NULL | Probable | As nectar-tap_saprotroph (Põlme et al. 2020); Decay sugar-rich substrates (Põlme et al. 2020) | Põlme S, et al. 2020. Fungal Diversity 105:1-16 ((https://doi.org/10.1007/s13225-020-00466-2)) |
| Otu58131 | 1 | k__Fungi(100);p__Ascomycota(100);c__Saccharomycetes(100);o__Saccharomycetales(100);f__Metschnikowiaceae(100);g__Metschnikowia(100);s__Metschnikowia_pulcherrima(100); | Metschnikowia | 13 | Saprotroph | \|Undefined Saprotroph\| | Yeast | NULL | Probable | As nectar-tap_saprotroph (Põlme et al. 2020); Decay sugar-rich substrates (Põlme et al. 2020) | Põlme S, et al. 2020. Fungal Diversity 105:1-16 ((https://doi.org/10.1007/s13225-020-00466-2)) |
| Otu58132 | 1 | k__Fungi(100);p__Ascomycota(100);c__Saccharomycetes(100);o__Saccharomycetales(100);f__Metschnikowiaceae(100);g__Metschnikowia(100);s__Metschnikowia_pulcherrima(100); | Metschnikowia | 13 | Saprotroph | \|Undefined Saprotroph\| | Yeast | NULL | Probable | As nectar-tap_saprotroph (Põlme et al. 2020); Decay sugar-rich substrates (Põlme et al. 2020) | Põlme S, et al. 2020. Fungal Diversity 105:1-16 ((https://doi.org/10.1007/s13225-020-00466-2)) |
| Otu58133 | 1 | k__Fungi(100);p__Ascomycota(100);c__Saccharomycetes(100);o__Saccharomycetales(100);f__Metschnikowiaceae(100);g__Metschnikowia(100);s__Metschnikowia_pulcherrima(100); | Metschnikowia | 13 | Saprotroph | \|Undefined Saprotroph\| | Yeast | NULL | Probable | As nectar-tap_saprotroph (Põlme et al. 2020); Decay sugar-rich substrates (Põlme et al. 2020) | Põlme S, et al. 2020. Fungal Diversity 105:1-16 ((https://doi.org/10.1007/s13225-020-00466-2)) |
| Otu58134 | 1 | k__Fungi(100);p__Ascomycota(100);c__Saccharomycetes(100);o__Saccharomycetales(100);f__Metschnikowiaceae(100);g__Metschnikowia(100);s__Metschnikowia_pulcherrima(100); | Metschnikowia | 13 | Saprotroph | \|Undefined Saprotroph\| | Yeast | NULL | Probable | As nectar-tap_saprotroph (Põlme et al. 2020); Decay sugar-rich substrates (Põlme et al. 2020) | Põlme S, et al. 2020. Fungal Diversity 105:1-16 ((https://doi.org/10.1007/s13225-020-00466-2)) |
| Otu58135 | 1 | k__Fungi(100);p__Ascomycota(100);c__Saccharomycetes(100);o__Saccharomycetales(100);f__Metschnikowiaceae(100);g__Metschnikowia(100);s__Metschnikowia_pulcherrima(100); | Metschnikowia | 13 | Saprotroph | \|Undefined Saprotroph\| | Yeast | NULL | Probable | As nectar-tap_saprotroph (Põlme et al. 2020); Decay sugar-rich substrates (Põlme et al. 2020) | Põlme S, et al. 2020. Fungal Diversity 105:1-16 ((https://doi.org/10.1007/s13225-020-00466-2)) |
| Otu58136 | 1 | k__Fungi(100);p__Ascomycota(100);c__Saccharomycetes(100);o__Saccharomycetales(100);f__Metschnikowiaceae(100);g__Metschnikowia(100);s__Metschnikowia_pulcherrima(100); | Metschnikowia | 13 | Saprotroph | \|Undefined Saprotroph\| | Yeast | NULL | Probable | As nectar-tap_saprotroph (Põlme et al. 2020); Decay sugar-rich substrates (Põlme et al. 2020) | Põlme S, et al. 2020. Fungal Diversity 105:1-16 ((https://doi.org/10.1007/s13225-020-00466-2)) |
| Otu58137 | 1 | k__Fungi(100);p__Ascomycota(100);c__Saccharomycetes(100);o__Saccharomycetales(100);f__Metschnikowiaceae(100);g__Metschnikowia(100);s__Metschnikowia_pulcherrima(100); | Metschnikowia | 13 | Saprotroph | \|Undefined Saprotroph\| | Yeast | NULL | Probable | As nectar-tap_saprotroph (Põlme et al. 2020); Decay sugar-rich substrates (Põlme et al. 2020) | Põlme S, et al. 2020. Fungal Diversity 105:1-16 ((https://doi.org/10.1007/s13225-020-00466-2)) |
| Otu58138 | 1 | k__Fungi(100);p__Ascomycota(100);c__Saccharomycetes(100);o__Saccharomycetales(100);f__Metschnikowiaceae(100);g__Metschnikowia(100);s__Metschnikowia_pulcherrima(100); | Metschnikowia | 13 | Saprotroph | \|Undefined Saprotroph\| | Yeast | NULL | Probable | As nectar-tap_saprotroph (Põlme et al. 2020); Decay sugar-rich substrates (Põlme et al. 2020) | Põlme S, et al. 2020. Fungal Diversity 105:1-16 ((https://doi.org/10.1007/s13225-020-00466-2)) |
| Otu58139 | 1 | k__Fungi(100);p__Ascomycota(100);c__Saccharomycetes(100);o__Saccharomycetales(100);f__Metschnikowiaceae(100);g__Metschnikowia(100);s__Metschnikowia_pulcherrima(100); | Metschnikowia | 13 | Saprotroph | \|Undefined Saprotroph\| | Yeast | NULL | Probable | As nectar-tap_saprotroph (Põlme et al. 2020); Decay sugar-rich substrates (Põlme et al. 2020) | Põlme S, et al. 2020. Fungal Diversity 105:1-16 ((https://doi.org/10.1007/s13225-020-00466-2)) |
| Otu58143 | 1 | k__Fungi(100);p__Ascomycota(100);c__Saccharomycetes(100);o__Saccharomycetales(100);f__Metschnikowiaceae(100);g__Metschnikowia(100);s__Metschnikowia_pulcherrima(100); | Metschnikowia | 13 | Saprotroph | \|Undefined Saprotroph\| | Yeast | NULL | Probable | As nectar-tap_saprotroph (Põlme et al. 2020); Decay sugar-rich substrates (Põlme et al. 2020) | Põlme S, et al. 2020. Fungal Diversity 105:1-16 ((https://doi.org/10.1007/s13225-020-00466-2)) |
| Otu58144 | 1 | k__Fungi(100);p__Ascomycota(100);c__Saccharomycetes(100);o__Saccharomycetales(100);f__Metschnikowiaceae(100);g__Metschnikowia(100);s__Metschnikowia_pulcherrima(100); | Metschnikowia | 13 | Saprotroph | \|Undefined Saprotroph\| | Yeast | NULL | Probable | As nectar-tap_saprotroph (Põlme et al. 2020); Decay sugar-rich substrates (Põlme et al. 2020) | Põlme S, et al. 2020. Fungal Diversity 105:1-16 ((https://doi.org/10.1007/s13225-020-00466-2)) |
| Otu58145 | 1 | k__Fungi(100);p__Ascomycota(100);c__Saccharomycetes(100);o__Saccharomycetales(100);f__Metschnikowiaceae(100);g__Metschnikowia(100);s__Metschnikowia_pulcherrima(100); | Metschnikowia | 13 | Saprotroph | \|Undefined Saprotroph\| | Yeast | NULL | Probable | As nectar-tap_saprotroph (Põlme et al. 2020); Decay sugar-rich substrates (Põlme et al. 2020) | Põlme S, et al. 2020. Fungal Diversity 105:1-16 ((https://doi.org/10.1007/s13225-020-00466-2)) |
| Otu58146 | 1 | k__Fungi(100);p__Ascomycota(100);c__Saccharomycetes(100);o__Saccharomycetales(100);f__Metschnikowiaceae(100);g__Metschnikowia(100);g__Metschnikowia_unclassified(100); | Metschnikowia | 13 | Saprotroph | \|Undefined Saprotroph\| | Yeast | NULL | Probable | As nectar-tap_saprotroph (Põlme et al. 2020); Decay sugar-rich substrates (Põlme et al. 2020) | Põlme S, et al. 2020. Fungal Diversity 105:1-16 ((https://doi.org/10.1007/s13225-020-00466-2)) |
| Otu58147 | 1 | k__Fungi(100);p__Ascomycota(100);c__Saccharomycetes(100);o__Saccharomycetales(100);f__Metschnikowiaceae(100);g__Metschnikowia(100);s__Metschnikowia_pulcherrima(100); | Metschnikowia | 13 | Saprotroph | \|Undefined Saprotroph\| | Yeast | NULL | Probable | As nectar-tap_saprotroph (Põlme et al. 2020); Decay sugar-rich substrates (Põlme et al. 2020) | Põlme S, et al. 2020. Fungal Diversity 105:1-16 ((https://doi.org/10.1007/s13225-020-00466-2)) |
| Otu58148 | 1 | k__Fungi(100);p__Ascomycota(100);c__Saccharomycetes(100);o__Saccharomycetales(100);f__Metschnikowiaceae(100);g__Metschnikowia(100);g__Metschnikowia_unclassified(100); | Metschnikowia | 13 | Saprotroph | \|Undefined Saprotroph\| | Yeast | NULL | Probable | As nectar-tap_saprotroph (Põlme et al. 2020); Decay sugar-rich substrates (Põlme et al. 2020) | Põlme S, et al. 2020. Fungal Diversity 105:1-16 ((https://doi.org/10.1007/s13225-020-00466-2)) |
| Otu58150 | 1 | k__Fungi(100);p__Ascomycota(100);c__Saccharomycetes(100);o__Saccharomycetales(100);f__Metschnikowiaceae(100);g__Metschnikowia(100);s__Metschnikowia_pulcherrima(100); | Metschnikowia | 13 | Saprotroph | \|Undefined Saprotroph\| | Yeast | NULL | Probable | As nectar-tap_saprotroph (Põlme et al. 2020); Decay sugar-rich substrates (Põlme et al. 2020) | Põlme S, et al. 2020. Fungal Diversity 105:1-16 ((https://doi.org/10.1007/s13225-020-00466-2)) |
| Otu58151 | 1 | k__Fungi(100);p__Ascomycota(100);c__Saccharomycetes(100);o__Saccharomycetales(100);f__Metschnikowiaceae(100);g__Metschnikowia(100);s__Metschnikowia_pulcherrima(100); | Metschnikowia | 13 | Saprotroph | \|Undefined Saprotroph\| | Yeast | NULL | Probable | As nectar-tap_saprotroph (Põlme et al. 2020); Decay sugar-rich substrates (Põlme et al. 2020) | Põlme S, et al. 2020. Fungal Diversity 105:1-16 ((https://doi.org/10.1007/s13225-020-00466-2)) |
| Otu58153 | 1 | k__Fungi(100);p__Ascomycota(100);c__Saccharomycetes(100);o__Saccharomycetales(100);f__Metschnikowiaceae(100);g__Metschnikowia(100);s__Metschnikowia_pulcherrima(100); | Metschnikowia | 13 | Saprotroph | \|Undefined Saprotroph\| | Yeast | NULL | Probable | As nectar-tap_saprotroph (Põlme et al. 2020); Decay sugar-rich substrates (Põlme et al. 2020) | Põlme S, et al. 2020. Fungal Diversity 105:1-16 ((https://doi.org/10.1007/s13225-020-00466-2)) |
| Otu58155 | 1 | k__Fungi(100);p__Ascomycota(100);c__Saccharomycetes(100);o__Saccharomycetales(100);f__Metschnikowiaceae(100);g__Metschnikowia(100);s__Metschnikowia_pulcherrima(100); | Metschnikowia | 13 | Saprotroph | \|Undefined Saprotroph\| | Yeast | NULL | Probable | As nectar-tap_saprotroph (Põlme et al. 2020); Decay sugar-rich substrates (Põlme et al. 2020) | Põlme S, et al. 2020. Fungal Diversity 105:1-16 ((https://doi.org/10.1007/s13225-020-00466-2)) |
| Otu58156 | 1 | k__Fungi(100);p__Ascomycota(100);c__Saccharomycetes(100);o__Saccharomycetales(100);f__Metschnikowiaceae(100);g__Metschnikowia(100);s__Metschnikowia_pulcherrima(100); | Metschnikowia | 13 | Saprotroph | \|Undefined Saprotroph\| | Yeast | NULL | Probable | As nectar-tap_saprotroph (Põlme et al. 2020); Decay sugar-rich substrates (Põlme et al. 2020) | Põlme S, et al. 2020. Fungal Diversity 105:1-16 ((https://doi.org/10.1007/s13225-020-00466-2)) |
| Otu58157 | 1 | k__Fungi(100);p__Ascomycota(100);c__Saccharomycetes(100);o__Saccharomycetales(100);f__Saccharomycetaceae(100);g__Kazachstania(100);s__Kazachstania_intestinalis(100); | Kazachstania | 13 | Saprotroph | \|Undefined Saprotroph\| | Yeast | NULL | Probable | As nectar-tap_saprotroph (Põlme et al. 2020); Decay sugar-rich substrates (Põlme et al. 2020) | Põlme S, et al. 2020. Fungal Diversity 105:1-16 ((https://doi.org/10.1007/s13225-020-00466-2)) |
| Otu58160 | 1 | k__Fungi(100);p__Ascomycota(100);c__Saccharomycetes(100);o__Saccharomycetales(100);f__Metschnikowiaceae(100);g__Metschnikowia(100);s__Metschnikowia_pulcherrima(100); | Metschnikowia | 13 | Saprotroph | \|Undefined Saprotroph\| | Yeast | NULL | Probable | As nectar-tap_saprotroph (Põlme et al. 2020); Decay sugar-rich substrates (Põlme et al. 2020) | Põlme S, et al. 2020. Fungal Diversity 105:1-16 ((https://doi.org/10.1007/s13225-020-00466-2)) |
| Otu58161 | 1 | k__Fungi(100);p__Ascomycota(100);c__Saccharomycetes(100);o__Saccharomycetales(100);f__Metschnikowiaceae(100);g__Metschnikowia(100);s__Metschnikowia_pulcherrima(100); | Metschnikowia | 13 | Saprotroph | \|Undefined Saprotroph\| | Yeast | NULL | Probable | As nectar-tap_saprotroph (Põlme et al. 2020); Decay sugar-rich substrates (Põlme et al. 2020) | Põlme S, et al. 2020. Fungal Diversity 105:1-16 ((https://doi.org/10.1007/s13225-020-00466-2)) |
| Otu58162 | 1 | k__Fungi(100);p__Ascomycota(100);c__Saccharomycetes(100);o__Saccharomycetales(100);f__Metschnikowiaceae(100);g__Metschnikowia(100);s__Metschnikowia_pulcherrima(100); | Metschnikowia | 13 | Saprotroph | \|Undefined Saprotroph\| | Yeast | NULL | Probable | As nectar-tap_saprotroph (Põlme et al. 2020); Decay sugar-rich substrates (Põlme et al. 2020) | Põlme S, et al. 2020. Fungal Diversity 105:1-16 ((https://doi.org/10.1007/s13225-020-00466-2)) |
| Otu58164 | 1 | k__Fungi(100);p__Ascomycota(100);c__Saccharomycetes(100);o__Saccharomycetales(100);f__Metschnikowiaceae(100);g__Metschnikowia(100);s__Metschnikowia_pulcherrima(100); | Metschnikowia | 13 | Saprotroph | \|Undefined Saprotroph\| | Yeast | NULL | Probable | As nectar-tap_saprotroph (Põlme et al. 2020); Decay sugar-rich substrates (Põlme et al. 2020) | Põlme S, et al. 2020. Fungal Diversity 105:1-16 ((https://doi.org/10.1007/s13225-020-00466-2)) |
| Otu58165 | 1 | k__Fungi(100);p__Ascomycota(100);c__Saccharomycetes(100);o__Saccharomycetales(100);f__Metschnikowiaceae(100);g__Metschnikowia(100);s__Metschnikowia_pulcherrima(100); | Metschnikowia | 13 | Saprotroph | \|Undefined Saprotroph\| | Yeast | NULL | Probable | As nectar-tap_saprotroph (Põlme et al. 2020); Decay sugar-rich substrates (Põlme et al. 2020) | Põlme S, et al. 2020. Fungal Diversity 105:1-16 ((https://doi.org/10.1007/s13225-020-00466-2)) |
| Otu58167 | 1 | k__Fungi(100);p__Ascomycota(100);c__Saccharomycetes(100);o__Saccharomycetales(100);f__Metschnikowiaceae(100);g__Metschnikowia(100);s__Metschnikowia_pulcherrima(100); | Metschnikowia | 13 | Saprotroph | \|Undefined Saprotroph\| | Yeast | NULL | Probable | As nectar-tap_saprotroph (Põlme et al. 2020); Decay sugar-rich substrates (Põlme et al. 2020) | Põlme S, et al. 2020. Fungal Diversity 105:1-16 ((https://doi.org/10.1007/s13225-020-00466-2)) |
| Otu58168 | 1 | k__Fungi(100);p__Ascomycota(100);c__Saccharomycetes(100);o__Saccharomycetales(100);f__Metschnikowiaceae(100);g__Metschnikowia(100);g__Metschnikowia_unclassified(100); | Metschnikowia | 13 | Saprotroph | \|Undefined Saprotroph\| | Yeast | NULL | Probable | As nectar-tap_saprotroph (Põlme et al. 2020); Decay sugar-rich substrates (Põlme et al. 2020) | Põlme S, et al. 2020. Fungal Diversity 105:1-16 ((https://doi.org/10.1007/s13225-020-00466-2)) |
| Otu58170 | 1 | k__Fungi(100);p__Ascomycota(100);c__Saccharomycetes(100);o__Saccharomycetales(100);f__Metschnikowiaceae(100);g__Metschnikowia(100);s__Metschnikowia_pulcherrima(100); | Metschnikowia | 13 | Saprotroph | \|Undefined Saprotroph\| | Yeast | NULL | Probable | As nectar-tap_saprotroph (Põlme et al. 2020); Decay sugar-rich substrates (Põlme et al. 2020) | Põlme S, et al. 2020. Fungal Diversity 105:1-16 ((https://doi.org/10.1007/s13225-020-00466-2)) |
| Otu58171 | 1 | k__Fungi(100);p__Ascomycota(100);c__Saccharomycetes(100);o__Saccharomycetales(100);f__Metschnikowiaceae(100);g__Metschnikowia(100);s__Metschnikowia_pulcherrima(100); | Metschnikowia | 13 | Saprotroph | \|Undefined Saprotroph\| | Yeast | NULL | Probable | As nectar-tap_saprotroph (Põlme et al. 2020); Decay sugar-rich substrates (Põlme et al. 2020) | Põlme S, et al. 2020. Fungal Diversity 105:1-16 ((https://doi.org/10.1007/s13225-020-00466-2)) |
| Otu58172 | 1 | k__Fungi(100);p__Ascomycota(100);c__Saccharomycetes(100);o__Saccharomycetales(100);f__Metschnikowiaceae(100);g__Metschnikowia(100);g__Metschnikowia_unclassified(100); | Metschnikowia | 13 | Saprotroph | \|Undefined Saprotroph\| | Yeast | NULL | Probable | As nectar-tap_saprotroph (Põlme et al. 2020); Decay sugar-rich substrates (Põlme et al. 2020) | Põlme S, et al. 2020. Fungal Diversity 105:1-16 ((https://doi.org/10.1007/s13225-020-00466-2)) |
| Otu58174 | 1 | k__Fungi(100);p__Ascomycota(100);c__Saccharomycetes(100);o__Saccharomycetales(100);f__Metschnikowiaceae(100);g__Metschnikowia(100);s__Metschnikowia_pulcherrima(100); | Metschnikowia | 13 | Saprotroph | \|Undefined Saprotroph\| | Yeast | NULL | Probable | As nectar-tap_saprotroph (Põlme et al. 2020); Decay sugar-rich substrates (Põlme et al. 2020) | Põlme S, et al. 2020. Fungal Diversity 105:1-16 ((https://doi.org/10.1007/s13225-020-00466-2)) |
| Otu58177 | 1 | k__Fungi(100);p__Ascomycota(100);c__Saccharomycetes(100);o__Saccharomycetales(100);f__Metschnikowiaceae(100);g__Metschnikowia(100);s__Metschnikowia_pulcherrima(100); | Metschnikowia | 13 | Saprotroph | \|Undefined Saprotroph\| | Yeast | NULL | Probable | As nectar-tap_saprotroph (Põlme et al. 2020); Decay sugar-rich substrates (Põlme et al. 2020) | Põlme S, et al. 2020. Fungal Diversity 105:1-16 ((https://doi.org/10.1007/s13225-020-00466-2)) |
| Otu58178 | 1 | k__Fungi(100);p__Ascomycota(100);c__Saccharomycetes(100);o__Saccharomycetales(100);f__Metschnikowiaceae(100);g__Metschnikowia(100);g__Metschnikowia_unclassified(100); | Metschnikowia | 13 | Saprotroph | \|Undefined Saprotroph\| | Yeast | NULL | Probable | As nectar-tap_saprotroph (Põlme et al. 2020); Decay sugar-rich substrates (Põlme et al. 2020) | Põlme S, et al. 2020. Fungal Diversity 105:1-16 ((https://doi.org/10.1007/s13225-020-00466-2)) |
| Otu58179 | 1 | k__Fungi(100);p__Ascomycota(100);c__Saccharomycetes(100);o__Saccharomycetales(100);f__Metschnikowiaceae(100);g__Metschnikowia(100);s__Metschnikowia_pulcherrima(100); | Metschnikowia | 13 | Saprotroph | \|Undefined Saprotroph\| | Yeast | NULL | Probable | As nectar-tap_saprotroph (Põlme et al. 2020); Decay sugar-rich substrates (Põlme et al. 2020) | Põlme S, et al. 2020. Fungal Diversity 105:1-16 ((https://doi.org/10.1007/s13225-020-00466-2)) |
| Otu58180 | 1 | k__Fungi(100);p__Ascomycota(100);c__Saccharomycetes(100);o__Saccharomycetales(100);f__Metschnikowiaceae(100);g__Metschnikowia(100);s__Metschnikowia_pulcherrima(100); | Metschnikowia | 13 | Saprotroph | \|Undefined Saprotroph\| | Yeast | NULL | Probable | As nectar-tap_saprotroph (Põlme et al. 2020); Decay sugar-rich substrates (Põlme et al. 2020) | Põlme S, et al. 2020. Fungal Diversity 105:1-16 ((https://doi.org/10.1007/s13225-020-00466-2)) |
| Otu58181 | 1 | k__Fungi(100);p__Ascomycota(100);c__Saccharomycetes(100);o__Saccharomycetales(100);f__Metschnikowiaceae(100);g__Metschnikowia(100);s__Metschnikowia_pulcherrima(100); | Metschnikowia | 13 | Saprotroph | \|Undefined Saprotroph\| | Yeast | NULL | Probable | As nectar-tap_saprotroph (Põlme et al. 2020); Decay sugar-rich substrates (Põlme et al. 2020) | Põlme S, et al. 2020. Fungal Diversity 105:1-16 ((https://doi.org/10.1007/s13225-020-00466-2)) |
| Otu58182 | 1 | k__Fungi(100);p__Ascomycota(100);c__Saccharomycetes(100);o__Saccharomycetales(100);f__Metschnikowiaceae(100);g__Metschnikowia(100);s__Metschnikowia_pulcherrima(100); | Metschnikowia | 13 | Saprotroph | \|Undefined Saprotroph\| | Yeast | NULL | Probable | As nectar-tap_saprotroph (Põlme et al. 2020); Decay sugar-rich substrates (Põlme et al. 2020) | Põlme S, et al. 2020. Fungal Diversity 105:1-16 ((https://doi.org/10.1007/s13225-020-00466-2)) |
| Otu58183 | 1 | k__Fungi(100);p__Ascomycota(100);c__Saccharomycetes(100);o__Saccharomycetales(100);f__Metschnikowiaceae(100);g__Metschnikowia(100);s__Metschnikowia_pulcherrima(100); | Metschnikowia | 13 | Saprotroph | \|Undefined Saprotroph\| | Yeast | NULL | Probable | As nectar-tap_saprotroph (Põlme et al. 2020); Decay sugar-rich substrates (Põlme et al. 2020) | Põlme S, et al. 2020. Fungal Diversity 105:1-16 ((https://doi.org/10.1007/s13225-020-00466-2)) |
| Otu58184 | 1 | k__Fungi(100);p__Ascomycota(100);c__Saccharomycetes(100);o__Saccharomycetales(100);f__Metschnikowiaceae(100);g__Metschnikowia(100);s__Metschnikowia_pulcherrima(100); | Metschnikowia | 13 | Saprotroph | \|Undefined Saprotroph\| | Yeast | NULL | Probable | As nectar-tap_saprotroph (Põlme et al. 2020); Decay sugar-rich substrates (Põlme et al. 2020) | Põlme S, et al. 2020. Fungal Diversity 105:1-16 ((https://doi.org/10.1007/s13225-020-00466-2)) |
| Otu58185 | 1 | k__Fungi(100);p__Ascomycota(100);c__Saccharomycetes(100);o__Saccharomycetales(100);f__Metschnikowiaceae(100);g__Metschnikowia(100);g__Metschnikowia_unclassified(100); | Metschnikowia | 13 | Saprotroph | \|Undefined Saprotroph\| | Yeast | NULL | Probable | As nectar-tap_saprotroph (Põlme et al. 2020); Decay sugar-rich substrates (Põlme et al. 2020) | Põlme S, et al. 2020. Fungal Diversity 105:1-16 ((https://doi.org/10.1007/s13225-020-00466-2)) |
| Otu58186 | 1 | k__Fungi(100);p__Ascomycota(100);c__Saccharomycetes(100);o__Saccharomycetales(100);f__Metschnikowiaceae(100);g__Metschnikowia(100);s__Metschnikowia_pulcherrima(100); | Metschnikowia | 13 | Saprotroph | \|Undefined Saprotroph\| | Yeast | NULL | Probable | As nectar-tap_saprotroph (Põlme et al. 2020); Decay sugar-rich substrates (Põlme et al. 2020) | Põlme S, et al. 2020. Fungal Diversity 105:1-16 ((https://doi.org/10.1007/s13225-020-00466-2)) |
| Otu58192 | 1 | k__Fungi(100);p__Ascomycota(100);c__Saccharomycetes(100);o__Saccharomycetales(100);f__Metschnikowiaceae(100);g__Metschnikowia(100);s__Metschnikowia_pulcherrima(100); | Metschnikowia | 13 | Saprotroph | \|Undefined Saprotroph\| | Yeast | NULL | Probable | As nectar-tap_saprotroph (Põlme et al. 2020); Decay sugar-rich substrates (Põlme et al. 2020) | Põlme S, et al. 2020. Fungal Diversity 105:1-16 ((https://doi.org/10.1007/s13225-020-00466-2)) |
| Otu58193 | 1 | k__Fungi(100);p__Ascomycota(100);c__Saccharomycetes(100);o__Saccharomycetales(100);f__Metschnikowiaceae(100);g__Metschnikowia(100);g__Metschnikowia_unclassified(100); | Metschnikowia | 13 | Saprotroph | \|Undefined Saprotroph\| | Yeast | NULL | Probable | As nectar-tap_saprotroph (Põlme et al. 2020); Decay sugar-rich substrates (Põlme et al. 2020) | Põlme S, et al. 2020. Fungal Diversity 105:1-16 ((https://doi.org/10.1007/s13225-020-00466-2)) |
| Otu58194 | 1 | k__Fungi(100);p__Ascomycota(100);c__Saccharomycetes(100);o__Saccharomycetales(100);f__Metschnikowiaceae(100);g__Metschnikowia(100);s__Metschnikowia_pulcherrima(100); | Metschnikowia | 13 | Saprotroph | \|Undefined Saprotroph\| | Yeast | NULL | Probable | As nectar-tap_saprotroph (Põlme et al. 2020); Decay sugar-rich substrates (Põlme et al. 2020) | Põlme S, et al. 2020. Fungal Diversity 105:1-16 ((https://doi.org/10.1007/s13225-020-00466-2)) |
| Otu58198 | 1 | k__Fungi(100);p__Ascomycota(100);c__Saccharomycetes(100);o__Saccharomycetales(100);f__Metschnikowiaceae(100);g__Metschnikowia(100);s__Metschnikowia_pulcherrima(100); | Metschnikowia | 13 | Saprotroph | \|Undefined Saprotroph\| | Yeast | NULL | Probable | As nectar-tap_saprotroph (Põlme et al. 2020); Decay sugar-rich substrates (Põlme et al. 2020) | Põlme S, et al. 2020. Fungal Diversity 105:1-16 ((https://doi.org/10.1007/s13225-020-00466-2)) |
| Otu58199 | 1 | k__Fungi(100);p__Ascomycota(100);c__Saccharomycetes(100);o__Saccharomycetales(100);f__Metschnikowiaceae(100);g__Metschnikowia(100);s__Metschnikowia_pulcherrima(100); | Metschnikowia | 13 | Saprotroph | \|Undefined Saprotroph\| | Yeast | NULL | Probable | As nectar-tap_saprotroph (Põlme et al. 2020); Decay sugar-rich substrates (Põlme et al. 2020) | Põlme S, et al. 2020. Fungal Diversity 105:1-16 ((https://doi.org/10.1007/s13225-020-00466-2)) |
| Otu58201 | 1 | k__Fungi(100);p__Ascomycota(100);c__Saccharomycetes(100);o__Saccharomycetales(100);f__Metschnikowiaceae(100);g__Metschnikowia(100);s__Metschnikowia_pulcherrima(100); | Metschnikowia | 13 | Saprotroph | \|Undefined Saprotroph\| | Yeast | NULL | Probable | As nectar-tap_saprotroph (Põlme et al. 2020); Decay sugar-rich substrates (Põlme et al. 2020) | Põlme S, et al. 2020. Fungal Diversity 105:1-16 ((https://doi.org/10.1007/s13225-020-00466-2)) |
| Otu58203 | 1 | k__Fungi(100);p__Ascomycota(100);c__Saccharomycetes(100);o__Saccharomycetales(100);f__Metschnikowiaceae(100);g__Metschnikowia(100);s__Metschnikowia_pulcherrima(100); | Metschnikowia | 13 | Saprotroph | \|Undefined Saprotroph\| | Yeast | NULL | Probable | As nectar-tap_saprotroph (Põlme et al. 2020); Decay sugar-rich substrates (Põlme et al. 2020) | Põlme S, et al. 2020. Fungal Diversity 105:1-16 ((https://doi.org/10.1007/s13225-020-00466-2)) |
| Otu58205 | 1 | k__Fungi(100);p__Ascomycota(100);c__Saccharomycetes(100);o__Saccharomycetales(100);f__Metschnikowiaceae(100);g__Metschnikowia(100);g__Metschnikowia_unclassified(100); | Metschnikowia | 13 | Saprotroph | \|Undefined Saprotroph\| | Yeast | NULL | Probable | As nectar-tap_saprotroph (Põlme et al. 2020); Decay sugar-rich substrates (Põlme et al. 2020) | Põlme S, et al. 2020. Fungal Diversity 105:1-16 ((https://doi.org/10.1007/s13225-020-00466-2)) |
| Otu58208 | 1 | k__Fungi(100);p__Ascomycota(100);c__Saccharomycetes(100);o__Saccharomycetales(100);f__Metschnikowiaceae(100);g__Metschnikowia(100);s__Metschnikowia_pulcherrima(100); | Metschnikowia | 13 | Saprotroph | \|Undefined Saprotroph\| | Yeast | NULL | Probable | As nectar-tap_saprotroph (Põlme et al. 2020); Decay sugar-rich substrates (Põlme et al. 2020) | Põlme S, et al. 2020. Fungal Diversity 105:1-16 ((https://doi.org/10.1007/s13225-020-00466-2)) |
| Otu58209 | 1 | k__Fungi(100);p__Ascomycota(100);c__Saccharomycetes(100);o__Saccharomycetales(100);f__Metschnikowiaceae(100);g__Metschnikowia(100);s__Metschnikowia_pulcherrima(100); | Metschnikowia | 13 | Saprotroph | \|Undefined Saprotroph\| | Yeast | NULL | Probable | As nectar-tap_saprotroph (Põlme et al. 2020); Decay sugar-rich substrates (Põlme et al. 2020) | Põlme S, et al. 2020. Fungal Diversity 105:1-16 ((https://doi.org/10.1007/s13225-020-00466-2)) |
| Otu58210 | 1 | k__Fungi(100);p__Ascomycota(100);c__Saccharomycetes(100);o__Saccharomycetales(100);f__Metschnikowiaceae(100);g__Metschnikowia(100);s__Metschnikowia_pulcherrima(100); | Metschnikowia | 13 | Saprotroph | \|Undefined Saprotroph\| | Yeast | NULL | Probable | As nectar-tap_saprotroph (Põlme et al. 2020); Decay sugar-rich substrates (Põlme et al. 2020) | Põlme S, et al. 2020. Fungal Diversity 105:1-16 ((https://doi.org/10.1007/s13225-020-00466-2)) |
| Otu58211 | 1 | k__Fungi(100);p__Ascomycota(100);c__Saccharomycetes(100);o__Saccharomycetales(100);f__Metschnikowiaceae(100);g__Metschnikowia(100);s__Metschnikowia_pulcherrima(100); | Metschnikowia | 13 | Saprotroph | \|Undefined Saprotroph\| | Yeast | NULL | Probable | As nectar-tap_saprotroph (Põlme et al. 2020); Decay sugar-rich substrates (Põlme et al. 2020) | Põlme S, et al. 2020. Fungal Diversity 105:1-16 ((https://doi.org/10.1007/s13225-020-00466-2)) |
| Otu58219 | 1 | k__Fungi(100);p__Ascomycota(100);c__Saccharomycetes(100);o__Saccharomycetales(100);f__Metschnikowiaceae(100);g__Metschnikowia(100);s__Metschnikowia_pulcherrima(100); | Metschnikowia | 13 | Saprotroph | \|Undefined Saprotroph\| | Yeast | NULL | Probable | As nectar-tap_saprotroph (Põlme et al. 2020); Decay sugar-rich substrates (Põlme et al. 2020) | Põlme S, et al. 2020. Fungal Diversity 105:1-16 ((https://doi.org/10.1007/s13225-020-00466-2)) |
| Otu58220 | 1 | k__Fungi(100);p__Ascomycota(100);c__Saccharomycetes(100);o__Saccharomycetales(100);f__Metschnikowiaceae(100);g__Metschnikowia(100);g__Metschnikowia_unclassified(100); | Metschnikowia | 13 | Saprotroph | \|Undefined Saprotroph\| | Yeast | NULL | Probable | As nectar-tap_saprotroph (Põlme et al. 2020); Decay sugar-rich substrates (Põlme et al. 2020) | Põlme S, et al. 2020. Fungal Diversity 105:1-16 ((https://doi.org/10.1007/s13225-020-00466-2)) |
| Otu58221 | 1 | k__Fungi(100);p__Ascomycota(100);c__Saccharomycetes(100);o__Saccharomycetales(100);f__Metschnikowiaceae(100);g__Metschnikowia(100);s__Metschnikowia_pulcherrima(100); | Metschnikowia | 13 | Saprotroph | \|Undefined Saprotroph\| | Yeast | NULL | Probable | As nectar-tap_saprotroph (Põlme et al. 2020); Decay sugar-rich substrates (Põlme et al. 2020) | Põlme S, et al. 2020. Fungal Diversity 105:1-16 ((https://doi.org/10.1007/s13225-020-00466-2)) |
| Otu58222 | 1 | k__Fungi(100);p__Ascomycota(100);c__Saccharomycetes(100);o__Saccharomycetales(100);f__Metschnikowiaceae(100);g__Metschnikowia(100);s__Metschnikowia_pulcherrima(100); | Metschnikowia | 13 | Saprotroph | \|Undefined Saprotroph\| | Yeast | NULL | Probable | As nectar-tap_saprotroph (Põlme et al. 2020); Decay sugar-rich substrates (Põlme et al. 2020) | Põlme S, et al. 2020. Fungal Diversity 105:1-16 ((https://doi.org/10.1007/s13225-020-00466-2)) |
| Otu58223 | 1 | k__Fungi(100);p__Ascomycota(100);c__Saccharomycetes(100);o__Saccharomycetales(100);f__Metschnikowiaceae(100);g__Metschnikowia(100);s__Metschnikowia_pulcherrima(100); | Metschnikowia | 13 | Saprotroph | \|Undefined Saprotroph\| | Yeast | NULL | Probable | As nectar-tap_saprotroph (Põlme et al. 2020); Decay sugar-rich substrates (Põlme et al. 2020) | Põlme S, et al. 2020. Fungal Diversity 105:1-16 ((https://doi.org/10.1007/s13225-020-00466-2)) |
| Otu58224 | 1 | k__Fungi(100);p__Ascomycota(100);c__Saccharomycetes(100);o__Saccharomycetales(100);f__Metschnikowiaceae(100);g__Metschnikowia(100);g__Metschnikowia_unclassified(100); | Metschnikowia | 13 | Saprotroph | \|Undefined Saprotroph\| | Yeast | NULL | Probable | As nectar-tap_saprotroph (Põlme et al. 2020); Decay sugar-rich substrates (Põlme et al. 2020) | Põlme S, et al. 2020. Fungal Diversity 105:1-16 ((https://doi.org/10.1007/s13225-020-00466-2)) |
| Otu58225 | 1 | k__Fungi(100);p__Ascomycota(100);c__Saccharomycetes(100);o__Saccharomycetales(100);f__Metschnikowiaceae(100);g__Metschnikowia(100);g__Metschnikowia_unclassified(100); | Metschnikowia | 13 | Saprotroph | \|Undefined Saprotroph\| | Yeast | NULL | Probable | As nectar-tap_saprotroph (Põlme et al. 2020); Decay sugar-rich substrates (Põlme et al. 2020) | Põlme S, et al. 2020. Fungal Diversity 105:1-16 ((https://doi.org/10.1007/s13225-020-00466-2)) |
| Otu58227 | 1 | k__Fungi(100);p__Ascomycota(100);c__Saccharomycetes(100);o__Saccharomycetales(100);f__Metschnikowiaceae(100);g__Metschnikowia(100);s__Metschnikowia_pulcherrima(100); | Metschnikowia | 13 | Saprotroph | \|Undefined Saprotroph\| | Yeast | NULL | Probable | As nectar-tap_saprotroph (Põlme et al. 2020); Decay sugar-rich substrates (Põlme et al. 2020) | Põlme S, et al. 2020. Fungal Diversity 105:1-16 ((https://doi.org/10.1007/s13225-020-00466-2)) |
| Otu58229 | 1 | k__Fungi(100);p__Ascomycota(100);c__Saccharomycetes(100);o__Saccharomycetales(100);f__Metschnikowiaceae(100);g__Metschnikowia(100);s__Metschnikowia_pulcherrima(100); | Metschnikowia | 13 | Saprotroph | \|Undefined Saprotroph\| | Yeast | NULL | Probable | As nectar-tap_saprotroph (Põlme et al. 2020); Decay sugar-rich substrates (Põlme et al. 2020) | Põlme S, et al. 2020. Fungal Diversity 105:1-16 ((https://doi.org/10.1007/s13225-020-00466-2)) |
| Otu58230 | 1 | k__Fungi(100);p__Ascomycota(100);c__Saccharomycetes(100);o__Saccharomycetales(100);f__Metschnikowiaceae(100);g__Metschnikowia(100);s__Metschnikowia_pulcherrima(100); | Metschnikowia | 13 | Saprotroph | \|Undefined Saprotroph\| | Yeast | NULL | Probable | As nectar-tap_saprotroph (Põlme et al. 2020); Decay sugar-rich substrates (Põlme et al. 2020) | Põlme S, et al. 2020. Fungal Diversity 105:1-16 ((https://doi.org/10.1007/s13225-020-00466-2)) |
| Otu58231 | 1 | k__Fungi(100);p__Ascomycota(100);c__Saccharomycetes(100);o__Saccharomycetales(100);f__Metschnikowiaceae(100);g__Metschnikowia(100);s__Metschnikowia_pulcherrima(100); | Metschnikowia | 13 | Saprotroph | \|Undefined Saprotroph\| | Yeast | NULL | Probable | As nectar-tap_saprotroph (Põlme et al. 2020); Decay sugar-rich substrates (Põlme et al. 2020) | Põlme S, et al. 2020. Fungal Diversity 105:1-16 ((https://doi.org/10.1007/s13225-020-00466-2)) |
| Otu58233 | 1 | k__Fungi(100);p__Ascomycota(100);c__Saccharomycetes(100);o__Saccharomycetales(100);f__Metschnikowiaceae(100);g__Metschnikowia(100);s__Metschnikowia_pulcherrima(100); | Metschnikowia | 13 | Saprotroph | \|Undefined Saprotroph\| | Yeast | NULL | Probable | As nectar-tap_saprotroph (Põlme et al. 2020); Decay sugar-rich substrates (Põlme et al. 2020) | Põlme S, et al. 2020. Fungal Diversity 105:1-16 ((https://doi.org/10.1007/s13225-020-00466-2)) |
| Otu58234 | 1 | k__Fungi(100);p__Ascomycota(100);c__Saccharomycetes(100);o__Saccharomycetales(100);f__Metschnikowiaceae(100);g__Metschnikowia(100);s__Metschnikowia_pulcherrima(100); | Metschnikowia | 13 | Saprotroph | \|Undefined Saprotroph\| | Yeast | NULL | Probable | As nectar-tap_saprotroph (Põlme et al. 2020); Decay sugar-rich substrates (Põlme et al. 2020) | Põlme S, et al. 2020. Fungal Diversity 105:1-16 ((https://doi.org/10.1007/s13225-020-00466-2)) |
| Otu58235 | 1 | k__Fungi(100);p__Ascomycota(100);c__Saccharomycetes(100);o__Saccharomycetales(100);f__Metschnikowiaceae(100);g__Metschnikowia(100);s__Metschnikowia_pulcherrima(100); | Metschnikowia | 13 | Saprotroph | \|Undefined Saprotroph\| | Yeast | NULL | Probable | As nectar-tap_saprotroph (Põlme et al. 2020); Decay sugar-rich substrates (Põlme et al. 2020) | Põlme S, et al. 2020. Fungal Diversity 105:1-16 ((https://doi.org/10.1007/s13225-020-00466-2)) |
| Otu58236 | 1 | k__Fungi(100);p__Ascomycota(100);c__Saccharomycetes(100);o__Saccharomycetales(100);f__Metschnikowiaceae(100);g__Metschnikowia(100);s__Metschnikowia_pulcherrima(100); | Metschnikowia | 13 | Saprotroph | \|Undefined Saprotroph\| | Yeast | NULL | Probable | As nectar-tap_saprotroph (Põlme et al. 2020); Decay sugar-rich substrates (Põlme et al. 2020) | Põlme S, et al. 2020. Fungal Diversity 105:1-16 ((https://doi.org/10.1007/s13225-020-00466-2)) |
| Otu58237 | 1 | k__Fungi(100);p__Ascomycota(100);c__Saccharomycetes(100);o__Saccharomycetales(100);f__Metschnikowiaceae(100);g__Metschnikowia(100);g__Metschnikowia_unclassified(100); | Metschnikowia | 13 | Saprotroph | \|Undefined Saprotroph\| | Yeast | NULL | Probable | As nectar-tap_saprotroph (Põlme et al. 2020); Decay sugar-rich substrates (Põlme et al. 2020) | Põlme S, et al. 2020. Fungal Diversity 105:1-16 ((https://doi.org/10.1007/s13225-020-00466-2)) |
| Otu58238 | 1 | k__Fungi(100);p__Ascomycota(100);c__Saccharomycetes(100);o__Saccharomycetales(100);f__Metschnikowiaceae(100);g__Metschnikowia(100);s__Metschnikowia_pulcherrima(100); | Metschnikowia | 13 | Saprotroph | \|Undefined Saprotroph\| | Yeast | NULL | Probable | As nectar-tap_saprotroph (Põlme et al. 2020); Decay sugar-rich substrates (Põlme et al. 2020) | Põlme S, et al. 2020. Fungal Diversity 105:1-16 ((https://doi.org/10.1007/s13225-020-00466-2)) |
| Otu58239 | 1 | k__Fungi(100);p__Ascomycota(100);c__Saccharomycetes(100);o__Saccharomycetales(100);f__Metschnikowiaceae(100);g__Metschnikowia(100);s__Metschnikowia_pulcherrima(100); | Metschnikowia | 13 | Saprotroph | \|Undefined Saprotroph\| | Yeast | NULL | Probable | As nectar-tap_saprotroph (Põlme et al. 2020); Decay sugar-rich substrates (Põlme et al. 2020) | Põlme S, et al. 2020. Fungal Diversity 105:1-16 ((https://doi.org/10.1007/s13225-020-00466-2)) |
| Otu58240 | 1 | k__Fungi(100);p__Ascomycota(100);c__Saccharomycetes(100);o__Saccharomycetales(100);f__Metschnikowiaceae(100);g__Metschnikowia(100);s__Metschnikowia_pulcherrima(100); | Metschnikowia | 13 | Saprotroph | \|Undefined Saprotroph\| | Yeast | NULL | Probable | As nectar-tap_saprotroph (Põlme et al. 2020); Decay sugar-rich substrates (Põlme et al. 2020) | Põlme S, et al. 2020. Fungal Diversity 105:1-16 ((https://doi.org/10.1007/s13225-020-00466-2)) |
| Otu58245 | 1 | k__Fungi(100);p__Ascomycota(100);c__Saccharomycetes(100);o__Saccharomycetales(100);f__Metschnikowiaceae(100);g__Metschnikowia(100);s__Metschnikowia_pulcherrima(100); | Metschnikowia | 13 | Saprotroph | \|Undefined Saprotroph\| | Yeast | NULL | Probable | As nectar-tap_saprotroph (Põlme et al. 2020); Decay sugar-rich substrates (Põlme et al. 2020) | Põlme S, et al. 2020. Fungal Diversity 105:1-16 ((https://doi.org/10.1007/s13225-020-00466-2)) |
| Otu58246 | 1 | k__Fungi(100);p__Ascomycota(100);c__Saccharomycetes(100);o__Saccharomycetales(100);f__Metschnikowiaceae(100);g__Metschnikowia(100);g__Metschnikowia_unclassified(100); | Metschnikowia | 13 | Saprotroph | \|Undefined Saprotroph\| | Yeast | NULL | Probable | As nectar-tap_saprotroph (Põlme et al. 2020); Decay sugar-rich substrates (Põlme et al. 2020) | Põlme S, et al. 2020. Fungal Diversity 105:1-16 ((https://doi.org/10.1007/s13225-020-00466-2)) |
| Otu58247 | 1 | k__Fungi(100);p__Ascomycota(100);c__Saccharomycetes(100);o__Saccharomycetales(100);f__Metschnikowiaceae(100);g__Metschnikowia(100);s__Metschnikowia_pulcherrima(100); | Metschnikowia | 13 | Saprotroph | \|Undefined Saprotroph\| | Yeast | NULL | Probable | As nectar-tap_saprotroph (Põlme et al. 2020); Decay sugar-rich substrates (Põlme et al. 2020) | Põlme S, et al. 2020. Fungal Diversity 105:1-16 ((https://doi.org/10.1007/s13225-020-00466-2)) |
| Otu58248 | 1 | k__Fungi(100);p__Ascomycota(100);c__Saccharomycetes(100);o__Saccharomycetales(100);f__Metschnikowiaceae(100);g__Metschnikowia(100);s__Metschnikowia_pulcherrima(100); | Metschnikowia | 13 | Saprotroph | \|Undefined Saprotroph\| | Yeast | NULL | Probable | As nectar-tap_saprotroph (Põlme et al. 2020); Decay sugar-rich substrates (Põlme et al. 2020) | Põlme S, et al. 2020. Fungal Diversity 105:1-16 ((https://doi.org/10.1007/s13225-020-00466-2)) |
| Otu58249 | 1 | k__Fungi(100);p__Ascomycota(100);c__Saccharomycetes(100);o__Saccharomycetales(100);f__Metschnikowiaceae(100);g__Metschnikowia(100);s__Metschnikowia_pulcherrima(100); | Metschnikowia | 13 | Saprotroph | \|Undefined Saprotroph\| | Yeast | NULL | Probable | As nectar-tap_saprotroph (Põlme et al. 2020); Decay sugar-rich substrates (Põlme et al. 2020) | Põlme S, et al. 2020. Fungal Diversity 105:1-16 ((https://doi.org/10.1007/s13225-020-00466-2)) |
| Otu58250 | 1 | k__Fungi(100);p__Ascomycota(100);c__Saccharomycetes(100);o__Saccharomycetales(100);f__Metschnikowiaceae(100);g__Metschnikowia(100);g__Metschnikowia_unclassified(100); | Metschnikowia | 13 | Saprotroph | \|Undefined Saprotroph\| | Yeast | NULL | Probable | As nectar-tap_saprotroph (Põlme et al. 2020); Decay sugar-rich substrates (Põlme et al. 2020) | Põlme S, et al. 2020. Fungal Diversity 105:1-16 ((https://doi.org/10.1007/s13225-020-00466-2)) |
| Otu58251 | 1 | k__Fungi(100);p__Ascomycota(100);c__Saccharomycetes(100);o__Saccharomycetales(100);f__Metschnikowiaceae(100);g__Metschnikowia(100);g__Metschnikowia_unclassified(100); | Metschnikowia | 13 | Saprotroph | \|Undefined Saprotroph\| | Yeast | NULL | Probable | As nectar-tap_saprotroph (Põlme et al. 2020); Decay sugar-rich substrates (Põlme et al. 2020) | Põlme S, et al. 2020. Fungal Diversity 105:1-16 ((https://doi.org/10.1007/s13225-020-00466-2)) |
| Otu58252 | 1 | k__Fungi(100);p__Ascomycota(100);c__Saccharomycetes(100);o__Saccharomycetales(100);f__Metschnikowiaceae(100);g__Metschnikowia(100);s__Metschnikowia_pulcherrima(100); | Metschnikowia | 13 | Saprotroph | \|Undefined Saprotroph\| | Yeast | NULL | Probable | As nectar-tap_saprotroph (Põlme et al. 2020); Decay sugar-rich substrates (Põlme et al. 2020) | Põlme S, et al. 2020. Fungal Diversity 105:1-16 ((https://doi.org/10.1007/s13225-020-00466-2)) |
| Otu58253 | 1 | k__Fungi(100);p__Ascomycota(100);c__Saccharomycetes(100);o__Saccharomycetales(100);f__Metschnikowiaceae(100);g__Metschnikowia(100);s__Metschnikowia_pulcherrima(100); | Metschnikowia | 13 | Saprotroph | \|Undefined Saprotroph\| | Yeast | NULL | Probable | As nectar-tap_saprotroph (Põlme et al. 2020); Decay sugar-rich substrates (Põlme et al. 2020) | Põlme S, et al. 2020. Fungal Diversity 105:1-16 ((https://doi.org/10.1007/s13225-020-00466-2)) |
| Otu58254 | 1 | k__Fungi(100);p__Ascomycota(100);c__Saccharomycetes(100);o__Saccharomycetales(100);f__Metschnikowiaceae(100);g__Metschnikowia(100);s__Metschnikowia_pulcherrima(100); | Metschnikowia | 13 | Saprotroph | \|Undefined Saprotroph\| | Yeast | NULL | Probable | As nectar-tap_saprotroph (Põlme et al. 2020); Decay sugar-rich substrates (Põlme et al. 2020) | Põlme S, et al. 2020. Fungal Diversity 105:1-16 ((https://doi.org/10.1007/s13225-020-00466-2)) |
| Otu58256 | 1 | k__Fungi(100);p__Ascomycota(100);c__Saccharomycetes(100);o__Saccharomycetales(100);f__Metschnikowiaceae(100);g__Metschnikowia(100);s__Metschnikowia_pulcherrima(100); | Metschnikowia | 13 | Saprotroph | \|Undefined Saprotroph\| | Yeast | NULL | Probable | As nectar-tap_saprotroph (Põlme et al. 2020); Decay sugar-rich substrates (Põlme et al. 2020) | Põlme S, et al. 2020. Fungal Diversity 105:1-16 ((https://doi.org/10.1007/s13225-020-00466-2)) |
| Otu58257 | 1 | k__Fungi(100);p__Ascomycota(100);c__Saccharomycetes(100);o__Saccharomycetales(100);f__Metschnikowiaceae(100);g__Metschnikowia(100);s__Metschnikowia_pulcherrima(100); | Metschnikowia | 13 | Saprotroph | \|Undefined Saprotroph\| | Yeast | NULL | Probable | As nectar-tap_saprotroph (Põlme et al. 2020); Decay sugar-rich substrates (Põlme et al. 2020) | Põlme S, et al. 2020. Fungal Diversity 105:1-16 ((https://doi.org/10.1007/s13225-020-00466-2)) |
| Otu58258 | 1 | k__Fungi(100);p__Ascomycota(100);c__Saccharomycetes(100);o__Saccharomycetales(100);f__Metschnikowiaceae(100);g__Metschnikowia(100);g__Metschnikowia_unclassified(100); | Metschnikowia | 13 | Saprotroph | \|Undefined Saprotroph\| | Yeast | NULL | Probable | As nectar-tap_saprotroph (Põlme et al. 2020); Decay sugar-rich substrates (Põlme et al. 2020) | Põlme S, et al. 2020. Fungal Diversity 105:1-16 ((https://doi.org/10.1007/s13225-020-00466-2)) |
| Otu58259 | 1 | k__Fungi(100);p__Ascomycota(100);c__Saccharomycetes(100);o__Saccharomycetales(100);f__Metschnikowiaceae(100);g__Metschnikowia(100);s__Metschnikowia_pulcherrima(100); | Metschnikowia | 13 | Saprotroph | \|Undefined Saprotroph\| | Yeast | NULL | Probable | As nectar-tap_saprotroph (Põlme et al. 2020); Decay sugar-rich substrates (Põlme et al. 2020) | Põlme S, et al. 2020. Fungal Diversity 105:1-16 ((https://doi.org/10.1007/s13225-020-00466-2)) |
| Otu58260 | 1 | k__Fungi(100);p__Ascomycota(100);c__Saccharomycetes(100);o__Saccharomycetales(100);f__Metschnikowiaceae(100);g__Metschnikowia(100);s__Metschnikowia_pulcherrima(100); | Metschnikowia | 13 | Saprotroph | \|Undefined Saprotroph\| | Yeast | NULL | Probable | As nectar-tap_saprotroph (Põlme et al. 2020); Decay sugar-rich substrates (Põlme et al. 2020) | Põlme S, et al. 2020. Fungal Diversity 105:1-16 ((https://doi.org/10.1007/s13225-020-00466-2)) |
| Otu58261 | 1 | k__Fungi(100);p__Ascomycota(100);c__Saccharomycetes(100);o__Saccharomycetales(100);f__Metschnikowiaceae(100);g__Metschnikowia(100);s__Metschnikowia_pulcherrima(100); | Metschnikowia | 13 | Saprotroph | \|Undefined Saprotroph\| | Yeast | NULL | Probable | As nectar-tap_saprotroph (Põlme et al. 2020); Decay sugar-rich substrates (Põlme et al. 2020) | Põlme S, et al. 2020. Fungal Diversity 105:1-16 ((https://doi.org/10.1007/s13225-020-00466-2)) |
| Otu58262 | 1 | k__Fungi(100);p__Ascomycota(100);c__Saccharomycetes(100);o__Saccharomycetales(100);f__Metschnikowiaceae(100);g__Metschnikowia(100);g__Metschnikowia_unclassified(100); | Metschnikowia | 13 | Saprotroph | \|Undefined Saprotroph\| | Yeast | NULL | Probable | As nectar-tap_saprotroph (Põlme et al. 2020); Decay sugar-rich substrates (Põlme et al. 2020) | Põlme S, et al. 2020. Fungal Diversity 105:1-16 ((https://doi.org/10.1007/s13225-020-00466-2)) |
| Otu58263 | 1 | k__Fungi(100);p__Ascomycota(100);c__Saccharomycetes(100);o__Saccharomycetales(100);f__Metschnikowiaceae(100);g__Metschnikowia(100);s__Metschnikowia_pulcherrima(100); | Metschnikowia | 13 | Saprotroph | \|Undefined Saprotroph\| | Yeast | NULL | Probable | As nectar-tap_saprotroph (Põlme et al. 2020); Decay sugar-rich substrates (Põlme et al. 2020) | Põlme S, et al. 2020. Fungal Diversity 105:1-16 ((https://doi.org/10.1007/s13225-020-00466-2)) |
| Otu58264 | 1 | k__Fungi(100);p__Ascomycota(100);c__Saccharomycetes(100);o__Saccharomycetales(100);f__Metschnikowiaceae(100);g__Metschnikowia(100);s__Metschnikowia_pulcherrima(100); | Metschnikowia | 13 | Saprotroph | \|Undefined Saprotroph\| | Yeast | NULL | Probable | As nectar-tap_saprotroph (Põlme et al. 2020); Decay sugar-rich substrates (Põlme et al. 2020) | Põlme S, et al. 2020. Fungal Diversity 105:1-16 ((https://doi.org/10.1007/s13225-020-00466-2)) |
| Otu58265 | 1 | k__Fungi(100);p__Ascomycota(100);c__Saccharomycetes(100);o__Saccharomycetales(100);f__Metschnikowiaceae(100);g__Metschnikowia(100);s__Metschnikowia_pulcherrima(100); | Metschnikowia | 13 | Saprotroph | \|Undefined Saprotroph\| | Yeast | NULL | Probable | As nectar-tap_saprotroph (Põlme et al. 2020); Decay sugar-rich substrates (Põlme et al. 2020) | Põlme S, et al. 2020. Fungal Diversity 105:1-16 ((https://doi.org/10.1007/s13225-020-00466-2)) |
| Otu58266 | 1 | k__Fungi(100);p__Ascomycota(100);c__Saccharomycetes(100);o__Saccharomycetales(100);f__Metschnikowiaceae(100);g__Metschnikowia(100);g__Metschnikowia_unclassified(100); | Metschnikowia | 13 | Saprotroph | \|Undefined Saprotroph\| | Yeast | NULL | Probable | As nectar-tap_saprotroph (Põlme et al. 2020); Decay sugar-rich substrates (Põlme et al. 2020) | Põlme S, et al. 2020. Fungal Diversity 105:1-16 ((https://doi.org/10.1007/s13225-020-00466-2)) |
| Otu58267 | 1 | k__Fungi(100);p__Ascomycota(100);c__Saccharomycetes(100);o__Saccharomycetales(100);f__Metschnikowiaceae(100);g__Metschnikowia(100);g__Metschnikowia_unclassified(100); | Metschnikowia | 13 | Saprotroph | \|Undefined Saprotroph\| | Yeast | NULL | Probable | As nectar-tap_saprotroph (Põlme et al. 2020); Decay sugar-rich substrates (Põlme et al. 2020) | Põlme S, et al. 2020. Fungal Diversity 105:1-16 ((https://doi.org/10.1007/s13225-020-00466-2)) |
| Otu58268 | 1 | k__Fungi(100);p__Ascomycota(100);c__Saccharomycetes(100);o__Saccharomycetales(100);f__Metschnikowiaceae(100);g__Metschnikowia(100);s__Metschnikowia_pulcherrima(100); | Metschnikowia | 13 | Saprotroph | \|Undefined Saprotroph\| | Yeast | NULL | Probable | As nectar-tap_saprotroph (Põlme et al. 2020); Decay sugar-rich substrates (Põlme et al. 2020) | Põlme S, et al. 2020. Fungal Diversity 105:1-16 ((https://doi.org/10.1007/s13225-020-00466-2)) |
| Otu58269 | 1 | k__Fungi(100);p__Ascomycota(100);c__Saccharomycetes(100);o__Saccharomycetales(100);f__Metschnikowiaceae(100);g__Metschnikowia(100);s__Metschnikowia_pulcherrima(100); | Metschnikowia | 13 | Saprotroph | \|Undefined Saprotroph\| | Yeast | NULL | Probable | As nectar-tap_saprotroph (Põlme et al. 2020); Decay sugar-rich substrates (Põlme et al. 2020) | Põlme S, et al. 2020. Fungal Diversity 105:1-16 ((https://doi.org/10.1007/s13225-020-00466-2)) |
| Otu58270 | 1 | k__Fungi(100);p__Ascomycota(100);c__Saccharomycetes(100);o__Saccharomycetales(100);f__Metschnikowiaceae(100);g__Metschnikowia(100);s__Metschnikowia_pulcherrima(100); | Metschnikowia | 13 | Saprotroph | \|Undefined Saprotroph\| | Yeast | NULL | Probable | As nectar-tap_saprotroph (Põlme et al. 2020); Decay sugar-rich substrates (Põlme et al. 2020) | Põlme S, et al. 2020. Fungal Diversity 105:1-16 ((https://doi.org/10.1007/s13225-020-00466-2)) |
| Otu58274 | 1 | k__Fungi(100);p__Ascomycota(100);c__Dothideomycetes(100);o__Pleosporales(100);f__Pleosporales_family_Incertae_sedis(100);g__Didymella(100);s__Didymella_sp(100); | Didymella | 13 | Pathotroph-Saprotroph | Animal Pathogen-\|Plant Pathogen\|-Plant Saprotroph-Undefined Saprotroph | Microfungus | NULL | Probable | As litter_saprotroph (Põlme et al. 2020); Decay leaf, fruit, seed, roots, soil, animal material (Põlme et al. 2020) | Tedersoo L, et al. 2014. Science 346:e1256688 ((https://doi.org/10.1126/science.1256688)); Põlme S, et al. 2020. Fungal Diversity 105:1-16 ((https://doi.org/10.1007/s13225-020-00466-2)) |
| Otu58282 | 1 | k__Fungi(100);p__Ascomycota(100);c__Saccharomycetes(100);o__Saccharomycetales(100);f__Metschnikowiaceae(100);g__Metschnikowia(100);g__Metschnikowia_unclassified(100); | Metschnikowia | 13 | Saprotroph | \|Undefined Saprotroph\| | Yeast | NULL | Probable | As nectar-tap_saprotroph (Põlme et al. 2020); Decay sugar-rich substrates (Põlme et al. 2020) | Põlme S, et al. 2020. Fungal Diversity 105:1-16 ((https://doi.org/10.1007/s13225-020-00466-2)) |
| Otu58283 | 1 | k__Fungi(100);p__Ascomycota(100);c__Saccharomycetes(100);o__Saccharomycetales(100);f__Metschnikowiaceae(100);g__Metschnikowia(100);g__Metschnikowia_unclassified(100); | Metschnikowia | 13 | Saprotroph | \|Undefined Saprotroph\| | Yeast | NULL | Probable | As nectar-tap_saprotroph (Põlme et al. 2020); Decay sugar-rich substrates (Põlme et al. 2020) | Põlme S, et al. 2020. Fungal Diversity 105:1-16 ((https://doi.org/10.1007/s13225-020-00466-2)) |
| Otu58284 | 1 | k__Fungi(100);p__Ascomycota(100);c__Saccharomycetes(100);o__Saccharomycetales(100);f__Metschnikowiaceae(100);g__Metschnikowia(100);s__Metschnikowia_pulcherrima(100); | Metschnikowia | 13 | Saprotroph | \|Undefined Saprotroph\| | Yeast | NULL | Probable | As nectar-tap_saprotroph (Põlme et al. 2020); Decay sugar-rich substrates (Põlme et al. 2020) | Põlme S, et al. 2020. Fungal Diversity 105:1-16 ((https://doi.org/10.1007/s13225-020-00466-2)) |
| Otu58285 | 1 | k__Fungi(100);p__Ascomycota(100);c__Saccharomycetes(100);o__Saccharomycetales(100);f__Metschnikowiaceae(100);g__Metschnikowia(100);g__Metschnikowia_unclassified(100); | Metschnikowia | 13 | Saprotroph | \|Undefined Saprotroph\| | Yeast | NULL | Probable | As nectar-tap_saprotroph (Põlme et al. 2020); Decay sugar-rich substrates (Põlme et al. 2020) | Põlme S, et al. 2020. Fungal Diversity 105:1-16 ((https://doi.org/10.1007/s13225-020-00466-2)) |
| Otu58286 | 1 | k__Fungi(100);p__Ascomycota(100);c__Saccharomycetes(100);o__Saccharomycetales(100);f__Metschnikowiaceae(100);g__Metschnikowia(100);s__Metschnikowia_pulcherrima(100); | Metschnikowia | 13 | Saprotroph | \|Undefined Saprotroph\| | Yeast | NULL | Probable | As nectar-tap_saprotroph (Põlme et al. 2020); Decay sugar-rich substrates (Põlme et al. 2020) | Põlme S, et al. 2020. Fungal Diversity 105:1-16 ((https://doi.org/10.1007/s13225-020-00466-2)) |
| Otu58289 | 1 | k__Fungi(100);p__Ascomycota(100);c__Saccharomycetes(100);o__Saccharomycetales(100);f__Metschnikowiaceae(100);g__Metschnikowia(100);g__Metschnikowia_unclassified(100); | Metschnikowia | 13 | Saprotroph | \|Undefined Saprotroph\| | Yeast | NULL | Probable | As nectar-tap_saprotroph (Põlme et al. 2020); Decay sugar-rich substrates (Põlme et al. 2020) | Põlme S, et al. 2020. Fungal Diversity 105:1-16 ((https://doi.org/10.1007/s13225-020-00466-2)) |
| Otu58290 | 1 | k__Fungi(100);p__Ascomycota(100);c__Saccharomycetes(100);o__Saccharomycetales(100);f__Metschnikowiaceae(100);g__Metschnikowia(100);g__Metschnikowia_unclassified(100); | Metschnikowia | 13 | Saprotroph | \|Undefined Saprotroph\| | Yeast | NULL | Probable | As nectar-tap_saprotroph (Põlme et al. 2020); Decay sugar-rich substrates (Põlme et al. 2020) | Põlme S, et al. 2020. Fungal Diversity 105:1-16 ((https://doi.org/10.1007/s13225-020-00466-2)) |
| Otu58291 | 1 | k__Fungi(100);p__Ascomycota(100);c__Saccharomycetes(100);o__Saccharomycetales(100);f__Metschnikowiaceae(100);g__Metschnikowia(100);s__Metschnikowia_pulcherrima(100); | Metschnikowia | 13 | Saprotroph | \|Undefined Saprotroph\| | Yeast | NULL | Probable | As nectar-tap_saprotroph (Põlme et al. 2020); Decay sugar-rich substrates (Põlme et al. 2020) | Põlme S, et al. 2020. Fungal Diversity 105:1-16 ((https://doi.org/10.1007/s13225-020-00466-2)) |
| Otu58292 | 1 | k__Fungi(100);p__Ascomycota(100);c__Saccharomycetes(100);o__Saccharomycetales(100);f__Metschnikowiaceae(100);g__Metschnikowia(100);s__Metschnikowia_pulcherrima(100); | Metschnikowia | 13 | Saprotroph | \|Undefined Saprotroph\| | Yeast | NULL | Probable | As nectar-tap_saprotroph (Põlme et al. 2020); Decay sugar-rich substrates (Põlme et al. 2020) | Põlme S, et al. 2020. Fungal Diversity 105:1-16 ((https://doi.org/10.1007/s13225-020-00466-2)) |
| Otu58294 | 1 | k__Fungi(100);p__Ascomycota(100);c__Saccharomycetes(100);o__Saccharomycetales(100);f__Metschnikowiaceae(100);g__Metschnikowia(100);g__Metschnikowia_unclassified(100); | Metschnikowia | 13 | Saprotroph | \|Undefined Saprotroph\| | Yeast | NULL | Probable | As nectar-tap_saprotroph (Põlme et al. 2020); Decay sugar-rich substrates (Põlme et al. 2020) | Põlme S, et al. 2020. Fungal Diversity 105:1-16 ((https://doi.org/10.1007/s13225-020-00466-2)) |
| Otu58295 | 1 | k__Fungi(100);p__Ascomycota(100);c__Saccharomycetes(100);o__Saccharomycetales(100);f__Metschnikowiaceae(100);g__Metschnikowia(100);s__Metschnikowia_pulcherrima(100); | Metschnikowia | 13 | Saprotroph | \|Undefined Saprotroph\| | Yeast | NULL | Probable | As nectar-tap_saprotroph (Põlme et al. 2020); Decay sugar-rich substrates (Põlme et al. 2020) | Põlme S, et al. 2020. Fungal Diversity 105:1-16 ((https://doi.org/10.1007/s13225-020-00466-2)) |
| Otu58296 | 1 | k__Fungi(100);p__Ascomycota(100);c__Saccharomycetes(100);o__Saccharomycetales(100);f__Metschnikowiaceae(100);g__Metschnikowia(100);g__Metschnikowia_unclassified(100); | Metschnikowia | 13 | Saprotroph | \|Undefined Saprotroph\| | Yeast | NULL | Probable | As nectar-tap_saprotroph (Põlme et al. 2020); Decay sugar-rich substrates (Põlme et al. 2020) | Põlme S, et al. 2020. Fungal Diversity 105:1-16 ((https://doi.org/10.1007/s13225-020-00466-2)) |
| Otu58297 | 1 | k__Fungi(100);p__Ascomycota(100);c__Saccharomycetes(100);o__Saccharomycetales(100);f__Metschnikowiaceae(100);g__Metschnikowia(100);s__Metschnikowia_pulcherrima(100); | Metschnikowia | 13 | Saprotroph | \|Undefined Saprotroph\| | Yeast | NULL | Probable | As nectar-tap_saprotroph (Põlme et al. 2020); Decay sugar-rich substrates (Põlme et al. 2020) | Põlme S, et al. 2020. Fungal Diversity 105:1-16 ((https://doi.org/10.1007/s13225-020-00466-2)) |
| Otu58298 | 1 | k__Fungi(100);p__Ascomycota(100);c__Saccharomycetes(100);o__Saccharomycetales(100);f__Metschnikowiaceae(100);g__Metschnikowia(100);s__Metschnikowia_pulcherrima(100); | Metschnikowia | 13 | Saprotroph | \|Undefined Saprotroph\| | Yeast | NULL | Probable | As nectar-tap_saprotroph (Põlme et al. 2020); Decay sugar-rich substrates (Põlme et al. 2020) | Põlme S, et al. 2020. Fungal Diversity 105:1-16 ((https://doi.org/10.1007/s13225-020-00466-2)) |
| Otu58299 | 1 | k__Fungi(100);p__Ascomycota(100);c__Saccharomycetes(100);o__Saccharomycetales(100);f__Metschnikowiaceae(100);g__Metschnikowia(100);s__Metschnikowia_pulcherrima(100); | Metschnikowia | 13 | Saprotroph | \|Undefined Saprotroph\| | Yeast | NULL | Probable | As nectar-tap_saprotroph (Põlme et al. 2020); Decay sugar-rich substrates (Põlme et al. 2020) | Põlme S, et al. 2020. Fungal Diversity 105:1-16 ((https://doi.org/10.1007/s13225-020-00466-2)) |
| Otu58300 | 1 | k__Fungi(100);p__Ascomycota(100);c__Saccharomycetes(100);o__Saccharomycetales(100);f__Metschnikowiaceae(100);g__Metschnikowia(100);s__Metschnikowia_pulcherrima(100); | Metschnikowia | 13 | Saprotroph | \|Undefined Saprotroph\| | Yeast | NULL | Probable | As nectar-tap_saprotroph (Põlme et al. 2020); Decay sugar-rich substrates (Põlme et al. 2020) | Põlme S, et al. 2020. Fungal Diversity 105:1-16 ((https://doi.org/10.1007/s13225-020-00466-2)) |
| Otu58301 | 1 | k__Fungi(100);p__Ascomycota(100);c__Saccharomycetes(100);o__Saccharomycetales(100);f__Metschnikowiaceae(100);g__Metschnikowia(100);s__Metschnikowia_pulcherrima(100); | Metschnikowia | 13 | Saprotroph | \|Undefined Saprotroph\| | Yeast | NULL | Probable | As nectar-tap_saprotroph (Põlme et al. 2020); Decay sugar-rich substrates (Põlme et al. 2020) | Põlme S, et al. 2020. Fungal Diversity 105:1-16 ((https://doi.org/10.1007/s13225-020-00466-2)) |
| Otu58302 | 1 | k__Fungi(100);p__Ascomycota(100);c__Saccharomycetes(100);o__Saccharomycetales(100);f__Metschnikowiaceae(100);g__Metschnikowia(100);s__Metschnikowia_pulcherrima(100); | Metschnikowia | 13 | Saprotroph | \|Undefined Saprotroph\| | Yeast | NULL | Probable | As nectar-tap_saprotroph (Põlme et al. 2020); Decay sugar-rich substrates (Põlme et al. 2020) | Põlme S, et al. 2020. Fungal Diversity 105:1-16 ((https://doi.org/10.1007/s13225-020-00466-2)) |
| Otu58303 | 1 | k__Fungi(100);p__Ascomycota(100);c__Saccharomycetes(100);o__Saccharomycetales(100);f__Metschnikowiaceae(100);g__Metschnikowia(100);g__Metschnikowia_unclassified(100); | Metschnikowia | 13 | Saprotroph | \|Undefined Saprotroph\| | Yeast | NULL | Probable | As nectar-tap_saprotroph (Põlme et al. 2020); Decay sugar-rich substrates (Põlme et al. 2020) | Põlme S, et al. 2020. Fungal Diversity 105:1-16 ((https://doi.org/10.1007/s13225-020-00466-2)) |
| Otu58304 | 1 | k__Fungi(100);p__Ascomycota(100);c__Saccharomycetes(100);o__Saccharomycetales(100);f__Metschnikowiaceae(100);g__Metschnikowia(100);s__Metschnikowia_pulcherrima(100); | Metschnikowia | 13 | Saprotroph | \|Undefined Saprotroph\| | Yeast | NULL | Probable | As nectar-tap_saprotroph (Põlme et al. 2020); Decay sugar-rich substrates (Põlme et al. 2020) | Põlme S, et al. 2020. Fungal Diversity 105:1-16 ((https://doi.org/10.1007/s13225-020-00466-2)) |
| Otu58305 | 1 | k__Fungi(100);p__Ascomycota(100);c__Saccharomycetes(100);o__Saccharomycetales(100);f__Metschnikowiaceae(100);g__Metschnikowia(100);s__Metschnikowia_pulcherrima(100); | Metschnikowia | 13 | Saprotroph | \|Undefined Saprotroph\| | Yeast | NULL | Probable | As nectar-tap_saprotroph (Põlme et al. 2020); Decay sugar-rich substrates (Põlme et al. 2020) | Põlme S, et al. 2020. Fungal Diversity 105:1-16 ((https://doi.org/10.1007/s13225-020-00466-2)) |
| Otu58306 | 1 | k__Fungi(100);p__Ascomycota(100);c__Saccharomycetes(100);o__Saccharomycetales(100);f__Metschnikowiaceae(100);g__Metschnikowia(100);s__Metschnikowia_pulcherrima(100); | Metschnikowia | 13 | Saprotroph | \|Undefined Saprotroph\| | Yeast | NULL | Probable | As nectar-tap_saprotroph (Põlme et al. 2020); Decay sugar-rich substrates (Põlme et al. 2020) | Põlme S, et al. 2020. Fungal Diversity 105:1-16 ((https://doi.org/10.1007/s13225-020-00466-2)) |
| Otu58307 | 1 | k__Fungi(100);p__Ascomycota(100);c__Saccharomycetes(100);o__Saccharomycetales(100);f__Metschnikowiaceae(100);g__Metschnikowia(100);g__Metschnikowia_unclassified(100); | Metschnikowia | 13 | Saprotroph | \|Undefined Saprotroph\| | Yeast | NULL | Probable | As nectar-tap_saprotroph (Põlme et al. 2020); Decay sugar-rich substrates (Põlme et al. 2020) | Põlme S, et al. 2020. Fungal Diversity 105:1-16 ((https://doi.org/10.1007/s13225-020-00466-2)) |
| Otu58308 | 1 | k__Fungi(100);p__Ascomycota(100);c__Saccharomycetes(100);o__Saccharomycetales(100);f__Metschnikowiaceae(100);g__Metschnikowia(100);s__Metschnikowia_pulcherrima(100); | Metschnikowia | 13 | Saprotroph | \|Undefined Saprotroph\| | Yeast | NULL | Probable | As nectar-tap_saprotroph (Põlme et al. 2020); Decay sugar-rich substrates (Põlme et al. 2020) | Põlme S, et al. 2020. Fungal Diversity 105:1-16 ((https://doi.org/10.1007/s13225-020-00466-2)) |
| Otu58309 | 1 | k__Fungi(100);p__Ascomycota(100);c__Saccharomycetes(100);o__Saccharomycetales(100);f__Metschnikowiaceae(100);g__Metschnikowia(100);s__Metschnikowia_pulcherrima(100); | Metschnikowia | 13 | Saprotroph | \|Undefined Saprotroph\| | Yeast | NULL | Probable | As nectar-tap_saprotroph (Põlme et al. 2020); Decay sugar-rich substrates (Põlme et al. 2020) | Põlme S, et al. 2020. Fungal Diversity 105:1-16 ((https://doi.org/10.1007/s13225-020-00466-2)) |
| Otu58310 | 1 | k__Fungi(100);p__Ascomycota(100);c__Saccharomycetes(100);o__Saccharomycetales(100);f__Metschnikowiaceae(100);g__Metschnikowia(100);s__Metschnikowia_pulcherrima(100); | Metschnikowia | 13 | Saprotroph | \|Undefined Saprotroph\| | Yeast | NULL | Probable | As nectar-tap_saprotroph (Põlme et al. 2020); Decay sugar-rich substrates (Põlme et al. 2020) | Põlme S, et al. 2020. Fungal Diversity 105:1-16 ((https://doi.org/10.1007/s13225-020-00466-2)) |
| Otu58311 | 1 | k__Fungi(100);p__Ascomycota(100);c__Saccharomycetes(100);o__Saccharomycetales(100);f__Metschnikowiaceae(100);g__Metschnikowia(100);s__Metschnikowia_pulcherrima(100); | Metschnikowia | 13 | Saprotroph | \|Undefined Saprotroph\| | Yeast | NULL | Probable | As nectar-tap_saprotroph (Põlme et al. 2020); Decay sugar-rich substrates (Põlme et al. 2020) | Põlme S, et al. 2020. Fungal Diversity 105:1-16 ((https://doi.org/10.1007/s13225-020-00466-2)) |
| Otu58312 | 1 | k__Fungi(100);p__Ascomycota(100);c__Saccharomycetes(100);o__Saccharomycetales(100);f__Metschnikowiaceae(100);g__Metschnikowia(100);s__Metschnikowia_pulcherrima(100); | Metschnikowia | 13 | Saprotroph | \|Undefined Saprotroph\| | Yeast | NULL | Probable | As nectar-tap_saprotroph (Põlme et al. 2020); Decay sugar-rich substrates (Põlme et al. 2020) | Põlme S, et al. 2020. Fungal Diversity 105:1-16 ((https://doi.org/10.1007/s13225-020-00466-2)) |
| Otu58313 | 1 | k__Fungi(100);p__Ascomycota(100);c__Saccharomycetes(100);o__Saccharomycetales(100);f__Metschnikowiaceae(100);g__Metschnikowia(100);s__Metschnikowia_pulcherrima(100); | Metschnikowia | 13 | Saprotroph | \|Undefined Saprotroph\| | Yeast | NULL | Probable | As nectar-tap_saprotroph (Põlme et al. 2020); Decay sugar-rich substrates (Põlme et al. 2020) | Põlme S, et al. 2020. Fungal Diversity 105:1-16 ((https://doi.org/10.1007/s13225-020-00466-2)) |
| Otu58314 | 1 | k__Fungi(100);p__Ascomycota(100);c__Saccharomycetes(100);o__Saccharomycetales(100);f__Metschnikowiaceae(100);g__Metschnikowia(100);g__Metschnikowia_unclassified(100); | Metschnikowia | 13 | Saprotroph | \|Undefined Saprotroph\| | Yeast | NULL | Probable | As nectar-tap_saprotroph (Põlme et al. 2020); Decay sugar-rich substrates (Põlme et al. 2020) | Põlme S, et al. 2020. Fungal Diversity 105:1-16 ((https://doi.org/10.1007/s13225-020-00466-2)) |
| Otu58315 | 1 | k__Fungi(100);p__Ascomycota(100);c__Saccharomycetes(100);o__Saccharomycetales(100);f__Metschnikowiaceae(100);g__Metschnikowia(100);g__Metschnikowia_unclassified(100); | Metschnikowia | 13 | Saprotroph | \|Undefined Saprotroph\| | Yeast | NULL | Probable | As nectar-tap_saprotroph (Põlme et al. 2020); Decay sugar-rich substrates (Põlme et al. 2020) | Põlme S, et al. 2020. Fungal Diversity 105:1-16 ((https://doi.org/10.1007/s13225-020-00466-2)) |
| Otu58316 | 1 | k__Fungi(100);p__Ascomycota(100);c__Saccharomycetes(100);o__Saccharomycetales(100);f__Metschnikowiaceae(100);g__Metschnikowia(100);s__Metschnikowia_pulcherrima(100); | Metschnikowia | 13 | Saprotroph | \|Undefined Saprotroph\| | Yeast | NULL | Probable | As nectar-tap_saprotroph (Põlme et al. 2020); Decay sugar-rich substrates (Põlme et al. 2020) | Põlme S, et al. 2020. Fungal Diversity 105:1-16 ((https://doi.org/10.1007/s13225-020-00466-2)) |
| Otu58317 | 1 | k__Fungi(100);p__Ascomycota(100);c__Saccharomycetes(100);o__Saccharomycetales(100);f__Metschnikowiaceae(100);g__Metschnikowia(100);g__Metschnikowia_unclassified(100); | Metschnikowia | 13 | Saprotroph | \|Undefined Saprotroph\| | Yeast | NULL | Probable | As nectar-tap_saprotroph (Põlme et al. 2020); Decay sugar-rich substrates (Põlme et al. 2020) | Põlme S, et al. 2020. Fungal Diversity 105:1-16 ((https://doi.org/10.1007/s13225-020-00466-2)) |
| Otu58318 | 1 | k__Fungi(100);p__Ascomycota(100);c__Saccharomycetes(100);o__Saccharomycetales(100);f__Metschnikowiaceae(100);g__Metschnikowia(100);g__Metschnikowia_unclassified(100); | Metschnikowia | 13 | Saprotroph | \|Undefined Saprotroph\| | Yeast | NULL | Probable | As nectar-tap_saprotroph (Põlme et al. 2020); Decay sugar-rich substrates (Põlme et al. 2020) | Põlme S, et al. 2020. Fungal Diversity 105:1-16 ((https://doi.org/10.1007/s13225-020-00466-2)) |
| Otu58319 | 1 | k__Fungi(100);p__Ascomycota(100);c__Saccharomycetes(100);o__Saccharomycetales(100);f__Metschnikowiaceae(100);g__Metschnikowia(100);s__Metschnikowia_pulcherrima(100); | Metschnikowia | 13 | Saprotroph | \|Undefined Saprotroph\| | Yeast | NULL | Probable | As nectar-tap_saprotroph (Põlme et al. 2020); Decay sugar-rich substrates (Põlme et al. 2020) | Põlme S, et al. 2020. Fungal Diversity 105:1-16 ((https://doi.org/10.1007/s13225-020-00466-2)) |
| Otu58320 | 1 | k__Fungi(100);p__Ascomycota(100);c__Saccharomycetes(100);o__Saccharomycetales(100);f__Metschnikowiaceae(100);g__Metschnikowia(100);s__Metschnikowia_pulcherrima(100); | Metschnikowia | 13 | Saprotroph | \|Undefined Saprotroph\| | Yeast | NULL | Probable | As nectar-tap_saprotroph (Põlme et al. 2020); Decay sugar-rich substrates (Põlme et al. 2020) | Põlme S, et al. 2020. Fungal Diversity 105:1-16 ((https://doi.org/10.1007/s13225-020-00466-2)) |
| Otu58321 | 1 | k__Fungi(100);p__Ascomycota(100);c__Saccharomycetes(100);o__Saccharomycetales(100);f__Metschnikowiaceae(100);g__Metschnikowia(100);s__Metschnikowia_pulcherrima(100); | Metschnikowia | 13 | Saprotroph | \|Undefined Saprotroph\| | Yeast | NULL | Probable | As nectar-tap_saprotroph (Põlme et al. 2020); Decay sugar-rich substrates (Põlme et al. 2020) | Põlme S, et al. 2020. Fungal Diversity 105:1-16 ((https://doi.org/10.1007/s13225-020-00466-2)) |
| Otu58322 | 1 | k__Fungi(100);p__Ascomycota(100);c__Saccharomycetes(100);o__Saccharomycetales(100);f__Metschnikowiaceae(100);g__Metschnikowia(100);s__Metschnikowia_pulcherrima(100); | Metschnikowia | 13 | Saprotroph | \|Undefined Saprotroph\| | Yeast | NULL | Probable | As nectar-tap_saprotroph (Põlme et al. 2020); Decay sugar-rich substrates (Põlme et al. 2020) | Põlme S, et al. 2020. Fungal Diversity 105:1-16 ((https://doi.org/10.1007/s13225-020-00466-2)) |
| Otu58323 | 1 | k__Fungi(100);p__Ascomycota(100);c__Saccharomycetes(100);o__Saccharomycetales(100);f__Metschnikowiaceae(100);g__Metschnikowia(100);g__Metschnikowia_unclassified(100); | Metschnikowia | 13 | Saprotroph | \|Undefined Saprotroph\| | Yeast | NULL | Probable | As nectar-tap_saprotroph (Põlme et al. 2020); Decay sugar-rich substrates (Põlme et al. 2020) | Põlme S, et al. 2020. Fungal Diversity 105:1-16 ((https://doi.org/10.1007/s13225-020-00466-2)) |
| Otu58324 | 1 | k__Fungi(100);p__Ascomycota(100);c__Saccharomycetes(100);o__Saccharomycetales(100);f__Metschnikowiaceae(100);g__Metschnikowia(100);s__Metschnikowia_pulcherrima(100); | Metschnikowia | 13 | Saprotroph | \|Undefined Saprotroph\| | Yeast | NULL | Probable | As nectar-tap_saprotroph (Põlme et al. 2020); Decay sugar-rich substrates (Põlme et al. 2020) | Põlme S, et al. 2020. Fungal Diversity 105:1-16 ((https://doi.org/10.1007/s13225-020-00466-2)) |
| Otu58325 | 1 | k__Fungi(100);p__Ascomycota(100);c__Saccharomycetes(100);o__Saccharomycetales(100);f__Metschnikowiaceae(100);g__Metschnikowia(100);s__Metschnikowia_pulcherrima(100); | Metschnikowia | 13 | Saprotroph | \|Undefined Saprotroph\| | Yeast | NULL | Probable | As nectar-tap_saprotroph (Põlme et al. 2020); Decay sugar-rich substrates (Põlme et al. 2020) | Põlme S, et al. 2020. Fungal Diversity 105:1-16 ((https://doi.org/10.1007/s13225-020-00466-2)) |
| Otu58326 | 1 | k__Fungi(100);p__Ascomycota(100);c__Saccharomycetes(100);o__Saccharomycetales(100);f__Metschnikowiaceae(100);g__Metschnikowia(100);s__Metschnikowia_pulcherrima(100); | Metschnikowia | 13 | Saprotroph | \|Undefined Saprotroph\| | Yeast | NULL | Probable | As nectar-tap_saprotroph (Põlme et al. 2020); Decay sugar-rich substrates (Põlme et al. 2020) | Põlme S, et al. 2020. Fungal Diversity 105:1-16 ((https://doi.org/10.1007/s13225-020-00466-2)) |
| Otu58327 | 1 | k__Fungi(100);p__Ascomycota(100);c__Saccharomycetes(100);o__Saccharomycetales(100);f__Metschnikowiaceae(100);g__Metschnikowia(100);g__Metschnikowia_unclassified(100); | Metschnikowia | 13 | Saprotroph | \|Undefined Saprotroph\| | Yeast | NULL | Probable | As nectar-tap_saprotroph (Põlme et al. 2020); Decay sugar-rich substrates (Põlme et al. 2020) | Põlme S, et al. 2020. Fungal Diversity 105:1-16 ((https://doi.org/10.1007/s13225-020-00466-2)) |
| Otu58328 | 1 | k__Fungi(100);p__Ascomycota(100);c__Saccharomycetes(100);o__Saccharomycetales(100);f__Metschnikowiaceae(100);g__Metschnikowia(100);s__Metschnikowia_pulcherrima(100); | Metschnikowia | 13 | Saprotroph | \|Undefined Saprotroph\| | Yeast | NULL | Probable | As nectar-tap_saprotroph (Põlme et al. 2020); Decay sugar-rich substrates (Põlme et al. 2020) | Põlme S, et al. 2020. Fungal Diversity 105:1-16 ((https://doi.org/10.1007/s13225-020-00466-2)) |
| Otu58329 | 1 | k__Fungi(100);p__Ascomycota(100);c__Saccharomycetes(100);o__Saccharomycetales(100);f__Metschnikowiaceae(100);g__Metschnikowia(100);s__Metschnikowia_pulcherrima(100); | Metschnikowia | 13 | Saprotroph | \|Undefined Saprotroph\| | Yeast | NULL | Probable | As nectar-tap_saprotroph (Põlme et al. 2020); Decay sugar-rich substrates (Põlme et al. 2020) | Põlme S, et al. 2020. Fungal Diversity 105:1-16 ((https://doi.org/10.1007/s13225-020-00466-2)) |
| Otu58330 | 1 | k__Fungi(100);p__Ascomycota(100);c__Saccharomycetes(100);o__Saccharomycetales(100);f__Metschnikowiaceae(100);g__Metschnikowia(100);g__Metschnikowia_unclassified(100); | Metschnikowia | 13 | Saprotroph | \|Undefined Saprotroph\| | Yeast | NULL | Probable | As nectar-tap_saprotroph (Põlme et al. 2020); Decay sugar-rich substrates (Põlme et al. 2020) | Põlme S, et al. 2020. Fungal Diversity 105:1-16 ((https://doi.org/10.1007/s13225-020-00466-2)) |
| Otu58331 | 1 | k__Fungi(100);p__Ascomycota(100);c__Saccharomycetes(100);o__Saccharomycetales(100);f__Metschnikowiaceae(100);g__Metschnikowia(100);s__Metschnikowia_pulcherrima(100); | Metschnikowia | 13 | Saprotroph | \|Undefined Saprotroph\| | Yeast | NULL | Probable | As nectar-tap_saprotroph (Põlme et al. 2020); Decay sugar-rich substrates (Põlme et al. 2020) | Põlme S, et al. 2020. Fungal Diversity 105:1-16 ((https://doi.org/10.1007/s13225-020-00466-2)) |
| Otu58333 | 1 | k__Fungi(100);p__Ascomycota(100);c__Saccharomycetes(100);o__Saccharomycetales(100);f__Metschnikowiaceae(100);g__Metschnikowia(100);s__Metschnikowia_pulcherrima(100); | Metschnikowia | 13 | Saprotroph | \|Undefined Saprotroph\| | Yeast | NULL | Probable | As nectar-tap_saprotroph (Põlme et al. 2020); Decay sugar-rich substrates (Põlme et al. 2020) | Põlme S, et al. 2020. Fungal Diversity 105:1-16 ((https://doi.org/10.1007/s13225-020-00466-2)) |
| Otu58335 | 1 | k__Fungi(100);p__Ascomycota(100);c__Saccharomycetes(100);o__Saccharomycetales(100);f__Metschnikowiaceae(100);g__Metschnikowia(100);s__Metschnikowia_pulcherrima(100); | Metschnikowia | 13 | Saprotroph | \|Undefined Saprotroph\| | Yeast | NULL | Probable | As nectar-tap_saprotroph (Põlme et al. 2020); Decay sugar-rich substrates (Põlme et al. 2020) | Põlme S, et al. 2020. Fungal Diversity 105:1-16 ((https://doi.org/10.1007/s13225-020-00466-2)) |
| Otu58336 | 1 | k__Fungi(100);p__Ascomycota(100);c__Saccharomycetes(100);o__Saccharomycetales(100);f__Metschnikowiaceae(100);g__Metschnikowia(100);s__Metschnikowia_pulcherrima(100); | Metschnikowia | 13 | Saprotroph | \|Undefined Saprotroph\| | Yeast | NULL | Probable | As nectar-tap_saprotroph (Põlme et al. 2020); Decay sugar-rich substrates (Põlme et al. 2020) | Põlme S, et al. 2020. Fungal Diversity 105:1-16 ((https://doi.org/10.1007/s13225-020-00466-2)) |
| Otu58337 | 1 | k__Fungi(100);p__Ascomycota(100);c__Saccharomycetes(100);o__Saccharomycetales(100);f__Metschnikowiaceae(100);g__Metschnikowia(100);g__Metschnikowia_unclassified(100); | Metschnikowia | 13 | Saprotroph | \|Undefined Saprotroph\| | Yeast | NULL | Probable | As nectar-tap_saprotroph (Põlme et al. 2020); Decay sugar-rich substrates (Põlme et al. 2020) | Põlme S, et al. 2020. Fungal Diversity 105:1-16 ((https://doi.org/10.1007/s13225-020-00466-2)) |
| Otu58338 | 1 | k__Fungi(100);p__Ascomycota(100);c__Saccharomycetes(100);o__Saccharomycetales(100);f__Metschnikowiaceae(100);g__Metschnikowia(100);s__Metschnikowia_pulcherrima(100); | Metschnikowia | 13 | Saprotroph | \|Undefined Saprotroph\| | Yeast | NULL | Probable | As nectar-tap_saprotroph (Põlme et al. 2020); Decay sugar-rich substrates (Põlme et al. 2020) | Põlme S, et al. 2020. Fungal Diversity 105:1-16 ((https://doi.org/10.1007/s13225-020-00466-2)) |
| Otu58339 | 1 | k__Fungi(100);p__Ascomycota(100);c__Saccharomycetes(100);o__Saccharomycetales(100);f__Metschnikowiaceae(100);g__Metschnikowia(100);s__Metschnikowia_pulcherrima(100); | Metschnikowia | 13 | Saprotroph | \|Undefined Saprotroph\| | Yeast | NULL | Probable | As nectar-tap_saprotroph (Põlme et al. 2020); Decay sugar-rich substrates (Põlme et al. 2020) | Põlme S, et al. 2020. Fungal Diversity 105:1-16 ((https://doi.org/10.1007/s13225-020-00466-2)) |
| Otu58340 | 1 | k__Fungi(100);p__Ascomycota(100);c__Saccharomycetes(100);o__Saccharomycetales(100);f__Metschnikowiaceae(100);g__Metschnikowia(100);s__Metschnikowia_pulcherrima(100); | Metschnikowia | 13 | Saprotroph | \|Undefined Saprotroph\| | Yeast | NULL | Probable | As nectar-tap_saprotroph (Põlme et al. 2020); Decay sugar-rich substrates (Põlme et al. 2020) | Põlme S, et al. 2020. Fungal Diversity 105:1-16 ((https://doi.org/10.1007/s13225-020-00466-2)) |
| Otu58341 | 1 | k__Fungi(100);p__Ascomycota(100);c__Saccharomycetes(100);o__Saccharomycetales(100);f__Metschnikowiaceae(100);g__Metschnikowia(100);g__Metschnikowia_unclassified(100); | Metschnikowia | 13 | Saprotroph | \|Undefined Saprotroph\| | Yeast | NULL | Probable | As nectar-tap_saprotroph (Põlme et al. 2020); Decay sugar-rich substrates (Põlme et al. 2020) | Põlme S, et al. 2020. Fungal Diversity 105:1-16 ((https://doi.org/10.1007/s13225-020-00466-2)) |
| Otu58342 | 1 | k__Fungi(100);p__Ascomycota(100);c__Saccharomycetes(100);o__Saccharomycetales(100);f__Metschnikowiaceae(100);g__Metschnikowia(100);g__Metschnikowia_unclassified(100); | Metschnikowia | 13 | Saprotroph | \|Undefined Saprotroph\| | Yeast | NULL | Probable | As nectar-tap_saprotroph (Põlme et al. 2020); Decay sugar-rich substrates (Põlme et al. 2020) | Põlme S, et al. 2020. Fungal Diversity 105:1-16 ((https://doi.org/10.1007/s13225-020-00466-2)) |
| Otu58343 | 1 | k__Fungi(100);p__Ascomycota(100);c__Saccharomycetes(100);o__Saccharomycetales(100);f__Metschnikowiaceae(100);g__Metschnikowia(100);g__Metschnikowia_unclassified(100); | Metschnikowia | 13 | Saprotroph | \|Undefined Saprotroph\| | Yeast | NULL | Probable | As nectar-tap_saprotroph (Põlme et al. 2020); Decay sugar-rich substrates (Põlme et al. 2020) | Põlme S, et al. 2020. Fungal Diversity 105:1-16 ((https://doi.org/10.1007/s13225-020-00466-2)) |
| Otu58344 | 1 | k__Fungi(100);p__Ascomycota(100);c__Saccharomycetes(100);o__Saccharomycetales(100);f__Metschnikowiaceae(100);g__Metschnikowia(100);g__Metschnikowia_unclassified(100); | Metschnikowia | 13 | Saprotroph | \|Undefined Saprotroph\| | Yeast | NULL | Probable | As nectar-tap_saprotroph (Põlme et al. 2020); Decay sugar-rich substrates (Põlme et al. 2020) | Põlme S, et al. 2020. Fungal Diversity 105:1-16 ((https://doi.org/10.1007/s13225-020-00466-2)) |
| Otu58345 | 1 | k__Fungi(100);p__Ascomycota(100);c__Saccharomycetes(100);o__Saccharomycetales(100);f__Metschnikowiaceae(100);g__Metschnikowia(100);s__Metschnikowia_pulcherrima(100); | Metschnikowia | 13 | Saprotroph | \|Undefined Saprotroph\| | Yeast | NULL | Probable | As nectar-tap_saprotroph (Põlme et al. 2020); Decay sugar-rich substrates (Põlme et al. 2020) | Põlme S, et al. 2020. Fungal Diversity 105:1-16 ((https://doi.org/10.1007/s13225-020-00466-2)) |

| | | | | | | | | | | | |
|---|---|---|---|---|---|---|---|---|---|---|---|
| Otu58346 | 1 | k__Fungi(100);p__Ascomycota(100);c__Saccharomycetes(100);o__Saccharomycetales(100);f__Metschnikowiaceae(100);g__Metschnikowia(100);g__Metschnikowia_unclassified(100); | Metschnikowia | 13 | Saprotroph | \|Undefined Saprotroph\| | Yeast | NULL | Probable | As nectar-tap_saprotroph (Põlme et al. 2020); Decay sugar-rich substrates (Põlme et al. 2020) | Põlme S, et al. 2020. Fungal Diversity 105:1-16 ((https://doi.org/10.1007/s13225-020-00466-2)) |
| Otu58347 | 1 | k__Fungi(100);p__Ascomycota(100);c__Saccharomycetes(100);o__Saccharomycetales(100);f__Metschnikowiaceae(100);g__Metschnikowia(100);g__Metschnikowia_unclassified(100); | Metschnikowia | 13 | Saprotroph | \|Undefined Saprotroph\| | Yeast | NULL | Probable | As nectar-tap_saprotroph (Põlme et al. 2020); Decay sugar-rich substrates (Põlme et al. 2020) | Põlme S, et al. 2020. Fungal Diversity 105:1-16 ((https://doi.org/10.1007/s13225-020-00466-2)) |
| Otu58349 | 1 | k__Fungi(100);p__Ascomycota(100);c__Saccharomycetes(100);o__Saccharomycetales(100);f__Metschnikowiaceae(100);g__Metschnikowia(100);s__Metschnikowia_pulcherrima(100); | Metschnikowia | 13 | Saprotroph | \|Undefined Saprotroph\| | Yeast | NULL | Probable | As nectar-tap_saprotroph (Põlme et al. 2020); Decay sugar-rich substrates (Põlme et al. 2020) | Põlme S, et al. 2020. Fungal Diversity 105:1-16 ((https://doi.org/10.1007/s13225-020-00466-2)) |
| Otu58350 | 1 | k__Fungi(100);p__Ascomycota(100);c__Saccharomycetes(100);o__Saccharomycetales(100);f__Metschnikowiaceae(100);g__Metschnikowia(100);s__Metschnikowia_pulcherrima(100); | Metschnikowia | 13 | Saprotroph | \|Undefined Saprotroph\| | Yeast | NULL | Probable | As nectar-tap_saprotroph (Põlme et al. 2020); Decay sugar-rich substrates (Põlme et al. 2020) | Põlme S, et al. 2020. Fungal Diversity 105:1-16 ((https://doi.org/10.1007/s13225-020-00466-2)) |
| Otu58351 | 1 | k__Fungi(100);p__Ascomycota(100);c__Saccharomycetes(100);o__Saccharomycetales(100);f__Metschnikowiaceae(100);g__Metschnikowia(100);g__Metschnikowia_unclassified(100); | Metschnikowia | 13 | Saprotroph | \|Undefined Saprotroph\| | Yeast | NULL | Probable | As nectar-tap_saprotroph (Põlme et al. 2020); Decay sugar-rich substrates (Põlme et al. 2020) | Põlme S, et al. 2020. Fungal Diversity 105:1-16 ((https://doi.org/10.1007/s13225-020-00466-2)) |
| Otu58352 | 1 | k__Fungi(100);p__Ascomycota(100);c__Saccharomycetes(100);o__Saccharomycetales(100);f__Metschnikowiaceae(100);g__Metschnikowia(100);s__Metschnikowia_pulcherrima(100); | Metschnikowia | 13 | Saprotroph | \|Undefined Saprotroph\| | Yeast | NULL | Probable | As nectar-tap_saprotroph (Põlme et al. 2020); Decay sugar-rich substrates (Põlme et al. 2020) | Põlme S, et al. 2020. Fungal Diversity 105:1-16 ((https://doi.org/10.1007/s13225-020-00466-2)) |
| Otu58354 | 1 | k__Fungi(100);p__Ascomycota(100);c__Saccharomycetes(100);o__Saccharomycetales(100);f__Metschnikowiaceae(100);g__Metschnikowia(100);g__Metschnikowia_unclassified(100); | Metschnikowia | 13 | Saprotroph | \|Undefined Saprotroph\| | Yeast | NULL | Probable | As nectar-tap_saprotroph (Põlme et al. 2020); Decay sugar-rich substrates (Põlme et al. 2020) | Põlme S, et al. 2020. Fungal Diversity 105:1-16 ((https://doi.org/10.1007/s13225-020-00466-2)) |
| Otu58355 | 1 | k__Fungi(100);p__Ascomycota(100);c__Saccharomycetes(100);o__Saccharomycetales(100);f__Metschnikowiaceae(100);g__Metschnikowia(100);s__Metschnikowia_pulcherrima(100); | Metschnikowia | 13 | Saprotroph | \|Undefined Saprotroph\| | Yeast | NULL | Probable | As nectar-tap_saprotroph (Põlme et al. 2020); Decay sugar-rich substrates (Põlme et al. 2020) | Põlme S, et al. 2020. Fungal Diversity 105:1-16 ((https://doi.org/10.1007/s13225-020-00466-2)) |
| Otu58361 | 1 | k__Fungi(100);p__Ascomycota(100);c__Saccharomycetes(100);o__Saccharomycetales(100);f__Metschnikowiaceae(100);g__Metschnikowia(100);s__Metschnikowia_pulcherrima(100); | Metschnikowia | 13 | Saprotroph | \|Undefined Saprotroph\| | Yeast | NULL | Probable | As nectar-tap_saprotroph (Põlme et al. 2020); Decay sugar-rich substrates (Põlme et al. 2020) | Põlme S, et al. 2020. Fungal Diversity 105:1-16 ((https://doi.org/10.1007/s13225-020-00466-2)) |
| Otu58362 | 1 | k__Fungi(100);p__Ascomycota(100);c__Saccharomycetes(100);o__Saccharomycetales(100);f__Metschnikowiaceae(100);g__Metschnikowia(100);s__Metschnikowia_pulcherrima(100); | Metschnikowia | 13 | Saprotroph | \|Undefined Saprotroph\| | Yeast | NULL | Probable | As nectar-tap_saprotroph (Põlme et al. 2020); Decay sugar-rich substrates (Põlme et al. 2020) | Põlme S, et al. 2020. Fungal Diversity 105:1-16 ((https://doi.org/10.1007/s13225-020-00466-2)) |
| Otu58363 | 1 | k__Fungi(100);p__Ascomycota(100);c__Saccharomycetes(100);o__Saccharomycetales(100);f__Metschnikowiaceae(100);g__Metschnikowia(100);g__Metschnikowia_unclassified(100); | Metschnikowia | 13 | Saprotroph | \|Undefined Saprotroph\| | Yeast | NULL | Probable | As nectar-tap_saprotroph (Põlme et al. 2020); Decay sugar-rich substrates (Põlme et al. 2020) | Põlme S, et al. 2020. Fungal Diversity 105:1-16 ((https://doi.org/10.1007/s13225-020-00466-2)) |
| Otu58366 | 1 | k__Fungi(100);p__Ascomycota(100);c__Saccharomycetes(100);o__Saccharomycetales(100);f__Metschnikowiaceae(100);g__Metschnikowia(100);g__Metschnikowia_unclassified(100); | Metschnikowia | 13 | Saprotroph | \|Undefined Saprotroph\| | Yeast | NULL | Probable | As nectar-tap_saprotroph (Põlme et al. 2020); Decay sugar-rich substrates (Põlme et al. 2020) | Põlme S, et al. 2020. Fungal Diversity 105:1-16 ((https://doi.org/10.1007/s13225-020-00466-2)) |
| Otu58367 | 1 | k__Fungi(100);p__Ascomycota(100);c__Saccharomycetes(100);o__Saccharomycetales(100);f__Metschnikowiaceae(100);g__Metschnikowia(100);s__Metschnikowia_pulcherrima(100); | Metschnikowia | 13 | Saprotroph | \|Undefined Saprotroph\| | Yeast | NULL | Probable | As nectar-tap_saprotroph (Põlme et al. 2020); Decay sugar-rich substrates (Põlme et al. 2020) | Põlme S, et al. 2020. Fungal Diversity 105:1-16 ((https://doi.org/10.1007/s13225-020-00466-2)) |
| Otu58368 | 1 | k__Fungi(100);p__Ascomycota(100);c__Saccharomycetes(100);o__Saccharomycetales(100);f__Metschnikowiaceae(100);g__Metschnikowia(100);g__Metschnikowia_unclassified(100); | Metschnikowia | 13 | Saprotroph | \|Undefined Saprotroph\| | Yeast | NULL | Probable | As nectar-tap_saprotroph (Põlme et al. 2020); Decay sugar-rich substrates (Põlme et al. 2020) | Põlme S, et al. 2020. Fungal Diversity 105:1-16 ((https://doi.org/10.1007/s13225-020-00466-2)) |
| Otu58369 | 1 | k__Fungi(100);p__Ascomycota(100);c__Saccharomycetes(100);o__Saccharomycetales(100);f__Metschnikowiaceae(100);g__Metschnikowia(100);s__Metschnikowia_pulcherrima(100); | Metschnikowia | 13 | Saprotroph | \|Undefined Saprotroph\| | Yeast | NULL | Probable | As nectar-tap_saprotroph (Põlme et al. 2020); Decay sugar-rich substrates (Põlme et al. 2020) | Põlme S, et al. 2020. Fungal Diversity 105:1-16 ((https://doi.org/10.1007/s13225-020-00466-2)) |
| Otu58370 | 1 | k__Fungi(100);p__Ascomycota(100);c__Saccharomycetes(100);o__Saccharomycetales(100);f__Metschnikowiaceae(100);g__Metschnikowia(100);s__Metschnikowia_pulcherrima(100); | Metschnikowia | 13 | Saprotroph | \|Undefined Saprotroph\| | Yeast | NULL | Probable | As nectar-tap_saprotroph (Põlme et al. 2020); Decay sugar-rich substrates (Põlme et al. 2020) | Põlme S, et al. 2020. Fungal Diversity 105:1-16 ((https://doi.org/10.1007/s13225-020-00466-2)) |
| Otu58371 | 1 | k__Fungi(100);p__Ascomycota(100);c__Saccharomycetes(100);o__Saccharomycetales(100);f__Metschnikowiaceae(100);g__Metschnikowia(100);s__Metschnikowia_pulcherrima(100); | Metschnikowia | 13 | Saprotroph | \|Undefined Saprotroph\| | Yeast | NULL | Probable | As nectar-tap_saprotroph (Põlme et al. 2020); Decay sugar-rich substrates (Põlme et al. 2020) | Põlme S, et al. 2020. Fungal Diversity 105:1-16 ((https://doi.org/10.1007/s13225-020-00466-2)) |
| Otu58372 | 1 | k__Fungi(100);p__Ascomycota(100);c__Saccharomycetes(100);o__Saccharomycetales(100);f__Metschnikowiaceae(100);g__Metschnikowia(100);s__Metschnikowia_pulcherrima(100); | Metschnikowia | 13 | Saprotroph | \|Undefined Saprotroph\| | Yeast | NULL | Probable | As nectar-tap_saprotroph (Põlme et al. 2020); Decay sugar-rich substrates (Põlme et al. 2020) | Põlme S, et al. 2020. Fungal Diversity 105:1-16 ((https://doi.org/10.1007/s13225-020-00466-2)) |
| Otu58373 | 1 | k__Fungi(100);p__Ascomycota(100);c__Saccharomycetes(100);o__Saccharomycetales(100);f__Metschnikowiaceae(100);g__Metschnikowia(100);s__Metschnikowia_pulcherrima(100); | Metschnikowia | 13 | Saprotroph | \|Undefined Saprotroph\| | Yeast | NULL | Probable | As nectar-tap_saprotroph (Põlme et al. 2020); Decay sugar-rich substrates (Põlme et al. 2020) | Põlme S, et al. 2020. Fungal Diversity 105:1-16 ((https://doi.org/10.1007/s13225-020-00466-2)) |
| Otu58374 | 1 | k__Fungi(100);p__Ascomycota(100);c__Saccharomycetes(100);o__Saccharomycetales(100);f__Metschnikowiaceae(100);g__Metschnikowia(100);s__Metschnikowia_pulcherrima(100); | Metschnikowia | 13 | Saprotroph | \|Undefined Saprotroph\| | Yeast | NULL | Probable | As nectar-tap_saprotroph (Põlme et al. 2020); Decay sugar-rich substrates (Põlme et al. 2020) | Põlme S, et al. 2020. Fungal Diversity 105:1-16 ((https://doi.org/10.1007/s13225-020-00466-2)) |
| Otu58375 | 1 | k__Fungi(100);p__Ascomycota(100);c__Saccharomycetes(100);o__Saccharomycetales(100);f__Metschnikowiaceae(100);g__Metschnikowia(100);s__Metschnikowia_pulcherrima(100); | Metschnikowia | 13 | Saprotroph | \|Undefined Saprotroph\| | Yeast | NULL | Probable | As nectar-tap_saprotroph (Põlme et al. 2020); Decay sugar-rich substrates (Põlme et al. 2020) | Põlme S, et al. 2020. Fungal Diversity 105:1-16 ((https://doi.org/10.1007/s13225-020-00466-2)) |
| Otu58376 | 1 | k__Fungi(100);p__Ascomycota(100);c__Saccharomycetes(100);o__Saccharomycetales(100);f__Metschnikowiaceae(100);g__Metschnikowia(100);g__Metschnikowia_unclassified(100); | Metschnikowia | 13 | Saprotroph | \|Undefined Saprotroph\| | Yeast | NULL | Probable | As nectar-tap_saprotroph (Põlme et al. 2020); Decay sugar-rich substrates (Põlme et al. 2020) | Põlme S, et al. 2020. Fungal Diversity 105:1-16 ((https://doi.org/10.1007/s13225-020-00466-2)) |
| Otu58377 | 1 | k__Fungi(100);p__Ascomycota(100);c__Saccharomycetes(100);o__Saccharomycetales(100);f__Metschnikowiaceae(100);g__Metschnikowia(100);s__Metschnikowia_pulcherrima(100); | Metschnikowia | 13 | Saprotroph | \|Undefined Saprotroph\| | Yeast | NULL | Probable | As nectar-tap_saprotroph (Põlme et al. 2020); Decay sugar-rich substrates (Põlme et al. 2020) | Põlme S, et al. 2020. Fungal Diversity 105:1-16 ((https://doi.org/10.1007/s13225-020-00466-2)) |
| Otu58378 | 1 | k__Fungi(100);p__Ascomycota(100);c__Saccharomycetes(100);o__Saccharomycetales(100);f__Metschnikowiaceae(100);g__Metschnikowia(100);s__Metschnikowia_pulcherrima(100); | Metschnikowia | 13 | Saprotroph | \|Undefined Saprotroph\| | Yeast | NULL | Probable | As nectar-tap_saprotroph (Põlme et al. 2020); Decay sugar-rich substrates (Põlme et al. 2020) | Põlme S, et al. 2020. Fungal Diversity 105:1-16 ((https://doi.org/10.1007/s13225-020-00466-2)) |
| Otu58379 | 1 | k__Fungi(100);p__Ascomycota(100);c__Saccharomycetes(100);o__Saccharomycetales(100);f__Metschnikowiaceae(100);g__Metschnikowia(100);s__Metschnikowia_pulcherrima(100); | Metschnikowia | 13 | Saprotroph | \|Undefined Saprotroph\| | Yeast | NULL | Probable | As nectar-tap_saprotroph (Põlme et al. 2020); Decay sugar-rich substrates (Põlme et al. 2020) | Põlme S, et al. 2020. Fungal Diversity 105:1-16 ((https://doi.org/10.1007/s13225-020-00466-2)) |
| Otu58380 | 1 | k__Fungi(100);p__Ascomycota(100);c__Saccharomycetes(100);o__Saccharomycetales(100);f__Metschnikowiaceae(100);g__Metschnikowia(100);s__Metschnikowia_pulcherrima(100); | Metschnikowia | 13 | Saprotroph | \|Undefined Saprotroph\| | Yeast | NULL | Probable | As nectar-tap_saprotroph (Põlme et al. 2020); Decay sugar-rich substrates (Põlme et al. 2020) | Põlme S, et al. 2020. Fungal Diversity 105:1-16 ((https://doi.org/10.1007/s13225-020-00466-2)) |
| Otu58381 | 1 | k__Fungi(100);p__Ascomycota(100);c__Saccharomycetes(100);o__Saccharomycetales(100);f__Metschnikowiaceae(100);g__Metschnikowia(100);g__Metschnikowia_unclassified(100); | Metschnikowia | 13 | Saprotroph | \|Undefined Saprotroph\| | Yeast | NULL | Probable | As nectar-tap_saprotroph (Põlme et al. 2020); Decay sugar-rich substrates (Põlme et al. 2020) | Põlme S, et al. 2020. Fungal Diversity 105:1-16 ((https://doi.org/10.1007/s13225-020-00466-2)) |
| Otu58382 | 1 | k__Fungi(100);p__Ascomycota(100);c__Saccharomycetes(100);o__Saccharomycetales(100);f__Metschnikowiaceae(100);g__Metschnikowia(100);s__Metschnikowia_pulcherrima(100); | Metschnikowia | 13 | Saprotroph | \|Undefined Saprotroph\| | Yeast | NULL | Probable | As nectar-tap_saprotroph (Põlme et al. 2020); Decay sugar-rich substrates (Põlme et al. 2020) | Põlme S, et al. 2020. Fungal Diversity 105:1-16 ((https://doi.org/10.1007/s13225-020-00466-2)) |
| Otu58383 | 1 | k__Fungi(100);p__Ascomycota(100);c__Saccharomycetes(100);o__Saccharomycetales(100);f__Metschnikowiaceae(100);g__Metschnikowia(100);s__Metschnikowia_pulcherrima(100); | Metschnikowia | 13 | Saprotroph | \|Undefined Saprotroph\| | Yeast | NULL | Probable | As nectar-tap_saprotroph (Põlme et al. 2020); Decay sugar-rich substrates (Põlme et al. 2020) | Põlme S, et al. 2020. Fungal Diversity 105:1-16 ((https://doi.org/10.1007/s13225-020-00466-2)) |
| Otu58384 | 1 | k__Fungi(100);p__Ascomycota(100);c__Saccharomycetes(100);o__Saccharomycetales(100);f__Metschnikowiaceae(100);g__Metschnikowia(100);s__Metschnikowia_pulcherrima(100); | Metschnikowia | 13 | Saprotroph | \|Undefined Saprotroph\| | Yeast | NULL | Probable | As nectar-tap_saprotroph (Põlme et al. 2020); Decay sugar-rich substrates (Põlme et al. 2020) | Põlme S, et al. 2020. Fungal Diversity 105:1-16 ((https://doi.org/10.1007/s13225-020-00466-2)) |
| Otu58385 | 1 | k__Fungi(100);p__Ascomycota(100);c__Saccharomycetes(100);o__Saccharomycetales(100);f__Metschnikowiaceae(100);g__Metschnikowia(100);s__Metschnikowia_pulcherrima(100); | Metschnikowia | 13 | Saprotroph | \|Undefined Saprotroph\| | Yeast | NULL | Probable | As nectar-tap_saprotroph (Põlme et al. 2020); Decay sugar-rich substrates (Põlme et al. 2020) | Põlme S, et al. 2020. Fungal Diversity 105:1-16 ((https://doi.org/10.1007/s13225-020-00466-2)) |
| Otu58386 | 1 | k__Fungi(100);p__Ascomycota(100);c__Saccharomycetes(100);o__Saccharomycetales(100);f__Metschnikowiaceae(100);g__Metschnikowia(100);s__Metschnikowia_pulcherrima(100); | Metschnikowia | 13 | Saprotroph | \|Undefined Saprotroph\| | Yeast | NULL | Probable | As nectar-tap_saprotroph (Põlme et al. 2020); Decay sugar-rich substrates (Põlme et al. 2020) | Põlme S, et al. 2020. Fungal Diversity 105:1-16 ((https://doi.org/10.1007/s13225-020-00466-2)) |
| Otu58387 | 1 | k__Fungi(100);p__Ascomycota(100);c__Saccharomycetes(100);o__Saccharomycetales(100);f__Metschnikowiaceae(100);g__Metschnikowia(100);s__Metschnikowia_pulcherrima(100); | Metschnikowia | 13 | Saprotroph | \|Undefined Saprotroph\| | Yeast | NULL | Probable | As nectar-tap_saprotroph (Põlme et al. 2020); Decay sugar-rich substrates (Põlme et al. 2020) | Põlme S, et al. 2020. Fungal Diversity 105:1-16 ((https://doi.org/10.1007/s13225-020-00466-2)) |
| Otu58388 | 1 | k__Fungi(100);p__Ascomycota(100);c__Saccharomycetes(100);o__Saccharomycetales(100);f__Metschnikowiaceae(100);g__Metschnikowia(100);s__Metschnikowia_pulcherrima(100); | Metschnikowia | 13 | Saprotroph | \|Undefined Saprotroph\| | Yeast | NULL | Probable | As nectar-tap_saprotroph (Põlme et al. 2020); Decay sugar-rich substrates (Põlme et al. 2020) | Põlme S, et al. 2020. Fungal Diversity 105:1-16 ((https://doi.org/10.1007/s13225-020-00466-2)) |
| Otu58389 | 1 | k__Fungi(100);p__Ascomycota(100);c__Saccharomycetes(100);o__Saccharomycetales(100);f__Metschnikowiaceae(100);g__Metschnikowia(100);s__Metschnikowia_pulcherrima(100); | Metschnikowia | 13 | Saprotroph | \|Undefined Saprotroph\| | Yeast | NULL | Probable | As nectar-tap_saprotroph (Põlme et al. 2020); Decay sugar-rich substrates (Põlme et al. 2020) | Põlme S, et al. 2020. Fungal Diversity 105:1-16 ((https://doi.org/10.1007/s13225-020-00466-2)) |
| Otu58390 | 1 | k__Fungi(100);p__Ascomycota(100);c__Saccharomycetes(100);o__Saccharomycetales(100);f__Metschnikowiaceae(100);g__Metschnikowia(100);s__Metschnikowia_pulcherrima(100); | Metschnikowia | 13 | Saprotroph | \|Undefined Saprotroph\| | Yeast | NULL | Probable | As nectar-tap_saprotroph (Põlme et al. 2020); Decay sugar-rich substrates (Põlme et al. 2020) | Põlme S, et al. 2020. Fungal Diversity 105:1-16 ((https://doi.org/10.1007/s13225-020-00466-2)) |
| Otu58391 | 1 | k__Fungi(100);p__Ascomycota(100);c__Saccharomycetes(100);o__Saccharomycetales(100);f__Metschnikowiaceae(100);g__Metschnikowia(100);g__Metschnikowia_unclassified(100); | Metschnikowia | 13 | Saprotroph | \|Undefined Saprotroph\| | Yeast | NULL | Probable | As nectar-tap_saprotroph (Põlme et al. 2020); Decay sugar-rich substrates (Põlme et al. 2020) | Põlme S, et al. 2020. Fungal Diversity 105:1-16 ((https://doi.org/10.1007/s13225-020-00466-2)) |
| Otu58392 | 1 | k__Fungi(100);p__Ascomycota(100);c__Saccharomycetes(100);o__Saccharomycetales(100);f__Metschnikowiaceae(100);g__Metschnikowia(100);s__Metschnikowia_pulcherrima(100); | Metschnikowia | 13 | Saprotroph | \|Undefined Saprotroph\| | Yeast | NULL | Probable | As nectar-tap_saprotroph (Põlme et al. 2020); Decay sugar-rich substrates (Põlme et al. 2020) | Põlme S, et al. 2020. Fungal Diversity 105:1-16 ((https://doi.org/10.1007/s13225-020-00466-2)) |
| Otu58393 | 1 | k__Fungi(100);p__Ascomycota(100);c__Saccharomycetes(100);o__Saccharomycetales(100);f__Metschnikowiaceae(100);g__Metschnikowia(100);s__Metschnikowia_pulcherrima(100); | Metschnikowia | 13 | Saprotroph | \|Undefined Saprotroph\| | Yeast | NULL | Probable | As nectar-tap_saprotroph (Põlme et al. 2020); Decay sugar-rich substrates (Põlme et al. 2020) | Põlme S, et al. 2020. Fungal Diversity 105:1-16 ((https://doi.org/10.1007/s13225-020-00466-2)) |
| Otu58394 | 1 | k__Fungi(100);p__Ascomycota(100);c__Saccharomycetes(100);o__Saccharomycetales(100);f__Metschnikowiaceae(100);g__Metschnikowia(100);g__Metschnikowia_unclassified(100); | Metschnikowia | 13 | Saprotroph | \|Undefined Saprotroph\| | Yeast | NULL | Probable | As nectar-tap_saprotroph (Põlme et al. 2020); Decay sugar-rich substrates (Põlme et al. 2020) | Põlme S, et al. 2020. Fungal Diversity 105:1-16 ((https://doi.org/10.1007/s13225-020-00466-2)) |
| Otu58395 | 1 | k__Fungi(100);p__Ascomycota(100);c__Saccharomycetes(100);o__Saccharomycetales(100);f__Metschnikowiaceae(100);g__Metschnikowia(100);s__Metschnikowia_pulcherrima(100); | Metschnikowia | 13 | Saprotroph | \|Undefined Saprotroph\| | Yeast | NULL | Probable | As nectar-tap_saprotroph (Põlme et al. 2020); Decay sugar-rich substrates (Põlme et al. 2020) | Põlme S, et al. 2020. Fungal Diversity 105:1-16 ((https://doi.org/10.1007/s13225-020-00466-2)) |
| Otu58400 | 1 | k__Fungi(100);p__Ascomycota(100);c__Saccharomycetes(100);o__Saccharomycetales(100);f__Metschnikowiaceae(100);g__Metschnikowia(100);g__Metschnikowia_unclassified(100); | Metschnikowia | 13 | Saprotroph | \|Undefined Saprotroph\| | Yeast | NULL | Probable | As nectar-tap_saprotroph (Põlme et al. 2020); Decay sugar-rich substrates (Põlme et al. 2020) | Põlme S, et al. 2020. Fungal Diversity 105:1-16 ((https://doi.org/10.1007/s13225-020-00466-2)) |
| Otu58401 | 1 | k__Fungi(100);p__Ascomycota(100);c__Saccharomycetes(100);o__Saccharomycetales(100);f__Metschnikowiaceae(100);g__Metschnikowia(100);s__Metschnikowia_pulcherrima(100); | Metschnikowia | 13 | Saprotroph | \|Undefined Saprotroph\| | Yeast | NULL | Probable | As nectar-tap_saprotroph (Põlme et al. 2020); Decay sugar-rich substrates (Põlme et al. 2020) | Põlme S, et al. 2020. Fungal Diversity 105:1-16 ((https://doi.org/10.1007/s13225-020-00466-2)) |
| Otu58402 | 1 | k__Fungi(100);p__Ascomycota(100);c__Saccharomycetes(100);o__Saccharomycetales(100);f__Metschnikowiaceae(100);g__Metschnikowia(100);s__Metschnikowia_pulcherrima(100); | Metschnikowia | 13 | Saprotroph | \|Undefined Saprotroph\| | Yeast | NULL | Probable | As nectar-tap_saprotroph (Põlme et al. 2020); Decay sugar-rich substrates (Põlme et al. 2020) | Põlme S, et al. 2020. Fungal Diversity 105:1-16 ((https://doi.org/10.1007/s13225-020-00466-2)) |
| Otu58403 | 1 | k__Fungi(100);p__Ascomycota(100);c__Saccharomycetes(100);o__Saccharomycetales(100);f__Metschnikowiaceae(100);g__Metschnikowia(100);s__Metschnikowia_pulcherrima(100); | Metschnikowia | 13 | Saprotroph | \|Undefined Saprotroph\| | Yeast | NULL | Probable | As nectar-tap_saprotroph (Põlme et al. 2020); Decay sugar-rich substrates (Põlme et al. 2020) | Põlme S, et al. 2020. Fungal Diversity 105:1-16 ((https://doi.org/10.1007/s13225-020-00466-2)) |
| Otu58404 | 1 | k__Fungi(100);p__Ascomycota(100);c__Saccharomycetes(100);o__Saccharomycetales(100);f__Metschnikowiaceae(100);g__Metschnikowia(100);s__Metschnikowia_pulcherrima(100); | Metschnikowia | 13 | Saprotroph | \|Undefined Saprotroph\| | Yeast | NULL | Probable | As nectar-tap_saprotroph (Põlme et al. 2020); Decay sugar-rich substrates (Põlme et al. 2020) | Põlme S, et al. 2020. Fungal Diversity 105:1-16 ((https://doi.org/10.1007/s13225-020-00466-2)) |
| Otu58405 | 1 | k__Fungi(100);p__Ascomycota(100);c__Saccharomycetes(100);o__Saccharomycetales(100);f__Metschnikowiaceae(100);g__Metschnikowia(100);s__Metschnikowia_pulcherrima(100); | Metschnikowia | 13 | Saprotroph | \|Undefined Saprotroph\| | Yeast | NULL | Probable | As nectar-tap_saprotroph (Põlme et al. 2020); Decay sugar-rich substrates (Põlme et al. 2020) | Põlme S, et al. 2020. Fungal Diversity 105:1-16 ((https://doi.org/10.1007/s13225-020-00466-2)) |
| Otu58406 | 1 | k__Fungi(100);p__Ascomycota(100);c__Saccharomycetes(100);o__Saccharomycetales(100);f__Metschnikowiaceae(100);g__Metschnikowia(100);g__Metschnikowia_unclassified(100); | Metschnikowia | 13 | Saprotroph | \|Undefined Saprotroph\| | Yeast | NULL | Probable | As nectar-tap_saprotroph (Põlme et al. 2020); Decay sugar-rich substrates (Põlme et al. 2020) | Põlme S, et al. 2020. Fungal Diversity 105:1-16 ((https://doi.org/10.1007/s13225-020-00466-2)) |
| Otu58407 | 1 | k__Fungi(100);p__Ascomycota(100);c__Saccharomycetes(100);o__Saccharomycetales(100);f__Metschnikowiaceae(100);g__Metschnikowia(100);g__Metschnikowia_unclassified(100); | Metschnikowia | 13 | Saprotroph | \|Undefined Saprotroph\| | Yeast | NULL | Probable | As nectar-tap_saprotroph (Põlme et al. 2020); Decay sugar-rich substrates (Põlme et al. 2020) | Põlme S, et al. 2020. Fungal Diversity 105:1-16 ((https://doi.org/10.1007/s13225-020-00466-2)) |
| Otu58408 | 1 | k__Fungi(100);p__Ascomycota(100);c__Saccharomycetes(100);o__Saccharomycetales(100);f__Metschnikowiaceae(100);g__Metschnikowia(100);g__Metschnikowia_unclassified(100); | Metschnikowia | 13 | Saprotroph | \|Undefined Saprotroph\| | Yeast | NULL | Probable | As nectar-tap_saprotroph (Põlme et al. 2020); Decay sugar-rich substrates (Põlme et al. 2020) | Põlme S, et al. 2020. Fungal Diversity 105:1-16 ((https://doi.org/10.1007/s13225-020-00466-2)) |
| Otu58409 | 1 | k__Fungi(100);p__Ascomycota(100);c__Saccharomycetes(100);o__Saccharomycetales(100);f__Metschnikowiaceae(100);g__Metschnikowia(100);g__Metschnikowia_unclassified(100); | Metschnikowia | 13 | Saprotroph | \|Undefined Saprotroph\| | Yeast | NULL | Probable | As nectar-tap_saprotroph (Põlme et al. 2020); Decay sugar-rich substrates (Põlme et al. 2020) | Põlme S, et al. 2020. Fungal Diversity 105:1-16 ((https://doi.org/10.1007/s13225-020-00466-2)) |
| Otu58410 | 1 | k__Fungi(100);p__Ascomycota(100);c__Saccharomycetes(100);o__Saccharomycetales(100);f__Metschnikowiaceae(100);g__Metschnikowia(100);s__Metschnikowia_pulcherrima(100); | Metschnikowia | 13 | Saprotroph | \|Undefined Saprotroph\| | Yeast | NULL | Probable | As nectar-tap_saprotroph (Põlme et al. 2020); Decay sugar-rich substrates (Põlme et al. 2020) | Põlme S, et al. 2020. Fungal Diversity 105:1-16 ((https://doi.org/10.1007/s13225-020-00466-2)) |
| Otu58411 | 1 | k__Fungi(100);p__Ascomycota(100);c__Saccharomycetes(100);o__Saccharomycetales(100);f__Metschnikowiaceae(100);g__Metschnikowia(100);g__Metschnikowia_unclassified(100); | Metschnikowia | 13 | Saprotroph | \|Undefined Saprotroph\| | Yeast | NULL | Probable | As nectar-tap_saprotroph (Põlme et al. 2020); Decay sugar-rich substrates (Põlme et al. 2020) | Põlme S, et al. 2020. Fungal Diversity 105:1-16 ((https://doi.org/10.1007/s13225-020-00466-2)) |
| Otu58412 | 1 | k__Fungi(100);p__Ascomycota(100);c__Saccharomycetes(100);o__Saccharomycetales(100);f__Metschnikowiaceae(100);g__Metschnikowia(100);s__Metschnikowia_pulcherrima(100); | Metschnikowia | 13 | Saprotroph | \|Undefined Saprotroph\| | Yeast | NULL | Probable | As nectar-tap_saprotroph (Põlme et al. 2020); Decay sugar-rich substrates (Põlme et al. 2020) | Põlme S, et al. 2020. Fungal Diversity 105:1-16 ((https://doi.org/10.1007/s13225-020-00466-2)) |
| Otu58414 | 1 | k__Fungi(100);p__Ascomycota(100);c__Saccharomycetes(100);o__Saccharomycetales(100);f__Metschnikowiaceae(100);g__Metschnikowia(100);s__Metschnikowia_pulcherrima(100); | Metschnikowia | 13 | Saprotroph | \|Undefined Saprotroph\| | Yeast | NULL | Probable | As nectar-tap_saprotroph (Põlme et al. 2020); Decay sugar-rich substrates (Põlme et al. 2020) | Põlme S, et al. 2020. Fungal Diversity 105:1-16 ((https://doi.org/10.1007/s13225-020-00466-2)) |
| Otu58415 | 1 | k__Fungi(100);p__Ascomycota(100);c__Saccharomycetes(100);o__Saccharomycetales(100);f__Metschnikowiaceae(100);g__Metschnikowia(100);g__Metschnikowia_unclassified(100); | Metschnikowia | 13 | Saprotroph | \|Undefined Saprotroph\| | Yeast | NULL | Probable | As nectar-tap_saprotroph (Põlme et al. 2020); Decay sugar-rich substrates (Põlme et al. 2020) | Põlme S, et al. 2020. Fungal Diversity 105:1-16 ((https://doi.org/10.1007/s13225-020-00466-2)) |
| Otu58416 | 1 | k__Fungi(100);p__Ascomycota(100);c__Saccharomycetes(100);o__Saccharomycetales(100);f__Metschnikowiaceae(100);g__Metschnikowia(100);s__Metschnikowia_pulcherrima(100); | Metschnikowia | 13 | Saprotroph | \|Undefined Saprotroph\| | Yeast | NULL | Probable | As nectar-tap_saprotroph (Põlme et al. 2020); Decay sugar-rich substrates (Põlme et al. 2020) | Põlme S, et al. 2020. Fungal Diversity 105:1-16 ((https://doi.org/10.1007/s13225-020-00466-2)) |
| Otu58417 | 1 | k__Fungi(100);p__Ascomycota(100);c__Saccharomycetes(100);o__Saccharomycetales(100);f__Metschnikowiaceae(100);g__Metschnikowia(100);s__Metschnikowia_pulcherrima(100); | Metschnikowia | 13 | Saprotroph | \|Undefined Saprotroph\| | Yeast | NULL | Probable | As nectar-tap_saprotroph (Põlme et al. 2020); Decay sugar-rich substrates (Põlme et al. 2020) | Põlme S, et al. 2020. Fungal Diversity 105:1-16 ((https://doi.org/10.1007/s13225-020-00466-2)) |
| Otu58418 | 1 | k__Fungi(100);p__Ascomycota(100);c__Saccharomycetes(100);o__Saccharomycetales(100);f__Metschnikowiaceae(100);g__Metschnikowia(100);s__Metschnikowia_pulcherrima(100); | Metschnikowia | 13 | Saprotroph | \|Undefined Saprotroph\| | Yeast | NULL | Probable | As nectar-tap_saprotroph (Põlme et al. 2020); Decay sugar-rich substrates (Põlme et al. 2020) | Põlme S, et al. 2020. Fungal Diversity 105:1-16 ((https://doi.org/10.1007/s13225-020-00466-2)) |
| Otu58419 | 1 | k__Fungi(100);p__Ascomycota(100);c__Saccharomycetes(100);o__Saccharomycetales(100);f__Metschnikowiaceae(100);g__Metschnikowia(100);s__Metschnikowia_pulcherrima(100); | Metschnikowia | 13 | Saprotroph | \|Undefined Saprotroph\| | Yeast | NULL | Probable | As nectar-tap_saprotroph (Põlme et al. 2020); Decay sugar-rich substrates (Põlme et al. 2020) | Põlme S, et al. 2020. Fungal Diversity 105:1-16 ((https://doi.org/10.1007/s13225-020-00466-2)) |
| Otu58420 | 1 | k__Fungi(100);p__Ascomycota(100);c__Saccharomycetes(100);o__Saccharomycetales(100);f__Metschnikowiaceae(100);g__Metschnikowia(100);s__Metschnikowia_pulcherrima(100); | Metschnikowia | 13 | Saprotroph | \|Undefined Saprotroph\| | Yeast | NULL | Probable | As nectar-tap_saprotroph (Põlme et al. 2020); Decay sugar-rich substrates (Põlme et al. 2020) | Põlme S, et al. 2020. Fungal Diversity 105:1-16 ((https://doi.org/10.1007/s13225-020-00466-2)) |
| Otu58421 | 1 | k__Fungi(100);p__Ascomycota(100);c__Saccharomycetes(100);o__Saccharomycetales(100);f__Metschnikowiaceae(100);g__Metschnikowia(100);s__Metschnikowia_pulcherrima(100); | Metschnikowia | 13 | Saprotroph | \|Undefined Saprotroph\| | Yeast | NULL | Probable | As nectar-tap_saprotroph (Põlme et al. 2020); Decay sugar-rich substrates (Põlme et al. 2020) | Põlme S, et al. 2020. Fungal Diversity 105:1-16 ((https://doi.org/10.1007/s13225-020-00466-2)) |
| Otu58422 | 1 | k__Fungi(100);p__Ascomycota(100);c__Saccharomycetes(100);o__Saccharomycetales(100);f__Metschnikowiaceae(100);g__Metschnikowia(100);s__Metschnikowia_pulcherrima(100); | Metschnikowia | 13 | Saprotroph | \|Undefined Saprotroph\| | Yeast | NULL | Probable | As nectar-tap_saprotroph (Põlme et al. 2020); Decay sugar-rich substrates (Põlme et al. 2020) | Põlme S, et al. 2020. Fungal Diversity 105:1-16 ((https://doi.org/10.1007/s13225-020-00466-2)) |
| Otu58423 | 1 | k__Fungi(100);p__Ascomycota(100);c__Saccharomycetes(100);o__Saccharomycetales(100);f__Metschnikowiaceae(100);g__Metschnikowia(100);g__Metschnikowia_unclassified(100); | Metschnikowia | 13 | Saprotroph | \|Undefined Saprotroph\| | Yeast | NULL | Probable | As nectar-tap_saprotroph (Põlme et al. 2020); Decay sugar-rich substrates (Põlme et al. 2020) | Põlme S, et al. 2020. Fungal Diversity 105:1-16 ((https://doi.org/10.1007/s13225-020-00466-2)) |
| Otu58424 | 1 | k__Fungi(100);p__Ascomycota(100);c__Saccharomycetes(100);o__Saccharomycetales(100);f__Metschnikowiaceae(100);g__Metschnikowia(100);g__Metschnikowia_unclassified(100); | Metschnikowia | 13 | Saprotroph | \|Undefined Saprotroph\| | Yeast | NULL | Probable | As nectar-tap_saprotroph (Põlme et al. 2020); Decay sugar-rich substrates (Põlme et al. 2020) | Põlme S, et al. 2020. Fungal Diversity 105:1-16 ((https://doi.org/10.1007/s13225-020-00466-2)) |
| Otu58425 | 1 | k__Fungi(100);p__Ascomycota(100);c__Saccharomycetes(100);o__Saccharomycetales(100);f__Metschnikowiaceae(100);g__Metschnikowia(100);s__Metschnikowia_pulcherrima(100); | Metschnikowia | 13 | Saprotroph | \|Undefined Saprotroph\| | Yeast | NULL | Probable | As nectar-tap_saprotroph (Põlme et al. 2020); Decay sugar-rich substrates (Põlme et al. 2020) | Põlme S, et al. 2020. Fungal Diversity 105:1-16 ((https://doi.org/10.1007/s13225-020-00466-2)) |
| Otu58426 | 1 | k__Fungi(100);p__Ascomycota(100);c__Saccharomycetes(100);o__Saccharomycetales(100);f__Metschnikowiaceae(100);g__Metschnikowia(100);s__Metschnikowia_pulcherrima(100); | Metschnikowia | 13 | Saprotroph | \|Undefined Saprotroph\| | Yeast | NULL | Probable | As nectar-tap_saprotroph (Põlme et al. 2020); Decay sugar-rich substrates (Põlme et al. 2020) | Põlme S, et al. 2020. Fungal Diversity 105:1-16 ((https://doi.org/10.1007/s13225-020-00466-2)) |
| Otu58427 | 1 | k__Fungi(100);p__Ascomycota(100);c__Saccharomycetes(100);o__Saccharomycetales(100);f__Metschnikowiaceae(100);g__Metschnikowia(100);s__Metschnikowia_pulcherrima(100); | Metschnikowia | 13 | Saprotroph | \|Undefined Saprotroph\| | Yeast | NULL | Probable | As nectar-tap_saprotroph (Põlme et al. 2020); Decay sugar-rich substrates (Põlme et al. 2020) | Põlme S, et al. 2020. Fungal Diversity 105:1-16 ((https://doi.org/10.1007/s13225-020-00466-2)) |
| Otu58428 | 1 | k__Fungi(100);p__Ascomycota(100);c__Saccharomycetes(100);o__Saccharomycetales(100);f__Metschnikowiaceae(100);g__Metschnikowia(100);g__Metschnikowia_unclassified(100); | Metschnikowia | 13 | Saprotroph | \|Undefined Saprotroph\| | Yeast | NULL | Probable | As nectar-tap_saprotroph (Põlme et al. 2020); Decay sugar-rich substrates (Põlme et al. 2020) | Põlme S, et al. 2020. Fungal Diversity 105:1-16 ((https://doi.org/10.1007/s13225-020-00466-2)) |
| Otu58429 | 1 | k__Fungi(100);p__Ascomycota(100);c__Saccharomycetes(100);o__Saccharomycetales(100);f__Metschnikowiaceae(100);g__Metschnikowia(100);s__Metschnikowia_pulcherrima(100); | Metschnikowia | 13 | Saprotroph | \|Undefined Saprotroph\| | Yeast | NULL | Probable | As nectar-tap_saprotroph (Põlme et al. 2020); Decay sugar-rich substrates (Põlme et al. 2020) | Põlme S, et al. 2020. Fungal Diversity 105:1-16 ((https://doi.org/10.1007/s13225-020-00466-2)) |
| Otu58430 | 1 | k__Fungi(100);p__Ascomycota(100);c__Saccharomycetes(100);o__Saccharomycetales(100);f__Metschnikowiaceae(100);g__Metschnikowia(100);s__Metschnikowia_pulcherrima(100); | Metschnikowia | 13 | Saprotroph | \|Undefined Saprotroph\| | Yeast | NULL | Probable | As nectar-tap_saprotroph (Põlme et al. 2020); Decay sugar-rich substrates (Põlme et al. 2020) | Põlme S, et al. 2020. Fungal Diversity 105:1-16 ((https://doi.org/10.1007/s13225-020-00466-2)) |
| Otu58433 | 1 | k__Fungi(100);p__Ascomycota(100);c__Saccharomycetes(100);o__Saccharomycetales(100);f__Metschnikowiaceae(100);g__Metschnikowia(100);g__Metschnikowia_unclassified(100); | Metschnikowia | 13 | Saprotroph | \|Undefined Saprotroph\| | Yeast | NULL | Probable | As nectar-tap_saprotroph (Põlme et al. 2020); Decay sugar-rich substrates (Põlme et al. 2020) | Põlme S, et al. 2020. Fungal Diversity 105:1-16 ((https://doi.org/10.1007/s13225-020-00466-2)) |
| Otu58434 | 1 | k__Fungi(100);p__Ascomycota(100);c__Saccharomycetes(100);o__Saccharomycetales(100);f__Metschnikowiaceae(100);g__Metschnikowia(100);s__Metschnikowia_pulcherrima(100); | Metschnikowia | 13 | Saprotroph | \|Undefined Saprotroph\| | Yeast | NULL | Probable | As nectar-tap_saprotroph (Põlme et al. 2020); Decay sugar-rich substrates (Põlme et al. 2020) | Põlme S, et al. 2020. Fungal Diversity 105:1-16 ((https://doi.org/10.1007/s13225-020-00466-2)) |
| Otu58435 | 1 | k__Fungi(100);p__Ascomycota(100);c__Saccharomycetes(100);o__Saccharomycetales(100);f__Metschnikowiaceae(100);g__Metschnikowia(100);s__Metschnikowia_pulcherrima(100); | Metschnikowia | 13 | Saprotroph | \|Undefined Saprotroph\| | Yeast | NULL | Probable | As nectar-tap_saprotroph (Põlme et al. 2020); Decay sugar-rich substrates (Põlme et al. 2020) | Põlme S, et al. 2020. Fungal Diversity 105:1-16 ((https://doi.org/10.1007/s13225-020-00466-2)) |
| Otu58436 | 1 | k__Fungi(100);p__Ascomycota(100);c__Saccharomycetes(100);o__Saccharomycetales(100);f__Metschnikowiaceae(100);g__Metschnikowia(100);g__Metschnikowia_unclassified(100); | Metschnikowia | 13 | Saprotroph | \|Undefined Saprotroph\| | Yeast | NULL | Probable | As nectar-tap_saprotroph (Põlme et al. 2020); Decay sugar-rich substrates (Põlme et al. 2020) | Põlme S, et al. 2020. Fungal Diversity 105:1-16 ((https://doi.org/10.1007/s13225-020-00466-2)) |
| Otu58437 | 1 | k__Fungi(100);p__Ascomycota(100);c__Saccharomycetes(100);o__Saccharomycetales(100);f__Metschnikowiaceae(100);g__Metschnikowia(100);s__Metschnikowia_pulcherrima(100); | Metschnikowia | 13 | Saprotroph | \|Undefined Saprotroph\| | Yeast | NULL | Probable | As nectar-tap_saprotroph (Põlme et al. 2020); Decay sugar-rich substrates (Põlme et al. 2020) | Põlme S, et al. 2020. Fungal Diversity 105:1-16 ((https://doi.org/10.1007/s13225-020-00466-2)) |
| Otu58438 | 1 | k__Fungi(100);p__Basidiomycota(100);c__Microbotryomycetes(100);o__Sporidiobolales(100);f__Sporidiobolales_family_Incertae_sedis(100);g__Rhodotorula(100);s__Rhodotorula_taiwanensis(100); | Rhodotorula | 13 | Pathotroph-Saprotroph-Symbiotroph | Animal Pathogen-Animal Symbiotroph-Endophyte-Plant Pathogen-\|Undefined Saprotroph\| | Yeast | NULL | Probable | As foliar_endophyte (Põlme et al. 2020); Opportunistic human pathogen | Põlme S, et al. 2020. Fungal Diversity 105:1-16 ((https://doi.org/10.1007/s13225-020-00466-2)); https://www.cdc.gov (https://www.cdc.gov/) |
| Otu58440 | 1 | k__Fungi(100);p__Ascomycota(100);c__Saccharomycetes(100);o__Saccharomycetales(100);f__Metschnikowiaceae(100);g__Metschnikowia(100);g__Metschnikowia_unclassified(100); | Metschnikowia | 13 | Saprotroph | \|Undefined Saprotroph\| | Yeast | NULL | Probable | As nectar-tap_saprotroph (Põlme et al. 2020); Decay sugar-rich substrates (Põlme et al. 2020) | Põlme S, et al. 2020. Fungal Diversity 105:1-16 ((https://doi.org/10.1007/s13225-020-00466-2)) |
| Otu58441 | 1 | k__Fungi(100);p__Ascomycota(100);c__Saccharomycetes(100);o__Saccharomycetales(100);f__Metschnikowiaceae(100);g__Metschnikowia(100);s__Metschnikowia_pulcherrima(100); | Metschnikowia | 13 | Saprotroph | \|Undefined Saprotroph\| | Yeast | NULL | Probable | As nectar-tap_saprotroph (Põlme et al. 2020); Decay sugar-rich substrates (Põlme et al. 2020) | Põlme S, et al. 2020. Fungal Diversity 105:1-16 ((https://doi.org/10.1007/s13225-020-00466-2)) |
| Otu58442 | 1 | k__Fungi(100);p__Ascomycota(100);c__Saccharomycetes(100);o__Saccharomycetales(100);f__Metschnikowiaceae(100);g__Metschnikowia(100);s__Metschnikowia_pulcherrima(100); | Metschnikowia | 13 | Saprotroph | \|Undefined Saprotroph\| | Yeast | NULL | Probable | As nectar-tap_saprotroph (Põlme et al. 2020); Decay sugar-rich substrates (Põlme et al. 2020) | Põlme S, et al. 2020. Fungal Diversity 105:1-16 ((https://doi.org/10.1007/s13225-020-00466-2)) |
| Otu58445 | 1 | k__Fungi(100);p__Ascomycota(100);c__Saccharomycetes(100);o__Saccharomycetales(100);f__Metschnikowiaceae(100);g__Metschnikowia(100);g__Metschnikowia_unclassified(100); | Metschnikowia | 13 | Saprotroph | \|Undefined Saprotroph\| | Yeast | NULL | Probable | As nectar-tap_saprotroph (Põlme et al. 2020); Decay sugar-rich substrates (Põlme et al. 2020) | Põlme S, et al. 2020. Fungal Diversity 105:1-16 ((https://doi.org/10.1007/s13225-020-00466-2)) |
| Otu58447 | 1 | k__Fungi(100);p__Ascomycota(100);c__Saccharomycetes(100);o__Saccharomycetales(100);f__Metschnikowiaceae(100);g__Metschnikowia(100);s__Metschnikowia_pulcherrima(100); | Metschnikowia | 13 | Saprotroph | \|Undefined Saprotroph\| | Yeast | NULL | Probable | As nectar-tap_saprotroph (Põlme et al. 2020); Decay sugar-rich substrates (Põlme et al. 2020) | Põlme S, et al. 2020. Fungal Diversity 105:1-16 ((https://doi.org/10.1007/s13225-020-00466-2)) |
| Otu58448 | 1 | k__Fungi(100);p__Ascomycota(100);c__Saccharomycetes(100);o__Saccharomycetales(100);f__Metschnikowiaceae(100);g__Metschnikowia(100);g__Metschnikowia_unclassified(100); | Metschnikowia | 13 | Saprotroph | \|Undefined Saprotroph\| | Yeast | NULL | Probable | As nectar-tap_saprotroph (Põlme et al. 2020); Decay sugar-rich substrates (Põlme et al. 2020) | Põlme S, et al. 2020. Fungal Diversity 105:1-16 ((https://doi.org/10.1007/s13225-020-00466-2)) |
| Otu58449 | 1 | k__Fungi(100);p__Ascomycota(100);c__Saccharomycetes(100);o__Saccharomycetales(100);f__Metschnikowiaceae(100);g__Metschnikowia(100);s__Metschnikowia_pulcherrima(100); | Metschnikowia | 13 | Saprotroph | \|Undefined Saprotroph\| | Yeast | NULL | Probable | As nectar-tap_saprotroph (Põlme et al. 2020); Decay sugar-rich substrates (Põlme et al. 2020) | Põlme S, et al. 2020. Fungal Diversity 105:1-16 ((https://doi.org/10.1007/s13225-020-00466-2)) |
| Otu58452 | 1 | k__Fungi(100);p__Ascomycota(100);c__Saccharomycetes(100);o__Saccharomycetales(100);f__Metschnikowiaceae(100);g__Metschnikowia(100);s__Metschnikowia_pulcherrima(100); | Metschnikowia | 13 | Saprotroph | \|Undefined Saprotroph\| | Yeast | NULL | Probable | As nectar-tap_saprotroph (Põlme et al. 2020); Decay sugar-rich substrates (Põlme et al. 2020) | Põlme S, et al. 2020. Fungal Diversity 105:1-16 ((https://doi.org/10.1007/s13225-020-00466-2)) |
| Otu58453 | 1 | k__Fungi(100);p__Ascomycota(100);c__Saccharomycetes(100);o__Saccharomycetales(100);f__Metschnikowiaceae(100);g__Metschnikowia(100);g__Metschnikowia_unclassified(100); | Metschnikowia | 13 | Saprotroph | \|Undefined Saprotroph\| | Yeast | NULL | Probable | As nectar-tap_saprotroph (Põlme et al. 2020); Decay sugar-rich substrates (Põlme et al. 2020) | Põlme S, et al. 2020. Fungal Diversity 105:1-16 ((https://doi.org/10.1007/s13225-020-00466-2)) |
| Otu58454 | 1 | k__Fungi(100);p__Ascomycota(100);c__Saccharomycetes(100);o__Saccharomycetales(100);f__Metschnikowiaceae(100);g__Metschnikowia(100);g__Metschnikowia_unclassified(100); | Metschnikowia | 13 | Saprotroph | \|Undefined Saprotroph\| | Yeast | NULL | Probable | As nectar-tap_saprotroph (Põlme et al. 2020); Decay sugar-rich substrates (Põlme et al. 2020) | Põlme S, et al. 2020. Fungal Diversity 105:1-16 ((https://doi.org/10.1007/s13225-020-00466-2)) |
| Otu58455 | 1 | k__Fungi(100);p__Ascomycota(100);c__Saccharomycetes(100);o__Saccharomycetales(100);f__Metschnikowiaceae(100);g__Metschnikowia(100);s__Metschnikowia_pulcherrima(100); | Metschnikowia | 13 | Saprotroph | \|Undefined Saprotroph\| | Yeast | NULL | Probable | As nectar-tap_saprotroph (Põlme et al. 2020); Decay sugar-rich substrates (Põlme et al. 2020) | Põlme S, et al. 2020. Fungal Diversity 105:1-16 ((https://doi.org/10.1007/s13225-020-00466-2)) |
| Otu58456 | 1 | k__Fungi(100);p__Ascomycota(100);c__Saccharomycetes(100);o__Saccharomycetales(100);f__Metschnikowiaceae(100);g__Metschnikowia(100);s__Metschnikowia_pulcherrima(100); | Metschnikowia | 13 | Saprotroph | \|Undefined Saprotroph\| | Yeast | NULL | Probable | As nectar-tap_saprotroph (Põlme et al. 2020); Decay sugar-rich substrates (Põlme et al. 2020) | Põlme S, et al. 2020. Fungal Diversity 105:1-16 ((https://doi.org/10.1007/s13225-020-00466-2)) |
| Otu58457 | 1 | k__Fungi(100);p__Ascomycota(100);c__Saccharomycetes(100);o__Saccharomycetales(100);f__Metschnikowiaceae(100);g__Metschnikowia(100);g__Metschnikowia_unclassified(100); | Metschnikowia | 13 | Saprotroph | \|Undefined Saprotroph\| | Yeast | NULL | Probable | As nectar-tap_saprotroph (Põlme et al. 2020); Decay sugar-rich substrates (Põlme et al. 2020) | Põlme S, et al. 2020. Fungal Diversity 105:1-16 ((https://doi.org/10.1007/s13225-020-00466-2)) |
| Otu58458 | 1 | k__Fungi(100);p__Ascomycota(100);c__Saccharomycetes(100);o__Saccharomycetales(100);f__Metschnikowiaceae(100);g__Metschnikowia(100);s__Metschnikowia_pulcherrima(100); | Metschnikowia | 13 | Saprotroph | \|Undefined Saprotroph\| | Yeast | NULL | Probable | As nectar-tap_saprotroph (Põlme et al. 2020); Decay sugar-rich substrates (Põlme et al. 2020) | Põlme S, et al. 2020. Fungal Diversity 105:1-16 ((https://doi.org/10.1007/s13225-020-00466-2)) |
| Otu58460 | 1 | k__Fungi(100);p__Ascomycota(100);c__Saccharomycetes(100);o__Saccharomycetales(100);f__Metschnikowiaceae(100);g__Metschnikowia(100);s__Metschnikowia_pulcherrima(100); | Metschnikowia | 13 | Saprotroph | \|Undefined Saprotroph\| | Yeast | NULL | Probable | As nectar-tap_saprotroph (Põlme et al. 2020); Decay sugar-rich substrates (Põlme et al. 2020) | Põlme S, et al. 2020. Fungal Diversity 105:1-16 ((https://doi.org/10.1007/s13225-020-00466-2)) |
| Otu58461 | 1 | k__Fungi(100);p__Ascomycota(100);c__Saccharomycetes(100);o__Saccharomycetales(100);f__Metschnikowiaceae(100);g__Metschnikowia(100);s__Metschnikowia_pulcherrima(100); | Metschnikowia | 13 | Saprotroph | \|Undefined Saprotroph\| | Yeast | NULL | Probable | As nectar-tap_saprotroph (Põlme et al. 2020); Decay sugar-rich substrates (Põlme et al. 2020) | Põlme S, et al. 2020. Fungal Diversity 105:1-16 ((https://doi.org/10.1007/s13225-020-00466-2)) |
| Otu58462 | 1 | k__Fungi(100);p__Ascomycota(100);c__Saccharomycetes(100);o__Saccharomycetales(100);f__Metschnikowiaceae(100);g__Metschnikowia(100);s__Metschnikowia_pulcherrima(100); | Metschnikowia | 13 | Saprotroph | \|Undefined Saprotroph\| | Yeast | NULL | Probable | As nectar-tap_saprotroph (Põlme et al. 2020); Decay sugar-rich substrates (Põlme et al. 2020) | Põlme S, et al. 2020. Fungal Diversity 105:1-16 ((https://doi.org/10.1007/s13225-020-00466-2)) |
| Otu58463 | 1 | k__Fungi(100);p__Ascomycota(100);c__Saccharomycetes(100);o__Saccharomycetales(100);f__Metschnikowiaceae(100);g__Metschnikowia(100);s__Metschnikowia_pulcherrima(100); | Metschnikowia | 13 | Saprotroph | \|Undefined Saprotroph\| | Yeast | NULL | Probable | As nectar-tap_saprotroph (Põlme et al. 2020); Decay sugar-rich substrates (Põlme et al. 2020) | Põlme S, et al. 2020. Fungal Diversity 105:1-16 ((https://doi.org/10.1007/s13225-020-00466-2)) |
| Otu58464 | 1 | k__Fungi(100);p__Ascomycota(100);c__Saccharomycetes(100);o__Saccharomycetales(100);f__Metschnikowiaceae(100);g__Metschnikowia(100);s__Metschnikowia_pulcherrima(100); | Metschnikowia | 13 | Saprotroph | \|Undefined Saprotroph\| | Yeast | NULL | Probable | As nectar-tap_saprotroph (Põlme et al. 2020); Decay sugar-rich substrates (Põlme et al. 2020) | Põlme S, et al. 2020. Fungal Diversity 105:1-16 ((https://doi.org/10.1007/s13225-020-00466-2)) |
| Otu58465 | 1 | k__Fungi(100);p__Ascomycota(100);c__Saccharomycetes(100);o__Saccharomycetales(100);f__Metschnikowiaceae(100);g__Metschnikowia(100);s__Metschnikowia_pulcherrima(100); | Metschnikowia | 13 | Saprotroph | \|Undefined Saprotroph\| | Yeast | NULL | Probable | As nectar-tap_saprotroph (Põlme et al. 2020); Decay sugar-rich substrates (Põlme et al. 2020) | Põlme S, et al. 2020. Fungal Diversity 105:1-16 ((https://doi.org/10.1007/s13225-020-00466-2)) |
| Otu58466 | 1 | k__Fungi(100);p__Ascomycota(100);c__Saccharomycetes(100);o__Saccharomycetales(100);f__Metschnikowiaceae(100);g__Metschnikowia(100);s__Metschnikowia_pulcherrima(100); | Metschnikowia | 13 | Saprotroph | \|Undefined Saprotroph\| | Yeast | NULL | Probable | As nectar-tap_saprotroph (Põlme et al. 2020); Decay sugar-rich substrates (Põlme et al. 2020) | Põlme S, et al. 2020. Fungal Diversity 105:1-16 ((https://doi.org/10.1007/s13225-020-00466-2)) |
| Otu58467 | 1 | k__Fungi(100);p__Ascomycota(100);c__Saccharomycetes(100);o__Saccharomycetales(100);f__Metschnikowiaceae(100);g__Metschnikowia(100);s__Metschnikowia_pulcherrima(100); | Metschnikowia | 13 | Saprotroph | \|Undefined Saprotroph\| | Yeast | NULL | Probable | As nectar-tap_saprotroph (Põlme et al. 2020); Decay sugar-rich substrates (Põlme et al. 2020) | Põlme S, et al. 2020. Fungal Diversity 105:1-16 ((https://doi.org/10.1007/s13225-020-00466-2)) |
| Otu58468 | 1 | k__Fungi(100);p__Ascomycota(100);c__Saccharomycetes(100);o__Saccharomycetales(100);f__Metschnikowiaceae(100);g__Metschnikowia(100);g__Metschnikowia_unclassified(100); | Metschnikowia | 13 | Saprotroph | \|Undefined Saprotroph\| | Yeast | NULL | Probable | As nectar-tap_saprotroph (Põlme et al. 2020); Decay sugar-rich substrates (Põlme et al. 2020) | Põlme S, et al. 2020. Fungal Diversity 105:1-16 ((https://doi.org/10.1007/s13225-020-00466-2)) |
| Otu58469 | 1 | k__Fungi(100);p__Ascomycota(100);c__Saccharomycetes(100);o__Saccharomycetales(100);f__Metschnikowiaceae(100);g__Metschnikowia(100);s__Metschnikowia_pulcherrima(100); | Metschnikowia | 13 | Saprotroph | \|Undefined Saprotroph\| | Yeast | NULL | Probable | As nectar-tap_saprotroph (Põlme et al. 2020); Decay sugar-rich substrates (Põlme et al. 2020) | Põlme S, et al. 2020. Fungal Diversity 105:1-16 ((https://doi.org/10.1007/s13225-020-00466-2)) |
| Otu58470 | 1 | k__Fungi(100);p__Ascomycota(100);c__Saccharomycetes(100);o__Saccharomycetales(100);f__Metschnikowiaceae(100);g__Metschnikowia(100);s__Metschnikowia_pulcherrima(100); | Metschnikowia | 13 | Saprotroph | \|Undefined Saprotroph\| | Yeast | NULL | Probable | As nectar-tap_saprotroph (Põlme et al. 2020); Decay sugar-rich substrates (Põlme et al. 2020) | Põlme S, et al. 2020. Fungal Diversity 105:1-16 ((https://doi.org/10.1007/s13225-020-00466-2)) |
| Otu58471 | 1 | k__Fungi(100);p__Ascomycota(100);c__Saccharomycetes(100);o__Saccharomycetales(100);f__Metschnikowiaceae(100);g__Metschnikowia(100);s__Metschnikowia_pulcherrima(100); | Metschnikowia | 13 | Saprotroph | \|Undefined Saprotroph\| | Yeast | NULL | Probable | As nectar-tap_saprotroph (Põlme et al. 2020); Decay sugar-rich substrates (Põlme et al. 2020) | Põlme S, et al. 2020. Fungal Diversity 105:1-16 ((https://doi.org/10.1007/s13225-020-00466-2)) |
| Otu58472 | 1 | k__Fungi(100);p__Ascomycota(100);c__Saccharomycetes(100);o__Saccharomycetales(100);f__Metschnikowiaceae(100);g__Metschnikowia(100);s__Metschnikowia_pulcherrima(100); | Metschnikowia | 13 | Saprotroph | \|Undefined Saprotroph\| | Yeast | NULL | Probable | As nectar-tap_saprotroph (Põlme et al. 2020); Decay sugar-rich substrates (Põlme et al. 2020) | Põlme S, et al. 2020. Fungal Diversity 105:1-16 ((https://doi.org/10.1007/s13225-020-00466-2)) |
| Otu58473 | 1 | k__Fungi(100);p__Ascomycota(100);c__Saccharomycetes(100);o__Saccharomycetales(100);f__Metschnikowiaceae(100);g__Metschnikowia(100);s__Metschnikowia_pulcherrima(100); | Metschnikowia | 13 | Saprotroph | \|Undefined Saprotroph\| | Yeast | NULL | Probable | As nectar-tap_saprotroph (Põlme et al. 2020); Decay sugar-rich substrates (Põlme et al. 2020) | Põlme S, et al. 2020. Fungal Diversity 105:1-16 ((https://doi.org/10.1007/s13225-020-00466-2)) |
| Otu58474 | 1 | k__Fungi(100);p__Ascomycota(100);c__Saccharomycetes(100);o__Saccharomycetales(100);f__Metschnikowiaceae(100);g__Metschnikowia(100);s__Metschnikowia_pulcherrima(100); | Metschnikowia | 13 | Saprotroph | \|Undefined Saprotroph\| | Yeast | NULL | Probable | As nectar-tap_saprotroph (Põlme et al. 2020); Decay sugar-rich substrates (Põlme et al. 2020) | Põlme S, et al. 2020. Fungal Diversity 105:1-16 ((https://doi.org/10.1007/s13225-020-00466-2)) |
| Otu58475 | 1 | k__Fungi(100);p__Ascomycota(100);c__Saccharomycetes(100);o__Saccharomycetales(100);f__Metschnikowiaceae(100);g__Metschnikowia(100);s__Metschnikowia_pulcherrima(100); | Metschnikowia | 13 | Saprotroph | \|Undefined Saprotroph\| | Yeast | NULL | Probable | As nectar-tap_saprotroph (Põlme et al. 2020); Decay sugar-rich substrates (Põlme et al. 2020) | Põlme S, et al. 2020. Fungal Diversity 105:1-16 ((https://doi.org/10.1007/s13225-020-00466-2)) |
| Otu58476 | 1 | k__Fungi(100);p__Ascomycota(100);c__Saccharomycetes(100);o__Saccharomycetales(100);f__Metschnikowiaceae(100);g__Metschnikowia(100);s__Metschnikowia_pulcherrima(100); | Metschnikowia | 13 | Saprotroph | \|Undefined Saprotroph\| | Yeast | NULL | Probable | As nectar-tap_saprotroph (Põlme et al. 2020); Decay sugar-rich substrates (Põlme et al. 2020) | Põlme S, et al. 2020. Fungal Diversity 105:1-16 ((https://doi.org/10.1007/s13225-020-00466-2)) |
| Otu58477 | 1 | k__Fungi(100);p__Ascomycota(100);c__Saccharomycetes(100);o__Saccharomycetales(100);f__Metschnikowiaceae(100);g__Metschnikowia(100);g__Metschnikowia_unclassified(100); | Metschnikowia | 13 | Saprotroph | \|Undefined Saprotroph\| | Yeast | NULL | Probable | As nectar-tap_saprotroph (Põlme et al. 2020); Decay sugar-rich substrates (Põlme et al. 2020) | Põlme S, et al. 2020. Fungal Diversity 105:1-16 ((https://doi.org/10.1007/s13225-020-00466-2)) |
| Otu58478 | 1 | k__Fungi(100);p__Ascomycota(100);c__Saccharomycetes(100);o__Saccharomycetales(100);f__Metschnikowiaceae(100);g__Metschnikowia(100);s__Metschnikowia_pulcherrima(100); | Metschnikowia | 13 | Saprotroph | \|Undefined Saprotroph\| | Yeast | NULL | Probable | As nectar-tap_saprotroph (Põlme et al. 2020); Decay sugar-rich substrates (Põlme et al. 2020) | Põlme S, et al. 2020. Fungal Diversity 105:1-16 ((https://doi.org/10.1007/s13225-020-00466-2)) |
| Otu58479 | 1 | k__Fungi(100);p__Ascomycota(100);c__Saccharomycetes(100);o__Saccharomycetales(100);f__Metschnikowiaceae(100);g__Metschnikowia(100);s__Metschnikowia_pulcherrima(100); | Metschnikowia | 13 | Saprotroph | \|Undefined Saprotroph\| | Yeast | NULL | Probable | As nectar-tap_saprotroph (Põlme et al. 2020); Decay sugar-rich substrates (Põlme et al. 2020) | Põlme S, et al. 2020. Fungal Diversity 105:1-16 ((https://doi.org/10.1007/s13225-020-00466-2)) |
| Otu58480 | 1 | k__Fungi(100);p__Ascomycota(100);c__Saccharomycetes(100);o__Saccharomycetales(100);f__Metschnikowiaceae(100);g__Metschnikowia(100);s__Metschnikowia_pulcherrima(100); | Metschnikowia | 13 | Saprotroph | \|Undefined Saprotroph\| | Yeast | NULL | Probable | As nectar-tap_saprotroph (Põlme et al. 2020); Decay sugar-rich substrates (Põlme et al. 2020) | Põlme S, et al. 2020. Fungal Diversity 105:1-16 ((https://doi.org/10.1007/s13225-020-00466-2)) |
| Otu58481 | 1 | k__Fungi(100);p__Ascomycota(100);c__Saccharomycetes(100);o__Saccharomycetales(100);f__Metschnikowiaceae(100);g__Metschnikowia(100);g__Metschnikowia_unclassified(100); | Metschnikowia | 13 | Saprotroph | \|Undefined Saprotroph\| | Yeast | NULL | Probable | As nectar-tap_saprotroph (Põlme et al. 2020); Decay sugar-rich substrates (Põlme et al. 2020) | Põlme S, et al. 2020. Fungal Diversity 105:1-16 ((https://doi.org/10.1007/s13225-020-00466-2)) |
| Otu58482 | 1 | k__Fungi(100);p__Ascomycota(100);c__Saccharomycetes(100);o__Saccharomycetales(100);f__Metschnikowiaceae(100);g__Metschnikowia(100);s__Metschnikowia_pulcherrima(100); | Metschnikowia | 13 | Saprotroph | \|Undefined Saprotroph\| | Yeast | NULL | Probable | As nectar-tap_saprotroph (Põlme et al. 2020); Decay sugar-rich substrates (Põlme et al. 2020) | Põlme S, et al. 2020. Fungal Diversity 105:1-16 ((https://doi.org/10.1007/s13225-020-00466-2)) |
| Otu58483 | 1 | k__Fungi(100);p__Ascomycota(100);c__Saccharomycetes(100);o__Saccharomycetales(100);f__Metschnikowiaceae(100);g__Metschnikowia(100);s__Metschnikowia_pulcherrima(100); | Metschnikowia | 13 | Saprotroph | \|Undefined Saprotroph\| | Yeast | NULL | Probable | As nectar-tap_saprotroph (Põlme et al. 2020); Decay sugar-rich substrates (Põlme et al. 2020) | Põlme S, et al. 2020. Fungal Diversity 105:1-16 ((https://doi.org/10.1007/s13225-020-00466-2)) |
| Otu58484 | 1 | k__Fungi(100);p__Ascomycota(100);c__Saccharomycetes(100);o__Saccharomycetales(100);f__Metschnikowiaceae(100);g__Metschnikowia(100);s__Metschnikowia_pulcherrima(100); | Metschnikowia | 13 | Saprotroph | \|Undefined Saprotroph\| | Yeast | NULL | Probable | As nectar-tap_saprotroph (Põlme et al. 2020); Decay sugar-rich substrates (Põlme et al. 2020) | Põlme S, et al. 2020. Fungal Diversity 105:1-16 ((https://doi.org/10.1007/s13225-020-00466-2)) |
| Otu58485 | 1 | k__Fungi(100);p__Ascomycota(100);c__Saccharomycetes(100);o__Saccharomycetales(100);f__Metschnikowiaceae(100);g__Metschnikowia(100);g__Metschnikowia_unclassified(100); | Metschnikowia | 13 | Saprotroph | \|Undefined Saprotroph\| | Yeast | NULL | Probable | As nectar-tap_saprotroph (Põlme et al. 2020); Decay sugar-rich substrates (Põlme et al. 2020) | Põlme S, et al. 2020. Fungal Diversity 105:1-16 ((https://doi.org/10.1007/s13225-020-00466-2)) |
| Otu58486 | 1 | k__Fungi(100);p__Ascomycota(100);c__Saccharomycetes(100);o__Saccharomycetales(100);f__Metschnikowiaceae(100);g__Metschnikowia(100);g__Metschnikowia_unclassified(100); | Metschnikowia | 13 | Saprotroph | \|Undefined Saprotroph\| | Yeast | NULL | Probable | As nectar-tap_saprotroph (Põlme et al. 2020); Decay sugar-rich substrates (Põlme et al. 2020) | Põlme S, et al. 2020. Fungal Diversity 105:1-16 ((https://doi.org/10.1007/s13225-020-00466-2)) |
| Otu58487 | 1 | k__Fungi(100);p__Ascomycota(100);c__Saccharomycetes(100);o__Saccharomycetales(100);f__Metschnikowiaceae(100);g__Metschnikowia(100);g__Metschnikowia_unclassified(100); | Metschnikowia | 13 | Saprotroph | \|Undefined Saprotroph\| | Yeast | NULL | Probable | As nectar-tap_saprotroph (Põlme et al. 2020); Decay sugar-rich substrates (Põlme et al. 2020) | Põlme S, et al. 2020. Fungal Diversity 105:1-16 ((https://doi.org/10.1007/s13225-020-00466-2)) |
| Otu58488 | 1 | k__Fungi(100);p__Ascomycota(100);c__Saccharomycetes(100);o__Saccharomycetales(100);f__Metschnikowiaceae(100);g__Metschnikowia(100);g__Metschnikowia_unclassified(100); | Metschnikowia | 13 | Saprotroph | \|Undefined Saprotroph\| | Yeast | NULL | Probable | As nectar-tap_saprotroph (Põlme et al. 2020); Decay sugar-rich substrates (Põlme et al. 2020) | Põlme S, et al. 2020. Fungal Diversity 105:1-16 ((https://doi.org/10.1007/s13225-020-00466-2)) |
| Otu58489 | 1 | k__Fungi(100);p__Ascomycota(100);c__Saccharomycetes(100);o__Saccharomycetales(100);f__Metschnikowiaceae(100);g__Metschnikowia(100);g__Metschnikowia_unclassified(100); | Metschnikowia | 13 | Saprotroph | \|Undefined Saprotroph\| | Yeast | NULL | Probable | As nectar-tap_saprotroph (Põlme et al. 2020); Decay sugar-rich substrates (Põlme et al. 2020) | Põlme S, et al. 2020. Fungal Diversity 105:1-16 ((https://doi.org/10.1007/s13225-020-00466-2)) |
| Otu58491 | 1 | k__Fungi(100);p__Ascomycota(100);c__Saccharomycetes(100);o__Saccharomycetales(100);f__Metschnikowiaceae(100);g__Metschnikowia(100);s__Metschnikowia_pulcherrima(100); | Metschnikowia | 13 | Saprotroph | \|Undefined Saprotroph\| | Yeast | NULL | Probable | As nectar-tap_saprotroph (Põlme et al. 2020); Decay sugar-rich substrates (Põlme et al. 2020) | Põlme S, et al. 2020. Fungal Diversity 105:1-16 ((https://doi.org/10.1007/s13225-020-00466-2)) |
| Otu58492 | 1 | k__Fungi(100);p__Ascomycota(100);c__Saccharomycetes(100);o__Saccharomycetales(100);f__Metschnikowiaceae(100);g__Metschnikowia(100);s__Metschnikowia_pulcherrima(100); | Metschnikowia | 13 | Saprotroph | \|Undefined Saprotroph\| | Yeast | NULL | Probable | As nectar-tap_saprotroph (Põlme et al. 2020); Decay sugar-rich substrates (Põlme et al. 2020) | Põlme S, et al. 2020. Fungal Diversity 105:1-16 ((https://doi.org/10.1007/s13225-020-00466-2)) |
| Otu58493 | 1 | k__Fungi(100);p__Ascomycota(100);c__Saccharomycetes(100);o__Saccharomycetales(100);f__Metschnikowiaceae(100);g__Metschnikowia(100);s__Metschnikowia_pulcherrima(100); | Metschnikowia | 13 | Saprotroph | \|Undefined Saprotroph\| | Yeast | NULL | Probable | As nectar-tap_saprotroph (Põlme et al. 2020); Decay sugar-rich substrates (Põlme et al. 2020) | Põlme S, et al. 2020. Fungal Diversity 105:1-16 ((https://doi.org/10.1007/s13225-020-00466-2)) |
| Otu58494 | 1 | k__Fungi(100);p__Ascomycota(100);c__Saccharomycetes(100);o__Saccharomycetales(100);f__Metschnikowiaceae(100);g__Metschnikowia(100);s__Metschnikowia_pulcherrima(100); | Metschnikowia | 13 | Saprotroph | \|Undefined Saprotroph\| | Yeast | NULL | Probable | As nectar-tap_saprotroph (Põlme et al. 2020); Decay sugar-rich substrates (Põlme et al. 2020) | Põlme S, et al. 2020. Fungal Diversity 105:1-16 ((https://doi.org/10.1007/s13225-020-00466-2)) |
| Otu58495 | 1 | k__Fungi(100);p__Ascomycota(100);c__Saccharomycetes(100);o__Saccharomycetales(100);f__Metschnikowiaceae(100);g__Metschnikowia(100);s__Metschnikowia_pulcherrima(100); | Metschnikowia | 13 | Saprotroph | \|Undefined Saprotroph\| | Yeast | NULL | Probable | As nectar-tap_saprotroph (Põlme et al. 2020); Decay sugar-rich substrates (Põlme et al. 2020) | Põlme S, et al. 2020. Fungal Diversity 105:1-16 ((https://doi.org/10.1007/s13225-020-00466-2)) |
| Otu58496 | 1 | k__Fungi(100);p__Ascomycota(100);c__Saccharomycetes(100);o__Saccharomycetales(100);f__Metschnikowiaceae(100);g__Metschnikowia(100);g__Metschnikowia_unclassified(100); | Metschnikowia | 13 | Saprotroph | \|Undefined Saprotroph\| | Yeast | NULL | Probable | As nectar-tap_saprotroph (Põlme et al. 2020); Decay sugar-rich substrates (Põlme et al. 2020) | Põlme S, et al. 2020. Fungal Diversity 105:1-16 ((https://doi.org/10.1007/s13225-020-00466-2)) |
| Otu58497 | 1 | k__Fungi(100);p__Ascomycota(100);c__Saccharomycetes(100);o__Saccharomycetales(100);f__Metschnikowiaceae(100);g__Metschnikowia(100);s__Metschnikowia_pulcherrima(100); | Metschnikowia | 13 | Saprotroph | \|Undefined Saprotroph\| | Yeast | NULL | Probable | As nectar-tap_saprotroph (Põlme et al. 2020); Decay sugar-rich substrates (Põlme et al. 2020) | Põlme S, et al. 2020. Fungal Diversity 105:1-16 ((https://doi.org/10.1007/s13225-020-00466-2)) |
| Otu58499 | 1 | k__Fungi(100);p__Ascomycota(100);c__Saccharomycetes(100);o__Saccharomycetales(100);f__Metschnikowiaceae(100);g__Metschnikowia(100);s__Metschnikowia_pulcherrima(100); | Metschnikowia | 13 | Saprotroph | \|Undefined Saprotroph\| | Yeast | NULL | Probable | As nectar-tap_saprotroph (Põlme et al. 2020); Decay sugar-rich substrates (Põlme et al. 2020) | Põlme S, et al. 2020. Fungal Diversity 105:1-16 ((https://doi.org/10.1007/s13225-020-00466-2)) |
| Otu58500 | 1 | k__Fungi(100);p__Ascomycota(100);c__Saccharomycetes(100);o__Saccharomycetales(100);f__Metschnikowiaceae(100);g__Metschnikowia(100);s__Metschnikowia_pulcherrima(100); | Metschnikowia | 13 | Saprotroph | \|Undefined Saprotroph\| | Yeast | NULL | Probable | As nectar-tap_saprotroph (Põlme et al. 2020); Decay sugar-rich substrates (Põlme et al. 2020) | Põlme S, et al. 2020. Fungal Diversity 105:1-16 ((https://doi.org/10.1007/s13225-020-00466-2)) |
| Otu58501 | 1 | k__Fungi(100);p__Ascomycota(100);c__Saccharomycetes(100);o__Saccharomycetales(100);f__Metschnikowiaceae(100);g__Metschnikowia(100);s__Metschnikowia_pulcherrima(100); | Metschnikowia | 13 | Saprotroph | \|Undefined Saprotroph\| | Yeast | NULL | Probable | As nectar-tap_saprotroph (Põlme et al. 2020); Decay sugar-rich substrates (Põlme et al. 2020) | Põlme S, et al. 2020. Fungal Diversity 105:1-16 ((https://doi.org/10.1007/s13225-020-00466-2)) |
| Otu58502 | 1 | k__Fungi(100);p__Ascomycota(100);c__Saccharomycetes(100);o__Saccharomycetales(100);f__Metschnikowiaceae(100);g__Metschnikowia(100);g__Metschnikowia_unclassified(100); | Metschnikowia | 13 | Saprotroph | \|Undefined Saprotroph\| | Yeast | NULL | Probable | As nectar-tap_saprotroph (Põlme et al. 2020); Decay sugar-rich substrates (Põlme et al. 2020) | Põlme S, et al. 2020. Fungal Diversity 105:1-16 ((https://doi.org/10.1007/s13225-020-00466-2)) |
| Otu58503 | 1 | k__Fungi(100);p__Ascomycota(100);c__Saccharomycetes(100);o__Saccharomycetales(100);f__Metschnikowiaceae(100);g__Metschnikowia(100);g__Metschnikowia_unclassified(100); | Metschnikowia | 13 | Saprotroph | \|Undefined Saprotroph\| | Yeast | NULL | Probable | As nectar-tap_saprotroph (Põlme et al. 2020); Decay sugar-rich substrates (Põlme et al. 2020) | Põlme S, et al. 2020. Fungal Diversity 105:1-16 ((https://doi.org/10.1007/s13225-020-00466-2)) |
| Otu58505 | 1 | k__Fungi(100);p__Ascomycota(100);c__Saccharomycetes(100);o__Saccharomycetales(100);f__Metschnikowiaceae(100);g__Metschnikowia(100);s__Metschnikowia_pulcherrima(100); | Metschnikowia | 13 | Saprotroph | \|Undefined Saprotroph\| | Yeast | NULL | Probable | As nectar-tap_saprotroph (Põlme et al. 2020); Decay sugar-rich substrates (Põlme et al. 2020) | Põlme S, et al. 2020. Fungal Diversity 105:1-16 ((https://doi.org/10.1007/s13225-020-00466-2)) |
| Otu58510 | 1 | k__Fungi(100);p__Ascomycota(100);c__Saccharomycetes(100);o__Saccharomycetales(100);f__Metschnikowiaceae(100);g__Metschnikowia(100);s__Metschnikowia_pulcherrima(100); | Metschnikowia | 13 | Saprotroph | \|Undefined Saprotroph\| | Yeast | NULL | Probable | As nectar-tap_saprotroph (Põlme et al. 2020); Decay sugar-rich substrates (Põlme et al. 2020) | Põlme S, et al. 2020. Fungal Diversity 105:1-16 ((https://doi.org/10.1007/s13225-020-00466-2)) |
| Otu58512 | 1 | k__Fungi(100);p__Ascomycota(100);c__Saccharomycetes(100);o__Saccharomycetales(100);f__Metschnikowiaceae(100);g__Metschnikowia(100);g__Metschnikowia_unclassified(100); | Metschnikowia | 13 | Saprotroph | \|Undefined Saprotroph\| | Yeast | NULL | Probable | As nectar-tap_saprotroph (Põlme et al. 2020); Decay sugar-rich substrates (Põlme et al. 2020) | Põlme S, et al. 2020. Fungal Diversity 105:1-16 ((https://doi.org/10.1007/s13225-020-00466-2)) |
| Otu58513 | 1 | k__Fungi(100);p__Ascomycota(100);c__Saccharomycetes(100);o__Saccharomycetales(100);f__Metschnikowiaceae(100);g__Metschnikowia(100);s__Metschnikowia_pulcherrima(100); | Metschnikowia | 13 | Saprotroph | \|Undefined Saprotroph\| | Yeast | NULL | Probable | As nectar-tap_saprotroph (Põlme et al. 2020); Decay sugar-rich substrates (Põlme et al. 2020) | Põlme S, et al. 2020. Fungal Diversity 105:1-16 ((https://doi.org/10.1007/s13225-020-00466-2)) |
| Otu58515 | 1 | k__Fungi(100);p__Ascomycota(100);c__Saccharomycetes(100);o__Saccharomycetales(100);f__Metschnikowiaceae(100);g__Metschnikowia(100);s__Metschnikowia_pulcherrima(100); | Metschnikowia | 13 | Saprotroph | \|Undefined Saprotroph\| | Yeast | NULL | Probable | As nectar-tap_saprotroph (Põlme et al. 2020); Decay sugar-rich substrates (Põlme et al. 2020) | Põlme S, et al. 2020. Fungal Diversity 105:1-16 ((https://doi.org/10.1007/s13225-020-00466-2)) |
| Otu58516 | 1 | k__Fungi(100);p__Ascomycota(100);c__Saccharomycetes(100);o__Saccharomycetales(100);f__Metschnikowiaceae(100);g__Metschnikowia(100);g__Metschnikowia_unclassified(100); | Metschnikowia | 13 | Saprotroph | \|Undefined Saprotroph\| | Yeast | NULL | Probable | As nectar-tap_saprotroph (Põlme et al. 2020); Decay sugar-rich substrates (Põlme et al. 2020) | Põlme S, et al. 2020. Fungal Diversity 105:1-16 ((https://doi.org/10.1007/s13225-020-00466-2)) |
| Otu58517 | 1 | k__Fungi(100);p__Ascomycota(100);c__Saccharomycetes(100);o__Saccharomycetales(100);f__Metschnikowiaceae(100);g__Metschnikowia(100);s__Metschnikowia_pulcherrima(100); | Metschnikowia | 13 | Saprotroph | \|Undefined Saprotroph\| | Yeast | NULL | Probable | As nectar-tap_saprotroph (Põlme et al. 2020); Decay sugar-rich substrates (Põlme et al. 2020) | Põlme S, et al. 2020. Fungal Diversity 105:1-16 ((https://doi.org/10.1007/s13225-020-00466-2)) |
| Otu58518 | 1 | k__Fungi(100);p__Ascomycota(100);c__Saccharomycetes(100);o__Saccharomycetales(100);f__Metschnikowiaceae(100);g__Metschnikowia(100);g__Metschnikowia_unclassified(100); | Metschnikowia | 13 | Saprotroph | \|Undefined Saprotroph\| | Yeast | NULL | Probable | As nectar-tap_saprotroph (Põlme et al. 2020); Decay sugar-rich substrates (Põlme et al. 2020) | Põlme S, et al. 2020. Fungal Diversity 105:1-16 ((https://doi.org/10.1007/s13225-020-00466-2)) |
| Otu58519 | 1 | k__Fungi(100);p__Ascomycota(100);c__Saccharomycetes(100);o__Saccharomycetales(100);f__Metschnikowiaceae(100);g__Metschnikowia(100);s__Metschnikowia_pulcherrima(100); | Metschnikowia | 13 | Saprotroph | \|Undefined Saprotroph\| | Yeast | NULL | Probable | As nectar-tap_saprotroph (Põlme et al. 2020); Decay sugar-rich substrates (Põlme et al. 2020) | Põlme S, et al. 2020. Fungal Diversity 105:1-16 ((https://doi.org/10.1007/s13225-020-00466-2)) |
| Otu58522 | 1 | k__Fungi(100);p__Ascomycota(100);c__Saccharomycetes(100);o__Saccharomycetales(100);f__Metschnikowiaceae(100);g__Metschnikowia(100);s__Metschnikowia_pulcherrima(100); | Metschnikowia | 13 | Saprotroph | \|Undefined Saprotroph\| | Yeast | NULL | Probable | As nectar-tap_saprotroph (Põlme et al. 2020); Decay sugar-rich substrates (Põlme et al. 2020) | Põlme S, et al. 2020. Fungal Diversity 105:1-16 ((https://doi.org/10.1007/s13225-020-00466-2)) |
| Otu58525 | 1 | k__Fungi(100);p__Ascomycota(100);c__Saccharomycetes(100);o__Saccharomycetales(100);f__Metschnikowiaceae(100);g__Metschnikowia(100);s__Metschnikowia_pulcherrima(100); | Metschnikowia | 13 | Saprotroph | \|Undefined Saprotroph\| | Yeast | NULL | Probable | As nectar-tap_saprotroph (Põlme et al. 2020); Decay sugar-rich substrates (Põlme et al. 2020) | Põlme S, et al. 2020. Fungal Diversity 105:1-16 ((https://doi.org/10.1007/s13225-020-00466-2)) |
| Otu58526 | 1 | k__Fungi(100);p__Ascomycota(100);c__Saccharomycetes(100);o__Saccharomycetales(100);f__Metschnikowiaceae(100);g__Metschnikowia(100);s__Metschnikowia_pulcherrima(100); | Metschnikowia | 13 | Saprotroph | \|Undefined Saprotroph\| | Yeast | NULL | Probable | As nectar-tap_saprotroph (Põlme et al. 2020); Decay sugar-rich substrates (Põlme et al. 2020) | Põlme S, et al. 2020. Fungal Diversity 105:1-16 ((https://doi.org/10.1007/s13225-020-00466-2)) |
| Otu58527 | 1 | k__Fungi(100);p__Ascomycota(100);c__Saccharomycetes(100);o__Saccharomycetales(100);f__Metschnikowiaceae(100);g__Metschnikowia(100);s__Metschnikowia_pulcherrima(100); | Metschnikowia | 13 | Saprotroph | \|Undefined Saprotroph\| | Yeast | NULL | Probable | As nectar-tap_saprotroph (Põlme et al. 2020); Decay sugar-rich substrates (Põlme et al. 2020) | Põlme S, et al. 2020. Fungal Diversity 105:1-16 ((https://doi.org/10.1007/s13225-020-00466-2)) |
| Otu58528 | 1 | k__Fungi(100);p__Ascomycota(100);c__Saccharomycetes(100);o__Saccharomycetales(100);f__Metschnikowiaceae(100);g__Metschnikowia(100);s__Metschnikowia_pulcherrima(100); | Metschnikowia | 13 | Saprotroph | \|Undefined Saprotroph\| | Yeast | NULL | Probable | As nectar-tap_saprotroph (Põlme et al. 2020); Decay sugar-rich substrates (Põlme et al. 2020) | Põlme S, et al. 2020. Fungal Diversity 105:1-16 ((https://doi.org/10.1007/s13225-020-00466-2)) |
| Otu58529 | 1 | k__Fungi(100);p__Ascomycota(100);c__Saccharomycetes(100);o__Saccharomycetales(100);f__Metschnikowiaceae(100);g__Metschnikowia(100);s__Metschnikowia_pulcherrima(100); | Metschnikowia | 13 | Saprotroph | \|Undefined Saprotroph\| | Yeast | NULL | Probable | As nectar-tap_saprotroph (Põlme et al. 2020); Decay sugar-rich substrates (Põlme et al. 2020) | Põlme S, et al. 2020. Fungal Diversity 105:1-16 ((https://doi.org/10.1007/s13225-020-00466-2)) |
| Otu58530 | 1 | k__Fungi(100);p__Ascomycota(100);c__Saccharomycetes(100);o__Saccharomycetales(100);f__Metschnikowiaceae(100);g__Metschnikowia(100);s__Metschnikowia_pulcherrima(100); | Metschnikowia | 13 | Saprotroph | \|Undefined Saprotroph\| | Yeast | NULL | Probable | As nectar-tap_saprotroph (Põlme et al. 2020); Decay sugar-rich substrates (Põlme et al. 2020) | Põlme S, et al. 2020. Fungal Diversity 105:1-16 ((https://doi.org/10.1007/s13225-020-00466-2)) |
| Otu58533 | 1 | k__Fungi(100);p__Ascomycota(100);c__Saccharomycetes(100);o__Saccharomycetales(100);f__Metschnikowiaceae(100);g__Metschnikowia(100);s__Metschnikowia_pulcherrima(100); | Metschnikowia | 13 | Saprotroph | \|Undefined Saprotroph\| | Yeast | NULL | Probable | As nectar-tap_saprotroph (Põlme et al. 2020); Decay sugar-rich substrates (Põlme et al. 2020) | Põlme S, et al. 2020. Fungal Diversity 105:1-16 ((https://doi.org/10.1007/s13225-020-00466-2)) |
| Otu58534 | 1 | k__Fungi(100);p__Ascomycota(100);c__Saccharomycetes(100);o__Saccharomycetales(100);f__Metschnikowiaceae(100);g__Metschnikowia(100);s__Metschnikowia_pulcherrima(100); | Metschnikowia | 13 | Saprotroph | \|Undefined Saprotroph\| | Yeast | NULL | Probable | As nectar-tap_saprotroph (Põlme et al. 2020); Decay sugar-rich substrates (Põlme et al. 2020) | Põlme S, et al. 2020. Fungal Diversity 105:1-16 ((https://doi.org/10.1007/s13225-020-00466-2)) |
| Otu58536 | 1 | k__Fungi(100);p__Ascomycota(100);c__Saccharomycetes(100);o__Saccharomycetales(100);f__Metschnikowiaceae(100);g__Metschnikowia(100);s__Metschnikowia_pulcherrima(100); | Metschnikowia | 13 | Saprotroph | \|Undefined Saprotroph\| | Yeast | NULL | Probable | As nectar-tap_saprotroph (Põlme et al. 2020); Decay sugar-rich substrates (Põlme et al. 2020) | Põlme S, et al. 2020. Fungal Diversity 105:1-16 ((https://doi.org/10.1007/s13225-020-00466-2)) |
| Otu58537 | 1 | k__Fungi(100);p__Ascomycota(100);c__Saccharomycetes(100);o__Saccharomycetales(100);f__Metschnikowiaceae(100);g__Metschnikowia(100);s__Metschnikowia_pulcherrima(100); | Metschnikowia | 13 | Saprotroph | \|Undefined Saprotroph\| | Yeast | NULL | Probable | As nectar-tap_saprotroph (Põlme et al. 2020); Decay sugar-rich substrates (Põlme et al. 2020) | Põlme S, et al. 2020. Fungal Diversity 105:1-16 ((https://doi.org/10.1007/s13225-020-00466-2)) |
| Otu58538 | 1 | k__Fungi(100);p__Ascomycota(100);c__Saccharomycetes(100);o__Saccharomycetales(100);f__Metschnikowiaceae(100);g__Metschnikowia(100);s__Metschnikowia_pulcherrima(100); | Metschnikowia | 13 | Saprotroph | \|Undefined Saprotroph\| | Yeast | NULL | Probable | As nectar-tap_saprotroph (Põlme et al. 2020); Decay sugar-rich substrates (Põlme et al. 2020) | Põlme S, et al. 2020. Fungal Diversity 105:1-16 ((https://doi.org/10.1007/s13225-020-00466-2)) |
| Otu58539 | 1 | k__Fungi(100);p__Ascomycota(100);c__Saccharomycetes(100);o__Saccharomycetales(100);f__Metschnikowiaceae(100);g__Metschnikowia(100);s__Metschnikowia_pulcherrima(100); | Metschnikowia | 13 | Saprotroph | \|Undefined Saprotroph\| | Yeast | NULL | Probable | As nectar-tap_saprotroph (Põlme et al. 2020); Decay sugar-rich substrates (Põlme et al. 2020) | Põlme S, et al. 2020. Fungal Diversity 105:1-16 ((https://doi.org/10.1007/s13225-020-00466-2)) |
| Otu58540 | 1 | k__Fungi(100);p__Ascomycota(100);c__Saccharomycetes(100);o__Saccharomycetales(100);f__Metschnikowiaceae(100);g__Metschnikowia(100);s__Metschnikowia_pulcherrima(100); | Metschnikowia | 13 | Saprotroph | \|Undefined Saprotroph\| | Yeast | NULL | Probable | As nectar-tap_saprotroph (Põlme et al. 2020); Decay sugar-rich substrates (Põlme et al. 2020) | Põlme S, et al. 2020. Fungal Diversity 105:1-16 ((https://doi.org/10.1007/s13225-020-00466-2)) |
| Otu58541 | 1 | k__Fungi(100);p__Ascomycota(100);c__Saccharomycetes(100);o__Saccharomycetales(100);f__Metschnikowiaceae(100);g__Metschnikowia(100);s__Metschnikowia_pulcherrima(100); | Metschnikowia | 13 | Saprotroph | \|Undefined Saprotroph\| | Yeast | NULL | Probable | As nectar-tap_saprotroph (Põlme et al. 2020); Decay sugar-rich substrates (Põlme et al. 2020) | Põlme S, et al. 2020. Fungal Diversity 105:1-16 ((https://doi.org/10.1007/s13225-020-00466-2)) |
| Otu58543 | 1 | k__Fungi(100);p__Ascomycota(100);c__Saccharomycetes(100);o__Saccharomycetales(100);f__Metschnikowiaceae(100);g__Metschnikowia(100);s__Metschnikowia_pulcherrima(100); | Metschnikowia | 13 | Saprotroph | \|Undefined Saprotroph\| | Yeast | NULL | Probable | As nectar-tap_saprotroph (Põlme et al. 2020); Decay sugar-rich substrates (Põlme et al. 2020) | Põlme S, et al. 2020. Fungal Diversity 105:1-16 ((https://doi.org/10.1007/s13225-020-00466-2)) |
| Otu58545 | 1 | k__Fungi(100);p__Ascomycota(100);c__Saccharomycetes(100);o__Saccharomycetales(100);f__Metschnikowiaceae(100);g__Metschnikowia(100);g__Metschnikowia_unclassified(100); | Metschnikowia | 13 | Saprotroph | \|Undefined Saprotroph\| | Yeast | NULL | Probable | As nectar-tap_saprotroph (Põlme et al. 2020); Decay sugar-rich substrates (Põlme et al. 2020) | Põlme S, et al. 2020. Fungal Diversity 105:1-16 ((https://doi.org/10.1007/s13225-020-00466-2)) |
| Otu58546 | 1 | k__Fungi(100);p__Ascomycota(100);c__Saccharomycetes(100);o__Saccharomycetales(100);f__Metschnikowiaceae(100);g__Metschnikowia(100);g__Metschnikowia_unclassified(100); | Metschnikowia | 13 | Saprotroph | \|Undefined Saprotroph\| | Yeast | NULL | Probable | As nectar-tap_saprotroph (Põlme et al. 2020); Decay sugar-rich substrates (Põlme et al. 2020) | Põlme S, et al. 2020. Fungal Diversity 105:1-16 ((https://doi.org/10.1007/s13225-020-00466-2)) |
| Otu58547 | 1 | k__Fungi(100);p__Ascomycota(100);c__Saccharomycetes(100);o__Saccharomycetales(100);f__Metschnikowiaceae(100);g__Metschnikowia(100);s__Metschnikowia_pulcherrima(100); | Metschnikowia | 13 | Saprotroph | \|Undefined Saprotroph\| | Yeast | NULL | Probable | As nectar-tap_saprotroph (Põlme et al. 2020); Decay sugar-rich substrates (Põlme et al. 2020) | Põlme S, et al. 2020. Fungal Diversity 105:1-16 ((https://doi.org/10.1007/s13225-020-00466-2)) |
| Otu58548 | 1 | k__Fungi(100);p__Ascomycota(100);c__Saccharomycetes(100);o__Saccharomycetales(100);f__Metschnikowiaceae(100);g__Metschnikowia(100);s__Metschnikowia_pulcherrima(100); | Metschnikowia | 13 | Saprotroph | \|Undefined Saprotroph\| | Yeast | NULL | Probable | As nectar-tap_saprotroph (Põlme et al. 2020); Decay sugar-rich substrates (Põlme et al. 2020) | Põlme S, et al. 2020. Fungal Diversity 105:1-16 ((https://doi.org/10.1007/s13225-020-00466-2)) |
| Otu58549 | 1 | k__Fungi(100);p__Ascomycota(100);c__Saccharomycetes(100);o__Saccharomycetales(100);f__Metschnikowiaceae(100);g__Metschnikowia(100);s__Metschnikowia_pulcherrima(100); | Metschnikowia | 13 | Saprotroph | \|Undefined Saprotroph\| | Yeast | NULL | Probable | As nectar-tap_saprotroph (Põlme et al. 2020); Decay sugar-rich substrates (Põlme et al. 2020) | Põlme S, et al. 2020. Fungal Diversity 105:1-16 ((https://doi.org/10.1007/s13225-020-00466-2)) |
| Otu58550 | 1 | k__Fungi(100);p__Ascomycota(100);c__Saccharomycetes(100);o__Saccharomycetales(100);f__Metschnikowiaceae(100);g__Metschnikowia(100);s__Metschnikowia_pulcherrima(100); | Metschnikowia | 13 | Saprotroph | \|Undefined Saprotroph\| | Yeast | NULL | Probable | As nectar-tap_saprotroph (Põlme et al. 2020); Decay sugar-rich substrates (Põlme et al. 2020) | Põlme S, et al. 2020. Fungal Diversity 105:1-16 ((https://doi.org/10.1007/s13225-020-00466-2)) |
| Otu58551 | 1 | k__Fungi(100);p__Ascomycota(100);c__Saccharomycetes(100);o__Saccharomycetales(100);f__Metschnikowiaceae(100);g__Metschnikowia(100);s__Metschnikowia_pulcherrima(100); | Metschnikowia | 13 | Saprotroph | \|Undefined Saprotroph\| | Yeast | NULL | Probable | As nectar-tap_saprotroph (Põlme et al. 2020); Decay sugar-rich substrates (Põlme et al. 2020) | Põlme S, et al. 2020. Fungal Diversity 105:1-16 ((https://doi.org/10.1007/s13225-020-00466-2)) |
| Otu58552 | 1 | k__Fungi(100);p__Ascomycota(100);c__Saccharomycetes(100);o__Saccharomycetales(100);f__Metschnikowiaceae(100);g__Metschnikowia(100);s__Metschnikowia_pulcherrima(100); | Metschnikowia | 13 | Saprotroph | \|Undefined Saprotroph\| | Yeast | NULL | Probable | As nectar-tap_saprotroph (Põlme et al. 2020); Decay sugar-rich substrates (Põlme et al. 2020) | Põlme S, et al. 2020. Fungal Diversity 105:1-16 ((https://doi.org/10.1007/s13225-020-00466-2)) |
| Otu58553 | 1 | k__Fungi(100);p__Ascomycota(100);c__Saccharomycetes(100);o__Saccharomycetales(100);f__Metschnikowiaceae(100);g__Metschnikowia(100);s__Metschnikowia_pulcherrima(100); | Metschnikowia | 13 | Saprotroph | \|Undefined Saprotroph\| | Yeast | NULL | Probable | As nectar-tap_saprotroph (Põlme et al. 2020); Decay sugar-rich substrates (Põlme et al. 2020) | Põlme S, et al. 2020. Fungal Diversity 105:1-16 ((https://doi.org/10.1007/s13225-020-00466-2)) |
| Otu58554 | 1 | k__Fungi(100);p__Ascomycota(100);c__Saccharomycetes(100);o__Saccharomycetales(100);f__Metschnikowiaceae(100);g__Metschnikowia(100);s__Metschnikowia_pulcherrima(100); | Metschnikowia | 13 | Saprotroph | \|Undefined Saprotroph\| | Yeast | NULL | Probable | As nectar-tap_saprotroph (Põlme et al. 2020); Decay sugar-rich substrates (Põlme et al. 2020) | Põlme S, et al. 2020. Fungal Diversity 105:1-16 ((https://doi.org/10.1007/s13225-020-00466-2)) |
| Otu58555 | 1 | k__Fungi(100);p__Ascomycota(100);c__Saccharomycetes(100);o__Saccharomycetales(100);f__Metschnikowiaceae(100);g__Metschnikowia(100);s__Metschnikowia_pulcherrima(100); | Metschnikowia | 13 | Saprotroph | \|Undefined Saprotroph\| | Yeast | NULL | Probable | As nectar-tap_saprotroph (Põlme et al. 2020); Decay sugar-rich substrates (Põlme et al. 2020) | Põlme S, et al. 2020. Fungal Diversity 105:1-16 ((https://doi.org/10.1007/s13225-020-00466-2)) |
| Otu58557 | 1 | k__Fungi(100);p__Ascomycota(100);c__Saccharomycetes(100);o__Saccharomycetales(100);f__Metschnikowiaceae(100);g__Metschnikowia(100);s__Metschnikowia_pulcherrima(100); | Metschnikowia | 13 | Saprotroph | \|Undefined Saprotroph\| | Yeast | NULL | Probable | As nectar-tap_saprotroph (Põlme et al. 2020); Decay sugar-rich substrates (Põlme et al. 2020) | Põlme S, et al. 2020. Fungal Diversity 105:1-16 ((https://doi.org/10.1007/s13225-020-00466-2)) |
| Otu58558 | 1 | k__Fungi(100);p__Ascomycota(100);c__Saccharomycetes(100);o__Saccharomycetales(100);f__Metschnikowiaceae(100);g__Metschnikowia(100);s__Metschnikowia_pulcherrima(100); | Metschnikowia | 13 | Saprotroph | \|Undefined Saprotroph\| | Yeast | NULL | Probable | As nectar-tap_saprotroph (Põlme et al. 2020); Decay sugar-rich substrates (Põlme et al. 2020) | Põlme S, et al. 2020. Fungal Diversity 105:1-16 ((https://doi.org/10.1007/s13225-020-00466-2)) |
| Otu58559 | 1 | k__Fungi(100);p__Ascomycota(100);c__Saccharomycetes(100);o__Saccharomycetales(100);f__Metschnikowiaceae(100);g__Metschnikowia(100);s__Metschnikowia_pulcherrima(100); | Metschnikowia | 13 | Saprotroph | \|Undefined Saprotroph\| | Yeast | NULL | Probable | As nectar-tap_saprotroph (Põlme et al. 2020); Decay sugar-rich substrates (Põlme et al. 2020) | Põlme S, et al. 2020. Fungal Diversity 105:1-16 ((https://doi.org/10.1007/s13225-020-00466-2)) |
| Otu58560 | 1 | k__Fungi(100);p__Ascomycota(100);c__Saccharomycetes(100);o__Saccharomycetales(100);f__Metschnikowiaceae(100);g__Metschnikowia(100);g__Metschnikowia_unclassified(100); | Metschnikowia | 13 | Saprotroph | \|Undefined Saprotroph\| | Yeast | NULL | Probable | As nectar-tap_saprotroph (Põlme et al. 2020); Decay sugar-rich substrates (Põlme et al. 2020) | Põlme S, et al. 2020. Fungal Diversity 105:1-16 ((https://doi.org/10.1007/s13225-020-00466-2)) |
| Otu58561 | 1 | k__Fungi(100);p__Ascomycota(100);c__Saccharomycetes(100);o__Saccharomycetales(100);f__Metschnikowiaceae(100);g__Metschnikowia(100);s__Metschnikowia_pulcherrima(100); | Metschnikowia | 13 | Saprotroph | \|Undefined Saprotroph\| | Yeast | NULL | Probable | As nectar-tap_saprotroph (Põlme et al. 2020); Decay sugar-rich substrates (Põlme et al. 2020) | Põlme S, et al. 2020. Fungal Diversity 105:1-16 ((https://doi.org/10.1007/s13225-020-00466-2)) |
| Otu58562 | 1 | k__Fungi(100);p__Ascomycota(100);c__Saccharomycetes(100);o__Saccharomycetales(100);f__Metschnikowiaceae(100);g__Metschnikowia(100);g__Metschnikowia_unclassified(100); | Metschnikowia | 13 | Saprotroph | \|Undefined Saprotroph\| | Yeast | NULL | Probable | As nectar-tap_saprotroph (Põlme et al. 2020); Decay sugar-rich substrates (Põlme et al. 2020) | Põlme S, et al. 2020. Fungal Diversity 105:1-16 ((https://doi.org/10.1007/s13225-020-00466-2)) |
| Otu58563 | 1 | k__Fungi(100);p__Ascomycota(100);c__Saccharomycetes(100);o__Saccharomycetales(100);f__Metschnikowiaceae(100);g__Metschnikowia(100);g__Metschnikowia_unclassified(100); | Metschnikowia | 13 | Saprotroph | \|Undefined Saprotroph\| | Yeast | NULL | Probable | As nectar-tap_saprotroph (Põlme et al. 2020); Decay sugar-rich substrates (Põlme et al. 2020) | Põlme S, et al. 2020. Fungal Diversity 105:1-16 ((https://doi.org/10.1007/s13225-020-00466-2)) |
| Otu58564 | 1 | k__Fungi(100);p__Ascomycota(100);c__Saccharomycetes(100);o__Saccharomycetales(100);f__Metschnikowiaceae(100);g__Metschnikowia(100);g__Metschnikowia_unclassified(100); | Metschnikowia | 13 | Saprotroph | \|Undefined Saprotroph\| | Yeast | NULL | Probable | As nectar-tap_saprotroph (Põlme et al. 2020); Decay sugar-rich substrates (Põlme et al. 2020) | Põlme S, et al. 2020. Fungal Diversity 105:1-16 ((https://doi.org/10.1007/s13225-020-00466-2)) |
| Otu58565 | 1 | k__Fungi(100);p__Ascomycota(100);c__Saccharomycetes(100);o__Saccharomycetales(100);f__Metschnikowiaceae(100);g__Metschnikowia(100);g__Metschnikowia_unclassified(100); | Metschnikowia | 13 | Saprotroph | \|Undefined Saprotroph\| | Yeast | NULL | Probable | As nectar-tap_saprotroph (Põlme et al. 2020); Decay sugar-rich substrates (Põlme et al. 2020) | Põlme S, et al. 2020. Fungal Diversity 105:1-16 ((https://doi.org/10.1007/s13225-020-00466-2)) |
| Otu58566 | 1 | k__Fungi(100);p__Ascomycota(100);c__Saccharomycetes(100);o__Saccharomycetales(100);f__Metschnikowiaceae(100);g__Metschnikowia(100);g__Metschnikowia_unclassified(100); | Metschnikowia | 13 | Saprotroph | \|Undefined Saprotroph\| | Yeast | NULL | Probable | As nectar-tap_saprotroph (Põlme et al. 2020); Decay sugar-rich substrates (Põlme et al. 2020) | Põlme S, et al. 2020. Fungal Diversity 105:1-16 ((https://doi.org/10.1007/s13225-020-00466-2)) |
| Otu58567 | 1 | k__Fungi(100);p__Ascomycota(100);c__Saccharomycetes(100);o__Saccharomycetales(100);f__Metschnikowiaceae(100);g__Metschnikowia(100);g__Metschnikowia_unclassified(100); | Metschnikowia | 13 | Saprotroph | \|Undefined Saprotroph\| | Yeast | NULL | Probable | As nectar-tap_saprotroph (Põlme et al. 2020); Decay sugar-rich substrates (Põlme et al. 2020) | Põlme S, et al. 2020. Fungal Diversity 105:1-16 ((https://doi.org/10.1007/s13225-020-00466-2)) |
| Otu58568 | 1 | k__Fungi(100);p__Ascomycota(100);c__Saccharomycetes(100);o__Saccharomycetales(100);f__Metschnikowiaceae(100);g__Metschnikowia(100);g__Metschnikowia_unclassified(100); | Metschnikowia | 13 | Saprotroph | \|Undefined Saprotroph\| | Yeast | NULL | Probable | As nectar-tap_saprotroph (Põlme et al. 2020); Decay sugar-rich substrates (Põlme et al. 2020) | Põlme S, et al. 2020. Fungal Diversity 105:1-16 ((https://doi.org/10.1007/s13225-020-00466-2)) |
| Otu58569 | 1 | k__Fungi(100);p__Ascomycota(100);c__Saccharomycetes(100);o__Saccharomycetales(100);f__Metschnikowiaceae(100);g__Metschnikowia(100);s__Metschnikowia_pulcherrima(100); | Metschnikowia | 13 | Saprotroph | \|Undefined Saprotroph\| | Yeast | NULL | Probable | As nectar-tap_saprotroph (Põlme et al. 2020); Decay sugar-rich substrates (Põlme et al. 2020) | Põlme S, et al. 2020. Fungal Diversity 105:1-16 ((https://doi.org/10.1007/s13225-020-00466-2)) |
| Otu58570 | 1 | k__Fungi(100);p__Ascomycota(100);c__Saccharomycetes(100);o__Saccharomycetales(100);f__Metschnikowiaceae(100);g__Metschnikowia(100);g__Metschnikowia_unclassified(100); | Metschnikowia | 13 | Saprotroph | \|Undefined Saprotroph\| | Yeast | NULL | Probable | As nectar-tap_saprotroph (Põlme et al. 2020); Decay sugar-rich substrates (Põlme et al. 2020) | Põlme S, et al. 2020. Fungal Diversity 105:1-16 ((https://doi.org/10.1007/s13225-020-00466-2)) |
| Otu58571 | 1 | k__Fungi(100);p__Ascomycota(100);c__Saccharomycetes(100);o__Saccharomycetales(100);f__Metschnikowiaceae(100);g__Metschnikowia(100);s__Metschnikowia_pulcherrima(100); | Metschnikowia | 13 | Saprotroph | \|Undefined Saprotroph\| | Yeast | NULL | Probable | As nectar-tap_saprotroph (Põlme et al. 2020); Decay sugar-rich substrates (Põlme et al. 2020) | Põlme S, et al. 2020. Fungal Diversity 105:1-16 ((https://doi.org/10.1007/s13225-020-00466-2)) |
| Otu58572 | 1 | k__Fungi(100);p__Ascomycota(100);c__Saccharomycetes(100);o__Saccharomycetales(100);f__Metschnikowiaceae(100);g__Metschnikowia(100);s__Metschnikowia_pulcherrima(100); | Metschnikowia | 13 | Saprotroph | \|Undefined Saprotroph\| | Yeast | NULL | Probable | As nectar-tap_saprotroph (Põlme et al. 2020); Decay sugar-rich substrates (Põlme et al. 2020) | Põlme S, et al. 2020. Fungal Diversity 105:1-16 ((https://doi.org/10.1007/s13225-020-00466-2)) |
| Otu58573 | 1 | k__Fungi(100);p__Ascomycota(100);c__Saccharomycetes(100);o__Saccharomycetales(100);f__Metschnikowiaceae(100);g__Metschnikowia(100);s__Metschnikowia_pulcherrima(100); | Metschnikowia | 13 | Saprotroph | \|Undefined Saprotroph\| | Yeast | NULL | Probable | As nectar-tap_saprotroph (Põlme et al. 2020); Decay sugar-rich substrates (Põlme et al. 2020) | Põlme S, et al. 2020. Fungal Diversity 105:1-16 ((https://doi.org/10.1007/s13225-020-00466-2)) |
| Otu58574 | 1 | k__Fungi(100);p__Ascomycota(100);c__Saccharomycetes(100);o__Saccharomycetales(100);f__Metschnikowiaceae(100);g__Metschnikowia(100);s__Metschnikowia_pulcherrima(100); | Metschnikowia | 13 | Saprotroph | \|Undefined Saprotroph\| | Yeast | NULL | Probable | As nectar-tap_saprotroph (Põlme et al. 2020); Decay sugar-rich substrates (Põlme et al. 2020) | Põlme S, et al. 2020. Fungal Diversity 105:1-16 ((https://doi.org/10.1007/s13225-020-00466-2)) |
| Otu58575 | 1 | k__Fungi(100);p__Ascomycota(100);c__Saccharomycetes(100);o__Saccharomycetales(100);f__Metschnikowiaceae(100);g__Metschnikowia(100);g__Metschnikowia_unclassified(100); | Metschnikowia | 13 | Saprotroph | \|Undefined Saprotroph\| | Yeast | NULL | Probable | As nectar-tap_saprotroph (Põlme et al. 2020); Decay sugar-rich substrates (Põlme et al. 2020) | Põlme S, et al. 2020. Fungal Diversity 105:1-16 ((https://doi.org/10.1007/s13225-020-00466-2)) |
| Otu58576 | 1 | k__Fungi(100);p__Ascomycota(100);c__Saccharomycetes(100);o__Saccharomycetales(100);f__Metschnikowiaceae(100);g__Metschnikowia(100);s__Metschnikowia_pulcherrima(100); | Metschnikowia | 13 | Saprotroph | \|Undefined Saprotroph\| | Yeast | NULL | Probable | As nectar-tap_saprotroph (Põlme et al. 2020); Decay sugar-rich substrates (Põlme et al. 2020) | Põlme S, et al. 2020. Fungal Diversity 105:1-16 ((https://doi.org/10.1007/s13225-020-00466-2)) |
| Otu58577 | 1 | k__Fungi(100);p__Ascomycota(100);c__Saccharomycetes(100);o__Saccharomycetales(100);f__Metschnikowiaceae(100);g__Metschnikowia(100);s__Metschnikowia_pulcherrima(100); | Metschnikowia | 13 | Saprotroph | \|Undefined Saprotroph\| | Yeast | NULL | Probable | As nectar-tap_saprotroph (Põlme et al. 2020); Decay sugar-rich substrates (Põlme et al. 2020) | Põlme S, et al. 2020. Fungal Diversity 105:1-16 ((https://doi.org/10.1007/s13225-020-00466-2)) |
| Otu58578 | 1 | k__Fungi(100);p__Ascomycota(100);c__Saccharomycetes(100);o__Saccharomycetales(100);f__Metschnikowiaceae(100);g__Metschnikowia(100);s__Metschnikowia_pulcherrima(100); | Metschnikowia | 13 | Saprotroph | \|Undefined Saprotroph\| | Yeast | NULL | Probable | As nectar-tap_saprotroph (Põlme et al. 2020); Decay sugar-rich substrates (Põlme et al. 2020) | Põlme S, et al. 2020. Fungal Diversity 105:1-16 ((https://doi.org/10.1007/s13225-020-00466-2)) |
| Otu58579 | 1 | k__Fungi(100);p__Ascomycota(100);c__Saccharomycetes(100);o__Saccharomycetales(100);f__Metschnikowiaceae(100);g__Metschnikowia(100);s__Metschnikowia_pulcherrima(100); | Metschnikowia | 13 | Saprotroph | \|Undefined Saprotroph\| | Yeast | NULL | Probable | As nectar-tap_saprotroph (Põlme et al. 2020); Decay sugar-rich substrates (Põlme et al. 2020) | Põlme S, et al. 2020. Fungal Diversity 105:1-16 ((https://doi.org/10.1007/s13225-020-00466-2)) |
| Otu58580 | 1 | k__Fungi(100);p__Ascomycota(100);c__Saccharomycetes(100);o__Saccharomycetales(100);f__Metschnikowiaceae(100);g__Metschnikowia(100);s__Metschnikowia_pulcherrima(100); | Metschnikowia | 13 | Saprotroph | \|Undefined Saprotroph\| | Yeast | NULL | Probable | As nectar-tap_saprotroph (Põlme et al. 2020); Decay sugar-rich substrates (Põlme et al. 2020) | Põlme S, et al. 2020. Fungal Diversity 105:1-16 ((https://doi.org/10.1007/s13225-020-00466-2)) |
| Otu58581 | 1 | k__Fungi(100);p__Ascomycota(100);c__Saccharomycetes(100);o__Saccharomycetales(100);f__Metschnikowiaceae(100);g__Metschnikowia(100);s__Metschnikowia_pulcherrima(100); | Metschnikowia | 13 | Saprotroph | \|Undefined Saprotroph\| | Yeast | NULL | Probable | As nectar-tap_saprotroph (Põlme et al. 2020); Decay sugar-rich substrates (Põlme et al. 2020) | Põlme S, et al. 2020. Fungal Diversity 105:1-16 ((https://doi.org/10.1007/s13225-020-00466-2)) |
| Otu58582 | 1 | k__Fungi(100);p__Ascomycota(100);c__Saccharomycetes(100);o__Saccharomycetales(100);f__Metschnikowiaceae(100);g__Metschnikowia(100);s__Metschnikowia_pulcherrima(100); | Metschnikowia | 13 | Saprotroph | \|Undefined Saprotroph\| | Yeast | NULL | Probable | As nectar-tap_saprotroph (Põlme et al. 2020); Decay sugar-rich substrates (Põlme et al. 2020) | Põlme S, et al. 2020. Fungal Diversity 105:1-16 ((https://doi.org/10.1007/s13225-020-00466-2)) |
| Otu58584 | 1 | k__Fungi(100);p__Ascomycota(100);c__Saccharomycetes(100);o__Saccharomycetales(100);f__Metschnikowiaceae(100);g__Metschnikowia(100);s__Metschnikowia_pulcherrima(100); | Metschnikowia | 13 | Saprotroph | \|Undefined Saprotroph\| | Yeast | NULL | Probable | As nectar-tap_saprotroph (Põlme et al. 2020); Decay sugar-rich substrates (Põlme et al. 2020) | Põlme S, et al. 2020. Fungal Diversity 105:1-16 ((https://doi.org/10.1007/s13225-020-00466-2)) |
| Otu58585 | 1 | k__Fungi(100);p__Ascomycota(100);c__Saccharomycetes(100);o__Saccharomycetales(100);f__Metschnikowiaceae(100);g__Metschnikowia(100);g__Metschnikowia_unclassified(100); | Metschnikowia | 13 | Saprotroph | \|Undefined Saprotroph\| | Yeast | NULL | Probable | As nectar-tap_saprotroph (Põlme et al. 2020); Decay sugar-rich substrates (Põlme et al. 2020) | Põlme S, et al. 2020. Fungal Diversity 105:1-16 ((https://doi.org/10.1007/s13225-020-00466-2)) |
| Otu58586 | 1 | k__Fungi(100);p__Ascomycota(100);c__Saccharomycetes(100);o__Saccharomycetales(100);f__Metschnikowiaceae(100);g__Metschnikowia(100);g__Metschnikowia_unclassified(100); | Metschnikowia | 13 | Saprotroph | \|Undefined Saprotroph\| | Yeast | NULL | Probable | As nectar-tap_saprotroph (Põlme et al. 2020); Decay sugar-rich substrates (Põlme et al. 2020) | Põlme S, et al. 2020. Fungal Diversity 105:1-16 ((https://doi.org/10.1007/s13225-020-00466-2)) |
| Otu58587 | 1 | k__Fungi(100);p__Ascomycota(100);c__Saccharomycetes(100);o__Saccharomycetales(100);f__Metschnikowiaceae(100);g__Metschnikowia(100);s__Metschnikowia_pulcherrima(100); | Metschnikowia | 13 | Saprotroph | \|Undefined Saprotroph\| | Yeast | NULL | Probable | As nectar-tap_saprotroph (Põlme et al. 2020); Decay sugar-rich substrates (Põlme et al. 2020) | Põlme S, et al. 2020. Fungal Diversity 105:1-16 ((https://doi.org/10.1007/s13225-020-00466-2)) |
| Otu58588 | 1 | k__Fungi(100);p__Ascomycota(100);c__Saccharomycetes(100);o__Saccharomycetales(100);f__Metschnikowiaceae(100);g__Metschnikowia(100);g__Metschnikowia_unclassified(100); | Metschnikowia | 13 | Saprotroph | \|Undefined Saprotroph\| | Yeast | NULL | Probable | As nectar-tap_saprotroph (Põlme et al. 2020); Decay sugar-rich substrates (Põlme et al. 2020) | Põlme S, et al. 2020. Fungal Diversity 105:1-16 ((https://doi.org/10.1007/s13225-020-00466-2)) |
| Otu58589 | 1 | k__Fungi(100);p__Ascomycota(100);c__Saccharomycetes(100);o__Saccharomycetales(100);f__Metschnikowiaceae(100);g__Metschnikowia(100);s__Metschnikowia_pulcherrima(100); | Metschnikowia | 13 | Saprotroph | \|Undefined Saprotroph\| | Yeast | NULL | Probable | As nectar-tap_saprotroph (Põlme et al. 2020); Decay sugar-rich substrates (Põlme et al. 2020) | Põlme S, et al. 2020. Fungal Diversity 105:1-16 ((https://doi.org/10.1007/s13225-020-00466-2)) |
| Otu58590 | 1 | k__Fungi(100);p__Ascomycota(100);c__Saccharomycetes(100);o__Saccharomycetales(100);f__Metschnikowiaceae(100);g__Metschnikowia(100);s__Metschnikowia_pulcherrima(100); | Metschnikowia | 13 | Saprotroph | \|Undefined Saprotroph\| | Yeast | NULL | Probable | As nectar-tap_saprotroph (Põlme et al. 2020); Decay sugar-rich substrates (Põlme et al. 2020) | Põlme S, et al. 2020. Fungal Diversity 105:1-16 ((https://doi.org/10.1007/s13225-020-00466-2)) |
| Otu58591 | 1 | k__Fungi(100);p__Ascomycota(100);c__Saccharomycetes(100);o__Saccharomycetales(100);f__Metschnikowiaceae(100);g__Metschnikowia(100);s__Metschnikowia_pulcherrima(100); | Metschnikowia | 13 | Saprotroph | \|Undefined Saprotroph\| | Yeast | NULL | Probable | As nectar-tap_saprotroph (Põlme et al. 2020); Decay sugar-rich substrates (Põlme et al. 2020) | Põlme S, et al. 2020. Fungal Diversity 105:1-16 ((https://doi.org/10.1007/s13225-020-00466-2)) |
| Otu58592 | 1 | k__Fungi(100);p__Ascomycota(100);c__Saccharomycetes(100);o__Saccharomycetales(100);f__Metschnikowiaceae(100);g__Metschnikowia(100);s__Metschnikowia_pulcherrima(100); | Metschnikowia | 13 | Saprotroph | \|Undefined Saprotroph\| | Yeast | NULL | Probable | As nectar-tap_saprotroph (Põlme et al. 2020); Decay sugar-rich substrates (Põlme et al. 2020) | Põlme S, et al. 2020. Fungal Diversity 105:1-16 ((https://doi.org/10.1007/s13225-020-00466-2)) |
| Otu58593 | 1 | k__Fungi(100);p__Ascomycota(100);c__Saccharomycetes(100);o__Saccharomycetales(100);f__Metschnikowiaceae(100);g__Metschnikowia(100);s__Metschnikowia_pulcherrima(100); | Metschnikowia | 13 | Saprotroph | \|Undefined Saprotroph\| | Yeast | NULL | Probable | As nectar-tap_saprotroph (Põlme et al. 2020); Decay sugar-rich substrates (Põlme et al. 2020) | Põlme S, et al. 2020. Fungal Diversity 105:1-16 ((https://doi.org/10.1007/s13225-020-00466-2)) |
| Otu58594 | 1 | k__Fungi(100);p__Ascomycota(100);c__Saccharomycetes(100);o__Saccharomycetales(100);f__Metschnikowiaceae(100);g__Metschnikowia(100);s__Metschnikowia_pulcherrima(100); | Metschnikowia | 13 | Saprotroph | \|Undefined Saprotroph\| | Yeast | NULL | Probable | As nectar-tap_saprotroph (Põlme et al. 2020); Decay sugar-rich substrates (Põlme et al. 2020) | Põlme S, et al. 2020. Fungal Diversity 105:1-16 ((https://doi.org/10.1007/s13225-020-00466-2)) |
| Otu58595 | 1 | k__Fungi(100);p__Ascomycota(100);c__Saccharomycetes(100);o__Saccharomycetales(100);f__Metschnikowiaceae(100);g__Metschnikowia(100);s__Metschnikowia_pulcherrima(100); | Metschnikowia | 13 | Saprotroph | \|Undefined Saprotroph\| | Yeast | NULL | Probable | As nectar-tap_saprotroph (Põlme et al. 2020); Decay sugar-rich substrates (Põlme et al. 2020) | Põlme S, et al. 2020. Fungal Diversity 105:1-16 ((https://doi.org/10.1007/s13225-020-00466-2)) |
| Otu58596 | 1 | k__Fungi(100);p__Ascomycota(100);c__Saccharomycetes(100);o__Saccharomycetales(100);f__Metschnikowiaceae(100);g__Metschnikowia(100);g__Metschnikowia_unclassified(100); | Metschnikowia | 13 | Saprotroph | \|Undefined Saprotroph\| | Yeast | NULL | Probable | As nectar-tap_saprotroph (Põlme et al. 2020); Decay sugar-rich substrates (Põlme et al. 2020) | Põlme S, et al. 2020. Fungal Diversity 105:1-16 ((https://doi.org/10.1007/s13225-020-00466-2)) |
| Otu58597 | 1 | k__Fungi(100);p__Ascomycota(100);c__Saccharomycetes(100);o__Saccharomycetales(100);f__Metschnikowiaceae(100);g__Metschnikowia(100);g__Metschnikowia_unclassified(100); | Metschnikowia | 13 | Saprotroph | \|Undefined Saprotroph\| | Yeast | NULL | Probable | As nectar-tap_saprotroph (Põlme et al. 2020); Decay sugar-rich substrates (Põlme et al. 2020) | Põlme S, et al. 2020. Fungal Diversity 105:1-16 ((https://doi.org/10.1007/s13225-020-00466-2)) |
| Otu58602 | 1 | k__Fungi(100);p__Ascomycota(100);c__Saccharomycetes(100);o__Saccharomycetales(100);f__Metschnikowiaceae(100);g__Metschnikowia(100);s__Metschnikowia_pulcherrima(100); | Metschnikowia | 13 | Saprotroph | \|Undefined Saprotroph\| | Yeast | NULL | Probable | As nectar-tap_saprotroph (Põlme et al. 2020); Decay sugar-rich substrates (Põlme et al. 2020) | Põlme S, et al. 2020. Fungal Diversity 105:1-16 ((https://doi.org/10.1007/s13225-020-00466-2)) |
| Otu58603 | 1 | k__Fungi(100);p__Ascomycota(100);c__Saccharomycetes(100);o__Saccharomycetales(100);f__Metschnikowiaceae(100);g__Metschnikowia(100);g__Metschnikowia_unclassified(100); | Metschnikowia | 13 | Saprotroph | \|Undefined Saprotroph\| | Yeast | NULL | Probable | As nectar-tap_saprotroph (Põlme et al. 2020); Decay sugar-rich substrates (Põlme et al. 2020) | Põlme S, et al. 2020. Fungal Diversity 105:1-16 ((https://doi.org/10.1007/s13225-020-00466-2)) |
| Otu58604 | 1 | k__Fungi(100);p__Ascomycota(100);c__Saccharomycetes(100);o__Saccharomycetales(100);f__Metschnikowiaceae(100);g__Metschnikowia(100);s__Metschnikowia_pulcherrima(100); | Metschnikowia | 13 | Saprotroph | \|Undefined Saprotroph\| | Yeast | NULL | Probable | As nectar-tap_saprotroph (Põlme et al. 2020); Decay sugar-rich substrates (Põlme et al. 2020) | Põlme S, et al. 2020. Fungal Diversity 105:1-16 ((https://doi.org/10.1007/s13225-020-00466-2)) |
| Otu58606 | 1 | k__Fungi(100);p__Ascomycota(100);c__Saccharomycetes(100);o__Saccharomycetales(100);f__Metschnikowiaceae(100);g__Metschnikowia(100);s__Metschnikowia_pulcherrima(100); | Metschnikowia | 13 | Saprotroph | \|Undefined Saprotroph\| | Yeast | NULL | Probable | As nectar-tap_saprotroph (Põlme et al. 2020); Decay sugar-rich substrates (Põlme et al. 2020) | Põlme S, et al. 2020. Fungal Diversity 105:1-16 ((https://doi.org/10.1007/s13225-020-00466-2)) |
| Otu58607 | 1 | k__Fungi(100);p__Ascomycota(100);c__Saccharomycetes(100);o__Saccharomycetales(100);f__Metschnikowiaceae(100);g__Metschnikowia(100);s__Metschnikowia_pulcherrima(100); | Metschnikowia | 13 | Saprotroph | \|Undefined Saprotroph\| | Yeast | NULL | Probable | As nectar-tap_saprotroph (Põlme et al. 2020); Decay sugar-rich substrates (Põlme et al. 2020) | Põlme S, et al. 2020. Fungal Diversity 105:1-16 ((https://doi.org/10.1007/s13225-020-00466-2)) |
| Otu58608 | 1 | k__Fungi(100);p__Ascomycota(100);c__Saccharomycetes(100);o__Saccharomycetales(100);f__Metschnikowiaceae(100);g__Metschnikowia(100);s__Metschnikowia_pulcherrima(100); | Metschnikowia | 13 | Saprotroph | \|Undefined Saprotroph\| | Yeast | NULL | Probable | As nectar-tap_saprotroph (Põlme et al. 2020); Decay sugar-rich substrates (Põlme et al. 2020) | Põlme S, et al. 2020. Fungal Diversity 105:1-16 ((https://doi.org/10.1007/s13225-020-00466-2)) |
| Otu58609 | 1 | k__Fungi(100);p__Ascomycota(100);c__Saccharomycetes(100);o__Saccharomycetales(100);f__Metschnikowiaceae(100);g__Metschnikowia(100);s__Metschnikowia_pulcherrima(100); | Metschnikowia | 13 | Saprotroph | \|Undefined Saprotroph\| | Yeast | NULL | Probable | As nectar-tap_saprotroph (Põlme et al. 2020); Decay sugar-rich substrates (Põlme et al. 2020) | Põlme S, et al. 2020. Fungal Diversity 105:1-16 ((https://doi.org/10.1007/s13225-020-00466-2)) |

| | | | | | | | | | | | |
|---|---|---|---|---|---|---|---|---|---|---|---|
| Otu58610 | 1 | k__Fungi(100);p__Ascomycota(100);c__Saccharomycetes(100);o__Saccharomycetales(100);f__Metschnikowiaceae(100);g__Metschnikowia(100);s__Metschnikowia_pulcherrima(100); | Metschnikowia | 13 | Saprotroph | \|Undefined Saprotroph\| | Yeast | NULL | Probable | As nectar-tap_saprotroph (Põlme et al. 2020); Decay sugar-rich substrates (Põlme et al. 2020) | Põlme S, et al. 2020. Fungal Diversity 105:1-16 ((https://doi.org/10.1007/s13225-020-00466-2)) |
| Otu58611 | 1 | k__Fungi(100);p__Ascomycota(100);c__Saccharomycetes(100);o__Saccharomycetales(100);f__Metschnikowiaceae(100);g__Metschnikowia(100);s__Metschnikowia_pulcherrima(100); | Metschnikowia | 13 | Saprotroph | \|Undefined Saprotroph\| | Yeast | NULL | Probable | As nectar-tap_saprotroph (Põlme et al. 2020); Decay sugar-rich substrates (Põlme et al. 2020) | Põlme S, et al. 2020. Fungal Diversity 105:1-16 ((https://doi.org/10.1007/s13225-020-00466-2)) |
| Otu58612 | 1 | k__Fungi(100);p__Ascomycota(100);c__Saccharomycetes(100);o__Saccharomycetales(100);f__Metschnikowiaceae(100);g__Metschnikowia(100);s__Metschnikowia_pulcherrima(100); | Metschnikowia | 13 | Saprotroph | \|Undefined Saprotroph\| | Yeast | NULL | Probable | As nectar-tap_saprotroph (Põlme et al. 2020); Decay sugar-rich substrates (Põlme et al. 2020) | Põlme S, et al. 2020. Fungal Diversity 105:1-16 ((https://doi.org/10.1007/s13225-020-00466-2)) |
| Otu58613 | 1 | k__Fungi(100);p__Ascomycota(100);c__Saccharomycetes(100);o__Saccharomycetales(100);f__Metschnikowiaceae(100);g__Metschnikowia(100);s__Metschnikowia_pulcherrima(100); | Metschnikowia | 13 | Saprotroph | \|Undefined Saprotroph\| | Yeast | NULL | Probable | As nectar-tap_saprotroph (Põlme et al. 2020); Decay sugar-rich substrates (Põlme et al. 2020) | Põlme S, et al. 2020. Fungal Diversity 105:1-16 ((https://doi.org/10.1007/s13225-020-00466-2)) |
| Otu58614 | 1 | k__Fungi(100);p__Ascomycota(100);c__Saccharomycetes(100);o__Saccharomycetales(100);f__Metschnikowiaceae(100);g__Metschnikowia(100);s__Metschnikowia_pulcherrima(100); | Metschnikowia | 13 | Saprotroph | \|Undefined Saprotroph\| | Yeast | NULL | Probable | As nectar-tap_saprotroph (Põlme et al. 2020); Decay sugar-rich substrates (Põlme et al. 2020) | Põlme S, et al. 2020. Fungal Diversity 105:1-16 ((https://doi.org/10.1007/s13225-020-00466-2)) |
| Otu58615 | 1 | k__Fungi(100);p__Ascomycota(100);c__Saccharomycetes(100);o__Saccharomycetales(100);f__Metschnikowiaceae(100);g__Metschnikowia(100);s__Metschnikowia_pulcherrima(100); | Metschnikowia | 13 | Saprotroph | \|Undefined Saprotroph\| | Yeast | NULL | Probable | As nectar-tap_saprotroph (Põlme et al. 2020); Decay sugar-rich substrates (Põlme et al. 2020) | Põlme S, et al. 2020. Fungal Diversity 105:1-16 ((https://doi.org/10.1007/s13225-020-00466-2)) |
| Otu58616 | 1 | k__Fungi(100);p__Ascomycota(100);c__Saccharomycetes(100);o__Saccharomycetales(100);f__Metschnikowiaceae(100);g__Metschnikowia(100);s__Metschnikowia_pulcherrima(100); | Metschnikowia | 13 | Saprotroph | \|Undefined Saprotroph\| | Yeast | NULL | Probable | As nectar-tap_saprotroph (Põlme et al. 2020); Decay sugar-rich substrates (Põlme et al. 2020) | Põlme S, et al. 2020. Fungal Diversity 105:1-16 ((https://doi.org/10.1007/s13225-020-00466-2)) |
| Otu58617 | 1 | k__Fungi(100);p__Ascomycota(100);c__Saccharomycetes(100);o__Saccharomycetales(100);f__Metschnikowiaceae(100);g__Metschnikowia(100);s__Metschnikowia_pulcherrima(100); | Metschnikowia | 13 | Saprotroph | \|Undefined Saprotroph\| | Yeast | NULL | Probable | As nectar-tap_saprotroph (Põlme et al. 2020); Decay sugar-rich substrates (Põlme et al. 2020) | Põlme S, et al. 2020. Fungal Diversity 105:1-16 ((https://doi.org/10.1007/s13225-020-00466-2)) |
| Otu58618 | 1 | k__Fungi(100);p__Ascomycota(100);c__Saccharomycetes(100);o__Saccharomycetales(100);f__Metschnikowiaceae(100);g__Metschnikowia(100);s__Metschnikowia_pulcherrima(100); | Metschnikowia | 13 | Saprotroph | \|Undefined Saprotroph\| | Yeast | NULL | Probable | As nectar-tap_saprotroph (Põlme et al. 2020); Decay sugar-rich substrates (Põlme et al. 2020) | Põlme S, et al. 2020. Fungal Diversity 105:1-16 ((https://doi.org/10.1007/s13225-020-00466-2)) |
| Otu58619 | 1 | k__Fungi(100);p__Ascomycota(100);c__Saccharomycetes(100);o__Saccharomycetales(100);f__Metschnikowiaceae(100);g__Metschnikowia(100);s__Metschnikowia_pulcherrima(100); | Metschnikowia | 13 | Saprotroph | \|Undefined Saprotroph\| | Yeast | NULL | Probable | As nectar-tap_saprotroph (Põlme et al. 2020); Decay sugar-rich substrates (Põlme et al. 2020) | Põlme S, et al. 2020. Fungal Diversity 105:1-16 ((https://doi.org/10.1007/s13225-020-00466-2)) |
| Otu58620 | 1 | k__Fungi(100);p__Ascomycota(100);c__Saccharomycetes(100);o__Saccharomycetales(100);f__Metschnikowiaceae(100);g__Metschnikowia(100);s__Metschnikowia_pulcherrima(100); | Metschnikowia | 13 | Saprotroph | \|Undefined Saprotroph\| | Yeast | NULL | Probable | As nectar-tap_saprotroph (Põlme et al. 2020); Decay sugar-rich substrates (Põlme et al. 2020) | Põlme S, et al. 2020. Fungal Diversity 105:1-16 ((https://doi.org/10.1007/s13225-020-00466-2)) |
| Otu58621 | 1 | k__Fungi(100);p__Ascomycota(100);c__Saccharomycetes(100);o__Saccharomycetales(100);f__Metschnikowiaceae(100);g__Metschnikowia(100);s__Metschnikowia_pulcherrima(100); | Metschnikowia | 13 | Saprotroph | \|Undefined Saprotroph\| | Yeast | NULL | Probable | As nectar-tap_saprotroph (Põlme et al. 2020); Decay sugar-rich substrates (Põlme et al. 2020) | Põlme S, et al. 2020. Fungal Diversity 105:1-16 ((https://doi.org/10.1007/s13225-020-00466-2)) |
| Otu58622 | 1 | k__Fungi(100);p__Ascomycota(100);c__Saccharomycetes(100);o__Saccharomycetales(100);f__Metschnikowiaceae(100);g__Metschnikowia(100);g__Metschnikowia_unclassified(100); | Metschnikowia | 13 | Saprotroph | \|Undefined Saprotroph\| | Yeast | NULL | Probable | As nectar-tap_saprotroph (Põlme et al. 2020); Decay sugar-rich substrates (Põlme et al. 2020) | Põlme S, et al. 2020. Fungal Diversity 105:1-16 ((https://doi.org/10.1007/s13225-020-00466-2)) |
| Otu58623 | 1 | k__Fungi(100);p__Ascomycota(100);c__Saccharomycetes(100);o__Saccharomycetales(100);f__Metschnikowiaceae(100);g__Metschnikowia(100);s__Metschnikowia_pulcherrima(100); | Metschnikowia | 13 | Saprotroph | \|Undefined Saprotroph\| | Yeast | NULL | Probable | As nectar-tap_saprotroph (Põlme et al. 2020); Decay sugar-rich substrates (Põlme et al. 2020) | Põlme S, et al. 2020. Fungal Diversity 105:1-16 ((https://doi.org/10.1007/s13225-020-00466-2)) |
| Otu58624 | 1 | k__Fungi(100);p__Ascomycota(100);c__Saccharomycetes(100);o__Saccharomycetales(100);f__Metschnikowiaceae(100);g__Metschnikowia(100);s__Metschnikowia_pulcherrima(100); | Metschnikowia | 13 | Saprotroph | \|Undefined Saprotroph\| | Yeast | NULL | Probable | As nectar-tap_saprotroph (Põlme et al. 2020); Decay sugar-rich substrates (Põlme et al. 2020) | Põlme S, et al. 2020. Fungal Diversity 105:1-16 ((https://doi.org/10.1007/s13225-020-00466-2)) |
| Otu58625 | 1 | k__Fungi(100);p__Ascomycota(100);c__Saccharomycetes(100);o__Saccharomycetales(100);f__Metschnikowiaceae(100);g__Metschnikowia(100);s__Metschnikowia_pulcherrima(100); | Metschnikowia | 13 | Saprotroph | \|Undefined Saprotroph\| | Yeast | NULL | Probable | As nectar-tap_saprotroph (Põlme et al. 2020); Decay sugar-rich substrates (Põlme et al. 2020) | Põlme S, et al. 2020. Fungal Diversity 105:1-16 ((https://doi.org/10.1007/s13225-020-00466-2)) |
| Otu58626 | 1 | k__Fungi(100);p__Ascomycota(100);c__Saccharomycetes(100);o__Saccharomycetales(100);f__Metschnikowiaceae(100);g__Metschnikowia(100);s__Metschnikowia_pulcherrima(100); | Metschnikowia | 13 | Saprotroph | \|Undefined Saprotroph\| | Yeast | NULL | Probable | As nectar-tap_saprotroph (Põlme et al. 2020); Decay sugar-rich substrates (Põlme et al. 2020) | Põlme S, et al. 2020. Fungal Diversity 105:1-16 ((https://doi.org/10.1007/s13225-020-00466-2)) |
| Otu58627 | 1 | k__Fungi(100);p__Ascomycota(100);c__Saccharomycetes(100);o__Saccharomycetales(100);f__Metschnikowiaceae(100);g__Metschnikowia(100);g__Metschnikowia_unclassified(100); | Metschnikowia | 13 | Saprotroph | \|Undefined Saprotroph\| | Yeast | NULL | Probable | As nectar-tap_saprotroph (Põlme et al. 2020); Decay sugar-rich substrates (Põlme et al. 2020) | Põlme S, et al. 2020. Fungal Diversity 105:1-16 ((https://doi.org/10.1007/s13225-020-00466-2)) |
| Otu58628 | 1 | k__Fungi(100);p__Ascomycota(100);c__Saccharomycetes(100);o__Saccharomycetales(100);f__Metschnikowiaceae(100);g__Metschnikowia(100);g__Metschnikowia_unclassified(100); | Metschnikowia | 13 | Saprotroph | \|Undefined Saprotroph\| | Yeast | NULL | Probable | As nectar-tap_saprotroph (Põlme et al. 2020); Decay sugar-rich substrates (Põlme et al. 2020) | Põlme S, et al. 2020. Fungal Diversity 105:1-16 ((https://doi.org/10.1007/s13225-020-00466-2)) |
| Otu58629 | 1 | k__Fungi(100);p__Ascomycota(100);c__Saccharomycetes(100);o__Saccharomycetales(100);f__Metschnikowiaceae(100);g__Metschnikowia(100);s__Metschnikowia_pulcherrima(100); | Metschnikowia | 13 | Saprotroph | \|Undefined Saprotroph\| | Yeast | NULL | Probable | As nectar-tap_saprotroph (Põlme et al. 2020); Decay sugar-rich substrates (Põlme et al. 2020) | Põlme S, et al. 2020. Fungal Diversity 105:1-16 ((https://doi.org/10.1007/s13225-020-00466-2)) |
| Otu58630 | 1 | k__Fungi(100);p__Ascomycota(100);c__Saccharomycetes(100);o__Saccharomycetales(100);f__Metschnikowiaceae(100);g__Metschnikowia(100);s__Metschnikowia_pulcherrima(100); | Metschnikowia | 13 | Saprotroph | \|Undefined Saprotroph\| | Yeast | NULL | Probable | As nectar-tap_saprotroph (Põlme et al. 2020); Decay sugar-rich substrates (Põlme et al. 2020) | Põlme S, et al. 2020. Fungal Diversity 105:1-16 ((https://doi.org/10.1007/s13225-020-00466-2)) |
| Otu58631 | 1 | k__Fungi(100);p__Ascomycota(100);c__Saccharomycetes(100);o__Saccharomycetales(100);f__Metschnikowiaceae(100);g__Metschnikowia(100);s__Metschnikowia_pulcherrima(100); | Metschnikowia | 13 | Saprotroph | \|Undefined Saprotroph\| | Yeast | NULL | Probable | As nectar-tap_saprotroph (Põlme et al. 2020); Decay sugar-rich substrates (Põlme et al. 2020) | Põlme S, et al. 2020. Fungal Diversity 105:1-16 ((https://doi.org/10.1007/s13225-020-00466-2)) |
| Otu58634 | 1 | k__Fungi(100);p__Ascomycota(100);c__Saccharomycetes(100);o__Saccharomycetales(100);f__Metschnikowiaceae(100);g__Metschnikowia(100);s__Metschnikowia_pulcherrima(100); | Metschnikowia | 13 | Saprotroph | \|Undefined Saprotroph\| | Yeast | NULL | Probable | As nectar-tap_saprotroph (Põlme et al. 2020); Decay sugar-rich substrates (Põlme et al. 2020) | Põlme S, et al. 2020. Fungal Diversity 105:1-16 ((https://doi.org/10.1007/s13225-020-00466-2)) |
| Otu58635 | 1 | k__Fungi(100);p__Ascomycota(100);c__Saccharomycetes(100);o__Saccharomycetales(100);f__Metschnikowiaceae(100);g__Metschnikowia(100);g__Metschnikowia_unclassified(100); | Metschnikowia | 13 | Saprotroph | \|Undefined Saprotroph\| | Yeast | NULL | Probable | As nectar-tap_saprotroph (Põlme et al. 2020); Decay sugar-rich substrates (Põlme et al. 2020) | Põlme S, et al. 2020. Fungal Diversity 105:1-16 ((https://doi.org/10.1007/s13225-020-00466-2)) |
| Otu58636 | 1 | k__Fungi(100);p__Ascomycota(100);c__Saccharomycetes(100);o__Saccharomycetales(100);f__Metschnikowiaceae(100);g__Metschnikowia(100);g__Metschnikowia_unclassified(100); | Metschnikowia | 13 | Saprotroph | \|Undefined Saprotroph\| | Yeast | NULL | Probable | As nectar-tap_saprotroph (Põlme et al. 2020); Decay sugar-rich substrates (Põlme et al. 2020) | Põlme S, et al. 2020. Fungal Diversity 105:1-16 ((https://doi.org/10.1007/s13225-020-00466-2)) |
| Otu58637 | 1 | k__Fungi(100);p__Ascomycota(100);c__Saccharomycetes(100);o__Saccharomycetales(100);f__Metschnikowiaceae(100);g__Metschnikowia(100);s__Metschnikowia_pulcherrima(100); | Metschnikowia | 13 | Saprotroph | \|Undefined Saprotroph\| | Yeast | NULL | Probable | As nectar-tap_saprotroph (Põlme et al. 2020); Decay sugar-rich substrates (Põlme et al. 2020) | Põlme S, et al. 2020. Fungal Diversity 105:1-16 ((https://doi.org/10.1007/s13225-020-00466-2)) |
| Otu58638 | 1 | k__Fungi(100);p__Ascomycota(100);c__Saccharomycetes(100);o__Saccharomycetales(100);f__Metschnikowiaceae(100);g__Metschnikowia(100);s__Metschnikowia_pulcherrima(100); | Metschnikowia | 13 | Saprotroph | \|Undefined Saprotroph\| | Yeast | NULL | Probable | As nectar-tap_saprotroph (Põlme et al. 2020); Decay sugar-rich substrates (Põlme et al. 2020) | Põlme S, et al. 2020. Fungal Diversity 105:1-16 ((https://doi.org/10.1007/s13225-020-00466-2)) |
| Otu58639 | 1 | k__Fungi(100);p__Ascomycota(100);c__Saccharomycetes(100);o__Saccharomycetales(100);f__Metschnikowiaceae(100);g__Metschnikowia(100);s__Metschnikowia_pulcherrima(100); | Metschnikowia | 13 | Saprotroph | \|Undefined Saprotroph\| | Yeast | NULL | Probable | As nectar-tap_saprotroph (Põlme et al. 2020); Decay sugar-rich substrates (Põlme et al. 2020) | Põlme S, et al. 2020. Fungal Diversity 105:1-16 ((https://doi.org/10.1007/s13225-020-00466-2)) |
| Otu58640 | 1 | k__Fungi(100);p__Ascomycota(100);c__Saccharomycetes(100);o__Saccharomycetales(100);f__Metschnikowiaceae(100);g__Metschnikowia(100);g__Metschnikowia_unclassified(100); | Metschnikowia | 13 | Saprotroph | \|Undefined Saprotroph\| | Yeast | NULL | Probable | As nectar-tap_saprotroph (Põlme et al. 2020); Decay sugar-rich substrates (Põlme et al. 2020) | Põlme S, et al. 2020. Fungal Diversity 105:1-16 ((https://doi.org/10.1007/s13225-020-00466-2)) |
| Otu58641 | 1 | k__Fungi(100);p__Ascomycota(100);c__Saccharomycetes(100);o__Saccharomycetales(100);f__Metschnikowiaceae(100);g__Metschnikowia(100);s__Metschnikowia_pulcherrima(100); | Metschnikowia | 13 | Saprotroph | \|Undefined Saprotroph\| | Yeast | NULL | Probable | As nectar-tap_saprotroph (Põlme et al. 2020); Decay sugar-rich substrates (Põlme et al. 2020) | Põlme S, et al. 2020. Fungal Diversity 105:1-16 ((https://doi.org/10.1007/s13225-020-00466-2)) |
| Otu58643 | 1 | k__Fungi(100);p__Ascomycota(100);c__Saccharomycetes(100);o__Saccharomycetales(100);f__Metschnikowiaceae(100);g__Metschnikowia(100);s__Metschnikowia_pulcherrima(100); | Metschnikowia | 13 | Saprotroph | \|Undefined Saprotroph\| | Yeast | NULL | Probable | As nectar-tap_saprotroph (Põlme et al. 2020); Decay sugar-rich substrates (Põlme et al. 2020) | Põlme S, et al. 2020. Fungal Diversity 105:1-16 ((https://doi.org/10.1007/s13225-020-00466-2)) |
| Otu58644 | 1 | k__Fungi(100);p__Ascomycota(100);c__Saccharomycetes(100);o__Saccharomycetales(100);f__Metschnikowiaceae(100);g__Metschnikowia(100);s__Metschnikowia_pulcherrima(100); | Metschnikowia | 13 | Saprotroph | \|Undefined Saprotroph\| | Yeast | NULL | Probable | As nectar-tap_saprotroph (Põlme et al. 2020); Decay sugar-rich substrates (Põlme et al. 2020) | Põlme S, et al. 2020. Fungal Diversity 105:1-16 ((https://doi.org/10.1007/s13225-020-00466-2)) |
| Otu58646 | 1 | k__Fungi(100);p__Ascomycota(100);c__Saccharomycetes(100);o__Saccharomycetales(100);f__Metschnikowiaceae(100);g__Metschnikowia(100);s__Metschnikowia_pulcherrima(100); | Metschnikowia | 13 | Saprotroph | \|Undefined Saprotroph\| | Yeast | NULL | Probable | As nectar-tap_saprotroph (Põlme et al. 2020); Decay sugar-rich substrates (Põlme et al. 2020) | Põlme S, et al. 2020. Fungal Diversity 105:1-16 ((https://doi.org/10.1007/s13225-020-00466-2)) |
| Otu58647 | 1 | k__Fungi(100);p__Ascomycota(100);c__Saccharomycetes(100);o__Saccharomycetales(100);f__Metschnikowiaceae(100);g__Metschnikowia(100);s__Metschnikowia_pulcherrima(100); | Metschnikowia | 13 | Saprotroph | \|Undefined Saprotroph\| | Yeast | NULL | Probable | As nectar-tap_saprotroph (Põlme et al. 2020); Decay sugar-rich substrates (Põlme et al. 2020) | Põlme S, et al. 2020. Fungal Diversity 105:1-16 ((https://doi.org/10.1007/s13225-020-00466-2)) |
| Otu58648 | 1 | k__Fungi(100);p__Ascomycota(100);c__Saccharomycetes(100);o__Saccharomycetales(100);f__Metschnikowiaceae(100);g__Metschnikowia(100);g__Metschnikowia_unclassified(100); | Metschnikowia | 13 | Saprotroph | \|Undefined Saprotroph\| | Yeast | NULL | Probable | As nectar-tap_saprotroph (Põlme et al. 2020); Decay sugar-rich substrates (Põlme et al. 2020) | Põlme S, et al. 2020. Fungal Diversity 105:1-16 ((https://doi.org/10.1007/s13225-020-00466-2)) |
| Otu58652 | 1 | k__Fungi(100);p__Ascomycota(100);c__Saccharomycetes(100);o__Saccharomycetales(100);f__Metschnikowiaceae(100);g__Metschnikowia(100);s__Metschnikowia_pulcherrima(100); | Metschnikowia | 13 | Saprotroph | \|Undefined Saprotroph\| | Yeast | NULL | Probable | As nectar-tap_saprotroph (Põlme et al. 2020); Decay sugar-rich substrates (Põlme et al. 2020) | Põlme S, et al. 2020. Fungal Diversity 105:1-16 ((https://doi.org/10.1007/s13225-020-00466-2)) |
| Otu58653 | 1 | k__Fungi(100);p__Ascomycota(100);c__Saccharomycetes(100);o__Saccharomycetales(100);f__Metschnikowiaceae(100);g__Metschnikowia(100);s__Metschnikowia_pulcherrima(100); | Metschnikowia | 13 | Saprotroph | \|Undefined Saprotroph\| | Yeast | NULL | Probable | As nectar-tap_saprotroph (Põlme et al. 2020); Decay sugar-rich substrates (Põlme et al. 2020) | Põlme S, et al. 2020. Fungal Diversity 105:1-16 ((https://doi.org/10.1007/s13225-020-00466-2)) |
| Otu58654 | 1 | k__Fungi(100);p__Ascomycota(100);c__Saccharomycetes(100);o__Saccharomycetales(100);f__Metschnikowiaceae(100);g__Metschnikowia(100);s__Metschnikowia_pulcherrima(100); | Metschnikowia | 13 | Saprotroph | \|Undefined Saprotroph\| | Yeast | NULL | Probable | As nectar-tap_saprotroph (Põlme et al. 2020); Decay sugar-rich substrates (Põlme et al. 2020) | Põlme S, et al. 2020. Fungal Diversity 105:1-16 ((https://doi.org/10.1007/s13225-020-00466-2)) |
| Otu58655 | 1 | k__Fungi(100);p__Ascomycota(100);c__Saccharomycetes(100);o__Saccharomycetales(100);f__Metschnikowiaceae(100);g__Metschnikowia(100);s__Metschnikowia_pulcherrima(100); | Metschnikowia | 13 | Saprotroph | \|Undefined Saprotroph\| | Yeast | NULL | Probable | As nectar-tap_saprotroph (Põlme et al. 2020); Decay sugar-rich substrates (Põlme et al. 2020) | Põlme S, et al. 2020. Fungal Diversity 105:1-16 ((https://doi.org/10.1007/s13225-020-00466-2)) |
| Otu58656 | 1 | k__Fungi(100);p__Ascomycota(100);c__Saccharomycetes(100);o__Saccharomycetales(100);f__Metschnikowiaceae(100);g__Metschnikowia(100);s__Metschnikowia_pulcherrima(100); | Metschnikowia | 13 | Saprotroph | \|Undefined Saprotroph\| | Yeast | NULL | Probable | As nectar-tap_saprotroph (Põlme et al. 2020); Decay sugar-rich substrates (Põlme et al. 2020) | Põlme S, et al. 2020. Fungal Diversity 105:1-16 ((https://doi.org/10.1007/s13225-020-00466-2)) |
| Otu58657 | 1 | k__Fungi(100);p__Ascomycota(100);c__Saccharomycetes(100);o__Saccharomycetales(100);f__Metschnikowiaceae(100);g__Metschnikowia(100);g__Metschnikowia_unclassified(100); | Metschnikowia | 13 | Saprotroph | \|Undefined Saprotroph\| | Yeast | NULL | Probable | As nectar-tap_saprotroph (Põlme et al. 2020); Decay sugar-rich substrates (Põlme et al. 2020) | Põlme S, et al. 2020. Fungal Diversity 105:1-16 ((https://doi.org/10.1007/s13225-020-00466-2)) |
| Otu58658 | 1 | k__Fungi(100);p__Ascomycota(100);c__Saccharomycetes(100);o__Saccharomycetales(100);f__Metschnikowiaceae(100);g__Metschnikowia(100);s__Metschnikowia_pulcherrima(100); | Metschnikowia | 13 | Saprotroph | \|Undefined Saprotroph\| | Yeast | NULL | Probable | As nectar-tap_saprotroph (Põlme et al. 2020); Decay sugar-rich substrates (Põlme et al. 2020) | Põlme S, et al. 2020. Fungal Diversity 105:1-16 ((https://doi.org/10.1007/s13225-020-00466-2)) |
| Otu58659 | 1 | k__Fungi(100);p__Ascomycota(100);c__Saccharomycetes(100);o__Saccharomycetales(100);f__Metschnikowiaceae(100);g__Metschnikowia(100);s__Metschnikowia_pulcherrima(100); | Metschnikowia | 13 | Saprotroph | \|Undefined Saprotroph\| | Yeast | NULL | Probable | As nectar-tap_saprotroph (Põlme et al. 2020); Decay sugar-rich substrates (Põlme et al. 2020) | Põlme S, et al. 2020. Fungal Diversity 105:1-16 ((https://doi.org/10.1007/s13225-020-00466-2)) |
| Otu58660 | 1 | k__Fungi(100);p__Ascomycota(100);c__Saccharomycetes(100);o__Saccharomycetales(100);f__Metschnikowiaceae(100);g__Metschnikowia(100);g__Metschnikowia_unclassified(100); | Metschnikowia | 13 | Saprotroph | \|Undefined Saprotroph\| | Yeast | NULL | Probable | As nectar-tap_saprotroph (Põlme et al. 2020); Decay sugar-rich substrates (Põlme et al. 2020) | Põlme S, et al. 2020. Fungal Diversity 105:1-16 ((https://doi.org/10.1007/s13225-020-00466-2)) |
| Otu58661 | 1 | k__Fungi(100);p__Ascomycota(100);c__Saccharomycetes(100);o__Saccharomycetales(100);f__Metschnikowiaceae(100);g__Metschnikowia(100);s__Metschnikowia_pulcherrima(100); | Metschnikowia | 13 | Saprotroph | \|Undefined Saprotroph\| | Yeast | NULL | Probable | As nectar-tap_saprotroph (Põlme et al. 2020); Decay sugar-rich substrates (Põlme et al. 2020) | Põlme S, et al. 2020. Fungal Diversity 105:1-16 ((https://doi.org/10.1007/s13225-020-00466-2)) |
| Otu58662 | 1 | k__Fungi(100);p__Ascomycota(100);c__Saccharomycetes(100);o__Saccharomycetales(100);f__Metschnikowiaceae(100);g__Metschnikowia(100);s__Metschnikowia_pulcherrima(100); | Metschnikowia | 13 | Saprotroph | \|Undefined Saprotroph\| | Yeast | NULL | Probable | As nectar-tap_saprotroph (Põlme et al. 2020); Decay sugar-rich substrates (Põlme et al. 2020) | Põlme S, et al. 2020. Fungal Diversity 105:1-16 ((https://doi.org/10.1007/s13225-020-00466-2)) |
| Otu58663 | 1 | k__Fungi(100);p__Ascomycota(100);c__Saccharomycetes(100);o__Saccharomycetales(100);f__Metschnikowiaceae(100);g__Metschnikowia(100);s__Metschnikowia_pulcherrima(100); | Metschnikowia | 13 | Saprotroph | \|Undefined Saprotroph\| | Yeast | NULL | Probable | As nectar-tap_saprotroph (Põlme et al. 2020); Decay sugar-rich substrates (Põlme et al. 2020) | Põlme S, et al. 2020. Fungal Diversity 105:1-16 ((https://doi.org/10.1007/s13225-020-00466-2)) |
| Otu58664 | 1 | k__Fungi(100);p__Ascomycota(100);c__Saccharomycetes(100);o__Saccharomycetales(100);f__Metschnikowiaceae(100);g__Metschnikowia(100);s__Metschnikowia_pulcherrima(100); | Metschnikowia | 13 | Saprotroph | \|Undefined Saprotroph\| | Yeast | NULL | Probable | As nectar-tap_saprotroph (Põlme et al. 2020); Decay sugar-rich substrates (Põlme et al. 2020) | Põlme S, et al. 2020. Fungal Diversity 105:1-16 ((https://doi.org/10.1007/s13225-020-00466-2)) |
| Otu58665 | 1 | k__Fungi(100);p__Ascomycota(100);c__Saccharomycetes(100);o__Saccharomycetales(100);f__Metschnikowiaceae(100);g__Metschnikowia(100);s__Metschnikowia_pulcherrima(100); | Metschnikowia | 13 | Saprotroph | \|Undefined Saprotroph\| | Yeast | NULL | Probable | As nectar-tap_saprotroph (Põlme et al. 2020); Decay sugar-rich substrates (Põlme et al. 2020) | Põlme S, et al. 2020. Fungal Diversity 105:1-16 ((https://doi.org/10.1007/s13225-020-00466-2)) |
| Otu58666 | 1 | k__Fungi(100);p__Ascomycota(100);c__Saccharomycetes(100);o__Saccharomycetales(100);f__Metschnikowiaceae(100);g__Metschnikowia(100);s__Metschnikowia_pulcherrima(100); | Metschnikowia | 13 | Saprotroph | \|Undefined Saprotroph\| | Yeast | NULL | Probable | As nectar-tap_saprotroph (Põlme et al. 2020); Decay sugar-rich substrates (Põlme et al. 2020) | Põlme S, et al. 2020. Fungal Diversity 105:1-16 ((https://doi.org/10.1007/s13225-020-00466-2)) |
| Otu58667 | 1 | k__Fungi(100);p__Ascomycota(100);c__Saccharomycetes(100);o__Saccharomycetales(100);f__Metschnikowiaceae(100);g__Metschnikowia(100);g__Metschnikowia_unclassified(100); | Metschnikowia | 13 | Saprotroph | \|Undefined Saprotroph\| | Yeast | NULL | Probable | As nectar-tap_saprotroph (Põlme et al. 2020); Decay sugar-rich substrates (Põlme et al. 2020) | Põlme S, et al. 2020. Fungal Diversity 105:1-16 ((https://doi.org/10.1007/s13225-020-00466-2)) |
| Otu58668 | 1 | k__Fungi(100);p__Ascomycota(100);c__Saccharomycetes(100);o__Saccharomycetales(100);f__Metschnikowiaceae(100);g__Metschnikowia(100);g__Metschnikowia_unclassified(100); | Metschnikowia | 13 | Saprotroph | \|Undefined Saprotroph\| | Yeast | NULL | Probable | As nectar-tap_saprotroph (Põlme et al. 2020); Decay sugar-rich substrates (Põlme et al. 2020) | Põlme S, et al. 2020. Fungal Diversity 105:1-16 ((https://doi.org/10.1007/s13225-020-00466-2)) |
| Otu58669 | 1 | k__Fungi(100);p__Ascomycota(100);c__Saccharomycetes(100);o__Saccharomycetales(100);f__Metschnikowiaceae(100);g__Metschnikowia(100);s__Metschnikowia_pulcherrima(100); | Metschnikowia | 13 | Saprotroph | \|Undefined Saprotroph\| | Yeast | NULL | Probable | As nectar-tap_saprotroph (Põlme et al. 2020); Decay sugar-rich substrates (Põlme et al. 2020) | Põlme S, et al. 2020. Fungal Diversity 105:1-16 ((https://doi.org/10.1007/s13225-020-00466-2)) |
| Otu58670 | 1 | k__Fungi(100);p__Ascomycota(100);c__Saccharomycetes(100);o__Saccharomycetales(100);f__Metschnikowiaceae(100);g__Metschnikowia(100);s__Metschnikowia_pulcherrima(100); | Metschnikowia | 13 | Saprotroph | \|Undefined Saprotroph\| | Yeast | NULL | Probable | As nectar-tap_saprotroph (Põlme et al. 2020); Decay sugar-rich substrates (Põlme et al. 2020) | Põlme S, et al. 2020. Fungal Diversity 105:1-16 ((https://doi.org/10.1007/s13225-020-00466-2)) |
| Otu58671 | 1 | k__Fungi(100);p__Ascomycota(100);c__Saccharomycetes(100);o__Saccharomycetales(100);f__Metschnikowiaceae(100);g__Metschnikowia(100);g__Metschnikowia_unclassified(100); | Metschnikowia | 13 | Saprotroph | \|Undefined Saprotroph\| | Yeast | NULL | Probable | As nectar-tap_saprotroph (Põlme et al. 2020); Decay sugar-rich substrates (Põlme et al. 2020) | Põlme S, et al. 2020. Fungal Diversity 105:1-16 ((https://doi.org/10.1007/s13225-020-00466-2)) |
| Otu58672 | 1 | k__Fungi(100);p__Ascomycota(100);c__Saccharomycetes(100);o__Saccharomycetales(100);f__Metschnikowiaceae(100);g__Metschnikowia(100);s__Metschnikowia_pulcherrima(100); | Metschnikowia | 13 | Saprotroph | \|Undefined Saprotroph\| | Yeast | NULL | Probable | As nectar-tap_saprotroph (Põlme et al. 2020); Decay sugar-rich substrates (Põlme et al. 2020) | Põlme S, et al. 2020. Fungal Diversity 105:1-16 ((https://doi.org/10.1007/s13225-020-00466-2)) |
| Otu58673 | 1 | k__Fungi(100);p__Ascomycota(100);c__Saccharomycetes(100);o__Saccharomycetales(100);f__Metschnikowiaceae(100);g__Metschnikowia(100);s__Metschnikowia_pulcherrima(100); | Metschnikowia | 13 | Saprotroph | \|Undefined Saprotroph\| | Yeast | NULL | Probable | As nectar-tap_saprotroph (Põlme et al. 2020); Decay sugar-rich substrates (Põlme et al. 2020) | Põlme S, et al. 2020. Fungal Diversity 105:1-16 ((https://doi.org/10.1007/s13225-020-00466-2)) |
| Otu58674 | 1 | k__Fungi(100);p__Ascomycota(100);c__Saccharomycetes(100);o__Saccharomycetales(100);f__Metschnikowiaceae(100);g__Metschnikowia(100);s__Metschnikowia_pulcherrima(100); | Metschnikowia | 13 | Saprotroph | \|Undefined Saprotroph\| | Yeast | NULL | Probable | As nectar-tap_saprotroph (Põlme et al. 2020); Decay sugar-rich substrates (Põlme et al. 2020) | Põlme S, et al. 2020. Fungal Diversity 105:1-16 ((https://doi.org/10.1007/s13225-020-00466-2)) |
| Otu58675 | 1 | k__Fungi(100);p__Ascomycota(100);c__Saccharomycetes(100);o__Saccharomycetales(100);f__Metschnikowiaceae(100);g__Metschnikowia(100);s__Metschnikowia_pulcherrima(100); | Metschnikowia | 13 | Saprotroph | \|Undefined Saprotroph\| | Yeast | NULL | Probable | As nectar-tap_saprotroph (Põlme et al. 2020); Decay sugar-rich substrates (Põlme et al. 2020) | Põlme S, et al. 2020. Fungal Diversity 105:1-16 ((https://doi.org/10.1007/s13225-020-00466-2)) |
| Otu58677 | 1 | k__Fungi(100);p__Ascomycota(100);c__Saccharomycetes(100);o__Saccharomycetales(100);f__Metschnikowiaceae(100);g__Metschnikowia(100);g__Metschnikowia_unclassified(100); | Metschnikowia | 13 | Saprotroph | \|Undefined Saprotroph\| | Yeast | NULL | Probable | As nectar-tap_saprotroph (Põlme et al. 2020); Decay sugar-rich substrates (Põlme et al. 2020) | Põlme S, et al. 2020. Fungal Diversity 105:1-16 ((https://doi.org/10.1007/s13225-020-00466-2)) |
| Otu58678 | 1 | k__Fungi(100);p__Ascomycota(100);c__Saccharomycetes(100);o__Saccharomycetales(100);f__Metschnikowiaceae(100);g__Metschnikowia(100);s__Metschnikowia_pulcherrima(100); | Metschnikowia | 13 | Saprotroph | \|Undefined Saprotroph\| | Yeast | NULL | Probable | As nectar-tap_saprotroph (Põlme et al. 2020); Decay sugar-rich substrates (Põlme et al. 2020) | Põlme S, et al. 2020. Fungal Diversity 105:1-16 ((https://doi.org/10.1007/s13225-020-00466-2)) |
| Otu58679 | 1 | k__Fungi(100);p__Ascomycota(100);c__Saccharomycetes(100);o__Saccharomycetales(100);f__Metschnikowiaceae(100);g__Metschnikowia(100);s__Metschnikowia_pulcherrima(100); | Metschnikowia | 13 | Saprotroph | \|Undefined Saprotroph\| | Yeast | NULL | Probable | As nectar-tap_saprotroph (Põlme et al. 2020); Decay sugar-rich substrates (Põlme et al. 2020) | Põlme S, et al. 2020. Fungal Diversity 105:1-16 ((https://doi.org/10.1007/s13225-020-00466-2)) |
| Otu58680 | 1 | k__Fungi(100);p__Ascomycota(100);c__Saccharomycetes(100);o__Saccharomycetales(100);f__Metschnikowiaceae(100);g__Metschnikowia(100);s__Metschnikowia_pulcherrima(100); | Metschnikowia | 13 | Saprotroph | \|Undefined Saprotroph\| | Yeast | NULL | Probable | As nectar-tap_saprotroph (Põlme et al. 2020); Decay sugar-rich substrates (Põlme et al. 2020) | Põlme S, et al. 2020. Fungal Diversity 105:1-16 ((https://doi.org/10.1007/s13225-020-00466-2)) |
| Otu58681 | 1 | k__Fungi(100);p__Ascomycota(100);c__Saccharomycetes(100);o__Saccharomycetales(100);f__Metschnikowiaceae(100);g__Metschnikowia(100);s__Metschnikowia_pulcherrima(100); | Metschnikowia | 13 | Saprotroph | \|Undefined Saprotroph\| | Yeast | NULL | Probable | As nectar-tap_saprotroph (Põlme et al. 2020); Decay sugar-rich substrates (Põlme et al. 2020) | Põlme S, et al. 2020. Fungal Diversity 105:1-16 ((https://doi.org/10.1007/s13225-020-00466-2)) |
| Otu58683 | 1 | k__Fungi(100);p__Ascomycota(100);c__Lecanoromycetes(100);o__Lecanorales(100);o__Lecanorales_unclassified(100);o__Lecanorales_unclassified(100);o__Lecanorales_unclassified(100); | Lecanorales | 7 | Pathotroph-Symbiotroph | Lichen Parasite-Lichenized | Microfungus-Thallus | NULL | Probable | Lichenized with green algae, occasionally lichenicolous, on rocks, mosses and wood (Cannon & Kirk 2007); ...an order of mostly lichen-forming fungi (Kirk et al. 2008) | Cannon PF, Kirk PM. 2007. Fungal Families of the World. CAB International, Cambridge (ISBN: 978-0851998275); Kirk PM et al. 2008. Dictionary of the Fungi. Tenth Edition. CAB International, Wallingford (ISBN: 978-0851998268); Esslinger TL. 2014. North Dakota State University ((http://www.ndsu.edu/pubweb/~esslinge/chcklst/chcklst7.htm)) |
| Otu58684 | 1 | k__Fungi(100);p__Ascomycota(100);c__Saccharomycetes(100);o__Saccharomycetales(100);f__Metschnikowiaceae(100);g__Metschnikowia(100);s__Metschnikowia_pulcherrima(100); | Metschnikowia | 13 | Saprotroph | \|Undefined Saprotroph\| | Yeast | NULL | Probable | As nectar-tap_saprotroph (Põlme et al. 2020); Decay sugar-rich substrates (Põlme et al. 2020) | Põlme S, et al. 2020. Fungal Diversity 105:1-16 ((https://doi.org/10.1007/s13225-020-00466-2)) |
| Otu58685 | 1 | k__Fungi(100);p__Ascomycota(100);c__Saccharomycetes(100);o__Saccharomycetales(100);f__Metschnikowiaceae(100);g__Metschnikowia(100);s__Metschnikowia_pulcherrima(100); | Metschnikowia | 13 | Saprotroph | \|Undefined Saprotroph\| | Yeast | NULL | Probable | As nectar-tap_saprotroph (Põlme et al. 2020); Decay sugar-rich substrates (Põlme et al. 2020) | Põlme S, et al. 2020. Fungal Diversity 105:1-16 ((https://doi.org/10.1007/s13225-020-00466-2)) |
| Otu58686 | 1 | k__Fungi(100);p__Ascomycota(100);c__Saccharomycetes(100);o__Saccharomycetales(100);f__Metschnikowiaceae(100);g__Metschnikowia(100);s__Metschnikowia_pulcherrima(100); | Metschnikowia | 13 | Saprotroph | \|Undefined Saprotroph\| | Yeast | NULL | Probable | As nectar-tap_saprotroph (Põlme et al. 2020); Decay sugar-rich substrates (Põlme et al. 2020) | Põlme S, et al. 2020. Fungal Diversity 105:1-16 ((https://doi.org/10.1007/s13225-020-00466-2)) |
| Otu58687 | 1 | k__Fungi(100);p__Ascomycota(100);c__Saccharomycetes(100);o__Saccharomycetales(100);f__Metschnikowiaceae(100);g__Metschnikowia(100);s__Metschnikowia_pulcherrima(100); | Metschnikowia | 13 | Saprotroph | \|Undefined Saprotroph\| | Yeast | NULL | Probable | As nectar-tap_saprotroph (Põlme et al. 2020); Decay sugar-rich substrates (Põlme et al. 2020) | Põlme S, et al. 2020. Fungal Diversity 105:1-16 ((https://doi.org/10.1007/s13225-020-00466-2)) |
| Otu58688 | 1 | k__Fungi(100);p__Ascomycota(100);c__Saccharomycetes(100);o__Saccharomycetales(100);f__Metschnikowiaceae(100);g__Metschnikowia(100);s__Metschnikowia_pulcherrima(100); | Metschnikowia | 13 | Saprotroph | \|Undefined Saprotroph\| | Yeast | NULL | Probable | As nectar-tap_saprotroph (Põlme et al. 2020); Decay sugar-rich substrates (Põlme et al. 2020) | Põlme S, et al. 2020. Fungal Diversity 105:1-16 ((https://doi.org/10.1007/s13225-020-00466-2)) |
| Otu58689 | 1 | k__Fungi(100);p__Ascomycota(100);c__Saccharomycetes(100);o__Saccharomycetales(100);f__Metschnikowiaceae(100);g__Metschnikowia(100);s__Metschnikowia_pulcherrima(100); | Metschnikowia | 13 | Saprotroph | \|Undefined Saprotroph\| | Yeast | NULL | Probable | As nectar-tap_saprotroph (Põlme et al. 2020); Decay sugar-rich substrates (Põlme et al. 2020) | Põlme S, et al. 2020. Fungal Diversity 105:1-16 ((https://doi.org/10.1007/s13225-020-00466-2)) |
| Otu58690 | 1 | k__Fungi(100);p__Ascomycota(100);c__Saccharomycetes(100);o__Saccharomycetales(100);f__Metschnikowiaceae(100);g__Metschnikowia(100);s__Metschnikowia_pulcherrima(100); | Metschnikowia | 13 | Saprotroph | \|Undefined Saprotroph\| | Yeast | NULL | Probable | As nectar-tap_saprotroph (Põlme et al. 2020); Decay sugar-rich substrates (Põlme et al. 2020) | Põlme S, et al. 2020. Fungal Diversity 105:1-16 ((https://doi.org/10.1007/s13225-020-00466-2)) |
| Otu58691 | 1 | k__Fungi(100);p__Ascomycota(100);c__Saccharomycetes(100);o__Saccharomycetales(100);f__Metschnikowiaceae(100);g__Metschnikowia(100);s__Metschnikowia_pulcherrima(100); | Metschnikowia | 13 | Saprotroph | \|Undefined Saprotroph\| | Yeast | NULL | Probable | As nectar-tap_saprotroph (Põlme et al. 2020); Decay sugar-rich substrates (Põlme et al. 2020) | Põlme S, et al. 2020. Fungal Diversity 105:1-16 ((https://doi.org/10.1007/s13225-020-00466-2)) |
| Otu58692 | 1 | k__Fungi(100);p__Ascomycota(100);c__Saccharomycetes(100);o__Saccharomycetales(100);f__Metschnikowiaceae(100);g__Metschnikowia(100);s__Metschnikowia_pulcherrima(100); | Metschnikowia | 13 | Saprotroph | \|Undefined Saprotroph\| | Yeast | NULL | Probable | As nectar-tap_saprotroph (Põlme et al. 2020); Decay sugar-rich substrates (Põlme et al. 2020) | Põlme S, et al. 2020. Fungal Diversity 105:1-16 ((https://doi.org/10.1007/s13225-020-00466-2)) |
| Otu58693 | 1 | k__Fungi(100);p__Ascomycota(100);c__Saccharomycetes(100);o__Saccharomycetales(100);f__Metschnikowiaceae(100);g__Metschnikowia(100);s__Metschnikowia_pulcherrima(100); | Metschnikowia | 13 | Saprotroph | \|Undefined Saprotroph\| | Yeast | NULL | Probable | As nectar-tap_saprotroph (Põlme et al. 2020); Decay sugar-rich substrates (Põlme et al. 2020) | Põlme S, et al. 2020. Fungal Diversity 105:1-16 ((https://doi.org/10.1007/s13225-020-00466-2)) |
| Otu58694 | 1 | k__Fungi(100);p__Ascomycota(100);c__Saccharomycetes(100);o__Saccharomycetales(100);f__Metschnikowiaceae(100);g__Metschnikowia(100);s__Metschnikowia_pulcherrima(100); | Metschnikowia | 13 | Saprotroph | \|Undefined Saprotroph\| | Yeast | NULL | Probable | As nectar-tap_saprotroph (Põlme et al. 2020); Decay sugar-rich substrates (Põlme et al. 2020) | Põlme S, et al. 2020. Fungal Diversity 105:1-16 ((https://doi.org/10.1007/s13225-020-00466-2)) |
| Otu58695 | 1 | k__Fungi(100);p__Ascomycota(100);c__Saccharomycetes(100);o__Saccharomycetales(100);f__Metschnikowiaceae(100);g__Metschnikowia(100);g__Metschnikowia_unclassified(100); | Metschnikowia | 13 | Saprotroph | \|Undefined Saprotroph\| | Yeast | NULL | Probable | As nectar-tap_saprotroph (Põlme et al. 2020); Decay sugar-rich substrates (Põlme et al. 2020) | Põlme S, et al. 2020. Fungal Diversity 105:1-16 ((https://doi.org/10.1007/s13225-020-00466-2)) |
| Otu58696 | 1 | k__Fungi(100);p__Ascomycota(100);c__Saccharomycetes(100);o__Saccharomycetales(100);f__Metschnikowiaceae(100);g__Metschnikowia(100);s__Metschnikowia_pulcherrima(100); | Metschnikowia | 13 | Saprotroph | \|Undefined Saprotroph\| | Yeast | NULL | Probable | As nectar-tap_saprotroph (Põlme et al. 2020); Decay sugar-rich substrates (Põlme et al. 2020) | Põlme S, et al. 2020. Fungal Diversity 105:1-16 ((https://doi.org/10.1007/s13225-020-00466-2)) |
| Otu58697 | 1 | k__Fungi(100);p__Ascomycota(100);c__Saccharomycetes(100);o__Saccharomycetales(100);f__Metschnikowiaceae(100);g__Metschnikowia(100);s__Metschnikowia_pulcherrima(100); | Metschnikowia | 13 | Saprotroph | \|Undefined Saprotroph\| | Yeast | NULL | Probable | As nectar-tap_saprotroph (Põlme et al. 2020); Decay sugar-rich substrates (Põlme et al. 2020) | Põlme S, et al. 2020. Fungal Diversity 105:1-16 ((https://doi.org/10.1007/s13225-020-00466-2)) |
| Otu58698 | 1 | k__Fungi(100);p__Ascomycota(100);c__Saccharomycetes(100);o__Saccharomycetales(100);f__Metschnikowiaceae(100);g__Metschnikowia(100);g__Metschnikowia_unclassified(100); | Metschnikowia | 13 | Saprotroph | \|Undefined Saprotroph\| | Yeast | NULL | Probable | As nectar-tap_saprotroph (Põlme et al. 2020); Decay sugar-rich substrates (Põlme et al. 2020) | Põlme S, et al. 2020. Fungal Diversity 105:1-16 ((https://doi.org/10.1007/s13225-020-00466-2)) |
| Otu58699 | 1 | k__Fungi(100);p__Ascomycota(100);c__Saccharomycetes(100);o__Saccharomycetales(100);f__Metschnikowiaceae(100);g__Metschnikowia(100);g__Metschnikowia_unclassified(100); | Metschnikowia | 13 | Saprotroph | \|Undefined Saprotroph\| | Yeast | NULL | Probable | As nectar-tap_saprotroph (Põlme et al. 2020); Decay sugar-rich substrates (Põlme et al. 2020) | Põlme S, et al. 2020. Fungal Diversity 105:1-16 ((https://doi.org/10.1007/s13225-020-00466-2)) |
| Otu58700 | 1 | k__Fungi(100);p__Ascomycota(100);c__Saccharomycetes(100);o__Saccharomycetales(100);f__Metschnikowiaceae(100);g__Metschnikowia(100);s__Metschnikowia_pulcherrima(100); | Metschnikowia | 13 | Saprotroph | \|Undefined Saprotroph\| | Yeast | NULL | Probable | As nectar-tap_saprotroph (Põlme et al. 2020); Decay sugar-rich substrates (Põlme et al. 2020) | Põlme S, et al. 2020. Fungal Diversity 105:1-16 ((https://doi.org/10.1007/s13225-020-00466-2)) |
| Otu58701 | 1 | k__Fungi(100);p__Ascomycota(100);c__Saccharomycetes(100);o__Saccharomycetales(100);f__Metschnikowiaceae(100);g__Metschnikowia(100);s__Metschnikowia_pulcherrima(100); | Metschnikowia | 13 | Saprotroph | \|Undefined Saprotroph\| | Yeast | NULL | Probable | As nectar-tap_saprotroph (Põlme et al. 2020); Decay sugar-rich substrates (Põlme et al. 2020) | Põlme S, et al. 2020. Fungal Diversity 105:1-16 ((https://doi.org/10.1007/s13225-020-00466-2)) |
| Otu58702 | 1 | k__Fungi(100);p__Ascomycota(100);c__Saccharomycetes(100);o__Saccharomycetales(100);f__Metschnikowiaceae(100);g__Metschnikowia(100);s__Metschnikowia_pulcherrima(100); | Metschnikowia | 13 | Saprotroph | \|Undefined Saprotroph\| | Yeast | NULL | Probable | As nectar-tap_saprotroph (Põlme et al. 2020); Decay sugar-rich substrates (Põlme et al. 2020) | Põlme S, et al. 2020. Fungal Diversity 105:1-16 ((https://doi.org/10.1007/s13225-020-00466-2)) |
| Otu58703 | 1 | k__Fungi(100);p__Ascomycota(100);c__Saccharomycetes(100);o__Saccharomycetales(100);f__Metschnikowiaceae(100);g__Metschnikowia(100);s__Metschnikowia_pulcherrima(100); | Metschnikowia | 13 | Saprotroph | \|Undefined Saprotroph\| | Yeast | NULL | Probable | As nectar-tap_saprotroph (Põlme et al. 2020); Decay sugar-rich substrates (Põlme et al. 2020) | Põlme S, et al. 2020. Fungal Diversity 105:1-16 ((https://doi.org/10.1007/s13225-020-00466-2)) |
| Otu58704 | 1 | k__Fungi(100);p__Ascomycota(100);c__Saccharomycetes(100);o__Saccharomycetales(100);f__Metschnikowiaceae(100);g__Metschnikowia(100);g__Metschnikowia_unclassified(100); | Metschnikowia | 13 | Saprotroph | \|Undefined Saprotroph\| | Yeast | NULL | Probable | As nectar-tap_saprotroph (Põlme et al. 2020); Decay sugar-rich substrates (Põlme et al. 2020) | Põlme S, et al. 2020. Fungal Diversity 105:1-16 ((https://doi.org/10.1007/s13225-020-00466-2)) |
| Otu58705 | 1 | k__Fungi(100);p__Ascomycota(100);c__Saccharomycetes(100);o__Saccharomycetales(100);f__Metschnikowiaceae(100);g__Metschnikowia(100);s__Metschnikowia_pulcherrima(100); | Metschnikowia | 13 | Saprotroph | \|Undefined Saprotroph\| | Yeast | NULL | Probable | As nectar-tap_saprotroph (Põlme et al. 2020); Decay sugar-rich substrates (Põlme et al. 2020) | Põlme S, et al. 2020. Fungal Diversity 105:1-16 ((https://doi.org/10.1007/s13225-020-00466-2)) |
| Otu58706 | 1 | k__Fungi(100);p__Ascomycota(100);c__Saccharomycetes(100);o__Saccharomycetales(100);f__Metschnikowiaceae(100);g__Metschnikowia(100);g__Metschnikowia_unclassified(100); | Metschnikowia | 13 | Saprotroph | \|Undefined Saprotroph\| | Yeast | NULL | Probable | As nectar-tap_saprotroph (Põlme et al. 2020); Decay sugar-rich substrates (Põlme et al. 2020) | Põlme S, et al. 2020. Fungal Diversity 105:1-16 ((https://doi.org/10.1007/s13225-020-00466-2)) |
| Otu58707 | 1 | k__Fungi(100);p__Ascomycota(100);c__Saccharomycetes(100);o__Saccharomycetales(100);f__Metschnikowiaceae(100);g__Metschnikowia(100);s__Metschnikowia_pulcherrima(100); | Metschnikowia | 13 | Saprotroph | \|Undefined Saprotroph\| | Yeast | NULL | Probable | As nectar-tap_saprotroph (Põlme et al. 2020); Decay sugar-rich substrates (Põlme et al. 2020) | Põlme S, et al. 2020. Fungal Diversity 105:1-16 ((https://doi.org/10.1007/s13225-020-00466-2)) |
| Otu58708 | 1 | k__Fungi(100);p__Ascomycota(100);c__Saccharomycetes(100);o__Saccharomycetales(100);f__Metschnikowiaceae(100);g__Metschnikowia(100);s__Metschnikowia_pulcherrima(100); | Metschnikowia | 13 | Saprotroph | \|Undefined Saprotroph\| | Yeast | NULL | Probable | As nectar-tap_saprotroph (Põlme et al. 2020); Decay sugar-rich substrates (Põlme et al. 2020) | Põlme S, et al. 2020. Fungal Diversity 105:1-16 ((https://doi.org/10.1007/s13225-020-00466-2)) |
| Otu58709 | 1 | k__Fungi(100);p__Ascomycota(100);c__Saccharomycetes(100);o__Saccharomycetales(100);f__Metschnikowiaceae(100);g__Metschnikowia(100);g__Metschnikowia_unclassified(100); | Metschnikowia | 13 | Saprotroph | \|Undefined Saprotroph\| | Yeast | NULL | Probable | As nectar-tap_saprotroph (Põlme et al. 2020); Decay sugar-rich substrates (Põlme et al. 2020) | Põlme S, et al. 2020. Fungal Diversity 105:1-16 ((https://doi.org/10.1007/s13225-020-00466-2)) |
| Otu58710 | 1 | k__Fungi(100);p__Ascomycota(100);c__Saccharomycetes(100);o__Saccharomycetales(100);f__Metschnikowiaceae(100);g__Metschnikowia(100);s__Metschnikowia_pulcherrima(100); | Metschnikowia | 13 | Saprotroph | \|Undefined Saprotroph\| | Yeast | NULL | Probable | As nectar-tap_saprotroph (Põlme et al. 2020); Decay sugar-rich substrates (Põlme et al. 2020) | Põlme S, et al. 2020. Fungal Diversity 105:1-16 ((https://doi.org/10.1007/s13225-020-00466-2)) |
| Otu58711 | 1 | k__Fungi(100);p__Ascomycota(100);c__Saccharomycetes(100);o__Saccharomycetales(100);f__Metschnikowiaceae(100);g__Metschnikowia(100);s__Metschnikowia_pulcherrima(100); | Metschnikowia | 13 | Saprotroph | \|Undefined Saprotroph\| | Yeast | NULL | Probable | As nectar-tap_saprotroph (Põlme et al. 2020); Decay sugar-rich substrates (Põlme et al. 2020) | Põlme S, et al. 2020. Fungal Diversity 105:1-16 ((https://doi.org/10.1007/s13225-020-00466-2)) |
| Otu58724 | 1 | k__Fungi(100);p__Ascomycota(100);c__Saccharomycetes(100);o__Saccharomycetales(100);f__Metschnikowiaceae(100);g__Metschnikowia(100);s__Metschnikowia_pulcherrima(100); | Metschnikowia | 13 | Saprotroph | \|Undefined Saprotroph\| | Yeast | NULL | Probable | As nectar-tap_saprotroph (Põlme et al. 2020); Decay sugar-rich substrates (Põlme et al. 2020) | Põlme S, et al. 2020. Fungal Diversity 105:1-16 ((https://doi.org/10.1007/s13225-020-00466-2)) |
| Otu58728 | 1 | k__Fungi(100);p__Ascomycota(100);c__Saccharomycetes(100);o__Saccharomycetales(100);f__Metschnikowiaceae(100);g__Metschnikowia(100);g__Metschnikowia_unclassified(100); | Metschnikowia | 13 | Saprotroph | \|Undefined Saprotroph\| | Yeast | NULL | Probable | As nectar-tap_saprotroph (Põlme et al. 2020); Decay sugar-rich substrates (Põlme et al. 2020) | Põlme S, et al. 2020. Fungal Diversity 105:1-16 ((https://doi.org/10.1007/s13225-020-00466-2)) |
| Otu58729 | 1 | k__Fungi(100);p__Ascomycota(100);c__Saccharomycetes(100);o__Saccharomycetales(100);f__Metschnikowiaceae(100);g__Metschnikowia(100);s__Metschnikowia_pulcherrima(100); | Metschnikowia | 13 | Saprotroph | \|Undefined Saprotroph\| | Yeast | NULL | Probable | As nectar-tap_saprotroph (Põlme et al. 2020); Decay sugar-rich substrates (Põlme et al. 2020) | Põlme S, et al. 2020. Fungal Diversity 105:1-16 ((https://doi.org/10.1007/s13225-020-00466-2)) |
| Otu58730 | 1 | k__Fungi(100);p__Ascomycota(100);c__Saccharomycetes(100);o__Saccharomycetales(100);f__Metschnikowiaceae(100);g__Metschnikowia(100);s__Metschnikowia_pulcherrima(100); | Metschnikowia | 13 | Saprotroph | \|Undefined Saprotroph\| | Yeast | NULL | Probable | As nectar-tap_saprotroph (Põlme et al. 2020); Decay sugar-rich substrates (Põlme et al. 2020) | Põlme S, et al. 2020. Fungal Diversity 105:1-16 ((https://doi.org/10.1007/s13225-020-00466-2)) |
| Otu58731 | 1 | k__Fungi(100);p__Ascomycota(100);c__Saccharomycetes(100);o__Saccharomycetales(100);f__Metschnikowiaceae(100);g__Metschnikowia(100);s__Metschnikowia_pulcherrima(100); | Metschnikowia | 13 | Saprotroph | \|Undefined Saprotroph\| | Yeast | NULL | Probable | As nectar-tap_saprotroph (Põlme et al. 2020); Decay sugar-rich substrates (Põlme et al. 2020) | Põlme S, et al. 2020. Fungal Diversity 105:1-16 ((https://doi.org/10.1007/s13225-020-00466-2)) |
| Otu58732 | 1 | k__Fungi(100);p__Ascomycota(100);c__Saccharomycetes(100);o__Saccharomycetales(100);f__Metschnikowiaceae(100);g__Metschnikowia(100);s__Metschnikowia_pulcherrima(100); | Metschnikowia | 13 | Saprotroph | \|Undefined Saprotroph\| | Yeast | NULL | Probable | As nectar-tap_saprotroph (Põlme et al. 2020); Decay sugar-rich substrates (Põlme et al. 2020) | Põlme S, et al. 2020. Fungal Diversity 105:1-16 ((https://doi.org/10.1007/s13225-020-00466-2)) |
| Otu58733 | 1 | k__Fungi(100);p__Ascomycota(100);c__Saccharomycetes(100);o__Saccharomycetales(100);f__Metschnikowiaceae(100);g__Metschnikowia(100);s__Metschnikowia_pulcherrima(100); | Metschnikowia | 13 | Saprotroph | \|Undefined Saprotroph\| | Yeast | NULL | Probable | As nectar-tap_saprotroph (Põlme et al. 2020); Decay sugar-rich substrates (Põlme et al. 2020) | Põlme S, et al. 2020. Fungal Diversity 105:1-16 ((https://doi.org/10.1007/s13225-020-00466-2)) |
| Otu58734 | 1 | k__Fungi(100);p__Ascomycota(100);c__Saccharomycetes(100);o__Saccharomycetales(100);f__Metschnikowiaceae(100);g__Metschnikowia(100);s__Metschnikowia_pulcherrima(100); | Metschnikowia | 13 | Saprotroph | \|Undefined Saprotroph\| | Yeast | NULL | Probable | As nectar-tap_saprotroph (Põlme et al. 2020); Decay sugar-rich substrates (Põlme et al. 2020) | Põlme S, et al. 2020. Fungal Diversity 105:1-16 ((https://doi.org/10.1007/s13225-020-00466-2)) |
| Otu58736 | 1 | k__Fungi(100);p__Ascomycota(100);c__Saccharomycetes(100);o__Saccharomycetales(100);f__Metschnikowiaceae(100);g__Metschnikowia(100);s__Metschnikowia_pulcherrima(100); | Metschnikowia | 13 | Saprotroph | \|Undefined Saprotroph\| | Yeast | NULL | Probable | As nectar-tap_saprotroph (Põlme et al. 2020); Decay sugar-rich substrates (Põlme et al. 2020) | Põlme S, et al. 2020. Fungal Diversity 105:1-16 ((https://doi.org/10.1007/s13225-020-00466-2)) |
| Otu58738 | 1 | k__Fungi(100);p__Ascomycota(100);c__Saccharomycetes(100);o__Saccharomycetales(100);f__Metschnikowiaceae(100);g__Metschnikowia(100);s__Metschnikowia_pulcherrima(100); | Metschnikowia | 13 | Saprotroph | \|Undefined Saprotroph\| | Yeast | NULL | Probable | As nectar-tap_saprotroph (Põlme et al. 2020); Decay sugar-rich substrates (Põlme et al. 2020) | Põlme S, et al. 2020. Fungal Diversity 105:1-16 ((https://doi.org/10.1007/s13225-020-00466-2)) |
| Otu58739 | 1 | k__Fungi(100);p__Ascomycota(100);c__Saccharomycetes(100);o__Saccharomycetales(100);f__Metschnikowiaceae(100);g__Metschnikowia(100);s__Metschnikowia_pulcherrima(100); | Metschnikowia | 13 | Saprotroph | \|Undefined Saprotroph\| | Yeast | NULL | Probable | As nectar-tap_saprotroph (Põlme et al. 2020); Decay sugar-rich substrates (Põlme et al. 2020) | Põlme S, et al. 2020. Fungal Diversity 105:1-16 ((https://doi.org/10.1007/s13225-020-00466-2)) |
| Otu58740 | 1 | k__Fungi(100);p__Ascomycota(100);c__Saccharomycetes(100);o__Saccharomycetales(100);f__Metschnikowiaceae(100);g__Metschnikowia(100);g__Metschnikowia_unclassified(100); | Metschnikowia | 13 | Saprotroph | \|Undefined Saprotroph\| | Yeast | NULL | Probable | As nectar-tap_saprotroph (Põlme et al. 2020); Decay sugar-rich substrates (Põlme et al. 2020) | Põlme S, et al. 2020. Fungal Diversity 105:1-16 ((https://doi.org/10.1007/s13225-020-00466-2)) |
| Otu58741 | 1 | k__Fungi(100);p__Ascomycota(100);c__Saccharomycetes(100);o__Saccharomycetales(100);f__Metschnikowiaceae(100);g__Metschnikowia(100);s__Metschnikowia_pulcherrima(100); | Metschnikowia | 13 | Saprotroph | \|Undefined Saprotroph\| | Yeast | NULL | Probable | As nectar-tap_saprotroph (Põlme et al. 2020); Decay sugar-rich substrates (Põlme et al. 2020) | Põlme S, et al. 2020. Fungal Diversity 105:1-16 ((https://doi.org/10.1007/s13225-020-00466-2)) |
| Otu58742 | 1 | k__Fungi(100);p__Ascomycota(100);c__Saccharomycetes(100);o__Saccharomycetales(100);f__Metschnikowiaceae(100);g__Metschnikowia(100);s__Metschnikowia_pulcherrima(100); | Metschnikowia | 13 | Saprotroph | \|Undefined Saprotroph\| | Yeast | NULL | Probable | As nectar-tap_saprotroph (Põlme et al. 2020); Decay sugar-rich substrates (Põlme et al. 2020) | Põlme S, et al. 2020. Fungal Diversity 105:1-16 ((https://doi.org/10.1007/s13225-020-00466-2)) |
| Otu58743 | 1 | k__Fungi(100);p__Ascomycota(100);c__Saccharomycetes(100);o__Saccharomycetales(100);f__Metschnikowiaceae(100);g__Metschnikowia(100);s__Metschnikowia_pulcherrima(100); | Metschnikowia | 13 | Saprotroph | \|Undefined Saprotroph\| | Yeast | NULL | Probable | As nectar-tap_saprotroph (Põlme et al. 2020); Decay sugar-rich substrates (Põlme et al. 2020) | Põlme S, et al. 2020. Fungal Diversity 105:1-16 ((https://doi.org/10.1007/s13225-020-00466-2)) |
| Otu58744 | 1 | k__Fungi(100);p__Ascomycota(100);c__Saccharomycetes(100);o__Saccharomycetales(100);f__Metschnikowiaceae(100);g__Metschnikowia(100);s__Metschnikowia_pulcherrima(100); | Metschnikowia | 13 | Saprotroph | \|Undefined Saprotroph\| | Yeast | NULL | Probable | As nectar-tap_saprotroph (Põlme et al. 2020); Decay sugar-rich substrates (Põlme et al. 2020) | Põlme S, et al. 2020. Fungal Diversity 105:1-16 ((https://doi.org/10.1007/s13225-020-00466-2)) |
| Otu58747 | 1 | k__Fungi(100);p__Ascomycota(100);c__Saccharomycetes(100);o__Saccharomycetales(100);f__Metschnikowiaceae(100);g__Metschnikowia(100);s__Metschnikowia_pulcherrima(100); | Metschnikowia | 13 | Saprotroph | \|Undefined Saprotroph\| | Yeast | NULL | Probable | As nectar-tap_saprotroph (Põlme et al. 2020); Decay sugar-rich substrates (Põlme et al. 2020) | Põlme S, et al. 2020. Fungal Diversity 105:1-16 ((https://doi.org/10.1007/s13225-020-00466-2)) |
| Otu58748 | 1 | k__Fungi(100);p__Ascomycota(100);c__Saccharomycetes(100);o__Saccharomycetales(100);f__Metschnikowiaceae(100);g__Metschnikowia(100);g__Metschnikowia_unclassified(100); | Metschnikowia | 13 | Saprotroph | \|Undefined Saprotroph\| | Yeast | NULL | Probable | As nectar-tap_saprotroph (Põlme et al. 2020); Decay sugar-rich substrates (Põlme et al. 2020) | Põlme S, et al. 2020. Fungal Diversity 105:1-16 ((https://doi.org/10.1007/s13225-020-00466-2)) |
| Otu58749 | 1 | k__Fungi(100);p__Ascomycota(100);c__Saccharomycetes(100);o__Saccharomycetales(100);f__Metschnikowiaceae(100);g__Metschnikowia(100);g__Metschnikowia_unclassified(100); | Metschnikowia | 13 | Saprotroph | \|Undefined Saprotroph\| | Yeast | NULL | Probable | As nectar-tap_saprotroph (Põlme et al. 2020); Decay sugar-rich substrates (Põlme et al. 2020) | Põlme S, et al. 2020. Fungal Diversity 105:1-16 ((https://doi.org/10.1007/s13225-020-00466-2)) |
| Otu58750 | 1 | k__Fungi(100);p__Ascomycota(100);c__Saccharomycetes(100);o__Saccharomycetales(100);f__Metschnikowiaceae(100);g__Metschnikowia(100);s__Metschnikowia_pulcherrima(100); | Metschnikowia | 13 | Saprotroph | \|Undefined Saprotroph\| | Yeast | NULL | Probable | As nectar-tap_saprotroph (Põlme et al. 2020); Decay sugar-rich substrates (Põlme et al. 2020) | Põlme S, et al. 2020. Fungal Diversity 105:1-16 ((https://doi.org/10.1007/s13225-020-00466-2)) |
| Otu58769 | 1 | k__Fungi(100);p__Ascomycota(100);c__Saccharomycetes(100);o__Saccharomycetales(100);f__Metschnikowiaceae(100);g__Metschnikowia(100);s__Metschnikowia_pulcherrima(100); | Metschnikowia | 13 | Saprotroph | \|Undefined Saprotroph\| | Yeast | NULL | Probable | As nectar-tap_saprotroph (Põlme et al. 2020); Decay sugar-rich substrates (Põlme et al. 2020) | Põlme S, et al. 2020. Fungal Diversity 105:1-16 ((https://doi.org/10.1007/s13225-020-00466-2)) |
| Otu58774 | 1 | k__Fungi(100);p__Ascomycota(100);c__Saccharomycetes(100);o__Saccharomycetales(100);f__Metschnikowiaceae(100);g__Metschnikowia(100);s__Metschnikowia_pulcherrima(100); | Metschnikowia | 13 | Saprotroph | \|Undefined Saprotroph\| | Yeast | NULL | Probable | As nectar-tap_saprotroph (Põlme et al. 2020); Decay sugar-rich substrates (Põlme et al. 2020) | Põlme S, et al. 2020. Fungal Diversity 105:1-16 ((https://doi.org/10.1007/s13225-020-00466-2)) |
| Otu58779 | 1 | k__Fungi(100);p__Ascomycota(100);c__Saccharomycetes(100);o__Saccharomycetales(100);f__Metschnikowiaceae(100);g__Metschnikowia(100);g__Metschnikowia_unclassified(100); | Metschnikowia | 13 | Saprotroph | \|Undefined Saprotroph\| | Yeast | NULL | Probable | As nectar-tap_saprotroph (Põlme et al. 2020); Decay sugar-rich substrates (Põlme et al. 2020) | Põlme S, et al. 2020. Fungal Diversity 105:1-16 ((https://doi.org/10.1007/s13225-020-00466-2)) |
| Otu58780 | 1 | k__Fungi(100);p__Ascomycota(100);c__Saccharomycetes(100);o__Saccharomycetales(100);f__Metschnikowiaceae(100);g__Metschnikowia(100);g__Metschnikowia_unclassified(100); | Metschnikowia | 13 | Saprotroph | \|Undefined Saprotroph\| | Yeast | NULL | Probable | As nectar-tap_saprotroph (Põlme et al. 2020); Decay sugar-rich substrates (Põlme et al. 2020) | Põlme S, et al. 2020. Fungal Diversity 105:1-16 ((https://doi.org/10.1007/s13225-020-00466-2)) |
| Otu58786 | 1 | k__Fungi(100);p__Ascomycota(100);c__Saccharomycetes(100);o__Saccharomycetales(100);f__Metschnikowiaceae(100);g__Metschnikowia(100);s__Metschnikowia_pulcherrima(100); | Metschnikowia | 13 | Saprotroph | \|Undefined Saprotroph\| | Yeast | NULL | Probable | As nectar-tap_saprotroph (Põlme et al. 2020); Decay sugar-rich substrates (Põlme et al. 2020) | Põlme S, et al. 2020. Fungal Diversity 105:1-16 ((https://doi.org/10.1007/s13225-020-00466-2)) |
| Otu58787 | 1 | k__Fungi(100);p__Ascomycota(100);c__Saccharomycetes(100);o__Saccharomycetales(100);f__Metschnikowiaceae(100);g__Metschnikowia(100);g__Metschnikowia_unclassified(100); | Metschnikowia | 13 | Saprotroph | \|Undefined Saprotroph\| | Yeast | NULL | Probable | As nectar-tap_saprotroph (Põlme et al. 2020); Decay sugar-rich substrates (Põlme et al. 2020) | Põlme S, et al. 2020. Fungal Diversity 105:1-16 ((https://doi.org/10.1007/s13225-020-00466-2)) |
| Otu58788 | 1 | k__Fungi(100);p__Ascomycota(100);c__Saccharomycetes(100);o__Saccharomycetales(100);f__Metschnikowiaceae(100);g__Metschnikowia(100);s__Metschnikowia_pulcherrima(100); | Metschnikowia | 13 | Saprotroph | \|Undefined Saprotroph\| | Yeast | NULL | Probable | As nectar-tap_saprotroph (Põlme et al. 2020); Decay sugar-rich substrates (Põlme et al. 2020) | Põlme S, et al. 2020. Fungal Diversity 105:1-16 ((https://doi.org/10.1007/s13225-020-00466-2)) |
| Otu58790 | 1 | k__Fungi(100);p__Ascomycota(100);c__Saccharomycetes(100);o__Saccharomycetales(100);f__Metschnikowiaceae(100);g__Metschnikowia(100);s__Metschnikowia_pulcherrima(100); | Metschnikowia | 13 | Saprotroph | \|Undefined Saprotroph\| | Yeast | NULL | Probable | As nectar-tap_saprotroph (Põlme et al. 2020); Decay sugar-rich substrates (Põlme et al. 2020) | Põlme S, et al. 2020. Fungal Diversity 105:1-16 ((https://doi.org/10.1007/s13225-020-00466-2)) |
| Otu58791 | 1 | k__Fungi(100);p__Ascomycota(100);c__Saccharomycetes(100);o__Saccharomycetales(100);f__Metschnikowiaceae(100);g__Metschnikowia(100);g__Metschnikowia_unclassified(100); | Metschnikowia | 13 | Saprotroph | \|Undefined Saprotroph\| | Yeast | NULL | Probable | As nectar-tap_saprotroph (Põlme et al. 2020); Decay sugar-rich substrates (Põlme et al. 2020) | Põlme S, et al. 2020. Fungal Diversity 105:1-16 ((https://doi.org/10.1007/s13225-020-00466-2)) |
| Otu58793 | 1 | k__Fungi(100);p__Ascomycota(100);c__Saccharomycetes(100);o__Saccharomycetales(100);f__Metschnikowiaceae(100);g__Metschnikowia(100);g__Metschnikowia_unclassified(100); | Metschnikowia | 13 | Saprotroph | \|Undefined Saprotroph\| | Yeast | NULL | Probable | As nectar-tap_saprotroph (Põlme et al. 2020); Decay sugar-rich substrates (Põlme et al. 2020) | Põlme S, et al. 2020. Fungal Diversity 105:1-16 ((https://doi.org/10.1007/s13225-020-00466-2)) |
| Otu58794 | 1 | k__Fungi(100);p__Ascomycota(100);c__Saccharomycetes(100);o__Saccharomycetales(100);f__Metschnikowiaceae(100);g__Metschnikowia(100);g__Metschnikowia_unclassified(100); | Metschnikowia | 13 | Saprotroph | \|Undefined Saprotroph\| | Yeast | NULL | Probable | As nectar-tap_saprotroph (Põlme et al. 2020); Decay sugar-rich substrates (Põlme et al. 2020) | Põlme S, et al. 2020. Fungal Diversity 105:1-16 ((https://doi.org/10.1007/s13225-020-00466-2)) |
| Otu58798 | 1 | k__Fungi(100);p__Ascomycota(100);c__Saccharomycetes(100);o__Saccharomycetales(100);f__Metschnikowiaceae(100);g__Metschnikowia(100);s__Metschnikowia_pulcherrima(100); | Metschnikowia | 13 | Saprotroph | \|Undefined Saprotroph\| | Yeast | NULL | Probable | As nectar-tap_saprotroph (Põlme et al. 2020); Decay sugar-rich substrates (Põlme et al. 2020) | Põlme S, et al. 2020. Fungal Diversity 105:1-16 ((https://doi.org/10.1007/s13225-020-00466-2)) |
| Otu58800 | 1 | k__Fungi(100);p__Ascomycota(100);c__Saccharomycetes(100);o__Saccharomycetales(100);f__Metschnikowiaceae(100);g__Metschnikowia(100);s__Metschnikowia_pulcherrima(100); | Metschnikowia | 13 | Saprotroph | \|Undefined Saprotroph\| | Yeast | NULL | Probable | As nectar-tap_saprotroph (Põlme et al. 2020); Decay sugar-rich substrates (Põlme et al. 2020) | Põlme S, et al. 2020. Fungal Diversity 105:1-16 ((https://doi.org/10.1007/s13225-020-00466-2)) |
| Otu58801 | 1 | k__Fungi(100);p__Ascomycota(100);c__Saccharomycetes(100);o__Saccharomycetales(100);f__Metschnikowiaceae(100);g__Metschnikowia(100);g__Metschnikowia_unclassified(100); | Metschnikowia | 13 | Saprotroph | \|Undefined Saprotroph\| | Yeast | NULL | Probable | As nectar-tap_saprotroph (Põlme et al. 2020); Decay sugar-rich substrates (Põlme et al. 2020) | Põlme S, et al. 2020. Fungal Diversity 105:1-16 ((https://doi.org/10.1007/s13225-020-00466-2)) |
| Otu58802 | 1 | k__Fungi(100);p__Ascomycota(100);c__Saccharomycetes(100);o__Saccharomycetales(100);f__Metschnikowiaceae(100);g__Metschnikowia(100);s__Metschnikowia_pulcherrima(100); | Metschnikowia | 13 | Saprotroph | \|Undefined Saprotroph\| | Yeast | NULL | Probable | As nectar-tap_saprotroph (Põlme et al. 2020); Decay sugar-rich substrates (Põlme et al. 2020) | Põlme S, et al. 2020. Fungal Diversity 105:1-16 ((https://doi.org/10.1007/s13225-020-00466-2)) |
| Otu58803 | 1 | k__Fungi(100);p__Ascomycota(100);c__Saccharomycetes(100);o__Saccharomycetales(100);f__Metschnikowiaceae(100);g__Metschnikowia(100);g__Metschnikowia_unclassified(100); | Metschnikowia | 13 | Saprotroph | \|Undefined Saprotroph\| | Yeast | NULL | Probable | As nectar-tap_saprotroph (Põlme et al. 2020); Decay sugar-rich substrates (Põlme et al. 2020) | Põlme S, et al. 2020. Fungal Diversity 105:1-16 ((https://doi.org/10.1007/s13225-020-00466-2)) |
| Otu58816 | 1 | k__Fungi(100);p__Ascomycota(100);c__Saccharomycetes(100);o__Saccharomycetales(100);f__Metschnikowiaceae(100);g__Metschnikowia(100);s__Metschnikowia_pulcherrima(100); | Metschnikowia | 13 | Saprotroph | \|Undefined Saprotroph\| | Yeast | NULL | Probable | As nectar-tap_saprotroph (Põlme et al. 2020); Decay sugar-rich substrates (Põlme et al. 2020) | Põlme S, et al. 2020. Fungal Diversity 105:1-16 ((https://doi.org/10.1007/s13225-020-00466-2)) |
| Otu58817 | 1 | k__Fungi(100);p__Ascomycota(100);c__Saccharomycetes(100);o__Saccharomycetales(100);f__Metschnikowiaceae(100);g__Metschnikowia(100);g__Metschnikowia_unclassified(100); | Metschnikowia | 13 | Saprotroph | \|Undefined Saprotroph\| | Yeast | NULL | Probable | As nectar-tap_saprotroph (Põlme et al. 2020); Decay sugar-rich substrates (Põlme et al. 2020) | Põlme S, et al. 2020. Fungal Diversity 105:1-16 ((https://doi.org/10.1007/s13225-020-00466-2)) |
| Otu58818 | 1 | k__Fungi(100);p__Ascomycota(100);c__Saccharomycetes(100);o__Saccharomycetales(100);f__Metschnikowiaceae(100);g__Metschnikowia(100);s__Metschnikowia_pulcherrima(100); | Metschnikowia | 13 | Saprotroph | \|Undefined Saprotroph\| | Yeast | NULL | Probable | As nectar-tap_saprotroph (Põlme et al. 2020); Decay sugar-rich substrates (Põlme et al. 2020) | Põlme S, et al. 2020. Fungal Diversity 105:1-16 ((https://doi.org/10.1007/s13225-020-00466-2)) |
| Otu58819 | 1 | k__Fungi(100);p__Ascomycota(100);c__Saccharomycetes(100);o__Saccharomycetales(100);f__Metschnikowiaceae(100);g__Metschnikowia(100);s__Metschnikowia_pulcherrima(100); | Metschnikowia | 13 | Saprotroph | \|Undefined Saprotroph\| | Yeast | NULL | Probable | As nectar-tap_saprotroph (Põlme et al. 2020); Decay sugar-rich substrates (Põlme et al. 2020) | Põlme S, et al. 2020. Fungal Diversity 105:1-16 ((https://doi.org/10.1007/s13225-020-00466-2)) |
| Otu58820 | 1 | k__Fungi(100);p__Ascomycota(100);c__Saccharomycetes(100);o__Saccharomycetales(100);f__Metschnikowiaceae(100);g__Metschnikowia(100);s__Metschnikowia_pulcherrima(100); | Metschnikowia | 13 | Saprotroph | \|Undefined Saprotroph\| | Yeast | NULL | Probable | As nectar-tap_saprotroph (Põlme et al. 2020); Decay sugar-rich substrates (Põlme et al. 2020) | Põlme S, et al. 2020. Fungal Diversity 105:1-16 ((https://doi.org/10.1007/s13225-020-00466-2)) |
| Otu58821 | 1 | k__Fungi(100);p__Ascomycota(100);c__Saccharomycetes(100);o__Saccharomycetales(100);f__Metschnikowiaceae(100);g__Metschnikowia(100);s__Metschnikowia_pulcherrima(100); | Metschnikowia | 13 | Saprotroph | \|Undefined Saprotroph\| | Yeast | NULL | Probable | As nectar-tap_saprotroph (Põlme et al. 2020); Decay sugar-rich substrates (Põlme et al. 2020) | Põlme S, et al. 2020. Fungal Diversity 105:1-16 ((https://doi.org/10.1007/s13225-020-00466-2)) |
| Otu58822 | 1 | k__Fungi(100);p__Ascomycota(100);c__Saccharomycetes(100);o__Saccharomycetales(100);f__Metschnikowiaceae(100);g__Metschnikowia(100);s__Metschnikowia_pulcherrima(100); | Metschnikowia | 13 | Saprotroph | \|Undefined Saprotroph\| | Yeast | NULL | Probable | As nectar-tap_saprotroph (Põlme et al. 2020); Decay sugar-rich substrates (Põlme et al. 2020) | Põlme S, et al. 2020. Fungal Diversity 105:1-16 ((https://doi.org/10.1007/s13225-020-00466-2)) |
| Otu58823 | 1 | k__Fungi(100);p__Ascomycota(100);c__Saccharomycetes(100);o__Saccharomycetales(100);f__Metschnikowiaceae(100);g__Metschnikowia(100);s__Metschnikowia_pulcherrima(100); | Metschnikowia | 13 | Saprotroph | \|Undefined Saprotroph\| | Yeast | NULL | Probable | As nectar-tap_saprotroph (Põlme et al. 2020); Decay sugar-rich substrates (Põlme et al. 2020) | Põlme S, et al. 2020. Fungal Diversity 105:1-16 ((https://doi.org/10.1007/s13225-020-00466-2)) |
| Otu58824 | 1 | k__Fungi(100);p__Ascomycota(100);c__Saccharomycetes(100);o__Saccharomycetales(100);f__Metschnikowiaceae(100);g__Metschnikowia(100);s__Metschnikowia_pulcherrima(100); | Metschnikowia | 13 | Saprotroph | \|Undefined Saprotroph\| | Yeast | NULL | Probable | As nectar-tap_saprotroph (Põlme et al. 2020); Decay sugar-rich substrates (Põlme et al. 2020) | Põlme S, et al. 2020. Fungal Diversity 105:1-16 ((https://doi.org/10.1007/s13225-020-00466-2)) |
| Otu58826 | 1 | k__Fungi(100);p__Ascomycota(100);c__Saccharomycetes(100);o__Saccharomycetales(100);f__Metschnikowiaceae(100);g__Metschnikowia(100);s__Metschnikowia_pulcherrima(100); | Metschnikowia | 13 | Saprotroph | \|Undefined Saprotroph\| | Yeast | NULL | Probable | As nectar-tap_saprotroph (Põlme et al. 2020); Decay sugar-rich substrates (Põlme et al. 2020) | Põlme S, et al. 2020. Fungal Diversity 105:1-16 ((https://doi.org/10.1007/s13225-020-00466-2)) |
| Otu58827 | 1 | k__Fungi(100);p__Ascomycota(100);c__Saccharomycetes(100);o__Saccharomycetales(100);f__Metschnikowiaceae(100);g__Metschnikowia(100);s__Metschnikowia_pulcherrima(100); | Metschnikowia | 13 | Saprotroph | \|Undefined Saprotroph\| | Yeast | NULL | Probable | As nectar-tap_saprotroph (Põlme et al. 2020); Decay sugar-rich substrates (Põlme et al. 2020) | Põlme S, et al. 2020. Fungal Diversity 105:1-16 ((https://doi.org/10.1007/s13225-020-00466-2)) |
| Otu58829 | 1 | k__Fungi(100);p__Ascomycota(100);c__Saccharomycetes(100);o__Saccharomycetales(100);f__Metschnikowiaceae(100);g__Metschnikowia(100);g__Metschnikowia_unclassified(100); | Metschnikowia | 13 | Saprotroph | \|Undefined Saprotroph\| | Yeast | NULL | Probable | As nectar-tap_saprotroph (Põlme et al. 2020); Decay sugar-rich substrates (Põlme et al. 2020) | Põlme S, et al. 2020. Fungal Diversity 105:1-16 ((https://doi.org/10.1007/s13225-020-00466-2)) |
| Otu58830 | 1 | k__Fungi(100);p__Ascomycota(100);c__Saccharomycetes(100);o__Saccharomycetales(100);f__Metschnikowiaceae(100);g__Metschnikowia(100);s__Metschnikowia_pulcherrima(100); | Metschnikowia | 13 | Saprotroph | \|Undefined Saprotroph\| | Yeast | NULL | Probable | As nectar-tap_saprotroph (Põlme et al. 2020); Decay sugar-rich substrates (Põlme et al. 2020) | Põlme S, et al. 2020. Fungal Diversity 105:1-16 ((https://doi.org/10.1007/s13225-020-00466-2)) |
| Otu58831 | 1 | k__Fungi(100);p__Ascomycota(100);c__Saccharomycetes(100);o__Saccharomycetales(100);f__Metschnikowiaceae(100);g__Metschnikowia(100);s__Metschnikowia_pulcherrima(100); | Metschnikowia | 13 | Saprotroph | \|Undefined Saprotroph\| | Yeast | NULL | Probable | As nectar-tap_saprotroph (Põlme et al. 2020); Decay sugar-rich substrates (Põlme et al. 2020) | Põlme S, et al. 2020. Fungal Diversity 105:1-16 ((https://doi.org/10.1007/s13225-020-00466-2)) |
| Otu58832 | 1 | k__Fungi(100);p__Ascomycota(100);c__Saccharomycetes(100);o__Saccharomycetales(100);f__Metschnikowiaceae(100);g__Metschnikowia(100);s__Metschnikowia_pulcherrima(100); | Metschnikowia | 13 | Saprotroph | \|Undefined Saprotroph\| | Yeast | NULL | Probable | As nectar-tap_saprotroph (Põlme et al. 2020); Decay sugar-rich substrates (Põlme et al. 2020) | Põlme S, et al. 2020. Fungal Diversity 105:1-16 ((https://doi.org/10.1007/s13225-020-00466-2)) |
| Otu58833 | 1 | k__Fungi(100);p__Ascomycota(100);c__Saccharomycetes(100);o__Saccharomycetales(100);f__Metschnikowiaceae(100);g__Metschnikowia(100);s__Metschnikowia_pulcherrima(100); | Metschnikowia | 13 | Saprotroph | \|Undefined Saprotroph\| | Yeast | NULL | Probable | As nectar-tap_saprotroph (Põlme et al. 2020); Decay sugar-rich substrates (Põlme et al. 2020) | Põlme S, et al. 2020. Fungal Diversity 105:1-16 ((https://doi.org/10.1007/s13225-020-00466-2)) |
| Otu58834 | 1 | k__Fungi(100);p__Ascomycota(100);c__Saccharomycetes(100);o__Saccharomycetales(100);f__Metschnikowiaceae(100);g__Metschnikowia(100);g__Metschnikowia_unclassified(100); | Metschnikowia | 13 | Saprotroph | \|Undefined Saprotroph\| | Yeast | NULL | Probable | As nectar-tap_saprotroph (Põlme et al. 2020); Decay sugar-rich substrates (Põlme et al. 2020) | Põlme S, et al. 2020. Fungal Diversity 105:1-16 ((https://doi.org/10.1007/s13225-020-00466-2)) |
| Otu58835 | 1 | k__Fungi(100);p__Ascomycota(100);c__Saccharomycetes(100);o__Saccharomycetales(100);f__Metschnikowiaceae(100);g__Metschnikowia(100);g__Metschnikowia_unclassified(100); | Metschnikowia | 13 | Saprotroph | \|Undefined Saprotroph\| | Yeast | NULL | Probable | As nectar-tap_saprotroph (Põlme et al. 2020); Decay sugar-rich substrates (Põlme et al. 2020) | Põlme S, et al. 2020. Fungal Diversity 105:1-16 ((https://doi.org/10.1007/s13225-020-00466-2)) |
| Otu58836 | 1 | k__Fungi(100);p__Ascomycota(100);c__Saccharomycetes(100);o__Saccharomycetales(100);f__Metschnikowiaceae(100);g__Metschnikowia(100);g__Metschnikowia_unclassified(100); | Metschnikowia | 13 | Saprotroph | \|Undefined Saprotroph\| | Yeast | NULL | Probable | As nectar-tap_saprotroph (Põlme et al. 2020); Decay sugar-rich substrates (Põlme et al. 2020) | Põlme S, et al. 2020. Fungal Diversity 105:1-16 ((https://doi.org/10.1007/s13225-020-00466-2)) |
| Otu58837 | 1 | k__Fungi(100);p__Ascomycota(100);c__Saccharomycetes(100);o__Saccharomycetales(100);f__Metschnikowiaceae(100);g__Metschnikowia(100);s__Metschnikowia_pulcherrima(100); | Metschnikowia | 13 | Saprotroph | \|Undefined Saprotroph\| | Yeast | NULL | Probable | As nectar-tap_saprotroph (Põlme et al. 2020); Decay sugar-rich substrates (Põlme et al. 2020) | Põlme S, et al. 2020. Fungal Diversity 105:1-16 ((https://doi.org/10.1007/s13225-020-00466-2)) |
| Otu58838 | 1 | k__Fungi(100);p__Ascomycota(100);c__Saccharomycetes(100);o__Saccharomycetales(100);f__Metschnikowiaceae(100);g__Metschnikowia(100);s__Metschnikowia_pulcherrima(100); | Metschnikowia | 13 | Saprotroph | \|Undefined Saprotroph\| | Yeast | NULL | Probable | As nectar-tap_saprotroph (Põlme et al. 2020); Decay sugar-rich substrates (Põlme et al. 2020) | Põlme S, et al. 2020. Fungal Diversity 105:1-16 ((https://doi.org/10.1007/s13225-020-00466-2)) |
| Otu58839 | 1 | k__Fungi(100);p__Ascomycota(100);c__Saccharomycetes(100);o__Saccharomycetales(100);f__Metschnikowiaceae(100);g__Metschnikowia(100);g__Metschnikowia_unclassified(100); | Metschnikowia | 13 | Saprotroph | \|Undefined Saprotroph\| | Yeast | NULL | Probable | As nectar-tap_saprotroph (Põlme et al. 2020); Decay sugar-rich substrates (Põlme et al. 2020) | Põlme S, et al. 2020. Fungal Diversity 105:1-16 ((https://doi.org/10.1007/s13225-020-00466-2)) |
| Otu58840 | 1 | k__Fungi(100);p__Ascomycota(100);c__Saccharomycetes(100);o__Saccharomycetales(100);f__Metschnikowiaceae(100);g__Metschnikowia(100);g__Metschnikowia_unclassified(100); | Metschnikowia | 13 | Saprotroph | \|Undefined Saprotroph\| | Yeast | NULL | Probable | As nectar-tap_saprotroph (Põlme et al. 2020); Decay sugar-rich substrates (Põlme et al. 2020) | Põlme S, et al. 2020. Fungal Diversity 105:1-16 ((https://doi.org/10.1007/s13225-020-00466-2)) |
| Otu58842 | 1 | k__Fungi(100);p__Ascomycota(100);c__Saccharomycetes(100);o__Saccharomycetales(100);f__Metschnikowiaceae(100);g__Metschnikowia(100);g__Metschnikowia_unclassified(100); | Metschnikowia | 13 | Saprotroph | \|Undefined Saprotroph\| | Yeast | NULL | Probable | As nectar-tap_saprotroph (Põlme et al. 2020); Decay sugar-rich substrates (Põlme et al. 2020) | Põlme S, et al. 2020. Fungal Diversity 105:1-16 ((https://doi.org/10.1007/s13225-020-00466-2)) |
| Otu58843 | 1 | k__Fungi(100);p__Ascomycota(100);c__Saccharomycetes(100);o__Saccharomycetales(100);f__Metschnikowiaceae(100);g__Metschnikowia(100);s__Metschnikowia_pulcherrima(100); | Metschnikowia | 13 | Saprotroph | \|Undefined Saprotroph\| | Yeast | NULL | Probable | As nectar-tap_saprotroph (Põlme et al. 2020); Decay sugar-rich substrates (Põlme et al. 2020) | Põlme S, et al. 2020. Fungal Diversity 105:1-16 ((https://doi.org/10.1007/s13225-020-00466-2)) |
| Otu58844 | 1 | k__Fungi(100);p__Ascomycota(100);c__Saccharomycetes(100);o__Saccharomycetales(100);f__Metschnikowiaceae(100);g__Metschnikowia(100);s__Metschnikowia_pulcherrima(100); | Metschnikowia | 13 | Saprotroph | \|Undefined Saprotroph\| | Yeast | NULL | Probable | As nectar-tap_saprotroph (Põlme et al. 2020); Decay sugar-rich substrates (Põlme et al. 2020) | Põlme S, et al. 2020. Fungal Diversity 105:1-16 ((https://doi.org/10.1007/s13225-020-00466-2)) |
| Otu58846 | 1 | k__Fungi(100);p__Ascomycota(100);c__Saccharomycetes(100);o__Saccharomycetales(100);f__Metschnikowiaceae(100);g__Metschnikowia(100);s__Metschnikowia_pulcherrima(100); | Metschnikowia | 13 | Saprotroph | \|Undefined Saprotroph\| | Yeast | NULL | Probable | As nectar-tap_saprotroph (Põlme et al. 2020); Decay sugar-rich substrates (Põlme et al. 2020) | Põlme S, et al. 2020. Fungal Diversity 105:1-16 ((https://doi.org/10.1007/s13225-020-00466-2)) |
| Otu58847 | 1 | k__Fungi(100);p__Ascomycota(100);c__Saccharomycetes(100);o__Saccharomycetales(100);f__Metschnikowiaceae(100);g__Metschnikowia(100);s__Metschnikowia_pulcherrima(100); | Metschnikowia | 13 | Saprotroph | \|Undefined Saprotroph\| | Yeast | NULL | Probable | As nectar-tap_saprotroph (Põlme et al. 2020); Decay sugar-rich substrates (Põlme et al. 2020) | Põlme S, et al. 2020. Fungal Diversity 105:1-16 ((https://doi.org/10.1007/s13225-020-00466-2)) |
| Otu58848 | 1 | k__Fungi(100);p__Ascomycota(100);c__Saccharomycetes(100);o__Saccharomycetales(100);f__Metschnikowiaceae(100);g__Metschnikowia(100);s__Metschnikowia_pulcherrima(100); | Metschnikowia | 13 | Saprotroph | \|Undefined Saprotroph\| | Yeast | NULL | Probable | As nectar-tap_saprotroph (Põlme et al. 2020); Decay sugar-rich substrates (Põlme et al. 2020) | Põlme S, et al. 2020. Fungal Diversity 105:1-16 ((https://doi.org/10.1007/s13225-020-00466-2)) |
| Otu58850 | 1 | k__Fungi(100);p__Ascomycota(100);c__Saccharomycetes(100);o__Saccharomycetales(100);f__Metschnikowiaceae(100);g__Metschnikowia(100);s__Metschnikowia_pulcherrima(100); | Metschnikowia | 13 | Saprotroph | \|Undefined Saprotroph\| | Yeast | NULL | Probable | As nectar-tap_saprotroph (Põlme et al. 2020); Decay sugar-rich substrates (Põlme et al. 2020) | Põlme S, et al. 2020. Fungal Diversity 105:1-16 ((https://doi.org/10.1007/s13225-020-00466-2)) |
| Otu58851 | 1 | k__Fungi(100);p__Ascomycota(100);c__Saccharomycetes(100);o__Saccharomycetales(100);f__Metschnikowiaceae(100);g__Metschnikowia(100);s__Metschnikowia_pulcherrima(100); | Metschnikowia | 13 | Saprotroph | \|Undefined Saprotroph\| | Yeast | NULL | Probable | As nectar-tap_saprotroph (Põlme et al. 2020); Decay sugar-rich substrates (Põlme et al. 2020) | Põlme S, et al. 2020. Fungal Diversity 105:1-16 ((https://doi.org/10.1007/s13225-020-00466-2)) |
| Otu58854 | 1 | k__Fungi(100);p__Ascomycota(100);c__Saccharomycetes(100);o__Saccharomycetales(100);f__Metschnikowiaceae(100);g__Metschnikowia(100);g__Metschnikowia_unclassified(100); | Metschnikowia | 13 | Saprotroph | \|Undefined Saprotroph\| | Yeast | NULL | Probable | As nectar-tap_saprotroph (Põlme et al. 2020); Decay sugar-rich substrates (Põlme et al. 2020) | Põlme S, et al. 2020. Fungal Diversity 105:1-16 ((https://doi.org/10.1007/s13225-020-00466-2)) |
| Otu58858 | 1 | k__Fungi(100);p__Ascomycota(100);c__Saccharomycetes(100);o__Saccharomycetales(100);f__Metschnikowiaceae(100);g__Metschnikowia(100);s__Metschnikowia_pulcherrima(100); | Metschnikowia | 13 | Saprotroph | \|Undefined Saprotroph\| | Yeast | NULL | Probable | As nectar-tap_saprotroph (Põlme et al. 2020); Decay sugar-rich substrates (Põlme et al. 2020) | Põlme S, et al. 2020. Fungal Diversity 105:1-16 ((https://doi.org/10.1007/s13225-020-00466-2)) |
| Otu58861 | 1 | k__Fungi(100);p__Ascomycota(100);c__Saccharomycetes(100);o__Saccharomycetales(100);f__Metschnikowiaceae(100);g__Metschnikowia(100);s__Metschnikowia_pulcherrima(100); | Metschnikowia | 13 | Saprotroph | \|Undefined Saprotroph\| | Yeast | NULL | Probable | As nectar-tap_saprotroph (Põlme et al. 2020); Decay sugar-rich substrates (Põlme et al. 2020) | Põlme S, et al. 2020. Fungal Diversity 105:1-16 ((https://doi.org/10.1007/s13225-020-00466-2)) |
| Otu58862 | 1 | k__Fungi(100);p__Ascomycota(100);c__Saccharomycetes(100);o__Saccharomycetales(100);f__Metschnikowiaceae(100);g__Metschnikowia(100);s__Metschnikowia_pulcherrima(100); | Metschnikowia | 13 | Saprotroph | \|Undefined Saprotroph\| | Yeast | NULL | Probable | As nectar-tap_saprotroph (Põlme et al. 2020); Decay sugar-rich substrates (Põlme et al. 2020) | Põlme S, et al. 2020. Fungal Diversity 105:1-16 ((https://doi.org/10.1007/s13225-020-00466-2)) |
| Otu58863 | 1 | k__Fungi(100);p__Ascomycota(100);c__Saccharomycetes(100);o__Saccharomycetales(100);f__Metschnikowiaceae(100);g__Metschnikowia(100);s__Metschnikowia_pulcherrima(100); | Metschnikowia | 13 | Saprotroph | \|Undefined Saprotroph\| | Yeast | NULL | Probable | As nectar-tap_saprotroph (Põlme et al. 2020); Decay sugar-rich substrates (Põlme et al. 2020) | Põlme S, et al. 2020. Fungal Diversity 105:1-16 ((https://doi.org/10.1007/s13225-020-00466-2)) |
| Otu58867 | 1 | k__Fungi(100);p__Ascomycota(100);c__Saccharomycetes(100);o__Saccharomycetales(100);f__Metschnikowiaceae(100);g__Metschnikowia(100);s__Metschnikowia_pulcherrima(100); | Metschnikowia | 13 | Saprotroph | \|Undefined Saprotroph\| | Yeast | NULL | Probable | As nectar-tap_saprotroph (Põlme et al. 2020); Decay sugar-rich substrates (Põlme et al. 2020) | Põlme S, et al. 2020. Fungal Diversity 105:1-16 ((https://doi.org/10.1007/s13225-020-00466-2)) |
| Otu58868 | 1 | k__Fungi(100);p__Ascomycota(100);c__Saccharomycetes(100);o__Saccharomycetales(100);f__Metschnikowiaceae(100);g__Metschnikowia(100);g__Metschnikowia_unclassified(100); | Metschnikowia | 13 | Saprotroph | \|Undefined Saprotroph\| | Yeast | NULL | Probable | As nectar-tap_saprotroph (Põlme et al. 2020); Decay sugar-rich substrates (Põlme et al. 2020) | Põlme S, et al. 2020. Fungal Diversity 105:1-16 ((https://doi.org/10.1007/s13225-020-00466-2)) |
| Otu58869 | 1 | k__Fungi(100);p__Ascomycota(100);c__Saccharomycetes(100);o__Saccharomycetales(100);f__Metschnikowiaceae(100);g__Metschnikowia(100);s__Metschnikowia_pulcherrima(100); | Metschnikowia | 13 | Saprotroph | \|Undefined Saprotroph\| | Yeast | NULL | Probable | As nectar-tap_saprotroph (Põlme et al. 2020); Decay sugar-rich substrates (Põlme et al. 2020) | Põlme S, et al. 2020. Fungal Diversity 105:1-16 ((https://doi.org/10.1007/s13225-020-00466-2)) |
| Otu58871 | 1 | k__Fungi(100);p__Ascomycota(100);c__Saccharomycetes(100);o__Saccharomycetales(100);f__Metschnikowiaceae(100);g__Metschnikowia(100);s__Metschnikowia_pulcherrima(100); | Metschnikowia | 13 | Saprotroph | \|Undefined Saprotroph\| | Yeast | NULL | Probable | As nectar-tap_saprotroph (Põlme et al. 2020); Decay sugar-rich substrates (Põlme et al. 2020) | Põlme S, et al. 2020. Fungal Diversity 105:1-16 ((https://doi.org/10.1007/s13225-020-00466-2)) |
| Otu58872 | 1 | k__Fungi(100);p__Ascomycota(100);c__Saccharomycetes(100);o__Saccharomycetales(100);f__Metschnikowiaceae(100);g__Metschnikowia(100);g__Metschnikowia_unclassified(100); | Metschnikowia | 13 | Saprotroph | \|Undefined Saprotroph\| | Yeast | NULL | Probable | As nectar-tap_saprotroph (Põlme et al. 2020); Decay sugar-rich substrates (Põlme et al. 2020) | Põlme S, et al. 2020. Fungal Diversity 105:1-16 ((https://doi.org/10.1007/s13225-020-00466-2)) |
| Otu58873 | 1 | k__Fungi(100);p__Ascomycota(100);c__Saccharomycetes(100);o__Saccharomycetales(100);f__Metschnikowiaceae(100);g__Metschnikowia(100);s__Metschnikowia_pulcherrima(100); | Metschnikowia | 13 | Saprotroph | \|Undefined Saprotroph\| | Yeast | NULL | Probable | As nectar-tap_saprotroph (Põlme et al. 2020); Decay sugar-rich substrates (Põlme et al. 2020) | Põlme S, et al. 2020. Fungal Diversity 105:1-16 ((https://doi.org/10.1007/s13225-020-00466-2)) |
| Otu58874 | 1 | k__Fungi(100);p__Ascomycota(100);c__Saccharomycetes(100);o__Saccharomycetales(100);f__Metschnikowiaceae(100);g__Metschnikowia(100);g__Metschnikowia_unclassified(100); | Metschnikowia | 13 | Saprotroph | \|Undefined Saprotroph\| | Yeast | NULL | Probable | As nectar-tap_saprotroph (Põlme et al. 2020); Decay sugar-rich substrates (Põlme et al. 2020) | Põlme S, et al. 2020. Fungal Diversity 105:1-16 ((https://doi.org/10.1007/s13225-020-00466-2)) |
| Otu58875 | 1 | k__Fungi(100);p__Ascomycota(100);c__Saccharomycetes(100);o__Saccharomycetales(100);f__Metschnikowiaceae(100);g__Metschnikowia(100);g__Metschnikowia_unclassified(100); | Metschnikowia | 13 | Saprotroph | \|Undefined Saprotroph\| | Yeast | NULL | Probable | As nectar-tap_saprotroph (Põlme et al. 2020); Decay sugar-rich substrates (Põlme et al. 2020) | Põlme S, et al. 2020. Fungal Diversity 105:1-16 ((https://doi.org/10.1007/s13225-020-00466-2)) |
| Otu58876 | 1 | k__Fungi(100);p__Ascomycota(100);c__Saccharomycetes(100);o__Saccharomycetales(100);f__Metschnikowiaceae(100);g__Metschnikowia(100);s__Metschnikowia_pulcherrima(100); | Metschnikowia | 13 | Saprotroph | \|Undefined Saprotroph\| | Yeast | NULL | Probable | As nectar-tap_saprotroph (Põlme et al. 2020); Decay sugar-rich substrates (Põlme et al. 2020) | Põlme S, et al. 2020. Fungal Diversity 105:1-16 ((https://doi.org/10.1007/s13225-020-00466-2)) |
| Otu58877 | 1 | k__Fungi(100);p__Ascomycota(100);c__Saccharomycetes(100);o__Saccharomycetales(100);f__Metschnikowiaceae(100);g__Metschnikowia(100);s__Metschnikowia_pulcherrima(100); | Metschnikowia | 13 | Saprotroph | \|Undefined Saprotroph\| | Yeast | NULL | Probable | As nectar-tap_saprotroph (Põlme et al. 2020); Decay sugar-rich substrates (Põlme et al. 2020) | Põlme S, et al. 2020. Fungal Diversity 105:1-16 ((https://doi.org/10.1007/s13225-020-00466-2)) |
| Otu58879 | 1 | k__Fungi(100);p__Ascomycota(100);c__Saccharomycetes(100);o__Saccharomycetales(100);f__Metschnikowiaceae(100);g__Metschnikowia(100);s__Metschnikowia_pulcherrima(100); | Metschnikowia | 13 | Saprotroph | \|Undefined Saprotroph\| | Yeast | NULL | Probable | As nectar-tap_saprotroph (Põlme et al. 2020); Decay sugar-rich substrates (Põlme et al. 2020) | Põlme S, et al. 2020. Fungal Diversity 105:1-16 ((https://doi.org/10.1007/s13225-020-00466-2)) |
| Otu58881 | 1 | k__Fungi(100);p__Ascomycota(100);c__Saccharomycetes(100);o__Saccharomycetales(100);f__Metschnikowiaceae(100);g__Metschnikowia(100);s__Metschnikowia_pulcherrima(100); | Metschnikowia | 13 | Saprotroph | \|Undefined Saprotroph\| | Yeast | NULL | Probable | As nectar-tap_saprotroph (Põlme et al. 2020); Decay sugar-rich substrates (Põlme et al. 2020) | Põlme S, et al. 2020. Fungal Diversity 105:1-16 ((https://doi.org/10.1007/s13225-020-00466-2)) |
| Otu58882 | 1 | k__Fungi(100);p__Ascomycota(100);c__Saccharomycetes(100);o__Saccharomycetales(100);f__Metschnikowiaceae(100);g__Metschnikowia(100);s__Metschnikowia_pulcherrima(100); | Metschnikowia | 13 | Saprotroph | \|Undefined Saprotroph\| | Yeast | NULL | Probable | As nectar-tap_saprotroph (Põlme et al. 2020); Decay sugar-rich substrates (Põlme et al. 2020) | Põlme S, et al. 2020. Fungal Diversity 105:1-16 ((https://doi.org/10.1007/s13225-020-00466-2)) |
| Otu58883 | 1 | k__Fungi(100);p__Ascomycota(100);c__Saccharomycetes(100);o__Saccharomycetales(100);f__Metschnikowiaceae(100);g__Metschnikowia(100);g__Metschnikowia_unclassified(100); | Metschnikowia | 13 | Saprotroph | \|Undefined Saprotroph\| | Yeast | NULL | Probable | As nectar-tap_saprotroph (Põlme et al. 2020); Decay sugar-rich substrates (Põlme et al. 2020) | Põlme S, et al. 2020. Fungal Diversity 105:1-16 ((https://doi.org/10.1007/s13225-020-00466-2)) |
| Otu58884 | 1 | k__Fungi(100);p__Ascomycota(100);c__Saccharomycetes(100);o__Saccharomycetales(100);f__Metschnikowiaceae(100);g__Metschnikowia(100);s__Metschnikowia_pulcherrima(100); | Metschnikowia | 13 | Saprotroph | \|Undefined Saprotroph\| | Yeast | NULL | Probable | As nectar-tap_saprotroph (Põlme et al. 2020); Decay sugar-rich substrates (Põlme et al. 2020) | Põlme S, et al. 2020. Fungal Diversity 105:1-16 ((https://doi.org/10.1007/s13225-020-00466-2)) |
| Otu58885 | 1 | k__Fungi(100);p__Ascomycota(100);c__Saccharomycetes(100);o__Saccharomycetales(100);f__Metschnikowiaceae(100);g__Metschnikowia(100);s__Metschnikowia_pulcherrima(100); | Metschnikowia | 13 | Saprotroph | \|Undefined Saprotroph\| | Yeast | NULL | Probable | As nectar-tap_saprotroph (Põlme et al. 2020); Decay sugar-rich substrates (Põlme et al. 2020) | Põlme S, et al. 2020. Fungal Diversity 105:1-16 ((https://doi.org/10.1007/s13225-020-00466-2)) |
| Otu58886 | 1 | k__Fungi(100);p__Ascomycota(100);c__Saccharomycetes(100);o__Saccharomycetales(100);f__Metschnikowiaceae(100);g__Metschnikowia(100);s__Metschnikowia_pulcherrima(100); | Metschnikowia | 13 | Saprotroph | \|Undefined Saprotroph\| | Yeast | NULL | Probable | As nectar-tap_saprotroph (Põlme et al. 2020); Decay sugar-rich substrates (Põlme et al. 2020) | Põlme S, et al. 2020. Fungal Diversity 105:1-16 ((https://doi.org/10.1007/s13225-020-00466-2)) |
| Otu58887 | 1 | k__Fungi(100);p__Ascomycota(100);c__Saccharomycetes(100);o__Saccharomycetales(100);f__Metschnikowiaceae(100);g__Metschnikowia(100);g__Metschnikowia_unclassified(100); | Metschnikowia | 13 | Saprotroph | \|Undefined Saprotroph\| | Yeast | NULL | Probable | As nectar-tap_saprotroph (Põlme et al. 2020); Decay sugar-rich substrates (Põlme et al. 2020) | Põlme S, et al. 2020. Fungal Diversity 105:1-16 ((https://doi.org/10.1007/s13225-020-00466-2)) |
| Otu58888 | 1 | k__Fungi(100);p__Ascomycota(100);c__Saccharomycetes(100);o__Saccharomycetales(100);f__Metschnikowiaceae(100);g__Metschnikowia(100);s__Metschnikowia_pulcherrima(100); | Metschnikowia | 13 | Saprotroph | \|Undefined Saprotroph\| | Yeast | NULL | Probable | As nectar-tap_saprotroph (Põlme et al. 2020); Decay sugar-rich substrates (Põlme et al. 2020) | Põlme S, et al. 2020. Fungal Diversity 105:1-16 ((https://doi.org/10.1007/s13225-020-00466-2)) |
| Otu58889 | 1 | k__Fungi(100);p__Ascomycota(100);c__Saccharomycetes(100);o__Saccharomycetales(100);f__Metschnikowiaceae(100);g__Metschnikowia(100);s__Metschnikowia_pulcherrima(100); | Metschnikowia | 13 | Saprotroph | \|Undefined Saprotroph\| | Yeast | NULL | Probable | As nectar-tap_saprotroph (Põlme et al. 2020); Decay sugar-rich substrates (Põlme et al. 2020) | Põlme S, et al. 2020. Fungal Diversity 105:1-16 ((https://doi.org/10.1007/s13225-020-00466-2)) |
| Otu58890 | 1 | k__Fungi(100);p__Ascomycota(100);c__Saccharomycetes(100);o__Saccharomycetales(100);f__Metschnikowiaceae(100);g__Metschnikowia(100);s__Metschnikowia_pulcherrima(100); | Metschnikowia | 13 | Saprotroph | \|Undefined Saprotroph\| | Yeast | NULL | Probable | As nectar-tap_saprotroph (Põlme et al. 2020); Decay sugar-rich substrates (Põlme et al. 2020) | Põlme S, et al. 2020. Fungal Diversity 105:1-16 ((https://doi.org/10.1007/s13225-020-00466-2)) |
| Otu58891 | 1 | k__Fungi(100);p__Ascomycota(100);c__Saccharomycetes(100);o__Saccharomycetales(100);f__Metschnikowiaceae(100);g__Metschnikowia(100);s__Metschnikowia_pulcherrima(100); | Metschnikowia | 13 | Saprotroph | \|Undefined Saprotroph\| | Yeast | NULL | Probable | As nectar-tap_saprotroph (Põlme et al. 2020); Decay sugar-rich substrates (Põlme et al. 2020) | Põlme S, et al. 2020. Fungal Diversity 105:1-16 ((https://doi.org/10.1007/s13225-020-00466-2)) |
| Otu58892 | 1 | k__Fungi(100);p__Ascomycota(100);c__Saccharomycetes(100);o__Saccharomycetales(100);f__Metschnikowiaceae(100);g__Metschnikowia(100);g__Metschnikowia_unclassified(100); | Metschnikowia | 13 | Saprotroph | \|Undefined Saprotroph\| | Yeast | NULL | Probable | As nectar-tap_saprotroph (Põlme et al. 2020); Decay sugar-rich substrates (Põlme et al. 2020) | Põlme S, et al. 2020. Fungal Diversity 105:1-16 ((https://doi.org/10.1007/s13225-020-00466-2)) |
| Otu58893 | 1 | k__Fungi(100);p__Ascomycota(100);c__Saccharomycetes(100);o__Saccharomycetales(100);f__Metschnikowiaceae(100);g__Metschnikowia(100);s__Metschnikowia_pulcherrima(100); | Metschnikowia | 13 | Saprotroph | \|Undefined Saprotroph\| | Yeast | NULL | Probable | As nectar-tap_saprotroph (Põlme et al. 2020); Decay sugar-rich substrates (Põlme et al. 2020) | Põlme S, et al. 2020. Fungal Diversity 105:1-16 ((https://doi.org/10.1007/s13225-020-00466-2)) |
| Otu58894 | 1 | k__Fungi(100);p__Ascomycota(100);c__Saccharomycetes(100);o__Saccharomycetales(100);f__Metschnikowiaceae(100);g__Metschnikowia(100);g__Metschnikowia_unclassified(100); | Metschnikowia | 13 | Saprotroph | \|Undefined Saprotroph\| | Yeast | NULL | Probable | As nectar-tap_saprotroph (Põlme et al. 2020); Decay sugar-rich substrates (Põlme et al. 2020) | Põlme S, et al. 2020. Fungal Diversity 105:1-16 ((https://doi.org/10.1007/s13225-020-00466-2)) |
| Otu58895 | 1 | k__Fungi(100);p__Ascomycota(100);c__Saccharomycetes(100);o__Saccharomycetales(100);f__Metschnikowiaceae(100);g__Metschnikowia(100);g__Metschnikowia_unclassified(100); | Metschnikowia | 13 | Saprotroph | \|Undefined Saprotroph\| | Yeast | NULL | Probable | As nectar-tap_saprotroph (Põlme et al. 2020); Decay sugar-rich substrates (Põlme et al. 2020) | Põlme S, et al. 2020. Fungal Diversity 105:1-16 ((https://doi.org/10.1007/s13225-020-00466-2)) |
| Otu58896 | 1 | k__Fungi(100);p__Ascomycota(100);c__Saccharomycetes(100);o__Saccharomycetales(100);f__Metschnikowiaceae(100);g__Metschnikowia(100);g__Metschnikowia_unclassified(100); | Metschnikowia | 13 | Saprotroph | \|Undefined Saprotroph\| | Yeast | NULL | Probable | As nectar-tap_saprotroph (Põlme et al. 2020); Decay sugar-rich substrates (Põlme et al. 2020) | Põlme S, et al. 2020. Fungal Diversity 105:1-16 ((https://doi.org/10.1007/s13225-020-00466-2)) |
| Otu58897 | 1 | k__Fungi(100);p__Ascomycota(100);c__Saccharomycetes(100);o__Saccharomycetales(100);f__Metschnikowiaceae(100);g__Metschnikowia(100);g__Metschnikowia_unclassified(100); | Metschnikowia | 13 | Saprotroph | \|Undefined Saprotroph\| | Yeast | NULL | Probable | As nectar-tap_saprotroph (Põlme et al. 2020); Decay sugar-rich substrates (Põlme et al. 2020) | Põlme S, et al. 2020. Fungal Diversity 105:1-16 ((https://doi.org/10.1007/s13225-020-00466-2)) |
| Otu58898 | 1 | k__Fungi(100);p__Ascomycota(100);c__Saccharomycetes(100);o__Saccharomycetales(100);f__Metschnikowiaceae(100);g__Metschnikowia(100);g__Metschnikowia_unclassified(100); | Metschnikowia | 13 | Saprotroph | \|Undefined Saprotroph\| | Yeast | NULL | Probable | As nectar-tap_saprotroph (Põlme et al. 2020); Decay sugar-rich substrates (Põlme et al. 2020) | Põlme S, et al. 2020. Fungal Diversity 105:1-16 ((https://doi.org/10.1007/s13225-020-00466-2)) |
| Otu58899 | 1 | k__Fungi(100);p__Ascomycota(100);c__Saccharomycetes(100);o__Saccharomycetales(100);f__Metschnikowiaceae(100);g__Metschnikowia(100);g__Metschnikowia_unclassified(100); | Metschnikowia | 13 | Saprotroph | \|Undefined Saprotroph\| | Yeast | NULL | Probable | As nectar-tap_saprotroph (Põlme et al. 2020); Decay sugar-rich substrates (Põlme et al. 2020) | Põlme S, et al. 2020. Fungal Diversity 105:1-16 ((https://doi.org/10.1007/s13225-020-00466-2)) |
| Otu58900 | 1 | k__Fungi(100);p__Ascomycota(100);c__Saccharomycetes(100);o__Saccharomycetales(100);f__Metschnikowiaceae(100);g__Metschnikowia(100);s__Metschnikowia_pulcherrima(100); | Metschnikowia | 13 | Saprotroph | \|Undefined Saprotroph\| | Yeast | NULL | Probable | As nectar-tap_saprotroph (Põlme et al. 2020); Decay sugar-rich substrates (Põlme et al. 2020) | Põlme S, et al. 2020. Fungal Diversity 105:1-16 ((https://doi.org/10.1007/s13225-020-00466-2)) |
| Otu58901 | 1 | k__Fungi(100);p__Ascomycota(100);c__Saccharomycetes(100);o__Saccharomycetales(100);f__Metschnikowiaceae(100);g__Metschnikowia(100);s__Metschnikowia_pulcherrima(100); | Metschnikowia | 13 | Saprotroph | \|Undefined Saprotroph\| | Yeast | NULL | Probable | As nectar-tap_saprotroph (Põlme et al. 2020); Decay sugar-rich substrates (Põlme et al. 2020) | Põlme S, et al. 2020. Fungal Diversity 105:1-16 ((https://doi.org/10.1007/s13225-020-00466-2)) |
| Otu58903 | 1 | k__Fungi(100);p__Ascomycota(100);c__Saccharomycetes(100);o__Saccharomycetales(100);f__Metschnikowiaceae(100);g__Metschnikowia(100);s__Metschnikowia_pulcherrima(100); | Metschnikowia | 13 | Saprotroph | \|Undefined Saprotroph\| | Yeast | NULL | Probable | As nectar-tap_saprotroph (Põlme et al. 2020); Decay sugar-rich substrates (Põlme et al. 2020) | Põlme S, et al. 2020. Fungal Diversity 105:1-16 ((https://doi.org/10.1007/s13225-020-00466-2)) |
| Otu58904 | 1 | k__Fungi(100);p__Ascomycota(100);c__Saccharomycetes(100);o__Saccharomycetales(100);f__Metschnikowiaceae(100);g__Metschnikowia(100);s__Metschnikowia_pulcherrima(100); | Metschnikowia | 13 | Saprotroph | \|Undefined Saprotroph\| | Yeast | NULL | Probable | As nectar-tap_saprotroph (Põlme et al. 2020); Decay sugar-rich substrates (Põlme et al. 2020) | Põlme S, et al. 2020. Fungal Diversity 105:1-16 ((https://doi.org/10.1007/s13225-020-00466-2)) |
| Otu58906 | 1 | k__Fungi(100);p__Ascomycota(100);c__Saccharomycetes(100);o__Saccharomycetales(100);f__Metschnikowiaceae(100);g__Metschnikowia(100);s__Metschnikowia_pulcherrima(100); | Metschnikowia | 13 | Saprotroph | \|Undefined Saprotroph\| | Yeast | NULL | Probable | As nectar-tap_saprotroph (Põlme et al. 2020); Decay sugar-rich substrates (Põlme et al. 2020) | Põlme S, et al. 2020. Fungal Diversity 105:1-16 ((https://doi.org/10.1007/s13225-020-00466-2)) |
| Otu58907 | 1 | k__Fungi(100);p__Ascomycota(100);c__Saccharomycetes(100);o__Saccharomycetales(100);f__Metschnikowiaceae(100);g__Metschnikowia(100);g__Metschnikowia_unclassified(100); | Metschnikowia | 13 | Saprotroph | \|Undefined Saprotroph\| | Yeast | NULL | Probable | As nectar-tap_saprotroph (Põlme et al. 2020); Decay sugar-rich substrates (Põlme et al. 2020) | Põlme S, et al. 2020. Fungal Diversity 105:1-16 ((https://doi.org/10.1007/s13225-020-00466-2)) |
| Otu58908 | 1 | k__Fungi(100);p__Ascomycota(100);c__Saccharomycetes(100);o__Saccharomycetales(100);f__Metschnikowiaceae(100);g__Metschnikowia(100);s__Metschnikowia_pulcherrima(100); | Metschnikowia | 13 | Saprotroph | \|Undefined Saprotroph\| | Yeast | NULL | Probable | As nectar-tap_saprotroph (Põlme et al. 2020); Decay sugar-rich substrates (Põlme et al. 2020) | Põlme S, et al. 2020. Fungal Diversity 105:1-16 ((https://doi.org/10.1007/s13225-020-00466-2)) |
| Otu58909 | 1 | k__Fungi(100);p__Ascomycota(100);c__Saccharomycetes(100);o__Saccharomycetales(100);f__Metschnikowiaceae(100);g__Metschnikowia(100);g__Metschnikowia_unclassified(100); | Metschnikowia | 13 | Saprotroph | \|Undefined Saprotroph\| | Yeast | NULL | Probable | As nectar-tap_saprotroph (Põlme et al. 2020); Decay sugar-rich substrates (Põlme et al. 2020) | Põlme S, et al. 2020. Fungal Diversity 105:1-16 ((https://doi.org/10.1007/s13225-020-00466-2)) |
| Otu58910 | 1 | k__Fungi(100);p__Ascomycota(100);c__Saccharomycetes(100);o__Saccharomycetales(100);f__Metschnikowiaceae(100);g__Metschnikowia(100);g__Metschnikowia_unclassified(100); | Metschnikowia | 13 | Saprotroph | \|Undefined Saprotroph\| | Yeast | NULL | Probable | As nectar-tap_saprotroph (Põlme et al. 2020); Decay sugar-rich substrates (Põlme et al. 2020) | Põlme S, et al. 2020. Fungal Diversity 105:1-16 ((https://doi.org/10.1007/s13225-020-00466-2)) |
| Otu58911 | 1 | k__Fungi(100);p__Ascomycota(100);c__Saccharomycetes(100);o__Saccharomycetales(100);f__Metschnikowiaceae(100);g__Metschnikowia(100);g__Metschnikowia_unclassified(100); | Metschnikowia | 13 | Saprotroph | \|Undefined Saprotroph\| | Yeast | NULL | Probable | As nectar-tap_saprotroph (Põlme et al. 2020); Decay sugar-rich substrates (Põlme et al. 2020) | Põlme S, et al. 2020. Fungal Diversity 105:1-16 ((https://doi.org/10.1007/s13225-020-00466-2)) |
| Otu58912 | 1 | k__Fungi(100);p__Ascomycota(100);c__Saccharomycetes(100);o__Saccharomycetales(100);f__Metschnikowiaceae(100);g__Metschnikowia(100);s__Metschnikowia_pulcherrima(100); | Metschnikowia | 13 | Saprotroph | \|Undefined Saprotroph\| | Yeast | NULL | Probable | As nectar-tap_saprotroph (Põlme et al. 2020); Decay sugar-rich substrates (Põlme et al. 2020) | Põlme S, et al. 2020. Fungal Diversity 105:1-16 ((https://doi.org/10.1007/s13225-020-00466-2)) |
| Otu58913 | 1 | k__Fungi(100);p__Ascomycota(100);c__Saccharomycetes(100);o__Saccharomycetales(100);f__Metschnikowiaceae(100);g__Metschnikowia(100);s__Metschnikowia_pulcherrima(100); | Metschnikowia | 13 | Saprotroph | \|Undefined Saprotroph\| | Yeast | NULL | Probable | As nectar-tap_saprotroph (Põlme et al. 2020); Decay sugar-rich substrates (Põlme et al. 2020) | Põlme S, et al. 2020. Fungal Diversity 105:1-16 ((https://doi.org/10.1007/s13225-020-00466-2)) |
| Otu58914 | 1 | k__Fungi(100);p__Ascomycota(100);c__Saccharomycetes(100);o__Saccharomycetales(100);f__Metschnikowiaceae(100);g__Metschnikowia(100);s__Metschnikowia_pulcherrima(100); | Metschnikowia | 13 | Saprotroph | \|Undefined Saprotroph\| | Yeast | NULL | Probable | As nectar-tap_saprotroph (Põlme et al. 2020); Decay sugar-rich substrates (Põlme et al. 2020) | Põlme S, et al. 2020. Fungal Diversity 105:1-16 ((https://doi.org/10.1007/s13225-020-00466-2)) |
| Otu58915 | 1 | k__Fungi(100);p__Ascomycota(100);c__Saccharomycetes(100);o__Saccharomycetales(100);f__Metschnikowiaceae(100);g__Metschnikowia(100);s__Metschnikowia_pulcherrima(100); | Metschnikowia | 13 | Saprotroph | \|Undefined Saprotroph\| | Yeast | NULL | Probable | As nectar-tap_saprotroph (Põlme et al. 2020); Decay sugar-rich substrates (Põlme et al. 2020) | Põlme S, et al. 2020. Fungal Diversity 105:1-16 ((https://doi.org/10.1007/s13225-020-00466-2)) |
| Otu58916 | 1 | k__Fungi(100);p__Ascomycota(100);c__Saccharomycetes(100);o__Saccharomycetales(100);f__Metschnikowiaceae(100);g__Metschnikowia(100);g__Metschnikowia_unclassified(100); | Metschnikowia | 13 | Saprotroph | \|Undefined Saprotroph\| | Yeast | NULL | Probable | As nectar-tap_saprotroph (Põlme et al. 2020); Decay sugar-rich substrates (Põlme et al. 2020) | Põlme S, et al. 2020. Fungal Diversity 105:1-16 ((https://doi.org/10.1007/s13225-020-00466-2)) |
| Otu58917 | 1 | k__Fungi(100);p__Ascomycota(100);c__Saccharomycetes(100);o__Saccharomycetales(100);f__Metschnikowiaceae(100);g__Metschnikowia(100);s__Metschnikowia_pulcherrima(100); | Metschnikowia | 13 | Saprotroph | \|Undefined Saprotroph\| | Yeast | NULL | Probable | As nectar-tap_saprotroph (Põlme et al. 2020); Decay sugar-rich substrates (Põlme et al. 2020) | Põlme S, et al. 2020. Fungal Diversity 105:1-16 ((https://doi.org/10.1007/s13225-020-00466-2)) |
| Otu58919 | 1 | k__Fungi(100);p__Ascomycota(100);c__Saccharomycetes(100);o__Saccharomycetales(100);f__Metschnikowiaceae(100);g__Metschnikowia(100);s__Metschnikowia_pulcherrima(100); | Metschnikowia | 13 | Saprotroph | \|Undefined Saprotroph\| | Yeast | NULL | Probable | As nectar-tap_saprotroph (Põlme et al. 2020); Decay sugar-rich substrates (Põlme et al. 2020) | Põlme S, et al. 2020. Fungal Diversity 105:1-16 ((https://doi.org/10.1007/s13225-020-00466-2)) |
| Otu58920 | 1 | k__Fungi(100);p__Ascomycota(100);c__Saccharomycetes(100);o__Saccharomycetales(100);f__Metschnikowiaceae(100);g__Metschnikowia(100);g__Metschnikowia_unclassified(100); | Metschnikowia | 13 | Saprotroph | \|Undefined Saprotroph\| | Yeast | NULL | Probable | As nectar-tap_saprotroph (Põlme et al. 2020); Decay sugar-rich substrates (Põlme et al. 2020) | Põlme S, et al. 2020. Fungal Diversity 105:1-16 ((https://doi.org/10.1007/s13225-020-00466-2)) |
| Otu58921 | 1 | k__Fungi(100);p__Ascomycota(100);c__Saccharomycetes(100);o__Saccharomycetales(100);f__Metschnikowiaceae(100);g__Metschnikowia(100);s__Metschnikowia_pulcherrima(100); | Metschnikowia | 13 | Saprotroph | \|Undefined Saprotroph\| | Yeast | NULL | Probable | As nectar-tap_saprotroph (Põlme et al. 2020); Decay sugar-rich substrates (Põlme et al. 2020) | Põlme S, et al. 2020. Fungal Diversity 105:1-16 ((https://doi.org/10.1007/s13225-020-00466-2)) |
| Otu58922 | 1 | k__Fungi(100);p__Ascomycota(100);c__Saccharomycetes(100);o__Saccharomycetales(100);f__Metschnikowiaceae(100);g__Metschnikowia(100);s__Metschnikowia_pulcherrima(100); | Metschnikowia | 13 | Saprotroph | \|Undefined Saprotroph\| | Yeast | NULL | Probable | As nectar-tap_saprotroph (Põlme et al. 2020); Decay sugar-rich substrates (Põlme et al. 2020) | Põlme S, et al. 2020. Fungal Diversity 105:1-16 ((https://doi.org/10.1007/s13225-020-00466-2)) |

| | | | | | | | | | | | |
|---|---|---|---|---|---|---|---|---|---|---|---|
| Otu58923 | 1 | k__Fungi(100);p__Ascomycota(100);c__Saccharomycetes(100);o__Saccharomycetales(100);f__Metschnikowiaceae(100);g__Metschnikowia(100);s__Metschnikowia_pulcherrima(100); | Metschnikowia | 13 | Saprotroph | \|Undefined Saprotroph\| | Yeast | NULL | Probable | As nectar-tap_saprotroph (Põlme et al. 2020); Decay sugar-rich substrates (Põlme et al. 2020) | Põlme S, et al. 2020. Fungal Diversity 105:1-16 ((https://doi.org/10.1007/s13225-020-00466-2)) |
| Otu58924 | 1 | k__Fungi(100);p__Ascomycota(100);c__Saccharomycetes(100);o__Saccharomycetales(100);f__Metschnikowiaceae(100);g__Metschnikowia(100);s__Metschnikowia_pulcherrima(100); | Metschnikowia | 13 | Saprotroph | \|Undefined Saprotroph\| | Yeast | NULL | Probable | As nectar-tap_saprotroph (Põlme et al. 2020); Decay sugar-rich substrates (Põlme et al. 2020) | Põlme S, et al. 2020. Fungal Diversity 105:1-16 ((https://doi.org/10.1007/s13225-020-00466-2)) |
| Otu58925 | 1 | k__Fungi(100);p__Ascomycota(100);c__Saccharomycetes(100);o__Saccharomycetales(100);f__Metschnikowiaceae(100);g__Metschnikowia(100);s__Metschnikowia_pulcherrima(100); | Metschnikowia | 13 | Saprotroph | \|Undefined Saprotroph\| | Yeast | NULL | Probable | As nectar-tap_saprotroph (Põlme et al. 2020); Decay sugar-rich substrates (Põlme et al. 2020) | Põlme S, et al. 2020. Fungal Diversity 105:1-16 ((https://doi.org/10.1007/s13225-020-00466-2)) |
| Otu58926 | 1 | k__Fungi(100);p__Ascomycota(100);c__Saccharomycetes(100);o__Saccharomycetales(100);f__Metschnikowiaceae(100);g__Metschnikowia(100);g__Metschnikowia_unclassified(100); | Metschnikowia | 13 | Saprotroph | \|Undefined Saprotroph\| | Yeast | NULL | Probable | As nectar-tap_saprotroph (Põlme et al. 2020); Decay sugar-rich substrates (Põlme et al. 2020) | Põlme S, et al. 2020. Fungal Diversity 105:1-16 ((https://doi.org/10.1007/s13225-020-00466-2)) |
| Otu58927 | 1 | k__Fungi(100);p__Ascomycota(100);c__Saccharomycetes(100);o__Saccharomycetales(100);f__Metschnikowiaceae(100);g__Metschnikowia(100);g__Metschnikowia_unclassified(100); | Metschnikowia | 13 | Saprotroph | \|Undefined Saprotroph\| | Yeast | NULL | Probable | As nectar-tap_saprotroph (Põlme et al. 2020); Decay sugar-rich substrates (Põlme et al. 2020) | Põlme S, et al. 2020. Fungal Diversity 105:1-16 ((https://doi.org/10.1007/s13225-020-00466-2)) |
| Otu58928 | 1 | k__Fungi(100);p__Ascomycota(100);c__Saccharomycetes(100);o__Saccharomycetales(100);f__Metschnikowiaceae(100);g__Metschnikowia(100);s__Metschnikowia_pulcherrima(100); | Metschnikowia | 13 | Saprotroph | \|Undefined Saprotroph\| | Yeast | NULL | Probable | As nectar-tap_saprotroph (Põlme et al. 2020); Decay sugar-rich substrates (Põlme et al. 2020) | Põlme S, et al. 2020. Fungal Diversity 105:1-16 ((https://doi.org/10.1007/s13225-020-00466-2)) |
| Otu58929 | 1 | k__Fungi(100);p__Ascomycota(100);c__Saccharomycetes(100);o__Saccharomycetales(100);f__Metschnikowiaceae(100);g__Metschnikowia(100);s__Metschnikowia_pulcherrima(100); | Metschnikowia | 13 | Saprotroph | \|Undefined Saprotroph\| | Yeast | NULL | Probable | As nectar-tap_saprotroph (Põlme et al. 2020); Decay sugar-rich substrates (Põlme et al. 2020) | Põlme S, et al. 2020. Fungal Diversity 105:1-16 ((https://doi.org/10.1007/s13225-020-00466-2)) |
| Otu58930 | 1 | k__Fungi(100);p__Ascomycota(100);c__Saccharomycetes(100);o__Saccharomycetales(100);f__Metschnikowiaceae(100);g__Metschnikowia(100);s__Metschnikowia_pulcherrima(100); | Metschnikowia | 13 | Saprotroph | \|Undefined Saprotroph\| | Yeast | NULL | Probable | As nectar-tap_saprotroph (Põlme et al. 2020); Decay sugar-rich substrates (Põlme et al. 2020) | Põlme S, et al. 2020. Fungal Diversity 105:1-16 ((https://doi.org/10.1007/s13225-020-00466-2)) |
| Otu58931 | 1 | k__Fungi(100);p__Ascomycota(100);c__Saccharomycetes(100);o__Saccharomycetales(100);f__Metschnikowiaceae(100);g__Metschnikowia(100);s__Metschnikowia_pulcherrima(100); | Metschnikowia | 13 | Saprotroph | \|Undefined Saprotroph\| | Yeast | NULL | Probable | As nectar-tap_saprotroph (Põlme et al. 2020); Decay sugar-rich substrates (Põlme et al. 2020) | Põlme S, et al. 2020. Fungal Diversity 105:1-16 ((https://doi.org/10.1007/s13225-020-00466-2)) |
| Otu58932 | 1 | k__Fungi(100);p__Ascomycota(100);c__Saccharomycetes(100);o__Saccharomycetales(100);f__Metschnikowiaceae(100);g__Metschnikowia(100);s__Metschnikowia_pulcherrima(100); | Metschnikowia | 13 | Saprotroph | \|Undefined Saprotroph\| | Yeast | NULL | Probable | As nectar-tap_saprotroph (Põlme et al. 2020); Decay sugar-rich substrates (Põlme et al. 2020) | Põlme S, et al. 2020. Fungal Diversity 105:1-16 ((https://doi.org/10.1007/s13225-020-00466-2)) |
| Otu58933 | 1 | k__Fungi(100);p__Ascomycota(100);c__Saccharomycetes(100);o__Saccharomycetales(100);f__Metschnikowiaceae(100);g__Metschnikowia(100);s__Metschnikowia_pulcherrima(100); | Metschnikowia | 13 | Saprotroph | \|Undefined Saprotroph\| | Yeast | NULL | Probable | As nectar-tap_saprotroph (Põlme et al. 2020); Decay sugar-rich substrates (Põlme et al. 2020) | Põlme S, et al. 2020. Fungal Diversity 105:1-16 ((https://doi.org/10.1007/s13225-020-00466-2)) |
| Otu58934 | 1 | k__Fungi(100);p__Ascomycota(100);c__Saccharomycetes(100);o__Saccharomycetales(100);f__Metschnikowiaceae(100);g__Metschnikowia(100);s__Metschnikowia_pulcherrima(100); | Metschnikowia | 13 | Saprotroph | \|Undefined Saprotroph\| | Yeast | NULL | Probable | As nectar-tap_saprotroph (Põlme et al. 2020); Decay sugar-rich substrates (Põlme et al. 2020) | Põlme S, et al. 2020. Fungal Diversity 105:1-16 ((https://doi.org/10.1007/s13225-020-00466-2)) |
| Otu58935 | 1 | k__Fungi(100);p__Ascomycota(100);c__Saccharomycetes(100);o__Saccharomycetales(100);f__Metschnikowiaceae(100);g__Metschnikowia(100);s__Metschnikowia_pulcherrima(100); | Metschnikowia | 13 | Saprotroph | \|Undefined Saprotroph\| | Yeast | NULL | Probable | As nectar-tap_saprotroph (Põlme et al. 2020); Decay sugar-rich substrates (Põlme et al. 2020) | Põlme S, et al. 2020. Fungal Diversity 105:1-16 ((https://doi.org/10.1007/s13225-020-00466-2)) |
| Otu58936 | 1 | k__Fungi(100);p__Ascomycota(100);c__Saccharomycetes(100);o__Saccharomycetales(100);f__Metschnikowiaceae(100);g__Metschnikowia(100);s__Metschnikowia_pulcherrima(100); | Metschnikowia | 13 | Saprotroph | \|Undefined Saprotroph\| | Yeast | NULL | Probable | As nectar-tap_saprotroph (Põlme et al. 2020); Decay sugar-rich substrates (Põlme et al. 2020) | Põlme S, et al. 2020. Fungal Diversity 105:1-16 ((https://doi.org/10.1007/s13225-020-00466-2)) |
| Otu58937 | 1 | k__Fungi(100);p__Ascomycota(100);c__Saccharomycetes(100);o__Saccharomycetales(100);f__Metschnikowiaceae(100);g__Metschnikowia(100);s__Metschnikowia_pulcherrima(100); | Metschnikowia | 13 | Saprotroph | \|Undefined Saprotroph\| | Yeast | NULL | Probable | As nectar-tap_saprotroph (Põlme et al. 2020); Decay sugar-rich substrates (Põlme et al. 2020) | Põlme S, et al. 2020. Fungal Diversity 105:1-16 ((https://doi.org/10.1007/s13225-020-00466-2)) |
| Otu58938 | 1 | k__Fungi(100);p__Ascomycota(100);c__Saccharomycetes(100);o__Saccharomycetales(100);f__Metschnikowiaceae(100);g__Metschnikowia(100);s__Metschnikowia_pulcherrima(100); | Metschnikowia | 13 | Saprotroph | \|Undefined Saprotroph\| | Yeast | NULL | Probable | As nectar-tap_saprotroph (Põlme et al. 2020); Decay sugar-rich substrates (Põlme et al. 2020) | Põlme S, et al. 2020. Fungal Diversity 105:1-16 ((https://doi.org/10.1007/s13225-020-00466-2)) |
| Otu58939 | 1 | k__Fungi(100);p__Ascomycota(100);c__Saccharomycetes(100);o__Saccharomycetales(100);f__Metschnikowiaceae(100);g__Metschnikowia(100);g__Metschnikowia_unclassified(100); | Metschnikowia | 13 | Saprotroph | \|Undefined Saprotroph\| | Yeast | NULL | Probable | As nectar-tap_saprotroph (Põlme et al. 2020); Decay sugar-rich substrates (Põlme et al. 2020) | Põlme S, et al. 2020. Fungal Diversity 105:1-16 ((https://doi.org/10.1007/s13225-020-00466-2)) |
| Otu58940 | 1 | k__Fungi(100);p__Ascomycota(100);c__Saccharomycetes(100);o__Saccharomycetales(100);f__Metschnikowiaceae(100);g__Metschnikowia(100);s__Metschnikowia_pulcherrima(100); | Metschnikowia | 13 | Saprotroph | \|Undefined Saprotroph\| | Yeast | NULL | Probable | As nectar-tap_saprotroph (Põlme et al. 2020); Decay sugar-rich substrates (Põlme et al. 2020) | Põlme S, et al. 2020. Fungal Diversity 105:1-16 ((https://doi.org/10.1007/s13225-020-00466-2)) |
| Otu58941 | 1 | k__Fungi(100);p__Ascomycota(100);c__Saccharomycetes(100);o__Saccharomycetales(100);f__Metschnikowiaceae(100);g__Metschnikowia(100);s__Metschnikowia_pulcherrima(100); | Metschnikowia | 13 | Saprotroph | \|Undefined Saprotroph\| | Yeast | NULL | Probable | As nectar-tap_saprotroph (Põlme et al. 2020); Decay sugar-rich substrates (Põlme et al. 2020) | Põlme S, et al. 2020. Fungal Diversity 105:1-16 ((https://doi.org/10.1007/s13225-020-00466-2)) |
| Otu58942 | 1 | k__Fungi(100);p__Ascomycota(100);c__Saccharomycetes(100);o__Saccharomycetales(100);f__Metschnikowiaceae(100);g__Metschnikowia(100);s__Metschnikowia_pulcherrima(100); | Metschnikowia | 13 | Saprotroph | \|Undefined Saprotroph\| | Yeast | NULL | Probable | As nectar-tap_saprotroph (Põlme et al. 2020); Decay sugar-rich substrates (Põlme et al. 2020) | Põlme S, et al. 2020. Fungal Diversity 105:1-16 ((https://doi.org/10.1007/s13225-020-00466-2)) |
| Otu58943 | 1 | k__Fungi(100);p__Ascomycota(100);c__Saccharomycetes(100);o__Saccharomycetales(100);f__Metschnikowiaceae(100);g__Metschnikowia(100);s__Metschnikowia_pulcherrima(100); | Metschnikowia | 13 | Saprotroph | \|Undefined Saprotroph\| | Yeast | NULL | Probable | As nectar-tap_saprotroph (Põlme et al. 2020); Decay sugar-rich substrates (Põlme et al. 2020) | Põlme S, et al. 2020. Fungal Diversity 105:1-16 ((https://doi.org/10.1007/s13225-020-00466-2)) |
| Otu58944 | 1 | k__Fungi(100);p__Ascomycota(100);c__Saccharomycetes(100);o__Saccharomycetales(100);f__Metschnikowiaceae(100);g__Metschnikowia(100);s__Metschnikowia_pulcherrima(100); | Metschnikowia | 13 | Saprotroph | \|Undefined Saprotroph\| | Yeast | NULL | Probable | As nectar-tap_saprotroph (Põlme et al. 2020); Decay sugar-rich substrates (Põlme et al. 2020) | Põlme S, et al. 2020. Fungal Diversity 105:1-16 ((https://doi.org/10.1007/s13225-020-00466-2)) |
| Otu58945 | 1 | k__Fungi(100);p__Ascomycota(100);c__Saccharomycetes(100);o__Saccharomycetales(100);f__Metschnikowiaceae(100);g__Metschnikowia(100);s__Metschnikowia_pulcherrima(100); | Metschnikowia | 13 | Saprotroph | \|Undefined Saprotroph\| | Yeast | NULL | Probable | As nectar-tap_saprotroph (Põlme et al. 2020); Decay sugar-rich substrates (Põlme et al. 2020) | Põlme S, et al. 2020. Fungal Diversity 105:1-16 ((https://doi.org/10.1007/s13225-020-00466-2)) |
| Otu58946 | 1 | k__Fungi(100);p__Ascomycota(100);c__Saccharomycetes(100);o__Saccharomycetales(100);f__Metschnikowiaceae(100);g__Metschnikowia(100);s__Metschnikowia_pulcherrima(100); | Metschnikowia | 13 | Saprotroph | \|Undefined Saprotroph\| | Yeast | NULL | Probable | As nectar-tap_saprotroph (Põlme et al. 2020); Decay sugar-rich substrates (Põlme et al. 2020) | Põlme S, et al. 2020. Fungal Diversity 105:1-16 ((https://doi.org/10.1007/s13225-020-00466-2)) |
| Otu58948 | 1 | k__Fungi(100);p__Ascomycota(100);c__Saccharomycetes(100);o__Saccharomycetales(100);f__Metschnikowiaceae(100);g__Metschnikowia(100);g__Metschnikowia_unclassified(100); | Metschnikowia | 13 | Saprotroph | \|Undefined Saprotroph\| | Yeast | NULL | Probable | As nectar-tap_saprotroph (Põlme et al. 2020); Decay sugar-rich substrates (Põlme et al. 2020) | Põlme S, et al. 2020. Fungal Diversity 105:1-16 ((https://doi.org/10.1007/s13225-020-00466-2)) |
| Otu58949 | 1 | k__Fungi(100);p__Ascomycota(100);c__Saccharomycetes(100);o__Saccharomycetales(100);f__Metschnikowiaceae(100);g__Metschnikowia(100);s__Metschnikowia_pulcherrima(100); | Metschnikowia | 13 | Saprotroph | \|Undefined Saprotroph\| | Yeast | NULL | Probable | As nectar-tap_saprotroph (Põlme et al. 2020); Decay sugar-rich substrates (Põlme et al. 2020) | Põlme S, et al. 2020. Fungal Diversity 105:1-16 ((https://doi.org/10.1007/s13225-020-00466-2)) |
| Otu58950 | 1 | k__Fungi(100);p__Ascomycota(100);c__Saccharomycetes(100);o__Saccharomycetales(100);f__Metschnikowiaceae(100);g__Metschnikowia(100);s__Metschnikowia_pulcherrima(100); | Metschnikowia | 13 | Saprotroph | \|Undefined Saprotroph\| | Yeast | NULL | Probable | As nectar-tap_saprotroph (Põlme et al. 2020); Decay sugar-rich substrates (Põlme et al. 2020) | Põlme S, et al. 2020. Fungal Diversity 105:1-16 ((https://doi.org/10.1007/s13225-020-00466-2)) |
| Otu58951 | 1 | k__Fungi(100);p__Ascomycota(100);c__Saccharomycetes(100);o__Saccharomycetales(100);f__Metschnikowiaceae(100);g__Metschnikowia(100);g__Metschnikowia_unclassified(100); | Metschnikowia | 13 | Saprotroph | \|Undefined Saprotroph\| | Yeast | NULL | Probable | As nectar-tap_saprotroph (Põlme et al. 2020); Decay sugar-rich substrates (Põlme et al. 2020) | Põlme S, et al. 2020. Fungal Diversity 105:1-16 ((https://doi.org/10.1007/s13225-020-00466-2)) |
| Otu58952 | 1 | k__Fungi(100);p__Ascomycota(100);c__Saccharomycetes(100);o__Saccharomycetales(100);f__Metschnikowiaceae(100);g__Metschnikowia(100);s__Metschnikowia_pulcherrima(100); | Metschnikowia | 13 | Saprotroph | \|Undefined Saprotroph\| | Yeast | NULL | Probable | As nectar-tap_saprotroph (Põlme et al. 2020); Decay sugar-rich substrates (Põlme et al. 2020) | Põlme S, et al. 2020. Fungal Diversity 105:1-16 ((https://doi.org/10.1007/s13225-020-00466-2)) |
| Otu58953 | 1 | k__Fungi(100);p__Ascomycota(100);c__Saccharomycetes(100);o__Saccharomycetales(100);f__Metschnikowiaceae(100);g__Metschnikowia(100);s__Metschnikowia_pulcherrima(100); | Metschnikowia | 13 | Saprotroph | \|Undefined Saprotroph\| | Yeast | NULL | Probable | As nectar-tap_saprotroph (Põlme et al. 2020); Decay sugar-rich substrates (Põlme et al. 2020) | Põlme S, et al. 2020. Fungal Diversity 105:1-16 ((https://doi.org/10.1007/s13225-020-00466-2)) |
| Otu58954 | 1 | k__Fungi(100);p__Ascomycota(100);c__Saccharomycetes(100);o__Saccharomycetales(100);f__Metschnikowiaceae(100);g__Metschnikowia(100);s__Metschnikowia_pulcherrima(100); | Metschnikowia | 13 | Saprotroph | \|Undefined Saprotroph\| | Yeast | NULL | Probable | As nectar-tap_saprotroph (Põlme et al. 2020); Decay sugar-rich substrates (Põlme et al. 2020) | Põlme S, et al. 2020. Fungal Diversity 105:1-16 ((https://doi.org/10.1007/s13225-020-00466-2)) |
| Otu58955 | 1 | k__Fungi(100);p__Ascomycota(100);c__Saccharomycetes(100);o__Saccharomycetales(100);f__Metschnikowiaceae(100);g__Metschnikowia(100);s__Metschnikowia_pulcherrima(100); | Metschnikowia | 13 | Saprotroph | \|Undefined Saprotroph\| | Yeast | NULL | Probable | As nectar-tap_saprotroph (Põlme et al. 2020); Decay sugar-rich substrates (Põlme et al. 2020) | Põlme S, et al. 2020. Fungal Diversity 105:1-16 ((https://doi.org/10.1007/s13225-020-00466-2)) |
| Otu58956 | 1 | k__Fungi(100);p__Ascomycota(100);c__Saccharomycetes(100);o__Saccharomycetales(100);f__Metschnikowiaceae(100);g__Metschnikowia(100);s__Metschnikowia_pulcherrima(100); | Metschnikowia | 13 | Saprotroph | \|Undefined Saprotroph\| | Yeast | NULL | Probable | As nectar-tap_saprotroph (Põlme et al. 2020); Decay sugar-rich substrates (Põlme et al. 2020) | Põlme S, et al. 2020. Fungal Diversity 105:1-16 ((https://doi.org/10.1007/s13225-020-00466-2)) |
| Otu58957 | 1 | k__Fungi(100);p__Ascomycota(100);c__Saccharomycetes(100);o__Saccharomycetales(100);f__Metschnikowiaceae(100);g__Metschnikowia(100);g__Metschnikowia_unclassified(100); | Metschnikowia | 13 | Saprotroph | \|Undefined Saprotroph\| | Yeast | NULL | Probable | As nectar-tap_saprotroph (Põlme et al. 2020); Decay sugar-rich substrates (Põlme et al. 2020) | Põlme S, et al. 2020. Fungal Diversity 105:1-16 ((https://doi.org/10.1007/s13225-020-00466-2)) |
| Otu58958 | 1 | k__Fungi(100);p__Ascomycota(100);c__Saccharomycetes(100);o__Saccharomycetales(100);f__Metschnikowiaceae(100);g__Metschnikowia(100);s__Metschnikowia_pulcherrima(100); | Metschnikowia | 13 | Saprotroph | \|Undefined Saprotroph\| | Yeast | NULL | Probable | As nectar-tap_saprotroph (Põlme et al. 2020); Decay sugar-rich substrates (Põlme et al. 2020) | Põlme S, et al. 2020. Fungal Diversity 105:1-16 ((https://doi.org/10.1007/s13225-020-00466-2)) |
| Otu58959 | 1 | k__Fungi(100);p__Ascomycota(100);c__Saccharomycetes(100);o__Saccharomycetales(100);f__Metschnikowiaceae(100);g__Metschnikowia(100);s__Metschnikowia_pulcherrima(100); | Metschnikowia | 13 | Saprotroph | \|Undefined Saprotroph\| | Yeast | NULL | Probable | As nectar-tap_saprotroph (Põlme et al. 2020); Decay sugar-rich substrates (Põlme et al. 2020) | Põlme S, et al. 2020. Fungal Diversity 105:1-16 ((https://doi.org/10.1007/s13225-020-00466-2)) |
| Otu58960 | 1 | k__Fungi(100);p__Ascomycota(100);c__Saccharomycetes(100);o__Saccharomycetales(100);f__Metschnikowiaceae(100);g__Metschnikowia(100);s__Metschnikowia_pulcherrima(100); | Metschnikowia | 13 | Saprotroph | \|Undefined Saprotroph\| | Yeast | NULL | Probable | As nectar-tap_saprotroph (Põlme et al. 2020); Decay sugar-rich substrates (Põlme et al. 2020) | Põlme S, et al. 2020. Fungal Diversity 105:1-16 ((https://doi.org/10.1007/s13225-020-00466-2)) |
| Otu58961 | 1 | k__Fungi(100);p__Ascomycota(100);c__Saccharomycetes(100);o__Saccharomycetales(100);f__Metschnikowiaceae(100);g__Metschnikowia(100);g__Metschnikowia_unclassified(100); | Metschnikowia | 13 | Saprotroph | \|Undefined Saprotroph\| | Yeast | NULL | Probable | As nectar-tap_saprotroph (Põlme et al. 2020); Decay sugar-rich substrates (Põlme et al. 2020) | Põlme S, et al. 2020. Fungal Diversity 105:1-16 ((https://doi.org/10.1007/s13225-020-00466-2)) |
| Otu58962 | 1 | k__Fungi(100);p__Ascomycota(100);c__Saccharomycetes(100);o__Saccharomycetales(100);f__Metschnikowiaceae(100);g__Metschnikowia(100);g__Metschnikowia_unclassified(100); | Metschnikowia | 13 | Saprotroph | \|Undefined Saprotroph\| | Yeast | NULL | Probable | As nectar-tap_saprotroph (Põlme et al. 2020); Decay sugar-rich substrates (Põlme et al. 2020) | Põlme S, et al. 2020. Fungal Diversity 105:1-16 ((https://doi.org/10.1007/s13225-020-00466-2)) |
| Otu58963 | 1 | k__Fungi(100);p__Ascomycota(100);c__Saccharomycetes(100);o__Saccharomycetales(100);f__Metschnikowiaceae(100);g__Metschnikowia(100);s__Metschnikowia_pulcherrima(100); | Metschnikowia | 13 | Saprotroph | \|Undefined Saprotroph\| | Yeast | NULL | Probable | As nectar-tap_saprotroph (Põlme et al. 2020); Decay sugar-rich substrates (Põlme et al. 2020) | Põlme S, et al. 2020. Fungal Diversity 105:1-16 ((https://doi.org/10.1007/s13225-020-00466-2)) |
| Otu58964 | 1 | k__Fungi(100);p__Ascomycota(100);c__Saccharomycetes(100);o__Saccharomycetales(100);f__Metschnikowiaceae(100);g__Metschnikowia(100);g__Metschnikowia_unclassified(100); | Metschnikowia | 13 | Saprotroph | \|Undefined Saprotroph\| | Yeast | NULL | Probable | As nectar-tap_saprotroph (Põlme et al. 2020); Decay sugar-rich substrates (Põlme et al. 2020) | Põlme S, et al. 2020. Fungal Diversity 105:1-16 ((https://doi.org/10.1007/s13225-020-00466-2)) |
| Otu58965 | 1 | k__Fungi(100);p__Ascomycota(100);c__Saccharomycetes(100);o__Saccharomycetales(100);f__Metschnikowiaceae(100);g__Metschnikowia(100);s__Metschnikowia_pulcherrima(100); | Metschnikowia | 13 | Saprotroph | \|Undefined Saprotroph\| | Yeast | NULL | Probable | As nectar-tap_saprotroph (Põlme et al. 2020); Decay sugar-rich substrates (Põlme et al. 2020) | Põlme S, et al. 2020. Fungal Diversity 105:1-16 ((https://doi.org/10.1007/s13225-020-00466-2)) |
| Otu58966 | 1 | k__Fungi(100);p__Ascomycota(100);c__Saccharomycetes(100);o__Saccharomycetales(100);f__Metschnikowiaceae(100);g__Metschnikowia(100);g__Metschnikowia_unclassified(100); | Metschnikowia | 13 | Saprotroph | \|Undefined Saprotroph\| | Yeast | NULL | Probable | As nectar-tap_saprotroph (Põlme et al. 2020); Decay sugar-rich substrates (Põlme et al. 2020) | Põlme S, et al. 2020. Fungal Diversity 105:1-16 ((https://doi.org/10.1007/s13225-020-00466-2)) |
| Otu58967 | 1 | k__Fungi(100);p__Ascomycota(100);c__Saccharomycetes(100);o__Saccharomycetales(100);f__Metschnikowiaceae(100);g__Metschnikowia(100);s__Metschnikowia_pulcherrima(100); | Metschnikowia | 13 | Saprotroph | \|Undefined Saprotroph\| | Yeast | NULL | Probable | As nectar-tap_saprotroph (Põlme et al. 2020); Decay sugar-rich substrates (Põlme et al. 2020) | Põlme S, et al. 2020. Fungal Diversity 105:1-16 ((https://doi.org/10.1007/s13225-020-00466-2)) |
| Otu58968 | 1 | k__Fungi(100);p__Ascomycota(100);c__Saccharomycetes(100);o__Saccharomycetales(100);f__Metschnikowiaceae(100);g__Metschnikowia(100);s__Metschnikowia_pulcherrima(100); | Metschnikowia | 13 | Saprotroph | \|Undefined Saprotroph\| | Yeast | NULL | Probable | As nectar-tap_saprotroph (Põlme et al. 2020); Decay sugar-rich substrates (Põlme et al. 2020) | Põlme S, et al. 2020. Fungal Diversity 105:1-16 ((https://doi.org/10.1007/s13225-020-00466-2)) |
| Otu58969 | 1 | k__Fungi(100);p__Ascomycota(100);c__Saccharomycetes(100);o__Saccharomycetales(100);f__Metschnikowiaceae(100);g__Metschnikowia(100);s__Metschnikowia_pulcherrima(100); | Metschnikowia | 13 | Saprotroph | \|Undefined Saprotroph\| | Yeast | NULL | Probable | As nectar-tap_saprotroph (Põlme et al. 2020); Decay sugar-rich substrates (Põlme et al. 2020) | Põlme S, et al. 2020. Fungal Diversity 105:1-16 ((https://doi.org/10.1007/s13225-020-00466-2)) |
| Otu58970 | 1 | k__Fungi(100);p__Ascomycota(100);c__Saccharomycetes(100);o__Saccharomycetales(100);f__Metschnikowiaceae(100);g__Metschnikowia(100);s__Metschnikowia_pulcherrima(100); | Metschnikowia | 13 | Saprotroph | \|Undefined Saprotroph\| | Yeast | NULL | Probable | As nectar-tap_saprotroph (Põlme et al. 2020); Decay sugar-rich substrates (Põlme et al. 2020) | Põlme S, et al. 2020. Fungal Diversity 105:1-16 ((https://doi.org/10.1007/s13225-020-00466-2)) |
| Otu58971 | 1 | k__Fungi(100);p__Ascomycota(100);c__Saccharomycetes(100);o__Saccharomycetales(100);f__Metschnikowiaceae(100);g__Metschnikowia(100);s__Metschnikowia_pulcherrima(100); | Metschnikowia | 13 | Saprotroph | \|Undefined Saprotroph\| | Yeast | NULL | Probable | As nectar-tap_saprotroph (Põlme et al. 2020); Decay sugar-rich substrates (Põlme et al. 2020) | Põlme S, et al. 2020. Fungal Diversity 105:1-16 ((https://doi.org/10.1007/s13225-020-00466-2)) |
| Otu58972 | 1 | k__Fungi(100);p__Ascomycota(100);c__Saccharomycetes(100);o__Saccharomycetales(100);f__Metschnikowiaceae(100);g__Metschnikowia(100);g__Metschnikowia_unclassified(100); | Metschnikowia | 13 | Saprotroph | \|Undefined Saprotroph\| | Yeast | NULL | Probable | As nectar-tap_saprotroph (Põlme et al. 2020); Decay sugar-rich substrates (Põlme et al. 2020) | Põlme S, et al. 2020. Fungal Diversity 105:1-16 ((https://doi.org/10.1007/s13225-020-00466-2)) |
| Otu58973 | 1 | k__Fungi(100);p__Ascomycota(100);c__Saccharomycetes(100);o__Saccharomycetales(100);f__Metschnikowiaceae(100);g__Metschnikowia(100);s__Metschnikowia_pulcherrima(100); | Metschnikowia | 13 | Saprotroph | \|Undefined Saprotroph\| | Yeast | NULL | Probable | As nectar-tap_saprotroph (Põlme et al. 2020); Decay sugar-rich substrates (Põlme et al. 2020) | Põlme S, et al. 2020. Fungal Diversity 105:1-16 ((https://doi.org/10.1007/s13225-020-00466-2)) |
| Otu58974 | 1 | k__Fungi(100);p__Ascomycota(100);c__Saccharomycetes(100);o__Saccharomycetales(100);f__Metschnikowiaceae(100);g__Metschnikowia(100);s__Metschnikowia_pulcherrima(100); | Metschnikowia | 13 | Saprotroph | \|Undefined Saprotroph\| | Yeast | NULL | Probable | As nectar-tap_saprotroph (Põlme et al. 2020); Decay sugar-rich substrates (Põlme et al. 2020) | Põlme S, et al. 2020. Fungal Diversity 105:1-16 ((https://doi.org/10.1007/s13225-020-00466-2)) |
| Otu58975 | 1 | k__Fungi(100);p__Ascomycota(100);c__Saccharomycetes(100);o__Saccharomycetales(100);f__Metschnikowiaceae(100);g__Metschnikowia(100);s__Metschnikowia_pulcherrima(100); | Metschnikowia | 13 | Saprotroph | \|Undefined Saprotroph\| | Yeast | NULL | Probable | As nectar-tap_saprotroph (Põlme et al. 2020); Decay sugar-rich substrates (Põlme et al. 2020) | Põlme S, et al. 2020. Fungal Diversity 105:1-16 ((https://doi.org/10.1007/s13225-020-00466-2)) |
| Otu58976 | 1 | k__Fungi(100);p__Ascomycota(100);c__Saccharomycetes(100);o__Saccharomycetales(100);f__Metschnikowiaceae(100);g__Metschnikowia(100);s__Metschnikowia_pulcherrima(100); | Metschnikowia | 13 | Saprotroph | \|Undefined Saprotroph\| | Yeast | NULL | Probable | As nectar-tap_saprotroph (Põlme et al. 2020); Decay sugar-rich substrates (Põlme et al. 2020) | Põlme S, et al. 2020. Fungal Diversity 105:1-16 ((https://doi.org/10.1007/s13225-020-00466-2)) |
| Otu58977 | 1 | k__Fungi(100);p__Ascomycota(100);c__Saccharomycetes(100);o__Saccharomycetales(100);f__Metschnikowiaceae(100);g__Metschnikowia(100);s__Metschnikowia_pulcherrima(100); | Metschnikowia | 13 | Saprotroph | \|Undefined Saprotroph\| | Yeast | NULL | Probable | As nectar-tap_saprotroph (Põlme et al. 2020); Decay sugar-rich substrates (Põlme et al. 2020) | Põlme S, et al. 2020. Fungal Diversity 105:1-16 ((https://doi.org/10.1007/s13225-020-00466-2)) |
| Otu58978 | 1 | k__Fungi(100);p__Ascomycota(100);c__Saccharomycetes(100);o__Saccharomycetales(100);f__Metschnikowiaceae(100);g__Metschnikowia(100);s__Metschnikowia_pulcherrima(100); | Metschnikowia | 13 | Saprotroph | \|Undefined Saprotroph\| | Yeast | NULL | Probable | As nectar-tap_saprotroph (Põlme et al. 2020); Decay sugar-rich substrates (Põlme et al. 2020) | Põlme S, et al. 2020. Fungal Diversity 105:1-16 ((https://doi.org/10.1007/s13225-020-00466-2)) |
| Otu58979 | 1 | k__Fungi(100);p__Ascomycota(100);c__Saccharomycetes(100);o__Saccharomycetales(100);f__Metschnikowiaceae(100);g__Metschnikowia(100);s__Metschnikowia_pulcherrima(100); | Metschnikowia | 13 | Saprotroph | \|Undefined Saprotroph\| | Yeast | NULL | Probable | As nectar-tap_saprotroph (Põlme et al. 2020); Decay sugar-rich substrates (Põlme et al. 2020) | Põlme S, et al. 2020. Fungal Diversity 105:1-16 ((https://doi.org/10.1007/s13225-020-00466-2)) |
| Otu58980 | 1 | k__Fungi(100);p__Ascomycota(100);c__Saccharomycetes(100);o__Saccharomycetales(100);f__Metschnikowiaceae(100);g__Metschnikowia(100);s__Metschnikowia_pulcherrima(100); | Metschnikowia | 13 | Saprotroph | \|Undefined Saprotroph\| | Yeast | NULL | Probable | As nectar-tap_saprotroph (Põlme et al. 2020); Decay sugar-rich substrates (Põlme et al. 2020) | Põlme S, et al. 2020. Fungal Diversity 105:1-16 ((https://doi.org/10.1007/s13225-020-00466-2)) |
| Otu58981 | 1 | k__Fungi(100);p__Ascomycota(100);c__Saccharomycetes(100);o__Saccharomycetales(100);f__Metschnikowiaceae(100);g__Metschnikowia(100);s__Metschnikowia_pulcherrima(100); | Metschnikowia | 13 | Saprotroph | \|Undefined Saprotroph\| | Yeast | NULL | Probable | As nectar-tap_saprotroph (Põlme et al. 2020); Decay sugar-rich substrates (Põlme et al. 2020) | Põlme S, et al. 2020. Fungal Diversity 105:1-16 ((https://doi.org/10.1007/s13225-020-00466-2)) |
| Otu58982 | 1 | k__Fungi(100);p__Ascomycota(100);c__Saccharomycetes(100);o__Saccharomycetales(100);f__Metschnikowiaceae(100);g__Metschnikowia(100);s__Metschnikowia_pulcherrima(100); | Metschnikowia | 13 | Saprotroph | \|Undefined Saprotroph\| | Yeast | NULL | Probable | As nectar-tap_saprotroph (Põlme et al. 2020); Decay sugar-rich substrates (Põlme et al. 2020) | Põlme S, et al. 2020. Fungal Diversity 105:1-16 ((https://doi.org/10.1007/s13225-020-00466-2)) |
| Otu58983 | 1 | k__Fungi(100);p__Ascomycota(100);c__Saccharomycetes(100);o__Saccharomycetales(100);f__Metschnikowiaceae(100);g__Metschnikowia(100);s__Metschnikowia_pulcherrima(100); | Metschnikowia | 13 | Saprotroph | \|Undefined Saprotroph\| | Yeast | NULL | Probable | As nectar-tap_saprotroph (Põlme et al. 2020); Decay sugar-rich substrates (Põlme et al. 2020) | Põlme S, et al. 2020. Fungal Diversity 105:1-16 ((https://doi.org/10.1007/s13225-020-00466-2)) |
| Otu58985 | 1 | k__Fungi(100);p__Ascomycota(100);c__Saccharomycetes(100);o__Saccharomycetales(100);f__Metschnikowiaceae(100);g__Metschnikowia(100);s__Metschnikowia_pulcherrima(100); | Metschnikowia | 13 | Saprotroph | \|Undefined Saprotroph\| | Yeast | NULL | Probable | As nectar-tap_saprotroph (Põlme et al. 2020); Decay sugar-rich substrates (Põlme et al. 2020) | Põlme S, et al. 2020. Fungal Diversity 105:1-16 ((https://doi.org/10.1007/s13225-020-00466-2)) |
| Otu58987 | 1 | k__Fungi(100);p__Ascomycota(100);c__Saccharomycetes(100);o__Saccharomycetales(100);f__Metschnikowiaceae(100);g__Metschnikowia(100);s__Metschnikowia_pulcherrima(100); | Metschnikowia | 13 | Saprotroph | \|Undefined Saprotroph\| | Yeast | NULL | Probable | As nectar-tap_saprotroph (Põlme et al. 2020); Decay sugar-rich substrates (Põlme et al. 2020) | Põlme S, et al. 2020. Fungal Diversity 105:1-16 ((https://doi.org/10.1007/s13225-020-00466-2)) |
| Otu58988 | 1 | k__Fungi(100);p__Ascomycota(100);c__Saccharomycetes(100);o__Saccharomycetales(100);f__Metschnikowiaceae(100);g__Metschnikowia(100);s__Metschnikowia_pulcherrima(100); | Metschnikowia | 13 | Saprotroph | \|Undefined Saprotroph\| | Yeast | NULL | Probable | As nectar-tap_saprotroph (Põlme et al. 2020); Decay sugar-rich substrates (Põlme et al. 2020) | Põlme S, et al. 2020. Fungal Diversity 105:1-16 ((https://doi.org/10.1007/s13225-020-00466-2)) |
| Otu58989 | 1 | k__Fungi(100);p__Ascomycota(100);c__Saccharomycetes(100);o__Saccharomycetales(100);f__Metschnikowiaceae(100);g__Metschnikowia(100);s__Metschnikowia_pulcherrima(100); | Metschnikowia | 13 | Saprotroph | \|Undefined Saprotroph\| | Yeast | NULL | Probable | As nectar-tap_saprotroph (Põlme et al. 2020); Decay sugar-rich substrates (Põlme et al. 2020) | Põlme S, et al. 2020. Fungal Diversity 105:1-16 ((https://doi.org/10.1007/s13225-020-00466-2)) |
| Otu58990 | 1 | k__Fungi(100);p__Ascomycota(100);c__Saccharomycetes(100);o__Saccharomycetales(100);f__Metschnikowiaceae(100);g__Metschnikowia(100);s__Metschnikowia_pulcherrima(100); | Metschnikowia | 13 | Saprotroph | \|Undefined Saprotroph\| | Yeast | NULL | Probable | As nectar-tap_saprotroph (Põlme et al. 2020); Decay sugar-rich substrates (Põlme et al. 2020) | Põlme S, et al. 2020. Fungal Diversity 105:1-16 ((https://doi.org/10.1007/s13225-020-00466-2)) |
| Otu58991 | 1 | k__Fungi(100);p__Ascomycota(100);c__Saccharomycetes(100);o__Saccharomycetales(100);f__Metschnikowiaceae(100);g__Metschnikowia(100);s__Metschnikowia_pulcherrima(100); | Metschnikowia | 13 | Saprotroph | \|Undefined Saprotroph\| | Yeast | NULL | Probable | As nectar-tap_saprotroph (Põlme et al. 2020); Decay sugar-rich substrates (Põlme et al. 2020) | Põlme S, et al. 2020. Fungal Diversity 105:1-16 ((https://doi.org/10.1007/s13225-020-00466-2)) |
| Otu58992 | 1 | k__Fungi(100);p__Ascomycota(100);c__Saccharomycetes(100);o__Saccharomycetales(100);f__Metschnikowiaceae(100);g__Metschnikowia(100);s__Metschnikowia_pulcherrima(100); | Metschnikowia | 13 | Saprotroph | \|Undefined Saprotroph\| | Yeast | NULL | Probable | As nectar-tap_saprotroph (Põlme et al. 2020); Decay sugar-rich substrates (Põlme et al. 2020) | Põlme S, et al. 2020. Fungal Diversity 105:1-16 ((https://doi.org/10.1007/s13225-020-00466-2)) |
| Otu58993 | 1 | k__Fungi(100);p__Ascomycota(100);c__Saccharomycetes(100);o__Saccharomycetales(100);f__Metschnikowiaceae(100);g__Metschnikowia(100);s__Metschnikowia_pulcherrima(100); | Metschnikowia | 13 | Saprotroph | \|Undefined Saprotroph\| | Yeast | NULL | Probable | As nectar-tap_saprotroph (Põlme et al. 2020); Decay sugar-rich substrates (Põlme et al. 2020) | Põlme S, et al. 2020. Fungal Diversity 105:1-16 ((https://doi.org/10.1007/s13225-020-00466-2)) |
| Otu58994 | 1 | k__Fungi(100);p__Ascomycota(100);c__Saccharomycetes(100);o__Saccharomycetales(100);f__Metschnikowiaceae(100);g__Metschnikowia(100);g__Metschnikowia_unclassified(100); | Metschnikowia | 13 | Saprotroph | \|Undefined Saprotroph\| | Yeast | NULL | Probable | As nectar-tap_saprotroph (Põlme et al. 2020); Decay sugar-rich substrates (Põlme et al. 2020) | Põlme S, et al. 2020. Fungal Diversity 105:1-16 ((https://doi.org/10.1007/s13225-020-00466-2)) |
| Otu58995 | 1 | k__Fungi(100);p__Ascomycota(100);c__Saccharomycetes(100);o__Saccharomycetales(100);f__Metschnikowiaceae(100);g__Metschnikowia(100);s__Metschnikowia_pulcherrima(100); | Metschnikowia | 13 | Saprotroph | \|Undefined Saprotroph\| | Yeast | NULL | Probable | As nectar-tap_saprotroph (Põlme et al. 2020); Decay sugar-rich substrates (Põlme et al. 2020) | Põlme S, et al. 2020. Fungal Diversity 105:1-16 ((https://doi.org/10.1007/s13225-020-00466-2)) |
| Otu58996 | 1 | k__Fungi(100);p__Ascomycota(100);c__Saccharomycetes(100);o__Saccharomycetales(100);f__Metschnikowiaceae(100);g__Metschnikowia(100);g__Metschnikowia_unclassified(100); | Metschnikowia | 13 | Saprotroph | \|Undefined Saprotroph\| | Yeast | NULL | Probable | As nectar-tap_saprotroph (Põlme et al. 2020); Decay sugar-rich substrates (Põlme et al. 2020) | Põlme S, et al. 2020. Fungal Diversity 105:1-16 ((https://doi.org/10.1007/s13225-020-00466-2)) |
| Otu58997 | 1 | k__Fungi(100);p__Ascomycota(100);c__Saccharomycetes(100);o__Saccharomycetales(100);f__Metschnikowiaceae(100);g__Metschnikowia(100);s__Metschnikowia_pulcherrima(100); | Metschnikowia | 13 | Saprotroph | \|Undefined Saprotroph\| | Yeast | NULL | Probable | As nectar-tap_saprotroph (Põlme et al. 2020); Decay sugar-rich substrates (Põlme et al. 2020) | Põlme S, et al. 2020. Fungal Diversity 105:1-16 ((https://doi.org/10.1007/s13225-020-00466-2)) |
| Otu58998 | 1 | k__Fungi(100);p__Ascomycota(100);c__Saccharomycetes(100);o__Saccharomycetales(100);f__Metschnikowiaceae(100);g__Metschnikowia(100);g__Metschnikowia_unclassified(100); | Metschnikowia | 13 | Saprotroph | \|Undefined Saprotroph\| | Yeast | NULL | Probable | As nectar-tap_saprotroph (Põlme et al. 2020); Decay sugar-rich substrates (Põlme et al. 2020) | Põlme S, et al. 2020. Fungal Diversity 105:1-16 ((https://doi.org/10.1007/s13225-020-00466-2)) |
| Otu58999 | 1 | k__Fungi(100);p__Ascomycota(100);c__Saccharomycetes(100);o__Saccharomycetales(100);f__Metschnikowiaceae(100);g__Metschnikowia(100);g__Metschnikowia_unclassified(100); | Metschnikowia | 13 | Saprotroph | \|Undefined Saprotroph\| | Yeast | NULL | Probable | As nectar-tap_saprotroph (Põlme et al. 2020); Decay sugar-rich substrates (Põlme et al. 2020) | Põlme S, et al. 2020. Fungal Diversity 105:1-16 ((https://doi.org/10.1007/s13225-020-00466-2)) |
| Otu59000 | 1 | k__Fungi(100);p__Ascomycota(100);c__Saccharomycetes(100);o__Saccharomycetales(100);f__Metschnikowiaceae(100);g__Metschnikowia(100);s__Metschnikowia_pulcherrima(100); | Metschnikowia | 13 | Saprotroph | \|Undefined Saprotroph\| | Yeast | NULL | Probable | As nectar-tap_saprotroph (Põlme et al. 2020); Decay sugar-rich substrates (Põlme et al. 2020) | Põlme S, et al. 2020. Fungal Diversity 105:1-16 ((https://doi.org/10.1007/s13225-020-00466-2)) |
| Otu59001 | 1 | k__Fungi(100);p__Ascomycota(100);c__Saccharomycetes(100);o__Saccharomycetales(100);f__Metschnikowiaceae(100);g__Metschnikowia(100);s__Metschnikowia_pulcherrima(100); | Metschnikowia | 13 | Saprotroph | \|Undefined Saprotroph\| | Yeast | NULL | Probable | As nectar-tap_saprotroph (Põlme et al. 2020); Decay sugar-rich substrates (Põlme et al. 2020) | Põlme S, et al. 2020. Fungal Diversity 105:1-16 ((https://doi.org/10.1007/s13225-020-00466-2)) |
| Otu59002 | 1 | k__Fungi(100);p__Ascomycota(100);c__Saccharomycetes(100);o__Saccharomycetales(100);f__Metschnikowiaceae(100);g__Metschnikowia(100);g__Metschnikowia_unclassified(100); | Metschnikowia | 13 | Saprotroph | \|Undefined Saprotroph\| | Yeast | NULL | Probable | As nectar-tap_saprotroph (Põlme et al. 2020); Decay sugar-rich substrates (Põlme et al. 2020) | Põlme S, et al. 2020. Fungal Diversity 105:1-16 ((https://doi.org/10.1007/s13225-020-00466-2)) |
| Otu59003 | 1 | k__Fungi(100);p__Ascomycota(100);c__Saccharomycetes(100);o__Saccharomycetales(100);f__Metschnikowiaceae(100);g__Metschnikowia(100);g__Metschnikowia_unclassified(100); | Metschnikowia | 13 | Saprotroph | \|Undefined Saprotroph\| | Yeast | NULL | Probable | As nectar-tap_saprotroph (Põlme et al. 2020); Decay sugar-rich substrates (Põlme et al. 2020) | Põlme S, et al. 2020. Fungal Diversity 105:1-16 ((https://doi.org/10.1007/s13225-020-00466-2)) |
| Otu59004 | 1 | k__Fungi(100);p__Ascomycota(100);c__Saccharomycetes(100);o__Saccharomycetales(100);f__Metschnikowiaceae(100);g__Metschnikowia(100);s__Metschnikowia_pulcherrima(100); | Metschnikowia | 13 | Saprotroph | \|Undefined Saprotroph\| | Yeast | NULL | Probable | As nectar-tap_saprotroph (Põlme et al. 2020); Decay sugar-rich substrates (Põlme et al. 2020) | Põlme S, et al. 2020. Fungal Diversity 105:1-16 ((https://doi.org/10.1007/s13225-020-00466-2)) |
| Otu59005 | 1 | k__Fungi(100);p__Ascomycota(100);c__Saccharomycetes(100);o__Saccharomycetales(100);f__Metschnikowiaceae(100);g__Metschnikowia(100);s__Metschnikowia_pulcherrima(100); | Metschnikowia | 13 | Saprotroph | \|Undefined Saprotroph\| | Yeast | NULL | Probable | As nectar-tap_saprotroph (Põlme et al. 2020); Decay sugar-rich substrates (Põlme et al. 2020) | Põlme S, et al. 2020. Fungal Diversity 105:1-16 ((https://doi.org/10.1007/s13225-020-00466-2)) |
| Otu59006 | 1 | k__Fungi(100);p__Ascomycota(100);c__Saccharomycetes(100);o__Saccharomycetales(100);f__Metschnikowiaceae(100);g__Metschnikowia(100);s__Metschnikowia_pulcherrima(100); | Metschnikowia | 13 | Saprotroph | \|Undefined Saprotroph\| | Yeast | NULL | Probable | As nectar-tap_saprotroph (Põlme et al. 2020); Decay sugar-rich substrates (Põlme et al. 2020) | Põlme S, et al. 2020. Fungal Diversity 105:1-16 ((https://doi.org/10.1007/s13225-020-00466-2)) |
| Otu59007 | 1 | k__Fungi(100);p__Ascomycota(100);c__Saccharomycetes(100);o__Saccharomycetales(100);f__Metschnikowiaceae(100);g__Metschnikowia(100);g__Metschnikowia_unclassified(100); | Metschnikowia | 13 | Saprotroph | \|Undefined Saprotroph\| | Yeast | NULL | Probable | As nectar-tap_saprotroph (Põlme et al. 2020); Decay sugar-rich substrates (Põlme et al. 2020) | Põlme S, et al. 2020. Fungal Diversity 105:1-16 ((https://doi.org/10.1007/s13225-020-00466-2)) |
| Otu59008 | 1 | k__Fungi(100);p__Basidiomycota(100);c__Agaricomycetes(100);o__Polyporales(100);f__Fomitopsidaceae(100);g__Piptoporus(100);s__Piptoporus_betulinus(100); | Piptoporus | 13 | Pathotroph-Saprotroph | \|Plant Pathogen\|-Wood Saprotroph | Polyporoid | Brown Rot | Highly Probable | As brown-rot (Põlme et al. 2020); As wood_pathogen (Põlme et al. 2020); Decay wood (Põlme et al. 2020) | Gilbertson RL, Ryvarden L. 1987-1987. North American Polypores. Fungiflora, Oslo (ISBN: 978-0945345060); Põlme S, et al. 2020. Fungal Diversity 105:1-16 ((https://doi.org/10.1007/s13225-020-00466-2)) |
| Otu59009 | 1 | k__Fungi(100);p__Ascomycota(100);c__Saccharomycetes(100);o__Saccharomycetales(100);f__Metschnikowiaceae(100);g__Metschnikowia(100);s__Metschnikowia_pulcherrima(100); | Metschnikowia | 13 | Saprotroph | \|Undefined Saprotroph\| | Yeast | NULL | Probable | As nectar-tap_saprotroph (Põlme et al. 2020); Decay sugar-rich substrates (Põlme et al. 2020) | Põlme S, et al. 2020. Fungal Diversity 105:1-16 ((https://doi.org/10.1007/s13225-020-00466-2)) |
| Otu59010 | 1 | k__Fungi(100);p__Ascomycota(100);c__Saccharomycetes(100);o__Saccharomycetales(100);f__Metschnikowiaceae(100);g__Metschnikowia(100);g__Metschnikowia_unclassified(100); | Metschnikowia | 13 | Saprotroph | \|Undefined Saprotroph\| | Yeast | NULL | Probable | As nectar-tap_saprotroph (Põlme et al. 2020); Decay sugar-rich substrates (Põlme et al. 2020) | Põlme S, et al. 2020. Fungal Diversity 105:1-16 ((https://doi.org/10.1007/s13225-020-00466-2)) |
| Otu59011 | 1 | k__Fungi(100);p__Ascomycota(100);c__Saccharomycetes(100);o__Saccharomycetales(100);f__Metschnikowiaceae(100);g__Metschnikowia(100);s__Metschnikowia_pulcherrima(100); | Metschnikowia | 13 | Saprotroph | \|Undefined Saprotroph\| | Yeast | NULL | Probable | As nectar-tap_saprotroph (Põlme et al. 2020); Decay sugar-rich substrates (Põlme et al. 2020) | Põlme S, et al. 2020. Fungal Diversity 105:1-16 ((https://doi.org/10.1007/s13225-020-00466-2)) |
| Otu59012 | 1 | k__Fungi(100);p__Ascomycota(100);c__Saccharomycetes(100);o__Saccharomycetales(100);f__Metschnikowiaceae(100);g__Metschnikowia(100);s__Metschnikowia_pulcherrima(100); | Metschnikowia | 13 | Saprotroph | \|Undefined Saprotroph\| | Yeast | NULL | Probable | As nectar-tap_saprotroph (Põlme et al. 2020); Decay sugar-rich substrates (Põlme et al. 2020) | Põlme S, et al. 2020. Fungal Diversity 105:1-16 ((https://doi.org/10.1007/s13225-020-00466-2)) |
| Otu59014 | 1 | k__Fungi(100);p__Ascomycota(100);c__Saccharomycetes(100);o__Saccharomycetales(100);f__Metschnikowiaceae(100);g__Metschnikowia(100);s__Metschnikowia_pulcherrima(100); | Metschnikowia | 13 | Saprotroph | \|Undefined Saprotroph\| | Yeast | NULL | Probable | As nectar-tap_saprotroph (Põlme et al. 2020); Decay sugar-rich substrates (Põlme et al. 2020) | Põlme S, et al. 2020. Fungal Diversity 105:1-16 ((https://doi.org/10.1007/s13225-020-00466-2)) |
| Otu59015 | 1 | k__Fungi(100);p__Ascomycota(100);c__Saccharomycetes(100);o__Saccharomycetales(100);f__Metschnikowiaceae(100);g__Metschnikowia(100);g__Metschnikowia_unclassified(100); | Metschnikowia | 13 | Saprotroph | \|Undefined Saprotroph\| | Yeast | NULL | Probable | As nectar-tap_saprotroph (Põlme et al. 2020); Decay sugar-rich substrates (Põlme et al. 2020) | Põlme S, et al. 2020. Fungal Diversity 105:1-16 ((https://doi.org/10.1007/s13225-020-00466-2)) |
| Otu59016 | 1 | k__Fungi(100);p__Ascomycota(100);c__Saccharomycetes(100);o__Saccharomycetales(100);f__Metschnikowiaceae(100);g__Metschnikowia(100);s__Metschnikowia_pulcherrima(100); | Metschnikowia | 13 | Saprotroph | \|Undefined Saprotroph\| | Yeast | NULL | Probable | As nectar-tap_saprotroph (Põlme et al. 2020); Decay sugar-rich substrates (Põlme et al. 2020) | Põlme S, et al. 2020. Fungal Diversity 105:1-16 ((https://doi.org/10.1007/s13225-020-00466-2)) |
| Otu59017 | 1 | k__Fungi(100);p__Ascomycota(100);c__Saccharomycetes(100);o__Saccharomycetales(100);f__Metschnikowiaceae(100);g__Metschnikowia(100);g__Metschnikowia_unclassified(100); | Metschnikowia | 13 | Saprotroph | \|Undefined Saprotroph\| | Yeast | NULL | Probable | As nectar-tap_saprotroph (Põlme et al. 2020); Decay sugar-rich substrates (Põlme et al. 2020) | Põlme S, et al. 2020. Fungal Diversity 105:1-16 ((https://doi.org/10.1007/s13225-020-00466-2)) |
| Otu59018 | 1 | k__Fungi(100);p__Ascomycota(100);c__Saccharomycetes(100);o__Saccharomycetales(100);f__Metschnikowiaceae(100);g__Metschnikowia(100);s__Metschnikowia_pulcherrima(100); | Metschnikowia | 13 | Saprotroph | \|Undefined Saprotroph\| | Yeast | NULL | Probable | As nectar-tap_saprotroph (Põlme et al. 2020); Decay sugar-rich substrates (Põlme et al. 2020) | Põlme S, et al. 2020. Fungal Diversity 105:1-16 ((https://doi.org/10.1007/s13225-020-00466-2)) |
| Otu59019 | 1 | k__Fungi(100);p__Ascomycota(100);c__Saccharomycetes(100);o__Saccharomycetales(100);f__Metschnikowiaceae(100);g__Metschnikowia(100);s__Metschnikowia_pulcherrima(100); | Metschnikowia | 13 | Saprotroph | \|Undefined Saprotroph\| | Yeast | NULL | Probable | As nectar-tap_saprotroph (Põlme et al. 2020); Decay sugar-rich substrates (Põlme et al. 2020) | Põlme S, et al. 2020. Fungal Diversity 105:1-16 ((https://doi.org/10.1007/s13225-020-00466-2)) |
| Otu59020 | 1 | k__Fungi(100);p__Ascomycota(100);c__Saccharomycetes(100);o__Saccharomycetales(100);f__Metschnikowiaceae(100);g__Metschnikowia(100);s__Metschnikowia_pulcherrima(100); | Metschnikowia | 13 | Saprotroph | \|Undefined Saprotroph\| | Yeast | NULL | Probable | As nectar-tap_saprotroph (Põlme et al. 2020); Decay sugar-rich substrates (Põlme et al. 2020) | Põlme S, et al. 2020. Fungal Diversity 105:1-16 ((https://doi.org/10.1007/s13225-020-00466-2)) |
| Otu59021 | 1 | k__Fungi(100);p__Ascomycota(100);c__Saccharomycetes(100);o__Saccharomycetales(100);f__Metschnikowiaceae(100);g__Metschnikowia(100);s__Metschnikowia_pulcherrima(100); | Metschnikowia | 13 | Saprotroph | \|Undefined Saprotroph\| | Yeast | NULL | Probable | As nectar-tap_saprotroph (Põlme et al. 2020); Decay sugar-rich substrates (Põlme et al. 2020) | Põlme S, et al. 2020. Fungal Diversity 105:1-16 ((https://doi.org/10.1007/s13225-020-00466-2)) |
| Otu59022 | 1 | k__Fungi(100);p__Ascomycota(100);c__Saccharomycetes(100);o__Saccharomycetales(100);f__Metschnikowiaceae(100);g__Metschnikowia(100);g__Metschnikowia_unclassified(100); | Metschnikowia | 13 | Saprotroph | \|Undefined Saprotroph\| | Yeast | NULL | Probable | As nectar-tap_saprotroph (Põlme et al. 2020); Decay sugar-rich substrates (Põlme et al. 2020) | Põlme S, et al. 2020. Fungal Diversity 105:1-16 ((https://doi.org/10.1007/s13225-020-00466-2)) |
| Otu59023 | 1 | k__Fungi(100);p__Ascomycota(100);c__Saccharomycetes(100);o__Saccharomycetales(100);f__Metschnikowiaceae(100);g__Metschnikowia(100);s__Metschnikowia_pulcherrima(100); | Metschnikowia | 13 | Saprotroph | \|Undefined Saprotroph\| | Yeast | NULL | Probable | As nectar-tap_saprotroph (Põlme et al. 2020); Decay sugar-rich substrates (Põlme et al. 2020) | Põlme S, et al. 2020. Fungal Diversity 105:1-16 ((https://doi.org/10.1007/s13225-020-00466-2)) |
| Otu59024 | 1 | k__Fungi(100);p__Ascomycota(100);c__Saccharomycetes(100);o__Saccharomycetales(100);f__Metschnikowiaceae(100);g__Metschnikowia(100);s__Metschnikowia_pulcherrima(100); | Metschnikowia | 13 | Saprotroph | \|Undefined Saprotroph\| | Yeast | NULL | Probable | As nectar-tap_saprotroph (Põlme et al. 2020); Decay sugar-rich substrates (Põlme et al. 2020) | Põlme S, et al. 2020. Fungal Diversity 105:1-16 ((https://doi.org/10.1007/s13225-020-00466-2)) |
| Otu59025 | 1 | k__Fungi(100);p__Ascomycota(100);c__Saccharomycetes(100);o__Saccharomycetales(100);f__Metschnikowiaceae(100);g__Metschnikowia(100);s__Metschnikowia_pulcherrima(100); | Metschnikowia | 13 | Saprotroph | \|Undefined Saprotroph\| | Yeast | NULL | Probable | As nectar-tap_saprotroph (Põlme et al. 2020); Decay sugar-rich substrates (Põlme et al. 2020) | Põlme S, et al. 2020. Fungal Diversity 105:1-16 ((https://doi.org/10.1007/s13225-020-00466-2)) |
| Otu59027 | 1 | k__Fungi(100);p__Ascomycota(100);c__Saccharomycetes(100);o__Saccharomycetales(100);f__Metschnikowiaceae(100);g__Metschnikowia(100);s__Metschnikowia_pulcherrima(100); | Metschnikowia | 13 | Saprotroph | \|Undefined Saprotroph\| | Yeast | NULL | Probable | As nectar-tap_saprotroph (Põlme et al. 2020); Decay sugar-rich substrates (Põlme et al. 2020) | Põlme S, et al. 2020. Fungal Diversity 105:1-16 ((https://doi.org/10.1007/s13225-020-00466-2)) |
| Otu59028 | 1 | k__Fungi(100);p__Ascomycota(100);c__Saccharomycetes(100);o__Saccharomycetales(100);f__Metschnikowiaceae(100);g__Metschnikowia(100);s__Metschnikowia_pulcherrima(100); | Metschnikowia | 13 | Saprotroph | \|Undefined Saprotroph\| | Yeast | NULL | Probable | As nectar-tap_saprotroph (Põlme et al. 2020); Decay sugar-rich substrates (Põlme et al. 2020) | Põlme S, et al. 2020. Fungal Diversity 105:1-16 ((https://doi.org/10.1007/s13225-020-00466-2)) |
| Otu59029 | 1 | k__Fungi(100);p__Ascomycota(100);c__Saccharomycetes(100);o__Saccharomycetales(100);f__Metschnikowiaceae(100);g__Metschnikowia(100);s__Metschnikowia_pulcherrima(100); | Metschnikowia | 13 | Saprotroph | \|Undefined Saprotroph\| | Yeast | NULL | Probable | As nectar-tap_saprotroph (Põlme et al. 2020); Decay sugar-rich substrates (Põlme et al. 2020) | Põlme S, et al. 2020. Fungal Diversity 105:1-16 ((https://doi.org/10.1007/s13225-020-00466-2)) |
| Otu59030 | 1 | k__Fungi(100);p__Ascomycota(100);c__Saccharomycetes(100);o__Saccharomycetales(100);f__Metschnikowiaceae(100);g__Metschnikowia(100);s__Metschnikowia_pulcherrima(100); | Metschnikowia | 13 | Saprotroph | \|Undefined Saprotroph\| | Yeast | NULL | Probable | As nectar-tap_saprotroph (Põlme et al. 2020); Decay sugar-rich substrates (Põlme et al. 2020) | Põlme S, et al. 2020. Fungal Diversity 105:1-16 ((https://doi.org/10.1007/s13225-020-00466-2)) |
| Otu59031 | 1 | k__Fungi(100);p__Ascomycota(100);c__Saccharomycetes(100);o__Saccharomycetales(100);f__Metschnikowiaceae(100);g__Metschnikowia(100);s__Metschnikowia_pulcherrima(100); | Metschnikowia | 13 | Saprotroph | \|Undefined Saprotroph\| | Yeast | NULL | Probable | As nectar-tap_saprotroph (Põlme et al. 2020); Decay sugar-rich substrates (Põlme et al. 2020) | Põlme S, et al. 2020. Fungal Diversity 105:1-16 ((https://doi.org/10.1007/s13225-020-00466-2)) |
| Otu59032 | 1 | k__Fungi(100);p__Ascomycota(100);c__Saccharomycetes(100);o__Saccharomycetales(100);f__Metschnikowiaceae(100);g__Metschnikowia(100);g__Metschnikowia_unclassified(100); | Metschnikowia | 13 | Saprotroph | \|Undefined Saprotroph\| | Yeast | NULL | Probable | As nectar-tap_saprotroph (Põlme et al. 2020); Decay sugar-rich substrates (Põlme et al. 2020) | Põlme S, et al. 2020. Fungal Diversity 105:1-16 ((https://doi.org/10.1007/s13225-020-00466-2)) |
| Otu59033 | 1 | k__Fungi(100);p__Ascomycota(100);c__Saccharomycetes(100);o__Saccharomycetales(100);f__Metschnikowiaceae(100);g__Metschnikowia(100);s__Metschnikowia_pulcherrima(100); | Metschnikowia | 13 | Saprotroph | \|Undefined Saprotroph\| | Yeast | NULL | Probable | As nectar-tap_saprotroph (Põlme et al. 2020); Decay sugar-rich substrates (Põlme et al. 2020) | Põlme S, et al. 2020. Fungal Diversity 105:1-16 ((https://doi.org/10.1007/s13225-020-00466-2)) |
| Otu59034 | 1 | k__Fungi(100);p__Ascomycota(100);c__Saccharomycetes(100);o__Saccharomycetales(100);f__Metschnikowiaceae(100);g__Metschnikowia(100);s__Metschnikowia_pulcherrima(100); | Metschnikowia | 13 | Saprotroph | \|Undefined Saprotroph\| | Yeast | NULL | Probable | As nectar-tap_saprotroph (Põlme et al. 2020); Decay sugar-rich substrates (Põlme et al. 2020) | Põlme S, et al. 2020. Fungal Diversity 105:1-16 ((https://doi.org/10.1007/s13225-020-00466-2)) |
| Otu59035 | 1 | k__Fungi(100);p__Ascomycota(100);c__Saccharomycetes(100);o__Saccharomycetales(100);f__Metschnikowiaceae(100);g__Metschnikowia(100);s__Metschnikowia_pulcherrima(100); | Metschnikowia | 13 | Saprotroph | \|Undefined Saprotroph\| | Yeast | NULL | Probable | As nectar-tap_saprotroph (Põlme et al. 2020); Decay sugar-rich substrates (Põlme et al. 2020) | Põlme S, et al. 2020. Fungal Diversity 105:1-16 ((https://doi.org/10.1007/s13225-020-00466-2)) |
| Otu59038 | 1 | k__Fungi(100);p__Ascomycota(100);c__Saccharomycetes(100);o__Saccharomycetales(100);f__Metschnikowiaceae(100);g__Metschnikowia(100);s__Metschnikowia_pulcherrima(100); | Metschnikowia | 13 | Saprotroph | \|Undefined Saprotroph\| | Yeast | NULL | Probable | As nectar-tap_saprotroph (Põlme et al. 2020); Decay sugar-rich substrates (Põlme et al. 2020) | Põlme S, et al. 2020. Fungal Diversity 105:1-16 ((https://doi.org/10.1007/s13225-020-00466-2)) |
| Otu59040 | 1 | k__Fungi(100);p__Ascomycota(100);c__Saccharomycetes(100);o__Saccharomycetales(100);f__Metschnikowiaceae(100);g__Metschnikowia(100);s__Metschnikowia_pulcherrima(100); | Metschnikowia | 13 | Saprotroph | \|Undefined Saprotroph\| | Yeast | NULL | Probable | As nectar-tap_saprotroph (Põlme et al. 2020); Decay sugar-rich substrates (Põlme et al. 2020) | Põlme S, et al. 2020. Fungal Diversity 105:1-16 ((https://doi.org/10.1007/s13225-020-00466-2)) |
| Otu59041 | 1 | k__Fungi(100);p__Ascomycota(100);c__Saccharomycetes(100);o__Saccharomycetales(100);f__Metschnikowiaceae(100);g__Metschnikowia(100);s__Metschnikowia_pulcherrima(100); | Metschnikowia | 13 | Saprotroph | \|Undefined Saprotroph\| | Yeast | NULL | Probable | As nectar-tap_saprotroph (Põlme et al. 2020); Decay sugar-rich substrates (Põlme et al. 2020) | Põlme S, et al. 2020. Fungal Diversity 105:1-16 ((https://doi.org/10.1007/s13225-020-00466-2)) |
| Otu59042 | 1 | k__Fungi(100);p__Ascomycota(100);c__Saccharomycetes(100);o__Saccharomycetales(100);f__Metschnikowiaceae(100);g__Metschnikowia(100);s__Metschnikowia_pulcherrima(100); | Metschnikowia | 13 | Saprotroph | \|Undefined Saprotroph\| | Yeast | NULL | Probable | As nectar-tap_saprotroph (Põlme et al. 2020); Decay sugar-rich substrates (Põlme et al. 2020) | Põlme S, et al. 2020. Fungal Diversity 105:1-16 ((https://doi.org/10.1007/s13225-020-00466-2)) |
| Otu59043 | 1 | k__Fungi(100);p__Ascomycota(100);c__Saccharomycetes(100);o__Saccharomycetales(100);f__Metschnikowiaceae(100);g__Metschnikowia(100);s__Metschnikowia_pulcherrima(100); | Metschnikowia | 13 | Saprotroph | \|Undefined Saprotroph\| | Yeast | NULL | Probable | As nectar-tap_saprotroph (Põlme et al. 2020); Decay sugar-rich substrates (Põlme et al. 2020) | Põlme S, et al. 2020. Fungal Diversity 105:1-16 ((https://doi.org/10.1007/s13225-020-00466-2)) |
| Otu59044 | 1 | k__Fungi(100);p__Ascomycota(100);c__Saccharomycetes(100);o__Saccharomycetales(100);f__Metschnikowiaceae(100);g__Metschnikowia(100);g__Metschnikowia_unclassified(100); | Metschnikowia | 13 | Saprotroph | \|Undefined Saprotroph\| | Yeast | NULL | Probable | As nectar-tap_saprotroph (Põlme et al. 2020); Decay sugar-rich substrates (Põlme et al. 2020) | Põlme S, et al. 2020. Fungal Diversity 105:1-16 ((https://doi.org/10.1007/s13225-020-00466-2)) |
| Otu59045 | 1 | k__Fungi(100);p__Ascomycota(100);c__Saccharomycetes(100);o__Saccharomycetales(100);f__Metschnikowiaceae(100);g__Metschnikowia(100);s__Metschnikowia_pulcherrima(100); | Metschnikowia | 13 | Saprotroph | \|Undefined Saprotroph\| | Yeast | NULL | Probable | As nectar-tap_saprotroph (Põlme et al. 2020); Decay sugar-rich substrates (Põlme et al. 2020) | Põlme S, et al. 2020. Fungal Diversity 105:1-16 ((https://doi.org/10.1007/s13225-020-00466-2)) |
| Otu59046 | 1 | k__Fungi(100);p__Ascomycota(100);c__Saccharomycetes(100);o__Saccharomycetales(100);f__Metschnikowiaceae(100);g__Metschnikowia(100);s__Metschnikowia_pulcherrima(100); | Metschnikowia | 13 | Saprotroph | \|Undefined Saprotroph\| | Yeast | NULL | Probable | As nectar-tap_saprotroph (Põlme et al. 2020); Decay sugar-rich substrates (Põlme et al. 2020) | Põlme S, et al. 2020. Fungal Diversity 105:1-16 ((https://doi.org/10.1007/s13225-020-00466-2)) |
| Otu59048 | 1 | k__Fungi(100);p__Ascomycota(100);c__Saccharomycetes(100);o__Saccharomycetales(100);f__Metschnikowiaceae(100);g__Metschnikowia(100);s__Metschnikowia_pulcherrima(100); | Metschnikowia | 13 | Saprotroph | \|Undefined Saprotroph\| | Yeast | NULL | Probable | As nectar-tap_saprotroph (Põlme et al. 2020); Decay sugar-rich substrates (Põlme et al. 2020) | Põlme S, et al. 2020. Fungal Diversity 105:1-16 ((https://doi.org/10.1007/s13225-020-00466-2)) |
| Otu59049 | 1 | k__Fungi(100);p__Ascomycota(100);c__Saccharomycetes(100);o__Saccharomycetales(100);f__Metschnikowiaceae(100);g__Metschnikowia(100);s__Metschnikowia_pulcherrima(100); | Metschnikowia | 13 | Saprotroph | \|Undefined Saprotroph\| | Yeast | NULL | Probable | As nectar-tap_saprotroph (Põlme et al. 2020); Decay sugar-rich substrates (Põlme et al. 2020) | Põlme S, et al. 2020. Fungal Diversity 105:1-16 ((https://doi.org/10.1007/s13225-020-00466-2)) |
| Otu59050 | 1 | k__Fungi(100);p__Ascomycota(100);c__Saccharomycetes(100);o__Saccharomycetales(100);f__Metschnikowiaceae(100);g__Metschnikowia(100);g__Metschnikowia_unclassified(100); | Metschnikowia | 13 | Saprotroph | \|Undefined Saprotroph\| | Yeast | NULL | Probable | As nectar-tap_saprotroph (Põlme et al. 2020); Decay sugar-rich substrates (Põlme et al. 2020) | Põlme S, et al. 2020. Fungal Diversity 105:1-16 ((https://doi.org/10.1007/s13225-020-00466-2)) |
| Otu59051 | 1 | k__Fungi(100);p__Ascomycota(100);c__Saccharomycetes(100);o__Saccharomycetales(100);f__Metschnikowiaceae(100);g__Metschnikowia(100);g__Metschnikowia_unclassified(100); | Metschnikowia | 13 | Saprotroph | \|Undefined Saprotroph\| | Yeast | NULL | Probable | As nectar-tap_saprotroph (Põlme et al. 2020); Decay sugar-rich substrates (Põlme et al. 2020) | Põlme S, et al. 2020. Fungal Diversity 105:1-16 ((https://doi.org/10.1007/s13225-020-00466-2)) |
| Otu59052 | 1 | k__Fungi(100);p__Ascomycota(100);c__Saccharomycetes(100);o__Saccharomycetales(100);f__Metschnikowiaceae(100);g__Metschnikowia(100);s__Metschnikowia_pulcherrima(100); | Metschnikowia | 13 | Saprotroph | \|Undefined Saprotroph\| | Yeast | NULL | Probable | As nectar-tap_saprotroph (Põlme et al. 2020); Decay sugar-rich substrates (Põlme et al. 2020) | Põlme S, et al. 2020. Fungal Diversity 105:1-16 ((https://doi.org/10.1007/s13225-020-00466-2)) |
| Otu59053 | 1 | k__Fungi(100);p__Ascomycota(100);c__Saccharomycetes(100);o__Saccharomycetales(100);f__Metschnikowiaceae(100);g__Metschnikowia(100);g__Metschnikowia_unclassified(100); | Metschnikowia | 13 | Saprotroph | \|Undefined Saprotroph\| | Yeast | NULL | Probable | As nectar-tap_saprotroph (Põlme et al. 2020); Decay sugar-rich substrates (Põlme et al. 2020) | Põlme S, et al. 2020. Fungal Diversity 105:1-16 ((https://doi.org/10.1007/s13225-020-00466-2)) |
| Otu59055 | 1 | k__Fungi(100);p__Ascomycota(100);c__Saccharomycetes(100);o__Saccharomycetales(100);f__Metschnikowiaceae(100);g__Metschnikowia(100);g__Metschnikowia_unclassified(100); | Metschnikowia | 13 | Saprotroph | \|Undefined Saprotroph\| | Yeast | NULL | Probable | As nectar-tap_saprotroph (Põlme et al. 2020); Decay sugar-rich substrates (Põlme et al. 2020) | Põlme S, et al. 2020. Fungal Diversity 105:1-16 ((https://doi.org/10.1007/s13225-020-00466-2)) |
| Otu59057 | 1 | k__Fungi(100);p__Ascomycota(100);c__Saccharomycetes(100);o__Saccharomycetales(100);f__Metschnikowiaceae(100);g__Metschnikowia(100);s__Metschnikowia_pulcherrima(100); | Metschnikowia | 13 | Saprotroph | \|Undefined Saprotroph\| | Yeast | NULL | Probable | As nectar-tap_saprotroph (Põlme et al. 2020); Decay sugar-rich substrates (Põlme et al. 2020) | Põlme S, et al. 2020. Fungal Diversity 105:1-16 ((https://doi.org/10.1007/s13225-020-00466-2)) |
| Otu59058 | 1 | k__Fungi(100);p__Ascomycota(100);c__Saccharomycetes(100);o__Saccharomycetales(100);f__Metschnikowiaceae(100);g__Metschnikowia(100);g__Metschnikowia_unclassified(100); | Metschnikowia | 13 | Saprotroph | \|Undefined Saprotroph\| | Yeast | NULL | Probable | As nectar-tap_saprotroph (Põlme et al. 2020); Decay sugar-rich substrates (Põlme et al. 2020) | Põlme S, et al. 2020. Fungal Diversity 105:1-16 ((https://doi.org/10.1007/s13225-020-00466-2)) |
| Otu59059 | 1 | k__Fungi(100);p__Ascomycota(100);c__Saccharomycetes(100);o__Saccharomycetales(100);f__Metschnikowiaceae(100);g__Metschnikowia(100);s__Metschnikowia_pulcherrima(100); | Metschnikowia | 13 | Saprotroph | \|Undefined Saprotroph\| | Yeast | NULL | Probable | As nectar-tap_saprotroph (Põlme et al. 2020); Decay sugar-rich substrates (Põlme et al. 2020) | Põlme S, et al. 2020. Fungal Diversity 105:1-16 ((https://doi.org/10.1007/s13225-020-00466-2)) |
| Otu59061 | 1 | k__Fungi(100);p__Ascomycota(100);c__Saccharomycetes(100);o__Saccharomycetales(100);f__Metschnikowiaceae(100);g__Metschnikowia(100);g__Metschnikowia_unclassified(100); | Metschnikowia | 13 | Saprotroph | \|Undefined Saprotroph\| | Yeast | NULL | Probable | As nectar-tap_saprotroph (Põlme et al. 2020); Decay sugar-rich substrates (Põlme et al. 2020) | Põlme S, et al. 2020. Fungal Diversity 105:1-16 ((https://doi.org/10.1007/s13225-020-00466-2)) |
| Otu59062 | 1 | k__Fungi(100);p__Ascomycota(100);c__Saccharomycetes(100);o__Saccharomycetales(100);f__Metschnikowiaceae(100);g__Metschnikowia(100);s__Metschnikowia_pulcherrima(100); | Metschnikowia | 13 | Saprotroph | \|Undefined Saprotroph\| | Yeast | NULL | Probable | As nectar-tap_saprotroph (Põlme et al. 2020); Decay sugar-rich substrates (Põlme et al. 2020) | Põlme S, et al. 2020. Fungal Diversity 105:1-16 ((https://doi.org/10.1007/s13225-020-00466-2)) |
| Otu59063 | 1 | k__Fungi(100);p__Ascomycota(100);c__Saccharomycetes(100);o__Saccharomycetales(100);f__Metschnikowiaceae(100);g__Metschnikowia(100);s__Metschnikowia_pulcherrima(100); | Metschnikowia | 13 | Saprotroph | \|Undefined Saprotroph\| | Yeast | NULL | Probable | As nectar-tap_saprotroph (Põlme et al. 2020); Decay sugar-rich substrates (Põlme et al. 2020) | Põlme S, et al. 2020. Fungal Diversity 105:1-16 ((https://doi.org/10.1007/s13225-020-00466-2)) |
| Otu59067 | 1 | k__Fungi(100);p__Ascomycota(100);c__Saccharomycetes(100);o__Saccharomycetales(100);f__Metschnikowiaceae(100);g__Metschnikowia(100);s__Metschnikowia_pulcherrima(100); | Metschnikowia | 13 | Saprotroph | \|Undefined Saprotroph\| | Yeast | NULL | Probable | As nectar-tap_saprotroph (Põlme et al. 2020); Decay sugar-rich substrates (Põlme et al. 2020) | Põlme S, et al. 2020. Fungal Diversity 105:1-16 ((https://doi.org/10.1007/s13225-020-00466-2)) |
| Otu59069 | 1 | k__Fungi(100);p__Ascomycota(100);c__Saccharomycetes(100);o__Saccharomycetales(100);f__Metschnikowiaceae(100);g__Metschnikowia(100);s__Metschnikowia_pulcherrima(100); | Metschnikowia | 13 | Saprotroph | \|Undefined Saprotroph\| | Yeast | NULL | Probable | As nectar-tap_saprotroph (Põlme et al. 2020); Decay sugar-rich substrates (Põlme et al. 2020) | Põlme S, et al. 2020. Fungal Diversity 105:1-16 ((https://doi.org/10.1007/s13225-020-00466-2)) |
| Otu59070 | 1 | k__Fungi(100);p__Ascomycota(100);c__Saccharomycetes(100);o__Saccharomycetales(100);f__Metschnikowiaceae(100);g__Metschnikowia(100);g__Metschnikowia_unclassified(100); | Metschnikowia | 13 | Saprotroph | \|Undefined Saprotroph\| | Yeast | NULL | Probable | As nectar-tap_saprotroph (Põlme et al. 2020); Decay sugar-rich substrates (Põlme et al. 2020) | Põlme S, et al. 2020. Fungal Diversity 105:1-16 ((https://doi.org/10.1007/s13225-020-00466-2)) |
| Otu59071 | 1 | k__Fungi(100);p__Ascomycota(100);c__Saccharomycetes(100);o__Saccharomycetales(100);f__Metschnikowiaceae(100);g__Metschnikowia(100);g__Metschnikowia_unclassified(100); | Metschnikowia | 13 | Saprotroph | \|Undefined Saprotroph\| | Yeast | NULL | Probable | As nectar-tap_saprotroph (Põlme et al. 2020); Decay sugar-rich substrates (Põlme et al. 2020) | Põlme S, et al. 2020. Fungal Diversity 105:1-16 ((https://doi.org/10.1007/s13225-020-00466-2)) |
| Otu59072 | 1 | k__Fungi(100);p__Ascomycota(100);c__Saccharomycetes(100);o__Saccharomycetales(100);f__Metschnikowiaceae(100);g__Metschnikowia(100);s__Metschnikowia_pulcherrima(100); | Metschnikowia | 13 | Saprotroph | \|Undefined Saprotroph\| | Yeast | NULL | Probable | As nectar-tap_saprotroph (Põlme et al. 2020); Decay sugar-rich substrates (Põlme et al. 2020) | Põlme S, et al. 2020. Fungal Diversity 105:1-16 ((https://doi.org/10.1007/s13225-020-00466-2)) |
| Otu59073 | 1 | k__Fungi(100);p__Ascomycota(100);c__Saccharomycetes(100);o__Saccharomycetales(100);f__Metschnikowiaceae(100);g__Metschnikowia(100);s__Metschnikowia_pulcherrima(100); | Metschnikowia | 13 | Saprotroph | \|Undefined Saprotroph\| | Yeast | NULL | Probable | As nectar-tap_saprotroph (Põlme et al. 2020); Decay sugar-rich substrates (Põlme et al. 2020) | Põlme S, et al. 2020. Fungal Diversity 105:1-16 ((https://doi.org/10.1007/s13225-020-00466-2)) |
| Otu59074 | 1 | k__Fungi(100);p__Ascomycota(100);c__Saccharomycetes(100);o__Saccharomycetales(100);f__Metschnikowiaceae(100);g__Metschnikowia(100);s__Metschnikowia_pulcherrima(100); | Metschnikowia | 13 | Saprotroph | \|Undefined Saprotroph\| | Yeast | NULL | Probable | As nectar-tap_saprotroph (Põlme et al. 2020); Decay sugar-rich substrates (Põlme et al. 2020) | Põlme S, et al. 2020. Fungal Diversity 105:1-16 ((https://doi.org/10.1007/s13225-020-00466-2)) |
| Otu59075 | 1 | k__Fungi(100);p__Ascomycota(100);c__Saccharomycetes(100);o__Saccharomycetales(100);f__Metschnikowiaceae(100);g__Metschnikowia(100);g__Metschnikowia_unclassified(100); | Metschnikowia | 13 | Saprotroph | \|Undefined Saprotroph\| | Yeast | NULL | Probable | As nectar-tap_saprotroph (Põlme et al. 2020); Decay sugar-rich substrates (Põlme et al. 2020) | Põlme S, et al. 2020. Fungal Diversity 105:1-16 ((https://doi.org/10.1007/s13225-020-00466-2)) |
| Otu59077 | 1 | k__Fungi(100);p__Ascomycota(100);c__Saccharomycetes(100);o__Saccharomycetales(100);f__Metschnikowiaceae(100);g__Metschnikowia(100);s__Metschnikowia_pulcherrima(100); | Metschnikowia | 13 | Saprotroph | \|Undefined Saprotroph\| | Yeast | NULL | Probable | As nectar-tap_saprotroph (Põlme et al. 2020); Decay sugar-rich substrates (Põlme et al. 2020) | Põlme S, et al. 2020. Fungal Diversity 105:1-16 ((https://doi.org/10.1007/s13225-020-00466-2)) |
| Otu59078 | 1 | k__Fungi(100);p__Ascomycota(100);c__Saccharomycetes(100);o__Saccharomycetales(100);f__Metschnikowiaceae(100);g__Metschnikowia(100);g__Metschnikowia_unclassified(100); | Metschnikowia | 13 | Saprotroph | \|Undefined Saprotroph\| | Yeast | NULL | Probable | As nectar-tap_saprotroph (Põlme et al. 2020); Decay sugar-rich substrates (Põlme et al. 2020) | Põlme S, et al. 2020. Fungal Diversity 105:1-16 ((https://doi.org/10.1007/s13225-020-00466-2)) |
| Otu59079 | 1 | k__Fungi(100);p__Ascomycota(100);c__Saccharomycetes(100);o__Saccharomycetales(100);f__Metschnikowiaceae(100);g__Metschnikowia(100);s__Metschnikowia_pulcherrima(100); | Metschnikowia | 13 | Saprotroph | \|Undefined Saprotroph\| | Yeast | NULL | Probable | As nectar-tap_saprotroph (Põlme et al. 2020); Decay sugar-rich substrates (Põlme et al. 2020) | Põlme S, et al. 2020. Fungal Diversity 105:1-16 ((https://doi.org/10.1007/s13225-020-00466-2)) |
| Otu59080 | 1 | k__Fungi(100);p__Ascomycota(100);c__Saccharomycetes(100);o__Saccharomycetales(100);f__Metschnikowiaceae(100);g__Metschnikowia(100);s__Metschnikowia_pulcherrima(100); | Metschnikowia | 13 | Saprotroph | \|Undefined Saprotroph\| | Yeast | NULL | Probable | As nectar-tap_saprotroph (Põlme et al. 2020); Decay sugar-rich substrates (Põlme et al. 2020) | Põlme S, et al. 2020. Fungal Diversity 105:1-16 ((https://doi.org/10.1007/s13225-020-00466-2)) |
| Otu59081 | 1 | k__Fungi(100);p__Ascomycota(100);c__Saccharomycetes(100);o__Saccharomycetales(100);f__Metschnikowiaceae(100);g__Metschnikowia(100);s__Metschnikowia_pulcherrima(100); | Metschnikowia | 13 | Saprotroph | \|Undefined Saprotroph\| | Yeast | NULL | Probable | As nectar-tap_saprotroph (Põlme et al. 2020); Decay sugar-rich substrates (Põlme et al. 2020) | Põlme S, et al. 2020. Fungal Diversity 105:1-16 ((https://doi.org/10.1007/s13225-020-00466-2)) |
| Otu59082 | 1 | k__Fungi(100);p__Ascomycota(100);c__Saccharomycetes(100);o__Saccharomycetales(100);f__Metschnikowiaceae(100);g__Metschnikowia(100);g__Metschnikowia_unclassified(100); | Metschnikowia | 13 | Saprotroph | \|Undefined Saprotroph\| | Yeast | NULL | Probable | As nectar-tap_saprotroph (Põlme et al. 2020); Decay sugar-rich substrates (Põlme et al. 2020) | Põlme S, et al. 2020. Fungal Diversity 105:1-16 ((https://doi.org/10.1007/s13225-020-00466-2)) |
| Otu59083 | 1 | k__Fungi(100);p__Ascomycota(100);c__Saccharomycetes(100);o__Saccharomycetales(100);f__Metschnikowiaceae(100);g__Metschnikowia(100);s__Metschnikowia_pulcherrima(100); | Metschnikowia | 13 | Saprotroph | \|Undefined Saprotroph\| | Yeast | NULL | Probable | As nectar-tap_saprotroph (Põlme et al. 2020); Decay sugar-rich substrates (Põlme et al. 2020) | Põlme S, et al. 2020. Fungal Diversity 105:1-16 ((https://doi.org/10.1007/s13225-020-00466-2)) |
| Otu59085 | 1 | k__Fungi(100);p__Ascomycota(100);c__Saccharomycetes(100);o__Saccharomycetales(100);f__Metschnikowiaceae(100);g__Metschnikowia(100);g__Metschnikowia_unclassified(100); | Metschnikowia | 13 | Saprotroph | \|Undefined Saprotroph\| | Yeast | NULL | Probable | As nectar-tap_saprotroph (Põlme et al. 2020); Decay sugar-rich substrates (Põlme et al. 2020) | Põlme S, et al. 2020. Fungal Diversity 105:1-16 ((https://doi.org/10.1007/s13225-020-00466-2)) |
| Otu59086 | 1 | k__Fungi(100);p__Ascomycota(100);c__Saccharomycetes(100);o__Saccharomycetales(100);f__Metschnikowiaceae(100);g__Metschnikowia(100);s__Metschnikowia_pulcherrima(100); | Metschnikowia | 13 | Saprotroph | \|Undefined Saprotroph\| | Yeast | NULL | Probable | As nectar-tap_saprotroph (Põlme et al. 2020); Decay sugar-rich substrates (Põlme et al. 2020) | Põlme S, et al. 2020. Fungal Diversity 105:1-16 ((https://doi.org/10.1007/s13225-020-00466-2)) |
| Otu59087 | 1 | k__Fungi(100);p__Ascomycota(100);c__Saccharomycetes(100);o__Saccharomycetales(100);f__Metschnikowiaceae(100);g__Metschnikowia(100);s__Metschnikowia_pulcherrima(100); | Metschnikowia | 13 | Saprotroph | \|Undefined Saprotroph\| | Yeast | NULL | Probable | As nectar-tap_saprotroph (Põlme et al. 2020); Decay sugar-rich substrates (Põlme et al. 2020) | Põlme S, et al. 2020. Fungal Diversity 105:1-16 ((https://doi.org/10.1007/s13225-020-00466-2)) |
| Otu59089 | 1 | k__Fungi(100);p__Ascomycota(100);c__Saccharomycetes(100);o__Saccharomycetales(100);f__Metschnikowiaceae(100);g__Metschnikowia(100);s__Metschnikowia_pulcherrima(100); | Metschnikowia | 13 | Saprotroph | \|Undefined Saprotroph\| | Yeast | NULL | Probable | As nectar-tap_saprotroph (Põlme et al. 2020); Decay sugar-rich substrates (Põlme et al. 2020) | Põlme S, et al. 2020. Fungal Diversity 105:1-16 ((https://doi.org/10.1007/s13225-020-00466-2)) |
| Otu59090 | 1 | k__Fungi(100);p__Ascomycota(100);c__Saccharomycetes(100);o__Saccharomycetales(100);f__Metschnikowiaceae(100);g__Metschnikowia(100);s__Metschnikowia_pulcherrima(100); | Metschnikowia | 13 | Saprotroph | \|Undefined Saprotroph\| | Yeast | NULL | Probable | As nectar-tap_saprotroph (Põlme et al. 2020); Decay sugar-rich substrates (Põlme et al. 2020) | Põlme S, et al. 2020. Fungal Diversity 105:1-16 ((https://doi.org/10.1007/s13225-020-00466-2)) |
| Otu59091 | 1 | k__Fungi(100);p__Ascomycota(100);c__Saccharomycetes(100);o__Saccharomycetales(100);f__Metschnikowiaceae(100);g__Metschnikowia(100);s__Metschnikowia_pulcherrima(100); | Metschnikowia | 13 | Saprotroph | \|Undefined Saprotroph\| | Yeast | NULL | Probable | As nectar-tap_saprotroph (Põlme et al. 2020); Decay sugar-rich substrates (Põlme et al. 2020) | Põlme S, et al. 2020. Fungal Diversity 105:1-16 ((https://doi.org/10.1007/s13225-020-00466-2)) |
| Otu59092 | 1 | k__Fungi(100);p__Ascomycota(100);c__Saccharomycetes(100);o__Saccharomycetales(100);f__Metschnikowiaceae(100);g__Metschnikowia(100);s__Metschnikowia_pulcherrima(100); | Metschnikowia | 13 | Saprotroph | \|Undefined Saprotroph\| | Yeast | NULL | Probable | As nectar-tap_saprotroph (Põlme et al. 2020); Decay sugar-rich substrates (Põlme et al. 2020) | Põlme S, et al. 2020. Fungal Diversity 105:1-16 ((https://doi.org/10.1007/s13225-020-00466-2)) |
| Otu59093 | 1 | k__Fungi(100);p__Ascomycota(100);c__Saccharomycetes(100);o__Saccharomycetales(100);f__Metschnikowiaceae(100);g__Metschnikowia(100);g__Metschnikowia_unclassified(100); | Metschnikowia | 13 | Saprotroph | \|Undefined Saprotroph\| | Yeast | NULL | Probable | As nectar-tap_saprotroph (Põlme et al. 2020); Decay sugar-rich substrates (Põlme et al. 2020) | Põlme S, et al. 2020. Fungal Diversity 105:1-16 ((https://doi.org/10.1007/s13225-020-00466-2)) |
| Otu59094 | 1 | k__Fungi(100);p__Ascomycota(100);c__Saccharomycetes(100);o__Saccharomycetales(100);f__Metschnikowiaceae(100);g__Metschnikowia(100);s__Metschnikowia_pulcherrima(100); | Metschnikowia | 13 | Saprotroph | \|Undefined Saprotroph\| | Yeast | NULL | Probable | As nectar-tap_saprotroph (Põlme et al. 2020); Decay sugar-rich substrates (Põlme et al. 2020) | Põlme S, et al. 2020. Fungal Diversity 105:1-16 ((https://doi.org/10.1007/s13225-020-00466-2)) |
| Otu59096 | 1 | k__Fungi(100);p__Ascomycota(100);c__Saccharomycetes(100);o__Saccharomycetales(100);f__Metschnikowiaceae(100);g__Metschnikowia(100);s__Metschnikowia_pulcherrima(100); | Metschnikowia | 13 | Saprotroph | \|Undefined Saprotroph\| | Yeast | NULL | Probable | As nectar-tap_saprotroph (Põlme et al. 2020); Decay sugar-rich substrates (Põlme et al. 2020) | Põlme S, et al. 2020. Fungal Diversity 105:1-16 ((https://doi.org/10.1007/s13225-020-00466-2)) |
| Otu59097 | 1 | k__Fungi(100);p__Ascomycota(100);c__Saccharomycetes(100);o__Saccharomycetales(100);f__Metschnikowiaceae(100);g__Metschnikowia(100);g__Metschnikowia_unclassified(100); | Metschnikowia | 13 | Saprotroph | \|Undefined Saprotroph\| | Yeast | NULL | Probable | As nectar-tap_saprotroph (Põlme et al. 2020); Decay sugar-rich substrates (Põlme et al. 2020) | Põlme S, et al. 2020. Fungal Diversity 105:1-16 ((https://doi.org/10.1007/s13225-020-00466-2)) |
| Otu59098 | 1 | k__Fungi(100);p__Ascomycota(100);c__Saccharomycetes(100);o__Saccharomycetales(100);f__Metschnikowiaceae(100);g__Metschnikowia(100);g__Metschnikowia_unclassified(100); | Metschnikowia | 13 | Saprotroph | \|Undefined Saprotroph\| | Yeast | NULL | Probable | As nectar-tap_saprotroph (Põlme et al. 2020); Decay sugar-rich substrates (Põlme et al. 2020) | Põlme S, et al. 2020. Fungal Diversity 105:1-16 ((https://doi.org/10.1007/s13225-020-00466-2)) |
| Otu59100 | 1 | k__Fungi(100);p__Ascomycota(100);c__Saccharomycetes(100);o__Saccharomycetales(100);f__Metschnikowiaceae(100);g__Metschnikowia(100);g__Metschnikowia_unclassified(100); | Metschnikowia | 13 | Saprotroph | \|Undefined Saprotroph\| | Yeast | NULL | Probable | As nectar-tap_saprotroph (Põlme et al. 2020); Decay sugar-rich substrates (Põlme et al. 2020) | Põlme S, et al. 2020. Fungal Diversity 105:1-16 ((https://doi.org/10.1007/s13225-020-00466-2)) |
| Otu59102 | 1 | k__Fungi(100);p__Ascomycota(100);c__Saccharomycetes(100);o__Saccharomycetales(100);f__Metschnikowiaceae(100);g__Metschnikowia(100);s__Metschnikowia_pulcherrima(100); | Metschnikowia | 13 | Saprotroph | \|Undefined Saprotroph\| | Yeast | NULL | Probable | As nectar-tap_saprotroph (Põlme et al. 2020); Decay sugar-rich substrates (Põlme et al. 2020) | Põlme S, et al. 2020. Fungal Diversity 105:1-16 ((https://doi.org/10.1007/s13225-020-00466-2)) |
| Otu59103 | 1 | k__Fungi(100);p__Ascomycota(100);c__Saccharomycetes(100);o__Saccharomycetales(100);f__Metschnikowiaceae(100);g__Metschnikowia(100);s__Metschnikowia_pulcherrima(100); | Metschnikowia | 13 | Saprotroph | \|Undefined Saprotroph\| | Yeast | NULL | Probable | As nectar-tap_saprotroph (Põlme et al. 2020); Decay sugar-rich substrates (Põlme et al. 2020) | Põlme S, et al. 2020. Fungal Diversity 105:1-16 ((https://doi.org/10.1007/s13225-020-00466-2)) |
| Otu59104 | 1 | k__Fungi(100);p__Ascomycota(100);c__Saccharomycetes(100);o__Saccharomycetales(100);f__Metschnikowiaceae(100);g__Metschnikowia(100);s__Metschnikowia_pulcherrima(100); | Metschnikowia | 13 | Saprotroph | \|Undefined Saprotroph\| | Yeast | NULL | Probable | As nectar-tap_saprotroph (Põlme et al. 2020); Decay sugar-rich substrates (Põlme et al. 2020) | Põlme S, et al. 2020. Fungal Diversity 105:1-16 ((https://doi.org/10.1007/s13225-020-00466-2)) |
| Otu59105 | 1 | k__Fungi(100);p__Ascomycota(100);c__Saccharomycetes(100);o__Saccharomycetales(100);f__Metschnikowiaceae(100);g__Metschnikowia(100);s__Metschnikowia_pulcherrima(100); | Metschnikowia | 13 | Saprotroph | \|Undefined Saprotroph\| | Yeast | NULL | Probable | As nectar-tap_saprotroph (Põlme et al. 2020); Decay sugar-rich substrates (Põlme et al. 2020) | Põlme S, et al. 2020. Fungal Diversity 105:1-16 ((https://doi.org/10.1007/s13225-020-00466-2)) |
| Otu59106 | 1 | k__Fungi(100);p__Ascomycota(100);c__Saccharomycetes(100);o__Saccharomycetales(100);f__Metschnikowiaceae(100);g__Metschnikowia(100);s__Metschnikowia_pulcherrima(100); | Metschnikowia | 13 | Saprotroph | \|Undefined Saprotroph\| | Yeast | NULL | Probable | As nectar-tap_saprotroph (Põlme et al. 2020); Decay sugar-rich substrates (Põlme et al. 2020) | Põlme S, et al. 2020. Fungal Diversity 105:1-16 ((https://doi.org/10.1007/s13225-020-00466-2)) |
| Otu59107 | 1 | k__Fungi(100);p__Ascomycota(100);c__Saccharomycetes(100);o__Saccharomycetales(100);f__Metschnikowiaceae(100);g__Metschnikowia(100);s__Metschnikowia_pulcherrima(100); | Metschnikowia | 13 | Saprotroph | \|Undefined Saprotroph\| | Yeast | NULL | Probable | As nectar-tap_saprotroph (Põlme et al. 2020); Decay sugar-rich substrates (Põlme et al. 2020) | Põlme S, et al. 2020. Fungal Diversity 105:1-16 ((https://doi.org/10.1007/s13225-020-00466-2)) |
| Otu59108 | 1 | k__Fungi(100);p__Ascomycota(100);c__Saccharomycetes(100);o__Saccharomycetales(100);f__Metschnikowiaceae(100);g__Metschnikowia(100);s__Metschnikowia_pulcherrima(100); | Metschnikowia | 13 | Saprotroph | \|Undefined Saprotroph\| | Yeast | NULL | Probable | As nectar-tap_saprotroph (Põlme et al. 2020); Decay sugar-rich substrates (Põlme et al. 2020) | Põlme S, et al. 2020. Fungal Diversity 105:1-16 ((https://doi.org/10.1007/s13225-020-00466-2)) |
| Otu59109 | 1 | k__Fungi(100);p__Ascomycota(100);c__Saccharomycetes(100);o__Saccharomycetales(100);f__Metschnikowiaceae(100);g__Metschnikowia(100);s__Metschnikowia_pulcherrima(100); | Metschnikowia | 13 | Saprotroph | \|Undefined Saprotroph\| | Yeast | NULL | Probable | As nectar-tap_saprotroph (Põlme et al. 2020); Decay sugar-rich substrates (Põlme et al. 2020) | Põlme S, et al. 2020. Fungal Diversity 105:1-16 ((https://doi.org/10.1007/s13225-020-00466-2)) |
| Otu59110 | 1 | k__Fungi(100);p__Ascomycota(100);c__Saccharomycetes(100);o__Saccharomycetales(100);f__Metschnikowiaceae(100);g__Metschnikowia(100);g__Metschnikowia_unclassified(100); | Metschnikowia | 13 | Saprotroph | \|Undefined Saprotroph\| | Yeast | NULL | Probable | As nectar-tap_saprotroph (Põlme et al. 2020); Decay sugar-rich substrates (Põlme et al. 2020) | Põlme S, et al. 2020. Fungal Diversity 105:1-16 ((https://doi.org/10.1007/s13225-020-00466-2)) |
| Otu59114 | 1 | k__Fungi(100);p__Ascomycota(100);c__Saccharomycetes(100);o__Saccharomycetales(100);f__Metschnikowiaceae(100);g__Metschnikowia(100);s__Metschnikowia_pulcherrima(100); | Metschnikowia | 13 | Saprotroph | \|Undefined Saprotroph\| | Yeast | NULL | Probable | As nectar-tap_saprotroph (Põlme et al. 2020); Decay sugar-rich substrates (Põlme et al. 2020) | Põlme S, et al. 2020. Fungal Diversity 105:1-16 ((https://doi.org/10.1007/s13225-020-00466-2)) |
| Otu59115 | 1 | k__Fungi(100);p__Ascomycota(100);c__Saccharomycetes(100);o__Saccharomycetales(100);f__Metschnikowiaceae(100);g__Metschnikowia(100);g__Metschnikowia_unclassified(100); | Metschnikowia | 13 | Saprotroph | \|Undefined Saprotroph\| | Yeast | NULL | Probable | As nectar-tap_saprotroph (Põlme et al. 2020); Decay sugar-rich substrates (Põlme et al. 2020) | Põlme S, et al. 2020. Fungal Diversity 105:1-16 ((https://doi.org/10.1007/s13225-020-00466-2)) |
| Otu59118 | 1 | k__Fungi(100);p__Ascomycota(100);c__Saccharomycetes(100);o__Saccharomycetales(100);f__Metschnikowiaceae(100);g__Metschnikowia(100);s__Metschnikowia_pulcherrima(100); | Metschnikowia | 13 | Saprotroph | \|Undefined Saprotroph\| | Yeast | NULL | Probable | As nectar-tap_saprotroph (Põlme et al. 2020); Decay sugar-rich substrates (Põlme et al. 2020) | Põlme S, et al. 2020. Fungal Diversity 105:1-16 ((https://doi.org/10.1007/s13225-020-00466-2)) |
| Otu59119 | 1 | k__Fungi(100);p__Ascomycota(100);c__Saccharomycetes(100);o__Saccharomycetales(100);f__Metschnikowiaceae(100);g__Metschnikowia(100);s__Metschnikowia_pulcherrima(100); | Metschnikowia | 13 | Saprotroph | \|Undefined Saprotroph\| | Yeast | NULL | Probable | As nectar-tap_saprotroph (Põlme et al. 2020); Decay sugar-rich substrates (Põlme et al. 2020) | Põlme S, et al. 2020. Fungal Diversity 105:1-16 ((https://doi.org/10.1007/s13225-020-00466-2)) |
| Otu59120 | 1 | k__Fungi(100);p__Ascomycota(100);c__Saccharomycetes(100);o__Saccharomycetales(100);f__Metschnikowiaceae(100);g__Metschnikowia(100);s__Metschnikowia_pulcherrima(100); | Metschnikowia | 13 | Saprotroph | \|Undefined Saprotroph\| | Yeast | NULL | Probable | As nectar-tap_saprotroph (Põlme et al. 2020); Decay sugar-rich substrates (Põlme et al. 2020) | Põlme S, et al. 2020. Fungal Diversity 105:1-16 ((https://doi.org/10.1007/s13225-020-00466-2)) |
| Otu59121 | 1 | k__Fungi(100);p__Ascomycota(100);c__Saccharomycetes(100);o__Saccharomycetales(100);f__Metschnikowiaceae(100);g__Metschnikowia(100);s__Metschnikowia_pulcherrima(100); | Metschnikowia | 13 | Saprotroph | \|Undefined Saprotroph\| | Yeast | NULL | Probable | As nectar-tap_saprotroph (Põlme et al. 2020); Decay sugar-rich substrates (Põlme et al. 2020) | Põlme S, et al. 2020. Fungal Diversity 105:1-16 ((https://doi.org/10.1007/s13225-020-00466-2)) |
| Otu59122 | 1 | k__Fungi(100);p__Ascomycota(100);c__Saccharomycetes(100);o__Saccharomycetales(100);f__Metschnikowiaceae(100);g__Metschnikowia(100);s__Metschnikowia_pulcherrima(100); | Metschnikowia | 13 | Saprotroph | \|Undefined Saprotroph\| | Yeast | NULL | Probable | As nectar-tap_saprotroph (Põlme et al. 2020); Decay sugar-rich substrates (Põlme et al. 2020) | Põlme S, et al. 2020. Fungal Diversity 105:1-16 ((https://doi.org/10.1007/s13225-020-00466-2)) |
| Otu59124 | 1 | k__Fungi(100);p__Ascomycota(100);c__Saccharomycetes(100);o__Saccharomycetales(100);f__Metschnikowiaceae(100);g__Metschnikowia(100);g__Metschnikowia_unclassified(100); | Metschnikowia | 13 | Saprotroph | \|Undefined Saprotroph\| | Yeast | NULL | Probable | As nectar-tap_saprotroph (Põlme et al. 2020); Decay sugar-rich substrates (Põlme et al. 2020) | Põlme S, et al. 2020. Fungal Diversity 105:1-16 ((https://doi.org/10.1007/s13225-020-00466-2)) |
| Otu59125 | 1 | k__Fungi(100);p__Ascomycota(100);c__Saccharomycetes(100);o__Saccharomycetales(100);f__Metschnikowiaceae(100);g__Metschnikowia(100);s__Metschnikowia_pulcherrima(100); | Metschnikowia | 13 | Saprotroph | \|Undefined Saprotroph\| | Yeast | NULL | Probable | As nectar-tap_saprotroph (Põlme et al. 2020); Decay sugar-rich substrates (Põlme et al. 2020) | Põlme S, et al. 2020. Fungal Diversity 105:1-16 ((https://doi.org/10.1007/s13225-020-00466-2)) |
| Otu59128 | 1 | k__Fungi(100);p__Ascomycota(100);c__Saccharomycetes(100);o__Saccharomycetales(100);f__Metschnikowiaceae(100);g__Metschnikowia(100);g__Metschnikowia_unclassified(100); | Metschnikowia | 13 | Saprotroph | \|Undefined Saprotroph\| | Yeast | NULL | Probable | As nectar-tap_saprotroph (Põlme et al. 2020); Decay sugar-rich substrates (Põlme et al. 2020) | Põlme S, et al. 2020. Fungal Diversity 105:1-16 ((https://doi.org/10.1007/s13225-020-00466-2)) |
| Otu59142 | 1 | k__Fungi(100);p__Ascomycota(100);c__Saccharomycetes(100);o__Saccharomycetales(100);o__Saccharomycetales_unclassified(100);o__Saccharomycetales_unclassified(100);o__Saccharomycetales_unclassified(100); | Saccharomycetales | 7 | Saprotroph | Undefined Saprotroph | Yeast | NULL | Possible | NULL | Sterkenburg E, et al. 2015. New Phytologist 207:1145-1158 ((https://doi.org/10.1111/nph.13426)) |
| Otu59161 | 1 | k__Fungi(100);p__Ascomycota(100);c__Saccharomycetes(100);o__Saccharomycetales(100);f__Metschnikowiaceae(100);g__Metschnikowia(100);s__Metschnikowia_pulcherrima(100); | Metschnikowia | 13 | Saprotroph | \|Undefined Saprotroph\| | Yeast | NULL | Probable | As nectar-tap_saprotroph (Põlme et al. 2020); Decay sugar-rich substrates (Põlme et al. 2020) | Põlme S, et al. 2020. Fungal Diversity 105:1-16 ((https://doi.org/10.1007/s13225-020-00466-2)) |
| Otu59162 | 1 | k__Fungi(100);p__Ascomycota(100);c__Saccharomycetes(100);o__Saccharomycetales(100);f__Metschnikowiaceae(100);g__Metschnikowia(100);s__Metschnikowia_pulcherrima(100); | Metschnikowia | 13 | Saprotroph | \|Undefined Saprotroph\| | Yeast | NULL | Probable | As nectar-tap_saprotroph (Põlme et al. 2020); Decay sugar-rich substrates (Põlme et al. 2020) | Põlme S, et al. 2020. Fungal Diversity 105:1-16 ((https://doi.org/10.1007/s13225-020-00466-2)) |
| Otu59163 | 1 | k__Fungi(100);p__Ascomycota(100);c__Saccharomycetes(100);o__Saccharomycetales(100);f__Metschnikowiaceae(100);g__Metschnikowia(100);s__Metschnikowia_pulcherrima(100); | Metschnikowia | 13 | Saprotroph | \|Undefined Saprotroph\| | Yeast | NULL | Probable | As nectar-tap_saprotroph (Põlme et al. 2020); Decay sugar-rich substrates (Põlme et al. 2020) | Põlme S, et al. 2020. Fungal Diversity 105:1-16 ((https://doi.org/10.1007/s13225-020-00466-2)) |
| Otu59164 | 1 | k__Fungi(100);p__Ascomycota(100);c__Saccharomycetes(100);o__Saccharomycetales(100);f__Metschnikowiaceae(100);g__Metschnikowia(100);s__Metschnikowia_pulcherrima(100); | Metschnikowia | 13 | Saprotroph | \|Undefined Saprotroph\| | Yeast | NULL | Probable | As nectar-tap_saprotroph (Põlme et al. 2020); Decay sugar-rich substrates (Põlme et al. 2020) | Põlme S, et al. 2020. Fungal Diversity 105:1-16 ((https://doi.org/10.1007/s13225-020-00466-2)) |
| Otu59166 | 1 | k__Fungi(100);p__Ascomycota(100);c__Saccharomycetes(100);o__Saccharomycetales(100);f__Metschnikowiaceae(100);g__Metschnikowia(100);s__Metschnikowia_pulcherrima(100); | Metschnikowia | 13 | Saprotroph | \|Undefined Saprotroph\| | Yeast | NULL | Probable | As nectar-tap_saprotroph (Põlme et al. 2020); Decay sugar-rich substrates (Põlme et al. 2020) | Põlme S, et al. 2020. Fungal Diversity 105:1-16 ((https://doi.org/10.1007/s13225-020-00466-2)) |
| Otu59167 | 1 | k__Fungi(100);p__Ascomycota(100);c__Saccharomycetes(100);o__Saccharomycetales(100);f__Metschnikowiaceae(100);g__Metschnikowia(100);s__Metschnikowia_pulcherrima(100); | Metschnikowia | 13 | Saprotroph | \|Undefined Saprotroph\| | Yeast | NULL | Probable | As nectar-tap_saprotroph (Põlme et al. 2020); Decay sugar-rich substrates (Põlme et al. 2020) | Põlme S, et al. 2020. Fungal Diversity 105:1-16 ((https://doi.org/10.1007/s13225-020-00466-2)) |
| Otu59168 | 1 | k__Fungi(100);p__Ascomycota(100);c__Saccharomycetes(100);o__Saccharomycetales(100);f__Metschnikowiaceae(100);g__Metschnikowia(100);s__Metschnikowia_pulcherrima(100); | Metschnikowia | 13 | Saprotroph | \|Undefined Saprotroph\| | Yeast | NULL | Probable | As nectar-tap_saprotroph (Põlme et al. 2020); Decay sugar-rich substrates (Põlme et al. 2020) | Põlme S, et al. 2020. Fungal Diversity 105:1-16 ((https://doi.org/10.1007/s13225-020-00466-2)) |
| Otu59170 | 1 | k__Fungi(100);p__Ascomycota(100);c__Saccharomycetes(100);o__Saccharomycetales(100);f__Metschnikowiaceae(100);g__Metschnikowia(100);s__Metschnikowia_pulcherrima(100); | Metschnikowia | 13 | Saprotroph | \|Undefined Saprotroph\| | Yeast | NULL | Probable | As nectar-tap_saprotroph (Põlme et al. 2020); Decay sugar-rich substrates (Põlme et al. 2020) | Põlme S, et al. 2020. Fungal Diversity 105:1-16 ((https://doi.org/10.1007/s13225-020-00466-2)) |
| Otu59171 | 1 | k__Fungi(100);p__Ascomycota(100);c__Saccharomycetes(100);o__Saccharomycetales(100);f__Metschnikowiaceae(100);g__Metschnikowia(100);g__Metschnikowia_unclassified(100); | Metschnikowia | 13 | Saprotroph | \|Undefined Saprotroph\| | Yeast | NULL | Probable | As nectar-tap_saprotroph (Põlme et al. 2020); Decay sugar-rich substrates (Põlme et al. 2020) | Põlme S, et al. 2020. Fungal Diversity 105:1-16 ((https://doi.org/10.1007/s13225-020-00466-2)) |
| Otu59172 | 1 | k__Fungi(100);p__Ascomycota(100);c__Saccharomycetes(100);o__Saccharomycetales(100);f__Metschnikowiaceae(100);g__Metschnikowia(100);s__Metschnikowia_pulcherrima(100); | Metschnikowia | 13 | Saprotroph | \|Undefined Saprotroph\| | Yeast | NULL | Probable | As nectar-tap_saprotroph (Põlme et al. 2020); Decay sugar-rich substrates (Põlme et al. 2020) | Põlme S, et al. 2020. Fungal Diversity 105:1-16 ((https://doi.org/10.1007/s13225-020-00466-2)) |
| Otu59175 | 1 | k__Fungi(100);p__Ascomycota(100);c__Saccharomycetes(100);o__Saccharomycetales(100);f__Metschnikowiaceae(100);g__Metschnikowia(100);s__Metschnikowia_pulcherrima(100); | Metschnikowia | 13 | Saprotroph | \|Undefined Saprotroph\| | Yeast | NULL | Probable | As nectar-tap_saprotroph (Põlme et al. 2020); Decay sugar-rich substrates (Põlme et al. 2020) | Põlme S, et al. 2020. Fungal Diversity 105:1-16 ((https://doi.org/10.1007/s13225-020-00466-2)) |
| Otu59178 | 1 | k__Fungi(100);p__Ascomycota(100);c__Saccharomycetes(100);o__Saccharomycetales(100);f__Metschnikowiaceae(100);g__Metschnikowia(100);g__Metschnikowia_unclassified(100); | Metschnikowia | 13 | Saprotroph | \|Undefined Saprotroph\| | Yeast | NULL | Probable | As nectar-tap_saprotroph (Põlme et al. 2020); Decay sugar-rich substrates (Põlme et al. 2020) | Põlme S, et al. 2020. Fungal Diversity 105:1-16 ((https://doi.org/10.1007/s13225-020-00466-2)) |
| Otu59179 | 1 | k__Fungi(100);p__Ascomycota(100);c__Saccharomycetes(100);o__Saccharomycetales(100);f__Metschnikowiaceae(100);g__Metschnikowia(100);s__Metschnikowia_pulcherrima(100); | Metschnikowia | 13 | Saprotroph | \|Undefined Saprotroph\| | Yeast | NULL | Probable | As nectar-tap_saprotroph (Põlme et al. 2020); Decay sugar-rich substrates (Põlme et al. 2020) | Põlme S, et al. 2020. Fungal Diversity 105:1-16 ((https://doi.org/10.1007/s13225-020-00466-2)) |
| Otu59180 | 1 | k__Fungi(100);p__Ascomycota(100);c__Saccharomycetes(100);o__Saccharomycetales(100);f__Metschnikowiaceae(100);g__Metschnikowia(100);g__Metschnikowia_unclassified(100); | Metschnikowia | 13 | Saprotroph | \|Undefined Saprotroph\| | Yeast | NULL | Probable | As nectar-tap_saprotroph (Põlme et al. 2020); Decay sugar-rich substrates (Põlme et al. 2020) | Põlme S, et al. 2020. Fungal Diversity 105:1-16 ((https://doi.org/10.1007/s13225-020-00466-2)) |
| Otu59181 | 1 | k__Fungi(100);p__Ascomycota(100);c__Saccharomycetes(100);o__Saccharomycetales(100);f__Metschnikowiaceae(100);g__Metschnikowia(100);g__Metschnikowia_unclassified(100); | Metschnikowia | 13 | Saprotroph | \|Undefined Saprotroph\| | Yeast | NULL | Probable | As nectar-tap_saprotroph (Põlme et al. 2020); Decay sugar-rich substrates (Põlme et al. 2020) | Põlme S, et al. 2020. Fungal Diversity 105:1-16 ((https://doi.org/10.1007/s13225-020-00466-2)) |
| Otu59182 | 1 | k__Fungi(100);p__Ascomycota(100);c__Saccharomycetes(100);o__Saccharomycetales(100);f__Metschnikowiaceae(100);g__Metschnikowia(100);s__Metschnikowia_pulcherrima(100); | Metschnikowia | 13 | Saprotroph | \|Undefined Saprotroph\| | Yeast | NULL | Probable | As nectar-tap_saprotroph (Põlme et al. 2020); Decay sugar-rich substrates (Põlme et al. 2020) | Põlme S, et al. 2020. Fungal Diversity 105:1-16 ((https://doi.org/10.1007/s13225-020-00466-2)) |
| Otu59183 | 1 | k__Fungi(100);p__Ascomycota(100);c__Saccharomycetes(100);o__Saccharomycetales(100);f__Metschnikowiaceae(100);g__Metschnikowia(100);s__Metschnikowia_pulcherrima(100); | Metschnikowia | 13 | Saprotroph | \|Undefined Saprotroph\| | Yeast | NULL | Probable | As nectar-tap_saprotroph (Põlme et al. 2020); Decay sugar-rich substrates (Põlme et al. 2020) | Põlme S, et al. 2020. Fungal Diversity 105:1-16 ((https://doi.org/10.1007/s13225-020-00466-2)) |
| Otu59184 | 1 | k__Fungi(100);p__Ascomycota(100);c__Saccharomycetes(100);o__Saccharomycetales(100);f__Metschnikowiaceae(100);g__Metschnikowia(100);g__Metschnikowia_unclassified(100); | Metschnikowia | 13 | Saprotroph | \|Undefined Saprotroph\| | Yeast | NULL | Probable | As nectar-tap_saprotroph (Põlme et al. 2020); Decay sugar-rich substrates (Põlme et al. 2020) | Põlme S, et al. 2020. Fungal Diversity 105:1-16 ((https://doi.org/10.1007/s13225-020-00466-2)) |
| Otu59185 | 1 | k__Fungi(100);p__Ascomycota(100);c__Saccharomycetes(100);o__Saccharomycetales(100);f__Metschnikowiaceae(100);g__Metschnikowia(100);s__Metschnikowia_pulcherrima(100); | Metschnikowia | 13 | Saprotroph | \|Undefined Saprotroph\| | Yeast | NULL | Probable | As nectar-tap_saprotroph (Põlme et al. 2020); Decay sugar-rich substrates (Põlme et al. 2020) | Põlme S, et al. 2020. Fungal Diversity 105:1-16 ((https://doi.org/10.1007/s13225-020-00466-2)) |
| Otu59186 | 1 | k__Fungi(100);p__Ascomycota(100);c__Saccharomycetes(100);o__Saccharomycetales(100);f__Metschnikowiaceae(100);g__Metschnikowia(100);s__Metschnikowia_pulcherrima(100); | Metschnikowia | 13 | Saprotroph | \|Undefined Saprotroph\| | Yeast | NULL | Probable | As nectar-tap_saprotroph (Põlme et al. 2020); Decay sugar-rich substrates (Põlme et al. 2020) | Põlme S, et al. 2020. Fungal Diversity 105:1-16 ((https://doi.org/10.1007/s13225-020-00466-2)) |
| Otu59187 | 1 | k__Fungi(100);p__Ascomycota(100);c__Saccharomycetes(100);o__Saccharomycetales(100);f__Metschnikowiaceae(100);g__Metschnikowia(100);g__Metschnikowia_unclassified(100); | Metschnikowia | 13 | Saprotroph | \|Undefined Saprotroph\| | Yeast | NULL | Probable | As nectar-tap_saprotroph (Põlme et al. 2020); Decay sugar-rich substrates (Põlme et al. 2020) | Põlme S, et al. 2020. Fungal Diversity 105:1-16 ((https://doi.org/10.1007/s13225-020-00466-2)) |
| Otu59191 | 1 | k__Fungi(100);p__Ascomycota(100);c__Saccharomycetes(100);o__Saccharomycetales(100);f__Metschnikowiaceae(100);g__Metschnikowia(100);s__Metschnikowia_pulcherrima(100); | Metschnikowia | 13 | Saprotroph | \|Undefined Saprotroph\| | Yeast | NULL | Probable | As nectar-tap_saprotroph (Põlme et al. 2020); Decay sugar-rich substrates (Põlme et al. 2020) | Põlme S, et al. 2020. Fungal Diversity 105:1-16 ((https://doi.org/10.1007/s13225-020-00466-2)) |
| Otu59192 | 1 | k__Fungi(100);p__Ascomycota(100);c__Saccharomycetes(100);o__Saccharomycetales(100);f__Metschnikowiaceae(100);g__Metschnikowia(100);s__Metschnikowia_pulcherrima(100); | Metschnikowia | 13 | Saprotroph | \|Undefined Saprotroph\| | Yeast | NULL | Probable | As nectar-tap_saprotroph (Põlme et al. 2020); Decay sugar-rich substrates (Põlme et al. 2020) | Põlme S, et al. 2020. Fungal Diversity 105:1-16 ((https://doi.org/10.1007/s13225-020-00466-2)) |
| Otu59193 | 1 | k__Fungi(100);p__Ascomycota(100);c__Saccharomycetes(100);o__Saccharomycetales(100);f__Metschnikowiaceae(100);g__Metschnikowia(100);s__Metschnikowia_pulcherrima(100); | Metschnikowia | 13 | Saprotroph | \|Undefined Saprotroph\| | Yeast | NULL | Probable | As nectar-tap_saprotroph (Põlme et al. 2020); Decay sugar-rich substrates (Põlme et al. 2020) | Põlme S, et al. 2020. Fungal Diversity 105:1-16 ((https://doi.org/10.1007/s13225-020-00466-2)) |
| Otu59195 | 1 | k__Fungi(100);p__Ascomycota(100);c__Saccharomycetes(100);o__Saccharomycetales(100);f__Metschnikowiaceae(100);g__Metschnikowia(100);s__Metschnikowia_pulcherrima(100); | Metschnikowia | 13 | Saprotroph | \|Undefined Saprotroph\| | Yeast | NULL | Probable | As nectar-tap_saprotroph (Põlme et al. 2020); Decay sugar-rich substrates (Põlme et al. 2020) | Põlme S, et al. 2020. Fungal Diversity 105:1-16 ((https://doi.org/10.1007/s13225-020-00466-2)) |
| Otu59196 | 1 | k__Fungi(100);p__Ascomycota(100);c__Saccharomycetes(100);o__Saccharomycetales(100);f__Metschnikowiaceae(100);g__Metschnikowia(100);s__Metschnikowia_pulcherrima(100); | Metschnikowia | 13 | Saprotroph | \|Undefined Saprotroph\| | Yeast | NULL | Probable | As nectar-tap_saprotroph (Põlme et al. 2020); Decay sugar-rich substrates (Põlme et al. 2020) | Põlme S, et al. 2020. Fungal Diversity 105:1-16 ((https://doi.org/10.1007/s13225-020-00466-2)) |
| Otu59199 | 1 | k__Fungi(100);p__Ascomycota(100);c__Saccharomycetes(100);o__Saccharomycetales(100);f__Saccharomycetales_family_Incertae_sedis(100);g__Candida(100);g__Candida_unclassified(100); | Candida | 13 | Pathotroph-Saprotroph-Symbiotroph | Animal Pathogen-Endophyte-Endosymbiont-Epiphyte-\|Undefined Saprotroph\| | Dimorphic Yeast-Yeast | NULL | Probable | As nectar-tap_saprotroph (Põlme et al. 2020); Decay sugar-rich substrates (Põlme et al. 2020); This asexual genus is polyphyletic and contain species that function very differently depend on what they are. Caution should be taken when interpreting guilds for Candida species. | Manolakaki D, et al. 2010. Virulence. 1: 367-375 ((https://doi.org/10.4161/viru.1.5.12796)); Mekbib SB, et al. 2011. Tropical Plant Pathology 36:233-240 ((https://doi.org/10.1590/S1982-56762011000400004)); Põlme S, et al. 2020. Fungal Diversity 105:1-16 ((https://doi.org/10.1007/s13225-020-00466-2)) |
| Otu59229 | 1 | k__Fungi(100);p__Ascomycota(100);c__Saccharomycetes(100);o__Saccharomycetales(100);f__Metschnikowiaceae(100);g__Metschnikowia(100);s__Metschnikowia_pulcherrima(100); | Metschnikowia | 13 | Saprotroph | \|Undefined Saprotroph\| | Yeast | NULL | Probable | As nectar-tap_saprotroph (Põlme et al. 2020); Decay sugar-rich substrates (Põlme et al. 2020) | Põlme S, et al. 2020. Fungal Diversity 105:1-16 ((https://doi.org/10.1007/s13225-020-00466-2)) |
| Otu59230 | 1 | k__Fungi(100);p__Ascomycota(100);c__Saccharomycetes(100);o__Saccharomycetales(100);f__Metschnikowiaceae(100);g__Metschnikowia(100);s__Metschnikowia_pulcherrima(100); | Metschnikowia | 13 | Saprotroph | \|Undefined Saprotroph\| | Yeast | NULL | Probable | As nectar-tap_saprotroph (Põlme et al. 2020); Decay sugar-rich substrates (Põlme et al. 2020) | Põlme S, et al. 2020. Fungal Diversity 105:1-16 ((https://doi.org/10.1007/s13225-020-00466-2)) |
| Otu59231 | 1 | k__Fungi(100);p__Ascomycota(100);c__Saccharomycetes(100);o__Saccharomycetales(100);f__Metschnikowiaceae(100);g__Metschnikowia(100);s__Metschnikowia_pulcherrima(100); | Metschnikowia | 13 | Saprotroph | \|Undefined Saprotroph\| | Yeast | NULL | Probable | As nectar-tap_saprotroph (Põlme et al. 2020); Decay sugar-rich substrates (Põlme et al. 2020) | Põlme S, et al. 2020. Fungal Diversity 105:1-16 ((https://doi.org/10.1007/s13225-020-00466-2)) |
| Otu59232 | 1 | k__Fungi(100);p__Ascomycota(100);c__Saccharomycetes(100);o__Saccharomycetales(100);f__Metschnikowiaceae(100);g__Metschnikowia(100);s__Metschnikowia_pulcherrima(100); | Metschnikowia | 13 | Saprotroph | \|Undefined Saprotroph\| | Yeast | NULL | Probable | As nectar-tap_saprotroph (Põlme et al. 2020); Decay sugar-rich substrates (Põlme et al. 2020) | Põlme S, et al. 2020. Fungal Diversity 105:1-16 ((https://doi.org/10.1007/s13225-020-00466-2)) |
| Otu59233 | 1 | k__Fungi(100);p__Ascomycota(100);c__Saccharomycetes(100);o__Saccharomycetales(100);f__Metschnikowiaceae(100);g__Metschnikowia(100);g__Metschnikowia_unclassified(100); | Metschnikowia | 13 | Saprotroph | \|Undefined Saprotroph\| | Yeast | NULL | Probable | As nectar-tap_saprotroph (Põlme et al. 2020); Decay sugar-rich substrates (Põlme et al. 2020) | Põlme S, et al. 2020. Fungal Diversity 105:1-16 ((https://doi.org/10.1007/s13225-020-00466-2)) |
| Otu59234 | 1 | k__Fungi(100);p__Ascomycota(100);c__Saccharomycetes(100);o__Saccharomycetales(100);f__Metschnikowiaceae(100);g__Metschnikowia(100);s__Metschnikowia_pulcherrima(100); | Metschnikowia | 13 | Saprotroph | \|Undefined Saprotroph\| | Yeast | NULL | Probable | As nectar-tap_saprotroph (Põlme et al. 2020); Decay sugar-rich substrates (Põlme et al. 2020) | Põlme S, et al. 2020. Fungal Diversity 105:1-16 ((https://doi.org/10.1007/s13225-020-00466-2)) |
| Otu59235 | 1 | k__Fungi(100);p__Ascomycota(100);c__Saccharomycetes(100);o__Saccharomycetales(100);f__Metschnikowiaceae(100);g__Metschnikowia(100);g__Metschnikowia_unclassified(100); | Metschnikowia | 13 | Saprotroph | \|Undefined Saprotroph\| | Yeast | NULL | Probable | As nectar-tap_saprotroph (Põlme et al. 2020); Decay sugar-rich substrates (Põlme et al. 2020) | Põlme S, et al. 2020. Fungal Diversity 105:1-16 ((https://doi.org/10.1007/s13225-020-00466-2)) |
| Otu59236 | 1 | k__Fungi(100);p__Ascomycota(100);c__Saccharomycetes(100);o__Saccharomycetales(100);f__Metschnikowiaceae(100);g__Metschnikowia(100);s__Metschnikowia_pulcherrima(100); | Metschnikowia | 13 | Saprotroph | \|Undefined Saprotroph\| | Yeast | NULL | Probable | As nectar-tap_saprotroph (Põlme et al. 2020); Decay sugar-rich substrates (Põlme et al. 2020) | Põlme S, et al. 2020. Fungal Diversity 105:1-16 ((https://doi.org/10.1007/s13225-020-00466-2)) |
| Otu59237 | 1 | k__Fungi(100);p__Ascomycota(100);c__Saccharomycetes(100);o__Saccharomycetales(100);f__Metschnikowiaceae(100);g__Metschnikowia(100);s__Metschnikowia_pulcherrima(100); | Metschnikowia | 13 | Saprotroph | \|Undefined Saprotroph\| | Yeast | NULL | Probable | As nectar-tap_saprotroph (Põlme et al. 2020); Decay sugar-rich substrates (Põlme et al. 2020) | Põlme S, et al. 2020. Fungal Diversity 105:1-16 ((https://doi.org/10.1007/s13225-020-00466-2)) |
| Otu59238 | 1 | k__Fungi(100);p__Ascomycota(100);c__Saccharomycetes(100);o__Saccharomycetales(100);f__Metschnikowiaceae(100);g__Metschnikowia(100);s__Metschnikowia_pulcherrima(100); | Metschnikowia | 13 | Saprotroph | \|Undefined Saprotroph\| | Yeast | NULL | Probable | As nectar-tap_saprotroph (Põlme et al. 2020); Decay sugar-rich substrates (Põlme et al. 2020) | Põlme S, et al. 2020. Fungal Diversity 105:1-16 ((https://doi.org/10.1007/s13225-020-00466-2)) |

| | | | | | | | | | | | |
|---|---|---|---|---|---|---|---|---|---|---|---|
| Otu59239 | 1 | k__Fungi(100);p__Ascomycota(100);c__Saccharomycetes(100);o__Saccharomycetales(100);f__Metschnikowiaceae(100);g__Metschnikowia(100);s__Metschnikowia_pulcherrima(100); | Metschnikowia | 13 | Saprotroph | \|Undefined Saprotroph\| | Yeast | NULL | Probable | As nectar-tap_saprotroph (Põlme et al. 2020); Decay sugar-rich substrates (Põlme et al. 2020) | Põlme S, et al. 2020. Fungal Diversity 105:1-16 ((https://doi.org/10.1007/s13225-020-00466-2)) |
| Otu59240 | 1 | k__Fungi(100);p__Ascomycota(100);c__Saccharomycetes(100);o__Saccharomycetales(100);f__Metschnikowiaceae(100);g__Metschnikowia(100);s__Metschnikowia_pulcherrima(100); | Metschnikowia | 13 | Saprotroph | \|Undefined Saprotroph\| | Yeast | NULL | Probable | As nectar-tap_saprotroph (Põlme et al. 2020); Decay sugar-rich substrates (Põlme et al. 2020) | Põlme S, et al. 2020. Fungal Diversity 105:1-16 ((https://doi.org/10.1007/s13225-020-00466-2)) |
| Otu59241 | 1 | k__Fungi(100);p__Ascomycota(100);c__Saccharomycetes(100);o__Saccharomycetales(100);f__Metschnikowiaceae(100);g__Metschnikowia(100);s__Metschnikowia_pulcherrima(100); | Metschnikowia | 13 | Saprotroph | \|Undefined Saprotroph\| | Yeast | NULL | Probable | As nectar-tap_saprotroph (Põlme et al. 2020); Decay sugar-rich substrates (Põlme et al. 2020) | Põlme S, et al. 2020. Fungal Diversity 105:1-16 ((https://doi.org/10.1007/s13225-020-00466-2)) |
| Otu59242 | 1 | k__Fungi(100);p__Ascomycota(100);c__Saccharomycetes(100);o__Saccharomycetales(100);f__Metschnikowiaceae(100);g__Metschnikowia(100);s__Metschnikowia_pulcherrima(100); | Metschnikowia | 13 | Saprotroph | \|Undefined Saprotroph\| | Yeast | NULL | Probable | As nectar-tap_saprotroph (Põlme et al. 2020); Decay sugar-rich substrates (Põlme et al. 2020) | Põlme S, et al. 2020. Fungal Diversity 105:1-16 ((https://doi.org/10.1007/s13225-020-00466-2)) |
| Otu59243 | 1 | k__Fungi(100);p__Ascomycota(100);c__Saccharomycetes(100);o__Saccharomycetales(100);f__Metschnikowiaceae(100);g__Metschnikowia(100);g__Metschnikowia_unclassified(100); | Metschnikowia | 13 | Saprotroph | \|Undefined Saprotroph\| | Yeast | NULL | Probable | As nectar-tap_saprotroph (Põlme et al. 2020); Decay sugar-rich substrates (Põlme et al. 2020) | Põlme S, et al. 2020. Fungal Diversity 105:1-16 ((https://doi.org/10.1007/s13225-020-00466-2)) |
| Otu59245 | 1 | k__Fungi(100);p__Ascomycota(100);c__Saccharomycetes(100);o__Saccharomycetales(100);f__Metschnikowiaceae(100);g__Metschnikowia(100);s__Metschnikowia_pulcherrima(100); | Metschnikowia | 13 | Saprotroph | \|Undefined Saprotroph\| | Yeast | NULL | Probable | As nectar-tap_saprotroph (Põlme et al. 2020); Decay sugar-rich substrates (Põlme et al. 2020) | Põlme S, et al. 2020. Fungal Diversity 105:1-16 ((https://doi.org/10.1007/s13225-020-00466-2)) |
| Otu59246 | 1 | k__Fungi(100);p__Ascomycota(100);c__Saccharomycetes(100);o__Saccharomycetales(100);f__Metschnikowiaceae(100);g__Metschnikowia(100);s__Metschnikowia_pulcherrima(100); | Metschnikowia | 13 | Saprotroph | \|Undefined Saprotroph\| | Yeast | NULL | Probable | As nectar-tap_saprotroph (Põlme et al. 2020); Decay sugar-rich substrates (Põlme et al. 2020) | Põlme S, et al. 2020. Fungal Diversity 105:1-16 ((https://doi.org/10.1007/s13225-020-00466-2)) |
| Otu59247 | 1 | k__Fungi(100);p__Ascomycota(100);c__Saccharomycetes(100);o__Saccharomycetales(100);f__Metschnikowiaceae(100);g__Metschnikowia(100);s__Metschnikowia_pulcherrima(100); | Metschnikowia | 13 | Saprotroph | \|Undefined Saprotroph\| | Yeast | NULL | Probable | As nectar-tap_saprotroph (Põlme et al. 2020); Decay sugar-rich substrates (Põlme et al. 2020) | Põlme S, et al. 2020. Fungal Diversity 105:1-16 ((https://doi.org/10.1007/s13225-020-00466-2)) |
| Otu59248 | 1 | k__Fungi(100);p__Ascomycota(100);c__Saccharomycetes(100);o__Saccharomycetales(100);f__Metschnikowiaceae(100);g__Metschnikowia(100);s__Metschnikowia_pulcherrima(100); | Metschnikowia | 13 | Saprotroph | \|Undefined Saprotroph\| | Yeast | NULL | Probable | As nectar-tap_saprotroph (Põlme et al. 2020); Decay sugar-rich substrates (Põlme et al. 2020) | Põlme S, et al. 2020. Fungal Diversity 105:1-16 ((https://doi.org/10.1007/s13225-020-00466-2)) |
| Otu59250 | 1 | k__Fungi(100);p__Ascomycota(100);c__Saccharomycetes(100);o__Saccharomycetales(100);f__Metschnikowiaceae(100);g__Metschnikowia(100);s__Metschnikowia_pulcherrima(100); | Metschnikowia | 13 | Saprotroph | \|Undefined Saprotroph\| | Yeast | NULL | Probable | As nectar-tap_saprotroph (Põlme et al. 2020); Decay sugar-rich substrates (Põlme et al. 2020) | Põlme S, et al. 2020. Fungal Diversity 105:1-16 ((https://doi.org/10.1007/s13225-020-00466-2)) |
| Otu59252 | 1 | k__Fungi(100);p__Ascomycota(100);c__Saccharomycetes(100);o__Saccharomycetales(100);f__Metschnikowiaceae(100);g__Metschnikowia(100);s__Metschnikowia_pulcherrima(100); | Metschnikowia | 13 | Saprotroph | \|Undefined Saprotroph\| | Yeast | NULL | Probable | As nectar-tap_saprotroph (Põlme et al. 2020); Decay sugar-rich substrates (Põlme et al. 2020) | Põlme S, et al. 2020. Fungal Diversity 105:1-16 ((https://doi.org/10.1007/s13225-020-00466-2)) |
| Otu59253 | 1 | k__Fungi(100);p__Ascomycota(100);c__Saccharomycetes(100);o__Saccharomycetales(100);f__Metschnikowiaceae(100);g__Metschnikowia(100);s__Metschnikowia_pulcherrima(100); | Metschnikowia | 13 | Saprotroph | \|Undefined Saprotroph\| | Yeast | NULL | Probable | As nectar-tap_saprotroph (Põlme et al. 2020); Decay sugar-rich substrates (Põlme et al. 2020) | Põlme S, et al. 2020. Fungal Diversity 105:1-16 ((https://doi.org/10.1007/s13225-020-00466-2)) |
| Otu59255 | 1 | k__Fungi(100);p__Ascomycota(100);c__Saccharomycetes(100);o__Saccharomycetales(100);f__Metschnikowiaceae(100);g__Metschnikowia(100);s__Metschnikowia_pulcherrima(100); | Metschnikowia | 13 | Saprotroph | \|Undefined Saprotroph\| | Yeast | NULL | Probable | As nectar-tap_saprotroph (Põlme et al. 2020); Decay sugar-rich substrates (Põlme et al. 2020) | Põlme S, et al. 2020. Fungal Diversity 105:1-16 ((https://doi.org/10.1007/s13225-020-00466-2)) |
| Otu59256 | 1 | k__Fungi(100);p__Ascomycota(100);c__Saccharomycetes(100);o__Saccharomycetales(100);f__Metschnikowiaceae(100);g__Metschnikowia(100);s__Metschnikowia_pulcherrima(100); | Metschnikowia | 13 | Saprotroph | \|Undefined Saprotroph\| | Yeast | NULL | Probable | As nectar-tap_saprotroph (Põlme et al. 2020); Decay sugar-rich substrates (Põlme et al. 2020) | Põlme S, et al. 2020. Fungal Diversity 105:1-16 ((https://doi.org/10.1007/s13225-020-00466-2)) |
| Otu59257 | 1 | k__Fungi(100);p__Ascomycota(100);c__Saccharomycetes(100);o__Saccharomycetales(100);f__Metschnikowiaceae(100);g__Metschnikowia(100);s__Metschnikowia_pulcherrima(100); | Metschnikowia | 13 | Saprotroph | \|Undefined Saprotroph\| | Yeast | NULL | Probable | As nectar-tap_saprotroph (Põlme et al. 2020); Decay sugar-rich substrates (Põlme et al. 2020) | Põlme S, et al. 2020. Fungal Diversity 105:1-16 ((https://doi.org/10.1007/s13225-020-00466-2)) |
| Otu59258 | 1 | k__Fungi(100);p__Ascomycota(100);c__Saccharomycetes(100);o__Saccharomycetales(100);f__Metschnikowiaceae(100);g__Metschnikowia(100);s__Metschnikowia_pulcherrima(100); | Metschnikowia | 13 | Saprotroph | \|Undefined Saprotroph\| | Yeast | NULL | Probable | As nectar-tap_saprotroph (Põlme et al. 2020); Decay sugar-rich substrates (Põlme et al. 2020) | Põlme S, et al. 2020. Fungal Diversity 105:1-16 ((https://doi.org/10.1007/s13225-020-00466-2)) |
| Otu59259 | 1 | k__Fungi(100);p__Ascomycota(100);c__Saccharomycetes(100);o__Saccharomycetales(100);f__Metschnikowiaceae(100);g__Metschnikowia(100);s__Metschnikowia_pulcherrima(100); | Metschnikowia | 13 | Saprotroph | \|Undefined Saprotroph\| | Yeast | NULL | Probable | As nectar-tap_saprotroph (Põlme et al. 2020); Decay sugar-rich substrates (Põlme et al. 2020) | Põlme S, et al. 2020. Fungal Diversity 105:1-16 ((https://doi.org/10.1007/s13225-020-00466-2)) |
| Otu59260 | 1 | k__Fungi(100);p__Ascomycota(100);c__Saccharomycetes(100);o__Saccharomycetales(100);f__Metschnikowiaceae(100);g__Metschnikowia(100);s__Metschnikowia_pulcherrima(100); | Metschnikowia | 13 | Saprotroph | \|Undefined Saprotroph\| | Yeast | NULL | Probable | As nectar-tap_saprotroph (Põlme et al. 2020); Decay sugar-rich substrates (Põlme et al. 2020) | Põlme S, et al. 2020. Fungal Diversity 105:1-16 ((https://doi.org/10.1007/s13225-020-00466-2)) |
| Otu59261 | 1 | k__Fungi(100);p__Ascomycota(100);c__Saccharomycetes(100);o__Saccharomycetales(100);f__Metschnikowiaceae(100);g__Metschnikowia(100);g__Metschnikowia_unclassified(100); | Metschnikowia | 13 | Saprotroph | \|Undefined Saprotroph\| | Yeast | NULL | Probable | As nectar-tap_saprotroph (Põlme et al. 2020); Decay sugar-rich substrates (Põlme et al. 2020) | Põlme S, et al. 2020. Fungal Diversity 105:1-16 ((https://doi.org/10.1007/s13225-020-00466-2)) |
| Otu59262 | 1 | k__Fungi(100);p__Ascomycota(100);c__Saccharomycetes(100);o__Saccharomycetales(100);f__Metschnikowiaceae(100);g__Metschnikowia(100);g__Metschnikowia_unclassified(100); | Metschnikowia | 13 | Saprotroph | \|Undefined Saprotroph\| | Yeast | NULL | Probable | As nectar-tap_saprotroph (Põlme et al. 2020); Decay sugar-rich substrates (Põlme et al. 2020) | Põlme S, et al. 2020. Fungal Diversity 105:1-16 ((https://doi.org/10.1007/s13225-020-00466-2)) |
| Otu59263 | 1 | k__Fungi(100);p__Ascomycota(100);c__Saccharomycetes(100);o__Saccharomycetales(100);f__Metschnikowiaceae(100);g__Metschnikowia(100);g__Metschnikowia_unclassified(100); | Metschnikowia | 13 | Saprotroph | \|Undefined Saprotroph\| | Yeast | NULL | Probable | As nectar-tap_saprotroph (Põlme et al. 2020); Decay sugar-rich substrates (Põlme et al. 2020) | Põlme S, et al. 2020. Fungal Diversity 105:1-16 ((https://doi.org/10.1007/s13225-020-00466-2)) |
| Otu59264 | 1 | k__Fungi(100);p__Ascomycota(100);c__Saccharomycetes(100);o__Saccharomycetales(100);f__Metschnikowiaceae(100);g__Metschnikowia(100);g__Metschnikowia_unclassified(100); | Metschnikowia | 13 | Saprotroph | \|Undefined Saprotroph\| | Yeast | NULL | Probable | As nectar-tap_saprotroph (Põlme et al. 2020); Decay sugar-rich substrates (Põlme et al. 2020) | Põlme S, et al. 2020. Fungal Diversity 105:1-16 ((https://doi.org/10.1007/s13225-020-00466-2)) |
| Otu59265 | 1 | k__Fungi(100);p__Ascomycota(100);c__Saccharomycetes(100);o__Saccharomycetales(100);f__Metschnikowiaceae(100);g__Metschnikowia(100);s__Metschnikowia_pulcherrima(100); | Metschnikowia | 13 | Saprotroph | \|Undefined Saprotroph\| | Yeast | NULL | Probable | As nectar-tap_saprotroph (Põlme et al. 2020); Decay sugar-rich substrates (Põlme et al. 2020) | Põlme S, et al. 2020. Fungal Diversity 105:1-16 ((https://doi.org/10.1007/s13225-020-00466-2)) |
| Otu59266 | 1 | k__Fungi(100);p__Ascomycota(100);c__Saccharomycetes(100);o__Saccharomycetales(100);f__Metschnikowiaceae(100);g__Metschnikowia(100);s__Metschnikowia_pulcherrima(100); | Metschnikowia | 13 | Saprotroph | \|Undefined Saprotroph\| | Yeast | NULL | Probable | As nectar-tap_saprotroph (Põlme et al. 2020); Decay sugar-rich substrates (Põlme et al. 2020) | Põlme S, et al. 2020. Fungal Diversity 105:1-16 ((https://doi.org/10.1007/s13225-020-00466-2)) |
| Otu59267 | 1 | k__Fungi(100);p__Ascomycota(100);c__Saccharomycetes(100);o__Saccharomycetales(100);f__Metschnikowiaceae(100);g__Metschnikowia(100);g__Metschnikowia_unclassified(100); | Metschnikowia | 13 | Saprotroph | \|Undefined Saprotroph\| | Yeast | NULL | Probable | As nectar-tap_saprotroph (Põlme et al. 2020); Decay sugar-rich substrates (Põlme et al. 2020) | Põlme S, et al. 2020. Fungal Diversity 105:1-16 ((https://doi.org/10.1007/s13225-020-00466-2)) |
| Otu59268 | 1 | k__Fungi(100);p__Ascomycota(100);c__Saccharomycetes(100);o__Saccharomycetales(100);f__Metschnikowiaceae(100);g__Metschnikowia(100);s__Metschnikowia_pulcherrima(100); | Metschnikowia | 13 | Saprotroph | \|Undefined Saprotroph\| | Yeast | NULL | Probable | As nectar-tap_saprotroph (Põlme et al. 2020); Decay sugar-rich substrates (Põlme et al. 2020) | Põlme S, et al. 2020. Fungal Diversity 105:1-16 ((https://doi.org/10.1007/s13225-020-00466-2)) |
| Otu59270 | 1 | k__Fungi(100);p__Ascomycota(100);c__Saccharomycetes(100);o__Saccharomycetales(100);f__Metschnikowiaceae(100);g__Metschnikowia(100);s__Metschnikowia_pulcherrima(100); | Metschnikowia | 13 | Saprotroph | \|Undefined Saprotroph\| | Yeast | NULL | Probable | As nectar-tap_saprotroph (Põlme et al. 2020); Decay sugar-rich substrates (Põlme et al. 2020) | Põlme S, et al. 2020. Fungal Diversity 105:1-16 ((https://doi.org/10.1007/s13225-020-00466-2)) |
| Otu59271 | 1 | k__Fungi(100);p__Ascomycota(100);c__Saccharomycetes(100);o__Saccharomycetales(100);f__Metschnikowiaceae(100);g__Metschnikowia(100);s__Metschnikowia_pulcherrima(100); | Metschnikowia | 13 | Saprotroph | \|Undefined Saprotroph\| | Yeast | NULL | Probable | As nectar-tap_saprotroph (Põlme et al. 2020); Decay sugar-rich substrates (Põlme et al. 2020) | Põlme S, et al. 2020. Fungal Diversity 105:1-16 ((https://doi.org/10.1007/s13225-020-00466-2)) |
| Otu59272 | 1 | k__Fungi(100);p__Ascomycota(100);c__Saccharomycetes(100);o__Saccharomycetales(100);f__Metschnikowiaceae(100);g__Metschnikowia(100);s__Metschnikowia_pulcherrima(100); | Metschnikowia | 13 | Saprotroph | \|Undefined Saprotroph\| | Yeast | NULL | Probable | As nectar-tap_saprotroph (Põlme et al. 2020); Decay sugar-rich substrates (Põlme et al. 2020) | Põlme S, et al. 2020. Fungal Diversity 105:1-16 ((https://doi.org/10.1007/s13225-020-00466-2)) |
| Otu59273 | 1 | k__Fungi(100);p__Ascomycota(100);c__Saccharomycetes(100);o__Saccharomycetales(100);f__Metschnikowiaceae(100);g__Metschnikowia(100);s__Metschnikowia_pulcherrima(100); | Metschnikowia | 13 | Saprotroph | \|Undefined Saprotroph\| | Yeast | NULL | Probable | As nectar-tap_saprotroph (Põlme et al. 2020); Decay sugar-rich substrates (Põlme et al. 2020) | Põlme S, et al. 2020. Fungal Diversity 105:1-16 ((https://doi.org/10.1007/s13225-020-00466-2)) |
| Otu59274 | 1 | k__Fungi(100);p__Ascomycota(100);c__Saccharomycetes(100);o__Saccharomycetales(100);f__Metschnikowiaceae(100);g__Metschnikowia(100);s__Metschnikowia_pulcherrima(100); | Metschnikowia | 13 | Saprotroph | \|Undefined Saprotroph\| | Yeast | NULL | Probable | As nectar-tap_saprotroph (Põlme et al. 2020); Decay sugar-rich substrates (Põlme et al. 2020) | Põlme S, et al. 2020. Fungal Diversity 105:1-16 ((https://doi.org/10.1007/s13225-020-00466-2)) |
| Otu59275 | 1 | k__Fungi(100);p__Ascomycota(100);c__Saccharomycetes(100);o__Saccharomycetales(100);f__Metschnikowiaceae(100);g__Metschnikowia(100);g__Metschnikowia_unclassified(100); | Metschnikowia | 13 | Saprotroph | \|Undefined Saprotroph\| | Yeast | NULL | Probable | As nectar-tap_saprotroph (Põlme et al. 2020); Decay sugar-rich substrates (Põlme et al. 2020) | Põlme S, et al. 2020. Fungal Diversity 105:1-16 ((https://doi.org/10.1007/s13225-020-00466-2)) |
| Otu59276 | 1 | k__Fungi(100);p__Ascomycota(100);c__Saccharomycetes(100);o__Saccharomycetales(100);f__Metschnikowiaceae(100);g__Metschnikowia(100);s__Metschnikowia_pulcherrima(100); | Metschnikowia | 13 | Saprotroph | \|Undefined Saprotroph\| | Yeast | NULL | Probable | As nectar-tap_saprotroph (Põlme et al. 2020); Decay sugar-rich substrates (Põlme et al. 2020) | Põlme S, et al. 2020. Fungal Diversity 105:1-16 ((https://doi.org/10.1007/s13225-020-00466-2)) |
| Otu59277 | 1 | k__Fungi(100);p__Ascomycota(100);c__Saccharomycetes(100);o__Saccharomycetales(100);f__Metschnikowiaceae(100);g__Metschnikowia(100);s__Metschnikowia_pulcherrima(100); | Metschnikowia | 13 | Saprotroph | \|Undefined Saprotroph\| | Yeast | NULL | Probable | As nectar-tap_saprotroph (Põlme et al. 2020); Decay sugar-rich substrates (Põlme et al. 2020) | Põlme S, et al. 2020. Fungal Diversity 105:1-16 ((https://doi.org/10.1007/s13225-020-00466-2)) |
| Otu59278 | 1 | k__Fungi(100);p__Ascomycota(100);c__Saccharomycetes(100);o__Saccharomycetales(100);f__Metschnikowiaceae(100);g__Metschnikowia(100);s__Metschnikowia_pulcherrima(100); | Metschnikowia | 13 | Saprotroph | \|Undefined Saprotroph\| | Yeast | NULL | Probable | As nectar-tap_saprotroph (Põlme et al. 2020); Decay sugar-rich substrates (Põlme et al. 2020) | Põlme S, et al. 2020. Fungal Diversity 105:1-16 ((https://doi.org/10.1007/s13225-020-00466-2)) |
| Otu59279 | 1 | k__Fungi(100);p__Ascomycota(100);c__Saccharomycetes(100);o__Saccharomycetales(100);f__Metschnikowiaceae(100);g__Metschnikowia(100);s__Metschnikowia_pulcherrima(100); | Metschnikowia | 13 | Saprotroph | \|Undefined Saprotroph\| | Yeast | NULL | Probable | As nectar-tap_saprotroph (Põlme et al. 2020); Decay sugar-rich substrates (Põlme et al. 2020) | Põlme S, et al. 2020. Fungal Diversity 105:1-16 ((https://doi.org/10.1007/s13225-020-00466-2)) |
| Otu59280 | 1 | k__Fungi(100);p__Ascomycota(100);c__Saccharomycetes(100);o__Saccharomycetales(100);f__Metschnikowiaceae(100);g__Metschnikowia(100);s__Metschnikowia_pulcherrima(100); | Metschnikowia | 13 | Saprotroph | \|Undefined Saprotroph\| | Yeast | NULL | Probable | As nectar-tap_saprotroph (Põlme et al. 2020); Decay sugar-rich substrates (Põlme et al. 2020) | Põlme S, et al. 2020. Fungal Diversity 105:1-16 ((https://doi.org/10.1007/s13225-020-00466-2)) |
| Otu59281 | 1 | k__Fungi(100);p__Ascomycota(100);c__Saccharomycetes(100);o__Saccharomycetales(100);f__Metschnikowiaceae(100);g__Metschnikowia(100);s__Metschnikowia_pulcherrima(100); | Metschnikowia | 13 | Saprotroph | \|Undefined Saprotroph\| | Yeast | NULL | Probable | As nectar-tap_saprotroph (Põlme et al. 2020); Decay sugar-rich substrates (Põlme et al. 2020) | Põlme S, et al. 2020. Fungal Diversity 105:1-16 ((https://doi.org/10.1007/s13225-020-00466-2)) |
| Otu59308 | 1 | k__Fungi(100);p__Ascomycota(100);c__Saccharomycetes(100);o__Saccharomycetales(100);f__Metschnikowiaceae(100);g__Metschnikowia(100);s__Metschnikowia_pulcherrima(100); | Metschnikowia | 13 | Saprotroph | \|Undefined Saprotroph\| | Yeast | NULL | Probable | As nectar-tap_saprotroph (Põlme et al. 2020); Decay sugar-rich substrates (Põlme et al. 2020) | Põlme S, et al. 2020. Fungal Diversity 105:1-16 ((https://doi.org/10.1007/s13225-020-00466-2)) |
| Otu59311 | 1 | k__Fungi(100);p__Ascomycota(100);c__Saccharomycetes(100);o__Saccharomycetales(100);f__Metschnikowiaceae(100);g__Metschnikowia(100);g__Metschnikowia_unclassified(100); | Metschnikowia | 13 | Saprotroph | \|Undefined Saprotroph\| | Yeast | NULL | Probable | As nectar-tap_saprotroph (Põlme et al. 2020); Decay sugar-rich substrates (Põlme et al. 2020) | Põlme S, et al. 2020. Fungal Diversity 105:1-16 ((https://doi.org/10.1007/s13225-020-00466-2)) |
| Otu59313 | 1 | k__Fungi(100);p__Ascomycota(100);c__Saccharomycetes(100);o__Saccharomycetales(100);f__Metschnikowiaceae(100);g__Metschnikowia(100);g__Metschnikowia_unclassified(100); | Metschnikowia | 13 | Saprotroph | \|Undefined Saprotroph\| | Yeast | NULL | Probable | As nectar-tap_saprotroph (Põlme et al. 2020); Decay sugar-rich substrates (Põlme et al. 2020) | Põlme S, et al. 2020. Fungal Diversity 105:1-16 ((https://doi.org/10.1007/s13225-020-00466-2)) |
| Otu59316 | 1 | k__Fungi(100);p__Ascomycota(100);c__Saccharomycetes(100);o__Saccharomycetales(100);f__Metschnikowiaceae(100);g__Metschnikowia(100);s__Metschnikowia_pulcherrima(100); | Metschnikowia | 13 | Saprotroph | \|Undefined Saprotroph\| | Yeast | NULL | Probable | As nectar-tap_saprotroph (Põlme et al. 2020); Decay sugar-rich substrates (Põlme et al. 2020) | Põlme S, et al. 2020. Fungal Diversity 105:1-16 ((https://doi.org/10.1007/s13225-020-00466-2)) |
| Otu59317 | 1 | k__Fungi(100);p__Ascomycota(100);c__Saccharomycetes(100);o__Saccharomycetales(100);f__Metschnikowiaceae(100);g__Metschnikowia(100);g__Metschnikowia_unclassified(100); | Metschnikowia | 13 | Saprotroph | \|Undefined Saprotroph\| | Yeast | NULL | Probable | As nectar-tap_saprotroph (Põlme et al. 2020); Decay sugar-rich substrates (Põlme et al. 2020) | Põlme S, et al. 2020. Fungal Diversity 105:1-16 ((https://doi.org/10.1007/s13225-020-00466-2)) |
| Otu59318 | 1 | k__Fungi(100);p__Ascomycota(100);c__Saccharomycetes(100);o__Saccharomycetales(100);f__Metschnikowiaceae(100);g__Metschnikowia(100);s__Metschnikowia_pulcherrima(100); | Metschnikowia | 13 | Saprotroph | \|Undefined Saprotroph\| | Yeast | NULL | Probable | As nectar-tap_saprotroph (Põlme et al. 2020); Decay sugar-rich substrates (Põlme et al. 2020) | Põlme S, et al. 2020. Fungal Diversity 105:1-16 ((https://doi.org/10.1007/s13225-020-00466-2)) |
| Otu59322 | 1 | k__Fungi(100);p__Ascomycota(100);c__Saccharomycetes(100);o__Saccharomycetales(100);f__Metschnikowiaceae(100);g__Metschnikowia(100);s__Metschnikowia_pulcherrima(100); | Metschnikowia | 13 | Saprotroph | \|Undefined Saprotroph\| | Yeast | NULL | Probable | As nectar-tap_saprotroph (Põlme et al. 2020); Decay sugar-rich substrates (Põlme et al. 2020) | Põlme S, et al. 2020. Fungal Diversity 105:1-16 ((https://doi.org/10.1007/s13225-020-00466-2)) |
| Otu59323 | 1 | k__Fungi(100);p__Ascomycota(100);c__Saccharomycetes(100);o__Saccharomycetales(100);f__Metschnikowiaceae(100);g__Metschnikowia(100);s__Metschnikowia_pulcherrima(100); | Metschnikowia | 13 | Saprotroph | \|Undefined Saprotroph\| | Yeast | NULL | Probable | As nectar-tap_saprotroph (Põlme et al. 2020); Decay sugar-rich substrates (Põlme et al. 2020) | Põlme S, et al. 2020. Fungal Diversity 105:1-16 ((https://doi.org/10.1007/s13225-020-00466-2)) |
| Otu59324 | 1 | k__Fungi(100);p__Ascomycota(100);c__Saccharomycetes(100);o__Saccharomycetales(100);f__Metschnikowiaceae(100);g__Metschnikowia(100);s__Metschnikowia_pulcherrima(100); | Metschnikowia | 13 | Saprotroph | \|Undefined Saprotroph\| | Yeast | NULL | Probable | As nectar-tap_saprotroph (Põlme et al. 2020); Decay sugar-rich substrates (Põlme et al. 2020) | Põlme S, et al. 2020. Fungal Diversity 105:1-16 ((https://doi.org/10.1007/s13225-020-00466-2)) |
| Otu59325 | 1 | k__Fungi(100);p__Ascomycota(100);c__Saccharomycetes(100);o__Saccharomycetales(100);f__Metschnikowiaceae(100);g__Metschnikowia(100);s__Metschnikowia_pulcherrima(100); | Metschnikowia | 13 | Saprotroph | \|Undefined Saprotroph\| | Yeast | NULL | Probable | As nectar-tap_saprotroph (Põlme et al. 2020); Decay sugar-rich substrates (Põlme et al. 2020) | Põlme S, et al. 2020. Fungal Diversity 105:1-16 ((https://doi.org/10.1007/s13225-020-00466-2)) |
| Otu59326 | 1 | k__Fungi(100);p__Ascomycota(100);c__Saccharomycetes(100);o__Saccharomycetales(100);f__Metschnikowiaceae(100);g__Metschnikowia(100);g__Metschnikowia_unclassified(100); | Metschnikowia | 13 | Saprotroph | \|Undefined Saprotroph\| | Yeast | NULL | Probable | As nectar-tap_saprotroph (Põlme et al. 2020); Decay sugar-rich substrates (Põlme et al. 2020) | Põlme S, et al. 2020. Fungal Diversity 105:1-16 ((https://doi.org/10.1007/s13225-020-00466-2)) |
| Otu59327 | 1 | k__Fungi(100);p__Ascomycota(100);c__Saccharomycetes(100);o__Saccharomycetales(100);f__Metschnikowiaceae(100);g__Metschnikowia(100);s__Metschnikowia_pulcherrima(100); | Metschnikowia | 13 | Saprotroph | \|Undefined Saprotroph\| | Yeast | NULL | Probable | As nectar-tap_saprotroph (Põlme et al. 2020); Decay sugar-rich substrates (Põlme et al. 2020) | Põlme S, et al. 2020. Fungal Diversity 105:1-16 ((https://doi.org/10.1007/s13225-020-00466-2)) |
| Otu59335 | 1 | k__Fungi(100);p__Ascomycota(100);c__Saccharomycetes(100);o__Saccharomycetales(100);f__Metschnikowiaceae(100);g__Metschnikowia(100);s__Metschnikowia_pulcherrima(100); | Metschnikowia | 13 | Saprotroph | \|Undefined Saprotroph\| | Yeast | NULL | Probable | As nectar-tap_saprotroph (Põlme et al. 2020); Decay sugar-rich substrates (Põlme et al. 2020) | Põlme S, et al. 2020. Fungal Diversity 105:1-16 ((https://doi.org/10.1007/s13225-020-00466-2)) |
| Otu59336 | 1 | k__Fungi(100);p__Ascomycota(100);c__Saccharomycetes(100);o__Saccharomycetales(100);f__Metschnikowiaceae(100);g__Metschnikowia(100);s__Metschnikowia_pulcherrima(100); | Metschnikowia | 13 | Saprotroph | \|Undefined Saprotroph\| | Yeast | NULL | Probable | As nectar-tap_saprotroph (Põlme et al. 2020); Decay sugar-rich substrates (Põlme et al. 2020) | Põlme S, et al. 2020. Fungal Diversity 105:1-16 ((https://doi.org/10.1007/s13225-020-00466-2)) |
| Otu59337 | 1 | k__Fungi(100);p__Ascomycota(100);c__Saccharomycetes(100);o__Saccharomycetales(100);f__Metschnikowiaceae(100);g__Metschnikowia(100);g__Metschnikowia_unclassified(100); | Metschnikowia | 13 | Saprotroph | \|Undefined Saprotroph\| | Yeast | NULL | Probable | As nectar-tap_saprotroph (Põlme et al. 2020); Decay sugar-rich substrates (Põlme et al. 2020) | Põlme S, et al. 2020. Fungal Diversity 105:1-16 ((https://doi.org/10.1007/s13225-020-00466-2)) |
| Otu59338 | 1 | k__Fungi(100);p__Ascomycota(100);c__Saccharomycetes(100);o__Saccharomycetales(100);f__Metschnikowiaceae(100);g__Metschnikowia(100);s__Metschnikowia_pulcherrima(100); | Metschnikowia | 13 | Saprotroph | \|Undefined Saprotroph\| | Yeast | NULL | Probable | As nectar-tap_saprotroph (Põlme et al. 2020); Decay sugar-rich substrates (Põlme et al. 2020) | Põlme S, et al. 2020. Fungal Diversity 105:1-16 ((https://doi.org/10.1007/s13225-020-00466-2)) |
| Otu59339 | 1 | k__Fungi(100);p__Ascomycota(100);c__Saccharomycetes(100);o__Saccharomycetales(100);f__Metschnikowiaceae(100);g__Metschnikowia(100);s__Metschnikowia_pulcherrima(100); | Metschnikowia | 13 | Saprotroph | \|Undefined Saprotroph\| | Yeast | NULL | Probable | As nectar-tap_saprotroph (Põlme et al. 2020); Decay sugar-rich substrates (Põlme et al. 2020) | Põlme S, et al. 2020. Fungal Diversity 105:1-16 ((https://doi.org/10.1007/s13225-020-00466-2)) |
| Otu59340 | 1 | k__Fungi(100);p__Ascomycota(100);c__Saccharomycetes(100);o__Saccharomycetales(100);f__Metschnikowiaceae(100);g__Metschnikowia(100);s__Metschnikowia_pulcherrima(100); | Metschnikowia | 13 | Saprotroph | \|Undefined Saprotroph\| | Yeast | NULL | Probable | As nectar-tap_saprotroph (Põlme et al. 2020); Decay sugar-rich substrates (Põlme et al. 2020) | Põlme S, et al. 2020. Fungal Diversity 105:1-16 ((https://doi.org/10.1007/s13225-020-00466-2)) |
| Otu59341 | 1 | k__Fungi(100);p__Ascomycota(100);c__Saccharomycetes(100);o__Saccharomycetales(100);f__Metschnikowiaceae(100);g__Metschnikowia(100);s__Metschnikowia_pulcherrima(100); | Metschnikowia | 13 | Saprotroph | \|Undefined Saprotroph\| | Yeast | NULL | Probable | As nectar-tap_saprotroph (Põlme et al. 2020); Decay sugar-rich substrates (Põlme et al. 2020) | Põlme S, et al. 2020. Fungal Diversity 105:1-16 ((https://doi.org/10.1007/s13225-020-00466-2)) |
| Otu59342 | 1 | k__Fungi(100);p__Ascomycota(100);c__Saccharomycetes(100);o__Saccharomycetales(100);f__Metschnikowiaceae(100);g__Metschnikowia(100);s__Metschnikowia_pulcherrima(100); | Metschnikowia | 13 | Saprotroph | \|Undefined Saprotroph\| | Yeast | NULL | Probable | As nectar-tap_saprotroph (Põlme et al. 2020); Decay sugar-rich substrates (Põlme et al. 2020) | Põlme S, et al. 2020. Fungal Diversity 105:1-16 ((https://doi.org/10.1007/s13225-020-00466-2)) |
| Otu59343 | 1 | k__Fungi(100);p__Ascomycota(100);c__Saccharomycetes(100);o__Saccharomycetales(100);f__Metschnikowiaceae(100);g__Metschnikowia(100);s__Metschnikowia_pulcherrima(100); | Metschnikowia | 13 | Saprotroph | \|Undefined Saprotroph\| | Yeast | NULL | Probable | As nectar-tap_saprotroph (Põlme et al. 2020); Decay sugar-rich substrates (Põlme et al. 2020) | Põlme S, et al. 2020. Fungal Diversity 105:1-16 ((https://doi.org/10.1007/s13225-020-00466-2)) |
| Otu59344 | 1 | k__Fungi(100);p__Ascomycota(100);c__Saccharomycetes(100);o__Saccharomycetales(100);f__Metschnikowiaceae(100);g__Metschnikowia(100);s__Metschnikowia_pulcherrima(100); | Metschnikowia | 13 | Saprotroph | \|Undefined Saprotroph\| | Yeast | NULL | Probable | As nectar-tap_saprotroph (Põlme et al. 2020); Decay sugar-rich substrates (Põlme et al. 2020) | Põlme S, et al. 2020. Fungal Diversity 105:1-16 ((https://doi.org/10.1007/s13225-020-00466-2)) |
| Otu59345 | 1 | k__Fungi(100);p__Ascomycota(100);c__Saccharomycetes(100);o__Saccharomycetales(100);f__Metschnikowiaceae(100);g__Metschnikowia(100);s__Metschnikowia_pulcherrima(100); | Metschnikowia | 13 | Saprotroph | \|Undefined Saprotroph\| | Yeast | NULL | Probable | As nectar-tap_saprotroph (Põlme et al. 2020); Decay sugar-rich substrates (Põlme et al. 2020) | Põlme S, et al. 2020. Fungal Diversity 105:1-16 ((https://doi.org/10.1007/s13225-020-00466-2)) |
| Otu59346 | 1 | k__Fungi(100);p__Ascomycota(100);c__Saccharomycetes(100);o__Saccharomycetales(100);f__Metschnikowiaceae(100);g__Metschnikowia(100);g__Metschnikowia_unclassified(100); | Metschnikowia | 13 | Saprotroph | \|Undefined Saprotroph\| | Yeast | NULL | Probable | As nectar-tap_saprotroph (Põlme et al. 2020); Decay sugar-rich substrates (Põlme et al. 2020) | Põlme S, et al. 2020. Fungal Diversity 105:1-16 ((https://doi.org/10.1007/s13225-020-00466-2)) |
| Otu59347 | 1 | k__Fungi(100);p__Ascomycota(100);c__Saccharomycetes(100);o__Saccharomycetales(100);f__Metschnikowiaceae(100);g__Metschnikowia(100);s__Metschnikowia_pulcherrima(100); | Metschnikowia | 13 | Saprotroph | \|Undefined Saprotroph\| | Yeast | NULL | Probable | As nectar-tap_saprotroph (Põlme et al. 2020); Decay sugar-rich substrates (Põlme et al. 2020) | Põlme S, et al. 2020. Fungal Diversity 105:1-16 ((https://doi.org/10.1007/s13225-020-00466-2)) |
| Otu59348 | 1 | k__Fungi(100);p__Ascomycota(100);c__Saccharomycetes(100);o__Saccharomycetales(100);f__Metschnikowiaceae(100);g__Metschnikowia(100);s__Metschnikowia_pulcherrima(100); | Metschnikowia | 13 | Saprotroph | \|Undefined Saprotroph\| | Yeast | NULL | Probable | As nectar-tap_saprotroph (Põlme et al. 2020); Decay sugar-rich substrates (Põlme et al. 2020) | Põlme S, et al. 2020. Fungal Diversity 105:1-16 ((https://doi.org/10.1007/s13225-020-00466-2)) |
| Otu59351 | 1 | k__Fungi(100);p__Ascomycota(100);c__Saccharomycetes(100);o__Saccharomycetales(100);f__Metschnikowiaceae(100);g__Metschnikowia(100);s__Metschnikowia_pulcherrima(100); | Metschnikowia | 13 | Saprotroph | \|Undefined Saprotroph\| | Yeast | NULL | Probable | As nectar-tap_saprotroph (Põlme et al. 2020); Decay sugar-rich substrates (Põlme et al. 2020) | Põlme S, et al. 2020. Fungal Diversity 105:1-16 ((https://doi.org/10.1007/s13225-020-00466-2)) |
| Otu59352 | 1 | k__Fungi(100);p__Ascomycota(100);c__Saccharomycetes(100);o__Saccharomycetales(100);f__Metschnikowiaceae(100);g__Metschnikowia(100);g__Metschnikowia_unclassified(100); | Metschnikowia | 13 | Saprotroph | \|Undefined Saprotroph\| | Yeast | NULL | Probable | As nectar-tap_saprotroph (Põlme et al. 2020); Decay sugar-rich substrates (Põlme et al. 2020) | Põlme S, et al. 2020. Fungal Diversity 105:1-16 ((https://doi.org/10.1007/s13225-020-00466-2)) |
| Otu59353 | 1 | k__Fungi(100);p__Ascomycota(100);c__Saccharomycetes(100);o__Saccharomycetales(100);f__Metschnikowiaceae(100);g__Metschnikowia(100);s__Metschnikowia_pulcherrima(100); | Metschnikowia | 13 | Saprotroph | \|Undefined Saprotroph\| | Yeast | NULL | Probable | As nectar-tap_saprotroph (Põlme et al. 2020); Decay sugar-rich substrates (Põlme et al. 2020) | Põlme S, et al. 2020. Fungal Diversity 105:1-16 ((https://doi.org/10.1007/s13225-020-00466-2)) |
| Otu59354 | 1 | k__Fungi(100);p__Ascomycota(100);c__Saccharomycetes(100);o__Saccharomycetales(100);f__Metschnikowiaceae(100);g__Metschnikowia(100);s__Metschnikowia_pulcherrima(100); | Metschnikowia | 13 | Saprotroph | \|Undefined Saprotroph\| | Yeast | NULL | Probable | As nectar-tap_saprotroph (Põlme et al. 2020); Decay sugar-rich substrates (Põlme et al. 2020) | Põlme S, et al. 2020. Fungal Diversity 105:1-16 ((https://doi.org/10.1007/s13225-020-00466-2)) |
| Otu59355 | 1 | k__Fungi(100);p__Ascomycota(100);c__Saccharomycetes(100);o__Saccharomycetales(100);f__Metschnikowiaceae(100);g__Metschnikowia(100);s__Metschnikowia_pulcherrima(100); | Metschnikowia | 13 | Saprotroph | \|Undefined Saprotroph\| | Yeast | NULL | Probable | As nectar-tap_saprotroph (Põlme et al. 2020); Decay sugar-rich substrates (Põlme et al. 2020) | Põlme S, et al. 2020. Fungal Diversity 105:1-16 ((https://doi.org/10.1007/s13225-020-00466-2)) |
| Otu59356 | 1 | k__Fungi(100);p__Ascomycota(100);c__Saccharomycetes(100);o__Saccharomycetales(100);f__Metschnikowiaceae(100);g__Metschnikowia(100);s__Metschnikowia_pulcherrima(100); | Metschnikowia | 13 | Saprotroph | \|Undefined Saprotroph\| | Yeast | NULL | Probable | As nectar-tap_saprotroph (Põlme et al. 2020); Decay sugar-rich substrates (Põlme et al. 2020) | Põlme S, et al. 2020. Fungal Diversity 105:1-16 ((https://doi.org/10.1007/s13225-020-00466-2)) |
| Otu59357 | 1 | k__Fungi(100);p__Ascomycota(100);c__Saccharomycetes(100);o__Saccharomycetales(100);f__Metschnikowiaceae(100);g__Metschnikowia(100);s__Metschnikowia_pulcherrima(100); | Metschnikowia | 13 | Saprotroph | \|Undefined Saprotroph\| | Yeast | NULL | Probable | As nectar-tap_saprotroph (Põlme et al. 2020); Decay sugar-rich substrates (Põlme et al. 2020) | Põlme S, et al. 2020. Fungal Diversity 105:1-16 ((https://doi.org/10.1007/s13225-020-00466-2)) |
| Otu59358 | 1 | k__Fungi(100);p__Ascomycota(100);c__Saccharomycetes(100);o__Saccharomycetales(100);f__Metschnikowiaceae(100);g__Metschnikowia(100);s__Metschnikowia_pulcherrima(100); | Metschnikowia | 13 | Saprotroph | \|Undefined Saprotroph\| | Yeast | NULL | Probable | As nectar-tap_saprotroph (Põlme et al. 2020); Decay sugar-rich substrates (Põlme et al. 2020) | Põlme S, et al. 2020. Fungal Diversity 105:1-16 ((https://doi.org/10.1007/s13225-020-00466-2)) |
| Otu59361 | 1 | k__Fungi(100);p__Ascomycota(100);c__Sordariomycetes(100);o__Hypocreales(100);f__Nectriaceae(100);g__Gibberella(100);g__Gibberella_unclassified(100); | Gibberella | 13 | Pathotroph-Saprotroph-Symbiotroph | Endophyte-\|Plant Pathogen\|-Plant Saprotroph-Undefined Saprotroph | Microfungus | Soft Rot | Probable | As foliar_endophyte (Põlme et al. 2020); As litter_saprotroph (Põlme et al. 2020); Causing damping-off disease (Põlme et al. 2020); Decay leaf, fruit, seed, roots, soil, animal material (Põlme et al. 2020) | Tedersoo L, et al. 2014. Science 346:e1256688 ((https://doi.org/10.1126/science.1256688)); Põlme S, et al. 2020. Fungal Diversity 105:1-16 ((https://doi.org/10.1007/s13225-020-00466-2)) |
| Otu59362 | 1 | k__Fungi(100);p__Ascomycota(100);c__Saccharomycetes(100);o__Saccharomycetales(100);f__Metschnikowiaceae(100);g__Metschnikowia(100);s__Metschnikowia_pulcherrima(100); | Metschnikowia | 13 | Saprotroph | \|Undefined Saprotroph\| | Yeast | NULL | Probable | As nectar-tap_saprotroph (Põlme et al. 2020); Decay sugar-rich substrates (Põlme et al. 2020) | Põlme S, et al. 2020. Fungal Diversity 105:1-16 ((https://doi.org/10.1007/s13225-020-00466-2)) |
| Otu59363 | 1 | k__Fungi(100);p__Ascomycota(100);c__Saccharomycetes(100);o__Saccharomycetales(100);f__Metschnikowiaceae(100);g__Metschnikowia(100);s__Metschnikowia_pulcherrima(100); | Metschnikowia | 13 | Saprotroph | \|Undefined Saprotroph\| | Yeast | NULL | Probable | As nectar-tap_saprotroph (Põlme et al. 2020); Decay sugar-rich substrates (Põlme et al. 2020) | Põlme S, et al. 2020. Fungal Diversity 105:1-16 ((https://doi.org/10.1007/s13225-020-00466-2)) |
| Otu59364 | 1 | k__Fungi(100);p__Ascomycota(100);c__Saccharomycetes(100);o__Saccharomycetales(100);f__Metschnikowiaceae(100);g__Metschnikowia(100);s__Metschnikowia_pulcherrima(100); | Metschnikowia | 13 | Saprotroph | \|Undefined Saprotroph\| | Yeast | NULL | Probable | As nectar-tap_saprotroph (Põlme et al. 2020); Decay sugar-rich substrates (Põlme et al. 2020) | Põlme S, et al. 2020. Fungal Diversity 105:1-16 ((https://doi.org/10.1007/s13225-020-00466-2)) |
| Otu59366 | 1 | k__Fungi(100);p__Ascomycota(100);c__Saccharomycetes(100);o__Saccharomycetales(100);f__Metschnikowiaceae(100);g__Metschnikowia(100);g__Metschnikowia_unclassified(100); | Metschnikowia | 13 | Saprotroph | \|Undefined Saprotroph\| | Yeast | NULL | Probable | As nectar-tap_saprotroph (Põlme et al. 2020); Decay sugar-rich substrates (Põlme et al. 2020) | Põlme S, et al. 2020. Fungal Diversity 105:1-16 ((https://doi.org/10.1007/s13225-020-00466-2)) |
| Otu59367 | 1 | k__Fungi(100);p__Ascomycota(100);c__Saccharomycetes(100);o__Saccharomycetales(100);f__Metschnikowiaceae(100);g__Metschnikowia(100);s__Metschnikowia_pulcherrima(100); | Metschnikowia | 13 | Saprotroph | \|Undefined Saprotroph\| | Yeast | NULL | Probable | As nectar-tap_saprotroph (Põlme et al. 2020); Decay sugar-rich substrates (Põlme et al. 2020) | Põlme S, et al. 2020. Fungal Diversity 105:1-16 ((https://doi.org/10.1007/s13225-020-00466-2)) |
| Otu59372 | 1 | k__Fungi(100);p__Ascomycota(100);c__Saccharomycetes(100);o__Saccharomycetales(100);f__Metschnikowiaceae(100);g__Metschnikowia(100);s__Metschnikowia_pulcherrima(100); | Metschnikowia | 13 | Saprotroph | \|Undefined Saprotroph\| | Yeast | NULL | Probable | As nectar-tap_saprotroph (Põlme et al. 2020); Decay sugar-rich substrates (Põlme et al. 2020) | Põlme S, et al. 2020. Fungal Diversity 105:1-16 ((https://doi.org/10.1007/s13225-020-00466-2)) |
| Otu59374 | 1 | k__Fungi(100);p__Ascomycota(100);c__Saccharomycetes(100);o__Saccharomycetales(100);f__Metschnikowiaceae(100);g__Metschnikowia(100);s__Metschnikowia_pulcherrima(100); | Metschnikowia | 13 | Saprotroph | \|Undefined Saprotroph\| | Yeast | NULL | Probable | As nectar-tap_saprotroph (Põlme et al. 2020); Decay sugar-rich substrates (Põlme et al. 2020) | Põlme S, et al. 2020. Fungal Diversity 105:1-16 ((https://doi.org/10.1007/s13225-020-00466-2)) |
| Otu59375 | 1 | k__Fungi(100);p__Ascomycota(100);c__Saccharomycetes(100);o__Saccharomycetales(100);f__Metschnikowiaceae(100);g__Metschnikowia(100);g__Metschnikowia_unclassified(100); | Metschnikowia | 13 | Saprotroph | \|Undefined Saprotroph\| | Yeast | NULL | Probable | As nectar-tap_saprotroph (Põlme et al. 2020); Decay sugar-rich substrates (Põlme et al. 2020) | Põlme S, et al. 2020. Fungal Diversity 105:1-16 ((https://doi.org/10.1007/s13225-020-00466-2)) |
| Otu59376 | 1 | k__Fungi(100);p__Ascomycota(100);c__Saccharomycetes(100);o__Saccharomycetales(100);f__Metschnikowiaceae(100);g__Metschnikowia(100);g__Metschnikowia_unclassified(100); | Metschnikowia | 13 | Saprotroph | \|Undefined Saprotroph\| | Yeast | NULL | Probable | As nectar-tap_saprotroph (Põlme et al. 2020); Decay sugar-rich substrates (Põlme et al. 2020) | Põlme S, et al. 2020. Fungal Diversity 105:1-16 ((https://doi.org/10.1007/s13225-020-00466-2)) |
| Otu59377 | 1 | k__Fungi(100);p__Ascomycota(100);c__Saccharomycetes(100);o__Saccharomycetales(100);f__Metschnikowiaceae(100);g__Metschnikowia(100);g__Metschnikowia_unclassified(100); | Metschnikowia | 13 | Saprotroph | \|Undefined Saprotroph\| | Yeast | NULL | Probable | As nectar-tap_saprotroph (Põlme et al. 2020); Decay sugar-rich substrates (Põlme et al. 2020) | Põlme S, et al. 2020. Fungal Diversity 105:1-16 ((https://doi.org/10.1007/s13225-020-00466-2)) |
| Otu59378 | 1 | k__Fungi(100);p__Ascomycota(100);c__Saccharomycetes(100);o__Saccharomycetales(100);f__Metschnikowiaceae(100);g__Metschnikowia(100);g__Metschnikowia_unclassified(100); | Metschnikowia | 13 | Saprotroph | \|Undefined Saprotroph\| | Yeast | NULL | Probable | As nectar-tap_saprotroph (Põlme et al. 2020); Decay sugar-rich substrates (Põlme et al. 2020) | Põlme S, et al. 2020. Fungal Diversity 105:1-16 ((https://doi.org/10.1007/s13225-020-00466-2)) |
| Otu59379 | 1 | k__Fungi(100);p__Ascomycota(100);c__Saccharomycetes(100);o__Saccharomycetales(100);f__Metschnikowiaceae(100);g__Metschnikowia(100);g__Metschnikowia_unclassified(100); | Metschnikowia | 13 | Saprotroph | \|Undefined Saprotroph\| | Yeast | NULL | Probable | As nectar-tap_saprotroph (Põlme et al. 2020); Decay sugar-rich substrates (Põlme et al. 2020) | Põlme S, et al. 2020. Fungal Diversity 105:1-16 ((https://doi.org/10.1007/s13225-020-00466-2)) |
| Otu59380 | 1 | k__Fungi(100);p__Ascomycota(100);c__Saccharomycetes(100);o__Saccharomycetales(100);f__Metschnikowiaceae(100);g__Metschnikowia(100);g__Metschnikowia_unclassified(100); | Metschnikowia | 13 | Saprotroph | \|Undefined Saprotroph\| | Yeast | NULL | Probable | As nectar-tap_saprotroph (Põlme et al. 2020); Decay sugar-rich substrates (Põlme et al. 2020) | Põlme S, et al. 2020. Fungal Diversity 105:1-16 ((https://doi.org/10.1007/s13225-020-00466-2)) |
| Otu59381 | 1 | k__Fungi(100);p__Ascomycota(100);c__Saccharomycetes(100);o__Saccharomycetales(100);f__Metschnikowiaceae(100);g__Metschnikowia(100);g__Metschnikowia_unclassified(100); | Metschnikowia | 13 | Saprotroph | \|Undefined Saprotroph\| | Yeast | NULL | Probable | As nectar-tap_saprotroph (Põlme et al. 2020); Decay sugar-rich substrates (Põlme et al. 2020) | Põlme S, et al. 2020. Fungal Diversity 105:1-16 ((https://doi.org/10.1007/s13225-020-00466-2)) |
| Otu59382 | 1 | k__Fungi(100);p__Ascomycota(100);c__Saccharomycetes(100);o__Saccharomycetales(100);f__Metschnikowiaceae(100);g__Metschnikowia(100);g__Metschnikowia_unclassified(100); | Metschnikowia | 13 | Saprotroph | \|Undefined Saprotroph\| | Yeast | NULL | Probable | As nectar-tap_saprotroph (Põlme et al. 2020); Decay sugar-rich substrates (Põlme et al. 2020) | Põlme S, et al. 2020. Fungal Diversity 105:1-16 ((https://doi.org/10.1007/s13225-020-00466-2)) |
| Otu59383 | 1 | k__Fungi(100);p__Ascomycota(100);c__Saccharomycetes(100);o__Saccharomycetales(100);f__Metschnikowiaceae(100);g__Metschnikowia(100);s__Metschnikowia_pulcherrima(100); | Metschnikowia | 13 | Saprotroph | \|Undefined Saprotroph\| | Yeast | NULL | Probable | As nectar-tap_saprotroph (Põlme et al. 2020); Decay sugar-rich substrates (Põlme et al. 2020) | Põlme S, et al. 2020. Fungal Diversity 105:1-16 ((https://doi.org/10.1007/s13225-020-00466-2)) |
| Otu59384 | 1 | k__Fungi(100);p__Ascomycota(100);c__Saccharomycetes(100);o__Saccharomycetales(100);f__Metschnikowiaceae(100);g__Metschnikowia(100);s__Metschnikowia_pulcherrima(100); | Metschnikowia | 13 | Saprotroph | \|Undefined Saprotroph\| | Yeast | NULL | Probable | As nectar-tap_saprotroph (Põlme et al. 2020); Decay sugar-rich substrates (Põlme et al. 2020) | Põlme S, et al. 2020. Fungal Diversity 105:1-16 ((https://doi.org/10.1007/s13225-020-00466-2)) |
| Otu59385 | 1 | k__Fungi(100);p__Ascomycota(100);c__Saccharomycetes(100);o__Saccharomycetales(100);f__Metschnikowiaceae(100);g__Metschnikowia(100);s__Metschnikowia_pulcherrima(100); | Metschnikowia | 13 | Saprotroph | \|Undefined Saprotroph\| | Yeast | NULL | Probable | As nectar-tap_saprotroph (Põlme et al. 2020); Decay sugar-rich substrates (Põlme et al. 2020) | Põlme S, et al. 2020. Fungal Diversity 105:1-16 ((https://doi.org/10.1007/s13225-020-00466-2)) |
| Otu59386 | 1 | k__Fungi(100);p__Ascomycota(100);c__Saccharomycetes(100);o__Saccharomycetales(100);f__Metschnikowiaceae(100);g__Metschnikowia(100);s__Metschnikowia_pulcherrima(100); | Metschnikowia | 13 | Saprotroph | \|Undefined Saprotroph\| | Yeast | NULL | Probable | As nectar-tap_saprotroph (Põlme et al. 2020); Decay sugar-rich substrates (Põlme et al. 2020) | Põlme S, et al. 2020. Fungal Diversity 105:1-16 ((https://doi.org/10.1007/s13225-020-00466-2)) |
| Otu59387 | 1 | k__Fungi(100);p__Ascomycota(100);c__Saccharomycetes(100);o__Saccharomycetales(100);f__Metschnikowiaceae(100);g__Metschnikowia(100);s__Metschnikowia_pulcherrima(100); | Metschnikowia | 13 | Saprotroph | \|Undefined Saprotroph\| | Yeast | NULL | Probable | As nectar-tap_saprotroph (Põlme et al. 2020); Decay sugar-rich substrates (Põlme et al. 2020) | Põlme S, et al. 2020. Fungal Diversity 105:1-16 ((https://doi.org/10.1007/s13225-020-00466-2)) |
| Otu59388 | 1 | k__Fungi(100);p__Ascomycota(100);c__Saccharomycetes(100);o__Saccharomycetales(100);f__Metschnikowiaceae(100);g__Metschnikowia(100);s__Metschnikowia_pulcherrima(100); | Metschnikowia | 13 | Saprotroph | \|Undefined Saprotroph\| | Yeast | NULL | Probable | As nectar-tap_saprotroph (Põlme et al. 2020); Decay sugar-rich substrates (Põlme et al. 2020) | Põlme S, et al. 2020. Fungal Diversity 105:1-16 ((https://doi.org/10.1007/s13225-020-00466-2)) |
| Otu59389 | 1 | k__Fungi(100);p__Ascomycota(100);c__Saccharomycetes(100);o__Saccharomycetales(100);f__Metschnikowiaceae(100);g__Metschnikowia(100);s__Metschnikowia_pulcherrima(100); | Metschnikowia | 13 | Saprotroph | \|Undefined Saprotroph\| | Yeast | NULL | Probable | As nectar-tap_saprotroph (Põlme et al. 2020); Decay sugar-rich substrates (Põlme et al. 2020) | Põlme S, et al. 2020. Fungal Diversity 105:1-16 ((https://doi.org/10.1007/s13225-020-00466-2)) |
| Otu59390 | 1 | k__Fungi(100);p__Ascomycota(100);c__Saccharomycetes(100);o__Saccharomycetales(100);f__Metschnikowiaceae(100);g__Metschnikowia(100);g__Metschnikowia_unclassified(100); | Metschnikowia | 13 | Saprotroph | \|Undefined Saprotroph\| | Yeast | NULL | Probable | As nectar-tap_saprotroph (Põlme et al. 2020); Decay sugar-rich substrates (Põlme et al. 2020) | Põlme S, et al. 2020. Fungal Diversity 105:1-16 ((https://doi.org/10.1007/s13225-020-00466-2)) |
| Otu59391 | 1 | k__Fungi(100);p__Ascomycota(100);c__Saccharomycetes(100);o__Saccharomycetales(100);f__Metschnikowiaceae(100);g__Metschnikowia(100);s__Metschnikowia_pulcherrima(100); | Metschnikowia | 13 | Saprotroph | \|Undefined Saprotroph\| | Yeast | NULL | Probable | As nectar-tap_saprotroph (Põlme et al. 2020); Decay sugar-rich substrates (Põlme et al. 2020) | Põlme S, et al. 2020. Fungal Diversity 105:1-16 ((https://doi.org/10.1007/s13225-020-00466-2)) |
| Otu59392 | 1 | k__Fungi(100);p__Ascomycota(100);c__Saccharomycetes(100);o__Saccharomycetales(100);f__Metschnikowiaceae(100);g__Metschnikowia(100);s__Metschnikowia_pulcherrima(100); | Metschnikowia | 13 | Saprotroph | \|Undefined Saprotroph\| | Yeast | NULL | Probable | As nectar-tap_saprotroph (Põlme et al. 2020); Decay sugar-rich substrates (Põlme et al. 2020) | Põlme S, et al. 2020. Fungal Diversity 105:1-16 ((https://doi.org/10.1007/s13225-020-00466-2)) |
| Otu59393 | 1 | k__Fungi(100);p__Ascomycota(100);c__Saccharomycetes(100);o__Saccharomycetales(100);f__Metschnikowiaceae(100);g__Metschnikowia(100);s__Metschnikowia_pulcherrima(100); | Metschnikowia | 13 | Saprotroph | \|Undefined Saprotroph\| | Yeast | NULL | Probable | As nectar-tap_saprotroph (Põlme et al. 2020); Decay sugar-rich substrates (Põlme et al. 2020) | Põlme S, et al. 2020. Fungal Diversity 105:1-16 ((https://doi.org/10.1007/s13225-020-00466-2)) |
| Otu59394 | 1 | k__Fungi(100);p__Ascomycota(100);c__Saccharomycetes(100);o__Saccharomycetales(100);f__Metschnikowiaceae(100);g__Metschnikowia(100);s__Metschnikowia_pulcherrima(100); | Metschnikowia | 13 | Saprotroph | \|Undefined Saprotroph\| | Yeast | NULL | Probable | As nectar-tap_saprotroph (Põlme et al. 2020); Decay sugar-rich substrates (Põlme et al. 2020) | Põlme S, et al. 2020. Fungal Diversity 105:1-16 ((https://doi.org/10.1007/s13225-020-00466-2)) |
| Otu59395 | 1 | k__Fungi(100);p__Ascomycota(100);c__Saccharomycetes(100);o__Saccharomycetales(100);f__Metschnikowiaceae(100);g__Metschnikowia(100);s__Metschnikowia_pulcherrima(100); | Metschnikowia | 13 | Saprotroph | \|Undefined Saprotroph\| | Yeast | NULL | Probable | As nectar-tap_saprotroph (Põlme et al. 2020); Decay sugar-rich substrates (Põlme et al. 2020) | Põlme S, et al. 2020. Fungal Diversity 105:1-16 ((https://doi.org/10.1007/s13225-020-00466-2)) |
| Otu59396 | 1 | k__Fungi(100);p__Ascomycota(100);c__Saccharomycetes(100);o__Saccharomycetales(100);f__Metschnikowiaceae(100);g__Metschnikowia(100);g__Metschnikowia_unclassified(100); | Metschnikowia | 13 | Saprotroph | \|Undefined Saprotroph\| | Yeast | NULL | Probable | As nectar-tap_saprotroph (Põlme et al. 2020); Decay sugar-rich substrates (Põlme et al. 2020) | Põlme S, et al. 2020. Fungal Diversity 105:1-16 ((https://doi.org/10.1007/s13225-020-00466-2)) |
| Otu59398 | 1 | k__Fungi(100);p__Ascomycota(100);c__Saccharomycetes(100);o__Saccharomycetales(100);f__Metschnikowiaceae(100);g__Metschnikowia(100);g__Metschnikowia_unclassified(100); | Metschnikowia | 13 | Saprotroph | \|Undefined Saprotroph\| | Yeast | NULL | Probable | As nectar-tap_saprotroph (Põlme et al. 2020); Decay sugar-rich substrates (Põlme et al. 2020) | Põlme S, et al. 2020. Fungal Diversity 105:1-16 ((https://doi.org/10.1007/s13225-020-00466-2)) |
| Otu59399 | 1 | k__Fungi(100);p__Ascomycota(100);c__Saccharomycetes(100);o__Saccharomycetales(100);f__Metschnikowiaceae(100);g__Metschnikowia(100);s__Metschnikowia_pulcherrima(100); | Metschnikowia | 13 | Saprotroph | \|Undefined Saprotroph\| | Yeast | NULL | Probable | As nectar-tap_saprotroph (Põlme et al. 2020); Decay sugar-rich substrates (Põlme et al. 2020) | Põlme S, et al. 2020. Fungal Diversity 105:1-16 ((https://doi.org/10.1007/s13225-020-00466-2)) |
| Otu59403 | 1 | k__Fungi(100);p__Ascomycota(100);c__Saccharomycetes(100);o__Saccharomycetales(100);f__Metschnikowiaceae(100);g__Metschnikowia(100);s__Metschnikowia_pulcherrima(100); | Metschnikowia | 13 | Saprotroph | \|Undefined Saprotroph\| | Yeast | NULL | Probable | As nectar-tap_saprotroph (Põlme et al. 2020); Decay sugar-rich substrates (Põlme et al. 2020) | Põlme S, et al. 2020. Fungal Diversity 105:1-16 ((https://doi.org/10.1007/s13225-020-00466-2)) |
| Otu59407 | 1 | k__Fungi(100);p__Ascomycota(100);c__Saccharomycetes(100);o__Saccharomycetales(100);f__Metschnikowiaceae(100);g__Metschnikowia(100);g__Metschnikowia_unclassified(100); | Metschnikowia | 13 | Saprotroph | \|Undefined Saprotroph\| | Yeast | NULL | Probable | As nectar-tap_saprotroph (Põlme et al. 2020); Decay sugar-rich substrates (Põlme et al. 2020) | Põlme S, et al. 2020. Fungal Diversity 105:1-16 ((https://doi.org/10.1007/s13225-020-00466-2)) |
| Otu59412 | 1 | k__Fungi(100);p__Ascomycota(100);c__Saccharomycetes(100);o__Saccharomycetales(100);f__Metschnikowiaceae(100);g__Metschnikowia(100);s__Metschnikowia_pulcherrima(100); | Metschnikowia | 13 | Saprotroph | \|Undefined Saprotroph\| | Yeast | NULL | Probable | As nectar-tap_saprotroph (Põlme et al. 2020); Decay sugar-rich substrates (Põlme et al. 2020) | Põlme S, et al. 2020. Fungal Diversity 105:1-16 ((https://doi.org/10.1007/s13225-020-00466-2)) |
| Otu59413 | 1 | k__Fungi(100);p__Ascomycota(100);c__Saccharomycetes(100);o__Saccharomycetales(100);f__Metschnikowiaceae(100);g__Metschnikowia(100);s__Metschnikowia_pulcherrima(100); | Metschnikowia | 13 | Saprotroph | \|Undefined Saprotroph\| | Yeast | NULL | Probable | As nectar-tap_saprotroph (Põlme et al. 2020); Decay sugar-rich substrates (Põlme et al. 2020) | Põlme S, et al. 2020. Fungal Diversity 105:1-16 ((https://doi.org/10.1007/s13225-020-00466-2)) |
| Otu59414 | 1 | k__Fungi(100);p__Ascomycota(100);c__Saccharomycetes(100);o__Saccharomycetales(100);f__Metschnikowiaceae(100);g__Metschnikowia(100);s__Metschnikowia_sp(100); | Metschnikowia | 13 | Saprotroph | \|Undefined Saprotroph\| | Yeast | NULL | Probable | As nectar-tap_saprotroph (Põlme et al. 2020); Decay sugar-rich substrates (Põlme et al. 2020) | Põlme S, et al. 2020. Fungal Diversity 105:1-16 ((https://doi.org/10.1007/s13225-020-00466-2)) |
| Otu59415 | 1 | k__Fungi(100);p__Ascomycota(100);c__Saccharomycetes(100);o__Saccharomycetales(100);f__Metschnikowiaceae(100);g__Metschnikowia(100);s__Metschnikowia_pulcherrima(100); | Metschnikowia | 13 | Saprotroph | \|Undefined Saprotroph\| | Yeast | NULL | Probable | As nectar-tap_saprotroph (Põlme et al. 2020); Decay sugar-rich substrates (Põlme et al. 2020) | Põlme S, et al. 2020. Fungal Diversity 105:1-16 ((https://doi.org/10.1007/s13225-020-00466-2)) |
| Otu59417 | 1 | k__Fungi(100);p__Ascomycota(100);c__Saccharomycetes(100);o__Saccharomycetales(100);f__Metschnikowiaceae(100);g__Metschnikowia(100);s__Metschnikowia_pulcherrima(100); | Metschnikowia | 13 | Saprotroph | \|Undefined Saprotroph\| | Yeast | NULL | Probable | As nectar-tap_saprotroph (Põlme et al. 2020); Decay sugar-rich substrates (Põlme et al. 2020) | Põlme S, et al. 2020. Fungal Diversity 105:1-16 ((https://doi.org/10.1007/s13225-020-00466-2)) |
| Otu59419 | 1 | k__Fungi(100);p__Ascomycota(100);c__Saccharomycetes(100);o__Saccharomycetales(100);f__Metschnikowiaceae(100);g__Metschnikowia(100);s__Metschnikowia_pulcherrima(100); | Metschnikowia | 13 | Saprotroph | \|Undefined Saprotroph\| | Yeast | NULL | Probable | As nectar-tap_saprotroph (Põlme et al. 2020); Decay sugar-rich substrates (Põlme et al. 2020) | Põlme S, et al. 2020. Fungal Diversity 105:1-16 ((https://doi.org/10.1007/s13225-020-00466-2)) |
| Otu59420 | 1 | k__Fungi(100);p__Ascomycota(100);c__Saccharomycetes(100);o__Saccharomycetales(100);f__Metschnikowiaceae(100);g__Metschnikowia(100);s__Metschnikowia_pulcherrima(100); | Metschnikowia | 13 | Saprotroph | \|Undefined Saprotroph\| | Yeast | NULL | Probable | As nectar-tap_saprotroph (Põlme et al. 2020); Decay sugar-rich substrates (Põlme et al. 2020) | Põlme S, et al. 2020. Fungal Diversity 105:1-16 ((https://doi.org/10.1007/s13225-020-00466-2)) |
| Otu59421 | 1 | k__Fungi(100);p__Ascomycota(100);c__Saccharomycetes(100);o__Saccharomycetales(100);f__Metschnikowiaceae(100);g__Metschnikowia(100);s__Metschnikowia_pulcherrima(100); | Metschnikowia | 13 | Saprotroph | \|Undefined Saprotroph\| | Yeast | NULL | Probable | As nectar-tap_saprotroph (Põlme et al. 2020); Decay sugar-rich substrates (Põlme et al. 2020) | Põlme S, et al. 2020. Fungal Diversity 105:1-16 ((https://doi.org/10.1007/s13225-020-00466-2)) |
| Otu59422 | 1 | k__Fungi(100);p__Ascomycota(100);c__Saccharomycetes(100);o__Saccharomycetales(100);f__Metschnikowiaceae(100);g__Metschnikowia(100);s__Metschnikowia_pulcherrima(100); | Metschnikowia | 13 | Saprotroph | \|Undefined Saprotroph\| | Yeast | NULL | Probable | As nectar-tap_saprotroph (Põlme et al. 2020); Decay sugar-rich substrates (Põlme et al. 2020) | Põlme S, et al. 2020. Fungal Diversity 105:1-16 ((https://doi.org/10.1007/s13225-020-00466-2)) |
| Otu59423 | 1 | k__Fungi(100);p__Ascomycota(100);c__Saccharomycetes(100);o__Saccharomycetales(100);f__Metschnikowiaceae(100);g__Metschnikowia(100);s__Metschnikowia_pulcherrima(100); | Metschnikowia | 13 | Saprotroph | \|Undefined Saprotroph\| | Yeast | NULL | Probable | As nectar-tap_saprotroph (Põlme et al. 2020); Decay sugar-rich substrates (Põlme et al. 2020) | Põlme S, et al. 2020. Fungal Diversity 105:1-16 ((https://doi.org/10.1007/s13225-020-00466-2)) |
| Otu59424 | 1 | k__Fungi(100);p__Ascomycota(100);c__Saccharomycetes(100);o__Saccharomycetales(100);f__Metschnikowiaceae(100);g__Metschnikowia(100);g__Metschnikowia_unclassified(100); | Metschnikowia | 13 | Saprotroph | \|Undefined Saprotroph\| | Yeast | NULL | Probable | As nectar-tap_saprotroph (Põlme et al. 2020); Decay sugar-rich substrates (Põlme et al. 2020) | Põlme S, et al. 2020. Fungal Diversity 105:1-16 ((https://doi.org/10.1007/s13225-020-00466-2)) |
| Otu59426 | 1 | k__Fungi(100);p__Ascomycota(100);c__Saccharomycetes(100);o__Saccharomycetales(100);f__Metschnikowiaceae(100);g__Metschnikowia(100);s__Metschnikowia_pulcherrima(100); | Metschnikowia | 13 | Saprotroph | \|Undefined Saprotroph\| | Yeast | NULL | Probable | As nectar-tap_saprotroph (Põlme et al. 2020); Decay sugar-rich substrates (Põlme et al. 2020) | Põlme S, et al. 2020. Fungal Diversity 105:1-16 ((https://doi.org/10.1007/s13225-020-00466-2)) |
| Otu59427 | 1 | k__Fungi(100);p__Ascomycota(100);c__Saccharomycetes(100);o__Saccharomycetales(100);f__Metschnikowiaceae(100);g__Metschnikowia(100);s__Metschnikowia_pulcherrima(100); | Metschnikowia | 13 | Saprotroph | \|Undefined Saprotroph\| | Yeast | NULL | Probable | As nectar-tap_saprotroph (Põlme et al. 2020); Decay sugar-rich substrates (Põlme et al. 2020) | Põlme S, et al. 2020. Fungal Diversity 105:1-16 ((https://doi.org/10.1007/s13225-020-00466-2)) |
| Otu59428 | 1 | k__Fungi(100);p__Ascomycota(100);c__Saccharomycetes(100);o__Saccharomycetales(100);f__Metschnikowiaceae(100);g__Metschnikowia(100);s__Metschnikowia_pulcherrima(100); | Metschnikowia | 13 | Saprotroph | \|Undefined Saprotroph\| | Yeast | NULL | Probable | As nectar-tap_saprotroph (Põlme et al. 2020); Decay sugar-rich substrates (Põlme et al. 2020) | Põlme S, et al. 2020. Fungal Diversity 105:1-16 ((https://doi.org/10.1007/s13225-020-00466-2)) |
| Otu59429 | 1 | k__Fungi(100);p__Ascomycota(100);c__Saccharomycetes(100);o__Saccharomycetales(100);f__Metschnikowiaceae(100);g__Metschnikowia(100);g__Metschnikowia_unclassified(100); | Metschnikowia | 13 | Saprotroph | \|Undefined Saprotroph\| | Yeast | NULL | Probable | As nectar-tap_saprotroph (Põlme et al. 2020); Decay sugar-rich substrates (Põlme et al. 2020) | Põlme S, et al. 2020. Fungal Diversity 105:1-16 ((https://doi.org/10.1007/s13225-020-00466-2)) |
| Otu59430 | 1 | k__Fungi(100);p__Ascomycota(100);c__Saccharomycetes(100);o__Saccharomycetales(100);f__Metschnikowiaceae(100);g__Metschnikowia(100);s__Metschnikowia_pulcherrima(100); | Metschnikowia | 13 | Saprotroph | \|Undefined Saprotroph\| | Yeast | NULL | Probable | As nectar-tap_saprotroph (Põlme et al. 2020); Decay sugar-rich substrates (Põlme et al. 2020) | Põlme S, et al. 2020. Fungal Diversity 105:1-16 ((https://doi.org/10.1007/s13225-020-00466-2)) |
| Otu59431 | 1 | k__Fungi(100);p__Ascomycota(100);c__Saccharomycetes(100);o__Saccharomycetales(100);f__Metschnikowiaceae(100);g__Metschnikowia(100);g__Metschnikowia_unclassified(100); | Metschnikowia | 13 | Saprotroph | \|Undefined Saprotroph\| | Yeast | NULL | Probable | As nectar-tap_saprotroph (Põlme et al. 2020); Decay sugar-rich substrates (Põlme et al. 2020) | Põlme S, et al. 2020. Fungal Diversity 105:1-16 ((https://doi.org/10.1007/s13225-020-00466-2)) |
| Otu59432 | 1 | k__Fungi(100);p__Ascomycota(100);c__Saccharomycetes(100);o__Saccharomycetales(100);f__Metschnikowiaceae(100);g__Metschnikowia(100);s__Metschnikowia_pulcherrima(100); | Metschnikowia | 13 | Saprotroph | \|Undefined Saprotroph\| | Yeast | NULL | Probable | As nectar-tap_saprotroph (Põlme et al. 2020); Decay sugar-rich substrates (Põlme et al. 2020) | Põlme S, et al. 2020. Fungal Diversity 105:1-16 ((https://doi.org/10.1007/s13225-020-00466-2)) |
| Otu59433 | 1 | k__Fungi(100);p__Ascomycota(100);c__Saccharomycetes(100);o__Saccharomycetales(100);f__Metschnikowiaceae(100);g__Metschnikowia(100);s__Metschnikowia_pulcherrima(100); | Metschnikowia | 13 | Saprotroph | \|Undefined Saprotroph\| | Yeast | NULL | Probable | As nectar-tap_saprotroph (Põlme et al. 2020); Decay sugar-rich substrates (Põlme et al. 2020) | Põlme S, et al. 2020. Fungal Diversity 105:1-16 ((https://doi.org/10.1007/s13225-020-00466-2)) |
| Otu59434 | 1 | k__Fungi(100);p__Ascomycota(100);c__Saccharomycetes(100);o__Saccharomycetales(100);f__Metschnikowiaceae(100);g__Metschnikowia(100);s__Metschnikowia_pulcherrima(100); | Metschnikowia | 13 | Saprotroph | \|Undefined Saprotroph\| | Yeast | NULL | Probable | As nectar-tap_saprotroph (Põlme et al. 2020); Decay sugar-rich substrates (Põlme et al. 2020) | Põlme S, et al. 2020. Fungal Diversity 105:1-16 ((https://doi.org/10.1007/s13225-020-00466-2)) |
| Otu59435 | 1 | k__Fungi(100);p__Ascomycota(100);c__Saccharomycetes(100);o__Saccharomycetales(100);f__Metschnikowiaceae(100);g__Metschnikowia(100);s__Metschnikowia_pulcherrima(100); | Metschnikowia | 13 | Saprotroph | \|Undefined Saprotroph\| | Yeast | NULL | Probable | As nectar-tap_saprotroph (Põlme et al. 2020); Decay sugar-rich substrates (Põlme et al. 2020) | Põlme S, et al. 2020. Fungal Diversity 105:1-16 ((https://doi.org/10.1007/s13225-020-00466-2)) |
| Otu59436 | 1 | k__Fungi(100);p__Ascomycota(100);c__Saccharomycetes(100);o__Saccharomycetales(100);f__Metschnikowiaceae(100);g__Metschnikowia(100);g__Metschnikowia_unclassified(100); | Metschnikowia | 13 | Saprotroph | \|Undefined Saprotroph\| | Yeast | NULL | Probable | As nectar-tap_saprotroph (Põlme et al. 2020); Decay sugar-rich substrates (Põlme et al. 2020) | Põlme S, et al. 2020. Fungal Diversity 105:1-16 ((https://doi.org/10.1007/s13225-020-00466-2)) |
| Otu59437 | 1 | k__Fungi(100);p__Ascomycota(100);c__Saccharomycetes(100);o__Saccharomycetales(100);f__Metschnikowiaceae(100);g__Metschnikowia(100);s__Metschnikowia_pulcherrima(100); | Metschnikowia | 13 | Saprotroph | \|Undefined Saprotroph\| | Yeast | NULL | Probable | As nectar-tap_saprotroph (Põlme et al. 2020); Decay sugar-rich substrates (Põlme et al. 2020) | Põlme S, et al. 2020. Fungal Diversity 105:1-16 ((https://doi.org/10.1007/s13225-020-00466-2)) |
| Otu59443 | 1 | k__Fungi(100);p__Ascomycota(100);c__Saccharomycetes(100);o__Saccharomycetales(100);f__Metschnikowiaceae(100);g__Metschnikowia(100);s__Metschnikowia_pulcherrima(100); | Metschnikowia | 13 | Saprotroph | \|Undefined Saprotroph\| | Yeast | NULL | Probable | As nectar-tap_saprotroph (Põlme et al. 2020); Decay sugar-rich substrates (Põlme et al. 2020) | Põlme S, et al. 2020. Fungal Diversity 105:1-16 ((https://doi.org/10.1007/s13225-020-00466-2)) |
| Otu59444 | 1 | k__Fungi(100);p__Ascomycota(100);c__Saccharomycetes(100);o__Saccharomycetales(100);f__Metschnikowiaceae(100);g__Metschnikowia(100);g__Metschnikowia_unclassified(100); | Metschnikowia | 13 | Saprotroph | \|Undefined Saprotroph\| | Yeast | NULL | Probable | As nectar-tap_saprotroph (Põlme et al. 2020); Decay sugar-rich substrates (Põlme et al. 2020) | Põlme S, et al. 2020. Fungal Diversity 105:1-16 ((https://doi.org/10.1007/s13225-020-00466-2)) |
| Otu59452 | 1 | k__Fungi(100);p__Ascomycota(100);c__Saccharomycetes(100);o__Saccharomycetales(100);f__Metschnikowiaceae(100);g__Metschnikowia(100);g__Metschnikowia_unclassified(100); | Metschnikowia | 13 | Saprotroph | \|Undefined Saprotroph\| | Yeast | NULL | Probable | As nectar-tap_saprotroph (Põlme et al. 2020); Decay sugar-rich substrates (Põlme et al. 2020) | Põlme S, et al. 2020. Fungal Diversity 105:1-16 ((https://doi.org/10.1007/s13225-020-00466-2)) |
| Otu59453 | 1 | k__Fungi(100);p__Ascomycota(100);c__Saccharomycetes(100);o__Saccharomycetales(100);f__Metschnikowiaceae(100);g__Metschnikowia(100);g__Metschnikowia_unclassified(100); | Metschnikowia | 13 | Saprotroph | \|Undefined Saprotroph\| | Yeast | NULL | Probable | As nectar-tap_saprotroph (Põlme et al. 2020); Decay sugar-rich substrates (Põlme et al. 2020) | Põlme S, et al. 2020. Fungal Diversity 105:1-16 ((https://doi.org/10.1007/s13225-020-00466-2)) |
| Otu59454 | 1 | k__Fungi(100);p__Ascomycota(100);c__Saccharomycetes(100);o__Saccharomycetales(100);f__Metschnikowiaceae(100);g__Metschnikowia(100);s__Metschnikowia_pulcherrima(100); | Metschnikowia | 13 | Saprotroph | \|Undefined Saprotroph\| | Yeast | NULL | Probable | As nectar-tap_saprotroph (Põlme et al. 2020); Decay sugar-rich substrates (Põlme et al. 2020) | Põlme S, et al. 2020. Fungal Diversity 105:1-16 ((https://doi.org/10.1007/s13225-020-00466-2)) |
| Otu59457 | 1 | k__Fungi(100);p__Ascomycota(100);c__Saccharomycetes(100);o__Saccharomycetales(100);f__Metschnikowiaceae(100);g__Metschnikowia(100);s__Metschnikowia_pulcherrima(100); | Metschnikowia | 13 | Saprotroph | \|Undefined Saprotroph\| | Yeast | NULL | Probable | As nectar-tap_saprotroph (Põlme et al. 2020); Decay sugar-rich substrates (Põlme et al. 2020) | Põlme S, et al. 2020. Fungal Diversity 105:1-16 ((https://doi.org/10.1007/s13225-020-00466-2)) |
| Otu59460 | 1 | k__Fungi(100);p__Ascomycota(100);c__Saccharomycetes(100);o__Saccharomycetales(100);f__Metschnikowiaceae(100);g__Metschnikowia(100);s__Metschnikowia_pulcherrima(100); | Metschnikowia | 13 | Saprotroph | \|Undefined Saprotroph\| | Yeast | NULL | Probable | As nectar-tap_saprotroph (Põlme et al. 2020); Decay sugar-rich substrates (Põlme et al. 2020) | Põlme S, et al. 2020. Fungal Diversity 105:1-16 ((https://doi.org/10.1007/s13225-020-00466-2)) |
| Otu59461 | 1 | k__Fungi(100);p__Ascomycota(100);c__Saccharomycetes(100);o__Saccharomycetales(100);f__Metschnikowiaceae(100);g__Metschnikowia(100);s__Metschnikowia_pulcherrima(100); | Metschnikowia | 13 | Saprotroph | \|Undefined Saprotroph\| | Yeast | NULL | Probable | As nectar-tap_saprotroph (Põlme et al. 2020); Decay sugar-rich substrates (Põlme et al. 2020) | Põlme S, et al. 2020. Fungal Diversity 105:1-16 ((https://doi.org/10.1007/s13225-020-00466-2)) |
| Otu59462 | 1 | k__Fungi(100);p__Ascomycota(100);c__Saccharomycetes(100);o__Saccharomycetales(100);f__Metschnikowiaceae(100);g__Metschnikowia(100);s__Metschnikowia_pulcherrima(100); | Metschnikowia | 13 | Saprotroph | \|Undefined Saprotroph\| | Yeast | NULL | Probable | As nectar-tap_saprotroph (Põlme et al. 2020); Decay sugar-rich substrates (Põlme et al. 2020) | Põlme S, et al. 2020. Fungal Diversity 105:1-16 ((https://doi.org/10.1007/s13225-020-00466-2)) |
| Otu59463 | 1 | k__Fungi(100);p__Ascomycota(100);c__Saccharomycetes(100);o__Saccharomycetales(100);f__Metschnikowiaceae(100);g__Metschnikowia(100);s__Metschnikowia_pulcherrima(100); | Metschnikowia | 13 | Saprotroph | \|Undefined Saprotroph\| | Yeast | NULL | Probable | As nectar-tap_saprotroph (Põlme et al. 2020); Decay sugar-rich substrates (Põlme et al. 2020) | Põlme S, et al. 2020. Fungal Diversity 105:1-16 ((https://doi.org/10.1007/s13225-020-00466-2)) |
| Otu59466 | 1 | k__Fungi(100);p__Ascomycota(100);c__Saccharomycetes(100);o__Saccharomycetales(100);f__Metschnikowiaceae(100);g__Metschnikowia(100);s__Metschnikowia_pulcherrima(100); | Metschnikowia | 13 | Saprotroph | \|Undefined Saprotroph\| | Yeast | NULL | Probable | As nectar-tap_saprotroph (Põlme et al. 2020); Decay sugar-rich substrates (Põlme et al. 2020) | Põlme S, et al. 2020. Fungal Diversity 105:1-16 ((https://doi.org/10.1007/s13225-020-00466-2)) |
| Otu59469 | 1 | k__Fungi(100);p__Ascomycota(100);c__Saccharomycetes(100);o__Saccharomycetales(100);f__Metschnikowiaceae(100);g__Metschnikowia(100);g__Metschnikowia_unclassified(100); | Metschnikowia | 13 | Saprotroph | \|Undefined Saprotroph\| | Yeast | NULL | Probable | As nectar-tap_saprotroph (Põlme et al. 2020); Decay sugar-rich substrates (Põlme et al. 2020) | Põlme S, et al. 2020. Fungal Diversity 105:1-16 ((https://doi.org/10.1007/s13225-020-00466-2)) |
| Otu59470 | 1 | k__Fungi(100);p__Ascomycota(100);c__Saccharomycetes(100);o__Saccharomycetales(100);f__Metschnikowiaceae(100);g__Metschnikowia(100);s__Metschnikowia_pulcherrima(100); | Metschnikowia | 13 | Saprotroph | \|Undefined Saprotroph\| | Yeast | NULL | Probable | As nectar-tap_saprotroph (Põlme et al. 2020); Decay sugar-rich substrates (Põlme et al. 2020) | Põlme S, et al. 2020. Fungal Diversity 105:1-16 ((https://doi.org/10.1007/s13225-020-00466-2)) |
| Otu59471 | 1 | k__Fungi(100);p__Ascomycota(100);c__Saccharomycetes(100);o__Saccharomycetales(100);f__Metschnikowiaceae(100);g__Metschnikowia(100);s__Metschnikowia_pulcherrima(100); | Metschnikowia | 13 | Saprotroph | \|Undefined Saprotroph\| | Yeast | NULL | Probable | As nectar-tap_saprotroph (Põlme et al. 2020); Decay sugar-rich substrates (Põlme et al. 2020) | Põlme S, et al. 2020. Fungal Diversity 105:1-16 ((https://doi.org/10.1007/s13225-020-00466-2)) |
| Otu59472 | 1 | k__Fungi(100);p__Ascomycota(100);c__Saccharomycetes(100);o__Saccharomycetales(100);f__Metschnikowiaceae(100);g__Metschnikowia(100);s__Metschnikowia_pulcherrima(100); | Metschnikowia | 13 | Saprotroph | \|Undefined Saprotroph\| | Yeast | NULL | Probable | As nectar-tap_saprotroph (Põlme et al. 2020); Decay sugar-rich substrates (Põlme et al. 2020) | Põlme S, et al. 2020. Fungal Diversity 105:1-16 ((https://doi.org/10.1007/s13225-020-00466-2)) |
| Otu59477 | 1 | k__Fungi(100);p__Ascomycota(100);c__Saccharomycetes(100);o__Saccharomycetales(100);f__Metschnikowiaceae(100);g__Metschnikowia(100);s__Metschnikowia_pulcherrima(100); | Metschnikowia | 13 | Saprotroph | \|Undefined Saprotroph\| | Yeast | NULL | Probable | As nectar-tap_saprotroph (Põlme et al. 2020); Decay sugar-rich substrates (Põlme et al. 2020) | Põlme S, et al. 2020. Fungal Diversity 105:1-16 ((https://doi.org/10.1007/s13225-020-00466-2)) |
| Otu59483 | 1 | k__Fungi(100);p__Ascomycota(100);c__Saccharomycetes(100);o__Saccharomycetales(100);f__Metschnikowiaceae(100);g__Metschnikowia(100);s__Metschnikowia_pulcherrima(100); | Metschnikowia | 13 | Saprotroph | \|Undefined Saprotroph\| | Yeast | NULL | Probable | As nectar-tap_saprotroph (Põlme et al. 2020); Decay sugar-rich substrates (Põlme et al. 2020) | Põlme S, et al. 2020. Fungal Diversity 105:1-16 ((https://doi.org/10.1007/s13225-020-00466-2)) |
| Otu59487 | 1 | k__Fungi(100);p__Ascomycota(100);c__Saccharomycetes(100);o__Saccharomycetales(100);f__Metschnikowiaceae(100);g__Metschnikowia(100);g__Metschnikowia_unclassified(100); | Metschnikowia | 13 | Saprotroph | \|Undefined Saprotroph\| | Yeast | NULL | Probable | As nectar-tap_saprotroph (Põlme et al. 2020); Decay sugar-rich substrates (Põlme et al. 2020) | Põlme S, et al. 2020. Fungal Diversity 105:1-16 ((https://doi.org/10.1007/s13225-020-00466-2)) |
| Otu59490 | 1 | k__Fungi(100);p__Ascomycota(100);c__Saccharomycetes(100);o__Saccharomycetales(100);f__Metschnikowiaceae(100);g__Metschnikowia(100);s__Metschnikowia_pulcherrima(100); | Metschnikowia | 13 | Saprotroph | \|Undefined Saprotroph\| | Yeast | NULL | Probable | As nectar-tap_saprotroph (Põlme et al. 2020); Decay sugar-rich substrates (Põlme et al. 2020) | Põlme S, et al. 2020. Fungal Diversity 105:1-16 ((https://doi.org/10.1007/s13225-020-00466-2)) |
| Otu59492 | 1 | k__Fungi(100);p__Ascomycota(100);c__Saccharomycetes(100);o__Saccharomycetales(100);f__Metschnikowiaceae(100);g__Metschnikowia(100);s__Metschnikowia_pulcherrima(100); | Metschnikowia | 13 | Saprotroph | \|Undefined Saprotroph\| | Yeast | NULL | Probable | As nectar-tap_saprotroph (Põlme et al. 2020); Decay sugar-rich substrates (Põlme et al. 2020) | Põlme S, et al. 2020. Fungal Diversity 105:1-16 ((https://doi.org/10.1007/s13225-020-00466-2)) |
| Otu59498 | 1 | k__Fungi(100);p__Ascomycota(100);c__Saccharomycetes(100);o__Saccharomycetales(100);f__Metschnikowiaceae(100);g__Metschnikowia(100);s__Metschnikowia_pulcherrima(100); | Metschnikowia | 13 | Saprotroph | \|Undefined Saprotroph\| | Yeast | NULL | Probable | As nectar-tap_saprotroph (Põlme et al. 2020); Decay sugar-rich substrates (Põlme et al. 2020) | Põlme S, et al. 2020. Fungal Diversity 105:1-16 ((https://doi.org/10.1007/s13225-020-00466-2)) |
| Otu59499 | 1 | k__Fungi(100);p__Ascomycota(100);c__Saccharomycetes(100);o__Saccharomycetales(100);f__Metschnikowiaceae(100);g__Metschnikowia(100);s__Metschnikowia_pulcherrima(100); | Metschnikowia | 13 | Saprotroph | \|Undefined Saprotroph\| | Yeast | NULL | Probable | As nectar-tap_saprotroph (Põlme et al. 2020); Decay sugar-rich substrates (Põlme et al. 2020) | Põlme S, et al. 2020. Fungal Diversity 105:1-16 ((https://doi.org/10.1007/s13225-020-00466-2)) |
| Otu59502 | 1 | k__Fungi(100);p__Ascomycota(100);c__Saccharomycetes(100);o__Saccharomycetales(100);f__Metschnikowiaceae(100);g__Metschnikowia(100);s__Metschnikowia_pulcherrima(100); | Metschnikowia | 13 | Saprotroph | \|Undefined Saprotroph\| | Yeast | NULL | Probable | As nectar-tap_saprotroph (Põlme et al. 2020); Decay sugar-rich substrates (Põlme et al. 2020) | Põlme S, et al. 2020. Fungal Diversity 105:1-16 ((https://doi.org/10.1007/s13225-020-00466-2)) |
| Otu59504 | 1 | k__Fungi(100);p__Ascomycota(100);c__Saccharomycetes(100);o__Saccharomycetales(100);f__Metschnikowiaceae(100);g__Metschnikowia(100);s__Metschnikowia_pulcherrima(100); | Metschnikowia | 13 | Saprotroph | \|Undefined Saprotroph\| | Yeast | NULL | Probable | As nectar-tap_saprotroph (Põlme et al. 2020); Decay sugar-rich substrates (Põlme et al. 2020) | Põlme S, et al. 2020. Fungal Diversity 105:1-16 ((https://doi.org/10.1007/s13225-020-00466-2)) |
| Otu59507 | 1 | k__Fungi(100);p__Ascomycota(100);c__Saccharomycetes(100);o__Saccharomycetales(100);f__Metschnikowiaceae(100);g__Metschnikowia(100);s__Metschnikowia_pulcherrima(100); | Metschnikowia | 13 | Saprotroph | \|Undefined Saprotroph\| | Yeast | NULL | Probable | As nectar-tap_saprotroph (Põlme et al. 2020); Decay sugar-rich substrates (Põlme et al. 2020) | Põlme S, et al. 2020. Fungal Diversity 105:1-16 ((https://doi.org/10.1007/s13225-020-00466-2)) |
| Otu59508 | 1 | k__Fungi(100);p__Ascomycota(100);c__Saccharomycetes(100);o__Saccharomycetales(100);f__Metschnikowiaceae(100);g__Metschnikowia(100);s__Metschnikowia_pulcherrima(100); | Metschnikowia | 13 | Saprotroph | \|Undefined Saprotroph\| | Yeast | NULL | Probable | As nectar-tap_saprotroph (Põlme et al. 2020); Decay sugar-rich substrates (Põlme et al. 2020) | Põlme S, et al. 2020. Fungal Diversity 105:1-16 ((https://doi.org/10.1007/s13225-020-00466-2)) |
| Otu59511 | 1 | k__Fungi(100);p__Ascomycota(100);c__Saccharomycetes(100);o__Saccharomycetales(100);f__Metschnikowiaceae(100);g__Metschnikowia(100);g__Metschnikowia_unclassified(100); | Metschnikowia | 13 | Saprotroph | \|Undefined Saprotroph\| | Yeast | NULL | Probable | As nectar-tap_saprotroph (Põlme et al. 2020); Decay sugar-rich substrates (Põlme et al. 2020) | Põlme S, et al. 2020. Fungal Diversity 105:1-16 ((https://doi.org/10.1007/s13225-020-00466-2)) |
| Otu59514 | 1 | k__Fungi(100);p__Ascomycota(100);c__Saccharomycetes(100);o__Saccharomycetales(100);f__Metschnikowiaceae(100);g__Metschnikowia(100);s__Metschnikowia_pulcherrima(100); | Metschnikowia | 13 | Saprotroph | \|Undefined Saprotroph\| | Yeast | NULL | Probable | As nectar-tap_saprotroph (Põlme et al. 2020); Decay sugar-rich substrates (Põlme et al. 2020) | Põlme S, et al. 2020. Fungal Diversity 105:1-16 ((https://doi.org/10.1007/s13225-020-00466-2)) |
| Otu59516 | 1 | k__Fungi(100);p__Ascomycota(100);c__Saccharomycetes(100);o__Saccharomycetales(100);f__Metschnikowiaceae(100);g__Metschnikowia(100);s__Metschnikowia_pulcherrima(100); | Metschnikowia | 13 | Saprotroph | \|Undefined Saprotroph\| | Yeast | NULL | Probable | As nectar-tap_saprotroph (Põlme et al. 2020); Decay sugar-rich substrates (Põlme et al. 2020) | Põlme S, et al. 2020. Fungal Diversity 105:1-16 ((https://doi.org/10.1007/s13225-020-00466-2)) |
| Otu59517 | 1 | k__Fungi(100);p__Ascomycota(100);c__Saccharomycetes(100);o__Saccharomycetales(100);f__Metschnikowiaceae(100);g__Metschnikowia(100);s__Metschnikowia_pulcherrima(100); | Metschnikowia | 13 | Saprotroph | \|Undefined Saprotroph\| | Yeast | NULL | Probable | As nectar-tap_saprotroph (Põlme et al. 2020); Decay sugar-rich substrates (Põlme et al. 2020) | Põlme S, et al. 2020. Fungal Diversity 105:1-16 ((https://doi.org/10.1007/s13225-020-00466-2)) |
| Otu59518 | 1 | k__Fungi(100);p__Ascomycota(100);c__Saccharomycetes(100);o__Saccharomycetales(100);f__Metschnikowiaceae(100);g__Metschnikowia(100);g__Metschnikowia_unclassified(100); | Metschnikowia | 13 | Saprotroph | \|Undefined Saprotroph\| | Yeast | NULL | Probable | As nectar-tap_saprotroph (Põlme et al. 2020); Decay sugar-rich substrates (Põlme et al. 2020) | Põlme S, et al. 2020. Fungal Diversity 105:1-16 ((https://doi.org/10.1007/s13225-020-00466-2)) |
| Otu59519 | 1 | k__Fungi(100);p__Ascomycota(100);c__Saccharomycetes(100);o__Saccharomycetales(100);f__Metschnikowiaceae(100);g__Metschnikowia(100);s__Metschnikowia_pulcherrima(100); | Metschnikowia | 13 | Saprotroph | \|Undefined Saprotroph\| | Yeast | NULL | Probable | As nectar-tap_saprotroph (Põlme et al. 2020); Decay sugar-rich substrates (Põlme et al. 2020) | Põlme S, et al. 2020. Fungal Diversity 105:1-16 ((https://doi.org/10.1007/s13225-020-00466-2)) |
| Otu59520 | 1 | k__Fungi(100);p__Ascomycota(100);c__Saccharomycetes(100);o__Saccharomycetales(100);f__Metschnikowiaceae(100);g__Metschnikowia(100);g__Metschnikowia_unclassified(100); | Metschnikowia | 13 | Saprotroph | \|Undefined Saprotroph\| | Yeast | NULL | Probable | As nectar-tap_saprotroph (Põlme et al. 2020); Decay sugar-rich substrates (Põlme et al. 2020) | Põlme S, et al. 2020. Fungal Diversity 105:1-16 ((https://doi.org/10.1007/s13225-020-00466-2)) |
| Otu59521 | 1 | k__Fungi(100);p__Ascomycota(100);c__Saccharomycetes(100);o__Saccharomycetales(100);f__Metschnikowiaceae(100);g__Metschnikowia(100);g__Metschnikowia_unclassified(100); | Metschnikowia | 13 | Saprotroph | \|Undefined Saprotroph\| | Yeast | NULL | Probable | As nectar-tap_saprotroph (Põlme et al. 2020); Decay sugar-rich substrates (Põlme et al. 2020) | Põlme S, et al. 2020. Fungal Diversity 105:1-16 ((https://doi.org/10.1007/s13225-020-00466-2)) |
| Otu59522 | 1 | k__Fungi(100);p__Ascomycota(100);c__Saccharomycetes(100);o__Saccharomycetales(100);f__Metschnikowiaceae(100);g__Metschnikowia(100);g__Metschnikowia_unclassified(100); | Metschnikowia | 13 | Saprotroph | \|Undefined Saprotroph\| | Yeast | NULL | Probable | As nectar-tap_saprotroph (Põlme et al. 2020); Decay sugar-rich substrates (Põlme et al. 2020) | Põlme S, et al. 2020. Fungal Diversity 105:1-16 ((https://doi.org/10.1007/s13225-020-00466-2)) |
| Otu59523 | 1 | k__Fungi(100);p__Ascomycota(100);c__Saccharomycetes(100);o__Saccharomycetales(100);f__Metschnikowiaceae(100);g__Metschnikowia(100);s__Metschnikowia_pulcherrima(100); | Metschnikowia | 13 | Saprotroph | \|Undefined Saprotroph\| | Yeast | NULL | Probable | As nectar-tap_saprotroph (Põlme et al. 2020); Decay sugar-rich substrates (Põlme et al. 2020) | Põlme S, et al. 2020. Fungal Diversity 105:1-16 ((https://doi.org/10.1007/s13225-020-00466-2)) |
| Otu59524 | 1 | k__Fungi(100);p__Ascomycota(100);c__Saccharomycetes(100);o__Saccharomycetales(100);f__Metschnikowiaceae(100);g__Metschnikowia(100);s__Metschnikowia_pulcherrima(100); | Metschnikowia | 13 | Saprotroph | \|Undefined Saprotroph\| | Yeast | NULL | Probable | As nectar-tap_saprotroph (Põlme et al. 2020); Decay sugar-rich substrates (Põlme et al. 2020) | Põlme S, et al. 2020. Fungal Diversity 105:1-16 ((https://doi.org/10.1007/s13225-020-00466-2)) |
| Otu59525 | 1 | k__Fungi(100);p__Ascomycota(100);c__Saccharomycetes(100);o__Saccharomycetales(100);f__Metschnikowiaceae(100);g__Metschnikowia(100);s__Metschnikowia_pulcherrima(100); | Metschnikowia | 13 | Saprotroph | \|Undefined Saprotroph\| | Yeast | NULL | Probable | As nectar-tap_saprotroph (Põlme et al. 2020); Decay sugar-rich substrates (Põlme et al. 2020) | Põlme S, et al. 2020. Fungal Diversity 105:1-16 ((https://doi.org/10.1007/s13225-020-00466-2)) |
| Otu59528 | 1 | k__Fungi(100);p__Ascomycota(100);c__Saccharomycetes(100);o__Saccharomycetales(100);f__Metschnikowiaceae(100);g__Metschnikowia(100);g__Metschnikowia_unclassified(100); | Metschnikowia | 13 | Saprotroph | \|Undefined Saprotroph\| | Yeast | NULL | Probable | As nectar-tap_saprotroph (Põlme et al. 2020); Decay sugar-rich substrates (Põlme et al. 2020) | Põlme S, et al. 2020. Fungal Diversity 105:1-16 ((https://doi.org/10.1007/s13225-020-00466-2)) |
| Otu59529 | 1 | k__Fungi(100);p__Ascomycota(100);c__Saccharomycetes(100);o__Saccharomycetales(100);f__Metschnikowiaceae(100);g__Metschnikowia(100);s__Metschnikowia_pulcherrima(100); | Metschnikowia | 13 | Saprotroph | \|Undefined Saprotroph\| | Yeast | NULL | Probable | As nectar-tap_saprotroph (Põlme et al. 2020); Decay sugar-rich substrates (Põlme et al. 2020) | Põlme S, et al. 2020. Fungal Diversity 105:1-16 ((https://doi.org/10.1007/s13225-020-00466-2)) |
| Otu59530 | 1 | k__Fungi(100);p__Ascomycota(100);c__Saccharomycetes(100);o__Saccharomycetales(100);f__Metschnikowiaceae(100);g__Metschnikowia(100);s__Metschnikowia_pulcherrima(100); | Metschnikowia | 13 | Saprotroph | \|Undefined Saprotroph\| | Yeast | NULL | Probable | As nectar-tap_saprotroph (Põlme et al. 2020); Decay sugar-rich substrates (Põlme et al. 2020) | Põlme S, et al. 2020. Fungal Diversity 105:1-16 ((https://doi.org/10.1007/s13225-020-00466-2)) |
| Otu59531 | 1 | k__Fungi(100);p__Ascomycota(100);c__Saccharomycetes(100);o__Saccharomycetales(100);f__Metschnikowiaceae(100);g__Metschnikowia(100);s__Metschnikowia_pulcherrima(100); | Metschnikowia | 13 | Saprotroph | \|Undefined Saprotroph\| | Yeast | NULL | Probable | As nectar-tap_saprotroph (Põlme et al. 2020); Decay sugar-rich substrates (Põlme et al. 2020) | Põlme S, et al. 2020. Fungal Diversity 105:1-16 ((https://doi.org/10.1007/s13225-020-00466-2)) |
| Otu59532 | 1 | k__Fungi(100);p__Ascomycota(100);c__Saccharomycetes(100);o__Saccharomycetales(100);f__Metschnikowiaceae(100);g__Metschnikowia(100);s__Metschnikowia_pulcherrima(100); | Metschnikowia | 13 | Saprotroph | \|Undefined Saprotroph\| | Yeast | NULL | Probable | As nectar-tap_saprotroph (Põlme et al. 2020); Decay sugar-rich substrates (Põlme et al. 2020) | Põlme S, et al. 2020. Fungal Diversity 105:1-16 ((https://doi.org/10.1007/s13225-020-00466-2)) |
| Otu59533 | 1 | k__Fungi(100);p__Ascomycota(100);c__Saccharomycetes(100);o__Saccharomycetales(100);f__Metschnikowiaceae(100);g__Metschnikowia(100);s__Metschnikowia_pulcherrima(100); | Metschnikowia | 13 | Saprotroph | \|Undefined Saprotroph\| | Yeast | NULL | Probable | As nectar-tap_saprotroph (Põlme et al. 2020); Decay sugar-rich substrates (Põlme et al. 2020) | Põlme S, et al. 2020. Fungal Diversity 105:1-16 ((https://doi.org/10.1007/s13225-020-00466-2)) |
| Otu59534 | 1 | k__Fungi(100);p__Ascomycota(100);c__Saccharomycetes(100);o__Saccharomycetales(100);f__Metschnikowiaceae(100);g__Metschnikowia(100);s__Metschnikowia_pulcherrima(100); | Metschnikowia | 13 | Saprotroph | \|Undefined Saprotroph\| | Yeast | NULL | Probable | As nectar-tap_saprotroph (Põlme et al. 2020); Decay sugar-rich substrates (Põlme et al. 2020) | Põlme S, et al. 2020. Fungal Diversity 105:1-16 ((https://doi.org/10.1007/s13225-020-00466-2)) |
| Otu59535 | 1 | k__Fungi(100);p__Ascomycota(100);c__Saccharomycetes(100);o__Saccharomycetales(100);f__Metschnikowiaceae(100);g__Metschnikowia(100);s__Metschnikowia_pulcherrima(100); | Metschnikowia | 13 | Saprotroph | \|Undefined Saprotroph\| | Yeast | NULL | Probable | As nectar-tap_saprotroph (Põlme et al. 2020); Decay sugar-rich substrates (Põlme et al. 2020) | Põlme S, et al. 2020. Fungal Diversity 105:1-16 ((https://doi.org/10.1007/s13225-020-00466-2)) |
| Otu59536 | 1 | k__Fungi(100);p__Ascomycota(100);c__Saccharomycetes(100);o__Saccharomycetales(100);f__Metschnikowiaceae(100);g__Metschnikowia(100);s__Metschnikowia_pulcherrima(100); | Metschnikowia | 13 | Saprotroph | \|Undefined Saprotroph\| | Yeast | NULL | Probable | As nectar-tap_saprotroph (Põlme et al. 2020); Decay sugar-rich substrates (Põlme et al. 2020) | Põlme S, et al. 2020. Fungal Diversity 105:1-16 ((https://doi.org/10.1007/s13225-020-00466-2)) |
| Otu59537 | 1 | k__Fungi(100);p__Ascomycota(100);c__Saccharomycetes(100);o__Saccharomycetales(100);f__Metschnikowiaceae(100);g__Metschnikowia(100);s__Metschnikowia_pulcherrima(100); | Metschnikowia | 13 | Saprotroph | \|Undefined Saprotroph\| | Yeast | NULL | Probable | As nectar-tap_saprotroph (Põlme et al. 2020); Decay sugar-rich substrates (Põlme et al. 2020) | Põlme S, et al. 2020. Fungal Diversity 105:1-16 ((https://doi.org/10.1007/s13225-020-00466-2)) |
| Otu59538 | 1 | k__Fungi(100);p__Ascomycota(100);c__Saccharomycetes(100);o__Saccharomycetales(100);f__Metschnikowiaceae(100);g__Metschnikowia(100);s__Metschnikowia_pulcherrima(100); | Metschnikowia | 13 | Saprotroph | \|Undefined Saprotroph\| | Yeast | NULL | Probable | As nectar-tap_saprotroph (Põlme et al. 2020); Decay sugar-rich substrates (Põlme et al. 2020) | Põlme S, et al. 2020. Fungal Diversity 105:1-16 ((https://doi.org/10.1007/s13225-020-00466-2)) |
| Otu59539 | 1 | k__Fungi(100);p__Ascomycota(100);c__Saccharomycetes(100);o__Saccharomycetales(100);f__Metschnikowiaceae(100);g__Metschnikowia(100);g__Metschnikowia_unclassified(100); | Metschnikowia | 13 | Saprotroph | \|Undefined Saprotroph\| | Yeast | NULL | Probable | As nectar-tap_saprotroph (Põlme et al. 2020); Decay sugar-rich substrates (Põlme et al. 2020) | Põlme S, et al. 2020. Fungal Diversity 105:1-16 ((https://doi.org/10.1007/s13225-020-00466-2)) |
| Otu59540 | 1 | k__Fungi(100);p__Ascomycota(100);c__Saccharomycetes(100);o__Saccharomycetales(100);f__Metschnikowiaceae(100);g__Metschnikowia(100);s__Metschnikowia_pulcherrima(100); | Metschnikowia | 13 | Saprotroph | \|Undefined Saprotroph\| | Yeast | NULL | Probable | As nectar-tap_saprotroph (Põlme et al. 2020); Decay sugar-rich substrates (Põlme et al. 2020) | Põlme S, et al. 2020. Fungal Diversity 105:1-16 ((https://doi.org/10.1007/s13225-020-00466-2)) |
| Otu59542 | 1 | k__Fungi(100);p__Ascomycota(100);c__Saccharomycetes(100);o__Saccharomycetales(100);f__Metschnikowiaceae(100);g__Metschnikowia(100);g__Metschnikowia_unclassified(100); | Metschnikowia | 13 | Saprotroph | \|Undefined Saprotroph\| | Yeast | NULL | Probable | As nectar-tap_saprotroph (Põlme et al. 2020); Decay sugar-rich substrates (Põlme et al. 2020) | Põlme S, et al. 2020. Fungal Diversity 105:1-16 ((https://doi.org/10.1007/s13225-020-00466-2)) |
| Otu59543 | 1 | k__Fungi(100);p__Ascomycota(100);c__Saccharomycetes(100);o__Saccharomycetales(100);f__Metschnikowiaceae(100);g__Metschnikowia(100);s__Metschnikowia_pulcherrima(100); | Metschnikowia | 13 | Saprotroph | \|Undefined Saprotroph\| | Yeast | NULL | Probable | As nectar-tap_saprotroph (Põlme et al. 2020); Decay sugar-rich substrates (Põlme et al. 2020) | Põlme S, et al. 2020. Fungal Diversity 105:1-16 ((https://doi.org/10.1007/s13225-020-00466-2)) |
| Otu59545 | 1 | k__Fungi(100);p__Ascomycota(100);c__Saccharomycetes(100);o__Saccharomycetales(100);f__Metschnikowiaceae(100);g__Metschnikowia(100);s__Metschnikowia_pulcherrima(100); | Metschnikowia | 13 | Saprotroph | \|Undefined Saprotroph\| | Yeast | NULL | Probable | As nectar-tap_saprotroph (Põlme et al. 2020); Decay sugar-rich substrates (Põlme et al. 2020) | Põlme S, et al. 2020. Fungal Diversity 105:1-16 ((https://doi.org/10.1007/s13225-020-00466-2)) |
| Otu59546 | 1 | k__Fungi(100);p__Ascomycota(100);c__Saccharomycetes(100);o__Saccharomycetales(100);f__Metschnikowiaceae(100);g__Metschnikowia(100);s__Metschnikowia_pulcherrima(100); | Metschnikowia | 13 | Saprotroph | \|Undefined Saprotroph\| | Yeast | NULL | Probable | As nectar-tap_saprotroph (Põlme et al. 2020); Decay sugar-rich substrates (Põlme et al. 2020) | Põlme S, et al. 2020. Fungal Diversity 105:1-16 ((https://doi.org/10.1007/s13225-020-00466-2)) |
| Otu59547 | 1 | k__Fungi(100);p__Ascomycota(100);c__Saccharomycetes(100);o__Saccharomycetales(100);f__Metschnikowiaceae(100);g__Metschnikowia(100);s__Metschnikowia_pulcherrima(100); | Metschnikowia | 13 | Saprotroph | \|Undefined Saprotroph\| | Yeast | NULL | Probable | As nectar-tap_saprotroph (Põlme et al. 2020); Decay sugar-rich substrates (Põlme et al. 2020) | Põlme S, et al. 2020. Fungal Diversity 105:1-16 ((https://doi.org/10.1007/s13225-020-00466-2)) |
| Otu59548 | 1 | k__Fungi(100);p__Ascomycota(100);c__Saccharomycetes(100);o__Saccharomycetales(100);f__Metschnikowiaceae(100);g__Metschnikowia(100);s__Metschnikowia_pulcherrima(100); | Metschnikowia | 13 | Saprotroph | \|Undefined Saprotroph\| | Yeast | NULL | Probable | As nectar-tap_saprotroph (Põlme et al. 2020); Decay sugar-rich substrates (Põlme et al. 2020) | Põlme S, et al. 2020. Fungal Diversity 105:1-16 ((https://doi.org/10.1007/s13225-020-00466-2)) |
| Otu59549 | 1 | k__Fungi(100);p__Ascomycota(100);c__Saccharomycetes(100);o__Saccharomycetales(100);f__Metschnikowiaceae(100);g__Metschnikowia(100);s__Metschnikowia_pulcherrima(100); | Metschnikowia | 13 | Saprotroph | \|Undefined Saprotroph\| | Yeast | NULL | Probable | As nectar-tap_saprotroph (Põlme et al. 2020); Decay sugar-rich substrates (Põlme et al. 2020) | Põlme S, et al. 2020. Fungal Diversity 105:1-16 ((https://doi.org/10.1007/s13225-020-00466-2)) |
| Otu59551 | 1 | k__Fungi(100);p__Ascomycota(100);c__Saccharomycetes(100);o__Saccharomycetales(100);f__Metschnikowiaceae(100);g__Metschnikowia(100);s__Metschnikowia_pulcherrima(100); | Metschnikowia | 13 | Saprotroph | \|Undefined Saprotroph\| | Yeast | NULL | Probable | As nectar-tap_saprotroph (Põlme et al. 2020); Decay sugar-rich substrates (Põlme et al. 2020) | Põlme S, et al. 2020. Fungal Diversity 105:1-16 ((https://doi.org/10.1007/s13225-020-00466-2)) |
| Otu59554 | 1 | k__Fungi(100);p__Ascomycota(100);c__Saccharomycetes(100);o__Saccharomycetales(100);f__Metschnikowiaceae(100);g__Metschnikowia(100);s__Metschnikowia_pulcherrima(100); | Metschnikowia | 13 | Saprotroph | \|Undefined Saprotroph\| | Yeast | NULL | Probable | As nectar-tap_saprotroph (Põlme et al. 2020); Decay sugar-rich substrates (Põlme et al. 2020) | Põlme S, et al. 2020. Fungal Diversity 105:1-16 ((https://doi.org/10.1007/s13225-020-00466-2)) |
| Otu59556 | 1 | k__Fungi(100);p__Ascomycota(100);c__Saccharomycetes(100);o__Saccharomycetales(100);f__Metschnikowiaceae(100);g__Metschnikowia(100);s__Metschnikowia_pulcherrima(100); | Metschnikowia | 13 | Saprotroph | \|Undefined Saprotroph\| | Yeast | NULL | Probable | As nectar-tap_saprotroph (Põlme et al. 2020); Decay sugar-rich substrates (Põlme et al. 2020) | Põlme S, et al. 2020. Fungal Diversity 105:1-16 ((https://doi.org/10.1007/s13225-020-00466-2)) |
| Otu59560 | 1 | k__Fungi(100);p__Ascomycota(100);c__Saccharomycetes(100);o__Saccharomycetales(100);f__Metschnikowiaceae(100);g__Metschnikowia(100);s__Metschnikowia_pulcherrima(100); | Metschnikowia | 13 | Saprotroph | \|Undefined Saprotroph\| | Yeast | NULL | Probable | As nectar-tap_saprotroph (Põlme et al. 2020); Decay sugar-rich substrates (Põlme et al. 2020) | Põlme S, et al. 2020. Fungal Diversity 105:1-16 ((https://doi.org/10.1007/s13225-020-00466-2)) |
| Otu59561 | 1 | k__Fungi(100);p__Ascomycota(100);c__Saccharomycetes(100);o__Saccharomycetales(100);f__Metschnikowiaceae(100);g__Metschnikowia(100);s__Metschnikowia_pulcherrima(100); | Metschnikowia | 13 | Saprotroph | \|Undefined Saprotroph\| | Yeast | NULL | Probable | As nectar-tap_saprotroph (Põlme et al. 2020); Decay sugar-rich substrates (Põlme et al. 2020) | Põlme S, et al. 2020. Fungal Diversity 105:1-16 ((https://doi.org/10.1007/s13225-020-00466-2)) |
| Otu59562 | 1 | k__Fungi(100);p__Ascomycota(100);c__Saccharomycetes(100);o__Saccharomycetales(100);f__Metschnikowiaceae(100);g__Metschnikowia(100);s__Metschnikowia_pulcherrima(100); | Metschnikowia | 13 | Saprotroph | \|Undefined Saprotroph\| | Yeast | NULL | Probable | As nectar-tap_saprotroph (Põlme et al. 2020); Decay sugar-rich substrates (Põlme et al. 2020) | Põlme S, et al. 2020. Fungal Diversity 105:1-16 ((https://doi.org/10.1007/s13225-020-00466-2)) |
| Otu59563 | 1 | k__Fungi(100);p__Ascomycota(100);c__Saccharomycetes(100);o__Saccharomycetales(100);f__Metschnikowiaceae(100);g__Metschnikowia(100);s__Metschnikowia_pulcherrima(100); | Metschnikowia | 13 | Saprotroph | \|Undefined Saprotroph\| | Yeast | NULL | Probable | As nectar-tap_saprotroph (Põlme et al. 2020); Decay sugar-rich substrates (Põlme et al. 2020) | Põlme S, et al. 2020. Fungal Diversity 105:1-16 ((https://doi.org/10.1007/s13225-020-00466-2)) |
| Otu59564 | 1 | k__Fungi(100);p__Ascomycota(100);c__Saccharomycetes(100);o__Saccharomycetales(100);f__Metschnikowiaceae(100);g__Metschnikowia(100);s__Metschnikowia_pulcherrima(100); | Metschnikowia | 13 | Saprotroph | \|Undefined Saprotroph\| | Yeast | NULL | Probable | As nectar-tap_saprotroph (Põlme et al. 2020); Decay sugar-rich substrates (Põlme et al. 2020) | Põlme S, et al. 2020. Fungal Diversity 105:1-16 ((https://doi.org/10.1007/s13225-020-00466-2)) |
| Otu59565 | 1 | k__Fungi(100);p__Ascomycota(100);c__Saccharomycetes(100);o__Saccharomycetales(100);f__Metschnikowiaceae(100);g__Metschnikowia(100);s__Metschnikowia_pulcherrima(100); | Metschnikowia | 13 | Saprotroph | \|Undefined Saprotroph\| | Yeast | NULL | Probable | As nectar-tap_saprotroph (Põlme et al. 2020); Decay sugar-rich substrates (Põlme et al. 2020) | Põlme S, et al. 2020. Fungal Diversity 105:1-16 ((https://doi.org/10.1007/s13225-020-00466-2)) |
| Otu59567 | 1 | k__Fungi(100);p__Ascomycota(100);c__Saccharomycetes(100);o__Saccharomycetales(100);f__Metschnikowiaceae(100);g__Metschnikowia(100);s__Metschnikowia_pulcherrima(100); | Metschnikowia | 13 | Saprotroph | \|Undefined Saprotroph\| | Yeast | NULL | Probable | As nectar-tap_saprotroph (Põlme et al. 2020); Decay sugar-rich substrates (Põlme et al. 2020) | Põlme S, et al. 2020. Fungal Diversity 105:1-16 ((https://doi.org/10.1007/s13225-020-00466-2)) |
| Otu59568 | 1 | k__Fungi(100);p__Ascomycota(100);c__Saccharomycetes(100);o__Saccharomycetales(100);f__Metschnikowiaceae(100);g__Metschnikowia(100);s__Metschnikowia_pulcherrima(100); | Metschnikowia | 13 | Saprotroph | \|Undefined Saprotroph\| | Yeast | NULL | Probable | As nectar-tap_saprotroph (Põlme et al. 2020); Decay sugar-rich substrates (Põlme et al. 2020) | Põlme S, et al. 2020. Fungal Diversity 105:1-16 ((https://doi.org/10.1007/s13225-020-00466-2)) |
| Otu59569 | 1 | k__Fungi(100);p__Ascomycota(100);c__Saccharomycetes(100);o__Saccharomycetales(100);f__Metschnikowiaceae(100);g__Metschnikowia(100);g__Metschnikowia_unclassified(100); | Metschnikowia | 13 | Saprotroph | \|Undefined Saprotroph\| | Yeast | NULL | Probable | As nectar-tap_saprotroph (Põlme et al. 2020); Decay sugar-rich substrates (Põlme et al. 2020) | Põlme S, et al. 2020. Fungal Diversity 105:1-16 ((https://doi.org/10.1007/s13225-020-00466-2)) |
| Otu59570 | 1 | k__Fungi(100);p__Ascomycota(100);c__Saccharomycetes(100);o__Saccharomycetales(100);f__Metschnikowiaceae(100);g__Metschnikowia(100);g__Metschnikowia_unclassified(100); | Metschnikowia | 13 | Saprotroph | \|Undefined Saprotroph\| | Yeast | NULL | Probable | As nectar-tap_saprotroph (Põlme et al. 2020); Decay sugar-rich substrates (Põlme et al. 2020) | Põlme S, et al. 2020. Fungal Diversity 105:1-16 ((https://doi.org/10.1007/s13225-020-00466-2)) |
| Otu59571 | 1 | k__Fungi(100);p__Ascomycota(100);c__Saccharomycetes(100);o__Saccharomycetales(100);f__Metschnikowiaceae(100);g__Metschnikowia(100);s__Metschnikowia_pulcherrima(100); | Metschnikowia | 13 | Saprotroph | \|Undefined Saprotroph\| | Yeast | NULL | Probable | As nectar-tap_saprotroph (Põlme et al. 2020); Decay sugar-rich substrates (Põlme et al. 2020) | Põlme S, et al. 2020. Fungal Diversity 105:1-16 ((https://doi.org/10.1007/s13225-020-00466-2)) |
| Otu59572 | 1 | k__Fungi(100);p__Ascomycota(100);c__Saccharomycetes(100);o__Saccharomycetales(100);f__Metschnikowiaceae(100);g__Metschnikowia(100);s__Metschnikowia_pulcherrima(100); | Metschnikowia | 13 | Saprotroph | \|Undefined Saprotroph\| | Yeast | NULL | Probable | As nectar-tap_saprotroph (Põlme et al. 2020); Decay sugar-rich substrates (Põlme et al. 2020) | Põlme S, et al. 2020. Fungal Diversity 105:1-16 ((https://doi.org/10.1007/s13225-020-00466-2)) |
| Otu59573 | 1 | k__Fungi(100);p__Ascomycota(100);c__Saccharomycetes(100);o__Saccharomycetales(100);f__Metschnikowiaceae(100);g__Metschnikowia(100);s__Metschnikowia_pulcherrima(100); | Metschnikowia | 13 | Saprotroph | \|Undefined Saprotroph\| | Yeast | NULL | Probable | As nectar-tap_saprotroph (Põlme et al. 2020); Decay sugar-rich substrates (Põlme et al. 2020) | Põlme S, et al. 2020. Fungal Diversity 105:1-16 ((https://doi.org/10.1007/s13225-020-00466-2)) |
| Otu59574 | 1 | k__Fungi(100);p__Ascomycota(100);c__Saccharomycetes(100);o__Saccharomycetales(100);f__Metschnikowiaceae(100);g__Metschnikowia(100);s__Metschnikowia_pulcherrima(100); | Metschnikowia | 13 | Saprotroph | \|Undefined Saprotroph\| | Yeast | NULL | Probable | As nectar-tap_saprotroph (Põlme et al. 2020); Decay sugar-rich substrates (Põlme et al. 2020) | Põlme S, et al. 2020. Fungal Diversity 105:1-16 ((https://doi.org/10.1007/s13225-020-00466-2)) |
| Otu59575 | 1 | k__Fungi(100);p__Ascomycota(100);c__Saccharomycetes(100);o__Saccharomycetales(100);f__Metschnikowiaceae(100);g__Metschnikowia(100);s__Metschnikowia_pulcherrima(100); | Metschnikowia | 13 | Saprotroph | \|Undefined Saprotroph\| | Yeast | NULL | Probable | As nectar-tap_saprotroph (Põlme et al. 2020); Decay sugar-rich substrates (Põlme et al. 2020) | Põlme S, et al. 2020. Fungal Diversity 105:1-16 ((https://doi.org/10.1007/s13225-020-00466-2)) |
| Otu59576 | 1 | k__Fungi(100);p__Ascomycota(100);c__Saccharomycetes(100);o__Saccharomycetales(100);f__Metschnikowiaceae(100);g__Metschnikowia(100);s__Metschnikowia_pulcherrima(100); | Metschnikowia | 13 | Saprotroph | \|Undefined Saprotroph\| | Yeast | NULL | Probable | As nectar-tap_saprotroph (Põlme et al. 2020); Decay sugar-rich substrates (Põlme et al. 2020) | Põlme S, et al. 2020. Fungal Diversity 105:1-16 ((https://doi.org/10.1007/s13225-020-00466-2)) |
| Otu59577 | 1 | k__Fungi(100);p__Ascomycota(100);c__Saccharomycetes(100);o__Saccharomycetales(100);f__Metschnikowiaceae(100);g__Metschnikowia(100);s__Metschnikowia_pulcherrima(100); | Metschnikowia | 13 | Saprotroph | \|Undefined Saprotroph\| | Yeast | NULL | Probable | As nectar-tap_saprotroph (Põlme et al. 2020); Decay sugar-rich substrates (Põlme et al. 2020) | Põlme S, et al. 2020. Fungal Diversity 105:1-16 ((https://doi.org/10.1007/s13225-020-00466-2)) |
| Otu59578 | 1 | k__Fungi(100);p__Ascomycota(100);c__Saccharomycetes(100);o__Saccharomycetales(100);f__Metschnikowiaceae(100);g__Metschnikowia(100);s__Metschnikowia_pulcherrima(100); | Metschnikowia | 13 | Saprotroph | \|Undefined Saprotroph\| | Yeast | NULL | Probable | As nectar-tap_saprotroph (Põlme et al. 2020); Decay sugar-rich substrates (Põlme et al. 2020) | Põlme S, et al. 2020. Fungal Diversity 105:1-16 ((https://doi.org/10.1007/s13225-020-00466-2)) |
| Otu59579 | 1 | k__Fungi(100);p__Ascomycota(100);c__Saccharomycetes(100);o__Saccharomycetales(100);f__Metschnikowiaceae(100);g__Metschnikowia(100);g__Metschnikowia_unclassified(100); | Metschnikowia | 13 | Saprotroph | \|Undefined Saprotroph\| | Yeast | NULL | Probable | As nectar-tap_saprotroph (Põlme et al. 2020); Decay sugar-rich substrates (Põlme et al. 2020) | Põlme S, et al. 2020. Fungal Diversity 105:1-16 ((https://doi.org/10.1007/s13225-020-00466-2)) |
| Otu59580 | 1 | k__Fungi(100);p__Ascomycota(100);c__Saccharomycetes(100);o__Saccharomycetales(100);f__Metschnikowiaceae(100);g__Metschnikowia(100);g__Metschnikowia_unclassified(100); | Metschnikowia | 13 | Saprotroph | \|Undefined Saprotroph\| | Yeast | NULL | Probable | As nectar-tap_saprotroph (Põlme et al. 2020); Decay sugar-rich substrates (Põlme et al. 2020) | Põlme S, et al. 2020. Fungal Diversity 105:1-16 ((https://doi.org/10.1007/s13225-020-00466-2)) |
| Otu59581 | 1 | k__Fungi(100);p__Ascomycota(100);c__Saccharomycetes(100);o__Saccharomycetales(100);f__Metschnikowiaceae(100);g__Metschnikowia(100);g__Metschnikowia_unclassified(100); | Metschnikowia | 13 | Saprotroph | \|Undefined Saprotroph\| | Yeast | NULL | Probable | As nectar-tap_saprotroph (Põlme et al. 2020); Decay sugar-rich substrates (Põlme et al. 2020) | Põlme S, et al. 2020. Fungal Diversity 105:1-16 ((https://doi.org/10.1007/s13225-020-00466-2)) |
| Otu59582 | 1 | k__Fungi(100);p__Ascomycota(100);c__Saccharomycetes(100);o__Saccharomycetales(100);f__Metschnikowiaceae(100);g__Metschnikowia(100);s__Metschnikowia_pulcherrima(100); | Metschnikowia | 13 | Saprotroph | \|Undefined Saprotroph\| | Yeast | NULL | Probable | As nectar-tap_saprotroph (Põlme et al. 2020); Decay sugar-rich substrates (Põlme et al. 2020) | Põlme S, et al. 2020. Fungal Diversity 105:1-16 ((https://doi.org/10.1007/s13225-020-00466-2)) |
| Otu59583 | 1 | k__Fungi(100);p__Ascomycota(100);c__Saccharomycetes(100);o__Saccharomycetales(100);f__Metschnikowiaceae(100);g__Metschnikowia(100);g__Metschnikowia_unclassified(100); | Metschnikowia | 13 | Saprotroph | \|Undefined Saprotroph\| | Yeast | NULL | Probable | As nectar-tap_saprotroph (Põlme et al. 2020); Decay sugar-rich substrates (Põlme et al. 2020) | Põlme S, et al. 2020. Fungal Diversity 105:1-16 ((https://doi.org/10.1007/s13225-020-00466-2)) |
| Otu59584 | 1 | k__Fungi(100);p__Ascomycota(100);c__Saccharomycetes(100);o__Saccharomycetales(100);f__Metschnikowiaceae(100);g__Metschnikowia(100);s__Metschnikowia_pulcherrima(100); | Metschnikowia | 13 | Saprotroph | \|Undefined Saprotroph\| | Yeast | NULL | Probable | As nectar-tap_saprotroph (Põlme et al. 2020); Decay sugar-rich substrates (Põlme et al. 2020) | Põlme S, et al. 2020. Fungal Diversity 105:1-16 ((https://doi.org/10.1007/s13225-020-00466-2)) |
| Otu59585 | 1 | k__Fungi(100);p__Ascomycota(100);c__Saccharomycetes(100);o__Saccharomycetales(100);f__Metschnikowiaceae(100);g__Metschnikowia(100);s__Metschnikowia_pulcherrima(100); | Metschnikowia | 13 | Saprotroph | \|Undefined Saprotroph\| | Yeast | NULL | Probable | As nectar-tap_saprotroph (Põlme et al. 2020); Decay sugar-rich substrates (Põlme et al. 2020) | Põlme S, et al. 2020. Fungal Diversity 105:1-16 ((https://doi.org/10.1007/s13225-020-00466-2)) |
| Otu59586 | 1 | k__Fungi(100);p__Ascomycota(100);c__Saccharomycetes(100);o__Saccharomycetales(100);f__Metschnikowiaceae(100);g__Metschnikowia(100);s__Metschnikowia_pulcherrima(100); | Metschnikowia | 13 | Saprotroph | \|Undefined Saprotroph\| | Yeast | NULL | Probable | As nectar-tap_saprotroph (Põlme et al. 2020); Decay sugar-rich substrates (Põlme et al. 2020) | Põlme S, et al. 2020. Fungal Diversity 105:1-16 ((https://doi.org/10.1007/s13225-020-00466-2)) |

| | | | | | | | | | | | |
|---|---|---|---|---|---|---|---|---|---|---|---|
| Otu59587 | 1 | k__Fungi(100);p__Ascomycota(100);c__Saccharomycetes(100);o__Saccharomycetales(100);f__Metschnikowiaceae(100);g__Metschnikowia(100);s__Metschnikowia_pulcherrima(100); | Metschnikowia | 13 | Saprotroph | \|Undefined Saprotroph\| | Yeast | NULL | Probable | As nectar-tap_saprotroph (Põlme et al. 2020); Decay sugar-rich substrates (Põlme et al. 2020) | Põlme S, et al. 2020. Fungal Diversity 105:1-16 ((https://doi.org/10.1007/s13225-020-00466-2)) |
| Otu59588 | 1 | k__Fungi(100);p__Ascomycota(100);c__Saccharomycetes(100);o__Saccharomycetales(100);f__Metschnikowiaceae(100);g__Metschnikowia(100);g__Metschnikowia_unclassified(100); | Metschnikowia | 13 | Saprotroph | \|Undefined Saprotroph\| | Yeast | NULL | Probable | As nectar-tap_saprotroph (Põlme et al. 2020); Decay sugar-rich substrates (Põlme et al. 2020) | Põlme S, et al. 2020. Fungal Diversity 105:1-16 ((https://doi.org/10.1007/s13225-020-00466-2)) |
| Otu59590 | 1 | k__Fungi(100);p__Ascomycota(100);c__Saccharomycetes(100);o__Saccharomycetales(100);f__Metschnikowiaceae(100);g__Metschnikowia(100);s__Metschnikowia_pulcherrima(100); | Metschnikowia | 13 | Saprotroph | \|Undefined Saprotroph\| | Yeast | NULL | Probable | As nectar-tap_saprotroph (Põlme et al. 2020); Decay sugar-rich substrates (Põlme et al. 2020) | Põlme S, et al. 2020. Fungal Diversity 105:1-16 ((https://doi.org/10.1007/s13225-020-00466-2)) |
| Otu59591 | 1 | k__Fungi(100);p__Ascomycota(100);c__Saccharomycetes(100);o__Saccharomycetales(100);f__Metschnikowiaceae(100);g__Metschnikowia(100);s__Metschnikowia_pulcherrima(100); | Metschnikowia | 13 | Saprotroph | \|Undefined Saprotroph\| | Yeast | NULL | Probable | As nectar-tap_saprotroph (Põlme et al. 2020); Decay sugar-rich substrates (Põlme et al. 2020) | Põlme S, et al. 2020. Fungal Diversity 105:1-16 ((https://doi.org/10.1007/s13225-020-00466-2)) |
| Otu59592 | 1 | k__Fungi(100);p__Ascomycota(100);c__Saccharomycetes(100);o__Saccharomycetales(100);f__Metschnikowiaceae(100);g__Metschnikowia(100);s__Metschnikowia_pulcherrima(100); | Metschnikowia | 13 | Saprotroph | \|Undefined Saprotroph\| | Yeast | NULL | Probable | As nectar-tap_saprotroph (Põlme et al. 2020); Decay sugar-rich substrates (Põlme et al. 2020) | Põlme S, et al. 2020. Fungal Diversity 105:1-16 ((https://doi.org/10.1007/s13225-020-00466-2)) |
| Otu59593 | 1 | k__Fungi(100);p__Ascomycota(100);c__Saccharomycetes(100);o__Saccharomycetales(100);f__Metschnikowiaceae(100);g__Metschnikowia(100);g__Metschnikowia_unclassified(100); | Metschnikowia | 13 | Saprotroph | \|Undefined Saprotroph\| | Yeast | NULL | Probable | As nectar-tap_saprotroph (Põlme et al. 2020); Decay sugar-rich substrates (Põlme et al. 2020) | Põlme S, et al. 2020. Fungal Diversity 105:1-16 ((https://doi.org/10.1007/s13225-020-00466-2)) |
| Otu59594 | 1 | k__Fungi(100);p__Ascomycota(100);c__Saccharomycetes(100);o__Saccharomycetales(100);f__Metschnikowiaceae(100);g__Metschnikowia(100);g__Metschnikowia_unclassified(100); | Metschnikowia | 13 | Saprotroph | \|Undefined Saprotroph\| | Yeast | NULL | Probable | As nectar-tap_saprotroph (Põlme et al. 2020); Decay sugar-rich substrates (Põlme et al. 2020) | Põlme S, et al. 2020. Fungal Diversity 105:1-16 ((https://doi.org/10.1007/s13225-020-00466-2)) |
| Otu59596 | 1 | k__Fungi(100);p__Ascomycota(100);c__Saccharomycetes(100);o__Saccharomycetales(100);f__Metschnikowiaceae(100);g__Metschnikowia(100);s__Metschnikowia_pulcherrima(100); | Metschnikowia | 13 | Saprotroph | \|Undefined Saprotroph\| | Yeast | NULL | Probable | As nectar-tap_saprotroph (Põlme et al. 2020); Decay sugar-rich substrates (Põlme et al. 2020) | Põlme S, et al. 2020. Fungal Diversity 105:1-16 ((https://doi.org/10.1007/s13225-020-00466-2)) |
| Otu59597 | 1 | k__Fungi(100);p__Ascomycota(100);c__Saccharomycetes(100);o__Saccharomycetales(100);f__Metschnikowiaceae(100);g__Metschnikowia(100);g__Metschnikowia_unclassified(100); | Metschnikowia | 13 | Saprotroph | \|Undefined Saprotroph\| | Yeast | NULL | Probable | As nectar-tap_saprotroph (Põlme et al. 2020); Decay sugar-rich substrates (Põlme et al. 2020) | Põlme S, et al. 2020. Fungal Diversity 105:1-16 ((https://doi.org/10.1007/s13225-020-00466-2)) |
| Otu59598 | 1 | k__Fungi(100);p__Ascomycota(100);c__Saccharomycetes(100);o__Saccharomycetales(100);f__Metschnikowiaceae(100);g__Metschnikowia(100);s__Metschnikowia_pulcherrima(100); | Metschnikowia | 13 | Saprotroph | \|Undefined Saprotroph\| | Yeast | NULL | Probable | As nectar-tap_saprotroph (Põlme et al. 2020); Decay sugar-rich substrates (Põlme et al. 2020) | Põlme S, et al. 2020. Fungal Diversity 105:1-16 ((https://doi.org/10.1007/s13225-020-00466-2)) |
| Otu59599 | 1 | k__Fungi(100);p__Ascomycota(100);c__Saccharomycetes(100);o__Saccharomycetales(100);f__Metschnikowiaceae(100);g__Metschnikowia(100);s__Metschnikowia_pulcherrima(100); | Metschnikowia | 13 | Saprotroph | \|Undefined Saprotroph\| | Yeast | NULL | Probable | As nectar-tap_saprotroph (Põlme et al. 2020); Decay sugar-rich substrates (Põlme et al. 2020) | Põlme S, et al. 2020. Fungal Diversity 105:1-16 ((https://doi.org/10.1007/s13225-020-00466-2)) |
| Otu59600 | 1 | k__Fungi(100);p__Ascomycota(100);c__Saccharomycetes(100);o__Saccharomycetales(100);f__Metschnikowiaceae(100);g__Metschnikowia(100);s__Metschnikowia_pulcherrima(100); | Metschnikowia | 13 | Saprotroph | \|Undefined Saprotroph\| | Yeast | NULL | Probable | As nectar-tap_saprotroph (Põlme et al. 2020); Decay sugar-rich substrates (Põlme et al. 2020) | Põlme S, et al. 2020. Fungal Diversity 105:1-16 ((https://doi.org/10.1007/s13225-020-00466-2)) |
| Otu59601 | 1 | k__Fungi(100);p__Ascomycota(100);c__Saccharomycetes(100);o__Saccharomycetales(100);f__Metschnikowiaceae(100);g__Metschnikowia(100);s__Metschnikowia_pulcherrima(100); | Metschnikowia | 13 | Saprotroph | \|Undefined Saprotroph\| | Yeast | NULL | Probable | As nectar-tap_saprotroph (Põlme et al. 2020); Decay sugar-rich substrates (Põlme et al. 2020) | Põlme S, et al. 2020. Fungal Diversity 105:1-16 ((https://doi.org/10.1007/s13225-020-00466-2)) |
| Otu59602 | 1 | k__Fungi(100);p__Ascomycota(100);c__Saccharomycetes(100);o__Saccharomycetales(100);f__Metschnikowiaceae(100);g__Metschnikowia(100);s__Metschnikowia_pulcherrima(100); | Metschnikowia | 13 | Saprotroph | \|Undefined Saprotroph\| | Yeast | NULL | Probable | As nectar-tap_saprotroph (Põlme et al. 2020); Decay sugar-rich substrates (Põlme et al. 2020) | Põlme S, et al. 2020. Fungal Diversity 105:1-16 ((https://doi.org/10.1007/s13225-020-00466-2)) |
| Otu59603 | 1 | k__Fungi(100);p__Ascomycota(100);c__Saccharomycetes(100);o__Saccharomycetales(100);f__Metschnikowiaceae(100);g__Metschnikowia(100);g__Metschnikowia_unclassified(100); | Metschnikowia | 13 | Saprotroph | \|Undefined Saprotroph\| | Yeast | NULL | Probable | As nectar-tap_saprotroph (Põlme et al. 2020); Decay sugar-rich substrates (Põlme et al. 2020) | Põlme S, et al. 2020. Fungal Diversity 105:1-16 ((https://doi.org/10.1007/s13225-020-00466-2)) |
| Otu59604 | 1 | k__Fungi(100);p__Ascomycota(100);c__Saccharomycetes(100);o__Saccharomycetales(100);f__Metschnikowiaceae(100);g__Metschnikowia(100);s__Metschnikowia_pulcherrima(100); | Metschnikowia | 13 | Saprotroph | \|Undefined Saprotroph\| | Yeast | NULL | Probable | As nectar-tap_saprotroph (Põlme et al. 2020); Decay sugar-rich substrates (Põlme et al. 2020) | Põlme S, et al. 2020. Fungal Diversity 105:1-16 ((https://doi.org/10.1007/s13225-020-00466-2)) |
| Otu59605 | 1 | k__Fungi(100);p__Ascomycota(100);c__Saccharomycetes(100);o__Saccharomycetales(100);f__Metschnikowiaceae(100);g__Metschnikowia(100);g__Metschnikowia_unclassified(100); | Metschnikowia | 13 | Saprotroph | \|Undefined Saprotroph\| | Yeast | NULL | Probable | As nectar-tap_saprotroph (Põlme et al. 2020); Decay sugar-rich substrates (Põlme et al. 2020) | Põlme S, et al. 2020. Fungal Diversity 105:1-16 ((https://doi.org/10.1007/s13225-020-00466-2)) |
| Otu59606 | 1 | k__Fungi(100);p__Ascomycota(100);c__Saccharomycetes(100);o__Saccharomycetales(100);f__Metschnikowiaceae(100);g__Metschnikowia(100);s__Metschnikowia_pulcherrima(100); | Metschnikowia | 13 | Saprotroph | \|Undefined Saprotroph\| | Yeast | NULL | Probable | As nectar-tap_saprotroph (Põlme et al. 2020); Decay sugar-rich substrates (Põlme et al. 2020) | Põlme S, et al. 2020. Fungal Diversity 105:1-16 ((https://doi.org/10.1007/s13225-020-00466-2)) |
| Otu59607 | 1 | k__Fungi(100);p__Ascomycota(100);c__Saccharomycetes(100);o__Saccharomycetales(100);f__Metschnikowiaceae(100);g__Metschnikowia(100);s__Metschnikowia_pulcherrima(100); | Metschnikowia | 13 | Saprotroph | \|Undefined Saprotroph\| | Yeast | NULL | Probable | As nectar-tap_saprotroph (Põlme et al. 2020); Decay sugar-rich substrates (Põlme et al. 2020) | Põlme S, et al. 2020. Fungal Diversity 105:1-16 ((https://doi.org/10.1007/s13225-020-00466-2)) |
| Otu59608 | 1 | k__Fungi(100);p__Ascomycota(100);c__Saccharomycetes(100);o__Saccharomycetales(100);f__Metschnikowiaceae(100);g__Metschnikowia(100);s__Metschnikowia_pulcherrima(100); | Metschnikowia | 13 | Saprotroph | \|Undefined Saprotroph\| | Yeast | NULL | Probable | As nectar-tap_saprotroph (Põlme et al. 2020); Decay sugar-rich substrates (Põlme et al. 2020) | Põlme S, et al. 2020. Fungal Diversity 105:1-16 ((https://doi.org/10.1007/s13225-020-00466-2)) |
| Otu59609 | 1 | k__Fungi(100);p__Ascomycota(100);c__Saccharomycetes(100);o__Saccharomycetales(100);f__Metschnikowiaceae(100);g__Metschnikowia(100);s__Metschnikowia_pulcherrima(100); | Metschnikowia | 13 | Saprotroph | \|Undefined Saprotroph\| | Yeast | NULL | Probable | As nectar-tap_saprotroph (Põlme et al. 2020); Decay sugar-rich substrates (Põlme et al. 2020) | Põlme S, et al. 2020. Fungal Diversity 105:1-16 ((https://doi.org/10.1007/s13225-020-00466-2)) |
| Otu59610 | 1 | k__Fungi(100);p__Ascomycota(100);c__Saccharomycetes(100);o__Saccharomycetales(100);f__Metschnikowiaceae(100);g__Metschnikowia(100);s__Metschnikowia_pulcherrima(100); | Metschnikowia | 13 | Saprotroph | \|Undefined Saprotroph\| | Yeast | NULL | Probable | As nectar-tap_saprotroph (Põlme et al. 2020); Decay sugar-rich substrates (Põlme et al. 2020) | Põlme S, et al. 2020. Fungal Diversity 105:1-16 ((https://doi.org/10.1007/s13225-020-00466-2)) |
| Otu59611 | 1 | k__Fungi(100);p__Ascomycota(100);c__Saccharomycetes(100);o__Saccharomycetales(100);f__Metschnikowiaceae(100);g__Metschnikowia(100);s__Metschnikowia_pulcherrima(100); | Metschnikowia | 13 | Saprotroph | \|Undefined Saprotroph\| | Yeast | NULL | Probable | As nectar-tap_saprotroph (Põlme et al. 2020); Decay sugar-rich substrates (Põlme et al. 2020) | Põlme S, et al. 2020. Fungal Diversity 105:1-16 ((https://doi.org/10.1007/s13225-020-00466-2)) |
| Otu59612 | 1 | k__Fungi(100);p__Ascomycota(100);c__Saccharomycetes(100);o__Saccharomycetales(100);f__Metschnikowiaceae(100);g__Metschnikowia(100);s__Metschnikowia_pulcherrima(100); | Metschnikowia | 13 | Saprotroph | \|Undefined Saprotroph\| | Yeast | NULL | Probable | As nectar-tap_saprotroph (Põlme et al. 2020); Decay sugar-rich substrates (Põlme et al. 2020) | Põlme S, et al. 2020. Fungal Diversity 105:1-16 ((https://doi.org/10.1007/s13225-020-00466-2)) |
| Otu59614 | 1 | k__Fungi(100);p__Ascomycota(100);c__Saccharomycetes(100);o__Saccharomycetales(100);f__Metschnikowiaceae(100);g__Metschnikowia(100);s__Metschnikowia_pulcherrima(100); | Metschnikowia | 13 | Saprotroph | \|Undefined Saprotroph\| | Yeast | NULL | Probable | As nectar-tap_saprotroph (Põlme et al. 2020); Decay sugar-rich substrates (Põlme et al. 2020) | Põlme S, et al. 2020. Fungal Diversity 105:1-16 ((https://doi.org/10.1007/s13225-020-00466-2)) |
| Otu59615 | 1 | k__Fungi(100);p__Ascomycota(100);c__Saccharomycetes(100);o__Saccharomycetales(100);f__Metschnikowiaceae(100);g__Metschnikowia(100);g__Metschnikowia_unclassified(100); | Metschnikowia | 13 | Saprotroph | \|Undefined Saprotroph\| | Yeast | NULL | Probable | As nectar-tap_saprotroph (Põlme et al. 2020); Decay sugar-rich substrates (Põlme et al. 2020) | Põlme S, et al. 2020. Fungal Diversity 105:1-16 ((https://doi.org/10.1007/s13225-020-00466-2)) |
| Otu59616 | 1 | k__Fungi(100);p__Ascomycota(100);c__Saccharomycetes(100);o__Saccharomycetales(100);f__Metschnikowiaceae(100);g__Metschnikowia(100);s__Metschnikowia_pulcherrima(100); | Metschnikowia | 13 | Saprotroph | \|Undefined Saprotroph\| | Yeast | NULL | Probable | As nectar-tap_saprotroph (Põlme et al. 2020); Decay sugar-rich substrates (Põlme et al. 2020) | Põlme S, et al. 2020. Fungal Diversity 105:1-16 ((https://doi.org/10.1007/s13225-020-00466-2)) |
| Otu59617 | 1 | k__Fungi(100);p__Ascomycota(100);c__Saccharomycetes(100);o__Saccharomycetales(100);f__Metschnikowiaceae(100);g__Metschnikowia(100);s__Metschnikowia_pulcherrima(100); | Metschnikowia | 13 | Saprotroph | \|Undefined Saprotroph\| | Yeast | NULL | Probable | As nectar-tap_saprotroph (Põlme et al. 2020); Decay sugar-rich substrates (Põlme et al. 2020) | Põlme S, et al. 2020. Fungal Diversity 105:1-16 ((https://doi.org/10.1007/s13225-020-00466-2)) |
| Otu59618 | 1 | k__Fungi(100);p__Ascomycota(100);c__Saccharomycetes(100);o__Saccharomycetales(100);f__Metschnikowiaceae(100);g__Metschnikowia(100);g__Metschnikowia_unclassified(100); | Metschnikowia | 13 | Saprotroph | \|Undefined Saprotroph\| | Yeast | NULL | Probable | As nectar-tap_saprotroph (Põlme et al. 2020); Decay sugar-rich substrates (Põlme et al. 2020) | Põlme S, et al. 2020. Fungal Diversity 105:1-16 ((https://doi.org/10.1007/s13225-020-00466-2)) |
| Otu59619 | 1 | k__Fungi(100);p__Ascomycota(100);c__Saccharomycetes(100);o__Saccharomycetales(100);f__Metschnikowiaceae(100);g__Metschnikowia(100);g__Metschnikowia_unclassified(100); | Metschnikowia | 13 | Saprotroph | \|Undefined Saprotroph\| | Yeast | NULL | Probable | As nectar-tap_saprotroph (Põlme et al. 2020); Decay sugar-rich substrates (Põlme et al. 2020) | Põlme S, et al. 2020. Fungal Diversity 105:1-16 ((https://doi.org/10.1007/s13225-020-00466-2)) |
| Otu59620 | 1 | k__Fungi(100);p__Ascomycota(100);c__Saccharomycetes(100);o__Saccharomycetales(100);f__Metschnikowiaceae(100);g__Metschnikowia(100);s__Metschnikowia_pulcherrima(100); | Metschnikowia | 13 | Saprotroph | \|Undefined Saprotroph\| | Yeast | NULL | Probable | As nectar-tap_saprotroph (Põlme et al. 2020); Decay sugar-rich substrates (Põlme et al. 2020) | Põlme S, et al. 2020. Fungal Diversity 105:1-16 ((https://doi.org/10.1007/s13225-020-00466-2)) |
| Otu59621 | 1 | k__Fungi(100);p__Ascomycota(100);c__Saccharomycetes(100);o__Saccharomycetales(100);f__Metschnikowiaceae(100);g__Metschnikowia(100);s__Metschnikowia_pulcherrima(100); | Metschnikowia | 13 | Saprotroph | \|Undefined Saprotroph\| | Yeast | NULL | Probable | As nectar-tap_saprotroph (Põlme et al. 2020); Decay sugar-rich substrates (Põlme et al. 2020) | Põlme S, et al. 2020. Fungal Diversity 105:1-16 ((https://doi.org/10.1007/s13225-020-00466-2)) |
| Otu59622 | 1 | k__Fungi(100);p__Ascomycota(100);c__Saccharomycetes(100);o__Saccharomycetales(100);f__Metschnikowiaceae(100);g__Metschnikowia(100);s__Metschnikowia_pulcherrima(100); | Metschnikowia | 13 | Saprotroph | \|Undefined Saprotroph\| | Yeast | NULL | Probable | As nectar-tap_saprotroph (Põlme et al. 2020); Decay sugar-rich substrates (Põlme et al. 2020) | Põlme S, et al. 2020. Fungal Diversity 105:1-16 ((https://doi.org/10.1007/s13225-020-00466-2)) |
| Otu59623 | 1 | k__Fungi(100);p__Ascomycota(100);c__Saccharomycetes(100);o__Saccharomycetales(100);f__Metschnikowiaceae(100);g__Metschnikowia(100);s__Metschnikowia_pulcherrima(100); | Metschnikowia | 13 | Saprotroph | \|Undefined Saprotroph\| | Yeast | NULL | Probable | As nectar-tap_saprotroph (Põlme et al. 2020); Decay sugar-rich substrates (Põlme et al. 2020) | Põlme S, et al. 2020. Fungal Diversity 105:1-16 ((https://doi.org/10.1007/s13225-020-00466-2)) |
| Otu59624 | 1 | k__Fungi(100);p__Ascomycota(100);c__Saccharomycetes(100);o__Saccharomycetales(100);f__Metschnikowiaceae(100);g__Metschnikowia(100);s__Metschnikowia_pulcherrima(100); | Metschnikowia | 13 | Saprotroph | \|Undefined Saprotroph\| | Yeast | NULL | Probable | As nectar-tap_saprotroph (Põlme et al. 2020); Decay sugar-rich substrates (Põlme et al. 2020) | Põlme S, et al. 2020. Fungal Diversity 105:1-16 ((https://doi.org/10.1007/s13225-020-00466-2)) |
| Otu59625 | 1 | k__Fungi(100);p__Ascomycota(100);c__Saccharomycetes(100);o__Saccharomycetales(100);f__Metschnikowiaceae(100);g__Metschnikowia(100);s__Metschnikowia_pulcherrima(100); | Metschnikowia | 13 | Saprotroph | \|Undefined Saprotroph\| | Yeast | NULL | Probable | As nectar-tap_saprotroph (Põlme et al. 2020); Decay sugar-rich substrates (Põlme et al. 2020) | Põlme S, et al. 2020. Fungal Diversity 105:1-16 ((https://doi.org/10.1007/s13225-020-00466-2)) |
| Otu59626 | 1 | k__Fungi(100);p__Ascomycota(100);c__Saccharomycetes(100);o__Saccharomycetales(100);f__Metschnikowiaceae(100);g__Metschnikowia(100);g__Metschnikowia_unclassified(100); | Metschnikowia | 13 | Saprotroph | \|Undefined Saprotroph\| | Yeast | NULL | Probable | As nectar-tap_saprotroph (Põlme et al. 2020); Decay sugar-rich substrates (Põlme et al. 2020) | Põlme S, et al. 2020. Fungal Diversity 105:1-16 ((https://doi.org/10.1007/s13225-020-00466-2)) |
| Otu59627 | 1 | k__Fungi(100);p__Ascomycota(100);c__Saccharomycetes(100);o__Saccharomycetales(100);f__Metschnikowiaceae(100);g__Metschnikowia(100);g__Metschnikowia_unclassified(100); | Metschnikowia | 13 | Saprotroph | \|Undefined Saprotroph\| | Yeast | NULL | Probable | As nectar-tap_saprotroph (Põlme et al. 2020); Decay sugar-rich substrates (Põlme et al. 2020) | Põlme S, et al. 2020. Fungal Diversity 105:1-16 ((https://doi.org/10.1007/s13225-020-00466-2)) |
| Otu59628 | 1 | k__Fungi(100);p__Ascomycota(100);c__Saccharomycetes(100);o__Saccharomycetales(100);f__Metschnikowiaceae(100);g__Metschnikowia(100);s__Metschnikowia_pulcherrima(100); | Metschnikowia | 13 | Saprotroph | \|Undefined Saprotroph\| | Yeast | NULL | Probable | As nectar-tap_saprotroph (Põlme et al. 2020); Decay sugar-rich substrates (Põlme et al. 2020) | Põlme S, et al. 2020. Fungal Diversity 105:1-16 ((https://doi.org/10.1007/s13225-020-00466-2)) |
| Otu59629 | 1 | k__Fungi(100);p__Ascomycota(100);c__Saccharomycetes(100);o__Saccharomycetales(100);f__Metschnikowiaceae(100);g__Metschnikowia(100);s__Metschnikowia_pulcherrima(100); | Metschnikowia | 13 | Saprotroph | \|Undefined Saprotroph\| | Yeast | NULL | Probable | As nectar-tap_saprotroph (Põlme et al. 2020); Decay sugar-rich substrates (Põlme et al. 2020) | Põlme S, et al. 2020. Fungal Diversity 105:1-16 ((https://doi.org/10.1007/s13225-020-00466-2)) |
| Otu59630 | 1 | k__Fungi(100);p__Ascomycota(100);c__Saccharomycetes(100);o__Saccharomycetales(100);f__Metschnikowiaceae(100);g__Metschnikowia(100);g__Metschnikowia_unclassified(100); | Metschnikowia | 13 | Saprotroph | \|Undefined Saprotroph\| | Yeast | NULL | Probable | As nectar-tap_saprotroph (Põlme et al. 2020); Decay sugar-rich substrates (Põlme et al. 2020) | Põlme S, et al. 2020. Fungal Diversity 105:1-16 ((https://doi.org/10.1007/s13225-020-00466-2)) |
| Otu59631 | 1 | k__Fungi(100);p__Ascomycota(100);c__Saccharomycetes(100);o__Saccharomycetales(100);f__Metschnikowiaceae(100);g__Metschnikowia(100);g__Metschnikowia_unclassified(100); | Metschnikowia | 13 | Saprotroph | \|Undefined Saprotroph\| | Yeast | NULL | Probable | As nectar-tap_saprotroph (Põlme et al. 2020); Decay sugar-rich substrates (Põlme et al. 2020) | Põlme S, et al. 2020. Fungal Diversity 105:1-16 ((https://doi.org/10.1007/s13225-020-00466-2)) |
| Otu59632 | 1 | k__Fungi(100);p__Ascomycota(100);c__Saccharomycetes(100);o__Saccharomycetales(100);f__Metschnikowiaceae(100);g__Metschnikowia(100);g__Metschnikowia_unclassified(100); | Metschnikowia | 13 | Saprotroph | \|Undefined Saprotroph\| | Yeast | NULL | Probable | As nectar-tap_saprotroph (Põlme et al. 2020); Decay sugar-rich substrates (Põlme et al. 2020) | Põlme S, et al. 2020. Fungal Diversity 105:1-16 ((https://doi.org/10.1007/s13225-020-00466-2)) |
| Otu59633 | 1 | k__Fungi(100);p__Ascomycota(100);c__Saccharomycetes(100);o__Saccharomycetales(100);f__Metschnikowiaceae(100);g__Metschnikowia(100);s__Metschnikowia_pulcherrima(100); | Metschnikowia | 13 | Saprotroph | \|Undefined Saprotroph\| | Yeast | NULL | Probable | As nectar-tap_saprotroph (Põlme et al. 2020); Decay sugar-rich substrates (Põlme et al. 2020) | Põlme S, et al. 2020. Fungal Diversity 105:1-16 ((https://doi.org/10.1007/s13225-020-00466-2)) |
| Otu59634 | 1 | k__Fungi(100);p__Ascomycota(100);c__Saccharomycetes(100);o__Saccharomycetales(100);f__Metschnikowiaceae(100);g__Metschnikowia(100);s__Metschnikowia_pulcherrima(100); | Metschnikowia | 13 | Saprotroph | \|Undefined Saprotroph\| | Yeast | NULL | Probable | As nectar-tap_saprotroph (Põlme et al. 2020); Decay sugar-rich substrates (Põlme et al. 2020) | Põlme S, et al. 2020. Fungal Diversity 105:1-16 ((https://doi.org/10.1007/s13225-020-00466-2)) |
| Otu59635 | 1 | k__Fungi(100);p__Ascomycota(100);c__Saccharomycetes(100);o__Saccharomycetales(100);f__Metschnikowiaceae(100);g__Metschnikowia(100);s__Metschnikowia_pulcherrima(100); | Metschnikowia | 13 | Saprotroph | \|Undefined Saprotroph\| | Yeast | NULL | Probable | As nectar-tap_saprotroph (Põlme et al. 2020); Decay sugar-rich substrates (Põlme et al. 2020) | Põlme S, et al. 2020. Fungal Diversity 105:1-16 ((https://doi.org/10.1007/s13225-020-00466-2)) |
| Otu59636 | 1 | k__Fungi(100);p__Ascomycota(100);c__Saccharomycetes(100);o__Saccharomycetales(100);f__Metschnikowiaceae(100);g__Metschnikowia(100);s__Metschnikowia_pulcherrima(100); | Metschnikowia | 13 | Saprotroph | \|Undefined Saprotroph\| | Yeast | NULL | Probable | As nectar-tap_saprotroph (Põlme et al. 2020); Decay sugar-rich substrates (Põlme et al. 2020) | Põlme S, et al. 2020. Fungal Diversity 105:1-16 ((https://doi.org/10.1007/s13225-020-00466-2)) |
| Otu59637 | 1 | k__Fungi(100);p__Ascomycota(100);c__Saccharomycetes(100);o__Saccharomycetales(100);f__Metschnikowiaceae(100);g__Metschnikowia(100);s__Metschnikowia_pulcherrima(100); | Metschnikowia | 13 | Saprotroph | \|Undefined Saprotroph\| | Yeast | NULL | Probable | As nectar-tap_saprotroph (Põlme et al. 2020); Decay sugar-rich substrates (Põlme et al. 2020) | Põlme S, et al. 2020. Fungal Diversity 105:1-16 ((https://doi.org/10.1007/s13225-020-00466-2)) |
| Otu59638 | 1 | k__Fungi(100);p__Ascomycota(100);c__Saccharomycetes(100);o__Saccharomycetales(100);f__Metschnikowiaceae(100);g__Metschnikowia(100);s__Metschnikowia_pulcherrima(100); | Metschnikowia | 13 | Saprotroph | \|Undefined Saprotroph\| | Yeast | NULL | Probable | As nectar-tap_saprotroph (Põlme et al. 2020); Decay sugar-rich substrates (Põlme et al. 2020) | Põlme S, et al. 2020. Fungal Diversity 105:1-16 ((https://doi.org/10.1007/s13225-020-00466-2)) |
| Otu59639 | 1 | k__Fungi(100);p__Ascomycota(100);c__Saccharomycetes(100);o__Saccharomycetales(100);f__Metschnikowiaceae(100);g__Metschnikowia(100);g__Metschnikowia_unclassified(100); | Metschnikowia | 13 | Saprotroph | \|Undefined Saprotroph\| | Yeast | NULL | Probable | As nectar-tap_saprotroph (Põlme et al. 2020); Decay sugar-rich substrates (Põlme et al. 2020) | Põlme S, et al. 2020. Fungal Diversity 105:1-16 ((https://doi.org/10.1007/s13225-020-00466-2)) |
| Otu59640 | 1 | k__Fungi(100);p__Ascomycota(100);c__Saccharomycetes(100);o__Saccharomycetales(100);f__Metschnikowiaceae(100);g__Metschnikowia(100);g__Metschnikowia_unclassified(100); | Metschnikowia | 13 | Saprotroph | \|Undefined Saprotroph\| | Yeast | NULL | Probable | As nectar-tap_saprotroph (Põlme et al. 2020); Decay sugar-rich substrates (Põlme et al. 2020) | Põlme S, et al. 2020. Fungal Diversity 105:1-16 ((https://doi.org/10.1007/s13225-020-00466-2)) |
| Otu59641 | 1 | k__Fungi(100);p__Ascomycota(100);c__Saccharomycetes(100);o__Saccharomycetales(100);f__Metschnikowiaceae(100);g__Metschnikowia(100);s__Metschnikowia_pulcherrima(100); | Metschnikowia | 13 | Saprotroph | \|Undefined Saprotroph\| | Yeast | NULL | Probable | As nectar-tap_saprotroph (Põlme et al. 2020); Decay sugar-rich substrates (Põlme et al. 2020) | Põlme S, et al. 2020. Fungal Diversity 105:1-16 ((https://doi.org/10.1007/s13225-020-00466-2)) |
| Otu59642 | 1 | k__Fungi(100);p__Ascomycota(100);c__Saccharomycetes(100);o__Saccharomycetales(100);f__Metschnikowiaceae(100);g__Metschnikowia(100);g__Metschnikowia_unclassified(100); | Metschnikowia | 13 | Saprotroph | \|Undefined Saprotroph\| | Yeast | NULL | Probable | As nectar-tap_saprotroph (Põlme et al. 2020); Decay sugar-rich substrates (Põlme et al. 2020) | Põlme S, et al. 2020. Fungal Diversity 105:1-16 ((https://doi.org/10.1007/s13225-020-00466-2)) |
| Otu59643 | 1 | k__Fungi(100);p__Ascomycota(100);c__Saccharomycetes(100);o__Saccharomycetales(100);f__Metschnikowiaceae(100);g__Metschnikowia(100);s__Metschnikowia_pulcherrima(100); | Metschnikowia | 13 | Saprotroph | \|Undefined Saprotroph\| | Yeast | NULL | Probable | As nectar-tap_saprotroph (Põlme et al. 2020); Decay sugar-rich substrates (Põlme et al. 2020) | Põlme S, et al. 2020. Fungal Diversity 105:1-16 ((https://doi.org/10.1007/s13225-020-00466-2)) |
| Otu59644 | 1 | k__Fungi(100);p__Ascomycota(100);c__Saccharomycetes(100);o__Saccharomycetales(100);f__Metschnikowiaceae(100);g__Metschnikowia(100);s__Metschnikowia_pulcherrima(100); | Metschnikowia | 13 | Saprotroph | \|Undefined Saprotroph\| | Yeast | NULL | Probable | As nectar-tap_saprotroph (Põlme et al. 2020); Decay sugar-rich substrates (Põlme et al. 2020) | Põlme S, et al. 2020. Fungal Diversity 105:1-16 ((https://doi.org/10.1007/s13225-020-00466-2)) |
| Otu59645 | 1 | k__Fungi(100);p__Ascomycota(100);c__Saccharomycetes(100);o__Saccharomycetales(100);f__Metschnikowiaceae(100);g__Metschnikowia(100);g__Metschnikowia_unclassified(100); | Metschnikowia | 13 | Saprotroph | \|Undefined Saprotroph\| | Yeast | NULL | Probable | As nectar-tap_saprotroph (Põlme et al. 2020); Decay sugar-rich substrates (Põlme et al. 2020) | Põlme S, et al. 2020. Fungal Diversity 105:1-16 ((https://doi.org/10.1007/s13225-020-00466-2)) |
| Otu59646 | 1 | k__Fungi(100);p__Ascomycota(100);c__Saccharomycetes(100);o__Saccharomycetales(100);f__Metschnikowiaceae(100);g__Metschnikowia(100);s__Metschnikowia_pulcherrima(100); | Metschnikowia | 13 | Saprotroph | \|Undefined Saprotroph\| | Yeast | NULL | Probable | As nectar-tap_saprotroph (Põlme et al. 2020); Decay sugar-rich substrates (Põlme et al. 2020) | Põlme S, et al. 2020. Fungal Diversity 105:1-16 ((https://doi.org/10.1007/s13225-020-00466-2)) |
| Otu59647 | 1 | k__Fungi(100);p__Ascomycota(100);c__Saccharomycetes(100);o__Saccharomycetales(100);f__Metschnikowiaceae(100);g__Metschnikowia(100);s__Metschnikowia_pulcherrima(100); | Metschnikowia | 13 | Saprotroph | \|Undefined Saprotroph\| | Yeast | NULL | Probable | As nectar-tap_saprotroph (Põlme et al. 2020); Decay sugar-rich substrates (Põlme et al. 2020) | Põlme S, et al. 2020. Fungal Diversity 105:1-16 ((https://doi.org/10.1007/s13225-020-00466-2)) |
| Otu59648 | 1 | k__Fungi(100);p__Ascomycota(100);c__Saccharomycetes(100);o__Saccharomycetales(100);f__Metschnikowiaceae(100);g__Metschnikowia(100);s__Metschnikowia_pulcherrima(100); | Metschnikowia | 13 | Saprotroph | \|Undefined Saprotroph\| | Yeast | NULL | Probable | As nectar-tap_saprotroph (Põlme et al. 2020); Decay sugar-rich substrates (Põlme et al. 2020) | Põlme S, et al. 2020. Fungal Diversity 105:1-16 ((https://doi.org/10.1007/s13225-020-00466-2)) |
| Otu59649 | 1 | k__Fungi(100);p__Ascomycota(100);c__Saccharomycetes(100);o__Saccharomycetales(100);f__Metschnikowiaceae(100);g__Metschnikowia(100);s__Metschnikowia_pulcherrima(100); | Metschnikowia | 13 | Saprotroph | \|Undefined Saprotroph\| | Yeast | NULL | Probable | As nectar-tap_saprotroph (Põlme et al. 2020); Decay sugar-rich substrates (Põlme et al. 2020) | Põlme S, et al. 2020. Fungal Diversity 105:1-16 ((https://doi.org/10.1007/s13225-020-00466-2)) |
| Otu59650 | 1 | k__Fungi(100);p__Ascomycota(100);c__Saccharomycetes(100);o__Saccharomycetales(100);f__Metschnikowiaceae(100);g__Metschnikowia(100);s__Metschnikowia_pulcherrima(100); | Metschnikowia | 13 | Saprotroph | \|Undefined Saprotroph\| | Yeast | NULL | Probable | As nectar-tap_saprotroph (Põlme et al. 2020); Decay sugar-rich substrates (Põlme et al. 2020) | Põlme S, et al. 2020. Fungal Diversity 105:1-16 ((https://doi.org/10.1007/s13225-020-00466-2)) |
| Otu59651 | 1 | k__Fungi(100);p__Ascomycota(100);c__Saccharomycetes(100);o__Saccharomycetales(100);f__Metschnikowiaceae(100);g__Metschnikowia(100);s__Metschnikowia_pulcherrima(100); | Metschnikowia | 13 | Saprotroph | \|Undefined Saprotroph\| | Yeast | NULL | Probable | As nectar-tap_saprotroph (Põlme et al. 2020); Decay sugar-rich substrates (Põlme et al. 2020) | Põlme S, et al. 2020. Fungal Diversity 105:1-16 ((https://doi.org/10.1007/s13225-020-00466-2)) |
| Otu59652 | 1 | k__Fungi(100);p__Ascomycota(100);c__Saccharomycetes(100);o__Saccharomycetales(100);f__Metschnikowiaceae(100);g__Metschnikowia(100);s__Metschnikowia_pulcherrima(100); | Metschnikowia | 13 | Saprotroph | \|Undefined Saprotroph\| | Yeast | NULL | Probable | As nectar-tap_saprotroph (Põlme et al. 2020); Decay sugar-rich substrates (Põlme et al. 2020) | Põlme S, et al. 2020. Fungal Diversity 105:1-16 ((https://doi.org/10.1007/s13225-020-00466-2)) |
| Otu59653 | 1 | k__Fungi(100);p__Ascomycota(100);c__Saccharomycetes(100);o__Saccharomycetales(100);f__Metschnikowiaceae(100);g__Metschnikowia(100);g__Metschnikowia_unclassified(100); | Metschnikowia | 13 | Saprotroph | \|Undefined Saprotroph\| | Yeast | NULL | Probable | As nectar-tap_saprotroph (Põlme et al. 2020); Decay sugar-rich substrates (Põlme et al. 2020) | Põlme S, et al. 2020. Fungal Diversity 105:1-16 ((https://doi.org/10.1007/s13225-020-00466-2)) |
| Otu59654 | 1 | k__Fungi(100);p__Ascomycota(100);c__Saccharomycetes(100);o__Saccharomycetales(100);f__Metschnikowiaceae(100);g__Metschnikowia(100);g__Metschnikowia_unclassified(100); | Metschnikowia | 13 | Saprotroph | \|Undefined Saprotroph\| | Yeast | NULL | Probable | As nectar-tap_saprotroph (Põlme et al. 2020); Decay sugar-rich substrates (Põlme et al. 2020) | Põlme S, et al. 2020. Fungal Diversity 105:1-16 ((https://doi.org/10.1007/s13225-020-00466-2)) |
| Otu59655 | 1 | k__Fungi(100);p__Ascomycota(100);c__Saccharomycetes(100);o__Saccharomycetales(100);f__Metschnikowiaceae(100);g__Metschnikowia(100);g__Metschnikowia_unclassified(100); | Metschnikowia | 13 | Saprotroph | \|Undefined Saprotroph\| | Yeast | NULL | Probable | As nectar-tap_saprotroph (Põlme et al. 2020); Decay sugar-rich substrates (Põlme et al. 2020) | Põlme S, et al. 2020. Fungal Diversity 105:1-16 ((https://doi.org/10.1007/s13225-020-00466-2)) |
| Otu59656 | 1 | k__Fungi(100);p__Ascomycota(100);c__Saccharomycetes(100);o__Saccharomycetales(100);f__Metschnikowiaceae(100);g__Metschnikowia(100);g__Metschnikowia_unclassified(100); | Metschnikowia | 13 | Saprotroph | \|Undefined Saprotroph\| | Yeast | NULL | Probable | As nectar-tap_saprotroph (Põlme et al. 2020); Decay sugar-rich substrates (Põlme et al. 2020) | Põlme S, et al. 2020. Fungal Diversity 105:1-16 ((https://doi.org/10.1007/s13225-020-00466-2)) |
| Otu59657 | 1 | k__Fungi(100);p__Ascomycota(100);c__Saccharomycetes(100);o__Saccharomycetales(100);f__Metschnikowiaceae(100);g__Metschnikowia(100);s__Metschnikowia_pulcherrima(100); | Metschnikowia | 13 | Saprotroph | \|Undefined Saprotroph\| | Yeast | NULL | Probable | As nectar-tap_saprotroph (Põlme et al. 2020); Decay sugar-rich substrates (Põlme et al. 2020) | Põlme S, et al. 2020. Fungal Diversity 105:1-16 ((https://doi.org/10.1007/s13225-020-00466-2)) |
| Otu59658 | 1 | k__Fungi(100);p__Ascomycota(100);c__Saccharomycetes(100);o__Saccharomycetales(100);f__Metschnikowiaceae(100);g__Metschnikowia(100);s__Metschnikowia_pulcherrima(100); | Metschnikowia | 13 | Saprotroph | \|Undefined Saprotroph\| | Yeast | NULL | Probable | As nectar-tap_saprotroph (Põlme et al. 2020); Decay sugar-rich substrates (Põlme et al. 2020) | Põlme S, et al. 2020. Fungal Diversity 105:1-16 ((https://doi.org/10.1007/s13225-020-00466-2)) |
| Otu59659 | 1 | k__Fungi(100);p__Ascomycota(100);c__Saccharomycetes(100);o__Saccharomycetales(100);f__Metschnikowiaceae(100);g__Metschnikowia(100);s__Metschnikowia_pulcherrima(100); | Metschnikowia | 13 | Saprotroph | \|Undefined Saprotroph\| | Yeast | NULL | Probable | As nectar-tap_saprotroph (Põlme et al. 2020); Decay sugar-rich substrates (Põlme et al. 2020) | Põlme S, et al. 2020. Fungal Diversity 105:1-16 ((https://doi.org/10.1007/s13225-020-00466-2)) |
| Otu59660 | 1 | k__Fungi(100);p__Ascomycota(100);c__Saccharomycetes(100);o__Saccharomycetales(100);f__Metschnikowiaceae(100);g__Metschnikowia(100);g__Metschnikowia_unclassified(100); | Metschnikowia | 13 | Saprotroph | \|Undefined Saprotroph\| | Yeast | NULL | Probable | As nectar-tap_saprotroph (Põlme et al. 2020); Decay sugar-rich substrates (Põlme et al. 2020) | Põlme S, et al. 2020. Fungal Diversity 105:1-16 ((https://doi.org/10.1007/s13225-020-00466-2)) |
| Otu59661 | 1 | k__Fungi(100);p__Ascomycota(100);c__Saccharomycetes(100);o__Saccharomycetales(100);f__Metschnikowiaceae(100);g__Metschnikowia(100);s__Metschnikowia_pulcherrima(100); | Metschnikowia | 13 | Saprotroph | \|Undefined Saprotroph\| | Yeast | NULL | Probable | As nectar-tap_saprotroph (Põlme et al. 2020); Decay sugar-rich substrates (Põlme et al. 2020) | Põlme S, et al. 2020. Fungal Diversity 105:1-16 ((https://doi.org/10.1007/s13225-020-00466-2)) |
| Otu59662 | 1 | k__Fungi(100);p__Ascomycota(100);c__Saccharomycetes(100);o__Saccharomycetales(100);f__Metschnikowiaceae(100);g__Metschnikowia(100);s__Metschnikowia_pulcherrima(100); | Metschnikowia | 13 | Saprotroph | \|Undefined Saprotroph\| | Yeast | NULL | Probable | As nectar-tap_saprotroph (Põlme et al. 2020); Decay sugar-rich substrates (Põlme et al. 2020) | Põlme S, et al. 2020. Fungal Diversity 105:1-16 ((https://doi.org/10.1007/s13225-020-00466-2)) |
| Otu59663 | 1 | k__Fungi(100);p__Ascomycota(100);c__Saccharomycetes(100);o__Saccharomycetales(100);f__Metschnikowiaceae(100);g__Metschnikowia(100);s__Metschnikowia_pulcherrima(100); | Metschnikowia | 13 | Saprotroph | \|Undefined Saprotroph\| | Yeast | NULL | Probable | As nectar-tap_saprotroph (Põlme et al. 2020); Decay sugar-rich substrates (Põlme et al. 2020) | Põlme S, et al. 2020. Fungal Diversity 105:1-16 ((https://doi.org/10.1007/s13225-020-00466-2)) |
| Otu59664 | 1 | k__Fungi(100);p__Ascomycota(100);c__Saccharomycetes(100);o__Saccharomycetales(100);f__Metschnikowiaceae(100);g__Metschnikowia(100);s__Metschnikowia_pulcherrima(100); | Metschnikowia | 13 | Saprotroph | \|Undefined Saprotroph\| | Yeast | NULL | Probable | As nectar-tap_saprotroph (Põlme et al. 2020); Decay sugar-rich substrates (Põlme et al. 2020) | Põlme S, et al. 2020. Fungal Diversity 105:1-16 ((https://doi.org/10.1007/s13225-020-00466-2)) |
| Otu59665 | 1 | k__Fungi(100);p__Ascomycota(100);c__Saccharomycetes(100);o__Saccharomycetales(100);f__Metschnikowiaceae(100);g__Metschnikowia(100);s__Metschnikowia_pulcherrima(100); | Metschnikowia | 13 | Saprotroph | \|Undefined Saprotroph\| | Yeast | NULL | Probable | As nectar-tap_saprotroph (Põlme et al. 2020); Decay sugar-rich substrates (Põlme et al. 2020) | Põlme S, et al. 2020. Fungal Diversity 105:1-16 ((https://doi.org/10.1007/s13225-020-00466-2)) |
| Otu59666 | 1 | k__Fungi(100);p__Ascomycota(100);c__Saccharomycetes(100);o__Saccharomycetales(100);f__Metschnikowiaceae(100);g__Metschnikowia(100);g__Metschnikowia_unclassified(100); | Metschnikowia | 13 | Saprotroph | \|Undefined Saprotroph\| | Yeast | NULL | Probable | As nectar-tap_saprotroph (Põlme et al. 2020); Decay sugar-rich substrates (Põlme et al. 2020) | Põlme S, et al. 2020. Fungal Diversity 105:1-16 ((https://doi.org/10.1007/s13225-020-00466-2)) |
| Otu59667 | 1 | k__Fungi(100);p__Ascomycota(100);c__Saccharomycetes(100);o__Saccharomycetales(100);f__Metschnikowiaceae(100);g__Metschnikowia(100);g__Metschnikowia_unclassified(100); | Metschnikowia | 13 | Saprotroph | \|Undefined Saprotroph\| | Yeast | NULL | Probable | As nectar-tap_saprotroph (Põlme et al. 2020); Decay sugar-rich substrates (Põlme et al. 2020) | Põlme S, et al. 2020. Fungal Diversity 105:1-16 ((https://doi.org/10.1007/s13225-020-00466-2)) |
| Otu59668 | 1 | k__Fungi(100);p__Ascomycota(100);c__Saccharomycetes(100);o__Saccharomycetales(100);f__Metschnikowiaceae(100);g__Metschnikowia(100);s__Metschnikowia_pulcherrima(100); | Metschnikowia | 13 | Saprotroph | \|Undefined Saprotroph\| | Yeast | NULL | Probable | As nectar-tap_saprotroph (Põlme et al. 2020); Decay sugar-rich substrates (Põlme et al. 2020) | Põlme S, et al. 2020. Fungal Diversity 105:1-16 ((https://doi.org/10.1007/s13225-020-00466-2)) |
| Otu59669 | 1 | k__Fungi(100);p__Ascomycota(100);c__Saccharomycetes(100);o__Saccharomycetales(100);f__Metschnikowiaceae(100);g__Metschnikowia(100);s__Metschnikowia_pulcherrima(100); | Metschnikowia | 13 | Saprotroph | \|Undefined Saprotroph\| | Yeast | NULL | Probable | As nectar-tap_saprotroph (Põlme et al. 2020); Decay sugar-rich substrates (Põlme et al. 2020) | Põlme S, et al. 2020. Fungal Diversity 105:1-16 ((https://doi.org/10.1007/s13225-020-00466-2)) |
| Otu59670 | 1 | k__Fungi(100);p__Ascomycota(100);c__Saccharomycetes(100);o__Saccharomycetales(100);f__Metschnikowiaceae(100);g__Metschnikowia(100);s__Metschnikowia_pulcherrima(100); | Metschnikowia | 13 | Saprotroph | \|Undefined Saprotroph\| | Yeast | NULL | Probable | As nectar-tap_saprotroph (Põlme et al. 2020); Decay sugar-rich substrates (Põlme et al. 2020) | Põlme S, et al. 2020. Fungal Diversity 105:1-16 ((https://doi.org/10.1007/s13225-020-00466-2)) |
| Otu59671 | 1 | k__Fungi(100);p__Ascomycota(100);c__Saccharomycetes(100);o__Saccharomycetales(100);f__Metschnikowiaceae(100);g__Metschnikowia(100);s__Metschnikowia_pulcherrima(100); | Metschnikowia | 13 | Saprotroph | \|Undefined Saprotroph\| | Yeast | NULL | Probable | As nectar-tap_saprotroph (Põlme et al. 2020); Decay sugar-rich substrates (Põlme et al. 2020) | Põlme S, et al. 2020. Fungal Diversity 105:1-16 ((https://doi.org/10.1007/s13225-020-00466-2)) |
| Otu59672 | 1 | k__Fungi(100);p__Ascomycota(100);c__Saccharomycetes(100);o__Saccharomycetales(100);f__Metschnikowiaceae(100);g__Metschnikowia(100);s__Metschnikowia_pulcherrima(100); | Metschnikowia | 13 | Saprotroph | \|Undefined Saprotroph\| | Yeast | NULL | Probable | As nectar-tap_saprotroph (Põlme et al. 2020); Decay sugar-rich substrates (Põlme et al. 2020) | Põlme S, et al. 2020. Fungal Diversity 105:1-16 ((https://doi.org/10.1007/s13225-020-00466-2)) |
| Otu59673 | 1 | k__Fungi(100);p__Ascomycota(100);c__Saccharomycetes(100);o__Saccharomycetales(100);f__Metschnikowiaceae(100);g__Metschnikowia(100);s__Metschnikowia_pulcherrima(100); | Metschnikowia | 13 | Saprotroph | \|Undefined Saprotroph\| | Yeast | NULL | Probable | As nectar-tap_saprotroph (Põlme et al. 2020); Decay sugar-rich substrates (Põlme et al. 2020) | Põlme S, et al. 2020. Fungal Diversity 105:1-16 ((https://doi.org/10.1007/s13225-020-00466-2)) |
| Otu59674 | 1 | k__Fungi(100);p__Ascomycota(100);c__Saccharomycetes(100);o__Saccharomycetales(100);f__Metschnikowiaceae(100);g__Metschnikowia(100);s__Metschnikowia_pulcherrima(100); | Metschnikowia | 13 | Saprotroph | \|Undefined Saprotroph\| | Yeast | NULL | Probable | As nectar-tap_saprotroph (Põlme et al. 2020); Decay sugar-rich substrates (Põlme et al. 2020) | Põlme S, et al. 2020. Fungal Diversity 105:1-16 ((https://doi.org/10.1007/s13225-020-00466-2)) |
| Otu59675 | 1 | k__Fungi(100);p__Ascomycota(100);c__Saccharomycetes(100);o__Saccharomycetales(100);f__Metschnikowiaceae(100);g__Metschnikowia(100);s__Metschnikowia_pulcherrima(100); | Metschnikowia | 13 | Saprotroph | \|Undefined Saprotroph\| | Yeast | NULL | Probable | As nectar-tap_saprotroph (Põlme et al. 2020); Decay sugar-rich substrates (Põlme et al. 2020) | Põlme S, et al. 2020. Fungal Diversity 105:1-16 ((https://doi.org/10.1007/s13225-020-00466-2)) |
| Otu59676 | 1 | k__Fungi(100);p__Ascomycota(100);c__Saccharomycetes(100);o__Saccharomycetales(100);f__Metschnikowiaceae(100);g__Metschnikowia(100);s__Metschnikowia_pulcherrima(100); | Metschnikowia | 13 | Saprotroph | \|Undefined Saprotroph\| | Yeast | NULL | Probable | As nectar-tap_saprotroph (Põlme et al. 2020); Decay sugar-rich substrates (Põlme et al. 2020) | Põlme S, et al. 2020. Fungal Diversity 105:1-16 ((https://doi.org/10.1007/s13225-020-00466-2)) |
| Otu59677 | 1 | k__Fungi(100);p__Ascomycota(100);c__Saccharomycetes(100);o__Saccharomycetales(100);f__Metschnikowiaceae(100);g__Metschnikowia(100);s__Metschnikowia_pulcherrima(100); | Metschnikowia | 13 | Saprotroph | \|Undefined Saprotroph\| | Yeast | NULL | Probable | As nectar-tap_saprotroph (Põlme et al. 2020); Decay sugar-rich substrates (Põlme et al. 2020) | Põlme S, et al. 2020. Fungal Diversity 105:1-16 ((https://doi.org/10.1007/s13225-020-00466-2)) |
| Otu59678 | 1 | k__Fungi(100);p__Ascomycota(100);c__Saccharomycetes(100);o__Saccharomycetales(100);f__Metschnikowiaceae(100);g__Metschnikowia(100);s__Metschnikowia_pulcherrima(100); | Metschnikowia | 13 | Saprotroph | \|Undefined Saprotroph\| | Yeast | NULL | Probable | As nectar-tap_saprotroph (Põlme et al. 2020); Decay sugar-rich substrates (Põlme et al. 2020) | Põlme S, et al. 2020. Fungal Diversity 105:1-16 ((https://doi.org/10.1007/s13225-020-00466-2)) |
| Otu59679 | 1 | k__Fungi(100);p__Ascomycota(100);c__Saccharomycetes(100);o__Saccharomycetales(100);f__Metschnikowiaceae(100);g__Metschnikowia(100);s__Metschnikowia_pulcherrima(100); | Metschnikowia | 13 | Saprotroph | \|Undefined Saprotroph\| | Yeast | NULL | Probable | As nectar-tap_saprotroph (Põlme et al. 2020); Decay sugar-rich substrates (Põlme et al. 2020) | Põlme S, et al. 2020. Fungal Diversity 105:1-16 ((https://doi.org/10.1007/s13225-020-00466-2)) |
| Otu59680 | 1 | k__Fungi(100);p__Ascomycota(100);c__Saccharomycetes(100);o__Saccharomycetales(100);f__Metschnikowiaceae(100);g__Metschnikowia(100);s__Metschnikowia_pulcherrima(100); | Metschnikowia | 13 | Saprotroph | \|Undefined Saprotroph\| | Yeast | NULL | Probable | As nectar-tap_saprotroph (Põlme et al. 2020); Decay sugar-rich substrates (Põlme et al. 2020) | Põlme S, et al. 2020. Fungal Diversity 105:1-16 ((https://doi.org/10.1007/s13225-020-00466-2)) |
| Otu59681 | 1 | k__Fungi(100);p__Ascomycota(100);c__Saccharomycetes(100);o__Saccharomycetales(100);f__Metschnikowiaceae(100);g__Metschnikowia(100);s__Metschnikowia_pulcherrima(100); | Metschnikowia | 13 | Saprotroph | \|Undefined Saprotroph\| | Yeast | NULL | Probable | As nectar-tap_saprotroph (Põlme et al. 2020); Decay sugar-rich substrates (Põlme et al. 2020) | Põlme S, et al. 2020. Fungal Diversity 105:1-16 ((https://doi.org/10.1007/s13225-020-00466-2)) |
| Otu59682 | 1 | k__Fungi(100);p__Ascomycota(100);c__Saccharomycetes(100);o__Saccharomycetales(100);f__Metschnikowiaceae(100);g__Metschnikowia(100);s__Metschnikowia_pulcherrima(100); | Metschnikowia | 13 | Saprotroph | \|Undefined Saprotroph\| | Yeast | NULL | Probable | As nectar-tap_saprotroph (Põlme et al. 2020); Decay sugar-rich substrates (Põlme et al. 2020) | Põlme S, et al. 2020. Fungal Diversity 105:1-16 ((https://doi.org/10.1007/s13225-020-00466-2)) |
| Otu59683 | 1 | k__Fungi(100);p__Ascomycota(100);c__Saccharomycetes(100);o__Saccharomycetales(100);f__Metschnikowiaceae(100);g__Metschnikowia(100);s__Metschnikowia_pulcherrima(100); | Metschnikowia | 13 | Saprotroph | \|Undefined Saprotroph\| | Yeast | NULL | Probable | As nectar-tap_saprotroph (Põlme et al. 2020); Decay sugar-rich substrates (Põlme et al. 2020) | Põlme S, et al. 2020. Fungal Diversity 105:1-16 ((https://doi.org/10.1007/s13225-020-00466-2)) |
| Otu59684 | 1 | k__Fungi(100);p__Ascomycota(100);c__Saccharomycetes(100);o__Saccharomycetales(100);f__Metschnikowiaceae(100);g__Metschnikowia(100);s__Metschnikowia_pulcherrima(100); | Metschnikowia | 13 | Saprotroph | \|Undefined Saprotroph\| | Yeast | NULL | Probable | As nectar-tap_saprotroph (Põlme et al. 2020); Decay sugar-rich substrates (Põlme et al. 2020) | Põlme S, et al. 2020. Fungal Diversity 105:1-16 ((https://doi.org/10.1007/s13225-020-00466-2)) |
| Otu59685 | 1 | k__Fungi(100);p__Ascomycota(100);c__Saccharomycetes(100);o__Saccharomycetales(100);f__Metschnikowiaceae(100);g__Metschnikowia(100);s__Metschnikowia_pulcherrima(100); | Metschnikowia | 13 | Saprotroph | \|Undefined Saprotroph\| | Yeast | NULL | Probable | As nectar-tap_saprotroph (Põlme et al. 2020); Decay sugar-rich substrates (Põlme et al. 2020) | Põlme S, et al. 2020. Fungal Diversity 105:1-16 ((https://doi.org/10.1007/s13225-020-00466-2)) |
| Otu59686 | 1 | k__Fungi(100);p__Ascomycota(100);c__Saccharomycetes(100);o__Saccharomycetales(100);f__Metschnikowiaceae(100);g__Metschnikowia(100);s__Metschnikowia_pulcherrima(100); | Metschnikowia | 13 | Saprotroph | \|Undefined Saprotroph\| | Yeast | NULL | Probable | As nectar-tap_saprotroph (Põlme et al. 2020); Decay sugar-rich substrates (Põlme et al. 2020) | Põlme S, et al. 2020. Fungal Diversity 105:1-16 ((https://doi.org/10.1007/s13225-020-00466-2)) |
| Otu59687 | 1 | k__Fungi(100);p__Ascomycota(100);c__Saccharomycetes(100);o__Saccharomycetales(100);f__Metschnikowiaceae(100);g__Metschnikowia(100);g__Metschnikowia_unclassified(100); | Metschnikowia | 13 | Saprotroph | \|Undefined Saprotroph\| | Yeast | NULL | Probable | As nectar-tap_saprotroph (Põlme et al. 2020); Decay sugar-rich substrates (Põlme et al. 2020) | Põlme S, et al. 2020. Fungal Diversity 105:1-16 ((https://doi.org/10.1007/s13225-020-00466-2)) |
| Otu59688 | 1 | k__Fungi(100);p__Ascomycota(100);c__Saccharomycetes(100);o__Saccharomycetales(100);f__Metschnikowiaceae(100);g__Metschnikowia(100);g__Metschnikowia_unclassified(100); | Metschnikowia | 13 | Saprotroph | \|Undefined Saprotroph\| | Yeast | NULL | Probable | As nectar-tap_saprotroph (Põlme et al. 2020); Decay sugar-rich substrates (Põlme et al. 2020) | Põlme S, et al. 2020. Fungal Diversity 105:1-16 ((https://doi.org/10.1007/s13225-020-00466-2)) |
| Otu59689 | 1 | k__Fungi(100);p__Ascomycota(100);c__Saccharomycetes(100);o__Saccharomycetales(100);f__Metschnikowiaceae(100);g__Metschnikowia(100);g__Metschnikowia_unclassified(100); | Metschnikowia | 13 | Saprotroph | \|Undefined Saprotroph\| | Yeast | NULL | Probable | As nectar-tap_saprotroph (Põlme et al. 2020); Decay sugar-rich substrates (Põlme et al. 2020) | Põlme S, et al. 2020. Fungal Diversity 105:1-16 ((https://doi.org/10.1007/s13225-020-00466-2)) |
| Otu59690 | 1 | k__Fungi(100);p__Ascomycota(100);c__Saccharomycetes(100);o__Saccharomycetales(100);f__Metschnikowiaceae(100);g__Metschnikowia(100);g__Metschnikowia_unclassified(100); | Metschnikowia | 13 | Saprotroph | \|Undefined Saprotroph\| | Yeast | NULL | Probable | As nectar-tap_saprotroph (Põlme et al. 2020); Decay sugar-rich substrates (Põlme et al. 2020) | Põlme S, et al. 2020. Fungal Diversity 105:1-16 ((https://doi.org/10.1007/s13225-020-00466-2)) |
| Otu59691 | 1 | k__Fungi(100);p__Ascomycota(100);c__Saccharomycetes(100);o__Saccharomycetales(100);f__Metschnikowiaceae(100);g__Metschnikowia(100);s__Metschnikowia_pulcherrima(100); | Metschnikowia | 13 | Saprotroph | \|Undefined Saprotroph\| | Yeast | NULL | Probable | As nectar-tap_saprotroph (Põlme et al. 2020); Decay sugar-rich substrates (Põlme et al. 2020) | Põlme S, et al. 2020. Fungal Diversity 105:1-16 ((https://doi.org/10.1007/s13225-020-00466-2)) |
| Otu59692 | 1 | k__Fungi(100);p__Ascomycota(100);c__Saccharomycetes(100);o__Saccharomycetales(100);f__Metschnikowiaceae(100);g__Metschnikowia(100);s__Metschnikowia_pulcherrima(100); | Metschnikowia | 13 | Saprotroph | \|Undefined Saprotroph\| | Yeast | NULL | Probable | As nectar-tap_saprotroph (Põlme et al. 2020); Decay sugar-rich substrates (Põlme et al. 2020) | Põlme S, et al. 2020. Fungal Diversity 105:1-16 ((https://doi.org/10.1007/s13225-020-00466-2)) |
| Otu59693 | 1 | k__Fungi(100);p__Ascomycota(100);c__Saccharomycetes(100);o__Saccharomycetales(100);f__Metschnikowiaceae(100);g__Metschnikowia(100);s__Metschnikowia_pulcherrima(100); | Metschnikowia | 13 | Saprotroph | \|Undefined Saprotroph\| | Yeast | NULL | Probable | As nectar-tap_saprotroph (Põlme et al. 2020); Decay sugar-rich substrates (Põlme et al. 2020) | Põlme S, et al. 2020. Fungal Diversity 105:1-16 ((https://doi.org/10.1007/s13225-020-00466-2)) |
| Otu59694 | 1 | k__Fungi(100);p__Ascomycota(100);c__Saccharomycetes(100);o__Saccharomycetales(100);f__Metschnikowiaceae(100);g__Metschnikowia(100);s__Metschnikowia_pulcherrima(100); | Metschnikowia | 13 | Saprotroph | \|Undefined Saprotroph\| | Yeast | NULL | Probable | As nectar-tap_saprotroph (Põlme et al. 2020); Decay sugar-rich substrates (Põlme et al. 2020) | Põlme S, et al. 2020. Fungal Diversity 105:1-16 ((https://doi.org/10.1007/s13225-020-00466-2)) |
| Otu59695 | 1 | k__Fungi(100);p__Ascomycota(100);c__Saccharomycetes(100);o__Saccharomycetales(100);f__Metschnikowiaceae(100);g__Metschnikowia(100);s__Metschnikowia_pulcherrima(100); | Metschnikowia | 13 | Saprotroph | \|Undefined Saprotroph\| | Yeast | NULL | Probable | As nectar-tap_saprotroph (Põlme et al. 2020); Decay sugar-rich substrates (Põlme et al. 2020) | Põlme S, et al. 2020. Fungal Diversity 105:1-16 ((https://doi.org/10.1007/s13225-020-00466-2)) |
| Otu59696 | 1 | k__Fungi(100);p__Ascomycota(100);c__Saccharomycetes(100);o__Saccharomycetales(100);f__Metschnikowiaceae(100);g__Metschnikowia(100);s__Metschnikowia_pulcherrima(100); | Metschnikowia | 13 | Saprotroph | \|Undefined Saprotroph\| | Yeast | NULL | Probable | As nectar-tap_saprotroph (Põlme et al. 2020); Decay sugar-rich substrates (Põlme et al. 2020) | Põlme S, et al. 2020. Fungal Diversity 105:1-16 ((https://doi.org/10.1007/s13225-020-00466-2)) |
| Otu59697 | 1 | k__Fungi(100);p__Ascomycota(100);c__Saccharomycetes(100);o__Saccharomycetales(100);f__Metschnikowiaceae(100);g__Metschnikowia(100);s__Metschnikowia_pulcherrima(100); | Metschnikowia | 13 | Saprotroph | \|Undefined Saprotroph\| | Yeast | NULL | Probable | As nectar-tap_saprotroph (Põlme et al. 2020); Decay sugar-rich substrates (Põlme et al. 2020) | Põlme S, et al. 2020. Fungal Diversity 105:1-16 ((https://doi.org/10.1007/s13225-020-00466-2)) |
| Otu59698 | 1 | k__Fungi(100);p__Ascomycota(100);c__Saccharomycetes(100);o__Saccharomycetales(100);f__Metschnikowiaceae(100);g__Metschnikowia(100);s__Metschnikowia_pulcherrima(100); | Metschnikowia | 13 | Saprotroph | \|Undefined Saprotroph\| | Yeast | NULL | Probable | As nectar-tap_saprotroph (Põlme et al. 2020); Decay sugar-rich substrates (Põlme et al. 2020) | Põlme S, et al. 2020. Fungal Diversity 105:1-16 ((https://doi.org/10.1007/s13225-020-00466-2)) |
| Otu59699 | 1 | k__Fungi(100);p__Ascomycota(100);c__Saccharomycetes(100);o__Saccharomycetales(100);f__Metschnikowiaceae(100);g__Metschnikowia(100);g__Metschnikowia_unclassified(100); | Metschnikowia | 13 | Saprotroph | \|Undefined Saprotroph\| | Yeast | NULL | Probable | As nectar-tap_saprotroph (Põlme et al. 2020); Decay sugar-rich substrates (Põlme et al. 2020) | Põlme S, et al. 2020. Fungal Diversity 105:1-16 ((https://doi.org/10.1007/s13225-020-00466-2)) |
| Otu59700 | 1 | k__Fungi(100);p__Ascomycota(100);c__Saccharomycetes(100);o__Saccharomycetales(100);f__Metschnikowiaceae(100);g__Metschnikowia(100);s__Metschnikowia_pulcherrima(100); | Metschnikowia | 13 | Saprotroph | \|Undefined Saprotroph\| | Yeast | NULL | Probable | As nectar-tap_saprotroph (Põlme et al. 2020); Decay sugar-rich substrates (Põlme et al. 2020) | Põlme S, et al. 2020. Fungal Diversity 105:1-16 ((https://doi.org/10.1007/s13225-020-00466-2)) |
| Otu59701 | 1 | k__Fungi(100);p__Ascomycota(100);c__Saccharomycetes(100);o__Saccharomycetales(100);f__Metschnikowiaceae(100);g__Metschnikowia(100);s__Metschnikowia_pulcherrima(100); | Metschnikowia | 13 | Saprotroph | \|Undefined Saprotroph\| | Yeast | NULL | Probable | As nectar-tap_saprotroph (Põlme et al. 2020); Decay sugar-rich substrates (Põlme et al. 2020) | Põlme S, et al. 2020. Fungal Diversity 105:1-16 ((https://doi.org/10.1007/s13225-020-00466-2)) |
| Otu59702 | 1 | k__Fungi(100);p__Ascomycota(100);c__Saccharomycetes(100);o__Saccharomycetales(100);f__Metschnikowiaceae(100);g__Metschnikowia(100);s__Metschnikowia_pulcherrima(100); | Metschnikowia | 13 | Saprotroph | \|Undefined Saprotroph\| | Yeast | NULL | Probable | As nectar-tap_saprotroph (Põlme et al. 2020); Decay sugar-rich substrates (Põlme et al. 2020) | Põlme S, et al. 2020. Fungal Diversity 105:1-16 ((https://doi.org/10.1007/s13225-020-00466-2)) |
| Otu59703 | 1 | k__Fungi(100);p__Ascomycota(100);c__Saccharomycetes(100);o__Saccharomycetales(100);f__Metschnikowiaceae(100);g__Metschnikowia(100);s__Metschnikowia_pulcherrima(100); | Metschnikowia | 13 | Saprotroph | \|Undefined Saprotroph\| | Yeast | NULL | Probable | As nectar-tap_saprotroph (Põlme et al. 2020); Decay sugar-rich substrates (Põlme et al. 2020) | Põlme S, et al. 2020. Fungal Diversity 105:1-16 ((https://doi.org/10.1007/s13225-020-00466-2)) |
| Otu59704 | 1 | k__Fungi(100);p__Ascomycota(100);c__Saccharomycetes(100);o__Saccharomycetales(100);f__Metschnikowiaceae(100);g__Metschnikowia(100);s__Metschnikowia_pulcherrima(100); | Metschnikowia | 13 | Saprotroph | \|Undefined Saprotroph\| | Yeast | NULL | Probable | As nectar-tap_saprotroph (Põlme et al. 2020); Decay sugar-rich substrates (Põlme et al. 2020) | Põlme S, et al. 2020. Fungal Diversity 105:1-16 ((https://doi.org/10.1007/s13225-020-00466-2)) |
| Otu59705 | 1 | k__Fungi(100);p__Ascomycota(100);c__Saccharomycetes(100);o__Saccharomycetales(100);f__Metschnikowiaceae(100);g__Metschnikowia(100);s__Metschnikowia_pulcherrima(100); | Metschnikowia | 13 | Saprotroph | \|Undefined Saprotroph\| | Yeast | NULL | Probable | As nectar-tap_saprotroph (Põlme et al. 2020); Decay sugar-rich substrates (Põlme et al. 2020) | Põlme S, et al. 2020. Fungal Diversity 105:1-16 ((https://doi.org/10.1007/s13225-020-00466-2)) |
| Otu59706 | 1 | k__Fungi(100);p__Ascomycota(100);c__Saccharomycetes(100);o__Saccharomycetales(100);f__Metschnikowiaceae(100);g__Metschnikowia(100);s__Metschnikowia_pulcherrima(100); | Metschnikowia | 13 | Saprotroph | \|Undefined Saprotroph\| | Yeast | NULL | Probable | As nectar-tap_saprotroph (Põlme et al. 2020); Decay sugar-rich substrates (Põlme et al. 2020) | Põlme S, et al. 2020. Fungal Diversity 105:1-16 ((https://doi.org/10.1007/s13225-020-00466-2)) |
| Otu59707 | 1 | k__Fungi(100);p__Ascomycota(100);c__Saccharomycetes(100);o__Saccharomycetales(100);f__Metschnikowiaceae(100);g__Metschnikowia(100);s__Metschnikowia_pulcherrima(100); | Metschnikowia | 13 | Saprotroph | \|Undefined Saprotroph\| | Yeast | NULL | Probable | As nectar-tap_saprotroph (Põlme et al. 2020); Decay sugar-rich substrates (Põlme et al. 2020) | Põlme S, et al. 2020. Fungal Diversity 105:1-16 ((https://doi.org/10.1007/s13225-020-00466-2)) |
| Otu59708 | 1 | k__Fungi(100);p__Ascomycota(100);c__Saccharomycetes(100);o__Saccharomycetales(100);f__Metschnikowiaceae(100);g__Metschnikowia(100);g__Metschnikowia_unclassified(100); | Metschnikowia | 13 | Saprotroph | \|Undefined Saprotroph\| | Yeast | NULL | Probable | As nectar-tap_saprotroph (Põlme et al. 2020); Decay sugar-rich substrates (Põlme et al. 2020) | Põlme S, et al. 2020. Fungal Diversity 105:1-16 ((https://doi.org/10.1007/s13225-020-00466-2)) |
| Otu59709 | 1 | k__Fungi(100);p__Ascomycota(100);c__Saccharomycetes(100);o__Saccharomycetales(100);f__Metschnikowiaceae(100);g__Metschnikowia(100);g__Metschnikowia_unclassified(100); | Metschnikowia | 13 | Saprotroph | \|Undefined Saprotroph\| | Yeast | NULL | Probable | As nectar-tap_saprotroph (Põlme et al. 2020); Decay sugar-rich substrates (Põlme et al. 2020) | Põlme S, et al. 2020. Fungal Diversity 105:1-16 ((https://doi.org/10.1007/s13225-020-00466-2)) |
| Otu59710 | 1 | k__Fungi(100);p__Ascomycota(100);c__Saccharomycetes(100);o__Saccharomycetales(100);f__Metschnikowiaceae(100);g__Metschnikowia(100);s__Metschnikowia_pulcherrima(100); | Metschnikowia | 13 | Saprotroph | \|Undefined Saprotroph\| | Yeast | NULL | Probable | As nectar-tap_saprotroph (Põlme et al. 2020); Decay sugar-rich substrates (Põlme et al. 2020) | Põlme S, et al. 2020. Fungal Diversity 105:1-16 ((https://doi.org/10.1007/s13225-020-00466-2)) |
| Otu59711 | 1 | k__Fungi(100);p__Ascomycota(100);c__Saccharomycetes(100);o__Saccharomycetales(100);f__Metschnikowiaceae(100);g__Metschnikowia(100);g__Metschnikowia_unclassified(100); | Metschnikowia | 13 | Saprotroph | \|Undefined Saprotroph\| | Yeast | NULL | Probable | As nectar-tap_saprotroph (Põlme et al. 2020); Decay sugar-rich substrates (Põlme et al. 2020) | Põlme S, et al. 2020. Fungal Diversity 105:1-16 ((https://doi.org/10.1007/s13225-020-00466-2)) |
| Otu59712 | 1 | k__Fungi(100);p__Ascomycota(100);c__Saccharomycetes(100);o__Saccharomycetales(100);f__Metschnikowiaceae(100);g__Metschnikowia(100);s__Metschnikowia_pulcherrima(100); | Metschnikowia | 13 | Saprotroph | \|Undefined Saprotroph\| | Yeast | NULL | Probable | As nectar-tap_saprotroph (Põlme et al. 2020); Decay sugar-rich substrates (Põlme et al. 2020) | Põlme S, et al. 2020. Fungal Diversity 105:1-16 ((https://doi.org/10.1007/s13225-020-00466-2)) |
| Otu59713 | 1 | k__Fungi(100);p__Ascomycota(100);c__Saccharomycetes(100);o__Saccharomycetales(100);f__Metschnikowiaceae(100);g__Metschnikowia(100);s__Metschnikowia_pulcherrima(100); | Metschnikowia | 13 | Saprotroph | \|Undefined Saprotroph\| | Yeast | NULL | Probable | As nectar-tap_saprotroph (Põlme et al. 2020); Decay sugar-rich substrates (Põlme et al. 2020) | Põlme S, et al. 2020. Fungal Diversity 105:1-16 ((https://doi.org/10.1007/s13225-020-00466-2)) |
| Otu59714 | 1 | k__Fungi(100);p__Ascomycota(100);c__Saccharomycetes(100);o__Saccharomycetales(100);f__Metschnikowiaceae(100);g__Metschnikowia(100);s__Metschnikowia_pulcherrima(100); | Metschnikowia | 13 | Saprotroph | \|Undefined Saprotroph\| | Yeast | NULL | Probable | As nectar-tap_saprotroph (Põlme et al. 2020); Decay sugar-rich substrates (Põlme et al. 2020) | Põlme S, et al. 2020. Fungal Diversity 105:1-16 ((https://doi.org/10.1007/s13225-020-00466-2)) |
| Otu59715 | 1 | k__Fungi(100);p__Ascomycota(100);c__Saccharomycetes(100);o__Saccharomycetales(100);f__Metschnikowiaceae(100);g__Metschnikowia(100);s__Metschnikowia_pulcherrima(100); | Metschnikowia | 13 | Saprotroph | \|Undefined Saprotroph\| | Yeast | NULL | Probable | As nectar-tap_saprotroph (Põlme et al. 2020); Decay sugar-rich substrates (Põlme et al. 2020) | Põlme S, et al. 2020. Fungal Diversity 105:1-16 ((https://doi.org/10.1007/s13225-020-00466-2)) |
| Otu59716 | 1 | k__Fungi(100);p__Ascomycota(100);c__Saccharomycetes(100);o__Saccharomycetales(100);f__Metschnikowiaceae(100);g__Metschnikowia(100);s__Metschnikowia_pulcherrima(100); | Metschnikowia | 13 | Saprotroph | \|Undefined Saprotroph\| | Yeast | NULL | Probable | As nectar-tap_saprotroph (Põlme et al. 2020); Decay sugar-rich substrates (Põlme et al. 2020) | Põlme S, et al. 2020. Fungal Diversity 105:1-16 ((https://doi.org/10.1007/s13225-020-00466-2)) |
| Otu59717 | 1 | k__Fungi(100);p__Ascomycota(100);c__Saccharomycetes(100);o__Saccharomycetales(100);f__Metschnikowiaceae(100);g__Metschnikowia(100);s__Metschnikowia_pulcherrima(100); | Metschnikowia | 13 | Saprotroph | \|Undefined Saprotroph\| | Yeast | NULL | Probable | As nectar-tap_saprotroph (Põlme et al. 2020); Decay sugar-rich substrates (Põlme et al. 2020) | Põlme S, et al. 2020. Fungal Diversity 105:1-16 ((https://doi.org/10.1007/s13225-020-00466-2)) |
| Otu59718 | 1 | k__Fungi(100);p__Ascomycota(100);c__Saccharomycetes(100);o__Saccharomycetales(100);f__Metschnikowiaceae(100);g__Metschnikowia(100);s__Metschnikowia_pulcherrima(100); | Metschnikowia | 13 | Saprotroph | \|Undefined Saprotroph\| | Yeast | NULL | Probable | As nectar-tap_saprotroph (Põlme et al. 2020); Decay sugar-rich substrates (Põlme et al. 2020) | Põlme S, et al. 2020. Fungal Diversity 105:1-16 ((https://doi.org/10.1007/s13225-020-00466-2)) |
| Otu59719 | 1 | k__Fungi(100);p__Ascomycota(100);c__Saccharomycetes(100);o__Saccharomycetales(100);f__Metschnikowiaceae(100);g__Metschnikowia(100);s__Metschnikowia_pulcherrima(100); | Metschnikowia | 13 | Saprotroph | \|Undefined Saprotroph\| | Yeast | NULL | Probable | As nectar-tap_saprotroph (Põlme et al. 2020); Decay sugar-rich substrates (Põlme et al. 2020) | Põlme S, et al. 2020. Fungal Diversity 105:1-16 ((https://doi.org/10.1007/s13225-020-00466-2)) |
| Otu59720 | 1 | k__Fungi(100);p__Ascomycota(100);c__Saccharomycetes(100);o__Saccharomycetales(100);f__Metschnikowiaceae(100);g__Metschnikowia(100);s__Metschnikowia_pulcherrima(100); | Metschnikowia | 13 | Saprotroph | \|Undefined Saprotroph\| | Yeast | NULL | Probable | As nectar-tap_saprotroph (Põlme et al. 2020); Decay sugar-rich substrates (Põlme et al. 2020) | Põlme S, et al. 2020. Fungal Diversity 105:1-16 ((https://doi.org/10.1007/s13225-020-00466-2)) |
| Otu59721 | 1 | k__Fungi(100);p__Ascomycota(100);c__Saccharomycetes(100);o__Saccharomycetales(100);f__Metschnikowiaceae(100);g__Metschnikowia(100);s__Metschnikowia_pulcherrima(100); | Metschnikowia | 13 | Saprotroph | \|Undefined Saprotroph\| | Yeast | NULL | Probable | As nectar-tap_saprotroph (Põlme et al. 2020); Decay sugar-rich substrates (Põlme et al. 2020) | Põlme S, et al. 2020. Fungal Diversity 105:1-16 ((https://doi.org/10.1007/s13225-020-00466-2)) |
| Otu59722 | 1 | k__Fungi(100);p__Ascomycota(100);c__Saccharomycetes(100);o__Saccharomycetales(100);f__Metschnikowiaceae(100);g__Metschnikowia(100);s__Metschnikowia_pulcherrima(100); | Metschnikowia | 13 | Saprotroph | \|Undefined Saprotroph\| | Yeast | NULL | Probable | As nectar-tap_saprotroph (Põlme et al. 2020); Decay sugar-rich substrates (Põlme et al. 2020) | Põlme S, et al. 2020. Fungal Diversity 105:1-16 ((https://doi.org/10.1007/s13225-020-00466-2)) |
| Otu59723 | 1 | k__Fungi(100);p__Ascomycota(100);c__Saccharomycetes(100);o__Saccharomycetales(100);f__Metschnikowiaceae(100);g__Metschnikowia(100);s__Metschnikowia_pulcherrima(100); | Metschnikowia | 13 | Saprotroph | \|Undefined Saprotroph\| | Yeast | NULL | Probable | As nectar-tap_saprotroph (Põlme et al. 2020); Decay sugar-rich substrates (Põlme et al. 2020) | Põlme S, et al. 2020. Fungal Diversity 105:1-16 ((https://doi.org/10.1007/s13225-020-00466-2)) |
| Otu59724 | 1 | k__Fungi(100);p__Ascomycota(100);c__Saccharomycetes(100);o__Saccharomycetales(100);f__Metschnikowiaceae(100);g__Metschnikowia(100);g__Metschnikowia_unclassified(100); | Metschnikowia | 13 | Saprotroph | \|Undefined Saprotroph\| | Yeast | NULL | Probable | As nectar-tap_saprotroph (Põlme et al. 2020); Decay sugar-rich substrates (Põlme et al. 2020) | Põlme S, et al. 2020. Fungal Diversity 105:1-16 ((https://doi.org/10.1007/s13225-020-00466-2)) |
| Otu59725 | 1 | k__Fungi(100);p__Ascomycota(100);c__Saccharomycetes(100);o__Saccharomycetales(100);f__Metschnikowiaceae(100);g__Metschnikowia(100);s__Metschnikowia_pulcherrima(100); | Metschnikowia | 13 | Saprotroph | \|Undefined Saprotroph\| | Yeast | NULL | Probable | As nectar-tap_saprotroph (Põlme et al. 2020); Decay sugar-rich substrates (Põlme et al. 2020) | Põlme S, et al. 2020. Fungal Diversity 105:1-16 ((https://doi.org/10.1007/s13225-020-00466-2)) |
| Otu59726 | 1 | k__Fungi(100);p__Ascomycota(100);c__Saccharomycetes(100);o__Saccharomycetales(100);f__Metschnikowiaceae(100);g__Metschnikowia(100);s__Metschnikowia_pulcherrima(100); | Metschnikowia | 13 | Saprotroph | \|Undefined Saprotroph\| | Yeast | NULL | Probable | As nectar-tap_saprotroph (Põlme et al. 2020); Decay sugar-rich substrates (Põlme et al. 2020) | Põlme S, et al. 2020. Fungal Diversity 105:1-16 ((https://doi.org/10.1007/s13225-020-00466-2)) |
| Otu59727 | 1 | k__Fungi(100);p__Ascomycota(100);c__Saccharomycetes(100);o__Saccharomycetales(100);f__Metschnikowiaceae(100);g__Metschnikowia(100);s__Metschnikowia_pulcherrima(100); | Metschnikowia | 13 | Saprotroph | \|Undefined Saprotroph\| | Yeast | NULL | Probable | As nectar-tap_saprotroph (Põlme et al. 2020); Decay sugar-rich substrates (Põlme et al. 2020) | Põlme S, et al. 2020. Fungal Diversity 105:1-16 ((https://doi.org/10.1007/s13225-020-00466-2)) |
| Otu59728 | 1 | k__Fungi(100);p__Ascomycota(100);c__Saccharomycetes(100);o__Saccharomycetales(100);f__Metschnikowiaceae(100);g__Metschnikowia(100);s__Metschnikowia_pulcherrima(100); | Metschnikowia | 13 | Saprotroph | \|Undefined Saprotroph\| | Yeast | NULL | Probable | As nectar-tap_saprotroph (Põlme et al. 2020); Decay sugar-rich substrates (Põlme et al. 2020) | Põlme S, et al. 2020. Fungal Diversity 105:1-16 ((https://doi.org/10.1007/s13225-020-00466-2)) |
| Otu59729 | 1 | k__Fungi(100);p__Ascomycota(100);c__Saccharomycetes(100);o__Saccharomycetales(100);f__Metschnikowiaceae(100);g__Metschnikowia(100);s__Metschnikowia_pulcherrima(100); | Metschnikowia | 13 | Saprotroph | \|Undefined Saprotroph\| | Yeast | NULL | Probable | As nectar-tap_saprotroph (Põlme et al. 2020); Decay sugar-rich substrates (Põlme et al. 2020) | Põlme S, et al. 2020. Fungal Diversity 105:1-16 ((https://doi.org/10.1007/s13225-020-00466-2)) |
| Otu59730 | 1 | k__Fungi(100);p__Ascomycota(100);c__Saccharomycetes(100);o__Saccharomycetales(100);f__Metschnikowiaceae(100);g__Metschnikowia(100);s__Metschnikowia_pulcherrima(100); | Metschnikowia | 13 | Saprotroph | \|Undefined Saprotroph\| | Yeast | NULL | Probable | As nectar-tap_saprotroph (Põlme et al. 2020); Decay sugar-rich substrates (Põlme et al. 2020) | Põlme S, et al. 2020. Fungal Diversity 105:1-16 ((https://doi.org/10.1007/s13225-020-00466-2)) |
| Otu59731 | 1 | k__Fungi(100);p__Ascomycota(100);c__Saccharomycetes(100);o__Saccharomycetales(100);f__Metschnikowiaceae(100);g__Metschnikowia(100);s__Metschnikowia_pulcherrima(100); | Metschnikowia | 13 | Saprotroph | \|Undefined Saprotroph\| | Yeast | NULL | Probable | As nectar-tap_saprotroph (Põlme et al. 2020); Decay sugar-rich substrates (Põlme et al. 2020) | Põlme S, et al. 2020. Fungal Diversity 105:1-16 ((https://doi.org/10.1007/s13225-020-00466-2)) |
| Otu59732 | 1 | k__Fungi(100);p__Ascomycota(100);c__Saccharomycetes(100);o__Saccharomycetales(100);f__Metschnikowiaceae(100);g__Metschnikowia(100);g__Metschnikowia_unclassified(100); | Metschnikowia | 13 | Saprotroph | \|Undefined Saprotroph\| | Yeast | NULL | Probable | As nectar-tap_saprotroph (Põlme et al. 2020); Decay sugar-rich substrates (Põlme et al. 2020) | Põlme S, et al. 2020. Fungal Diversity 105:1-16 ((https://doi.org/10.1007/s13225-020-00466-2)) |
| Otu59733 | 1 | k__Fungi(100);p__Ascomycota(100);c__Saccharomycetes(100);o__Saccharomycetales(100);f__Metschnikowiaceae(100);g__Metschnikowia(100);g__Metschnikowia_unclassified(100); | Metschnikowia | 13 | Saprotroph | \|Undefined Saprotroph\| | Yeast | NULL | Probable | As nectar-tap_saprotroph (Põlme et al. 2020); Decay sugar-rich substrates (Põlme et al. 2020) | Põlme S, et al. 2020. Fungal Diversity 105:1-16 ((https://doi.org/10.1007/s13225-020-00466-2)) |
| Otu59735 | 1 | k__Fungi(100);p__Ascomycota(100);c__Saccharomycetes(100);o__Saccharomycetales(100);f__Metschnikowiaceae(100);g__Metschnikowia(100);g__Metschnikowia_unclassified(100); | Metschnikowia | 13 | Saprotroph | \|Undefined Saprotroph\| | Yeast | NULL | Probable | As nectar-tap_saprotroph (Põlme et al. 2020); Decay sugar-rich substrates (Põlme et al. 2020) | Põlme S, et al. 2020. Fungal Diversity 105:1-16 ((https://doi.org/10.1007/s13225-020-00466-2)) |
| Otu59736 | 1 | k__Fungi(100);p__Ascomycota(100);c__Saccharomycetes(100);o__Saccharomycetales(100);f__Metschnikowiaceae(100);g__Metschnikowia(100);s__Metschnikowia_pulcherrima(100); | Metschnikowia | 13 | Saprotroph | \|Undefined Saprotroph\| | Yeast | NULL | Probable | As nectar-tap_saprotroph (Põlme et al. 2020); Decay sugar-rich substrates (Põlme et al. 2020) | Põlme S, et al. 2020. Fungal Diversity 105:1-16 ((https://doi.org/10.1007/s13225-020-00466-2)) |
| Otu59737 | 1 | k__Fungi(100);p__Ascomycota(100);c__Saccharomycetes(100);o__Saccharomycetales(100);f__Metschnikowiaceae(100);g__Metschnikowia(100);g__Metschnikowia_unclassified(100); | Metschnikowia | 13 | Saprotroph | \|Undefined Saprotroph\| | Yeast | NULL | Probable | As nectar-tap_saprotroph (Põlme et al. 2020); Decay sugar-rich substrates (Põlme et al. 2020) | Põlme S, et al. 2020. Fungal Diversity 105:1-16 ((https://doi.org/10.1007/s13225-020-00466-2)) |
| Otu59738 | 1 | k__Fungi(100);p__Ascomycota(100);c__Saccharomycetes(100);o__Saccharomycetales(100);f__Metschnikowiaceae(100);g__Metschnikowia(100);s__Metschnikowia_pulcherrima(100); | Metschnikowia | 13 | Saprotroph | \|Undefined Saprotroph\| | Yeast | NULL | Probable | As nectar-tap_saprotroph (Põlme et al. 2020); Decay sugar-rich substrates (Põlme et al. 2020) | Põlme S, et al. 2020. Fungal Diversity 105:1-16 ((https://doi.org/10.1007/s13225-020-00466-2)) |
| Otu59739 | 1 | k__Fungi(100);p__Ascomycota(100);c__Saccharomycetes(100);o__Saccharomycetales(100);f__Metschnikowiaceae(100);g__Metschnikowia(100);g__Metschnikowia_unclassified(100); | Metschnikowia | 13 | Saprotroph | \|Undefined Saprotroph\| | Yeast | NULL | Probable | As nectar-tap_saprotroph (Põlme et al. 2020); Decay sugar-rich substrates (Põlme et al. 2020) | Põlme S, et al. 2020. Fungal Diversity 105:1-16 ((https://doi.org/10.1007/s13225-020-00466-2)) |
| Otu59740 | 1 | k__Fungi(100);p__Ascomycota(100);c__Saccharomycetes(100);o__Saccharomycetales(100);f__Metschnikowiaceae(100);g__Metschnikowia(100);s__Metschnikowia_pulcherrima(100); | Metschnikowia | 13 | Saprotroph | \|Undefined Saprotroph\| | Yeast | NULL | Probable | As nectar-tap_saprotroph (Põlme et al. 2020); Decay sugar-rich substrates (Põlme et al. 2020) | Põlme S, et al. 2020. Fungal Diversity 105:1-16 ((https://doi.org/10.1007/s13225-020-00466-2)) |
| Otu59741 | 1 | k__Fungi(100);p__Ascomycota(100);c__Saccharomycetes(100);o__Saccharomycetales(100);f__Metschnikowiaceae(100);g__Metschnikowia(100);g__Metschnikowia_unclassified(100); | Metschnikowia | 13 | Saprotroph | \|Undefined Saprotroph\| | Yeast | NULL | Probable | As nectar-tap_saprotroph (Põlme et al. 2020); Decay sugar-rich substrates (Põlme et al. 2020) | Põlme S, et al. 2020. Fungal Diversity 105:1-16 ((https://doi.org/10.1007/s13225-020-00466-2)) |
| Otu59742 | 1 | k__Fungi(100);p__Ascomycota(100);c__Saccharomycetes(100);o__Saccharomycetales(100);f__Metschnikowiaceae(100);g__Metschnikowia(100);s__Metschnikowia_pulcherrima(100); | Metschnikowia | 13 | Saprotroph | \|Undefined Saprotroph\| | Yeast | NULL | Probable | As nectar-tap_saprotroph (Põlme et al. 2020); Decay sugar-rich substrates (Põlme et al. 2020) | Põlme S, et al. 2020. Fungal Diversity 105:1-16 ((https://doi.org/10.1007/s13225-020-00466-2)) |
| Otu59743 | 1 | k__Fungi(100);p__Ascomycota(100);c__Saccharomycetes(100);o__Saccharomycetales(100);f__Metschnikowiaceae(100);g__Metschnikowia(100);s__Metschnikowia_pulcherrima(100); | Metschnikowia | 13 | Saprotroph | \|Undefined Saprotroph\| | Yeast | NULL | Probable | As nectar-tap_saprotroph (Põlme et al. 2020); Decay sugar-rich substrates (Põlme et al. 2020) | Põlme S, et al. 2020. Fungal Diversity 105:1-16 ((https://doi.org/10.1007/s13225-020-00466-2)) |
| Otu59744 | 1 | k__Fungi(100);p__Ascomycota(100);c__Saccharomycetes(100);o__Saccharomycetales(100);f__Metschnikowiaceae(100);g__Metschnikowia(100);s__Metschnikowia_pulcherrima(100); | Metschnikowia | 13 | Saprotroph | \|Undefined Saprotroph\| | Yeast | NULL | Probable | As nectar-tap_saprotroph (Põlme et al. 2020); Decay sugar-rich substrates (Põlme et al. 2020) | Põlme S, et al. 2020. Fungal Diversity 105:1-16 ((https://doi.org/10.1007/s13225-020-00466-2)) |
| Otu59745 | 1 | k__Fungi(100);p__Ascomycota(100);c__Saccharomycetes(100);o__Saccharomycetales(100);f__Metschnikowiaceae(100);g__Metschnikowia(100);s__Metschnikowia_pulcherrima(100); | Metschnikowia | 13 | Saprotroph | \|Undefined Saprotroph\| | Yeast | NULL | Probable | As nectar-tap_saprotroph (Põlme et al. 2020); Decay sugar-rich substrates (Põlme et al. 2020) | Põlme S, et al. 2020. Fungal Diversity 105:1-16 ((https://doi.org/10.1007/s13225-020-00466-2)) |
| Otu59746 | 1 | k__Fungi(100);p__Ascomycota(100);c__Saccharomycetes(100);o__Saccharomycetales(100);f__Metschnikowiaceae(100);g__Metschnikowia(100);s__Metschnikowia_pulcherrima(100); | Metschnikowia | 13 | Saprotroph | \|Undefined Saprotroph\| | Yeast | NULL | Probable | As nectar-tap_saprotroph (Põlme et al. 2020); Decay sugar-rich substrates (Põlme et al. 2020) | Põlme S, et al. 2020. Fungal Diversity 105:1-16 ((https://doi.org/10.1007/s13225-020-00466-2)) |
| Otu59747 | 1 | k__Fungi(100);p__Ascomycota(100);c__Saccharomycetes(100);o__Saccharomycetales(100);f__Metschnikowiaceae(100);g__Metschnikowia(100);g__Metschnikowia_unclassified(100); | Metschnikowia | 13 | Saprotroph | \|Undefined Saprotroph\| | Yeast | NULL | Probable | As nectar-tap_saprotroph (Põlme et al. 2020); Decay sugar-rich substrates (Põlme et al. 2020) | Põlme S, et al. 2020. Fungal Diversity 105:1-16 ((https://doi.org/10.1007/s13225-020-00466-2)) |
| Otu59748 | 1 | k__Fungi(100);p__Ascomycota(100);c__Saccharomycetes(100);o__Saccharomycetales(100);f__Metschnikowiaceae(100);g__Metschnikowia(100);g__Metschnikowia_unclassified(100); | Metschnikowia | 13 | Saprotroph | \|Undefined Saprotroph\| | Yeast | NULL | Probable | As nectar-tap_saprotroph (Põlme et al. 2020); Decay sugar-rich substrates (Põlme et al. 2020) | Põlme S, et al. 2020. Fungal Diversity 105:1-16 ((https://doi.org/10.1007/s13225-020-00466-2)) |
| Otu59749 | 1 | k__Fungi(100);p__Ascomycota(100);c__Saccharomycetes(100);o__Saccharomycetales(100);f__Metschnikowiaceae(100);g__Metschnikowia(100);g__Metschnikowia_unclassified(100); | Metschnikowia | 13 | Saprotroph | \|Undefined Saprotroph\| | Yeast | NULL | Probable | As nectar-tap_saprotroph (Põlme et al. 2020); Decay sugar-rich substrates (Põlme et al. 2020) | Põlme S, et al. 2020. Fungal Diversity 105:1-16 ((https://doi.org/10.1007/s13225-020-00466-2)) |
| Otu59750 | 1 | k__Fungi(100);p__Ascomycota(100);c__Saccharomycetes(100);o__Saccharomycetales(100);f__Metschnikowiaceae(100);g__Metschnikowia(100);s__Metschnikowia_pulcherrima(100); | Metschnikowia | 13 | Saprotroph | \|Undefined Saprotroph\| | Yeast | NULL | Probable | As nectar-tap_saprotroph (Põlme et al. 2020); Decay sugar-rich substrates (Põlme et al. 2020) | Põlme S, et al. 2020. Fungal Diversity 105:1-16 ((https://doi.org/10.1007/s13225-020-00466-2)) |
| Otu59751 | 1 | k__Fungi(100);p__Ascomycota(100);c__Saccharomycetes(100);o__Saccharomycetales(100);f__Metschnikowiaceae(100);g__Metschnikowia(100);s__Metschnikowia_pulcherrima(100); | Metschnikowia | 13 | Saprotroph | \|Undefined Saprotroph\| | Yeast | NULL | Probable | As nectar-tap_saprotroph (Põlme et al. 2020); Decay sugar-rich substrates (Põlme et al. 2020) | Põlme S, et al. 2020. Fungal Diversity 105:1-16 ((https://doi.org/10.1007/s13225-020-00466-2)) |
| Otu59752 | 1 | k__Fungi(100);p__Ascomycota(100);c__Saccharomycetes(100);o__Saccharomycetales(100);f__Metschnikowiaceae(100);g__Metschnikowia(100);s__Metschnikowia_pulcherrima(100); | Metschnikowia | 13 | Saprotroph | \|Undefined Saprotroph\| | Yeast | NULL | Probable | As nectar-tap_saprotroph (Põlme et al. 2020); Decay sugar-rich substrates (Põlme et al. 2020) | Põlme S, et al. 2020. Fungal Diversity 105:1-16 ((https://doi.org/10.1007/s13225-020-00466-2)) |
| Otu59753 | 1 | k__Fungi(100);p__Ascomycota(100);c__Saccharomycetes(100);o__Saccharomycetales(100);f__Metschnikowiaceae(100);g__Metschnikowia(100);g__Metschnikowia_unclassified(100); | Metschnikowia | 13 | Saprotroph | \|Undefined Saprotroph\| | Yeast | NULL | Probable | As nectar-tap_saprotroph (Põlme et al. 2020); Decay sugar-rich substrates (Põlme et al. 2020) | Põlme S, et al. 2020. Fungal Diversity 105:1-16 ((https://doi.org/10.1007/s13225-020-00466-2)) |
| Otu59754 | 1 | k__Fungi(100);p__Ascomycota(100);c__Saccharomycetes(100);o__Saccharomycetales(100);f__Metschnikowiaceae(100);g__Metschnikowia(100);s__Metschnikowia_pulcherrima(100); | Metschnikowia | 13 | Saprotroph | \|Undefined Saprotroph\| | Yeast | NULL | Probable | As nectar-tap_saprotroph (Põlme et al. 2020); Decay sugar-rich substrates (Põlme et al. 2020) | Põlme S, et al. 2020. Fungal Diversity 105:1-16 ((https://doi.org/10.1007/s13225-020-00466-2)) |
| Otu59755 | 1 | k__Fungi(100);p__Ascomycota(100);c__Saccharomycetes(100);o__Saccharomycetales(100);f__Metschnikowiaceae(100);g__Metschnikowia(100);s__Metschnikowia_pulcherrima(100); | Metschnikowia | 13 | Saprotroph | \|Undefined Saprotroph\| | Yeast | NULL | Probable | As nectar-tap_saprotroph (Põlme et al. 2020); Decay sugar-rich substrates (Põlme et al. 2020) | Põlme S, et al. 2020. Fungal Diversity 105:1-16 ((https://doi.org/10.1007/s13225-020-00466-2)) |
| Otu59756 | 1 | k__Fungi(100);p__Ascomycota(100);c__Saccharomycetes(100);o__Saccharomycetales(100);f__Metschnikowiaceae(100);g__Metschnikowia(100);s__Metschnikowia_pulcherrima(100); | Metschnikowia | 13 | Saprotroph | \|Undefined Saprotroph\| | Yeast | NULL | Probable | As nectar-tap_saprotroph (Põlme et al. 2020); Decay sugar-rich substrates (Põlme et al. 2020) | Põlme S, et al. 2020. Fungal Diversity 105:1-16 ((https://doi.org/10.1007/s13225-020-00466-2)) |
| Otu59757 | 1 | k__Fungi(100);p__Ascomycota(100);c__Saccharomycetes(100);o__Saccharomycetales(100);f__Metschnikowiaceae(100);g__Metschnikowia(100);s__Metschnikowia_pulcherrima(100); | Metschnikowia | 13 | Saprotroph | \|Undefined Saprotroph\| | Yeast | NULL | Probable | As nectar-tap_saprotroph (Põlme et al. 2020); Decay sugar-rich substrates (Põlme et al. 2020) | Põlme S, et al. 2020. Fungal Diversity 105:1-16 ((https://doi.org/10.1007/s13225-020-00466-2)) |
| Otu59758 | 1 | k__Fungi(100);p__Ascomycota(100);c__Saccharomycetes(100);o__Saccharomycetales(100);f__Metschnikowiaceae(100);g__Metschnikowia(100);s__Metschnikowia_pulcherrima(100); | Metschnikowia | 13 | Saprotroph | \|Undefined Saprotroph\| | Yeast | NULL | Probable | As nectar-tap_saprotroph (Põlme et al. 2020); Decay sugar-rich substrates (Põlme et al. 2020) | Põlme S, et al. 2020. Fungal Diversity 105:1-16 ((https://doi.org/10.1007/s13225-020-00466-2)) |
| Otu59759 | 1 | k__Fungi(100);p__Ascomycota(100);c__Saccharomycetes(100);o__Saccharomycetales(100);f__Metschnikowiaceae(100);g__Metschnikowia(100);g__Metschnikowia_unclassified(100); | Metschnikowia | 13 | Saprotroph | \|Undefined Saprotroph\| | Yeast | NULL | Probable | As nectar-tap_saprotroph (Põlme et al. 2020); Decay sugar-rich substrates (Põlme et al. 2020) | Põlme S, et al. 2020. Fungal Diversity 105:1-16 ((https://doi.org/10.1007/s13225-020-00466-2)) |
| Otu59760 | 1 | k__Fungi(100);p__Ascomycota(100);c__Saccharomycetes(100);o__Saccharomycetales(100);f__Metschnikowiaceae(100);g__Metschnikowia(100);g__Metschnikowia_unclassified(100); | Metschnikowia | 13 | Saprotroph | \|Undefined Saprotroph\| | Yeast | NULL | Probable | As nectar-tap_saprotroph (Põlme et al. 2020); Decay sugar-rich substrates (Põlme et al. 2020) | Põlme S, et al. 2020. Fungal Diversity 105:1-16 ((https://doi.org/10.1007/s13225-020-00466-2)) |
| Otu59761 | 1 | k__Fungi(100);p__Ascomycota(100);c__Saccharomycetes(100);o__Saccharomycetales(100);f__Metschnikowiaceae(100);g__Metschnikowia(100);s__Metschnikowia_pulcherrima(100); | Metschnikowia | 13 | Saprotroph | \|Undefined Saprotroph\| | Yeast | NULL | Probable | As nectar-tap_saprotroph (Põlme et al. 2020); Decay sugar-rich substrates (Põlme et al. 2020) | Põlme S, et al. 2020. Fungal Diversity 105:1-16 ((https://doi.org/10.1007/s13225-020-00466-2)) |
| Otu59762 | 1 | k__Fungi(100);p__Ascomycota(100);c__Saccharomycetes(100);o__Saccharomycetales(100);f__Metschnikowiaceae(100);g__Metschnikowia(100);s__Metschnikowia_pulcherrima(100); | Metschnikowia | 13 | Saprotroph | \|Undefined Saprotroph\| | Yeast | NULL | Probable | As nectar-tap_saprotroph (Põlme et al. 2020); Decay sugar-rich substrates (Põlme et al. 2020) | Põlme S, et al. 2020. Fungal Diversity 105:1-16 ((https://doi.org/10.1007/s13225-020-00466-2)) |
| Otu59763 | 1 | k__Fungi(100);p__Ascomycota(100);c__Saccharomycetes(100);o__Saccharomycetales(100);f__Metschnikowiaceae(100);g__Metschnikowia(100);s__Metschnikowia_pulcherrima(100); | Metschnikowia | 13 | Saprotroph | \|Undefined Saprotroph\| | Yeast | NULL | Probable | As nectar-tap_saprotroph (Põlme et al. 2020); Decay sugar-rich substrates (Põlme et al. 2020) | Põlme S, et al. 2020. Fungal Diversity 105:1-16 ((https://doi.org/10.1007/s13225-020-00466-2)) |
| Otu59764 | 1 | k__Fungi(100);p__Ascomycota(100);c__Saccharomycetes(100);o__Saccharomycetales(100);f__Metschnikowiaceae(100);g__Metschnikowia(100);s__Metschnikowia_pulcherrima(100); | Metschnikowia | 13 | Saprotroph | \|Undefined Saprotroph\| | Yeast | NULL | Probable | As nectar-tap_saprotroph (Põlme et al. 2020); Decay sugar-rich substrates (Põlme et al. 2020) | Põlme S, et al. 2020. Fungal Diversity 105:1-16 ((https://doi.org/10.1007/s13225-020-00466-2)) |
| Otu59765 | 1 | k__Fungi(100);p__Ascomycota(100);c__Saccharomycetes(100);o__Saccharomycetales(100);f__Metschnikowiaceae(100);g__Metschnikowia(100);s__Metschnikowia_sp(100); | Metschnikowia | 13 | Saprotroph | \|Undefined Saprotroph\| | Yeast | NULL | Probable | As nectar-tap_saprotroph (Põlme et al. 2020); Decay sugar-rich substrates (Põlme et al. 2020) | Põlme S, et al. 2020. Fungal Diversity 105:1-16 ((https://doi.org/10.1007/s13225-020-00466-2)) |
| Otu59766 | 1 | k__Fungi(100);p__Ascomycota(100);c__Saccharomycetes(100);o__Saccharomycetales(100);f__Metschnikowiaceae(100);g__Metschnikowia(100);g__Metschnikowia_unclassified(100); | Metschnikowia | 13 | Saprotroph | \|Undefined Saprotroph\| | Yeast | NULL | Probable | As nectar-tap_saprotroph (Põlme et al. 2020); Decay sugar-rich substrates (Põlme et al. 2020) | Põlme S, et al. 2020. Fungal Diversity 105:1-16 ((https://doi.org/10.1007/s13225-020-00466-2)) |
| Otu59767 | 1 | k__Fungi(100);p__Ascomycota(100);c__Saccharomycetes(100);o__Saccharomycetales(100);f__Metschnikowiaceae(100);g__Metschnikowia(100);s__Metschnikowia_pulcherrima(100); | Metschnikowia | 13 | Saprotroph | \|Undefined Saprotroph\| | Yeast | NULL | Probable | As nectar-tap_saprotroph (Põlme et al. 2020); Decay sugar-rich substrates (Põlme et al. 2020) | Põlme S, et al. 2020. Fungal Diversity 105:1-16 ((https://doi.org/10.1007/s13225-020-00466-2)) |
| Otu59768 | 1 | k__Fungi(100);p__Ascomycota(100);c__Saccharomycetes(100);o__Saccharomycetales(100);f__Metschnikowiaceae(100);g__Metschnikowia(100);s__Metschnikowia_pulcherrima(100); | Metschnikowia | 13 | Saprotroph | \|Undefined Saprotroph\| | Yeast | NULL | Probable | As nectar-tap_saprotroph (Põlme et al. 2020); Decay sugar-rich substrates (Põlme et al. 2020) | Põlme S, et al. 2020. Fungal Diversity 105:1-16 ((https://doi.org/10.1007/s13225-020-00466-2)) |
| Otu59769 | 1 | k__Fungi(100);p__Ascomycota(100);c__Saccharomycetes(100);o__Saccharomycetales(100);f__Metschnikowiaceae(100);g__Metschnikowia(100);s__Metschnikowia_pulcherrima(100); | Metschnikowia | 13 | Saprotroph | \|Undefined Saprotroph\| | Yeast | NULL | Probable | As nectar-tap_saprotroph (Põlme et al. 2020); Decay sugar-rich substrates (Põlme et al. 2020) | Põlme S, et al. 2020. Fungal Diversity 105:1-16 ((https://doi.org/10.1007/s13225-020-00466-2)) |
| Otu59770 | 1 | k__Fungi(100);p__Ascomycota(100);c__Saccharomycetes(100);o__Saccharomycetales(100);f__Metschnikowiaceae(100);g__Metschnikowia(100);g__Metschnikowia_unclassified(100); | Metschnikowia | 13 | Saprotroph | \|Undefined Saprotroph\| | Yeast | NULL | Probable | As nectar-tap_saprotroph (Põlme et al. 2020); Decay sugar-rich substrates (Põlme et al. 2020) | Põlme S, et al. 2020. Fungal Diversity 105:1-16 ((https://doi.org/10.1007/s13225-020-00466-2)) |
| Otu59771 | 1 | k__Fungi(100);p__Ascomycota(100);c__Saccharomycetes(100);o__Saccharomycetales(100);f__Metschnikowiaceae(100);g__Metschnikowia(100);s__Metschnikowia_pulcherrima(100); | Metschnikowia | 13 | Saprotroph | \|Undefined Saprotroph\| | Yeast | NULL | Probable | As nectar-tap_saprotroph (Põlme et al. 2020); Decay sugar-rich substrates (Põlme et al. 2020) | Põlme S, et al. 2020. Fungal Diversity 105:1-16 ((https://doi.org/10.1007/s13225-020-00466-2)) |
| Otu59772 | 1 | k__Fungi(100);p__Ascomycota(100);c__Saccharomycetes(100);o__Saccharomycetales(100);f__Metschnikowiaceae(100);g__Metschnikowia(100);s__Metschnikowia_pulcherrima(100); | Metschnikowia | 13 | Saprotroph | \|Undefined Saprotroph\| | Yeast | NULL | Probable | As nectar-tap_saprotroph (Põlme et al. 2020); Decay sugar-rich substrates (Põlme et al. 2020) | Põlme S, et al. 2020. Fungal Diversity 105:1-16 ((https://doi.org/10.1007/s13225-020-00466-2)) |
| Otu59773 | 1 | k__Fungi(100);p__Ascomycota(100);c__Saccharomycetes(100);o__Saccharomycetales(100);f__Metschnikowiaceae(100);g__Metschnikowia(100);s__Metschnikowia_pulcherrima(100); | Metschnikowia | 13 | Saprotroph | \|Undefined Saprotroph\| | Yeast | NULL | Probable | As nectar-tap_saprotroph (Põlme et al. 2020); Decay sugar-rich substrates (Põlme et al. 2020) | Põlme S, et al. 2020. Fungal Diversity 105:1-16 ((https://doi.org/10.1007/s13225-020-00466-2)) |
| Otu59774 | 1 | k__Fungi(100);p__Ascomycota(100);c__Saccharomycetes(100);o__Saccharomycetales(100);f__Metschnikowiaceae(100);g__Metschnikowia(100);s__Metschnikowia_pulcherrima(100); | Metschnikowia | 13 | Saprotroph | \|Undefined Saprotroph\| | Yeast | NULL | Probable | As nectar-tap_saprotroph (Põlme et al. 2020); Decay sugar-rich substrates (Põlme et al. 2020) | Põlme S, et al. 2020. Fungal Diversity 105:1-16 ((https://doi.org/10.1007/s13225-020-00466-2)) |
| Otu59775 | 1 | k__Fungi(100);p__Ascomycota(100);c__Saccharomycetes(100);o__Saccharomycetales(100);f__Metschnikowiaceae(100);g__Metschnikowia(100);g__Metschnikowia_unclassified(100); | Metschnikowia | 13 | Saprotroph | \|Undefined Saprotroph\| | Yeast | NULL | Probable | As nectar-tap_saprotroph (Põlme et al. 2020); Decay sugar-rich substrates (Põlme et al. 2020) | Põlme S, et al. 2020. Fungal Diversity 105:1-16 ((https://doi.org/10.1007/s13225-020-00466-2)) |
| Otu59776 | 1 | k__Fungi(100);p__Ascomycota(100);c__Saccharomycetes(100);o__Saccharomycetales(100);f__Metschnikowiaceae(100);g__Metschnikowia(100);g__Metschnikowia_unclassified(100); | Metschnikowia | 13 | Saprotroph | \|Undefined Saprotroph\| | Yeast | NULL | Probable | As nectar-tap_saprotroph (Põlme et al. 2020); Decay sugar-rich substrates (Põlme et al. 2020) | Põlme S, et al. 2020. Fungal Diversity 105:1-16 ((https://doi.org/10.1007/s13225-020-00466-2)) |
| Otu59777 | 1 | k__Fungi(100);p__Ascomycota(100);c__Saccharomycetes(100);o__Saccharomycetales(100);f__Metschnikowiaceae(100);g__Metschnikowia(100);s__Metschnikowia_pulcherrima(100); | Metschnikowia | 13 | Saprotroph | \|Undefined Saprotroph\| | Yeast | NULL | Probable | As nectar-tap_saprotroph (Põlme et al. 2020); Decay sugar-rich substrates (Põlme et al. 2020) | Põlme S, et al. 2020. Fungal Diversity 105:1-16 ((https://doi.org/10.1007/s13225-020-00466-2)) |
| Otu59778 | 1 | k__Fungi(100);p__Ascomycota(100);c__Saccharomycetes(100);o__Saccharomycetales(100);f__Metschnikowiaceae(100);g__Metschnikowia(100);s__Metschnikowia_pulcherrima(100); | Metschnikowia | 13 | Saprotroph | \|Undefined Saprotroph\| | Yeast | NULL | Probable | As nectar-tap_saprotroph (Põlme et al. 2020); Decay sugar-rich substrates (Põlme et al. 2020) | Põlme S, et al. 2020. Fungal Diversity 105:1-16 ((https://doi.org/10.1007/s13225-020-00466-2)) |
| Otu59779 | 1 | k__Fungi(100);p__Ascomycota(100);c__Saccharomycetes(100);o__Saccharomycetales(100);f__Metschnikowiaceae(100);g__Metschnikowia(100);s__Metschnikowia_pulcherrima(100); | Metschnikowia | 13 | Saprotroph | \|Undefined Saprotroph\| | Yeast | NULL | Probable | As nectar-tap_saprotroph (Põlme et al. 2020); Decay sugar-rich substrates (Põlme et al. 2020) | Põlme S, et al. 2020. Fungal Diversity 105:1-16 ((https://doi.org/10.1007/s13225-020-00466-2)) |
| Otu59780 | 1 | k__Fungi(100);p__Ascomycota(100);c__Saccharomycetes(100);o__Saccharomycetales(100);f__Metschnikowiaceae(100);g__Metschnikowia(100);s__Metschnikowia_pulcherrima(100); | Metschnikowia | 13 | Saprotroph | \|Undefined Saprotroph\| | Yeast | NULL | Probable | As nectar-tap_saprotroph (Põlme et al. 2020); Decay sugar-rich substrates (Põlme et al. 2020) | Põlme S, et al. 2020. Fungal Diversity 105:1-16 ((https://doi.org/10.1007/s13225-020-00466-2)) |
| Otu59781 | 1 | k__Fungi(100);p__Ascomycota(100);c__Saccharomycetes(100);o__Saccharomycetales(100);f__Metschnikowiaceae(100);g__Metschnikowia(100);g__Metschnikowia_unclassified(100); | Metschnikowia | 13 | Saprotroph | \|Undefined Saprotroph\| | Yeast | NULL | Probable | As nectar-tap_saprotroph (Põlme et al. 2020); Decay sugar-rich substrates (Põlme et al. 2020) | Põlme S, et al. 2020. Fungal Diversity 105:1-16 ((https://doi.org/10.1007/s13225-020-00466-2)) |
| Otu59782 | 1 | k__Fungi(100);p__Ascomycota(100);c__Saccharomycetes(100);o__Saccharomycetales(100);f__Metschnikowiaceae(100);g__Metschnikowia(100);s__Metschnikowia_pulcherrima(100); | Metschnikowia | 13 | Saprotroph | \|Undefined Saprotroph\| | Yeast | NULL | Probable | As nectar-tap_saprotroph (Põlme et al. 2020); Decay sugar-rich substrates (Põlme et al. 2020) | Põlme S, et al. 2020. Fungal Diversity 105:1-16 ((https://doi.org/10.1007/s13225-020-00466-2)) |
| Otu59783 | 1 | k__Fungi(100);p__Ascomycota(100);c__Saccharomycetes(100);o__Saccharomycetales(100);f__Metschnikowiaceae(100);g__Metschnikowia(100);s__Metschnikowia_pulcherrima(100); | Metschnikowia | 13 | Saprotroph | \|Undefined Saprotroph\| | Yeast | NULL | Probable | As nectar-tap_saprotroph (Põlme et al. 2020); Decay sugar-rich substrates (Põlme et al. 2020) | Põlme S, et al. 2020. Fungal Diversity 105:1-16 ((https://doi.org/10.1007/s13225-020-00466-2)) |
| Otu59784 | 1 | k__Fungi(100);p__Ascomycota(100);c__Saccharomycetes(100);o__Saccharomycetales(100);f__Metschnikowiaceae(100);g__Metschnikowia(100);s__Metschnikowia_pulcherrima(100); | Metschnikowia | 13 | Saprotroph | \|Undefined Saprotroph\| | Yeast | NULL | Probable | As nectar-tap_saprotroph (Põlme et al. 2020); Decay sugar-rich substrates (Põlme et al. 2020) | Põlme S, et al. 2020. Fungal Diversity 105:1-16 ((https://doi.org/10.1007/s13225-020-00466-2)) |
| Otu59785 | 1 | k__Fungi(100);p__Ascomycota(100);c__Saccharomycetes(100);o__Saccharomycetales(100);f__Metschnikowiaceae(100);g__Metschnikowia(100);s__Metschnikowia_pulcherrima(100); | Metschnikowia | 13 | Saprotroph | \|Undefined Saprotroph\| | Yeast | NULL | Probable | As nectar-tap_saprotroph (Põlme et al. 2020); Decay sugar-rich substrates (Põlme et al. 2020) | Põlme S, et al. 2020. Fungal Diversity 105:1-16 ((https://doi.org/10.1007/s13225-020-00466-2)) |
| Otu59786 | 1 | k__Fungi(100);p__Ascomycota(100);c__Saccharomycetes(100);o__Saccharomycetales(100);f__Metschnikowiaceae(100);g__Metschnikowia(100);g__Metschnikowia_unclassified(100); | Metschnikowia | 13 | Saprotroph | \|Undefined Saprotroph\| | Yeast | NULL | Probable | As nectar-tap_saprotroph (Põlme et al. 2020); Decay sugar-rich substrates (Põlme et al. 2020) | Põlme S, et al. 2020. Fungal Diversity 105:1-16 ((https://doi.org/10.1007/s13225-020-00466-2)) |
| Otu59787 | 1 | k__Fungi(100);p__Ascomycota(100);c__Saccharomycetes(100);o__Saccharomycetales(100);f__Metschnikowiaceae(100);g__Metschnikowia(100);s__Metschnikowia_pulcherrima(100); | Metschnikowia | 13 | Saprotroph | \|Undefined Saprotroph\| | Yeast | NULL | Probable | As nectar-tap_saprotroph (Põlme et al. 2020); Decay sugar-rich substrates (Põlme et al. 2020) | Põlme S, et al. 2020. Fungal Diversity 105:1-16 ((https://doi.org/10.1007/s13225-020-00466-2)) |
| Otu59788 | 1 | k__Fungi(100);p__Ascomycota(100);c__Saccharomycetes(100);o__Saccharomycetales(100);f__Metschnikowiaceae(100);g__Metschnikowia(100);s__Metschnikowia_pulcherrima(100); | Metschnikowia | 13 | Saprotroph | \|Undefined Saprotroph\| | Yeast | NULL | Probable | As nectar-tap_saprotroph (Põlme et al. 2020); Decay sugar-rich substrates (Põlme et al. 2020) | Põlme S, et al. 2020. Fungal Diversity 105:1-16 ((https://doi.org/10.1007/s13225-020-00466-2)) |
| Otu59789 | 1 | k__Fungi(100);p__Ascomycota(100);c__Saccharomycetes(100);o__Saccharomycetales(100);f__Metschnikowiaceae(100);g__Metschnikowia(100);g__Metschnikowia_unclassified(100); | Metschnikowia | 13 | Saprotroph | \|Undefined Saprotroph\| | Yeast | NULL | Probable | As nectar-tap_saprotroph (Põlme et al. 2020); Decay sugar-rich substrates (Põlme et al. 2020) | Põlme S, et al. 2020. Fungal Diversity 105:1-16 ((https://doi.org/10.1007/s13225-020-00466-2)) |
| Otu59790 | 1 | k__Fungi(100);p__Ascomycota(100);c__Saccharomycetes(100);o__Saccharomycetales(100);f__Metschnikowiaceae(100);g__Metschnikowia(100);g__Metschnikowia_unclassified(100); | Metschnikowia | 13 | Saprotroph | \|Undefined Saprotroph\| | Yeast | NULL | Probable | As nectar-tap_saprotroph (Põlme et al. 2020); Decay sugar-rich substrates (Põlme et al. 2020) | Põlme S, et al. 2020. Fungal Diversity 105:1-16 ((https://doi.org/10.1007/s13225-020-00466-2)) |
| Otu59791 | 1 | k__Fungi(100);p__Ascomycota(100);c__Saccharomycetes(100);o__Saccharomycetales(100);f__Metschnikowiaceae(100);g__Metschnikowia(100);s__Metschnikowia_pulcherrima(100); | Metschnikowia | 13 | Saprotroph | \|Undefined Saprotroph\| | Yeast | NULL | Probable | As nectar-tap_saprotroph (Põlme et al. 2020); Decay sugar-rich substrates (Põlme et al. 2020) | Põlme S, et al. 2020. Fungal Diversity 105:1-16 ((https://doi.org/10.1007/s13225-020-00466-2)) |
| Otu59792 | 1 | k__Fungi(100);p__Ascomycota(100);c__Saccharomycetes(100);o__Saccharomycetales(100);f__Metschnikowiaceae(100);g__Metschnikowia(100);s__Metschnikowia_pulcherrima(100); | Metschnikowia | 13 | Saprotroph | \|Undefined Saprotroph\| | Yeast | NULL | Probable | As nectar-tap_saprotroph (Põlme et al. 2020); Decay sugar-rich substrates (Põlme et al. 2020) | Põlme S, et al. 2020. Fungal Diversity 105:1-16 ((https://doi.org/10.1007/s13225-020-00466-2)) |
| Otu59793 | 1 | k__Fungi(100);p__Ascomycota(100);c__Saccharomycetes(100);o__Saccharomycetales(100);f__Metschnikowiaceae(100);g__Metschnikowia(100);g__Metschnikowia_unclassified(100); | Metschnikowia | 13 | Saprotroph | \|Undefined Saprotroph\| | Yeast | NULL | Probable | As nectar-tap_saprotroph (Põlme et al. 2020); Decay sugar-rich substrates (Põlme et al. 2020) | Põlme S, et al. 2020. Fungal Diversity 105:1-16 ((https://doi.org/10.1007/s13225-020-00466-2)) |
| Otu59794 | 1 | k__Fungi(100);p__Ascomycota(100);c__Saccharomycetes(100);o__Saccharomycetales(100);f__Metschnikowiaceae(100);g__Metschnikowia(100);s__Metschnikowia_pulcherrima(100); | Metschnikowia | 13 | Saprotroph | \|Undefined Saprotroph\| | Yeast | NULL | Probable | As nectar-tap_saprotroph (Põlme et al. 2020); Decay sugar-rich substrates (Põlme et al. 2020) | Põlme S, et al. 2020. Fungal Diversity 105:1-16 ((https://doi.org/10.1007/s13225-020-00466-2)) |
| Otu59795 | 1 | k__Fungi(100);p__Ascomycota(100);c__Saccharomycetes(100);o__Saccharomycetales(100);f__Metschnikowiaceae(100);g__Metschnikowia(100);s__Metschnikowia_pulcherrima(100); | Metschnikowia | 13 | Saprotroph | \|Undefined Saprotroph\| | Yeast | NULL | Probable | As nectar-tap_saprotroph (Põlme et al. 2020); Decay sugar-rich substrates (Põlme et al. 2020) | Põlme S, et al. 2020. Fungal Diversity 105:1-16 ((https://doi.org/10.1007/s13225-020-00466-2)) |
| Otu59796 | 1 | k__Fungi(100);p__Ascomycota(100);c__Saccharomycetes(100);o__Saccharomycetales(100);f__Metschnikowiaceae(100);g__Metschnikowia(100);s__Metschnikowia_pulcherrima(100); | Metschnikowia | 13 | Saprotroph | \|Undefined Saprotroph\| | Yeast | NULL | Probable | As nectar-tap_saprotroph (Põlme et al. 2020); Decay sugar-rich substrates (Põlme et al. 2020) | Põlme S, et al. 2020. Fungal Diversity 105:1-16 ((https://doi.org/10.1007/s13225-020-00466-2)) |
| Otu59797 | 1 | k__Fungi(100);p__Ascomycota(100);c__Saccharomycetes(100);o__Saccharomycetales(100);f__Metschnikowiaceae(100);g__Metschnikowia(100);s__Metschnikowia_pulcherrima(100); | Metschnikowia | 13 | Saprotroph | \|Undefined Saprotroph\| | Yeast | NULL | Probable | As nectar-tap_saprotroph (Põlme et al. 2020); Decay sugar-rich substrates (Põlme et al. 2020) | Põlme S, et al. 2020. Fungal Diversity 105:1-16 ((https://doi.org/10.1007/s13225-020-00466-2)) |
| Otu59798 | 1 | k__Fungi(100);p__Ascomycota(100);c__Saccharomycetes(100);o__Saccharomycetales(100);f__Metschnikowiaceae(100);g__Metschnikowia(100);s__Metschnikowia_pulcherrima(100); | Metschnikowia | 13 | Saprotroph | \|Undefined Saprotroph\| | Yeast | NULL | Probable | As nectar-tap_saprotroph (Põlme et al. 2020); Decay sugar-rich substrates (Põlme et al. 2020) | Põlme S, et al. 2020. Fungal Diversity 105:1-16 ((https://doi.org/10.1007/s13225-020-00466-2)) |
| Otu59799 | 1 | k__Fungi(100);p__Ascomycota(100);c__Saccharomycetes(100);o__Saccharomycetales(100);f__Metschnikowiaceae(100);g__Metschnikowia(100);s__Metschnikowia_pulcherrima(100); | Metschnikowia | 13 | Saprotroph | \|Undefined Saprotroph\| | Yeast | NULL | Probable | As nectar-tap_saprotroph (Põlme et al. 2020); Decay sugar-rich substrates (Põlme et al. 2020) | Põlme S, et al. 2020. Fungal Diversity 105:1-16 ((https://doi.org/10.1007/s13225-020-00466-2)) |
| Otu59800 | 1 | k__Fungi(100);p__Ascomycota(100);c__Saccharomycetes(100);o__Saccharomycetales(100);f__Metschnikowiaceae(100);g__Metschnikowia(100);s__Metschnikowia_pulcherrima(100); | Metschnikowia | 13 | Saprotroph | \|Undefined Saprotroph\| | Yeast | NULL | Probable | As nectar-tap_saprotroph (Põlme et al. 2020); Decay sugar-rich substrates (Põlme et al. 2020) | Põlme S, et al. 2020. Fungal Diversity 105:1-16 ((https://doi.org/10.1007/s13225-020-00466-2)) |
| Otu59801 | 1 | k__Fungi(100);p__Ascomycota(100);c__Saccharomycetes(100);o__Saccharomycetales(100);f__Metschnikowiaceae(100);g__Metschnikowia(100);s__Metschnikowia_pulcherrima(100); | Metschnikowia | 13 | Saprotroph | \|Undefined Saprotroph\| | Yeast | NULL | Probable | As nectar-tap_saprotroph (Põlme et al. 2020); Decay sugar-rich substrates (Põlme et al. 2020) | Põlme S, et al. 2020. Fungal Diversity 105:1-16 ((https://doi.org/10.1007/s13225-020-00466-2)) |
| Otu59802 | 1 | k__Fungi(100);p__Ascomycota(100);c__Saccharomycetes(100);o__Saccharomycetales(100);f__Metschnikowiaceae(100);g__Metschnikowia(100);s__Metschnikowia_pulcherrima(100); | Metschnikowia | 13 | Saprotroph | \|Undefined Saprotroph\| | Yeast | NULL | Probable | As nectar-tap_saprotroph (Põlme et al. 2020); Decay sugar-rich substrates (Põlme et al. 2020) | Põlme S, et al. 2020. Fungal Diversity 105:1-16 ((https://doi.org/10.1007/s13225-020-00466-2)) |
| Otu59803 | 1 | k__Fungi(100);p__Ascomycota(100);c__Saccharomycetes(100);o__Saccharomycetales(100);f__Metschnikowiaceae(100);g__Metschnikowia(100);s__Metschnikowia_pulcherrima(100); | Metschnikowia | 13 | Saprotroph | \|Undefined Saprotroph\| | Yeast | NULL | Probable | As nectar-tap_saprotroph (Põlme et al. 2020); Decay sugar-rich substrates (Põlme et al. 2020) | Põlme S, et al. 2020. Fungal Diversity 105:1-16 ((https://doi.org/10.1007/s13225-020-00466-2)) |
| Otu59804 | 1 | k__Fungi(100);p__Ascomycota(100);c__Saccharomycetes(100);o__Saccharomycetales(100);f__Metschnikowiaceae(100);g__Metschnikowia(100);s__Metschnikowia_pulcherrima(100); | Metschnikowia | 13 | Saprotroph | \|Undefined Saprotroph\| | Yeast | NULL | Probable | As nectar-tap_saprotroph (Põlme et al. 2020); Decay sugar-rich substrates (Põlme et al. 2020) | Põlme S, et al. 2020. Fungal Diversity 105:1-16 ((https://doi.org/10.1007/s13225-020-00466-2)) |
| Otu59805 | 1 | k__Fungi(100);p__Ascomycota(100);c__Saccharomycetes(100);o__Saccharomycetales(100);f__Metschnikowiaceae(100);g__Metschnikowia(100);s__Metschnikowia_pulcherrima(100); | Metschnikowia | 13 | Saprotroph | \|Undefined Saprotroph\| | Yeast | NULL | Probable | As nectar-tap_saprotroph (Põlme et al. 2020); Decay sugar-rich substrates (Põlme et al. 2020) | Põlme S, et al. 2020. Fungal Diversity 105:1-16 ((https://doi.org/10.1007/s13225-020-00466-2)) |
| Otu59806 | 1 | k__Fungi(100);p__Ascomycota(100);c__Saccharomycetes(100);o__Saccharomycetales(100);f__Metschnikowiaceae(100);g__Metschnikowia(100);s__Metschnikowia_pulcherrima(100); | Metschnikowia | 13 | Saprotroph | \|Undefined Saprotroph\| | Yeast | NULL | Probable | As nectar-tap_saprotroph (Põlme et al. 2020); Decay sugar-rich substrates (Põlme et al. 2020) | Põlme S, et al. 2020. Fungal Diversity 105:1-16 ((https://doi.org/10.1007/s13225-020-00466-2)) |

| | | | | | | | | | | | |
|---|---|---|---|---|---|---|---|---|---|---|---|
| Otu59809 | 1 | k__Fungi(100);p__Ascomycota(100);c__Saccharomycetes(100);o__Saccharomycetales(100);f__Metschnikowiaceae(100);g__Metschnikowia(100);s__Metschnikowia_pulcherrima(100); | Metschnikowia | 13 | Saprotroph | \|Undefined Saprotroph\| | Yeast | NULL | Probable | As nectar-tap_saprotroph (Põlme et al. 2020); Decay sugar-rich substrates (Põlme et al. 2020) | Põlme S, et al. 2020. Fungal Diversity 105:1-16 ((https://doi.org/10.1007/s13225-020-00466-2)) |
| Otu59810 | 1 | k__Fungi(100);p__Ascomycota(100);c__Saccharomycetes(100);o__Saccharomycetales(100);f__Metschnikowiaceae(100);g__Metschnikowia(100);s__Metschnikowia_pulcherrima(100); | Metschnikowia | 13 | Saprotroph | \|Undefined Saprotroph\| | Yeast | NULL | Probable | As nectar-tap_saprotroph (Põlme et al. 2020); Decay sugar-rich substrates (Põlme et al. 2020) | Põlme S, et al. 2020. Fungal Diversity 105:1-16 ((https://doi.org/10.1007/s13225-020-00466-2)) |
| Otu59811 | 1 | k__Fungi(100);p__Ascomycota(100);c__Saccharomycetes(100);o__Saccharomycetales(100);f__Metschnikowiaceae(100);g__Metschnikowia(100);s__Metschnikowia_pulcherrima(100); | Metschnikowia | 13 | Saprotroph | \|Undefined Saprotroph\| | Yeast | NULL | Probable | As nectar-tap_saprotroph (Põlme et al. 2020); Decay sugar-rich substrates (Põlme et al. 2020) | Põlme S, et al. 2020. Fungal Diversity 105:1-16 ((https://doi.org/10.1007/s13225-020-00466-2)) |
| Otu59812 | 1 | k__Fungi(100);p__Ascomycota(100);c__Saccharomycetes(100);o__Saccharomycetales(100);f__Metschnikowiaceae(100);g__Metschnikowia(100);g__Metschnikowia_unclassified(100); | Metschnikowia | 13 | Saprotroph | \|Undefined Saprotroph\| | Yeast | NULL | Probable | As nectar-tap_saprotroph (Põlme et al. 2020); Decay sugar-rich substrates (Põlme et al. 2020) | Põlme S, et al. 2020. Fungal Diversity 105:1-16 ((https://doi.org/10.1007/s13225-020-00466-2)) |
| Otu59813 | 1 | k__Fungi(100);p__Ascomycota(100);c__Saccharomycetes(100);o__Saccharomycetales(100);f__Metschnikowiaceae(100);g__Metschnikowia(100);s__Metschnikowia_pulcherrima(100); | Metschnikowia | 13 | Saprotroph | \|Undefined Saprotroph\| | Yeast | NULL | Probable | As nectar-tap_saprotroph (Põlme et al. 2020); Decay sugar-rich substrates (Põlme et al. 2020) | Põlme S, et al. 2020. Fungal Diversity 105:1-16 ((https://doi.org/10.1007/s13225-020-00466-2)) |
| Otu59814 | 1 | k__Fungi(100);p__Ascomycota(100);c__Saccharomycetes(100);o__Saccharomycetales(100);f__Metschnikowiaceae(100);g__Metschnikowia(100);s__Metschnikowia_pulcherrima(100); | Metschnikowia | 13 | Saprotroph | \|Undefined Saprotroph\| | Yeast | NULL | Probable | As nectar-tap_saprotroph (Põlme et al. 2020); Decay sugar-rich substrates (Põlme et al. 2020) | Põlme S, et al. 2020. Fungal Diversity 105:1-16 ((https://doi.org/10.1007/s13225-020-00466-2)) |
| Otu59815 | 1 | k__Fungi(100);p__Ascomycota(100);c__Saccharomycetes(100);o__Saccharomycetales(100);f__Metschnikowiaceae(100);g__Metschnikowia(100);g__Metschnikowia_unclassified(100); | Metschnikowia | 13 | Saprotroph | \|Undefined Saprotroph\| | Yeast | NULL | Probable | As nectar-tap_saprotroph (Põlme et al. 2020); Decay sugar-rich substrates (Põlme et al. 2020) | Põlme S, et al. 2020. Fungal Diversity 105:1-16 ((https://doi.org/10.1007/s13225-020-00466-2)) |
| Otu59816 | 1 | k__Fungi(100);p__Ascomycota(100);c__Saccharomycetes(100);o__Saccharomycetales(100);f__Metschnikowiaceae(100);g__Metschnikowia(100);g__Metschnikowia_unclassified(100); | Metschnikowia | 13 | Saprotroph | \|Undefined Saprotroph\| | Yeast | NULL | Probable | As nectar-tap_saprotroph (Põlme et al. 2020); Decay sugar-rich substrates (Põlme et al. 2020) | Põlme S, et al. 2020. Fungal Diversity 105:1-16 ((https://doi.org/10.1007/s13225-020-00466-2)) |
| Otu59817 | 1 | k__Fungi(100);p__Ascomycota(100);c__Saccharomycetes(100);o__Saccharomycetales(100);f__Metschnikowiaceae(100);g__Metschnikowia(100);g__Metschnikowia_unclassified(100); | Metschnikowia | 13 | Saprotroph | \|Undefined Saprotroph\| | Yeast | NULL | Probable | As nectar-tap_saprotroph (Põlme et al. 2020); Decay sugar-rich substrates (Põlme et al. 2020) | Põlme S, et al. 2020. Fungal Diversity 105:1-16 ((https://doi.org/10.1007/s13225-020-00466-2)) |
| Otu59818 | 1 | k__Fungi(100);p__Ascomycota(100);c__Saccharomycetes(100);o__Saccharomycetales(100);f__Metschnikowiaceae(100);g__Metschnikowia(100);s__Metschnikowia_pulcherrima(100); | Metschnikowia | 13 | Saprotroph | \|Undefined Saprotroph\| | Yeast | NULL | Probable | As nectar-tap_saprotroph (Põlme et al. 2020); Decay sugar-rich substrates (Põlme et al. 2020) | Põlme S, et al. 2020. Fungal Diversity 105:1-16 ((https://doi.org/10.1007/s13225-020-00466-2)) |
| Otu59819 | 1 | k__Fungi(100);p__Ascomycota(100);c__Saccharomycetes(100);o__Saccharomycetales(100);f__Metschnikowiaceae(100);g__Metschnikowia(100);s__Metschnikowia_pulcherrima(100); | Metschnikowia | 13 | Saprotroph | \|Undefined Saprotroph\| | Yeast | NULL | Probable | As nectar-tap_saprotroph (Põlme et al. 2020); Decay sugar-rich substrates (Põlme et al. 2020) | Põlme S, et al. 2020. Fungal Diversity 105:1-16 ((https://doi.org/10.1007/s13225-020-00466-2)) |
| Otu59820 | 1 | k__Fungi(100);p__Ascomycota(100);c__Saccharomycetes(100);o__Saccharomycetales(100);f__Metschnikowiaceae(100);g__Metschnikowia(100);s__Metschnikowia_pulcherrima(100); | Metschnikowia | 13 | Saprotroph | \|Undefined Saprotroph\| | Yeast | NULL | Probable | As nectar-tap_saprotroph (Põlme et al. 2020); Decay sugar-rich substrates (Põlme et al. 2020) | Põlme S, et al. 2020. Fungal Diversity 105:1-16 ((https://doi.org/10.1007/s13225-020-00466-2)) |
| Otu59821 | 1 | k__Fungi(100);p__Ascomycota(100);c__Saccharomycetes(100);o__Saccharomycetales(100);f__Metschnikowiaceae(100);g__Metschnikowia(100);s__Metschnikowia_pulcherrima(100); | Metschnikowia | 13 | Saprotroph | \|Undefined Saprotroph\| | Yeast | NULL | Probable | As nectar-tap_saprotroph (Põlme et al. 2020); Decay sugar-rich substrates (Põlme et al. 2020) | Põlme S, et al. 2020. Fungal Diversity 105:1-16 ((https://doi.org/10.1007/s13225-020-00466-2)) |
| Otu59822 | 1 | k__Fungi(100);p__Ascomycota(100);c__Saccharomycetes(100);o__Saccharomycetales(100);f__Metschnikowiaceae(100);g__Metschnikowia(100);s__Metschnikowia_pulcherrima(100); | Metschnikowia | 13 | Saprotroph | \|Undefined Saprotroph\| | Yeast | NULL | Probable | As nectar-tap_saprotroph (Põlme et al. 2020); Decay sugar-rich substrates (Põlme et al. 2020) | Põlme S, et al. 2020. Fungal Diversity 105:1-16 ((https://doi.org/10.1007/s13225-020-00466-2)) |
| Otu59823 | 1 | k__Fungi(100);p__Ascomycota(100);c__Saccharomycetes(100);o__Saccharomycetales(100);f__Metschnikowiaceae(100);g__Metschnikowia(100);s__Metschnikowia_pulcherrima(100); | Metschnikowia | 13 | Saprotroph | \|Undefined Saprotroph\| | Yeast | NULL | Probable | As nectar-tap_saprotroph (Põlme et al. 2020); Decay sugar-rich substrates (Põlme et al. 2020) | Põlme S, et al. 2020. Fungal Diversity 105:1-16 ((https://doi.org/10.1007/s13225-020-00466-2)) |
| Otu59824 | 1 | k__Fungi(100);p__Ascomycota(100);c__Saccharomycetes(100);o__Saccharomycetales(100);f__Metschnikowiaceae(100);g__Metschnikowia(100);s__Metschnikowia_pulcherrima(100); | Metschnikowia | 13 | Saprotroph | \|Undefined Saprotroph\| | Yeast | NULL | Probable | As nectar-tap_saprotroph (Põlme et al. 2020); Decay sugar-rich substrates (Põlme et al. 2020) | Põlme S, et al. 2020. Fungal Diversity 105:1-16 ((https://doi.org/10.1007/s13225-020-00466-2)) |
| Otu59825 | 1 | k__Fungi(100);p__Ascomycota(100);c__Saccharomycetes(100);o__Saccharomycetales(100);f__Metschnikowiaceae(100);g__Metschnikowia(100);s__Metschnikowia_pulcherrima(100); | Metschnikowia | 13 | Saprotroph | \|Undefined Saprotroph\| | Yeast | NULL | Probable | As nectar-tap_saprotroph (Põlme et al. 2020); Decay sugar-rich substrates (Põlme et al. 2020) | Põlme S, et al. 2020. Fungal Diversity 105:1-16 ((https://doi.org/10.1007/s13225-020-00466-2)) |
| Otu59826 | 1 | k__Fungi(100);p__Ascomycota(100);c__Saccharomycetes(100);o__Saccharomycetales(100);f__Metschnikowiaceae(100);g__Metschnikowia(100);s__Metschnikowia_pulcherrima(100); | Metschnikowia | 13 | Saprotroph | \|Undefined Saprotroph\| | Yeast | NULL | Probable | As nectar-tap_saprotroph (Põlme et al. 2020); Decay sugar-rich substrates (Põlme et al. 2020) | Põlme S, et al. 2020. Fungal Diversity 105:1-16 ((https://doi.org/10.1007/s13225-020-00466-2)) |
| Otu59827 | 1 | k__Fungi(100);p__Ascomycota(100);c__Saccharomycetes(100);o__Saccharomycetales(100);f__Metschnikowiaceae(100);g__Metschnikowia(100);s__Metschnikowia_pulcherrima(100); | Metschnikowia | 13 | Saprotroph | \|Undefined Saprotroph\| | Yeast | NULL | Probable | As nectar-tap_saprotroph (Põlme et al. 2020); Decay sugar-rich substrates (Põlme et al. 2020) | Põlme S, et al. 2020. Fungal Diversity 105:1-16 ((https://doi.org/10.1007/s13225-020-00466-2)) |
| Otu59828 | 1 | k__Fungi(100);p__Ascomycota(100);c__Saccharomycetes(100);o__Saccharomycetales(100);f__Metschnikowiaceae(100);g__Metschnikowia(100);s__Metschnikowia_pulcherrima(100); | Metschnikowia | 13 | Saprotroph | \|Undefined Saprotroph\| | Yeast | NULL | Probable | As nectar-tap_saprotroph (Põlme et al. 2020); Decay sugar-rich substrates (Põlme et al. 2020) | Põlme S, et al. 2020. Fungal Diversity 105:1-16 ((https://doi.org/10.1007/s13225-020-00466-2)) |
| Otu59829 | 1 | k__Fungi(100);p__Ascomycota(100);c__Saccharomycetes(100);o__Saccharomycetales(100);f__Metschnikowiaceae(100);g__Metschnikowia(100);s__Metschnikowia_pulcherrima(100); | Metschnikowia | 13 | Saprotroph | \|Undefined Saprotroph\| | Yeast | NULL | Probable | As nectar-tap_saprotroph (Põlme et al. 2020); Decay sugar-rich substrates (Põlme et al. 2020) | Põlme S, et al. 2020. Fungal Diversity 105:1-16 ((https://doi.org/10.1007/s13225-020-00466-2)) |
| Otu59830 | 1 | k__Fungi(100);p__Ascomycota(100);c__Saccharomycetes(100);o__Saccharomycetales(100);f__Metschnikowiaceae(100);g__Metschnikowia(100);s__Metschnikowia_pulcherrima(100); | Metschnikowia | 13 | Saprotroph | \|Undefined Saprotroph\| | Yeast | NULL | Probable | As nectar-tap_saprotroph (Põlme et al. 2020); Decay sugar-rich substrates (Põlme et al. 2020) | Põlme S, et al. 2020. Fungal Diversity 105:1-16 ((https://doi.org/10.1007/s13225-020-00466-2)) |
| Otu59831 | 1 | k__Fungi(100);p__Ascomycota(100);c__Saccharomycetes(100);o__Saccharomycetales(100);f__Metschnikowiaceae(100);g__Metschnikowia(100);s__Metschnikowia_pulcherrima(100); | Metschnikowia | 13 | Saprotroph | \|Undefined Saprotroph\| | Yeast | NULL | Probable | As nectar-tap_saprotroph (Põlme et al. 2020); Decay sugar-rich substrates (Põlme et al. 2020) | Põlme S, et al. 2020. Fungal Diversity 105:1-16 ((https://doi.org/10.1007/s13225-020-00466-2)) |
| Otu59833 | 1 | k__Fungi(100);p__Ascomycota(100);c__Saccharomycetes(100);o__Saccharomycetales(100);f__Metschnikowiaceae(100);g__Metschnikowia(100);s__Metschnikowia_pulcherrima(100); | Metschnikowia | 13 | Saprotroph | \|Undefined Saprotroph\| | Yeast | NULL | Probable | As nectar-tap_saprotroph (Põlme et al. 2020); Decay sugar-rich substrates (Põlme et al. 2020) | Põlme S, et al. 2020. Fungal Diversity 105:1-16 ((https://doi.org/10.1007/s13225-020-00466-2)) |
| Otu59834 | 1 | k__Fungi(100);p__Ascomycota(100);c__Saccharomycetes(100);o__Saccharomycetales(100);f__Metschnikowiaceae(100);g__Metschnikowia(100);s__Metschnikowia_pulcherrima(100); | Metschnikowia | 13 | Saprotroph | \|Undefined Saprotroph\| | Yeast | NULL | Probable | As nectar-tap_saprotroph (Põlme et al. 2020); Decay sugar-rich substrates (Põlme et al. 2020) | Põlme S, et al. 2020. Fungal Diversity 105:1-16 ((https://doi.org/10.1007/s13225-020-00466-2)) |
| Otu59835 | 1 | k__Fungi(100);p__Ascomycota(100);c__Saccharomycetes(100);o__Saccharomycetales(100);f__Metschnikowiaceae(100);g__Metschnikowia(100);s__Metschnikowia_pulcherrima(100); | Metschnikowia | 13 | Saprotroph | \|Undefined Saprotroph\| | Yeast | NULL | Probable | As nectar-tap_saprotroph (Põlme et al. 2020); Decay sugar-rich substrates (Põlme et al. 2020) | Põlme S, et al. 2020. Fungal Diversity 105:1-16 ((https://doi.org/10.1007/s13225-020-00466-2)) |
| Otu59836 | 1 | k__Fungi(100);p__Ascomycota(100);c__Saccharomycetes(100);o__Saccharomycetales(100);f__Metschnikowiaceae(100);g__Metschnikowia(100);s__Metschnikowia_pulcherrima(100); | Metschnikowia | 13 | Saprotroph | \|Undefined Saprotroph\| | Yeast | NULL | Probable | As nectar-tap_saprotroph (Põlme et al. 2020); Decay sugar-rich substrates (Põlme et al. 2020) | Põlme S, et al. 2020. Fungal Diversity 105:1-16 ((https://doi.org/10.1007/s13225-020-00466-2)) |
| Otu59837 | 1 | k__Fungi(100);p__Ascomycota(100);c__Saccharomycetes(100);o__Saccharomycetales(100);f__Metschnikowiaceae(100);g__Metschnikowia(100);s__Metschnikowia_pulcherrima(100); | Metschnikowia | 13 | Saprotroph | \|Undefined Saprotroph\| | Yeast | NULL | Probable | As nectar-tap_saprotroph (Põlme et al. 2020); Decay sugar-rich substrates (Põlme et al. 2020) | Põlme S, et al. 2020. Fungal Diversity 105:1-16 ((https://doi.org/10.1007/s13225-020-00466-2)) |
| Otu59838 | 1 | k__Fungi(100);p__Ascomycota(100);c__Saccharomycetes(100);o__Saccharomycetales(100);f__Metschnikowiaceae(100);g__Metschnikowia(100);s__Metschnikowia_pulcherrima(100); | Metschnikowia | 13 | Saprotroph | \|Undefined Saprotroph\| | Yeast | NULL | Probable | As nectar-tap_saprotroph (Põlme et al. 2020); Decay sugar-rich substrates (Põlme et al. 2020) | Põlme S, et al. 2020. Fungal Diversity 105:1-16 ((https://doi.org/10.1007/s13225-020-00466-2)) |
| Otu59839 | 1 | k__Fungi(100);p__Ascomycota(100);c__Saccharomycetes(100);o__Saccharomycetales(100);f__Metschnikowiaceae(100);g__Metschnikowia(100);g__Metschnikowia_unclassified(100); | Metschnikowia | 13 | Saprotroph | \|Undefined Saprotroph\| | Yeast | NULL | Probable | As nectar-tap_saprotroph (Põlme et al. 2020); Decay sugar-rich substrates (Põlme et al. 2020) | Põlme S, et al. 2020. Fungal Diversity 105:1-16 ((https://doi.org/10.1007/s13225-020-00466-2)) |
| Otu59840 | 1 | k__Fungi(100);p__Ascomycota(100);c__Saccharomycetes(100);o__Saccharomycetales(100);f__Metschnikowiaceae(100);g__Metschnikowia(100);s__Metschnikowia_pulcherrima(100); | Metschnikowia | 13 | Saprotroph | \|Undefined Saprotroph\| | Yeast | NULL | Probable | As nectar-tap_saprotroph (Põlme et al. 2020); Decay sugar-rich substrates (Põlme et al. 2020) | Põlme S, et al. 2020. Fungal Diversity 105:1-16 ((https://doi.org/10.1007/s13225-020-00466-2)) |
| Otu59841 | 1 | k__Fungi(100);p__Ascomycota(100);c__Saccharomycetes(100);o__Saccharomycetales(100);f__Metschnikowiaceae(100);g__Metschnikowia(100);s__Metschnikowia_pulcherrima(100); | Metschnikowia | 13 | Saprotroph | \|Undefined Saprotroph\| | Yeast | NULL | Probable | As nectar-tap_saprotroph (Põlme et al. 2020); Decay sugar-rich substrates (Põlme et al. 2020) | Põlme S, et al. 2020. Fungal Diversity 105:1-16 ((https://doi.org/10.1007/s13225-020-00466-2)) |
| Otu59844 | 1 | k__Fungi(100);p__Ascomycota(100);c__Saccharomycetes(100);o__Saccharomycetales(100);f__Metschnikowiaceae(100);g__Metschnikowia(100);s__Metschnikowia_pulcherrima(100); | Metschnikowia | 13 | Saprotroph | \|Undefined Saprotroph\| | Yeast | NULL | Probable | As nectar-tap_saprotroph (Põlme et al. 2020); Decay sugar-rich substrates (Põlme et al. 2020) | Põlme S, et al. 2020. Fungal Diversity 105:1-16 ((https://doi.org/10.1007/s13225-020-00466-2)) |
| Otu59845 | 1 | k__Fungi(100);p__Ascomycota(100);c__Saccharomycetes(100);o__Saccharomycetales(100);f__Metschnikowiaceae(100);g__Metschnikowia(100);s__Metschnikowia_pulcherrima(100); | Metschnikowia | 13 | Saprotroph | \|Undefined Saprotroph\| | Yeast | NULL | Probable | As nectar-tap_saprotroph (Põlme et al. 2020); Decay sugar-rich substrates (Põlme et al. 2020) | Põlme S, et al. 2020. Fungal Diversity 105:1-16 ((https://doi.org/10.1007/s13225-020-00466-2)) |
| Otu59846 | 1 | k__Fungi(100);p__Ascomycota(100);c__Saccharomycetes(100);o__Saccharomycetales(100);f__Metschnikowiaceae(100);g__Metschnikowia(100);g__Metschnikowia_unclassified(100); | Metschnikowia | 13 | Saprotroph | \|Undefined Saprotroph\| | Yeast | NULL | Probable | As nectar-tap_saprotroph (Põlme et al. 2020); Decay sugar-rich substrates (Põlme et al. 2020) | Põlme S, et al. 2020. Fungal Diversity 105:1-16 ((https://doi.org/10.1007/s13225-020-00466-2)) |
| Otu59847 | 1 | k__Fungi(100);p__Ascomycota(100);c__Saccharomycetes(100);o__Saccharomycetales(100);f__Metschnikowiaceae(100);g__Metschnikowia(100);s__Metschnikowia_pulcherrima(100); | Metschnikowia | 13 | Saprotroph | \|Undefined Saprotroph\| | Yeast | NULL | Probable | As nectar-tap_saprotroph (Põlme et al. 2020); Decay sugar-rich substrates (Põlme et al. 2020) | Põlme S, et al. 2020. Fungal Diversity 105:1-16 ((https://doi.org/10.1007/s13225-020-00466-2)) |
| Otu59848 | 1 | k__Fungi(100);p__Ascomycota(100);c__Saccharomycetes(100);o__Saccharomycetales(100);f__Metschnikowiaceae(100);g__Metschnikowia(100);s__Metschnikowia_pulcherrima(100); | Metschnikowia | 13 | Saprotroph | \|Undefined Saprotroph\| | Yeast | NULL | Probable | As nectar-tap_saprotroph (Põlme et al. 2020); Decay sugar-rich substrates (Põlme et al. 2020) | Põlme S, et al. 2020. Fungal Diversity 105:1-16 ((https://doi.org/10.1007/s13225-020-00466-2)) |
| Otu59849 | 1 | k__Fungi(100);p__Ascomycota(100);c__Saccharomycetes(100);o__Saccharomycetales(100);f__Metschnikowiaceae(100);g__Metschnikowia(100);s__Metschnikowia_pulcherrima(100); | Metschnikowia | 13 | Saprotroph | \|Undefined Saprotroph\| | Yeast | NULL | Probable | As nectar-tap_saprotroph (Põlme et al. 2020); Decay sugar-rich substrates (Põlme et al. 2020) | Põlme S, et al. 2020. Fungal Diversity 105:1-16 ((https://doi.org/10.1007/s13225-020-00466-2)) |
| Otu59850 | 1 | k__Fungi(100);p__Ascomycota(100);c__Saccharomycetes(100);o__Saccharomycetales(100);f__Metschnikowiaceae(100);g__Metschnikowia(100);g__Metschnikowia_unclassified(100); | Metschnikowia | 13 | Saprotroph | \|Undefined Saprotroph\| | Yeast | NULL | Probable | As nectar-tap_saprotroph (Põlme et al. 2020); Decay sugar-rich substrates (Põlme et al. 2020) | Põlme S, et al. 2020. Fungal Diversity 105:1-16 ((https://doi.org/10.1007/s13225-020-00466-2)) |
| Otu59851 | 1 | k__Fungi(100);p__Ascomycota(100);c__Saccharomycetes(100);o__Saccharomycetales(100);f__Metschnikowiaceae(100);g__Metschnikowia(100);s__Metschnikowia_pulcherrima(100); | Metschnikowia | 13 | Saprotroph | \|Undefined Saprotroph\| | Yeast | NULL | Probable | As nectar-tap_saprotroph (Põlme et al. 2020); Decay sugar-rich substrates (Põlme et al. 2020) | Põlme S, et al. 2020. Fungal Diversity 105:1-16 ((https://doi.org/10.1007/s13225-020-00466-2)) |
| Otu59853 | 1 | k__Fungi(100);p__Ascomycota(100);c__Saccharomycetes(100);o__Saccharomycetales(100);f__Metschnikowiaceae(100);g__Metschnikowia(100);s__Metschnikowia_pulcherrima(100); | Metschnikowia | 13 | Saprotroph | \|Undefined Saprotroph\| | Yeast | NULL | Probable | As nectar-tap_saprotroph (Põlme et al. 2020); Decay sugar-rich substrates (Põlme et al. 2020) | Põlme S, et al. 2020. Fungal Diversity 105:1-16 ((https://doi.org/10.1007/s13225-020-00466-2)) |
| Otu59854 | 1 | k__Fungi(100);p__Ascomycota(100);c__Saccharomycetes(100);o__Saccharomycetales(100);f__Metschnikowiaceae(100);g__Metschnikowia(100);s__Metschnikowia_pulcherrima(100); | Metschnikowia | 13 | Saprotroph | \|Undefined Saprotroph\| | Yeast | NULL | Probable | As nectar-tap_saprotroph (Põlme et al. 2020); Decay sugar-rich substrates (Põlme et al. 2020) | Põlme S, et al. 2020. Fungal Diversity 105:1-16 ((https://doi.org/10.1007/s13225-020-00466-2)) |
| Otu59855 | 1 | k__Fungi(100);p__Ascomycota(100);c__Saccharomycetes(100);o__Saccharomycetales(100);f__Metschnikowiaceae(100);g__Metschnikowia(100);s__Metschnikowia_pulcherrima(100); | Metschnikowia | 13 | Saprotroph | \|Undefined Saprotroph\| | Yeast | NULL | Probable | As nectar-tap_saprotroph (Põlme et al. 2020); Decay sugar-rich substrates (Põlme et al. 2020) | Põlme S, et al. 2020. Fungal Diversity 105:1-16 ((https://doi.org/10.1007/s13225-020-00466-2)) |
| Otu59856 | 1 | k__Fungi(100);p__Ascomycota(100);c__Saccharomycetes(100);o__Saccharomycetales(100);f__Metschnikowiaceae(100);g__Metschnikowia(100);s__Metschnikowia_pulcherrima(100); | Metschnikowia | 13 | Saprotroph | \|Undefined Saprotroph\| | Yeast | NULL | Probable | As nectar-tap_saprotroph (Põlme et al. 2020); Decay sugar-rich substrates (Põlme et al. 2020) | Põlme S, et al. 2020. Fungal Diversity 105:1-16 ((https://doi.org/10.1007/s13225-020-00466-2)) |
| Otu59858 | 1 | k__Fungi(100);p__Ascomycota(100);c__Saccharomycetes(100);o__Saccharomycetales(100);f__Metschnikowiaceae(100);g__Metschnikowia(100);s__Metschnikowia_pulcherrima(100); | Metschnikowia | 13 | Saprotroph | \|Undefined Saprotroph\| | Yeast | NULL | Probable | As nectar-tap_saprotroph (Põlme et al. 2020); Decay sugar-rich substrates (Põlme et al. 2020) | Põlme S, et al. 2020. Fungal Diversity 105:1-16 ((https://doi.org/10.1007/s13225-020-00466-2)) |
| Otu59859 | 1 | k__Fungi(100);p__Ascomycota(100);c__Saccharomycetes(100);o__Saccharomycetales(100);f__Metschnikowiaceae(100);g__Metschnikowia(100);s__Metschnikowia_pulcherrima(100); | Metschnikowia | 13 | Saprotroph | \|Undefined Saprotroph\| | Yeast | NULL | Probable | As nectar-tap_saprotroph (Põlme et al. 2020); Decay sugar-rich substrates (Põlme et al. 2020) | Põlme S, et al. 2020. Fungal Diversity 105:1-16 ((https://doi.org/10.1007/s13225-020-00466-2)) |
| Otu59860 | 1 | k__Fungi(100);p__Ascomycota(100);c__Saccharomycetes(100);o__Saccharomycetales(100);f__Metschnikowiaceae(100);g__Metschnikowia(100);s__Metschnikowia_pulcherrima(100); | Metschnikowia | 13 | Saprotroph | \|Undefined Saprotroph\| | Yeast | NULL | Probable | As nectar-tap_saprotroph (Põlme et al. 2020); Decay sugar-rich substrates (Põlme et al. 2020) | Põlme S, et al. 2020. Fungal Diversity 105:1-16 ((https://doi.org/10.1007/s13225-020-00466-2)) |
| Otu59861 | 1 | k__Fungi(100);p__Ascomycota(100);c__Saccharomycetes(100);o__Saccharomycetales(100);f__Metschnikowiaceae(100);g__Metschnikowia(100);s__Metschnikowia_pulcherrima(100); | Metschnikowia | 13 | Saprotroph | \|Undefined Saprotroph\| | Yeast | NULL | Probable | As nectar-tap_saprotroph (Põlme et al. 2020); Decay sugar-rich substrates (Põlme et al. 2020) | Põlme S, et al. 2020. Fungal Diversity 105:1-16 ((https://doi.org/10.1007/s13225-020-00466-2)) |
| Otu59863 | 1 | k__Fungi(100);p__Ascomycota(100);c__Saccharomycetes(100);o__Saccharomycetales(100);f__Metschnikowiaceae(100);g__Metschnikowia(100);s__Metschnikowia_pulcherrima(100); | Metschnikowia | 13 | Saprotroph | \|Undefined Saprotroph\| | Yeast | NULL | Probable | As nectar-tap_saprotroph (Põlme et al. 2020); Decay sugar-rich substrates (Põlme et al. 2020) | Põlme S, et al. 2020. Fungal Diversity 105:1-16 ((https://doi.org/10.1007/s13225-020-00466-2)) |
| Otu59865 | 1 | k__Fungi(100);p__Ascomycota(100);c__Saccharomycetes(100);o__Saccharomycetales(100);f__Metschnikowiaceae(100);g__Metschnikowia(100);s__Metschnikowia_pulcherrima(100); | Metschnikowia | 13 | Saprotroph | \|Undefined Saprotroph\| | Yeast | NULL | Probable | As nectar-tap_saprotroph (Põlme et al. 2020); Decay sugar-rich substrates (Põlme et al. 2020) | Põlme S, et al. 2020. Fungal Diversity 105:1-16 ((https://doi.org/10.1007/s13225-020-00466-2)) |
| Otu59868 | 1 | k__Fungi(100);p__Ascomycota(100);c__Saccharomycetes(100);o__Saccharomycetales(100);f__Metschnikowiaceae(100);g__Metschnikowia(100);g__Metschnikowia_unclassified(100); | Metschnikowia | 13 | Saprotroph | \|Undefined Saprotroph\| | Yeast | NULL | Probable | As nectar-tap_saprotroph (Põlme et al. 2020); Decay sugar-rich substrates (Põlme et al. 2020) | Põlme S, et al. 2020. Fungal Diversity 105:1-16 ((https://doi.org/10.1007/s13225-020-00466-2)) |
| Otu59869 | 1 | k__Fungi(100);p__Ascomycota(100);c__Saccharomycetes(100);o__Saccharomycetales(100);f__Metschnikowiaceae(100);g__Metschnikowia(100);s__Metschnikowia_pulcherrima(100); | Metschnikowia | 13 | Saprotroph | \|Undefined Saprotroph\| | Yeast | NULL | Probable | As nectar-tap_saprotroph (Põlme et al. 2020); Decay sugar-rich substrates (Põlme et al. 2020) | Põlme S, et al. 2020. Fungal Diversity 105:1-16 ((https://doi.org/10.1007/s13225-020-00466-2)) |
| Otu59870 | 1 | k__Fungi(100);p__Ascomycota(100);c__Saccharomycetes(100);o__Saccharomycetales(100);f__Metschnikowiaceae(100);g__Metschnikowia(100);s__Metschnikowia_pulcherrima(100); | Metschnikowia | 13 | Saprotroph | \|Undefined Saprotroph\| | Yeast | NULL | Probable | As nectar-tap_saprotroph (Põlme et al. 2020); Decay sugar-rich substrates (Põlme et al. 2020) | Põlme S, et al. 2020. Fungal Diversity 105:1-16 ((https://doi.org/10.1007/s13225-020-00466-2)) |
| Otu59872 | 1 | k__Fungi(100);p__Ascomycota(100);c__Saccharomycetes(100);o__Saccharomycetales(100);f__Metschnikowiaceae(100);g__Metschnikowia(100);s__Metschnikowia_pulcherrima(100); | Metschnikowia | 13 | Saprotroph | \|Undefined Saprotroph\| | Yeast | NULL | Probable | As nectar-tap_saprotroph (Põlme et al. 2020); Decay sugar-rich substrates (Põlme et al. 2020) | Põlme S, et al. 2020. Fungal Diversity 105:1-16 ((https://doi.org/10.1007/s13225-020-00466-2)) |
| Otu59874 | 1 | k__Fungi(100);p__Ascomycota(100);c__Saccharomycetes(100);o__Saccharomycetales(100);f__Metschnikowiaceae(100);g__Metschnikowia(100);s__Metschnikowia_pulcherrima(100); | Metschnikowia | 13 | Saprotroph | \|Undefined Saprotroph\| | Yeast | NULL | Probable | As nectar-tap_saprotroph (Põlme et al. 2020); Decay sugar-rich substrates (Põlme et al. 2020) | Põlme S, et al. 2020. Fungal Diversity 105:1-16 ((https://doi.org/10.1007/s13225-020-00466-2)) |
| Otu59875 | 1 | k__Fungi(100);p__Ascomycota(100);c__Saccharomycetes(100);o__Saccharomycetales(100);f__Metschnikowiaceae(100);g__Metschnikowia(100);s__Metschnikowia_pulcherrima(100); | Metschnikowia | 13 | Saprotroph | \|Undefined Saprotroph\| | Yeast | NULL | Probable | As nectar-tap_saprotroph (Põlme et al. 2020); Decay sugar-rich substrates (Põlme et al. 2020) | Põlme S, et al. 2020. Fungal Diversity 105:1-16 ((https://doi.org/10.1007/s13225-020-00466-2)) |
| Otu59877 | 1 | k__Fungi(100);p__Ascomycota(100);c__Saccharomycetes(100);o__Saccharomycetales(100);f__Metschnikowiaceae(100);g__Metschnikowia(100);s__Metschnikowia_pulcherrima(100); | Metschnikowia | 13 | Saprotroph | \|Undefined Saprotroph\| | Yeast | NULL | Probable | As nectar-tap_saprotroph (Põlme et al. 2020); Decay sugar-rich substrates (Põlme et al. 2020) | Põlme S, et al. 2020. Fungal Diversity 105:1-16 ((https://doi.org/10.1007/s13225-020-00466-2)) |
| Otu59878 | 1 | k__Fungi(100);p__Ascomycota(100);c__Saccharomycetes(100);o__Saccharomycetales(100);f__Metschnikowiaceae(100);g__Metschnikowia(100);s__Metschnikowia_pulcherrima(100); | Metschnikowia | 13 | Saprotroph | \|Undefined Saprotroph\| | Yeast | NULL | Probable | As nectar-tap_saprotroph (Põlme et al. 2020); Decay sugar-rich substrates (Põlme et al. 2020) | Põlme S, et al. 2020. Fungal Diversity 105:1-16 ((https://doi.org/10.1007/s13225-020-00466-2)) |
| Otu59879 | 1 | k__Fungi(100);p__Ascomycota(100);c__Saccharomycetes(100);o__Saccharomycetales(100);f__Metschnikowiaceae(100);g__Metschnikowia(100);g__Metschnikowia_unclassified(100); | Metschnikowia | 13 | Saprotroph | \|Undefined Saprotroph\| | Yeast | NULL | Probable | As nectar-tap_saprotroph (Põlme et al. 2020); Decay sugar-rich substrates (Põlme et al. 2020) | Põlme S, et al. 2020. Fungal Diversity 105:1-16 ((https://doi.org/10.1007/s13225-020-00466-2)) |
| Otu59880 | 1 | k__Fungi(100);p__Ascomycota(100);c__Saccharomycetes(100);o__Saccharomycetales(100);f__Metschnikowiaceae(100);g__Metschnikowia(100);s__Metschnikowia_pulcherrima(100); | Metschnikowia | 13 | Saprotroph | \|Undefined Saprotroph\| | Yeast | NULL | Probable | As nectar-tap_saprotroph (Põlme et al. 2020); Decay sugar-rich substrates (Põlme et al. 2020) | Põlme S, et al. 2020. Fungal Diversity 105:1-16 ((https://doi.org/10.1007/s13225-020-00466-2)) |
| Otu59881 | 1 | k__Fungi(100);p__Ascomycota(100);c__Saccharomycetes(100);o__Saccharomycetales(100);f__Metschnikowiaceae(100);g__Metschnikowia(100);s__Metschnikowia_pulcherrima(100); | Metschnikowia | 13 | Saprotroph | \|Undefined Saprotroph\| | Yeast | NULL | Probable | As nectar-tap_saprotroph (Põlme et al. 2020); Decay sugar-rich substrates (Põlme et al. 2020) | Põlme S, et al. 2020. Fungal Diversity 105:1-16 ((https://doi.org/10.1007/s13225-020-00466-2)) |
| Otu59882 | 1 | k__Fungi(100);p__Ascomycota(100);c__Saccharomycetes(100);o__Saccharomycetales(100);f__Metschnikowiaceae(100);g__Metschnikowia(100);s__Metschnikowia_pulcherrima(100); | Metschnikowia | 13 | Saprotroph | \|Undefined Saprotroph\| | Yeast | NULL | Probable | As nectar-tap_saprotroph (Põlme et al. 2020); Decay sugar-rich substrates (Põlme et al. 2020) | Põlme S, et al. 2020. Fungal Diversity 105:1-16 ((https://doi.org/10.1007/s13225-020-00466-2)) |
| Otu59883 | 1 | k__Fungi(100);p__Ascomycota(100);c__Saccharomycetes(100);o__Saccharomycetales(100);f__Metschnikowiaceae(100);g__Metschnikowia(100);s__Metschnikowia_pulcherrima(100); | Metschnikowia | 13 | Saprotroph | \|Undefined Saprotroph\| | Yeast | NULL | Probable | As nectar-tap_saprotroph (Põlme et al. 2020); Decay sugar-rich substrates (Põlme et al. 2020) | Põlme S, et al. 2020. Fungal Diversity 105:1-16 ((https://doi.org/10.1007/s13225-020-00466-2)) |
| Otu59884 | 1 | k__Fungi(100);p__Ascomycota(100);c__Saccharomycetes(100);o__Saccharomycetales(100);f__Metschnikowiaceae(100);g__Metschnikowia(100);g__Metschnikowia_unclassified(100); | Metschnikowia | 13 | Saprotroph | \|Undefined Saprotroph\| | Yeast | NULL | Probable | As nectar-tap_saprotroph (Põlme et al. 2020); Decay sugar-rich substrates (Põlme et al. 2020) | Põlme S, et al. 2020. Fungal Diversity 105:1-16 ((https://doi.org/10.1007/s13225-020-00466-2)) |
| Otu59885 | 1 | k__Fungi(100);p__Ascomycota(100);c__Saccharomycetes(100);o__Saccharomycetales(100);f__Metschnikowiaceae(100);g__Metschnikowia(100);s__Metschnikowia_pulcherrima(100); | Metschnikowia | 13 | Saprotroph | \|Undefined Saprotroph\| | Yeast | NULL | Probable | As nectar-tap_saprotroph (Põlme et al. 2020); Decay sugar-rich substrates (Põlme et al. 2020) | Põlme S, et al. 2020. Fungal Diversity 105:1-16 ((https://doi.org/10.1007/s13225-020-00466-2)) |
| Otu59886 | 1 | k__Fungi(100);p__Ascomycota(100);c__Saccharomycetes(100);o__Saccharomycetales(100);f__Metschnikowiaceae(100);g__Metschnikowia(100);g__Metschnikowia_unclassified(100); | Metschnikowia | 13 | Saprotroph | \|Undefined Saprotroph\| | Yeast | NULL | Probable | As nectar-tap_saprotroph (Põlme et al. 2020); Decay sugar-rich substrates (Põlme et al. 2020) | Põlme S, et al. 2020. Fungal Diversity 105:1-16 ((https://doi.org/10.1007/s13225-020-00466-2)) |
| Otu59887 | 1 | k__Fungi(100);p__Ascomycota(100);c__Saccharomycetes(100);o__Saccharomycetales(100);f__Metschnikowiaceae(100);g__Metschnikowia(100);g__Metschnikowia_unclassified(100); | Metschnikowia | 13 | Saprotroph | \|Undefined Saprotroph\| | Yeast | NULL | Probable | As nectar-tap_saprotroph (Põlme et al. 2020); Decay sugar-rich substrates (Põlme et al. 2020) | Põlme S, et al. 2020. Fungal Diversity 105:1-16 ((https://doi.org/10.1007/s13225-020-00466-2)) |
| Otu59888 | 1 | k__Fungi(100);p__Ascomycota(100);c__Saccharomycetes(100);o__Saccharomycetales(100);f__Metschnikowiaceae(100);g__Metschnikowia(100);s__Metschnikowia_pulcherrima(100); | Metschnikowia | 13 | Saprotroph | \|Undefined Saprotroph\| | Yeast | NULL | Probable | As nectar-tap_saprotroph (Põlme et al. 2020); Decay sugar-rich substrates (Põlme et al. 2020) | Põlme S, et al. 2020. Fungal Diversity 105:1-16 ((https://doi.org/10.1007/s13225-020-00466-2)) |
| Otu59889 | 1 | k__Fungi(100);p__Ascomycota(100);c__Saccharomycetes(100);o__Saccharomycetales(100);f__Metschnikowiaceae(100);g__Metschnikowia(100);s__Metschnikowia_pulcherrima(100); | Metschnikowia | 13 | Saprotroph | \|Undefined Saprotroph\| | Yeast | NULL | Probable | As nectar-tap_saprotroph (Põlme et al. 2020); Decay sugar-rich substrates (Põlme et al. 2020) | Põlme S, et al. 2020. Fungal Diversity 105:1-16 ((https://doi.org/10.1007/s13225-020-00466-2)) |
| Otu59890 | 1 | k__Fungi(100);p__Ascomycota(100);c__Saccharomycetes(100);o__Saccharomycetales(100);f__Metschnikowiaceae(100);g__Metschnikowia(100);g__Metschnikowia_unclassified(100); | Metschnikowia | 13 | Saprotroph | \|Undefined Saprotroph\| | Yeast | NULL | Probable | As nectar-tap_saprotroph (Põlme et al. 2020); Decay sugar-rich substrates (Põlme et al. 2020) | Põlme S, et al. 2020. Fungal Diversity 105:1-16 ((https://doi.org/10.1007/s13225-020-00466-2)) |
| Otu59891 | 1 | k__Fungi(100);p__Ascomycota(100);c__Saccharomycetes(100);o__Saccharomycetales(100);f__Metschnikowiaceae(100);g__Metschnikowia(100);s__Metschnikowia_pulcherrima(100); | Metschnikowia | 13 | Saprotroph | \|Undefined Saprotroph\| | Yeast | NULL | Probable | As nectar-tap_saprotroph (Põlme et al. 2020); Decay sugar-rich substrates (Põlme et al. 2020) | Põlme S, et al. 2020. Fungal Diversity 105:1-16 ((https://doi.org/10.1007/s13225-020-00466-2)) |
| Otu59893 | 1 | k__Fungi(100);p__Ascomycota(100);c__Saccharomycetes(100);o__Saccharomycetales(100);f__Metschnikowiaceae(100);g__Metschnikowia(100);s__Metschnikowia_pulcherrima(100); | Metschnikowia | 13 | Saprotroph | \|Undefined Saprotroph\| | Yeast | NULL | Probable | As nectar-tap_saprotroph (Põlme et al. 2020); Decay sugar-rich substrates (Põlme et al. 2020) | Põlme S, et al. 2020. Fungal Diversity 105:1-16 ((https://doi.org/10.1007/s13225-020-00466-2)) |
| Otu59894 | 1 | k__Fungi(100);p__Ascomycota(100);c__Saccharomycetes(100);o__Saccharomycetales(100);f__Metschnikowiaceae(100);g__Metschnikowia(100);g__Metschnikowia_unclassified(100); | Metschnikowia | 13 | Saprotroph | \|Undefined Saprotroph\| | Yeast | NULL | Probable | As nectar-tap_saprotroph (Põlme et al. 2020); Decay sugar-rich substrates (Põlme et al. 2020) | Põlme S, et al. 2020. Fungal Diversity 105:1-16 ((https://doi.org/10.1007/s13225-020-00466-2)) |
| Otu59895 | 1 | k__Fungi(100);p__Ascomycota(100);c__Saccharomycetes(100);o__Saccharomycetales(100);f__Metschnikowiaceae(100);g__Metschnikowia(100);g__Metschnikowia_unclassified(100); | Metschnikowia | 13 | Saprotroph | \|Undefined Saprotroph\| | Yeast | NULL | Probable | As nectar-tap_saprotroph (Põlme et al. 2020); Decay sugar-rich substrates (Põlme et al. 2020) | Põlme S, et al. 2020. Fungal Diversity 105:1-16 ((https://doi.org/10.1007/s13225-020-00466-2)) |
| Otu59902 | 1 | k__Fungi(100);p__Ascomycota(100);c__Saccharomycetes(100);o__Saccharomycetales(100);f__Metschnikowiaceae(100);g__Metschnikowia(100);g__Metschnikowia_unclassified(100); | Metschnikowia | 13 | Saprotroph | \|Undefined Saprotroph\| | Yeast | NULL | Probable | As nectar-tap_saprotroph (Põlme et al. 2020); Decay sugar-rich substrates (Põlme et al. 2020) | Põlme S, et al. 2020. Fungal Diversity 105:1-16 ((https://doi.org/10.1007/s13225-020-00466-2)) |
| Otu59903 | 1 | k__Fungi(100);p__Ascomycota(100);c__Saccharomycetes(100);o__Saccharomycetales(100);f__Metschnikowiaceae(100);g__Metschnikowia(100);s__Metschnikowia_pulcherrima(100); | Metschnikowia | 13 | Saprotroph | \|Undefined Saprotroph\| | Yeast | NULL | Probable | As nectar-tap_saprotroph (Põlme et al. 2020); Decay sugar-rich substrates (Põlme et al. 2020) | Põlme S, et al. 2020. Fungal Diversity 105:1-16 ((https://doi.org/10.1007/s13225-020-00466-2)) |
| Otu59904 | 1 | k__Fungi(100);p__Ascomycota(100);c__Saccharomycetes(100);o__Saccharomycetales(100);f__Metschnikowiaceae(100);g__Metschnikowia(100);s__Metschnikowia_pulcherrima(100); | Metschnikowia | 13 | Saprotroph | \|Undefined Saprotroph\| | Yeast | NULL | Probable | As nectar-tap_saprotroph (Põlme et al. 2020); Decay sugar-rich substrates (Põlme et al. 2020) | Põlme S, et al. 2020. Fungal Diversity 105:1-16 ((https://doi.org/10.1007/s13225-020-00466-2)) |
| Otu59906 | 1 | k__Fungi(100);p__Ascomycota(100);c__Saccharomycetes(100);o__Saccharomycetales(100);f__Metschnikowiaceae(100);g__Metschnikowia(100);s__Metschnikowia_pulcherrima(100); | Metschnikowia | 13 | Saprotroph | \|Undefined Saprotroph\| | Yeast | NULL | Probable | As nectar-tap_saprotroph (Põlme et al. 2020); Decay sugar-rich substrates (Põlme et al. 2020) | Põlme S, et al. 2020. Fungal Diversity 105:1-16 ((https://doi.org/10.1007/s13225-020-00466-2)) |
| Otu59907 | 1 | k__Fungi(100);p__Ascomycota(100);c__Saccharomycetes(100);o__Saccharomycetales(100);f__Metschnikowiaceae(100);g__Metschnikowia(100);s__Metschnikowia_pulcherrima(100); | Metschnikowia | 13 | Saprotroph | \|Undefined Saprotroph\| | Yeast | NULL | Probable | As nectar-tap_saprotroph (Põlme et al. 2020); Decay sugar-rich substrates (Põlme et al. 2020) | Põlme S, et al. 2020. Fungal Diversity 105:1-16 ((https://doi.org/10.1007/s13225-020-00466-2)) |
| Otu59908 | 1 | k__Fungi(100);p__Ascomycota(100);c__Saccharomycetes(100);o__Saccharomycetales(100);f__Metschnikowiaceae(100);g__Metschnikowia(100);s__Metschnikowia_pulcherrima(100); | Metschnikowia | 13 | Saprotroph | \|Undefined Saprotroph\| | Yeast | NULL | Probable | As nectar-tap_saprotroph (Põlme et al. 2020); Decay sugar-rich substrates (Põlme et al. 2020) | Põlme S, et al. 2020. Fungal Diversity 105:1-16 ((https://doi.org/10.1007/s13225-020-00466-2)) |
| Otu59909 | 1 | k__Fungi(100);p__Ascomycota(100);c__Saccharomycetes(100);o__Saccharomycetales(100);f__Metschnikowiaceae(100);g__Metschnikowia(100);s__Metschnikowia_pulcherrima(100); | Metschnikowia | 13 | Saprotroph | \|Undefined Saprotroph\| | Yeast | NULL | Probable | As nectar-tap_saprotroph (Põlme et al. 2020); Decay sugar-rich substrates (Põlme et al. 2020) | Põlme S, et al. 2020. Fungal Diversity 105:1-16 ((https://doi.org/10.1007/s13225-020-00466-2)) |
| Otu59910 | 1 | k__Fungi(100);p__Ascomycota(100);c__Saccharomycetes(100);o__Saccharomycetales(100);f__Metschnikowiaceae(100);g__Metschnikowia(100);s__Metschnikowia_pulcherrima(100); | Metschnikowia | 13 | Saprotroph | \|Undefined Saprotroph\| | Yeast | NULL | Probable | As nectar-tap_saprotroph (Põlme et al. 2020); Decay sugar-rich substrates (Põlme et al. 2020) | Põlme S, et al. 2020. Fungal Diversity 105:1-16 ((https://doi.org/10.1007/s13225-020-00466-2)) |
| Otu59911 | 1 | k__Fungi(100);p__Ascomycota(100);c__Saccharomycetes(100);o__Saccharomycetales(100);f__Metschnikowiaceae(100);g__Metschnikowia(100);s__Metschnikowia_pulcherrima(100); | Metschnikowia | 13 | Saprotroph | \|Undefined Saprotroph\| | Yeast | NULL | Probable | As nectar-tap_saprotroph (Põlme et al. 2020); Decay sugar-rich substrates (Põlme et al. 2020) | Põlme S, et al. 2020. Fungal Diversity 105:1-16 ((https://doi.org/10.1007/s13225-020-00466-2)) |
| Otu59912 | 1 | k__Fungi(100);p__Ascomycota(100);c__Saccharomycetes(100);o__Saccharomycetales(100);f__Metschnikowiaceae(100);g__Metschnikowia(100);s__Metschnikowia_pulcherrima(100); | Metschnikowia | 13 | Saprotroph | \|Undefined Saprotroph\| | Yeast | NULL | Probable | As nectar-tap_saprotroph (Põlme et al. 2020); Decay sugar-rich substrates (Põlme et al. 2020) | Põlme S, et al. 2020. Fungal Diversity 105:1-16 ((https://doi.org/10.1007/s13225-020-00466-2)) |
| Otu59913 | 1 | k__Fungi(100);p__Ascomycota(100);c__Saccharomycetes(100);o__Saccharomycetales(100);f__Metschnikowiaceae(100);g__Metschnikowia(100);s__Metschnikowia_pulcherrima(100); | Metschnikowia | 13 | Saprotroph | \|Undefined Saprotroph\| | Yeast | NULL | Probable | As nectar-tap_saprotroph (Põlme et al. 2020); Decay sugar-rich substrates (Põlme et al. 2020) | Põlme S, et al. 2020. Fungal Diversity 105:1-16 ((https://doi.org/10.1007/s13225-020-00466-2)) |
| Otu59914 | 1 | k__Fungi(100);p__Ascomycota(100);c__Saccharomycetes(100);o__Saccharomycetales(100);f__Metschnikowiaceae(100);g__Metschnikowia(100);g__Metschnikowia_unclassified(100); | Metschnikowia | 13 | Saprotroph | \|Undefined Saprotroph\| | Yeast | NULL | Probable | As nectar-tap_saprotroph (Põlme et al. 2020); Decay sugar-rich substrates (Põlme et al. 2020) | Põlme S, et al. 2020. Fungal Diversity 105:1-16 ((https://doi.org/10.1007/s13225-020-00466-2)) |
| Otu59915 | 1 | k__Fungi(100);p__Ascomycota(100);c__Saccharomycetes(100);o__Saccharomycetales(100);f__Metschnikowiaceae(100);g__Metschnikowia(100);s__Metschnikowia_pulcherrima(100); | Metschnikowia | 13 | Saprotroph | \|Undefined Saprotroph\| | Yeast | NULL | Probable | As nectar-tap_saprotroph (Põlme et al. 2020); Decay sugar-rich substrates (Põlme et al. 2020) | Põlme S, et al. 2020. Fungal Diversity 105:1-16 ((https://doi.org/10.1007/s13225-020-00466-2)) |
| Otu59916 | 1 | k__Fungi(100);p__Ascomycota(100);c__Saccharomycetes(100);o__Saccharomycetales(100);f__Metschnikowiaceae(100);g__Metschnikowia(100);s__Metschnikowia_pulcherrima(100); | Metschnikowia | 13 | Saprotroph | \|Undefined Saprotroph\| | Yeast | NULL | Probable | As nectar-tap_saprotroph (Põlme et al. 2020); Decay sugar-rich substrates (Põlme et al. 2020) | Põlme S, et al. 2020. Fungal Diversity 105:1-16 ((https://doi.org/10.1007/s13225-020-00466-2)) |
| Otu59918 | 1 | k__Fungi(100);p__Ascomycota(100);c__Saccharomycetes(100);o__Saccharomycetales(100);f__Metschnikowiaceae(100);g__Metschnikowia(100);g__Metschnikowia_unclassified(100); | Metschnikowia | 13 | Saprotroph | \|Undefined Saprotroph\| | Yeast | NULL | Probable | As nectar-tap_saprotroph (Põlme et al. 2020); Decay sugar-rich substrates (Põlme et al. 2020) | Põlme S, et al. 2020. Fungal Diversity 105:1-16 ((https://doi.org/10.1007/s13225-020-00466-2)) |
| Otu59919 | 1 | k__Fungi(100);p__Ascomycota(100);c__Saccharomycetes(100);o__Saccharomycetales(100);f__Metschnikowiaceae(100);g__Metschnikowia(100);s__Metschnikowia_pulcherrima(100); | Metschnikowia | 13 | Saprotroph | \|Undefined Saprotroph\| | Yeast | NULL | Probable | As nectar-tap_saprotroph (Põlme et al. 2020); Decay sugar-rich substrates (Põlme et al. 2020) | Põlme S, et al. 2020. Fungal Diversity 105:1-16 ((https://doi.org/10.1007/s13225-020-00466-2)) |
| Otu59920 | 1 | k__Fungi(100);p__Ascomycota(100);c__Saccharomycetes(100);o__Saccharomycetales(100);f__Metschnikowiaceae(100);g__Metschnikowia(100);s__Metschnikowia_pulcherrima(100); | Metschnikowia | 13 | Saprotroph | \|Undefined Saprotroph\| | Yeast | NULL | Probable | As nectar-tap_saprotroph (Põlme et al. 2020); Decay sugar-rich substrates (Põlme et al. 2020) | Põlme S, et al. 2020. Fungal Diversity 105:1-16 ((https://doi.org/10.1007/s13225-020-00466-2)) |
| Otu59921 | 1 | k__Fungi(100);p__Ascomycota(100);c__Saccharomycetes(100);o__Saccharomycetales(100);f__Metschnikowiaceae(100);g__Metschnikowia(100);s__Metschnikowia_pulcherrima(100); | Metschnikowia | 13 | Saprotroph | \|Undefined Saprotroph\| | Yeast | NULL | Probable | As nectar-tap_saprotroph (Põlme et al. 2020); Decay sugar-rich substrates (Põlme et al. 2020) | Põlme S, et al. 2020. Fungal Diversity 105:1-16 ((https://doi.org/10.1007/s13225-020-00466-2)) |
| Otu59922 | 1 | k__Fungi(100);p__Ascomycota(100);c__Saccharomycetes(100);o__Saccharomycetales(100);f__Metschnikowiaceae(100);g__Metschnikowia(100);s__Metschnikowia_pulcherrima(100); | Metschnikowia | 13 | Saprotroph | \|Undefined Saprotroph\| | Yeast | NULL | Probable | As nectar-tap_saprotroph (Põlme et al. 2020); Decay sugar-rich substrates (Põlme et al. 2020) | Põlme S, et al. 2020. Fungal Diversity 105:1-16 ((https://doi.org/10.1007/s13225-020-00466-2)) |
| Otu59923 | 1 | k__Fungi(100);p__Ascomycota(100);c__Saccharomycetes(100);o__Saccharomycetales(100);f__Metschnikowiaceae(100);g__Metschnikowia(100);s__Metschnikowia_pulcherrima(100); | Metschnikowia | 13 | Saprotroph | \|Undefined Saprotroph\| | Yeast | NULL | Probable | As nectar-tap_saprotroph (Põlme et al. 2020); Decay sugar-rich substrates (Põlme et al. 2020) | Põlme S, et al. 2020. Fungal Diversity 105:1-16 ((https://doi.org/10.1007/s13225-020-00466-2)) |
| Otu59924 | 1 | k__Fungi(100);p__Ascomycota(100);c__Saccharomycetes(100);o__Saccharomycetales(100);f__Metschnikowiaceae(100);g__Metschnikowia(100);s__Metschnikowia_pulcherrima(100); | Metschnikowia | 13 | Saprotroph | \|Undefined Saprotroph\| | Yeast | NULL | Probable | As nectar-tap_saprotroph (Põlme et al. 2020); Decay sugar-rich substrates (Põlme et al. 2020) | Põlme S, et al. 2020. Fungal Diversity 105:1-16 ((https://doi.org/10.1007/s13225-020-00466-2)) |
| Otu59925 | 1 | k__Fungi(100);p__Ascomycota(100);c__Saccharomycetes(100);o__Saccharomycetales(100);f__Metschnikowiaceae(100);g__Metschnikowia(100);s__Metschnikowia_pulcherrima(100); | Metschnikowia | 13 | Saprotroph | \|Undefined Saprotroph\| | Yeast | NULL | Probable | As nectar-tap_saprotroph (Põlme et al. 2020); Decay sugar-rich substrates (Põlme et al. 2020) | Põlme S, et al. 2020. Fungal Diversity 105:1-16 ((https://doi.org/10.1007/s13225-020-00466-2)) |
| Otu59926 | 1 | k__Fungi(100);p__Ascomycota(100);c__Saccharomycetes(100);o__Saccharomycetales(100);f__Metschnikowiaceae(100);g__Metschnikowia(100);s__Metschnikowia_pulcherrima(100); | Metschnikowia | 13 | Saprotroph | \|Undefined Saprotroph\| | Yeast | NULL | Probable | As nectar-tap_saprotroph (Põlme et al. 2020); Decay sugar-rich substrates (Põlme et al. 2020) | Põlme S, et al. 2020. Fungal Diversity 105:1-16 ((https://doi.org/10.1007/s13225-020-00466-2)) |
| Otu59927 | 1 | k__Fungi(100);p__Ascomycota(100);c__Saccharomycetes(100);o__Saccharomycetales(100);f__Metschnikowiaceae(100);g__Metschnikowia(100);s__Metschnikowia_pulcherrima(100); | Metschnikowia | 13 | Saprotroph | \|Undefined Saprotroph\| | Yeast | NULL | Probable | As nectar-tap_saprotroph (Põlme et al. 2020); Decay sugar-rich substrates (Põlme et al. 2020) | Põlme S, et al. 2020. Fungal Diversity 105:1-16 ((https://doi.org/10.1007/s13225-020-00466-2)) |
| Otu59928 | 1 | k__Fungi(100);p__Ascomycota(100);c__Saccharomycetes(100);o__Saccharomycetales(100);f__Metschnikowiaceae(100);g__Metschnikowia(100);s__Metschnikowia_pulcherrima(100); | Metschnikowia | 13 | Saprotroph | \|Undefined Saprotroph\| | Yeast | NULL | Probable | As nectar-tap_saprotroph (Põlme et al. 2020); Decay sugar-rich substrates (Põlme et al. 2020) | Põlme S, et al. 2020. Fungal Diversity 105:1-16 ((https://doi.org/10.1007/s13225-020-00466-2)) |
| Otu59929 | 1 | k__Fungi(100);p__Ascomycota(100);c__Saccharomycetes(100);o__Saccharomycetales(100);f__Metschnikowiaceae(100);g__Metschnikowia(100);g__Metschnikowia_unclassified(100); | Metschnikowia | 13 | Saprotroph | \|Undefined Saprotroph\| | Yeast | NULL | Probable | As nectar-tap_saprotroph (Põlme et al. 2020); Decay sugar-rich substrates (Põlme et al. 2020) | Põlme S, et al. 2020. Fungal Diversity 105:1-16 ((https://doi.org/10.1007/s13225-020-00466-2)) |
| Otu59930 | 1 | k__Fungi(100);p__Ascomycota(100);c__Saccharomycetes(100);o__Saccharomycetales(100);f__Metschnikowiaceae(100);g__Metschnikowia(100);s__Metschnikowia_pulcherrima(100); | Metschnikowia | 13 | Saprotroph | \|Undefined Saprotroph\| | Yeast | NULL | Probable | As nectar-tap_saprotroph (Põlme et al. 2020); Decay sugar-rich substrates (Põlme et al. 2020) | Põlme S, et al. 2020. Fungal Diversity 105:1-16 ((https://doi.org/10.1007/s13225-020-00466-2)) |
| Otu59931 | 1 | k__Fungi(100);p__Ascomycota(100);c__Saccharomycetes(100);o__Saccharomycetales(100);f__Metschnikowiaceae(100);g__Metschnikowia(100);s__Metschnikowia_pulcherrima(100); | Metschnikowia | 13 | Saprotroph | \|Undefined Saprotroph\| | Yeast | NULL | Probable | As nectar-tap_saprotroph (Põlme et al. 2020); Decay sugar-rich substrates (Põlme et al. 2020) | Põlme S, et al. 2020. Fungal Diversity 105:1-16 ((https://doi.org/10.1007/s13225-020-00466-2)) |
| Otu59932 | 1 | k__Fungi(100);p__Ascomycota(100);c__Saccharomycetes(100);o__Saccharomycetales(100);f__Metschnikowiaceae(100);g__Metschnikowia(100);s__Metschnikowia_pulcherrima(100); | Metschnikowia | 13 | Saprotroph | \|Undefined Saprotroph\| | Yeast | NULL | Probable | As nectar-tap_saprotroph (Põlme et al. 2020); Decay sugar-rich substrates (Põlme et al. 2020) | Põlme S, et al. 2020. Fungal Diversity 105:1-16 ((https://doi.org/10.1007/s13225-020-00466-2)) |
| Otu59933 | 1 | k__Fungi(100);p__Ascomycota(100);c__Saccharomycetes(100);o__Saccharomycetales(100);f__Metschnikowiaceae(100);g__Metschnikowia(100);g__Metschnikowia_unclassified(100); | Metschnikowia | 13 | Saprotroph | \|Undefined Saprotroph\| | Yeast | NULL | Probable | As nectar-tap_saprotroph (Põlme et al. 2020); Decay sugar-rich substrates (Põlme et al. 2020) | Põlme S, et al. 2020. Fungal Diversity 105:1-16 ((https://doi.org/10.1007/s13225-020-00466-2)) |
| Otu59934 | 1 | k__Fungi(100);p__Ascomycota(100);c__Saccharomycetes(100);o__Saccharomycetales(100);f__Metschnikowiaceae(100);g__Metschnikowia(100);s__Metschnikowia_pulcherrima(100); | Metschnikowia | 13 | Saprotroph | \|Undefined Saprotroph\| | Yeast | NULL | Probable | As nectar-tap_saprotroph (Põlme et al. 2020); Decay sugar-rich substrates (Põlme et al. 2020) | Põlme S, et al. 2020. Fungal Diversity 105:1-16 ((https://doi.org/10.1007/s13225-020-00466-2)) |
| Otu59935 | 1 | k__Fungi(100);p__Ascomycota(100);c__Saccharomycetes(100);o__Saccharomycetales(100);f__Metschnikowiaceae(100);g__Metschnikowia(100);s__Metschnikowia_pulcherrima(100); | Metschnikowia | 13 | Saprotroph | \|Undefined Saprotroph\| | Yeast | NULL | Probable | As nectar-tap_saprotroph (Põlme et al. 2020); Decay sugar-rich substrates (Põlme et al. 2020) | Põlme S, et al. 2020. Fungal Diversity 105:1-16 ((https://doi.org/10.1007/s13225-020-00466-2)) |
| Otu59936 | 1 | k__Fungi(100);p__Ascomycota(100);c__Saccharomycetes(100);o__Saccharomycetales(100);f__Metschnikowiaceae(100);g__Metschnikowia(100);s__Metschnikowia_pulcherrima(100); | Metschnikowia | 13 | Saprotroph | \|Undefined Saprotroph\| | Yeast | NULL | Probable | As nectar-tap_saprotroph (Põlme et al. 2020); Decay sugar-rich substrates (Põlme et al. 2020) | Põlme S, et al. 2020. Fungal Diversity 105:1-16 ((https://doi.org/10.1007/s13225-020-00466-2)) |
| Otu59937 | 1 | k__Fungi(100);p__Ascomycota(100);c__Saccharomycetes(100);o__Saccharomycetales(100);f__Metschnikowiaceae(100);g__Metschnikowia(100);g__Metschnikowia_unclassified(100); | Metschnikowia | 13 | Saprotroph | \|Undefined Saprotroph\| | Yeast | NULL | Probable | As nectar-tap_saprotroph (Põlme et al. 2020); Decay sugar-rich substrates (Põlme et al. 2020) | Põlme S, et al. 2020. Fungal Diversity 105:1-16 ((https://doi.org/10.1007/s13225-020-00466-2)) |
| Otu59938 | 1 | k__Fungi(100);p__Ascomycota(100);c__Saccharomycetes(100);o__Saccharomycetales(100);f__Metschnikowiaceae(100);g__Metschnikowia(100);s__Metschnikowia_pulcherrima(100); | Metschnikowia | 13 | Saprotroph | \|Undefined Saprotroph\| | Yeast | NULL | Probable | As nectar-tap_saprotroph (Põlme et al. 2020); Decay sugar-rich substrates (Põlme et al. 2020) | Põlme S, et al. 2020. Fungal Diversity 105:1-16 ((https://doi.org/10.1007/s13225-020-00466-2)) |
| Otu59939 | 1 | k__Fungi(100);p__Ascomycota(100);c__Saccharomycetes(100);o__Saccharomycetales(100);f__Metschnikowiaceae(100);g__Metschnikowia(100);g__Metschnikowia_unclassified(100); | Metschnikowia | 13 | Saprotroph | \|Undefined Saprotroph\| | Yeast | NULL | Probable | As nectar-tap_saprotroph (Põlme et al. 2020); Decay sugar-rich substrates (Põlme et al. 2020) | Põlme S, et al. 2020. Fungal Diversity 105:1-16 ((https://doi.org/10.1007/s13225-020-00466-2)) |
| Otu59940 | 1 | k__Fungi(100);p__Ascomycota(100);c__Saccharomycetes(100);o__Saccharomycetales(100);f__Metschnikowiaceae(100);g__Metschnikowia(100);g__Metschnikowia_unclassified(100); | Metschnikowia | 13 | Saprotroph | \|Undefined Saprotroph\| | Yeast | NULL | Probable | As nectar-tap_saprotroph (Põlme et al. 2020); Decay sugar-rich substrates (Põlme et al. 2020) | Põlme S, et al. 2020. Fungal Diversity 105:1-16 ((https://doi.org/10.1007/s13225-020-00466-2)) |
| Otu59941 | 1 | k__Fungi(100);p__Ascomycota(100);c__Saccharomycetes(100);o__Saccharomycetales(100);f__Metschnikowiaceae(100);g__Metschnikowia(100);s__Metschnikowia_pulcherrima(100); | Metschnikowia | 13 | Saprotroph | \|Undefined Saprotroph\| | Yeast | NULL | Probable | As nectar-tap_saprotroph (Põlme et al. 2020); Decay sugar-rich substrates (Põlme et al. 2020) | Põlme S, et al. 2020. Fungal Diversity 105:1-16 ((https://doi.org/10.1007/s13225-020-00466-2)) |
| Otu59942 | 1 | k__Fungi(100);p__Ascomycota(100);c__Saccharomycetes(100);o__Saccharomycetales(100);f__Metschnikowiaceae(100);g__Metschnikowia(100);s__Metschnikowia_pulcherrima(100); | Metschnikowia | 13 | Saprotroph | \|Undefined Saprotroph\| | Yeast | NULL | Probable | As nectar-tap_saprotroph (Põlme et al. 2020); Decay sugar-rich substrates (Põlme et al. 2020) | Põlme S, et al. 2020. Fungal Diversity 105:1-16 ((https://doi.org/10.1007/s13225-020-00466-2)) |
| Otu59944 | 1 | k__Fungi(100);p__Ascomycota(100);c__Saccharomycetes(100);o__Saccharomycetales(100);f__Metschnikowiaceae(100);g__Metschnikowia(100);s__Metschnikowia_pulcherrima(100); | Metschnikowia | 13 | Saprotroph | \|Undefined Saprotroph\| | Yeast | NULL | Probable | As nectar-tap_saprotroph (Põlme et al. 2020); Decay sugar-rich substrates (Põlme et al. 2020) | Põlme S, et al. 2020. Fungal Diversity 105:1-16 ((https://doi.org/10.1007/s13225-020-00466-2)) |
| Otu59945 | 1 | k__Fungi(100);p__Ascomycota(100);c__Saccharomycetes(100);o__Saccharomycetales(100);f__Metschnikowiaceae(100);g__Metschnikowia(100);s__Metschnikowia_pulcherrima(100); | Metschnikowia | 13 | Saprotroph | \|Undefined Saprotroph\| | Yeast | NULL | Probable | As nectar-tap_saprotroph (Põlme et al. 2020); Decay sugar-rich substrates (Põlme et al. 2020) | Põlme S, et al. 2020. Fungal Diversity 105:1-16 ((https://doi.org/10.1007/s13225-020-00466-2)) |
| Otu59946 | 1 | k__Fungi(100);p__Ascomycota(100);c__Saccharomycetes(100);o__Saccharomycetales(100);f__Metschnikowiaceae(100);g__Metschnikowia(100);s__Metschnikowia_pulcherrima(100); | Metschnikowia | 13 | Saprotroph | \|Undefined Saprotroph\| | Yeast | NULL | Probable | As nectar-tap_saprotroph (Põlme et al. 2020); Decay sugar-rich substrates (Põlme et al. 2020) | Põlme S, et al. 2020. Fungal Diversity 105:1-16 ((https://doi.org/10.1007/s13225-020-00466-2)) |
| Otu59947 | 1 | k__Fungi(100);p__Ascomycota(100);c__Saccharomycetes(100);o__Saccharomycetales(100);f__Metschnikowiaceae(100);g__Metschnikowia(100);g__Metschnikowia_unclassified(100); | Metschnikowia | 13 | Saprotroph | \|Undefined Saprotroph\| | Yeast | NULL | Probable | As nectar-tap_saprotroph (Põlme et al. 2020); Decay sugar-rich substrates (Põlme et al. 2020) | Põlme S, et al. 2020. Fungal Diversity 105:1-16 ((https://doi.org/10.1007/s13225-020-00466-2)) |
| Otu59948 | 1 | k__Fungi(100);p__Ascomycota(100);c__Saccharomycetes(100);o__Saccharomycetales(100);f__Metschnikowiaceae(100);g__Metschnikowia(100);s__Metschnikowia_pulcherrima(100); | Metschnikowia | 13 | Saprotroph | \|Undefined Saprotroph\| | Yeast | NULL | Probable | As nectar-tap_saprotroph (Põlme et al. 2020); Decay sugar-rich substrates (Põlme et al. 2020) | Põlme S, et al. 2020. Fungal Diversity 105:1-16 ((https://doi.org/10.1007/s13225-020-00466-2)) |
| Otu59949 | 1 | k__Fungi(100);p__Ascomycota(100);c__Saccharomycetes(100);o__Saccharomycetales(100);f__Metschnikowiaceae(100);g__Metschnikowia(100);s__Metschnikowia_pulcherrima(100); | Metschnikowia | 13 | Saprotroph | \|Undefined Saprotroph\| | Yeast | NULL | Probable | As nectar-tap_saprotroph (Põlme et al. 2020); Decay sugar-rich substrates (Põlme et al. 2020) | Põlme S, et al. 2020. Fungal Diversity 105:1-16 ((https://doi.org/10.1007/s13225-020-00466-2)) |
| Otu59950 | 1 | k__Fungi(100);p__Ascomycota(100);c__Saccharomycetes(100);o__Saccharomycetales(100);f__Metschnikowiaceae(100);g__Metschnikowia(100);g__Metschnikowia_unclassified(100); | Metschnikowia | 13 | Saprotroph | \|Undefined Saprotroph\| | Yeast | NULL | Probable | As nectar-tap_saprotroph (Põlme et al. 2020); Decay sugar-rich substrates (Põlme et al. 2020) | Põlme S, et al. 2020. Fungal Diversity 105:1-16 ((https://doi.org/10.1007/s13225-020-00466-2)) |
| Otu59951 | 1 | k__Fungi(100);p__Ascomycota(100);c__Saccharomycetes(100);o__Saccharomycetales(100);f__Metschnikowiaceae(100);g__Metschnikowia(100);s__Metschnikowia_pulcherrima(100); | Metschnikowia | 13 | Saprotroph | \|Undefined Saprotroph\| | Yeast | NULL | Probable | As nectar-tap_saprotroph (Põlme et al. 2020); Decay sugar-rich substrates (Põlme et al. 2020) | Põlme S, et al. 2020. Fungal Diversity 105:1-16 ((https://doi.org/10.1007/s13225-020-00466-2)) |
| Otu59952 | 1 | k__Fungi(100);p__Ascomycota(100);c__Saccharomycetes(100);o__Saccharomycetales(100);f__Metschnikowiaceae(100);g__Metschnikowia(100);s__Metschnikowia_pulcherrima(100); | Metschnikowia | 13 | Saprotroph | \|Undefined Saprotroph\| | Yeast | NULL | Probable | As nectar-tap_saprotroph (Põlme et al. 2020); Decay sugar-rich substrates (Põlme et al. 2020) | Põlme S, et al. 2020. Fungal Diversity 105:1-16 ((https://doi.org/10.1007/s13225-020-00466-2)) |
| Otu59953 | 1 | k__Fungi(100);p__Ascomycota(100);c__Saccharomycetes(100);o__Saccharomycetales(100);f__Metschnikowiaceae(100);g__Metschnikowia(100);s__Metschnikowia_pulcherrima(100); | Metschnikowia | 13 | Saprotroph | \|Undefined Saprotroph\| | Yeast | NULL | Probable | As nectar-tap_saprotroph (Põlme et al. 2020); Decay sugar-rich substrates (Põlme et al. 2020) | Põlme S, et al. 2020. Fungal Diversity 105:1-16 ((https://doi.org/10.1007/s13225-020-00466-2)) |
| Otu59954 | 1 | k__Fungi(100);p__Ascomycota(100);c__Saccharomycetes(100);o__Saccharomycetales(100);f__Metschnikowiaceae(100);g__Metschnikowia(100);s__Metschnikowia_pulcherrima(100); | Metschnikowia | 13 | Saprotroph | \|Undefined Saprotroph\| | Yeast | NULL | Probable | As nectar-tap_saprotroph (Põlme et al. 2020); Decay sugar-rich substrates (Põlme et al. 2020) | Põlme S, et al. 2020. Fungal Diversity 105:1-16 ((https://doi.org/10.1007/s13225-020-00466-2)) |
| Otu59957 | 1 | k__Fungi(100);p__Ascomycota(100);c__Saccharomycetes(100);o__Saccharomycetales(100);f__Saccharomycetales_family_Incertae_sedis(100);f__Saccharomycetales_family_Incertae_sedis_unclassified(100);f__Saccharomycetales_family_Incertae_sedis_unclassified(100); | Saccharomycetales | 7 | Saprotroph | Undefined Saprotroph | Yeast | NULL | Possible | NULL | Sterkenburg E, et al. 2015. New Phytologist 207:1145-1158 ((https://doi.org/10.1111/nph.13426)) |
| Otu59962 | 1 | k__Fungi(100);p__Ascomycota(100);c__Saccharomycetes(100);o__Saccharomycetales(100);f__Metschnikowiaceae(100);g__Metschnikowia(100);g__Metschnikowia_unclassified(100); | Metschnikowia | 13 | Saprotroph | \|Undefined Saprotroph\| | Yeast | NULL | Probable | As nectar-tap_saprotroph (Põlme et al. 2020); Decay sugar-rich substrates (Põlme et al. 2020) | Põlme S, et al. 2020. Fungal Diversity 105:1-16 ((https://doi.org/10.1007/s13225-020-00466-2)) |
| Otu59963 | 1 | k__Fungi(100);p__Ascomycota(100);c__Saccharomycetes(100);o__Saccharomycetales(100);f__Saccharomycetales_family_Incertae_sedis(100);f__Saccharomycetales_family_Incertae_sedis_unclassified(100);f__Saccharomycetales_family_Incertae_sedis_unclassified(100); | Saccharomycetales | 7 | Saprotroph | Undefined Saprotroph | Yeast | NULL | Possible | NULL | Sterkenburg E, et al. 2015. New Phytologist 207:1145-1158 ((https://doi.org/10.1111/nph.13426)) |
| Otu59964 | 1 | k__Fungi(100);p__Ascomycota(100);c__Saccharomycetes(100);o__Saccharomycetales(100);f__Metschnikowiaceae(100);g__Metschnikowia(100);s__Metschnikowia_pulcherrima(100); | Metschnikowia | 13 | Saprotroph | \|Undefined Saprotroph\| | Yeast | NULL | Probable | As nectar-tap_saprotroph (Põlme et al. 2020); Decay sugar-rich substrates (Põlme et al. 2020) | Põlme S, et al. 2020. Fungal Diversity 105:1-16 ((https://doi.org/10.1007/s13225-020-00466-2)) |
| Otu59970 | 1 | k__Fungi(100);p__Ascomycota(100);c__Saccharomycetes(100);o__Saccharomycetales(100);f__Saccharomycetales_family_Incertae_sedis(100);g__Candida(100);g__Candida_unclassified(100); | Candida | 13 | Pathotroph-Saprotroph-Symbiotroph | Animal Pathogen-Endophyte-Endosymbiont-Epiphyte-\|Undefined Saprotroph\| | Dimorphic Yeast-Yeast | NULL | Probable | As nectar-tap_saprotroph (Põlme et al. 2020); Decay sugar-rich substrates (Põlme et al. 2020); This asexual genus is polyphyletic and contain species that function very differently depend on what they are. Caution should be taken when interpreting guilds for Candida species. | Manolakaki D, et al. 2010. Virulence. 1: 367-375 ((https://doi.org/10.4161/viru.1.5.12796)); Mekbib SB, et al. 2011. Tropical Plant Pathology 36:233-240 ((https://doi.org/10.1590/S1982-56762011000400004)); Põlme S, et al. 2020. Fungal Diversity 105:1-16 ((https://doi.org/10.1007/s13225-020-00466-2)) |
| Otu59971 | 1 | k__Fungi(100);p__Ascomycota(100);c__Saccharomycetes(100);o__Saccharomycetales(100);f__Metschnikowiaceae(100);g__Metschnikowia(100);s__Metschnikowia_pulcherrima(100); | Metschnikowia | 13 | Saprotroph | \|Undefined Saprotroph\| | Yeast | NULL | Probable | As nectar-tap_saprotroph (Põlme et al. 2020); Decay sugar-rich substrates (Põlme et al. 2020) | Põlme S, et al. 2020. Fungal Diversity 105:1-16 ((https://doi.org/10.1007/s13225-020-00466-2)) |
| Otu59972 | 1 | k__Fungi(100);p__Ascomycota(100);c__Saccharomycetes(100);o__Saccharomycetales(100);f__Metschnikowiaceae(100);g__Metschnikowia(100);g__Metschnikowia_unclassified(100); | Metschnikowia | 13 | Saprotroph | \|Undefined Saprotroph\| | Yeast | NULL | Probable | As nectar-tap_saprotroph (Põlme et al. 2020); Decay sugar-rich substrates (Põlme et al. 2020) | Põlme S, et al. 2020. Fungal Diversity 105:1-16 ((https://doi.org/10.1007/s13225-020-00466-2)) |
| Otu59973 | 1 | k__Fungi(100);p__Ascomycota(100);c__Saccharomycetes(100);o__Saccharomycetales(100);f__Metschnikowiaceae(100);g__Metschnikowia(100);s__Metschnikowia_pulcherrima(100); | Metschnikowia | 13 | Saprotroph | \|Undefined Saprotroph\| | Yeast | NULL | Probable | As nectar-tap_saprotroph (Põlme et al. 2020); Decay sugar-rich substrates (Põlme et al. 2020) | Põlme S, et al. 2020. Fungal Diversity 105:1-16 ((https://doi.org/10.1007/s13225-020-00466-2)) |
| Otu59974 | 1 | k__Fungi(100);p__Ascomycota(100);c__Saccharomycetes(100);o__Saccharomycetales(100);f__Metschnikowiaceae(100);g__Metschnikowia(100);s__Metschnikowia_pulcherrima(100); | Metschnikowia | 13 | Saprotroph | \|Undefined Saprotroph\| | Yeast | NULL | Probable | As nectar-tap_saprotroph (Põlme et al. 2020); Decay sugar-rich substrates (Põlme et al. 2020) | Põlme S, et al. 2020. Fungal Diversity 105:1-16 ((https://doi.org/10.1007/s13225-020-00466-2)) |
| Otu59975 | 1 | k__Fungi(100);p__Ascomycota(100);c__Saccharomycetes(100);o__Saccharomycetales(100);f__Metschnikowiaceae(100);g__Metschnikowia(100);s__Metschnikowia_pulcherrima(100); | Metschnikowia | 13 | Saprotroph | \|Undefined Saprotroph\| | Yeast | NULL | Probable | As nectar-tap_saprotroph (Põlme et al. 2020); Decay sugar-rich substrates (Põlme et al. 2020) | Põlme S, et al. 2020. Fungal Diversity 105:1-16 ((https://doi.org/10.1007/s13225-020-00466-2)) |
| Otu59978 | 1 | k__Fungi(100);p__Ascomycota(100);c__Saccharomycetes(100);o__Saccharomycetales(100);f__Metschnikowiaceae(100);g__Metschnikowia(100);s__Metschnikowia_pulcherrima(100); | Metschnikowia | 13 | Saprotroph | \|Undefined Saprotroph\| | Yeast | NULL | Probable | As nectar-tap_saprotroph (Põlme et al. 2020); Decay sugar-rich substrates (Põlme et al. 2020) | Põlme S, et al. 2020. Fungal Diversity 105:1-16 ((https://doi.org/10.1007/s13225-020-00466-2)) |
| Otu59979 | 1 | k__Fungi(100);p__Ascomycota(100);c__Saccharomycetes(100);o__Saccharomycetales(100);f__Metschnikowiaceae(100);g__Metschnikowia(100);s__Metschnikowia_pulcherrima(100); | Metschnikowia | 13 | Saprotroph | \|Undefined Saprotroph\| | Yeast | NULL | Probable | As nectar-tap_saprotroph (Põlme et al. 2020); Decay sugar-rich substrates (Põlme et al. 2020) | Põlme S, et al. 2020. Fungal Diversity 105:1-16 ((https://doi.org/10.1007/s13225-020-00466-2)) |
| Otu59980 | 1 | k__Fungi(100);p__Ascomycota(100);c__Saccharomycetes(100);o__Saccharomycetales(100);f__Metschnikowiaceae(100);g__Metschnikowia(100);s__Metschnikowia_pulcherrima(100); | Metschnikowia | 13 | Saprotroph | \|Undefined Saprotroph\| | Yeast | NULL | Probable | As nectar-tap_saprotroph (Põlme et al. 2020); Decay sugar-rich substrates (Põlme et al. 2020) | Põlme S, et al. 2020. Fungal Diversity 105:1-16 ((https://doi.org/10.1007/s13225-020-00466-2)) |
| Otu59981 | 1 | k__Fungi(100);p__Ascomycota(100);c__Saccharomycetes(100);o__Saccharomycetales(100);f__Metschnikowiaceae(100);g__Metschnikowia(100);s__Metschnikowia_pulcherrima(100); | Metschnikowia | 13 | Saprotroph | \|Undefined Saprotroph\| | Yeast | NULL | Probable | As nectar-tap_saprotroph (Põlme et al. 2020); Decay sugar-rich substrates (Põlme et al. 2020) | Põlme S, et al. 2020. Fungal Diversity 105:1-16 ((https://doi.org/10.1007/s13225-020-00466-2)) |
| Otu59985 | 1 | k__Fungi(100);p__Ascomycota(100);c__Saccharomycetes(100);o__Saccharomycetales(100);f__Metschnikowiaceae(100);g__Metschnikowia(100);s__Metschnikowia_pulcherrima(100); | Metschnikowia | 13 | Saprotroph | \|Undefined Saprotroph\| | Yeast | NULL | Probable | As nectar-tap_saprotroph (Põlme et al. 2020); Decay sugar-rich substrates (Põlme et al. 2020) | Põlme S, et al. 2020. Fungal Diversity 105:1-16 ((https://doi.org/10.1007/s13225-020-00466-2)) |
| Otu59990 | 1 | k__Fungi(100);p__Ascomycota(100);c__Saccharomycetes(100);o__Saccharomycetales(100);f__Metschnikowiaceae(100);g__Metschnikowia(100);s__Metschnikowia_pulcherrima(100); | Metschnikowia | 13 | Saprotroph | \|Undefined Saprotroph\| | Yeast | NULL | Probable | As nectar-tap_saprotroph (Põlme et al. 2020); Decay sugar-rich substrates (Põlme et al. 2020) | Põlme S, et al. 2020. Fungal Diversity 105:1-16 ((https://doi.org/10.1007/s13225-020-00466-2)) |
| Otu59992 | 1 | k__Fungi(100);p__Ascomycota(100);c__Saccharomycetes(100);o__Saccharomycetales(100);f__Metschnikowiaceae(100);g__Metschnikowia(100);s__Metschnikowia_pulcherrima(100); | Metschnikowia | 13 | Saprotroph | \|Undefined Saprotroph\| | Yeast | NULL | Probable | As nectar-tap_saprotroph (Põlme et al. 2020); Decay sugar-rich substrates (Põlme et al. 2020) | Põlme S, et al. 2020. Fungal Diversity 105:1-16 ((https://doi.org/10.1007/s13225-020-00466-2)) |
| Otu59994 | 1 | k__Fungi(100);p__Ascomycota(100);c__Saccharomycetes(100);o__Saccharomycetales(100);f__Metschnikowiaceae(100);g__Metschnikowia(100);s__Metschnikowia_pulcherrima(100); | Metschnikowia | 13 | Saprotroph | \|Undefined Saprotroph\| | Yeast | NULL | Probable | As nectar-tap_saprotroph (Põlme et al. 2020); Decay sugar-rich substrates (Põlme et al. 2020) | Põlme S, et al. 2020. Fungal Diversity 105:1-16 ((https://doi.org/10.1007/s13225-020-00466-2)) |
| Otu59995 | 1 | k__Fungi(100);p__Ascomycota(100);c__Saccharomycetes(100);o__Saccharomycetales(100);f__Metschnikowiaceae(100);g__Metschnikowia(100);s__Metschnikowia_pulcherrima(100); | Metschnikowia | 13 | Saprotroph | \|Undefined Saprotroph\| | Yeast | NULL | Probable | As nectar-tap_saprotroph (Põlme et al. 2020); Decay sugar-rich substrates (Põlme et al. 2020) | Põlme S, et al. 2020. Fungal Diversity 105:1-16 ((https://doi.org/10.1007/s13225-020-00466-2)) |
| Otu59996 | 1 | k__Fungi(100);p__Ascomycota(100);c__Saccharomycetes(100);o__Saccharomycetales(100);f__Metschnikowiaceae(100);g__Metschnikowia(100);g__Metschnikowia_unclassified(100); | Metschnikowia | 13 | Saprotroph | \|Undefined Saprotroph\| | Yeast | NULL | Probable | As nectar-tap_saprotroph (Põlme et al. 2020); Decay sugar-rich substrates (Põlme et al. 2020) | Põlme S, et al. 2020. Fungal Diversity 105:1-16 ((https://doi.org/10.1007/s13225-020-00466-2)) |
| Otu59997 | 1 | k__Fungi(100);p__Ascomycota(100);c__Saccharomycetes(100);o__Saccharomycetales(100);f__Saccharomycetales_family_Incertae_sedis(100);g__Candida(100);g__Candida_unclassified(100); | Candida | 13 | Pathotroph-Saprotroph-Symbiotroph | Animal Pathogen-Endophyte-Endosymbiont-Epiphyte-\|Undefined Saprotroph\| | Dimorphic Yeast-Yeast | NULL | Probable | As nectar-tap_saprotroph (Põlme et al. 2020); Decay sugar-rich substrates (Põlme et al. 2020); This asexual genus is polyphyletic and contain species that function very differently depend on what they are. Caution should be taken when interpreting guilds for Candida species. | Manolakaki D, et al. 2010. Virulence. 1: 367-375 ((https://doi.org/10.4161/viru.1.5.12796)); Mekbib SB, et al. 2011. Tropical Plant Pathology 36:233-240 ((https://doi.org/10.1590/S1982-56762011000400004)); Põlme S, et al. 2020. Fungal Diversity 105:1-16 ((https://doi.org/10.1007/s13225-020-00466-2)) |
| Otu59998 | 1 | k__Fungi(100);p__Ascomycota(100);c__Saccharomycetes(100);o__Saccharomycetales(100);f__Metschnikowiaceae(100);g__Metschnikowia(100);s__Metschnikowia_pulcherrima(100); | Metschnikowia | 13 | Saprotroph | \|Undefined Saprotroph\| | Yeast | NULL | Probable | As nectar-tap_saprotroph (Põlme et al. 2020); Decay sugar-rich substrates (Põlme et al. 2020) | Põlme S, et al. 2020. Fungal Diversity 105:1-16 ((https://doi.org/10.1007/s13225-020-00466-2)) |
| Otu60007 | 1 | k__Fungi(100);p__Ascomycota(100);c__Saccharomycetes(100);o__Saccharomycetales(100);f__Saccharomycetales_family_Incertae_sedis(100);g__Candida(100);g__Candida_unclassified(100); | Candida | 13 | Pathotroph-Saprotroph-Symbiotroph | Animal Pathogen-Endophyte-Endosymbiont-Epiphyte-\|Undefined Saprotroph\| | Dimorphic Yeast-Yeast | NULL | Probable | As nectar-tap_saprotroph (Põlme et al. 2020); Decay sugar-rich substrates (Põlme et al. 2020); This asexual genus is polyphyletic and contain species that function very differently depend on what they are. Caution should be taken when interpreting guilds for Candida species. | Manolakaki D, et al. 2010. Virulence. 1: 367-375 ((https://doi.org/10.4161/viru.1.5.12796)); Mekbib SB, et al. 2011. Tropical Plant Pathology 36:233-240 ((https://doi.org/10.1590/S1982-56762011000400004)); Põlme S, et al. 2020. Fungal Diversity 105:1-16 ((https://doi.org/10.1007/s13225-020-00466-2)) |
| Otu60024 | 1 | k__Fungi(100);p__Ascomycota(100);c__Saccharomycetes(100);o__Saccharomycetales(100);f__Metschnikowiaceae(100);g__Metschnikowia(100);g__Metschnikowia_unclassified(100); | Metschnikowia | 13 | Saprotroph | \|Undefined Saprotroph\| | Yeast | NULL | Probable | As nectar-tap_saprotroph (Põlme et al. 2020); Decay sugar-rich substrates (Põlme et al. 2020) | Põlme S, et al. 2020. Fungal Diversity 105:1-16 ((https://doi.org/10.1007/s13225-020-00466-2)) |
| Otu60026 | 1 | k__Fungi(100);p__Ascomycota(100);c__Saccharomycetes(100);o__Saccharomycetales(100);f__Metschnikowiaceae(100);g__Metschnikowia(100);s__Metschnikowia_pulcherrima(100); | Metschnikowia | 13 | Saprotroph | \|Undefined Saprotroph\| | Yeast | NULL | Probable | As nectar-tap_saprotroph (Põlme et al. 2020); Decay sugar-rich substrates (Põlme et al. 2020) | Põlme S, et al. 2020. Fungal Diversity 105:1-16 ((https://doi.org/10.1007/s13225-020-00466-2)) |
| Otu60027 | 1 | k__Fungi(100);p__Ascomycota(100);c__Saccharomycetes(100);o__Saccharomycetales(100);f__Metschnikowiaceae(100);g__Metschnikowia(100);s__Metschnikowia_pulcherrima(100); | Metschnikowia | 13 | Saprotroph | \|Undefined Saprotroph\| | Yeast | NULL | Probable | As nectar-tap_saprotroph (Põlme et al. 2020); Decay sugar-rich substrates (Põlme et al. 2020) | Põlme S, et al. 2020. Fungal Diversity 105:1-16 ((https://doi.org/10.1007/s13225-020-00466-2)) |
| Otu60030 | 1 | k__Fungi(100);p__Ascomycota(100);c__Saccharomycetes(100);o__Saccharomycetales(100);f__Metschnikowiaceae(100);g__Metschnikowia(100);s__Metschnikowia_pulcherrima(100); | Metschnikowia | 13 | Saprotroph | \|Undefined Saprotroph\| | Yeast | NULL | Probable | As nectar-tap_saprotroph (Põlme et al. 2020); Decay sugar-rich substrates (Põlme et al. 2020) | Põlme S, et al. 2020. Fungal Diversity 105:1-16 ((https://doi.org/10.1007/s13225-020-00466-2)) |
| Otu60034 | 1 | k__Fungi(100);p__Ascomycota(100);c__Saccharomycetes(100);o__Saccharomycetales(100);f__Metschnikowiaceae(100);g__Metschnikowia(100);s__Metschnikowia_pulcherrima(100); | Metschnikowia | 13 | Saprotroph | \|Undefined Saprotroph\| | Yeast | NULL | Probable | As nectar-tap_saprotroph (Põlme et al. 2020); Decay sugar-rich substrates (Põlme et al. 2020) | Põlme S, et al. 2020. Fungal Diversity 105:1-16 ((https://doi.org/10.1007/s13225-020-00466-2)) |
| Otu60035 | 1 | k__Fungi(100);p__Ascomycota(100);c__Saccharomycetes(100);o__Saccharomycetales(100);f__Metschnikowiaceae(100);g__Metschnikowia(100);s__Metschnikowia_pulcherrima(100); | Metschnikowia | 13 | Saprotroph | \|Undefined Saprotroph\| | Yeast | NULL | Probable | As nectar-tap_saprotroph (Põlme et al. 2020); Decay sugar-rich substrates (Põlme et al. 2020) | Põlme S, et al. 2020. Fungal Diversity 105:1-16 ((https://doi.org/10.1007/s13225-020-00466-2)) |
| Otu60037 | 1 | k__Fungi(100);p__Ascomycota(100);c__Saccharomycetes(100);o__Saccharomycetales(100);f__Metschnikowiaceae(100);g__Metschnikowia(100);s__Metschnikowia_pulcherrima(100); | Metschnikowia | 13 | Saprotroph | \|Undefined Saprotroph\| | Yeast | NULL | Probable | As nectar-tap_saprotroph (Põlme et al. 2020); Decay sugar-rich substrates (Põlme et al. 2020) | Põlme S, et al. 2020. Fungal Diversity 105:1-16 ((https://doi.org/10.1007/s13225-020-00466-2)) |
| Otu60046 | 1 | k__Fungi(100);p__Ascomycota(100);c__Saccharomycetes(100);o__Saccharomycetales(100);f__Metschnikowiaceae(100);g__Metschnikowia(100);g__Metschnikowia_unclassified(100); | Metschnikowia | 13 | Saprotroph | \|Undefined Saprotroph\| | Yeast | NULL | Probable | As nectar-tap_saprotroph (Põlme et al. 2020); Decay sugar-rich substrates (Põlme et al. 2020) | Põlme S, et al. 2020. Fungal Diversity 105:1-16 ((https://doi.org/10.1007/s13225-020-00466-2)) |
| Otu60047 | 1 | k__Fungi(100);p__Ascomycota(100);c__Saccharomycetes(100);o__Saccharomycetales(100);f__Metschnikowiaceae(100);g__Metschnikowia(100);s__Metschnikowia_pulcherrima(100); | Metschnikowia | 13 | Saprotroph | \|Undefined Saprotroph\| | Yeast | NULL | Probable | As nectar-tap_saprotroph (Põlme et al. 2020); Decay sugar-rich substrates (Põlme et al. 2020) | Põlme S, et al. 2020. Fungal Diversity 105:1-16 ((https://doi.org/10.1007/s13225-020-00466-2)) |
| Otu60048 | 1 | k__Fungi(100);p__Ascomycota(100);c__Saccharomycetes(100);o__Saccharomycetales(100);f__Metschnikowiaceae(100);g__Metschnikowia(100);s__Metschnikowia_pulcherrima(100); | Metschnikowia | 13 | Saprotroph | \|Undefined Saprotroph\| | Yeast | NULL | Probable | As nectar-tap_saprotroph (Põlme et al. 2020); Decay sugar-rich substrates (Põlme et al. 2020) | Põlme S, et al. 2020. Fungal Diversity 105:1-16 ((https://doi.org/10.1007/s13225-020-00466-2)) |
| Otu60064 | 1 | k__Fungi(100);p__Ascomycota(100);c__Saccharomycetes(100);o__Saccharomycetales(100);f__Metschnikowiaceae(100);g__Metschnikowia(100);s__Metschnikowia_pulcherrima(100); | Metschnikowia | 13 | Saprotroph | \|Undefined Saprotroph\| | Yeast | NULL | Probable | As nectar-tap_saprotroph (Põlme et al. 2020); Decay sugar-rich substrates (Põlme et al. 2020) | Põlme S, et al. 2020. Fungal Diversity 105:1-16 ((https://doi.org/10.1007/s13225-020-00466-2)) |
| Otu60072 | 1 | k__Fungi(100);p__Ascomycota(100);c__Saccharomycetes(100);o__Saccharomycetales(100);f__Metschnikowiaceae(100);g__Metschnikowia(100);g__Metschnikowia_unclassified(100); | Metschnikowia | 13 | Saprotroph | \|Undefined Saprotroph\| | Yeast | NULL | Probable | As nectar-tap_saprotroph (Põlme et al. 2020); Decay sugar-rich substrates (Põlme et al. 2020) | Põlme S, et al. 2020. Fungal Diversity 105:1-16 ((https://doi.org/10.1007/s13225-020-00466-2)) |
| Otu60074 | 1 | k__Fungi(100);p__Ascomycota(100);c__Saccharomycetes(100);o__Saccharomycetales(100);f__Metschnikowiaceae(100);g__Metschnikowia(100);s__Metschnikowia_pulcherrima(100); | Metschnikowia | 13 | Saprotroph | \|Undefined Saprotroph\| | Yeast | NULL | Probable | As nectar-tap_saprotroph (Põlme et al. 2020); Decay sugar-rich substrates (Põlme et al. 2020) | Põlme S, et al. 2020. Fungal Diversity 105:1-16 ((https://doi.org/10.1007/s13225-020-00466-2)) |
| Otu60075 | 1 | k__Fungi(100);p__Ascomycota(100);c__Saccharomycetes(100);o__Saccharomycetales(100);f__Metschnikowiaceae(100);g__Metschnikowia(100);s__Metschnikowia_pulcherrima(100); | Metschnikowia | 13 | Saprotroph | \|Undefined Saprotroph\| | Yeast | NULL | Probable | As nectar-tap_saprotroph (Põlme et al. 2020); Decay sugar-rich substrates (Põlme et al. 2020) | Põlme S, et al. 2020. Fungal Diversity 105:1-16 ((https://doi.org/10.1007/s13225-020-00466-2)) |
| Otu60076 | 1 | k__Fungi(100);p__Ascomycota(100);c__Saccharomycetes(100);o__Saccharomycetales(100);f__Metschnikowiaceae(100);g__Metschnikowia(100);s__Metschnikowia_pulcherrima(100); | Metschnikowia | 13 | Saprotroph | \|Undefined Saprotroph\| | Yeast | NULL | Probable | As nectar-tap_saprotroph (Põlme et al. 2020); Decay sugar-rich substrates (Põlme et al. 2020) | Põlme S, et al. 2020. Fungal Diversity 105:1-16 ((https://doi.org/10.1007/s13225-020-00466-2)) |
| Otu60077 | 1 | k__Fungi(100);p__Ascomycota(100);c__Saccharomycetes(100);o__Saccharomycetales(100);f__Metschnikowiaceae(100);g__Metschnikowia(100);g__Metschnikowia_unclassified(100); | Metschnikowia | 13 | Saprotroph | \|Undefined Saprotroph\| | Yeast | NULL | Probable | As nectar-tap_saprotroph (Põlme et al. 2020); Decay sugar-rich substrates (Põlme et al. 2020) | Põlme S, et al. 2020. Fungal Diversity 105:1-16 ((https://doi.org/10.1007/s13225-020-00466-2)) |
| Otu60078 | 1 | k__Fungi(100);p__Ascomycota(100);c__Saccharomycetes(100);o__Saccharomycetales(100);f__Metschnikowiaceae(100);g__Metschnikowia(100);s__Metschnikowia_pulcherrima(100); | Metschnikowia | 13 | Saprotroph | \|Undefined Saprotroph\| | Yeast | NULL | Probable | As nectar-tap_saprotroph (Põlme et al. 2020); Decay sugar-rich substrates (Põlme et al. 2020) | Põlme S, et al. 2020. Fungal Diversity 105:1-16 ((https://doi.org/10.1007/s13225-020-00466-2)) |
| Otu60079 | 1 | k__Fungi(100);p__Ascomycota(100);c__Saccharomycetes(100);o__Saccharomycetales(100);f__Metschnikowiaceae(100);g__Metschnikowia(100);s__Metschnikowia_pulcherrima(100); | Metschnikowia | 13 | Saprotroph | \|Undefined Saprotroph\| | Yeast | NULL | Probable | As nectar-tap_saprotroph (Põlme et al. 2020); Decay sugar-rich substrates (Põlme et al. 2020) | Põlme S, et al. 2020. Fungal Diversity 105:1-16 ((https://doi.org/10.1007/s13225-020-00466-2)) |
| Otu60084 | 1 | k__Fungi(100);p__Ascomycota(100);c__Saccharomycetes(100);o__Saccharomycetales(100);f__Metschnikowiaceae(100);g__Metschnikowia(100);s__Metschnikowia_pulcherrima(100); | Metschnikowia | 13 | Saprotroph | \|Undefined Saprotroph\| | Yeast | NULL | Probable | As nectar-tap_saprotroph (Põlme et al. 2020); Decay sugar-rich substrates (Põlme et al. 2020) | Põlme S, et al. 2020. Fungal Diversity 105:1-16 ((https://doi.org/10.1007/s13225-020-00466-2)) |
| Otu60085 | 1 | k__Fungi(100);p__Ascomycota(100);c__Saccharomycetes(100);o__Saccharomycetales(100);f__Metschnikowiaceae(100);g__Metschnikowia(100);g__Metschnikowia_unclassified(100); | Metschnikowia | 13 | Saprotroph | \|Undefined Saprotroph\| | Yeast | NULL | Probable | As nectar-tap_saprotroph (Põlme et al. 2020); Decay sugar-rich substrates (Põlme et al. 2020) | Põlme S, et al. 2020. Fungal Diversity 105:1-16 ((https://doi.org/10.1007/s13225-020-00466-2)) |
| Otu60091 | 1 | k__Fungi(100);p__Ascomycota(100);c__Saccharomycetes(100);o__Saccharomycetales(100);f__Metschnikowiaceae(100);g__Metschnikowia(100);s__Metschnikowia_pulcherrima(100); | Metschnikowia | 13 | Saprotroph | \|Undefined Saprotroph\| | Yeast | NULL | Probable | As nectar-tap_saprotroph (Põlme et al. 2020); Decay sugar-rich substrates (Põlme et al. 2020) | Põlme S, et al. 2020. Fungal Diversity 105:1-16 ((https://doi.org/10.1007/s13225-020-00466-2)) |
| Otu60093 | 1 | k__Fungi(100);p__Ascomycota(100);c__Saccharomycetes(100);o__Saccharomycetales(100);f__Metschnikowiaceae(100);g__Metschnikowia(100);s__Metschnikowia_pulcherrima(100); | Metschnikowia | 13 | Saprotroph | \|Undefined Saprotroph\| | Yeast | NULL | Probable | As nectar-tap_saprotroph (Põlme et al. 2020); Decay sugar-rich substrates (Põlme et al. 2020) | Põlme S, et al. 2020. Fungal Diversity 105:1-16 ((https://doi.org/10.1007/s13225-020-00466-2)) |
| Otu60094 | 1 | k__Fungi(100);p__Ascomycota(100);c__Saccharomycetes(100);o__Saccharomycetales(100);f__Metschnikowiaceae(100);g__Metschnikowia(100);s__Metschnikowia_pulcherrima(100); | Metschnikowia | 13 | Saprotroph | \|Undefined Saprotroph\| | Yeast | NULL | Probable | As nectar-tap_saprotroph (Põlme et al. 2020); Decay sugar-rich substrates (Põlme et al. 2020) | Põlme S, et al. 2020. Fungal Diversity 105:1-16 ((https://doi.org/10.1007/s13225-020-00466-2)) |
| Otu60095 | 1 | k__Fungi(100);p__Ascomycota(100);c__Saccharomycetes(100);o__Saccharomycetales(100);f__Metschnikowiaceae(100);g__Metschnikowia(100);s__Metschnikowia_pulcherrima(100); | Metschnikowia | 13 | Saprotroph | \|Undefined Saprotroph\| | Yeast | NULL | Probable | As nectar-tap_saprotroph (Põlme et al. 2020); Decay sugar-rich substrates (Põlme et al. 2020) | Põlme S, et al. 2020. Fungal Diversity 105:1-16 ((https://doi.org/10.1007/s13225-020-00466-2)) |
| Otu60096 | 1 | k__Fungi(100);p__Ascomycota(100);c__Saccharomycetes(100);o__Saccharomycetales(100);f__Metschnikowiaceae(100);g__Metschnikowia(100);g__Metschnikowia_unclassified(100); | Metschnikowia | 13 | Saprotroph | \|Undefined Saprotroph\| | Yeast | NULL | Probable | As nectar-tap_saprotroph (Põlme et al. 2020); Decay sugar-rich substrates (Põlme et al. 2020) | Põlme S, et al. 2020. Fungal Diversity 105:1-16 ((https://doi.org/10.1007/s13225-020-00466-2)) |
| Otu60097 | 1 | k__Fungi(100);p__Ascomycota(100);c__Saccharomycetes(100);o__Saccharomycetales(100);f__Metschnikowiaceae(100);g__Metschnikowia(100);g__Metschnikowia_unclassified(100); | Metschnikowia | 13 | Saprotroph | \|Undefined Saprotroph\| | Yeast | NULL | Probable | As nectar-tap_saprotroph (Põlme et al. 2020); Decay sugar-rich substrates (Põlme et al. 2020) | Põlme S, et al. 2020. Fungal Diversity 105:1-16 ((https://doi.org/10.1007/s13225-020-00466-2)) |
| Otu60098 | 1 | k__Fungi(100);p__Ascomycota(100);c__Saccharomycetes(100);o__Saccharomycetales(100);f__Metschnikowiaceae(100);g__Metschnikowia(100);g__Metschnikowia_unclassified(100); | Metschnikowia | 13 | Saprotroph | \|Undefined Saprotroph\| | Yeast | NULL | Probable | As nectar-tap_saprotroph (Põlme et al. 2020); Decay sugar-rich substrates (Põlme et al. 2020) | Põlme S, et al. 2020. Fungal Diversity 105:1-16 ((https://doi.org/10.1007/s13225-020-00466-2)) |
| Otu60099 | 1 | k__Fungi(100);p__Ascomycota(100);c__Saccharomycetes(100);o__Saccharomycetales(100);f__Metschnikowiaceae(100);g__Metschnikowia(100);g__Metschnikowia_unclassified(100); | Metschnikowia | 13 | Saprotroph | \|Undefined Saprotroph\| | Yeast | NULL | Probable | As nectar-tap_saprotroph (Põlme et al. 2020); Decay sugar-rich substrates (Põlme et al. 2020) | Põlme S, et al. 2020. Fungal Diversity 105:1-16 ((https://doi.org/10.1007/s13225-020-00466-2)) |
| Otu60108 | 1 | k__Fungi(100);p__Ascomycota(100);c__Saccharomycetes(100);o__Saccharomycetales(100);f__Metschnikowiaceae(100);g__Metschnikowia(100);s__Metschnikowia_pulcherrima(100); | Metschnikowia | 13 | Saprotroph | \|Undefined Saprotroph\| | Yeast | NULL | Probable | As nectar-tap_saprotroph (Põlme et al. 2020); Decay sugar-rich substrates (Põlme et al. 2020) | Põlme S, et al. 2020. Fungal Diversity 105:1-16 ((https://doi.org/10.1007/s13225-020-00466-2)) |
| Otu60109 | 1 | k__Fungi(100);p__Ascomycota(100);c__Saccharomycetes(100);o__Saccharomycetales(100);f__Saccharomycetales_family_Incertae_sedis(100);g__Candida(100);g__Candida_unclassified(100); | Candida | 13 | Pathotroph-Saprotroph-Symbiotroph | Animal Pathogen-Endophyte-Endosymbiont-Epiphyte-\|Undefined Saprotroph\| | Dimorphic Yeast-Yeast | NULL | Probable | As nectar-tap_saprotroph (Põlme et al. 2020); Decay sugar-rich substrates (Põlme et al. 2020); This asexual genus is polyphyletic and contain species that function very differently depend on what they are. Caution should be taken when interpreting guilds for Candida species. | Manolakaki D, et al. 2010. Virulence. 1: 367-375 ((https://doi.org/10.4161/viru.1.5.12796)); Mekbib SB, et al. 2011. Tropical Plant Pathology 36:233-240 ((https://doi.org/10.1590/S1982-56762011000400004)); Põlme S, et al. 2020. Fungal Diversity 105:1-16 ((https://doi.org/10.1007/s13225-020-00466-2)) |
| Otu60110 | 1 | k__Fungi(100);p__Ascomycota(100);c__Saccharomycetes(100);o__Saccharomycetales(100);f__Metschnikowiaceae(100);g__Metschnikowia(100);s__Metschnikowia_pulcherrima(100); | Metschnikowia | 13 | Saprotroph | \|Undefined Saprotroph\| | Yeast | NULL | Probable | As nectar-tap_saprotroph (Põlme et al. 2020); Decay sugar-rich substrates (Põlme et al. 2020) | Põlme S, et al. 2020. Fungal Diversity 105:1-16 ((https://doi.org/10.1007/s13225-020-00466-2)) |
| Otu60116 | 1 | k__Fungi(100);p__Ascomycota(100);c__Saccharomycetes(100);o__Saccharomycetales(100);f__Saccharomycetales_family_Incertae_sedis(100);f__Saccharomycetales_family_Incertae_sedis_unclassified(100);f__Saccharomycetales_family_Incertae_sedis_unclassified(100); | Saccharomycetales | 7 | Saprotroph | Undefined Saprotroph | Yeast | NULL | Possible | NULL | Sterkenburg E, et al. 2015. New Phytologist 207:1145-1158 ((https://doi.org/10.1111/nph.13426)) |
| Otu60120 | 1 | k__Fungi(100);p__Ascomycota(100);c__Saccharomycetes(100);o__Saccharomycetales(100);f__Metschnikowiaceae(100);g__Metschnikowia(100);s__Metschnikowia_pulcherrima(100); | Metschnikowia | 13 | Saprotroph | \|Undefined Saprotroph\| | Yeast | NULL | Probable | As nectar-tap_saprotroph (Põlme et al. 2020); Decay sugar-rich substrates (Põlme et al. 2020) | Põlme S, et al. 2020. Fungal Diversity 105:1-16 ((https://doi.org/10.1007/s13225-020-00466-2)) |
| Otu60133 | 1 | k__Fungi(100);p__Ascomycota(100);c__Saccharomycetes(100);o__Saccharomycetales(100);f__Metschnikowiaceae(100);g__Metschnikowia(100);s__Metschnikowia_pulcherrima(100); | Metschnikowia | 13 | Saprotroph | \|Undefined Saprotroph\| | Yeast | NULL | Probable | As nectar-tap_saprotroph (Põlme et al. 2020); Decay sugar-rich substrates (Põlme et al. 2020) | Põlme S, et al. 2020. Fungal Diversity 105:1-16 ((https://doi.org/10.1007/s13225-020-00466-2)) |
| Otu60134 | 1 | k__Fungi(100);p__Basidiomycota(100);c__Agaricomycetes(100);o__Agaricales(100);f__Cortinariaceae(100);g__Cortinarius(100);s__Cortinarius_schlerophyllarum(100); | Cortinarius | 13 | Symbiotroph | \|Ectomycorrhizal\| | Agaricoid | NULL | Highly Probable | NULL | Rinaldi AC, et al. 2008. Fungal Diversity 33:1-45 ((http://www.fungaldiversity.org/fdp/sfdp/33-1.pdf)); Tedersoo L, et al. 2010. Mycorrhiza 20:217-263 ((https://doi.org/10.1007/s00572-009-0274-x)); Põlme S, et al. 2020. Fungal Diversity 105:1-16 ((https://doi.org/10.1007/s13225-020-00466-2)); http://mycorrhizas.info/ecmf.html ((http://mycorrhizas.info/ecmf.html)) |
| Otu60135 | 1 | k__Fungi(100);p__Basidiomycota(100);c__Exobasidiomycetes(100);o__Microstromatales(100);f__Microstromataceae(100);g__Microstroma(100);s__Microstroma_juglandis(100); | Microstroma | 13 | Pathotroph | \|Plant Pathogen\| | Dimorphic | NULL | Probable | NULL | Tedersoo L, et al. 2014. Science 346:e1256688 ((https://doi.org/10.1126/science.1256688)); Põlme S, et al. 2020. Fungal Diversity 105:1-16 ((https://doi.org/10.1007/s13225-020-00466-2)) |
| Otu60138 | 1 | k__Fungi(100);p__Ascomycota(100);c__Saccharomycetes(100);o__Saccharomycetales(100);f__Metschnikowiaceae(100);g__Metschnikowia(100);s__Metschnikowia_pulcherrima(100); | Metschnikowia | 13 | Saprotroph | \|Undefined Saprotroph\| | Yeast | NULL | Probable | As nectar-tap_saprotroph (Põlme et al. 2020); Decay sugar-rich substrates (Põlme et al. 2020) | Põlme S, et al. 2020. Fungal Diversity 105:1-16 ((https://doi.org/10.1007/s13225-020-00466-2)) |
| Otu60139 | 1 | k__Fungi(100);p__Ascomycota(100);c__Saccharomycetes(100);o__Saccharomycetales(100);f__Metschnikowiaceae(100);g__Metschnikowia(100);s__Metschnikowia_pulcherrima(100); | Metschnikowia | 13 | Saprotroph | \|Undefined Saprotroph\| | Yeast | NULL | Probable | As nectar-tap_saprotroph (Põlme et al. 2020); Decay sugar-rich substrates (Põlme et al. 2020) | Põlme S, et al. 2020. Fungal Diversity 105:1-16 ((https://doi.org/10.1007/s13225-020-00466-2)) |
| Otu60141 | 1 | k__Fungi(100);p__Ascomycota(100);c__Saccharomycetes(100);o__Saccharomycetales(100);f__Metschnikowiaceae(100);g__Metschnikowia(100);s__Metschnikowia_pulcherrima(100); | Metschnikowia | 13 | Saprotroph | \|Undefined Saprotroph\| | Yeast | NULL | Probable | As nectar-tap_saprotroph (Põlme et al. 2020); Decay sugar-rich substrates (Põlme et al. 2020) | Põlme S, et al. 2020. Fungal Diversity 105:1-16 ((https://doi.org/10.1007/s13225-020-00466-2)) |
| Otu60143 | 1 | k__Fungi(100);p__Ascomycota(100);c__Saccharomycetes(100);o__Saccharomycetales(100);f__Metschnikowiaceae(100);g__Metschnikowia(100);s__Metschnikowia_pulcherrima(100); | Metschnikowia | 13 | Saprotroph | \|Undefined Saprotroph\| | Yeast | NULL | Probable | As nectar-tap_saprotroph (Põlme et al. 2020); Decay sugar-rich substrates (Põlme et al. 2020) | Põlme S, et al. 2020. Fungal Diversity 105:1-16 ((https://doi.org/10.1007/s13225-020-00466-2)) |
| Otu60144 | 1 | k__Fungi(100);p__Ascomycota(100);c__Saccharomycetes(100);o__Saccharomycetales(100);f__Metschnikowiaceae(100);g__Metschnikowia(100);g__Metschnikowia_unclassified(100); | Metschnikowia | 13 | Saprotroph | \|Undefined Saprotroph\| | Yeast | NULL | Probable | As nectar-tap_saprotroph (Põlme et al. 2020); Decay sugar-rich substrates (Põlme et al. 2020) | Põlme S, et al. 2020. Fungal Diversity 105:1-16 ((https://doi.org/10.1007/s13225-020-00466-2)) |
| Otu60146 | 1 | k__Fungi(100);p__Ascomycota(100);c__Saccharomycetes(100);o__Saccharomycetales(100);f__Metschnikowiaceae(100);g__Metschnikowia(100);s__Metschnikowia_pulcherrima(100); | Metschnikowia | 13 | Saprotroph | \|Undefined Saprotroph\| | Yeast | NULL | Probable | As nectar-tap_saprotroph (Põlme et al. 2020); Decay sugar-rich substrates (Põlme et al. 2020) | Põlme S, et al. 2020. Fungal Diversity 105:1-16 ((https://doi.org/10.1007/s13225-020-00466-2)) |
| Otu60147 | 1 | k__Fungi(100);p__Ascomycota(100);c__Saccharomycetes(100);o__Saccharomycetales(100);f__Metschnikowiaceae(100);g__Metschnikowia(100);s__Metschnikowia_pulcherrima(100); | Metschnikowia | 13 | Saprotroph | \|Undefined Saprotroph\| | Yeast | NULL | Probable | As nectar-tap_saprotroph (Põlme et al. 2020); Decay sugar-rich substrates (Põlme et al. 2020) | Põlme S, et al. 2020. Fungal Diversity 105:1-16 ((https://doi.org/10.1007/s13225-020-00466-2)) |
| Otu60148 | 1 | k__Fungi(100);p__Ascomycota(100);c__Saccharomycetes(100);o__Saccharomycetales(100);f__Metschnikowiaceae(100);g__Metschnikowia(100);s__Metschnikowia_pulcherrima(100); | Metschnikowia | 13 | Saprotroph | \|Undefined Saprotroph\| | Yeast | NULL | Probable | As nectar-tap_saprotroph (Põlme et al. 2020); Decay sugar-rich substrates (Põlme et al. 2020) | Põlme S, et al. 2020. Fungal Diversity 105:1-16 ((https://doi.org/10.1007/s13225-020-00466-2)) |
| Otu60149 | 1 | k__Fungi(100);p__Ascomycota(100);c__Saccharomycetes(100);o__Saccharomycetales(100);f__Metschnikowiaceae(100);g__Metschnikowia(100);s__Metschnikowia_pulcherrima(100); | Metschnikowia | 13 | Saprotroph | \|Undefined Saprotroph\| | Yeast | NULL | Probable | As nectar-tap_saprotroph (Põlme et al. 2020); Decay sugar-rich substrates (Põlme et al. 2020) | Põlme S, et al. 2020. Fungal Diversity 105:1-16 ((https://doi.org/10.1007/s13225-020-00466-2)) |
| Otu60150 | 1 | k__Fungi(100);p__Ascomycota(100);c__Saccharomycetes(100);o__Saccharomycetales(100);f__Metschnikowiaceae(100);g__Metschnikowia(100);s__Metschnikowia_pulcherrima(100); | Metschnikowia | 13 | Saprotroph | \|Undefined Saprotroph\| | Yeast | NULL | Probable | As nectar-tap_saprotroph (Põlme et al. 2020); Decay sugar-rich substrates (Põlme et al. 2020) | Põlme S, et al. 2020. Fungal Diversity 105:1-16 ((https://doi.org/10.1007/s13225-020-00466-2)) |
| Otu60152 | 1 | k__Fungi(100);p__Ascomycota(100);c__Saccharomycetes(100);o__Saccharomycetales(100);f__Metschnikowiaceae(100);g__Metschnikowia(100);s__Metschnikowia_pulcherrima(100); | Metschnikowia | 13 | Saprotroph | \|Undefined Saprotroph\| | Yeast | NULL | Probable | As nectar-tap_saprotroph (Põlme et al. 2020); Decay sugar-rich substrates (Põlme et al. 2020) | Põlme S, et al. 2020. Fungal Diversity 105:1-16 ((https://doi.org/10.1007/s13225-020-00466-2)) |
| Otu60153 | 1 | k__Fungi(100);p__Ascomycota(100);c__Saccharomycetes(100);o__Saccharomycetales(100);f__Metschnikowiaceae(100);g__Metschnikowia(100);s__Metschnikowia_pulcherrima(100); | Metschnikowia | 13 | Saprotroph | \|Undefined Saprotroph\| | Yeast | NULL | Probable | As nectar-tap_saprotroph (Põlme et al. 2020); Decay sugar-rich substrates (Põlme et al. 2020) | Põlme S, et al. 2020. Fungal Diversity 105:1-16 ((https://doi.org/10.1007/s13225-020-00466-2)) |
| Otu60156 | 1 | k__Fungi(100);p__Basidiomycota(100);c__Dacrymycetes(100);o__Dacrymycetales(100);f__Dacrymycetaceae(100);g__Cerinosterus(100);s__Cerinosterus_sp(100); | Cerinosterus | 13 | Saprotroph | \|Undefined Saprotroph\| | Microfungus | Brown Rot | Probable | NULL | Tedersoo L, et al. 2014. Science 346:e1256688 ((https://doi.org/10.1126/science.1256688)); Põlme S, et al. 2020. Fungal Diversity 105:1-16 ((https://doi.org/10.1007/s13225-020-00466-2)) |
| Otu60164 | 1 | k__Fungi(100);p__Ascomycota(100);c__Saccharomycetes(100);o__Saccharomycetales(100);f__Trichomonascaceae(100);g__Blastobotrys(100);g__Blastobotrys_unclassified(100); | Blastobotrys | 13 | Saprotroph | \|Undefined Saprotroph\| | Yeast | Its unusual metal | Highly Probable | As nectar-tap_saprotroph (Põlme et al. 2020); Decay sugar-rich substrates (Põlme et al. 2020); In the course of identifying alternative microorganisms for biotechnological interest, A. adeninivorans has already proved its strengthened competitiveness as a promising cell factory for many more applications. | Kunze G, et al. 2014. Biotechnology for Biofuels 7:66 ((https://doi.org/10.1186/1754-6834-7-66)); Põlme S, et al. 2020. Fungal Diversity 105:1-16 ((https://doi.org/10.1007/s13225-020-00466-2)) |
| Otu60166 | 1 | k__Fungi(100);p__Ascomycota(100);c__Leotiomycetes(100);o__Helotiales(100);f__Helotiales_family_Incertae_sedis(100);g__Hyphodiscus(100);s__Hyphodiscus_hymeniophilus(100); | Hyphodiscus | 13 | Pathotroph-Saprotroph | \|Fungal Parasite\|-Wood Saprotroph | Microfungus | NULL | Highly Probable | As mycoparasite (Põlme et al. 2020); Decay fungal material (Põlme et al. 2020) | Põlme S, et al. 2020. Fungal Diversity 105:1-16 ((https://doi.org/10.1007/s13225-020-00466-2)); List compiled by John Plischke III ((https://goodmushroom.weebly.com/uploads/5/7/1/7/57176535/fungi_on_fungi_pdf_lq.pdf)) |
| Otu60168 | 1 | k__Fungi(100);p__Ascomycota(100);c__Saccharomycetes(100);o__Saccharomycetales(100);f__Metschnikowiaceae(100);g__Metschnikowia(100);s__Metschnikowia_pulcherrima(100); | Metschnikowia | 13 | Saprotroph | \|Undefined Saprotroph\| | Yeast | NULL | Probable | As nectar-tap_saprotroph (Põlme et al. 2020); Decay sugar-rich substrates (Põlme et al. 2020) | Põlme S, et al. 2020. Fungal Diversity 105:1-16 ((https://doi.org/10.1007/s13225-020-00466-2)) |
| Otu60176 | 1 | k__Fungi(100);p__Ascomycota(100);c__Lecanoromycetes(100);o__Lecanorales(100);f__Parmeliaceae(100);g__Tuckermannopsis(100);s__Tuckermannopsis_orbata(100); | Tuckermannopsis | 13 | Symbiotroph | \|Lichenized\| | Thallus | NULL | Probable | NULL | Põlme S, et al. 2020. Fungal Diversity 105:1-16 ((https://doi.org/10.1007/s13225-020-00466-2)) |
| Otu60184 | 1 | k__Fungi(100);p__Basidiomycota(100);c__Tremellomycetes(100);o__Tremellales(100);f__Tremellales_family_Incertae_sedis(100);g__Cryptococcus(100);s__Cryptococcus_sp_JCM_11356(100); | Cryptococcus | 13 | Pathotroph-Saprotroph-Symbiotroph | Animal Pathogen-Endophyte-Epiphyte-\|Undefined Saprotroph\| | Dimorphic Yeast-Yeast | NULL | Possible | As foliar_endophyte (Põlme et al. 2020); Decay leaf, fruit, seed, soil, dung, animal material (Põlme et al. 2020) | Andrews JH, et al. 1983. Phytopathology 73:228-234 ((https://doi.org/10.1094/Phyto-73-228)); Kurtzman CP, et al. (eds.) 2011. The Yeasts, a Taxonomic Study. Fifth Edition. Vols 1-3. Elsevier, San Diego (ISBN: 9780444521491); Põlme S, et al. 2020. Fungal Diversity 105:1-16 ((https://doi.org/10.1007/s13225-020-00466-2)) |
| Otu60185 | 1 | k__Fungi(100);p__Ascomycota(100);c__Saccharomycetes(100);o__Saccharomycetales(100);f__Metschnikowiaceae(100);g__Metschnikowia(100);s__Metschnikowia_pulcherrima(100); | Metschnikowia | 13 | Saprotroph | \|Undefined Saprotroph\| | Yeast | NULL | Probable | As nectar-tap_saprotroph (Põlme et al. 2020); Decay sugar-rich substrates (Põlme et al. 2020) | Põlme S, et al. 2020. Fungal Diversity 105:1-16 ((https://doi.org/10.1007/s13225-020-00466-2)) |
| Otu60186 | 1 | k__Fungi(100);p__Ascomycota(100);c__Saccharomycetes(100);o__Saccharomycetales(100);f__Metschnikowiaceae(100);g__Metschnikowia(100);s__Metschnikowia_pulcherrima(100); | Metschnikowia | 13 | Saprotroph | \|Undefined Saprotroph\| | Yeast | NULL | Probable | As nectar-tap_saprotroph (Põlme et al. 2020); Decay sugar-rich substrates (Põlme et al. 2020) | Põlme S, et al. 2020. Fungal Diversity 105:1-16 ((https://doi.org/10.1007/s13225-020-00466-2)) |
| Otu60189 | 1 | k__Fungi(100);p__Ascomycota(100);c__Saccharomycetes(100);o__Saccharomycetales(100);f__Metschnikowiaceae(100);g__Metschnikowia(100);s__Metschnikowia_pulcherrima(100); | Metschnikowia | 13 | Saprotroph | \|Undefined Saprotroph\| | Yeast | NULL | Probable | As nectar-tap_saprotroph (Põlme et al. 2020); Decay sugar-rich substrates (Põlme et al. 2020) | Põlme S, et al. 2020. Fungal Diversity 105:1-16 ((https://doi.org/10.1007/s13225-020-00466-2)) |

| | | | | | | | | | | | |
|---|---|---|---|---|---|---|---|---|---|---|---|
| Otu60191 | 1 | k__Fungi(100);p__Ascomycota(100);c__Sordariomycetes(100);o__Hypocreales(100);f__Nectriaceae(100);g__Fusarium(100);g__Fusarium_unclassified(100); | Fusarium | 13 | Pathotroph-Saprotroph-Symbiotroph | Animal Pathogen-Endophyte-Lichen Parasite-\|Plant Pathogen\|-Plant Saprotroph-Undefined Saprotroph-Wood Saprotroph | Microfungus | Soft Rot | Possible | As foliar_endophyte (Põlme et al. 2020); As litter_saprotroph (Põlme et al. 2020); Causing damping-off disease (Põlme et al. 2020); Decay leaf, fruit, seed, roots, soil, animal material (Põlme et al. 2020); Likely opportunistic human pathogen (Irinyi et al. 2015) | Duncan CG, Eslyn WE. 1966. Mycologia 58:642-664 ((https://doi.org/10.2307/3757045)); Seehann G, et al. 1975. List of Fungi in Soft-Rot Tissue Tests. Institute fur Holzbiologie und Holzschutz, Bundesforschungsanstalt fur Forst - und Holzwirtshaft, Hamburg - Lohbrugge, Germany; Costa IPMW, et al. 2012. Mycotaxon 119:e26 ((http://mycotaxon.com/resources/checklists/costa_v119_checklist.pdf)); Tedersoo L, et al. 2014. Science 346:e1256688 ((https://doi.org/10.1126/science.1256688)); Irinyi L, et al. 2015. Medical Mycology 53:313-337 ((https://doi.org/10.1093/mmy/myv008)); Schroers, et al. 2016. Mycologia 108:806-819 ((https://doi.org/10.3852/15-255)); Põlme S, et al. 2020. Fungal Diversity 105:1-16 ((https://doi.org/10.1007/s13225-020-00466-2)); The American Phytopathological Society Common Names of Plant Diseases ((http://www.apsnet.org/publications/commonnames/Pages/default.aspx)); Lawrey JD, Diederich P. 2016. http://www.lichenicolous.net (accessed 24 May 2017) ((http://www.lichenicolous.net)) |
| Otu60193 | 1 | k__Fungi(100);p__Ascomycota(100);c__Saccharomycetes(100);o__Saccharomycetales(100);f__Metschnikowiaceae(100);g__Metschnikowia(100);s__Metschnikowia_pulcherrima(100); | Metschnikowia | 13 | Saprotroph | \|Undefined Saprotroph\| | Yeast | NULL | Probable | As nectar-tap_saprotroph (Põlme et al. 2020); Decay sugar-rich substrates (Põlme et al. 2020) | Põlme S, et al. 2020. Fungal Diversity 105:1-16 ((https://doi.org/10.1007/s13225-020-00466-2)) |
| Otu60194 | 1 | k__Fungi(100);p__Ascomycota(100);c__Saccharomycetes(100);o__Saccharomycetales(100);f__Metschnikowiaceae(100);g__Metschnikowia(100);s__Metschnikowia_pulcherrima(100); | Metschnikowia | 13 | Saprotroph | \|Undefined Saprotroph\| | Yeast | NULL | Probable | As nectar-tap_saprotroph (Põlme et al. 2020); Decay sugar-rich substrates (Põlme et al. 2020) | Põlme S, et al. 2020. Fungal Diversity 105:1-16 ((https://doi.org/10.1007/s13225-020-00466-2)) |
| Otu60195 | 1 | k__Fungi(100);p__Ascomycota(100);c__Saccharomycetes(100);o__Saccharomycetales(100);f__Metschnikowiaceae(100);g__Metschnikowia(100);s__Metschnikowia_pulcherrima(100); | Metschnikowia | 13 | Saprotroph | \|Undefined Saprotroph\| | Yeast | NULL | Probable | As nectar-tap_saprotroph (Põlme et al. 2020); Decay sugar-rich substrates (Põlme et al. 2020) | Põlme S, et al. 2020. Fungal Diversity 105:1-16 ((https://doi.org/10.1007/s13225-020-00466-2)) |
| Otu60196 | 1 | k__Fungi(100);p__Ascomycota(100);c__Saccharomycetes(100);o__Saccharomycetales(100);f__Metschnikowiaceae(100);g__Metschnikowia(100);s__Metschnikowia_pulcherrima(100); | Metschnikowia | 13 | Saprotroph | \|Undefined Saprotroph\| | Yeast | NULL | Probable | As nectar-tap_saprotroph (Põlme et al. 2020); Decay sugar-rich substrates (Põlme et al. 2020) | Põlme S, et al. 2020. Fungal Diversity 105:1-16 ((https://doi.org/10.1007/s13225-020-00466-2)) |
| Otu60198 | 1 | k__Fungi(100);p__Ascomycota(100);c__Saccharomycetes(100);o__Saccharomycetales(100);f__Metschnikowiaceae(100);g__Metschnikowia(100);s__Metschnikowia_pulcherrima(100); | Metschnikowia | 13 | Saprotroph | \|Undefined Saprotroph\| | Yeast | NULL | Probable | As nectar-tap_saprotroph (Põlme et al. 2020); Decay sugar-rich substrates (Põlme et al. 2020) | Põlme S, et al. 2020. Fungal Diversity 105:1-16 ((https://doi.org/10.1007/s13225-020-00466-2)) |
| Otu60200 | 1 | k__Fungi(100);p__Ascomycota(100);c__Saccharomycetes(100);o__Saccharomycetales(100);f__Metschnikowiaceae(100);g__Metschnikowia(100);s__Metschnikowia_pulcherrima(100); | Metschnikowia | 13 | Saprotroph | \|Undefined Saprotroph\| | Yeast | NULL | Probable | As nectar-tap_saprotroph (Põlme et al. 2020); Decay sugar-rich substrates (Põlme et al. 2020) | Põlme S, et al. 2020. Fungal Diversity 105:1-16 ((https://doi.org/10.1007/s13225-020-00466-2)) |
| Otu60201 | 1 | k__Fungi(100);p__Ascomycota(100);c__Saccharomycetes(100);o__Saccharomycetales(100);f__Metschnikowiaceae(100);g__Metschnikowia(100);g__Metschnikowia_unclassified(100); | Metschnikowia | 13 | Saprotroph | \|Undefined Saprotroph\| | Yeast | NULL | Probable | As nectar-tap_saprotroph (Põlme et al. 2020); Decay sugar-rich substrates (Põlme et al. 2020) | Põlme S, et al. 2020. Fungal Diversity 105:1-16 ((https://doi.org/10.1007/s13225-020-00466-2)) |
| Otu60210 | 1 | k__Fungi(100);p__Ascomycota(100);c__Leotiomycetes(100);o__Helotiales(100);f__Helotiales_family_Incertae_sedis(100);g__Rhexocercosporidium(100);s__Rhexocercosporidium_panacis(100); | Rhexocercosporidium | 13 | Pathotroph-Saprotroph | \|Plant Pathogen\|-Plant Saprotroph | Microfungus | NULL | Probable | As litter_saprotroph (Põlme et al. 2020); As root_pathogen (Põlme et al. 2020); Decay leaf, fruit, seed, roots (Põlme et al. 2020) | Tedersoo L, et al. 2014. Science 346:e1256688 ((https://doi.org/10.1126/science.1256688)); Põlme S, et al. 2020. Fungal Diversity 105:1-16 ((https://doi.org/10.1007/s13225-020-00466-2)) |
| Otu60212 | 1 | k__Fungi(100);p__Ascomycota(100);c__Ascomycota_class_Incertae_sedis(100);o__Ascomycota_order_Incertae_sedis(100);f__Ascomycota_family_Incertae_sedis(100);g__Miniancora(100);s__Miniancora_allisoniensis(100); | Miniancora | 13 | Saprotroph | \|Plant Saprotroph\| | Microfungus | NULL | Probable | As litter_saprotroph (Põlme et al. 2020); Decay leaf, fruit, seed (Põlme et al. 2020) | Põlme S, et al. 2020. Fungal Diversity 105:1-16 ((https://doi.org/10.1007/s13225-020-00466-2)) |
| Otu60214 | 1 | k__Fungi(100);p__Ascomycota(100);c__Leotiomycetes(100);o__Helotiales(100);f__Helotiaceae(100);g__Hymenoscyphus(100);g__Hymenoscyphus_unclassified(100); | Hymenoscyphus | 13 | Pathotroph-Saprotroph-Symbiotroph | Bryophyte Parasite-Ectomycorrhizal-Ericoid Mycorrhizal-Plant Pathogen-\|Plant Saprotroph\|-Undefined Saprotroph | Microfungus | NULL | Possible | As litter_saprotroph (Põlme et al. 2020); Dark Septate Microfungus; Syn. Pezoloma among other (see e.g., Pezoloma ericae); Decay leaf, fruit, seed (Põlme et al. 2020) | Baral HO, Krieglsteiner L. 2006. Acta Mycologica 41:11-20 ((https://doi.org/10.5586/am.2006.003)) (as a bryophyte parasite); Huhtinen S, et al. 2010. Nova Hedwigia 90:413-431 ((https://doi.org/10.1127/0029-5035/2010/0090-0413)); Bent E, et al. 2011. Applied and Environmental Microbiology 77:3351-3359 ((https://doi.org/10.1128/AEM.02575-10)) (as ectomycorrhizae); Tedersoo L, et al. 2014. Science 346:1256688 ((https://doi.org/10.1126/science.1256688)) (as saprotroph); Põlme S, et al. 2020. Fungal Diversity 105:1-16 ((https://doi.org/10.1007/s13225-020-00466-2)) |
| Otu60221 | 1 | k__Fungi(100);p__Ascomycota(100);c__Saccharomycetes(100);o__Saccharomycetales(100);f__Metschnikowiaceae(100);g__Metschnikowia(100);g__Metschnikowia_unclassified(100); | Metschnikowia | 13 | Saprotroph | \|Undefined Saprotroph\| | Yeast | NULL | Probable | As nectar-tap_saprotroph (Põlme et al. 2020); Decay sugar-rich substrates (Põlme et al. 2020) | Põlme S, et al. 2020. Fungal Diversity 105:1-16 ((https://doi.org/10.1007/s13225-020-00466-2)) |
| Otu60227 | 1 | k__Fungi(100);p__Ascomycota(100);c__Saccharomycetes(100);o__Saccharomycetales(100);f__Metschnikowiaceae(100);g__Metschnikowia(100);s__Metschnikowia_pulcherrima(100); | Metschnikowia | 13 | Saprotroph | \|Undefined Saprotroph\| | Yeast | NULL | Probable | As nectar-tap_saprotroph (Põlme et al. 2020); Decay sugar-rich substrates (Põlme et al. 2020) | Põlme S, et al. 2020. Fungal Diversity 105:1-16 ((https://doi.org/10.1007/s13225-020-00466-2)) |
| Otu60231 | 1 | k__Fungi(100);p__Ascomycota(100);c__Saccharomycetes(100);o__Saccharomycetales(100);f__Saccharomycetales_family_Incertae_sedis(100);g__Candida(100);g__Candida_unclassified(100); | Candida | 13 | Pathotroph-Saprotroph-Symbiotroph | Animal Pathogen-Endophyte-Endosymbiont-Epiphyte-\|Undefined Saprotroph\| | Dimorphic Yeast-Yeast | NULL | Probable | As nectar-tap_saprotroph (Põlme et al. 2020); Decay sugar-rich substrates (Põlme et al. 2020); This asexual genus is polyphyletic and contain species that function very differently depend on what they are. Caution should be taken when interpreting guilds for Candida species. | Manolakaki D, et al. 2010. Virulence. 1: 367-375 ((https://doi.org/10.4161/viru.1.5.12796)); Mekbib SB, et al. 2011. Tropical Plant Pathology 36:233-240 ((https://doi.org/10.1590/S1982-56762011000400004)); Põlme S, et al. 2020. Fungal Diversity 105:1-16 ((https://doi.org/10.1007/s13225-020-00466-2)) |
| Otu60232 | 1 | k__Fungi(100);p__Ascomycota(100);c__Saccharomycetes(100);o__Saccharomycetales(100);f__Metschnikowiaceae(100);g__Metschnikowia(100);s__Metschnikowia_pulcherrima(100); | Metschnikowia | 13 | Saprotroph | \|Undefined Saprotroph\| | Yeast | NULL | Probable | As nectar-tap_saprotroph (Põlme et al. 2020); Decay sugar-rich substrates (Põlme et al. 2020) | Põlme S, et al. 2020. Fungal Diversity 105:1-16 ((https://doi.org/10.1007/s13225-020-00466-2)) |
| Otu60241 | 1 | k__Fungi(100);p__Ascomycota(100);c__Saccharomycetes(100);o__Saccharomycetales(100);f__Metschnikowiaceae(100);g__Metschnikowia(100);s__Metschnikowia_pulcherrima(100); | Metschnikowia | 13 | Saprotroph | \|Undefined Saprotroph\| | Yeast | NULL | Probable | As nectar-tap_saprotroph (Põlme et al. 2020); Decay sugar-rich substrates (Põlme et al. 2020) | Põlme S, et al. 2020. Fungal Diversity 105:1-16 ((https://doi.org/10.1007/s13225-020-00466-2)) |
| Otu60243 | 1 | k__Fungi(100);p__Ascomycota(100);c__Saccharomycetes(100);o__Saccharomycetales(100);f__Metschnikowiaceae(100);g__Metschnikowia(100);s__Metschnikowia_pulcherrima(100); | Metschnikowia | 13 | Saprotroph | \|Undefined Saprotroph\| | Yeast | NULL | Probable | As nectar-tap_saprotroph (Põlme et al. 2020); Decay sugar-rich substrates (Põlme et al. 2020) | Põlme S, et al. 2020. Fungal Diversity 105:1-16 ((https://doi.org/10.1007/s13225-020-00466-2)) |
| Otu60247 | 1 | k__Fungi(100);p__Ascomycota(100);c__Saccharomycetes(100);o__Saccharomycetales(100);f__Metschnikowiaceae(100);g__Metschnikowia(100);g__Metschnikowia_unclassified(100); | Metschnikowia | 13 | Saprotroph | \|Undefined Saprotroph\| | Yeast | NULL | Probable | As nectar-tap_saprotroph (Põlme et al. 2020); Decay sugar-rich substrates (Põlme et al. 2020) | Põlme S, et al. 2020. Fungal Diversity 105:1-16 ((https://doi.org/10.1007/s13225-020-00466-2)) |
| Otu60248 | 1 | k__Fungi(100);p__Ascomycota(100);c__Saccharomycetes(100);o__Saccharomycetales(100);f__Metschnikowiaceae(100);g__Metschnikowia(100);s__Metschnikowia_pulcherrima(100); | Metschnikowia | 13 | Saprotroph | \|Undefined Saprotroph\| | Yeast | NULL | Probable | As nectar-tap_saprotroph (Põlme et al. 2020); Decay sugar-rich substrates (Põlme et al. 2020) | Põlme S, et al. 2020. Fungal Diversity 105:1-16 ((https://doi.org/10.1007/s13225-020-00466-2)) |
| Otu60249 | 1 | k__Fungi(100);p__Ascomycota(100);c__Saccharomycetes(100);o__Saccharomycetales(100);f__Metschnikowiaceae(100);g__Metschnikowia(100);s__Metschnikowia_pulcherrima(100); | Metschnikowia | 13 | Saprotroph | \|Undefined Saprotroph\| | Yeast | NULL | Probable | As nectar-tap_saprotroph (Põlme et al. 2020); Decay sugar-rich substrates (Põlme et al. 2020) | Põlme S, et al. 2020. Fungal Diversity 105:1-16 ((https://doi.org/10.1007/s13225-020-00466-2)) |
| Otu60253 | 1 | k__Fungi(100);p__Ascomycota(100);c__Saccharomycetes(100);o__Saccharomycetales(100);f__Metschnikowiaceae(100);g__Metschnikowia(100);s__Metschnikowia_pulcherrima(100); | Metschnikowia | 13 | Saprotroph | \|Undefined Saprotroph\| | Yeast | NULL | Probable | As nectar-tap_saprotroph (Põlme et al. 2020); Decay sugar-rich substrates (Põlme et al. 2020) | Põlme S, et al. 2020. Fungal Diversity 105:1-16 ((https://doi.org/10.1007/s13225-020-00466-2)) |
| Otu60256 | 1 | k__Fungi(100);p__Ascomycota(100);c__Saccharomycetes(100);o__Saccharomycetales(100);f__Metschnikowiaceae(100);g__Metschnikowia(100);s__Metschnikowia_pulcherrima(100); | Metschnikowia | 13 | Saprotroph | \|Undefined Saprotroph\| | Yeast | NULL | Probable | As nectar-tap_saprotroph (Põlme et al. 2020); Decay sugar-rich substrates (Põlme et al. 2020) | Põlme S, et al. 2020. Fungal Diversity 105:1-16 ((https://doi.org/10.1007/s13225-020-00466-2)) |
| Otu60257 | 1 | k__Fungi(100);p__Ascomycota(100);c__Saccharomycetes(100);o__Saccharomycetales(100);f__Metschnikowiaceae(100);g__Metschnikowia(100);s__Metschnikowia_pulcherrima(100); | Metschnikowia | 13 | Saprotroph | \|Undefined Saprotroph\| | Yeast | NULL | Probable | As nectar-tap_saprotroph (Põlme et al. 2020); Decay sugar-rich substrates (Põlme et al. 2020) | Põlme S, et al. 2020. Fungal Diversity 105:1-16 ((https://doi.org/10.1007/s13225-020-00466-2)) |
| Otu60258 | 1 | k__Fungi(100);p__Ascomycota(100);c__Saccharomycetes(100);o__Saccharomycetales(100);f__Metschnikowiaceae(100);g__Metschnikowia(100);s__Metschnikowia_pulcherrima(100); | Metschnikowia | 13 | Saprotroph | \|Undefined Saprotroph\| | Yeast | NULL | Probable | As nectar-tap_saprotroph (Põlme et al. 2020); Decay sugar-rich substrates (Põlme et al. 2020) | Põlme S, et al. 2020. Fungal Diversity 105:1-16 ((https://doi.org/10.1007/s13225-020-00466-2)) |
| Otu60259 | 1 | k__Fungi(100);p__Ascomycota(100);c__Saccharomycetes(100);o__Saccharomycetales(100);f__Metschnikowiaceae(100);g__Metschnikowia(100);s__Metschnikowia_pulcherrima(100); | Metschnikowia | 13 | Saprotroph | \|Undefined Saprotroph\| | Yeast | NULL | Probable | As nectar-tap_saprotroph (Põlme et al. 2020); Decay sugar-rich substrates (Põlme et al. 2020) | Põlme S, et al. 2020. Fungal Diversity 105:1-16 ((https://doi.org/10.1007/s13225-020-00466-2)) |
| Otu60260 | 1 | k__Fungi(100);p__Ascomycota(100);c__Saccharomycetes(100);o__Saccharomycetales(100);f__Metschnikowiaceae(100);g__Metschnikowia(100);s__Metschnikowia_pulcherrima(100); | Metschnikowia | 13 | Saprotroph | \|Undefined Saprotroph\| | Yeast | NULL | Probable | As nectar-tap_saprotroph (Põlme et al. 2020); Decay sugar-rich substrates (Põlme et al. 2020) | Põlme S, et al. 2020. Fungal Diversity 105:1-16 ((https://doi.org/10.1007/s13225-020-00466-2)) |
| Otu60261 | 1 | k__Fungi(100);p__Ascomycota(100);c__Saccharomycetes(100);o__Saccharomycetales(100);f__Metschnikowiaceae(100);g__Metschnikowia(100);s__Metschnikowia_pulcherrima(100); | Metschnikowia | 13 | Saprotroph | \|Undefined Saprotroph\| | Yeast | NULL | Probable | As nectar-tap_saprotroph (Põlme et al. 2020); Decay sugar-rich substrates (Põlme et al. 2020) | Põlme S, et al. 2020. Fungal Diversity 105:1-16 ((https://doi.org/10.1007/s13225-020-00466-2)) |
| Otu60262 | 1 | k__Fungi(100);p__Ascomycota(100);c__Saccharomycetes(100);o__Saccharomycetales(100);f__Metschnikowiaceae(100);g__Metschnikowia(100);s__Metschnikowia_pulcherrima(100); | Metschnikowia | 13 | Saprotroph | \|Undefined Saprotroph\| | Yeast | NULL | Probable | As nectar-tap_saprotroph (Põlme et al. 2020); Decay sugar-rich substrates (Põlme et al. 2020) | Põlme S, et al. 2020. Fungal Diversity 105:1-16 ((https://doi.org/10.1007/s13225-020-00466-2)) |
| Otu60265 | 1 | k__Fungi(100);p__Ascomycota(100);c__Saccharomycetes(100);o__Saccharomycetales(100);f__Metschnikowiaceae(100);g__Metschnikowia(100);s__Metschnikowia_pulcherrima(100); | Metschnikowia | 13 | Saprotroph | \|Undefined Saprotroph\| | Yeast | NULL | Probable | As nectar-tap_saprotroph (Põlme et al. 2020); Decay sugar-rich substrates (Põlme et al. 2020) | Põlme S, et al. 2020. Fungal Diversity 105:1-16 ((https://doi.org/10.1007/s13225-020-00466-2)) |
| Otu60296 | 1 | k__Fungi(100);p__Ascomycota(100);c__Saccharomycetes(100);o__Saccharomycetales(100);f__Metschnikowiaceae(100);g__Metschnikowia(100);g__Metschnikowia_unclassified(100); | Metschnikowia | 13 | Saprotroph | \|Undefined Saprotroph\| | Yeast | NULL | Probable | As nectar-tap_saprotroph (Põlme et al. 2020); Decay sugar-rich substrates (Põlme et al. 2020) | Põlme S, et al. 2020. Fungal Diversity 105:1-16 ((https://doi.org/10.1007/s13225-020-00466-2)) |
| Otu60302 | 1 | k__Fungi(100);p__Ascomycota(100);c__Saccharomycetes(100);o__Saccharomycetales(100);f__Metschnikowiaceae(100);g__Metschnikowia(100);g__Metschnikowia_unclassified(100); | Metschnikowia | 13 | Saprotroph | \|Undefined Saprotroph\| | Yeast | NULL | Probable | As nectar-tap_saprotroph (Põlme et al. 2020); Decay sugar-rich substrates (Põlme et al. 2020) | Põlme S, et al. 2020. Fungal Diversity 105:1-16 ((https://doi.org/10.1007/s13225-020-00466-2)) |
| Otu60319 | 1 | k__Fungi(100);p__Ascomycota(100);c__Saccharomycetes(100);o__Saccharomycetales(100);o__Saccharomycetales_unclassified(100);o__Saccharomycetales_unclassified(100);o__Saccharomycetales_unclassified(100); | Saccharomycetales | 7 | Saprotroph | Undefined Saprotroph | Yeast | NULL | Possible | NULL | Sterkenburg E, et al. 2015. New Phytologist 207:1145-1158 ((https://doi.org/10.1111/nph.13426)) |
| Otu60329 | 1 | k__Fungi(100);p__Ascomycota(100);c__Saccharomycetes(100);o__Saccharomycetales(100);f__Saccharomycetales_family_Incertae_sedis(100);f__Saccharomycetales_family_Incertae_sedis_unclassified(100);f__Saccharomycetales_family_Incertae_sedis_unclassified(100); | Saccharomycetales | 7 | Saprotroph | Undefined Saprotroph | Yeast | NULL | Possible | NULL | Sterkenburg E, et al. 2015. New Phytologist 207:1145-1158 ((https://doi.org/10.1111/nph.13426)) |
| Otu60333 | 1 | k__Fungi(100);p__Ascomycota(100);c__Saccharomycetes(100);o__Saccharomycetales(100);o__Saccharomycetales_unclassified(100);o__Saccharomycetales_unclassified(100);o__Saccharomycetales_unclassified(100); | Saccharomycetales | 7 | Saprotroph | Undefined Saprotroph | Yeast | NULL | Possible | NULL | Sterkenburg E, et al. 2015. New Phytologist 207:1145-1158 ((https://doi.org/10.1111/nph.13426)) |
| Otu60338 | 1 | k__Fungi(100);p__Ascomycota(100);c__Saccharomycetes(100);o__Saccharomycetales(100);f__Metschnikowiaceae(100);g__Metschnikowia(100);g__Metschnikowia_unclassified(100); | Metschnikowia | 13 | Saprotroph | \|Undefined Saprotroph\| | Yeast | NULL | Probable | As nectar-tap_saprotroph (Põlme et al. 2020); Decay sugar-rich substrates (Põlme et al. 2020) | Põlme S, et al. 2020. Fungal Diversity 105:1-16 ((https://doi.org/10.1007/s13225-020-00466-2)) |
| Otu60339 | 1 | k__Fungi(100);p__Ascomycota(100);c__Saccharomycetes(100);o__Saccharomycetales(100);f__Metschnikowiaceae(100);g__Metschnikowia(100);s__Metschnikowia_pulcherrima(100); | Metschnikowia | 13 | Saprotroph | \|Undefined Saprotroph\| | Yeast | NULL | Probable | As nectar-tap_saprotroph (Põlme et al. 2020); Decay sugar-rich substrates (Põlme et al. 2020) | Põlme S, et al. 2020. Fungal Diversity 105:1-16 ((https://doi.org/10.1007/s13225-020-00466-2)) |
| Otu60348 | 1 | k__Fungi(100);p__Ascomycota(100);c__Saccharomycetes(100);o__Saccharomycetales(100);f__Saccharomycetales_family_Incertae_sedis(100);g__Candida(100);g__Candida_unclassified(100); | Candida | 13 | Pathotroph-Saprotroph-Symbiotroph | Animal Pathogen-Endophyte-Endosymbiont-Epiphyte-\|Undefined Saprotroph\| | Dimorphic Yeast-Yeast | NULL | Probable | As nectar-tap_saprotroph (Põlme et al. 2020); Decay sugar-rich substrates (Põlme et al. 2020); This asexual genus is polyphyletic and contain species that function very differently depend on what they are. Caution should be taken when interpreting guilds for Candida species. | Manolakaki D, et al. 2010. Virulence. 1: 367-375 ((https://doi.org/10.4161/viru.1.5.12796)); Mekbib SB, et al. 2011. Tropical Plant Pathology 36:233-240 ((https://doi.org/10.1590/S1982-56762011000400004)); Põlme S, et al. 2020. Fungal Diversity 105:1-16 ((https://doi.org/10.1007/s13225-020-00466-2)) |
| Otu60362 | 1 | k__Fungi(100);p__Ascomycota(100);c__Saccharomycetes(100);o__Saccharomycetales(100);f__Metschnikowiaceae(100);g__Metschnikowia(100);s__Metschnikowia_pulcherrima(100); | Metschnikowia | 13 | Saprotroph | \|Undefined Saprotroph\| | Yeast | NULL | Probable | As nectar-tap_saprotroph (Põlme et al. 2020); Decay sugar-rich substrates (Põlme et al. 2020) | Põlme S, et al. 2020. Fungal Diversity 105:1-16 ((https://doi.org/10.1007/s13225-020-00466-2)) |
| Otu60368 | 1 | k__Fungi(100);p__Ascomycota(100);c__Saccharomycetes(100);o__Saccharomycetales(100);f__Metschnikowiaceae(100);g__Metschnikowia(100);s__Metschnikowia_pulcherrima(100); | Metschnikowia | 13 | Saprotroph | \|Undefined Saprotroph\| | Yeast | NULL | Probable | As nectar-tap_saprotroph (Põlme et al. 2020); Decay sugar-rich substrates (Põlme et al. 2020) | Põlme S, et al. 2020. Fungal Diversity 105:1-16 ((https://doi.org/10.1007/s13225-020-00466-2)) |
| Otu60369 | 1 | k__Fungi(100);p__Ascomycota(100);c__Saccharomycetes(100);o__Saccharomycetales(100);f__Saccharomycetales_family_Incertae_sedis(100);f__Saccharomycetales_family_Incertae_sedis_unclassified(100);f__Saccharomycetales_family_Incertae_sedis_unclassified(100); | Saccharomycetales | 7 | Saprotroph | Undefined Saprotroph | Yeast | NULL | Possible | NULL | Sterkenburg E, et al. 2015. New Phytologist 207:1145-1158 ((https://doi.org/10.1111/nph.13426)) |
| Otu60378 | 1 | k__Fungi(100);p__Ascomycota(100);c__Saccharomycetes(100);o__Saccharomycetales(100);f__Metschnikowiaceae(100);g__Metschnikowia(100);s__Metschnikowia_pulcherrima(100); | Metschnikowia | 13 | Saprotroph | \|Undefined Saprotroph\| | Yeast | NULL | Probable | As nectar-tap_saprotroph (Põlme et al. 2020); Decay sugar-rich substrates (Põlme et al. 2020) | Põlme S, et al. 2020. Fungal Diversity 105:1-16 ((https://doi.org/10.1007/s13225-020-00466-2)) |
| Otu60383 | 1 | k__Fungi(100);p__Ascomycota(100);c__Saccharomycetes(100);o__Saccharomycetales(100);f__Metschnikowiaceae(100);g__Metschnikowia(100);g__Metschnikowia_unclassified(100); | Metschnikowia | 13 | Saprotroph | \|Undefined Saprotroph\| | Yeast | NULL | Probable | As nectar-tap_saprotroph (Põlme et al. 2020); Decay sugar-rich substrates (Põlme et al. 2020) | Põlme S, et al. 2020. Fungal Diversity 105:1-16 ((https://doi.org/10.1007/s13225-020-00466-2)) |
| Otu60394 | 1 | k__Fungi(100);p__Ascomycota(100);c__Saccharomycetes(100);o__Saccharomycetales(100);f__Metschnikowiaceae(100);g__Metschnikowia(100);s__Metschnikowia_pulcherrima(100); | Metschnikowia | 13 | Saprotroph | \|Undefined Saprotroph\| | Yeast | NULL | Probable | As nectar-tap_saprotroph (Põlme et al. 2020); Decay sugar-rich substrates (Põlme et al. 2020) | Põlme S, et al. 2020. Fungal Diversity 105:1-16 ((https://doi.org/10.1007/s13225-020-00466-2)) |
| Otu60401 | 1 | k__Fungi(100);p__Ascomycota(100);c__Saccharomycetes(100);o__Saccharomycetales(100);f__Metschnikowiaceae(100);g__Metschnikowia(100);s__Metschnikowia_pulcherrima(100); | Metschnikowia | 13 | Saprotroph | \|Undefined Saprotroph\| | Yeast | NULL | Probable | As nectar-tap_saprotroph (Põlme et al. 2020); Decay sugar-rich substrates (Põlme et al. 2020) | Põlme S, et al. 2020. Fungal Diversity 105:1-16 ((https://doi.org/10.1007/s13225-020-00466-2)) |
| Otu60409 | 1 | k__Fungi(100);p__Ascomycota(100);c__Saccharomycetes(100);o__Saccharomycetales(100);f__Metschnikowiaceae(100);g__Metschnikowia(100);s__Metschnikowia_pulcherrima(100); | Metschnikowia | 13 | Saprotroph | \|Undefined Saprotroph\| | Yeast | NULL | Probable | As nectar-tap_saprotroph (Põlme et al. 2020); Decay sugar-rich substrates (Põlme et al. 2020) | Põlme S, et al. 2020. Fungal Diversity 105:1-16 ((https://doi.org/10.1007/s13225-020-00466-2)) |
| Otu60415 | 1 | k__Fungi(100);p__Ascomycota(100);c__Saccharomycetes(100);o__Saccharomycetales(100);f__Metschnikowiaceae(100);g__Metschnikowia(100);s__Metschnikowia_pulcherrima(100); | Metschnikowia | 13 | Saprotroph | \|Undefined Saprotroph\| | Yeast | NULL | Probable | As nectar-tap_saprotroph (Põlme et al. 2020); Decay sugar-rich substrates (Põlme et al. 2020) | Põlme S, et al. 2020. Fungal Diversity 105:1-16 ((https://doi.org/10.1007/s13225-020-00466-2)) |
| Otu60418 | 1 | k__Fungi(100);p__Ascomycota(100);c__Saccharomycetes(100);o__Saccharomycetales(100);f__Metschnikowiaceae(100);g__Metschnikowia(100);s__Metschnikowia_pulcherrima(100); | Metschnikowia | 13 | Saprotroph | \|Undefined Saprotroph\| | Yeast | NULL | Probable | As nectar-tap_saprotroph (Põlme et al. 2020); Decay sugar-rich substrates (Põlme et al. 2020) | Põlme S, et al. 2020. Fungal Diversity 105:1-16 ((https://doi.org/10.1007/s13225-020-00466-2)) |
| Otu60424 | 1 | k__Fungi(100);p__Ascomycota(100);c__Saccharomycetes(100);o__Saccharomycetales(100);f__Metschnikowiaceae(100);g__Metschnikowia(100);g__Metschnikowia_unclassified(100); | Metschnikowia | 13 | Saprotroph | \|Undefined Saprotroph\| | Yeast | NULL | Probable | As nectar-tap_saprotroph (Põlme et al. 2020); Decay sugar-rich substrates (Põlme et al. 2020) | Põlme S, et al. 2020. Fungal Diversity 105:1-16 ((https://doi.org/10.1007/s13225-020-00466-2)) |
| Otu60425 | 1 | k__Fungi(100);p__Ascomycota(100);c__Saccharomycetes(100);o__Saccharomycetales(100);f__Metschnikowiaceae(100);g__Metschnikowia(100);g__Metschnikowia_unclassified(100); | Metschnikowia | 13 | Saprotroph | \|Undefined Saprotroph\| | Yeast | NULL | Probable | As nectar-tap_saprotroph (Põlme et al. 2020); Decay sugar-rich substrates (Põlme et al. 2020) | Põlme S, et al. 2020. Fungal Diversity 105:1-16 ((https://doi.org/10.1007/s13225-020-00466-2)) |
| Otu60426 | 1 | k__Fungi(100);p__Ascomycota(100);c__Saccharomycetes(100);o__Saccharomycetales(100);f__Metschnikowiaceae(100);g__Metschnikowia(100);s__Metschnikowia_pulcherrima(100); | Metschnikowia | 13 | Saprotroph | \|Undefined Saprotroph\| | Yeast | NULL | Probable | As nectar-tap_saprotroph (Põlme et al. 2020); Decay sugar-rich substrates (Põlme et al. 2020) | Põlme S, et al. 2020. Fungal Diversity 105:1-16 ((https://doi.org/10.1007/s13225-020-00466-2)) |
| Otu60428 | 1 | k__Fungi(100);p__Ascomycota(100);c__Saccharomycetes(100);o__Saccharomycetales(100);f__Metschnikowiaceae(100);g__Metschnikowia(100);s__Metschnikowia_pulcherrima(100); | Metschnikowia | 13 | Saprotroph | \|Undefined Saprotroph\| | Yeast | NULL | Probable | As nectar-tap_saprotroph (Põlme et al. 2020); Decay sugar-rich substrates (Põlme et al. 2020) | Põlme S, et al. 2020. Fungal Diversity 105:1-16 ((https://doi.org/10.1007/s13225-020-00466-2)) |
| Otu60430 | 1 | k__Fungi(100);p__Ascomycota(100);c__Saccharomycetes(100);o__Saccharomycetales(100);f__Metschnikowiaceae(100);g__Metschnikowia(100);s__Metschnikowia_pulcherrima(100); | Metschnikowia | 13 | Saprotroph | \|Undefined Saprotroph\| | Yeast | NULL | Probable | As nectar-tap_saprotroph (Põlme et al. 2020); Decay sugar-rich substrates (Põlme et al. 2020) | Põlme S, et al. 2020. Fungal Diversity 105:1-16 ((https://doi.org/10.1007/s13225-020-00466-2)) |
| Otu60438 | 1 | k__Fungi(100);p__Ascomycota(100);c__Saccharomycetes(100);o__Saccharomycetales(100);f__Metschnikowiaceae(100);g__Metschnikowia(100);s__Metschnikowia_pulcherrima(100); | Metschnikowia | 13 | Saprotroph | \|Undefined Saprotroph\| | Yeast | NULL | Probable | As nectar-tap_saprotroph (Põlme et al. 2020); Decay sugar-rich substrates (Põlme et al. 2020) | Põlme S, et al. 2020. Fungal Diversity 105:1-16 ((https://doi.org/10.1007/s13225-020-00466-2)) |
| Otu60442 | 1 | k__Fungi(100);p__Ascomycota(100);c__Saccharomycetes(100);o__Saccharomycetales(100);o__Saccharomycetales_unclassified(100);o__Saccharomycetales_unclassified(100);o__Saccharomycetales_unclassified(100); | Saccharomycetales | 7 | Saprotroph | Undefined Saprotroph | Yeast | NULL | Possible | NULL | Sterkenburg E, et al. 2015. New Phytologist 207:1145-1158 ((https://doi.org/10.1111/nph.13426)) |
| Otu60443 | 1 | k__Fungi(100);p__Ascomycota(100);c__Saccharomycetes(100);o__Saccharomycetales(100);f__Metschnikowiaceae(100);g__Metschnikowia(100);s__Metschnikowia_pulcherrima(100); | Metschnikowia | 13 | Saprotroph | \|Undefined Saprotroph\| | Yeast | NULL | Probable | As nectar-tap_saprotroph (Põlme et al. 2020); Decay sugar-rich substrates (Põlme et al. 2020) | Põlme S, et al. 2020. Fungal Diversity 105:1-16 ((https://doi.org/10.1007/s13225-020-00466-2)) |
| Otu60446 | 1 | k__Fungi(100);p__Ascomycota(100);c__Saccharomycetes(100);o__Saccharomycetales(100);f__Metschnikowiaceae(100);g__Metschnikowia(100);s__Metschnikowia_pulcherrima(100); | Metschnikowia | 13 | Saprotroph | \|Undefined Saprotroph\| | Yeast | NULL | Probable | As nectar-tap_saprotroph (Põlme et al. 2020); Decay sugar-rich substrates (Põlme et al. 2020) | Põlme S, et al. 2020. Fungal Diversity 105:1-16 ((https://doi.org/10.1007/s13225-020-00466-2)) |
| Otu60459 | 1 | k__Fungi(100);p__Ascomycota(100);c__Saccharomycetes(100);o__Saccharomycetales(100);f__Metschnikowiaceae(100);g__Metschnikowia(100);s__Metschnikowia_pulcherrima(100); | Metschnikowia | 13 | Saprotroph | \|Undefined Saprotroph\| | Yeast | NULL | Probable | As nectar-tap_saprotroph (Põlme et al. 2020); Decay sugar-rich substrates (Põlme et al. 2020) | Põlme S, et al. 2020. Fungal Diversity 105:1-16 ((https://doi.org/10.1007/s13225-020-00466-2)) |
| Otu60464 | 1 | k__Fungi(100);p__Basidiomycota(100);c__Tremellomycetes(100);o__Tremellales(100);f__Tremellales_family_Incertae_sedis(100);g__Cryptococcus(100);s__Cryptococcus_sp_JCM_11356(100); | Cryptococcus | 13 | Pathotroph-Saprotroph-Symbiotroph | Animal Pathogen-Endophyte-Epiphyte-\|Undefined Saprotroph\| | Dimorphic Yeast-Yeast | NULL | Possible | As foliar_endophyte (Põlme et al. 2020); Decay leaf, fruit, seed, soil, dung, animal material (Põlme et al. 2020) | Andrews JH, et al. 1983. Phytopathology 73:228-234 ((https://doi.org/10.1094/Phyto-73-228)); Kurtzman CP, et al. (eds.) 2011. The Yeasts, a Taxonomic Study. Fifth Edition. Vols 1-3. Elsevier, San Diego (ISBN: 9780444521491); Põlme S, et al. 2020. Fungal Diversity 105:1-16 ((https://doi.org/10.1007/s13225-020-00466-2)) |
| Otu60465 | 1 | k__Fungi(100);p__Ascomycota(100);c__Saccharomycetes(100);o__Saccharomycetales(100);f__Metschnikowiaceae(100);g__Metschnikowia(100);s__Metschnikowia_pulcherrima(100); | Metschnikowia | 13 | Saprotroph | \|Undefined Saprotroph\| | Yeast | NULL | Probable | As nectar-tap_saprotroph (Põlme et al. 2020); Decay sugar-rich substrates (Põlme et al. 2020) | Põlme S, et al. 2020. Fungal Diversity 105:1-16 ((https://doi.org/10.1007/s13225-020-00466-2)) |
| Otu60468 | 1 | k__Fungi(100);p__Ascomycota(100);c__Saccharomycetes(100);o__Saccharomycetales(100);f__Metschnikowiaceae(100);g__Metschnikowia(100);g__Metschnikowia_unclassified(100); | Metschnikowia | 13 | Saprotroph | \|Undefined Saprotroph\| | Yeast | NULL | Probable | As nectar-tap_saprotroph (Põlme et al. 2020); Decay sugar-rich substrates (Põlme et al. 2020) | Põlme S, et al. 2020. Fungal Diversity 105:1-16 ((https://doi.org/10.1007/s13225-020-00466-2)) |
| Otu60470 | 1 | k__Fungi(100);p__Basidiomycota(100);c__Tremellomycetes(100);o__Tremellales(100);f__Tremellales_family_Incertae_sedis(100);g__Cryptococcus(100);s__Cryptococcus_sp_JCM_11356(100); | Cryptococcus | 13 | Pathotroph-Saprotroph-Symbiotroph | Animal Pathogen-Endophyte-Epiphyte-\|Undefined Saprotroph\| | Dimorphic Yeast-Yeast | NULL | Possible | As foliar_endophyte (Põlme et al. 2020); Decay leaf, fruit, seed, soil, dung, animal material (Põlme et al. 2020) | Andrews JH, et al. 1983. Phytopathology 73:228-234 ((https://doi.org/10.1094/Phyto-73-228)); Kurtzman CP, et al. (eds.) 2011. The Yeasts, a Taxonomic Study. Fifth Edition. Vols 1-3. Elsevier, San Diego (ISBN: 9780444521491); Põlme S, et al. 2020. Fungal Diversity 105:1-16 ((https://doi.org/10.1007/s13225-020-00466-2)) |
| Otu60471 | 1 | k__Fungi(100);p__Ascomycota(100);c__Saccharomycetes(100);o__Saccharomycetales(100);f__Metschnikowiaceae(100);g__Metschnikowia(100);s__Metschnikowia_pulcherrima(100); | Metschnikowia | 13 | Saprotroph | \|Undefined Saprotroph\| | Yeast | NULL | Probable | As nectar-tap_saprotroph (Põlme et al. 2020); Decay sugar-rich substrates (Põlme et al. 2020) | Põlme S, et al. 2020. Fungal Diversity 105:1-16 ((https://doi.org/10.1007/s13225-020-00466-2)) |
| Otu60472 | 1 | k__Fungi(100);p__Ascomycota(100);c__Saccharomycetes(100);o__Saccharomycetales(100);f__Metschnikowiaceae(100);g__Metschnikowia(100);s__Metschnikowia_pulcherrima(100); | Metschnikowia | 13 | Saprotroph | \|Undefined Saprotroph\| | Yeast | NULL | Probable | As nectar-tap_saprotroph (Põlme et al. 2020); Decay sugar-rich substrates (Põlme et al. 2020) | Põlme S, et al. 2020. Fungal Diversity 105:1-16 ((https://doi.org/10.1007/s13225-020-00466-2)) |
| Otu60473 | 1 | k__Fungi(100);p__Ascomycota(100);c__Saccharomycetes(100);o__Saccharomycetales(100);f__Metschnikowiaceae(100);g__Metschnikowia(100);s__Metschnikowia_pulcherrima(100); | Metschnikowia | 13 | Saprotroph | \|Undefined Saprotroph\| | Yeast | NULL | Probable | As nectar-tap_saprotroph (Põlme et al. 2020); Decay sugar-rich substrates (Põlme et al. 2020) | Põlme S, et al. 2020. Fungal Diversity 105:1-16 ((https://doi.org/10.1007/s13225-020-00466-2)) |
| Otu60528 | 1 | k__Fungi(100);p__Ascomycota(100);c__Saccharomycetes(100);o__Saccharomycetales(100);f__Metschnikowiaceae(100);g__Metschnikowia(100);g__Metschnikowia_unclassified(100); | Metschnikowia | 13 | Saprotroph | \|Undefined Saprotroph\| | Yeast | NULL | Probable | As nectar-tap_saprotroph (Põlme et al. 2020); Decay sugar-rich substrates (Põlme et al. 2020) | Põlme S, et al. 2020. Fungal Diversity 105:1-16 ((https://doi.org/10.1007/s13225-020-00466-2)) |
| Otu60568 | 1 | k__Fungi(100);p__Ascomycota(100);c__Saccharomycetes(100);o__Saccharomycetales(100);f__Metschnikowiaceae(100);g__Metschnikowia(100);s__Metschnikowia_pulcherrima(100); | Metschnikowia | 13 | Saprotroph | \|Undefined Saprotroph\| | Yeast | NULL | Probable | As nectar-tap_saprotroph (Põlme et al. 2020); Decay sugar-rich substrates (Põlme et al. 2020) | Põlme S, et al. 2020. Fungal Diversity 105:1-16 ((https://doi.org/10.1007/s13225-020-00466-2)) |
| Otu60581 | 1 | k__Fungi(100);p__Ascomycota(100);c__Saccharomycetes(100);o__Saccharomycetales(100);f__Metschnikowiaceae(100);g__Metschnikowia(100);s__Metschnikowia_pulcherrima(100); | Metschnikowia | 13 | Saprotroph | \|Undefined Saprotroph\| | Yeast | NULL | Probable | As nectar-tap_saprotroph (Põlme et al. 2020); Decay sugar-rich substrates (Põlme et al. 2020) | Põlme S, et al. 2020. Fungal Diversity 105:1-16 ((https://doi.org/10.1007/s13225-020-00466-2)) |
| Otu60582 | 1 | k__Fungi(100);p__Ascomycota(100);c__Saccharomycetes(100);o__Saccharomycetales(100);f__Metschnikowiaceae(100);g__Metschnikowia(100);s__Metschnikowia_pulcherrima(100); | Metschnikowia | 13 | Saprotroph | \|Undefined Saprotroph\| | Yeast | NULL | Probable | As nectar-tap_saprotroph (Põlme et al. 2020); Decay sugar-rich substrates (Põlme et al. 2020) | Põlme S, et al. 2020. Fungal Diversity 105:1-16 ((https://doi.org/10.1007/s13225-020-00466-2)) |
| Otu60597 | 1 | k__Fungi(100);p__Ascomycota(100);c__Saccharomycetes(100);o__Saccharomycetales(100);f__Metschnikowiaceae(100);g__Metschnikowia(100);s__Metschnikowia_pulcherrima(100); | Metschnikowia | 13 | Saprotroph | \|Undefined Saprotroph\| | Yeast | NULL | Probable | As nectar-tap_saprotroph (Põlme et al. 2020); Decay sugar-rich substrates (Põlme et al. 2020) | Põlme S, et al. 2020. Fungal Diversity 105:1-16 ((https://doi.org/10.1007/s13225-020-00466-2)) |
| Otu60598 | 1 | k__Fungi(100);p__Ascomycota(100);c__Saccharomycetes(100);o__Saccharomycetales(100);f__Metschnikowiaceae(100);g__Metschnikowia(100);s__Metschnikowia_pulcherrima(100); | Metschnikowia | 13 | Saprotroph | \|Undefined Saprotroph\| | Yeast | NULL | Probable | As nectar-tap_saprotroph (Põlme et al. 2020); Decay sugar-rich substrates (Põlme et al. 2020) | Põlme S, et al. 2020. Fungal Diversity 105:1-16 ((https://doi.org/10.1007/s13225-020-00466-2)) |
| Otu60599 | 1 | k__Fungi(100);p__Ascomycota(100);c__Saccharomycetes(100);o__Saccharomycetales(100);f__Metschnikowiaceae(100);g__Metschnikowia(100);g__Metschnikowia_unclassified(100); | Metschnikowia | 13 | Saprotroph | \|Undefined Saprotroph\| | Yeast | NULL | Probable | As nectar-tap_saprotroph (Põlme et al. 2020); Decay sugar-rich substrates (Põlme et al. 2020) | Põlme S, et al. 2020. Fungal Diversity 105:1-16 ((https://doi.org/10.1007/s13225-020-00466-2)) |
| Otu60600 | 1 | k__Fungi(100);p__Ascomycota(100);c__Saccharomycetes(100);o__Saccharomycetales(100);f__Metschnikowiaceae(100);g__Metschnikowia(100);s__Metschnikowia_pulcherrima(100); | Metschnikowia | 13 | Saprotroph | \|Undefined Saprotroph\| | Yeast | NULL | Probable | As nectar-tap_saprotroph (Põlme et al. 2020); Decay sugar-rich substrates (Põlme et al. 2020) | Põlme S, et al. 2020. Fungal Diversity 105:1-16 ((https://doi.org/10.1007/s13225-020-00466-2)) |
| Otu60601 | 1 | k__Fungi(100);p__Ascomycota(100);c__Saccharomycetes(100);o__Saccharomycetales(100);f__Metschnikowiaceae(100);g__Metschnikowia(100);s__Metschnikowia_pulcherrima(100); | Metschnikowia | 13 | Saprotroph | \|Undefined Saprotroph\| | Yeast | NULL | Probable | As nectar-tap_saprotroph (Põlme et al. 2020); Decay sugar-rich substrates (Põlme et al. 2020) | Põlme S, et al. 2020. Fungal Diversity 105:1-16 ((https://doi.org/10.1007/s13225-020-00466-2)) |
| Otu60604 | 1 | k__Fungi(100);p__Ascomycota(100);c__Saccharomycetes(100);o__Saccharomycetales(100);f__Metschnikowiaceae(100);g__Metschnikowia(100);s__Metschnikowia_pulcherrima(100); | Metschnikowia | 13 | Saprotroph | \|Undefined Saprotroph\| | Yeast | NULL | Probable | As nectar-tap_saprotroph (Põlme et al. 2020); Decay sugar-rich substrates (Põlme et al. 2020) | Põlme S, et al. 2020. Fungal Diversity 105:1-16 ((https://doi.org/10.1007/s13225-020-00466-2)) |
| Otu60606 | 1 | k__Fungi(100);p__Ascomycota(100);c__Saccharomycetes(100);o__Saccharomycetales(100);f__Metschnikowiaceae(100);g__Metschnikowia(100);s__Metschnikowia_pulcherrima(100); | Metschnikowia | 13 | Saprotroph | \|Undefined Saprotroph\| | Yeast | NULL | Probable | As nectar-tap_saprotroph (Põlme et al. 2020); Decay sugar-rich substrates (Põlme et al. 2020) | Põlme S, et al. 2020. Fungal Diversity 105:1-16 ((https://doi.org/10.1007/s13225-020-00466-2)) |
| Otu60607 | 1 | k__Fungi(100);p__Ascomycota(100);c__Saccharomycetes(100);o__Saccharomycetales(100);f__Metschnikowiaceae(100);g__Metschnikowia(100);s__Metschnikowia_pulcherrima(100); | Metschnikowia | 13 | Saprotroph | \|Undefined Saprotroph\| | Yeast | NULL | Probable | As nectar-tap_saprotroph (Põlme et al. 2020); Decay sugar-rich substrates (Põlme et al. 2020) | Põlme S, et al. 2020. Fungal Diversity 105:1-16 ((https://doi.org/10.1007/s13225-020-00466-2)) |
| Otu60608 | 1 | k__Fungi(100);p__Ascomycota(100);c__Saccharomycetes(100);o__Saccharomycetales(100);f__Metschnikowiaceae(100);g__Metschnikowia(100);g__Metschnikowia_unclassified(100); | Metschnikowia | 13 | Saprotroph | \|Undefined Saprotroph\| | Yeast | NULL | Probable | As nectar-tap_saprotroph (Põlme et al. 2020); Decay sugar-rich substrates (Põlme et al. 2020) | Põlme S, et al. 2020. Fungal Diversity 105:1-16 ((https://doi.org/10.1007/s13225-020-00466-2)) |
| Otu60609 | 1 | k__Fungi(100);p__Ascomycota(100);c__Saccharomycetes(100);o__Saccharomycetales(100);f__Metschnikowiaceae(100);g__Metschnikowia(100);s__Metschnikowia_pulcherrima(100); | Metschnikowia | 13 | Saprotroph | \|Undefined Saprotroph\| | Yeast | NULL | Probable | As nectar-tap_saprotroph (Põlme et al. 2020); Decay sugar-rich substrates (Põlme et al. 2020) | Põlme S, et al. 2020. Fungal Diversity 105:1-16 ((https://doi.org/10.1007/s13225-020-00466-2)) |
| Otu60610 | 1 | k__Fungi(100);p__Ascomycota(100);c__Saccharomycetes(100);o__Saccharomycetales(100);f__Metschnikowiaceae(100);g__Metschnikowia(100);s__Metschnikowia_pulcherrima(100); | Metschnikowia | 13 | Saprotroph | \|Undefined Saprotroph\| | Yeast | NULL | Probable | As nectar-tap_saprotroph (Põlme et al. 2020); Decay sugar-rich substrates (Põlme et al. 2020) | Põlme S, et al. 2020. Fungal Diversity 105:1-16 ((https://doi.org/10.1007/s13225-020-00466-2)) |
| Otu60611 | 1 | k__Fungi(100);p__Ascomycota(100);c__Saccharomycetes(100);o__Saccharomycetales(100);f__Metschnikowiaceae(100);g__Metschnikowia(100);s__Metschnikowia_pulcherrima(100); | Metschnikowia | 13 | Saprotroph | \|Undefined Saprotroph\| | Yeast | NULL | Probable | As nectar-tap_saprotroph (Põlme et al. 2020); Decay sugar-rich substrates (Põlme et al. 2020) | Põlme S, et al. 2020. Fungal Diversity 105:1-16 ((https://doi.org/10.1007/s13225-020-00466-2)) |
| Otu60612 | 1 | k__Fungi(100);p__Ascomycota(100);c__Saccharomycetes(100);o__Saccharomycetales(100);f__Metschnikowiaceae(100);g__Metschnikowia(100);s__Metschnikowia_pulcherrima(100); | Metschnikowia | 13 | Saprotroph | \|Undefined Saprotroph\| | Yeast | NULL | Probable | As nectar-tap_saprotroph (Põlme et al. 2020); Decay sugar-rich substrates (Põlme et al. 2020) | Põlme S, et al. 2020. Fungal Diversity 105:1-16 ((https://doi.org/10.1007/s13225-020-00466-2)) |
| Otu60614 | 1 | k__Fungi(100);p__Ascomycota(100);c__Saccharomycetes(100);o__Saccharomycetales(100);f__Metschnikowiaceae(100);g__Metschnikowia(100);g__Metschnikowia_unclassified(100); | Metschnikowia | 13 | Saprotroph | \|Undefined Saprotroph\| | Yeast | NULL | Probable | As nectar-tap_saprotroph (Põlme et al. 2020); Decay sugar-rich substrates (Põlme et al. 2020) | Põlme S, et al. 2020. Fungal Diversity 105:1-16 ((https://doi.org/10.1007/s13225-020-00466-2)) |
| Otu60615 | 1 | k__Fungi(100);p__Ascomycota(100);c__Saccharomycetes(100);o__Saccharomycetales(100);f__Metschnikowiaceae(100);g__Metschnikowia(100);s__Metschnikowia_pulcherrima(100); | Metschnikowia | 13 | Saprotroph | \|Undefined Saprotroph\| | Yeast | NULL | Probable | As nectar-tap_saprotroph (Põlme et al. 2020); Decay sugar-rich substrates (Põlme et al. 2020) | Põlme S, et al. 2020. Fungal Diversity 105:1-16 ((https://doi.org/10.1007/s13225-020-00466-2)) |
| Otu60616 | 1 | k__Fungi(100);p__Ascomycota(100);c__Saccharomycetes(100);o__Saccharomycetales(100);f__Metschnikowiaceae(100);g__Metschnikowia(100);s__Metschnikowia_pulcherrima(100); | Metschnikowia | 13 | Saprotroph | \|Undefined Saprotroph\| | Yeast | NULL | Probable | As nectar-tap_saprotroph (Põlme et al. 2020); Decay sugar-rich substrates (Põlme et al. 2020) | Põlme S, et al. 2020. Fungal Diversity 105:1-16 ((https://doi.org/10.1007/s13225-020-00466-2)) |
| Otu60619 | 1 | k__Fungi(100);p__Ascomycota(100);c__Saccharomycetes(100);o__Saccharomycetales(100);f__Metschnikowiaceae(100);g__Metschnikowia(100);s__Metschnikowia_pulcherrima(100); | Metschnikowia | 13 | Saprotroph | \|Undefined Saprotroph\| | Yeast | NULL | Probable | As nectar-tap_saprotroph (Põlme et al. 2020); Decay sugar-rich substrates (Põlme et al. 2020) | Põlme S, et al. 2020. Fungal Diversity 105:1-16 ((https://doi.org/10.1007/s13225-020-00466-2)) |
| Otu60639 | 1 | k__Fungi(100);p__Ascomycota(100);c__Saccharomycetes(100);o__Saccharomycetales(100);f__Saccharomycetales_family_Incertae_sedis(100);g__Candida(100);s__Candida_cretensis(100); | Candida | 13 | Pathotroph-Saprotroph-Symbiotroph | Animal Pathogen-Endophyte-Endosymbiont-Epiphyte-\|Undefined Saprotroph\| | Dimorphic Yeast-Yeast | NULL | Probable | As nectar-tap_saprotroph (Põlme et al. 2020); Decay sugar-rich substrates (Põlme et al. 2020); This asexual genus is polyphyletic and contain species that function very differently depend on what they are. Caution should be taken when interpreting guilds for Candida species. | Manolakaki D, et al. 2010. Virulence. 1: 367-375 ((https://doi.org/10.4161/viru.1.5.12796)); Mekbib SB, et al. 2011. Tropical Plant Pathology 36:233-240 ((https://doi.org/10.1590/S1982-56762011000400004)); Põlme S, et al. 2020. Fungal Diversity 105:1-16 ((https://doi.org/10.1007/s13225-020-00466-2)) |
| Otu60646 | 1 | k__Fungi(100);p__Ascomycota(100);c__Saccharomycetes(100);o__Saccharomycetales(100);o__Saccharomycetales_unclassified(100);o__Saccharomycetales_unclassified(100);o__Saccharomycetales_unclassified(100); | Saccharomycetales | 7 | Saprotroph | Undefined Saprotroph | Yeast | NULL | Possible | NULL | Sterkenburg E, et al. 2015. New Phytologist 207:1145-1158 ((https://doi.org/10.1111/nph.13426)) |
| Otu60648 | 1 | k__Fungi(100);p__Ascomycota(100);c__Saccharomycetes(100);o__Saccharomycetales(100);f__Metschnikowiaceae(100);g__Metschnikowia(100);g__Metschnikowia_unclassified(100); | Metschnikowia | 13 | Saprotroph | \|Undefined Saprotroph\| | Yeast | NULL | Probable | As nectar-tap_saprotroph (Põlme et al. 2020); Decay sugar-rich substrates (Põlme et al. 2020) | Põlme S, et al. 2020. Fungal Diversity 105:1-16 ((https://doi.org/10.1007/s13225-020-00466-2)) |
| Otu60649 | 1 | k__Fungi(100);p__Ascomycota(100);c__Saccharomycetes(100);o__Saccharomycetales(100);f__Metschnikowiaceae(100);g__Metschnikowia(100);s__Metschnikowia_pulcherrima(100); | Metschnikowia | 13 | Saprotroph | \|Undefined Saprotroph\| | Yeast | NULL | Probable | As nectar-tap_saprotroph (Põlme et al. 2020); Decay sugar-rich substrates (Põlme et al. 2020) | Põlme S, et al. 2020. Fungal Diversity 105:1-16 ((https://doi.org/10.1007/s13225-020-00466-2)) |
| Otu60650 | 1 | k__Fungi(100);p__Ascomycota(100);c__Sordariomycetes(100);o__Chaetosphaeriales(100);f__Chaetosphaeriaceae(100);f__Chaetosphaeriaceae_unclassified(100);f__Chaetosphaeriaceae_unclassified(100); | Chaetosphaeriaceae | 9 | Saprotroph | Plant Saprotroph-Wood Saprotroph | Microfungus | NULL | Probable | Saprobes associated with decaying woody and herbaceous plant material, including timber and timber products in damp buildings (Cannon & Kirk 2007) | Cannon PF, Kirk PM. 2007. Fungal Families of the World. CAB International, Cambridge (ISBN: 978-0851998275) |
| Otu60679 | 1 | k__Fungi(100);p__Ascomycota(100);c__Saccharomycetes(100);o__Saccharomycetales(100);f__Metschnikowiaceae(100);g__Metschnikowia(100);s__Metschnikowia_pulcherrima(100); | Metschnikowia | 13 | Saprotroph | \|Undefined Saprotroph\| | Yeast | NULL | Probable | As nectar-tap_saprotroph (Põlme et al. 2020); Decay sugar-rich substrates (Põlme et al. 2020) | Põlme S, et al. 2020. Fungal Diversity 105:1-16 ((https://doi.org/10.1007/s13225-020-00466-2)) |
| Otu60683 | 1 | k__Fungi(100);p__Ascomycota(100);c__Saccharomycetes(100);o__Saccharomycetales(100);f__Metschnikowiaceae(100);g__Metschnikowia(100);s__Metschnikowia_pulcherrima(100); | Metschnikowia | 13 | Saprotroph | \|Undefined Saprotroph\| | Yeast | NULL | Probable | As nectar-tap_saprotroph (Põlme et al. 2020); Decay sugar-rich substrates (Põlme et al. 2020) | Põlme S, et al. 2020. Fungal Diversity 105:1-16 ((https://doi.org/10.1007/s13225-020-00466-2)) |
| Otu60684 | 1 | k__Fungi(100);p__Ascomycota(100);c__Saccharomycetes(100);o__Saccharomycetales(100);f__Metschnikowiaceae(100);g__Metschnikowia(100);g__Metschnikowia_unclassified(100); | Metschnikowia | 13 | Saprotroph | \|Undefined Saprotroph\| | Yeast | NULL | Probable | As nectar-tap_saprotroph (Põlme et al. 2020); Decay sugar-rich substrates (Põlme et al. 2020) | Põlme S, et al. 2020. Fungal Diversity 105:1-16 ((https://doi.org/10.1007/s13225-020-00466-2)) |
| Otu60686 | 1 | k__Fungi(100);p__Ascomycota(100);c__Saccharomycetes(100);o__Saccharomycetales(100);f__Metschnikowiaceae(100);g__Metschnikowia(100);s__Metschnikowia_pulcherrima(100); | Metschnikowia | 13 | Saprotroph | \|Undefined Saprotroph\| | Yeast | NULL | Probable | As nectar-tap_saprotroph (Põlme et al. 2020); Decay sugar-rich substrates (Põlme et al. 2020) | Põlme S, et al. 2020. Fungal Diversity 105:1-16 ((https://doi.org/10.1007/s13225-020-00466-2)) |
| Otu60687 | 1 | k__Fungi(100);p__Ascomycota(100);c__Saccharomycetes(100);o__Saccharomycetales(100);f__Metschnikowiaceae(100);g__Metschnikowia(100);s__Metschnikowia_pulcherrima(100); | Metschnikowia | 13 | Saprotroph | \|Undefined Saprotroph\| | Yeast | NULL | Probable | As nectar-tap_saprotroph (Põlme et al. 2020); Decay sugar-rich substrates (Põlme et al. 2020) | Põlme S, et al. 2020. Fungal Diversity 105:1-16 ((https://doi.org/10.1007/s13225-020-00466-2)) |
| Otu60689 | 1 | k__Fungi(100);p__Ascomycota(100);c__Saccharomycetes(100);o__Saccharomycetales(100);f__Metschnikowiaceae(100);g__Metschnikowia(100);g__Metschnikowia_unclassified(100); | Metschnikowia | 13 | Saprotroph | \|Undefined Saprotroph\| | Yeast | NULL | Probable | As nectar-tap_saprotroph (Põlme et al. 2020); Decay sugar-rich substrates (Põlme et al. 2020) | Põlme S, et al. 2020. Fungal Diversity 105:1-16 ((https://doi.org/10.1007/s13225-020-00466-2)) |
| Otu60690 | 1 | k__Fungi(100);p__Ascomycota(100);c__Saccharomycetes(100);o__Saccharomycetales(100);f__Metschnikowiaceae(100);g__Metschnikowia(100);s__Metschnikowia_pulcherrima(100); | Metschnikowia | 13 | Saprotroph | \|Undefined Saprotroph\| | Yeast | NULL | Probable | As nectar-tap_saprotroph (Põlme et al. 2020); Decay sugar-rich substrates (Põlme et al. 2020) | Põlme S, et al. 2020. Fungal Diversity 105:1-16 ((https://doi.org/10.1007/s13225-020-00466-2)) |
| Otu60705 | 1 | k__Fungi(100);p__Ascomycota(100);c__Saccharomycetes(100);o__Saccharomycetales(100);f__Metschnikowiaceae(100);g__Metschnikowia(100);s__Metschnikowia_pulcherrima(100); | Metschnikowia | 13 | Saprotroph | \|Undefined Saprotroph\| | Yeast | NULL | Probable | As nectar-tap_saprotroph (Põlme et al. 2020); Decay sugar-rich substrates (Põlme et al. 2020) | Põlme S, et al. 2020. Fungal Diversity 105:1-16 ((https://doi.org/10.1007/s13225-020-00466-2)) |
| Otu60706 | 1 | k__Fungi(100);p__Ascomycota(100);c__Sordariomycetes(100);o__Chaetosphaeriales(100);f__Chaetosphaeriaceae(100);g__Menispora(100);s__Menispora_ciliata(100); | Menispora | 13 | Saprotroph-Symbiotroph | Endophyte-Plant Saprotroph-Undefined Saprotroph-\|Wood Saprotroph\| | Microfungus | NULL | Probable | As foliar_endophyte (Põlme et al. 2020); Decay wood, leaf, fruit, seed (Põlme et al. 2020) | Tedersoo L, et al. 2014. Science 346:e1256688 ((https://doi.org/10.1126/science.1256688)); Christian N, et al. 2016. Microbial Ecology 72: 621-632 ((https://doi.org/10.1007/s00248-016-0804-y)) (as endophyte); Põlme S, et al. 2020. Fungal Diversity 105:1-16 ((https://doi.org/10.1007/s13225-020-00466-2)) |
| Otu60710 | 1 | k__Fungi(100);p__Ascomycota(100);c__Saccharomycetes(100);o__Saccharomycetales(100);f__Metschnikowiaceae(100);g__Metschnikowia(100);s__Metschnikowia_pulcherrima(100); | Metschnikowia | 13 | Saprotroph | \|Undefined Saprotroph\| | Yeast | NULL | Probable | As nectar-tap_saprotroph (Põlme et al. 2020); Decay sugar-rich substrates (Põlme et al. 2020) | Põlme S, et al. 2020. Fungal Diversity 105:1-16 ((https://doi.org/10.1007/s13225-020-00466-2)) |
| Otu60715 | 1 | k__Fungi(100);p__Ascomycota(100);c__Saccharomycetes(100);o__Saccharomycetales(100);f__Metschnikowiaceae(100);g__Metschnikowia(100);g__Metschnikowia_unclassified(100); | Metschnikowia | 13 | Saprotroph | \|Undefined Saprotroph\| | Yeast | NULL | Probable | As nectar-tap_saprotroph (Põlme et al. 2020); Decay sugar-rich substrates (Põlme et al. 2020) | Põlme S, et al. 2020. Fungal Diversity 105:1-16 ((https://doi.org/10.1007/s13225-020-00466-2)) |
| Otu60716 | 1 | k__Fungi(100);p__Ascomycota(100);c__Saccharomycetes(100);o__Saccharomycetales(100);f__Metschnikowiaceae(100);g__Metschnikowia(100);s__Metschnikowia_pulcherrima(100); | Metschnikowia | 13 | Saprotroph | \|Undefined Saprotroph\| | Yeast | NULL | Probable | As nectar-tap_saprotroph (Põlme et al. 2020); Decay sugar-rich substrates (Põlme et al. 2020) | Põlme S, et al. 2020. Fungal Diversity 105:1-16 ((https://doi.org/10.1007/s13225-020-00466-2)) |
| Otu60723 | 1 | k__Fungi(100);p__Ascomycota(100);c__Saccharomycetes(100);o__Saccharomycetales(100);f__Metschnikowiaceae(100);g__Metschnikowia(100);s__Metschnikowia_pulcherrima(100); | Metschnikowia | 13 | Saprotroph | \|Undefined Saprotroph\| | Yeast | NULL | Probable | As nectar-tap_saprotroph (Põlme et al. 2020); Decay sugar-rich substrates (Põlme et al. 2020) | Põlme S, et al. 2020. Fungal Diversity 105:1-16 ((https://doi.org/10.1007/s13225-020-00466-2)) |
| Otu60724 | 1 | k__Fungi(100);p__Ascomycota(100);c__Saccharomycetes(100);o__Saccharomycetales(100);f__Metschnikowiaceae(100);g__Metschnikowia(100);s__Metschnikowia_pulcherrima(100); | Metschnikowia | 13 | Saprotroph | \|Undefined Saprotroph\| | Yeast | NULL | Probable | As nectar-tap_saprotroph (Põlme et al. 2020); Decay sugar-rich substrates (Põlme et al. 2020) | Põlme S, et al. 2020. Fungal Diversity 105:1-16 ((https://doi.org/10.1007/s13225-020-00466-2)) |
| Otu60725 | 1 | k__Fungi(100);p__Ascomycota(100);c__Saccharomycetes(100);o__Saccharomycetales(100);f__Metschnikowiaceae(100);g__Metschnikowia(100);g__Metschnikowia_unclassified(100); | Metschnikowia | 13 | Saprotroph | \|Undefined Saprotroph\| | Yeast | NULL | Probable | As nectar-tap_saprotroph (Põlme et al. 2020); Decay sugar-rich substrates (Põlme et al. 2020) | Põlme S, et al. 2020. Fungal Diversity 105:1-16 ((https://doi.org/10.1007/s13225-020-00466-2)) |
| Otu60726 | 1 | k__Fungi(100);p__Ascomycota(100);c__Saccharomycetes(100);o__Saccharomycetales(100);f__Metschnikowiaceae(100);g__Metschnikowia(100);s__Metschnikowia_pulcherrima(100); | Metschnikowia | 13 | Saprotroph | \|Undefined Saprotroph\| | Yeast | NULL | Probable | As nectar-tap_saprotroph (Põlme et al. 2020); Decay sugar-rich substrates (Põlme et al. 2020) | Põlme S, et al. 2020. Fungal Diversity 105:1-16 ((https://doi.org/10.1007/s13225-020-00466-2)) |
| Otu60731 | 1 | k__Fungi(100);p__Ascomycota(100);c__Saccharomycetes(100);o__Saccharomycetales(100);f__Metschnikowiaceae(100);g__Metschnikowia(100);s__Metschnikowia_pulcherrima(100); | Metschnikowia | 13 | Saprotroph | \|Undefined Saprotroph\| | Yeast | NULL | Probable | As nectar-tap_saprotroph (Põlme et al. 2020); Decay sugar-rich substrates (Põlme et al. 2020) | Põlme S, et al. 2020. Fungal Diversity 105:1-16 ((https://doi.org/10.1007/s13225-020-00466-2)) |
| Otu60738 | 1 | k__Fungi(100);p__Ascomycota(100);c__Saccharomycetes(100);o__Saccharomycetales(100);f__Metschnikowiaceae(100);g__Metschnikowia(100);s__Metschnikowia_pulcherrima(100); | Metschnikowia | 13 | Saprotroph | \|Undefined Saprotroph\| | Yeast | NULL | Probable | As nectar-tap_saprotroph (Põlme et al. 2020); Decay sugar-rich substrates (Põlme et al. 2020) | Põlme S, et al. 2020. Fungal Diversity 105:1-16 ((https://doi.org/10.1007/s13225-020-00466-2)) |
| Otu60747 | 1 | k__Fungi(100);p__Basidiomycota(100);c__Tremellomycetes(100);o__Tremellales(100);f__Tremellales_family_Incertae_sedis(100);g__Dioszegia(100);g__Dioszegia_unclassified(100); | Dioszegia | 13 | Saprotroph | \|Plant Saprotroph\|-Undefined Saprotroph | Yeast | NULL | Probable | As litter_saprotroph (Põlme et al. 2020); Decay leaf, fruit, seed (Põlme et al. 2020); The Dioszegia subclade sensu Inacio (Inacio et al. 20015) is characterized by orange; yellow or orange to deep orange colonies. Isolated from plant leaves and soil. | Põlme S, et al. 2020. Fungal Diversity 105:1-16 ((https://doi.org/10.1007/s13225-020-00466-2)); Takashima M, et al. 2011 International Journal of Systematic and Evolutionary Microbiology 61:1736-1739 ((https://doi.org/10.1099/ijs.0.025254-0)) |
| Otu60750 | 1 | k__Fungi(100);p__Ascomycota(100);c__Saccharomycetes(100);o__Saccharomycetales(100);f__Saccharomycetales_family_Incertae_sedis(100);g__Candida(100);g__Candida_unclassified(100); | Candida | 13 | Pathotroph-Saprotroph-Symbiotroph | Animal Pathogen-Endophyte-Endosymbiont-Epiphyte-\|Undefined Saprotroph\| | Dimorphic Yeast-Yeast | NULL | Probable | As nectar-tap_saprotroph (Põlme et al. 2020); Decay sugar-rich substrates (Põlme et al. 2020); This asexual genus is polyphyletic and contain species that function very differently depend on what they are. Caution should be taken when interpreting guilds for Candida species. | Manolakaki D, et al. 2010. Virulence. 1: 367-375 ((https://doi.org/10.4161/viru.1.5.12796)); Mekbib SB, et al. 2011. Tropical Plant Pathology 36:233-240 ((https://doi.org/10.1590/S1982-56762011000400004)); Põlme S, et al. 2020. Fungal Diversity 105:1-16 ((https://doi.org/10.1007/s13225-020-00466-2)) |
| Otu60760 | 1 | k__Fungi(100);p__Ascomycota(100);c__Saccharomycetes(100);o__Saccharomycetales(100);f__Metschnikowiaceae(100);g__Metschnikowia(100);s__Metschnikowia_pulcherrima(100); | Metschnikowia | 13 | Saprotroph | \|Undefined Saprotroph\| | Yeast | NULL | Probable | As nectar-tap_saprotroph (Põlme et al. 2020); Decay sugar-rich substrates (Põlme et al. 2020) | Põlme S, et al. 2020. Fungal Diversity 105:1-16 ((https://doi.org/10.1007/s13225-020-00466-2)) |
| Otu60762 | 1 | k__Fungi(100);p__Ascomycota(100);c__Saccharomycetes(100);o__Saccharomycetales(100);f__Saccharomycetales_family_Incertae_sedis(100);g__Candida(100);g__Candida_unclassified(100); | Candida | 13 | Pathotroph-Saprotroph-Symbiotroph | Animal Pathogen-Endophyte-Endosymbiont-Epiphyte-\|Undefined Saprotroph\| | Dimorphic Yeast-Yeast | NULL | Probable | As nectar-tap_saprotroph (Põlme et al. 2020); Decay sugar-rich substrates (Põlme et al. 2020); This asexual genus is polyphyletic and contain species that function very differently depend on what they are. Caution should be taken when interpreting guilds for Candida species. | Manolakaki D, et al. 2010. Virulence. 1: 367-375 ((https://doi.org/10.4161/viru.1.5.12796)); Mekbib SB, et al. 2011. Tropical Plant Pathology 36:233-240 ((https://doi.org/10.1590/S1982-56762011000400004)); Põlme S, et al. 2020. Fungal Diversity 105:1-16 ((https://doi.org/10.1007/s13225-020-00466-2)) |
| Otu60763 | 1 | k__Fungi(100);p__Basidiomycota(100);c__Tremellomycetes(100);o__Tremellales(100);f__Tremellales_family_Incertae_sedis(100);g__Cryptococcus(100);s__Cryptococcus_sp_JCM_11356(100); | Cryptococcus | 13 | Pathotroph-Saprotroph-Symbiotroph | Animal Pathogen-Endophyte-Epiphyte-\|Undefined Saprotroph\| | Dimorphic Yeast-Yeast | NULL | Possible | As foliar_endophyte (Põlme et al. 2020); Decay leaf, fruit, seed, soil, dung, animal material (Põlme et al. 2020) | Andrews JH, et al. 1983. Phytopathology 73:228-234 ((https://doi.org/10.1094/Phyto-73-228)); Kurtzman CP, et al. (eds.) 2011. The Yeasts, a Taxonomic Study. Fifth Edition. Vols 1-3. Elsevier, San Diego (ISBN: 9780444521491); Põlme S, et al. 2020. Fungal Diversity 105:1-16 ((https://doi.org/10.1007/s13225-020-00466-2)) |
| Otu60764 | 1 | k__Fungi(100);p__Ascomycota(100);c__Saccharomycetes(100);o__Saccharomycetales(100);f__Metschnikowiaceae(100);g__Metschnikowia(100);s__Metschnikowia_pulcherrima(100); | Metschnikowia | 13 | Saprotroph | \|Undefined Saprotroph\| | Yeast | NULL | Probable | As nectar-tap_saprotroph (Põlme et al. 2020); Decay sugar-rich substrates (Põlme et al. 2020) | Põlme S, et al. 2020. Fungal Diversity 105:1-16 ((https://doi.org/10.1007/s13225-020-00466-2)) |
| Otu60765 | 1 | k__Fungi(100);p__Ascomycota(100);c__Saccharomycetes(100);o__Saccharomycetales(100);f__Metschnikowiaceae(100);g__Metschnikowia(100);s__Metschnikowia_pulcherrima(100); | Metschnikowia | 13 | Saprotroph | \|Undefined Saprotroph\| | Yeast | NULL | Probable | As nectar-tap_saprotroph (Põlme et al. 2020); Decay sugar-rich substrates (Põlme et al. 2020) | Põlme S, et al. 2020. Fungal Diversity 105:1-16 ((https://doi.org/10.1007/s13225-020-00466-2)) |
| Otu60768 | 1 | k__Fungi(100);p__Ascomycota(100);c__Saccharomycetes(100);o__Saccharomycetales(100);f__Metschnikowiaceae(100);g__Metschnikowia(100);s__Metschnikowia_pulcherrima(100); | Metschnikowia | 13 | Saprotroph | \|Undefined Saprotroph\| | Yeast | NULL | Probable | As nectar-tap_saprotroph (Põlme et al. 2020); Decay sugar-rich substrates (Põlme et al. 2020) | Põlme S, et al. 2020. Fungal Diversity 105:1-16 ((https://doi.org/10.1007/s13225-020-00466-2)) |
| Otu60771 | 1 | k__Fungi(100);p__Ascomycota(100);c__Saccharomycetes(100);o__Saccharomycetales(100);f__Metschnikowiaceae(100);g__Metschnikowia(100);s__Metschnikowia_pulcherrima(100); | Metschnikowia | 13 | Saprotroph | \|Undefined Saprotroph\| | Yeast | NULL | Probable | As nectar-tap_saprotroph (Põlme et al. 2020); Decay sugar-rich substrates (Põlme et al. 2020) | Põlme S, et al. 2020. Fungal Diversity 105:1-16 ((https://doi.org/10.1007/s13225-020-00466-2)) |
| Otu60776 | 1 | k__Fungi(100);p__Ascomycota(100);c__Saccharomycetes(100);o__Saccharomycetales(100);f__Metschnikowiaceae(100);g__Metschnikowia(100);g__Metschnikowia_unclassified(100); | Metschnikowia | 13 | Saprotroph | \|Undefined Saprotroph\| | Yeast | NULL | Probable | As nectar-tap_saprotroph (Põlme et al. 2020); Decay sugar-rich substrates (Põlme et al. 2020) | Põlme S, et al. 2020. Fungal Diversity 105:1-16 ((https://doi.org/10.1007/s13225-020-00466-2)) |
| Otu60777 | 1 | k__Fungi(100);p__Ascomycota(100);c__Saccharomycetes(100);o__Saccharomycetales(100);f__Saccharomycetales_family_Incertae_sedis(100);g__Candida(100);g__Candida_unclassified(100); | Candida | 13 | Pathotroph-Saprotroph-Symbiotroph | Animal Pathogen-Endophyte-Endosymbiont-Epiphyte-\|Undefined Saprotroph\| | Dimorphic Yeast-Yeast | NULL | Probable | As nectar-tap_saprotroph (Põlme et al. 2020); Decay sugar-rich substrates (Põlme et al. 2020); This asexual genus is polyphyletic and contain species that function very differently depend on what they are. Caution should be taken when interpreting guilds for Candida species. | Manolakaki D, et al. 2010. Virulence. 1: 367-375 ((https://doi.org/10.4161/viru.1.5.12796)); Mekbib SB, et al. 2011. Tropical Plant Pathology 36:233-240 ((https://doi.org/10.1590/S1982-56762011000400004)); Põlme S, et al. 2020. Fungal Diversity 105:1-16 ((https://doi.org/10.1007/s13225-020-00466-2)) |
| Otu60780 | 1 | k__Fungi(100);p__Ascomycota(100);c__Saccharomycetes(100);o__Saccharomycetales(100);f__Metschnikowiaceae(100);g__Metschnikowia(100);s__Metschnikowia_pulcherrima(100); | Metschnikowia | 13 | Saprotroph | \|Undefined Saprotroph\| | Yeast | NULL | Probable | As nectar-tap_saprotroph (Põlme et al. 2020); Decay sugar-rich substrates (Põlme et al. 2020) | Põlme S, et al. 2020. Fungal Diversity 105:1-16 ((https://doi.org/10.1007/s13225-020-00466-2)) |
| Otu60781 | 1 | k__Fungi(100);p__Ascomycota(100);c__Saccharomycetes(100);o__Saccharomycetales(100);f__Saccharomycetales_family_Incertae_sedis(100);g__Candida(100);g__Candida_unclassified(100); | Candida | 13 | Pathotroph-Saprotroph-Symbiotroph | Animal Pathogen-Endophyte-Endosymbiont-Epiphyte-\|Undefined Saprotroph\| | Dimorphic Yeast-Yeast | NULL | Probable | As nectar-tap_saprotroph (Põlme et al. 2020); Decay sugar-rich substrates (Põlme et al. 2020); This asexual genus is polyphyletic and contain species that function very differently depend on what they are. Caution should be taken when interpreting guilds for Candida species. | Manolakaki D, et al. 2010. Virulence. 1: 367-375 ((https://doi.org/10.4161/viru.1.5.12796)); Mekbib SB, et al. 2011. Tropical Plant Pathology 36:233-240 ((https://doi.org/10.1590/S1982-56762011000400004)); Põlme S, et al. 2020. Fungal Diversity 105:1-16 ((https://doi.org/10.1007/s13225-020-00466-2)) |
| Otu60785 | 1 | k__Fungi(100);p__Ascomycota(100);c__Saccharomycetes(100);o__Saccharomycetales(100);f__Metschnikowiaceae(100);g__Metschnikowia(100);s__Metschnikowia_pulcherrima(100); | Metschnikowia | 13 | Saprotroph | \|Undefined Saprotroph\| | Yeast | NULL | Probable | As nectar-tap_saprotroph (Põlme et al. 2020); Decay sugar-rich substrates (Põlme et al. 2020) | Põlme S, et al. 2020. Fungal Diversity 105:1-16 ((https://doi.org/10.1007/s13225-020-00466-2)) |
| Otu60786 | 1 | k__Fungi(100);p__Ascomycota(100);c__Saccharomycetes(100);o__Saccharomycetales(100);f__Metschnikowiaceae(100);g__Metschnikowia(100);g__Metschnikowia_unclassified(100); | Metschnikowia | 13 | Saprotroph | \|Undefined Saprotroph\| | Yeast | NULL | Probable | As nectar-tap_saprotroph (Põlme et al. 2020); Decay sugar-rich substrates (Põlme et al. 2020) | Põlme S, et al. 2020. Fungal Diversity 105:1-16 ((https://doi.org/10.1007/s13225-020-00466-2)) |
| Otu60787 | 1 | k__Fungi(100);p__Ascomycota(100);c__Saccharomycetes(100);o__Saccharomycetales(100);f__Metschnikowiaceae(100);g__Metschnikowia(100);s__Metschnikowia_pulcherrima(100); | Metschnikowia | 13 | Saprotroph | \|Undefined Saprotroph\| | Yeast | NULL | Probable | As nectar-tap_saprotroph (Põlme et al. 2020); Decay sugar-rich substrates (Põlme et al. 2020) | Põlme S, et al. 2020. Fungal Diversity 105:1-16 ((https://doi.org/10.1007/s13225-020-00466-2)) |
| Otu60788 | 1 | k__Fungi(100);p__Ascomycota(100);c__Saccharomycetes(100);o__Saccharomycetales(100);f__Metschnikowiaceae(100);g__Metschnikowia(100);s__Metschnikowia_pulcherrima(100); | Metschnikowia | 13 | Saprotroph | \|Undefined Saprotroph\| | Yeast | NULL | Probable | As nectar-tap_saprotroph (Põlme et al. 2020); Decay sugar-rich substrates (Põlme et al. 2020) | Põlme S, et al. 2020. Fungal Diversity 105:1-16 ((https://doi.org/10.1007/s13225-020-00466-2)) |
| Otu60789 | 1 | k__Fungi(100);p__Ascomycota(100);c__Saccharomycetes(100);o__Saccharomycetales(100);f__Metschnikowiaceae(100);g__Metschnikowia(100);g__Metschnikowia_unclassified(100); | Metschnikowia | 13 | Saprotroph | \|Undefined Saprotroph\| | Yeast | NULL | Probable | As nectar-tap_saprotroph (Põlme et al. 2020); Decay sugar-rich substrates (Põlme et al. 2020) | Põlme S, et al. 2020. Fungal Diversity 105:1-16 ((https://doi.org/10.1007/s13225-020-00466-2)) |
| Otu60790 | 1 | k__Fungi(100);p__Ascomycota(100);c__Saccharomycetes(100);o__Saccharomycetales(100);f__Metschnikowiaceae(100);g__Metschnikowia(100);s__Metschnikowia_pulcherrima(100); | Metschnikowia | 13 | Saprotroph | \|Undefined Saprotroph\| | Yeast | NULL | Probable | As nectar-tap_saprotroph (Põlme et al. 2020); Decay sugar-rich substrates (Põlme et al. 2020) | Põlme S, et al. 2020. Fungal Diversity 105:1-16 ((https://doi.org/10.1007/s13225-020-00466-2)) |
| Otu60791 | 1 | k__Fungi(100);p__Ascomycota(100);c__Saccharomycetes(100);o__Saccharomycetales(100);f__Metschnikowiaceae(100);g__Metschnikowia(100);g__Metschnikowia_unclassified(100); | Metschnikowia | 13 | Saprotroph | \|Undefined Saprotroph\| | Yeast | NULL | Probable | As nectar-tap_saprotroph (Põlme et al. 2020); Decay sugar-rich substrates (Põlme et al. 2020) | Põlme S, et al. 2020. Fungal Diversity 105:1-16 ((https://doi.org/10.1007/s13225-020-00466-2)) |
| Otu60792 | 1 | k__Fungi(100);p__Ascomycota(100);c__Saccharomycetes(100);o__Saccharomycetales(100);f__Metschnikowiaceae(100);g__Metschnikowia(100);s__Metschnikowia_pulcherrima(100); | Metschnikowia | 13 | Saprotroph | \|Undefined Saprotroph\| | Yeast | NULL | Probable | As nectar-tap_saprotroph (Põlme et al. 2020); Decay sugar-rich substrates (Põlme et al. 2020) | Põlme S, et al. 2020. Fungal Diversity 105:1-16 ((https://doi.org/10.1007/s13225-020-00466-2)) |
| Otu60797 | 1 | k__Fungi(100);p__Ascomycota(100);c__Saccharomycetes(100);o__Saccharomycetales(100);f__Metschnikowiaceae(100);g__Metschnikowia(100);s__Metschnikowia_pulcherrima(100); | Metschnikowia | 13 | Saprotroph | \|Undefined Saprotroph\| | Yeast | NULL | Probable | As nectar-tap_saprotroph (Põlme et al. 2020); Decay sugar-rich substrates (Põlme et al. 2020) | Põlme S, et al. 2020. Fungal Diversity 105:1-16 ((https://doi.org/10.1007/s13225-020-00466-2)) |
| Otu60798 | 1 | k__Fungi(100);p__Ascomycota(100);c__Saccharomycetes(100);o__Saccharomycetales(100);f__Metschnikowiaceae(100);g__Metschnikowia(100);s__Metschnikowia_pulcherrima(100); | Metschnikowia | 13 | Saprotroph | \|Undefined Saprotroph\| | Yeast | NULL | Probable | As nectar-tap_saprotroph (Põlme et al. 2020); Decay sugar-rich substrates (Põlme et al. 2020) | Põlme S, et al. 2020. Fungal Diversity 105:1-16 ((https://doi.org/10.1007/s13225-020-00466-2)) |
| Otu60799 | 1 | k__Fungi(100);p__Ascomycota(100);c__Saccharomycetes(100);o__Saccharomycetales(100);f__Metschnikowiaceae(100);g__Metschnikowia(100);s__Metschnikowia_pulcherrima(100); | Metschnikowia | 13 | Saprotroph | \|Undefined Saprotroph\| | Yeast | NULL | Probable | As nectar-tap_saprotroph (Põlme et al. 2020); Decay sugar-rich substrates (Põlme et al. 2020) | Põlme S, et al. 2020. Fungal Diversity 105:1-16 ((https://doi.org/10.1007/s13225-020-00466-2)) |
| Otu60800 | 1 | k__Fungi(100);p__Ascomycota(100);c__Saccharomycetes(100);o__Saccharomycetales(100);f__Metschnikowiaceae(100);g__Metschnikowia(100);s__Metschnikowia_pulcherrima(100); | Metschnikowia | 13 | Saprotroph | \|Undefined Saprotroph\| | Yeast | NULL | Probable | As nectar-tap_saprotroph (Põlme et al. 2020); Decay sugar-rich substrates (Põlme et al. 2020) | Põlme S, et al. 2020. Fungal Diversity 105:1-16 ((https://doi.org/10.1007/s13225-020-00466-2)) |
| Otu60801 | 1 | k__Fungi(100);p__Ascomycota(100);c__Saccharomycetes(100);o__Saccharomycetales(100);f__Metschnikowiaceae(100);g__Metschnikowia(100);s__Metschnikowia_pulcherrima(100); | Metschnikowia | 13 | Saprotroph | \|Undefined Saprotroph\| | Yeast | NULL | Probable | As nectar-tap_saprotroph (Põlme et al. 2020); Decay sugar-rich substrates (Põlme et al. 2020) | Põlme S, et al. 2020. Fungal Diversity 105:1-16 ((https://doi.org/10.1007/s13225-020-00466-2)) |
| Otu60802 | 1 | k__Fungi(100);p__Ascomycota(100);c__Saccharomycetes(100);o__Saccharomycetales(100);f__Metschnikowiaceae(100);g__Metschnikowia(100);s__Metschnikowia_pulcherrima(100); | Metschnikowia | 13 | Saprotroph | \|Undefined Saprotroph\| | Yeast | NULL | Probable | As nectar-tap_saprotroph (Põlme et al. 2020); Decay sugar-rich substrates (Põlme et al. 2020) | Põlme S, et al. 2020. Fungal Diversity 105:1-16 ((https://doi.org/10.1007/s13225-020-00466-2)) |
| Otu60803 | 1 | k__Fungi(100);p__Ascomycota(100);c__Saccharomycetes(100);o__Saccharomycetales(100);f__Metschnikowiaceae(100);g__Metschnikowia(100);s__Metschnikowia_pulcherrima(100); | Metschnikowia | 13 | Saprotroph | \|Undefined Saprotroph\| | Yeast | NULL | Probable | As nectar-tap_saprotroph (Põlme et al. 2020); Decay sugar-rich substrates (Põlme et al. 2020) | Põlme S, et al. 2020. Fungal Diversity 105:1-16 ((https://doi.org/10.1007/s13225-020-00466-2)) |
| Otu60804 | 1 | k__Fungi(100);p__Ascomycota(100);c__Saccharomycetes(100);o__Saccharomycetales(100);f__Metschnikowiaceae(100);g__Metschnikowia(100);s__Metschnikowia_pulcherrima(100); | Metschnikowia | 13 | Saprotroph | \|Undefined Saprotroph\| | Yeast | NULL | Probable | As nectar-tap_saprotroph (Põlme et al. 2020); Decay sugar-rich substrates (Põlme et al. 2020) | Põlme S, et al. 2020. Fungal Diversity 105:1-16 ((https://doi.org/10.1007/s13225-020-00466-2)) |
| Otu60805 | 1 | k__Fungi(100);p__Ascomycota(100);c__Saccharomycetes(100);o__Saccharomycetales(100);f__Metschnikowiaceae(100);g__Metschnikowia(100);s__Metschnikowia_pulcherrima(100); | Metschnikowia | 13 | Saprotroph | \|Undefined Saprotroph\| | Yeast | NULL | Probable | As nectar-tap_saprotroph (Põlme et al. 2020); Decay sugar-rich substrates (Põlme et al. 2020) | Põlme S, et al. 2020. Fungal Diversity 105:1-16 ((https://doi.org/10.1007/s13225-020-00466-2)) |
| Otu60806 | 1 | k__Fungi(100);p__Ascomycota(100);c__Saccharomycetes(100);o__Saccharomycetales(100);f__Metschnikowiaceae(100);g__Metschnikowia(100);s__Metschnikowia_pulcherrima(100); | Metschnikowia | 13 | Saprotroph | \|Undefined Saprotroph\| | Yeast | NULL | Probable | As nectar-tap_saprotroph (Põlme et al. 2020); Decay sugar-rich substrates (Põlme et al. 2020) | Põlme S, et al. 2020. Fungal Diversity 105:1-16 ((https://doi.org/10.1007/s13225-020-00466-2)) |
| Otu60807 | 1 | k__Fungi(100);p__Ascomycota(100);c__Saccharomycetes(100);o__Saccharomycetales(100);f__Metschnikowiaceae(100);g__Metschnikowia(100);s__Metschnikowia_pulcherrima(100); | Metschnikowia | 13 | Saprotroph | \|Undefined Saprotroph\| | Yeast | NULL | Probable | As nectar-tap_saprotroph (Põlme et al. 2020); Decay sugar-rich substrates (Põlme et al. 2020) | Põlme S, et al. 2020. Fungal Diversity 105:1-16 ((https://doi.org/10.1007/s13225-020-00466-2)) |
| Otu60808 | 1 | k__Fungi(100);p__Ascomycota(100);c__Saccharomycetes(100);o__Saccharomycetales(100);f__Metschnikowiaceae(100);g__Metschnikowia(100);s__Metschnikowia_pulcherrima(100); | Metschnikowia | 13 | Saprotroph | \|Undefined Saprotroph\| | Yeast | NULL | Probable | As nectar-tap_saprotroph (Põlme et al. 2020); Decay sugar-rich substrates (Põlme et al. 2020) | Põlme S, et al. 2020. Fungal Diversity 105:1-16 ((https://doi.org/10.1007/s13225-020-00466-2)) |
| Otu60809 | 1 | k__Fungi(100);p__Ascomycota(100);c__Saccharomycetes(100);o__Saccharomycetales(100);f__Metschnikowiaceae(100);g__Metschnikowia(100);s__Metschnikowia_pulcherrima(100); | Metschnikowia | 13 | Saprotroph | \|Undefined Saprotroph\| | Yeast | NULL | Probable | As nectar-tap_saprotroph (Põlme et al. 2020); Decay sugar-rich substrates (Põlme et al. 2020) | Põlme S, et al. 2020. Fungal Diversity 105:1-16 ((https://doi.org/10.1007/s13225-020-00466-2)) |
| Otu60810 | 1 | k__Fungi(100);p__Ascomycota(100);c__Saccharomycetes(100);o__Saccharomycetales(100);f__Metschnikowiaceae(100);g__Metschnikowia(100);s__Metschnikowia_pulcherrima(100); | Metschnikowia | 13 | Saprotroph | \|Undefined Saprotroph\| | Yeast | NULL | Probable | As nectar-tap_saprotroph (Põlme et al. 2020); Decay sugar-rich substrates (Põlme et al. 2020) | Põlme S, et al. 2020. Fungal Diversity 105:1-16 ((https://doi.org/10.1007/s13225-020-00466-2)) |
| Otu60811 | 1 | k__Fungi(100);p__Ascomycota(100);c__Saccharomycetes(100);o__Saccharomycetales(100);f__Metschnikowiaceae(100);g__Metschnikowia(100);s__Metschnikowia_pulcherrima(100); | Metschnikowia | 13 | Saprotroph | \|Undefined Saprotroph\| | Yeast | NULL | Probable | As nectar-tap_saprotroph (Põlme et al. 2020); Decay sugar-rich substrates (Põlme et al. 2020) | Põlme S, et al. 2020. Fungal Diversity 105:1-16 ((https://doi.org/10.1007/s13225-020-00466-2)) |
| Otu60812 | 1 | k__Fungi(100);p__Ascomycota(100);c__Saccharomycetes(100);o__Saccharomycetales(100);f__Metschnikowiaceae(100);g__Metschnikowia(100);s__Metschnikowia_pulcherrima(100); | Metschnikowia | 13 | Saprotroph | \|Undefined Saprotroph\| | Yeast | NULL | Probable | As nectar-tap_saprotroph (Põlme et al. 2020); Decay sugar-rich substrates (Põlme et al. 2020) | Põlme S, et al. 2020. Fungal Diversity 105:1-16 ((https://doi.org/10.1007/s13225-020-00466-2)) |
| Otu60813 | 1 | k__Fungi(100);p__Ascomycota(100);c__Saccharomycetes(100);o__Saccharomycetales(100);f__Metschnikowiaceae(100);g__Metschnikowia(100);s__Metschnikowia_pulcherrima(100); | Metschnikowia | 13 | Saprotroph | \|Undefined Saprotroph\| | Yeast | NULL | Probable | As nectar-tap_saprotroph (Põlme et al. 2020); Decay sugar-rich substrates (Põlme et al. 2020) | Põlme S, et al. 2020. Fungal Diversity 105:1-16 ((https://doi.org/10.1007/s13225-020-00466-2)) |
| Otu60814 | 1 | k__Fungi(100);p__Ascomycota(100);c__Saccharomycetes(100);o__Saccharomycetales(100);f__Metschnikowiaceae(100);g__Metschnikowia(100);g__Metschnikowia_unclassified(100); | Metschnikowia | 13 | Saprotroph | \|Undefined Saprotroph\| | Yeast | NULL | Probable | As nectar-tap_saprotroph (Põlme et al. 2020); Decay sugar-rich substrates (Põlme et al. 2020) | Põlme S, et al. 2020. Fungal Diversity 105:1-16 ((https://doi.org/10.1007/s13225-020-00466-2)) |
| Otu60815 | 1 | k__Fungi(100);p__Ascomycota(100);c__Saccharomycetes(100);o__Saccharomycetales(100);f__Metschnikowiaceae(100);g__Metschnikowia(100);s__Metschnikowia_pulcherrima(100); | Metschnikowia | 13 | Saprotroph | \|Undefined Saprotroph\| | Yeast | NULL | Probable | As nectar-tap_saprotroph (Põlme et al. 2020); Decay sugar-rich substrates (Põlme et al. 2020) | Põlme S, et al. 2020. Fungal Diversity 105:1-16 ((https://doi.org/10.1007/s13225-020-00466-2)) |
| Otu60816 | 1 | k__Fungi(100);p__Ascomycota(100);c__Saccharomycetes(100);o__Saccharomycetales(100);f__Metschnikowiaceae(100);g__Metschnikowia(100);g__Metschnikowia_unclassified(100); | Metschnikowia | 13 | Saprotroph | \|Undefined Saprotroph\| | Yeast | NULL | Probable | As nectar-tap_saprotroph (Põlme et al. 2020); Decay sugar-rich substrates (Põlme et al. 2020) | Põlme S, et al. 2020. Fungal Diversity 105:1-16 ((https://doi.org/10.1007/s13225-020-00466-2)) |
| Otu60817 | 1 | k__Fungi(100);p__Ascomycota(100);c__Saccharomycetes(100);o__Saccharomycetales(100);f__Metschnikowiaceae(100);g__Metschnikowia(100);g__Metschnikowia_unclassified(100); | Metschnikowia | 13 | Saprotroph | \|Undefined Saprotroph\| | Yeast | NULL | Probable | As nectar-tap_saprotroph (Põlme et al. 2020); Decay sugar-rich substrates (Põlme et al. 2020) | Põlme S, et al. 2020. Fungal Diversity 105:1-16 ((https://doi.org/10.1007/s13225-020-00466-2)) |
| Otu60818 | 1 | k__Fungi(100);p__Ascomycota(100);c__Saccharomycetes(100);o__Saccharomycetales(100);f__Metschnikowiaceae(100);g__Metschnikowia(100);g__Metschnikowia_unclassified(100); | Metschnikowia | 13 | Saprotroph | \|Undefined Saprotroph\| | Yeast | NULL | Probable | As nectar-tap_saprotroph (Põlme et al. 2020); Decay sugar-rich substrates (Põlme et al. 2020) | Põlme S, et al. 2020. Fungal Diversity 105:1-16 ((https://doi.org/10.1007/s13225-020-00466-2)) |
| Otu60819 | 1 | k__Fungi(100);p__Ascomycota(100);c__Saccharomycetes(100);o__Saccharomycetales(100);f__Metschnikowiaceae(100);g__Metschnikowia(100);s__Metschnikowia_pulcherrima(100); | Metschnikowia | 13 | Saprotroph | \|Undefined Saprotroph\| | Yeast | NULL | Probable | As nectar-tap_saprotroph (Põlme et al. 2020); Decay sugar-rich substrates (Põlme et al. 2020) | Põlme S, et al. 2020. Fungal Diversity 105:1-16 ((https://doi.org/10.1007/s13225-020-00466-2)) |
| Otu60820 | 1 | k__Fungi(100);p__Ascomycota(100);c__Saccharomycetes(100);o__Saccharomycetales(100);f__Metschnikowiaceae(100);g__Metschnikowia(100);s__Metschnikowia_pulcherrima(100); | Metschnikowia | 13 | Saprotroph | \|Undefined Saprotroph\| | Yeast | NULL | Probable | As nectar-tap_saprotroph (Põlme et al. 2020); Decay sugar-rich substrates (Põlme et al. 2020) | Põlme S, et al. 2020. Fungal Diversity 105:1-16 ((https://doi.org/10.1007/s13225-020-00466-2)) |
| Otu60821 | 1 | k__Fungi(100);p__Ascomycota(100);c__Saccharomycetes(100);o__Saccharomycetales(100);f__Metschnikowiaceae(100);g__Metschnikowia(100);g__Metschnikowia_unclassified(100); | Metschnikowia | 13 | Saprotroph | \|Undefined Saprotroph\| | Yeast | NULL | Probable | As nectar-tap_saprotroph (Põlme et al. 2020); Decay sugar-rich substrates (Põlme et al. 2020) | Põlme S, et al. 2020. Fungal Diversity 105:1-16 ((https://doi.org/10.1007/s13225-020-00466-2)) |
| Otu60822 | 1 | k__Fungi(100);p__Ascomycota(100);c__Saccharomycetes(100);o__Saccharomycetales(100);f__Metschnikowiaceae(100);g__Metschnikowia(100);s__Metschnikowia_pulcherrima(100); | Metschnikowia | 13 | Saprotroph | \|Undefined Saprotroph\| | Yeast | NULL | Probable | As nectar-tap_saprotroph (Põlme et al. 2020); Decay sugar-rich substrates (Põlme et al. 2020) | Põlme S, et al. 2020. Fungal Diversity 105:1-16 ((https://doi.org/10.1007/s13225-020-00466-2)) |
| Otu60823 | 1 | k__Fungi(100);p__Ascomycota(100);c__Saccharomycetes(100);o__Saccharomycetales(100);f__Metschnikowiaceae(100);g__Metschnikowia(100);s__Metschnikowia_pulcherrima(100); | Metschnikowia | 13 | Saprotroph | \|Undefined Saprotroph\| | Yeast | NULL | Probable | As nectar-tap_saprotroph (Põlme et al. 2020); Decay sugar-rich substrates (Põlme et al. 2020) | Põlme S, et al. 2020. Fungal Diversity 105:1-16 ((https://doi.org/10.1007/s13225-020-00466-2)) |
| Otu60824 | 1 | k__Fungi(100);p__Ascomycota(100);c__Saccharomycetes(100);o__Saccharomycetales(100);f__Metschnikowiaceae(100);g__Metschnikowia(100);s__Metschnikowia_pulcherrima(100); | Metschnikowia | 13 | Saprotroph | \|Undefined Saprotroph\| | Yeast | NULL | Probable | As nectar-tap_saprotroph (Põlme et al. 2020); Decay sugar-rich substrates (Põlme et al. 2020) | Põlme S, et al. 2020. Fungal Diversity 105:1-16 ((https://doi.org/10.1007/s13225-020-00466-2)) |
| Otu60825 | 1 | k__Fungi(100);p__Ascomycota(100);c__Saccharomycetes(100);o__Saccharomycetales(100);f__Metschnikowiaceae(100);g__Metschnikowia(100);g__Metschnikowia_unclassified(100); | Metschnikowia | 13 | Saprotroph | \|Undefined Saprotroph\| | Yeast | NULL | Probable | As nectar-tap_saprotroph (Põlme et al. 2020); Decay sugar-rich substrates (Põlme et al. 2020) | Põlme S, et al. 2020. Fungal Diversity 105:1-16 ((https://doi.org/10.1007/s13225-020-00466-2)) |
| Otu60826 | 1 | k__Fungi(100);p__Ascomycota(100);c__Saccharomycetes(100);o__Saccharomycetales(100);f__Metschnikowiaceae(100);g__Metschnikowia(100);s__Metschnikowia_pulcherrima(100); | Metschnikowia | 13 | Saprotroph | \|Undefined Saprotroph\| | Yeast | NULL | Probable | As nectar-tap_saprotroph (Põlme et al. 2020); Decay sugar-rich substrates (Põlme et al. 2020) | Põlme S, et al. 2020. Fungal Diversity 105:1-16 ((https://doi.org/10.1007/s13225-020-00466-2)) |
| Otu60827 | 1 | k__Fungi(100);p__Ascomycota(100);c__Saccharomycetes(100);o__Saccharomycetales(100);f__Metschnikowiaceae(100);g__Metschnikowia(100);s__Metschnikowia_pulcherrima(100); | Metschnikowia | 13 | Saprotroph | \|Undefined Saprotroph\| | Yeast | NULL | Probable | As nectar-tap_saprotroph (Põlme et al. 2020); Decay sugar-rich substrates (Põlme et al. 2020) | Põlme S, et al. 2020. Fungal Diversity 105:1-16 ((https://doi.org/10.1007/s13225-020-00466-2)) |
| Otu60828 | 1 | k__Fungi(100);p__Ascomycota(100);c__Saccharomycetes(100);o__Saccharomycetales(100);f__Metschnikowiaceae(100);g__Metschnikowia(100);g__Metschnikowia_unclassified(100); | Metschnikowia | 13 | Saprotroph | \|Undefined Saprotroph\| | Yeast | NULL | Probable | As nectar-tap_saprotroph (Põlme et al. 2020); Decay sugar-rich substrates (Põlme et al. 2020) | Põlme S, et al. 2020. Fungal Diversity 105:1-16 ((https://doi.org/10.1007/s13225-020-00466-2)) |
| Otu60829 | 1 | k__Fungi(100);p__Ascomycota(100);c__Saccharomycetes(100);o__Saccharomycetales(100);f__Metschnikowiaceae(100);g__Metschnikowia(100);g__Metschnikowia_unclassified(100); | Metschnikowia | 13 | Saprotroph | \|Undefined Saprotroph\| | Yeast | NULL | Probable | As nectar-tap_saprotroph (Põlme et al. 2020); Decay sugar-rich substrates (Põlme et al. 2020) | Põlme S, et al. 2020. Fungal Diversity 105:1-16 ((https://doi.org/10.1007/s13225-020-00466-2)) |
| Otu60830 | 1 | k__Fungi(100);p__Ascomycota(100);c__Saccharomycetes(100);o__Saccharomycetales(100);f__Metschnikowiaceae(100);g__Metschnikowia(100);s__Metschnikowia_pulcherrima(100); | Metschnikowia | 13 | Saprotroph | \|Undefined Saprotroph\| | Yeast | NULL | Probable | As nectar-tap_saprotroph (Põlme et al. 2020); Decay sugar-rich substrates (Põlme et al. 2020) | Põlme S, et al. 2020. Fungal Diversity 105:1-16 ((https://doi.org/10.1007/s13225-020-00466-2)) |
| Otu60831 | 1 | k__Fungi(100);p__Ascomycota(100);c__Saccharomycetes(100);o__Saccharomycetales(100);f__Metschnikowiaceae(100);g__Metschnikowia(100);s__Metschnikowia_pulcherrima(100); | Metschnikowia | 13 | Saprotroph | \|Undefined Saprotroph\| | Yeast | NULL | Probable | As nectar-tap_saprotroph (Põlme et al. 2020); Decay sugar-rich substrates (Põlme et al. 2020) | Põlme S, et al. 2020. Fungal Diversity 105:1-16 ((https://doi.org/10.1007/s13225-020-00466-2)) |
| Otu60832 | 1 | k__Fungi(100);p__Ascomycota(100);c__Saccharomycetes(100);o__Saccharomycetales(100);f__Metschnikowiaceae(100);g__Metschnikowia(100);s__Metschnikowia_pulcherrima(100); | Metschnikowia | 13 | Saprotroph | \|Undefined Saprotroph\| | Yeast | NULL | Probable | As nectar-tap_saprotroph (Põlme et al. 2020); Decay sugar-rich substrates (Põlme et al. 2020) | Põlme S, et al. 2020. Fungal Diversity 105:1-16 ((https://doi.org/10.1007/s13225-020-00466-2)) |
| Otu60833 | 1 | k__Fungi(100);p__Ascomycota(100);c__Saccharomycetes(100);o__Saccharomycetales(100);f__Metschnikowiaceae(100);g__Metschnikowia(100);s__Metschnikowia_pulcherrima(100); | Metschnikowia | 13 | Saprotroph | \|Undefined Saprotroph\| | Yeast | NULL | Probable | As nectar-tap_saprotroph (Põlme et al. 2020); Decay sugar-rich substrates (Põlme et al. 2020) | Põlme S, et al. 2020. Fungal Diversity 105:1-16 ((https://doi.org/10.1007/s13225-020-00466-2)) |
| Otu60834 | 1 | k__Fungi(100);p__Ascomycota(100);c__Saccharomycetes(100);o__Saccharomycetales(100);f__Metschnikowiaceae(100);g__Metschnikowia(100);s__Metschnikowia_pulcherrima(100); | Metschnikowia | 13 | Saprotroph | \|Undefined Saprotroph\| | Yeast | NULL | Probable | As nectar-tap_saprotroph (Põlme et al. 2020); Decay sugar-rich substrates (Põlme et al. 2020) | Põlme S, et al. 2020. Fungal Diversity 105:1-16 ((https://doi.org/10.1007/s13225-020-00466-2)) |
| Otu60835 | 1 | k__Fungi(100);p__Ascomycota(100);c__Saccharomycetes(100);o__Saccharomycetales(100);f__Metschnikowiaceae(100);g__Metschnikowia(100);s__Metschnikowia_pulcherrima(100); | Metschnikowia | 13 | Saprotroph | \|Undefined Saprotroph\| | Yeast | NULL | Probable | As nectar-tap_saprotroph (Põlme et al. 2020); Decay sugar-rich substrates (Põlme et al. 2020) | Põlme S, et al. 2020. Fungal Diversity 105:1-16 ((https://doi.org/10.1007/s13225-020-00466-2)) |
| Otu60836 | 1 | k__Fungi(100);p__Ascomycota(100);c__Saccharomycetes(100);o__Saccharomycetales(100);f__Metschnikowiaceae(100);g__Metschnikowia(100);g__Metschnikowia_unclassified(100); | Metschnikowia | 13 | Saprotroph | \|Undefined Saprotroph\| | Yeast | NULL | Probable | As nectar-tap_saprotroph (Põlme et al. 2020); Decay sugar-rich substrates (Põlme et al. 2020) | Põlme S, et al. 2020. Fungal Diversity 105:1-16 ((https://doi.org/10.1007/s13225-020-00466-2)) |
| Otu60837 | 1 | k__Fungi(100);p__Ascomycota(100);c__Saccharomycetes(100);o__Saccharomycetales(100);f__Metschnikowiaceae(100);g__Metschnikowia(100);s__Metschnikowia_pulcherrima(100); | Metschnikowia | 13 | Saprotroph | \|Undefined Saprotroph\| | Yeast | NULL | Probable | As nectar-tap_saprotroph (Põlme et al. 2020); Decay sugar-rich substrates (Põlme et al. 2020) | Põlme S, et al. 2020. Fungal Diversity 105:1-16 ((https://doi.org/10.1007/s13225-020-00466-2)) |
| Otu60838 | 1 | k__Fungi(100);p__Ascomycota(100);c__Saccharomycetes(100);o__Saccharomycetales(100);f__Metschnikowiaceae(100);g__Metschnikowia(100);s__Metschnikowia_pulcherrima(100); | Metschnikowia | 13 | Saprotroph | \|Undefined Saprotroph\| | Yeast | NULL | Probable | As nectar-tap_saprotroph (Põlme et al. 2020); Decay sugar-rich substrates (Põlme et al. 2020) | Põlme S, et al. 2020. Fungal Diversity 105:1-16 ((https://doi.org/10.1007/s13225-020-00466-2)) |
| Otu60839 | 1 | k__Fungi(100);p__Basidiomycota(100);c__Microbotryomycetes(100);o__Sporidiobolales(100);f__Sporidiobolales_family_Incertae_sedis(100);g__Sporobolomyces(100);s__Sporobolomyces_lactosus(100); | Sporobolomyces | 13 | Pathotroph-Saprotroph | \|Fungal Parasite\|-Plant Saprotroph | Yeast | NULL | Probable | As litter_saprotroph (Põlme et al. 2020); As mycoparasite (Põlme et al. 2020); Decay fungal material (Põlme et al. 2020) | Tedersoo L, et al. 2014. Science 346:e1256688 ((https://doi.org/10.1126/science.1256688)); Punzhong W, et al. 2016. Molecular Ecology 25:4059-4074 ((https://doi.org/10.1111/mec.13735)); Põlme S, et al. 2020. Fungal Diversity 105:1-16 ((https://doi.org/10.1007/s13225-020-00466-2)) |
| Otu60840 | 1 | k__Fungi(100);p__Ascomycota(100);c__Saccharomycetes(100);o__Saccharomycetales(100);f__Metschnikowiaceae(100);g__Metschnikowia(100);s__Metschnikowia_pulcherrima(100); | Metschnikowia | 13 | Saprotroph | \|Undefined Saprotroph\| | Yeast | NULL | Probable | As nectar-tap_saprotroph (Põlme et al. 2020); Decay sugar-rich substrates (Põlme et al. 2020) | Põlme S, et al. 2020. Fungal Diversity 105:1-16 ((https://doi.org/10.1007/s13225-020-00466-2)) |
| Otu60841 | 1 | k__Fungi(100);p__Ascomycota(100);c__Saccharomycetes(100);o__Saccharomycetales(100);f__Metschnikowiaceae(100);g__Metschnikowia(100);g__Metschnikowia_unclassified(100); | Metschnikowia | 13 | Saprotroph | \|Undefined Saprotroph\| | Yeast | NULL | Probable | As nectar-tap_saprotroph (Põlme et al. 2020); Decay sugar-rich substrates (Põlme et al. 2020) | Põlme S, et al. 2020. Fungal Diversity 105:1-16 ((https://doi.org/10.1007/s13225-020-00466-2)) |
| Otu60842 | 1 | k__Fungi(100);p__Ascomycota(100);c__Saccharomycetes(100);o__Saccharomycetales(100);f__Metschnikowiaceae(100);g__Metschnikowia(100);s__Metschnikowia_pulcherrima(100); | Metschnikowia | 13 | Saprotroph | \|Undefined Saprotroph\| | Yeast | NULL | Probable | As nectar-tap_saprotroph (Põlme et al. 2020); Decay sugar-rich substrates (Põlme et al. 2020) | Põlme S, et al. 2020. Fungal Diversity 105:1-16 ((https://doi.org/10.1007/s13225-020-00466-2)) |
| Otu60843 | 1 | k__Fungi(100);p__Ascomycota(100);c__Saccharomycetes(100);o__Saccharomycetales(100);f__Metschnikowiaceae(100);g__Metschnikowia(100);s__Metschnikowia_pulcherrima(100); | Metschnikowia | 13 | Saprotroph | \|Undefined Saprotroph\| | Yeast | NULL | Probable | As nectar-tap_saprotroph (Põlme et al. 2020); Decay sugar-rich substrates (Põlme et al. 2020) | Põlme S, et al. 2020. Fungal Diversity 105:1-16 ((https://doi.org/10.1007/s13225-020-00466-2)) |
| Otu60844 | 1 | k__Fungi(100);p__Ascomycota(100);c__Saccharomycetes(100);o__Saccharomycetales(100);f__Metschnikowiaceae(100);g__Metschnikowia(100);g__Metschnikowia_unclassified(100); | Metschnikowia | 13 | Saprotroph | \|Undefined Saprotroph\| | Yeast | NULL | Probable | As nectar-tap_saprotroph (Põlme et al. 2020); Decay sugar-rich substrates (Põlme et al. 2020) | Põlme S, et al. 2020. Fungal Diversity 105:1-16 ((https://doi.org/10.1007/s13225-020-00466-2)) |
| Otu60845 | 1 | k__Fungi(100);p__Ascomycota(100);c__Saccharomycetes(100);o__Saccharomycetales(100);f__Metschnikowiaceae(100);g__Metschnikowia(100);s__Metschnikowia_pulcherrima(100); | Metschnikowia | 13 | Saprotroph | \|Undefined Saprotroph\| | Yeast | NULL | Probable | As nectar-tap_saprotroph (Põlme et al. 2020); Decay sugar-rich substrates (Põlme et al. 2020) | Põlme S, et al. 2020. Fungal Diversity 105:1-16 ((https://doi.org/10.1007/s13225-020-00466-2)) |
| Otu60846 | 1 | k__Fungi(100);p__Ascomycota(100);c__Saccharomycetes(100);o__Saccharomycetales(100);f__Metschnikowiaceae(100);g__Metschnikowia(100);g__Metschnikowia_unclassified(100); | Metschnikowia | 13 | Saprotroph | \|Undefined Saprotroph\| | Yeast | NULL | Probable | As nectar-tap_saprotroph (Põlme et al. 2020); Decay sugar-rich substrates (Põlme et al. 2020) | Põlme S, et al. 2020. Fungal Diversity 105:1-16 ((https://doi.org/10.1007/s13225-020-00466-2)) |
| Otu60847 | 1 | k__Fungi(100);p__Ascomycota(100);c__Saccharomycetes(100);o__Saccharomycetales(100);f__Metschnikowiaceae(100);g__Metschnikowia(100);s__Metschnikowia_pulcherrima(100); | Metschnikowia | 13 | Saprotroph | \|Undefined Saprotroph\| | Yeast | NULL | Probable | As nectar-tap_saprotroph (Põlme et al. 2020); Decay sugar-rich substrates (Põlme et al. 2020) | Põlme S, et al. 2020. Fungal Diversity 105:1-16 ((https://doi.org/10.1007/s13225-020-00466-2)) |
| Otu60850 | 1 | k__Fungi(100);p__Ascomycota(100);c__Saccharomycetes(100);o__Saccharomycetales(100);f__Metschnikowiaceae(100);g__Metschnikowia(100);s__Metschnikowia_pulcherrima(100); | Metschnikowia | 13 | Saprotroph | \|Undefined Saprotroph\| | Yeast | NULL | Probable | As nectar-tap_saprotroph (Põlme et al. 2020); Decay sugar-rich substrates (Põlme et al. 2020) | Põlme S, et al. 2020. Fungal Diversity 105:1-16 ((https://doi.org/10.1007/s13225-020-00466-2)) |
| Otu60851 | 1 | k__Fungi(100);p__Ascomycota(100);c__Saccharomycetes(100);o__Saccharomycetales(100);f__Metschnikowiaceae(100);g__Metschnikowia(100);s__Metschnikowia_pulcherrima(100); | Metschnikowia | 13 | Saprotroph | \|Undefined Saprotroph\| | Yeast | NULL | Probable | As nectar-tap_saprotroph (Põlme et al. 2020); Decay sugar-rich substrates (Põlme et al. 2020) | Põlme S, et al. 2020. Fungal Diversity 105:1-16 ((https://doi.org/10.1007/s13225-020-00466-2)) |
| Otu60852 | 1 | k__Fungi(100);p__Ascomycota(100);c__Saccharomycetes(100);o__Saccharomycetales(100);f__Metschnikowiaceae(100);g__Metschnikowia(100);s__Metschnikowia_pulcherrima(100); | Metschnikowia | 13 | Saprotroph | \|Undefined Saprotroph\| | Yeast | NULL | Probable | As nectar-tap_saprotroph (Põlme et al. 2020); Decay sugar-rich substrates (Põlme et al. 2020) | Põlme S, et al. 2020. Fungal Diversity 105:1-16 ((https://doi.org/10.1007/s13225-020-00466-2)) |
| Otu60853 | 1 | k__Fungi(100);p__Ascomycota(100);c__Saccharomycetes(100);o__Saccharomycetales(100);f__Metschnikowiaceae(100);g__Metschnikowia(100);g__Metschnikowia_unclassified(100); | Metschnikowia | 13 | Saprotroph | \|Undefined Saprotroph\| | Yeast | NULL | Probable | As nectar-tap_saprotroph (Põlme et al. 2020); Decay sugar-rich substrates (Põlme et al. 2020) | Põlme S, et al. 2020. Fungal Diversity 105:1-16 ((https://doi.org/10.1007/s13225-020-00466-2)) |
| Otu60854 | 1 | k__Fungi(100);p__Ascomycota(100);c__Saccharomycetes(100);o__Saccharomycetales(100);f__Metschnikowiaceae(100);g__Metschnikowia(100);s__Metschnikowia_pulcherrima(100); | Metschnikowia | 13 | Saprotroph | \|Undefined Saprotroph\| | Yeast | NULL | Probable | As nectar-tap_saprotroph (Põlme et al. 2020); Decay sugar-rich substrates (Põlme et al. 2020) | Põlme S, et al. 2020. Fungal Diversity 105:1-16 ((https://doi.org/10.1007/s13225-020-00466-2)) |
| Otu60855 | 1 | k__Fungi(100);p__Ascomycota(100);c__Saccharomycetes(100);o__Saccharomycetales(100);f__Metschnikowiaceae(100);g__Metschnikowia(100);s__Metschnikowia_pulcherrima(100); | Metschnikowia | 13 | Saprotroph | \|Undefined Saprotroph\| | Yeast | NULL | Probable | As nectar-tap_saprotroph (Põlme et al. 2020); Decay sugar-rich substrates (Põlme et al. 2020) | Põlme S, et al. 2020. Fungal Diversity 105:1-16 ((https://doi.org/10.1007/s13225-020-00466-2)) |
| Otu60856 | 1 | k__Fungi(100);p__Ascomycota(100);c__Saccharomycetes(100);o__Saccharomycetales(100);f__Metschnikowiaceae(100);g__Metschnikowia(100);g__Metschnikowia_unclassified(100); | Metschnikowia | 13 | Saprotroph | \|Undefined Saprotroph\| | Yeast | NULL | Probable | As nectar-tap_saprotroph (Põlme et al. 2020); Decay sugar-rich substrates (Põlme et al. 2020) | Põlme S, et al. 2020. Fungal Diversity 105:1-16 ((https://doi.org/10.1007/s13225-020-00466-2)) |
| Otu60857 | 1 | k__Fungi(100);p__Ascomycota(100);c__Saccharomycetes(100);o__Saccharomycetales(100);f__Metschnikowiaceae(100);g__Metschnikowia(100);s__Metschnikowia_pulcherrima(100); | Metschnikowia | 13 | Saprotroph | \|Undefined Saprotroph\| | Yeast | NULL | Probable | As nectar-tap_saprotroph (Põlme et al. 2020); Decay sugar-rich substrates (Põlme et al. 2020) | Põlme S, et al. 2020. Fungal Diversity 105:1-16 ((https://doi.org/10.1007/s13225-020-00466-2)) |
| Otu60861 | 1 | k__Fungi(100);p__Ascomycota(100);c__Saccharomycetes(100);o__Saccharomycetales(100);f__Metschnikowiaceae(100);g__Metschnikowia(100);s__Metschnikowia_pulcherrima(100); | Metschnikowia | 13 | Saprotroph | \|Undefined Saprotroph\| | Yeast | NULL | Probable | As nectar-tap_saprotroph (Põlme et al. 2020); Decay sugar-rich substrates (Põlme et al. 2020) | Põlme S, et al. 2020. Fungal Diversity 105:1-16 ((https://doi.org/10.1007/s13225-020-00466-2)) |
| Otu60862 | 1 | k__Fungi(100);p__Ascomycota(100);c__Saccharomycetes(100);o__Saccharomycetales(100);f__Metschnikowiaceae(100);g__Metschnikowia(100);s__Metschnikowia_pulcherrima(100); | Metschnikowia | 13 | Saprotroph | \|Undefined Saprotroph\| | Yeast | NULL | Probable | As nectar-tap_saprotroph (Põlme et al. 2020); Decay sugar-rich substrates (Põlme et al. 2020) | Põlme S, et al. 2020. Fungal Diversity 105:1-16 ((https://doi.org/10.1007/s13225-020-00466-2)) |
| Otu60863 | 1 | k__Fungi(100);p__Ascomycota(100);c__Saccharomycetes(100);o__Saccharomycetales(100);f__Metschnikowiaceae(100);g__Metschnikowia(100);s__Metschnikowia_pulcherrima(100); | Metschnikowia | 13 | Saprotroph | \|Undefined Saprotroph\| | Yeast | NULL | Probable | As nectar-tap_saprotroph (Põlme et al. 2020); Decay sugar-rich substrates (Põlme et al. 2020) | Põlme S, et al. 2020. Fungal Diversity 105:1-16 ((https://doi.org/10.1007/s13225-020-00466-2)) |

| | | | | | | | | | | | |
|---|---|---|---|---|---|---|---|---|---|---|---|
| Otu60864 | 1 | k__Fungi(100);p__Ascomycota(100);c__Saccharomycetes(100);o__Saccharomycetales(100);f__Metschnikowiaceae(100);g__Metschnikowia(100);s__Metschnikowia_pulcherrima(100); | Metschnikowia | 13 | Saprotroph | \|Undefined Saprotroph\| | Yeast | NULL | Probable | As nectar-tap_saprotroph (Põlme et al. 2020); Decay sugar-rich substrates (Põlme et al. 2020) | Põlme S, et al. 2020. Fungal Diversity 105:1-16 ((https://doi.org/10.1007/s13225-020-00466-2)) |
| Otu60865 | 1 | k__Fungi(100);p__Ascomycota(100);c__Saccharomycetes(100);o__Saccharomycetales(100);f__Metschnikowiaceae(100);g__Metschnikowia(100);s__Metschnikowia_pulcherrima(100); | Metschnikowia | 13 | Saprotroph | \|Undefined Saprotroph\| | Yeast | NULL | Probable | As nectar-tap_saprotroph (Põlme et al. 2020); Decay sugar-rich substrates (Põlme et al. 2020) | Põlme S, et al. 2020. Fungal Diversity 105:1-16 ((https://doi.org/10.1007/s13225-020-00466-2)) |
| Otu60866 | 1 | k__Fungi(100);p__Ascomycota(100);c__Saccharomycetes(100);o__Saccharomycetales(100);f__Metschnikowiaceae(100);g__Metschnikowia(100);s__Metschnikowia_pulcherrima(100); | Metschnikowia | 13 | Saprotroph | \|Undefined Saprotroph\| | Yeast | NULL | Probable | As nectar-tap_saprotroph (Põlme et al. 2020); Decay sugar-rich substrates (Põlme et al. 2020) | Põlme S, et al. 2020. Fungal Diversity 105:1-16 ((https://doi.org/10.1007/s13225-020-00466-2)) |
| Otu60867 | 1 | k__Fungi(100);p__Ascomycota(100);c__Saccharomycetes(100);o__Saccharomycetales(100);f__Metschnikowiaceae(100);g__Metschnikowia(100);s__Metschnikowia_pulcherrima(100); | Metschnikowia | 13 | Saprotroph | \|Undefined Saprotroph\| | Yeast | NULL | Probable | As nectar-tap_saprotroph (Põlme et al. 2020); Decay sugar-rich substrates (Põlme et al. 2020) | Põlme S, et al. 2020. Fungal Diversity 105:1-16 ((https://doi.org/10.1007/s13225-020-00466-2)) |
| Otu60868 | 1 | k__Fungi(100);p__Ascomycota(100);c__Saccharomycetes(100);o__Saccharomycetales(100);f__Metschnikowiaceae(100);g__Metschnikowia(100);s__Metschnikowia_pulcherrima(100); | Metschnikowia | 13 | Saprotroph | \|Undefined Saprotroph\| | Yeast | NULL | Probable | As nectar-tap_saprotroph (Põlme et al. 2020); Decay sugar-rich substrates (Põlme et al. 2020) | Põlme S, et al. 2020. Fungal Diversity 105:1-16 ((https://doi.org/10.1007/s13225-020-00466-2)) |
| Otu60869 | 1 | k__Fungi(100);p__Ascomycota(100);c__Saccharomycetes(100);o__Saccharomycetales(100);f__Metschnikowiaceae(100);g__Metschnikowia(100);g__Metschnikowia_unclassified(100); | Metschnikowia | 13 | Saprotroph | \|Undefined Saprotroph\| | Yeast | NULL | Probable | As nectar-tap_saprotroph (Põlme et al. 2020); Decay sugar-rich substrates (Põlme et al. 2020) | Põlme S, et al. 2020. Fungal Diversity 105:1-16 ((https://doi.org/10.1007/s13225-020-00466-2)) |
| Otu60870 | 1 | k__Fungi(100);p__Ascomycota(100);c__Saccharomycetes(100);o__Saccharomycetales(100);f__Metschnikowiaceae(100);g__Metschnikowia(100);s__Metschnikowia_pulcherrima(100); | Metschnikowia | 13 | Saprotroph | \|Undefined Saprotroph\| | Yeast | NULL | Probable | As nectar-tap_saprotroph (Põlme et al. 2020); Decay sugar-rich substrates (Põlme et al. 2020) | Põlme S, et al. 2020. Fungal Diversity 105:1-16 ((https://doi.org/10.1007/s13225-020-00466-2)) |
| Otu60871 | 1 | k__Fungi(100);p__Ascomycota(100);c__Saccharomycetes(100);o__Saccharomycetales(100);f__Metschnikowiaceae(100);g__Metschnikowia(100);s__Metschnikowia_pulcherrima(100); | Metschnikowia | 13 | Saprotroph | \|Undefined Saprotroph\| | Yeast | NULL | Probable | As nectar-tap_saprotroph (Põlme et al. 2020); Decay sugar-rich substrates (Põlme et al. 2020) | Põlme S, et al. 2020. Fungal Diversity 105:1-16 ((https://doi.org/10.1007/s13225-020-00466-2)) |
| Otu60872 | 1 | k__Fungi(100);p__Ascomycota(100);c__Saccharomycetes(100);o__Saccharomycetales(100);f__Metschnikowiaceae(100);g__Metschnikowia(100);s__Metschnikowia_pulcherrima(100); | Metschnikowia | 13 | Saprotroph | \|Undefined Saprotroph\| | Yeast | NULL | Probable | As nectar-tap_saprotroph (Põlme et al. 2020); Decay sugar-rich substrates (Põlme et al. 2020) | Põlme S, et al. 2020. Fungal Diversity 105:1-16 ((https://doi.org/10.1007/s13225-020-00466-2)) |
| Otu60873 | 1 | k__Fungi(100);p__Ascomycota(100);c__Saccharomycetes(100);o__Saccharomycetales(100);f__Metschnikowiaceae(100);g__Metschnikowia(100);g__Metschnikowia_unclassified(100); | Metschnikowia | 13 | Saprotroph | \|Undefined Saprotroph\| | Yeast | NULL | Probable | As nectar-tap_saprotroph (Põlme et al. 2020); Decay sugar-rich substrates (Põlme et al. 2020) | Põlme S, et al. 2020. Fungal Diversity 105:1-16 ((https://doi.org/10.1007/s13225-020-00466-2)) |
| Otu60874 | 1 | k__Fungi(100);p__Ascomycota(100);c__Saccharomycetes(100);o__Saccharomycetales(100);f__Metschnikowiaceae(100);g__Metschnikowia(100);s__Metschnikowia_pulcherrima(100); | Metschnikowia | 13 | Saprotroph | \|Undefined Saprotroph\| | Yeast | NULL | Probable | As nectar-tap_saprotroph (Põlme et al. 2020); Decay sugar-rich substrates (Põlme et al. 2020) | Põlme S, et al. 2020. Fungal Diversity 105:1-16 ((https://doi.org/10.1007/s13225-020-00466-2)) |
| Otu60875 | 1 | k__Fungi(100);p__Ascomycota(100);c__Saccharomycetes(100);o__Saccharomycetales(100);f__Metschnikowiaceae(100);g__Metschnikowia(100);s__Metschnikowia_pulcherrima(100); | Metschnikowia | 13 | Saprotroph | \|Undefined Saprotroph\| | Yeast | NULL | Probable | As nectar-tap_saprotroph (Põlme et al. 2020); Decay sugar-rich substrates (Põlme et al. 2020) | Põlme S, et al. 2020. Fungal Diversity 105:1-16 ((https://doi.org/10.1007/s13225-020-00466-2)) |
| Otu60876 | 1 | k__Fungi(100);p__Ascomycota(100);c__Saccharomycetes(100);o__Saccharomycetales(100);f__Metschnikowiaceae(100);g__Metschnikowia(100);s__Metschnikowia_pulcherrima(100); | Metschnikowia | 13 | Saprotroph | \|Undefined Saprotroph\| | Yeast | NULL | Probable | As nectar-tap_saprotroph (Põlme et al. 2020); Decay sugar-rich substrates (Põlme et al. 2020) | Põlme S, et al. 2020. Fungal Diversity 105:1-16 ((https://doi.org/10.1007/s13225-020-00466-2)) |
| Otu60877 | 1 | k__Fungi(100);p__Ascomycota(100);c__Saccharomycetes(100);o__Saccharomycetales(100);f__Metschnikowiaceae(100);g__Metschnikowia(100);s__Metschnikowia_pulcherrima(100); | Metschnikowia | 13 | Saprotroph | \|Undefined Saprotroph\| | Yeast | NULL | Probable | As nectar-tap_saprotroph (Põlme et al. 2020); Decay sugar-rich substrates (Põlme et al. 2020) | Põlme S, et al. 2020. Fungal Diversity 105:1-16 ((https://doi.org/10.1007/s13225-020-00466-2)) |
| Otu60878 | 1 | k__Fungi(100);p__Ascomycota(100);c__Saccharomycetes(100);o__Saccharomycetales(100);f__Metschnikowiaceae(100);g__Metschnikowia(100);s__Metschnikowia_pulcherrima(100); | Metschnikowia | 13 | Saprotroph | \|Undefined Saprotroph\| | Yeast | NULL | Probable | As nectar-tap_saprotroph (Põlme et al. 2020); Decay sugar-rich substrates (Põlme et al. 2020) | Põlme S, et al. 2020. Fungal Diversity 105:1-16 ((https://doi.org/10.1007/s13225-020-00466-2)) |
| Otu60879 | 1 | k__Fungi(100);p__Ascomycota(100);c__Saccharomycetes(100);o__Saccharomycetales(100);f__Metschnikowiaceae(100);g__Metschnikowia(100);s__Metschnikowia_pulcherrima(100); | Metschnikowia | 13 | Saprotroph | \|Undefined Saprotroph\| | Yeast | NULL | Probable | As nectar-tap_saprotroph (Põlme et al. 2020); Decay sugar-rich substrates (Põlme et al. 2020) | Põlme S, et al. 2020. Fungal Diversity 105:1-16 ((https://doi.org/10.1007/s13225-020-00466-2)) |
| Otu60880 | 1 | k__Fungi(100);p__Ascomycota(100);c__Saccharomycetes(100);o__Saccharomycetales(100);f__Metschnikowiaceae(100);g__Metschnikowia(100);s__Metschnikowia_pulcherrima(100); | Metschnikowia | 13 | Saprotroph | \|Undefined Saprotroph\| | Yeast | NULL | Probable | As nectar-tap_saprotroph (Põlme et al. 2020); Decay sugar-rich substrates (Põlme et al. 2020) | Põlme S, et al. 2020. Fungal Diversity 105:1-16 ((https://doi.org/10.1007/s13225-020-00466-2)) |
| Otu60882 | 1 | k__Fungi(100);p__Ascomycota(100);c__Saccharomycetes(100);o__Saccharomycetales(100);f__Metschnikowiaceae(100);g__Metschnikowia(100);s__Metschnikowia_pulcherrima(100); | Metschnikowia | 13 | Saprotroph | \|Undefined Saprotroph\| | Yeast | NULL | Probable | As nectar-tap_saprotroph (Põlme et al. 2020); Decay sugar-rich substrates (Põlme et al. 2020) | Põlme S, et al. 2020. Fungal Diversity 105:1-16 ((https://doi.org/10.1007/s13225-020-00466-2)) |
| Otu60884 | 1 | k__Fungi(100);p__Ascomycota(100);c__Saccharomycetes(100);o__Saccharomycetales(100);f__Metschnikowiaceae(100);g__Metschnikowia(100);s__Metschnikowia_pulcherrima(100); | Metschnikowia | 13 | Saprotroph | \|Undefined Saprotroph\| | Yeast | NULL | Probable | As nectar-tap_saprotroph (Põlme et al. 2020); Decay sugar-rich substrates (Põlme et al. 2020) | Põlme S, et al. 2020. Fungal Diversity 105:1-16 ((https://doi.org/10.1007/s13225-020-00466-2)) |
| Otu60885 | 1 | k__Fungi(100);p__Ascomycota(100);c__Saccharomycetes(100);o__Saccharomycetales(100);f__Metschnikowiaceae(100);g__Metschnikowia(100);s__Metschnikowia_pulcherrima(100); | Metschnikowia | 13 | Saprotroph | \|Undefined Saprotroph\| | Yeast | NULL | Probable | As nectar-tap_saprotroph (Põlme et al. 2020); Decay sugar-rich substrates (Põlme et al. 2020) | Põlme S, et al. 2020. Fungal Diversity 105:1-16 ((https://doi.org/10.1007/s13225-020-00466-2)) |
| Otu60888 | 1 | k__Fungi(100);p__Ascomycota(100);c__Saccharomycetes(100);o__Saccharomycetales(100);f__Metschnikowiaceae(100);g__Metschnikowia(100);g__Metschnikowia_unclassified(100); | Metschnikowia | 13 | Saprotroph | \|Undefined Saprotroph\| | Yeast | NULL | Probable | As nectar-tap_saprotroph (Põlme et al. 2020); Decay sugar-rich substrates (Põlme et al. 2020) | Põlme S, et al. 2020. Fungal Diversity 105:1-16 ((https://doi.org/10.1007/s13225-020-00466-2)) |
| Otu60889 | 1 | k__Fungi(100);p__Ascomycota(100);c__Saccharomycetes(100);o__Saccharomycetales(100);f__Metschnikowiaceae(100);g__Metschnikowia(100);s__Metschnikowia_pulcherrima(100); | Metschnikowia | 13 | Saprotroph | \|Undefined Saprotroph\| | Yeast | NULL | Probable | As nectar-tap_saprotroph (Põlme et al. 2020); Decay sugar-rich substrates (Põlme et al. 2020) | Põlme S, et al. 2020. Fungal Diversity 105:1-16 ((https://doi.org/10.1007/s13225-020-00466-2)) |
| Otu60890 | 1 | k__Fungi(100);p__Ascomycota(100);c__Saccharomycetes(100);o__Saccharomycetales(100);f__Metschnikowiaceae(100);g__Metschnikowia(100);s__Metschnikowia_pulcherrima(100); | Metschnikowia | 13 | Saprotroph | \|Undefined Saprotroph\| | Yeast | NULL | Probable | As nectar-tap_saprotroph (Põlme et al. 2020); Decay sugar-rich substrates (Põlme et al. 2020) | Põlme S, et al. 2020. Fungal Diversity 105:1-16 ((https://doi.org/10.1007/s13225-020-00466-2)) |
| Otu60891 | 1 | k__Fungi(100);p__Ascomycota(100);c__Saccharomycetes(100);o__Saccharomycetales(100);f__Metschnikowiaceae(100);g__Metschnikowia(100);s__Metschnikowia_pulcherrima(100); | Metschnikowia | 13 | Saprotroph | \|Undefined Saprotroph\| | Yeast | NULL | Probable | As nectar-tap_saprotroph (Põlme et al. 2020); Decay sugar-rich substrates (Põlme et al. 2020) | Põlme S, et al. 2020. Fungal Diversity 105:1-16 ((https://doi.org/10.1007/s13225-020-00466-2)) |
| Otu60892 | 1 | k__Fungi(100);p__Ascomycota(100);c__Saccharomycetes(100);o__Saccharomycetales(100);f__Metschnikowiaceae(100);g__Metschnikowia(100);g__Metschnikowia_unclassified(100); | Metschnikowia | 13 | Saprotroph | \|Undefined Saprotroph\| | Yeast | NULL | Probable | As nectar-tap_saprotroph (Põlme et al. 2020); Decay sugar-rich substrates (Põlme et al. 2020) | Põlme S, et al. 2020. Fungal Diversity 105:1-16 ((https://doi.org/10.1007/s13225-020-00466-2)) |
| Otu60893 | 1 | k__Fungi(100);p__Ascomycota(100);c__Saccharomycetes(100);o__Saccharomycetales(100);f__Metschnikowiaceae(100);g__Metschnikowia(100);s__Metschnikowia_pulcherrima(100); | Metschnikowia | 13 | Saprotroph | \|Undefined Saprotroph\| | Yeast | NULL | Probable | As nectar-tap_saprotroph (Põlme et al. 2020); Decay sugar-rich substrates (Põlme et al. 2020) | Põlme S, et al. 2020. Fungal Diversity 105:1-16 ((https://doi.org/10.1007/s13225-020-00466-2)) |
| Otu60894 | 1 | k__Fungi(100);p__Ascomycota(100);c__Saccharomycetes(100);o__Saccharomycetales(100);f__Metschnikowiaceae(100);g__Metschnikowia(100);s__Metschnikowia_pulcherrima(100); | Metschnikowia | 13 | Saprotroph | \|Undefined Saprotroph\| | Yeast | NULL | Probable | As nectar-tap_saprotroph (Põlme et al. 2020); Decay sugar-rich substrates (Põlme et al. 2020) | Põlme S, et al. 2020. Fungal Diversity 105:1-16 ((https://doi.org/10.1007/s13225-020-00466-2)) |
| Otu60895 | 1 | k__Fungi(100);p__Ascomycota(100);c__Saccharomycetes(100);o__Saccharomycetales(100);f__Metschnikowiaceae(100);g__Metschnikowia(100);g__Metschnikowia_unclassified(100); | Metschnikowia | 13 | Saprotroph | \|Undefined Saprotroph\| | Yeast | NULL | Probable | As nectar-tap_saprotroph (Põlme et al. 2020); Decay sugar-rich substrates (Põlme et al. 2020) | Põlme S, et al. 2020. Fungal Diversity 105:1-16 ((https://doi.org/10.1007/s13225-020-00466-2)) |
| Otu60896 | 1 | k__Fungi(100);p__Ascomycota(100);c__Saccharomycetes(100);o__Saccharomycetales(100);f__Metschnikowiaceae(100);g__Metschnikowia(100);s__Metschnikowia_pulcherrima(100); | Metschnikowia | 13 | Saprotroph | \|Undefined Saprotroph\| | Yeast | NULL | Probable | As nectar-tap_saprotroph (Põlme et al. 2020); Decay sugar-rich substrates (Põlme et al. 2020) | Põlme S, et al. 2020. Fungal Diversity 105:1-16 ((https://doi.org/10.1007/s13225-020-00466-2)) |
| Otu60897 | 1 | k__Fungi(100);p__Ascomycota(100);c__Saccharomycetes(100);o__Saccharomycetales(100);f__Metschnikowiaceae(100);g__Metschnikowia(100);s__Metschnikowia_pulcherrima(100); | Metschnikowia | 13 | Saprotroph | \|Undefined Saprotroph\| | Yeast | NULL | Probable | As nectar-tap_saprotroph (Põlme et al. 2020); Decay sugar-rich substrates (Põlme et al. 2020) | Põlme S, et al. 2020. Fungal Diversity 105:1-16 ((https://doi.org/10.1007/s13225-020-00466-2)) |
| Otu60898 | 1 | k__Fungi(100);p__Ascomycota(100);c__Saccharomycetes(100);o__Saccharomycetales(100);f__Metschnikowiaceae(100);g__Metschnikowia(100);s__Metschnikowia_pulcherrima(100); | Metschnikowia | 13 | Saprotroph | \|Undefined Saprotroph\| | Yeast | NULL | Probable | As nectar-tap_saprotroph (Põlme et al. 2020); Decay sugar-rich substrates (Põlme et al. 2020) | Põlme S, et al. 2020. Fungal Diversity 105:1-16 ((https://doi.org/10.1007/s13225-020-00466-2)) |
| Otu60899 | 1 | k__Fungi(100);p__Ascomycota(100);c__Saccharomycetes(100);o__Saccharomycetales(100);f__Metschnikowiaceae(100);g__Metschnikowia(100);s__Metschnikowia_pulcherrima(100); | Metschnikowia | 13 | Saprotroph | \|Undefined Saprotroph\| | Yeast | NULL | Probable | As nectar-tap_saprotroph (Põlme et al. 2020); Decay sugar-rich substrates (Põlme et al. 2020) | Põlme S, et al. 2020. Fungal Diversity 105:1-16 ((https://doi.org/10.1007/s13225-020-00466-2)) |
| Otu60900 | 1 | k__Fungi(100);p__Ascomycota(100);c__Saccharomycetes(100);o__Saccharomycetales(100);f__Metschnikowiaceae(100);g__Metschnikowia(100);g__Metschnikowia_unclassified(100); | Metschnikowia | 13 | Saprotroph | \|Undefined Saprotroph\| | Yeast | NULL | Probable | As nectar-tap_saprotroph (Põlme et al. 2020); Decay sugar-rich substrates (Põlme et al. 2020) | Põlme S, et al. 2020. Fungal Diversity 105:1-16 ((https://doi.org/10.1007/s13225-020-00466-2)) |
| Otu60901 | 1 | k__Fungi(100);p__Ascomycota(100);c__Saccharomycetes(100);o__Saccharomycetales(100);f__Metschnikowiaceae(100);g__Metschnikowia(100);s__Metschnikowia_pulcherrima(100); | Metschnikowia | 13 | Saprotroph | \|Undefined Saprotroph\| | Yeast | NULL | Probable | As nectar-tap_saprotroph (Põlme et al. 2020); Decay sugar-rich substrates (Põlme et al. 2020) | Põlme S, et al. 2020. Fungal Diversity 105:1-16 ((https://doi.org/10.1007/s13225-020-00466-2)) |
| Otu60902 | 1 | k__Fungi(100);p__Ascomycota(100);c__Saccharomycetes(100);o__Saccharomycetales(100);f__Metschnikowiaceae(100);g__Metschnikowia(100);g__Metschnikowia_unclassified(100); | Metschnikowia | 13 | Saprotroph | \|Undefined Saprotroph\| | Yeast | NULL | Probable | As nectar-tap_saprotroph (Põlme et al. 2020); Decay sugar-rich substrates (Põlme et al. 2020) | Põlme S, et al. 2020. Fungal Diversity 105:1-16 ((https://doi.org/10.1007/s13225-020-00466-2)) |
| Otu60903 | 1 | k__Fungi(100);p__Ascomycota(100);c__Saccharomycetes(100);o__Saccharomycetales(100);f__Metschnikowiaceae(100);g__Metschnikowia(100);g__Metschnikowia_unclassified(100); | Metschnikowia | 13 | Saprotroph | \|Undefined Saprotroph\| | Yeast | NULL | Probable | As nectar-tap_saprotroph (Põlme et al. 2020); Decay sugar-rich substrates (Põlme et al. 2020) | Põlme S, et al. 2020. Fungal Diversity 105:1-16 ((https://doi.org/10.1007/s13225-020-00466-2)) |
| Otu60904 | 1 | k__Fungi(100);p__Ascomycota(100);c__Saccharomycetes(100);o__Saccharomycetales(100);f__Metschnikowiaceae(100);g__Metschnikowia(100);g__Metschnikowia_unclassified(100); | Metschnikowia | 13 | Saprotroph | \|Undefined Saprotroph\| | Yeast | NULL | Probable | As nectar-tap_saprotroph (Põlme et al. 2020); Decay sugar-rich substrates (Põlme et al. 2020) | Põlme S, et al. 2020. Fungal Diversity 105:1-16 ((https://doi.org/10.1007/s13225-020-00466-2)) |
| Otu60905 | 1 | k__Fungi(100);p__Ascomycota(100);c__Saccharomycetes(100);o__Saccharomycetales(100);f__Metschnikowiaceae(100);g__Metschnikowia(100);g__Metschnikowia_unclassified(100); | Metschnikowia | 13 | Saprotroph | \|Undefined Saprotroph\| | Yeast | NULL | Probable | As nectar-tap_saprotroph (Põlme et al. 2020); Decay sugar-rich substrates (Põlme et al. 2020) | Põlme S, et al. 2020. Fungal Diversity 105:1-16 ((https://doi.org/10.1007/s13225-020-00466-2)) |
| Otu60906 | 1 | k__Fungi(100);p__Ascomycota(100);c__Saccharomycetes(100);o__Saccharomycetales(100);f__Metschnikowiaceae(100);g__Metschnikowia(100);s__Metschnikowia_pulcherrima(100); | Metschnikowia | 13 | Saprotroph | \|Undefined Saprotroph\| | Yeast | NULL | Probable | As nectar-tap_saprotroph (Põlme et al. 2020); Decay sugar-rich substrates (Põlme et al. 2020) | Põlme S, et al. 2020. Fungal Diversity 105:1-16 ((https://doi.org/10.1007/s13225-020-00466-2)) |
| Otu60907 | 1 | k__Fungi(100);p__Ascomycota(100);c__Saccharomycetes(100);o__Saccharomycetales(100);f__Metschnikowiaceae(100);g__Metschnikowia(100);g__Metschnikowia_unclassified(100); | Metschnikowia | 13 | Saprotroph | \|Undefined Saprotroph\| | Yeast | NULL | Probable | As nectar-tap_saprotroph (Põlme et al. 2020); Decay sugar-rich substrates (Põlme et al. 2020) | Põlme S, et al. 2020. Fungal Diversity 105:1-16 ((https://doi.org/10.1007/s13225-020-00466-2)) |
| Otu60908 | 1 | k__Fungi(100);p__Ascomycota(100);c__Saccharomycetes(100);o__Saccharomycetales(100);f__Metschnikowiaceae(100);g__Metschnikowia(100);g__Metschnikowia_unclassified(100); | Metschnikowia | 13 | Saprotroph | \|Undefined Saprotroph\| | Yeast | NULL | Probable | As nectar-tap_saprotroph (Põlme et al. 2020); Decay sugar-rich substrates (Põlme et al. 2020) | Põlme S, et al. 2020. Fungal Diversity 105:1-16 ((https://doi.org/10.1007/s13225-020-00466-2)) |
| Otu60909 | 1 | k__Fungi(100);p__Ascomycota(100);c__Saccharomycetes(100);o__Saccharomycetales(100);f__Metschnikowiaceae(100);g__Metschnikowia(100);g__Metschnikowia_unclassified(100); | Metschnikowia | 13 | Saprotroph | \|Undefined Saprotroph\| | Yeast | NULL | Probable | As nectar-tap_saprotroph (Põlme et al. 2020); Decay sugar-rich substrates (Põlme et al. 2020) | Põlme S, et al. 2020. Fungal Diversity 105:1-16 ((https://doi.org/10.1007/s13225-020-00466-2)) |
| Otu60910 | 1 | k__Fungi(100);p__Ascomycota(100);c__Saccharomycetes(100);o__Saccharomycetales(100);f__Metschnikowiaceae(100);g__Metschnikowia(100);g__Metschnikowia_unclassified(100); | Metschnikowia | 13 | Saprotroph | \|Undefined Saprotroph\| | Yeast | NULL | Probable | As nectar-tap_saprotroph (Põlme et al. 2020); Decay sugar-rich substrates (Põlme et al. 2020) | Põlme S, et al. 2020. Fungal Diversity 105:1-16 ((https://doi.org/10.1007/s13225-020-00466-2)) |
| Otu60911 | 1 | k__Fungi(100);p__Ascomycota(100);c__Saccharomycetes(100);o__Saccharomycetales(100);f__Metschnikowiaceae(100);g__Metschnikowia(100);g__Metschnikowia_unclassified(100); | Metschnikowia | 13 | Saprotroph | \|Undefined Saprotroph\| | Yeast | NULL | Probable | As nectar-tap_saprotroph (Põlme et al. 2020); Decay sugar-rich substrates (Põlme et al. 2020) | Põlme S, et al. 2020. Fungal Diversity 105:1-16 ((https://doi.org/10.1007/s13225-020-00466-2)) |
| Otu60912 | 1 | k__Fungi(100);p__Ascomycota(100);c__Saccharomycetes(100);o__Saccharomycetales(100);f__Metschnikowiaceae(100);g__Metschnikowia(100);g__Metschnikowia_unclassified(100); | Metschnikowia | 13 | Saprotroph | \|Undefined Saprotroph\| | Yeast | NULL | Probable | As nectar-tap_saprotroph (Põlme et al. 2020); Decay sugar-rich substrates (Põlme et al. 2020) | Põlme S, et al. 2020. Fungal Diversity 105:1-16 ((https://doi.org/10.1007/s13225-020-00466-2)) |
| Otu60913 | 1 | k__Fungi(100);p__Ascomycota(100);c__Saccharomycetes(100);o__Saccharomycetales(100);f__Metschnikowiaceae(100);g__Metschnikowia(100);s__Metschnikowia_pulcherrima(100); | Metschnikowia | 13 | Saprotroph | \|Undefined Saprotroph\| | Yeast | NULL | Probable | As nectar-tap_saprotroph (Põlme et al. 2020); Decay sugar-rich substrates (Põlme et al. 2020) | Põlme S, et al. 2020. Fungal Diversity 105:1-16 ((https://doi.org/10.1007/s13225-020-00466-2)) |
| Otu60914 | 1 | k__Fungi(100);p__Ascomycota(100);c__Saccharomycetes(100);o__Saccharomycetales(100);f__Metschnikowiaceae(100);g__Metschnikowia(100);s__Metschnikowia_pulcherrima(100); | Metschnikowia | 13 | Saprotroph | \|Undefined Saprotroph\| | Yeast | NULL | Probable | As nectar-tap_saprotroph (Põlme et al. 2020); Decay sugar-rich substrates (Põlme et al. 2020) | Põlme S, et al. 2020. Fungal Diversity 105:1-16 ((https://doi.org/10.1007/s13225-020-00466-2)) |
| Otu60915 | 1 | k__Fungi(100);p__Ascomycota(100);c__Saccharomycetes(100);o__Saccharomycetales(100);f__Metschnikowiaceae(100);g__Metschnikowia(100);g__Metschnikowia_unclassified(100); | Metschnikowia | 13 | Saprotroph | \|Undefined Saprotroph\| | Yeast | NULL | Probable | As nectar-tap_saprotroph (Põlme et al. 2020); Decay sugar-rich substrates (Põlme et al. 2020) | Põlme S, et al. 2020. Fungal Diversity 105:1-16 ((https://doi.org/10.1007/s13225-020-00466-2)) |
| Otu60916 | 1 | k__Fungi(100);p__Ascomycota(100);c__Saccharomycetes(100);o__Saccharomycetales(100);f__Metschnikowiaceae(100);g__Metschnikowia(100);s__Metschnikowia_pulcherrima(100); | Metschnikowia | 13 | Saprotroph | \|Undefined Saprotroph\| | Yeast | NULL | Probable | As nectar-tap_saprotroph (Põlme et al. 2020); Decay sugar-rich substrates (Põlme et al. 2020) | Põlme S, et al. 2020. Fungal Diversity 105:1-16 ((https://doi.org/10.1007/s13225-020-00466-2)) |
| Otu60917 | 1 | k__Fungi(100);p__Ascomycota(100);c__Saccharomycetes(100);o__Saccharomycetales(100);f__Metschnikowiaceae(100);g__Metschnikowia(100);s__Metschnikowia_pulcherrima(100); | Metschnikowia | 13 | Saprotroph | \|Undefined Saprotroph\| | Yeast | NULL | Probable | As nectar-tap_saprotroph (Põlme et al. 2020); Decay sugar-rich substrates (Põlme et al. 2020) | Põlme S, et al. 2020. Fungal Diversity 105:1-16 ((https://doi.org/10.1007/s13225-020-00466-2)) |
| Otu60920 | 1 | k__Fungi(100);p__Ascomycota(100);c__Saccharomycetes(100);o__Saccharomycetales(100);f__Metschnikowiaceae(100);g__Metschnikowia(100);s__Metschnikowia_pulcherrima(100); | Metschnikowia | 13 | Saprotroph | \|Undefined Saprotroph\| | Yeast | NULL | Probable | As nectar-tap_saprotroph (Põlme et al. 2020); Decay sugar-rich substrates (Põlme et al. 2020) | Põlme S, et al. 2020. Fungal Diversity 105:1-16 ((https://doi.org/10.1007/s13225-020-00466-2)) |
| Otu60922 | 1 | k__Fungi(100);p__Ascomycota(100);c__Saccharomycetes(100);o__Saccharomycetales(100);f__Metschnikowiaceae(100);g__Metschnikowia(100);s__Metschnikowia_pulcherrima(100); | Metschnikowia | 13 | Saprotroph | \|Undefined Saprotroph\| | Yeast | NULL | Probable | As nectar-tap_saprotroph (Põlme et al. 2020); Decay sugar-rich substrates (Põlme et al. 2020) | Põlme S, et al. 2020. Fungal Diversity 105:1-16 ((https://doi.org/10.1007/s13225-020-00466-2)) |
| Otu60923 | 1 | k__Fungi(100);p__Ascomycota(100);c__Saccharomycetes(100);o__Saccharomycetales(100);f__Metschnikowiaceae(100);g__Metschnikowia(100);s__Metschnikowia_pulcherrima(100); | Metschnikowia | 13 | Saprotroph | \|Undefined Saprotroph\| | Yeast | NULL | Probable | As nectar-tap_saprotroph (Põlme et al. 2020); Decay sugar-rich substrates (Põlme et al. 2020) | Põlme S, et al. 2020. Fungal Diversity 105:1-16 ((https://doi.org/10.1007/s13225-020-00466-2)) |
| Otu60924 | 1 | k__Fungi(100);p__Ascomycota(100);c__Saccharomycetes(100);o__Saccharomycetales(100);f__Metschnikowiaceae(100);g__Metschnikowia(100);s__Metschnikowia_pulcherrima(100); | Metschnikowia | 13 | Saprotroph | \|Undefined Saprotroph\| | Yeast | NULL | Probable | As nectar-tap_saprotroph (Põlme et al. 2020); Decay sugar-rich substrates (Põlme et al. 2020) | Põlme S, et al. 2020. Fungal Diversity 105:1-16 ((https://doi.org/10.1007/s13225-020-00466-2)) |
| Otu60925 | 1 | k__Fungi(100);p__Ascomycota(100);c__Saccharomycetes(100);o__Saccharomycetales(100);f__Metschnikowiaceae(100);g__Metschnikowia(100);s__Metschnikowia_pulcherrima(100); | Metschnikowia | 13 | Saprotroph | \|Undefined Saprotroph\| | Yeast | NULL | Probable | As nectar-tap_saprotroph (Põlme et al. 2020); Decay sugar-rich substrates (Põlme et al. 2020) | Põlme S, et al. 2020. Fungal Diversity 105:1-16 ((https://doi.org/10.1007/s13225-020-00466-2)) |
| Otu60926 | 1 | k__Fungi(100);p__Ascomycota(100);c__Saccharomycetes(100);o__Saccharomycetales(100);f__Metschnikowiaceae(100);g__Metschnikowia(100);s__Metschnikowia_pulcherrima(100); | Metschnikowia | 13 | Saprotroph | \|Undefined Saprotroph\| | Yeast | NULL | Probable | As nectar-tap_saprotroph (Põlme et al. 2020); Decay sugar-rich substrates (Põlme et al. 2020) | Põlme S, et al. 2020. Fungal Diversity 105:1-16 ((https://doi.org/10.1007/s13225-020-00466-2)) |
| Otu60927 | 1 | k__Fungi(100);p__Ascomycota(100);c__Saccharomycetes(100);o__Saccharomycetales(100);f__Metschnikowiaceae(100);g__Metschnikowia(100);s__Metschnikowia_pulcherrima(100); | Metschnikowia | 13 | Saprotroph | \|Undefined Saprotroph\| | Yeast | NULL | Probable | As nectar-tap_saprotroph (Põlme et al. 2020); Decay sugar-rich substrates (Põlme et al. 2020) | Põlme S, et al. 2020. Fungal Diversity 105:1-16 ((https://doi.org/10.1007/s13225-020-00466-2)) |
| Otu60928 | 1 | k__Fungi(100);p__Ascomycota(100);c__Saccharomycetes(100);o__Saccharomycetales(100);f__Metschnikowiaceae(100);g__Metschnikowia(100);s__Metschnikowia_pulcherrima(100); | Metschnikowia | 13 | Saprotroph | \|Undefined Saprotroph\| | Yeast | NULL | Probable | As nectar-tap_saprotroph (Põlme et al. 2020); Decay sugar-rich substrates (Põlme et al. 2020) | Põlme S, et al. 2020. Fungal Diversity 105:1-16 ((https://doi.org/10.1007/s13225-020-00466-2)) |
| Otu60929 | 1 | k__Fungi(100);p__Ascomycota(100);c__Saccharomycetes(100);o__Saccharomycetales(100);f__Metschnikowiaceae(100);g__Metschnikowia(100);s__Metschnikowia_pulcherrima(100); | Metschnikowia | 13 | Saprotroph | \|Undefined Saprotroph\| | Yeast | NULL | Probable | As nectar-tap_saprotroph (Põlme et al. 2020); Decay sugar-rich substrates (Põlme et al. 2020) | Põlme S, et al. 2020. Fungal Diversity 105:1-16 ((https://doi.org/10.1007/s13225-020-00466-2)) |
| Otu60930 | 1 | k__Fungi(100);p__Ascomycota(100);c__Saccharomycetes(100);o__Saccharomycetales(100);f__Metschnikowiaceae(100);g__Metschnikowia(100);s__Metschnikowia_pulcherrima(100); | Metschnikowia | 13 | Saprotroph | \|Undefined Saprotroph\| | Yeast | NULL | Probable | As nectar-tap_saprotroph (Põlme et al. 2020); Decay sugar-rich substrates (Põlme et al. 2020) | Põlme S, et al. 2020. Fungal Diversity 105:1-16 ((https://doi.org/10.1007/s13225-020-00466-2)) |
| Otu60931 | 1 | k__Fungi(100);p__Ascomycota(100);c__Saccharomycetes(100);o__Saccharomycetales(100);f__Metschnikowiaceae(100);g__Metschnikowia(100);s__Metschnikowia_pulcherrima(100); | Metschnikowia | 13 | Saprotroph | \|Undefined Saprotroph\| | Yeast | NULL | Probable | As nectar-tap_saprotroph (Põlme et al. 2020); Decay sugar-rich substrates (Põlme et al. 2020) | Põlme S, et al. 2020. Fungal Diversity 105:1-16 ((https://doi.org/10.1007/s13225-020-00466-2)) |
| Otu60934 | 1 | k__Fungi(100);p__Ascomycota(100);c__Saccharomycetes(100);o__Saccharomycetales(100);f__Metschnikowiaceae(100);g__Metschnikowia(100);s__Metschnikowia_pulcherrima(100); | Metschnikowia | 13 | Saprotroph | \|Undefined Saprotroph\| | Yeast | NULL | Probable | As nectar-tap_saprotroph (Põlme et al. 2020); Decay sugar-rich substrates (Põlme et al. 2020) | Põlme S, et al. 2020. Fungal Diversity 105:1-16 ((https://doi.org/10.1007/s13225-020-00466-2)) |
| Otu60935 | 1 | k__Fungi(100);p__Ascomycota(100);c__Saccharomycetes(100);o__Saccharomycetales(100);f__Metschnikowiaceae(100);g__Metschnikowia(100);s__Metschnikowia_pulcherrima(100); | Metschnikowia | 13 | Saprotroph | \|Undefined Saprotroph\| | Yeast | NULL | Probable | As nectar-tap_saprotroph (Põlme et al. 2020); Decay sugar-rich substrates (Põlme et al. 2020) | Põlme S, et al. 2020. Fungal Diversity 105:1-16 ((https://doi.org/10.1007/s13225-020-00466-2)) |
| Otu60936 | 1 | k__Fungi(100);p__Ascomycota(100);c__Saccharomycetes(100);o__Saccharomycetales(100);f__Metschnikowiaceae(100);g__Metschnikowia(100);s__Metschnikowia_pulcherrima(100); | Metschnikowia | 13 | Saprotroph | \|Undefined Saprotroph\| | Yeast | NULL | Probable | As nectar-tap_saprotroph (Põlme et al. 2020); Decay sugar-rich substrates (Põlme et al. 2020) | Põlme S, et al. 2020. Fungal Diversity 105:1-16 ((https://doi.org/10.1007/s13225-020-00466-2)) |
| Otu60937 | 1 | k__Fungi(100);p__Ascomycota(100);c__Saccharomycetes(100);o__Saccharomycetales(100);f__Metschnikowiaceae(100);g__Metschnikowia(100);g__Metschnikowia_unclassified(100); | Metschnikowia | 13 | Saprotroph | \|Undefined Saprotroph\| | Yeast | NULL | Probable | As nectar-tap_saprotroph (Põlme et al. 2020); Decay sugar-rich substrates (Põlme et al. 2020) | Põlme S, et al. 2020. Fungal Diversity 105:1-16 ((https://doi.org/10.1007/s13225-020-00466-2)) |
| Otu60938 | 1 | k__Fungi(100);p__Ascomycota(100);c__Saccharomycetes(100);o__Saccharomycetales(100);f__Metschnikowiaceae(100);g__Metschnikowia(100);s__Metschnikowia_pulcherrima(100); | Metschnikowia | 13 | Saprotroph | \|Undefined Saprotroph\| | Yeast | NULL | Probable | As nectar-tap_saprotroph (Põlme et al. 2020); Decay sugar-rich substrates (Põlme et al. 2020) | Põlme S, et al. 2020. Fungal Diversity 105:1-16 ((https://doi.org/10.1007/s13225-020-00466-2)) |
| Otu60939 | 1 | k__Fungi(100);p__Ascomycota(100);c__Saccharomycetes(100);o__Saccharomycetales(100);f__Metschnikowiaceae(100);g__Metschnikowia(100);s__Metschnikowia_pulcherrima(100); | Metschnikowia | 13 | Saprotroph | \|Undefined Saprotroph\| | Yeast | NULL | Probable | As nectar-tap_saprotroph (Põlme et al. 2020); Decay sugar-rich substrates (Põlme et al. 2020) | Põlme S, et al. 2020. Fungal Diversity 105:1-16 ((https://doi.org/10.1007/s13225-020-00466-2)) |
| Otu60940 | 1 | k__Fungi(100);p__Ascomycota(100);c__Saccharomycetes(100);o__Saccharomycetales(100);f__Metschnikowiaceae(100);g__Metschnikowia(100);s__Metschnikowia_pulcherrima(100); | Metschnikowia | 13 | Saprotroph | \|Undefined Saprotroph\| | Yeast | NULL | Probable | As nectar-tap_saprotroph (Põlme et al. 2020); Decay sugar-rich substrates (Põlme et al. 2020) | Põlme S, et al. 2020. Fungal Diversity 105:1-16 ((https://doi.org/10.1007/s13225-020-00466-2)) |
| Otu60941 | 1 | k__Fungi(100);p__Ascomycota(100);c__Saccharomycetes(100);o__Saccharomycetales(100);f__Metschnikowiaceae(100);g__Metschnikowia(100);s__Metschnikowia_pulcherrima(100); | Metschnikowia | 13 | Saprotroph | \|Undefined Saprotroph\| | Yeast | NULL | Probable | As nectar-tap_saprotroph (Põlme et al. 2020); Decay sugar-rich substrates (Põlme et al. 2020) | Põlme S, et al. 2020. Fungal Diversity 105:1-16 ((https://doi.org/10.1007/s13225-020-00466-2)) |
| Otu60942 | 1 | k__Fungi(100);p__Ascomycota(100);c__Saccharomycetes(100);o__Saccharomycetales(100);f__Metschnikowiaceae(100);g__Metschnikowia(100);g__Metschnikowia_unclassified(100); | Metschnikowia | 13 | Saprotroph | \|Undefined Saprotroph\| | Yeast | NULL | Probable | As nectar-tap_saprotroph (Põlme et al. 2020); Decay sugar-rich substrates (Põlme et al. 2020) | Põlme S, et al. 2020. Fungal Diversity 105:1-16 ((https://doi.org/10.1007/s13225-020-00466-2)) |
| Otu60943 | 1 | k__Fungi(100);p__Ascomycota(100);c__Saccharomycetes(100);o__Saccharomycetales(100);f__Metschnikowiaceae(100);g__Metschnikowia(100);s__Metschnikowia_pulcherrima(100); | Metschnikowia | 13 | Saprotroph | \|Undefined Saprotroph\| | Yeast | NULL | Probable | As nectar-tap_saprotroph (Põlme et al. 2020); Decay sugar-rich substrates (Põlme et al. 2020) | Põlme S, et al. 2020. Fungal Diversity 105:1-16 ((https://doi.org/10.1007/s13225-020-00466-2)) |
| Otu60944 | 1 | k__Fungi(100);p__Ascomycota(100);c__Saccharomycetes(100);o__Saccharomycetales(100);f__Metschnikowiaceae(100);g__Metschnikowia(100);s__Metschnikowia_pulcherrima(100); | Metschnikowia | 13 | Saprotroph | \|Undefined Saprotroph\| | Yeast | NULL | Probable | As nectar-tap_saprotroph (Põlme et al. 2020); Decay sugar-rich substrates (Põlme et al. 2020) | Põlme S, et al. 2020. Fungal Diversity 105:1-16 ((https://doi.org/10.1007/s13225-020-00466-2)) |
| Otu60945 | 1 | k__Fungi(100);p__Ascomycota(100);c__Saccharomycetes(100);o__Saccharomycetales(100);f__Metschnikowiaceae(100);g__Metschnikowia(100);s__Metschnikowia_pulcherrima(100); | Metschnikowia | 13 | Saprotroph | \|Undefined Saprotroph\| | Yeast | NULL | Probable | As nectar-tap_saprotroph (Põlme et al. 2020); Decay sugar-rich substrates (Põlme et al. 2020) | Põlme S, et al. 2020. Fungal Diversity 105:1-16 ((https://doi.org/10.1007/s13225-020-00466-2)) |
| Otu60946 | 1 | k__Fungi(100);p__Ascomycota(100);c__Saccharomycetes(100);o__Saccharomycetales(100);f__Metschnikowiaceae(100);g__Metschnikowia(100);s__Metschnikowia_pulcherrima(100); | Metschnikowia | 13 | Saprotroph | \|Undefined Saprotroph\| | Yeast | NULL | Probable | As nectar-tap_saprotroph (Põlme et al. 2020); Decay sugar-rich substrates (Põlme et al. 2020) | Põlme S, et al. 2020. Fungal Diversity 105:1-16 ((https://doi.org/10.1007/s13225-020-00466-2)) |
| Otu60947 | 1 | k__Fungi(100);p__Ascomycota(100);c__Saccharomycetes(100);o__Saccharomycetales(100);f__Metschnikowiaceae(100);g__Metschnikowia(100);s__Metschnikowia_pulcherrima(100); | Metschnikowia | 13 | Saprotroph | \|Undefined Saprotroph\| | Yeast | NULL | Probable | As nectar-tap_saprotroph (Põlme et al. 2020); Decay sugar-rich substrates (Põlme et al. 2020) | Põlme S, et al. 2020. Fungal Diversity 105:1-16 ((https://doi.org/10.1007/s13225-020-00466-2)) |
| Otu60948 | 1 | k__Fungi(100);p__Ascomycota(100);c__Saccharomycetes(100);o__Saccharomycetales(100);f__Metschnikowiaceae(100);g__Metschnikowia(100);s__Metschnikowia_pulcherrima(100); | Metschnikowia | 13 | Saprotroph | \|Undefined Saprotroph\| | Yeast | NULL | Probable | As nectar-tap_saprotroph (Põlme et al. 2020); Decay sugar-rich substrates (Põlme et al. 2020) | Põlme S, et al. 2020. Fungal Diversity 105:1-16 ((https://doi.org/10.1007/s13225-020-00466-2)) |
| Otu60949 | 1 | k__Fungi(100);p__Ascomycota(100);c__Saccharomycetes(100);o__Saccharomycetales(100);f__Metschnikowiaceae(100);g__Metschnikowia(100);s__Metschnikowia_pulcherrima(100); | Metschnikowia | 13 | Saprotroph | \|Undefined Saprotroph\| | Yeast | NULL | Probable | As nectar-tap_saprotroph (Põlme et al. 2020); Decay sugar-rich substrates (Põlme et al. 2020) | Põlme S, et al. 2020. Fungal Diversity 105:1-16 ((https://doi.org/10.1007/s13225-020-00466-2)) |
| Otu60950 | 1 | k__Fungi(100);p__Ascomycota(100);c__Saccharomycetes(100);o__Saccharomycetales(100);f__Metschnikowiaceae(100);g__Metschnikowia(100);s__Metschnikowia_pulcherrima(100); | Metschnikowia | 13 | Saprotroph | \|Undefined Saprotroph\| | Yeast | NULL | Probable | As nectar-tap_saprotroph (Põlme et al. 2020); Decay sugar-rich substrates (Põlme et al. 2020) | Põlme S, et al. 2020. Fungal Diversity 105:1-16 ((https://doi.org/10.1007/s13225-020-00466-2)) |
| Otu60951 | 1 | k__Fungi(100);p__Ascomycota(100);c__Saccharomycetes(100);o__Saccharomycetales(100);f__Metschnikowiaceae(100);g__Metschnikowia(100);g__Metschnikowia_unclassified(100); | Metschnikowia | 13 | Saprotroph | \|Undefined Saprotroph\| | Yeast | NULL | Probable | As nectar-tap_saprotroph (Põlme et al. 2020); Decay sugar-rich substrates (Põlme et al. 2020) | Põlme S, et al. 2020. Fungal Diversity 105:1-16 ((https://doi.org/10.1007/s13225-020-00466-2)) |
| Otu60952 | 1 | k__Fungi(100);p__Ascomycota(100);c__Saccharomycetes(100);o__Saccharomycetales(100);f__Metschnikowiaceae(100);g__Metschnikowia(100);g__Metschnikowia_unclassified(100); | Metschnikowia | 13 | Saprotroph | \|Undefined Saprotroph\| | Yeast | NULL | Probable | As nectar-tap_saprotroph (Põlme et al. 2020); Decay sugar-rich substrates (Põlme et al. 2020) | Põlme S, et al. 2020. Fungal Diversity 105:1-16 ((https://doi.org/10.1007/s13225-020-00466-2)) |
| Otu60953 | 1 | k__Fungi(100);p__Ascomycota(100);c__Saccharomycetes(100);o__Saccharomycetales(100);f__Metschnikowiaceae(100);g__Metschnikowia(100);s__Metschnikowia_pulcherrima(100); | Metschnikowia | 13 | Saprotroph | \|Undefined Saprotroph\| | Yeast | NULL | Probable | As nectar-tap_saprotroph (Põlme et al. 2020); Decay sugar-rich substrates (Põlme et al. 2020) | Põlme S, et al. 2020. Fungal Diversity 105:1-16 ((https://doi.org/10.1007/s13225-020-00466-2)) |
| Otu60954 | 1 | k__Fungi(100);p__Ascomycota(100);c__Saccharomycetes(100);o__Saccharomycetales(100);f__Metschnikowiaceae(100);g__Metschnikowia(100);s__Metschnikowia_pulcherrima(100); | Metschnikowia | 13 | Saprotroph | \|Undefined Saprotroph\| | Yeast | NULL | Probable | As nectar-tap_saprotroph (Põlme et al. 2020); Decay sugar-rich substrates (Põlme et al. 2020) | Põlme S, et al. 2020. Fungal Diversity 105:1-16 ((https://doi.org/10.1007/s13225-020-00466-2)) |
| Otu60955 | 1 | k__Fungi(100);p__Ascomycota(100);c__Saccharomycetes(100);o__Saccharomycetales(100);f__Metschnikowiaceae(100);g__Metschnikowia(100);s__Metschnikowia_pulcherrima(100); | Metschnikowia | 13 | Saprotroph | \|Undefined Saprotroph\| | Yeast | NULL | Probable | As nectar-tap_saprotroph (Põlme et al. 2020); Decay sugar-rich substrates (Põlme et al. 2020) | Põlme S, et al. 2020. Fungal Diversity 105:1-16 ((https://doi.org/10.1007/s13225-020-00466-2)) |
| Otu60956 | 1 | k__Fungi(100);p__Ascomycota(100);c__Saccharomycetes(100);o__Saccharomycetales(100);f__Metschnikowiaceae(100);g__Metschnikowia(100);s__Metschnikowia_pulcherrima(100); | Metschnikowia | 13 | Saprotroph | \|Undefined Saprotroph\| | Yeast | NULL | Probable | As nectar-tap_saprotroph (Põlme et al. 2020); Decay sugar-rich substrates (Põlme et al. 2020) | Põlme S, et al. 2020. Fungal Diversity 105:1-16 ((https://doi.org/10.1007/s13225-020-00466-2)) |
| Otu60957 | 1 | k__Fungi(100);p__Ascomycota(100);c__Saccharomycetes(100);o__Saccharomycetales(100);f__Metschnikowiaceae(100);g__Metschnikowia(100);s__Metschnikowia_pulcherrima(100); | Metschnikowia | 13 | Saprotroph | \|Undefined Saprotroph\| | Yeast | NULL | Probable | As nectar-tap_saprotroph (Põlme et al. 2020); Decay sugar-rich substrates (Põlme et al. 2020) | Põlme S, et al. 2020. Fungal Diversity 105:1-16 ((https://doi.org/10.1007/s13225-020-00466-2)) |
| Otu60958 | 1 | k__Fungi(100);p__Ascomycota(100);c__Saccharomycetes(100);o__Saccharomycetales(100);f__Metschnikowiaceae(100);g__Metschnikowia(100);g__Metschnikowia_unclassified(100); | Metschnikowia | 13 | Saprotroph | \|Undefined Saprotroph\| | Yeast | NULL | Probable | As nectar-tap_saprotroph (Põlme et al. 2020); Decay sugar-rich substrates (Põlme et al. 2020) | Põlme S, et al. 2020. Fungal Diversity 105:1-16 ((https://doi.org/10.1007/s13225-020-00466-2)) |
| Otu60959 | 1 | k__Fungi(100);p__Ascomycota(100);c__Saccharomycetes(100);o__Saccharomycetales(100);f__Metschnikowiaceae(100);g__Metschnikowia(100);s__Metschnikowia_pulcherrima(100); | Metschnikowia | 13 | Saprotroph | \|Undefined Saprotroph\| | Yeast | NULL | Probable | As nectar-tap_saprotroph (Põlme et al. 2020); Decay sugar-rich substrates (Põlme et al. 2020) | Põlme S, et al. 2020. Fungal Diversity 105:1-16 ((https://doi.org/10.1007/s13225-020-00466-2)) |
| Otu60960 | 1 | k__Fungi(100);p__Ascomycota(100);c__Saccharomycetes(100);o__Saccharomycetales(100);f__Metschnikowiaceae(100);g__Metschnikowia(100);g__Metschnikowia_unclassified(100); | Metschnikowia | 13 | Saprotroph | \|Undefined Saprotroph\| | Yeast | NULL | Probable | As nectar-tap_saprotroph (Põlme et al. 2020); Decay sugar-rich substrates (Põlme et al. 2020) | Põlme S, et al. 2020. Fungal Diversity 105:1-16 ((https://doi.org/10.1007/s13225-020-00466-2)) |
| Otu60961 | 1 | k__Fungi(100);p__Ascomycota(100);c__Saccharomycetes(100);o__Saccharomycetales(100);f__Metschnikowiaceae(100);g__Metschnikowia(100);s__Metschnikowia_pulcherrima(100); | Metschnikowia | 13 | Saprotroph | \|Undefined Saprotroph\| | Yeast | NULL | Probable | As nectar-tap_saprotroph (Põlme et al. 2020); Decay sugar-rich substrates (Põlme et al. 2020) | Põlme S, et al. 2020. Fungal Diversity 105:1-16 ((https://doi.org/10.1007/s13225-020-00466-2)) |
| Otu60963 | 1 | k__Fungi(100);p__Ascomycota(100);c__Saccharomycetes(100);o__Saccharomycetales(100);f__Metschnikowiaceae(100);g__Metschnikowia(100);s__Metschnikowia_pulcherrima(100); | Metschnikowia | 13 | Saprotroph | \|Undefined Saprotroph\| | Yeast | NULL | Probable | As nectar-tap_saprotroph (Põlme et al. 2020); Decay sugar-rich substrates (Põlme et al. 2020) | Põlme S, et al. 2020. Fungal Diversity 105:1-16 ((https://doi.org/10.1007/s13225-020-00466-2)) |
| Otu60964 | 1 | k__Fungi(100);p__Ascomycota(100);c__Saccharomycetes(100);o__Saccharomycetales(100);f__Metschnikowiaceae(100);g__Metschnikowia(100);s__Metschnikowia_pulcherrima(100); | Metschnikowia | 13 | Saprotroph | \|Undefined Saprotroph\| | Yeast | NULL | Probable | As nectar-tap_saprotroph (Põlme et al. 2020); Decay sugar-rich substrates (Põlme et al. 2020) | Põlme S, et al. 2020. Fungal Diversity 105:1-16 ((https://doi.org/10.1007/s13225-020-00466-2)) |
| Otu60965 | 1 | k__Fungi(100);p__Ascomycota(100);c__Saccharomycetes(100);o__Saccharomycetales(100);f__Metschnikowiaceae(100);g__Metschnikowia(100);s__Metschnikowia_pulcherrima(100); | Metschnikowia | 13 | Saprotroph | \|Undefined Saprotroph\| | Yeast | NULL | Probable | As nectar-tap_saprotroph (Põlme et al. 2020); Decay sugar-rich substrates (Põlme et al. 2020) | Põlme S, et al. 2020. Fungal Diversity 105:1-16 ((https://doi.org/10.1007/s13225-020-00466-2)) |
| Otu60966 | 1 | k__Fungi(100);p__Ascomycota(100);c__Saccharomycetes(100);o__Saccharomycetales(100);f__Metschnikowiaceae(100);g__Metschnikowia(100);s__Metschnikowia_pulcherrima(100); | Metschnikowia | 13 | Saprotroph | \|Undefined Saprotroph\| | Yeast | NULL | Probable | As nectar-tap_saprotroph (Põlme et al. 2020); Decay sugar-rich substrates (Põlme et al. 2020) | Põlme S, et al. 2020. Fungal Diversity 105:1-16 ((https://doi.org/10.1007/s13225-020-00466-2)) |
| Otu60967 | 1 | k__Fungi(100);p__Ascomycota(100);c__Saccharomycetes(100);o__Saccharomycetales(100);f__Metschnikowiaceae(100);g__Metschnikowia(100);s__Metschnikowia_pulcherrima(100); | Metschnikowia | 13 | Saprotroph | \|Undefined Saprotroph\| | Yeast | NULL | Probable | As nectar-tap_saprotroph (Põlme et al. 2020); Decay sugar-rich substrates (Põlme et al. 2020) | Põlme S, et al. 2020. Fungal Diversity 105:1-16 ((https://doi.org/10.1007/s13225-020-00466-2)) |
| Otu60968 | 1 | k__Fungi(100);p__Ascomycota(100);c__Saccharomycetes(100);o__Saccharomycetales(100);f__Metschnikowiaceae(100);g__Metschnikowia(100);s__Metschnikowia_pulcherrima(100); | Metschnikowia | 13 | Saprotroph | \|Undefined Saprotroph\| | Yeast | NULL | Probable | As nectar-tap_saprotroph (Põlme et al. 2020); Decay sugar-rich substrates (Põlme et al. 2020) | Põlme S, et al. 2020. Fungal Diversity 105:1-16 ((https://doi.org/10.1007/s13225-020-00466-2)) |
| Otu60969 | 1 | k__Fungi(100);p__Ascomycota(100);c__Saccharomycetes(100);o__Saccharomycetales(100);f__Metschnikowiaceae(100);g__Metschnikowia(100);g__Metschnikowia_unclassified(100); | Metschnikowia | 13 | Saprotroph | \|Undefined Saprotroph\| | Yeast | NULL | Probable | As nectar-tap_saprotroph (Põlme et al. 2020); Decay sugar-rich substrates (Põlme et al. 2020) | Põlme S, et al. 2020. Fungal Diversity 105:1-16 ((https://doi.org/10.1007/s13225-020-00466-2)) |
| Otu60970 | 1 | k__Fungi(100);p__Ascomycota(100);c__Saccharomycetes(100);o__Saccharomycetales(100);f__Metschnikowiaceae(100);g__Metschnikowia(100);s__Metschnikowia_pulcherrima(100); | Metschnikowia | 13 | Saprotroph | \|Undefined Saprotroph\| | Yeast | NULL | Probable | As nectar-tap_saprotroph (Põlme et al. 2020); Decay sugar-rich substrates (Põlme et al. 2020) | Põlme S, et al. 2020. Fungal Diversity 105:1-16 ((https://doi.org/10.1007/s13225-020-00466-2)) |
| Otu60971 | 1 | k__Fungi(100);p__Ascomycota(100);c__Saccharomycetes(100);o__Saccharomycetales(100);f__Metschnikowiaceae(100);g__Metschnikowia(100);g__Metschnikowia_unclassified(100); | Metschnikowia | 13 | Saprotroph | \|Undefined Saprotroph\| | Yeast | NULL | Probable | As nectar-tap_saprotroph (Põlme et al. 2020); Decay sugar-rich substrates (Põlme et al. 2020) | Põlme S, et al. 2020. Fungal Diversity 105:1-16 ((https://doi.org/10.1007/s13225-020-00466-2)) |
| Otu60972 | 1 | k__Fungi(100);p__Ascomycota(100);c__Saccharomycetes(100);o__Saccharomycetales(100);f__Metschnikowiaceae(100);g__Metschnikowia(100);s__Metschnikowia_pulcherrima(100); | Metschnikowia | 13 | Saprotroph | \|Undefined Saprotroph\| | Yeast | NULL | Probable | As nectar-tap_saprotroph (Põlme et al. 2020); Decay sugar-rich substrates (Põlme et al. 2020) | Põlme S, et al. 2020. Fungal Diversity 105:1-16 ((https://doi.org/10.1007/s13225-020-00466-2)) |
| Otu60973 | 1 | k__Fungi(100);p__Ascomycota(100);c__Saccharomycetes(100);o__Saccharomycetales(100);f__Metschnikowiaceae(100);g__Metschnikowia(100);s__Metschnikowia_pulcherrima(100); | Metschnikowia | 13 | Saprotroph | \|Undefined Saprotroph\| | Yeast | NULL | Probable | As nectar-tap_saprotroph (Põlme et al. 2020); Decay sugar-rich substrates (Põlme et al. 2020) | Põlme S, et al. 2020. Fungal Diversity 105:1-16 ((https://doi.org/10.1007/s13225-020-00466-2)) |
| Otu60974 | 1 | k__Fungi(100);p__Ascomycota(100);c__Saccharomycetes(100);o__Saccharomycetales(100);f__Metschnikowiaceae(100);g__Metschnikowia(100);s__Metschnikowia_pulcherrima(100); | Metschnikowia | 13 | Saprotroph | \|Undefined Saprotroph\| | Yeast | NULL | Probable | As nectar-tap_saprotroph (Põlme et al. 2020); Decay sugar-rich substrates (Põlme et al. 2020) | Põlme S, et al. 2020. Fungal Diversity 105:1-16 ((https://doi.org/10.1007/s13225-020-00466-2)) |
| Otu60975 | 1 | k__Fungi(100);p__Ascomycota(100);c__Saccharomycetes(100);o__Saccharomycetales(100);f__Metschnikowiaceae(100);g__Metschnikowia(100);s__Metschnikowia_pulcherrima(100); | Metschnikowia | 13 | Saprotroph | \|Undefined Saprotroph\| | Yeast | NULL | Probable | As nectar-tap_saprotroph (Põlme et al. 2020); Decay sugar-rich substrates (Põlme et al. 2020) | Põlme S, et al. 2020. Fungal Diversity 105:1-16 ((https://doi.org/10.1007/s13225-020-00466-2)) |
| Otu60976 | 1 | k__Fungi(100);p__Ascomycota(100);c__Saccharomycetes(100);o__Saccharomycetales(100);f__Metschnikowiaceae(100);g__Metschnikowia(100);s__Metschnikowia_pulcherrima(100); | Metschnikowia | 13 | Saprotroph | \|Undefined Saprotroph\| | Yeast | NULL | Probable | As nectar-tap_saprotroph (Põlme et al. 2020); Decay sugar-rich substrates (Põlme et al. 2020) | Põlme S, et al. 2020. Fungal Diversity 105:1-16 ((https://doi.org/10.1007/s13225-020-00466-2)) |
| Otu60977 | 1 | k__Fungi(100);p__Ascomycota(100);c__Saccharomycetes(100);o__Saccharomycetales(100);f__Metschnikowiaceae(100);g__Metschnikowia(100);s__Metschnikowia_pulcherrima(100); | Metschnikowia | 13 | Saprotroph | \|Undefined Saprotroph\| | Yeast | NULL | Probable | As nectar-tap_saprotroph (Põlme et al. 2020); Decay sugar-rich substrates (Põlme et al. 2020) | Põlme S, et al. 2020. Fungal Diversity 105:1-16 ((https://doi.org/10.1007/s13225-020-00466-2)) |
| Otu60978 | 1 | k__Fungi(100);p__Ascomycota(100);c__Saccharomycetes(100);o__Saccharomycetales(100);f__Metschnikowiaceae(100);g__Metschnikowia(100);s__Metschnikowia_pulcherrima(100); | Metschnikowia | 13 | Saprotroph | \|Undefined Saprotroph\| | Yeast | NULL | Probable | As nectar-tap_saprotroph (Põlme et al. 2020); Decay sugar-rich substrates (Põlme et al. 2020) | Põlme S, et al. 2020. Fungal Diversity 105:1-16 ((https://doi.org/10.1007/s13225-020-00466-2)) |
| Otu60979 | 1 | k__Fungi(100);p__Ascomycota(100);c__Saccharomycetes(100);o__Saccharomycetales(100);f__Metschnikowiaceae(100);g__Metschnikowia(100);s__Metschnikowia_pulcherrima(100); | Metschnikowia | 13 | Saprotroph | \|Undefined Saprotroph\| | Yeast | NULL | Probable | As nectar-tap_saprotroph (Põlme et al. 2020); Decay sugar-rich substrates (Põlme et al. 2020) | Põlme S, et al. 2020. Fungal Diversity 105:1-16 ((https://doi.org/10.1007/s13225-020-00466-2)) |
| Otu60981 | 1 | k__Fungi(100);p__Ascomycota(100);c__Saccharomycetes(100);o__Saccharomycetales(100);f__Metschnikowiaceae(100);g__Metschnikowia(100);g__Metschnikowia_unclassified(100); | Metschnikowia | 13 | Saprotroph | \|Undefined Saprotroph\| | Yeast | NULL | Probable | As nectar-tap_saprotroph (Põlme et al. 2020); Decay sugar-rich substrates (Põlme et al. 2020) | Põlme S, et al. 2020. Fungal Diversity 105:1-16 ((https://doi.org/10.1007/s13225-020-00466-2)) |
| Otu60982 | 1 | k__Fungi(100);p__Ascomycota(100);c__Saccharomycetes(100);o__Saccharomycetales(100);f__Metschnikowiaceae(100);g__Metschnikowia(100);g__Metschnikowia_unclassified(100); | Metschnikowia | 13 | Saprotroph | \|Undefined Saprotroph\| | Yeast | NULL | Probable | As nectar-tap_saprotroph (Põlme et al. 2020); Decay sugar-rich substrates (Põlme et al. 2020) | Põlme S, et al. 2020. Fungal Diversity 105:1-16 ((https://doi.org/10.1007/s13225-020-00466-2)) |
| Otu60983 | 1 | k__Fungi(100);p__Ascomycota(100);c__Saccharomycetes(100);o__Saccharomycetales(100);f__Metschnikowiaceae(100);g__Metschnikowia(100);s__Metschnikowia_pulcherrima(100); | Metschnikowia | 13 | Saprotroph | \|Undefined Saprotroph\| | Yeast | NULL | Probable | As nectar-tap_saprotroph (Põlme et al. 2020); Decay sugar-rich substrates (Põlme et al. 2020) | Põlme S, et al. 2020. Fungal Diversity 105:1-16 ((https://doi.org/10.1007/s13225-020-00466-2)) |
| Otu60984 | 1 | k__Fungi(100);p__Ascomycota(100);c__Saccharomycetes(100);o__Saccharomycetales(100);f__Metschnikowiaceae(100);g__Metschnikowia(100);s__Metschnikowia_pulcherrima(100); | Metschnikowia | 13 | Saprotroph | \|Undefined Saprotroph\| | Yeast | NULL | Probable | As nectar-tap_saprotroph (Põlme et al. 2020); Decay sugar-rich substrates (Põlme et al. 2020) | Põlme S, et al. 2020. Fungal Diversity 105:1-16 ((https://doi.org/10.1007/s13225-020-00466-2)) |
| Otu60985 | 1 | k__Fungi(100);p__Ascomycota(100);c__Saccharomycetes(100);o__Saccharomycetales(100);f__Metschnikowiaceae(100);g__Metschnikowia(100);s__Metschnikowia_pulcherrima(100); | Metschnikowia | 13 | Saprotroph | \|Undefined Saprotroph\| | Yeast | NULL | Probable | As nectar-tap_saprotroph (Põlme et al. 2020); Decay sugar-rich substrates (Põlme et al. 2020) | Põlme S, et al. 2020. Fungal Diversity 105:1-16 ((https://doi.org/10.1007/s13225-020-00466-2)) |
| Otu60986 | 1 | k__Fungi(100);p__Ascomycota(100);c__Saccharomycetes(100);o__Saccharomycetales(100);f__Metschnikowiaceae(100);g__Metschnikowia(100);s__Metschnikowia_pulcherrima(100); | Metschnikowia | 13 | Saprotroph | \|Undefined Saprotroph\| | Yeast | NULL | Probable | As nectar-tap_saprotroph (Põlme et al. 2020); Decay sugar-rich substrates (Põlme et al. 2020) | Põlme S, et al. 2020. Fungal Diversity 105:1-16 ((https://doi.org/10.1007/s13225-020-00466-2)) |
| Otu60987 | 1 | k__Fungi(100);p__Ascomycota(100);c__Saccharomycetes(100);o__Saccharomycetales(100);f__Metschnikowiaceae(100);g__Metschnikowia(100);s__Metschnikowia_pulcherrima(100); | Metschnikowia | 13 | Saprotroph | \|Undefined Saprotroph\| | Yeast | NULL | Probable | As nectar-tap_saprotroph (Põlme et al. 2020); Decay sugar-rich substrates (Põlme et al. 2020) | Põlme S, et al. 2020. Fungal Diversity 105:1-16 ((https://doi.org/10.1007/s13225-020-00466-2)) |
| Otu60988 | 1 | k__Fungi(100);p__Ascomycota(100);c__Saccharomycetes(100);o__Saccharomycetales(100);f__Metschnikowiaceae(100);g__Metschnikowia(100);s__Metschnikowia_pulcherrima(100); | Metschnikowia | 13 | Saprotroph | \|Undefined Saprotroph\| | Yeast | NULL | Probable | As nectar-tap_saprotroph (Põlme et al. 2020); Decay sugar-rich substrates (Põlme et al. 2020) | Põlme S, et al. 2020. Fungal Diversity 105:1-16 ((https://doi.org/10.1007/s13225-020-00466-2)) |
| Otu60989 | 1 | k__Fungi(100);p__Ascomycota(100);c__Saccharomycetes(100);o__Saccharomycetales(100);f__Metschnikowiaceae(100);g__Metschnikowia(100);s__Metschnikowia_pulcherrima(100); | Metschnikowia | 13 | Saprotroph | \|Undefined Saprotroph\| | Yeast | NULL | Probable | As nectar-tap_saprotroph (Põlme et al. 2020); Decay sugar-rich substrates (Põlme et al. 2020) | Põlme S, et al. 2020. Fungal Diversity 105:1-16 ((https://doi.org/10.1007/s13225-020-00466-2)) |
| Otu60990 | 1 | k__Fungi(100);p__Ascomycota(100);c__Saccharomycetes(100);o__Saccharomycetales(100);f__Metschnikowiaceae(100);g__Metschnikowia(100);s__Metschnikowia_pulcherrima(100); | Metschnikowia | 13 | Saprotroph | \|Undefined Saprotroph\| | Yeast | NULL | Probable | As nectar-tap_saprotroph (Põlme et al. 2020); Decay sugar-rich substrates (Põlme et al. 2020) | Põlme S, et al. 2020. Fungal Diversity 105:1-16 ((https://doi.org/10.1007/s13225-020-00466-2)) |
| Otu60991 | 1 | k__Fungi(100);p__Ascomycota(100);c__Saccharomycetes(100);o__Saccharomycetales(100);f__Metschnikowiaceae(100);g__Metschnikowia(100);g__Metschnikowia_unclassified(100); | Metschnikowia | 13 | Saprotroph | \|Undefined Saprotroph\| | Yeast | NULL | Probable | As nectar-tap_saprotroph (Põlme et al. 2020); Decay sugar-rich substrates (Põlme et al. 2020) | Põlme S, et al. 2020. Fungal Diversity 105:1-16 ((https://doi.org/10.1007/s13225-020-00466-2)) |
| Otu60992 | 1 | k__Fungi(100);p__Ascomycota(100);c__Saccharomycetes(100);o__Saccharomycetales(100);f__Metschnikowiaceae(100);g__Metschnikowia(100);s__Metschnikowia_pulcherrima(100); | Metschnikowia | 13 | Saprotroph | \|Undefined Saprotroph\| | Yeast | NULL | Probable | As nectar-tap_saprotroph (Põlme et al. 2020); Decay sugar-rich substrates (Põlme et al. 2020) | Põlme S, et al. 2020. Fungal Diversity 105:1-16 ((https://doi.org/10.1007/s13225-020-00466-2)) |
| Otu60993 | 1 | k__Fungi(100);p__Basidiomycota(100);c__Microbotryomycetes(100);o__Sporidiobolales(100);f__Sporidiobolales_family_Incertae_sedis(100);g__Sporobolomyces(100);s__Sporobolomyces_lactosus(100); | Sporobolomyces | 13 | Pathotroph-Saprotroph | \|Fungal Parasite\|-Plant Saprotroph | Yeast | NULL | Probable | As litter_saprotroph (Põlme et al. 2020); As mycoparasite (Põlme et al. 2020); Decay fungal material (Põlme et al. 2020) | Tedersoo L, et al. 2014. Science 346:e1256688 ((https://doi.org/10.1126/science.1256688)); Purahong W, et al. 2016. Molecular Ecology 25:4059-4074 ((https://doi.org/10.1111/mec.13739)); Põlme S, et al. 2020. Fungal Diversity 105:1-16 ((https://doi.org/10.1007/s13225-020-00466-2)) |
| Otu60994 | 1 | k__Fungi(100);p__Ascomycota(100);c__Saccharomycetes(100);o__Saccharomycetales(100);f__Metschnikowiaceae(100);g__Metschnikowia(100);s__Metschnikowia_pulcherrima(100); | Metschnikowia | 13 | Saprotroph | \|Undefined Saprotroph\| | Yeast | NULL | Probable | As nectar-tap_saprotroph (Põlme et al. 2020); Decay sugar-rich substrates (Põlme et al. 2020) | Põlme S, et al. 2020. Fungal Diversity 105:1-16 ((https://doi.org/10.1007/s13225-020-00466-2)) |
| Otu60995 | 1 | k__Fungi(100);p__Ascomycota(100);c__Saccharomycetes(100);o__Saccharomycetales(100);f__Metschnikowiaceae(100);g__Metschnikowia(100);g__Metschnikowia_unclassified(100); | Metschnikowia | 13 | Saprotroph | \|Undefined Saprotroph\| | Yeast | NULL | Probable | As nectar-tap_saprotroph (Põlme et al. 2020); Decay sugar-rich substrates (Põlme et al. 2020) | Põlme S, et al. 2020. Fungal Diversity 105:1-16 ((https://doi.org/10.1007/s13225-020-00466-2)) |
| Otu60996 | 1 | k__Fungi(100);p__Ascomycota(100);c__Saccharomycetes(100);o__Saccharomycetales(100);f__Metschnikowiaceae(100);g__Metschnikowia(100);s__Metschnikowia_pulcherrima(100); | Metschnikowia | 13 | Saprotroph | \|Undefined Saprotroph\| | Yeast | NULL | Probable | As nectar-tap_saprotroph (Põlme et al. 2020); Decay sugar-rich substrates (Põlme et al. 2020) | Põlme S, et al. 2020. Fungal Diversity 105:1-16 ((https://doi.org/10.1007/s13225-020-00466-2)) |
| Otu60997 | 1 | k__Fungi(100);p__Ascomycota(100);c__Saccharomycetes(100);o__Saccharomycetales(100);f__Metschnikowiaceae(100);g__Metschnikowia(100);g__Metschnikowia_unclassified(100); | Metschnikowia | 13 | Saprotroph | \|Undefined Saprotroph\| | Yeast | NULL | Probable | As nectar-tap_saprotroph (Põlme et al. 2020); Decay sugar-rich substrates (Põlme et al. 2020) | Põlme S, et al. 2020. Fungal Diversity 105:1-16 ((https://doi.org/10.1007/s13225-020-00466-2)) |
| Otu60998 | 1 | k__Fungi(100);p__Ascomycota(100);c__Saccharomycetes(100);o__Saccharomycetales(100);f__Metschnikowiaceae(100);g__Metschnikowia(100);s__Metschnikowia_pulcherrima(100); | Metschnikowia | 13 | Saprotroph | \|Undefined Saprotroph\| | Yeast | NULL | Probable | As nectar-tap_saprotroph (Põlme et al. 2020); Decay sugar-rich substrates (Põlme et al. 2020) | Põlme S, et al. 2020. Fungal Diversity 105:1-16 ((https://doi.org/10.1007/s13225-020-00466-2)) |
| Otu60999 | 1 | k__Fungi(100);p__Ascomycota(100);c__Saccharomycetes(100);o__Saccharomycetales(100);f__Metschnikowiaceae(100);g__Metschnikowia(100);s__Metschnikowia_pulcherrima(100); | Metschnikowia | 13 | Saprotroph | \|Undefined Saprotroph\| | Yeast | NULL | Probable | As nectar-tap_saprotroph (Põlme et al. 2020); Decay sugar-rich substrates (Põlme et al. 2020) | Põlme S, et al. 2020. Fungal Diversity 105:1-16 ((https://doi.org/10.1007/s13225-020-00466-2)) |
| Otu61000 | 1 | k__Fungi(100);p__Ascomycota(100);c__Saccharomycetes(100);o__Saccharomycetales(100);f__Metschnikowiaceae(100);g__Metschnikowia(100);s__Metschnikowia_pulcherrima(100); | Metschnikowia | 13 | Saprotroph | \|Undefined Saprotroph\| | Yeast | NULL | Probable | As nectar-tap_saprotroph (Põlme et al. 2020); Decay sugar-rich substrates (Põlme et al. 2020) | Põlme S, et al. 2020. Fungal Diversity 105:1-16 ((https://doi.org/10.1007/s13225-020-00466-2)) |
| Otu61001 | 1 | k__Fungi(100);p__Ascomycota(100);c__Saccharomycetes(100);o__Saccharomycetales(100);f__Metschnikowiaceae(100);g__Metschnikowia(100);s__Metschnikowia_pulcherrima(100); | Metschnikowia | 13 | Saprotroph | \|Undefined Saprotroph\| | Yeast | NULL | Probable | As nectar-tap_saprotroph (Põlme et al. 2020); Decay sugar-rich substrates (Põlme et al. 2020) | Põlme S, et al. 2020. Fungal Diversity 105:1-16 ((https://doi.org/10.1007/s13225-020-00466-2)) |
| Otu61002 | 1 | k__Fungi(100);p__Ascomycota(100);c__Saccharomycetes(100);o__Saccharomycetales(100);f__Metschnikowiaceae(100);g__Metschnikowia(100);s__Metschnikowia_pulcherrima(100); | Metschnikowia | 13 | Saprotroph | \|Undefined Saprotroph\| | Yeast | NULL | Probable | As nectar-tap_saprotroph (Põlme et al. 2020); Decay sugar-rich substrates (Põlme et al. 2020) | Põlme S, et al. 2020. Fungal Diversity 105:1-16 ((https://doi.org/10.1007/s13225-020-00466-2)) |
| Otu61003 | 1 | k__Fungi(100);p__Ascomycota(100);c__Saccharomycetes(100);o__Saccharomycetales(100);f__Metschnikowiaceae(100);g__Metschnikowia(100);g__Metschnikowia_unclassified(100); | Metschnikowia | 13 | Saprotroph | \|Undefined Saprotroph\| | Yeast | NULL | Probable | As nectar-tap_saprotroph (Põlme et al. 2020); Decay sugar-rich substrates (Põlme et al. 2020) | Põlme S, et al. 2020. Fungal Diversity 105:1-16 ((https://doi.org/10.1007/s13225-020-00466-2)) |
| Otu61004 | 1 | k__Fungi(100);p__Ascomycota(100);c__Saccharomycetes(100);o__Saccharomycetales(100);f__Metschnikowiaceae(100);g__Metschnikowia(100);s__Metschnikowia_pulcherrima(100); | Metschnikowia | 13 | Saprotroph | \|Undefined Saprotroph\| | Yeast | NULL | Probable | As nectar-tap_saprotroph (Põlme et al. 2020); Decay sugar-rich substrates (Põlme et al. 2020) | Põlme S, et al. 2020. Fungal Diversity 105:1-16 ((https://doi.org/10.1007/s13225-020-00466-2)) |
| Otu61005 | 1 | k__Fungi(100);p__Ascomycota(100);c__Saccharomycetes(100);o__Saccharomycetales(100);f__Metschnikowiaceae(100);g__Metschnikowia(100);s__Metschnikowia_pulcherrima(100); | Metschnikowia | 13 | Saprotroph | \|Undefined Saprotroph\| | Yeast | NULL | Probable | As nectar-tap_saprotroph (Põlme et al. 2020); Decay sugar-rich substrates (Põlme et al. 2020) | Põlme S, et al. 2020. Fungal Diversity 105:1-16 ((https://doi.org/10.1007/s13225-020-00466-2)) |
| Otu61006 | 1 | k__Fungi(100);p__Ascomycota(100);c__Saccharomycetes(100);o__Saccharomycetales(100);f__Metschnikowiaceae(100);g__Metschnikowia(100);g__Metschnikowia_unclassified(100); | Metschnikowia | 13 | Saprotroph | \|Undefined Saprotroph\| | Yeast | NULL | Probable | As nectar-tap_saprotroph (Põlme et al. 2020); Decay sugar-rich substrates (Põlme et al. 2020) | Põlme S, et al. 2020. Fungal Diversity 105:1-16 ((https://doi.org/10.1007/s13225-020-00466-2)) |
| Otu61007 | 1 | k__Fungi(100);p__Ascomycota(100);c__Saccharomycetes(100);o__Saccharomycetales(100);f__Metschnikowiaceae(100);g__Metschnikowia(100);s__Metschnikowia_pulcherrima(100); | Metschnikowia | 13 | Saprotroph | \|Undefined Saprotroph\| | Yeast | NULL | Probable | As nectar-tap_saprotroph (Põlme et al. 2020); Decay sugar-rich substrates (Põlme et al. 2020) | Põlme S, et al. 2020. Fungal Diversity 105:1-16 ((https://doi.org/10.1007/s13225-020-00466-2)) |
| Otu61008 | 1 | k__Fungi(100);p__Ascomycota(100);c__Saccharomycetes(100);o__Saccharomycetales(100);f__Metschnikowiaceae(100);g__Metschnikowia(100);s__Metschnikowia_pulcherrima(100); | Metschnikowia | 13 | Saprotroph | \|Undefined Saprotroph\| | Yeast | NULL | Probable | As nectar-tap_saprotroph (Põlme et al. 2020); Decay sugar-rich substrates (Põlme et al. 2020) | Põlme S, et al. 2020. Fungal Diversity 105:1-16 ((https://doi.org/10.1007/s13225-020-00466-2)) |
| Otu61009 | 1 | k__Fungi(100);p__Ascomycota(100);c__Saccharomycetes(100);o__Saccharomycetales(100);f__Metschnikowiaceae(100);g__Metschnikowia(100);s__Metschnikowia_pulcherrima(100); | Metschnikowia | 13 | Saprotroph | \|Undefined Saprotroph\| | Yeast | NULL | Probable | As nectar-tap_saprotroph (Põlme et al. 2020); Decay sugar-rich substrates (Põlme et al. 2020) | Põlme S, et al. 2020. Fungal Diversity 105:1-16 ((https://doi.org/10.1007/s13225-020-00466-2)) |
| Otu61010 | 1 | k__Fungi(100);p__Ascomycota(100);c__Saccharomycetes(100);o__Saccharomycetales(100);f__Metschnikowiaceae(100);g__Metschnikowia(100);s__Metschnikowia_pulcherrima(100); | Metschnikowia | 13 | Saprotroph | \|Undefined Saprotroph\| | Yeast | NULL | Probable | As nectar-tap_saprotroph (Põlme et al. 2020); Decay sugar-rich substrates (Põlme et al. 2020) | Põlme S, et al. 2020. Fungal Diversity 105:1-16 ((https://doi.org/10.1007/s13225-020-00466-2)) |
| Otu61011 | 1 | k__Fungi(100);p__Ascomycota(100);c__Saccharomycetes(100);o__Saccharomycetales(100);f__Metschnikowiaceae(100);g__Metschnikowia(100);s__Metschnikowia_pulcherrima(100); | Metschnikowia | 13 | Saprotroph | \|Undefined Saprotroph\| | Yeast | NULL | Probable | As nectar-tap_saprotroph (Põlme et al. 2020); Decay sugar-rich substrates (Põlme et al. 2020) | Põlme S, et al. 2020. Fungal Diversity 105:1-16 ((https://doi.org/10.1007/s13225-020-00466-2)) |
| Otu61012 | 1 | k__Fungi(100);p__Ascomycota(100);c__Saccharomycetes(100);o__Saccharomycetales(100);f__Metschnikowiaceae(100);g__Metschnikowia(100);s__Metschnikowia_pulcherrima(100); | Metschnikowia | 13 | Saprotroph | \|Undefined Saprotroph\| | Yeast | NULL | Probable | As nectar-tap_saprotroph (Põlme et al. 2020); Decay sugar-rich substrates (Põlme et al. 2020) | Põlme S, et al. 2020. Fungal Diversity 105:1-16 ((https://doi.org/10.1007/s13225-020-00466-2)) |
| Otu61013 | 1 | k__Fungi(100);p__Ascomycota(100);c__Saccharomycetes(100);o__Saccharomycetales(100);f__Metschnikowiaceae(100);g__Metschnikowia(100);s__Metschnikowia_pulcherrima(100); | Metschnikowia | 13 | Saprotroph | \|Undefined Saprotroph\| | Yeast | NULL | Probable | As nectar-tap_saprotroph (Põlme et al. 2020); Decay sugar-rich substrates (Põlme et al. 2020) | Põlme S, et al. 2020. Fungal Diversity 105:1-16 ((https://doi.org/10.1007/s13225-020-00466-2)) |
| Otu61014 | 1 | k__Fungi(100);p__Ascomycota(100);c__Saccharomycetes(100);o__Saccharomycetales(100);f__Metschnikowiaceae(100);g__Metschnikowia(100);s__Metschnikowia_pulcherrima(100); | Metschnikowia | 13 | Saprotroph | \|Undefined Saprotroph\| | Yeast | NULL | Probable | As nectar-tap_saprotroph (Põlme et al. 2020); Decay sugar-rich substrates (Põlme et al. 2020) | Põlme S, et al. 2020. Fungal Diversity 105:1-16 ((https://doi.org/10.1007/s13225-020-00466-2)) |
| Otu61015 | 1 | k__Fungi(100);p__Ascomycota(100);c__Saccharomycetes(100);o__Saccharomycetales(100);f__Metschnikowiaceae(100);g__Metschnikowia(100);g__Metschnikowia_unclassified(100); | Metschnikowia | 13 | Saprotroph | \|Undefined Saprotroph\| | Yeast | NULL | Probable | As nectar-tap_saprotroph (Põlme et al. 2020); Decay sugar-rich substrates (Põlme et al. 2020) | Põlme S, et al. 2020. Fungal Diversity 105:1-16 ((https://doi.org/10.1007/s13225-020-00466-2)) |
| Otu61016 | 1 | k__Fungi(100);p__Ascomycota(100);c__Saccharomycetes(100);o__Saccharomycetales(100);f__Metschnikowiaceae(100);g__Metschnikowia(100);g__Metschnikowia_unclassified(100); | Metschnikowia | 13 | Saprotroph | \|Undefined Saprotroph\| | Yeast | NULL | Probable | As nectar-tap_saprotroph (Põlme et al. 2020); Decay sugar-rich substrates (Põlme et al. 2020) | Põlme S, et al. 2020. Fungal Diversity 105:1-16 ((https://doi.org/10.1007/s13225-020-00466-2)) |
| Otu61017 | 1 | k__Fungi(100);p__Ascomycota(100);c__Saccharomycetes(100);o__Saccharomycetales(100);f__Metschnikowiaceae(100);g__Metschnikowia(100);g__Metschnikowia_unclassified(100); | Metschnikowia | 13 | Saprotroph | \|Undefined Saprotroph\| | Yeast | NULL | Probable | As nectar-tap_saprotroph (Põlme et al. 2020); Decay sugar-rich substrates (Põlme et al. 2020) | Põlme S, et al. 2020. Fungal Diversity 105:1-16 ((https://doi.org/10.1007/s13225-020-00466-2)) |
| Otu61018 | 1 | k__Fungi(100);p__Ascomycota(100);c__Saccharomycetes(100);o__Saccharomycetales(100);f__Metschnikowiaceae(100);g__Metschnikowia(100);s__Metschnikowia_pulcherrima(100); | Metschnikowia | 13 | Saprotroph | \|Undefined Saprotroph\| | Yeast | NULL | Probable | As nectar-tap_saprotroph (Põlme et al. 2020); Decay sugar-rich substrates (Põlme et al. 2020) | Põlme S, et al. 2020. Fungal Diversity 105:1-16 ((https://doi.org/10.1007/s13225-020-00466-2)) |
| Otu61019 | 1 | k__Fungi(100);p__Ascomycota(100);c__Saccharomycetes(100);o__Saccharomycetales(100);f__Metschnikowiaceae(100);g__Metschnikowia(100);g__Metschnikowia_unclassified(100); | Metschnikowia | 13 | Saprotroph | \|Undefined Saprotroph\| | Yeast | NULL | Probable | As nectar-tap_saprotroph (Põlme et al. 2020); Decay sugar-rich substrates (Põlme et al. 2020) | Põlme S, et al. 2020. Fungal Diversity 105:1-16 ((https://doi.org/10.1007/s13225-020-00466-2)) |
| Otu61020 | 1 | k__Fungi(100);p__Ascomycota(100);c__Saccharomycetes(100);o__Saccharomycetales(100);f__Metschnikowiaceae(100);g__Metschnikowia(100);s__Metschnikowia_pulcherrima(100); | Metschnikowia | 13 | Saprotroph | \|Undefined Saprotroph\| | Yeast | NULL | Probable | As nectar-tap_saprotroph (Põlme et al. 2020); Decay sugar-rich substrates (Põlme et al. 2020) | Põlme S, et al. 2020. Fungal Diversity 105:1-16 ((https://doi.org/10.1007/s13225-020-00466-2)) |
| Otu61022 | 1 | k__Fungi(100);p__Ascomycota(100);c__Saccharomycetes(100);o__Saccharomycetales(100);f__Metschnikowiaceae(100);g__Metschnikowia(100);s__Metschnikowia_pulcherrima(100); | Metschnikowia | 13 | Saprotroph | \|Undefined Saprotroph\| | Yeast | NULL | Probable | As nectar-tap_saprotroph (Põlme et al. 2020); Decay sugar-rich substrates (Põlme et al. 2020) | Põlme S, et al. 2020. Fungal Diversity 105:1-16 ((https://doi.org/10.1007/s13225-020-00466-2)) |
| Otu61023 | 1 | k__Fungi(100);p__Ascomycota(100);c__Saccharomycetes(100);o__Saccharomycetales(100);f__Metschnikowiaceae(100);g__Metschnikowia(100);g__Metschnikowia_unclassified(100); | Metschnikowia | 13 | Saprotroph | \|Undefined Saprotroph\| | Yeast | NULL | Probable | As nectar-tap_saprotroph (Põlme et al. 2020); Decay sugar-rich substrates (Põlme et al. 2020) | Põlme S, et al. 2020. Fungal Diversity 105:1-16 ((https://doi.org/10.1007/s13225-020-00466-2)) |
| Otu61024 | 1 | k__Fungi(100);p__Ascomycota(100);c__Saccharomycetes(100);o__Saccharomycetales(100);f__Metschnikowiaceae(100);g__Metschnikowia(100);s__Metschnikowia_pulcherrima(100); | Metschnikowia | 13 | Saprotroph | \|Undefined Saprotroph\| | Yeast | NULL | Probable | As nectar-tap_saprotroph (Põlme et al. 2020); Decay sugar-rich substrates (Põlme et al. 2020) | Põlme S, et al. 2020. Fungal Diversity 105:1-16 ((https://doi.org/10.1007/s13225-020-00466-2)) |
| Otu61025 | 1 | k__Fungi(100);p__Ascomycota(100);c__Saccharomycetes(100);o__Saccharomycetales(100);f__Metschnikowiaceae(100);g__Metschnikowia(100);s__Metschnikowia_pulcherrima(100); | Metschnikowia | 13 | Saprotroph | \|Undefined Saprotroph\| | Yeast | NULL | Probable | As nectar-tap_saprotroph (Põlme et al. 2020); Decay sugar-rich substrates (Põlme et al. 2020) | Põlme S, et al. 2020. Fungal Diversity 105:1-16 ((https://doi.org/10.1007/s13225-020-00466-2)) |
| Otu61026 | 1 | k__Fungi(100);p__Ascomycota(100);c__Saccharomycetes(100);o__Saccharomycetales(100);f__Metschnikowiaceae(100);g__Metschnikowia(100);s__Metschnikowia_pulcherrima(100); | Metschnikowia | 13 | Saprotroph | \|Undefined Saprotroph\| | Yeast | NULL | Probable | As nectar-tap_saprotroph (Põlme et al. 2020); Decay sugar-rich substrates (Põlme et al. 2020) | Põlme S, et al. 2020. Fungal Diversity 105:1-16 ((https://doi.org/10.1007/s13225-020-00466-2)) |
| Otu61027 | 1 | k__Fungi(100);p__Ascomycota(100);c__Saccharomycetes(100);o__Saccharomycetales(100);f__Metschnikowiaceae(100);g__Metschnikowia(100);s__Metschnikowia_pulcherrima(100); | Metschnikowia | 13 | Saprotroph | \|Undefined Saprotroph\| | Yeast | NULL | Probable | As nectar-tap_saprotroph (Põlme et al. 2020); Decay sugar-rich substrates (Põlme et al. 2020) | Põlme S, et al. 2020. Fungal Diversity 105:1-16 ((https://doi.org/10.1007/s13225-020-00466-2)) |
| Otu61028 | 1 | k__Fungi(100);p__Ascomycota(100);c__Saccharomycetes(100);o__Saccharomycetales(100);f__Metschnikowiaceae(100);g__Metschnikowia(100);s__Metschnikowia_pulcherrima(100); | Metschnikowia | 13 | Saprotroph | \|Undefined Saprotroph\| | Yeast | NULL | Probable | As nectar-tap_saprotroph (Põlme et al. 2020); Decay sugar-rich substrates (Põlme et al. 2020) | Põlme S, et al. 2020. Fungal Diversity 105:1-16 ((https://doi.org/10.1007/s13225-020-00466-2)) |
| Otu61029 | 1 | k__Fungi(100);p__Ascomycota(100);c__Saccharomycetes(100);o__Saccharomycetales(100);f__Metschnikowiaceae(100);g__Metschnikowia(100);s__Metschnikowia_pulcherrima(100); | Metschnikowia | 13 | Saprotroph | \|Undefined Saprotroph\| | Yeast | NULL | Probable | As nectar-tap_saprotroph (Põlme et al. 2020); Decay sugar-rich substrates (Põlme et al. 2020) | Põlme S, et al. 2020. Fungal Diversity 105:1-16 ((https://doi.org/10.1007/s13225-020-00466-2)) |
| Otu61030 | 1 | k__Fungi(100);p__Ascomycota(100);c__Saccharomycetes(100);o__Saccharomycetales(100);f__Metschnikowiaceae(100);g__Metschnikowia(100);s__Metschnikowia_pulcherrima(100); | Metschnikowia | 13 | Saprotroph | \|Undefined Saprotroph\| | Yeast | NULL | Probable | As nectar-tap_saprotroph (Põlme et al. 2020); Decay sugar-rich substrates (Põlme et al. 2020) | Põlme S, et al. 2020. Fungal Diversity 105:1-16 ((https://doi.org/10.1007/s13225-020-00466-2)) |
| Otu61031 | 1 | k__Fungi(100);p__Ascomycota(100);c__Saccharomycetes(100);o__Saccharomycetales(100);f__Metschnikowiaceae(100);g__Metschnikowia(100);g__Metschnikowia_unclassified(100); | Metschnikowia | 13 | Saprotroph | \|Undefined Saprotroph\| | Yeast | NULL | Probable | As nectar-tap_saprotroph (Põlme et al. 2020); Decay sugar-rich substrates (Põlme et al. 2020) | Põlme S, et al. 2020. Fungal Diversity 105:1-16 ((https://doi.org/10.1007/s13225-020-00466-2)) |
| Otu61032 | 1 | k__Fungi(100);p__Ascomycota(100);c__Saccharomycetes(100);o__Saccharomycetales(100);f__Metschnikowiaceae(100);g__Metschnikowia(100);s__Metschnikowia_pulcherrima(100); | Metschnikowia | 13 | Saprotroph | \|Undefined Saprotroph\| | Yeast | NULL | Probable | As nectar-tap_saprotroph (Põlme et al. 2020); Decay sugar-rich substrates (Põlme et al. 2020) | Põlme S, et al. 2020. Fungal Diversity 105:1-16 ((https://doi.org/10.1007/s13225-020-00466-2)) |
| Otu61033 | 1 | k__Fungi(100);p__Ascomycota(100);c__Saccharomycetes(100);o__Saccharomycetales(100);f__Metschnikowiaceae(100);g__Metschnikowia(100);s__Metschnikowia_pulcherrima(100); | Metschnikowia | 13 | Saprotroph | \|Undefined Saprotroph\| | Yeast | NULL | Probable | As nectar-tap_saprotroph (Põlme et al. 2020); Decay sugar-rich substrates (Põlme et al. 2020) | Põlme S, et al. 2020. Fungal Diversity 105:1-16 ((https://doi.org/10.1007/s13225-020-00466-2)) |
| Otu61034 | 1 | k__Fungi(100);p__Ascomycota(100);c__Saccharomycetes(100);o__Saccharomycetales(100);f__Metschnikowiaceae(100);g__Metschnikowia(100);s__Metschnikowia_pulcherrima(100); | Metschnikowia | 13 | Saprotroph | \|Undefined Saprotroph\| | Yeast | NULL | Probable | As nectar-tap_saprotroph (Põlme et al. 2020); Decay sugar-rich substrates (Põlme et al. 2020) | Põlme S, et al. 2020. Fungal Diversity 105:1-16 ((https://doi.org/10.1007/s13225-020-00466-2)) |
| Otu61035 | 1 | k__Fungi(100);p__Ascomycota(100);c__Saccharomycetes(100);o__Saccharomycetales(100);f__Metschnikowiaceae(100);g__Metschnikowia(100);s__Metschnikowia_pulcherrima(100); | Metschnikowia | 13 | Saprotroph | \|Undefined Saprotroph\| | Yeast | NULL | Probable | As nectar-tap_saprotroph (Põlme et al. 2020); Decay sugar-rich substrates (Põlme et al. 2020) | Põlme S, et al. 2020. Fungal Diversity 105:1-16 ((https://doi.org/10.1007/s13225-020-00466-2)) |
| Otu61036 | 1 | k__Fungi(100);p__Ascomycota(100);c__Saccharomycetes(100);o__Saccharomycetales(100);f__Metschnikowiaceae(100);g__Metschnikowia(100);g__Metschnikowia_unclassified(100); | Metschnikowia | 13 | Saprotroph | \|Undefined Saprotroph\| | Yeast | NULL | Probable | As nectar-tap_saprotroph (Põlme et al. 2020); Decay sugar-rich substrates (Põlme et al. 2020) | Põlme S, et al. 2020. Fungal Diversity 105:1-16 ((https://doi.org/10.1007/s13225-020-00466-2)) |
| Otu61037 | 1 | k__Fungi(100);p__Ascomycota(100);c__Saccharomycetes(100);o__Saccharomycetales(100);f__Metschnikowiaceae(100);g__Metschnikowia(100);s__Metschnikowia_pulcherrima(100); | Metschnikowia | 13 | Saprotroph | \|Undefined Saprotroph\| | Yeast | NULL | Probable | As nectar-tap_saprotroph (Põlme et al. 2020); Decay sugar-rich substrates (Põlme et al. 2020) | Põlme S, et al. 2020. Fungal Diversity 105:1-16 ((https://doi.org/10.1007/s13225-020-00466-2)) |
| Otu61038 | 1 | k__Fungi(100);p__Ascomycota(100);c__Saccharomycetes(100);o__Saccharomycetales(100);f__Metschnikowiaceae(100);g__Metschnikowia(100);g__Metschnikowia_unclassified(100); | Metschnikowia | 13 | Saprotroph | \|Undefined Saprotroph\| | Yeast | NULL | Probable | As nectar-tap_saprotroph (Põlme et al. 2020); Decay sugar-rich substrates (Põlme et al. 2020) | Põlme S, et al. 2020. Fungal Diversity 105:1-16 ((https://doi.org/10.1007/s13225-020-00466-2)) |
| Otu61039 | 1 | k__Fungi(100);p__Ascomycota(100);c__Saccharomycetes(100);o__Saccharomycetales(100);f__Metschnikowiaceae(100);g__Metschnikowia(100);s__Metschnikowia_pulcherrima(100); | Metschnikowia | 13 | Saprotroph | \|Undefined Saprotroph\| | Yeast | NULL | Probable | As nectar-tap_saprotroph (Põlme et al. 2020); Decay sugar-rich substrates (Põlme et al. 2020) | Põlme S, et al. 2020. Fungal Diversity 105:1-16 ((https://doi.org/10.1007/s13225-020-00466-2)) |
| Otu61040 | 1 | k__Fungi(100);p__Ascomycota(100);c__Saccharomycetes(100);o__Saccharomycetales(100);f__Metschnikowiaceae(100);g__Metschnikowia(100);s__Metschnikowia_pulcherrima(100); | Metschnikowia | 13 | Saprotroph | \|Undefined Saprotroph\| | Yeast | NULL | Probable | As nectar-tap_saprotroph (Põlme et al. 2020); Decay sugar-rich substrates (Põlme et al. 2020) | Põlme S, et al. 2020. Fungal Diversity 105:1-16 ((https://doi.org/10.1007/s13225-020-00466-2)) |
| Otu61041 | 1 | k__Fungi(100);p__Ascomycota(100);c__Saccharomycetes(100);o__Saccharomycetales(100);f__Metschnikowiaceae(100);g__Metschnikowia(100);s__Metschnikowia_pulcherrima(100); | Metschnikowia | 13 | Saprotroph | \|Undefined Saprotroph\| | Yeast | NULL | Probable | As nectar-tap_saprotroph (Põlme et al. 2020); Decay sugar-rich substrates (Põlme et al. 2020) | Põlme S, et al. 2020. Fungal Diversity 105:1-16 ((https://doi.org/10.1007/s13225-020-00466-2)) |
| Otu61042 | 1 | k__Fungi(100);p__Ascomycota(100);c__Saccharomycetes(100);o__Saccharomycetales(100);f__Metschnikowiaceae(100);g__Metschnikowia(100);g__Metschnikowia_unclassified(100); | Metschnikowia | 13 | Saprotroph | \|Undefined Saprotroph\| | Yeast | NULL | Probable | As nectar-tap_saprotroph (Põlme et al. 2020); Decay sugar-rich substrates (Põlme et al. 2020) | Põlme S, et al. 2020. Fungal Diversity 105:1-16 ((https://doi.org/10.1007/s13225-020-00466-2)) |
| Otu61043 | 1 | k__Fungi(100);p__Ascomycota(100);c__Saccharomycetes(100);o__Saccharomycetales(100);f__Metschnikowiaceae(100);g__Metschnikowia(100);s__Metschnikowia_pulcherrima(100); | Metschnikowia | 13 | Saprotroph | \|Undefined Saprotroph\| | Yeast | NULL | Probable | As nectar-tap_saprotroph (Põlme et al. 2020); Decay sugar-rich substrates (Põlme et al. 2020) | Põlme S, et al. 2020. Fungal Diversity 105:1-16 ((https://doi.org/10.1007/s13225-020-00466-2)) |
| Otu61044 | 1 | k__Fungi(100);p__Ascomycota(100);c__Saccharomycetes(100);o__Saccharomycetales(100);f__Metschnikowiaceae(100);g__Metschnikowia(100);s__Metschnikowia_pulcherrima(100); | Metschnikowia | 13 | Saprotroph | \|Undefined Saprotroph\| | Yeast | NULL | Probable | As nectar-tap_saprotroph (Põlme et al. 2020); Decay sugar-rich substrates (Põlme et al. 2020) | Põlme S, et al. 2020. Fungal Diversity 105:1-16 ((https://doi.org/10.1007/s13225-020-00466-2)) |
| Otu61046 | 1 | k__Fungi(100);p__Ascomycota(100);c__Saccharomycetes(100);o__Saccharomycetales(100);f__Metschnikowiaceae(100);g__Metschnikowia(100);g__Metschnikowia_unclassified(100); | Metschnikowia | 13 | Saprotroph | \|Undefined Saprotroph\| | Yeast | NULL | Probable | As nectar-tap_saprotroph (Põlme et al. 2020); Decay sugar-rich substrates (Põlme et al. 2020) | Põlme S, et al. 2020. Fungal Diversity 105:1-16 ((https://doi.org/10.1007/s13225-020-00466-2)) |
| Otu61047 | 1 | k__Fungi(100);p__Ascomycota(100);c__Saccharomycetes(100);o__Saccharomycetales(100);f__Metschnikowiaceae(100);g__Metschnikowia(100);s__Metschnikowia_pulcherrima(100); | Metschnikowia | 13 | Saprotroph | \|Undefined Saprotroph\| | Yeast | NULL | Probable | As nectar-tap_saprotroph (Põlme et al. 2020); Decay sugar-rich substrates (Põlme et al. 2020) | Põlme S, et al. 2020. Fungal Diversity 105:1-16 ((https://doi.org/10.1007/s13225-020-00466-2)) |
| Otu61048 | 1 | k__Fungi(100);p__Ascomycota(100);c__Saccharomycetes(100);o__Saccharomycetales(100);f__Metschnikowiaceae(100);g__Metschnikowia(100);g__Metschnikowia_unclassified(100); | Metschnikowia | 13 | Saprotroph | \|Undefined Saprotroph\| | Yeast | NULL | Probable | As nectar-tap_saprotroph (Põlme et al. 2020); Decay sugar-rich substrates (Põlme et al. 2020) | Põlme S, et al. 2020. Fungal Diversity 105:1-16 ((https://doi.org/10.1007/s13225-020-00466-2)) |
| Otu61049 | 1 | k__Fungi(100);p__Ascomycota(100);c__Saccharomycetes(100);o__Saccharomycetales(100);f__Metschnikowiaceae(100);g__Metschnikowia(100);s__Metschnikowia_pulcherrima(100); | Metschnikowia | 13 | Saprotroph | \|Undefined Saprotroph\| | Yeast | NULL | Probable | As nectar-tap_saprotroph (Põlme et al. 2020); Decay sugar-rich substrates (Põlme et al. 2020) | Põlme S, et al. 2020. Fungal Diversity 105:1-16 ((https://doi.org/10.1007/s13225-020-00466-2)) |
| Otu61050 | 1 | k__Fungi(100);p__Ascomycota(100);c__Saccharomycetes(100);o__Saccharomycetales(100);f__Metschnikowiaceae(100);g__Metschnikowia(100);s__Metschnikowia_pulcherrima(100); | Metschnikowia | 13 | Saprotroph | \|Undefined Saprotroph\| | Yeast | NULL | Probable | As nectar-tap_saprotroph (Põlme et al. 2020); Decay sugar-rich substrates (Põlme et al. 2020) | Põlme S, et al. 2020. Fungal Diversity 105:1-16 ((https://doi.org/10.1007/s13225-020-00466-2)) |
| Otu61051 | 1 | k__Fungi(100);p__Ascomycota(100);c__Saccharomycetes(100);o__Saccharomycetales(100);f__Metschnikowiaceae(100);g__Metschnikowia(100);s__Metschnikowia_pulcherrima(100); | Metschnikowia | 13 | Saprotroph | \|Undefined Saprotroph\| | Yeast | NULL | Probable | As nectar-tap_saprotroph (Põlme et al. 2020); Decay sugar-rich substrates (Põlme et al. 2020) | Põlme S, et al. 2020. Fungal Diversity 105:1-16 ((https://doi.org/10.1007/s13225-020-00466-2)) |
| Otu61052 | 1 | k__Fungi(100);p__Ascomycota(100);c__Saccharomycetes(100);o__Saccharomycetales(100);f__Metschnikowiaceae(100);g__Metschnikowia(100);s__Metschnikowia_pulcherrima(100); | Metschnikowia | 13 | Saprotroph | \|Undefined Saprotroph\| | Yeast | NULL | Probable | As nectar-tap_saprotroph (Põlme et al. 2020); Decay sugar-rich substrates (Põlme et al. 2020) | Põlme S, et al. 2020. Fungal Diversity 105:1-16 ((https://doi.org/10.1007/s13225-020-00466-2)) |
| Otu61053 | 1 | k__Fungi(100);p__Ascomycota(100);c__Saccharomycetes(100);o__Saccharomycetales(100);f__Metschnikowiaceae(100);g__Metschnikowia(100);s__Metschnikowia_pulcherrima(100); | Metschnikowia | 13 | Saprotroph | \|Undefined Saprotroph\| | Yeast | NULL | Probable | As nectar-tap_saprotroph (Põlme et al. 2020); Decay sugar-rich substrates (Põlme et al. 2020) | Põlme S, et al. 2020. Fungal Diversity 105:1-16 ((https://doi.org/10.1007/s13225-020-00466-2)) |
| Otu61054 | 1 | k__Fungi(100);p__Ascomycota(100);c__Saccharomycetes(100);o__Saccharomycetales(100);f__Metschnikowiaceae(100);g__Metschnikowia(100);s__Metschnikowia_pulcherrima(100); | Metschnikowia | 13 | Saprotroph | \|Undefined Saprotroph\| | Yeast | NULL | Probable | As nectar-tap_saprotroph (Põlme et al. 2020); Decay sugar-rich substrates (Põlme et al. 2020) | Põlme S, et al. 2020. Fungal Diversity 105:1-16 ((https://doi.org/10.1007/s13225-020-00466-2)) |
| Otu61055 | 1 | k__Fungi(100);p__Ascomycota(100);c__Saccharomycetes(100);o__Saccharomycetales(100);f__Metschnikowiaceae(100);g__Metschnikowia(100);g__Metschnikowia_unclassified(100); | Metschnikowia | 13 | Saprotroph | \|Undefined Saprotroph\| | Yeast | NULL | Probable | As nectar-tap_saprotroph (Põlme et al. 2020); Decay sugar-rich substrates (Põlme et al. 2020) | Põlme S, et al. 2020. Fungal Diversity 105:1-16 ((https://doi.org/10.1007/s13225-020-00466-2)) |
| Otu61056 | 1 | k__Fungi(100);p__Ascomycota(100);c__Saccharomycetes(100);o__Saccharomycetales(100);f__Metschnikowiaceae(100);g__Metschnikowia(100);s__Metschnikowia_pulcherrima(100); | Metschnikowia | 13 | Saprotroph | \|Undefined Saprotroph\| | Yeast | NULL | Probable | As nectar-tap_saprotroph (Põlme et al. 2020); Decay sugar-rich substrates (Põlme et al. 2020) | Põlme S, et al. 2020. Fungal Diversity 105:1-16 ((https://doi.org/10.1007/s13225-020-00466-2)) |
| Otu61058 | 1 | k__Fungi(100);p__Ascomycota(100);c__Saccharomycetes(100);o__Saccharomycetales(100);f__Metschnikowiaceae(100);g__Metschnikowia(100);g__Metschnikowia_unclassified(100); | Metschnikowia | 13 | Saprotroph | \|Undefined Saprotroph\| | Yeast | NULL | Probable | As nectar-tap_saprotroph (Põlme et al. 2020); Decay sugar-rich substrates (Põlme et al. 2020) | Põlme S, et al. 2020. Fungal Diversity 105:1-16 ((https://doi.org/10.1007/s13225-020-00466-2)) |
| Otu61059 | 1 | k__Fungi(100);p__Ascomycota(100);c__Saccharomycetes(100);o__Saccharomycetales(100);f__Metschnikowiaceae(100);g__Metschnikowia(100);s__Metschnikowia_pulcherrima(100); | Metschnikowia | 13 | Saprotroph | \|Undefined Saprotroph\| | Yeast | NULL | Probable | As nectar-tap_saprotroph (Põlme et al. 2020); Decay sugar-rich substrates (Põlme et al. 2020) | Põlme S, et al. 2020. Fungal Diversity 105:1-16 ((https://doi.org/10.1007/s13225-020-00466-2)) |
| Otu61060 | 1 | k__Fungi(100);p__Ascomycota(100);c__Saccharomycetes(100);o__Saccharomycetales(100);f__Metschnikowiaceae(100);g__Metschnikowia(100);s__Metschnikowia_pulcherrima(100); | Metschnikowia | 13 | Saprotroph | \|Undefined Saprotroph\| | Yeast | NULL | Probable | As nectar-tap_saprotroph (Põlme et al. 2020); Decay sugar-rich substrates (Põlme et al. 2020) | Põlme S, et al. 2020. Fungal Diversity 105:1-16 ((https://doi.org/10.1007/s13225-020-00466-2)) |
| Otu61061 | 1 | k__Fungi(100);p__Ascomycota(100);c__Saccharomycetes(100);o__Saccharomycetales(100);f__Metschnikowiaceae(100);g__Metschnikowia(100);s__Metschnikowia_pulcherrima(100); | Metschnikowia | 13 | Saprotroph | \|Undefined Saprotroph\| | Yeast | NULL | Probable | As nectar-tap_saprotroph (Põlme et al. 2020); Decay sugar-rich substrates (Põlme et al. 2020) | Põlme S, et al. 2020. Fungal Diversity 105:1-16 ((https://doi.org/10.1007/s13225-020-00466-2)) |
| Otu61062 | 1 | k__Fungi(100);p__Ascomycota(100);c__Saccharomycetes(100);o__Saccharomycetales(100);f__Metschnikowiaceae(100);g__Metschnikowia(100);g__Metschnikowia_unclassified(100); | Metschnikowia | 13 | Saprotroph | \|Undefined Saprotroph\| | Yeast | NULL | Probable | As nectar-tap_saprotroph (Põlme et al. 2020); Decay sugar-rich substrates (Põlme et al. 2020) | Põlme S, et al. 2020. Fungal Diversity 105:1-16 ((https://doi.org/10.1007/s13225-020-00466-2)) |
| Otu61063 | 1 | k__Fungi(100);p__Ascomycota(100);c__Saccharomycetes(100);o__Saccharomycetales(100);f__Metschnikowiaceae(100);g__Metschnikowia(100);s__Metschnikowia_pulcherrima(100); | Metschnikowia | 13 | Saprotroph | \|Undefined Saprotroph\| | Yeast | NULL | Probable | As nectar-tap_saprotroph (Põlme et al. 2020); Decay sugar-rich substrates (Põlme et al. 2020) | Põlme S, et al. 2020. Fungal Diversity 105:1-16 ((https://doi.org/10.1007/s13225-020-00466-2)) |
| Otu61064 | 1 | k__Fungi(100);p__Ascomycota(100);c__Saccharomycetes(100);o__Saccharomycetales(100);f__Metschnikowiaceae(100);g__Metschnikowia(100);s__Metschnikowia_pulcherrima(100); | Metschnikowia | 13 | Saprotroph | \|Undefined Saprotroph\| | Yeast | NULL | Probable | As nectar-tap_saprotroph (Põlme et al. 2020); Decay sugar-rich substrates (Põlme et al. 2020) | Põlme S, et al. 2020. Fungal Diversity 105:1-16 ((https://doi.org/10.1007/s13225-020-00466-2)) |
| Otu61065 | 1 | k__Fungi(100);p__Ascomycota(100);c__Saccharomycetes(100);o__Saccharomycetales(100);f__Metschnikowiaceae(100);g__Metschnikowia(100);s__Metschnikowia_pulcherrima(100); | Metschnikowia | 13 | Saprotroph | \|Undefined Saprotroph\| | Yeast | NULL | Probable | As nectar-tap_saprotroph (Põlme et al. 2020); Decay sugar-rich substrates (Põlme et al. 2020) | Põlme S, et al. 2020. Fungal Diversity 105:1-16 ((https://doi.org/10.1007/s13225-020-00466-2)) |
| Otu61066 | 1 | k__Fungi(100);p__Ascomycota(100);c__Saccharomycetes(100);o__Saccharomycetales(100);f__Metschnikowiaceae(100);g__Metschnikowia(100);s__Metschnikowia_pulcherrima(100); | Metschnikowia | 13 | Saprotroph | \|Undefined Saprotroph\| | Yeast | NULL | Probable | As nectar-tap_saprotroph (Põlme et al. 2020); Decay sugar-rich substrates (Põlme et al. 2020) | Põlme S, et al. 2020. Fungal Diversity 105:1-16 ((https://doi.org/10.1007/s13225-020-00466-2)) |
| Otu61067 | 1 | k__Fungi(100);p__Ascomycota(100);c__Saccharomycetes(100);o__Saccharomycetales(100);f__Metschnikowiaceae(100);g__Metschnikowia(100);g__Metschnikowia_unclassified(100); | Metschnikowia | 13 | Saprotroph | \|Undefined Saprotroph\| | Yeast | NULL | Probable | As nectar-tap_saprotroph (Põlme et al. 2020); Decay sugar-rich substrates (Põlme et al. 2020) | Põlme S, et al. 2020. Fungal Diversity 105:1-16 ((https://doi.org/10.1007/s13225-020-00466-2)) |
| Otu61069 | 1 | k__Fungi(100);p__Ascomycota(100);c__Saccharomycetes(100);o__Saccharomycetales(100);f__Metschnikowiaceae(100);g__Metschnikowia(100);g__Metschnikowia_unclassified(100); | Metschnikowia | 13 | Saprotroph | \|Undefined Saprotroph\| | Yeast | NULL | Probable | As nectar-tap_saprotroph (Põlme et al. 2020); Decay sugar-rich substrates (Põlme et al. 2020) | Põlme S, et al. 2020. Fungal Diversity 105:1-16 ((https://doi.org/10.1007/s13225-020-00466-2)) |
| Otu61070 | 1 | k__Fungi(100);p__Ascomycota(100);c__Saccharomycetes(100);o__Saccharomycetales(100);f__Metschnikowiaceae(100);g__Metschnikowia(100);s__Metschnikowia_pulcherrima(100); | Metschnikowia | 13 | Saprotroph | \|Undefined Saprotroph\| | Yeast | NULL | Probable | As nectar-tap_saprotroph (Põlme et al. 2020); Decay sugar-rich substrates (Põlme et al. 2020) | Põlme S, et al. 2020. Fungal Diversity 105:1-16 ((https://doi.org/10.1007/s13225-020-00466-2)) |
| Otu61071 | 1 | k__Fungi(100);p__Ascomycota(100);c__Saccharomycetes(100);o__Saccharomycetales(100);f__Metschnikowiaceae(100);g__Metschnikowia(100);s__Metschnikowia_pulcherrima(100); | Metschnikowia | 13 | Saprotroph | \|Undefined Saprotroph\| | Yeast | NULL | Probable | As nectar-tap_saprotroph (Põlme et al. 2020); Decay sugar-rich substrates (Põlme et al. 2020) | Põlme S, et al. 2020. Fungal Diversity 105:1-16 ((https://doi.org/10.1007/s13225-020-00466-2)) |
| Otu61072 | 1 | k__Fungi(100);p__Ascomycota(100);c__Saccharomycetes(100);o__Saccharomycetales(100);f__Metschnikowiaceae(100);g__Metschnikowia(100);s__Metschnikowia_pulcherrima(100); | Metschnikowia | 13 | Saprotroph | \|Undefined Saprotroph\| | Yeast | NULL | Probable | As nectar-tap_saprotroph (Põlme et al. 2020); Decay sugar-rich substrates (Põlme et al. 2020) | Põlme S, et al. 2020. Fungal Diversity 105:1-16 ((https://doi.org/10.1007/s13225-020-00466-2)) |
| Otu61074 | 1 | k__Fungi(100);p__Ascomycota(100);c__Saccharomycetes(100);o__Saccharomycetales(100);f__Metschnikowiaceae(100);g__Metschnikowia(100);s__Metschnikowia_pulcherrima(100); | Metschnikowia | 13 | Saprotroph | \|Undefined Saprotroph\| | Yeast | NULL | Probable | As nectar-tap_saprotroph (Põlme et al. 2020); Decay sugar-rich substrates (Põlme et al. 2020) | Põlme S, et al. 2020. Fungal Diversity 105:1-16 ((https://doi.org/10.1007/s13225-020-00466-2)) |
| Otu61075 | 1 | k__Fungi(100);p__Ascomycota(100);c__Saccharomycetes(100);o__Saccharomycetales(100);f__Metschnikowiaceae(100);g__Metschnikowia(100);s__Metschnikowia_pulcherrima(100); | Metschnikowia | 13 | Saprotroph | \|Undefined Saprotroph\| | Yeast | NULL | Probable | As nectar-tap_saprotroph (Põlme et al. 2020); Decay sugar-rich substrates (Põlme et al. 2020) | Põlme S, et al. 2020. Fungal Diversity 105:1-16 ((https://doi.org/10.1007/s13225-020-00466-2)) |
| Otu61076 | 1 | k__Fungi(100);p__Ascomycota(100);c__Saccharomycetes(100);o__Saccharomycetales(100);f__Metschnikowiaceae(100);g__Metschnikowia(100);s__Metschnikowia_pulcherrima(100); | Metschnikowia | 13 | Saprotroph | \|Undefined Saprotroph\| | Yeast | NULL | Probable | As nectar-tap_saprotroph (Põlme et al. 2020); Decay sugar-rich substrates (Põlme et al. 2020) | Põlme S, et al. 2020. Fungal Diversity 105:1-16 ((https://doi.org/10.1007/s13225-020-00466-2)) |
| Otu61077 | 1 | k__Fungi(100);p__Ascomycota(100);c__Saccharomycetes(100);o__Saccharomycetales(100);f__Metschnikowiaceae(100);g__Metschnikowia(100);s__Metschnikowia_pulcherrima(100); | Metschnikowia | 13 | Saprotroph | \|Undefined Saprotroph\| | Yeast | NULL | Probable | As nectar-tap_saprotroph (Põlme et al. 2020); Decay sugar-rich substrates (Põlme et al. 2020) | Põlme S, et al. 2020. Fungal Diversity 105:1-16 ((https://doi.org/10.1007/s13225-020-00466-2)) |
| Otu61078 | 1 | k__Fungi(100);p__Ascomycota(100);c__Saccharomycetes(100);o__Saccharomycetales(100);f__Metschnikowiaceae(100);g__Metschnikowia(100);g__Metschnikowia_unclassified(100); | Metschnikowia | 13 | Saprotroph | \|Undefined Saprotroph\| | Yeast | NULL | Probable | As nectar-tap_saprotroph (Põlme et al. 2020); Decay sugar-rich substrates (Põlme et al. 2020) | Põlme S, et al. 2020. Fungal Diversity 105:1-16 ((https://doi.org/10.1007/s13225-020-00466-2)) |
| Otu61079 | 1 | k__Fungi(100);p__Ascomycota(100);c__Saccharomycetes(100);o__Saccharomycetales(100);f__Metschnikowiaceae(100);g__Metschnikowia(100);g__Metschnikowia_unclassified(100); | Metschnikowia | 13 | Saprotroph | \|Undefined Saprotroph\| | Yeast | NULL | Probable | As nectar-tap_saprotroph (Põlme et al. 2020); Decay sugar-rich substrates (Põlme et al. 2020) | Põlme S, et al. 2020. Fungal Diversity 105:1-16 ((https://doi.org/10.1007/s13225-020-00466-2)) |
| Otu61080 | 1 | k__Fungi(100);p__Ascomycota(100);c__Saccharomycetes(100);o__Saccharomycetales(100);f__Metschnikowiaceae(100);g__Metschnikowia(100);g__Metschnikowia_unclassified(100); | Metschnikowia | 13 | Saprotroph | \|Undefined Saprotroph\| | Yeast | NULL | Probable | As nectar-tap_saprotroph (Põlme et al. 2020); Decay sugar-rich substrates (Põlme et al. 2020) | Põlme S, et al. 2020. Fungal Diversity 105:1-16 ((https://doi.org/10.1007/s13225-020-00466-2)) |
| Otu61081 | 1 | k__Fungi(100);p__Ascomycota(100);c__Saccharomycetes(100);o__Saccharomycetales(100);f__Metschnikowiaceae(100);g__Metschnikowia(100);g__Metschnikowia_unclassified(100); | Metschnikowia | 13 | Saprotroph | \|Undefined Saprotroph\| | Yeast | NULL | Probable | As nectar-tap_saprotroph (Põlme et al. 2020); Decay sugar-rich substrates (Põlme et al. 2020) | Põlme S, et al. 2020. Fungal Diversity 105:1-16 ((https://doi.org/10.1007/s13225-020-00466-2)) |
| Otu61082 | 1 | k__Fungi(100);p__Ascomycota(100);c__Saccharomycetes(100);o__Saccharomycetales(100);f__Metschnikowiaceae(100);g__Metschnikowia(100);s__Metschnikowia_pulcherrima(100); | Metschnikowia | 13 | Saprotroph | \|Undefined Saprotroph\| | Yeast | NULL | Probable | As nectar-tap_saprotroph (Põlme et al. 2020); Decay sugar-rich substrates (Põlme et al. 2020) | Põlme S, et al. 2020. Fungal Diversity 105:1-16 ((https://doi.org/10.1007/s13225-020-00466-2)) |
| Otu61083 | 1 | k__Fungi(100);p__Ascomycota(100);c__Saccharomycetes(100);o__Saccharomycetales(100);f__Metschnikowiaceae(100);g__Metschnikowia(100);s__Metschnikowia_pulcherrima(100); | Metschnikowia | 13 | Saprotroph | \|Undefined Saprotroph\| | Yeast | NULL | Probable | As nectar-tap_saprotroph (Põlme et al. 2020); Decay sugar-rich substrates (Põlme et al. 2020) | Põlme S, et al. 2020. Fungal Diversity 105:1-16 ((https://doi.org/10.1007/s13225-020-00466-2)) |
| Otu61084 | 1 | k__Fungi(100);p__Ascomycota(100);c__Saccharomycetes(100);o__Saccharomycetales(100);f__Metschnikowiaceae(100);g__Metschnikowia(100);g__Metschnikowia_unclassified(100); | Metschnikowia | 13 | Saprotroph | \|Undefined Saprotroph\| | Yeast | NULL | Probable | As nectar-tap_saprotroph (Põlme et al. 2020); Decay sugar-rich substrates (Põlme et al. 2020) | Põlme S, et al. 2020. Fungal Diversity 105:1-16 ((https://doi.org/10.1007/s13225-020-00466-2)) |
| Otu61085 | 1 | k__Fungi(100);p__Ascomycota(100);c__Saccharomycetes(100);o__Saccharomycetales(100);f__Metschnikowiaceae(100);g__Metschnikowia(100);s__Metschnikowia_pulcherrima(100); | Metschnikowia | 13 | Saprotroph | \|Undefined Saprotroph\| | Yeast | NULL | Probable | As nectar-tap_saprotroph (Põlme et al. 2020); Decay sugar-rich substrates (Põlme et al. 2020) | Põlme S, et al. 2020. Fungal Diversity 105:1-16 ((https://doi.org/10.1007/s13225-020-00466-2)) |
| Otu61086 | 1 | k__Fungi(100);p__Ascomycota(100);c__Saccharomycetes(100);o__Saccharomycetales(100);f__Metschnikowiaceae(100);g__Metschnikowia(100);s__Metschnikowia_pulcherrima(100); | Metschnikowia | 13 | Saprotroph | \|Undefined Saprotroph\| | Yeast | NULL | Probable | As nectar-tap_saprotroph (Põlme et al. 2020); Decay sugar-rich substrates (Põlme et al. 2020) | Põlme S, et al. 2020. Fungal Diversity 105:1-16 ((https://doi.org/10.1007/s13225-020-00466-2)) |
| Otu61087 | 1 | k__Fungi(100);p__Ascomycota(100);c__Saccharomycetes(100);o__Saccharomycetales(100);f__Metschnikowiaceae(100);g__Metschnikowia(100);s__Metschnikowia_pulcherrima(100); | Metschnikowia | 13 | Saprotroph | \|Undefined Saprotroph\| | Yeast | NULL | Probable | As nectar-tap_saprotroph (Põlme et al. 2020); Decay sugar-rich substrates (Põlme et al. 2020) | Põlme S, et al. 2020. Fungal Diversity 105:1-16 ((https://doi.org/10.1007/s13225-020-00466-2)) |
| Otu61088 | 1 | k__Fungi(100);p__Ascomycota(100);c__Saccharomycetes(100);o__Saccharomycetales(100);f__Metschnikowiaceae(100);g__Metschnikowia(100);s__Metschnikowia_pulcherrima(100); | Metschnikowia | 13 | Saprotroph | \|Undefined Saprotroph\| | Yeast | NULL | Probable | As nectar-tap_saprotroph (Põlme et al. 2020); Decay sugar-rich substrates (Põlme et al. 2020) | Põlme S, et al. 2020. Fungal Diversity 105:1-16 ((https://doi.org/10.1007/s13225-020-00466-2)) |
| Otu61089 | 1 | k__Fungi(100);p__Ascomycota(100);c__Saccharomycetes(100);o__Saccharomycetales(100);f__Metschnikowiaceae(100);g__Metschnikowia(100);s__Metschnikowia_pulcherrima(100); | Metschnikowia | 13 | Saprotroph | \|Undefined Saprotroph\| | Yeast | NULL | Probable | As nectar-tap_saprotroph (Põlme et al. 2020); Decay sugar-rich substrates (Põlme et al. 2020) | Põlme S, et al. 2020. Fungal Diversity 105:1-16 ((https://doi.org/10.1007/s13225-020-00466-2)) |
| Otu61090 | 1 | k__Fungi(100);p__Ascomycota(100);c__Saccharomycetes(100);o__Saccharomycetales(100);f__Metschnikowiaceae(100);g__Metschnikowia(100);s__Metschnikowia_pulcherrima(100); | Metschnikowia | 13 | Saprotroph | \|Undefined Saprotroph\| | Yeast | NULL | Probable | As nectar-tap_saprotroph (Põlme et al. 2020); Decay sugar-rich substrates (Põlme et al. 2020) | Põlme S, et al. 2020. Fungal Diversity 105:1-16 ((https://doi.org/10.1007/s13225-020-00466-2)) |
| Otu61091 | 1 | k__Fungi(100);p__Ascomycota(100);c__Saccharomycetes(100);o__Saccharomycetales(100);f__Metschnikowiaceae(100);g__Metschnikowia(100);s__Metschnikowia_pulcherrima(100); | Metschnikowia | 13 | Saprotroph | \|Undefined Saprotroph\| | Yeast | NULL | Probable | As nectar-tap_saprotroph (Põlme et al. 2020); Decay sugar-rich substrates (Põlme et al. 2020) | Põlme S, et al. 2020. Fungal Diversity 105:1-16 ((https://doi.org/10.1007/s13225-020-00466-2)) |
| Otu61092 | 1 | k__Fungi(100);p__Ascomycota(100);c__Saccharomycetes(100);o__Saccharomycetales(100);f__Metschnikowiaceae(100);g__Metschnikowia(100);g__Metschnikowia_unclassified(100); | Metschnikowia | 13 | Saprotroph | \|Undefined Saprotroph\| | Yeast | NULL | Probable | As nectar-tap_saprotroph (Põlme et al. 2020); Decay sugar-rich substrates (Põlme et al. 2020) | Põlme S, et al. 2020. Fungal Diversity 105:1-16 ((https://doi.org/10.1007/s13225-020-00466-2)) |
| Otu61093 | 1 | k__Fungi(100);p__Ascomycota(100);c__Saccharomycetes(100);o__Saccharomycetales(100);f__Metschnikowiaceae(100);g__Metschnikowia(100);s__Metschnikowia_pulcherrima(100); | Metschnikowia | 13 | Saprotroph | \|Undefined Saprotroph\| | Yeast | NULL | Probable | As nectar-tap_saprotroph (Põlme et al. 2020); Decay sugar-rich substrates (Põlme et al. 2020) | Põlme S, et al. 2020. Fungal Diversity 105:1-16 ((https://doi.org/10.1007/s13225-020-00466-2)) |
| Otu61094 | 1 | k__Fungi(100);p__Ascomycota(100);c__Saccharomycetes(100);o__Saccharomycetales(100);f__Metschnikowiaceae(100);g__Metschnikowia(100);s__Metschnikowia_pulcherrima(100); | Metschnikowia | 13 | Saprotroph | \|Undefined Saprotroph\| | Yeast | NULL | Probable | As nectar-tap_saprotroph (Põlme et al. 2020); Decay sugar-rich substrates (Põlme et al. 2020) | Põlme S, et al. 2020. Fungal Diversity 105:1-16 ((https://doi.org/10.1007/s13225-020-00466-2)) |

| | | | | | | | | | | | |
|---|---|---|---|---|---|---|---|---|---|---|---|
| Otu61095 | 1 | k__Fungi(100);p__Ascomycota(100);c__Saccharomycetes(100);o__Saccharomycetales(100);f__Metschnikowiaceae(100);g__Metschnikowia(100);g__Metschnikowia_unclassified(100); | Metschnikowia | 13 | Saprotroph | \|Undefined Saprotroph\| | Yeast | NULL | Probable | As nectar-tap_saprotroph (Põlme et al. 2020); Decay sugar-rich substrates (Põlme et al. 2020) | Põlme S, et al. 2020. Fungal Diversity 105:1-16 ((https://doi.org/10.1007/s13225-020-00466-2)) |
| Otu61096 | 1 | k__Fungi(100);p__Ascomycota(100);c__Saccharomycetes(100);o__Saccharomycetales(100);f__Metschnikowiaceae(100);g__Metschnikowia(100);g__Metschnikowia_unclassified(100); | Metschnikowia | 13 | Saprotroph | \|Undefined Saprotroph\| | Yeast | NULL | Probable | As nectar-tap_saprotroph (Põlme et al. 2020); Decay sugar-rich substrates (Põlme et al. 2020) | Põlme S, et al. 2020. Fungal Diversity 105:1-16 ((https://doi.org/10.1007/s13225-020-00466-2)) |
| Otu61097 | 1 | k__Fungi(100);p__Ascomycota(100);c__Saccharomycetes(100);o__Saccharomycetales(100);f__Metschnikowiaceae(100);g__Metschnikowia(100);s__Metschnikowia_pulcherrima(100); | Metschnikowia | 13 | Saprotroph | \|Undefined Saprotroph\| | Yeast | NULL | Probable | As nectar-tap_saprotroph (Põlme et al. 2020); Decay sugar-rich substrates (Põlme et al. 2020) | Põlme S, et al. 2020. Fungal Diversity 105:1-16 ((https://doi.org/10.1007/s13225-020-00466-2)) |
| Otu61098 | 1 | k__Fungi(100);p__Ascomycota(100);c__Saccharomycetes(100);o__Saccharomycetales(100);f__Metschnikowiaceae(100);g__Metschnikowia(100);g__Metschnikowia_unclassified(100); | Metschnikowia | 13 | Saprotroph | \|Undefined Saprotroph\| | Yeast | NULL | Probable | As nectar-tap_saprotroph (Põlme et al. 2020); Decay sugar-rich substrates (Põlme et al. 2020) | Põlme S, et al. 2020. Fungal Diversity 105:1-16 ((https://doi.org/10.1007/s13225-020-00466-2)) |
| Otu61099 | 1 | k__Fungi(100);p__Ascomycota(100);c__Saccharomycetes(100);o__Saccharomycetales(100);f__Metschnikowiaceae(100);g__Metschnikowia(100);g__Metschnikowia_unclassified(100); | Metschnikowia | 13 | Saprotroph | \|Undefined Saprotroph\| | Yeast | NULL | Probable | As nectar-tap_saprotroph (Põlme et al. 2020); Decay sugar-rich substrates (Põlme et al. 2020) | Põlme S, et al. 2020. Fungal Diversity 105:1-16 ((https://doi.org/10.1007/s13225-020-00466-2)) |
| Otu61100 | 1 | k__Fungi(100);p__Ascomycota(100);c__Saccharomycetes(100);o__Saccharomycetales(100);f__Metschnikowiaceae(100);g__Metschnikowia(100);g__Metschnikowia_unclassified(100); | Metschnikowia | 13 | Saprotroph | \|Undefined Saprotroph\| | Yeast | NULL | Probable | As nectar-tap_saprotroph (Põlme et al. 2020); Decay sugar-rich substrates (Põlme et al. 2020) | Põlme S, et al. 2020. Fungal Diversity 105:1-16 ((https://doi.org/10.1007/s13225-020-00466-2)) |
| Otu61101 | 1 | k__Fungi(100);p__Ascomycota(100);c__Saccharomycetes(100);o__Saccharomycetales(100);f__Metschnikowiaceae(100);g__Metschnikowia(100);s__Metschnikowia_pulcherrima(100); | Metschnikowia | 13 | Saprotroph | \|Undefined Saprotroph\| | Yeast | NULL | Probable | As nectar-tap_saprotroph (Põlme et al. 2020); Decay sugar-rich substrates (Põlme et al. 2020) | Põlme S, et al. 2020. Fungal Diversity 105:1-16 ((https://doi.org/10.1007/s13225-020-00466-2)) |
| Otu61102 | 1 | k__Fungi(100);p__Ascomycota(100);c__Saccharomycetes(100);o__Saccharomycetales(100);f__Metschnikowiaceae(100);g__Metschnikowia(100);s__Metschnikowia_pulcherrima(100); | Metschnikowia | 13 | Saprotroph | \|Undefined Saprotroph\| | Yeast | NULL | Probable | As nectar-tap_saprotroph (Põlme et al. 2020); Decay sugar-rich substrates (Põlme et al. 2020) | Põlme S, et al. 2020. Fungal Diversity 105:1-16 ((https://doi.org/10.1007/s13225-020-00466-2)) |
| Otu61103 | 1 | k__Fungi(100);p__Ascomycota(100);c__Saccharomycetes(100);o__Saccharomycetales(100);f__Metschnikowiaceae(100);g__Metschnikowia(100);g__Metschnikowia_unclassified(100); | Metschnikowia | 13 | Saprotroph | \|Undefined Saprotroph\| | Yeast | NULL | Probable | As nectar-tap_saprotroph (Põlme et al. 2020); Decay sugar-rich substrates (Põlme et al. 2020) | Põlme S, et al. 2020. Fungal Diversity 105:1-16 ((https://doi.org/10.1007/s13225-020-00466-2)) |
| Otu61104 | 1 | k__Fungi(100);p__Ascomycota(100);c__Saccharomycetes(100);o__Saccharomycetales(100);f__Metschnikowiaceae(100);g__Metschnikowia(100);s__Metschnikowia_pulcherrima(100); | Metschnikowia | 13 | Saprotroph | \|Undefined Saprotroph\| | Yeast | NULL | Probable | As nectar-tap_saprotroph (Põlme et al. 2020); Decay sugar-rich substrates (Põlme et al. 2020) | Põlme S, et al. 2020. Fungal Diversity 105:1-16 ((https://doi.org/10.1007/s13225-020-00466-2)) |
| Otu61105 | 1 | k__Fungi(100);p__Ascomycota(100);c__Saccharomycetes(100);o__Saccharomycetales(100);f__Metschnikowiaceae(100);g__Metschnikowia(100);s__Metschnikowia_pulcherrima(100); | Metschnikowia | 13 | Saprotroph | \|Undefined Saprotroph\| | Yeast | NULL | Probable | As nectar-tap_saprotroph (Põlme et al. 2020); Decay sugar-rich substrates (Põlme et al. 2020) | Põlme S, et al. 2020. Fungal Diversity 105:1-16 ((https://doi.org/10.1007/s13225-020-00466-2)) |
| Otu61106 | 1 | k__Fungi(100);p__Ascomycota(100);c__Saccharomycetes(100);o__Saccharomycetales(100);f__Metschnikowiaceae(100);g__Metschnikowia(100);s__Metschnikowia_pulcherrima(100); | Metschnikowia | 13 | Saprotroph | \|Undefined Saprotroph\| | Yeast | NULL | Probable | As nectar-tap_saprotroph (Põlme et al. 2020); Decay sugar-rich substrates (Põlme et al. 2020) | Põlme S, et al. 2020. Fungal Diversity 105:1-16 ((https://doi.org/10.1007/s13225-020-00466-2)) |
| Otu61107 | 1 | k__Fungi(100);p__Ascomycota(100);c__Saccharomycetes(100);o__Saccharomycetales(100);f__Metschnikowiaceae(100);g__Metschnikowia(100);g__Metschnikowia_unclassified(100); | Metschnikowia | 13 | Saprotroph | \|Undefined Saprotroph\| | Yeast | NULL | Probable | As nectar-tap_saprotroph (Põlme et al. 2020); Decay sugar-rich substrates (Põlme et al. 2020) | Põlme S, et al. 2020. Fungal Diversity 105:1-16 ((https://doi.org/10.1007/s13225-020-00466-2)) |
| Otu61108 | 1 | k__Fungi(100);p__Ascomycota(100);c__Saccharomycetes(100);o__Saccharomycetales(100);f__Metschnikowiaceae(100);g__Metschnikowia(100);s__Metschnikowia_pulcherrima(100); | Metschnikowia | 13 | Saprotroph | \|Undefined Saprotroph\| | Yeast | NULL | Probable | As nectar-tap_saprotroph (Põlme et al. 2020); Decay sugar-rich substrates (Põlme et al. 2020) | Põlme S, et al. 2020. Fungal Diversity 105:1-16 ((https://doi.org/10.1007/s13225-020-00466-2)) |
| Otu61109 | 1 | k__Fungi(100);p__Ascomycota(100);c__Saccharomycetes(100);o__Saccharomycetales(100);f__Metschnikowiaceae(100);g__Metschnikowia(100);g__Metschnikowia_unclassified(100); | Metschnikowia | 13 | Saprotroph | \|Undefined Saprotroph\| | Yeast | NULL | Probable | As nectar-tap_saprotroph (Põlme et al. 2020); Decay sugar-rich substrates (Põlme et al. 2020) | Põlme S, et al. 2020. Fungal Diversity 105:1-16 ((https://doi.org/10.1007/s13225-020-00466-2)) |
| Otu61110 | 1 | k__Fungi(100);p__Ascomycota(100);c__Saccharomycetes(100);o__Saccharomycetales(100);f__Metschnikowiaceae(100);g__Metschnikowia(100);s__Metschnikowia_pulcherrima(100); | Metschnikowia | 13 | Saprotroph | \|Undefined Saprotroph\| | Yeast | NULL | Probable | As nectar-tap_saprotroph (Põlme et al. 2020); Decay sugar-rich substrates (Põlme et al. 2020) | Põlme S, et al. 2020. Fungal Diversity 105:1-16 ((https://doi.org/10.1007/s13225-020-00466-2)) |
| Otu61111 | 1 | k__Fungi(100);p__Ascomycota(100);c__Saccharomycetes(100);o__Saccharomycetales(100);f__Metschnikowiaceae(100);g__Metschnikowia(100);s__Metschnikowia_pulcherrima(100); | Metschnikowia | 13 | Saprotroph | \|Undefined Saprotroph\| | Yeast | NULL | Probable | As nectar-tap_saprotroph (Põlme et al. 2020); Decay sugar-rich substrates (Põlme et al. 2020) | Põlme S, et al. 2020. Fungal Diversity 105:1-16 ((https://doi.org/10.1007/s13225-020-00466-2)) |
| Otu61112 | 1 | k__Fungi(100);p__Ascomycota(100);c__Saccharomycetes(100);o__Saccharomycetales(100);f__Metschnikowiaceae(100);g__Metschnikowia(100);g__Metschnikowia_unclassified(100); | Metschnikowia | 13 | Saprotroph | \|Undefined Saprotroph\| | Yeast | NULL | Probable | As nectar-tap_saprotroph (Põlme et al. 2020); Decay sugar-rich substrates (Põlme et al. 2020) | Põlme S, et al. 2020. Fungal Diversity 105:1-16 ((https://doi.org/10.1007/s13225-020-00466-2)) |
| Otu61113 | 1 | k__Fungi(100);p__Ascomycota(100);c__Saccharomycetes(100);o__Saccharomycetales(100);f__Metschnikowiaceae(100);g__Metschnikowia(100);s__Metschnikowia_pulcherrima(100); | Metschnikowia | 13 | Saprotroph | \|Undefined Saprotroph\| | Yeast | NULL | Probable | As nectar-tap_saprotroph (Põlme et al. 2020); Decay sugar-rich substrates (Põlme et al. 2020) | Põlme S, et al. 2020. Fungal Diversity 105:1-16 ((https://doi.org/10.1007/s13225-020-00466-2)) |
| Otu61114 | 1 | k__Fungi(100);p__Ascomycota(100);c__Saccharomycetes(100);o__Saccharomycetales(100);f__Metschnikowiaceae(100);g__Metschnikowia(100);s__Metschnikowia_pulcherrima(100); | Metschnikowia | 13 | Saprotroph | \|Undefined Saprotroph\| | Yeast | NULL | Probable | As nectar-tap_saprotroph (Põlme et al. 2020); Decay sugar-rich substrates (Põlme et al. 2020) | Põlme S, et al. 2020. Fungal Diversity 105:1-16 ((https://doi.org/10.1007/s13225-020-00466-2)) |
| Otu61116 | 1 | k__Fungi(100);p__Ascomycota(100);c__Saccharomycetes(100);o__Saccharomycetales(100);f__Metschnikowiaceae(100);g__Metschnikowia(100);s__Metschnikowia_pulcherrima(100); | Metschnikowia | 13 | Saprotroph | \|Undefined Saprotroph\| | Yeast | NULL | Probable | As nectar-tap_saprotroph (Põlme et al. 2020); Decay sugar-rich substrates (Põlme et al. 2020) | Põlme S, et al. 2020. Fungal Diversity 105:1-16 ((https://doi.org/10.1007/s13225-020-00466-2)) |
| Otu61117 | 1 | k__Fungi(100);p__Ascomycota(100);c__Saccharomycetes(100);o__Saccharomycetales(100);f__Metschnikowiaceae(100);g__Metschnikowia(100);s__Metschnikowia_pulcherrima(100); | Metschnikowia | 13 | Saprotroph | \|Undefined Saprotroph\| | Yeast | NULL | Probable | As nectar-tap_saprotroph (Põlme et al. 2020); Decay sugar-rich substrates (Põlme et al. 2020) | Põlme S, et al. 2020. Fungal Diversity 105:1-16 ((https://doi.org/10.1007/s13225-020-00466-2)) |
| Otu61118 | 1 | k__Fungi(100);p__Ascomycota(100);c__Saccharomycetes(100);o__Saccharomycetales(100);f__Metschnikowiaceae(100);g__Metschnikowia(100);s__Metschnikowia_pulcherrima(100); | Metschnikowia | 13 | Saprotroph | \|Undefined Saprotroph\| | Yeast | NULL | Probable | As nectar-tap_saprotroph (Põlme et al. 2020); Decay sugar-rich substrates (Põlme et al. 2020) | Põlme S, et al. 2020. Fungal Diversity 105:1-16 ((https://doi.org/10.1007/s13225-020-00466-2)) |
| Otu61119 | 1 | k__Fungi(100);p__Ascomycota(100);c__Saccharomycetes(100);o__Saccharomycetales(100);f__Metschnikowiaceae(100);g__Metschnikowia(100);s__Metschnikowia_pulcherrima(100); | Metschnikowia | 13 | Saprotroph | \|Undefined Saprotroph\| | Yeast | NULL | Probable | As nectar-tap_saprotroph (Põlme et al. 2020); Decay sugar-rich substrates (Põlme et al. 2020) | Põlme S, et al. 2020. Fungal Diversity 105:1-16 ((https://doi.org/10.1007/s13225-020-00466-2)) |
| Otu61120 | 1 | k__Fungi(100);p__Ascomycota(100);c__Saccharomycetes(100);o__Saccharomycetales(100);f__Metschnikowiaceae(100);g__Metschnikowia(100);g__Metschnikowia_unclassified(100); | Metschnikowia | 13 | Saprotroph | \|Undefined Saprotroph\| | Yeast | NULL | Probable | As nectar-tap_saprotroph (Põlme et al. 2020); Decay sugar-rich substrates (Põlme et al. 2020) | Põlme S, et al. 2020. Fungal Diversity 105:1-16 ((https://doi.org/10.1007/s13225-020-00466-2)) |
| Otu61121 | 1 | k__Fungi(100);p__Ascomycota(100);c__Saccharomycetes(100);o__Saccharomycetales(100);f__Metschnikowiaceae(100);g__Metschnikowia(100);s__Metschnikowia_pulcherrima(100); | Metschnikowia | 13 | Saprotroph | \|Undefined Saprotroph\| | Yeast | NULL | Probable | As nectar-tap_saprotroph (Põlme et al. 2020); Decay sugar-rich substrates (Põlme et al. 2020) | Põlme S, et al. 2020. Fungal Diversity 105:1-16 ((https://doi.org/10.1007/s13225-020-00466-2)) |
| Otu61122 | 1 | k__Fungi(100);p__Ascomycota(100);c__Saccharomycetes(100);o__Saccharomycetales(100);f__Metschnikowiaceae(100);g__Metschnikowia(100);g__Metschnikowia_unclassified(100); | Metschnikowia | 13 | Saprotroph | \|Undefined Saprotroph\| | Yeast | NULL | Probable | As nectar-tap_saprotroph (Põlme et al. 2020); Decay sugar-rich substrates (Põlme et al. 2020) | Põlme S, et al. 2020. Fungal Diversity 105:1-16 ((https://doi.org/10.1007/s13225-020-00466-2)) |
| Otu61123 | 1 | k__Fungi(100);p__Ascomycota(100);c__Saccharomycetes(100);o__Saccharomycetales(100);f__Metschnikowiaceae(100);g__Metschnikowia(100);s__Metschnikowia_pulcherrima(100); | Metschnikowia | 13 | Saprotroph | \|Undefined Saprotroph\| | Yeast | NULL | Probable | As nectar-tap_saprotroph (Põlme et al. 2020); Decay sugar-rich substrates (Põlme et al. 2020) | Põlme S, et al. 2020. Fungal Diversity 105:1-16 ((https://doi.org/10.1007/s13225-020-00466-2)) |
| Otu61126 | 1 | k__Fungi(100);p__Ascomycota(100);c__Saccharomycetes(100);o__Saccharomycetales(100);f__Metschnikowiaceae(100);g__Metschnikowia(100);s__Metschnikowia_pulcherrima(100); | Metschnikowia | 13 | Saprotroph | \|Undefined Saprotroph\| | Yeast | NULL | Probable | As nectar-tap_saprotroph (Põlme et al. 2020); Decay sugar-rich substrates (Põlme et al. 2020) | Põlme S, et al. 2020. Fungal Diversity 105:1-16 ((https://doi.org/10.1007/s13225-020-00466-2)) |
| Otu61127 | 1 | k__Fungi(100);p__Ascomycota(100);c__Saccharomycetes(100);o__Saccharomycetales(100);f__Metschnikowiaceae(100);g__Metschnikowia(100);g__Metschnikowia_unclassified(100); | Metschnikowia | 13 | Saprotroph | \|Undefined Saprotroph\| | Yeast | NULL | Probable | As nectar-tap_saprotroph (Põlme et al. 2020); Decay sugar-rich substrates (Põlme et al. 2020) | Põlme S, et al. 2020. Fungal Diversity 105:1-16 ((https://doi.org/10.1007/s13225-020-00466-2)) |
| Otu61128 | 1 | k__Fungi(100);p__Ascomycota(100);c__Saccharomycetes(100);o__Saccharomycetales(100);f__Metschnikowiaceae(100);g__Metschnikowia(100);s__Metschnikowia_pulcherrima(100); | Metschnikowia | 13 | Saprotroph | \|Undefined Saprotroph\| | Yeast | NULL | Probable | As nectar-tap_saprotroph (Põlme et al. 2020); Decay sugar-rich substrates (Põlme et al. 2020) | Põlme S, et al. 2020. Fungal Diversity 105:1-16 ((https://doi.org/10.1007/s13225-020-00466-2)) |
| Otu61129 | 1 | k__Fungi(100);p__Ascomycota(100);c__Saccharomycetes(100);o__Saccharomycetales(100);f__Metschnikowiaceae(100);g__Metschnikowia(100);g__Metschnikowia_unclassified(100); | Metschnikowia | 13 | Saprotroph | \|Undefined Saprotroph\| | Yeast | NULL | Probable | As nectar-tap_saprotroph (Põlme et al. 2020); Decay sugar-rich substrates (Põlme et al. 2020) | Põlme S, et al. 2020. Fungal Diversity 105:1-16 ((https://doi.org/10.1007/s13225-020-00466-2)) |
| Otu61130 | 1 | k__Fungi(100);p__Ascomycota(100);c__Saccharomycetes(100);o__Saccharomycetales(100);f__Metschnikowiaceae(100);g__Metschnikowia(100);g__Metschnikowia_unclassified(100); | Metschnikowia | 13 | Saprotroph | \|Undefined Saprotroph\| | Yeast | NULL | Probable | As nectar-tap_saprotroph (Põlme et al. 2020); Decay sugar-rich substrates (Põlme et al. 2020) | Põlme S, et al. 2020. Fungal Diversity 105:1-16 ((https://doi.org/10.1007/s13225-020-00466-2)) |
| Otu61131 | 1 | k__Fungi(100);p__Ascomycota(100);c__Saccharomycetes(100);o__Saccharomycetales(100);f__Metschnikowiaceae(100);g__Metschnikowia(100);g__Metschnikowia_unclassified(100); | Metschnikowia | 13 | Saprotroph | \|Undefined Saprotroph\| | Yeast | NULL | Probable | As nectar-tap_saprotroph (Põlme et al. 2020); Decay sugar-rich substrates (Põlme et al. 2020) | Põlme S, et al. 2020. Fungal Diversity 105:1-16 ((https://doi.org/10.1007/s13225-020-00466-2)) |
| Otu61132 | 1 | k__Fungi(100);p__Ascomycota(100);c__Saccharomycetes(100);o__Saccharomycetales(100);f__Metschnikowiaceae(100);g__Metschnikowia(100);s__Metschnikowia_pulcherrima(100); | Metschnikowia | 13 | Saprotroph | \|Undefined Saprotroph\| | Yeast | NULL | Probable | As nectar-tap_saprotroph (Põlme et al. 2020); Decay sugar-rich substrates (Põlme et al. 2020) | Põlme S, et al. 2020. Fungal Diversity 105:1-16 ((https://doi.org/10.1007/s13225-020-00466-2)) |
| Otu61133 | 1 | k__Fungi(100);p__Ascomycota(100);c__Saccharomycetes(100);o__Saccharomycetales(100);f__Metschnikowiaceae(100);g__Metschnikowia(100);s__Metschnikowia_pulcherrima(100); | Metschnikowia | 13 | Saprotroph | \|Undefined Saprotroph\| | Yeast | NULL | Probable | As nectar-tap_saprotroph (Põlme et al. 2020); Decay sugar-rich substrates (Põlme et al. 2020) | Põlme S, et al. 2020. Fungal Diversity 105:1-16 ((https://doi.org/10.1007/s13225-020-00466-2)) |
| Otu61134 | 1 | k__Fungi(100);p__Ascomycota(100);c__Saccharomycetes(100);o__Saccharomycetales(100);f__Metschnikowiaceae(100);g__Metschnikowia(100);s__Metschnikowia_pulcherrima(100); | Metschnikowia | 13 | Saprotroph | \|Undefined Saprotroph\| | Yeast | NULL | Probable | As nectar-tap_saprotroph (Põlme et al. 2020); Decay sugar-rich substrates (Põlme et al. 2020) | Põlme S, et al. 2020. Fungal Diversity 105:1-16 ((https://doi.org/10.1007/s13225-020-00466-2)) |
| Otu61135 | 1 | k__Fungi(100);p__Ascomycota(100);c__Saccharomycetes(100);o__Saccharomycetales(100);f__Metschnikowiaceae(100);g__Metschnikowia(100);g__Metschnikowia_unclassified(100); | Metschnikowia | 13 | Saprotroph | \|Undefined Saprotroph\| | Yeast | NULL | Probable | As nectar-tap_saprotroph (Põlme et al. 2020); Decay sugar-rich substrates (Põlme et al. 2020) | Põlme S, et al. 2020. Fungal Diversity 105:1-16 ((https://doi.org/10.1007/s13225-020-00466-2)) |
| Otu61136 | 1 | k__Fungi(100);p__Ascomycota(100);c__Saccharomycetes(100);o__Saccharomycetales(100);f__Metschnikowiaceae(100);g__Metschnikowia(100);s__Metschnikowia_pulcherrima(100); | Metschnikowia | 13 | Saprotroph | \|Undefined Saprotroph\| | Yeast | NULL | Probable | As nectar-tap_saprotroph (Põlme et al. 2020); Decay sugar-rich substrates (Põlme et al. 2020) | Põlme S, et al. 2020. Fungal Diversity 105:1-16 ((https://doi.org/10.1007/s13225-020-00466-2)) |
| Otu61137 | 1 | k__Fungi(100);p__Ascomycota(100);c__Saccharomycetes(100);o__Saccharomycetales(100);f__Metschnikowiaceae(100);g__Metschnikowia(100);g__Metschnikowia_unclassified(100); | Metschnikowia | 13 | Saprotroph | \|Undefined Saprotroph\| | Yeast | NULL | Probable | As nectar-tap_saprotroph (Põlme et al. 2020); Decay sugar-rich substrates (Põlme et al. 2020) | Põlme S, et al. 2020. Fungal Diversity 105:1-16 ((https://doi.org/10.1007/s13225-020-00466-2)) |
| Otu61138 | 1 | k__Fungi(100);p__Ascomycota(100);c__Saccharomycetes(100);o__Saccharomycetales(100);f__Metschnikowiaceae(100);g__Metschnikowia(100);s__Metschnikowia_pulcherrima(100); | Metschnikowia | 13 | Saprotroph | \|Undefined Saprotroph\| | Yeast | NULL | Probable | As nectar-tap_saprotroph (Põlme et al. 2020); Decay sugar-rich substrates (Põlme et al. 2020) | Põlme S, et al. 2020. Fungal Diversity 105:1-16 ((https://doi.org/10.1007/s13225-020-00466-2)) |
| Otu61139 | 1 | k__Fungi(100);p__Ascomycota(100);c__Saccharomycetes(100);o__Saccharomycetales(100);f__Metschnikowiaceae(100);g__Metschnikowia(100);s__Metschnikowia_pulcherrima(100); | Metschnikowia | 13 | Saprotroph | \|Undefined Saprotroph\| | Yeast | NULL | Probable | As nectar-tap_saprotroph (Põlme et al. 2020); Decay sugar-rich substrates (Põlme et al. 2020) | Põlme S, et al. 2020. Fungal Diversity 105:1-16 ((https://doi.org/10.1007/s13225-020-00466-2)) |
| Otu61140 | 1 | k__Fungi(100);p__Ascomycota(100);c__Saccharomycetes(100);o__Saccharomycetales(100);f__Metschnikowiaceae(100);g__Metschnikowia(100);s__Metschnikowia_pulcherrima(100); | Metschnikowia | 13 | Saprotroph | \|Undefined Saprotroph\| | Yeast | NULL | Probable | As nectar-tap_saprotroph (Põlme et al. 2020); Decay sugar-rich substrates (Põlme et al. 2020) | Põlme S, et al. 2020. Fungal Diversity 105:1-16 ((https://doi.org/10.1007/s13225-020-00466-2)) |
| Otu61141 | 1 | k__Fungi(100);p__Ascomycota(100);c__Saccharomycetes(100);o__Saccharomycetales(100);f__Metschnikowiaceae(100);g__Metschnikowia(100);g__Metschnikowia_unclassified(100); | Metschnikowia | 13 | Saprotroph | \|Undefined Saprotroph\| | Yeast | NULL | Probable | As nectar-tap_saprotroph (Põlme et al. 2020); Decay sugar-rich substrates (Põlme et al. 2020) | Põlme S, et al. 2020. Fungal Diversity 105:1-16 ((https://doi.org/10.1007/s13225-020-00466-2)) |
| Otu61142 | 1 | k__Fungi(100);p__Ascomycota(100);c__Saccharomycetes(100);o__Saccharomycetales(100);f__Metschnikowiaceae(100);g__Metschnikowia(100);s__Metschnikowia_pulcherrima(100); | Metschnikowia | 13 | Saprotroph | \|Undefined Saprotroph\| | Yeast | NULL | Probable | As nectar-tap_saprotroph (Põlme et al. 2020); Decay sugar-rich substrates (Põlme et al. 2020) | Põlme S, et al. 2020. Fungal Diversity 105:1-16 ((https://doi.org/10.1007/s13225-020-00466-2)) |
| Otu61143 | 1 | k__Fungi(100);p__Ascomycota(100);c__Saccharomycetes(100);o__Saccharomycetales(100);f__Metschnikowiaceae(100);g__Metschnikowia(100);s__Metschnikowia_pulcherrima(100); | Metschnikowia | 13 | Saprotroph | \|Undefined Saprotroph\| | Yeast | NULL | Probable | As nectar-tap_saprotroph (Põlme et al. 2020); Decay sugar-rich substrates (Põlme et al. 2020) | Põlme S, et al. 2020. Fungal Diversity 105:1-16 ((https://doi.org/10.1007/s13225-020-00466-2)) |
| Otu61144 | 1 | k__Fungi(100);p__Ascomycota(100);c__Saccharomycetes(100);o__Saccharomycetales(100);f__Metschnikowiaceae(100);g__Metschnikowia(100);s__Metschnikowia_pulcherrima(100); | Metschnikowia | 13 | Saprotroph | \|Undefined Saprotroph\| | Yeast | NULL | Probable | As nectar-tap_saprotroph (Põlme et al. 2020); Decay sugar-rich substrates (Põlme et al. 2020) | Põlme S, et al. 2020. Fungal Diversity 105:1-16 ((https://doi.org/10.1007/s13225-020-00466-2)) |
| Otu61145 | 1 | k__Fungi(100);p__Ascomycota(100);c__Saccharomycetes(100);o__Saccharomycetales(100);f__Metschnikowiaceae(100);g__Metschnikowia(100);s__Metschnikowia_pulcherrima(100); | Metschnikowia | 13 | Saprotroph | \|Undefined Saprotroph\| | Yeast | NULL | Probable | As nectar-tap_saprotroph (Põlme et al. 2020); Decay sugar-rich substrates (Põlme et al. 2020) | Põlme S, et al. 2020. Fungal Diversity 105:1-16 ((https://doi.org/10.1007/s13225-020-00466-2)) |
| Otu61146 | 1 | k__Fungi(100);p__Ascomycota(100);c__Saccharomycetes(100);o__Saccharomycetales(100);f__Metschnikowiaceae(100);g__Metschnikowia(100);g__Metschnikowia_unclassified(100); | Metschnikowia | 13 | Saprotroph | \|Undefined Saprotroph\| | Yeast | NULL | Probable | As nectar-tap_saprotroph (Põlme et al. 2020); Decay sugar-rich substrates (Põlme et al. 2020) | Põlme S, et al. 2020. Fungal Diversity 105:1-16 ((https://doi.org/10.1007/s13225-020-00466-2)) |
| Otu61147 | 1 | k__Fungi(100);p__Ascomycota(100);c__Saccharomycetes(100);o__Saccharomycetales(100);f__Metschnikowiaceae(100);g__Metschnikowia(100);g__Metschnikowia_unclassified(100); | Metschnikowia | 13 | Saprotroph | \|Undefined Saprotroph\| | Yeast | NULL | Probable | As nectar-tap_saprotroph (Põlme et al. 2020); Decay sugar-rich substrates (Põlme et al. 2020) | Põlme S, et al. 2020. Fungal Diversity 105:1-16 ((https://doi.org/10.1007/s13225-020-00466-2)) |
| Otu61149 | 1 | k__Fungi(100);p__Ascomycota(100);c__Saccharomycetes(100);o__Saccharomycetales(100);f__Metschnikowiaceae(100);g__Metschnikowia(100);g__Metschnikowia_unclassified(100); | Metschnikowia | 13 | Saprotroph | \|Undefined Saprotroph\| | Yeast | NULL | Probable | As nectar-tap_saprotroph (Põlme et al. 2020); Decay sugar-rich substrates (Põlme et al. 2020) | Põlme S, et al. 2020. Fungal Diversity 105:1-16 ((https://doi.org/10.1007/s13225-020-00466-2)) |
| Otu61150 | 1 | k__Fungi(100);p__Ascomycota(100);c__Saccharomycetes(100);o__Saccharomycetales(100);f__Metschnikowiaceae(100);g__Metschnikowia(100);g__Metschnikowia_unclassified(100); | Metschnikowia | 13 | Saprotroph | \|Undefined Saprotroph\| | Yeast | NULL | Probable | As nectar-tap_saprotroph (Põlme et al. 2020); Decay sugar-rich substrates (Põlme et al. 2020) | Põlme S, et al. 2020. Fungal Diversity 105:1-16 ((https://doi.org/10.1007/s13225-020-00466-2)) |
| Otu61151 | 1 | k__Fungi(100);p__Ascomycota(100);c__Saccharomycetes(100);o__Saccharomycetales(100);f__Saccharomycetales_family_Incertae_sedis(100);g__Candida(100);g__Candida_unclassified(100); | Candida | 13 | Pathotroph-Saprotroph-Symbiotroph | Animal Pathogen-Endophyte-Endosymbiont-Epiphyte-\|Undefined Saprotroph\| | Dimorphic Yeast-Yeast | NULL | Probable | As nectar-tap_saprotroph (Põlme et al. 2020); Decay sugar-rich substrates (Põlme et al. 2020); This asexual genus is polyphyletic and contain species that function very differently depend on what they are. Caution should be taken when interpreting guilds for Candida species. | Manolakaki D, et al. 2010. Virulence. 1: 367-375 ((https://doi.org/10.4161/viru.1.5.12796)); Mekbib SB, et al. 2011. Tropical Plant Pathology 36:233-240 ((https://doi.org/10.1590/S1982-56762011000400004)); Põlme S, et al. 2020. Fungal Diversity 105:1-16 ((https://doi.org/10.1007/s13225-020-00466-2)) |
| Otu61153 | 1 | k__Fungi(100);p__Ascomycota(100);c__Saccharomycetes(100);o__Saccharomycetales(100);f__Metschnikowiaceae(100);g__Metschnikowia(100);g__Metschnikowia_unclassified(100); | Metschnikowia | 13 | Saprotroph | \|Undefined Saprotroph\| | Yeast | NULL | Probable | As nectar-tap_saprotroph (Põlme et al. 2020); Decay sugar-rich substrates (Põlme et al. 2020) | Põlme S, et al. 2020. Fungal Diversity 105:1-16 ((https://doi.org/10.1007/s13225-020-00466-2)) |
| Otu61154 | 1 | k__Fungi(100);p__Ascomycota(100);c__Saccharomycetes(100);o__Saccharomycetales(100);f__Metschnikowiaceae(100);g__Metschnikowia(100);g__Metschnikowia_unclassified(100); | Metschnikowia | 13 | Saprotroph | \|Undefined Saprotroph\| | Yeast | NULL | Probable | As nectar-tap_saprotroph (Põlme et al. 2020); Decay sugar-rich substrates (Põlme et al. 2020) | Põlme S, et al. 2020. Fungal Diversity 105:1-16 ((https://doi.org/10.1007/s13225-020-00466-2)) |
| Otu61156 | 1 | k__Fungi(100);p__Ascomycota(100);c__Saccharomycetes(100);o__Saccharomycetales(100);f__Metschnikowiaceae(100);g__Metschnikowia(100);g__Metschnikowia_unclassified(100); | Metschnikowia | 13 | Saprotroph | \|Undefined Saprotroph\| | Yeast | NULL | Probable | As nectar-tap_saprotroph (Põlme et al. 2020); Decay sugar-rich substrates (Põlme et al. 2020) | Põlme S, et al. 2020. Fungal Diversity 105:1-16 ((https://doi.org/10.1007/s13225-020-00466-2)) |
| Otu61157 | 1 | k__Fungi(100);p__Ascomycota(100);c__Saccharomycetes(100);o__Saccharomycetales(100);f__Metschnikowiaceae(100);g__Metschnikowia(100);s__Metschnikowia_pulcherrima(100); | Metschnikowia | 13 | Saprotroph | \|Undefined Saprotroph\| | Yeast | NULL | Probable | As nectar-tap_saprotroph (Põlme et al. 2020); Decay sugar-rich substrates (Põlme et al. 2020) | Põlme S, et al. 2020. Fungal Diversity 105:1-16 ((https://doi.org/10.1007/s13225-020-00466-2)) |
| Otu61158 | 1 | k__Fungi(100);p__Ascomycota(100);c__Saccharomycetes(100);o__Saccharomycetales(100);f__Metschnikowiaceae(100);g__Metschnikowia(100);s__Metschnikowia_pulcherrima(100); | Metschnikowia | 13 | Saprotroph | \|Undefined Saprotroph\| | Yeast | NULL | Probable | As nectar-tap_saprotroph (Põlme et al. 2020); Decay sugar-rich substrates (Põlme et al. 2020) | Põlme S, et al. 2020. Fungal Diversity 105:1-16 ((https://doi.org/10.1007/s13225-020-00466-2)) |
| Otu61159 | 1 | k__Fungi(100);p__Ascomycota(100);c__Saccharomycetes(100);o__Saccharomycetales(100);f__Metschnikowiaceae(100);g__Metschnikowia(100);s__Metschnikowia_pulcherrima(100); | Metschnikowia | 13 | Saprotroph | \|Undefined Saprotroph\| | Yeast | NULL | Probable | As nectar-tap_saprotroph (Põlme et al. 2020); Decay sugar-rich substrates (Põlme et al. 2020) | Põlme S, et al. 2020. Fungal Diversity 105:1-16 ((https://doi.org/10.1007/s13225-020-00466-2)) |
| Otu61160 | 1 | k__Fungi(100);p__Ascomycota(100);c__Saccharomycetes(100);o__Saccharomycetales(100);f__Metschnikowiaceae(100);g__Metschnikowia(100);s__Metschnikowia_pulcherrima(100); | Metschnikowia | 13 | Saprotroph | \|Undefined Saprotroph\| | Yeast | NULL | Probable | As nectar-tap_saprotroph (Põlme et al. 2020); Decay sugar-rich substrates (Põlme et al. 2020) | Põlme S, et al. 2020. Fungal Diversity 105:1-16 ((https://doi.org/10.1007/s13225-020-00466-2)) |
| Otu61161 | 1 | k__Fungi(100);p__Ascomycota(100);c__Saccharomycetes(100);o__Saccharomycetales(100);f__Metschnikowiaceae(100);g__Metschnikowia(100);s__Metschnikowia_pulcherrima(100); | Metschnikowia | 13 | Saprotroph | \|Undefined Saprotroph\| | Yeast | NULL | Probable | As nectar-tap_saprotroph (Põlme et al. 2020); Decay sugar-rich substrates (Põlme et al. 2020) | Põlme S, et al. 2020. Fungal Diversity 105:1-16 ((https://doi.org/10.1007/s13225-020-00466-2)) |
| Otu61162 | 1 | k__Fungi(100);p__Ascomycota(100);c__Saccharomycetes(100);o__Saccharomycetales(100);f__Metschnikowiaceae(100);g__Metschnikowia(100);s__Metschnikowia_pulcherrima(100); | Metschnikowia | 13 | Saprotroph | \|Undefined Saprotroph\| | Yeast | NULL | Probable | As nectar-tap_saprotroph (Põlme et al. 2020); Decay sugar-rich substrates (Põlme et al. 2020) | Põlme S, et al. 2020. Fungal Diversity 105:1-16 ((https://doi.org/10.1007/s13225-020-00466-2)) |
| Otu61163 | 1 | k__Fungi(100);p__Ascomycota(100);c__Saccharomycetes(100);o__Saccharomycetales(100);f__Metschnikowiaceae(100);g__Metschnikowia(100);s__Metschnikowia_pulcherrima(100); | Metschnikowia | 13 | Saprotroph | \|Undefined Saprotroph\| | Yeast | NULL | Probable | As nectar-tap_saprotroph (Põlme et al. 2020); Decay sugar-rich substrates (Põlme et al. 2020) | Põlme S, et al. 2020. Fungal Diversity 105:1-16 ((https://doi.org/10.1007/s13225-020-00466-2)) |
| Otu61164 | 1 | k__Fungi(100);p__Ascomycota(100);c__Saccharomycetes(100);o__Saccharomycetales(100);f__Metschnikowiaceae(100);g__Metschnikowia(100);s__Metschnikowia_pulcherrima(100); | Metschnikowia | 13 | Saprotroph | \|Undefined Saprotroph\| | Yeast | NULL | Probable | As nectar-tap_saprotroph (Põlme et al. 2020); Decay sugar-rich substrates (Põlme et al. 2020) | Põlme S, et al. 2020. Fungal Diversity 105:1-16 ((https://doi.org/10.1007/s13225-020-00466-2)) |
| Otu61165 | 1 | k__Fungi(100);p__Ascomycota(100);c__Saccharomycetes(100);o__Saccharomycetales(100);f__Metschnikowiaceae(100);g__Metschnikowia(100);s__Metschnikowia_pulcherrima(100); | Metschnikowia | 13 | Saprotroph | \|Undefined Saprotroph\| | Yeast | NULL | Probable | As nectar-tap_saprotroph (Põlme et al. 2020); Decay sugar-rich substrates (Põlme et al. 2020) | Põlme S, et al. 2020. Fungal Diversity 105:1-16 ((https://doi.org/10.1007/s13225-020-00466-2)) |
| Otu61166 | 1 | k__Fungi(100);p__Ascomycota(100);c__Saccharomycetes(100);o__Saccharomycetales(100);f__Metschnikowiaceae(100);g__Metschnikowia(100);s__Metschnikowia_pulcherrima(100); | Metschnikowia | 13 | Saprotroph | \|Undefined Saprotroph\| | Yeast | NULL | Probable | As nectar-tap_saprotroph (Põlme et al. 2020); Decay sugar-rich substrates (Põlme et al. 2020) | Põlme S, et al. 2020. Fungal Diversity 105:1-16 ((https://doi.org/10.1007/s13225-020-00466-2)) |
| Otu61167 | 1 | k__Fungi(100);p__Ascomycota(100);c__Saccharomycetes(100);o__Saccharomycetales(100);f__Metschnikowiaceae(100);g__Metschnikowia(100);s__Metschnikowia_pulcherrima(100); | Metschnikowia | 13 | Saprotroph | \|Undefined Saprotroph\| | Yeast | NULL | Probable | As nectar-tap_saprotroph (Põlme et al. 2020); Decay sugar-rich substrates (Põlme et al. 2020) | Põlme S, et al. 2020. Fungal Diversity 105:1-16 ((https://doi.org/10.1007/s13225-020-00466-2)) |
| Otu61168 | 1 | k__Fungi(100);p__Ascomycota(100);c__Saccharomycetes(100);o__Saccharomycetales(100);f__Metschnikowiaceae(100);g__Metschnikowia(100);s__Metschnikowia_pulcherrima(100); | Metschnikowia | 13 | Saprotroph | \|Undefined Saprotroph\| | Yeast | NULL | Probable | As nectar-tap_saprotroph (Põlme et al. 2020); Decay sugar-rich substrates (Põlme et al. 2020) | Põlme S, et al. 2020. Fungal Diversity 105:1-16 ((https://doi.org/10.1007/s13225-020-00466-2)) |
| Otu61169 | 1 | k__Fungi(100);p__Ascomycota(100);c__Saccharomycetes(100);o__Saccharomycetales(100);f__Metschnikowiaceae(100);g__Metschnikowia(100);s__Metschnikowia_pulcherrima(100); | Metschnikowia | 13 | Saprotroph | \|Undefined Saprotroph\| | Yeast | NULL | Probable | As nectar-tap_saprotroph (Põlme et al. 2020); Decay sugar-rich substrates (Põlme et al. 2020) | Põlme S, et al. 2020. Fungal Diversity 105:1-16 ((https://doi.org/10.1007/s13225-020-00466-2)) |
| Otu61170 | 1 | k__Fungi(100);p__Ascomycota(100);c__Saccharomycetes(100);o__Saccharomycetales(100);f__Metschnikowiaceae(100);g__Metschnikowia(100);s__Metschnikowia_pulcherrima(100); | Metschnikowia | 13 | Saprotroph | \|Undefined Saprotroph\| | Yeast | NULL | Probable | As nectar-tap_saprotroph (Põlme et al. 2020); Decay sugar-rich substrates (Põlme et al. 2020) | Põlme S, et al. 2020. Fungal Diversity 105:1-16 ((https://doi.org/10.1007/s13225-020-00466-2)) |
| Otu61171 | 1 | k__Fungi(100);p__Ascomycota(100);c__Saccharomycetes(100);o__Saccharomycetales(100);f__Metschnikowiaceae(100);g__Metschnikowia(100);s__Metschnikowia_pulcherrima(100); | Metschnikowia | 13 | Saprotroph | \|Undefined Saprotroph\| | Yeast | NULL | Probable | As nectar-tap_saprotroph (Põlme et al. 2020); Decay sugar-rich substrates (Põlme et al. 2020) | Põlme S, et al. 2020. Fungal Diversity 105:1-16 ((https://doi.org/10.1007/s13225-020-00466-2)) |
| Otu61172 | 1 | k__Fungi(100);p__Ascomycota(100);c__Saccharomycetes(100);o__Saccharomycetales(100);f__Metschnikowiaceae(100);g__Metschnikowia(100);s__Metschnikowia_pulcherrima(100); | Metschnikowia | 13 | Saprotroph | \|Undefined Saprotroph\| | Yeast | NULL | Probable | As nectar-tap_saprotroph (Põlme et al. 2020); Decay sugar-rich substrates (Põlme et al. 2020) | Põlme S, et al. 2020. Fungal Diversity 105:1-16 ((https://doi.org/10.1007/s13225-020-00466-2)) |
| Otu61173 | 1 | k__Fungi(100);p__Ascomycota(100);c__Saccharomycetes(100);o__Saccharomycetales(100);f__Metschnikowiaceae(100);g__Metschnikowia(100);s__Metschnikowia_pulcherrima(100); | Metschnikowia | 13 | Saprotroph | \|Undefined Saprotroph\| | Yeast | NULL | Probable | As nectar-tap_saprotroph (Põlme et al. 2020); Decay sugar-rich substrates (Põlme et al. 2020) | Põlme S, et al. 2020. Fungal Diversity 105:1-16 ((https://doi.org/10.1007/s13225-020-00466-2)) |
| Otu61174 | 1 | k__Fungi(100);p__Ascomycota(100);c__Saccharomycetes(100);o__Saccharomycetales(100);f__Metschnikowiaceae(100);g__Metschnikowia(100);s__Metschnikowia_pulcherrima(100); | Metschnikowia | 13 | Saprotroph | \|Undefined Saprotroph\| | Yeast | NULL | Probable | As nectar-tap_saprotroph (Põlme et al. 2020); Decay sugar-rich substrates (Põlme et al. 2020) | Põlme S, et al. 2020. Fungal Diversity 105:1-16 ((https://doi.org/10.1007/s13225-020-00466-2)) |
| Otu61175 | 1 | k__Fungi(100);p__Ascomycota(100);c__Saccharomycetes(100);o__Saccharomycetales(100);f__Metschnikowiaceae(100);g__Metschnikowia(100);g__Metschnikowia_unclassified(100); | Metschnikowia | 13 | Saprotroph | \|Undefined Saprotroph\| | Yeast | NULL | Probable | As nectar-tap_saprotroph (Põlme et al. 2020); Decay sugar-rich substrates (Põlme et al. 2020) | Põlme S, et al. 2020. Fungal Diversity 105:1-16 ((https://doi.org/10.1007/s13225-020-00466-2)) |
| Otu61176 | 1 | k__Fungi(100);p__Ascomycota(100);c__Saccharomycetes(100);o__Saccharomycetales(100);f__Metschnikowiaceae(100);g__Metschnikowia(100);s__Metschnikowia_pulcherrima(100); | Metschnikowia | 13 | Saprotroph | \|Undefined Saprotroph\| | Yeast | NULL | Probable | As nectar-tap_saprotroph (Põlme et al. 2020); Decay sugar-rich substrates (Põlme et al. 2020) | Põlme S, et al. 2020. Fungal Diversity 105:1-16 ((https://doi.org/10.1007/s13225-020-00466-2)) |
| Otu61177 | 1 | k__Fungi(100);p__Ascomycota(100);c__Saccharomycetes(100);o__Saccharomycetales(100);f__Metschnikowiaceae(100);g__Metschnikowia(100);s__Metschnikowia_pulcherrima(100); | Metschnikowia | 13 | Saprotroph | \|Undefined Saprotroph\| | Yeast | NULL | Probable | As nectar-tap_saprotroph (Põlme et al. 2020); Decay sugar-rich substrates (Põlme et al. 2020) | Põlme S, et al. 2020. Fungal Diversity 105:1-16 ((https://doi.org/10.1007/s13225-020-00466-2)) |
| Otu61178 | 1 | k__Fungi(100);p__Ascomycota(100);c__Saccharomycetes(100);o__Saccharomycetales(100);f__Metschnikowiaceae(100);g__Metschnikowia(100);s__Metschnikowia_pulcherrima(100); | Metschnikowia | 13 | Saprotroph | \|Undefined Saprotroph\| | Yeast | NULL | Probable | As nectar-tap_saprotroph (Põlme et al. 2020); Decay sugar-rich substrates (Põlme et al. 2020) | Põlme S, et al. 2020. Fungal Diversity 105:1-16 ((https://doi.org/10.1007/s13225-020-00466-2)) |
| Otu61179 | 1 | k__Fungi(100);p__Ascomycota(100);c__Saccharomycetes(100);o__Saccharomycetales(100);f__Metschnikowiaceae(100);g__Metschnikowia(100);g__Metschnikowia_unclassified(100); | Metschnikowia | 13 | Saprotroph | \|Undefined Saprotroph\| | Yeast | NULL | Probable | As nectar-tap_saprotroph (Põlme et al. 2020); Decay sugar-rich substrates (Põlme et al. 2020) | Põlme S, et al. 2020. Fungal Diversity 105:1-16 ((https://doi.org/10.1007/s13225-020-00466-2)) |
| Otu61180 | 1 | k__Fungi(100);p__Ascomycota(100);c__Saccharomycetes(100);o__Saccharomycetales(100);f__Metschnikowiaceae(100);g__Metschnikowia(100);s__Metschnikowia_pulcherrima(100); | Metschnikowia | 13 | Saprotroph | \|Undefined Saprotroph\| | Yeast | NULL | Probable | As nectar-tap_saprotroph (Põlme et al. 2020); Decay sugar-rich substrates (Põlme et al. 2020) | Põlme S, et al. 2020. Fungal Diversity 105:1-16 ((https://doi.org/10.1007/s13225-020-00466-2)) |
| Otu61181 | 1 | k__Fungi(100);p__Ascomycota(100);c__Saccharomycetes(100);o__Saccharomycetales(100);f__Metschnikowiaceae(100);g__Metschnikowia(100);s__Metschnikowia_pulcherrima(100); | Metschnikowia | 13 | Saprotroph | \|Undefined Saprotroph\| | Yeast | NULL | Probable | As nectar-tap_saprotroph (Põlme et al. 2020); Decay sugar-rich substrates (Põlme et al. 2020) | Põlme S, et al. 2020. Fungal Diversity 105:1-16 ((https://doi.org/10.1007/s13225-020-00466-2)) |
| Otu61182 | 1 | k__Fungi(100);p__Ascomycota(100);c__Saccharomycetes(100);o__Saccharomycetales(100);f__Metschnikowiaceae(100);g__Metschnikowia(100);s__Metschnikowia_pulcherrima(100); | Metschnikowia | 13 | Saprotroph | \|Undefined Saprotroph\| | Yeast | NULL | Probable | As nectar-tap_saprotroph (Põlme et al. 2020); Decay sugar-rich substrates (Põlme et al. 2020) | Põlme S, et al. 2020. Fungal Diversity 105:1-16 ((https://doi.org/10.1007/s13225-020-00466-2)) |
| Otu61183 | 1 | k__Fungi(100);p__Ascomycota(100);c__Saccharomycetes(100);o__Saccharomycetales(100);f__Metschnikowiaceae(100);g__Metschnikowia(100);s__Metschnikowia_pulcherrima(100); | Metschnikowia | 13 | Saprotroph | \|Undefined Saprotroph\| | Yeast | NULL | Probable | As nectar-tap_saprotroph (Põlme et al. 2020); Decay sugar-rich substrates (Põlme et al. 2020) | Põlme S, et al. 2020. Fungal Diversity 105:1-16 ((https://doi.org/10.1007/s13225-020-00466-2)) |
| Otu61184 | 1 | k__Fungi(100);p__Ascomycota(100);c__Saccharomycetes(100);o__Saccharomycetales(100);f__Metschnikowiaceae(100);g__Metschnikowia(100);s__Metschnikowia_pulcherrima(100); | Metschnikowia | 13 | Saprotroph | \|Undefined Saprotroph\| | Yeast | NULL | Probable | As nectar-tap_saprotroph (Põlme et al. 2020); Decay sugar-rich substrates (Põlme et al. 2020) | Põlme S, et al. 2020. Fungal Diversity 105:1-16 ((https://doi.org/10.1007/s13225-020-00466-2)) |
| Otu61185 | 1 | k__Fungi(100);p__Ascomycota(100);c__Saccharomycetes(100);o__Saccharomycetales(100);f__Metschnikowiaceae(100);g__Metschnikowia(100);g__Metschnikowia_unclassified(100); | Metschnikowia | 13 | Saprotroph | \|Undefined Saprotroph\| | Yeast | NULL | Probable | As nectar-tap_saprotroph (Põlme et al. 2020); Decay sugar-rich substrates (Põlme et al. 2020) | Põlme S, et al. 2020. Fungal Diversity 105:1-16 ((https://doi.org/10.1007/s13225-020-00466-2)) |
| Otu61186 | 1 | k__Fungi(100);p__Ascomycota(100);c__Saccharomycetes(100);o__Saccharomycetales(100);f__Metschnikowiaceae(100);g__Metschnikowia(100);s__Metschnikowia_pulcherrima(100); | Metschnikowia | 13 | Saprotroph | \|Undefined Saprotroph\| | Yeast | NULL | Probable | As nectar-tap_saprotroph (Põlme et al. 2020); Decay sugar-rich substrates (Põlme et al. 2020) | Põlme S, et al. 2020. Fungal Diversity 105:1-16 ((https://doi.org/10.1007/s13225-020-00466-2)) |
| Otu61187 | 1 | k__Fungi(100);p__Ascomycota(100);c__Saccharomycetes(100);o__Saccharomycetales(100);f__Metschnikowiaceae(100);g__Metschnikowia(100);s__Metschnikowia_pulcherrima(100); | Metschnikowia | 13 | Saprotroph | \|Undefined Saprotroph\| | Yeast | NULL | Probable | As nectar-tap_saprotroph (Põlme et al. 2020); Decay sugar-rich substrates (Põlme et al. 2020) | Põlme S, et al. 2020. Fungal Diversity 105:1-16 ((https://doi.org/10.1007/s13225-020-00466-2)) |
| Otu61188 | 1 | k__Fungi(100);p__Ascomycota(100);c__Saccharomycetes(100);o__Saccharomycetales(100);f__Metschnikowiaceae(100);g__Metschnikowia(100);s__Metschnikowia_pulcherrima(100); | Metschnikowia | 13 | Saprotroph | \|Undefined Saprotroph\| | Yeast | NULL | Probable | As nectar-tap_saprotroph (Põlme et al. 2020); Decay sugar-rich substrates (Põlme et al. 2020) | Põlme S, et al. 2020. Fungal Diversity 105:1-16 ((https://doi.org/10.1007/s13225-020-00466-2)) |
| Otu61189 | 1 | k__Fungi(100);p__Ascomycota(100);c__Saccharomycetes(100);o__Saccharomycetales(100);f__Metschnikowiaceae(100);g__Metschnikowia(100);s__Metschnikowia_pulcherrima(100); | Metschnikowia | 13 | Saprotroph | \|Undefined Saprotroph\| | Yeast | NULL | Probable | As nectar-tap_saprotroph (Põlme et al. 2020); Decay sugar-rich substrates (Põlme et al. 2020) | Põlme S, et al. 2020. Fungal Diversity 105:1-16 ((https://doi.org/10.1007/s13225-020-00466-2)) |
| Otu61190 | 1 | k__Fungi(100);p__Ascomycota(100);c__Saccharomycetes(100);o__Saccharomycetales(100);f__Metschnikowiaceae(100);g__Metschnikowia(100);g__Metschnikowia_unclassified(100); | Metschnikowia | 13 | Saprotroph | \|Undefined Saprotroph\| | Yeast | NULL | Probable | As nectar-tap_saprotroph (Põlme et al. 2020); Decay sugar-rich substrates (Põlme et al. 2020) | Põlme S, et al. 2020. Fungal Diversity 105:1-16 ((https://doi.org/10.1007/s13225-020-00466-2)) |
| Otu61191 | 1 | k__Fungi(100);p__Ascomycota(100);c__Saccharomycetes(100);o__Saccharomycetales(100);f__Metschnikowiaceae(100);g__Metschnikowia(100);s__Metschnikowia_pulcherrima(100); | Metschnikowia | 13 | Saprotroph | \|Undefined Saprotroph\| | Yeast | NULL | Probable | As nectar-tap_saprotroph (Põlme et al. 2020); Decay sugar-rich substrates (Põlme et al. 2020) | Põlme S, et al. 2020. Fungal Diversity 105:1-16 ((https://doi.org/10.1007/s13225-020-00466-2)) |
| Otu61192 | 1 | k__Fungi(100);p__Ascomycota(100);c__Saccharomycetes(100);o__Saccharomycetales(100);f__Metschnikowiaceae(100);g__Metschnikowia(100);s__Metschnikowia_pulcherrima(100); | Metschnikowia | 13 | Saprotroph | \|Undefined Saprotroph\| | Yeast | NULL | Probable | As nectar-tap_saprotroph (Põlme et al. 2020); Decay sugar-rich substrates (Põlme et al. 2020) | Põlme S, et al. 2020. Fungal Diversity 105:1-16 ((https://doi.org/10.1007/s13225-020-00466-2)) |
| Otu61193 | 1 | k__Fungi(100);p__Ascomycota(100);c__Saccharomycetes(100);o__Saccharomycetales(100);f__Metschnikowiaceae(100);g__Metschnikowia(100);s__Metschnikowia_pulcherrima(100); | Metschnikowia | 13 | Saprotroph | \|Undefined Saprotroph\| | Yeast | NULL | Probable | As nectar-tap_saprotroph (Põlme et al. 2020); Decay sugar-rich substrates (Põlme et al. 2020) | Põlme S, et al. 2020. Fungal Diversity 105:1-16 ((https://doi.org/10.1007/s13225-020-00466-2)) |
| Otu61194 | 1 | k__Fungi(100);p__Ascomycota(100);c__Saccharomycetes(100);o__Saccharomycetales(100);f__Metschnikowiaceae(100);g__Metschnikowia(100);s__Metschnikowia_pulcherrima(100); | Metschnikowia | 13 | Saprotroph | \|Undefined Saprotroph\| | Yeast | NULL | Probable | As nectar-tap_saprotroph (Põlme et al. 2020); Decay sugar-rich substrates (Põlme et al. 2020) | Põlme S, et al. 2020. Fungal Diversity 105:1-16 ((https://doi.org/10.1007/s13225-020-00466-2)) |
| Otu61195 | 1 | k__Fungi(100);p__Ascomycota(100);c__Saccharomycetes(100);o__Saccharomycetales(100);f__Metschnikowiaceae(100);g__Metschnikowia(100);s__Metschnikowia_pulcherrima(100); | Metschnikowia | 13 | Saprotroph | \|Undefined Saprotroph\| | Yeast | NULL | Probable | As nectar-tap_saprotroph (Põlme et al. 2020); Decay sugar-rich substrates (Põlme et al. 2020) | Põlme S, et al. 2020. Fungal Diversity 105:1-16 ((https://doi.org/10.1007/s13225-020-00466-2)) |
| Otu61196 | 1 | k__Fungi(100);p__Ascomycota(100);c__Saccharomycetes(100);o__Saccharomycetales(100);f__Metschnikowiaceae(100);g__Metschnikowia(100);s__Metschnikowia_pulcherrima(100); | Metschnikowia | 13 | Saprotroph | \|Undefined Saprotroph\| | Yeast | NULL | Probable | As nectar-tap_saprotroph (Põlme et al. 2020); Decay sugar-rich substrates (Põlme et al. 2020) | Põlme S, et al. 2020. Fungal Diversity 105:1-16 ((https://doi.org/10.1007/s13225-020-00466-2)) |
| Otu61197 | 1 | k__Fungi(100);p__Ascomycota(100);c__Saccharomycetes(100);o__Saccharomycetales(100);f__Metschnikowiaceae(100);g__Metschnikowia(100);s__Metschnikowia_pulcherrima(100); | Metschnikowia | 13 | Saprotroph | \|Undefined Saprotroph\| | Yeast | NULL | Probable | As nectar-tap_saprotroph (Põlme et al. 2020); Decay sugar-rich substrates (Põlme et al. 2020) | Põlme S, et al. 2020. Fungal Diversity 105:1-16 ((https://doi.org/10.1007/s13225-020-00466-2)) |
| Otu61199 | 1 | k__Fungi(100);p__Ascomycota(100);c__Saccharomycetes(100);o__Saccharomycetales(100);f__Metschnikowiaceae(100);g__Metschnikowia(100);s__Metschnikowia_pulcherrima(100); | Metschnikowia | 13 | Saprotroph | \|Undefined Saprotroph\| | Yeast | NULL | Probable | As nectar-tap_saprotroph (Põlme et al. 2020); Decay sugar-rich substrates (Põlme et al. 2020) | Põlme S, et al. 2020. Fungal Diversity 105:1-16 ((https://doi.org/10.1007/s13225-020-00466-2)) |
| Otu61214 | 1 | k__Fungi(100);p__Ascomycota(100);c__Saccharomycetes(100);o__Saccharomycetales(100);f__Saccharomycetales_family_Incertae_sedis(100);g__Candida(100);s__Candida_sp_AS23486(100); | Candida | 13 | Pathotroph-Saprotroph-Symbiotroph | Animal Pathogen-Endophyte-Endosymbiont-Epiphyte-\|Undefined Saprotroph\| | Dimorphic Yeast-Yeast | NULL | Probable | As nectar-tap_saprotroph (Põlme et al. 2020); Decay sugar-rich substrates (Põlme et al. 2020); This asexual genus is polyphyletic and contain species that function very differently depend on what they are. Caution should be taken when interpreting guilds for Candida species. | Manolakaki D, et al. 2010. Virulence. 1: 367-375 ((https://doi.org/10.4161/viru.1.5.12796)); Mekbib SB, et al. 2011. Tropical Plant Pathology 36:233-240 ((https://doi.org/10.1590/S1982-56762011000400004)); Põlme S, et al. 2020. Fungal Diversity 105:1-16 ((https://doi.org/10.1007/s13225-020-00466-2)) |
| Otu61216 | 1 | k__Fungi(100);p__Ascomycota(100);c__Saccharomycetes(100);o__Saccharomycetales(100);f__Metschnikowiaceae(100);g__Metschnikowia(100);s__Metschnikowia_pulcherrima(100); | Metschnikowia | 13 | Saprotroph | \|Undefined Saprotroph\| | Yeast | NULL | Probable | As nectar-tap_saprotroph (Põlme et al. 2020); Decay sugar-rich substrates (Põlme et al. 2020) | Põlme S, et al. 2020. Fungal Diversity 105:1-16 ((https://doi.org/10.1007/s13225-020-00466-2)) |
| Otu61227 | 1 | k__Fungi(100);p__Ascomycota(100);c__Saccharomycetes(100);o__Saccharomycetales(100);f__Metschnikowiaceae(100);g__Metschnikowia(100);s__Metschnikowia_pulcherrima(100); | Metschnikowia | 13 | Saprotroph | \|Undefined Saprotroph\| | Yeast | NULL | Probable | As nectar-tap_saprotroph (Põlme et al. 2020); Decay sugar-rich substrates (Põlme et al. 2020) | Põlme S, et al. 2020. Fungal Diversity 105:1-16 ((https://doi.org/10.1007/s13225-020-00466-2)) |
| Otu61230 | 1 | k__Fungi(100);p__Ascomycota(100);c__Saccharomycetes(100);o__Saccharomycetales(100);f__Saccharomycetales_family_Incertae_sedis(100);g__Candida(100);s__Candida_cretensis(100); | Candida | 13 | Pathotroph-Saprotroph-Symbiotroph | Animal Pathogen-Endophyte-Endosymbiont-Epiphyte-\|Undefined Saprotroph\| | Dimorphic Yeast-Yeast | NULL | Probable | As nectar-tap_saprotroph (Põlme et al. 2020); Decay sugar-rich substrates (Põlme et al. 2020); This asexual genus is polyphyletic and contain species that function very differently depend on what they are. Caution should be taken when interpreting guilds for Candida species. | Manolakaki D, et al. 2010. Virulence. 1: 367-375 ((https://doi.org/10.4161/viru.1.5.12796)); Mekbib SB, et al. 2011. Tropical Plant Pathology 36:233-240 ((https://doi.org/10.1590/S1982-56762011000400004)); Põlme S, et al. 2020. Fungal Diversity 105:1-16 ((https://doi.org/10.1007/s13225-020-00466-2)) |
| Otu61231 | 1 | k__Fungi(100);p__Ascomycota(100);c__Saccharomycetes(100);o__Saccharomycetales(100);f__Saccharomycetales_family_Incertae_sedis(100);f__Saccharomycetales_family_Incertae_sedis_unclassified(100);f__Saccharomycetales_family_Incertae_sedis_unclassified(100); | Saccharomycetales | 7 | Saprotroph | Undefined Saprotroph | Yeast | NULL | Possible | NULL | Sterkenburg E, et al. 2015. New Phytologist 207:1145-1158 ((https://doi.org/10.1111/nph.13426)) |
| Otu61236 | 1 | k__Fungi(100);p__Ascomycota(100);c__Saccharomycetes(100);o__Saccharomycetales(100);f__Metschnikowiaceae(100);g__Metschnikowia(100);g__Metschnikowia_unclassified(100); | Metschnikowia | 13 | Saprotroph | \|Undefined Saprotroph\| | Yeast | NULL | Probable | As nectar-tap_saprotroph (Põlme et al. 2020); Decay sugar-rich substrates (Põlme et al. 2020) | Põlme S, et al. 2020. Fungal Diversity 105:1-16 ((https://doi.org/10.1007/s13225-020-00466-2)) |
| Otu61237 | 1 | k__Fungi(100);p__Ascomycota(100);c__Saccharomycetes(100);o__Saccharomycetales(100);f__Metschnikowiaceae(100);g__Metschnikowia(100);s__Metschnikowia_pulcherrima(100); | Metschnikowia | 13 | Saprotroph | \|Undefined Saprotroph\| | Yeast | NULL | Probable | As nectar-tap_saprotroph (Põlme et al. 2020); Decay sugar-rich substrates (Põlme et al. 2020) | Põlme S, et al. 2020. Fungal Diversity 105:1-16 ((https://doi.org/10.1007/s13225-020-00466-2)) |
| Otu61238 | 1 | k__Fungi(100);p__Ascomycota(100);c__Saccharomycetes(100);o__Saccharomycetales(100);f__Metschnikowiaceae(100);g__Metschnikowia(100);s__Metschnikowia_pulcherrima(100); | Metschnikowia | 13 | Saprotroph | \|Undefined Saprotroph\| | Yeast | NULL | Probable | As nectar-tap_saprotroph (Põlme et al. 2020); Decay sugar-rich substrates (Põlme et al. 2020) | Põlme S, et al. 2020. Fungal Diversity 105:1-16 ((https://doi.org/10.1007/s13225-020-00466-2)) |
| Otu61243 | 1 | k__Fungi(100);p__Ascomycota(100);c__Saccharomycetes(100);o__Saccharomycetales(100);f__Metschnikowiaceae(100);g__Metschnikowia(100);g__Metschnikowia_unclassified(100); | Metschnikowia | 13 | Saprotroph | \|Undefined Saprotroph\| | Yeast | NULL | Probable | As nectar-tap_saprotroph (Põlme et al. 2020); Decay sugar-rich substrates (Põlme et al. 2020) | Põlme S, et al. 2020. Fungal Diversity 105:1-16 ((https://doi.org/10.1007/s13225-020-00466-2)) |
| Otu61244 | 1 | k__Fungi(100);p__Ascomycota(100);c__Dothideomycetes(100);o__Dothideomycetes_order_Incertae_sedis(100);f__Pseudeurotiaceae(100);g__Pseudeurotium(100);g__Pseudeurotium_unclassified(100); | Pseudeurotium | 13 | Saprotroph | \|Dung Saprotroph\|-Undefined Saprotroph | Microfungus | NULL | Probable | As soil_saprotroph (Põlme et al. 2020); Decay soil, dung (Põlme et al. 2020) | Tedersoo L, et al. 2014. Science 346:e1256688 ((https://doi.org/10.1126/science.1256688)); Põlme S, et al. 2020. Fungal Diversity 105:1-16 ((https://doi.org/10.1007/s13225-020-00466-2)) |
| Otu61247 | 1 | k__Fungi(100);p__Ascomycota(100);c__Saccharomycetes(100);o__Saccharomycetales(100);f__Metschnikowiaceae(100);g__Metschnikowia(100);g__Metschnikowia_unclassified(100); | Metschnikowia | 13 | Saprotroph | \|Undefined Saprotroph\| | Yeast | NULL | Probable | As nectar-tap_saprotroph (Põlme et al. 2020); Decay sugar-rich substrates (Põlme et al. 2020) | Põlme S, et al. 2020. Fungal Diversity 105:1-16 ((https://doi.org/10.1007/s13225-020-00466-2)) |
| Otu61248 | 1 | k__Fungi(100);p__Ascomycota(100);c__Saccharomycetes(100);o__Saccharomycetales(100);f__Metschnikowiaceae(100);g__Metschnikowia(100);g__Metschnikowia_unclassified(100); | Metschnikowia | 13 | Saprotroph | \|Undefined Saprotroph\| | Yeast | NULL | Probable | As nectar-tap_saprotroph (Põlme et al. 2020); Decay sugar-rich substrates (Põlme et al. 2020) | Põlme S, et al. 2020. Fungal Diversity 105:1-16 ((https://doi.org/10.1007/s13225-020-00466-2)) |
| Otu61254 | 1 | k__Fungi(100);p__Ascomycota(100);c__Saccharomycetes(100);o__Saccharomycetales(100);f__Saccharomycetales_family_Incertae_sedis(100);f__Saccharomycetales_family_Incertae_sedis_unclassified(100);f__Saccharomycetales_family_Incertae_sedis_unclassified(100); | Saccharomycetales | 7 | Saprotroph | Undefined Saprotroph | Yeast | NULL | Possible | NULL | Sterkenburg E, et al. 2015. New Phytologist 207:1145-1158 ((https://doi.org/10.1111/nph.13426)) |
| Otu61258 | 1 | k__Fungi(100);p__Ascomycota(100);c__Saccharomycetes(100);o__Saccharomycetales(100);f__Metschnikowiaceae(100);g__Metschnikowia(100);s__Metschnikowia_pulcherrima(100); | Metschnikowia | 13 | Saprotroph | \|Undefined Saprotroph\| | Yeast | NULL | Probable | As nectar-tap_saprotroph (Põlme et al. 2020); Decay sugar-rich substrates (Põlme et al. 2020) | Põlme S, et al. 2020. Fungal Diversity 105:1-16 ((https://doi.org/10.1007/s13225-020-00466-2)) |
| Otu61261 | 1 | k__Fungi(100);p__Basidiomycota(100);c__Tremellomycetes(100);o__Tremellales(100);f__Tremellales_family_Incertae_sedis(100);g__Cryptococcus(100);s__Cryptococcus_sp_JCM_11356(100); | Cryptococcus | 13 | Pathotroph-Saprotroph-Symbiotroph | Animal Pathogen-Endophyte-Epiphyte-\|Undefined Saprotroph\| | Dimorphic Yeast-Yeast | NULL | Possible | As foliar_endophyte (Põlme et al. 2020); Decay leaf, fruit, seed, soil, dung, animal material (Põlme et al. 2020) | Andrews JH, et al. 1983. Phytopathology 73:228-234 ((https://doi.org/10.1094/Phyto-73-228)); Kurtzman CP, et al. (eds.) 2011. The Yeasts, a Taxonomic Study. Fifth Edition. Vols 1-3. Elsevier, San Diego (ISBN: 9780444521491); Põlme S, et al. 2020. Fungal Diversity 105:1-16 ((https://doi.org/10.1007/s13225-020-00466-2)) |
| Otu61269 | 1 | k__Fungi(100);p__Ascomycota(100);c__Saccharomycetes(100);o__Saccharomycetales(100);f__Saccharomycetales_family_Incertae_sedis(100);g__Candida(100);g__Candida_unclassified(100); | Candida | 13 | Pathotroph-Saprotroph-Symbiotroph | Animal Pathogen-Endophyte-Endosymbiont-Epiphyte-\|Undefined Saprotroph\| | Dimorphic Yeast-Yeast | NULL | Probable | As nectar-tap_saprotroph (Põlme et al. 2020); Decay sugar-rich substrates (Põlme et al. 2020); This asexual genus is polyphyletic and contain species that function very differently depend on what they are. Caution should be taken when interpreting guilds for Candida species. | Manolakaki D, et al. 2010. Virulence. 1: 367-375 ((https://doi.org/10.4161/viru.1.5.12796)); Mekbib SB, et al. 2011. Tropical Plant Pathology 36:233-240 ((https://doi.org/10.1590/S1982-56762011000400004)); Põlme S, et al. 2020. Fungal Diversity 105:1-16 ((https://doi.org/10.1007/s13225-020-00466-2)) |
| Otu61272 | 1 | k__Fungi(100);p__Ascomycota(100);c__Saccharomycetes(100);o__Saccharomycetales(100);f__Metschnikowiaceae(100);g__Metschnikowia(100);g__Metschnikowia_unclassified(100); | Metschnikowia | 13 | Saprotroph | \|Undefined Saprotroph\| | Yeast | NULL | Probable | As nectar-tap_saprotroph (Põlme et al. 2020); Decay sugar-rich substrates (Põlme et al. 2020) | Põlme S, et al. 2020. Fungal Diversity 105:1-16 ((https://doi.org/10.1007/s13225-020-00466-2)) |
| Otu61291 | 1 | k__Fungi(100);p__Ascomycota(100);c__Saccharomycetes(100);o__Saccharomycetales(100);f__Metschnikowiaceae(100);g__Metschnikowia(100);s__Metschnikowia_pulcherrima(100); | Metschnikowia | 13 | Saprotroph | \|Undefined Saprotroph\| | Yeast | NULL | Probable | As nectar-tap_saprotroph (Põlme et al. 2020); Decay sugar-rich substrates (Põlme et al. 2020) | Põlme S, et al. 2020. Fungal Diversity 105:1-16 ((https://doi.org/10.1007/s13225-020-00466-2)) |
| Otu61304 | 1 | k__Fungi(100);p__Ascomycota(100);c__Sordariomycetes(100);o__Coniochaetales(100);f__Coniochaetaceae(100);g__Lecythophora(100);s__Lecythophora_sp(100); | Lecythophora | 13 | Pathotroph-Saprotroph-Symbiotroph | Endophyte-Plant Pathogen-\|Undefined Saprotroph\| | Microfungus | NULL | Highly Probable | As foliar_endophyte (Põlme et al. 2020); Dark Septate Endophyte, known from plant roots.; Decay leaf, fruit, seed, soil, dung, animal material (Põlme et al. 2020) | Taylor DL, et al. 2014. Ecological Monographs 84:3-20 ((https://esajournals.onlinelibrary.wiley.com/doi/10.1890/12-1693.1)); Põlme S, et al. 2020. Fungal Diversity 105:1-16 ((https://doi.org/10.1007/s13225-020-00466-2)) |
| Otu61305 | 1 | k__Fungi(100);p__Ascomycota(100);c__Saccharomycetes(100);o__Saccharomycetales(100);f__Metschnikowiaceae(100);g__Metschnikowia(100);s__Metschnikowia_pulcherrima(100); | Metschnikowia | 13 | Saprotroph | \|Undefined Saprotroph\| | Yeast | NULL | Probable | As nectar-tap_saprotroph (Põlme et al. 2020); Decay sugar-rich substrates (Põlme et al. 2020) | Põlme S, et al. 2020. Fungal Diversity 105:1-16 ((https://doi.org/10.1007/s13225-020-00466-2)) |
| Otu61310 | 1 | k__Fungi(100);p__Ascomycota(100);c__Dothideomycetes(100);o__Capnodiales(100);f__Capnodiales_family_Incertae_sedis(100);g__Capnobotryella(100);s__Capnobotryella_sp_MA_4624(100); | Capnobotryella | 13 | Saprotroph | \|Plant Saprotroph\|-Undefined Saprotroph | Microfungus | NULL | Probable | As litter_saprotroph (Põlme et al. 2020); Decay leaf, fruit, seed, soil (Põlme et al. 2020) | Tedersoo L, et al. 2014. Science 346:e1256688 ((https://doi.org/10.1126/science.1256688)); Põlme S, et al. 2020. Fungal Diversity 105:1-16 ((https://doi.org/10.1007/s13225-020-00466-2)) |
| Otu61312 | 1 | k__Fungi(100);p__Ascomycota(100);c__Lecanoromycetes(100);o__Teloschistales(100);f__Caliciaceae(100);g__Amandinea(100);s__Amandinea_punctata(100); | Amandinea | 13 | Symbiotroph | \|Lichenized\| | Thallus | NULL | Highly Probable | NULL | Esslinger TL. 2014. North Dakota State University ((http://www.ndsu.edu/pubweb/~esslinge/chcklst/chcklst7.htm)); Põlme S, et al. 2020. Fungal Diversity 105:1-16 ((https://doi.org/10.1007/s13225-020-00466-2)) |
| Otu61314 | 1 | k__Fungi(100);p__Ascomycota(100);c__Dothideomycetes(100);o__Pleosporales(100);f__Leptosphaeriaceae(100);g__Leptosphaeria(100);g__Leptosphaeria_unclassified(100); | Leptosphaeria | 13 | Pathotroph-Saprotroph | \|Plant Pathogen\|-Wood Saprotroph | Microfungus | NULL | Probable | As wood_pathogen (Põlme et al. 2020); Decay wood (Põlme et al. 2020) | Tedersoo L, et al. 2014. Science 346:e1256688 ((https://doi.org/10.1126/science.1256688)); Põlme S, et al. 2020. Fungal Diversity 105:1-16 ((https://doi.org/10.1007/s13225-020-00466-2)) |
| Otu61319 | 1 | k__Fungi(100);p__Ascomycota(100);c__Dothideomycetes(100);o__Capnodiales(100);f__Mycosphaerellaceae(100);g__Ramichloridium(100);s__Ramichloridium_luteum(100); | Ramichloridium | 13 | Pathotroph-Pathotroph-Saprotroph | \|Plant Pathogen\|-Plant Saprotroph | Microfungus | NULL | Probable | As litter_saprotroph (Põlme et al. 2020); Decay leaf, fruit, seed (Põlme et al. 2020) | Tedersoo L, et al. 2014. Science 346:e1256688 ((https://doi.org/10.1126/science.1256688)); Põlme S, et al. 2020. Fungal Diversity 105:1-16 ((https://doi.org/10.1007/s13225-020-00466-2)) |
| Otu61321 | 1 | k__Fungi(100);p__Ascomycota(100);c__Saccharomycetes(100);o__Saccharomycetales(100);f__Metschnikowiaceae(100);g__Metschnikowia(100);s__Metschnikowia_pulcherrima(100); | Metschnikowia | 13 | Saprotroph | \|Undefined Saprotroph\| | Yeast | NULL | Probable | As nectar-tap_saprotroph (Põlme et al. 2020); Decay sugar-rich substrates (Põlme et al. 2020) | Põlme S, et al. 2020. Fungal Diversity 105:1-16 ((https://doi.org/10.1007/s13225-020-00466-2)) |
| Otu61332 | 1 | k__Fungi(100);p__Ascomycota(100);c__Dothideomycetes(100);o__Pleosporales(100);f__Pleosporales_family_Incertae_sedis(100);g__Phoma(100);g__Phoma_unclassified(100); | Phoma | 13 | Pathotroph-Saprotroph-Symbiotroph | Dung Saprotroph-Endophyte-Lichen Parasite-\|Plant Pathogen\|-Plant Saprotroph-Undefined Saprotroph-Wood Saprotroph | Microfungus | Soft Rot | Possible | As foliar_endophyte (Põlme et al. 2020); As litter_saprotroph (Põlme et al. 2020); Decay leaf, fruit, seed, roots, soil (Põlme et al. 2020) | Richardson MJ. 1972. Transactions of the British Mycological Society 58:37-48 ((https://doi.org/10.1016/S0007-1536(72)80069-X)); Seehann G, et al. 1975. List of Fungi in Soft-Rot Tissue Tests. Institute fur Holzbiologie und Holzschutz, Bundesforschungsanstalt fur Forst - und Holzwirtshaft, Hamburg - Lohbrugge, Germany; de Gruyter J, et al. 2010. Mycologia 102:1066-1081 ((https://doi.org/10.3852/09-240)); Costa IPMW, et al. 2012. Mycotaxon 119:e26 ((http://mycotaxon.com/resources/checklists/costa_v119_checklist.pdf)); Tedersoo L, et al. 2014. Science 346:e1256688 ((https://doi.org/10.1126/science.1256688)); Lawrey JD, Diederich P. 2016. http://www.lichenicolous.net (accessed 24 May 2017) ((http://www.lichenicolous.net)); Põlme S, et al. 2020. Fungal Diversity 105:1-16 ((https://doi.org/10.1007/s13225-020-00466-2)) |
| Otu61333 | 1 | k__Fungi(100);p__Ascomycota(100);c__Saccharomycetes(100);o__Saccharomycetales(100);f__Metschnikowiaceae(100);g__Metschnikowia(100);s__Metschnikowia_pulcherrima(100); | Metschnikowia | 13 | Saprotroph | \|Undefined Saprotroph\| | Yeast | NULL | Probable | As nectar-tap_saprotroph (Põlme et al. 2020); Decay sugar-rich substrates (Põlme et al. 2020) | Põlme S, et al. 2020. Fungal Diversity 105:1-16 ((https://doi.org/10.1007/s13225-020-00466-2)) |
| Otu61355 | 1 | k__Fungi(100);p__Ascomycota(100);c__Saccharomycetes(100);o__Saccharomycetales(100);f__Metschnikowiaceae(100);g__Metschnikowia(100);s__Metschnikowia_pulcherrima(100); | Metschnikowia | 13 | Saprotroph | \|Undefined Saprotroph\| | Yeast | NULL | Probable | As nectar-tap_saprotroph (Põlme et al. 2020); Decay sugar-rich substrates (Põlme et al. 2020) | Põlme S, et al. 2020. Fungal Diversity 105:1-16 ((https://doi.org/10.1007/s13225-020-00466-2)) |
| Otu61360 | 1 | k__Fungi(100);p__Ascomycota(100);c__Saccharomycetes(100);o__Saccharomycetales(100);f__Saccharomycetales_family_Incertae_sedis(100);f__Saccharomycetales_family_Incertae_sedis_unclassified(100);f__Saccharomycetales_family_Incertae_sedis_unclassified(100); | Saccharomycetales | 7 | Saprotroph | Undefined Saprotroph | Yeast | NULL | Possible | NULL | Sterkenburg E, et al. 2015. New Phytologist 207:1145-1158 ((https://doi.org/10.1111/nph.13426)) |
| Otu61361 | 1 | k__Fungi(100);p__Ascomycota(100);c__Saccharomycetes(100);o__Saccharomycetales(100);f__Saccharomycetales_family_Incertae_sedis(100);g__Candida(100);g__Candida_unclassified(100); | Candida | 13 | Pathotroph-Saprotroph-Symbiotroph | Animal Pathogen-Endophyte-Endosymbiont-Epiphyte-\|Undefined Saprotroph\| | Dimorphic Yeast-Yeast | NULL | Probable | As nectar-tap_saprotroph (Põlme et al. 2020); Decay sugar-rich substrates (Põlme et al. 2020); This asexual genus is polyphyletic and contain species that function very differently depend on what they are. Caution should be taken when interpreting guilds for Candida species. | Manolakaki D, et al. 2010. Virulence. 1: 367-375 ((https://doi.org/10.4161/viru.1.5.12796)); Mekbib SB, et al. 2011. Tropical Plant Pathology 36:233-240 ((https://doi.org/10.1590/S1982-56762011000400004)); Põlme S, et al. 2020. Fungal Diversity 105:1-16 ((https://doi.org/10.1007/s13225-020-00466-2)) |
| Otu61362 | 1 | k__Fungi(100);p__Ascomycota(100);c__Saccharomycetes(100);o__Saccharomycetales(100);f__Saccharomycetales_family_Incertae_sedis(100);g__Candida(100);g__Candida_unclassified(100); | Candida | 13 | Pathotroph-Saprotroph-Symbiotroph | Animal Pathogen-Endophyte-Endosymbiont-Epiphyte-\|Undefined Saprotroph\| | Dimorphic Yeast-Yeast | NULL | Probable | As nectar-tap_saprotroph (Põlme et al. 2020); Decay sugar-rich substrates (Põlme et al. 2020); This asexual genus is polyphyletic and contain species that function very differently depend on what they are. Caution should be taken when interpreting guilds for Candida species. | Manolakaki D, et al. 2010. Virulence. 1: 367-375 ((https://doi.org/10.4161/viru.1.5.12796)); Mekbib SB, et al. 2011. Tropical Plant Pathology 36:233-240 ((https://doi.org/10.1590/S1982-56762011000400004)); Põlme S, et al. 2020. Fungal Diversity 105:1-16 ((https://doi.org/10.1007/s13225-020-00466-2)) |
| Otu61397 | 1 | k__Fungi(100);p__Ascomycota(100);c__Saccharomycetes(100);o__Saccharomycetales(100);f__Metschnikowiaceae(100);g__Metschnikowia(100);g__Metschnikowia_unclassified(100); | Metschnikowia | 13 | Saprotroph | \|Undefined Saprotroph\| | Yeast | NULL | Probable | As nectar-tap_saprotroph (Põlme et al. 2020); Decay sugar-rich substrates (Põlme et al. 2020) | Põlme S, et al. 2020. Fungal Diversity 105:1-16 ((https://doi.org/10.1007/s13225-020-00466-2)) |
| Otu61408 | 1 | k__Fungi(100);p__Ascomycota(100);c__Saccharomycetes(100);o__Saccharomycetales(100);f__Metschnikowiaceae(100);g__Metschnikowia(100);s__Metschnikowia_pulcherrima(100); | Metschnikowia | 13 | Saprotroph | \|Undefined Saprotroph\| | Yeast | NULL | Probable | As nectar-tap_saprotroph (Põlme et al. 2020); Decay sugar-rich substrates (Põlme et al. 2020) | Põlme S, et al. 2020. Fungal Diversity 105:1-16 ((https://doi.org/10.1007/s13225-020-00466-2)) |
| Otu61409 | 1 | k__Fungi(100);p__Ascomycota(100);c__Saccharomycetes(100);o__Saccharomycetales(100);f__Metschnikowiaceae(100);g__Metschnikowia(100);g__Metschnikowia_unclassified(100); | Metschnikowia | 13 | Saprotroph | \|Undefined Saprotroph\| | Yeast | NULL | Probable | As nectar-tap_saprotroph (Põlme et al. 2020); Decay sugar-rich substrates (Põlme et al. 2020) | Põlme S, et al. 2020. Fungal Diversity 105:1-16 ((https://doi.org/10.1007/s13225-020-00466-2)) |
| Otu61410 | 1 | k__Fungi(100);p__Ascomycota(100);c__Saccharomycetes(100);o__Saccharomycetales(100);f__Metschnikowiaceae(100);g__Metschnikowia(100);s__Metschnikowia_pulcherrima(100); | Metschnikowia | 13 | Saprotroph | \|Undefined Saprotroph\| | Yeast | NULL | Probable | As nectar-tap_saprotroph (Põlme et al. 2020); Decay sugar-rich substrates (Põlme et al. 2020) | Põlme S, et al. 2020. Fungal Diversity 105:1-16 ((https://doi.org/10.1007/s13225-020-00466-2)) |
| Otu61411 | 1 | k__Fungi(100);p__Ascomycota(100);c__Saccharomycetes(100);o__Saccharomycetales(100);f__Metschnikowiaceae(100);g__Metschnikowia(100);g__Metschnikowia_unclassified(100); | Metschnikowia | 13 | Saprotroph | \|Undefined Saprotroph\| | Yeast | NULL | Probable | As nectar-tap_saprotroph (Põlme et al. 2020); Decay sugar-rich substrates (Põlme et al. 2020) | Põlme S, et al. 2020. Fungal Diversity 105:1-16 ((https://doi.org/10.1007/s13225-020-00466-2)) |
| Otu61412 | 1 | k__Fungi(100);p__Ascomycota(100);c__Saccharomycetes(100);o__Saccharomycetales(100);f__Metschnikowiaceae(100);g__Metschnikowia(100);g__Metschnikowia_unclassified(100); | Metschnikowia | 13 | Saprotroph | \|Undefined Saprotroph\| | Yeast | NULL | Probable | As nectar-tap_saprotroph (Põlme et al. 2020); Decay sugar-rich substrates (Põlme et al. 2020) | Põlme S, et al. 2020. Fungal Diversity 105:1-16 ((https://doi.org/10.1007/s13225-020-00466-2)) |
| Otu61413 | 1 | k__Fungi(100);p__Ascomycota(100);c__Saccharomycetes(100);o__Saccharomycetales(100);f__Metschnikowiaceae(100);g__Metschnikowia(100);g__Metschnikowia_unclassified(100); | Metschnikowia | 13 | Saprotroph | \|Undefined Saprotroph\| | Yeast | NULL | Probable | As nectar-tap_saprotroph (Põlme et al. 2020); Decay sugar-rich substrates (Põlme et al. 2020) | Põlme S, et al. 2020. Fungal Diversity 105:1-16 ((https://doi.org/10.1007/s13225-020-00466-2)) |
| Otu61414 | 1 | k__Fungi(100);p__Ascomycota(100);c__Saccharomycetes(100);o__Saccharomycetales(100);f__Metschnikowiaceae(100);g__Metschnikowia(100);g__Metschnikowia_unclassified(100); | Metschnikowia | 13 | Saprotroph | \|Undefined Saprotroph\| | Yeast | NULL | Probable | As nectar-tap_saprotroph (Põlme et al. 2020); Decay sugar-rich substrates (Põlme et al. 2020) | Põlme S, et al. 2020. Fungal Diversity 105:1-16 ((https://doi.org/10.1007/s13225-020-00466-2)) |
| Otu61415 | 1 | k__Fungi(100);p__Ascomycota(100);c__Sordariomycetes(100);o__Sordariales(100);f__Cephalothecaceae(100);g__Phialemonium(100);s__Phialemonium_atrogriseum(100); | Phialemonium | 13 | Pathotroph-Saprotroph | \|Animal Parasite\|-Undefined Saprotroph | Microfungus | NULL | Probable | As animal_decomposer (Põlme et al. 2020); Decay animal material (Põlme et al. 2020) | Tedersoo L, et al. 2014. Science 346:e1256688 ((https://doi.org/10.1126/science.1256688)); Põlme S, et al. 2020. Fungal Diversity 105:1-16 ((https://doi.org/10.1007/s13225-020-00466-2)) |
| Otu61416 | 1 | k__Fungi(100);p__Ascomycota(100);c__Lecanoromycetes(100);o__Lecanorales(100);o__Lecanorales_unclassified(100);o__Lecanorales_unclassified(100);o__Lecanorales_unclassified(100); | Lecanorales | 7 | Pathotroph-Symbiotroph | Lichen Parasite-Lichenized | Microfungus-Thallus | NULL | Probable | Lichenized with green algae, occasionally lichenicolous, on rocks, mosses and wood (Cannon & Kirk 2007); ...an order of mostly lichen-forming fungi (Kirk et al. 2008) | Cannon PF, Kirk PM. 2007. Fungal Families of the World. CAB International, Cambridge (ISBN: 978-0851998275); Kirk PM et al. 2008. Dictionary of the Fungi. Tenth Edition. CAB International, Wallingford (ISBN: 978-0851998268); Esslinger TL. 2014. North Dakota State University ((http://www.ndsu.edu/pubweb/~esslinge/chcklst/chcklst7.htm)) |
| Otu61417 | 1 | k__Fungi(100);p__Ascomycota(100);c__Saccharomycetes(100);o__Saccharomycetales(100);f__Metschnikowiaceae(100);g__Metschnikowia(100);g__Metschnikowia_unclassified(100); | Metschnikowia | 13 | Saprotroph | \|Undefined Saprotroph\| | Yeast | NULL | Probable | As nectar-tap_saprotroph (Põlme et al. 2020); Decay sugar-rich substrates (Põlme et al. 2020) | Põlme S, et al. 2020. Fungal Diversity 105:1-16 ((https://doi.org/10.1007/s13225-020-00466-2)) |
| Otu61418 | 1 | k__Fungi(100);p__Ascomycota(100);c__Saccharomycetes(100);o__Saccharomycetales(100);f__Metschnikowiaceae(100);g__Metschnikowia(100);g__Metschnikowia_unclassified(100); | Metschnikowia | 13 | Saprotroph | \|Undefined Saprotroph\| | Yeast | NULL | Probable | As nectar-tap_saprotroph (Põlme et al. 2020); Decay sugar-rich substrates (Põlme et al. 2020) | Põlme S, et al. 2020. Fungal Diversity 105:1-16 ((https://doi.org/10.1007/s13225-020-00466-2)) |
| Otu61419 | 1 | k__Fungi(100);p__Ascomycota(100);c__Saccharomycetes(100);o__Saccharomycetales(100);f__Metschnikowiaceae(100);g__Metschnikowia(100);s__Metschnikowia_pulcherrima(100); | Metschnikowia | 13 | Saprotroph | \|Undefined Saprotroph\| | Yeast | NULL | Probable | As nectar-tap_saprotroph (Põlme et al. 2020); Decay sugar-rich substrates (Põlme et al. 2020) | Põlme S, et al. 2020. Fungal Diversity 105:1-16 ((https://doi.org/10.1007/s13225-020-00466-2)) |
| Otu61420 | 1 | k__Fungi(100);p__Ascomycota(100);c__Saccharomycetes(100);o__Saccharomycetales(100);f__Metschnikowiaceae(100);g__Metschnikowia(100);s__Metschnikowia_pulcherrima(100); | Metschnikowia | 13 | Saprotroph | \|Undefined Saprotroph\| | Yeast | NULL | Probable | As nectar-tap_saprotroph (Põlme et al. 2020); Decay sugar-rich substrates (Põlme et al. 2020) | Põlme S, et al. 2020. Fungal Diversity 105:1-16 ((https://doi.org/10.1007/s13225-020-00466-2)) |
| Otu61421 | 1 | k__Fungi(100);p__Ascomycota(100);c__Saccharomycetes(100);o__Saccharomycetales(100);f__Metschnikowiaceae(100);g__Metschnikowia(100);s__Metschnikowia_pulcherrima(100); | Metschnikowia | 13 | Saprotroph | \|Undefined Saprotroph\| | Yeast | NULL | Probable | As nectar-tap_saprotroph (Põlme et al. 2020); Decay sugar-rich substrates (Põlme et al. 2020) | Põlme S, et al. 2020. Fungal Diversity 105:1-16 ((https://doi.org/10.1007/s13225-020-00466-2)) |
| Otu61422 | 1 | k__Fungi(100);p__Ascomycota(100);c__Saccharomycetes(100);o__Saccharomycetales(100);f__Metschnikowiaceae(100);g__Metschnikowia(100);s__Metschnikowia_pulcherrima(100); | Metschnikowia | 13 | Saprotroph | \|Undefined Saprotroph\| | Yeast | NULL | Probable | As nectar-tap_saprotroph (Põlme et al. 2020); Decay sugar-rich substrates (Põlme et al. 2020) | Põlme S, et al. 2020. Fungal Diversity 105:1-16 ((https://doi.org/10.1007/s13225-020-00466-2)) |
| Otu61423 | 1 | k__Fungi(100);p__Ascomycota(100);c__Saccharomycetes(100);o__Saccharomycetales(100);f__Metschnikowiaceae(100);g__Metschnikowia(100);s__Metschnikowia_pulcherrima(100); | Metschnikowia | 13 | Saprotroph | \|Undefined Saprotroph\| | Yeast | NULL | Probable | As nectar-tap_saprotroph (Põlme et al. 2020); Decay sugar-rich substrates (Põlme et al. 2020) | Põlme S, et al. 2020. Fungal Diversity 105:1-16 ((https://doi.org/10.1007/s13225-020-00466-2)) |
| Otu61424 | 1 | k__Fungi(100);p__Ascomycota(100);c__Saccharomycetes(100);o__Saccharomycetales(100);f__Metschnikowiaceae(100);g__Metschnikowia(100);s__Metschnikowia_pulcherrima(100); | Metschnikowia | 13 | Saprotroph | \|Undefined Saprotroph\| | Yeast | NULL | Probable | As nectar-tap_saprotroph (Põlme et al. 2020); Decay sugar-rich substrates (Põlme et al. 2020) | Põlme S, et al. 2020. Fungal Diversity 105:1-16 ((https://doi.org/10.1007/s13225-020-00466-2)) |
| Otu61425 | 1 | k__Fungi(100);p__Ascomycota(100);c__Saccharomycetes(100);o__Saccharomycetales(100);f__Metschnikowiaceae(100);g__Metschnikowia(100);g__Metschnikowia_unclassified(100); | Metschnikowia | 13 | Saprotroph | \|Undefined Saprotroph\| | Yeast | NULL | Probable | As nectar-tap_saprotroph (Põlme et al. 2020); Decay sugar-rich substrates (Põlme et al. 2020) | Põlme S, et al. 2020. Fungal Diversity 105:1-16 ((https://doi.org/10.1007/s13225-020-00466-2)) |
| Otu61426 | 1 | k__Fungi(100);p__Ascomycota(100);c__Saccharomycetes(100);o__Saccharomycetales(100);f__Metschnikowiaceae(100);g__Metschnikowia(100);s__Metschnikowia_pulcherrima(100); | Metschnikowia | 13 | Saprotroph | \|Undefined Saprotroph\| | Yeast | NULL | Probable | As nectar-tap_saprotroph (Põlme et al. 2020); Decay sugar-rich substrates (Põlme et al. 2020) | Põlme S, et al. 2020. Fungal Diversity 105:1-16 ((https://doi.org/10.1007/s13225-020-00466-2)) |
| Otu61427 | 1 | k__Fungi(100);p__Ascomycota(100);c__Saccharomycetes(100);o__Saccharomycetales(100);f__Metschnikowiaceae(100);g__Metschnikowia(100);g__Metschnikowia_unclassified(100); | Metschnikowia | 13 | Saprotroph | \|Undefined Saprotroph\| | Yeast | NULL | Probable | As nectar-tap_saprotroph (Põlme et al. 2020); Decay sugar-rich substrates (Põlme et al. 2020) | Põlme S, et al. 2020. Fungal Diversity 105:1-16 ((https://doi.org/10.1007/s13225-020-00466-2)) |
| Otu61428 | 1 | k__Fungi(100);p__Ascomycota(100);c__Saccharomycetes(100);o__Saccharomycetales(100);f__Metschnikowiaceae(100);g__Metschnikowia(100);s__Metschnikowia_pulcherrima(100); | Metschnikowia | 13 | Saprotroph | \|Undefined Saprotroph\| | Yeast | NULL | Probable | As nectar-tap_saprotroph (Põlme et al. 2020); Decay sugar-rich substrates (Põlme et al. 2020) | Põlme S, et al. 2020. Fungal Diversity 105:1-16 ((https://doi.org/10.1007/s13225-020-00466-2)) |
| Otu61429 | 1 | k__Fungi(100);p__Ascomycota(100);c__Saccharomycetes(100);o__Saccharomycetales(100);f__Metschnikowiaceae(100);g__Metschnikowia(100);g__Metschnikowia_unclassified(100); | Metschnikowia | 13 | Saprotroph | \|Undefined Saprotroph\| | Yeast | NULL | Probable | As nectar-tap_saprotroph (Põlme et al. 2020); Decay sugar-rich substrates (Põlme et al. 2020) | Põlme S, et al. 2020. Fungal Diversity 105:1-16 ((https://doi.org/10.1007/s13225-020-00466-2)) |
| Otu61430 | 1 | k__Fungi(100);p__Ascomycota(100);c__Saccharomycetes(100);o__Saccharomycetales(100);f__Metschnikowiaceae(100);g__Metschnikowia(100);g__Metschnikowia_unclassified(100); | Metschnikowia | 13 | Saprotroph | \|Undefined Saprotroph\| | Yeast | NULL | Probable | As nectar-tap_saprotroph (Põlme et al. 2020); Decay sugar-rich substrates (Põlme et al. 2020) | Põlme S, et al. 2020. Fungal Diversity 105:1-16 ((https://doi.org/10.1007/s13225-020-00466-2)) |
| Otu61431 | 1 | k__Fungi(100);p__Ascomycota(100);c__Saccharomycetes(100);o__Saccharomycetales(100);f__Metschnikowiaceae(100);g__Metschnikowia(100);s__Metschnikowia_pulcherrima(100); | Metschnikowia | 13 | Saprotroph | \|Undefined Saprotroph\| | Yeast | NULL | Probable | As nectar-tap_saprotroph (Põlme et al. 2020); Decay sugar-rich substrates (Põlme et al. 2020) | Põlme S, et al. 2020. Fungal Diversity 105:1-16 ((https://doi.org/10.1007/s13225-020-00466-2)) |
| Otu61432 | 1 | k__Fungi(100);p__Ascomycota(100);c__Saccharomycetes(100);o__Saccharomycetales(100);f__Metschnikowiaceae(100);g__Metschnikowia(100);s__Metschnikowia_pulcherrima(100); | Metschnikowia | 13 | Saprotroph | \|Undefined Saprotroph\| | Yeast | NULL | Probable | As nectar-tap_saprotroph (Põlme et al. 2020); Decay sugar-rich substrates (Põlme et al. 2020) | Põlme S, et al. 2020. Fungal Diversity 105:1-16 ((https://doi.org/10.1007/s13225-020-00466-2)) |
| Otu61433 | 1 | k__Fungi(100);p__Ascomycota(100);c__Saccharomycetes(100);o__Saccharomycetales(100);f__Metschnikowiaceae(100);g__Metschnikowia(100);s__Metschnikowia_pulcherrima(100); | Metschnikowia | 13 | Saprotroph | \|Undefined Saprotroph\| | Yeast | NULL | Probable | As nectar-tap_saprotroph (Põlme et al. 2020); Decay sugar-rich substrates (Põlme et al. 2020) | Põlme S, et al. 2020. Fungal Diversity 105:1-16 ((https://doi.org/10.1007/s13225-020-00466-2)) |
| Otu61434 | 1 | k__Fungi(100);p__Ascomycota(100);c__Saccharomycetes(100);o__Saccharomycetales(100);f__Metschnikowiaceae(100);g__Metschnikowia(100);s__Metschnikowia_pulcherrima(100); | Metschnikowia | 13 | Saprotroph | \|Undefined Saprotroph\| | Yeast | NULL | Probable | As nectar-tap_saprotroph (Põlme et al. 2020); Decay sugar-rich substrates (Põlme et al. 2020) | Põlme S, et al. 2020. Fungal Diversity 105:1-16 ((https://doi.org/10.1007/s13225-020-00466-2)) |
| Otu61435 | 1 | k__Fungi(100);p__Ascomycota(100);c__Saccharomycetes(100);o__Saccharomycetales(100);f__Metschnikowiaceae(100);g__Metschnikowia(100);g__Metschnikowia_unclassified(100); | Metschnikowia | 13 | Saprotroph | \|Undefined Saprotroph\| | Yeast | NULL | Probable | As nectar-tap_saprotroph (Põlme et al. 2020); Decay sugar-rich substrates (Põlme et al. 2020) | Põlme S, et al. 2020. Fungal Diversity 105:1-16 ((https://doi.org/10.1007/s13225-020-00466-2)) |
| Otu61436 | 1 | k__Fungi(100);p__Ascomycota(100);c__Saccharomycetes(100);o__Saccharomycetales(100);f__Metschnikowiaceae(100);g__Metschnikowia(100);s__Metschnikowia_pulcherrima(100); | Metschnikowia | 13 | Saprotroph | \|Undefined Saprotroph\| | Yeast | NULL | Probable | As nectar-tap_saprotroph (Põlme et al. 2020); Decay sugar-rich substrates (Põlme et al. 2020) | Põlme S, et al. 2020. Fungal Diversity 105:1-16 ((https://doi.org/10.1007/s13225-020-00466-2)) |
| Otu61437 | 1 | k__Fungi(100);p__Ascomycota(100);c__Saccharomycetes(100);o__Saccharomycetales(100);f__Metschnikowiaceae(100);g__Metschnikowia(100);s__Metschnikowia_pulcherrima(100); | Metschnikowia | 13 | Saprotroph | \|Undefined Saprotroph\| | Yeast | NULL | Probable | As nectar-tap_saprotroph (Põlme et al. 2020); Decay sugar-rich substrates (Põlme et al. 2020) | Põlme S, et al. 2020. Fungal Diversity 105:1-16 ((https://doi.org/10.1007/s13225-020-00466-2)) |
| Otu61438 | 1 | k__Fungi(100);p__Ascomycota(100);c__Saccharomycetes(100);o__Saccharomycetales(100);f__Metschnikowiaceae(100);g__Metschnikowia(100);s__Metschnikowia_pulcherrima(100); | Metschnikowia | 13 | Saprotroph | \|Undefined Saprotroph\| | Yeast | NULL | Probable | As nectar-tap_saprotroph (Põlme et al. 2020); Decay sugar-rich substrates (Põlme et al. 2020) | Põlme S, et al. 2020. Fungal Diversity 105:1-16 ((https://doi.org/10.1007/s13225-020-00466-2)) |
| Otu61439 | 1 | k__Fungi(100);p__Ascomycota(100);c__Saccharomycetes(100);o__Saccharomycetales(100);f__Metschnikowiaceae(100);g__Metschnikowia(100);s__Metschnikowia_pulcherrima(100); | Metschnikowia | 13 | Saprotroph | \|Undefined Saprotroph\| | Yeast | NULL | Probable | As nectar-tap_saprotroph (Põlme et al. 2020); Decay sugar-rich substrates (Põlme et al. 2020) | Põlme S, et al. 2020. Fungal Diversity 105:1-16 ((https://doi.org/10.1007/s13225-020-00466-2)) |
| Otu61440 | 1 | k__Fungi(100);p__Ascomycota(100);c__Saccharomycetes(100);o__Saccharomycetales(100);f__Metschnikowiaceae(100);g__Metschnikowia(100);s__Metschnikowia_pulcherrima(100); | Metschnikowia | 13 | Saprotroph | \|Undefined Saprotroph\| | Yeast | NULL | Probable | As nectar-tap_saprotroph (Põlme et al. 2020); Decay sugar-rich substrates (Põlme et al. 2020) | Põlme S, et al. 2020. Fungal Diversity 105:1-16 ((https://doi.org/10.1007/s13225-020-00466-2)) |
| Otu61441 | 1 | k__Fungi(100);p__Ascomycota(100);c__Saccharomycetes(100);o__Saccharomycetales(100);f__Metschnikowiaceae(100);g__Metschnikowia(100);s__Metschnikowia_pulcherrima(100); | Metschnikowia | 13 | Saprotroph | \|Undefined Saprotroph\| | Yeast | NULL | Probable | As nectar-tap_saprotroph (Põlme et al. 2020); Decay sugar-rich substrates (Põlme et al. 2020) | Põlme S, et al. 2020. Fungal Diversity 105:1-16 ((https://doi.org/10.1007/s13225-020-00466-2)) |
| Otu61442 | 1 | k__Fungi(100);p__Ascomycota(100);c__Saccharomycetes(100);o__Saccharomycetales(100);f__Metschnikowiaceae(100);g__Metschnikowia(100);s__Metschnikowia_pulcherrima(100); | Metschnikowia | 13 | Saprotroph | \|Undefined Saprotroph\| | Yeast | NULL | Probable | As nectar-tap_saprotroph (Põlme et al. 2020); Decay sugar-rich substrates (Põlme et al. 2020) | Põlme S, et al. 2020. Fungal Diversity 105:1-16 ((https://doi.org/10.1007/s13225-020-00466-2)) |
| Otu61443 | 1 | k__Fungi(100);p__Ascomycota(100);c__Saccharomycetes(100);o__Saccharomycetales(100);f__Metschnikowiaceae(100);g__Metschnikowia(100);g__Metschnikowia_unclassified(100); | Metschnikowia | 13 | Saprotroph | \|Undefined Saprotroph\| | Yeast | NULL | Probable | As nectar-tap_saprotroph (Põlme et al. 2020); Decay sugar-rich substrates (Põlme et al. 2020) | Põlme S, et al. 2020. Fungal Diversity 105:1-16 ((https://doi.org/10.1007/s13225-020-00466-2)) |
| Otu61444 | 1 | k__Fungi(100);p__Ascomycota(100);c__Saccharomycetes(100);o__Saccharomycetales(100);f__Metschnikowiaceae(100);g__Metschnikowia(100);s__Metschnikowia_pulcherrima(100); | Metschnikowia | 13 | Saprotroph | \|Undefined Saprotroph\| | Yeast | NULL | Probable | As nectar-tap_saprotroph (Põlme et al. 2020); Decay sugar-rich substrates (Põlme et al. 2020) | Põlme S, et al. 2020. Fungal Diversity 105:1-16 ((https://doi.org/10.1007/s13225-020-00466-2)) |
| Otu61445 | 1 | k__Fungi(100);p__Ascomycota(100);c__Saccharomycetes(100);o__Saccharomycetales(100);f__Metschnikowiaceae(100);g__Metschnikowia(100);s__Metschnikowia_pulcherrima(100); | Metschnikowia | 13 | Saprotroph | \|Undefined Saprotroph\| | Yeast | NULL | Probable | As nectar-tap_saprotroph (Põlme et al. 2020); Decay sugar-rich substrates (Põlme et al. 2020) | Põlme S, et al. 2020. Fungal Diversity 105:1-16 ((https://doi.org/10.1007/s13225-020-00466-2)) |
| Otu61446 | 1 | k__Fungi(100);p__Ascomycota(100);c__Saccharomycetes(100);o__Saccharomycetales(100);f__Metschnikowiaceae(100);g__Metschnikowia(100);s__Metschnikowia_pulcherrima(100); | Metschnikowia | 13 | Saprotroph | \|Undefined Saprotroph\| | Yeast | NULL | Probable | As nectar-tap_saprotroph (Põlme et al. 2020); Decay sugar-rich substrates (Põlme et al. 2020) | Põlme S, et al. 2020. Fungal Diversity 105:1-16 ((https://doi.org/10.1007/s13225-020-00466-2)) |
| Otu61447 | 1 | k__Fungi(100);p__Ascomycota(100);c__Saccharomycetes(100);o__Saccharomycetales(100);f__Metschnikowiaceae(100);g__Metschnikowia(100);s__Metschnikowia_pulcherrima(100); | Metschnikowia | 13 | Saprotroph | \|Undefined Saprotroph\| | Yeast | NULL | Probable | As nectar-tap_saprotroph (Põlme et al. 2020); Decay sugar-rich substrates (Põlme et al. 2020) | Põlme S, et al. 2020. Fungal Diversity 105:1-16 ((https://doi.org/10.1007/s13225-020-00466-2)) |
| Otu61448 | 1 | k__Fungi(100);p__Ascomycota(100);c__Saccharomycetes(100);o__Saccharomycetales(100);f__Metschnikowiaceae(100);g__Metschnikowia(100);g__Metschnikowia_unclassified(100); | Metschnikowia | 13 | Saprotroph | \|Undefined Saprotroph\| | Yeast | NULL | Probable | As nectar-tap_saprotroph (Põlme et al. 2020); Decay sugar-rich substrates (Põlme et al. 2020) | Põlme S, et al. 2020. Fungal Diversity 105:1-16 ((https://doi.org/10.1007/s13225-020-00466-2)) |
| Otu61449 | 1 | k__Fungi(100);p__Ascomycota(100);c__Saccharomycetes(100);o__Saccharomycetales(100);f__Metschnikowiaceae(100);g__Metschnikowia(100);s__Metschnikowia_pulcherrima(100); | Metschnikowia | 13 | Saprotroph | \|Undefined Saprotroph\| | Yeast | NULL | Probable | As nectar-tap_saprotroph (Põlme et al. 2020); Decay sugar-rich substrates (Põlme et al. 2020) | Põlme S, et al. 2020. Fungal Diversity 105:1-16 ((https://doi.org/10.1007/s13225-020-00466-2)) |
| Otu61451 | 1 | k__Fungi(100);p__Ascomycota(100);c__Saccharomycetes(100);o__Saccharomycetales(100);f__Metschnikowiaceae(100);g__Metschnikowia(100);s__Metschnikowia_pulcherrima(100); | Metschnikowia | 13 | Saprotroph | \|Undefined Saprotroph\| | Yeast | NULL | Probable | As nectar-tap_saprotroph (Põlme et al. 2020); Decay sugar-rich substrates (Põlme et al. 2020) | Põlme S, et al. 2020. Fungal Diversity 105:1-16 ((https://doi.org/10.1007/s13225-020-00466-2)) |
| Otu61452 | 1 | k__Fungi(100);p__Ascomycota(100);c__Saccharomycetes(100);o__Saccharomycetales(100);f__Metschnikowiaceae(100);g__Metschnikowia(100);s__Metschnikowia_pulcherrima(100); | Metschnikowia | 13 | Saprotroph | \|Undefined Saprotroph\| | Yeast | NULL | Probable | As nectar-tap_saprotroph (Põlme et al. 2020); Decay sugar-rich substrates (Põlme et al. 2020) | Põlme S, et al. 2020. Fungal Diversity 105:1-16 ((https://doi.org/10.1007/s13225-020-00466-2)) |
| Otu61453 | 1 | k__Fungi(100);p__Ascomycota(100);c__Saccharomycetes(100);o__Saccharomycetales(100);f__Metschnikowiaceae(100);g__Metschnikowia(100);g__Metschnikowia_unclassified(100); | Metschnikowia | 13 | Saprotroph | \|Undefined Saprotroph\| | Yeast | NULL | Probable | As nectar-tap_saprotroph (Põlme et al. 2020); Decay sugar-rich substrates (Põlme et al. 2020) | Põlme S, et al. 2020. Fungal Diversity 105:1-16 ((https://doi.org/10.1007/s13225-020-00466-2)) |
| Otu61454 | 1 | k__Fungi(100);p__Ascomycota(100);c__Saccharomycetes(100);o__Saccharomycetales(100);f__Metschnikowiaceae(100);g__Metschnikowia(100);s__Metschnikowia_pulcherrima(100); | Metschnikowia | 13 | Saprotroph | \|Undefined Saprotroph\| | Yeast | NULL | Probable | As nectar-tap_saprotroph (Põlme et al. 2020); Decay sugar-rich substrates (Põlme et al. 2020) | Põlme S, et al. 2020. Fungal Diversity 105:1-16 ((https://doi.org/10.1007/s13225-020-00466-2)) |
| Otu61455 | 1 | k__Fungi(100);p__Ascomycota(100);c__Saccharomycetes(100);o__Saccharomycetales(100);f__Metschnikowiaceae(100);g__Metschnikowia(100);s__Metschnikowia_pulcherrima(100); | Metschnikowia | 13 | Saprotroph | \|Undefined Saprotroph\| | Yeast | NULL | Probable | As nectar-tap_saprotroph (Põlme et al. 2020); Decay sugar-rich substrates (Põlme et al. 2020) | Põlme S, et al. 2020. Fungal Diversity 105:1-16 ((https://doi.org/10.1007/s13225-020-00466-2)) |
| Otu61456 | 1 | k__Fungi(100);p__Ascomycota(100);c__Saccharomycetes(100);o__Saccharomycetales(100);f__Metschnikowiaceae(100);g__Metschnikowia(100);g__Metschnikowia_unclassified(100); | Metschnikowia | 13 | Saprotroph | \|Undefined Saprotroph\| | Yeast | NULL | Probable | As nectar-tap_saprotroph (Põlme et al. 2020); Decay sugar-rich substrates (Põlme et al. 2020) | Põlme S, et al. 2020. Fungal Diversity 105:1-16 ((https://doi.org/10.1007/s13225-020-00466-2)) |
| Otu61457 | 1 | k__Fungi(100);p__Ascomycota(100);c__Saccharomycetes(100);o__Saccharomycetales(100);f__Metschnikowiaceae(100);g__Metschnikowia(100);g__Metschnikowia_unclassified(100); | Metschnikowia | 13 | Saprotroph | \|Undefined Saprotroph\| | Yeast | NULL | Probable | As nectar-tap_saprotroph (Põlme et al. 2020); Decay sugar-rich substrates (Põlme et al. 2020) | Põlme S, et al. 2020. Fungal Diversity 105:1-16 ((https://doi.org/10.1007/s13225-020-00466-2)) |
| Otu61458 | 1 | k__Fungi(100);p__Ascomycota(100);c__Saccharomycetes(100);o__Saccharomycetales(100);f__Metschnikowiaceae(100);g__Metschnikowia(100);s__Metschnikowia_pulcherrima(100); | Metschnikowia | 13 | Saprotroph | \|Undefined Saprotroph\| | Yeast | NULL | Probable | As nectar-tap_saprotroph (Põlme et al. 2020); Decay sugar-rich substrates (Põlme et al. 2020) | Põlme S, et al. 2020. Fungal Diversity 105:1-16 ((https://doi.org/10.1007/s13225-020-00466-2)) |
| Otu61459 | 1 | k__Fungi(100);p__Ascomycota(100);c__Saccharomycetes(100);o__Saccharomycetales(100);f__Metschnikowiaceae(100);g__Metschnikowia(100);s__Metschnikowia_pulcherrima(100); | Metschnikowia | 13 | Saprotroph | \|Undefined Saprotroph\| | Yeast | NULL | Probable | As nectar-tap_saprotroph (Põlme et al. 2020); Decay sugar-rich substrates (Põlme et al. 2020) | Põlme S, et al. 2020. Fungal Diversity 105:1-16 ((https://doi.org/10.1007/s13225-020-00466-2)) |
| Otu61460 | 1 | k__Fungi(100);p__Ascomycota(100);c__Saccharomycetes(100);o__Saccharomycetales(100);f__Metschnikowiaceae(100);g__Metschnikowia(100);s__Metschnikowia_pulcherrima(100); | Metschnikowia | 13 | Saprotroph | \|Undefined Saprotroph\| | Yeast | NULL | Probable | As nectar-tap_saprotroph (Põlme et al. 2020); Decay sugar-rich substrates (Põlme et al. 2020) | Põlme S, et al. 2020. Fungal Diversity 105:1-16 ((https://doi.org/10.1007/s13225-020-00466-2)) |
| Otu61461 | 1 | k__Fungi(100);p__Ascomycota(100);c__Saccharomycetes(100);o__Saccharomycetales(100);f__Metschnikowiaceae(100);g__Metschnikowia(100);g__Metschnikowia_unclassified(100); | Metschnikowia | 13 | Saprotroph | \|Undefined Saprotroph\| | Yeast | NULL | Probable | As nectar-tap_saprotroph (Põlme et al. 2020); Decay sugar-rich substrates (Põlme et al. 2020) | Põlme S, et al. 2020. Fungal Diversity 105:1-16 ((https://doi.org/10.1007/s13225-020-00466-2)) |
| Otu61462 | 1 | k__Fungi(100);p__Ascomycota(100);c__Saccharomycetes(100);o__Saccharomycetales(100);f__Metschnikowiaceae(100);g__Metschnikowia(100);g__Metschnikowia_unclassified(100); | Metschnikowia | 13 | Saprotroph | \|Undefined Saprotroph\| | Yeast | NULL | Probable | As nectar-tap_saprotroph (Põlme et al. 2020); Decay sugar-rich substrates (Põlme et al. 2020) | Põlme S, et al. 2020. Fungal Diversity 105:1-16 ((https://doi.org/10.1007/s13225-020-00466-2)) |
| Otu61463 | 1 | k__Fungi(100);p__Ascomycota(100);c__Saccharomycetes(100);o__Saccharomycetales(100);f__Metschnikowiaceae(100);g__Metschnikowia(100);g__Metschnikowia_unclassified(100); | Metschnikowia | 13 | Saprotroph | \|Undefined Saprotroph\| | Yeast | NULL | Probable | As nectar-tap_saprotroph (Põlme et al. 2020); Decay sugar-rich substrates (Põlme et al. 2020) | Põlme S, et al. 2020. Fungal Diversity 105:1-16 ((https://doi.org/10.1007/s13225-020-00466-2)) |
| Otu61464 | 1 | k__Fungi(100);p__Ascomycota(100);c__Saccharomycetes(100);o__Saccharomycetales(100);f__Metschnikowiaceae(100);g__Metschnikowia(100);s__Metschnikowia_pulcherrima(100); | Metschnikowia | 13 | Saprotroph | \|Undefined Saprotroph\| | Yeast | NULL | Probable | As nectar-tap_saprotroph (Põlme et al. 2020); Decay sugar-rich substrates (Põlme et al. 2020) | Põlme S, et al. 2020. Fungal Diversity 105:1-16 ((https://doi.org/10.1007/s13225-020-00466-2)) |
| Otu61465 | 1 | k__Fungi(100);p__Ascomycota(100);c__Saccharomycetes(100);o__Saccharomycetales(100);f__Metschnikowiaceae(100);g__Metschnikowia(100);g__Metschnikowia_unclassified(100); | Metschnikowia | 13 | Saprotroph | \|Undefined Saprotroph\| | Yeast | NULL | Probable | As nectar-tap_saprotroph (Põlme et al. 2020); Decay sugar-rich substrates (Põlme et al. 2020) | Põlme S, et al. 2020. Fungal Diversity 105:1-16 ((https://doi.org/10.1007/s13225-020-00466-2)) |
| Otu61466 | 1 | k__Fungi(100);p__Ascomycota(100);c__Saccharomycetes(100);o__Saccharomycetales(100);f__Metschnikowiaceae(100);g__Metschnikowia(100);s__Metschnikowia_pulcherrima(100); | Metschnikowia | 13 | Saprotroph | \|Undefined Saprotroph\| | Yeast | NULL | Probable | As nectar-tap_saprotroph (Põlme et al. 2020); Decay sugar-rich substrates (Põlme et al. 2020) | Põlme S, et al. 2020. Fungal Diversity 105:1-16 ((https://doi.org/10.1007/s13225-020-00466-2)) |
| Otu61467 | 1 | k__Fungi(100);p__Ascomycota(100);c__Saccharomycetes(100);o__Saccharomycetales(100);f__Metschnikowiaceae(100);g__Metschnikowia(100);s__Metschnikowia_pulcherrima(100); | Metschnikowia | 13 | Saprotroph | \|Undefined Saprotroph\| | Yeast | NULL | Probable | As nectar-tap_saprotroph (Põlme et al. 2020); Decay sugar-rich substrates (Põlme et al. 2020) | Põlme S, et al. 2020. Fungal Diversity 105:1-16 ((https://doi.org/10.1007/s13225-020-00466-2)) |
| Otu61468 | 1 | k__Fungi(100);p__Ascomycota(100);c__Saccharomycetes(100);o__Saccharomycetales(100);f__Metschnikowiaceae(100);g__Metschnikowia(100);s__Metschnikowia_pulcherrima(100); | Metschnikowia | 13 | Saprotroph | \|Undefined Saprotroph\| | Yeast | NULL | Probable | As nectar-tap_saprotroph (Põlme et al. 2020); Decay sugar-rich substrates (Põlme et al. 2020) | Põlme S, et al. 2020. Fungal Diversity 105:1-16 ((https://doi.org/10.1007/s13225-020-00466-2)) |
| Otu61469 | 1 | k__Fungi(100);p__Ascomycota(100);c__Saccharomycetes(100);o__Saccharomycetales(100);f__Metschnikowiaceae(100);g__Metschnikowia(100);s__Metschnikowia_pulcherrima(100); | Metschnikowia | 13 | Saprotroph | \|Undefined Saprotroph\| | Yeast | NULL | Probable | As nectar-tap_saprotroph (Põlme et al. 2020); Decay sugar-rich substrates (Põlme et al. 2020) | Põlme S, et al. 2020. Fungal Diversity 105:1-16 ((https://doi.org/10.1007/s13225-020-00466-2)) |
| Otu61470 | 1 | k__Fungi(100);p__Ascomycota(100);c__Saccharomycetes(100);o__Saccharomycetales(100);f__Metschnikowiaceae(100);g__Metschnikowia(100);s__Metschnikowia_pulcherrima(100); | Metschnikowia | 13 | Saprotroph | \|Undefined Saprotroph\| | Yeast | NULL | Probable | As nectar-tap_saprotroph (Põlme et al. 2020); Decay sugar-rich substrates (Põlme et al. 2020) | Põlme S, et al. 2020. Fungal Diversity 105:1-16 ((https://doi.org/10.1007/s13225-020-00466-2)) |
| Otu61471 | 1 | k__Fungi(100);p__Ascomycota(100);c__Saccharomycetes(100);o__Saccharomycetales(100);f__Metschnikowiaceae(100);g__Metschnikowia(100);s__Metschnikowia_pulcherrima(100); | Metschnikowia | 13 | Saprotroph | \|Undefined Saprotroph\| | Yeast | NULL | Probable | As nectar-tap_saprotroph (Põlme et al. 2020); Decay sugar-rich substrates (Põlme et al. 2020) | Põlme S, et al. 2020. Fungal Diversity 105:1-16 ((https://doi.org/10.1007/s13225-020-00466-2)) |
| Otu61472 | 1 | k__Fungi(100);p__Ascomycota(100);c__Saccharomycetes(100);o__Saccharomycetales(100);f__Metschnikowiaceae(100);g__Metschnikowia(100);s__Metschnikowia_pulcherrima(100); | Metschnikowia | 13 | Saprotroph | \|Undefined Saprotroph\| | Yeast | NULL | Probable | As nectar-tap_saprotroph (Põlme et al. 2020); Decay sugar-rich substrates (Põlme et al. 2020) | Põlme S, et al. 2020. Fungal Diversity 105:1-16 ((https://doi.org/10.1007/s13225-020-00466-2)) |
| Otu61473 | 1 | k__Fungi(100);p__Ascomycota(100);c__Saccharomycetes(100);o__Saccharomycetales(100);f__Metschnikowiaceae(100);g__Metschnikowia(100);g__Metschnikowia_unclassified(100); | Metschnikowia | 13 | Saprotroph | \|Undefined Saprotroph\| | Yeast | NULL | Probable | As nectar-tap_saprotroph (Põlme et al. 2020); Decay sugar-rich substrates (Põlme et al. 2020) | Põlme S, et al. 2020. Fungal Diversity 105:1-16 ((https://doi.org/10.1007/s13225-020-00466-2)) |
| Otu61474 | 1 | k__Fungi(100);p__Ascomycota(100);c__Saccharomycetes(100);o__Saccharomycetales(100);f__Metschnikowiaceae(100);g__Metschnikowia(100);s__Metschnikowia_pulcherrima(100); | Metschnikowia | 13 | Saprotroph | \|Undefined Saprotroph\| | Yeast | NULL | Probable | As nectar-tap_saprotroph (Põlme et al. 2020); Decay sugar-rich substrates (Põlme et al. 2020) | Põlme S, et al. 2020. Fungal Diversity 105:1-16 ((https://doi.org/10.1007/s13225-020-00466-2)) |

| | | | | | | | | | | | |
|---|---|---|---|---|---|---|---|---|---|---|---|
| Otu61475 | 1 | k__Fungi(100);p__Ascomycota(100);c__Saccharomycetes(100);o__Saccharomycetales(100);f__Metschnikowiaceae(100);g__Metschnikowia(100);s__Metschnikowia_pulcherrima(100); | Metschnikowia | 13 | Saprotroph | \|Undefined Saprotroph\| | Yeast | NULL | Probable | As nectar-tap_saprotroph (Põlme et al. 2020); Decay sugar-rich substrates (Põlme et al. 2020) | Põlme S, et al. 2020. Fungal Diversity 105:1-16 ((https://doi.org/10.1007/s13225-020-00466-2)) |
| Otu61476 | 1 | k__Fungi(100);p__Ascomycota(100);c__Saccharomycetes(100);o__Saccharomycetales(100);f__Metschnikowiaceae(100);g__Metschnikowia(100);s__Metschnikowia_pulcherrima(100); | Metschnikowia | 13 | Saprotroph | \|Undefined Saprotroph\| | Yeast | NULL | Probable | As nectar-tap_saprotroph (Põlme et al. 2020); Decay sugar-rich substrates (Põlme et al. 2020) | Põlme S, et al. 2020. Fungal Diversity 105:1-16 ((https://doi.org/10.1007/s13225-020-00466-2)) |
| Otu61477 | 1 | k__Fungi(100);p__Ascomycota(100);c__Saccharomycetes(100);o__Saccharomycetales(100);f__Metschnikowiaceae(100);g__Metschnikowia(100);g__Metschnikowia_unclassified(100); | Metschnikowia | 13 | Saprotroph | \|Undefined Saprotroph\| | Yeast | NULL | Probable | As nectar-tap_saprotroph (Põlme et al. 2020); Decay sugar-rich substrates (Põlme et al. 2020) | Põlme S, et al. 2020. Fungal Diversity 105:1-16 ((https://doi.org/10.1007/s13225-020-00466-2)) |
| Otu61478 | 1 | k__Fungi(100);p__Ascomycota(100);c__Saccharomycetes(100);o__Saccharomycetales(100);f__Metschnikowiaceae(100);g__Metschnikowia(100);g__Metschnikowia_unclassified(100); | Metschnikowia | 13 | Saprotroph | \|Undefined Saprotroph\| | Yeast | NULL | Probable | As nectar-tap_saprotroph (Põlme et al. 2020); Decay sugar-rich substrates (Põlme et al. 2020) | Põlme S, et al. 2020. Fungal Diversity 105:1-16 ((https://doi.org/10.1007/s13225-020-00466-2)) |
| Otu61479 | 1 | k__Fungi(100);p__Ascomycota(100);c__Saccharomycetes(100);o__Saccharomycetales(100);f__Metschnikowiaceae(100);g__Metschnikowia(100);s__Metschnikowia_pulcherrima(100); | Metschnikowia | 13 | Saprotroph | \|Undefined Saprotroph\| | Yeast | NULL | Probable | As nectar-tap_saprotroph (Põlme et al. 2020); Decay sugar-rich substrates (Põlme et al. 2020) | Põlme S, et al. 2020. Fungal Diversity 105:1-16 ((https://doi.org/10.1007/s13225-020-00466-2)) |
| Otu61480 | 1 | k__Fungi(100);p__Ascomycota(100);c__Saccharomycetes(100);o__Saccharomycetales(100);f__Metschnikowiaceae(100);g__Metschnikowia(100);s__Metschnikowia_pulcherrima(100); | Metschnikowia | 13 | Saprotroph | \|Undefined Saprotroph\| | Yeast | NULL | Probable | As nectar-tap_saprotroph (Põlme et al. 2020); Decay sugar-rich substrates (Põlme et al. 2020) | Põlme S, et al. 2020. Fungal Diversity 105:1-16 ((https://doi.org/10.1007/s13225-020-00466-2)) |
| Otu61481 | 1 | k__Fungi(100);p__Ascomycota(100);c__Saccharomycetes(100);o__Saccharomycetales(100);f__Metschnikowiaceae(100);g__Metschnikowia(100);s__Metschnikowia_pulcherrima(100); | Metschnikowia | 13 | Saprotroph | \|Undefined Saprotroph\| | Yeast | NULL | Probable | As nectar-tap_saprotroph (Põlme et al. 2020); Decay sugar-rich substrates (Põlme et al. 2020) | Põlme S, et al. 2020. Fungal Diversity 105:1-16 ((https://doi.org/10.1007/s13225-020-00466-2)) |
| Otu61482 | 1 | k__Fungi(100);p__Ascomycota(100);c__Saccharomycetes(100);o__Saccharomycetales(100);f__Metschnikowiaceae(100);g__Metschnikowia(100);g__Metschnikowia_unclassified(100); | Metschnikowia | 13 | Saprotroph | \|Undefined Saprotroph\| | Yeast | NULL | Probable | As nectar-tap_saprotroph (Põlme et al. 2020); Decay sugar-rich substrates (Põlme et al. 2020) | Põlme S, et al. 2020. Fungal Diversity 105:1-16 ((https://doi.org/10.1007/s13225-020-00466-2)) |
| Otu61483 | 1 | k__Fungi(100);p__Ascomycota(100);c__Saccharomycetes(100);o__Saccharomycetales(100);f__Metschnikowiaceae(100);g__Metschnikowia(100);s__Metschnikowia_pulcherrima(100); | Metschnikowia | 13 | Saprotroph | \|Undefined Saprotroph\| | Yeast | NULL | Probable | As nectar-tap_saprotroph (Põlme et al. 2020); Decay sugar-rich substrates (Põlme et al. 2020) | Põlme S, et al. 2020. Fungal Diversity 105:1-16 ((https://doi.org/10.1007/s13225-020-00466-2)) |
| Otu61484 | 1 | k__Fungi(100);p__Ascomycota(100);c__Saccharomycetes(100);o__Saccharomycetales(100);f__Metschnikowiaceae(100);g__Metschnikowia(100);s__Metschnikowia_pulcherrima(100); | Metschnikowia | 13 | Saprotroph | \|Undefined Saprotroph\| | Yeast | NULL | Probable | As nectar-tap_saprotroph (Põlme et al. 2020); Decay sugar-rich substrates (Põlme et al. 2020) | Põlme S, et al. 2020. Fungal Diversity 105:1-16 ((https://doi.org/10.1007/s13225-020-00466-2)) |
| Otu61485 | 1 | k__Fungi(100);p__Ascomycota(100);c__Saccharomycetes(100);o__Saccharomycetales(100);f__Metschnikowiaceae(100);g__Metschnikowia(100);s__Metschnikowia_pulcherrima(100); | Metschnikowia | 13 | Saprotroph | \|Undefined Saprotroph\| | Yeast | NULL | Probable | As nectar-tap_saprotroph (Põlme et al. 2020); Decay sugar-rich substrates (Põlme et al. 2020) | Põlme S, et al. 2020. Fungal Diversity 105:1-16 ((https://doi.org/10.1007/s13225-020-00466-2)) |
| Otu61486 | 1 | k__Fungi(100);p__Ascomycota(100);c__Saccharomycetes(100);o__Saccharomycetales(100);f__Metschnikowiaceae(100);g__Metschnikowia(100);g__Metschnikowia_unclassified(100); | Metschnikowia | 13 | Saprotroph | \|Undefined Saprotroph\| | Yeast | NULL | Probable | As nectar-tap_saprotroph (Põlme et al. 2020); Decay sugar-rich substrates (Põlme et al. 2020) | Põlme S, et al. 2020. Fungal Diversity 105:1-16 ((https://doi.org/10.1007/s13225-020-00466-2)) |
| Otu61487 | 1 | k__Fungi(100);p__Ascomycota(100);c__Saccharomycetes(100);o__Saccharomycetales(100);f__Metschnikowiaceae(100);g__Metschnikowia(100);s__Metschnikowia_pulcherrima(100); | Metschnikowia | 13 | Saprotroph | \|Undefined Saprotroph\| | Yeast | NULL | Probable | As nectar-tap_saprotroph (Põlme et al. 2020); Decay sugar-rich substrates (Põlme et al. 2020) | Põlme S, et al. 2020. Fungal Diversity 105:1-16 ((https://doi.org/10.1007/s13225-020-00466-2)) |
| Otu61488 | 1 | k__Fungi(100);p__Ascomycota(100);c__Saccharomycetes(100);o__Saccharomycetales(100);f__Metschnikowiaceae(100);g__Metschnikowia(100);s__Metschnikowia_pulcherrima(100); | Metschnikowia | 13 | Saprotroph | \|Undefined Saprotroph\| | Yeast | NULL | Probable | As nectar-tap_saprotroph (Põlme et al. 2020); Decay sugar-rich substrates (Põlme et al. 2020) | Põlme S, et al. 2020. Fungal Diversity 105:1-16 ((https://doi.org/10.1007/s13225-020-00466-2)) |
| Otu61489 | 1 | k__Fungi(100);p__Ascomycota(100);c__Saccharomycetes(100);o__Saccharomycetales(100);f__Metschnikowiaceae(100);g__Metschnikowia(100);s__Metschnikowia_pulcherrima(100); | Metschnikowia | 13 | Saprotroph | \|Undefined Saprotroph\| | Yeast | NULL | Probable | As nectar-tap_saprotroph (Põlme et al. 2020); Decay sugar-rich substrates (Põlme et al. 2020) | Põlme S, et al. 2020. Fungal Diversity 105:1-16 ((https://doi.org/10.1007/s13225-020-00466-2)) |
| Otu61490 | 1 | k__Fungi(100);p__Ascomycota(100);c__Saccharomycetes(100);o__Saccharomycetales(100);f__Metschnikowiaceae(100);g__Metschnikowia(100);s__Metschnikowia_pulcherrima(100); | Metschnikowia | 13 | Saprotroph | \|Undefined Saprotroph\| | Yeast | NULL | Probable | As nectar-tap_saprotroph (Põlme et al. 2020); Decay sugar-rich substrates (Põlme et al. 2020) | Põlme S, et al. 2020. Fungal Diversity 105:1-16 ((https://doi.org/10.1007/s13225-020-00466-2)) |
| Otu61491 | 1 | k__Fungi(100);p__Ascomycota(100);c__Saccharomycetes(100);o__Saccharomycetales(100);f__Metschnikowiaceae(100);g__Metschnikowia(100);s__Metschnikowia_pulcherrima(100); | Metschnikowia | 13 | Saprotroph | \|Undefined Saprotroph\| | Yeast | NULL | Probable | As nectar-tap_saprotroph (Põlme et al. 2020); Decay sugar-rich substrates (Põlme et al. 2020) | Põlme S, et al. 2020. Fungal Diversity 105:1-16 ((https://doi.org/10.1007/s13225-020-00466-2)) |
| Otu61492 | 1 | k__Fungi(100);p__Ascomycota(100);c__Saccharomycetes(100);o__Saccharomycetales(100);f__Metschnikowiaceae(100);g__Metschnikowia(100);s__Metschnikowia_pulcherrima(100); | Metschnikowia | 13 | Saprotroph | \|Undefined Saprotroph\| | Yeast | NULL | Probable | As nectar-tap_saprotroph (Põlme et al. 2020); Decay sugar-rich substrates (Põlme et al. 2020) | Põlme S, et al. 2020. Fungal Diversity 105:1-16 ((https://doi.org/10.1007/s13225-020-00466-2)) |
| Otu61493 | 1 | k__Fungi(100);p__Ascomycota(100);c__Saccharomycetes(100);o__Saccharomycetales(100);f__Metschnikowiaceae(100);g__Metschnikowia(100);s__Metschnikowia_pulcherrima(100); | Metschnikowia | 13 | Saprotroph | \|Undefined Saprotroph\| | Yeast | NULL | Probable | As nectar-tap_saprotroph (Põlme et al. 2020); Decay sugar-rich substrates (Põlme et al. 2020) | Põlme S, et al. 2020. Fungal Diversity 105:1-16 ((https://doi.org/10.1007/s13225-020-00466-2)) |
| Otu61494 | 1 | k__Fungi(100);p__Ascomycota(100);c__Saccharomycetes(100);o__Saccharomycetales(100);f__Metschnikowiaceae(100);g__Metschnikowia(100);s__Metschnikowia_pulcherrima(100); | Metschnikowia | 13 | Saprotroph | \|Undefined Saprotroph\| | Yeast | NULL | Probable | As nectar-tap_saprotroph (Põlme et al. 2020); Decay sugar-rich substrates (Põlme et al. 2020) | Põlme S, et al. 2020. Fungal Diversity 105:1-16 ((https://doi.org/10.1007/s13225-020-00466-2)) |
| Otu61495 | 1 | k__Fungi(100);p__Ascomycota(100);c__Saccharomycetes(100);o__Saccharomycetales(100);f__Metschnikowiaceae(100);g__Metschnikowia(100);s__Metschnikowia_pulcherrima(100); | Metschnikowia | 13 | Saprotroph | \|Undefined Saprotroph\| | Yeast | NULL | Probable | As nectar-tap_saprotroph (Põlme et al. 2020); Decay sugar-rich substrates (Põlme et al. 2020) | Põlme S, et al. 2020. Fungal Diversity 105:1-16 ((https://doi.org/10.1007/s13225-020-00466-2)) |
| Otu61496 | 1 | k__Fungi(100);p__Ascomycota(100);c__Saccharomycetes(100);o__Saccharomycetales(100);f__Metschnikowiaceae(100);g__Metschnikowia(100);s__Metschnikowia_pulcherrima(100); | Metschnikowia | 13 | Saprotroph | \|Undefined Saprotroph\| | Yeast | NULL | Probable | As nectar-tap_saprotroph (Põlme et al. 2020); Decay sugar-rich substrates (Põlme et al. 2020) | Põlme S, et al. 2020. Fungal Diversity 105:1-16 ((https://doi.org/10.1007/s13225-020-00466-2)) |
| Otu61497 | 1 | k__Fungi(100);p__Ascomycota(100);c__Saccharomycetes(100);o__Saccharomycetales(100);f__Metschnikowiaceae(100);g__Metschnikowia(100);s__Metschnikowia_pulcherrima(100); | Metschnikowia | 13 | Saprotroph | \|Undefined Saprotroph\| | Yeast | NULL | Probable | As nectar-tap_saprotroph (Põlme et al. 2020); Decay sugar-rich substrates (Põlme et al. 2020) | Põlme S, et al. 2020. Fungal Diversity 105:1-16 ((https://doi.org/10.1007/s13225-020-00466-2)) |
| Otu61498 | 1 | k__Fungi(100);p__Ascomycota(100);c__Saccharomycetes(100);o__Saccharomycetales(100);f__Metschnikowiaceae(100);g__Metschnikowia(100);s__Metschnikowia_pulcherrima(100); | Metschnikowia | 13 | Saprotroph | \|Undefined Saprotroph\| | Yeast | NULL | Probable | As nectar-tap_saprotroph (Põlme et al. 2020); Decay sugar-rich substrates (Põlme et al. 2020) | Põlme S, et al. 2020. Fungal Diversity 105:1-16 ((https://doi.org/10.1007/s13225-020-00466-2)) |
| Otu61499 | 1 | k__Fungi(100);p__Ascomycota(100);c__Saccharomycetes(100);o__Saccharomycetales(100);f__Metschnikowiaceae(100);g__Metschnikowia(100);s__Metschnikowia_pulcherrima(100); | Metschnikowia | 13 | Saprotroph | \|Undefined Saprotroph\| | Yeast | NULL | Probable | As nectar-tap_saprotroph (Põlme et al. 2020); Decay sugar-rich substrates (Põlme et al. 2020) | Põlme S, et al. 2020. Fungal Diversity 105:1-16 ((https://doi.org/10.1007/s13225-020-00466-2)) |
| Otu61500 | 1 | k__Fungi(100);p__Ascomycota(100);c__Saccharomycetes(100);o__Saccharomycetales(100);f__Metschnikowiaceae(100);g__Metschnikowia(100);g__Metschnikowia_unclassified(100); | Metschnikowia | 13 | Saprotroph | \|Undefined Saprotroph\| | Yeast | NULL | Probable | As nectar-tap_saprotroph (Põlme et al. 2020); Decay sugar-rich substrates (Põlme et al. 2020) | Põlme S, et al. 2020. Fungal Diversity 105:1-16 ((https://doi.org/10.1007/s13225-020-00466-2)) |
| Otu61501 | 1 | k__Fungi(100);p__Ascomycota(100);c__Saccharomycetes(100);o__Saccharomycetales(100);f__Metschnikowiaceae(100);g__Metschnikowia(100);g__Metschnikowia_unclassified(100); | Metschnikowia | 13 | Saprotroph | \|Undefined Saprotroph\| | Yeast | NULL | Probable | As nectar-tap_saprotroph (Põlme et al. 2020); Decay sugar-rich substrates (Põlme et al. 2020) | Põlme S, et al. 2020. Fungal Diversity 105:1-16 ((https://doi.org/10.1007/s13225-020-00466-2)) |
| Otu61502 | 1 | k__Fungi(100);p__Ascomycota(100);c__Saccharomycetes(100);o__Saccharomycetales(100);f__Metschnikowiaceae(100);g__Metschnikowia(100);s__Metschnikowia_pulcherrima(100); | Metschnikowia | 13 | Saprotroph | \|Undefined Saprotroph\| | Yeast | NULL | Probable | As nectar-tap_saprotroph (Põlme et al. 2020); Decay sugar-rich substrates (Põlme et al. 2020) | Põlme S, et al. 2020. Fungal Diversity 105:1-16 ((https://doi.org/10.1007/s13225-020-00466-2)) |
| Otu61503 | 1 | k__Fungi(100);p__Ascomycota(100);c__Saccharomycetes(100);o__Saccharomycetales(100);f__Metschnikowiaceae(100);g__Metschnikowia(100);g__Metschnikowia_unclassified(100); | Metschnikowia | 13 | Saprotroph | \|Undefined Saprotroph\| | Yeast | NULL | Probable | As nectar-tap_saprotroph (Põlme et al. 2020); Decay sugar-rich substrates (Põlme et al. 2020) | Põlme S, et al. 2020. Fungal Diversity 105:1-16 ((https://doi.org/10.1007/s13225-020-00466-2)) |
| Otu61504 | 1 | k__Fungi(100);p__Ascomycota(100);c__Saccharomycetes(100);o__Saccharomycetales(100);f__Metschnikowiaceae(100);g__Metschnikowia(100);s__Metschnikowia_pulcherrima(100); | Metschnikowia | 13 | Saprotroph | \|Undefined Saprotroph\| | Yeast | NULL | Probable | As nectar-tap_saprotroph (Põlme et al. 2020); Decay sugar-rich substrates (Põlme et al. 2020) | Põlme S, et al. 2020. Fungal Diversity 105:1-16 ((https://doi.org/10.1007/s13225-020-00466-2)) |
| Otu61506 | 1 | k__Fungi(100);p__Ascomycota(100);c__Saccharomycetes(100);o__Saccharomycetales(100);f__Metschnikowiaceae(100);g__Metschnikowia(100);s__Metschnikowia_pulcherrima(100); | Metschnikowia | 13 | Saprotroph | \|Undefined Saprotroph\| | Yeast | NULL | Probable | As nectar-tap_saprotroph (Põlme et al. 2020); Decay sugar-rich substrates (Põlme et al. 2020) | Põlme S, et al. 2020. Fungal Diversity 105:1-16 ((https://doi.org/10.1007/s13225-020-00466-2)) |
| Otu61507 | 1 | k__Fungi(100);p__Ascomycota(100);c__Saccharomycetes(100);o__Saccharomycetales(100);f__Metschnikowiaceae(100);g__Metschnikowia(100);s__Metschnikowia_pulcherrima(100); | Metschnikowia | 13 | Saprotroph | \|Undefined Saprotroph\| | Yeast | NULL | Probable | As nectar-tap_saprotroph (Põlme et al. 2020); Decay sugar-rich substrates (Põlme et al. 2020) | Põlme S, et al. 2020. Fungal Diversity 105:1-16 ((https://doi.org/10.1007/s13225-020-00466-2)) |
| Otu61508 | 1 | k__Fungi(100);p__Ascomycota(100);c__Saccharomycetes(100);o__Saccharomycetales(100);f__Metschnikowiaceae(100);g__Metschnikowia(100);s__Metschnikowia_pulcherrima(100); | Metschnikowia | 13 | Saprotroph | \|Undefined Saprotroph\| | Yeast | NULL | Probable | As nectar-tap_saprotroph (Põlme et al. 2020); Decay sugar-rich substrates (Põlme et al. 2020) | Põlme S, et al. 2020. Fungal Diversity 105:1-16 ((https://doi.org/10.1007/s13225-020-00466-2)) |
| Otu61509 | 1 | k__Fungi(100);p__Ascomycota(100);c__Saccharomycetes(100);o__Saccharomycetales(100);f__Metschnikowiaceae(100);g__Metschnikowia(100);s__Metschnikowia_pulcherrima(100); | Metschnikowia | 13 | Saprotroph | \|Undefined Saprotroph\| | Yeast | NULL | Probable | As nectar-tap_saprotroph (Põlme et al. 2020); Decay sugar-rich substrates (Põlme et al. 2020) | Põlme S, et al. 2020. Fungal Diversity 105:1-16 ((https://doi.org/10.1007/s13225-020-00466-2)) |
| Otu61511 | 1 | k__Fungi(100);p__Ascomycota(100);c__Saccharomycetes(100);o__Saccharomycetales(100);f__Metschnikowiaceae(100);g__Metschnikowia(100);s__Metschnikowia_pulcherrima(100); | Metschnikowia | 13 | Saprotroph | \|Undefined Saprotroph\| | Yeast | NULL | Probable | As nectar-tap_saprotroph (Põlme et al. 2020); Decay sugar-rich substrates (Põlme et al. 2020) | Põlme S, et al. 2020. Fungal Diversity 105:1-16 ((https://doi.org/10.1007/s13225-020-00466-2)) |
| Otu61512 | 1 | k__Fungi(100);p__Ascomycota(100);c__Saccharomycetes(100);o__Saccharomycetales(100);f__Metschnikowiaceae(100);g__Metschnikowia(100);g__Metschnikowia_unclassified(100); | Metschnikowia | 13 | Saprotroph | \|Undefined Saprotroph\| | Yeast | NULL | Probable | As nectar-tap_saprotroph (Põlme et al. 2020); Decay sugar-rich substrates (Põlme et al. 2020) | Põlme S, et al. 2020. Fungal Diversity 105:1-16 ((https://doi.org/10.1007/s13225-020-00466-2)) |
| Otu61513 | 1 | k__Fungi(100);p__Ascomycota(100);c__Saccharomycetes(100);o__Saccharomycetales(100);f__Metschnikowiaceae(100);g__Metschnikowia(100);s__Metschnikowia_pulcherrima(100); | Metschnikowia | 13 | Saprotroph | \|Undefined Saprotroph\| | Yeast | NULL | Probable | As nectar-tap_saprotroph (Põlme et al. 2020); Decay sugar-rich substrates (Põlme et al. 2020) | Põlme S, et al. 2020. Fungal Diversity 105:1-16 ((https://doi.org/10.1007/s13225-020-00466-2)) |
| Otu61514 | 1 | k__Fungi(100);p__Ascomycota(100);c__Saccharomycetes(100);o__Saccharomycetales(100);f__Metschnikowiaceae(100);g__Metschnikowia(100);g__Metschnikowia_unclassified(100); | Metschnikowia | 13 | Saprotroph | \|Undefined Saprotroph\| | Yeast | NULL | Probable | As nectar-tap_saprotroph (Põlme et al. 2020); Decay sugar-rich substrates (Põlme et al. 2020) | Põlme S, et al. 2020. Fungal Diversity 105:1-16 ((https://doi.org/10.1007/s13225-020-00466-2)) |
| Otu61515 | 1 | k__Fungi(100);p__Ascomycota(100);c__Saccharomycetes(100);o__Saccharomycetales(100);f__Metschnikowiaceae(100);g__Metschnikowia(100);s__Metschnikowia_pulcherrima(100); | Metschnikowia | 13 | Saprotroph | \|Undefined Saprotroph\| | Yeast | NULL | Probable | As nectar-tap_saprotroph (Põlme et al. 2020); Decay sugar-rich substrates (Põlme et al. 2020) | Põlme S, et al. 2020. Fungal Diversity 105:1-16 ((https://doi.org/10.1007/s13225-020-00466-2)) |
| Otu61516 | 1 | k__Fungi(100);p__Ascomycota(100);c__Saccharomycetes(100);o__Saccharomycetales(100);f__Metschnikowiaceae(100);g__Metschnikowia(100);s__Metschnikowia_pulcherrima(100); | Metschnikowia | 13 | Saprotroph | \|Undefined Saprotroph\| | Yeast | NULL | Probable | As nectar-tap_saprotroph (Põlme et al. 2020); Decay sugar-rich substrates (Põlme et al. 2020) | Põlme S, et al. 2020. Fungal Diversity 105:1-16 ((https://doi.org/10.1007/s13225-020-00466-2)) |
| Otu61517 | 1 | k__Fungi(100);p__Ascomycota(100);c__Saccharomycetes(100);o__Saccharomycetales(100);f__Metschnikowiaceae(100);g__Metschnikowia(100);g__Metschnikowia_unclassified(100); | Metschnikowia | 13 | Saprotroph | \|Undefined Saprotroph\| | Yeast | NULL | Probable | As nectar-tap_saprotroph (Põlme et al. 2020); Decay sugar-rich substrates (Põlme et al. 2020) | Põlme S, et al. 2020. Fungal Diversity 105:1-16 ((https://doi.org/10.1007/s13225-020-00466-2)) |
| Otu61518 | 1 | k__Fungi(100);p__Ascomycota(100);c__Saccharomycetes(100);o__Saccharomycetales(100);f__Metschnikowiaceae(100);g__Metschnikowia(100);s__Metschnikowia_pulcherrima(100); | Metschnikowia | 13 | Saprotroph | \|Undefined Saprotroph\| | Yeast | NULL | Probable | As nectar-tap_saprotroph (Põlme et al. 2020); Decay sugar-rich substrates (Põlme et al. 2020) | Põlme S, et al. 2020. Fungal Diversity 105:1-16 ((https://doi.org/10.1007/s13225-020-00466-2)) |
| Otu61519 | 1 | k__Fungi(100);p__Ascomycota(100);c__Sordariomycetes(100);o__Coniochaetales(100);f__Coniochaetaceae(100);g__Coniochaeta(100);s__Coniochaeta_sp(100); | Coniochaeta | 13 | Pathotroph-Saprotroph-Symbiotroph | Animal Pathogen-Dung Saprotroph-Endophyte-Lichen Parasite-Plant Pathogen-\|Undefined Saprotroph\| | Microfungus | NULL | Possible | As foliar_endophyte (Põlme et al. 2020); Decay leaf, fruit, seed, soil, dung, animal material (Põlme et al. 2020) | Bell A. 1983. Dung Fungi: An Illustrated Guide to Coprophilous Fungi in New Zealand. Victoria University Press, Wellington (ISBN: 978-0864730015); Damm U, et al. 2010. Persoonia 24:60-80 ((http://doi.org/10.3767/003158510X500705)); Costa IPMW, et al. 2012. Mycotaxon 119:e26 ((http://mycotaxon.com/resources/checklists/costa_v119_checklist.pdf)); Tedersoo L, et al. 2014. Science 346: 1256688 ((https://doi.org/10.1126/science.1256688)); Irinyi L, et al. 2015. Medical Mycology 53:313-337 ((https://doi.org/10.1093/mmy/myv008)); Lawrey JD, Diederich P. 2016. http://www.lichenicolous.net (accessed 24 May 2017) ((http://www.lichenicolous.net)); Põlme S, et al. 2020. Fungal Diversity 105:1-16 ((https://doi.org/10.1007/s13225-020-00466-2)) |
| Otu61520 | 1 | k__Fungi(100);p__Ascomycota(100);c__Saccharomycetes(100);o__Saccharomycetales(100);f__Metschnikowiaceae(100);g__Metschnikowia(100);s__Metschnikowia_pulcherrima(100); | Metschnikowia | 13 | Saprotroph | \|Undefined Saprotroph\| | Yeast | NULL | Probable | As nectar-tap_saprotroph (Põlme et al. 2020); Decay sugar-rich substrates (Põlme et al. 2020) | Põlme S, et al. 2020. Fungal Diversity 105:1-16 ((https://doi.org/10.1007/s13225-020-00466-2)) |
| Otu61521 | 1 | k__Fungi(100);p__Ascomycota(100);c__Saccharomycetes(100);o__Saccharomycetales(100);f__Metschnikowiaceae(100);g__Metschnikowia(100);s__Metschnikowia_pulcherrima(100); | Metschnikowia | 13 | Saprotroph | \|Undefined Saprotroph\| | Yeast | NULL | Probable | As nectar-tap_saprotroph (Põlme et al. 2020); Decay sugar-rich substrates (Põlme et al. 2020) | Põlme S, et al. 2020. Fungal Diversity 105:1-16 ((https://doi.org/10.1007/s13225-020-00466-2)) |
| Otu61522 | 1 | k__Fungi(100);p__Ascomycota(100);c__Saccharomycetes(100);o__Saccharomycetales(100);f__Metschnikowiaceae(100);g__Metschnikowia(100);s__Metschnikowia_pulcherrima(100); | Metschnikowia | 13 | Saprotroph | \|Undefined Saprotroph\| | Yeast | NULL | Probable | As nectar-tap_saprotroph (Põlme et al. 2020); Decay sugar-rich substrates (Põlme et al. 2020) | Põlme S, et al. 2020. Fungal Diversity 105:1-16 ((https://doi.org/10.1007/s13225-020-00466-2)) |
| Otu61523 | 1 | k__Fungi(100);p__Ascomycota(100);c__Saccharomycetes(100);o__Saccharomycetales(100);f__Metschnikowiaceae(100);g__Metschnikowia(100);g__Metschnikowia_unclassified(100); | Metschnikowia | 13 | Saprotroph | \|Undefined Saprotroph\| | Yeast | NULL | Probable | As nectar-tap_saprotroph (Põlme et al. 2020); Decay sugar-rich substrates (Põlme et al. 2020) | Põlme S, et al. 2020. Fungal Diversity 105:1-16 ((https://doi.org/10.1007/s13225-020-00466-2)) |
| Otu61524 | 1 | k__Fungi(100);p__Ascomycota(100);c__Saccharomycetes(100);o__Saccharomycetales(100);f__Metschnikowiaceae(100);g__Metschnikowia(100);s__Metschnikowia_pulcherrima(100); | Metschnikowia | 13 | Saprotroph | \|Undefined Saprotroph\| | Yeast | NULL | Probable | As nectar-tap_saprotroph (Põlme et al. 2020); Decay sugar-rich substrates (Põlme et al. 2020) | Põlme S, et al. 2020. Fungal Diversity 105:1-16 ((https://doi.org/10.1007/s13225-020-00466-2)) |
| Otu61525 | 1 | k__Fungi(100);p__Ascomycota(100);c__Saccharomycetes(100);o__Saccharomycetales(100);f__Metschnikowiaceae(100);g__Metschnikowia(100);g__Metschnikowia_unclassified(100); | Metschnikowia | 13 | Saprotroph | \|Undefined Saprotroph\| | Yeast | NULL | Probable | As nectar-tap_saprotroph (Põlme et al. 2020); Decay sugar-rich substrates (Põlme et al. 2020) | Põlme S, et al. 2020. Fungal Diversity 105:1-16 ((https://doi.org/10.1007/s13225-020-00466-2)) |
| Otu61526 | 1 | k__Fungi(100);p__Ascomycota(100);c__Saccharomycetes(100);o__Saccharomycetales(100);f__Metschnikowiaceae(100);g__Metschnikowia(100);g__Metschnikowia_unclassified(100); | Metschnikowia | 13 | Saprotroph | \|Undefined Saprotroph\| | Yeast | NULL | Probable | As nectar-tap_saprotroph (Põlme et al. 2020); Decay sugar-rich substrates (Põlme et al. 2020) | Põlme S, et al. 2020. Fungal Diversity 105:1-16 ((https://doi.org/10.1007/s13225-020-00466-2)) |
| Otu61527 | 1 | k__Fungi(100);p__Ascomycota(100);c__Saccharomycetes(100);o__Saccharomycetales(100);f__Metschnikowiaceae(100);g__Metschnikowia(100);s__Metschnikowia_pulcherrima(100); | Metschnikowia | 13 | Saprotroph | \|Undefined Saprotroph\| | Yeast | NULL | Probable | As nectar-tap_saprotroph (Põlme et al. 2020); Decay sugar-rich substrates (Põlme et al. 2020) | Põlme S, et al. 2020. Fungal Diversity 105:1-16 ((https://doi.org/10.1007/s13225-020-00466-2)) |
| Otu61528 | 1 | k__Fungi(100);p__Ascomycota(100);c__Saccharomycetes(100);o__Saccharomycetales(100);f__Metschnikowiaceae(100);g__Metschnikowia(100);s__Metschnikowia_pulcherrima(100); | Metschnikowia | 13 | Saprotroph | \|Undefined Saprotroph\| | Yeast | NULL | Probable | As nectar-tap_saprotroph (Põlme et al. 2020); Decay sugar-rich substrates (Põlme et al. 2020) | Põlme S, et al. 2020. Fungal Diversity 105:1-16 ((https://doi.org/10.1007/s13225-020-00466-2)) |
| Otu61529 | 1 | k__Fungi(100);p__Ascomycota(100);c__Saccharomycetes(100);o__Saccharomycetales(100);f__Metschnikowiaceae(100);g__Metschnikowia(100);s__Metschnikowia_pulcherrima(100); | Metschnikowia | 13 | Saprotroph | \|Undefined Saprotroph\| | Yeast | NULL | Probable | As nectar-tap_saprotroph (Põlme et al. 2020); Decay sugar-rich substrates (Põlme et al. 2020) | Põlme S, et al. 2020. Fungal Diversity 105:1-16 ((https://doi.org/10.1007/s13225-020-00466-2)) |
| Otu61530 | 1 | k__Fungi(100);p__Ascomycota(100);c__Saccharomycetes(100);o__Saccharomycetales(100);f__Metschnikowiaceae(100);g__Metschnikowia(100);s__Metschnikowia_pulcherrima(100); | Metschnikowia | 13 | Saprotroph | \|Undefined Saprotroph\| | Yeast | NULL | Probable | As nectar-tap_saprotroph (Põlme et al. 2020); Decay sugar-rich substrates (Põlme et al. 2020) | Põlme S, et al. 2020. Fungal Diversity 105:1-16 ((https://doi.org/10.1007/s13225-020-00466-2)) |
| Otu61531 | 1 | k__Fungi(100);p__Ascomycota(100);c__Saccharomycetes(100);o__Saccharomycetales(100);f__Metschnikowiaceae(100);g__Metschnikowia(100);g__Metschnikowia_unclassified(100); | Metschnikowia | 13 | Saprotroph | \|Undefined Saprotroph\| | Yeast | NULL | Probable | As nectar-tap_saprotroph (Põlme et al. 2020); Decay sugar-rich substrates (Põlme et al. 2020) | Põlme S, et al. 2020. Fungal Diversity 105:1-16 ((https://doi.org/10.1007/s13225-020-00466-2)) |
| Otu61532 | 1 | k__Fungi(100);p__Ascomycota(100);c__Saccharomycetes(100);o__Saccharomycetales(100);f__Metschnikowiaceae(100);g__Metschnikowia(100);g__Metschnikowia_unclassified(100); | Metschnikowia | 13 | Saprotroph | \|Undefined Saprotroph\| | Yeast | NULL | Probable | As nectar-tap_saprotroph (Põlme et al. 2020); Decay sugar-rich substrates (Põlme et al. 2020) | Põlme S, et al. 2020. Fungal Diversity 105:1-16 ((https://doi.org/10.1007/s13225-020-00466-2)) |
| Otu61533 | 1 | k__Fungi(100);p__Ascomycota(100);c__Saccharomycetes(100);o__Saccharomycetales(100);f__Metschnikowiaceae(100);g__Metschnikowia(100);s__Metschnikowia_pulcherrima(100); | Metschnikowia | 13 | Saprotroph | \|Undefined Saprotroph\| | Yeast | NULL | Probable | As nectar-tap_saprotroph (Põlme et al. 2020); Decay sugar-rich substrates (Põlme et al. 2020) | Põlme S, et al. 2020. Fungal Diversity 105:1-16 ((https://doi.org/10.1007/s13225-020-00466-2)) |
| Otu61534 | 1 | k__Fungi(100);p__Ascomycota(100);c__Saccharomycetes(100);o__Saccharomycetales(100);f__Metschnikowiaceae(100);g__Metschnikowia(100);s__Metschnikowia_pulcherrima(100); | Metschnikowia | 13 | Saprotroph | \|Undefined Saprotroph\| | Yeast | NULL | Probable | As nectar-tap_saprotroph (Põlme et al. 2020); Decay sugar-rich substrates (Põlme et al. 2020) | Põlme S, et al. 2020. Fungal Diversity 105:1-16 ((https://doi.org/10.1007/s13225-020-00466-2)) |
| Otu61535 | 1 | k__Fungi(100);p__Ascomycota(100);c__Saccharomycetes(100);o__Saccharomycetales(100);f__Metschnikowiaceae(100);g__Metschnikowia(100);s__Metschnikowia_pulcherrima(100); | Metschnikowia | 13 | Saprotroph | \|Undefined Saprotroph\| | Yeast | NULL | Probable | As nectar-tap_saprotroph (Põlme et al. 2020); Decay sugar-rich substrates (Põlme et al. 2020) | Põlme S, et al. 2020. Fungal Diversity 105:1-16 ((https://doi.org/10.1007/s13225-020-00466-2)) |
| Otu61536 | 1 | k__Fungi(100);p__Ascomycota(100);c__Saccharomycetes(100);o__Saccharomycetales(100);f__Metschnikowiaceae(100);g__Metschnikowia(100);s__Metschnikowia_pulcherrima(100); | Metschnikowia | 13 | Saprotroph | \|Undefined Saprotroph\| | Yeast | NULL | Probable | As nectar-tap_saprotroph (Põlme et al. 2020); Decay sugar-rich substrates (Põlme et al. 2020) | Põlme S, et al. 2020. Fungal Diversity 105:1-16 ((https://doi.org/10.1007/s13225-020-00466-2)) |
| Otu61537 | 1 | k__Fungi(100);p__Ascomycota(100);c__Saccharomycetes(100);o__Saccharomycetales(100);f__Metschnikowiaceae(100);g__Metschnikowia(100);s__Metschnikowia_pulcherrima(100); | Metschnikowia | 13 | Saprotroph | \|Undefined Saprotroph\| | Yeast | NULL | Probable | As nectar-tap_saprotroph (Põlme et al. 2020); Decay sugar-rich substrates (Põlme et al. 2020) | Põlme S, et al. 2020. Fungal Diversity 105:1-16 ((https://doi.org/10.1007/s13225-020-00466-2)) |
| Otu61538 | 1 | k__Fungi(100);p__Ascomycota(100);c__Saccharomycetes(100);o__Saccharomycetales(100);f__Metschnikowiaceae(100);g__Metschnikowia(100);g__Metschnikowia_unclassified(100); | Metschnikowia | 13 | Saprotroph | \|Undefined Saprotroph\| | Yeast | NULL | Probable | As nectar-tap_saprotroph (Põlme et al. 2020); Decay sugar-rich substrates (Põlme et al. 2020) | Põlme S, et al. 2020. Fungal Diversity 105:1-16 ((https://doi.org/10.1007/s13225-020-00466-2)) |
| Otu61546 | 1 | k__Fungi(100);p__Ascomycota(100);c__Saccharomycetes(100);o__Saccharomycetales(100);f__Metschnikowiaceae(100);g__Metschnikowia(100);s__Metschnikowia_pulcherrima(100); | Metschnikowia | 13 | Saprotroph | \|Undefined Saprotroph\| | Yeast | NULL | Probable | As nectar-tap_saprotroph (Põlme et al. 2020); Decay sugar-rich substrates (Põlme et al. 2020) | Põlme S, et al. 2020. Fungal Diversity 105:1-16 ((https://doi.org/10.1007/s13225-020-00466-2)) |
| Otu61548 | 1 | k__Fungi(100);p__Ascomycota(100);c__Saccharomycetes(100);o__Saccharomycetales(100);f__Saccharomycetales_family_Incertae_sedis(100);g__Candida(100);g__Candida_unclassified(100); | Candida | 13 | Pathotroph-Saprotroph-Symbiotroph | Animal Pathogen-Endophyte-Endosymbiont-Epiphyte-\|Undefined Saprotroph\| | Dimorphic Yeast-Yeast | NULL | Probable | As nectar-tap_saprotroph (Põlme et al. 2020); Decay sugar-rich substrates (Põlme et al. 2020); This asexual genus is polyphyletic and contain species that function very differently depend on what they are. Caution should be taken when interpreting guilds for Candida species. | Manolakaki D, et al. 2010. Virulence. 1: 367-375 ((https://doi.org/10.4161/viru.1.5.12796)); Mekbib SB, et al. 2011. Tropical Plant Pathology 36:233-240 ((https://doi.org/10.1590/S1982-56762011000400004)); Põlme S, et al. 2020. Fungal Diversity 105:1-16 ((https://doi.org/10.1007/s13225-020-00466-2)) |
| Otu61549 | 1 | k__Fungi(100);p__Ascomycota(100);c__Saccharomycetes(100);o__Saccharomycetales(100);f__Metschnikowiaceae(100);g__Metschnikowia(100);g__Metschnikowia_unclassified(100); | Metschnikowia | 13 | Saprotroph | \|Undefined Saprotroph\| | Yeast | NULL | Probable | As nectar-tap_saprotroph (Põlme et al. 2020); Decay sugar-rich substrates (Põlme et al. 2020) | Põlme S, et al. 2020. Fungal Diversity 105:1-16 ((https://doi.org/10.1007/s13225-020-00466-2)) |
| Otu61551 | 1 | k__Fungi(100);p__Ascomycota(100);c__Saccharomycetes(100);o__Saccharomycetales(100);f__Metschnikowiaceae(100);g__Metschnikowia(100);s__Metschnikowia_pulcherrima(100); | Metschnikowia | 13 | Saprotroph | \|Undefined Saprotroph\| | Yeast | NULL | Probable | As nectar-tap_saprotroph (Põlme et al. 2020); Decay sugar-rich substrates (Põlme et al. 2020) | Põlme S, et al. 2020. Fungal Diversity 105:1-16 ((https://doi.org/10.1007/s13225-020-00466-2)) |
| Otu61552 | 1 | k__Fungi(100);p__Ascomycota(100);c__Saccharomycetes(100);o__Saccharomycetales(100);f__Metschnikowiaceae(100);g__Metschnikowia(100);s__Metschnikowia_pulcherrima(100); | Metschnikowia | 13 | Saprotroph | \|Undefined Saprotroph\| | Yeast | NULL | Probable | As nectar-tap_saprotroph (Põlme et al. 2020); Decay sugar-rich substrates (Põlme et al. 2020) | Põlme S, et al. 2020. Fungal Diversity 105:1-16 ((https://doi.org/10.1007/s13225-020-00466-2)) |
| Otu61553 | 1 | k__Fungi(100);p__Ascomycota(100);c__Saccharomycetes(100);o__Saccharomycetales(100);f__Metschnikowiaceae(100);g__Metschnikowia(100);s__Metschnikowia_pulcherrima(100); | Metschnikowia | 13 | Saprotroph | \|Undefined Saprotroph\| | Yeast | NULL | Probable | As nectar-tap_saprotroph (Põlme et al. 2020); Decay sugar-rich substrates (Põlme et al. 2020) | Põlme S, et al. 2020. Fungal Diversity 105:1-16 ((https://doi.org/10.1007/s13225-020-00466-2)) |
| Otu61554 | 1 | k__Fungi(100);p__Ascomycota(100);c__Saccharomycetes(100);o__Saccharomycetales(100);f__Metschnikowiaceae(100);g__Metschnikowia(100);s__Metschnikowia_pulcherrima(100); | Metschnikowia | 13 | Saprotroph | \|Undefined Saprotroph\| | Yeast | NULL | Probable | As nectar-tap_saprotroph (Põlme et al. 2020); Decay sugar-rich substrates (Põlme et al. 2020) | Põlme S, et al. 2020. Fungal Diversity 105:1-16 ((https://doi.org/10.1007/s13225-020-00466-2)) |
| Otu61556 | 1 | k__Fungi(100);p__Ascomycota(100);c__Saccharomycetes(100);o__Saccharomycetales(100);f__Metschnikowiaceae(100);g__Metschnikowia(100);g__Metschnikowia_unclassified(100); | Metschnikowia | 13 | Saprotroph | \|Undefined Saprotroph\| | Yeast | NULL | Probable | As nectar-tap_saprotroph (Põlme et al. 2020); Decay sugar-rich substrates (Põlme et al. 2020) | Põlme S, et al. 2020. Fungal Diversity 105:1-16 ((https://doi.org/10.1007/s13225-020-00466-2)) |
| Otu61557 | 1 | k__Fungi(100);p__Ascomycota(100);c__Saccharomycetes(100);o__Saccharomycetales(100);f__Metschnikowiaceae(100);g__Metschnikowia(100);g__Metschnikowia_unclassified(100); | Metschnikowia | 13 | Saprotroph | \|Undefined Saprotroph\| | Yeast | NULL | Probable | As nectar-tap_saprotroph (Põlme et al. 2020); Decay sugar-rich substrates (Põlme et al. 2020) | Põlme S, et al. 2020. Fungal Diversity 105:1-16 ((https://doi.org/10.1007/s13225-020-00466-2)) |
| Otu61558 | 1 | k__Fungi(100);p__Ascomycota(100);c__Saccharomycetes(100);o__Saccharomycetales(100);f__Metschnikowiaceae(100);g__Metschnikowia(100);s__Metschnikowia_pulcherrima(100); | Metschnikowia | 13 | Saprotroph | \|Undefined Saprotroph\| | Yeast | NULL | Probable | As nectar-tap_saprotroph (Põlme et al. 2020); Decay sugar-rich substrates (Põlme et al. 2020) | Põlme S, et al. 2020. Fungal Diversity 105:1-16 ((https://doi.org/10.1007/s13225-020-00466-2)) |
| Otu61559 | 1 | k__Fungi(100);p__Ascomycota(100);c__Saccharomycetes(100);o__Saccharomycetales(100);f__Metschnikowiaceae(100);g__Metschnikowia(100);s__Metschnikowia_pulcherrima(100); | Metschnikowia | 13 | Saprotroph | \|Undefined Saprotroph\| | Yeast | NULL | Probable | As nectar-tap_saprotroph (Põlme et al. 2020); Decay sugar-rich substrates (Põlme et al. 2020) | Põlme S, et al. 2020. Fungal Diversity 105:1-16 ((https://doi.org/10.1007/s13225-020-00466-2)) |
| Otu61560 | 1 | k__Fungi(100);p__Ascomycota(100);c__Saccharomycetes(100);o__Saccharomycetales(100);f__Metschnikowiaceae(100);g__Metschnikowia(100);s__Metschnikowia_pulcherrima(100); | Metschnikowia | 13 | Saprotroph | \|Undefined Saprotroph\| | Yeast | NULL | Probable | As nectar-tap_saprotroph (Põlme et al. 2020); Decay sugar-rich substrates (Põlme et al. 2020) | Põlme S, et al. 2020. Fungal Diversity 105:1-16 ((https://doi.org/10.1007/s13225-020-00466-2)) |
| Otu61561 | 1 | k__Fungi(100);p__Ascomycota(100);c__Saccharomycetes(100);o__Saccharomycetales(100);f__Metschnikowiaceae(100);g__Metschnikowia(100);g__Metschnikowia_unclassified(100); | Metschnikowia | 13 | Saprotroph | \|Undefined Saprotroph\| | Yeast | NULL | Probable | As nectar-tap_saprotroph (Põlme et al. 2020); Decay sugar-rich substrates (Põlme et al. 2020) | Põlme S, et al. 2020. Fungal Diversity 105:1-16 ((https://doi.org/10.1007/s13225-020-00466-2)) |
| Otu61562 | 1 | k__Fungi(100);p__Ascomycota(100);c__Saccharomycetes(100);o__Saccharomycetales(100);f__Metschnikowiaceae(100);g__Metschnikowia(100);s__Metschnikowia_pulcherrima(100); | Metschnikowia | 13 | Saprotroph | \|Undefined Saprotroph\| | Yeast | NULL | Probable | As nectar-tap_saprotroph (Põlme et al. 2020); Decay sugar-rich substrates (Põlme et al. 2020) | Põlme S, et al. 2020. Fungal Diversity 105:1-16 ((https://doi.org/10.1007/s13225-020-00466-2)) |
| Otu61563 | 1 | k__Fungi(100);p__Ascomycota(100);c__Saccharomycetes(100);o__Saccharomycetales(100);f__Metschnikowiaceae(100);g__Metschnikowia(100);g__Metschnikowia_unclassified(100); | Metschnikowia | 13 | Saprotroph | \|Undefined Saprotroph\| | Yeast | NULL | Probable | As nectar-tap_saprotroph (Põlme et al. 2020); Decay sugar-rich substrates (Põlme et al. 2020) | Põlme S, et al. 2020. Fungal Diversity 105:1-16 ((https://doi.org/10.1007/s13225-020-00466-2)) |
| Otu61564 | 1 | k__Fungi(100);p__Ascomycota(100);c__Saccharomycetes(100);o__Saccharomycetales(100);f__Metschnikowiaceae(100);g__Metschnikowia(100);s__Metschnikowia_pulcherrima(100); | Metschnikowia | 13 | Saprotroph | \|Undefined Saprotroph\| | Yeast | NULL | Probable | As nectar-tap_saprotroph (Põlme et al. 2020); Decay sugar-rich substrates (Põlme et al. 2020) | Põlme S, et al. 2020. Fungal Diversity 105:1-16 ((https://doi.org/10.1007/s13225-020-00466-2)) |
| Otu61565 | 1 | k__Fungi(100);p__Ascomycota(100);c__Saccharomycetes(100);o__Saccharomycetales(100);f__Metschnikowiaceae(100);g__Metschnikowia(100);g__Metschnikowia_unclassified(100); | Metschnikowia | 13 | Saprotroph | \|Undefined Saprotroph\| | Yeast | NULL | Probable | As nectar-tap_saprotroph (Põlme et al. 2020); Decay sugar-rich substrates (Põlme et al. 2020) | Põlme S, et al. 2020. Fungal Diversity 105:1-16 ((https://doi.org/10.1007/s13225-020-00466-2)) |
| Otu61566 | 1 | k__Fungi(100);p__Ascomycota(100);c__Saccharomycetes(100);o__Saccharomycetales(100);f__Metschnikowiaceae(100);g__Metschnikowia(100);s__Metschnikowia_pulcherrima(100); | Metschnikowia | 13 | Saprotroph | \|Undefined Saprotroph\| | Yeast | NULL | Probable | As nectar-tap_saprotroph (Põlme et al. 2020); Decay sugar-rich substrates (Põlme et al. 2020) | Põlme S, et al. 2020. Fungal Diversity 105:1-16 ((https://doi.org/10.1007/s13225-020-00466-2)) |
| Otu61567 | 1 | k__Fungi(100);p__Ascomycota(100);c__Saccharomycetes(100);o__Saccharomycetales(100);f__Metschnikowiaceae(100);g__Metschnikowia(100);s__Metschnikowia_pulcherrima(100); | Metschnikowia | 13 | Saprotroph | \|Undefined Saprotroph\| | Yeast | NULL | Probable | As nectar-tap_saprotroph (Põlme et al. 2020); Decay sugar-rich substrates (Põlme et al. 2020) | Põlme S, et al. 2020. Fungal Diversity 105:1-16 ((https://doi.org/10.1007/s13225-020-00466-2)) |
| Otu61568 | 1 | k__Fungi(100);p__Ascomycota(100);c__Saccharomycetes(100);o__Saccharomycetales(100);f__Metschnikowiaceae(100);g__Metschnikowia(100);g__Metschnikowia_unclassified(100); | Metschnikowia | 13 | Saprotroph | \|Undefined Saprotroph\| | Yeast | NULL | Probable | As nectar-tap_saprotroph (Põlme et al. 2020); Decay sugar-rich substrates (Põlme et al. 2020) | Põlme S, et al. 2020. Fungal Diversity 105:1-16 ((https://doi.org/10.1007/s13225-020-00466-2)) |
| Otu61569 | 1 | k__Fungi(100);p__Ascomycota(100);c__Saccharomycetes(100);o__Saccharomycetales(100);f__Metschnikowiaceae(100);g__Metschnikowia(100);g__Metschnikowia_unclassified(100); | Metschnikowia | 13 | Saprotroph | \|Undefined Saprotroph\| | Yeast | NULL | Probable | As nectar-tap_saprotroph (Põlme et al. 2020); Decay sugar-rich substrates (Põlme et al. 2020) | Põlme S, et al. 2020. Fungal Diversity 105:1-16 ((https://doi.org/10.1007/s13225-020-00466-2)) |
| Otu61570 | 1 | k__Fungi(100);p__Ascomycota(100);c__Saccharomycetes(100);o__Saccharomycetales(100);f__Metschnikowiaceae(100);g__Metschnikowia(100);s__Metschnikowia_pulcherrima(100); | Metschnikowia | 13 | Saprotroph | \|Undefined Saprotroph\| | Yeast | NULL | Probable | As nectar-tap_saprotroph (Põlme et al. 2020); Decay sugar-rich substrates (Põlme et al. 2020) | Põlme S, et al. 2020. Fungal Diversity 105:1-16 ((https://doi.org/10.1007/s13225-020-00466-2)) |
| Otu61571 | 1 | k__Fungi(100);p__Ascomycota(100);c__Saccharomycetes(100);o__Saccharomycetales(100);f__Metschnikowiaceae(100);g__Metschnikowia(100);s__Metschnikowia_pulcherrima(100); | Metschnikowia | 13 | Saprotroph | \|Undefined Saprotroph\| | Yeast | NULL | Probable | As nectar-tap_saprotroph (Põlme et al. 2020); Decay sugar-rich substrates (Põlme et al. 2020) | Põlme S, et al. 2020. Fungal Diversity 105:1-16 ((https://doi.org/10.1007/s13225-020-00466-2)) |
| Otu61572 | 1 | k__Fungi(100);p__Ascomycota(100);c__Saccharomycetes(100);o__Saccharomycetales(100);f__Metschnikowiaceae(100);g__Metschnikowia(100);s__Metschnikowia_pulcherrima(100); | Metschnikowia | 13 | Saprotroph | \|Undefined Saprotroph\| | Yeast | NULL | Probable | As nectar-tap_saprotroph (Põlme et al. 2020); Decay sugar-rich substrates (Põlme et al. 2020) | Põlme S, et al. 2020. Fungal Diversity 105:1-16 ((https://doi.org/10.1007/s13225-020-00466-2)) |
| Otu61573 | 1 | k__Fungi(100);p__Ascomycota(100);c__Saccharomycetes(100);o__Saccharomycetales(100);f__Metschnikowiaceae(100);g__Metschnikowia(100);s__Metschnikowia_pulcherrima(100); | Metschnikowia | 13 | Saprotroph | \|Undefined Saprotroph\| | Yeast | NULL | Probable | As nectar-tap_saprotroph (Põlme et al. 2020); Decay sugar-rich substrates (Põlme et al. 2020) | Põlme S, et al. 2020. Fungal Diversity 105:1-16 ((https://doi.org/10.1007/s13225-020-00466-2)) |
| Otu61574 | 1 | k__Fungi(100);p__Ascomycota(100);c__Saccharomycetes(100);o__Saccharomycetales(100);f__Metschnikowiaceae(100);g__Metschnikowia(100);s__Metschnikowia_pulcherrima(100); | Metschnikowia | 13 | Saprotroph | \|Undefined Saprotroph\| | Yeast | NULL | Probable | As nectar-tap_saprotroph (Põlme et al. 2020); Decay sugar-rich substrates (Põlme et al. 2020) | Põlme S, et al. 2020. Fungal Diversity 105:1-16 ((https://doi.org/10.1007/s13225-020-00466-2)) |
| Otu61575 | 1 | k__Fungi(100);p__Ascomycota(100);c__Saccharomycetes(100);o__Saccharomycetales(100);f__Metschnikowiaceae(100);g__Metschnikowia(100);s__Metschnikowia_pulcherrima(100); | Metschnikowia | 13 | Saprotroph | \|Undefined Saprotroph\| | Yeast | NULL | Probable | As nectar-tap_saprotroph (Põlme et al. 2020); Decay sugar-rich substrates (Põlme et al. 2020) | Põlme S, et al. 2020. Fungal Diversity 105:1-16 ((https://doi.org/10.1007/s13225-020-00466-2)) |
| Otu61576 | 1 | k__Fungi(100);p__Ascomycota(100);c__Saccharomycetes(100);o__Saccharomycetales(100);f__Metschnikowiaceae(100);g__Metschnikowia(100);s__Metschnikowia_pulcherrima(100); | Metschnikowia | 13 | Saprotroph | \|Undefined Saprotroph\| | Yeast | NULL | Probable | As nectar-tap_saprotroph (Põlme et al. 2020); Decay sugar-rich substrates (Põlme et al. 2020) | Põlme S, et al. 2020. Fungal Diversity 105:1-16 ((https://doi.org/10.1007/s13225-020-00466-2)) |
| Otu61577 | 1 | k__Fungi(100);p__Ascomycota(100);c__Saccharomycetes(100);o__Saccharomycetales(100);f__Metschnikowiaceae(100);g__Metschnikowia(100);s__Metschnikowia_pulcherrima(100); | Metschnikowia | 13 | Saprotroph | \|Undefined Saprotroph\| | Yeast | NULL | Probable | As nectar-tap_saprotroph (Põlme et al. 2020); Decay sugar-rich substrates (Põlme et al. 2020) | Põlme S, et al. 2020. Fungal Diversity 105:1-16 ((https://doi.org/10.1007/s13225-020-00466-2)) |
| Otu61579 | 1 | k__Fungi(100);p__Ascomycota(100);c__Saccharomycetes(100);o__Saccharomycetales(100);f__Metschnikowiaceae(100);g__Metschnikowia(100);s__Metschnikowia_pulcherrima(100); | Metschnikowia | 13 | Saprotroph | \|Undefined Saprotroph\| | Yeast | NULL | Probable | As nectar-tap_saprotroph (Põlme et al. 2020); Decay sugar-rich substrates (Põlme et al. 2020) | Põlme S, et al. 2020. Fungal Diversity 105:1-16 ((https://doi.org/10.1007/s13225-020-00466-2)) |
| Otu61580 | 1 | k__Fungi(100);p__Ascomycota(100);c__Saccharomycetes(100);o__Saccharomycetales(100);f__Metschnikowiaceae(100);g__Metschnikowia(100);s__Metschnikowia_pulcherrima(100); | Metschnikowia | 13 | Saprotroph | \|Undefined Saprotroph\| | Yeast | NULL | Probable | As nectar-tap_saprotroph (Põlme et al. 2020); Decay sugar-rich substrates (Põlme et al. 2020) | Põlme S, et al. 2020. Fungal Diversity 105:1-16 ((https://doi.org/10.1007/s13225-020-00466-2)) |
| Otu61581 | 1 | k__Fungi(100);p__Ascomycota(100);c__Saccharomycetes(100);o__Saccharomycetales(100);f__Metschnikowiaceae(100);g__Metschnikowia(100);s__Metschnikowia_pulcherrima(100); | Metschnikowia | 13 | Saprotroph | \|Undefined Saprotroph\| | Yeast | NULL | Probable | As nectar-tap_saprotroph (Põlme et al. 2020); Decay sugar-rich substrates (Põlme et al. 2020) | Põlme S, et al. 2020. Fungal Diversity 105:1-16 ((https://doi.org/10.1007/s13225-020-00466-2)) |
| Otu61584 | 1 | k__Fungi(100);p__Ascomycota(100);c__Saccharomycetes(100);o__Saccharomycetales(100);f__Metschnikowiaceae(100);g__Metschnikowia(100);s__Metschnikowia_pulcherrima(100); | Metschnikowia | 13 | Saprotroph | \|Undefined Saprotroph\| | Yeast | NULL | Probable | As nectar-tap_saprotroph (Põlme et al. 2020); Decay sugar-rich substrates (Põlme et al. 2020) | Põlme S, et al. 2020. Fungal Diversity 105:1-16 ((https://doi.org/10.1007/s13225-020-00466-2)) |
| Otu61585 | 1 | k__Fungi(100);p__Ascomycota(100);c__Saccharomycetes(100);o__Saccharomycetales(100);f__Metschnikowiaceae(100);g__Metschnikowia(100);s__Metschnikowia_pulcherrima(100); | Metschnikowia | 13 | Saprotroph | \|Undefined Saprotroph\| | Yeast | NULL | Probable | As nectar-tap_saprotroph (Põlme et al. 2020); Decay sugar-rich substrates (Põlme et al. 2020) | Põlme S, et al. 2020. Fungal Diversity 105:1-16 ((https://doi.org/10.1007/s13225-020-00466-2)) |
| Otu61586 | 1 | k__Fungi(100);p__Ascomycota(100);c__Saccharomycetes(100);o__Saccharomycetales(100);f__Metschnikowiaceae(100);g__Metschnikowia(100);s__Metschnikowia_pulcherrima(100); | Metschnikowia | 13 | Saprotroph | \|Undefined Saprotroph\| | Yeast | NULL | Probable | As nectar-tap_saprotroph (Põlme et al. 2020); Decay sugar-rich substrates (Põlme et al. 2020) | Põlme S, et al. 2020. Fungal Diversity 105:1-16 ((https://doi.org/10.1007/s13225-020-00466-2)) |
| Otu61587 | 1 | k__Fungi(100);p__Ascomycota(100);c__Saccharomycetes(100);o__Saccharomycetales(100);f__Metschnikowiaceae(100);g__Metschnikowia(100);s__Metschnikowia_pulcherrima(100); | Metschnikowia | 13 | Saprotroph | \|Undefined Saprotroph\| | Yeast | NULL | Probable | As nectar-tap_saprotroph (Põlme et al. 2020); Decay sugar-rich substrates (Põlme et al. 2020) | Põlme S, et al. 2020. Fungal Diversity 105:1-16 ((https://doi.org/10.1007/s13225-020-00466-2)) |
| Otu61588 | 1 | k__Fungi(100);p__Ascomycota(100);c__Saccharomycetes(100);o__Saccharomycetales(100);f__Metschnikowiaceae(100);g__Metschnikowia(100);g__Metschnikowia_unclassified(100); | Metschnikowia | 13 | Saprotroph | \|Undefined Saprotroph\| | Yeast | NULL | Probable | As nectar-tap_saprotroph (Põlme et al. 2020); Decay sugar-rich substrates (Põlme et al. 2020) | Põlme S, et al. 2020. Fungal Diversity 105:1-16 ((https://doi.org/10.1007/s13225-020-00466-2)) |
| Otu61589 | 1 | k__Fungi(100);p__Ascomycota(100);c__Saccharomycetes(100);o__Saccharomycetales(100);f__Metschnikowiaceae(100);g__Metschnikowia(100);g__Metschnikowia_unclassified(100); | Metschnikowia | 13 | Saprotroph | \|Undefined Saprotroph\| | Yeast | NULL | Probable | As nectar-tap_saprotroph (Põlme et al. 2020); Decay sugar-rich substrates (Põlme et al. 2020) | Põlme S, et al. 2020. Fungal Diversity 105:1-16 ((https://doi.org/10.1007/s13225-020-00466-2)) |
| Otu61591 | 1 | k__Fungi(100);p__Ascomycota(100);c__Saccharomycetes(100);o__Saccharomycetales(100);f__Metschnikowiaceae(100);g__Metschnikowia(100);g__Metschnikowia_unclassified(100); | Metschnikowia | 13 | Saprotroph | \|Undefined Saprotroph\| | Yeast | NULL | Probable | As nectar-tap_saprotroph (Põlme et al. 2020); Decay sugar-rich substrates (Põlme et al. 2020) | Põlme S, et al. 2020. Fungal Diversity 105:1-16 ((https://doi.org/10.1007/s13225-020-00466-2)) |
| Otu61592 | 1 | k__Fungi(100);p__Ascomycota(100);c__Saccharomycetes(100);o__Saccharomycetales(100);f__Metschnikowiaceae(100);g__Metschnikowia(100);s__Metschnikowia_pulcherrima(100); | Metschnikowia | 13 | Saprotroph | \|Undefined Saprotroph\| | Yeast | NULL | Probable | As nectar-tap_saprotroph (Põlme et al. 2020); Decay sugar-rich substrates (Põlme et al. 2020) | Põlme S, et al. 2020. Fungal Diversity 105:1-16 ((https://doi.org/10.1007/s13225-020-00466-2)) |
| Otu61593 | 1 | k__Fungi(100);p__Ascomycota(100);c__Saccharomycetes(100);o__Saccharomycetales(100);f__Metschnikowiaceae(100);g__Metschnikowia(100);s__Metschnikowia_pulcherrima(100); | Metschnikowia | 13 | Saprotroph | \|Undefined Saprotroph\| | Yeast | NULL | Probable | As nectar-tap_saprotroph (Põlme et al. 2020); Decay sugar-rich substrates (Põlme et al. 2020) | Põlme S, et al. 2020. Fungal Diversity 105:1-16 ((https://doi.org/10.1007/s13225-020-00466-2)) |
| Otu61594 | 1 | k__Fungi(100);p__Ascomycota(100);c__Saccharomycetes(100);o__Saccharomycetales(100);f__Metschnikowiaceae(100);g__Metschnikowia(100);s__Metschnikowia_pulcherrima(100); | Metschnikowia | 13 | Saprotroph | \|Undefined Saprotroph\| | Yeast | NULL | Probable | As nectar-tap_saprotroph (Põlme et al. 2020); Decay sugar-rich substrates (Põlme et al. 2020) | Põlme S, et al. 2020. Fungal Diversity 105:1-16 ((https://doi.org/10.1007/s13225-020-00466-2)) |
| Otu61595 | 1 | k__Fungi(100);p__Ascomycota(100);c__Saccharomycetes(100);o__Saccharomycetales(100);f__Metschnikowiaceae(100);g__Metschnikowia(100);s__Metschnikowia_pulcherrima(100); | Metschnikowia | 13 | Saprotroph | \|Undefined Saprotroph\| | Yeast | NULL | Probable | As nectar-tap_saprotroph (Põlme et al. 2020); Decay sugar-rich substrates (Põlme et al. 2020) | Põlme S, et al. 2020. Fungal Diversity 105:1-16 ((https://doi.org/10.1007/s13225-020-00466-2)) |
| Otu61596 | 1 | k__Fungi(100);p__Ascomycota(100);c__Saccharomycetes(100);o__Saccharomycetales(100);f__Metschnikowiaceae(100);g__Metschnikowia(100);g__Metschnikowia_unclassified(100); | Metschnikowia | 13 | Saprotroph | \|Undefined Saprotroph\| | Yeast | NULL | Probable | As nectar-tap_saprotroph (Põlme et al. 2020); Decay sugar-rich substrates (Põlme et al. 2020) | Põlme S, et al. 2020. Fungal Diversity 105:1-16 ((https://doi.org/10.1007/s13225-020-00466-2)) |
| Otu61597 | 1 | k__Fungi(100);p__Ascomycota(100);c__Saccharomycetes(100);o__Saccharomycetales(100);f__Metschnikowiaceae(100);g__Metschnikowia(100);g__Metschnikowia_unclassified(100); | Metschnikowia | 13 | Saprotroph | \|Undefined Saprotroph\| | Yeast | NULL | Probable | As nectar-tap_saprotroph (Põlme et al. 2020); Decay sugar-rich substrates (Põlme et al. 2020) | Põlme S, et al. 2020. Fungal Diversity 105:1-16 ((https://doi.org/10.1007/s13225-020-00466-2)) |
| Otu61598 | 1 | k__Fungi(100);p__Ascomycota(100);c__Saccharomycetes(100);o__Saccharomycetales(100);f__Metschnikowiaceae(100);g__Metschnikowia(100);s__Metschnikowia_pulcherrima(100); | Metschnikowia | 13 | Saprotroph | \|Undefined Saprotroph\| | Yeast | NULL | Probable | As nectar-tap_saprotroph (Põlme et al. 2020); Decay sugar-rich substrates (Põlme et al. 2020) | Põlme S, et al. 2020. Fungal Diversity 105:1-16 ((https://doi.org/10.1007/s13225-020-00466-2)) |
| Otu61599 | 1 | k__Fungi(100);p__Ascomycota(100);c__Saccharomycetes(100);o__Saccharomycetales(100);f__Metschnikowiaceae(100);g__Metschnikowia(100);s__Metschnikowia_pulcherrima(100); | Metschnikowia | 13 | Saprotroph | \|Undefined Saprotroph\| | Yeast | NULL | Probable | As nectar-tap_saprotroph (Põlme et al. 2020); Decay sugar-rich substrates (Põlme et al. 2020) | Põlme S, et al. 2020. Fungal Diversity 105:1-16 ((https://doi.org/10.1007/s13225-020-00466-2)) |
| Otu61600 | 1 | k__Fungi(100);p__Ascomycota(100);c__Saccharomycetes(100);o__Saccharomycetales(100);f__Metschnikowiaceae(100);g__Metschnikowia(100);g__Metschnikowia_unclassified(100); | Metschnikowia | 13 | Saprotroph | \|Undefined Saprotroph\| | Yeast | NULL | Probable | As nectar-tap_saprotroph (Põlme et al. 2020); Decay sugar-rich substrates (Põlme et al. 2020) | Põlme S, et al. 2020. Fungal Diversity 105:1-16 ((https://doi.org/10.1007/s13225-020-00466-2)) |
| Otu61601 | 1 | k__Fungi(100);p__Ascomycota(100);c__Saccharomycetes(100);o__Saccharomycetales(100);f__Metschnikowiaceae(100);g__Metschnikowia(100);s__Metschnikowia_pulcherrima(100); | Metschnikowia | 13 | Saprotroph | \|Undefined Saprotroph\| | Yeast | NULL | Probable | As nectar-tap_saprotroph (Põlme et al. 2020); Decay sugar-rich substrates (Põlme et al. 2020) | Põlme S, et al. 2020. Fungal Diversity 105:1-16 ((https://doi.org/10.1007/s13225-020-00466-2)) |
| Otu61605 | 1 | k__Fungi(100);p__Ascomycota(100);c__Sordariomycetes(100);o__Sordariales(100);f__Chaetomiaceae(100);g__Humicola(100);s__Humicola_grisea_var._grisea(100); | Humicola | 13 | Pathotroph-Saprotroph | Plant Pathogen-Undefined Saprotroph-\|Wood Saprotroph\| | Microfungus | Soft Rot | Probable | Decay wood (Põlme et al. 2020) | Seehann G, et al. 1975. List of Fungi in Soft-Rot Tissue Tests. Institute fur Holzbiologie und Holzschutz, Bundesforschungsanstalt fur Forst - und Holzwirtschaft, Hamburg - Lohbrugge, Germany; Costa IPMW, et al. 2012. Mycotaxon 119:e26 ((http://mycotaxon.com/resources/checklists/costa_v119_checklist.pdf)); Tedersoo L, et al. 2014. Science 346:e1256688 ((https://doi.org/10.1126/science.1256688)); Põlme S, et al. 2020. Fungal Diversity 105:1-16 ((https://doi.org/10.1007/s13225-020-00466-2)) |
| Otu61606 | 1 | k__Fungi(100);p__Ascomycota(100);c__Dothideomycetes(100);o__Pleosporales(100);f__Didymosphaeriaceae(100);g__Roussoella(100);s__Roussoella_japanensis(100); | Roussoella | 13 | Saprotroph | Undefined Saprotroph-\|Wood Saprotroph\| | Microfungus | NULL | Probable | Decay wood (Põlme et al. 2020) | Tedersoo L, et al. 2014. Science 346:e1256688 ((https://doi.org/10.1126/science.1256688)); Põlme S, et al. 2020. Fungal Diversity 105:1-16 ((https://doi.org/10.1007/s13225-020-00466-2)) |
| Otu61608 | 1 | k__Fungi(100);p__Ascomycota(100);c__Saccharomycetes(100);o__Saccharomycetales(100);f__Metschnikowiaceae(100);g__Metschnikowia(100);g__Metschnikowia_unclassified(100); | Metschnikowia | 13 | Saprotroph | \|Undefined Saprotroph\| | Yeast | NULL | Probable | As nectar-tap_saprotroph (Põlme et al. 2020); Decay sugar-rich substrates (Põlme et al. 2020) | Põlme S, et al. 2020. Fungal Diversity 105:1-16 ((https://doi.org/10.1007/s13225-020-00466-2)) |
| Otu61609 | 1 | k__Fungi(100);p__Ascomycota(100);c__Saccharomycetes(100);o__Saccharomycetales(100);f__Metschnikowiaceae(100);g__Metschnikowia(100);s__Metschnikowia_pulcherrima(100); | Metschnikowia | 13 | Saprotroph | \|Undefined Saprotroph\| | Yeast | NULL | Probable | As nectar-tap_saprotroph (Põlme et al. 2020); Decay sugar-rich substrates (Põlme et al. 2020) | Põlme S, et al. 2020. Fungal Diversity 105:1-16 ((https://doi.org/10.1007/s13225-020-00466-2)) |
| Otu61610 | 1 | k__Fungi(100);p__Ascomycota(100);c__Saccharomycetes(100);o__Saccharomycetales(100);f__Metschnikowiaceae(100);g__Metschnikowia(100);s__Metschnikowia_pulcherrima(100); | Metschnikowia | 13 | Saprotroph | \|Undefined Saprotroph\| | Yeast | NULL | Probable | As nectar-tap_saprotroph (Põlme et al. 2020); Decay sugar-rich substrates (Põlme et al. 2020) | Põlme S, et al. 2020. Fungal Diversity 105:1-16 ((https://doi.org/10.1007/s13225-020-00466-2)) |
| Otu61611 | 1 | k__Fungi(100);p__Ascomycota(100);c__Saccharomycetes(100);o__Saccharomycetales(100);f__Metschnikowiaceae(100);g__Metschnikowia(100);s__Metschnikowia_pulcherrima(100); | Metschnikowia | 13 | Saprotroph | \|Undefined Saprotroph\| | Yeast | NULL | Probable | As nectar-tap_saprotroph (Põlme et al. 2020); Decay sugar-rich substrates (Põlme et al. 2020) | Põlme S, et al. 2020. Fungal Diversity 105:1-16 ((https://doi.org/10.1007/s13225-020-00466-2)) |
| Otu61612 | 1 | k__Fungi(100);p__Ascomycota(100);c__Saccharomycetes(100);o__Saccharomycetales(100);f__Metschnikowiaceae(100);g__Metschnikowia(100);s__Metschnikowia_pulcherrima(100); | Metschnikowia | 13 | Saprotroph | \|Undefined Saprotroph\| | Yeast | NULL | Probable | As nectar-tap_saprotroph (Põlme et al. 2020); Decay sugar-rich substrates (Põlme et al. 2020) | Põlme S, et al. 2020. Fungal Diversity 105:1-16 ((https://doi.org/10.1007/s13225-020-00466-2)) |
| Otu61613 | 1 | k__Fungi(100);p__Ascomycota(100);c__Saccharomycetes(100);o__Saccharomycetales(100);f__Metschnikowiaceae(100);g__Metschnikowia(100);g__Metschnikowia_unclassified(100); | Metschnikowia | 13 | Saprotroph | \|Undefined Saprotroph\| | Yeast | NULL | Probable | As nectar-tap_saprotroph (Põlme et al. 2020); Decay sugar-rich substrates (Põlme et al. 2020) | Põlme S, et al. 2020. Fungal Diversity 105:1-16 ((https://doi.org/10.1007/s13225-020-00466-2)) |
| Otu61614 | 1 | k__Fungi(100);p__Ascomycota(100);c__Saccharomycetes(100);o__Saccharomycetales(100);f__Metschnikowiaceae(100);g__Metschnikowia(100);s__Metschnikowia_pulcherrima(100); | Metschnikowia | 13 | Saprotroph | \|Undefined Saprotroph\| | Yeast | NULL | Probable | As nectar-tap_saprotroph (Põlme et al. 2020); Decay sugar-rich substrates (Põlme et al. 2020) | Põlme S, et al. 2020. Fungal Diversity 105:1-16 ((https://doi.org/10.1007/s13225-020-00466-2)) |
| Otu61615 | 1 | k__Fungi(100);p__Ascomycota(100);c__Saccharomycetes(100);o__Saccharomycetales(100);f__Saccharomycetales_family_Incertae_sedis(100);g__Candida(100);g__Candida_unclassified(100); | Candida | 13 | Pathotroph-Saprotroph-Symbiotroph | Animal Pathogen-Endophyte-Endosymbiont-Epiphyte-\|Undefined Saprotroph\| | Dimorphic Yeast-Yeast | NULL | Probable | As nectar-tap_saprotroph (Põlme et al. 2020); Decay sugar-rich substrates (Põlme et al. 2020); This asexual genus is polyphyletic and contain species that function very differently depend on what they are. Caution should be taken when interpreting guilds for Candida species. | Manolakaki D, et al. 2010. Virulence. 1: 367-375 ((https://doi.org/10.4161/viru.1.5.12796)); Mekbib SB, et al. 2011. Tropical Plant Pathology 36:233-240 ((https://doi.org/10.1590/S1982-56762011000400004)); Põlme S, et al. 2020. Fungal Diversity 105:1-16 ((https://doi.org/10.1007/s13225-020-00466-2)) |
| Otu61616 | 1 | k__Fungi(100);p__Ascomycota(100);c__Saccharomycetes(100);o__Saccharomycetales(100);f__Metschnikowiaceae(100);g__Metschnikowia(100);s__Metschnikowia_pulcherrima(100); | Metschnikowia | 13 | Saprotroph | \|Undefined Saprotroph\| | Yeast | NULL | Probable | As nectar-tap_saprotroph (Põlme et al. 2020); Decay sugar-rich substrates (Põlme et al. 2020) | Põlme S, et al. 2020. Fungal Diversity 105:1-16 ((https://doi.org/10.1007/s13225-020-00466-2)) |
| Otu61618 | 1 | k__Fungi(100);p__Ascomycota(100);c__Saccharomycetes(100);o__Saccharomycetales(100);f__Metschnikowiaceae(100);g__Metschnikowia(100);s__Metschnikowia_pulcherrima(100); | Metschnikowia | 13 | Saprotroph | \|Undefined Saprotroph\| | Yeast | NULL | Probable | As nectar-tap_saprotroph (Põlme et al. 2020); Decay sugar-rich substrates (Põlme et al. 2020) | Põlme S, et al. 2020. Fungal Diversity 105:1-16 ((https://doi.org/10.1007/s13225-020-00466-2)) |
| Otu61620 | 1 | k__Fungi(100);p__Ascomycota(100);c__Saccharomycetes(100);o__Saccharomycetales(100);f__Saccharomycetales_family_Incertae_sedis(100);f__Saccharomycetales_family_Incertae_sedis_unclassified(100);f__Saccharomycetales_family_Incertae_sedis_unclassified(100); | Saccharomycetales | 7 | Saprotroph | Undefined Saprotroph | Yeast | NULL | Possible | NULL | Sterkenburg E, et al. 2015. New Phytologist 207:1145-1158 ((https://doi.org/10.1111/nph.13426)) |
| Otu61621 | 1 | k__Fungi(100);p__Ascomycota(100);c__Saccharomycetes(100);o__Saccharomycetales(100);f__Saccharomycetales_family_Incertae_sedis(100);g__Candida(100);g__Candida_unclassified(100); | Candida | 13 | Pathotroph-Saprotroph-Symbiotroph | Animal Pathogen-Endophyte-Endosymbiont-Epiphyte-\|Undefined Saprotroph\| | Dimorphic Yeast-Yeast | NULL | Probable | As nectar-tap_saprotroph (Põlme et al. 2020); Decay sugar-rich substrates (Põlme et al. 2020); This asexual genus is polyphyletic and contain species that function very differently depend on what they are. Caution should be taken when interpreting guilds for Candida species. | Manolakaki D, et al. 2010. Virulence. 1: 367-375 ((https://doi.org/10.4161/viru.1.5.12796)); Mekbib SB, et al. 2011. Tropical Plant Pathology 36:233-240 ((https://doi.org/10.1590/S1982-56762011000400004)); Põlme S, et al. 2020. Fungal Diversity 105:1-16 ((https://doi.org/10.1007/s13225-020-00466-2)) |
| Otu61622 | 1 | k__Fungi(100);p__Ascomycota(100);c__Saccharomycetes(100);o__Saccharomycetales(100);f__Saccharomycetales_family_Incertae_sedis(100);g__Candida(100);g__Candida_unclassified(100); | Candida | 13 | Pathotroph-Saprotroph-Symbiotroph | Animal Pathogen-Endophyte-Endosymbiont-Epiphyte-\|Undefined Saprotroph\| | Dimorphic Yeast-Yeast | NULL | Probable | As nectar-tap_saprotroph (Põlme et al. 2020); Decay sugar-rich substrates (Põlme et al. 2020); This asexual genus is polyphyletic and contain species that function very differently depend on what they are. Caution should be taken when interpreting guilds for Candida species. | Manolakaki D, et al. 2010. Virulence. 1: 367-375 ((https://doi.org/10.4161/viru.1.5.12796)); Mekbib SB, et al. 2011. Tropical Plant Pathology 36:233-240 ((https://doi.org/10.1590/S1982-56762011000400004)); Põlme S, et al. 2020. Fungal Diversity 105:1-16 ((https://doi.org/10.1007/s13225-020-00466-2)) |
| Otu61623 | 1 | k__Fungi(100);p__Ascomycota(100);c__Saccharomycetes(100);o__Saccharomycetales(100);f__Metschnikowiaceae(100);g__Metschnikowia(100);s__Metschnikowia_pulcherrima(100); | Metschnikowia | 13 | Saprotroph | \|Undefined Saprotroph\| | Yeast | NULL | Probable | As nectar-tap_saprotroph (Põlme et al. 2020); Decay sugar-rich substrates (Põlme et al. 2020) | Põlme S, et al. 2020. Fungal Diversity 105:1-16 ((https://doi.org/10.1007/s13225-020-00466-2)) |
| Otu61626 | 1 | k__Fungi(100);p__Ascomycota(100);c__Saccharomycetes(100);o__Saccharomycetales(100);f__Metschnikowiaceae(100);g__Metschnikowia(100);s__Metschnikowia_pulcherrima(100); | Metschnikowia | 13 | Saprotroph | \|Undefined Saprotroph\| | Yeast | NULL | Probable | As nectar-tap_saprotroph (Põlme et al. 2020); Decay sugar-rich substrates (Põlme et al. 2020) | Põlme S, et al. 2020. Fungal Diversity 105:1-16 ((https://doi.org/10.1007/s13225-020-00466-2)) |
| Otu61627 | 1 | k__Fungi(100);p__Ascomycota(100);c__Saccharomycetes(100);o__Saccharomycetales(100);f__Metschnikowiaceae(100);g__Metschnikowia(100);s__Metschnikowia_pulcherrima(100); | Metschnikowia | 13 | Saprotroph | \|Undefined Saprotroph\| | Yeast | NULL | Probable | As nectar-tap_saprotroph (Põlme et al. 2020); Decay sugar-rich substrates (Põlme et al. 2020) | Põlme S, et al. 2020. Fungal Diversity 105:1-16 ((https://doi.org/10.1007/s13225-020-00466-2)) |
| Otu61628 | 1 | k__Fungi(100);p__Ascomycota(100);c__Saccharomycetes(100);o__Saccharomycetales(100);f__Metschnikowiaceae(100);g__Metschnikowia(100);g__Metschnikowia_unclassified(100); | Metschnikowia | 13 | Saprotroph | \|Undefined Saprotroph\| | Yeast | NULL | Probable | As nectar-tap_saprotroph (Põlme et al. 2020); Decay sugar-rich substrates (Põlme et al. 2020) | Põlme S, et al. 2020. Fungal Diversity 105:1-16 ((https://doi.org/10.1007/s13225-020-00466-2)) |
| Otu61630 | 1 | k__Fungi(100);p__Ascomycota(100);c__Saccharomycetes(100);o__Saccharomycetales(100);f__Metschnikowiaceae(100);g__Metschnikowia(100);g__Metschnikowia_unclassified(100); | Metschnikowia | 13 | Saprotroph | \|Undefined Saprotroph\| | Yeast | NULL | Probable | As nectar-tap_saprotroph (Põlme et al. 2020); Decay sugar-rich substrates (Põlme et al. 2020) | Põlme S, et al. 2020. Fungal Diversity 105:1-16 ((https://doi.org/10.1007/s13225-020-00466-2)) |
| Otu61632 | 1 | k__Fungi(100);p__Ascomycota(100);c__Saccharomycetes(100);o__Saccharomycetales(100);f__Saccharomycetales_family_Incertae_sedis(100);g__Candida(100);g__Candida_unclassified(100); | Candida | 13 | Pathotroph-Saprotroph-Symbiotroph | Animal Pathogen-Endophyte-Endosymbiont-Epiphyte-\|Undefined Saprotroph\| | Dimorphic Yeast-Yeast | NULL | Probable | As nectar-tap_saprotroph (Põlme et al. 2020); Decay sugar-rich substrates (Põlme et al. 2020); This asexual genus is polyphyletic and contain species that function very differently depend on what they are. Caution should be taken when interpreting guilds for Candida species. | Manolakaki D, et al. 2010. Virulence. 1: 367-375 ((https://doi.org/10.4161/viru.1.5.12796)); Mekbib SB, et al. 2011. Tropical Plant Pathology 36:233-240 ((https://doi.org/10.1590/S1982-56762011000400004)); Põlme S, et al. 2020. Fungal Diversity 105:1-16 ((https://doi.org/10.1007/s13225-020-00466-2)) |
| Otu61641 | 1 | k__Fungi(100);p__Ascomycota(100);c__Saccharomycetes(100);o__Saccharomycetales(100);f__Metschnikowiaceae(100);g__Metschnikowia(100);g__Metschnikowia_unclassified(100); | Metschnikowia | 13 | Saprotroph | \|Undefined Saprotroph\| | Yeast | NULL | Probable | As nectar-tap_saprotroph (Põlme et al. 2020); Decay sugar-rich substrates (Põlme et al. 2020) | Põlme S, et al. 2020. Fungal Diversity 105:1-16 ((https://doi.org/10.1007/s13225-020-00466-2)) |
| Otu61644 | 1 | k__Fungi(100);p__Ascomycota(100);c__Saccharomycetes(100);o__Saccharomycetales(100);f__Metschnikowiaceae(100);g__Metschnikowia(100);s__Metschnikowia_pulcherrima(100); | Metschnikowia | 13 | Saprotroph | \|Undefined Saprotroph\| | Yeast | NULL | Probable | As nectar-tap_saprotroph (Põlme et al. 2020); Decay sugar-rich substrates (Põlme et al. 2020) | Põlme S, et al. 2020. Fungal Diversity 105:1-16 ((https://doi.org/10.1007/s13225-020-00466-2)) |
| Otu61648 | 1 | k__Fungi(100);p__Ascomycota(100);c__Saccharomycetes(100);o__Saccharomycetales(100);f__Metschnikowiaceae(100);g__Metschnikowia(100);s__Metschnikowia_pulcherrima(100); | Metschnikowia | 13 | Saprotroph | \|Undefined Saprotroph\| | Yeast | NULL | Probable | As nectar-tap_saprotroph (Põlme et al. 2020); Decay sugar-rich substrates (Põlme et al. 2020) | Põlme S, et al. 2020. Fungal Diversity 105:1-16 ((https://doi.org/10.1007/s13225-020-00466-2)) |
| Otu61649 | 1 | k__Fungi(100);p__Ascomycota(100);c__Saccharomycetes(100);o__Saccharomycetales(100);f__Metschnikowiaceae(100);g__Metschnikowia(100);s__Metschnikowia_pulcherrima(100); | Metschnikowia | 13 | Saprotroph | \|Undefined Saprotroph\| | Yeast | NULL | Probable | As nectar-tap_saprotroph (Põlme et al. 2020); Decay sugar-rich substrates (Põlme et al. 2020) | Põlme S, et al. 2020. Fungal Diversity 105:1-16 ((https://doi.org/10.1007/s13225-020-00466-2)) |
| Otu61651 | 1 | k__Fungi(100);p__Basidiomycota(100);c__Microbotryomycetes(100);o__Sporidiobolales(100);f__Sporidiobolales_family_Incertae_sedis(100);g__Sporobolomyces(100);s__Sporobolomyces_lactosus(100); | Sporobolomyces | 13 | Pathotroph-Saprotroph | \|Fungal Parasite\|-Plant Saprotroph | Yeast | NULL | Probable | As litter_saprotroph (Põlme et al. 2020); As mycoparasite (Põlme et al. 2020); Decay fungal material (Põlme et al. 2020) | Tedersoo L, et al. 2014. Science 346:e1256688 ((https://doi.org/10.1126/science.1256688)); Purahong W, et al. 2016. Molecular Ecology 25:4059-4074 ((https://doi.org/10.1111/mec.13739)); Põlme S, et al. 2020. Fungal Diversity 105:1-16 ((https://doi.org/10.1007/s13225-020-00466-2)) |
| Otu61653 | 1 | k__Fungi(100);p__Ascomycota(100);c__Saccharomycetes(100);o__Saccharomycetales(100);f__Metschnikowiaceae(100);g__Metschnikowia(100);s__Metschnikowia_pulcherrima(100); | Metschnikowia | 13 | Saprotroph | \|Undefined Saprotroph\| | Yeast | NULL | Probable | As nectar-tap_saprotroph (Põlme et al. 2020); Decay sugar-rich substrates (Põlme et al. 2020) | Põlme S, et al. 2020. Fungal Diversity 105:1-16 ((https://doi.org/10.1007/s13225-020-00466-2)) |
| Otu61654 | 1 | k__Fungi(100);p__Ascomycota(100);c__Saccharomycetes(100);o__Saccharomycetales(100);f__Metschnikowiaceae(100);g__Metschnikowia(100);s__Metschnikowia_pulcherrima(100); | Metschnikowia | 13 | Saprotroph | \|Undefined Saprotroph\| | Yeast | NULL | Probable | As nectar-tap_saprotroph (Põlme et al. 2020); Decay sugar-rich substrates (Põlme et al. 2020) | Põlme S, et al. 2020. Fungal Diversity 105:1-16 ((https://doi.org/10.1007/s13225-020-00466-2)) |
| Otu61656 | 1 | k__Fungi(100);p__Ascomycota(100);c__Saccharomycetes(100);o__Saccharomycetales(100);f__Metschnikowiaceae(100);g__Metschnikowia(100);s__Metschnikowia_pulcherrima(100); | Metschnikowia | 13 | Saprotroph | \|Undefined Saprotroph\| | Yeast | NULL | Probable | As nectar-tap_saprotroph (Põlme et al. 2020); Decay sugar-rich substrates (Põlme et al. 2020) | Põlme S, et al. 2020. Fungal Diversity 105:1-16 ((https://doi.org/10.1007/s13225-020-00466-2)) |
| Otu61657 | 1 | k__Fungi(100);p__Ascomycota(100);c__Saccharomycetes(100);o__Saccharomycetales(100);f__Metschnikowiaceae(100);g__Metschnikowia(100);s__Metschnikowia_pulcherrima(100); | Metschnikowia | 13 | Saprotroph | \|Undefined Saprotroph\| | Yeast | NULL | Probable | As nectar-tap_saprotroph (Põlme et al. 2020); Decay sugar-rich substrates (Põlme et al. 2020) | Põlme S, et al. 2020. Fungal Diversity 105:1-16 ((https://doi.org/10.1007/s13225-020-00466-2)) |
| Otu61658 | 1 | k__Fungi(100);p__Ascomycota(100);c__Saccharomycetes(100);o__Saccharomycetales(100);f__Metschnikowiaceae(100);g__Metschnikowia(100);s__Metschnikowia_pulcherrima(100); | Metschnikowia | 13 | Saprotroph | \|Undefined Saprotroph\| | Yeast | NULL | Probable | As nectar-tap_saprotroph (Põlme et al. 2020); Decay sugar-rich substrates (Põlme et al. 2020) | Põlme S, et al. 2020. Fungal Diversity 105:1-16 ((https://doi.org/10.1007/s13225-020-00466-2)) |
| Otu61659 | 1 | k__Fungi(100);p__Ascomycota(100);c__Saccharomycetes(100);o__Saccharomycetales(100);f__Metschnikowiaceae(100);g__Metschnikowia(100);s__Metschnikowia_pulcherrima(100); | Metschnikowia | 13 | Saprotroph | \|Undefined Saprotroph\| | Yeast | NULL | Probable | As nectar-tap_saprotroph (Põlme et al. 2020); Decay sugar-rich substrates (Põlme et al. 2020) | Põlme S, et al. 2020. Fungal Diversity 105:1-16 ((https://doi.org/10.1007/s13225-020-00466-2)) |
| Otu61660 | 1 | k__Fungi(100);p__Ascomycota(100);c__Saccharomycetes(100);o__Saccharomycetales(100);f__Metschnikowiaceae(100);g__Metschnikowia(100);s__Metschnikowia_pulcherrima(100); | Metschnikowia | 13 | Saprotroph | \|Undefined Saprotroph\| | Yeast | NULL | Probable | As nectar-tap_saprotroph (Põlme et al. 2020); Decay sugar-rich substrates (Põlme et al. 2020) | Põlme S, et al. 2020. Fungal Diversity 105:1-16 ((https://doi.org/10.1007/s13225-020-00466-2)) |
| Otu61661 | 1 | k__Fungi(100);p__Ascomycota(100);c__Saccharomycetes(100);o__Saccharomycetales(100);f__Metschnikowiaceae(100);g__Metschnikowia(100);s__Metschnikowia_pulcherrima(100); | Metschnikowia | 13 | Saprotroph | \|Undefined Saprotroph\| | Yeast | NULL | Probable | As nectar-tap_saprotroph (Põlme et al. 2020); Decay sugar-rich substrates (Põlme et al. 2020) | Põlme S, et al. 2020. Fungal Diversity 105:1-16 ((https://doi.org/10.1007/s13225-020-00466-2)) |
| Otu61662 | 1 | k__Fungi(100);p__Ascomycota(100);c__Saccharomycetes(100);o__Saccharomycetales(100);f__Metschnikowiaceae(100);g__Metschnikowia(100);g__Metschnikowia_unclassified(100); | Metschnikowia | 13 | Saprotroph | \|Undefined Saprotroph\| | Yeast | NULL | Probable | As nectar-tap_saprotroph (Põlme et al. 2020); Decay sugar-rich substrates (Põlme et al. 2020) | Põlme S, et al. 2020. Fungal Diversity 105:1-16 ((https://doi.org/10.1007/s13225-020-00466-2)) |
| Otu61663 | 1 | k__Fungi(100);p__Ascomycota(100);c__Saccharomycetes(100);o__Saccharomycetales(100);f__Metschnikowiaceae(100);g__Metschnikowia(100);s__Metschnikowia_pulcherrima(100); | Metschnikowia | 13 | Saprotroph | \|Undefined Saprotroph\| | Yeast | NULL | Probable | As nectar-tap_saprotroph (Põlme et al. 2020); Decay sugar-rich substrates (Põlme et al. 2020) | Põlme S, et al. 2020. Fungal Diversity 105:1-16 ((https://doi.org/10.1007/s13225-020-00466-2)) |
| Otu61665 | 1 | k__Fungi(100);p__Ascomycota(100);c__Saccharomycetes(100);o__Saccharomycetales(100);f__Metschnikowiaceae(100);g__Metschnikowia(100);s__Metschnikowia_pulcherrima(100); | Metschnikowia | 13 | Saprotroph | \|Undefined Saprotroph\| | Yeast | NULL | Probable | As nectar-tap_saprotroph (Põlme et al. 2020); Decay sugar-rich substrates (Põlme et al. 2020) | Põlme S, et al. 2020. Fungal Diversity 105:1-16 ((https://doi.org/10.1007/s13225-020-00466-2)) |
| Otu61666 | 1 | k__Fungi(100);p__Ascomycota(100);c__Saccharomycetes(100);o__Saccharomycetales(100);f__Metschnikowiaceae(100);g__Metschnikowia(100);s__Metschnikowia_pulcherrima(100); | Metschnikowia | 13 | Saprotroph | \|Undefined Saprotroph\| | Yeast | NULL | Probable | As nectar-tap_saprotroph (Põlme et al. 2020); Decay sugar-rich substrates (Põlme et al. 2020) | Põlme S, et al. 2020. Fungal Diversity 105:1-16 ((https://doi.org/10.1007/s13225-020-00466-2)) |
| Otu61668 | 1 | k__Fungi(100);p__Ascomycota(100);c__Saccharomycetes(100);o__Saccharomycetales(100);f__Metschnikowiaceae(100);g__Metschnikowia(100);s__Metschnikowia_pulcherrima(100); | Metschnikowia | 13 | Saprotroph | \|Undefined Saprotroph\| | Yeast | NULL | Probable | As nectar-tap_saprotroph (Põlme et al. 2020); Decay sugar-rich substrates (Põlme et al. 2020) | Põlme S, et al. 2020. Fungal Diversity 105:1-16 ((https://doi.org/10.1007/s13225-020-00466-2)) |
| Otu61669 | 1 | k__Fungi(100);p__Ascomycota(100);c__Saccharomycetes(100);o__Saccharomycetales(100);f__Metschnikowiaceae(100);g__Metschnikowia(100);s__Metschnikowia_pulcherrima(100); | Metschnikowia | 13 | Saprotroph | \|Undefined Saprotroph\| | Yeast | NULL | Probable | As nectar-tap_saprotroph (Põlme et al. 2020); Decay sugar-rich substrates (Põlme et al. 2020) | Põlme S, et al. 2020. Fungal Diversity 105:1-16 ((https://doi.org/10.1007/s13225-020-00466-2)) |
| Otu61680 | 1 | k__Fungi(100);p__Ascomycota(100);c__Saccharomycetes(100);o__Saccharomycetales(100);f__Metschnikowiaceae(100);g__Metschnikowia(100);g__Metschnikowia_unclassified(100); | Metschnikowia | 13 | Saprotroph | \|Undefined Saprotroph\| | Yeast | NULL | Probable | As nectar-tap_saprotroph (Põlme et al. 2020); Decay sugar-rich substrates (Põlme et al. 2020) | Põlme S, et al. 2020. Fungal Diversity 105:1-16 ((https://doi.org/10.1007/s13225-020-00466-2)) |
| Otu61681 | 1 | k__Fungi(100);p__Ascomycota(100);c__Saccharomycetes(100);o__Saccharomycetales(100);f__Metschnikowiaceae(100);g__Metschnikowia(100);s__Metschnikowia_pulcherrima(100); | Metschnikowia | 13 | Saprotroph | \|Undefined Saprotroph\| | Yeast | NULL | Probable | As nectar-tap_saprotroph (Põlme et al. 2020); Decay sugar-rich substrates (Põlme et al. 2020) | Põlme S, et al. 2020. Fungal Diversity 105:1-16 ((https://doi.org/10.1007/s13225-020-00466-2)) |
| Otu61683 | 1 | k__Fungi(100);p__Ascomycota(100);c__Saccharomycetes(100);o__Saccharomycetales(100);f__Metschnikowiaceae(100);g__Metschnikowia(100);s__Metschnikowia_pulcherrima(100); | Metschnikowia | 13 | Saprotroph | \|Undefined Saprotroph\| | Yeast | NULL | Probable | As nectar-tap_saprotroph (Põlme et al. 2020); Decay sugar-rich substrates (Põlme et al. 2020) | Põlme S, et al. 2020. Fungal Diversity 105:1-16 ((https://doi.org/10.1007/s13225-020-00466-2)) |
| Otu61687 | 1 | k__Fungi(100);p__Ascomycota(100);c__Saccharomycetes(100);o__Saccharomycetales(100);f__Metschnikowiaceae(100);g__Metschnikowia(100);g__Metschnikowia_unclassified(100); | Metschnikowia | 13 | Saprotroph | \|Undefined Saprotroph\| | Yeast | NULL | Probable | As nectar-tap_saprotroph (Põlme et al. 2020); Decay sugar-rich substrates (Põlme et al. 2020) | Põlme S, et al. 2020. Fungal Diversity 105:1-16 ((https://doi.org/10.1007/s13225-020-00466-2)) |
| Otu61688 | 1 | k__Fungi(100);p__Ascomycota(100);c__Saccharomycetes(100);o__Saccharomycetales(100);f__Metschnikowiaceae(100);g__Metschnikowia(100);s__Metschnikowia_pulcherrima(100); | Metschnikowia | 13 | Saprotroph | \|Undefined Saprotroph\| | Yeast | NULL | Probable | As nectar-tap_saprotroph (Põlme et al. 2020); Decay sugar-rich substrates (Põlme et al. 2020) | Põlme S, et al. 2020. Fungal Diversity 105:1-16 ((https://doi.org/10.1007/s13225-020-00466-2)) |
| Otu61690 | 1 | k__Fungi(100);p__Ascomycota(100);c__Saccharomycetes(100);o__Saccharomycetales(100);f__Metschnikowiaceae(100);g__Metschnikowia(100);s__Metschnikowia_pulcherrima(100); | Metschnikowia | 13 | Saprotroph | \|Undefined Saprotroph\| | Yeast | NULL | Probable | As nectar-tap_saprotroph (Põlme et al. 2020); Decay sugar-rich substrates (Põlme et al. 2020) | Põlme S, et al. 2020. Fungal Diversity 105:1-16 ((https://doi.org/10.1007/s13225-020-00466-2)) |
| Otu61694 | 1 | k__Fungi(100);p__Ascomycota(100);c__Saccharomycetes(100);o__Saccharomycetales(100);f__Saccharomycetales_family_Incertae_sedis(100);g__Candida(100);g__Candida_unclassified(100); | Candida | 13 | Pathotroph-Saprotroph-Symbiotroph | Animal Pathogen-Endophyte-Endosymbiont-Epiphyte-\|Undefined Saprotroph\| | Dimorphic Yeast-Yeast | NULL | Probable | As nectar-tap_saprotroph (Põlme et al. 2020); Decay sugar-rich substrates (Põlme et al. 2020); This asexual genus is polyphyletic and contain species that function very differently depend on what they are. Caution should be taken when interpreting guilds for Candida species. | Manolakaki D, et al. 2010. Virulence. 1: 367-375 ((https://doi.org/10.4161/viru.1.5.12796)); Mekbib SB, et al. 2011. Tropical Plant Pathology 36:233-240 ((https://doi.org/10.1590/S1982-56762011000400004)); Põlme S, et al. 2020. Fungal Diversity 105:1-16 ((https://doi.org/10.1007/s13225-020-00466-2)) |
| Otu61696 | 1 | k__Fungi(100);p__Ascomycota(100);c__Saccharomycetes(100);o__Saccharomycetales(100);f__Saccharomycetales_family_Incertae_sedis(100);g__Candida(100);g__Candida_unclassified(100); | Candida | 13 | Pathotroph-Saprotroph-Symbiotroph | Animal Pathogen-Endophyte-Endosymbiont-Epiphyte-\|Undefined Saprotroph\| | Dimorphic Yeast-Yeast | NULL | Probable | As nectar-tap_saprotroph (Põlme et al. 2020); Decay sugar-rich substrates (Põlme et al. 2020); This asexual genus is polyphyletic and contain species that function very differently depend on what they are. Caution should be taken when interpreting guilds for Candida species. | Manolakaki D, et al. 2010. Virulence. 1: 367-375 ((https://doi.org/10.4161/viru.1.5.12796)); Mekbib SB, et al. 2011. Tropical Plant Pathology 36:233-240 ((https://doi.org/10.1590/S1982-56762011000400004)); Põlme S, et al. 2020. Fungal Diversity 105:1-16 ((https://doi.org/10.1007/s13225-020-00466-2)) |
| Otu61698 | 1 | k__Fungi(100);p__Ascomycota(100);c__Saccharomycetes(100);o__Saccharomycetales(100);f__Metschnikowiaceae(100);g__Metschnikowia(100);s__Metschnikowia_pulcherrima(100); | Metschnikowia | 13 | Saprotroph | \|Undefined Saprotroph\| | Yeast | NULL | Probable | As nectar-tap_saprotroph (Põlme et al. 2020); Decay sugar-rich substrates (Põlme et al. 2020) | Põlme S, et al. 2020. Fungal Diversity 105:1-16 ((https://doi.org/10.1007/s13225-020-00466-2)) |
| Otu61699 | 1 | k__Fungi(100);p__Ascomycota(100);c__Saccharomycetes(100);o__Saccharomycetales(100);f__Metschnikowiaceae(100);g__Metschnikowia(100);s__Metschnikowia_pulcherrima(100); | Metschnikowia | 13 | Saprotroph | \|Undefined Saprotroph\| | Yeast | NULL | Probable | As nectar-tap_saprotroph (Põlme et al. 2020); Decay sugar-rich substrates (Põlme et al. 2020) | Põlme S, et al. 2020. Fungal Diversity 105:1-16 ((https://doi.org/10.1007/s13225-020-00466-2)) |
| Otu61703 | 1 | k__Fungi(100);p__Ascomycota(100);c__Saccharomycetes(100);o__Saccharomycetales(100);f__Metschnikowiaceae(100);g__Metschnikowia(100);s__Metschnikowia_pulcherrima(100); | Metschnikowia | 13 | Saprotroph | \|Undefined Saprotroph\| | Yeast | NULL | Probable | As nectar-tap_saprotroph (Põlme et al. 2020); Decay sugar-rich substrates (Põlme et al. 2020) | Põlme S, et al. 2020. Fungal Diversity 105:1-16 ((https://doi.org/10.1007/s13225-020-00466-2)) |
| Otu61705 | 1 | k__Fungi(100);p__Ascomycota(100);c__Saccharomycetes(100);o__Saccharomycetales(100);f__Metschnikowiaceae(100);g__Metschnikowia(100);s__Metschnikowia_pulcherrima(100); | Metschnikowia | 13 | Saprotroph | \|Undefined Saprotroph\| | Yeast | NULL | Probable | As nectar-tap_saprotroph (Põlme et al. 2020); Decay sugar-rich substrates (Põlme et al. 2020) | Põlme S, et al. 2020. Fungal Diversity 105:1-16 ((https://doi.org/10.1007/s13225-020-00466-2)) |
| Otu61706 | 1 | k__Fungi(100);p__Ascomycota(100);c__Saccharomycetes(100);o__Saccharomycetales(100);f__Metschnikowiaceae(100);g__Metschnikowia(100);s__Metschnikowia_pulcherrima(100); | Metschnikowia | 13 | Saprotroph | \|Undefined Saprotroph\| | Yeast | NULL | Probable | As nectar-tap_saprotroph (Põlme et al. 2020); Decay sugar-rich substrates (Põlme et al. 2020) | Põlme S, et al. 2020. Fungal Diversity 105:1-16 ((https://doi.org/10.1007/s13225-020-00466-2)) |
| Otu61709 | 1 | k__Fungi(100);p__Ascomycota(100);c__Saccharomycetes(100);o__Saccharomycetales(100);f__Metschnikowiaceae(100);g__Metschnikowia(100);s__Metschnikowia_pulcherrima(100); | Metschnikowia | 13 | Saprotroph | \|Undefined Saprotroph\| | Yeast | NULL | Probable | As nectar-tap_saprotroph (Põlme et al. 2020); Decay sugar-rich substrates (Põlme et al. 2020) | Põlme S, et al. 2020. Fungal Diversity 105:1-16 ((https://doi.org/10.1007/s13225-020-00466-2)) |
| Otu61711 | 1 | k__Fungi(100);p__Ascomycota(100);c__Saccharomycetes(100);o__Saccharomycetales(100);f__Saccharomycetales_family_Incertae_sedis(100);g__Candida(100);g__Candida_unclassified(100); | Candida | 13 | Pathotroph-Saprotroph-Symbiotroph | Animal Pathogen-Endophyte-Endosymbiont-Epiphyte-\|Undefined Saprotroph\| | Dimorphic Yeast-Yeast | NULL | Probable | As nectar-tap_saprotroph (Põlme et al. 2020); Decay sugar-rich substrates (Põlme et al. 2020); This asexual genus is polyphyletic and contain species that function very differently depend on what they are. Caution should be taken when interpreting guilds for Candida species. | Manolakaki D, et al. 2010. Virulence. 1: 367-375 ((https://doi.org/10.4161/viru.1.5.12796)); Mekbib SB, et al. 2011. Tropical Plant Pathology 36:233-240 ((https://doi.org/10.1590/S1982-56762011000400004)); Põlme S, et al. 2020. Fungal Diversity 105:1-16 ((https://doi.org/10.1007/s13225-020-00466-2)) |
| Otu61713 | 1 | k__Fungi(100);p__Ascomycota(100);c__Saccharomycetes(100);o__Saccharomycetales(100);f__Metschnikowiaceae(100);g__Metschnikowia(100);s__Metschnikowia_pulcherrima(100); | Metschnikowia | 13 | Saprotroph | \|Undefined Saprotroph\| | Yeast | NULL | Probable | As nectar-tap_saprotroph (Põlme et al. 2020); Decay sugar-rich substrates (Põlme et al. 2020) | Põlme S, et al. 2020. Fungal Diversity 105:1-16 ((https://doi.org/10.1007/s13225-020-00466-2)) |
| Otu61715 | 1 | k__Fungi(100);p__Ascomycota(100);c__Saccharomycetes(100);o__Saccharomycetales(100);f__Metschnikowiaceae(100);g__Metschnikowia(100);s__Metschnikowia_pulcherrima(100); | Metschnikowia | 13 | Saprotroph | \|Undefined Saprotroph\| | Yeast | NULL | Probable | As nectar-tap_saprotroph (Põlme et al. 2020); Decay sugar-rich substrates (Põlme et al. 2020) | Põlme S, et al. 2020. Fungal Diversity 105:1-16 ((https://doi.org/10.1007/s13225-020-00466-2)) |
| Otu61717 | 1 | k__Fungi(100);p__Ascomycota(100);c__Saccharomycetes(100);o__Saccharomycetales(100);f__Metschnikowiaceae(100);g__Metschnikowia(100);g__Metschnikowia_unclassified(100); | Metschnikowia | 13 | Saprotroph | \|Undefined Saprotroph\| | Yeast | NULL | Probable | As nectar-tap_saprotroph (Põlme et al. 2020); Decay sugar-rich substrates (Põlme et al. 2020) | Põlme S, et al. 2020. Fungal Diversity 105:1-16 ((https://doi.org/10.1007/s13225-020-00466-2)) |
| Otu61721 | 1 | k__Fungi(100);p__Ascomycota(100);c__Saccharomycetes(100);o__Saccharomycetales(100);f__Metschnikowiaceae(100);g__Metschnikowia(100);s__Metschnikowia_pulcherrima(100); | Metschnikowia | 13 | Saprotroph | \|Undefined Saprotroph\| | Yeast | NULL | Probable | As nectar-tap_saprotroph (Põlme et al. 2020); Decay sugar-rich substrates (Põlme et al. 2020) | Põlme S, et al. 2020. Fungal Diversity 105:1-16 ((https://doi.org/10.1007/s13225-020-00466-2)) |
| Otu61722 | 1 | k__Fungi(100);p__Ascomycota(100);c__Saccharomycetes(100);o__Saccharomycetales(100);f__Metschnikowiaceae(100);g__Metschnikowia(100);s__Metschnikowia_pulcherrima(100); | Metschnikowia | 13 | Saprotroph | \|Undefined Saprotroph\| | Yeast | NULL | Probable | As nectar-tap_saprotroph (Põlme et al. 2020); Decay sugar-rich substrates (Põlme et al. 2020) | Põlme S, et al. 2020. Fungal Diversity 105:1-16 ((https://doi.org/10.1007/s13225-020-00466-2)) |
| Otu61723 | 1 | k__Fungi(100);p__Ascomycota(100);c__Saccharomycetes(100);o__Saccharomycetales(100);f__Metschnikowiaceae(100);g__Metschnikowia(100);s__Metschnikowia_pulcherrima(100); | Metschnikowia | 13 | Saprotroph | \|Undefined Saprotroph\| | Yeast | NULL | Probable | As nectar-tap_saprotroph (Põlme et al. 2020); Decay sugar-rich substrates (Põlme et al. 2020) | Põlme S, et al. 2020. Fungal Diversity 105:1-16 ((https://doi.org/10.1007/s13225-020-00466-2)) |
| Otu61724 | 1 | k__Fungi(100);p__Ascomycota(100);c__Saccharomycetes(100);o__Saccharomycetales(100);f__Metschnikowiaceae(100);g__Metschnikowia(100);s__Metschnikowia_pulcherrima(100); | Metschnikowia | 13 | Saprotroph | \|Undefined Saprotroph\| | Yeast | NULL | Probable | As nectar-tap_saprotroph (Põlme et al. 2020); Decay sugar-rich substrates (Põlme et al. 2020) | Põlme S, et al. 2020. Fungal Diversity 105:1-16 ((https://doi.org/10.1007/s13225-020-00466-2)) |
| Otu61729 | 1 | k__Fungi(100);p__Ascomycota(100);c__Saccharomycetes(100);o__Saccharomycetales(100);f__Metschnikowiaceae(100);g__Metschnikowia(100);s__Metschnikowia_pulcherrima(100); | Metschnikowia | 13 | Saprotroph | \|Undefined Saprotroph\| | Yeast | NULL | Probable | As nectar-tap_saprotroph (Põlme et al. 2020); Decay sugar-rich substrates (Põlme et al. 2020) | Põlme S, et al. 2020. Fungal Diversity 105:1-16 ((https://doi.org/10.1007/s13225-020-00466-2)) |
| Otu61732 | 1 | k__Fungi(100);p__Ascomycota(100);c__Saccharomycetes(100);o__Saccharomycetales(100);f__Metschnikowiaceae(100);g__Metschnikowia(100);s__Metschnikowia_pulcherrima(100); | Metschnikowia | 13 | Saprotroph | \|Undefined Saprotroph\| | Yeast | NULL | Probable | As nectar-tap_saprotroph (Põlme et al. 2020); Decay sugar-rich substrates (Põlme et al. 2020) | Põlme S, et al. 2020. Fungal Diversity 105:1-16 ((https://doi.org/10.1007/s13225-020-00466-2)) |
| Otu61733 | 1 | k__Fungi(100);p__Ascomycota(100);c__Saccharomycetes(100);o__Saccharomycetales(100);f__Metschnikowiaceae(100);g__Metschnikowia(100);s__Metschnikowia_pulcherrima(100); | Metschnikowia | 13 | Saprotroph | \|Undefined Saprotroph\| | Yeast | NULL | Probable | As nectar-tap_saprotroph (Põlme et al. 2020); Decay sugar-rich substrates (Põlme et al. 2020) | Põlme S, et al. 2020. Fungal Diversity 105:1-16 ((https://doi.org/10.1007/s13225-020-00466-2)) |
| Otu61748 | 1 | k__Fungi(100);p__Ascomycota(100);c__Saccharomycetes(100);o__Saccharomycetales(100);f__Metschnikowiaceae(100);g__Metschnikowia(100);s__Metschnikowia_pulcherrima(100); | Metschnikowia | 13 | Saprotroph | \|Undefined Saprotroph\| | Yeast | NULL | Probable | As nectar-tap_saprotroph (Põlme et al. 2020); Decay sugar-rich substrates (Põlme et al. 2020) | Põlme S, et al. 2020. Fungal Diversity 105:1-16 ((https://doi.org/10.1007/s13225-020-00466-2)) |
| Otu61751 | 1 | k__Fungi(100);p__Ascomycota(100);c__Saccharomycetes(100);o__Saccharomycetales(100);f__Saccharomycetales_family_Incertae_sedis(100);g__Candida(100);g__Candida_unclassified(100); | Candida | 13 | Pathotroph-Saprotroph-Symbiotroph | Animal Pathogen-Endophyte-Endosymbiont-Epiphyte-\|Undefined Saprotroph\| | Dimorphic Yeast-Yeast | NULL | Probable | As nectar-tap_saprotroph (Põlme et al. 2020); Decay sugar-rich substrates (Põlme et al. 2020); This asexual genus is polyphyletic and contain species that function very differently depend on what they are. Caution should be taken when interpreting guilds for Candida species. | Manolakaki D, et al. 2010. Virulence. 1: 367-375 ((https://doi.org/10.4161/viru.1.5.12796)); Mekbib SB, et al. 2011. Tropical Plant Pathology 36:233-240 ((https://doi.org/10.1590/S1982-56762011000400004)); Põlme S, et al. 2020. Fungal Diversity 105:1-16 ((https://doi.org/10.1007/s13225-020-00466-2)) |
| Otu61753 | 1 | k__Fungi(100);p__Ascomycota(100);c__Saccharomycetes(100);o__Saccharomycetales(100);f__Metschnikowiaceae(100);g__Metschnikowia(100);g__Metschnikowia_unclassified(100); | Metschnikowia | 13 | Saprotroph | \|Undefined Saprotroph\| | Yeast | NULL | Probable | As nectar-tap_saprotroph (Põlme et al. 2020); Decay sugar-rich substrates (Põlme et al. 2020) | Põlme S, et al. 2020. Fungal Diversity 105:1-16 ((https://doi.org/10.1007/s13225-020-00466-2)) |
| Otu61755 | 1 | k__Fungi(100);p__Ascomycota(100);c__Saccharomycetes(100);o__Saccharomycetales(100);f__Metschnikowiaceae(100);g__Metschnikowia(100);s__Metschnikowia_pulcherrima(100); | Metschnikowia | 13 | Saprotroph | \|Undefined Saprotroph\| | Yeast | NULL | Probable | As nectar-tap_saprotroph (Põlme et al. 2020); Decay sugar-rich substrates (Põlme et al. 2020) | Põlme S, et al. 2020. Fungal Diversity 105:1-16 ((https://doi.org/10.1007/s13225-020-00466-2)) |
| Otu61758 | 1 | k__Fungi(100);p__Ascomycota(100);c__Saccharomycetes(100);o__Saccharomycetales(100);o__Saccharomycetales_unclassified(100);o__Saccharomycetales_unclassified(100);o__Saccharomycetales_unclassified(100); | Saccharomycetales | 7 | Saprotroph | Undefined Saprotroph | Yeast | NULL | Possible | NULL | Sterkenburg E, et al. 2015. New Phytologist 207:1145-1158 ((https://doi.org/10.1111/nph.13426)) |
| Otu61762 | 1 | k__Fungi(100);p__Ascomycota(100);c__Saccharomycetes(100);o__Saccharomycetales(100);f__Metschnikowiaceae(100);g__Metschnikowia(100);s__Metschnikowia_pulcherrima(100); | Metschnikowia | 13 | Saprotroph | \|Undefined Saprotroph\| | Yeast | NULL | Probable | As nectar-tap_saprotroph (Põlme et al. 2020); Decay sugar-rich substrates (Põlme et al. 2020) | Põlme S, et al. 2020. Fungal Diversity 105:1-16 ((https://doi.org/10.1007/s13225-020-00466-2)) |
| Otu61768 | 1 | k__Fungi(100);p__Ascomycota(100);c__Saccharomycetes(100);o__Saccharomycetales(100);f__Metschnikowiaceae(100);g__Metschnikowia(100);g__Metschnikowia_unclassified(100); | Metschnikowia | 13 | Saprotroph | \|Undefined Saprotroph\| | Yeast | NULL | Probable | As nectar-tap_saprotroph (Põlme et al. 2020); Decay sugar-rich substrates (Põlme et al. 2020) | Põlme S, et al. 2020. Fungal Diversity 105:1-16 ((https://doi.org/10.1007/s13225-020-00466-2)) |
| Otu61769 | 1 | k__Fungi(100);p__Ascomycota(100);c__Saccharomycetes(100);o__Saccharomycetales(100);f__Metschnikowiaceae(100);g__Metschnikowia(100);s__Metschnikowia_pulcherrima(100); | Metschnikowia | 13 | Saprotroph | \|Undefined Saprotroph\| | Yeast | NULL | Probable | As nectar-tap_saprotroph (Põlme et al. 2020); Decay sugar-rich substrates (Põlme et al. 2020) | Põlme S, et al. 2020. Fungal Diversity 105:1-16 ((https://doi.org/10.1007/s13225-020-00466-2)) |
| Otu61770 | 1 | k__Fungi(100);p__Ascomycota(100);c__Saccharomycetes(100);o__Saccharomycetales(100);f__Metschnikowiaceae(100);g__Metschnikowia(100);s__Metschnikowia_pulcherrima(100); | Metschnikowia | 13 | Saprotroph | \|Undefined Saprotroph\| | Yeast | NULL | Probable | As nectar-tap_saprotroph (Põlme et al. 2020); Decay sugar-rich substrates (Põlme et al. 2020) | Põlme S, et al. 2020. Fungal Diversity 105:1-16 ((https://doi.org/10.1007/s13225-020-00466-2)) |
| Otu61771 | 1 | k__Fungi(100);p__Ascomycota(100);c__Saccharomycetes(100);o__Saccharomycetales(100);f__Metschnikowiaceae(100);g__Metschnikowia(100);s__Metschnikowia_pulcherrima(100); | Metschnikowia | 13 | Saprotroph | \|Undefined Saprotroph\| | Yeast | NULL | Probable | As nectar-tap_saprotroph (Põlme et al. 2020); Decay sugar-rich substrates (Põlme et al. 2020) | Põlme S, et al. 2020. Fungal Diversity 105:1-16 ((https://doi.org/10.1007/s13225-020-00466-2)) |
| Otu61772 | 1 | k__Fungi(100);p__Ascomycota(100);c__Saccharomycetes(100);o__Saccharomycetales(100);f__Metschnikowiaceae(100);g__Metschnikowia(100);s__Metschnikowia_pulcherrima(100); | Metschnikowia | 13 | Saprotroph | \|Undefined Saprotroph\| | Yeast | NULL | Probable | As nectar-tap_saprotroph (Põlme et al. 2020); Decay sugar-rich substrates (Põlme et al. 2020) | Põlme S, et al. 2020. Fungal Diversity 105:1-16 ((https://doi.org/10.1007/s13225-020-00466-2)) |
| Otu61776 | 1 | k__Fungi(100);p__Ascomycota(100);c__Saccharomycetes(100);o__Saccharomycetales(100);f__Metschnikowiaceae(100);g__Metschnikowia(100);s__Metschnikowia_pulcherrima(100); | Metschnikowia | 13 | Saprotroph | \|Undefined Saprotroph\| | Yeast | NULL | Probable | As nectar-tap_saprotroph (Põlme et al. 2020); Decay sugar-rich substrates (Põlme et al. 2020) | Põlme S, et al. 2020. Fungal Diversity 105:1-16 ((https://doi.org/10.1007/s13225-020-00466-2)) |
| Otu61777 | 1 | k__Fungi(100);p__Ascomycota(100);c__Saccharomycetes(100);o__Saccharomycetales(100);f__Metschnikowiaceae(100);g__Metschnikowia(100);s__Metschnikowia_pulcherrima(100); | Metschnikowia | 13 | Saprotroph | \|Undefined Saprotroph\| | Yeast | NULL | Probable | As nectar-tap_saprotroph (Põlme et al. 2020); Decay sugar-rich substrates (Põlme et al. 2020) | Põlme S, et al. 2020. Fungal Diversity 105:1-16 ((https://doi.org/10.1007/s13225-020-00466-2)) |
| Otu61778 | 1 | k__Fungi(100);p__Ascomycota(100);c__Saccharomycetes(100);o__Saccharomycetales(100);f__Metschnikowiaceae(100);g__Metschnikowia(100);s__Metschnikowia_pulcherrima(100); | Metschnikowia | 13 | Saprotroph | \|Undefined Saprotroph\| | Yeast | NULL | Probable | As nectar-tap_saprotroph (Põlme et al. 2020); Decay sugar-rich substrates (Põlme et al. 2020) | Põlme S, et al. 2020. Fungal Diversity 105:1-16 ((https://doi.org/10.1007/s13225-020-00466-2)) |
| Otu61779 | 1 | k__Fungi(100);p__Ascomycota(100);c__Saccharomycetes(100);o__Saccharomycetales(100);f__Metschnikowiaceae(100);g__Metschnikowia(100);s__Metschnikowia_pulcherrima(100); | Metschnikowia | 13 | Saprotroph | \|Undefined Saprotroph\| | Yeast | NULL | Probable | As nectar-tap_saprotroph (Põlme et al. 2020); Decay sugar-rich substrates (Põlme et al. 2020) | Põlme S, et al. 2020. Fungal Diversity 105:1-16 ((https://doi.org/10.1007/s13225-020-00466-2)) |
| Otu61780 | 1 | k__Fungi(100);p__Ascomycota(100);c__Saccharomycetes(100);o__Saccharomycetales(100);f__Metschnikowiaceae(100);g__Metschnikowia(100);s__Metschnikowia_pulcherrima(100); | Metschnikowia | 13 | Saprotroph | \|Undefined Saprotroph\| | Yeast | NULL | Probable | As nectar-tap_saprotroph (Põlme et al. 2020); Decay sugar-rich substrates (Põlme et al. 2020) | Põlme S, et al. 2020. Fungal Diversity 105:1-16 ((https://doi.org/10.1007/s13225-020-00466-2)) |
| Otu61781 | 1 | k__Fungi(100);p__Ascomycota(100);c__Saccharomycetes(100);o__Saccharomycetales(100);f__Metschnikowiaceae(100);g__Metschnikowia(100);s__Metschnikowia_pulcherrima(100); | Metschnikowia | 13 | Saprotroph | \|Undefined Saprotroph\| | Yeast | NULL | Probable | As nectar-tap_saprotroph (Põlme et al. 2020); Decay sugar-rich substrates (Põlme et al. 2020) | Põlme S, et al. 2020. Fungal Diversity 105:1-16 ((https://doi.org/10.1007/s13225-020-00466-2)) |
| Otu61782 | 1 | k__Fungi(100);p__Ascomycota(100);c__Saccharomycetes(100);o__Saccharomycetales(100);f__Metschnikowiaceae(100);g__Metschnikowia(100);s__Metschnikowia_pulcherrima(100); | Metschnikowia | 13 | Saprotroph | \|Undefined Saprotroph\| | Yeast | NULL | Probable | As nectar-tap_saprotroph (Põlme et al. 2020); Decay sugar-rich substrates (Põlme et al. 2020) | Põlme S, et al. 2020. Fungal Diversity 105:1-16 ((https://doi.org/10.1007/s13225-020-00466-2)) |
| Otu61783 | 1 | k__Fungi(100);p__Ascomycota(100);c__Saccharomycetes(100);o__Saccharomycetales(100);f__Metschnikowiaceae(100);g__Metschnikowia(100);s__Metschnikowia_pulcherrima(100); | Metschnikowia | 13 | Saprotroph | \|Undefined Saprotroph\| | Yeast | NULL | Probable | As nectar-tap_saprotroph (Põlme et al. 2020); Decay sugar-rich substrates (Põlme et al. 2020) | Põlme S, et al. 2020. Fungal Diversity 105:1-16 ((https://doi.org/10.1007/s13225-020-00466-2)) |
| Otu61784 | 1 | k__Fungi(100);p__Ascomycota(100);c__Saccharomycetes(100);o__Saccharomycetales(100);f__Metschnikowiaceae(100);g__Metschnikowia(100);s__Metschnikowia_pulcherrima(100); | Metschnikowia | 13 | Saprotroph | \|Undefined Saprotroph\| | Yeast | NULL | Probable | As nectar-tap_saprotroph (Põlme et al. 2020); Decay sugar-rich substrates (Põlme et al. 2020) | Põlme S, et al. 2020. Fungal Diversity 105:1-16 ((https://doi.org/10.1007/s13225-020-00466-2)) |

| | | | | | | | | | | | |
|---|---|---|---|---|---|---|---|---|---|---|---|
| Otu61785 | 1 | k__Fungi(100);p__Ascomycota(100);c__Saccharomycetes(100);o__Saccharomycetales(100);f__Metschnikowiaceae(100);g__Metschnikowia(100);s__Metschnikowia_pulcherrima(100); | Metschnikowia | 13 | Saprotroph | \|Undefined Saprotroph\| | Yeast | NULL | Probable | As nectar-tap_saprotroph (Põlme et al. 2020); Decay sugar-rich substrates (Põlme et al. 2020) | Põlme S, et al. 2020. Fungal Diversity 105:1-16 ((https://doi.org/10.1007/s13225-020-00466-2)) |
| Otu61786 | 1 | k__Fungi(100);p__Ascomycota(100);c__Saccharomycetes(100);o__Saccharomycetales(100);f__Metschnikowiaceae(100);g__Metschnikowia(100);s__Metschnikowia_pulcherrima(100); | Metschnikowia | 13 | Saprotroph | \|Undefined Saprotroph\| | Yeast | NULL | Probable | As nectar-tap_saprotroph (Põlme et al. 2020); Decay sugar-rich substrates (Põlme et al. 2020) | Põlme S, et al. 2020. Fungal Diversity 105:1-16 ((https://doi.org/10.1007/s13225-020-00466-2)) |
| Otu61788 | 1 | k__Fungi(100);p__Ascomycota(100);c__Saccharomycetes(100);o__Saccharomycetales(100);f__Metschnikowiaceae(100);g__Metschnikowia(100);s__Metschnikowia_pulcherrima(100); | Metschnikowia | 13 | Saprotroph | \|Undefined Saprotroph\| | Yeast | NULL | Probable | As nectar-tap_saprotroph (Põlme et al. 2020); Decay sugar-rich substrates (Põlme et al. 2020) | Põlme S, et al. 2020. Fungal Diversity 105:1-16 ((https://doi.org/10.1007/s13225-020-00466-2)) |
| Otu61796 | 1 | k__Fungi(100);p__Ascomycota(100);c__Saccharomycetes(100);o__Saccharomycetales(100);f__Saccharomycetales_family_Incertae_sedis(100);g__Candida(100);s__Candida_cretensis(100); | Candida | 13 | Pathotroph-Saprotroph-Symbiotroph | Animal Pathogen-Endophyte-Endosymbiont-Epiphyte-\|Undefined Saprotroph\| | Dimorphic Yeast-Yeast | NULL | Probable | As nectar-tap_saprotroph (Põlme et al. 2020); Decay sugar-rich substrates (Põlme et al. 2020); This asexual genus is polyphyletic and contain species that function very differently depend on what they are. Caution should be taken when interpreting guilds for Candida species. | Manolakaki D, et al. 2010. Virulence. 1: 367-375 ((https://doi.org/10.4161/viru.1.5.12796)); Mekbib SB, et al. 2011. Tropical Plant Pathology 36:233-240 ((https://doi.org/10.1590/S1982-56762011000400004)); Põlme S, et al. 2020. Fungal Diversity 105:1-16 ((https://doi.org/10.1007/s13225-020-00466-2)) |
| Otu61797 | 1 | k__Fungi(100);p__Ascomycota(100);c__Saccharomycetes(100);o__Saccharomycetales(100);f__Saccharomycetales_family_Incertae_sedis(100);g__Candida(100);g__Candida_unclassified(100); | Candida | 13 | Pathotroph-Saprotroph-Symbiotroph | Animal Pathogen-Endophyte-Endosymbiont-Epiphyte-\|Undefined Saprotroph\| | Dimorphic Yeast-Yeast | NULL | Probable | As nectar-tap_saprotroph (Põlme et al. 2020); Decay sugar-rich substrates (Põlme et al. 2020); This asexual genus is polyphyletic and contain species that function very differently depend on what they are. Caution should be taken when interpreting guilds for Candida species. | Manolakaki D, et al. 2010. Virulence. 1: 367-375 ((https://doi.org/10.4161/viru.1.5.12796)); Mekbib SB, et al. 2011. Tropical Plant Pathology 36:233-240 ((https://doi.org/10.1590/S1982-56762011000400004)); Põlme S, et al. 2020. Fungal Diversity 105:1-16 ((https://doi.org/10.1007/s13225-020-00466-2)) |
| Otu61798 | 1 | k__Fungi(100);p__Ascomycota(100);c__Saccharomycetes(100);o__Saccharomycetales(100);f__Metschnikowiaceae(100);g__Metschnikowia(100);s__Metschnikowia_pulcherrima(100); | Metschnikowia | 13 | Saprotroph | \|Undefined Saprotroph\| | Yeast | NULL | Probable | As nectar-tap_saprotroph (Põlme et al. 2020); Decay sugar-rich substrates (Põlme et al. 2020) | Põlme S, et al. 2020. Fungal Diversity 105:1-16 ((https://doi.org/10.1007/s13225-020-00466-2)) |
| Otu61806 | 1 | k__Fungi(100);p__Ascomycota(100);c__Saccharomycetes(100);o__Saccharomycetales(100);f__Metschnikowiaceae(100);g__Metschnikowia(100);s__Metschnikowia_pulcherrima(100); | Metschnikowia | 13 | Saprotroph | \|Undefined Saprotroph\| | Yeast | NULL | Probable | As nectar-tap_saprotroph (Põlme et al. 2020); Decay sugar-rich substrates (Põlme et al. 2020) | Põlme S, et al. 2020. Fungal Diversity 105:1-16 ((https://doi.org/10.1007/s13225-020-00466-2)) |
| Otu61808 | 1 | k__Fungi(100);p__Ascomycota(100);c__Sordariomycetes(100);o__Chaetosphaeriales(100);f__Chaetosphaeriaceae(100);g__Menispora(100);s__Menispora_ciliata(100); | Menispora | 13 | Saprotroph-Symbiotroph | Endophyte-Plant Saprotroph-Undefined Saprotroph-\|Wood Saprotroph\| | Microfungus | NULL | Probable | As foliar_endophyte (Põlme et al. 2020); Decay wood, leaf, fruit, seed (Põlme et al. 2020) | Tedersoo L, et al. 2014. Science 346:e1256688 ((https://doi.org/10.1126/science.1256688)); Christian N, et al. 2016. Microbial Ecology 72:621-632 ((https://doi.org/10.1007/s00248-016-0804-y)) (as endophyte); Põlme S, et al. 2020. Fungal Diversity 105:1-16 ((https://doi.org/10.1007/s13225-020-00466-2)) |
| Otu61819 | 1 | k__Fungi(100);p__Ascomycota(100);c__Saccharomycetes(100);o__Saccharomycetales(100);f__Metschnikowiaceae(100);g__Metschnikowia(100);s__Metschnikowia_pulcherrima(100); | Metschnikowia | 13 | Saprotroph | \|Undefined Saprotroph\| | Yeast | NULL | Probable | As nectar-tap_saprotroph (Põlme et al. 2020); Decay sugar-rich substrates (Põlme et al. 2020) | Põlme S, et al. 2020. Fungal Diversity 105:1-16 ((https://doi.org/10.1007/s13225-020-00466-2)) |
| Otu61820 | 1 | k__Fungi(100);p__Ascomycota(100);c__Saccharomycetes(100);o__Saccharomycetales(100);f__Metschnikowiaceae(100);g__Metschnikowia(100);s__Metschnikowia_pulcherrima(100); | Metschnikowia | 13 | Saprotroph | \|Undefined Saprotroph\| | Yeast | NULL | Probable | As nectar-tap_saprotroph (Põlme et al. 2020); Decay sugar-rich substrates (Põlme et al. 2020) | Põlme S, et al. 2020. Fungal Diversity 105:1-16 ((https://doi.org/10.1007/s13225-020-00466-2)) |
| Otu61821 | 1 | k__Fungi(100);p__Ascomycota(100);c__Saccharomycetes(100);o__Saccharomycetales(100);f__Metschnikowiaceae(100);g__Metschnikowia(100);s__Metschnikowia_pulcherrima(100); | Metschnikowia | 13 | Saprotroph | \|Undefined Saprotroph\| | Yeast | NULL | Probable | As nectar-tap_saprotroph (Põlme et al. 2020); Decay sugar-rich substrates (Põlme et al. 2020) | Põlme S, et al. 2020. Fungal Diversity 105:1-16 ((https://doi.org/10.1007/s13225-020-00466-2)) |
| Otu61822 | 1 | k__Fungi(100);p__Ascomycota(100);c__Saccharomycetes(100);o__Saccharomycetales(100);f__Metschnikowiaceae(100);g__Metschnikowia(100);s__Metschnikowia_pulcherrima(100); | Metschnikowia | 13 | Saprotroph | \|Undefined Saprotroph\| | Yeast | NULL | Probable | As nectar-tap_saprotroph (Põlme et al. 2020); Decay sugar-rich substrates (Põlme et al. 2020) | Põlme S, et al. 2020. Fungal Diversity 105:1-16 ((https://doi.org/10.1007/s13225-020-00466-2)) |
| Otu61823 | 1 | k__Fungi(100);p__Ascomycota(100);c__Sordariomycetes(100);o__Chaetosphaeriales(100);f__Chaetosphaeriaceae(100);f__Chaetosphaeriaceae_unclassified(100);f__Chaetosphaeriaceae_unclassified(100); | Chaetosphaeriaceae | 9 | Saprotroph | Plant Saprotroph-Wood Saprotroph | Microfungus | NULL | Probable | Saprobes associated with decaying woody and herbaceous plant material, including timber and timber products in damp buildings (Cannon & Kirk 2007) | Cannon PF, Kirk PM. 2007. Fungal Families of the World. CAB International, Cambridge (ISBN: 978-0851998275) |
| Otu61832 | 1 | k__Fungi(100);p__Ascomycota(100);c__Saccharomycetes(100);o__Saccharomycetales(100);f__Metschnikowiaceae(100);g__Metschnikowia(100);s__Metschnikowia_pulcherrima(100); | Metschnikowia | 13 | Saprotroph | \|Undefined Saprotroph\| | Yeast | NULL | Probable | As nectar-tap_saprotroph (Põlme et al. 2020); Decay sugar-rich substrates (Põlme et al. 2020) | Põlme S, et al. 2020. Fungal Diversity 105:1-16 ((https://doi.org/10.1007/s13225-020-00466-2)) |
| Otu61835 | 1 | k__Fungi(100);p__Basidiomycota(100);c__Microbotryomycetes(100);o__Sporidiobolales(100);f__Sporidiobolales_family_Incertae_sedis(100);g__Sporobolomyces(100);s__Sporobolomyces_lactosus(100); | Sporobolomyces | 13 | Pathotroph-Saprotroph | \|Fungal Parasite\|-Plant Saprotroph | Yeast | NULL | Probable | As litter_saprotroph (Põlme et al. 2020); As mycoparasite (Põlme et al. 2020); Decay fungal material (Põlme et al. 2020) | Tedersoo L, et al. 2014. Science 346:e1256688 ((https://doi.org/10.1126/science.1256688)); Purahong W, et al. 2016. Molecular Ecology 25:4059-4074 ((https://doi.org/10.1111/mec.13739)); Põlme S, et al. 2020. Fungal Diversity 105:1-16 ((https://doi.org/10.1007/s13225-020-00466-2)) |
| Otu61837 | 1 | k__Fungi(100);p__Ascomycota(100);c__Saccharomycetes(100);o__Saccharomycetales(100);f__Metschnikowiaceae(100);g__Metschnikowia(100);s__Metschnikowia_pulcherrima(100); | Metschnikowia | 13 | Saprotroph | \|Undefined Saprotroph\| | Yeast | NULL | Probable | As nectar-tap_saprotroph (Põlme et al. 2020); Decay sugar-rich substrates (Põlme et al. 2020) | Põlme S, et al. 2020. Fungal Diversity 105:1-16 ((https://doi.org/10.1007/s13225-020-00466-2)) |
| Otu61838 | 1 | k__Fungi(100);p__Ascomycota(100);c__Saccharomycetes(100);o__Saccharomycetales(100);f__Metschnikowiaceae(100);g__Metschnikowia(100);s__Metschnikowia_pulcherrima(100); | Metschnikowia | 13 | Saprotroph | \|Undefined Saprotroph\| | Yeast | NULL | Probable | As nectar-tap_saprotroph (Põlme et al. 2020); Decay sugar-rich substrates (Põlme et al. 2020) | Põlme S, et al. 2020. Fungal Diversity 105:1-16 ((https://doi.org/10.1007/s13225-020-00466-2)) |
| Otu61840 | 1 | k__Fungi(100);p__Ascomycota(100);c__Saccharomycetes(100);o__Saccharomycetales(100);f__Metschnikowiaceae(100);g__Metschnikowia(100);s__Metschnikowia_pulcherrima(100); | Metschnikowia | 13 | Saprotroph | \|Undefined Saprotroph\| | Yeast | NULL | Probable | As nectar-tap_saprotroph (Põlme et al. 2020); Decay sugar-rich substrates (Põlme et al. 2020) | Põlme S, et al. 2020. Fungal Diversity 105:1-16 ((https://doi.org/10.1007/s13225-020-00466-2)) |
| Otu61843 | 1 | k__Fungi(100);p__Ascomycota(100);c__Saccharomycetes(100);o__Saccharomycetales(100);f__Metschnikowiaceae(100);g__Metschnikowia(100);s__Metschnikowia_pulcherrima(100); | Metschnikowia | 13 | Saprotroph | \|Undefined Saprotroph\| | Yeast | NULL | Probable | As nectar-tap_saprotroph (Põlme et al. 2020); Decay sugar-rich substrates (Põlme et al. 2020) | Põlme S, et al. 2020. Fungal Diversity 105:1-16 ((https://doi.org/10.1007/s13225-020-00466-2)) |
| Otu61846 | 1 | k__Fungi(100);p__Ascomycota(100);c__Saccharomycetes(100);o__Saccharomycetales(100);f__Metschnikowiaceae(100);g__Metschnikowia(100);g__Metschnikowia_unclassified(100); | Metschnikowia | 13 | Saprotroph | \|Undefined Saprotroph\| | Yeast | NULL | Probable | As nectar-tap_saprotroph (Põlme et al. 2020); Decay sugar-rich substrates (Põlme et al. 2020) | Põlme S, et al. 2020. Fungal Diversity 105:1-16 ((https://doi.org/10.1007/s13225-020-00466-2)) |
| Otu61847 | 1 | k__Fungi(100);p__Ascomycota(100);c__Saccharomycetes(100);o__Saccharomycetales(100);f__Metschnikowiaceae(100);g__Metschnikowia(100);g__Metschnikowia_unclassified(100); | Metschnikowia | 13 | Saprotroph | \|Undefined Saprotroph\| | Yeast | NULL | Probable | As nectar-tap_saprotroph (Põlme et al. 2020); Decay sugar-rich substrates (Põlme et al. 2020) | Põlme S, et al. 2020. Fungal Diversity 105:1-16 ((https://doi.org/10.1007/s13225-020-00466-2)) |
| Otu61853 | 1 | k__Fungi(100);p__Ascomycota(100);c__Saccharomycetes(100);o__Saccharomycetales(100);f__Saccharomycetales_family_Incertae_sedis(100);g__Candida(100);g__Candida_unclassified(100); | Candida | 13 | Pathotroph-Saprotroph-Symbiotroph | Animal Pathogen-Endophyte-Endosymbiont-Epiphyte-\|Undefined Saprotroph\| | Dimorphic Yeast-Yeast | NULL | Probable | As nectar-tap_saprotroph (Põlme et al. 2020); Decay sugar-rich substrates (Põlme et al. 2020); This asexual genus is polyphyletic and contain species that function very differently depend on what they are. Caution should be taken when interpreting guilds for Candida species. | Manolakaki D, et al. 2010. Virulence. 1: 367-375 ((https://doi.org/10.4161/viru.1.5.12796)); Mekbib SB, et al. 2011. Tropical Plant Pathology 36:233-240 ((https://doi.org/10.1590/S1982-56762011000400004)); Põlme S, et al. 2020. Fungal Diversity 105:1-16 ((https://doi.org/10.1007/s13225-020-00466-2)) |
| Otu61855 | 1 | k__Fungi(100);p__Ascomycota(100);c__Saccharomycetes(100);o__Saccharomycetales(100);f__Saccharomycetales_family_Incertae_sedis(100);g__Candida(100);g__Candida_unclassified(100); | Candida | 13 | Pathotroph-Saprotroph-Symbiotroph | Animal Pathogen-Endophyte-Endosymbiont-Epiphyte-\|Undefined Saprotroph\| | Dimorphic Yeast-Yeast | NULL | Probable | As nectar-tap_saprotroph (Põlme et al. 2020); Decay sugar-rich substrates (Põlme et al. 2020); This asexual genus is polyphyletic and contain species that function very differently depend on what they are. Caution should be taken when interpreting guilds for Candida species. | Manolakaki D, et al. 2010. Virulence. 1: 367-375 ((https://doi.org/10.4161/viru.1.5.12796)); Mekbib SB, et al. 2011. Tropical Plant Pathology 36:233-240 ((https://doi.org/10.1590/S1982-56762011000400004)); Põlme S, et al. 2020. Fungal Diversity 105:1-16 ((https://doi.org/10.1007/s13225-020-00466-2)) |
| Otu61857 | 1 | k__Fungi(100);p__Ascomycota(100);c__Saccharomycetes(100);o__Saccharomycetales(100);f__Saccharomycetales_family_Incertae_sedis(100);g__Candida(100);g__Candida_unclassified(100); | Candida | 13 | Pathotroph-Saprotroph-Symbiotroph | Animal Pathogen-Endophyte-Endosymbiont-Epiphyte-\|Undefined Saprotroph\| | Dimorphic Yeast-Yeast | NULL | Probable | As nectar-tap_saprotroph (Põlme et al. 2020); Decay sugar-rich substrates (Põlme et al. 2020); This asexual genus is polyphyletic and contain species that function very differently depend on what they are. Caution should be taken when interpreting guilds for Candida species. | Manolakaki D, et al. 2010. Virulence. 1: 367-375 ((https://doi.org/10.4161/viru.1.5.12796)); Mekbib SB, et al. 2011. Tropical Plant Pathology 36:233-240 ((https://doi.org/10.1590/S1982-56762011000400004)); Põlme S, et al. 2020. Fungal Diversity 105:1-16 ((https://doi.org/10.1007/s13225-020-00466-2)) |
| Otu61858 | 1 | k__Fungi(100);p__Ascomycota(100);c__Saccharomycetes(100);o__Saccharomycetales(100);f__Metschnikowiaceae(100);g__Metschnikowia(100);s__Metschnikowia_pulcherrima(100); | Metschnikowia | 13 | Saprotroph | \|Undefined Saprotroph\| | Yeast | NULL | Probable | As nectar-tap_saprotroph (Põlme et al. 2020); Decay sugar-rich substrates (Põlme et al. 2020) | Põlme S, et al. 2020. Fungal Diversity 105:1-16 ((https://doi.org/10.1007/s13225-020-00466-2)) |
| Otu61863 | 1 | k__Fungi(100);p__Ascomycota(100);c__Saccharomycetes(100);o__Saccharomycetales(100);f__Metschnikowiaceae(100);g__Metschnikowia(100);s__Metschnikowia_pulcherrima(100); | Metschnikowia | 13 | Saprotroph | \|Undefined Saprotroph\| | Yeast | NULL | Probable | As nectar-tap_saprotroph (Põlme et al. 2020); Decay sugar-rich substrates (Põlme et al. 2020) | Põlme S, et al. 2020. Fungal Diversity 105:1-16 ((https://doi.org/10.1007/s13225-020-00466-2)) |
| Otu61864 | 1 | k__Fungi(100);p__Ascomycota(100);c__Dothideomycetes(100);o__Capnodiales(100);f__Mycosphaerellaceae(100);g__Pseudocercospora(100);g__Pseudocercospora_unclassified(100); | Pseudocercospora | 13 | Pathotroph | \|Plant Pathogen\| | Microfungus | NULL | Probable | NULL | Tedersoo L, et al. 2014. Science 346:e1256688 ((https://doi.org/10.1126/science.1256688)); Põlme S, et al. 2020. Fungal Diversity 105:1-16 ((https://doi.org/10.1007/s13225-020-00466-2)) |
| Otu61866 | 1 | k__Fungi(100);p__Ascomycota(100);c__Saccharomycetes(100);o__Saccharomycetales(100);f__Saccharomycetales_family_Incertae_sedis(100);g__Candida(100);g__Candida_unclassified(100); | Candida | 13 | Pathotroph-Saprotroph-Symbiotroph | Animal Pathogen-Endophyte-Endosymbiont-Epiphyte-\|Undefined Saprotroph\| | Dimorphic Yeast-Yeast | NULL | Probable | As nectar-tap_saprotroph (Põlme et al. 2020); Decay sugar-rich substrates (Põlme et al. 2020); This asexual genus is polyphyletic and contain species that function very differently depend on what they are. Caution should be taken when interpreting guilds for Candida species. | Manolakaki D, et al. 2010. Virulence. 1: 367-375 ((https://doi.org/10.4161/viru.1.5.12796)); Mekbib SB, et al. 2011. Tropical Plant Pathology 36:233-240 ((https://doi.org/10.1590/S1982-56762011000400004)); Põlme S, et al. 2020. Fungal Diversity 105:1-16 ((https://doi.org/10.1007/s13225-020-00466-2)) |
| Otu61870 | 1 | k__Fungi(100);p__Ascomycota(100);c__Saccharomycetes(100);o__Saccharomycetales(100);o__Saccharomycetales_unclassified(100);o__Saccharomycetales_unclassified(100);o__Saccharomycetales_unclassified(100); | Saccharomycetales | 7 | Saprotroph | Undefined Saprotroph | Yeast | NULL | Possible | NULL | Sterkenburg E, et al. 2015. New Phytologist 207:1145-1158 ((https://doi.org/10.1111/nph.13426)) |
| Otu61871 | 1 | k__Fungi(100);p__Ascomycota(100);c__Saccharomycetes(100);o__Saccharomycetales(100);f__Saccharomycetales_family_Incertae_sedis(100);g__Candida(100);g__Candida_unclassified(100); | Candida | 13 | Pathotroph-Saprotroph-Symbiotroph | Animal Pathogen-Endophyte-Endosymbiont-Epiphyte-\|Undefined Saprotroph\| | Dimorphic Yeast-Yeast | NULL | Probable | As nectar-tap_saprotroph (Põlme et al. 2020); Decay sugar-rich substrates (Põlme et al. 2020); This asexual genus is polyphyletic and contain species that function very differently depend on what they are. Caution should be taken when interpreting guilds for Candida species. | Manolakaki D, et al. 2010. Virulence. 1: 367-375 ((https://doi.org/10.4161/viru.1.5.12796)); Mekbib SB, et al. 2011. Tropical Plant Pathology 36:233-240 ((https://doi.org/10.1590/S1982-56762011000400004)); Põlme S, et al. 2020. Fungal Diversity 105:1-16 ((https://doi.org/10.1007/s13225-020-00466-2)) |
| Otu61872 | 1 | k__Fungi(100);p__Ascomycota(100);c__Saccharomycetes(100);o__Saccharomycetales(100);f__Metschnikowiaceae(100);g__Metschnikowia(100);s__Metschnikowia_pulcherrima(100); | Metschnikowia | 13 | Saprotroph | \|Undefined Saprotroph\| | Yeast | NULL | Probable | As nectar-tap_saprotroph (Põlme et al. 2020); Decay sugar-rich substrates (Põlme et al. 2020) | Põlme S, et al. 2020. Fungal Diversity 105:1-16 ((https://doi.org/10.1007/s13225-020-00466-2)) |
| Otu61874 | 1 | k__Fungi(100);p__Ascomycota(100);c__Saccharomycetes(100);o__Saccharomycetales(100);f__Metschnikowiaceae(100);g__Metschnikowia(100);s__Metschnikowia_pulcherrima(100); | Metschnikowia | 13 | Saprotroph | \|Undefined Saprotroph\| | Yeast | NULL | Probable | As nectar-tap_saprotroph (Põlme et al. 2020); Decay sugar-rich substrates (Põlme et al. 2020) | Põlme S, et al. 2020. Fungal Diversity 105:1-16 ((https://doi.org/10.1007/s13225-020-00466-2)) |
| Otu61879 | 1 | k__Fungi(100);p__Ascomycota(100);c__Saccharomycetes(100);o__Saccharomycetales(100);f__Metschnikowiaceae(100);g__Metschnikowia(100);s__Metschnikowia_pulcherrima(100); | Metschnikowia | 13 | Saprotroph | \|Undefined Saprotroph\| | Yeast | NULL | Probable | As nectar-tap_saprotroph (Põlme et al. 2020); Decay sugar-rich substrates (Põlme et al. 2020) | Põlme S, et al. 2020. Fungal Diversity 105:1-16 ((https://doi.org/10.1007/s13225-020-00466-2)) |
| Otu61881 | 1 | k__Fungi(100);p__Ascomycota(100);c__Saccharomycetes(100);o__Saccharomycetales(100);f__Metschnikowiaceae(100);g__Metschnikowia(100);g__Metschnikowia_unclassified(100); | Metschnikowia | 13 | Saprotroph | \|Undefined Saprotroph\| | Yeast | NULL | Probable | As nectar-tap_saprotroph (Põlme et al. 2020); Decay sugar-rich substrates (Põlme et al. 2020) | Põlme S, et al. 2020. Fungal Diversity 105:1-16 ((https://doi.org/10.1007/s13225-020-00466-2)) |
| Otu61883 | 1 | k__Fungi(100);p__Ascomycota(100);c__Saccharomycetes(100);o__Saccharomycetales(100);f__Metschnikowiaceae(100);g__Metschnikowia(100);g__Metschnikowia_unclassified(100); | Metschnikowia | 13 | Saprotroph | \|Undefined Saprotroph\| | Yeast | NULL | Probable | As nectar-tap_saprotroph (Põlme et al. 2020); Decay sugar-rich substrates (Põlme et al. 2020) | Põlme S, et al. 2020. Fungal Diversity 105:1-16 ((https://doi.org/10.1007/s13225-020-00466-2)) |
| Otu61884 | 1 | k__Fungi(100);p__Ascomycota(100);c__Saccharomycetes(100);o__Saccharomycetales(100);f__Metschnikowiaceae(100);g__Metschnikowia(100);g__Metschnikowia_unclassified(100); | Metschnikowia | 13 | Saprotroph | \|Undefined Saprotroph\| | Yeast | NULL | Probable | As nectar-tap_saprotroph (Põlme et al. 2020); Decay sugar-rich substrates (Põlme et al. 2020) | Põlme S, et al. 2020. Fungal Diversity 105:1-16 ((https://doi.org/10.1007/s13225-020-00466-2)) |
| Otu61885 | 1 | k__Fungi(100);p__Ascomycota(100);c__Saccharomycetes(100);o__Saccharomycetales(100);f__Metschnikowiaceae(100);g__Metschnikowia(100);g__Metschnikowia_unclassified(100); | Metschnikowia | 13 | Saprotroph | \|Undefined Saprotroph\| | Yeast | NULL | Probable | As nectar-tap_saprotroph (Põlme et al. 2020); Decay sugar-rich substrates (Põlme et al. 2020) | Põlme S, et al. 2020. Fungal Diversity 105:1-16 ((https://doi.org/10.1007/s13225-020-00466-2)) |
| Otu61886 | 1 | k__Fungi(100);p__Ascomycota(100);c__Saccharomycetes(100);o__Saccharomycetales(100);f__Metschnikowiaceae(100);g__Metschnikowia(100);s__Metschnikowia_pulcherrima(100); | Metschnikowia | 13 | Saprotroph | \|Undefined Saprotroph\| | Yeast | NULL | Probable | As nectar-tap_saprotroph (Põlme et al. 2020); Decay sugar-rich substrates (Põlme et al. 2020) | Põlme S, et al. 2020. Fungal Diversity 105:1-16 ((https://doi.org/10.1007/s13225-020-00466-2)) |
| Otu61887 | 1 | k__Fungi(100);p__Ascomycota(100);c__Saccharomycetes(100);o__Saccharomycetales(100);f__Metschnikowiaceae(100);g__Metschnikowia(100);s__Metschnikowia_pulcherrima(100); | Metschnikowia | 13 | Saprotroph | \|Undefined Saprotroph\| | Yeast | NULL | Probable | As nectar-tap_saprotroph (Põlme et al. 2020); Decay sugar-rich substrates (Põlme et al. 2020) | Põlme S, et al. 2020. Fungal Diversity 105:1-16 ((https://doi.org/10.1007/s13225-020-00466-2)) |
| Otu61888 | 1 | k__Fungi(100);p__Ascomycota(100);c__Saccharomycetes(100);o__Saccharomycetales(100);f__Metschnikowiaceae(100);g__Metschnikowia(100);s__Metschnikowia_pulcherrima(100); | Metschnikowia | 13 | Saprotroph | \|Undefined Saprotroph\| | Yeast | NULL | Probable | As nectar-tap_saprotroph (Põlme et al. 2020); Decay sugar-rich substrates (Põlme et al. 2020) | Põlme S, et al. 2020. Fungal Diversity 105:1-16 ((https://doi.org/10.1007/s13225-020-00466-2)) |
| Otu61889 | 1 | k__Fungi(100);p__Ascomycota(100);c__Saccharomycetes(100);o__Saccharomycetales(100);f__Metschnikowiaceae(100);g__Metschnikowia(100);s__Metschnikowia_pulcherrima(100); | Metschnikowia | 13 | Saprotroph | \|Undefined Saprotroph\| | Yeast | NULL | Probable | As nectar-tap_saprotroph (Põlme et al. 2020); Decay sugar-rich substrates (Põlme et al. 2020) | Põlme S, et al. 2020. Fungal Diversity 105:1-16 ((https://doi.org/10.1007/s13225-020-00466-2)) |
| Otu61890 | 1 | k__Fungi(100);p__Ascomycota(100);c__Saccharomycetes(100);o__Saccharomycetales(100);f__Metschnikowiaceae(100);g__Metschnikowia(100);s__Metschnikowia_pulcherrima(100); | Metschnikowia | 13 | Saprotroph | \|Undefined Saprotroph\| | Yeast | NULL | Probable | As nectar-tap_saprotroph (Põlme et al. 2020); Decay sugar-rich substrates (Põlme et al. 2020) | Põlme S, et al. 2020. Fungal Diversity 105:1-16 ((https://doi.org/10.1007/s13225-020-00466-2)) |
| Otu61891 | 1 | k__Fungi(100);p__Ascomycota(100);c__Saccharomycetes(100);o__Saccharomycetales(100);f__Metschnikowiaceae(100);g__Metschnikowia(100);s__Metschnikowia_pulcherrima(100); | Metschnikowia | 13 | Saprotroph | \|Undefined Saprotroph\| | Yeast | NULL | Probable | As nectar-tap_saprotroph (Põlme et al. 2020); Decay sugar-rich substrates (Põlme et al. 2020) | Põlme S, et al. 2020. Fungal Diversity 105:1-16 ((https://doi.org/10.1007/s13225-020-00466-2)) |
| Otu61892 | 1 | k__Fungi(100);p__Ascomycota(100);c__Saccharomycetes(100);o__Saccharomycetales(100);f__Metschnikowiaceae(100);g__Metschnikowia(100);s__Metschnikowia_pulcherrima(100); | Metschnikowia | 13 | Saprotroph | \|Undefined Saprotroph\| | Yeast | NULL | Probable | As nectar-tap_saprotroph (Põlme et al. 2020); Decay sugar-rich substrates (Põlme et al. 2020) | Põlme S, et al. 2020. Fungal Diversity 105:1-16 ((https://doi.org/10.1007/s13225-020-00466-2)) |
| Otu61893 | 1 | k__Fungi(100);p__Ascomycota(100);c__Saccharomycetes(100);o__Saccharomycetales(100);f__Metschnikowiaceae(100);g__Metschnikowia(100);s__Metschnikowia_pulcherrima(100); | Metschnikowia | 13 | Saprotroph | \|Undefined Saprotroph\| | Yeast | NULL | Probable | As nectar-tap_saprotroph (Põlme et al. 2020); Decay sugar-rich substrates (Põlme et al. 2020) | Põlme S, et al. 2020. Fungal Diversity 105:1-16 ((https://doi.org/10.1007/s13225-020-00466-2)) |
| Otu61894 | 1 | k__Fungi(100);p__Ascomycota(100);c__Saccharomycetes(100);o__Saccharomycetales(100);f__Metschnikowiaceae(100);g__Metschnikowia(100);s__Metschnikowia_pulcherrima(100); | Metschnikowia | 13 | Saprotroph | \|Undefined Saprotroph\| | Yeast | NULL | Probable | As nectar-tap_saprotroph (Põlme et al. 2020); Decay sugar-rich substrates (Põlme et al. 2020) | Põlme S, et al. 2020. Fungal Diversity 105:1-16 ((https://doi.org/10.1007/s13225-020-00466-2)) |
| Otu61897 | 1 | k__Fungi(100);p__Ascomycota(100);c__Saccharomycetes(100);o__Saccharomycetales(100);f__Metschnikowiaceae(100);g__Metschnikowia(100);s__Metschnikowia_pulcherrima(100); | Metschnikowia | 13 | Saprotroph | \|Undefined Saprotroph\| | Yeast | NULL | Probable | As nectar-tap_saprotroph (Põlme et al. 2020); Decay sugar-rich substrates (Põlme et al. 2020) | Põlme S, et al. 2020. Fungal Diversity 105:1-16 ((https://doi.org/10.1007/s13225-020-00466-2)) |
| Otu61899 | 1 | k__Fungi(100);p__Ascomycota(100);c__Saccharomycetes(100);o__Saccharomycetales(100);f__Metschnikowiaceae(100);g__Metschnikowia(100);s__Metschnikowia_pulcherrima(100); | Metschnikowia | 13 | Saprotroph | \|Undefined Saprotroph\| | Yeast | NULL | Probable | As nectar-tap_saprotroph (Põlme et al. 2020); Decay sugar-rich substrates (Põlme et al. 2020) | Põlme S, et al. 2020. Fungal Diversity 105:1-16 ((https://doi.org/10.1007/s13225-020-00466-2)) |
| Otu61900 | 1 | k__Fungi(100);p__Ascomycota(100);c__Saccharomycetes(100);o__Saccharomycetales(100);f__Metschnikowiaceae(100);g__Metschnikowia(100);s__Metschnikowia_pulcherrima(100); | Metschnikowia | 13 | Saprotroph | \|Undefined Saprotroph\| | Yeast | NULL | Probable | As nectar-tap_saprotroph (Põlme et al. 2020); Decay sugar-rich substrates (Põlme et al. 2020) | Põlme S, et al. 2020. Fungal Diversity 105:1-16 ((https://doi.org/10.1007/s13225-020-00466-2)) |
| Otu61901 | 1 | k__Fungi(100);p__Ascomycota(100);c__Saccharomycetes(100);o__Saccharomycetales(100);f__Metschnikowiaceae(100);g__Metschnikowia(100);s__Metschnikowia_pulcherrima(100); | Metschnikowia | 13 | Saprotroph | \|Undefined Saprotroph\| | Yeast | NULL | Probable | As nectar-tap_saprotroph (Põlme et al. 2020); Decay sugar-rich substrates (Põlme et al. 2020) | Põlme S, et al. 2020. Fungal Diversity 105:1-16 ((https://doi.org/10.1007/s13225-020-00466-2)) |
| Otu61904 | 1 | k__Fungi(100);p__Ascomycota(100);c__Saccharomycetes(100);o__Saccharomycetales(100);f__Metschnikowiaceae(100);g__Metschnikowia(100);s__Metschnikowia_pulcherrima(100); | Metschnikowia | 13 | Saprotroph | \|Undefined Saprotroph\| | Yeast | NULL | Probable | As nectar-tap_saprotroph (Põlme et al. 2020); Decay sugar-rich substrates (Põlme et al. 2020) | Põlme S, et al. 2020. Fungal Diversity 105:1-16 ((https://doi.org/10.1007/s13225-020-00466-2)) |
| Otu61907 | 1 | k__Fungi(100);p__Ascomycota(100);c__Saccharomycetes(100);o__Saccharomycetales(100);f__Metschnikowiaceae(100);g__Metschnikowia(100);s__Metschnikowia_pulcherrima(100); | Metschnikowia | 13 | Saprotroph | \|Undefined Saprotroph\| | Yeast | NULL | Probable | As nectar-tap_saprotroph (Põlme et al. 2020); Decay sugar-rich substrates (Põlme et al. 2020) | Põlme S, et al. 2020. Fungal Diversity 105:1-16 ((https://doi.org/10.1007/s13225-020-00466-2)) |
| Otu61908 | 1 | k__Fungi(100);p__Ascomycota(100);c__Saccharomycetes(100);o__Saccharomycetales(100);f__Metschnikowiaceae(100);g__Metschnikowia(100);s__Metschnikowia_pulcherrima(100); | Metschnikowia | 13 | Saprotroph | \|Undefined Saprotroph\| | Yeast | NULL | Probable | As nectar-tap_saprotroph (Põlme et al. 2020); Decay sugar-rich substrates (Põlme et al. 2020) | Põlme S, et al. 2020. Fungal Diversity 105:1-16 ((https://doi.org/10.1007/s13225-020-00466-2)) |
| Otu61909 | 1 | k__Fungi(100);p__Ascomycota(100);c__Saccharomycetes(100);o__Saccharomycetales(100);f__Metschnikowiaceae(100);g__Metschnikowia(100);s__Metschnikowia_pulcherrima(100); | Metschnikowia | 13 | Saprotroph | \|Undefined Saprotroph\| | Yeast | NULL | Probable | As nectar-tap_saprotroph (Põlme et al. 2020); Decay sugar-rich substrates (Põlme et al. 2020) | Põlme S, et al. 2020. Fungal Diversity 105:1-16 ((https://doi.org/10.1007/s13225-020-00466-2)) |
| Otu61910 | 1 | k__Fungi(100);p__Ascomycota(100);c__Saccharomycetes(100);o__Saccharomycetales(100);f__Metschnikowiaceae(100);g__Metschnikowia(100);g__Metschnikowia_unclassified(100); | Metschnikowia | 13 | Saprotroph | \|Undefined Saprotroph\| | Yeast | NULL | Probable | As nectar-tap_saprotroph (Põlme et al. 2020); Decay sugar-rich substrates (Põlme et al. 2020) | Põlme S, et al. 2020. Fungal Diversity 105:1-16 ((https://doi.org/10.1007/s13225-020-00466-2)) |
| Otu61911 | 1 | k__Fungi(100);p__Ascomycota(100);c__Saccharomycetes(100);o__Saccharomycetales(100);f__Metschnikowiaceae(100);g__Metschnikowia(100);g__Metschnikowia_unclassified(100); | Metschnikowia | 13 | Saprotroph | \|Undefined Saprotroph\| | Yeast | NULL | Probable | As nectar-tap_saprotroph (Põlme et al. 2020); Decay sugar-rich substrates (Põlme et al. 2020) | Põlme S, et al. 2020. Fungal Diversity 105:1-16 ((https://doi.org/10.1007/s13225-020-00466-2)) |
| Otu61912 | 1 | k__Fungi(100);p__Ascomycota(100);c__Saccharomycetes(100);o__Saccharomycetales(100);f__Metschnikowiaceae(100);g__Metschnikowia(100);s__Metschnikowia_pulcherrima(100); | Metschnikowia | 13 | Saprotroph | \|Undefined Saprotroph\| | Yeast | NULL | Probable | As nectar-tap_saprotroph (Põlme et al. 2020); Decay sugar-rich substrates (Põlme et al. 2020) | Põlme S, et al. 2020. Fungal Diversity 105:1-16 ((https://doi.org/10.1007/s13225-020-00466-2)) |
| Otu61916 | 1 | k__Fungi(100);p__Ascomycota(100);c__Saccharomycetes(100);o__Saccharomycetales(100);f__Metschnikowiaceae(100);g__Metschnikowia(100);g__Metschnikowia_unclassified(100); | Metschnikowia | 13 | Saprotroph | \|Undefined Saprotroph\| | Yeast | NULL | Probable | As nectar-tap_saprotroph (Põlme et al. 2020); Decay sugar-rich substrates (Põlme et al. 2020) | Põlme S, et al. 2020. Fungal Diversity 105:1-16 ((https://doi.org/10.1007/s13225-020-00466-2)) |
| Otu61917 | 1 | k__Fungi(100);p__Ascomycota(100);c__Saccharomycetes(100);o__Saccharomycetales(100);f__Metschnikowiaceae(100);g__Metschnikowia(100);s__Metschnikowia_pulcherrima(100); | Metschnikowia | 13 | Saprotroph | \|Undefined Saprotroph\| | Yeast | NULL | Probable | As nectar-tap_saprotroph (Põlme et al. 2020); Decay sugar-rich substrates (Põlme et al. 2020) | Põlme S, et al. 2020. Fungal Diversity 105:1-16 ((https://doi.org/10.1007/s13225-020-00466-2)) |
| Otu61923 | 1 | k__Fungi(100);p__Ascomycota(100);c__Saccharomycetes(100);o__Saccharomycetales(100);f__Metschnikowiaceae(100);g__Metschnikowia(100);s__Metschnikowia_pulcherrima(100); | Metschnikowia | 13 | Saprotroph | \|Undefined Saprotroph\| | Yeast | NULL | Probable | As nectar-tap_saprotroph (Põlme et al. 2020); Decay sugar-rich substrates (Põlme et al. 2020) | Põlme S, et al. 2020. Fungal Diversity 105:1-16 ((https://doi.org/10.1007/s13225-020-00466-2)) |
| Otu61925 | 1 | k__Fungi(100);p__Ascomycota(100);c__Saccharomycetes(100);o__Saccharomycetales(100);f__Metschnikowiaceae(100);g__Metschnikowia(100);s__Metschnikowia_pulcherrima(100); | Metschnikowia | 13 | Saprotroph | \|Undefined Saprotroph\| | Yeast | NULL | Probable | As nectar-tap_saprotroph (Põlme et al. 2020); Decay sugar-rich substrates (Põlme et al. 2020) | Põlme S, et al. 2020. Fungal Diversity 105:1-16 ((https://doi.org/10.1007/s13225-020-00466-2)) |
| Otu61926 | 1 | k__Fungi(100);p__Ascomycota(100);c__Saccharomycetes(100);o__Saccharomycetales(100);f__Metschnikowiaceae(100);g__Metschnikowia(100);s__Metschnikowia_pulcherrima(100); | Metschnikowia | 13 | Saprotroph | \|Undefined Saprotroph\| | Yeast | NULL | Probable | As nectar-tap_saprotroph (Põlme et al. 2020); Decay sugar-rich substrates (Põlme et al. 2020) | Põlme S, et al. 2020. Fungal Diversity 105:1-16 ((https://doi.org/10.1007/s13225-020-00466-2)) |
| Otu61928 | 1 | k__Fungi(100);p__Ascomycota(100);c__Saccharomycetes(100);o__Saccharomycetales(100);f__Metschnikowiaceae(100);g__Metschnikowia(100);s__Metschnikowia_pulcherrima(100); | Metschnikowia | 13 | Saprotroph | \|Undefined Saprotroph\| | Yeast | NULL | Probable | As nectar-tap_saprotroph (Põlme et al. 2020); Decay sugar-rich substrates (Põlme et al. 2020) | Põlme S, et al. 2020. Fungal Diversity 105:1-16 ((https://doi.org/10.1007/s13225-020-00466-2)) |
| Otu61933 | 1 | k__Fungi(100);p__Ascomycota(100);c__Saccharomycetes(100);o__Saccharomycetales(100);f__Metschnikowiaceae(100);g__Metschnikowia(100);s__Metschnikowia_pulcherrima(100); | Metschnikowia | 13 | Saprotroph | \|Undefined Saprotroph\| | Yeast | NULL | Probable | As nectar-tap_saprotroph (Põlme et al. 2020); Decay sugar-rich substrates (Põlme et al. 2020) | Põlme S, et al. 2020. Fungal Diversity 105:1-16 ((https://doi.org/10.1007/s13225-020-00466-2)) |
| Otu61936 | 1 | k__Fungi(100);p__Ascomycota(100);c__Saccharomycetes(100);o__Saccharomycetales(100);f__Saccharomycetales_family_Incertae_sedis(100);g__Candida(100);g__Candida_unclassified(100); | Candida | 13 | Pathotroph-Saprotroph-Symbiotroph | Animal Pathogen-Endophyte-Endosymbiont-Epiphyte-\|Undefined Saprotroph\| | Dimorphic Yeast-Yeast | NULL | Probable | As nectar-tap_saprotroph (Põlme et al. 2020); Decay sugar-rich substrates (Põlme et al. 2020); This asexual genus is polyphyletic and contain species that function very differently depend on what they are. Caution should be taken when interpreting guilds for Candida species. | Manolakaki D, et al. 2010. Virulence. 1: 367-375 ((https://doi.org/10.4161/viru.1.5.12796)); Mekbib SB, et al. 2011. Tropical Plant Pathology 36:233-240 ((https://doi.org/10.1590/S1982-56762011000400004)); Põlme S, et al. 2020. Fungal Diversity 105:1-16 ((https://doi.org/10.1007/s13225-020-00466-2)) |
| Otu61947 | 1 | k__Fungi(100);p__Ascomycota(100);c__Saccharomycetes(100);o__Saccharomycetales(100);f__Saccharomycetales_family_Incertae_sedis(100);g__Candida(100);s__Candida_cretensis(100); | Candida | 13 | Pathotroph-Saprotroph-Symbiotroph | Animal Pathogen-Endophyte-Endosymbiont-Epiphyte-\|Undefined Saprotroph\| | Dimorphic Yeast-Yeast | NULL | Probable | As nectar-tap_saprotroph (Põlme et al. 2020); Decay sugar-rich substrates (Põlme et al. 2020); This asexual genus is polyphyletic and contain species that function very differently depend on what they are. Caution should be taken when interpreting guilds for Candida species. | Manolakaki D, et al. 2010. Virulence. 1: 367-375 ((https://doi.org/10.4161/viru.1.5.12796)); Mekbib SB, et al. 2011. Tropical Plant Pathology 36:233-240 ((https://doi.org/10.1590/S1982-56762011000400004)); Põlme S, et al. 2020. Fungal Diversity 105:1-16 ((https://doi.org/10.1007/s13225-020-00466-2)) |
| Otu61948 | 1 | k__Fungi(100);p__Ascomycota(100);c__Saccharomycetes(100);o__Saccharomycetales(100);o__Saccharomycetales_unclassified(100);o__Saccharomycetales_unclassified(100);o__Saccharomycetales_unclassified(100); | Saccharomycetales | 7 | Saprotroph | Undefined Saprotroph | Yeast | NULL | Possible | NULL | Sterkenburg E, et al. 2015. New Phytologist 207:1145-1158 ((https://doi.org/10.1111/nph.13426)) |
| Otu61954 | 1 | k__Fungi(100);p__Ascomycota(100);c__Saccharomycetes(100);o__Saccharomycetales(100);f__Metschnikowiaceae(100);g__Metschnikowia(100);s__Metschnikowia_pulcherrima(100); | Metschnikowia | 13 | Saprotroph | \|Undefined Saprotroph\| | Yeast | NULL | Probable | As nectar-tap_saprotroph (Põlme et al. 2020); Decay sugar-rich substrates (Põlme et al. 2020) | Põlme S, et al. 2020. Fungal Diversity 105:1-16 ((https://doi.org/10.1007/s13225-020-00466-2)) |
| Otu61956 | 1 | k__Fungi(100);p__Ascomycota(100);c__Saccharomycetes(100);o__Saccharomycetales(100);f__Metschnikowiaceae(100);g__Metschnikowia(100);s__Metschnikowia_pulcherrima(100); | Metschnikowia | 13 | Saprotroph | \|Undefined Saprotroph\| | Yeast | NULL | Probable | As nectar-tap_saprotroph (Põlme et al. 2020); Decay sugar-rich substrates (Põlme et al. 2020) | Põlme S, et al. 2020. Fungal Diversity 105:1-16 ((https://doi.org/10.1007/s13225-020-00466-2)) |
| Otu61967 | 1 | k__Fungi(100);p__Ascomycota(100);c__Saccharomycetes(100);o__Saccharomycetales(100);f__Saccharomycetales_family_Incertae_sedis(100);g__Candida(100);g__Candida_unclassified(100); | Candida | 13 | Pathotroph-Saprotroph-Symbiotroph | Animal Pathogen-Endophyte-Endosymbiont-Epiphyte-\|Undefined Saprotroph\| | Dimorphic Yeast-Yeast | NULL | Probable | As nectar-tap_saprotroph (Põlme et al. 2020); Decay sugar-rich substrates (Põlme et al. 2020); This asexual genus is polyphyletic and contain species that function very differently depend on what they are. Caution should be taken when interpreting guilds for Candida species. | Manolakaki D, et al. 2010. Virulence. 1: 367-375 ((https://doi.org/10.4161/viru.1.5.12796)); Mekbib SB, et al. 2011. Tropical Plant Pathology 36:233-240 ((https://doi.org/10.1590/S1982-56762011000400004)); Põlme S, et al. 2020. Fungal Diversity 105:1-16 ((https://doi.org/10.1007/s13225-020-00466-2)) |
| Otu61969 | 1 | k__Fungi(100);p__Ascomycota(100);c__Saccharomycetes(100);o__Saccharomycetales(100);f__Saccharomycetales_family_Incertae_sedis(100);f__Saccharomycetales_family_Incertae_sedis_unclassified(100);f__Saccharomycetales_family_Incertae_sedis_unclassified(100); | Saccharomycetales | 7 | Saprotroph | Undefined Saprotroph | Yeast | NULL | Possible | NULL | Sterkenburg E, et al. 2015. New Phytologist 207:1145-1158 ((https://doi.org/10.1111/nph.13426)) |
| Otu61970 | 1 | k__Fungi(100);p__Ascomycota(100);c__Saccharomycetes(100);o__Saccharomycetales(100);f__Saccharomycetales_family_Incertae_sedis(100);g__Candida(100);g__Candida_unclassified(100); | Candida | 13 | Pathotroph-Saprotroph-Symbiotroph | Animal Pathogen-Endophyte-Endosymbiont-Epiphyte-\|Undefined Saprotroph\| | Dimorphic Yeast-Yeast | NULL | Probable | As nectar-tap_saprotroph (Põlme et al. 2020); Decay sugar-rich substrates (Põlme et al. 2020); This asexual genus is polyphyletic and contain species that function very differently depend on what they are. Caution should be taken when interpreting guilds for Candida species. | Manolakaki D, et al. 2010. Virulence. 1: 367-375 ((https://doi.org/10.4161/viru.1.5.12796)); Mekbib SB, et al. 2011. Tropical Plant Pathology 36:233-240 ((https://doi.org/10.1590/S1982-56762011000400004)); Põlme S, et al. 2020. Fungal Diversity 105:1-16 ((https://doi.org/10.1007/s13225-020-00466-2)) |
| Otu61971 | 1 | k__Fungi(100);p__Ascomycota(100);c__Saccharomycetes(100);o__Saccharomycetales(100);f__Saccharomycetales_family_Incertae_sedis(100);g__Candida(100);g__Candida_unclassified(100); | Candida | 13 | Pathotroph-Saprotroph-Symbiotroph | Animal Pathogen-Endophyte-Endosymbiont-Epiphyte-\|Undefined Saprotroph\| | Dimorphic Yeast-Yeast | NULL | Probable | As nectar-tap_saprotroph (Põlme et al. 2020); Decay sugar-rich substrates (Põlme et al. 2020); This asexual genus is polyphyletic and contain species that function very differently depend on what they are. Caution should be taken when interpreting guilds for Candida species. | Manolakaki D, et al. 2010. Virulence. 1: 367-375 ((https://doi.org/10.4161/viru.1.5.12796)); Mekbib SB, et al. 2011. Tropical Plant Pathology 36:233-240 ((https://doi.org/10.1590/S1982-56762011000400004)); Põlme S, et al. 2020. Fungal Diversity 105:1-16 ((https://doi.org/10.1007/s13225-020-00466-2)) |
| Otu61974 | 1 | k__Fungi(100);p__Ascomycota(100);c__Saccharomycetes(100);o__Saccharomycetales(100);f__Saccharomycetales_family_Incertae_sedis(100);f__Saccharomycetales_family_Incertae_sedis_unclassified(100);f__Saccharomycetales_family_Incertae_sedis_unclassified(100); | Saccharomycetales | 7 | Saprotroph | Undefined Saprotroph | Yeast | NULL | Possible | NULL | Sterkenburg E, et al. 2015. New Phytologist 207:1145-1158 ((https://doi.org/10.1111/nph.13426)) |
| Otu61975 | 1 | k__Fungi(100);p__Ascomycota(100);c__Saccharomycetes(100);o__Saccharomycetales(100);o__Saccharomycetales_unclassified(100);o__Saccharomycetales_unclassified(100);o__Saccharomycetales_unclassified(100); | Saccharomycetales | 7 | Saprotroph | Undefined Saprotroph | Yeast | NULL | Possible | NULL | Sterkenburg E, et al. 2015. New Phytologist 207:1145-1158 ((https://doi.org/10.1111/nph.13426)) |
| Otu61976 | 1 | k__Fungi(100);p__Ascomycota(100);c__Saccharomycetes(100);o__Saccharomycetales(100);f__Saccharomycetales_family_Incertae_sedis(100);g__Candida(100);g__Candida_unclassified(100); | Candida | 13 | Pathotroph-Saprotroph-Symbiotroph | Animal Pathogen-Endophyte-Endosymbiont-Epiphyte-\|Undefined Saprotroph\| | Dimorphic Yeast-Yeast | NULL | Probable | As nectar-tap_saprotroph (Põlme et al. 2020); Decay sugar-rich substrates (Põlme et al. 2020); This asexual genus is polyphyletic and contain species that function very differently depend on what they are. Caution should be taken when interpreting guilds for Candida species. | Manolakaki D, et al. 2010. Virulence. 1: 367-375 ((https://doi.org/10.4161/viru.1.5.12796)); Mekbib SB, et al. 2011. Tropical Plant Pathology 36:233-240 ((https://doi.org/10.1590/S1982-56762011000400004)); Põlme S, et al. 2020. Fungal Diversity 105:1-16 ((https://doi.org/10.1007/s13225-020-00466-2)) |
| Otu61977 | 1 | k__Fungi(100);p__Ascomycota(100);c__Saccharomycetes(100);o__Saccharomycetales(100);f__Saccharomycetales_family_Incertae_sedis(100);g__Candida(100);g__Candida_unclassified(100); | Candida | 13 | Pathotroph-Saprotroph-Symbiotroph | Animal Pathogen-Endophyte-Endosymbiont-Epiphyte-\|Undefined Saprotroph\| | Dimorphic Yeast-Yeast | NULL | Probable | As nectar-tap_saprotroph (Põlme et al. 2020); Decay sugar-rich substrates (Põlme et al. 2020); This asexual genus is polyphyletic and contain species that function very differently depend on what they are. Caution should be taken when interpreting guilds for Candida species. | Manolakaki D, et al. 2010. Virulence. 1: 367-375 ((https://doi.org/10.4161/viru.1.5.12796)); Mekbib SB, et al. 2011. Tropical Plant Pathology 36:233-240 ((https://doi.org/10.1590/S1982-56762011000400004)); Põlme S, et al. 2020. Fungal Diversity 105:1-16 ((https://doi.org/10.1007/s13225-020-00466-2)) |
| Otu61979 | 1 | k__Fungi(100);p__Ascomycota(100);c__Saccharomycetes(100);o__Saccharomycetales(100);f__Metschnikowiaceae(100);g__Metschnikowia(100);s__Metschnikowia_pulcherrima(100); | Metschnikowia | 13 | Saprotroph | \|Undefined Saprotroph\| | Yeast | NULL | Probable | As nectar-tap_saprotroph (Põlme et al. 2020); Decay sugar-rich substrates (Põlme et al. 2020) | Põlme S, et al. 2020. Fungal Diversity 105:1-16 ((https://doi.org/10.1007/s13225-020-00466-2)) |
| Otu61982 | 1 | k__Fungi(100);p__Ascomycota(100);c__Saccharomycetes(100);o__Saccharomycetales(100);f__Metschnikowiaceae(100);g__Metschnikowia(100);s__Metschnikowia_pulcherrima(100); | Metschnikowia | 13 | Saprotroph | \|Undefined Saprotroph\| | Yeast | NULL | Probable | As nectar-tap_saprotroph (Põlme et al. 2020); Decay sugar-rich substrates (Põlme et al. 2020) | Põlme S, et al. 2020. Fungal Diversity 105:1-16 ((https://doi.org/10.1007/s13225-020-00466-2)) |
| Otu61986 | 1 | k__Fungi(100);p__Ascomycota(100);c__Saccharomycetes(100);o__Saccharomycetales(100);f__Metschnikowiaceae(100);g__Metschnikowia(100);g__Metschnikowia_unclassified(100); | Metschnikowia | 13 | Saprotroph | \|Undefined Saprotroph\| | Yeast | NULL | Probable | As nectar-tap_saprotroph (Põlme et al. 2020); Decay sugar-rich substrates (Põlme et al. 2020) | Põlme S, et al. 2020. Fungal Diversity 105:1-16 ((https://doi.org/10.1007/s13225-020-00466-2)) |
| Otu61987 | 1 | k__Fungi(100);p__Ascomycota(100);c__Saccharomycetes(100);o__Saccharomycetales(100);f__Metschnikowiaceae(100);g__Metschnikowia(100);s__Metschnikowia_pulcherrima(100); | Metschnikowia | 13 | Saprotroph | \|Undefined Saprotroph\| | Yeast | NULL | Probable | As nectar-tap_saprotroph (Põlme et al. 2020); Decay sugar-rich substrates (Põlme et al. 2020) | Põlme S, et al. 2020. Fungal Diversity 105:1-16 ((https://doi.org/10.1007/s13225-020-00466-2)) |
| Otu61994 | 1 | k__Fungi(100);p__Ascomycota(100);c__Dothideomycetes(100);o__Capnodiales(100);f__Capnodiales_family_Incertae_sedis(100);g__Hispidoconidioma(100);s__Hispidoconidioma_sp(100); | Hispidoconidioma | 13 | Saprotroph | \|Wood Saprotroph\| | Microfungus | NULL | Probable | Decay wood (Põlme et al. 2020) | Põlme S, et al. 2020. Fungal Diversity 105:1-16 ((https://doi.org/10.1007/s13225-020-00466-2)) |
| Otu62000 | 1 | k__Fungi(100);p__Ascomycota(100);c__Saccharomycetes(100);o__Saccharomycetales(100);f__Metschnikowiaceae(100);g__Metschnikowia(100);s__Metschnikowia_pulcherrima(100); | Metschnikowia | 13 | Saprotroph | \|Undefined Saprotroph\| | Yeast | NULL | Probable | As nectar-tap_saprotroph (Põlme et al. 2020); Decay sugar-rich substrates (Põlme et al. 2020) | Põlme S, et al. 2020. Fungal Diversity 105:1-16 ((https://doi.org/10.1007/s13225-020-00466-2)) |
| Otu62005 | 1 | k__Fungi(100);p__Basidiomycota(100);c__Microbotryomycetes(100);o__Sporidiobolales(100);f__Sporidiobolales_family_Incertae_sedis(100);g__Sporobolomyces(100);s__Sporobolomyces_lactosus(100); | Sporobolomyces | 13 | Pathotroph-Saprotroph | \|Fungal Parasite\|-Plant Saprotroph | Yeast | NULL | Probable | As litter_saprotroph (Põlme et al. 2020); As mycoparasite (Põlme et al. 2020); Decay fungal material (Põlme et al. 2020) | Tedersoo L, et al. 2014. Science 346:e1256688 ((https://doi.org/10.1126/science.1256688)); Purahong W, et al. 2016. Molecular Ecology 25:4059-4074 ((https://doi.org/10.1111/mec.13739)); Põlme S, et al. 2020. Fungal Diversity 105:1-16 ((https://doi.org/10.1007/s13225-020-00466-2)) |
| Otu62007 | 1 | k__Fungi(100);p__Ascomycota(100);c__Saccharomycetes(100);o__Saccharomycetales(100);f__Metschnikowiaceae(100);g__Metschnikowia(100);s__Metschnikowia_pulcherrima(100); | Metschnikowia | 13 | Saprotroph | \|Undefined Saprotroph\| | Yeast | NULL | Probable | As nectar-tap_saprotroph (Põlme et al. 2020); Decay sugar-rich substrates (Põlme et al. 2020) | Põlme S, et al. 2020. Fungal Diversity 105:1-16 ((https://doi.org/10.1007/s13225-020-00466-2)) |
| Otu62008 | 1 | k__Fungi(100);p__Ascomycota(100);c__Saccharomycetes(100);o__Saccharomycetales(100);f__Metschnikowiaceae(100);g__Metschnikowia(100);g__Metschnikowia_unclassified(100); | Metschnikowia | 13 | Saprotroph | \|Undefined Saprotroph\| | Yeast | NULL | Probable | As nectar-tap_saprotroph (Põlme et al. 2020); Decay sugar-rich substrates (Põlme et al. 2020) | Põlme S, et al. 2020. Fungal Diversity 105:1-16 ((https://doi.org/10.1007/s13225-020-00466-2)) |
| Otu62014 | 1 | k__Fungi(100);p__Basidiomycota(100);c__Microbotryomycetes(100);o__Sporidiobolales(100);f__Sporidiobolales_family_Incertae_sedis(100);g__Sporobolomyces(100);s__Sporobolomyces_lactosus(100); | Sporobolomyces | 13 | Pathotroph-Saprotroph | \|Fungal Parasite\|-Plant Saprotroph | Yeast | NULL | Probable | As litter_saprotroph (Põlme et al. 2020); As mycoparasite (Põlme et al. 2020); Decay fungal material (Põlme et al. 2020) | Tedersoo L, et al. 2014. Science 346:e1256688 ((https://doi.org/10.1126/science.1256688)); Purahong W, et al. 2016. Molecular Ecology 25:4059-4074 ((https://doi.org/10.1111/mec.13739)); Põlme S, et al. 2020. Fungal Diversity 105:1-16 ((https://doi.org/10.1007/s13225-020-00466-2)) |
| Otu62015 | 1 | k__Fungi(100);p__Ascomycota(100);c__Saccharomycetes(100);o__Saccharomycetales(100);f__Metschnikowiaceae(100);g__Metschnikowia(100);g__Metschnikowia_unclassified(100); | Metschnikowia | 13 | Saprotroph | \|Undefined Saprotroph\| | Yeast | NULL | Probable | As nectar-tap_saprotroph (Põlme et al. 2020); Decay sugar-rich substrates (Põlme et al. 2020) | Põlme S, et al. 2020. Fungal Diversity 105:1-16 ((https://doi.org/10.1007/s13225-020-00466-2)) |
| Otu62016 | 1 | k__Fungi(100);p__Ascomycota(100);c__Dothideomycetes(100);o__Capnodiales(100);f__Mycosphaerellaceae(100);f__Mycosphaerellaceae_unclassified(100);f__Mycosphaerellaceae_unclassified(100); | Mycosphaerellaceae | 9 | Pathotroph-Saprotroph | Plant Pathogen-Undefined Saprotroph | Microfungus | NULL | Probable | Biotrophic, necrotrophic or saprobic on various plant tissues (Cannon & Kirk 2007) | Cannon PF, Kirk PM. 2007. Fungal Families of the World. CAB International, Cambridge (ISBN: 978-0851998275) |
| Otu62017 | 1 | k__Fungi(100);p__Ascomycota(100);c__Saccharomycetes(100);o__Saccharomycetales(100);f__Metschnikowiaceae(100);g__Metschnikowia(100);g__Metschnikowia_unclassified(100); | Metschnikowia | 13 | Saprotroph | \|Undefined Saprotroph\| | Yeast | NULL | Probable | As nectar-tap_saprotroph (Põlme et al. 2020); Decay sugar-rich substrates (Põlme et al. 2020) | Põlme S, et al. 2020. Fungal Diversity 105:1-16 ((https://doi.org/10.1007/s13225-020-00466-2)) |
| Otu62019 | 1 | k__Fungi(100);p__Ascomycota(100);c__Saccharomycetes(100);o__Saccharomycetales(100);f__Metschnikowiaceae(100);g__Metschnikowia(100);s__Metschnikowia_pulcherrima(100); | Metschnikowia | 13 | Saprotroph | \|Undefined Saprotroph\| | Yeast | NULL | Probable | As nectar-tap_saprotroph (Põlme et al. 2020); Decay sugar-rich substrates (Põlme et al. 2020) | Põlme S, et al. 2020. Fungal Diversity 105:1-16 ((https://doi.org/10.1007/s13225-020-00466-2)) |
| Otu62022 | 1 | k__Fungi(100);p__Ascomycota(100);c__Saccharomycetes(100);o__Saccharomycetales(100);f__Metschnikowiaceae(100);g__Metschnikowia(100);g__Metschnikowia_unclassified(100); | Metschnikowia | 13 | Saprotroph | \|Undefined Saprotroph\| | Yeast | NULL | Probable | As nectar-tap_saprotroph (Põlme et al. 2020); Decay sugar-rich substrates (Põlme et al. 2020) | Põlme S, et al. 2020. Fungal Diversity 105:1-16 ((https://doi.org/10.1007/s13225-020-00466-2)) |
| Otu62023 | 1 | k__Fungi(100);p__Ascomycota(100);c__Saccharomycetes(100);o__Saccharomycetales(100);f__Metschnikowiaceae(100);g__Metschnikowia(100);s__Metschnikowia_pulcherrima(100); | Metschnikowia | 13 | Saprotroph | \|Undefined Saprotroph\| | Yeast | NULL | Probable | As nectar-tap_saprotroph (Põlme et al. 2020); Decay sugar-rich substrates (Põlme et al. 2020) | Põlme S, et al. 2020. Fungal Diversity 105:1-16 ((https://doi.org/10.1007/s13225-020-00466-2)) |
| Otu62029 | 1 | k__Fungi(100);p__Ascomycota(100);c__Saccharomycetes(100);o__Saccharomycetales(100);f__Metschnikowiaceae(100);g__Metschnikowia(100);s__Metschnikowia_pulcherrima(100); | Metschnikowia | 13 | Saprotroph | \|Undefined Saprotroph\| | Yeast | NULL | Probable | As nectar-tap_saprotroph (Põlme et al. 2020); Decay sugar-rich substrates (Põlme et al. 2020) | Põlme S, et al. 2020. Fungal Diversity 105:1-16 ((https://doi.org/10.1007/s13225-020-00466-2)) |
| Otu62032 | 1 | k__Fungi(100);p__Ascomycota(100);c__Saccharomycetes(100);o__Saccharomycetales(100);f__Metschnikowiaceae(100);g__Metschnikowia(100);s__Metschnikowia_pulcherrima(100); | Metschnikowia | 13 | Saprotroph | \|Undefined Saprotroph\| | Yeast | NULL | Probable | As nectar-tap_saprotroph (Põlme et al. 2020); Decay sugar-rich substrates (Põlme et al. 2020) | Põlme S, et al. 2020. Fungal Diversity 105:1-16 ((https://doi.org/10.1007/s13225-020-00466-2)) |
| Otu62037 | 1 | k__Fungi(100);p__Ascomycota(100);c__Saccharomycetes(100);o__Saccharomycetales(100);f__Metschnikowiaceae(100);g__Metschnikowia(100);s__Metschnikowia_pulcherrima(100); | Metschnikowia | 13 | Saprotroph | \|Undefined Saprotroph\| | Yeast | NULL | Probable | As nectar-tap_saprotroph (Põlme et al. 2020); Decay sugar-rich substrates (Põlme et al. 2020) | Põlme S, et al. 2020. Fungal Diversity 105:1-16 ((https://doi.org/10.1007/s13225-020-00466-2)) |
| Otu62038 | 1 | k__Fungi(100);p__Ascomycota(100);c__Saccharomycetes(100);o__Saccharomycetales(100);f__Metschnikowiaceae(100);g__Metschnikowia(100);s__Metschnikowia_pulcherrima(100); | Metschnikowia | 13 | Saprotroph | \|Undefined Saprotroph\| | Yeast | NULL | Probable | As nectar-tap_saprotroph (Põlme et al. 2020); Decay sugar-rich substrates (Põlme et al. 2020) | Põlme S, et al. 2020. Fungal Diversity 105:1-16 ((https://doi.org/10.1007/s13225-020-00466-2)) |
| Otu62039 | 1 | k__Fungi(100);p__Ascomycota(100);c__Saccharomycetes(100);o__Saccharomycetales(100);f__Metschnikowiaceae(100);g__Metschnikowia(100);g__Metschnikowia_unclassified(100); | Metschnikowia | 13 | Saprotroph | \|Undefined Saprotroph\| | Yeast | NULL | Probable | As nectar-tap_saprotroph (Põlme et al. 2020); Decay sugar-rich substrates (Põlme et al. 2020) | Põlme S, et al. 2020. Fungal Diversity 105:1-16 ((https://doi.org/10.1007/s13225-020-00466-2)) |
| Otu62040 | 1 | k__Fungi(100);p__Ascomycota(100);c__Saccharomycetes(100);o__Saccharomycetales(100);f__Metschnikowiaceae(100);g__Metschnikowia(100);s__Metschnikowia_pulcherrima(100); | Metschnikowia | 13 | Saprotroph | \|Undefined Saprotroph\| | Yeast | NULL | Probable | As nectar-tap_saprotroph (Põlme et al. 2020); Decay sugar-rich substrates (Põlme et al. 2020) | Põlme S, et al. 2020. Fungal Diversity 105:1-16 ((https://doi.org/10.1007/s13225-020-00466-2)) |
| Otu62041 | 1 | k__Fungi(100);p__Ascomycota(100);c__Saccharomycetes(100);o__Saccharomycetales(100);f__Metschnikowiaceae(100);g__Metschnikowia(100);s__Metschnikowia_pulcherrima(100); | Metschnikowia | 13 | Saprotroph | \|Undefined Saprotroph\| | Yeast | NULL | Probable | As nectar-tap_saprotroph (Põlme et al. 2020); Decay sugar-rich substrates (Põlme et al. 2020) | Põlme S, et al. 2020. Fungal Diversity 105:1-16 ((https://doi.org/10.1007/s13225-020-00466-2)) |
| Otu62042 | 1 | k__Fungi(100);p__Ascomycota(100);c__Saccharomycetes(100);o__Saccharomycetales(100);f__Metschnikowiaceae(100);g__Metschnikowia(100);g__Metschnikowia_unclassified(100); | Metschnikowia | 13 | Saprotroph | \|Undefined Saprotroph\| | Yeast | NULL | Probable | As nectar-tap_saprotroph (Põlme et al. 2020); Decay sugar-rich substrates (Põlme et al. 2020) | Põlme S, et al. 2020. Fungal Diversity 105:1-16 ((https://doi.org/10.1007/s13225-020-00466-2)) |
| Otu62045 | 1 | k__Fungi(100);p__Ascomycota(100);c__Saccharomycetes(100);o__Saccharomycetales(100);f__Metschnikowiaceae(100);g__Metschnikowia(100);s__Metschnikowia_pulcherrima(100); | Metschnikowia | 13 | Saprotroph | \|Undefined Saprotroph\| | Yeast | NULL | Probable | As nectar-tap_saprotroph (Põlme et al. 2020); Decay sugar-rich substrates (Põlme et al. 2020) | Põlme S, et al. 2020. Fungal Diversity 105:1-16 ((https://doi.org/10.1007/s13225-020-00466-2)) |
| Otu62050 | 1 | k__Fungi(100);p__Ascomycota(100);c__Saccharomycetes(100);o__Saccharomycetales(100);f__Metschnikowiaceae(100);g__Metschnikowia(100);s__Metschnikowia_pulcherrima(100); | Metschnikowia | 13 | Saprotroph | \|Undefined Saprotroph\| | Yeast | NULL | Probable | As nectar-tap_saprotroph (Põlme et al. 2020); Decay sugar-rich substrates (Põlme et al. 2020) | Põlme S, et al. 2020. Fungal Diversity 105:1-16 ((https://doi.org/10.1007/s13225-020-00466-2)) |
| Otu62054 | 1 | k__Fungi(100);p__Ascomycota(100);c__Saccharomycetes(100);o__Saccharomycetales(100);f__Metschnikowiaceae(100);g__Metschnikowia(100);s__Metschnikowia_pulcherrima(100); | Metschnikowia | 13 | Saprotroph | \|Undefined Saprotroph\| | Yeast | NULL | Probable | As nectar-tap_saprotroph (Põlme et al. 2020); Decay sugar-rich substrates (Põlme et al. 2020) | Põlme S, et al. 2020. Fungal Diversity 105:1-16 ((https://doi.org/10.1007/s13225-020-00466-2)) |
| Otu62055 | 1 | k__Fungi(100);p__Ascomycota(100);c__Saccharomycetes(100);o__Saccharomycetales(100);f__Metschnikowiaceae(100);g__Metschnikowia(100);s__Metschnikowia_pulcherrima(100); | Metschnikowia | 13 | Saprotroph | \|Undefined Saprotroph\| | Yeast | NULL | Probable | As nectar-tap_saprotroph (Põlme et al. 2020); Decay sugar-rich substrates (Põlme et al. 2020) | Põlme S, et al. 2020. Fungal Diversity 105:1-16 ((https://doi.org/10.1007/s13225-020-00466-2)) |
| Otu62056 | 1 | k__Fungi(100);p__Ascomycota(100);c__Saccharomycetes(100);o__Saccharomycetales(100);f__Metschnikowiaceae(100);g__Metschnikowia(100);s__Metschnikowia_pulcherrima(100); | Metschnikowia | 13 | Saprotroph | \|Undefined Saprotroph\| | Yeast | NULL | Probable | As nectar-tap_saprotroph (Põlme et al. 2020); Decay sugar-rich substrates (Põlme et al. 2020) | Põlme S, et al. 2020. Fungal Diversity 105:1-16 ((https://doi.org/10.1007/s13225-020-00466-2)) |
| Otu62057 | 1 | k__Fungi(100);p__Ascomycota(100);c__Saccharomycetes(100);o__Saccharomycetales(100);f__Metschnikowiaceae(100);g__Metschnikowia(100);s__Metschnikowia_pulcherrima(100); | Metschnikowia | 13 | Saprotroph | \|Undefined Saprotroph\| | Yeast | NULL | Probable | As nectar-tap_saprotroph (Põlme et al. 2020); Decay sugar-rich substrates (Põlme et al. 2020) | Põlme S, et al. 2020. Fungal Diversity 105:1-16 ((https://doi.org/10.1007/s13225-020-00466-2)) |
| Otu62058 | 1 | k__Fungi(100);p__Ascomycota(100);c__Saccharomycetes(100);o__Saccharomycetales(100);f__Metschnikowiaceae(100);g__Metschnikowia(100);s__Metschnikowia_pulcherrima(100); | Metschnikowia | 13 | Saprotroph | \|Undefined Saprotroph\| | Yeast | NULL | Probable | As nectar-tap_saprotroph (Põlme et al. 2020); Decay sugar-rich substrates (Põlme et al. 2020) | Põlme S, et al. 2020. Fungal Diversity 105:1-16 ((https://doi.org/10.1007/s13225-020-00466-2)) |
| Otu62059 | 1 | k__Fungi(100);p__Ascomycota(100);c__Saccharomycetes(100);o__Saccharomycetales(100);f__Metschnikowiaceae(100);g__Metschnikowia(100);s__Metschnikowia_pulcherrima(100); | Metschnikowia | 13 | Saprotroph | \|Undefined Saprotroph\| | Yeast | NULL | Probable | As nectar-tap_saprotroph (Põlme et al. 2020); Decay sugar-rich substrates (Põlme et al. 2020) | Põlme S, et al. 2020. Fungal Diversity 105:1-16 ((https://doi.org/10.1007/s13225-020-00466-2)) |
| Otu62060 | 1 | k__Fungi(100);p__Ascomycota(100);c__Saccharomycetes(100);o__Saccharomycetales(100);f__Metschnikowiaceae(100);g__Metschnikowia(100);s__Metschnikowia_pulcherrima(100); | Metschnikowia | 13 | Saprotroph | \|Undefined Saprotroph\| | Yeast | NULL | Probable | As nectar-tap_saprotroph (Põlme et al. 2020); Decay sugar-rich substrates (Põlme et al. 2020) | Põlme S, et al. 2020. Fungal Diversity 105:1-16 ((https://doi.org/10.1007/s13225-020-00466-2)) |
| Otu62061 | 1 | k__Fungi(100);p__Ascomycota(100);c__Saccharomycetes(100);o__Saccharomycetales(100);f__Metschnikowiaceae(100);g__Metschnikowia(100);g__Metschnikowia_unclassified(100); | Metschnikowia | 13 | Saprotroph | \|Undefined Saprotroph\| | Yeast | NULL | Probable | As nectar-tap_saprotroph (Põlme et al. 2020); Decay sugar-rich substrates (Põlme et al. 2020) | Põlme S, et al. 2020. Fungal Diversity 105:1-16 ((https://doi.org/10.1007/s13225-020-00466-2)) |
| Otu62062 | 1 | k__Fungi(100);p__Ascomycota(100);c__Saccharomycetes(100);o__Saccharomycetales(100);f__Metschnikowiaceae(100);g__Metschnikowia(100);s__Metschnikowia_pulcherrima(100); | Metschnikowia | 13 | Saprotroph | \|Undefined Saprotroph\| | Yeast | NULL | Probable | As nectar-tap_saprotroph (Põlme et al. 2020); Decay sugar-rich substrates (Põlme et al. 2020) | Põlme S, et al. 2020. Fungal Diversity 105:1-16 ((https://doi.org/10.1007/s13225-020-00466-2)) |
| Otu62063 | 1 | k__Fungi(100);p__Ascomycota(100);c__Saccharomycetes(100);o__Saccharomycetales(100);f__Metschnikowiaceae(100);g__Metschnikowia(100);g__Metschnikowia_unclassified(100); | Metschnikowia | 13 | Saprotroph | \|Undefined Saprotroph\| | Yeast | NULL | Probable | As nectar-tap_saprotroph (Põlme et al. 2020); Decay sugar-rich substrates (Põlme et al. 2020) | Põlme S, et al. 2020. Fungal Diversity 105:1-16 ((https://doi.org/10.1007/s13225-020-00466-2)) |
| Otu62064 | 1 | k__Fungi(100);p__Ascomycota(100);c__Saccharomycetes(100);o__Saccharomycetales(100);f__Metschnikowiaceae(100);g__Metschnikowia(100);s__Metschnikowia_pulcherrima(100); | Metschnikowia | 13 | Saprotroph | \|Undefined Saprotroph\| | Yeast | NULL | Probable | As nectar-tap_saprotroph (Põlme et al. 2020); Decay sugar-rich substrates (Põlme et al. 2020) | Põlme S, et al. 2020. Fungal Diversity 105:1-16 ((https://doi.org/10.1007/s13225-020-00466-2)) |
| Otu62065 | 1 | k__Fungi(100);p__Ascomycota(100);c__Saccharomycetes(100);o__Saccharomycetales(100);f__Metschnikowiaceae(100);g__Metschnikowia(100);g__Metschnikowia_unclassified(100); | Metschnikowia | 13 | Saprotroph | \|Undefined Saprotroph\| | Yeast | NULL | Probable | As nectar-tap_saprotroph (Põlme et al. 2020); Decay sugar-rich substrates (Põlme et al. 2020) | Põlme S, et al. 2020. Fungal Diversity 105:1-16 ((https://doi.org/10.1007/s13225-020-00466-2)) |
| Otu62068 | 1 | k__Fungi(100);p__Ascomycota(100);c__Saccharomycetes(100);o__Saccharomycetales(100);f__Metschnikowiaceae(100);g__Metschnikowia(100);s__Metschnikowia_pulcherrima(100); | Metschnikowia | 13 | Saprotroph | \|Undefined Saprotroph\| | Yeast | NULL | Probable | As nectar-tap_saprotroph (Põlme et al. 2020); Decay sugar-rich substrates (Põlme et al. 2020) | Põlme S, et al. 2020. Fungal Diversity 105:1-16 ((https://doi.org/10.1007/s13225-020-00466-2)) |
| Otu62069 | 1 | k__Fungi(100);p__Ascomycota(100);c__Saccharomycetes(100);o__Saccharomycetales(100);f__Metschnikowiaceae(100);g__Metschnikowia(100);s__Metschnikowia_pulcherrima(100); | Metschnikowia | 13 | Saprotroph | \|Undefined Saprotroph\| | Yeast | NULL | Probable | As nectar-tap_saprotroph (Põlme et al. 2020); Decay sugar-rich substrates (Põlme et al. 2020) | Põlme S, et al. 2020. Fungal Diversity 105:1-16 ((https://doi.org/10.1007/s13225-020-00466-2)) |
| Otu62070 | 1 | k__Fungi(100);p__Ascomycota(100);c__Saccharomycetes(100);o__Saccharomycetales(100);f__Metschnikowiaceae(100);g__Metschnikowia(100);g__Metschnikowia_unclassified(100); | Metschnikowia | 13 | Saprotroph | \|Undefined Saprotroph\| | Yeast | NULL | Probable | As nectar-tap_saprotroph (Põlme et al. 2020); Decay sugar-rich substrates (Põlme et al. 2020) | Põlme S, et al. 2020. Fungal Diversity 105:1-16 ((https://doi.org/10.1007/s13225-020-00466-2)) |
| Otu62071 | 1 | k__Fungi(100);p__Ascomycota(100);c__Saccharomycetes(100);o__Saccharomycetales(100);f__Metschnikowiaceae(100);g__Metschnikowia(100);s__Metschnikowia_pulcherrima(100); | Metschnikowia | 13 | Saprotroph | \|Undefined Saprotroph\| | Yeast | NULL | Probable | As nectar-tap_saprotroph (Põlme et al. 2020); Decay sugar-rich substrates (Põlme et al. 2020) | Põlme S, et al. 2020. Fungal Diversity 105:1-16 ((https://doi.org/10.1007/s13225-020-00466-2)) |
| Otu62072 | 1 | k__Fungi(100);p__Ascomycota(100);c__Saccharomycetes(100);o__Saccharomycetales(100);f__Metschnikowiaceae(100);g__Metschnikowia(100);s__Metschnikowia_pulcherrima(100); | Metschnikowia | 13 | Saprotroph | \|Undefined Saprotroph\| | Yeast | NULL | Probable | As nectar-tap_saprotroph (Põlme et al. 2020); Decay sugar-rich substrates (Põlme et al. 2020) | Põlme S, et al. 2020. Fungal Diversity 105:1-16 ((https://doi.org/10.1007/s13225-020-00466-2)) |
| Otu62073 | 1 | k__Fungi(100);p__Ascomycota(100);c__Saccharomycetes(100);o__Saccharomycetales(100);f__Metschnikowiaceae(100);g__Metschnikowia(100);g__Metschnikowia_unclassified(100); | Metschnikowia | 13 | Saprotroph | \|Undefined Saprotroph\| | Yeast | NULL | Probable | As nectar-tap_saprotroph (Põlme et al. 2020); Decay sugar-rich substrates (Põlme et al. 2020) | Põlme S, et al. 2020. Fungal Diversity 105:1-16 ((https://doi.org/10.1007/s13225-020-00466-2)) |
| Otu62076 | 1 | k__Fungi(100);p__Ascomycota(100);c__Saccharomycetes(100);o__Saccharomycetales(100);f__Metschnikowiaceae(100);g__Metschnikowia(100);s__Metschnikowia_pulcherrima(100); | Metschnikowia | 13 | Saprotroph | \|Undefined Saprotroph\| | Yeast | NULL | Probable | As nectar-tap_saprotroph (Põlme et al. 2020); Decay sugar-rich substrates (Põlme et al. 2020) | Põlme S, et al. 2020. Fungal Diversity 105:1-16 ((https://doi.org/10.1007/s13225-020-00466-2)) |
| Otu62077 | 1 | k__Fungi(100);p__Ascomycota(100);c__Saccharomycetes(100);o__Saccharomycetales(100);f__Metschnikowiaceae(100);g__Metschnikowia(100);s__Metschnikowia_pulcherrima(100); | Metschnikowia | 13 | Saprotroph | \|Undefined Saprotroph\| | Yeast | NULL | Probable | As nectar-tap_saprotroph (Põlme et al. 2020); Decay sugar-rich substrates (Põlme et al. 2020) | Põlme S, et al. 2020. Fungal Diversity 105:1-16 ((https://doi.org/10.1007/s13225-020-00466-2)) |
| Otu62078 | 1 | k__Fungi(100);p__Ascomycota(100);c__Saccharomycetes(100);o__Saccharomycetales(100);f__Metschnikowiaceae(100);g__Metschnikowia(100);s__Metschnikowia_pulcherrima(100); | Metschnikowia | 13 | Saprotroph | \|Undefined Saprotroph\| | Yeast | NULL | Probable | As nectar-tap_saprotroph (Põlme et al. 2020); Decay sugar-rich substrates (Põlme et al. 2020) | Põlme S, et al. 2020. Fungal Diversity 105:1-16 ((https://doi.org/10.1007/s13225-020-00466-2)) |
| Otu62079 | 1 | k__Fungi(100);p__Ascomycota(100);c__Saccharomycetes(100);o__Saccharomycetales(100);f__Metschnikowiaceae(100);g__Metschnikowia(100);s__Metschnikowia_pulcherrima(100); | Metschnikowia | 13 | Saprotroph | \|Undefined Saprotroph\| | Yeast | NULL | Probable | As nectar-tap_saprotroph (Põlme et al. 2020); Decay sugar-rich substrates (Põlme et al. 2020) | Põlme S, et al. 2020. Fungal Diversity 105:1-16 ((https://doi.org/10.1007/s13225-020-00466-2)) |
| Otu62080 | 1 | k__Fungi(100);p__Ascomycota(100);c__Saccharomycetes(100);o__Saccharomycetales(100);f__Metschnikowiaceae(100);g__Metschnikowia(100);g__Metschnikowia_unclassified(100); | Metschnikowia | 13 | Saprotroph | \|Undefined Saprotroph\| | Yeast | NULL | Probable | As nectar-tap_saprotroph (Põlme et al. 2020); Decay sugar-rich substrates (Põlme et al. 2020) | Põlme S, et al. 2020. Fungal Diversity 105:1-16 ((https://doi.org/10.1007/s13225-020-00466-2)) |
| Otu62083 | 1 | k__Fungi(100);p__Ascomycota(100);c__Saccharomycetes(100);o__Saccharomycetales(100);f__Saccharomycetales_family_Incertae_sedis(100);f__Saccharomycetales_family_Incertae_sedis_unclassified(100);f__Saccharomycetales_family_Incertae_sedis_unclassified(100); | Saccharomycetales | 7 | Saprotroph | Undefined Saprotroph | Yeast | NULL | Possible | NULL | Sterkenburg E, et al. 2015. New Phytologist 207:1145-1158 ((https://doi.org/10.1111/nph.13426)) |
| Otu62087 | 1 | k__Fungi(100);p__Ascomycota(100);c__Saccharomycetes(100);o__Saccharomycetales(100);f__Metschnikowiaceae(100);g__Metschnikowia(100);g__Metschnikowia_unclassified(100); | Metschnikowia | 13 | Saprotroph | \|Undefined Saprotroph\| | Yeast | NULL | Probable | As nectar-tap_saprotroph (Põlme et al. 2020); Decay sugar-rich substrates (Põlme et al. 2020) | Põlme S, et al. 2020. Fungal Diversity 105:1-16 ((https://doi.org/10.1007/s13225-020-00466-2)) |
| Otu62093 | 1 | k__Fungi(100);p__Ascomycota(100);c__Saccharomycetes(100);o__Saccharomycetales(100);f__Metschnikowiaceae(100);g__Metschnikowia(100);s__Metschnikowia_pulcherrima(100); | Metschnikowia | 13 | Saprotroph | \|Undefined Saprotroph\| | Yeast | NULL | Probable | As nectar-tap_saprotroph (Põlme et al. 2020); Decay sugar-rich substrates (Põlme et al. 2020) | Põlme S, et al. 2020. Fungal Diversity 105:1-16 ((https://doi.org/10.1007/s13225-020-00466-2)) |
| Otu62096 | 1 | k__Fungi(100);p__Ascomycota(100);c__Saccharomycetes(100);o__Saccharomycetales(100);f__Metschnikowiaceae(100);g__Metschnikowia(100);s__Metschnikowia_pulcherrima(100); | Metschnikowia | 13 | Saprotroph | \|Undefined Saprotroph\| | Yeast | NULL | Probable | As nectar-tap_saprotroph (Põlme et al. 2020); Decay sugar-rich substrates (Põlme et al. 2020) | Põlme S, et al. 2020. Fungal Diversity 105:1-16 ((https://doi.org/10.1007/s13225-020-00466-2)) |
| Otu62099 | 1 | k__Fungi(100);p__Ascomycota(100);c__Saccharomycetes(100);o__Saccharomycetales(100);f__Metschnikowiaceae(100);g__Metschnikowia(100);s__Metschnikowia_pulcherrima(100); | Metschnikowia | 13 | Saprotroph | \|Undefined Saprotroph\| | Yeast | NULL | Probable | As nectar-tap_saprotroph (Põlme et al. 2020); Decay sugar-rich substrates (Põlme et al. 2020) | Põlme S, et al. 2020. Fungal Diversity 105:1-16 ((https://doi.org/10.1007/s13225-020-00466-2)) |
| Otu62100 | 1 | k__Fungi(100);p__Ascomycota(100);c__Saccharomycetes(100);o__Saccharomycetales(100);f__Metschnikowiaceae(100);g__Metschnikowia(100);g__Metschnikowia_unclassified(100); | Metschnikowia | 13 | Saprotroph | \|Undefined Saprotroph\| | Yeast | NULL | Probable | As nectar-tap_saprotroph (Põlme et al. 2020); Decay sugar-rich substrates (Põlme et al. 2020) | Põlme S, et al. 2020. Fungal Diversity 105:1-16 ((https://doi.org/10.1007/s13225-020-00466-2)) |
| Otu62101 | 1 | k__Fungi(100);p__Ascomycota(100);c__Saccharomycetes(100);o__Saccharomycetales(100);f__Metschnikowiaceae(100);g__Metschnikowia(100);s__Metschnikowia_pulcherrima(100); | Metschnikowia | 13 | Saprotroph | \|Undefined Saprotroph\| | Yeast | NULL | Probable | As nectar-tap_saprotroph (Põlme et al. 2020); Decay sugar-rich substrates (Põlme et al. 2020) | Põlme S, et al. 2020. Fungal Diversity 105:1-16 ((https://doi.org/10.1007/s13225-020-00466-2)) |
| Otu62102 | 1 | k__Fungi(100);p__Ascomycota(100);c__Saccharomycetes(100);o__Saccharomycetales(100);f__Metschnikowiaceae(100);g__Metschnikowia(100);s__Metschnikowia_pulcherrima(100); | Metschnikowia | 13 | Saprotroph | \|Undefined Saprotroph\| | Yeast | NULL | Probable | As nectar-tap_saprotroph (Põlme et al. 2020); Decay sugar-rich substrates (Põlme et al. 2020) | Põlme S, et al. 2020. Fungal Diversity 105:1-16 ((https://doi.org/10.1007/s13225-020-00466-2)) |
| Otu62103 | 1 | k__Fungi(100);p__Ascomycota(100);c__Saccharomycetes(100);o__Saccharomycetales(100);f__Metschnikowiaceae(100);g__Metschnikowia(100);s__Metschnikowia_pulcherrima(100); | Metschnikowia | 13 | Saprotroph | \|Undefined Saprotroph\| | Yeast | NULL | Probable | As nectar-tap_saprotroph (Põlme et al. 2020); Decay sugar-rich substrates (Põlme et al. 2020) | Põlme S, et al. 2020. Fungal Diversity 105:1-16 ((https://doi.org/10.1007/s13225-020-00466-2)) |
| Otu62104 | 1 | k__Fungi(100);p__Ascomycota(100);c__Saccharomycetes(100);o__Saccharomycetales(100);f__Metschnikowiaceae(100);g__Metschnikowia(100);s__Metschnikowia_pulcherrima(100); | Metschnikowia | 13 | Saprotroph | \|Undefined Saprotroph\| | Yeast | NULL | Probable | As nectar-tap_saprotroph (Põlme et al. 2020); Decay sugar-rich substrates (Põlme et al. 2020) | Põlme S, et al. 2020. Fungal Diversity 105:1-16 ((https://doi.org/10.1007/s13225-020-00466-2)) |
| Otu62105 | 1 | k__Fungi(100);p__Ascomycota(100);c__Saccharomycetes(100);o__Saccharomycetales(100);f__Metschnikowiaceae(100);g__Metschnikowia(100);s__Metschnikowia_pulcherrima(100); | Metschnikowia | 13 | Saprotroph | \|Undefined Saprotroph\| | Yeast | NULL | Probable | As nectar-tap_saprotroph (Põlme et al. 2020); Decay sugar-rich substrates (Põlme et al. 2020) | Põlme S, et al. 2020. Fungal Diversity 105:1-16 ((https://doi.org/10.1007/s13225-020-00466-2)) |
| Otu62108 | 1 | k__Fungi(100);p__Ascomycota(100);c__Saccharomycetes(100);o__Saccharomycetales(100);f__Metschnikowiaceae(100);g__Metschnikowia(100);s__Metschnikowia_pulcherrima(100); | Metschnikowia | 13 | Saprotroph | \|Undefined Saprotroph\| | Yeast | NULL | Probable | As nectar-tap_saprotroph (Põlme et al. 2020); Decay sugar-rich substrates (Põlme et al. 2020) | Põlme S, et al. 2020. Fungal Diversity 105:1-16 ((https://doi.org/10.1007/s13225-020-00466-2)) |
| Otu62110 | 1 | k__Fungi(100);p__Ascomycota(100);c__Saccharomycetes(100);o__Saccharomycetales(100);f__Metschnikowiaceae(100);g__Metschnikowia(100);s__Metschnikowia_pulcherrima(100); | Metschnikowia | 13 | Saprotroph | \|Undefined Saprotroph\| | Yeast | NULL | Probable | As nectar-tap_saprotroph (Põlme et al. 2020); Decay sugar-rich substrates (Põlme et al. 2020) | Põlme S, et al. 2020. Fungal Diversity 105:1-16 ((https://doi.org/10.1007/s13225-020-00466-2)) |
| Otu62113 | 1 | k__Fungi(100);p__Ascomycota(100);c__Saccharomycetes(100);o__Saccharomycetales(100);f__Metschnikowiaceae(100);g__Metschnikowia(100);g__Metschnikowia_unclassified(100); | Metschnikowia | 13 | Saprotroph | \|Undefined Saprotroph\| | Yeast | NULL | Probable | As nectar-tap_saprotroph (Põlme et al. 2020); Decay sugar-rich substrates (Põlme et al. 2020) | Põlme S, et al. 2020. Fungal Diversity 105:1-16 ((https://doi.org/10.1007/s13225-020-00466-2)) |
| Otu62121 | 1 | k__Fungi(100);p__Ascomycota(100);c__Saccharomycetes(100);o__Saccharomycetales(100);f__Metschnikowiaceae(100);g__Metschnikowia(100);s__Metschnikowia_pulcherrima(100); | Metschnikowia | 13 | Saprotroph | \|Undefined Saprotroph\| | Yeast | NULL | Probable | As nectar-tap_saprotroph (Põlme et al. 2020); Decay sugar-rich substrates (Põlme et al. 2020) | Põlme S, et al. 2020. Fungal Diversity 105:1-16 ((https://doi.org/10.1007/s13225-020-00466-2)) |
| Otu62127 | 1 | k__Fungi(100);p__Basidiomycota(100);c__Tremellomycetes(100);o__Tremellales(100);f__Tremellales_family_Incertae_sedis(100);g__Hannaella(100);s__Hannaella_sp(100); | Hannaella | 13 | Pathotroph-Saprotroph | \|Animal Parasite\|-Undefined Saprotroph | Yeast | NULL | Probable | As animal_decomposer (Põlme et al. 2020); Decay animal material (Põlme et al. 2020) | Põlme S, et al. 2020. Fungal Diversity 105:1-16 ((https://doi.org/10.1007/s13225-020-00466-2)) |
| Otu62129 | 1 | k__Fungi(100);p__Ascomycota(100);c__Dothideomycetes(100);o__Pleosporales(100);f__Montagnulaceae(100);f__Montagnulaceae_unclassified(100);f__Montagnulaceae_unclassified(100); | Montagnulaceae | 9 | Pathotroph-Saprotroph | Fungal Parasite-Plant Pathogen-Undefined Saprotroph | Microfungus | NULL | Probable | Species in this clade can be saprobic, plant pathogenic, or mycoparasitic (Zhang et al. 2009) | Zhang Y, et al. 2009. Studies in Mycology 64:85-102 ((https://doi.org/10.3114/sim.2009.64.04)) |
| Otu62131 | 1 | k__Fungi(100);p__Ascomycota(100);c__Saccharomycetes(100);o__Saccharomycetales(100);f__Metschnikowiaceae(100);g__Metschnikowia(100);s__Metschnikowia_pulcherrima(100); | Metschnikowia | 13 | Saprotroph | \|Undefined Saprotroph\| | Yeast | NULL | Probable | As nectar-tap_saprotroph (Põlme et al. 2020); Decay sugar-rich substrates (Põlme et al. 2020) | Põlme S, et al. 2020. Fungal Diversity 105:1-16 ((https://doi.org/10.1007/s13225-020-00466-2)) |
| Otu62136 | 1 | k__Fungi(100);p__Ascomycota(100);c__Saccharomycetes(100);o__Saccharomycetales(100);f__Metschnikowiaceae(100);g__Metschnikowia(100);s__Metschnikowia_pulcherrima(100); | Metschnikowia | 13 | Saprotroph | \|Undefined Saprotroph\| | Yeast | NULL | Probable | As nectar-tap_saprotroph (Põlme et al. 2020); Decay sugar-rich substrates (Põlme et al. 2020) | Põlme S, et al. 2020. Fungal Diversity 105:1-16 ((https://doi.org/10.1007/s13225-020-00466-2)) |
| Otu62137 | 1 | k__Fungi(100);p__Ascomycota(100);c__Saccharomycetes(100);o__Saccharomycetales(100);f__Metschnikowiaceae(100);g__Metschnikowia(100);s__Metschnikowia_pulcherrima(100); | Metschnikowia | 13 | Saprotroph | \|Undefined Saprotroph\| | Yeast | NULL | Probable | As nectar-tap_saprotroph (Põlme et al. 2020); Decay sugar-rich substrates (Põlme et al. 2020) | Põlme S, et al. 2020. Fungal Diversity 105:1-16 ((https://doi.org/10.1007/s13225-020-00466-2)) |
| Otu62139 | 1 | k__Fungi(100);p__Ascomycota(100);c__Saccharomycetes(100);o__Saccharomycetales(100);f__Metschnikowiaceae(100);g__Metschnikowia(100);s__Metschnikowia_pulcherrima(100); | Metschnikowia | 13 | Saprotroph | \|Undefined Saprotroph\| | Yeast | NULL | Probable | As nectar-tap_saprotroph (Põlme et al. 2020); Decay sugar-rich substrates (Põlme et al. 2020) | Põlme S, et al. 2020. Fungal Diversity 105:1-16 ((https://doi.org/10.1007/s13225-020-00466-2)) |
| Otu62140 | 1 | k__Fungi(100);p__Ascomycota(100);c__Saccharomycetes(100);o__Saccharomycetales(100);f__Metschnikowiaceae(100);g__Metschnikowia(100);s__Metschnikowia_pulcherrima(100); | Metschnikowia | 13 | Saprotroph | \|Undefined Saprotroph\| | Yeast | NULL | Probable | As nectar-tap_saprotroph (Põlme et al. 2020); Decay sugar-rich substrates (Põlme et al. 2020) | Põlme S, et al. 2020. Fungal Diversity 105:1-16 ((https://doi.org/10.1007/s13225-020-00466-2)) |
| Otu62145 | 1 | k__Fungi(100);p__Ascomycota(100);c__Saccharomycetes(100);o__Saccharomycetales(100);f__Metschnikowiaceae(100);g__Metschnikowia(100);s__Metschnikowia_pulcherrima(100); | Metschnikowia | 13 | Saprotroph | \|Undefined Saprotroph\| | Yeast | NULL | Probable | As nectar-tap_saprotroph (Põlme et al. 2020); Decay sugar-rich substrates (Põlme et al. 2020) | Põlme S, et al. 2020. Fungal Diversity 105:1-16 ((https://doi.org/10.1007/s13225-020-00466-2)) |
| Otu62147 | 1 | k__Fungi(100);p__Ascomycota(100);c__Saccharomycetes(100);o__Saccharomycetales(100);f__Metschnikowiaceae(100);g__Metschnikowia(100);g__Metschnikowia_unclassified(100); | Metschnikowia | 13 | Saprotroph | \|Undefined Saprotroph\| | Yeast | NULL | Probable | As nectar-tap_saprotroph (Põlme et al. 2020); Decay sugar-rich substrates (Põlme et al. 2020) | Põlme S, et al. 2020. Fungal Diversity 105:1-16 ((https://doi.org/10.1007/s13225-020-00466-2)) |
| Otu62149 | 1 | k__Fungi(100);p__Ascomycota(100);c__Saccharomycetes(100);o__Saccharomycetales(100);f__Metschnikowiaceae(100);g__Metschnikowia(100);g__Metschnikowia_unclassified(100); | Metschnikowia | 13 | Saprotroph | \|Undefined Saprotroph\| | Yeast | NULL | Probable | As nectar-tap_saprotroph (Põlme et al. 2020); Decay sugar-rich substrates (Põlme et al. 2020) | Põlme S, et al. 2020. Fungal Diversity 105:1-16 ((https://doi.org/10.1007/s13225-020-00466-2)) |
| Otu62151 | 1 | k__Fungi(100);p__Ascomycota(100);c__Saccharomycetes(100);o__Saccharomycetales(100);f__Metschnikowiaceae(100);g__Metschnikowia(100);s__Metschnikowia_pulcherrima(100); | Metschnikowia | 13 | Saprotroph | \|Undefined Saprotroph\| | Yeast | NULL | Probable | As nectar-tap_saprotroph (Põlme et al. 2020); Decay sugar-rich substrates (Põlme et al. 2020) | Põlme S, et al. 2020. Fungal Diversity 105:1-16 ((https://doi.org/10.1007/s13225-020-00466-2)) |
| Otu62154 | 1 | k__Fungi(100);p__Ascomycota(100);c__Saccharomycetes(100);o__Saccharomycetales(100);f__Metschnikowiaceae(100);g__Metschnikowia(100);s__Metschnikowia_pulcherrima(100); | Metschnikowia | 13 | Saprotroph | \|Undefined Saprotroph\| | Yeast | NULL | Probable | As nectar-tap_saprotroph (Põlme et al. 2020); Decay sugar-rich substrates (Põlme et al. 2020) | Põlme S, et al. 2020. Fungal Diversity 105:1-16 ((https://doi.org/10.1007/s13225-020-00466-2)) |
| Otu62158 | 1 | k__Fungi(100);p__Ascomycota(100);c__Saccharomycetes(100);o__Saccharomycetales(100);f__Metschnikowiaceae(100);g__Metschnikowia(100);g__Metschnikowia_unclassified(100); | Metschnikowia | 13 | Saprotroph | \|Undefined Saprotroph\| | Yeast | NULL | Probable | As nectar-tap_saprotroph (Põlme et al. 2020); Decay sugar-rich substrates (Põlme et al. 2020) | Põlme S, et al. 2020. Fungal Diversity 105:1-16 ((https://doi.org/10.1007/s13225-020-00466-2)) |
| Otu62159 | 1 | k__Fungi(100);p__Ascomycota(100);c__Saccharomycetes(100);o__Saccharomycetales(100);f__Metschnikowiaceae(100);g__Metschnikowia(100);g__Metschnikowia_unclassified(100); | Metschnikowia | 13 | Saprotroph | \|Undefined Saprotroph\| | Yeast | NULL | Probable | As nectar-tap_saprotroph (Põlme et al. 2020); Decay sugar-rich substrates (Põlme et al. 2020) | Põlme S, et al. 2020. Fungal Diversity 105:1-16 ((https://doi.org/10.1007/s13225-020-00466-2)) |
| Otu62161 | 1 | k__Fungi(100);p__Ascomycota(100);c__Saccharomycetes(100);o__Saccharomycetales(100);f__Metschnikowiaceae(100);g__Metschnikowia(100);s__Metschnikowia_pulcherrima(100); | Metschnikowia | 13 | Saprotroph | \|Undefined Saprotroph\| | Yeast | NULL | Probable | As nectar-tap_saprotroph (Põlme et al. 2020); Decay sugar-rich substrates (Põlme et al. 2020) | Põlme S, et al. 2020. Fungal Diversity 105:1-16 ((https://doi.org/10.1007/s13225-020-00466-2)) |
| Otu62162 | 1 | k__Fungi(100);p__Ascomycota(100);c__Saccharomycetes(100);o__Saccharomycetales(100);f__Metschnikowiaceae(100);g__Metschnikowia(100);s__Metschnikowia_pulcherrima(100); | Metschnikowia | 13 | Saprotroph | \|Undefined Saprotroph\| | Yeast | NULL | Probable | As nectar-tap_saprotroph (Põlme et al. 2020); Decay sugar-rich substrates (Põlme et al. 2020) | Põlme S, et al. 2020. Fungal Diversity 105:1-16 ((https://doi.org/10.1007/s13225-020-00466-2)) |
| Otu62163 | 1 | k__Fungi(100);p__Ascomycota(100);c__Saccharomycetes(100);o__Saccharomycetales(100);f__Metschnikowiaceae(100);g__Metschnikowia(100);s__Metschnikowia_pulcherrima(100); | Metschnikowia | 13 | Saprotroph | \|Undefined Saprotroph\| | Yeast | NULL | Probable | As nectar-tap_saprotroph (Põlme et al. 2020); Decay sugar-rich substrates (Põlme et al. 2020) | Põlme S, et al. 2020. Fungal Diversity 105:1-16 ((https://doi.org/10.1007/s13225-020-00466-2)) |
| Otu62164 | 1 | k__Fungi(100);p__Ascomycota(100);c__Saccharomycetes(100);o__Saccharomycetales(100);f__Metschnikowiaceae(100);g__Metschnikowia(100);s__Metschnikowia_pulcherrima(100); | Metschnikowia | 13 | Saprotroph | \|Undefined Saprotroph\| | Yeast | NULL | Probable | As nectar-tap_saprotroph (Põlme et al. 2020); Decay sugar-rich substrates (Põlme et al. 2020) | Põlme S, et al. 2020. Fungal Diversity 105:1-16 ((https://doi.org/10.1007/s13225-020-00466-2)) |
| Otu62165 | 1 | k__Fungi(100);p__Ascomycota(100);c__Saccharomycetes(100);o__Saccharomycetales(100);f__Metschnikowiaceae(100);g__Metschnikowia(100);s__Metschnikowia_pulcherrima(100); | Metschnikowia | 13 | Saprotroph | \|Undefined Saprotroph\| | Yeast | NULL | Probable | As nectar-tap_saprotroph (Põlme et al. 2020); Decay sugar-rich substrates (Põlme et al. 2020) | Põlme S, et al. 2020. Fungal Diversity 105:1-16 ((https://doi.org/10.1007/s13225-020-00466-2)) |
| Otu62166 | 1 | k__Fungi(100);p__Ascomycota(100);c__Saccharomycetes(100);o__Saccharomycetales(100);f__Metschnikowiaceae(100);g__Metschnikowia(100);s__Metschnikowia_pulcherrima(100); | Metschnikowia | 13 | Saprotroph | \|Undefined Saprotroph\| | Yeast | NULL | Probable | As nectar-tap_saprotroph (Põlme et al. 2020); Decay sugar-rich substrates (Põlme et al. 2020) | Põlme S, et al. 2020. Fungal Diversity 105:1-16 ((https://doi.org/10.1007/s13225-020-00466-2)) |
| Otu62167 | 1 | k__Fungi(100);p__Ascomycota(100);c__Saccharomycetes(100);o__Saccharomycetales(100);f__Metschnikowiaceae(100);g__Metschnikowia(100);s__Metschnikowia_pulcherrima(100); | Metschnikowia | 13 | Saprotroph | \|Undefined Saprotroph\| | Yeast | NULL | Probable | As nectar-tap_saprotroph (Põlme et al. 2020); Decay sugar-rich substrates (Põlme et al. 2020) | Põlme S, et al. 2020. Fungal Diversity 105:1-16 ((https://doi.org/10.1007/s13225-020-00466-2)) |
| Otu62168 | 1 | k__Fungi(100);p__Ascomycota(100);c__Saccharomycetes(100);o__Saccharomycetales(100);f__Metschnikowiaceae(100);g__Metschnikowia(100);s__Metschnikowia_pulcherrima(100); | Metschnikowia | 13 | Saprotroph | \|Undefined Saprotroph\| | Yeast | NULL | Probable | As nectar-tap_saprotroph (Põlme et al. 2020); Decay sugar-rich substrates (Põlme et al. 2020) | Põlme S, et al. 2020. Fungal Diversity 105:1-16 ((https://doi.org/10.1007/s13225-020-00466-2)) |
| Otu62169 | 1 | k__Fungi(100);p__Ascomycota(100);c__Saccharomycetes(100);o__Saccharomycetales(100);f__Metschnikowiaceae(100);g__Metschnikowia(100);s__Metschnikowia_pulcherrima(100); | Metschnikowia | 13 | Saprotroph | \|Undefined Saprotroph\| | Yeast | NULL | Probable | As nectar-tap_saprotroph (Põlme et al. 2020); Decay sugar-rich substrates (Põlme et al. 2020) | Põlme S, et al. 2020. Fungal Diversity 105:1-16 ((https://doi.org/10.1007/s13225-020-00466-2)) |
| Otu62170 | 1 | k__Fungi(100);p__Ascomycota(100);c__Saccharomycetes(100);o__Saccharomycetales(100);f__Metschnikowiaceae(100);g__Metschnikowia(100);s__Metschnikowia_pulcherrima(100); | Metschnikowia | 13 | Saprotroph | \|Undefined Saprotroph\| | Yeast | NULL | Probable | As nectar-tap_saprotroph (Põlme et al. 2020); Decay sugar-rich substrates (Põlme et al. 2020) | Põlme S, et al. 2020. Fungal Diversity 105:1-16 ((https://doi.org/10.1007/s13225-020-00466-2)) |
| Otu62172 | 1 | k__Fungi(100);p__Ascomycota(100);c__Dothideomycetes(100);o__Dothideomycetes_order_Incertae_sedis(100);f__Myxotrichaceae(100);g__Oidiodendron(100);s__Oidiodendron_sp_22MS12(100); | Oidiodendron | 13 | Pathotroph-Saprotroph-Symbiotroph | Endophyte-Ericoid Mycorrhizal-\|Undefined Saprotroph\| | Microfungus | NULL | Probable | As root_endophyte (Põlme et al. 2020); As soil_saprotroph (Põlme et al. 2020); Dark Septate Endophyte; Decay soil (Põlme et al. 2020) | Newsham KK. 2011. New Phytologist 190:783-793 ((https://doi.org/10.1111/j.1469-8137.2010.03611.x)) (as dark septate root endophytes); Põlme S, et al. 2020. Fungal Diversity 105:1-16 ((https://doi.org/10.1007/s13225-020-00466-2)) |
| Otu62173 | 1 | k__Fungi(100);p__Ascomycota(100);c__Saccharomycetes(100);o__Saccharomycetales(100);f__Metschnikowiaceae(100);g__Metschnikowia(100);g__Metschnikowia_unclassified(100); | Metschnikowia | 13 | Saprotroph | \|Undefined Saprotroph\| | Yeast | NULL | Probable | As nectar-tap_saprotroph (Põlme et al. 2020); Decay sugar-rich substrates (Põlme et al. 2020) | Põlme S, et al. 2020. Fungal Diversity 105:1-16 ((https://doi.org/10.1007/s13225-020-00466-2)) |
| Otu62174 | 1 | k__Fungi(100);p__Ascomycota(100);c__Saccharomycetes(100);o__Saccharomycetales(100);f__Metschnikowiaceae(100);g__Metschnikowia(100);g__Metschnikowia_unclassified(100); | Metschnikowia | 13 | Saprotroph | \|Undefined Saprotroph\| | Yeast | NULL | Probable | As nectar-tap_saprotroph (Põlme et al. 2020); Decay sugar-rich substrates (Põlme et al. 2020) | Põlme S, et al. 2020. Fungal Diversity 105:1-16 ((https://doi.org/10.1007/s13225-020-00466-2)) |
| Otu62175 | 1 | k__Fungi(100);p__Ascomycota(100);c__Saccharomycetes(100);o__Saccharomycetales(100);f__Metschnikowiaceae(100);g__Metschnikowia(100);s__Metschnikowia_pulcherrima(100); | Metschnikowia | 13 | Saprotroph | \|Undefined Saprotroph\| | Yeast | NULL | Probable | As nectar-tap_saprotroph (Põlme et al. 2020); Decay sugar-rich substrates (Põlme et al. 2020) | Põlme S, et al. 2020. Fungal Diversity 105:1-16 ((https://doi.org/10.1007/s13225-020-00466-2)) |
| Otu62176 | 1 | k__Fungi(100);p__Ascomycota(100);c__Saccharomycetes(100);o__Saccharomycetales(100);f__Metschnikowiaceae(100);g__Metschnikowia(100);s__Metschnikowia_pulcherrima(100); | Metschnikowia | 13 | Saprotroph | \|Undefined Saprotroph\| | Yeast | NULL | Probable | As nectar-tap_saprotroph (Põlme et al. 2020); Decay sugar-rich substrates (Põlme et al. 2020) | Põlme S, et al. 2020. Fungal Diversity 105:1-16 ((https://doi.org/10.1007/s13225-020-00466-2)) |
| Otu62177 | 1 | k__Fungi(100);p__Ascomycota(100);c__Saccharomycetes(100);o__Saccharomycetales(100);f__Metschnikowiaceae(100);g__Metschnikowia(100);s__Metschnikowia_pulcherrima(100); | Metschnikowia | 13 | Saprotroph | \|Undefined Saprotroph\| | Yeast | NULL | Probable | As nectar-tap_saprotroph (Põlme et al. 2020); Decay sugar-rich substrates (Põlme et al. 2020) | Põlme S, et al. 2020. Fungal Diversity 105:1-16 ((https://doi.org/10.1007/s13225-020-00466-2)) |
| Otu62178 | 1 | k__Fungi(100);p__Ascomycota(100);c__Saccharomycetes(100);o__Saccharomycetales(100);f__Metschnikowiaceae(100);g__Metschnikowia(100);s__Metschnikowia_pulcherrima(100); | Metschnikowia | 13 | Saprotroph | \|Undefined Saprotroph\| | Yeast | NULL | Probable | As nectar-tap_saprotroph (Põlme et al. 2020); Decay sugar-rich substrates (Põlme et al. 2020) | Põlme S, et al. 2020. Fungal Diversity 105:1-16 ((https://doi.org/10.1007/s13225-020-00466-2)) |
| Otu62179 | 1 | k__Fungi(100);p__Ascomycota(100);c__Saccharomycetes(100);o__Saccharomycetales(100);f__Metschnikowiaceae(100);g__Metschnikowia(100);s__Metschnikowia_pulcherrima(100); | Metschnikowia | 13 | Saprotroph | \|Undefined Saprotroph\| | Yeast | NULL | Probable | As nectar-tap_saprotroph (Põlme et al. 2020); Decay sugar-rich substrates (Põlme et al. 2020) | Põlme S, et al. 2020. Fungal Diversity 105:1-16 ((https://doi.org/10.1007/s13225-020-00466-2)) |
| Otu62180 | 1 | k__Fungi(100);p__Ascomycota(100);c__Saccharomycetes(100);o__Saccharomycetales(100);f__Metschnikowiaceae(100);g__Metschnikowia(100);s__Metschnikowia_pulcherrima(100); | Metschnikowia | 13 | Saprotroph | \|Undefined Saprotroph\| | Yeast | NULL | Probable | As nectar-tap_saprotroph (Põlme et al. 2020); Decay sugar-rich substrates (Põlme et al. 2020) | Põlme S, et al. 2020. Fungal Diversity 105:1-16 ((https://doi.org/10.1007/s13225-020-00466-2)) |
| Otu62181 | 1 | k__Fungi(100);p__Ascomycota(100);c__Saccharomycetes(100);o__Saccharomycetales(100);f__Metschnikowiaceae(100);g__Metschnikowia(100);g__Metschnikowia_unclassified(100); | Metschnikowia | 13 | Saprotroph | \|Undefined Saprotroph\| | Yeast | NULL | Probable | As nectar-tap_saprotroph (Põlme et al. 2020); Decay sugar-rich substrates (Põlme et al. 2020) | Põlme S, et al. 2020. Fungal Diversity 105:1-16 ((https://doi.org/10.1007/s13225-020-00466-2)) |
| Otu62182 | 1 | k__Fungi(100);p__Ascomycota(100);c__Saccharomycetes(100);o__Saccharomycetales(100);f__Metschnikowiaceae(100);g__Metschnikowia(100);s__Metschnikowia_pulcherrima(100); | Metschnikowia | 13 | Saprotroph | \|Undefined Saprotroph\| | Yeast | NULL | Probable | As nectar-tap_saprotroph (Põlme et al. 2020); Decay sugar-rich substrates (Põlme et al. 2020) | Põlme S, et al. 2020. Fungal Diversity 105:1-16 ((https://doi.org/10.1007/s13225-020-00466-2)) |
| Otu62184 | 1 | k__Fungi(100);p__Ascomycota(100);c__Saccharomycetes(100);o__Saccharomycetales(100);f__Metschnikowiaceae(100);g__Metschnikowia(100);s__Metschnikowia_pulcherrima(100); | Metschnikowia | 13 | Saprotroph | \|Undefined Saprotroph\| | Yeast | NULL | Probable | As nectar-tap_saprotroph (Põlme et al. 2020); Decay sugar-rich substrates (Põlme et al. 2020) | Põlme S, et al. 2020. Fungal Diversity 105:1-16 ((https://doi.org/10.1007/s13225-020-00466-2)) |
| Otu62185 | 1 | k__Fungi(100);p__Ascomycota(100);c__Saccharomycetes(100);o__Saccharomycetales(100);f__Metschnikowiaceae(100);g__Metschnikowia(100);s__Metschnikowia_pulcherrima(100); | Metschnikowia | 13 | Saprotroph | \|Undefined Saprotroph\| | Yeast | NULL | Probable | As nectar-tap_saprotroph (Põlme et al. 2020); Decay sugar-rich substrates (Põlme et al. 2020) | Põlme S, et al. 2020. Fungal Diversity 105:1-16 ((https://doi.org/10.1007/s13225-020-00466-2)) |
| Otu62186 | 1 | k__Fungi(100);p__Ascomycota(100);c__Saccharomycetes(100);o__Saccharomycetales(100);f__Metschnikowiaceae(100);g__Metschnikowia(100);s__Metschnikowia_pulcherrima(100); | Metschnikowia | 13 | Saprotroph | \|Undefined Saprotroph\| | Yeast | NULL | Probable | As nectar-tap_saprotroph (Põlme et al. 2020); Decay sugar-rich substrates (Põlme et al. 2020) | Põlme S, et al. 2020. Fungal Diversity 105:1-16 ((https://doi.org/10.1007/s13225-020-00466-2)) |
| Otu62187 | 1 | k__Fungi(100);p__Ascomycota(100);c__Saccharomycetes(100);o__Saccharomycetales(100);f__Metschnikowiaceae(100);g__Metschnikowia(100);s__Metschnikowia_pulcherrima(100); | Metschnikowia | 13 | Saprotroph | \|Undefined Saprotroph\| | Yeast | NULL | Probable | As nectar-tap_saprotroph (Põlme et al. 2020); Decay sugar-rich substrates (Põlme et al. 2020) | Põlme S, et al. 2020. Fungal Diversity 105:1-16 ((https://doi.org/10.1007/s13225-020-00466-2)) |
| Otu62189 | 1 | k__Fungi(100);p__Ascomycota(100);c__Saccharomycetes(100);o__Saccharomycetales(100);f__Metschnikowiaceae(100);g__Metschnikowia(100);s__Metschnikowia_pulcherrima(100); | Metschnikowia | 13 | Saprotroph | \|Undefined Saprotroph\| | Yeast | NULL | Probable | As nectar-tap_saprotroph (Põlme et al. 2020); Decay sugar-rich substrates (Põlme et al. 2020) | Põlme S, et al. 2020. Fungal Diversity 105:1-16 ((https://doi.org/10.1007/s13225-020-00466-2)) |
| Otu62192 | 1 | k__Fungi(100);p__Ascomycota(100);c__Saccharomycetes(100);o__Saccharomycetales(100);f__Metschnikowiaceae(100);g__Metschnikowia(100);s__Metschnikowia_pulcherrima(100); | Metschnikowia | 13 | Saprotroph | \|Undefined Saprotroph\| | Yeast | NULL | Probable | As nectar-tap_saprotroph (Põlme et al. 2020); Decay sugar-rich substrates (Põlme et al. 2020) | Põlme S, et al. 2020. Fungal Diversity 105:1-16 ((https://doi.org/10.1007/s13225-020-00466-2)) |
| Otu62195 | 1 | k__Fungi(100);p__Ascomycota(100);c__Saccharomycetes(100);o__Saccharomycetales(100);f__Metschnikowiaceae(100);g__Metschnikowia(100);s__Metschnikowia_pulcherrima(100); | Metschnikowia | 13 | Saprotroph | \|Undefined Saprotroph\| | Yeast | NULL | Probable | As nectar-tap_saprotroph (Põlme et al. 2020); Decay sugar-rich substrates (Põlme et al. 2020) | Põlme S, et al. 2020. Fungal Diversity 105:1-16 ((https://doi.org/10.1007/s13225-020-00466-2)) |
| Otu62196 | 1 | k__Fungi(100);p__Ascomycota(100);c__Saccharomycetes(100);o__Saccharomycetales(100);f__Metschnikowiaceae(100);g__Metschnikowia(100);g__Metschnikowia_unclassified(100); | Metschnikowia | 13 | Saprotroph | \|Undefined Saprotroph\| | Yeast | NULL | Probable | As nectar-tap_saprotroph (Põlme et al. 2020); Decay sugar-rich substrates (Põlme et al. 2020) | Põlme S, et al. 2020. Fungal Diversity 105:1-16 ((https://doi.org/10.1007/s13225-020-00466-2)) |
| Otu62197 | 1 | k__Fungi(100);p__Ascomycota(100);c__Saccharomycetes(100);o__Saccharomycetales(100);f__Metschnikowiaceae(100);g__Metschnikowia(100);s__Metschnikowia_pulcherrima(100); | Metschnikowia | 13 | Saprotroph | \|Undefined Saprotroph\| | Yeast | NULL | Probable | As nectar-tap_saprotroph (Põlme et al. 2020); Decay sugar-rich substrates (Põlme et al. 2020) | Põlme S, et al. 2020. Fungal Diversity 105:1-16 ((https://doi.org/10.1007/s13225-020-00466-2)) |
| Otu62198 | 1 | k__Fungi(100);p__Ascomycota(100);c__Saccharomycetes(100);o__Saccharomycetales(100);f__Metschnikowiaceae(100);g__Metschnikowia(100);g__Metschnikowia_unclassified(100); | Metschnikowia | 13 | Saprotroph | \|Undefined Saprotroph\| | Yeast | NULL | Probable | As nectar-tap_saprotroph (Põlme et al. 2020); Decay sugar-rich substrates (Põlme et al. 2020) | Põlme S, et al. 2020. Fungal Diversity 105:1-16 ((https://doi.org/10.1007/s13225-020-00466-2)) |
| Otu62200 | 1 | k__Fungi(100);p__Ascomycota(100);c__Saccharomycetes(100);o__Saccharomycetales(100);f__Metschnikowiaceae(100);g__Metschnikowia(100);s__Metschnikowia_pulcherrima(100); | Metschnikowia | 13 | Saprotroph | \|Undefined Saprotroph\| | Yeast | NULL | Probable | As nectar-tap_saprotroph (Põlme et al. 2020); Decay sugar-rich substrates (Põlme et al. 2020) | Põlme S, et al. 2020. Fungal Diversity 105:1-16 ((https://doi.org/10.1007/s13225-020-00466-2)) |
| Otu62201 | 1 | k__Fungi(100);p__Ascomycota(100);c__Saccharomycetes(100);o__Saccharomycetales(100);f__Metschnikowiaceae(100);g__Metschnikowia(100);s__Metschnikowia_pulcherrima(100); | Metschnikowia | 13 | Saprotroph | \|Undefined Saprotroph\| | Yeast | NULL | Probable | As nectar-tap_saprotroph (Põlme et al. 2020); Decay sugar-rich substrates (Põlme et al. 2020) | Põlme S, et al. 2020. Fungal Diversity 105:1-16 ((https://doi.org/10.1007/s13225-020-00466-2)) |
| Otu62202 | 1 | k__Fungi(100);p__Ascomycota(100);c__Saccharomycetes(100);o__Saccharomycetales(100);f__Metschnikowiaceae(100);g__Metschnikowia(100);g__Metschnikowia_unclassified(100); | Metschnikowia | 13 | Saprotroph | \|Undefined Saprotroph\| | Yeast | NULL | Probable | As nectar-tap_saprotroph (Põlme et al. 2020); Decay sugar-rich substrates (Põlme et al. 2020) | Põlme S, et al. 2020. Fungal Diversity 105:1-16 ((https://doi.org/10.1007/s13225-020-00466-2)) |
| Otu62203 | 1 | k__Fungi(100);p__Ascomycota(100);c__Saccharomycetes(100);o__Saccharomycetales(100);f__Metschnikowiaceae(100);g__Metschnikowia(100);s__Metschnikowia_pulcherrima(100); | Metschnikowia | 13 | Saprotroph | \|Undefined Saprotroph\| | Yeast | NULL | Probable | As nectar-tap_saprotroph (Põlme et al. 2020); Decay sugar-rich substrates (Põlme et al. 2020) | Põlme S, et al. 2020. Fungal Diversity 105:1-16 ((https://doi.org/10.1007/s13225-020-00466-2)) |

| | | | | | | | | | | | |
|---|---|---|---|---|---|---|---|---|---|---|---|
| Otu62803 | 1 | k__Fungi(100);p__Ascomycota(100);c__Saccharomycetes(100);o__Saccharomycetales(100);f__Metschnikowiaceae(100);g__Metschnikowia(100);s__Metschnikowia_pulcherrima(100); | Metschnikowia | 13 | Saprotroph | \|Undefined Saprotroph\| | Yeast | NULL | Probable | As nectar-tap_saprotroph (Põlme et al. 2020); Decay sugar-rich substrates (Põlme et al. 2020) | Põlme S, et al. 2020. Fungal Diversity 105:1-16 ((https://doi.org/10.1007/s13225-020-00466-2)) |
| Otu62804 | 1 | k__Fungi(100);p__Ascomycota(100);c__Saccharomycetes(100);o__Saccharomycetales(100);f__Metschnikowiaceae(100);g__Metschnikowia(100);g__Metschnikowia_unclassified(100); | Metschnikowia | 13 | Saprotroph | \|Undefined Saprotroph\| | Yeast | NULL | Probable | As nectar-tap_saprotroph (Põlme et al. 2020); Decay sugar-rich substrates (Põlme et al. 2020) | Põlme S, et al. 2020. Fungal Diversity 105:1-16 ((https://doi.org/10.1007/s13225-020-00466-2)) |
| Otu62805 | 1 | k__Fungi(100);p__Ascomycota(100);c__Saccharomycetes(100);o__Saccharomycetales(100);f__Metschnikowiaceae(100);g__Metschnikowia(100);s__Metschnikowia_pulcherrima(100); | Metschnikowia | 13 | Saprotroph | \|Undefined Saprotroph\| | Yeast | NULL | Probable | As nectar-tap_saprotroph (Põlme et al. 2020); Decay sugar-rich substrates (Põlme et al. 2020) | Põlme S, et al. 2020. Fungal Diversity 105:1-16 ((https://doi.org/10.1007/s13225-020-00466-2)) |
| Otu62809 | 1 | k__Fungi(100);p__Ascomycota(100);c__Saccharomycetes(100);o__Saccharomycetales(100);f__Metschnikowiaceae(100);g__Metschnikowia(100);s__Metschnikowia_pulcherrima(100); | Metschnikowia | 13 | Saprotroph | \|Undefined Saprotroph\| | Yeast | NULL | Probable | As nectar-tap_saprotroph (Põlme et al. 2020); Decay sugar-rich substrates (Põlme et al. 2020) | Põlme S, et al. 2020. Fungal Diversity 105:1-16 ((https://doi.org/10.1007/s13225-020-00466-2)) |
| Otu62810 | 1 | k__Fungi(100);p__Ascomycota(100);c__Saccharomycetes(100);o__Saccharomycetales(100);f__Metschnikowiaceae(100);g__Metschnikowia(100);s__Metschnikowia_pulcherrima(100); | Metschnikowia | 13 | Saprotroph | \|Undefined Saprotroph\| | Yeast | NULL | Probable | As nectar-tap_saprotroph (Põlme et al. 2020); Decay sugar-rich substrates (Põlme et al. 2020) | Põlme S, et al. 2020. Fungal Diversity 105:1-16 ((https://doi.org/10.1007/s13225-020-00466-2)) |
| Otu62813 | 1 | k__Fungi(100);p__Ascomycota(100);c__Saccharomycetes(100);o__Saccharomycetales(100);f__Metschnikowiaceae(100);g__Metschnikowia(100);s__Metschnikowia_pulcherrima(100); | Metschnikowia | 13 | Saprotroph | \|Undefined Saprotroph\| | Yeast | NULL | Probable | As nectar-tap_saprotroph (Põlme et al. 2020); Decay sugar-rich substrates (Põlme et al. 2020) | Põlme S, et al. 2020. Fungal Diversity 105:1-16 ((https://doi.org/10.1007/s13225-020-00466-2)) |
| Otu62825 | 1 | k__Fungi(100);p__Ascomycota(100);c__Saccharomycetes(100);o__Saccharomycetales(100);f__Metschnikowiaceae(100);g__Metschnikowia(100);s__Metschnikowia_pulcherrima(100); | Metschnikowia | 13 | Saprotroph | \|Undefined Saprotroph\| | Yeast | NULL | Probable | As nectar-tap_saprotroph (Põlme et al. 2020); Decay sugar-rich substrates (Põlme et al. 2020) | Põlme S, et al. 2020. Fungal Diversity 105:1-16 ((https://doi.org/10.1007/s13225-020-00466-2)) |
| Otu62827 | 1 | k__Fungi(100);p__Ascomycota(100);c__Saccharomycetes(100);o__Saccharomycetales(100);f__Metschnikowiaceae(100);g__Metschnikowia(100);s__Metschnikowia_pulcherrima(100); | Metschnikowia | 13 | Saprotroph | \|Undefined Saprotroph\| | Yeast | NULL | Probable | As nectar-tap_saprotroph (Põlme et al. 2020); Decay sugar-rich substrates (Põlme et al. 2020) | Põlme S, et al. 2020. Fungal Diversity 105:1-16 ((https://doi.org/10.1007/s13225-020-00466-2)) |
| Otu62828 | 1 | k__Fungi(100);p__Ascomycota(100);c__Saccharomycetes(100);o__Saccharomycetales(100);f__Metschnikowiaceae(100);g__Metschnikowia(100);s__Metschnikowia_pulcherrima(100); | Metschnikowia | 13 | Saprotroph | \|Undefined Saprotroph\| | Yeast | NULL | Probable | As nectar-tap_saprotroph (Põlme et al. 2020); Decay sugar-rich substrates (Põlme et al. 2020) | Põlme S, et al. 2020. Fungal Diversity 105:1-16 ((https://doi.org/10.1007/s13225-020-00466-2)) |
| Otu62831 | 1 | k__Fungi(100);p__Ascomycota(100);c__Saccharomycetes(100);o__Saccharomycetales(100);f__Metschnikowiaceae(100);g__Metschnikowia(100);s__Metschnikowia_pulcherrima(100); | Metschnikowia | 13 | Saprotroph | \|Undefined Saprotroph\| | Yeast | NULL | Probable | As nectar-tap_saprotroph (Põlme et al. 2020); Decay sugar-rich substrates (Põlme et al. 2020) | Põlme S, et al. 2020. Fungal Diversity 105:1-16 ((https://doi.org/10.1007/s13225-020-00466-2)) |
| Otu62832 | 1 | k__Fungi(100);p__Ascomycota(100);c__Saccharomycetes(100);o__Saccharomycetales(100);f__Metschnikowiaceae(100);g__Metschnikowia(100);s__Metschnikowia_pulcherrima(100); | Metschnikowia | 13 | Saprotroph | \|Undefined Saprotroph\| | Yeast | NULL | Probable | As nectar-tap_saprotroph (Põlme et al. 2020); Decay sugar-rich substrates (Põlme et al. 2020) | Põlme S, et al. 2020. Fungal Diversity 105:1-16 ((https://doi.org/10.1007/s13225-020-00466-2)) |
| Otu62842 | 1 | k__Fungi(100);p__Ascomycota(100);c__Saccharomycetes(100);o__Saccharomycetales(100);f__Metschnikowiaceae(100);g__Metschnikowia(100);g__Metschnikowia_unclassified(100); | Metschnikowia | 13 | Saprotroph | \|Undefined Saprotroph\| | Yeast | NULL | Probable | As nectar-tap_saprotroph (Põlme et al. 2020); Decay sugar-rich substrates (Põlme et al. 2020) | Põlme S, et al. 2020. Fungal Diversity 105:1-16 ((https://doi.org/10.1007/s13225-020-00466-2)) |
| Otu62846 | 1 | k__Fungi(100);p__Ascomycota(100);c__Saccharomycetes(100);o__Saccharomycetales(100);f__Metschnikowiaceae(100);g__Metschnikowia(100);s__Metschnikowia_pulcherrima(100); | Metschnikowia | 13 | Saprotroph | \|Undefined Saprotroph\| | Yeast | NULL | Probable | As nectar-tap_saprotroph (Põlme et al. 2020); Decay sugar-rich substrates (Põlme et al. 2020) | Põlme S, et al. 2020. Fungal Diversity 105:1-16 ((https://doi.org/10.1007/s13225-020-00466-2)) |
| Otu62862 | 1 | k__Fungi(100);p__Ascomycota(100);c__Saccharomycetes(100);o__Saccharomycetales(100);f__Metschnikowiaceae(100);g__Metschnikowia(100);s__Metschnikowia_pulcherrima(100); | Metschnikowia | 13 | Saprotroph | \|Undefined Saprotroph\| | Yeast | NULL | Probable | As nectar-tap_saprotroph (Põlme et al. 2020); Decay sugar-rich substrates (Põlme et al. 2020) | Põlme S, et al. 2020. Fungal Diversity 105:1-16 ((https://doi.org/10.1007/s13225-020-00466-2)) |
| Otu62863 | 1 | k__Fungi(100);p__Ascomycota(100);c__Saccharomycetes(100);o__Saccharomycetales(100);f__Metschnikowiaceae(100);g__Metschnikowia(100);s__Metschnikowia_pulcherrima(100); | Metschnikowia | 13 | Saprotroph | \|Undefined Saprotroph\| | Yeast | NULL | Probable | As nectar-tap_saprotroph (Põlme et al. 2020); Decay sugar-rich substrates (Põlme et al. 2020) | Põlme S, et al. 2020. Fungal Diversity 105:1-16 ((https://doi.org/10.1007/s13225-020-00466-2)) |
| Otu62866 | 1 | k__Fungi(100);p__Ascomycota(100);c__Saccharomycetes(100);o__Saccharomycetales(100);f__Metschnikowiaceae(100);g__Metschnikowia(100);s__Metschnikowia_pulcherrima(100); | Metschnikowia | 13 | Saprotroph | \|Undefined Saprotroph\| | Yeast | NULL | Probable | As nectar-tap_saprotroph (Põlme et al. 2020); Decay sugar-rich substrates (Põlme et al. 2020) | Põlme S, et al. 2020. Fungal Diversity 105:1-16 ((https://doi.org/10.1007/s13225-020-00466-2)) |
| Otu62867 | 1 | k__Fungi(100);p__Ascomycota(100);c__Saccharomycetes(100);o__Saccharomycetales(100);f__Metschnikowiaceae(100);g__Metschnikowia(100);s__Metschnikowia_pulcherrima(100); | Metschnikowia | 13 | Saprotroph | \|Undefined Saprotroph\| | Yeast | NULL | Probable | As nectar-tap_saprotroph (Põlme et al. 2020); Decay sugar-rich substrates (Põlme et al. 2020) | Põlme S, et al. 2020. Fungal Diversity 105:1-16 ((https://doi.org/10.1007/s13225-020-00466-2)) |
| Otu62868 | 1 | k__Fungi(100);p__Ascomycota(100);c__Saccharomycetes(100);o__Saccharomycetales(100);f__Metschnikowiaceae(100);g__Metschnikowia(100);s__Metschnikowia_pulcherrima(100); | Metschnikowia | 13 | Saprotroph | \|Undefined Saprotroph\| | Yeast | NULL | Probable | As nectar-tap_saprotroph (Põlme et al. 2020); Decay sugar-rich substrates (Põlme et al. 2020) | Põlme S, et al. 2020. Fungal Diversity 105:1-16 ((https://doi.org/10.1007/s13225-020-00466-2)) |
| Otu62870 | 1 | k__Fungi(100);p__Ascomycota(100);c__Saccharomycetes(100);o__Saccharomycetales(100);f__Metschnikowiaceae(100);g__Metschnikowia(100);g__Metschnikowia_unclassified(100); | Metschnikowia | 13 | Saprotroph | \|Undefined Saprotroph\| | Yeast | NULL | Probable | As nectar-tap_saprotroph (Põlme et al. 2020); Decay sugar-rich substrates (Põlme et al. 2020) | Põlme S, et al. 2020. Fungal Diversity 105:1-16 ((https://doi.org/10.1007/s13225-020-00466-2)) |
| Otu62872 | 1 | k__Fungi(100);p__Ascomycota(100);c__Saccharomycetes(100);o__Saccharomycetales(100);f__Metschnikowiaceae(100);g__Metschnikowia(100);s__Metschnikowia_pulcherrima(100); | Metschnikowia | 13 | Saprotroph | \|Undefined Saprotroph\| | Yeast | NULL | Probable | As nectar-tap_saprotroph (Põlme et al. 2020); Decay sugar-rich substrates (Põlme et al. 2020) | Põlme S, et al. 2020. Fungal Diversity 105:1-16 ((https://doi.org/10.1007/s13225-020-00466-2)) |
| Otu62873 | 1 | k__Fungi(100);p__Ascomycota(100);c__Saccharomycetes(100);o__Saccharomycetales(100);f__Metschnikowiaceae(100);g__Metschnikowia(100);g__Metschnikowia_unclassified(100); | Metschnikowia | 13 | Saprotroph | \|Undefined Saprotroph\| | Yeast | NULL | Probable | As nectar-tap_saprotroph (Põlme et al. 2020); Decay sugar-rich substrates (Põlme et al. 2020) | Põlme S, et al. 2020. Fungal Diversity 105:1-16 ((https://doi.org/10.1007/s13225-020-00466-2)) |
| Otu62875 | 1 | k__Fungi(100);p__Ascomycota(100);c__Saccharomycetes(100);o__Saccharomycetales(100);f__Metschnikowiaceae(100);g__Metschnikowia(100);s__Metschnikowia_pulcherrima(100); | Metschnikowia | 13 | Saprotroph | \|Undefined Saprotroph\| | Yeast | NULL | Probable | As nectar-tap_saprotroph (Põlme et al. 2020); Decay sugar-rich substrates (Põlme et al. 2020) | Põlme S, et al. 2020. Fungal Diversity 105:1-16 ((https://doi.org/10.1007/s13225-020-00466-2)) |
| Otu62878 | 1 | k__Fungi(100);p__Ascomycota(100);c__Saccharomycetes(100);o__Saccharomycetales(100);f__Metschnikowiaceae(100);g__Metschnikowia(100);s__Metschnikowia_pulcherrima(100); | Metschnikowia | 13 | Saprotroph | \|Undefined Saprotroph\| | Yeast | NULL | Probable | As nectar-tap_saprotroph (Põlme et al. 2020); Decay sugar-rich substrates (Põlme et al. 2020) | Põlme S, et al. 2020. Fungal Diversity 105:1-16 ((https://doi.org/10.1007/s13225-020-00466-2)) |
| Otu62879 | 1 | k__Fungi(100);p__Ascomycota(100);c__Saccharomycetes(100);o__Saccharomycetales(100);f__Metschnikowiaceae(100);g__Metschnikowia(100);s__Metschnikowia_pulcherrima(100); | Metschnikowia | 13 | Saprotroph | \|Undefined Saprotroph\| | Yeast | NULL | Probable | As nectar-tap_saprotroph (Põlme et al. 2020); Decay sugar-rich substrates (Põlme et al. 2020) | Põlme S, et al. 2020. Fungal Diversity 105:1-16 ((https://doi.org/10.1007/s13225-020-00466-2)) |
| Otu62881 | 1 | k__Fungi(100);p__Ascomycota(100);c__Saccharomycetes(100);o__Saccharomycetales(100);f__Metschnikowiaceae(100);g__Metschnikowia(100);s__Metschnikowia_pulcherrima(100); | Metschnikowia | 13 | Saprotroph | \|Undefined Saprotroph\| | Yeast | NULL | Probable | As nectar-tap_saprotroph (Põlme et al. 2020); Decay sugar-rich substrates (Põlme et al. 2020) | Põlme S, et al. 2020. Fungal Diversity 105:1-16 ((https://doi.org/10.1007/s13225-020-00466-2)) |
| Otu62888 | 1 | k__Fungi(100);p__Ascomycota(100);c__Saccharomycetes(100);o__Saccharomycetales(100);f__Metschnikowiaceae(100);g__Metschnikowia(100);g__Metschnikowia_unclassified(100); | Metschnikowia | 13 | Saprotroph | \|Undefined Saprotroph\| | Yeast | NULL | Probable | As nectar-tap_saprotroph (Põlme et al. 2020); Decay sugar-rich substrates (Põlme et al. 2020) | Põlme S, et al. 2020. Fungal Diversity 105:1-16 ((https://doi.org/10.1007/s13225-020-00466-2)) |
| Otu62889 | 1 | k__Fungi(100);p__Ascomycota(100);c__Saccharomycetes(100);o__Saccharomycetales(100);f__Metschnikowiaceae(100);g__Metschnikowia(100);g__Metschnikowia_unclassified(100); | Metschnikowia | 13 | Saprotroph | \|Undefined Saprotroph\| | Yeast | NULL | Probable | As nectar-tap_saprotroph (Põlme et al. 2020); Decay sugar-rich substrates (Põlme et al. 2020) | Põlme S, et al. 2020. Fungal Diversity 105:1-16 ((https://doi.org/10.1007/s13225-020-00466-2)) |
| Otu62890 | 1 | k__Fungi(100);p__Ascomycota(100);c__Saccharomycetes(100);o__Saccharomycetales(100);f__Metschnikowiaceae(100);g__Metschnikowia(100);g__Metschnikowia_unclassified(100); | Metschnikowia | 13 | Saprotroph | \|Undefined Saprotroph\| | Yeast | NULL | Probable | As nectar-tap_saprotroph (Põlme et al. 2020); Decay sugar-rich substrates (Põlme et al. 2020) | Põlme S, et al. 2020. Fungal Diversity 105:1-16 ((https://doi.org/10.1007/s13225-020-00466-2)) |
| Otu62891 | 1 | k__Fungi(100);p__Ascomycota(100);c__Saccharomycetes(100);o__Saccharomycetales(100);f__Metschnikowiaceae(100);g__Metschnikowia(100);s__Metschnikowia_pulcherrima(100); | Metschnikowia | 13 | Saprotroph | \|Undefined Saprotroph\| | Yeast | NULL | Probable | As nectar-tap_saprotroph (Põlme et al. 2020); Decay sugar-rich substrates (Põlme et al. 2020) | Põlme S, et al. 2020. Fungal Diversity 105:1-16 ((https://doi.org/10.1007/s13225-020-00466-2)) |
| Otu62892 | 1 | k__Fungi(100);p__Ascomycota(100);c__Saccharomycetes(100);o__Saccharomycetales(100);f__Metschnikowiaceae(100);g__Metschnikowia(100);s__Metschnikowia_pulcherrima(100); | Metschnikowia | 13 | Saprotroph | \|Undefined Saprotroph\| | Yeast | NULL | Probable | As nectar-tap_saprotroph (Põlme et al. 2020); Decay sugar-rich substrates (Põlme et al. 2020) | Põlme S, et al. 2020. Fungal Diversity 105:1-16 ((https://doi.org/10.1007/s13225-020-00466-2)) |
| Otu62895 | 1 | k__Fungi(100);p__Ascomycota(100);c__Saccharomycetes(100);o__Saccharomycetales(100);f__Metschnikowiaceae(100);g__Metschnikowia(100);s__Metschnikowia_pulcherrima(100); | Metschnikowia | 13 | Saprotroph | \|Undefined Saprotroph\| | Yeast | NULL | Probable | As nectar-tap_saprotroph (Põlme et al. 2020); Decay sugar-rich substrates (Põlme et al. 2020) | Põlme S, et al. 2020. Fungal Diversity 105:1-16 ((https://doi.org/10.1007/s13225-020-00466-2)) |
| Otu62897 | 1 | k__Fungi(100);p__Ascomycota(100);c__Saccharomycetes(100);o__Saccharomycetales(100);f__Metschnikowiaceae(100);g__Metschnikowia(100);s__Metschnikowia_pulcherrima(100); | Metschnikowia | 13 | Saprotroph | \|Undefined Saprotroph\| | Yeast | NULL | Probable | As nectar-tap_saprotroph (Põlme et al. 2020); Decay sugar-rich substrates (Põlme et al. 2020) | Põlme S, et al. 2020. Fungal Diversity 105:1-16 ((https://doi.org/10.1007/s13225-020-00466-2)) |
| Otu62898 | 1 | k__Fungi(100);p__Ascomycota(100);c__Saccharomycetes(100);o__Saccharomycetales(100);f__Metschnikowiaceae(100);g__Metschnikowia(100);s__Metschnikowia_pulcherrima(100); | Metschnikowia | 13 | Saprotroph | \|Undefined Saprotroph\| | Yeast | NULL | Probable | As nectar-tap_saprotroph (Põlme et al. 2020); Decay sugar-rich substrates (Põlme et al. 2020) | Põlme S, et al. 2020. Fungal Diversity 105:1-16 ((https://doi.org/10.1007/s13225-020-00466-2)) |
| Otu62900 | 1 | k__Fungi(100);p__Ascomycota(100);c__Saccharomycetes(100);o__Saccharomycetales(100);f__Metschnikowiaceae(100);g__Metschnikowia(100);s__Metschnikowia_pulcherrima(100); | Metschnikowia | 13 | Saprotroph | \|Undefined Saprotroph\| | Yeast | NULL | Probable | As nectar-tap_saprotroph (Põlme et al. 2020); Decay sugar-rich substrates (Põlme et al. 2020) | Põlme S, et al. 2020. Fungal Diversity 105:1-16 ((https://doi.org/10.1007/s13225-020-00466-2)) |
| Otu62904 | 1 | k__Fungi(100);p__Ascomycota(100);c__Saccharomycetes(100);o__Saccharomycetales(100);f__Metschnikowiaceae(100);g__Metschnikowia(100);g__Metschnikowia_unclassified(100); | Metschnikowia | 13 | Saprotroph | \|Undefined Saprotroph\| | Yeast | NULL | Probable | As nectar-tap_saprotroph (Põlme et al. 2020); Decay sugar-rich substrates (Põlme et al. 2020) | Põlme S, et al. 2020. Fungal Diversity 105:1-16 ((https://doi.org/10.1007/s13225-020-00466-2)) |
| Otu62905 | 1 | k__Fungi(100);p__Ascomycota(100);c__Saccharomycetes(100);o__Saccharomycetales(100);f__Metschnikowiaceae(100);g__Metschnikowia(100);s__Metschnikowia_pulcherrima(100); | Metschnikowia | 13 | Saprotroph | \|Undefined Saprotroph\| | Yeast | NULL | Probable | As nectar-tap_saprotroph (Põlme et al. 2020); Decay sugar-rich substrates (Põlme et al. 2020) | Põlme S, et al. 2020. Fungal Diversity 105:1-16 ((https://doi.org/10.1007/s13225-020-00466-2)) |
| Otu62907 | 1 | k__Fungi(100);p__Ascomycota(100);c__Saccharomycetes(100);o__Saccharomycetales(100);f__Metschnikowiaceae(100);g__Metschnikowia(100);s__Metschnikowia_pulcherrima(100); | Metschnikowia | 13 | Saprotroph | \|Undefined Saprotroph\| | Yeast | NULL | Probable | As nectar-tap_saprotroph (Põlme et al. 2020); Decay sugar-rich substrates (Põlme et al. 2020) | Põlme S, et al. 2020. Fungal Diversity 105:1-16 ((https://doi.org/10.1007/s13225-020-00466-2)) |
| Otu62908 | 1 | k__Fungi(100);p__Ascomycota(100);c__Saccharomycetes(100);o__Saccharomycetales(100);f__Metschnikowiaceae(100);g__Metschnikowia(100);g__Metschnikowia_unclassified(100); | Metschnikowia | 13 | Saprotroph | \|Undefined Saprotroph\| | Yeast | NULL | Probable | As nectar-tap_saprotroph (Põlme et al. 2020); Decay sugar-rich substrates (Põlme et al. 2020) | Põlme S, et al. 2020. Fungal Diversity 105:1-16 ((https://doi.org/10.1007/s13225-020-00466-2)) |
| Otu62909 | 1 | k__Fungi(100);p__Ascomycota(100);c__Saccharomycetes(100);o__Saccharomycetales(100);f__Metschnikowiaceae(100);g__Metschnikowia(100);s__Metschnikowia_pulcherrima(100); | Metschnikowia | 13 | Saprotroph | \|Undefined Saprotroph\| | Yeast | NULL | Probable | As nectar-tap_saprotroph (Põlme et al. 2020); Decay sugar-rich substrates (Põlme et al. 2020) | Põlme S, et al. 2020. Fungal Diversity 105:1-16 ((https://doi.org/10.1007/s13225-020-00466-2)) |
| Otu62911 | 1 | k__Fungi(100);p__Ascomycota(100);c__Saccharomycetes(100);o__Saccharomycetales(100);f__Metschnikowiaceae(100);g__Metschnikowia(100);s__Metschnikowia_pulcherrima(100); | Metschnikowia | 13 | Saprotroph | \|Undefined Saprotroph\| | Yeast | NULL | Probable | As nectar-tap_saprotroph (Põlme et al. 2020); Decay sugar-rich substrates (Põlme et al. 2020) | Põlme S, et al. 2020. Fungal Diversity 105:1-16 ((https://doi.org/10.1007/s13225-020-00466-2)) |
| Otu62912 | 1 | k__Fungi(100);p__Ascomycota(100);c__Saccharomycetes(100);o__Saccharomycetales(100);f__Metschnikowiaceae(100);g__Metschnikowia(100);s__Metschnikowia_pulcherrima(100); | Metschnikowia | 13 | Saprotroph | \|Undefined Saprotroph\| | Yeast | NULL | Probable | As nectar-tap_saprotroph (Põlme et al. 2020); Decay sugar-rich substrates (Põlme et al. 2020) | Põlme S, et al. 2020. Fungal Diversity 105:1-16 ((https://doi.org/10.1007/s13225-020-00466-2)) |
| Otu62925 | 1 | k__Fungi(100);p__Ascomycota(100);c__Saccharomycetes(100);o__Saccharomycetales(100);f__Metschnikowiaceae(100);g__Metschnikowia(100);s__Metschnikowia_pulcherrima(100); | Metschnikowia | 13 | Saprotroph | \|Undefined Saprotroph\| | Yeast | NULL | Probable | As nectar-tap_saprotroph (Põlme et al. 2020); Decay sugar-rich substrates (Põlme et al. 2020) | Põlme S, et al. 2020. Fungal Diversity 105:1-16 ((https://doi.org/10.1007/s13225-020-00466-2)) |
| Otu62931 | 1 | k__Fungi(100);p__Ascomycota(100);c__Saccharomycetes(100);o__Saccharomycetales(100);f__Saccharomycetales_family_Incertae_sedis(100);g__Candida(100);g__Candida_unclassified(100); | Candida | 13 | Pathotroph-Saprotroph-Symbiotroph | Animal Pathogen-Endophyte-Endosymbiont-Epiphyte-\|Undefined Saprotroph\| | Dimorphic Yeast-Yeast | NULL | Probable | As nectar-tap_saprotroph (Põlme et al. 2020); Decay sugar-rich substrates (Põlme et al. 2020); This asexual genus is polyphyletic and contain species that function very differently depend on what they are. Caution should be taken when interpreting guilds for Candida species. | Manolakaki D, et al. 2010. Virulence. 1: 367-375 ((https://doi.org/10.4161/viru.1.5.12796)); Mekbib SB, et al. 2011. Tropical Plant Pathology 36:233-240 ((https://doi.org/10.1590/S1982-56762011000400004)); Põlme S, et al. 2020. Fungal Diversity 105:1-16 ((https://doi.org/10.1007/s13225-020-00466-2)) |
| Otu62932 | 1 | k__Fungi(100);p__Ascomycota(100);c__Saccharomycetes(100);o__Saccharomycetales(100);f__Metschnikowiaceae(100);g__Metschnikowia(100);s__Metschnikowia_pulcherrima(100); | Metschnikowia | 13 | Saprotroph | \|Undefined Saprotroph\| | Yeast | NULL | Probable | As nectar-tap_saprotroph (Põlme et al. 2020); Decay sugar-rich substrates (Põlme et al. 2020) | Põlme S, et al. 2020. Fungal Diversity 105:1-16 ((https://doi.org/10.1007/s13225-020-00466-2)) |
| Otu62933 | 1 | k__Fungi(100);p__Ascomycota(100);c__Saccharomycetes(100);o__Saccharomycetales(100);f__Metschnikowiaceae(100);g__Metschnikowia(100);s__Metschnikowia_pulcherrima(100); | Metschnikowia | 13 | Saprotroph | \|Undefined Saprotroph\| | Yeast | NULL | Probable | As nectar-tap_saprotroph (Põlme et al. 2020); Decay sugar-rich substrates (Põlme et al. 2020) | Põlme S, et al. 2020. Fungal Diversity 105:1-16 ((https://doi.org/10.1007/s13225-020-00466-2)) |
| Otu62938 | 1 | k__Fungi(100);p__Ascomycota(100);c__Saccharomycetes(100);o__Saccharomycetales(100);f__Metschnikowiaceae(100);g__Metschnikowia(100);s__Metschnikowia_pulcherrima(100); | Metschnikowia | 13 | Saprotroph | \|Undefined Saprotroph\| | Yeast | NULL | Probable | As nectar-tap_saprotroph (Põlme et al. 2020); Decay sugar-rich substrates (Põlme et al. 2020) | Põlme S, et al. 2020. Fungal Diversity 105:1-16 ((https://doi.org/10.1007/s13225-020-00466-2)) |
| Otu62939 | 1 | k__Fungi(100);p__Ascomycota(100);c__Saccharomycetes(100);o__Saccharomycetales(100);f__Metschnikowiaceae(100);g__Metschnikowia(100);g__Metschnikowia_unclassified(100); | Metschnikowia | 13 | Saprotroph | \|Undefined Saprotroph\| | Yeast | NULL | Probable | As nectar-tap_saprotroph (Põlme et al. 2020); Decay sugar-rich substrates (Põlme et al. 2020) | Põlme S, et al. 2020. Fungal Diversity 105:1-16 ((https://doi.org/10.1007/s13225-020-00466-2)) |
| Otu62942 | 1 | k__Fungi(100);p__Ascomycota(100);c__Saccharomycetes(100);o__Saccharomycetales(100);f__Saccharomycetales_family_Incertae_sedis(100);g__Candida(100);g__Candida_unclassified(100); | Candida | 13 | Pathotroph-Saprotroph-Symbiotroph | Animal Pathogen-Endophyte-Endosymbiont-Epiphyte-\|Undefined Saprotroph\| | Dimorphic Yeast-Yeast | NULL | Probable | As nectar-tap_saprotroph (Põlme et al. 2020); Decay sugar-rich substrates (Põlme et al. 2020); This asexual genus is polyphyletic and contain species that function very differently depend on what they are. Caution should be taken when interpreting guilds for Candida species. | Manolakaki D, et al. 2010. Virulence. 1: 367-375 ((https://doi.org/10.4161/viru.1.5.12796)); Mekbib SB, et al. 2011. Tropical Plant Pathology 36:233-240 ((https://doi.org/10.1590/S1982-56762011000400004)); Põlme S, et al. 2020. Fungal Diversity 105:1-16 ((https://doi.org/10.1007/s13225-020-00466-2)) |
| Otu62943 | 1 | k__Fungi(100);p__Ascomycota(100);c__Saccharomycetes(100);o__Saccharomycetales(100);f__Metschnikowiaceae(100);g__Metschnikowia(100);s__Metschnikowia_pulcherrima(100); | Metschnikowia | 13 | Saprotroph | \|Undefined Saprotroph\| | Yeast | NULL | Probable | As nectar-tap_saprotroph (Põlme et al. 2020); Decay sugar-rich substrates (Põlme et al. 2020) | Põlme S, et al. 2020. Fungal Diversity 105:1-16 ((https://doi.org/10.1007/s13225-020-00466-2)) |
| Otu62945 | 1 | k__Fungi(100);p__Ascomycota(100);c__Saccharomycetes(100);o__Saccharomycetales(100);f__Metschnikowiaceae(100);g__Metschnikowia(100);g__Metschnikowia_unclassified(100); | Metschnikowia | 13 | Saprotroph | \|Undefined Saprotroph\| | Yeast | NULL | Probable | As nectar-tap_saprotroph (Põlme et al. 2020); Decay sugar-rich substrates (Põlme et al. 2020) | Põlme S, et al. 2020. Fungal Diversity 105:1-16 ((https://doi.org/10.1007/s13225-020-00466-2)) |
| Otu62950 | 1 | k__Fungi(100);p__Ascomycota(100);c__Saccharomycetes(100);o__Saccharomycetales(100);f__Metschnikowiaceae(100);g__Metschnikowia(100);s__Metschnikowia_pulcherrima(100); | Metschnikowia | 13 | Saprotroph | \|Undefined Saprotroph\| | Yeast | NULL | Probable | As nectar-tap_saprotroph (Põlme et al. 2020); Decay sugar-rich substrates (Põlme et al. 2020) | Põlme S, et al. 2020. Fungal Diversity 105:1-16 ((https://doi.org/10.1007/s13225-020-00466-2)) |
| Otu62951 | 1 | k__Fungi(100);p__Ascomycota(100);c__Saccharomycetes(100);o__Saccharomycetales(100);f__Metschnikowiaceae(100);g__Metschnikowia(100);s__Metschnikowia_pulcherrima(100); | Metschnikowia | 13 | Saprotroph | \|Undefined Saprotroph\| | Yeast | NULL | Probable | As nectar-tap_saprotroph (Põlme et al. 2020); Decay sugar-rich substrates (Põlme et al. 2020) | Põlme S, et al. 2020. Fungal Diversity 105:1-16 ((https://doi.org/10.1007/s13225-020-00466-2)) |
| Otu62953 | 1 | k__Fungi(100);p__Ascomycota(100);c__Saccharomycetes(100);o__Saccharomycetales(100);f__Metschnikowiaceae(100);g__Metschnikowia(100);s__Metschnikowia_pulcherrima(100); | Metschnikowia | 13 | Saprotroph | \|Undefined Saprotroph\| | Yeast | NULL | Probable | As nectar-tap_saprotroph (Põlme et al. 2020); Decay sugar-rich substrates (Põlme et al. 2020) | Põlme S, et al. 2020. Fungal Diversity 105:1-16 ((https://doi.org/10.1007/s13225-020-00466-2)) |
| Otu62954 | 1 | k__Fungi(100);p__Ascomycota(100);c__Saccharomycetes(100);o__Saccharomycetales(100);f__Metschnikowiaceae(100);g__Metschnikowia(100);s__Metschnikowia_pulcherrima(100); | Metschnikowia | 13 | Saprotroph | \|Undefined Saprotroph\| | Yeast | NULL | Probable | As nectar-tap_saprotroph (Põlme et al. 2020); Decay sugar-rich substrates (Põlme et al. 2020) | Põlme S, et al. 2020. Fungal Diversity 105:1-16 ((https://doi.org/10.1007/s13225-020-00466-2)) |
| Otu62955 | 1 | k__Fungi(100);p__Ascomycota(100);c__Saccharomycetes(100);o__Saccharomycetales(100);f__Metschnikowiaceae(100);g__Metschnikowia(100);s__Metschnikowia_pulcherrima(100); | Metschnikowia | 13 | Saprotroph | \|Undefined Saprotroph\| | Yeast | NULL | Probable | As nectar-tap_saprotroph (Põlme et al. 2020); Decay sugar-rich substrates (Põlme et al. 2020) | Põlme S, et al. 2020. Fungal Diversity 105:1-16 ((https://doi.org/10.1007/s13225-020-00466-2)) |
| Otu62956 | 1 | k__Fungi(100);p__Ascomycota(100);c__Saccharomycetes(100);o__Saccharomycetales(100);f__Metschnikowiaceae(100);g__Metschnikowia(100);s__Metschnikowia_pulcherrima(100); | Metschnikowia | 13 | Saprotroph | \|Undefined Saprotroph\| | Yeast | NULL | Probable | As nectar-tap_saprotroph (Põlme et al. 2020); Decay sugar-rich substrates (Põlme et al. 2020) | Põlme S, et al. 2020. Fungal Diversity 105:1-16 ((https://doi.org/10.1007/s13225-020-00466-2)) |
| Otu62959 | 1 | k__Fungi(100);p__Ascomycota(100);c__Saccharomycetes(100);o__Saccharomycetales(100);f__Metschnikowiaceae(100);g__Metschnikowia(100);g__Metschnikowia_unclassified(100); | Metschnikowia | 13 | Saprotroph | \|Undefined Saprotroph\| | Yeast | NULL | Probable | As nectar-tap_saprotroph (Põlme et al. 2020); Decay sugar-rich substrates (Põlme et al. 2020) | Põlme S, et al. 2020. Fungal Diversity 105:1-16 ((https://doi.org/10.1007/s13225-020-00466-2)) |
| Otu62960 | 1 | k__Fungi(100);p__Ascomycota(100);c__Saccharomycetes(100);o__Saccharomycetales(100);f__Metschnikowiaceae(100);g__Metschnikowia(100);s__Metschnikowia_pulcherrima(100); | Metschnikowia | 13 | Saprotroph | \|Undefined Saprotroph\| | Yeast | NULL | Probable | As nectar-tap_saprotroph (Põlme et al. 2020); Decay sugar-rich substrates (Põlme et al. 2020) | Põlme S, et al. 2020. Fungal Diversity 105:1-16 ((https://doi.org/10.1007/s13225-020-00466-2)) |
| Otu62961 | 1 | k__Fungi(100);p__Ascomycota(100);c__Saccharomycetes(100);o__Saccharomycetales(100);f__Metschnikowiaceae(100);g__Metschnikowia(100);s__Metschnikowia_pulcherrima(100); | Metschnikowia | 13 | Saprotroph | \|Undefined Saprotroph\| | Yeast | NULL | Probable | As nectar-tap_saprotroph (Põlme et al. 2020); Decay sugar-rich substrates (Põlme et al. 2020) | Põlme S, et al. 2020. Fungal Diversity 105:1-16 ((https://doi.org/10.1007/s13225-020-00466-2)) |
| Otu62965 | 1 | k__Fungi(100);p__Ascomycota(100);c__Sordariomycetes(100);o__Hypocreales(100);f__Nectriaceae(100);f__Nectriaceae_unclassified(100);f__Nectriaceae_unclassified(100); | Nectriaceae | 9 | Pathotroph-Saprotroph-Symbiotroph | Animal Pathogen-Endophyte-Fungal Parasite-Lichen Parasite-Plant Pathogen-Wood Saprotroph | Microfungus | NULL | Possible | Fusarium species are amongst the economically significant plant pathogens (Cannon & Kirk 2007) | Cannon PF, Kirk PM. 2007. Fungal Families of the World. CAB International, Cambridge (ISBN: 978-0851998275); Hirooka Y, et al. 2012. Studies in Mycology 71-210 ((https://doi.org/10.3114/sim0001)); Tedersoo L, et al. 2014. Science 346:e1256688 ((https://doi.org/10.1126/science.1256688)); The American Phytopathological Society Common Names of Plant Diseases ((http://www.apsnet. [illegible] |
| Otu62966 | 1 | k__Fungi(100);p__Ascomycota(100);c__Saccharomycetes(100);o__Saccharomycetales(100);f__Metschnikowiaceae(100);g__Metschnikowia(100);s__Metschnikowia_pulcherrima(100); | Metschnikowia | 13 | Saprotroph | \|Undefined Saprotroph\| | Yeast | NULL | Probable | As nectar-tap_saprotroph (Põlme et al. 2020); Decay sugar-rich substrates (Põlme et al. 2020) | Põlme S, et al. 2020. Fungal Diversity 105:1-16 ((https://doi.org/10.1007/s13225-020-00466-2)) |
| Otu62967 | 1 | k__Fungi(100);p__Ascomycota(100);c__Saccharomycetes(100);o__Saccharomycetales(100);f__Metschnikowiaceae(100);g__Metschnikowia(100);g__Metschnikowia_unclassified(100); | Metschnikowia | 13 | Saprotroph | \|Undefined Saprotroph\| | Yeast | NULL | Probable | As nectar-tap_saprotroph (Põlme et al. 2020); Decay sugar-rich substrates (Põlme et al. 2020) | Põlme S, et al. 2020. Fungal Diversity 105:1-16 ((https://doi.org/10.1007/s13225-020-00466-2)) |
| Otu62968 | 1 | k__Fungi(100);p__Ascomycota(100);c__Saccharomycetes(100);o__Saccharomycetales(100);f__Metschnikowiaceae(100);g__Metschnikowia(100);s__Metschnikowia_pulcherrima(100); | Metschnikowia | 13 | Saprotroph | \|Undefined Saprotroph\| | Yeast | NULL | Probable | As nectar-tap_saprotroph (Põlme et al. 2020); Decay sugar-rich substrates (Põlme et al. 2020) | Põlme S, et al. 2020. Fungal Diversity 105:1-16 ((https://doi.org/10.1007/s13225-020-00466-2)) |
| Otu62969 | 1 | k__Fungi(100);p__Ascomycota(100);c__Saccharomycetes(100);o__Saccharomycetales(100);f__Metschnikowiaceae(100);g__Metschnikowia(100);s__Metschnikowia_pulcherrima(100); | Metschnikowia | 13 | Saprotroph | \|Undefined Saprotroph\| | Yeast | NULL | Probable | As nectar-tap_saprotroph (Põlme et al. 2020); Decay sugar-rich substrates (Põlme et al. 2020) | Põlme S, et al. 2020. Fungal Diversity 105:1-16 ((https://doi.org/10.1007/s13225-020-00466-2)) |
| Otu62970 | 1 | k__Fungi(100);p__Ascomycota(100);c__Saccharomycetes(100);o__Saccharomycetales(100);f__Metschnikowiaceae(100);g__Metschnikowia(100);s__Metschnikowia_pulcherrima(100); | Metschnikowia | 13 | Saprotroph | \|Undefined Saprotroph\| | Yeast | NULL | Probable | As nectar-tap_saprotroph (Põlme et al. 2020); Decay sugar-rich substrates (Põlme et al. 2020) | Põlme S, et al. 2020. Fungal Diversity 105:1-16 ((https://doi.org/10.1007/s13225-020-00466-2)) |
| Otu62971 | 1 | k__Fungi(100);p__Ascomycota(100);c__Saccharomycetes(100);o__Saccharomycetales(100);f__Metschnikowiaceae(100);g__Metschnikowia(100);s__Metschnikowia_pulcherrima(100); | Metschnikowia | 13 | Saprotroph | \|Undefined Saprotroph\| | Yeast | NULL | Probable | As nectar-tap_saprotroph (Põlme et al. 2020); Decay sugar-rich substrates (Põlme et al. 2020) | Põlme S, et al. 2020. Fungal Diversity 105:1-16 ((https://doi.org/10.1007/s13225-020-00466-2)) |
| Otu62972 | 1 | k__Fungi(100);p__Ascomycota(100);c__Saccharomycetes(100);o__Saccharomycetales(100);f__Metschnikowiaceae(100);g__Metschnikowia(100);g__Metschnikowia_unclassified(100); | Metschnikowia | 13 | Saprotroph | \|Undefined Saprotroph\| | Yeast | NULL | Probable | As nectar-tap_saprotroph (Põlme et al. 2020); Decay sugar-rich substrates (Põlme et al. 2020) | Põlme S, et al. 2020. Fungal Diversity 105:1-16 ((https://doi.org/10.1007/s13225-020-00466-2)) |
| Otu62973 | 1 | k__Fungi(100);p__Ascomycota(100);c__Saccharomycetes(100);o__Saccharomycetales(100);f__Metschnikowiaceae(100);g__Metschnikowia(100);s__Metschnikowia_pulcherrima(100); | Metschnikowia | 13 | Saprotroph | \|Undefined Saprotroph\| | Yeast | NULL | Probable | As nectar-tap_saprotroph (Põlme et al. 2020); Decay sugar-rich substrates (Põlme et al. 2020) | Põlme S, et al. 2020. Fungal Diversity 105:1-16 ((https://doi.org/10.1007/s13225-020-00466-2)) |
| Otu62974 | 1 | k__Fungi(100);p__Ascomycota(100);c__Saccharomycetes(100);o__Saccharomycetales(100);f__Metschnikowiaceae(100);g__Metschnikowia(100);g__Metschnikowia_unclassified(100); | Metschnikowia | 13 | Saprotroph | \|Undefined Saprotroph\| | Yeast | NULL | Probable | As nectar-tap_saprotroph (Põlme et al. 2020); Decay sugar-rich substrates (Põlme et al. 2020) | Põlme S, et al. 2020. Fungal Diversity 105:1-16 ((https://doi.org/10.1007/s13225-020-00466-2)) |
| Otu62975 | 1 | k__Fungi(100);p__Ascomycota(100);c__Saccharomycetes(100);o__Saccharomycetales(100);f__Metschnikowiaceae(100);g__Metschnikowia(100);s__Metschnikowia_pulcherrima(100); | Metschnikowia | 13 | Saprotroph | \|Undefined Saprotroph\| | Yeast | NULL | Probable | As nectar-tap_saprotroph (Põlme et al. 2020); Decay sugar-rich substrates (Põlme et al. 2020) | Põlme S, et al. 2020. Fungal Diversity 105:1-16 ((https://doi.org/10.1007/s13225-020-00466-2)) |
| Otu62976 | 1 | k__Fungi(100);p__Ascomycota(100);c__Saccharomycetes(100);o__Saccharomycetales(100);f__Metschnikowiaceae(100);g__Metschnikowia(100);s__Metschnikowia_pulcherrima(100); | Metschnikowia | 13 | Saprotroph | \|Undefined Saprotroph\| | Yeast | NULL | Probable | As nectar-tap_saprotroph (Põlme et al. 2020); Decay sugar-rich substrates (Põlme et al. 2020) | Põlme S, et al. 2020. Fungal Diversity 105:1-16 ((https://doi.org/10.1007/s13225-020-00466-2)) |
| Otu62977 | 1 | k__Fungi(100);p__Ascomycota(100);c__Saccharomycetes(100);o__Saccharomycetales(100);f__Metschnikowiaceae(100);g__Metschnikowia(100);s__Metschnikowia_pulcherrima(100); | Metschnikowia | 13 | Saprotroph | \|Undefined Saprotroph\| | Yeast | NULL | Probable | As nectar-tap_saprotroph (Põlme et al. 2020); Decay sugar-rich substrates (Põlme et al. 2020) | Põlme S, et al. 2020. Fungal Diversity 105:1-16 ((https://doi.org/10.1007/s13225-020-00466-2)) |
| Otu62978 | 1 | k__Fungi(100);p__Ascomycota(100);c__Saccharomycetes(100);o__Saccharomycetales(100);f__Metschnikowiaceae(100);g__Metschnikowia(100);s__Metschnikowia_pulcherrima(100); | Metschnikowia | 13 | Saprotroph | \|Undefined Saprotroph\| | Yeast | NULL | Probable | As nectar-tap_saprotroph (Põlme et al. 2020); Decay sugar-rich substrates (Põlme et al. 2020) | Põlme S, et al. 2020. Fungal Diversity 105:1-16 ((https://doi.org/10.1007/s13225-020-00466-2)) |
| Otu62979 | 1 | k__Fungi(100);p__Ascomycota(100);c__Saccharomycetes(100);o__Saccharomycetales(100);f__Metschnikowiaceae(100);g__Metschnikowia(100);s__Metschnikowia_pulcherrima(100); | Metschnikowia | 13 | Saprotroph | \|Undefined Saprotroph\| | Yeast | NULL | Probable | As nectar-tap_saprotroph (Põlme et al. 2020); Decay sugar-rich substrates (Põlme et al. 2020) | Põlme S, et al. 2020. Fungal Diversity 105:1-16 ((https://doi.org/10.1007/s13225-020-00466-2)) |
| Otu62980 | 1 | k__Fungi(100);p__Ascomycota(100);c__Saccharomycetes(100);o__Saccharomycetales(100);f__Metschnikowiaceae(100);g__Metschnikowia(100);s__Metschnikowia_pulcherrima(100); | Metschnikowia | 13 | Saprotroph | \|Undefined Saprotroph\| | Yeast | NULL | Probable | As nectar-tap_saprotroph (Põlme et al. 2020); Decay sugar-rich substrates (Põlme et al. 2020) | Põlme S, et al. 2020. Fungal Diversity 105:1-16 ((https://doi.org/10.1007/s13225-020-00466-2)) |
| Otu62982 | 1 | k__Fungi(100);p__Ascomycota(100);c__Saccharomycetes(100);o__Saccharomycetales(100);f__Metschnikowiaceae(100);g__Metschnikowia(100);g__Metschnikowia_unclassified(100); | Metschnikowia | 13 | Saprotroph | \|Undefined Saprotroph\| | Yeast | NULL | Probable | As nectar-tap_saprotroph (Põlme et al. 2020); Decay sugar-rich substrates (Põlme et al. 2020) | Põlme S, et al. 2020. Fungal Diversity 105:1-16 ((https://doi.org/10.1007/s13225-020-00466-2)) |
| Otu62985 | 1 | k__Fungi(100);p__Ascomycota(100);c__Saccharomycetes(100);o__Saccharomycetales(100);f__Metschnikowiaceae(100);g__Metschnikowia(100);s__Metschnikowia_pulcherrima(100); | Metschnikowia | 13 | Saprotroph | \|Undefined Saprotroph\| | Yeast | NULL | Probable | As nectar-tap_saprotroph (Põlme et al. 2020); Decay sugar-rich substrates (Põlme et al. 2020) | Põlme S, et al. 2020. Fungal Diversity 105:1-16 ((https://doi.org/10.1007/s13225-020-00466-2)) |
| Otu62986 | 1 | k__Fungi(100);p__Ascomycota(100);c__Saccharomycetes(100);o__Saccharomycetales(100);f__Metschnikowiaceae(100);g__Metschnikowia(100);s__Metschnikowia_pulcherrima(100); | Metschnikowia | 13 | Saprotroph | \|Undefined Saprotroph\| | Yeast | NULL | Probable | As nectar-tap_saprotroph (Põlme et al. 2020); Decay sugar-rich substrates (Põlme et al. 2020) | Põlme S, et al. 2020. Fungal Diversity 105:1-16 ((https://doi.org/10.1007/s13225-020-00466-2)) |
| Otu62989 | 1 | k__Fungi(100);p__Ascomycota(100);c__Saccharomycetes(100);o__Saccharomycetales(100);f__Metschnikowiaceae(100);g__Metschnikowia(100);s__Metschnikowia_pulcherrima(100); | Metschnikowia | 13 | Saprotroph | \|Undefined Saprotroph\| | Yeast | NULL | Probable | As nectar-tap_saprotroph (Põlme et al. 2020); Decay sugar-rich substrates (Põlme et al. 2020) | Põlme S, et al. 2020. Fungal Diversity 105:1-16 ((https://doi.org/10.1007/s13225-020-00466-2)) |
| Otu62991 | 1 | k__Fungi(100);p__Ascomycota(100);c__Saccharomycetes(100);o__Saccharomycetales(100);f__Saccharomycetales_family_Incertae_sedis(100);g__Candida(100);s__Candida_cretensis(100); | Candida | 13 | Pathotroph-Saprotroph-Symbiotroph | Animal Pathogen-Endophyte-Endosymbiont-Epiphyte-\|Undefined Saprotroph\| | Dimorphic Yeast-Yeast | NULL | Probable | As nectar-tap_saprotroph (Põlme et al. 2020); Decay sugar-rich substrates (Põlme et al. 2020); This asexual genus is polyphyletic and contain species that function very differently depend on what they are. Caution should be taken when interpreting guilds for Candida species. | Manolakaki D, et al. 2010. Virulence. 1: 367-375 ((https://doi.org/10.4161/viru.1.5.12796)); Mekbib SB, et al. 2011. Tropical Plant Pathology 36:233-240 ((https://doi.org/10.1590/S1982-56762011000400004)); Põlme S, et al. 2020. Fungal Diversity 105:1-16 ((https://doi.org/10.1007/s13225-020-00466-2)) |
| Otu62993 | 1 | k__Fungi(100);p__Ascomycota(100);c__Saccharomycetes(100);o__Saccharomycetales(100);f__Metschnikowiaceae(100);g__Metschnikowia(100);s__Metschnikowia_pulcherrima(100); | Metschnikowia | 13 | Saprotroph | \|Undefined Saprotroph\| | Yeast | NULL | Probable | As nectar-tap_saprotroph (Põlme et al. 2020); Decay sugar-rich substrates (Põlme et al. 2020) | Põlme S, et al. 2020. Fungal Diversity 105:1-16 ((https://doi.org/10.1007/s13225-020-00466-2)) |
| Otu62996 | 1 | k__Fungi(100);p__Ascomycota(100);c__Saccharomycetes(100);o__Saccharomycetales(100);f__Metschnikowiaceae(100);g__Metschnikowia(100);s__Metschnikowia_pulcherrima(100); | Metschnikowia | 13 | Saprotroph | \|Undefined Saprotroph\| | Yeast | NULL | Probable | As nectar-tap_saprotroph (Põlme et al. 2020); Decay sugar-rich substrates (Põlme et al. 2020) | Põlme S, et al. 2020. Fungal Diversity 105:1-16 ((https://doi.org/10.1007/s13225-020-00466-2)) |
| Otu63003 | 1 | k__Fungi(100);p__Ascomycota(100);c__Saccharomycetes(100);o__Saccharomycetales(100);f__Metschnikowiaceae(100);g__Metschnikowia(100);s__Metschnikowia_pulcherrima(100); | Metschnikowia | 13 | Saprotroph | \|Undefined Saprotroph\| | Yeast | NULL | Probable | As nectar-tap_saprotroph (Põlme et al. 2020); Decay sugar-rich substrates (Põlme et al. 2020) | Põlme S, et al. 2020. Fungal Diversity 105:1-16 ((https://doi.org/10.1007/s13225-020-00466-2)) |
| Otu63007 | 1 | k__Fungi(100);p__Ascomycota(100);c__Saccharomycetes(100);o__Saccharomycetales(100);f__Metschnikowiaceae(100);g__Metschnikowia(100);s__Metschnikowia_pulcherrima(100); | Metschnikowia | 13 | Saprotroph | \|Undefined Saprotroph\| | Yeast | NULL | Probable | As nectar-tap_saprotroph (Põlme et al. 2020); Decay sugar-rich substrates (Põlme et al. 2020) | Põlme S, et al. 2020. Fungal Diversity 105:1-16 ((https://doi.org/10.1007/s13225-020-00466-2)) |
| Otu63008 | 1 | k__Fungi(100);p__Ascomycota(100);c__Saccharomycetes(100);o__Saccharomycetales(100);f__Metschnikowiaceae(100);g__Metschnikowia(100);s__Metschnikowia_pulcherrima(100); | Metschnikowia | 13 | Saprotroph | \|Undefined Saprotroph\| | Yeast | NULL | Probable | As nectar-tap_saprotroph (Põlme et al. 2020); Decay sugar-rich substrates (Põlme et al. 2020) | Põlme S, et al. 2020. Fungal Diversity 105:1-16 ((https://doi.org/10.1007/s13225-020-00466-2)) |
| Otu63012 | 1 | k__Fungi(100);p__Ascomycota(100);c__Saccharomycetes(100);o__Saccharomycetales(100);f__Metschnikowiaceae(100);g__Metschnikowia(100);s__Metschnikowia_pulcherrima(100); | Metschnikowia | 13 | Saprotroph | \|Undefined Saprotroph\| | Yeast | NULL | Probable | As nectar-tap_saprotroph (Põlme et al. 2020); Decay sugar-rich substrates (Põlme et al. 2020) | Põlme S, et al. 2020. Fungal Diversity 105:1-16 ((https://doi.org/10.1007/s13225-020-00466-2)) |
| Otu63014 | 1 | k__Fungi(100);p__Ascomycota(100);c__Saccharomycetes(100);o__Saccharomycetales(100);f__Metschnikowiaceae(100);g__Metschnikowia(100);s__Metschnikowia_pulcherrima(100); | Metschnikowia | 13 | Saprotroph | \|Undefined Saprotroph\| | Yeast | NULL | Probable | As nectar-tap_saprotroph (Põlme et al. 2020); Decay sugar-rich substrates (Põlme et al. 2020) | Põlme S, et al. 2020. Fungal Diversity 105:1-16 ((https://doi.org/10.1007/s13225-020-00466-2)) |
| Otu63015 | 1 | k__Fungi(100);p__Ascomycota(100);c__Saccharomycetes(100);o__Saccharomycetales(100);f__Metschnikowiaceae(100);g__Metschnikowia(100);s__Metschnikowia_pulcherrima(100); | Metschnikowia | 13 | Saprotroph | \|Undefined Saprotroph\| | Yeast | NULL | Probable | As nectar-tap_saprotroph (Põlme et al. 2020); Decay sugar-rich substrates (Põlme et al. 2020) | Põlme S, et al. 2020. Fungal Diversity 105:1-16 ((https://doi.org/10.1007/s13225-020-00466-2)) |
| Otu63027 | 1 | k__Fungi(100);p__Ascomycota(100);c__Saccharomycetes(100);o__Saccharomycetales(100);f__Metschnikowiaceae(100);g__Metschnikowia(100);s__Metschnikowia_pulcherrima(100); | Metschnikowia | 13 | Saprotroph | \|Undefined Saprotroph\| | Yeast | NULL | Probable | As nectar-tap_saprotroph (Põlme et al. 2020); Decay sugar-rich substrates (Põlme et al. 2020) | Põlme S, et al. 2020. Fungal Diversity 105:1-16 ((https://doi.org/10.1007/s13225-020-00466-2)) |
| Otu63035 | 1 | k__Fungi(100);p__Ascomycota(100);c__Saccharomycetes(100);o__Saccharomycetales(100);f__Metschnikowiaceae(100);g__Metschnikowia(100);s__Metschnikowia_pulcherrima(100); | Metschnikowia | 13 | Saprotroph | \|Undefined Saprotroph\| | Yeast | NULL | Probable | As nectar-tap_saprotroph (Põlme et al. 2020); Decay sugar-rich substrates (Põlme et al. 2020) | Põlme S, et al. 2020. Fungal Diversity 105:1-16 ((https://doi.org/10.1007/s13225-020-00466-2)) |
| Otu63038 | 1 | k__Fungi(100);p__Ascomycota(100);c__Saccharomycetes(100);o__Saccharomycetales(100);f__Metschnikowiaceae(100);g__Metschnikowia(100);s__Metschnikowia_pulcherrima(100); | Metschnikowia | 13 | Saprotroph | \|Undefined Saprotroph\| | Yeast | NULL | Probable | As nectar-tap_saprotroph (Põlme et al. 2020); Decay sugar-rich substrates (Põlme et al. 2020) | Põlme S, et al. 2020. Fungal Diversity 105:1-16 ((https://doi.org/10.1007/s13225-020-00466-2)) |
| Otu63039 | 1 | k__Fungi(100);p__Ascomycota(100);c__Saccharomycetes(100);o__Saccharomycetales(100);f__Metschnikowiaceae(100);g__Metschnikowia(100);g__Metschnikowia_unclassified(100); | Metschnikowia | 13 | Saprotroph | \|Undefined Saprotroph\| | Yeast | NULL | Probable | As nectar-tap_saprotroph (Põlme et al. 2020); Decay sugar-rich substrates (Põlme et al. 2020) | Põlme S, et al. 2020. Fungal Diversity 105:1-16 ((https://doi.org/10.1007/s13225-020-00466-2)) |
| Otu63041 | 1 | k__Fungi(100);p__Ascomycota(100);c__Saccharomycetes(100);o__Saccharomycetales(100);f__Metschnikowiaceae(100);g__Metschnikowia(100);g__Metschnikowia_unclassified(100); | Metschnikowia | 13 | Saprotroph | \|Undefined Saprotroph\| | Yeast | NULL | Probable | As nectar-tap_saprotroph (Põlme et al. 2020); Decay sugar-rich substrates (Põlme et al. 2020) | Põlme S, et al. 2020. Fungal Diversity 105:1-16 ((https://doi.org/10.1007/s13225-020-00466-2)) |
| Otu63043 | 1 | k__Fungi(100);p__Ascomycota(100);c__Saccharomycetes(100);o__Saccharomycetales(100);f__Metschnikowiaceae(100);g__Metschnikowia(100);s__Metschnikowia_pulcherrima(100); | Metschnikowia | 13 | Saprotroph | \|Undefined Saprotroph\| | Yeast | NULL | Probable | As nectar-tap_saprotroph (Põlme et al. 2020); Decay sugar-rich substrates (Põlme et al. 2020) | Põlme S, et al. 2020. Fungal Diversity 105:1-16 ((https://doi.org/10.1007/s13225-020-00466-2)) |
| Otu63045 | 1 | k__Fungi(100);p__Ascomycota(100);c__Saccharomycetes(100);o__Saccharomycetales(100);f__Metschnikowiaceae(100);g__Metschnikowia(100);s__Metschnikowia_pulcherrima(100); | Metschnikowia | 13 | Saprotroph | \|Undefined Saprotroph\| | Yeast | NULL | Probable | As nectar-tap_saprotroph (Põlme et al. 2020); Decay sugar-rich substrates (Põlme et al. 2020) | Põlme S, et al. 2020. Fungal Diversity 105:1-16 ((https://doi.org/10.1007/s13225-020-00466-2)) |
| Otu63046 | 1 | k__Fungi(100);p__Ascomycota(100);c__Saccharomycetes(100);o__Saccharomycetales(100);f__Metschnikowiaceae(100);g__Metschnikowia(100);g__Metschnikowia_unclassified(100); | Metschnikowia | 13 | Saprotroph | \|Undefined Saprotroph\| | Yeast | NULL | Probable | As nectar-tap_saprotroph (Põlme et al. 2020); Decay sugar-rich substrates (Põlme et al. 2020) | Põlme S, et al. 2020. Fungal Diversity 105:1-16 ((https://doi.org/10.1007/s13225-020-00466-2)) |
| Otu63055 | 1 | k__Fungi(100);p__Ascomycota(100);c__Saccharomycetes(100);o__Saccharomycetales(100);f__Saccharomycetales_family_Incertae_sedis(100);g__Candida(100);g__Candida_unclassified(100); | Candida | 13 | Pathotroph-Saprotroph-Symbiotroph | Animal Pathogen-Endophyte-Endosymbiont-Epiphyte-\|Undefined Saprotroph\| | Dimorphic Yeast-Yeast | NULL | Probable | As nectar-tap_saprotroph (Põlme et al. 2020); Decay sugar-rich substrates (Põlme et al. 2020); This asexual genus is polyphyletic and contain species that function very differently depend on what they are. Caution should be taken when interpreting guilds for Candida species. | Manolakaki D, et al. 2010. Virulence. 1: 367-375 ((https://doi.org/10.4161/viru.1.5.12796)); Mekbib SB, et al. 2011. Tropical Plant Pathology 36:233-240 ((https://doi.org/10.1590/S1982-56762011000400004)); Põlme S, et al. 2020. Fungal Diversity 105:1-16 ((https://doi.org/10.1007/s13225-020-00466-2)) |
| Otu63068 | 1 | k__Fungi(100);p__Ascomycota(100);c__Saccharomycetes(100);o__Saccharomycetales(100);f__Metschnikowiaceae(100);g__Metschnikowia(100);s__Metschnikowia_pulcherrima(100); | Metschnikowia | 13 | Saprotroph | \|Undefined Saprotroph\| | Yeast | NULL | Probable | As nectar-tap_saprotroph (Põlme et al. 2020); Decay sugar-rich substrates (Põlme et al. 2020) | Põlme S, et al. 2020. Fungal Diversity 105:1-16 ((https://doi.org/10.1007/s13225-020-00466-2)) |
| Otu63070 | 1 | k__Fungi(100);p__Ascomycota(100);c__Saccharomycetes(100);o__Saccharomycetales(100);f__Metschnikowiaceae(100);g__Metschnikowia(100);g__Metschnikowia_unclassified(100); | Metschnikowia | 13 | Saprotroph | \|Undefined Saprotroph\| | Yeast | NULL | Probable | As nectar-tap_saprotroph (Põlme et al. 2020); Decay sugar-rich substrates (Põlme et al. 2020) | Põlme S, et al. 2020. Fungal Diversity 105:1-16 ((https://doi.org/10.1007/s13225-020-00466-2)) |
| Otu63075 | 1 | k__Fungi(100);p__Ascomycota(100);c__Saccharomycetes(100);o__Saccharomycetales(100);f__Metschnikowiaceae(100);g__Metschnikowia(100);s__Metschnikowia_pulcherrima(100); | Metschnikowia | 13 | Saprotroph | \|Undefined Saprotroph\| | Yeast | NULL | Probable | As nectar-tap_saprotroph (Põlme et al. 2020); Decay sugar-rich substrates (Põlme et al. 2020) | Põlme S, et al. 2020. Fungal Diversity 105:1-16 ((https://doi.org/10.1007/s13225-020-00466-2)) |
| Otu63077 | 1 | k__Fungi(100);p__Ascomycota(100);c__Saccharomycetes(100);o__Saccharomycetales(100);f__Metschnikowiaceae(100);g__Metschnikowia(100);g__Metschnikowia_unclassified(100); | Metschnikowia | 13 | Saprotroph | \|Undefined Saprotroph\| | Yeast | NULL | Probable | As nectar-tap_saprotroph (Põlme et al. 2020); Decay sugar-rich substrates (Põlme et al. 2020) | Põlme S, et al. 2020. Fungal Diversity 105:1-16 ((https://doi.org/10.1007/s13225-020-00466-2)) |
| Otu63080 | 1 | k__Fungi(100);p__Ascomycota(100);c__Saccharomycetes(100);o__Saccharomycetales(100);f__Metschnikowiaceae(100);g__Metschnikowia(100);s__Metschnikowia_pulcherrima(100); | Metschnikowia | 13 | Saprotroph | \|Undefined Saprotroph\| | Yeast | NULL | Probable | As nectar-tap_saprotroph (Põlme et al. 2020); Decay sugar-rich substrates (Põlme et al. 2020) | Põlme S, et al. 2020. Fungal Diversity 105:1-16 ((https://doi.org/10.1007/s13225-020-00466-2)) |
| Otu63084 | 1 | k__Fungi(100);p__Ascomycota(100);c__Saccharomycetes(100);o__Saccharomycetales(100);f__Metschnikowiaceae(100);g__Metschnikowia(100);s__Metschnikowia_pulcherrima(100); | Metschnikowia | 13 | Saprotroph | \|Undefined Saprotroph\| | Yeast | NULL | Probable | As nectar-tap_saprotroph (Põlme et al. 2020); Decay sugar-rich substrates (Põlme et al. 2020) | Põlme S, et al. 2020. Fungal Diversity 105:1-16 ((https://doi.org/10.1007/s13225-020-00466-2)) |
| Otu63086 | 1 | k__Fungi(100);p__Ascomycota(100);c__Saccharomycetes(100);o__Saccharomycetales(100);f__Saccharomycetales_family_Incertae_sedis(100);g__Candida(100);g__Candida_unclassified(100); | Candida | 13 | Pathotroph-Saprotroph-Symbiotroph | Animal Pathogen-Endophyte-Endosymbiont-Epiphyte-\|Undefined Saprotroph\| | Dimorphic Yeast-Yeast | NULL | Probable | As nectar-tap_saprotroph (Põlme et al. 2020); Decay sugar-rich substrates (Põlme et al. 2020); This asexual genus is polyphyletic and contain species that function very differently depend on what they are. Caution should be taken when interpreting guilds for Candida species. | Manolakaki D, et al. 2010. Virulence. 1: 367-375 ((https://doi.org/10.4161/viru.1.5.12796)); Mekbib SB, et al. 2011. Tropical Plant Pathology 36:233-240 ((https://doi.org/10.1590/S1982-56762011000400004)); Põlme S, et al. 2020. Fungal Diversity 105:1-16 ((https://doi.org/10.1007/s13225-020-00466-2)) |
| Otu63088 | 1 | k__Fungi(100);p__Ascomycota(100);c__Saccharomycetes(100);o__Saccharomycetales(100);f__Metschnikowiaceae(100);g__Metschnikowia(100);s__Metschnikowia_pulcherrima(100); | Metschnikowia | 13 | Saprotroph | \|Undefined Saprotroph\| | Yeast | NULL | Probable | As nectar-tap_saprotroph (Põlme et al. 2020); Decay sugar-rich substrates (Põlme et al. 2020) | Põlme S, et al. 2020. Fungal Diversity 105:1-16 ((https://doi.org/10.1007/s13225-020-00466-2)) |
| Otu63090 | 1 | k__Fungi(100);p__Ascomycota(100);c__Saccharomycetes(100);o__Saccharomycetales(100);f__Saccharomycetales_family_Incertae_sedis(100);g__Candida(100);s__Candida_sp_AS23486(100); | Candida | 13 | Pathotroph-Saprotroph-Symbiotroph | Animal Pathogen-Endophyte-Endosymbiont-Epiphyte-\|Undefined Saprotroph\| | Dimorphic Yeast-Yeast | NULL | Probable | As nectar-tap_saprotroph (Põlme et al. 2020); Decay sugar-rich substrates (Põlme et al. 2020); This asexual genus is polyphyletic and contain species that function very differently depend on what they are. Caution should be taken when interpreting guilds for Candida species. | Manolakaki D, et al. 2010. Virulence. 1: 367-375 ((https://doi.org/10.4161/viru.1.5.12796)); Mekbib SB, et al. 2011. Tropical Plant Pathology 36:233-240 ((https://doi.org/10.1590/S1982-56762011000400004)); Põlme S, et al. 2020. Fungal Diversity 105:1-16 ((https://doi.org/10.1007/s13225-020-00466-2)) |
| Otu63093 | 1 | k__Fungi(100);p__Ascomycota(100);c__Saccharomycetes(100);o__Saccharomycetales(100);f__Metschnikowiaceae(100);g__Metschnikowia(100);s__Metschnikowia_pulcherrima(100); | Metschnikowia | 13 | Saprotroph | \|Undefined Saprotroph\| | Yeast | NULL | Probable | As nectar-tap_saprotroph (Põlme et al. 2020); Decay sugar-rich substrates (Põlme et al. 2020) | Põlme S, et al. 2020. Fungal Diversity 105:1-16 ((https://doi.org/10.1007/s13225-020-00466-2)) |
| Otu63094 | 1 | k__Fungi(100);p__Ascomycota(100);c__Saccharomycetes(100);o__Saccharomycetales(100);f__Metschnikowiaceae(100);g__Metschnikowia(100);s__Metschnikowia_pulcherrima(100); | Metschnikowia | 13 | Saprotroph | \|Undefined Saprotroph\| | Yeast | NULL | Probable | As nectar-tap_saprotroph (Põlme et al. 2020); Decay sugar-rich substrates (Põlme et al. 2020) | Põlme S, et al. 2020. Fungal Diversity 105:1-16 ((https://doi.org/10.1007/s13225-020-00466-2)) |
| Otu63104 | 1 | k__Fungi(100);p__Ascomycota(100);c__Saccharomycetes(100);o__Saccharomycetales(100);f__Metschnikowiaceae(100);g__Metschnikowia(100);s__Metschnikowia_pulcherrima(100); | Metschnikowia | 13 | Saprotroph | \|Undefined Saprotroph\| | Yeast | NULL | Probable | As nectar-tap_saprotroph (Põlme et al. 2020); Decay sugar-rich substrates (Põlme et al. 2020) | Põlme S, et al. 2020. Fungal Diversity 105:1-16 ((https://doi.org/10.1007/s13225-020-00466-2)) |
| Otu63105 | 1 | k__Fungi(100);p__Ascomycota(100);c__Saccharomycetes(100);o__Saccharomycetales(100);f__Metschnikowiaceae(100);g__Metschnikowia(100);s__Metschnikowia_pulcherrima(100); | Metschnikowia | 13 | Saprotroph | \|Undefined Saprotroph\| | Yeast | NULL | Probable | As nectar-tap_saprotroph (Põlme et al. 2020); Decay sugar-rich substrates (Põlme et al. 2020) | Põlme S, et al. 2020. Fungal Diversity 105:1-16 ((https://doi.org/10.1007/s13225-020-00466-2)) |
| Otu63107 | 1 | k__Fungi(100);p__Ascomycota(100);c__Saccharomycetes(100);o__Saccharomycetales(100);f__Metschnikowiaceae(100);g__Metschnikowia(100);s__Metschnikowia_pulcherrima(100); | Metschnikowia | 13 | Saprotroph | \|Undefined Saprotroph\| | Yeast | NULL | Probable | As nectar-tap_saprotroph (Põlme et al. 2020); Decay sugar-rich substrates (Põlme et al. 2020) | Põlme S, et al. 2020. Fungal Diversity 105:1-16 ((https://doi.org/10.1007/s13225-020-00466-2)) |
| Otu63108 | 1 | k__Fungi(100);p__Ascomycota(100);c__Saccharomycetes(100);o__Saccharomycetales(100);f__Metschnikowiaceae(100);g__Metschnikowia(100);s__Metschnikowia_pulcherrima(100); | Metschnikowia | 13 | Saprotroph | \|Undefined Saprotroph\| | Yeast | NULL | Probable | As nectar-tap_saprotroph (Põlme et al. 2020); Decay sugar-rich substrates (Põlme et al. 2020) | Põlme S, et al. 2020. Fungal Diversity 105:1-16 ((https://doi.org/10.1007/s13225-020-00466-2)) |
| Otu63109 | 1 | k__Fungi(100);p__Ascomycota(100);c__Saccharomycetes(100);o__Saccharomycetales(100);f__Metschnikowiaceae(100);g__Metschnikowia(100);s__Metschnikowia_pulcherrima(100); | Metschnikowia | 13 | Saprotroph | \|Undefined Saprotroph\| | Yeast | NULL | Probable | As nectar-tap_saprotroph (Põlme et al. 2020); Decay sugar-rich substrates (Põlme et al. 2020) | Põlme S, et al. 2020. Fungal Diversity 105:1-16 ((https://doi.org/10.1007/s13225-020-00466-2)) |
| Otu63110 | 1 | k__Fungi(100);p__Ascomycota(100);c__Saccharomycetes(100);o__Saccharomycetales(100);f__Metschnikowiaceae(100);g__Metschnikowia(100);s__Metschnikowia_pulcherrima(100); | Metschnikowia | 13 | Saprotroph | \|Undefined Saprotroph\| | Yeast | NULL | Probable | As nectar-tap_saprotroph (Põlme et al. 2020); Decay sugar-rich substrates (Põlme et al. 2020) | Põlme S, et al. 2020. Fungal Diversity 105:1-16 ((https://doi.org/10.1007/s13225-020-00466-2)) |
| Otu63111 | 1 | k__Fungi(100);p__Ascomycota(100);c__Saccharomycetes(100);o__Saccharomycetales(100);f__Metschnikowiaceae(100);g__Metschnikowia(100);s__Metschnikowia_pulcherrima(100); | Metschnikowia | 13 | Saprotroph | \|Undefined Saprotroph\| | Yeast | NULL | Probable | As nectar-tap_saprotroph (Põlme et al. 2020); Decay sugar-rich substrates (Põlme et al. 2020) | Põlme S, et al. 2020. Fungal Diversity 105:1-16 ((https://doi.org/10.1007/s13225-020-00466-2)) |
| Otu63112 | 1 | k__Fungi(100);p__Ascomycota(100);c__Saccharomycetes(100);o__Saccharomycetales(100);f__Metschnikowiaceae(100);g__Metschnikowia(100);s__Metschnikowia_pulcherrima(100); | Metschnikowia | 13 | Saprotroph | \|Undefined Saprotroph\| | Yeast | NULL | Probable | As nectar-tap_saprotroph (Põlme et al. 2020); Decay sugar-rich substrates (Põlme et al. 2020) | Põlme S, et al. 2020. Fungal Diversity 105:1-16 ((https://doi.org/10.1007/s13225-020-00466-2)) |
| Otu63113 | 1 | k__Fungi(100);p__Ascomycota(100);c__Saccharomycetes(100);o__Saccharomycetales(100);f__Metschnikowiaceae(100);g__Metschnikowia(100);g__Metschnikowia_unclassified(100); | Metschnikowia | 13 | Saprotroph | \|Undefined Saprotroph\| | Yeast | NULL | Probable | As nectar-tap_saprotroph (Põlme et al. 2020); Decay sugar-rich substrates (Põlme et al. 2020) | Põlme S, et al. 2020. Fungal Diversity 105:1-16 ((https://doi.org/10.1007/s13225-020-00466-2)) |
| Otu63114 | 1 | k__Fungi(100);p__Ascomycota(100);c__Saccharomycetes(100);o__Saccharomycetales(100);f__Metschnikowiaceae(100);g__Metschnikowia(100);s__Metschnikowia_pulcherrima(100); | Metschnikowia | 13 | Saprotroph | \|Undefined Saprotroph\| | Yeast | NULL | Probable | As nectar-tap_saprotroph (Põlme et al. 2020); Decay sugar-rich substrates (Põlme et al. 2020) | Põlme S, et al. 2020. Fungal Diversity 105:1-16 ((https://doi.org/10.1007/s13225-020-00466-2)) |
| Otu63115 | 1 | k__Fungi(100);p__Basidiomycota(100);c__Tremellomycetes(100);o__Tremellales(100);f__Tremellales_family_Incertae_sedis(100);g__Dioszegia(100);s__Dioszegia_changbaiensis(100); | Dioszegia | 13 | Saprotroph | \|Plant Saprotroph\|-Undefined Saprotroph | Yeast | NULL | Probable | As litter_saprotroph (Põlme et al. 2020); Decay leaf, fruit, seed (Põlme et al. 2020); The Dioszegia subclade sensu Inacio (Inacio et al. 20015) is characterized by orange; yellow or orange to deep orange colonies. Isolated from plant leaves and soil. | Põlme S, et al. 2020. Fungal Diversity 105:1-16 ((https://doi.org/10.1007/s13225-020-00466-2)); Takashima M, et al. 2011 International Journal of Systematic and Evolutionary Microbiology 61:1736-1739 (https://doi.org/10.1099/ijs.0.025254-0)) |
| Otu63116 | 1 | k__Fungi(100);p__Ascomycota(100);c__Saccharomycetes(100);o__Saccharomycetales(100);f__Metschnikowiaceae(100);g__Metschnikowia(100);s__Metschnikowia_pulcherrima(100); | Metschnikowia | 13 | Saprotroph | \|Undefined Saprotroph\| | Yeast | NULL | Probable | As nectar-tap_saprotroph (Põlme et al. 2020); Decay sugar-rich substrates (Põlme et al. 2020) | Põlme S, et al. 2020. Fungal Diversity 105:1-16 ((https://doi.org/10.1007/s13225-020-00466-2)) |
| Otu63117 | 1 | k__Fungi(100);p__Ascomycota(100);c__Saccharomycetes(100);o__Saccharomycetales(100);f__Metschnikowiaceae(100);g__Metschnikowia(100);s__Metschnikowia_pulcherrima(100); | Metschnikowia | 13 | Saprotroph | \|Undefined Saprotroph\| | Yeast | NULL | Probable | As nectar-tap_saprotroph (Põlme et al. 2020); Decay sugar-rich substrates (Põlme et al. 2020) | Põlme S, et al. 2020. Fungal Diversity 105:1-16 ((https://doi.org/10.1007/s13225-020-00466-2)) |
| Otu63136 | 1 | k__Fungi(100);p__Ascomycota(100);c__Saccharomycetes(100);o__Saccharomycetales(100);f__Metschnikowiaceae(100);g__Metschnikowia(100);s__Metschnikowia_pulcherrima(100); | Metschnikowia | 13 | Saprotroph | \|Undefined Saprotroph\| | Yeast | NULL | Probable | As nectar-tap_saprotroph (Põlme et al. 2020); Decay sugar-rich substrates (Põlme et al. 2020) | Põlme S, et al. 2020. Fungal Diversity 105:1-16 ((https://doi.org/10.1007/s13225-020-00466-2)) |
| Otu63140 | 1 | k__Fungi(100);p__Ascomycota(100);c__Saccharomycetes(100);o__Saccharomycetales(100);f__Saccharomycetales_family_Incertae_sedis(100);g__Candida(100);g__Candida_unclassified(100); | Candida | 13 | Pathotroph-Saprotroph-Symbiotroph | Animal Pathogen-Endophyte-Endosymbiont-Epiphyte-\|Undefined Saprotroph\| | Dimorphic Yeast-Yeast | NULL | Probable | As nectar-tap_saprotroph (Põlme et al. 2020); Decay sugar-rich substrates (Põlme et al. 2020); This asexual genus is polyphyletic and contain species that function very differently depend on what they are. Caution should be taken when interpreting guilds for Candida species. | Manolakaki D, et al. 2010. Virulence. 1: 367-375 ((https://doi.org/10.4161/viru.1.5.12796)); Mekbib SB, et al. 2011. Tropical Plant Pathology 36:233-240 ((https://doi.org/10.1590/S1982-56762011000400004)); Põlme S, et al. 2020. Fungal Diversity 105:1-16 ((https://doi.org/10.1007/s13225-020-00466-2)) |
| Otu63152 | 1 | k__Fungi(100);p__Ascomycota(100);c__Saccharomycetes(100);o__Saccharomycetales(100);f__Metschnikowiaceae(100);g__Metschnikowia(100);s__Metschnikowia_pulcherrima(100); | Metschnikowia | 13 | Saprotroph | \|Undefined Saprotroph\| | Yeast | NULL | Probable | As nectar-tap_saprotroph (Põlme et al. 2020); Decay sugar-rich substrates (Põlme et al. 2020) | Põlme S, et al. 2020. Fungal Diversity 105:1-16 ((https://doi.org/10.1007/s13225-020-00466-2)) |
| Otu63154 | 1 | k__Fungi(100);p__Ascomycota(100);c__Saccharomycetes(100);o__Saccharomycetales(100);f__Metschnikowiaceae(100);g__Metschnikowia(100);s__Metschnikowia_pulcherrima(100); | Metschnikowia | 13 | Saprotroph | \|Undefined Saprotroph\| | Yeast | NULL | Probable | As nectar-tap_saprotroph (Põlme et al. 2020); Decay sugar-rich substrates (Põlme et al. 2020) | Põlme S, et al. 2020. Fungal Diversity 105:1-16 ((https://doi.org/10.1007/s13225-020-00466-2)) |
| Otu63155 | 1 | k__Fungi(100);p__Ascomycota(100);c__Saccharomycetes(100);o__Saccharomycetales(100);f__Metschnikowiaceae(100);g__Metschnikowia(100);s__Metschnikowia_pulcherrima(100); | Metschnikowia | 13 | Saprotroph | \|Undefined Saprotroph\| | Yeast | NULL | Probable | As nectar-tap_saprotroph (Põlme et al. 2020); Decay sugar-rich substrates (Põlme et al. 2020) | Põlme S, et al. 2020. Fungal Diversity 105:1-16 ((https://doi.org/10.1007/s13225-020-00466-2)) |
| Otu63156 | 1 | k__Fungi(100);p__Ascomycota(100);c__Saccharomycetes(100);o__Saccharomycetales(100);f__Metschnikowiaceae(100);g__Metschnikowia(100);g__Metschnikowia_unclassified(100); | Metschnikowia | 13 | Saprotroph | \|Undefined Saprotroph\| | Yeast | NULL | Probable | As nectar-tap_saprotroph (Põlme et al. 2020); Decay sugar-rich substrates (Põlme et al. 2020) | Põlme S, et al. 2020. Fungal Diversity 105:1-16 ((https://doi.org/10.1007/s13225-020-00466-2)) |
| Otu63157 | 1 | k__Fungi(100);p__Ascomycota(100);c__Saccharomycetes(100);o__Saccharomycetales(100);f__Metschnikowiaceae(100);g__Metschnikowia(100);s__Metschnikowia_pulcherrima(100); | Metschnikowia | 13 | Saprotroph | \|Undefined Saprotroph\| | Yeast | NULL | Probable | As nectar-tap_saprotroph (Põlme et al. 2020); Decay sugar-rich substrates (Põlme et al. 2020) | Põlme S, et al. 2020. Fungal Diversity 105:1-16 ((https://doi.org/10.1007/s13225-020-00466-2)) |
| Otu63158 | 1 | k__Fungi(100);p__Ascomycota(100);c__Saccharomycetes(100);o__Saccharomycetales(100);f__Metschnikowiaceae(100);g__Metschnikowia(100);s__Metschnikowia_pulcherrima(100); | Metschnikowia | 13 | Saprotroph | \|Undefined Saprotroph\| | Yeast | NULL | Probable | As nectar-tap_saprotroph (Põlme et al. 2020); Decay sugar-rich substrates (Põlme et al. 2020) | Põlme S, et al. 2020. Fungal Diversity 105:1-16 ((https://doi.org/10.1007/s13225-020-00466-2)) |
| Otu63160 | 1 | k__Fungi(100);p__Ascomycota(100);c__Saccharomycetes(100);o__Saccharomycetales(100);f__Saccharomycetales_family_Incertae_sedis(100);g__Candida(100);g__Candida_unclassified(100); | Candida | 13 | Pathotroph-Saprotroph-Symbiotroph | Animal Pathogen-Endophyte-Endosymbiont-Epiphyte-\|Undefined Saprotroph\| | Dimorphic Yeast-Yeast | NULL | Probable | As nectar-tap_saprotroph (Põlme et al. 2020); Decay sugar-rich substrates (Põlme et al. 2020); This asexual genus is polyphyletic and contain species that function very differently depend on what they are. Caution should be taken when interpreting guilds for Candida species. | Manolakaki D, et al. 2010. Virulence. 1: 367-375 ((https://doi.org/10.4161/viru.1.5.12796)); Mekbib SB, et al. 2011. Tropical Plant Pathology 36:233-240 ((https://doi.org/10.1590/S1982-56762011000400004)); Põlme S, et al. 2020. Fungal Diversity 105:1-16 ((https://doi.org/10.1007/s13225-020-00466-2)) |
| Otu63176 | 1 | k__Fungi(100);p__Ascomycota(100);c__Saccharomycetes(100);o__Saccharomycetales(100);o__Saccharomycetales_unclassified(100);o__Saccharomycetales_unclassified(100);o__Saccharomycetales_unclassified(100); | Saccharomycetales | 7 | Saprotroph | Undefined Saprotroph | Yeast | NULL | Possible | NULL | Sterkenburg E, et al. 2015. New Phytologist 207:1145-1158 ((https://doi.org/10.1111/nph.13426)) |
| Otu63179 | 1 | k__Fungi(100);p__Ascomycota(100);c__Saccharomycetes(100);o__Saccharomycetales(100);f__Metschnikowiaceae(100);g__Metschnikowia(100);g__Metschnikowia_unclassified(100); | Metschnikowia | 13 | Saprotroph | \|Undefined Saprotroph\| | Yeast | NULL | Probable | As nectar-tap_saprotroph (Põlme et al. 2020); Decay sugar-rich substrates (Põlme et al. 2020) | Põlme S, et al. 2020. Fungal Diversity 105:1-16 ((https://doi.org/10.1007/s13225-020-00466-2)) |
| Otu63190 | 1 | k__Fungi(100);p__Ascomycota(100);c__Saccharomycetes(100);o__Saccharomycetales(100);f__Metschnikowiaceae(100);g__Metschnikowia(100);s__Metschnikowia_pulcherrima(100); | Metschnikowia | 13 | Saprotroph | \|Undefined Saprotroph\| | Yeast | NULL | Probable | As nectar-tap_saprotroph (Põlme et al. 2020); Decay sugar-rich substrates (Põlme et al. 2020) | Põlme S, et al. 2020. Fungal Diversity 105:1-16 ((https://doi.org/10.1007/s13225-020-00466-2)) |
| Otu63191 | 1 | k__Fungi(100);p__Ascomycota(100);c__Saccharomycetes(100);o__Saccharomycetales(100);f__Metschnikowiaceae(100);g__Metschnikowia(100);s__Metschnikowia_pulcherrima(100); | Metschnikowia | 13 | Saprotroph | \|Undefined Saprotroph\| | Yeast | NULL | Probable | As nectar-tap_saprotroph (Põlme et al. 2020); Decay sugar-rich substrates (Põlme et al. 2020) | Põlme S, et al. 2020. Fungal Diversity 105:1-16 ((https://doi.org/10.1007/s13225-020-00466-2)) |
| Otu63192 | 1 | k__Fungi(100);p__Ascomycota(100);c__Saccharomycetes(100);o__Saccharomycetales(100);f__Metschnikowiaceae(100);g__Metschnikowia(100);s__Metschnikowia_pulcherrima(100); | Metschnikowia | 13 | Saprotroph | \|Undefined Saprotroph\| | Yeast | NULL | Probable | As nectar-tap_saprotroph (Põlme et al. 2020); Decay sugar-rich substrates (Põlme et al. 2020) | Põlme S, et al. 2020. Fungal Diversity 105:1-16 ((https://doi.org/10.1007/s13225-020-00466-2)) |
| Otu63195 | 1 | k__Fungi(100);p__Ascomycota(100);c__Saccharomycetes(100);o__Saccharomycetales(100);f__Metschnikowiaceae(100);g__Metschnikowia(100);s__Metschnikowia_pulcherrima(100); | Metschnikowia | 13 | Saprotroph | \|Undefined Saprotroph\| | Yeast | NULL | Probable | As nectar-tap_saprotroph (Põlme et al. 2020); Decay sugar-rich substrates (Põlme et al. 2020) | Põlme S, et al. 2020. Fungal Diversity 105:1-16 ((https://doi.org/10.1007/s13225-020-00466-2)) |
| Otu63196 | 1 | k__Fungi(100);p__Ascomycota(100);c__Saccharomycetes(100);o__Saccharomycetales(100);f__Metschnikowiaceae(100);g__Metschnikowia(100);s__Metschnikowia_pulcherrima(100); | Metschnikowia | 13 | Saprotroph | \|Undefined Saprotroph\| | Yeast | NULL | Probable | As nectar-tap_saprotroph (Põlme et al. 2020); Decay sugar-rich substrates (Põlme et al. 2020) | Põlme S, et al. 2020. Fungal Diversity 105:1-16 ((https://doi.org/10.1007/s13225-020-00466-2)) |
| Otu63197 | 1 | k__Fungi(100);p__Ascomycota(100);c__Saccharomycetes(100);o__Saccharomycetales(100);f__Metschnikowiaceae(100);g__Metschnikowia(100);s__Metschnikowia_pulcherrima(100); | Metschnikowia | 13 | Saprotroph | \|Undefined Saprotroph\| | Yeast | NULL | Probable | As nectar-tap_saprotroph (Põlme et al. 2020); Decay sugar-rich substrates (Põlme et al. 2020) | Põlme S, et al. 2020. Fungal Diversity 105:1-16 ((https://doi.org/10.1007/s13225-020-00466-2)) |
| Otu63198 | 1 | k__Fungi(100);p__Ascomycota(100);c__Saccharomycetes(100);o__Saccharomycetales(100);f__Metschnikowiaceae(100);g__Metschnikowia(100);s__Metschnikowia_pulcherrima(100); | Metschnikowia | 13 | Saprotroph | \|Undefined Saprotroph\| | Yeast | NULL | Probable | As nectar-tap_saprotroph (Põlme et al. 2020); Decay sugar-rich substrates (Põlme et al. 2020) | Põlme S, et al. 2020. Fungal Diversity 105:1-16 ((https://doi.org/10.1007/s13225-020-00466-2)) |
| Otu63199 | 1 | k__Fungi(100);p__Ascomycota(100);c__Saccharomycetes(100);o__Saccharomycetales(100);f__Saccharomycetales_family_Incertae_sedis(100);g__Candida(100);g__Candida_unclassified(100); | Candida | 13 | Pathotroph-Saprotroph-Symbiotroph | Animal Pathogen-Endophyte-Endosymbiont-Epiphyte-\|Undefined Saprotroph\| | Dimorphic Yeast-Yeast | NULL | Probable | As nectar-tap_saprotroph (Põlme et al. 2020); Decay sugar-rich substrates (Põlme et al. 2020); This asexual genus is polyphyletic and contain species that function very differently depend on what they are. Caution should be taken when interpreting guilds for Candida species. | Manolakaki D, et al. 2010. Virulence. 1: 367-375 ((https://doi.org/10.4161/viru.1.5.12796)); Mekbib SB, et al. 2011. Tropical Plant Pathology 36:233-240 ((https://doi.org/10.1590/S1982-56762011000400004)); Põlme S, et al. 2020. Fungal Diversity 105:1-16 ((https://doi.org/10.1007/s13225-020-00466-2)) |
| Otu63200 | 1 | k__Fungi(100);p__Ascomycota(100);c__Saccharomycetes(100);o__Saccharomycetales(100);f__Metschnikowiaceae(100);g__Metschnikowia(100);g__Metschnikowia_unclassified(100); | Metschnikowia | 13 | Saprotroph | \|Undefined Saprotroph\| | Yeast | NULL | Probable | As nectar-tap_saprotroph (Põlme et al. 2020); Decay sugar-rich substrates (Põlme et al. 2020) | Põlme S, et al. 2020. Fungal Diversity 105:1-16 ((https://doi.org/10.1007/s13225-020-00466-2)) |
| Otu63201 | 1 | k__Fungi(100);p__Ascomycota(100);c__Saccharomycetes(100);o__Saccharomycetales(100);f__Metschnikowiaceae(100);g__Metschnikowia(100);s__Metschnikowia_pulcherrima(100); | Metschnikowia | 13 | Saprotroph | \|Undefined Saprotroph\| | Yeast | NULL | Probable | As nectar-tap_saprotroph (Põlme et al. 2020); Decay sugar-rich substrates (Põlme et al. 2020) | Põlme S, et al. 2020. Fungal Diversity 105:1-16 ((https://doi.org/10.1007/s13225-020-00466-2)) |
| Otu63202 | 1 | k__Fungi(100);p__Ascomycota(100);c__Saccharomycetes(100);o__Saccharomycetales(100);f__Metschnikowiaceae(100);g__Metschnikowia(100);s__Metschnikowia_pulcherrima(100); | Metschnikowia | 13 | Saprotroph | \|Undefined Saprotroph\| | Yeast | NULL | Probable | As nectar-tap_saprotroph (Põlme et al. 2020); Decay sugar-rich substrates (Põlme et al. 2020) | Põlme S, et al. 2020. Fungal Diversity 105:1-16 ((https://doi.org/10.1007/s13225-020-00466-2)) |
| Otu63205 | 1 | k__Fungi(100);p__Ascomycota(100);c__Saccharomycetes(100);o__Saccharomycetales(100);f__Metschnikowiaceae(100);g__Metschnikowia(100);g__Metschnikowia_unclassified(100); | Metschnikowia | 13 | Saprotroph | \|Undefined Saprotroph\| | Yeast | NULL | Probable | As nectar-tap_saprotroph (Põlme et al. 2020); Decay sugar-rich substrates (Põlme et al. 2020) | Põlme S, et al. 2020. Fungal Diversity 105:1-16 ((https://doi.org/10.1007/s13225-020-00466-2)) |
| Otu63208 | 1 | k__Fungi(100);p__Ascomycota(100);c__Dothideomycetes(100);o__Dothideomycetes_order_Incertae_sedis(100);f__Myxotrichaceae(100);g__Oidiodendron(100);s__Oidiodendron_maius(100); | Oidiodendron | 13 | Pathotroph-Saprotroph-Symbiotroph | Endophyte-Ericoid Mycorrhizal-\|Undefined Saprotroph\| | Microfungus | NULL | Probable | As root_endophyte (Põlme et al. 2020); As soil_saprotroph (Põlme et al. 2020); Dark Septate Endophyte; Decay soil (Põlme et al. 2020) | Newsham KK. 2011. New Phytologist 190:783-793 ((https://doi.org/10.1111/j.1469-8137.2010.03611.x)) (as dark septate root endophytes); Põlme S, et al. 2020. Fungal Diversity 105:1-16 ((https://doi.org/10.1007/s13225-020-00466-2)) |
| Otu63209 | 1 | k__Fungi(100);p__Ascomycota(100);c__Saccharomycetes(100);o__Saccharomycetales(100);f__Metschnikowiaceae(100);g__Metschnikowia(100);s__Metschnikowia_pulcherrima(100); | Metschnikowia | 13 | Saprotroph | \|Undefined Saprotroph\| | Yeast | NULL | Probable | As nectar-tap_saprotroph (Põlme et al. 2020); Decay sugar-rich substrates (Põlme et al. 2020) | Põlme S, et al. 2020. Fungal Diversity 105:1-16 ((https://doi.org/10.1007/s13225-020-00466-2)) |
| Otu63210 | 1 | k__Fungi(100);p__Ascomycota(100);c__Saccharomycetes(100);o__Saccharomycetales(100);f__Metschnikowiaceae(100);g__Metschnikowia(100);s__Metschnikowia_pulcherrima(100); | Metschnikowia | 13 | Saprotroph | \|Undefined Saprotroph\| | Yeast | NULL | Probable | As nectar-tap_saprotroph (Põlme et al. 2020); Decay sugar-rich substrates (Põlme et al. 2020) | Põlme S, et al. 2020. Fungal Diversity 105:1-16 ((https://doi.org/10.1007/s13225-020-00466-2)) |
| Otu63211 | 1 | k__Fungi(100);p__Ascomycota(100);c__Saccharomycetes(100);o__Saccharomycetales(100);f__Metschnikowiaceae(100);g__Metschnikowia(100);s__Metschnikowia_pulcherrima(100); | Metschnikowia | 13 | Saprotroph | \|Undefined Saprotroph\| | Yeast | NULL | Probable | As nectar-tap_saprotroph (Põlme et al. 2020); Decay sugar-rich substrates (Põlme et al. 2020) | Põlme S, et al. 2020. Fungal Diversity 105:1-16 ((https://doi.org/10.1007/s13225-020-00466-2)) |
| Otu63212 | 1 | k__Fungi(100);p__Ascomycota(100);c__Saccharomycetes(100);o__Saccharomycetales(100);f__Metschnikowiaceae(100);g__Metschnikowia(100);s__Metschnikowia_pulcherrima(100); | Metschnikowia | 13 | Saprotroph | \|Undefined Saprotroph\| | Yeast | NULL | Probable | As nectar-tap_saprotroph (Põlme et al. 2020); Decay sugar-rich substrates (Põlme et al. 2020) | Põlme S, et al. 2020. Fungal Diversity 105:1-16 ((https://doi.org/10.1007/s13225-020-00466-2)) |
| Otu63213 | 1 | k__Fungi(100);p__Ascomycota(100);c__Saccharomycetes(100);o__Saccharomycetales(100);f__Metschnikowiaceae(100);g__Metschnikowia(100);s__Metschnikowia_pulcherrima(100); | Metschnikowia | 13 | Saprotroph | \|Undefined Saprotroph\| | Yeast | NULL | Probable | As nectar-tap_saprotroph (Põlme et al. 2020); Decay sugar-rich substrates (Põlme et al. 2020) | Põlme S, et al. 2020. Fungal Diversity 105:1-16 ((https://doi.org/10.1007/s13225-020-00466-2)) |
| Otu63214 | 1 | k__Fungi(100);p__Ascomycota(100);c__Saccharomycetes(100);o__Saccharomycetales(100);f__Metschnikowiaceae(100);g__Metschnikowia(100);s__Metschnikowia_pulcherrima(100); | Metschnikowia | 13 | Saprotroph | \|Undefined Saprotroph\| | Yeast | NULL | Probable | As nectar-tap_saprotroph (Põlme et al. 2020); Decay sugar-rich substrates (Põlme et al. 2020) | Põlme S, et al. 2020. Fungal Diversity 105:1-16 ((https://doi.org/10.1007/s13225-020-00466-2)) |
| Otu63215 | 1 | k__Fungi(100);p__Ascomycota(100);c__Saccharomycetes(100);o__Saccharomycetales(100);f__Metschnikowiaceae(100);g__Metschnikowia(100);s__Metschnikowia_pulcherrima(100); | Metschnikowia | 13 | Saprotroph | \|Undefined Saprotroph\| | Yeast | NULL | Probable | As nectar-tap_saprotroph (Põlme et al. 2020); Decay sugar-rich substrates (Põlme et al. 2020) | Põlme S, et al. 2020. Fungal Diversity 105:1-16 ((https://doi.org/10.1007/s13225-020-00466-2)) |
| Otu63220 | 1 | k__Fungi(100);p__Ascomycota(100);c__Saccharomycetes(100);o__Saccharomycetales(100);f__Metschnikowiaceae(100);g__Metschnikowia(100);s__Metschnikowia_pulcherrima(100); | Metschnikowia | 13 | Saprotroph | \|Undefined Saprotroph\| | Yeast | NULL | Probable | As nectar-tap_saprotroph (Põlme et al. 2020); Decay sugar-rich substrates (Põlme et al. 2020) | Põlme S, et al. 2020. Fungal Diversity 105:1-16 ((https://doi.org/10.1007/s13225-020-00466-2)) |
| Otu63225 | 1 | k__Fungi(100);p__Ascomycota(100);c__Saccharomycetes(100);o__Saccharomycetales(100);f__Metschnikowiaceae(100);g__Metschnikowia(100);g__Metschnikowia_unclassified(100); | Metschnikowia | 13 | Saprotroph | \|Undefined Saprotroph\| | Yeast | NULL | Probable | As nectar-tap_saprotroph (Põlme et al. 2020); Decay sugar-rich substrates (Põlme et al. 2020) | Põlme S, et al. 2020. Fungal Diversity 105:1-16 ((https://doi.org/10.1007/s13225-020-00466-2)) |
| Otu63226 | 1 | k__Fungi(100);p__Ascomycota(100);c__Saccharomycetes(100);o__Saccharomycetales(100);f__Metschnikowiaceae(100);g__Metschnikowia(100);s__Metschnikowia_pulcherrima(100); | Metschnikowia | 13 | Saprotroph | \|Undefined Saprotroph\| | Yeast | NULL | Probable | As nectar-tap_saprotroph (Põlme et al. 2020); Decay sugar-rich substrates (Põlme et al. 2020) | Põlme S, et al. 2020. Fungal Diversity 105:1-16 ((https://doi.org/10.1007/s13225-020-00466-2)) |
| Otu63228 | 1 | k__Fungi(100);p__Ascomycota(100);c__Saccharomycetes(100);o__Saccharomycetales(100);f__Metschnikowiaceae(100);g__Metschnikowia(100);s__Metschnikowia_pulcherrima(100); | Metschnikowia | 13 | Saprotroph | \|Undefined Saprotroph\| | Yeast | NULL | Probable | As nectar-tap_saprotroph (Põlme et al. 2020); Decay sugar-rich substrates (Põlme et al. 2020) | Põlme S, et al. 2020. Fungal Diversity 105:1-16 ((https://doi.org/10.1007/s13225-020-00466-2)) |
| Otu63230 | 1 | k__Fungi(100);p__Ascomycota(100);c__Saccharomycetes(100);o__Saccharomycetales(100);f__Metschnikowiaceae(100);g__Metschnikowia(100);g__Metschnikowia_unclassified(100); | Metschnikowia | 13 | Saprotroph | \|Undefined Saprotroph\| | Yeast | NULL | Probable | As nectar-tap_saprotroph (Põlme et al. 2020); Decay sugar-rich substrates (Põlme et al. 2020) | Põlme S, et al. 2020. Fungal Diversity 105:1-16 ((https://doi.org/10.1007/s13225-020-00466-2)) |
| Otu63231 | 1 | k__Fungi(100);p__Ascomycota(100);c__Saccharomycetes(100);o__Saccharomycetales(100);f__Metschnikowiaceae(100);g__Metschnikowia(100);s__Metschnikowia_pulcherrima(100); | Metschnikowia | 13 | Saprotroph | \|Undefined Saprotroph\| | Yeast | NULL | Probable | As nectar-tap_saprotroph (Põlme et al. 2020); Decay sugar-rich substrates (Põlme et al. 2020) | Põlme S, et al. 2020. Fungal Diversity 105:1-16 ((https://doi.org/10.1007/s13225-020-00466-2)) |
| Otu63232 | 1 | k__Fungi(100);p__Ascomycota(100);c__Saccharomycetes(100);o__Saccharomycetales(100);f__Metschnikowiaceae(100);g__Metschnikowia(100);s__Metschnikowia_pulcherrima(100); | Metschnikowia | 13 | Saprotroph | \|Undefined Saprotroph\| | Yeast | NULL | Probable | As nectar-tap_saprotroph (Põlme et al. 2020); Decay sugar-rich substrates (Põlme et al. 2020) | Põlme S, et al. 2020. Fungal Diversity 105:1-16 ((https://doi.org/10.1007/s13225-020-00466-2)) |
| Otu63236 | 1 | k__Fungi(100);p__Ascomycota(100);c__Saccharomycetes(100);o__Saccharomycetales(100);f__Metschnikowiaceae(100);g__Metschnikowia(100);s__Metschnikowia_pulcherrima(100); | Metschnikowia | 13 | Saprotroph | \|Undefined Saprotroph\| | Yeast | NULL | Probable | As nectar-tap_saprotroph (Põlme et al. 2020); Decay sugar-rich substrates (Põlme et al. 2020) | Põlme S, et al. 2020. Fungal Diversity 105:1-16 ((https://doi.org/10.1007/s13225-020-00466-2)) |
| Otu63237 | 1 | k__Fungi(100);p__Ascomycota(100);c__Saccharomycetes(100);o__Saccharomycetales(100);f__Metschnikowiaceae(100);g__Metschnikowia(100);s__Metschnikowia_pulcherrima(100); | Metschnikowia | 13 | Saprotroph | \|Undefined Saprotroph\| | Yeast | NULL | Probable | As nectar-tap_saprotroph (Põlme et al. 2020); Decay sugar-rich substrates (Põlme et al. 2020) | Põlme S, et al. 2020. Fungal Diversity 105:1-16 ((https://doi.org/10.1007/s13225-020-00466-2)) |
| Otu63238 | 1 | k__Fungi(100);p__Ascomycota(100);c__Saccharomycetes(100);o__Saccharomycetales(100);f__Metschnikowiaceae(100);g__Metschnikowia(100);s__Metschnikowia_pulcherrima(100); | Metschnikowia | 13 | Saprotroph | \|Undefined Saprotroph\| | Yeast | NULL | Probable | As nectar-tap_saprotroph (Põlme et al. 2020); Decay sugar-rich substrates (Põlme et al. 2020) | Põlme S, et al. 2020. Fungal Diversity 105:1-16 ((https://doi.org/10.1007/s13225-020-00466-2)) |
| Otu63239 | 1 | k__Fungi(100);p__Ascomycota(100);c__Saccharomycetes(100);o__Saccharomycetales(100);f__Metschnikowiaceae(100);g__Metschnikowia(100);s__Metschnikowia_pulcherrima(100); | Metschnikowia | 13 | Saprotroph | \|Undefined Saprotroph\| | Yeast | NULL | Probable | As nectar-tap_saprotroph (Põlme et al. 2020); Decay sugar-rich substrates (Põlme et al. 2020) | Põlme S, et al. 2020. Fungal Diversity 105:1-16 ((https://doi.org/10.1007/s13225-020-00466-2)) |
| Otu63248 | 1 | k__Fungi(100);p__Ascomycota(100);c__Saccharomycetes(100);o__Saccharomycetales(100);f__Metschnikowiaceae(100);g__Metschnikowia(100);s__Metschnikowia_pulcherrima(100); | Metschnikowia | 13 | Saprotroph | \|Undefined Saprotroph\| | Yeast | NULL | Probable | As nectar-tap_saprotroph (Põlme et al. 2020); Decay sugar-rich substrates (Põlme et al. 2020) | Põlme S, et al. 2020. Fungal Diversity 105:1-16 ((https://doi.org/10.1007/s13225-020-00466-2)) |
| Otu63249 | 1 | k__Fungi(100);p__Ascomycota(100);c__Saccharomycetes(100);o__Saccharomycetales(100);f__Metschnikowiaceae(100);g__Metschnikowia(100);s__Metschnikowia_pulcherrima(100); | Metschnikowia | 13 | Saprotroph | \|Undefined Saprotroph\| | Yeast | NULL | Probable | As nectar-tap_saprotroph (Põlme et al. 2020); Decay sugar-rich substrates (Põlme et al. 2020) | Põlme S, et al. 2020. Fungal Diversity 105:1-16 ((https://doi.org/10.1007/s13225-020-00466-2)) |
| Otu63256 | 1 | k__Fungi(100);p__Ascomycota(100);c__Saccharomycetes(100);o__Saccharomycetales(100);f__Saccharomycetales_family_Incertae_sedis(100);g__Candida(100);g__Candida_unclassified(100); | Candida | 13 | Pathotroph-Saprotroph-Symbiotroph | Animal Pathogen-Endophyte-Endosymbiont-Epiphyte-\|Undefined Saprotroph\| | Dimorphic Yeast-Yeast | NULL | Probable | As nectar-tap_saprotroph (Põlme et al. 2020); Decay sugar-rich substrates (Põlme et al. 2020); This asexual genus is polyphyletic and contain species that function very differently depend on what they are. Caution should be taken when interpreting guilds for Candida species. | Manolakaki D, et al. 2010. Virulence. 1: 367-375 ((https://doi.org/10.4161/viru.1.5.12796)); Mekbib SB, et al. 2011. Tropical Plant Pathology 36:233-240 ((https://doi.org/10.1590/S1982-56762011000400004)); Põlme S, et al. 2020. Fungal Diversity 105:1-16 ((https://doi.org/10.1007/s13225-020-00466-2)) |
| Otu63257 | 1 | k__Fungi(100);p__Ascomycota(100);c__Saccharomycetes(100);o__Saccharomycetales(100);f__Saccharomycetales_family_Incertae_sedis(100);g__Candida(100);g__Candida_unclassified(100); | Candida | 13 | Pathotroph-Saprotroph-Symbiotroph | Animal Pathogen-Endophyte-Endosymbiont-Epiphyte-\|Undefined Saprotroph\| | Dimorphic Yeast-Yeast | NULL | Probable | As nectar-tap_saprotroph (Põlme et al. 2020); Decay sugar-rich substrates (Põlme et al. 2020); This asexual genus is polyphyletic and contain species that function very differently depend on what they are. Caution should be taken when interpreting guilds for Candida species. | Manolakaki D, et al. 2010. Virulence. 1: 367-375 ((https://doi.org/10.4161/viru.1.5.12796)); Mekbib SB, et al. 2011. Tropical Plant Pathology 36:233-240 ((https://doi.org/10.1590/S1982-56762011000400004)); Põlme S, et al. 2020. Fungal Diversity 105:1-16 ((https://doi.org/10.1007/s13225-020-00466-2)) |
| Otu63262 | 1 | k__Fungi(100);p__Ascomycota(100);c__Saccharomycetes(100);o__Saccharomycetales(100);f__Metschnikowiaceae(100);g__Metschnikowia(100);s__Metschnikowia_pulcherrima(100); | Metschnikowia | 13 | Saprotroph | \|Undefined Saprotroph\| | Yeast | NULL | Probable | As nectar-tap_saprotroph (Põlme et al. 2020); Decay sugar-rich substrates (Põlme et al. 2020) | Põlme S, et al. 2020. Fungal Diversity 105:1-16 ((https://doi.org/10.1007/s13225-020-00466-2)) |
| Otu63268 | 1 | k__Fungi(100);p__Ascomycota(100);c__Saccharomycetes(100);o__Saccharomycetales(100);f__Saccharomycetales_family_Incertae_sedis(100);g__Candida(100);g__Candida_unclassified(100); | Candida | 13 | Pathotroph-Saprotroph-Symbiotroph | Animal Pathogen-Endophyte-Endosymbiont-Epiphyte-\|Undefined Saprotroph\| | Dimorphic Yeast-Yeast | NULL | Probable | As nectar-tap_saprotroph (Põlme et al. 2020); Decay sugar-rich substrates (Põlme et al. 2020); This asexual genus is polyphyletic and contain species that function very differently depend on what they are. Caution should be taken when interpreting guilds for Candida species. | Manolakaki D, et al. 2010. Virulence. 1: 367-375 ((https://doi.org/10.4161/viru.1.5.12796)); Mekbib SB, et al. 2011. Tropical Plant Pathology 36:233-240 ((https://doi.org/10.1590/S1982-56762011000400004)); Põlme S, et al. 2020. Fungal Diversity 105:1-16 ((https://doi.org/10.1007/s13225-020-00466-2)) |
| Otu63269 | 1 | k__Fungi(100);p__Ascomycota(100);c__Sordariomycetes(100);o__Hypocreales(100);f__Nectriaceae(100);g__Gibberella(100);g__Gibberella_unclassified(100); | Gibberella | 13 | Pathotroph-Saprotroph-Symbiotroph | Endophyte-\|Plant Pathogen\|-Plant Saprotroph-Undefined Saprotroph | Microfungus | Soft Rot | Probable | As foliar_endophyte (Põlme et al. 2020); As litter_saprotroph (Põlme et al. 2020); Causing damping-off disease (Põlme et al. 2020); Decay leaf, fruit, seed, roots, soil, animal material (Põlme et al. 2020) | Tedersoo L, et al. 2014. Science 346:e1256688 ((https://doi.org/10.1126/science.1256688)); Põlme S, et al. 2020. Fungal Diversity 105:1-16 ((https://doi.org/10.1007/s13225-020-00466-2)) |
| Otu63270 | 1 | k__Fungi(100);p__Ascomycota(100);c__Saccharomycetes(100);o__Saccharomycetales(100);f__Metschnikowiaceae(100);g__Metschnikowia(100);s__Metschnikowia_pulcherrima(100); | Metschnikowia | 13 | Saprotroph | \|Undefined Saprotroph\| | Yeast | NULL | Probable | As nectar-tap_saprotroph (Põlme et al. 2020); Decay sugar-rich substrates (Põlme et al. 2020) | Põlme S, et al. 2020. Fungal Diversity 105:1-16 ((https://doi.org/10.1007/s13225-020-00466-2)) |
| Otu63271 | 1 | k__Fungi(100);p__Ascomycota(100);c__Saccharomycetes(100);o__Saccharomycetales(100);f__Saccharomycetales_family_Incertae_sedis(100);g__Candida(100);s__Candida_sp_AS23486(100); | Candida | 13 | Pathotroph-Saprotroph-Symbiotroph | Animal Pathogen-Endophyte-Endosymbiont-Epiphyte-\|Undefined Saprotroph\| | Dimorphic Yeast-Yeast | NULL | Probable | As nectar-tap_saprotroph (Põlme et al. 2020); Decay sugar-rich substrates (Põlme et al. 2020); This asexual genus is polyphyletic and contain species that function very differently depend on what they are. Caution should be taken when interpreting guilds for Candida species. | Manolakaki D, et al. 2010. Virulence. 1: 367-375 ((https://doi.org/10.4161/viru.1.5.12796)); Mekbib SB, et al. 2011. Tropical Plant Pathology 36:233-240 ((https://doi.org/10.1590/S1982-56762011000400004)); Põlme S, et al. 2020. Fungal Diversity 105:1-16 ((https://doi.org/10.1007/s13225-020-00466-2)) |
| Otu63272 | 1 | k__Fungi(100);p__Ascomycota(100);c__Saccharomycetes(100);o__Saccharomycetales(100);f__Metschnikowiaceae(100);g__Metschnikowia(100);s__Metschnikowia_pulcherrima(100); | Metschnikowia | 13 | Saprotroph | \|Undefined Saprotroph\| | Yeast | NULL | Probable | As nectar-tap_saprotroph (Põlme et al. 2020); Decay sugar-rich substrates (Põlme et al. 2020) | Põlme S, et al. 2020. Fungal Diversity 105:1-16 ((https://doi.org/10.1007/s13225-020-00466-2)) |
| Otu12968 | 1 | k__Fungi(100);p__Ascomycota(100);p__Ascomycota_unclassified(100);p__Ascomycota_unclassified(100);p__Ascomycota_unclassified(100);p__Ascomycota_unclassified(100);p__Ascomycota_unclassified(100); | - | - | - | - | - | - | - | Unassigned | - |
| Otu12969 | 1 | k__Fungi(100);p__Ascomycota(100);p__Ascomycota_unclassified(100);p__Ascomycota_unclassified(100);p__Ascomycota_unclassified(100);p__Ascomycota_unclassified(100);p__Ascomycota_unclassified(100); | - | - | - | - | - | - | - | Unassigned | - |
| Otu12971 | 1 | k__Fungi(100);p__Ascomycota(100);p__Ascomycota_unclassified(100);p__Ascomycota_unclassified(100);p__Ascomycota_unclassified(100);p__Ascomycota_unclassified(100);p__Ascomycota_unclassified(100); | - | - | - | - | - | - | - | Unassigned | - |
| Otu12974 | 1 | k__Fungi(100);k__Fungi_unclassified(100);k__Fungi_unclassified(100);k__Fungi_unclassified(100);k__Fungi_unclassified(100);k__Fungi_unclassified(100);k__Fungi_unclassified(100); | - | - | - | - | - | - | - | Unassigned | - |
| Otu12975 | 1 | k__Fungi(100);p__Ascomycota(100);p__Ascomycota_unclassified(100);p__Ascomycota_unclassified(100);p__Ascomycota_unclassified(100);p__Ascomycota_unclassified(100);p__Ascomycota_unclassified(100); | - | - | - | - | - | - | - | Unassigned | - |
| Otu12979 | 1 | k__Fungi(100);p__Ascomycota(100);p__Ascomycota_unclassified(100);p__Ascomycota_unclassified(100);p__Ascomycota_unclassified(100);p__Ascomycota_unclassified(100);p__Ascomycota_unclassified(100); | - | - | - | - | - | - | - | Unassigned | - |
| Otu12980 | 1 | k__Fungi(100);p__Ascomycota(100);p__Ascomycota_unclassified(100);p__Ascomycota_unclassified(100);p__Ascomycota_unclassified(100);p__Ascomycota_unclassified(100);p__Ascomycota_unclassified(100); | - | - | - | - | - | - | - | Unassigned | - |
| Otu12981 | 1 | k__Fungi(100);k__Fungi_unclassified(100);k__Fungi_unclassified(100);k__Fungi_unclassified(100);k__Fungi_unclassified(100);k__Fungi_unclassified(100);k__Fungi_unclassified(100); | - | - | - | - | - | - | - | Unassigned | - |
| Otu12982 | 1 | k__Fungi(100);p__Ascomycota(100);p__Ascomycota_unclassified(100);p__Ascomycota_unclassified(100);p__Ascomycota_unclassified(100);p__Ascomycota_unclassified(100);p__Ascomycota_unclassified(100); | - | - | - | - | - | - | - | Unassigned | - |
| Otu12985 | 1 | k__Fungi(100);k__Fungi_unclassified(100);k__Fungi_unclassified(100);k__Fungi_unclassified(100);k__Fungi_unclassified(100);k__Fungi_unclassified(100);k__Fungi_unclassified(100); | - | - | - | - | - | - | - | Unassigned | - |
| Otu12986 | 1 | k__Fungi(100);p__Ascomycota(100);p__Ascomycota_unclassified(100);p__Ascomycota_unclassified(100);p__Ascomycota_unclassified(100);p__Ascomycota_unclassified(100);p__Ascomycota_unclassified(100); | - | - | - | - | - | - | - | Unassigned | - |
| Otu12988 | 1 | k__Fungi(100);k__Fungi_unclassified(100);k__Fungi_unclassified(100);k__Fungi_unclassified(100);k__Fungi_unclassified(100);k__Fungi_unclassified(100);k__Fungi_unclassified(100); | - | - | - | - | - | - | - | Unassigned | - |
| Otu12989 | 1 | k__Fungi(100);k__Fungi_unclassified(100);k__Fungi_unclassified(100);k__Fungi_unclassified(100);k__Fungi_unclassified(100);k__Fungi_unclassified(100);k__Fungi_unclassified(100); | - | - | - | - | - | - | - | Unassigned | - |
| Otu12990 | 1 | k__Fungi(100);p__Ascomycota(100);p__Ascomycota_unclassified(100);p__Ascomycota_unclassified(100);p__Ascomycota_unclassified(100);p__Ascomycota_unclassified(100);p__Ascomycota_unclassified(100); | - | - | - | - | - | - | - | Unassigned | - |
| Otu12992 | 1 | k__Fungi(100);k__Fungi_unclassified(100);k__Fungi_unclassified(100);k__Fungi_unclassified(100);k__Fungi_unclassified(100);k__Fungi_unclassified(100);k__Fungi_unclassified(100); | - | - | - | - | - | - | - | Unassigned | - |
| Otu12996 | 1 | k__Fungi(100);k__Fungi_unclassified(100);k__Fungi_unclassified(100);k__Fungi_unclassified(100);k__Fungi_unclassified(100);k__Fungi_unclassified(100);k__Fungi_unclassified(100); | - | - | - | - | - | - | - | Unassigned | - |
| Otu13004 | 1 | k__Fungi(100);k__Fungi_unclassified(100);k__Fungi_unclassified(100);k__Fungi_unclassified(100);k__Fungi_unclassified(100);k__Fungi_unclassified(100);k__Fungi_unclassified(100); | - | - | - | - | - | - | - | Unassigned | - |
| Otu13007 | 1 | k__Fungi(100);p__Ascomycota(100);p__Ascomycota_unclassified(100);p__Ascomycota_unclassified(100);p__Ascomycota_unclassified(100);p__Ascomycota_unclassified(100);p__Ascomycota_unclassified(100); | - | - | - | - | - | - | - | Unassigned | - |
| Otu13009 | 1 | k__Fungi(100);p__Ascomycota(100);p__Ascomycota_unclassified(100);p__Ascomycota_unclassified(100);p__Ascomycota_unclassified(100);p__Ascomycota_unclassified(100);p__Ascomycota_unclassified(100); | - | - | - | - | - | - | - | Unassigned | - |
| Otu13010 | 1 | k__Fungi(100);k__Fungi_unclassified(100);k__Fungi_unclassified(100);k__Fungi_unclassified(100);k__Fungi_unclassified(100);k__Fungi_unclassified(100);k__Fungi_unclassified(100); | - | - | - | - | - | - | - | Unassigned | - |
| Otu13013 | 1 | k__Fungi(100);p__Ascomycota(100);p__Ascomycota_unclassified(100);p__Ascomycota_unclassified(100);p__Ascomycota_unclassified(100);p__Ascomycota_unclassified(100);p__Ascomycota_unclassified(100); | - | - | - | - | - | - | - | Unassigned | - |

| | | | | | | | | | | | |
|---|---|---|---|---|---|---|---|---|---|---|---|
| Otu13014 | 1 | k__Fungi(100);k__Fungi_unclassified(100);k__Fungi_unclassified(100);k__Fungi_unclassified(100);k__Fungi_unclassified(100);k__Fungi_unclassified(100);k__Fungi_unclassified(100); | - | - | - | - | - | - | - | Unassigned | - |
| Otu13016 | 1 | k__Fungi(100);p__Ascomycota(100);p__Ascomycota_unclassified(100);p__Ascomycota_unclassified(100);p__Ascomycota_unclassified(100);p__Ascomycota_unclassified(100);p__Ascomycota_unclassified(100); | - | - | - | - | - | - | - | Unassigned | - |
| Otu13017 | 1 | k__Fungi(100);p__Ascomycota(100);p__Ascomycota_unclassified(100);p__Ascomycota_unclassified(100);p__Ascomycota_unclassified(100);p__Ascomycota_unclassified(100);p__Ascomycota_unclassified(100); | - | - | - | - | - | - | - | Unassigned | - |
| Otu13019 | 1 | k__Fungi(100);p__Ascomycota(100);p__Ascomycota_unclassified(100);p__Ascomycota_unclassified(100);p__Ascomycota_unclassified(100);p__Ascomycota_unclassified(100);p__Ascomycota_unclassified(100); | - | - | - | - | - | - | - | Unassigned | - |
| Otu13022 | 1 | k__Fungi(100);p__Ascomycota(100);p__Ascomycota_unclassified(100);p__Ascomycota_unclassified(100);p__Ascomycota_unclassified(100);p__Ascomycota_unclassified(100);p__Ascomycota_unclassified(100); | - | - | - | - | - | - | - | Unassigned | - |
| Otu13023 | 1 | k__Fungi(100);k__Fungi_unclassified(100);k__Fungi_unclassified(100);k__Fungi_unclassified(100);k__Fungi_unclassified(100);k__Fungi_unclassified(100);k__Fungi_unclassified(100); | - | - | - | - | - | - | - | Unassigned | - |
| Otu13025 | 1 | k__Fungi(100);p__Ascomycota(100);p__Ascomycota_unclassified(100);p__Ascomycota_unclassified(100);p__Ascomycota_unclassified(100);p__Ascomycota_unclassified(100);p__Ascomycota_unclassified(100); | - | - | - | - | - | - | - | Unassigned | - |
| Otu13026 | 1 | k__Fungi(100);p__Ascomycota(100);p__Ascomycota_unclassified(100);p__Ascomycota_unclassified(100);p__Ascomycota_unclassified(100);p__Ascomycota_unclassified(100);p__Ascomycota_unclassified(100); | - | - | - | - | - | - | - | Unassigned | - |
| Otu13027 | 1 | k__Fungi(100);p__Ascomycota(100);p__Ascomycota_unclassified(100);p__Ascomycota_unclassified(100);p__Ascomycota_unclassified(100);p__Ascomycota_unclassified(100);p__Ascomycota_unclassified(100); | - | - | - | - | - | - | - | Unassigned | - |
| Otu13028 | 1 | k__Fungi(100);p__Ascomycota(100);p__Ascomycota_unclassified(100);p__Ascomycota_unclassified(100);p__Ascomycota_unclassified(100);p__Ascomycota_unclassified(100);p__Ascomycota_unclassified(100); | - | - | - | - | - | - | - | Unassigned | - |
| Otu13029 | 1 | k__Fungi(100);p__Ascomycota(100);p__Ascomycota_unclassified(100);p__Ascomycota_unclassified(100);p__Ascomycota_unclassified(100);p__Ascomycota_unclassified(100);p__Ascomycota_unclassified(100); | - | - | - | - | - | - | - | Unassigned | - |
| Otu13030 | 1 | k__Fungi(100);p__Ascomycota(100);p__Ascomycota_unclassified(100);p__Ascomycota_unclassified(100);p__Ascomycota_unclassified(100);p__Ascomycota_unclassified(100);p__Ascomycota_unclassified(100); | - | - | - | - | - | - | - | Unassigned | - |
| Otu13031 | 1 | k__Fungi(100);k__Fungi_unclassified(100);k__Fungi_unclassified(100);k__Fungi_unclassified(100);k__Fungi_unclassified(100);k__Fungi_unclassified(100);k__Fungi_unclassified(100); | - | - | - | - | - | - | - | Unassigned | - |
| Otu13032 | 1 | k__Fungi(100);p__Ascomycota(100);p__Ascomycota_unclassified(100);p__Ascomycota_unclassified(100);p__Ascomycota_unclassified(100);p__Ascomycota_unclassified(100);p__Ascomycota_unclassified(100); | - | - | - | - | - | - | - | Unassigned | - |
| Otu13038 | 1 | k__Fungi(100);p__Ascomycota(100);p__Ascomycota_unclassified(100);p__Ascomycota_unclassified(100);p__Ascomycota_unclassified(100);p__Ascomycota_unclassified(100);p__Ascomycota_unclassified(100); | - | - | - | - | - | - | - | Unassigned | - |
| Otu13039 | 1 | k__Fungi(100);p__Ascomycota(100);p__Ascomycota_unclassified(100);p__Ascomycota_unclassified(100);p__Ascomycota_unclassified(100);p__Ascomycota_unclassified(100);p__Ascomycota_unclassified(100); | - | - | - | - | - | - | - | Unassigned | - |
| Otu13042 | 1 | k__Fungi(100);p__Ascomycota(100);p__Ascomycota_unclassified(100);p__Ascomycota_unclassified(100);p__Ascomycota_unclassified(100);p__Ascomycota_unclassified(100);p__Ascomycota_unclassified(100); | - | - | - | - | - | - | - | Unassigned | - |
| Otu13043 | 1 | k__Fungi(100);p__Ascomycota(100);p__Ascomycota_unclassified(100);p__Ascomycota_unclassified(100);p__Ascomycota_unclassified(100);p__Ascomycota_unclassified(100);p__Ascomycota_unclassified(100); | - | - | - | - | - | - | - | Unassigned | - |
| Otu13045 | 1 | k__Fungi(100);p__Ascomycota(100);p__Ascomycota_unclassified(100);p__Ascomycota_unclassified(100);p__Ascomycota_unclassified(100);p__Ascomycota_unclassified(100);p__Ascomycota_unclassified(100); | - | - | - | - | - | - | - | Unassigned | - |
| Otu13046 | 1 | k__Fungi(100);p__Ascomycota(100);p__Ascomycota_unclassified(100);p__Ascomycota_unclassified(100);p__Ascomycota_unclassified(100);p__Ascomycota_unclassified(100);p__Ascomycota_unclassified(100); | - | - | - | - | - | - | - | Unassigned | - |
| Otu13047 | 1 | k__Fungi(100);k__Fungi_unclassified(100);k__Fungi_unclassified(100);k__Fungi_unclassified(100);k__Fungi_unclassified(100);k__Fungi_unclassified(100);k__Fungi_unclassified(100); | - | - | - | - | - | - | - | Unassigned | - |
| Otu13053 | 1 | k__Fungi(100);p__Ascomycota(100);p__Ascomycota_unclassified(100);p__Ascomycota_unclassified(100);p__Ascomycota_unclassified(100);p__Ascomycota_unclassified(100);p__Ascomycota_unclassified(100); | - | - | - | - | - | - | - | Unassigned | - |
| Otu13054 | 1 | k__Fungi(100);p__Ascomycota(100);p__Ascomycota_unclassified(100);p__Ascomycota_unclassified(100);p__Ascomycota_unclassified(100);p__Ascomycota_unclassified(100);p__Ascomycota_unclassified(100); | - | - | - | - | - | - | - | Unassigned | - |
| Otu13057 | 1 | k__Fungi(100);k__Fungi_unclassified(100);k__Fungi_unclassified(100);k__Fungi_unclassified(100);k__Fungi_unclassified(100);k__Fungi_unclassified(100);k__Fungi_unclassified(100); | - | - | - | - | - | - | - | Unassigned | - |
| Otu13060 | 1 | k__Fungi(100);k__Fungi_unclassified(100);k__Fungi_unclassified(100);k__Fungi_unclassified(100);k__Fungi_unclassified(100);k__Fungi_unclassified(100);k__Fungi_unclassified(100); | - | - | - | - | - | - | - | Unassigned | - |
| Otu13067 | 1 | k__Fungi(100);p__Ascomycota(100);p__Ascomycota_unclassified(100);p__Ascomycota_unclassified(100);p__Ascomycota_unclassified(100);p__Ascomycota_unclassified(100);p__Ascomycota_unclassified(100); | - | - | - | - | - | - | - | Unassigned | - |
| Otu13068 | 1 | k__Fungi(100);p__Ascomycota(100);p__Ascomycota_unclassified(100);p__Ascomycota_unclassified(100);p__Ascomycota_unclassified(100);p__Ascomycota_unclassified(100);p__Ascomycota_unclassified(100); | - | - | - | - | - | - | - | Unassigned | - |
| Otu13072 | 1 | k__Fungi(100);p__Ascomycota(100);p__Ascomycota_unclassified(100);p__Ascomycota_unclassified(100);p__Ascomycota_unclassified(100);p__Ascomycota_unclassified(100);p__Ascomycota_unclassified(100); | - | - | - | - | - | - | - | Unassigned | - |
| Otu13073 | 1 | k__Fungi(100);p__Ascomycota(100);p__Ascomycota_unclassified(100);p__Ascomycota_unclassified(100);p__Ascomycota_unclassified(100);p__Ascomycota_unclassified(100);p__Ascomycota_unclassified(100); | - | - | - | - | - | - | - | Unassigned | - |
| Otu13075 | 1 | k__Fungi(100);p__Ascomycota(100);p__Ascomycota_unclassified(100);p__Ascomycota_unclassified(100);p__Ascomycota_unclassified(100);p__Ascomycota_unclassified(100);p__Ascomycota_unclassified(100); | - | - | - | - | - | - | - | Unassigned | - |
| Otu13078 | 1 | k__Fungi(100);p__Ascomycota(100);p__Ascomycota_unclassified(100);p__Ascomycota_unclassified(100);p__Ascomycota_unclassified(100);p__Ascomycota_unclassified(100);p__Ascomycota_unclassified(100); | - | - | - | - | - | - | - | Unassigned | - |
| Otu13083 | 1 | k__Fungi(100);k__Fungi_unclassified(100);k__Fungi_unclassified(100);k__Fungi_unclassified(100);k__Fungi_unclassified(100);k__Fungi_unclassified(100);k__Fungi_unclassified(100); | - | - | - | - | - | - | - | Unassigned | - |
| Otu13086 | 1 | k__Fungi(100);p__Ascomycota(100);p__Ascomycota_unclassified(100);p__Ascomycota_unclassified(100);p__Ascomycota_unclassified(100);p__Ascomycota_unclassified(100);p__Ascomycota_unclassified(100); | - | - | - | - | - | - | - | Unassigned | - |
| Otu13092 | 1 | k__Fungi(100);k__Fungi_unclassified(100);k__Fungi_unclassified(100);k__Fungi_unclassified(100);k__Fungi_unclassified(100);k__Fungi_unclassified(100);k__Fungi_unclassified(100); | - | - | - | - | - | - | - | Unassigned | - |
| Otu13093 | 1 | k__Fungi(100);k__Fungi_unclassified(100);k__Fungi_unclassified(100);k__Fungi_unclassified(100);k__Fungi_unclassified(100);k__Fungi_unclassified(100);k__Fungi_unclassified(100); | - | - | - | - | - | - | - | Unassigned | - |
| Otu13094 | 1 | k__Fungi(100);p__Ascomycota(100);p__Ascomycota_unclassified(100);p__Ascomycota_unclassified(100);p__Ascomycota_unclassified(100);p__Ascomycota_unclassified(100);p__Ascomycota_unclassified(100); | - | - | - | - | - | - | - | Unassigned | - |
| Otu13097 | 1 | k__Fungi(100);p__Ascomycota(100);p__Ascomycota_unclassified(100);p__Ascomycota_unclassified(100);p__Ascomycota_unclassified(100);p__Ascomycota_unclassified(100);p__Ascomycota_unclassified(100); | - | - | - | - | - | - | - | Unassigned | - |
| Otu13100 | 1 | k__Fungi(100);k__Fungi_unclassified(100);k__Fungi_unclassified(100);k__Fungi_unclassified(100);k__Fungi_unclassified(100);k__Fungi_unclassified(100);k__Fungi_unclassified(100); | - | - | - | - | - | - | - | Unassigned | - |
| Otu13102 | 1 | k__Fungi(100);k__Fungi_unclassified(100);k__Fungi_unclassified(100);k__Fungi_unclassified(100);k__Fungi_unclassified(100);k__Fungi_unclassified(100);k__Fungi_unclassified(100); | - | - | - | - | - | - | - | Unassigned | - |
| Otu13104 | 1 | k__Fungi(100);k__Fungi_unclassified(100);k__Fungi_unclassified(100);k__Fungi_unclassified(100);k__Fungi_unclassified(100);k__Fungi_unclassified(100);k__Fungi_unclassified(100); | - | - | - | - | - | - | - | Unassigned | - |
| Otu13105 | 1 | k__Fungi(100);p__Ascomycota(100);p__Ascomycota_unclassified(100);p__Ascomycota_unclassified(100);p__Ascomycota_unclassified(100);p__Ascomycota_unclassified(100);p__Ascomycota_unclassified(100); | - | - | - | - | - | - | - | Unassigned | - |
| Otu13110 | 1 | k__Fungi(100);k__Fungi_unclassified(100);k__Fungi_unclassified(100);k__Fungi_unclassified(100);k__Fungi_unclassified(100);k__Fungi_unclassified(100);k__Fungi_unclassified(100); | - | - | - | - | - | - | - | Unassigned | - |
| Otu13113 | 1 | k__Fungi(100);p__Ascomycota(100);p__Ascomycota_unclassified(100);p__Ascomycota_unclassified(100);p__Ascomycota_unclassified(100);p__Ascomycota_unclassified(100);p__Ascomycota_unclassified(100); | - | - | - | - | - | - | - | Unassigned | - |
| Otu13115 | 1 | k__Fungi(100);p__Ascomycota(100);p__Ascomycota_unclassified(100);p__Ascomycota_unclassified(100);p__Ascomycota_unclassified(100);p__Ascomycota_unclassified(100);p__Ascomycota_unclassified(100); | - | - | - | - | - | - | - | Unassigned | - |
| Otu13117 | 1 | k__Fungi(100);p__Ascomycota(100);p__Ascomycota_unclassified(100);p__Ascomycota_unclassified(100);p__Ascomycota_unclassified(100);p__Ascomycota_unclassified(100);p__Ascomycota_unclassified(100); | - | - | - | - | - | - | - | Unassigned | - |
| Otu13119 | 1 | k__Fungi(100);p__Ascomycota(100);p__Ascomycota_unclassified(100);p__Ascomycota_unclassified(100);p__Ascomycota_unclassified(100);p__Ascomycota_unclassified(100);p__Ascomycota_unclassified(100); | - | - | - | - | - | - | - | Unassigned | - |
| Otu13124 | 1 | k__Fungi(100);k__Fungi_unclassified(100);k__Fungi_unclassified(100);k__Fungi_unclassified(100);k__Fungi_unclassified(100);k__Fungi_unclassified(100);k__Fungi_unclassified(100); | - | - | - | - | - | - | - | Unassigned | - |
| Otu13132 | 1 | k__Fungi(100);k__Fungi_unclassified(100);k__Fungi_unclassified(100);k__Fungi_unclassified(100);k__Fungi_unclassified(100);k__Fungi_unclassified(100);k__Fungi_unclassified(100); | - | - | - | - | - | - | - | Unassigned | - |
| Otu13135 | 1 | k__Fungi(100);p__Ascomycota(100);p__Ascomycota_unclassified(100);p__Ascomycota_unclassified(100);p__Ascomycota_unclassified(100);p__Ascomycota_unclassified(100);p__Ascomycota_unclassified(100); | - | - | - | - | - | - | - | Unassigned | - |
| Otu13141 | 1 | k__Fungi(100);k__Fungi_unclassified(100);k__Fungi_unclassified(100);k__Fungi_unclassified(100);k__Fungi_unclassified(100);k__Fungi_unclassified(100);k__Fungi_unclassified(100); | - | - | - | - | - | - | - | Unassigned | - |
| Otu13142 | 1 | k__Fungi(100);p__Ascomycota(100);p__Ascomycota_unclassified(100);p__Ascomycota_unclassified(100);p__Ascomycota_unclassified(100);p__Ascomycota_unclassified(100);p__Ascomycota_unclassified(100); | - | - | - | - | - | - | - | Unassigned | - |
| Otu13145 | 1 | k__Fungi(100);p__Ascomycota(100);p__Ascomycota_unclassified(100);p__Ascomycota_unclassified(100);p__Ascomycota_unclassified(100);p__Ascomycota_unclassified(100);p__Ascomycota_unclassified(100); | - | - | - | - | - | - | - | Unassigned | - |
| Otu13150 | 1 | k__Fungi(100);p__Ascomycota(100);p__Ascomycota_unclassified(100);p__Ascomycota_unclassified(100);p__Ascomycota_unclassified(100);p__Ascomycota_unclassified(100);p__Ascomycota_unclassified(100); | - | - | - | - | - | - | - | Unassigned | - |
| Otu13151 | 1 | k__Fungi(100);p__Ascomycota(100);p__Ascomycota_unclassified(100);p__Ascomycota_unclassified(100);p__Ascomycota_unclassified(100);p__Ascomycota_unclassified(100);p__Ascomycota_unclassified(100); | - | - | - | - | - | - | - | Unassigned | - |
| Otu13154 | 1 | k__Fungi(100);k__Fungi_unclassified(100);k__Fungi_unclassified(100);k__Fungi_unclassified(100);k__Fungi_unclassified(100);k__Fungi_unclassified(100);k__Fungi_unclassified(100); | - | - | - | - | - | - | - | Unassigned | - |
| Otu13155 | 1 | k__Fungi(100);k__Fungi_unclassified(100);k__Fungi_unclassified(100);k__Fungi_unclassified(100);k__Fungi_unclassified(100);k__Fungi_unclassified(100);k__Fungi_unclassified(100); | - | - | - | - | - | - | - | Unassigned | - |
| Otu13159 | 1 | k__Fungi(100);k__Fungi_unclassified(100);k__Fungi_unclassified(100);k__Fungi_unclassified(100);k__Fungi_unclassified(100);k__Fungi_unclassified(100);k__Fungi_unclassified(100); | - | - | - | - | - | - | - | Unassigned | - |
| Otu13162 | 1 | k__Fungi(100);k__Fungi_unclassified(100);k__Fungi_unclassified(100);k__Fungi_unclassified(100);k__Fungi_unclassified(100);k__Fungi_unclassified(100);k__Fungi_unclassified(100); | - | - | - | - | - | - | - | Unassigned | - |
| Otu13163 | 1 | k__Fungi(100);p__Ascomycota(100);p__Ascomycota_unclassified(100);p__Ascomycota_unclassified(100);p__Ascomycota_unclassified(100);p__Ascomycota_unclassified(100);p__Ascomycota_unclassified(100); | - | - | - | - | - | - | - | Unassigned | - |
| Otu13164 | 1 | k__Fungi(100);k__Fungi_unclassified(100);k__Fungi_unclassified(100);k__Fungi_unclassified(100);k__Fungi_unclassified(100);k__Fungi_unclassified(100);k__Fungi_unclassified(100); | - | - | - | - | - | - | - | Unassigned | - |
| Otu13166 | 1 | k__Fungi(100);k__Fungi_unclassified(100);k__Fungi_unclassified(100);k__Fungi_unclassified(100);k__Fungi_unclassified(100);k__Fungi_unclassified(100);k__Fungi_unclassified(100); | - | - | - | - | - | - | - | Unassigned | - |
| Otu13168 | 1 | k__Fungi(100);p__Ascomycota(100);p__Ascomycota_unclassified(100);p__Ascomycota_unclassified(100);p__Ascomycota_unclassified(100);p__Ascomycota_unclassified(100);p__Ascomycota_unclassified(100); | - | - | - | - | - | - | - | Unassigned | - |
| Otu13170 | 1 | k__Fungi(100);p__Ascomycota(100);p__Ascomycota_unclassified(100);p__Ascomycota_unclassified(100);p__Ascomycota_unclassified(100);p__Ascomycota_unclassified(100);p__Ascomycota_unclassified(100); | - | - | - | - | - | - | - | Unassigned | - |
| Otu13178 | 1 | k__Fungi(100);k__Fungi_unclassified(100);k__Fungi_unclassified(100);k__Fungi_unclassified(100);k__Fungi_unclassified(100);k__Fungi_unclassified(100);k__Fungi_unclassified(100); | - | - | - | - | - | - | - | Unassigned | - |
| Otu13183 | 1 | k__Fungi(100);k__Fungi_unclassified(100);k__Fungi_unclassified(100);k__Fungi_unclassified(100);k__Fungi_unclassified(100);k__Fungi_unclassified(100);k__Fungi_unclassified(100); | - | - | - | - | - | - | - | Unassigned | - |
| Otu13187 | 1 | k__Fungi(100);k__Fungi_unclassified(100);k__Fungi_unclassified(100);k__Fungi_unclassified(100);k__Fungi_unclassified(100);k__Fungi_unclassified(100);k__Fungi_unclassified(100); | - | - | - | - | - | - | - | Unassigned | - |
| Otu13190 | 1 | k__Fungi(100);p__Ascomycota(100);p__Ascomycota_unclassified(100);p__Ascomycota_unclassified(100);p__Ascomycota_unclassified(100);p__Ascomycota_unclassified(100);p__Ascomycota_unclassified(100); | - | - | - | - | - | - | - | Unassigned | - |
| Otu13192 | 1 | k__Fungi(100);p__Ascomycota(100);p__Ascomycota_unclassified(100);p__Ascomycota_unclassified(100);p__Ascomycota_unclassified(100);p__Ascomycota_unclassified(100);p__Ascomycota_unclassified(100); | - | - | - | - | - | - | - | Unassigned | - |
| Otu13193 | 1 | k__Fungi(100);p__Ascomycota(100);p__Ascomycota_unclassified(100);p__Ascomycota_unclassified(100);p__Ascomycota_unclassified(100);p__Ascomycota_unclassified(100);p__Ascomycota_unclassified(100); | - | - | - | - | - | - | - | Unassigned | - |
| Otu13198 | 1 | k__Fungi(100);p__Ascomycota(100);p__Ascomycota_unclassified(100);p__Ascomycota_unclassified(100);p__Ascomycota_unclassified(100);p__Ascomycota_unclassified(100);p__Ascomycota_unclassified(100); | - | - | - | - | - | - | - | Unassigned | - |
| Otu13200 | 1 | k__Fungi(100);k__Fungi_unclassified(100);k__Fungi_unclassified(100);k__Fungi_unclassified(100);k__Fungi_unclassified(100);k__Fungi_unclassified(100);k__Fungi_unclassified(100); | - | - | - | - | - | - | - | Unassigned | - |
| Otu13201 | 1 | k__Fungi(100);k__Fungi_unclassified(100);k__Fungi_unclassified(100);k__Fungi_unclassified(100);k__Fungi_unclassified(100);k__Fungi_unclassified(100);k__Fungi_unclassified(100); | - | - | - | - | - | - | - | Unassigned | - |
| Otu13202 | 1 | k__Fungi(100);k__Fungi_unclassified(100);k__Fungi_unclassified(100);k__Fungi_unclassified(100);k__Fungi_unclassified(100);k__Fungi_unclassified(100);k__Fungi_unclassified(100); | - | - | - | - | - | - | - | Unassigned | - |
| Otu13206 | 1 | k__Fungi(100);k__Fungi_unclassified(100);k__Fungi_unclassified(100);k__Fungi_unclassified(100);k__Fungi_unclassified(100);k__Fungi_unclassified(100);k__Fungi_unclassified(100); | - | - | - | - | - | - | - | Unassigned | - |
| Otu13208 | 1 | k__Fungi(100);k__Fungi_unclassified(100);k__Fungi_unclassified(100);k__Fungi_unclassified(100);k__Fungi_unclassified(100);k__Fungi_unclassified(100);k__Fungi_unclassified(100); | - | - | - | - | - | - | - | Unassigned | - |
| Otu13215 | 1 | k__Fungi(100);k__Fungi_unclassified(100);k__Fungi_unclassified(100);k__Fungi_unclassified(100);k__Fungi_unclassified(100);k__Fungi_unclassified(100);k__Fungi_unclassified(100); | - | - | - | - | - | - | - | Unassigned | - |
| Otu13216 | 1 | k__Fungi(100);p__Ascomycota(100);p__Ascomycota_unclassified(100);p__Ascomycota_unclassified(100);p__Ascomycota_unclassified(100);p__Ascomycota_unclassified(100);p__Ascomycota_unclassified(100); | - | - | - | - | - | - | - | Unassigned | - |
| Otu13217 | 1 | k__Fungi(100);p__Ascomycota(100);p__Ascomycota_unclassified(100);p__Ascomycota_unclassified(100);p__Ascomycota_unclassified(100);p__Ascomycota_unclassified(100);p__Ascomycota_unclassified(100); | - | - | - | - | - | - | - | Unassigned | - |
| Otu13218 | 1 | k__Fungi(100);p__Ascomycota(100);p__Ascomycota_unclassified(100);p__Ascomycota_unclassified(100);p__Ascomycota_unclassified(100);p__Ascomycota_unclassified(100);p__Ascomycota_unclassified(100); | - | - | - | - | - | - | - | Unassigned | - |
| Otu13219 | 1 | k__Fungi(100);k__Fungi_unclassified(100);k__Fungi_unclassified(100);k__Fungi_unclassified(100);k__Fungi_unclassified(100);k__Fungi_unclassified(100);k__Fungi_unclassified(100); | - | - | - | - | - | - | - | Unassigned | - |
| Otu13221 | 1 | k__Fungi(100);p__Ascomycota(100);p__Ascomycota_unclassified(100);p__Ascomycota_unclassified(100);p__Ascomycota_unclassified(100);p__Ascomycota_unclassified(100);p__Ascomycota_unclassified(100); | - | - | - | - | - | - | - | Unassigned | - |
| Otu13223 | 1 | k__Fungi(100);p__Ascomycota(100);p__Ascomycota_unclassified(100);p__Ascomycota_unclassified(100);p__Ascomycota_unclassified(100);p__Ascomycota_unclassified(100);p__Ascomycota_unclassified(100); | - | - | - | - | - | - | - | Unassigned | - |
| Otu13230 | 1 | k__Fungi(100);p__Ascomycota(100);p__Ascomycota_unclassified(100);p__Ascomycota_unclassified(100);p__Ascomycota_unclassified(100);p__Ascomycota_unclassified(100);p__Ascomycota_unclassified(100); | - | - | - | - | - | - | - | Unassigned | - |
| Otu13233 | 1 | k__Fungi(100);k__Fungi_unclassified(100);k__Fungi_unclassified(100);k__Fungi_unclassified(100);k__Fungi_unclassified(100);k__Fungi_unclassified(100);k__Fungi_unclassified(100); | - | - | - | - | - | - | - | Unassigned | - |
| Otu13236 | 1 | k__Fungi(100);k__Fungi_unclassified(100);k__Fungi_unclassified(100);k__Fungi_unclassified(100);k__Fungi_unclassified(100);k__Fungi_unclassified(100);k__Fungi_unclassified(100); | - | - | - | - | - | - | - | Unassigned | - |
| Otu13245 | 1 | k__Fungi(100);k__Fungi_unclassified(100);k__Fungi_unclassified(100);k__Fungi_unclassified(100);k__Fungi_unclassified(100);k__Fungi_unclassified(100);k__Fungi_unclassified(100); | - | - | - | - | - | - | - | Unassigned | - |
| Otu13249 | 1 | k__Fungi(100);p__Ascomycota(100);p__Ascomycota_unclassified(100);p__Ascomycota_unclassified(100);p__Ascomycota_unclassified(100);p__Ascomycota_unclassified(100);p__Ascomycota_unclassified(100); | - | - | - | - | - | - | - | Unassigned | - |
| Otu13250 | 1 | k__Fungi(100);p__Ascomycota(100);p__Ascomycota_unclassified(100);p__Ascomycota_unclassified(100);p__Ascomycota_unclassified(100);p__Ascomycota_unclassified(100);p__Ascomycota_unclassified(100); | - | - | - | - | - | - | - | Unassigned | - |
| Otu13252 | 1 | k__Fungi(100);k__Fungi_unclassified(100);k__Fungi_unclassified(100);k__Fungi_unclassified(100);k__Fungi_unclassified(100);k__Fungi_unclassified(100);k__Fungi_unclassified(100); | - | - | - | - | - | - | - | Unassigned | - |
| Otu13253 | 1 | k__Fungi(100);k__Fungi_unclassified(100);k__Fungi_unclassified(100);k__Fungi_unclassified(100);k__Fungi_unclassified(100);k__Fungi_unclassified(100);k__Fungi_unclassified(100); | - | - | - | - | - | - | - | Unassigned | - |
| Otu13256 | 1 | k__Fungi(100);p__Ascomycota(100);p__Ascomycota_unclassified(100);p__Ascomycota_unclassified(100);p__Ascomycota_unclassified(100);p__Ascomycota_unclassified(100);p__Ascomycota_unclassified(100); | - | - | - | - | - | - | - | Unassigned | - |
| Otu13258 | 1 | k__Fungi(100);k__Fungi_unclassified(100);k__Fungi_unclassified(100);k__Fungi_unclassified(100);k__Fungi_unclassified(100);k__Fungi_unclassified(100);k__Fungi_unclassified(100); | - | - | - | - | - | - | - | Unassigned | - |
| Otu13259 | 1 | k__Fungi(100);k__Fungi_unclassified(100);k__Fungi_unclassified(100);k__Fungi_unclassified(100);k__Fungi_unclassified(100);k__Fungi_unclassified(100);k__Fungi_unclassified(100); | - | - | - | - | - | - | - | Unassigned | - |
| Otu13263 | 1 | k__Fungi(100);p__Ascomycota(100);p__Ascomycota_unclassified(100);p__Ascomycota_unclassified(100);p__Ascomycota_unclassified(100);p__Ascomycota_unclassified(100);p__Ascomycota_unclassified(100); | - | - | - | - | - | - | - | Unassigned | - |
| Otu13264 | 1 | k__Fungi(100);p__Ascomycota(100);p__Ascomycota_unclassified(100);p__Ascomycota_unclassified(100);p__Ascomycota_unclassified(100);p__Ascomycota_unclassified(100);p__Ascomycota_unclassified(100); | - | - | - | - | - | - | - | Unassigned | - |
| Otu13268 | 1 | k__Fungi(100);p__Ascomycota(100);p__Ascomycota_unclassified(100);p__Ascomycota_unclassified(100);p__Ascomycota_unclassified(100);p__Ascomycota_unclassified(100);p__Ascomycota_unclassified(100); | - | - | - | - | - | - | - | Unassigned | - |
| Otu13274 | 1 | k__Fungi(100);p__Ascomycota(100);p__Ascomycota_unclassified(100);p__Ascomycota_unclassified(100);p__Ascomycota_unclassified(100);p__Ascomycota_unclassified(100);p__Ascomycota_unclassified(100); | - | - | - | - | - | - | - | Unassigned | - |
| Otu13278 | 1 | k__Fungi(100);p__Ascomycota(100);p__Ascomycota_unclassified(100);p__Ascomycota_unclassified(100);p__Ascomycota_unclassified(100);p__Ascomycota_unclassified(100);p__Ascomycota_unclassified(100); | - | - | - | - | - | - | - | Unassigned | - |
| Otu13279 | 1 | k__Fungi(100);p__Ascomycota(100);p__Ascomycota_unclassified(100);p__Ascomycota_unclassified(100);p__Ascomycota_unclassified(100);p__Ascomycota_unclassified(100);p__Ascomycota_unclassified(100); | - | - | - | - | - | - | - | Unassigned | - |
| Otu13281 | 1 | k__Fungi(100);k__Fungi_unclassified(100);k__Fungi_unclassified(100);k__Fungi_unclassified(100);k__Fungi_unclassified(100);k__Fungi_unclassified(100);k__Fungi_unclassified(100); | - | - | - | - | - | - | - | Unassigned | - |
| Otu13282 | 1 | k__Fungi(100);p__Ascomycota(100);p__Ascomycota_unclassified(100);p__Ascomycota_unclassified(100);p__Ascomycota_unclassified(100);p__Ascomycota_unclassified(100);p__Ascomycota_unclassified(100); | - | - | - | - | - | - | - | Unassigned | - |
| Otu13284 | 1 | k__Fungi(100);k__Fungi_unclassified(100);k__Fungi_unclassified(100);k__Fungi_unclassified(100);k__Fungi_unclassified(100);k__Fungi_unclassified(100);k__Fungi_unclassified(100); | - | - | - | - | - | - | - | Unassigned | - |
| Otu13286 | 1 | k__Fungi(100);k__Fungi_unclassified(100);k__Fungi_unclassified(100);k__Fungi_unclassified(100);k__Fungi_unclassified(100);k__Fungi_unclassified(100);k__Fungi_unclassified(100); | - | - | - | - | - | - | - | Unassigned | - |
| Otu13287 | 1 | k__Fungi(100);p__Ascomycota(100);p__Ascomycota_unclassified(100);p__Ascomycota_unclassified(100);p__Ascomycota_unclassified(100);p__Ascomycota_unclassified(100);p__Ascomycota_unclassified(100); | - | - | - | - | - | - | - | Unassigned | - |
| Otu13290 | 1 | k__Fungi(100);k__Fungi_unclassified(100);k__Fungi_unclassified(100);k__Fungi_unclassified(100);k__Fungi_unclassified(100);k__Fungi_unclassified(100);k__Fungi_unclassified(100); | - | - | - | - | - | - | - | Unassigned | - |
| Otu13293 | 1 | k__Fungi(100);k__Fungi_unclassified(100);k__Fungi_unclassified(100);k__Fungi_unclassified(100);k__Fungi_unclassified(100);k__Fungi_unclassified(100);k__Fungi_unclassified(100); | - | - | - | - | - | - | - | Unassigned | - |
| Otu13294 | 1 | k__Fungi(100);p__Ascomycota(100);p__Ascomycota_unclassified(100);p__Ascomycota_unclassified(100);p__Ascomycota_unclassified(100);p__Ascomycota_unclassified(100);p__Ascomycota_unclassified(100); | - | - | - | - | - | - | - | Unassigned | - |
| Otu13300 | 1 | k__Fungi(100);k__Fungi_unclassified(100);k__Fungi_unclassified(100);k__Fungi_unclassified(100);k__Fungi_unclassified(100);k__Fungi_unclassified(100);k__Fungi_unclassified(100); | - | - | - | - | - | - | - | Unassigned | - |
| Otu13301 | 1 | k__Fungi(100);k__Fungi_unclassified(100);k__Fungi_unclassified(100);k__Fungi_unclassified(100);k__Fungi_unclassified(100);k__Fungi_unclassified(100);k__Fungi_unclassified(100); | - | - | - | - | - | - | - | Unassigned | - |
| Otu13302 | 1 | k__Fungi(100);k__Fungi_unclassified(100);k__Fungi_unclassified(100);k__Fungi_unclassified(100);k__Fungi_unclassified(100);k__Fungi_unclassified(100);k__Fungi_unclassified(100); | - | - | - | - | - | - | - | Unassigned | - |
| Otu13311 | 1 | k__Fungi(100);k__Fungi_unclassified(100);k__Fungi_unclassified(100);k__Fungi_unclassified(100);k__Fungi_unclassified(100);k__Fungi_unclassified(100);k__Fungi_unclassified(100); | - | - | - | - | - | - | - | Unassigned | - |
| Otu13315 | 1 | k__Fungi(100);p__Ascomycota(100);p__Ascomycota_unclassified(100);p__Ascomycota_unclassified(100);p__Ascomycota_unclassified(100);p__Ascomycota_unclassified(100);p__Ascomycota_unclassified(100); | - | - | - | - | - | - | - | Unassigned | - |
| Otu13316 | 1 | k__Fungi(100);p__Ascomycota(100);p__Ascomycota_unclassified(100);p__Ascomycota_unclassified(100);p__Ascomycota_unclassified(100);p__Ascomycota_unclassified(100);p__Ascomycota_unclassified(100); | - | - | - | - | - | - | - | Unassigned | - |
| Otu13324 | 1 | k__Fungi(100);k__Fungi_unclassified(100);k__Fungi_unclassified(100);k__Fungi_unclassified(100);k__Fungi_unclassified(100);k__Fungi_unclassified(100);k__Fungi_unclassified(100); | - | - | - | - | - | - | - | Unassigned | - |
| Otu13329 | 1 | k__Fungi(100);k__Fungi_unclassified(100);k__Fungi_unclassified(100);k__Fungi_unclassified(100);k__Fungi_unclassified(100);k__Fungi_unclassified(100);k__Fungi_unclassified(100); | - | - | - | - | - | - | - | Unassigned | - |
| Otu13330 | 1 | k__Fungi(100);k__Fungi_unclassified(100);k__Fungi_unclassified(100);k__Fungi_unclassified(100);k__Fungi_unclassified(100);k__Fungi_unclassified(100);k__Fungi_unclassified(100); | - | - | - | - | - | - | - | Unassigned | - |
| Otu13332 | 1 | k__Fungi(100);k__Fungi_unclassified(100);k__Fungi_unclassified(100);k__Fungi_unclassified(100);k__Fungi_unclassified(100);k__Fungi_unclassified(100);k__Fungi_unclassified(100); | - | - | - | - | - | - | - | Unassigned | - |
| Otu13335 | 1 | k__Fungi(100);k__Fungi_unclassified(100);k__Fungi_unclassified(100);k__Fungi_unclassified(100);k__Fungi_unclassified(100);k__Fungi_unclassified(100);k__Fungi_unclassified(100); | - | - | - | - | - | - | - | Unassigned | - |
| Otu13346 | 1 | k__Fungi(100);k__Fungi_unclassified(100);k__Fungi_unclassified(100);k__Fungi_unclassified(100);k__Fungi_unclassified(100);k__Fungi_unclassified(100);k__Fungi_unclassified(100); | - | - | - | - | - | - | - | Unassigned | - |
| Otu13347 | 1 | k__Fungi(100);p__Ascomycota(100);p__Ascomycota_unclassified(100);p__Ascomycota_unclassified(100);p__Ascomycota_unclassified(100);p__Ascomycota_unclassified(100);p__Ascomycota_unclassified(100); | - | - | - | - | - | - | - | Unassigned | - |
| Otu13351 | 1 | k__Fungi(100);k__Fungi_unclassified(100);k__Fungi_unclassified(100);k__Fungi_unclassified(100);k__Fungi_unclassified(100);k__Fungi_unclassified(100);k__Fungi_unclassified(100); | - | - | - | - | - | - | - | Unassigned | - |
| Otu13353 | 1 | k__Fungi(100);k__Fungi_unclassified(100);k__Fungi_unclassified(100);k__Fungi_unclassified(100);k__Fungi_unclassified(100);k__Fungi_unclassified(100);k__Fungi_unclassified(100); | - | - | - | - | - | - | - | Unassigned | - |
| Otu13354 | 1 | k__Fungi(100);p__Ascomycota(100);p__Ascomycota_unclassified(100);p__Ascomycota_unclassified(100);p__Ascomycota_unclassified(100);p__Ascomycota_unclassified(100);p__Ascomycota_unclassified(100); | - | - | - | - | - | - | - | Unassigned | - |
| Otu13355 | 1 | k__Fungi(100);p__Ascomycota(100);p__Ascomycota_unclassified(100);p__Ascomycota_unclassified(100);p__Ascomycota_unclassified(100);p__Ascomycota_unclassified(100);p__Ascomycota_unclassified(100); | - | - | - | - | - | - | - | Unassigned | - |
| Otu13357 | 1 | k__Fungi(100);k__Fungi_unclassified(100);k__Fungi_unclassified(100);k__Fungi_unclassified(100);k__Fungi_unclassified(100);k__Fungi_unclassified(100);k__Fungi_unclassified(100); | - | - | - | - | - | - | - | Unassigned | - |
| Otu13363 | 1 | k__Fungi(100);k__Fungi_unclassified(100);k__Fungi_unclassified(100);k__Fungi_unclassified(100);k__Fungi_unclassified(100);k__Fungi_unclassified(100);k__Fungi_unclassified(100); | - | - | - | - | - | - | - | Unassigned | - |
| Otu13365 | 1 | k__Fungi(100);k__Fungi_unclassified(100);k__Fungi_unclassified(100);k__Fungi_unclassified(100);k__Fungi_unclassified(100);k__Fungi_unclassified(100);k__Fungi_unclassified(100); | - | - | - | - | - | - | - | Unassigned | - |
| Otu13368 | 1 | k__Fungi(100);p__Ascomycota(100);p__Ascomycota_unclassified(100);p__Ascomycota_unclassified(100);p__Ascomycota_unclassified(100);p__Ascomycota_unclassified(100);p__Ascomycota_unclassified(100); | - | - | - | - | - | - | - | Unassigned | - |
| Otu13373 | 1 | k__Fungi(100);p__Ascomycota(100);p__Ascomycota_unclassified(100);p__Ascomycota_unclassified(100);p__Ascomycota_unclassified(100);p__Ascomycota_unclassified(100);p__Ascomycota_unclassified(100); | - | - | - | - | - | - | - | Unassigned | - |
| Otu13379 | 1 | k__Fungi(100);k__Fungi_unclassified(100);k__Fungi_unclassified(100);k__Fungi_unclassified(100);k__Fungi_unclassified(100);k__Fungi_unclassified(100);k__Fungi_unclassified(100); | - | - | - | - | - | - | - | Unassigned | - |
| Otu13381 | 1 | k__Fungi(100);p__Ascomycota(100);p__Ascomycota_unclassified(100);p__Ascomycota_unclassified(100);p__Ascomycota_unclassified(100);p__Ascomycota_unclassified(100);p__Ascomycota_unclassified(100); | - | - | - | - | - | - | - | Unassigned | - |
| Otu13387 | 1 | k__Fungi(100);k__Fungi_unclassified(100);k__Fungi_unclassified(100);k__Fungi_unclassified(100);k__Fungi_unclassified(100);k__Fungi_unclassified(100);k__Fungi_unclassified(100); | - | - | - | - | - | - | - | Unassigned | - |
| Otu13390 | 1 | k__Fungi(100);p__Ascomycota(100);p__Ascomycota_unclassified(100);p__Ascomycota_unclassified(100);p__Ascomycota_unclassified(100);p__Ascomycota_unclassified(100);p__Ascomycota_unclassified(100); | - | - | - | - | - | - | - | Unassigned | - |
| Otu13398 | 1 | k__Fungi(100);p__Ascomycota(100);p__Ascomycota_unclassified(100);p__Ascomycota_unclassified(100);p__Ascomycota_unclassified(100);p__Ascomycota_unclassified(100);p__Ascomycota_unclassified(100); | - | - | - | - | - | - | - | Unassigned | - |
| Otu13402 | 1 | k__Fungi(100);k__Fungi_unclassified(100);k__Fungi_unclassified(100);k__Fungi_unclassified(100);k__Fungi_unclassified(100);k__Fungi_unclassified(100);k__Fungi_unclassified(100); | - | - | - | - | - | - | - | Unassigned | - |
| Otu13407 | 1 | k__Fungi(100);k__Fungi_unclassified(100);k__Fungi_unclassified(100);k__Fungi_unclassified(100);k__Fungi_unclassified(100);k__Fungi_unclassified(100);k__Fungi_unclassified(100); | - | - | - | - | - | - | - | Unassigned | - |
| Otu13410 | 1 | k__Fungi(100);p__Ascomycota(100);p__Ascomycota_unclassified(100);p__Ascomycota_unclassified(100);p__Ascomycota_unclassified(100);p__Ascomycota_unclassified(100);p__Ascomycota_unclassified(100); | - | - | - | - | - | - | - | Unassigned | - |
| Otu13412 | 1 | k__Fungi(100);k__Fungi_unclassified(100);k__Fungi_unclassified(100);k__Fungi_unclassified(100);k__Fungi_unclassified(100);k__Fungi_unclassified(100);k__Fungi_unclassified(100); | - | - | - | - | - | - | - | Unassigned | - |
| Otu13421 | 1 | k__Fungi(100);p__Ascomycota(100);p__Ascomycota_unclassified(100);p__Ascomycota_unclassified(100);p__Ascomycota_unclassified(100);p__Ascomycota_unclassified(100);p__Ascomycota_unclassified(100); | - | - | - | - | - | - | - | Unassigned | - |
| Otu13428 | 1 | k__Fungi(100);p__Ascomycota(100);p__Ascomycota_unclassified(100);p__Ascomycota_unclassified(100);p__Ascomycota_unclassified(100);p__Ascomycota_unclassified(100);p__Ascomycota_unclassified(100); | - | - | - | - | - | - | - | Unassigned | - |
| Otu13429 | 1 | k__Fungi(100);p__Ascomycota(100);p__Ascomycota_unclassified(100);p__Ascomycota_unclassified(100);p__Ascomycota_unclassified(100);p__Ascomycota_unclassified(100);p__Ascomycota_unclassified(100); | - | - | - | - | - | - | - | Unassigned | - |
| Otu13430 | 1 | k__Fungi(100);k__Fungi_unclassified(100);k__Fungi_unclassified(100);k__Fungi_unclassified(100);k__Fungi_unclassified(100);k__Fungi_unclassified(100);k__Fungi_unclassified(100); | - | - | - | - | - | - | - | Unassigned | - |
| Otu13431 | 1 | k__Fungi(100);p__Ascomycota(100);p__Ascomycota_unclassified(100);p__Ascomycota_unclassified(100);p__Ascomycota_unclassified(100);p__Ascomycota_unclassified(100);p__Ascomycota_unclassified(100); | - | - | - | - | - | - | - | Unassigned | - |
| Otu13434 | 1 | k__Fungi(100);k__Fungi_unclassified(100);k__Fungi_unclassified(100);k__Fungi_unclassified(100);k__Fungi_unclassified(100);k__Fungi_unclassified(100);k__Fungi_unclassified(100); | - | - | - | - | - | - | - | Unassigned | - |
| Otu13437 | 1 | k__Fungi(100);k__Fungi_unclassified(100);k__Fungi_unclassified(100);k__Fungi_unclassified(100);k__Fungi_unclassified(100);k__Fungi_unclassified(100);k__Fungi_unclassified(100); | - | - | - | - | - | - | - | Unassigned | - |
| Otu13438 | 1 | k__Fungi(100);k__Fungi_unclassified(100);k__Fungi_unclassified(100);k__Fungi_unclassified(100);k__Fungi_unclassified(100);k__Fungi_unclassified(100);k__Fungi_unclassified(100); | - | - | - | - | - | - | - | Unassigned | - |
| Otu13439 | 1 | k__Fungi(100);k__Fungi_unclassified(100);k__Fungi_unclassified(100);k__Fungi_unclassified(100);k__Fungi_unclassified(100);k__Fungi_unclassified(100);k__Fungi_unclassified(100); | - | - | - | - | - | - | - | Unassigned | - |
| Otu13440 | 1 | k__Fungi(100);p__Ascomycota(100);p__Ascomycota_unclassified(100);p__Ascomycota_unclassified(100);p__Ascomycota_unclassified(100);p__Ascomycota_unclassified(100);p__Ascomycota_unclassified(100); | - | - | - | - | - | - | - | Unassigned | - |
| Otu13443 | 1 | k__Fungi(100);p__Ascomycota(100);p__Ascomycota_unclassified(100);p__Ascomycota_unclassified(100);p__Ascomycota_unclassified(100);p__Ascomycota_unclassified(100);p__Ascomycota_unclassified(100); | - | - | - | - | - | - | - | Unassigned | - |
| Otu13445 | 1 | k__Fungi(100);k__Fungi_unclassified(100);k__Fungi_unclassified(100);k__Fungi_unclassified(100);k__Fungi_unclassified(100);k__Fungi_unclassified(100);k__Fungi_unclassified(100); | - | - | - | - | - | - | - | Unassigned | - |
| Otu13446 | 1 | k__Fungi(100);p__Ascomycota(100);p__Ascomycota_unclassified(100);p__Ascomycota_unclassified(100);p__Ascomycota_unclassified(100);p__Ascomycota_unclassified(100);p__Ascomycota_unclassified(100); | - | - | - | - | - | - | - | Unassigned | - |
| Otu13447 | 1 | k__Fungi(100);p__Ascomycota(100);p__Ascomycota_unclassified(100);p__Ascomycota_unclassified(100);p__Ascomycota_unclassified(100);p__Ascomycota_unclassified(100);p__Ascomycota_unclassified(100); | - | - | - | - | - | - | - | Unassigned | - |
| Otu13448 | 1 | k__Fungi(100);p__Ascomycota(100);p__Ascomycota_unclassified(100);p__Ascomycota_unclassified(100);p__Ascomycota_unclassified(100);p__Ascomycota_unclassified(100);p__Ascomycota_unclassified(100); | - | - | - | - | - | - | - | Unassigned | - |
| Otu13450 | 1 | k__Fungi(100);p__Ascomycota(100);p__Ascomycota_unclassified(100);p__Ascomycota_unclassified(100);p__Ascomycota_unclassified(100);p__Ascomycota_unclassified(100);p__Ascomycota_unclassified(100); | - | - | - | - | - | - | - | Unassigned | - |
| Otu13452 | 1 | k__Fungi(100);p__Ascomycota(100);p__Ascomycota_unclassified(100);p__Ascomycota_unclassified(100);p__Ascomycota_unclassified(100);p__Ascomycota_unclassified(100);p__Ascomycota_unclassified(100); | - | - | - | - | - | - | - | Unassigned | - |
| Otu13453 | 1 | k__Fungi(100);k__Fungi_unclassified(100);k__Fungi_unclassified(100);k__Fungi_unclassified(100);k__Fungi_unclassified(100);k__Fungi_unclassified(100);k__Fungi_unclassified(100); | - | - | - | - | - | - | - | Unassigned | - |
| Otu13455 | 1 | k__Fungi(100);k__Fungi_unclassified(100);k__Fungi_unclassified(100);k__Fungi_unclassified(100);k__Fungi_unclassified(100);k__Fungi_unclassified(100);k__Fungi_unclassified(100); | - | - | - | - | - | - | - | Unassigned | - |
| Otu13456 | 1 | k__Fungi(100);p__Ascomycota(100);p__Ascomycota_unclassified(100);p__Ascomycota_unclassified(100);p__Ascomycota_unclassified(100);p__Ascomycota_unclassified(100);p__Ascomycota_unclassified(100); | - | - | - | - | - | - | - | Unassigned | - |
| Otu13458 | 1 | k__Fungi(100);k__Fungi_unclassified(100);k__Fungi_unclassified(100);k__Fungi_unclassified(100);k__Fungi_unclassified(100);k__Fungi_unclassified(100);k__Fungi_unclassified(100); | - | - | - | - | - | - | - | Unassigned | - |
| Otu13461 | 1 | k__Fungi(100);p__Ascomycota(100);p__Ascomycota_unclassified(100);p__Ascomycota_unclassified(100);p__Ascomycota_unclassified(100);p__Ascomycota_unclassified(100);p__Ascomycota_unclassified(100); | - | - | - | - | - | - | - | Unassigned | - |
| Otu13465 | 1 | k__Fungi(100);k__Fungi_unclassified(100);k__Fungi_unclassified(100);k__Fungi_unclassified(100);k__Fungi_unclassified(100);k__Fungi_unclassified(100);k__Fungi_unclassified(100); | - | - | - | - | - | - | - | Unassigned | - |
| Otu13466 | 1 | k__Fungi(100);p__Ascomycota(100);p__Ascomycota_unclassified(100);p__Ascomycota_unclassified(100);p__Ascomycota_unclassified(100);p__Ascomycota_unclassified(100);p__Ascomycota_unclassified(100); | - | - | - | - | - | - | - | Unassigned | - |
| Otu13467 | 1 | k__Fungi(100);p__Ascomycota(100);p__Ascomycota_unclassified(100);p__Ascomycota_unclassified(100);p__Ascomycota_unclassified(100);p__Ascomycota_unclassified(100);p__Ascomycota_unclassified(100); | - | - | - | - | - | - | - | Unassigned | - |
| Otu13468 | 1 | k__Fungi(100);p__Ascomycota(100);p__Ascomycota_unclassified(100);p__Ascomycota_unclassified(100);p__Ascomycota_unclassified(100);p__Ascomycota_unclassified(100);p__Ascomycota_unclassified(100); | - | - | - | - | - | - | - | Unassigned | - |
| Otu13470 | 1 | k__Fungi(100);p__Ascomycota(100);p__Ascomycota_unclassified(100);p__Ascomycota_unclassified(100);p__Ascomycota_unclassified(100);p__Ascomycota_unclassified(100);p__Ascomycota_unclassified(100); | - | - | - | - | - | - | - | Unassigned | - |
| Otu13471 | 1 | k__Fungi(100);k__Fungi_unclassified(100);k__Fungi_unclassified(100);k__Fungi_unclassified(100);k__Fungi_unclassified(100);k__Fungi_unclassified(100);k__Fungi_unclassified(100); | - | - | - | - | - | - | - | Unassigned | - |
| Otu13472 | 1 | k__Fungi(100);k__Fungi_unclassified(100);k__Fungi_unclassified(100);k__Fungi_unclassified(100);k__Fungi_unclassified(100);k__Fungi_unclassified(100);k__Fungi_unclassified(100); | - | - | - | - | - | - | - | Unassigned | - |
| Otu13473 | 1 | k__Fungi(100);p__Ascomycota(100);p__Ascomycota_unclassified(100);p__Ascomycota_unclassified(100);p__Ascomycota_unclassified(100);p__Ascomycota_unclassified(100);p__Ascomycota_unclassified(100); | - | - | - | - | - | - | - | Unassigned | - |
| Otu13474 | 1 | k__Fungi(100);p__Ascomycota(100);p__Ascomycota_unclassified(100);p__Ascomycota_unclassified(100);p__Ascomycota_unclassified(100);p__Ascomycota_unclassified(100);p__Ascomycota_unclassified(100); | - | - | - | - | - | - | - | Unassigned | - |
| Otu13475 | 1 | k__Fungi(100);k__Fungi_unclassified(100);k__Fungi_unclassified(100);k__Fungi_unclassified(100);k__Fungi_unclassified(100);k__Fungi_unclassified(100);k__Fungi_unclassified(100); | - | - | - | - | - | - | - | Unassigned | - |
| Otu13476 | 1 | k__Fungi(100);k__Fungi_unclassified(100);k__Fungi_unclassified(100);k__Fungi_unclassified(100);k__Fungi_unclassified(100);k__Fungi_unclassified(100);k__Fungi_unclassified(100); | - | - | - | - | - | - | - | Unassigned | - |
| Otu13479 | 1 | k__Fungi(100);k__Fungi_unclassified(100);k__Fungi_unclassified(100);k__Fungi_unclassified(100);k__Fungi_unclassified(100);k__Fungi_unclassified(100);k__Fungi_unclassified(100); | - | - | - | - | - | - | - | Unassigned | - |
| Otu13480 | 1 | k__Fungi(100);p__Ascomycota(100);p__Ascomycota_unclassified(100);p__Ascomycota_unclassified(100);p__Ascomycota_unclassified(100);p__Ascomycota_unclassified(100);p__Ascomycota_unclassified(100); | - | - | - | - | - | - | - | Unassigned | - |
| Otu13482 | 1 | k__Fungi(100);p__Ascomycota(100);p__Ascomycota_unclassified(100);p__Ascomycota_unclassified(100);p__Ascomycota_unclassified(100);p__Ascomycota_unclassified(100);p__Ascomycota_unclassified(100); | - | - | - | - | - | - | - | Unassigned | - |
| Otu13483 | 1 | k__Fungi(100);p__Ascomycota(100);p__Ascomycota_unclassified(100);p__Ascomycota_unclassified(100);p__Ascomycota_unclassified(100);p__Ascomycota_unclassified(100);p__Ascomycota_unclassified(100); | - | - | - | - | - | - | - | Unassigned | - |
| Otu13486 | 1 | k__Fungi(100);k__Fungi_unclassified(100);k__Fungi_unclassified(100);k__Fungi_unclassified(100);k__Fungi_unclassified(100);k__Fungi_unclassified(100);k__Fungi_unclassified(100); | - | - | - | - | - | - | - | Unassigned | - |
| Otu13489 | 1 | k__Fungi(100);k__Fungi_unclassified(100);k__Fungi_unclassified(100);k__Fungi_unclassified(100);k__Fungi_unclassified(100);k__Fungi_unclassified(100);k__Fungi_unclassified(100); | - | - | - | - | - | - | - | Unassigned | - |
| Otu13492 | 1 | k__Fungi(100);p__Ascomycota(100);p__Ascomycota_unclassified(100);p__Ascomycota_unclassified(100);p__Ascomycota_unclassified(100);p__Ascomycota_unclassified(100);p__Ascomycota_unclassified(100); | - | - | - | - | - | - | - | Unassigned | - |
| Otu13493 | 1 | k__Fungi(100);p__Ascomycota(100);p__Ascomycota_unclassified(100);p__Ascomycota_unclassified(100);p__Ascomycota_unclassified(100);p__Ascomycota_unclassified(100);p__Ascomycota_unclassified(100); | - | - | - | - | - | - | - | Unassigned | - |
| Otu13494 | 1 | k__Fungi(100);k__Fungi_unclassified(100);k__Fungi_unclassified(100);k__Fungi_unclassified(100);k__Fungi_unclassified(100);k__Fungi_unclassified(100);k__Fungi_unclassified(100); | - | - | - | - | - | - | - | Unassigned | - |
| Otu13495 | 1 | k__Fungi(100);p__Ascomycota(100);p__Ascomycota_unclassified(100);p__Ascomycota_unclassified(100);p__Ascomycota_unclassified(100);p__Ascomycota_unclassified(100);p__Ascomycota_unclassified(100); | - | - | - | - | - | - | - | Unassigned | - |
| Otu13496 | 1 | k__Fungi(100);k__Fungi_unclassified(100);k__Fungi_unclassified(100);k__Fungi_unclassified(100);k__Fungi_unclassified(100);k__Fungi_unclassified(100);k__Fungi_unclassified(100); | - | - | - | - | - | - | - | Unassigned | - |
| Otu13497 | 1 | k__Fungi(100);p__Ascomycota(100);p__Ascomycota_unclassified(100);p__Ascomycota_unclassified(100);p__Ascomycota_unclassified(100);p__Ascomycota_unclassified(100);p__Ascomycota_unclassified(100); | - | - | - | - | - | - | - | Unassigned | - |
| Otu13499 | 1 | k__Fungi(100);k__Fungi_unclassified(100);k__Fungi_unclassified(100);k__Fungi_unclassified(100);k__Fungi_unclassified(100);k__Fungi_unclassified(100);k__Fungi_unclassified(100); | - | - | - | - | - | - | - | Unassigned | - |
| Otu13500 | 1 | k__Fungi(100);k__Fungi_unclassified(100);k__Fungi_unclassified(100);k__Fungi_unclassified(100);k__Fungi_unclassified(100);k__Fungi_unclassified(100);k__Fungi_unclassified(100); | - | - | - | - | - | - | - | Unassigned | - |
| Otu13502 | 1 | k__Fungi(100);p__Ascomycota(100);p__Ascomycota_unclassified(100);p__Ascomycota_unclassified(100);p__Ascomycota_unclassified(100);p__Ascomycota_unclassified(100);p__Ascomycota_unclassified(100); | - | - | - | - | - | - | - | Unassigned | - |
| Otu13504 | 1 | k__Fungi(100);p__Ascomycota(100);p__Ascomycota_unclassified(100);p__Ascomycota_unclassified(100);p__Ascomycota_unclassified(100);p__Ascomycota_unclassified(100);p__Ascomycota_unclassified(100); | - | - | - | - | - | - | - | Unassigned | - |
| Otu13505 | 1 | k__Fungi(100);k__Fungi_unclassified(100);k__Fungi_unclassified(100);k__Fungi_unclassified(100);k__Fungi_unclassified(100);k__Fungi_unclassified(100);k__Fungi_unclassified(100); | - | - | - | - | - | - | - | Unassigned | - |
| Otu13506 | 1 | k__Fungi(100);p__Ascomycota(100);p__Ascomycota_unclassified(100);p__Ascomycota_unclassified(100);p__Ascomycota_unclassified(100);p__Ascomycota_unclassified(100);p__Ascomycota_unclassified(100); | - | - | - | - | - | - | - | Unassigned | - |
| Otu13510 | 1 | k__Fungi(100);p__Ascomycota(100);p__Ascomycota_unclassified(100);p__Ascomycota_unclassified(100);p__Ascomycota_unclassified(100);p__Ascomycota_unclassified(100);p__Ascomycota_unclassified(100); | - | - | - | - | - | - | - | Unassigned | - |
| Otu13511 | 1 | k__Fungi(100);p__Ascomycota(100);p__Ascomycota_unclassified(100);p__Ascomycota_unclassified(100);p__Ascomycota_unclassified(100);p__Ascomycota_unclassified(100);p__Ascomycota_unclassified(100); | - | - | - | - | - | - | - | Unassigned | - |
| Otu13514 | 1 | k__Fungi(100);p__Ascomycota(100);p__Ascomycota_unclassified(100);p__Ascomycota_unclassified(100);p__Ascomycota_unclassified(100);p__Ascomycota_unclassified(100);p__Ascomycota_unclassified(100); | - | - | - | - | - | - | - | Unassigned | - |
| Otu13516 | 1 | k__Fungi(100);k__Fungi_unclassified(100);k__Fungi_unclassified(100);k__Fungi_unclassified(100);k__Fungi_unclassified(100);k__Fungi_unclassified(100);k__Fungi_unclassified(100); | - | - | - | - | - | - | - | Unassigned | - |
| Otu13518 | 1 | k__Fungi(100);p__Ascomycota(100);p__Ascomycota_unclassified(100);p__Ascomycota_unclassified(100);p__Ascomycota_unclassified(100);p__Ascomycota_unclassified(100);p__Ascomycota_unclassified(100); | - | - | - | - | - | - | - | Unassigned | - |
| Otu13521 | 1 | k__Fungi(100);p__Ascomycota(100);p__Ascomycota_unclassified(100);p__Ascomycota_unclassified(100);p__Ascomycota_unclassified(100);p__Ascomycota_unclassified(100);p__Ascomycota_unclassified(100); | - | - | - | - | - | - | - | Unassigned | - |
| Otu13522 | 1 | k__Fungi(100);p__Ascomycota(100);p__Ascomycota_unclassified(100);p__Ascomycota_unclassified(100);p__Ascomycota_unclassified(100);p__Ascomycota_unclassified(100);p__Ascomycota_unclassified(100); | - | - | - | - | - | - | - | Unassigned | - |
| Otu13524 | 1 | k__Fungi(100);k__Fungi_unclassified(100);k__Fungi_unclassified(100);k__Fungi_unclassified(100);k__Fungi_unclassified(100);k__Fungi_unclassified(100);k__Fungi_unclassified(100); | - | - | - | - | - | - | - | Unassigned | - |
| Otu13526 | 1 | k__Fungi(100);k__Fungi_unclassified(100);k__Fungi_unclassified(100);k__Fungi_unclassified(100);k__Fungi_unclassified(100);k__Fungi_unclassified(100);k__Fungi_unclassified(100); | - | - | - | - | - | - | - | Unassigned | - |
| Otu13528 | 1 | k__Fungi(100);p__Ascomycota(100);p__Ascomycota_unclassified(100);p__Ascomycota_unclassified(100);p__Ascomycota_unclassified(100);p__Ascomycota_unclassified(100);p__Ascomycota_unclassified(100); | - | - | - | - | - | - | - | Unassigned | - |
| Otu13533 | 1 | k__Fungi(100);k__Fungi_unclassified(100);k__Fungi_unclassified(100);k__Fungi_unclassified(100);k__Fungi_unclassified(100);k__Fungi_unclassified(100);k__Fungi_unclassified(100); | - | - | - | - | - | - | - | Unassigned | - |
| Otu13535 | 1 | k__Fungi(100);k__Fungi_unclassified(100);k__Fungi_unclassified(100);k__Fungi_unclassified(100);k__Fungi_unclassified(100);k__Fungi_unclassified(100);k__Fungi_unclassified(100); | - | - | - | - | - | - | - | Unassigned | - |
| Otu13536 | 1 | k__Fungi(100);p__Ascomycota(100);p__Ascomycota_unclassified(100);p__Ascomycota_unclassified(100);p__Ascomycota_unclassified(100);p__Ascomycota_unclassified(100);p__Ascomycota_unclassified(100); | - | - | - | - | - | - | - | Unassigned | - |
| Otu13537 | 1 | k__Fungi(100);p__Ascomycota(100);p__Ascomycota_unclassified(100);p__Ascomycota_unclassified(100);p__Ascomycota_unclassified(100);p__Ascomycota_unclassified(100);p__Ascomycota_unclassified(100); | - | - | - | - | - | - | - | Unassigned | - |
| Otu13539 | 1 | k__Fungi(100);p__Ascomycota(100);p__Ascomycota_unclassified(100);p__Ascomycota_unclassified(100);p__Ascomycota_unclassified(100);p__Ascomycota_unclassified(100);p__Ascomycota_unclassified(100); | - | - | - | - | - | - | - | Unassigned | - |
| Otu13540 | 1 | k__Fungi(100);p__Ascomycota(100);p__Ascomycota_unclassified(100);p__Ascomycota_unclassified(100);p__Ascomycota_unclassified(100);p__Ascomycota_unclassified(100);p__Ascomycota_unclassified(100); | - | - | - | - | - | - | - | Unassigned | - |
| Otu13541 | 1 | k__Fungi(100);p__Ascomycota(100);p__Ascomycota_unclassified(100);p__Ascomycota_unclassified(100);p__Ascomycota_unclassified(100);p__Ascomycota_unclassified(100);p__Ascomycota_unclassified(100); | - | - | - | - | - | - | - | Unassigned | - |
| Otu13542 | 1 | k__Fungi(100);p__Ascomycota(100);p__Ascomycota_unclassified(100);p__Ascomycota_unclassified(100);p__Ascomycota_unclassified(100);p__Ascomycota_unclassified(100);p__Ascomycota_unclassified(100); | - | - | - | - | - | - | - | Unassigned | - |
| Otu13543 | 1 | k__Fungi(100);p__Ascomycota(100);p__Ascomycota_unclassified(100);p__Ascomycota_unclassified(100);p__Ascomycota_unclassified(100);p__Ascomycota_unclassified(100);p__Ascomycota_unclassified(100); | - | - | - | - | - | - | - | Unassigned | - |
| Otu13544 | 1 | k__Fungi(100);p__Ascomycota(100);p__Ascomycota_unclassified(100);p__Ascomycota_unclassified(100);p__Ascomycota_unclassified(100);p__Ascomycota_unclassified(100);p__Ascomycota_unclassified(100); | - | - | - | - | - | - | - | Unassigned | - |
| Otu13547 | 1 | k__Fungi(100);p__Ascomycota(100);p__Ascomycota_unclassified(100);p__Ascomycota_unclassified(100);p__Ascomycota_unclassified(100);p__Ascomycota_unclassified(100);p__Ascomycota_unclassified(100); | - | - | - | - | - | - | - | Unassigned | - |
| Otu13551 | 1 | k__Fungi(100);p__Ascomycota(100);p__Ascomycota_unclassified(100);p__Ascomycota_unclassified(100);p__Ascomycota_unclassified(100);p__Ascomycota_unclassified(100);p__Ascomycota_unclassified(100); | - | - | - | - | - | - | - | Unassigned | - |
| Otu13552 | 1 | k__Fungi(100);p__Ascomycota(100);p__Ascomycota_unclassified(100);p__Ascomycota_unclassified(100);p__Ascomycota_unclassified(100);p__Ascomycota_unclassified(100);p__Ascomycota_unclassified(100); | - | - | - | - | - | - | - | Unassigned | - |
| Otu13554 | 1 | k__Fungi(100);p__Ascomycota(100);p__Ascomycota_unclassified(100);p__Ascomycota_unclassified(100);p__Ascomycota_unclassified(100);p__Ascomycota_unclassified(100);p__Ascomycota_unclassified(100); | - | - | - | - | - | - | - | Unassigned | - |
| Otu13555 | 1 | k__Fungi(100);p__Ascomycota(100);p__Ascomycota_unclassified(100);p__Ascomycota_unclassified(100);p__Ascomycota_unclassified(100);p__Ascomycota_unclassified(100);p__Ascomycota_unclassified(100); | - | - | - | - | - | - | - | Unassigned | - |
| Otu13556 | 1 | k__Fungi(100);p__Ascomycota(100);p__Ascomycota_unclassified(100);p__Ascomycota_unclassified(100);p__Ascomycota_unclassified(100);p__Ascomycota_unclassified(100);p__Ascomycota_unclassified(100); | - | - | - | - | - | - | - | Unassigned | - |
| Otu13558 | 1 | k__Fungi(100);k__Fungi_unclassified(100);k__Fungi_unclassified(100);k__Fungi_unclassified(100);k__Fungi_unclassified(100);k__Fungi_unclassified(100);k__Fungi_unclassified(100); | - | - | - | - | - | - | - | Unassigned | - |

| | | | | | | | | | | | |
|---|---|---|---|---|---|---|---|---|---|---|---|
| Otu13559 | 1 | k__Fungi(100);p__Ascomycota(100);p__Ascomycota_unclassified(100);p__Ascomycota_unclassified(100);p__Ascomycota_unclassified(100);p__Ascomycota_unclassified(100);p__Ascomycota_unclassified(100); | - | - | - | - | - | - | - | Unassigned | - |
| Otu13562 | 1 | k__Fungi(100);p__Ascomycota(100);p__Ascomycota_unclassified(100);p__Ascomycota_unclassified(100);p__Ascomycota_unclassified(100);p__Ascomycota_unclassified(100);p__Ascomycota_unclassified(100); | - | - | - | - | - | - | - | Unassigned | - |
| Otu13567 | 1 | k__Fungi(100);k__Fungi_unclassified(100);k__Fungi_unclassified(100);k__Fungi_unclassified(100);k__Fungi_unclassified(100);k__Fungi_unclassified(100);k__Fungi_unclassified(100); | - | - | - | - | - | - | - | Unassigned | - |
| Otu13572 | 1 | k__Fungi(100);k__Fungi_unclassified(100);k__Fungi_unclassified(100);k__Fungi_unclassified(100);k__Fungi_unclassified(100);k__Fungi_unclassified(100);k__Fungi_unclassified(100); | - | - | - | - | - | - | - | Unassigned | - |
| Otu13573 | 1 | k__Fungi(100);k__Fungi_unclassified(100);k__Fungi_unclassified(100);k__Fungi_unclassified(100);k__Fungi_unclassified(100);k__Fungi_unclassified(100);k__Fungi_unclassified(100); | - | - | - | - | - | - | - | Unassigned | - |
| Otu13574 | 1 | k__Fungi(100);p__Ascomycota(100);p__Ascomycota_unclassified(100);p__Ascomycota_unclassified(100);p__Ascomycota_unclassified(100);p__Ascomycota_unclassified(100);p__Ascomycota_unclassified(100); | - | - | - | - | - | - | - | Unassigned | - |
| Otu13575 | 1 | k__Fungi(100);k__Fungi_unclassified(100);k__Fungi_unclassified(100);k__Fungi_unclassified(100);k__Fungi_unclassified(100);k__Fungi_unclassified(100);k__Fungi_unclassified(100); | - | - | - | - | - | - | - | Unassigned | - |
| Otu13579 | 1 | k__Fungi(100);k__Fungi_unclassified(100);k__Fungi_unclassified(100);k__Fungi_unclassified(100);k__Fungi_unclassified(100);k__Fungi_unclassified(100);k__Fungi_unclassified(100); | - | - | - | - | - | - | - | Unassigned | - |
| Otu13580 | 1 | k__Fungi(100);k__Fungi_unclassified(100);k__Fungi_unclassified(100);k__Fungi_unclassified(100);k__Fungi_unclassified(100);k__Fungi_unclassified(100);k__Fungi_unclassified(100); | - | - | - | - | - | - | - | Unassigned | - |
| Otu13583 | 1 | k__Fungi(100);k__Fungi_unclassified(100);k__Fungi_unclassified(100);k__Fungi_unclassified(100);k__Fungi_unclassified(100);k__Fungi_unclassified(100);k__Fungi_unclassified(100); | - | - | - | - | - | - | - | Unassigned | - |
| Otu13585 | 1 | k__Fungi(100);k__Fungi_unclassified(100);k__Fungi_unclassified(100);k__Fungi_unclassified(100);k__Fungi_unclassified(100);k__Fungi_unclassified(100);k__Fungi_unclassified(100); | - | - | - | - | - | - | - | Unassigned | - |
| Otu13586 | 1 | k__Fungi(100);p__Ascomycota(100);p__Ascomycota_unclassified(100);p__Ascomycota_unclassified(100);p__Ascomycota_unclassified(100);p__Ascomycota_unclassified(100);p__Ascomycota_unclassified(100); | - | - | - | - | - | - | - | Unassigned | - |
| Otu13589 | 1 | k__Fungi(100);k__Fungi_unclassified(100);k__Fungi_unclassified(100);k__Fungi_unclassified(100);k__Fungi_unclassified(100);k__Fungi_unclassified(100);k__Fungi_unclassified(100); | - | - | - | - | - | - | - | Unassigned | - |
| Otu13596 | 1 | k__Fungi(100);p__Ascomycota(100);p__Ascomycota_unclassified(100);p__Ascomycota_unclassified(100);p__Ascomycota_unclassified(100);p__Ascomycota_unclassified(100);p__Ascomycota_unclassified(100); | - | - | - | - | - | - | - | Unassigned | - |
| Otu13599 | 1 | k__Fungi(100);k__Fungi_unclassified(100);k__Fungi_unclassified(100);k__Fungi_unclassified(100);k__Fungi_unclassified(100);k__Fungi_unclassified(100);k__Fungi_unclassified(100); | - | - | - | - | - | - | - | Unassigned | - |
| Otu13605 | 1 | k__Fungi(100);k__Fungi_unclassified(100);k__Fungi_unclassified(100);k__Fungi_unclassified(100);k__Fungi_unclassified(100);k__Fungi_unclassified(100);k__Fungi_unclassified(100); | - | - | - | - | - | - | - | Unassigned | - |
| Otu13608 | 1 | k__Fungi(100);p__Ascomycota(100);p__Ascomycota_unclassified(100);p__Ascomycota_unclassified(100);p__Ascomycota_unclassified(100);p__Ascomycota_unclassified(100);p__Ascomycota_unclassified(100); | - | - | - | - | - | - | - | Unassigned | - |
| Otu13609 | 1 | k__Fungi(100);k__Fungi_unclassified(100);k__Fungi_unclassified(100);k__Fungi_unclassified(100);k__Fungi_unclassified(100);k__Fungi_unclassified(100);k__Fungi_unclassified(100); | - | - | - | - | - | - | - | Unassigned | - |
| Otu13611 | 1 | k__Fungi(100);p__Ascomycota(100);p__Ascomycota_unclassified(100);p__Ascomycota_unclassified(100);p__Ascomycota_unclassified(100);p__Ascomycota_unclassified(100);p__Ascomycota_unclassified(100); | - | - | - | - | - | - | - | Unassigned | - |
| Otu13612 | 1 | k__Fungi(100);k__Fungi_unclassified(100);k__Fungi_unclassified(100);k__Fungi_unclassified(100);k__Fungi_unclassified(100);k__Fungi_unclassified(100);k__Fungi_unclassified(100); | - | - | - | - | - | - | - | Unassigned | - |
| Otu13613 | 1 | k__Fungi(100);p__Ascomycota(100);p__Ascomycota_unclassified(100);p__Ascomycota_unclassified(100);p__Ascomycota_unclassified(100);p__Ascomycota_unclassified(100);p__Ascomycota_unclassified(100); | - | - | - | - | - | - | - | Unassigned | - |
| Otu13614 | 1 | k__Fungi(100);k__Fungi_unclassified(100);k__Fungi_unclassified(100);k__Fungi_unclassified(100);k__Fungi_unclassified(100);k__Fungi_unclassified(100);k__Fungi_unclassified(100); | - | - | - | - | - | - | - | Unassigned | - |
| Otu13615 | 1 | k__Fungi(100);k__Fungi_unclassified(100);k__Fungi_unclassified(100);k__Fungi_unclassified(100);k__Fungi_unclassified(100);k__Fungi_unclassified(100);k__Fungi_unclassified(100); | - | - | - | - | - | - | - | Unassigned | - |
| Otu13620 | 1 | k__Fungi(100);k__Fungi_unclassified(100);k__Fungi_unclassified(100);k__Fungi_unclassified(100);k__Fungi_unclassified(100);k__Fungi_unclassified(100);k__Fungi_unclassified(100); | - | - | - | - | - | - | - | Unassigned | - |
| Otu13622 | 1 | k__Fungi(100);k__Fungi_unclassified(100);k__Fungi_unclassified(100);k__Fungi_unclassified(100);k__Fungi_unclassified(100);k__Fungi_unclassified(100);k__Fungi_unclassified(100); | - | - | - | - | - | - | - | Unassigned | - |
| Otu13623 | 1 | k__Fungi(100);k__Fungi_unclassified(100);k__Fungi_unclassified(100);k__Fungi_unclassified(100);k__Fungi_unclassified(100);k__Fungi_unclassified(100);k__Fungi_unclassified(100); | - | - | - | - | - | - | - | Unassigned | - |
| Otu13627 | 1 | k__Fungi(100);k__Fungi_unclassified(100);k__Fungi_unclassified(100);k__Fungi_unclassified(100);k__Fungi_unclassified(100);k__Fungi_unclassified(100);k__Fungi_unclassified(100); | - | - | - | - | - | - | - | Unassigned | - |
| Otu13629 | 1 | k__Fungi(100);p__Ascomycota(100);p__Ascomycota_unclassified(100);p__Ascomycota_unclassified(100);p__Ascomycota_unclassified(100);p__Ascomycota_unclassified(100);p__Ascomycota_unclassified(100); | - | - | - | - | - | - | - | Unassigned | - |
| Otu13632 | 1 | k__Fungi(100);k__Fungi_unclassified(100);k__Fungi_unclassified(100);k__Fungi_unclassified(100);k__Fungi_unclassified(100);k__Fungi_unclassified(100);k__Fungi_unclassified(100); | - | - | - | - | - | - | - | Unassigned | - |
| Otu13633 | 1 | k__Fungi(100);k__Fungi_unclassified(100);k__Fungi_unclassified(100);k__Fungi_unclassified(100);k__Fungi_unclassified(100);k__Fungi_unclassified(100);k__Fungi_unclassified(100); | - | - | - | - | - | - | - | Unassigned | - |
| Otu13634 | 1 | k__Fungi(100);k__Fungi_unclassified(100);k__Fungi_unclassified(100);k__Fungi_unclassified(100);k__Fungi_unclassified(100);k__Fungi_unclassified(100);k__Fungi_unclassified(100); | - | - | - | - | - | - | - | Unassigned | - |
| Otu13635 | 1 | k__Fungi(100);k__Fungi_unclassified(100);k__Fungi_unclassified(100);k__Fungi_unclassified(100);k__Fungi_unclassified(100);k__Fungi_unclassified(100);k__Fungi_unclassified(100); | - | - | - | - | - | - | - | Unassigned | - |
| Otu13636 | 1 | k__Fungi(100);p__Ascomycota(100);p__Ascomycota_unclassified(100);p__Ascomycota_unclassified(100);p__Ascomycota_unclassified(100);p__Ascomycota_unclassified(100);p__Ascomycota_unclassified(100); | - | - | - | - | - | - | - | Unassigned | - |
| Otu13639 | 1 | k__Fungi(100);p__Ascomycota(100);p__Ascomycota_unclassified(100);p__Ascomycota_unclassified(100);p__Ascomycota_unclassified(100);p__Ascomycota_unclassified(100);p__Ascomycota_unclassified(100); | - | - | - | - | - | - | - | Unassigned | - |
| Otu13640 | 1 | k__Fungi(100);p__Ascomycota(100);p__Ascomycota_unclassified(100);p__Ascomycota_unclassified(100);p__Ascomycota_unclassified(100);p__Ascomycota_unclassified(100);p__Ascomycota_unclassified(100); | - | - | - | - | - | - | - | Unassigned | - |
| Otu13641 | 1 | k__Fungi(100);p__Ascomycota(100);p__Ascomycota_unclassified(100);p__Ascomycota_unclassified(100);p__Ascomycota_unclassified(100);p__Ascomycota_unclassified(100);p__Ascomycota_unclassified(100); | - | - | - | - | - | - | - | Unassigned | - |
| Otu13642 | 1 | k__Fungi(100);p__Ascomycota(100);p__Ascomycota_unclassified(100);p__Ascomycota_unclassified(100);p__Ascomycota_unclassified(100);p__Ascomycota_unclassified(100);p__Ascomycota_unclassified(100); | - | - | - | - | - | - | - | Unassigned | - |
| Otu13643 | 1 | k__Fungi(100);p__Ascomycota(100);p__Ascomycota_unclassified(100);p__Ascomycota_unclassified(100);p__Ascomycota_unclassified(100);p__Ascomycota_unclassified(100);p__Ascomycota_unclassified(100); | - | - | - | - | - | - | - | Unassigned | - |
| Otu13644 | 1 | k__Fungi(100);p__Ascomycota(100);p__Ascomycota_unclassified(100);p__Ascomycota_unclassified(100);p__Ascomycota_unclassified(100);p__Ascomycota_unclassified(100);p__Ascomycota_unclassified(100); | - | - | - | - | - | - | - | Unassigned | - |
| Otu13647 | 1 | k__Fungi(100);k__Fungi_unclassified(100);k__Fungi_unclassified(100);k__Fungi_unclassified(100);k__Fungi_unclassified(100);k__Fungi_unclassified(100);k__Fungi_unclassified(100); | - | - | - | - | - | - | - | Unassigned | - |
| Otu13653 | 1 | k__Fungi(100);k__Fungi_unclassified(100);k__Fungi_unclassified(100);k__Fungi_unclassified(100);k__Fungi_unclassified(100);k__Fungi_unclassified(100);k__Fungi_unclassified(100); | - | - | - | - | - | - | - | Unassigned | - |
| Otu13654 | 1 | k__Fungi(100);k__Fungi_unclassified(100);k__Fungi_unclassified(100);k__Fungi_unclassified(100);k__Fungi_unclassified(100);k__Fungi_unclassified(100);k__Fungi_unclassified(100); | - | - | - | - | - | - | - | Unassigned | - |
| Otu13655 | 1 | k__Fungi(100);k__Fungi_unclassified(100);k__Fungi_unclassified(100);k__Fungi_unclassified(100);k__Fungi_unclassified(100);k__Fungi_unclassified(100);k__Fungi_unclassified(100); | - | - | - | - | - | - | - | Unassigned | - |
| Otu13656 | 1 | k__Fungi(100);p__Ascomycota(100);p__Ascomycota_unclassified(100);p__Ascomycota_unclassified(100);p__Ascomycota_unclassified(100);p__Ascomycota_unclassified(100);p__Ascomycota_unclassified(100); | - | - | - | - | - | - | - | Unassigned | - |
| Otu13657 | 1 | k__Fungi(100);k__Fungi_unclassified(100);k__Fungi_unclassified(100);k__Fungi_unclassified(100);k__Fungi_unclassified(100);k__Fungi_unclassified(100);k__Fungi_unclassified(100); | - | - | - | - | - | - | - | Unassigned | - |
| Otu13658 | 1 | k__Fungi(100);k__Fungi_unclassified(100);k__Fungi_unclassified(100);k__Fungi_unclassified(100);k__Fungi_unclassified(100);k__Fungi_unclassified(100);k__Fungi_unclassified(100); | - | - | - | - | - | - | - | Unassigned | - |
| Otu13659 | 1 | k__Fungi(100);k__Fungi_unclassified(100);k__Fungi_unclassified(100);k__Fungi_unclassified(100);k__Fungi_unclassified(100);k__Fungi_unclassified(100);k__Fungi_unclassified(100); | - | - | - | - | - | - | - | Unassigned | - |
| Otu13662 | 1 | k__Fungi(100);k__Fungi_unclassified(100);k__Fungi_unclassified(100);k__Fungi_unclassified(100);k__Fungi_unclassified(100);k__Fungi_unclassified(100);k__Fungi_unclassified(100); | - | - | - | - | - | - | - | Unassigned | - |
| Otu13663 | 1 | k__Fungi(100);k__Fungi_unclassified(100);k__Fungi_unclassified(100);k__Fungi_unclassified(100);k__Fungi_unclassified(100);k__Fungi_unclassified(100);k__Fungi_unclassified(100); | - | - | - | - | - | - | - | Unassigned | - |
| Otu13664 | 1 | k__Fungi(100);p__Ascomycota(100);p__Ascomycota_unclassified(100);p__Ascomycota_unclassified(100);p__Ascomycota_unclassified(100);p__Ascomycota_unclassified(100);p__Ascomycota_unclassified(100); | - | - | - | - | - | - | - | Unassigned | - |
| Otu13669 | 1 | k__Fungi(100);k__Fungi_unclassified(100);k__Fungi_unclassified(100);k__Fungi_unclassified(100);k__Fungi_unclassified(100);k__Fungi_unclassified(100);k__Fungi_unclassified(100); | - | - | - | - | - | - | - | Unassigned | - |
| Otu13675 | 1 | k__Fungi(100);k__Fungi_unclassified(100);k__Fungi_unclassified(100);k__Fungi_unclassified(100);k__Fungi_unclassified(100);k__Fungi_unclassified(100);k__Fungi_unclassified(100); | - | - | - | - | - | - | - | Unassigned | - |
| Otu13678 | 1 | k__Fungi(100);k__Fungi_unclassified(100);k__Fungi_unclassified(100);k__Fungi_unclassified(100);k__Fungi_unclassified(100);k__Fungi_unclassified(100);k__Fungi_unclassified(100); | - | - | - | - | - | - | - | Unassigned | - |
| Otu13679 | 1 | k__Fungi(100);k__Fungi_unclassified(100);k__Fungi_unclassified(100);k__Fungi_unclassified(100);k__Fungi_unclassified(100);k__Fungi_unclassified(100);k__Fungi_unclassified(100); | - | - | - | - | - | - | - | Unassigned | - |
| Otu13680 | 1 | k__Fungi(100);k__Fungi_unclassified(100);k__Fungi_unclassified(100);k__Fungi_unclassified(100);k__Fungi_unclassified(100);k__Fungi_unclassified(100);k__Fungi_unclassified(100); | - | - | - | - | - | - | - | Unassigned | - |
| Otu13686 | 1 | k__Fungi(100);k__Fungi_unclassified(100);k__Fungi_unclassified(100);k__Fungi_unclassified(100);k__Fungi_unclassified(100);k__Fungi_unclassified(100);k__Fungi_unclassified(100); | - | - | - | - | - | - | - | Unassigned | - |
| Otu13687 | 1 | k__Fungi(100);p__Ascomycota(100);p__Ascomycota_unclassified(100);p__Ascomycota_unclassified(100);p__Ascomycota_unclassified(100);p__Ascomycota_unclassified(100);p__Ascomycota_unclassified(100); | - | - | - | - | - | - | - | Unassigned | - |
| Otu13690 | 1 | k__Fungi(100);p__Ascomycota(100);p__Ascomycota_unclassified(100);p__Ascomycota_unclassified(100);p__Ascomycota_unclassified(100);p__Ascomycota_unclassified(100);p__Ascomycota_unclassified(100); | - | - | - | - | - | - | - | Unassigned | - |
| Otu13692 | 1 | k__Fungi(100);k__Fungi_unclassified(100);k__Fungi_unclassified(100);k__Fungi_unclassified(100);k__Fungi_unclassified(100);k__Fungi_unclassified(100);k__Fungi_unclassified(100); | - | - | - | - | - | - | - | Unassigned | - |
| Otu13694 | 1 | k__Fungi(100);p__Ascomycota(100);p__Ascomycota_unclassified(100);p__Ascomycota_unclassified(100);p__Ascomycota_unclassified(100);p__Ascomycota_unclassified(100);p__Ascomycota_unclassified(100); | - | - | - | - | - | - | - | Unassigned | - |
| Otu13700 | 1 | k__Fungi(100);p__Ascomycota(100);p__Ascomycota_unclassified(100);p__Ascomycota_unclassified(100);p__Ascomycota_unclassified(100);p__Ascomycota_unclassified(100);p__Ascomycota_unclassified(100); | - | - | - | - | - | - | - | Unassigned | - |
| Otu13703 | 1 | k__Fungi(100);k__Fungi_unclassified(100);k__Fungi_unclassified(100);k__Fungi_unclassified(100);k__Fungi_unclassified(100);k__Fungi_unclassified(100);k__Fungi_unclassified(100); | - | - | - | - | - | - | - | Unassigned | - |
| Otu13709 | 1 | k__Fungi(100);p__Ascomycota(100);p__Ascomycota_unclassified(100);p__Ascomycota_unclassified(100);p__Ascomycota_unclassified(100);p__Ascomycota_unclassified(100);p__Ascomycota_unclassified(100); | - | - | - | - | - | - | - | Unassigned | - |
| Otu13718 | 1 | k__Fungi(100);k__Fungi_unclassified(100);k__Fungi_unclassified(100);k__Fungi_unclassified(100);k__Fungi_unclassified(100);k__Fungi_unclassified(100);k__Fungi_unclassified(100); | - | - | - | - | - | - | - | Unassigned | - |
| Otu13720 | 1 | k__Fungi(100);p__Ascomycota(100);p__Ascomycota_unclassified(100);p__Ascomycota_unclassified(100);p__Ascomycota_unclassified(100);p__Ascomycota_unclassified(100);p__Ascomycota_unclassified(100); | - | - | - | - | - | - | - | Unassigned | - |
| Otu13723 | 1 | k__Fungi(100);k__Fungi_unclassified(100);k__Fungi_unclassified(100);k__Fungi_unclassified(100);k__Fungi_unclassified(100);k__Fungi_unclassified(100);k__Fungi_unclassified(100); | - | - | - | - | - | - | - | Unassigned | - |
| Otu13725 | 1 | k__Fungi(100);k__Fungi_unclassified(100);k__Fungi_unclassified(100);k__Fungi_unclassified(100);k__Fungi_unclassified(100);k__Fungi_unclassified(100);k__Fungi_unclassified(100); | - | - | - | - | - | - | - | Unassigned | - |
| Otu13729 | 1 | k__Fungi(100);p__Ascomycota(100);p__Ascomycota_unclassified(100);p__Ascomycota_unclassified(100);p__Ascomycota_unclassified(100);p__Ascomycota_unclassified(100);p__Ascomycota_unclassified(100); | - | - | - | - | - | - | - | Unassigned | - |
| Otu13730 | 1 | k__Fungi(100);k__Fungi_unclassified(100);k__Fungi_unclassified(100);k__Fungi_unclassified(100);k__Fungi_unclassified(100);k__Fungi_unclassified(100);k__Fungi_unclassified(100); | - | - | - | - | - | - | - | Unassigned | - |
| Otu13732 | 1 | k__Fungi(100);p__Ascomycota(100);p__Ascomycota_unclassified(100);p__Ascomycota_unclassified(100);p__Ascomycota_unclassified(100);p__Ascomycota_unclassified(100);p__Ascomycota_unclassified(100); | - | - | - | - | - | - | - | Unassigned | - |
| Otu13733 | 1 | k__Fungi(100);p__Ascomycota(100);p__Ascomycota_unclassified(100);p__Ascomycota_unclassified(100);p__Ascomycota_unclassified(100);p__Ascomycota_unclassified(100);p__Ascomycota_unclassified(100); | - | - | - | - | - | - | - | Unassigned | - |
| Otu13736 | 1 | k__Fungi(100);k__Fungi_unclassified(100);k__Fungi_unclassified(100);k__Fungi_unclassified(100);k__Fungi_unclassified(100);k__Fungi_unclassified(100);k__Fungi_unclassified(100); | - | - | - | - | - | - | - | Unassigned | - |
| Otu13742 | 1 | k__Fungi(100);p__Ascomycota(100);p__Ascomycota_unclassified(100);p__Ascomycota_unclassified(100);p__Ascomycota_unclassified(100);p__Ascomycota_unclassified(100);p__Ascomycota_unclassified(100); | - | - | - | - | - | - | - | Unassigned | - |
| Otu13747 | 1 | k__Fungi(100);p__Ascomycota(100);p__Ascomycota_unclassified(100);p__Ascomycota_unclassified(100);p__Ascomycota_unclassified(100);p__Ascomycota_unclassified(100);p__Ascomycota_unclassified(100); | - | - | - | - | - | - | - | Unassigned | - |
| Otu13749 | 1 | k__Fungi(100);k__Fungi_unclassified(100);k__Fungi_unclassified(100);k__Fungi_unclassified(100);k__Fungi_unclassified(100);k__Fungi_unclassified(100);k__Fungi_unclassified(100); | - | - | - | - | - | - | - | Unassigned | - |
| Otu13750 | 1 | k__Fungi(100);k__Fungi_unclassified(100);k__Fungi_unclassified(100);k__Fungi_unclassified(100);k__Fungi_unclassified(100);k__Fungi_unclassified(100);k__Fungi_unclassified(100); | - | - | - | - | - | - | - | Unassigned | - |
| Otu13751 | 1 | k__Fungi(100);k__Fungi_unclassified(100);k__Fungi_unclassified(100);k__Fungi_unclassified(100);k__Fungi_unclassified(100);k__Fungi_unclassified(100);k__Fungi_unclassified(100); | - | - | - | - | - | - | - | Unassigned | - |
| Otu13754 | 1 | k__Fungi(100);k__Fungi_unclassified(100);k__Fungi_unclassified(100);k__Fungi_unclassified(100);k__Fungi_unclassified(100);k__Fungi_unclassified(100);k__Fungi_unclassified(100); | - | - | - | - | - | - | - | Unassigned | - |
| Otu13758 | 1 | k__Fungi(100);k__Fungi_unclassified(100);k__Fungi_unclassified(100);k__Fungi_unclassified(100);k__Fungi_unclassified(100);k__Fungi_unclassified(100);k__Fungi_unclassified(100); | - | - | - | - | - | - | - | Unassigned | - |
| Otu13759 | 1 | k__Fungi(100);p__Ascomycota(100);p__Ascomycota_unclassified(100);p__Ascomycota_unclassified(100);p__Ascomycota_unclassified(100);p__Ascomycota_unclassified(100);p__Ascomycota_unclassified(100); | - | - | - | - | - | - | - | Unassigned | - |
| Otu13763 | 1 | k__Fungi(100);k__Fungi_unclassified(100);k__Fungi_unclassified(100);k__Fungi_unclassified(100);k__Fungi_unclassified(100);k__Fungi_unclassified(100);k__Fungi_unclassified(100); | - | - | - | - | - | - | - | Unassigned | - |
| Otu13765 | 1 | k__Fungi(100);p__Ascomycota(100);p__Ascomycota_unclassified(100);p__Ascomycota_unclassified(100);p__Ascomycota_unclassified(100);p__Ascomycota_unclassified(100);p__Ascomycota_unclassified(100); | - | - | - | - | - | - | - | Unassigned | - |
| Otu13772 | 1 | k__Fungi(100);k__Fungi_unclassified(100);k__Fungi_unclassified(100);k__Fungi_unclassified(100);k__Fungi_unclassified(100);k__Fungi_unclassified(100);k__Fungi_unclassified(100); | - | - | - | - | - | - | - | Unassigned | - |
| Otu13778 | 1 | k__Fungi(100);p__Ascomycota(100);p__Ascomycota_unclassified(100);p__Ascomycota_unclassified(100);p__Ascomycota_unclassified(100);p__Ascomycota_unclassified(100);p__Ascomycota_unclassified(100); | - | - | - | - | - | - | - | Unassigned | - |
| Otu13792 | 1 | k__Fungi(100);p__Ascomycota(100);p__Ascomycota_unclassified(100);p__Ascomycota_unclassified(100);p__Ascomycota_unclassified(100);p__Ascomycota_unclassified(100);p__Ascomycota_unclassified(100); | - | - | - | - | - | - | - | Unassigned | - |
| Otu13806 | 1 | k__Fungi(100);p__Ascomycota(100);p__Ascomycota_unclassified(100);p__Ascomycota_unclassified(100);p__Ascomycota_unclassified(100);p__Ascomycota_unclassified(100);p__Ascomycota_unclassified(100); | - | - | - | - | - | - | - | Unassigned | - |
| Otu13807 | 1 | k__Fungi(100);p__Ascomycota(100);p__Ascomycota_unclassified(100);p__Ascomycota_unclassified(100);p__Ascomycota_unclassified(100);p__Ascomycota_unclassified(100);p__Ascomycota_unclassified(100); | - | - | - | - | - | - | - | Unassigned | - |
| Otu13826 | 1 | k__Fungi(100);p__Ascomycota(100);p__Ascomycota_unclassified(100);p__Ascomycota_unclassified(100);p__Ascomycota_unclassified(100);p__Ascomycota_unclassified(100);p__Ascomycota_unclassified(100); | - | - | - | - | - | - | - | Unassigned | - |
| Otu13831 | 1 | k__Fungi(100);p__Ascomycota(100);p__Ascomycota_unclassified(100);p__Ascomycota_unclassified(100);p__Ascomycota_unclassified(100);p__Ascomycota_unclassified(100);p__Ascomycota_unclassified(100); | - | - | - | - | - | - | - | Unassigned | - |
| Otu13839 | 1 | k__Fungi(100);k__Fungi_unclassified(100);k__Fungi_unclassified(100);k__Fungi_unclassified(100);k__Fungi_unclassified(100);k__Fungi_unclassified(100);k__Fungi_unclassified(100); | - | - | - | - | - | - | - | Unassigned | - |
| Otu13841 | 1 | k__Fungi(100);p__Ascomycota(100);p__Ascomycota_unclassified(100);p__Ascomycota_unclassified(100);p__Ascomycota_unclassified(100);p__Ascomycota_unclassified(100);p__Ascomycota_unclassified(100); | - | - | - | - | - | - | - | Unassigned | - |
| Otu13845 | 1 | k__Fungi(100);p__Ascomycota(100);p__Ascomycota_unclassified(100);p__Ascomycota_unclassified(100);p__Ascomycota_unclassified(100);p__Ascomycota_unclassified(100);p__Ascomycota_unclassified(100); | - | - | - | - | - | - | - | Unassigned | - |
| Otu13848 | 1 | k__Fungi(100);p__Ascomycota(100);p__Ascomycota_unclassified(100);p__Ascomycota_unclassified(100);p__Ascomycota_unclassified(100);p__Ascomycota_unclassified(100);p__Ascomycota_unclassified(100); | - | - | - | - | - | - | - | Unassigned | - |
| Otu13850 | 1 | k__Fungi(100);k__Fungi_unclassified(100);k__Fungi_unclassified(100);k__Fungi_unclassified(100);k__Fungi_unclassified(100);k__Fungi_unclassified(100);k__Fungi_unclassified(100); | - | - | - | - | - | - | - | Unassigned | - |
| Otu13853 | 1 | k__Fungi(100);k__Fungi_unclassified(100);k__Fungi_unclassified(100);k__Fungi_unclassified(100);k__Fungi_unclassified(100);k__Fungi_unclassified(100);k__Fungi_unclassified(100); | - | - | - | - | - | - | - | Unassigned | - |
| Otu13856 | 1 | k__Fungi(100);p__Ascomycota(100);p__Ascomycota_unclassified(100);p__Ascomycota_unclassified(100);p__Ascomycota_unclassified(100);p__Ascomycota_unclassified(100);p__Ascomycota_unclassified(100); | - | - | - | - | - | - | - | Unassigned | - |
| Otu13857 | 1 | k__Fungi(100);k__Fungi_unclassified(100);k__Fungi_unclassified(100);k__Fungi_unclassified(100);k__Fungi_unclassified(100);k__Fungi_unclassified(100);k__Fungi_unclassified(100); | - | - | - | - | - | - | - | Unassigned | - |
| Otu13858 | 1 | k__Fungi(100);p__Ascomycota(100);c__Sordariomycetes(100);c__Sordariomycetes_unclassified(100);c__Sordariomycetes_unclassified(100);c__Sordariomycetes_unclassified(100);c__Sordariomycetes_unclassified(100); | - | - | - | - | - | - | - | Unassigned | - |
| Otu13870 | 1 | k__Fungi(100);k__Fungi_unclassified(100);k__Fungi_unclassified(100);k__Fungi_unclassified(100);k__Fungi_unclassified(100);k__Fungi_unclassified(100);k__Fungi_unclassified(100); | - | - | - | - | - | - | - | Unassigned | - |
| Otu13877 | 1 | k__Fungi(100);k__Fungi_unclassified(100);k__Fungi_unclassified(100);k__Fungi_unclassified(100);k__Fungi_unclassified(100);k__Fungi_unclassified(100);k__Fungi_unclassified(100); | - | - | - | - | - | - | - | Unassigned | - |
| Otu13889 | 1 | k__Fungi(100);k__Fungi_unclassified(100);k__Fungi_unclassified(100);k__Fungi_unclassified(100);k__Fungi_unclassified(100);k__Fungi_unclassified(100);k__Fungi_unclassified(100); | - | - | - | - | - | - | - | Unassigned | - |
| Otu13894 | 1 | k__Fungi(100);p__Ascomycota(100);p__Ascomycota_unclassified(100);p__Ascomycota_unclassified(100);p__Ascomycota_unclassified(100);p__Ascomycota_unclassified(100);p__Ascomycota_unclassified(100); | - | - | - | - | - | - | - | Unassigned | - |
| Otu13897 | 1 | k__Fungi(100);p__Ascomycota(100);p__Ascomycota_unclassified(100);p__Ascomycota_unclassified(100);p__Ascomycota_unclassified(100);p__Ascomycota_unclassified(100);p__Ascomycota_unclassified(100); | - | - | - | - | - | - | - | Unassigned | - |
| Otu13898 | 1 | k__Fungi(100);k__Fungi_unclassified(100);k__Fungi_unclassified(100);k__Fungi_unclassified(100);k__Fungi_unclassified(100);k__Fungi_unclassified(100);k__Fungi_unclassified(100); | - | - | - | - | - | - | - | Unassigned | - |
| Otu13900 | 1 | k__Fungi(100);p__Ascomycota(100);p__Ascomycota_unclassified(100);p__Ascomycota_unclassified(100);p__Ascomycota_unclassified(100);p__Ascomycota_unclassified(100);p__Ascomycota_unclassified(100); | - | - | - | - | - | - | - | Unassigned | - |
| Otu13903 | 1 | k__Fungi(100);k__Fungi_unclassified(100);k__Fungi_unclassified(100);k__Fungi_unclassified(100);k__Fungi_unclassified(100);k__Fungi_unclassified(100);k__Fungi_unclassified(100); | - | - | - | - | - | - | - | Unassigned | - |
| Otu13914 | 1 | k__Fungi(100);k__Fungi_unclassified(100);k__Fungi_unclassified(100);k__Fungi_unclassified(100);k__Fungi_unclassified(100);k__Fungi_unclassified(100);k__Fungi_unclassified(100); | - | - | - | - | - | - | - | Unassigned | - |
| Otu13917 | 1 | k__Fungi(100);k__Fungi_unclassified(100);k__Fungi_unclassified(100);k__Fungi_unclassified(100);k__Fungi_unclassified(100);k__Fungi_unclassified(100);k__Fungi_unclassified(100); | - | - | - | - | - | - | - | Unassigned | - |
| Otu13922 | 1 | k__Fungi(100);p__Ascomycota(100);p__Ascomycota_unclassified(100);p__Ascomycota_unclassified(100);p__Ascomycota_unclassified(100);p__Ascomycota_unclassified(100);p__Ascomycota_unclassified(100); | - | - | - | - | - | - | - | Unassigned | - |
| Otu13923 | 1 | k__Fungi(100);k__Fungi_unclassified(100);k__Fungi_unclassified(100);k__Fungi_unclassified(100);k__Fungi_unclassified(100);k__Fungi_unclassified(100);k__Fungi_unclassified(100); | - | - | - | - | - | - | - | Unassigned | - |
| Otu13924 | 1 | k__Fungi(100);p__Ascomycota(100);p__Ascomycota_unclassified(100);p__Ascomycota_unclassified(100);p__Ascomycota_unclassified(100);p__Ascomycota_unclassified(100);p__Ascomycota_unclassified(100); | - | - | - | - | - | - | - | Unassigned | - |
| Otu13928 | 1 | k__Fungi(100);p__Ascomycota(100);p__Ascomycota_unclassified(100);p__Ascomycota_unclassified(100);p__Ascomycota_unclassified(100);p__Ascomycota_unclassified(100);p__Ascomycota_unclassified(100); | - | - | - | - | - | - | - | Unassigned | - |
| Otu13929 | 1 | k__Fungi(100);p__Ascomycota(100);p__Ascomycota_unclassified(100);p__Ascomycota_unclassified(100);p__Ascomycota_unclassified(100);p__Ascomycota_unclassified(100);p__Ascomycota_unclassified(100); | - | - | - | - | - | - | - | Unassigned | - |
| Otu13932 | 1 | k__Fungi(100);k__Fungi_unclassified(100);k__Fungi_unclassified(100);k__Fungi_unclassified(100);k__Fungi_unclassified(100);k__Fungi_unclassified(100);k__Fungi_unclassified(100); | - | - | - | - | - | - | - | Unassigned | - |
| Otu13943 | 1 | k__Fungi(100);p__Ascomycota(100);p__Ascomycota_unclassified(100);p__Ascomycota_unclassified(100);p__Ascomycota_unclassified(100);p__Ascomycota_unclassified(100);p__Ascomycota_unclassified(100); | - | - | - | - | - | - | - | Unassigned | - |
| Otu13946 | 1 | k__Fungi(100);k__Fungi_unclassified(100);k__Fungi_unclassified(100);k__Fungi_unclassified(100);k__Fungi_unclassified(100);k__Fungi_unclassified(100);k__Fungi_unclassified(100); | - | - | - | - | - | - | - | Unassigned | - |
| Otu13951 | 1 | k__Fungi(100);p__Ascomycota(100);p__Ascomycota_unclassified(100);p__Ascomycota_unclassified(100);p__Ascomycota_unclassified(100);p__Ascomycota_unclassified(100);p__Ascomycota_unclassified(100); | - | - | - | - | - | - | - | Unassigned | - |
| Otu13952 | 1 | k__Fungi(100);k__Fungi_unclassified(100);k__Fungi_unclassified(100);k__Fungi_unclassified(100);k__Fungi_unclassified(100);k__Fungi_unclassified(100);k__Fungi_unclassified(100); | - | - | - | - | - | - | - | Unassigned | - |
| Otu13953 | 1 | k__Fungi(100);p__Ascomycota(100);p__Ascomycota_unclassified(100);p__Ascomycota_unclassified(100);p__Ascomycota_unclassified(100);p__Ascomycota_unclassified(100);p__Ascomycota_unclassified(100); | - | - | - | - | - | - | - | Unassigned | - |
| Otu13954 | 1 | k__Fungi(100);p__Ascomycota(100);p__Ascomycota_unclassified(100);p__Ascomycota_unclassified(100);p__Ascomycota_unclassified(100);p__Ascomycota_unclassified(100);p__Ascomycota_unclassified(100); | - | - | - | - | - | - | - | Unassigned | - |
| Otu13955 | 1 | k__Fungi(100);k__Fungi_unclassified(100);k__Fungi_unclassified(100);k__Fungi_unclassified(100);k__Fungi_unclassified(100);k__Fungi_unclassified(100);k__Fungi_unclassified(100); | - | - | - | - | - | - | - | Unassigned | - |
| Otu13958 | 1 | k__Fungi(100);p__Ascomycota(100);p__Ascomycota_unclassified(100);p__Ascomycota_unclassified(100);p__Ascomycota_unclassified(100);p__Ascomycota_unclassified(100);p__Ascomycota_unclassified(100); | - | - | - | - | - | - | - | Unassigned | - |
| Otu13960 | 1 | k__Fungi(100);k__Fungi_unclassified(100);k__Fungi_unclassified(100);k__Fungi_unclassified(100);k__Fungi_unclassified(100);k__Fungi_unclassified(100);k__Fungi_unclassified(100); | - | - | - | - | - | - | - | Unassigned | - |
| Otu13961 | 1 | k__Fungi(100);k__Fungi_unclassified(100);k__Fungi_unclassified(100);k__Fungi_unclassified(100);k__Fungi_unclassified(100);k__Fungi_unclassified(100);k__Fungi_unclassified(100); | - | - | - | - | - | - | - | Unassigned | - |
| Otu13962 | 1 | k__Fungi(100);k__Fungi_unclassified(100);k__Fungi_unclassified(100);k__Fungi_unclassified(100);k__Fungi_unclassified(100);k__Fungi_unclassified(100);k__Fungi_unclassified(100); | - | - | - | - | - | - | - | Unassigned | - |
| Otu13963 | 1 | k__Fungi(100);k__Fungi_unclassified(100);k__Fungi_unclassified(100);k__Fungi_unclassified(100);k__Fungi_unclassified(100);k__Fungi_unclassified(100);k__Fungi_unclassified(100); | - | - | - | - | - | - | - | Unassigned | - |
| Otu13964 | 1 | k__Fungi(100);p__Ascomycota(100);p__Ascomycota_unclassified(100);p__Ascomycota_unclassified(100);p__Ascomycota_unclassified(100);p__Ascomycota_unclassified(100);p__Ascomycota_unclassified(100); | - | - | - | - | - | - | - | Unassigned | - |
| Otu13966 | 1 | k__Fungi(100);p__Ascomycota(100);p__Ascomycota_unclassified(100);p__Ascomycota_unclassified(100);p__Ascomycota_unclassified(100);p__Ascomycota_unclassified(100);p__Ascomycota_unclassified(100); | - | - | - | - | - | - | - | Unassigned | - |
| Otu13968 | 1 | k__Fungi(100);k__Fungi_unclassified(100);k__Fungi_unclassified(100);k__Fungi_unclassified(100);k__Fungi_unclassified(100);k__Fungi_unclassified(100);k__Fungi_unclassified(100); | - | - | - | - | - | - | - | Unassigned | - |
| Otu13969 | 1 | k__Fungi(100);k__Fungi_unclassified(100);k__Fungi_unclassified(100);k__Fungi_unclassified(100);k__Fungi_unclassified(100);k__Fungi_unclassified(100);k__Fungi_unclassified(100); | - | - | - | - | - | - | - | Unassigned | - |
| Otu13971 | 1 | k__Fungi(100);k__Fungi_unclassified(100);k__Fungi_unclassified(100);k__Fungi_unclassified(100);k__Fungi_unclassified(100);k__Fungi_unclassified(100);k__Fungi_unclassified(100); | - | - | - | - | - | - | - | Unassigned | - |
| Otu13972 | 1 | k__Fungi(100);k__Fungi_unclassified(100);k__Fungi_unclassified(100);k__Fungi_unclassified(100);k__Fungi_unclassified(100);k__Fungi_unclassified(100);k__Fungi_unclassified(100); | - | - | - | - | - | - | - | Unassigned | - |
| Otu13973 | 1 | k__Fungi(100);k__Fungi_unclassified(100);k__Fungi_unclassified(100);k__Fungi_unclassified(100);k__Fungi_unclassified(100);k__Fungi_unclassified(100);k__Fungi_unclassified(100); | - | - | - | - | - | - | - | Unassigned | - |
| Otu13975 | 1 | k__Fungi(100);p__Ascomycota(100);p__Ascomycota_unclassified(100);p__Ascomycota_unclassified(100);p__Ascomycota_unclassified(100);p__Ascomycota_unclassified(100);p__Ascomycota_unclassified(100); | - | - | - | - | - | - | - | Unassigned | - |
| Otu13977 | 1 | k__Fungi(100);k__Fungi_unclassified(100);k__Fungi_unclassified(100);k__Fungi_unclassified(100);k__Fungi_unclassified(100);k__Fungi_unclassified(100);k__Fungi_unclassified(100); | - | - | - | - | - | - | - | Unassigned | - |
| Otu13981 | 1 | k__Fungi(100);p__Ascomycota(100);p__Ascomycota_unclassified(100);p__Ascomycota_unclassified(100);p__Ascomycota_unclassified(100);p__Ascomycota_unclassified(100);p__Ascomycota_unclassified(100); | - | - | - | - | - | - | - | Unassigned | - |
| Otu13983 | 1 | k__Fungi(100);k__Fungi_unclassified(100);k__Fungi_unclassified(100);k__Fungi_unclassified(100);k__Fungi_unclassified(100);k__Fungi_unclassified(100);k__Fungi_unclassified(100); | - | - | - | - | - | - | - | Unassigned | - |
| Otu13984 | 1 | k__Fungi(100);p__Ascomycota(100);p__Ascomycota_unclassified(100);p__Ascomycota_unclassified(100);p__Ascomycota_unclassified(100);p__Ascomycota_unclassified(100);p__Ascomycota_unclassified(100); | - | - | - | - | - | - | - | Unassigned | - |
| Otu13989 | 1 | k__Fungi(100);k__Fungi_unclassified(100);k__Fungi_unclassified(100);k__Fungi_unclassified(100);k__Fungi_unclassified(100);k__Fungi_unclassified(100);k__Fungi_unclassified(100); | - | - | - | - | - | - | - | Unassigned | - |
| Otu13990 | 1 | k__Fungi(100);p__Ascomycota(100);p__Ascomycota_unclassified(100);p__Ascomycota_unclassified(100);p__Ascomycota_unclassified(100);p__Ascomycota_unclassified(100);p__Ascomycota_unclassified(100); | - | - | - | - | - | - | - | Unassigned | - |
| Otu13992 | 1 | k__Fungi(100);p__Ascomycota(100);p__Ascomycota_unclassified(100);p__Ascomycota_unclassified(100);p__Ascomycota_unclassified(100);p__Ascomycota_unclassified(100);p__Ascomycota_unclassified(100); | - | - | - | - | - | - | - | Unassigned | - |
| Otu13993 | 1 | k__Fungi(100);k__Fungi_unclassified(100);k__Fungi_unclassified(100);k__Fungi_unclassified(100);k__Fungi_unclassified(100);k__Fungi_unclassified(100);k__Fungi_unclassified(100); | - | - | - | - | - | - | - | Unassigned | - |
| Otu13994 | 1 | k__Fungi(100);k__Fungi_unclassified(100);k__Fungi_unclassified(100);k__Fungi_unclassified(100);k__Fungi_unclassified(100);k__Fungi_unclassified(100);k__Fungi_unclassified(100); | - | - | - | - | - | - | - | Unassigned | - |
| Otu13995 | 1 | k__Fungi(100);k__Fungi_unclassified(100);k__Fungi_unclassified(100);k__Fungi_unclassified(100);k__Fungi_unclassified(100);k__Fungi_unclassified(100);k__Fungi_unclassified(100); | - | - | - | - | - | - | - | Unassigned | - |
| Otu14000 | 1 | k__Fungi(100);k__Fungi_unclassified(100);k__Fungi_unclassified(100);k__Fungi_unclassified(100);k__Fungi_unclassified(100);k__Fungi_unclassified(100);k__Fungi_unclassified(100); | - | - | - | - | - | - | - | Unassigned | - |
| Otu14001 | 1 | k__Fungi(100);k__Fungi_unclassified(100);k__Fungi_unclassified(100);k__Fungi_unclassified(100);k__Fungi_unclassified(100);k__Fungi_unclassified(100);k__Fungi_unclassified(100); | - | - | - | - | - | - | - | Unassigned | - |
| Otu14011 | 1 | k__Fungi(100);k__Fungi_unclassified(100);k__Fungi_unclassified(100);k__Fungi_unclassified(100);k__Fungi_unclassified(100);k__Fungi_unclassified(100);k__Fungi_unclassified(100); | - | - | - | - | - | - | - | Unassigned | - |
| Otu14016 | 1 | k__Fungi(100);k__Fungi_unclassified(100);k__Fungi_unclassified(100);k__Fungi_unclassified(100);k__Fungi_unclassified(100);k__Fungi_unclassified(100);k__Fungi_unclassified(100); | - | - | - | - | - | - | - | Unassigned | - |
| Otu14018 | 1 | k__Fungi(100);k__Fungi_unclassified(100);k__Fungi_unclassified(100);k__Fungi_unclassified(100);k__Fungi_unclassified(100);k__Fungi_unclassified(100);k__Fungi_unclassified(100); | - | - | - | - | - | - | - | Unassigned | - |
| Otu14020 | 1 | k__Fungi(100);p__Ascomycota(100);p__Ascomycota_unclassified(100);p__Ascomycota_unclassified(100);p__Ascomycota_unclassified(100);p__Ascomycota_unclassified(100);p__Ascomycota_unclassified(100); | - | - | - | - | - | - | - | Unassigned | - |
| Otu14030 | 1 | k__Fungi(100);k__Fungi_unclassified(100);k__Fungi_unclassified(100);k__Fungi_unclassified(100);k__Fungi_unclassified(100);k__Fungi_unclassified(100);k__Fungi_unclassified(100); | - | - | - | - | - | - | - | Unassigned | - |
| Otu14034 | 1 | k__Fungi(100);k__Fungi_unclassified(100);k__Fungi_unclassified(100);k__Fungi_unclassified(100);k__Fungi_unclassified(100);k__Fungi_unclassified(100);k__Fungi_unclassified(100); | - | - | - | - | - | - | - | Unassigned | - |
| Otu14046 | 1 | k__Fungi(100);k__Fungi_unclassified(100);k__Fungi_unclassified(100);k__Fungi_unclassified(100);k__Fungi_unclassified(100);k__Fungi_unclassified(100);k__Fungi_unclassified(100); | - | - | - | - | - | - | - | Unassigned | - |
| Otu14047 | 1 | k__Fungi(100);k__Fungi_unclassified(100);k__Fungi_unclassified(100);k__Fungi_unclassified(100);k__Fungi_unclassified(100);k__Fungi_unclassified(100);k__Fungi_unclassified(100); | - | - | - | - | - | - | - | Unassigned | - |
| Otu14048 | 1 | k__Fungi(100);k__Fungi_unclassified(100);k__Fungi_unclassified(100);k__Fungi_unclassified(100);k__Fungi_unclassified(100);k__Fungi_unclassified(100);k__Fungi_unclassified(100); | - | - | - | - | - | - | - | Unassigned | - |
| Otu14061 | 1 | k__Fungi(100);k__Fungi_unclassified(100);k__Fungi_unclassified(100);k__Fungi_unclassified(100);k__Fungi_unclassified(100);k__Fungi_unclassified(100);k__Fungi_unclassified(100); | - | - | - | - | - | - | - | Unassigned | - |
| Otu14063 | 1 | k__Fungi(100);k__Fungi_unclassified(100);k__Fungi_unclassified(100);k__Fungi_unclassified(100);k__Fungi_unclassified(100);k__Fungi_unclassified(100);k__Fungi_unclassified(100); | - | - | - | - | - | - | - | Unassigned | - |
| Otu14064 | 1 | k__Fungi(100);k__Fungi_unclassified(100);k__Fungi_unclassified(100);k__Fungi_unclassified(100);k__Fungi_unclassified(100);k__Fungi_unclassified(100);k__Fungi_unclassified(100); | - | - | - | - | - | - | - | Unassigned | - |
| Otu14069 | 1 | k__Fungi(100);p__Ascomycota(100);p__Ascomycota_unclassified(100);p__Ascomycota_unclassified(100);p__Ascomycota_unclassified(100);p__Ascomycota_unclassified(100);p__Ascomycota_unclassified(100); | - | - | - | - | - | - | - | Unassigned | - |
| Otu14075 | 1 | k__Fungi(100);k__Fungi_unclassified(100);k__Fungi_unclassified(100);k__Fungi_unclassified(100);k__Fungi_unclassified(100);k__Fungi_unclassified(100);k__Fungi_unclassified(100); | - | - | - | - | - | - | - | Unassigned | - |
| Otu14076 | 1 | k__Fungi(100);p__Ascomycota(100);p__Ascomycota_unclassified(100);p__Ascomycota_unclassified(100);p__Ascomycota_unclassified(100);p__Ascomycota_unclassified(100);p__Ascomycota_unclassified(100); | - | - | - | - | - | - | - | Unassigned | - |
| Otu14078 | 1 | k__Fungi(100);k__Fungi_unclassified(100);k__Fungi_unclassified(100);k__Fungi_unclassified(100);k__Fungi_unclassified(100);k__Fungi_unclassified(100);k__Fungi_unclassified(100); | - | - | - | - | - | - | - | Unassigned | - |
| Otu14080 | 1 | k__Fungi(100);k__Fungi_unclassified(100);k__Fungi_unclassified(100);k__Fungi_unclassified(100);k__Fungi_unclassified(100);k__Fungi_unclassified(100);k__Fungi_unclassified(100); | - | - | - | - | - | - | - | Unassigned | - |
| Otu14083 | 1 | k__Fungi(100);k__Fungi_unclassified(100);k__Fungi_unclassified(100);k__Fungi_unclassified(100);k__Fungi_unclassified(100);k__Fungi_unclassified(100);k__Fungi_unclassified(100); | - | - | - | - | - | - | - | Unassigned | - |
| Otu14089 | 1 | k__Fungi(100);p__Ascomycota(100);p__Ascomycota_unclassified(100);p__Ascomycota_unclassified(100);p__Ascomycota_unclassified(100);p__Ascomycota_unclassified(100);p__Ascomycota_unclassified(100); | - | - | - | - | - | - | - | Unassigned | - |
| Otu14094 | 1 | k__Fungi(100);k__Fungi_unclassified(100);k__Fungi_unclassified(100);k__Fungi_unclassified(100);k__Fungi_unclassified(100);k__Fungi_unclassified(100);k__Fungi_unclassified(100); | - | - | - | - | - | - | - | Unassigned | - |
| Otu14096 | 1 | k__Fungi(100);p__Ascomycota(100);p__Ascomycota_unclassified(100);p__Ascomycota_unclassified(100);p__Ascomycota_unclassified(100);p__Ascomycota_unclassified(100);p__Ascomycota_unclassified(100); | - | - | - | - | - | - | - | Unassigned | - |
| Otu14100 | 1 | k__Fungi(100);k__Fungi_unclassified(100);k__Fungi_unclassified(100);k__Fungi_unclassified(100);k__Fungi_unclassified(100);k__Fungi_unclassified(100);k__Fungi_unclassified(100); | - | - | - | - | - | - | - | Unassigned | - |
| Otu14103 | 1 | k__Fungi(100);k__Fungi_unclassified(100);k__Fungi_unclassified(100);k__Fungi_unclassified(100);k__Fungi_unclassified(100);k__Fungi_unclassified(100);k__Fungi_unclassified(100); | - | - | - | - | - | - | - | Unassigned | - |
| Otu14104 | 1 | k__Fungi(100);k__Fungi_unclassified(100);k__Fungi_unclassified(100);k__Fungi_unclassified(100);k__Fungi_unclassified(100);k__Fungi_unclassified(100);k__Fungi_unclassified(100); | - | - | - | - | - | - | - | Unassigned | - |
| Otu14105 | 1 | k__Fungi(100);k__Fungi_unclassified(100);k__Fungi_unclassified(100);k__Fungi_unclassified(100);k__Fungi_unclassified(100);k__Fungi_unclassified(100);k__Fungi_unclassified(100); | - | - | - | - | - | - | - | Unassigned | - |
| Otu14108 | 1 | k__Fungi(100);k__Fungi_unclassified(100);k__Fungi_unclassified(100);k__Fungi_unclassified(100);k__Fungi_unclassified(100);k__Fungi_unclassified(100);k__Fungi_unclassified(100); | - | - | - | - | - | - | - | Unassigned | - |
| Otu14109 | 1 | k__Fungi(100);k__Fungi_unclassified(100);k__Fungi_unclassified(100);k__Fungi_unclassified(100);k__Fungi_unclassified(100);k__Fungi_unclassified(100);k__Fungi_unclassified(100); | - | - | - | - | - | - | - | Unassigned | - |
| Otu14110 | 1 | k__Fungi(100);p__Ascomycota(100);p__Ascomycota_unclassified(100);p__Ascomycota_unclassified(100);p__Ascomycota_unclassified(100);p__Ascomycota_unclassified(100);p__Ascomycota_unclassified(100); | - | - | - | - | - | - | - | Unassigned | - |
| Otu14111 | 1 | k__Fungi(100);k__Fungi_unclassified(100);k__Fungi_unclassified(100);k__Fungi_unclassified(100);k__Fungi_unclassified(100);k__Fungi_unclassified(100);k__Fungi_unclassified(100); | - | - | - | - | - | - | - | Unassigned | - |
| Otu14112 | 1 | k__Fungi(100);k__Fungi_unclassified(100);k__Fungi_unclassified(100);k__Fungi_unclassified(100);k__Fungi_unclassified(100);k__Fungi_unclassified(100);k__Fungi_unclassified(100); | - | - | - | - | - | - | - | Unassigned | - |
| Otu14113 | 1 | k__Fungi(100);k__Fungi_unclassified(100);k__Fungi_unclassified(100);k__Fungi_unclassified(100);k__Fungi_unclassified(100);k__Fungi_unclassified(100);k__Fungi_unclassified(100); | - | - | - | - | - | - | - | Unassigned | - |
| Otu14114 | 1 | k__Fungi(100);k__Fungi_unclassified(100);k__Fungi_unclassified(100);k__Fungi_unclassified(100);k__Fungi_unclassified(100);k__Fungi_unclassified(100);k__Fungi_unclassified(100); | - | - | - | - | - | - | - | Unassigned | - |
| Otu14115 | 1 | k__Fungi(100);k__Fungi_unclassified(100);k__Fungi_unclassified(100);k__Fungi_unclassified(100);k__Fungi_unclassified(100);k__Fungi_unclassified(100);k__Fungi_unclassified(100); | - | - | - | - | - | - | - | Unassigned | - |
| Otu14118 | 1 | k__Fungi(100);k__Fungi_unclassified(100);k__Fungi_unclassified(100);k__Fungi_unclassified(100);k__Fungi_unclassified(100);k__Fungi_unclassified(100);k__Fungi_unclassified(100); | - | - | - | - | - | - | - | Unassigned | - |
| Otu14119 | 1 | k__Fungi(100);p__Ascomycota(100);p__Ascomycota_unclassified(100);p__Ascomycota_unclassified(100);p__Ascomycota_unclassified(100);p__Ascomycota_unclassified(100);p__Ascomycota_unclassified(100); | - | - | - | - | - | - | - | Unassigned | - |
| Otu14120 | 1 | k__Fungi(100);p__Ascomycota(100);p__Ascomycota_unclassified(100);p__Ascomycota_unclassified(100);p__Ascomycota_unclassified(100);p__Ascomycota_unclassified(100);p__Ascomycota_unclassified(100); | - | - | - | - | - | - | - | Unassigned | - |
| Otu14121 | 1 | k__Fungi(100);p__Ascomycota(100);p__Ascomycota_unclassified(100);p__Ascomycota_unclassified(100);p__Ascomycota_unclassified(100);p__Ascomycota_unclassified(100);p__Ascomycota_unclassified(100); | - | - | - | - | - | - | - | Unassigned | - |
| Otu14123 | 1 | k__Fungi(100);k__Fungi_unclassified(100);k__Fungi_unclassified(100);k__Fungi_unclassified(100);k__Fungi_unclassified(100);k__Fungi_unclassified(100);k__Fungi_unclassified(100); | - | - | - | - | - | - | - | Unassigned | - |
| Otu14124 | 1 | k__Fungi(100);k__Fungi_unclassified(100);k__Fungi_unclassified(100);k__Fungi_unclassified(100);k__Fungi_unclassified(100);k__Fungi_unclassified(100);k__Fungi_unclassified(100); | - | - | - | - | - | - | - | Unassigned | - |
| Otu14126 | 1 | k__Fungi(100);k__Fungi_unclassified(100);k__Fungi_unclassified(100);k__Fungi_unclassified(100);k__Fungi_unclassified(100);k__Fungi_unclassified(100);k__Fungi_unclassified(100); | - | - | - | - | - | - | - | Unassigned | - |
| Otu14128 | 1 | k__Fungi(100);k__Fungi_unclassified(100);k__Fungi_unclassified(100);k__Fungi_unclassified(100);k__Fungi_unclassified(100);k__Fungi_unclassified(100);k__Fungi_unclassified(100); | - | - | - | - | - | - | - | Unassigned | - |
| Otu14130 | 1 | k__Fungi(100);p__Ascomycota(100);p__Ascomycota_unclassified(100);p__Ascomycota_unclassified(100);p__Ascomycota_unclassified(100);p__Ascomycota_unclassified(100);p__Ascomycota_unclassified(100); | - | - | - | - | - | - | - | Unassigned | - |
| Otu14131 | 1 | k__Fungi(100);k__Fungi_unclassified(100);k__Fungi_unclassified(100);k__Fungi_unclassified(100);k__Fungi_unclassified(100);k__Fungi_unclassified(100);k__Fungi_unclassified(100); | - | - | - | - | - | - | - | Unassigned | - |
| Otu14132 | 1 | k__Fungi(100);p__Ascomycota(100);p__Ascomycota_unclassified(100);p__Ascomycota_unclassified(100);p__Ascomycota_unclassified(100);p__Ascomycota_unclassified(100);p__Ascomycota_unclassified(100); | - | - | - | - | - | - | - | Unassigned | - |
| Otu14133 | 1 | k__Fungi(100);p__Ascomycota(100);p__Ascomycota_unclassified(100);p__Ascomycota_unclassified(100);p__Ascomycota_unclassified(100);p__Ascomycota_unclassified(100);p__Ascomycota_unclassified(100); | - | - | - | - | - | - | - | Unassigned | - |
| Otu14134 | 1 | k__Fungi(100);k__Fungi_unclassified(100);k__Fungi_unclassified(100);k__Fungi_unclassified(100);k__Fungi_unclassified(100);k__Fungi_unclassified(100);k__Fungi_unclassified(100); | - | - | - | - | - | - | - | Unassigned | - |
| Otu14136 | 1 | k__Fungi(100);p__Ascomycota(100);p__Ascomycota_unclassified(100);p__Ascomycota_unclassified(100);p__Ascomycota_unclassified(100);p__Ascomycota_unclassified(100);p__Ascomycota_unclassified(100); | - | - | - | - | - | - | - | Unassigned | - |
| Otu14142 | 1 | k__Fungi(100);p__Ascomycota(100);p__Ascomycota_unclassified(100);p__Ascomycota_unclassified(100);p__Ascomycota_unclassified(100);p__Ascomycota_unclassified(100);p__Ascomycota_unclassified(100); | - | - | - | - | - | - | - | Unassigned | - |
| Otu14143 | 1 | k__Fungi(100);p__Ascomycota(100);p__Ascomycota_unclassified(100);p__Ascomycota_unclassified(100);p__Ascomycota_unclassified(100);p__Ascomycota_unclassified(100);p__Ascomycota_unclassified(100); | - | - | - | - | - | - | - | Unassigned | - |
| Otu14144 | 1 | k__Fungi(100);k__Fungi_unclassified(100);k__Fungi_unclassified(100);k__Fungi_unclassified(100);k__Fungi_unclassified(100);k__Fungi_unclassified(100);k__Fungi_unclassified(100); | - | - | - | - | - | - | - | Unassigned | - |
| Otu14145 | 1 | k__Fungi(100);p__Ascomycota(100);p__Ascomycota_unclassified(100);p__Ascomycota_unclassified(100);p__Ascomycota_unclassified(100);p__Ascomycota_unclassified(100);p__Ascomycota_unclassified(100); | - | - | - | - | - | - | - | Unassigned | - |
| Otu14148 | 1 | k__Fungi(100);k__Fungi_unclassified(100);k__Fungi_unclassified(100);k__Fungi_unclassified(100);k__Fungi_unclassified(100);k__Fungi_unclassified(100);k__Fungi_unclassified(100); | - | - | - | - | - | - | - | Unassigned | - |
| Otu14149 | 1 | k__Fungi(100);k__Fungi_unclassified(100);k__Fungi_unclassified(100);k__Fungi_unclassified(100);k__Fungi_unclassified(100);k__Fungi_unclassified(100);k__Fungi_unclassified(100); | - | - | - | - | - | - | - | Unassigned | - |
| Otu14152 | 1 | k__Fungi(100);k__Fungi_unclassified(100);k__Fungi_unclassified(100);k__Fungi_unclassified(100);k__Fungi_unclassified(100);k__Fungi_unclassified(100);k__Fungi_unclassified(100); | - | - | - | - | - | - | - | Unassigned | - |
| Otu14154 | 1 | k__Fungi(100);k__Fungi_unclassified(100);k__Fungi_unclassified(100);k__Fungi_unclassified(100);k__Fungi_unclassified(100);k__Fungi_unclassified(100);k__Fungi_unclassified(100); | - | - | - | - | - | - | - | Unassigned | - |
| Otu14156 | 1 | k__Fungi(100);p__Ascomycota(100);p__Ascomycota_unclassified(100);p__Ascomycota_unclassified(100);p__Ascomycota_unclassified(100);p__Ascomycota_unclassified(100);p__Ascomycota_unclassified(100); | - | - | - | - | - | - | - | Unassigned | - |
| Otu14157 | 1 | k__Fungi(100);p__Ascomycota(100);p__Ascomycota_unclassified(100);p__Ascomycota_unclassified(100);p__Ascomycota_unclassified(100);p__Ascomycota_unclassified(100);p__Ascomycota_unclassified(100); | - | - | - | - | - | - | - | Unassigned | - |
| Otu14158 | 1 | k__Fungi(100);p__Ascomycota(100);p__Ascomycota_unclassified(100);p__Ascomycota_unclassified(100);p__Ascomycota_unclassified(100);p__Ascomycota_unclassified(100);p__Ascomycota_unclassified(100); | - | - | - | - | - | - | - | Unassigned | - |
| Otu14160 | 1 | k__Fungi(100);p__Ascomycota(100);p__Ascomycota_unclassified(100);p__Ascomycota_unclassified(100);p__Ascomycota_unclassified(100);p__Ascomycota_unclassified(100);p__Ascomycota_unclassified(100); | - | - | - | - | - | - | - | Unassigned | - |
| Otu14168 | 1 | k__Fungi(100);p__Ascomycota(100);p__Ascomycota_unclassified(100);p__Ascomycota_unclassified(100);p__Ascomycota_unclassified(100);p__Ascomycota_unclassified(100);p__Ascomycota_unclassified(100); | - | - | - | - | - | - | - | Unassigned | - |
| Otu14171 | 1 | k__Fungi(100);p__Ascomycota(100);p__Ascomycota_unclassified(100);p__Ascomycota_unclassified(100);p__Ascomycota_unclassified(100);p__Ascomycota_unclassified(100);p__Ascomycota_unclassified(100); | - | - | - | - | - | - | - | Unassigned | - |
| Otu14172 | 1 | k__Fungi(100);p__Ascomycota(100);p__Ascomycota_unclassified(100);p__Ascomycota_unclassified(100);p__Ascomycota_unclassified(100);p__Ascomycota_unclassified(100);p__Ascomycota_unclassified(100); | - | - | - | - | - | - | - | Unassigned | - |
| Otu14173 | 1 | k__Fungi(100);p__Ascomycota(100);p__Ascomycota_unclassified(100);p__Ascomycota_unclassified(100);p__Ascomycota_unclassified(100);p__Ascomycota_unclassified(100);p__Ascomycota_unclassified(100); | - | - | - | - | - | - | - | Unassigned | - |
| Otu14176 | 1 | k__Fungi(100);k__Fungi_unclassified(100);k__Fungi_unclassified(100);k__Fungi_unclassified(100);k__Fungi_unclassified(100);k__Fungi_unclassified(100);k__Fungi_unclassified(100); | - | - | - | - | - | - | - | Unassigned | - |
| Otu14177 | 1 | k__Fungi(100);p__Ascomycota(100);p__Ascomycota_unclassified(100);p__Ascomycota_unclassified(100);p__Ascomycota_unclassified(100);p__Ascomycota_unclassified(100);p__Ascomycota_unclassified(100); | - | - | - | - | - | - | - | Unassigned | - |
| Otu14178 | 1 | k__Fungi(100);k__Fungi_unclassified(100);k__Fungi_unclassified(100);k__Fungi_unclassified(100);k__Fungi_unclassified(100);k__Fungi_unclassified(100);k__Fungi_unclassified(100); | - | - | - | - | - | - | - | Unassigned | - |
| Otu14179 | 1 | k__Fungi(100);k__Fungi_unclassified(100);k__Fungi_unclassified(100);k__Fungi_unclassified(100);k__Fungi_unclassified(100);k__Fungi_unclassified(100);k__Fungi_unclassified(100); | - | - | - | - | - | - | - | Unassigned | - |
| Otu14182 | 1 | k__Fungi(100);p__Ascomycota(100);p__Ascomycota_unclassified(100);p__Ascomycota_unclassified(100);p__Ascomycota_unclassified(100);p__Ascomycota_unclassified(100);p__Ascomycota_unclassified(100); | - | - | - | - | - | - | - | Unassigned | - |
| Otu14185 | 1 | k__Fungi(100);p__Ascomycota(100);p__Ascomycota_unclassified(100);p__Ascomycota_unclassified(100);p__Ascomycota_unclassified(100);p__Ascomycota_unclassified(100);p__Ascomycota_unclassified(100); | - | - | - | - | - | - | - | Unassigned | - |
| Otu14186 | 1 | k__Fungi(100);p__Ascomycota(100);p__Ascomycota_unclassified(100);p__Ascomycota_unclassified(100);p__Ascomycota_unclassified(100);p__Ascomycota_unclassified(100);p__Ascomycota_unclassified(100); | - | - | - | - | - | - | - | Unassigned | - |

| | | | | | | | | | | | |
|---|---|---|---|---|---|---|---|---|---|---|---|
| Otu14187 | 1 | k__Fungi(100);p__Ascomycota(100);p__Ascomycota_unclassified(100);p__Ascomycota_unclassified(100);p__Ascomycota_unclassified(100);p__Ascomycota_unclassified(100);p__Ascomycota_unclassified(100); | - | - | - | - | - | - | - | Unassigned | - |
| Otu14188 | 1 | k__Fungi(100);k__Fungi_unclassified(100);k__Fungi_unclassified(100);k__Fungi_unclassified(100);k__Fungi_unclassified(100);k__Fungi_unclassified(100);k__Fungi_unclassified(100); | - | - | - | - | - | - | - | Unassigned | - |
| Otu14189 | 1 | k__Fungi(100);p__Ascomycota(100);p__Ascomycota_unclassified(100);p__Ascomycota_unclassified(100);p__Ascomycota_unclassified(100);p__Ascomycota_unclassified(100);p__Ascomycota_unclassified(100); | - | - | - | - | - | - | - | Unassigned | - |
| Otu14190 | 1 | k__Fungi(100);k__Fungi_unclassified(100);k__Fungi_unclassified(100);k__Fungi_unclassified(100);k__Fungi_unclassified(100);k__Fungi_unclassified(100);k__Fungi_unclassified(100); | - | - | - | - | - | - | - | Unassigned | - |
| Otu14192 | 1 | k__Fungi(100);p__Ascomycota(100);p__Ascomycota_unclassified(100);p__Ascomycota_unclassified(100);p__Ascomycota_unclassified(100);p__Ascomycota_unclassified(100);p__Ascomycota_unclassified(100); | - | - | - | - | - | - | - | Unassigned | - |
| Otu14193 | 1 | k__Fungi(100);k__Fungi_unclassified(100);k__Fungi_unclassified(100);k__Fungi_unclassified(100);k__Fungi_unclassified(100);k__Fungi_unclassified(100);k__Fungi_unclassified(100); | - | - | - | - | - | - | - | Unassigned | - |
| Otu14194 | 1 | k__Fungi(100);k__Fungi_unclassified(100);k__Fungi_unclassified(100);k__Fungi_unclassified(100);k__Fungi_unclassified(100);k__Fungi_unclassified(100);k__Fungi_unclassified(100); | - | - | - | - | - | - | - | Unassigned | - |
| Otu14197 | 1 | k__Fungi(100);k__Fungi_unclassified(100);k__Fungi_unclassified(100);k__Fungi_unclassified(100);k__Fungi_unclassified(100);k__Fungi_unclassified(100);k__Fungi_unclassified(100); | - | - | - | - | - | - | - | Unassigned | - |
| Otu14200 | 1 | k__Fungi(100);p__Ascomycota(100);p__Ascomycota_unclassified(100);p__Ascomycota_unclassified(100);p__Ascomycota_unclassified(100);p__Ascomycota_unclassified(100);p__Ascomycota_unclassified(100); | - | - | - | - | - | - | - | Unassigned | - |
| Otu14203 | 1 | k__Fungi(100);k__Fungi_unclassified(100);k__Fungi_unclassified(100);k__Fungi_unclassified(100);k__Fungi_unclassified(100);k__Fungi_unclassified(100);k__Fungi_unclassified(100); | - | - | - | - | - | - | - | Unassigned | - |
| Otu14207 | 1 | k__Fungi(100);k__Fungi_unclassified(100);k__Fungi_unclassified(100);k__Fungi_unclassified(100);k__Fungi_unclassified(100);k__Fungi_unclassified(100);k__Fungi_unclassified(100); | - | - | - | - | - | - | - | Unassigned | - |
| Otu14210 | 1 | k__Fungi(100);k__Fungi_unclassified(100);k__Fungi_unclassified(100);k__Fungi_unclassified(100);k__Fungi_unclassified(100);k__Fungi_unclassified(100);k__Fungi_unclassified(100); | - | - | - | - | - | - | - | Unassigned | - |
| Otu14213 | 1 | k__Fungi(100);k__Fungi_unclassified(100);k__Fungi_unclassified(100);k__Fungi_unclassified(100);k__Fungi_unclassified(100);k__Fungi_unclassified(100);k__Fungi_unclassified(100); | - | - | - | - | - | - | - | Unassigned | - |
| Otu14215 | 1 | k__Fungi(100);p__Ascomycota(100);p__Ascomycota_unclassified(100);p__Ascomycota_unclassified(100);p__Ascomycota_unclassified(100);p__Ascomycota_unclassified(100);p__Ascomycota_unclassified(100); | - | - | - | - | - | - | - | Unassigned | - |
| Otu14216 | 1 | k__Fungi(100);p__Ascomycota(100);p__Ascomycota_unclassified(100);p__Ascomycota_unclassified(100);p__Ascomycota_unclassified(100);p__Ascomycota_unclassified(100);p__Ascomycota_unclassified(100); | - | - | - | - | - | - | - | Unassigned | - |
| Otu14217 | 1 | k__Fungi(100);k__Fungi_unclassified(100);k__Fungi_unclassified(100);k__Fungi_unclassified(100);k__Fungi_unclassified(100);k__Fungi_unclassified(100);k__Fungi_unclassified(100); | - | - | - | - | - | - | - | Unassigned | - |
| Otu14218 | 1 | k__Fungi(100);k__Fungi_unclassified(100);k__Fungi_unclassified(100);k__Fungi_unclassified(100);k__Fungi_unclassified(100);k__Fungi_unclassified(100);k__Fungi_unclassified(100); | - | - | - | - | - | - | - | Unassigned | - |
| Otu14219 | 1 | k__Fungi(100);p__Ascomycota(100);p__Ascomycota_unclassified(100);p__Ascomycota_unclassified(100);p__Ascomycota_unclassified(100);p__Ascomycota_unclassified(100);p__Ascomycota_unclassified(100); | - | - | - | - | - | - | - | Unassigned | - |
| Otu14220 | 1 | k__Fungi(100);p__Ascomycota(100);p__Ascomycota_unclassified(100);p__Ascomycota_unclassified(100);p__Ascomycota_unclassified(100);p__Ascomycota_unclassified(100);p__Ascomycota_unclassified(100); | - | - | - | - | - | - | - | Unassigned | - |
| Otu14222 | 1 | k__Fungi(100);k__Fungi_unclassified(100);k__Fungi_unclassified(100);k__Fungi_unclassified(100);k__Fungi_unclassified(100);k__Fungi_unclassified(100);k__Fungi_unclassified(100); | - | - | - | - | - | - | - | Unassigned | - |
| Otu14224 | 1 | k__Fungi(100);k__Fungi_unclassified(100);k__Fungi_unclassified(100);k__Fungi_unclassified(100);k__Fungi_unclassified(100);k__Fungi_unclassified(100);k__Fungi_unclassified(100); | - | - | - | - | - | - | - | Unassigned | - |
| Otu14227 | 1 | k__Fungi(100);p__Ascomycota(100);p__Ascomycota_unclassified(100);p__Ascomycota_unclassified(100);p__Ascomycota_unclassified(100);p__Ascomycota_unclassified(100);p__Ascomycota_unclassified(100); | - | - | - | - | - | - | - | Unassigned | - |
| Otu14228 | 1 | k__Fungi(100);k__Fungi_unclassified(100);k__Fungi_unclassified(100);k__Fungi_unclassified(100);k__Fungi_unclassified(100);k__Fungi_unclassified(100);k__Fungi_unclassified(100); | - | - | - | - | - | - | - | Unassigned | - |
| Otu14229 | 1 | k__Fungi(100);p__Ascomycota(100);p__Ascomycota_unclassified(100);p__Ascomycota_unclassified(100);p__Ascomycota_unclassified(100);p__Ascomycota_unclassified(100);p__Ascomycota_unclassified(100); | - | - | - | - | - | - | - | Unassigned | - |
| Otu14230 | 1 | k__Fungi(100);k__Fungi_unclassified(100);k__Fungi_unclassified(100);k__Fungi_unclassified(100);k__Fungi_unclassified(100);k__Fungi_unclassified(100);k__Fungi_unclassified(100); | - | - | - | - | - | - | - | Unassigned | - |
| Otu14231 | 1 | k__Fungi(100);p__Ascomycota(100);p__Ascomycota_unclassified(100);p__Ascomycota_unclassified(100);p__Ascomycota_unclassified(100);p__Ascomycota_unclassified(100);p__Ascomycota_unclassified(100); | - | - | - | - | - | - | - | Unassigned | - |
| Otu14237 | 1 | k__Fungi(100);k__Fungi_unclassified(100);k__Fungi_unclassified(100);k__Fungi_unclassified(100);k__Fungi_unclassified(100);k__Fungi_unclassified(100);k__Fungi_unclassified(100); | - | - | - | - | - | - | - | Unassigned | - |
| Otu14238 | 1 | k__Fungi(100);p__Ascomycota(100);p__Ascomycota_unclassified(100);p__Ascomycota_unclassified(100);p__Ascomycota_unclassified(100);p__Ascomycota_unclassified(100);p__Ascomycota_unclassified(100); | - | - | - | - | - | - | - | Unassigned | - |
| Otu14241 | 1 | k__Fungi(100);k__Fungi_unclassified(100);k__Fungi_unclassified(100);k__Fungi_unclassified(100);k__Fungi_unclassified(100);k__Fungi_unclassified(100);k__Fungi_unclassified(100); | - | - | - | - | - | - | - | Unassigned | - |
| Otu14242 | 1 | k__Fungi(100);k__Fungi_unclassified(100);k__Fungi_unclassified(100);k__Fungi_unclassified(100);k__Fungi_unclassified(100);k__Fungi_unclassified(100);k__Fungi_unclassified(100); | - | - | - | - | - | - | - | Unassigned | - |
| Otu14244 | 1 | k__Fungi(100);p__Ascomycota(100);p__Ascomycota_unclassified(100);p__Ascomycota_unclassified(100);p__Ascomycota_unclassified(100);p__Ascomycota_unclassified(100);p__Ascomycota_unclassified(100); | - | - | - | - | - | - | - | Unassigned | - |
| Otu14246 | 1 | k__Fungi(100);k__Fungi_unclassified(100);k__Fungi_unclassified(100);k__Fungi_unclassified(100);k__Fungi_unclassified(100);k__Fungi_unclassified(100);k__Fungi_unclassified(100); | - | - | - | - | - | - | - | Unassigned | - |
| Otu14252 | 1 | k__Fungi(100);p__Ascomycota(100);p__Ascomycota_unclassified(100);p__Ascomycota_unclassified(100);p__Ascomycota_unclassified(100);p__Ascomycota_unclassified(100);p__Ascomycota_unclassified(100); | - | - | - | - | - | - | - | Unassigned | - |
| Otu14253 | 1 | k__Fungi(100);k__Fungi_unclassified(100);k__Fungi_unclassified(100);k__Fungi_unclassified(100);k__Fungi_unclassified(100);k__Fungi_unclassified(100);k__Fungi_unclassified(100); | - | - | - | - | - | - | - | Unassigned | - |
| Otu14254 | 1 | k__Fungi(100);p__Ascomycota(100);p__Ascomycota_unclassified(100);p__Ascomycota_unclassified(100);p__Ascomycota_unclassified(100);p__Ascomycota_unclassified(100);p__Ascomycota_unclassified(100); | - | - | - | - | - | - | - | Unassigned | - |
| Otu14262 | 1 | k__Fungi(100);k__Fungi_unclassified(100);k__Fungi_unclassified(100);k__Fungi_unclassified(100);k__Fungi_unclassified(100);k__Fungi_unclassified(100);k__Fungi_unclassified(100); | - | - | - | - | - | - | - | Unassigned | - |
| Otu14266 | 1 | k__Fungi(100);p__Ascomycota(100);p__Ascomycota_unclassified(100);p__Ascomycota_unclassified(100);p__Ascomycota_unclassified(100);p__Ascomycota_unclassified(100);p__Ascomycota_unclassified(100); | - | - | - | - | - | - | - | Unassigned | - |
| Otu14268 | 1 | k__Fungi(100);k__Fungi_unclassified(100);k__Fungi_unclassified(100);k__Fungi_unclassified(100);k__Fungi_unclassified(100);k__Fungi_unclassified(100);k__Fungi_unclassified(100); | - | - | - | - | - | - | - | Unassigned | - |
| Otu14269 | 1 | k__Fungi(100);k__Fungi_unclassified(100);k__Fungi_unclassified(100);k__Fungi_unclassified(100);k__Fungi_unclassified(100);k__Fungi_unclassified(100);k__Fungi_unclassified(100); | - | - | - | - | - | - | - | Unassigned | - |
| Otu14272 | 1 | k__Fungi(100);k__Fungi_unclassified(100);k__Fungi_unclassified(100);k__Fungi_unclassified(100);k__Fungi_unclassified(100);k__Fungi_unclassified(100);k__Fungi_unclassified(100); | - | - | - | - | - | - | - | Unassigned | - |
| Otu14275 | 1 | k__Fungi(100);k__Fungi_unclassified(100);k__Fungi_unclassified(100);k__Fungi_unclassified(100);k__Fungi_unclassified(100);k__Fungi_unclassified(100);k__Fungi_unclassified(100); | - | - | - | - | - | - | - | Unassigned | - |
| Otu14277 | 1 | k__Fungi(100);p__Ascomycota(100);p__Ascomycota_unclassified(100);p__Ascomycota_unclassified(100);p__Ascomycota_unclassified(100);p__Ascomycota_unclassified(100);p__Ascomycota_unclassified(100); | - | - | - | - | - | - | - | Unassigned | - |
| Otu14278 | 1 | k__Fungi(100);p__Ascomycota(100);p__Ascomycota_unclassified(100);p__Ascomycota_unclassified(100);p__Ascomycota_unclassified(100);p__Ascomycota_unclassified(100);p__Ascomycota_unclassified(100); | - | - | - | - | - | - | - | Unassigned | - |
| Otu14280 | 1 | k__Fungi(100);p__Ascomycota(100);p__Ascomycota_unclassified(100);p__Ascomycota_unclassified(100);p__Ascomycota_unclassified(100);p__Ascomycota_unclassified(100);p__Ascomycota_unclassified(100); | - | - | - | - | - | - | - | Unassigned | - |
| Otu14283 | 1 | k__Fungi(100);k__Fungi_unclassified(100);k__Fungi_unclassified(100);k__Fungi_unclassified(100);k__Fungi_unclassified(100);k__Fungi_unclassified(100);k__Fungi_unclassified(100); | - | - | - | - | - | - | - | Unassigned | - |
| Otu14285 | 1 | k__Fungi(100);k__Fungi_unclassified(100);k__Fungi_unclassified(100);k__Fungi_unclassified(100);k__Fungi_unclassified(100);k__Fungi_unclassified(100);k__Fungi_unclassified(100); | - | - | - | - | - | - | - | Unassigned | - |
| Otu14292 | 1 | k__Fungi(100);p__Ascomycota(100);p__Ascomycota_unclassified(100);p__Ascomycota_unclassified(100);p__Ascomycota_unclassified(100);p__Ascomycota_unclassified(100);p__Ascomycota_unclassified(100); | - | - | - | - | - | - | - | Unassigned | - |
| Otu14306 | 1 | k__Fungi(100);k__Fungi_unclassified(100);k__Fungi_unclassified(100);k__Fungi_unclassified(100);k__Fungi_unclassified(100);k__Fungi_unclassified(100);k__Fungi_unclassified(100); | - | - | - | - | - | - | - | Unassigned | - |
| Otu14308 | 1 | k__Fungi(100);k__Fungi_unclassified(100);k__Fungi_unclassified(100);k__Fungi_unclassified(100);k__Fungi_unclassified(100);k__Fungi_unclassified(100);k__Fungi_unclassified(100); | - | - | - | - | - | - | - | Unassigned | - |
| Otu14310 | 1 | k__Fungi(100);p__Ascomycota(100);p__Ascomycota_unclassified(100);p__Ascomycota_unclassified(100);p__Ascomycota_unclassified(100);p__Ascomycota_unclassified(100);p__Ascomycota_unclassified(100); | - | - | - | - | - | - | - | Unassigned | - |
| Otu14311 | 1 | k__Fungi(100);k__Fungi_unclassified(100);k__Fungi_unclassified(100);k__Fungi_unclassified(100);k__Fungi_unclassified(100);k__Fungi_unclassified(100);k__Fungi_unclassified(100); | - | - | - | - | - | - | - | Unassigned | - |
| Otu14312 | 1 | k__Fungi(100);p__Ascomycota(100);p__Ascomycota_unclassified(100);p__Ascomycota_unclassified(100);p__Ascomycota_unclassified(100);p__Ascomycota_unclassified(100);p__Ascomycota_unclassified(100); | - | - | - | - | - | - | - | Unassigned | - |
| Otu14313 | 1 | k__Fungi(100);p__Ascomycota(100);p__Ascomycota_unclassified(100);p__Ascomycota_unclassified(100);p__Ascomycota_unclassified(100);p__Ascomycota_unclassified(100);p__Ascomycota_unclassified(100); | - | - | - | - | - | - | - | Unassigned | - |
| Otu14314 | 1 | k__Fungi(100);p__Ascomycota(100);p__Ascomycota_unclassified(100);p__Ascomycota_unclassified(100);p__Ascomycota_unclassified(100);p__Ascomycota_unclassified(100);p__Ascomycota_unclassified(100); | - | - | - | - | - | - | - | Unassigned | - |
| Otu14315 | 1 | k__Fungi(100);k__Fungi_unclassified(100);k__Fungi_unclassified(100);k__Fungi_unclassified(100);k__Fungi_unclassified(100);k__Fungi_unclassified(100);k__Fungi_unclassified(100); | - | - | - | - | - | - | - | Unassigned | - |
| Otu14316 | 1 | k__Fungi(100);k__Fungi_unclassified(100);k__Fungi_unclassified(100);k__Fungi_unclassified(100);k__Fungi_unclassified(100);k__Fungi_unclassified(100);k__Fungi_unclassified(100); | - | - | - | - | - | - | - | Unassigned | - |
| Otu14318 | 1 | k__Fungi(100);p__Ascomycota(100);p__Ascomycota_unclassified(100);p__Ascomycota_unclassified(100);p__Ascomycota_unclassified(100);p__Ascomycota_unclassified(100);p__Ascomycota_unclassified(100); | - | - | - | - | - | - | - | Unassigned | - |
| Otu14319 | 1 | k__Fungi(100);p__Ascomycota(100);p__Ascomycota_unclassified(100);p__Ascomycota_unclassified(100);p__Ascomycota_unclassified(100);p__Ascomycota_unclassified(100);p__Ascomycota_unclassified(100); | - | - | - | - | - | - | - | Unassigned | - |
| Otu14320 | 1 | k__Fungi(100);p__Ascomycota(100);p__Ascomycota_unclassified(100);p__Ascomycota_unclassified(100);p__Ascomycota_unclassified(100);p__Ascomycota_unclassified(100);p__Ascomycota_unclassified(100); | - | - | - | - | - | - | - | Unassigned | - |
| Otu14324 | 1 | k__Fungi(100);p__Ascomycota(100);p__Ascomycota_unclassified(100);p__Ascomycota_unclassified(100);p__Ascomycota_unclassified(100);p__Ascomycota_unclassified(100);p__Ascomycota_unclassified(100); | - | - | - | - | - | - | - | Unassigned | - |
| Otu14325 | 1 | k__Fungi(100);p__Ascomycota(100);p__Ascomycota_unclassified(100);p__Ascomycota_unclassified(100);p__Ascomycota_unclassified(100);p__Ascomycota_unclassified(100);p__Ascomycota_unclassified(100); | - | - | - | - | - | - | - | Unassigned | - |
| Otu14326 | 1 | k__Fungi(100);k__Fungi_unclassified(100);k__Fungi_unclassified(100);k__Fungi_unclassified(100);k__Fungi_unclassified(100);k__Fungi_unclassified(100);k__Fungi_unclassified(100); | - | - | - | - | - | - | - | Unassigned | - |
| Otu14327 | 1 | k__Fungi(100);p__Ascomycota(100);p__Ascomycota_unclassified(100);p__Ascomycota_unclassified(100);p__Ascomycota_unclassified(100);p__Ascomycota_unclassified(100);p__Ascomycota_unclassified(100); | - | - | - | - | - | - | - | Unassigned | - |
| Otu14328 | 1 | k__Fungi(100);p__Ascomycota(100);p__Ascomycota_unclassified(100);p__Ascomycota_unclassified(100);p__Ascomycota_unclassified(100);p__Ascomycota_unclassified(100);p__Ascomycota_unclassified(100); | - | - | - | - | - | - | - | Unassigned | - |
| Otu14329 | 1 | k__Fungi(100);p__Ascomycota(100);p__Ascomycota_unclassified(100);p__Ascomycota_unclassified(100);p__Ascomycota_unclassified(100);p__Ascomycota_unclassified(100);p__Ascomycota_unclassified(100); | - | - | - | - | - | - | - | Unassigned | - |
| Otu14332 | 1 | k__Fungi(100);p__Ascomycota(100);p__Ascomycota_unclassified(100);p__Ascomycota_unclassified(100);p__Ascomycota_unclassified(100);p__Ascomycota_unclassified(100);p__Ascomycota_unclassified(100); | - | - | - | - | - | - | - | Unassigned | - |
| Otu14334 | 1 | k__Fungi(100);p__Ascomycota(100);p__Ascomycota_unclassified(100);p__Ascomycota_unclassified(100);p__Ascomycota_unclassified(100);p__Ascomycota_unclassified(100);p__Ascomycota_unclassified(100); | - | - | - | - | - | - | - | Unassigned | - |
| Otu14336 | 1 | k__Fungi(100);k__Fungi_unclassified(100);k__Fungi_unclassified(100);k__Fungi_unclassified(100);k__Fungi_unclassified(100);k__Fungi_unclassified(100);k__Fungi_unclassified(100); | - | - | - | - | - | - | - | Unassigned | - |
| Otu14349 | 1 | k__Fungi(100);k__Fungi_unclassified(100);k__Fungi_unclassified(100);k__Fungi_unclassified(100);k__Fungi_unclassified(100);k__Fungi_unclassified(100);k__Fungi_unclassified(100); | - | - | - | - | - | - | - | Unassigned | - |
| Otu14354 | 1 | k__Fungi(100);k__Fungi_unclassified(100);k__Fungi_unclassified(100);k__Fungi_unclassified(100);k__Fungi_unclassified(100);k__Fungi_unclassified(100);k__Fungi_unclassified(100); | - | - | - | - | - | - | - | Unassigned | - |
| Otu14357 | 1 | k__Fungi(100);p__Ascomycota(100);p__Ascomycota_unclassified(100);p__Ascomycota_unclassified(100);p__Ascomycota_unclassified(100);p__Ascomycota_unclassified(100);p__Ascomycota_unclassified(100); | - | - | - | - | - | - | - | Unassigned | - |
| Otu14361 | 1 | k__Fungi(100);p__Ascomycota(100);p__Ascomycota_unclassified(100);p__Ascomycota_unclassified(100);p__Ascomycota_unclassified(100);p__Ascomycota_unclassified(100);p__Ascomycota_unclassified(100); | - | - | - | - | - | - | - | Unassigned | - |
| Otu14367 | 1 | k__Fungi(100);p__Ascomycota(100);p__Ascomycota_unclassified(100);p__Ascomycota_unclassified(100);p__Ascomycota_unclassified(100);p__Ascomycota_unclassified(100);p__Ascomycota_unclassified(100); | - | - | - | - | - | - | - | Unassigned | - |
| Otu14374 | 1 | k__Fungi(100);k__Fungi_unclassified(100);k__Fungi_unclassified(100);k__Fungi_unclassified(100);k__Fungi_unclassified(100);k__Fungi_unclassified(100);k__Fungi_unclassified(100); | - | - | - | - | - | - | - | Unassigned | - |
| Otu14376 | 1 | k__Fungi(100);k__Fungi_unclassified(100);k__Fungi_unclassified(100);k__Fungi_unclassified(100);k__Fungi_unclassified(100);k__Fungi_unclassified(100);k__Fungi_unclassified(100); | - | - | - | - | - | - | - | Unassigned | - |
| Otu14378 | 1 | k__Fungi(100);k__Fungi_unclassified(100);k__Fungi_unclassified(100);k__Fungi_unclassified(100);k__Fungi_unclassified(100);k__Fungi_unclassified(100);k__Fungi_unclassified(100); | - | - | - | - | - | - | - | Unassigned | - |
| Otu14380 | 1 | k__Fungi(100);k__Fungi_unclassified(100);k__Fungi_unclassified(100);k__Fungi_unclassified(100);k__Fungi_unclassified(100);k__Fungi_unclassified(100);k__Fungi_unclassified(100); | - | - | - | - | - | - | - | Unassigned | - |
| Otu14381 | 1 | k__Fungi(100);p__Ascomycota(100);p__Ascomycota_unclassified(100);p__Ascomycota_unclassified(100);p__Ascomycota_unclassified(100);p__Ascomycota_unclassified(100);p__Ascomycota_unclassified(100); | - | - | - | - | - | - | - | Unassigned | - |
| Otu14383 | 1 | k__Fungi(100);k__Fungi_unclassified(100);k__Fungi_unclassified(100);k__Fungi_unclassified(100);k__Fungi_unclassified(100);k__Fungi_unclassified(100);k__Fungi_unclassified(100); | - | - | - | - | - | - | - | Unassigned | - |
| Otu14392 | 1 | k__Fungi(100);p__Ascomycota(100);p__Ascomycota_unclassified(100);p__Ascomycota_unclassified(100);p__Ascomycota_unclassified(100);p__Ascomycota_unclassified(100);p__Ascomycota_unclassified(100); | - | - | - | - | - | - | - | Unassigned | - |
| Otu14394 | 1 | k__Fungi(100);p__Ascomycota(100);p__Ascomycota_unclassified(100);p__Ascomycota_unclassified(100);p__Ascomycota_unclassified(100);p__Ascomycota_unclassified(100);p__Ascomycota_unclassified(100); | - | - | - | - | - | - | - | Unassigned | - |
| Otu14395 | 1 | k__Fungi(100);k__Fungi_unclassified(100);k__Fungi_unclassified(100);k__Fungi_unclassified(100);k__Fungi_unclassified(100);k__Fungi_unclassified(100);k__Fungi_unclassified(100); | - | - | - | - | - | - | - | Unassigned | - |
| Otu14396 | 1 | k__Fungi(100);p__Ascomycota(100);p__Ascomycota_unclassified(100);p__Ascomycota_unclassified(100);p__Ascomycota_unclassified(100);p__Ascomycota_unclassified(100);p__Ascomycota_unclassified(100); | - | - | - | - | - | - | - | Unassigned | - |
| Otu14397 | 1 | k__Fungi(100);k__Fungi_unclassified(100);k__Fungi_unclassified(100);k__Fungi_unclassified(100);k__Fungi_unclassified(100);k__Fungi_unclassified(100);k__Fungi_unclassified(100); | - | - | - | - | - | - | - | Unassigned | - |
| Otu14403 | 1 | k__Fungi(100);p__Ascomycota(100);p__Ascomycota_unclassified(100);p__Ascomycota_unclassified(100);p__Ascomycota_unclassified(100);p__Ascomycota_unclassified(100);p__Ascomycota_unclassified(100); | - | - | - | - | - | - | - | Unassigned | - |
| Otu14407 | 1 | k__Fungi(100);p__Ascomycota(100);p__Ascomycota_unclassified(100);p__Ascomycota_unclassified(100);p__Ascomycota_unclassified(100);p__Ascomycota_unclassified(100);p__Ascomycota_unclassified(100); | - | - | - | - | - | - | - | Unassigned | - |
| Otu14413 | 1 | k__Fungi(100);k__Fungi_unclassified(100);k__Fungi_unclassified(100);k__Fungi_unclassified(100);k__Fungi_unclassified(100);k__Fungi_unclassified(100);k__Fungi_unclassified(100); | - | - | - | - | - | - | - | Unassigned | - |
| Otu14419 | 1 | k__Fungi(100);p__Ascomycota(100);p__Ascomycota_unclassified(100);p__Ascomycota_unclassified(100);p__Ascomycota_unclassified(100);p__Ascomycota_unclassified(100);p__Ascomycota_unclassified(100); | - | - | - | - | - | - | - | Unassigned | - |
| Otu14428 | 1 | k__Fungi(100);k__Fungi_unclassified(100);k__Fungi_unclassified(100);k__Fungi_unclassified(100);k__Fungi_unclassified(100);k__Fungi_unclassified(100);k__Fungi_unclassified(100); | - | - | - | - | - | - | - | Unassigned | - |
| Otu14429 | 1 | k__Fungi(100);k__Fungi_unclassified(100);k__Fungi_unclassified(100);k__Fungi_unclassified(100);k__Fungi_unclassified(100);k__Fungi_unclassified(100);k__Fungi_unclassified(100); | - | - | - | - | - | - | - | Unassigned | - |
| Otu14430 | 1 | k__Fungi(100);p__Ascomycota(100);p__Ascomycota_unclassified(100);p__Ascomycota_unclassified(100);p__Ascomycota_unclassified(100);p__Ascomycota_unclassified(100);p__Ascomycota_unclassified(100); | - | - | - | - | - | - | - | Unassigned | - |
| Otu14432 | 1 | k__Fungi(100);p__Ascomycota(100);p__Ascomycota_unclassified(100);p__Ascomycota_unclassified(100);p__Ascomycota_unclassified(100);p__Ascomycota_unclassified(100);p__Ascomycota_unclassified(100); | - | - | - | - | - | - | - | Unassigned | - |
| Otu14433 | 1 | k__Fungi(100);k__Fungi_unclassified(100);k__Fungi_unclassified(100);k__Fungi_unclassified(100);k__Fungi_unclassified(100);k__Fungi_unclassified(100);k__Fungi_unclassified(100); | - | - | - | - | - | - | - | Unassigned | - |
| Otu14434 | 1 | k__Fungi(100);p__Ascomycota(100);p__Ascomycota_unclassified(100);p__Ascomycota_unclassified(100);p__Ascomycota_unclassified(100);p__Ascomycota_unclassified(100);p__Ascomycota_unclassified(100); | - | - | - | - | - | - | - | Unassigned | - |
| Otu14435 | 1 | k__Fungi(100);k__Fungi_unclassified(100);k__Fungi_unclassified(100);k__Fungi_unclassified(100);k__Fungi_unclassified(100);k__Fungi_unclassified(100);k__Fungi_unclassified(100); | - | - | - | - | - | - | - | Unassigned | - |
| Otu14436 | 1 | k__Fungi(100);p__Ascomycota(100);p__Ascomycota_unclassified(100);p__Ascomycota_unclassified(100);p__Ascomycota_unclassified(100);p__Ascomycota_unclassified(100);p__Ascomycota_unclassified(100); | - | - | - | - | - | - | - | Unassigned | - |
| Otu14437 | 1 | k__Fungi(100);p__Ascomycota(100);p__Ascomycota_unclassified(100);p__Ascomycota_unclassified(100);p__Ascomycota_unclassified(100);p__Ascomycota_unclassified(100);p__Ascomycota_unclassified(100); | - | - | - | - | - | - | - | Unassigned | - |
| Otu14439 | 1 | k__Fungi(100);k__Fungi_unclassified(100);k__Fungi_unclassified(100);k__Fungi_unclassified(100);k__Fungi_unclassified(100);k__Fungi_unclassified(100);k__Fungi_unclassified(100); | - | - | - | - | - | - | - | Unassigned | - |
| Otu14441 | 1 | k__Fungi(100);k__Fungi_unclassified(100);k__Fungi_unclassified(100);k__Fungi_unclassified(100);k__Fungi_unclassified(100);k__Fungi_unclassified(100);k__Fungi_unclassified(100); | - | - | - | - | - | - | - | Unassigned | - |
| Otu14442 | 1 | k__Fungi(100);k__Fungi_unclassified(100);k__Fungi_unclassified(100);k__Fungi_unclassified(100);k__Fungi_unclassified(100);k__Fungi_unclassified(100);k__Fungi_unclassified(100); | - | - | - | - | - | - | - | Unassigned | - |
| Otu14445 | 1 | k__Fungi(100);k__Fungi_unclassified(100);k__Fungi_unclassified(100);k__Fungi_unclassified(100);k__Fungi_unclassified(100);k__Fungi_unclassified(100);k__Fungi_unclassified(100); | - | - | - | - | - | - | - | Unassigned | - |
| Otu14449 | 1 | k__Fungi(100);k__Fungi_unclassified(100);k__Fungi_unclassified(100);k__Fungi_unclassified(100);k__Fungi_unclassified(100);k__Fungi_unclassified(100);k__Fungi_unclassified(100); | - | - | - | - | - | - | - | Unassigned | - |
| Otu14450 | 1 | k__Fungi(100);k__Fungi_unclassified(100);k__Fungi_unclassified(100);k__Fungi_unclassified(100);k__Fungi_unclassified(100);k__Fungi_unclassified(100);k__Fungi_unclassified(100); | - | - | - | - | - | - | - | Unassigned | - |
| Otu14451 | 1 | k__Fungi(100);p__Ascomycota(100);p__Ascomycota_unclassified(100);p__Ascomycota_unclassified(100);p__Ascomycota_unclassified(100);p__Ascomycota_unclassified(100);p__Ascomycota_unclassified(100); | - | - | - | - | - | - | - | Unassigned | - |
| Otu14457 | 1 | k__Fungi(100);k__Fungi_unclassified(100);k__Fungi_unclassified(100);k__Fungi_unclassified(100);k__Fungi_unclassified(100);k__Fungi_unclassified(100);k__Fungi_unclassified(100); | - | - | - | - | - | - | - | Unassigned | - |
| Otu14458 | 1 | k__Fungi(100);k__Fungi_unclassified(100);k__Fungi_unclassified(100);k__Fungi_unclassified(100);k__Fungi_unclassified(100);k__Fungi_unclassified(100);k__Fungi_unclassified(100); | - | - | - | - | - | - | - | Unassigned | - |
| Otu14459 | 1 | k__Fungi(100);p__Ascomycota(100);p__Ascomycota_unclassified(100);p__Ascomycota_unclassified(100);p__Ascomycota_unclassified(100);p__Ascomycota_unclassified(100);p__Ascomycota_unclassified(100); | - | - | - | - | - | - | - | Unassigned | - |
| Otu14461 | 1 | k__Fungi(100);p__Ascomycota(100);p__Ascomycota_unclassified(100);p__Ascomycota_unclassified(100);p__Ascomycota_unclassified(100);p__Ascomycota_unclassified(100);p__Ascomycota_unclassified(100); | - | - | - | - | - | - | - | Unassigned | - |
| Otu14462 | 1 | k__Fungi(100);k__Fungi_unclassified(100);k__Fungi_unclassified(100);k__Fungi_unclassified(100);k__Fungi_unclassified(100);k__Fungi_unclassified(100);k__Fungi_unclassified(100); | - | - | - | - | - | - | - | Unassigned | - |
| Otu14463 | 1 | k__Fungi(100);k__Fungi_unclassified(100);k__Fungi_unclassified(100);k__Fungi_unclassified(100);k__Fungi_unclassified(100);k__Fungi_unclassified(100);k__Fungi_unclassified(100); | - | - | - | - | - | - | - | Unassigned | - |
| Otu14466 | 1 | k__Fungi(100);p__Ascomycota(100);p__Ascomycota_unclassified(100);p__Ascomycota_unclassified(100);p__Ascomycota_unclassified(100);p__Ascomycota_unclassified(100);p__Ascomycota_unclassified(100); | - | - | - | - | - | - | - | Unassigned | - |
| Otu14468 | 1 | k__Fungi(100);p__Ascomycota(100);p__Ascomycota_unclassified(100);p__Ascomycota_unclassified(100);p__Ascomycota_unclassified(100);p__Ascomycota_unclassified(100);p__Ascomycota_unclassified(100); | - | - | - | - | - | - | - | Unassigned | - |
| Otu14469 | 1 | k__Fungi(100);k__Fungi_unclassified(100);k__Fungi_unclassified(100);k__Fungi_unclassified(100);k__Fungi_unclassified(100);k__Fungi_unclassified(100);k__Fungi_unclassified(100); | - | - | - | - | - | - | - | Unassigned | - |
| Otu14470 | 1 | k__Fungi(100);k__Fungi_unclassified(100);k__Fungi_unclassified(100);k__Fungi_unclassified(100);k__Fungi_unclassified(100);k__Fungi_unclassified(100);k__Fungi_unclassified(100); | - | - | - | - | - | - | - | Unassigned | - |
| Otu14477 | 1 | k__Fungi(100);k__Fungi_unclassified(100);k__Fungi_unclassified(100);k__Fungi_unclassified(100);k__Fungi_unclassified(100);k__Fungi_unclassified(100);k__Fungi_unclassified(100); | - | - | - | - | - | - | - | Unassigned | - |
| Otu14478 | 1 | k__Fungi(100);k__Fungi_unclassified(100);k__Fungi_unclassified(100);k__Fungi_unclassified(100);k__Fungi_unclassified(100);k__Fungi_unclassified(100);k__Fungi_unclassified(100); | - | - | - | - | - | - | - | Unassigned | - |
| Otu14479 | 1 | k__Fungi(100);k__Fungi_unclassified(100);k__Fungi_unclassified(100);k__Fungi_unclassified(100);k__Fungi_unclassified(100);k__Fungi_unclassified(100);k__Fungi_unclassified(100); | - | - | - | - | - | - | - | Unassigned | - |
| Otu14480 | 1 | k__Fungi(100);p__Ascomycota(100);p__Ascomycota_unclassified(100);p__Ascomycota_unclassified(100);p__Ascomycota_unclassified(100);p__Ascomycota_unclassified(100);p__Ascomycota_unclassified(100); | - | - | - | - | - | - | - | Unassigned | - |
| Otu14481 | 1 | k__Fungi(100);p__Ascomycota(100);p__Ascomycota_unclassified(100);p__Ascomycota_unclassified(100);p__Ascomycota_unclassified(100);p__Ascomycota_unclassified(100);p__Ascomycota_unclassified(100); | - | - | - | - | - | - | - | Unassigned | - |
| Otu14490 | 1 | k__Fungi(100);k__Fungi_unclassified(100);k__Fungi_unclassified(100);k__Fungi_unclassified(100);k__Fungi_unclassified(100);k__Fungi_unclassified(100);k__Fungi_unclassified(100); | - | - | - | - | - | - | - | Unassigned | - |
| Otu14494 | 1 | k__Fungi(100);p__Ascomycota(100);p__Ascomycota_unclassified(100);p__Ascomycota_unclassified(100);p__Ascomycota_unclassified(100);p__Ascomycota_unclassified(100);p__Ascomycota_unclassified(100); | - | - | - | - | - | - | - | Unassigned | - |
| Otu14498 | 1 | k__Fungi(100);k__Fungi_unclassified(100);k__Fungi_unclassified(100);k__Fungi_unclassified(100);k__Fungi_unclassified(100);k__Fungi_unclassified(100);k__Fungi_unclassified(100); | - | - | - | - | - | - | - | Unassigned | - |
| Otu14510 | 1 | k__Fungi(100);k__Fungi_unclassified(100);k__Fungi_unclassified(100);k__Fungi_unclassified(100);k__Fungi_unclassified(100);k__Fungi_unclassified(100);k__Fungi_unclassified(100); | - | - | - | - | - | - | - | Unassigned | - |
| Otu14523 | 1 | k__Fungi(100);p__Ascomycota(100);p__Ascomycota_unclassified(100);p__Ascomycota_unclassified(100);p__Ascomycota_unclassified(100);p__Ascomycota_unclassified(100);p__Ascomycota_unclassified(100); | - | - | - | - | - | - | - | Unassigned | - |
| Otu14524 | 1 | k__Fungi(100);k__Fungi_unclassified(100);k__Fungi_unclassified(100);k__Fungi_unclassified(100);k__Fungi_unclassified(100);k__Fungi_unclassified(100);k__Fungi_unclassified(100); | - | - | - | - | - | - | - | Unassigned | - |
| Otu14527 | 1 | k__Fungi(100);p__Ascomycota(100);p__Ascomycota_unclassified(100);p__Ascomycota_unclassified(100);p__Ascomycota_unclassified(100);p__Ascomycota_unclassified(100);p__Ascomycota_unclassified(100); | - | - | - | - | - | - | - | Unassigned | - |
| Otu14529 | 1 | k__Fungi(100);k__Fungi_unclassified(100);k__Fungi_unclassified(100);k__Fungi_unclassified(100);k__Fungi_unclassified(100);k__Fungi_unclassified(100);k__Fungi_unclassified(100); | - | - | - | - | - | - | - | Unassigned | - |
| Otu14535 | 1 | k__Fungi(100);k__Fungi_unclassified(100);k__Fungi_unclassified(100);k__Fungi_unclassified(100);k__Fungi_unclassified(100);k__Fungi_unclassified(100);k__Fungi_unclassified(100); | - | - | - | - | - | - | - | Unassigned | - |
| Otu14539 | 1 | k__Fungi(100);k__Fungi_unclassified(100);k__Fungi_unclassified(100);k__Fungi_unclassified(100);k__Fungi_unclassified(100);k__Fungi_unclassified(100);k__Fungi_unclassified(100); | - | - | - | - | - | - | - | Unassigned | - |
| Otu14542 | 1 | k__Fungi(100);p__Ascomycota(100);p__Ascomycota_unclassified(100);p__Ascomycota_unclassified(100);p__Ascomycota_unclassified(100);p__Ascomycota_unclassified(100);p__Ascomycota_unclassified(100); | - | - | - | - | - | - | - | Unassigned | - |
| Otu14544 | 1 | k__Fungi(100);p__Ascomycota(100);p__Ascomycota_unclassified(100);p__Ascomycota_unclassified(100);p__Ascomycota_unclassified(100);p__Ascomycota_unclassified(100);p__Ascomycota_unclassified(100); | - | - | - | - | - | - | - | Unassigned | - |
| Otu14545 | 1 | k__Fungi(100);p__Ascomycota(100);p__Ascomycota_unclassified(100);p__Ascomycota_unclassified(100);p__Ascomycota_unclassified(100);p__Ascomycota_unclassified(100);p__Ascomycota_unclassified(100); | - | - | - | - | - | - | - | Unassigned | - |
| Otu14547 | 1 | k__Fungi(100);k__Fungi_unclassified(100);k__Fungi_unclassified(100);k__Fungi_unclassified(100);k__Fungi_unclassified(100);k__Fungi_unclassified(100);k__Fungi_unclassified(100); | - | - | - | - | - | - | - | Unassigned | - |
| Otu14548 | 1 | k__Fungi(100);k__Fungi_unclassified(100);k__Fungi_unclassified(100);k__Fungi_unclassified(100);k__Fungi_unclassified(100);k__Fungi_unclassified(100);k__Fungi_unclassified(100); | - | - | - | - | - | - | - | Unassigned | - |
| Otu14549 | 1 | k__Fungi(100);k__Fungi_unclassified(100);k__Fungi_unclassified(100);k__Fungi_unclassified(100);k__Fungi_unclassified(100);k__Fungi_unclassified(100);k__Fungi_unclassified(100); | - | - | - | - | - | - | - | Unassigned | - |
| Otu14551 | 1 | k__Fungi(100);p__Ascomycota(100);p__Ascomycota_unclassified(100);p__Ascomycota_unclassified(100);p__Ascomycota_unclassified(100);p__Ascomycota_unclassified(100);p__Ascomycota_unclassified(100); | - | - | - | - | - | - | - | Unassigned | - |
| Otu14552 | 1 | k__Fungi(100);p__Ascomycota(100);p__Ascomycota_unclassified(100);p__Ascomycota_unclassified(100);p__Ascomycota_unclassified(100);p__Ascomycota_unclassified(100);p__Ascomycota_unclassified(100); | - | - | - | - | - | - | - | Unassigned | - |
| Otu14553 | 1 | k__Fungi(100);p__Ascomycota(100);p__Ascomycota_unclassified(100);p__Ascomycota_unclassified(100);p__Ascomycota_unclassified(100);p__Ascomycota_unclassified(100);p__Ascomycota_unclassified(100); | - | - | - | - | - | - | - | Unassigned | - |
| Otu14554 | 1 | k__Fungi(100);k__Fungi_unclassified(100);k__Fungi_unclassified(100);k__Fungi_unclassified(100);k__Fungi_unclassified(100);k__Fungi_unclassified(100);k__Fungi_unclassified(100); | - | - | - | - | - | - | - | Unassigned | - |
| Otu14556 | 1 | k__Fungi(100);k__Fungi_unclassified(100);k__Fungi_unclassified(100);k__Fungi_unclassified(100);k__Fungi_unclassified(100);k__Fungi_unclassified(100);k__Fungi_unclassified(100); | - | - | - | - | - | - | - | Unassigned | - |
| Otu14559 | 1 | k__Fungi(100);p__Ascomycota(100);p__Ascomycota_unclassified(100);p__Ascomycota_unclassified(100);p__Ascomycota_unclassified(100);p__Ascomycota_unclassified(100);p__Ascomycota_unclassified(100); | - | - | - | - | - | - | - | Unassigned | - |
| Otu14571 | 1 | k__Fungi(100);p__Ascomycota(100);p__Ascomycota_unclassified(100);p__Ascomycota_unclassified(100);p__Ascomycota_unclassified(100);p__Ascomycota_unclassified(100);p__Ascomycota_unclassified(100); | - | - | - | - | - | - | - | Unassigned | - |
| Otu14579 | 1 | k__Fungi(100);k__Fungi_unclassified(100);k__Fungi_unclassified(100);k__Fungi_unclassified(100);k__Fungi_unclassified(100);k__Fungi_unclassified(100);k__Fungi_unclassified(100); | - | - | - | - | - | - | - | Unassigned | - |
| Otu14580 | 1 | k__Fungi(100);p__Ascomycota(100);p__Ascomycota_unclassified(100);p__Ascomycota_unclassified(100);p__Ascomycota_unclassified(100);p__Ascomycota_unclassified(100);p__Ascomycota_unclassified(100); | - | - | - | - | - | - | - | Unassigned | - |
| Otu14581 | 1 | k__Fungi(100);k__Fungi_unclassified(100);k__Fungi_unclassified(100);k__Fungi_unclassified(100);k__Fungi_unclassified(100);k__Fungi_unclassified(100);k__Fungi_unclassified(100); | - | - | - | - | - | - | - | Unassigned | - |
| Otu14584 | 1 | k__Fungi(100);k__Fungi_unclassified(100);k__Fungi_unclassified(100);k__Fungi_unclassified(100);k__Fungi_unclassified(100);k__Fungi_unclassified(100);k__Fungi_unclassified(100); | - | - | - | - | - | - | - | Unassigned | - |
| Otu14589 | 1 | k__Fungi(100);k__Fungi_unclassified(100);k__Fungi_unclassified(100);k__Fungi_unclassified(100);k__Fungi_unclassified(100);k__Fungi_unclassified(100);k__Fungi_unclassified(100); | - | - | - | - | - | - | - | Unassigned | - |
| Otu14594 | 1 | k__Fungi(100);k__Fungi_unclassified(100);k__Fungi_unclassified(100);k__Fungi_unclassified(100);k__Fungi_unclassified(100);k__Fungi_unclassified(100);k__Fungi_unclassified(100); | - | - | - | - | - | - | - | Unassigned | - |
| Otu14596 | 1 | k__Fungi(100);k__Fungi_unclassified(100);k__Fungi_unclassified(100);k__Fungi_unclassified(100);k__Fungi_unclassified(100);k__Fungi_unclassified(100);k__Fungi_unclassified(100); | - | - | - | - | - | - | - | Unassigned | - |
| Otu14601 | 1 | k__Fungi(100);k__Fungi_unclassified(100);k__Fungi_unclassified(100);k__Fungi_unclassified(100);k__Fungi_unclassified(100);k__Fungi_unclassified(100);k__Fungi_unclassified(100); | - | - | - | - | - | - | - | Unassigned | - |
| Otu14602 | 1 | k__Fungi(100);k__Fungi_unclassified(100);k__Fungi_unclassified(100);k__Fungi_unclassified(100);k__Fungi_unclassified(100);k__Fungi_unclassified(100);k__Fungi_unclassified(100); | - | - | - | - | - | - | - | Unassigned | - |
| Otu14608 | 1 | k__Fungi(100);k__Fungi_unclassified(100);k__Fungi_unclassified(100);k__Fungi_unclassified(100);k__Fungi_unclassified(100);k__Fungi_unclassified(100);k__Fungi_unclassified(100); | - | - | - | - | - | - | - | Unassigned | - |
| Otu14609 | 1 | k__Fungi(100);k__Fungi_unclassified(100);k__Fungi_unclassified(100);k__Fungi_unclassified(100);k__Fungi_unclassified(100);k__Fungi_unclassified(100);k__Fungi_unclassified(100); | - | - | - | - | - | - | - | Unassigned | - |
| Otu14617 | 1 | k__Fungi(100);k__Fungi_unclassified(100);k__Fungi_unclassified(100);k__Fungi_unclassified(100);k__Fungi_unclassified(100);k__Fungi_unclassified(100);k__Fungi_unclassified(100); | - | - | - | - | - | - | - | Unassigned | - |
| Otu14620 | 1 | k__Fungi(100);k__Fungi_unclassified(100);k__Fungi_unclassified(100);k__Fungi_unclassified(100);k__Fungi_unclassified(100);k__Fungi_unclassified(100);k__Fungi_unclassified(100); | - | - | - | - | - | - | - | Unassigned | - |
| Otu14633 | 1 | k__Fungi(100);k__Fungi_unclassified(100);k__Fungi_unclassified(100);k__Fungi_unclassified(100);k__Fungi_unclassified(100);k__Fungi_unclassified(100);k__Fungi_unclassified(100); | - | - | - | - | - | - | - | Unassigned | - |
| Otu14634 | 1 | k__Fungi(100);k__Fungi_unclassified(100);k__Fungi_unclassified(100);k__Fungi_unclassified(100);k__Fungi_unclassified(100);k__Fungi_unclassified(100);k__Fungi_unclassified(100); | - | - | - | - | - | - | - | Unassigned | - |
| Otu14635 | 1 | k__Fungi(100);k__Fungi_unclassified(100);k__Fungi_unclassified(100);k__Fungi_unclassified(100);k__Fungi_unclassified(100);k__Fungi_unclassified(100);k__Fungi_unclassified(100); | - | - | - | - | - | - | - | Unassigned | - |
| Otu14648 | 1 | k__Fungi(100);p__Ascomycota(100);p__Ascomycota_unclassified(100);p__Ascomycota_unclassified(100);p__Ascomycota_unclassified(100);p__Ascomycota_unclassified(100);p__Ascomycota_unclassified(100); | - | - | - | - | - | - | - | Unassigned | - |
| Otu14661 | 1 | k__Fungi(100);k__Fungi_unclassified(100);k__Fungi_unclassified(100);k__Fungi_unclassified(100);k__Fungi_unclassified(100);k__Fungi_unclassified(100);k__Fungi_unclassified(100); | - | - | - | - | - | - | - | Unassigned | - |
| Otu14662 | 1 | k__Fungi(100);k__Fungi_unclassified(100);k__Fungi_unclassified(100);k__Fungi_unclassified(100);k__Fungi_unclassified(100);k__Fungi_unclassified(100);k__Fungi_unclassified(100); | - | - | - | - | - | - | - | Unassigned | - |
| Otu14667 | 1 | k__Fungi(100);k__Fungi_unclassified(100);k__Fungi_unclassified(100);k__Fungi_unclassified(100);k__Fungi_unclassified(100);k__Fungi_unclassified(100);k__Fungi_unclassified(100); | - | - | - | - | - | - | - | Unassigned | - |
| Otu14680 | 1 | k__Fungi(100);p__Ascomycota(100);p__Ascomycota_unclassified(100);p__Ascomycota_unclassified(100);p__Ascomycota_unclassified(100);p__Ascomycota_unclassified(100);p__Ascomycota_unclassified(100); | - | - | - | - | - | - | - | Unassigned | - |
| Otu14681 | 1 | k__Fungi(100);k__Fungi_unclassified(100);k__Fungi_unclassified(100);k__Fungi_unclassified(100);k__Fungi_unclassified(100);k__Fungi_unclassified(100);k__Fungi_unclassified(100); | - | - | - | - | - | - | - | Unassigned | - |
| Otu14683 | 1 | k__Fungi(100);k__Fungi_unclassified(100);k__Fungi_unclassified(100);k__Fungi_unclassified(100);k__Fungi_unclassified(100);k__Fungi_unclassified(100);k__Fungi_unclassified(100); | - | - | - | - | - | - | - | Unassigned | - |
| Otu14685 | 1 | k__Fungi(100);k__Fungi_unclassified(100);k__Fungi_unclassified(100);k__Fungi_unclassified(100);k__Fungi_unclassified(100);k__Fungi_unclassified(100);k__Fungi_unclassified(100); | - | - | - | - | - | - | - | Unassigned | - |
| Otu14690 | 1 | k__Fungi(100);k__Fungi_unclassified(100);k__Fungi_unclassified(100);k__Fungi_unclassified(100);k__Fungi_unclassified(100);k__Fungi_unclassified(100);k__Fungi_unclassified(100); | - | - | - | - | - | - | - | Unassigned | - |
| Otu14692 | 1 | k__Fungi(100);k__Fungi_unclassified(100);k__Fungi_unclassified(100);k__Fungi_unclassified(100);k__Fungi_unclassified(100);k__Fungi_unclassified(100);k__Fungi_unclassified(100); | - | - | - | - | - | - | - | Unassigned | - |
| Otu14696 | 1 | k__Fungi(100);k__Fungi_unclassified(100);k__Fungi_unclassified(100);k__Fungi_unclassified(100);k__Fungi_unclassified(100);k__Fungi_unclassified(100);k__Fungi_unclassified(100); | - | - | - | - | - | - | - | Unassigned | - |
| Otu14697 | 1 | k__Fungi(100);p__Ascomycota(100);p__Ascomycota_unclassified(100);p__Ascomycota_unclassified(100);p__Ascomycota_unclassified(100);p__Ascomycota_unclassified(100);p__Ascomycota_unclassified(100); | - | - | - | - | - | - | - | Unassigned | - |
| Otu14699 | 1 | k__Fungi(100);k__Fungi_unclassified(100);k__Fungi_unclassified(100);k__Fungi_unclassified(100);k__Fungi_unclassified(100);k__Fungi_unclassified(100);k__Fungi_unclassified(100); | - | - | - | - | - | - | - | Unassigned | - |
| Otu14700 | 1 | k__Fungi(100);p__Ascomycota(100);p__Ascomycota_unclassified(100);p__Ascomycota_unclassified(100);p__Ascomycota_unclassified(100);p__Ascomycota_unclassified(100);p__Ascomycota_unclassified(100); | - | - | - | - | - | - | - | Unassigned | - |
| Otu14703 | 1 | k__Fungi(100);k__Fungi_unclassified(100);k__Fungi_unclassified(100);k__Fungi_unclassified(100);k__Fungi_unclassified(100);k__Fungi_unclassified(100);k__Fungi_unclassified(100); | - | - | - | - | - | - | - | Unassigned | - |
| Otu14705 | 1 | k__Fungi(100);k__Fungi_unclassified(100);k__Fungi_unclassified(100);k__Fungi_unclassified(100);k__Fungi_unclassified(100);k__Fungi_unclassified(100);k__Fungi_unclassified(100); | - | - | - | - | - | - | - | Unassigned | - |
| Otu14706 | 1 | k__Fungi(100);k__Fungi_unclassified(100);k__Fungi_unclassified(100);k__Fungi_unclassified(100);k__Fungi_unclassified(100);k__Fungi_unclassified(100);k__Fungi_unclassified(100); | - | - | - | - | - | - | - | Unassigned | - |
| Otu14709 | 1 | k__Fungi(100);p__Ascomycota(100);p__Ascomycota_unclassified(100);p__Ascomycota_unclassified(100);p__Ascomycota_unclassified(100);p__Ascomycota_unclassified(100);p__Ascomycota_unclassified(100); | - | - | - | - | - | - | - | Unassigned | - |
| Otu14710 | 1 | k__Fungi(100);p__Ascomycota(100);p__Ascomycota_unclassified(100);p__Ascomycota_unclassified(100);p__Ascomycota_unclassified(100);p__Ascomycota_unclassified(100);p__Ascomycota_unclassified(100); | - | - | - | - | - | - | - | Unassigned | - |
| Otu14711 | 1 | k__Fungi(100);p__Ascomycota(100);p__Ascomycota_unclassified(100);p__Ascomycota_unclassified(100);p__Ascomycota_unclassified(100);p__Ascomycota_unclassified(100);p__Ascomycota_unclassified(100); | - | - | - | - | - | - | - | Unassigned | - |
| Otu14712 | 1 | k__Fungi(100);p__Ascomycota(100);p__Ascomycota_unclassified(100);p__Ascomycota_unclassified(100);p__Ascomycota_unclassified(100);p__Ascomycota_unclassified(100);p__Ascomycota_unclassified(100); | - | - | - | - | - | - | - | Unassigned | - |
| Otu14718 | 1 | k__Fungi(100);p__Ascomycota(100);p__Ascomycota_unclassified(100);p__Ascomycota_unclassified(100);p__Ascomycota_unclassified(100);p__Ascomycota_unclassified(100);p__Ascomycota_unclassified(100); | - | - | - | - | - | - | - | Unassigned | - |
| Otu14724 | 1 | k__Fungi(100);k__Fungi_unclassified(100);k__Fungi_unclassified(100);k__Fungi_unclassified(100);k__Fungi_unclassified(100);k__Fungi_unclassified(100);k__Fungi_unclassified(100); | - | - | - | - | - | - | - | Unassigned | - |
| Otu14729 | 1 | k__Fungi(100);k__Fungi_unclassified(100);k__Fungi_unclassified(100);k__Fungi_unclassified(100);k__Fungi_unclassified(100);k__Fungi_unclassified(100);k__Fungi_unclassified(100); | - | - | - | - | - | - | - | Unassigned | - |
| Otu14730 | 1 | k__Fungi(100);k__Fungi_unclassified(100);k__Fungi_unclassified(100);k__Fungi_unclassified(100);k__Fungi_unclassified(100);k__Fungi_unclassified(100);k__Fungi_unclassified(100); | - | - | - | - | - | - | - | Unassigned | - |
| Otu14731 | 1 | k__Fungi(100);p__Ascomycota(100);p__Ascomycota_unclassified(100);p__Ascomycota_unclassified(100);p__Ascomycota_unclassified(100);p__Ascomycota_unclassified(100);p__Ascomycota_unclassified(100); | - | - | - | - | - | - | - | Unassigned | - |
| Otu14732 | 1 | k__Fungi(100);p__Ascomycota(100);p__Ascomycota_unclassified(100);p__Ascomycota_unclassified(100);p__Ascomycota_unclassified(100);p__Ascomycota_unclassified(100);p__Ascomycota_unclassified(100); | - | - | - | - | - | - | - | Unassigned | - |
| Otu14733 | 1 | k__Fungi(100);k__Fungi_unclassified(100);k__Fungi_unclassified(100);k__Fungi_unclassified(100);k__Fungi_unclassified(100);k__Fungi_unclassified(100);k__Fungi_unclassified(100); | - | - | - | - | - | - | - | Unassigned | - |
| Otu14736 | 1 | k__Fungi(100);k__Fungi_unclassified(100);k__Fungi_unclassified(100);k__Fungi_unclassified(100);k__Fungi_unclassified(100);k__Fungi_unclassified(100);k__Fungi_unclassified(100); | - | - | - | - | - | - | - | Unassigned | - |
| Otu14738 | 1 | k__Fungi(100);p__Ascomycota(100);p__Ascomycota_unclassified(100);p__Ascomycota_unclassified(100);p__Ascomycota_unclassified(100);p__Ascomycota_unclassified(100);p__Ascomycota_unclassified(100); | - | - | - | - | - | - | - | Unassigned | - |
| Otu14739 | 1 | k__Fungi(100);k__Fungi_unclassified(100);k__Fungi_unclassified(100);k__Fungi_unclassified(100);k__Fungi_unclassified(100);k__Fungi_unclassified(100);k__Fungi_unclassified(100); | - | - | - | - | - | - | - | Unassigned | - |
| Otu14742 | 1 | k__Fungi(100);p__Ascomycota(100);p__Ascomycota_unclassified(100);p__Ascomycota_unclassified(100);p__Ascomycota_unclassified(100);p__Ascomycota_unclassified(100);p__Ascomycota_unclassified(100); | - | - | - | - | - | - | - | Unassigned | - |
| Otu14744 | 1 | k__Fungi(100);k__Fungi_unclassified(100);k__Fungi_unclassified(100);k__Fungi_unclassified(100);k__Fungi_unclassified(100);k__Fungi_unclassified(100);k__Fungi_unclassified(100); | - | - | - | - | - | - | - | Unassigned | - |
| Otu14749 | 1 | k__Fungi(100);k__Fungi_unclassified(100);k__Fungi_unclassified(100);k__Fungi_unclassified(100);k__Fungi_unclassified(100);k__Fungi_unclassified(100);k__Fungi_unclassified(100); | - | - | - | - | - | - | - | Unassigned | - |
| Otu14750 | 1 | k__Fungi(100);p__Ascomycota(100);p__Ascomycota_unclassified(100);p__Ascomycota_unclassified(100);p__Ascomycota_unclassified(100);p__Ascomycota_unclassified(100);p__Ascomycota_unclassified(100); | - | - | - | - | - | - | - | Unassigned | - |
| Otu14751 | 1 | k__Fungi(100);k__Fungi_unclassified(100);k__Fungi_unclassified(100);k__Fungi_unclassified(100);k__Fungi_unclassified(100);k__Fungi_unclassified(100);k__Fungi_unclassified(100); | - | - | - | - | - | - | - | Unassigned | - |
| Otu14754 | 1 | k__Fungi(100);k__Fungi_unclassified(100);k__Fungi_unclassified(100);k__Fungi_unclassified(100);k__Fungi_unclassified(100);k__Fungi_unclassified(100);k__Fungi_unclassified(100); | - | - | - | - | - | - | - | Unassigned | - |
| Otu14757 | 1 | k__Fungi(100);k__Fungi_unclassified(100);k__Fungi_unclassified(100);k__Fungi_unclassified(100);k__Fungi_unclassified(100);k__Fungi_unclassified(100);k__Fungi_unclassified(100); | - | - | - | - | - | - | - | Unassigned | - |
| Otu14758 | 1 | k__Fungi(100);k__Fungi_unclassified(100);k__Fungi_unclassified(100);k__Fungi_unclassified(100);k__Fungi_unclassified(100);k__Fungi_unclassified(100);k__Fungi_unclassified(100); | - | - | - | - | - | - | - | Unassigned | - |
| Otu14760 | 1 | k__Fungi(100);k__Fungi_unclassified(100);k__Fungi_unclassified(100);k__Fungi_unclassified(100);k__Fungi_unclassified(100);k__Fungi_unclassified(100);k__Fungi_unclassified(100); | - | - | - | - | - | - | - | Unassigned | - |
| Otu14761 | 1 | k__Fungi(100);k__Fungi_unclassified(100);k__Fungi_unclassified(100);k__Fungi_unclassified(100);k__Fungi_unclassified(100);k__Fungi_unclassified(100);k__Fungi_unclassified(100); | - | - | - | - | - | - | - | Unassigned | - |
| Otu14762 | 1 | k__Fungi(100);p__Ascomycota(100);p__Ascomycota_unclassified(100);p__Ascomycota_unclassified(100);p__Ascomycota_unclassified(100);p__Ascomycota_unclassified(100);p__Ascomycota_unclassified(100); | - | - | - | - | - | - | - | Unassigned | - |
| Otu14763 | 1 | k__Fungi(100);k__Fungi_unclassified(100);k__Fungi_unclassified(100);k__Fungi_unclassified(100);k__Fungi_unclassified(100);k__Fungi_unclassified(100);k__Fungi_unclassified(100); | - | - | - | - | - | - | - | Unassigned | - |
| Otu14766 | 1 | k__Fungi(100);k__Fungi_unclassified(100);k__Fungi_unclassified(100);k__Fungi_unclassified(100);k__Fungi_unclassified(100);k__Fungi_unclassified(100);k__Fungi_unclassified(100); | - | - | - | - | - | - | - | Unassigned | - |
| Otu14767 | 1 | k__Fungi(100);p__Ascomycota(100);p__Ascomycota_unclassified(100);p__Ascomycota_unclassified(100);p__Ascomycota_unclassified(100);p__Ascomycota_unclassified(100);p__Ascomycota_unclassified(100); | - | - | - | - | - | - | - | Unassigned | - |
| Otu14771 | 1 | k__Fungi(100);p__Ascomycota(100);p__Ascomycota_unclassified(100);p__Ascomycota_unclassified(100);p__Ascomycota_unclassified(100);p__Ascomycota_unclassified(100);p__Ascomycota_unclassified(100); | - | - | - | - | - | - | - | Unassigned | - |
| Otu14772 | 1 | k__Fungi(100);k__Fungi_unclassified(100);k__Fungi_unclassified(100);k__Fungi_unclassified(100);k__Fungi_unclassified(100);k__Fungi_unclassified(100);k__Fungi_unclassified(100); | - | - | - | - | - | - | - | Unassigned | - |
| Otu14778 | 1 | k__Fungi(100);p__Ascomycota(100);p__Ascomycota_unclassified(100);p__Ascomycota_unclassified(100);p__Ascomycota_unclassified(100);p__Ascomycota_unclassified(100);p__Ascomycota_unclassified(100); | - | - | - | - | - | - | - | Unassigned | - |
| Otu14779 | 1 | k__Fungi(100);p__Ascomycota(100);p__Ascomycota_unclassified(100);p__Ascomycota_unclassified(100);p__Ascomycota_unclassified(100);p__Ascomycota_unclassified(100);p__Ascomycota_unclassified(100); | - | - | - | - | - | - | - | Unassigned | - |
| Otu14782 | 1 | k__Fungi(100);p__Ascomycota(100);p__Ascomycota_unclassified(100);p__Ascomycota_unclassified(100);p__Ascomycota_unclassified(100);p__Ascomycota_unclassified(100);p__Ascomycota_unclassified(100); | - | - | - | - | - | - | - | Unassigned | - |
| Otu14785 | 1 | k__Fungi(100);p__Ascomycota(100);p__Ascomycota_unclassified(100);p__Ascomycota_unclassified(100);p__Ascomycota_unclassified(100);p__Ascomycota_unclassified(100);p__Ascomycota_unclassified(100); | - | - | - | - | - | - | - | Unassigned | - |
| Otu14791 | 1 | k__Fungi(100);k__Fungi_unclassified(100);k__Fungi_unclassified(100);k__Fungi_unclassified(100);k__Fungi_unclassified(100);k__Fungi_unclassified(100);k__Fungi_unclassified(100); | - | - | - | - | - | - | - | Unassigned | - |
| Otu14792 | 1 | k__Fungi(100);k__Fungi_unclassified(100);k__Fungi_unclassified(100);k__Fungi_unclassified(100);k__Fungi_unclassified(100);k__Fungi_unclassified(100);k__Fungi_unclassified(100); | - | - | - | - | - | - | - | Unassigned | - |
| Otu14793 | 1 | k__Fungi(100);p__Ascomycota(100);p__Ascomycota_unclassified(100);p__Ascomycota_unclassified(100);p__Ascomycota_unclassified(100);p__Ascomycota_unclassified(100);p__Ascomycota_unclassified(100); | - | - | - | - | - | - | - | Unassigned | - |
| Otu14796 | 1 | k__Fungi(100);k__Fungi_unclassified(100);k__Fungi_unclassified(100);k__Fungi_unclassified(100);k__Fungi_unclassified(100);k__Fungi_unclassified(100);k__Fungi_unclassified(100); | - | - | - | - | - | - | - | Unassigned | - |
| Otu14797 | 1 | k__Fungi(100);k__Fungi_unclassified(100);k__Fungi_unclassified(100);k__Fungi_unclassified(100);k__Fungi_unclassified(100);k__Fungi_unclassified(100);k__Fungi_unclassified(100); | - | - | - | - | - | - | - | Unassigned | - |
| Otu14798 | 1 | k__Fungi(100);k__Fungi_unclassified(100);k__Fungi_unclassified(100);k__Fungi_unclassified(100);k__Fungi_unclassified(100);k__Fungi_unclassified(100);k__Fungi_unclassified(100); | - | - | - | - | - | - | - | Unassigned | - |
| Otu14804 | 1 | k__Fungi(100);k__Fungi_unclassified(100);k__Fungi_unclassified(100);k__Fungi_unclassified(100);k__Fungi_unclassified(100);k__Fungi_unclassified(100);k__Fungi_unclassified(100); | - | - | - | - | - | - | - | Unassigned | - |

| | | | | | | | | | | | |
|---|---|---|---|---|---|---|---|---|---|---|---|
| Otu14805 | 1 | k__Fungi(100);p__Ascomycota(100);p__Ascomycota_unclassified(100);p__Ascomycota_unclassified(100);p__Ascomycota_unclassified(100);p__Ascomycota_unclassified(100);p__Ascomycota_unclassified(100); | - | - | - | - | - | - | - | Unassigned | - |
| Otu14806 | 1 | k__Fungi(100);k__Fungi_unclassified(100);k__Fungi_unclassified(100);k__Fungi_unclassified(100);k__Fungi_unclassified(100);k__Fungi_unclassified(100);k__Fungi_unclassified(100); | - | - | - | - | - | - | - | Unassigned | - |
| Otu14808 | 1 | k__Fungi(100);k__Fungi_unclassified(100);k__Fungi_unclassified(100);k__Fungi_unclassified(100);k__Fungi_unclassified(100);k__Fungi_unclassified(100);k__Fungi_unclassified(100); | - | - | - | - | - | - | - | Unassigned | - |
| Otu14809 | 1 | k__Fungi(100);k__Fungi_unclassified(100);k__Fungi_unclassified(100);k__Fungi_unclassified(100);k__Fungi_unclassified(100);k__Fungi_unclassified(100);k__Fungi_unclassified(100); | - | - | - | - | - | - | - | Unassigned | - |
| Otu14810 | 1 | k__Fungi(100);k__Fungi_unclassified(100);k__Fungi_unclassified(100);k__Fungi_unclassified(100);k__Fungi_unclassified(100);k__Fungi_unclassified(100);k__Fungi_unclassified(100); | - | - | - | - | - | - | - | Unassigned | - |
| Otu14811 | 1 | k__Fungi(100);p__Ascomycota(100);p__Ascomycota_unclassified(100);p__Ascomycota_unclassified(100);p__Ascomycota_unclassified(100);p__Ascomycota_unclassified(100);p__Ascomycota_unclassified(100); | - | - | - | - | - | - | - | Unassigned | - |
| Otu14816 | 1 | k__Fungi(100);p__Ascomycota(100);p__Ascomycota_unclassified(100);p__Ascomycota_unclassified(100);p__Ascomycota_unclassified(100);p__Ascomycota_unclassified(100);p__Ascomycota_unclassified(100); | - | - | - | - | - | - | - | Unassigned | - |
| Otu14817 | 1 | k__Fungi(100);p__Ascomycota(100);p__Ascomycota_unclassified(100);p__Ascomycota_unclassified(100);p__Ascomycota_unclassified(100);p__Ascomycota_unclassified(100);p__Ascomycota_unclassified(100); | - | - | - | - | - | - | - | Unassigned | - |
| Otu14818 | 1 | k__Fungi(100);k__Fungi_unclassified(100);k__Fungi_unclassified(100);k__Fungi_unclassified(100);k__Fungi_unclassified(100);k__Fungi_unclassified(100);k__Fungi_unclassified(100); | - | - | - | - | - | - | - | Unassigned | - |
| Otu14819 | 1 | k__Fungi(100);k__Fungi_unclassified(100);k__Fungi_unclassified(100);k__Fungi_unclassified(100);k__Fungi_unclassified(100);k__Fungi_unclassified(100);k__Fungi_unclassified(100); | - | - | - | - | - | - | - | Unassigned | - |
| Otu14821 | 1 | k__Fungi(100);k__Fungi_unclassified(100);k__Fungi_unclassified(100);k__Fungi_unclassified(100);k__Fungi_unclassified(100);k__Fungi_unclassified(100);k__Fungi_unclassified(100); | - | - | - | - | - | - | - | Unassigned | - |
| Otu14822 | 1 | k__Fungi(100);k__Fungi_unclassified(100);k__Fungi_unclassified(100);k__Fungi_unclassified(100);k__Fungi_unclassified(100);k__Fungi_unclassified(100);k__Fungi_unclassified(100); | - | - | - | - | - | - | - | Unassigned | - |
| Otu14824 | 1 | k__Fungi(100);k__Fungi_unclassified(100);k__Fungi_unclassified(100);k__Fungi_unclassified(100);k__Fungi_unclassified(100);k__Fungi_unclassified(100);k__Fungi_unclassified(100); | - | - | - | - | - | - | - | Unassigned | - |
| Otu14835 | 1 | k__Fungi(100);k__Fungi_unclassified(100);k__Fungi_unclassified(100);k__Fungi_unclassified(100);k__Fungi_unclassified(100);k__Fungi_unclassified(100);k__Fungi_unclassified(100); | - | - | - | - | - | - | - | Unassigned | - |
| Otu14836 | 1 | k__Fungi(100);k__Fungi_unclassified(100);k__Fungi_unclassified(100);k__Fungi_unclassified(100);k__Fungi_unclassified(100);k__Fungi_unclassified(100);k__Fungi_unclassified(100); | - | - | - | - | - | - | - | Unassigned | - |
| Otu14838 | 1 | k__Fungi(100);k__Fungi_unclassified(100);k__Fungi_unclassified(100);k__Fungi_unclassified(100);k__Fungi_unclassified(100);k__Fungi_unclassified(100);k__Fungi_unclassified(100); | - | - | - | - | - | - | - | Unassigned | - |
| Otu14840 | 1 | k__Fungi(100);k__Fungi_unclassified(100);k__Fungi_unclassified(100);k__Fungi_unclassified(100);k__Fungi_unclassified(100);k__Fungi_unclassified(100);k__Fungi_unclassified(100); | - | - | - | - | - | - | - | Unassigned | - |
| Otu14842 | 1 | k__Fungi(100);k__Fungi_unclassified(100);k__Fungi_unclassified(100);k__Fungi_unclassified(100);k__Fungi_unclassified(100);k__Fungi_unclassified(100);k__Fungi_unclassified(100); | - | - | - | - | - | - | - | Unassigned | - |
| Otu14844 | 1 | k__Fungi(100);p__Ascomycota(100);p__Ascomycota_unclassified(100);p__Ascomycota_unclassified(100);p__Ascomycota_unclassified(100);p__Ascomycota_unclassified(100);p__Ascomycota_unclassified(100); | - | - | - | - | - | - | - | Unassigned | - |
| Otu14845 | 1 | k__Fungi(100);k__Fungi_unclassified(100);k__Fungi_unclassified(100);k__Fungi_unclassified(100);k__Fungi_unclassified(100);k__Fungi_unclassified(100);k__Fungi_unclassified(100); | - | - | - | - | - | - | - | Unassigned | - |
| Otu14847 | 1 | k__Fungi(100);k__Fungi_unclassified(100);k__Fungi_unclassified(100);k__Fungi_unclassified(100);k__Fungi_unclassified(100);k__Fungi_unclassified(100);k__Fungi_unclassified(100); | - | - | - | - | - | - | - | Unassigned | - |
| Otu14853 | 1 | k__Fungi(100);k__Fungi_unclassified(100);k__Fungi_unclassified(100);k__Fungi_unclassified(100);k__Fungi_unclassified(100);k__Fungi_unclassified(100);k__Fungi_unclassified(100); | - | - | - | - | - | - | - | Unassigned | - |
| Otu14855 | 1 | k__Fungi(100);k__Fungi_unclassified(100);k__Fungi_unclassified(100);k__Fungi_unclassified(100);k__Fungi_unclassified(100);k__Fungi_unclassified(100);k__Fungi_unclassified(100); | - | - | - | - | - | - | - | Unassigned | - |
| Otu14856 | 1 | k__Fungi(100);k__Fungi_unclassified(100);k__Fungi_unclassified(100);k__Fungi_unclassified(100);k__Fungi_unclassified(100);k__Fungi_unclassified(100);k__Fungi_unclassified(100); | - | - | - | - | - | - | - | Unassigned | - |
| Otu14865 | 1 | k__Fungi(100);k__Fungi_unclassified(100);k__Fungi_unclassified(100);k__Fungi_unclassified(100);k__Fungi_unclassified(100);k__Fungi_unclassified(100);k__Fungi_unclassified(100); | - | - | - | - | - | - | - | Unassigned | - |
| Otu14873 | 1 | k__Fungi(100);p__Basidiomycota(100);p__Basidiomycota_unclassified(100);p__Basidiomycota_unclassified(100);p__Basidiomycota_unclassified(100);p__Basidiomycota_unclassified(100);p__Basidiomycota_unclassified(100); | - | - | - | - | - | - | - | Unassigned | - |
| Otu14876 | 1 | k__Fungi(100);p__Ascomycota(100);p__Ascomycota_unclassified(100);p__Ascomycota_unclassified(100);p__Ascomycota_unclassified(100);p__Ascomycota_unclassified(100);p__Ascomycota_unclassified(100); | - | - | - | - | - | - | - | Unassigned | - |
| Otu14878 | 1 | k__Fungi(100);k__Fungi_unclassified(100);k__Fungi_unclassified(100);k__Fungi_unclassified(100);k__Fungi_unclassified(100);k__Fungi_unclassified(100);k__Fungi_unclassified(100); | - | - | - | - | - | - | - | Unassigned | - |
| Otu14880 | 1 | k__Fungi(100);p__Ascomycota(100);p__Ascomycota_unclassified(100);p__Ascomycota_unclassified(100);p__Ascomycota_unclassified(100);p__Ascomycota_unclassified(100);p__Ascomycota_unclassified(100); | - | - | - | - | - | - | - | Unassigned | - |
| Otu14885 | 1 | k__Fungi(100);k__Fungi_unclassified(100);k__Fungi_unclassified(100);k__Fungi_unclassified(100);k__Fungi_unclassified(100);k__Fungi_unclassified(100);k__Fungi_unclassified(100); | - | - | - | - | - | - | - | Unassigned | - |
| Otu14887 | 1 | k__Fungi(100);p__Ascomycota(100);p__Ascomycota_unclassified(100);p__Ascomycota_unclassified(100);p__Ascomycota_unclassified(100);p__Ascomycota_unclassified(100);p__Ascomycota_unclassified(100); | - | - | - | - | - | - | - | Unassigned | - |
| Otu14893 | 1 | k__Fungi(100);p__Ascomycota(100);p__Ascomycota_unclassified(100);p__Ascomycota_unclassified(100);p__Ascomycota_unclassified(100);p__Ascomycota_unclassified(100);p__Ascomycota_unclassified(100); | - | - | - | - | - | - | - | Unassigned | - |
| Otu14895 | 1 | k__Fungi(100);k__Fungi_unclassified(100);k__Fungi_unclassified(100);k__Fungi_unclassified(100);k__Fungi_unclassified(100);k__Fungi_unclassified(100);k__Fungi_unclassified(100); | - | - | - | - | - | - | - | Unassigned | - |
| Otu14897 | 1 | k__Fungi(100);k__Fungi_unclassified(100);k__Fungi_unclassified(100);k__Fungi_unclassified(100);k__Fungi_unclassified(100);k__Fungi_unclassified(100);k__Fungi_unclassified(100); | - | - | - | - | - | - | - | Unassigned | - |
| Otu14909 | 1 | k__Fungi(100);k__Fungi_unclassified(100);k__Fungi_unclassified(100);k__Fungi_unclassified(100);k__Fungi_unclassified(100);k__Fungi_unclassified(100);k__Fungi_unclassified(100); | - | - | - | - | - | - | - | Unassigned | - |
| Otu14910 | 1 | k__Fungi(100);k__Fungi_unclassified(100);k__Fungi_unclassified(100);k__Fungi_unclassified(100);k__Fungi_unclassified(100);k__Fungi_unclassified(100);k__Fungi_unclassified(100); | - | - | - | - | - | - | - | Unassigned | - |
| Otu14914 | 1 | k__Fungi(100);k__Fungi_unclassified(100);k__Fungi_unclassified(100);k__Fungi_unclassified(100);k__Fungi_unclassified(100);k__Fungi_unclassified(100);k__Fungi_unclassified(100); | - | - | - | - | - | - | - | Unassigned | - |
| Otu14915 | 1 | k__Fungi(100);p__Ascomycota(100);p__Ascomycota_unclassified(100);p__Ascomycota_unclassified(100);p__Ascomycota_unclassified(100);p__Ascomycota_unclassified(100);p__Ascomycota_unclassified(100); | - | - | - | - | - | - | - | Unassigned | - |
| Otu14916 | 1 | k__Fungi(100);p__Ascomycota(100);p__Ascomycota_unclassified(100);p__Ascomycota_unclassified(100);p__Ascomycota_unclassified(100);p__Ascomycota_unclassified(100);p__Ascomycota_unclassified(100); | - | - | - | - | - | - | - | Unassigned | - |
| Otu14917 | 1 | k__Fungi(100);k__Fungi_unclassified(100);k__Fungi_unclassified(100);k__Fungi_unclassified(100);k__Fungi_unclassified(100);k__Fungi_unclassified(100);k__Fungi_unclassified(100); | - | - | - | - | - | - | - | Unassigned | - |
| Otu14920 | 1 | k__Fungi(100);k__Fungi_unclassified(100);k__Fungi_unclassified(100);k__Fungi_unclassified(100);k__Fungi_unclassified(100);k__Fungi_unclassified(100);k__Fungi_unclassified(100); | - | - | - | - | - | - | - | Unassigned | - |
| Otu14921 | 1 | k__Fungi(100);k__Fungi_unclassified(100);k__Fungi_unclassified(100);k__Fungi_unclassified(100);k__Fungi_unclassified(100);k__Fungi_unclassified(100);k__Fungi_unclassified(100); | - | - | - | - | - | - | - | Unassigned | - |
| Otu14922 | 1 | k__Fungi(100);p__Ascomycota(100);p__Ascomycota_unclassified(100);p__Ascomycota_unclassified(100);p__Ascomycota_unclassified(100);p__Ascomycota_unclassified(100);p__Ascomycota_unclassified(100); | - | - | - | - | - | - | - | Unassigned | - |
| Otu14926 | 1 | k__Fungi(100);k__Fungi_unclassified(100);k__Fungi_unclassified(100);k__Fungi_unclassified(100);k__Fungi_unclassified(100);k__Fungi_unclassified(100);k__Fungi_unclassified(100); | - | - | - | - | - | - | - | Unassigned | - |
| Otu14928 | 1 | k__Fungi(100);k__Fungi_unclassified(100);k__Fungi_unclassified(100);k__Fungi_unclassified(100);k__Fungi_unclassified(100);k__Fungi_unclassified(100);k__Fungi_unclassified(100); | - | - | - | - | - | - | - | Unassigned | - |
| Otu14931 | 1 | k__Fungi(100);k__Fungi_unclassified(100);k__Fungi_unclassified(100);k__Fungi_unclassified(100);k__Fungi_unclassified(100);k__Fungi_unclassified(100);k__Fungi_unclassified(100); | - | - | - | - | - | - | - | Unassigned | - |
| Otu14932 | 1 | k__Fungi(100);p__Ascomycota(100);p__Ascomycota_unclassified(100);p__Ascomycota_unclassified(100);p__Ascomycota_unclassified(100);p__Ascomycota_unclassified(100);p__Ascomycota_unclassified(100); | - | - | - | - | - | - | - | Unassigned | - |
| Otu14934 | 1 | k__Fungi(100);p__Ascomycota(100);p__Ascomycota_unclassified(100);p__Ascomycota_unclassified(100);p__Ascomycota_unclassified(100);p__Ascomycota_unclassified(100);p__Ascomycota_unclassified(100); | - | - | - | - | - | - | - | Unassigned | - |
| Otu14936 | 1 | k__Fungi(100);k__Fungi_unclassified(100);k__Fungi_unclassified(100);k__Fungi_unclassified(100);k__Fungi_unclassified(100);k__Fungi_unclassified(100);k__Fungi_unclassified(100); | - | - | - | - | - | - | - | Unassigned | - |
| Otu14941 | 1 | k__Fungi(100);p__Ascomycota(100);p__Ascomycota_unclassified(100);p__Ascomycota_unclassified(100);p__Ascomycota_unclassified(100);p__Ascomycota_unclassified(100);p__Ascomycota_unclassified(100); | - | - | - | - | - | - | - | Unassigned | - |
| Otu14942 | 1 | k__Fungi(100);k__Fungi_unclassified(100);k__Fungi_unclassified(100);k__Fungi_unclassified(100);k__Fungi_unclassified(100);k__Fungi_unclassified(100);k__Fungi_unclassified(100); | - | - | - | - | - | - | - | Unassigned | - |
| Otu14944 | 1 | k__Fungi(100);k__Fungi_unclassified(100);k__Fungi_unclassified(100);k__Fungi_unclassified(100);k__Fungi_unclassified(100);k__Fungi_unclassified(100);k__Fungi_unclassified(100); | - | - | - | - | - | - | - | Unassigned | - |
| Otu14945 | 1 | k__Fungi(100);k__Fungi_unclassified(100);k__Fungi_unclassified(100);k__Fungi_unclassified(100);k__Fungi_unclassified(100);k__Fungi_unclassified(100);k__Fungi_unclassified(100); | - | - | - | - | - | - | - | Unassigned | - |
| Otu14947 | 1 | k__Fungi(100);k__Fungi_unclassified(100);k__Fungi_unclassified(100);k__Fungi_unclassified(100);k__Fungi_unclassified(100);k__Fungi_unclassified(100);k__Fungi_unclassified(100); | - | - | - | - | - | - | - | Unassigned | - |
| Otu14950 | 1 | k__Fungi(100);k__Fungi_unclassified(100);k__Fungi_unclassified(100);k__Fungi_unclassified(100);k__Fungi_unclassified(100);k__Fungi_unclassified(100);k__Fungi_unclassified(100); | - | - | - | - | - | - | - | Unassigned | - |
| Otu14954 | 1 | k__Fungi(100);k__Fungi_unclassified(100);k__Fungi_unclassified(100);k__Fungi_unclassified(100);k__Fungi_unclassified(100);k__Fungi_unclassified(100);k__Fungi_unclassified(100); | - | - | - | - | - | - | - | Unassigned | - |
| Otu14955 | 1 | k__Fungi(100);k__Fungi_unclassified(100);k__Fungi_unclassified(100);k__Fungi_unclassified(100);k__Fungi_unclassified(100);k__Fungi_unclassified(100);k__Fungi_unclassified(100); | - | - | - | - | - | - | - | Unassigned | - |
| Otu14956 | 1 | k__Fungi(100);p__Ascomycota(100);p__Ascomycota_unclassified(100);p__Ascomycota_unclassified(100);p__Ascomycota_unclassified(100);p__Ascomycota_unclassified(100);p__Ascomycota_unclassified(100); | - | - | - | - | - | - | - | Unassigned | - |
| Otu14958 | 1 | k__Fungi(100);k__Fungi_unclassified(100);k__Fungi_unclassified(100);k__Fungi_unclassified(100);k__Fungi_unclassified(100);k__Fungi_unclassified(100);k__Fungi_unclassified(100); | - | - | - | - | - | - | - | Unassigned | - |
| Otu14961 | 1 | k__Fungi(100);k__Fungi_unclassified(100);k__Fungi_unclassified(100);k__Fungi_unclassified(100);k__Fungi_unclassified(100);k__Fungi_unclassified(100);k__Fungi_unclassified(100); | - | - | - | - | - | - | - | Unassigned | - |
| Otu14962 | 1 | k__Fungi(100);k__Fungi_unclassified(100);k__Fungi_unclassified(100);k__Fungi_unclassified(100);k__Fungi_unclassified(100);k__Fungi_unclassified(100);k__Fungi_unclassified(100); | - | - | - | - | - | - | - | Unassigned | - |
| Otu14963 | 1 | k__Fungi(100);k__Fungi_unclassified(100);k__Fungi_unclassified(100);k__Fungi_unclassified(100);k__Fungi_unclassified(100);k__Fungi_unclassified(100);k__Fungi_unclassified(100); | - | - | - | - | - | - | - | Unassigned | - |
| Otu14966 | 1 | k__Fungi(100);k__Fungi_unclassified(100);k__Fungi_unclassified(100);k__Fungi_unclassified(100);k__Fungi_unclassified(100);k__Fungi_unclassified(100);k__Fungi_unclassified(100); | - | - | - | - | - | - | - | Unassigned | - |
| Otu14972 | 1 | k__Fungi(100);k__Fungi_unclassified(100);k__Fungi_unclassified(100);k__Fungi_unclassified(100);k__Fungi_unclassified(100);k__Fungi_unclassified(100);k__Fungi_unclassified(100); | - | - | - | - | - | - | - | Unassigned | - |
| Otu14978 | 1 | k__Fungi(100);k__Fungi_unclassified(100);k__Fungi_unclassified(100);k__Fungi_unclassified(100);k__Fungi_unclassified(100);k__Fungi_unclassified(100);k__Fungi_unclassified(100); | - | - | - | - | - | - | - | Unassigned | - |
| Otu14981 | 1 | k__Fungi(100);k__Fungi_unclassified(100);k__Fungi_unclassified(100);k__Fungi_unclassified(100);k__Fungi_unclassified(100);k__Fungi_unclassified(100);k__Fungi_unclassified(100); | - | - | - | - | - | - | - | Unassigned | - |
| Otu14983 | 1 | k__Fungi(100);k__Fungi_unclassified(100);k__Fungi_unclassified(100);k__Fungi_unclassified(100);k__Fungi_unclassified(100);k__Fungi_unclassified(100);k__Fungi_unclassified(100); | - | - | - | - | - | - | - | Unassigned | - |
| Otu14988 | 1 | k__Fungi(100);p__Ascomycota(100);p__Ascomycota_unclassified(100);p__Ascomycota_unclassified(100);p__Ascomycota_unclassified(100);p__Ascomycota_unclassified(100);p__Ascomycota_unclassified(100); | - | - | - | - | - | - | - | Unassigned | - |
| Otu14990 | 1 | k__Fungi(100);k__Fungi_unclassified(100);k__Fungi_unclassified(100);k__Fungi_unclassified(100);k__Fungi_unclassified(100);k__Fungi_unclassified(100);k__Fungi_unclassified(100); | - | - | - | - | - | - | - | Unassigned | - |
| Otu14991 | 1 | k__Fungi(100);k__Fungi_unclassified(100);k__Fungi_unclassified(100);k__Fungi_unclassified(100);k__Fungi_unclassified(100);k__Fungi_unclassified(100);k__Fungi_unclassified(100); | - | - | - | - | - | - | - | Unassigned | - |
| Otu14992 | 1 | k__Fungi(100);p__Ascomycota(100);p__Ascomycota_unclassified(100);p__Ascomycota_unclassified(100);p__Ascomycota_unclassified(100);p__Ascomycota_unclassified(100);p__Ascomycota_unclassified(100); | - | - | - | - | - | - | - | Unassigned | - |
| Otu14994 | 1 | k__Fungi(100);k__Fungi_unclassified(100);k__Fungi_unclassified(100);k__Fungi_unclassified(100);k__Fungi_unclassified(100);k__Fungi_unclassified(100);k__Fungi_unclassified(100); | - | - | - | - | - | - | - | Unassigned | - |
| Otu14996 | 1 | k__Fungi(100);k__Fungi_unclassified(100);k__Fungi_unclassified(100);k__Fungi_unclassified(100);k__Fungi_unclassified(100);k__Fungi_unclassified(100);k__Fungi_unclassified(100); | - | - | - | - | - | - | - | Unassigned | - |
| Otu14998 | 1 | k__Fungi(100);k__Fungi_unclassified(100);k__Fungi_unclassified(100);k__Fungi_unclassified(100);k__Fungi_unclassified(100);k__Fungi_unclassified(100);k__Fungi_unclassified(100); | - | - | - | - | - | - | - | Unassigned | - |
| Otu15000 | 1 | k__Fungi(100);p__Ascomycota(100);p__Ascomycota_unclassified(100);p__Ascomycota_unclassified(100);p__Ascomycota_unclassified(100);p__Ascomycota_unclassified(100);p__Ascomycota_unclassified(100); | - | - | - | - | - | - | - | Unassigned | - |
| Otu15001 | 1 | k__Fungi(100);k__Fungi_unclassified(100);k__Fungi_unclassified(100);k__Fungi_unclassified(100);k__Fungi_unclassified(100);k__Fungi_unclassified(100);k__Fungi_unclassified(100); | - | - | - | - | - | - | - | Unassigned | - |
| Otu15003 | 1 | k__Fungi(100);p__Ascomycota(100);p__Ascomycota_unclassified(100);p__Ascomycota_unclassified(100);p__Ascomycota_unclassified(100);p__Ascomycota_unclassified(100);p__Ascomycota_unclassified(100); | - | - | - | - | - | - | - | Unassigned | - |
| Otu15004 | 1 | k__Fungi(100);k__Fungi_unclassified(100);k__Fungi_unclassified(100);k__Fungi_unclassified(100);k__Fungi_unclassified(100);k__Fungi_unclassified(100);k__Fungi_unclassified(100); | - | - | - | - | - | - | - | Unassigned | - |
| Otu15006 | 1 | k__Fungi(100);k__Fungi_unclassified(100);k__Fungi_unclassified(100);k__Fungi_unclassified(100);k__Fungi_unclassified(100);k__Fungi_unclassified(100);k__Fungi_unclassified(100); | - | - | - | - | - | - | - | Unassigned | - |
| Otu15008 | 1 | k__Fungi(100);p__Ascomycota(100);p__Ascomycota_unclassified(100);p__Ascomycota_unclassified(100);p__Ascomycota_unclassified(100);p__Ascomycota_unclassified(100);p__Ascomycota_unclassified(100); | - | - | - | - | - | - | - | Unassigned | - |
| Otu15009 | 1 | k__Fungi(100);p__Ascomycota(100);p__Ascomycota_unclassified(100);p__Ascomycota_unclassified(100);p__Ascomycota_unclassified(100);p__Ascomycota_unclassified(100);p__Ascomycota_unclassified(100); | - | - | - | - | - | - | - | Unassigned | - |
| Otu15010 | 1 | k__Fungi(100);p__Ascomycota(100);p__Ascomycota_unclassified(100);p__Ascomycota_unclassified(100);p__Ascomycota_unclassified(100);p__Ascomycota_unclassified(100);p__Ascomycota_unclassified(100); | - | - | - | - | - | - | - | Unassigned | - |
| Otu15011 | 1 | k__Fungi(100);k__Fungi_unclassified(100);k__Fungi_unclassified(100);k__Fungi_unclassified(100);k__Fungi_unclassified(100);k__Fungi_unclassified(100);k__Fungi_unclassified(100); | - | - | - | - | - | - | - | Unassigned | - |
| Otu15015 | 1 | k__Fungi(100);k__Fungi_unclassified(100);k__Fungi_unclassified(100);k__Fungi_unclassified(100);k__Fungi_unclassified(100);k__Fungi_unclassified(100);k__Fungi_unclassified(100); | - | - | - | - | - | - | - | Unassigned | - |
| Otu15017 | 1 | k__Fungi(100);k__Fungi_unclassified(100);k__Fungi_unclassified(100);k__Fungi_unclassified(100);k__Fungi_unclassified(100);k__Fungi_unclassified(100);k__Fungi_unclassified(100); | - | - | - | - | - | - | - | Unassigned | - |
| Otu15018 | 1 | k__Fungi(100);p__Ascomycota(100);p__Ascomycota_unclassified(100);p__Ascomycota_unclassified(100);p__Ascomycota_unclassified(100);p__Ascomycota_unclassified(100);p__Ascomycota_unclassified(100); | - | - | - | - | - | - | - | Unassigned | - |
| Otu15019 | 1 | k__Fungi(100);k__Fungi_unclassified(100);k__Fungi_unclassified(100);k__Fungi_unclassified(100);k__Fungi_unclassified(100);k__Fungi_unclassified(100);k__Fungi_unclassified(100); | - | - | - | - | - | - | - | Unassigned | - |
| Otu15022 | 1 | k__Fungi(100);k__Fungi_unclassified(100);k__Fungi_unclassified(100);k__Fungi_unclassified(100);k__Fungi_unclassified(100);k__Fungi_unclassified(100);k__Fungi_unclassified(100); | - | - | - | - | - | - | - | Unassigned | - |
| Otu15023 | 1 | k__Fungi(100);k__Fungi_unclassified(100);k__Fungi_unclassified(100);k__Fungi_unclassified(100);k__Fungi_unclassified(100);k__Fungi_unclassified(100);k__Fungi_unclassified(100); | - | - | - | - | - | - | - | Unassigned | - |
| Otu15024 | 1 | k__Fungi(100);k__Fungi_unclassified(100);k__Fungi_unclassified(100);k__Fungi_unclassified(100);k__Fungi_unclassified(100);k__Fungi_unclassified(100);k__Fungi_unclassified(100); | - | - | - | - | - | - | - | Unassigned | - |
| Otu15029 | 1 | k__Fungi(100);k__Fungi_unclassified(100);k__Fungi_unclassified(100);k__Fungi_unclassified(100);k__Fungi_unclassified(100);k__Fungi_unclassified(100);k__Fungi_unclassified(100); | - | - | - | - | - | - | - | Unassigned | - |
| Otu15030 | 1 | k__Fungi(100);k__Fungi_unclassified(100);k__Fungi_unclassified(100);k__Fungi_unclassified(100);k__Fungi_unclassified(100);k__Fungi_unclassified(100);k__Fungi_unclassified(100); | - | - | - | - | - | - | - | Unassigned | - |
| Otu15044 | 1 | k__Fungi(100);k__Fungi_unclassified(100);k__Fungi_unclassified(100);k__Fungi_unclassified(100);k__Fungi_unclassified(100);k__Fungi_unclassified(100);k__Fungi_unclassified(100); | - | - | - | - | - | - | - | Unassigned | - |
| Otu15045 | 1 | k__Fungi(100);k__Fungi_unclassified(100);k__Fungi_unclassified(100);k__Fungi_unclassified(100);k__Fungi_unclassified(100);k__Fungi_unclassified(100);k__Fungi_unclassified(100); | - | - | - | - | - | - | - | Unassigned | - |
| Otu15047 | 1 | k__Fungi(100);k__Fungi_unclassified(100);k__Fungi_unclassified(100);k__Fungi_unclassified(100);k__Fungi_unclassified(100);k__Fungi_unclassified(100);k__Fungi_unclassified(100); | - | - | - | - | - | - | - | Unassigned | - |
| Otu15056 | 1 | k__Fungi(100);p__Ascomycota(100);p__Ascomycota_unclassified(100);p__Ascomycota_unclassified(100);p__Ascomycota_unclassified(100);p__Ascomycota_unclassified(100);p__Ascomycota_unclassified(100); | - | - | - | - | - | - | - | Unassigned | - |
| Otu15059 | 1 | k__Fungi(100);k__Fungi_unclassified(100);k__Fungi_unclassified(100);k__Fungi_unclassified(100);k__Fungi_unclassified(100);k__Fungi_unclassified(100);k__Fungi_unclassified(100); | - | - | - | - | - | - | - | Unassigned | - |
| Otu15060 | 1 | k__Fungi(100);k__Fungi_unclassified(100);k__Fungi_unclassified(100);k__Fungi_unclassified(100);k__Fungi_unclassified(100);k__Fungi_unclassified(100);k__Fungi_unclassified(100); | - | - | - | - | - | - | - | Unassigned | - |
| Otu15064 | 1 | k__Fungi(100);k__Fungi_unclassified(100);k__Fungi_unclassified(100);k__Fungi_unclassified(100);k__Fungi_unclassified(100);k__Fungi_unclassified(100);k__Fungi_unclassified(100); | - | - | - | - | - | - | - | Unassigned | - |
| Otu15067 | 1 | k__Fungi(100);k__Fungi_unclassified(100);k__Fungi_unclassified(100);k__Fungi_unclassified(100);k__Fungi_unclassified(100);k__Fungi_unclassified(100);k__Fungi_unclassified(100); | - | - | - | - | - | - | - | Unassigned | - |
| Otu15071 | 1 | k__Fungi(100);k__Fungi_unclassified(100);k__Fungi_unclassified(100);k__Fungi_unclassified(100);k__Fungi_unclassified(100);k__Fungi_unclassified(100);k__Fungi_unclassified(100); | - | - | - | - | - | - | - | Unassigned | - |
| Otu15072 | 1 | k__Fungi(100);k__Fungi_unclassified(100);k__Fungi_unclassified(100);k__Fungi_unclassified(100);k__Fungi_unclassified(100);k__Fungi_unclassified(100);k__Fungi_unclassified(100); | - | - | - | - | - | - | - | Unassigned | - |
| Otu15073 | 1 | k__Fungi(100);p__Ascomycota(100);p__Ascomycota_unclassified(100);p__Ascomycota_unclassified(100);p__Ascomycota_unclassified(100);p__Ascomycota_unclassified(100);p__Ascomycota_unclassified(100); | - | - | - | - | - | - | - | Unassigned | - |
| Otu15074 | 1 | k__Fungi(100);p__Basidiomycota(100);p__Basidiomycota_unclassified(100);p__Basidiomycota_unclassified(100);p__Basidiomycota_unclassified(100);p__Basidiomycota_unclassified(100);p__Basidiomycota_unclassified(100); | - | - | - | - | - | - | - | Unassigned | - |
| Otu15075 | 1 | k__Fungi(100);k__Fungi_unclassified(100);k__Fungi_unclassified(100);k__Fungi_unclassified(100);k__Fungi_unclassified(100);k__Fungi_unclassified(100);k__Fungi_unclassified(100); | - | - | - | - | - | - | - | Unassigned | - |
| Otu15079 | 1 | k__Fungi(100);k__Fungi_unclassified(100);k__Fungi_unclassified(100);k__Fungi_unclassified(100);k__Fungi_unclassified(100);k__Fungi_unclassified(100);k__Fungi_unclassified(100); | - | - | - | - | - | - | - | Unassigned | - |
| Otu15082 | 1 | k__Fungi(100);k__Fungi_unclassified(100);k__Fungi_unclassified(100);k__Fungi_unclassified(100);k__Fungi_unclassified(100);k__Fungi_unclassified(100);k__Fungi_unclassified(100); | - | - | - | - | - | - | - | Unassigned | - |
| Otu15085 | 1 | k__Fungi(100);k__Fungi_unclassified(100);k__Fungi_unclassified(100);k__Fungi_unclassified(100);k__Fungi_unclassified(100);k__Fungi_unclassified(100);k__Fungi_unclassified(100); | - | - | - | - | - | - | - | Unassigned | - |
| Otu15090 | 1 | k__Fungi(100);k__Fungi_unclassified(100);k__Fungi_unclassified(100);k__Fungi_unclassified(100);k__Fungi_unclassified(100);k__Fungi_unclassified(100);k__Fungi_unclassified(100); | - | - | - | - | - | - | - | Unassigned | - |
| Otu15108 | 1 | k__Fungi(100);k__Fungi_unclassified(100);k__Fungi_unclassified(100);k__Fungi_unclassified(100);k__Fungi_unclassified(100);k__Fungi_unclassified(100);k__Fungi_unclassified(100); | - | - | - | - | - | - | - | Unassigned | - |
| Otu15113 | 1 | k__Fungi(100);k__Fungi_unclassified(100);k__Fungi_unclassified(100);k__Fungi_unclassified(100);k__Fungi_unclassified(100);k__Fungi_unclassified(100);k__Fungi_unclassified(100); | - | - | - | - | - | - | - | Unassigned | - |
| Otu15115 | 1 | k__Fungi(100);k__Fungi_unclassified(100);k__Fungi_unclassified(100);k__Fungi_unclassified(100);k__Fungi_unclassified(100);k__Fungi_unclassified(100);k__Fungi_unclassified(100); | - | - | - | - | - | - | - | Unassigned | - |
| Otu15119 | 1 | k__Fungi(100);p__Ascomycota(100);p__Ascomycota_unclassified(100);p__Ascomycota_unclassified(100);p__Ascomycota_unclassified(100);p__Ascomycota_unclassified(100);p__Ascomycota_unclassified(100); | - | - | - | - | - | - | - | Unassigned | - |
| Otu15125 | 1 | k__Fungi(100);k__Fungi_unclassified(100);k__Fungi_unclassified(100);k__Fungi_unclassified(100);k__Fungi_unclassified(100);k__Fungi_unclassified(100);k__Fungi_unclassified(100); | - | - | - | - | - | - | - | Unassigned | - |
| Otu15130 | 1 | k__Fungi(100);k__Fungi_unclassified(100);k__Fungi_unclassified(100);k__Fungi_unclassified(100);k__Fungi_unclassified(100);k__Fungi_unclassified(100);k__Fungi_unclassified(100); | - | - | - | - | - | - | - | Unassigned | - |
| Otu15145 | 1 | k__Fungi(100);p__Ascomycota(100);p__Ascomycota_unclassified(100);p__Ascomycota_unclassified(100);p__Ascomycota_unclassified(100);p__Ascomycota_unclassified(100);p__Ascomycota_unclassified(100); | - | - | - | - | - | - | - | Unassigned | - |
| Otu15146 | 1 | k__Fungi(100);k__Fungi_unclassified(100);k__Fungi_unclassified(100);k__Fungi_unclassified(100);k__Fungi_unclassified(100);k__Fungi_unclassified(100);k__Fungi_unclassified(100); | - | - | - | - | - | - | - | Unassigned | - |
| Otu15151 | 1 | k__Fungi(100);k__Fungi_unclassified(100);k__Fungi_unclassified(100);k__Fungi_unclassified(100);k__Fungi_unclassified(100);k__Fungi_unclassified(100);k__Fungi_unclassified(100); | - | - | - | - | - | - | - | Unassigned | - |
| Otu15155 | 1 | k__Fungi(100);k__Fungi_unclassified(100);k__Fungi_unclassified(100);k__Fungi_unclassified(100);k__Fungi_unclassified(100);k__Fungi_unclassified(100);k__Fungi_unclassified(100); | - | - | - | - | - | - | - | Unassigned | - |
| Otu15163 | 1 | k__Fungi(100);k__Fungi_unclassified(100);k__Fungi_unclassified(100);k__Fungi_unclassified(100);k__Fungi_unclassified(100);k__Fungi_unclassified(100);k__Fungi_unclassified(100); | - | - | - | - | - | - | - | Unassigned | - |
| Otu15179 | 1 | k__Fungi(100);p__Ascomycota(100);p__Ascomycota_unclassified(100);p__Ascomycota_unclassified(100);p__Ascomycota_unclassified(100);p__Ascomycota_unclassified(100);p__Ascomycota_unclassified(100); | - | - | - | - | - | - | - | Unassigned | - |
| Otu15209 | 1 | k__Fungi(100);k__Fungi_unclassified(100);k__Fungi_unclassified(100);k__Fungi_unclassified(100);k__Fungi_unclassified(100);k__Fungi_unclassified(100);k__Fungi_unclassified(100); | - | - | - | - | - | - | - | Unassigned | - |
| Otu15214 | 1 | k__Fungi(100);p__Ascomycota(100);p__Ascomycota_unclassified(100);p__Ascomycota_unclassified(100);p__Ascomycota_unclassified(100);p__Ascomycota_unclassified(100);p__Ascomycota_unclassified(100); | - | - | - | - | - | - | - | Unassigned | - |
| Otu15222 | 1 | k__Fungi(100);k__Fungi_unclassified(100);k__Fungi_unclassified(100);k__Fungi_unclassified(100);k__Fungi_unclassified(100);k__Fungi_unclassified(100);k__Fungi_unclassified(100); | - | - | - | - | - | - | - | Unassigned | - |
| Otu15243 | 1 | k__Fungi(100);k__Fungi_unclassified(100);k__Fungi_unclassified(100);k__Fungi_unclassified(100);k__Fungi_unclassified(100);k__Fungi_unclassified(100);k__Fungi_unclassified(100); | - | - | - | - | - | - | - | Unassigned | - |
| Otu15244 | 1 | k__Fungi(100);k__Fungi_unclassified(100);k__Fungi_unclassified(100);k__Fungi_unclassified(100);k__Fungi_unclassified(100);k__Fungi_unclassified(100);k__Fungi_unclassified(100); | - | - | - | - | - | - | - | Unassigned | - |
| Otu15251 | 1 | k__Fungi(100);k__Fungi_unclassified(100);k__Fungi_unclassified(100);k__Fungi_unclassified(100);k__Fungi_unclassified(100);k__Fungi_unclassified(100);k__Fungi_unclassified(100); | - | - | - | - | - | - | - | Unassigned | - |
| Otu15253 | 1 | k__Fungi(100);k__Fungi_unclassified(100);k__Fungi_unclassified(100);k__Fungi_unclassified(100);k__Fungi_unclassified(100);k__Fungi_unclassified(100);k__Fungi_unclassified(100); | - | - | - | - | - | - | - | Unassigned | - |
| Otu15256 | 1 | k__Fungi(100);p__Ascomycota(100);p__Ascomycota_unclassified(100);p__Ascomycota_unclassified(100);p__Ascomycota_unclassified(100);p__Ascomycota_unclassified(100);p__Ascomycota_unclassified(100); | - | - | - | - | - | - | - | Unassigned | - |
| Otu15261 | 1 | k__Fungi(100);p__Ascomycota(100);p__Ascomycota_unclassified(100);p__Ascomycota_unclassified(100);p__Ascomycota_unclassified(100);p__Ascomycota_unclassified(100);p__Ascomycota_unclassified(100); | - | - | - | - | - | - | - | Unassigned | - |
| Otu15263 | 1 | k__Fungi(100);k__Fungi_unclassified(100);k__Fungi_unclassified(100);k__Fungi_unclassified(100);k__Fungi_unclassified(100);k__Fungi_unclassified(100);k__Fungi_unclassified(100); | - | - | - | - | - | - | - | Unassigned | - |
| Otu15283 | 1 | k__Fungi(100);k__Fungi_unclassified(100);k__Fungi_unclassified(100);k__Fungi_unclassified(100);k__Fungi_unclassified(100);k__Fungi_unclassified(100);k__Fungi_unclassified(100); | - | - | - | - | - | - | - | Unassigned | - |
| Otu15293 | 1 | k__Fungi(100);p__Ascomycota(100);p__Ascomycota_unclassified(100);p__Ascomycota_unclassified(100);p__Ascomycota_unclassified(100);p__Ascomycota_unclassified(100);p__Ascomycota_unclassified(100); | - | - | - | - | - | - | - | Unassigned | - |
| Otu15306 | 1 | k__Fungi(100);k__Fungi_unclassified(100);k__Fungi_unclassified(100);k__Fungi_unclassified(100);k__Fungi_unclassified(100);k__Fungi_unclassified(100);k__Fungi_unclassified(100); | - | - | - | - | - | - | - | Unassigned | - |
| Otu15311 | 1 | k__Fungi(100);k__Fungi_unclassified(100);k__Fungi_unclassified(100);k__Fungi_unclassified(100);k__Fungi_unclassified(100);k__Fungi_unclassified(100);k__Fungi_unclassified(100); | - | - | - | - | - | - | - | Unassigned | - |
| Otu15312 | 1 | k__Fungi(100);k__Fungi_unclassified(100);k__Fungi_unclassified(100);k__Fungi_unclassified(100);k__Fungi_unclassified(100);k__Fungi_unclassified(100);k__Fungi_unclassified(100); | - | - | - | - | - | - | - | Unassigned | - |
| Otu15316 | 1 | k__Fungi(100);k__Fungi_unclassified(100);k__Fungi_unclassified(100);k__Fungi_unclassified(100);k__Fungi_unclassified(100);k__Fungi_unclassified(100);k__Fungi_unclassified(100); | - | - | - | - | - | - | - | Unassigned | - |
| Otu15318 | 1 | k__Fungi(100);k__Fungi_unclassified(100);k__Fungi_unclassified(100);k__Fungi_unclassified(100);k__Fungi_unclassified(100);k__Fungi_unclassified(100);k__Fungi_unclassified(100); | - | - | - | - | - | - | - | Unassigned | - |
| Otu15319 | 1 | k__Fungi(100);k__Fungi_unclassified(100);k__Fungi_unclassified(100);k__Fungi_unclassified(100);k__Fungi_unclassified(100);k__Fungi_unclassified(100);k__Fungi_unclassified(100); | - | - | - | - | - | - | - | Unassigned | - |
| Otu15326 | 1 | k__Fungi(100);p__Ascomycota(100);p__Ascomycota_unclassified(100);p__Ascomycota_unclassified(100);p__Ascomycota_unclassified(100);p__Ascomycota_unclassified(100);p__Ascomycota_unclassified(100); | - | - | - | - | - | - | - | Unassigned | - |
| Otu15328 | 1 | k__Fungi(100);k__Fungi_unclassified(100);k__Fungi_unclassified(100);k__Fungi_unclassified(100);k__Fungi_unclassified(100);k__Fungi_unclassified(100);k__Fungi_unclassified(100); | - | - | - | - | - | - | - | Unassigned | - |
| Otu15329 | 1 | k__Fungi(100);k__Fungi_unclassified(100);k__Fungi_unclassified(100);k__Fungi_unclassified(100);k__Fungi_unclassified(100);k__Fungi_unclassified(100);k__Fungi_unclassified(100); | - | - | - | - | - | - | - | Unassigned | - |
| Otu15331 | 1 | k__Fungi(100);p__Ascomycota(100);p__Ascomycota_unclassified(100);p__Ascomycota_unclassified(100);p__Ascomycota_unclassified(100);p__Ascomycota_unclassified(100);p__Ascomycota_unclassified(100); | - | - | - | - | - | - | - | Unassigned | - |
| Otu15332 | 1 | k__Fungi(100);k__Fungi_unclassified(100);k__Fungi_unclassified(100);k__Fungi_unclassified(100);k__Fungi_unclassified(100);k__Fungi_unclassified(100);k__Fungi_unclassified(100); | - | - | - | - | - | - | - | Unassigned | - |
| Otu15333 | 1 | k__Fungi(100);k__Fungi_unclassified(100);k__Fungi_unclassified(100);k__Fungi_unclassified(100);k__Fungi_unclassified(100);k__Fungi_unclassified(100);k__Fungi_unclassified(100); | - | - | - | - | - | - | - | Unassigned | - |
| Otu15335 | 1 | k__Fungi(100);k__Fungi_unclassified(100);k__Fungi_unclassified(100);k__Fungi_unclassified(100);k__Fungi_unclassified(100);k__Fungi_unclassified(100);k__Fungi_unclassified(100); | - | - | - | - | - | - | - | Unassigned | - |
| Otu15339 | 1 | k__Fungi(100);p__Ascomycota(100);p__Ascomycota_unclassified(100);p__Ascomycota_unclassified(100);p__Ascomycota_unclassified(100);p__Ascomycota_unclassified(100);p__Ascomycota_unclassified(100); | - | - | - | - | - | - | - | Unassigned | - |
| Otu15346 | 1 | k__Fungi(100);k__Fungi_unclassified(100);k__Fungi_unclassified(100);k__Fungi_unclassified(100);k__Fungi_unclassified(100);k__Fungi_unclassified(100);k__Fungi_unclassified(100); | - | - | - | - | - | - | - | Unassigned | - |
| Otu15350 | 1 | k__Fungi(100);p__Ascomycota(100);p__Ascomycota_unclassified(100);p__Ascomycota_unclassified(100);p__Ascomycota_unclassified(100);p__Ascomycota_unclassified(100);p__Ascomycota_unclassified(100); | - | - | - | - | - | - | - | Unassigned | - |
| Otu15354 | 1 | k__Fungi(100);k__Fungi_unclassified(100);k__Fungi_unclassified(100);k__Fungi_unclassified(100);k__Fungi_unclassified(100);k__Fungi_unclassified(100);k__Fungi_unclassified(100); | - | - | - | - | - | - | - | Unassigned | - |
| Otu15357 | 1 | k__Fungi(100);p__Ascomycota(100);p__Ascomycota_unclassified(100);p__Ascomycota_unclassified(100);p__Ascomycota_unclassified(100);p__Ascomycota_unclassified(100);p__Ascomycota_unclassified(100); | - | - | - | - | - | - | - | Unassigned | - |
| Otu15358 | 1 | k__Fungi(100);p__Ascomycota(100);p__Ascomycota_unclassified(100);p__Ascomycota_unclassified(100);p__Ascomycota_unclassified(100);p__Ascomycota_unclassified(100);p__Ascomycota_unclassified(100); | - | - | - | - | - | - | - | Unassigned | - |
| Otu15363 | 1 | k__Fungi(100);p__Ascomycota(100);p__Ascomycota_unclassified(100);p__Ascomycota_unclassified(100);p__Ascomycota_unclassified(100);p__Ascomycota_unclassified(100);p__Ascomycota_unclassified(100); | - | - | - | - | - | - | - | Unassigned | - |
| Otu15365 | 1 | k__Fungi(100);p__Ascomycota(100);p__Ascomycota_unclassified(100);p__Ascomycota_unclassified(100);p__Ascomycota_unclassified(100);p__Ascomycota_unclassified(100);p__Ascomycota_unclassified(100); | - | - | - | - | - | - | - | Unassigned | - |
| Otu15371 | 1 | k__Fungi(100);p__Ascomycota(100);p__Ascomycota_unclassified(100);p__Ascomycota_unclassified(100);p__Ascomycota_unclassified(100);p__Ascomycota_unclassified(100);p__Ascomycota_unclassified(100); | - | - | - | - | - | - | - | Unassigned | - |
| Otu15372 | 1 | k__Fungi(100);k__Fungi_unclassified(100);k__Fungi_unclassified(100);k__Fungi_unclassified(100);k__Fungi_unclassified(100);k__Fungi_unclassified(100);k__Fungi_unclassified(100); | - | - | - | - | - | - | - | Unassigned | - |
| Otu15376 | 1 | k__Fungi(100);k__Fungi_unclassified(100);k__Fungi_unclassified(100);k__Fungi_unclassified(100);k__Fungi_unclassified(100);k__Fungi_unclassified(100);k__Fungi_unclassified(100); | - | - | - | - | - | - | - | Unassigned | - |
| Otu15378 | 1 | k__Fungi(100);k__Fungi_unclassified(100);k__Fungi_unclassified(100);k__Fungi_unclassified(100);k__Fungi_unclassified(100);k__Fungi_unclassified(100);k__Fungi_unclassified(100); | - | - | - | - | - | - | - | Unassigned | - |
| Otu15379 | 1 | k__Fungi(100);k__Fungi_unclassified(100);k__Fungi_unclassified(100);k__Fungi_unclassified(100);k__Fungi_unclassified(100);k__Fungi_unclassified(100);k__Fungi_unclassified(100); | - | - | - | - | - | - | - | Unassigned | - |
| Otu15380 | 1 | k__Fungi(100);k__Fungi_unclassified(100);k__Fungi_unclassified(100);k__Fungi_unclassified(100);k__Fungi_unclassified(100);k__Fungi_unclassified(100);k__Fungi_unclassified(100); | - | - | - | - | - | - | - | Unassigned | - |
| Otu15385 | 1 | k__Fungi(100);k__Fungi_unclassified(100);k__Fungi_unclassified(100);k__Fungi_unclassified(100);k__Fungi_unclassified(100);k__Fungi_unclassified(100);k__Fungi_unclassified(100); | - | - | - | - | - | - | - | Unassigned | - |
| Otu15386 | 1 | k__Fungi(100);k__Fungi_unclassified(100);k__Fungi_unclassified(100);k__Fungi_unclassified(100);k__Fungi_unclassified(100);k__Fungi_unclassified(100);k__Fungi_unclassified(100); | - | - | - | - | - | - | - | Unassigned | - |
| Otu15387 | 1 | k__Fungi(100);k__Fungi_unclassified(100);k__Fungi_unclassified(100);k__Fungi_unclassified(100);k__Fungi_unclassified(100);k__Fungi_unclassified(100);k__Fungi_unclassified(100); | - | - | - | - | - | - | - | Unassigned | - |
| Otu15393 | 1 | k__Fungi(100);p__Ascomycota(100);p__Ascomycota_unclassified(100);p__Ascomycota_unclassified(100);p__Ascomycota_unclassified(100);p__Ascomycota_unclassified(100);p__Ascomycota_unclassified(100); | - | - | - | - | - | - | - | Unassigned | - |
| Otu15397 | 1 | k__Fungi(100);k__Fungi_unclassified(100);k__Fungi_unclassified(100);k__Fungi_unclassified(100);k__Fungi_unclassified(100);k__Fungi_unclassified(100);k__Fungi_unclassified(100); | - | - | - | - | - | - | - | Unassigned | - |
| Otu15407 | 1 | k__Fungi(100);p__Ascomycota(100);p__Ascomycota_unclassified(100);p__Ascomycota_unclassified(100);p__Ascomycota_unclassified(100);p__Ascomycota_unclassified(100);p__Ascomycota_unclassified(100); | - | - | - | - | - | - | - | Unassigned | - |
| Otu15408 | 1 | k__Fungi(100);k__Fungi_unclassified(100);k__Fungi_unclassified(100);k__Fungi_unclassified(100);k__Fungi_unclassified(100);k__Fungi_unclassified(100);k__Fungi_unclassified(100); | - | - | - | - | - | - | - | Unassigned | - |
| Otu15422 | 1 | k__Fungi(100);k__Fungi_unclassified(100);k__Fungi_unclassified(100);k__Fungi_unclassified(100);k__Fungi_unclassified(100);k__Fungi_unclassified(100);k__Fungi_unclassified(100); | - | - | - | - | - | - | - | Unassigned | - |
| Otu15426 | 1 | k__Fungi(100);p__Ascomycota(100);c__Leotiomycetes(100);c__Leotiomycetes_unclassified(100);c__Leotiomycetes_unclassified(100);c__Leotiomycetes_unclassified(100);c__Leotiomycetes_unclassified(100); | - | - | - | - | - | - | - | Unassigned | - |
| Otu15427 | 1 | k__Fungi(100);k__Fungi_unclassified(100);k__Fungi_unclassified(100);k__Fungi_unclassified(100);k__Fungi_unclassified(100);k__Fungi_unclassified(100);k__Fungi_unclassified(100); | - | - | - | - | - | - | - | Unassigned | - |
| Otu15428 | 1 | k__Fungi(100);k__Fungi_unclassified(100);k__Fungi_unclassified(100);k__Fungi_unclassified(100);k__Fungi_unclassified(100);k__Fungi_unclassified(100);k__Fungi_unclassified(100); | - | - | - | - | - | - | - | Unassigned | - |
| Otu15432 | 1 | k__Fungi(100);k__Fungi_unclassified(100);k__Fungi_unclassified(100);k__Fungi_unclassified(100);k__Fungi_unclassified(100);k__Fungi_unclassified(100);k__Fungi_unclassified(100); | - | - | - | - | - | - | - | Unassigned | - |
| Otu15434 | 1 | k__Fungi(100);p__Ascomycota(100);p__Ascomycota_unclassified(100);p__Ascomycota_unclassified(100);p__Ascomycota_unclassified(100);p__Ascomycota_unclassified(100);p__Ascomycota_unclassified(100); | - | - | - | - | - | - | - | Unassigned | - |
| Otu15436 | 1 | k__Fungi(100);k__Fungi_unclassified(100);k__Fungi_unclassified(100);k__Fungi_unclassified(100);k__Fungi_unclassified(100);k__Fungi_unclassified(100);k__Fungi_unclassified(100); | - | - | - | - | - | - | - | Unassigned | - |
| Otu15440 | 1 | k__Fungi(100);p__Ascomycota(100);p__Ascomycota_unclassified(100);p__Ascomycota_unclassified(100);p__Ascomycota_unclassified(100);p__Ascomycota_unclassified(100);p__Ascomycota_unclassified(100); | - | - | - | - | - | - | - | Unassigned | - |
| Otu15441 | 1 | k__Fungi(100);k__Fungi_unclassified(100);k__Fungi_unclassified(100);k__Fungi_unclassified(100);k__Fungi_unclassified(100);k__Fungi_unclassified(100);k__Fungi_unclassified(100); | - | - | - | - | - | - | - | Unassigned | - |
| Otu15455 | 1 | k__Fungi(100);k__Fungi_unclassified(100);k__Fungi_unclassified(100);k__Fungi_unclassified(100);k__Fungi_unclassified(100);k__Fungi_unclassified(100);k__Fungi_unclassified(100); | - | - | - | - | - | - | - | Unassigned | - |
| Otu15459 | 1 | k__Fungi(100);p__Ascomycota(100);p__Ascomycota_unclassified(100);p__Ascomycota_unclassified(100);p__Ascomycota_unclassified(100);p__Ascomycota_unclassified(100);p__Ascomycota_unclassified(100); | - | - | - | - | - | - | - | Unassigned | - |
| Otu15464 | 1 | k__Fungi(100);p__Ascomycota(100);p__Ascomycota_unclassified(100);p__Ascomycota_unclassified(100);p__Ascomycota_unclassified(100);p__Ascomycota_unclassified(100);p__Ascomycota_unclassified(100); | - | - | - | - | - | - | - | Unassigned | - |
| Otu15467 | 1 | k__Fungi(100);p__Ascomycota(100);p__Ascomycota_unclassified(100);p__Ascomycota_unclassified(100);p__Ascomycota_unclassified(100);p__Ascomycota_unclassified(100);p__Ascomycota_unclassified(100); | - | - | - | - | - | - | - | Unassigned | - |
| Otu15481 | 1 | k__Fungi(100);k__Fungi_unclassified(100);k__Fungi_unclassified(100);k__Fungi_unclassified(100);k__Fungi_unclassified(100);k__Fungi_unclassified(100);k__Fungi_unclassified(100); | - | - | - | - | - | - | - | Unassigned | - |
| Otu15489 | 1 | k__Fungi(100);k__Fungi_unclassified(100);k__Fungi_unclassified(100);k__Fungi_unclassified(100);k__Fungi_unclassified(100);k__Fungi_unclassified(100);k__Fungi_unclassified(100); | - | - | - | - | - | - | - | Unassigned | - |
| Otu15501 | 1 | k__Fungi(100);k__Fungi_unclassified(100);k__Fungi_unclassified(100);k__Fungi_unclassified(100);k__Fungi_unclassified(100);k__Fungi_unclassified(100);k__Fungi_unclassified(100); | - | - | - | - | - | - | - | Unassigned | - |
| Otu15505 | 1 | k__Fungi(100);p__Ascomycota(100);p__Ascomycota_unclassified(100);p__Ascomycota_unclassified(100);p__Ascomycota_unclassified(100);p__Ascomycota_unclassified(100);p__Ascomycota_unclassified(100); | - | - | - | - | - | - | - | Unassigned | - |
| Otu15509 | 1 | k__Fungi(100);p__Ascomycota(100);p__Ascomycota_unclassified(100);p__Ascomycota_unclassified(100);p__Ascomycota_unclassified(100);p__Ascomycota_unclassified(100);p__Ascomycota_unclassified(100); | - | - | - | - | - | - | - | Unassigned | - |
| Otu15512 | 1 | k__Fungi(100);k__Fungi_unclassified(100);k__Fungi_unclassified(100);k__Fungi_unclassified(100);k__Fungi_unclassified(100);k__Fungi_unclassified(100);k__Fungi_unclassified(100); | - | - | - | - | - | - | - | Unassigned | - |
| Otu15517 | 1 | k__Fungi(100);k__Fungi_unclassified(100);k__Fungi_unclassified(100);k__Fungi_unclassified(100);k__Fungi_unclassified(100);k__Fungi_unclassified(100);k__Fungi_unclassified(100); | - | - | - | - | - | - | - | Unassigned | - |
| Otu15522 | 1 | k__Fungi(100);k__Fungi_unclassified(100);k__Fungi_unclassified(100);k__Fungi_unclassified(100);k__Fungi_unclassified(100);k__Fungi_unclassified(100);k__Fungi_unclassified(100); | - | - | - | - | - | - | - | Unassigned | - |
| Otu15531 | 1 | k__Fungi(100);p__Ascomycota(100);p__Ascomycota_unclassified(100);p__Ascomycota_unclassified(100);p__Ascomycota_unclassified(100);p__Ascomycota_unclassified(100);p__Ascomycota_unclassified(100); | - | - | - | - | - | - | - | Unassigned | - |
| Otu15534 | 1 | k__Fungi(100);k__Fungi_unclassified(100);k__Fungi_unclassified(100);k__Fungi_unclassified(100);k__Fungi_unclassified(100);k__Fungi_unclassified(100);k__Fungi_unclassified(100); | - | - | - | - | - | - | - | Unassigned | - |
| Otu15557 | 1 | k__Fungi(100);k__Fungi_unclassified(100);k__Fungi_unclassified(100);k__Fungi_unclassified(100);k__Fungi_unclassified(100);k__Fungi_unclassified(100);k__Fungi_unclassified(100); | - | - | - | - | - | - | - | Unassigned | - |
| Otu15558 | 1 | k__Fungi(100);k__Fungi_unclassified(100);k__Fungi_unclassified(100);k__Fungi_unclassified(100);k__Fungi_unclassified(100);k__Fungi_unclassified(100);k__Fungi_unclassified(100); | - | - | - | - | - | - | - | Unassigned | - |
| Otu15565 | 1 | k__Fungi(100);k__Fungi_unclassified(100);k__Fungi_unclassified(100);k__Fungi_unclassified(100);k__Fungi_unclassified(100);k__Fungi_unclassified(100);k__Fungi_unclassified(100); | - | - | - | - | - | - | - | Unassigned | - |
| Otu15566 | 1 | k__Fungi(100);k__Fungi_unclassified(100);k__Fungi_unclassified(100);k__Fungi_unclassified(100);k__Fungi_unclassified(100);k__Fungi_unclassified(100);k__Fungi_unclassified(100); | - | - | - | - | - | - | - | Unassigned | - |
| Otu15571 | 1 | k__Fungi(100);p__Ascomycota(100);p__Ascomycota_unclassified(100);p__Ascomycota_unclassified(100);p__Ascomycota_unclassified(100);p__Ascomycota_unclassified(100);p__Ascomycota_unclassified(100); | - | - | - | - | - | - | - | Unassigned | - |
| Otu15573 | 1 | k__Fungi(100);p__Ascomycota(100);p__Ascomycota_unclassified(100);p__Ascomycota_unclassified(100);p__Ascomycota_unclassified(100);p__Ascomycota_unclassified(100);p__Ascomycota_unclassified(100); | - | - | - | - | - | - | - | Unassigned | - |
| Otu15575 | 1 | k__Fungi(100);k__Fungi_unclassified(100);k__Fungi_unclassified(100);k__Fungi_unclassified(100);k__Fungi_unclassified(100);k__Fungi_unclassified(100);k__Fungi_unclassified(100); | - | - | - | - | - | - | - | Unassigned | - |
| Otu15592 | 1 | k__Fungi(100);k__Fungi_unclassified(100);k__Fungi_unclassified(100);k__Fungi_unclassified(100);k__Fungi_unclassified(100);k__Fungi_unclassified(100);k__Fungi_unclassified(100); | - | - | - | - | - | - | - | Unassigned | - |
| Otu15598 | 1 | k__Fungi(100);k__Fungi_unclassified(100);k__Fungi_unclassified(100);k__Fungi_unclassified(100);k__Fungi_unclassified(100);k__Fungi_unclassified(100);k__Fungi_unclassified(100); | - | - | - | - | - | - | - | Unassigned | - |
| Otu15605 | 1 | k__Fungi(100);k__Fungi_unclassified(100);k__Fungi_unclassified(100);k__Fungi_unclassified(100);k__Fungi_unclassified(100);k__Fungi_unclassified(100);k__Fungi_unclassified(100); | - | - | - | - | - | - | - | Unassigned | - |
| Otu15611 | 1 | k__Fungi(100);k__Fungi_unclassified(100);k__Fungi_unclassified(100);k__Fungi_unclassified(100);k__Fungi_unclassified(100);k__Fungi_unclassified(100);k__Fungi_unclassified(100); | - | - | - | - | - | - | - | Unassigned | - |
| Otu15617 | 1 | k__Fungi(100);k__Fungi_unclassified(100);k__Fungi_unclassified(100);k__Fungi_unclassified(100);k__Fungi_unclassified(100);k__Fungi_unclassified(100);k__Fungi_unclassified(100); | - | - | - | - | - | - | - | Unassigned | - |
| Otu15620 | 1 | k__Fungi(100);k__Fungi_unclassified(100);k__Fungi_unclassified(100);k__Fungi_unclassified(100);k__Fungi_unclassified(100);k__Fungi_unclassified(100);k__Fungi_unclassified(100); | - | - | - | - | - | - | - | Unassigned | - |
| Otu15639 | 1 | k__Fungi(100);p__Ascomycota(100);c__Dothideomycetes(100);c__Dothideomycetes_unclassified(100);c__Dothideomycetes_unclassified(100);c__Dothideomycetes_unclassified(100);c__Dothideomycetes_unclassified(100); | - | - | - | - | - | - | - | Unassigned | - |
| Otu15640 | 1 | k__Fungi(100);k__Fungi_unclassified(100);k__Fungi_unclassified(100);k__Fungi_unclassified(100);k__Fungi_unclassified(100);k__Fungi_unclassified(100);k__Fungi_unclassified(100); | - | - | - | - | - | - | - | Unassigned | - |
| Otu15641 | 1 | k__Fungi(100);p__Ascomycota(100);p__Ascomycota_unclassified(100);p__Ascomycota_unclassified(100);p__Ascomycota_unclassified(100);p__Ascomycota_unclassified(100);p__Ascomycota_unclassified(100); | - | - | - | - | - | - | - | Unassigned | - |
| Otu15642 | 1 | k__Fungi(100);k__Fungi_unclassified(100);k__Fungi_unclassified(100);k__Fungi_unclassified(100);k__Fungi_unclassified(100);k__Fungi_unclassified(100);k__Fungi_unclassified(100); | - | - | - | - | - | - | - | Unassigned | - |
| Otu15645 | 1 | k__Fungi(100);k__Fungi_unclassified(100);k__Fungi_unclassified(100);k__Fungi_unclassified(100);k__Fungi_unclassified(100);k__Fungi_unclassified(100);k__Fungi_unclassified(100); | - | - | - | - | - | - | - | Unassigned | - |
| Otu15647 | 1 | k__Fungi(100);k__Fungi_unclassified(100);k__Fungi_unclassified(100);k__Fungi_unclassified(100);k__Fungi_unclassified(100);k__Fungi_unclassified(100);k__Fungi_unclassified(100); | - | - | - | - | - | - | - | Unassigned | - |
| Otu15650 | 1 | k__Fungi(100);k__Fungi_unclassified(100);k__Fungi_unclassified(100);k__Fungi_unclassified(100);k__Fungi_unclassified(100);k__Fungi_unclassified(100);k__Fungi_unclassified(100); | - | - | - | - | - | - | - | Unassigned | - |
| Otu15651 | 1 | k__Fungi(100);k__Fungi_unclassified(100);k__Fungi_unclassified(100);k__Fungi_unclassified(100);k__Fungi_unclassified(100);k__Fungi_unclassified(100);k__Fungi_unclassified(100); | - | - | - | - | - | - | - | Unassigned | - |
| Otu15653 | 1 | k__Fungi(100);k__Fungi_unclassified(100);k__Fungi_unclassified(100);k__Fungi_unclassified(100);k__Fungi_unclassified(100);k__Fungi_unclassified(100);k__Fungi_unclassified(100); | - | - | - | - | - | - | - | Unassigned | - |
| Otu15654 | 1 | k__Fungi(100);k__Fungi_unclassified(100);k__Fungi_unclassified(100);k__Fungi_unclassified(100);k__Fungi_unclassified(100);k__Fungi_unclassified(100);k__Fungi_unclassified(100); | - | - | - | - | - | - | - | Unassigned | - |
| Otu15655 | 1 | k__Fungi(100);k__Fungi_unclassified(100);k__Fungi_unclassified(100);k__Fungi_unclassified(100);k__Fungi_unclassified(100);k__Fungi_unclassified(100);k__Fungi_unclassified(100); | - | - | - | - | - | - | - | Unassigned | - |
| Otu15658 | 1 | k__Fungi(100);k__Fungi_unclassified(100);k__Fungi_unclassified(100);k__Fungi_unclassified(100);k__Fungi_unclassified(100);k__Fungi_unclassified(100);k__Fungi_unclassified(100); | - | - | - | - | - | - | - | Unassigned | - |
| Otu15659 | 1 | k__Fungi(100);k__Fungi_unclassified(100);k__Fungi_unclassified(100);k__Fungi_unclassified(100);k__Fungi_unclassified(100);k__Fungi_unclassified(100);k__Fungi_unclassified(100); | - | - | - | - | - | - | - | Unassigned | - |

| | | | | | | | | | | | |
|---|---|---|---|---|---|---|---|---|---|---|---|
| Otu15661 | 1 | k__Fungi(100);p__Ascomycota(100);p__Ascomycota_unclassified(100);p__Ascomycota_unclassified(100);p__Ascomycota_unclassified(100);p__Ascomycota_unclassified(100);p__Ascomycota_unclassified(100); | - | - | - | - | - | - | - | Unassigned | - |
| Otu15662 | 1 | k__Fungi(100);k__Fungi_unclassified(100);k__Fungi_unclassified(100);k__Fungi_unclassified(100);k__Fungi_unclassified(100);k__Fungi_unclassified(100);k__Fungi_unclassified(100); | - | - | - | - | - | - | - | Unassigned | - |
| Otu15665 | 1 | k__Fungi(100);p__Ascomycota(100);p__Ascomycota_unclassified(100);p__Ascomycota_unclassified(100);p__Ascomycota_unclassified(100);p__Ascomycota_unclassified(100);p__Ascomycota_unclassified(100); | - | - | - | - | - | - | - | Unassigned | - |
| Otu15675 | 1 | k__Fungi(100);p__Ascomycota(100);p__Ascomycota_unclassified(100);p__Ascomycota_unclassified(100);p__Ascomycota_unclassified(100);p__Ascomycota_unclassified(100);p__Ascomycota_unclassified(100); | - | - | - | - | - | - | - | Unassigned | - |
| Otu15676 | 1 | k__Fungi(100);k__Fungi_unclassified(100);k__Fungi_unclassified(100);k__Fungi_unclassified(100);k__Fungi_unclassified(100);k__Fungi_unclassified(100);k__Fungi_unclassified(100); | - | - | - | - | - | - | - | Unassigned | - |
| Otu15678 | 1 | k__Fungi(100);p__Ascomycota(100);p__Ascomycota_unclassified(100);p__Ascomycota_unclassified(100);p__Ascomycota_unclassified(100);p__Ascomycota_unclassified(100);p__Ascomycota_unclassified(100); | - | - | - | - | - | - | - | Unassigned | - |
| Otu15684 | 1 | k__Fungi(100);k__Fungi_unclassified(100);k__Fungi_unclassified(100);k__Fungi_unclassified(100);k__Fungi_unclassified(100);k__Fungi_unclassified(100);k__Fungi_unclassified(100); | - | - | - | - | - | - | - | Unassigned | - |
| Otu15685 | 1 | k__Fungi(100);p__Ascomycota(100);p__Ascomycota_unclassified(100);p__Ascomycota_unclassified(100);p__Ascomycota_unclassified(100);p__Ascomycota_unclassified(100);p__Ascomycota_unclassified(100); | - | - | - | - | - | - | - | Unassigned | - |
| Otu15688 | 1 | k__Fungi(100);k__Fungi_unclassified(100);k__Fungi_unclassified(100);k__Fungi_unclassified(100);k__Fungi_unclassified(100);k__Fungi_unclassified(100);k__Fungi_unclassified(100); | - | - | - | - | - | - | - | Unassigned | - |
| Otu15689 | 1 | k__Fungi(100);k__Fungi_unclassified(100);k__Fungi_unclassified(100);k__Fungi_unclassified(100);k__Fungi_unclassified(100);k__Fungi_unclassified(100);k__Fungi_unclassified(100); | - | - | - | - | - | - | - | Unassigned | - |
| Otu15690 | 1 | k__Fungi(100);k__Fungi_unclassified(100);k__Fungi_unclassified(100);k__Fungi_unclassified(100);k__Fungi_unclassified(100);k__Fungi_unclassified(100);k__Fungi_unclassified(100); | - | - | - | - | - | - | - | Unassigned | - |
| Otu15696 | 1 | k__Fungi(100);k__Fungi_unclassified(100);k__Fungi_unclassified(100);k__Fungi_unclassified(100);k__Fungi_unclassified(100);k__Fungi_unclassified(100);k__Fungi_unclassified(100); | - | - | - | - | - | - | - | Unassigned | - |
| Otu15704 | 1 | k__Fungi(100);p__Ascomycota(100);p__Ascomycota_unclassified(100);p__Ascomycota_unclassified(100);p__Ascomycota_unclassified(100);p__Ascomycota_unclassified(100);p__Ascomycota_unclassified(100); | - | - | - | - | - | - | - | Unassigned | - |
| Otu15705 | 1 | k__Fungi(100);k__Fungi_unclassified(100);k__Fungi_unclassified(100);k__Fungi_unclassified(100);k__Fungi_unclassified(100);k__Fungi_unclassified(100);k__Fungi_unclassified(100); | - | - | - | - | - | - | - | Unassigned | - |
| Otu15710 | 1 | k__Fungi(100);k__Fungi_unclassified(100);k__Fungi_unclassified(100);k__Fungi_unclassified(100);k__Fungi_unclassified(100);k__Fungi_unclassified(100);k__Fungi_unclassified(100); | - | - | - | - | - | - | - | Unassigned | - |
| Otu15712 | 1 | k__Fungi(100);k__Fungi_unclassified(100);k__Fungi_unclassified(100);k__Fungi_unclassified(100);k__Fungi_unclassified(100);k__Fungi_unclassified(100);k__Fungi_unclassified(100); | - | - | - | - | - | - | - | Unassigned | - |
| Otu15714 | 1 | k__Fungi(100);p__Ascomycota(100);p__Ascomycota_unclassified(100);p__Ascomycota_unclassified(100);p__Ascomycota_unclassified(100);p__Ascomycota_unclassified(100);p__Ascomycota_unclassified(100); | - | - | - | - | - | - | - | Unassigned | - |
| Otu15715 | 1 | k__Fungi(100);p__Ascomycota(100);p__Ascomycota_unclassified(100);p__Ascomycota_unclassified(100);p__Ascomycota_unclassified(100);p__Ascomycota_unclassified(100);p__Ascomycota_unclassified(100); | - | - | - | - | - | - | - | Unassigned | - |
| Otu15717 | 1 | k__Fungi(100);k__Fungi_unclassified(100);k__Fungi_unclassified(100);k__Fungi_unclassified(100);k__Fungi_unclassified(100);k__Fungi_unclassified(100);k__Fungi_unclassified(100); | - | - | - | - | - | - | - | Unassigned | - |
| Otu15718 | 1 | k__Fungi(100);p__Ascomycota(100);p__Ascomycota_unclassified(100);p__Ascomycota_unclassified(100);p__Ascomycota_unclassified(100);p__Ascomycota_unclassified(100);p__Ascomycota_unclassified(100); | - | - | - | - | - | - | - | Unassigned | - |
| Otu15720 | 1 | k__Fungi(100);k__Fungi_unclassified(100);k__Fungi_unclassified(100);k__Fungi_unclassified(100);k__Fungi_unclassified(100);k__Fungi_unclassified(100);k__Fungi_unclassified(100); | - | - | - | - | - | - | - | Unassigned | - |
| Otu15724 | 1 | k__Fungi(100);p__Ascomycota(100);p__Ascomycota_unclassified(100);p__Ascomycota_unclassified(100);p__Ascomycota_unclassified(100);p__Ascomycota_unclassified(100);p__Ascomycota_unclassified(100); | - | - | - | - | - | - | - | Unassigned | - |
| Otu15725 | 1 | k__Fungi(100);k__Fungi_unclassified(100);k__Fungi_unclassified(100);k__Fungi_unclassified(100);k__Fungi_unclassified(100);k__Fungi_unclassified(100);k__Fungi_unclassified(100); | - | - | - | - | - | - | - | Unassigned | - |
| Otu15727 | 1 | k__Fungi(100);k__Fungi_unclassified(100);k__Fungi_unclassified(100);k__Fungi_unclassified(100);k__Fungi_unclassified(100);k__Fungi_unclassified(100);k__Fungi_unclassified(100); | - | - | - | - | - | - | - | Unassigned | - |
| Otu15729 | 1 | k__Fungi(100);k__Fungi_unclassified(100);k__Fungi_unclassified(100);k__Fungi_unclassified(100);k__Fungi_unclassified(100);k__Fungi_unclassified(100);k__Fungi_unclassified(100); | - | - | - | - | - | - | - | Unassigned | - |
| Otu15742 | 1 | k__Fungi(100);p__Ascomycota(100);p__Ascomycota_unclassified(100);p__Ascomycota_unclassified(100);p__Ascomycota_unclassified(100);p__Ascomycota_unclassified(100);p__Ascomycota_unclassified(100); | - | - | - | - | - | - | - | Unassigned | - |
| Otu15744 | 1 | k__Fungi(100);k__Fungi_unclassified(100);k__Fungi_unclassified(100);k__Fungi_unclassified(100);k__Fungi_unclassified(100);k__Fungi_unclassified(100);k__Fungi_unclassified(100); | - | - | - | - | - | - | - | Unassigned | - |
| Otu15746 | 1 | k__Fungi(100);k__Fungi_unclassified(100);k__Fungi_unclassified(100);k__Fungi_unclassified(100);k__Fungi_unclassified(100);k__Fungi_unclassified(100);k__Fungi_unclassified(100); | - | - | - | - | - | - | - | Unassigned | - |
| Otu15752 | 1 | k__Fungi(100);k__Fungi_unclassified(100);k__Fungi_unclassified(100);k__Fungi_unclassified(100);k__Fungi_unclassified(100);k__Fungi_unclassified(100);k__Fungi_unclassified(100); | - | - | - | - | - | - | - | Unassigned | - |
| Otu15759 | 1 | k__Fungi(100);p__Ascomycota(100);p__Ascomycota_unclassified(100);p__Ascomycota_unclassified(100);p__Ascomycota_unclassified(100);p__Ascomycota_unclassified(100);p__Ascomycota_unclassified(100); | - | - | - | - | - | - | - | Unassigned | - |
| Otu15762 | 1 | k__Fungi(100);k__Fungi_unclassified(100);k__Fungi_unclassified(100);k__Fungi_unclassified(100);k__Fungi_unclassified(100);k__Fungi_unclassified(100);k__Fungi_unclassified(100); | - | - | - | - | - | - | - | Unassigned | - |
| Otu15767 | 1 | k__Fungi(100);k__Fungi_unclassified(100);k__Fungi_unclassified(100);k__Fungi_unclassified(100);k__Fungi_unclassified(100);k__Fungi_unclassified(100);k__Fungi_unclassified(100); | - | - | - | - | - | - | - | Unassigned | - |
| Otu15769 | 1 | k__Fungi(100);k__Fungi_unclassified(100);k__Fungi_unclassified(100);k__Fungi_unclassified(100);k__Fungi_unclassified(100);k__Fungi_unclassified(100);k__Fungi_unclassified(100); | - | - | - | - | - | - | - | Unassigned | - |
| Otu15787 | 1 | k__Fungi(100);p__Ascomycota(100);p__Ascomycota_unclassified(100);p__Ascomycota_unclassified(100);p__Ascomycota_unclassified(100);p__Ascomycota_unclassified(100);p__Ascomycota_unclassified(100); | - | - | - | - | - | - | - | Unassigned | - |
| Otu15798 | 1 | k__Fungi(100);p__Ascomycota(100);p__Ascomycota_unclassified(100);p__Ascomycota_unclassified(100);p__Ascomycota_unclassified(100);p__Ascomycota_unclassified(100);p__Ascomycota_unclassified(100); | - | - | - | - | - | - | - | Unassigned | - |
| Otu15800 | 1 | k__Fungi(100);k__Fungi_unclassified(100);k__Fungi_unclassified(100);k__Fungi_unclassified(100);k__Fungi_unclassified(100);k__Fungi_unclassified(100);k__Fungi_unclassified(100); | - | - | - | - | - | - | - | Unassigned | - |
| Otu15805 | 1 | k__Fungi(100);p__Ascomycota(100);p__Ascomycota_unclassified(100);p__Ascomycota_unclassified(100);p__Ascomycota_unclassified(100);p__Ascomycota_unclassified(100);p__Ascomycota_unclassified(100); | - | - | - | - | - | - | - | Unassigned | - |
| Otu15823 | 1 | k__Fungi(100);k__Fungi_unclassified(100);k__Fungi_unclassified(100);k__Fungi_unclassified(100);k__Fungi_unclassified(100);k__Fungi_unclassified(100);k__Fungi_unclassified(100); | - | - | - | - | - | - | - | Unassigned | - |
| Otu15824 | 1 | k__Fungi(100);p__Ascomycota(100);p__Ascomycota_unclassified(100);p__Ascomycota_unclassified(100);p__Ascomycota_unclassified(100);p__Ascomycota_unclassified(100);p__Ascomycota_unclassified(100); | - | - | - | - | - | - | - | Unassigned | - |
| Otu15832 | 1 | k__Fungi(100);p__Ascomycota(100);p__Ascomycota_unclassified(100);p__Ascomycota_unclassified(100);p__Ascomycota_unclassified(100);p__Ascomycota_unclassified(100);p__Ascomycota_unclassified(100); | - | - | - | - | - | - | - | Unassigned | - |
| Otu15835 | 1 | k__Fungi(100);p__Ascomycota(100);p__Ascomycota_unclassified(100);p__Ascomycota_unclassified(100);p__Ascomycota_unclassified(100);p__Ascomycota_unclassified(100);p__Ascomycota_unclassified(100); | - | - | - | - | - | - | - | Unassigned | - |
| Otu15837 | 1 | k__Fungi(100);p__Ascomycota(100);p__Ascomycota_unclassified(100);p__Ascomycota_unclassified(100);p__Ascomycota_unclassified(100);p__Ascomycota_unclassified(100);p__Ascomycota_unclassified(100); | - | - | - | - | - | - | - | Unassigned | - |
| Otu15847 | 1 | k__Fungi(100);k__Fungi_unclassified(100);k__Fungi_unclassified(100);k__Fungi_unclassified(100);k__Fungi_unclassified(100);k__Fungi_unclassified(100);k__Fungi_unclassified(100); | - | - | - | - | - | - | - | Unassigned | - |
| Otu15896 | 1 | k__Fungi(100);k__Fungi_unclassified(100);k__Fungi_unclassified(100);k__Fungi_unclassified(100);k__Fungi_unclassified(100);k__Fungi_unclassified(100);k__Fungi_unclassified(100); | - | - | - | - | - | - | - | Unassigned | - |
| Otu15907 | 1 | k__Fungi(100);k__Fungi_unclassified(100);k__Fungi_unclassified(100);k__Fungi_unclassified(100);k__Fungi_unclassified(100);k__Fungi_unclassified(100);k__Fungi_unclassified(100); | - | - | - | - | - | - | - | Unassigned | - |
| Otu15908 | 1 | k__Fungi(100);k__Fungi_unclassified(100);k__Fungi_unclassified(100);k__Fungi_unclassified(100);k__Fungi_unclassified(100);k__Fungi_unclassified(100);k__Fungi_unclassified(100); | - | - | - | - | - | - | - | Unassigned | - |
| Otu15926 | 1 | k__Fungi(100);k__Fungi_unclassified(100);k__Fungi_unclassified(100);k__Fungi_unclassified(100);k__Fungi_unclassified(100);k__Fungi_unclassified(100);k__Fungi_unclassified(100); | - | - | - | - | - | - | - | Unassigned | - |
| Otu15928 | 1 | k__Fungi(100);k__Fungi_unclassified(100);k__Fungi_unclassified(100);k__Fungi_unclassified(100);k__Fungi_unclassified(100);k__Fungi_unclassified(100);k__Fungi_unclassified(100); | - | - | - | - | - | - | - | Unassigned | - |
| Otu15929 | 1 | k__Fungi(100);k__Fungi_unclassified(100);k__Fungi_unclassified(100);k__Fungi_unclassified(100);k__Fungi_unclassified(100);k__Fungi_unclassified(100);k__Fungi_unclassified(100); | - | - | - | - | - | - | - | Unassigned | - |
| Otu15931 | 1 | k__Fungi(100);k__Fungi_unclassified(100);k__Fungi_unclassified(100);k__Fungi_unclassified(100);k__Fungi_unclassified(100);k__Fungi_unclassified(100);k__Fungi_unclassified(100); | - | - | - | - | - | - | - | Unassigned | - |
| Otu15936 | 1 | k__Fungi(100);p__Ascomycota(100);p__Ascomycota_unclassified(100);p__Ascomycota_unclassified(100);p__Ascomycota_unclassified(100);p__Ascomycota_unclassified(100);p__Ascomycota_unclassified(100); | - | - | - | - | - | - | - | Unassigned | - |
| Otu15939 | 1 | k__Fungi(100);k__Fungi_unclassified(100);k__Fungi_unclassified(100);k__Fungi_unclassified(100);k__Fungi_unclassified(100);k__Fungi_unclassified(100);k__Fungi_unclassified(100); | - | - | - | - | - | - | - | Unassigned | - |
| Otu15963 | 1 | k__Fungi(100);k__Fungi_unclassified(100);k__Fungi_unclassified(100);k__Fungi_unclassified(100);k__Fungi_unclassified(100);k__Fungi_unclassified(100);k__Fungi_unclassified(100); | - | - | - | - | - | - | - | Unassigned | - |
| Otu15980 | 1 | k__Fungi(100);k__Fungi_unclassified(100);k__Fungi_unclassified(100);k__Fungi_unclassified(100);k__Fungi_unclassified(100);k__Fungi_unclassified(100);k__Fungi_unclassified(100); | - | - | - | - | - | - | - | Unassigned | - |
| Otu15984 | 1 | k__Fungi(100);p__Ascomycota(100);p__Ascomycota_unclassified(100);p__Ascomycota_unclassified(100);p__Ascomycota_unclassified(100);p__Ascomycota_unclassified(100);p__Ascomycota_unclassified(100); | - | - | - | - | - | - | - | Unassigned | - |
| Otu15986 | 1 | k__Fungi(100);k__Fungi_unclassified(100);k__Fungi_unclassified(100);k__Fungi_unclassified(100);k__Fungi_unclassified(100);k__Fungi_unclassified(100);k__Fungi_unclassified(100); | - | - | - | - | - | - | - | Unassigned | - |
| Otu15989 | 1 | k__Fungi(100);p__Ascomycota(100);p__Ascomycota_unclassified(100);p__Ascomycota_unclassified(100);p__Ascomycota_unclassified(100);p__Ascomycota_unclassified(100);p__Ascomycota_unclassified(100); | - | - | - | - | - | - | - | Unassigned | - |
| Otu15990 | 1 | k__Fungi(100);k__Fungi_unclassified(100);k__Fungi_unclassified(100);k__Fungi_unclassified(100);k__Fungi_unclassified(100);k__Fungi_unclassified(100);k__Fungi_unclassified(100); | - | - | - | - | - | - | - | Unassigned | - |
| Otu15998 | 1 | k__Fungi(100);k__Fungi_unclassified(100);k__Fungi_unclassified(100);k__Fungi_unclassified(100);k__Fungi_unclassified(100);k__Fungi_unclassified(100);k__Fungi_unclassified(100); | - | - | - | - | - | - | - | Unassigned | - |
| Otu16004 | 1 | k__Fungi(100);k__Fungi_unclassified(100);k__Fungi_unclassified(100);k__Fungi_unclassified(100);k__Fungi_unclassified(100);k__Fungi_unclassified(100);k__Fungi_unclassified(100); | - | - | - | - | - | - | - | Unassigned | - |
| Otu16005 | 1 | k__Fungi(100);p__Ascomycota(100);p__Ascomycota_unclassified(100);p__Ascomycota_unclassified(100);p__Ascomycota_unclassified(100);p__Ascomycota_unclassified(100);p__Ascomycota_unclassified(100); | - | - | - | - | - | - | - | Unassigned | - |
| Otu16006 | 1 | k__Fungi(100);p__Ascomycota(100);p__Ascomycota_unclassified(100);p__Ascomycota_unclassified(100);p__Ascomycota_unclassified(100);p__Ascomycota_unclassified(100);p__Ascomycota_unclassified(100); | - | - | - | - | - | - | - | Unassigned | - |
| Otu16009 | 1 | k__Fungi(100);k__Fungi_unclassified(100);k__Fungi_unclassified(100);k__Fungi_unclassified(100);k__Fungi_unclassified(100);k__Fungi_unclassified(100);k__Fungi_unclassified(100); | - | - | - | - | - | - | - | Unassigned | - |
| Otu16010 | 1 | k__Fungi(100);p__Ascomycota(100);p__Ascomycota_unclassified(100);p__Ascomycota_unclassified(100);p__Ascomycota_unclassified(100);p__Ascomycota_unclassified(100);p__Ascomycota_unclassified(100); | - | - | - | - | - | - | - | Unassigned | - |
| Otu16011 | 1 | k__Fungi(100);p__Ascomycota(100);p__Ascomycota_unclassified(100);p__Ascomycota_unclassified(100);p__Ascomycota_unclassified(100);p__Ascomycota_unclassified(100);p__Ascomycota_unclassified(100); | - | - | - | - | - | - | - | Unassigned | - |
| Otu16012 | 1 | k__Fungi(100);p__Ascomycota(100);p__Ascomycota_unclassified(100);p__Ascomycota_unclassified(100);p__Ascomycota_unclassified(100);p__Ascomycota_unclassified(100);p__Ascomycota_unclassified(100); | - | - | - | - | - | - | - | Unassigned | - |
| Otu16014 | 1 | k__Fungi(100);p__Ascomycota(100);p__Ascomycota_unclassified(100);p__Ascomycota_unclassified(100);p__Ascomycota_unclassified(100);p__Ascomycota_unclassified(100);p__Ascomycota_unclassified(100); | - | - | - | - | - | - | - | Unassigned | - |
| Otu16015 | 1 | k__Fungi(100);p__Ascomycota(100);p__Ascomycota_unclassified(100);p__Ascomycota_unclassified(100);p__Ascomycota_unclassified(100);p__Ascomycota_unclassified(100);p__Ascomycota_unclassified(100); | - | - | - | - | - | - | - | Unassigned | - |
| Otu16016 | 1 | k__Fungi(100);p__Ascomycota(100);p__Ascomycota_unclassified(100);p__Ascomycota_unclassified(100);p__Ascomycota_unclassified(100);p__Ascomycota_unclassified(100);p__Ascomycota_unclassified(100); | - | - | - | - | - | - | - | Unassigned | - |
| Otu16019 | 1 | k__Fungi(100);k__Fungi_unclassified(100);k__Fungi_unclassified(100);k__Fungi_unclassified(100);k__Fungi_unclassified(100);k__Fungi_unclassified(100);k__Fungi_unclassified(100); | - | - | - | - | - | - | - | Unassigned | - |
| Otu16021 | 1 | k__Fungi(100);k__Fungi_unclassified(100);k__Fungi_unclassified(100);k__Fungi_unclassified(100);k__Fungi_unclassified(100);k__Fungi_unclassified(100);k__Fungi_unclassified(100); | - | - | - | - | - | - | - | Unassigned | - |
| Otu16024 | 1 | k__Fungi(100);p__Ascomycota(100);p__Ascomycota_unclassified(100);p__Ascomycota_unclassified(100);p__Ascomycota_unclassified(100);p__Ascomycota_unclassified(100);p__Ascomycota_unclassified(100); | - | - | - | - | - | - | - | Unassigned | - |
| Otu16027 | 1 | k__Fungi(100);p__Ascomycota(100);p__Ascomycota_unclassified(100);p__Ascomycota_unclassified(100);p__Ascomycota_unclassified(100);p__Ascomycota_unclassified(100);p__Ascomycota_unclassified(100); | - | - | - | - | - | - | - | Unassigned | - |
| Otu16029 | 1 | k__Fungi(100);k__Fungi_unclassified(100);k__Fungi_unclassified(100);k__Fungi_unclassified(100);k__Fungi_unclassified(100);k__Fungi_unclassified(100);k__Fungi_unclassified(100); | - | - | - | - | - | - | - | Unassigned | - |
| Otu16030 | 1 | k__Fungi(100);k__Fungi_unclassified(100);k__Fungi_unclassified(100);k__Fungi_unclassified(100);k__Fungi_unclassified(100);k__Fungi_unclassified(100);k__Fungi_unclassified(100); | - | - | - | - | - | - | - | Unassigned | - |
| Otu16031 | 1 | k__Fungi(100);k__Fungi_unclassified(100);k__Fungi_unclassified(100);k__Fungi_unclassified(100);k__Fungi_unclassified(100);k__Fungi_unclassified(100);k__Fungi_unclassified(100); | - | - | - | - | - | - | - | Unassigned | - |
| Otu16032 | 1 | k__Fungi(100);k__Fungi_unclassified(100);k__Fungi_unclassified(100);k__Fungi_unclassified(100);k__Fungi_unclassified(100);k__Fungi_unclassified(100);k__Fungi_unclassified(100); | - | - | - | - | - | - | - | Unassigned | - |
| Otu16037 | 1 | k__Fungi(100);p__Ascomycota(100);p__Ascomycota_unclassified(100);p__Ascomycota_unclassified(100);p__Ascomycota_unclassified(100);p__Ascomycota_unclassified(100);p__Ascomycota_unclassified(100); | - | - | - | - | - | - | - | Unassigned | - |
| Otu16041 | 1 | k__Fungi(100);k__Fungi_unclassified(100);k__Fungi_unclassified(100);k__Fungi_unclassified(100);k__Fungi_unclassified(100);k__Fungi_unclassified(100);k__Fungi_unclassified(100); | - | - | - | - | - | - | - | Unassigned | - |
| Otu16045 | 1 | k__Fungi(100);k__Fungi_unclassified(100);k__Fungi_unclassified(100);k__Fungi_unclassified(100);k__Fungi_unclassified(100);k__Fungi_unclassified(100);k__Fungi_unclassified(100); | - | - | - | - | - | - | - | Unassigned | - |
| Otu16046 | 1 | k__Fungi(100);k__Fungi_unclassified(100);k__Fungi_unclassified(100);k__Fungi_unclassified(100);k__Fungi_unclassified(100);k__Fungi_unclassified(100);k__Fungi_unclassified(100); | - | - | - | - | - | - | - | Unassigned | - |
| Otu16048 | 1 | k__Fungi(100);p__Ascomycota(100);p__Ascomycota_unclassified(100);p__Ascomycota_unclassified(100);p__Ascomycota_unclassified(100);p__Ascomycota_unclassified(100);p__Ascomycota_unclassified(100); | - | - | - | - | - | - | - | Unassigned | - |
| Otu16052 | 1 | k__Fungi(100);k__Fungi_unclassified(100);k__Fungi_unclassified(100);k__Fungi_unclassified(100);k__Fungi_unclassified(100);k__Fungi_unclassified(100);k__Fungi_unclassified(100); | - | - | - | - | - | - | - | Unassigned | - |
| Otu16056 | 1 | k__Fungi(100);k__Fungi_unclassified(100);k__Fungi_unclassified(100);k__Fungi_unclassified(100);k__Fungi_unclassified(100);k__Fungi_unclassified(100);k__Fungi_unclassified(100); | - | - | - | - | - | - | - | Unassigned | - |
| Otu16057 | 1 | k__Fungi(100);p__Ascomycota(100);p__Ascomycota_unclassified(100);p__Ascomycota_unclassified(100);p__Ascomycota_unclassified(100);p__Ascomycota_unclassified(100);p__Ascomycota_unclassified(100); | - | - | - | - | - | - | - | Unassigned | - |
| Otu16060 | 1 | k__Fungi(100);p__Ascomycota(100);p__Ascomycota_unclassified(100);p__Ascomycota_unclassified(100);p__Ascomycota_unclassified(100);p__Ascomycota_unclassified(100);p__Ascomycota_unclassified(100); | - | - | - | - | - | - | - | Unassigned | - |
| Otu16063 | 1 | k__Fungi(100);k__Fungi_unclassified(100);k__Fungi_unclassified(100);k__Fungi_unclassified(100);k__Fungi_unclassified(100);k__Fungi_unclassified(100);k__Fungi_unclassified(100); | - | - | - | - | - | - | - | Unassigned | - |
| Otu16067 | 1 | k__Fungi(100);p__Ascomycota(100);p__Ascomycota_unclassified(100);p__Ascomycota_unclassified(100);p__Ascomycota_unclassified(100);p__Ascomycota_unclassified(100);p__Ascomycota_unclassified(100); | - | - | - | - | - | - | - | Unassigned | - |
| Otu16068 | 1 | k__Fungi(100);k__Fungi_unclassified(100);k__Fungi_unclassified(100);k__Fungi_unclassified(100);k__Fungi_unclassified(100);k__Fungi_unclassified(100);k__Fungi_unclassified(100); | - | - | - | - | - | - | - | Unassigned | - |
| Otu16069 | 1 | k__Fungi(100);k__Fungi_unclassified(100);k__Fungi_unclassified(100);k__Fungi_unclassified(100);k__Fungi_unclassified(100);k__Fungi_unclassified(100);k__Fungi_unclassified(100); | - | - | - | - | - | - | - | Unassigned | - |
| Otu16072 | 1 | k__Fungi(100);k__Fungi_unclassified(100);k__Fungi_unclassified(100);k__Fungi_unclassified(100);k__Fungi_unclassified(100);k__Fungi_unclassified(100);k__Fungi_unclassified(100); | - | - | - | - | - | - | - | Unassigned | - |
| Otu16073 | 1 | k__Fungi(100);k__Fungi_unclassified(100);k__Fungi_unclassified(100);k__Fungi_unclassified(100);k__Fungi_unclassified(100);k__Fungi_unclassified(100);k__Fungi_unclassified(100); | - | - | - | - | - | - | - | Unassigned | - |
| Otu16075 | 1 | k__Fungi(100);k__Fungi_unclassified(100);k__Fungi_unclassified(100);k__Fungi_unclassified(100);k__Fungi_unclassified(100);k__Fungi_unclassified(100);k__Fungi_unclassified(100); | - | - | - | - | - | - | - | Unassigned | - |
| Otu16077 | 1 | k__Fungi(100);k__Fungi_unclassified(100);k__Fungi_unclassified(100);k__Fungi_unclassified(100);k__Fungi_unclassified(100);k__Fungi_unclassified(100);k__Fungi_unclassified(100); | - | - | - | - | - | - | - | Unassigned | - |
| Otu16080 | 1 | k__Fungi(100);p__Ascomycota(100);p__Ascomycota_unclassified(100);p__Ascomycota_unclassified(100);p__Ascomycota_unclassified(100);p__Ascomycota_unclassified(100);p__Ascomycota_unclassified(100); | - | - | - | - | - | - | - | Unassigned | - |
| Otu16085 | 1 | k__Fungi(100);k__Fungi_unclassified(100);k__Fungi_unclassified(100);k__Fungi_unclassified(100);k__Fungi_unclassified(100);k__Fungi_unclassified(100);k__Fungi_unclassified(100); | - | - | - | - | - | - | - | Unassigned | - |
| Otu16087 | 1 | k__Fungi(100);k__Fungi_unclassified(100);k__Fungi_unclassified(100);k__Fungi_unclassified(100);k__Fungi_unclassified(100);k__Fungi_unclassified(100);k__Fungi_unclassified(100); | - | - | - | - | - | - | - | Unassigned | - |
| Otu16088 | 1 | k__Fungi(100);p__Ascomycota(100);p__Ascomycota_unclassified(100);p__Ascomycota_unclassified(100);p__Ascomycota_unclassified(100);p__Ascomycota_unclassified(100);p__Ascomycota_unclassified(100); | - | - | - | - | - | - | - | Unassigned | - |
| Otu16090 | 1 | k__Fungi(100);k__Fungi_unclassified(100);k__Fungi_unclassified(100);k__Fungi_unclassified(100);k__Fungi_unclassified(100);k__Fungi_unclassified(100);k__Fungi_unclassified(100); | - | - | - | - | - | - | - | Unassigned | - |
| Otu16091 | 1 | k__Fungi(100);k__Fungi_unclassified(100);k__Fungi_unclassified(100);k__Fungi_unclassified(100);k__Fungi_unclassified(100);k__Fungi_unclassified(100);k__Fungi_unclassified(100); | - | - | - | - | - | - | - | Unassigned | - |
| Otu16092 | 1 | k__Fungi(100);k__Fungi_unclassified(100);k__Fungi_unclassified(100);k__Fungi_unclassified(100);k__Fungi_unclassified(100);k__Fungi_unclassified(100);k__Fungi_unclassified(100); | - | - | - | - | - | - | - | Unassigned | - |
| Otu16094 | 1 | k__Fungi(100);k__Fungi_unclassified(100);k__Fungi_unclassified(100);k__Fungi_unclassified(100);k__Fungi_unclassified(100);k__Fungi_unclassified(100);k__Fungi_unclassified(100); | - | - | - | - | - | - | - | Unassigned | - |
| Otu16095 | 1 | k__Fungi(100);p__Ascomycota(100);p__Ascomycota_unclassified(100);p__Ascomycota_unclassified(100);p__Ascomycota_unclassified(100);p__Ascomycota_unclassified(100);p__Ascomycota_unclassified(100); | - | - | - | - | - | - | - | Unassigned | - |
| Otu16097 | 1 | k__Fungi(100);p__Ascomycota(100);p__Ascomycota_unclassified(100);p__Ascomycota_unclassified(100);p__Ascomycota_unclassified(100);p__Ascomycota_unclassified(100);p__Ascomycota_unclassified(100); | - | - | - | - | - | - | - | Unassigned | - |
| Otu16098 | 1 | k__Fungi(100);k__Fungi_unclassified(100);k__Fungi_unclassified(100);k__Fungi_unclassified(100);k__Fungi_unclassified(100);k__Fungi_unclassified(100);k__Fungi_unclassified(100); | - | - | - | - | - | - | - | Unassigned | - |
| Otu16100 | 1 | k__Fungi(100);p__Ascomycota(100);p__Ascomycota_unclassified(100);p__Ascomycota_unclassified(100);p__Ascomycota_unclassified(100);p__Ascomycota_unclassified(100);p__Ascomycota_unclassified(100); | - | - | - | - | - | - | - | Unassigned | - |
| Otu16101 | 1 | k__Fungi(100);k__Fungi_unclassified(100);k__Fungi_unclassified(100);k__Fungi_unclassified(100);k__Fungi_unclassified(100);k__Fungi_unclassified(100);k__Fungi_unclassified(100); | - | - | - | - | - | - | - | Unassigned | - |
| Otu16104 | 1 | k__Fungi(100);p__Ascomycota(100);p__Ascomycota_unclassified(100);p__Ascomycota_unclassified(100);p__Ascomycota_unclassified(100);p__Ascomycota_unclassified(100);p__Ascomycota_unclassified(100); | - | - | - | - | - | - | - | Unassigned | - |
| Otu16105 | 1 | k__Fungi(100);k__Fungi_unclassified(100);k__Fungi_unclassified(100);k__Fungi_unclassified(100);k__Fungi_unclassified(100);k__Fungi_unclassified(100);k__Fungi_unclassified(100); | - | - | - | - | - | - | - | Unassigned | - |
| Otu16106 | 1 | k__Fungi(100);p__Ascomycota(100);p__Ascomycota_unclassified(100);p__Ascomycota_unclassified(100);p__Ascomycota_unclassified(100);p__Ascomycota_unclassified(100);p__Ascomycota_unclassified(100); | - | - | - | - | - | - | - | Unassigned | - |
| Otu16107 | 1 | k__Fungi(100);k__Fungi_unclassified(100);k__Fungi_unclassified(100);k__Fungi_unclassified(100);k__Fungi_unclassified(100);k__Fungi_unclassified(100);k__Fungi_unclassified(100); | - | - | - | - | - | - | - | Unassigned | - |
| Otu16118 | 1 | k__Fungi(100);p__Ascomycota(100);p__Ascomycota_unclassified(100);p__Ascomycota_unclassified(100);p__Ascomycota_unclassified(100);p__Ascomycota_unclassified(100);p__Ascomycota_unclassified(100); | - | - | - | - | - | - | - | Unassigned | - |
| Otu16119 | 1 | k__Fungi(100);k__Fungi_unclassified(100);k__Fungi_unclassified(100);k__Fungi_unclassified(100);k__Fungi_unclassified(100);k__Fungi_unclassified(100);k__Fungi_unclassified(100); | - | - | - | - | - | - | - | Unassigned | - |
| Otu16121 | 1 | k__Fungi(100);k__Fungi_unclassified(100);k__Fungi_unclassified(100);k__Fungi_unclassified(100);k__Fungi_unclassified(100);k__Fungi_unclassified(100);k__Fungi_unclassified(100); | - | - | - | - | - | - | - | Unassigned | - |
| Otu16124 | 1 | k__Fungi(100);k__Fungi_unclassified(100);k__Fungi_unclassified(100);k__Fungi_unclassified(100);k__Fungi_unclassified(100);k__Fungi_unclassified(100);k__Fungi_unclassified(100); | - | - | - | - | - | - | - | Unassigned | - |
| Otu16125 | 1 | k__Fungi(100);k__Fungi_unclassified(100);k__Fungi_unclassified(100);k__Fungi_unclassified(100);k__Fungi_unclassified(100);k__Fungi_unclassified(100);k__Fungi_unclassified(100); | - | - | - | - | - | - | - | Unassigned | - |
| Otu16126 | 1 | k__Fungi(100);k__Fungi_unclassified(100);k__Fungi_unclassified(100);k__Fungi_unclassified(100);k__Fungi_unclassified(100);k__Fungi_unclassified(100);k__Fungi_unclassified(100); | - | - | - | - | - | - | - | Unassigned | - |
| Otu16127 | 1 | k__Fungi(100);k__Fungi_unclassified(100);k__Fungi_unclassified(100);k__Fungi_unclassified(100);k__Fungi_unclassified(100);k__Fungi_unclassified(100);k__Fungi_unclassified(100); | - | - | - | - | - | - | - | Unassigned | - |
| Otu16129 | 1 | k__Fungi(100);k__Fungi_unclassified(100);k__Fungi_unclassified(100);k__Fungi_unclassified(100);k__Fungi_unclassified(100);k__Fungi_unclassified(100);k__Fungi_unclassified(100); | - | - | - | - | - | - | - | Unassigned | - |
| Otu16130 | 1 | k__Fungi(100);p__Ascomycota(100);p__Ascomycota_unclassified(100);p__Ascomycota_unclassified(100);p__Ascomycota_unclassified(100);p__Ascomycota_unclassified(100);p__Ascomycota_unclassified(100); | - | - | - | - | - | - | - | Unassigned | - |
| Otu16135 | 1 | k__Fungi(100);k__Fungi_unclassified(100);k__Fungi_unclassified(100);k__Fungi_unclassified(100);k__Fungi_unclassified(100);k__Fungi_unclassified(100);k__Fungi_unclassified(100); | - | - | - | - | - | - | - | Unassigned | - |
| Otu16138 | 1 | k__Fungi(100);k__Fungi_unclassified(100);k__Fungi_unclassified(100);k__Fungi_unclassified(100);k__Fungi_unclassified(100);k__Fungi_unclassified(100);k__Fungi_unclassified(100); | - | - | - | - | - | - | - | Unassigned | - |
| Otu16140 | 1 | k__Fungi(100);k__Fungi_unclassified(100);k__Fungi_unclassified(100);k__Fungi_unclassified(100);k__Fungi_unclassified(100);k__Fungi_unclassified(100);k__Fungi_unclassified(100); | - | - | - | - | - | - | - | Unassigned | - |
| Otu16146 | 1 | k__Fungi(100);p__Ascomycota(100);p__Ascomycota_unclassified(100);p__Ascomycota_unclassified(100);p__Ascomycota_unclassified(100);p__Ascomycota_unclassified(100);p__Ascomycota_unclassified(100); | - | - | - | - | - | - | - | Unassigned | - |
| Otu16149 | 1 | k__Fungi(100);p__Ascomycota(100);p__Ascomycota_unclassified(100);p__Ascomycota_unclassified(100);p__Ascomycota_unclassified(100);p__Ascomycota_unclassified(100);p__Ascomycota_unclassified(100); | - | - | - | - | - | - | - | Unassigned | - |
| Otu16152 | 1 | k__Fungi(100);k__Fungi_unclassified(100);k__Fungi_unclassified(100);k__Fungi_unclassified(100);k__Fungi_unclassified(100);k__Fungi_unclassified(100);k__Fungi_unclassified(100); | - | - | - | - | - | - | - | Unassigned | - |
| Otu16160 | 1 | k__Fungi(100);k__Fungi_unclassified(100);k__Fungi_unclassified(100);k__Fungi_unclassified(100);k__Fungi_unclassified(100);k__Fungi_unclassified(100);k__Fungi_unclassified(100); | - | - | - | - | - | - | - | Unassigned | - |
| Otu16161 | 1 | k__Fungi(100);k__Fungi_unclassified(100);k__Fungi_unclassified(100);k__Fungi_unclassified(100);k__Fungi_unclassified(100);k__Fungi_unclassified(100);k__Fungi_unclassified(100); | - | - | - | - | - | - | - | Unassigned | - |
| Otu16163 | 1 | k__Fungi(100);k__Fungi_unclassified(100);k__Fungi_unclassified(100);k__Fungi_unclassified(100);k__Fungi_unclassified(100);k__Fungi_unclassified(100);k__Fungi_unclassified(100); | - | - | - | - | - | - | - | Unassigned | - |
| Otu16164 | 1 | k__Fungi(100);k__Fungi_unclassified(100);k__Fungi_unclassified(100);k__Fungi_unclassified(100);k__Fungi_unclassified(100);k__Fungi_unclassified(100);k__Fungi_unclassified(100); | - | - | - | - | - | - | - | Unassigned | - |
| Otu16166 | 1 | k__Fungi(100);p__Ascomycota(100);p__Ascomycota_unclassified(100);p__Ascomycota_unclassified(100);p__Ascomycota_unclassified(100);p__Ascomycota_unclassified(100);p__Ascomycota_unclassified(100); | - | - | - | - | - | - | - | Unassigned | - |
| Otu16169 | 1 | k__Fungi(100);k__Fungi_unclassified(100);k__Fungi_unclassified(100);k__Fungi_unclassified(100);k__Fungi_unclassified(100);k__Fungi_unclassified(100);k__Fungi_unclassified(100); | - | - | - | - | - | - | - | Unassigned | - |
| Otu16173 | 1 | k__Fungi(100);k__Fungi_unclassified(100);k__Fungi_unclassified(100);k__Fungi_unclassified(100);k__Fungi_unclassified(100);k__Fungi_unclassified(100);k__Fungi_unclassified(100); | - | - | - | - | - | - | - | Unassigned | - |
| Otu16176 | 1 | k__Fungi(100);k__Fungi_unclassified(100);k__Fungi_unclassified(100);k__Fungi_unclassified(100);k__Fungi_unclassified(100);k__Fungi_unclassified(100);k__Fungi_unclassified(100); | - | - | - | - | - | - | - | Unassigned | - |
| Otu16177 | 1 | k__Fungi(100);p__Ascomycota(100);p__Ascomycota_unclassified(100);p__Ascomycota_unclassified(100);p__Ascomycota_unclassified(100);p__Ascomycota_unclassified(100);p__Ascomycota_unclassified(100); | - | - | - | - | - | - | - | Unassigned | - |
| Otu16181 | 1 | k__Fungi(100);p__Ascomycota(100);p__Ascomycota_unclassified(100);p__Ascomycota_unclassified(100);p__Ascomycota_unclassified(100);p__Ascomycota_unclassified(100);p__Ascomycota_unclassified(100); | - | - | - | - | - | - | - | Unassigned | - |
| Otu16185 | 1 | k__Fungi(100);p__Ascomycota(100);p__Ascomycota_unclassified(100);p__Ascomycota_unclassified(100);p__Ascomycota_unclassified(100);p__Ascomycota_unclassified(100);p__Ascomycota_unclassified(100); | - | - | - | - | - | - | - | Unassigned | - |
| Otu16188 | 1 | k__Fungi(100);k__Fungi_unclassified(100);k__Fungi_unclassified(100);k__Fungi_unclassified(100);k__Fungi_unclassified(100);k__Fungi_unclassified(100);k__Fungi_unclassified(100); | - | - | - | - | - | - | - | Unassigned | - |
| Otu16189 | 1 | k__Fungi(100);k__Fungi_unclassified(100);k__Fungi_unclassified(100);k__Fungi_unclassified(100);k__Fungi_unclassified(100);k__Fungi_unclassified(100);k__Fungi_unclassified(100); | - | - | - | - | - | - | - | Unassigned | - |
| Otu16190 | 1 | k__Fungi(100);k__Fungi_unclassified(100);k__Fungi_unclassified(100);k__Fungi_unclassified(100);k__Fungi_unclassified(100);k__Fungi_unclassified(100);k__Fungi_unclassified(100); | - | - | - | - | - | - | - | Unassigned | - |
| Otu16191 | 1 | k__Fungi(100);k__Fungi_unclassified(100);k__Fungi_unclassified(100);k__Fungi_unclassified(100);k__Fungi_unclassified(100);k__Fungi_unclassified(100);k__Fungi_unclassified(100); | - | - | - | - | - | - | - | Unassigned | - |
| Otu16192 | 1 | k__Fungi(100);k__Fungi_unclassified(100);k__Fungi_unclassified(100);k__Fungi_unclassified(100);k__Fungi_unclassified(100);k__Fungi_unclassified(100);k__Fungi_unclassified(100); | - | - | - | - | - | - | - | Unassigned | - |
| Otu16193 | 1 | k__Fungi(100);p__Ascomycota(100);p__Ascomycota_unclassified(100);p__Ascomycota_unclassified(100);p__Ascomycota_unclassified(100);p__Ascomycota_unclassified(100);p__Ascomycota_unclassified(100); | - | - | - | - | - | - | - | Unassigned | - |
| Otu16195 | 1 | k__Fungi(100);p__Ascomycota(100);p__Ascomycota_unclassified(100);p__Ascomycota_unclassified(100);p__Ascomycota_unclassified(100);p__Ascomycota_unclassified(100);p__Ascomycota_unclassified(100); | - | - | - | - | - | - | - | Unassigned | - |
| Otu16196 | 1 | k__Fungi(100);k__Fungi_unclassified(100);k__Fungi_unclassified(100);k__Fungi_unclassified(100);k__Fungi_unclassified(100);k__Fungi_unclassified(100);k__Fungi_unclassified(100); | - | - | - | - | - | - | - | Unassigned | - |
| Otu16197 | 1 | k__Fungi(100);k__Fungi_unclassified(100);k__Fungi_unclassified(100);k__Fungi_unclassified(100);k__Fungi_unclassified(100);k__Fungi_unclassified(100);k__Fungi_unclassified(100); | - | - | - | - | - | - | - | Unassigned | - |
| Otu16199 | 1 | k__Fungi(100);k__Fungi_unclassified(100);k__Fungi_unclassified(100);k__Fungi_unclassified(100);k__Fungi_unclassified(100);k__Fungi_unclassified(100);k__Fungi_unclassified(100); | - | - | - | - | - | - | - | Unassigned | - |
| Otu16201 | 1 | k__Fungi(100);k__Fungi_unclassified(100);k__Fungi_unclassified(100);k__Fungi_unclassified(100);k__Fungi_unclassified(100);k__Fungi_unclassified(100);k__Fungi_unclassified(100); | - | - | - | - | - | - | - | Unassigned | - |
| Otu16202 | 1 | k__Fungi(100);k__Fungi_unclassified(100);k__Fungi_unclassified(100);k__Fungi_unclassified(100);k__Fungi_unclassified(100);k__Fungi_unclassified(100);k__Fungi_unclassified(100); | - | - | - | - | - | - | - | Unassigned | - |
| Otu16203 | 1 | k__Fungi(100);k__Fungi_unclassified(100);k__Fungi_unclassified(100);k__Fungi_unclassified(100);k__Fungi_unclassified(100);k__Fungi_unclassified(100);k__Fungi_unclassified(100); | - | - | - | - | - | - | - | Unassigned | - |
| Otu16205 | 1 | k__Fungi(100);k__Fungi_unclassified(100);k__Fungi_unclassified(100);k__Fungi_unclassified(100);k__Fungi_unclassified(100);k__Fungi_unclassified(100);k__Fungi_unclassified(100); | - | - | - | - | - | - | - | Unassigned | - |
| Otu16206 | 1 | k__Fungi(100);k__Fungi_unclassified(100);k__Fungi_unclassified(100);k__Fungi_unclassified(100);k__Fungi_unclassified(100);k__Fungi_unclassified(100);k__Fungi_unclassified(100); | - | - | - | - | - | - | - | Unassigned | - |
| Otu16208 | 1 | k__Fungi(100);k__Fungi_unclassified(100);k__Fungi_unclassified(100);k__Fungi_unclassified(100);k__Fungi_unclassified(100);k__Fungi_unclassified(100);k__Fungi_unclassified(100); | - | - | - | - | - | - | - | Unassigned | - |
| Otu16219 | 1 | k__Fungi(100);k__Fungi_unclassified(100);k__Fungi_unclassified(100);k__Fungi_unclassified(100);k__Fungi_unclassified(100);k__Fungi_unclassified(100);k__Fungi_unclassified(100); | - | - | - | - | - | - | - | Unassigned | - |
| Otu16235 | 1 | k__Fungi(100);p__Ascomycota(100);p__Ascomycota_unclassified(100);p__Ascomycota_unclassified(100);p__Ascomycota_unclassified(100);p__Ascomycota_unclassified(100);p__Ascomycota_unclassified(100); | - | - | - | - | - | - | - | Unassigned | - |
| Otu16238 | 1 | k__Fungi(100);p__Ascomycota(100);p__Ascomycota_unclassified(100);p__Ascomycota_unclassified(100);p__Ascomycota_unclassified(100);p__Ascomycota_unclassified(100);p__Ascomycota_unclassified(100); | - | - | - | - | - | - | - | Unassigned | - |
| Otu16239 | 1 | k__Fungi(100);p__Ascomycota(100);p__Ascomycota_unclassified(100);p__Ascomycota_unclassified(100);p__Ascomycota_unclassified(100);p__Ascomycota_unclassified(100);p__Ascomycota_unclassified(100); | - | - | - | - | - | - | - | Unassigned | - |
| Otu16240 | 1 | k__Fungi(100);k__Fungi_unclassified(100);k__Fungi_unclassified(100);k__Fungi_unclassified(100);k__Fungi_unclassified(100);k__Fungi_unclassified(100);k__Fungi_unclassified(100); | - | - | - | - | - | - | - | Unassigned | - |
| Otu16241 | 1 | k__Fungi(100);k__Fungi_unclassified(100);k__Fungi_unclassified(100);k__Fungi_unclassified(100);k__Fungi_unclassified(100);k__Fungi_unclassified(100);k__Fungi_unclassified(100); | - | - | - | - | - | - | - | Unassigned | - |
| Otu16248 | 1 | k__Fungi(100);k__Fungi_unclassified(100);k__Fungi_unclassified(100);k__Fungi_unclassified(100);k__Fungi_unclassified(100);k__Fungi_unclassified(100);k__Fungi_unclassified(100); | - | - | - | - | - | - | - | Unassigned | - |
| Otu16253 | 1 | k__Fungi(100);k__Fungi_unclassified(100);k__Fungi_unclassified(100);k__Fungi_unclassified(100);k__Fungi_unclassified(100);k__Fungi_unclassified(100);k__Fungi_unclassified(100); | - | - | - | - | - | - | - | Unassigned | - |
| Otu16256 | 1 | k__Fungi(100);p__Ascomycota(100);p__Ascomycota_unclassified(100);p__Ascomycota_unclassified(100);p__Ascomycota_unclassified(100);p__Ascomycota_unclassified(100);p__Ascomycota_unclassified(100); | - | - | - | - | - | - | - | Unassigned | - |
| Otu16258 | 1 | k__Fungi(100);k__Fungi_unclassified(100);k__Fungi_unclassified(100);k__Fungi_unclassified(100);k__Fungi_unclassified(100);k__Fungi_unclassified(100);k__Fungi_unclassified(100); | - | - | - | - | - | - | - | Unassigned | - |
| Otu16259 | 1 | k__Fungi(100);p__Ascomycota(100);p__Ascomycota_unclassified(100);p__Ascomycota_unclassified(100);p__Ascomycota_unclassified(100);p__Ascomycota_unclassified(100);p__Ascomycota_unclassified(100); | - | - | - | - | - | - | - | Unassigned | - |
| Otu16260 | 1 | k__Fungi(100);k__Fungi_unclassified(100);k__Fungi_unclassified(100);k__Fungi_unclassified(100);k__Fungi_unclassified(100);k__Fungi_unclassified(100);k__Fungi_unclassified(100); | - | - | - | - | - | - | - | Unassigned | - |
| Otu16261 | 1 | k__Fungi(100);k__Fungi_unclassified(100);k__Fungi_unclassified(100);k__Fungi_unclassified(100);k__Fungi_unclassified(100);k__Fungi_unclassified(100);k__Fungi_unclassified(100); | - | - | - | - | - | - | - | Unassigned | - |
| Otu16263 | 1 | k__Fungi(100);p__Ascomycota(100);p__Ascomycota_unclassified(100);p__Ascomycota_unclassified(100);p__Ascomycota_unclassified(100);p__Ascomycota_unclassified(100);p__Ascomycota_unclassified(100); | - | - | - | - | - | - | - | Unassigned | - |
| Otu16264 | 1 | k__Fungi(100);p__Ascomycota(100);p__Ascomycota_unclassified(100);p__Ascomycota_unclassified(100);p__Ascomycota_unclassified(100);p__Ascomycota_unclassified(100);p__Ascomycota_unclassified(100); | - | - | - | - | - | - | - | Unassigned | - |
| Otu16265 | 1 | k__Fungi(100);k__Fungi_unclassified(100);k__Fungi_unclassified(100);k__Fungi_unclassified(100);k__Fungi_unclassified(100);k__Fungi_unclassified(100);k__Fungi_unclassified(100); | - | - | - | - | - | - | - | Unassigned | - |
| Otu16270 | 1 | k__Fungi(100);k__Fungi_unclassified(100);k__Fungi_unclassified(100);k__Fungi_unclassified(100);k__Fungi_unclassified(100);k__Fungi_unclassified(100);k__Fungi_unclassified(100); | - | - | - | - | - | - | - | Unassigned | - |
| Otu16273 | 1 | k__Fungi(100);p__Ascomycota(100);p__Ascomycota_unclassified(100);p__Ascomycota_unclassified(100);p__Ascomycota_unclassified(100);p__Ascomycota_unclassified(100);p__Ascomycota_unclassified(100); | - | - | - | - | - | - | - | Unassigned | - |
| Otu16274 | 1 | k__Fungi(100);k__Fungi_unclassified(100);k__Fungi_unclassified(100);k__Fungi_unclassified(100);k__Fungi_unclassified(100);k__Fungi_unclassified(100);k__Fungi_unclassified(100); | - | - | - | - | - | - | - | Unassigned | - |
| Otu16279 | 1 | k__Fungi(100);p__Ascomycota(100);c__Leotiomycetes(100);o__Helotiales(100);o__Helotiales_unclassified(100);o__Helotiales_unclassified(100);o__Helotiales_unclassified(100); | - | - | - | - | - | - | - | Unassigned | - |
| Otu16291 | 1 | k__Fungi(100);p__Ascomycota(100);p__Ascomycota_unclassified(100);p__Ascomycota_unclassified(100);p__Ascomycota_unclassified(100);p__Ascomycota_unclassified(100);p__Ascomycota_unclassified(100); | - | - | - | - | - | - | - | Unassigned | - |
| Otu16293 | 1 | k__Fungi(100);k__Fungi_unclassified(100);k__Fungi_unclassified(100);k__Fungi_unclassified(100);k__Fungi_unclassified(100);k__Fungi_unclassified(100);k__Fungi_unclassified(100); | - | - | - | - | - | - | - | Unassigned | - |
| Otu16297 | 1 | k__Fungi(100);p__Ascomycota(100);p__Ascomycota_unclassified(100);p__Ascomycota_unclassified(100);p__Ascomycota_unclassified(100);p__Ascomycota_unclassified(100);p__Ascomycota_unclassified(100); | - | - | - | - | - | - | - | Unassigned | - |
| Otu16303 | 1 | k__Fungi(100);k__Fungi_unclassified(100);k__Fungi_unclassified(100);k__Fungi_unclassified(100);k__Fungi_unclassified(100);k__Fungi_unclassified(100);k__Fungi_unclassified(100); | - | - | - | - | - | - | - | Unassigned | - |
| Otu16310 | 1 | k__Fungi(100);p__Ascomycota(100);p__Ascomycota_unclassified(100);p__Ascomycota_unclassified(100);p__Ascomycota_unclassified(100);p__Ascomycota_unclassified(100);p__Ascomycota_unclassified(100); | - | - | - | - | - | - | - | Unassigned | - |
| Otu16321 | 1 | k__Fungi(100);p__Ascomycota(100);p__Ascomycota_unclassified(100);p__Ascomycota_unclassified(100);p__Ascomycota_unclassified(100);p__Ascomycota_unclassified(100);p__Ascomycota_unclassified(100); | - | - | - | - | - | - | - | Unassigned | - |
| Otu16322 | 1 | k__Fungi(100);p__Ascomycota(100);p__Ascomycota_unclassified(100);p__Ascomycota_unclassified(100);p__Ascomycota_unclassified(100);p__Ascomycota_unclassified(100);p__Ascomycota_unclassified(100); | - | - | - | - | - | - | - | Unassigned | - |
| Otu16324 | 1 | k__Fungi(100);p__Ascomycota(100);p__Ascomycota_unclassified(100);p__Ascomycota_unclassified(100);p__Ascomycota_unclassified(100);p__Ascomycota_unclassified(100);p__Ascomycota_unclassified(100); | - | - | - | - | - | - | - | Unassigned | - |
| Otu16330 | 1 | k__Fungi(100);k__Fungi_unclassified(100);k__Fungi_unclassified(100);k__Fungi_unclassified(100);k__Fungi_unclassified(100);k__Fungi_unclassified(100);k__Fungi_unclassified(100); | - | - | - | - | - | - | - | Unassigned | - |
| Otu16331 | 1 | k__Fungi(100);k__Fungi_unclassified(100);k__Fungi_unclassified(100);k__Fungi_unclassified(100);k__Fungi_unclassified(100);k__Fungi_unclassified(100);k__Fungi_unclassified(100); | - | - | - | - | - | - | - | Unassigned | - |
| Otu16333 | 1 | k__Fungi(100);k__Fungi_unclassified(100);k__Fungi_unclassified(100);k__Fungi_unclassified(100);k__Fungi_unclassified(100);k__Fungi_unclassified(100);k__Fungi_unclassified(100); | - | - | - | - | - | - | - | Unassigned | - |
| Otu16334 | 1 | k__Fungi(100);p__Ascomycota(100);p__Ascomycota_unclassified(100);p__Ascomycota_unclassified(100);p__Ascomycota_unclassified(100);p__Ascomycota_unclassified(100);p__Ascomycota_unclassified(100); | - | - | - | - | - | - | - | Unassigned | - |
| Otu16336 | 1 | k__Fungi(100);p__Ascomycota(100);p__Ascomycota_unclassified(100);p__Ascomycota_unclassified(100);p__Ascomycota_unclassified(100);p__Ascomycota_unclassified(100);p__Ascomycota_unclassified(100); | - | - | - | - | - | - | - | Unassigned | - |
| Otu16337 | 1 | k__Fungi(100);k__Fungi_unclassified(100);k__Fungi_unclassified(100);k__Fungi_unclassified(100);k__Fungi_unclassified(100);k__Fungi_unclassified(100);k__Fungi_unclassified(100); | - | - | - | - | - | - | - | Unassigned | - |
| Otu16338 | 1 | k__Fungi(100);p__Ascomycota(100);p__Ascomycota_unclassified(100);p__Ascomycota_unclassified(100);p__Ascomycota_unclassified(100);p__Ascomycota_unclassified(100);p__Ascomycota_unclassified(100); | - | - | - | - | - | - | - | Unassigned | - |
| Otu16339 | 1 | k__Fungi(100);k__Fungi_unclassified(100);k__Fungi_unclassified(100);k__Fungi_unclassified(100);k__Fungi_unclassified(100);k__Fungi_unclassified(100);k__Fungi_unclassified(100); | - | - | - | - | - | - | - | Unassigned | - |
| Otu16343 | 1 | k__Fungi(100);k__Fungi_unclassified(100);k__Fungi_unclassified(100);k__Fungi_unclassified(100);k__Fungi_unclassified(100);k__Fungi_unclassified(100);k__Fungi_unclassified(100); | - | - | - | - | - | - | - | Unassigned | - |
| Otu16348 | 1 | k__Fungi(100);k__Fungi_unclassified(100);k__Fungi_unclassified(100);k__Fungi_unclassified(100);k__Fungi_unclassified(100);k__Fungi_unclassified(100);k__Fungi_unclassified(100); | - | - | - | - | - | - | - | Unassigned | - |
| Otu16351 | 1 | k__Fungi(100);k__Fungi_unclassified(100);k__Fungi_unclassified(100);k__Fungi_unclassified(100);k__Fungi_unclassified(100);k__Fungi_unclassified(100);k__Fungi_unclassified(100); | - | - | - | - | - | - | - | Unassigned | - |
| Otu16358 | 1 | k__Fungi(100);p__Ascomycota(100);p__Ascomycota_unclassified(100);p__Ascomycota_unclassified(100);p__Ascomycota_unclassified(100);p__Ascomycota_unclassified(100);p__Ascomycota_unclassified(100); | - | - | - | - | - | - | - | Unassigned | - |
| Otu16361 | 1 | k__Fungi(100);k__Fungi_unclassified(100);k__Fungi_unclassified(100);k__Fungi_unclassified(100);k__Fungi_unclassified(100);k__Fungi_unclassified(100);k__Fungi_unclassified(100); | - | - | - | - | - | - | - | Unassigned | - |
| Otu16363 | 1 | k__Fungi(100);p__Ascomycota(100);p__Ascomycota_unclassified(100);p__Ascomycota_unclassified(100);p__Ascomycota_unclassified(100);p__Ascomycota_unclassified(100);p__Ascomycota_unclassified(100); | - | - | - | - | - | - | - | Unassigned | - |
| Otu16373 | 1 | k__Fungi(100);k__Fungi_unclassified(100);k__Fungi_unclassified(100);k__Fungi_unclassified(100);k__Fungi_unclassified(100);k__Fungi_unclassified(100);k__Fungi_unclassified(100); | - | - | - | - | - | - | - | Unassigned | - |
| Otu16381 | 1 | k__Fungi(100);p__Ascomycota(100);p__Ascomycota_unclassified(100);p__Ascomycota_unclassified(100);p__Ascomycota_unclassified(100);p__Ascomycota_unclassified(100);p__Ascomycota_unclassified(100); | - | - | - | - | - | - | - | Unassigned | - |
| Otu16388 | 1 | k__Fungi(100);k__Fungi_unclassified(100);k__Fungi_unclassified(100);k__Fungi_unclassified(100);k__Fungi_unclassified(100);k__Fungi_unclassified(100);k__Fungi_unclassified(100); | - | - | - | - | - | - | - | Unassigned | - |
| Otu16390 | 1 | k__Fungi(100);p__Ascomycota(100);p__Ascomycota_unclassified(100);p__Ascomycota_unclassified(100);p__Ascomycota_unclassified(100);p__Ascomycota_unclassified(100);p__Ascomycota_unclassified(100); | - | - | - | - | - | - | - | Unassigned | - |
| Otu16394 | 1 | k__Fungi(100);p__Ascomycota(100);p__Ascomycota_unclassified(100);p__Ascomycota_unclassified(100);p__Ascomycota_unclassified(100);p__Ascomycota_unclassified(100);p__Ascomycota_unclassified(100); | - | - | - | - | - | - | - | Unassigned | - |
| Otu16407 | 1 | k__Fungi(100);k__Fungi_unclassified(100);k__Fungi_unclassified(100);k__Fungi_unclassified(100);k__Fungi_unclassified(100);k__Fungi_unclassified(100);k__Fungi_unclassified(100); | - | - | - | - | - | - | - | Unassigned | - |
| Otu16410 | 1 | k__Fungi(100);k__Fungi_unclassified(100);k__Fungi_unclassified(100);k__Fungi_unclassified(100);k__Fungi_unclassified(100);k__Fungi_unclassified(100);k__Fungi_unclassified(100); | - | - | - | - | - | - | - | Unassigned | - |
| Otu16413 | 1 | k__Fungi(100);k__Fungi_unclassified(100);k__Fungi_unclassified(100);k__Fungi_unclassified(100);k__Fungi_unclassified(100);k__Fungi_unclassified(100);k__Fungi_unclassified(100); | - | - | - | - | - | - | - | Unassigned | - |
| Otu16415 | 1 | k__Fungi(100);p__Ascomycota(100);p__Ascomycota_unclassified(100);p__Ascomycota_unclassified(100);p__Ascomycota_unclassified(100);p__Ascomycota_unclassified(100);p__Ascomycota_unclassified(100); | - | - | - | - | - | - | - | Unassigned | - |
| Otu16427 | 1 | k__Fungi(100);k__Fungi_unclassified(100);k__Fungi_unclassified(100);k__Fungi_unclassified(100);k__Fungi_unclassified(100);k__Fungi_unclassified(100);k__Fungi_unclassified(100); | - | - | - | - | - | - | - | Unassigned | - |
| Otu16430 | 1 | k__Fungi(100);p__Ascomycota(100);p__Ascomycota_unclassified(100);p__Ascomycota_unclassified(100);p__Ascomycota_unclassified(100);p__Ascomycota_unclassified(100);p__Ascomycota_unclassified(100); | - | - | - | - | - | - | - | Unassigned | - |
| Otu16431 | 1 | k__Fungi(100);k__Fungi_unclassified(100);k__Fungi_unclassified(100);k__Fungi_unclassified(100);k__Fungi_unclassified(100);k__Fungi_unclassified(100);k__Fungi_unclassified(100); | - | - | - | - | - | - | - | Unassigned | - |
| Otu16432 | 1 | k__Fungi(100);p__Ascomycota(100);p__Ascomycota_unclassified(100);p__Ascomycota_unclassified(100);p__Ascomycota_unclassified(100);p__Ascomycota_unclassified(100);p__Ascomycota_unclassified(100); | - | - | - | - | - | - | - | Unassigned | - |
| Otu16434 | 1 | k__Fungi(100);k__Fungi_unclassified(100);k__Fungi_unclassified(100);k__Fungi_unclassified(100);k__Fungi_unclassified(100);k__Fungi_unclassified(100);k__Fungi_unclassified(100); | - | - | - | - | - | - | - | Unassigned | - |
| Otu16441 | 1 | k__Fungi(100);k__Fungi_unclassified(100);k__Fungi_unclassified(100);k__Fungi_unclassified(100);k__Fungi_unclassified(100);k__Fungi_unclassified(100);k__Fungi_unclassified(100); | - | - | - | - | - | - | - | Unassigned | - |
| Otu16456 | 1 | k__Fungi(100);k__Fungi_unclassified(100);k__Fungi_unclassified(100);k__Fungi_unclassified(100);k__Fungi_unclassified(100);k__Fungi_unclassified(100);k__Fungi_unclassified(100); | - | - | - | - | - | - | - | Unassigned | - |
| Otu16474 | 1 | k__Fungi(100);p__Ascomycota(100);p__Ascomycota_unclassified(100);p__Ascomycota_unclassified(100);p__Ascomycota_unclassified(100);p__Ascomycota_unclassified(100);p__Ascomycota_unclassified(100); | - | - | - | - | - | - | - | Unassigned | - |
| Otu16478 | 1 | k__Fungi(100);k__Fungi_unclassified(100);k__Fungi_unclassified(100);k__Fungi_unclassified(100);k__Fungi_unclassified(100);k__Fungi_unclassified(100);k__Fungi_unclassified(100); | - | - | - | - | - | - | - | Unassigned | - |
| Otu16479 | 1 | k__Fungi(100);p__Ascomycota(100);p__Ascomycota_unclassified(100);p__Ascomycota_unclassified(100);p__Ascomycota_unclassified(100);p__Ascomycota_unclassified(100);p__Ascomycota_unclassified(100); | - | - | - | - | - | - | - | Unassigned | - |
| Otu16480 | 1 | k__Fungi(100);k__Fungi_unclassified(100);k__Fungi_unclassified(100);k__Fungi_unclassified(100);k__Fungi_unclassified(100);k__Fungi_unclassified(100);k__Fungi_unclassified(100); | - | - | - | - | - | - | - | Unassigned | - |
| Otu16484 | 1 | k__Fungi(100);p__Ascomycota(100);p__Ascomycota_unclassified(100);p__Ascomycota_unclassified(100);p__Ascomycota_unclassified(100);p__Ascomycota_unclassified(100);p__Ascomycota_unclassified(100); | - | - | - | - | - | - | - | Unassigned | - |

| | | | | | | | | | | | |
|---|---|---|---|---|---|---|---|---|---|---|---|
| Otu16486 | 1 | k__Fungi(100);p__Ascomycota(100);p__Ascomycota_unclassified(100);p__Ascomycota_unclassified(100);p__Ascomycota_unclassified(100);p__Ascomycota_unclassified(100);p__Ascomycota_unclassified(100); | - | - | - | - | - | - | - | Unassigned | - |
| Otu16492 | 1 | k__Fungi(100);p__Ascomycota(100);p__Ascomycota_unclassified(100);p__Ascomycota_unclassified(100);p__Ascomycota_unclassified(100);p__Ascomycota_unclassified(100);p__Ascomycota_unclassified(100); | - | - | - | - | - | - | - | Unassigned | - |
| Otu16500 | 1 | k__Fungi(100);p__Ascomycota(100);p__Ascomycota_unclassified(100);p__Ascomycota_unclassified(100);p__Ascomycota_unclassified(100);p__Ascomycota_unclassified(100);p__Ascomycota_unclassified(100); | - | - | - | - | - | - | - | Unassigned | - |
| Otu16504 | 1 | k__Fungi(100);k__Fungi_unclassified(100);k__Fungi_unclassified(100);k__Fungi_unclassified(100);k__Fungi_unclassified(100);k__Fungi_unclassified(100);k__Fungi_unclassified(100); | - | - | - | - | - | - | - | Unassigned | - |
| Otu16508 | 1 | k__Fungi(100);p__Ascomycota(100);p__Ascomycota_unclassified(100);p__Ascomycota_unclassified(100);p__Ascomycota_unclassified(100);p__Ascomycota_unclassified(100);p__Ascomycota_unclassified(100); | - | - | - | - | - | - | - | Unassigned | - |
| Otu16509 | 1 | k__Fungi(100);p__Ascomycota(100);p__Ascomycota_unclassified(100);p__Ascomycota_unclassified(100);p__Ascomycota_unclassified(100);p__Ascomycota_unclassified(100);p__Ascomycota_unclassified(100); | - | - | - | - | - | - | - | Unassigned | - |
| Otu16511 | 1 | k__Fungi(100);p__Ascomycota(100);p__Ascomycota_unclassified(100);p__Ascomycota_unclassified(100);p__Ascomycota_unclassified(100);p__Ascomycota_unclassified(100);p__Ascomycota_unclassified(100); | - | - | - | - | - | - | - | Unassigned | - |
| Otu16514 | 1 | k__Fungi(100);p__Ascomycota(100);p__Ascomycota_unclassified(100);p__Ascomycota_unclassified(100);p__Ascomycota_unclassified(100);p__Ascomycota_unclassified(100);p__Ascomycota_unclassified(100); | - | - | - | - | - | - | - | Unassigned | - |
| Otu16515 | 1 | k__Fungi(100);k__Fungi_unclassified(100);k__Fungi_unclassified(100);k__Fungi_unclassified(100);k__Fungi_unclassified(100);k__Fungi_unclassified(100);k__Fungi_unclassified(100); | - | - | - | - | - | - | - | Unassigned | - |
| Otu16516 | 1 | k__Fungi(100);p__Ascomycota(100);p__Ascomycota_unclassified(100);p__Ascomycota_unclassified(100);p__Ascomycota_unclassified(100);p__Ascomycota_unclassified(100);p__Ascomycota_unclassified(100); | - | - | - | - | - | - | - | Unassigned | - |
| Otu16517 | 1 | k__Fungi(100);p__Ascomycota(100);p__Ascomycota_unclassified(100);p__Ascomycota_unclassified(100);p__Ascomycota_unclassified(100);p__Ascomycota_unclassified(100);p__Ascomycota_unclassified(100); | - | - | - | - | - | - | - | Unassigned | - |
| Otu16518 | 1 | k__Fungi(100);k__Fungi_unclassified(100);k__Fungi_unclassified(100);k__Fungi_unclassified(100);k__Fungi_unclassified(100);k__Fungi_unclassified(100);k__Fungi_unclassified(100); | - | - | - | - | - | - | - | Unassigned | - |
| Otu16521 | 1 | k__Fungi(100);p__Ascomycota(100);p__Ascomycota_unclassified(100);p__Ascomycota_unclassified(100);p__Ascomycota_unclassified(100);p__Ascomycota_unclassified(100);p__Ascomycota_unclassified(100); | - | - | - | - | - | - | - | Unassigned | - |
| Otu16522 | 1 | k__Fungi(100);k__Fungi_unclassified(100);k__Fungi_unclassified(100);k__Fungi_unclassified(100);k__Fungi_unclassified(100);k__Fungi_unclassified(100);k__Fungi_unclassified(100); | - | - | - | - | - | - | - | Unassigned | - |
| Otu16523 | 1 | k__Fungi(100);p__Ascomycota(100);p__Ascomycota_unclassified(100);p__Ascomycota_unclassified(100);p__Ascomycota_unclassified(100);p__Ascomycota_unclassified(100);p__Ascomycota_unclassified(100); | - | - | - | - | - | - | - | Unassigned | - |
| Otu16524 | 1 | k__Fungi(100);k__Fungi_unclassified(100);k__Fungi_unclassified(100);k__Fungi_unclassified(100);k__Fungi_unclassified(100);k__Fungi_unclassified(100);k__Fungi_unclassified(100); | - | - | - | - | - | - | - | Unassigned | - |
| Otu16525 | 1 | k__Fungi(100);p__Ascomycota(100);p__Ascomycota_unclassified(100);p__Ascomycota_unclassified(100);p__Ascomycota_unclassified(100);p__Ascomycota_unclassified(100);p__Ascomycota_unclassified(100); | - | - | - | - | - | - | - | Unassigned | - |
| Otu16526 | 1 | k__Fungi(100);k__Fungi_unclassified(100);k__Fungi_unclassified(100);k__Fungi_unclassified(100);k__Fungi_unclassified(100);k__Fungi_unclassified(100);k__Fungi_unclassified(100); | - | - | - | - | - | - | - | Unassigned | - |
| Otu16528 | 1 | k__Fungi(100);k__Fungi_unclassified(100);k__Fungi_unclassified(100);k__Fungi_unclassified(100);k__Fungi_unclassified(100);k__Fungi_unclassified(100);k__Fungi_unclassified(100); | - | - | - | - | - | - | - | Unassigned | - |
| Otu16529 | 1 | k__Fungi(100);p__Ascomycota(100);c__Dothideomycetes(100);o__Pleosporales(100);o__Pleosporales_unclassified(100);o__Pleosporales_unclassified(100);o__Pleosporales_unclassified(100); | - | - | - | - | - | - | - | Unassigned | - |
| Otu16531 | 1 | k__Fungi(100);k__Fungi_unclassified(100);k__Fungi_unclassified(100);k__Fungi_unclassified(100);k__Fungi_unclassified(100);k__Fungi_unclassified(100);k__Fungi_unclassified(100); | - | - | - | - | - | - | - | Unassigned | - |
| Otu16537 | 1 | k__Fungi(100);k__Fungi_unclassified(100);k__Fungi_unclassified(100);k__Fungi_unclassified(100);k__Fungi_unclassified(100);k__Fungi_unclassified(100);k__Fungi_unclassified(100); | - | - | - | - | - | - | - | Unassigned | - |
| Otu16538 | 1 | k__Fungi(100);k__Fungi_unclassified(100);k__Fungi_unclassified(100);k__Fungi_unclassified(100);k__Fungi_unclassified(100);k__Fungi_unclassified(100);k__Fungi_unclassified(100); | - | - | - | - | - | - | - | Unassigned | - |
| Otu16540 | 1 | k__Fungi(100);k__Fungi_unclassified(100);k__Fungi_unclassified(100);k__Fungi_unclassified(100);k__Fungi_unclassified(100);k__Fungi_unclassified(100);k__Fungi_unclassified(100); | - | - | - | - | - | - | - | Unassigned | - |
| Otu16541 | 1 | k__Fungi(100);k__Fungi_unclassified(100);k__Fungi_unclassified(100);k__Fungi_unclassified(100);k__Fungi_unclassified(100);k__Fungi_unclassified(100);k__Fungi_unclassified(100); | - | - | - | - | - | - | - | Unassigned | - |
| Otu16543 | 1 | k__Fungi(100);p__Ascomycota(100);p__Ascomycota_unclassified(100);p__Ascomycota_unclassified(100);p__Ascomycota_unclassified(100);p__Ascomycota_unclassified(100);p__Ascomycota_unclassified(100); | - | - | - | - | - | - | - | Unassigned | - |
| Otu16544 | 1 | k__Fungi(100);k__Fungi_unclassified(100);k__Fungi_unclassified(100);k__Fungi_unclassified(100);k__Fungi_unclassified(100);k__Fungi_unclassified(100);k__Fungi_unclassified(100); | - | - | - | - | - | - | - | Unassigned | - |
| Otu16546 | 1 | k__Fungi(100);p__Ascomycota(100);p__Ascomycota_unclassified(100);p__Ascomycota_unclassified(100);p__Ascomycota_unclassified(100);p__Ascomycota_unclassified(100);p__Ascomycota_unclassified(100); | - | - | - | - | - | - | - | Unassigned | - |
| Otu16547 | 1 | k__Fungi(100);k__Fungi_unclassified(100);k__Fungi_unclassified(100);k__Fungi_unclassified(100);k__Fungi_unclassified(100);k__Fungi_unclassified(100);k__Fungi_unclassified(100); | - | - | - | - | - | - | - | Unassigned | - |
| Otu16558 | 1 | k__Fungi(100);k__Fungi_unclassified(100);k__Fungi_unclassified(100);k__Fungi_unclassified(100);k__Fungi_unclassified(100);k__Fungi_unclassified(100);k__Fungi_unclassified(100); | - | - | - | - | - | - | - | Unassigned | - |
| Otu16561 | 1 | k__Fungi(100);k__Fungi_unclassified(100);k__Fungi_unclassified(100);k__Fungi_unclassified(100);k__Fungi_unclassified(100);k__Fungi_unclassified(100);k__Fungi_unclassified(100); | - | - | - | - | - | - | - | Unassigned | - |
| Otu16567 | 1 | k__Fungi(100);p__Ascomycota(100);p__Ascomycota_unclassified(100);p__Ascomycota_unclassified(100);p__Ascomycota_unclassified(100);p__Ascomycota_unclassified(100);p__Ascomycota_unclassified(100); | - | - | - | - | - | - | - | Unassigned | - |
| Otu16568 | 1 | k__Fungi(100);k__Fungi_unclassified(100);k__Fungi_unclassified(100);k__Fungi_unclassified(100);k__Fungi_unclassified(100);k__Fungi_unclassified(100);k__Fungi_unclassified(100); | - | - | - | - | - | - | - | Unassigned | - |
| Otu16569 | 1 | k__Fungi(100);k__Fungi_unclassified(100);k__Fungi_unclassified(100);k__Fungi_unclassified(100);k__Fungi_unclassified(100);k__Fungi_unclassified(100);k__Fungi_unclassified(100); | - | - | - | - | - | - | - | Unassigned | - |
| Otu16583 | 1 | k__Fungi(100);k__Fungi_unclassified(100);k__Fungi_unclassified(100);k__Fungi_unclassified(100);k__Fungi_unclassified(100);k__Fungi_unclassified(100);k__Fungi_unclassified(100); | - | - | - | - | - | - | - | Unassigned | - |
| Otu16584 | 1 | k__Fungi(100);p__Ascomycota(100);p__Ascomycota_unclassified(100);p__Ascomycota_unclassified(100);p__Ascomycota_unclassified(100);p__Ascomycota_unclassified(100);p__Ascomycota_unclassified(100); | - | - | - | - | - | - | - | Unassigned | - |
| Otu16613 | 1 | k__Fungi(100);k__Fungi_unclassified(100);k__Fungi_unclassified(100);k__Fungi_unclassified(100);k__Fungi_unclassified(100);k__Fungi_unclassified(100);k__Fungi_unclassified(100); | - | - | - | - | - | - | - | Unassigned | - |
| Otu16614 | 1 | k__Fungi(100);p__Ascomycota(100);p__Ascomycota_unclassified(100);p__Ascomycota_unclassified(100);p__Ascomycota_unclassified(100);p__Ascomycota_unclassified(100);p__Ascomycota_unclassified(100); | - | - | - | - | - | - | - | Unassigned | - |
| Otu16620 | 1 | k__Fungi(100);k__Fungi_unclassified(100);k__Fungi_unclassified(100);k__Fungi_unclassified(100);k__Fungi_unclassified(100);k__Fungi_unclassified(100);k__Fungi_unclassified(100); | - | - | - | - | - | - | - | Unassigned | - |
| Otu16621 | 1 | k__Fungi(100);p__Ascomycota(100);p__Ascomycota_unclassified(100);p__Ascomycota_unclassified(100);p__Ascomycota_unclassified(100);p__Ascomycota_unclassified(100);p__Ascomycota_unclassified(100); | - | - | - | - | - | - | - | Unassigned | - |
| Otu16640 | 1 | k__Fungi(100);k__Fungi_unclassified(100);k__Fungi_unclassified(100);k__Fungi_unclassified(100);k__Fungi_unclassified(100);k__Fungi_unclassified(100);k__Fungi_unclassified(100); | - | - | - | - | - | - | - | Unassigned | - |
| Otu16666 | 1 | k__Fungi(100);p__Ascomycota(100);p__Ascomycota_unclassified(100);p__Ascomycota_unclassified(100);p__Ascomycota_unclassified(100);p__Ascomycota_unclassified(100);p__Ascomycota_unclassified(100); | - | - | - | - | - | - | - | Unassigned | - |
| Otu16709 | 1 | k__Fungi(100);k__Fungi_unclassified(100);k__Fungi_unclassified(100);k__Fungi_unclassified(100);k__Fungi_unclassified(100);k__Fungi_unclassified(100);k__Fungi_unclassified(100); | - | - | - | - | - | - | - | Unassigned | - |
| Otu16710 | 1 | k__Fungi(100);k__Fungi_unclassified(100);k__Fungi_unclassified(100);k__Fungi_unclassified(100);k__Fungi_unclassified(100);k__Fungi_unclassified(100);k__Fungi_unclassified(100); | - | - | - | - | - | - | - | Unassigned | - |
| Otu16712 | 1 | k__Fungi(100);k__Fungi_unclassified(100);k__Fungi_unclassified(100);k__Fungi_unclassified(100);k__Fungi_unclassified(100);k__Fungi_unclassified(100);k__Fungi_unclassified(100); | - | - | - | - | - | - | - | Unassigned | - |
| Otu16713 | 1 | k__Fungi(100);k__Fungi_unclassified(100);k__Fungi_unclassified(100);k__Fungi_unclassified(100);k__Fungi_unclassified(100);k__Fungi_unclassified(100);k__Fungi_unclassified(100); | - | - | - | - | - | - | - | Unassigned | - |
| Otu16734 | 1 | k__Fungi(100);p__Ascomycota(100);p__Ascomycota_unclassified(100);p__Ascomycota_unclassified(100);p__Ascomycota_unclassified(100);p__Ascomycota_unclassified(100);p__Ascomycota_unclassified(100); | - | - | - | - | - | - | - | Unassigned | - |
| Otu16735 | 1 | k__Fungi(100);p__Ascomycota(100);p__Ascomycota_unclassified(100);p__Ascomycota_unclassified(100);p__Ascomycota_unclassified(100);p__Ascomycota_unclassified(100);p__Ascomycota_unclassified(100); | - | - | - | - | - | - | - | Unassigned | - |
| Otu16741 | 1 | k__Fungi(100);p__Ascomycota(100);p__Ascomycota_unclassified(100);p__Ascomycota_unclassified(100);p__Ascomycota_unclassified(100);p__Ascomycota_unclassified(100);p__Ascomycota_unclassified(100); | - | - | - | - | - | - | - | Unassigned | - |
| Otu16743 | 1 | k__Fungi(100);p__Ascomycota(100);p__Ascomycota_unclassified(100);p__Ascomycota_unclassified(100);p__Ascomycota_unclassified(100);p__Ascomycota_unclassified(100);p__Ascomycota_unclassified(100); | - | - | - | - | - | - | - | Unassigned | - |
| Otu16744 | 1 | k__Fungi(100);p__Ascomycota(100);p__Ascomycota_unclassified(100);p__Ascomycota_unclassified(100);p__Ascomycota_unclassified(100);p__Ascomycota_unclassified(100);p__Ascomycota_unclassified(100); | - | - | - | - | - | - | - | Unassigned | - |
| Otu16745 | 1 | k__Fungi(100);p__Ascomycota(100);p__Ascomycota_unclassified(100);p__Ascomycota_unclassified(100);p__Ascomycota_unclassified(100);p__Ascomycota_unclassified(100);p__Ascomycota_unclassified(100); | - | - | - | - | - | - | - | Unassigned | - |
| Otu16746 | 1 | k__Fungi(100);p__Ascomycota(100);p__Ascomycota_unclassified(100);p__Ascomycota_unclassified(100);p__Ascomycota_unclassified(100);p__Ascomycota_unclassified(100);p__Ascomycota_unclassified(100); | - | - | - | - | - | - | - | Unassigned | - |
| Otu16747 | 1 | k__Fungi(100);p__Ascomycota(100);p__Ascomycota_unclassified(100);p__Ascomycota_unclassified(100);p__Ascomycota_unclassified(100);p__Ascomycota_unclassified(100);p__Ascomycota_unclassified(100); | - | - | - | - | - | - | - | Unassigned | - |
| Otu16748 | 1 | k__Fungi(100);p__Ascomycota(100);p__Ascomycota_unclassified(100);p__Ascomycota_unclassified(100);p__Ascomycota_unclassified(100);p__Ascomycota_unclassified(100);p__Ascomycota_unclassified(100); | - | - | - | - | - | - | - | Unassigned | - |
| Otu16749 | 1 | k__Fungi(100);p__Ascomycota(100);p__Ascomycota_unclassified(100);p__Ascomycota_unclassified(100);p__Ascomycota_unclassified(100);p__Ascomycota_unclassified(100);p__Ascomycota_unclassified(100); | - | - | - | - | - | - | - | Unassigned | - |
| Otu16750 | 1 | k__Fungi(100);p__Ascomycota(100);p__Ascomycota_unclassified(100);p__Ascomycota_unclassified(100);p__Ascomycota_unclassified(100);p__Ascomycota_unclassified(100);p__Ascomycota_unclassified(100); | - | - | - | - | - | - | - | Unassigned | - |
| Otu16751 | 1 | k__Fungi(100);p__Ascomycota(100);p__Ascomycota_unclassified(100);p__Ascomycota_unclassified(100);p__Ascomycota_unclassified(100);p__Ascomycota_unclassified(100);p__Ascomycota_unclassified(100); | - | - | - | - | - | - | - | Unassigned | - |
| Otu16752 | 1 | k__Fungi(100);k__Fungi_unclassified(100);k__Fungi_unclassified(100);k__Fungi_unclassified(100);k__Fungi_unclassified(100);k__Fungi_unclassified(100);k__Fungi_unclassified(100); | - | - | - | - | - | - | - | Unassigned | - |
| Otu16755 | 1 | k__Fungi(100);p__Ascomycota(100);p__Ascomycota_unclassified(100);p__Ascomycota_unclassified(100);p__Ascomycota_unclassified(100);p__Ascomycota_unclassified(100);p__Ascomycota_unclassified(100); | - | - | - | - | - | - | - | Unassigned | - |
| Otu16757 | 1 | k__Fungi(100);p__Ascomycota(100);p__Ascomycota_unclassified(100);p__Ascomycota_unclassified(100);p__Ascomycota_unclassified(100);p__Ascomycota_unclassified(100);p__Ascomycota_unclassified(100); | - | - | - | - | - | - | - | Unassigned | - |
| Otu16759 | 1 | k__Fungi(100);p__Ascomycota(100);p__Ascomycota_unclassified(100);p__Ascomycota_unclassified(100);p__Ascomycota_unclassified(100);p__Ascomycota_unclassified(100);p__Ascomycota_unclassified(100); | - | - | - | - | - | - | - | Unassigned | - |
| Otu16773 | 1 | k__Fungi(100);p__Ascomycota(100);p__Ascomycota_unclassified(100);p__Ascomycota_unclassified(100);p__Ascomycota_unclassified(100);p__Ascomycota_unclassified(100);p__Ascomycota_unclassified(100); | - | - | - | - | - | - | - | Unassigned | - |
| Otu16777 | 1 | k__Fungi(100);k__Fungi_unclassified(100);k__Fungi_unclassified(100);k__Fungi_unclassified(100);k__Fungi_unclassified(100);k__Fungi_unclassified(100);k__Fungi_unclassified(100); | - | - | - | - | - | - | - | Unassigned | - |
| Otu16783 | 1 | k__Fungi(100);p__Ascomycota(100);p__Ascomycota_unclassified(100);p__Ascomycota_unclassified(100);p__Ascomycota_unclassified(100);p__Ascomycota_unclassified(100);p__Ascomycota_unclassified(100); | - | - | - | - | - | - | - | Unassigned | - |
| Otu16788 | 1 | k__Fungi(100);k__Fungi_unclassified(100);k__Fungi_unclassified(100);k__Fungi_unclassified(100);k__Fungi_unclassified(100);k__Fungi_unclassified(100);k__Fungi_unclassified(100); | - | - | - | - | - | - | - | Unassigned | - |
| Otu16792 | 1 | k__Fungi(100);k__Fungi_unclassified(100);k__Fungi_unclassified(100);k__Fungi_unclassified(100);k__Fungi_unclassified(100);k__Fungi_unclassified(100);k__Fungi_unclassified(100); | - | - | - | - | - | - | - | Unassigned | - |
| Otu16793 | 1 | k__Fungi(100);k__Fungi_unclassified(100);k__Fungi_unclassified(100);k__Fungi_unclassified(100);k__Fungi_unclassified(100);k__Fungi_unclassified(100);k__Fungi_unclassified(100); | - | - | - | - | - | - | - | Unassigned | - |
| Otu16794 | 1 | k__Fungi(100);p__Ascomycota(100);p__Ascomycota_unclassified(100);p__Ascomycota_unclassified(100);p__Ascomycota_unclassified(100);p__Ascomycota_unclassified(100);p__Ascomycota_unclassified(100); | - | - | - | - | - | - | - | Unassigned | - |
| Otu16795 | 1 | k__Fungi(100);k__Fungi_unclassified(100);k__Fungi_unclassified(100);k__Fungi_unclassified(100);k__Fungi_unclassified(100);k__Fungi_unclassified(100);k__Fungi_unclassified(100); | - | - | - | - | - | - | - | Unassigned | - |
| Otu16796 | 1 | k__Fungi(100);k__Fungi_unclassified(100);k__Fungi_unclassified(100);k__Fungi_unclassified(100);k__Fungi_unclassified(100);k__Fungi_unclassified(100);k__Fungi_unclassified(100); | - | - | - | - | - | - | - | Unassigned | - |
| Otu16797 | 1 | k__Fungi(100);p__Ascomycota(100);p__Ascomycota_unclassified(100);p__Ascomycota_unclassified(100);p__Ascomycota_unclassified(100);p__Ascomycota_unclassified(100);p__Ascomycota_unclassified(100); | - | - | - | - | - | - | - | Unassigned | - |
| Otu16798 | 1 | k__Fungi(100);k__Fungi_unclassified(100);k__Fungi_unclassified(100);k__Fungi_unclassified(100);k__Fungi_unclassified(100);k__Fungi_unclassified(100);k__Fungi_unclassified(100); | - | - | - | - | - | - | - | Unassigned | - |
| Otu16804 | 1 | k__Fungi(100);k__Fungi_unclassified(100);k__Fungi_unclassified(100);k__Fungi_unclassified(100);k__Fungi_unclassified(100);k__Fungi_unclassified(100);k__Fungi_unclassified(100); | - | - | - | - | - | - | - | Unassigned | - |
| Otu16809 | 1 | k__Fungi(100);p__Ascomycota(100);p__Ascomycota_unclassified(100);p__Ascomycota_unclassified(100);p__Ascomycota_unclassified(100);p__Ascomycota_unclassified(100);p__Ascomycota_unclassified(100); | - | - | - | - | - | - | - | Unassigned | - |
| Otu16814 | 1 | k__Fungi(100);k__Fungi_unclassified(100);k__Fungi_unclassified(100);k__Fungi_unclassified(100);k__Fungi_unclassified(100);k__Fungi_unclassified(100);k__Fungi_unclassified(100); | - | - | - | - | - | - | - | Unassigned | - |
| Otu16815 | 1 | k__Fungi(100);k__Fungi_unclassified(100);k__Fungi_unclassified(100);k__Fungi_unclassified(100);k__Fungi_unclassified(100);k__Fungi_unclassified(100);k__Fungi_unclassified(100); | - | - | - | - | - | - | - | Unassigned | - |
| Otu16819 | 1 | k__Fungi(100);p__Ascomycota(100);p__Ascomycota_unclassified(100);p__Ascomycota_unclassified(100);p__Ascomycota_unclassified(100);p__Ascomycota_unclassified(100);p__Ascomycota_unclassified(100); | - | - | - | - | - | - | - | Unassigned | - |
| Otu16821 | 1 | k__Fungi(100);k__Fungi_unclassified(100);k__Fungi_unclassified(100);k__Fungi_unclassified(100);k__Fungi_unclassified(100);k__Fungi_unclassified(100);k__Fungi_unclassified(100); | - | - | - | - | - | - | - | Unassigned | - |
| Otu16822 | 1 | k__Fungi(100);k__Fungi_unclassified(100);k__Fungi_unclassified(100);k__Fungi_unclassified(100);k__Fungi_unclassified(100);k__Fungi_unclassified(100);k__Fungi_unclassified(100); | - | - | - | - | - | - | - | Unassigned | - |
| Otu16825 | 1 | k__Fungi(100);p__Ascomycota(100);p__Ascomycota_unclassified(100);p__Ascomycota_unclassified(100);p__Ascomycota_unclassified(100);p__Ascomycota_unclassified(100);p__Ascomycota_unclassified(100); | - | - | - | - | - | - | - | Unassigned | - |
| Otu16827 | 1 | k__Fungi(100);k__Fungi_unclassified(100);k__Fungi_unclassified(100);k__Fungi_unclassified(100);k__Fungi_unclassified(100);k__Fungi_unclassified(100);k__Fungi_unclassified(100); | - | - | - | - | - | - | - | Unassigned | - |
| Otu16828 | 1 | k__Fungi(100);k__Fungi_unclassified(100);k__Fungi_unclassified(100);k__Fungi_unclassified(100);k__Fungi_unclassified(100);k__Fungi_unclassified(100);k__Fungi_unclassified(100); | - | - | - | - | - | - | - | Unassigned | - |
| Otu16829 | 1 | k__Fungi(100);k__Fungi_unclassified(100);k__Fungi_unclassified(100);k__Fungi_unclassified(100);k__Fungi_unclassified(100);k__Fungi_unclassified(100);k__Fungi_unclassified(100); | - | - | - | - | - | - | - | Unassigned | - |
| Otu16831 | 1 | k__Fungi(100);p__Ascomycota(100);p__Ascomycota_unclassified(100);p__Ascomycota_unclassified(100);p__Ascomycota_unclassified(100);p__Ascomycota_unclassified(100);p__Ascomycota_unclassified(100); | - | - | - | - | - | - | - | Unassigned | - |
| Otu16839 | 1 | k__Fungi(100);p__Ascomycota(100);c__Leotiomycetes(100);o__Helotiales(100);f__Helotiales_family_Incertae_sedis(100);f__Helotiales_family_Incertae_sedis_unclassified(100);f__Helotiales_family_Incertae_sedis_unclassified(100); | - | - | - | - | - | - | - | Unassigned | - |
| Otu16841 | 1 | k__Fungi(100);p__Ascomycota(100);p__Ascomycota_unclassified(100);p__Ascomycota_unclassified(100);p__Ascomycota_unclassified(100);p__Ascomycota_unclassified(100);p__Ascomycota_unclassified(100); | - | - | - | - | - | - | - | Unassigned | - |
| Otu16842 | 1 | k__Fungi(100);p__Ascomycota(100);p__Ascomycota_unclassified(100);p__Ascomycota_unclassified(100);p__Ascomycota_unclassified(100);p__Ascomycota_unclassified(100);p__Ascomycota_unclassified(100); | - | - | - | - | - | - | - | Unassigned | - |
| Otu16851 | 1 | k__Fungi(100);p__Ascomycota(100);c__Dothideomycetes(100);o__Capnodiales(100);o__Capnodiales_unclassified(100);o__Capnodiales_unclassified(100);o__Capnodiales_unclassified(100); | - | - | - | - | - | - | - | Unassigned | - |
| Otu16852 | 1 | k__Fungi(100);k__Fungi_unclassified(100);k__Fungi_unclassified(100);k__Fungi_unclassified(100);k__Fungi_unclassified(100);k__Fungi_unclassified(100);k__Fungi_unclassified(100); | - | - | - | - | - | - | - | Unassigned | - |
| Otu16854 | 1 | k__Fungi(100);p__Ascomycota(100);p__Ascomycota_unclassified(100);p__Ascomycota_unclassified(100);p__Ascomycota_unclassified(100);p__Ascomycota_unclassified(100);p__Ascomycota_unclassified(100); | - | - | - | - | - | - | - | Unassigned | - |
| Otu16858 | 1 | k__Fungi(100);p__Ascomycota(100);p__Ascomycota_unclassified(100);p__Ascomycota_unclassified(100);p__Ascomycota_unclassified(100);p__Ascomycota_unclassified(100);p__Ascomycota_unclassified(100); | - | - | - | - | - | - | - | Unassigned | - |
| Otu16861 | 1 | k__Fungi(100);p__Ascomycota(100);c__Sordariomycetes(100);c__Sordariomycetes_unclassified(100);c__Sordariomycetes_unclassified(100);c__Sordariomycetes_unclassified(100);c__Sordariomycetes_unclassified(100); | - | - | - | - | - | - | - | Unassigned | - |
| Otu16862 | 1 | k__Fungi(100);k__Fungi_unclassified(100);k__Fungi_unclassified(100);k__Fungi_unclassified(100);k__Fungi_unclassified(100);k__Fungi_unclassified(100);k__Fungi_unclassified(100); | - | - | - | - | - | - | - | Unassigned | - |
| Otu16869 | 1 | k__Fungi(100);p__Ascomycota(100);p__Ascomycota_unclassified(100);p__Ascomycota_unclassified(100);p__Ascomycota_unclassified(100);p__Ascomycota_unclassified(100);p__Ascomycota_unclassified(100); | - | - | - | - | - | - | - | Unassigned | - |
| Otu16876 | 1 | k__Fungi(100);k__Fungi_unclassified(100);k__Fungi_unclassified(100);k__Fungi_unclassified(100);k__Fungi_unclassified(100);k__Fungi_unclassified(100);k__Fungi_unclassified(100); | - | - | - | - | - | - | - | Unassigned | - |
| Otu16880 | 1 | k__Fungi(100);k__Fungi_unclassified(100);k__Fungi_unclassified(100);k__Fungi_unclassified(100);k__Fungi_unclassified(100);k__Fungi_unclassified(100);k__Fungi_unclassified(100); | - | - | - | - | - | - | - | Unassigned | - |
| Otu16881 | 1 | k__Fungi(100);p__Ascomycota(100);p__Ascomycota_unclassified(100);p__Ascomycota_unclassified(100);p__Ascomycota_unclassified(100);p__Ascomycota_unclassified(100);p__Ascomycota_unclassified(100); | - | - | - | - | - | - | - | Unassigned | - |
| Otu16900 | 1 | k__Fungi(100);p__Ascomycota(100);p__Ascomycota_unclassified(100);p__Ascomycota_unclassified(100);p__Ascomycota_unclassified(100);p__Ascomycota_unclassified(100);p__Ascomycota_unclassified(100); | - | - | - | - | - | - | - | Unassigned | - |
| Otu16914 | 1 | k__Fungi(100);k__Fungi_unclassified(100);k__Fungi_unclassified(100);k__Fungi_unclassified(100);k__Fungi_unclassified(100);k__Fungi_unclassified(100);k__Fungi_unclassified(100); | - | - | - | - | - | - | - | Unassigned | - |
| Otu16935 | 1 | k__Fungi(100);k__Fungi_unclassified(100);k__Fungi_unclassified(100);k__Fungi_unclassified(100);k__Fungi_unclassified(100);k__Fungi_unclassified(100);k__Fungi_unclassified(100); | - | - | - | - | - | - | - | Unassigned | - |
| Otu16936 | 1 | k__Fungi(100);k__Fungi_unclassified(100);k__Fungi_unclassified(100);k__Fungi_unclassified(100);k__Fungi_unclassified(100);k__Fungi_unclassified(100);k__Fungi_unclassified(100); | - | - | - | - | - | - | - | Unassigned | - |
| Otu16938 | 1 | k__Fungi(100);k__Fungi_unclassified(100);k__Fungi_unclassified(100);k__Fungi_unclassified(100);k__Fungi_unclassified(100);k__Fungi_unclassified(100);k__Fungi_unclassified(100); | - | - | - | - | - | - | - | Unassigned | - |
| Otu16943 | 1 | k__Fungi(100);p__Ascomycota(100);p__Ascomycota_unclassified(100);p__Ascomycota_unclassified(100);p__Ascomycota_unclassified(100);p__Ascomycota_unclassified(100);p__Ascomycota_unclassified(100); | - | - | - | - | - | - | - | Unassigned | - |
| Otu16944 | 1 | k__Fungi(100);k__Fungi_unclassified(100);k__Fungi_unclassified(100);k__Fungi_unclassified(100);k__Fungi_unclassified(100);k__Fungi_unclassified(100);k__Fungi_unclassified(100); | - | - | - | - | - | - | - | Unassigned | - |
| Otu16945 | 1 | k__Fungi(100);p__Ascomycota(100);p__Ascomycota_unclassified(100);p__Ascomycota_unclassified(100);p__Ascomycota_unclassified(100);p__Ascomycota_unclassified(100);p__Ascomycota_unclassified(100); | - | - | - | - | - | - | - | Unassigned | - |
| Otu16950 | 1 | k__Fungi(100);k__Fungi_unclassified(100);k__Fungi_unclassified(100);k__Fungi_unclassified(100);k__Fungi_unclassified(100);k__Fungi_unclassified(100);k__Fungi_unclassified(100); | - | - | - | - | - | - | - | Unassigned | - |
| Otu16951 | 1 | k__Fungi(100);p__Ascomycota(100);p__Ascomycota_unclassified(100);p__Ascomycota_unclassified(100);p__Ascomycota_unclassified(100);p__Ascomycota_unclassified(100);p__Ascomycota_unclassified(100); | - | - | - | - | - | - | - | Unassigned | - |
| Otu16955 | 1 | k__Fungi(100);k__Fungi_unclassified(100);k__Fungi_unclassified(100);k__Fungi_unclassified(100);k__Fungi_unclassified(100);k__Fungi_unclassified(100);k__Fungi_unclassified(100); | - | - | - | - | - | - | - | Unassigned | - |
| Otu16956 | 1 | k__Fungi(100);p__Ascomycota(100);p__Ascomycota_unclassified(100);p__Ascomycota_unclassified(100);p__Ascomycota_unclassified(100);p__Ascomycota_unclassified(100);p__Ascomycota_unclassified(100); | - | - | - | - | - | - | - | Unassigned | - |
| Otu16968 | 1 | k__Fungi(100);k__Fungi_unclassified(100);k__Fungi_unclassified(100);k__Fungi_unclassified(100);k__Fungi_unclassified(100);k__Fungi_unclassified(100);k__Fungi_unclassified(100); | - | - | - | - | - | - | - | Unassigned | - |
| Otu16975 | 1 | k__Fungi(100);p__Ascomycota(100);p__Ascomycota_unclassified(100);p__Ascomycota_unclassified(100);p__Ascomycota_unclassified(100);p__Ascomycota_unclassified(100);p__Ascomycota_unclassified(100); | - | - | - | - | - | - | - | Unassigned | - |
| Otu16976 | 1 | k__Fungi(100);p__Ascomycota(100);p__Ascomycota_unclassified(100);p__Ascomycota_unclassified(100);p__Ascomycota_unclassified(100);p__Ascomycota_unclassified(100);p__Ascomycota_unclassified(100); | - | - | - | - | - | - | - | Unassigned | - |
| Otu16977 | 1 | k__Fungi(100);p__Ascomycota(100);p__Ascomycota_unclassified(100);p__Ascomycota_unclassified(100);p__Ascomycota_unclassified(100);p__Ascomycota_unclassified(100);p__Ascomycota_unclassified(100); | - | - | - | - | - | - | - | Unassigned | - |
| Otu16978 | 1 | k__Fungi(100);k__Fungi_unclassified(100);k__Fungi_unclassified(100);k__Fungi_unclassified(100);k__Fungi_unclassified(100);k__Fungi_unclassified(100);k__Fungi_unclassified(100); | - | - | - | - | - | - | - | Unassigned | - |
| Otu16979 | 1 | k__Fungi(100);k__Fungi_unclassified(100);k__Fungi_unclassified(100);k__Fungi_unclassified(100);k__Fungi_unclassified(100);k__Fungi_unclassified(100);k__Fungi_unclassified(100); | - | - | - | - | - | - | - | Unassigned | - |
| Otu16980 | 1 | k__Fungi(100);p__Ascomycota(100);p__Ascomycota_unclassified(100);p__Ascomycota_unclassified(100);p__Ascomycota_unclassified(100);p__Ascomycota_unclassified(100);p__Ascomycota_unclassified(100); | - | - | - | - | - | - | - | Unassigned | - |
| Otu16982 | 1 | k__Fungi(100);k__Fungi_unclassified(100);k__Fungi_unclassified(100);k__Fungi_unclassified(100);k__Fungi_unclassified(100);k__Fungi_unclassified(100);k__Fungi_unclassified(100); | - | - | - | - | - | - | - | Unassigned | - |
| Otu16983 | 1 | k__Fungi(100);k__Fungi_unclassified(100);k__Fungi_unclassified(100);k__Fungi_unclassified(100);k__Fungi_unclassified(100);k__Fungi_unclassified(100);k__Fungi_unclassified(100); | - | - | - | - | - | - | - | Unassigned | - |
| Otu16984 | 1 | k__Fungi(100);k__Fungi_unclassified(100);k__Fungi_unclassified(100);k__Fungi_unclassified(100);k__Fungi_unclassified(100);k__Fungi_unclassified(100);k__Fungi_unclassified(100); | - | - | - | - | - | - | - | Unassigned | - |
| Otu16985 | 1 | k__Fungi(100);p__Ascomycota(100);p__Ascomycota_unclassified(100);p__Ascomycota_unclassified(100);p__Ascomycota_unclassified(100);p__Ascomycota_unclassified(100);p__Ascomycota_unclassified(100); | - | - | - | - | - | - | - | Unassigned | - |
| Otu16990 | 1 | k__Fungi(100);k__Fungi_unclassified(100);k__Fungi_unclassified(100);k__Fungi_unclassified(100);k__Fungi_unclassified(100);k__Fungi_unclassified(100);k__Fungi_unclassified(100); | - | - | - | - | - | - | - | Unassigned | - |
| Otu16991 | 1 | k__Fungi(100);p__Ascomycota(100);p__Ascomycota_unclassified(100);p__Ascomycota_unclassified(100);p__Ascomycota_unclassified(100);p__Ascomycota_unclassified(100);p__Ascomycota_unclassified(100); | - | - | - | - | - | - | - | Unassigned | - |
| Otu16992 | 1 | k__Fungi(100);p__Ascomycota(100);p__Ascomycota_unclassified(100);p__Ascomycota_unclassified(100);p__Ascomycota_unclassified(100);p__Ascomycota_unclassified(100);p__Ascomycota_unclassified(100); | - | - | - | - | - | - | - | Unassigned | - |
| Otu16993 | 1 | k__Fungi(100);k__Fungi_unclassified(100);k__Fungi_unclassified(100);k__Fungi_unclassified(100);k__Fungi_unclassified(100);k__Fungi_unclassified(100);k__Fungi_unclassified(100); | - | - | - | - | - | - | - | Unassigned | - |
| Otu17003 | 1 | k__Fungi(100);p__Ascomycota(100);p__Ascomycota_unclassified(100);p__Ascomycota_unclassified(100);p__Ascomycota_unclassified(100);p__Ascomycota_unclassified(100);p__Ascomycota_unclassified(100); | - | - | - | - | - | - | - | Unassigned | - |
| Otu17014 | 1 | k__Fungi(100);k__Fungi_unclassified(100);k__Fungi_unclassified(100);k__Fungi_unclassified(100);k__Fungi_unclassified(100);k__Fungi_unclassified(100);k__Fungi_unclassified(100); | - | - | - | - | - | - | - | Unassigned | - |
| Otu17016 | 1 | k__Fungi(100);p__Ascomycota(100);p__Ascomycota_unclassified(100);p__Ascomycota_unclassified(100);p__Ascomycota_unclassified(100);p__Ascomycota_unclassified(100);p__Ascomycota_unclassified(100); | - | - | - | - | - | - | - | Unassigned | - |
| Otu17017 | 1 | k__Fungi(100);k__Fungi_unclassified(100);k__Fungi_unclassified(100);k__Fungi_unclassified(100);k__Fungi_unclassified(100);k__Fungi_unclassified(100);k__Fungi_unclassified(100); | - | - | - | - | - | - | - | Unassigned | - |
| Otu17019 | 1 | k__Fungi(100);p__Ascomycota(100);p__Ascomycota_unclassified(100);p__Ascomycota_unclassified(100);p__Ascomycota_unclassified(100);p__Ascomycota_unclassified(100);p__Ascomycota_unclassified(100); | - | - | - | - | - | - | - | Unassigned | - |
| Otu17021 | 1 | k__Fungi(100);k__Fungi_unclassified(100);k__Fungi_unclassified(100);k__Fungi_unclassified(100);k__Fungi_unclassified(100);k__Fungi_unclassified(100);k__Fungi_unclassified(100); | - | - | - | - | - | - | - | Unassigned | - |
| Otu17022 | 1 | k__Fungi(100);p__Ascomycota(100);p__Ascomycota_unclassified(100);p__Ascomycota_unclassified(100);p__Ascomycota_unclassified(100);p__Ascomycota_unclassified(100);p__Ascomycota_unclassified(100); | - | - | - | - | - | - | - | Unassigned | - |
| Otu17027 | 1 | k__Fungi(100);p__Ascomycota(100);p__Ascomycota_unclassified(100);p__Ascomycota_unclassified(100);p__Ascomycota_unclassified(100);p__Ascomycota_unclassified(100);p__Ascomycota_unclassified(100); | - | - | - | - | - | - | - | Unassigned | - |
| Otu17031 | 1 | k__Fungi(100);p__Ascomycota(100);p__Ascomycota_unclassified(100);p__Ascomycota_unclassified(100);p__Ascomycota_unclassified(100);p__Ascomycota_unclassified(100);p__Ascomycota_unclassified(100); | - | - | - | - | - | - | - | Unassigned | - |
| Otu17033 | 1 | k__Fungi(100);k__Fungi_unclassified(100);k__Fungi_unclassified(100);k__Fungi_unclassified(100);k__Fungi_unclassified(100);k__Fungi_unclassified(100);k__Fungi_unclassified(100); | - | - | - | - | - | - | - | Unassigned | - |
| Otu17058 | 1 | k__Fungi(100);k__Fungi_unclassified(100);k__Fungi_unclassified(100);k__Fungi_unclassified(100);k__Fungi_unclassified(100);k__Fungi_unclassified(100);k__Fungi_unclassified(100); | - | - | - | - | - | - | - | Unassigned | - |
| Otu17065 | 1 | k__Fungi(100);p__Ascomycota(100);p__Ascomycota_unclassified(100);p__Ascomycota_unclassified(100);p__Ascomycota_unclassified(100);p__Ascomycota_unclassified(100);p__Ascomycota_unclassified(100); | - | - | - | - | - | - | - | Unassigned | - |
| Otu17072 | 1 | k__Fungi(100);k__Fungi_unclassified(100);k__Fungi_unclassified(100);k__Fungi_unclassified(100);k__Fungi_unclassified(100);k__Fungi_unclassified(100);k__Fungi_unclassified(100); | - | - | - | - | - | - | - | Unassigned | - |
| Otu17080 | 1 | k__Fungi(100);p__Ascomycota(100);p__Ascomycota_unclassified(100);p__Ascomycota_unclassified(100);p__Ascomycota_unclassified(100);p__Ascomycota_unclassified(100);p__Ascomycota_unclassified(100); | - | - | - | - | - | - | - | Unassigned | - |
| Otu17081 | 1 | k__Fungi(100);p__Ascomycota(100);p__Ascomycota_unclassified(100);p__Ascomycota_unclassified(100);p__Ascomycota_unclassified(100);p__Ascomycota_unclassified(100);p__Ascomycota_unclassified(100); | - | - | - | - | - | - | - | Unassigned | - |
| Otu17083 | 1 | k__Fungi(100);k__Fungi_unclassified(100);k__Fungi_unclassified(100);k__Fungi_unclassified(100);k__Fungi_unclassified(100);k__Fungi_unclassified(100);k__Fungi_unclassified(100); | - | - | - | - | - | - | - | Unassigned | - |
| Otu17084 | 1 | k__Fungi(100);k__Fungi_unclassified(100);k__Fungi_unclassified(100);k__Fungi_unclassified(100);k__Fungi_unclassified(100);k__Fungi_unclassified(100);k__Fungi_unclassified(100); | - | - | - | - | - | - | - | Unassigned | - |
| Otu17089 | 1 | k__Fungi(100);k__Fungi_unclassified(100);k__Fungi_unclassified(100);k__Fungi_unclassified(100);k__Fungi_unclassified(100);k__Fungi_unclassified(100);k__Fungi_unclassified(100); | - | - | - | - | - | - | - | Unassigned | - |
| Otu17095 | 1 | k__Fungi(100);k__Fungi_unclassified(100);k__Fungi_unclassified(100);k__Fungi_unclassified(100);k__Fungi_unclassified(100);k__Fungi_unclassified(100);k__Fungi_unclassified(100); | - | - | - | - | - | - | - | Unassigned | - |
| Otu17099 | 1 | k__Fungi(100);k__Fungi_unclassified(100);k__Fungi_unclassified(100);k__Fungi_unclassified(100);k__Fungi_unclassified(100);k__Fungi_unclassified(100);k__Fungi_unclassified(100); | - | - | - | - | - | - | - | Unassigned | - |
| Otu17100 | 1 | k__Fungi(100);k__Fungi_unclassified(100);k__Fungi_unclassified(100);k__Fungi_unclassified(100);k__Fungi_unclassified(100);k__Fungi_unclassified(100);k__Fungi_unclassified(100); | - | - | - | - | - | - | - | Unassigned | - |
| Otu17102 | 1 | k__Fungi(100);k__Fungi_unclassified(100);k__Fungi_unclassified(100);k__Fungi_unclassified(100);k__Fungi_unclassified(100);k__Fungi_unclassified(100);k__Fungi_unclassified(100); | - | - | - | - | - | - | - | Unassigned | - |
| Otu17103 | 1 | k__Fungi(100);p__Ascomycota(100);p__Ascomycota_unclassified(100);p__Ascomycota_unclassified(100);p__Ascomycota_unclassified(100);p__Ascomycota_unclassified(100);p__Ascomycota_unclassified(100); | - | - | - | - | - | - | - | Unassigned | - |
| Otu17105 | 1 | k__Fungi(100);k__Fungi_unclassified(100);k__Fungi_unclassified(100);k__Fungi_unclassified(100);k__Fungi_unclassified(100);k__Fungi_unclassified(100);k__Fungi_unclassified(100); | - | - | - | - | - | - | - | Unassigned | - |
| Otu17106 | 1 | k__Fungi(100);k__Fungi_unclassified(100);k__Fungi_unclassified(100);k__Fungi_unclassified(100);k__Fungi_unclassified(100);k__Fungi_unclassified(100);k__Fungi_unclassified(100); | - | - | - | - | - | - | - | Unassigned | - |
| Otu17110 | 1 | k__Fungi(100);p__Ascomycota(100);p__Ascomycota_unclassified(100);p__Ascomycota_unclassified(100);p__Ascomycota_unclassified(100);p__Ascomycota_unclassified(100);p__Ascomycota_unclassified(100); | - | - | - | - | - | - | - | Unassigned | - |
| Otu17111 | 1 | k__Fungi(100);k__Fungi_unclassified(100);k__Fungi_unclassified(100);k__Fungi_unclassified(100);k__Fungi_unclassified(100);k__Fungi_unclassified(100);k__Fungi_unclassified(100); | - | - | - | - | - | - | - | Unassigned | - |
| Otu17116 | 1 | k__Fungi(100);k__Fungi_unclassified(100);k__Fungi_unclassified(100);k__Fungi_unclassified(100);k__Fungi_unclassified(100);k__Fungi_unclassified(100);k__Fungi_unclassified(100); | - | - | - | - | - | - | - | Unassigned | - |
| Otu17117 | 1 | k__Fungi(100);p__Ascomycota(100);p__Ascomycota_unclassified(100);p__Ascomycota_unclassified(100);p__Ascomycota_unclassified(100);p__Ascomycota_unclassified(100);p__Ascomycota_unclassified(100); | - | - | - | - | - | - | - | Unassigned | - |
| Otu17118 | 1 | k__Fungi(100);k__Fungi_unclassified(100);k__Fungi_unclassified(100);k__Fungi_unclassified(100);k__Fungi_unclassified(100);k__Fungi_unclassified(100);k__Fungi_unclassified(100); | - | - | - | - | - | - | - | Unassigned | - |
| Otu17119 | 1 | k__Fungi(100);k__Fungi_unclassified(100);k__Fungi_unclassified(100);k__Fungi_unclassified(100);k__Fungi_unclassified(100);k__Fungi_unclassified(100);k__Fungi_unclassified(100); | - | - | - | - | - | - | - | Unassigned | - |
| Otu17120 | 1 | k__Fungi(100);k__Fungi_unclassified(100);k__Fungi_unclassified(100);k__Fungi_unclassified(100);k__Fungi_unclassified(100);k__Fungi_unclassified(100);k__Fungi_unclassified(100); | - | - | - | - | - | - | - | Unassigned | - |
| Otu17131 | 1 | k__Fungi(100);p__Ascomycota(100);p__Ascomycota_unclassified(100);p__Ascomycota_unclassified(100);p__Ascomycota_unclassified(100);p__Ascomycota_unclassified(100);p__Ascomycota_unclassified(100); | - | - | - | - | - | - | - | Unassigned | - |
| Otu17134 | 1 | k__Fungi(100);p__Ascomycota(100);p__Ascomycota_unclassified(100);p__Ascomycota_unclassified(100);p__Ascomycota_unclassified(100);p__Ascomycota_unclassified(100);p__Ascomycota_unclassified(100); | - | - | - | - | - | - | - | Unassigned | - |
| Otu17136 | 1 | k__Fungi(100);p__Ascomycota(100);p__Ascomycota_unclassified(100);p__Ascomycota_unclassified(100);p__Ascomycota_unclassified(100);p__Ascomycota_unclassified(100);p__Ascomycota_unclassified(100); | - | - | - | - | - | - | - | Unassigned | - |
| Otu17139 | 1 | k__Fungi(100);k__Fungi_unclassified(100);k__Fungi_unclassified(100);k__Fungi_unclassified(100);k__Fungi_unclassified(100);k__Fungi_unclassified(100);k__Fungi_unclassified(100); | - | - | - | - | - | - | - | Unassigned | - |
| Otu17140 | 1 | k__Fungi(100);k__Fungi_unclassified(100);k__Fungi_unclassified(100);k__Fungi_unclassified(100);k__Fungi_unclassified(100);k__Fungi_unclassified(100);k__Fungi_unclassified(100); | - | - | - | - | - | - | - | Unassigned | - |
| Otu17141 | 1 | k__Fungi(100);p__Ascomycota(100);p__Ascomycota_unclassified(100);p__Ascomycota_unclassified(100);p__Ascomycota_unclassified(100);p__Ascomycota_unclassified(100);p__Ascomycota_unclassified(100); | - | - | - | - | - | - | - | Unassigned | - |
| Otu17142 | 1 | k__Fungi(100);p__Ascomycota(100);p__Ascomycota_unclassified(100);p__Ascomycota_unclassified(100);p__Ascomycota_unclassified(100);p__Ascomycota_unclassified(100);p__Ascomycota_unclassified(100); | - | - | - | - | - | - | - | Unassigned | - |
| Otu17144 | 1 | k__Fungi(100);k__Fungi_unclassified(100);k__Fungi_unclassified(100);k__Fungi_unclassified(100);k__Fungi_unclassified(100);k__Fungi_unclassified(100);k__Fungi_unclassified(100); | - | - | - | - | - | - | - | Unassigned | - |
| Otu17146 | 1 | k__Fungi(100);k__Fungi_unclassified(100);k__Fungi_unclassified(100);k__Fungi_unclassified(100);k__Fungi_unclassified(100);k__Fungi_unclassified(100);k__Fungi_unclassified(100); | - | - | - | - | - | - | - | Unassigned | - |
| Otu17147 | 1 | k__Fungi(100);k__Fungi_unclassified(100);k__Fungi_unclassified(100);k__Fungi_unclassified(100);k__Fungi_unclassified(100);k__Fungi_unclassified(100);k__Fungi_unclassified(100); | - | - | - | - | - | - | - | Unassigned | - |
| Otu17149 | 1 | k__Fungi(100);p__Ascomycota(100);p__Ascomycota_unclassified(100);p__Ascomycota_unclassified(100);p__Ascomycota_unclassified(100);p__Ascomycota_unclassified(100);p__Ascomycota_unclassified(100); | - | - | - | - | - | - | - | Unassigned | - |
| Otu17158 | 1 | k__Fungi(100);p__Ascomycota(100);p__Ascomycota_unclassified(100);p__Ascomycota_unclassified(100);p__Ascomycota_unclassified(100);p__Ascomycota_unclassified(100);p__Ascomycota_unclassified(100); | - | - | - | - | - | - | - | Unassigned | - |
| Otu17165 | 1 | k__Fungi(100);p__Ascomycota(100);p__Ascomycota_unclassified(100);p__Ascomycota_unclassified(100);p__Ascomycota_unclassified(100);p__Ascomycota_unclassified(100);p__Ascomycota_unclassified(100); | - | - | - | - | - | - | - | Unassigned | - |
| Otu17167 | 1 | k__Fungi(100);p__Ascomycota(100);p__Ascomycota_unclassified(100);p__Ascomycota_unclassified(100);p__Ascomycota_unclassified(100);p__Ascomycota_unclassified(100);p__Ascomycota_unclassified(100); | - | - | - | - | - | - | - | Unassigned | - |
| Otu17172 | 1 | k__Fungi(100);p__Ascomycota(100);p__Ascomycota_unclassified(100);p__Ascomycota_unclassified(100);p__Ascomycota_unclassified(100);p__Ascomycota_unclassified(100);p__Ascomycota_unclassified(100); | - | - | - | - | - | - | - | Unassigned | - |
| Otu17174 | 1 | k__Fungi(100);p__Ascomycota(100);p__Ascomycota_unclassified(100);p__Ascomycota_unclassified(100);p__Ascomycota_unclassified(100);p__Ascomycota_unclassified(100);p__Ascomycota_unclassified(100); | - | - | - | - | - | - | - | Unassigned | - |
| Otu17178 | 1 | k__Fungi(100);p__Ascomycota(100);p__Ascomycota_unclassified(100);p__Ascomycota_unclassified(100);p__Ascomycota_unclassified(100);p__Ascomycota_unclassified(100);p__Ascomycota_unclassified(100); | - | - | - | - | - | - | - | Unassigned | - |
| Otu17179 | 1 | k__Fungi(100);k__Fungi_unclassified(100);k__Fungi_unclassified(100);k__Fungi_unclassified(100);k__Fungi_unclassified(100);k__Fungi_unclassified(100);k__Fungi_unclassified(100); | - | - | - | - | - | - | - | Unassigned | - |
| Otu17183 | 1 | k__Fungi(100);p__Ascomycota(100);p__Ascomycota_unclassified(100);p__Ascomycota_unclassified(100);p__Ascomycota_unclassified(100);p__Ascomycota_unclassified(100);p__Ascomycota_unclassified(100); | - | - | - | - | - | - | - | Unassigned | - |
| Otu17185 | 1 | k__Fungi(100);k__Fungi_unclassified(100);k__Fungi_unclassified(100);k__Fungi_unclassified(100);k__Fungi_unclassified(100);k__Fungi_unclassified(100);k__Fungi_unclassified(100); | - | - | - | - | - | - | - | Unassigned | - |
| Otu17186 | 1 | k__Fungi(100);k__Fungi_unclassified(100);k__Fungi_unclassified(100);k__Fungi_unclassified(100);k__Fungi_unclassified(100);k__Fungi_unclassified(100);k__Fungi_unclassified(100); | - | - | - | - | - | - | - | Unassigned | - |
| Otu17189 | 1 | k__Fungi(100);k__Fungi_unclassified(100);k__Fungi_unclassified(100);k__Fungi_unclassified(100);k__Fungi_unclassified(100);k__Fungi_unclassified(100);k__Fungi_unclassified(100); | - | - | - | - | - | - | - | Unassigned | - |
| Otu17226 | 1 | k__Fungi(100);p__Ascomycota(100);p__Ascomycota_unclassified(100);p__Ascomycota_unclassified(100);p__Ascomycota_unclassified(100);p__Ascomycota_unclassified(100);p__Ascomycota_unclassified(100); | - | - | - | - | - | - | - | Unassigned | - |
| Otu17236 | 1 | k__Fungi(100);k__Fungi_unclassified(100);k__Fungi_unclassified(100);k__Fungi_unclassified(100);k__Fungi_unclassified(100);k__Fungi_unclassified(100);k__Fungi_unclassified(100); | - | - | - | - | - | - | - | Unassigned | - |
| Otu17237 | 1 | k__Fungi(100);k__Fungi_unclassified(100);k__Fungi_unclassified(100);k__Fungi_unclassified(100);k__Fungi_unclassified(100);k__Fungi_unclassified(100);k__Fungi_unclassified(100); | - | - | - | - | - | - | - | Unassigned | - |
| Otu17246 | 1 | k__Fungi(100);k__Fungi_unclassified(100);k__Fungi_unclassified(100);k__Fungi_unclassified(100);k__Fungi_unclassified(100);k__Fungi_unclassified(100);k__Fungi_unclassified(100); | - | - | - | - | - | - | - | Unassigned | - |
| Otu17258 | 1 | k__Fungi(100);p__Ascomycota(100);p__Ascomycota_unclassified(100);p__Ascomycota_unclassified(100);p__Ascomycota_unclassified(100);p__Ascomycota_unclassified(100);p__Ascomycota_unclassified(100); | - | - | - | - | - | - | - | Unassigned | - |
| Otu17259 | 1 | k__Fungi(100);k__Fungi_unclassified(100);k__Fungi_unclassified(100);k__Fungi_unclassified(100);k__Fungi_unclassified(100);k__Fungi_unclassified(100);k__Fungi_unclassified(100); | - | - | - | - | - | - | - | Unassigned | - |
| Otu17261 | 1 | k__Fungi(100);k__Fungi_unclassified(100);k__Fungi_unclassified(100);k__Fungi_unclassified(100);k__Fungi_unclassified(100);k__Fungi_unclassified(100);k__Fungi_unclassified(100); | - | - | - | - | - | - | - | Unassigned | - |
| Otu17262 | 1 | k__Fungi(100);k__Fungi_unclassified(100);k__Fungi_unclassified(100);k__Fungi_unclassified(100);k__Fungi_unclassified(100);k__Fungi_unclassified(100);k__Fungi_unclassified(100); | - | - | - | - | - | - | - | Unassigned | - |
| Otu17269 | 1 | k__Fungi(100);k__Fungi_unclassified(100);k__Fungi_unclassified(100);k__Fungi_unclassified(100);k__Fungi_unclassified(100);k__Fungi_unclassified(100);k__Fungi_unclassified(100); | - | - | - | - | - | - | - | Unassigned | - |
| Otu17280 | 1 | k__Fungi(100);k__Fungi_unclassified(100);k__Fungi_unclassified(100);k__Fungi_unclassified(100);k__Fungi_unclassified(100);k__Fungi_unclassified(100);k__Fungi_unclassified(100); | - | - | - | - | - | - | - | Unassigned | - |
| Otu17282 | 1 | k__Fungi(100);p__Ascomycota(100);p__Ascomycota_unclassified(100);p__Ascomycota_unclassified(100);p__Ascomycota_unclassified(100);p__Ascomycota_unclassified(100);p__Ascomycota_unclassified(100); | - | - | - | - | - | - | - | Unassigned | - |
| Otu17283 | 1 | k__Fungi(100);k__Fungi_unclassified(100);k__Fungi_unclassified(100);k__Fungi_unclassified(100);k__Fungi_unclassified(100);k__Fungi_unclassified(100);k__Fungi_unclassified(100); | - | - | - | - | - | - | - | Unassigned | - |
| Otu17286 | 1 | k__Fungi(100);k__Fungi_unclassified(100);k__Fungi_unclassified(100);k__Fungi_unclassified(100);k__Fungi_unclassified(100);k__Fungi_unclassified(100);k__Fungi_unclassified(100); | - | - | - | - | - | - | - | Unassigned | - |
| Otu17287 | 1 | k__Fungi(100);p__Ascomycota(100);p__Ascomycota_unclassified(100);p__Ascomycota_unclassified(100);p__Ascomycota_unclassified(100);p__Ascomycota_unclassified(100);p__Ascomycota_unclassified(100); | - | - | - | - | - | - | - | Unassigned | - |
| Otu17288 | 1 | k__Fungi(100);p__Ascomycota(100);p__Ascomycota_unclassified(100);p__Ascomycota_unclassified(100);p__Ascomycota_unclassified(100);p__Ascomycota_unclassified(100);p__Ascomycota_unclassified(100); | - | - | - | - | - | - | - | Unassigned | - |
| Otu17289 | 1 | k__Fungi(100);p__Ascomycota(100);p__Ascomycota_unclassified(100);p__Ascomycota_unclassified(100);p__Ascomycota_unclassified(100);p__Ascomycota_unclassified(100);p__Ascomycota_unclassified(100); | - | - | - | - | - | - | - | Unassigned | - |
| Otu17290 | 1 | k__Fungi(100);k__Fungi_unclassified(100);k__Fungi_unclassified(100);k__Fungi_unclassified(100);k__Fungi_unclassified(100);k__Fungi_unclassified(100);k__Fungi_unclassified(100); | - | - | - | - | - | - | - | Unassigned | - |
| Otu17291 | 1 | k__Fungi(100);k__Fungi_unclassified(100);k__Fungi_unclassified(100);k__Fungi_unclassified(100);k__Fungi_unclassified(100);k__Fungi_unclassified(100);k__Fungi_unclassified(100); | - | - | - | - | - | - | - | Unassigned | - |
| Otu17292 | 1 | k__Fungi(100);k__Fungi_unclassified(100);k__Fungi_unclassified(100);k__Fungi_unclassified(100);k__Fungi_unclassified(100);k__Fungi_unclassified(100);k__Fungi_unclassified(100); | - | - | - | - | - | - | - | Unassigned | - |
| Otu17293 | 1 | k__Fungi(100);p__Ascomycota(100);p__Ascomycota_unclassified(100);p__Ascomycota_unclassified(100);p__Ascomycota_unclassified(100);p__Ascomycota_unclassified(100);p__Ascomycota_unclassified(100); | - | - | - | - | - | - | - | Unassigned | - |
| Otu17294 | 1 | k__Fungi(100);p__Ascomycota(100);p__Ascomycota_unclassified(100);p__Ascomycota_unclassified(100);p__Ascomycota_unclassified(100);p__Ascomycota_unclassified(100);p__Ascomycota_unclassified(100); | - | - | - | - | - | - | - | Unassigned | - |
| Otu17296 | 1 | k__Fungi(100);k__Fungi_unclassified(100);k__Fungi_unclassified(100);k__Fungi_unclassified(100);k__Fungi_unclassified(100);k__Fungi_unclassified(100);k__Fungi_unclassified(100); | - | - | - | - | - | - | - | Unassigned | - |
| Otu17297 | 1 | k__Fungi(100);p__Ascomycota(100);p__Ascomycota_unclassified(100);p__Ascomycota_unclassified(100);p__Ascomycota_unclassified(100);p__Ascomycota_unclassified(100);p__Ascomycota_unclassified(100); | - | - | - | - | - | - | - | Unassigned | - |
| Otu17303 | 1 | k__Fungi(100);p__Ascomycota(100);p__Ascomycota_unclassified(100);p__Ascomycota_unclassified(100);p__Ascomycota_unclassified(100);p__Ascomycota_unclassified(100);p__Ascomycota_unclassified(100); | - | - | - | - | - | - | - | Unassigned | - |
| Otu17304 | 1 | k__Fungi(100);p__Ascomycota(100);p__Ascomycota_unclassified(100);p__Ascomycota_unclassified(100);p__Ascomycota_unclassified(100);p__Ascomycota_unclassified(100);p__Ascomycota_unclassified(100); | - | - | - | - | - | - | - | Unassigned | - |
| Otu17307 | 1 | k__Fungi(100);p__Ascomycota(100);p__Ascomycota_unclassified(100);p__Ascomycota_unclassified(100);p__Ascomycota_unclassified(100);p__Ascomycota_unclassified(100);p__Ascomycota_unclassified(100); | - | - | - | - | - | - | - | Unassigned | - |
| Otu17308 | 1 | k__Fungi(100);k__Fungi_unclassified(100);k__Fungi_unclassified(100);k__Fungi_unclassified(100);k__Fungi_unclassified(100);k__Fungi_unclassified(100);k__Fungi_unclassified(100); | - | - | - | - | - | - | - | Unassigned | - |
| Otu17309 | 1 | k__Fungi(100);p__Ascomycota(100);p__Ascomycota_unclassified(100);p__Ascomycota_unclassified(100);p__Ascomycota_unclassified(100);p__Ascomycota_unclassified(100);p__Ascomycota_unclassified(100); | - | - | - | - | - | - | - | Unassigned | - |
| Otu17310 | 1 | k__Fungi(100);p__Ascomycota(100);p__Ascomycota_unclassified(100);p__Ascomycota_unclassified(100);p__Ascomycota_unclassified(100);p__Ascomycota_unclassified(100);p__Ascomycota_unclassified(100); | - | - | - | - | - | - | - | Unassigned | - |
| Otu17311 | 1 | k__Fungi(100);k__Fungi_unclassified(100);k__Fungi_unclassified(100);k__Fungi_unclassified(100);k__Fungi_unclassified(100);k__Fungi_unclassified(100);k__Fungi_unclassified(100); | - | - | - | - | - | - | - | Unassigned | - |
| Otu17312 | 1 | k__Fungi(100);p__Ascomycota(100);p__Ascomycota_unclassified(100);p__Ascomycota_unclassified(100);p__Ascomycota_unclassified(100);p__Ascomycota_unclassified(100);p__Ascomycota_unclassified(100); | - | - | - | - | - | - | - | Unassigned | - |
| Otu17319 | 1 | k__Fungi(100);p__Ascomycota(100);p__Ascomycota_unclassified(100);p__Ascomycota_unclassified(100);p__Ascomycota_unclassified(100);p__Ascomycota_unclassified(100);p__Ascomycota_unclassified(100); | - | - | - | - | - | - | - | Unassigned | - |
| Otu17322 | 1 | k__Fungi(100);p__Ascomycota(100);p__Ascomycota_unclassified(100);p__Ascomycota_unclassified(100);p__Ascomycota_unclassified(100);p__Ascomycota_unclassified(100);p__Ascomycota_unclassified(100); | - | - | - | - | - | - | - | Unassigned | - |
| Otu17329 | 1 | k__Fungi(100);k__Fungi_unclassified(100);k__Fungi_unclassified(100);k__Fungi_unclassified(100);k__Fungi_unclassified(100);k__Fungi_unclassified(100);k__Fungi_unclassified(100); | - | - | - | - | - | - | - | Unassigned | - |
| Otu17331 | 1 | k__Fungi(100);k__Fungi_unclassified(100);k__Fungi_unclassified(100);k__Fungi_unclassified(100);k__Fungi_unclassified(100);k__Fungi_unclassified(100);k__Fungi_unclassified(100); | - | - | - | - | - | - | - | Unassigned | - |
| Otu17332 | 1 | k__Fungi(100);p__Ascomycota(100);p__Ascomycota_unclassified(100);p__Ascomycota_unclassified(100);p__Ascomycota_unclassified(100);p__Ascomycota_unclassified(100);p__Ascomycota_unclassified(100); | - | - | - | - | - | - | - | Unassigned | - |
| Otu17333 | 1 | k__Fungi(100);p__Ascomycota(100);p__Ascomycota_unclassified(100);p__Ascomycota_unclassified(100);p__Ascomycota_unclassified(100);p__Ascomycota_unclassified(100);p__Ascomycota_unclassified(100); | - | - | - | - | - | - | - | Unassigned | - |

| | | | | | | | | | | | |
|---|---|---|---|---|---|---|---|---|---|---|---|
| Otu17336 | 1 | k__Fungi(100);p__Ascomycota(100);p__Ascomycota_unclassified(100);p__Ascomycota_unclassified(100);p__Ascomycota_unclassified(100);p__Ascomycota_unclassified(100);p__Ascomycota_unclassified(100); | - | - | - | - | - | - | - | Unassigned | - |
| Otu17337 | 1 | k__Fungi(100);k__Fungi_unclassified(100);k__Fungi_unclassified(100);k__Fungi_unclassified(100);k__Fungi_unclassified(100);k__Fungi_unclassified(100);k__Fungi_unclassified(100); | - | - | - | - | - | - | - | Unassigned | - |
| Otu17338 | 1 | k__Fungi(100);p__Ascomycota(100);p__Ascomycota_unclassified(100);p__Ascomycota_unclassified(100);p__Ascomycota_unclassified(100);p__Ascomycota_unclassified(100);p__Ascomycota_unclassified(100); | - | - | - | - | - | - | - | Unassigned | - |
| Otu17339 | 1 | k__Fungi(100);k__Fungi_unclassified(100);k__Fungi_unclassified(100);k__Fungi_unclassified(100);k__Fungi_unclassified(100);k__Fungi_unclassified(100);k__Fungi_unclassified(100); | - | - | - | - | - | - | - | Unassigned | - |
| Otu17340 | 1 | k__Fungi(100);p__Ascomycota(100);p__Ascomycota_unclassified(100);p__Ascomycota_unclassified(100);p__Ascomycota_unclassified(100);p__Ascomycota_unclassified(100);p__Ascomycota_unclassified(100); | - | - | - | - | - | - | - | Unassigned | - |
| Otu17341 | 1 | k__Fungi(100);k__Fungi_unclassified(100);k__Fungi_unclassified(100);k__Fungi_unclassified(100);k__Fungi_unclassified(100);k__Fungi_unclassified(100);k__Fungi_unclassified(100); | - | - | - | - | - | - | - | Unassigned | - |
| Otu17342 | 1 | k__Fungi(100);p__Ascomycota(100);p__Ascomycota_unclassified(100);p__Ascomycota_unclassified(100);p__Ascomycota_unclassified(100);p__Ascomycota_unclassified(100);p__Ascomycota_unclassified(100); | - | - | - | - | - | - | - | Unassigned | - |
| Otu17343 | 1 | k__Fungi(100);k__Fungi_unclassified(100);k__Fungi_unclassified(100);k__Fungi_unclassified(100);k__Fungi_unclassified(100);k__Fungi_unclassified(100);k__Fungi_unclassified(100); | - | - | - | - | - | - | - | Unassigned | - |
| Otu17346 | 1 | k__Fungi(100);k__Fungi_unclassified(100);k__Fungi_unclassified(100);k__Fungi_unclassified(100);k__Fungi_unclassified(100);k__Fungi_unclassified(100);k__Fungi_unclassified(100); | - | - | - | - | - | - | - | Unassigned | - |
| Otu17347 | 1 | k__Fungi(100);p__Ascomycota(100);p__Ascomycota_unclassified(100);p__Ascomycota_unclassified(100);p__Ascomycota_unclassified(100);p__Ascomycota_unclassified(100);p__Ascomycota_unclassified(100); | - | - | - | - | - | - | - | Unassigned | - |
| Otu17348 | 1 | k__Fungi(100);p__Ascomycota(100);p__Ascomycota_unclassified(100);p__Ascomycota_unclassified(100);p__Ascomycota_unclassified(100);p__Ascomycota_unclassified(100);p__Ascomycota_unclassified(100); | - | - | - | - | - | - | - | Unassigned | - |
| Otu17349 | 1 | k__Fungi(100);p__Ascomycota(100);p__Ascomycota_unclassified(100);p__Ascomycota_unclassified(100);p__Ascomycota_unclassified(100);p__Ascomycota_unclassified(100);p__Ascomycota_unclassified(100); | - | - | - | - | - | - | - | Unassigned | - |
| Otu17350 | 1 | k__Fungi(100);k__Fungi_unclassified(100);k__Fungi_unclassified(100);k__Fungi_unclassified(100);k__Fungi_unclassified(100);k__Fungi_unclassified(100);k__Fungi_unclassified(100); | - | - | - | - | - | - | - | Unassigned | - |
| Otu17351 | 1 | k__Fungi(100);k__Fungi_unclassified(100);k__Fungi_unclassified(100);k__Fungi_unclassified(100);k__Fungi_unclassified(100);k__Fungi_unclassified(100);k__Fungi_unclassified(100); | - | - | - | - | - | - | - | Unassigned | - |
| Otu17353 | 1 | k__Fungi(100);p__Ascomycota(100);p__Ascomycota_unclassified(100);p__Ascomycota_unclassified(100);p__Ascomycota_unclassified(100);p__Ascomycota_unclassified(100);p__Ascomycota_unclassified(100); | - | - | - | - | - | - | - | Unassigned | - |
| Otu17354 | 1 | k__Fungi(100);p__Ascomycota(100);p__Ascomycota_unclassified(100);p__Ascomycota_unclassified(100);p__Ascomycota_unclassified(100);p__Ascomycota_unclassified(100);p__Ascomycota_unclassified(100); | - | - | - | - | - | - | - | Unassigned | - |
| Otu17358 | 1 | k__Fungi(100);k__Fungi_unclassified(100);k__Fungi_unclassified(100);k__Fungi_unclassified(100);k__Fungi_unclassified(100);k__Fungi_unclassified(100);k__Fungi_unclassified(100); | - | - | - | - | - | - | - | Unassigned | - |
| Otu17359 | 1 | k__Fungi(100);k__Fungi_unclassified(100);k__Fungi_unclassified(100);k__Fungi_unclassified(100);k__Fungi_unclassified(100);k__Fungi_unclassified(100);k__Fungi_unclassified(100); | - | - | - | - | - | - | - | Unassigned | - |
| Otu17360 | 1 | k__Fungi(100);p__Ascomycota(100);p__Ascomycota_unclassified(100);p__Ascomycota_unclassified(100);p__Ascomycota_unclassified(100);p__Ascomycota_unclassified(100);p__Ascomycota_unclassified(100); | - | - | - | - | - | - | - | Unassigned | - |
| Otu17361 | 1 | k__Fungi(100);k__Fungi_unclassified(100);k__Fungi_unclassified(100);k__Fungi_unclassified(100);k__Fungi_unclassified(100);k__Fungi_unclassified(100);k__Fungi_unclassified(100); | - | - | - | - | - | - | - | Unassigned | - |
| Otu17363 | 1 | k__Fungi(100);k__Fungi_unclassified(100);k__Fungi_unclassified(100);k__Fungi_unclassified(100);k__Fungi_unclassified(100);k__Fungi_unclassified(100);k__Fungi_unclassified(100); | - | - | - | - | - | - | - | Unassigned | - |
| Otu17364 | 1 | k__Fungi(100);k__Fungi_unclassified(100);k__Fungi_unclassified(100);k__Fungi_unclassified(100);k__Fungi_unclassified(100);k__Fungi_unclassified(100);k__Fungi_unclassified(100); | - | - | - | - | - | - | - | Unassigned | - |
| Otu17365 | 1 | k__Fungi(100);p__Ascomycota(100);p__Ascomycota_unclassified(100);p__Ascomycota_unclassified(100);p__Ascomycota_unclassified(100);p__Ascomycota_unclassified(100);p__Ascomycota_unclassified(100); | - | - | - | - | - | - | - | Unassigned | - |
| Otu17367 | 1 | k__Fungi(100);p__Ascomycota(100);p__Ascomycota_unclassified(100);p__Ascomycota_unclassified(100);p__Ascomycota_unclassified(100);p__Ascomycota_unclassified(100);p__Ascomycota_unclassified(100); | - | - | - | - | - | - | - | Unassigned | - |
| Otu17368 | 1 | k__Fungi(100);p__Ascomycota(100);p__Ascomycota_unclassified(100);p__Ascomycota_unclassified(100);p__Ascomycota_unclassified(100);p__Ascomycota_unclassified(100);p__Ascomycota_unclassified(100); | - | - | - | - | - | - | - | Unassigned | - |
| Otu17369 | 1 | k__Fungi(100);p__Ascomycota(100);p__Ascomycota_unclassified(100);p__Ascomycota_unclassified(100);p__Ascomycota_unclassified(100);p__Ascomycota_unclassified(100);p__Ascomycota_unclassified(100); | - | - | - | - | - | - | - | Unassigned | - |
| Otu17370 | 1 | k__Fungi(100);k__Fungi_unclassified(100);k__Fungi_unclassified(100);k__Fungi_unclassified(100);k__Fungi_unclassified(100);k__Fungi_unclassified(100);k__Fungi_unclassified(100); | - | - | - | - | - | - | - | Unassigned | - |
| Otu17372 | 1 | k__Fungi(100);p__Ascomycota(100);p__Ascomycota_unclassified(100);p__Ascomycota_unclassified(100);p__Ascomycota_unclassified(100);p__Ascomycota_unclassified(100);p__Ascomycota_unclassified(100); | - | - | - | - | - | - | - | Unassigned | - |
| Otu17374 | 1 | k__Fungi(100);p__Ascomycota(100);p__Ascomycota_unclassified(100);p__Ascomycota_unclassified(100);p__Ascomycota_unclassified(100);p__Ascomycota_unclassified(100);p__Ascomycota_unclassified(100); | - | - | - | - | - | - | - | Unassigned | - |
| Otu17375 | 1 | k__Fungi(100);p__Ascomycota(100);p__Ascomycota_unclassified(100);p__Ascomycota_unclassified(100);p__Ascomycota_unclassified(100);p__Ascomycota_unclassified(100);p__Ascomycota_unclassified(100); | - | - | - | - | - | - | - | Unassigned | - |
| Otu17377 | 1 | k__Fungi(100);p__Ascomycota(100);p__Ascomycota_unclassified(100);p__Ascomycota_unclassified(100);p__Ascomycota_unclassified(100);p__Ascomycota_unclassified(100);p__Ascomycota_unclassified(100); | - | - | - | - | - | - | - | Unassigned | - |
| Otu17378 | 1 | k__Fungi(100);p__Ascomycota(100);p__Ascomycota_unclassified(100);p__Ascomycota_unclassified(100);p__Ascomycota_unclassified(100);p__Ascomycota_unclassified(100);p__Ascomycota_unclassified(100); | - | - | - | - | - | - | - | Unassigned | - |
| Otu17379 | 1 | k__Fungi(100);p__Ascomycota(100);p__Ascomycota_unclassified(100);p__Ascomycota_unclassified(100);p__Ascomycota_unclassified(100);p__Ascomycota_unclassified(100);p__Ascomycota_unclassified(100); | - | - | - | - | - | - | - | Unassigned | - |
| Otu17380 | 1 | k__Fungi(100);k__Fungi_unclassified(100);k__Fungi_unclassified(100);k__Fungi_unclassified(100);k__Fungi_unclassified(100);k__Fungi_unclassified(100);k__Fungi_unclassified(100); | - | - | - | - | - | - | - | Unassigned | - |
| Otu17386 | 1 | k__Fungi(100);p__Ascomycota(100);p__Ascomycota_unclassified(100);p__Ascomycota_unclassified(100);p__Ascomycota_unclassified(100);p__Ascomycota_unclassified(100);p__Ascomycota_unclassified(100); | - | - | - | - | - | - | - | Unassigned | - |
| Otu17391 | 1 | k__Fungi(100);p__Ascomycota(100);p__Ascomycota_unclassified(100);p__Ascomycota_unclassified(100);p__Ascomycota_unclassified(100);p__Ascomycota_unclassified(100);p__Ascomycota_unclassified(100); | - | - | - | - | - | - | - | Unassigned | - |
| Otu17403 | 1 | k__Fungi(100);p__Ascomycota(100);p__Ascomycota_unclassified(100);p__Ascomycota_unclassified(100);p__Ascomycota_unclassified(100);p__Ascomycota_unclassified(100);p__Ascomycota_unclassified(100); | - | - | - | - | - | - | - | Unassigned | - |
| Otu17405 | 1 | k__Fungi(100);k__Fungi_unclassified(100);k__Fungi_unclassified(100);k__Fungi_unclassified(100);k__Fungi_unclassified(100);k__Fungi_unclassified(100);k__Fungi_unclassified(100); | - | - | - | - | - | - | - | Unassigned | - |
| Otu17406 | 1 | k__Fungi(100);k__Fungi_unclassified(100);k__Fungi_unclassified(100);k__Fungi_unclassified(100);k__Fungi_unclassified(100);k__Fungi_unclassified(100);k__Fungi_unclassified(100); | - | - | - | - | - | - | - | Unassigned | - |
| Otu17410 | 1 | k__Fungi(100);p__Ascomycota(100);p__Ascomycota_unclassified(100);p__Ascomycota_unclassified(100);p__Ascomycota_unclassified(100);p__Ascomycota_unclassified(100);p__Ascomycota_unclassified(100); | - | - | - | - | - | - | - | Unassigned | - |
| Otu17411 | 1 | k__Fungi(100);p__Ascomycota(100);p__Ascomycota_unclassified(100);p__Ascomycota_unclassified(100);p__Ascomycota_unclassified(100);p__Ascomycota_unclassified(100);p__Ascomycota_unclassified(100); | - | - | - | - | - | - | - | Unassigned | - |
| Otu17412 | 1 | k__Fungi(100);k__Fungi_unclassified(100);k__Fungi_unclassified(100);k__Fungi_unclassified(100);k__Fungi_unclassified(100);k__Fungi_unclassified(100);k__Fungi_unclassified(100); | - | - | - | - | - | - | - | Unassigned | - |
| Otu17413 | 1 | k__Fungi(100);k__Fungi_unclassified(100);k__Fungi_unclassified(100);k__Fungi_unclassified(100);k__Fungi_unclassified(100);k__Fungi_unclassified(100);k__Fungi_unclassified(100); | - | - | - | - | - | - | - | Unassigned | - |
| Otu17414 | 1 | k__Fungi(100);p__Ascomycota(100);p__Ascomycota_unclassified(100);p__Ascomycota_unclassified(100);p__Ascomycota_unclassified(100);p__Ascomycota_unclassified(100);p__Ascomycota_unclassified(100); | - | - | - | - | - | - | - | Unassigned | - |
| Otu17415 | 1 | k__Fungi(100);k__Fungi_unclassified(100);k__Fungi_unclassified(100);k__Fungi_unclassified(100);k__Fungi_unclassified(100);k__Fungi_unclassified(100);k__Fungi_unclassified(100); | - | - | - | - | - | - | - | Unassigned | - |
| Otu17416 | 1 | k__Fungi(100);p__Ascomycota(100);p__Ascomycota_unclassified(100);p__Ascomycota_unclassified(100);p__Ascomycota_unclassified(100);p__Ascomycota_unclassified(100);p__Ascomycota_unclassified(100); | - | - | - | - | - | - | - | Unassigned | - |
| Otu17418 | 1 | k__Fungi(100);p__Ascomycota(100);p__Ascomycota_unclassified(100);p__Ascomycota_unclassified(100);p__Ascomycota_unclassified(100);p__Ascomycota_unclassified(100);p__Ascomycota_unclassified(100); | - | - | - | - | - | - | - | Unassigned | - |
| Otu17419 | 1 | k__Fungi(100);k__Fungi_unclassified(100);k__Fungi_unclassified(100);k__Fungi_unclassified(100);k__Fungi_unclassified(100);k__Fungi_unclassified(100);k__Fungi_unclassified(100); | - | - | - | - | - | - | - | Unassigned | - |
| Otu17420 | 1 | k__Fungi(100);p__Ascomycota(100);p__Ascomycota_unclassified(100);p__Ascomycota_unclassified(100);p__Ascomycota_unclassified(100);p__Ascomycota_unclassified(100);p__Ascomycota_unclassified(100); | - | - | - | - | - | - | - | Unassigned | - |
| Otu17421 | 1 | k__Fungi(100);k__Fungi_unclassified(100);k__Fungi_unclassified(100);k__Fungi_unclassified(100);k__Fungi_unclassified(100);k__Fungi_unclassified(100);k__Fungi_unclassified(100); | - | - | - | - | - | - | - | Unassigned | - |
| Otu17422 | 1 | k__Fungi(100);p__Ascomycota(100);p__Ascomycota_unclassified(100);p__Ascomycota_unclassified(100);p__Ascomycota_unclassified(100);p__Ascomycota_unclassified(100);p__Ascomycota_unclassified(100); | - | - | - | - | - | - | - | Unassigned | - |
| Otu17423 | 1 | k__Fungi(100);k__Fungi_unclassified(100);k__Fungi_unclassified(100);k__Fungi_unclassified(100);k__Fungi_unclassified(100);k__Fungi_unclassified(100);k__Fungi_unclassified(100); | - | - | - | - | - | - | - | Unassigned | - |
| Otu17424 | 1 | k__Fungi(100);p__Ascomycota(100);p__Ascomycota_unclassified(100);p__Ascomycota_unclassified(100);p__Ascomycota_unclassified(100);p__Ascomycota_unclassified(100);p__Ascomycota_unclassified(100); | - | - | - | - | - | - | - | Unassigned | - |
| Otu17425 | 1 | k__Fungi(100);p__Ascomycota(100);p__Ascomycota_unclassified(100);p__Ascomycota_unclassified(100);p__Ascomycota_unclassified(100);p__Ascomycota_unclassified(100);p__Ascomycota_unclassified(100); | - | - | - | - | - | - | - | Unassigned | - |
| Otu17426 | 1 | k__Fungi(100);p__Ascomycota(100);p__Ascomycota_unclassified(100);p__Ascomycota_unclassified(100);p__Ascomycota_unclassified(100);p__Ascomycota_unclassified(100);p__Ascomycota_unclassified(100); | - | - | - | - | - | - | - | Unassigned | - |
| Otu17427 | 1 | k__Fungi(100);p__Ascomycota(100);p__Ascomycota_unclassified(100);p__Ascomycota_unclassified(100);p__Ascomycota_unclassified(100);p__Ascomycota_unclassified(100);p__Ascomycota_unclassified(100); | - | - | - | - | - | - | - | Unassigned | - |
| Otu17428 | 1 | k__Fungi(100);p__Ascomycota(100);p__Ascomycota_unclassified(100);p__Ascomycota_unclassified(100);p__Ascomycota_unclassified(100);p__Ascomycota_unclassified(100);p__Ascomycota_unclassified(100); | - | - | - | - | - | - | - | Unassigned | - |
| Otu17429 | 1 | k__Fungi(100);p__Ascomycota(100);p__Ascomycota_unclassified(100);p__Ascomycota_unclassified(100);p__Ascomycota_unclassified(100);p__Ascomycota_unclassified(100);p__Ascomycota_unclassified(100); | - | - | - | - | - | - | - | Unassigned | - |
| Otu17430 | 1 | k__Fungi(100);p__Ascomycota(100);p__Ascomycota_unclassified(100);p__Ascomycota_unclassified(100);p__Ascomycota_unclassified(100);p__Ascomycota_unclassified(100);p__Ascomycota_unclassified(100); | - | - | - | - | - | - | - | Unassigned | - |
| Otu17432 | 1 | k__Fungi(100);k__Fungi_unclassified(100);k__Fungi_unclassified(100);k__Fungi_unclassified(100);k__Fungi_unclassified(100);k__Fungi_unclassified(100);k__Fungi_unclassified(100); | - | - | - | - | - | - | - | Unassigned | - |
| Otu17436 | 1 | k__Fungi(100);p__Ascomycota(100);p__Ascomycota_unclassified(100);p__Ascomycota_unclassified(100);p__Ascomycota_unclassified(100);p__Ascomycota_unclassified(100);p__Ascomycota_unclassified(100); | - | - | - | - | - | - | - | Unassigned | - |
| Otu17438 | 1 | k__Fungi(100);k__Fungi_unclassified(100);k__Fungi_unclassified(100);k__Fungi_unclassified(100);k__Fungi_unclassified(100);k__Fungi_unclassified(100);k__Fungi_unclassified(100); | - | - | - | - | - | - | - | Unassigned | - |
| Otu17441 | 1 | k__Fungi(100);k__Fungi_unclassified(100);k__Fungi_unclassified(100);k__Fungi_unclassified(100);k__Fungi_unclassified(100);k__Fungi_unclassified(100);k__Fungi_unclassified(100); | - | - | - | - | - | - | - | Unassigned | - |
| Otu17444 | 1 | k__Fungi(100);p__Ascomycota(100);p__Ascomycota_unclassified(100);p__Ascomycota_unclassified(100);p__Ascomycota_unclassified(100);p__Ascomycota_unclassified(100);p__Ascomycota_unclassified(100); | - | - | - | - | - | - | - | Unassigned | - |
| Otu17447 | 1 | k__Fungi(100);k__Fungi_unclassified(100);k__Fungi_unclassified(100);k__Fungi_unclassified(100);k__Fungi_unclassified(100);k__Fungi_unclassified(100);k__Fungi_unclassified(100); | - | - | - | - | - | - | - | Unassigned | - |
| Otu17448 | 1 | k__Fungi(100);k__Fungi_unclassified(100);k__Fungi_unclassified(100);k__Fungi_unclassified(100);k__Fungi_unclassified(100);k__Fungi_unclassified(100);k__Fungi_unclassified(100); | - | - | - | - | - | - | - | Unassigned | - |
| Otu17451 | 1 | k__Fungi(100);p__Ascomycota(100);p__Ascomycota_unclassified(100);p__Ascomycota_unclassified(100);p__Ascomycota_unclassified(100);p__Ascomycota_unclassified(100);p__Ascomycota_unclassified(100); | - | - | - | - | - | - | - | Unassigned | - |
| Otu17452 | 1 | k__Fungi(100);p__Ascomycota(100);p__Ascomycota_unclassified(100);p__Ascomycota_unclassified(100);p__Ascomycota_unclassified(100);p__Ascomycota_unclassified(100);p__Ascomycota_unclassified(100); | - | - | - | - | - | - | - | Unassigned | - |
| Otu17453 | 1 | k__Fungi(100);k__Fungi_unclassified(100);k__Fungi_unclassified(100);k__Fungi_unclassified(100);k__Fungi_unclassified(100);k__Fungi_unclassified(100);k__Fungi_unclassified(100); | - | - | - | - | - | - | - | Unassigned | - |
| Otu17454 | 1 | k__Fungi(100);p__Ascomycota(100);p__Ascomycota_unclassified(100);p__Ascomycota_unclassified(100);p__Ascomycota_unclassified(100);p__Ascomycota_unclassified(100);p__Ascomycota_unclassified(100); | - | - | - | - | - | - | - | Unassigned | - |
| Otu17455 | 1 | k__Fungi(100);p__Ascomycota(100);p__Ascomycota_unclassified(100);p__Ascomycota_unclassified(100);p__Ascomycota_unclassified(100);p__Ascomycota_unclassified(100);p__Ascomycota_unclassified(100); | - | - | - | - | - | - | - | Unassigned | - |
| Otu17456 | 1 | k__Fungi(100);p__Ascomycota(100);p__Ascomycota_unclassified(100);p__Ascomycota_unclassified(100);p__Ascomycota_unclassified(100);p__Ascomycota_unclassified(100);p__Ascomycota_unclassified(100); | - | - | - | - | - | - | - | Unassigned | - |
| Otu17458 | 1 | k__Fungi(100);k__Fungi_unclassified(100);k__Fungi_unclassified(100);k__Fungi_unclassified(100);k__Fungi_unclassified(100);k__Fungi_unclassified(100);k__Fungi_unclassified(100); | - | - | - | - | - | - | - | Unassigned | - |
| Otu17459 | 1 | k__Fungi(100);k__Fungi_unclassified(100);k__Fungi_unclassified(100);k__Fungi_unclassified(100);k__Fungi_unclassified(100);k__Fungi_unclassified(100);k__Fungi_unclassified(100); | - | - | - | - | - | - | - | Unassigned | - |
| Otu17460 | 1 | k__Fungi(100);p__Ascomycota(100);p__Ascomycota_unclassified(100);p__Ascomycota_unclassified(100);p__Ascomycota_unclassified(100);p__Ascomycota_unclassified(100);p__Ascomycota_unclassified(100); | - | - | - | - | - | - | - | Unassigned | - |
| Otu17461 | 1 | k__Fungi(100);p__Ascomycota(100);p__Ascomycota_unclassified(100);p__Ascomycota_unclassified(100);p__Ascomycota_unclassified(100);p__Ascomycota_unclassified(100);p__Ascomycota_unclassified(100); | - | - | - | - | - | - | - | Unassigned | - |
| Otu17462 | 1 | k__Fungi(100);p__Ascomycota(100);p__Ascomycota_unclassified(100);p__Ascomycota_unclassified(100);p__Ascomycota_unclassified(100);p__Ascomycota_unclassified(100);p__Ascomycota_unclassified(100); | - | - | - | - | - | - | - | Unassigned | - |
| Otu17468 | 1 | k__Fungi(100);p__Ascomycota(100);p__Ascomycota_unclassified(100);p__Ascomycota_unclassified(100);p__Ascomycota_unclassified(100);p__Ascomycota_unclassified(100);p__Ascomycota_unclassified(100); | - | - | - | - | - | - | - | Unassigned | - |
| Otu17469 | 1 | k__Fungi(100);p__Ascomycota(100);p__Ascomycota_unclassified(100);p__Ascomycota_unclassified(100);p__Ascomycota_unclassified(100);p__Ascomycota_unclassified(100);p__Ascomycota_unclassified(100); | - | - | - | - | - | - | - | Unassigned | - |
| Otu17470 | 1 | k__Fungi(100);p__Ascomycota(100);p__Ascomycota_unclassified(100);p__Ascomycota_unclassified(100);p__Ascomycota_unclassified(100);p__Ascomycota_unclassified(100);p__Ascomycota_unclassified(100); | - | - | - | - | - | - | - | Unassigned | - |
| Otu17471 | 1 | k__Fungi(100);p__Ascomycota(100);p__Ascomycota_unclassified(100);p__Ascomycota_unclassified(100);p__Ascomycota_unclassified(100);p__Ascomycota_unclassified(100);p__Ascomycota_unclassified(100); | - | - | - | - | - | - | - | Unassigned | - |
| Otu17472 | 1 | k__Fungi(100);p__Ascomycota(100);p__Ascomycota_unclassified(100);p__Ascomycota_unclassified(100);p__Ascomycota_unclassified(100);p__Ascomycota_unclassified(100);p__Ascomycota_unclassified(100); | - | - | - | - | - | - | - | Unassigned | - |
| Otu17473 | 1 | k__Fungi(100);k__Fungi_unclassified(100);k__Fungi_unclassified(100);k__Fungi_unclassified(100);k__Fungi_unclassified(100);k__Fungi_unclassified(100);k__Fungi_unclassified(100); | - | - | - | - | - | - | - | Unassigned | - |
| Otu17477 | 1 | k__Fungi(100);k__Fungi_unclassified(100);k__Fungi_unclassified(100);k__Fungi_unclassified(100);k__Fungi_unclassified(100);k__Fungi_unclassified(100);k__Fungi_unclassified(100); | - | - | - | - | - | - | - | Unassigned | - |
| Otu17478 | 1 | k__Fungi(100);p__Ascomycota(100);p__Ascomycota_unclassified(100);p__Ascomycota_unclassified(100);p__Ascomycota_unclassified(100);p__Ascomycota_unclassified(100);p__Ascomycota_unclassified(100); | - | - | - | - | - | - | - | Unassigned | - |
| Otu17479 | 1 | k__Fungi(100);p__Ascomycota(100);p__Ascomycota_unclassified(100);p__Ascomycota_unclassified(100);p__Ascomycota_unclassified(100);p__Ascomycota_unclassified(100);p__Ascomycota_unclassified(100); | - | - | - | - | - | - | - | Unassigned | - |
| Otu17480 | 1 | k__Fungi(100);k__Fungi_unclassified(100);k__Fungi_unclassified(100);k__Fungi_unclassified(100);k__Fungi_unclassified(100);k__Fungi_unclassified(100);k__Fungi_unclassified(100); | - | - | - | - | - | - | - | Unassigned | - |
| Otu17481 | 1 | k__Fungi(100);p__Ascomycota(100);p__Ascomycota_unclassified(100);p__Ascomycota_unclassified(100);p__Ascomycota_unclassified(100);p__Ascomycota_unclassified(100);p__Ascomycota_unclassified(100); | - | - | - | - | - | - | - | Unassigned | - |
| Otu17482 | 1 | k__Fungi(100);k__Fungi_unclassified(100);k__Fungi_unclassified(100);k__Fungi_unclassified(100);k__Fungi_unclassified(100);k__Fungi_unclassified(100);k__Fungi_unclassified(100); | - | - | - | - | - | - | - | Unassigned | - |
| Otu17483 | 1 | k__Fungi(100);k__Fungi_unclassified(100);k__Fungi_unclassified(100);k__Fungi_unclassified(100);k__Fungi_unclassified(100);k__Fungi_unclassified(100);k__Fungi_unclassified(100); | - | - | - | - | - | - | - | Unassigned | - |
| Otu17484 | 1 | k__Fungi(100);p__Ascomycota(100);p__Ascomycota_unclassified(100);p__Ascomycota_unclassified(100);p__Ascomycota_unclassified(100);p__Ascomycota_unclassified(100);p__Ascomycota_unclassified(100); | - | - | - | - | - | - | - | Unassigned | - |
| Otu17485 | 1 | k__Fungi(100);k__Fungi_unclassified(100);k__Fungi_unclassified(100);k__Fungi_unclassified(100);k__Fungi_unclassified(100);k__Fungi_unclassified(100);k__Fungi_unclassified(100); | - | - | - | - | - | - | - | Unassigned | - |
| Otu17486 | 1 | k__Fungi(100);k__Fungi_unclassified(100);k__Fungi_unclassified(100);k__Fungi_unclassified(100);k__Fungi_unclassified(100);k__Fungi_unclassified(100);k__Fungi_unclassified(100); | - | - | - | - | - | - | - | Unassigned | - |
| Otu17487 | 1 | k__Fungi(100);k__Fungi_unclassified(100);k__Fungi_unclassified(100);k__Fungi_unclassified(100);k__Fungi_unclassified(100);k__Fungi_unclassified(100);k__Fungi_unclassified(100); | - | - | - | - | - | - | - | Unassigned | - |
| Otu17490 | 1 | k__Fungi(100);k__Fungi_unclassified(100);k__Fungi_unclassified(100);k__Fungi_unclassified(100);k__Fungi_unclassified(100);k__Fungi_unclassified(100);k__Fungi_unclassified(100); | - | - | - | - | - | - | - | Unassigned | - |
| Otu17491 | 1 | k__Fungi(100);k__Fungi_unclassified(100);k__Fungi_unclassified(100);k__Fungi_unclassified(100);k__Fungi_unclassified(100);k__Fungi_unclassified(100);k__Fungi_unclassified(100); | - | - | - | - | - | - | - | Unassigned | - |
| Otu17492 | 1 | k__Fungi(100);k__Fungi_unclassified(100);k__Fungi_unclassified(100);k__Fungi_unclassified(100);k__Fungi_unclassified(100);k__Fungi_unclassified(100);k__Fungi_unclassified(100); | - | - | - | - | - | - | - | Unassigned | - |
| Otu17494 | 1 | k__Fungi(100);p__Ascomycota(100);p__Ascomycota_unclassified(100);p__Ascomycota_unclassified(100);p__Ascomycota_unclassified(100);p__Ascomycota_unclassified(100);p__Ascomycota_unclassified(100); | - | - | - | - | - | - | - | Unassigned | - |
| Otu17499 | 1 | k__Fungi(100);k__Fungi_unclassified(100);k__Fungi_unclassified(100);k__Fungi_unclassified(100);k__Fungi_unclassified(100);k__Fungi_unclassified(100);k__Fungi_unclassified(100); | - | - | - | - | - | - | - | Unassigned | - |
| Otu17501 | 1 | k__Fungi(100);p__Ascomycota(100);p__Ascomycota_unclassified(100);p__Ascomycota_unclassified(100);p__Ascomycota_unclassified(100);p__Ascomycota_unclassified(100);p__Ascomycota_unclassified(100); | - | - | - | - | - | - | - | Unassigned | - |
| Otu17502 | 1 | k__Fungi(100);p__Ascomycota(100);p__Ascomycota_unclassified(100);p__Ascomycota_unclassified(100);p__Ascomycota_unclassified(100);p__Ascomycota_unclassified(100);p__Ascomycota_unclassified(100); | - | - | - | - | - | - | - | Unassigned | - |
| Otu17503 | 1 | k__Fungi(100);k__Fungi_unclassified(100);k__Fungi_unclassified(100);k__Fungi_unclassified(100);k__Fungi_unclassified(100);k__Fungi_unclassified(100);k__Fungi_unclassified(100); | - | - | - | - | - | - | - | Unassigned | - |
| Otu17505 | 1 | k__Fungi(100);k__Fungi_unclassified(100);k__Fungi_unclassified(100);k__Fungi_unclassified(100);k__Fungi_unclassified(100);k__Fungi_unclassified(100);k__Fungi_unclassified(100); | - | - | - | - | - | - | - | Unassigned | - |
| Otu17506 | 1 | k__Fungi(100);k__Fungi_unclassified(100);k__Fungi_unclassified(100);k__Fungi_unclassified(100);k__Fungi_unclassified(100);k__Fungi_unclassified(100);k__Fungi_unclassified(100); | - | - | - | - | - | - | - | Unassigned | - |
| Otu17508 | 1 | k__Fungi(100);k__Fungi_unclassified(100);k__Fungi_unclassified(100);k__Fungi_unclassified(100);k__Fungi_unclassified(100);k__Fungi_unclassified(100);k__Fungi_unclassified(100); | - | - | - | - | - | - | - | Unassigned | - |
| Otu17510 | 1 | k__Fungi(100);p__Ascomycota(100);p__Ascomycota_unclassified(100);p__Ascomycota_unclassified(100);p__Ascomycota_unclassified(100);p__Ascomycota_unclassified(100);p__Ascomycota_unclassified(100); | - | - | - | - | - | - | - | Unassigned | - |
| Otu17511 | 1 | k__Fungi(100);p__Ascomycota(100);p__Ascomycota_unclassified(100);p__Ascomycota_unclassified(100);p__Ascomycota_unclassified(100);p__Ascomycota_unclassified(100);p__Ascomycota_unclassified(100); | - | - | - | - | - | - | - | Unassigned | - |
| Otu17512 | 1 | k__Fungi(100);k__Fungi_unclassified(100);k__Fungi_unclassified(100);k__Fungi_unclassified(100);k__Fungi_unclassified(100);k__Fungi_unclassified(100);k__Fungi_unclassified(100); | - | - | - | - | - | - | - | Unassigned | - |
| Otu17513 | 1 | k__Fungi(100);p__Ascomycota(100);p__Ascomycota_unclassified(100);p__Ascomycota_unclassified(100);p__Ascomycota_unclassified(100);p__Ascomycota_unclassified(100);p__Ascomycota_unclassified(100); | - | - | - | - | - | - | - | Unassigned | - |
| Otu17514 | 1 | k__Fungi(100);p__Ascomycota(100);p__Ascomycota_unclassified(100);p__Ascomycota_unclassified(100);p__Ascomycota_unclassified(100);p__Ascomycota_unclassified(100);p__Ascomycota_unclassified(100); | - | - | - | - | - | - | - | Unassigned | - |
| Otu17515 | 1 | k__Fungi(100);k__Fungi_unclassified(100);k__Fungi_unclassified(100);k__Fungi_unclassified(100);k__Fungi_unclassified(100);k__Fungi_unclassified(100);k__Fungi_unclassified(100); | - | - | - | - | - | - | - | Unassigned | - |
| Otu17516 | 1 | k__Fungi(100);p__Ascomycota(100);p__Ascomycota_unclassified(100);p__Ascomycota_unclassified(100);p__Ascomycota_unclassified(100);p__Ascomycota_unclassified(100);p__Ascomycota_unclassified(100); | - | - | - | - | - | - | - | Unassigned | - |
| Otu17519 | 1 | k__Fungi(100);k__Fungi_unclassified(100);k__Fungi_unclassified(100);k__Fungi_unclassified(100);k__Fungi_unclassified(100);k__Fungi_unclassified(100);k__Fungi_unclassified(100); | - | - | - | - | - | - | - | Unassigned | - |
| Otu17520 | 1 | k__Fungi(100);k__Fungi_unclassified(100);k__Fungi_unclassified(100);k__Fungi_unclassified(100);k__Fungi_unclassified(100);k__Fungi_unclassified(100);k__Fungi_unclassified(100); | - | - | - | - | - | - | - | Unassigned | - |
| Otu17521 | 1 | k__Fungi(100);k__Fungi_unclassified(100);k__Fungi_unclassified(100);k__Fungi_unclassified(100);k__Fungi_unclassified(100);k__Fungi_unclassified(100);k__Fungi_unclassified(100); | - | - | - | - | - | - | - | Unassigned | - |
| Otu17523 | 1 | k__Fungi(100);k__Fungi_unclassified(100);k__Fungi_unclassified(100);k__Fungi_unclassified(100);k__Fungi_unclassified(100);k__Fungi_unclassified(100);k__Fungi_unclassified(100); | - | - | - | - | - | - | - | Unassigned | - |
| Otu17524 | 1 | k__Fungi(100);p__Ascomycota(100);p__Ascomycota_unclassified(100);p__Ascomycota_unclassified(100);p__Ascomycota_unclassified(100);p__Ascomycota_unclassified(100);p__Ascomycota_unclassified(100); | - | - | - | - | - | - | - | Unassigned | - |
| Otu17525 | 1 | k__Fungi(100);k__Fungi_unclassified(100);k__Fungi_unclassified(100);k__Fungi_unclassified(100);k__Fungi_unclassified(100);k__Fungi_unclassified(100);k__Fungi_unclassified(100); | - | - | - | - | - | - | - | Unassigned | - |
| Otu17526 | 1 | k__Fungi(100);p__Ascomycota(100);p__Ascomycota_unclassified(100);p__Ascomycota_unclassified(100);p__Ascomycota_unclassified(100);p__Ascomycota_unclassified(100);p__Ascomycota_unclassified(100); | - | - | - | - | - | - | - | Unassigned | - |
| Otu17527 | 1 | k__Fungi(100);k__Fungi_unclassified(100);k__Fungi_unclassified(100);k__Fungi_unclassified(100);k__Fungi_unclassified(100);k__Fungi_unclassified(100);k__Fungi_unclassified(100); | - | - | - | - | - | - | - | Unassigned | - |
| Otu17528 | 1 | k__Fungi(100);p__Ascomycota(100);p__Ascomycota_unclassified(100);p__Ascomycota_unclassified(100);p__Ascomycota_unclassified(100);p__Ascomycota_unclassified(100);p__Ascomycota_unclassified(100); | - | - | - | - | - | - | - | Unassigned | - |
| Otu17529 | 1 | k__Fungi(100);p__Ascomycota(100);p__Ascomycota_unclassified(100);p__Ascomycota_unclassified(100);p__Ascomycota_unclassified(100);p__Ascomycota_unclassified(100);p__Ascomycota_unclassified(100); | - | - | - | - | - | - | - | Unassigned | - |
| Otu17530 | 1 | k__Fungi(100);k__Fungi_unclassified(100);k__Fungi_unclassified(100);k__Fungi_unclassified(100);k__Fungi_unclassified(100);k__Fungi_unclassified(100);k__Fungi_unclassified(100); | - | - | - | - | - | - | - | Unassigned | - |
| Otu17531 | 1 | k__Fungi(100);k__Fungi_unclassified(100);k__Fungi_unclassified(100);k__Fungi_unclassified(100);k__Fungi_unclassified(100);k__Fungi_unclassified(100);k__Fungi_unclassified(100); | - | - | - | - | - | - | - | Unassigned | - |
| Otu17532 | 1 | k__Fungi(100);p__Ascomycota(100);p__Ascomycota_unclassified(100);p__Ascomycota_unclassified(100);p__Ascomycota_unclassified(100);p__Ascomycota_unclassified(100);p__Ascomycota_unclassified(100); | - | - | - | - | - | - | - | Unassigned | - |
| Otu17533 | 1 | k__Fungi(100);p__Ascomycota(100);p__Ascomycota_unclassified(100);p__Ascomycota_unclassified(100);p__Ascomycota_unclassified(100);p__Ascomycota_unclassified(100);p__Ascomycota_unclassified(100); | - | - | - | - | - | - | - | Unassigned | - |
| Otu17534 | 1 | k__Fungi(100);p__Ascomycota(100);p__Ascomycota_unclassified(100);p__Ascomycota_unclassified(100);p__Ascomycota_unclassified(100);p__Ascomycota_unclassified(100);p__Ascomycota_unclassified(100); | - | - | - | - | - | - | - | Unassigned | - |
| Otu17535 | 1 | k__Fungi(100);k__Fungi_unclassified(100);k__Fungi_unclassified(100);k__Fungi_unclassified(100);k__Fungi_unclassified(100);k__Fungi_unclassified(100);k__Fungi_unclassified(100); | - | - | - | - | - | - | - | Unassigned | - |
| Otu17536 | 1 | k__Fungi(100);p__Ascomycota(100);p__Ascomycota_unclassified(100);p__Ascomycota_unclassified(100);p__Ascomycota_unclassified(100);p__Ascomycota_unclassified(100);p__Ascomycota_unclassified(100); | - | - | - | - | - | - | - | Unassigned | - |
| Otu17537 | 1 | k__Fungi(100);p__Ascomycota(100);p__Ascomycota_unclassified(100);p__Ascomycota_unclassified(100);p__Ascomycota_unclassified(100);p__Ascomycota_unclassified(100);p__Ascomycota_unclassified(100); | - | - | - | - | - | - | - | Unassigned | - |
| Otu17538 | 1 | k__Fungi(100);p__Ascomycota(100);p__Ascomycota_unclassified(100);p__Ascomycota_unclassified(100);p__Ascomycota_unclassified(100);p__Ascomycota_unclassified(100);p__Ascomycota_unclassified(100); | - | - | - | - | - | - | - | Unassigned | - |
| Otu17540 | 1 | k__Fungi(100);p__Ascomycota(100);p__Ascomycota_unclassified(100);p__Ascomycota_unclassified(100);p__Ascomycota_unclassified(100);p__Ascomycota_unclassified(100);p__Ascomycota_unclassified(100); | - | - | - | - | - | - | - | Unassigned | - |
| Otu17552 | 1 | k__Fungi(100);k__Fungi_unclassified(100);k__Fungi_unclassified(100);k__Fungi_unclassified(100);k__Fungi_unclassified(100);k__Fungi_unclassified(100);k__Fungi_unclassified(100); | - | - | - | - | - | - | - | Unassigned | - |
| Otu17553 | 1 | k__Fungi(100);k__Fungi_unclassified(100);k__Fungi_unclassified(100);k__Fungi_unclassified(100);k__Fungi_unclassified(100);k__Fungi_unclassified(100);k__Fungi_unclassified(100); | - | - | - | - | - | - | - | Unassigned | - |
| Otu17554 | 1 | k__Fungi(100);p__Ascomycota(100);p__Ascomycota_unclassified(100);p__Ascomycota_unclassified(100);p__Ascomycota_unclassified(100);p__Ascomycota_unclassified(100);p__Ascomycota_unclassified(100); | - | - | - | - | - | - | - | Unassigned | - |
| Otu17555 | 1 | k__Fungi(100);p__Ascomycota(100);p__Ascomycota_unclassified(100);p__Ascomycota_unclassified(100);p__Ascomycota_unclassified(100);p__Ascomycota_unclassified(100);p__Ascomycota_unclassified(100); | - | - | - | - | - | - | - | Unassigned | - |
| Otu17556 | 1 | k__Fungi(100);p__Ascomycota(100);p__Ascomycota_unclassified(100);p__Ascomycota_unclassified(100);p__Ascomycota_unclassified(100);p__Ascomycota_unclassified(100);p__Ascomycota_unclassified(100); | - | - | - | - | - | - | - | Unassigned | - |
| Otu17557 | 1 | k__Fungi(100);p__Ascomycota(100);p__Ascomycota_unclassified(100);p__Ascomycota_unclassified(100);p__Ascomycota_unclassified(100);p__Ascomycota_unclassified(100);p__Ascomycota_unclassified(100); | - | - | - | - | - | - | - | Unassigned | - |
| Otu17558 | 1 | k__Fungi(100);p__Ascomycota(100);p__Ascomycota_unclassified(100);p__Ascomycota_unclassified(100);p__Ascomycota_unclassified(100);p__Ascomycota_unclassified(100);p__Ascomycota_unclassified(100); | - | - | - | - | - | - | - | Unassigned | - |
| Otu17559 | 1 | k__Fungi(100);p__Ascomycota(100);p__Ascomycota_unclassified(100);p__Ascomycota_unclassified(100);p__Ascomycota_unclassified(100);p__Ascomycota_unclassified(100);p__Ascomycota_unclassified(100); | - | - | - | - | - | - | - | Unassigned | - |
| Otu17560 | 1 | k__Fungi(100);k__Fungi_unclassified(100);k__Fungi_unclassified(100);k__Fungi_unclassified(100);k__Fungi_unclassified(100);k__Fungi_unclassified(100);k__Fungi_unclassified(100); | - | - | - | - | - | - | - | Unassigned | - |
| Otu17561 | 1 | k__Fungi(100);k__Fungi_unclassified(100);k__Fungi_unclassified(100);k__Fungi_unclassified(100);k__Fungi_unclassified(100);k__Fungi_unclassified(100);k__Fungi_unclassified(100); | - | - | - | - | - | - | - | Unassigned | - |
| Otu17562 | 1 | k__Fungi(100);p__Ascomycota(100);p__Ascomycota_unclassified(100);p__Ascomycota_unclassified(100);p__Ascomycota_unclassified(100);p__Ascomycota_unclassified(100);p__Ascomycota_unclassified(100); | - | - | - | - | - | - | - | Unassigned | - |
| Otu17563 | 1 | k__Fungi(100);p__Ascomycota(100);p__Ascomycota_unclassified(100);p__Ascomycota_unclassified(100);p__Ascomycota_unclassified(100);p__Ascomycota_unclassified(100);p__Ascomycota_unclassified(100); | - | - | - | - | - | - | - | Unassigned | - |
| Otu17564 | 1 | k__Fungi(100);p__Ascomycota(100);p__Ascomycota_unclassified(100);p__Ascomycota_unclassified(100);p__Ascomycota_unclassified(100);p__Ascomycota_unclassified(100);p__Ascomycota_unclassified(100); | - | - | - | - | - | - | - | Unassigned | - |
| Otu17565 | 1 | k__Fungi(100);p__Ascomycota(100);p__Ascomycota_unclassified(100);p__Ascomycota_unclassified(100);p__Ascomycota_unclassified(100);p__Ascomycota_unclassified(100);p__Ascomycota_unclassified(100); | - | - | - | - | - | - | - | Unassigned | - |
| Otu17567 | 1 | k__Fungi(100);p__Ascomycota(100);p__Ascomycota_unclassified(100);p__Ascomycota_unclassified(100);p__Ascomycota_unclassified(100);p__Ascomycota_unclassified(100);p__Ascomycota_unclassified(100); | - | - | - | - | - | - | - | Unassigned | - |
| Otu17568 | 1 | k__Fungi(100);p__Ascomycota(100);p__Ascomycota_unclassified(100);p__Ascomycota_unclassified(100);p__Ascomycota_unclassified(100);p__Ascomycota_unclassified(100);p__Ascomycota_unclassified(100); | - | - | - | - | - | - | - | Unassigned | - |
| Otu17569 | 1 | k__Fungi(100);k__Fungi_unclassified(100);k__Fungi_unclassified(100);k__Fungi_unclassified(100);k__Fungi_unclassified(100);k__Fungi_unclassified(100);k__Fungi_unclassified(100); | - | - | - | - | - | - | - | Unassigned | - |
| Otu17570 | 1 | k__Fungi(100);p__Ascomycota(100);p__Ascomycota_unclassified(100);p__Ascomycota_unclassified(100);p__Ascomycota_unclassified(100);p__Ascomycota_unclassified(100);p__Ascomycota_unclassified(100); | - | - | - | - | - | - | - | Unassigned | - |
| Otu17572 | 1 | k__Fungi(100);p__Ascomycota(100);p__Ascomycota_unclassified(100);p__Ascomycota_unclassified(100);p__Ascomycota_unclassified(100);p__Ascomycota_unclassified(100);p__Ascomycota_unclassified(100); | - | - | - | - | - | - | - | Unassigned | - |
| Otu17573 | 1 | k__Fungi(100);k__Fungi_unclassified(100);k__Fungi_unclassified(100);k__Fungi_unclassified(100);k__Fungi_unclassified(100);k__Fungi_unclassified(100);k__Fungi_unclassified(100); | - | - | - | - | - | - | - | Unassigned | - |
| Otu17574 | 1 | k__Fungi(100);k__Fungi_unclassified(100);k__Fungi_unclassified(100);k__Fungi_unclassified(100);k__Fungi_unclassified(100);k__Fungi_unclassified(100);k__Fungi_unclassified(100); | - | - | - | - | - | - | - | Unassigned | - |
| Otu17575 | 1 | k__Fungi(100);p__Ascomycota(100);p__Ascomycota_unclassified(100);p__Ascomycota_unclassified(100);p__Ascomycota_unclassified(100);p__Ascomycota_unclassified(100);p__Ascomycota_unclassified(100); | - | - | - | - | - | - | - | Unassigned | - |
| Otu17576 | 1 | k__Fungi(100);p__Ascomycota(100);p__Ascomycota_unclassified(100);p__Ascomycota_unclassified(100);p__Ascomycota_unclassified(100);p__Ascomycota_unclassified(100);p__Ascomycota_unclassified(100); | - | - | - | - | - | - | - | Unassigned | - |
| Otu17577 | 1 | k__Fungi(100);p__Ascomycota(100);p__Ascomycota_unclassified(100);p__Ascomycota_unclassified(100);p__Ascomycota_unclassified(100);p__Ascomycota_unclassified(100);p__Ascomycota_unclassified(100); | - | - | - | - | - | - | - | Unassigned | - |
| Otu17578 | 1 | k__Fungi(100);p__Ascomycota(100);p__Ascomycota_unclassified(100);p__Ascomycota_unclassified(100);p__Ascomycota_unclassified(100);p__Ascomycota_unclassified(100);p__Ascomycota_unclassified(100); | - | - | - | - | - | - | - | Unassigned | - |
| Otu17579 | 1 | k__Fungi(100);p__Ascomycota(100);p__Ascomycota_unclassified(100);p__Ascomycota_unclassified(100);p__Ascomycota_unclassified(100);p__Ascomycota_unclassified(100);p__Ascomycota_unclassified(100); | - | - | - | - | - | - | - | Unassigned | - |
| Otu17592 | 1 | k__Fungi(100);k__Fungi_unclassified(100);k__Fungi_unclassified(100);k__Fungi_unclassified(100);k__Fungi_unclassified(100);k__Fungi_unclassified(100);k__Fungi_unclassified(100); | - | - | - | - | - | - | - | Unassigned | - |
| Otu17593 | 1 | k__Fungi(100);p__Ascomycota(100);p__Ascomycota_unclassified(100);p__Ascomycota_unclassified(100);p__Ascomycota_unclassified(100);p__Ascomycota_unclassified(100);p__Ascomycota_unclassified(100); | - | - | - | - | - | - | - | Unassigned | - |
| Otu17594 | 1 | k__Fungi(100);p__Ascomycota(100);p__Ascomycota_unclassified(100);p__Ascomycota_unclassified(100);p__Ascomycota_unclassified(100);p__Ascomycota_unclassified(100);p__Ascomycota_unclassified(100); | - | - | - | - | - | - | - | Unassigned | - |
| Otu17596 | 1 | k__Fungi(100);p__Ascomycota(100);p__Ascomycota_unclassified(100);p__Ascomycota_unclassified(100);p__Ascomycota_unclassified(100);p__Ascomycota_unclassified(100);p__Ascomycota_unclassified(100); | - | - | - | - | - | - | - | Unassigned | - |
| Otu17597 | 1 | k__Fungi(100);p__Ascomycota(100);p__Ascomycota_unclassified(100);p__Ascomycota_unclassified(100);p__Ascomycota_unclassified(100);p__Ascomycota_unclassified(100);p__Ascomycota_unclassified(100); | - | - | - | - | - | - | - | Unassigned | - |
| Otu17598 | 1 | k__Fungi(100);p__Ascomycota(100);p__Ascomycota_unclassified(100);p__Ascomycota_unclassified(100);p__Ascomycota_unclassified(100);p__Ascomycota_unclassified(100);p__Ascomycota_unclassified(100); | - | - | - | - | - | - | - | Unassigned | - |
| Otu17599 | 1 | k__Fungi(100);k__Fungi_unclassified(100);k__Fungi_unclassified(100);k__Fungi_unclassified(100);k__Fungi_unclassified(100);k__Fungi_unclassified(100);k__Fungi_unclassified(100); | - | - | - | - | - | - | - | Unassigned | - |
| Otu17600 | 1 | k__Fungi(100);p__Ascomycota(100);p__Ascomycota_unclassified(100);p__Ascomycota_unclassified(100);p__Ascomycota_unclassified(100);p__Ascomycota_unclassified(100);p__Ascomycota_unclassified(100); | - | - | - | - | - | - | - | Unassigned | - |
| Otu17601 | 1 | k__Fungi(100);k__Fungi_unclassified(100);k__Fungi_unclassified(100);k__Fungi_unclassified(100);k__Fungi_unclassified(100);k__Fungi_unclassified(100);k__Fungi_unclassified(100); | - | - | - | - | - | - | - | Unassigned | - |
| Otu17603 | 1 | k__Fungi(100);p__Ascomycota(100);p__Ascomycota_unclassified(100);p__Ascomycota_unclassified(100);p__Ascomycota_unclassified(100);p__Ascomycota_unclassified(100);p__Ascomycota_unclassified(100); | - | - | - | - | - | - | - | Unassigned | - |
| Otu17606 | 1 | k__Fungi(100);k__Fungi_unclassified(100);k__Fungi_unclassified(100);k__Fungi_unclassified(100);k__Fungi_unclassified(100);k__Fungi_unclassified(100);k__Fungi_unclassified(100); | - | - | - | - | - | - | - | Unassigned | - |
| Otu17607 | 1 | k__Fungi(100);k__Fungi_unclassified(100);k__Fungi_unclassified(100);k__Fungi_unclassified(100);k__Fungi_unclassified(100);k__Fungi_unclassified(100);k__Fungi_unclassified(100); | - | - | - | - | - | - | - | Unassigned | - |
| Otu17608 | 1 | k__Fungi(100);p__Ascomycota(100);p__Ascomycota_unclassified(100);p__Ascomycota_unclassified(100);p__Ascomycota_unclassified(100);p__Ascomycota_unclassified(100);p__Ascomycota_unclassified(100); | - | - | - | - | - | - | - | Unassigned | - |
| Otu17609 | 1 | k__Fungi(100);p__Ascomycota(100);p__Ascomycota_unclassified(100);p__Ascomycota_unclassified(100);p__Ascomycota_unclassified(100);p__Ascomycota_unclassified(100);p__Ascomycota_unclassified(100); | - | - | - | - | - | - | - | Unassigned | - |
| Otu17611 | 1 | k__Fungi(100);p__Ascomycota(100);p__Ascomycota_unclassified(100);p__Ascomycota_unclassified(100);p__Ascomycota_unclassified(100);p__Ascomycota_unclassified(100);p__Ascomycota_unclassified(100); | - | - | - | - | - | - | - | Unassigned | - |
| Otu17612 | 1 | k__Fungi(100);p__Ascomycota(100);p__Ascomycota_unclassified(100);p__Ascomycota_unclassified(100);p__Ascomycota_unclassified(100);p__Ascomycota_unclassified(100);p__Ascomycota_unclassified(100); | - | - | - | - | - | - | - | Unassigned | - |
| Otu17613 | 1 | k__Fungi(100);k__Fungi_unclassified(100);k__Fungi_unclassified(100);k__Fungi_unclassified(100);k__Fungi_unclassified(100);k__Fungi_unclassified(100);k__Fungi_unclassified(100); | - | - | - | - | - | - | - | Unassigned | - |
| Otu17614 | 1 | k__Fungi(100);k__Fungi_unclassified(100);k__Fungi_unclassified(100);k__Fungi_unclassified(100);k__Fungi_unclassified(100);k__Fungi_unclassified(100);k__Fungi_unclassified(100); | - | - | - | - | - | - | - | Unassigned | - |
| Otu17615 | 1 | k__Fungi(100);p__Ascomycota(100);p__Ascomycota_unclassified(100);p__Ascomycota_unclassified(100);p__Ascomycota_unclassified(100);p__Ascomycota_unclassified(100);p__Ascomycota_unclassified(100); | - | - | - | - | - | - | - | Unassigned | - |
| Otu17617 | 1 | k__Fungi(100);p__Ascomycota(100);p__Ascomycota_unclassified(100);p__Ascomycota_unclassified(100);p__Ascomycota_unclassified(100);p__Ascomycota_unclassified(100);p__Ascomycota_unclassified(100); | - | - | - | - | - | - | - | Unassigned | - |
| Otu17639 | 1 | k__Fungi(100);k__Fungi_unclassified(100);k__Fungi_unclassified(100);k__Fungi_unclassified(100);k__Fungi_unclassified(100);k__Fungi_unclassified(100);k__Fungi_unclassified(100); | - | - | - | - | - | - | - | Unassigned | - |
| Otu17641 | 1 | k__Fungi(100);k__Fungi_unclassified(100);k__Fungi_unclassified(100);k__Fungi_unclassified(100);k__Fungi_unclassified(100);k__Fungi_unclassified(100);k__Fungi_unclassified(100); | - | - | - | - | - | - | - | Unassigned | - |
| Otu17645 | 1 | k__Fungi(100);k__Fungi_unclassified(100);k__Fungi_unclassified(100);k__Fungi_unclassified(100);k__Fungi_unclassified(100);k__Fungi_unclassified(100);k__Fungi_unclassified(100); | - | - | - | - | - | - | - | Unassigned | - |
| Otu17648 | 1 | k__Fungi(100);p__Ascomycota(100);p__Ascomycota_unclassified(100);p__Ascomycota_unclassified(100);p__Ascomycota_unclassified(100);p__Ascomycota_unclassified(100);p__Ascomycota_unclassified(100); | - | - | - | - | - | - | - | Unassigned | - |
| Otu17649 | 1 | k__Fungi(100);p__Ascomycota(100);p__Ascomycota_unclassified(100);p__Ascomycota_unclassified(100);p__Ascomycota_unclassified(100);p__Ascomycota_unclassified(100);p__Ascomycota_unclassified(100); | - | - | - | - | - | - | - | Unassigned | - |
| Otu17650 | 1 | k__Fungi(100);p__Ascomycota(100);p__Ascomycota_unclassified(100);p__Ascomycota_unclassified(100);p__Ascomycota_unclassified(100);p__Ascomycota_unclassified(100);p__Ascomycota_unclassified(100); | - | - | - | - | - | - | - | Unassigned | - |
| Otu17651 | 1 | k__Fungi(100);p__Ascomycota(100);p__Ascomycota_unclassified(100);p__Ascomycota_unclassified(100);p__Ascomycota_unclassified(100);p__Ascomycota_unclassified(100);p__Ascomycota_unclassified(100); | - | - | - | - | - | - | - | Unassigned | - |
| Otu17656 | 1 | k__Fungi(100);p__Ascomycota(100);p__Ascomycota_unclassified(100);p__Ascomycota_unclassified(100);p__Ascomycota_unclassified(100);p__Ascomycota_unclassified(100);p__Ascomycota_unclassified(100); | - | - | - | - | - | - | - | Unassigned | - |
| Otu17671 | 1 | k__Fungi(100);p__Ascomycota(100);p__Ascomycota_unclassified(100);p__Ascomycota_unclassified(100);p__Ascomycota_unclassified(100);p__Ascomycota_unclassified(100);p__Ascomycota_unclassified(100); | - | - | - | - | - | - | - | Unassigned | - |
| Otu17672 | 1 | k__Fungi(100);p__Ascomycota(100);p__Ascomycota_unclassified(100);p__Ascomycota_unclassified(100);p__Ascomycota_unclassified(100);p__Ascomycota_unclassified(100);p__Ascomycota_unclassified(100); | - | - | - | - | - | - | - | Unassigned | - |
| Otu17710 | 1 | k__Fungi(100);k__Fungi_unclassified(100);k__Fungi_unclassified(100);k__Fungi_unclassified(100);k__Fungi_unclassified(100);k__Fungi_unclassified(100);k__Fungi_unclassified(100); | - | - | - | - | - | - | - | Unassigned | - |
| Otu17725 | 1 | k__Fungi(100);k__Fungi_unclassified(100);k__Fungi_unclassified(100);k__Fungi_unclassified(100);k__Fungi_unclassified(100);k__Fungi_unclassified(100);k__Fungi_unclassified(100); | - | - | - | - | - | - | - | Unassigned | - |
| Otu17758 | 1 | k__Fungi(100);k__Fungi_unclassified(100);k__Fungi_unclassified(100);k__Fungi_unclassified(100);k__Fungi_unclassified(100);k__Fungi_unclassified(100);k__Fungi_unclassified(100); | - | - | - | - | - | - | - | Unassigned | - |
| Otu17781 | 1 | k__Fungi(100);k__Fungi_unclassified(100);k__Fungi_unclassified(100);k__Fungi_unclassified(100);k__Fungi_unclassified(100);k__Fungi_unclassified(100);k__Fungi_unclassified(100); | - | - | - | - | - | - | - | Unassigned | - |
| Otu17783 | 1 | k__Fungi(100);p__Ascomycota(100);c__Eurotiomycetes(100);o__Eurotiales(100);f__Trichocomaceae(100);f__Trichocomaceae_unclassified(100);f__Trichocomaceae_unclassified(100); | - | - | - | - | - | - | - | Unassigned | - |
| Otu17784 | 1 | k__Fungi(100);p__Basidiomycota(100);c__Agaricomycetes(100);o__Polyporales(100);o__Polyporales_unclassified(100);o__Polyporales_unclassified(100);o__Polyporales_unclassified(100); | - | - | - | - | - | - | - | Unassigned | - |
| Otu17787 | 1 | k__Plantae(100);p__unclassified_Plantae(100);c__unclassified_Plantae(100);o__unclassified_Plantae(100);f__unclassified_Plantae(100);g__unclassified_Plantae(100);s__Plantae_sp(100); | - | - | - | - | - | - | - | Unassigned | - |
| Otu17796 | 1 | k__Fungi(100);p__Ascomycota(100);p__Ascomycota_unclassified(100);p__Ascomycota_unclassified(100);p__Ascomycota_unclassified(100);p__Ascomycota_unclassified(100);p__Ascomycota_unclassified(100); | - | - | - | - | - | - | - | Unassigned | - |
| Otu17797 | 1 | k__Fungi(100);p__Ascomycota(100);p__Ascomycota_unclassified(100);p__Ascomycota_unclassified(100);p__Ascomycota_unclassified(100);p__Ascomycota_unclassified(100);p__Ascomycota_unclassified(100); | - | - | - | - | - | - | - | Unassigned | - |
| Otu17798 | 1 | k__Fungi(100);p__Ascomycota(100);p__Ascomycota_unclassified(100);p__Ascomycota_unclassified(100);p__Ascomycota_unclassified(100);p__Ascomycota_unclassified(100);p__Ascomycota_unclassified(100); | - | - | - | - | - | - | - | Unassigned | - |
| Otu17802 | 1 | k__Fungi(100);p__Ascomycota(100);p__Ascomycota_unclassified(100);p__Ascomycota_unclassified(100);p__Ascomycota_unclassified(100);p__Ascomycota_unclassified(100);p__Ascomycota_unclassified(100); | - | - | - | - | - | - | - | Unassigned | - |
| Otu17815 | 1 | k__Fungi(100);p__Ascomycota(100);p__Ascomycota_unclassified(100);p__Ascomycota_unclassified(100);p__Ascomycota_unclassified(100);p__Ascomycota_unclassified(100);p__Ascomycota_unclassified(100); | - | - | - | - | - | - | - | Unassigned | - |
| Otu17817 | 1 | k__Fungi(100);p__Ascomycota(100);p__Ascomycota_unclassified(100);p__Ascomycota_unclassified(100);p__Ascomycota_unclassified(100);p__Ascomycota_unclassified(100);p__Ascomycota_unclassified(100); | - | - | - | - | - | - | - | Unassigned | - |
| Otu17819 | 1 | k__Fungi(100);k__Fungi_unclassified(100);k__Fungi_unclassified(100);k__Fungi_unclassified(100);k__Fungi_unclassified(100);k__Fungi_unclassified(100);k__Fungi_unclassified(100); | - | - | - | - | - | - | - | Unassigned | - |
| Otu17820 | 1 | k__Fungi(100);k__Fungi_unclassified(100);k__Fungi_unclassified(100);k__Fungi_unclassified(100);k__Fungi_unclassified(100);k__Fungi_unclassified(100);k__Fungi_unclassified(100); | - | - | - | - | - | - | - | Unassigned | - |
| Otu17821 | 1 | k__Fungi(100);p__Ascomycota(100);p__Ascomycota_unclassified(100);p__Ascomycota_unclassified(100);p__Ascomycota_unclassified(100);p__Ascomycota_unclassified(100);p__Ascomycota_unclassified(100); | - | - | - | - | - | - | - | Unassigned | - |
| Otu17825 | 1 | k__Plantae(100);p__unclassified_Plantae(100);c__unclassified_Plantae(100);o__unclassified_Plantae(100);f__unclassified_Plantae(100);g__unclassified_Plantae(100);s__Plantae_sp(100); | - | - | - | - | - | - | - | Unassigned | - |
| Otu17857 | 1 | k__Fungi(100);k__Fungi_unclassified(100);k__Fungi_unclassified(100);k__Fungi_unclassified(100);k__Fungi_unclassified(100);k__Fungi_unclassified(100);k__Fungi_unclassified(100); | - | - | - | - | - | - | - | Unassigned | - |
| Otu17858 | 1 | k__Fungi(100);k__Fungi_unclassified(100);k__Fungi_unclassified(100);k__Fungi_unclassified(100);k__Fungi_unclassified(100);k__Fungi_unclassified(100);k__Fungi_unclassified(100); | - | - | - | - | - | - | - | Unassigned | - |
| Otu17859 | 1 | k__Fungi(100);k__Fungi_unclassified(100);k__Fungi_unclassified(100);k__Fungi_unclassified(100);k__Fungi_unclassified(100);k__Fungi_unclassified(100);k__Fungi_unclassified(100); | - | - | - | - | - | - | - | Unassigned | - |
| Otu17863 | 1 | k__Fungi(100);p__Ascomycota(100);p__Ascomycota_unclassified(100);p__Ascomycota_unclassified(100);p__Ascomycota_unclassified(100);p__Ascomycota_unclassified(100);p__Ascomycota_unclassified(100); | - | - | - | - | - | - | - | Unassigned | - |
| Otu17964 | 1 | k__Fungi(100);k__Fungi_unclassified(100);k__Fungi_unclassified(100);k__Fungi_unclassified(100);k__Fungi_unclassified(100);k__Fungi_unclassified(100);k__Fungi_unclassified(100); | - | - | - | - | - | - | - | Unassigned | - |
| Otu17994 | 1 | k__Fungi(100);p__Ascomycota(100);p__Ascomycota_unclassified(100);p__Ascomycota_unclassified(100);p__Ascomycota_unclassified(100);p__Ascomycota_unclassified(100);p__Ascomycota_unclassified(100); | - | - | - | - | - | - | - | Unassigned | - |
| Otu17999 | 1 | k__Fungi(100);p__Ascomycota(100);p__Ascomycota_unclassified(100);p__Ascomycota_unclassified(100);p__Ascomycota_unclassified(100);p__Ascomycota_unclassified(100);p__Ascomycota_unclassified(100); | - | - | - | - | - | - | - | Unassigned | - |
| Otu18000 | 1 | k__Fungi(100);p__Ascomycota(100);p__Ascomycota_unclassified(100);p__Ascomycota_unclassified(100);p__Ascomycota_unclassified(100);p__Ascomycota_unclassified(100);p__Ascomycota_unclassified(100); | - | - | - | - | - | - | - | Unassigned | - |
| Otu18001 | 1 | k__Fungi(100);p__Ascomycota(100);p__Ascomycota_unclassified(100);p__Ascomycota_unclassified(100);p__Ascomycota_unclassified(100);p__Ascomycota_unclassified(100);p__Ascomycota_unclassified(100); | - | - | - | - | - | - | - | Unassigned | - |
| Otu18004 | 1 | k__Fungi(100);p__Ascomycota(100);p__Ascomycota_unclassified(100);p__Ascomycota_unclassified(100);p__Ascomycota_unclassified(100);p__Ascomycota_unclassified(100);p__Ascomycota_unclassified(100); | - | - | - | - | - | - | - | Unassigned | - |
| Otu18006 | 1 | k__Fungi(100);k__Fungi_unclassified(100);k__Fungi_unclassified(100);k__Fungi_unclassified(100);k__Fungi_unclassified(100);k__Fungi_unclassified(100);k__Fungi_unclassified(100); | - | - | - | - | - | - | - | Unassigned | - |

| | | | | | | | | | | | |
|---|---|---|---|---|---|---|---|---|---|---|---|
| Otu18007 | 1 | k__Fungi(100);k__Fungi_unclassified(100);k__Fungi_unclassified(100);k__Fungi_unclassified(100);k__Fungi_unclassified(100);k__Fungi_unclassified(100); | - | - | - | - | - | - | - | Unassigned | - |
| Otu18009 | 1 | k__Fungi(100);p__Ascomycota(100);p__Ascomycota_unclassified(100);p__Ascomycota_unclassified(100);p__Ascomycota_unclassified(100);p__Ascomycota_unclassified(100);p__Ascomycota_unclassified(100); | - | - | - | - | - | - | - | Unassigned | - |
| Otu18043 | 1 | k__Fungi(100);p__Ascomycota(100);p__Ascomycota_unclassified(100);p__Ascomycota_unclassified(100);p__Ascomycota_unclassified(100);p__Ascomycota_unclassified(100);p__Ascomycota_unclassified(100); | - | - | - | - | - | - | - | Unassigned | - |
| Otu18048 | 1 | k__Fungi(100);k__Fungi_unclassified(100);k__Fungi_unclassified(100);k__Fungi_unclassified(100);k__Fungi_unclassified(100);k__Fungi_unclassified(100); | - | - | - | - | - | - | - | Unassigned | - |
| Otu18051 | 1 | k__Fungi(100);p__Ascomycota(100);p__Ascomycota_unclassified(100);p__Ascomycota_unclassified(100);p__Ascomycota_unclassified(100);p__Ascomycota_unclassified(100);p__Ascomycota_unclassified(100); | - | - | - | - | - | - | - | Unassigned | - |
| Otu18052 | 1 | k__Fungi(100);p__Ascomycota(100);p__Ascomycota_unclassified(100);p__Ascomycota_unclassified(100);p__Ascomycota_unclassified(100);p__Ascomycota_unclassified(100);p__Ascomycota_unclassified(100); | - | - | - | - | - | - | - | Unassigned | - |
| Otu18054 | 1 | k__Fungi(100);k__Fungi_unclassified(100);k__Fungi_unclassified(100);k__Fungi_unclassified(100);k__Fungi_unclassified(100);k__Fungi_unclassified(100); | - | - | - | - | - | - | - | Unassigned | - |
| Otu18055 | 1 | k__Fungi(100);k__Fungi_unclassified(100);k__Fungi_unclassified(100);k__Fungi_unclassified(100);k__Fungi_unclassified(100);k__Fungi_unclassified(100); | - | - | - | - | - | - | - | Unassigned | - |
| Otu18056 | 1 | k__Fungi(100);p__Ascomycota(100);p__Ascomycota_unclassified(100);p__Ascomycota_unclassified(100);p__Ascomycota_unclassified(100);p__Ascomycota_unclassified(100);p__Ascomycota_unclassified(100); | - | - | - | - | - | - | - | Unassigned | - |
| Otu18077 | 1 | k__Fungi(100);p__Ascomycota(100);p__Ascomycota_unclassified(100);p__Ascomycota_unclassified(100);p__Ascomycota_unclassified(100);p__Ascomycota_unclassified(100);p__Ascomycota_unclassified(100); | - | - | - | - | - | - | - | Unassigned | - |
| Otu18078 | 1 | k__Fungi(100);k__Fungi_unclassified(100);k__Fungi_unclassified(100);k__Fungi_unclassified(100);k__Fungi_unclassified(100);k__Fungi_unclassified(100); | - | - | - | - | - | - | - | Unassigned | - |
| Otu18079 | 1 | k__Fungi(100);k__Fungi_unclassified(100);k__Fungi_unclassified(100);k__Fungi_unclassified(100);k__Fungi_unclassified(100);k__Fungi_unclassified(100); | - | - | - | - | - | - | - | Unassigned | - |
| Otu18081 | 1 | k__Fungi(100);p__Ascomycota(100);p__Ascomycota_unclassified(100);p__Ascomycota_unclassified(100);p__Ascomycota_unclassified(100);p__Ascomycota_unclassified(100);p__Ascomycota_unclassified(100); | - | - | - | - | - | - | - | Unassigned | - |
| Otu18116 | 1 | k__Fungi(100);p__Ascomycota(100);p__Ascomycota_unclassified(100);p__Ascomycota_unclassified(100);p__Ascomycota_unclassified(100);p__Ascomycota_unclassified(100);p__Ascomycota_unclassified(100); | - | - | - | - | - | - | - | Unassigned | - |
| Otu18125 | 1 | k__Fungi(100);k__Fungi_unclassified(100);k__Fungi_unclassified(100);k__Fungi_unclassified(100);k__Fungi_unclassified(100);k__Fungi_unclassified(100); | - | - | - | - | - | - | - | Unassigned | - |
| Otu18137 | 1 | k__Fungi(100);p__Ascomycota(100);p__Ascomycota_unclassified(100);p__Ascomycota_unclassified(100);p__Ascomycota_unclassified(100);p__Ascomycota_unclassified(100);p__Ascomycota_unclassified(100); | - | - | - | - | - | - | - | Unassigned | - |
| Otu18144 | 1 | k__Fungi(100);k__Fungi_unclassified(100);k__Fungi_unclassified(100);k__Fungi_unclassified(100);k__Fungi_unclassified(100);k__Fungi_unclassified(100); | - | - | - | - | - | - | - | Unassigned | - |
| Otu18146 | 1 | k__Fungi(100);p__Ascomycota(100);p__Ascomycota_unclassified(100);p__Ascomycota_unclassified(100);p__Ascomycota_unclassified(100);p__Ascomycota_unclassified(100);p__Ascomycota_unclassified(100); | - | - | - | - | - | - | - | Unassigned | - |
| Otu18149 | 1 | k__Fungi(100);p__Ascomycota(100);p__Ascomycota_unclassified(100);p__Ascomycota_unclassified(100);p__Ascomycota_unclassified(100);p__Ascomycota_unclassified(100);p__Ascomycota_unclassified(100); | - | - | - | - | - | - | - | Unassigned | - |
| Otu18151 | 1 | k__Fungi(100);k__Fungi_unclassified(100);k__Fungi_unclassified(100);k__Fungi_unclassified(100);k__Fungi_unclassified(100);k__Fungi_unclassified(100); | - | - | - | - | - | - | - | Unassigned | - |
| Otu18152 | 1 | k__Fungi(100);k__Fungi_unclassified(100);k__Fungi_unclassified(100);k__Fungi_unclassified(100);k__Fungi_unclassified(100);k__Fungi_unclassified(100); | - | - | - | - | - | - | - | Unassigned | - |
| Otu18154 | 1 | k__Fungi(100);k__Fungi_unclassified(100);k__Fungi_unclassified(100);k__Fungi_unclassified(100);k__Fungi_unclassified(100);k__Fungi_unclassified(100); | - | - | - | - | - | - | - | Unassigned | - |
| Otu18156 | 1 | k__Fungi(100);k__Fungi_unclassified(100);k__Fungi_unclassified(100);k__Fungi_unclassified(100);k__Fungi_unclassified(100);k__Fungi_unclassified(100); | - | - | - | - | - | - | - | Unassigned | - |
| Otu18157 | 1 | k__Fungi(100);p__Ascomycota(100);p__Ascomycota_unclassified(100);p__Ascomycota_unclassified(100);p__Ascomycota_unclassified(100);p__Ascomycota_unclassified(100);p__Ascomycota_unclassified(100); | - | - | - | - | - | - | - | Unassigned | - |
| Otu18158 | 1 | k__Fungi(100);k__Fungi_unclassified(100);k__Fungi_unclassified(100);k__Fungi_unclassified(100);k__Fungi_unclassified(100);k__Fungi_unclassified(100); | - | - | - | - | - | - | - | Unassigned | - |
| Otu18159 | 1 | k__Fungi(100);p__Ascomycota(100);p__Ascomycota_unclassified(100);p__Ascomycota_unclassified(100);p__Ascomycota_unclassified(100);p__Ascomycota_unclassified(100);p__Ascomycota_unclassified(100); | - | - | - | - | - | - | - | Unassigned | - |
| Otu18160 | 1 | k__Fungi(100);k__Fungi_unclassified(100);k__Fungi_unclassified(100);k__Fungi_unclassified(100);k__Fungi_unclassified(100);k__Fungi_unclassified(100); | - | - | - | - | - | - | - | Unassigned | - |
| Otu18161 | 1 | k__Fungi(100);k__Fungi_unclassified(100);k__Fungi_unclassified(100);k__Fungi_unclassified(100);k__Fungi_unclassified(100);k__Fungi_unclassified(100); | - | - | - | - | - | - | - | Unassigned | - |
| Otu18162 | 1 | k__Fungi(100);k__Fungi_unclassified(100);k__Fungi_unclassified(100);k__Fungi_unclassified(100);k__Fungi_unclassified(100);k__Fungi_unclassified(100); | - | - | - | - | - | - | - | Unassigned | - |
| Otu18173 | 1 | k__Fungi(100);p__Ascomycota(100);p__Ascomycota_unclassified(100);p__Ascomycota_unclassified(100);p__Ascomycota_unclassified(100);p__Ascomycota_unclassified(100);p__Ascomycota_unclassified(100); | - | - | - | - | - | - | - | Unassigned | - |
| Otu18174 | 1 | k__Fungi(100);p__Ascomycota(100);p__Ascomycota_unclassified(100);p__Ascomycota_unclassified(100);p__Ascomycota_unclassified(100);p__Ascomycota_unclassified(100);p__Ascomycota_unclassified(100); | - | - | - | - | - | - | - | Unassigned | - |
| Otu18184 | 1 | k__Fungi(100);k__Fungi_unclassified(100);k__Fungi_unclassified(100);k__Fungi_unclassified(100);k__Fungi_unclassified(100);k__Fungi_unclassified(100); | - | - | - | - | - | - | - | Unassigned | - |
| Otu18195 | 1 | k__Fungi(100);k__Fungi_unclassified(100);k__Fungi_unclassified(100);k__Fungi_unclassified(100);k__Fungi_unclassified(100);k__Fungi_unclassified(100); | - | - | - | - | - | - | - | Unassigned | - |
| Otu18196 | 1 | k__Fungi(100);k__Fungi_unclassified(100);k__Fungi_unclassified(100);k__Fungi_unclassified(100);k__Fungi_unclassified(100);k__Fungi_unclassified(100); | - | - | - | - | - | - | - | Unassigned | - |
| Otu18214 | 1 | k__Fungi(100);k__Fungi_unclassified(100);k__Fungi_unclassified(100);k__Fungi_unclassified(100);k__Fungi_unclassified(100);k__Fungi_unclassified(100); | - | - | - | - | - | - | - | Unassigned | - |
| Otu18216 | 1 | k__Fungi(100);p__Ascomycota(100);p__Ascomycota_unclassified(100);p__Ascomycota_unclassified(100);p__Ascomycota_unclassified(100);p__Ascomycota_unclassified(100);p__Ascomycota_unclassified(100); | - | - | - | - | - | - | - | Unassigned | - |
| Otu18235 | 1 | k__Fungi(100);k__Fungi_unclassified(100);k__Fungi_unclassified(100);k__Fungi_unclassified(100);k__Fungi_unclassified(100);k__Fungi_unclassified(100); | - | - | - | - | - | - | - | Unassigned | - |
| Otu18236 | 1 | k__Fungi(100);k__Fungi_unclassified(100);k__Fungi_unclassified(100);k__Fungi_unclassified(100);k__Fungi_unclassified(100);k__Fungi_unclassified(100); | - | - | - | - | - | - | - | Unassigned | - |
| Otu18242 | 1 | k__Fungi(100);p__Ascomycota(100);p__Ascomycota_unclassified(100);p__Ascomycota_unclassified(100);p__Ascomycota_unclassified(100);p__Ascomycota_unclassified(100);p__Ascomycota_unclassified(100); | - | - | - | - | - | - | - | Unassigned | - |
| Otu18249 | 1 | k__Fungi(100);k__Fungi_unclassified(100);k__Fungi_unclassified(100);k__Fungi_unclassified(100);k__Fungi_unclassified(100);k__Fungi_unclassified(100); | - | - | - | - | - | - | - | Unassigned | - |
| Otu18250 | 1 | k__Fungi(100);p__Ascomycota(100);c__Dothideomycetes(100);o__Capnodiales(100);o__Capnodiales_unclassified(100);o__Capnodiales_unclassified(100);o__Capnodiales_unclassified(100); | - | - | - | - | - | - | - | Unassigned | - |
| Otu18268 | 1 | k__Fungi(100);p__Ascomycota(100);p__Ascomycota_unclassified(100);p__Ascomycota_unclassified(100);p__Ascomycota_unclassified(100);p__Ascomycota_unclassified(100);p__Ascomycota_unclassified(100); | - | - | - | - | - | - | - | Unassigned | - |
| Otu18269 | 1 | k__Fungi(100);p__Ascomycota(100);p__Ascomycota_unclassified(100);p__Ascomycota_unclassified(100);p__Ascomycota_unclassified(100);p__Ascomycota_unclassified(100);p__Ascomycota_unclassified(100); | - | - | - | - | - | - | - | Unassigned | - |
| Otu18273 | 1 | k__Fungi(100);p__Ascomycota(100);p__Ascomycota_unclassified(100);p__Ascomycota_unclassified(100);p__Ascomycota_unclassified(100);p__Ascomycota_unclassified(100);p__Ascomycota_unclassified(100); | - | - | - | - | - | - | - | Unassigned | - |
| Otu18275 | 1 | k__Fungi(100);p__Ascomycota(100);p__Ascomycota_unclassified(100);p__Ascomycota_unclassified(100);p__Ascomycota_unclassified(100);p__Ascomycota_unclassified(100);p__Ascomycota_unclassified(100); | - | - | - | - | - | - | - | Unassigned | - |
| Otu18277 | 1 | k__Fungi(100);p__Ascomycota(100);p__Ascomycota_unclassified(100);p__Ascomycota_unclassified(100);p__Ascomycota_unclassified(100);p__Ascomycota_unclassified(100);p__Ascomycota_unclassified(100); | - | - | - | - | - | - | - | Unassigned | - |
| Otu18278 | 1 | k__Fungi(100);k__Fungi_unclassified(100);k__Fungi_unclassified(100);k__Fungi_unclassified(100);k__Fungi_unclassified(100);k__Fungi_unclassified(100); | - | - | - | - | - | - | - | Unassigned | - |
| Otu18279 | 1 | k__Fungi(100);p__Ascomycota(100);p__Ascomycota_unclassified(100);p__Ascomycota_unclassified(100);p__Ascomycota_unclassified(100);p__Ascomycota_unclassified(100);p__Ascomycota_unclassified(100); | - | - | - | - | - | - | - | Unassigned | - |
| Otu18288 | 1 | k__Fungi(100);p__Ascomycota(100);p__Ascomycota_unclassified(100);p__Ascomycota_unclassified(100);p__Ascomycota_unclassified(100);p__Ascomycota_unclassified(100);p__Ascomycota_unclassified(100); | - | - | - | - | - | - | - | Unassigned | - |
| Otu18289 | 1 | k__Fungi(100);p__Ascomycota(100);p__Ascomycota_unclassified(100);p__Ascomycota_unclassified(100);p__Ascomycota_unclassified(100);p__Ascomycota_unclassified(100);p__Ascomycota_unclassified(100); | - | - | - | - | - | - | - | Unassigned | - |
| Otu18332 | 1 | k__Fungi(100);p__Ascomycota(100);p__Ascomycota_unclassified(100);p__Ascomycota_unclassified(100);p__Ascomycota_unclassified(100);p__Ascomycota_unclassified(100);p__Ascomycota_unclassified(100); | - | - | - | - | - | - | - | Unassigned | - |
| Otu18344 | 1 | k__Fungi(100);p__Ascomycota(100);p__Ascomycota_unclassified(100);p__Ascomycota_unclassified(100);p__Ascomycota_unclassified(100);p__Ascomycota_unclassified(100);p__Ascomycota_unclassified(100); | - | - | - | - | - | - | - | Unassigned | - |
| Otu18358 | 1 | k__Fungi(100);k__Fungi_unclassified(100);k__Fungi_unclassified(100);k__Fungi_unclassified(100);k__Fungi_unclassified(100);k__Fungi_unclassified(100); | - | - | - | - | - | - | - | Unassigned | - |
| Otu18398 | 1 | k__Fungi(100);p__Basidiomycota(100);p__Basidiomycota_unclassified(100);p__Basidiomycota_unclassified(100);p__Basidiomycota_unclassified(100);p__Basidiomycota_unclassified(100);p__Basidiomycota_unclassified(100); | - | - | - | - | - | - | - | Unassigned | - |
| Otu18399 | 1 | k__Fungi(100);k__Fungi_unclassified(100);k__Fungi_unclassified(100);k__Fungi_unclassified(100);k__Fungi_unclassified(100);k__Fungi_unclassified(100); | - | - | - | - | - | - | - | Unassigned | - |
| Otu18400 | 1 | k__Fungi(100);p__Ascomycota(100);p__Ascomycota_unclassified(100);p__Ascomycota_unclassified(100);p__Ascomycota_unclassified(100);p__Ascomycota_unclassified(100);p__Ascomycota_unclassified(100); | - | - | - | - | - | - | - | Unassigned | - |
| Otu18423 | 1 | k__Fungi(100);k__Fungi_unclassified(100);k__Fungi_unclassified(100);k__Fungi_unclassified(100);k__Fungi_unclassified(100);k__Fungi_unclassified(100); | - | - | - | - | - | - | - | Unassigned | - |
| Otu18431 | 1 | k__Fungi(100);k__Fungi_unclassified(100);k__Fungi_unclassified(100);k__Fungi_unclassified(100);k__Fungi_unclassified(100);k__Fungi_unclassified(100); | - | - | - | - | - | - | - | Unassigned | - |
| Otu18435 | 1 | k__Fungi(100);p__Ascomycota(100);p__Ascomycota_unclassified(100);p__Ascomycota_unclassified(100);p__Ascomycota_unclassified(100);p__Ascomycota_unclassified(100);p__Ascomycota_unclassified(100); | - | - | - | - | - | - | - | Unassigned | - |
| Otu18438 | 1 | k__Fungi(100);k__Fungi_unclassified(100);k__Fungi_unclassified(100);k__Fungi_unclassified(100);k__Fungi_unclassified(100);k__Fungi_unclassified(100); | - | - | - | - | - | - | - | Unassigned | - |
| Otu18442 | 1 | k__Fungi(100);k__Fungi_unclassified(100);k__Fungi_unclassified(100);k__Fungi_unclassified(100);k__Fungi_unclassified(100);k__Fungi_unclassified(100); | - | - | - | - | - | - | - | Unassigned | - |
| Otu18447 | 1 | k__Fungi(100);k__Fungi_unclassified(100);k__Fungi_unclassified(100);k__Fungi_unclassified(100);k__Fungi_unclassified(100);k__Fungi_unclassified(100); | - | - | - | - | - | - | - | Unassigned | - |
| Otu18448 | 1 | k__Fungi(100);k__Fungi_unclassified(100);k__Fungi_unclassified(100);k__Fungi_unclassified(100);k__Fungi_unclassified(100);k__Fungi_unclassified(100); | - | - | - | - | - | - | - | Unassigned | - |
| Otu18449 | 1 | k__Fungi(100);p__Ascomycota(100);p__Ascomycota_unclassified(100);p__Ascomycota_unclassified(100);p__Ascomycota_unclassified(100);p__Ascomycota_unclassified(100);p__Ascomycota_unclassified(100); | - | - | - | - | - | - | - | Unassigned | - |
| Otu18460 | 1 | k__Fungi(100);p__Ascomycota(100);p__Ascomycota_unclassified(100);p__Ascomycota_unclassified(100);p__Ascomycota_unclassified(100);p__Ascomycota_unclassified(100);p__Ascomycota_unclassified(100); | - | - | - | - | - | - | - | Unassigned | - |
| Otu18473 | 1 | k__Fungi(100);k__Fungi_unclassified(100);k__Fungi_unclassified(100);k__Fungi_unclassified(100);k__Fungi_unclassified(100);k__Fungi_unclassified(100); | - | - | - | - | - | - | - | Unassigned | - |
| Otu18474 | 1 | k__Fungi(100);k__Fungi_unclassified(100);k__Fungi_unclassified(100);k__Fungi_unclassified(100);k__Fungi_unclassified(100);k__Fungi_unclassified(100); | - | - | - | - | - | - | - | Unassigned | - |
| Otu18475 | 1 | k__Fungi(100);k__Fungi_unclassified(100);k__Fungi_unclassified(100);k__Fungi_unclassified(100);k__Fungi_unclassified(100);k__Fungi_unclassified(100); | - | - | - | - | - | - | - | Unassigned | - |
| Otu18477 | 1 | k__Fungi(100);k__Fungi_unclassified(100);k__Fungi_unclassified(100);k__Fungi_unclassified(100);k__Fungi_unclassified(100);k__Fungi_unclassified(100); | - | - | - | - | - | - | - | Unassigned | - |
| Otu18481 | 1 | k__Fungi(100);k__Fungi_unclassified(100);k__Fungi_unclassified(100);k__Fungi_unclassified(100);k__Fungi_unclassified(100);k__Fungi_unclassified(100); | - | - | - | - | - | - | - | Unassigned | - |
| Otu18491 | 1 | k__Fungi(100);k__Fungi_unclassified(100);k__Fungi_unclassified(100);k__Fungi_unclassified(100);k__Fungi_unclassified(100);k__Fungi_unclassified(100); | - | - | - | - | - | - | - | Unassigned | - |
| Otu18493 | 1 | k__Fungi(100);k__Fungi_unclassified(100);k__Fungi_unclassified(100);k__Fungi_unclassified(100);k__Fungi_unclassified(100);k__Fungi_unclassified(100); | - | - | - | - | - | - | - | Unassigned | - |
| Otu18504 | 1 | k__Fungi(100);k__Fungi_unclassified(100);k__Fungi_unclassified(100);k__Fungi_unclassified(100);k__Fungi_unclassified(100);k__Fungi_unclassified(100); | - | - | - | - | - | - | - | Unassigned | - |
| Otu18527 | 1 | k__Fungi(100);k__Fungi_unclassified(100);k__Fungi_unclassified(100);k__Fungi_unclassified(100);k__Fungi_unclassified(100);k__Fungi_unclassified(100); | - | - | - | - | - | - | - | Unassigned | - |
| Otu18531 | 1 | k__Fungi(100);p__Ascomycota(100);p__Ascomycota_unclassified(100);p__Ascomycota_unclassified(100);p__Ascomycota_unclassified(100);p__Ascomycota_unclassified(100);p__Ascomycota_unclassified(100); | - | - | - | - | - | - | - | Unassigned | - |
| Otu18532 | 1 | k__Fungi(100);p__Ascomycota(100);p__Ascomycota_unclassified(100);p__Ascomycota_unclassified(100);p__Ascomycota_unclassified(100);p__Ascomycota_unclassified(100);p__Ascomycota_unclassified(100); | - | - | - | - | - | - | - | Unassigned | - |
| Otu18534 | 1 | k__Fungi(100);k__Fungi_unclassified(100);k__Fungi_unclassified(100);k__Fungi_unclassified(100);k__Fungi_unclassified(100);k__Fungi_unclassified(100); | - | - | - | - | - | - | - | Unassigned | - |
| Otu18537 | 1 | k__Fungi(100);k__Fungi_unclassified(100);k__Fungi_unclassified(100);k__Fungi_unclassified(100);k__Fungi_unclassified(100);k__Fungi_unclassified(100); | - | - | - | - | - | - | - | Unassigned | - |
| Otu18539 | 1 | k__Fungi(100);p__Ascomycota(100);p__Ascomycota_unclassified(100);p__Ascomycota_unclassified(100);p__Ascomycota_unclassified(100);p__Ascomycota_unclassified(100);p__Ascomycota_unclassified(100); | - | - | - | - | - | - | - | Unassigned | - |
| Otu18542 | 1 | k__Fungi(100);k__Fungi_unclassified(100);k__Fungi_unclassified(100);k__Fungi_unclassified(100);k__Fungi_unclassified(100);k__Fungi_unclassified(100); | - | - | - | - | - | - | - | Unassigned | - |
| Otu18543 | 1 | k__Fungi(100);k__Fungi_unclassified(100);k__Fungi_unclassified(100);k__Fungi_unclassified(100);k__Fungi_unclassified(100);k__Fungi_unclassified(100); | - | - | - | - | - | - | - | Unassigned | - |
| Otu18548 | 1 | k__Fungi(100);p__Ascomycota(100);p__Ascomycota_unclassified(100);p__Ascomycota_unclassified(100);p__Ascomycota_unclassified(100);p__Ascomycota_unclassified(100);p__Ascomycota_unclassified(100); | - | - | - | - | - | - | - | Unassigned | - |
| Otu18558 | 1 | k__Fungi(100);p__Ascomycota(100);p__Ascomycota_unclassified(100);p__Ascomycota_unclassified(100);p__Ascomycota_unclassified(100);p__Ascomycota_unclassified(100);p__Ascomycota_unclassified(100); | - | - | - | - | - | - | - | Unassigned | - |
| Otu18563 | 1 | k__Fungi(100);p__Ascomycota(100);p__Ascomycota_unclassified(100);p__Ascomycota_unclassified(100);p__Ascomycota_unclassified(100);p__Ascomycota_unclassified(100);p__Ascomycota_unclassified(100); | - | - | - | - | - | - | - | Unassigned | - |
| Otu18572 | 1 | k__Fungi(100);k__Fungi_unclassified(100);k__Fungi_unclassified(100);k__Fungi_unclassified(100);k__Fungi_unclassified(100);k__Fungi_unclassified(100); | - | - | - | - | - | - | - | Unassigned | - |
| Otu18633 | 1 | k__Fungi(100);k__Fungi_unclassified(100);k__Fungi_unclassified(100);k__Fungi_unclassified(100);k__Fungi_unclassified(100);k__Fungi_unclassified(100); | - | - | - | - | - | - | - | Unassigned | - |
| Otu18634 | 1 | k__Fungi(100);p__Ascomycota(100);p__Ascomycota_unclassified(100);p__Ascomycota_unclassified(100);p__Ascomycota_unclassified(100);p__Ascomycota_unclassified(100);p__Ascomycota_unclassified(100); | - | - | - | - | - | - | - | Unassigned | - |
| Otu18638 | 1 | k__Fungi(100);k__Fungi_unclassified(100);k__Fungi_unclassified(100);k__Fungi_unclassified(100);k__Fungi_unclassified(100);k__Fungi_unclassified(100); | - | - | - | - | - | - | - | Unassigned | - |
| Otu18639 | 1 | k__Fungi(100);k__Fungi_unclassified(100);k__Fungi_unclassified(100);k__Fungi_unclassified(100);k__Fungi_unclassified(100);k__Fungi_unclassified(100); | - | - | - | - | - | - | - | Unassigned | - |
| Otu18659 | 1 | k__Fungi(100);p__Ascomycota(100);p__Ascomycota_unclassified(100);p__Ascomycota_unclassified(100);p__Ascomycota_unclassified(100);p__Ascomycota_unclassified(100);p__Ascomycota_unclassified(100); | - | - | - | - | - | - | - | Unassigned | - |
| Otu18660 | 1 | k__Fungi(100);k__Fungi_unclassified(100);k__Fungi_unclassified(100);k__Fungi_unclassified(100);k__Fungi_unclassified(100);k__Fungi_unclassified(100); | - | - | - | - | - | - | - | Unassigned | - |
| Otu18661 | 1 | k__Fungi(100);k__Fungi_unclassified(100);k__Fungi_unclassified(100);k__Fungi_unclassified(100);k__Fungi_unclassified(100);k__Fungi_unclassified(100); | - | - | - | - | - | - | - | Unassigned | - |
| Otu18670 | 1 | k__Fungi(100);p__Ascomycota(100);p__Ascomycota_unclassified(100);p__Ascomycota_unclassified(100);p__Ascomycota_unclassified(100);p__Ascomycota_unclassified(100);p__Ascomycota_unclassified(100); | - | - | - | - | - | - | - | Unassigned | - |
| Otu18671 | 1 | k__Fungi(100);p__Ascomycota(100);p__Ascomycota_unclassified(100);p__Ascomycota_unclassified(100);p__Ascomycota_unclassified(100);p__Ascomycota_unclassified(100);p__Ascomycota_unclassified(100); | - | - | - | - | - | - | - | Unassigned | - |
| Otu18672 | 1 | k__Fungi(100);p__Ascomycota(100);p__Ascomycota_unclassified(100);p__Ascomycota_unclassified(100);p__Ascomycota_unclassified(100);p__Ascomycota_unclassified(100);p__Ascomycota_unclassified(100); | - | - | - | - | - | - | - | Unassigned | - |
| Otu18673 | 1 | k__Fungi(100);k__Fungi_unclassified(100);k__Fungi_unclassified(100);k__Fungi_unclassified(100);k__Fungi_unclassified(100);k__Fungi_unclassified(100); | - | - | - | - | - | - | - | Unassigned | - |
| Otu18677 | 1 | k__Fungi(100);k__Fungi_unclassified(100);k__Fungi_unclassified(100);k__Fungi_unclassified(100);k__Fungi_unclassified(100);k__Fungi_unclassified(100); | - | - | - | - | - | - | - | Unassigned | - |
| Otu18678 | 1 | k__Fungi(100);p__Basidiomycota(100);p__Basidiomycota_unclassified(100);p__Basidiomycota_unclassified(100);p__Basidiomycota_unclassified(100);p__Basidiomycota_unclassified(100);p__Basidiomycota_unclassified(100); | - | - | - | - | - | - | - | Unassigned | - |
| Otu18679 | 1 | k__Fungi(100);k__Fungi_unclassified(100);k__Fungi_unclassified(100);k__Fungi_unclassified(100);k__Fungi_unclassified(100);k__Fungi_unclassified(100); | - | - | - | - | - | - | - | Unassigned | - |
| Otu18680 | 1 | k__Fungi(100);p__Ascomycota(100);p__Ascomycota_unclassified(100);p__Ascomycota_unclassified(100);p__Ascomycota_unclassified(100);p__Ascomycota_unclassified(100);p__Ascomycota_unclassified(100); | - | - | - | - | - | - | - | Unassigned | - |
| Otu18681 | 1 | k__Fungi(100);p__Ascomycota(100);p__Ascomycota_unclassified(100);p__Ascomycota_unclassified(100);p__Ascomycota_unclassified(100);p__Ascomycota_unclassified(100);p__Ascomycota_unclassified(100); | - | - | - | - | - | - | - | Unassigned | - |
| Otu18696 | 1 | k__Fungi(100);p__Ascomycota(100);p__Ascomycota_unclassified(100);p__Ascomycota_unclassified(100);p__Ascomycota_unclassified(100);p__Ascomycota_unclassified(100);p__Ascomycota_unclassified(100); | - | - | - | - | - | - | - | Unassigned | - |
| Otu18697 | 1 | k__Fungi(100);p__Ascomycota(100);p__Ascomycota_unclassified(100);p__Ascomycota_unclassified(100);p__Ascomycota_unclassified(100);p__Ascomycota_unclassified(100);p__Ascomycota_unclassified(100); | - | - | - | - | - | - | - | Unassigned | - |
| Otu18698 | 1 | k__Fungi(100);k__Fungi_unclassified(100);k__Fungi_unclassified(100);k__Fungi_unclassified(100);k__Fungi_unclassified(100);k__Fungi_unclassified(100); | - | - | - | - | - | - | - | Unassigned | - |
| Otu18699 | 1 | k__Fungi(100);p__Ascomycota(100);p__Ascomycota_unclassified(100);p__Ascomycota_unclassified(100);p__Ascomycota_unclassified(100);p__Ascomycota_unclassified(100);p__Ascomycota_unclassified(100); | - | - | - | - | - | - | - | Unassigned | - |
| Otu18701 | 1 | k__Fungi(100);p__Ascomycota(100);p__Ascomycota_unclassified(100);p__Ascomycota_unclassified(100);p__Ascomycota_unclassified(100);p__Ascomycota_unclassified(100);p__Ascomycota_unclassified(100); | - | - | - | - | - | - | - | Unassigned | - |
| Otu18704 | 1 | k__Fungi(100);k__Fungi_unclassified(100);k__Fungi_unclassified(100);k__Fungi_unclassified(100);k__Fungi_unclassified(100);k__Fungi_unclassified(100); | - | - | - | - | - | - | - | Unassigned | - |
| Otu18712 | 1 | k__Fungi(100);k__Fungi_unclassified(100);k__Fungi_unclassified(100);k__Fungi_unclassified(100);k__Fungi_unclassified(100);k__Fungi_unclassified(100); | - | - | - | - | - | - | - | Unassigned | - |
| Otu18713 | 1 | k__Fungi(100);p__Ascomycota(100);p__Ascomycota_unclassified(100);p__Ascomycota_unclassified(100);p__Ascomycota_unclassified(100);p__Ascomycota_unclassified(100);p__Ascomycota_unclassified(100); | - | - | - | - | - | - | - | Unassigned | - |
| Otu18717 | 1 | k__Fungi(100);p__Ascomycota(100);p__Ascomycota_unclassified(100);p__Ascomycota_unclassified(100);p__Ascomycota_unclassified(100);p__Ascomycota_unclassified(100);p__Ascomycota_unclassified(100); | - | - | - | - | - | - | - | Unassigned | - |
| Otu18733 | 1 | k__Fungi(100);k__Fungi_unclassified(100);k__Fungi_unclassified(100);k__Fungi_unclassified(100);k__Fungi_unclassified(100);k__Fungi_unclassified(100); | - | - | - | - | - | - | - | Unassigned | - |
| Otu18742 | 1 | k__Fungi(100);p__Ascomycota(100);p__Ascomycota_unclassified(100);p__Ascomycota_unclassified(100);p__Ascomycota_unclassified(100);p__Ascomycota_unclassified(100);p__Ascomycota_unclassified(100); | - | - | - | - | - | - | - | Unassigned | - |
| Otu18745 | 1 | k__Fungi(100);k__Fungi_unclassified(100);k__Fungi_unclassified(100);k__Fungi_unclassified(100);k__Fungi_unclassified(100);k__Fungi_unclassified(100); | - | - | - | - | - | - | - | Unassigned | - |
| Otu18746 | 1 | k__Fungi(100);k__Fungi_unclassified(100);k__Fungi_unclassified(100);k__Fungi_unclassified(100);k__Fungi_unclassified(100);k__Fungi_unclassified(100); | - | - | - | - | - | - | - | Unassigned | - |
| Otu18747 | 1 | k__Fungi(100);k__Fungi_unclassified(100);k__Fungi_unclassified(100);k__Fungi_unclassified(100);k__Fungi_unclassified(100);k__Fungi_unclassified(100); | - | - | - | - | - | - | - | Unassigned | - |
| Otu18755 | 1 | k__Fungi(100);p__Ascomycota(100);p__Ascomycota_unclassified(100);p__Ascomycota_unclassified(100);p__Ascomycota_unclassified(100);p__Ascomycota_unclassified(100);p__Ascomycota_unclassified(100); | - | - | - | - | - | - | - | Unassigned | - |
| Otu18789 | 1 | k__Fungi(100);k__Fungi_unclassified(100);k__Fungi_unclassified(100);k__Fungi_unclassified(100);k__Fungi_unclassified(100);k__Fungi_unclassified(100); | - | - | - | - | - | - | - | Unassigned | - |
| Otu18792 | 1 | k__Fungi(100);p__Ascomycota(100);p__Ascomycota_unclassified(100);p__Ascomycota_unclassified(100);p__Ascomycota_unclassified(100);p__Ascomycota_unclassified(100);p__Ascomycota_unclassified(100); | - | - | - | - | - | - | - | Unassigned | - |
| Otu18810 | 1 | k__Fungi(100);p__Ascomycota(100);p__Ascomycota_unclassified(100);p__Ascomycota_unclassified(100);p__Ascomycota_unclassified(100);p__Ascomycota_unclassified(100);p__Ascomycota_unclassified(100); | - | - | - | - | - | - | - | Unassigned | - |
| Otu18836 | 1 | k__Fungi(100);k__Fungi_unclassified(100);k__Fungi_unclassified(100);k__Fungi_unclassified(100);k__Fungi_unclassified(100);k__Fungi_unclassified(100); | - | - | - | - | - | - | - | Unassigned | - |
| Otu18837 | 1 | k__Fungi(100);p__Ascomycota(100);p__Ascomycota_unclassified(100);p__Ascomycota_unclassified(100);p__Ascomycota_unclassified(100);p__Ascomycota_unclassified(100);p__Ascomycota_unclassified(100); | - | - | - | - | - | - | - | Unassigned | - |
| Otu18887 | 1 | k__Fungi(100);p__Ascomycota(100);p__Ascomycota_unclassified(100);p__Ascomycota_unclassified(100);p__Ascomycota_unclassified(100);p__Ascomycota_unclassified(100);p__Ascomycota_unclassified(100); | - | - | - | - | - | - | - | Unassigned | - |
| Otu18888 | 1 | k__Fungi(100);p__Ascomycota(100);p__Ascomycota_unclassified(100);p__Ascomycota_unclassified(100);p__Ascomycota_unclassified(100);p__Ascomycota_unclassified(100);p__Ascomycota_unclassified(100); | - | - | - | - | - | - | - | Unassigned | - |
| Otu18912 | 1 | k__Fungi(100);p__Ascomycota(100);p__Ascomycota_unclassified(100);p__Ascomycota_unclassified(100);p__Ascomycota_unclassified(100);p__Ascomycota_unclassified(100);p__Ascomycota_unclassified(100); | - | - | - | - | - | - | - | Unassigned | - |
| Otu18932 | 1 | k__Fungi(100);k__Fungi_unclassified(100);k__Fungi_unclassified(100);k__Fungi_unclassified(100);k__Fungi_unclassified(100);k__Fungi_unclassified(100); | - | - | - | - | - | - | - | Unassigned | - |
| Otu18933 | 1 | k__Fungi(100);k__Fungi_unclassified(100);k__Fungi_unclassified(100);k__Fungi_unclassified(100);k__Fungi_unclassified(100);k__Fungi_unclassified(100); | - | - | - | - | - | - | - | Unassigned | - |
| Otu18946 | 1 | k__Fungi(100);k__Fungi_unclassified(100);k__Fungi_unclassified(100);k__Fungi_unclassified(100);k__Fungi_unclassified(100);k__Fungi_unclassified(100); | - | - | - | - | - | - | - | Unassigned | - |
| Otu18947 | 1 | k__Fungi(100);k__Fungi_unclassified(100);k__Fungi_unclassified(100);k__Fungi_unclassified(100);k__Fungi_unclassified(100);k__Fungi_unclassified(100); | - | - | - | - | - | - | - | Unassigned | - |
| Otu18954 | 1 | k__Fungi(100);p__Ascomycota(100);p__Ascomycota_unclassified(100);p__Ascomycota_unclassified(100);p__Ascomycota_unclassified(100);p__Ascomycota_unclassified(100);p__Ascomycota_unclassified(100); | - | - | - | - | - | - | - | Unassigned | - |
| Otu18971 | 1 | k__Fungi(100);k__Fungi_unclassified(100);k__Fungi_unclassified(100);k__Fungi_unclassified(100);k__Fungi_unclassified(100);k__Fungi_unclassified(100); | - | - | - | - | - | - | - | Unassigned | - |
| Otu18979 | 1 | k__Fungi(100);p__Ascomycota(100);p__Ascomycota_unclassified(100);p__Ascomycota_unclassified(100);p__Ascomycota_unclassified(100);p__Ascomycota_unclassified(100);p__Ascomycota_unclassified(100); | - | - | - | - | - | - | - | Unassigned | - |
| Otu18984 | 1 | k__Fungi(100);k__Fungi_unclassified(100);k__Fungi_unclassified(100);k__Fungi_unclassified(100);k__Fungi_unclassified(100);k__Fungi_unclassified(100); | - | - | - | - | - | - | - | Unassigned | - |
| Otu18985 | 1 | k__Fungi(100);p__Ascomycota(100);p__Ascomycota_unclassified(100);p__Ascomycota_unclassified(100);p__Ascomycota_unclassified(100);p__Ascomycota_unclassified(100);p__Ascomycota_unclassified(100); | - | - | - | - | - | - | - | Unassigned | - |
| Otu18986 | 1 | k__Fungi(100);p__Ascomycota(100);p__Ascomycota_unclassified(100);p__Ascomycota_unclassified(100);p__Ascomycota_unclassified(100);p__Ascomycota_unclassified(100);p__Ascomycota_unclassified(100); | - | - | - | - | - | - | - | Unassigned | - |
| Otu18987 | 1 | k__Fungi(100);k__Fungi_unclassified(100);k__Fungi_unclassified(100);k__Fungi_unclassified(100);k__Fungi_unclassified(100);k__Fungi_unclassified(100); | - | - | - | - | - | - | - | Unassigned | - |
| Otu18988 | 1 | k__Fungi(100);k__Fungi_unclassified(100);k__Fungi_unclassified(100);k__Fungi_unclassified(100);k__Fungi_unclassified(100);k__Fungi_unclassified(100); | - | - | - | - | - | - | - | Unassigned | - |
| Otu18989 | 1 | k__Fungi(100);k__Fungi_unclassified(100);k__Fungi_unclassified(100);k__Fungi_unclassified(100);k__Fungi_unclassified(100);k__Fungi_unclassified(100); | - | - | - | - | - | - | - | Unassigned | - |
| Otu18991 | 1 | k__Fungi(100);k__Fungi_unclassified(100);k__Fungi_unclassified(100);k__Fungi_unclassified(100);k__Fungi_unclassified(100);k__Fungi_unclassified(100); | - | - | - | - | - | - | - | Unassigned | - |
| Otu18992 | 1 | k__Fungi(100);k__Fungi_unclassified(100);k__Fungi_unclassified(100);k__Fungi_unclassified(100);k__Fungi_unclassified(100);k__Fungi_unclassified(100); | - | - | - | - | - | - | - | Unassigned | - |
| Otu19017 | 1 | k__Fungi(100);k__Fungi_unclassified(100);k__Fungi_unclassified(100);k__Fungi_unclassified(100);k__Fungi_unclassified(100);k__Fungi_unclassified(100); | - | - | - | - | - | - | - | Unassigned | - |
| Otu19018 | 1 | k__Fungi(100);k__Fungi_unclassified(100);k__Fungi_unclassified(100);k__Fungi_unclassified(100);k__Fungi_unclassified(100);k__Fungi_unclassified(100); | - | - | - | - | - | - | - | Unassigned | - |
| Otu19022 | 1 | k__Fungi(100);k__Fungi_unclassified(100);k__Fungi_unclassified(100);k__Fungi_unclassified(100);k__Fungi_unclassified(100);k__Fungi_unclassified(100); | - | - | - | - | - | - | - | Unassigned | - |
| Otu19035 | 1 | k__Fungi(100);k__Fungi_unclassified(100);k__Fungi_unclassified(100);k__Fungi_unclassified(100);k__Fungi_unclassified(100);k__Fungi_unclassified(100); | - | - | - | - | - | - | - | Unassigned | - |
| Otu19036 | 1 | k__Fungi(100);k__Fungi_unclassified(100);k__Fungi_unclassified(100);k__Fungi_unclassified(100);k__Fungi_unclassified(100);k__Fungi_unclassified(100); | - | - | - | - | - | - | - | Unassigned | - |
| Otu19037 | 1 | k__Fungi(100);k__Fungi_unclassified(100);k__Fungi_unclassified(100);k__Fungi_unclassified(100);k__Fungi_unclassified(100);k__Fungi_unclassified(100); | - | - | - | - | - | - | - | Unassigned | - |
| Otu19038 | 1 | k__Fungi(100);k__Fungi_unclassified(100);k__Fungi_unclassified(100);k__Fungi_unclassified(100);k__Fungi_unclassified(100);k__Fungi_unclassified(100); | - | - | - | - | - | - | - | Unassigned | - |
| Otu19039 | 1 | k__Fungi(100);p__Ascomycota(100);p__Ascomycota_unclassified(100);p__Ascomycota_unclassified(100);p__Ascomycota_unclassified(100);p__Ascomycota_unclassified(100);p__Ascomycota_unclassified(100); | - | - | - | - | - | - | - | Unassigned | - |
| Otu19041 | 1 | k__Fungi(100);k__Fungi_unclassified(100);k__Fungi_unclassified(100);k__Fungi_unclassified(100);k__Fungi_unclassified(100);k__Fungi_unclassified(100); | - | - | - | - | - | - | - | Unassigned | - |
| Otu19043 | 1 | k__Fungi(100);p__Ascomycota(100);p__Ascomycota_unclassified(100);p__Ascomycota_unclassified(100);p__Ascomycota_unclassified(100);p__Ascomycota_unclassified(100);p__Ascomycota_unclassified(100); | - | - | - | - | - | - | - | Unassigned | - |
| Otu19046 | 1 | k__Fungi(100);k__Fungi_unclassified(100);k__Fungi_unclassified(100);k__Fungi_unclassified(100);k__Fungi_unclassified(100);k__Fungi_unclassified(100); | - | - | - | - | - | - | - | Unassigned | - |
| Otu19047 | 1 | k__Fungi(100);k__Fungi_unclassified(100);k__Fungi_unclassified(100);k__Fungi_unclassified(100);k__Fungi_unclassified(100);k__Fungi_unclassified(100); | - | - | - | - | - | - | - | Unassigned | - |
| Otu19048 | 1 | k__Fungi(100);k__Fungi_unclassified(100);k__Fungi_unclassified(100);k__Fungi_unclassified(100);k__Fungi_unclassified(100);k__Fungi_unclassified(100); | - | - | - | - | - | - | - | Unassigned | - |
| Otu19049 | 1 | k__Fungi(100);p__Ascomycota(100);p__Ascomycota_unclassified(100);p__Ascomycota_unclassified(100);p__Ascomycota_unclassified(100);p__Ascomycota_unclassified(100);p__Ascomycota_unclassified(100); | - | - | - | - | - | - | - | Unassigned | - |
| Otu19051 | 1 | k__Fungi(100);p__Ascomycota(100);p__Ascomycota_unclassified(100);p__Ascomycota_unclassified(100);p__Ascomycota_unclassified(100);p__Ascomycota_unclassified(100);p__Ascomycota_unclassified(100); | - | - | - | - | - | - | - | Unassigned | - |
| Otu19056 | 1 | k__Fungi(100);p__Ascomycota(100);p__Ascomycota_unclassified(100);p__Ascomycota_unclassified(100);p__Ascomycota_unclassified(100);p__Ascomycota_unclassified(100);p__Ascomycota_unclassified(100); | - | - | - | - | - | - | - | Unassigned | - |
| Otu19060 | 1 | k__Fungi(100);k__Fungi_unclassified(100);k__Fungi_unclassified(100);k__Fungi_unclassified(100);k__Fungi_unclassified(100);k__Fungi_unclassified(100); | - | - | - | - | - | - | - | Unassigned | - |
| Otu19064 | 1 | k__Fungi(100);k__Fungi_unclassified(100);k__Fungi_unclassified(100);k__Fungi_unclassified(100);k__Fungi_unclassified(100);k__Fungi_unclassified(100); | - | - | - | - | - | - | - | Unassigned | - |
| Otu19066 | 1 | k__Fungi(100);p__Ascomycota(100);p__Ascomycota_unclassified(100);p__Ascomycota_unclassified(100);p__Ascomycota_unclassified(100);p__Ascomycota_unclassified(100);p__Ascomycota_unclassified(100); | - | - | - | - | - | - | - | Unassigned | - |
| Otu19067 | 1 | k__Fungi(100);p__Ascomycota(100);p__Ascomycota_unclassified(100);p__Ascomycota_unclassified(100);p__Ascomycota_unclassified(100);p__Ascomycota_unclassified(100);p__Ascomycota_unclassified(100); | - | - | - | - | - | - | - | Unassigned | - |
| Otu19068 | 1 | k__Fungi(100);p__Ascomycota(100);p__Ascomycota_unclassified(100);p__Ascomycota_unclassified(100);p__Ascomycota_unclassified(100);p__Ascomycota_unclassified(100);p__Ascomycota_unclassified(100); | - | - | - | - | - | - | - | Unassigned | - |
| Otu19080 | 1 | k__Fungi(100);p__Ascomycota(100);p__Ascomycota_unclassified(100);p__Ascomycota_unclassified(100);p__Ascomycota_unclassified(100);p__Ascomycota_unclassified(100);p__Ascomycota_unclassified(100); | - | - | - | - | - | - | - | Unassigned | - |
| Otu19093 | 1 | k__Fungi(100);k__Fungi_unclassified(100);k__Fungi_unclassified(100);k__Fungi_unclassified(100);k__Fungi_unclassified(100);k__Fungi_unclassified(100); | - | - | - | - | - | - | - | Unassigned | - |
| Otu19099 | 1 | k__Fungi(100);k__Fungi_unclassified(100);k__Fungi_unclassified(100);k__Fungi_unclassified(100);k__Fungi_unclassified(100);k__Fungi_unclassified(100); | - | - | - | - | - | - | - | Unassigned | - |
| Otu19111 | 1 | k__Fungi(100);k__Fungi_unclassified(100);k__Fungi_unclassified(100);k__Fungi_unclassified(100);k__Fungi_unclassified(100);k__Fungi_unclassified(100); | - | - | - | - | - | - | - | Unassigned | - |
| Otu19125 | 1 | k__Fungi(100);p__Ascomycota(100);p__Ascomycota_unclassified(100);p__Ascomycota_unclassified(100);p__Ascomycota_unclassified(100);p__Ascomycota_unclassified(100);p__Ascomycota_unclassified(100); | - | - | - | - | - | - | - | Unassigned | - |
| Otu19126 | 1 | k__Fungi(100);k__Fungi_unclassified(100);k__Fungi_unclassified(100);k__Fungi_unclassified(100);k__Fungi_unclassified(100);k__Fungi_unclassified(100); | - | - | - | - | - | - | - | Unassigned | - |
| Otu19132 | 1 | k__Fungi(100);k__Fungi_unclassified(100);k__Fungi_unclassified(100);k__Fungi_unclassified(100);k__Fungi_unclassified(100);k__Fungi_unclassified(100); | - | - | - | - | - | - | - | Unassigned | - |
| Otu19133 | 1 | k__Fungi(100);k__Fungi_unclassified(100);k__Fungi_unclassified(100);k__Fungi_unclassified(100);k__Fungi_unclassified(100);k__Fungi_unclassified(100); | - | - | - | - | - | - | - | Unassigned | - |
| Otu19137 | 1 | k__Fungi(100);k__Fungi_unclassified(100);k__Fungi_unclassified(100);k__Fungi_unclassified(100);k__Fungi_unclassified(100);k__Fungi_unclassified(100); | - | - | - | - | - | - | - | Unassigned | - |
| Otu19142 | 1 | k__Fungi(100);k__Fungi_unclassified(100);k__Fungi_unclassified(100);k__Fungi_unclassified(100);k__Fungi_unclassified(100);k__Fungi_unclassified(100); | - | - | - | - | - | - | - | Unassigned | - |
| Otu19144 | 1 | k__Fungi(100);p__Ascomycota(100);p__Ascomycota_unclassified(100);p__Ascomycota_unclassified(100);p__Ascomycota_unclassified(100);p__Ascomycota_unclassified(100);p__Ascomycota_unclassified(100); | - | - | - | - | - | - | - | Unassigned | - |
| Otu19145 | 1 | k__Fungi(100);k__Fungi_unclassified(100);k__Fungi_unclassified(100);k__Fungi_unclassified(100);k__Fungi_unclassified(100);k__Fungi_unclassified(100); | - | - | - | - | - | - | - | Unassigned | - |
| Otu19146 | 1 | k__Fungi(100);k__Fungi_unclassified(100);k__Fungi_unclassified(100);k__Fungi_unclassified(100);k__Fungi_unclassified(100);k__Fungi_unclassified(100); | - | - | - | - | - | - | - | Unassigned | - |
| Otu19148 | 1 | k__Fungi(100);p__Ascomycota(100);p__Ascomycota_unclassified(100);p__Ascomycota_unclassified(100);p__Ascomycota_unclassified(100);p__Ascomycota_unclassified(100);p__Ascomycota_unclassified(100); | - | - | - | - | - | - | - | Unassigned | - |
| Otu19151 | 1 | k__Fungi(100);k__Fungi_unclassified(100);k__Fungi_unclassified(100);k__Fungi_unclassified(100);k__Fungi_unclassified(100);k__Fungi_unclassified(100); | - | - | - | - | - | - | - | Unassigned | - |
| Otu19152 | 1 | k__Fungi(100);k__Fungi_unclassified(100);k__Fungi_unclassified(100);k__Fungi_unclassified(100);k__Fungi_unclassified(100);k__Fungi_unclassified(100); | - | - | - | - | - | - | - | Unassigned | - |
| Otu19155 | 1 | k__Fungi(100);k__Fungi_unclassified(100);k__Fungi_unclassified(100);k__Fungi_unclassified(100);k__Fungi_unclassified(100);k__Fungi_unclassified(100); | - | - | - | - | - | - | - | Unassigned | - |
| Otu19156 | 1 | k__Fungi(100);k__Fungi_unclassified(100);k__Fungi_unclassified(100);k__Fungi_unclassified(100);k__Fungi_unclassified(100);k__Fungi_unclassified(100); | - | - | - | - | - | - | - | Unassigned | - |
| Otu19157 | 1 | k__Fungi(100);k__Fungi_unclassified(100);k__Fungi_unclassified(100);k__Fungi_unclassified(100);k__Fungi_unclassified(100);k__Fungi_unclassified(100); | - | - | - | - | - | - | - | Unassigned | - |
| Otu19158 | 1 | k__Fungi(100);p__Ascomycota(100);p__Ascomycota_unclassified(100);p__Ascomycota_unclassified(100);p__Ascomycota_unclassified(100);p__Ascomycota_unclassified(100);p__Ascomycota_unclassified(100); | - | - | - | - | - | - | - | Unassigned | - |
| Otu19159 | 1 | k__Fungi(100);p__Ascomycota(100);p__Ascomycota_unclassified(100);p__Ascomycota_unclassified(100);p__Ascomycota_unclassified(100);p__Ascomycota_unclassified(100);p__Ascomycota_unclassified(100); | - | - | - | - | - | - | - | Unassigned | - |
| Otu19166 | 1 | k__Fungi(100);k__Fungi_unclassified(100);k__Fungi_unclassified(100);k__Fungi_unclassified(100);k__Fungi_unclassified(100);k__Fungi_unclassified(100); | - | - | - | - | - | - | - | Unassigned | - |
| Otu19167 | 1 | k__Fungi(100);p__Ascomycota(100);p__Ascomycota_unclassified(100);p__Ascomycota_unclassified(100);p__Ascomycota_unclassified(100);p__Ascomycota_unclassified(100);p__Ascomycota_unclassified(100); | - | - | - | - | - | - | - | Unassigned | - |
| Otu19168 | 1 | k__Fungi(100);p__Ascomycota(100);p__Ascomycota_unclassified(100);p__Ascomycota_unclassified(100);p__Ascomycota_unclassified(100);p__Ascomycota_unclassified(100);p__Ascomycota_unclassified(100); | - | - | - | - | - | - | - | Unassigned | - |
| Otu19169 | 1 | k__Fungi(100);p__Ascomycota(100);p__Ascomycota_unclassified(100);p__Ascomycota_unclassified(100);p__Ascomycota_unclassified(100);p__Ascomycota_unclassified(100);p__Ascomycota_unclassified(100); | - | - | - | - | - | - | - | Unassigned | - |
| Otu19170 | 1 | k__Fungi(100);p__Ascomycota(100);p__Ascomycota_unclassified(100);p__Ascomycota_unclassified(100);p__Ascomycota_unclassified(100);p__Ascomycota_unclassified(100);p__Ascomycota_unclassified(100); | - | - | - | - | - | - | - | Unassigned | - |
| Otu19171 | 1 | k__Fungi(100);p__Ascomycota(100);p__Ascomycota_unclassified(100);p__Ascomycota_unclassified(100);p__Ascomycota_unclassified(100);p__Ascomycota_unclassified(100);p__Ascomycota_unclassified(100); | - | - | - | - | - | - | - | Unassigned | - |
| Otu19172 | 1 | k__Fungi(100);k__Fungi_unclassified(100);k__Fungi_unclassified(100);k__Fungi_unclassified(100);k__Fungi_unclassified(100);k__Fungi_unclassified(100); | - | - | - | - | - | - | - | Unassigned | - |
| Otu19173 | 1 | k__Fungi(100);p__Ascomycota(100);p__Ascomycota_unclassified(100);p__Ascomycota_unclassified(100);p__Ascomycota_unclassified(100);p__Ascomycota_unclassified(100);p__Ascomycota_unclassified(100); | - | - | - | - | - | - | - | Unassigned | - |
| Otu19174 | 1 | k__Fungi(100);k__Fungi_unclassified(100);k__Fungi_unclassified(100);k__Fungi_unclassified(100);k__Fungi_unclassified(100);k__Fungi_unclassified(100); | - | - | - | - | - | - | - | Unassigned | - |
| Otu19175 | 1 | k__Fungi(100);k__Fungi_unclassified(100);k__Fungi_unclassified(100);k__Fungi_unclassified(100);k__Fungi_unclassified(100);k__Fungi_unclassified(100); | - | - | - | - | - | - | - | Unassigned | - |
| Otu19176 | 1 | k__Fungi(100);k__Fungi_unclassified(100);k__Fungi_unclassified(100);k__Fungi_unclassified(100);k__Fungi_unclassified(100);k__Fungi_unclassified(100); | - | - | - | - | - | - | - | Unassigned | - |
| Otu19218 | 1 | k__Fungi(100);k__Fungi_unclassified(100);k__Fungi_unclassified(100);k__Fungi_unclassified(100);k__Fungi_unclassified(100);k__Fungi_unclassified(100); | - | - | - | - | - | - | - | Unassigned | - |
| Otu19219 | 1 | k__Fungi(100);p__Ascomycota(100);p__Ascomycota_unclassified(100);p__Ascomycota_unclassified(100);p__Ascomycota_unclassified(100);p__Ascomycota_unclassified(100);p__Ascomycota_unclassified(100); | - | - | - | - | - | - | - | Unassigned | - |
| Otu19220 | 1 | k__Fungi(100);p__Ascomycota(100);p__Ascomycota_unclassified(100);p__Ascomycota_unclassified(100);p__Ascomycota_unclassified(100);p__Ascomycota_unclassified(100);p__Ascomycota_unclassified(100); | - | - | - | - | - | - | - | Unassigned | - |
| Otu19246 | 1 | k__Fungi(100);p__Ascomycota(100);p__Ascomycota_unclassified(100);p__Ascomycota_unclassified(100);p__Ascomycota_unclassified(100);p__Ascomycota_unclassified(100);p__Ascomycota_unclassified(100); | - | - | - | - | - | - | - | Unassigned | - |
| Otu19248 | 1 | k__Fungi(100);p__Ascomycota(100);p__Ascomycota_unclassified(100);p__Ascomycota_unclassified(100);p__Ascomycota_unclassified(100);p__Ascomycota_unclassified(100);p__Ascomycota_unclassified(100); | - | - | - | - | - | - | - | Unassigned | - |
| Otu19251 | 1 | k__Fungi(100);p__Ascomycota(100);p__Ascomycota_unclassified(100);p__Ascomycota_unclassified(100);p__Ascomycota_unclassified(100);p__Ascomycota_unclassified(100);p__Ascomycota_unclassified(100); | - | - | - | - | - | - | - | Unassigned | - |
| Otu19252 | 1 | k__Fungi(100);p__Ascomycota(100);p__Ascomycota_unclassified(100);p__Ascomycota_unclassified(100);p__Ascomycota_unclassified(100);p__Ascomycota_unclassified(100);p__Ascomycota_unclassified(100); | - | - | - | - | - | - | - | Unassigned | - |
| Otu19253 | 1 | k__Fungi(100);p__Ascomycota(100);p__Ascomycota_unclassified(100);p__Ascomycota_unclassified(100);p__Ascomycota_unclassified(100);p__Ascomycota_unclassified(100);p__Ascomycota_unclassified(100); | - | - | - | - | - | - | - | Unassigned | - |
| Otu19288 | 1 | k__Fungi(100);k__Fungi_unclassified(100);k__Fungi_unclassified(100);k__Fungi_unclassified(100);k__Fungi_unclassified(100);k__Fungi_unclassified(100); | - | - | - | - | - | - | - | Unassigned | - |
| Otu19289 | 1 | k__Fungi(100);k__Fungi_unclassified(100);k__Fungi_unclassified(100);k__Fungi_unclassified(100);k__Fungi_unclassified(100);k__Fungi_unclassified(100); | - | - | - | - | - | - | - | Unassigned | - |
| Otu19290 | 1 | k__Fungi(100);p__Ascomycota(100);p__Ascomycota_unclassified(100);p__Ascomycota_unclassified(100);p__Ascomycota_unclassified(100);p__Ascomycota_unclassified(100);p__Ascomycota_unclassified(100); | - | - | - | - | - | - | - | Unassigned | - |
| Otu19291 | 1 | k__Fungi(100);p__Ascomycota(100);p__Ascomycota_unclassified(100);p__Ascomycota_unclassified(100);p__Ascomycota_unclassified(100);p__Ascomycota_unclassified(100);p__Ascomycota_unclassified(100); | - | - | - | - | - | - | - | Unassigned | - |
| Otu19293 | 1 | k__Fungi(100);k__Fungi_unclassified(100);k__Fungi_unclassified(100);k__Fungi_unclassified(100);k__Fungi_unclassified(100);k__Fungi_unclassified(100); | - | - | - | - | - | - | - | Unassigned | - |
| Otu19295 | 1 | k__Fungi(100);k__Fungi_unclassified(100);k__Fungi_unclassified(100);k__Fungi_unclassified(100);k__Fungi_unclassified(100);k__Fungi_unclassified(100); | - | - | - | - | - | - | - | Unassigned | - |
| Otu19319 | 1 | k__Fungi(100);p__Ascomycota(100);p__Ascomycota_unclassified(100);p__Ascomycota_unclassified(100);p__Ascomycota_unclassified(100);p__Ascomycota_unclassified(100);p__Ascomycota_unclassified(100); | - | - | - | - | - | - | - | Unassigned | - |
| Otu19334 | 1 | k__Fungi(100);p__Ascomycota(100);p__Ascomycota_unclassified(100);p__Ascomycota_unclassified(100);p__Ascomycota_unclassified(100);p__Ascomycota_unclassified(100);p__Ascomycota_unclassified(100); | - | - | - | - | - | - | - | Unassigned | - |
| Otu19335 | 1 | k__Fungi(100);p__Ascomycota(100);p__Ascomycota_unclassified(100);p__Ascomycota_unclassified(100);p__Ascomycota_unclassified(100);p__Ascomycota_unclassified(100);p__Ascomycota_unclassified(100); | - | - | - | - | - | - | - | Unassigned | - |
| Otu19353 | 1 | k__Fungi(100);k__Fungi_unclassified(100);k__Fungi_unclassified(100);k__Fungi_unclassified(100);k__Fungi_unclassified(100);k__Fungi_unclassified(100); | - | - | - | - | - | - | - | Unassigned | - |
| Otu19358 | 1 | k__Fungi(100);k__Fungi_unclassified(100);k__Fungi_unclassified(100);k__Fungi_unclassified(100);k__Fungi_unclassified(100);k__Fungi_unclassified(100); | - | - | - | - | - | - | - | Unassigned | - |
| Otu19372 | 1 | k__Fungi(100);k__Fungi_unclassified(100);k__Fungi_unclassified(100);k__Fungi_unclassified(100);k__Fungi_unclassified(100);k__Fungi_unclassified(100); | - | - | - | - | - | - | - | Unassigned | - |
| Otu19374 | 1 | k__Fungi(100);k__Fungi_unclassified(100);k__Fungi_unclassified(100);k__Fungi_unclassified(100);k__Fungi_unclassified(100);k__Fungi_unclassified(100); | - | - | - | - | - | - | - | Unassigned | - |
| Otu19375 | 1 | k__Fungi(100);k__Fungi_unclassified(100);k__Fungi_unclassified(100);k__Fungi_unclassified(100);k__Fungi_unclassified(100);k__Fungi_unclassified(100); | - | - | - | - | - | - | - | Unassigned | - |
| Otu19377 | 1 | k__Fungi(100);p__Ascomycota(100);p__Ascomycota_unclassified(100);p__Ascomycota_unclassified(100);p__Ascomycota_unclassified(100);p__Ascomycota_unclassified(100);p__Ascomycota_unclassified(100); | - | - | - | - | - | - | - | Unassigned | - |
| Otu19379 | 1 | k__Fungi(100);k__Fungi_unclassified(100);k__Fungi_unclassified(100);k__Fungi_unclassified(100);k__Fungi_unclassified(100);k__Fungi_unclassified(100); | - | - | - | - | - | - | - | Unassigned | - |

| | | | | | | | | | | | |
|---|---|---|---|---|---|---|---|---|---|---|---|
| Otu19380 | 1 | k__Fungi(100);k__Fungi_unclassified(100);k__Fungi_unclassified(100);k__Fungi_unclassified(100);k__Fungi_unclassified(100);k__Fungi_unclassified(100);k__Fungi_unclassified(100); | - | - | - | - | - | - | - | Unassigned | - |
| Otu19501 | 1 | k__Fungi(100);p__Ascomycota(100);p__Ascomycota_unclassified(100);p__Ascomycota_unclassified(100);p__Ascomycota_unclassified(100);p__Ascomycota_unclassified(100);p__Ascomycota_unclassified(100); | - | - | - | - | - | - | - | Unassigned | - |
| Otu19508 | 1 | k__Fungi(100);k__Fungi_unclassified(100);k__Fungi_unclassified(100);k__Fungi_unclassified(100);k__Fungi_unclassified(100);k__Fungi_unclassified(100);k__Fungi_unclassified(100); | - | - | - | - | - | - | - | Unassigned | - |
| Otu19524 | 1 | k__Fungi(100);p__Ascomycota(100);p__Ascomycota_unclassified(100);p__Ascomycota_unclassified(100);p__Ascomycota_unclassified(100);p__Ascomycota_unclassified(100);p__Ascomycota_unclassified(100); | - | - | - | - | - | - | - | Unassigned | - |
| Otu19596 | 1 | k__Fungi(100);p__Ascomycota(100);p__Ascomycota_unclassified(100);p__Ascomycota_unclassified(100);p__Ascomycota_unclassified(100);p__Ascomycota_unclassified(100);p__Ascomycota_unclassified(100); | - | - | - | - | - | - | - | Unassigned | - |
| Otu19618 | 1 | k__Fungi(100);k__Fungi_unclassified(100);k__Fungi_unclassified(100);k__Fungi_unclassified(100);k__Fungi_unclassified(100);k__Fungi_unclassified(100);k__Fungi_unclassified(100); | - | - | - | - | - | - | - | Unassigned | - |
| Otu19626 | 1 | k__Fungi(100);p__Ascomycota(100);p__Ascomycota_unclassified(100);p__Ascomycota_unclassified(100);p__Ascomycota_unclassified(100);p__Ascomycota_unclassified(100);p__Ascomycota_unclassified(100); | - | - | - | - | - | - | - | Unassigned | - |
| Otu19628 | 1 | k__Fungi(100);p__Ascomycota(100);p__Ascomycota_unclassified(100);p__Ascomycota_unclassified(100);p__Ascomycota_unclassified(100);p__Ascomycota_unclassified(100);p__Ascomycota_unclassified(100); | - | - | - | - | - | - | - | Unassigned | - |
| Otu19629 | 1 | k__Fungi(100);p__Ascomycota(100);p__Ascomycota_unclassified(100);p__Ascomycota_unclassified(100);p__Ascomycota_unclassified(100);p__Ascomycota_unclassified(100);p__Ascomycota_unclassified(100); | - | - | - | - | - | - | - | Unassigned | - |
| Otu19632 | 1 | k__Fungi(100);p__Ascomycota(100);p__Ascomycota_unclassified(100);p__Ascomycota_unclassified(100);p__Ascomycota_unclassified(100);p__Ascomycota_unclassified(100);p__Ascomycota_unclassified(100); | - | - | - | - | - | - | - | Unassigned | - |
| Otu19635 | 1 | k__Fungi(100);k__Fungi_unclassified(100);k__Fungi_unclassified(100);k__Fungi_unclassified(100);k__Fungi_unclassified(100);k__Fungi_unclassified(100);k__Fungi_unclassified(100); | - | - | - | - | - | - | - | Unassigned | - |
| Otu19654 | 1 | k__Fungi(100);k__Fungi_unclassified(100);k__Fungi_unclassified(100);k__Fungi_unclassified(100);k__Fungi_unclassified(100);k__Fungi_unclassified(100);k__Fungi_unclassified(100); | - | - | - | - | - | - | - | Unassigned | - |
| Otu19724 | 1 | k__Fungi(100);p__Ascomycota(100);p__Ascomycota_unclassified(100);p__Ascomycota_unclassified(100);p__Ascomycota_unclassified(100);p__Ascomycota_unclassified(100);p__Ascomycota_unclassified(100); | - | - | - | - | - | - | - | Unassigned | - |
| Otu19733 | 1 | k__Fungi(100);k__Fungi_unclassified(100);k__Fungi_unclassified(100);k__Fungi_unclassified(100);k__Fungi_unclassified(100);k__Fungi_unclassified(100);k__Fungi_unclassified(100); | - | - | - | - | - | - | - | Unassigned | - |
| Otu19747 | 1 | k__Fungi(100);k__Fungi_unclassified(100);k__Fungi_unclassified(100);k__Fungi_unclassified(100);k__Fungi_unclassified(100);k__Fungi_unclassified(100);k__Fungi_unclassified(100); | - | - | - | - | - | - | - | Unassigned | - |
| Otu19748 | 1 | k__Fungi(100);p__Ascomycota(100);p__Ascomycota_unclassified(100);p__Ascomycota_unclassified(100);p__Ascomycota_unclassified(100);p__Ascomycota_unclassified(100);p__Ascomycota_unclassified(100); | - | - | - | - | - | - | - | Unassigned | - |
| Otu19758 | 1 | k__Fungi(100);k__Fungi_unclassified(100);k__Fungi_unclassified(100);k__Fungi_unclassified(100);k__Fungi_unclassified(100);k__Fungi_unclassified(100);k__Fungi_unclassified(100); | - | - | - | - | - | - | - | Unassigned | - |
| Otu19763 | 1 | k__Fungi(100);p__Ascomycota(100);p__Ascomycota_unclassified(100);p__Ascomycota_unclassified(100);p__Ascomycota_unclassified(100);p__Ascomycota_unclassified(100);p__Ascomycota_unclassified(100); | - | - | - | - | - | - | - | Unassigned | - |
| Otu19765 | 1 | k__Fungi(100);k__Fungi_unclassified(100);k__Fungi_unclassified(100);k__Fungi_unclassified(100);k__Fungi_unclassified(100);k__Fungi_unclassified(100);k__Fungi_unclassified(100); | - | - | - | - | - | - | - | Unassigned | - |
| Otu19789 | 1 | k__Fungi(100);p__Ascomycota(100);p__Ascomycota_unclassified(100);p__Ascomycota_unclassified(100);p__Ascomycota_unclassified(100);p__Ascomycota_unclassified(100);p__Ascomycota_unclassified(100); | - | - | - | - | - | - | - | Unassigned | - |
| Otu19817 | 1 | k__Fungi(100);k__Fungi_unclassified(100);k__Fungi_unclassified(100);k__Fungi_unclassified(100);k__Fungi_unclassified(100);k__Fungi_unclassified(100);k__Fungi_unclassified(100); | - | - | - | - | - | - | - | Unassigned | - |
| Otu19850 | 1 | k__Fungi(100);k__Fungi_unclassified(100);k__Fungi_unclassified(100);k__Fungi_unclassified(100);k__Fungi_unclassified(100);k__Fungi_unclassified(100);k__Fungi_unclassified(100); | - | - | - | - | - | - | - | Unassigned | - |
| Otu19858 | 1 | k__Fungi(100);k__Fungi_unclassified(100);k__Fungi_unclassified(100);k__Fungi_unclassified(100);k__Fungi_unclassified(100);k__Fungi_unclassified(100);k__Fungi_unclassified(100); | - | - | - | - | - | - | - | Unassigned | - |
| Otu19870 | 1 | k__Fungi(100);k__Fungi_unclassified(100);k__Fungi_unclassified(100);k__Fungi_unclassified(100);k__Fungi_unclassified(100);k__Fungi_unclassified(100);k__Fungi_unclassified(100); | - | - | - | - | - | - | - | Unassigned | - |
| Otu19877 | 1 | k__Fungi(100);p__Ascomycota(100);p__Ascomycota_unclassified(100);p__Ascomycota_unclassified(100);p__Ascomycota_unclassified(100);p__Ascomycota_unclassified(100);p__Ascomycota_unclassified(100); | - | - | - | - | - | - | - | Unassigned | - |
| Otu19878 | 1 | k__Fungi(100);p__Ascomycota(100);p__Ascomycota_unclassified(100);p__Ascomycota_unclassified(100);p__Ascomycota_unclassified(100);p__Ascomycota_unclassified(100);p__Ascomycota_unclassified(100); | - | - | - | - | - | - | - | Unassigned | - |
| Otu19898 | 1 | k__Fungi(100);k__Fungi_unclassified(100);k__Fungi_unclassified(100);k__Fungi_unclassified(100);k__Fungi_unclassified(100);k__Fungi_unclassified(100);k__Fungi_unclassified(100); | - | - | - | - | - | - | - | Unassigned | - |
| Otu19905 | 1 | k__Fungi(100);k__Fungi_unclassified(100);k__Fungi_unclassified(100);k__Fungi_unclassified(100);k__Fungi_unclassified(100);k__Fungi_unclassified(100);k__Fungi_unclassified(100); | - | - | - | - | - | - | - | Unassigned | - |
| Otu19906 | 1 | k__Fungi(100);p__Ascomycota(100);p__Ascomycota_unclassified(100);p__Ascomycota_unclassified(100);p__Ascomycota_unclassified(100);p__Ascomycota_unclassified(100);p__Ascomycota_unclassified(100); | - | - | - | - | - | - | - | Unassigned | - |
| Otu19907 | 1 | k__Fungi(100);p__Ascomycota(100);p__Ascomycota_unclassified(100);p__Ascomycota_unclassified(100);p__Ascomycota_unclassified(100);p__Ascomycota_unclassified(100);p__Ascomycota_unclassified(100); | - | - | - | - | - | - | - | Unassigned | - |
| Otu19908 | 1 | k__Fungi(100);k__Fungi_unclassified(100);k__Fungi_unclassified(100);k__Fungi_unclassified(100);k__Fungi_unclassified(100);k__Fungi_unclassified(100);k__Fungi_unclassified(100); | - | - | - | - | - | - | - | Unassigned | - |
| Otu19910 | 1 | k__Fungi(100);p__Ascomycota(100);p__Ascomycota_unclassified(100);p__Ascomycota_unclassified(100);p__Ascomycota_unclassified(100);p__Ascomycota_unclassified(100);p__Ascomycota_unclassified(100); | - | - | - | - | - | - | - | Unassigned | - |
| Otu19911 | 1 | k__Fungi(100);p__Ascomycota(100);p__Ascomycota_unclassified(100);p__Ascomycota_unclassified(100);p__Ascomycota_unclassified(100);p__Ascomycota_unclassified(100);p__Ascomycota_unclassified(100); | - | - | - | - | - | - | - | Unassigned | - |
| Otu19912 | 1 | k__Fungi(100);k__Fungi_unclassified(100);k__Fungi_unclassified(100);k__Fungi_unclassified(100);k__Fungi_unclassified(100);k__Fungi_unclassified(100);k__Fungi_unclassified(100); | - | - | - | - | - | - | - | Unassigned | - |
| Otu19913 | 1 | k__Fungi(100);k__Fungi_unclassified(100);k__Fungi_unclassified(100);k__Fungi_unclassified(100);k__Fungi_unclassified(100);k__Fungi_unclassified(100);k__Fungi_unclassified(100); | - | - | - | - | - | - | - | Unassigned | - |
| Otu19914 | 1 | k__Fungi(100);p__Ascomycota(100);p__Ascomycota_unclassified(100);p__Ascomycota_unclassified(100);p__Ascomycota_unclassified(100);p__Ascomycota_unclassified(100);p__Ascomycota_unclassified(100); | - | - | - | - | - | - | - | Unassigned | - |
| Otu19915 | 1 | k__Fungi(100);k__Fungi_unclassified(100);k__Fungi_unclassified(100);k__Fungi_unclassified(100);k__Fungi_unclassified(100);k__Fungi_unclassified(100);k__Fungi_unclassified(100); | - | - | - | - | - | - | - | Unassigned | - |
| Otu19917 | 1 | k__Fungi(100);p__Ascomycota(100);p__Ascomycota_unclassified(100);p__Ascomycota_unclassified(100);p__Ascomycota_unclassified(100);p__Ascomycota_unclassified(100);p__Ascomycota_unclassified(100); | - | - | - | - | - | - | - | Unassigned | - |
| Otu19918 | 1 | k__Fungi(100);k__Fungi_unclassified(100);k__Fungi_unclassified(100);k__Fungi_unclassified(100);k__Fungi_unclassified(100);k__Fungi_unclassified(100);k__Fungi_unclassified(100); | - | - | - | - | - | - | - | Unassigned | - |
| Otu19919 | 1 | k__Fungi(100);p__Ascomycota(100);p__Ascomycota_unclassified(100);p__Ascomycota_unclassified(100);p__Ascomycota_unclassified(100);p__Ascomycota_unclassified(100);p__Ascomycota_unclassified(100); | - | - | - | - | - | - | - | Unassigned | - |
| Otu19920 | 1 | k__Fungi(100);p__Ascomycota(100);p__Ascomycota_unclassified(100);p__Ascomycota_unclassified(100);p__Ascomycota_unclassified(100);p__Ascomycota_unclassified(100);p__Ascomycota_unclassified(100); | - | - | - | - | - | - | - | Unassigned | - |
| Otu19930 | 1 | k__Fungi(100);p__Ascomycota(100);p__Ascomycota_unclassified(100);p__Ascomycota_unclassified(100);p__Ascomycota_unclassified(100);p__Ascomycota_unclassified(100);p__Ascomycota_unclassified(100); | - | - | - | - | - | - | - | Unassigned | - |
| Otu19932 | 1 | k__Fungi(100);p__Ascomycota(100);p__Ascomycota_unclassified(100);p__Ascomycota_unclassified(100);p__Ascomycota_unclassified(100);p__Ascomycota_unclassified(100);p__Ascomycota_unclassified(100); | - | - | - | - | - | - | - | Unassigned | - |
| Otu19933 | 1 | k__Fungi(100);p__Ascomycota(100);p__Ascomycota_unclassified(100);p__Ascomycota_unclassified(100);p__Ascomycota_unclassified(100);p__Ascomycota_unclassified(100);p__Ascomycota_unclassified(100); | - | - | - | - | - | - | - | Unassigned | - |
| Otu19934 | 1 | k__Fungi(100);p__Ascomycota(100);p__Ascomycota_unclassified(100);p__Ascomycota_unclassified(100);p__Ascomycota_unclassified(100);p__Ascomycota_unclassified(100);p__Ascomycota_unclassified(100); | - | - | - | - | - | - | - | Unassigned | - |
| Otu19935 | 1 | k__Fungi(100);k__Fungi_unclassified(100);k__Fungi_unclassified(100);k__Fungi_unclassified(100);k__Fungi_unclassified(100);k__Fungi_unclassified(100);k__Fungi_unclassified(100); | - | - | - | - | - | - | - | Unassigned | - |
| Otu19936 | 1 | k__Fungi(100);p__Ascomycota(100);p__Ascomycota_unclassified(100);p__Ascomycota_unclassified(100);p__Ascomycota_unclassified(100);p__Ascomycota_unclassified(100);p__Ascomycota_unclassified(100); | - | - | - | - | - | - | - | Unassigned | - |
| Otu19938 | 1 | k__Fungi(100);k__Fungi_unclassified(100);k__Fungi_unclassified(100);k__Fungi_unclassified(100);k__Fungi_unclassified(100);k__Fungi_unclassified(100);k__Fungi_unclassified(100); | - | - | - | - | - | - | - | Unassigned | - |
| Otu19941 | 1 | k__Fungi(100);k__Fungi_unclassified(100);k__Fungi_unclassified(100);k__Fungi_unclassified(100);k__Fungi_unclassified(100);k__Fungi_unclassified(100);k__Fungi_unclassified(100); | - | - | - | - | - | - | - | Unassigned | - |
| Otu19943 | 1 | k__Fungi(100);p__Ascomycota(100);p__Ascomycota_unclassified(100);p__Ascomycota_unclassified(100);p__Ascomycota_unclassified(100);p__Ascomycota_unclassified(100);p__Ascomycota_unclassified(100); | - | - | - | - | - | - | - | Unassigned | - |
| Otu19944 | 1 | k__Fungi(100);p__Ascomycota(100);p__Ascomycota_unclassified(100);p__Ascomycota_unclassified(100);p__Ascomycota_unclassified(100);p__Ascomycota_unclassified(100);p__Ascomycota_unclassified(100); | - | - | - | - | - | - | - | Unassigned | - |
| Otu19945 | 1 | k__Fungi(100);p__Ascomycota(100);p__Ascomycota_unclassified(100);p__Ascomycota_unclassified(100);p__Ascomycota_unclassified(100);p__Ascomycota_unclassified(100);p__Ascomycota_unclassified(100); | - | - | - | - | - | - | - | Unassigned | - |
| Otu19946 | 1 | k__Fungi(100);p__Ascomycota(100);p__Ascomycota_unclassified(100);p__Ascomycota_unclassified(100);p__Ascomycota_unclassified(100);p__Ascomycota_unclassified(100);p__Ascomycota_unclassified(100); | - | - | - | - | - | - | - | Unassigned | - |
| Otu19948 | 1 | k__Fungi(100);k__Fungi_unclassified(100);k__Fungi_unclassified(100);k__Fungi_unclassified(100);k__Fungi_unclassified(100);k__Fungi_unclassified(100);k__Fungi_unclassified(100); | - | - | - | - | - | - | - | Unassigned | - |
| Otu19949 | 1 | k__Fungi(100);k__Fungi_unclassified(100);k__Fungi_unclassified(100);k__Fungi_unclassified(100);k__Fungi_unclassified(100);k__Fungi_unclassified(100);k__Fungi_unclassified(100); | - | - | - | - | - | - | - | Unassigned | - |
| Otu19951 | 1 | k__Fungi(100);k__Fungi_unclassified(100);k__Fungi_unclassified(100);k__Fungi_unclassified(100);k__Fungi_unclassified(100);k__Fungi_unclassified(100);k__Fungi_unclassified(100); | - | - | - | - | - | - | - | Unassigned | - |
| Otu19952 | 1 | k__Fungi(100);k__Fungi_unclassified(100);k__Fungi_unclassified(100);k__Fungi_unclassified(100);k__Fungi_unclassified(100);k__Fungi_unclassified(100);k__Fungi_unclassified(100); | - | - | - | - | - | - | - | Unassigned | - |
| Otu19953 | 1 | k__Fungi(100);p__Ascomycota(100);p__Ascomycota_unclassified(100);p__Ascomycota_unclassified(100);p__Ascomycota_unclassified(100);p__Ascomycota_unclassified(100);p__Ascomycota_unclassified(100); | - | - | - | - | - | - | - | Unassigned | - |
| Otu19954 | 1 | k__Fungi(100);p__Ascomycota(100);p__Ascomycota_unclassified(100);p__Ascomycota_unclassified(100);p__Ascomycota_unclassified(100);p__Ascomycota_unclassified(100);p__Ascomycota_unclassified(100); | - | - | - | - | - | - | - | Unassigned | - |
| Otu19955 | 1 | k__Fungi(100);p__Ascomycota(100);p__Ascomycota_unclassified(100);p__Ascomycota_unclassified(100);p__Ascomycota_unclassified(100);p__Ascomycota_unclassified(100);p__Ascomycota_unclassified(100); | - | - | - | - | - | - | - | Unassigned | - |
| Otu19956 | 1 | k__Fungi(100);k__Fungi_unclassified(100);k__Fungi_unclassified(100);k__Fungi_unclassified(100);k__Fungi_unclassified(100);k__Fungi_unclassified(100);k__Fungi_unclassified(100); | - | - | - | - | - | - | - | Unassigned | - |
| Otu19958 | 1 | k__Fungi(100);p__Ascomycota(100);p__Ascomycota_unclassified(100);p__Ascomycota_unclassified(100);p__Ascomycota_unclassified(100);p__Ascomycota_unclassified(100);p__Ascomycota_unclassified(100); | - | - | - | - | - | - | - | Unassigned | - |
| Otu19959 | 1 | k__Fungi(100);p__Ascomycota(100);p__Ascomycota_unclassified(100);p__Ascomycota_unclassified(100);p__Ascomycota_unclassified(100);p__Ascomycota_unclassified(100);p__Ascomycota_unclassified(100); | - | - | - | - | - | - | - | Unassigned | - |
| Otu19960 | 1 | k__Fungi(100);p__Ascomycota(100);p__Ascomycota_unclassified(100);p__Ascomycota_unclassified(100);p__Ascomycota_unclassified(100);p__Ascomycota_unclassified(100);p__Ascomycota_unclassified(100); | - | - | - | - | - | - | - | Unassigned | - |
| Otu19961 | 1 | k__Fungi(100);p__Ascomycota(100);p__Ascomycota_unclassified(100);p__Ascomycota_unclassified(100);p__Ascomycota_unclassified(100);p__Ascomycota_unclassified(100);p__Ascomycota_unclassified(100); | - | - | - | - | - | - | - | Unassigned | - |
| Otu19962 | 1 | k__Fungi(100);p__Ascomycota(100);p__Ascomycota_unclassified(100);p__Ascomycota_unclassified(100);p__Ascomycota_unclassified(100);p__Ascomycota_unclassified(100);p__Ascomycota_unclassified(100); | - | - | - | - | - | - | - | Unassigned | - |
| Otu19963 | 1 | k__Fungi(100);p__Ascomycota(100);p__Ascomycota_unclassified(100);p__Ascomycota_unclassified(100);p__Ascomycota_unclassified(100);p__Ascomycota_unclassified(100);p__Ascomycota_unclassified(100); | - | - | - | - | - | - | - | Unassigned | - |
| Otu19964 | 1 | k__Fungi(100);k__Fungi_unclassified(100);k__Fungi_unclassified(100);k__Fungi_unclassified(100);k__Fungi_unclassified(100);k__Fungi_unclassified(100);k__Fungi_unclassified(100); | - | - | - | - | - | - | - | Unassigned | - |
| Otu19968 | 1 | k__Fungi(100);k__Fungi_unclassified(100);k__Fungi_unclassified(100);k__Fungi_unclassified(100);k__Fungi_unclassified(100);k__Fungi_unclassified(100);k__Fungi_unclassified(100); | - | - | - | - | - | - | - | Unassigned | - |
| Otu19969 | 1 | k__Fungi(100);p__Ascomycota(100);p__Ascomycota_unclassified(100);p__Ascomycota_unclassified(100);p__Ascomycota_unclassified(100);p__Ascomycota_unclassified(100);p__Ascomycota_unclassified(100); | - | - | - | - | - | - | - | Unassigned | - |
| Otu19970 | 1 | k__Fungi(100);k__Fungi_unclassified(100);k__Fungi_unclassified(100);k__Fungi_unclassified(100);k__Fungi_unclassified(100);k__Fungi_unclassified(100);k__Fungi_unclassified(100); | - | - | - | - | - | - | - | Unassigned | - |
| Otu19971 | 1 | k__Fungi(100);p__Ascomycota(100);p__Ascomycota_unclassified(100);p__Ascomycota_unclassified(100);p__Ascomycota_unclassified(100);p__Ascomycota_unclassified(100);p__Ascomycota_unclassified(100); | - | - | - | - | - | - | - | Unassigned | - |
| Otu19974 | 1 | k__Fungi(100);k__Fungi_unclassified(100);k__Fungi_unclassified(100);k__Fungi_unclassified(100);k__Fungi_unclassified(100);k__Fungi_unclassified(100);k__Fungi_unclassified(100); | - | - | - | - | - | - | - | Unassigned | - |
| Otu19977 | 1 | k__Fungi(100);k__Fungi_unclassified(100);k__Fungi_unclassified(100);k__Fungi_unclassified(100);k__Fungi_unclassified(100);k__Fungi_unclassified(100);k__Fungi_unclassified(100); | - | - | - | - | - | - | - | Unassigned | - |
| Otu19981 | 1 | k__Fungi(100);k__Fungi_unclassified(100);k__Fungi_unclassified(100);k__Fungi_unclassified(100);k__Fungi_unclassified(100);k__Fungi_unclassified(100);k__Fungi_unclassified(100); | - | - | - | - | - | - | - | Unassigned | - |
| Otu19986 | 1 | k__Fungi(100);p__Ascomycota(100);p__Ascomycota_unclassified(100);p__Ascomycota_unclassified(100);p__Ascomycota_unclassified(100);p__Ascomycota_unclassified(100);p__Ascomycota_unclassified(100); | - | - | - | - | - | - | - | Unassigned | - |
| Otu19995 | 1 | k__Fungi(100);k__Fungi_unclassified(100);k__Fungi_unclassified(100);k__Fungi_unclassified(100);k__Fungi_unclassified(100);k__Fungi_unclassified(100);k__Fungi_unclassified(100); | - | - | - | - | - | - | - | Unassigned | - |
| Otu19996 | 1 | k__Fungi(100);p__Ascomycota(100);p__Ascomycota_unclassified(100);p__Ascomycota_unclassified(100);p__Ascomycota_unclassified(100);p__Ascomycota_unclassified(100);p__Ascomycota_unclassified(100); | - | - | - | - | - | - | - | Unassigned | - |
| Otu19998 | 1 | k__Fungi(100);p__Ascomycota(100);p__Ascomycota_unclassified(100);p__Ascomycota_unclassified(100);p__Ascomycota_unclassified(100);p__Ascomycota_unclassified(100);p__Ascomycota_unclassified(100); | - | - | - | - | - | - | - | Unassigned | - |
| Otu19999 | 1 | k__Fungi(100);k__Fungi_unclassified(100);k__Fungi_unclassified(100);k__Fungi_unclassified(100);k__Fungi_unclassified(100);k__Fungi_unclassified(100);k__Fungi_unclassified(100); | - | - | - | - | - | - | - | Unassigned | - |
| Otu20005 | 1 | k__Fungi(100);p__Ascomycota(100);p__Ascomycota_unclassified(100);p__Ascomycota_unclassified(100);p__Ascomycota_unclassified(100);p__Ascomycota_unclassified(100);p__Ascomycota_unclassified(100); | - | - | - | - | - | - | - | Unassigned | - |
| Otu20006 | 1 | k__Fungi(100);p__Ascomycota(100);p__Ascomycota_unclassified(100);p__Ascomycota_unclassified(100);p__Ascomycota_unclassified(100);p__Ascomycota_unclassified(100);p__Ascomycota_unclassified(100); | - | - | - | - | - | - | - | Unassigned | - |
| Otu20007 | 1 | k__Fungi(100);p__Ascomycota(100);p__Ascomycota_unclassified(100);p__Ascomycota_unclassified(100);p__Ascomycota_unclassified(100);p__Ascomycota_unclassified(100);p__Ascomycota_unclassified(100); | - | - | - | - | - | - | - | Unassigned | - |
| Otu20013 | 1 | k__Fungi(100);k__Fungi_unclassified(100);k__Fungi_unclassified(100);k__Fungi_unclassified(100);k__Fungi_unclassified(100);k__Fungi_unclassified(100);k__Fungi_unclassified(100); | - | - | - | - | - | - | - | Unassigned | - |
| Otu20016 | 1 | k__Fungi(100);k__Fungi_unclassified(100);k__Fungi_unclassified(100);k__Fungi_unclassified(100);k__Fungi_unclassified(100);k__Fungi_unclassified(100);k__Fungi_unclassified(100); | - | - | - | - | - | - | - | Unassigned | - |
| Otu20017 | 1 | k__Fungi(100);k__Fungi_unclassified(100);k__Fungi_unclassified(100);k__Fungi_unclassified(100);k__Fungi_unclassified(100);k__Fungi_unclassified(100);k__Fungi_unclassified(100); | - | - | - | - | - | - | - | Unassigned | - |
| Otu20018 | 1 | k__Fungi(100);p__Ascomycota(100);p__Ascomycota_unclassified(100);p__Ascomycota_unclassified(100);p__Ascomycota_unclassified(100);p__Ascomycota_unclassified(100);p__Ascomycota_unclassified(100); | - | - | - | - | - | - | - | Unassigned | - |
| Otu20019 | 1 | k__Fungi(100);k__Fungi_unclassified(100);k__Fungi_unclassified(100);k__Fungi_unclassified(100);k__Fungi_unclassified(100);k__Fungi_unclassified(100);k__Fungi_unclassified(100); | - | - | - | - | - | - | - | Unassigned | - |
| Otu20020 | 1 | k__Fungi(100);p__Ascomycota(100);p__Ascomycota_unclassified(100);p__Ascomycota_unclassified(100);p__Ascomycota_unclassified(100);p__Ascomycota_unclassified(100);p__Ascomycota_unclassified(100); | - | - | - | - | - | - | - | Unassigned | - |
| Otu20021 | 1 | k__Fungi(100);p__Ascomycota(100);p__Ascomycota_unclassified(100);p__Ascomycota_unclassified(100);p__Ascomycota_unclassified(100);p__Ascomycota_unclassified(100);p__Ascomycota_unclassified(100); | - | - | - | - | - | - | - | Unassigned | - |
| Otu20022 | 1 | k__Fungi(100);k__Fungi_unclassified(100);k__Fungi_unclassified(100);k__Fungi_unclassified(100);k__Fungi_unclassified(100);k__Fungi_unclassified(100);k__Fungi_unclassified(100); | - | - | - | - | - | - | - | Unassigned | - |
| Otu20023 | 1 | k__Fungi(100);p__Ascomycota(100);p__Ascomycota_unclassified(100);p__Ascomycota_unclassified(100);p__Ascomycota_unclassified(100);p__Ascomycota_unclassified(100);p__Ascomycota_unclassified(100); | - | - | - | - | - | - | - | Unassigned | - |
| Otu20027 | 1 | k__Fungi(100);k__Fungi_unclassified(100);k__Fungi_unclassified(100);k__Fungi_unclassified(100);k__Fungi_unclassified(100);k__Fungi_unclassified(100);k__Fungi_unclassified(100); | - | - | - | - | - | - | - | Unassigned | - |
| Otu20028 | 1 | k__Fungi(100);k__Fungi_unclassified(100);k__Fungi_unclassified(100);k__Fungi_unclassified(100);k__Fungi_unclassified(100);k__Fungi_unclassified(100);k__Fungi_unclassified(100); | - | - | - | - | - | - | - | Unassigned | - |
| Otu20029 | 1 | k__Fungi(100);k__Fungi_unclassified(100);k__Fungi_unclassified(100);k__Fungi_unclassified(100);k__Fungi_unclassified(100);k__Fungi_unclassified(100);k__Fungi_unclassified(100); | - | - | - | - | - | - | - | Unassigned | - |
| Otu20030 | 1 | k__Fungi(100);k__Fungi_unclassified(100);k__Fungi_unclassified(100);k__Fungi_unclassified(100);k__Fungi_unclassified(100);k__Fungi_unclassified(100);k__Fungi_unclassified(100); | - | - | - | - | - | - | - | Unassigned | - |
| Otu20031 | 1 | k__Fungi(100);p__Ascomycota(100);p__Ascomycota_unclassified(100);p__Ascomycota_unclassified(100);p__Ascomycota_unclassified(100);p__Ascomycota_unclassified(100);p__Ascomycota_unclassified(100); | - | - | - | - | - | - | - | Unassigned | - |
| Otu20033 | 1 | k__Fungi(100);k__Fungi_unclassified(100);k__Fungi_unclassified(100);k__Fungi_unclassified(100);k__Fungi_unclassified(100);k__Fungi_unclassified(100);k__Fungi_unclassified(100); | - | - | - | - | - | - | - | Unassigned | - |
| Otu20044 | 1 | k__Fungi(100);p__Ascomycota(100);p__Ascomycota_unclassified(100);p__Ascomycota_unclassified(100);p__Ascomycota_unclassified(100);p__Ascomycota_unclassified(100);p__Ascomycota_unclassified(100); | - | - | - | - | - | - | - | Unassigned | - |
| Otu20045 | 1 | k__Fungi(100);k__Fungi_unclassified(100);k__Fungi_unclassified(100);k__Fungi_unclassified(100);k__Fungi_unclassified(100);k__Fungi_unclassified(100);k__Fungi_unclassified(100); | - | - | - | - | - | - | - | Unassigned | - |
| Otu20053 | 1 | k__Fungi(100);k__Fungi_unclassified(100);k__Fungi_unclassified(100);k__Fungi_unclassified(100);k__Fungi_unclassified(100);k__Fungi_unclassified(100);k__Fungi_unclassified(100); | - | - | - | - | - | - | - | Unassigned | - |
| Otu20060 | 1 | k__Fungi(100);k__Fungi_unclassified(100);k__Fungi_unclassified(100);k__Fungi_unclassified(100);k__Fungi_unclassified(100);k__Fungi_unclassified(100);k__Fungi_unclassified(100); | - | - | - | - | - | - | - | Unassigned | - |
| Otu20061 | 1 | k__Fungi(100);k__Fungi_unclassified(100);k__Fungi_unclassified(100);k__Fungi_unclassified(100);k__Fungi_unclassified(100);k__Fungi_unclassified(100);k__Fungi_unclassified(100); | - | - | - | - | - | - | - | Unassigned | - |
| Otu20063 | 1 | k__Fungi(100);p__Ascomycota(100);p__Ascomycota_unclassified(100);p__Ascomycota_unclassified(100);p__Ascomycota_unclassified(100);p__Ascomycota_unclassified(100);p__Ascomycota_unclassified(100); | - | - | - | - | - | - | - | Unassigned | - |
| Otu20064 | 1 | k__Fungi(100);k__Fungi_unclassified(100);k__Fungi_unclassified(100);k__Fungi_unclassified(100);k__Fungi_unclassified(100);k__Fungi_unclassified(100);k__Fungi_unclassified(100); | - | - | - | - | - | - | - | Unassigned | - |
| Otu20065 | 1 | k__Fungi(100);p__Ascomycota(100);p__Ascomycota_unclassified(100);p__Ascomycota_unclassified(100);p__Ascomycota_unclassified(100);p__Ascomycota_unclassified(100);p__Ascomycota_unclassified(100); | - | - | - | - | - | - | - | Unassigned | - |
| Otu20066 | 1 | k__Fungi(100);k__Fungi_unclassified(100);k__Fungi_unclassified(100);k__Fungi_unclassified(100);k__Fungi_unclassified(100);k__Fungi_unclassified(100);k__Fungi_unclassified(100); | - | - | - | - | - | - | - | Unassigned | - |
| Otu20067 | 1 | k__Fungi(100);k__Fungi_unclassified(100);k__Fungi_unclassified(100);k__Fungi_unclassified(100);k__Fungi_unclassified(100);k__Fungi_unclassified(100);k__Fungi_unclassified(100); | - | - | - | - | - | - | - | Unassigned | - |
| Otu20074 | 1 | k__Fungi(100);k__Fungi_unclassified(100);k__Fungi_unclassified(100);k__Fungi_unclassified(100);k__Fungi_unclassified(100);k__Fungi_unclassified(100);k__Fungi_unclassified(100); | - | - | - | - | - | - | - | Unassigned | - |
| Otu20075 | 1 | k__Fungi(100);k__Fungi_unclassified(100);k__Fungi_unclassified(100);k__Fungi_unclassified(100);k__Fungi_unclassified(100);k__Fungi_unclassified(100);k__Fungi_unclassified(100); | - | - | - | - | - | - | - | Unassigned | - |
| Otu20076 | 1 | k__Fungi(100);p__Ascomycota(100);p__Ascomycota_unclassified(100);p__Ascomycota_unclassified(100);p__Ascomycota_unclassified(100);p__Ascomycota_unclassified(100);p__Ascomycota_unclassified(100); | - | - | - | - | - | - | - | Unassigned | - |
| Otu20078 | 1 | k__Fungi(100);k__Fungi_unclassified(100);k__Fungi_unclassified(100);k__Fungi_unclassified(100);k__Fungi_unclassified(100);k__Fungi_unclassified(100);k__Fungi_unclassified(100); | - | - | - | - | - | - | - | Unassigned | - |
| Otu20079 | 1 | k__Fungi(100);k__Fungi_unclassified(100);k__Fungi_unclassified(100);k__Fungi_unclassified(100);k__Fungi_unclassified(100);k__Fungi_unclassified(100);k__Fungi_unclassified(100); | - | - | - | - | - | - | - | Unassigned | - |
| Otu20080 | 1 | k__Fungi(100);k__Fungi_unclassified(100);k__Fungi_unclassified(100);k__Fungi_unclassified(100);k__Fungi_unclassified(100);k__Fungi_unclassified(100);k__Fungi_unclassified(100); | - | - | - | - | - | - | - | Unassigned | - |
| Otu20086 | 1 | k__Fungi(100);k__Fungi_unclassified(100);k__Fungi_unclassified(100);k__Fungi_unclassified(100);k__Fungi_unclassified(100);k__Fungi_unclassified(100);k__Fungi_unclassified(100); | - | - | - | - | - | - | - | Unassigned | - |
| Otu20087 | 1 | k__Fungi(100);k__Fungi_unclassified(100);k__Fungi_unclassified(100);k__Fungi_unclassified(100);k__Fungi_unclassified(100);k__Fungi_unclassified(100);k__Fungi_unclassified(100); | - | - | - | - | - | - | - | Unassigned | - |
| Otu20090 | 1 | k__Fungi(100);k__Fungi_unclassified(100);k__Fungi_unclassified(100);k__Fungi_unclassified(100);k__Fungi_unclassified(100);k__Fungi_unclassified(100);k__Fungi_unclassified(100); | - | - | - | - | - | - | - | Unassigned | - |
| Otu20092 | 1 | k__Fungi(100);p__Ascomycota(100);p__Ascomycota_unclassified(100);p__Ascomycota_unclassified(100);p__Ascomycota_unclassified(100);p__Ascomycota_unclassified(100);p__Ascomycota_unclassified(100); | - | - | - | - | - | - | - | Unassigned | - |
| Otu20094 | 1 | k__Fungi(100);k__Fungi_unclassified(100);k__Fungi_unclassified(100);k__Fungi_unclassified(100);k__Fungi_unclassified(100);k__Fungi_unclassified(100);k__Fungi_unclassified(100); | - | - | - | - | - | - | - | Unassigned | - |
| Otu20097 | 1 | k__Fungi(100);k__Fungi_unclassified(100);k__Fungi_unclassified(100);k__Fungi_unclassified(100);k__Fungi_unclassified(100);k__Fungi_unclassified(100);k__Fungi_unclassified(100); | - | - | - | - | - | - | - | Unassigned | - |
| Otu20099 | 1 | k__Fungi(100);k__Fungi_unclassified(100);k__Fungi_unclassified(100);k__Fungi_unclassified(100);k__Fungi_unclassified(100);k__Fungi_unclassified(100);k__Fungi_unclassified(100); | - | - | - | - | - | - | - | Unassigned | - |
| Otu20100 | 1 | k__Fungi(100);k__Fungi_unclassified(100);k__Fungi_unclassified(100);k__Fungi_unclassified(100);k__Fungi_unclassified(100);k__Fungi_unclassified(100);k__Fungi_unclassified(100); | - | - | - | - | - | - | - | Unassigned | - |
| Otu20102 | 1 | k__Fungi(100);k__Fungi_unclassified(100);k__Fungi_unclassified(100);k__Fungi_unclassified(100);k__Fungi_unclassified(100);k__Fungi_unclassified(100);k__Fungi_unclassified(100); | - | - | - | - | - | - | - | Unassigned | - |
| Otu20104 | 1 | k__Fungi(100);p__Ascomycota(100);p__Ascomycota_unclassified(100);p__Ascomycota_unclassified(100);p__Ascomycota_unclassified(100);p__Ascomycota_unclassified(100);p__Ascomycota_unclassified(100); | - | - | - | - | - | - | - | Unassigned | - |
| Otu20106 | 1 | k__Fungi(100);p__Ascomycota(100);p__Ascomycota_unclassified(100);p__Ascomycota_unclassified(100);p__Ascomycota_unclassified(100);p__Ascomycota_unclassified(100);p__Ascomycota_unclassified(100); | - | - | - | - | - | - | - | Unassigned | - |
| Otu20110 | 1 | k__Fungi(100);k__Fungi_unclassified(100);k__Fungi_unclassified(100);k__Fungi_unclassified(100);k__Fungi_unclassified(100);k__Fungi_unclassified(100);k__Fungi_unclassified(100); | - | - | - | - | - | - | - | Unassigned | - |
| Otu20112 | 1 | k__Fungi(100);p__Ascomycota(100);p__Ascomycota_unclassified(100);p__Ascomycota_unclassified(100);p__Ascomycota_unclassified(100);p__Ascomycota_unclassified(100);p__Ascomycota_unclassified(100); | - | - | - | - | - | - | - | Unassigned | - |
| Otu20113 | 1 | k__Fungi(100);p__Ascomycota(100);p__Ascomycota_unclassified(100);p__Ascomycota_unclassified(100);p__Ascomycota_unclassified(100);p__Ascomycota_unclassified(100);p__Ascomycota_unclassified(100); | - | - | - | - | - | - | - | Unassigned | - |
| Otu20114 | 1 | k__Fungi(100);k__Fungi_unclassified(100);k__Fungi_unclassified(100);k__Fungi_unclassified(100);k__Fungi_unclassified(100);k__Fungi_unclassified(100);k__Fungi_unclassified(100); | - | - | - | - | - | - | - | Unassigned | - |
| Otu20115 | 1 | k__Fungi(100);k__Fungi_unclassified(100);k__Fungi_unclassified(100);k__Fungi_unclassified(100);k__Fungi_unclassified(100);k__Fungi_unclassified(100);k__Fungi_unclassified(100); | - | - | - | - | - | - | - | Unassigned | - |
| Otu20118 | 1 | k__Fungi(100);p__Ascomycota(100);p__Ascomycota_unclassified(100);p__Ascomycota_unclassified(100);p__Ascomycota_unclassified(100);p__Ascomycota_unclassified(100);p__Ascomycota_unclassified(100); | - | - | - | - | - | - | - | Unassigned | - |
| Otu20120 | 1 | k__Fungi(100);k__Fungi_unclassified(100);k__Fungi_unclassified(100);k__Fungi_unclassified(100);k__Fungi_unclassified(100);k__Fungi_unclassified(100);k__Fungi_unclassified(100); | - | - | - | - | - | - | - | Unassigned | - |
| Otu20121 | 1 | k__Fungi(100);p__Ascomycota(100);p__Ascomycota_unclassified(100);p__Ascomycota_unclassified(100);p__Ascomycota_unclassified(100);p__Ascomycota_unclassified(100);p__Ascomycota_unclassified(100); | - | - | - | - | - | - | - | Unassigned | - |
| Otu20122 | 1 | k__Fungi(100);k__Fungi_unclassified(100);k__Fungi_unclassified(100);k__Fungi_unclassified(100);k__Fungi_unclassified(100);k__Fungi_unclassified(100);k__Fungi_unclassified(100); | - | - | - | - | - | - | - | Unassigned | - |
| Otu20123 | 1 | k__Fungi(100);p__Ascomycota(100);p__Ascomycota_unclassified(100);p__Ascomycota_unclassified(100);p__Ascomycota_unclassified(100);p__Ascomycota_unclassified(100);p__Ascomycota_unclassified(100); | - | - | - | - | - | - | - | Unassigned | - |
| Otu20125 | 1 | k__Fungi(100);k__Fungi_unclassified(100);k__Fungi_unclassified(100);k__Fungi_unclassified(100);k__Fungi_unclassified(100);k__Fungi_unclassified(100);k__Fungi_unclassified(100); | - | - | - | - | - | - | - | Unassigned | - |
| Otu20126 | 1 | k__Fungi(100);p__Ascomycota(100);p__Ascomycota_unclassified(100);p__Ascomycota_unclassified(100);p__Ascomycota_unclassified(100);p__Ascomycota_unclassified(100);p__Ascomycota_unclassified(100); | - | - | - | - | - | - | - | Unassigned | - |
| Otu20127 | 1 | k__Fungi(100);p__Ascomycota(100);p__Ascomycota_unclassified(100);p__Ascomycota_unclassified(100);p__Ascomycota_unclassified(100);p__Ascomycota_unclassified(100);p__Ascomycota_unclassified(100); | - | - | - | - | - | - | - | Unassigned | - |
| Otu20128 | 1 | k__Fungi(100);k__Fungi_unclassified(100);k__Fungi_unclassified(100);k__Fungi_unclassified(100);k__Fungi_unclassified(100);k__Fungi_unclassified(100);k__Fungi_unclassified(100); | - | - | - | - | - | - | - | Unassigned | - |
| Otu20132 | 1 | k__Fungi(100);k__Fungi_unclassified(100);k__Fungi_unclassified(100);k__Fungi_unclassified(100);k__Fungi_unclassified(100);k__Fungi_unclassified(100);k__Fungi_unclassified(100); | - | - | - | - | - | - | - | Unassigned | - |
| Otu20133 | 1 | k__Fungi(100);k__Fungi_unclassified(100);k__Fungi_unclassified(100);k__Fungi_unclassified(100);k__Fungi_unclassified(100);k__Fungi_unclassified(100);k__Fungi_unclassified(100); | - | - | - | - | - | - | - | Unassigned | - |
| Otu20135 | 1 | k__Fungi(100);k__Fungi_unclassified(100);k__Fungi_unclassified(100);k__Fungi_unclassified(100);k__Fungi_unclassified(100);k__Fungi_unclassified(100);k__Fungi_unclassified(100); | - | - | - | - | - | - | - | Unassigned | - |
| Otu20137 | 1 | k__Fungi(100);k__Fungi_unclassified(100);k__Fungi_unclassified(100);k__Fungi_unclassified(100);k__Fungi_unclassified(100);k__Fungi_unclassified(100);k__Fungi_unclassified(100); | - | - | - | - | - | - | - | Unassigned | - |
| Otu20141 | 1 | k__Fungi(100);k__Fungi_unclassified(100);k__Fungi_unclassified(100);k__Fungi_unclassified(100);k__Fungi_unclassified(100);k__Fungi_unclassified(100);k__Fungi_unclassified(100); | - | - | - | - | - | - | - | Unassigned | - |
| Otu20142 | 1 | k__Fungi(100);k__Fungi_unclassified(100);k__Fungi_unclassified(100);k__Fungi_unclassified(100);k__Fungi_unclassified(100);k__Fungi_unclassified(100);k__Fungi_unclassified(100); | - | - | - | - | - | - | - | Unassigned | - |
| Otu20143 | 1 | k__Fungi(100);p__Ascomycota(100);p__Ascomycota_unclassified(100);p__Ascomycota_unclassified(100);p__Ascomycota_unclassified(100);p__Ascomycota_unclassified(100);p__Ascomycota_unclassified(100); | - | - | - | - | - | - | - | Unassigned | - |
| Otu20146 | 1 | k__Fungi(100);k__Fungi_unclassified(100);k__Fungi_unclassified(100);k__Fungi_unclassified(100);k__Fungi_unclassified(100);k__Fungi_unclassified(100);k__Fungi_unclassified(100); | - | - | - | - | - | - | - | Unassigned | - |
| Otu20147 | 1 | k__Fungi(100);k__Fungi_unclassified(100);k__Fungi_unclassified(100);k__Fungi_unclassified(100);k__Fungi_unclassified(100);k__Fungi_unclassified(100);k__Fungi_unclassified(100); | - | - | - | - | - | - | - | Unassigned | - |
| Otu20148 | 1 | k__Fungi(100);p__Ascomycota(100);p__Ascomycota_unclassified(100);p__Ascomycota_unclassified(100);p__Ascomycota_unclassified(100);p__Ascomycota_unclassified(100);p__Ascomycota_unclassified(100); | - | - | - | - | - | - | - | Unassigned | - |
| Otu20149 | 1 | k__Fungi(100);p__Ascomycota(100);p__Ascomycota_unclassified(100);p__Ascomycota_unclassified(100);p__Ascomycota_unclassified(100);p__Ascomycota_unclassified(100);p__Ascomycota_unclassified(100); | - | - | - | - | - | - | - | Unassigned | - |
| Otu20151 | 1 | k__Fungi(100);p__Ascomycota(100);p__Ascomycota_unclassified(100);p__Ascomycota_unclassified(100);p__Ascomycota_unclassified(100);p__Ascomycota_unclassified(100);p__Ascomycota_unclassified(100); | - | - | - | - | - | - | - | Unassigned | - |
| Otu20152 | 1 | k__Fungi(100);k__Fungi_unclassified(100);k__Fungi_unclassified(100);k__Fungi_unclassified(100);k__Fungi_unclassified(100);k__Fungi_unclassified(100);k__Fungi_unclassified(100); | - | - | - | - | - | - | - | Unassigned | - |
| Otu20153 | 1 | k__Fungi(100);k__Fungi_unclassified(100);k__Fungi_unclassified(100);k__Fungi_unclassified(100);k__Fungi_unclassified(100);k__Fungi_unclassified(100);k__Fungi_unclassified(100); | - | - | - | - | - | - | - | Unassigned | - |
| Otu20154 | 1 | k__Fungi(100);p__Ascomycota(100);p__Ascomycota_unclassified(100);p__Ascomycota_unclassified(100);p__Ascomycota_unclassified(100);p__Ascomycota_unclassified(100);p__Ascomycota_unclassified(100); | - | - | - | - | - | - | - | Unassigned | - |
| Otu20156 | 1 | k__Fungi(100);p__Ascomycota(100);p__Ascomycota_unclassified(100);p__Ascomycota_unclassified(100);p__Ascomycota_unclassified(100);p__Ascomycota_unclassified(100);p__Ascomycota_unclassified(100); | - | - | - | - | - | - | - | Unassigned | - |
| Otu20157 | 1 | k__Fungi(100);k__Fungi_unclassified(100);k__Fungi_unclassified(100);k__Fungi_unclassified(100);k__Fungi_unclassified(100);k__Fungi_unclassified(100);k__Fungi_unclassified(100); | - | - | - | - | - | - | - | Unassigned | - |
| Otu20158 | 1 | k__Fungi(100);k__Fungi_unclassified(100);k__Fungi_unclassified(100);k__Fungi_unclassified(100);k__Fungi_unclassified(100);k__Fungi_unclassified(100);k__Fungi_unclassified(100); | - | - | - | - | - | - | - | Unassigned | - |
| Otu20160 | 1 | k__Fungi(100);k__Fungi_unclassified(100);k__Fungi_unclassified(100);k__Fungi_unclassified(100);k__Fungi_unclassified(100);k__Fungi_unclassified(100);k__Fungi_unclassified(100); | - | - | - | - | - | - | - | Unassigned | - |
| Otu20163 | 1 | k__Fungi(100);k__Fungi_unclassified(100);k__Fungi_unclassified(100);k__Fungi_unclassified(100);k__Fungi_unclassified(100);k__Fungi_unclassified(100);k__Fungi_unclassified(100); | - | - | - | - | - | - | - | Unassigned | - |
| Otu20164 | 1 | k__Fungi(100);k__Fungi_unclassified(100);k__Fungi_unclassified(100);k__Fungi_unclassified(100);k__Fungi_unclassified(100);k__Fungi_unclassified(100);k__Fungi_unclassified(100); | - | - | - | - | - | - | - | Unassigned | - |
| Otu20166 | 1 | k__Fungi(100);p__Ascomycota(100);p__Ascomycota_unclassified(100);p__Ascomycota_unclassified(100);p__Ascomycota_unclassified(100);p__Ascomycota_unclassified(100);p__Ascomycota_unclassified(100); | - | - | - | - | - | - | - | Unassigned | - |
| Otu20171 | 1 | k__Fungi(100);k__Fungi_unclassified(100);k__Fungi_unclassified(100);k__Fungi_unclassified(100);k__Fungi_unclassified(100);k__Fungi_unclassified(100);k__Fungi_unclassified(100); | - | - | - | - | - | - | - | Unassigned | - |
| Otu20172 | 1 | k__Fungi(100);p__Ascomycota(100);p__Ascomycota_unclassified(100);p__Ascomycota_unclassified(100);p__Ascomycota_unclassified(100);p__Ascomycota_unclassified(100);p__Ascomycota_unclassified(100); | - | - | - | - | - | - | - | Unassigned | - |
| Otu20173 | 1 | k__Fungi(100);k__Fungi_unclassified(100);k__Fungi_unclassified(100);k__Fungi_unclassified(100);k__Fungi_unclassified(100);k__Fungi_unclassified(100);k__Fungi_unclassified(100); | - | - | - | - | - | - | - | Unassigned | - |
| Otu20174 | 1 | k__Fungi(100);p__Ascomycota(100);p__Ascomycota_unclassified(100);p__Ascomycota_unclassified(100);p__Ascomycota_unclassified(100);p__Ascomycota_unclassified(100);p__Ascomycota_unclassified(100); | - | - | - | - | - | - | - | Unassigned | - |
| Otu20175 | 1 | k__Fungi(100);p__Ascomycota(100);p__Ascomycota_unclassified(100);p__Ascomycota_unclassified(100);p__Ascomycota_unclassified(100);p__Ascomycota_unclassified(100);p__Ascomycota_unclassified(100); | - | - | - | - | - | - | - | Unassigned | - |
| Otu20178 | 1 | k__Fungi(100);k__Fungi_unclassified(100);k__Fungi_unclassified(100);k__Fungi_unclassified(100);k__Fungi_unclassified(100);k__Fungi_unclassified(100);k__Fungi_unclassified(100); | - | - | - | - | - | - | - | Unassigned | - |
| Otu20179 | 1 | k__Fungi(100);k__Fungi_unclassified(100);k__Fungi_unclassified(100);k__Fungi_unclassified(100);k__Fungi_unclassified(100);k__Fungi_unclassified(100);k__Fungi_unclassified(100); | - | - | - | - | - | - | - | Unassigned | - |
| Otu20185 | 1 | k__Fungi(100);p__Ascomycota(100);p__Ascomycota_unclassified(100);p__Ascomycota_unclassified(100);p__Ascomycota_unclassified(100);p__Ascomycota_unclassified(100);p__Ascomycota_unclassified(100); | - | - | - | - | - | - | - | Unassigned | - |
| Otu20188 | 1 | k__Fungi(100);p__Ascomycota(100);p__Ascomycota_unclassified(100);p__Ascomycota_unclassified(100);p__Ascomycota_unclassified(100);p__Ascomycota_unclassified(100);p__Ascomycota_unclassified(100); | - | - | - | - | - | - | - | Unassigned | - |
| Otu20208 | 1 | k__Fungi(100);k__Fungi_unclassified(100);k__Fungi_unclassified(100);k__Fungi_unclassified(100);k__Fungi_unclassified(100);k__Fungi_unclassified(100);k__Fungi_unclassified(100); | - | - | - | - | - | - | - | Unassigned | - |
| Otu20209 | 1 | k__Fungi(100);k__Fungi_unclassified(100);k__Fungi_unclassified(100);k__Fungi_unclassified(100);k__Fungi_unclassified(100);k__Fungi_unclassified(100);k__Fungi_unclassified(100); | - | - | - | - | - | - | - | Unassigned | - |
| Otu20210 | 1 | k__Fungi(100);k__Fungi_unclassified(100);k__Fungi_unclassified(100);k__Fungi_unclassified(100);k__Fungi_unclassified(100);k__Fungi_unclassified(100);k__Fungi_unclassified(100); | - | - | - | - | - | - | - | Unassigned | - |
| Otu20222 | 1 | k__Fungi(100);k__Fungi_unclassified(100);k__Fungi_unclassified(100);k__Fungi_unclassified(100);k__Fungi_unclassified(100);k__Fungi_unclassified(100);k__Fungi_unclassified(100); | - | - | - | - | - | - | - | Unassigned | - |
| Otu20244 | 1 | k__Fungi(100);k__Fungi_unclassified(100);k__Fungi_unclassified(100);k__Fungi_unclassified(100);k__Fungi_unclassified(100);k__Fungi_unclassified(100);k__Fungi_unclassified(100); | - | - | - | - | - | - | - | Unassigned | - |
| Otu20246 | 1 | k__Fungi(100);p__Ascomycota(100);p__Ascomycota_unclassified(100);p__Ascomycota_unclassified(100);p__Ascomycota_unclassified(100);p__Ascomycota_unclassified(100);p__Ascomycota_unclassified(100); | - | - | - | - | - | - | - | Unassigned | - |
| Otu20247 | 1 | k__Fungi(100);p__Ascomycota(100);p__Ascomycota_unclassified(100);p__Ascomycota_unclassified(100);p__Ascomycota_unclassified(100);p__Ascomycota_unclassified(100);p__Ascomycota_unclassified(100); | - | - | - | - | - | - | - | Unassigned | - |
| Otu20248 | 1 | k__Fungi(100);k__Fungi_unclassified(100);k__Fungi_unclassified(100);k__Fungi_unclassified(100);k__Fungi_unclassified(100);k__Fungi_unclassified(100);k__Fungi_unclassified(100); | - | - | - | - | - | - | - | Unassigned | - |
| Otu20250 | 1 | k__Fungi(100);p__Ascomycota(100);c__Dothideomycetes(100);o__Capnodiales(100);o__Capnodiales_unclassified(100);o__Capnodiales_unclassified(100);o__Capnodiales_unclassified(100); | - | - | - | - | - | - | - | Unassigned | - |
| Otu20253 | 1 | k__Fungi(100);k__Fungi_unclassified(100);k__Fungi_unclassified(100);k__Fungi_unclassified(100);k__Fungi_unclassified(100);k__Fungi_unclassified(100);k__Fungi_unclassified(100); | - | - | - | - | - | - | - | Unassigned | - |
| Otu20282 | 1 | k__Fungi(100);k__Fungi_unclassified(100);k__Fungi_unclassified(100);k__Fungi_unclassified(100);k__Fungi_unclassified(100);k__Fungi_unclassified(100);k__Fungi_unclassified(100); | - | - | - | - | - | - | - | Unassigned | - |
| Otu20287 | 1 | k__Fungi(100);p__Ascomycota(100);p__Ascomycota_unclassified(100);p__Ascomycota_unclassified(100);p__Ascomycota_unclassified(100);p__Ascomycota_unclassified(100);p__Ascomycota_unclassified(100); | - | - | - | - | - | - | - | Unassigned | - |
| Otu20290 | 1 | k__Fungi(100);k__Fungi_unclassified(100);k__Fungi_unclassified(100);k__Fungi_unclassified(100);k__Fungi_unclassified(100);k__Fungi_unclassified(100);k__Fungi_unclassified(100); | - | - | - | - | - | - | - | Unassigned | - |
| Otu20292 | 1 | k__Fungi(100);k__Fungi_unclassified(100);k__Fungi_unclassified(100);k__Fungi_unclassified(100);k__Fungi_unclassified(100);k__Fungi_unclassified(100);k__Fungi_unclassified(100); | - | - | - | - | - | - | - | Unassigned | - |
| Otu20315 | 1 | k__Fungi(100);p__Ascomycota(100);p__Ascomycota_unclassified(100);p__Ascomycota_unclassified(100);p__Ascomycota_unclassified(100);p__Ascomycota_unclassified(100);p__Ascomycota_unclassified(100); | - | - | - | - | - | - | - | Unassigned | - |
| Otu20320 | 1 | k__Fungi(100);p__Ascomycota(100);p__Ascomycota_unclassified(100);p__Ascomycota_unclassified(100);p__Ascomycota_unclassified(100);p__Ascomycota_unclassified(100);p__Ascomycota_unclassified(100); | - | - | - | - | - | - | - | Unassigned | - |
| Otu20323 | 1 | k__Fungi(100);p__Ascomycota(100);p__Ascomycota_unclassified(100);p__Ascomycota_unclassified(100);p__Ascomycota_unclassified(100);p__Ascomycota_unclassified(100);p__Ascomycota_unclassified(100); | - | - | - | - | - | - | - | Unassigned | - |
| Otu20330 | 1 | k__Fungi(100);p__Ascomycota(100);p__Ascomycota_unclassified(100);p__Ascomycota_unclassified(100);p__Ascomycota_unclassified(100);p__Ascomycota_unclassified(100);p__Ascomycota_unclassified(100); | - | - | - | - | - | - | - | Unassigned | - |
| Otu20332 | 1 | k__Fungi(100);k__Fungi_unclassified(100);k__Fungi_unclassified(100);k__Fungi_unclassified(100);k__Fungi_unclassified(100);k__Fungi_unclassified(100);k__Fungi_unclassified(100); | - | - | - | - | - | - | - | Unassigned | - |
| Otu20333 | 1 | k__Fungi(100);k__Fungi_unclassified(100);k__Fungi_unclassified(100);k__Fungi_unclassified(100);k__Fungi_unclassified(100);k__Fungi_unclassified(100);k__Fungi_unclassified(100); | - | - | - | - | - | - | - | Unassigned | - |
| Otu20334 | 1 | k__Fungi(100);p__Ascomycota(100);p__Ascomycota_unclassified(100);p__Ascomycota_unclassified(100);p__Ascomycota_unclassified(100);p__Ascomycota_unclassified(100);p__Ascomycota_unclassified(100); | - | - | - | - | - | - | - | Unassigned | - |
| Otu20335 | 1 | k__Fungi(100);k__Fungi_unclassified(100);k__Fungi_unclassified(100);k__Fungi_unclassified(100);k__Fungi_unclassified(100);k__Fungi_unclassified(100);k__Fungi_unclassified(100); | - | - | - | - | - | - | - | Unassigned | - |
| Otu20336 | 1 | k__Fungi(100);p__Ascomycota(100);p__Ascomycota_unclassified(100);p__Ascomycota_unclassified(100);p__Ascomycota_unclassified(100);p__Ascomycota_unclassified(100);p__Ascomycota_unclassified(100); | - | - | - | - | - | - | - | Unassigned | - |
| Otu20337 | 1 | k__Fungi(100);p__Ascomycota(100);p__Ascomycota_unclassified(100);p__Ascomycota_unclassified(100);p__Ascomycota_unclassified(100);p__Ascomycota_unclassified(100);p__Ascomycota_unclassified(100); | - | - | - | - | - | - | - | Unassigned | - |
| Otu20339 | 1 | k__Fungi(100);k__Fungi_unclassified(100);k__Fungi_unclassified(100);k__Fungi_unclassified(100);k__Fungi_unclassified(100);k__Fungi_unclassified(100);k__Fungi_unclassified(100); | - | - | - | - | - | - | - | Unassigned | - |
| Otu20340 | 1 | k__Fungi(100);p__Ascomycota(100);p__Ascomycota_unclassified(100);p__Ascomycota_unclassified(100);p__Ascomycota_unclassified(100);p__Ascomycota_unclassified(100);p__Ascomycota_unclassified(100); | - | - | - | - | - | - | - | Unassigned | - |
| Otu20341 | 1 | k__Fungi(100);p__Ascomycota(100);p__Ascomycota_unclassified(100);p__Ascomycota_unclassified(100);p__Ascomycota_unclassified(100);p__Ascomycota_unclassified(100);p__Ascomycota_unclassified(100); | - | - | - | - | - | - | - | Unassigned | - |
| Otu20343 | 1 | k__Fungi(100);k__Fungi_unclassified(100);k__Fungi_unclassified(100);k__Fungi_unclassified(100);k__Fungi_unclassified(100);k__Fungi_unclassified(100);k__Fungi_unclassified(100); | - | - | - | - | - | - | - | Unassigned | - |
| Otu20345 | 1 | k__Fungi(100);p__Ascomycota(100);p__Ascomycota_unclassified(100);p__Ascomycota_unclassified(100);p__Ascomycota_unclassified(100);p__Ascomycota_unclassified(100);p__Ascomycota_unclassified(100); | - | - | - | - | - | - | - | Unassigned | - |
| Otu20348 | 1 | k__Fungi(100);k__Fungi_unclassified(100);k__Fungi_unclassified(100);k__Fungi_unclassified(100);k__Fungi_unclassified(100);k__Fungi_unclassified(100);k__Fungi_unclassified(100); | - | - | - | - | - | - | - | Unassigned | - |
| Otu20349 | 1 | k__Fungi(100);p__Ascomycota(100);p__Ascomycota_unclassified(100);p__Ascomycota_unclassified(100);p__Ascomycota_unclassified(100);p__Ascomycota_unclassified(100);p__Ascomycota_unclassified(100); | - | - | - | - | - | - | - | Unassigned | - |
| Otu20350 | 1 | k__Fungi(100);p__Ascomycota(100);p__Ascomycota_unclassified(100);p__Ascomycota_unclassified(100);p__Ascomycota_unclassified(100);p__Ascomycota_unclassified(100);p__Ascomycota_unclassified(100); | - | - | - | - | - | - | - | Unassigned | - |
| Otu20351 | 1 | k__Fungi(100);k__Fungi_unclassified(100);k__Fungi_unclassified(100);k__Fungi_unclassified(100);k__Fungi_unclassified(100);k__Fungi_unclassified(100);k__Fungi_unclassified(100); | - | - | - | - | - | - | - | Unassigned | - |
| Otu20354 | 1 | k__Fungi(100);k__Fungi_unclassified(100);k__Fungi_unclassified(100);k__Fungi_unclassified(100);k__Fungi_unclassified(100);k__Fungi_unclassified(100);k__Fungi_unclassified(100); | - | - | - | - | - | - | - | Unassigned | - |
| Otu20357 | 1 | k__Fungi(100);p__Ascomycota(100);p__Ascomycota_unclassified(100);p__Ascomycota_unclassified(100);p__Ascomycota_unclassified(100);p__Ascomycota_unclassified(100);p__Ascomycota_unclassified(100); | - | - | - | - | - | - | - | Unassigned | - |
| Otu20358 | 1 | k__Fungi(100);k__Fungi_unclassified(100);k__Fungi_unclassified(100);k__Fungi_unclassified(100);k__Fungi_unclassified(100);k__Fungi_unclassified(100);k__Fungi_unclassified(100); | - | - | - | - | - | - | - | Unassigned | - |
| Otu20374 | 1 | k__Fungi(100);k__Fungi_unclassified(100);k__Fungi_unclassified(100);k__Fungi_unclassified(100);k__Fungi_unclassified(100);k__Fungi_unclassified(100);k__Fungi_unclassified(100); | - | - | - | - | - | - | - | Unassigned | - |
| Otu20376 | 1 | k__Fungi(100);p__Ascomycota(100);p__Ascomycota_unclassified(100);p__Ascomycota_unclassified(100);p__Ascomycota_unclassified(100);p__Ascomycota_unclassified(100);p__Ascomycota_unclassified(100); | - | - | - | - | - | - | - | Unassigned | - |
| Otu20381 | 1 | k__Fungi(100);p__Ascomycota(100);p__Ascomycota_unclassified(100);p__Ascomycota_unclassified(100);p__Ascomycota_unclassified(100);p__Ascomycota_unclassified(100);p__Ascomycota_unclassified(100); | - | - | - | - | - | - | - | Unassigned | - |
| Otu20386 | 1 | k__Fungi(100);p__Ascomycota(100);p__Ascomycota_unclassified(100);p__Ascomycota_unclassified(100);p__Ascomycota_unclassified(100);p__Ascomycota_unclassified(100);p__Ascomycota_unclassified(100); | - | - | - | - | - | - | - | Unassigned | - |
| Otu20387 | 1 | k__Fungi(100);p__Ascomycota(100);p__Ascomycota_unclassified(100);p__Ascomycota_unclassified(100);p__Ascomycota_unclassified(100);p__Ascomycota_unclassified(100);p__Ascomycota_unclassified(100); | - | - | - | - | - | - | - | Unassigned | - |
| Otu20388 | 1 | k__Fungi(100);p__Ascomycota(100);p__Ascomycota_unclassified(100);p__Ascomycota_unclassified(100);p__Ascomycota_unclassified(100);p__Ascomycota_unclassified(100);p__Ascomycota_unclassified(100); | - | - | - | - | - | - | - | Unassigned | - |
| Otu20389 | 1 | k__Fungi(100);k__Fungi_unclassified(100);k__Fungi_unclassified(100);k__Fungi_unclassified(100);k__Fungi_unclassified(100);k__Fungi_unclassified(100);k__Fungi_unclassified(100); | - | - | - | - | - | - | - | Unassigned | - |
| Otu20390 | 1 | k__Fungi(100);p__Ascomycota(100);p__Ascomycota_unclassified(100);p__Ascomycota_unclassified(100);p__Ascomycota_unclassified(100);p__Ascomycota_unclassified(100);p__Ascomycota_unclassified(100); | - | - | - | - | - | - | - | Unassigned | - |
| Otu20392 | 1 | k__Fungi(100);k__Fungi_unclassified(100);k__Fungi_unclassified(100);k__Fungi_unclassified(100);k__Fungi_unclassified(100);k__Fungi_unclassified(100);k__Fungi_unclassified(100); | - | - | - | - | - | - | - | Unassigned | - |
| Otu20393 | 1 | k__Fungi(100);k__Fungi_unclassified(100);k__Fungi_unclassified(100);k__Fungi_unclassified(100);k__Fungi_unclassified(100);k__Fungi_unclassified(100);k__Fungi_unclassified(100); | - | - | - | - | - | - | - | Unassigned | - |

| | | | | | | | | | | | |
|---|---|---|---|---|---|---|---|---|---|---|---|
| Otu20394 | 1 | k__Fungi(100);p__Ascomycota(100);p__Ascomycota_unclassified(100);p__Ascomycota_unclassified(100);p__Ascomycota_unclassified(100);p__Ascomycota_unclassified(100);p__Ascomycota_unclassified(100); | - | - | - | - | - | - | - | Unassigned | - |
| Otu20395 | 1 | k__Fungi(100);p__Ascomycota(100);p__Ascomycota_unclassified(100);p__Ascomycota_unclassified(100);p__Ascomycota_unclassified(100);p__Ascomycota_unclassified(100);p__Ascomycota_unclassified(100); | - | - | - | - | - | - | - | Unassigned | - |
| Otu20399 | 1 | k__Fungi(100);k__Fungi_unclassified(100);k__Fungi_unclassified(100);k__Fungi_unclassified(100);k__Fungi_unclassified(100);k__Fungi_unclassified(100);k__Fungi_unclassified(100); | - | - | - | - | - | - | - | Unassigned | - |
| Otu20410 | 1 | k__Fungi(100);p__Ascomycota(100);p__Ascomycota_unclassified(100);p__Ascomycota_unclassified(100);p__Ascomycota_unclassified(100);p__Ascomycota_unclassified(100);p__Ascomycota_unclassified(100); | - | - | - | - | - | - | - | Unassigned | - |
| Otu20440 | 1 | k__Fungi(100);p__Ascomycota(100);p__Ascomycota_unclassified(100);p__Ascomycota_unclassified(100);p__Ascomycota_unclassified(100);p__Ascomycota_unclassified(100);p__Ascomycota_unclassified(100); | - | - | - | - | - | - | - | Unassigned | - |
| Otu20445 | 1 | k__Fungi(100);p__Ascomycota(100);p__Ascomycota_unclassified(100);p__Ascomycota_unclassified(100);p__Ascomycota_unclassified(100);p__Ascomycota_unclassified(100);p__Ascomycota_unclassified(100); | - | - | - | - | - | - | - | Unassigned | - |
| Otu20446 | 1 | k__Fungi(100);k__Fungi_unclassified(100);k__Fungi_unclassified(100);k__Fungi_unclassified(100);k__Fungi_unclassified(100);k__Fungi_unclassified(100);k__Fungi_unclassified(100); | - | - | - | - | - | - | - | Unassigned | - |
| Otu20452 | 1 | k__Fungi(100);k__Fungi_unclassified(100);k__Fungi_unclassified(100);k__Fungi_unclassified(100);k__Fungi_unclassified(100);k__Fungi_unclassified(100);k__Fungi_unclassified(100); | - | - | - | - | - | - | - | Unassigned | - |
| Otu20466 | 1 | k__Fungi(100);p__Ascomycota(100);p__Ascomycota_unclassified(100);p__Ascomycota_unclassified(100);p__Ascomycota_unclassified(100);p__Ascomycota_unclassified(100);p__Ascomycota_unclassified(100); | - | - | - | - | - | - | - | Unassigned | - |
| Otu20472 | 1 | k__Fungi(100);k__Fungi_unclassified(100);k__Fungi_unclassified(100);k__Fungi_unclassified(100);k__Fungi_unclassified(100);k__Fungi_unclassified(100);k__Fungi_unclassified(100); | - | - | - | - | - | - | - | Unassigned | - |
| Otu20475 | 1 | k__Fungi(100);k__Fungi_unclassified(100);k__Fungi_unclassified(100);k__Fungi_unclassified(100);k__Fungi_unclassified(100);k__Fungi_unclassified(100);k__Fungi_unclassified(100); | - | - | - | - | - | - | - | Unassigned | - |
| Otu20477 | 1 | k__Fungi(100);k__Fungi_unclassified(100);k__Fungi_unclassified(100);k__Fungi_unclassified(100);k__Fungi_unclassified(100);k__Fungi_unclassified(100);k__Fungi_unclassified(100); | - | - | - | - | - | - | - | Unassigned | - |
| Otu20479 | 1 | k__Fungi(100);k__Fungi_unclassified(100);k__Fungi_unclassified(100);k__Fungi_unclassified(100);k__Fungi_unclassified(100);k__Fungi_unclassified(100);k__Fungi_unclassified(100); | - | - | - | - | - | - | - | Unassigned | - |
| Otu20480 | 1 | k__Fungi(100);p__Ascomycota(100);p__Ascomycota_unclassified(100);p__Ascomycota_unclassified(100);p__Ascomycota_unclassified(100);p__Ascomycota_unclassified(100);p__Ascomycota_unclassified(100); | - | - | - | - | - | - | - | Unassigned | - |
| Otu20481 | 1 | k__Fungi(100);k__Fungi_unclassified(100);k__Fungi_unclassified(100);k__Fungi_unclassified(100);k__Fungi_unclassified(100);k__Fungi_unclassified(100);k__Fungi_unclassified(100); | - | - | - | - | - | - | - | Unassigned | - |
| Otu20482 | 1 | k__Fungi(100);p__Ascomycota(100);p__Ascomycota_unclassified(100);p__Ascomycota_unclassified(100);p__Ascomycota_unclassified(100);p__Ascomycota_unclassified(100);p__Ascomycota_unclassified(100); | - | - | - | - | - | - | - | Unassigned | - |
| Otu20484 | 1 | k__Fungi(100);k__Fungi_unclassified(100);k__Fungi_unclassified(100);k__Fungi_unclassified(100);k__Fungi_unclassified(100);k__Fungi_unclassified(100);k__Fungi_unclassified(100); | - | - | - | - | - | - | - | Unassigned | - |
| Otu20485 | 1 | k__Fungi(100);p__Ascomycota(100);p__Ascomycota_unclassified(100);p__Ascomycota_unclassified(100);p__Ascomycota_unclassified(100);p__Ascomycota_unclassified(100);p__Ascomycota_unclassified(100); | - | - | - | - | - | - | - | Unassigned | - |
| Otu20486 | 1 | k__Fungi(100);k__Fungi_unclassified(100);k__Fungi_unclassified(100);k__Fungi_unclassified(100);k__Fungi_unclassified(100);k__Fungi_unclassified(100);k__Fungi_unclassified(100); | - | - | - | - | - | - | - | Unassigned | - |
| Otu20487 | 1 | k__Fungi(100);p__Ascomycota(100);p__Ascomycota_unclassified(100);p__Ascomycota_unclassified(100);p__Ascomycota_unclassified(100);p__Ascomycota_unclassified(100);p__Ascomycota_unclassified(100); | - | - | - | - | - | - | - | Unassigned | - |
| Otu20488 | 1 | k__Fungi(100);p__Ascomycota(100);p__Ascomycota_unclassified(100);p__Ascomycota_unclassified(100);p__Ascomycota_unclassified(100);p__Ascomycota_unclassified(100);p__Ascomycota_unclassified(100); | - | - | - | - | - | - | - | Unassigned | - |
| Otu20490 | 1 | k__Fungi(100);p__Ascomycota(100);p__Ascomycota_unclassified(100);p__Ascomycota_unclassified(100);p__Ascomycota_unclassified(100);p__Ascomycota_unclassified(100);p__Ascomycota_unclassified(100); | - | - | - | - | - | - | - | Unassigned | - |
| Otu20491 | 1 | k__Fungi(100);k__Fungi_unclassified(100);k__Fungi_unclassified(100);k__Fungi_unclassified(100);k__Fungi_unclassified(100);k__Fungi_unclassified(100);k__Fungi_unclassified(100); | - | - | - | - | - | - | - | Unassigned | - |
| Otu20496 | 1 | k__Fungi(100);k__Fungi_unclassified(100);k__Fungi_unclassified(100);k__Fungi_unclassified(100);k__Fungi_unclassified(100);k__Fungi_unclassified(100);k__Fungi_unclassified(100); | - | - | - | - | - | - | - | Unassigned | - |
| Otu20498 | 1 | k__Fungi(100);p__Ascomycota(100);p__Ascomycota_unclassified(100);p__Ascomycota_unclassified(100);p__Ascomycota_unclassified(100);p__Ascomycota_unclassified(100);p__Ascomycota_unclassified(100); | - | - | - | - | - | - | - | Unassigned | - |
| Otu20507 | 1 | k__Fungi(100);p__Ascomycota(100);p__Ascomycota_unclassified(100);p__Ascomycota_unclassified(100);p__Ascomycota_unclassified(100);p__Ascomycota_unclassified(100);p__Ascomycota_unclassified(100); | - | - | - | - | - | - | - | Unassigned | - |
| Otu20519 | 1 | k__Fungi(100);p__Ascomycota(100);p__Ascomycota_unclassified(100);p__Ascomycota_unclassified(100);p__Ascomycota_unclassified(100);p__Ascomycota_unclassified(100);p__Ascomycota_unclassified(100); | - | - | - | - | - | - | - | Unassigned | - |
| Otu20520 | 1 | k__Fungi(100);p__Ascomycota(100);p__Ascomycota_unclassified(100);p__Ascomycota_unclassified(100);p__Ascomycota_unclassified(100);p__Ascomycota_unclassified(100);p__Ascomycota_unclassified(100); | - | - | - | - | - | - | - | Unassigned | - |
| Otu20521 | 1 | k__Fungi(100);k__Fungi_unclassified(100);k__Fungi_unclassified(100);k__Fungi_unclassified(100);k__Fungi_unclassified(100);k__Fungi_unclassified(100);k__Fungi_unclassified(100); | - | - | - | - | - | - | - | Unassigned | - |
| Otu20522 | 1 | k__Fungi(100);k__Fungi_unclassified(100);k__Fungi_unclassified(100);k__Fungi_unclassified(100);k__Fungi_unclassified(100);k__Fungi_unclassified(100);k__Fungi_unclassified(100); | - | - | - | - | - | - | - | Unassigned | - |
| Otu20523 | 1 | k__Fungi(100);k__Fungi_unclassified(100);k__Fungi_unclassified(100);k__Fungi_unclassified(100);k__Fungi_unclassified(100);k__Fungi_unclassified(100);k__Fungi_unclassified(100); | - | - | - | - | - | - | - | Unassigned | - |
| Otu20524 | 1 | k__Fungi(100);k__Fungi_unclassified(100);k__Fungi_unclassified(100);k__Fungi_unclassified(100);k__Fungi_unclassified(100);k__Fungi_unclassified(100);k__Fungi_unclassified(100); | - | - | - | - | - | - | - | Unassigned | - |
| Otu20525 | 1 | k__Fungi(100);p__Ascomycota(100);p__Ascomycota_unclassified(100);p__Ascomycota_unclassified(100);p__Ascomycota_unclassified(100);p__Ascomycota_unclassified(100);p__Ascomycota_unclassified(100); | - | - | - | - | - | - | - | Unassigned | - |
| Otu20526 | 1 | k__Fungi(100);p__Ascomycota(100);p__Ascomycota_unclassified(100);p__Ascomycota_unclassified(100);p__Ascomycota_unclassified(100);p__Ascomycota_unclassified(100);p__Ascomycota_unclassified(100); | - | - | - | - | - | - | - | Unassigned | - |
| Otu20537 | 1 | k__Fungi(100);k__Fungi_unclassified(100);k__Fungi_unclassified(100);k__Fungi_unclassified(100);k__Fungi_unclassified(100);k__Fungi_unclassified(100);k__Fungi_unclassified(100); | - | - | - | - | - | - | - | Unassigned | - |
| Otu20538 | 1 | k__Fungi(100);p__Ascomycota(100);p__Ascomycota_unclassified(100);p__Ascomycota_unclassified(100);p__Ascomycota_unclassified(100);p__Ascomycota_unclassified(100);p__Ascomycota_unclassified(100); | - | - | - | - | - | - | - | Unassigned | - |
| Otu20572 | 1 | k__Fungi(100);k__Fungi_unclassified(100);k__Fungi_unclassified(100);k__Fungi_unclassified(100);k__Fungi_unclassified(100);k__Fungi_unclassified(100);k__Fungi_unclassified(100); | - | - | - | - | - | - | - | Unassigned | - |
| Otu20581 | 1 | k__Fungi(100);k__Fungi_unclassified(100);k__Fungi_unclassified(100);k__Fungi_unclassified(100);k__Fungi_unclassified(100);k__Fungi_unclassified(100);k__Fungi_unclassified(100); | - | - | - | - | - | - | - | Unassigned | - |
| Otu20588 | 1 | k__Fungi(100);k__Fungi_unclassified(100);k__Fungi_unclassified(100);k__Fungi_unclassified(100);k__Fungi_unclassified(100);k__Fungi_unclassified(100);k__Fungi_unclassified(100); | - | - | - | - | - | - | - | Unassigned | - |
| Otu20592 | 1 | k__Fungi(100);k__Fungi_unclassified(100);k__Fungi_unclassified(100);k__Fungi_unclassified(100);k__Fungi_unclassified(100);k__Fungi_unclassified(100);k__Fungi_unclassified(100); | - | - | - | - | - | - | - | Unassigned | - |
| Otu20598 | 1 | k__Fungi(100);p__Ascomycota(100);p__Ascomycota_unclassified(100);p__Ascomycota_unclassified(100);p__Ascomycota_unclassified(100);p__Ascomycota_unclassified(100);p__Ascomycota_unclassified(100); | - | - | - | - | - | - | - | Unassigned | - |
| Otu20601 | 1 | k__Fungi(100);p__Ascomycota(100);p__Ascomycota_unclassified(100);p__Ascomycota_unclassified(100);p__Ascomycota_unclassified(100);p__Ascomycota_unclassified(100);p__Ascomycota_unclassified(100); | - | - | - | - | - | - | - | Unassigned | - |
| Otu20610 | 1 | k__Fungi(100);p__Ascomycota(100);p__Ascomycota_unclassified(100);p__Ascomycota_unclassified(100);p__Ascomycota_unclassified(100);p__Ascomycota_unclassified(100);p__Ascomycota_unclassified(100); | - | - | - | - | - | - | - | Unassigned | - |
| Otu20626 | 1 | k__Fungi(100);k__Fungi_unclassified(100);k__Fungi_unclassified(100);k__Fungi_unclassified(100);k__Fungi_unclassified(100);k__Fungi_unclassified(100);k__Fungi_unclassified(100); | - | - | - | - | - | - | - | Unassigned | - |
| Otu20628 | 1 | k__Fungi(100);p__Ascomycota(100);p__Ascomycota_unclassified(100);p__Ascomycota_unclassified(100);p__Ascomycota_unclassified(100);p__Ascomycota_unclassified(100);p__Ascomycota_unclassified(100); | - | - | - | - | - | - | - | Unassigned | - |
| Otu20639 | 1 | k__Fungi(100);k__Fungi_unclassified(100);k__Fungi_unclassified(100);k__Fungi_unclassified(100);k__Fungi_unclassified(100);k__Fungi_unclassified(100);k__Fungi_unclassified(100); | - | - | - | - | - | - | - | Unassigned | - |
| Otu20640 | 1 | k__Fungi(100);k__Fungi_unclassified(100);k__Fungi_unclassified(100);k__Fungi_unclassified(100);k__Fungi_unclassified(100);k__Fungi_unclassified(100);k__Fungi_unclassified(100); | - | - | - | - | - | - | - | Unassigned | - |
| Otu20643 | 1 | k__Fungi(100);k__Fungi_unclassified(100);k__Fungi_unclassified(100);k__Fungi_unclassified(100);k__Fungi_unclassified(100);k__Fungi_unclassified(100);k__Fungi_unclassified(100); | - | - | - | - | - | - | - | Unassigned | - |
| Otu20644 | 1 | k__Fungi(100);k__Fungi_unclassified(100);k__Fungi_unclassified(100);k__Fungi_unclassified(100);k__Fungi_unclassified(100);k__Fungi_unclassified(100);k__Fungi_unclassified(100); | - | - | - | - | - | - | - | Unassigned | - |
| Otu20646 | 1 | k__Fungi(100);k__Fungi_unclassified(100);k__Fungi_unclassified(100);k__Fungi_unclassified(100);k__Fungi_unclassified(100);k__Fungi_unclassified(100);k__Fungi_unclassified(100); | - | - | - | - | - | - | - | Unassigned | - |
| Otu20647 | 1 | k__Fungi(100);k__Fungi_unclassified(100);k__Fungi_unclassified(100);k__Fungi_unclassified(100);k__Fungi_unclassified(100);k__Fungi_unclassified(100);k__Fungi_unclassified(100); | - | - | - | - | - | - | - | Unassigned | - |
| Otu20648 | 1 | k__Fungi(100);p__Ascomycota(100);p__Ascomycota_unclassified(100);p__Ascomycota_unclassified(100);p__Ascomycota_unclassified(100);p__Ascomycota_unclassified(100);p__Ascomycota_unclassified(100); | - | - | - | - | - | - | - | Unassigned | - |
| Otu20649 | 1 | k__Fungi(100);k__Fungi_unclassified(100);k__Fungi_unclassified(100);k__Fungi_unclassified(100);k__Fungi_unclassified(100);k__Fungi_unclassified(100);k__Fungi_unclassified(100); | - | - | - | - | - | - | - | Unassigned | - |
| Otu20651 | 1 | k__Fungi(100);k__Fungi_unclassified(100);k__Fungi_unclassified(100);k__Fungi_unclassified(100);k__Fungi_unclassified(100);k__Fungi_unclassified(100);k__Fungi_unclassified(100); | - | - | - | - | - | - | - | Unassigned | - |
| Otu20652 | 1 | k__Fungi(100);k__Fungi_unclassified(100);k__Fungi_unclassified(100);k__Fungi_unclassified(100);k__Fungi_unclassified(100);k__Fungi_unclassified(100);k__Fungi_unclassified(100); | - | - | - | - | - | - | - | Unassigned | - |
| Otu20657 | 1 | k__Fungi(100);k__Fungi_unclassified(100);k__Fungi_unclassified(100);k__Fungi_unclassified(100);k__Fungi_unclassified(100);k__Fungi_unclassified(100);k__Fungi_unclassified(100); | - | - | - | - | - | - | - | Unassigned | - |
| Otu20658 | 1 | k__Fungi(100);p__Ascomycota(100);p__Ascomycota_unclassified(100);p__Ascomycota_unclassified(100);p__Ascomycota_unclassified(100);p__Ascomycota_unclassified(100);p__Ascomycota_unclassified(100); | - | - | - | - | - | - | - | Unassigned | - |
| Otu20669 | 1 | k__Fungi(100);k__Fungi_unclassified(100);k__Fungi_unclassified(100);k__Fungi_unclassified(100);k__Fungi_unclassified(100);k__Fungi_unclassified(100);k__Fungi_unclassified(100); | - | - | - | - | - | - | - | Unassigned | - |
| Otu20675 | 1 | k__Fungi(100);p__Ascomycota(100);p__Ascomycota_unclassified(100);p__Ascomycota_unclassified(100);p__Ascomycota_unclassified(100);p__Ascomycota_unclassified(100);p__Ascomycota_unclassified(100); | - | - | - | - | - | - | - | Unassigned | - |
| Otu20680 | 1 | k__Fungi(100);k__Fungi_unclassified(100);k__Fungi_unclassified(100);k__Fungi_unclassified(100);k__Fungi_unclassified(100);k__Fungi_unclassified(100);k__Fungi_unclassified(100); | - | - | - | - | - | - | - | Unassigned | - |
| Otu20681 | 1 | k__Fungi(100);k__Fungi_unclassified(100);k__Fungi_unclassified(100);k__Fungi_unclassified(100);k__Fungi_unclassified(100);k__Fungi_unclassified(100);k__Fungi_unclassified(100); | - | - | - | - | - | - | - | Unassigned | - |
| Otu20685 | 1 | k__Fungi(100);k__Fungi_unclassified(100);k__Fungi_unclassified(100);k__Fungi_unclassified(100);k__Fungi_unclassified(100);k__Fungi_unclassified(100);k__Fungi_unclassified(100); | - | - | - | - | - | - | - | Unassigned | - |
| Otu20686 | 1 | k__Fungi(100);p__Ascomycota(100);p__Ascomycota_unclassified(100);p__Ascomycota_unclassified(100);p__Ascomycota_unclassified(100);p__Ascomycota_unclassified(100);p__Ascomycota_unclassified(100); | - | - | - | - | - | - | - | Unassigned | - |
| Otu20709 | 1 | k__Fungi(100);k__Fungi_unclassified(100);k__Fungi_unclassified(100);k__Fungi_unclassified(100);k__Fungi_unclassified(100);k__Fungi_unclassified(100);k__Fungi_unclassified(100); | - | - | - | - | - | - | - | Unassigned | - |
| Otu20711 | 1 | k__Fungi(100);k__Fungi_unclassified(100);k__Fungi_unclassified(100);k__Fungi_unclassified(100);k__Fungi_unclassified(100);k__Fungi_unclassified(100);k__Fungi_unclassified(100); | - | - | - | - | - | - | - | Unassigned | - |
| Otu20717 | 1 | k__Fungi(100);k__Fungi_unclassified(100);k__Fungi_unclassified(100);k__Fungi_unclassified(100);k__Fungi_unclassified(100);k__Fungi_unclassified(100);k__Fungi_unclassified(100); | - | - | - | - | - | - | - | Unassigned | - |
| Otu20718 | 1 | k__Fungi(100);p__Ascomycota(100);p__Ascomycota_unclassified(100);p__Ascomycota_unclassified(100);p__Ascomycota_unclassified(100);p__Ascomycota_unclassified(100);p__Ascomycota_unclassified(100); | - | - | - | - | - | - | - | Unassigned | - |
| Otu20741 | 1 | k__Fungi(100);k__Fungi_unclassified(100);k__Fungi_unclassified(100);k__Fungi_unclassified(100);k__Fungi_unclassified(100);k__Fungi_unclassified(100);k__Fungi_unclassified(100); | - | - | - | - | - | - | - | Unassigned | - |
| Otu20746 | 1 | k__Fungi(100);k__Fungi_unclassified(100);k__Fungi_unclassified(100);k__Fungi_unclassified(100);k__Fungi_unclassified(100);k__Fungi_unclassified(100);k__Fungi_unclassified(100); | - | - | - | - | - | - | - | Unassigned | - |
| Otu20751 | 1 | k__Fungi(100);p__Ascomycota(100);p__Ascomycota_unclassified(100);p__Ascomycota_unclassified(100);p__Ascomycota_unclassified(100);p__Ascomycota_unclassified(100);p__Ascomycota_unclassified(100); | - | - | - | - | - | - | - | Unassigned | - |
| Otu20760 | 1 | k__Fungi(100);k__Fungi_unclassified(100);k__Fungi_unclassified(100);k__Fungi_unclassified(100);k__Fungi_unclassified(100);k__Fungi_unclassified(100);k__Fungi_unclassified(100); | - | - | - | - | - | - | - | Unassigned | - |
| Otu20769 | 1 | k__Fungi(100);p__Ascomycota(100);p__Ascomycota_unclassified(100);p__Ascomycota_unclassified(100);p__Ascomycota_unclassified(100);p__Ascomycota_unclassified(100);p__Ascomycota_unclassified(100); | - | - | - | - | - | - | - | Unassigned | - |
| Otu20771 | 1 | k__Fungi(100);k__Fungi_unclassified(100);k__Fungi_unclassified(100);k__Fungi_unclassified(100);k__Fungi_unclassified(100);k__Fungi_unclassified(100);k__Fungi_unclassified(100); | - | - | - | - | - | - | - | Unassigned | - |
| Otu20772 | 1 | k__Fungi(100);k__Fungi_unclassified(100);k__Fungi_unclassified(100);k__Fungi_unclassified(100);k__Fungi_unclassified(100);k__Fungi_unclassified(100);k__Fungi_unclassified(100); | - | - | - | - | - | - | - | Unassigned | - |
| Otu20773 | 1 | k__Fungi(100);p__Ascomycota(100);p__Ascomycota_unclassified(100);p__Ascomycota_unclassified(100);p__Ascomycota_unclassified(100);p__Ascomycota_unclassified(100);p__Ascomycota_unclassified(100); | - | - | - | - | - | - | - | Unassigned | - |
| Otu20774 | 1 | k__Fungi(100);p__Ascomycota(100);p__Ascomycota_unclassified(100);p__Ascomycota_unclassified(100);p__Ascomycota_unclassified(100);p__Ascomycota_unclassified(100);p__Ascomycota_unclassified(100); | - | - | - | - | - | - | - | Unassigned | - |
| Otu20775 | 1 | k__Fungi(100);k__Fungi_unclassified(100);k__Fungi_unclassified(100);k__Fungi_unclassified(100);k__Fungi_unclassified(100);k__Fungi_unclassified(100);k__Fungi_unclassified(100); | - | - | - | - | - | - | - | Unassigned | - |
| Otu20776 | 1 | k__Fungi(100);k__Fungi_unclassified(100);k__Fungi_unclassified(100);k__Fungi_unclassified(100);k__Fungi_unclassified(100);k__Fungi_unclassified(100);k__Fungi_unclassified(100); | - | - | - | - | - | - | - | Unassigned | - |
| Otu20779 | 1 | k__Fungi(100);p__Ascomycota(100);p__Ascomycota_unclassified(100);p__Ascomycota_unclassified(100);p__Ascomycota_unclassified(100);p__Ascomycota_unclassified(100);p__Ascomycota_unclassified(100); | - | - | - | - | - | - | - | Unassigned | - |
| Otu20780 | 1 | k__Fungi(100);p__Ascomycota(100);p__Ascomycota_unclassified(100);p__Ascomycota_unclassified(100);p__Ascomycota_unclassified(100);p__Ascomycota_unclassified(100);p__Ascomycota_unclassified(100); | - | - | - | - | - | - | - | Unassigned | - |
| Otu20782 | 1 | k__Fungi(100);p__Ascomycota(100);p__Ascomycota_unclassified(100);p__Ascomycota_unclassified(100);p__Ascomycota_unclassified(100);p__Ascomycota_unclassified(100);p__Ascomycota_unclassified(100); | - | - | - | - | - | - | - | Unassigned | - |
| Otu20783 | 1 | k__Fungi(100);p__Ascomycota(100);p__Ascomycota_unclassified(100);p__Ascomycota_unclassified(100);p__Ascomycota_unclassified(100);p__Ascomycota_unclassified(100);p__Ascomycota_unclassified(100); | - | - | - | - | - | - | - | Unassigned | - |
| Otu20803 | 1 | k__Fungi(100);k__Fungi_unclassified(100);k__Fungi_unclassified(100);k__Fungi_unclassified(100);k__Fungi_unclassified(100);k__Fungi_unclassified(100);k__Fungi_unclassified(100); | - | - | - | - | - | - | - | Unassigned | - |
| Otu20805 | 1 | k__Fungi(100);k__Fungi_unclassified(100);k__Fungi_unclassified(100);k__Fungi_unclassified(100);k__Fungi_unclassified(100);k__Fungi_unclassified(100);k__Fungi_unclassified(100); | - | - | - | - | - | - | - | Unassigned | - |
| Otu20807 | 1 | k__Fungi(100);p__Ascomycota(100);p__Ascomycota_unclassified(100);p__Ascomycota_unclassified(100);p__Ascomycota_unclassified(100);p__Ascomycota_unclassified(100);p__Ascomycota_unclassified(100); | - | - | - | - | - | - | - | Unassigned | - |
| Otu20809 | 1 | k__Fungi(100);k__Fungi_unclassified(100);k__Fungi_unclassified(100);k__Fungi_unclassified(100);k__Fungi_unclassified(100);k__Fungi_unclassified(100);k__Fungi_unclassified(100); | - | - | - | - | - | - | - | Unassigned | - |
| Otu20810 | 1 | k__Fungi(100);k__Fungi_unclassified(100);k__Fungi_unclassified(100);k__Fungi_unclassified(100);k__Fungi_unclassified(100);k__Fungi_unclassified(100);k__Fungi_unclassified(100); | - | - | - | - | - | - | - | Unassigned | - |
| Otu20811 | 1 | k__Fungi(100);p__Ascomycota(100);p__Ascomycota_unclassified(100);p__Ascomycota_unclassified(100);p__Ascomycota_unclassified(100);p__Ascomycota_unclassified(100);p__Ascomycota_unclassified(100); | - | - | - | - | - | - | - | Unassigned | - |
| Otu20812 | 1 | k__Fungi(100);k__Fungi_unclassified(100);k__Fungi_unclassified(100);k__Fungi_unclassified(100);k__Fungi_unclassified(100);k__Fungi_unclassified(100);k__Fungi_unclassified(100); | - | - | - | - | - | - | - | Unassigned | - |
| Otu20821 | 1 | k__Fungi(100);k__Fungi_unclassified(100);k__Fungi_unclassified(100);k__Fungi_unclassified(100);k__Fungi_unclassified(100);k__Fungi_unclassified(100);k__Fungi_unclassified(100); | - | - | - | - | - | - | - | Unassigned | - |
| Otu20826 | 1 | k__Fungi(100);k__Fungi_unclassified(100);k__Fungi_unclassified(100);k__Fungi_unclassified(100);k__Fungi_unclassified(100);k__Fungi_unclassified(100);k__Fungi_unclassified(100); | - | - | - | - | - | - | - | Unassigned | - |
| Otu20827 | 1 | k__Fungi(100);p__Ascomycota(100);p__Ascomycota_unclassified(100);p__Ascomycota_unclassified(100);p__Ascomycota_unclassified(100);p__Ascomycota_unclassified(100);p__Ascomycota_unclassified(100); | - | - | - | - | - | - | - | Unassigned | - |
| Otu20828 | 1 | k__Fungi(100);k__Fungi_unclassified(100);k__Fungi_unclassified(100);k__Fungi_unclassified(100);k__Fungi_unclassified(100);k__Fungi_unclassified(100);k__Fungi_unclassified(100); | - | - | - | - | - | - | - | Unassigned | - |
| Otu20830 | 1 | k__Fungi(100);k__Fungi_unclassified(100);k__Fungi_unclassified(100);k__Fungi_unclassified(100);k__Fungi_unclassified(100);k__Fungi_unclassified(100);k__Fungi_unclassified(100); | - | - | - | - | - | - | - | Unassigned | - |
| Otu20845 | 1 | k__Fungi(100);k__Fungi_unclassified(100);k__Fungi_unclassified(100);k__Fungi_unclassified(100);k__Fungi_unclassified(100);k__Fungi_unclassified(100);k__Fungi_unclassified(100); | - | - | - | - | - | - | - | Unassigned | - |
| Otu20853 | 1 | k__Fungi(100);p__Ascomycota(100);p__Ascomycota_unclassified(100);p__Ascomycota_unclassified(100);p__Ascomycota_unclassified(100);p__Ascomycota_unclassified(100);p__Ascomycota_unclassified(100); | - | - | - | - | - | - | - | Unassigned | - |
| Otu20854 | 1 | k__Fungi(100);k__Fungi_unclassified(100);k__Fungi_unclassified(100);k__Fungi_unclassified(100);k__Fungi_unclassified(100);k__Fungi_unclassified(100);k__Fungi_unclassified(100); | - | - | - | - | - | - | - | Unassigned | - |
| Otu20855 | 1 | k__Fungi(100);p__Ascomycota(100);p__Ascomycota_unclassified(100);p__Ascomycota_unclassified(100);p__Ascomycota_unclassified(100);p__Ascomycota_unclassified(100);p__Ascomycota_unclassified(100); | - | - | - | - | - | - | - | Unassigned | - |
| Otu20856 | 1 | k__Fungi(100);p__Ascomycota(100);p__Ascomycota_unclassified(100);p__Ascomycota_unclassified(100);p__Ascomycota_unclassified(100);p__Ascomycota_unclassified(100);p__Ascomycota_unclassified(100); | - | - | - | - | - | - | - | Unassigned | - |
| Otu20857 | 1 | k__Fungi(100);p__Ascomycota(100);p__Ascomycota_unclassified(100);p__Ascomycota_unclassified(100);p__Ascomycota_unclassified(100);p__Ascomycota_unclassified(100);p__Ascomycota_unclassified(100); | - | - | - | - | - | - | - | Unassigned | - |
| Otu20858 | 1 | k__Fungi(100);k__Fungi_unclassified(100);k__Fungi_unclassified(100);k__Fungi_unclassified(100);k__Fungi_unclassified(100);k__Fungi_unclassified(100);k__Fungi_unclassified(100); | - | - | - | - | - | - | - | Unassigned | - |
| Otu20859 | 1 | k__Fungi(100);p__Ascomycota(100);p__Ascomycota_unclassified(100);p__Ascomycota_unclassified(100);p__Ascomycota_unclassified(100);p__Ascomycota_unclassified(100);p__Ascomycota_unclassified(100); | - | - | - | - | - | - | - | Unassigned | - |
| Otu20862 | 1 | k__Fungi(100);p__Ascomycota(100);p__Ascomycota_unclassified(100);p__Ascomycota_unclassified(100);p__Ascomycota_unclassified(100);p__Ascomycota_unclassified(100);p__Ascomycota_unclassified(100); | - | - | - | - | - | - | - | Unassigned | - |
| Otu20863 | 1 | k__Fungi(100);p__Ascomycota(100);p__Ascomycota_unclassified(100);p__Ascomycota_unclassified(100);p__Ascomycota_unclassified(100);p__Ascomycota_unclassified(100);p__Ascomycota_unclassified(100); | - | - | - | - | - | - | - | Unassigned | - |
| Otu20870 | 1 | k__Fungi(100);k__Fungi_unclassified(100);k__Fungi_unclassified(100);k__Fungi_unclassified(100);k__Fungi_unclassified(100);k__Fungi_unclassified(100);k__Fungi_unclassified(100); | - | - | - | - | - | - | - | Unassigned | - |
| Otu20877 | 1 | k__Fungi(100);k__Fungi_unclassified(100);k__Fungi_unclassified(100);k__Fungi_unclassified(100);k__Fungi_unclassified(100);k__Fungi_unclassified(100);k__Fungi_unclassified(100); | - | - | - | - | - | - | - | Unassigned | - |
| Otu20884 | 1 | k__Fungi(100);k__Fungi_unclassified(100);k__Fungi_unclassified(100);k__Fungi_unclassified(100);k__Fungi_unclassified(100);k__Fungi_unclassified(100);k__Fungi_unclassified(100); | - | - | - | - | - | - | - | Unassigned | - |
| Otu20893 | 1 | k__Fungi(100);k__Fungi_unclassified(100);k__Fungi_unclassified(100);k__Fungi_unclassified(100);k__Fungi_unclassified(100);k__Fungi_unclassified(100);k__Fungi_unclassified(100); | - | - | - | - | - | - | - | Unassigned | - |
| Otu20894 | 1 | k__Fungi(100);k__Fungi_unclassified(100);k__Fungi_unclassified(100);k__Fungi_unclassified(100);k__Fungi_unclassified(100);k__Fungi_unclassified(100);k__Fungi_unclassified(100); | - | - | - | - | - | - | - | Unassigned | - |
| Otu20895 | 1 | k__Fungi(100);k__Fungi_unclassified(100);k__Fungi_unclassified(100);k__Fungi_unclassified(100);k__Fungi_unclassified(100);k__Fungi_unclassified(100);k__Fungi_unclassified(100); | - | - | - | - | - | - | - | Unassigned | - |
| Otu20896 | 1 | k__Fungi(100);k__Fungi_unclassified(100);k__Fungi_unclassified(100);k__Fungi_unclassified(100);k__Fungi_unclassified(100);k__Fungi_unclassified(100);k__Fungi_unclassified(100); | - | - | - | - | - | - | - | Unassigned | - |
| Otu20897 | 1 | k__Fungi(100);k__Fungi_unclassified(100);k__Fungi_unclassified(100);k__Fungi_unclassified(100);k__Fungi_unclassified(100);k__Fungi_unclassified(100);k__Fungi_unclassified(100); | - | - | - | - | - | - | - | Unassigned | - |
| Otu20898 | 1 | k__Fungi(100);p__Ascomycota(100);p__Ascomycota_unclassified(100);p__Ascomycota_unclassified(100);p__Ascomycota_unclassified(100);p__Ascomycota_unclassified(100);p__Ascomycota_unclassified(100); | - | - | - | - | - | - | - | Unassigned | - |
| Otu20899 | 1 | k__Fungi(100);p__Ascomycota(100);p__Ascomycota_unclassified(100);p__Ascomycota_unclassified(100);p__Ascomycota_unclassified(100);p__Ascomycota_unclassified(100);p__Ascomycota_unclassified(100); | - | - | - | - | - | - | - | Unassigned | - |
| Otu20900 | 1 | k__Fungi(100);p__Ascomycota(100);p__Ascomycota_unclassified(100);p__Ascomycota_unclassified(100);p__Ascomycota_unclassified(100);p__Ascomycota_unclassified(100);p__Ascomycota_unclassified(100); | - | - | - | - | - | - | - | Unassigned | - |
| Otu20908 | 1 | k__Fungi(100);k__Fungi_unclassified(100);k__Fungi_unclassified(100);k__Fungi_unclassified(100);k__Fungi_unclassified(100);k__Fungi_unclassified(100);k__Fungi_unclassified(100); | - | - | - | - | - | - | - | Unassigned | - |
| Otu20909 | 1 | k__Fungi(100);k__Fungi_unclassified(100);k__Fungi_unclassified(100);k__Fungi_unclassified(100);k__Fungi_unclassified(100);k__Fungi_unclassified(100);k__Fungi_unclassified(100); | - | - | - | - | - | - | - | Unassigned | - |
| Otu20915 | 1 | k__Fungi(100);k__Fungi_unclassified(100);k__Fungi_unclassified(100);k__Fungi_unclassified(100);k__Fungi_unclassified(100);k__Fungi_unclassified(100);k__Fungi_unclassified(100); | - | - | - | - | - | - | - | Unassigned | - |
| Otu20920 | 1 | k__Fungi(100);k__Fungi_unclassified(100);k__Fungi_unclassified(100);k__Fungi_unclassified(100);k__Fungi_unclassified(100);k__Fungi_unclassified(100);k__Fungi_unclassified(100); | - | - | - | - | - | - | - | Unassigned | - |
| Otu20922 | 1 | k__Fungi(100);p__Ascomycota(100);p__Ascomycota_unclassified(100);p__Ascomycota_unclassified(100);p__Ascomycota_unclassified(100);p__Ascomycota_unclassified(100);p__Ascomycota_unclassified(100); | - | - | - | - | - | - | - | Unassigned | - |
| Otu20923 | 1 | k__Fungi(100);k__Fungi_unclassified(100);k__Fungi_unclassified(100);k__Fungi_unclassified(100);k__Fungi_unclassified(100);k__Fungi_unclassified(100);k__Fungi_unclassified(100); | - | - | - | - | - | - | - | Unassigned | - |
| Otu20924 | 1 | k__Fungi(100);p__Ascomycota(100);p__Ascomycota_unclassified(100);p__Ascomycota_unclassified(100);p__Ascomycota_unclassified(100);p__Ascomycota_unclassified(100);p__Ascomycota_unclassified(100); | - | - | - | - | - | - | - | Unassigned | - |
| Otu20925 | 1 | k__Fungi(100);p__Ascomycota(100);p__Ascomycota_unclassified(100);p__Ascomycota_unclassified(100);p__Ascomycota_unclassified(100);p__Ascomycota_unclassified(100);p__Ascomycota_unclassified(100); | - | - | - | - | - | - | - | Unassigned | - |
| Otu20926 | 1 | k__Fungi(100);k__Fungi_unclassified(100);k__Fungi_unclassified(100);k__Fungi_unclassified(100);k__Fungi_unclassified(100);k__Fungi_unclassified(100);k__Fungi_unclassified(100); | - | - | - | - | - | - | - | Unassigned | - |
| Otu20927 | 1 | k__Fungi(100);k__Fungi_unclassified(100);k__Fungi_unclassified(100);k__Fungi_unclassified(100);k__Fungi_unclassified(100);k__Fungi_unclassified(100);k__Fungi_unclassified(100); | - | - | - | - | - | - | - | Unassigned | - |
| Otu20935 | 1 | k__Fungi(100);p__Ascomycota(100);p__Ascomycota_unclassified(100);p__Ascomycota_unclassified(100);p__Ascomycota_unclassified(100);p__Ascomycota_unclassified(100);p__Ascomycota_unclassified(100); | - | - | - | - | - | - | - | Unassigned | - |
| Otu20936 | 1 | k__Fungi(100);p__Ascomycota(100);p__Ascomycota_unclassified(100);p__Ascomycota_unclassified(100);p__Ascomycota_unclassified(100);p__Ascomycota_unclassified(100);p__Ascomycota_unclassified(100); | - | - | - | - | - | - | - | Unassigned | - |
| Otu20958 | 1 | k__Fungi(100);p__Ascomycota(100);p__Ascomycota_unclassified(100);p__Ascomycota_unclassified(100);p__Ascomycota_unclassified(100);p__Ascomycota_unclassified(100);p__Ascomycota_unclassified(100); | - | - | - | - | - | - | - | Unassigned | - |
| Otu20966 | 1 | k__Fungi(100);k__Fungi_unclassified(100);k__Fungi_unclassified(100);k__Fungi_unclassified(100);k__Fungi_unclassified(100);k__Fungi_unclassified(100);k__Fungi_unclassified(100); | - | - | - | - | - | - | - | Unassigned | - |
| Otu20967 | 1 | k__Fungi(100);p__Ascomycota(100);p__Ascomycota_unclassified(100);p__Ascomycota_unclassified(100);p__Ascomycota_unclassified(100);p__Ascomycota_unclassified(100);p__Ascomycota_unclassified(100); | - | - | - | - | - | - | - | Unassigned | - |
| Otu20973 | 1 | k__Fungi(100);k__Fungi_unclassified(100);k__Fungi_unclassified(100);k__Fungi_unclassified(100);k__Fungi_unclassified(100);k__Fungi_unclassified(100);k__Fungi_unclassified(100); | - | - | - | - | - | - | - | Unassigned | - |
| Otu20978 | 1 | k__Fungi(100);k__Fungi_unclassified(100);k__Fungi_unclassified(100);k__Fungi_unclassified(100);k__Fungi_unclassified(100);k__Fungi_unclassified(100);k__Fungi_unclassified(100); | - | - | - | - | - | - | - | Unassigned | - |
| Otu20980 | 1 | k__Fungi(100);k__Fungi_unclassified(100);k__Fungi_unclassified(100);k__Fungi_unclassified(100);k__Fungi_unclassified(100);k__Fungi_unclassified(100);k__Fungi_unclassified(100); | - | - | - | - | - | - | - | Unassigned | - |
| Otu20981 | 1 | k__Fungi(100);k__Fungi_unclassified(100);k__Fungi_unclassified(100);k__Fungi_unclassified(100);k__Fungi_unclassified(100);k__Fungi_unclassified(100);k__Fungi_unclassified(100); | - | - | - | - | - | - | - | Unassigned | - |
| Otu20990 | 1 | k__Fungi(100);p__Ascomycota(100);p__Ascomycota_unclassified(100);p__Ascomycota_unclassified(100);p__Ascomycota_unclassified(100);p__Ascomycota_unclassified(100);p__Ascomycota_unclassified(100); | - | - | - | - | - | - | - | Unassigned | - |
| Otu20995 | 1 | k__Fungi(100);k__Fungi_unclassified(100);k__Fungi_unclassified(100);k__Fungi_unclassified(100);k__Fungi_unclassified(100);k__Fungi_unclassified(100);k__Fungi_unclassified(100); | - | - | - | - | - | - | - | Unassigned | - |
| Otu20999 | 1 | k__Fungi(100);p__Ascomycota(100);p__Ascomycota_unclassified(100);p__Ascomycota_unclassified(100);p__Ascomycota_unclassified(100);p__Ascomycota_unclassified(100);p__Ascomycota_unclassified(100); | - | - | - | - | - | - | - | Unassigned | - |
| Otu21020 | 1 | k__Fungi(100);k__Fungi_unclassified(100);k__Fungi_unclassified(100);k__Fungi_unclassified(100);k__Fungi_unclassified(100);k__Fungi_unclassified(100);k__Fungi_unclassified(100); | - | - | - | - | - | - | - | Unassigned | - |
| Otu21021 | 1 | k__Fungi(100);p__Ascomycota(100);p__Ascomycota_unclassified(100);p__Ascomycota_unclassified(100);p__Ascomycota_unclassified(100);p__Ascomycota_unclassified(100);p__Ascomycota_unclassified(100); | - | - | - | - | - | - | - | Unassigned | - |
| Otu21024 | 1 | k__Fungi(100);k__Fungi_unclassified(100);k__Fungi_unclassified(100);k__Fungi_unclassified(100);k__Fungi_unclassified(100);k__Fungi_unclassified(100);k__Fungi_unclassified(100); | - | - | - | - | - | - | - | Unassigned | - |
| Otu21029 | 1 | k__Fungi(100);p__Ascomycota(100);p__Ascomycota_unclassified(100);p__Ascomycota_unclassified(100);p__Ascomycota_unclassified(100);p__Ascomycota_unclassified(100);p__Ascomycota_unclassified(100); | - | - | - | - | - | - | - | Unassigned | - |
| Otu21031 | 1 | k__Fungi(100);p__Ascomycota(100);p__Ascomycota_unclassified(100);p__Ascomycota_unclassified(100);p__Ascomycota_unclassified(100);p__Ascomycota_unclassified(100);p__Ascomycota_unclassified(100); | - | - | - | - | - | - | - | Unassigned | - |
| Otu21032 | 1 | k__Fungi(100);k__Fungi_unclassified(100);k__Fungi_unclassified(100);k__Fungi_unclassified(100);k__Fungi_unclassified(100);k__Fungi_unclassified(100);k__Fungi_unclassified(100); | - | - | - | - | - | - | - | Unassigned | - |
| Otu21033 | 1 | k__Fungi(100);k__Fungi_unclassified(100);k__Fungi_unclassified(100);k__Fungi_unclassified(100);k__Fungi_unclassified(100);k__Fungi_unclassified(100);k__Fungi_unclassified(100); | - | - | - | - | - | - | - | Unassigned | - |
| Otu21038 | 1 | k__Fungi(100);p__Ascomycota(100);p__Ascomycota_unclassified(100);p__Ascomycota_unclassified(100);p__Ascomycota_unclassified(100);p__Ascomycota_unclassified(100);p__Ascomycota_unclassified(100); | - | - | - | - | - | - | - | Unassigned | - |
| Otu21040 | 1 | k__Fungi(100);k__Fungi_unclassified(100);k__Fungi_unclassified(100);k__Fungi_unclassified(100);k__Fungi_unclassified(100);k__Fungi_unclassified(100);k__Fungi_unclassified(100); | - | - | - | - | - | - | - | Unassigned | - |
| Otu21041 | 1 | k__Fungi(100);k__Fungi_unclassified(100);k__Fungi_unclassified(100);k__Fungi_unclassified(100);k__Fungi_unclassified(100);k__Fungi_unclassified(100);k__Fungi_unclassified(100); | - | - | - | - | - | - | - | Unassigned | - |
| Otu21043 | 1 | k__Fungi(100);k__Fungi_unclassified(100);k__Fungi_unclassified(100);k__Fungi_unclassified(100);k__Fungi_unclassified(100);k__Fungi_unclassified(100);k__Fungi_unclassified(100); | - | - | - | - | - | - | - | Unassigned | - |
| Otu21047 | 1 | k__Fungi(100);k__Fungi_unclassified(100);k__Fungi_unclassified(100);k__Fungi_unclassified(100);k__Fungi_unclassified(100);k__Fungi_unclassified(100);k__Fungi_unclassified(100); | - | - | - | - | - | - | - | Unassigned | - |
| Otu21049 | 1 | k__Fungi(100);p__Ascomycota(100);p__Ascomycota_unclassified(100);p__Ascomycota_unclassified(100);p__Ascomycota_unclassified(100);p__Ascomycota_unclassified(100);p__Ascomycota_unclassified(100); | - | - | - | - | - | - | - | Unassigned | - |
| Otu21053 | 1 | k__Fungi(100);k__Fungi_unclassified(100);k__Fungi_unclassified(100);k__Fungi_unclassified(100);k__Fungi_unclassified(100);k__Fungi_unclassified(100);k__Fungi_unclassified(100); | - | - | - | - | - | - | - | Unassigned | - |
| Otu21054 | 1 | k__Fungi(100);k__Fungi_unclassified(100);k__Fungi_unclassified(100);k__Fungi_unclassified(100);k__Fungi_unclassified(100);k__Fungi_unclassified(100);k__Fungi_unclassified(100); | - | - | - | - | - | - | - | Unassigned | - |
| Otu21055 | 1 | k__Fungi(100);k__Fungi_unclassified(100);k__Fungi_unclassified(100);k__Fungi_unclassified(100);k__Fungi_unclassified(100);k__Fungi_unclassified(100);k__Fungi_unclassified(100); | - | - | - | - | - | - | - | Unassigned | - |
| Otu21056 | 1 | k__Fungi(100);p__Ascomycota(100);p__Ascomycota_unclassified(100);p__Ascomycota_unclassified(100);p__Ascomycota_unclassified(100);p__Ascomycota_unclassified(100);p__Ascomycota_unclassified(100); | - | - | - | - | - | - | - | Unassigned | - |
| Otu21057 | 1 | k__Fungi(100);k__Fungi_unclassified(100);k__Fungi_unclassified(100);k__Fungi_unclassified(100);k__Fungi_unclassified(100);k__Fungi_unclassified(100);k__Fungi_unclassified(100); | - | - | - | - | - | - | - | Unassigned | - |
| Otu21058 | 1 | k__Fungi(100);k__Fungi_unclassified(100);k__Fungi_unclassified(100);k__Fungi_unclassified(100);k__Fungi_unclassified(100);k__Fungi_unclassified(100);k__Fungi_unclassified(100); | - | - | - | - | - | - | - | Unassigned | - |
| Otu21061 | 1 | k__Fungi(100);k__Fungi_unclassified(100);k__Fungi_unclassified(100);k__Fungi_unclassified(100);k__Fungi_unclassified(100);k__Fungi_unclassified(100);k__Fungi_unclassified(100); | - | - | - | - | - | - | - | Unassigned | - |
| Otu21063 | 1 | k__Fungi(100);p__Ascomycota(100);p__Ascomycota_unclassified(100);p__Ascomycota_unclassified(100);p__Ascomycota_unclassified(100);p__Ascomycota_unclassified(100);p__Ascomycota_unclassified(100); | - | - | - | - | - | - | - | Unassigned | - |
| Otu21067 | 1 | k__Fungi(100);p__Ascomycota(100);p__Ascomycota_unclassified(100);p__Ascomycota_unclassified(100);p__Ascomycota_unclassified(100);p__Ascomycota_unclassified(100);p__Ascomycota_unclassified(100); | - | - | - | - | - | - | - | Unassigned | - |
| Otu21071 | 1 | k__Fungi(100);k__Fungi_unclassified(100);k__Fungi_unclassified(100);k__Fungi_unclassified(100);k__Fungi_unclassified(100);k__Fungi_unclassified(100);k__Fungi_unclassified(100); | - | - | - | - | - | - | - | Unassigned | - |
| Otu21072 | 1 | k__Fungi(100);k__Fungi_unclassified(100);k__Fungi_unclassified(100);k__Fungi_unclassified(100);k__Fungi_unclassified(100);k__Fungi_unclassified(100);k__Fungi_unclassified(100); | - | - | - | - | - | - | - | Unassigned | - |
| Otu21085 | 1 | k__Fungi(100);p__Ascomycota(100);p__Ascomycota_unclassified(100);p__Ascomycota_unclassified(100);p__Ascomycota_unclassified(100);p__Ascomycota_unclassified(100);p__Ascomycota_unclassified(100); | - | - | - | - | - | - | - | Unassigned | - |
| Otu21086 | 1 | k__Fungi(100);p__Ascomycota(100);p__Ascomycota_unclassified(100);p__Ascomycota_unclassified(100);p__Ascomycota_unclassified(100);p__Ascomycota_unclassified(100);p__Ascomycota_unclassified(100); | - | - | - | - | - | - | - | Unassigned | - |
| Otu21094 | 1 | k__Fungi(100);k__Fungi_unclassified(100);k__Fungi_unclassified(100);k__Fungi_unclassified(100);k__Fungi_unclassified(100);k__Fungi_unclassified(100);k__Fungi_unclassified(100); | - | - | - | - | - | - | - | Unassigned | - |
| Otu21095 | 1 | k__Fungi(100);k__Fungi_unclassified(100);k__Fungi_unclassified(100);k__Fungi_unclassified(100);k__Fungi_unclassified(100);k__Fungi_unclassified(100);k__Fungi_unclassified(100); | - | - | - | - | - | - | - | Unassigned | - |
| Otu21099 | 1 | k__Fungi(100);k__Fungi_unclassified(100);k__Fungi_unclassified(100);k__Fungi_unclassified(100);k__Fungi_unclassified(100);k__Fungi_unclassified(100);k__Fungi_unclassified(100); | - | - | - | - | - | - | - | Unassigned | - |
| Otu21100 | 1 | k__Fungi(100);k__Fungi_unclassified(100);k__Fungi_unclassified(100);k__Fungi_unclassified(100);k__Fungi_unclassified(100);k__Fungi_unclassified(100);k__Fungi_unclassified(100); | - | - | - | - | - | - | - | Unassigned | - |
| Otu21101 | 1 | k__Fungi(100);p__Ascomycota(100);p__Ascomycota_unclassified(100);p__Ascomycota_unclassified(100);p__Ascomycota_unclassified(100);p__Ascomycota_unclassified(100);p__Ascomycota_unclassified(100); | - | - | - | - | - | - | - | Unassigned | - |
| Otu21102 | 1 | k__Fungi(100);k__Fungi_unclassified(100);k__Fungi_unclassified(100);k__Fungi_unclassified(100);k__Fungi_unclassified(100);k__Fungi_unclassified(100);k__Fungi_unclassified(100); | - | - | - | - | - | - | - | Unassigned | - |
| Otu21103 | 1 | k__Fungi(100);k__Fungi_unclassified(100);k__Fungi_unclassified(100);k__Fungi_unclassified(100);k__Fungi_unclassified(100);k__Fungi_unclassified(100);k__Fungi_unclassified(100); | - | - | - | - | - | - | - | Unassigned | - |
| Otu21111 | 1 | k__Fungi(100);k__Fungi_unclassified(100);k__Fungi_unclassified(100);k__Fungi_unclassified(100);k__Fungi_unclassified(100);k__Fungi_unclassified(100);k__Fungi_unclassified(100); | - | - | - | - | - | - | - | Unassigned | - |
| Otu21112 | 1 | k__Fungi(100);k__Fungi_unclassified(100);k__Fungi_unclassified(100);k__Fungi_unclassified(100);k__Fungi_unclassified(100);k__Fungi_unclassified(100);k__Fungi_unclassified(100); | - | - | - | - | - | - | - | Unassigned | - |
| Otu21116 | 1 | k__Fungi(100);k__Fungi_unclassified(100);k__Fungi_unclassified(100);k__Fungi_unclassified(100);k__Fungi_unclassified(100);k__Fungi_unclassified(100);k__Fungi_unclassified(100); | - | - | - | - | - | - | - | Unassigned | - |
| Otu21117 | 1 | k__Fungi(100);p__Ascomycota(100);p__Ascomycota_unclassified(100);p__Ascomycota_unclassified(100);p__Ascomycota_unclassified(100);p__Ascomycota_unclassified(100);p__Ascomycota_unclassified(100); | - | - | - | - | - | - | - | Unassigned | - |
| Otu21119 | 1 | k__Fungi(100);k__Fungi_unclassified(100);k__Fungi_unclassified(100);k__Fungi_unclassified(100);k__Fungi_unclassified(100);k__Fungi_unclassified(100);k__Fungi_unclassified(100); | - | - | - | - | - | - | - | Unassigned | - |
| Otu21121 | 1 | k__Fungi(100);k__Fungi_unclassified(100);k__Fungi_unclassified(100);k__Fungi_unclassified(100);k__Fungi_unclassified(100);k__Fungi_unclassified(100);k__Fungi_unclassified(100); | - | - | - | - | - | - | - | Unassigned | - |
| Otu21126 | 1 | k__Fungi(100);p__Ascomycota(100);p__Ascomycota_unclassified(100);p__Ascomycota_unclassified(100);p__Ascomycota_unclassified(100);p__Ascomycota_unclassified(100);p__Ascomycota_unclassified(100); | - | - | - | - | - | - | - | Unassigned | - |
| Otu21127 | 1 | k__Fungi(100);k__Fungi_unclassified(100);k__Fungi_unclassified(100);k__Fungi_unclassified(100);k__Fungi_unclassified(100);k__Fungi_unclassified(100);k__Fungi_unclassified(100); | - | - | - | - | - | - | - | Unassigned | - |
| Otu21128 | 1 | k__Fungi(100);k__Fungi_unclassified(100);k__Fungi_unclassified(100);k__Fungi_unclassified(100);k__Fungi_unclassified(100);k__Fungi_unclassified(100);k__Fungi_unclassified(100); | - | - | - | - | - | - | - | Unassigned | - |
| Otu21131 | 1 | k__Fungi(100);k__Fungi_unclassified(100);k__Fungi_unclassified(100);k__Fungi_unclassified(100);k__Fungi_unclassified(100);k__Fungi_unclassified(100);k__Fungi_unclassified(100); | - | - | - | - | - | - | - | Unassigned | - |
| Otu21132 | 1 | k__Fungi(100);k__Fungi_unclassified(100);k__Fungi_unclassified(100);k__Fungi_unclassified(100);k__Fungi_unclassified(100);k__Fungi_unclassified(100);k__Fungi_unclassified(100); | - | - | - | - | - | - | - | Unassigned | - |
| Otu21133 | 1 | k__Fungi(100);p__Ascomycota(100);p__Ascomycota_unclassified(100);p__Ascomycota_unclassified(100);p__Ascomycota_unclassified(100);p__Ascomycota_unclassified(100);p__Ascomycota_unclassified(100); | - | - | - | - | - | - | - | Unassigned | - |
| Otu21137 | 1 | k__Fungi(100);p__Ascomycota(100);p__Ascomycota_unclassified(100);p__Ascomycota_unclassified(100);p__Ascomycota_unclassified(100);p__Ascomycota_unclassified(100);p__Ascomycota_unclassified(100); | - | - | - | - | - | - | - | Unassigned | - |
| Otu21139 | 1 | k__Fungi(100);k__Fungi_unclassified(100);k__Fungi_unclassified(100);k__Fungi_unclassified(100);k__Fungi_unclassified(100);k__Fungi_unclassified(100);k__Fungi_unclassified(100); | - | - | - | - | - | - | - | Unassigned | - |
| Otu21140 | 1 | k__Fungi(100);p__Ascomycota(100);p__Ascomycota_unclassified(100);p__Ascomycota_unclassified(100);p__Ascomycota_unclassified(100);p__Ascomycota_unclassified(100);p__Ascomycota_unclassified(100); | - | - | - | - | - | - | - | Unassigned | - |
| Otu21142 | 1 | k__Fungi(100);k__Fungi_unclassified(100);k__Fungi_unclassified(100);k__Fungi_unclassified(100);k__Fungi_unclassified(100);k__Fungi_unclassified(100);k__Fungi_unclassified(100); | - | - | - | - | - | - | - | Unassigned | - |
| Otu21143 | 1 | k__Fungi(100);k__Fungi_unclassified(100);k__Fungi_unclassified(100);k__Fungi_unclassified(100);k__Fungi_unclassified(100);k__Fungi_unclassified(100);k__Fungi_unclassified(100); | - | - | - | - | - | - | - | Unassigned | - |
| Otu21144 | 1 | k__Fungi(100);p__Ascomycota(100);p__Ascomycota_unclassified(100);p__Ascomycota_unclassified(100);p__Ascomycota_unclassified(100);p__Ascomycota_unclassified(100);p__Ascomycota_unclassified(100); | - | - | - | - | - | - | - | Unassigned | - |
| Otu21145 | 1 | k__Fungi(100);k__Fungi_unclassified(100);k__Fungi_unclassified(100);k__Fungi_unclassified(100);k__Fungi_unclassified(100);k__Fungi_unclassified(100);k__Fungi_unclassified(100); | - | - | - | - | - | - | - | Unassigned | - |
| Otu21149 | 1 | k__Fungi(100);k__Fungi_unclassified(100);k__Fungi_unclassified(100);k__Fungi_unclassified(100);k__Fungi_unclassified(100);k__Fungi_unclassified(100);k__Fungi_unclassified(100); | - | - | - | - | - | - | - | Unassigned | - |
| Otu21151 | 1 | k__Fungi(100);k__Fungi_unclassified(100);k__Fungi_unclassified(100);k__Fungi_unclassified(100);k__Fungi_unclassified(100);k__Fungi_unclassified(100);k__Fungi_unclassified(100); | - | - | - | - | - | - | - | Unassigned | - |
| Otu21152 | 1 | k__Fungi(100);p__Ascomycota(100);p__Ascomycota_unclassified(100);p__Ascomycota_unclassified(100);p__Ascomycota_unclassified(100);p__Ascomycota_unclassified(100);p__Ascomycota_unclassified(100); | - | - | - | - | - | - | - | Unassigned | - |
| Otu21154 | 1 | k__Fungi(100);k__Fungi_unclassified(100);k__Fungi_unclassified(100);k__Fungi_unclassified(100);k__Fungi_unclassified(100);k__Fungi_unclassified(100);k__Fungi_unclassified(100); | - | - | - | - | - | - | - | Unassigned | - |
| Otu21155 | 1 | k__Fungi(100);k__Fungi_unclassified(100);k__Fungi_unclassified(100);k__Fungi_unclassified(100);k__Fungi_unclassified(100);k__Fungi_unclassified(100);k__Fungi_unclassified(100); | - | - | - | - | - | - | - | Unassigned | - |
| Otu21156 | 1 | k__Fungi(100);p__Ascomycota(100);p__Ascomycota_unclassified(100);p__Ascomycota_unclassified(100);p__Ascomycota_unclassified(100);p__Ascomycota_unclassified(100);p__Ascomycota_unclassified(100); | - | - | - | - | - | - | - | Unassigned | - |
| Otu21157 | 1 | k__Fungi(100);k__Fungi_unclassified(100);k__Fungi_unclassified(100);k__Fungi_unclassified(100);k__Fungi_unclassified(100);k__Fungi_unclassified(100);k__Fungi_unclassified(100); | - | - | - | - | - | - | - | Unassigned | - |
| Otu21158 | 1 | k__Fungi(100);k__Fungi_unclassified(100);k__Fungi_unclassified(100);k__Fungi_unclassified(100);k__Fungi_unclassified(100);k__Fungi_unclassified(100);k__Fungi_unclassified(100); | - | - | - | - | - | - | - | Unassigned | - |
| Otu21159 | 1 | k__Fungi(100);k__Fungi_unclassified(100);k__Fungi_unclassified(100);k__Fungi_unclassified(100);k__Fungi_unclassified(100);k__Fungi_unclassified(100);k__Fungi_unclassified(100); | - | - | - | - | - | - | - | Unassigned | - |
| Otu21161 | 1 | k__Fungi(100);p__Ascomycota(100);p__Ascomycota_unclassified(100);p__Ascomycota_unclassified(100);p__Ascomycota_unclassified(100);p__Ascomycota_unclassified(100);p__Ascomycota_unclassified(100); | - | - | - | - | - | - | - | Unassigned | - |
| Otu21162 | 1 | k__Fungi(100);k__Fungi_unclassified(100);k__Fungi_unclassified(100);k__Fungi_unclassified(100);k__Fungi_unclassified(100);k__Fungi_unclassified(100);k__Fungi_unclassified(100); | - | - | - | - | - | - | - | Unassigned | - |
| Otu21163 | 1 | k__Fungi(100);k__Fungi_unclassified(100);k__Fungi_unclassified(100);k__Fungi_unclassified(100);k__Fungi_unclassified(100);k__Fungi_unclassified(100);k__Fungi_unclassified(100); | - | - | - | - | - | - | - | Unassigned | - |
| Otu21164 | 1 | k__Fungi(100);k__Fungi_unclassified(100);k__Fungi_unclassified(100);k__Fungi_unclassified(100);k__Fungi_unclassified(100);k__Fungi_unclassified(100);k__Fungi_unclassified(100); | - | - | - | - | - | - | - | Unassigned | - |
| Otu21165 | 1 | k__Fungi(100);k__Fungi_unclassified(100);k__Fungi_unclassified(100);k__Fungi_unclassified(100);k__Fungi_unclassified(100);k__Fungi_unclassified(100);k__Fungi_unclassified(100); | - | - | - | - | - | - | - | Unassigned | - |
| Otu21166 | 1 | k__Fungi(100);k__Fungi_unclassified(100);k__Fungi_unclassified(100);k__Fungi_unclassified(100);k__Fungi_unclassified(100);k__Fungi_unclassified(100);k__Fungi_unclassified(100); | - | - | - | - | - | - | - | Unassigned | - |
| Otu21169 | 1 | k__Fungi(100);p__Ascomycota(100);p__Ascomycota_unclassified(100);p__Ascomycota_unclassified(100);p__Ascomycota_unclassified(100);p__Ascomycota_unclassified(100);p__Ascomycota_unclassified(100); | - | - | - | - | - | - | - | Unassigned | - |
| Otu21177 | 1 | k__Fungi(100);k__Fungi_unclassified(100);k__Fungi_unclassified(100);k__Fungi_unclassified(100);k__Fungi_unclassified(100);k__Fungi_unclassified(100);k__Fungi_unclassified(100); | - | - | - | - | - | - | - | Unassigned | - |
| Otu21182 | 1 | k__Fungi(100);k__Fungi_unclassified(100);k__Fungi_unclassified(100);k__Fungi_unclassified(100);k__Fungi_unclassified(100);k__Fungi_unclassified(100);k__Fungi_unclassified(100); | - | - | - | - | - | - | - | Unassigned | - |
| Otu21183 | 1 | k__Fungi(100);k__Fungi_unclassified(100);k__Fungi_unclassified(100);k__Fungi_unclassified(100);k__Fungi_unclassified(100);k__Fungi_unclassified(100);k__Fungi_unclassified(100); | - | - | - | - | - | - | - | Unassigned | - |
| Otu21191 | 1 | k__Fungi(100);k__Fungi_unclassified(100);k__Fungi_unclassified(100);k__Fungi_unclassified(100);k__Fungi_unclassified(100);k__Fungi_unclassified(100);k__Fungi_unclassified(100); | - | - | - | - | - | - | - | Unassigned | - |
| Otu21192 | 1 | k__Fungi(100);p__Ascomycota(100);p__Ascomycota_unclassified(100);p__Ascomycota_unclassified(100);p__Ascomycota_unclassified(100);p__Ascomycota_unclassified(100);p__Ascomycota_unclassified(100); | - | - | - | - | - | - | - | Unassigned | - |
| Otu21196 | 1 | k__Fungi(100);k__Fungi_unclassified(100);k__Fungi_unclassified(100);k__Fungi_unclassified(100);k__Fungi_unclassified(100);k__Fungi_unclassified(100);k__Fungi_unclassified(100); | - | - | - | - | - | - | - | Unassigned | - |
| Otu21199 | 1 | k__Fungi(100);k__Fungi_unclassified(100);k__Fungi_unclassified(100);k__Fungi_unclassified(100);k__Fungi_unclassified(100);k__Fungi_unclassified(100);k__Fungi_unclassified(100); | - | - | - | - | - | - | - | Unassigned | - |
| Otu21200 | 1 | k__Fungi(100);k__Fungi_unclassified(100);k__Fungi_unclassified(100);k__Fungi_unclassified(100);k__Fungi_unclassified(100);k__Fungi_unclassified(100);k__Fungi_unclassified(100); | - | - | - | - | - | - | - | Unassigned | - |
| Otu21201 | 1 | k__Fungi(100);k__Fungi_unclassified(100);k__Fungi_unclassified(100);k__Fungi_unclassified(100);k__Fungi_unclassified(100);k__Fungi_unclassified(100);k__Fungi_unclassified(100); | - | - | - | - | - | - | - | Unassigned | - |
| Otu21202 | 1 | k__Fungi(100);k__Fungi_unclassified(100);k__Fungi_unclassified(100);k__Fungi_unclassified(100);k__Fungi_unclassified(100);k__Fungi_unclassified(100);k__Fungi_unclassified(100); | - | - | - | - | - | - | - | Unassigned | - |
| Otu21207 | 1 | k__Fungi(100);k__Fungi_unclassified(100);k__Fungi_unclassified(100);k__Fungi_unclassified(100);k__Fungi_unclassified(100);k__Fungi_unclassified(100);k__Fungi_unclassified(100); | - | - | - | - | - | - | - | Unassigned | - |

| | | | | | | | | | | | |
|---|---|---|---|---|---|---|---|---|---|---|---|
| Otu21214 | 1 | k__Fungi(100);k__Fungi_unclassified(100);k__Fungi_unclassified(100);k__Fungi_unclassified(100);k__Fungi_unclassified(100);k__Fungi_unclassified(100);k__Fungi_unclassified(100); | - | - | - | - | - | - | - | Unassigned | - |
| Otu21227 | 1 | k__Fungi(100);p__Ascomycota(100);p__Ascomycota_unclassified(100);p__Ascomycota_unclassified(100);p__Ascomycota_unclassified(100);p__Ascomycota_unclassified(100);p__Ascomycota_unclassified(100); | - | - | - | - | - | - | - | Unassigned | - |
| Otu21242 | 1 | k__Fungi(100);k__Fungi_unclassified(100);k__Fungi_unclassified(100);k__Fungi_unclassified(100);k__Fungi_unclassified(100);k__Fungi_unclassified(100);k__Fungi_unclassified(100); | - | - | - | - | - | - | - | Unassigned | - |
| Otu21244 | 1 | k__Fungi(100);p__Ascomycota(100);p__Ascomycota_unclassified(100);p__Ascomycota_unclassified(100);p__Ascomycota_unclassified(100);p__Ascomycota_unclassified(100);p__Ascomycota_unclassified(100); | - | - | - | - | - | - | - | Unassigned | - |
| Otu21250 | 1 | k__Fungi(100);p__Ascomycota(100);p__Ascomycota_unclassified(100);p__Ascomycota_unclassified(100);p__Ascomycota_unclassified(100);p__Ascomycota_unclassified(100);p__Ascomycota_unclassified(100); | - | - | - | - | - | - | - | Unassigned | - |
| Otu21270 | 1 | k__Fungi(100);p__Ascomycota(100);p__Ascomycota_unclassified(100);p__Ascomycota_unclassified(100);p__Ascomycota_unclassified(100);p__Ascomycota_unclassified(100);p__Ascomycota_unclassified(100); | - | - | - | - | - | - | - | Unassigned | - |
| Otu21274 | 1 | k__Fungi(100);k__Fungi_unclassified(100);k__Fungi_unclassified(100);k__Fungi_unclassified(100);k__Fungi_unclassified(100);k__Fungi_unclassified(100);k__Fungi_unclassified(100); | - | - | - | - | - | - | - | Unassigned | - |
| Otu21275 | 1 | k__Fungi(100);k__Fungi_unclassified(100);k__Fungi_unclassified(100);k__Fungi_unclassified(100);k__Fungi_unclassified(100);k__Fungi_unclassified(100);k__Fungi_unclassified(100); | - | - | - | - | - | - | - | Unassigned | - |
| Otu21276 | 1 | k__Fungi(100);k__Fungi_unclassified(100);k__Fungi_unclassified(100);k__Fungi_unclassified(100);k__Fungi_unclassified(100);k__Fungi_unclassified(100);k__Fungi_unclassified(100); | - | - | - | - | - | - | - | Unassigned | - |
| Otu21278 | 1 | k__Fungi(100);k__Fungi_unclassified(100);k__Fungi_unclassified(100);k__Fungi_unclassified(100);k__Fungi_unclassified(100);k__Fungi_unclassified(100);k__Fungi_unclassified(100); | - | - | - | - | - | - | - | Unassigned | - |
| Otu21279 | 1 | k__Fungi(100);k__Fungi_unclassified(100);k__Fungi_unclassified(100);k__Fungi_unclassified(100);k__Fungi_unclassified(100);k__Fungi_unclassified(100);k__Fungi_unclassified(100); | - | - | - | - | - | - | - | Unassigned | - |
| Otu21281 | 1 | k__Fungi(100);p__Ascomycota(100);p__Ascomycota_unclassified(100);p__Ascomycota_unclassified(100);p__Ascomycota_unclassified(100);p__Ascomycota_unclassified(100);p__Ascomycota_unclassified(100); | - | - | - | - | - | - | - | Unassigned | - |
| Otu21284 | 1 | k__Fungi(100);k__Fungi_unclassified(100);k__Fungi_unclassified(100);k__Fungi_unclassified(100);k__Fungi_unclassified(100);k__Fungi_unclassified(100);k__Fungi_unclassified(100); | - | - | - | - | - | - | - | Unassigned | - |
| Otu21320 | 1 | k__Fungi(100);p__Ascomycota(100);p__Ascomycota_unclassified(100);p__Ascomycota_unclassified(100);p__Ascomycota_unclassified(100);p__Ascomycota_unclassified(100);p__Ascomycota_unclassified(100); | - | - | - | - | - | - | - | Unassigned | - |
| Otu21322 | 1 | k__Fungi(100);k__Fungi_unclassified(100);k__Fungi_unclassified(100);k__Fungi_unclassified(100);k__Fungi_unclassified(100);k__Fungi_unclassified(100);k__Fungi_unclassified(100); | - | - | - | - | - | - | - | Unassigned | - |
| Otu21326 | 1 | k__Fungi(100);p__Ascomycota(100);p__Ascomycota_unclassified(100);p__Ascomycota_unclassified(100);p__Ascomycota_unclassified(100);p__Ascomycota_unclassified(100);p__Ascomycota_unclassified(100); | - | - | - | - | - | - | - | Unassigned | - |
| Otu21339 | 1 | k__Fungi(100);k__Fungi_unclassified(100);k__Fungi_unclassified(100);k__Fungi_unclassified(100);k__Fungi_unclassified(100);k__Fungi_unclassified(100);k__Fungi_unclassified(100); | - | - | - | - | - | - | - | Unassigned | - |
| Otu21340 | 1 | k__Fungi(100);k__Fungi_unclassified(100);k__Fungi_unclassified(100);k__Fungi_unclassified(100);k__Fungi_unclassified(100);k__Fungi_unclassified(100);k__Fungi_unclassified(100); | - | - | - | - | - | - | - | Unassigned | - |
| Otu21342 | 1 | k__Fungi(100);k__Fungi_unclassified(100);k__Fungi_unclassified(100);k__Fungi_unclassified(100);k__Fungi_unclassified(100);k__Fungi_unclassified(100);k__Fungi_unclassified(100); | - | - | - | - | - | - | - | Unassigned | - |
| Otu21359 | 1 | k__Fungi(100);k__Fungi_unclassified(100);k__Fungi_unclassified(100);k__Fungi_unclassified(100);k__Fungi_unclassified(100);k__Fungi_unclassified(100);k__Fungi_unclassified(100); | - | - | - | - | - | - | - | Unassigned | - |
| Otu21360 | 1 | k__Fungi(100);p__Ascomycota(100);p__Ascomycota_unclassified(100);p__Ascomycota_unclassified(100);p__Ascomycota_unclassified(100);p__Ascomycota_unclassified(100);p__Ascomycota_unclassified(100); | - | - | - | - | - | - | - | Unassigned | - |
| Otu21362 | 1 | k__Fungi(100);k__Fungi_unclassified(100);k__Fungi_unclassified(100);k__Fungi_unclassified(100);k__Fungi_unclassified(100);k__Fungi_unclassified(100);k__Fungi_unclassified(100); | - | - | - | - | - | - | - | Unassigned | - |
| Otu21371 | 1 | k__Fungi(100);p__Ascomycota(100);p__Ascomycota_unclassified(100);p__Ascomycota_unclassified(100);p__Ascomycota_unclassified(100);p__Ascomycota_unclassified(100);p__Ascomycota_unclassified(100); | - | - | - | - | - | - | - | Unassigned | - |
| Otu21385 | 1 | k__Fungi(100);k__Fungi_unclassified(100);k__Fungi_unclassified(100);k__Fungi_unclassified(100);k__Fungi_unclassified(100);k__Fungi_unclassified(100);k__Fungi_unclassified(100); | - | - | - | - | - | - | - | Unassigned | - |
| Otu21386 | 1 | k__Fungi(100);k__Fungi_unclassified(100);k__Fungi_unclassified(100);k__Fungi_unclassified(100);k__Fungi_unclassified(100);k__Fungi_unclassified(100);k__Fungi_unclassified(100); | - | - | - | - | - | - | - | Unassigned | - |
| Otu21389 | 1 | k__Fungi(100);p__Ascomycota(100);p__Ascomycota_unclassified(100);p__Ascomycota_unclassified(100);p__Ascomycota_unclassified(100);p__Ascomycota_unclassified(100);p__Ascomycota_unclassified(100); | - | - | - | - | - | - | - | Unassigned | - |
| Otu21391 | 1 | k__Fungi(100);p__Ascomycota(100);p__Ascomycota_unclassified(100);p__Ascomycota_unclassified(100);p__Ascomycota_unclassified(100);p__Ascomycota_unclassified(100);p__Ascomycota_unclassified(100); | - | - | - | - | - | - | - | Unassigned | - |
| Otu21394 | 1 | k__Fungi(100);k__Fungi_unclassified(100);k__Fungi_unclassified(100);k__Fungi_unclassified(100);k__Fungi_unclassified(100);k__Fungi_unclassified(100);k__Fungi_unclassified(100); | - | - | - | - | - | - | - | Unassigned | - |
| Otu21395 | 1 | k__Fungi(100);k__Fungi_unclassified(100);k__Fungi_unclassified(100);k__Fungi_unclassified(100);k__Fungi_unclassified(100);k__Fungi_unclassified(100);k__Fungi_unclassified(100); | - | - | - | - | - | - | - | Unassigned | - |
| Otu21396 | 1 | k__Fungi(100);k__Fungi_unclassified(100);k__Fungi_unclassified(100);k__Fungi_unclassified(100);k__Fungi_unclassified(100);k__Fungi_unclassified(100);k__Fungi_unclassified(100); | - | - | - | - | - | - | - | Unassigned | - |
| Otu21402 | 1 | k__Fungi(100);p__Ascomycota(100);p__Ascomycota_unclassified(100);p__Ascomycota_unclassified(100);p__Ascomycota_unclassified(100);p__Ascomycota_unclassified(100);p__Ascomycota_unclassified(100); | - | - | - | - | - | - | - | Unassigned | - |
| Otu21404 | 1 | k__Fungi(100);k__Fungi_unclassified(100);k__Fungi_unclassified(100);k__Fungi_unclassified(100);k__Fungi_unclassified(100);k__Fungi_unclassified(100);k__Fungi_unclassified(100); | - | - | - | - | - | - | - | Unassigned | - |
| Otu21405 | 1 | k__Fungi(100);k__Fungi_unclassified(100);k__Fungi_unclassified(100);k__Fungi_unclassified(100);k__Fungi_unclassified(100);k__Fungi_unclassified(100);k__Fungi_unclassified(100); | - | - | - | - | - | - | - | Unassigned | - |
| Otu21406 | 1 | k__Fungi(100);k__Fungi_unclassified(100);k__Fungi_unclassified(100);k__Fungi_unclassified(100);k__Fungi_unclassified(100);k__Fungi_unclassified(100);k__Fungi_unclassified(100); | - | - | - | - | - | - | - | Unassigned | - |
| Otu21409 | 1 | k__Fungi(100);k__Fungi_unclassified(100);k__Fungi_unclassified(100);k__Fungi_unclassified(100);k__Fungi_unclassified(100);k__Fungi_unclassified(100);k__Fungi_unclassified(100); | - | - | - | - | - | - | - | Unassigned | - |
| Otu21410 | 1 | k__Fungi(100);k__Fungi_unclassified(100);k__Fungi_unclassified(100);k__Fungi_unclassified(100);k__Fungi_unclassified(100);k__Fungi_unclassified(100);k__Fungi_unclassified(100); | - | - | - | - | - | - | - | Unassigned | - |
| Otu21420 | 1 | k__Fungi(100);k__Fungi_unclassified(100);k__Fungi_unclassified(100);k__Fungi_unclassified(100);k__Fungi_unclassified(100);k__Fungi_unclassified(100);k__Fungi_unclassified(100); | - | - | - | - | - | - | - | Unassigned | - |
| Otu21429 | 1 | k__Fungi(100);p__Ascomycota(100);p__Ascomycota_unclassified(100);p__Ascomycota_unclassified(100);p__Ascomycota_unclassified(100);p__Ascomycota_unclassified(100);p__Ascomycota_unclassified(100); | - | - | - | - | - | - | - | Unassigned | - |
| Otu21434 | 1 | k__Fungi(100);k__Fungi_unclassified(100);k__Fungi_unclassified(100);k__Fungi_unclassified(100);k__Fungi_unclassified(100);k__Fungi_unclassified(100);k__Fungi_unclassified(100); | - | - | - | - | - | - | - | Unassigned | - |
| Otu21435 | 1 | k__Fungi(100);k__Fungi_unclassified(100);k__Fungi_unclassified(100);k__Fungi_unclassified(100);k__Fungi_unclassified(100);k__Fungi_unclassified(100);k__Fungi_unclassified(100); | - | - | - | - | - | - | - | Unassigned | - |
| Otu21436 | 1 | k__Fungi(100);k__Fungi_unclassified(100);k__Fungi_unclassified(100);k__Fungi_unclassified(100);k__Fungi_unclassified(100);k__Fungi_unclassified(100);k__Fungi_unclassified(100); | - | - | - | - | - | - | - | Unassigned | - |
| Otu21448 | 1 | k__Fungi(100);k__Fungi_unclassified(100);k__Fungi_unclassified(100);k__Fungi_unclassified(100);k__Fungi_unclassified(100);k__Fungi_unclassified(100);k__Fungi_unclassified(100); | - | - | - | - | - | - | - | Unassigned | - |
| Otu21451 | 1 | k__Fungi(100);p__Ascomycota(100);p__Ascomycota_unclassified(100);p__Ascomycota_unclassified(100);p__Ascomycota_unclassified(100);p__Ascomycota_unclassified(100);p__Ascomycota_unclassified(100); | - | - | - | - | - | - | - | Unassigned | - |
| Otu21462 | 1 | k__Fungi(100);k__Fungi_unclassified(100);k__Fungi_unclassified(100);k__Fungi_unclassified(100);k__Fungi_unclassified(100);k__Fungi_unclassified(100);k__Fungi_unclassified(100); | - | - | - | - | - | - | - | Unassigned | - |
| Otu21463 | 1 | k__Fungi(100);k__Fungi_unclassified(100);k__Fungi_unclassified(100);k__Fungi_unclassified(100);k__Fungi_unclassified(100);k__Fungi_unclassified(100);k__Fungi_unclassified(100); | - | - | - | - | - | - | - | Unassigned | - |
| Otu21464 | 1 | k__Fungi(100);p__Ascomycota(100);p__Ascomycota_unclassified(100);p__Ascomycota_unclassified(100);p__Ascomycota_unclassified(100);p__Ascomycota_unclassified(100);p__Ascomycota_unclassified(100); | - | - | - | - | - | - | - | Unassigned | - |
| Otu21468 | 1 | k__Fungi(100);k__Fungi_unclassified(100);k__Fungi_unclassified(100);k__Fungi_unclassified(100);k__Fungi_unclassified(100);k__Fungi_unclassified(100);k__Fungi_unclassified(100); | - | - | - | - | - | - | - | Unassigned | - |
| Otu21469 | 1 | k__Fungi(100);p__Ascomycota(100);p__Ascomycota_unclassified(100);p__Ascomycota_unclassified(100);p__Ascomycota_unclassified(100);p__Ascomycota_unclassified(100);p__Ascomycota_unclassified(100); | - | - | - | - | - | - | - | Unassigned | - |
| Otu21470 | 1 | k__Fungi(100);k__Fungi_unclassified(100);k__Fungi_unclassified(100);k__Fungi_unclassified(100);k__Fungi_unclassified(100);k__Fungi_unclassified(100);k__Fungi_unclassified(100); | - | - | - | - | - | - | - | Unassigned | - |
| Otu21471 | 1 | k__Fungi(100);p__Ascomycota(100);p__Ascomycota_unclassified(100);p__Ascomycota_unclassified(100);p__Ascomycota_unclassified(100);p__Ascomycota_unclassified(100);p__Ascomycota_unclassified(100); | - | - | - | - | - | - | - | Unassigned | - |
| Otu21472 | 1 | k__Fungi(100);k__Fungi_unclassified(100);k__Fungi_unclassified(100);k__Fungi_unclassified(100);k__Fungi_unclassified(100);k__Fungi_unclassified(100);k__Fungi_unclassified(100); | - | - | - | - | - | - | - | Unassigned | - |
| Otu21474 | 1 | k__Fungi(100);p__Ascomycota(100);p__Ascomycota_unclassified(100);p__Ascomycota_unclassified(100);p__Ascomycota_unclassified(100);p__Ascomycota_unclassified(100);p__Ascomycota_unclassified(100); | - | - | - | - | - | - | - | Unassigned | - |
| Otu21494 | 1 | k__Fungi(100);k__Fungi_unclassified(100);k__Fungi_unclassified(100);k__Fungi_unclassified(100);k__Fungi_unclassified(100);k__Fungi_unclassified(100);k__Fungi_unclassified(100); | - | - | - | - | - | - | - | Unassigned | - |
| Otu21495 | 1 | k__Fungi(100);k__Fungi_unclassified(100);k__Fungi_unclassified(100);k__Fungi_unclassified(100);k__Fungi_unclassified(100);k__Fungi_unclassified(100);k__Fungi_unclassified(100); | - | - | - | - | - | - | - | Unassigned | - |
| Otu21496 | 1 | k__Fungi(100);k__Fungi_unclassified(100);k__Fungi_unclassified(100);k__Fungi_unclassified(100);k__Fungi_unclassified(100);k__Fungi_unclassified(100);k__Fungi_unclassified(100); | - | - | - | - | - | - | - | Unassigned | - |
| Otu21502 | 1 | k__Fungi(100);k__Fungi_unclassified(100);k__Fungi_unclassified(100);k__Fungi_unclassified(100);k__Fungi_unclassified(100);k__Fungi_unclassified(100);k__Fungi_unclassified(100); | - | - | - | - | - | - | - | Unassigned | - |
| Otu21507 | 1 | k__Fungi(100);k__Fungi_unclassified(100);k__Fungi_unclassified(100);k__Fungi_unclassified(100);k__Fungi_unclassified(100);k__Fungi_unclassified(100);k__Fungi_unclassified(100); | - | - | - | - | - | - | - | Unassigned | - |
| Otu21508 | 1 | k__Fungi(100);p__Ascomycota(100);p__Ascomycota_unclassified(100);p__Ascomycota_unclassified(100);p__Ascomycota_unclassified(100);p__Ascomycota_unclassified(100);p__Ascomycota_unclassified(100); | - | - | - | - | - | - | - | Unassigned | - |
| Otu21509 | 1 | k__Fungi(100);p__Ascomycota(100);p__Ascomycota_unclassified(100);p__Ascomycota_unclassified(100);p__Ascomycota_unclassified(100);p__Ascomycota_unclassified(100);p__Ascomycota_unclassified(100); | - | - | - | - | - | - | - | Unassigned | - |
| Otu21515 | 1 | k__Fungi(100);k__Fungi_unclassified(100);k__Fungi_unclassified(100);k__Fungi_unclassified(100);k__Fungi_unclassified(100);k__Fungi_unclassified(100);k__Fungi_unclassified(100); | - | - | - | - | - | - | - | Unassigned | - |
| Otu21518 | 1 | k__Fungi(100);k__Fungi_unclassified(100);k__Fungi_unclassified(100);k__Fungi_unclassified(100);k__Fungi_unclassified(100);k__Fungi_unclassified(100);k__Fungi_unclassified(100); | - | - | - | - | - | - | - | Unassigned | - |
| Otu21520 | 1 | k__Fungi(100);p__Ascomycota(100);p__Ascomycota_unclassified(100);p__Ascomycota_unclassified(100);p__Ascomycota_unclassified(100);p__Ascomycota_unclassified(100);p__Ascomycota_unclassified(100); | - | - | - | - | - | - | - | Unassigned | - |
| Otu21523 | 1 | k__Fungi(100);k__Fungi_unclassified(100);k__Fungi_unclassified(100);k__Fungi_unclassified(100);k__Fungi_unclassified(100);k__Fungi_unclassified(100);k__Fungi_unclassified(100); | - | - | - | - | - | - | - | Unassigned | - |
| Otu21524 | 1 | k__Fungi(100);p__Ascomycota(100);p__Ascomycota_unclassified(100);p__Ascomycota_unclassified(100);p__Ascomycota_unclassified(100);p__Ascomycota_unclassified(100);p__Ascomycota_unclassified(100); | - | - | - | - | - | - | - | Unassigned | - |
| Otu21525 | 1 | k__Fungi(100);k__Fungi_unclassified(100);k__Fungi_unclassified(100);k__Fungi_unclassified(100);k__Fungi_unclassified(100);k__Fungi_unclassified(100);k__Fungi_unclassified(100); | - | - | - | - | - | - | - | Unassigned | - |
| Otu21526 | 1 | k__Fungi(100);p__Ascomycota(100);p__Ascomycota_unclassified(100);p__Ascomycota_unclassified(100);p__Ascomycota_unclassified(100);p__Ascomycota_unclassified(100);p__Ascomycota_unclassified(100); | - | - | - | - | - | - | - | Unassigned | - |
| Otu21536 | 1 | k__Fungi(100);k__Fungi_unclassified(100);k__Fungi_unclassified(100);k__Fungi_unclassified(100);k__Fungi_unclassified(100);k__Fungi_unclassified(100);k__Fungi_unclassified(100); | - | - | - | - | - | - | - | Unassigned | - |
| Otu21539 | 1 | k__Fungi(100);p__Ascomycota(100);p__Ascomycota_unclassified(100);p__Ascomycota_unclassified(100);p__Ascomycota_unclassified(100);p__Ascomycota_unclassified(100);p__Ascomycota_unclassified(100); | - | - | - | - | - | - | - | Unassigned | - |
| Otu21545 | 1 | k__Fungi(100);k__Fungi_unclassified(100);k__Fungi_unclassified(100);k__Fungi_unclassified(100);k__Fungi_unclassified(100);k__Fungi_unclassified(100);k__Fungi_unclassified(100); | - | - | - | - | - | - | - | Unassigned | - |
| Otu21547 | 1 | k__Fungi(100);k__Fungi_unclassified(100);k__Fungi_unclassified(100);k__Fungi_unclassified(100);k__Fungi_unclassified(100);k__Fungi_unclassified(100);k__Fungi_unclassified(100); | - | - | - | - | - | - | - | Unassigned | - |
| Otu21556 | 1 | k__Fungi(100);k__Fungi_unclassified(100);k__Fungi_unclassified(100);k__Fungi_unclassified(100);k__Fungi_unclassified(100);k__Fungi_unclassified(100);k__Fungi_unclassified(100); | - | - | - | - | - | - | - | Unassigned | - |
| Otu21578 | 1 | k__Fungi(100);k__Fungi_unclassified(100);k__Fungi_unclassified(100);k__Fungi_unclassified(100);k__Fungi_unclassified(100);k__Fungi_unclassified(100);k__Fungi_unclassified(100); | - | - | - | - | - | - | - | Unassigned | - |
| Otu21591 | 1 | k__Fungi(100);p__Ascomycota(100);p__Ascomycota_unclassified(100);p__Ascomycota_unclassified(100);p__Ascomycota_unclassified(100);p__Ascomycota_unclassified(100);p__Ascomycota_unclassified(100); | - | - | - | - | - | - | - | Unassigned | - |
| Otu21592 | 1 | k__Fungi(100);p__Ascomycota(100);p__Ascomycota_unclassified(100);p__Ascomycota_unclassified(100);p__Ascomycota_unclassified(100);p__Ascomycota_unclassified(100);p__Ascomycota_unclassified(100); | - | - | - | - | - | - | - | Unassigned | - |
| Otu21593 | 1 | k__Fungi(100);p__Ascomycota(100);p__Ascomycota_unclassified(100);p__Ascomycota_unclassified(100);p__Ascomycota_unclassified(100);p__Ascomycota_unclassified(100);p__Ascomycota_unclassified(100); | - | - | - | - | - | - | - | Unassigned | - |
| Otu21594 | 1 | k__Fungi(100);k__Fungi_unclassified(100);k__Fungi_unclassified(100);k__Fungi_unclassified(100);k__Fungi_unclassified(100);k__Fungi_unclassified(100);k__Fungi_unclassified(100); | - | - | - | - | - | - | - | Unassigned | - |
| Otu21595 | 1 | k__Fungi(100);p__Ascomycota(100);p__Ascomycota_unclassified(100);p__Ascomycota_unclassified(100);p__Ascomycota_unclassified(100);p__Ascomycota_unclassified(100);p__Ascomycota_unclassified(100); | - | - | - | - | - | - | - | Unassigned | - |
| Otu21596 | 1 | k__Fungi(100);p__Ascomycota(100);p__Ascomycota_unclassified(100);p__Ascomycota_unclassified(100);p__Ascomycota_unclassified(100);p__Ascomycota_unclassified(100);p__Ascomycota_unclassified(100); | - | - | - | - | - | - | - | Unassigned | - |
| Otu21597 | 1 | k__Fungi(100);p__Ascomycota(100);p__Ascomycota_unclassified(100);p__Ascomycota_unclassified(100);p__Ascomycota_unclassified(100);p__Ascomycota_unclassified(100);p__Ascomycota_unclassified(100); | - | - | - | - | - | - | - | Unassigned | - |
| Otu21598 | 1 | k__Fungi(100);p__Ascomycota(100);p__Ascomycota_unclassified(100);p__Ascomycota_unclassified(100);p__Ascomycota_unclassified(100);p__Ascomycota_unclassified(100);p__Ascomycota_unclassified(100); | - | - | - | - | - | - | - | Unassigned | - |
| Otu21600 | 1 | k__Fungi(100);k__Fungi_unclassified(100);k__Fungi_unclassified(100);k__Fungi_unclassified(100);k__Fungi_unclassified(100);k__Fungi_unclassified(100);k__Fungi_unclassified(100); | - | - | - | - | - | - | - | Unassigned | - |
| Otu21605 | 1 | k__Fungi(100);p__Ascomycota(100);p__Ascomycota_unclassified(100);p__Ascomycota_unclassified(100);p__Ascomycota_unclassified(100);p__Ascomycota_unclassified(100);p__Ascomycota_unclassified(100); | - | - | - | - | - | - | - | Unassigned | - |
| Otu21606 | 1 | k__Fungi(100);k__Fungi_unclassified(100);k__Fungi_unclassified(100);k__Fungi_unclassified(100);k__Fungi_unclassified(100);k__Fungi_unclassified(100);k__Fungi_unclassified(100); | - | - | - | - | - | - | - | Unassigned | - |
| Otu21607 | 1 | k__Fungi(100);p__Ascomycota(100);p__Ascomycota_unclassified(100);p__Ascomycota_unclassified(100);p__Ascomycota_unclassified(100);p__Ascomycota_unclassified(100);p__Ascomycota_unclassified(100); | - | - | - | - | - | - | - | Unassigned | - |
| Otu21608 | 1 | k__Fungi(100);p__Ascomycota(100);p__Ascomycota_unclassified(100);p__Ascomycota_unclassified(100);p__Ascomycota_unclassified(100);p__Ascomycota_unclassified(100);p__Ascomycota_unclassified(100); | - | - | - | - | - | - | - | Unassigned | - |
| Otu21609 | 1 | k__Fungi(100);p__Ascomycota(100);p__Ascomycota_unclassified(100);p__Ascomycota_unclassified(100);p__Ascomycota_unclassified(100);p__Ascomycota_unclassified(100);p__Ascomycota_unclassified(100); | - | - | - | - | - | - | - | Unassigned | - |
| Otu21615 | 1 | k__Fungi(100);k__Fungi_unclassified(100);k__Fungi_unclassified(100);k__Fungi_unclassified(100);k__Fungi_unclassified(100);k__Fungi_unclassified(100);k__Fungi_unclassified(100); | - | - | - | - | - | - | - | Unassigned | - |
| Otu21648 | 1 | k__Fungi(100);k__Fungi_unclassified(100);k__Fungi_unclassified(100);k__Fungi_unclassified(100);k__Fungi_unclassified(100);k__Fungi_unclassified(100);k__Fungi_unclassified(100); | - | - | - | - | - | - | - | Unassigned | - |
| Otu21670 | 1 | k__Fungi(100);p__Ascomycota(100);p__Ascomycota_unclassified(100);p__Ascomycota_unclassified(100);p__Ascomycota_unclassified(100);p__Ascomycota_unclassified(100);p__Ascomycota_unclassified(100); | - | - | - | - | - | - | - | Unassigned | - |
| Otu21672 | 1 | k__Fungi(100);k__Fungi_unclassified(100);k__Fungi_unclassified(100);k__Fungi_unclassified(100);k__Fungi_unclassified(100);k__Fungi_unclassified(100);k__Fungi_unclassified(100); | - | - | - | - | - | - | - | Unassigned | - |
| Otu21677 | 1 | k__Fungi(100);k__Fungi_unclassified(100);k__Fungi_unclassified(100);k__Fungi_unclassified(100);k__Fungi_unclassified(100);k__Fungi_unclassified(100);k__Fungi_unclassified(100); | - | - | - | - | - | - | - | Unassigned | - |
| Otu21679 | 1 | k__Fungi(100);p__Ascomycota(100);p__Ascomycota_unclassified(100);p__Ascomycota_unclassified(100);p__Ascomycota_unclassified(100);p__Ascomycota_unclassified(100);p__Ascomycota_unclassified(100); | - | - | - | - | - | - | - | Unassigned | - |
| Otu21682 | 1 | k__Fungi(100);p__Ascomycota(100);p__Ascomycota_unclassified(100);p__Ascomycota_unclassified(100);p__Ascomycota_unclassified(100);p__Ascomycota_unclassified(100);p__Ascomycota_unclassified(100); | - | - | - | - | - | - | - | Unassigned | - |
| Otu21695 | 1 | k__Fungi(100);k__Fungi_unclassified(100);k__Fungi_unclassified(100);k__Fungi_unclassified(100);k__Fungi_unclassified(100);k__Fungi_unclassified(100);k__Fungi_unclassified(100); | - | - | - | - | - | - | - | Unassigned | - |
| Otu21705 | 1 | k__Fungi(100);k__Fungi_unclassified(100);k__Fungi_unclassified(100);k__Fungi_unclassified(100);k__Fungi_unclassified(100);k__Fungi_unclassified(100);k__Fungi_unclassified(100); | - | - | - | - | - | - | - | Unassigned | - |
| Otu21706 | 1 | k__Fungi(100);p__Ascomycota(100);p__Ascomycota_unclassified(100);p__Ascomycota_unclassified(100);p__Ascomycota_unclassified(100);p__Ascomycota_unclassified(100);p__Ascomycota_unclassified(100); | - | - | - | - | - | - | - | Unassigned | - |
| Otu21710 | 1 | k__Fungi(100);k__Fungi_unclassified(100);k__Fungi_unclassified(100);k__Fungi_unclassified(100);k__Fungi_unclassified(100);k__Fungi_unclassified(100);k__Fungi_unclassified(100); | - | - | - | - | - | - | - | Unassigned | - |
| Otu21711 | 1 | k__Fungi(100);p__Ascomycota(100);p__Ascomycota_unclassified(100);p__Ascomycota_unclassified(100);p__Ascomycota_unclassified(100);p__Ascomycota_unclassified(100);p__Ascomycota_unclassified(100); | - | - | - | - | - | - | - | Unassigned | - |
| Otu21712 | 1 | k__Fungi(100);k__Fungi_unclassified(100);k__Fungi_unclassified(100);k__Fungi_unclassified(100);k__Fungi_unclassified(100);k__Fungi_unclassified(100);k__Fungi_unclassified(100); | - | - | - | - | - | - | - | Unassigned | - |
| Otu21713 | 1 | k__Fungi(100);k__Fungi_unclassified(100);k__Fungi_unclassified(100);k__Fungi_unclassified(100);k__Fungi_unclassified(100);k__Fungi_unclassified(100);k__Fungi_unclassified(100); | - | - | - | - | - | - | - | Unassigned | - |
| Otu21720 | 1 | k__Fungi(100);k__Fungi_unclassified(100);k__Fungi_unclassified(100);k__Fungi_unclassified(100);k__Fungi_unclassified(100);k__Fungi_unclassified(100);k__Fungi_unclassified(100); | - | - | - | - | - | - | - | Unassigned | - |
| Otu21722 | 1 | k__Fungi(100);k__Fungi_unclassified(100);k__Fungi_unclassified(100);k__Fungi_unclassified(100);k__Fungi_unclassified(100);k__Fungi_unclassified(100);k__Fungi_unclassified(100); | - | - | - | - | - | - | - | Unassigned | - |
| Otu21723 | 1 | k__Fungi(100);p__Ascomycota(100);p__Ascomycota_unclassified(100);p__Ascomycota_unclassified(100);p__Ascomycota_unclassified(100);p__Ascomycota_unclassified(100);p__Ascomycota_unclassified(100); | - | - | - | - | - | - | - | Unassigned | - |
| Otu21731 | 1 | k__Fungi(100);k__Fungi_unclassified(100);k__Fungi_unclassified(100);k__Fungi_unclassified(100);k__Fungi_unclassified(100);k__Fungi_unclassified(100);k__Fungi_unclassified(100); | - | - | - | - | - | - | - | Unassigned | - |
| Otu21733 | 1 | k__Fungi(100);p__Ascomycota(100);c__Sordariomycetes(100);c__Sordariomycetes_unclassified(100);c__Sordariomycetes_unclassified(100);c__Sordariomycetes_unclassified(100);c__Sordariomycetes_unclassified(100); | - | - | - | - | - | - | - | Unassigned | - |
| Otu21740 | 1 | k__Fungi(100);k__Fungi_unclassified(100);k__Fungi_unclassified(100);k__Fungi_unclassified(100);k__Fungi_unclassified(100);k__Fungi_unclassified(100);k__Fungi_unclassified(100); | - | - | - | - | - | - | - | Unassigned | - |
| Otu21749 | 1 | k__Fungi(100);k__Fungi_unclassified(100);k__Fungi_unclassified(100);k__Fungi_unclassified(100);k__Fungi_unclassified(100);k__Fungi_unclassified(100);k__Fungi_unclassified(100); | - | - | - | - | - | - | - | Unassigned | - |
| Otu21750 | 1 | k__Fungi(100);k__Fungi_unclassified(100);k__Fungi_unclassified(100);k__Fungi_unclassified(100);k__Fungi_unclassified(100);k__Fungi_unclassified(100);k__Fungi_unclassified(100); | - | - | - | - | - | - | - | Unassigned | - |
| Otu21756 | 1 | k__Fungi(100);p__Ascomycota(100);p__Ascomycota_unclassified(100);p__Ascomycota_unclassified(100);p__Ascomycota_unclassified(100);p__Ascomycota_unclassified(100);p__Ascomycota_unclassified(100); | - | - | - | - | - | - | - | Unassigned | - |
| Otu21765 | 1 | k__Fungi(100);p__Ascomycota(100);p__Ascomycota_unclassified(100);p__Ascomycota_unclassified(100);p__Ascomycota_unclassified(100);p__Ascomycota_unclassified(100);p__Ascomycota_unclassified(100); | - | - | - | - | - | - | - | Unassigned | - |
| Otu21769 | 1 | k__Fungi(100);p__Ascomycota(100);p__Ascomycota_unclassified(100);p__Ascomycota_unclassified(100);p__Ascomycota_unclassified(100);p__Ascomycota_unclassified(100);p__Ascomycota_unclassified(100); | - | - | - | - | - | - | - | Unassigned | - |
| Otu21775 | 1 | k__Fungi(100);k__Fungi_unclassified(100);k__Fungi_unclassified(100);k__Fungi_unclassified(100);k__Fungi_unclassified(100);k__Fungi_unclassified(100);k__Fungi_unclassified(100); | - | - | - | - | - | - | - | Unassigned | - |
| Otu21777 | 1 | k__Fungi(100);p__Ascomycota(100);p__Ascomycota_unclassified(100);p__Ascomycota_unclassified(100);p__Ascomycota_unclassified(100);p__Ascomycota_unclassified(100);p__Ascomycota_unclassified(100); | - | - | - | - | - | - | - | Unassigned | - |
| Otu21778 | 1 | k__Fungi(100);k__Fungi_unclassified(100);k__Fungi_unclassified(100);k__Fungi_unclassified(100);k__Fungi_unclassified(100);k__Fungi_unclassified(100);k__Fungi_unclassified(100); | - | - | - | - | - | - | - | Unassigned | - |
| Otu21779 | 1 | k__Fungi(100);p__Ascomycota(100);p__Ascomycota_unclassified(100);p__Ascomycota_unclassified(100);p__Ascomycota_unclassified(100);p__Ascomycota_unclassified(100);p__Ascomycota_unclassified(100); | - | - | - | - | - | - | - | Unassigned | - |
| Otu21780 | 1 | k__Fungi(100);p__Ascomycota(100);p__Ascomycota_unclassified(100);p__Ascomycota_unclassified(100);p__Ascomycota_unclassified(100);p__Ascomycota_unclassified(100);p__Ascomycota_unclassified(100); | - | - | - | - | - | - | - | Unassigned | - |
| Otu21781 | 1 | k__Fungi(100);k__Fungi_unclassified(100);k__Fungi_unclassified(100);k__Fungi_unclassified(100);k__Fungi_unclassified(100);k__Fungi_unclassified(100);k__Fungi_unclassified(100); | - | - | - | - | - | - | - | Unassigned | - |
| Otu21782 | 1 | k__Fungi(100);p__Ascomycota(100);p__Ascomycota_unclassified(100);p__Ascomycota_unclassified(100);p__Ascomycota_unclassified(100);p__Ascomycota_unclassified(100);p__Ascomycota_unclassified(100); | - | - | - | - | - | - | - | Unassigned | - |
| Otu21783 | 1 | k__Fungi(100);k__Fungi_unclassified(100);k__Fungi_unclassified(100);k__Fungi_unclassified(100);k__Fungi_unclassified(100);k__Fungi_unclassified(100);k__Fungi_unclassified(100); | - | - | - | - | - | - | - | Unassigned | - |
| Otu21784 | 1 | k__Fungi(100);p__Ascomycota(100);p__Ascomycota_unclassified(100);p__Ascomycota_unclassified(100);p__Ascomycota_unclassified(100);p__Ascomycota_unclassified(100);p__Ascomycota_unclassified(100); | - | - | - | - | - | - | - | Unassigned | - |
| Otu21785 | 1 | k__Fungi(100);p__Ascomycota(100);p__Ascomycota_unclassified(100);p__Ascomycota_unclassified(100);p__Ascomycota_unclassified(100);p__Ascomycota_unclassified(100);p__Ascomycota_unclassified(100); | - | - | - | - | - | - | - | Unassigned | - |
| Otu21792 | 1 | k__Fungi(100);p__Ascomycota(100);p__Ascomycota_unclassified(100);p__Ascomycota_unclassified(100);p__Ascomycota_unclassified(100);p__Ascomycota_unclassified(100);p__Ascomycota_unclassified(100); | - | - | - | - | - | - | - | Unassigned | - |
| Otu21805 | 1 | k__Fungi(100);k__Fungi_unclassified(100);k__Fungi_unclassified(100);k__Fungi_unclassified(100);k__Fungi_unclassified(100);k__Fungi_unclassified(100);k__Fungi_unclassified(100); | - | - | - | - | - | - | - | Unassigned | - |
| Otu21809 | 1 | k__Fungi(100);k__Fungi_unclassified(100);k__Fungi_unclassified(100);k__Fungi_unclassified(100);k__Fungi_unclassified(100);k__Fungi_unclassified(100);k__Fungi_unclassified(100); | - | - | - | - | - | - | - | Unassigned | - |
| Otu21810 | 1 | k__Fungi(100);k__Fungi_unclassified(100);k__Fungi_unclassified(100);k__Fungi_unclassified(100);k__Fungi_unclassified(100);k__Fungi_unclassified(100);k__Fungi_unclassified(100); | - | - | - | - | - | - | - | Unassigned | - |
| Otu21812 | 1 | k__Fungi(100);p__Ascomycota(100);p__Ascomycota_unclassified(100);p__Ascomycota_unclassified(100);p__Ascomycota_unclassified(100);p__Ascomycota_unclassified(100);p__Ascomycota_unclassified(100); | - | - | - | - | - | - | - | Unassigned | - |
| Otu21813 | 1 | k__Fungi(100);p__Ascomycota(100);p__Ascomycota_unclassified(100);p__Ascomycota_unclassified(100);p__Ascomycota_unclassified(100);p__Ascomycota_unclassified(100);p__Ascomycota_unclassified(100); | - | - | - | - | - | - | - | Unassigned | - |
| Otu21815 | 1 | k__Fungi(100);p__Ascomycota(100);p__Ascomycota_unclassified(100);p__Ascomycota_unclassified(100);p__Ascomycota_unclassified(100);p__Ascomycota_unclassified(100);p__Ascomycota_unclassified(100); | - | - | - | - | - | - | - | Unassigned | - |
| Otu21817 | 1 | k__Fungi(100);k__Fungi_unclassified(100);k__Fungi_unclassified(100);k__Fungi_unclassified(100);k__Fungi_unclassified(100);k__Fungi_unclassified(100);k__Fungi_unclassified(100); | - | - | - | - | - | - | - | Unassigned | - |
| Otu21822 | 1 | k__Fungi(100);p__Ascomycota(100);p__Ascomycota_unclassified(100);p__Ascomycota_unclassified(100);p__Ascomycota_unclassified(100);p__Ascomycota_unclassified(100);p__Ascomycota_unclassified(100); | - | - | - | - | - | - | - | Unassigned | - |
| Otu21824 | 1 | k__Fungi(100);p__Ascomycota(100);p__Ascomycota_unclassified(100);p__Ascomycota_unclassified(100);p__Ascomycota_unclassified(100);p__Ascomycota_unclassified(100);p__Ascomycota_unclassified(100); | - | - | - | - | - | - | - | Unassigned | - |
| Otu21840 | 1 | k__Fungi(100);k__Fungi_unclassified(100);k__Fungi_unclassified(100);k__Fungi_unclassified(100);k__Fungi_unclassified(100);k__Fungi_unclassified(100);k__Fungi_unclassified(100); | - | - | - | - | - | - | - | Unassigned | - |
| Otu21841 | 1 | k__Fungi(100);p__Ascomycota(100);p__Ascomycota_unclassified(100);p__Ascomycota_unclassified(100);p__Ascomycota_unclassified(100);p__Ascomycota_unclassified(100);p__Ascomycota_unclassified(100); | - | - | - | - | - | - | - | Unassigned | - |
| Otu21842 | 1 | k__Fungi(100);p__Ascomycota(100);p__Ascomycota_unclassified(100);p__Ascomycota_unclassified(100);p__Ascomycota_unclassified(100);p__Ascomycota_unclassified(100);p__Ascomycota_unclassified(100); | - | - | - | - | - | - | - | Unassigned | - |
| Otu21843 | 1 | k__Fungi(100);k__Fungi_unclassified(100);k__Fungi_unclassified(100);k__Fungi_unclassified(100);k__Fungi_unclassified(100);k__Fungi_unclassified(100);k__Fungi_unclassified(100); | - | - | - | - | - | - | - | Unassigned | - |
| Otu21844 | 1 | k__Fungi(100);k__Fungi_unclassified(100);k__Fungi_unclassified(100);k__Fungi_unclassified(100);k__Fungi_unclassified(100);k__Fungi_unclassified(100);k__Fungi_unclassified(100); | - | - | - | - | - | - | - | Unassigned | - |
| Otu21851 | 1 | k__Fungi(100);k__Fungi_unclassified(100);k__Fungi_unclassified(100);k__Fungi_unclassified(100);k__Fungi_unclassified(100);k__Fungi_unclassified(100);k__Fungi_unclassified(100); | - | - | - | - | - | - | - | Unassigned | - |
| Otu21853 | 1 | k__Fungi(100);p__Ascomycota(100);p__Ascomycota_unclassified(100);p__Ascomycota_unclassified(100);p__Ascomycota_unclassified(100);p__Ascomycota_unclassified(100);p__Ascomycota_unclassified(100); | - | - | - | - | - | - | - | Unassigned | - |
| Otu21854 | 1 | k__Fungi(100);p__Ascomycota(100);p__Ascomycota_unclassified(100);p__Ascomycota_unclassified(100);p__Ascomycota_unclassified(100);p__Ascomycota_unclassified(100);p__Ascomycota_unclassified(100); | - | - | - | - | - | - | - | Unassigned | - |
| Otu21855 | 1 | k__Fungi(100);p__Ascomycota(100);p__Ascomycota_unclassified(100);p__Ascomycota_unclassified(100);p__Ascomycota_unclassified(100);p__Ascomycota_unclassified(100);p__Ascomycota_unclassified(100); | - | - | - | - | - | - | - | Unassigned | - |
| Otu21856 | 1 | k__Fungi(100);k__Fungi_unclassified(100);k__Fungi_unclassified(100);k__Fungi_unclassified(100);k__Fungi_unclassified(100);k__Fungi_unclassified(100);k__Fungi_unclassified(100); | - | - | - | - | - | - | - | Unassigned | - |
| Otu21857 | 1 | k__Fungi(100);p__Ascomycota(100);p__Ascomycota_unclassified(100);p__Ascomycota_unclassified(100);p__Ascomycota_unclassified(100);p__Ascomycota_unclassified(100);p__Ascomycota_unclassified(100); | - | - | - | - | - | - | - | Unassigned | - |
| Otu21858 | 1 | k__Fungi(100);k__Fungi_unclassified(100);k__Fungi_unclassified(100);k__Fungi_unclassified(100);k__Fungi_unclassified(100);k__Fungi_unclassified(100);k__Fungi_unclassified(100); | - | - | - | - | - | - | - | Unassigned | - |
| Otu21859 | 1 | k__Fungi(100);k__Fungi_unclassified(100);k__Fungi_unclassified(100);k__Fungi_unclassified(100);k__Fungi_unclassified(100);k__Fungi_unclassified(100);k__Fungi_unclassified(100); | - | - | - | - | - | - | - | Unassigned | - |
| Otu21860 | 1 | k__Fungi(100);p__Ascomycota(100);p__Ascomycota_unclassified(100);p__Ascomycota_unclassified(100);p__Ascomycota_unclassified(100);p__Ascomycota_unclassified(100);p__Ascomycota_unclassified(100); | - | - | - | - | - | - | - | Unassigned | - |
| Otu21861 | 1 | k__Fungi(100);k__Fungi_unclassified(100);k__Fungi_unclassified(100);k__Fungi_unclassified(100);k__Fungi_unclassified(100);k__Fungi_unclassified(100);k__Fungi_unclassified(100); | - | - | - | - | - | - | - | Unassigned | - |
| Otu21862 | 1 | k__Fungi(100);k__Fungi_unclassified(100);k__Fungi_unclassified(100);k__Fungi_unclassified(100);k__Fungi_unclassified(100);k__Fungi_unclassified(100);k__Fungi_unclassified(100); | - | - | - | - | - | - | - | Unassigned | - |
| Otu21865 | 1 | k__Fungi(100);p__Ascomycota(100);p__Ascomycota_unclassified(100);p__Ascomycota_unclassified(100);p__Ascomycota_unclassified(100);p__Ascomycota_unclassified(100);p__Ascomycota_unclassified(100); | - | - | - | - | - | - | - | Unassigned | - |
| Otu21866 | 1 | k__Fungi(100);k__Fungi_unclassified(100);k__Fungi_unclassified(100);k__Fungi_unclassified(100);k__Fungi_unclassified(100);k__Fungi_unclassified(100);k__Fungi_unclassified(100); | - | - | - | - | - | - | - | Unassigned | - |
| Otu21877 | 1 | k__Fungi(100);p__Basidiomycota(100);p__Basidiomycota_unclassified(100);p__Basidiomycota_unclassified(100);p__Basidiomycota_unclassified(100);p__Basidiomycota_unclassified(100);p__Basidiomycota_unclassified(100); | - | - | - | - | - | - | - | Unassigned | - |
| Otu21878 | 1 | k__Fungi(100);k__Fungi_unclassified(100);k__Fungi_unclassified(100);k__Fungi_unclassified(100);k__Fungi_unclassified(100);k__Fungi_unclassified(100);k__Fungi_unclassified(100); | - | - | - | - | - | - | - | Unassigned | - |
| Otu21879 | 1 | k__Fungi(100);k__Fungi_unclassified(100);k__Fungi_unclassified(100);k__Fungi_unclassified(100);k__Fungi_unclassified(100);k__Fungi_unclassified(100);k__Fungi_unclassified(100); | - | - | - | - | - | - | - | Unassigned | - |
| Otu21881 | 1 | k__Fungi(100);k__Fungi_unclassified(100);k__Fungi_unclassified(100);k__Fungi_unclassified(100);k__Fungi_unclassified(100);k__Fungi_unclassified(100);k__Fungi_unclassified(100); | - | - | - | - | - | - | - | Unassigned | - |
| Otu21883 | 1 | k__Fungi(100);p__Ascomycota(100);p__Ascomycota_unclassified(100);p__Ascomycota_unclassified(100);p__Ascomycota_unclassified(100);p__Ascomycota_unclassified(100);p__Ascomycota_unclassified(100); | - | - | - | - | - | - | - | Unassigned | - |
| Otu21884 | 1 | k__Fungi(100);p__Ascomycota(100);p__Ascomycota_unclassified(100);p__Ascomycota_unclassified(100);p__Ascomycota_unclassified(100);p__Ascomycota_unclassified(100);p__Ascomycota_unclassified(100); | - | - | - | - | - | - | - | Unassigned | - |
| Otu21885 | 1 | k__Fungi(100);p__Ascomycota(100);p__Ascomycota_unclassified(100);p__Ascomycota_unclassified(100);p__Ascomycota_unclassified(100);p__Ascomycota_unclassified(100);p__Ascomycota_unclassified(100); | - | - | - | - | - | - | - | Unassigned | - |
| Otu21888 | 1 | k__Fungi(100);k__Fungi_unclassified(100);k__Fungi_unclassified(100);k__Fungi_unclassified(100);k__Fungi_unclassified(100);k__Fungi_unclassified(100);k__Fungi_unclassified(100); | - | - | - | - | - | - | - | Unassigned | - |
| Otu21890 | 1 | k__Fungi(100);p__Ascomycota(100);p__Ascomycota_unclassified(100);p__Ascomycota_unclassified(100);p__Ascomycota_unclassified(100);p__Ascomycota_unclassified(100);p__Ascomycota_unclassified(100); | - | - | - | - | - | - | - | Unassigned | - |
| Otu21892 | 1 | k__Fungi(100);p__Ascomycota(100);p__Ascomycota_unclassified(100);p__Ascomycota_unclassified(100);p__Ascomycota_unclassified(100);p__Ascomycota_unclassified(100);p__Ascomycota_unclassified(100); | - | - | - | - | - | - | - | Unassigned | - |
| Otu21893 | 1 | k__Fungi(100);k__Fungi_unclassified(100);k__Fungi_unclassified(100);k__Fungi_unclassified(100);k__Fungi_unclassified(100);k__Fungi_unclassified(100);k__Fungi_unclassified(100); | - | - | - | - | - | - | - | Unassigned | - |
| Otu21895 | 1 | k__Fungi(100);k__Fungi_unclassified(100);k__Fungi_unclassified(100);k__Fungi_unclassified(100);k__Fungi_unclassified(100);k__Fungi_unclassified(100);k__Fungi_unclassified(100); | - | - | - | - | - | - | - | Unassigned | - |
| Otu21899 | 1 | k__Fungi(100);k__Fungi_unclassified(100);k__Fungi_unclassified(100);k__Fungi_unclassified(100);k__Fungi_unclassified(100);k__Fungi_unclassified(100);k__Fungi_unclassified(100); | - | - | - | - | - | - | - | Unassigned | - |
| Otu21900 | 1 | k__Fungi(100);k__Fungi_unclassified(100);k__Fungi_unclassified(100);k__Fungi_unclassified(100);k__Fungi_unclassified(100);k__Fungi_unclassified(100);k__Fungi_unclassified(100); | - | - | - | - | - | - | - | Unassigned | - |
| Otu21905 | 1 | k__Fungi(100);k__Fungi_unclassified(100);k__Fungi_unclassified(100);k__Fungi_unclassified(100);k__Fungi_unclassified(100);k__Fungi_unclassified(100);k__Fungi_unclassified(100); | - | - | - | - | - | - | - | Unassigned | - |
| Otu21921 | 1 | k__Fungi(100);k__Fungi_unclassified(100);k__Fungi_unclassified(100);k__Fungi_unclassified(100);k__Fungi_unclassified(100);k__Fungi_unclassified(100);k__Fungi_unclassified(100); | - | - | - | - | - | - | - | Unassigned | - |
| Otu21930 | 1 | k__Fungi(100);p__Ascomycota(100);p__Ascomycota_unclassified(100);p__Ascomycota_unclassified(100);p__Ascomycota_unclassified(100);p__Ascomycota_unclassified(100);p__Ascomycota_unclassified(100); | - | - | - | - | - | - | - | Unassigned | - |
| Otu21931 | 1 | k__Fungi(100);p__Ascomycota(100);p__Ascomycota_unclassified(100);p__Ascomycota_unclassified(100);p__Ascomycota_unclassified(100);p__Ascomycota_unclassified(100);p__Ascomycota_unclassified(100); | - | - | - | - | - | - | - | Unassigned | - |
| Otu21932 | 1 | k__Fungi(100);p__Ascomycota(100);p__Ascomycota_unclassified(100);p__Ascomycota_unclassified(100);p__Ascomycota_unclassified(100);p__Ascomycota_unclassified(100);p__Ascomycota_unclassified(100); | - | - | - | - | - | - | - | Unassigned | - |
| Otu21933 | 1 | k__Fungi(100);p__Ascomycota(100);p__Ascomycota_unclassified(100);p__Ascomycota_unclassified(100);p__Ascomycota_unclassified(100);p__Ascomycota_unclassified(100);p__Ascomycota_unclassified(100); | - | - | - | - | - | - | - | Unassigned | - |
| Otu21934 | 1 | k__Fungi(100);p__Ascomycota(100);p__Ascomycota_unclassified(100);p__Ascomycota_unclassified(100);p__Ascomycota_unclassified(100);p__Ascomycota_unclassified(100);p__Ascomycota_unclassified(100); | - | - | - | - | - | - | - | Unassigned | - |
| Otu21935 | 1 | k__Fungi(100);p__Ascomycota(100);p__Ascomycota_unclassified(100);p__Ascomycota_unclassified(100);p__Ascomycota_unclassified(100);p__Ascomycota_unclassified(100);p__Ascomycota_unclassified(100); | - | - | - | - | - | - | - | Unassigned | - |
| Otu21940 | 1 | k__Fungi(100);p__Ascomycota(100);p__Ascomycota_unclassified(100);p__Ascomycota_unclassified(100);p__Ascomycota_unclassified(100);p__Ascomycota_unclassified(100);p__Ascomycota_unclassified(100); | - | - | - | - | - | - | - | Unassigned | - |
| Otu21945 | 1 | k__Fungi(100);k__Fungi_unclassified(100);k__Fungi_unclassified(100);k__Fungi_unclassified(100);k__Fungi_unclassified(100);k__Fungi_unclassified(100);k__Fungi_unclassified(100); | - | - | - | - | - | - | - | Unassigned | - |
| Otu21950 | 1 | k__Fungi(100);p__Ascomycota(100);p__Ascomycota_unclassified(100);p__Ascomycota_unclassified(100);p__Ascomycota_unclassified(100);p__Ascomycota_unclassified(100);p__Ascomycota_unclassified(100); | - | - | - | - | - | - | - | Unassigned | - |
| Otu21954 | 1 | k__Fungi(100);p__Ascomycota(100);p__Ascomycota_unclassified(100);p__Ascomycota_unclassified(100);p__Ascomycota_unclassified(100);p__Ascomycota_unclassified(100);p__Ascomycota_unclassified(100); | - | - | - | - | - | - | - | Unassigned | - |
| Otu21981 | 1 | k__Fungi(100);k__Fungi_unclassified(100);k__Fungi_unclassified(100);k__Fungi_unclassified(100);k__Fungi_unclassified(100);k__Fungi_unclassified(100);k__Fungi_unclassified(100); | - | - | - | - | - | - | - | Unassigned | - |
| Otu21989 | 1 | k__Fungi(100);p__Ascomycota(100);p__Ascomycota_unclassified(100);p__Ascomycota_unclassified(100);p__Ascomycota_unclassified(100);p__Ascomycota_unclassified(100);p__Ascomycota_unclassified(100); | - | - | - | - | - | - | - | Unassigned | - |
| Otu21993 | 1 | k__Fungi(100);p__Ascomycota(100);p__Ascomycota_unclassified(100);p__Ascomycota_unclassified(100);p__Ascomycota_unclassified(100);p__Ascomycota_unclassified(100);p__Ascomycota_unclassified(100); | - | - | - | - | - | - | - | Unassigned | - |
| Otu21999 | 1 | k__Fungi(100);p__Ascomycota(100);p__Ascomycota_unclassified(100);p__Ascomycota_unclassified(100);p__Ascomycota_unclassified(100);p__Ascomycota_unclassified(100);p__Ascomycota_unclassified(100); | - | - | - | - | - | - | - | Unassigned | - |
| Otu22000 | 1 | k__Fungi(100);k__Fungi_unclassified(100);k__Fungi_unclassified(100);k__Fungi_unclassified(100);k__Fungi_unclassified(100);k__Fungi_unclassified(100);k__Fungi_unclassified(100); | - | - | - | - | - | - | - | Unassigned | - |
| Otu22001 | 1 | k__Fungi(100);p__Ascomycota(100);p__Ascomycota_unclassified(100);p__Ascomycota_unclassified(100);p__Ascomycota_unclassified(100);p__Ascomycota_unclassified(100);p__Ascomycota_unclassified(100); | - | - | - | - | - | - | - | Unassigned | - |
| Otu22002 | 1 | k__Fungi(100);p__Ascomycota(100);p__Ascomycota_unclassified(100);p__Ascomycota_unclassified(100);p__Ascomycota_unclassified(100);p__Ascomycota_unclassified(100);p__Ascomycota_unclassified(100); | - | - | - | - | - | - | - | Unassigned | - |
| Otu22004 | 1 | k__Fungi(100);p__Ascomycota(100);p__Ascomycota_unclassified(100);p__Ascomycota_unclassified(100);p__Ascomycota_unclassified(100);p__Ascomycota_unclassified(100);p__Ascomycota_unclassified(100); | - | - | - | - | - | - | - | Unassigned | - |
| Otu22006 | 1 | k__Fungi(100);p__Ascomycota(100);p__Ascomycota_unclassified(100);p__Ascomycota_unclassified(100);p__Ascomycota_unclassified(100);p__Ascomycota_unclassified(100);p__Ascomycota_unclassified(100); | - | - | - | - | - | - | - | Unassigned | - |
| Otu22007 | 1 | k__Fungi(100);k__Fungi_unclassified(100);k__Fungi_unclassified(100);k__Fungi_unclassified(100);k__Fungi_unclassified(100);k__Fungi_unclassified(100);k__Fungi_unclassified(100); | - | - | - | - | - | - | - | Unassigned | - |
| Otu22023 | 1 | k__Fungi(100);k__Fungi_unclassified(100);k__Fungi_unclassified(100);k__Fungi_unclassified(100);k__Fungi_unclassified(100);k__Fungi_unclassified(100);k__Fungi_unclassified(100); | - | - | - | - | - | - | - | Unassigned | - |
| Otu22024 | 1 | k__Fungi(100);k__Fungi_unclassified(100);k__Fungi_unclassified(100);k__Fungi_unclassified(100);k__Fungi_unclassified(100);k__Fungi_unclassified(100);k__Fungi_unclassified(100); | - | - | - | - | - | - | - | Unassigned | - |
| Otu22025 | 1 | k__Fungi(100);k__Fungi_unclassified(100);k__Fungi_unclassified(100);k__Fungi_unclassified(100);k__Fungi_unclassified(100);k__Fungi_unclassified(100);k__Fungi_unclassified(100); | - | - | - | - | - | - | - | Unassigned | - |
| Otu22026 | 1 | k__Fungi(100);k__Fungi_unclassified(100);k__Fungi_unclassified(100);k__Fungi_unclassified(100);k__Fungi_unclassified(100);k__Fungi_unclassified(100);k__Fungi_unclassified(100); | - | - | - | - | - | - | - | Unassigned | - |
| Otu22027 | 1 | k__Fungi(100);p__Ascomycota(100);p__Ascomycota_unclassified(100);p__Ascomycota_unclassified(100);p__Ascomycota_unclassified(100);p__Ascomycota_unclassified(100);p__Ascomycota_unclassified(100); | - | - | - | - | - | - | - | Unassigned | - |
| Otu22028 | 1 | k__Fungi(100);k__Fungi_unclassified(100);k__Fungi_unclassified(100);k__Fungi_unclassified(100);k__Fungi_unclassified(100);k__Fungi_unclassified(100);k__Fungi_unclassified(100); | - | - | - | - | - | - | - | Unassigned | - |
| Otu22029 | 1 | k__Fungi(100);p__Ascomycota(100);p__Ascomycota_unclassified(100);p__Ascomycota_unclassified(100);p__Ascomycota_unclassified(100);p__Ascomycota_unclassified(100);p__Ascomycota_unclassified(100); | - | - | - | - | - | - | - | Unassigned | - |
| Otu22033 | 1 | k__Fungi(100);k__Fungi_unclassified(100);k__Fungi_unclassified(100);k__Fungi_unclassified(100);k__Fungi_unclassified(100);k__Fungi_unclassified(100);k__Fungi_unclassified(100); | - | - | - | - | - | - | - | Unassigned | - |
| Otu22040 | 1 | k__Fungi(100);p__Ascomycota(100);p__Ascomycota_unclassified(100);p__Ascomycota_unclassified(100);p__Ascomycota_unclassified(100);p__Ascomycota_unclassified(100);p__Ascomycota_unclassified(100); | - | - | - | - | - | - | - | Unassigned | - |
| Otu22041 | 1 | k__Fungi(100);p__Ascomycota(100);p__Ascomycota_unclassified(100);p__Ascomycota_unclassified(100);p__Ascomycota_unclassified(100);p__Ascomycota_unclassified(100);p__Ascomycota_unclassified(100); | - | - | - | - | - | - | - | Unassigned | - |
| Otu22042 | 1 | k__Fungi(100);k__Fungi_unclassified(100);k__Fungi_unclassified(100);k__Fungi_unclassified(100);k__Fungi_unclassified(100);k__Fungi_unclassified(100);k__Fungi_unclassified(100); | - | - | - | - | - | - | - | Unassigned | - |
| Otu22043 | 1 | k__Fungi(100);p__Ascomycota(100);p__Ascomycota_unclassified(100);p__Ascomycota_unclassified(100);p__Ascomycota_unclassified(100);p__Ascomycota_unclassified(100);p__Ascomycota_unclassified(100); | - | - | - | - | - | - | - | Unassigned | - |
| Otu22045 | 1 | k__Fungi(100);k__Fungi_unclassified(100);k__Fungi_unclassified(100);k__Fungi_unclassified(100);k__Fungi_unclassified(100);k__Fungi_unclassified(100);k__Fungi_unclassified(100); | - | - | - | - | - | - | - | Unassigned | - |
| Otu22047 | 1 | k__Fungi(100);p__Ascomycota(100);p__Ascomycota_unclassified(100);p__Ascomycota_unclassified(100);p__Ascomycota_unclassified(100);p__Ascomycota_unclassified(100);p__Ascomycota_unclassified(100); | - | - | - | - | - | - | - | Unassigned | - |
| Otu22049 | 1 | k__Fungi(100);k__Fungi_unclassified(100);k__Fungi_unclassified(100);k__Fungi_unclassified(100);k__Fungi_unclassified(100);k__Fungi_unclassified(100);k__Fungi_unclassified(100); | - | - | - | - | - | - | - | Unassigned | - |
| Otu22050 | 1 | k__Fungi(100);p__Ascomycota(100);p__Ascomycota_unclassified(100);p__Ascomycota_unclassified(100);p__Ascomycota_unclassified(100);p__Ascomycota_unclassified(100);p__Ascomycota_unclassified(100); | - | - | - | - | - | - | - | Unassigned | - |
| Otu22051 | 1 | k__Fungi(100);p__Ascomycota(100);p__Ascomycota_unclassified(100);p__Ascomycota_unclassified(100);p__Ascomycota_unclassified(100);p__Ascomycota_unclassified(100);p__Ascomycota_unclassified(100); | - | - | - | - | - | - | - | Unassigned | - |
| Otu22052 | 1 | k__Fungi(100);k__Fungi_unclassified(100);k__Fungi_unclassified(100);k__Fungi_unclassified(100);k__Fungi_unclassified(100);k__Fungi_unclassified(100);k__Fungi_unclassified(100); | - | - | - | - | - | - | - | Unassigned | - |
| Otu22059 | 1 | k__Fungi(100);k__Fungi_unclassified(100);k__Fungi_unclassified(100);k__Fungi_unclassified(100);k__Fungi_unclassified(100);k__Fungi_unclassified(100);k__Fungi_unclassified(100); | - | - | - | - | - | - | - | Unassigned | - |
| Otu22062 | 1 | k__Fungi(100);k__Fungi_unclassified(100);k__Fungi_unclassified(100);k__Fungi_unclassified(100);k__Fungi_unclassified(100);k__Fungi_unclassified(100);k__Fungi_unclassified(100); | - | - | - | - | - | - | - | Unassigned | - |
| Otu22064 | 1 | k__Fungi(100);k__Fungi_unclassified(100);k__Fungi_unclassified(100);k__Fungi_unclassified(100);k__Fungi_unclassified(100);k__Fungi_unclassified(100);k__Fungi_unclassified(100); | - | - | - | - | - | - | - | Unassigned | - |
| Otu22066 | 1 | k__Fungi(100);k__Fungi_unclassified(100);k__Fungi_unclassified(100);k__Fungi_unclassified(100);k__Fungi_unclassified(100);k__Fungi_unclassified(100);k__Fungi_unclassified(100); | - | - | - | - | - | - | - | Unassigned | - |
| Otu22067 | 1 | k__Fungi(100);p__Ascomycota(100);p__Ascomycota_unclassified(100);p__Ascomycota_unclassified(100);p__Ascomycota_unclassified(100);p__Ascomycota_unclassified(100);p__Ascomycota_unclassified(100); | - | - | - | - | - | - | - | Unassigned | - |
| Otu22068 | 1 | k__Fungi(100);p__Ascomycota(100);p__Ascomycota_unclassified(100);p__Ascomycota_unclassified(100);p__Ascomycota_unclassified(100);p__Ascomycota_unclassified(100);p__Ascomycota_unclassified(100); | - | - | - | - | - | - | - | Unassigned | - |
| Otu22069 | 1 | k__Fungi(100);p__Ascomycota(100);p__Ascomycota_unclassified(100);p__Ascomycota_unclassified(100);p__Ascomycota_unclassified(100);p__Ascomycota_unclassified(100);p__Ascomycota_unclassified(100); | - | - | - | - | - | - | - | Unassigned | - |
| Otu22070 | 1 | k__Fungi(100);k__Fungi_unclassified(100);k__Fungi_unclassified(100);k__Fungi_unclassified(100);k__Fungi_unclassified(100);k__Fungi_unclassified(100);k__Fungi_unclassified(100); | - | - | - | - | - | - | - | Unassigned | - |
| Otu22073 | 1 | k__Fungi(100);k__Fungi_unclassified(100);k__Fungi_unclassified(100);k__Fungi_unclassified(100);k__Fungi_unclassified(100);k__Fungi_unclassified(100);k__Fungi_unclassified(100); | - | - | - | - | - | - | - | Unassigned | - |
| Otu22074 | 1 | k__Fungi(100);k__Fungi_unclassified(100);k__Fungi_unclassified(100);k__Fungi_unclassified(100);k__Fungi_unclassified(100);k__Fungi_unclassified(100);k__Fungi_unclassified(100); | - | - | - | - | - | - | - | Unassigned | - |
| Otu22075 | 1 | k__Fungi(100);k__Fungi_unclassified(100);k__Fungi_unclassified(100);k__Fungi_unclassified(100);k__Fungi_unclassified(100);k__Fungi_unclassified(100);k__Fungi_unclassified(100); | - | - | - | - | - | - | - | Unassigned | - |
| Otu22076 | 1 | k__Fungi(100);k__Fungi_unclassified(100);k__Fungi_unclassified(100);k__Fungi_unclassified(100);k__Fungi_unclassified(100);k__Fungi_unclassified(100);k__Fungi_unclassified(100); | - | - | - | - | - | - | - | Unassigned | - |
| Otu22077 | 1 | k__Fungi(100);k__Fungi_unclassified(100);k__Fungi_unclassified(100);k__Fungi_unclassified(100);k__Fungi_unclassified(100);k__Fungi_unclassified(100);k__Fungi_unclassified(100); | - | - | - | - | - | - | - | Unassigned | - |

| | | | | | | | | | | | |
|---|---|---|---|---|---|---|---|---|---|---|---|
| Otu22094 | 1 | k__Fungi(100);p__Ascomycota(100);p__Ascomycota_unclassified(100);p__Ascomycota_unclassified(100);p__Ascomycota_unclassified(100);p__Ascomycota_unclassified(100);p__Ascomycota_unclassified(100); | - | - | - | - | - | - | - | Unassigned | - |
| Otu22097 | 1 | k__Fungi(100);k__Fungi_unclassified(100);k__Fungi_unclassified(100);k__Fungi_unclassified(100);k__Fungi_unclassified(100);k__Fungi_unclassified(100);k__Fungi_unclassified(100); | - | - | - | - | - | - | - | Unassigned | - |
| Otu22113 | 1 | k__Fungi(100);k__Fungi_unclassified(100);k__Fungi_unclassified(100);k__Fungi_unclassified(100);k__Fungi_unclassified(100);k__Fungi_unclassified(100);k__Fungi_unclassified(100); | - | - | - | - | - | - | - | Unassigned | - |
| Otu22128 | 1 | k__Fungi(100);k__Fungi_unclassified(100);k__Fungi_unclassified(100);k__Fungi_unclassified(100);k__Fungi_unclassified(100);k__Fungi_unclassified(100);k__Fungi_unclassified(100); | - | - | - | - | - | - | - | Unassigned | - |
| Otu22130 | 1 | k__Fungi(100);p__Ascomycota(100);p__Ascomycota_unclassified(100);p__Ascomycota_unclassified(100);p__Ascomycota_unclassified(100);p__Ascomycota_unclassified(100);p__Ascomycota_unclassified(100); | - | - | - | - | - | - | - | Unassigned | - |
| Otu22137 | 1 | k__Fungi(100);k__Fungi_unclassified(100);k__Fungi_unclassified(100);k__Fungi_unclassified(100);k__Fungi_unclassified(100);k__Fungi_unclassified(100);k__Fungi_unclassified(100); | - | - | - | - | - | - | - | Unassigned | - |
| Otu22139 | 1 | k__Fungi(100);p__Ascomycota(100);p__Ascomycota_unclassified(100);p__Ascomycota_unclassified(100);p__Ascomycota_unclassified(100);p__Ascomycota_unclassified(100);p__Ascomycota_unclassified(100); | - | - | - | - | - | - | - | Unassigned | - |
| Otu22142 | 1 | k__Fungi(100);p__Ascomycota(100);p__Ascomycota_unclassified(100);p__Ascomycota_unclassified(100);p__Ascomycota_unclassified(100);p__Ascomycota_unclassified(100);p__Ascomycota_unclassified(100); | - | - | - | - | - | - | - | Unassigned | - |
| Otu22143 | 1 | k__Fungi(100);p__Ascomycota(100);p__Ascomycota_unclassified(100);p__Ascomycota_unclassified(100);p__Ascomycota_unclassified(100);p__Ascomycota_unclassified(100);p__Ascomycota_unclassified(100); | - | - | - | - | - | - | - | Unassigned | - |
| Otu22144 | 1 | k__Fungi(100);p__Ascomycota(100);p__Ascomycota_unclassified(100);p__Ascomycota_unclassified(100);p__Ascomycota_unclassified(100);p__Ascomycota_unclassified(100);p__Ascomycota_unclassified(100); | - | - | - | - | - | - | - | Unassigned | - |
| Otu22146 | 1 | k__Fungi(100);k__Fungi_unclassified(100);k__Fungi_unclassified(100);k__Fungi_unclassified(100);k__Fungi_unclassified(100);k__Fungi_unclassified(100);k__Fungi_unclassified(100); | - | - | - | - | - | - | - | Unassigned | - |
| Otu22147 | 1 | k__Fungi(100);p__Ascomycota(100);p__Ascomycota_unclassified(100);p__Ascomycota_unclassified(100);p__Ascomycota_unclassified(100);p__Ascomycota_unclassified(100);p__Ascomycota_unclassified(100); | - | - | - | - | - | - | - | Unassigned | - |
| Otu22148 | 1 | k__Fungi(100);k__Fungi_unclassified(100);k__Fungi_unclassified(100);k__Fungi_unclassified(100);k__Fungi_unclassified(100);k__Fungi_unclassified(100);k__Fungi_unclassified(100); | - | - | - | - | - | - | - | Unassigned | - |
| Otu22149 | 1 | k__Fungi(100);p__Ascomycota(100);p__Ascomycota_unclassified(100);p__Ascomycota_unclassified(100);p__Ascomycota_unclassified(100);p__Ascomycota_unclassified(100);p__Ascomycota_unclassified(100); | - | - | - | - | - | - | - | Unassigned | - |
| Otu22151 | 1 | k__Fungi(100);p__Ascomycota(100);p__Ascomycota_unclassified(100);p__Ascomycota_unclassified(100);p__Ascomycota_unclassified(100);p__Ascomycota_unclassified(100);p__Ascomycota_unclassified(100); | - | - | - | - | - | - | - | Unassigned | - |
| Otu22152 | 1 | k__Fungi(100);p__Ascomycota(100);p__Ascomycota_unclassified(100);p__Ascomycota_unclassified(100);p__Ascomycota_unclassified(100);p__Ascomycota_unclassified(100);p__Ascomycota_unclassified(100); | - | - | - | - | - | - | - | Unassigned | - |
| Otu22153 | 1 | k__Fungi(100);k__Fungi_unclassified(100);k__Fungi_unclassified(100);k__Fungi_unclassified(100);k__Fungi_unclassified(100);k__Fungi_unclassified(100);k__Fungi_unclassified(100); | - | - | - | - | - | - | - | Unassigned | - |
| Otu22154 | 1 | k__Fungi(100);p__Ascomycota(100);p__Ascomycota_unclassified(100);p__Ascomycota_unclassified(100);p__Ascomycota_unclassified(100);p__Ascomycota_unclassified(100);p__Ascomycota_unclassified(100); | - | - | - | - | - | - | - | Unassigned | - |
| Otu22155 | 1 | k__Fungi(100);p__Ascomycota(100);p__Ascomycota_unclassified(100);p__Ascomycota_unclassified(100);p__Ascomycota_unclassified(100);p__Ascomycota_unclassified(100);p__Ascomycota_unclassified(100); | - | - | - | - | - | - | - | Unassigned | - |
| Otu22156 | 1 | k__Fungi(100);p__Ascomycota(100);p__Ascomycota_unclassified(100);p__Ascomycota_unclassified(100);p__Ascomycota_unclassified(100);p__Ascomycota_unclassified(100);p__Ascomycota_unclassified(100); | - | - | - | - | - | - | - | Unassigned | - |
| Otu22157 | 1 | k__Fungi(100);k__Fungi_unclassified(100);k__Fungi_unclassified(100);k__Fungi_unclassified(100);k__Fungi_unclassified(100);k__Fungi_unclassified(100);k__Fungi_unclassified(100); | - | - | - | - | - | - | - | Unassigned | - |
| Otu22158 | 1 | k__Fungi(100);p__Ascomycota(100);p__Ascomycota_unclassified(100);p__Ascomycota_unclassified(100);p__Ascomycota_unclassified(100);p__Ascomycota_unclassified(100);p__Ascomycota_unclassified(100); | - | - | - | - | - | - | - | Unassigned | - |
| Otu22159 | 1 | k__Fungi(100);p__Ascomycota(100);p__Ascomycota_unclassified(100);p__Ascomycota_unclassified(100);p__Ascomycota_unclassified(100);p__Ascomycota_unclassified(100);p__Ascomycota_unclassified(100); | - | - | - | - | - | - | - | Unassigned | - |
| Otu22160 | 1 | k__Fungi(100);k__Fungi_unclassified(100);k__Fungi_unclassified(100);k__Fungi_unclassified(100);k__Fungi_unclassified(100);k__Fungi_unclassified(100);k__Fungi_unclassified(100); | - | - | - | - | - | - | - | Unassigned | - |
| Otu22162 | 1 | k__Fungi(100);k__Fungi_unclassified(100);k__Fungi_unclassified(100);k__Fungi_unclassified(100);k__Fungi_unclassified(100);k__Fungi_unclassified(100);k__Fungi_unclassified(100); | - | - | - | - | - | - | - | Unassigned | - |
| Otu22174 | 1 | k__Fungi(100);k__Fungi_unclassified(100);k__Fungi_unclassified(100);k__Fungi_unclassified(100);k__Fungi_unclassified(100);k__Fungi_unclassified(100);k__Fungi_unclassified(100); | - | - | - | - | - | - | - | Unassigned | - |
| Otu22175 | 1 | k__Fungi(100);k__Fungi_unclassified(100);k__Fungi_unclassified(100);k__Fungi_unclassified(100);k__Fungi_unclassified(100);k__Fungi_unclassified(100);k__Fungi_unclassified(100); | - | - | - | - | - | - | - | Unassigned | - |
| Otu22179 | 1 | k__Fungi(100);k__Fungi_unclassified(100);k__Fungi_unclassified(100);k__Fungi_unclassified(100);k__Fungi_unclassified(100);k__Fungi_unclassified(100);k__Fungi_unclassified(100); | - | - | - | - | - | - | - | Unassigned | - |
| Otu22180 | 1 | k__Fungi(100);k__Fungi_unclassified(100);k__Fungi_unclassified(100);k__Fungi_unclassified(100);k__Fungi_unclassified(100);k__Fungi_unclassified(100);k__Fungi_unclassified(100); | - | - | - | - | - | - | - | Unassigned | - |
| Otu22181 | 1 | k__Fungi(100);p__Ascomycota(100);p__Ascomycota_unclassified(100);p__Ascomycota_unclassified(100);p__Ascomycota_unclassified(100);p__Ascomycota_unclassified(100);p__Ascomycota_unclassified(100); | - | - | - | - | - | - | - | Unassigned | - |
| Otu22182 | 1 | k__Fungi(100);p__Ascomycota(100);p__Ascomycota_unclassified(100);p__Ascomycota_unclassified(100);p__Ascomycota_unclassified(100);p__Ascomycota_unclassified(100);p__Ascomycota_unclassified(100); | - | - | - | - | - | - | - | Unassigned | - |
| Otu22184 | 1 | k__Fungi(100);p__Ascomycota(100);p__Ascomycota_unclassified(100);p__Ascomycota_unclassified(100);p__Ascomycota_unclassified(100);p__Ascomycota_unclassified(100);p__Ascomycota_unclassified(100); | - | - | - | - | - | - | - | Unassigned | - |
| Otu22185 | 1 | k__Fungi(100);p__Ascomycota(100);p__Ascomycota_unclassified(100);p__Ascomycota_unclassified(100);p__Ascomycota_unclassified(100);p__Ascomycota_unclassified(100);p__Ascomycota_unclassified(100); | - | - | - | - | - | - | - | Unassigned | - |
| Otu22186 | 1 | k__Fungi(100);k__Fungi_unclassified(100);k__Fungi_unclassified(100);k__Fungi_unclassified(100);k__Fungi_unclassified(100);k__Fungi_unclassified(100);k__Fungi_unclassified(100); | - | - | - | - | - | - | - | Unassigned | - |
| Otu22188 | 1 | k__Fungi(100);p__Ascomycota(100);p__Ascomycota_unclassified(100);p__Ascomycota_unclassified(100);p__Ascomycota_unclassified(100);p__Ascomycota_unclassified(100);p__Ascomycota_unclassified(100); | - | - | - | - | - | - | - | Unassigned | - |
| Otu22191 | 1 | k__Fungi(100);k__Fungi_unclassified(100);k__Fungi_unclassified(100);k__Fungi_unclassified(100);k__Fungi_unclassified(100);k__Fungi_unclassified(100);k__Fungi_unclassified(100); | - | - | - | - | - | - | - | Unassigned | - |
| Otu22193 | 1 | k__Fungi(100);k__Fungi_unclassified(100);k__Fungi_unclassified(100);k__Fungi_unclassified(100);k__Fungi_unclassified(100);k__Fungi_unclassified(100);k__Fungi_unclassified(100); | - | - | - | - | - | - | - | Unassigned | - |
| Otu22194 | 1 | k__Fungi(100);k__Fungi_unclassified(100);k__Fungi_unclassified(100);k__Fungi_unclassified(100);k__Fungi_unclassified(100);k__Fungi_unclassified(100);k__Fungi_unclassified(100); | - | - | - | - | - | - | - | Unassigned | - |
| Otu22198 | 1 | k__Fungi(100);p__Ascomycota(100);p__Ascomycota_unclassified(100);p__Ascomycota_unclassified(100);p__Ascomycota_unclassified(100);p__Ascomycota_unclassified(100);p__Ascomycota_unclassified(100); | - | - | - | - | - | - | - | Unassigned | - |
| Otu22200 | 1 | k__Fungi(100);p__Ascomycota(100);p__Ascomycota_unclassified(100);p__Ascomycota_unclassified(100);p__Ascomycota_unclassified(100);p__Ascomycota_unclassified(100);p__Ascomycota_unclassified(100); | - | - | - | - | - | - | - | Unassigned | - |
| Otu22202 | 1 | k__Fungi(100);p__Ascomycota(100);p__Ascomycota_unclassified(100);p__Ascomycota_unclassified(100);p__Ascomycota_unclassified(100);p__Ascomycota_unclassified(100);p__Ascomycota_unclassified(100); | - | - | - | - | - | - | - | Unassigned | - |
| Otu22206 | 1 | k__Fungi(100);p__Ascomycota(100);p__Ascomycota_unclassified(100);p__Ascomycota_unclassified(100);p__Ascomycota_unclassified(100);p__Ascomycota_unclassified(100);p__Ascomycota_unclassified(100); | - | - | - | - | - | - | - | Unassigned | - |
| Otu22215 | 1 | k__Fungi(100);p__Ascomycota(100);p__Ascomycota_unclassified(100);p__Ascomycota_unclassified(100);p__Ascomycota_unclassified(100);p__Ascomycota_unclassified(100);p__Ascomycota_unclassified(100); | - | - | - | - | - | - | - | Unassigned | - |
| Otu22216 | 1 | k__Fungi(100);p__Ascomycota(100);p__Ascomycota_unclassified(100);p__Ascomycota_unclassified(100);p__Ascomycota_unclassified(100);p__Ascomycota_unclassified(100);p__Ascomycota_unclassified(100); | - | - | - | - | - | - | - | Unassigned | - |
| Otu22223 | 1 | k__Fungi(100);p__Ascomycota(100);p__Ascomycota_unclassified(100);p__Ascomycota_unclassified(100);p__Ascomycota_unclassified(100);p__Ascomycota_unclassified(100);p__Ascomycota_unclassified(100); | - | - | - | - | - | - | - | Unassigned | - |
| Otu22225 | 1 | k__Fungi(100);p__Ascomycota(100);p__Ascomycota_unclassified(100);p__Ascomycota_unclassified(100);p__Ascomycota_unclassified(100);p__Ascomycota_unclassified(100);p__Ascomycota_unclassified(100); | - | - | - | - | - | - | - | Unassigned | - |
| Otu22226 | 1 | k__Fungi(100);k__Fungi_unclassified(100);k__Fungi_unclassified(100);k__Fungi_unclassified(100);k__Fungi_unclassified(100);k__Fungi_unclassified(100);k__Fungi_unclassified(100); | - | - | - | - | - | - | - | Unassigned | - |
| Otu22229 | 1 | k__Fungi(100);k__Fungi_unclassified(100);k__Fungi_unclassified(100);k__Fungi_unclassified(100);k__Fungi_unclassified(100);k__Fungi_unclassified(100);k__Fungi_unclassified(100); | - | - | - | - | - | - | - | Unassigned | - |
| Otu22234 | 1 | k__Fungi(100);p__Ascomycota(100);p__Ascomycota_unclassified(100);p__Ascomycota_unclassified(100);p__Ascomycota_unclassified(100);p__Ascomycota_unclassified(100);p__Ascomycota_unclassified(100); | - | - | - | - | - | - | - | Unassigned | - |
| Otu22238 | 1 | k__Fungi(100);p__Ascomycota(100);p__Ascomycota_unclassified(100);p__Ascomycota_unclassified(100);p__Ascomycota_unclassified(100);p__Ascomycota_unclassified(100);p__Ascomycota_unclassified(100); | - | - | - | - | - | - | - | Unassigned | - |
| Otu22239 | 1 | k__Fungi(100);k__Fungi_unclassified(100);k__Fungi_unclassified(100);k__Fungi_unclassified(100);k__Fungi_unclassified(100);k__Fungi_unclassified(100);k__Fungi_unclassified(100); | - | - | - | - | - | - | - | Unassigned | - |
| Otu22240 | 1 | k__Fungi(100);k__Fungi_unclassified(100);k__Fungi_unclassified(100);k__Fungi_unclassified(100);k__Fungi_unclassified(100);k__Fungi_unclassified(100);k__Fungi_unclassified(100); | - | - | - | - | - | - | - | Unassigned | - |
| Otu22241 | 1 | k__Fungi(100);p__Ascomycota(100);p__Ascomycota_unclassified(100);p__Ascomycota_unclassified(100);p__Ascomycota_unclassified(100);p__Ascomycota_unclassified(100);p__Ascomycota_unclassified(100); | - | - | - | - | - | - | - | Unassigned | - |
| Otu22250 | 1 | k__Fungi(100);p__Ascomycota(100);p__Ascomycota_unclassified(100);p__Ascomycota_unclassified(100);p__Ascomycota_unclassified(100);p__Ascomycota_unclassified(100);p__Ascomycota_unclassified(100); | - | - | - | - | - | - | - | Unassigned | - |
| Otu22251 | 1 | k__Fungi(100);p__Ascomycota(100);p__Ascomycota_unclassified(100);p__Ascomycota_unclassified(100);p__Ascomycota_unclassified(100);p__Ascomycota_unclassified(100);p__Ascomycota_unclassified(100); | - | - | - | - | - | - | - | Unassigned | - |
| Otu22256 | 1 | k__Fungi(100);p__Ascomycota(100);p__Ascomycota_unclassified(100);p__Ascomycota_unclassified(100);p__Ascomycota_unclassified(100);p__Ascomycota_unclassified(100);p__Ascomycota_unclassified(100); | - | - | - | - | - | - | - | Unassigned | - |
| Otu22293 | 1 | k__Fungi(100);k__Fungi_unclassified(100);k__Fungi_unclassified(100);k__Fungi_unclassified(100);k__Fungi_unclassified(100);k__Fungi_unclassified(100);k__Fungi_unclassified(100); | - | - | - | - | - | - | - | Unassigned | - |
| Otu22295 | 1 | k__Fungi(100);k__Fungi_unclassified(100);k__Fungi_unclassified(100);k__Fungi_unclassified(100);k__Fungi_unclassified(100);k__Fungi_unclassified(100);k__Fungi_unclassified(100); | - | - | - | - | - | - | - | Unassigned | - |
| Otu22304 | 1 | k__Fungi(100);p__Ascomycota(100);p__Ascomycota_unclassified(100);p__Ascomycota_unclassified(100);p__Ascomycota_unclassified(100);p__Ascomycota_unclassified(100);p__Ascomycota_unclassified(100); | - | - | - | - | - | - | - | Unassigned | - |
| Otu22356 | 1 | k__Fungi(100);k__Fungi_unclassified(100);k__Fungi_unclassified(100);k__Fungi_unclassified(100);k__Fungi_unclassified(100);k__Fungi_unclassified(100);k__Fungi_unclassified(100); | - | - | - | - | - | - | - | Unassigned | - |
| Otu22390 | 1 | k__Fungi(100);k__Fungi_unclassified(100);k__Fungi_unclassified(100);k__Fungi_unclassified(100);k__Fungi_unclassified(100);k__Fungi_unclassified(100);k__Fungi_unclassified(100); | - | - | - | - | - | - | - | Unassigned | - |
| Otu22402 | 1 | k__Fungi(100);k__Fungi_unclassified(100);k__Fungi_unclassified(100);k__Fungi_unclassified(100);k__Fungi_unclassified(100);k__Fungi_unclassified(100);k__Fungi_unclassified(100); | - | - | - | - | - | - | - | Unassigned | - |
| Otu22405 | 1 | k__Fungi(100);p__Ascomycota(100);p__Ascomycota_unclassified(100);p__Ascomycota_unclassified(100);p__Ascomycota_unclassified(100);p__Ascomycota_unclassified(100);p__Ascomycota_unclassified(100); | - | - | - | - | - | - | - | Unassigned | - |
| Otu22410 | 1 | k__Fungi(100);k__Fungi_unclassified(100);k__Fungi_unclassified(100);k__Fungi_unclassified(100);k__Fungi_unclassified(100);k__Fungi_unclassified(100);k__Fungi_unclassified(100); | - | - | - | - | - | - | - | Unassigned | - |
| Otu22411 | 1 | k__Fungi(100);k__Fungi_unclassified(100);k__Fungi_unclassified(100);k__Fungi_unclassified(100);k__Fungi_unclassified(100);k__Fungi_unclassified(100);k__Fungi_unclassified(100); | - | - | - | - | - | - | - | Unassigned | - |
| Otu22425 | 1 | k__Fungi(100);p__Ascomycota(100);p__Ascomycota_unclassified(100);p__Ascomycota_unclassified(100);p__Ascomycota_unclassified(100);p__Ascomycota_unclassified(100);p__Ascomycota_unclassified(100); | - | - | - | - | - | - | - | Unassigned | - |
| Otu22426 | 1 | k__Fungi(100);k__Fungi_unclassified(100);k__Fungi_unclassified(100);k__Fungi_unclassified(100);k__Fungi_unclassified(100);k__Fungi_unclassified(100);k__Fungi_unclassified(100); | - | - | - | - | - | - | - | Unassigned | - |
| Otu22427 | 1 | k__Fungi(100);k__Fungi_unclassified(100);k__Fungi_unclassified(100);k__Fungi_unclassified(100);k__Fungi_unclassified(100);k__Fungi_unclassified(100);k__Fungi_unclassified(100); | - | - | - | - | - | - | - | Unassigned | - |
| Otu22441 | 1 | k__Fungi(100);k__Fungi_unclassified(100);k__Fungi_unclassified(100);k__Fungi_unclassified(100);k__Fungi_unclassified(100);k__Fungi_unclassified(100);k__Fungi_unclassified(100); | - | - | - | - | - | - | - | Unassigned | - |
| Otu22446 | 1 | k__Fungi(100);k__Fungi_unclassified(100);k__Fungi_unclassified(100);k__Fungi_unclassified(100);k__Fungi_unclassified(100);k__Fungi_unclassified(100);k__Fungi_unclassified(100); | - | - | - | - | - | - | - | Unassigned | - |
| Otu22460 | 1 | k__Fungi(100);k__Fungi_unclassified(100);k__Fungi_unclassified(100);k__Fungi_unclassified(100);k__Fungi_unclassified(100);k__Fungi_unclassified(100);k__Fungi_unclassified(100); | - | - | - | - | - | - | - | Unassigned | - |
| Otu22467 | 1 | k__Fungi(100);p__Ascomycota(100);p__Ascomycota_unclassified(100);p__Ascomycota_unclassified(100);p__Ascomycota_unclassified(100);p__Ascomycota_unclassified(100);p__Ascomycota_unclassified(100); | - | - | - | - | - | - | - | Unassigned | - |
| Otu22468 | 1 | k__Fungi(100);k__Fungi_unclassified(100);k__Fungi_unclassified(100);k__Fungi_unclassified(100);k__Fungi_unclassified(100);k__Fungi_unclassified(100);k__Fungi_unclassified(100); | - | - | - | - | - | - | - | Unassigned | - |
| Otu22494 | 1 | k__Fungi(100);p__Ascomycota(100);p__Ascomycota_unclassified(100);p__Ascomycota_unclassified(100);p__Ascomycota_unclassified(100);p__Ascomycota_unclassified(100);p__Ascomycota_unclassified(100); | - | - | - | - | - | - | - | Unassigned | - |
| Otu22495 | 1 | k__Fungi(100);k__Fungi_unclassified(100);k__Fungi_unclassified(100);k__Fungi_unclassified(100);k__Fungi_unclassified(100);k__Fungi_unclassified(100);k__Fungi_unclassified(100); | - | - | - | - | - | - | - | Unassigned | - |
| Otu22500 | 1 | k__Fungi(100);k__Fungi_unclassified(100);k__Fungi_unclassified(100);k__Fungi_unclassified(100);k__Fungi_unclassified(100);k__Fungi_unclassified(100);k__Fungi_unclassified(100); | - | - | - | - | - | - | - | Unassigned | - |
| Otu22510 | 1 | k__Fungi(100);k__Fungi_unclassified(100);k__Fungi_unclassified(100);k__Fungi_unclassified(100);k__Fungi_unclassified(100);k__Fungi_unclassified(100);k__Fungi_unclassified(100); | - | - | - | - | - | - | - | Unassigned | - |
| Otu22514 | 1 | k__Fungi(100);k__Fungi_unclassified(100);k__Fungi_unclassified(100);k__Fungi_unclassified(100);k__Fungi_unclassified(100);k__Fungi_unclassified(100);k__Fungi_unclassified(100); | - | - | - | - | - | - | - | Unassigned | - |
| Otu22515 | 1 | k__Fungi(100);k__Fungi_unclassified(100);k__Fungi_unclassified(100);k__Fungi_unclassified(100);k__Fungi_unclassified(100);k__Fungi_unclassified(100);k__Fungi_unclassified(100); | - | - | - | - | - | - | - | Unassigned | - |
| Otu22516 | 1 | k__Fungi(100);k__Fungi_unclassified(100);k__Fungi_unclassified(100);k__Fungi_unclassified(100);k__Fungi_unclassified(100);k__Fungi_unclassified(100);k__Fungi_unclassified(100); | - | - | - | - | - | - | - | Unassigned | - |
| Otu22517 | 1 | k__Fungi(100);k__Fungi_unclassified(100);k__Fungi_unclassified(100);k__Fungi_unclassified(100);k__Fungi_unclassified(100);k__Fungi_unclassified(100);k__Fungi_unclassified(100); | - | - | - | - | - | - | - | Unassigned | - |
| Otu22518 | 1 | k__Fungi(100);p__Ascomycota(100);p__Ascomycota_unclassified(100);p__Ascomycota_unclassified(100);p__Ascomycota_unclassified(100);p__Ascomycota_unclassified(100);p__Ascomycota_unclassified(100); | - | - | - | - | - | - | - | Unassigned | - |
| Otu22519 | 1 | k__Fungi(100);p__Ascomycota(100);p__Ascomycota_unclassified(100);p__Ascomycota_unclassified(100);p__Ascomycota_unclassified(100);p__Ascomycota_unclassified(100);p__Ascomycota_unclassified(100); | - | - | - | - | - | - | - | Unassigned | - |
| Otu22520 | 1 | k__Fungi(100);p__Ascomycota(100);p__Ascomycota_unclassified(100);p__Ascomycota_unclassified(100);p__Ascomycota_unclassified(100);p__Ascomycota_unclassified(100);p__Ascomycota_unclassified(100); | - | - | - | - | - | - | - | Unassigned | - |
| Otu22522 | 1 | k__Fungi(100);p__Ascomycota(100);p__Ascomycota_unclassified(100);p__Ascomycota_unclassified(100);p__Ascomycota_unclassified(100);p__Ascomycota_unclassified(100);p__Ascomycota_unclassified(100); | - | - | - | - | - | - | - | Unassigned | - |
| Otu22523 | 1 | k__Fungi(100);p__Ascomycota(100);p__Ascomycota_unclassified(100);p__Ascomycota_unclassified(100);p__Ascomycota_unclassified(100);p__Ascomycota_unclassified(100);p__Ascomycota_unclassified(100); | - | - | - | - | - | - | - | Unassigned | - |
| Otu22524 | 1 | k__Fungi(100);p__Ascomycota(100);p__Ascomycota_unclassified(100);p__Ascomycota_unclassified(100);p__Ascomycota_unclassified(100);p__Ascomycota_unclassified(100);p__Ascomycota_unclassified(100); | - | - | - | - | - | - | - | Unassigned | - |
| Otu22525 | 1 | k__Fungi(100);k__Fungi_unclassified(100);k__Fungi_unclassified(100);k__Fungi_unclassified(100);k__Fungi_unclassified(100);k__Fungi_unclassified(100);k__Fungi_unclassified(100); | - | - | - | - | - | - | - | Unassigned | - |
| Otu22535 | 1 | k__Fungi(100);k__Fungi_unclassified(100);k__Fungi_unclassified(100);k__Fungi_unclassified(100);k__Fungi_unclassified(100);k__Fungi_unclassified(100);k__Fungi_unclassified(100); | - | - | - | - | - | - | - | Unassigned | - |
| Otu22539 | 1 | k__Fungi(100);p__Ascomycota(100);p__Ascomycota_unclassified(100);p__Ascomycota_unclassified(100);p__Ascomycota_unclassified(100);p__Ascomycota_unclassified(100);p__Ascomycota_unclassified(100); | - | - | - | - | - | - | - | Unassigned | - |
| Otu22555 | 1 | k__Fungi(100);p__Ascomycota(100);p__Ascomycota_unclassified(100);p__Ascomycota_unclassified(100);p__Ascomycota_unclassified(100);p__Ascomycota_unclassified(100);p__Ascomycota_unclassified(100); | - | - | - | - | - | - | - | Unassigned | - |
| Otu22557 | 1 | k__Fungi(100);k__Fungi_unclassified(100);k__Fungi_unclassified(100);k__Fungi_unclassified(100);k__Fungi_unclassified(100);k__Fungi_unclassified(100);k__Fungi_unclassified(100); | - | - | - | - | - | - | - | Unassigned | - |
| Otu22558 | 1 | k__Fungi(100);k__Fungi_unclassified(100);k__Fungi_unclassified(100);k__Fungi_unclassified(100);k__Fungi_unclassified(100);k__Fungi_unclassified(100);k__Fungi_unclassified(100); | - | - | - | - | - | - | - | Unassigned | - |
| Otu22559 | 1 | k__Fungi(100);k__Fungi_unclassified(100);k__Fungi_unclassified(100);k__Fungi_unclassified(100);k__Fungi_unclassified(100);k__Fungi_unclassified(100);k__Fungi_unclassified(100); | - | - | - | - | - | - | - | Unassigned | - |
| Otu22560 | 1 | k__Fungi(100);k__Fungi_unclassified(100);k__Fungi_unclassified(100);k__Fungi_unclassified(100);k__Fungi_unclassified(100);k__Fungi_unclassified(100);k__Fungi_unclassified(100); | - | - | - | - | - | - | - | Unassigned | - |
| Otu22562 | 1 | k__Fungi(100);k__Fungi_unclassified(100);k__Fungi_unclassified(100);k__Fungi_unclassified(100);k__Fungi_unclassified(100);k__Fungi_unclassified(100);k__Fungi_unclassified(100); | - | - | - | - | - | - | - | Unassigned | - |
| Otu22563 | 1 | k__Fungi(100);k__Fungi_unclassified(100);k__Fungi_unclassified(100);k__Fungi_unclassified(100);k__Fungi_unclassified(100);k__Fungi_unclassified(100);k__Fungi_unclassified(100); | - | - | - | - | - | - | - | Unassigned | - |
| Otu22569 | 1 | k__Fungi(100);p__Ascomycota(100);p__Ascomycota_unclassified(100);p__Ascomycota_unclassified(100);p__Ascomycota_unclassified(100);p__Ascomycota_unclassified(100);p__Ascomycota_unclassified(100); | - | - | - | - | - | - | - | Unassigned | - |
| Otu22570 | 1 | k__Fungi(100);p__Ascomycota(100);p__Ascomycota_unclassified(100);p__Ascomycota_unclassified(100);p__Ascomycota_unclassified(100);p__Ascomycota_unclassified(100);p__Ascomycota_unclassified(100); | - | - | - | - | - | - | - | Unassigned | - |
| Otu22571 | 1 | k__Fungi(100);k__Fungi_unclassified(100);k__Fungi_unclassified(100);k__Fungi_unclassified(100);k__Fungi_unclassified(100);k__Fungi_unclassified(100);k__Fungi_unclassified(100); | - | - | - | - | - | - | - | Unassigned | - |
| Otu22572 | 1 | k__Fungi(100);k__Fungi_unclassified(100);k__Fungi_unclassified(100);k__Fungi_unclassified(100);k__Fungi_unclassified(100);k__Fungi_unclassified(100);k__Fungi_unclassified(100); | - | - | - | - | - | - | - | Unassigned | - |
| Otu22573 | 1 | k__Fungi(100);k__Fungi_unclassified(100);k__Fungi_unclassified(100);k__Fungi_unclassified(100);k__Fungi_unclassified(100);k__Fungi_unclassified(100);k__Fungi_unclassified(100); | - | - | - | - | - | - | - | Unassigned | - |
| Otu22579 | 1 | k__Fungi(100);p__Ascomycota(100);p__Ascomycota_unclassified(100);p__Ascomycota_unclassified(100);p__Ascomycota_unclassified(100);p__Ascomycota_unclassified(100);p__Ascomycota_unclassified(100); | - | - | - | - | - | - | - | Unassigned | - |
| Otu22580 | 1 | k__Fungi(100);k__Fungi_unclassified(100);k__Fungi_unclassified(100);k__Fungi_unclassified(100);k__Fungi_unclassified(100);k__Fungi_unclassified(100);k__Fungi_unclassified(100); | - | - | - | - | - | - | - | Unassigned | - |
| Otu22581 | 1 | k__Fungi(100);k__Fungi_unclassified(100);k__Fungi_unclassified(100);k__Fungi_unclassified(100);k__Fungi_unclassified(100);k__Fungi_unclassified(100);k__Fungi_unclassified(100); | - | - | - | - | - | - | - | Unassigned | - |
| Otu22591 | 1 | k__Fungi(100);k__Fungi_unclassified(100);k__Fungi_unclassified(100);k__Fungi_unclassified(100);k__Fungi_unclassified(100);k__Fungi_unclassified(100);k__Fungi_unclassified(100); | - | - | - | - | - | - | - | Unassigned | - |
| Otu22592 | 1 | k__Fungi(100);k__Fungi_unclassified(100);k__Fungi_unclassified(100);k__Fungi_unclassified(100);k__Fungi_unclassified(100);k__Fungi_unclassified(100);k__Fungi_unclassified(100); | - | - | - | - | - | - | - | Unassigned | - |
| Otu22593 | 1 | k__Fungi(100);k__Fungi_unclassified(100);k__Fungi_unclassified(100);k__Fungi_unclassified(100);k__Fungi_unclassified(100);k__Fungi_unclassified(100);k__Fungi_unclassified(100); | - | - | - | - | - | - | - | Unassigned | - |
| Otu22594 | 1 | k__Fungi(100);p__Ascomycota(100);p__Ascomycota_unclassified(100);p__Ascomycota_unclassified(100);p__Ascomycota_unclassified(100);p__Ascomycota_unclassified(100);p__Ascomycota_unclassified(100); | - | - | - | - | - | - | - | Unassigned | - |
| Otu22595 | 1 | k__Fungi(100);p__Ascomycota(100);p__Ascomycota_unclassified(100);p__Ascomycota_unclassified(100);p__Ascomycota_unclassified(100);p__Ascomycota_unclassified(100);p__Ascomycota_unclassified(100); | - | - | - | - | - | - | - | Unassigned | - |
| Otu22599 | 1 | k__Fungi(100);p__Ascomycota(100);p__Ascomycota_unclassified(100);p__Ascomycota_unclassified(100);p__Ascomycota_unclassified(100);p__Ascomycota_unclassified(100);p__Ascomycota_unclassified(100); | - | - | - | - | - | - | - | Unassigned | - |
| Otu22600 | 1 | k__Fungi(100);p__Ascomycota(100);p__Ascomycota_unclassified(100);p__Ascomycota_unclassified(100);p__Ascomycota_unclassified(100);p__Ascomycota_unclassified(100);p__Ascomycota_unclassified(100); | - | - | - | - | - | - | - | Unassigned | - |
| Otu22601 | 1 | k__Fungi(100);p__Ascomycota(100);p__Ascomycota_unclassified(100);p__Ascomycota_unclassified(100);p__Ascomycota_unclassified(100);p__Ascomycota_unclassified(100);p__Ascomycota_unclassified(100); | - | - | - | - | - | - | - | Unassigned | - |
| Otu22602 | 1 | k__Fungi(100);p__Ascomycota(100);p__Ascomycota_unclassified(100);p__Ascomycota_unclassified(100);p__Ascomycota_unclassified(100);p__Ascomycota_unclassified(100);p__Ascomycota_unclassified(100); | - | - | - | - | - | - | - | Unassigned | - |
| Otu22615 | 1 | k__Fungi(100);k__Fungi_unclassified(100);k__Fungi_unclassified(100);k__Fungi_unclassified(100);k__Fungi_unclassified(100);k__Fungi_unclassified(100);k__Fungi_unclassified(100); | - | - | - | - | - | - | - | Unassigned | - |
| Otu22617 | 1 | k__Fungi(100);p__Ascomycota(100);c__Leotiomycetes(100);o__Helotiales(100);o__Helotiales_unclassified(100);o__Helotiales_unclassified(100);o__Helotiales_unclassified(100); | - | - | - | - | - | - | - | Unassigned | - |
| Otu22622 | 1 | k__Fungi(100);p__Ascomycota(100);p__Ascomycota_unclassified(100);p__Ascomycota_unclassified(100);p__Ascomycota_unclassified(100);p__Ascomycota_unclassified(100);p__Ascomycota_unclassified(100); | - | - | - | - | - | - | - | Unassigned | - |
| Otu22623 | 1 | k__Fungi(100);p__Ascomycota(100);p__Ascomycota_unclassified(100);p__Ascomycota_unclassified(100);p__Ascomycota_unclassified(100);p__Ascomycota_unclassified(100);p__Ascomycota_unclassified(100); | - | - | - | - | - | - | - | Unassigned | - |
| Otu22649 | 1 | k__Fungi(100);k__Fungi_unclassified(100);k__Fungi_unclassified(100);k__Fungi_unclassified(100);k__Fungi_unclassified(100);k__Fungi_unclassified(100);k__Fungi_unclassified(100); | - | - | - | - | - | - | - | Unassigned | - |
| Otu22652 | 1 | k__Fungi(100);p__Ascomycota(100);p__Ascomycota_unclassified(100);p__Ascomycota_unclassified(100);p__Ascomycota_unclassified(100);p__Ascomycota_unclassified(100);p__Ascomycota_unclassified(100); | - | - | - | - | - | - | - | Unassigned | - |
| Otu22657 | 1 | k__Fungi(100);p__Ascomycota(100);p__Ascomycota_unclassified(100);p__Ascomycota_unclassified(100);p__Ascomycota_unclassified(100);p__Ascomycota_unclassified(100);p__Ascomycota_unclassified(100); | - | - | - | - | - | - | - | Unassigned | - |
| Otu22658 | 1 | k__Fungi(100);p__Ascomycota(100);p__Ascomycota_unclassified(100);p__Ascomycota_unclassified(100);p__Ascomycota_unclassified(100);p__Ascomycota_unclassified(100);p__Ascomycota_unclassified(100); | - | - | - | - | - | - | - | Unassigned | - |
| Otu22659 | 1 | k__Fungi(100);k__Fungi_unclassified(100);k__Fungi_unclassified(100);k__Fungi_unclassified(100);k__Fungi_unclassified(100);k__Fungi_unclassified(100);k__Fungi_unclassified(100); | - | - | - | - | - | - | - | Unassigned | - |
| Otu22660 | 1 | k__Fungi(100);p__Ascomycota(100);p__Ascomycota_unclassified(100);p__Ascomycota_unclassified(100);p__Ascomycota_unclassified(100);p__Ascomycota_unclassified(100);p__Ascomycota_unclassified(100); | - | - | - | - | - | - | - | Unassigned | - |
| Otu22661 | 1 | k__Fungi(100);p__Ascomycota(100);p__Ascomycota_unclassified(100);p__Ascomycota_unclassified(100);p__Ascomycota_unclassified(100);p__Ascomycota_unclassified(100);p__Ascomycota_unclassified(100); | - | - | - | - | - | - | - | Unassigned | - |
| Otu22662 | 1 | k__Fungi(100);p__Ascomycota(100);p__Ascomycota_unclassified(100);p__Ascomycota_unclassified(100);p__Ascomycota_unclassified(100);p__Ascomycota_unclassified(100);p__Ascomycota_unclassified(100); | - | - | - | - | - | - | - | Unassigned | - |
| Otu22663 | 1 | k__Fungi(100);k__Fungi_unclassified(100);k__Fungi_unclassified(100);k__Fungi_unclassified(100);k__Fungi_unclassified(100);k__Fungi_unclassified(100);k__Fungi_unclassified(100); | - | - | - | - | - | - | - | Unassigned | - |
| Otu22664 | 1 | k__Fungi(100);p__Ascomycota(100);p__Ascomycota_unclassified(100);p__Ascomycota_unclassified(100);p__Ascomycota_unclassified(100);p__Ascomycota_unclassified(100);p__Ascomycota_unclassified(100); | - | - | - | - | - | - | - | Unassigned | - |
| Otu22665 | 1 | k__Fungi(100);k__Fungi_unclassified(100);k__Fungi_unclassified(100);k__Fungi_unclassified(100);k__Fungi_unclassified(100);k__Fungi_unclassified(100);k__Fungi_unclassified(100); | - | - | - | - | - | - | - | Unassigned | - |
| Otu22666 | 1 | k__Fungi(100);p__Ascomycota(100);p__Ascomycota_unclassified(100);p__Ascomycota_unclassified(100);p__Ascomycota_unclassified(100);p__Ascomycota_unclassified(100);p__Ascomycota_unclassified(100); | - | - | - | - | - | - | - | Unassigned | - |
| Otu22667 | 1 | k__Fungi(100);p__Ascomycota(100);p__Ascomycota_unclassified(100);p__Ascomycota_unclassified(100);p__Ascomycota_unclassified(100);p__Ascomycota_unclassified(100);p__Ascomycota_unclassified(100); | - | - | - | - | - | - | - | Unassigned | - |
| Otu22668 | 1 | k__Fungi(100);k__Fungi_unclassified(100);k__Fungi_unclassified(100);k__Fungi_unclassified(100);k__Fungi_unclassified(100);k__Fungi_unclassified(100);k__Fungi_unclassified(100); | - | - | - | - | - | - | - | Unassigned | - |
| Otu22669 | 1 | k__Fungi(100);k__Fungi_unclassified(100);k__Fungi_unclassified(100);k__Fungi_unclassified(100);k__Fungi_unclassified(100);k__Fungi_unclassified(100);k__Fungi_unclassified(100); | - | - | - | - | - | - | - | Unassigned | - |
| Otu22670 | 1 | k__Fungi(100);p__Ascomycota(100);p__Ascomycota_unclassified(100);p__Ascomycota_unclassified(100);p__Ascomycota_unclassified(100);p__Ascomycota_unclassified(100);p__Ascomycota_unclassified(100); | - | - | - | - | - | - | - | Unassigned | - |
| Otu22671 | 1 | k__Fungi(100);p__Ascomycota(100);p__Ascomycota_unclassified(100);p__Ascomycota_unclassified(100);p__Ascomycota_unclassified(100);p__Ascomycota_unclassified(100);p__Ascomycota_unclassified(100); | - | - | - | - | - | - | - | Unassigned | - |
| Otu22672 | 1 | k__Fungi(100);k__Fungi_unclassified(100);k__Fungi_unclassified(100);k__Fungi_unclassified(100);k__Fungi_unclassified(100);k__Fungi_unclassified(100);k__Fungi_unclassified(100); | - | - | - | - | - | - | - | Unassigned | - |
| Otu22673 | 1 | k__Fungi(100);k__Fungi_unclassified(100);k__Fungi_unclassified(100);k__Fungi_unclassified(100);k__Fungi_unclassified(100);k__Fungi_unclassified(100);k__Fungi_unclassified(100); | - | - | - | - | - | - | - | Unassigned | - |
| Otu22674 | 1 | k__Fungi(100);p__Ascomycota(100);p__Ascomycota_unclassified(100);p__Ascomycota_unclassified(100);p__Ascomycota_unclassified(100);p__Ascomycota_unclassified(100);p__Ascomycota_unclassified(100); | - | - | - | - | - | - | - | Unassigned | - |
| Otu22675 | 1 | k__Fungi(100);p__Ascomycota(100);p__Ascomycota_unclassified(100);p__Ascomycota_unclassified(100);p__Ascomycota_unclassified(100);p__Ascomycota_unclassified(100);p__Ascomycota_unclassified(100); | - | - | - | - | - | - | - | Unassigned | - |
| Otu22676 | 1 | k__Fungi(100);k__Fungi_unclassified(100);k__Fungi_unclassified(100);k__Fungi_unclassified(100);k__Fungi_unclassified(100);k__Fungi_unclassified(100);k__Fungi_unclassified(100); | - | - | - | - | - | - | - | Unassigned | - |
| Otu22677 | 1 | k__Fungi(100);k__Fungi_unclassified(100);k__Fungi_unclassified(100);k__Fungi_unclassified(100);k__Fungi_unclassified(100);k__Fungi_unclassified(100);k__Fungi_unclassified(100); | - | - | - | - | - | - | - | Unassigned | - |
| Otu22678 | 1 | k__Fungi(100);p__Ascomycota(100);p__Ascomycota_unclassified(100);p__Ascomycota_unclassified(100);p__Ascomycota_unclassified(100);p__Ascomycota_unclassified(100);p__Ascomycota_unclassified(100); | - | - | - | - | - | - | - | Unassigned | - |
| Otu22679 | 1 | k__Fungi(100);p__Ascomycota(100);p__Ascomycota_unclassified(100);p__Ascomycota_unclassified(100);p__Ascomycota_unclassified(100);p__Ascomycota_unclassified(100);p__Ascomycota_unclassified(100); | - | - | - | - | - | - | - | Unassigned | - |
| Otu22680 | 1 | k__Fungi(100);k__Fungi_unclassified(100);k__Fungi_unclassified(100);k__Fungi_unclassified(100);k__Fungi_unclassified(100);k__Fungi_unclassified(100);k__Fungi_unclassified(100); | - | - | - | - | - | - | - | Unassigned | - |
| Otu22681 | 1 | k__Fungi(100);k__Fungi_unclassified(100);k__Fungi_unclassified(100);k__Fungi_unclassified(100);k__Fungi_unclassified(100);k__Fungi_unclassified(100);k__Fungi_unclassified(100); | - | - | - | - | - | - | - | Unassigned | - |
| Otu22704 | 1 | k__Fungi(100);k__Fungi_unclassified(100);k__Fungi_unclassified(100);k__Fungi_unclassified(100);k__Fungi_unclassified(100);k__Fungi_unclassified(100);k__Fungi_unclassified(100); | - | - | - | - | - | - | - | Unassigned | - |
| Otu22706 | 1 | k__Fungi(100);p__Ascomycota(100);p__Ascomycota_unclassified(100);p__Ascomycota_unclassified(100);p__Ascomycota_unclassified(100);p__Ascomycota_unclassified(100);p__Ascomycota_unclassified(100); | - | - | - | - | - | - | - | Unassigned | - |
| Otu22707 | 1 | k__Fungi(100);p__Ascomycota(100);p__Ascomycota_unclassified(100);p__Ascomycota_unclassified(100);p__Ascomycota_unclassified(100);p__Ascomycota_unclassified(100);p__Ascomycota_unclassified(100); | - | - | - | - | - | - | - | Unassigned | - |
| Otu22716 | 1 | k__Fungi(100);k__Fungi_unclassified(100);k__Fungi_unclassified(100);k__Fungi_unclassified(100);k__Fungi_unclassified(100);k__Fungi_unclassified(100);k__Fungi_unclassified(100); | - | - | - | - | - | - | - | Unassigned | - |
| Otu22720 | 1 | k__Fungi(100);p__Ascomycota(100);p__Ascomycota_unclassified(100);p__Ascomycota_unclassified(100);p__Ascomycota_unclassified(100);p__Ascomycota_unclassified(100);p__Ascomycota_unclassified(100); | - | - | - | - | - | - | - | Unassigned | - |
| Otu22725 | 1 | k__Fungi(100);p__Ascomycota(100);p__Ascomycota_unclassified(100);p__Ascomycota_unclassified(100);p__Ascomycota_unclassified(100);p__Ascomycota_unclassified(100);p__Ascomycota_unclassified(100); | - | - | - | - | - | - | - | Unassigned | - |
| Otu22749 | 1 | k__Fungi(100);p__Ascomycota(100);p__Ascomycota_unclassified(100);p__Ascomycota_unclassified(100);p__Ascomycota_unclassified(100);p__Ascomycota_unclassified(100);p__Ascomycota_unclassified(100); | - | - | - | - | - | - | - | Unassigned | - |
| Otu22752 | 1 | k__Fungi(100);p__Ascomycota(100);p__Ascomycota_unclassified(100);p__Ascomycota_unclassified(100);p__Ascomycota_unclassified(100);p__Ascomycota_unclassified(100);p__Ascomycota_unclassified(100); | - | - | - | - | - | - | - | Unassigned | - |
| Otu22755 | 1 | k__Fungi(100);p__Ascomycota(100);c__Sordariomycetes(100);c__Sordariomycetes_unclassified(100);c__Sordariomycetes_unclassified(100);c__Sordariomycetes_unclassified(100);c__Sordariomycetes_unclassified(100); | - | - | - | - | - | - | - | Unassigned | - |
| Otu22791 | 1 | k__Fungi(100);p__Ascomycota(100);p__Ascomycota_unclassified(100);p__Ascomycota_unclassified(100);p__Ascomycota_unclassified(100);p__Ascomycota_unclassified(100);p__Ascomycota_unclassified(100); | - | - | - | - | - | - | - | Unassigned | - |
| Otu22792 | 1 | k__Fungi(100);k__Fungi_unclassified(100);k__Fungi_unclassified(100);k__Fungi_unclassified(100);k__Fungi_unclassified(100);k__Fungi_unclassified(100);k__Fungi_unclassified(100); | - | - | - | - | - | - | - | Unassigned | - |
| Otu22793 | 1 | k__Fungi(100);p__Ascomycota(100);p__Ascomycota_unclassified(100);p__Ascomycota_unclassified(100);p__Ascomycota_unclassified(100);p__Ascomycota_unclassified(100);p__Ascomycota_unclassified(100); | - | - | - | - | - | - | - | Unassigned | - |
| Otu22800 | 1 | k__Fungi(100);k__Fungi_unclassified(100);k__Fungi_unclassified(100);k__Fungi_unclassified(100);k__Fungi_unclassified(100);k__Fungi_unclassified(100);k__Fungi_unclassified(100); | - | - | - | - | - | - | - | Unassigned | - |
| Otu22803 | 1 | k__Fungi(100);k__Fungi_unclassified(100);k__Fungi_unclassified(100);k__Fungi_unclassified(100);k__Fungi_unclassified(100);k__Fungi_unclassified(100);k__Fungi_unclassified(100); | - | - | - | - | - | - | - | Unassigned | - |
| Otu22815 | 1 | k__Fungi(100);k__Fungi_unclassified(100);k__Fungi_unclassified(100);k__Fungi_unclassified(100);k__Fungi_unclassified(100);k__Fungi_unclassified(100);k__Fungi_unclassified(100); | - | - | - | - | - | - | - | Unassigned | - |
| Otu22916 | 1 | k__Fungi(100);p__Ascomycota(100);p__Ascomycota_unclassified(100);p__Ascomycota_unclassified(100);p__Ascomycota_unclassified(100);p__Ascomycota_unclassified(100);p__Ascomycota_unclassified(100); | - | - | - | - | - | - | - | Unassigned | - |
| Otu22924 | 1 | k__Fungi(100);k__Fungi_unclassified(100);k__Fungi_unclassified(100);k__Fungi_unclassified(100);k__Fungi_unclassified(100);k__Fungi_unclassified(100);k__Fungi_unclassified(100); | - | - | - | - | - | - | - | Unassigned | - |
| Otu22926 | 1 | k__Fungi(100);k__Fungi_unclassified(100);k__Fungi_unclassified(100);k__Fungi_unclassified(100);k__Fungi_unclassified(100);k__Fungi_unclassified(100);k__Fungi_unclassified(100); | - | - | - | - | - | - | - | Unassigned | - |
| Otu22927 | 1 | k__Fungi(100);p__Ascomycota(100);p__Ascomycota_unclassified(100);p__Ascomycota_unclassified(100);p__Ascomycota_unclassified(100);p__Ascomycota_unclassified(100);p__Ascomycota_unclassified(100); | - | - | - | - | - | - | - | Unassigned | - |
| Otu22928 | 1 | k__Fungi(100);k__Fungi_unclassified(100);k__Fungi_unclassified(100);k__Fungi_unclassified(100);k__Fungi_unclassified(100);k__Fungi_unclassified(100);k__Fungi_unclassified(100); | - | - | - | - | - | - | - | Unassigned | - |
| Otu22939 | 1 | k__Fungi(100);k__Fungi_unclassified(100);k__Fungi_unclassified(100);k__Fungi_unclassified(100);k__Fungi_unclassified(100);k__Fungi_unclassified(100);k__Fungi_unclassified(100); | - | - | - | - | - | - | - | Unassigned | - |
| Otu22946 | 1 | k__Fungi(100);k__Fungi_unclassified(100);k__Fungi_unclassified(100);k__Fungi_unclassified(100);k__Fungi_unclassified(100);k__Fungi_unclassified(100);k__Fungi_unclassified(100); | - | - | - | - | - | - | - | Unassigned | - |
| Otu22958 | 1 | k__Fungi(100);p__Ascomycota(100);p__Ascomycota_unclassified(100);p__Ascomycota_unclassified(100);p__Ascomycota_unclassified(100);p__Ascomycota_unclassified(100);p__Ascomycota_unclassified(100); | - | - | - | - | - | - | - | Unassigned | - |
| Otu22959 | 1 | k__Fungi(100);k__Fungi_unclassified(100);k__Fungi_unclassified(100);k__Fungi_unclassified(100);k__Fungi_unclassified(100);k__Fungi_unclassified(100);k__Fungi_unclassified(100); | - | - | - | - | - | - | - | Unassigned | - |
| Otu22960 | 1 | k__Fungi(100);p__Ascomycota(100);p__Ascomycota_unclassified(100);p__Ascomycota_unclassified(100);p__Ascomycota_unclassified(100);p__Ascomycota_unclassified(100);p__Ascomycota_unclassified(100); | - | - | - | - | - | - | - | Unassigned | - |
| Otu22968 | 1 | k__Fungi(100);k__Fungi_unclassified(100);k__Fungi_unclassified(100);k__Fungi_unclassified(100);k__Fungi_unclassified(100);k__Fungi_unclassified(100);k__Fungi_unclassified(100); | - | - | - | - | - | - | - | Unassigned | - |
| Otu23001 | 1 | k__Fungi(100);k__Fungi_unclassified(100);k__Fungi_unclassified(100);k__Fungi_unclassified(100);k__Fungi_unclassified(100);k__Fungi_unclassified(100);k__Fungi_unclassified(100); | - | - | - | - | - | - | - | Unassigned | - |
| Otu23003 | 1 | k__Fungi(100);k__Fungi_unclassified(100);k__Fungi_unclassified(100);k__Fungi_unclassified(100);k__Fungi_unclassified(100);k__Fungi_unclassified(100);k__Fungi_unclassified(100); | - | - | - | - | - | - | - | Unassigned | - |
| Otu23005 | 1 | k__Fungi(100);k__Fungi_unclassified(100);k__Fungi_unclassified(100);k__Fungi_unclassified(100);k__Fungi_unclassified(100);k__Fungi_unclassified(100);k__Fungi_unclassified(100); | - | - | - | - | - | - | - | Unassigned | - |
| Otu23006 | 1 | k__Fungi(100);p__Ascomycota(100);p__Ascomycota_unclassified(100);p__Ascomycota_unclassified(100);p__Ascomycota_unclassified(100);p__Ascomycota_unclassified(100);p__Ascomycota_unclassified(100); | - | - | - | - | - | - | - | Unassigned | - |
| Otu23038 | 1 | k__Fungi(100);p__Ascomycota(100);p__Ascomycota_unclassified(100);p__Ascomycota_unclassified(100);p__Ascomycota_unclassified(100);p__Ascomycota_unclassified(100);p__Ascomycota_unclassified(100); | - | - | - | - | - | - | - | Unassigned | - |
| Otu23041 | 1 | k__Fungi(100);k__Fungi_unclassified(100);k__Fungi_unclassified(100);k__Fungi_unclassified(100);k__Fungi_unclassified(100);k__Fungi_unclassified(100);k__Fungi_unclassified(100); | - | - | - | - | - | - | - | Unassigned | - |
| Otu23073 | 1 | k__Fungi(100);k__Fungi_unclassified(100);k__Fungi_unclassified(100);k__Fungi_unclassified(100);k__Fungi_unclassified(100);k__Fungi_unclassified(100);k__Fungi_unclassified(100); | - | - | - | - | - | - | - | Unassigned | - |
| Otu23074 | 1 | k__Fungi(100);p__Ascomycota(100);p__Ascomycota_unclassified(100);p__Ascomycota_unclassified(100);p__Ascomycota_unclassified(100);p__Ascomycota_unclassified(100);p__Ascomycota_unclassified(100); | - | - | - | - | - | - | - | Unassigned | - |
| Otu23076 | 1 | k__Fungi(100);k__Fungi_unclassified(100);k__Fungi_unclassified(100);k__Fungi_unclassified(100);k__Fungi_unclassified(100);k__Fungi_unclassified(100);k__Fungi_unclassified(100); | - | - | - | - | - | - | - | Unassigned | - |
| Otu23077 | 1 | k__Fungi(100);p__Ascomycota(100);p__Ascomycota_unclassified(100);p__Ascomycota_unclassified(100);p__Ascomycota_unclassified(100);p__Ascomycota_unclassified(100);p__Ascomycota_unclassified(100); | - | - | - | - | - | - | - | Unassigned | - |
| Otu23081 | 1 | k__Fungi(100);k__Fungi_unclassified(100);k__Fungi_unclassified(100);k__Fungi_unclassified(100);k__Fungi_unclassified(100);k__Fungi_unclassified(100);k__Fungi_unclassified(100); | - | - | - | - | - | - | - | Unassigned | - |
| Otu23085 | 1 | k__Fungi(100);k__Fungi_unclassified(100);k__Fungi_unclassified(100);k__Fungi_unclassified(100);k__Fungi_unclassified(100);k__Fungi_unclassified(100);k__Fungi_unclassified(100); | - | - | - | - | - | - | - | Unassigned | - |
| Otu23120 | 1 | k__Fungi(100);k__Fungi_unclassified(100);k__Fungi_unclassified(100);k__Fungi_unclassified(100);k__Fungi_unclassified(100);k__Fungi_unclassified(100);k__Fungi_unclassified(100); | - | - | - | - | - | - | - | Unassigned | - |
| Otu23121 | 1 | k__Fungi(100);k__Fungi_unclassified(100);k__Fungi_unclassified(100);k__Fungi_unclassified(100);k__Fungi_unclassified(100);k__Fungi_unclassified(100);k__Fungi_unclassified(100); | - | - | - | - | - | - | - | Unassigned | - |
| Otu23122 | 1 | k__Fungi(100);k__Fungi_unclassified(100);k__Fungi_unclassified(100);k__Fungi_unclassified(100);k__Fungi_unclassified(100);k__Fungi_unclassified(100);k__Fungi_unclassified(100); | - | - | - | - | - | - | - | Unassigned | - |
| Otu23123 | 1 | k__Fungi(100);k__Fungi_unclassified(100);k__Fungi_unclassified(100);k__Fungi_unclassified(100);k__Fungi_unclassified(100);k__Fungi_unclassified(100);k__Fungi_unclassified(100); | - | - | - | - | - | - | - | Unassigned | - |
| Otu23125 | 1 | k__Fungi(100);k__Fungi_unclassified(100);k__Fungi_unclassified(100);k__Fungi_unclassified(100);k__Fungi_unclassified(100);k__Fungi_unclassified(100);k__Fungi_unclassified(100); | - | - | - | - | - | - | - | Unassigned | - |
| Otu23126 | 1 | k__Fungi(100);p__Ascomycota(100);p__Ascomycota_unclassified(100);p__Ascomycota_unclassified(100);p__Ascomycota_unclassified(100);p__Ascomycota_unclassified(100);p__Ascomycota_unclassified(100); | - | - | - | - | - | - | - | Unassigned | - |
| Otu23127 | 1 | k__Fungi(100);k__Fungi_unclassified(100);k__Fungi_unclassified(100);k__Fungi_unclassified(100);k__Fungi_unclassified(100);k__Fungi_unclassified(100);k__Fungi_unclassified(100); | - | - | - | - | - | - | - | Unassigned | - |
| Otu23136 | 1 | k__Fungi(100);k__Fungi_unclassified(100);k__Fungi_unclassified(100);k__Fungi_unclassified(100);k__Fungi_unclassified(100);k__Fungi_unclassified(100);k__Fungi_unclassified(100); | - | - | - | - | - | - | - | Unassigned | - |
| Otu23149 | 1 | k__Fungi(100);k__Fungi_unclassified(100);k__Fungi_unclassified(100);k__Fungi_unclassified(100);k__Fungi_unclassified(100);k__Fungi_unclassified(100);k__Fungi_unclassified(100); | - | - | - | - | - | - | - | Unassigned | - |
| Otu23150 | 1 | k__Fungi(100);k__Fungi_unclassified(100);k__Fungi_unclassified(100);k__Fungi_unclassified(100);k__Fungi_unclassified(100);k__Fungi_unclassified(100);k__Fungi_unclassified(100); | - | - | - | - | - | - | - | Unassigned | - |
| Otu23151 | 1 | k__Fungi(100);k__Fungi_unclassified(100);k__Fungi_unclassified(100);k__Fungi_unclassified(100);k__Fungi_unclassified(100);k__Fungi_unclassified(100);k__Fungi_unclassified(100); | - | - | - | - | - | - | - | Unassigned | - |
| Otu23152 | 1 | k__Fungi(100);k__Fungi_unclassified(100);k__Fungi_unclassified(100);k__Fungi_unclassified(100);k__Fungi_unclassified(100);k__Fungi_unclassified(100);k__Fungi_unclassified(100); | - | - | - | - | - | - | - | Unassigned | - |
| Otu23153 | 1 | k__Fungi(100);k__Fungi_unclassified(100);k__Fungi_unclassified(100);k__Fungi_unclassified(100);k__Fungi_unclassified(100);k__Fungi_unclassified(100);k__Fungi_unclassified(100); | - | - | - | - | - | - | - | Unassigned | - |
| Otu23154 | 1 | k__Fungi(100);k__Fungi_unclassified(100);k__Fungi_unclassified(100);k__Fungi_unclassified(100);k__Fungi_unclassified(100);k__Fungi_unclassified(100);k__Fungi_unclassified(100); | - | - | - | - | - | - | - | Unassigned | - |
| Otu23155 | 1 | k__Fungi(100);k__Fungi_unclassified(100);k__Fungi_unclassified(100);k__Fungi_unclassified(100);k__Fungi_unclassified(100);k__Fungi_unclassified(100);k__Fungi_unclassified(100); | - | - | - | - | - | - | - | Unassigned | - |
| Otu23156 | 1 | k__Fungi(100);k__Fungi_unclassified(100);k__Fungi_unclassified(100);k__Fungi_unclassified(100);k__Fungi_unclassified(100);k__Fungi_unclassified(100);k__Fungi_unclassified(100); | - | - | - | - | - | - | - | Unassigned | - |
| Otu23157 | 1 | k__Fungi(100);p__Ascomycota(100);p__Ascomycota_unclassified(100);p__Ascomycota_unclassified(100);p__Ascomycota_unclassified(100);p__Ascomycota_unclassified(100);p__Ascomycota_unclassified(100); | - | - | - | - | - | - | - | Unassigned | - |
| Otu23158 | 1 | k__Fungi(100);k__Fungi_unclassified(100);k__Fungi_unclassified(100);k__Fungi_unclassified(100);k__Fungi_unclassified(100);k__Fungi_unclassified(100);k__Fungi_unclassified(100); | - | - | - | - | - | - | - | Unassigned | - |
| Otu23159 | 1 | k__Fungi(100);k__Fungi_unclassified(100);k__Fungi_unclassified(100);k__Fungi_unclassified(100);k__Fungi_unclassified(100);k__Fungi_unclassified(100);k__Fungi_unclassified(100); | - | - | - | - | - | - | - | Unassigned | - |
| Otu23160 | 1 | k__Fungi(100);k__Fungi_unclassified(100);k__Fungi_unclassified(100);k__Fungi_unclassified(100);k__Fungi_unclassified(100);k__Fungi_unclassified(100);k__Fungi_unclassified(100); | - | - | - | - | - | - | - | Unassigned | - |
| Otu23161 | 1 | k__Fungi(100);p__Ascomycota(100);p__Ascomycota_unclassified(100);p__Ascomycota_unclassified(100);p__Ascomycota_unclassified(100);p__Ascomycota_unclassified(100);p__Ascomycota_unclassified(100); | - | - | - | - | - | - | - | Unassigned | - |
| Otu23162 | 1 | k__Fungi(100);p__Ascomycota(100);p__Ascomycota_unclassified(100);p__Ascomycota_unclassified(100);p__Ascomycota_unclassified(100);p__Ascomycota_unclassified(100);p__Ascomycota_unclassified(100); | - | - | - | - | - | - | - | Unassigned | - |
| Otu23163 | 1 | k__Fungi(100);k__Fungi_unclassified(100);k__Fungi_unclassified(100);k__Fungi_unclassified(100);k__Fungi_unclassified(100);k__Fungi_unclassified(100);k__Fungi_unclassified(100); | - | - | - | - | - | - | - | Unassigned | - |
| Otu23165 | 1 | k__Fungi(100);k__Fungi_unclassified(100);k__Fungi_unclassified(100);k__Fungi_unclassified(100);k__Fungi_unclassified(100);k__Fungi_unclassified(100);k__Fungi_unclassified(100); | - | - | - | - | - | - | - | Unassigned | - |
| Otu23170 | 1 | k__Fungi(100);p__Ascomycota(100);c__Leotiomycetes(100);c__Leotiomycetes_unclassified(100);c__Leotiomycetes_unclassified(100);c__Leotiomycetes_unclassified(100);c__Leotiomycetes_unclassified(100); | - | - | - | - | - | - | - | Unassigned | - |
| Otu23171 | 1 | k__Fungi(100);k__Fungi_unclassified(100);k__Fungi_unclassified(100);k__Fungi_unclassified(100);k__Fungi_unclassified(100);k__Fungi_unclassified(100);k__Fungi_unclassified(100); | - | - | - | - | - | - | - | Unassigned | - |
| Otu23175 | 1 | k__Fungi(100);k__Fungi_unclassified(100);k__Fungi_unclassified(100);k__Fungi_unclassified(100);k__Fungi_unclassified(100);k__Fungi_unclassified(100);k__Fungi_unclassified(100); | - | - | - | - | - | - | - | Unassigned | - |
| Otu23183 | 1 | k__Fungi(100);k__Fungi_unclassified(100);k__Fungi_unclassified(100);k__Fungi_unclassified(100);k__Fungi_unclassified(100);k__Fungi_unclassified(100);k__Fungi_unclassified(100); | - | - | - | - | - | - | - | Unassigned | - |
| Otu23185 | 1 | k__Fungi(100);p__Ascomycota(100);p__Ascomycota_unclassified(100);p__Ascomycota_unclassified(100);p__Ascomycota_unclassified(100);p__Ascomycota_unclassified(100);p__Ascomycota_unclassified(100); | - | - | - | - | - | - | - | Unassigned | - |
| Otu23189 | 1 | k__Fungi(100);k__Fungi_unclassified(100);k__Fungi_unclassified(100);k__Fungi_unclassified(100);k__Fungi_unclassified(100);k__Fungi_unclassified(100);k__Fungi_unclassified(100); | - | - | - | - | - | - | - | Unassigned | - |
| Otu23202 | 1 | k__Fungi(100);p__Ascomycota(100);p__Ascomycota_unclassified(100);p__Ascomycota_unclassified(100);p__Ascomycota_unclassified(100);p__Ascomycota_unclassified(100);p__Ascomycota_unclassified(100); | - | - | - | - | - | - | - | Unassigned | - |
| Otu23230 | 1 | k__Fungi(100);k__Fungi_unclassified(100);k__Fungi_unclassified(100);k__Fungi_unclassified(100);k__Fungi_unclassified(100);k__Fungi_unclassified(100);k__Fungi_unclassified(100); | - | - | - | - | - | - | - | Unassigned | - |
| Otu23231 | 1 | k__Fungi(100);k__Fungi_unclassified(100);k__Fungi_unclassified(100);k__Fungi_unclassified(100);k__Fungi_unclassified(100);k__Fungi_unclassified(100);k__Fungi_unclassified(100); | - | - | - | - | - | - | - | Unassigned | - |

| | | | | | | | | | | | |
|---|---|---|---|---|---|---|---|---|---|---|---|
| Otu23233 | 1 | k__Fungi(100);k__Fungi_unclassified(100);k__Fungi_unclassified(100);k__Fungi_unclassified(100);k__Fungi_unclassified(100);k__Fungi_unclassified(100);k__Fungi_unclassified(100); | - | - | - | - | - | - | - | Unassigned | - |
| Otu23234 | 1 | k__Fungi(100);p__Ascomycota(100);p__Ascomycota_unclassified(100);p__Ascomycota_unclassified(100);p__Ascomycota_unclassified(100);p__Ascomycota_unclassified(100);p__Ascomycota_unclassified(100); | - | - | - | - | - | - | - | Unassigned | - |
| Otu23236 | 1 | k__Fungi(100);p__Ascomycota(100);p__Ascomycota_unclassified(100);p__Ascomycota_unclassified(100);p__Ascomycota_unclassified(100);p__Ascomycota_unclassified(100);p__Ascomycota_unclassified(100); | - | - | - | - | - | - | - | Unassigned | - |
| Otu23237 | 1 | k__Fungi(100);k__Fungi_unclassified(100);k__Fungi_unclassified(100);k__Fungi_unclassified(100);k__Fungi_unclassified(100);k__Fungi_unclassified(100);k__Fungi_unclassified(100); | - | - | - | - | - | - | - | Unassigned | - |
| Otu23238 | 1 | k__Fungi(100);k__Fungi_unclassified(100);k__Fungi_unclassified(100);k__Fungi_unclassified(100);k__Fungi_unclassified(100);k__Fungi_unclassified(100);k__Fungi_unclassified(100); | - | - | - | - | - | - | - | Unassigned | - |
| Otu23239 | 1 | k__Fungi(100);p__Ascomycota(100);p__Ascomycota_unclassified(100);p__Ascomycota_unclassified(100);p__Ascomycota_unclassified(100);p__Ascomycota_unclassified(100);p__Ascomycota_unclassified(100); | - | - | - | - | - | - | - | Unassigned | - |
| Otu23240 | 1 | k__Fungi(100);k__Fungi_unclassified(100);k__Fungi_unclassified(100);k__Fungi_unclassified(100);k__Fungi_unclassified(100);k__Fungi_unclassified(100);k__Fungi_unclassified(100); | - | - | - | - | - | - | - | Unassigned | - |
| Otu23241 | 1 | k__Fungi(100);k__Fungi_unclassified(100);k__Fungi_unclassified(100);k__Fungi_unclassified(100);k__Fungi_unclassified(100);k__Fungi_unclassified(100);k__Fungi_unclassified(100); | - | - | - | - | - | - | - | Unassigned | - |
| Otu23242 | 1 | k__Fungi(100);p__Ascomycota(100);p__Ascomycota_unclassified(100);p__Ascomycota_unclassified(100);p__Ascomycota_unclassified(100);p__Ascomycota_unclassified(100);p__Ascomycota_unclassified(100); | - | - | - | - | - | - | - | Unassigned | - |
| Otu23243 | 1 | k__Fungi(100);k__Fungi_unclassified(100);k__Fungi_unclassified(100);k__Fungi_unclassified(100);k__Fungi_unclassified(100);k__Fungi_unclassified(100);k__Fungi_unclassified(100); | - | - | - | - | - | - | - | Unassigned | - |
| Otu23244 | 1 | k__Fungi(100);k__Fungi_unclassified(100);k__Fungi_unclassified(100);k__Fungi_unclassified(100);k__Fungi_unclassified(100);k__Fungi_unclassified(100);k__Fungi_unclassified(100); | - | - | - | - | - | - | - | Unassigned | - |
| Otu23245 | 1 | k__Fungi(100);k__Fungi_unclassified(100);k__Fungi_unclassified(100);k__Fungi_unclassified(100);k__Fungi_unclassified(100);k__Fungi_unclassified(100);k__Fungi_unclassified(100); | - | - | - | - | - | - | - | Unassigned | - |
| Otu23247 | 1 | k__Fungi(100);p__Ascomycota(100);p__Ascomycota_unclassified(100);p__Ascomycota_unclassified(100);p__Ascomycota_unclassified(100);p__Ascomycota_unclassified(100);p__Ascomycota_unclassified(100); | - | - | - | - | - | - | - | Unassigned | - |
| Otu23248 | 1 | k__Fungi(100);p__Ascomycota(100);p__Ascomycota_unclassified(100);p__Ascomycota_unclassified(100);p__Ascomycota_unclassified(100);p__Ascomycota_unclassified(100);p__Ascomycota_unclassified(100); | - | - | - | - | - | - | - | Unassigned | - |
| Otu23249 | 1 | k__Fungi(100);k__Fungi_unclassified(100);k__Fungi_unclassified(100);k__Fungi_unclassified(100);k__Fungi_unclassified(100);k__Fungi_unclassified(100);k__Fungi_unclassified(100); | - | - | - | - | - | - | - | Unassigned | - |
| Otu23250 | 1 | k__Fungi(100);k__Fungi_unclassified(100);k__Fungi_unclassified(100);k__Fungi_unclassified(100);k__Fungi_unclassified(100);k__Fungi_unclassified(100);k__Fungi_unclassified(100); | - | - | - | - | - | - | - | Unassigned | - |
| Otu23253 | 1 | k__Fungi(100);k__Fungi_unclassified(100);k__Fungi_unclassified(100);k__Fungi_unclassified(100);k__Fungi_unclassified(100);k__Fungi_unclassified(100);k__Fungi_unclassified(100); | - | - | - | - | - | - | - | Unassigned | - |
| Otu23254 | 1 | k__Fungi(100);k__Fungi_unclassified(100);k__Fungi_unclassified(100);k__Fungi_unclassified(100);k__Fungi_unclassified(100);k__Fungi_unclassified(100);k__Fungi_unclassified(100); | - | - | - | - | - | - | - | Unassigned | - |
| Otu23255 | 1 | k__Fungi(100);k__Fungi_unclassified(100);k__Fungi_unclassified(100);k__Fungi_unclassified(100);k__Fungi_unclassified(100);k__Fungi_unclassified(100);k__Fungi_unclassified(100); | - | - | - | - | - | - | - | Unassigned | - |
| Otu23256 | 1 | k__Fungi(100);k__Fungi_unclassified(100);k__Fungi_unclassified(100);k__Fungi_unclassified(100);k__Fungi_unclassified(100);k__Fungi_unclassified(100);k__Fungi_unclassified(100); | - | - | - | - | - | - | - | Unassigned | - |
| Otu23257 | 1 | k__Fungi(100);p__Ascomycota(100);p__Ascomycota_unclassified(100);p__Ascomycota_unclassified(100);p__Ascomycota_unclassified(100);p__Ascomycota_unclassified(100);p__Ascomycota_unclassified(100); | - | - | - | - | - | - | - | Unassigned | - |
| Otu23258 | 1 | k__Fungi(100);k__Fungi_unclassified(100);k__Fungi_unclassified(100);k__Fungi_unclassified(100);k__Fungi_unclassified(100);k__Fungi_unclassified(100);k__Fungi_unclassified(100); | - | - | - | - | - | - | - | Unassigned | - |
| Otu23259 | 1 | k__Fungi(100);k__Fungi_unclassified(100);k__Fungi_unclassified(100);k__Fungi_unclassified(100);k__Fungi_unclassified(100);k__Fungi_unclassified(100);k__Fungi_unclassified(100); | - | - | - | - | - | - | - | Unassigned | - |
| Otu23260 | 1 | k__Fungi(100);p__Ascomycota(100);p__Ascomycota_unclassified(100);p__Ascomycota_unclassified(100);p__Ascomycota_unclassified(100);p__Ascomycota_unclassified(100);p__Ascomycota_unclassified(100); | - | - | - | - | - | - | - | Unassigned | - |
| Otu23261 | 1 | k__Fungi(100);k__Fungi_unclassified(100);k__Fungi_unclassified(100);k__Fungi_unclassified(100);k__Fungi_unclassified(100);k__Fungi_unclassified(100);k__Fungi_unclassified(100); | - | - | - | - | - | - | - | Unassigned | - |
| Otu23262 | 1 | k__Fungi(100);k__Fungi_unclassified(100);k__Fungi_unclassified(100);k__Fungi_unclassified(100);k__Fungi_unclassified(100);k__Fungi_unclassified(100);k__Fungi_unclassified(100); | - | - | - | - | - | - | - | Unassigned | - |
| Otu23263 | 1 | k__Fungi(100);k__Fungi_unclassified(100);k__Fungi_unclassified(100);k__Fungi_unclassified(100);k__Fungi_unclassified(100);k__Fungi_unclassified(100);k__Fungi_unclassified(100); | - | - | - | - | - | - | - | Unassigned | - |
| Otu23264 | 1 | k__Fungi(100);p__Ascomycota(100);p__Ascomycota_unclassified(100);p__Ascomycota_unclassified(100);p__Ascomycota_unclassified(100);p__Ascomycota_unclassified(100);p__Ascomycota_unclassified(100); | - | - | - | - | - | - | - | Unassigned | - |
| Otu23265 | 1 | k__Fungi(100);p__Ascomycota(100);p__Ascomycota_unclassified(100);p__Ascomycota_unclassified(100);p__Ascomycota_unclassified(100);p__Ascomycota_unclassified(100);p__Ascomycota_unclassified(100); | - | - | - | - | - | - | - | Unassigned | - |
| Otu23266 | 1 | k__Fungi(100);k__Fungi_unclassified(100);k__Fungi_unclassified(100);k__Fungi_unclassified(100);k__Fungi_unclassified(100);k__Fungi_unclassified(100);k__Fungi_unclassified(100); | - | - | - | - | - | - | - | Unassigned | - |
| Otu23267 | 1 | k__Fungi(100);p__Ascomycota(100);p__Ascomycota_unclassified(100);p__Ascomycota_unclassified(100);p__Ascomycota_unclassified(100);p__Ascomycota_unclassified(100);p__Ascomycota_unclassified(100); | - | - | - | - | - | - | - | Unassigned | - |
| Otu23268 | 1 | k__Fungi(100);p__Ascomycota(100);p__Ascomycota_unclassified(100);p__Ascomycota_unclassified(100);p__Ascomycota_unclassified(100);p__Ascomycota_unclassified(100);p__Ascomycota_unclassified(100); | - | - | - | - | - | - | - | Unassigned | - |
| Otu23269 | 1 | k__Fungi(100);k__Fungi_unclassified(100);k__Fungi_unclassified(100);k__Fungi_unclassified(100);k__Fungi_unclassified(100);k__Fungi_unclassified(100);k__Fungi_unclassified(100); | - | - | - | - | - | - | - | Unassigned | - |
| Otu23270 | 1 | k__Fungi(100);k__Fungi_unclassified(100);k__Fungi_unclassified(100);k__Fungi_unclassified(100);k__Fungi_unclassified(100);k__Fungi_unclassified(100);k__Fungi_unclassified(100); | - | - | - | - | - | - | - | Unassigned | - |
| Otu23271 | 1 | k__Fungi(100);k__Fungi_unclassified(100);k__Fungi_unclassified(100);k__Fungi_unclassified(100);k__Fungi_unclassified(100);k__Fungi_unclassified(100);k__Fungi_unclassified(100); | - | - | - | - | - | - | - | Unassigned | - |
| Otu23272 | 1 | k__Fungi(100);k__Fungi_unclassified(100);k__Fungi_unclassified(100);k__Fungi_unclassified(100);k__Fungi_unclassified(100);k__Fungi_unclassified(100);k__Fungi_unclassified(100); | - | - | - | - | - | - | - | Unassigned | - |
| Otu23273 | 1 | k__Fungi(100);k__Fungi_unclassified(100);k__Fungi_unclassified(100);k__Fungi_unclassified(100);k__Fungi_unclassified(100);k__Fungi_unclassified(100);k__Fungi_unclassified(100); | - | - | - | - | - | - | - | Unassigned | - |
| Otu23274 | 1 | k__Fungi(100);k__Fungi_unclassified(100);k__Fungi_unclassified(100);k__Fungi_unclassified(100);k__Fungi_unclassified(100);k__Fungi_unclassified(100);k__Fungi_unclassified(100); | - | - | - | - | - | - | - | Unassigned | - |
| Otu23275 | 1 | k__Fungi(100);k__Fungi_unclassified(100);k__Fungi_unclassified(100);k__Fungi_unclassified(100);k__Fungi_unclassified(100);k__Fungi_unclassified(100);k__Fungi_unclassified(100); | - | - | - | - | - | - | - | Unassigned | - |
| Otu23276 | 1 | k__Fungi(100);k__Fungi_unclassified(100);k__Fungi_unclassified(100);k__Fungi_unclassified(100);k__Fungi_unclassified(100);k__Fungi_unclassified(100);k__Fungi_unclassified(100); | - | - | - | - | - | - | - | Unassigned | - |
| Otu23277 | 1 | k__Fungi(100);p__Ascomycota(100);p__Ascomycota_unclassified(100);p__Ascomycota_unclassified(100);p__Ascomycota_unclassified(100);p__Ascomycota_unclassified(100);p__Ascomycota_unclassified(100); | - | - | - | - | - | - | - | Unassigned | - |
| Otu23278 | 1 | k__Fungi(100);k__Fungi_unclassified(100);k__Fungi_unclassified(100);k__Fungi_unclassified(100);k__Fungi_unclassified(100);k__Fungi_unclassified(100);k__Fungi_unclassified(100); | - | - | - | - | - | - | - | Unassigned | - |
| Otu23279 | 1 | k__Fungi(100);k__Fungi_unclassified(100);k__Fungi_unclassified(100);k__Fungi_unclassified(100);k__Fungi_unclassified(100);k__Fungi_unclassified(100);k__Fungi_unclassified(100); | - | - | - | - | - | - | - | Unassigned | - |
| Otu23280 | 1 | k__Fungi(100);k__Fungi_unclassified(100);k__Fungi_unclassified(100);k__Fungi_unclassified(100);k__Fungi_unclassified(100);k__Fungi_unclassified(100);k__Fungi_unclassified(100); | - | - | - | - | - | - | - | Unassigned | - |
| Otu23281 | 1 | k__Fungi(100);p__Ascomycota(100);p__Ascomycota_unclassified(100);p__Ascomycota_unclassified(100);p__Ascomycota_unclassified(100);p__Ascomycota_unclassified(100);p__Ascomycota_unclassified(100); | - | - | - | - | - | - | - | Unassigned | - |
| Otu23282 | 1 | k__Fungi(100);k__Fungi_unclassified(100);k__Fungi_unclassified(100);k__Fungi_unclassified(100);k__Fungi_unclassified(100);k__Fungi_unclassified(100);k__Fungi_unclassified(100); | - | - | - | - | - | - | - | Unassigned | - |
| Otu23283 | 1 | k__Fungi(100);k__Fungi_unclassified(100);k__Fungi_unclassified(100);k__Fungi_unclassified(100);k__Fungi_unclassified(100);k__Fungi_unclassified(100);k__Fungi_unclassified(100); | - | - | - | - | - | - | - | Unassigned | - |
| Otu23285 | 1 | k__Fungi(100);k__Fungi_unclassified(100);k__Fungi_unclassified(100);k__Fungi_unclassified(100);k__Fungi_unclassified(100);k__Fungi_unclassified(100);k__Fungi_unclassified(100); | - | - | - | - | - | - | - | Unassigned | - |
| Otu23286 | 1 | k__Fungi(100);k__Fungi_unclassified(100);k__Fungi_unclassified(100);k__Fungi_unclassified(100);k__Fungi_unclassified(100);k__Fungi_unclassified(100);k__Fungi_unclassified(100); | - | - | - | - | - | - | - | Unassigned | - |
| Otu23287 | 1 | k__Fungi(100);k__Fungi_unclassified(100);k__Fungi_unclassified(100);k__Fungi_unclassified(100);k__Fungi_unclassified(100);k__Fungi_unclassified(100);k__Fungi_unclassified(100); | - | - | - | - | - | - | - | Unassigned | - |
| Otu23288 | 1 | k__Fungi(100);k__Fungi_unclassified(100);k__Fungi_unclassified(100);k__Fungi_unclassified(100);k__Fungi_unclassified(100);k__Fungi_unclassified(100);k__Fungi_unclassified(100); | - | - | - | - | - | - | - | Unassigned | - |
| Otu23289 | 1 | k__Fungi(100);p__Ascomycota(100);p__Ascomycota_unclassified(100);p__Ascomycota_unclassified(100);p__Ascomycota_unclassified(100);p__Ascomycota_unclassified(100);p__Ascomycota_unclassified(100); | - | - | - | - | - | - | - | Unassigned | - |
| Otu23290 | 1 | k__Fungi(100);p__Ascomycota(100);p__Ascomycota_unclassified(100);p__Ascomycota_unclassified(100);p__Ascomycota_unclassified(100);p__Ascomycota_unclassified(100);p__Ascomycota_unclassified(100); | - | - | - | - | - | - | - | Unassigned | - |
| Otu23291 | 1 | k__Fungi(100);p__Ascomycota(100);p__Ascomycota_unclassified(100);p__Ascomycota_unclassified(100);p__Ascomycota_unclassified(100);p__Ascomycota_unclassified(100);p__Ascomycota_unclassified(100); | - | - | - | - | - | - | - | Unassigned | - |
| Otu23292 | 1 | k__Fungi(100);p__Ascomycota(100);p__Ascomycota_unclassified(100);p__Ascomycota_unclassified(100);p__Ascomycota_unclassified(100);p__Ascomycota_unclassified(100);p__Ascomycota_unclassified(100); | - | - | - | - | - | - | - | Unassigned | - |
| Otu23293 | 1 | k__Fungi(100);p__Ascomycota(100);p__Ascomycota_unclassified(100);p__Ascomycota_unclassified(100);p__Ascomycota_unclassified(100);p__Ascomycota_unclassified(100);p__Ascomycota_unclassified(100); | - | - | - | - | - | - | - | Unassigned | - |
| Otu23294 | 1 | k__Fungi(100);k__Fungi_unclassified(100);k__Fungi_unclassified(100);k__Fungi_unclassified(100);k__Fungi_unclassified(100);k__Fungi_unclassified(100);k__Fungi_unclassified(100); | - | - | - | - | - | - | - | Unassigned | - |
| Otu23295 | 1 | k__Fungi(100);k__Fungi_unclassified(100);k__Fungi_unclassified(100);k__Fungi_unclassified(100);k__Fungi_unclassified(100);k__Fungi_unclassified(100);k__Fungi_unclassified(100); | - | - | - | - | - | - | - | Unassigned | - |
| Otu23296 | 1 | k__Fungi(100);k__Fungi_unclassified(100);k__Fungi_unclassified(100);k__Fungi_unclassified(100);k__Fungi_unclassified(100);k__Fungi_unclassified(100);k__Fungi_unclassified(100); | - | - | - | - | - | - | - | Unassigned | - |
| Otu23298 | 1 | k__Fungi(100);k__Fungi_unclassified(100);k__Fungi_unclassified(100);k__Fungi_unclassified(100);k__Fungi_unclassified(100);k__Fungi_unclassified(100);k__Fungi_unclassified(100); | - | - | - | - | - | - | - | Unassigned | - |
| Otu23299 | 1 | k__Fungi(100);k__Fungi_unclassified(100);k__Fungi_unclassified(100);k__Fungi_unclassified(100);k__Fungi_unclassified(100);k__Fungi_unclassified(100);k__Fungi_unclassified(100); | - | - | - | - | - | - | - | Unassigned | - |
| Otu23300 | 1 | k__Fungi(100);p__Ascomycota(100);p__Ascomycota_unclassified(100);p__Ascomycota_unclassified(100);p__Ascomycota_unclassified(100);p__Ascomycota_unclassified(100);p__Ascomycota_unclassified(100); | - | - | - | - | - | - | - | Unassigned | - |
| Otu23301 | 1 | k__Fungi(100);k__Fungi_unclassified(100);k__Fungi_unclassified(100);k__Fungi_unclassified(100);k__Fungi_unclassified(100);k__Fungi_unclassified(100);k__Fungi_unclassified(100); | - | - | - | - | - | - | - | Unassigned | - |
| Otu23302 | 1 | k__Fungi(100);k__Fungi_unclassified(100);k__Fungi_unclassified(100);k__Fungi_unclassified(100);k__Fungi_unclassified(100);k__Fungi_unclassified(100);k__Fungi_unclassified(100); | - | - | - | - | - | - | - | Unassigned | - |
| Otu23303 | 1 | k__Fungi(100);p__Ascomycota(100);p__Ascomycota_unclassified(100);p__Ascomycota_unclassified(100);p__Ascomycota_unclassified(100);p__Ascomycota_unclassified(100);p__Ascomycota_unclassified(100); | - | - | - | - | - | - | - | Unassigned | - |
| Otu23304 | 1 | k__Fungi(100);p__Ascomycota(100);p__Ascomycota_unclassified(100);p__Ascomycota_unclassified(100);p__Ascomycota_unclassified(100);p__Ascomycota_unclassified(100);p__Ascomycota_unclassified(100); | - | - | - | - | - | - | - | Unassigned | - |
| Otu23305 | 1 | k__Fungi(100);k__Fungi_unclassified(100);k__Fungi_unclassified(100);k__Fungi_unclassified(100);k__Fungi_unclassified(100);k__Fungi_unclassified(100);k__Fungi_unclassified(100); | - | - | - | - | - | - | - | Unassigned | - |
| Otu23308 | 1 | k__Fungi(100);k__Fungi_unclassified(100);k__Fungi_unclassified(100);k__Fungi_unclassified(100);k__Fungi_unclassified(100);k__Fungi_unclassified(100);k__Fungi_unclassified(100); | - | - | - | - | - | - | - | Unassigned | - |
| Otu23309 | 1 | k__Fungi(100);k__Fungi_unclassified(100);k__Fungi_unclassified(100);k__Fungi_unclassified(100);k__Fungi_unclassified(100);k__Fungi_unclassified(100);k__Fungi_unclassified(100); | - | - | - | - | - | - | - | Unassigned | - |
| Otu23311 | 1 | k__Fungi(100);k__Fungi_unclassified(100);k__Fungi_unclassified(100);k__Fungi_unclassified(100);k__Fungi_unclassified(100);k__Fungi_unclassified(100);k__Fungi_unclassified(100); | - | - | - | - | - | - | - | Unassigned | - |
| Otu23312 | 1 | k__Fungi(100);k__Fungi_unclassified(100);k__Fungi_unclassified(100);k__Fungi_unclassified(100);k__Fungi_unclassified(100);k__Fungi_unclassified(100);k__Fungi_unclassified(100); | - | - | - | - | - | - | - | Unassigned | - |
| Otu23313 | 1 | k__Fungi(100);p__Ascomycota(100);p__Ascomycota_unclassified(100);p__Ascomycota_unclassified(100);p__Ascomycota_unclassified(100);p__Ascomycota_unclassified(100);p__Ascomycota_unclassified(100); | - | - | - | - | - | - | - | Unassigned | - |
| Otu23314 | 1 | k__Fungi(100);k__Fungi_unclassified(100);k__Fungi_unclassified(100);k__Fungi_unclassified(100);k__Fungi_unclassified(100);k__Fungi_unclassified(100);k__Fungi_unclassified(100); | - | - | - | - | - | - | - | Unassigned | - |
| Otu23315 | 1 | k__Fungi(100);k__Fungi_unclassified(100);k__Fungi_unclassified(100);k__Fungi_unclassified(100);k__Fungi_unclassified(100);k__Fungi_unclassified(100);k__Fungi_unclassified(100); | - | - | - | - | - | - | - | Unassigned | - |
| Otu23316 | 1 | k__Fungi(100);k__Fungi_unclassified(100);k__Fungi_unclassified(100);k__Fungi_unclassified(100);k__Fungi_unclassified(100);k__Fungi_unclassified(100);k__Fungi_unclassified(100); | - | - | - | - | - | - | - | Unassigned | - |
| Otu23317 | 1 | k__Fungi(100);p__Ascomycota(100);p__Ascomycota_unclassified(100);p__Ascomycota_unclassified(100);p__Ascomycota_unclassified(100);p__Ascomycota_unclassified(100);p__Ascomycota_unclassified(100); | - | - | - | - | - | - | - | Unassigned | - |
| Otu23318 | 1 | k__Fungi(100);k__Fungi_unclassified(100);k__Fungi_unclassified(100);k__Fungi_unclassified(100);k__Fungi_unclassified(100);k__Fungi_unclassified(100);k__Fungi_unclassified(100); | - | - | - | - | - | - | - | Unassigned | - |
| Otu23319 | 1 | k__Fungi(100);p__Ascomycota(100);p__Ascomycota_unclassified(100);p__Ascomycota_unclassified(100);p__Ascomycota_unclassified(100);p__Ascomycota_unclassified(100);p__Ascomycota_unclassified(100); | - | - | - | - | - | - | - | Unassigned | - |
| Otu23327 | 1 | k__Fungi(100);p__Ascomycota(100);p__Ascomycota_unclassified(100);p__Ascomycota_unclassified(100);p__Ascomycota_unclassified(100);p__Ascomycota_unclassified(100);p__Ascomycota_unclassified(100); | - | - | - | - | - | - | - | Unassigned | - |
| Otu23331 | 1 | k__Fungi(100);k__Fungi_unclassified(100);k__Fungi_unclassified(100);k__Fungi_unclassified(100);k__Fungi_unclassified(100);k__Fungi_unclassified(100);k__Fungi_unclassified(100); | - | - | - | - | - | - | - | Unassigned | - |
| Otu23334 | 1 | k__Fungi(100);k__Fungi_unclassified(100);k__Fungi_unclassified(100);k__Fungi_unclassified(100);k__Fungi_unclassified(100);k__Fungi_unclassified(100);k__Fungi_unclassified(100); | - | - | - | - | - | - | - | Unassigned | - |
| Otu23336 | 1 | k__Fungi(100);k__Fungi_unclassified(100);k__Fungi_unclassified(100);k__Fungi_unclassified(100);k__Fungi_unclassified(100);k__Fungi_unclassified(100);k__Fungi_unclassified(100); | - | - | - | - | - | - | - | Unassigned | - |
| Otu23337 | 1 | k__Fungi(100);k__Fungi_unclassified(100);k__Fungi_unclassified(100);k__Fungi_unclassified(100);k__Fungi_unclassified(100);k__Fungi_unclassified(100);k__Fungi_unclassified(100); | - | - | - | - | - | - | - | Unassigned | - |
| Otu23338 | 1 | k__Fungi(100);p__Ascomycota(100);p__Ascomycota_unclassified(100);p__Ascomycota_unclassified(100);p__Ascomycota_unclassified(100);p__Ascomycota_unclassified(100);p__Ascomycota_unclassified(100); | - | - | - | - | - | - | - | Unassigned | - |
| Otu23339 | 1 | k__Fungi(100);p__Ascomycota(100);p__Ascomycota_unclassified(100);p__Ascomycota_unclassified(100);p__Ascomycota_unclassified(100);p__Ascomycota_unclassified(100);p__Ascomycota_unclassified(100); | - | - | - | - | - | - | - | Unassigned | - |
| Otu23340 | 1 | k__Fungi(100);k__Fungi_unclassified(100);k__Fungi_unclassified(100);k__Fungi_unclassified(100);k__Fungi_unclassified(100);k__Fungi_unclassified(100);k__Fungi_unclassified(100); | - | - | - | - | - | - | - | Unassigned | - |
| Otu23341 | 1 | k__Fungi(100);k__Fungi_unclassified(100);k__Fungi_unclassified(100);k__Fungi_unclassified(100);k__Fungi_unclassified(100);k__Fungi_unclassified(100);k__Fungi_unclassified(100); | - | - | - | - | - | - | - | Unassigned | - |
| Otu23342 | 1 | k__Fungi(100);k__Fungi_unclassified(100);k__Fungi_unclassified(100);k__Fungi_unclassified(100);k__Fungi_unclassified(100);k__Fungi_unclassified(100);k__Fungi_unclassified(100); | - | - | - | - | - | - | - | Unassigned | - |
| Otu23343 | 1 | k__Fungi(100);p__Ascomycota(100);p__Ascomycota_unclassified(100);p__Ascomycota_unclassified(100);p__Ascomycota_unclassified(100);p__Ascomycota_unclassified(100);p__Ascomycota_unclassified(100); | - | - | - | - | - | - | - | Unassigned | - |
| Otu23344 | 1 | k__Fungi(100);p__Ascomycota(100);p__Ascomycota_unclassified(100);p__Ascomycota_unclassified(100);p__Ascomycota_unclassified(100);p__Ascomycota_unclassified(100);p__Ascomycota_unclassified(100); | - | - | - | - | - | - | - | Unassigned | - |
| Otu23345 | 1 | k__Fungi(100);k__Fungi_unclassified(100);k__Fungi_unclassified(100);k__Fungi_unclassified(100);k__Fungi_unclassified(100);k__Fungi_unclassified(100);k__Fungi_unclassified(100); | - | - | - | - | - | - | - | Unassigned | - |
| Otu23346 | 1 | k__Fungi(100);p__Ascomycota(100);p__Ascomycota_unclassified(100);p__Ascomycota_unclassified(100);p__Ascomycota_unclassified(100);p__Ascomycota_unclassified(100);p__Ascomycota_unclassified(100); | - | - | - | - | - | - | - | Unassigned | - |
| Otu23347 | 1 | k__Fungi(100);p__Ascomycota(100);p__Ascomycota_unclassified(100);p__Ascomycota_unclassified(100);p__Ascomycota_unclassified(100);p__Ascomycota_unclassified(100);p__Ascomycota_unclassified(100); | - | - | - | - | - | - | - | Unassigned | - |
| Otu23348 | 1 | k__Fungi(100);k__Fungi_unclassified(100);k__Fungi_unclassified(100);k__Fungi_unclassified(100);k__Fungi_unclassified(100);k__Fungi_unclassified(100);k__Fungi_unclassified(100); | - | - | - | - | - | - | - | Unassigned | - |
| Otu23349 | 1 | k__Fungi(100);p__Ascomycota(100);p__Ascomycota_unclassified(100);p__Ascomycota_unclassified(100);p__Ascomycota_unclassified(100);p__Ascomycota_unclassified(100);p__Ascomycota_unclassified(100); | - | - | - | - | - | - | - | Unassigned | - |
| Otu23350 | 1 | k__Fungi(100);k__Fungi_unclassified(100);k__Fungi_unclassified(100);k__Fungi_unclassified(100);k__Fungi_unclassified(100);k__Fungi_unclassified(100);k__Fungi_unclassified(100); | - | - | - | - | - | - | - | Unassigned | - |
| Otu23351 | 1 | k__Fungi(100);k__Fungi_unclassified(100);k__Fungi_unclassified(100);k__Fungi_unclassified(100);k__Fungi_unclassified(100);k__Fungi_unclassified(100);k__Fungi_unclassified(100); | - | - | - | - | - | - | - | Unassigned | - |
| Otu23352 | 1 | k__Fungi(100);p__Ascomycota(100);p__Ascomycota_unclassified(100);p__Ascomycota_unclassified(100);p__Ascomycota_unclassified(100);p__Ascomycota_unclassified(100);p__Ascomycota_unclassified(100); | - | - | - | - | - | - | - | Unassigned | - |
| Otu23353 | 1 | k__Fungi(100);p__Ascomycota(100);p__Ascomycota_unclassified(100);p__Ascomycota_unclassified(100);p__Ascomycota_unclassified(100);p__Ascomycota_unclassified(100);p__Ascomycota_unclassified(100); | - | - | - | - | - | - | - | Unassigned | - |
| Otu23354 | 1 | k__Fungi(100);p__Ascomycota(100);p__Ascomycota_unclassified(100);p__Ascomycota_unclassified(100);p__Ascomycota_unclassified(100);p__Ascomycota_unclassified(100);p__Ascomycota_unclassified(100); | - | - | - | - | - | - | - | Unassigned | - |
| Otu23355 | 1 | k__Fungi(100);k__Fungi_unclassified(100);k__Fungi_unclassified(100);k__Fungi_unclassified(100);k__Fungi_unclassified(100);k__Fungi_unclassified(100);k__Fungi_unclassified(100); | - | - | - | - | - | - | - | Unassigned | - |
| Otu23356 | 1 | k__Fungi(100);p__Ascomycota(100);p__Ascomycota_unclassified(100);p__Ascomycota_unclassified(100);p__Ascomycota_unclassified(100);p__Ascomycota_unclassified(100);p__Ascomycota_unclassified(100); | - | - | - | - | - | - | - | Unassigned | - |
| Otu23357 | 1 | k__Fungi(100);p__Ascomycota(100);p__Ascomycota_unclassified(100);p__Ascomycota_unclassified(100);p__Ascomycota_unclassified(100);p__Ascomycota_unclassified(100);p__Ascomycota_unclassified(100); | - | - | - | - | - | - | - | Unassigned | - |
| Otu23358 | 1 | k__Fungi(100);k__Fungi_unclassified(100);k__Fungi_unclassified(100);k__Fungi_unclassified(100);k__Fungi_unclassified(100);k__Fungi_unclassified(100);k__Fungi_unclassified(100); | - | - | - | - | - | - | - | Unassigned | - |
| Otu23359 | 1 | k__Fungi(100);p__Ascomycota(100);p__Ascomycota_unclassified(100);p__Ascomycota_unclassified(100);p__Ascomycota_unclassified(100);p__Ascomycota_unclassified(100);p__Ascomycota_unclassified(100); | - | - | - | - | - | - | - | Unassigned | - |
| Otu23360 | 1 | k__Fungi(100);p__Ascomycota(100);p__Ascomycota_unclassified(100);p__Ascomycota_unclassified(100);p__Ascomycota_unclassified(100);p__Ascomycota_unclassified(100);p__Ascomycota_unclassified(100); | - | - | - | - | - | - | - | Unassigned | - |
| Otu23361 | 1 | k__Fungi(100);k__Fungi_unclassified(100);k__Fungi_unclassified(100);k__Fungi_unclassified(100);k__Fungi_unclassified(100);k__Fungi_unclassified(100);k__Fungi_unclassified(100); | - | - | - | - | - | - | - | Unassigned | - |
| Otu23362 | 1 | k__Fungi(100);p__Ascomycota(100);p__Ascomycota_unclassified(100);p__Ascomycota_unclassified(100);p__Ascomycota_unclassified(100);p__Ascomycota_unclassified(100);p__Ascomycota_unclassified(100); | - | - | - | - | - | - | - | Unassigned | - |
| Otu23363 | 1 | k__Fungi(100);p__Ascomycota(100);p__Ascomycota_unclassified(100);p__Ascomycota_unclassified(100);p__Ascomycota_unclassified(100);p__Ascomycota_unclassified(100);p__Ascomycota_unclassified(100); | - | - | - | - | - | - | - | Unassigned | - |
| Otu23365 | 1 | k__Fungi(100);k__Fungi_unclassified(100);k__Fungi_unclassified(100);k__Fungi_unclassified(100);k__Fungi_unclassified(100);k__Fungi_unclassified(100);k__Fungi_unclassified(100); | - | - | - | - | - | - | - | Unassigned | - |
| Otu23366 | 1 | k__Fungi(100);p__Ascomycota(100);p__Ascomycota_unclassified(100);p__Ascomycota_unclassified(100);p__Ascomycota_unclassified(100);p__Ascomycota_unclassified(100);p__Ascomycota_unclassified(100); | - | - | - | - | - | - | - | Unassigned | - |
| Otu23367 | 1 | k__Fungi(100);p__Ascomycota(100);p__Ascomycota_unclassified(100);p__Ascomycota_unclassified(100);p__Ascomycota_unclassified(100);p__Ascomycota_unclassified(100);p__Ascomycota_unclassified(100); | - | - | - | - | - | - | - | Unassigned | - |
| Otu23368 | 1 | k__Fungi(100);k__Fungi_unclassified(100);k__Fungi_unclassified(100);k__Fungi_unclassified(100);k__Fungi_unclassified(100);k__Fungi_unclassified(100);k__Fungi_unclassified(100); | - | - | - | - | - | - | - | Unassigned | - |
| Otu23369 | 1 | k__Fungi(100);k__Fungi_unclassified(100);k__Fungi_unclassified(100);k__Fungi_unclassified(100);k__Fungi_unclassified(100);k__Fungi_unclassified(100);k__Fungi_unclassified(100); | - | - | - | - | - | - | - | Unassigned | - |
| Otu23370 | 1 | k__Fungi(100);k__Fungi_unclassified(100);k__Fungi_unclassified(100);k__Fungi_unclassified(100);k__Fungi_unclassified(100);k__Fungi_unclassified(100);k__Fungi_unclassified(100); | - | - | - | - | - | - | - | Unassigned | - |
| Otu23371 | 1 | k__Fungi(100);k__Fungi_unclassified(100);k__Fungi_unclassified(100);k__Fungi_unclassified(100);k__Fungi_unclassified(100);k__Fungi_unclassified(100);k__Fungi_unclassified(100); | - | - | - | - | - | - | - | Unassigned | - |
| Otu23372 | 1 | k__Fungi(100);k__Fungi_unclassified(100);k__Fungi_unclassified(100);k__Fungi_unclassified(100);k__Fungi_unclassified(100);k__Fungi_unclassified(100);k__Fungi_unclassified(100); | - | - | - | - | - | - | - | Unassigned | - |
| Otu23373 | 1 | k__Fungi(100);k__Fungi_unclassified(100);k__Fungi_unclassified(100);k__Fungi_unclassified(100);k__Fungi_unclassified(100);k__Fungi_unclassified(100);k__Fungi_unclassified(100); | - | - | - | - | - | - | - | Unassigned | - |
| Otu23374 | 1 | k__Fungi(100);k__Fungi_unclassified(100);k__Fungi_unclassified(100);k__Fungi_unclassified(100);k__Fungi_unclassified(100);k__Fungi_unclassified(100);k__Fungi_unclassified(100); | - | - | - | - | - | - | - | Unassigned | - |
| Otu23375 | 1 | k__Fungi(100);k__Fungi_unclassified(100);k__Fungi_unclassified(100);k__Fungi_unclassified(100);k__Fungi_unclassified(100);k__Fungi_unclassified(100);k__Fungi_unclassified(100); | - | - | - | - | - | - | - | Unassigned | - |
| Otu23376 | 1 | k__Fungi(100);p__Ascomycota(100);p__Ascomycota_unclassified(100);p__Ascomycota_unclassified(100);p__Ascomycota_unclassified(100);p__Ascomycota_unclassified(100);p__Ascomycota_unclassified(100); | - | - | - | - | - | - | - | Unassigned | - |
| Otu23377 | 1 | k__Fungi(100);k__Fungi_unclassified(100);k__Fungi_unclassified(100);k__Fungi_unclassified(100);k__Fungi_unclassified(100);k__Fungi_unclassified(100);k__Fungi_unclassified(100); | - | - | - | - | - | - | - | Unassigned | - |
| Otu23378 | 1 | k__Fungi(100);k__Fungi_unclassified(100);k__Fungi_unclassified(100);k__Fungi_unclassified(100);k__Fungi_unclassified(100);k__Fungi_unclassified(100);k__Fungi_unclassified(100); | - | - | - | - | - | - | - | Unassigned | - |
| Otu23379 | 1 | k__Fungi(100);k__Fungi_unclassified(100);k__Fungi_unclassified(100);k__Fungi_unclassified(100);k__Fungi_unclassified(100);k__Fungi_unclassified(100);k__Fungi_unclassified(100); | - | - | - | - | - | - | - | Unassigned | - |
| Otu23380 | 1 | k__Fungi(100);k__Fungi_unclassified(100);k__Fungi_unclassified(100);k__Fungi_unclassified(100);k__Fungi_unclassified(100);k__Fungi_unclassified(100);k__Fungi_unclassified(100); | - | - | - | - | - | - | - | Unassigned | - |
| Otu23382 | 1 | k__Fungi(100);k__Fungi_unclassified(100);k__Fungi_unclassified(100);k__Fungi_unclassified(100);k__Fungi_unclassified(100);k__Fungi_unclassified(100);k__Fungi_unclassified(100); | - | - | - | - | - | - | - | Unassigned | - |
| Otu23384 | 1 | k__Fungi(100);k__Fungi_unclassified(100);k__Fungi_unclassified(100);k__Fungi_unclassified(100);k__Fungi_unclassified(100);k__Fungi_unclassified(100);k__Fungi_unclassified(100); | - | - | - | - | - | - | - | Unassigned | - |
| Otu23385 | 1 | k__Fungi(100);k__Fungi_unclassified(100);k__Fungi_unclassified(100);k__Fungi_unclassified(100);k__Fungi_unclassified(100);k__Fungi_unclassified(100);k__Fungi_unclassified(100); | - | - | - | - | - | - | - | Unassigned | - |
| Otu23386 | 1 | k__Fungi(100);k__Fungi_unclassified(100);k__Fungi_unclassified(100);k__Fungi_unclassified(100);k__Fungi_unclassified(100);k__Fungi_unclassified(100);k__Fungi_unclassified(100); | - | - | - | - | - | - | - | Unassigned | - |
| Otu23387 | 1 | k__Fungi(100);k__Fungi_unclassified(100);k__Fungi_unclassified(100);k__Fungi_unclassified(100);k__Fungi_unclassified(100);k__Fungi_unclassified(100);k__Fungi_unclassified(100); | - | - | - | - | - | - | - | Unassigned | - |
| Otu23388 | 1 | k__Fungi(100);k__Fungi_unclassified(100);k__Fungi_unclassified(100);k__Fungi_unclassified(100);k__Fungi_unclassified(100);k__Fungi_unclassified(100);k__Fungi_unclassified(100); | - | - | - | - | - | - | - | Unassigned | - |
| Otu23389 | 1 | k__Fungi(100);k__Fungi_unclassified(100);k__Fungi_unclassified(100);k__Fungi_unclassified(100);k__Fungi_unclassified(100);k__Fungi_unclassified(100);k__Fungi_unclassified(100); | - | - | - | - | - | - | - | Unassigned | - |
| Otu23390 | 1 | k__Fungi(100);p__Ascomycota(100);p__Ascomycota_unclassified(100);p__Ascomycota_unclassified(100);p__Ascomycota_unclassified(100);p__Ascomycota_unclassified(100);p__Ascomycota_unclassified(100); | - | - | - | - | - | - | - | Unassigned | - |
| Otu23391 | 1 | k__Fungi(100);k__Fungi_unclassified(100);k__Fungi_unclassified(100);k__Fungi_unclassified(100);k__Fungi_unclassified(100);k__Fungi_unclassified(100);k__Fungi_unclassified(100); | - | - | - | - | - | - | - | Unassigned | - |
| Otu23392 | 1 | k__Fungi(100);k__Fungi_unclassified(100);k__Fungi_unclassified(100);k__Fungi_unclassified(100);k__Fungi_unclassified(100);k__Fungi_unclassified(100);k__Fungi_unclassified(100); | - | - | - | - | - | - | - | Unassigned | - |
| Otu23393 | 1 | k__Fungi(100);k__Fungi_unclassified(100);k__Fungi_unclassified(100);k__Fungi_unclassified(100);k__Fungi_unclassified(100);k__Fungi_unclassified(100);k__Fungi_unclassified(100); | - | - | - | - | - | - | - | Unassigned | - |
| Otu23394 | 1 | k__Fungi(100);k__Fungi_unclassified(100);k__Fungi_unclassified(100);k__Fungi_unclassified(100);k__Fungi_unclassified(100);k__Fungi_unclassified(100);k__Fungi_unclassified(100); | - | - | - | - | - | - | - | Unassigned | - |
| Otu23395 | 1 | k__Fungi(100);k__Fungi_unclassified(100);k__Fungi_unclassified(100);k__Fungi_unclassified(100);k__Fungi_unclassified(100);k__Fungi_unclassified(100);k__Fungi_unclassified(100); | - | - | - | - | - | - | - | Unassigned | - |
| Otu23396 | 1 | k__Fungi(100);p__Ascomycota(100);p__Ascomycota_unclassified(100);p__Ascomycota_unclassified(100);p__Ascomycota_unclassified(100);p__Ascomycota_unclassified(100);p__Ascomycota_unclassified(100); | - | - | - | - | - | - | - | Unassigned | - |
| Otu23397 | 1 | k__Fungi(100);k__Fungi_unclassified(100);k__Fungi_unclassified(100);k__Fungi_unclassified(100);k__Fungi_unclassified(100);k__Fungi_unclassified(100);k__Fungi_unclassified(100); | - | - | - | - | - | - | - | Unassigned | - |
| Otu23398 | 1 | k__Fungi(100);k__Fungi_unclassified(100);k__Fungi_unclassified(100);k__Fungi_unclassified(100);k__Fungi_unclassified(100);k__Fungi_unclassified(100);k__Fungi_unclassified(100); | - | - | - | - | - | - | - | Unassigned | - |
| Otu23399 | 1 | k__Fungi(100);p__Ascomycota(100);p__Ascomycota_unclassified(100);p__Ascomycota_unclassified(100);p__Ascomycota_unclassified(100);p__Ascomycota_unclassified(100);p__Ascomycota_unclassified(100); | - | - | - | - | - | - | - | Unassigned | - |
| Otu23400 | 1 | k__Fungi(100);k__Fungi_unclassified(100);k__Fungi_unclassified(100);k__Fungi_unclassified(100);k__Fungi_unclassified(100);k__Fungi_unclassified(100);k__Fungi_unclassified(100); | - | - | - | - | - | - | - | Unassigned | - |
| Otu23401 | 1 | k__Fungi(100);k__Fungi_unclassified(100);k__Fungi_unclassified(100);k__Fungi_unclassified(100);k__Fungi_unclassified(100);k__Fungi_unclassified(100);k__Fungi_unclassified(100); | - | - | - | - | - | - | - | Unassigned | - |
| Otu23402 | 1 | k__Fungi(100);k__Fungi_unclassified(100);k__Fungi_unclassified(100);k__Fungi_unclassified(100);k__Fungi_unclassified(100);k__Fungi_unclassified(100);k__Fungi_unclassified(100); | - | - | - | - | - | - | - | Unassigned | - |
| Otu23403 | 1 | k__Fungi(100);k__Fungi_unclassified(100);k__Fungi_unclassified(100);k__Fungi_unclassified(100);k__Fungi_unclassified(100);k__Fungi_unclassified(100);k__Fungi_unclassified(100); | - | - | - | - | - | - | - | Unassigned | - |
| Otu23404 | 1 | k__Fungi(100);k__Fungi_unclassified(100);k__Fungi_unclassified(100);k__Fungi_unclassified(100);k__Fungi_unclassified(100);k__Fungi_unclassified(100);k__Fungi_unclassified(100); | - | - | - | - | - | - | - | Unassigned | - |
| Otu23405 | 1 | k__Fungi(100);k__Fungi_unclassified(100);k__Fungi_unclassified(100);k__Fungi_unclassified(100);k__Fungi_unclassified(100);k__Fungi_unclassified(100);k__Fungi_unclassified(100); | - | - | - | - | - | - | - | Unassigned | - |
| Otu23406 | 1 | k__Fungi(100);k__Fungi_unclassified(100);k__Fungi_unclassified(100);k__Fungi_unclassified(100);k__Fungi_unclassified(100);k__Fungi_unclassified(100);k__Fungi_unclassified(100); | - | - | - | - | - | - | - | Unassigned | - |
| Otu23408 | 1 | k__Fungi(100);p__Ascomycota(100);p__Ascomycota_unclassified(100);p__Ascomycota_unclassified(100);p__Ascomycota_unclassified(100);p__Ascomycota_unclassified(100);p__Ascomycota_unclassified(100); | - | - | - | - | - | - | - | Unassigned | - |
| Otu23409 | 1 | k__Fungi(100);k__Fungi_unclassified(100);k__Fungi_unclassified(100);k__Fungi_unclassified(100);k__Fungi_unclassified(100);k__Fungi_unclassified(100);k__Fungi_unclassified(100); | - | - | - | - | - | - | - | Unassigned | - |
| Otu23410 | 1 | k__Fungi(100);p__Ascomycota(100);p__Ascomycota_unclassified(100);p__Ascomycota_unclassified(100);p__Ascomycota_unclassified(100);p__Ascomycota_unclassified(100);p__Ascomycota_unclassified(100); | - | - | - | - | - | - | - | Unassigned | - |
| Otu23411 | 1 | k__Fungi(100);p__Ascomycota(100);p__Ascomycota_unclassified(100);p__Ascomycota_unclassified(100);p__Ascomycota_unclassified(100);p__Ascomycota_unclassified(100);p__Ascomycota_unclassified(100); | - | - | - | - | - | - | - | Unassigned | - |
| Otu23412 | 1 | k__Fungi(100);p__Ascomycota(100);p__Ascomycota_unclassified(100);p__Ascomycota_unclassified(100);p__Ascomycota_unclassified(100);p__Ascomycota_unclassified(100);p__Ascomycota_unclassified(100); | - | - | - | - | - | - | - | Unassigned | - |
| Otu23413 | 1 | k__Fungi(100);k__Fungi_unclassified(100);k__Fungi_unclassified(100);k__Fungi_unclassified(100);k__Fungi_unclassified(100);k__Fungi_unclassified(100);k__Fungi_unclassified(100); | - | - | - | - | - | - | - | Unassigned | - |
| Otu23414 | 1 | k__Fungi(100);k__Fungi_unclassified(100);k__Fungi_unclassified(100);k__Fungi_unclassified(100);k__Fungi_unclassified(100);k__Fungi_unclassified(100);k__Fungi_unclassified(100); | - | - | - | - | - | - | - | Unassigned | - |
| Otu23415 | 1 | k__Fungi(100);k__Fungi_unclassified(100);k__Fungi_unclassified(100);k__Fungi_unclassified(100);k__Fungi_unclassified(100);k__Fungi_unclassified(100);k__Fungi_unclassified(100); | - | - | - | - | - | - | - | Unassigned | - |
| Otu23416 | 1 | k__Fungi(100);k__Fungi_unclassified(100);k__Fungi_unclassified(100);k__Fungi_unclassified(100);k__Fungi_unclassified(100);k__Fungi_unclassified(100);k__Fungi_unclassified(100); | - | - | - | - | - | - | - | Unassigned | - |
| Otu23417 | 1 | k__Fungi(100);k__Fungi_unclassified(100);k__Fungi_unclassified(100);k__Fungi_unclassified(100);k__Fungi_unclassified(100);k__Fungi_unclassified(100);k__Fungi_unclassified(100); | - | - | - | - | - | - | - | Unassigned | - |
| Otu23418 | 1 | k__Fungi(100);p__Ascomycota(100);p__Ascomycota_unclassified(100);p__Ascomycota_unclassified(100);p__Ascomycota_unclassified(100);p__Ascomycota_unclassified(100);p__Ascomycota_unclassified(100); | - | - | - | - | - | - | - | Unassigned | - |
| Otu23419 | 1 | k__Fungi(100);k__Fungi_unclassified(100);k__Fungi_unclassified(100);k__Fungi_unclassified(100);k__Fungi_unclassified(100);k__Fungi_unclassified(100);k__Fungi_unclassified(100); | - | - | - | - | - | - | - | Unassigned | - |
| Otu23420 | 1 | k__Fungi(100);k__Fungi_unclassified(100);k__Fungi_unclassified(100);k__Fungi_unclassified(100);k__Fungi_unclassified(100);k__Fungi_unclassified(100);k__Fungi_unclassified(100); | - | - | - | - | - | - | - | Unassigned | - |
| Otu23421 | 1 | k__Fungi(100);p__Ascomycota(100);p__Ascomycota_unclassified(100);p__Ascomycota_unclassified(100);p__Ascomycota_unclassified(100);p__Ascomycota_unclassified(100);p__Ascomycota_unclassified(100); | - | - | - | - | - | - | - | Unassigned | - |
| Otu23422 | 1 | k__Fungi(100);k__Fungi_unclassified(100);k__Fungi_unclassified(100);k__Fungi_unclassified(100);k__Fungi_unclassified(100);k__Fungi_unclassified(100);k__Fungi_unclassified(100); | - | - | - | - | - | - | - | Unassigned | - |
| Otu23423 | 1 | k__Fungi(100);k__Fungi_unclassified(100);k__Fungi_unclassified(100);k__Fungi_unclassified(100);k__Fungi_unclassified(100);k__Fungi_unclassified(100);k__Fungi_unclassified(100); | - | - | - | - | - | - | - | Unassigned | - |
| Otu23424 | 1 | k__Fungi(100);k__Fungi_unclassified(100);k__Fungi_unclassified(100);k__Fungi_unclassified(100);k__Fungi_unclassified(100);k__Fungi_unclassified(100);k__Fungi_unclassified(100); | - | - | - | - | - | - | - | Unassigned | - |
| Otu23425 | 1 | k__Fungi(100);p__Ascomycota(100);p__Ascomycota_unclassified(100);p__Ascomycota_unclassified(100);p__Ascomycota_unclassified(100);p__Ascomycota_unclassified(100);p__Ascomycota_unclassified(100); | - | - | - | - | - | - | - | Unassigned | - |
| Otu23426 | 1 | k__Fungi(100);k__Fungi_unclassified(100);k__Fungi_unclassified(100);k__Fungi_unclassified(100);k__Fungi_unclassified(100);k__Fungi_unclassified(100);k__Fungi_unclassified(100); | - | - | - | - | - | - | - | Unassigned | - |
| Otu23427 | 1 | k__Fungi(100);k__Fungi_unclassified(100);k__Fungi_unclassified(100);k__Fungi_unclassified(100);k__Fungi_unclassified(100);k__Fungi_unclassified(100);k__Fungi_unclassified(100); | - | - | - | - | - | - | - | Unassigned | - |
| Otu23428 | 1 | k__Fungi(100);k__Fungi_unclassified(100);k__Fungi_unclassified(100);k__Fungi_unclassified(100);k__Fungi_unclassified(100);k__Fungi_unclassified(100);k__Fungi_unclassified(100); | - | - | - | - | - | - | - | Unassigned | - |
| Otu23430 | 1 | k__Fungi(100);k__Fungi_unclassified(100);k__Fungi_unclassified(100);k__Fungi_unclassified(100);k__Fungi_unclassified(100);k__Fungi_unclassified(100);k__Fungi_unclassified(100); | - | - | - | - | - | - | - | Unassigned | - |
| Otu23431 | 1 | k__Fungi(100);k__Fungi_unclassified(100);k__Fungi_unclassified(100);k__Fungi_unclassified(100);k__Fungi_unclassified(100);k__Fungi_unclassified(100);k__Fungi_unclassified(100); | - | - | - | - | - | - | - | Unassigned | - |
| Otu23432 | 1 | k__Fungi(100);p__Ascomycota(100);p__Ascomycota_unclassified(100);p__Ascomycota_unclassified(100);p__Ascomycota_unclassified(100);p__Ascomycota_unclassified(100);p__Ascomycota_unclassified(100); | - | - | - | - | - | - | - | Unassigned | - |
| Otu23433 | 1 | k__Fungi(100);k__Fungi_unclassified(100);k__Fungi_unclassified(100);k__Fungi_unclassified(100);k__Fungi_unclassified(100);k__Fungi_unclassified(100);k__Fungi_unclassified(100); | - | - | - | - | - | - | - | Unassigned | - |
| Otu23434 | 1 | k__Fungi(100);k__Fungi_unclassified(100);k__Fungi_unclassified(100);k__Fungi_unclassified(100);k__Fungi_unclassified(100);k__Fungi_unclassified(100);k__Fungi_unclassified(100); | - | - | - | - | - | - | - | Unassigned | - |
| Otu23435 | 1 | k__Fungi(100);k__Fungi_unclassified(100);k__Fungi_unclassified(100);k__Fungi_unclassified(100);k__Fungi_unclassified(100);k__Fungi_unclassified(100);k__Fungi_unclassified(100); | - | - | - | - | - | - | - | Unassigned | - |
| Otu23436 | 1 | k__Fungi(100);p__Ascomycota(100);p__Ascomycota_unclassified(100);p__Ascomycota_unclassified(100);p__Ascomycota_unclassified(100);p__Ascomycota_unclassified(100);p__Ascomycota_unclassified(100); | - | - | - | - | - | - | - | Unassigned | - |
| Otu23437 | 1 | k__Fungi(100);p__Ascomycota(100);p__Ascomycota_unclassified(100);p__Ascomycota_unclassified(100);p__Ascomycota_unclassified(100);p__Ascomycota_unclassified(100);p__Ascomycota_unclassified(100); | - | - | - | - | - | - | - | Unassigned | - |
| Otu23438 | 1 | k__Fungi(100);k__Fungi_unclassified(100);k__Fungi_unclassified(100);k__Fungi_unclassified(100);k__Fungi_unclassified(100);k__Fungi_unclassified(100);k__Fungi_unclassified(100); | - | - | - | - | - | - | - | Unassigned | - |
| Otu23439 | 1 | k__Fungi(100);k__Fungi_unclassified(100);k__Fungi_unclassified(100);k__Fungi_unclassified(100);k__Fungi_unclassified(100);k__Fungi_unclassified(100);k__Fungi_unclassified(100); | - | - | - | - | - | - | - | Unassigned | - |
| Otu23440 | 1 | k__Fungi(100);p__Ascomycota(100);p__Ascomycota_unclassified(100);p__Ascomycota_unclassified(100);p__Ascomycota_unclassified(100);p__Ascomycota_unclassified(100);p__Ascomycota_unclassified(100); | - | - | - | - | - | - | - | Unassigned | - |
| Otu23441 | 1 | k__Fungi(100);p__Ascomycota(100);p__Ascomycota_unclassified(100);p__Ascomycota_unclassified(100);p__Ascomycota_unclassified(100);p__Ascomycota_unclassified(100);p__Ascomycota_unclassified(100); | - | - | - | - | - | - | - | Unassigned | - |
| Otu23442 | 1 | k__Fungi(100);p__Ascomycota(100);p__Ascomycota_unclassified(100);p__Ascomycota_unclassified(100);p__Ascomycota_unclassified(100);p__Ascomycota_unclassified(100);p__Ascomycota_unclassified(100); | - | - | - | - | - | - | - | Unassigned | - |
| Otu23443 | 1 | k__Fungi(100);k__Fungi_unclassified(100);k__Fungi_unclassified(100);k__Fungi_unclassified(100);k__Fungi_unclassified(100);k__Fungi_unclassified(100);k__Fungi_unclassified(100); | - | - | - | - | - | - | - | Unassigned | - |
| Otu23444 | 1 | k__Fungi(100);k__Fungi_unclassified(100);k__Fungi_unclassified(100);k__Fungi_unclassified(100);k__Fungi_unclassified(100);k__Fungi_unclassified(100);k__Fungi_unclassified(100); | - | - | - | - | - | - | - | Unassigned | - |
| Otu23445 | 1 | k__Fungi(100);k__Fungi_unclassified(100);k__Fungi_unclassified(100);k__Fungi_unclassified(100);k__Fungi_unclassified(100);k__Fungi_unclassified(100);k__Fungi_unclassified(100); | - | - | - | - | - | - | - | Unassigned | - |
| Otu23446 | 1 | k__Fungi(100);k__Fungi_unclassified(100);k__Fungi_unclassified(100);k__Fungi_unclassified(100);k__Fungi_unclassified(100);k__Fungi_unclassified(100);k__Fungi_unclassified(100); | - | - | - | - | - | - | - | Unassigned | - |
| Otu23447 | 1 | k__Fungi(100);p__Ascomycota(100);p__Ascomycota_unclassified(100);p__Ascomycota_unclassified(100);p__Ascomycota_unclassified(100);p__Ascomycota_unclassified(100);p__Ascomycota_unclassified(100); | - | - | - | - | - | - | - | Unassigned | - |
| Otu23448 | 1 | k__Fungi(100);k__Fungi_unclassified(100);k__Fungi_unclassified(100);k__Fungi_unclassified(100);k__Fungi_unclassified(100);k__Fungi_unclassified(100);k__Fungi_unclassified(100); | - | - | - | - | - | - | - | Unassigned | - |
| Otu23449 | 1 | k__Fungi(100);p__Ascomycota(100);p__Ascomycota_unclassified(100);p__Ascomycota_unclassified(100);p__Ascomycota_unclassified(100);p__Ascomycota_unclassified(100);p__Ascomycota_unclassified(100); | - | - | - | - | - | - | - | Unassigned | - |
| Otu23450 | 1 | k__Fungi(100);k__Fungi_unclassified(100);k__Fungi_unclassified(100);k__Fungi_unclassified(100);k__Fungi_unclassified(100);k__Fungi_unclassified(100);k__Fungi_unclassified(100); | - | - | - | - | - | - | - | Unassigned | - |
| Otu23451 | 1 | k__Fungi(100);p__Ascomycota(100);p__Ascomycota_unclassified(100);p__Ascomycota_unclassified(100);p__Ascomycota_unclassified(100);p__Ascomycota_unclassified(100);p__Ascomycota_unclassified(100); | - | - | - | - | - | - | - | Unassigned | - |
| Otu23452 | 1 | k__Fungi(100);k__Fungi_unclassified(100);k__Fungi_unclassified(100);k__Fungi_unclassified(100);k__Fungi_unclassified(100);k__Fungi_unclassified(100);k__Fungi_unclassified(100); | - | - | - | - | - | - | - | Unassigned | - |
| Otu23453 | 1 | k__Fungi(100);k__Fungi_unclassified(100);k__Fungi_unclassified(100);k__Fungi_unclassified(100);k__Fungi_unclassified(100);k__Fungi_unclassified(100);k__Fungi_unclassified(100); | - | - | - | - | - | - | - | Unassigned | - |
| Otu23454 | 1 | k__Fungi(100);k__Fungi_unclassified(100);k__Fungi_unclassified(100);k__Fungi_unclassified(100);k__Fungi_unclassified(100);k__Fungi_unclassified(100);k__Fungi_unclassified(100); | - | - | - | - | - | - | - | Unassigned | - |
| Otu23455 | 1 | k__Fungi(100);k__Fungi_unclassified(100);k__Fungi_unclassified(100);k__Fungi_unclassified(100);k__Fungi_unclassified(100);k__Fungi_unclassified(100);k__Fungi_unclassified(100); | - | - | - | - | - | - | - | Unassigned | - |
| Otu23456 | 1 | k__Fungi(100);k__Fungi_unclassified(100);k__Fungi_unclassified(100);k__Fungi_unclassified(100);k__Fungi_unclassified(100);k__Fungi_unclassified(100);k__Fungi_unclassified(100); | - | - | - | - | - | - | - | Unassigned | - |
| Otu23457 | 1 | k__Fungi(100);k__Fungi_unclassified(100);k__Fungi_unclassified(100);k__Fungi_unclassified(100);k__Fungi_unclassified(100);k__Fungi_unclassified(100);k__Fungi_unclassified(100); | - | - | - | - | - | - | - | Unassigned | - |
| Otu23460 | 1 | k__Fungi(100);k__Fungi_unclassified(100);k__Fungi_unclassified(100);k__Fungi_unclassified(100);k__Fungi_unclassified(100);k__Fungi_unclassified(100);k__Fungi_unclassified(100); | - | - | - | - | - | - | - | Unassigned | - |
| Otu23461 | 1 | k__Fungi(100);k__Fungi_unclassified(100);k__Fungi_unclassified(100);k__Fungi_unclassified(100);k__Fungi_unclassified(100);k__Fungi_unclassified(100);k__Fungi_unclassified(100); | - | - | - | - | - | - | - | Unassigned | - |
| Otu23462 | 1 | k__Fungi(100);p__Ascomycota(100);p__Ascomycota_unclassified(100);p__Ascomycota_unclassified(100);p__Ascomycota_unclassified(100);p__Ascomycota_unclassified(100);p__Ascomycota_unclassified(100); | - | - | - | - | - | - | - | Unassigned | - |
| Otu23463 | 1 | k__Fungi(100);k__Fungi_unclassified(100);k__Fungi_unclassified(100);k__Fungi_unclassified(100);k__Fungi_unclassified(100);k__Fungi_unclassified(100);k__Fungi_unclassified(100); | - | - | - | - | - | - | - | Unassigned | - |
| Otu23464 | 1 | k__Fungi(100);p__Ascomycota(100);p__Ascomycota_unclassified(100);p__Ascomycota_unclassified(100);p__Ascomycota_unclassified(100);p__Ascomycota_unclassified(100);p__Ascomycota_unclassified(100); | - | - | - | - | - | - | - | Unassigned | - |
| Otu23465 | 1 | k__Fungi(100);p__Ascomycota(100);p__Ascomycota_unclassified(100);p__Ascomycota_unclassified(100);p__Ascomycota_unclassified(100);p__Ascomycota_unclassified(100);p__Ascomycota_unclassified(100); | - | - | - | - | - | - | - | Unassigned | - |
| Otu23466 | 1 | k__Fungi(100);p__Ascomycota(100);p__Ascomycota_unclassified(100);p__Ascomycota_unclassified(100);p__Ascomycota_unclassified(100);p__Ascomycota_unclassified(100);p__Ascomycota_unclassified(100); | - | - | - | - | - | - | - | Unassigned | - |
| Otu23467 | 1 | k__Fungi(100);p__Ascomycota(100);p__Ascomycota_unclassified(100);p__Ascomycota_unclassified(100);p__Ascomycota_unclassified(100);p__Ascomycota_unclassified(100);p__Ascomycota_unclassified(100); | - | - | - | - | - | - | - | Unassigned | - |
| Otu23468 | 1 | k__Fungi(100);k__Fungi_unclassified(100);k__Fungi_unclassified(100);k__Fungi_unclassified(100);k__Fungi_unclassified(100);k__Fungi_unclassified(100);k__Fungi_unclassified(100); | - | - | - | - | - | - | - | Unassigned | - |
| Otu23474 | 1 | k__Fungi(100);p__Ascomycota(100);p__Ascomycota_unclassified(100);p__Ascomycota_unclassified(100);p__Ascomycota_unclassified(100);p__Ascomycota_unclassified(100);p__Ascomycota_unclassified(100); | - | - | - | - | - | - | - | Unassigned | - |
| Otu23478 | 1 | k__Fungi(100);p__Ascomycota(100);p__Ascomycota_unclassified(100);p__Ascomycota_unclassified(100);p__Ascomycota_unclassified(100);p__Ascomycota_unclassified(100);p__Ascomycota_unclassified(100); | - | - | - | - | - | - | - | Unassigned | - |
| Otu23482 | 1 | k__Fungi(100);k__Fungi_unclassified(100);k__Fungi_unclassified(100);k__Fungi_unclassified(100);k__Fungi_unclassified(100);k__Fungi_unclassified(100);k__Fungi_unclassified(100); | - | - | - | - | - | - | - | Unassigned | - |
| Otu23484 | 1 | k__Fungi(100);k__Fungi_unclassified(100);k__Fungi_unclassified(100);k__Fungi_unclassified(100);k__Fungi_unclassified(100);k__Fungi_unclassified(100);k__Fungi_unclassified(100); | - | - | - | - | - | - | - | Unassigned | - |
| Otu23491 | 1 | k__Fungi(100);k__Fungi_unclassified(100);k__Fungi_unclassified(100);k__Fungi_unclassified(100);k__Fungi_unclassified(100);k__Fungi_unclassified(100);k__Fungi_unclassified(100); | - | - | - | - | - | - | - | Unassigned | - |
| Otu23492 | 1 | k__Fungi(100);p__Ascomycota(100);p__Ascomycota_unclassified(100);p__Ascomycota_unclassified(100);p__Ascomycota_unclassified(100);p__Ascomycota_unclassified(100);p__Ascomycota_unclassified(100); | - | - | - | - | - | - | - | Unassigned | - |
| Otu23503 | 1 | k__Fungi(100);p__Ascomycota(100);p__Ascomycota_unclassified(100);p__Ascomycota_unclassified(100);p__Ascomycota_unclassified(100);p__Ascomycota_unclassified(100);p__Ascomycota_unclassified(100); | - | - | - | - | - | - | - | Unassigned | - |
| Otu23504 | 1 | k__Fungi(100);k__Fungi_unclassified(100);k__Fungi_unclassified(100);k__Fungi_unclassified(100);k__Fungi_unclassified(100);k__Fungi_unclassified(100);k__Fungi_unclassified(100); | - | - | - | - | - | - | - | Unassigned | - |
| Otu23505 | 1 | k__Fungi(100);k__Fungi_unclassified(100);k__Fungi_unclassified(100);k__Fungi_unclassified(100);k__Fungi_unclassified(100);k__Fungi_unclassified(100);k__Fungi_unclassified(100); | - | - | - | - | - | - | - | Unassigned | - |

| | | | | | | | | | | | |
|---|---|---|---|---|---|---|---|---|---|---|---|
| Otu23506 | 1 | k__Fungi(100);k__Fungi_unclassified(100);k__Fungi_unclassified(100);k__Fungi_unclassified(100);k__Fungi_unclassified(100);k__Fungi_unclassified(100);k__Fungi_unclassified(100); | - | - | - | - | - | - | - | Unassigned | - |
| Otu23507 | 1 | k__Fungi(100);k__Fungi_unclassified(100);k__Fungi_unclassified(100);k__Fungi_unclassified(100);k__Fungi_unclassified(100);k__Fungi_unclassified(100);k__Fungi_unclassified(100); | - | - | - | - | - | - | - | Unassigned | - |
| Otu23511 | 1 | k__Fungi(100);k__Fungi_unclassified(100);k__Fungi_unclassified(100);k__Fungi_unclassified(100);k__Fungi_unclassified(100);k__Fungi_unclassified(100);k__Fungi_unclassified(100); | - | - | - | - | - | - | - | Unassigned | - |
| Otu23512 | 1 | k__Fungi(100);p__Ascomycota(100);p__Ascomycota_unclassified(100);p__Ascomycota_unclassified(100);p__Ascomycota_unclassified(100);p__Ascomycota_unclassified(100);p__Ascomycota_unclassified(100); | - | - | - | - | - | - | - | Unassigned | - |
| Otu23517 | 1 | k__Fungi(100);p__Ascomycota(100);p__Ascomycota_unclassified(100);p__Ascomycota_unclassified(100);p__Ascomycota_unclassified(100);p__Ascomycota_unclassified(100);p__Ascomycota_unclassified(100); | - | - | - | - | - | - | - | Unassigned | - |
| Otu23518 | 1 | k__Fungi(100);p__Ascomycota(100);p__Ascomycota_unclassified(100);p__Ascomycota_unclassified(100);p__Ascomycota_unclassified(100);p__Ascomycota_unclassified(100);p__Ascomycota_unclassified(100); | - | - | - | - | - | - | - | Unassigned | - |
| Otu23519 | 1 | k__Fungi(100);k__Fungi_unclassified(100);k__Fungi_unclassified(100);k__Fungi_unclassified(100);k__Fungi_unclassified(100);k__Fungi_unclassified(100);k__Fungi_unclassified(100); | - | - | - | - | - | - | - | Unassigned | - |
| Otu23522 | 1 | k__Fungi(100);k__Fungi_unclassified(100);k__Fungi_unclassified(100);k__Fungi_unclassified(100);k__Fungi_unclassified(100);k__Fungi_unclassified(100);k__Fungi_unclassified(100); | - | - | - | - | - | - | - | Unassigned | - |
| Otu23527 | 1 | k__Fungi(100);p__Ascomycota(100);p__Ascomycota_unclassified(100);p__Ascomycota_unclassified(100);p__Ascomycota_unclassified(100);p__Ascomycota_unclassified(100);p__Ascomycota_unclassified(100); | - | - | - | - | - | - | - | Unassigned | - |
| Otu23532 | 1 | k__Fungi(100);k__Fungi_unclassified(100);k__Fungi_unclassified(100);k__Fungi_unclassified(100);k__Fungi_unclassified(100);k__Fungi_unclassified(100);k__Fungi_unclassified(100); | - | - | - | - | - | - | - | Unassigned | - |
| Otu23534 | 1 | k__Fungi(100);k__Fungi_unclassified(100);k__Fungi_unclassified(100);k__Fungi_unclassified(100);k__Fungi_unclassified(100);k__Fungi_unclassified(100);k__Fungi_unclassified(100); | - | - | - | - | - | - | - | Unassigned | - |
| Otu23535 | 1 | k__Fungi(100);k__Fungi_unclassified(100);k__Fungi_unclassified(100);k__Fungi_unclassified(100);k__Fungi_unclassified(100);k__Fungi_unclassified(100);k__Fungi_unclassified(100); | - | - | - | - | - | - | - | Unassigned | - |
| Otu23536 | 1 | k__Fungi(100);k__Fungi_unclassified(100);k__Fungi_unclassified(100);k__Fungi_unclassified(100);k__Fungi_unclassified(100);k__Fungi_unclassified(100);k__Fungi_unclassified(100); | - | - | - | - | - | - | - | Unassigned | - |
| Otu23537 | 1 | k__Fungi(100);k__Fungi_unclassified(100);k__Fungi_unclassified(100);k__Fungi_unclassified(100);k__Fungi_unclassified(100);k__Fungi_unclassified(100);k__Fungi_unclassified(100); | - | - | - | - | - | - | - | Unassigned | - |
| Otu23539 | 1 | k__Fungi(100);p__Ascomycota(100);p__Ascomycota_unclassified(100);p__Ascomycota_unclassified(100);p__Ascomycota_unclassified(100);p__Ascomycota_unclassified(100);p__Ascomycota_unclassified(100); | - | - | - | - | - | - | - | Unassigned | - |
| Otu23540 | 1 | k__Fungi(100);k__Fungi_unclassified(100);k__Fungi_unclassified(100);k__Fungi_unclassified(100);k__Fungi_unclassified(100);k__Fungi_unclassified(100);k__Fungi_unclassified(100); | - | - | - | - | - | - | - | Unassigned | - |
| Otu23541 | 1 | k__Fungi(100);k__Fungi_unclassified(100);k__Fungi_unclassified(100);k__Fungi_unclassified(100);k__Fungi_unclassified(100);k__Fungi_unclassified(100);k__Fungi_unclassified(100); | - | - | - | - | - | - | - | Unassigned | - |
| Otu23543 | 1 | k__Fungi(100);k__Fungi_unclassified(100);k__Fungi_unclassified(100);k__Fungi_unclassified(100);k__Fungi_unclassified(100);k__Fungi_unclassified(100);k__Fungi_unclassified(100); | - | - | - | - | - | - | - | Unassigned | - |
| Otu23556 | 1 | k__Fungi(100);k__Fungi_unclassified(100);k__Fungi_unclassified(100);k__Fungi_unclassified(100);k__Fungi_unclassified(100);k__Fungi_unclassified(100);k__Fungi_unclassified(100); | - | - | - | - | - | - | - | Unassigned | - |
| Otu23557 | 1 | k__Fungi(100);k__Fungi_unclassified(100);k__Fungi_unclassified(100);k__Fungi_unclassified(100);k__Fungi_unclassified(100);k__Fungi_unclassified(100);k__Fungi_unclassified(100); | - | - | - | - | - | - | - | Unassigned | - |
| Otu23567 | 1 | k__Fungi(100);k__Fungi_unclassified(100);k__Fungi_unclassified(100);k__Fungi_unclassified(100);k__Fungi_unclassified(100);k__Fungi_unclassified(100);k__Fungi_unclassified(100); | - | - | - | - | - | - | - | Unassigned | - |
| Otu23582 | 1 | k__Fungi(100);k__Fungi_unclassified(100);k__Fungi_unclassified(100);k__Fungi_unclassified(100);k__Fungi_unclassified(100);k__Fungi_unclassified(100);k__Fungi_unclassified(100); | - | - | - | - | - | - | - | Unassigned | - |
| Otu23583 | 1 | k__Fungi(100);k__Fungi_unclassified(100);k__Fungi_unclassified(100);k__Fungi_unclassified(100);k__Fungi_unclassified(100);k__Fungi_unclassified(100);k__Fungi_unclassified(100); | - | - | - | - | - | - | - | Unassigned | - |
| Otu23584 | 1 | k__Fungi(100);k__Fungi_unclassified(100);k__Fungi_unclassified(100);k__Fungi_unclassified(100);k__Fungi_unclassified(100);k__Fungi_unclassified(100);k__Fungi_unclassified(100); | - | - | - | - | - | - | - | Unassigned | - |
| Otu23587 | 1 | k__Fungi(100);k__Fungi_unclassified(100);k__Fungi_unclassified(100);k__Fungi_unclassified(100);k__Fungi_unclassified(100);k__Fungi_unclassified(100);k__Fungi_unclassified(100); | - | - | - | - | - | - | - | Unassigned | - |
| Otu23600 | 1 | k__Fungi(100);p__Ascomycota(100);p__Ascomycota_unclassified(100);p__Ascomycota_unclassified(100);p__Ascomycota_unclassified(100);p__Ascomycota_unclassified(100);p__Ascomycota_unclassified(100); | - | - | - | - | - | - | - | Unassigned | - |
| Otu23602 | 1 | k__Fungi(100);k__Fungi_unclassified(100);k__Fungi_unclassified(100);k__Fungi_unclassified(100);k__Fungi_unclassified(100);k__Fungi_unclassified(100);k__Fungi_unclassified(100); | - | - | - | - | - | - | - | Unassigned | - |
| Otu23603 | 1 | k__Fungi(100);k__Fungi_unclassified(100);k__Fungi_unclassified(100);k__Fungi_unclassified(100);k__Fungi_unclassified(100);k__Fungi_unclassified(100);k__Fungi_unclassified(100); | - | - | - | - | - | - | - | Unassigned | - |
| Otu23605 | 1 | k__Fungi(100);k__Fungi_unclassified(100);k__Fungi_unclassified(100);k__Fungi_unclassified(100);k__Fungi_unclassified(100);k__Fungi_unclassified(100);k__Fungi_unclassified(100); | - | - | - | - | - | - | - | Unassigned | - |
| Otu23606 | 1 | k__Fungi(100);p__Ascomycota(100);p__Ascomycota_unclassified(100);p__Ascomycota_unclassified(100);p__Ascomycota_unclassified(100);p__Ascomycota_unclassified(100);p__Ascomycota_unclassified(100); | - | - | - | - | - | - | - | Unassigned | - |
| Otu23607 | 1 | k__Fungi(100);k__Fungi_unclassified(100);k__Fungi_unclassified(100);k__Fungi_unclassified(100);k__Fungi_unclassified(100);k__Fungi_unclassified(100);k__Fungi_unclassified(100); | - | - | - | - | - | - | - | Unassigned | - |
| Otu23610 | 1 | k__Fungi(100);k__Fungi_unclassified(100);k__Fungi_unclassified(100);k__Fungi_unclassified(100);k__Fungi_unclassified(100);k__Fungi_unclassified(100);k__Fungi_unclassified(100); | - | - | - | - | - | - | - | Unassigned | - |
| Otu23611 | 1 | k__Fungi(100);k__Fungi_unclassified(100);k__Fungi_unclassified(100);k__Fungi_unclassified(100);k__Fungi_unclassified(100);k__Fungi_unclassified(100);k__Fungi_unclassified(100); | - | - | - | - | - | - | - | Unassigned | - |
| Otu23615 | 1 | k__Fungi(100);k__Fungi_unclassified(100);k__Fungi_unclassified(100);k__Fungi_unclassified(100);k__Fungi_unclassified(100);k__Fungi_unclassified(100);k__Fungi_unclassified(100); | - | - | - | - | - | - | - | Unassigned | - |
| Otu23616 | 1 | k__Fungi(100);p__Ascomycota(100);p__Ascomycota_unclassified(100);p__Ascomycota_unclassified(100);p__Ascomycota_unclassified(100);p__Ascomycota_unclassified(100);p__Ascomycota_unclassified(100); | - | - | - | - | - | - | - | Unassigned | - |
| Otu23617 | 1 | k__Fungi(100);k__Fungi_unclassified(100);k__Fungi_unclassified(100);k__Fungi_unclassified(100);k__Fungi_unclassified(100);k__Fungi_unclassified(100);k__Fungi_unclassified(100); | - | - | - | - | - | - | - | Unassigned | - |
| Otu23618 | 1 | k__Fungi(100);p__Ascomycota(100);p__Ascomycota_unclassified(100);p__Ascomycota_unclassified(100);p__Ascomycota_unclassified(100);p__Ascomycota_unclassified(100);p__Ascomycota_unclassified(100); | - | - | - | - | - | - | - | Unassigned | - |
| Otu23619 | 1 | k__Fungi(100);k__Fungi_unclassified(100);k__Fungi_unclassified(100);k__Fungi_unclassified(100);k__Fungi_unclassified(100);k__Fungi_unclassified(100);k__Fungi_unclassified(100); | - | - | - | - | - | - | - | Unassigned | - |
| Otu23620 | 1 | k__Fungi(100);p__Ascomycota(100);p__Ascomycota_unclassified(100);p__Ascomycota_unclassified(100);p__Ascomycota_unclassified(100);p__Ascomycota_unclassified(100);p__Ascomycota_unclassified(100); | - | - | - | - | - | - | - | Unassigned | - |
| Otu23621 | 1 | k__Fungi(100);k__Fungi_unclassified(100);k__Fungi_unclassified(100);k__Fungi_unclassified(100);k__Fungi_unclassified(100);k__Fungi_unclassified(100);k__Fungi_unclassified(100); | - | - | - | - | - | - | - | Unassigned | - |
| Otu23622 | 1 | k__Fungi(100);k__Fungi_unclassified(100);k__Fungi_unclassified(100);k__Fungi_unclassified(100);k__Fungi_unclassified(100);k__Fungi_unclassified(100);k__Fungi_unclassified(100); | - | - | - | - | - | - | - | Unassigned | - |
| Otu23623 | 1 | k__Fungi(100);k__Fungi_unclassified(100);k__Fungi_unclassified(100);k__Fungi_unclassified(100);k__Fungi_unclassified(100);k__Fungi_unclassified(100);k__Fungi_unclassified(100); | - | - | - | - | - | - | - | Unassigned | - |
| Otu23624 | 1 | k__Fungi(100);p__Ascomycota(100);p__Ascomycota_unclassified(100);p__Ascomycota_unclassified(100);p__Ascomycota_unclassified(100);p__Ascomycota_unclassified(100);p__Ascomycota_unclassified(100); | - | - | - | - | - | - | - | Unassigned | - |
| Otu23625 | 1 | k__Fungi(100);p__Ascomycota(100);p__Ascomycota_unclassified(100);p__Ascomycota_unclassified(100);p__Ascomycota_unclassified(100);p__Ascomycota_unclassified(100);p__Ascomycota_unclassified(100); | - | - | - | - | - | - | - | Unassigned | - |
| Otu23626 | 1 | k__Fungi(100);p__Ascomycota(100);p__Ascomycota_unclassified(100);p__Ascomycota_unclassified(100);p__Ascomycota_unclassified(100);p__Ascomycota_unclassified(100);p__Ascomycota_unclassified(100); | - | - | - | - | - | - | - | Unassigned | - |
| Otu23627 | 1 | k__Fungi(100);k__Fungi_unclassified(100);k__Fungi_unclassified(100);k__Fungi_unclassified(100);k__Fungi_unclassified(100);k__Fungi_unclassified(100);k__Fungi_unclassified(100); | - | - | - | - | - | - | - | Unassigned | - |
| Otu23628 | 1 | k__Fungi(100);p__Ascomycota(100);p__Ascomycota_unclassified(100);p__Ascomycota_unclassified(100);p__Ascomycota_unclassified(100);p__Ascomycota_unclassified(100);p__Ascomycota_unclassified(100); | - | - | - | - | - | - | - | Unassigned | - |
| Otu23640 | 1 | k__Fungi(100);k__Fungi_unclassified(100);k__Fungi_unclassified(100);k__Fungi_unclassified(100);k__Fungi_unclassified(100);k__Fungi_unclassified(100);k__Fungi_unclassified(100); | - | - | - | - | - | - | - | Unassigned | - |
| Otu23647 | 1 | k__Fungi(100);k__Fungi_unclassified(100);k__Fungi_unclassified(100);k__Fungi_unclassified(100);k__Fungi_unclassified(100);k__Fungi_unclassified(100);k__Fungi_unclassified(100); | - | - | - | - | - | - | - | Unassigned | - |
| Otu23648 | 1 | k__Fungi(100);k__Fungi_unclassified(100);k__Fungi_unclassified(100);k__Fungi_unclassified(100);k__Fungi_unclassified(100);k__Fungi_unclassified(100);k__Fungi_unclassified(100); | - | - | - | - | - | - | - | Unassigned | - |
| Otu23650 | 1 | k__Fungi(100);k__Fungi_unclassified(100);k__Fungi_unclassified(100);k__Fungi_unclassified(100);k__Fungi_unclassified(100);k__Fungi_unclassified(100);k__Fungi_unclassified(100); | - | - | - | - | - | - | - | Unassigned | - |
| Otu23652 | 1 | k__Fungi(100);k__Fungi_unclassified(100);k__Fungi_unclassified(100);k__Fungi_unclassified(100);k__Fungi_unclassified(100);k__Fungi_unclassified(100);k__Fungi_unclassified(100); | - | - | - | - | - | - | - | Unassigned | - |
| Otu23653 | 1 | k__Fungi(100);p__Ascomycota(100);p__Ascomycota_unclassified(100);p__Ascomycota_unclassified(100);p__Ascomycota_unclassified(100);p__Ascomycota_unclassified(100);p__Ascomycota_unclassified(100); | - | - | - | - | - | - | - | Unassigned | - |
| Otu23654 | 1 | k__Fungi(100);k__Fungi_unclassified(100);k__Fungi_unclassified(100);k__Fungi_unclassified(100);k__Fungi_unclassified(100);k__Fungi_unclassified(100);k__Fungi_unclassified(100); | - | - | - | - | - | - | - | Unassigned | - |
| Otu23655 | 1 | k__Fungi(100);p__Ascomycota(100);p__Ascomycota_unclassified(100);p__Ascomycota_unclassified(100);p__Ascomycota_unclassified(100);p__Ascomycota_unclassified(100);p__Ascomycota_unclassified(100); | - | - | - | - | - | - | - | Unassigned | - |
| Otu23656 | 1 | k__Fungi(100);p__Ascomycota(100);p__Ascomycota_unclassified(100);p__Ascomycota_unclassified(100);p__Ascomycota_unclassified(100);p__Ascomycota_unclassified(100);p__Ascomycota_unclassified(100); | - | - | - | - | - | - | - | Unassigned | - |
| Otu23659 | 1 | k__Fungi(100);k__Fungi_unclassified(100);k__Fungi_unclassified(100);k__Fungi_unclassified(100);k__Fungi_unclassified(100);k__Fungi_unclassified(100);k__Fungi_unclassified(100); | - | - | - | - | - | - | - | Unassigned | - |
| Otu23660 | 1 | k__Fungi(100);p__Ascomycota(100);p__Ascomycota_unclassified(100);p__Ascomycota_unclassified(100);p__Ascomycota_unclassified(100);p__Ascomycota_unclassified(100);p__Ascomycota_unclassified(100); | - | - | - | - | - | - | - | Unassigned | - |
| Otu23663 | 1 | k__Fungi(100);k__Fungi_unclassified(100);k__Fungi_unclassified(100);k__Fungi_unclassified(100);k__Fungi_unclassified(100);k__Fungi_unclassified(100);k__Fungi_unclassified(100); | - | - | - | - | - | - | - | Unassigned | - |
| Otu23664 | 1 | k__Fungi(100);p__Ascomycota(100);p__Ascomycota_unclassified(100);p__Ascomycota_unclassified(100);p__Ascomycota_unclassified(100);p__Ascomycota_unclassified(100);p__Ascomycota_unclassified(100); | - | - | - | - | - | - | - | Unassigned | - |
| Otu23667 | 1 | k__Fungi(100);k__Fungi_unclassified(100);k__Fungi_unclassified(100);k__Fungi_unclassified(100);k__Fungi_unclassified(100);k__Fungi_unclassified(100);k__Fungi_unclassified(100); | - | - | - | - | - | - | - | Unassigned | - |
| Otu23668 | 1 | k__Fungi(100);k__Fungi_unclassified(100);k__Fungi_unclassified(100);k__Fungi_unclassified(100);k__Fungi_unclassified(100);k__Fungi_unclassified(100);k__Fungi_unclassified(100); | - | - | - | - | - | - | - | Unassigned | - |
| Otu23669 | 1 | k__Fungi(100);p__Ascomycota(100);p__Ascomycota_unclassified(100);p__Ascomycota_unclassified(100);p__Ascomycota_unclassified(100);p__Ascomycota_unclassified(100);p__Ascomycota_unclassified(100); | - | - | - | - | - | - | - | Unassigned | - |
| Otu23670 | 1 | k__Fungi(100);p__Ascomycota(100);p__Ascomycota_unclassified(100);p__Ascomycota_unclassified(100);p__Ascomycota_unclassified(100);p__Ascomycota_unclassified(100);p__Ascomycota_unclassified(100); | - | - | - | - | - | - | - | Unassigned | - |
| Otu23682 | 1 | k__Fungi(100);k__Fungi_unclassified(100);k__Fungi_unclassified(100);k__Fungi_unclassified(100);k__Fungi_unclassified(100);k__Fungi_unclassified(100);k__Fungi_unclassified(100); | - | - | - | - | - | - | - | Unassigned | - |
| Otu23683 | 1 | k__Fungi(100);p__Ascomycota(100);p__Ascomycota_unclassified(100);p__Ascomycota_unclassified(100);p__Ascomycota_unclassified(100);p__Ascomycota_unclassified(100);p__Ascomycota_unclassified(100); | - | - | - | - | - | - | - | Unassigned | - |
| Otu23684 | 1 | k__Fungi(100);k__Fungi_unclassified(100);k__Fungi_unclassified(100);k__Fungi_unclassified(100);k__Fungi_unclassified(100);k__Fungi_unclassified(100);k__Fungi_unclassified(100); | - | - | - | - | - | - | - | Unassigned | - |
| Otu23685 | 1 | k__Fungi(100);k__Fungi_unclassified(100);k__Fungi_unclassified(100);k__Fungi_unclassified(100);k__Fungi_unclassified(100);k__Fungi_unclassified(100);k__Fungi_unclassified(100); | - | - | - | - | - | - | - | Unassigned | - |
| Otu23688 | 1 | k__Fungi(100);k__Fungi_unclassified(100);k__Fungi_unclassified(100);k__Fungi_unclassified(100);k__Fungi_unclassified(100);k__Fungi_unclassified(100);k__Fungi_unclassified(100); | - | - | - | - | - | - | - | Unassigned | - |
| Otu23690 | 1 | k__Fungi(100);p__Ascomycota(100);p__Ascomycota_unclassified(100);p__Ascomycota_unclassified(100);p__Ascomycota_unclassified(100);p__Ascomycota_unclassified(100);p__Ascomycota_unclassified(100); | - | - | - | - | - | - | - | Unassigned | - |
| Otu23692 | 1 | k__Fungi(100);k__Fungi_unclassified(100);k__Fungi_unclassified(100);k__Fungi_unclassified(100);k__Fungi_unclassified(100);k__Fungi_unclassified(100);k__Fungi_unclassified(100); | - | - | - | - | - | - | - | Unassigned | - |
| Otu23695 | 1 | k__Fungi(100);p__Basidiomycota(100);p__Basidiomycota_unclassified(100);p__Basidiomycota_unclassified(100);p__Basidiomycota_unclassified(100);p__Basidiomycota_unclassified(100);p__Basidiomycota_unclassified(100); | - | - | - | - | - | - | - | Unassigned | - |
| Otu23697 | 1 | k__Fungi(100);p__Ascomycota(100);p__Ascomycota_unclassified(100);p__Ascomycota_unclassified(100);p__Ascomycota_unclassified(100);p__Ascomycota_unclassified(100);p__Ascomycota_unclassified(100); | - | - | - | - | - | - | - | Unassigned | - |
| Otu23698 | 1 | k__Fungi(100);k__Fungi_unclassified(100);k__Fungi_unclassified(100);k__Fungi_unclassified(100);k__Fungi_unclassified(100);k__Fungi_unclassified(100);k__Fungi_unclassified(100); | - | - | - | - | - | - | - | Unassigned | - |
| Otu23702 | 1 | k__Fungi(100);k__Fungi_unclassified(100);k__Fungi_unclassified(100);k__Fungi_unclassified(100);k__Fungi_unclassified(100);k__Fungi_unclassified(100);k__Fungi_unclassified(100); | - | - | - | - | - | - | - | Unassigned | - |
| Otu23703 | 1 | k__Fungi(100);p__Ascomycota(100);p__Ascomycota_unclassified(100);p__Ascomycota_unclassified(100);p__Ascomycota_unclassified(100);p__Ascomycota_unclassified(100);p__Ascomycota_unclassified(100); | - | - | - | - | - | - | - | Unassigned | - |
| Otu23706 | 1 | k__Fungi(100);k__Fungi_unclassified(100);k__Fungi_unclassified(100);k__Fungi_unclassified(100);k__Fungi_unclassified(100);k__Fungi_unclassified(100);k__Fungi_unclassified(100); | - | - | - | - | - | - | - | Unassigned | - |
| Otu23708 | 1 | k__Fungi(100);k__Fungi_unclassified(100);k__Fungi_unclassified(100);k__Fungi_unclassified(100);k__Fungi_unclassified(100);k__Fungi_unclassified(100);k__Fungi_unclassified(100); | - | - | - | - | - | - | - | Unassigned | - |
| Otu23709 | 1 | k__Fungi(100);p__Ascomycota(100);p__Ascomycota_unclassified(100);p__Ascomycota_unclassified(100);p__Ascomycota_unclassified(100);p__Ascomycota_unclassified(100);p__Ascomycota_unclassified(100); | - | - | - | - | - | - | - | Unassigned | - |
| Otu23710 | 1 | k__Fungi(100);k__Fungi_unclassified(100);k__Fungi_unclassified(100);k__Fungi_unclassified(100);k__Fungi_unclassified(100);k__Fungi_unclassified(100);k__Fungi_unclassified(100); | - | - | - | - | - | - | - | Unassigned | - |
| Otu23711 | 1 | k__Fungi(100);k__Fungi_unclassified(100);k__Fungi_unclassified(100);k__Fungi_unclassified(100);k__Fungi_unclassified(100);k__Fungi_unclassified(100);k__Fungi_unclassified(100); | - | - | - | - | - | - | - | Unassigned | - |
| Otu23712 | 1 | k__Fungi(100);k__Fungi_unclassified(100);k__Fungi_unclassified(100);k__Fungi_unclassified(100);k__Fungi_unclassified(100);k__Fungi_unclassified(100);k__Fungi_unclassified(100); | - | - | - | - | - | - | - | Unassigned | - |
| Otu23714 | 1 | k__Fungi(100);p__Ascomycota(100);p__Ascomycota_unclassified(100);p__Ascomycota_unclassified(100);p__Ascomycota_unclassified(100);p__Ascomycota_unclassified(100);p__Ascomycota_unclassified(100); | - | - | - | - | - | - | - | Unassigned | - |
| Otu23716 | 1 | k__Fungi(100);k__Fungi_unclassified(100);k__Fungi_unclassified(100);k__Fungi_unclassified(100);k__Fungi_unclassified(100);k__Fungi_unclassified(100);k__Fungi_unclassified(100); | - | - | - | - | - | - | - | Unassigned | - |
| Otu23718 | 1 | k__Fungi(100);k__Fungi_unclassified(100);k__Fungi_unclassified(100);k__Fungi_unclassified(100);k__Fungi_unclassified(100);k__Fungi_unclassified(100);k__Fungi_unclassified(100); | - | - | - | - | - | - | - | Unassigned | - |
| Otu23722 | 1 | k__Fungi(100);k__Fungi_unclassified(100);k__Fungi_unclassified(100);k__Fungi_unclassified(100);k__Fungi_unclassified(100);k__Fungi_unclassified(100);k__Fungi_unclassified(100); | - | - | - | - | - | - | - | Unassigned | - |
| Otu23723 | 1 | k__Fungi(100);p__Ascomycota(100);p__Ascomycota_unclassified(100);p__Ascomycota_unclassified(100);p__Ascomycota_unclassified(100);p__Ascomycota_unclassified(100);p__Ascomycota_unclassified(100); | - | - | - | - | - | - | - | Unassigned | - |
| Otu23725 | 1 | k__Fungi(100);p__Ascomycota(100);p__Ascomycota_unclassified(100);p__Ascomycota_unclassified(100);p__Ascomycota_unclassified(100);p__Ascomycota_unclassified(100);p__Ascomycota_unclassified(100); | - | - | - | - | - | - | - | Unassigned | - |
| Otu23728 | 1 | k__Fungi(100);p__Ascomycota(100);p__Ascomycota_unclassified(100);p__Ascomycota_unclassified(100);p__Ascomycota_unclassified(100);p__Ascomycota_unclassified(100);p__Ascomycota_unclassified(100); | - | - | - | - | - | - | - | Unassigned | - |
| Otu23740 | 1 | k__Fungi(100);k__Fungi_unclassified(100);k__Fungi_unclassified(100);k__Fungi_unclassified(100);k__Fungi_unclassified(100);k__Fungi_unclassified(100);k__Fungi_unclassified(100); | - | - | - | - | - | - | - | Unassigned | - |
| Otu23741 | 1 | k__Fungi(100);p__Ascomycota(100);p__Ascomycota_unclassified(100);p__Ascomycota_unclassified(100);p__Ascomycota_unclassified(100);p__Ascomycota_unclassified(100);p__Ascomycota_unclassified(100); | - | - | - | - | - | - | - | Unassigned | - |
| Otu23742 | 1 | k__Fungi(100);p__Ascomycota(100);p__Ascomycota_unclassified(100);p__Ascomycota_unclassified(100);p__Ascomycota_unclassified(100);p__Ascomycota_unclassified(100);p__Ascomycota_unclassified(100); | - | - | - | - | - | - | - | Unassigned | - |
| Otu23743 | 1 | k__Fungi(100);k__Fungi_unclassified(100);k__Fungi_unclassified(100);k__Fungi_unclassified(100);k__Fungi_unclassified(100);k__Fungi_unclassified(100);k__Fungi_unclassified(100); | - | - | - | - | - | - | - | Unassigned | - |
| Otu23748 | 1 | k__Fungi(100);k__Fungi_unclassified(100);k__Fungi_unclassified(100);k__Fungi_unclassified(100);k__Fungi_unclassified(100);k__Fungi_unclassified(100);k__Fungi_unclassified(100); | - | - | - | - | - | - | - | Unassigned | - |
| Otu23755 | 1 | k__Fungi(100);k__Fungi_unclassified(100);k__Fungi_unclassified(100);k__Fungi_unclassified(100);k__Fungi_unclassified(100);k__Fungi_unclassified(100);k__Fungi_unclassified(100); | - | - | - | - | - | - | - | Unassigned | - |
| Otu23759 | 1 | k__Fungi(100);k__Fungi_unclassified(100);k__Fungi_unclassified(100);k__Fungi_unclassified(100);k__Fungi_unclassified(100);k__Fungi_unclassified(100);k__Fungi_unclassified(100); | - | - | - | - | - | - | - | Unassigned | - |
| Otu23760 | 1 | k__Fungi(100);p__Ascomycota(100);p__Ascomycota_unclassified(100);p__Ascomycota_unclassified(100);p__Ascomycota_unclassified(100);p__Ascomycota_unclassified(100);p__Ascomycota_unclassified(100); | - | - | - | - | - | - | - | Unassigned | - |
| Otu23764 | 1 | k__Fungi(100);p__Ascomycota(100);p__Ascomycota_unclassified(100);p__Ascomycota_unclassified(100);p__Ascomycota_unclassified(100);p__Ascomycota_unclassified(100);p__Ascomycota_unclassified(100); | - | - | - | - | - | - | - | Unassigned | - |
| Otu23771 | 1 | k__Fungi(100);k__Fungi_unclassified(100);k__Fungi_unclassified(100);k__Fungi_unclassified(100);k__Fungi_unclassified(100);k__Fungi_unclassified(100);k__Fungi_unclassified(100); | - | - | - | - | - | - | - | Unassigned | - |
| Otu23782 | 1 | k__Fungi(100);p__Ascomycota(100);p__Ascomycota_unclassified(100);p__Ascomycota_unclassified(100);p__Ascomycota_unclassified(100);p__Ascomycota_unclassified(100);p__Ascomycota_unclassified(100); | - | - | - | - | - | - | - | Unassigned | - |
| Otu23785 | 1 | k__Fungi(100);p__Ascomycota(100);p__Ascomycota_unclassified(100);p__Ascomycota_unclassified(100);p__Ascomycota_unclassified(100);p__Ascomycota_unclassified(100);p__Ascomycota_unclassified(100); | - | - | - | - | - | - | - | Unassigned | - |
| Otu23786 | 1 | k__Fungi(100);p__Ascomycota(100);p__Ascomycota_unclassified(100);p__Ascomycota_unclassified(100);p__Ascomycota_unclassified(100);p__Ascomycota_unclassified(100);p__Ascomycota_unclassified(100); | - | - | - | - | - | - | - | Unassigned | - |
| Otu23788 | 1 | k__Fungi(100);k__Fungi_unclassified(100);k__Fungi_unclassified(100);k__Fungi_unclassified(100);k__Fungi_unclassified(100);k__Fungi_unclassified(100);k__Fungi_unclassified(100); | - | - | - | - | - | - | - | Unassigned | - |
| Otu23791 | 1 | k__Fungi(100);p__Ascomycota(100);p__Ascomycota_unclassified(100);p__Ascomycota_unclassified(100);p__Ascomycota_unclassified(100);p__Ascomycota_unclassified(100);p__Ascomycota_unclassified(100); | - | - | - | - | - | - | - | Unassigned | - |
| Otu23792 | 1 | k__Fungi(100);k__Fungi_unclassified(100);k__Fungi_unclassified(100);k__Fungi_unclassified(100);k__Fungi_unclassified(100);k__Fungi_unclassified(100);k__Fungi_unclassified(100); | - | - | - | - | - | - | - | Unassigned | - |
| Otu23793 | 1 | k__Fungi(100);k__Fungi_unclassified(100);k__Fungi_unclassified(100);k__Fungi_unclassified(100);k__Fungi_unclassified(100);k__Fungi_unclassified(100);k__Fungi_unclassified(100); | - | - | - | - | - | - | - | Unassigned | - |
| Otu23798 | 1 | k__Fungi(100);p__Ascomycota(100);p__Ascomycota_unclassified(100);p__Ascomycota_unclassified(100);p__Ascomycota_unclassified(100);p__Ascomycota_unclassified(100);p__Ascomycota_unclassified(100); | - | - | - | - | - | - | - | Unassigned | - |
| Otu23800 | 1 | k__Fungi(100);p__Ascomycota(100);c__Dothideomycetes(100);o__Capnodiales(100);f__Davidiellaceae(100);f__Davidiellaceae_unclassified(100);f__Davidiellaceae_unclassified(100); | - | - | - | - | - | - | - | Unassigned | - |
| Otu23801 | 1 | k__Fungi(100);k__Fungi_unclassified(100);k__Fungi_unclassified(100);k__Fungi_unclassified(100);k__Fungi_unclassified(100);k__Fungi_unclassified(100);k__Fungi_unclassified(100); | - | - | - | - | - | - | - | Unassigned | - |
| Otu23814 | 1 | k__Fungi(100);k__Fungi_unclassified(100);k__Fungi_unclassified(100);k__Fungi_unclassified(100);k__Fungi_unclassified(100);k__Fungi_unclassified(100);k__Fungi_unclassified(100); | - | - | - | - | - | - | - | Unassigned | - |
| Otu23818 | 1 | k__Fungi(100);k__Fungi_unclassified(100);k__Fungi_unclassified(100);k__Fungi_unclassified(100);k__Fungi_unclassified(100);k__Fungi_unclassified(100);k__Fungi_unclassified(100); | - | - | - | - | - | - | - | Unassigned | - |
| Otu23819 | 1 | k__Fungi(100);p__Ascomycota(100);p__Ascomycota_unclassified(100);p__Ascomycota_unclassified(100);p__Ascomycota_unclassified(100);p__Ascomycota_unclassified(100);p__Ascomycota_unclassified(100); | - | - | - | - | - | - | - | Unassigned | - |
| Otu23825 | 1 | k__Fungi(100);k__Fungi_unclassified(100);k__Fungi_unclassified(100);k__Fungi_unclassified(100);k__Fungi_unclassified(100);k__Fungi_unclassified(100);k__Fungi_unclassified(100); | - | - | - | - | - | - | - | Unassigned | - |
| Otu23826 | 1 | k__Fungi(100);p__Ascomycota(100);p__Ascomycota_unclassified(100);p__Ascomycota_unclassified(100);p__Ascomycota_unclassified(100);p__Ascomycota_unclassified(100);p__Ascomycota_unclassified(100); | - | - | - | - | - | - | - | Unassigned | - |
| Otu23827 | 1 | k__Fungi(100);k__Fungi_unclassified(100);k__Fungi_unclassified(100);k__Fungi_unclassified(100);k__Fungi_unclassified(100);k__Fungi_unclassified(100);k__Fungi_unclassified(100); | - | - | - | - | - | - | - | Unassigned | - |
| Otu23828 | 1 | k__Fungi(100);k__Fungi_unclassified(100);k__Fungi_unclassified(100);k__Fungi_unclassified(100);k__Fungi_unclassified(100);k__Fungi_unclassified(100);k__Fungi_unclassified(100); | - | - | - | - | - | - | - | Unassigned | - |
| Otu23829 | 1 | k__Fungi(100);k__Fungi_unclassified(100);k__Fungi_unclassified(100);k__Fungi_unclassified(100);k__Fungi_unclassified(100);k__Fungi_unclassified(100);k__Fungi_unclassified(100); | - | - | - | - | - | - | - | Unassigned | - |
| Otu23831 | 1 | k__Fungi(100);k__Fungi_unclassified(100);k__Fungi_unclassified(100);k__Fungi_unclassified(100);k__Fungi_unclassified(100);k__Fungi_unclassified(100);k__Fungi_unclassified(100); | - | - | - | - | - | - | - | Unassigned | - |
| Otu23833 | 1 | k__Fungi(100);k__Fungi_unclassified(100);k__Fungi_unclassified(100);k__Fungi_unclassified(100);k__Fungi_unclassified(100);k__Fungi_unclassified(100);k__Fungi_unclassified(100); | - | - | - | - | - | - | - | Unassigned | - |
| Otu23834 | 1 | k__Fungi(100);p__Ascomycota(100);c__Dothideomycetes(100);o__Capnodiales(100);o__Capnodiales_unclassified(100);o__Capnodiales_unclassified(100);o__Capnodiales_unclassified(100); | - | - | - | - | - | - | - | Unassigned | - |
| Otu23836 | 1 | k__Fungi(100);p__Ascomycota(100);p__Ascomycota_unclassified(100);p__Ascomycota_unclassified(100);p__Ascomycota_unclassified(100);p__Ascomycota_unclassified(100);p__Ascomycota_unclassified(100); | - | - | - | - | - | - | - | Unassigned | - |
| Otu23837 | 1 | k__Fungi(100);k__Fungi_unclassified(100);k__Fungi_unclassified(100);k__Fungi_unclassified(100);k__Fungi_unclassified(100);k__Fungi_unclassified(100);k__Fungi_unclassified(100); | - | - | - | - | - | - | - | Unassigned | - |
| Otu23838 | 1 | k__Fungi(100);p__Ascomycota(100);p__Ascomycota_unclassified(100);p__Ascomycota_unclassified(100);p__Ascomycota_unclassified(100);p__Ascomycota_unclassified(100);p__Ascomycota_unclassified(100); | - | - | - | - | - | - | - | Unassigned | - |
| Otu23839 | 1 | k__Fungi(100);k__Fungi_unclassified(100);k__Fungi_unclassified(100);k__Fungi_unclassified(100);k__Fungi_unclassified(100);k__Fungi_unclassified(100);k__Fungi_unclassified(100); | - | - | - | - | - | - | - | Unassigned | - |
| Otu23840 | 1 | k__Fungi(100);k__Fungi_unclassified(100);k__Fungi_unclassified(100);k__Fungi_unclassified(100);k__Fungi_unclassified(100);k__Fungi_unclassified(100);k__Fungi_unclassified(100); | - | - | - | - | - | - | - | Unassigned | - |
| Otu23841 | 1 | k__Fungi(100);p__Ascomycota(100);p__Ascomycota_unclassified(100);p__Ascomycota_unclassified(100);p__Ascomycota_unclassified(100);p__Ascomycota_unclassified(100);p__Ascomycota_unclassified(100); | - | - | - | - | - | - | - | Unassigned | - |
| Otu23843 | 1 | k__Fungi(100);k__Fungi_unclassified(100);k__Fungi_unclassified(100);k__Fungi_unclassified(100);k__Fungi_unclassified(100);k__Fungi_unclassified(100);k__Fungi_unclassified(100); | - | - | - | - | - | - | - | Unassigned | - |
| Otu23844 | 1 | k__Fungi(100);p__Ascomycota(100);p__Ascomycota_unclassified(100);p__Ascomycota_unclassified(100);p__Ascomycota_unclassified(100);p__Ascomycota_unclassified(100);p__Ascomycota_unclassified(100); | - | - | - | - | - | - | - | Unassigned | - |
| Otu23845 | 1 | k__Fungi(100);k__Fungi_unclassified(100);k__Fungi_unclassified(100);k__Fungi_unclassified(100);k__Fungi_unclassified(100);k__Fungi_unclassified(100);k__Fungi_unclassified(100); | - | - | - | - | - | - | - | Unassigned | - |
| Otu23846 | 1 | k__Fungi(100);p__Ascomycota(100);p__Ascomycota_unclassified(100);p__Ascomycota_unclassified(100);p__Ascomycota_unclassified(100);p__Ascomycota_unclassified(100);p__Ascomycota_unclassified(100); | - | - | - | - | - | - | - | Unassigned | - |
| Otu23847 | 1 | k__Fungi(100);p__Ascomycota(100);p__Ascomycota_unclassified(100);p__Ascomycota_unclassified(100);p__Ascomycota_unclassified(100);p__Ascomycota_unclassified(100);p__Ascomycota_unclassified(100); | - | - | - | - | - | - | - | Unassigned | - |
| Otu23848 | 1 | k__Fungi(100);k__Fungi_unclassified(100);k__Fungi_unclassified(100);k__Fungi_unclassified(100);k__Fungi_unclassified(100);k__Fungi_unclassified(100);k__Fungi_unclassified(100); | - | - | - | - | - | - | - | Unassigned | - |
| Otu23851 | 1 | k__Fungi(100);k__Fungi_unclassified(100);k__Fungi_unclassified(100);k__Fungi_unclassified(100);k__Fungi_unclassified(100);k__Fungi_unclassified(100);k__Fungi_unclassified(100); | - | - | - | - | - | - | - | Unassigned | - |
| Otu23852 | 1 | k__Fungi(100);p__Ascomycota(100);p__Ascomycota_unclassified(100);p__Ascomycota_unclassified(100);p__Ascomycota_unclassified(100);p__Ascomycota_unclassified(100);p__Ascomycota_unclassified(100); | - | - | - | - | - | - | - | Unassigned | - |
| Otu23853 | 1 | k__Fungi(100);p__Ascomycota(100);p__Ascomycota_unclassified(100);p__Ascomycota_unclassified(100);p__Ascomycota_unclassified(100);p__Ascomycota_unclassified(100);p__Ascomycota_unclassified(100); | - | - | - | - | - | - | - | Unassigned | - |
| Otu23855 | 1 | k__Fungi(100);p__Ascomycota(100);p__Ascomycota_unclassified(100);p__Ascomycota_unclassified(100);p__Ascomycota_unclassified(100);p__Ascomycota_unclassified(100);p__Ascomycota_unclassified(100); | - | - | - | - | - | - | - | Unassigned | - |
| Otu23857 | 1 | k__Fungi(100);k__Fungi_unclassified(100);k__Fungi_unclassified(100);k__Fungi_unclassified(100);k__Fungi_unclassified(100);k__Fungi_unclassified(100);k__Fungi_unclassified(100); | - | - | - | - | - | - | - | Unassigned | - |
| Otu23858 | 1 | k__Fungi(100);k__Fungi_unclassified(100);k__Fungi_unclassified(100);k__Fungi_unclassified(100);k__Fungi_unclassified(100);k__Fungi_unclassified(100);k__Fungi_unclassified(100); | - | - | - | - | - | - | - | Unassigned | - |
| Otu23860 | 1 | k__Fungi(100);k__Fungi_unclassified(100);k__Fungi_unclassified(100);k__Fungi_unclassified(100);k__Fungi_unclassified(100);k__Fungi_unclassified(100);k__Fungi_unclassified(100); | - | - | - | - | - | - | - | Unassigned | - |
| Otu23861 | 1 | k__Fungi(100);k__Fungi_unclassified(100);k__Fungi_unclassified(100);k__Fungi_unclassified(100);k__Fungi_unclassified(100);k__Fungi_unclassified(100);k__Fungi_unclassified(100); | - | - | - | - | - | - | - | Unassigned | - |
| Otu23862 | 1 | k__Fungi(100);k__Fungi_unclassified(100);k__Fungi_unclassified(100);k__Fungi_unclassified(100);k__Fungi_unclassified(100);k__Fungi_unclassified(100);k__Fungi_unclassified(100); | - | - | - | - | - | - | - | Unassigned | - |
| Otu23863 | 1 | k__Fungi(100);k__Fungi_unclassified(100);k__Fungi_unclassified(100);k__Fungi_unclassified(100);k__Fungi_unclassified(100);k__Fungi_unclassified(100);k__Fungi_unclassified(100); | - | - | - | - | - | - | - | Unassigned | - |
| Otu23864 | 1 | k__Fungi(100);k__Fungi_unclassified(100);k__Fungi_unclassified(100);k__Fungi_unclassified(100);k__Fungi_unclassified(100);k__Fungi_unclassified(100);k__Fungi_unclassified(100); | - | - | - | - | - | - | - | Unassigned | - |
| Otu23865 | 1 | k__Fungi(100);p__Ascomycota(100);p__Ascomycota_unclassified(100);p__Ascomycota_unclassified(100);p__Ascomycota_unclassified(100);p__Ascomycota_unclassified(100);p__Ascomycota_unclassified(100); | - | - | - | - | - | - | - | Unassigned | - |
| Otu23866 | 1 | k__Fungi(100);k__Fungi_unclassified(100);k__Fungi_unclassified(100);k__Fungi_unclassified(100);k__Fungi_unclassified(100);k__Fungi_unclassified(100);k__Fungi_unclassified(100); | - | - | - | - | - | - | - | Unassigned | - |
| Otu23869 | 1 | k__Fungi(100);k__Fungi_unclassified(100);k__Fungi_unclassified(100);k__Fungi_unclassified(100);k__Fungi_unclassified(100);k__Fungi_unclassified(100);k__Fungi_unclassified(100); | - | - | - | - | - | - | - | Unassigned | - |
| Otu23870 | 1 | k__Fungi(100);k__Fungi_unclassified(100);k__Fungi_unclassified(100);k__Fungi_unclassified(100);k__Fungi_unclassified(100);k__Fungi_unclassified(100);k__Fungi_unclassified(100); | - | - | - | - | - | - | - | Unassigned | - |
| Otu23871 | 1 | k__Fungi(100);k__Fungi_unclassified(100);k__Fungi_unclassified(100);k__Fungi_unclassified(100);k__Fungi_unclassified(100);k__Fungi_unclassified(100);k__Fungi_unclassified(100); | - | - | - | - | - | - | - | Unassigned | - |
| Otu23873 | 1 | k__Fungi(100);k__Fungi_unclassified(100);k__Fungi_unclassified(100);k__Fungi_unclassified(100);k__Fungi_unclassified(100);k__Fungi_unclassified(100);k__Fungi_unclassified(100); | - | - | - | - | - | - | - | Unassigned | - |
| Otu23876 | 1 | k__Fungi(100);k__Fungi_unclassified(100);k__Fungi_unclassified(100);k__Fungi_unclassified(100);k__Fungi_unclassified(100);k__Fungi_unclassified(100);k__Fungi_unclassified(100); | - | - | - | - | - | - | - | Unassigned | - |
| Otu23879 | 1 | k__Fungi(100);k__Fungi_unclassified(100);k__Fungi_unclassified(100);k__Fungi_unclassified(100);k__Fungi_unclassified(100);k__Fungi_unclassified(100);k__Fungi_unclassified(100); | - | - | - | - | - | - | - | Unassigned | - |
| Otu23881 | 1 | k__Fungi(100);p__Ascomycota(100);p__Ascomycota_unclassified(100);p__Ascomycota_unclassified(100);p__Ascomycota_unclassified(100);p__Ascomycota_unclassified(100);p__Ascomycota_unclassified(100); | - | - | - | - | - | - | - | Unassigned | - |
| Otu23887 | 1 | k__Fungi(100);k__Fungi_unclassified(100);k__Fungi_unclassified(100);k__Fungi_unclassified(100);k__Fungi_unclassified(100);k__Fungi_unclassified(100);k__Fungi_unclassified(100); | - | - | - | - | - | - | - | Unassigned | - |
| Otu23888 | 1 | k__Fungi(100);k__Fungi_unclassified(100);k__Fungi_unclassified(100);k__Fungi_unclassified(100);k__Fungi_unclassified(100);k__Fungi_unclassified(100);k__Fungi_unclassified(100); | - | - | - | - | - | - | - | Unassigned | - |
| Otu23889 | 1 | k__Fungi(100);k__Fungi_unclassified(100);k__Fungi_unclassified(100);k__Fungi_unclassified(100);k__Fungi_unclassified(100);k__Fungi_unclassified(100);k__Fungi_unclassified(100); | - | - | - | - | - | - | - | Unassigned | - |
| Otu23890 | 1 | k__Fungi(100);k__Fungi_unclassified(100);k__Fungi_unclassified(100);k__Fungi_unclassified(100);k__Fungi_unclassified(100);k__Fungi_unclassified(100);k__Fungi_unclassified(100); | - | - | - | - | - | - | - | Unassigned | - |
| Otu23891 | 1 | k__Fungi(100);k__Fungi_unclassified(100);k__Fungi_unclassified(100);k__Fungi_unclassified(100);k__Fungi_unclassified(100);k__Fungi_unclassified(100);k__Fungi_unclassified(100); | - | - | - | - | - | - | - | Unassigned | - |
| Otu23892 | 1 | k__Fungi(100);k__Fungi_unclassified(100);k__Fungi_unclassified(100);k__Fungi_unclassified(100);k__Fungi_unclassified(100);k__Fungi_unclassified(100);k__Fungi_unclassified(100); | - | - | - | - | - | - | - | Unassigned | - |
| Otu23893 | 1 | k__Fungi(100);k__Fungi_unclassified(100);k__Fungi_unclassified(100);k__Fungi_unclassified(100);k__Fungi_unclassified(100);k__Fungi_unclassified(100);k__Fungi_unclassified(100); | - | - | - | - | - | - | - | Unassigned | - |
| Otu23894 | 1 | k__Fungi(100);k__Fungi_unclassified(100);k__Fungi_unclassified(100);k__Fungi_unclassified(100);k__Fungi_unclassified(100);k__Fungi_unclassified(100);k__Fungi_unclassified(100); | - | - | - | - | - | - | - | Unassigned | - |
| Otu23895 | 1 | k__Fungi(100);p__Ascomycota(100);p__Ascomycota_unclassified(100);p__Ascomycota_unclassified(100);p__Ascomycota_unclassified(100);p__Ascomycota_unclassified(100);p__Ascomycota_unclassified(100); | - | - | - | - | - | - | - | Unassigned | - |
| Otu23896 | 1 | k__Fungi(100);k__Fungi_unclassified(100);k__Fungi_unclassified(100);k__Fungi_unclassified(100);k__Fungi_unclassified(100);k__Fungi_unclassified(100);k__Fungi_unclassified(100); | - | - | - | - | - | - | - | Unassigned | - |
| Otu23898 | 1 | k__Fungi(100);k__Fungi_unclassified(100);k__Fungi_unclassified(100);k__Fungi_unclassified(100);k__Fungi_unclassified(100);k__Fungi_unclassified(100);k__Fungi_unclassified(100); | - | - | - | - | - | - | - | Unassigned | - |
| Otu23901 | 1 | k__Fungi(100);k__Fungi_unclassified(100);k__Fungi_unclassified(100);k__Fungi_unclassified(100);k__Fungi_unclassified(100);k__Fungi_unclassified(100);k__Fungi_unclassified(100); | - | - | - | - | - | - | - | Unassigned | - |
| Otu23902 | 1 | k__Fungi(100);p__Ascomycota(100);p__Ascomycota_unclassified(100);p__Ascomycota_unclassified(100);p__Ascomycota_unclassified(100);p__Ascomycota_unclassified(100);p__Ascomycota_unclassified(100); | - | - | - | - | - | - | - | Unassigned | - |
| Otu23903 | 1 | k__Fungi(100);p__Ascomycota(100);p__Ascomycota_unclassified(100);p__Ascomycota_unclassified(100);p__Ascomycota_unclassified(100);p__Ascomycota_unclassified(100);p__Ascomycota_unclassified(100); | - | - | - | - | - | - | - | Unassigned | - |
| Otu23904 | 1 | k__Fungi(100);k__Fungi_unclassified(100);k__Fungi_unclassified(100);k__Fungi_unclassified(100);k__Fungi_unclassified(100);k__Fungi_unclassified(100);k__Fungi_unclassified(100); | - | - | - | - | - | - | - | Unassigned | - |
| Otu23905 | 1 | k__Fungi(100);p__Ascomycota(100);p__Ascomycota_unclassified(100);p__Ascomycota_unclassified(100);p__Ascomycota_unclassified(100);p__Ascomycota_unclassified(100);p__Ascomycota_unclassified(100); | - | - | - | - | - | - | - | Unassigned | - |
| Otu23907 | 1 | k__Fungi(100);k__Fungi_unclassified(100);k__Fungi_unclassified(100);k__Fungi_unclassified(100);k__Fungi_unclassified(100);k__Fungi_unclassified(100);k__Fungi_unclassified(100); | - | - | - | - | - | - | - | Unassigned | - |
| Otu23908 | 1 | k__Fungi(100);k__Fungi_unclassified(100);k__Fungi_unclassified(100);k__Fungi_unclassified(100);k__Fungi_unclassified(100);k__Fungi_unclassified(100);k__Fungi_unclassified(100); | - | - | - | - | - | - | - | Unassigned | - |
| Otu23913 | 1 | k__Fungi(100);p__Ascomycota(100);p__Ascomycota_unclassified(100);p__Ascomycota_unclassified(100);p__Ascomycota_unclassified(100);p__Ascomycota_unclassified(100);p__Ascomycota_unclassified(100); | - | - | - | - | - | - | - | Unassigned | - |
| Otu23918 | 1 | k__Fungi(100);k__Fungi_unclassified(100);k__Fungi_unclassified(100);k__Fungi_unclassified(100);k__Fungi_unclassified(100);k__Fungi_unclassified(100);k__Fungi_unclassified(100); | - | - | - | - | - | - | - | Unassigned | - |
| Otu23919 | 1 | k__Fungi(100);k__Fungi_unclassified(100);k__Fungi_unclassified(100);k__Fungi_unclassified(100);k__Fungi_unclassified(100);k__Fungi_unclassified(100);k__Fungi_unclassified(100); | - | - | - | - | - | - | - | Unassigned | - |
| Otu23922 | 1 | k__Fungi(100);k__Fungi_unclassified(100);k__Fungi_unclassified(100);k__Fungi_unclassified(100);k__Fungi_unclassified(100);k__Fungi_unclassified(100);k__Fungi_unclassified(100); | - | - | - | - | - | - | - | Unassigned | - |
| Otu23936 | 1 | k__Fungi(100);p__Ascomycota(100);c__Dothideomycetes(100);o__Capnodiales(100);f__Davidiellaceae(100);f__Davidiellaceae_unclassified(100);f__Davidiellaceae_unclassified(100); | - | - | - | - | - | - | - | Unassigned | - |
| Otu23941 | 1 | k__Fungi(100);p__Ascomycota(100);p__Ascomycota_unclassified(100);p__Ascomycota_unclassified(100);p__Ascomycota_unclassified(100);p__Ascomycota_unclassified(100);p__Ascomycota_unclassified(100); | - | - | - | - | - | - | - | Unassigned | - |
| Otu23942 | 1 | k__Fungi(100);p__Ascomycota(100);p__Ascomycota_unclassified(100);p__Ascomycota_unclassified(100);p__Ascomycota_unclassified(100);p__Ascomycota_unclassified(100);p__Ascomycota_unclassified(100); | - | - | - | - | - | - | - | Unassigned | - |
| Otu23943 | 1 | k__Fungi(100);p__Ascomycota(100);c__Leotiomycetes(100);c__Leotiomycetes_unclassified(100);c__Leotiomycetes_unclassified(100);c__Leotiomycetes_unclassified(100);c__Leotiomycetes_unclassified(100); | - | - | - | - | - | - | - | Unassigned | - |
| Otu23944 | 1 | k__Fungi(100);p__Ascomycota(100);c__Sordariomycetes(100);c__Sordariomycetes_unclassified(100);c__Sordariomycetes_unclassified(100);c__Sordariomycetes_unclassified(100);c__Sordariomycetes_unclassified(100); | - | - | - | - | - | - | - | Unassigned | - |
| Otu23947 | 1 | k__Fungi(100);p__Ascomycota(100);p__Ascomycota_unclassified(100);p__Ascomycota_unclassified(100);p__Ascomycota_unclassified(100);p__Ascomycota_unclassified(100);p__Ascomycota_unclassified(100); | - | - | - | - | - | - | - | Unassigned | - |
| Otu23948 | 1 | k__Fungi(100);k__Fungi_unclassified(100);k__Fungi_unclassified(100);k__Fungi_unclassified(100);k__Fungi_unclassified(100);k__Fungi_unclassified(100);k__Fungi_unclassified(100); | - | - | - | - | - | - | - | Unassigned | - |
| Otu23949 | 1 | k__Fungi(100);p__Ascomycota(100);p__Ascomycota_unclassified(100);p__Ascomycota_unclassified(100);p__Ascomycota_unclassified(100);p__Ascomycota_unclassified(100);p__Ascomycota_unclassified(100); | - | - | - | - | - | - | - | Unassigned | - |
| Otu23954 | 1 | k__Fungi(100);p__Ascomycota(100);p__Ascomycota_unclassified(100);p__Ascomycota_unclassified(100);p__Ascomycota_unclassified(100);p__Ascomycota_unclassified(100);p__Ascomycota_unclassified(100); | - | - | - | - | - | - | - | Unassigned | - |
| Otu23955 | 1 | k__Fungi(100);k__Fungi_unclassified(100);k__Fungi_unclassified(100);k__Fungi_unclassified(100);k__Fungi_unclassified(100);k__Fungi_unclassified(100);k__Fungi_unclassified(100); | - | - | - | - | - | - | - | Unassigned | - |
| Otu23959 | 1 | k__Fungi(100);k__Fungi_unclassified(100);k__Fungi_unclassified(100);k__Fungi_unclassified(100);k__Fungi_unclassified(100);k__Fungi_unclassified(100);k__Fungi_unclassified(100); | - | - | - | - | - | - | - | Unassigned | - |
| Otu23960 | 1 | k__Fungi(100);k__Fungi_unclassified(100);k__Fungi_unclassified(100);k__Fungi_unclassified(100);k__Fungi_unclassified(100);k__Fungi_unclassified(100);k__Fungi_unclassified(100); | - | - | - | - | - | - | - | Unassigned | - |
| Otu23961 | 1 | k__Fungi(100);k__Fungi_unclassified(100);k__Fungi_unclassified(100);k__Fungi_unclassified(100);k__Fungi_unclassified(100);k__Fungi_unclassified(100);k__Fungi_unclassified(100); | - | - | - | - | - | - | - | Unassigned | - |
| Otu23962 | 1 | k__Fungi(100);p__Ascomycota(100);p__Ascomycota_unclassified(100);p__Ascomycota_unclassified(100);p__Ascomycota_unclassified(100);p__Ascomycota_unclassified(100);p__Ascomycota_unclassified(100); | - | - | - | - | - | - | - | Unassigned | - |
| Otu23963 | 1 | k__Fungi(100);k__Fungi_unclassified(100);k__Fungi_unclassified(100);k__Fungi_unclassified(100);k__Fungi_unclassified(100);k__Fungi_unclassified(100);k__Fungi_unclassified(100); | - | - | - | - | - | - | - | Unassigned | - |
| Otu23964 | 1 | k__Fungi(100);k__Fungi_unclassified(100);k__Fungi_unclassified(100);k__Fungi_unclassified(100);k__Fungi_unclassified(100);k__Fungi_unclassified(100);k__Fungi_unclassified(100); | - | - | - | - | - | - | - | Unassigned | - |
| Otu23968 | 1 | k__Fungi(100);p__Ascomycota(100);p__Ascomycota_unclassified(100);p__Ascomycota_unclassified(100);p__Ascomycota_unclassified(100);p__Ascomycota_unclassified(100);p__Ascomycota_unclassified(100); | - | - | - | - | - | - | - | Unassigned | - |
| Otu23969 | 1 | k__Fungi(100);p__Ascomycota(100);p__Ascomycota_unclassified(100);p__Ascomycota_unclassified(100);p__Ascomycota_unclassified(100);p__Ascomycota_unclassified(100);p__Ascomycota_unclassified(100); | - | - | - | - | - | - | - | Unassigned | - |
| Otu23970 | 1 | k__Fungi(100);p__Ascomycota(100);c__unclassified_Ascomycota(100);o__unclassified_Ascomycota(100);f__unclassified_Ascomycota(100);g__unclassified_Ascomycota(100);s__uncultured_sordariomyceta(100); | - | - | - | - | - | - | - | Unassigned | - |
| Otu23973 | 1 | k__Fungi(100);k__Fungi_unclassified(100);k__Fungi_unclassified(100);k__Fungi_unclassified(100);k__Fungi_unclassified(100);k__Fungi_unclassified(100);k__Fungi_unclassified(100); | - | - | - | - | - | - | - | Unassigned | - |
| Otu23974 | 1 | k__Fungi(100);k__Fungi_unclassified(100);k__Fungi_unclassified(100);k__Fungi_unclassified(100);k__Fungi_unclassified(100);k__Fungi_unclassified(100);k__Fungi_unclassified(100); | - | - | - | - | - | - | - | Unassigned | - |
| Otu23975 | 1 | k__Fungi(100);p__Ascomycota(100);p__Ascomycota_unclassified(100);p__Ascomycota_unclassified(100);p__Ascomycota_unclassified(100);p__Ascomycota_unclassified(100);p__Ascomycota_unclassified(100); | - | - | - | - | - | - | - | Unassigned | - |
| Otu23976 | 1 | k__Fungi(100);p__Ascomycota(100);p__Ascomycota_unclassified(100);p__Ascomycota_unclassified(100);p__Ascomycota_unclassified(100);p__Ascomycota_unclassified(100);p__Ascomycota_unclassified(100); | - | - | - | - | - | - | - | Unassigned | - |
| Otu23977 | 1 | k__Fungi(100);k__Fungi_unclassified(100);k__Fungi_unclassified(100);k__Fungi_unclassified(100);k__Fungi_unclassified(100);k__Fungi_unclassified(100);k__Fungi_unclassified(100); | - | - | - | - | - | - | - | Unassigned | - |
| Otu23978 | 1 | k__Fungi(100);p__Ascomycota(100);p__Ascomycota_unclassified(100);p__Ascomycota_unclassified(100);p__Ascomycota_unclassified(100);p__Ascomycota_unclassified(100);p__Ascomycota_unclassified(100); | - | - | - | - | - | - | - | Unassigned | - |
| Otu23979 | 1 | k__Fungi(100);p__Ascomycota(100);p__Ascomycota_unclassified(100);p__Ascomycota_unclassified(100);p__Ascomycota_unclassified(100);p__Ascomycota_unclassified(100);p__Ascomycota_unclassified(100); | - | - | - | - | - | - | - | Unassigned | - |
| Otu23980 | 1 | k__Fungi(100);p__Ascomycota(100);p__Ascomycota_unclassified(100);p__Ascomycota_unclassified(100);p__Ascomycota_unclassified(100);p__Ascomycota_unclassified(100);p__Ascomycota_unclassified(100); | - | - | - | - | - | - | - | Unassigned | - |
| Otu23982 | 1 | k__Fungi(100);p__Ascomycota(100);p__Ascomycota_unclassified(100);p__Ascomycota_unclassified(100);p__Ascomycota_unclassified(100);p__Ascomycota_unclassified(100);p__Ascomycota_unclassified(100); | - | - | - | - | - | - | - | Unassigned | - |
| Otu23983 | 1 | k__Fungi(100);k__Fungi_unclassified(100);k__Fungi_unclassified(100);k__Fungi_unclassified(100);k__Fungi_unclassified(100);k__Fungi_unclassified(100);k__Fungi_unclassified(100); | - | - | - | - | - | - | - | Unassigned | - |
| Otu23985 | 1 | k__Fungi(100);p__Ascomycota(100);p__Ascomycota_unclassified(100);p__Ascomycota_unclassified(100);p__Ascomycota_unclassified(100);p__Ascomycota_unclassified(100);p__Ascomycota_unclassified(100); | - | - | - | - | - | - | - | Unassigned | - |
| Otu23986 | 1 | k__Fungi(100);k__Fungi_unclassified(100);k__Fungi_unclassified(100);k__Fungi_unclassified(100);k__Fungi_unclassified(100);k__Fungi_unclassified(100);k__Fungi_unclassified(100); | - | - | - | - | - | - | - | Unassigned | - |
| Otu23987 | 1 | k__Fungi(100);p__Ascomycota(100);p__Ascomycota_unclassified(100);p__Ascomycota_unclassified(100);p__Ascomycota_unclassified(100);p__Ascomycota_unclassified(100);p__Ascomycota_unclassified(100); | - | - | - | - | - | - | - | Unassigned | - |
| Otu23988 | 1 | k__Fungi(100);k__Fungi_unclassified(100);k__Fungi_unclassified(100);k__Fungi_unclassified(100);k__Fungi_unclassified(100);k__Fungi_unclassified(100);k__Fungi_unclassified(100); | - | - | - | - | - | - | - | Unassigned | - |
| Otu23989 | 1 | k__Fungi(100);k__Fungi_unclassified(100);k__Fungi_unclassified(100);k__Fungi_unclassified(100);k__Fungi_unclassified(100);k__Fungi_unclassified(100);k__Fungi_unclassified(100); | - | - | - | - | - | - | - | Unassigned | - |
| Otu23990 | 1 | k__Fungi(100);k__Fungi_unclassified(100);k__Fungi_unclassified(100);k__Fungi_unclassified(100);k__Fungi_unclassified(100);k__Fungi_unclassified(100);k__Fungi_unclassified(100); | - | - | - | - | - | - | - | Unassigned | - |
| Otu23991 | 1 | k__Fungi(100);k__Fungi_unclassified(100);k__Fungi_unclassified(100);k__Fungi_unclassified(100);k__Fungi_unclassified(100);k__Fungi_unclassified(100);k__Fungi_unclassified(100); | - | - | - | - | - | - | - | Unassigned | - |
| Otu23992 | 1 | k__Fungi(100);k__Fungi_unclassified(100);k__Fungi_unclassified(100);k__Fungi_unclassified(100);k__Fungi_unclassified(100);k__Fungi_unclassified(100);k__Fungi_unclassified(100); | - | - | - | - | - | - | - | Unassigned | - |
| Otu23993 | 1 | k__Fungi(100);k__Fungi_unclassified(100);k__Fungi_unclassified(100);k__Fungi_unclassified(100);k__Fungi_unclassified(100);k__Fungi_unclassified(100);k__Fungi_unclassified(100); | - | - | - | - | - | - | - | Unassigned | - |
| Otu23994 | 1 | k__Fungi(100);p__Ascomycota(100);p__Ascomycota_unclassified(100);p__Ascomycota_unclassified(100);p__Ascomycota_unclassified(100);p__Ascomycota_unclassified(100);p__Ascomycota_unclassified(100); | - | - | - | - | - | - | - | Unassigned | - |
| Otu23995 | 1 | k__Fungi(100);k__Fungi_unclassified(100);k__Fungi_unclassified(100);k__Fungi_unclassified(100);k__Fungi_unclassified(100);k__Fungi_unclassified(100);k__Fungi_unclassified(100); | - | - | - | - | - | - | - | Unassigned | - |
| Otu23996 | 1 | k__Fungi(100);p__Ascomycota(100);p__Ascomycota_unclassified(100);p__Ascomycota_unclassified(100);p__Ascomycota_unclassified(100);p__Ascomycota_unclassified(100);p__Ascomycota_unclassified(100); | - | - | - | - | - | - | - | Unassigned | - |
| Otu23999 | 1 | k__Fungi(100);p__Ascomycota(100);p__Ascomycota_unclassified(100);p__Ascomycota_unclassified(100);p__Ascomycota_unclassified(100);p__Ascomycota_unclassified(100);p__Ascomycota_unclassified(100); | - | - | - | - | - | - | - | Unassigned | - |

| | | | | | | | | | | | |
|---|---|---|---|---|---|---|---|---|---|---|---|
| Otu24011 | 1 | k__Fungi(100);k__Fungi_unclassified(100);k__Fungi_unclassified(100);k__Fungi_unclassified(100);k__Fungi_unclassified(100);k__Fungi_unclassified(100);k__Fungi_unclassified(100); | - | - | - | - | - | - | - | Unassigned | - |
| Otu24014 | 1 | k__Fungi(100);k__Fungi_unclassified(100);k__Fungi_unclassified(100);k__Fungi_unclassified(100);k__Fungi_unclassified(100);k__Fungi_unclassified(100);k__Fungi_unclassified(100); | - | - | - | - | - | - | - | Unassigned | - |
| Otu24027 | 1 | k__Fungi(100);k__Fungi_unclassified(100);k__Fungi_unclassified(100);k__Fungi_unclassified(100);k__Fungi_unclassified(100);k__Fungi_unclassified(100);k__Fungi_unclassified(100); | - | - | - | - | - | - | - | Unassigned | - |
| Otu24028 | 1 | k__Fungi(100);p__Ascomycota(100);p__Ascomycota_unclassified(100);p__Ascomycota_unclassified(100);p__Ascomycota_unclassified(100);p__Ascomycota_unclassified(100);p__Ascomycota_unclassified(100); | - | - | - | - | - | - | - | Unassigned | - |
| Otu24030 | 1 | k__Fungi(100);p__Ascomycota(100);p__Ascomycota_unclassified(100);p__Ascomycota_unclassified(100);p__Ascomycota_unclassified(100);p__Ascomycota_unclassified(100);p__Ascomycota_unclassified(100); | - | - | - | - | - | - | - | Unassigned | - |
| Otu24031 | 1 | k__Fungi(100);k__Fungi_unclassified(100);k__Fungi_unclassified(100);k__Fungi_unclassified(100);k__Fungi_unclassified(100);k__Fungi_unclassified(100);k__Fungi_unclassified(100); | - | - | - | - | - | - | - | Unassigned | - |
| Otu24032 | 1 | k__Fungi(100);p__Ascomycota(100);p__Ascomycota_unclassified(100);p__Ascomycota_unclassified(100);p__Ascomycota_unclassified(100);p__Ascomycota_unclassified(100);p__Ascomycota_unclassified(100); | - | - | - | - | - | - | - | Unassigned | - |
| Otu24033 | 1 | k__Fungi(100);k__Fungi_unclassified(100);k__Fungi_unclassified(100);k__Fungi_unclassified(100);k__Fungi_unclassified(100);k__Fungi_unclassified(100);k__Fungi_unclassified(100); | - | - | - | - | - | - | - | Unassigned | - |
| Otu24034 | 1 | k__Fungi(100);k__Fungi_unclassified(100);k__Fungi_unclassified(100);k__Fungi_unclassified(100);k__Fungi_unclassified(100);k__Fungi_unclassified(100);k__Fungi_unclassified(100); | - | - | - | - | - | - | - | Unassigned | - |
| Otu24035 | 1 | k__Fungi(100);k__Fungi_unclassified(100);k__Fungi_unclassified(100);k__Fungi_unclassified(100);k__Fungi_unclassified(100);k__Fungi_unclassified(100);k__Fungi_unclassified(100); | - | - | - | - | - | - | - | Unassigned | - |
| Otu24036 | 1 | k__Fungi(100);k__Fungi_unclassified(100);k__Fungi_unclassified(100);k__Fungi_unclassified(100);k__Fungi_unclassified(100);k__Fungi_unclassified(100);k__Fungi_unclassified(100); | - | - | - | - | - | - | - | Unassigned | - |
| Otu24037 | 1 | k__Fungi(100);k__Fungi_unclassified(100);k__Fungi_unclassified(100);k__Fungi_unclassified(100);k__Fungi_unclassified(100);k__Fungi_unclassified(100);k__Fungi_unclassified(100); | - | - | - | - | - | - | - | Unassigned | - |
| Otu24038 | 1 | k__Fungi(100);k__Fungi_unclassified(100);k__Fungi_unclassified(100);k__Fungi_unclassified(100);k__Fungi_unclassified(100);k__Fungi_unclassified(100);k__Fungi_unclassified(100); | - | - | - | - | - | - | - | Unassigned | - |
| Otu24039 | 1 | k__Fungi(100);p__Ascomycota(100);p__Ascomycota_unclassified(100);p__Ascomycota_unclassified(100);p__Ascomycota_unclassified(100);p__Ascomycota_unclassified(100);p__Ascomycota_unclassified(100); | - | - | - | - | - | - | - | Unassigned | - |
| Otu24040 | 1 | k__Fungi(100);p__Ascomycota(100);p__Ascomycota_unclassified(100);p__Ascomycota_unclassified(100);p__Ascomycota_unclassified(100);p__Ascomycota_unclassified(100);p__Ascomycota_unclassified(100); | - | - | - | - | - | - | - | Unassigned | - |
| Otu24042 | 1 | k__Fungi(100);k__Fungi_unclassified(100);k__Fungi_unclassified(100);k__Fungi_unclassified(100);k__Fungi_unclassified(100);k__Fungi_unclassified(100);k__Fungi_unclassified(100); | - | - | - | - | - | - | - | Unassigned | - |
| Otu24043 | 1 | k__Fungi(100);k__Fungi_unclassified(100);k__Fungi_unclassified(100);k__Fungi_unclassified(100);k__Fungi_unclassified(100);k__Fungi_unclassified(100);k__Fungi_unclassified(100); | - | - | - | - | - | - | - | Unassigned | - |
| Otu24044 | 1 | k__Fungi(100);k__Fungi_unclassified(100);k__Fungi_unclassified(100);k__Fungi_unclassified(100);k__Fungi_unclassified(100);k__Fungi_unclassified(100);k__Fungi_unclassified(100); | - | - | - | - | - | - | - | Unassigned | - |
| Otu24045 | 1 | k__Fungi(100);p__Ascomycota(100);p__Ascomycota_unclassified(100);p__Ascomycota_unclassified(100);p__Ascomycota_unclassified(100);p__Ascomycota_unclassified(100);p__Ascomycota_unclassified(100); | - | - | - | - | - | - | - | Unassigned | - |
| Otu24046 | 1 | k__Fungi(100);k__Fungi_unclassified(100);k__Fungi_unclassified(100);k__Fungi_unclassified(100);k__Fungi_unclassified(100);k__Fungi_unclassified(100);k__Fungi_unclassified(100); | - | - | - | - | - | - | - | Unassigned | - |
| Otu24047 | 1 | k__Fungi(100);k__Fungi_unclassified(100);k__Fungi_unclassified(100);k__Fungi_unclassified(100);k__Fungi_unclassified(100);k__Fungi_unclassified(100);k__Fungi_unclassified(100); | - | - | - | - | - | - | - | Unassigned | - |
| Otu24048 | 1 | k__Fungi(100);k__Fungi_unclassified(100);k__Fungi_unclassified(100);k__Fungi_unclassified(100);k__Fungi_unclassified(100);k__Fungi_unclassified(100);k__Fungi_unclassified(100); | - | - | - | - | - | - | - | Unassigned | - |
| Otu24049 | 1 | k__Fungi(100);k__Fungi_unclassified(100);k__Fungi_unclassified(100);k__Fungi_unclassified(100);k__Fungi_unclassified(100);k__Fungi_unclassified(100);k__Fungi_unclassified(100); | - | - | - | - | - | - | - | Unassigned | - |
| Otu24050 | 1 | k__Fungi(100);k__Fungi_unclassified(100);k__Fungi_unclassified(100);k__Fungi_unclassified(100);k__Fungi_unclassified(100);k__Fungi_unclassified(100);k__Fungi_unclassified(100); | - | - | - | - | - | - | - | Unassigned | - |
| Otu24051 | 1 | k__Fungi(100);p__Ascomycota(100);p__Ascomycota_unclassified(100);p__Ascomycota_unclassified(100);p__Ascomycota_unclassified(100);p__Ascomycota_unclassified(100);p__Ascomycota_unclassified(100); | - | - | - | - | - | - | - | Unassigned | - |
| Otu24052 | 1 | k__Fungi(100);k__Fungi_unclassified(100);k__Fungi_unclassified(100);k__Fungi_unclassified(100);k__Fungi_unclassified(100);k__Fungi_unclassified(100);k__Fungi_unclassified(100); | - | - | - | - | - | - | - | Unassigned | - |
| Otu24053 | 1 | k__Fungi(100);k__Fungi_unclassified(100);k__Fungi_unclassified(100);k__Fungi_unclassified(100);k__Fungi_unclassified(100);k__Fungi_unclassified(100);k__Fungi_unclassified(100); | - | - | - | - | - | - | - | Unassigned | - |
| Otu24054 | 1 | k__Fungi(100);p__Ascomycota(100);p__Ascomycota_unclassified(100);p__Ascomycota_unclassified(100);p__Ascomycota_unclassified(100);p__Ascomycota_unclassified(100);p__Ascomycota_unclassified(100); | - | - | - | - | - | - | - | Unassigned | - |
| Otu24055 | 1 | k__Fungi(100);k__Fungi_unclassified(100);k__Fungi_unclassified(100);k__Fungi_unclassified(100);k__Fungi_unclassified(100);k__Fungi_unclassified(100);k__Fungi_unclassified(100); | - | - | - | - | - | - | - | Unassigned | - |
| Otu24056 | 1 | k__Fungi(100);p__Ascomycota(100);p__Ascomycota_unclassified(100);p__Ascomycota_unclassified(100);p__Ascomycota_unclassified(100);p__Ascomycota_unclassified(100);p__Ascomycota_unclassified(100); | - | - | - | - | - | - | - | Unassigned | - |
| Otu24057 | 1 | k__Fungi(100);p__Ascomycota(100);p__Ascomycota_unclassified(100);p__Ascomycota_unclassified(100);p__Ascomycota_unclassified(100);p__Ascomycota_unclassified(100);p__Ascomycota_unclassified(100); | - | - | - | - | - | - | - | Unassigned | - |
| Otu24058 | 1 | k__Fungi(100);p__Ascomycota(100);p__Ascomycota_unclassified(100);p__Ascomycota_unclassified(100);p__Ascomycota_unclassified(100);p__Ascomycota_unclassified(100);p__Ascomycota_unclassified(100); | - | - | - | - | - | - | - | Unassigned | - |
| Otu24059 | 1 | k__Fungi(100);k__Fungi_unclassified(100);k__Fungi_unclassified(100);k__Fungi_unclassified(100);k__Fungi_unclassified(100);k__Fungi_unclassified(100);k__Fungi_unclassified(100); | - | - | - | - | - | - | - | Unassigned | - |
| Otu24060 | 1 | k__Fungi(100);p__Ascomycota(100);p__Ascomycota_unclassified(100);p__Ascomycota_unclassified(100);p__Ascomycota_unclassified(100);p__Ascomycota_unclassified(100);p__Ascomycota_unclassified(100); | - | - | - | - | - | - | - | Unassigned | - |
| Otu24062 | 1 | k__Fungi(100);p__Ascomycota(100);p__Ascomycota_unclassified(100);p__Ascomycota_unclassified(100);p__Ascomycota_unclassified(100);p__Ascomycota_unclassified(100);p__Ascomycota_unclassified(100); | - | - | - | - | - | - | - | Unassigned | - |
| Otu24063 | 1 | k__Fungi(100);p__Ascomycota(100);p__Ascomycota_unclassified(100);p__Ascomycota_unclassified(100);p__Ascomycota_unclassified(100);p__Ascomycota_unclassified(100);p__Ascomycota_unclassified(100); | - | - | - | - | - | - | - | Unassigned | - |
| Otu24065 | 1 | k__Fungi(100);k__Fungi_unclassified(100);k__Fungi_unclassified(100);k__Fungi_unclassified(100);k__Fungi_unclassified(100);k__Fungi_unclassified(100);k__Fungi_unclassified(100); | - | - | - | - | - | - | - | Unassigned | - |
| Otu24066 | 1 | k__Fungi(100);k__Fungi_unclassified(100);k__Fungi_unclassified(100);k__Fungi_unclassified(100);k__Fungi_unclassified(100);k__Fungi_unclassified(100);k__Fungi_unclassified(100); | - | - | - | - | - | - | - | Unassigned | - |
| Otu24068 | 1 | k__Fungi(100);k__Fungi_unclassified(100);k__Fungi_unclassified(100);k__Fungi_unclassified(100);k__Fungi_unclassified(100);k__Fungi_unclassified(100);k__Fungi_unclassified(100); | - | - | - | - | - | - | - | Unassigned | - |
| Otu24069 | 1 | k__Fungi(100);p__Ascomycota(100);p__Ascomycota_unclassified(100);p__Ascomycota_unclassified(100);p__Ascomycota_unclassified(100);p__Ascomycota_unclassified(100);p__Ascomycota_unclassified(100); | - | - | - | - | - | - | - | Unassigned | - |
| Otu24071 | 1 | k__Fungi(100);p__Ascomycota(100);p__Ascomycota_unclassified(100);p__Ascomycota_unclassified(100);p__Ascomycota_unclassified(100);p__Ascomycota_unclassified(100);p__Ascomycota_unclassified(100); | - | - | - | - | - | - | - | Unassigned | - |
| Otu24073 | 1 | k__Fungi(100);k__Fungi_unclassified(100);k__Fungi_unclassified(100);k__Fungi_unclassified(100);k__Fungi_unclassified(100);k__Fungi_unclassified(100);k__Fungi_unclassified(100); | - | - | - | - | - | - | - | Unassigned | - |
| Otu24074 | 1 | k__Fungi(100);p__Ascomycota(100);p__Ascomycota_unclassified(100);p__Ascomycota_unclassified(100);p__Ascomycota_unclassified(100);p__Ascomycota_unclassified(100);p__Ascomycota_unclassified(100); | - | - | - | - | - | - | - | Unassigned | - |
| Otu24076 | 1 | k__Fungi(100);k__Fungi_unclassified(100);k__Fungi_unclassified(100);k__Fungi_unclassified(100);k__Fungi_unclassified(100);k__Fungi_unclassified(100);k__Fungi_unclassified(100); | - | - | - | - | - | - | - | Unassigned | - |
| Otu24080 | 1 | k__Fungi(100);k__Fungi_unclassified(100);k__Fungi_unclassified(100);k__Fungi_unclassified(100);k__Fungi_unclassified(100);k__Fungi_unclassified(100);k__Fungi_unclassified(100); | - | - | - | - | - | - | - | Unassigned | - |
| Otu24081 | 1 | k__Fungi(100);k__Fungi_unclassified(100);k__Fungi_unclassified(100);k__Fungi_unclassified(100);k__Fungi_unclassified(100);k__Fungi_unclassified(100);k__Fungi_unclassified(100); | - | - | - | - | - | - | - | Unassigned | - |
| Otu24082 | 1 | k__Fungi(100);p__Ascomycota(100);p__Ascomycota_unclassified(100);p__Ascomycota_unclassified(100);p__Ascomycota_unclassified(100);p__Ascomycota_unclassified(100);p__Ascomycota_unclassified(100); | - | - | - | - | - | - | - | Unassigned | - |
| Otu24083 | 1 | k__Fungi(100);p__Ascomycota(100);p__Ascomycota_unclassified(100);p__Ascomycota_unclassified(100);p__Ascomycota_unclassified(100);p__Ascomycota_unclassified(100);p__Ascomycota_unclassified(100); | - | - | - | - | - | - | - | Unassigned | - |
| Otu24084 | 1 | k__Fungi(100);k__Fungi_unclassified(100);k__Fungi_unclassified(100);k__Fungi_unclassified(100);k__Fungi_unclassified(100);k__Fungi_unclassified(100);k__Fungi_unclassified(100); | - | - | - | - | - | - | - | Unassigned | - |
| Otu24085 | 1 | k__Fungi(100);k__Fungi_unclassified(100);k__Fungi_unclassified(100);k__Fungi_unclassified(100);k__Fungi_unclassified(100);k__Fungi_unclassified(100);k__Fungi_unclassified(100); | - | - | - | - | - | - | - | Unassigned | - |
| Otu24086 | 1 | k__Fungi(100);k__Fungi_unclassified(100);k__Fungi_unclassified(100);k__Fungi_unclassified(100);k__Fungi_unclassified(100);k__Fungi_unclassified(100);k__Fungi_unclassified(100); | - | - | - | - | - | - | - | Unassigned | - |
| Otu24087 | 1 | k__Fungi(100);k__Fungi_unclassified(100);k__Fungi_unclassified(100);k__Fungi_unclassified(100);k__Fungi_unclassified(100);k__Fungi_unclassified(100);k__Fungi_unclassified(100); | - | - | - | - | - | - | - | Unassigned | - |
| Otu24088 | 1 | k__Fungi(100);k__Fungi_unclassified(100);k__Fungi_unclassified(100);k__Fungi_unclassified(100);k__Fungi_unclassified(100);k__Fungi_unclassified(100);k__Fungi_unclassified(100); | - | - | - | - | - | - | - | Unassigned | - |
| Otu24090 | 1 | k__Fungi(100);p__Ascomycota(100);p__Ascomycota_unclassified(100);p__Ascomycota_unclassified(100);p__Ascomycota_unclassified(100);p__Ascomycota_unclassified(100);p__Ascomycota_unclassified(100); | - | - | - | - | - | - | - | Unassigned | - |
| Otu24092 | 1 | k__Fungi(100);p__Ascomycota(100);p__Ascomycota_unclassified(100);p__Ascomycota_unclassified(100);p__Ascomycota_unclassified(100);p__Ascomycota_unclassified(100);p__Ascomycota_unclassified(100); | - | - | - | - | - | - | - | Unassigned | - |
| Otu24093 | 1 | k__Fungi(100);k__Fungi_unclassified(100);k__Fungi_unclassified(100);k__Fungi_unclassified(100);k__Fungi_unclassified(100);k__Fungi_unclassified(100);k__Fungi_unclassified(100); | - | - | - | - | - | - | - | Unassigned | - |
| Otu24094 | 1 | k__Fungi(100);p__Ascomycota(100);p__Ascomycota_unclassified(100);p__Ascomycota_unclassified(100);p__Ascomycota_unclassified(100);p__Ascomycota_unclassified(100);p__Ascomycota_unclassified(100); | - | - | - | - | - | - | - | Unassigned | - |
| Otu24096 | 1 | k__Fungi(100);k__Fungi_unclassified(100);k__Fungi_unclassified(100);k__Fungi_unclassified(100);k__Fungi_unclassified(100);k__Fungi_unclassified(100);k__Fungi_unclassified(100); | - | - | - | - | - | - | - | Unassigned | - |
| Otu24097 | 1 | k__Fungi(100);p__Ascomycota(100);p__Ascomycota_unclassified(100);p__Ascomycota_unclassified(100);p__Ascomycota_unclassified(100);p__Ascomycota_unclassified(100);p__Ascomycota_unclassified(100); | - | - | - | - | - | - | - | Unassigned | - |
| Otu24098 | 1 | k__Fungi(100);p__Ascomycota(100);p__Ascomycota_unclassified(100);p__Ascomycota_unclassified(100);p__Ascomycota_unclassified(100);p__Ascomycota_unclassified(100);p__Ascomycota_unclassified(100); | - | - | - | - | - | - | - | Unassigned | - |
| Otu24099 | 1 | k__Fungi(100);k__Fungi_unclassified(100);k__Fungi_unclassified(100);k__Fungi_unclassified(100);k__Fungi_unclassified(100);k__Fungi_unclassified(100);k__Fungi_unclassified(100); | - | - | - | - | - | - | - | Unassigned | - |
| Otu24105 | 1 | k__Fungi(100);k__Fungi_unclassified(100);k__Fungi_unclassified(100);k__Fungi_unclassified(100);k__Fungi_unclassified(100);k__Fungi_unclassified(100);k__Fungi_unclassified(100); | - | - | - | - | - | - | - | Unassigned | - |
| Otu24108 | 1 | k__Fungi(100);p__Ascomycota(100);p__Ascomycota_unclassified(100);p__Ascomycota_unclassified(100);p__Ascomycota_unclassified(100);p__Ascomycota_unclassified(100);p__Ascomycota_unclassified(100); | - | - | - | - | - | - | - | Unassigned | - |
| Otu24110 | 1 | k__Fungi(100);p__Ascomycota(100);p__Ascomycota_unclassified(100);p__Ascomycota_unclassified(100);p__Ascomycota_unclassified(100);p__Ascomycota_unclassified(100);p__Ascomycota_unclassified(100); | - | - | - | - | - | - | - | Unassigned | - |
| Otu24111 | 1 | k__Fungi(100);p__Ascomycota(100);p__Ascomycota_unclassified(100);p__Ascomycota_unclassified(100);p__Ascomycota_unclassified(100);p__Ascomycota_unclassified(100);p__Ascomycota_unclassified(100); | - | - | - | - | - | - | - | Unassigned | - |
| Otu24113 | 1 | k__Fungi(100);p__Ascomycota(100);p__Ascomycota_unclassified(100);p__Ascomycota_unclassified(100);p__Ascomycota_unclassified(100);p__Ascomycota_unclassified(100);p__Ascomycota_unclassified(100); | - | - | - | - | - | - | - | Unassigned | - |
| Otu24114 | 1 | k__Fungi(100);k__Fungi_unclassified(100);k__Fungi_unclassified(100);k__Fungi_unclassified(100);k__Fungi_unclassified(100);k__Fungi_unclassified(100);k__Fungi_unclassified(100); | - | - | - | - | - | - | - | Unassigned | - |
| Otu24115 | 1 | k__Fungi(100);p__Ascomycota(100);p__Ascomycota_unclassified(100);p__Ascomycota_unclassified(100);p__Ascomycota_unclassified(100);p__Ascomycota_unclassified(100);p__Ascomycota_unclassified(100); | - | - | - | - | - | - | - | Unassigned | - |
| Otu24116 | 1 | k__Fungi(100);k__Fungi_unclassified(100);k__Fungi_unclassified(100);k__Fungi_unclassified(100);k__Fungi_unclassified(100);k__Fungi_unclassified(100);k__Fungi_unclassified(100); | - | - | - | - | - | - | - | Unassigned | - |
| Otu24117 | 1 | k__Fungi(100);p__Ascomycota(100);p__Ascomycota_unclassified(100);p__Ascomycota_unclassified(100);p__Ascomycota_unclassified(100);p__Ascomycota_unclassified(100);p__Ascomycota_unclassified(100); | - | - | - | - | - | - | - | Unassigned | - |
| Otu24119 | 1 | k__Fungi(100);k__Fungi_unclassified(100);k__Fungi_unclassified(100);k__Fungi_unclassified(100);k__Fungi_unclassified(100);k__Fungi_unclassified(100);k__Fungi_unclassified(100); | - | - | - | - | - | - | - | Unassigned | - |
| Otu24120 | 1 | k__Fungi(100);k__Fungi_unclassified(100);k__Fungi_unclassified(100);k__Fungi_unclassified(100);k__Fungi_unclassified(100);k__Fungi_unclassified(100);k__Fungi_unclassified(100); | - | - | - | - | - | - | - | Unassigned | - |
| Otu24125 | 1 | k__Fungi(100);k__Fungi_unclassified(100);k__Fungi_unclassified(100);k__Fungi_unclassified(100);k__Fungi_unclassified(100);k__Fungi_unclassified(100);k__Fungi_unclassified(100); | - | - | - | - | - | - | - | Unassigned | - |
| Otu24126 | 1 | k__Fungi(100);k__Fungi_unclassified(100);k__Fungi_unclassified(100);k__Fungi_unclassified(100);k__Fungi_unclassified(100);k__Fungi_unclassified(100);k__Fungi_unclassified(100); | - | - | - | - | - | - | - | Unassigned | - |
| Otu24127 | 1 | k__Fungi(100);k__Fungi_unclassified(100);k__Fungi_unclassified(100);k__Fungi_unclassified(100);k__Fungi_unclassified(100);k__Fungi_unclassified(100);k__Fungi_unclassified(100); | - | - | - | - | - | - | - | Unassigned | - |
| Otu24128 | 1 | k__Fungi(100);k__Fungi_unclassified(100);k__Fungi_unclassified(100);k__Fungi_unclassified(100);k__Fungi_unclassified(100);k__Fungi_unclassified(100);k__Fungi_unclassified(100); | - | - | - | - | - | - | - | Unassigned | - |
| Otu24129 | 1 | k__Fungi(100);k__Fungi_unclassified(100);k__Fungi_unclassified(100);k__Fungi_unclassified(100);k__Fungi_unclassified(100);k__Fungi_unclassified(100);k__Fungi_unclassified(100); | - | - | - | - | - | - | - | Unassigned | - |
| Otu24130 | 1 | k__Fungi(100);p__Ascomycota(100);p__Ascomycota_unclassified(100);p__Ascomycota_unclassified(100);p__Ascomycota_unclassified(100);p__Ascomycota_unclassified(100);p__Ascomycota_unclassified(100); | - | - | - | - | - | - | - | Unassigned | - |
| Otu24131 | 1 | k__Fungi(100);k__Fungi_unclassified(100);k__Fungi_unclassified(100);k__Fungi_unclassified(100);k__Fungi_unclassified(100);k__Fungi_unclassified(100);k__Fungi_unclassified(100); | - | - | - | - | - | - | - | Unassigned | - |
| Otu24132 | 1 | k__Fungi(100);k__Fungi_unclassified(100);k__Fungi_unclassified(100);k__Fungi_unclassified(100);k__Fungi_unclassified(100);k__Fungi_unclassified(100);k__Fungi_unclassified(100); | - | - | - | - | - | - | - | Unassigned | - |
| Otu24134 | 1 | k__Fungi(100);k__Fungi_unclassified(100);k__Fungi_unclassified(100);k__Fungi_unclassified(100);k__Fungi_unclassified(100);k__Fungi_unclassified(100);k__Fungi_unclassified(100); | - | - | - | - | - | - | - | Unassigned | - |
| Otu24135 | 1 | k__Fungi(100);p__Ascomycota(100);p__Ascomycota_unclassified(100);p__Ascomycota_unclassified(100);p__Ascomycota_unclassified(100);p__Ascomycota_unclassified(100);p__Ascomycota_unclassified(100); | - | - | - | - | - | - | - | Unassigned | - |
| Otu24136 | 1 | k__Fungi(100);k__Fungi_unclassified(100);k__Fungi_unclassified(100);k__Fungi_unclassified(100);k__Fungi_unclassified(100);k__Fungi_unclassified(100);k__Fungi_unclassified(100); | - | - | - | - | - | - | - | Unassigned | - |
| Otu24138 | 1 | k__Fungi(100);k__Fungi_unclassified(100);k__Fungi_unclassified(100);k__Fungi_unclassified(100);k__Fungi_unclassified(100);k__Fungi_unclassified(100);k__Fungi_unclassified(100); | - | - | - | - | - | - | - | Unassigned | - |
| Otu24148 | 1 | k__Fungi(100);k__Fungi_unclassified(100);k__Fungi_unclassified(100);k__Fungi_unclassified(100);k__Fungi_unclassified(100);k__Fungi_unclassified(100);k__Fungi_unclassified(100); | - | - | - | - | - | - | - | Unassigned | - |
| Otu24157 | 1 | k__Fungi(100);k__Fungi_unclassified(100);k__Fungi_unclassified(100);k__Fungi_unclassified(100);k__Fungi_unclassified(100);k__Fungi_unclassified(100);k__Fungi_unclassified(100); | - | - | - | - | - | - | - | Unassigned | - |
| Otu24158 | 1 | k__Fungi(100);k__Fungi_unclassified(100);k__Fungi_unclassified(100);k__Fungi_unclassified(100);k__Fungi_unclassified(100);k__Fungi_unclassified(100);k__Fungi_unclassified(100); | - | - | - | - | - | - | - | Unassigned | - |
| Otu24159 | 1 | k__Fungi(100);k__Fungi_unclassified(100);k__Fungi_unclassified(100);k__Fungi_unclassified(100);k__Fungi_unclassified(100);k__Fungi_unclassified(100);k__Fungi_unclassified(100); | - | - | - | - | - | - | - | Unassigned | - |
| Otu24161 | 1 | k__Fungi(100);p__Ascomycota(100);p__Ascomycota_unclassified(100);p__Ascomycota_unclassified(100);p__Ascomycota_unclassified(100);p__Ascomycota_unclassified(100);p__Ascomycota_unclassified(100); | - | - | - | - | - | - | - | Unassigned | - |
| Otu24162 | 1 | k__Fungi(100);p__Ascomycota(100);p__Ascomycota_unclassified(100);p__Ascomycota_unclassified(100);p__Ascomycota_unclassified(100);p__Ascomycota_unclassified(100);p__Ascomycota_unclassified(100); | - | - | - | - | - | - | - | Unassigned | - |
| Otu24163 | 1 | k__Fungi(100);p__Ascomycota(100);p__Ascomycota_unclassified(100);p__Ascomycota_unclassified(100);p__Ascomycota_unclassified(100);p__Ascomycota_unclassified(100);p__Ascomycota_unclassified(100); | - | - | - | - | - | - | - | Unassigned | - |
| Otu24164 | 1 | k__Fungi(100);p__Ascomycota(100);p__Ascomycota_unclassified(100);p__Ascomycota_unclassified(100);p__Ascomycota_unclassified(100);p__Ascomycota_unclassified(100);p__Ascomycota_unclassified(100); | - | - | - | - | - | - | - | Unassigned | - |
| Otu24166 | 1 | k__Fungi(100);p__Ascomycota(100);p__Ascomycota_unclassified(100);p__Ascomycota_unclassified(100);p__Ascomycota_unclassified(100);p__Ascomycota_unclassified(100);p__Ascomycota_unclassified(100); | - | - | - | - | - | - | - | Unassigned | - |
| Otu24167 | 1 | k__Fungi(100);k__Fungi_unclassified(100);k__Fungi_unclassified(100);k__Fungi_unclassified(100);k__Fungi_unclassified(100);k__Fungi_unclassified(100);k__Fungi_unclassified(100); | - | - | - | - | - | - | - | Unassigned | - |
| Otu24168 | 1 | k__Fungi(100);p__Ascomycota(100);p__Ascomycota_unclassified(100);p__Ascomycota_unclassified(100);p__Ascomycota_unclassified(100);p__Ascomycota_unclassified(100);p__Ascomycota_unclassified(100); | - | - | - | - | - | - | - | Unassigned | - |
| Otu24170 | 1 | k__Fungi(100);k__Fungi_unclassified(100);k__Fungi_unclassified(100);k__Fungi_unclassified(100);k__Fungi_unclassified(100);k__Fungi_unclassified(100);k__Fungi_unclassified(100); | - | - | - | - | - | - | - | Unassigned | - |
| Otu24172 | 1 | k__Fungi(100);p__Ascomycota(100);p__Ascomycota_unclassified(100);p__Ascomycota_unclassified(100);p__Ascomycota_unclassified(100);p__Ascomycota_unclassified(100);p__Ascomycota_unclassified(100); | - | - | - | - | - | - | - | Unassigned | - |
| Otu24173 | 1 | k__Fungi(100);k__Fungi_unclassified(100);k__Fungi_unclassified(100);k__Fungi_unclassified(100);k__Fungi_unclassified(100);k__Fungi_unclassified(100);k__Fungi_unclassified(100); | - | - | - | - | - | - | - | Unassigned | - |
| Otu24175 | 1 | k__Fungi(100);k__Fungi_unclassified(100);k__Fungi_unclassified(100);k__Fungi_unclassified(100);k__Fungi_unclassified(100);k__Fungi_unclassified(100);k__Fungi_unclassified(100); | - | - | - | - | - | - | - | Unassigned | - |
| Otu24176 | 1 | k__Fungi(100);k__Fungi_unclassified(100);k__Fungi_unclassified(100);k__Fungi_unclassified(100);k__Fungi_unclassified(100);k__Fungi_unclassified(100);k__Fungi_unclassified(100); | - | - | - | - | - | - | - | Unassigned | - |
| Otu24177 | 1 | k__Fungi(100);k__Fungi_unclassified(100);k__Fungi_unclassified(100);k__Fungi_unclassified(100);k__Fungi_unclassified(100);k__Fungi_unclassified(100);k__Fungi_unclassified(100); | - | - | - | - | - | - | - | Unassigned | - |
| Otu24178 | 1 | k__Fungi(100);k__Fungi_unclassified(100);k__Fungi_unclassified(100);k__Fungi_unclassified(100);k__Fungi_unclassified(100);k__Fungi_unclassified(100);k__Fungi_unclassified(100); | - | - | - | - | - | - | - | Unassigned | - |
| Otu24179 | 1 | k__Fungi(100);k__Fungi_unclassified(100);k__Fungi_unclassified(100);k__Fungi_unclassified(100);k__Fungi_unclassified(100);k__Fungi_unclassified(100);k__Fungi_unclassified(100); | - | - | - | - | - | - | - | Unassigned | - |
| Otu24180 | 1 | k__Fungi(100);k__Fungi_unclassified(100);k__Fungi_unclassified(100);k__Fungi_unclassified(100);k__Fungi_unclassified(100);k__Fungi_unclassified(100);k__Fungi_unclassified(100); | - | - | - | - | - | - | - | Unassigned | - |
| Otu24182 | 1 | k__Fungi(100);k__Fungi_unclassified(100);k__Fungi_unclassified(100);k__Fungi_unclassified(100);k__Fungi_unclassified(100);k__Fungi_unclassified(100);k__Fungi_unclassified(100); | - | - | - | - | - | - | - | Unassigned | - |
| Otu24183 | 1 | k__Fungi(100);p__Ascomycota(100);p__Ascomycota_unclassified(100);p__Ascomycota_unclassified(100);p__Ascomycota_unclassified(100);p__Ascomycota_unclassified(100);p__Ascomycota_unclassified(100); | - | - | - | - | - | - | - | Unassigned | - |
| Otu24187 | 1 | k__Fungi(100);k__Fungi_unclassified(100);k__Fungi_unclassified(100);k__Fungi_unclassified(100);k__Fungi_unclassified(100);k__Fungi_unclassified(100);k__Fungi_unclassified(100); | - | - | - | - | - | - | - | Unassigned | - |
| Otu24207 | 1 | k__Fungi(100);p__Ascomycota(100);p__Ascomycota_unclassified(100);p__Ascomycota_unclassified(100);p__Ascomycota_unclassified(100);p__Ascomycota_unclassified(100);p__Ascomycota_unclassified(100); | - | - | - | - | - | - | - | Unassigned | - |
| Otu24208 | 1 | k__Fungi(100);p__Ascomycota(100);p__Ascomycota_unclassified(100);p__Ascomycota_unclassified(100);p__Ascomycota_unclassified(100);p__Ascomycota_unclassified(100);p__Ascomycota_unclassified(100); | - | - | - | - | - | - | - | Unassigned | - |
| Otu24209 | 1 | k__Fungi(100);p__Ascomycota(100);p__Ascomycota_unclassified(100);p__Ascomycota_unclassified(100);p__Ascomycota_unclassified(100);p__Ascomycota_unclassified(100);p__Ascomycota_unclassified(100); | - | - | - | - | - | - | - | Unassigned | - |
| Otu24212 | 1 | k__Fungi(100);p__Ascomycota(100);p__Ascomycota_unclassified(100);p__Ascomycota_unclassified(100);p__Ascomycota_unclassified(100);p__Ascomycota_unclassified(100);p__Ascomycota_unclassified(100); | - | - | - | - | - | - | - | Unassigned | - |
| Otu24214 | 1 | k__Fungi(100);k__Fungi_unclassified(100);k__Fungi_unclassified(100);k__Fungi_unclassified(100);k__Fungi_unclassified(100);k__Fungi_unclassified(100);k__Fungi_unclassified(100); | - | - | - | - | - | - | - | Unassigned | - |
| Otu24215 | 1 | k__Fungi(100);k__Fungi_unclassified(100);k__Fungi_unclassified(100);k__Fungi_unclassified(100);k__Fungi_unclassified(100);k__Fungi_unclassified(100);k__Fungi_unclassified(100); | - | - | - | - | - | - | - | Unassigned | - |
| Otu24216 | 1 | k__Fungi(100);k__Fungi_unclassified(100);k__Fungi_unclassified(100);k__Fungi_unclassified(100);k__Fungi_unclassified(100);k__Fungi_unclassified(100);k__Fungi_unclassified(100); | - | - | - | - | - | - | - | Unassigned | - |
| Otu24217 | 1 | k__Fungi(100);p__Ascomycota(100);p__Ascomycota_unclassified(100);p__Ascomycota_unclassified(100);p__Ascomycota_unclassified(100);p__Ascomycota_unclassified(100);p__Ascomycota_unclassified(100); | - | - | - | - | - | - | - | Unassigned | - |
| Otu24220 | 1 | k__Fungi(100);p__Ascomycota(100);p__Ascomycota_unclassified(100);p__Ascomycota_unclassified(100);p__Ascomycota_unclassified(100);p__Ascomycota_unclassified(100);p__Ascomycota_unclassified(100); | - | - | - | - | - | - | - | Unassigned | - |
| Otu24222 | 1 | k__Fungi(100);k__Fungi_unclassified(100);k__Fungi_unclassified(100);k__Fungi_unclassified(100);k__Fungi_unclassified(100);k__Fungi_unclassified(100);k__Fungi_unclassified(100); | - | - | - | - | - | - | - | Unassigned | - |
| Otu24226 | 1 | k__Fungi(100);p__Ascomycota(100);p__Ascomycota_unclassified(100);p__Ascomycota_unclassified(100);p__Ascomycota_unclassified(100);p__Ascomycota_unclassified(100);p__Ascomycota_unclassified(100); | - | - | - | - | - | - | - | Unassigned | - |
| Otu24241 | 1 | k__Fungi(100);k__Fungi_unclassified(100);k__Fungi_unclassified(100);k__Fungi_unclassified(100);k__Fungi_unclassified(100);k__Fungi_unclassified(100);k__Fungi_unclassified(100); | - | - | - | - | - | - | - | Unassigned | - |
| Otu24242 | 1 | k__Fungi(100);p__Ascomycota(100);p__Ascomycota_unclassified(100);p__Ascomycota_unclassified(100);p__Ascomycota_unclassified(100);p__Ascomycota_unclassified(100);p__Ascomycota_unclassified(100); | - | - | - | - | - | - | - | Unassigned | - |
| Otu24243 | 1 | k__Fungi(100);k__Fungi_unclassified(100);k__Fungi_unclassified(100);k__Fungi_unclassified(100);k__Fungi_unclassified(100);k__Fungi_unclassified(100);k__Fungi_unclassified(100); | - | - | - | - | - | - | - | Unassigned | - |
| Otu24244 | 1 | k__Fungi(100);k__Fungi_unclassified(100);k__Fungi_unclassified(100);k__Fungi_unclassified(100);k__Fungi_unclassified(100);k__Fungi_unclassified(100);k__Fungi_unclassified(100); | - | - | - | - | - | - | - | Unassigned | - |
| Otu24245 | 1 | k__Fungi(100);k__Fungi_unclassified(100);k__Fungi_unclassified(100);k__Fungi_unclassified(100);k__Fungi_unclassified(100);k__Fungi_unclassified(100);k__Fungi_unclassified(100); | - | - | - | - | - | - | - | Unassigned | - |
| Otu24246 | 1 | k__Fungi(100);k__Fungi_unclassified(100);k__Fungi_unclassified(100);k__Fungi_unclassified(100);k__Fungi_unclassified(100);k__Fungi_unclassified(100);k__Fungi_unclassified(100); | - | - | - | - | - | - | - | Unassigned | - |
| Otu24247 | 1 | k__Fungi(100);k__Fungi_unclassified(100);k__Fungi_unclassified(100);k__Fungi_unclassified(100);k__Fungi_unclassified(100);k__Fungi_unclassified(100);k__Fungi_unclassified(100); | - | - | - | - | - | - | - | Unassigned | - |
| Otu24248 | 1 | k__Fungi(100);k__Fungi_unclassified(100);k__Fungi_unclassified(100);k__Fungi_unclassified(100);k__Fungi_unclassified(100);k__Fungi_unclassified(100);k__Fungi_unclassified(100); | - | - | - | - | - | - | - | Unassigned | - |
| Otu24249 | 1 | k__Fungi(100);p__Ascomycota(100);p__Ascomycota_unclassified(100);p__Ascomycota_unclassified(100);p__Ascomycota_unclassified(100);p__Ascomycota_unclassified(100);p__Ascomycota_unclassified(100); | - | - | - | - | - | - | - | Unassigned | - |
| Otu24250 | 1 | k__Fungi(100);k__Fungi_unclassified(100);k__Fungi_unclassified(100);k__Fungi_unclassified(100);k__Fungi_unclassified(100);k__Fungi_unclassified(100);k__Fungi_unclassified(100); | - | - | - | - | - | - | - | Unassigned | - |
| Otu24253 | 1 | k__Fungi(100);k__Fungi_unclassified(100);k__Fungi_unclassified(100);k__Fungi_unclassified(100);k__Fungi_unclassified(100);k__Fungi_unclassified(100);k__Fungi_unclassified(100); | - | - | - | - | - | - | - | Unassigned | - |
| Otu24254 | 1 | k__Fungi(100);k__Fungi_unclassified(100);k__Fungi_unclassified(100);k__Fungi_unclassified(100);k__Fungi_unclassified(100);k__Fungi_unclassified(100);k__Fungi_unclassified(100); | - | - | - | - | - | - | - | Unassigned | - |
| Otu24256 | 1 | k__Fungi(100);k__Fungi_unclassified(100);k__Fungi_unclassified(100);k__Fungi_unclassified(100);k__Fungi_unclassified(100);k__Fungi_unclassified(100);k__Fungi_unclassified(100); | - | - | - | - | - | - | - | Unassigned | - |
| Otu24257 | 1 | k__Fungi(100);k__Fungi_unclassified(100);k__Fungi_unclassified(100);k__Fungi_unclassified(100);k__Fungi_unclassified(100);k__Fungi_unclassified(100);k__Fungi_unclassified(100); | - | - | - | - | - | - | - | Unassigned | - |
| Otu24258 | 1 | k__Plantae(100);p__unclassified_Plantae(100);c__unclassified_Plantae(100);o__unclassified_Plantae(100);f__unclassified_Plantae(100);g__unclassified_Plantae(100);s__Plantae_sp(100); | - | - | - | - | - | - | - | Unassigned | - |
| Otu24261 | 1 | k__Fungi(100);p__Basidiomycota(100);c__Agaricomycetes(100);c__Agaricomycetes_unclassified(100);c__Agaricomycetes_unclassified(100);c__Agaricomycetes_unclassified(100);c__Agaricomycetes_unclassified(100); | - | - | - | - | - | - | - | Unassigned | - |
| Otu24262 | 1 | k__Fungi(100);p__Basidiomycota(100);c__Microbotryomycetes(100);o__Sporidiobolales(100);f__Sporidiobolales_family_Incertae_sedis(100);g__unclassified_Sporidiobolales(100);s__Sporidiobolales_sp(100); | - | - | - | - | - | - | - | Unassigned | - |
| Otu24265 | 1 | k__Fungi(100);p__Ascomycota(100);p__Ascomycota_unclassified(100);p__Ascomycota_unclassified(100);p__Ascomycota_unclassified(100);p__Ascomycota_unclassified(100);p__Ascomycota_unclassified(100); | - | - | - | - | - | - | - | Unassigned | - |
| Otu24266 | 1 | k__Fungi(100);k__Fungi_unclassified(100);k__Fungi_unclassified(100);k__Fungi_unclassified(100);k__Fungi_unclassified(100);k__Fungi_unclassified(100);k__Fungi_unclassified(100); | - | - | - | - | - | - | - | Unassigned | - |
| Otu24267 | 1 | k__Fungi(100);k__Fungi_unclassified(100);k__Fungi_unclassified(100);k__Fungi_unclassified(100);k__Fungi_unclassified(100);k__Fungi_unclassified(100);k__Fungi_unclassified(100); | - | - | - | - | - | - | - | Unassigned | - |
| Otu24268 | 1 | k__Fungi(100);p__Ascomycota(100);p__Ascomycota_unclassified(100);p__Ascomycota_unclassified(100);p__Ascomycota_unclassified(100);p__Ascomycota_unclassified(100);p__Ascomycota_unclassified(100); | - | - | - | - | - | - | - | Unassigned | - |
| Otu24269 | 1 | k__Fungi(100);k__Fungi_unclassified(100);k__Fungi_unclassified(100);k__Fungi_unclassified(100);k__Fungi_unclassified(100);k__Fungi_unclassified(100);k__Fungi_unclassified(100); | - | - | - | - | - | - | - | Unassigned | - |
| Otu24271 | 1 | k__Fungi(100);p__Ascomycota(100);p__Ascomycota_unclassified(100);p__Ascomycota_unclassified(100);p__Ascomycota_unclassified(100);p__Ascomycota_unclassified(100);p__Ascomycota_unclassified(100); | - | - | - | - | - | - | - | Unassigned | - |
| Otu24273 | 1 | k__Fungi(100);k__Fungi_unclassified(100);k__Fungi_unclassified(100);k__Fungi_unclassified(100);k__Fungi_unclassified(100);k__Fungi_unclassified(100);k__Fungi_unclassified(100); | - | - | - | - | - | - | - | Unassigned | - |
| Otu24274 | 1 | k__Fungi(100);p__Ascomycota(100);p__Ascomycota_unclassified(100);p__Ascomycota_unclassified(100);p__Ascomycota_unclassified(100);p__Ascomycota_unclassified(100);p__Ascomycota_unclassified(100); | - | - | - | - | - | - | - | Unassigned | - |
| Otu24275 | 1 | k__Fungi(100);p__Ascomycota(100);p__Ascomycota_unclassified(100);p__Ascomycota_unclassified(100);p__Ascomycota_unclassified(100);p__Ascomycota_unclassified(100);p__Ascomycota_unclassified(100); | - | - | - | - | - | - | - | Unassigned | - |
| Otu24276 | 1 | k__Fungi(100);p__Ascomycota(100);p__Ascomycota_unclassified(100);p__Ascomycota_unclassified(100);p__Ascomycota_unclassified(100);p__Ascomycota_unclassified(100);p__Ascomycota_unclassified(100); | - | - | - | - | - | - | - | Unassigned | - |
| Otu24277 | 1 | k__Fungi(100);k__Fungi_unclassified(100);k__Fungi_unclassified(100);k__Fungi_unclassified(100);k__Fungi_unclassified(100);k__Fungi_unclassified(100);k__Fungi_unclassified(100); | - | - | - | - | - | - | - | Unassigned | - |
| Otu24278 | 1 | k__Fungi(100);p__Ascomycota(100);p__Ascomycota_unclassified(100);p__Ascomycota_unclassified(100);p__Ascomycota_unclassified(100);p__Ascomycota_unclassified(100);p__Ascomycota_unclassified(100); | - | - | - | - | - | - | - | Unassigned | - |
| Otu24279 | 1 | k__Fungi(100);k__Fungi_unclassified(100);k__Fungi_unclassified(100);k__Fungi_unclassified(100);k__Fungi_unclassified(100);k__Fungi_unclassified(100);k__Fungi_unclassified(100); | - | - | - | - | - | - | - | Unassigned | - |
| Otu24281 | 1 | k__Fungi(100);p__Ascomycota(100);p__Ascomycota_unclassified(100);p__Ascomycota_unclassified(100);p__Ascomycota_unclassified(100);p__Ascomycota_unclassified(100);p__Ascomycota_unclassified(100); | - | - | - | - | - | - | - | Unassigned | - |
| Otu24282 | 1 | k__Fungi(100);p__Ascomycota(100);p__Ascomycota_unclassified(100);p__Ascomycota_unclassified(100);p__Ascomycota_unclassified(100);p__Ascomycota_unclassified(100);p__Ascomycota_unclassified(100); | - | - | - | - | - | - | - | Unassigned | - |
| Otu24283 | 1 | k__Fungi(100);k__Fungi_unclassified(100);k__Fungi_unclassified(100);k__Fungi_unclassified(100);k__Fungi_unclassified(100);k__Fungi_unclassified(100);k__Fungi_unclassified(100); | - | - | - | - | - | - | - | Unassigned | - |
| Otu24284 | 1 | k__Fungi(100);p__Ascomycota(100);p__Ascomycota_unclassified(100);p__Ascomycota_unclassified(100);p__Ascomycota_unclassified(100);p__Ascomycota_unclassified(100);p__Ascomycota_unclassified(100); | - | - | - | - | - | - | - | Unassigned | - |
| Otu24286 | 1 | k__Fungi(100);k__Fungi_unclassified(100);k__Fungi_unclassified(100);k__Fungi_unclassified(100);k__Fungi_unclassified(100);k__Fungi_unclassified(100);k__Fungi_unclassified(100); | - | - | - | - | - | - | - | Unassigned | - |
| Otu24290 | 1 | k__Fungi(100);k__Fungi_unclassified(100);k__Fungi_unclassified(100);k__Fungi_unclassified(100);k__Fungi_unclassified(100);k__Fungi_unclassified(100);k__Fungi_unclassified(100); | - | - | - | - | - | - | - | Unassigned | - |
| Otu24292 | 1 | k__Fungi(100);k__Fungi_unclassified(100);k__Fungi_unclassified(100);k__Fungi_unclassified(100);k__Fungi_unclassified(100);k__Fungi_unclassified(100);k__Fungi_unclassified(100); | - | - | - | - | - | - | - | Unassigned | - |
| Otu24293 | 1 | k__Fungi(100);k__Fungi_unclassified(100);k__Fungi_unclassified(100);k__Fungi_unclassified(100);k__Fungi_unclassified(100);k__Fungi_unclassified(100);k__Fungi_unclassified(100); | - | - | - | - | - | - | - | Unassigned | - |
| Otu24294 | 1 | k__Fungi(100);p__Ascomycota(100);p__Ascomycota_unclassified(100);p__Ascomycota_unclassified(100);p__Ascomycota_unclassified(100);p__Ascomycota_unclassified(100);p__Ascomycota_unclassified(100); | - | - | - | - | - | - | - | Unassigned | - |
| Otu24295 | 1 | k__Fungi(100);k__Fungi_unclassified(100);k__Fungi_unclassified(100);k__Fungi_unclassified(100);k__Fungi_unclassified(100);k__Fungi_unclassified(100);k__Fungi_unclassified(100); | - | - | - | - | - | - | - | Unassigned | - |
| Otu24296 | 1 | k__Fungi(100);k__Fungi_unclassified(100);k__Fungi_unclassified(100);k__Fungi_unclassified(100);k__Fungi_unclassified(100);k__Fungi_unclassified(100);k__Fungi_unclassified(100); | - | - | - | - | - | - | - | Unassigned | - |
| Otu24297 | 1 | k__Fungi(100);p__Ascomycota(100);p__Ascomycota_unclassified(100);p__Ascomycota_unclassified(100);p__Ascomycota_unclassified(100);p__Ascomycota_unclassified(100);p__Ascomycota_unclassified(100); | - | - | - | - | - | - | - | Unassigned | - |
| Otu24298 | 1 | k__Fungi(100);p__Ascomycota(100);p__Ascomycota_unclassified(100);p__Ascomycota_unclassified(100);p__Ascomycota_unclassified(100);p__Ascomycota_unclassified(100);p__Ascomycota_unclassified(100); | - | - | - | - | - | - | - | Unassigned | - |
| Otu24299 | 1 | k__Fungi(100);k__Fungi_unclassified(100);k__Fungi_unclassified(100);k__Fungi_unclassified(100);k__Fungi_unclassified(100);k__Fungi_unclassified(100);k__Fungi_unclassified(100); | - | - | - | - | - | - | - | Unassigned | - |
| Otu24300 | 1 | k__Fungi(100);k__Fungi_unclassified(100);k__Fungi_unclassified(100);k__Fungi_unclassified(100);k__Fungi_unclassified(100);k__Fungi_unclassified(100);k__Fungi_unclassified(100); | - | - | - | - | - | - | - | Unassigned | - |
| Otu24301 | 1 | k__Fungi(100);k__Fungi_unclassified(100);k__Fungi_unclassified(100);k__Fungi_unclassified(100);k__Fungi_unclassified(100);k__Fungi_unclassified(100);k__Fungi_unclassified(100); | - | - | - | - | - | - | - | Unassigned | - |
| Otu24302 | 1 | k__Fungi(100);p__Ascomycota(100);p__Ascomycota_unclassified(100);p__Ascomycota_unclassified(100);p__Ascomycota_unclassified(100);p__Ascomycota_unclassified(100);p__Ascomycota_unclassified(100); | - | - | - | - | - | - | - | Unassigned | - |
| Otu24303 | 1 | k__Fungi(100);p__Ascomycota(100);p__Ascomycota_unclassified(100);p__Ascomycota_unclassified(100);p__Ascomycota_unclassified(100);p__Ascomycota_unclassified(100);p__Ascomycota_unclassified(100); | - | - | - | - | - | - | - | Unassigned | - |
| Otu24304 | 1 | k__Fungi(100);p__Ascomycota(100);p__Ascomycota_unclassified(100);p__Ascomycota_unclassified(100);p__Ascomycota_unclassified(100);p__Ascomycota_unclassified(100);p__Ascomycota_unclassified(100); | - | - | - | - | - | - | - | Unassigned | - |
| Otu24306 | 1 | k__Fungi(100);k__Fungi_unclassified(100);k__Fungi_unclassified(100);k__Fungi_unclassified(100);k__Fungi_unclassified(100);k__Fungi_unclassified(100);k__Fungi_unclassified(100); | - | - | - | - | - | - | - | Unassigned | - |
| Otu24307 | 1 | k__Fungi(100);p__Ascomycota(100);p__Ascomycota_unclassified(100);p__Ascomycota_unclassified(100);p__Ascomycota_unclassified(100);p__Ascomycota_unclassified(100);p__Ascomycota_unclassified(100); | - | - | - | - | - | - | - | Unassigned | - |
| Otu24308 | 1 | k__Fungi(100);k__Fungi_unclassified(100);k__Fungi_unclassified(100);k__Fungi_unclassified(100);k__Fungi_unclassified(100);k__Fungi_unclassified(100);k__Fungi_unclassified(100); | - | - | - | - | - | - | - | Unassigned | - |
| Otu24309 | 1 | k__Fungi(100);k__Fungi_unclassified(100);k__Fungi_unclassified(100);k__Fungi_unclassified(100);k__Fungi_unclassified(100);k__Fungi_unclassified(100);k__Fungi_unclassified(100); | - | - | - | - | - | - | - | Unassigned | - |
| Otu24310 | 1 | k__Fungi(100);k__Fungi_unclassified(100);k__Fungi_unclassified(100);k__Fungi_unclassified(100);k__Fungi_unclassified(100);k__Fungi_unclassified(100);k__Fungi_unclassified(100); | - | - | - | - | - | - | - | Unassigned | - |
| Otu24312 | 1 | k__Fungi(100);p__Ascomycota(100);p__Ascomycota_unclassified(100);p__Ascomycota_unclassified(100);p__Ascomycota_unclassified(100);p__Ascomycota_unclassified(100);p__Ascomycota_unclassified(100); | - | - | - | - | - | - | - | Unassigned | - |
| Otu24313 | 1 | k__Fungi(100);p__Ascomycota(100);p__Ascomycota_unclassified(100);p__Ascomycota_unclassified(100);p__Ascomycota_unclassified(100);p__Ascomycota_unclassified(100);p__Ascomycota_unclassified(100); | - | - | - | - | - | - | - | Unassigned | - |
| Otu24315 | 1 | k__Fungi(100);p__Ascomycota(100);p__Ascomycota_unclassified(100);p__Ascomycota_unclassified(100);p__Ascomycota_unclassified(100);p__Ascomycota_unclassified(100);p__Ascomycota_unclassified(100); | - | - | - | - | - | - | - | Unassigned | - |
| Otu24316 | 1 | k__Fungi(100);k__Fungi_unclassified(100);k__Fungi_unclassified(100);k__Fungi_unclassified(100);k__Fungi_unclassified(100);k__Fungi_unclassified(100);k__Fungi_unclassified(100); | - | - | - | - | - | - | - | Unassigned | - |
| Otu24317 | 1 | k__Fungi(100);k__Fungi_unclassified(100);k__Fungi_unclassified(100);k__Fungi_unclassified(100);k__Fungi_unclassified(100);k__Fungi_unclassified(100);k__Fungi_unclassified(100); | - | - | - | - | - | - | - | Unassigned | - |
| Otu24318 | 1 | k__Fungi(100);k__Fungi_unclassified(100);k__Fungi_unclassified(100);k__Fungi_unclassified(100);k__Fungi_unclassified(100);k__Fungi_unclassified(100);k__Fungi_unclassified(100); | - | - | - | - | - | - | - | Unassigned | - |
| Otu24319 | 1 | k__Fungi(100);k__Fungi_unclassified(100);k__Fungi_unclassified(100);k__Fungi_unclassified(100);k__Fungi_unclassified(100);k__Fungi_unclassified(100);k__Fungi_unclassified(100); | - | - | - | - | - | - | - | Unassigned | - |
| Otu24320 | 1 | k__Fungi(100);k__Fungi_unclassified(100);k__Fungi_unclassified(100);k__Fungi_unclassified(100);k__Fungi_unclassified(100);k__Fungi_unclassified(100);k__Fungi_unclassified(100); | - | - | - | - | - | - | - | Unassigned | - |
| Otu24321 | 1 | k__Fungi(100);k__Fungi_unclassified(100);k__Fungi_unclassified(100);k__Fungi_unclassified(100);k__Fungi_unclassified(100);k__Fungi_unclassified(100);k__Fungi_unclassified(100); | - | - | - | - | - | - | - | Unassigned | - |
| Otu24322 | 1 | k__Fungi(100);k__Fungi_unclassified(100);k__Fungi_unclassified(100);k__Fungi_unclassified(100);k__Fungi_unclassified(100);k__Fungi_unclassified(100);k__Fungi_unclassified(100); | - | - | - | - | - | - | - | Unassigned | - |
| Otu24323 | 1 | k__Fungi(100);k__Fungi_unclassified(100);k__Fungi_unclassified(100);k__Fungi_unclassified(100);k__Fungi_unclassified(100);k__Fungi_unclassified(100);k__Fungi_unclassified(100); | - | - | - | - | - | - | - | Unassigned | - |
| Otu24324 | 1 | k__Fungi(100);p__Ascomycota(100);p__Ascomycota_unclassified(100);p__Ascomycota_unclassified(100);p__Ascomycota_unclassified(100);p__Ascomycota_unclassified(100);p__Ascomycota_unclassified(100); | - | - | - | - | - | - | - | Unassigned | - |
| Otu24325 | 1 | k__Fungi(100);k__Fungi_unclassified(100);k__Fungi_unclassified(100);k__Fungi_unclassified(100);k__Fungi_unclassified(100);k__Fungi_unclassified(100);k__Fungi_unclassified(100); | - | - | - | - | - | - | - | Unassigned | - |
| Otu24326 | 1 | k__Fungi(100);p__Ascomycota(100);p__Ascomycota_unclassified(100);p__Ascomycota_unclassified(100);p__Ascomycota_unclassified(100);p__Ascomycota_unclassified(100);p__Ascomycota_unclassified(100); | - | - | - | - | - | - | - | Unassigned | - |
| Otu24327 | 1 | k__Fungi(100);p__Ascomycota(100);p__Ascomycota_unclassified(100);p__Ascomycota_unclassified(100);p__Ascomycota_unclassified(100);p__Ascomycota_unclassified(100);p__Ascomycota_unclassified(100); | - | - | - | - | - | - | - | Unassigned | - |
| Otu24328 | 1 | k__Fungi(100);k__Fungi_unclassified(100);k__Fungi_unclassified(100);k__Fungi_unclassified(100);k__Fungi_unclassified(100);k__Fungi_unclassified(100);k__Fungi_unclassified(100); | - | - | - | - | - | - | - | Unassigned | - |
| Otu24329 | 1 | k__Fungi(100);k__Fungi_unclassified(100);k__Fungi_unclassified(100);k__Fungi_unclassified(100);k__Fungi_unclassified(100);k__Fungi_unclassified(100);k__Fungi_unclassified(100); | - | - | - | - | - | - | - | Unassigned | - |
| Otu24330 | 1 | k__Fungi(100);k__Fungi_unclassified(100);k__Fungi_unclassified(100);k__Fungi_unclassified(100);k__Fungi_unclassified(100);k__Fungi_unclassified(100);k__Fungi_unclassified(100); | - | - | - | - | - | - | - | Unassigned | - |
| Otu24333 | 1 | k__Fungi(100);p__Ascomycota(100);p__Ascomycota_unclassified(100);p__Ascomycota_unclassified(100);p__Ascomycota_unclassified(100);p__Ascomycota_unclassified(100);p__Ascomycota_unclassified(100); | - | - | - | - | - | - | - | Unassigned | - |
| Otu24334 | 1 | k__Fungi(100);p__Ascomycota(100);p__Ascomycota_unclassified(100);p__Ascomycota_unclassified(100);p__Ascomycota_unclassified(100);p__Ascomycota_unclassified(100);p__Ascomycota_unclassified(100); | - | - | - | - | - | - | - | Unassigned | - |
| Otu24335 | 1 | k__Fungi(100);k__Fungi_unclassified(100);k__Fungi_unclassified(100);k__Fungi_unclassified(100);k__Fungi_unclassified(100);k__Fungi_unclassified(100);k__Fungi_unclassified(100); | - | - | - | - | - | - | - | Unassigned | - |
| Otu24336 | 1 | k__Fungi(100);p__Ascomycota(100);p__Ascomycota_unclassified(100);p__Ascomycota_unclassified(100);p__Ascomycota_unclassified(100);p__Ascomycota_unclassified(100);p__Ascomycota_unclassified(100); | - | - | - | - | - | - | - | Unassigned | - |
| Otu24337 | 1 | k__Fungi(100);k__Fungi_unclassified(100);k__Fungi_unclassified(100);k__Fungi_unclassified(100);k__Fungi_unclassified(100);k__Fungi_unclassified(100);k__Fungi_unclassified(100); | - | - | - | - | - | - | - | Unassigned | - |
| Otu24338 | 1 | k__Fungi(100);k__Fungi_unclassified(100);k__Fungi_unclassified(100);k__Fungi_unclassified(100);k__Fungi_unclassified(100);k__Fungi_unclassified(100);k__Fungi_unclassified(100); | - | - | - | - | - | - | - | Unassigned | - |
| Otu24339 | 1 | k__Fungi(100);k__Fungi_unclassified(100);k__Fungi_unclassified(100);k__Fungi_unclassified(100);k__Fungi_unclassified(100);k__Fungi_unclassified(100);k__Fungi_unclassified(100); | - | - | - | - | - | - | - | Unassigned | - |
| Otu24340 | 1 | k__Fungi(100);k__Fungi_unclassified(100);k__Fungi_unclassified(100);k__Fungi_unclassified(100);k__Fungi_unclassified(100);k__Fungi_unclassified(100);k__Fungi_unclassified(100); | - | - | - | - | - | - | - | Unassigned | - |
| Otu24341 | 1 | k__Fungi(100);k__Fungi_unclassified(100);k__Fungi_unclassified(100);k__Fungi_unclassified(100);k__Fungi_unclassified(100);k__Fungi_unclassified(100);k__Fungi_unclassified(100); | - | - | - | - | - | - | - | Unassigned | - |
| Otu24344 | 1 | k__Fungi(100);p__Ascomycota(100);p__Ascomycota_unclassified(100);p__Ascomycota_unclassified(100);p__Ascomycota_unclassified(100);p__Ascomycota_unclassified(100);p__Ascomycota_unclassified(100); | - | - | - | - | - | - | - | Unassigned | - |
| Otu24346 | 1 | k__Fungi(100);k__Fungi_unclassified(100);k__Fungi_unclassified(100);k__Fungi_unclassified(100);k__Fungi_unclassified(100);k__Fungi_unclassified(100);k__Fungi_unclassified(100); | - | - | - | - | - | - | - | Unassigned | - |
| Otu24347 | 1 | k__Fungi(100);p__Ascomycota(100);p__Ascomycota_unclassified(100);p__Ascomycota_unclassified(100);p__Ascomycota_unclassified(100);p__Ascomycota_unclassified(100);p__Ascomycota_unclassified(100); | - | - | - | - | - | - | - | Unassigned | - |
| Otu24348 | 1 | k__Fungi(100);k__Fungi_unclassified(100);k__Fungi_unclassified(100);k__Fungi_unclassified(100);k__Fungi_unclassified(100);k__Fungi_unclassified(100);k__Fungi_unclassified(100); | - | - | - | - | - | - | - | Unassigned | - |
| Otu24349 | 1 | k__Fungi(100);k__Fungi_unclassified(100);k__Fungi_unclassified(100);k__Fungi_unclassified(100);k__Fungi_unclassified(100);k__Fungi_unclassified(100);k__Fungi_unclassified(100); | - | - | - | - | - | - | - | Unassigned | - |
| Otu24353 | 1 | k__Fungi(100);k__Fungi_unclassified(100);k__Fungi_unclassified(100);k__Fungi_unclassified(100);k__Fungi_unclassified(100);k__Fungi_unclassified(100);k__Fungi_unclassified(100); | - | - | - | - | - | - | - | Unassigned | - |
| Otu24355 | 1 | k__Fungi(100);k__Fungi_unclassified(100);k__Fungi_unclassified(100);k__Fungi_unclassified(100);k__Fungi_unclassified(100);k__Fungi_unclassified(100);k__Fungi_unclassified(100); | - | - | - | - | - | - | - | Unassigned | - |
| Otu24356 | 1 | k__Fungi(100);p__Ascomycota(100);p__Ascomycota_unclassified(100);p__Ascomycota_unclassified(100);p__Ascomycota_unclassified(100);p__Ascomycota_unclassified(100);p__Ascomycota_unclassified(100); | - | - | - | - | - | - | - | Unassigned | - |
| Otu24357 | 1 | k__Fungi(100);k__Fungi_unclassified(100);k__Fungi_unclassified(100);k__Fungi_unclassified(100);k__Fungi_unclassified(100);k__Fungi_unclassified(100);k__Fungi_unclassified(100); | - | - | - | - | - | - | - | Unassigned | - |
| Otu24358 | 1 | k__Fungi(100);k__Fungi_unclassified(100);k__Fungi_unclassified(100);k__Fungi_unclassified(100);k__Fungi_unclassified(100);k__Fungi_unclassified(100);k__Fungi_unclassified(100); | - | - | - | - | - | - | - | Unassigned | - |
| Otu24359 | 1 | k__Fungi(100);k__Fungi_unclassified(100);k__Fungi_unclassified(100);k__Fungi_unclassified(100);k__Fungi_unclassified(100);k__Fungi_unclassified(100);k__Fungi_unclassified(100); | - | - | - | - | - | - | - | Unassigned | - |
| Otu24361 | 1 | k__Fungi(100);k__Fungi_unclassified(100);k__Fungi_unclassified(100);k__Fungi_unclassified(100);k__Fungi_unclassified(100);k__Fungi_unclassified(100);k__Fungi_unclassified(100); | - | - | - | - | - | - | - | Unassigned | - |
| Otu24363 | 1 | k__Fungi(100);k__Fungi_unclassified(100);k__Fungi_unclassified(100);k__Fungi_unclassified(100);k__Fungi_unclassified(100);k__Fungi_unclassified(100);k__Fungi_unclassified(100); | - | - | - | - | - | - | - | Unassigned | - |
| Otu24364 | 1 | k__Fungi(100);k__Fungi_unclassified(100);k__Fungi_unclassified(100);k__Fungi_unclassified(100);k__Fungi_unclassified(100);k__Fungi_unclassified(100);k__Fungi_unclassified(100); | - | - | - | - | - | - | - | Unassigned | - |
| Otu24365 | 1 | k__Fungi(100);p__Ascomycota(100);p__Ascomycota_unclassified(100);p__Ascomycota_unclassified(100);p__Ascomycota_unclassified(100);p__Ascomycota_unclassified(100);p__Ascomycota_unclassified(100); | - | - | - | - | - | - | - | Unassigned | - |
| Otu24366 | 1 | k__Fungi(100);k__Fungi_unclassified(100);k__Fungi_unclassified(100);k__Fungi_unclassified(100);k__Fungi_unclassified(100);k__Fungi_unclassified(100);k__Fungi_unclassified(100); | - | - | - | - | - | - | - | Unassigned | - |
| Otu24367 | 1 | k__Fungi(100);p__Ascomycota(100);p__Ascomycota_unclassified(100);p__Ascomycota_unclassified(100);p__Ascomycota_unclassified(100);p__Ascomycota_unclassified(100);p__Ascomycota_unclassified(100); | - | - | - | - | - | - | - | Unassigned | - |

| | | | | | | | | | | |
|---|---|---|---|---|---|---|---|---|---|---|
| Otu24368 | 1 | k__Fungi(100);p__Ascomycota(100);p__Ascomycota_unclassified(100);p__Ascomycota_unclassified(100);p__Ascomycota_unclassified(100);p__Ascomycota_unclassified(100);p__Ascomycota_unclassified(100); | - | - | - | - | - | - | - | Unassigned | - |
| Otu24372 | 1 | k__Fungi(100);p__Ascomycota(100);p__Ascomycota_unclassified(100);p__Ascomycota_unclassified(100);p__Ascomycota_unclassified(100);p__Ascomycota_unclassified(100);p__Ascomycota_unclassified(100); | - | - | - | - | - | - | - | Unassigned | - |
| Otu24375 | 1 | k__Fungi(100);k__Fungi_unclassified(100);k__Fungi_unclassified(100);k__Fungi_unclassified(100);k__Fungi_unclassified(100);k__Fungi_unclassified(100);k__Fungi_unclassified(100); | - | - | - | - | - | - | - | Unassigned | - |
| Otu24376 | 1 | k__Fungi(100);k__Fungi_unclassified(100);k__Fungi_unclassified(100);k__Fungi_unclassified(100);k__Fungi_unclassified(100);k__Fungi_unclassified(100);k__Fungi_unclassified(100); | - | - | - | - | - | - | - | Unassigned | - |
| Otu24377 | 1 | k__Fungi(100);k__Fungi_unclassified(100);k__Fungi_unclassified(100);k__Fungi_unclassified(100);k__Fungi_unclassified(100);k__Fungi_unclassified(100);k__Fungi_unclassified(100); | - | - | - | - | - | - | - | Unassigned | - |
| Otu24379 | 1 | k__Fungi(100);p__Ascomycota(100);p__Ascomycota_unclassified(100);p__Ascomycota_unclassified(100);p__Ascomycota_unclassified(100);p__Ascomycota_unclassified(100);p__Ascomycota_unclassified(100); | - | - | - | - | - | - | - | Unassigned | - |
| Otu24380 | 1 | k__Fungi(100);k__Fungi_unclassified(100);k__Fungi_unclassified(100);k__Fungi_unclassified(100);k__Fungi_unclassified(100);k__Fungi_unclassified(100);k__Fungi_unclassified(100); | - | - | - | - | - | - | - | Unassigned | - |
| Otu24381 | 1 | k__Fungi(100);k__Fungi_unclassified(100);k__Fungi_unclassified(100);k__Fungi_unclassified(100);k__Fungi_unclassified(100);k__Fungi_unclassified(100);k__Fungi_unclassified(100); | - | - | - | - | - | - | - | Unassigned | - |
| Otu24384 | 1 | k__Fungi(100);p__Ascomycota(100);p__Ascomycota_unclassified(100);p__Ascomycota_unclassified(100);p__Ascomycota_unclassified(100);p__Ascomycota_unclassified(100);p__Ascomycota_unclassified(100); | - | - | - | - | - | - | - | Unassigned | - |
| Otu24385 | 1 | k__Fungi(100);p__Ascomycota(100);p__Ascomycota_unclassified(100);p__Ascomycota_unclassified(100);p__Ascomycota_unclassified(100);p__Ascomycota_unclassified(100);p__Ascomycota_unclassified(100); | - | - | - | - | - | - | - | Unassigned | - |
| Otu24392 | 1 | k__Fungi(100);k__Fungi_unclassified(100);k__Fungi_unclassified(100);k__Fungi_unclassified(100);k__Fungi_unclassified(100);k__Fungi_unclassified(100);k__Fungi_unclassified(100); | - | - | - | - | - | - | - | Unassigned | - |
| Otu24393 | 1 | k__Fungi(100);k__Fungi_unclassified(100);k__Fungi_unclassified(100);k__Fungi_unclassified(100);k__Fungi_unclassified(100);k__Fungi_unclassified(100);k__Fungi_unclassified(100); | - | - | - | - | - | - | - | Unassigned | - |
| Otu24394 | 1 | k__Fungi(100);k__Fungi_unclassified(100);k__Fungi_unclassified(100);k__Fungi_unclassified(100);k__Fungi_unclassified(100);k__Fungi_unclassified(100);k__Fungi_unclassified(100); | - | - | - | - | - | - | - | Unassigned | - |
| Otu24395 | 1 | k__Fungi(100);k__Fungi_unclassified(100);k__Fungi_unclassified(100);k__Fungi_unclassified(100);k__Fungi_unclassified(100);k__Fungi_unclassified(100);k__Fungi_unclassified(100); | - | - | - | - | - | - | - | Unassigned | - |
| Otu24396 | 1 | k__Fungi(100);k__Fungi_unclassified(100);k__Fungi_unclassified(100);k__Fungi_unclassified(100);k__Fungi_unclassified(100);k__Fungi_unclassified(100);k__Fungi_unclassified(100); | - | - | - | - | - | - | - | Unassigned | - |
| Otu24397 | 1 | k__Fungi(100);p__Ascomycota(100);p__Ascomycota_unclassified(100);p__Ascomycota_unclassified(100);p__Ascomycota_unclassified(100);p__Ascomycota_unclassified(100);p__Ascomycota_unclassified(100); | - | - | - | - | - | - | - | Unassigned | - |
| Otu24398 | 1 | k__Fungi(100);k__Fungi_unclassified(100);k__Fungi_unclassified(100);k__Fungi_unclassified(100);k__Fungi_unclassified(100);k__Fungi_unclassified(100);k__Fungi_unclassified(100); | - | - | - | - | - | - | - | Unassigned | - |
| Otu24399 | 1 | k__Fungi(100);p__Ascomycota(100);p__Ascomycota_unclassified(100);p__Ascomycota_unclassified(100);p__Ascomycota_unclassified(100);p__Ascomycota_unclassified(100);p__Ascomycota_unclassified(100); | - | - | - | - | - | - | - | Unassigned | - |
| Otu24401 | 1 | k__Fungi(100);k__Fungi_unclassified(100);k__Fungi_unclassified(100);k__Fungi_unclassified(100);k__Fungi_unclassified(100);k__Fungi_unclassified(100);k__Fungi_unclassified(100); | - | - | - | - | - | - | - | Unassigned | - |
| Otu24402 | 1 | k__Fungi(100);k__Fungi_unclassified(100);k__Fungi_unclassified(100);k__Fungi_unclassified(100);k__Fungi_unclassified(100);k__Fungi_unclassified(100);k__Fungi_unclassified(100); | - | - | - | - | - | - | - | Unassigned | - |
| Otu24406 | 1 | k__Fungi(100);k__Fungi_unclassified(100);k__Fungi_unclassified(100);k__Fungi_unclassified(100);k__Fungi_unclassified(100);k__Fungi_unclassified(100);k__Fungi_unclassified(100); | - | - | - | - | - | - | - | Unassigned | - |
| Otu24407 | 1 | k__Fungi(100);k__Fungi_unclassified(100);k__Fungi_unclassified(100);k__Fungi_unclassified(100);k__Fungi_unclassified(100);k__Fungi_unclassified(100);k__Fungi_unclassified(100); | - | - | - | - | - | - | - | Unassigned | - |
| Otu24408 | 1 | k__Fungi(100);p__Ascomycota(100);p__Ascomycota_unclassified(100);p__Ascomycota_unclassified(100);p__Ascomycota_unclassified(100);p__Ascomycota_unclassified(100);p__Ascomycota_unclassified(100); | - | - | - | - | - | - | - | Unassigned | - |
| Otu24409 | 1 | k__Fungi(100);k__Fungi_unclassified(100);k__Fungi_unclassified(100);k__Fungi_unclassified(100);k__Fungi_unclassified(100);k__Fungi_unclassified(100);k__Fungi_unclassified(100); | - | - | - | - | - | - | - | Unassigned | - |
| Otu24410 | 1 | k__Fungi(100);k__Fungi_unclassified(100);k__Fungi_unclassified(100);k__Fungi_unclassified(100);k__Fungi_unclassified(100);k__Fungi_unclassified(100);k__Fungi_unclassified(100); | - | - | - | - | - | - | - | Unassigned | - |
| Otu24411 | 1 | k__Fungi(100);k__Fungi_unclassified(100);k__Fungi_unclassified(100);k__Fungi_unclassified(100);k__Fungi_unclassified(100);k__Fungi_unclassified(100);k__Fungi_unclassified(100); | - | - | - | - | - | - | - | Unassigned | - |
| Otu24413 | 1 | k__Fungi(100);p__Ascomycota(100);p__Ascomycota_unclassified(100);p__Ascomycota_unclassified(100);p__Ascomycota_unclassified(100);p__Ascomycota_unclassified(100);p__Ascomycota_unclassified(100); | - | - | - | - | - | - | - | Unassigned | - |
| Otu24418 | 1 | k__Fungi(100);k__Fungi_unclassified(100);k__Fungi_unclassified(100);k__Fungi_unclassified(100);k__Fungi_unclassified(100);k__Fungi_unclassified(100);k__Fungi_unclassified(100); | - | - | - | - | - | - | - | Unassigned | - |
| Otu24419 | 1 | k__Fungi(100);k__Fungi_unclassified(100);k__Fungi_unclassified(100);k__Fungi_unclassified(100);k__Fungi_unclassified(100);k__Fungi_unclassified(100);k__Fungi_unclassified(100); | - | - | - | - | - | - | - | Unassigned | - |
| Otu24425 | 1 | k__Fungi(100);p__Ascomycota(100);p__Ascomycota_unclassified(100);p__Ascomycota_unclassified(100);p__Ascomycota_unclassified(100);p__Ascomycota_unclassified(100);p__Ascomycota_unclassified(100); | - | - | - | - | - | - | - | Unassigned | - |
| Otu24426 | 1 | k__Fungi(100);k__Fungi_unclassified(100);k__Fungi_unclassified(100);k__Fungi_unclassified(100);k__Fungi_unclassified(100);k__Fungi_unclassified(100);k__Fungi_unclassified(100); | - | - | - | - | - | - | - | Unassigned | - |
| Otu24427 | 1 | k__Fungi(100);k__Fungi_unclassified(100);k__Fungi_unclassified(100);k__Fungi_unclassified(100);k__Fungi_unclassified(100);k__Fungi_unclassified(100);k__Fungi_unclassified(100); | - | - | - | - | - | - | - | Unassigned | - |
| Otu24428 | 1 | k__Fungi(100);k__Fungi_unclassified(100);k__Fungi_unclassified(100);k__Fungi_unclassified(100);k__Fungi_unclassified(100);k__Fungi_unclassified(100);k__Fungi_unclassified(100); | - | - | - | - | - | - | - | Unassigned | - |
| Otu24429 | 1 | k__Fungi(100);k__Fungi_unclassified(100);k__Fungi_unclassified(100);k__Fungi_unclassified(100);k__Fungi_unclassified(100);k__Fungi_unclassified(100);k__Fungi_unclassified(100); | - | - | - | - | - | - | - | Unassigned | - |
| Otu24430 | 1 | k__Fungi(100);k__Fungi_unclassified(100);k__Fungi_unclassified(100);k__Fungi_unclassified(100);k__Fungi_unclassified(100);k__Fungi_unclassified(100);k__Fungi_unclassified(100); | - | - | - | - | - | - | - | Unassigned | - |
| Otu24431 | 1 | k__Fungi(100);p__Ascomycota(100);p__Ascomycota_unclassified(100);p__Ascomycota_unclassified(100);p__Ascomycota_unclassified(100);p__Ascomycota_unclassified(100);p__Ascomycota_unclassified(100); | - | - | - | - | - | - | - | Unassigned | - |
| Otu24432 | 1 | k__Fungi(100);k__Fungi_unclassified(100);k__Fungi_unclassified(100);k__Fungi_unclassified(100);k__Fungi_unclassified(100);k__Fungi_unclassified(100);k__Fungi_unclassified(100); | - | - | - | - | - | - | - | Unassigned | - |
| Otu24433 | 1 | k__Fungi(100);p__Ascomycota(100);p__Ascomycota_unclassified(100);p__Ascomycota_unclassified(100);p__Ascomycota_unclassified(100);p__Ascomycota_unclassified(100);p__Ascomycota_unclassified(100); | - | - | - | - | - | - | - | Unassigned | - |
| Otu24434 | 1 | k__Fungi(100);k__Fungi_unclassified(100);k__Fungi_unclassified(100);k__Fungi_unclassified(100);k__Fungi_unclassified(100);k__Fungi_unclassified(100);k__Fungi_unclassified(100); | - | - | - | - | - | - | - | Unassigned | - |
| Otu24435 | 1 | k__Fungi(100);p__Ascomycota(100);p__Ascomycota_unclassified(100);p__Ascomycota_unclassified(100);p__Ascomycota_unclassified(100);p__Ascomycota_unclassified(100);p__Ascomycota_unclassified(100); | - | - | - | - | - | - | - | Unassigned | - |
| Otu24436 | 1 | k__Fungi(100);p__Ascomycota(100);p__Ascomycota_unclassified(100);p__Ascomycota_unclassified(100);p__Ascomycota_unclassified(100);p__Ascomycota_unclassified(100);p__Ascomycota_unclassified(100); | - | - | - | - | - | - | - | Unassigned | - |
| Otu24437 | 1 | k__Fungi(100);k__Fungi_unclassified(100);k__Fungi_unclassified(100);k__Fungi_unclassified(100);k__Fungi_unclassified(100);k__Fungi_unclassified(100);k__Fungi_unclassified(100); | - | - | - | - | - | - | - | Unassigned | - |
| Otu24438 | 1 | k__Fungi(100);p__Ascomycota(100);p__Ascomycota_unclassified(100);p__Ascomycota_unclassified(100);p__Ascomycota_unclassified(100);p__Ascomycota_unclassified(100);p__Ascomycota_unclassified(100); | - | - | - | - | - | - | - | Unassigned | - |
| Otu24440 | 1 | k__Fungi(100);k__Fungi_unclassified(100);k__Fungi_unclassified(100);k__Fungi_unclassified(100);k__Fungi_unclassified(100);k__Fungi_unclassified(100);k__Fungi_unclassified(100); | - | - | - | - | - | - | - | Unassigned | - |
| Otu24441 | 1 | k__Fungi(100);k__Fungi_unclassified(100);k__Fungi_unclassified(100);k__Fungi_unclassified(100);k__Fungi_unclassified(100);k__Fungi_unclassified(100);k__Fungi_unclassified(100); | - | - | - | - | - | - | - | Unassigned | - |
| Otu24442 | 1 | k__Fungi(100);k__Fungi_unclassified(100);k__Fungi_unclassified(100);k__Fungi_unclassified(100);k__Fungi_unclassified(100);k__Fungi_unclassified(100);k__Fungi_unclassified(100); | - | - | - | - | - | - | - | Unassigned | - |
| Otu24443 | 1 | k__Fungi(100);p__Ascomycota(100);p__Ascomycota_unclassified(100);p__Ascomycota_unclassified(100);p__Ascomycota_unclassified(100);p__Ascomycota_unclassified(100);p__Ascomycota_unclassified(100); | - | - | - | - | - | - | - | Unassigned | - |
| Otu24444 | 1 | k__Fungi(100);p__Ascomycota(100);p__Ascomycota_unclassified(100);p__Ascomycota_unclassified(100);p__Ascomycota_unclassified(100);p__Ascomycota_unclassified(100);p__Ascomycota_unclassified(100); | - | - | - | - | - | - | - | Unassigned | - |
| Otu24445 | 1 | k__Fungi(100);k__Fungi_unclassified(100);k__Fungi_unclassified(100);k__Fungi_unclassified(100);k__Fungi_unclassified(100);k__Fungi_unclassified(100);k__Fungi_unclassified(100); | - | - | - | - | - | - | - | Unassigned | - |
| Otu24446 | 1 | k__Fungi(100);k__Fungi_unclassified(100);k__Fungi_unclassified(100);k__Fungi_unclassified(100);k__Fungi_unclassified(100);k__Fungi_unclassified(100);k__Fungi_unclassified(100); | - | - | - | - | - | - | - | Unassigned | - |
| Otu24447 | 1 | k__Fungi(100);k__Fungi_unclassified(100);k__Fungi_unclassified(100);k__Fungi_unclassified(100);k__Fungi_unclassified(100);k__Fungi_unclassified(100);k__Fungi_unclassified(100); | - | - | - | - | - | - | - | Unassigned | - |
| Otu24448 | 1 | k__Fungi(100);k__Fungi_unclassified(100);k__Fungi_unclassified(100);k__Fungi_unclassified(100);k__Fungi_unclassified(100);k__Fungi_unclassified(100);k__Fungi_unclassified(100); | - | - | - | - | - | - | - | Unassigned | - |
| Otu24449 | 1 | k__Fungi(100);p__Ascomycota(100);p__Ascomycota_unclassified(100);p__Ascomycota_unclassified(100);p__Ascomycota_unclassified(100);p__Ascomycota_unclassified(100);p__Ascomycota_unclassified(100); | - | - | - | - | - | - | - | Unassigned | - |
| Otu24450 | 1 | k__Fungi(100);k__Fungi_unclassified(100);k__Fungi_unclassified(100);k__Fungi_unclassified(100);k__Fungi_unclassified(100);k__Fungi_unclassified(100);k__Fungi_unclassified(100); | - | - | - | - | - | - | - | Unassigned | - |
| Otu24451 | 1 | k__Fungi(100);k__Fungi_unclassified(100);k__Fungi_unclassified(100);k__Fungi_unclassified(100);k__Fungi_unclassified(100);k__Fungi_unclassified(100);k__Fungi_unclassified(100); | - | - | - | - | - | - | - | Unassigned | - |
| Otu24452 | 1 | k__Fungi(100);k__Fungi_unclassified(100);k__Fungi_unclassified(100);k__Fungi_unclassified(100);k__Fungi_unclassified(100);k__Fungi_unclassified(100);k__Fungi_unclassified(100); | - | - | - | - | - | - | - | Unassigned | - |
| Otu24453 | 1 | k__Fungi(100);k__Fungi_unclassified(100);k__Fungi_unclassified(100);k__Fungi_unclassified(100);k__Fungi_unclassified(100);k__Fungi_unclassified(100);k__Fungi_unclassified(100); | - | - | - | - | - | - | - | Unassigned | - |
| Otu24454 | 1 | k__Fungi(100);k__Fungi_unclassified(100);k__Fungi_unclassified(100);k__Fungi_unclassified(100);k__Fungi_unclassified(100);k__Fungi_unclassified(100);k__Fungi_unclassified(100); | - | - | - | - | - | - | - | Unassigned | - |
| Otu24455 | 1 | k__Fungi(100);k__Fungi_unclassified(100);k__Fungi_unclassified(100);k__Fungi_unclassified(100);k__Fungi_unclassified(100);k__Fungi_unclassified(100);k__Fungi_unclassified(100); | - | - | - | - | - | - | - | Unassigned | - |
| Otu24456 | 1 | k__Fungi(100);p__Ascomycota(100);p__Ascomycota_unclassified(100);p__Ascomycota_unclassified(100);p__Ascomycota_unclassified(100);p__Ascomycota_unclassified(100);p__Ascomycota_unclassified(100); | - | - | - | - | - | - | - | Unassigned | - |
| Otu24457 | 1 | k__Fungi(100);p__Ascomycota(100);p__Ascomycota_unclassified(100);p__Ascomycota_unclassified(100);p__Ascomycota_unclassified(100);p__Ascomycota_unclassified(100);p__Ascomycota_unclassified(100); | - | - | - | - | - | - | - | Unassigned | - |
| Otu24458 | 1 | k__Fungi(100);p__Ascomycota(100);p__Ascomycota_unclassified(100);p__Ascomycota_unclassified(100);p__Ascomycota_unclassified(100);p__Ascomycota_unclassified(100);p__Ascomycota_unclassified(100); | - | - | - | - | - | - | - | Unassigned | - |
| Otu24459 | 1 | k__Fungi(100);k__Fungi_unclassified(100);k__Fungi_unclassified(100);k__Fungi_unclassified(100);k__Fungi_unclassified(100);k__Fungi_unclassified(100);k__Fungi_unclassified(100); | - | - | - | - | - | - | - | Unassigned | - |
| Otu24460 | 1 | k__Fungi(100);k__Fungi_unclassified(100);k__Fungi_unclassified(100);k__Fungi_unclassified(100);k__Fungi_unclassified(100);k__Fungi_unclassified(100);k__Fungi_unclassified(100); | - | - | - | - | - | - | - | Unassigned | - |
| Otu24461 | 1 | k__Fungi(100);k__Fungi_unclassified(100);k__Fungi_unclassified(100);k__Fungi_unclassified(100);k__Fungi_unclassified(100);k__Fungi_unclassified(100);k__Fungi_unclassified(100); | - | - | - | - | - | - | - | Unassigned | - |
| Otu24462 | 1 | k__Fungi(100);k__Fungi_unclassified(100);k__Fungi_unclassified(100);k__Fungi_unclassified(100);k__Fungi_unclassified(100);k__Fungi_unclassified(100);k__Fungi_unclassified(100); | - | - | - | - | - | - | - | Unassigned | - |
| Otu24463 | 1 | k__Fungi(100);k__Fungi_unclassified(100);k__Fungi_unclassified(100);k__Fungi_unclassified(100);k__Fungi_unclassified(100);k__Fungi_unclassified(100);k__Fungi_unclassified(100); | - | - | - | - | - | - | - | Unassigned | - |
| Otu24464 | 1 | k__Fungi(100);k__Fungi_unclassified(100);k__Fungi_unclassified(100);k__Fungi_unclassified(100);k__Fungi_unclassified(100);k__Fungi_unclassified(100);k__Fungi_unclassified(100); | - | - | - | - | - | - | - | Unassigned | - |
| Otu24465 | 1 | k__Fungi(100);k__Fungi_unclassified(100);k__Fungi_unclassified(100);k__Fungi_unclassified(100);k__Fungi_unclassified(100);k__Fungi_unclassified(100);k__Fungi_unclassified(100); | - | - | - | - | - | - | - | Unassigned | - |
| Otu24466 | 1 | k__Fungi(100);k__Fungi_unclassified(100);k__Fungi_unclassified(100);k__Fungi_unclassified(100);k__Fungi_unclassified(100);k__Fungi_unclassified(100);k__Fungi_unclassified(100); | - | - | - | - | - | - | - | Unassigned | - |
| Otu24467 | 1 | k__Fungi(100);k__Fungi_unclassified(100);k__Fungi_unclassified(100);k__Fungi_unclassified(100);k__Fungi_unclassified(100);k__Fungi_unclassified(100);k__Fungi_unclassified(100); | - | - | - | - | - | - | - | Unassigned | - |
| Otu24468 | 1 | k__Fungi(100);p__Ascomycota(100);p__Ascomycota_unclassified(100);p__Ascomycota_unclassified(100);p__Ascomycota_unclassified(100);p__Ascomycota_unclassified(100);p__Ascomycota_unclassified(100); | - | - | - | - | - | - | - | Unassigned | - |
| Otu24469 | 1 | k__Fungi(100);k__Fungi_unclassified(100);k__Fungi_unclassified(100);k__Fungi_unclassified(100);k__Fungi_unclassified(100);k__Fungi_unclassified(100);k__Fungi_unclassified(100); | - | - | - | - | - | - | - | Unassigned | - |
| Otu24470 | 1 | k__Fungi(100);k__Fungi_unclassified(100);k__Fungi_unclassified(100);k__Fungi_unclassified(100);k__Fungi_unclassified(100);k__Fungi_unclassified(100);k__Fungi_unclassified(100); | - | - | - | - | - | - | - | Unassigned | - |
| Otu24471 | 1 | k__Fungi(100);k__Fungi_unclassified(100);k__Fungi_unclassified(100);k__Fungi_unclassified(100);k__Fungi_unclassified(100);k__Fungi_unclassified(100);k__Fungi_unclassified(100); | - | - | - | - | - | - | - | Unassigned | - |
| Otu24472 | 1 | k__Fungi(100);p__Ascomycota(100);p__Ascomycota_unclassified(100);p__Ascomycota_unclassified(100);p__Ascomycota_unclassified(100);p__Ascomycota_unclassified(100);p__Ascomycota_unclassified(100); | - | - | - | - | - | - | - | Unassigned | - |
| Otu24475 | 1 | k__Fungi(100);k__Fungi_unclassified(100);k__Fungi_unclassified(100);k__Fungi_unclassified(100);k__Fungi_unclassified(100);k__Fungi_unclassified(100);k__Fungi_unclassified(100); | - | - | - | - | - | - | - | Unassigned | - |
| Otu24476 | 1 | k__Fungi(100);k__Fungi_unclassified(100);k__Fungi_unclassified(100);k__Fungi_unclassified(100);k__Fungi_unclassified(100);k__Fungi_unclassified(100);k__Fungi_unclassified(100); | - | - | - | - | - | - | - | Unassigned | - |
| Otu24478 | 1 | k__Fungi(100);k__Fungi_unclassified(100);k__Fungi_unclassified(100);k__Fungi_unclassified(100);k__Fungi_unclassified(100);k__Fungi_unclassified(100);k__Fungi_unclassified(100); | - | - | - | - | - | - | - | Unassigned | - |
| Otu24481 | 1 | k__Fungi(100);k__Fungi_unclassified(100);k__Fungi_unclassified(100);k__Fungi_unclassified(100);k__Fungi_unclassified(100);k__Fungi_unclassified(100);k__Fungi_unclassified(100); | - | - | - | - | - | - | - | Unassigned | - |
| Otu24482 | 1 | k__Fungi(100);k__Fungi_unclassified(100);k__Fungi_unclassified(100);k__Fungi_unclassified(100);k__Fungi_unclassified(100);k__Fungi_unclassified(100);k__Fungi_unclassified(100); | - | - | - | - | - | - | - | Unassigned | - |
| Otu24490 | 1 | k__Fungi(100);k__Fungi_unclassified(100);k__Fungi_unclassified(100);k__Fungi_unclassified(100);k__Fungi_unclassified(100);k__Fungi_unclassified(100);k__Fungi_unclassified(100); | - | - | - | - | - | - | - | Unassigned | - |
| Otu24493 | 1 | k__Fungi(100);p__Ascomycota(100);p__Ascomycota_unclassified(100);p__Ascomycota_unclassified(100);p__Ascomycota_unclassified(100);p__Ascomycota_unclassified(100);p__Ascomycota_unclassified(100); | - | - | - | - | - | - | - | Unassigned | - |
| Otu24494 | 1 | k__Fungi(100);k__Fungi_unclassified(100);k__Fungi_unclassified(100);k__Fungi_unclassified(100);k__Fungi_unclassified(100);k__Fungi_unclassified(100);k__Fungi_unclassified(100); | - | - | - | - | - | - | - | Unassigned | - |
| Otu24497 | 1 | k__Fungi(100);k__Fungi_unclassified(100);k__Fungi_unclassified(100);k__Fungi_unclassified(100);k__Fungi_unclassified(100);k__Fungi_unclassified(100);k__Fungi_unclassified(100); | - | - | - | - | - | - | - | Unassigned | - |
| Otu24499 | 1 | k__Fungi(100);k__Fungi_unclassified(100);k__Fungi_unclassified(100);k__Fungi_unclassified(100);k__Fungi_unclassified(100);k__Fungi_unclassified(100);k__Fungi_unclassified(100); | - | - | - | - | - | - | - | Unassigned | - |
| Otu24500 | 1 | k__Fungi(100);k__Fungi_unclassified(100);k__Fungi_unclassified(100);k__Fungi_unclassified(100);k__Fungi_unclassified(100);k__Fungi_unclassified(100);k__Fungi_unclassified(100); | - | - | - | - | - | - | - | Unassigned | - |
| Otu24501 | 1 | k__Fungi(100);k__Fungi_unclassified(100);k__Fungi_unclassified(100);k__Fungi_unclassified(100);k__Fungi_unclassified(100);k__Fungi_unclassified(100);k__Fungi_unclassified(100); | - | - | - | - | - | - | - | Unassigned | - |
| Otu24502 | 1 | k__Fungi(100);k__Fungi_unclassified(100);k__Fungi_unclassified(100);k__Fungi_unclassified(100);k__Fungi_unclassified(100);k__Fungi_unclassified(100);k__Fungi_unclassified(100); | - | - | - | - | - | - | - | Unassigned | - |
| Otu24503 | 1 | k__Fungi(100);p__Ascomycota(100);p__Ascomycota_unclassified(100);p__Ascomycota_unclassified(100);p__Ascomycota_unclassified(100);p__Ascomycota_unclassified(100);p__Ascomycota_unclassified(100); | - | - | - | - | - | - | - | Unassigned | - |
| Otu24504 | 1 | k__Fungi(100);k__Fungi_unclassified(100);k__Fungi_unclassified(100);k__Fungi_unclassified(100);k__Fungi_unclassified(100);k__Fungi_unclassified(100);k__Fungi_unclassified(100); | - | - | - | - | - | - | - | Unassigned | - |
| Otu24505 | 1 | k__Fungi(100);p__Ascomycota(100);p__Ascomycota_unclassified(100);p__Ascomycota_unclassified(100);p__Ascomycota_unclassified(100);p__Ascomycota_unclassified(100);p__Ascomycota_unclassified(100); | - | - | - | - | - | - | - | Unassigned | - |
| Otu24506 | 1 | k__Fungi(100);k__Fungi_unclassified(100);k__Fungi_unclassified(100);k__Fungi_unclassified(100);k__Fungi_unclassified(100);k__Fungi_unclassified(100);k__Fungi_unclassified(100); | - | - | - | - | - | - | - | Unassigned | - |
| Otu24507 | 1 | k__Fungi(100);p__Ascomycota(100);p__Ascomycota_unclassified(100);p__Ascomycota_unclassified(100);p__Ascomycota_unclassified(100);p__Ascomycota_unclassified(100);p__Ascomycota_unclassified(100); | - | - | - | - | - | - | - | Unassigned | - |
| Otu24508 | 1 | k__Fungi(100);p__Ascomycota(100);p__Ascomycota_unclassified(100);p__Ascomycota_unclassified(100);p__Ascomycota_unclassified(100);p__Ascomycota_unclassified(100);p__Ascomycota_unclassified(100); | - | - | - | - | - | - | - | Unassigned | - |
| Otu24509 | 1 | k__Fungi(100);k__Fungi_unclassified(100);k__Fungi_unclassified(100);k__Fungi_unclassified(100);k__Fungi_unclassified(100);k__Fungi_unclassified(100);k__Fungi_unclassified(100); | - | - | - | - | - | - | - | Unassigned | - |
| Otu24510 | 1 | k__Fungi(100);p__Ascomycota(100);p__Ascomycota_unclassified(100);p__Ascomycota_unclassified(100);p__Ascomycota_unclassified(100);p__Ascomycota_unclassified(100);p__Ascomycota_unclassified(100); | - | - | - | - | - | - | - | Unassigned | - |
| Otu24511 | 1 | k__Fungi(100);k__Fungi_unclassified(100);k__Fungi_unclassified(100);k__Fungi_unclassified(100);k__Fungi_unclassified(100);k__Fungi_unclassified(100);k__Fungi_unclassified(100); | - | - | - | - | - | - | - | Unassigned | - |
| Otu24512 | 1 | k__Fungi(100);p__Ascomycota(100);p__Ascomycota_unclassified(100);p__Ascomycota_unclassified(100);p__Ascomycota_unclassified(100);p__Ascomycota_unclassified(100);p__Ascomycota_unclassified(100); | - | - | - | - | - | - | - | Unassigned | - |
| Otu24513 | 1 | k__Fungi(100);p__Ascomycota(100);p__Ascomycota_unclassified(100);p__Ascomycota_unclassified(100);p__Ascomycota_unclassified(100);p__Ascomycota_unclassified(100);p__Ascomycota_unclassified(100); | - | - | - | - | - | - | - | Unassigned | - |
| Otu24514 | 1 | k__Fungi(100);k__Fungi_unclassified(100);k__Fungi_unclassified(100);k__Fungi_unclassified(100);k__Fungi_unclassified(100);k__Fungi_unclassified(100);k__Fungi_unclassified(100); | - | - | - | - | - | - | - | Unassigned | - |
| Otu24515 | 1 | k__Fungi(100);k__Fungi_unclassified(100);k__Fungi_unclassified(100);k__Fungi_unclassified(100);k__Fungi_unclassified(100);k__Fungi_unclassified(100);k__Fungi_unclassified(100); | - | - | - | - | - | - | - | Unassigned | - |
| Otu24516 | 1 | k__Fungi(100);k__Fungi_unclassified(100);k__Fungi_unclassified(100);k__Fungi_unclassified(100);k__Fungi_unclassified(100);k__Fungi_unclassified(100);k__Fungi_unclassified(100); | - | - | - | - | - | - | - | Unassigned | - |
| Otu24517 | 1 | k__Fungi(100);p__Ascomycota(100);p__Ascomycota_unclassified(100);p__Ascomycota_unclassified(100);p__Ascomycota_unclassified(100);p__Ascomycota_unclassified(100);p__Ascomycota_unclassified(100); | - | - | - | - | - | - | - | Unassigned | - |
| Otu24518 | 1 | k__Fungi(100);k__Fungi_unclassified(100);k__Fungi_unclassified(100);k__Fungi_unclassified(100);k__Fungi_unclassified(100);k__Fungi_unclassified(100);k__Fungi_unclassified(100); | - | - | - | - | - | - | - | Unassigned | - |
| Otu24519 | 1 | k__Fungi(100);k__Fungi_unclassified(100);k__Fungi_unclassified(100);k__Fungi_unclassified(100);k__Fungi_unclassified(100);k__Fungi_unclassified(100);k__Fungi_unclassified(100); | - | - | - | - | - | - | - | Unassigned | - |
| Otu24520 | 1 | k__Fungi(100);k__Fungi_unclassified(100);k__Fungi_unclassified(100);k__Fungi_unclassified(100);k__Fungi_unclassified(100);k__Fungi_unclassified(100);k__Fungi_unclassified(100); | - | - | - | - | - | - | - | Unassigned | - |
| Otu24521 | 1 | k__Fungi(100);k__Fungi_unclassified(100);k__Fungi_unclassified(100);k__Fungi_unclassified(100);k__Fungi_unclassified(100);k__Fungi_unclassified(100);k__Fungi_unclassified(100); | - | - | - | - | - | - | - | Unassigned | - |
| Otu24522 | 1 | k__Fungi(100);k__Fungi_unclassified(100);k__Fungi_unclassified(100);k__Fungi_unclassified(100);k__Fungi_unclassified(100);k__Fungi_unclassified(100);k__Fungi_unclassified(100); | - | - | - | - | - | - | - | Unassigned | - |
| Otu24523 | 1 | k__Fungi(100);p__Ascomycota(100);p__Ascomycota_unclassified(100);p__Ascomycota_unclassified(100);p__Ascomycota_unclassified(100);p__Ascomycota_unclassified(100);p__Ascomycota_unclassified(100); | - | - | - | - | - | - | - | Unassigned | - |
| Otu24524 | 1 | k__Fungi(100);p__Ascomycota(100);p__Ascomycota_unclassified(100);p__Ascomycota_unclassified(100);p__Ascomycota_unclassified(100);p__Ascomycota_unclassified(100);p__Ascomycota_unclassified(100); | - | - | - | - | - | - | - | Unassigned | - |
| Otu24525 | 1 | k__Fungi(100);k__Fungi_unclassified(100);k__Fungi_unclassified(100);k__Fungi_unclassified(100);k__Fungi_unclassified(100);k__Fungi_unclassified(100);k__Fungi_unclassified(100); | - | - | - | - | - | - | - | Unassigned | - |
| Otu24526 | 1 | k__Fungi(100);k__Fungi_unclassified(100);k__Fungi_unclassified(100);k__Fungi_unclassified(100);k__Fungi_unclassified(100);k__Fungi_unclassified(100);k__Fungi_unclassified(100); | - | - | - | - | - | - | - | Unassigned | - |
| Otu24527 | 1 | k__Fungi(100);k__Fungi_unclassified(100);k__Fungi_unclassified(100);k__Fungi_unclassified(100);k__Fungi_unclassified(100);k__Fungi_unclassified(100);k__Fungi_unclassified(100); | - | - | - | - | - | - | - | Unassigned | - |
| Otu24528 | 1 | k__Fungi(100);p__Ascomycota(100);p__Ascomycota_unclassified(100);p__Ascomycota_unclassified(100);p__Ascomycota_unclassified(100);p__Ascomycota_unclassified(100);p__Ascomycota_unclassified(100); | - | - | - | - | - | - | - | Unassigned | - |
| Otu24529 | 1 | k__Fungi(100);p__Ascomycota(100);p__Ascomycota_unclassified(100);p__Ascomycota_unclassified(100);p__Ascomycota_unclassified(100);p__Ascomycota_unclassified(100);p__Ascomycota_unclassified(100); | - | - | - | - | - | - | - | Unassigned | - |
| Otu24530 | 1 | k__Fungi(100);k__Fungi_unclassified(100);k__Fungi_unclassified(100);k__Fungi_unclassified(100);k__Fungi_unclassified(100);k__Fungi_unclassified(100);k__Fungi_unclassified(100); | - | - | - | - | - | - | - | Unassigned | - |
| Otu24532 | 1 | k__Fungi(100);k__Fungi_unclassified(100);k__Fungi_unclassified(100);k__Fungi_unclassified(100);k__Fungi_unclassified(100);k__Fungi_unclassified(100);k__Fungi_unclassified(100); | - | - | - | - | - | - | - | Unassigned | - |
| Otu24533 | 1 | k__Fungi(100);k__Fungi_unclassified(100);k__Fungi_unclassified(100);k__Fungi_unclassified(100);k__Fungi_unclassified(100);k__Fungi_unclassified(100);k__Fungi_unclassified(100); | - | - | - | - | - | - | - | Unassigned | - |
| Otu24534 | 1 | k__Fungi(100);k__Fungi_unclassified(100);k__Fungi_unclassified(100);k__Fungi_unclassified(100);k__Fungi_unclassified(100);k__Fungi_unclassified(100);k__Fungi_unclassified(100); | - | - | - | - | - | - | - | Unassigned | - |
| Otu24535 | 1 | k__Fungi(100);k__Fungi_unclassified(100);k__Fungi_unclassified(100);k__Fungi_unclassified(100);k__Fungi_unclassified(100);k__Fungi_unclassified(100);k__Fungi_unclassified(100); | - | - | - | - | - | - | - | Unassigned | - |
| Otu24536 | 1 | k__Fungi(100);p__Ascomycota(100);p__Ascomycota_unclassified(100);p__Ascomycota_unclassified(100);p__Ascomycota_unclassified(100);p__Ascomycota_unclassified(100);p__Ascomycota_unclassified(100); | - | - | - | - | - | - | - | Unassigned | - |
| Otu24537 | 1 | k__Fungi(100);p__Ascomycota(100);p__Ascomycota_unclassified(100);p__Ascomycota_unclassified(100);p__Ascomycota_unclassified(100);p__Ascomycota_unclassified(100);p__Ascomycota_unclassified(100); | - | - | - | - | - | - | - | Unassigned | - |
| Otu24538 | 1 | k__Fungi(100);k__Fungi_unclassified(100);k__Fungi_unclassified(100);k__Fungi_unclassified(100);k__Fungi_unclassified(100);k__Fungi_unclassified(100);k__Fungi_unclassified(100); | - | - | - | - | - | - | - | Unassigned | - |
| Otu24539 | 1 | k__Fungi(100);k__Fungi_unclassified(100);k__Fungi_unclassified(100);k__Fungi_unclassified(100);k__Fungi_unclassified(100);k__Fungi_unclassified(100);k__Fungi_unclassified(100); | - | - | - | - | - | - | - | Unassigned | - |
| Otu24540 | 1 | k__Fungi(100);k__Fungi_unclassified(100);k__Fungi_unclassified(100);k__Fungi_unclassified(100);k__Fungi_unclassified(100);k__Fungi_unclassified(100);k__Fungi_unclassified(100); | - | - | - | - | - | - | - | Unassigned | - |
| Otu24541 | 1 | k__Fungi(100);p__Ascomycota(100);p__Ascomycota_unclassified(100);p__Ascomycota_unclassified(100);p__Ascomycota_unclassified(100);p__Ascomycota_unclassified(100);p__Ascomycota_unclassified(100); | - | - | - | - | - | - | - | Unassigned | - |
| Otu24542 | 1 | k__Fungi(100);p__Ascomycota(100);p__Ascomycota_unclassified(100);p__Ascomycota_unclassified(100);p__Ascomycota_unclassified(100);p__Ascomycota_unclassified(100);p__Ascomycota_unclassified(100); | - | - | - | - | - | - | - | Unassigned | - |
| Otu24543 | 1 | k__Fungi(100);k__Fungi_unclassified(100);k__Fungi_unclassified(100);k__Fungi_unclassified(100);k__Fungi_unclassified(100);k__Fungi_unclassified(100);k__Fungi_unclassified(100); | - | - | - | - | - | - | - | Unassigned | - |
| Otu24544 | 1 | k__Fungi(100);k__Fungi_unclassified(100);k__Fungi_unclassified(100);k__Fungi_unclassified(100);k__Fungi_unclassified(100);k__Fungi_unclassified(100);k__Fungi_unclassified(100); | - | - | - | - | - | - | - | Unassigned | - |
| Otu24545 | 1 | k__Fungi(100);k__Fungi_unclassified(100);k__Fungi_unclassified(100);k__Fungi_unclassified(100);k__Fungi_unclassified(100);k__Fungi_unclassified(100);k__Fungi_unclassified(100); | - | - | - | - | - | - | - | Unassigned | - |
| Otu24546 | 1 | k__Fungi(100);p__Ascomycota(100);p__Ascomycota_unclassified(100);p__Ascomycota_unclassified(100);p__Ascomycota_unclassified(100);p__Ascomycota_unclassified(100);p__Ascomycota_unclassified(100); | - | - | - | - | - | - | - | Unassigned | - |
| Otu24547 | 1 | k__Fungi(100);k__Fungi_unclassified(100);k__Fungi_unclassified(100);k__Fungi_unclassified(100);k__Fungi_unclassified(100);k__Fungi_unclassified(100);k__Fungi_unclassified(100); | - | - | - | - | - | - | - | Unassigned | - |
| Otu24548 | 1 | k__Fungi(100);k__Fungi_unclassified(100);k__Fungi_unclassified(100);k__Fungi_unclassified(100);k__Fungi_unclassified(100);k__Fungi_unclassified(100);k__Fungi_unclassified(100); | - | - | - | - | - | - | - | Unassigned | - |
| Otu24549 | 1 | k__Fungi(100);k__Fungi_unclassified(100);k__Fungi_unclassified(100);k__Fungi_unclassified(100);k__Fungi_unclassified(100);k__Fungi_unclassified(100);k__Fungi_unclassified(100); | - | - | - | - | - | - | - | Unassigned | - |
| Otu24551 | 1 | k__Fungi(100);k__Fungi_unclassified(100);k__Fungi_unclassified(100);k__Fungi_unclassified(100);k__Fungi_unclassified(100);k__Fungi_unclassified(100);k__Fungi_unclassified(100); | - | - | - | - | - | - | - | Unassigned | - |
| Otu24552 | 1 | k__Fungi(100);k__Fungi_unclassified(100);k__Fungi_unclassified(100);k__Fungi_unclassified(100);k__Fungi_unclassified(100);k__Fungi_unclassified(100);k__Fungi_unclassified(100); | - | - | - | - | - | - | - | Unassigned | - |
| Otu24553 | 1 | k__Fungi(100);k__Fungi_unclassified(100);k__Fungi_unclassified(100);k__Fungi_unclassified(100);k__Fungi_unclassified(100);k__Fungi_unclassified(100);k__Fungi_unclassified(100); | - | - | - | - | - | - | - | Unassigned | - |
| Otu24554 | 1 | k__Fungi(100);k__Fungi_unclassified(100);k__Fungi_unclassified(100);k__Fungi_unclassified(100);k__Fungi_unclassified(100);k__Fungi_unclassified(100);k__Fungi_unclassified(100); | - | - | - | - | - | - | - | Unassigned | - |
| Otu24555 | 1 | k__Fungi(100);k__Fungi_unclassified(100);k__Fungi_unclassified(100);k__Fungi_unclassified(100);k__Fungi_unclassified(100);k__Fungi_unclassified(100);k__Fungi_unclassified(100); | - | - | - | - | - | - | - | Unassigned | - |
| Otu24556 | 1 | k__Fungi(100);p__Ascomycota(100);p__Ascomycota_unclassified(100);p__Ascomycota_unclassified(100);p__Ascomycota_unclassified(100);p__Ascomycota_unclassified(100);p__Ascomycota_unclassified(100); | - | - | - | - | - | - | - | Unassigned | - |
| Otu24557 | 1 | k__Fungi(100);k__Fungi_unclassified(100);k__Fungi_unclassified(100);k__Fungi_unclassified(100);k__Fungi_unclassified(100);k__Fungi_unclassified(100);k__Fungi_unclassified(100); | - | - | - | - | - | - | - | Unassigned | - |
| Otu24558 | 1 | k__Fungi(100);k__Fungi_unclassified(100);k__Fungi_unclassified(100);k__Fungi_unclassified(100);k__Fungi_unclassified(100);k__Fungi_unclassified(100);k__Fungi_unclassified(100); | - | - | - | - | - | - | - | Unassigned | - |
| Otu24560 | 1 | k__Fungi(100);k__Fungi_unclassified(100);k__Fungi_unclassified(100);k__Fungi_unclassified(100);k__Fungi_unclassified(100);k__Fungi_unclassified(100);k__Fungi_unclassified(100); | - | - | - | - | - | - | - | Unassigned | - |
| Otu24562 | 1 | k__Fungi(100);p__Ascomycota(100);p__Ascomycota_unclassified(100);p__Ascomycota_unclassified(100);p__Ascomycota_unclassified(100);p__Ascomycota_unclassified(100);p__Ascomycota_unclassified(100); | - | - | - | - | - | - | - | Unassigned | - |
| Otu24563 | 1 | k__Fungi(100);k__Fungi_unclassified(100);k__Fungi_unclassified(100);k__Fungi_unclassified(100);k__Fungi_unclassified(100);k__Fungi_unclassified(100);k__Fungi_unclassified(100); | - | - | - | - | - | - | - | Unassigned | - |
| Otu24565 | 1 | k__Fungi(100);p__Ascomycota(100);p__Ascomycota_unclassified(100);p__Ascomycota_unclassified(100);p__Ascomycota_unclassified(100);p__Ascomycota_unclassified(100);p__Ascomycota_unclassified(100); | - | - | - | - | - | - | - | Unassigned | - |
| Otu24566 | 1 | k__Fungi(100);k__Fungi_unclassified(100);k__Fungi_unclassified(100);k__Fungi_unclassified(100);k__Fungi_unclassified(100);k__Fungi_unclassified(100);k__Fungi_unclassified(100); | - | - | - | - | - | - | - | Unassigned | - |
| Otu24567 | 1 | k__Fungi(100);k__Fungi_unclassified(100);k__Fungi_unclassified(100);k__Fungi_unclassified(100);k__Fungi_unclassified(100);k__Fungi_unclassified(100);k__Fungi_unclassified(100); | - | - | - | - | - | - | - | Unassigned | - |
| Otu24568 | 1 | k__Fungi(100);k__Fungi_unclassified(100);k__Fungi_unclassified(100);k__Fungi_unclassified(100);k__Fungi_unclassified(100);k__Fungi_unclassified(100);k__Fungi_unclassified(100); | - | - | - | - | - | - | - | Unassigned | - |
| Otu24569 | 1 | k__Fungi(100);k__Fungi_unclassified(100);k__Fungi_unclassified(100);k__Fungi_unclassified(100);k__Fungi_unclassified(100);k__Fungi_unclassified(100);k__Fungi_unclassified(100); | - | - | - | - | - | - | - | Unassigned | - |
| Otu24570 | 1 | k__Fungi(100);k__Fungi_unclassified(100);k__Fungi_unclassified(100);k__Fungi_unclassified(100);k__Fungi_unclassified(100);k__Fungi_unclassified(100);k__Fungi_unclassified(100); | - | - | - | - | - | - | - | Unassigned | - |
| Otu24571 | 1 | k__Fungi(100);k__Fungi_unclassified(100);k__Fungi_unclassified(100);k__Fungi_unclassified(100);k__Fungi_unclassified(100);k__Fungi_unclassified(100);k__Fungi_unclassified(100); | - | - | - | - | - | - | - | Unassigned | - |
| Otu24572 | 1 | k__Fungi(100);p__Ascomycota(100);p__Ascomycota_unclassified(100);p__Ascomycota_unclassified(100);p__Ascomycota_unclassified(100);p__Ascomycota_unclassified(100);p__Ascomycota_unclassified(100); | - | - | - | - | - | - | - | Unassigned | - |
| Otu24573 | 1 | k__Fungi(100);k__Fungi_unclassified(100);k__Fungi_unclassified(100);k__Fungi_unclassified(100);k__Fungi_unclassified(100);k__Fungi_unclassified(100);k__Fungi_unclassified(100); | - | - | - | - | - | - | - | Unassigned | - |
| Otu24574 | 1 | k__Fungi(100);k__Fungi_unclassified(100);k__Fungi_unclassified(100);k__Fungi_unclassified(100);k__Fungi_unclassified(100);k__Fungi_unclassified(100);k__Fungi_unclassified(100); | - | - | - | - | - | - | - | Unassigned | - |
| Otu24575 | 1 | k__Fungi(100);k__Fungi_unclassified(100);k__Fungi_unclassified(100);k__Fungi_unclassified(100);k__Fungi_unclassified(100);k__Fungi_unclassified(100);k__Fungi_unclassified(100); | - | - | - | - | - | - | - | Unassigned | - |
| Otu24577 | 1 | k__Fungi(100);k__Fungi_unclassified(100);k__Fungi_unclassified(100);k__Fungi_unclassified(100);k__Fungi_unclassified(100);k__Fungi_unclassified(100);k__Fungi_unclassified(100); | - | - | - | - | - | - | - | Unassigned | - |
| Otu24578 | 1 | k__Fungi(100);k__Fungi_unclassified(100);k__Fungi_unclassified(100);k__Fungi_unclassified(100);k__Fungi_unclassified(100);k__Fungi_unclassified(100);k__Fungi_unclassified(100); | - | - | - | - | - | - | - | Unassigned | - |
| Otu24580 | 1 | k__Fungi(100);p__Ascomycota(100);p__Ascomycota_unclassified(100);p__Ascomycota_unclassified(100);p__Ascomycota_unclassified(100);p__Ascomycota_unclassified(100);p__Ascomycota_unclassified(100); | - | - | - | - | - | - | - | Unassigned | - |
| Otu24585 | 1 | k__Fungi(100);k__Fungi_unclassified(100);k__Fungi_unclassified(100);k__Fungi_unclassified(100);k__Fungi_unclassified(100);k__Fungi_unclassified(100);k__Fungi_unclassified(100); | - | - | - | - | - | - | - | Unassigned | - |
| Otu24588 | 1 | k__Fungi(100);k__Fungi_unclassified(100);k__Fungi_unclassified(100);k__Fungi_unclassified(100);k__Fungi_unclassified(100);k__Fungi_unclassified(100);k__Fungi_unclassified(100); | - | - | - | - | - | - | - | Unassigned | - |
| Otu24590 | 1 | k__Fungi(100);k__Fungi_unclassified(100);k__Fungi_unclassified(100);k__Fungi_unclassified(100);k__Fungi_unclassified(100);k__Fungi_unclassified(100);k__Fungi_unclassified(100); | - | - | - | - | - | - | - | Unassigned | - |
| Otu24591 | 1 | k__Fungi(100);k__Fungi_unclassified(100);k__Fungi_unclassified(100);k__Fungi_unclassified(100);k__Fungi_unclassified(100);k__Fungi_unclassified(100);k__Fungi_unclassified(100); | - | - | - | - | - | - | - | Unassigned | - |
| Otu24592 | 1 | k__Fungi(100);k__Fungi_unclassified(100);k__Fungi_unclassified(100);k__Fungi_unclassified(100);k__Fungi_unclassified(100);k__Fungi_unclassified(100);k__Fungi_unclassified(100); | - | - | - | - | - | - | - | Unassigned | - |
| Otu24593 | 1 | k__Fungi(100);k__Fungi_unclassified(100);k__Fungi_unclassified(100);k__Fungi_unclassified(100);k__Fungi_unclassified(100);k__Fungi_unclassified(100);k__Fungi_unclassified(100); | - | - | - | - | - | - | - | Unassigned | - |
| Otu24594 | 1 | k__Fungi(100);k__Fungi_unclassified(100);k__Fungi_unclassified(100);k__Fungi_unclassified(100);k__Fungi_unclassified(100);k__Fungi_unclassified(100);k__Fungi_unclassified(100); | - | - | - | - | - | - | - | Unassigned | - |
| Otu24595 | 1 | k__Fungi(100);k__Fungi_unclassified(100);k__Fungi_unclassified(100);k__Fungi_unclassified(100);k__Fungi_unclassified(100);k__Fungi_unclassified(100);k__Fungi_unclassified(100); | - | - | - | - | - | - | - | Unassigned | - |
| Otu24596 | 1 | k__Fungi(100);k__Fungi_unclassified(100);k__Fungi_unclassified(100);k__Fungi_unclassified(100);k__Fungi_unclassified(100);k__Fungi_unclassified(100);k__Fungi_unclassified(100); | - | - | - | - | - | - | - | Unassigned | - |
| Otu24597 | 1 | k__Fungi(100);k__Fungi_unclassified(100);k__Fungi_unclassified(100);k__Fungi_unclassified(100);k__Fungi_unclassified(100);k__Fungi_unclassified(100);k__Fungi_unclassified(100); | - | - | - | - | - | - | - | Unassigned | - |
| Otu24599 | 1 | k__Fungi(100);k__Fungi_unclassified(100);k__Fungi_unclassified(100);k__Fungi_unclassified(100);k__Fungi_unclassified(100);k__Fungi_unclassified(100);k__Fungi_unclassified(100); | - | - | - | - | - | - | - | Unassigned | - |
| Otu24600 | 1 | k__Fungi(100);k__Fungi_unclassified(100);k__Fungi_unclassified(100);k__Fungi_unclassified(100);k__Fungi_unclassified(100);k__Fungi_unclassified(100);k__Fungi_unclassified(100); | - | - | - | - | - | - | - | Unassigned | - |
| Otu24601 | 1 | k__Fungi(100);p__Ascomycota(100);p__Ascomycota_unclassified(100);p__Ascomycota_unclassified(100);p__Ascomycota_unclassified(100);p__Ascomycota_unclassified(100);p__Ascomycota_unclassified(100); | - | - | - | - | - | - | - | Unassigned | - |
| Otu24602 | 1 | k__Fungi(100);k__Fungi_unclassified(100);k__Fungi_unclassified(100);k__Fungi_unclassified(100);k__Fungi_unclassified(100);k__Fungi_unclassified(100);k__Fungi_unclassified(100); | - | - | - | - | - | - | - | Unassigned | - |
| Otu24603 | 1 | k__Fungi(100);k__Fungi_unclassified(100);k__Fungi_unclassified(100);k__Fungi_unclassified(100);k__Fungi_unclassified(100);k__Fungi_unclassified(100);k__Fungi_unclassified(100); | - | - | - | - | - | - | - | Unassigned | - |
| Otu24604 | 1 | k__Fungi(100);k__Fungi_unclassified(100);k__Fungi_unclassified(100);k__Fungi_unclassified(100);k__Fungi_unclassified(100);k__Fungi_unclassified(100);k__Fungi_unclassified(100); | - | - | - | - | - | - | - | Unassigned | - |
| Otu24605 | 1 | k__Fungi(100);k__Fungi_unclassified(100);k__Fungi_unclassified(100);k__Fungi_unclassified(100);k__Fungi_unclassified(100);k__Fungi_unclassified(100);k__Fungi_unclassified(100); | - | - | - | - | - | - | - | Unassigned | - |
| Otu24606 | 1 | k__Fungi(100);k__Fungi_unclassified(100);k__Fungi_unclassified(100);k__Fungi_unclassified(100);k__Fungi_unclassified(100);k__Fungi_unclassified(100);k__Fungi_unclassified(100); | - | - | - | - | - | - | - | Unassigned | - |
| Otu24607 | 1 | k__Fungi(100);p__Basidiomycota(100);p__Basidiomycota_unclassified(100);p__Basidiomycota_unclassified(100);p__Basidiomycota_unclassified(100);p__Basidiomycota_unclassified(100);p__Basidiomycota_unclassified(100); | - | - | - | - | - | - | - | Unassigned | - |
| Otu24608 | 1 | k__Fungi(100);k__Fungi_unclassified(100);k__Fungi_unclassified(100);k__Fungi_unclassified(100);k__Fungi_unclassified(100);k__Fungi_unclassified(100);k__Fungi_unclassified(100); | - | - | - | - | - | - | - | Unassigned | - |
| Otu24609 | 1 | k__Fungi(100);k__Fungi_unclassified(100);k__Fungi_unclassified(100);k__Fungi_unclassified(100);k__Fungi_unclassified(100);k__Fungi_unclassified(100);k__Fungi_unclassified(100); | - | - | - | - | - | - | - | Unassigned | - |
| Otu24610 | 1 | k__Fungi(100);p__Ascomycota(100);p__Ascomycota_unclassified(100);p__Ascomycota_unclassified(100);p__Ascomycota_unclassified(100);p__Ascomycota_unclassified(100);p__Ascomycota_unclassified(100); | - | - | - | - | - | - | - | Unassigned | - |
| Otu24611 | 1 | k__Fungi(100);k__Fungi_unclassified(100);k__Fungi_unclassified(100);k__Fungi_unclassified(100);k__Fungi_unclassified(100);k__Fungi_unclassified(100);k__Fungi_unclassified(100); | - | - | - | - | - | - | - | Unassigned | - |
| Otu24617 | 1 | k__Fungi(100);k__Fungi_unclassified(100);k__Fungi_unclassified(100);k__Fungi_unclassified(100);k__Fungi_unclassified(100);k__Fungi_unclassified(100);k__Fungi_unclassified(100); | - | - | - | - | - | - | - | Unassigned | - |
| Otu24625 | 1 | k__Fungi(100);p__Ascomycota(100);p__Ascomycota_unclassified(100);p__Ascomycota_unclassified(100);p__Ascomycota_unclassified(100);p__Ascomycota_unclassified(100);p__Ascomycota_unclassified(100); | - | - | - | - | - | - | - | Unassigned | - |
| Otu24630 | 1 | k__Fungi(100);k__Fungi_unclassified(100);k__Fungi_unclassified(100);k__Fungi_unclassified(100);k__Fungi_unclassified(100);k__Fungi_unclassified(100);k__Fungi_unclassified(100); | - | - | - | - | - | - | - | Unassigned | - |
| Otu24631 | 1 | k__Fungi(100);k__Fungi_unclassified(100);k__Fungi_unclassified(100);k__Fungi_unclassified(100);k__Fungi_unclassified(100);k__Fungi_unclassified(100);k__Fungi_unclassified(100); | - | - | - | - | - | - | - | Unassigned | - |
| Otu24632 | 1 | k__Fungi(100);p__Ascomycota(100);p__Ascomycota_unclassified(100);p__Ascomycota_unclassified(100);p__Ascomycota_unclassified(100);p__Ascomycota_unclassified(100);p__Ascomycota_unclassified(100); | - | - | - | - | - | - | - | Unassigned | - |
| Otu24634 | 1 | k__Fungi(100);p__Ascomycota(100);p__Ascomycota_unclassified(100);p__Ascomycota_unclassified(100);p__Ascomycota_unclassified(100);p__Ascomycota_unclassified(100);p__Ascomycota_unclassified(100); | - | - | - | - | - | - | - | Unassigned | - |
| Otu24635 | 1 | k__Fungi(100);k__Fungi_unclassified(100);k__Fungi_unclassified(100);k__Fungi_unclassified(100);k__Fungi_unclassified(100);k__Fungi_unclassified(100);k__Fungi_unclassified(100); | - | - | - | - | - | - | - | Unassigned | - |
| Otu24636 | 1 | k__Fungi(100);k__Fungi_unclassified(100);k__Fungi_unclassified(100);k__Fungi_unclassified(100);k__Fungi_unclassified(100);k__Fungi_unclassified(100);k__Fungi_unclassified(100); | - | - | - | - | - | - | - | Unassigned | - |
| Otu24637 | 1 | k__Fungi(100);p__Ascomycota(100);p__Ascomycota_unclassified(100);p__Ascomycota_unclassified(100);p__Ascomycota_unclassified(100);p__Ascomycota_unclassified(100);p__Ascomycota_unclassified(100); | - | - | - | - | - | - | - | Unassigned | - |
| Otu24644 | 1 | k__Fungi(100);k__Fungi_unclassified(100);k__Fungi_unclassified(100);k__Fungi_unclassified(100);k__Fungi_unclassified(100);k__Fungi_unclassified(100);k__Fungi_unclassified(100); | - | - | - | - | - | - | - | Unassigned | - |
| Otu24645 | 1 | k__Fungi(100);k__Fungi_unclassified(100);k__Fungi_unclassified(100);k__Fungi_unclassified(100);k__Fungi_unclassified(100);k__Fungi_unclassified(100);k__Fungi_unclassified(100); | - | - | - | - | - | - | - | Unassigned | - |
| Otu24646 | 1 | k__Fungi(100);p__Ascomycota(100);p__Ascomycota_unclassified(100);p__Ascomycota_unclassified(100);p__Ascomycota_unclassified(100);p__Ascomycota_unclassified(100);p__Ascomycota_unclassified(100); | - | - | - | - | - | - | - | Unassigned | - |
| Otu24647 | 1 | k__Fungi(100);k__Fungi_unclassified(100);k__Fungi_unclassified(100);k__Fungi_unclassified(100);k__Fungi_unclassified(100);k__Fungi_unclassified(100);k__Fungi_unclassified(100); | - | - | - | - | - | - | - | Unassigned | - |
| Otu24651 | 1 | k__Fungi(100);p__Ascomycota(100);p__Ascomycota_unclassified(100);p__Ascomycota_unclassified(100);p__Ascomycota_unclassified(100);p__Ascomycota_unclassified(100);p__Ascomycota_unclassified(100); | - | - | - | - | - | - | - | Unassigned | - |
| Otu24655 | 1 | k__Fungi(100);k__Fungi_unclassified(100);k__Fungi_unclassified(100);k__Fungi_unclassified(100);k__Fungi_unclassified(100);k__Fungi_unclassified(100);k__Fungi_unclassified(100); | - | - | - | - | - | - | - | Unassigned | - |
| Otu24658 | 1 | k__Fungi(100);p__Ascomycota(100);p__Ascomycota_unclassified(100);p__Ascomycota_unclassified(100);p__Ascomycota_unclassified(100);p__Ascomycota_unclassified(100);p__Ascomycota_unclassified(100); | - | - | - | - | - | - | - | Unassigned | - |
| Otu24664 | 1 | k__Fungi(100);k__Fungi_unclassified(100);k__Fungi_unclassified(100);k__Fungi_unclassified(100);k__Fungi_unclassified(100);k__Fungi_unclassified(100);k__Fungi_unclassified(100); | - | - | - | - | - | - | - | Unassigned | - |
| Otu24672 | 1 | k__Fungi(100);p__Ascomycota(100);p__Ascomycota_unclassified(100);p__Ascomycota_unclassified(100);p__Ascomycota_unclassified(100);p__Ascomycota_unclassified(100);p__Ascomycota_unclassified(100); | - | - | - | - | - | - | - | Unassigned | - |
| Otu24673 | 1 | k__Fungi(100);p__Ascomycota(100);p__Ascomycota_unclassified(100);p__Ascomycota_unclassified(100);p__Ascomycota_unclassified(100);p__Ascomycota_unclassified(100);p__Ascomycota_unclassified(100); | - | - | - | - | - | - | - | Unassigned | - |
| Otu24674 | 1 | k__Fungi(100);p__Ascomycota(100);p__Ascomycota_unclassified(100);p__Ascomycota_unclassified(100);p__Ascomycota_unclassified(100);p__Ascomycota_unclassified(100);p__Ascomycota_unclassified(100); | - | - | - | - | - | - | - | Unassigned | - |
| Otu24675 | 1 | k__Fungi(100);p__Ascomycota(100);p__Ascomycota_unclassified(100);p__Ascomycota_unclassified(100);p__Ascomycota_unclassified(100);p__Ascomycota_unclassified(100);p__Ascomycota_unclassified(100); | - | - | - | - | - | - | - | Unassigned | - |
| Otu24676 | 1 | k__Fungi(100);k__Fungi_unclassified(100);k__Fungi_unclassified(100);k__Fungi_unclassified(100);k__Fungi_unclassified(100);k__Fungi_unclassified(100);k__Fungi_unclassified(100); | - | - | - | - | - | - | - | Unassigned | - |
| Otu24677 | 1 | k__Fungi(100);k__Fungi_unclassified(100);k__Fungi_unclassified(100);k__Fungi_unclassified(100);k__Fungi_unclassified(100);k__Fungi_unclassified(100);k__Fungi_unclassified(100); | - | - | - | - | - | - | - | Unassigned | - |
| Otu24679 | 1 | k__Fungi(100);k__Fungi_unclassified(100);k__Fungi_unclassified(100);k__Fungi_unclassified(100);k__Fungi_unclassified(100);k__Fungi_unclassified(100);k__Fungi_unclassified(100); | - | - | - | - | - | - | - | Unassigned | - |
| Otu24680 | 1 | k__Fungi(100);k__Fungi_unclassified(100);k__Fungi_unclassified(100);k__Fungi_unclassified(100);k__Fungi_unclassified(100);k__Fungi_unclassified(100);k__Fungi_unclassified(100); | - | - | - | - | - | - | - | Unassigned | - |
| Otu24681 | 1 | k__Fungi(100);p__Ascomycota(100);p__Ascomycota_unclassified(100);p__Ascomycota_unclassified(100);p__Ascomycota_unclassified(100);p__Ascomycota_unclassified(100);p__Ascomycota_unclassified(100); | - | - | - | - | - | - | - | Unassigned | - |
| Otu24683 | 1 | k__Fungi(100);k__Fungi_unclassified(100);k__Fungi_unclassified(100);k__Fungi_unclassified(100);k__Fungi_unclassified(100);k__Fungi_unclassified(100);k__Fungi_unclassified(100); | - | - | - | - | - | - | - | Unassigned | - |
| Otu24684 | 1 | k__Fungi(100);p__Ascomycota(100);p__Ascomycota_unclassified(100);p__Ascomycota_unclassified(100);p__Ascomycota_unclassified(100);p__Ascomycota_unclassified(100);p__Ascomycota_unclassified(100); | - | - | - | - | - | - | - | Unassigned | - |
| Otu24685 | 1 | k__Fungi(100);k__Fungi_unclassified(100);k__Fungi_unclassified(100);k__Fungi_unclassified(100);k__Fungi_unclassified(100);k__Fungi_unclassified(100);k__Fungi_unclassified(100); | - | - | - | - | - | - | - | Unassigned | - |
| Otu24686 | 1 | k__Fungi(100);p__Ascomycota(100);p__Ascomycota_unclassified(100);p__Ascomycota_unclassified(100);p__Ascomycota_unclassified(100);p__Ascomycota_unclassified(100);p__Ascomycota_unclassified(100); | - | - | - | - | - | - | - | Unassigned | - |
| Otu24687 | 1 | k__Fungi(100);k__Fungi_unclassified(100);k__Fungi_unclassified(100);k__Fungi_unclassified(100);k__Fungi_unclassified(100);k__Fungi_unclassified(100);k__Fungi_unclassified(100); | - | - | - | - | - | - | - | Unassigned | - |
| Otu24688 | 1 | k__Fungi(100);k__Fungi_unclassified(100);k__Fungi_unclassified(100);k__Fungi_unclassified(100);k__Fungi_unclassified(100);k__Fungi_unclassified(100);k__Fungi_unclassified(100); | - | - | - | - | - | - | - | Unassigned | - |
| Otu24689 | 1 | k__Fungi(100);k__Fungi_unclassified(100);k__Fungi_unclassified(100);k__Fungi_unclassified(100);k__Fungi_unclassified(100);k__Fungi_unclassified(100);k__Fungi_unclassified(100); | - | - | - | - | - | - | - | Unassigned | - |

| | | | | | | | | | | | |
|---|---|---|---|---|---|---|---|---|---|---|---|
| Otu24690 | 1 | k__Fungi(100);p__Ascomycota(100);p__Ascomycota_unclassified(100);p__Ascomycota_unclassified(100);p__Ascomycota_unclassified(100);p__Ascomycota_unclassified(100);p__Ascomycota_unclassified(100); | - | - | - | - | - | - | - | Unassigned | - |
| Otu24691 | 1 | k__Fungi(100);p__Ascomycota(100);p__Ascomycota_unclassified(100);p__Ascomycota_unclassified(100);p__Ascomycota_unclassified(100);p__Ascomycota_unclassified(100);p__Ascomycota_unclassified(100); | - | - | - | - | - | - | - | Unassigned | - |
| Otu24692 | 1 | k__Fungi(100);k__Fungi_unclassified(100);k__Fungi_unclassified(100);k__Fungi_unclassified(100);k__Fungi_unclassified(100);k__Fungi_unclassified(100);k__Fungi_unclassified(100); | - | - | - | - | - | - | - | Unassigned | - |
| Otu24693 | 1 | k__Fungi(100);k__Fungi_unclassified(100);k__Fungi_unclassified(100);k__Fungi_unclassified(100);k__Fungi_unclassified(100);k__Fungi_unclassified(100);k__Fungi_unclassified(100); | - | - | - | - | - | - | - | Unassigned | - |
| Otu24694 | 1 | k__Fungi(100);p__Ascomycota(100);p__Ascomycota_unclassified(100);p__Ascomycota_unclassified(100);p__Ascomycota_unclassified(100);p__Ascomycota_unclassified(100);p__Ascomycota_unclassified(100); | - | - | - | - | - | - | - | Unassigned | - |
| Otu24695 | 1 | k__Fungi(100);k__Fungi_unclassified(100);k__Fungi_unclassified(100);k__Fungi_unclassified(100);k__Fungi_unclassified(100);k__Fungi_unclassified(100);k__Fungi_unclassified(100); | - | - | - | - | - | - | - | Unassigned | - |
| Otu24696 | 1 | k__Fungi(100);p__Ascomycota(100);p__Ascomycota_unclassified(100);p__Ascomycota_unclassified(100);p__Ascomycota_unclassified(100);p__Ascomycota_unclassified(100);p__Ascomycota_unclassified(100); | - | - | - | - | - | - | - | Unassigned | - |
| Otu24699 | 1 | k__Fungi(100);k__Fungi_unclassified(100);k__Fungi_unclassified(100);k__Fungi_unclassified(100);k__Fungi_unclassified(100);k__Fungi_unclassified(100);k__Fungi_unclassified(100); | - | - | - | - | - | - | - | Unassigned | - |
| Otu24700 | 1 | k__Fungi(100);p__Ascomycota(100);p__Ascomycota_unclassified(100);p__Ascomycota_unclassified(100);p__Ascomycota_unclassified(100);p__Ascomycota_unclassified(100);p__Ascomycota_unclassified(100); | - | - | - | - | - | - | - | Unassigned | - |
| Otu24701 | 1 | k__Fungi(100);p__Ascomycota(100);p__Ascomycota_unclassified(100);p__Ascomycota_unclassified(100);p__Ascomycota_unclassified(100);p__Ascomycota_unclassified(100);p__Ascomycota_unclassified(100); | - | - | - | - | - | - | - | Unassigned | - |
| Otu24702 | 1 | k__Fungi(100);p__Ascomycota(100);p__Ascomycota_unclassified(100);p__Ascomycota_unclassified(100);p__Ascomycota_unclassified(100);p__Ascomycota_unclassified(100);p__Ascomycota_unclassified(100); | - | - | - | - | - | - | - | Unassigned | - |
| Otu24703 | 1 | k__Fungi(100);p__Ascomycota(100);p__Ascomycota_unclassified(100);p__Ascomycota_unclassified(100);p__Ascomycota_unclassified(100);p__Ascomycota_unclassified(100);p__Ascomycota_unclassified(100); | - | - | - | - | - | - | - | Unassigned | - |
| Otu24705 | 1 | k__Fungi(100);k__Fungi_unclassified(100);k__Fungi_unclassified(100);k__Fungi_unclassified(100);k__Fungi_unclassified(100);k__Fungi_unclassified(100);k__Fungi_unclassified(100); | - | - | - | - | - | - | - | Unassigned | - |
| Otu24706 | 1 | k__Fungi(100);k__Fungi_unclassified(100);k__Fungi_unclassified(100);k__Fungi_unclassified(100);k__Fungi_unclassified(100);k__Fungi_unclassified(100);k__Fungi_unclassified(100); | - | - | - | - | - | - | - | Unassigned | - |
| Otu24713 | 1 | k__Fungi(100);p__Ascomycota(100);c__Dothideomycetes(100);o__Capnodiales(100);f__Davidiellaceae(100);f__Davidiellaceae_unclassified(100);f__Davidiellaceae_unclassified(100); | - | - | - | - | - | - | - | Unassigned | - |
| Otu24714 | 1 | k__Fungi(100);k__Fungi_unclassified(100);k__Fungi_unclassified(100);k__Fungi_unclassified(100);k__Fungi_unclassified(100);k__Fungi_unclassified(100);k__Fungi_unclassified(100); | - | - | - | - | - | - | - | Unassigned | - |
| Otu24715 | 1 | k__Fungi(100);k__Fungi_unclassified(100);k__Fungi_unclassified(100);k__Fungi_unclassified(100);k__Fungi_unclassified(100);k__Fungi_unclassified(100);k__Fungi_unclassified(100); | - | - | - | - | - | - | - | Unassigned | - |
| Otu24718 | 1 | k__Fungi(100);k__Fungi_unclassified(100);k__Fungi_unclassified(100);k__Fungi_unclassified(100);k__Fungi_unclassified(100);k__Fungi_unclassified(100);k__Fungi_unclassified(100); | - | - | - | - | - | - | - | Unassigned | - |
| Otu24720 | 1 | k__Fungi(100);k__Fungi_unclassified(100);k__Fungi_unclassified(100);k__Fungi_unclassified(100);k__Fungi_unclassified(100);k__Fungi_unclassified(100);k__Fungi_unclassified(100); | - | - | - | - | - | - | - | Unassigned | - |
| Otu24732 | 1 | k__Fungi(100);k__Fungi_unclassified(100);k__Fungi_unclassified(100);k__Fungi_unclassified(100);k__Fungi_unclassified(100);k__Fungi_unclassified(100);k__Fungi_unclassified(100); | - | - | - | - | - | - | - | Unassigned | - |
| Otu24744 | 1 | k__Fungi(100);k__Fungi_unclassified(100);k__Fungi_unclassified(100);k__Fungi_unclassified(100);k__Fungi_unclassified(100);k__Fungi_unclassified(100);k__Fungi_unclassified(100); | - | - | - | - | - | - | - | Unassigned | - |
| Otu24745 | 1 | k__Fungi(100);k__Fungi_unclassified(100);k__Fungi_unclassified(100);k__Fungi_unclassified(100);k__Fungi_unclassified(100);k__Fungi_unclassified(100);k__Fungi_unclassified(100); | - | - | - | - | - | - | - | Unassigned | - |
| Otu24746 | 1 | k__Fungi(100);p__Ascomycota(100);p__Ascomycota_unclassified(100);p__Ascomycota_unclassified(100);p__Ascomycota_unclassified(100);p__Ascomycota_unclassified(100);p__Ascomycota_unclassified(100); | - | - | - | - | - | - | - | Unassigned | - |
| Otu24747 | 1 | k__Fungi(100);k__Fungi_unclassified(100);k__Fungi_unclassified(100);k__Fungi_unclassified(100);k__Fungi_unclassified(100);k__Fungi_unclassified(100);k__Fungi_unclassified(100); | - | - | - | - | - | - | - | Unassigned | - |
| Otu24748 | 1 | k__Fungi(100);k__Fungi_unclassified(100);k__Fungi_unclassified(100);k__Fungi_unclassified(100);k__Fungi_unclassified(100);k__Fungi_unclassified(100);k__Fungi_unclassified(100); | - | - | - | - | - | - | - | Unassigned | - |
| Otu24751 | 1 | k__Fungi(100);k__Fungi_unclassified(100);k__Fungi_unclassified(100);k__Fungi_unclassified(100);k__Fungi_unclassified(100);k__Fungi_unclassified(100);k__Fungi_unclassified(100); | - | - | - | - | - | - | - | Unassigned | - |
| Otu24752 | 1 | k__Fungi(100);k__Fungi_unclassified(100);k__Fungi_unclassified(100);k__Fungi_unclassified(100);k__Fungi_unclassified(100);k__Fungi_unclassified(100);k__Fungi_unclassified(100); | - | - | - | - | - | - | - | Unassigned | - |
| Otu24753 | 1 | k__Fungi(100);p__Ascomycota(100);p__Ascomycota_unclassified(100);p__Ascomycota_unclassified(100);p__Ascomycota_unclassified(100);p__Ascomycota_unclassified(100);p__Ascomycota_unclassified(100); | - | - | - | - | - | - | - | Unassigned | - |
| Otu24754 | 1 | k__Fungi(100);k__Fungi_unclassified(100);k__Fungi_unclassified(100);k__Fungi_unclassified(100);k__Fungi_unclassified(100);k__Fungi_unclassified(100);k__Fungi_unclassified(100); | - | - | - | - | - | - | - | Unassigned | - |
| Otu24755 | 1 | k__Fungi(100);k__Fungi_unclassified(100);k__Fungi_unclassified(100);k__Fungi_unclassified(100);k__Fungi_unclassified(100);k__Fungi_unclassified(100);k__Fungi_unclassified(100); | - | - | - | - | - | - | - | Unassigned | - |
| Otu24759 | 1 | k__Fungi(100);k__Fungi_unclassified(100);k__Fungi_unclassified(100);k__Fungi_unclassified(100);k__Fungi_unclassified(100);k__Fungi_unclassified(100);k__Fungi_unclassified(100); | - | - | - | - | - | - | - | Unassigned | - |
| Otu24761 | 1 | k__Fungi(100);k__Fungi_unclassified(100);k__Fungi_unclassified(100);k__Fungi_unclassified(100);k__Fungi_unclassified(100);k__Fungi_unclassified(100);k__Fungi_unclassified(100); | - | - | - | - | - | - | - | Unassigned | - |
| Otu24763 | 1 | k__Fungi(100);k__Fungi_unclassified(100);k__Fungi_unclassified(100);k__Fungi_unclassified(100);k__Fungi_unclassified(100);k__Fungi_unclassified(100);k__Fungi_unclassified(100); | - | - | - | - | - | - | - | Unassigned | - |
| Otu24769 | 1 | k__Fungi(100);k__Fungi_unclassified(100);k__Fungi_unclassified(100);k__Fungi_unclassified(100);k__Fungi_unclassified(100);k__Fungi_unclassified(100);k__Fungi_unclassified(100); | - | - | - | - | - | - | - | Unassigned | - |
| Otu24770 | 1 | k__Fungi(100);p__Ascomycota(100);p__Ascomycota_unclassified(100);p__Ascomycota_unclassified(100);p__Ascomycota_unclassified(100);p__Ascomycota_unclassified(100);p__Ascomycota_unclassified(100); | - | - | - | - | - | - | - | Unassigned | - |
| Otu24777 | 1 | k__Fungi(100);p__Ascomycota(100);p__Ascomycota_unclassified(100);p__Ascomycota_unclassified(100);p__Ascomycota_unclassified(100);p__Ascomycota_unclassified(100);p__Ascomycota_unclassified(100); | - | - | - | - | - | - | - | Unassigned | - |
| Otu24778 | 1 | k__Fungi(100);k__Fungi_unclassified(100);k__Fungi_unclassified(100);k__Fungi_unclassified(100);k__Fungi_unclassified(100);k__Fungi_unclassified(100);k__Fungi_unclassified(100); | - | - | - | - | - | - | - | Unassigned | - |
| Otu24779 | 1 | k__Fungi(100);k__Fungi_unclassified(100);k__Fungi_unclassified(100);k__Fungi_unclassified(100);k__Fungi_unclassified(100);k__Fungi_unclassified(100);k__Fungi_unclassified(100); | - | - | - | - | - | - | - | Unassigned | - |
| Otu24780 | 1 | k__Fungi(100);k__Fungi_unclassified(100);k__Fungi_unclassified(100);k__Fungi_unclassified(100);k__Fungi_unclassified(100);k__Fungi_unclassified(100);k__Fungi_unclassified(100); | - | - | - | - | - | - | - | Unassigned | - |
| Otu24781 | 1 | k__Fungi(100);k__Fungi_unclassified(100);k__Fungi_unclassified(100);k__Fungi_unclassified(100);k__Fungi_unclassified(100);k__Fungi_unclassified(100);k__Fungi_unclassified(100); | - | - | - | - | - | - | - | Unassigned | - |
| Otu24782 | 1 | k__Fungi(100);k__Fungi_unclassified(100);k__Fungi_unclassified(100);k__Fungi_unclassified(100);k__Fungi_unclassified(100);k__Fungi_unclassified(100);k__Fungi_unclassified(100); | - | - | - | - | - | - | - | Unassigned | - |
| Otu24785 | 1 | k__Fungi(100);k__Fungi_unclassified(100);k__Fungi_unclassified(100);k__Fungi_unclassified(100);k__Fungi_unclassified(100);k__Fungi_unclassified(100);k__Fungi_unclassified(100); | - | - | - | - | - | - | - | Unassigned | - |
| Otu24787 | 1 | k__Fungi(100);p__Ascomycota(100);p__Ascomycota_unclassified(100);p__Ascomycota_unclassified(100);p__Ascomycota_unclassified(100);p__Ascomycota_unclassified(100);p__Ascomycota_unclassified(100); | - | - | - | - | - | - | - | Unassigned | - |
| Otu24788 | 1 | k__Fungi(100);k__Fungi_unclassified(100);k__Fungi_unclassified(100);k__Fungi_unclassified(100);k__Fungi_unclassified(100);k__Fungi_unclassified(100);k__Fungi_unclassified(100); | - | - | - | - | - | - | - | Unassigned | - |
| Otu24789 | 1 | k__Fungi(100);p__Ascomycota(100);p__Ascomycota_unclassified(100);p__Ascomycota_unclassified(100);p__Ascomycota_unclassified(100);p__Ascomycota_unclassified(100);p__Ascomycota_unclassified(100); | - | - | - | - | - | - | - | Unassigned | - |
| Otu24791 | 1 | k__Fungi(100);k__Fungi_unclassified(100);k__Fungi_unclassified(100);k__Fungi_unclassified(100);k__Fungi_unclassified(100);k__Fungi_unclassified(100);k__Fungi_unclassified(100); | - | - | - | - | - | - | - | Unassigned | - |
| Otu24799 | 1 | k__Fungi(100);k__Fungi_unclassified(100);k__Fungi_unclassified(100);k__Fungi_unclassified(100);k__Fungi_unclassified(100);k__Fungi_unclassified(100);k__Fungi_unclassified(100); | - | - | - | - | - | - | - | Unassigned | - |
| Otu24802 | 1 | k__Fungi(100);k__Fungi_unclassified(100);k__Fungi_unclassified(100);k__Fungi_unclassified(100);k__Fungi_unclassified(100);k__Fungi_unclassified(100);k__Fungi_unclassified(100); | - | - | - | - | - | - | - | Unassigned | - |
| Otu24807 | 1 | k__Fungi(100);k__Fungi_unclassified(100);k__Fungi_unclassified(100);k__Fungi_unclassified(100);k__Fungi_unclassified(100);k__Fungi_unclassified(100);k__Fungi_unclassified(100); | - | - | - | - | - | - | - | Unassigned | - |
| Otu24809 | 1 | k__Fungi(100);k__Fungi_unclassified(100);k__Fungi_unclassified(100);k__Fungi_unclassified(100);k__Fungi_unclassified(100);k__Fungi_unclassified(100);k__Fungi_unclassified(100); | - | - | - | - | - | - | - | Unassigned | - |
| Otu24810 | 1 | k__Fungi(100);p__Ascomycota(100);p__Ascomycota_unclassified(100);p__Ascomycota_unclassified(100);p__Ascomycota_unclassified(100);p__Ascomycota_unclassified(100);p__Ascomycota_unclassified(100); | - | - | - | - | - | - | - | Unassigned | - |
| Otu24811 | 1 | k__Fungi(100);p__Ascomycota(100);p__Ascomycota_unclassified(100);p__Ascomycota_unclassified(100);p__Ascomycota_unclassified(100);p__Ascomycota_unclassified(100);p__Ascomycota_unclassified(100); | - | - | - | - | - | - | - | Unassigned | - |
| Otu24812 | 1 | k__Fungi(100);k__Fungi_unclassified(100);k__Fungi_unclassified(100);k__Fungi_unclassified(100);k__Fungi_unclassified(100);k__Fungi_unclassified(100);k__Fungi_unclassified(100); | - | - | - | - | - | - | - | Unassigned | - |
| Otu24817 | 1 | k__Fungi(100);k__Fungi_unclassified(100);k__Fungi_unclassified(100);k__Fungi_unclassified(100);k__Fungi_unclassified(100);k__Fungi_unclassified(100);k__Fungi_unclassified(100); | - | - | - | - | - | - | - | Unassigned | - |
| Otu24818 | 1 | k__Fungi(100);p__Ascomycota(100);p__Ascomycota_unclassified(100);p__Ascomycota_unclassified(100);p__Ascomycota_unclassified(100);p__Ascomycota_unclassified(100);p__Ascomycota_unclassified(100); | - | - | - | - | - | - | - | Unassigned | - |
| Otu24823 | 1 | k__Fungi(100);p__Ascomycota(100);p__Ascomycota_unclassified(100);p__Ascomycota_unclassified(100);p__Ascomycota_unclassified(100);p__Ascomycota_unclassified(100);p__Ascomycota_unclassified(100); | - | - | - | - | - | - | - | Unassigned | - |
| Otu24825 | 1 | k__Fungi(100);k__Fungi_unclassified(100);k__Fungi_unclassified(100);k__Fungi_unclassified(100);k__Fungi_unclassified(100);k__Fungi_unclassified(100);k__Fungi_unclassified(100); | - | - | - | - | - | - | - | Unassigned | - |
| Otu24826 | 1 | k__Fungi(100);k__Fungi_unclassified(100);k__Fungi_unclassified(100);k__Fungi_unclassified(100);k__Fungi_unclassified(100);k__Fungi_unclassified(100);k__Fungi_unclassified(100); | - | - | - | - | - | - | - | Unassigned | - |
| Otu24828 | 1 | k__Fungi(100);p__Ascomycota(100);p__Ascomycota_unclassified(100);p__Ascomycota_unclassified(100);p__Ascomycota_unclassified(100);p__Ascomycota_unclassified(100);p__Ascomycota_unclassified(100); | - | - | - | - | - | - | - | Unassigned | - |
| Otu24833 | 1 | k__Fungi(100);p__Ascomycota(100);p__Ascomycota_unclassified(100);p__Ascomycota_unclassified(100);p__Ascomycota_unclassified(100);p__Ascomycota_unclassified(100);p__Ascomycota_unclassified(100); | - | - | - | - | - | - | - | Unassigned | - |
| Otu24836 | 1 | k__Fungi(100);k__Fungi_unclassified(100);k__Fungi_unclassified(100);k__Fungi_unclassified(100);k__Fungi_unclassified(100);k__Fungi_unclassified(100);k__Fungi_unclassified(100); | - | - | - | - | - | - | - | Unassigned | - |
| Otu24837 | 1 | k__Fungi(100);k__Fungi_unclassified(100);k__Fungi_unclassified(100);k__Fungi_unclassified(100);k__Fungi_unclassified(100);k__Fungi_unclassified(100);k__Fungi_unclassified(100); | - | - | - | - | - | - | - | Unassigned | - |
| Otu24839 | 1 | k__Fungi(100);k__Fungi_unclassified(100);k__Fungi_unclassified(100);k__Fungi_unclassified(100);k__Fungi_unclassified(100);k__Fungi_unclassified(100);k__Fungi_unclassified(100); | - | - | - | - | - | - | - | Unassigned | - |
| Otu24840 | 1 | k__Fungi(100);k__Fungi_unclassified(100);k__Fungi_unclassified(100);k__Fungi_unclassified(100);k__Fungi_unclassified(100);k__Fungi_unclassified(100);k__Fungi_unclassified(100); | - | - | - | - | - | - | - | Unassigned | - |
| Otu24841 | 1 | k__Fungi(100);k__Fungi_unclassified(100);k__Fungi_unclassified(100);k__Fungi_unclassified(100);k__Fungi_unclassified(100);k__Fungi_unclassified(100);k__Fungi_unclassified(100); | - | - | - | - | - | - | - | Unassigned | - |
| Otu24843 | 1 | k__Fungi(100);k__Fungi_unclassified(100);k__Fungi_unclassified(100);k__Fungi_unclassified(100);k__Fungi_unclassified(100);k__Fungi_unclassified(100);k__Fungi_unclassified(100); | - | - | - | - | - | - | - | Unassigned | - |
| Otu24846 | 1 | k__Fungi(100);k__Fungi_unclassified(100);k__Fungi_unclassified(100);k__Fungi_unclassified(100);k__Fungi_unclassified(100);k__Fungi_unclassified(100);k__Fungi_unclassified(100); | - | - | - | - | - | - | - | Unassigned | - |
| Otu24850 | 1 | k__Fungi(100);p__Ascomycota(100);p__Ascomycota_unclassified(100);p__Ascomycota_unclassified(100);p__Ascomycota_unclassified(100);p__Ascomycota_unclassified(100);p__Ascomycota_unclassified(100); | - | - | - | - | - | - | - | Unassigned | - |
| Otu24855 | 1 | k__Fungi(100);k__Fungi_unclassified(100);k__Fungi_unclassified(100);k__Fungi_unclassified(100);k__Fungi_unclassified(100);k__Fungi_unclassified(100);k__Fungi_unclassified(100); | - | - | - | - | - | - | - | Unassigned | - |
| Otu24856 | 1 | k__Fungi(100);k__Fungi_unclassified(100);k__Fungi_unclassified(100);k__Fungi_unclassified(100);k__Fungi_unclassified(100);k__Fungi_unclassified(100);k__Fungi_unclassified(100); | - | - | - | - | - | - | - | Unassigned | - |
| Otu24857 | 1 | k__Fungi(100);k__Fungi_unclassified(100);k__Fungi_unclassified(100);k__Fungi_unclassified(100);k__Fungi_unclassified(100);k__Fungi_unclassified(100);k__Fungi_unclassified(100); | - | - | - | - | - | - | - | Unassigned | - |
| Otu24858 | 1 | k__Fungi(100);k__Fungi_unclassified(100);k__Fungi_unclassified(100);k__Fungi_unclassified(100);k__Fungi_unclassified(100);k__Fungi_unclassified(100);k__Fungi_unclassified(100); | - | - | - | - | - | - | - | Unassigned | - |
| Otu24859 | 1 | k__Fungi(100);p__Ascomycota(100);p__Ascomycota_unclassified(100);p__Ascomycota_unclassified(100);p__Ascomycota_unclassified(100);p__Ascomycota_unclassified(100);p__Ascomycota_unclassified(100); | - | - | - | - | - | - | - | Unassigned | - |
| Otu24860 | 1 | k__Fungi(100);p__Ascomycota(100);p__Ascomycota_unclassified(100);p__Ascomycota_unclassified(100);p__Ascomycota_unclassified(100);p__Ascomycota_unclassified(100);p__Ascomycota_unclassified(100); | - | - | - | - | - | - | - | Unassigned | - |
| Otu24861 | 1 | k__Fungi(100);p__Ascomycota(100);p__Ascomycota_unclassified(100);p__Ascomycota_unclassified(100);p__Ascomycota_unclassified(100);p__Ascomycota_unclassified(100);p__Ascomycota_unclassified(100); | - | - | - | - | - | - | - | Unassigned | - |
| Otu24862 | 1 | k__Fungi(100);k__Fungi_unclassified(100);k__Fungi_unclassified(100);k__Fungi_unclassified(100);k__Fungi_unclassified(100);k__Fungi_unclassified(100);k__Fungi_unclassified(100); | - | - | - | - | - | - | - | Unassigned | - |
| Otu24864 | 1 | k__Fungi(100);k__Fungi_unclassified(100);k__Fungi_unclassified(100);k__Fungi_unclassified(100);k__Fungi_unclassified(100);k__Fungi_unclassified(100);k__Fungi_unclassified(100); | - | - | - | - | - | - | - | Unassigned | - |
| Otu24871 | 1 | k__Fungi(100);p__Ascomycota(100);p__Ascomycota_unclassified(100);p__Ascomycota_unclassified(100);p__Ascomycota_unclassified(100);p__Ascomycota_unclassified(100);p__Ascomycota_unclassified(100); | - | - | - | - | - | - | - | Unassigned | - |
| Otu24873 | 1 | k__Fungi(100);k__Fungi_unclassified(100);k__Fungi_unclassified(100);k__Fungi_unclassified(100);k__Fungi_unclassified(100);k__Fungi_unclassified(100);k__Fungi_unclassified(100); | - | - | - | - | - | - | - | Unassigned | - |
| Otu24874 | 1 | k__Fungi(100);p__Ascomycota(100);p__Ascomycota_unclassified(100);p__Ascomycota_unclassified(100);p__Ascomycota_unclassified(100);p__Ascomycota_unclassified(100);p__Ascomycota_unclassified(100); | - | - | - | - | - | - | - | Unassigned | - |
| Otu24875 | 1 | k__Fungi(100);p__Ascomycota(100);p__Ascomycota_unclassified(100);p__Ascomycota_unclassified(100);p__Ascomycota_unclassified(100);p__Ascomycota_unclassified(100);p__Ascomycota_unclassified(100); | - | - | - | - | - | - | - | Unassigned | - |
| Otu24883 | 1 | k__Fungi(100);k__Fungi_unclassified(100);k__Fungi_unclassified(100);k__Fungi_unclassified(100);k__Fungi_unclassified(100);k__Fungi_unclassified(100);k__Fungi_unclassified(100); | - | - | - | - | - | - | - | Unassigned | - |
| Otu24888 | 1 | k__Fungi(100);k__Fungi_unclassified(100);k__Fungi_unclassified(100);k__Fungi_unclassified(100);k__Fungi_unclassified(100);k__Fungi_unclassified(100);k__Fungi_unclassified(100); | - | - | - | - | - | - | - | Unassigned | - |
| Otu24895 | 1 | k__Fungi(100);p__Ascomycota(100);p__Ascomycota_unclassified(100);p__Ascomycota_unclassified(100);p__Ascomycota_unclassified(100);p__Ascomycota_unclassified(100);p__Ascomycota_unclassified(100); | - | - | - | - | - | - | - | Unassigned | - |
| Otu24900 | 1 | k__Fungi(100);k__Fungi_unclassified(100);k__Fungi_unclassified(100);k__Fungi_unclassified(100);k__Fungi_unclassified(100);k__Fungi_unclassified(100);k__Fungi_unclassified(100); | - | - | - | - | - | - | - | Unassigned | - |
| Otu24901 | 1 | k__Fungi(100);p__Ascomycota(100);p__Ascomycota_unclassified(100);p__Ascomycota_unclassified(100);p__Ascomycota_unclassified(100);p__Ascomycota_unclassified(100);p__Ascomycota_unclassified(100); | - | - | - | - | - | - | - | Unassigned | - |
| Otu24902 | 1 | k__Fungi(100);p__Ascomycota(100);p__Ascomycota_unclassified(100);p__Ascomycota_unclassified(100);p__Ascomycota_unclassified(100);p__Ascomycota_unclassified(100);p__Ascomycota_unclassified(100); | - | - | - | - | - | - | - | Unassigned | - |
| Otu24903 | 1 | k__Fungi(100);k__Fungi_unclassified(100);k__Fungi_unclassified(100);k__Fungi_unclassified(100);k__Fungi_unclassified(100);k__Fungi_unclassified(100);k__Fungi_unclassified(100); | - | - | - | - | - | - | - | Unassigned | - |
| Otu24905 | 1 | k__Fungi(100);p__Ascomycota(100);p__Ascomycota_unclassified(100);p__Ascomycota_unclassified(100);p__Ascomycota_unclassified(100);p__Ascomycota_unclassified(100);p__Ascomycota_unclassified(100); | - | - | - | - | - | - | - | Unassigned | - |
| Otu24906 | 1 | k__Fungi(100);p__Ascomycota(100);p__Ascomycota_unclassified(100);p__Ascomycota_unclassified(100);p__Ascomycota_unclassified(100);p__Ascomycota_unclassified(100);p__Ascomycota_unclassified(100); | - | - | - | - | - | - | - | Unassigned | - |
| Otu24930 | 1 | k__Fungi(100);k__Fungi_unclassified(100);k__Fungi_unclassified(100);k__Fungi_unclassified(100);k__Fungi_unclassified(100);k__Fungi_unclassified(100);k__Fungi_unclassified(100); | - | - | - | - | - | - | - | Unassigned | - |
| Otu24931 | 1 | k__Fungi(100);p__Ascomycota(100);p__Ascomycota_unclassified(100);p__Ascomycota_unclassified(100);p__Ascomycota_unclassified(100);p__Ascomycota_unclassified(100);p__Ascomycota_unclassified(100); | - | - | - | - | - | - | - | Unassigned | - |
| Otu24932 | 1 | k__Fungi(100);p__Ascomycota(100);p__Ascomycota_unclassified(100);p__Ascomycota_unclassified(100);p__Ascomycota_unclassified(100);p__Ascomycota_unclassified(100);p__Ascomycota_unclassified(100); | - | - | - | - | - | - | - | Unassigned | - |
| Otu24933 | 1 | k__Fungi(100);p__Ascomycota(100);p__Ascomycota_unclassified(100);p__Ascomycota_unclassified(100);p__Ascomycota_unclassified(100);p__Ascomycota_unclassified(100);p__Ascomycota_unclassified(100); | - | - | - | - | - | - | - | Unassigned | - |
| Otu24934 | 1 | k__Fungi(100);p__Ascomycota(100);p__Ascomycota_unclassified(100);p__Ascomycota_unclassified(100);p__Ascomycota_unclassified(100);p__Ascomycota_unclassified(100);p__Ascomycota_unclassified(100); | - | - | - | - | - | - | - | Unassigned | - |
| Otu24935 | 1 | k__Fungi(100);k__Fungi_unclassified(100);k__Fungi_unclassified(100);k__Fungi_unclassified(100);k__Fungi_unclassified(100);k__Fungi_unclassified(100);k__Fungi_unclassified(100); | - | - | - | - | - | - | - | Unassigned | - |
| Otu24936 | 1 | k__Fungi(100);k__Fungi_unclassified(100);k__Fungi_unclassified(100);k__Fungi_unclassified(100);k__Fungi_unclassified(100);k__Fungi_unclassified(100);k__Fungi_unclassified(100); | - | - | - | - | - | - | - | Unassigned | - |
| Otu24937 | 1 | k__Fungi(100);p__Ascomycota(100);p__Ascomycota_unclassified(100);p__Ascomycota_unclassified(100);p__Ascomycota_unclassified(100);p__Ascomycota_unclassified(100);p__Ascomycota_unclassified(100); | - | - | - | - | - | - | - | Unassigned | - |
| Otu24938 | 1 | k__Fungi(100);p__Ascomycota(100);p__Ascomycota_unclassified(100);p__Ascomycota_unclassified(100);p__Ascomycota_unclassified(100);p__Ascomycota_unclassified(100);p__Ascomycota_unclassified(100); | - | - | - | - | - | - | - | Unassigned | - |
| Otu24939 | 1 | k__Fungi(100);p__Ascomycota(100);p__Ascomycota_unclassified(100);p__Ascomycota_unclassified(100);p__Ascomycota_unclassified(100);p__Ascomycota_unclassified(100);p__Ascomycota_unclassified(100); | - | - | - | - | - | - | - | Unassigned | - |
| Otu24940 | 1 | k__Fungi(100);p__Ascomycota(100);p__Ascomycota_unclassified(100);p__Ascomycota_unclassified(100);p__Ascomycota_unclassified(100);p__Ascomycota_unclassified(100);p__Ascomycota_unclassified(100); | - | - | - | - | - | - | - | Unassigned | - |
| Otu24941 | 1 | k__Fungi(100);p__Ascomycota(100);p__Ascomycota_unclassified(100);p__Ascomycota_unclassified(100);p__Ascomycota_unclassified(100);p__Ascomycota_unclassified(100);p__Ascomycota_unclassified(100); | - | - | - | - | - | - | - | Unassigned | - |
| Otu24942 | 1 | k__Fungi(100);p__Ascomycota(100);p__Ascomycota_unclassified(100);p__Ascomycota_unclassified(100);p__Ascomycota_unclassified(100);p__Ascomycota_unclassified(100);p__Ascomycota_unclassified(100); | - | - | - | - | - | - | - | Unassigned | - |
| Otu24943 | 1 | k__Fungi(100);p__Ascomycota(100);p__Ascomycota_unclassified(100);p__Ascomycota_unclassified(100);p__Ascomycota_unclassified(100);p__Ascomycota_unclassified(100);p__Ascomycota_unclassified(100); | - | - | - | - | - | - | - | Unassigned | - |
| Otu24944 | 1 | k__Fungi(100);p__Ascomycota(100);p__Ascomycota_unclassified(100);p__Ascomycota_unclassified(100);p__Ascomycota_unclassified(100);p__Ascomycota_unclassified(100);p__Ascomycota_unclassified(100); | - | - | - | - | - | - | - | Unassigned | - |
| Otu24945 | 1 | k__Fungi(100);p__Ascomycota(100);p__Ascomycota_unclassified(100);p__Ascomycota_unclassified(100);p__Ascomycota_unclassified(100);p__Ascomycota_unclassified(100);p__Ascomycota_unclassified(100); | - | - | - | - | - | - | - | Unassigned | - |
| Otu24946 | 1 | k__Fungi(100);k__Fungi_unclassified(100);k__Fungi_unclassified(100);k__Fungi_unclassified(100);k__Fungi_unclassified(100);k__Fungi_unclassified(100);k__Fungi_unclassified(100); | - | - | - | - | - | - | - | Unassigned | - |
| Otu24947 | 1 | k__Fungi(100);p__Ascomycota(100);c__Sordariomycetes(100);c__Sordariomycetes_unclassified(100);c__Sordariomycetes_unclassified(100);c__Sordariomycetes_unclassified(100);c__Sordariomycetes_unclassified(100); | - | - | - | - | - | - | - | Unassigned | - |
| Otu24948 | 1 | k__Fungi(100);p__Ascomycota(100);p__Ascomycota_unclassified(100);p__Ascomycota_unclassified(100);p__Ascomycota_unclassified(100);p__Ascomycota_unclassified(100);p__Ascomycota_unclassified(100); | - | - | - | - | - | - | - | Unassigned | - |
| Otu24949 | 1 | k__Fungi(100);k__Fungi_unclassified(100);k__Fungi_unclassified(100);k__Fungi_unclassified(100);k__Fungi_unclassified(100);k__Fungi_unclassified(100);k__Fungi_unclassified(100); | - | - | - | - | - | - | - | Unassigned | - |
| Otu24951 | 1 | k__Fungi(100);p__Ascomycota(100);p__Ascomycota_unclassified(100);p__Ascomycota_unclassified(100);p__Ascomycota_unclassified(100);p__Ascomycota_unclassified(100);p__Ascomycota_unclassified(100); | - | - | - | - | - | - | - | Unassigned | - |
| Otu24952 | 1 | k__Fungi(100);p__Ascomycota(100);p__Ascomycota_unclassified(100);p__Ascomycota_unclassified(100);p__Ascomycota_unclassified(100);p__Ascomycota_unclassified(100);p__Ascomycota_unclassified(100); | - | - | - | - | - | - | - | Unassigned | - |
| Otu24953 | 1 | k__Fungi(100);p__Ascomycota(100);c__Leotiomycetes(100);c__Leotiomycetes_unclassified(100);c__Leotiomycetes_unclassified(100);c__Leotiomycetes_unclassified(100);c__Leotiomycetes_unclassified(100); | - | - | - | - | - | - | - | Unassigned | - |
| Otu24955 | 1 | k__Fungi(100);k__Fungi_unclassified(100);k__Fungi_unclassified(100);k__Fungi_unclassified(100);k__Fungi_unclassified(100);k__Fungi_unclassified(100);k__Fungi_unclassified(100); | - | - | - | - | - | - | - | Unassigned | - |
| Otu24956 | 1 | k__Fungi(100);p__Ascomycota(100);p__Ascomycota_unclassified(100);p__Ascomycota_unclassified(100);p__Ascomycota_unclassified(100);p__Ascomycota_unclassified(100);p__Ascomycota_unclassified(100); | - | - | - | - | - | - | - | Unassigned | - |
| Otu24957 | 1 | k__Fungi(100);p__Ascomycota(100);p__Ascomycota_unclassified(100);p__Ascomycota_unclassified(100);p__Ascomycota_unclassified(100);p__Ascomycota_unclassified(100);p__Ascomycota_unclassified(100); | - | - | - | - | - | - | - | Unassigned | - |
| Otu24968 | 1 | k__Fungi(100);k__Fungi_unclassified(100);k__Fungi_unclassified(100);k__Fungi_unclassified(100);k__Fungi_unclassified(100);k__Fungi_unclassified(100);k__Fungi_unclassified(100); | - | - | - | - | - | - | - | Unassigned | - |
| Otu24969 | 1 | k__Fungi(100);k__Fungi_unclassified(100);k__Fungi_unclassified(100);k__Fungi_unclassified(100);k__Fungi_unclassified(100);k__Fungi_unclassified(100);k__Fungi_unclassified(100); | - | - | - | - | - | - | - | Unassigned | - |
| Otu24972 | 1 | k__Fungi(100);p__Ascomycota(100);p__Ascomycota_unclassified(100);p__Ascomycota_unclassified(100);p__Ascomycota_unclassified(100);p__Ascomycota_unclassified(100);p__Ascomycota_unclassified(100); | - | - | - | - | - | - | - | Unassigned | - |
| Otu24973 | 1 | k__Fungi(100);k__Fungi_unclassified(100);k__Fungi_unclassified(100);k__Fungi_unclassified(100);k__Fungi_unclassified(100);k__Fungi_unclassified(100);k__Fungi_unclassified(100); | - | - | - | - | - | - | - | Unassigned | - |
| Otu24974 | 1 | k__Fungi(100);p__Ascomycota(100);p__Ascomycota_unclassified(100);p__Ascomycota_unclassified(100);p__Ascomycota_unclassified(100);p__Ascomycota_unclassified(100);p__Ascomycota_unclassified(100); | - | - | - | - | - | - | - | Unassigned | - |
| Otu24975 | 1 | k__Fungi(100);p__Ascomycota(100);p__Ascomycota_unclassified(100);p__Ascomycota_unclassified(100);p__Ascomycota_unclassified(100);p__Ascomycota_unclassified(100);p__Ascomycota_unclassified(100); | - | - | - | - | - | - | - | Unassigned | - |
| Otu24976 | 1 | k__Fungi(100);k__Fungi_unclassified(100);k__Fungi_unclassified(100);k__Fungi_unclassified(100);k__Fungi_unclassified(100);k__Fungi_unclassified(100);k__Fungi_unclassified(100); | - | - | - | - | - | - | - | Unassigned | - |
| Otu24977 | 1 | k__Fungi(100);p__Ascomycota(100);p__Ascomycota_unclassified(100);p__Ascomycota_unclassified(100);p__Ascomycota_unclassified(100);p__Ascomycota_unclassified(100);p__Ascomycota_unclassified(100); | - | - | - | - | - | - | - | Unassigned | - |
| Otu24978 | 1 | k__Fungi(100);k__Fungi_unclassified(100);k__Fungi_unclassified(100);k__Fungi_unclassified(100);k__Fungi_unclassified(100);k__Fungi_unclassified(100);k__Fungi_unclassified(100); | - | - | - | - | - | - | - | Unassigned | - |
| Otu24979 | 1 | k__Fungi(100);p__Ascomycota(100);p__Ascomycota_unclassified(100);p__Ascomycota_unclassified(100);p__Ascomycota_unclassified(100);p__Ascomycota_unclassified(100);p__Ascomycota_unclassified(100); | - | - | - | - | - | - | - | Unassigned | - |
| Otu24980 | 1 | k__Fungi(100);p__Ascomycota(100);p__Ascomycota_unclassified(100);p__Ascomycota_unclassified(100);p__Ascomycota_unclassified(100);p__Ascomycota_unclassified(100);p__Ascomycota_unclassified(100); | - | - | - | - | - | - | - | Unassigned | - |
| Otu24981 | 1 | k__Fungi(100);p__Ascomycota(100);p__Ascomycota_unclassified(100);p__Ascomycota_unclassified(100);p__Ascomycota_unclassified(100);p__Ascomycota_unclassified(100);p__Ascomycota_unclassified(100); | - | - | - | - | - | - | - | Unassigned | - |
| Otu24982 | 1 | k__Fungi(100);p__Ascomycota(100);p__Ascomycota_unclassified(100);p__Ascomycota_unclassified(100);p__Ascomycota_unclassified(100);p__Ascomycota_unclassified(100);p__Ascomycota_unclassified(100); | - | - | - | - | - | - | - | Unassigned | - |
| Otu24983 | 1 | k__Fungi(100);k__Fungi_unclassified(100);k__Fungi_unclassified(100);k__Fungi_unclassified(100);k__Fungi_unclassified(100);k__Fungi_unclassified(100);k__Fungi_unclassified(100); | - | - | - | - | - | - | - | Unassigned | - |
| Otu24984 | 1 | k__Fungi(100);k__Fungi_unclassified(100);k__Fungi_unclassified(100);k__Fungi_unclassified(100);k__Fungi_unclassified(100);k__Fungi_unclassified(100);k__Fungi_unclassified(100); | - | - | - | - | - | - | - | Unassigned | - |
| Otu24985 | 1 | k__Fungi(100);k__Fungi_unclassified(100);k__Fungi_unclassified(100);k__Fungi_unclassified(100);k__Fungi_unclassified(100);k__Fungi_unclassified(100);k__Fungi_unclassified(100); | - | - | - | - | - | - | - | Unassigned | - |
| Otu24987 | 1 | k__Fungi(100);p__Ascomycota(100);p__Ascomycota_unclassified(100);p__Ascomycota_unclassified(100);p__Ascomycota_unclassified(100);p__Ascomycota_unclassified(100);p__Ascomycota_unclassified(100); | - | - | - | - | - | - | - | Unassigned | - |
| Otu24989 | 1 | k__Fungi(100);k__Fungi_unclassified(100);k__Fungi_unclassified(100);k__Fungi_unclassified(100);k__Fungi_unclassified(100);k__Fungi_unclassified(100);k__Fungi_unclassified(100); | - | - | - | - | - | - | - | Unassigned | - |
| Otu24994 | 1 | k__Fungi(100);k__Fungi_unclassified(100);k__Fungi_unclassified(100);k__Fungi_unclassified(100);k__Fungi_unclassified(100);k__Fungi_unclassified(100);k__Fungi_unclassified(100); | - | - | - | - | - | - | - | Unassigned | - |
| Otu24995 | 1 | k__Fungi(100);p__Ascomycota(100);p__Ascomycota_unclassified(100);p__Ascomycota_unclassified(100);p__Ascomycota_unclassified(100);p__Ascomycota_unclassified(100);p__Ascomycota_unclassified(100); | - | - | - | - | - | - | - | Unassigned | - |
| Otu24996 | 1 | k__Fungi(100);k__Fungi_unclassified(100);k__Fungi_unclassified(100);k__Fungi_unclassified(100);k__Fungi_unclassified(100);k__Fungi_unclassified(100);k__Fungi_unclassified(100); | - | - | - | - | - | - | - | Unassigned | - |
| Otu24997 | 1 | k__Fungi(100);p__Ascomycota(100);p__Ascomycota_unclassified(100);p__Ascomycota_unclassified(100);p__Ascomycota_unclassified(100);p__Ascomycota_unclassified(100);p__Ascomycota_unclassified(100); | - | - | - | - | - | - | - | Unassigned | - |
| Otu24998 | 1 | k__Fungi(100);k__Fungi_unclassified(100);k__Fungi_unclassified(100);k__Fungi_unclassified(100);k__Fungi_unclassified(100);k__Fungi_unclassified(100);k__Fungi_unclassified(100); | - | - | - | - | - | - | - | Unassigned | - |
| Otu25000 | 1 | k__Fungi(100);p__Ascomycota(100);p__Ascomycota_unclassified(100);p__Ascomycota_unclassified(100);p__Ascomycota_unclassified(100);p__Ascomycota_unclassified(100);p__Ascomycota_unclassified(100); | - | - | - | - | - | - | - | Unassigned | - |
| Otu25003 | 1 | k__Fungi(100);p__Ascomycota(100);p__Ascomycota_unclassified(100);p__Ascomycota_unclassified(100);p__Ascomycota_unclassified(100);p__Ascomycota_unclassified(100);p__Ascomycota_unclassified(100); | - | - | - | - | - | - | - | Unassigned | - |
| Otu25005 | 1 | k__Fungi(100);k__Fungi_unclassified(100);k__Fungi_unclassified(100);k__Fungi_unclassified(100);k__Fungi_unclassified(100);k__Fungi_unclassified(100);k__Fungi_unclassified(100); | - | - | - | - | - | - | - | Unassigned | - |
| Otu25007 | 1 | k__Fungi(100);p__Ascomycota(100);p__Ascomycota_unclassified(100);p__Ascomycota_unclassified(100);p__Ascomycota_unclassified(100);p__Ascomycota_unclassified(100);p__Ascomycota_unclassified(100); | - | - | - | - | - | - | - | Unassigned | - |
| Otu25008 | 1 | k__Fungi(100);k__Fungi_unclassified(100);k__Fungi_unclassified(100);k__Fungi_unclassified(100);k__Fungi_unclassified(100);k__Fungi_unclassified(100);k__Fungi_unclassified(100); | - | - | - | - | - | - | - | Unassigned | - |
| Otu25009 | 1 | k__Fungi(100);k__Fungi_unclassified(100);k__Fungi_unclassified(100);k__Fungi_unclassified(100);k__Fungi_unclassified(100);k__Fungi_unclassified(100);k__Fungi_unclassified(100); | - | - | - | - | - | - | - | Unassigned | - |
| Otu25010 | 1 | k__Fungi(100);k__Fungi_unclassified(100);k__Fungi_unclassified(100);k__Fungi_unclassified(100);k__Fungi_unclassified(100);k__Fungi_unclassified(100);k__Fungi_unclassified(100); | - | - | - | - | - | - | - | Unassigned | - |
| Otu25026 | 1 | k__Fungi(100);k__Fungi_unclassified(100);k__Fungi_unclassified(100);k__Fungi_unclassified(100);k__Fungi_unclassified(100);k__Fungi_unclassified(100);k__Fungi_unclassified(100); | - | - | - | - | - | - | - | Unassigned | - |
| Otu25028 | 1 | k__Fungi(100);k__Fungi_unclassified(100);k__Fungi_unclassified(100);k__Fungi_unclassified(100);k__Fungi_unclassified(100);k__Fungi_unclassified(100);k__Fungi_unclassified(100); | - | - | - | - | - | - | - | Unassigned | - |
| Otu25029 | 1 | k__Fungi(100);p__Ascomycota(100);p__Ascomycota_unclassified(100);p__Ascomycota_unclassified(100);p__Ascomycota_unclassified(100);p__Ascomycota_unclassified(100);p__Ascomycota_unclassified(100); | - | - | - | - | - | - | - | Unassigned | - |
| Otu25030 | 1 | k__Fungi(100);p__Ascomycota(100);p__Ascomycota_unclassified(100);p__Ascomycota_unclassified(100);p__Ascomycota_unclassified(100);p__Ascomycota_unclassified(100);p__Ascomycota_unclassified(100); | - | - | - | - | - | - | - | Unassigned | - |
| Otu25031 | 1 | k__Fungi(100);k__Fungi_unclassified(100);k__Fungi_unclassified(100);k__Fungi_unclassified(100);k__Fungi_unclassified(100);k__Fungi_unclassified(100);k__Fungi_unclassified(100); | - | - | - | - | - | - | - | Unassigned | - |
| Otu25032 | 1 | k__Fungi(100);p__Ascomycota(100);p__Ascomycota_unclassified(100);p__Ascomycota_unclassified(100);p__Ascomycota_unclassified(100);p__Ascomycota_unclassified(100);p__Ascomycota_unclassified(100); | - | - | - | - | - | - | - | Unassigned | - |
| Otu25033 | 1 | k__Fungi(100);p__Ascomycota(100);p__Ascomycota_unclassified(100);p__Ascomycota_unclassified(100);p__Ascomycota_unclassified(100);p__Ascomycota_unclassified(100);p__Ascomycota_unclassified(100); | - | - | - | - | - | - | - | Unassigned | - |
| Otu25035 | 1 | k__Fungi(100);p__Ascomycota(100);p__Ascomycota_unclassified(100);p__Ascomycota_unclassified(100);p__Ascomycota_unclassified(100);p__Ascomycota_unclassified(100);p__Ascomycota_unclassified(100); | - | - | - | - | - | - | - | Unassigned | - |
| Otu25051 | 1 | k__Fungi(100);p__Ascomycota(100);p__Ascomycota_unclassified(100);p__Ascomycota_unclassified(100);p__Ascomycota_unclassified(100);p__Ascomycota_unclassified(100);p__Ascomycota_unclassified(100); | - | - | - | - | - | - | - | Unassigned | - |
| Otu25054 | 1 | k__Fungi(100);k__Fungi_unclassified(100);k__Fungi_unclassified(100);k__Fungi_unclassified(100);k__Fungi_unclassified(100);k__Fungi_unclassified(100);k__Fungi_unclassified(100); | - | - | - | - | - | - | - | Unassigned | - |
| Otu25055 | 1 | k__Fungi(100);k__Fungi_unclassified(100);k__Fungi_unclassified(100);k__Fungi_unclassified(100);k__Fungi_unclassified(100);k__Fungi_unclassified(100);k__Fungi_unclassified(100); | - | - | - | - | - | - | - | Unassigned | - |
| Otu25056 | 1 | k__Fungi(100);k__Fungi_unclassified(100);k__Fungi_unclassified(100);k__Fungi_unclassified(100);k__Fungi_unclassified(100);k__Fungi_unclassified(100);k__Fungi_unclassified(100); | - | - | - | - | - | - | - | Unassigned | - |
| Otu25058 | 1 | k__Fungi(100);p__Ascomycota(100);p__Ascomycota_unclassified(100);p__Ascomycota_unclassified(100);p__Ascomycota_unclassified(100);p__Ascomycota_unclassified(100);p__Ascomycota_unclassified(100); | - | - | - | - | - | - | - | Unassigned | - |
| Otu25059 | 1 | k__Fungi(100);k__Fungi_unclassified(100);k__Fungi_unclassified(100);k__Fungi_unclassified(100);k__Fungi_unclassified(100);k__Fungi_unclassified(100);k__Fungi_unclassified(100); | - | - | - | - | - | - | - | Unassigned | - |
| Otu25062 | 1 | k__Fungi(100);k__Fungi_unclassified(100);k__Fungi_unclassified(100);k__Fungi_unclassified(100);k__Fungi_unclassified(100);k__Fungi_unclassified(100);k__Fungi_unclassified(100); | - | - | - | - | - | - | - | Unassigned | - |
| Otu25064 | 1 | k__Fungi(100);k__Fungi_unclassified(100);k__Fungi_unclassified(100);k__Fungi_unclassified(100);k__Fungi_unclassified(100);k__Fungi_unclassified(100);k__Fungi_unclassified(100); | - | - | - | - | - | - | - | Unassigned | - |
| Otu25066 | 1 | k__Fungi(100);k__Fungi_unclassified(100);k__Fungi_unclassified(100);k__Fungi_unclassified(100);k__Fungi_unclassified(100);k__Fungi_unclassified(100);k__Fungi_unclassified(100); | - | - | - | - | - | - | - | Unassigned | - |
| Otu25068 | 1 | k__Fungi(100);p__Ascomycota(100);p__Ascomycota_unclassified(100);p__Ascomycota_unclassified(100);p__Ascomycota_unclassified(100);p__Ascomycota_unclassified(100);p__Ascomycota_unclassified(100); | - | - | - | - | - | - | - | Unassigned | - |
| Otu25069 | 1 | k__Fungi(100);k__Fungi_unclassified(100);k__Fungi_unclassified(100);k__Fungi_unclassified(100);k__Fungi_unclassified(100);k__Fungi_unclassified(100);k__Fungi_unclassified(100); | - | - | - | - | - | - | - | Unassigned | - |
| Otu25071 | 1 | k__Fungi(100);k__Fungi_unclassified(100);k__Fungi_unclassified(100);k__Fungi_unclassified(100);k__Fungi_unclassified(100);k__Fungi_unclassified(100);k__Fungi_unclassified(100); | - | - | - | - | - | - | - | Unassigned | - |
| Otu25072 | 1 | k__Fungi(100);k__Fungi_unclassified(100);k__Fungi_unclassified(100);k__Fungi_unclassified(100);k__Fungi_unclassified(100);k__Fungi_unclassified(100);k__Fungi_unclassified(100); | - | - | - | - | - | - | - | Unassigned | - |
| Otu25081 | 1 | k__Fungi(100);p__Ascomycota(100);c__Dothideomycetes(100);c__Dothideomycetes_unclassified(100);c__Dothideomycetes_unclassified(100);c__Dothideomycetes_unclassified(100);c__Dothideomycetes_unclassified(100); | - | - | - | - | - | - | - | Unassigned | - |
| Otu25096 | 1 | k__Fungi(100);p__Ascomycota(100);p__Ascomycota_unclassified(100);p__Ascomycota_unclassified(100);p__Ascomycota_unclassified(100);p__Ascomycota_unclassified(100);p__Ascomycota_unclassified(100); | - | - | - | - | - | - | - | Unassigned | - |
| Otu25097 | 1 | k__Fungi(100);k__Fungi_unclassified(100);k__Fungi_unclassified(100);k__Fungi_unclassified(100);k__Fungi_unclassified(100);k__Fungi_unclassified(100);k__Fungi_unclassified(100); | - | - | - | - | - | - | - | Unassigned | - |
| Otu25099 | 1 | k__Fungi(100);p__Ascomycota(100);p__Ascomycota_unclassified(100);p__Ascomycota_unclassified(100);p__Ascomycota_unclassified(100);p__Ascomycota_unclassified(100);p__Ascomycota_unclassified(100); | - | - | - | - | - | - | - | Unassigned | - |
| Otu25123 | 1 | k__Fungi(100);k__Fungi_unclassified(100);k__Fungi_unclassified(100);k__Fungi_unclassified(100);k__Fungi_unclassified(100);k__Fungi_unclassified(100);k__Fungi_unclassified(100); | - | - | - | - | - | - | - | Unassigned | - |
| Otu25125 | 1 | k__Fungi(100);k__Fungi_unclassified(100);k__Fungi_unclassified(100);k__Fungi_unclassified(100);k__Fungi_unclassified(100);k__Fungi_unclassified(100);k__Fungi_unclassified(100); | - | - | - | - | - | - | - | Unassigned | - |
| Otu25126 | 1 | k__Fungi(100);p__Ascomycota(100);p__Ascomycota_unclassified(100);p__Ascomycota_unclassified(100);p__Ascomycota_unclassified(100);p__Ascomycota_unclassified(100);p__Ascomycota_unclassified(100); | - | - | - | - | - | - | - | Unassigned | - |
| Otu25135 | 1 | k__Fungi(100);p__Ascomycota(100);p__Ascomycota_unclassified(100);p__Ascomycota_unclassified(100);p__Ascomycota_unclassified(100);p__Ascomycota_unclassified(100);p__Ascomycota_unclassified(100); | - | - | - | - | - | - | - | Unassigned | - |
| Otu25141 | 1 | k__Fungi(100);k__Fungi_unclassified(100);k__Fungi_unclassified(100);k__Fungi_unclassified(100);k__Fungi_unclassified(100);k__Fungi_unclassified(100);k__Fungi_unclassified(100); | - | - | - | - | - | - | - | Unassigned | - |
| Otu25143 | 1 | k__Fungi(100);k__Fungi_unclassified(100);k__Fungi_unclassified(100);k__Fungi_unclassified(100);k__Fungi_unclassified(100);k__Fungi_unclassified(100);k__Fungi_unclassified(100); | - | - | - | - | - | - | - | Unassigned | - |
| Otu25144 | 1 | k__Fungi(100);k__Fungi_unclassified(100);k__Fungi_unclassified(100);k__Fungi_unclassified(100);k__Fungi_unclassified(100);k__Fungi_unclassified(100);k__Fungi_unclassified(100); | - | - | - | - | - | - | - | Unassigned | - |
| Otu25160 | 1 | k__Fungi(100);k__Fungi_unclassified(100);k__Fungi_unclassified(100);k__Fungi_unclassified(100);k__Fungi_unclassified(100);k__Fungi_unclassified(100);k__Fungi_unclassified(100); | - | - | - | - | - | - | - | Unassigned | - |
| Otu25162 | 1 | k__Fungi(100);k__Fungi_unclassified(100);k__Fungi_unclassified(100);k__Fungi_unclassified(100);k__Fungi_unclassified(100);k__Fungi_unclassified(100);k__Fungi_unclassified(100); | - | - | - | - | - | - | - | Unassigned | - |
| Otu25165 | 1 | k__Fungi(100);k__Fungi_unclassified(100);k__Fungi_unclassified(100);k__Fungi_unclassified(100);k__Fungi_unclassified(100);k__Fungi_unclassified(100);k__Fungi_unclassified(100); | - | - | - | - | - | - | - | Unassigned | - |
| Otu25167 | 1 | k__Fungi(100);k__Fungi_unclassified(100);k__Fungi_unclassified(100);k__Fungi_unclassified(100);k__Fungi_unclassified(100);k__Fungi_unclassified(100);k__Fungi_unclassified(100); | - | - | - | - | - | - | - | Unassigned | - |
| Otu25168 | 1 | k__Fungi(100);k__Fungi_unclassified(100);k__Fungi_unclassified(100);k__Fungi_unclassified(100);k__Fungi_unclassified(100);k__Fungi_unclassified(100);k__Fungi_unclassified(100); | - | - | - | - | - | - | - | Unassigned | - |
| Otu25169 | 1 | k__Fungi(100);k__Fungi_unclassified(100);k__Fungi_unclassified(100);k__Fungi_unclassified(100);k__Fungi_unclassified(100);k__Fungi_unclassified(100);k__Fungi_unclassified(100); | - | - | - | - | - | - | - | Unassigned | - |
| Otu25172 | 1 | k__Fungi(100);p__Ascomycota(100);p__Ascomycota_unclassified(100);p__Ascomycota_unclassified(100);p__Ascomycota_unclassified(100);p__Ascomycota_unclassified(100);p__Ascomycota_unclassified(100); | - | - | - | - | - | - | - | Unassigned | - |
| Otu25174 | 1 | k__Fungi(100);k__Fungi_unclassified(100);k__Fungi_unclassified(100);k__Fungi_unclassified(100);k__Fungi_unclassified(100);k__Fungi_unclassified(100);k__Fungi_unclassified(100); | - | - | - | - | - | - | - | Unassigned | - |
| Otu25177 | 1 | k__Fungi(100);p__Ascomycota(100);p__Ascomycota_unclassified(100);p__Ascomycota_unclassified(100);p__Ascomycota_unclassified(100);p__Ascomycota_unclassified(100);p__Ascomycota_unclassified(100); | - | - | - | - | - | - | - | Unassigned | - |
| Otu25178 | 1 | k__Fungi(100);p__Ascomycota(100);p__Ascomycota_unclassified(100);p__Ascomycota_unclassified(100);p__Ascomycota_unclassified(100);p__Ascomycota_unclassified(100);p__Ascomycota_unclassified(100); | - | - | - | - | - | - | - | Unassigned | - |
| Otu25179 | 1 | k__Fungi(100);k__Fungi_unclassified(100);k__Fungi_unclassified(100);k__Fungi_unclassified(100);k__Fungi_unclassified(100);k__Fungi_unclassified(100);k__Fungi_unclassified(100); | - | - | - | - | - | - | - | Unassigned | - |
| Otu25181 | 1 | k__Fungi(100);k__Fungi_unclassified(100);k__Fungi_unclassified(100);k__Fungi_unclassified(100);k__Fungi_unclassified(100);k__Fungi_unclassified(100);k__Fungi_unclassified(100); | - | - | - | - | - | - | - | Unassigned | - |
| Otu25183 | 1 | k__Fungi(100);k__Fungi_unclassified(100);k__Fungi_unclassified(100);k__Fungi_unclassified(100);k__Fungi_unclassified(100);k__Fungi_unclassified(100);k__Fungi_unclassified(100); | - | - | - | - | - | - | - | Unassigned | - |
| Otu25185 | 1 | k__Fungi(100);p__Ascomycota(100);p__Ascomycota_unclassified(100);p__Ascomycota_unclassified(100);p__Ascomycota_unclassified(100);p__Ascomycota_unclassified(100);p__Ascomycota_unclassified(100); | - | - | - | - | - | - | - | Unassigned | - |
| Otu25196 | 1 | k__Fungi(100);k__Fungi_unclassified(100);k__Fungi_unclassified(100);k__Fungi_unclassified(100);k__Fungi_unclassified(100);k__Fungi_unclassified(100);k__Fungi_unclassified(100); | - | - | - | - | - | - | - | Unassigned | - |
| Otu25197 | 1 | k__Fungi(100);p__Ascomycota(100);p__Ascomycota_unclassified(100);p__Ascomycota_unclassified(100);p__Ascomycota_unclassified(100);p__Ascomycota_unclassified(100);p__Ascomycota_unclassified(100); | - | - | - | - | - | - | - | Unassigned | - |
| Otu25199 | 1 | k__Fungi(100);p__Ascomycota(100);p__Ascomycota_unclassified(100);p__Ascomycota_unclassified(100);p__Ascomycota_unclassified(100);p__Ascomycota_unclassified(100);p__Ascomycota_unclassified(100); | - | - | - | - | - | - | - | Unassigned | - |
| Otu25201 | 1 | k__Fungi(100);p__Ascomycota(100);p__Ascomycota_unclassified(100);p__Ascomycota_unclassified(100);p__Ascomycota_unclassified(100);p__Ascomycota_unclassified(100);p__Ascomycota_unclassified(100); | - | - | - | - | - | - | - | Unassigned | - |
| Otu25202 | 1 | k__Fungi(100);p__Ascomycota(100);p__Ascomycota_unclassified(100);p__Ascomycota_unclassified(100);p__Ascomycota_unclassified(100);p__Ascomycota_unclassified(100);p__Ascomycota_unclassified(100); | - | - | - | - | - | - | - | Unassigned | - |
| Otu25216 | 1 | k__Fungi(100);k__Fungi_unclassified(100);k__Fungi_unclassified(100);k__Fungi_unclassified(100);k__Fungi_unclassified(100);k__Fungi_unclassified(100);k__Fungi_unclassified(100); | - | - | - | - | - | - | - | Unassigned | - |
| Otu25217 | 1 | k__Fungi(100);k__Fungi_unclassified(100);k__Fungi_unclassified(100);k__Fungi_unclassified(100);k__Fungi_unclassified(100);k__Fungi_unclassified(100);k__Fungi_unclassified(100); | - | - | - | - | - | - | - | Unassigned | - |
| Otu25222 | 1 | k__Fungi(100);k__Fungi_unclassified(100);k__Fungi_unclassified(100);k__Fungi_unclassified(100);k__Fungi_unclassified(100);k__Fungi_unclassified(100);k__Fungi_unclassified(100); | - | - | - | - | - | - | - | Unassigned | - |
| Otu25225 | 1 | k__Fungi(100);k__Fungi_unclassified(100);k__Fungi_unclassified(100);k__Fungi_unclassified(100);k__Fungi_unclassified(100);k__Fungi_unclassified(100);k__Fungi_unclassified(100); | - | - | - | - | - | - | - | Unassigned | - |
| Otu25226 | 1 | k__Fungi(100);p__Ascomycota(100);p__Ascomycota_unclassified(100);p__Ascomycota_unclassified(100);p__Ascomycota_unclassified(100);p__Ascomycota_unclassified(100);p__Ascomycota_unclassified(100); | - | - | - | - | - | - | - | Unassigned | - |
| Otu25228 | 1 | k__Fungi(100);k__Fungi_unclassified(100);k__Fungi_unclassified(100);k__Fungi_unclassified(100);k__Fungi_unclassified(100);k__Fungi_unclassified(100);k__Fungi_unclassified(100); | - | - | - | - | - | - | - | Unassigned | - |
| Otu25231 | 1 | k__Fungi(100);k__Fungi_unclassified(100);k__Fungi_unclassified(100);k__Fungi_unclassified(100);k__Fungi_unclassified(100);k__Fungi_unclassified(100);k__Fungi_unclassified(100); | - | - | - | - | - | - | - | Unassigned | - |
| Otu25233 | 1 | k__Fungi(100);p__Ascomycota(100);p__Ascomycota_unclassified(100);p__Ascomycota_unclassified(100);p__Ascomycota_unclassified(100);p__Ascomycota_unclassified(100);p__Ascomycota_unclassified(100); | - | - | - | - | - | - | - | Unassigned | - |
| Otu25236 | 1 | k__Fungi(100);k__Fungi_unclassified(100);k__Fungi_unclassified(100);k__Fungi_unclassified(100);k__Fungi_unclassified(100);k__Fungi_unclassified(100);k__Fungi_unclassified(100); | - | - | - | - | - | - | - | Unassigned | - |
| Otu25237 | 1 | k__Fungi(100);k__Fungi_unclassified(100);k__Fungi_unclassified(100);k__Fungi_unclassified(100);k__Fungi_unclassified(100);k__Fungi_unclassified(100);k__Fungi_unclassified(100); | - | - | - | - | - | - | - | Unassigned | - |
| Otu25238 | 1 | k__Fungi(100);k__Fungi_unclassified(100);k__Fungi_unclassified(100);k__Fungi_unclassified(100);k__Fungi_unclassified(100);k__Fungi_unclassified(100);k__Fungi_unclassified(100); | - | - | - | - | - | - | - | Unassigned | - |
| Otu25241 | 1 | k__Fungi(100);k__Fungi_unclassified(100);k__Fungi_unclassified(100);k__Fungi_unclassified(100);k__Fungi_unclassified(100);k__Fungi_unclassified(100);k__Fungi_unclassified(100); | - | - | - | - | - | - | - | Unassigned | - |
| Otu25242 | 1 | k__Fungi(100);k__Fungi_unclassified(100);k__Fungi_unclassified(100);k__Fungi_unclassified(100);k__Fungi_unclassified(100);k__Fungi_unclassified(100);k__Fungi_unclassified(100); | - | - | - | - | - | - | - | Unassigned | - |
| Otu25244 | 1 | k__Fungi(100);k__Fungi_unclassified(100);k__Fungi_unclassified(100);k__Fungi_unclassified(100);k__Fungi_unclassified(100);k__Fungi_unclassified(100);k__Fungi_unclassified(100); | - | - | - | - | - | - | - | Unassigned | - |
| Otu25267 | 1 | k__Fungi(100);p__Ascomycota(100);p__Ascomycota_unclassified(100);p__Ascomycota_unclassified(100);p__Ascomycota_unclassified(100);p__Ascomycota_unclassified(100);p__Ascomycota_unclassified(100); | - | - | - | - | - | - | - | Unassigned | - |
| Otu25268 | 1 | k__Fungi(100);p__Ascomycota(100);p__Ascomycota_unclassified(100);p__Ascomycota_unclassified(100);p__Ascomycota_unclassified(100);p__Ascomycota_unclassified(100);p__Ascomycota_unclassified(100); | - | - | - | - | - | - | - | Unassigned | - |
| Otu25275 | 1 | k__Fungi(100);k__Fungi_unclassified(100);k__Fungi_unclassified(100);k__Fungi_unclassified(100);k__Fungi_unclassified(100);k__Fungi_unclassified(100);k__Fungi_unclassified(100); | - | - | - | - | - | - | - | Unassigned | - |
| Otu25280 | 1 | k__Fungi(100);k__Fungi_unclassified(100);k__Fungi_unclassified(100);k__Fungi_unclassified(100);k__Fungi_unclassified(100);k__Fungi_unclassified(100);k__Fungi_unclassified(100); | - | - | - | - | - | - | - | Unassigned | - |
| Otu25281 | 1 | k__Fungi(100);p__Ascomycota(100);p__Ascomycota_unclassified(100);p__Ascomycota_unclassified(100);p__Ascomycota_unclassified(100);p__Ascomycota_unclassified(100);p__Ascomycota_unclassified(100); | - | - | - | - | - | - | - | Unassigned | - |

| | | | | | | | | | | | |
|---|---|---|---|---|---|---|---|---|---|---|---|
| Otu25299 | 1 | k__Plantae(100);p__unclassified_Plantae(100);c__unclassified_Plantae(100);o__unclassified_Plantae(100);f__unclassified_Plantae(100);g__unclassified_Plantae(100);s__Plantae_sp(100); | - | - | - | - | - | - | - | Unassigned | - |
| Otu25302 | 1 | k__Fungi(100);p__Ascomycota(100);p__Ascomycota_unclassified(100);p__Ascomycota_unclassified(100);p__Ascomycota_unclassified(100);p__Ascomycota_unclassified(100);p__Ascomycota_unclassified(100); | - | - | - | - | - | - | - | Unassigned | - |
| Otu25307 | 1 | k__Fungi(100);p__Ascomycota(100);p__Ascomycota_unclassified(100);p__Ascomycota_unclassified(100);p__Ascomycota_unclassified(100);p__Ascomycota_unclassified(100);p__Ascomycota_unclassified(100); | - | - | - | - | - | - | - | Unassigned | - |
| Otu25308 | 1 | k__Fungi(100);k__Fungi_unclassified(100);k__Fungi_unclassified(100);k__Fungi_unclassified(100);k__Fungi_unclassified(100);k__Fungi_unclassified(100);k__Fungi_unclassified(100); | - | - | - | - | - | - | - | Unassigned | - |
| Otu25309 | 1 | k__Fungi(100);k__Fungi_unclassified(100);k__Fungi_unclassified(100);k__Fungi_unclassified(100);k__Fungi_unclassified(100);k__Fungi_unclassified(100);k__Fungi_unclassified(100); | - | - | - | - | - | - | - | Unassigned | - |
| Otu25310 | 1 | k__Fungi(100);p__Ascomycota(100);p__Ascomycota_unclassified(100);p__Ascomycota_unclassified(100);p__Ascomycota_unclassified(100);p__Ascomycota_unclassified(100);p__Ascomycota_unclassified(100); | - | - | - | - | - | - | - | Unassigned | - |
| Otu25311 | 1 | k__Fungi(100);p__Ascomycota(100);p__Ascomycota_unclassified(100);p__Ascomycota_unclassified(100);p__Ascomycota_unclassified(100);p__Ascomycota_unclassified(100);p__Ascomycota_unclassified(100); | - | - | - | - | - | - | - | Unassigned | - |
| Otu25312 | 1 | k__Fungi(100);k__Fungi_unclassified(100);k__Fungi_unclassified(100);k__Fungi_unclassified(100);k__Fungi_unclassified(100);k__Fungi_unclassified(100);k__Fungi_unclassified(100); | - | - | - | - | - | - | - | Unassigned | - |
| Otu25313 | 1 | k__Fungi(100);k__Fungi_unclassified(100);k__Fungi_unclassified(100);k__Fungi_unclassified(100);k__Fungi_unclassified(100);k__Fungi_unclassified(100);k__Fungi_unclassified(100); | - | - | - | - | - | - | - | Unassigned | - |
| Otu25314 | 1 | k__Fungi(100);p__Ascomycota(100);p__Ascomycota_unclassified(100);p__Ascomycota_unclassified(100);p__Ascomycota_unclassified(100);p__Ascomycota_unclassified(100);p__Ascomycota_unclassified(100); | - | - | - | - | - | - | - | Unassigned | - |
| Otu25315 | 1 | k__Fungi(100);p__Ascomycota(100);p__Ascomycota_unclassified(100);p__Ascomycota_unclassified(100);p__Ascomycota_unclassified(100);p__Ascomycota_unclassified(100);p__Ascomycota_unclassified(100); | - | - | - | - | - | - | - | Unassigned | - |
| Otu25316 | 1 | k__Fungi(100);p__Ascomycota(100);p__Ascomycota_unclassified(100);p__Ascomycota_unclassified(100);p__Ascomycota_unclassified(100);p__Ascomycota_unclassified(100);p__Ascomycota_unclassified(100); | - | - | - | - | - | - | - | Unassigned | - |
| Otu25317 | 1 | k__Fungi(100);k__Fungi_unclassified(100);k__Fungi_unclassified(100);k__Fungi_unclassified(100);k__Fungi_unclassified(100);k__Fungi_unclassified(100);k__Fungi_unclassified(100); | - | - | - | - | - | - | - | Unassigned | - |
| Otu25318 | 1 | k__Fungi(100);p__Ascomycota(100);p__Ascomycota_unclassified(100);p__Ascomycota_unclassified(100);p__Ascomycota_unclassified(100);p__Ascomycota_unclassified(100);p__Ascomycota_unclassified(100); | - | - | - | - | - | - | - | Unassigned | - |
| Otu25319 | 1 | k__Fungi(100);p__Ascomycota(100);p__Ascomycota_unclassified(100);p__Ascomycota_unclassified(100);p__Ascomycota_unclassified(100);p__Ascomycota_unclassified(100);p__Ascomycota_unclassified(100); | - | - | - | - | - | - | - | Unassigned | - |
| Otu25320 | 1 | k__Fungi(100);k__Fungi_unclassified(100);k__Fungi_unclassified(100);k__Fungi_unclassified(100);k__Fungi_unclassified(100);k__Fungi_unclassified(100);k__Fungi_unclassified(100); | - | - | - | - | - | - | - | Unassigned | - |
| Otu25321 | 1 | k__Fungi(100);p__Ascomycota(100);p__Ascomycota_unclassified(100);p__Ascomycota_unclassified(100);p__Ascomycota_unclassified(100);p__Ascomycota_unclassified(100);p__Ascomycota_unclassified(100); | - | - | - | - | - | - | - | Unassigned | - |
| Otu25322 | 1 | k__Fungi(100);p__Ascomycota(100);p__Ascomycota_unclassified(100);p__Ascomycota_unclassified(100);p__Ascomycota_unclassified(100);p__Ascomycota_unclassified(100);p__Ascomycota_unclassified(100); | - | - | - | - | - | - | - | Unassigned | - |
| Otu25323 | 1 | k__Fungi(100);p__Ascomycota(100);p__Ascomycota_unclassified(100);p__Ascomycota_unclassified(100);p__Ascomycota_unclassified(100);p__Ascomycota_unclassified(100);p__Ascomycota_unclassified(100); | - | - | - | - | - | - | - | Unassigned | - |
| Otu25324 | 1 | k__Fungi(100);p__Ascomycota(100);p__Ascomycota_unclassified(100);p__Ascomycota_unclassified(100);p__Ascomycota_unclassified(100);p__Ascomycota_unclassified(100);p__Ascomycota_unclassified(100); | - | - | - | - | - | - | - | Unassigned | - |
| Otu25325 | 1 | k__Fungi(100);p__Ascomycota(100);p__Ascomycota_unclassified(100);p__Ascomycota_unclassified(100);p__Ascomycota_unclassified(100);p__Ascomycota_unclassified(100);p__Ascomycota_unclassified(100); | - | - | - | - | - | - | - | Unassigned | - |
| Otu25326 | 1 | k__Fungi(100);k__Fungi_unclassified(100);k__Fungi_unclassified(100);k__Fungi_unclassified(100);k__Fungi_unclassified(100);k__Fungi_unclassified(100);k__Fungi_unclassified(100); | - | - | - | - | - | - | - | Unassigned | - |
| Otu25327 | 1 | k__Fungi(100);p__Ascomycota(100);p__Ascomycota_unclassified(100);p__Ascomycota_unclassified(100);p__Ascomycota_unclassified(100);p__Ascomycota_unclassified(100);p__Ascomycota_unclassified(100); | - | - | - | - | - | - | - | Unassigned | - |
| Otu25328 | 1 | k__Fungi(100);p__Ascomycota(100);p__Ascomycota_unclassified(100);p__Ascomycota_unclassified(100);p__Ascomycota_unclassified(100);p__Ascomycota_unclassified(100);p__Ascomycota_unclassified(100); | - | - | - | - | - | - | - | Unassigned | - |
| Otu25332 | 1 | k__Fungi(100);k__Fungi_unclassified(100);k__Fungi_unclassified(100);k__Fungi_unclassified(100);k__Fungi_unclassified(100);k__Fungi_unclassified(100);k__Fungi_unclassified(100); | - | - | - | - | - | - | - | Unassigned | - |
| Otu25335 | 1 | k__Fungi(100);p__Ascomycota(100);p__Ascomycota_unclassified(100);p__Ascomycota_unclassified(100);p__Ascomycota_unclassified(100);p__Ascomycota_unclassified(100);p__Ascomycota_unclassified(100); | - | - | - | - | - | - | - | Unassigned | - |
| Otu25343 | 1 | k__Fungi(100);p__Ascomycota(100);p__Ascomycota_unclassified(100);p__Ascomycota_unclassified(100);p__Ascomycota_unclassified(100);p__Ascomycota_unclassified(100);p__Ascomycota_unclassified(100); | - | - | - | - | - | - | - | Unassigned | - |
| Otu25347 | 1 | k__Fungi(100);p__Ascomycota(100);p__Ascomycota_unclassified(100);p__Ascomycota_unclassified(100);p__Ascomycota_unclassified(100);p__Ascomycota_unclassified(100);p__Ascomycota_unclassified(100); | - | - | - | - | - | - | - | Unassigned | - |
| Otu25348 | 1 | k__Fungi(100);p__Ascomycota(100);p__Ascomycota_unclassified(100);p__Ascomycota_unclassified(100);p__Ascomycota_unclassified(100);p__Ascomycota_unclassified(100);p__Ascomycota_unclassified(100); | - | - | - | - | - | - | - | Unassigned | - |
| Otu25355 | 1 | k__Fungi(100);k__Fungi_unclassified(100);k__Fungi_unclassified(100);k__Fungi_unclassified(100);k__Fungi_unclassified(100);k__Fungi_unclassified(100);k__Fungi_unclassified(100); | - | - | - | - | - | - | - | Unassigned | - |
| Otu25356 | 1 | k__Fungi(100);k__Fungi_unclassified(100);k__Fungi_unclassified(100);k__Fungi_unclassified(100);k__Fungi_unclassified(100);k__Fungi_unclassified(100);k__Fungi_unclassified(100); | - | - | - | - | - | - | - | Unassigned | - |
| Otu25357 | 1 | k__Fungi(100);p__Ascomycota(100);p__Ascomycota_unclassified(100);p__Ascomycota_unclassified(100);p__Ascomycota_unclassified(100);p__Ascomycota_unclassified(100);p__Ascomycota_unclassified(100); | - | - | - | - | - | - | - | Unassigned | - |
| Otu25361 | 1 | k__Fungi(100);k__Fungi_unclassified(100);k__Fungi_unclassified(100);k__Fungi_unclassified(100);k__Fungi_unclassified(100);k__Fungi_unclassified(100);k__Fungi_unclassified(100); | - | - | - | - | - | - | - | Unassigned | - |
| Otu25367 | 1 | k__Fungi(100);p__Ascomycota(100);p__Ascomycota_unclassified(100);p__Ascomycota_unclassified(100);p__Ascomycota_unclassified(100);p__Ascomycota_unclassified(100);p__Ascomycota_unclassified(100); | - | - | - | - | - | - | - | Unassigned | - |
| Otu25368 | 1 | k__Fungi(100);p__Ascomycota(100);p__Ascomycota_unclassified(100);p__Ascomycota_unclassified(100);p__Ascomycota_unclassified(100);p__Ascomycota_unclassified(100);p__Ascomycota_unclassified(100); | - | - | - | - | - | - | - | Unassigned | - |
| Otu25369 | 1 | k__Fungi(100);k__Fungi_unclassified(100);k__Fungi_unclassified(100);k__Fungi_unclassified(100);k__Fungi_unclassified(100);k__Fungi_unclassified(100);k__Fungi_unclassified(100); | - | - | - | - | - | - | - | Unassigned | - |
| Otu25371 | 1 | k__Fungi(100);k__Fungi_unclassified(100);k__Fungi_unclassified(100);k__Fungi_unclassified(100);k__Fungi_unclassified(100);k__Fungi_unclassified(100);k__Fungi_unclassified(100); | - | - | - | - | - | - | - | Unassigned | - |
| Otu25372 | 1 | k__Fungi(100);k__Fungi_unclassified(100);k__Fungi_unclassified(100);k__Fungi_unclassified(100);k__Fungi_unclassified(100);k__Fungi_unclassified(100);k__Fungi_unclassified(100); | - | - | - | - | - | - | - | Unassigned | - |
| Otu25379 | 1 | k__Fungi(100);k__Fungi_unclassified(100);k__Fungi_unclassified(100);k__Fungi_unclassified(100);k__Fungi_unclassified(100);k__Fungi_unclassified(100);k__Fungi_unclassified(100); | - | - | - | - | - | - | - | Unassigned | - |
| Otu25382 | 1 | k__Fungi(100);k__Fungi_unclassified(100);k__Fungi_unclassified(100);k__Fungi_unclassified(100);k__Fungi_unclassified(100);k__Fungi_unclassified(100);k__Fungi_unclassified(100); | - | - | - | - | - | - | - | Unassigned | - |
| Otu25383 | 1 | k__Fungi(100);k__Fungi_unclassified(100);k__Fungi_unclassified(100);k__Fungi_unclassified(100);k__Fungi_unclassified(100);k__Fungi_unclassified(100);k__Fungi_unclassified(100); | - | - | - | - | - | - | - | Unassigned | - |
| Otu25384 | 1 | k__Fungi(100);k__Fungi_unclassified(100);k__Fungi_unclassified(100);k__Fungi_unclassified(100);k__Fungi_unclassified(100);k__Fungi_unclassified(100);k__Fungi_unclassified(100); | - | - | - | - | - | - | - | Unassigned | - |
| Otu25386 | 1 | k__Fungi(100);k__Fungi_unclassified(100);k__Fungi_unclassified(100);k__Fungi_unclassified(100);k__Fungi_unclassified(100);k__Fungi_unclassified(100);k__Fungi_unclassified(100); | - | - | - | - | - | - | - | Unassigned | - |
| Otu25387 | 1 | k__Fungi(100);k__Fungi_unclassified(100);k__Fungi_unclassified(100);k__Fungi_unclassified(100);k__Fungi_unclassified(100);k__Fungi_unclassified(100);k__Fungi_unclassified(100); | - | - | - | - | - | - | - | Unassigned | - |
| Otu25388 | 1 | k__Fungi(100);p__Ascomycota(100);p__Ascomycota_unclassified(100);p__Ascomycota_unclassified(100);p__Ascomycota_unclassified(100);p__Ascomycota_unclassified(100);p__Ascomycota_unclassified(100); | - | - | - | - | - | - | - | Unassigned | - |
| Otu25389 | 1 | k__Fungi(100);k__Fungi_unclassified(100);k__Fungi_unclassified(100);k__Fungi_unclassified(100);k__Fungi_unclassified(100);k__Fungi_unclassified(100);k__Fungi_unclassified(100); | - | - | - | - | - | - | - | Unassigned | - |
| Otu25390 | 1 | k__Fungi(100);p__Ascomycota(100);p__Ascomycota_unclassified(100);p__Ascomycota_unclassified(100);p__Ascomycota_unclassified(100);p__Ascomycota_unclassified(100);p__Ascomycota_unclassified(100); | - | - | - | - | - | - | - | Unassigned | - |
| Otu25396 | 1 | k__Fungi(100);k__Fungi_unclassified(100);k__Fungi_unclassified(100);k__Fungi_unclassified(100);k__Fungi_unclassified(100);k__Fungi_unclassified(100);k__Fungi_unclassified(100); | - | - | - | - | - | - | - | Unassigned | - |
| Otu25397 | 1 | k__Fungi(100);p__Ascomycota(100);p__Ascomycota_unclassified(100);p__Ascomycota_unclassified(100);p__Ascomycota_unclassified(100);p__Ascomycota_unclassified(100);p__Ascomycota_unclassified(100); | - | - | - | - | - | - | - | Unassigned | - |
| Otu25398 | 1 | k__Fungi(100);k__Fungi_unclassified(100);k__Fungi_unclassified(100);k__Fungi_unclassified(100);k__Fungi_unclassified(100);k__Fungi_unclassified(100);k__Fungi_unclassified(100); | - | - | - | - | - | - | - | Unassigned | - |
| Otu25405 | 1 | k__Fungi(100);k__Fungi_unclassified(100);k__Fungi_unclassified(100);k__Fungi_unclassified(100);k__Fungi_unclassified(100);k__Fungi_unclassified(100);k__Fungi_unclassified(100); | - | - | - | - | - | - | - | Unassigned | - |
| Otu25406 | 1 | k__Fungi(100);k__Fungi_unclassified(100);k__Fungi_unclassified(100);k__Fungi_unclassified(100);k__Fungi_unclassified(100);k__Fungi_unclassified(100);k__Fungi_unclassified(100); | - | - | - | - | - | - | - | Unassigned | - |
| Otu25407 | 1 | k__Fungi(100);k__Fungi_unclassified(100);k__Fungi_unclassified(100);k__Fungi_unclassified(100);k__Fungi_unclassified(100);k__Fungi_unclassified(100);k__Fungi_unclassified(100); | - | - | - | - | - | - | - | Unassigned | - |
| Otu25412 | 1 | k__Fungi(100);p__Ascomycota(100);p__Ascomycota_unclassified(100);p__Ascomycota_unclassified(100);p__Ascomycota_unclassified(100);p__Ascomycota_unclassified(100);p__Ascomycota_unclassified(100); | - | - | - | - | - | - | - | Unassigned | - |
| Otu25413 | 1 | k__Fungi(100);p__Ascomycota(100);p__Ascomycota_unclassified(100);p__Ascomycota_unclassified(100);p__Ascomycota_unclassified(100);p__Ascomycota_unclassified(100);p__Ascomycota_unclassified(100); | - | - | - | - | - | - | - | Unassigned | - |
| Otu25422 | 1 | k__Fungi(100);k__Fungi_unclassified(100);k__Fungi_unclassified(100);k__Fungi_unclassified(100);k__Fungi_unclassified(100);k__Fungi_unclassified(100);k__Fungi_unclassified(100); | - | - | - | - | - | - | - | Unassigned | - |
| Otu25448 | 1 | k__Fungi(100);p__Ascomycota(100);p__Ascomycota_unclassified(100);p__Ascomycota_unclassified(100);p__Ascomycota_unclassified(100);p__Ascomycota_unclassified(100);p__Ascomycota_unclassified(100); | - | - | - | - | - | - | - | Unassigned | - |
| Otu25453 | 1 | k__Fungi(100);p__Basidiomycota(100);p__Basidiomycota_unclassified(100);p__Basidiomycota_unclassified(100);p__Basidiomycota_unclassified(100);p__Basidiomycota_unclassified(100);p__Basidiomycota_unclassified(100); | - | - | - | - | - | - | - | Unassigned | - |
| Otu25454 | 1 | k__Fungi(100);p__Ascomycota(100);p__Ascomycota_unclassified(100);p__Ascomycota_unclassified(100);p__Ascomycota_unclassified(100);p__Ascomycota_unclassified(100);p__Ascomycota_unclassified(100); | - | - | - | - | - | - | - | Unassigned | - |
| Otu25455 | 1 | k__Fungi(100);p__Ascomycota(100);p__Ascomycota_unclassified(100);p__Ascomycota_unclassified(100);p__Ascomycota_unclassified(100);p__Ascomycota_unclassified(100);p__Ascomycota_unclassified(100); | - | - | - | - | - | - | - | Unassigned | - |
| Otu25456 | 1 | k__Fungi(100);p__Ascomycota(100);p__Ascomycota_unclassified(100);p__Ascomycota_unclassified(100);p__Ascomycota_unclassified(100);p__Ascomycota_unclassified(100);p__Ascomycota_unclassified(100); | - | - | - | - | - | - | - | Unassigned | - |
| Otu25459 | 1 | k__Fungi(100);k__Fungi_unclassified(100);k__Fungi_unclassified(100);k__Fungi_unclassified(100);k__Fungi_unclassified(100);k__Fungi_unclassified(100);k__Fungi_unclassified(100); | - | - | - | - | - | - | - | Unassigned | - |
| Otu25460 | 1 | k__Fungi(100);p__Ascomycota(100);p__Ascomycota_unclassified(100);p__Ascomycota_unclassified(100);p__Ascomycota_unclassified(100);p__Ascomycota_unclassified(100);p__Ascomycota_unclassified(100); | - | - | - | - | - | - | - | Unassigned | - |
| Otu25461 | 1 | k__Fungi(100);p__Ascomycota(100);p__Ascomycota_unclassified(100);p__Ascomycota_unclassified(100);p__Ascomycota_unclassified(100);p__Ascomycota_unclassified(100);p__Ascomycota_unclassified(100); | - | - | - | - | - | - | - | Unassigned | - |
| Otu25463 | 1 | k__Fungi(100);p__Ascomycota(100);p__Ascomycota_unclassified(100);p__Ascomycota_unclassified(100);p__Ascomycota_unclassified(100);p__Ascomycota_unclassified(100);p__Ascomycota_unclassified(100); | - | - | - | - | - | - | - | Unassigned | - |
| Otu25464 | 1 | k__Fungi(100);p__Ascomycota(100);p__Ascomycota_unclassified(100);p__Ascomycota_unclassified(100);p__Ascomycota_unclassified(100);p__Ascomycota_unclassified(100);p__Ascomycota_unclassified(100); | - | - | - | - | - | - | - | Unassigned | - |
| Otu25465 | 1 | k__Fungi(100);p__Ascomycota(100);p__Ascomycota_unclassified(100);p__Ascomycota_unclassified(100);p__Ascomycota_unclassified(100);p__Ascomycota_unclassified(100);p__Ascomycota_unclassified(100); | - | - | - | - | - | - | - | Unassigned | - |
| Otu25466 | 1 | k__Fungi(100);k__Fungi_unclassified(100);k__Fungi_unclassified(100);k__Fungi_unclassified(100);k__Fungi_unclassified(100);k__Fungi_unclassified(100);k__Fungi_unclassified(100); | - | - | - | - | - | - | - | Unassigned | - |
| Otu25467 | 1 | k__Fungi(100);k__Fungi_unclassified(100);k__Fungi_unclassified(100);k__Fungi_unclassified(100);k__Fungi_unclassified(100);k__Fungi_unclassified(100);k__Fungi_unclassified(100); | - | - | - | - | - | - | - | Unassigned | - |
| Otu25471 | 1 | k__Fungi(100);p__Ascomycota(100);p__Ascomycota_unclassified(100);p__Ascomycota_unclassified(100);p__Ascomycota_unclassified(100);p__Ascomycota_unclassified(100);p__Ascomycota_unclassified(100); | - | - | - | - | - | - | - | Unassigned | - |
| Otu25472 | 1 | k__Fungi(100);k__Fungi_unclassified(100);k__Fungi_unclassified(100);k__Fungi_unclassified(100);k__Fungi_unclassified(100);k__Fungi_unclassified(100);k__Fungi_unclassified(100); | - | - | - | - | - | - | - | Unassigned | - |
| Otu25473 | 1 | k__Fungi(100);k__Fungi_unclassified(100);k__Fungi_unclassified(100);k__Fungi_unclassified(100);k__Fungi_unclassified(100);k__Fungi_unclassified(100);k__Fungi_unclassified(100); | - | - | - | - | - | - | - | Unassigned | - |
| Otu25474 | 1 | k__Fungi(100);k__Fungi_unclassified(100);k__Fungi_unclassified(100);k__Fungi_unclassified(100);k__Fungi_unclassified(100);k__Fungi_unclassified(100);k__Fungi_unclassified(100); | - | - | - | - | - | - | - | Unassigned | - |
| Otu25475 | 1 | k__Fungi(100);p__Ascomycota(100);p__Ascomycota_unclassified(100);p__Ascomycota_unclassified(100);p__Ascomycota_unclassified(100);p__Ascomycota_unclassified(100);p__Ascomycota_unclassified(100); | - | - | - | - | - | - | - | Unassigned | - |
| Otu25476 | 1 | k__Fungi(100);k__Fungi_unclassified(100);k__Fungi_unclassified(100);k__Fungi_unclassified(100);k__Fungi_unclassified(100);k__Fungi_unclassified(100);k__Fungi_unclassified(100); | - | - | - | - | - | - | - | Unassigned | - |
| Otu25477 | 1 | k__Fungi(100);p__Ascomycota(100);p__Ascomycota_unclassified(100);p__Ascomycota_unclassified(100);p__Ascomycota_unclassified(100);p__Ascomycota_unclassified(100);p__Ascomycota_unclassified(100); | - | - | - | - | - | - | - | Unassigned | - |
| Otu25478 | 1 | k__Fungi(100);k__Fungi_unclassified(100);k__Fungi_unclassified(100);k__Fungi_unclassified(100);k__Fungi_unclassified(100);k__Fungi_unclassified(100);k__Fungi_unclassified(100); | - | - | - | - | - | - | - | Unassigned | - |
| Otu25479 | 1 | k__Fungi(100);k__Fungi_unclassified(100);k__Fungi_unclassified(100);k__Fungi_unclassified(100);k__Fungi_unclassified(100);k__Fungi_unclassified(100);k__Fungi_unclassified(100); | - | - | - | - | - | - | - | Unassigned | - |
| Otu25480 | 1 | k__Fungi(100);k__Fungi_unclassified(100);k__Fungi_unclassified(100);k__Fungi_unclassified(100);k__Fungi_unclassified(100);k__Fungi_unclassified(100);k__Fungi_unclassified(100); | - | - | - | - | - | - | - | Unassigned | - |
| Otu25481 | 1 | k__Fungi(100);p__Ascomycota(100);p__Ascomycota_unclassified(100);p__Ascomycota_unclassified(100);p__Ascomycota_unclassified(100);p__Ascomycota_unclassified(100);p__Ascomycota_unclassified(100); | - | - | - | - | - | - | - | Unassigned | - |
| Otu25482 | 1 | k__Fungi(100);p__Ascomycota(100);p__Ascomycota_unclassified(100);p__Ascomycota_unclassified(100);p__Ascomycota_unclassified(100);p__Ascomycota_unclassified(100);p__Ascomycota_unclassified(100); | - | - | - | - | - | - | - | Unassigned | - |
| Otu25484 | 1 | k__Fungi(100);k__Fungi_unclassified(100);k__Fungi_unclassified(100);k__Fungi_unclassified(100);k__Fungi_unclassified(100);k__Fungi_unclassified(100);k__Fungi_unclassified(100); | - | - | - | - | - | - | - | Unassigned | - |
| Otu25485 | 1 | k__Fungi(100);k__Fungi_unclassified(100);k__Fungi_unclassified(100);k__Fungi_unclassified(100);k__Fungi_unclassified(100);k__Fungi_unclassified(100);k__Fungi_unclassified(100); | - | - | - | - | - | - | - | Unassigned | - |
| Otu25486 | 1 | k__Fungi(100);p__Ascomycota(100);p__Ascomycota_unclassified(100);p__Ascomycota_unclassified(100);p__Ascomycota_unclassified(100);p__Ascomycota_unclassified(100);p__Ascomycota_unclassified(100); | - | - | - | - | - | - | - | Unassigned | - |
| Otu25494 | 1 | k__Fungi(100);p__Ascomycota(100);p__Ascomycota_unclassified(100);p__Ascomycota_unclassified(100);p__Ascomycota_unclassified(100);p__Ascomycota_unclassified(100);p__Ascomycota_unclassified(100); | - | - | - | - | - | - | - | Unassigned | - |
| Otu25495 | 1 | k__Fungi(100);k__Fungi_unclassified(100);k__Fungi_unclassified(100);k__Fungi_unclassified(100);k__Fungi_unclassified(100);k__Fungi_unclassified(100);k__Fungi_unclassified(100); | - | - | - | - | - | - | - | Unassigned | - |
| Otu25496 | 1 | k__Fungi(100);p__Ascomycota(100);p__Ascomycota_unclassified(100);p__Ascomycota_unclassified(100);p__Ascomycota_unclassified(100);p__Ascomycota_unclassified(100);p__Ascomycota_unclassified(100); | - | - | - | - | - | - | - | Unassigned | - |
| Otu25497 | 1 | k__Fungi(100);p__Ascomycota(100);p__Ascomycota_unclassified(100);p__Ascomycota_unclassified(100);p__Ascomycota_unclassified(100);p__Ascomycota_unclassified(100);p__Ascomycota_unclassified(100); | - | - | - | - | - | - | - | Unassigned | - |
| Otu25498 | 1 | k__Fungi(100);k__Fungi_unclassified(100);k__Fungi_unclassified(100);k__Fungi_unclassified(100);k__Fungi_unclassified(100);k__Fungi_unclassified(100);k__Fungi_unclassified(100); | - | - | - | - | - | - | - | Unassigned | - |
| Otu25499 | 1 | k__Fungi(100);k__Fungi_unclassified(100);k__Fungi_unclassified(100);k__Fungi_unclassified(100);k__Fungi_unclassified(100);k__Fungi_unclassified(100);k__Fungi_unclassified(100); | - | - | - | - | - | - | - | Unassigned | - |
| Otu25500 | 1 | k__Fungi(100);p__Ascomycota(100);p__Ascomycota_unclassified(100);p__Ascomycota_unclassified(100);p__Ascomycota_unclassified(100);p__Ascomycota_unclassified(100);p__Ascomycota_unclassified(100); | - | - | - | - | - | - | - | Unassigned | - |
| Otu25502 | 1 | k__Fungi(100);p__Ascomycota(100);p__Ascomycota_unclassified(100);p__Ascomycota_unclassified(100);p__Ascomycota_unclassified(100);p__Ascomycota_unclassified(100);p__Ascomycota_unclassified(100); | - | - | - | - | - | - | - | Unassigned | - |
| Otu25503 | 1 | k__Fungi(100);p__Ascomycota(100);p__Ascomycota_unclassified(100);p__Ascomycota_unclassified(100);p__Ascomycota_unclassified(100);p__Ascomycota_unclassified(100);p__Ascomycota_unclassified(100); | - | - | - | - | - | - | - | Unassigned | - |
| Otu25504 | 1 | k__Fungi(100);p__Ascomycota(100);p__Ascomycota_unclassified(100);p__Ascomycota_unclassified(100);p__Ascomycota_unclassified(100);p__Ascomycota_unclassified(100);p__Ascomycota_unclassified(100); | - | - | - | - | - | - | - | Unassigned | - |
| Otu25505 | 1 | k__Fungi(100);k__Fungi_unclassified(100);k__Fungi_unclassified(100);k__Fungi_unclassified(100);k__Fungi_unclassified(100);k__Fungi_unclassified(100);k__Fungi_unclassified(100); | - | - | - | - | - | - | - | Unassigned | - |
| Otu25506 | 1 | k__Fungi(100);p__Ascomycota(100);p__Ascomycota_unclassified(100);p__Ascomycota_unclassified(100);p__Ascomycota_unclassified(100);p__Ascomycota_unclassified(100);p__Ascomycota_unclassified(100); | - | - | - | - | - | - | - | Unassigned | - |
| Otu25507 | 1 | k__Fungi(100);p__Ascomycota(100);p__Ascomycota_unclassified(100);p__Ascomycota_unclassified(100);p__Ascomycota_unclassified(100);p__Ascomycota_unclassified(100);p__Ascomycota_unclassified(100); | - | - | - | - | - | - | - | Unassigned | - |
| Otu25509 | 1 | k__Fungi(100);p__Ascomycota(100);p__Ascomycota_unclassified(100);p__Ascomycota_unclassified(100);p__Ascomycota_unclassified(100);p__Ascomycota_unclassified(100);p__Ascomycota_unclassified(100); | - | - | - | - | - | - | - | Unassigned | - |
| Otu25510 | 1 | k__Fungi(100);k__Fungi_unclassified(100);k__Fungi_unclassified(100);k__Fungi_unclassified(100);k__Fungi_unclassified(100);k__Fungi_unclassified(100);k__Fungi_unclassified(100); | - | - | - | - | - | - | - | Unassigned | - |
| Otu25511 | 1 | k__Fungi(100);p__Ascomycota(100);p__Ascomycota_unclassified(100);p__Ascomycota_unclassified(100);p__Ascomycota_unclassified(100);p__Ascomycota_unclassified(100);p__Ascomycota_unclassified(100); | - | - | - | - | - | - | - | Unassigned | - |
| Otu25512 | 1 | k__Fungi(100);k__Fungi_unclassified(100);k__Fungi_unclassified(100);k__Fungi_unclassified(100);k__Fungi_unclassified(100);k__Fungi_unclassified(100);k__Fungi_unclassified(100); | - | - | - | - | - | - | - | Unassigned | - |
| Otu25519 | 1 | k__Fungi(100);k__Fungi_unclassified(100);k__Fungi_unclassified(100);k__Fungi_unclassified(100);k__Fungi_unclassified(100);k__Fungi_unclassified(100);k__Fungi_unclassified(100); | - | - | - | - | - | - | - | Unassigned | - |
| Otu25523 | 1 | k__Fungi(100);k__Fungi_unclassified(100);k__Fungi_unclassified(100);k__Fungi_unclassified(100);k__Fungi_unclassified(100);k__Fungi_unclassified(100);k__Fungi_unclassified(100); | - | - | - | - | - | - | - | Unassigned | - |
| Otu25529 | 1 | k__Fungi(100);k__Fungi_unclassified(100);k__Fungi_unclassified(100);k__Fungi_unclassified(100);k__Fungi_unclassified(100);k__Fungi_unclassified(100);k__Fungi_unclassified(100); | - | - | - | - | - | - | - | Unassigned | - |
| Otu25533 | 1 | k__Fungi(100);p__Ascomycota(100);p__Ascomycota_unclassified(100);p__Ascomycota_unclassified(100);p__Ascomycota_unclassified(100);p__Ascomycota_unclassified(100);p__Ascomycota_unclassified(100); | - | - | - | - | - | - | - | Unassigned | - |
| Otu25541 | 1 | k__Fungi(100);p__Basidiomycota(100);p__Basidiomycota_unclassified(100);p__Basidiomycota_unclassified(100);p__Basidiomycota_unclassified(100);p__Basidiomycota_unclassified(100);p__Basidiomycota_unclassified(100); | - | - | - | - | - | - | - | Unassigned | - |
| Otu25544 | 1 | k__Fungi(100);p__Ascomycota(100);p__Ascomycota_unclassified(100);p__Ascomycota_unclassified(100);p__Ascomycota_unclassified(100);p__Ascomycota_unclassified(100);p__Ascomycota_unclassified(100); | - | - | - | - | - | - | - | Unassigned | - |
| Otu25548 | 1 | k__Fungi(100);p__Ascomycota(100);p__Ascomycota_unclassified(100);p__Ascomycota_unclassified(100);p__Ascomycota_unclassified(100);p__Ascomycota_unclassified(100);p__Ascomycota_unclassified(100); | - | - | - | - | - | - | - | Unassigned | - |
| Otu25550 | 1 | k__Fungi(100);k__Fungi_unclassified(100);k__Fungi_unclassified(100);k__Fungi_unclassified(100);k__Fungi_unclassified(100);k__Fungi_unclassified(100);k__Fungi_unclassified(100); | - | - | - | - | - | - | - | Unassigned | - |
| Otu25552 | 1 | k__Fungi(100);k__Fungi_unclassified(100);k__Fungi_unclassified(100);k__Fungi_unclassified(100);k__Fungi_unclassified(100);k__Fungi_unclassified(100);k__Fungi_unclassified(100); | - | - | - | - | - | - | - | Unassigned | - |
| Otu25553 | 1 | k__Fungi(100);p__Ascomycota(100);p__Ascomycota_unclassified(100);p__Ascomycota_unclassified(100);p__Ascomycota_unclassified(100);p__Ascomycota_unclassified(100);p__Ascomycota_unclassified(100); | - | - | - | - | - | - | - | Unassigned | - |
| Otu25558 | 1 | k__Fungi(100);p__Ascomycota(100);p__Ascomycota_unclassified(100);p__Ascomycota_unclassified(100);p__Ascomycota_unclassified(100);p__Ascomycota_unclassified(100);p__Ascomycota_unclassified(100); | - | - | - | - | - | - | - | Unassigned | - |
| Otu25559 | 1 | k__Fungi(100);p__Ascomycota(100);p__Ascomycota_unclassified(100);p__Ascomycota_unclassified(100);p__Ascomycota_unclassified(100);p__Ascomycota_unclassified(100);p__Ascomycota_unclassified(100); | - | - | - | - | - | - | - | Unassigned | - |
| Otu25568 | 1 | k__Fungi(100);p__Ascomycota(100);p__Ascomycota_unclassified(100);p__Ascomycota_unclassified(100);p__Ascomycota_unclassified(100);p__Ascomycota_unclassified(100);p__Ascomycota_unclassified(100); | - | - | - | - | - | - | - | Unassigned | - |
| Otu25569 | 1 | k__Fungi(100);p__Ascomycota(100);p__Ascomycota_unclassified(100);p__Ascomycota_unclassified(100);p__Ascomycota_unclassified(100);p__Ascomycota_unclassified(100);p__Ascomycota_unclassified(100); | - | - | - | - | - | - | - | Unassigned | - |
| Otu25570 | 1 | k__Fungi(100);p__Ascomycota(100);p__Ascomycota_unclassified(100);p__Ascomycota_unclassified(100);p__Ascomycota_unclassified(100);p__Ascomycota_unclassified(100);p__Ascomycota_unclassified(100); | - | - | - | - | - | - | - | Unassigned | - |
| Otu25571 | 1 | k__Fungi(100);p__Ascomycota(100);p__Ascomycota_unclassified(100);p__Ascomycota_unclassified(100);p__Ascomycota_unclassified(100);p__Ascomycota_unclassified(100);p__Ascomycota_unclassified(100); | - | - | - | - | - | - | - | Unassigned | - |
| Otu25572 | 1 | k__Fungi(100);k__Fungi_unclassified(100);k__Fungi_unclassified(100);k__Fungi_unclassified(100);k__Fungi_unclassified(100);k__Fungi_unclassified(100);k__Fungi_unclassified(100); | - | - | - | - | - | - | - | Unassigned | - |
| Otu25573 | 1 | k__Fungi(100);p__Ascomycota(100);p__Ascomycota_unclassified(100);p__Ascomycota_unclassified(100);p__Ascomycota_unclassified(100);p__Ascomycota_unclassified(100);p__Ascomycota_unclassified(100); | - | - | - | - | - | - | - | Unassigned | - |
| Otu25574 | 1 | k__Fungi(100);p__Ascomycota(100);p__Ascomycota_unclassified(100);p__Ascomycota_unclassified(100);p__Ascomycota_unclassified(100);p__Ascomycota_unclassified(100);p__Ascomycota_unclassified(100); | - | - | - | - | - | - | - | Unassigned | - |
| Otu25575 | 1 | k__Fungi(100);k__Fungi_unclassified(100);k__Fungi_unclassified(100);k__Fungi_unclassified(100);k__Fungi_unclassified(100);k__Fungi_unclassified(100);k__Fungi_unclassified(100); | - | - | - | - | - | - | - | Unassigned | - |
| Otu25576 | 1 | k__Fungi(100);k__Fungi_unclassified(100);k__Fungi_unclassified(100);k__Fungi_unclassified(100);k__Fungi_unclassified(100);k__Fungi_unclassified(100);k__Fungi_unclassified(100); | - | - | - | - | - | - | - | Unassigned | - |
| Otu25577 | 1 | k__Fungi(100);p__Ascomycota(100);p__Ascomycota_unclassified(100);p__Ascomycota_unclassified(100);p__Ascomycota_unclassified(100);p__Ascomycota_unclassified(100);p__Ascomycota_unclassified(100); | - | - | - | - | - | - | - | Unassigned | - |
| Otu25578 | 1 | k__Fungi(100);p__Ascomycota(100);p__Ascomycota_unclassified(100);p__Ascomycota_unclassified(100);p__Ascomycota_unclassified(100);p__Ascomycota_unclassified(100);p__Ascomycota_unclassified(100); | - | - | - | - | - | - | - | Unassigned | - |
| Otu25579 | 1 | k__Fungi(100);p__Ascomycota(100);p__Ascomycota_unclassified(100);p__Ascomycota_unclassified(100);p__Ascomycota_unclassified(100);p__Ascomycota_unclassified(100);p__Ascomycota_unclassified(100); | - | - | - | - | - | - | - | Unassigned | - |
| Otu25580 | 1 | k__Fungi(100);k__Fungi_unclassified(100);k__Fungi_unclassified(100);k__Fungi_unclassified(100);k__Fungi_unclassified(100);k__Fungi_unclassified(100);k__Fungi_unclassified(100); | - | - | - | - | - | - | - | Unassigned | - |
| Otu25581 | 1 | k__Fungi(100);p__Ascomycota(100);p__Ascomycota_unclassified(100);p__Ascomycota_unclassified(100);p__Ascomycota_unclassified(100);p__Ascomycota_unclassified(100);p__Ascomycota_unclassified(100); | - | - | - | - | - | - | - | Unassigned | - |
| Otu25582 | 1 | k__Fungi(100);p__Ascomycota(100);p__Ascomycota_unclassified(100);p__Ascomycota_unclassified(100);p__Ascomycota_unclassified(100);p__Ascomycota_unclassified(100);p__Ascomycota_unclassified(100); | - | - | - | - | - | - | - | Unassigned | - |
| Otu25583 | 1 | k__Fungi(100);p__Ascomycota(100);p__Ascomycota_unclassified(100);p__Ascomycota_unclassified(100);p__Ascomycota_unclassified(100);p__Ascomycota_unclassified(100);p__Ascomycota_unclassified(100); | - | - | - | - | - | - | - | Unassigned | - |
| Otu25584 | 1 | k__Fungi(100);k__Fungi_unclassified(100);k__Fungi_unclassified(100);k__Fungi_unclassified(100);k__Fungi_unclassified(100);k__Fungi_unclassified(100);k__Fungi_unclassified(100); | - | - | - | - | - | - | - | Unassigned | - |
| Otu25585 | 1 | k__Fungi(100);p__Ascomycota(100);p__Ascomycota_unclassified(100);p__Ascomycota_unclassified(100);p__Ascomycota_unclassified(100);p__Ascomycota_unclassified(100);p__Ascomycota_unclassified(100); | - | - | - | - | - | - | - | Unassigned | - |
| Otu25586 | 1 | k__Fungi(100);p__Ascomycota(100);p__Ascomycota_unclassified(100);p__Ascomycota_unclassified(100);p__Ascomycota_unclassified(100);p__Ascomycota_unclassified(100);p__Ascomycota_unclassified(100); | - | - | - | - | - | - | - | Unassigned | - |
| Otu25588 | 1 | k__Fungi(100);p__Ascomycota(100);p__Ascomycota_unclassified(100);p__Ascomycota_unclassified(100);p__Ascomycota_unclassified(100);p__Ascomycota_unclassified(100);p__Ascomycota_unclassified(100); | - | - | - | - | - | - | - | Unassigned | - |
| Otu25589 | 1 | k__Fungi(100);k__Fungi_unclassified(100);k__Fungi_unclassified(100);k__Fungi_unclassified(100);k__Fungi_unclassified(100);k__Fungi_unclassified(100);k__Fungi_unclassified(100); | - | - | - | - | - | - | - | Unassigned | - |
| Otu25590 | 1 | k__Fungi(100);p__Ascomycota(100);p__Ascomycota_unclassified(100);p__Ascomycota_unclassified(100);p__Ascomycota_unclassified(100);p__Ascomycota_unclassified(100);p__Ascomycota_unclassified(100); | - | - | - | - | - | - | - | Unassigned | - |
| Otu25592 | 1 | k__Fungi(100);p__Ascomycota(100);p__Ascomycota_unclassified(100);p__Ascomycota_unclassified(100);p__Ascomycota_unclassified(100);p__Ascomycota_unclassified(100);p__Ascomycota_unclassified(100); | - | - | - | - | - | - | - | Unassigned | - |
| Otu25595 | 1 | k__Fungi(100);k__Fungi_unclassified(100);k__Fungi_unclassified(100);k__Fungi_unclassified(100);k__Fungi_unclassified(100);k__Fungi_unclassified(100);k__Fungi_unclassified(100); | - | - | - | - | - | - | - | Unassigned | - |
| Otu25596 | 1 | k__Fungi(100);k__Fungi_unclassified(100);k__Fungi_unclassified(100);k__Fungi_unclassified(100);k__Fungi_unclassified(100);k__Fungi_unclassified(100);k__Fungi_unclassified(100); | - | - | - | - | - | - | - | Unassigned | - |
| Otu25597 | 1 | k__Fungi(100);p__Ascomycota(100);p__Ascomycota_unclassified(100);p__Ascomycota_unclassified(100);p__Ascomycota_unclassified(100);p__Ascomycota_unclassified(100);p__Ascomycota_unclassified(100); | - | - | - | - | - | - | - | Unassigned | - |
| Otu25598 | 1 | k__Fungi(100);p__Ascomycota(100);p__Ascomycota_unclassified(100);p__Ascomycota_unclassified(100);p__Ascomycota_unclassified(100);p__Ascomycota_unclassified(100);p__Ascomycota_unclassified(100); | - | - | - | - | - | - | - | Unassigned | - |
| Otu25599 | 1 | k__Fungi(100);p__Ascomycota(100);p__Ascomycota_unclassified(100);p__Ascomycota_unclassified(100);p__Ascomycota_unclassified(100);p__Ascomycota_unclassified(100);p__Ascomycota_unclassified(100); | - | - | - | - | - | - | - | Unassigned | - |
| Otu25600 | 1 | k__Fungi(100);k__Fungi_unclassified(100);k__Fungi_unclassified(100);k__Fungi_unclassified(100);k__Fungi_unclassified(100);k__Fungi_unclassified(100);k__Fungi_unclassified(100); | - | - | - | - | - | - | - | Unassigned | - |
| Otu25601 | 1 | k__Fungi(100);k__Fungi_unclassified(100);k__Fungi_unclassified(100);k__Fungi_unclassified(100);k__Fungi_unclassified(100);k__Fungi_unclassified(100);k__Fungi_unclassified(100); | - | - | - | - | - | - | - | Unassigned | - |
| Otu25602 | 1 | k__Fungi(100);p__Ascomycota(100);p__Ascomycota_unclassified(100);p__Ascomycota_unclassified(100);p__Ascomycota_unclassified(100);p__Ascomycota_unclassified(100);p__Ascomycota_unclassified(100); | - | - | - | - | - | - | - | Unassigned | - |
| Otu25603 | 1 | k__Fungi(100);p__Ascomycota(100);p__Ascomycota_unclassified(100);p__Ascomycota_unclassified(100);p__Ascomycota_unclassified(100);p__Ascomycota_unclassified(100);p__Ascomycota_unclassified(100); | - | - | - | - | - | - | - | Unassigned | - |
| Otu25604 | 1 | k__Fungi(100);k__Fungi_unclassified(100);k__Fungi_unclassified(100);k__Fungi_unclassified(100);k__Fungi_unclassified(100);k__Fungi_unclassified(100);k__Fungi_unclassified(100); | - | - | - | - | - | - | - | Unassigned | - |
| Otu25605 | 1 | k__Fungi(100);p__Ascomycota(100);p__Ascomycota_unclassified(100);p__Ascomycota_unclassified(100);p__Ascomycota_unclassified(100);p__Ascomycota_unclassified(100);p__Ascomycota_unclassified(100); | - | - | - | - | - | - | - | Unassigned | - |
| Otu25606 | 1 | k__Fungi(100);k__Fungi_unclassified(100);k__Fungi_unclassified(100);k__Fungi_unclassified(100);k__Fungi_unclassified(100);k__Fungi_unclassified(100);k__Fungi_unclassified(100); | - | - | - | - | - | - | - | Unassigned | - |
| Otu25607 | 1 | k__Fungi(100);k__Fungi_unclassified(100);k__Fungi_unclassified(100);k__Fungi_unclassified(100);k__Fungi_unclassified(100);k__Fungi_unclassified(100);k__Fungi_unclassified(100); | - | - | - | - | - | - | - | Unassigned | - |
| Otu25612 | 1 | k__Fungi(100);p__Ascomycota(100);p__Ascomycota_unclassified(100);p__Ascomycota_unclassified(100);p__Ascomycota_unclassified(100);p__Ascomycota_unclassified(100);p__Ascomycota_unclassified(100); | - | - | - | - | - | - | - | Unassigned | - |
| Otu25613 | 1 | k__Fungi(100);p__Ascomycota(100);p__Ascomycota_unclassified(100);p__Ascomycota_unclassified(100);p__Ascomycota_unclassified(100);p__Ascomycota_unclassified(100);p__Ascomycota_unclassified(100); | - | - | - | - | - | - | - | Unassigned | - |
| Otu25615 | 1 | k__Fungi(100);p__Ascomycota(100);p__Ascomycota_unclassified(100);p__Ascomycota_unclassified(100);p__Ascomycota_unclassified(100);p__Ascomycota_unclassified(100);p__Ascomycota_unclassified(100); | - | - | - | - | - | - | - | Unassigned | - |
| Otu25616 | 1 | k__Fungi(100);p__Ascomycota(100);p__Ascomycota_unclassified(100);p__Ascomycota_unclassified(100);p__Ascomycota_unclassified(100);p__Ascomycota_unclassified(100);p__Ascomycota_unclassified(100); | - | - | - | - | - | - | - | Unassigned | - |
| Otu25617 | 1 | k__Fungi(100);k__Fungi_unclassified(100);k__Fungi_unclassified(100);k__Fungi_unclassified(100);k__Fungi_unclassified(100);k__Fungi_unclassified(100);k__Fungi_unclassified(100); | - | - | - | - | - | - | - | Unassigned | - |
| Otu25618 | 1 | k__Fungi(100);p__Ascomycota(100);p__Ascomycota_unclassified(100);p__Ascomycota_unclassified(100);p__Ascomycota_unclassified(100);p__Ascomycota_unclassified(100);p__Ascomycota_unclassified(100); | - | - | - | - | - | - | - | Unassigned | - |
| Otu25619 | 1 | k__Fungi(100);p__Ascomycota(100);p__Ascomycota_unclassified(100);p__Ascomycota_unclassified(100);p__Ascomycota_unclassified(100);p__Ascomycota_unclassified(100);p__Ascomycota_unclassified(100); | - | - | - | - | - | - | - | Unassigned | - |
| Otu25621 | 1 | k__Fungi(100);p__Ascomycota(100);p__Ascomycota_unclassified(100);p__Ascomycota_unclassified(100);p__Ascomycota_unclassified(100);p__Ascomycota_unclassified(100);p__Ascomycota_unclassified(100); | - | - | - | - | - | - | - | Unassigned | - |
| Otu25623 | 1 | k__Fungi(100);k__Fungi_unclassified(100);k__Fungi_unclassified(100);k__Fungi_unclassified(100);k__Fungi_unclassified(100);k__Fungi_unclassified(100);k__Fungi_unclassified(100); | - | - | - | - | - | - | - | Unassigned | - |
| Otu25624 | 1 | k__Fungi(100);k__Fungi_unclassified(100);k__Fungi_unclassified(100);k__Fungi_unclassified(100);k__Fungi_unclassified(100);k__Fungi_unclassified(100);k__Fungi_unclassified(100); | - | - | - | - | - | - | - | Unassigned | - |
| Otu25625 | 1 | k__Fungi(100);p__Ascomycota(100);p__Ascomycota_unclassified(100);p__Ascomycota_unclassified(100);p__Ascomycota_unclassified(100);p__Ascomycota_unclassified(100);p__Ascomycota_unclassified(100); | - | - | - | - | - | - | - | Unassigned | - |
| Otu25632 | 1 | k__Fungi(100);p__Ascomycota(100);p__Ascomycota_unclassified(100);p__Ascomycota_unclassified(100);p__Ascomycota_unclassified(100);p__Ascomycota_unclassified(100);p__Ascomycota_unclassified(100); | - | - | - | - | - | - | - | Unassigned | - |
| Otu25634 | 1 | k__Fungi(100);k__Fungi_unclassified(100);k__Fungi_unclassified(100);k__Fungi_unclassified(100);k__Fungi_unclassified(100);k__Fungi_unclassified(100);k__Fungi_unclassified(100); | - | - | - | - | - | - | - | Unassigned | - |
| Otu25639 | 1 | k__Fungi(100);p__Ascomycota(100);p__Ascomycota_unclassified(100);p__Ascomycota_unclassified(100);p__Ascomycota_unclassified(100);p__Ascomycota_unclassified(100);p__Ascomycota_unclassified(100); | - | - | - | - | - | - | - | Unassigned | - |
| Otu25647 | 1 | k__Fungi(100);p__Ascomycota(100);p__Ascomycota_unclassified(100);p__Ascomycota_unclassified(100);p__Ascomycota_unclassified(100);p__Ascomycota_unclassified(100);p__Ascomycota_unclassified(100); | - | - | - | - | - | - | - | Unassigned | - |
| Otu25648 | 1 | k__Fungi(100);k__Fungi_unclassified(100);k__Fungi_unclassified(100);k__Fungi_unclassified(100);k__Fungi_unclassified(100);k__Fungi_unclassified(100);k__Fungi_unclassified(100); | - | - | - | - | - | - | - | Unassigned | - |
| Otu25649 | 1 | k__Fungi(100);k__Fungi_unclassified(100);k__Fungi_unclassified(100);k__Fungi_unclassified(100);k__Fungi_unclassified(100);k__Fungi_unclassified(100);k__Fungi_unclassified(100); | - | - | - | - | - | - | - | Unassigned | - |
| Otu25650 | 1 | k__Fungi(100);k__Fungi_unclassified(100);k__Fungi_unclassified(100);k__Fungi_unclassified(100);k__Fungi_unclassified(100);k__Fungi_unclassified(100);k__Fungi_unclassified(100); | - | - | - | - | - | - | - | Unassigned | - |
| Otu25652 | 1 | k__Fungi(100);p__Ascomycota(100);p__Ascomycota_unclassified(100);p__Ascomycota_unclassified(100);p__Ascomycota_unclassified(100);p__Ascomycota_unclassified(100);p__Ascomycota_unclassified(100); | - | - | - | - | - | - | - | Unassigned | - |
| Otu25658 | 1 | k__Fungi(100);k__Fungi_unclassified(100);k__Fungi_unclassified(100);k__Fungi_unclassified(100);k__Fungi_unclassified(100);k__Fungi_unclassified(100);k__Fungi_unclassified(100); | - | - | - | - | - | - | - | Unassigned | - |
| Otu25661 | 1 | k__Fungi(100);k__Fungi_unclassified(100);k__Fungi_unclassified(100);k__Fungi_unclassified(100);k__Fungi_unclassified(100);k__Fungi_unclassified(100);k__Fungi_unclassified(100); | - | - | - | - | - | - | - | Unassigned | - |
| Otu25662 | 1 | k__Fungi(100);p__Ascomycota(100);p__Ascomycota_unclassified(100);p__Ascomycota_unclassified(100);p__Ascomycota_unclassified(100);p__Ascomycota_unclassified(100);p__Ascomycota_unclassified(100); | - | - | - | - | - | - | - | Unassigned | - |
| Otu25663 | 1 | k__Fungi(100);p__Ascomycota(100);p__Ascomycota_unclassified(100);p__Ascomycota_unclassified(100);p__Ascomycota_unclassified(100);p__Ascomycota_unclassified(100);p__Ascomycota_unclassified(100); | - | - | - | - | - | - | - | Unassigned | - |
| Otu25664 | 1 | k__Fungi(100);p__Ascomycota(100);p__Ascomycota_unclassified(100);p__Ascomycota_unclassified(100);p__Ascomycota_unclassified(100);p__Ascomycota_unclassified(100);p__Ascomycota_unclassified(100); | - | - | - | - | - | - | - | Unassigned | - |
| Otu25665 | 1 | k__Fungi(100);p__Ascomycota(100);p__Ascomycota_unclassified(100);p__Ascomycota_unclassified(100);p__Ascomycota_unclassified(100);p__Ascomycota_unclassified(100);p__Ascomycota_unclassified(100); | - | - | - | - | - | - | - | Unassigned | - |
| Otu25667 | 1 | k__Fungi(100);p__Ascomycota(100);p__Ascomycota_unclassified(100);p__Ascomycota_unclassified(100);p__Ascomycota_unclassified(100);p__Ascomycota_unclassified(100);p__Ascomycota_unclassified(100); | - | - | - | - | - | - | - | Unassigned | - |
| Otu25668 | 1 | k__Fungi(100);k__Fungi_unclassified(100);k__Fungi_unclassified(100);k__Fungi_unclassified(100);k__Fungi_unclassified(100);k__Fungi_unclassified(100);k__Fungi_unclassified(100); | - | - | - | - | - | - | - | Unassigned | - |
| Otu25670 | 1 | k__Fungi(100);p__Ascomycota(100);p__Ascomycota_unclassified(100);p__Ascomycota_unclassified(100);p__Ascomycota_unclassified(100);p__Ascomycota_unclassified(100);p__Ascomycota_unclassified(100); | - | - | - | - | - | - | - | Unassigned | - |
| Otu25671 | 1 | k__Fungi(100);p__Ascomycota(100);p__Ascomycota_unclassified(100);p__Ascomycota_unclassified(100);p__Ascomycota_unclassified(100);p__Ascomycota_unclassified(100);p__Ascomycota_unclassified(100); | - | - | - | - | - | - | - | Unassigned | - |
| Otu25672 | 1 | k__Fungi(100);k__Fungi_unclassified(100);k__Fungi_unclassified(100);k__Fungi_unclassified(100);k__Fungi_unclassified(100);k__Fungi_unclassified(100);k__Fungi_unclassified(100); | - | - | - | - | - | - | - | Unassigned | - |
| Otu25673 | 1 | k__Fungi(100);k__Fungi_unclassified(100);k__Fungi_unclassified(100);k__Fungi_unclassified(100);k__Fungi_unclassified(100);k__Fungi_unclassified(100);k__Fungi_unclassified(100); | - | - | - | - | - | - | - | Unassigned | - |
| Otu25674 | 1 | k__Fungi(100);k__Fungi_unclassified(100);k__Fungi_unclassified(100);k__Fungi_unclassified(100);k__Fungi_unclassified(100);k__Fungi_unclassified(100);k__Fungi_unclassified(100); | - | - | - | - | - | - | - | Unassigned | - |
| Otu25676 | 1 | k__Fungi(100);k__Fungi_unclassified(100);k__Fungi_unclassified(100);k__Fungi_unclassified(100);k__Fungi_unclassified(100);k__Fungi_unclassified(100);k__Fungi_unclassified(100); | - | - | - | - | - | - | - | Unassigned | - |
| Otu25677 | 1 | k__Fungi(100);k__Fungi_unclassified(100);k__Fungi_unclassified(100);k__Fungi_unclassified(100);k__Fungi_unclassified(100);k__Fungi_unclassified(100);k__Fungi_unclassified(100); | - | - | - | - | - | - | - | Unassigned | - |
| Otu25678 | 1 | k__Fungi(100);k__Fungi_unclassified(100);k__Fungi_unclassified(100);k__Fungi_unclassified(100);k__Fungi_unclassified(100);k__Fungi_unclassified(100);k__Fungi_unclassified(100); | - | - | - | - | - | - | - | Unassigned | - |
| Otu25679 | 1 | k__Fungi(100);k__Fungi_unclassified(100);k__Fungi_unclassified(100);k__Fungi_unclassified(100);k__Fungi_unclassified(100);k__Fungi_unclassified(100);k__Fungi_unclassified(100); | - | - | - | - | - | - | - | Unassigned | - |
| Otu25680 | 1 | k__Fungi(100);p__Ascomycota(100);p__Ascomycota_unclassified(100);p__Ascomycota_unclassified(100);p__Ascomycota_unclassified(100);p__Ascomycota_unclassified(100);p__Ascomycota_unclassified(100); | - | - | - | - | - | - | - | Unassigned | - |
| Otu25681 | 1 | k__Fungi(100);k__Fungi_unclassified(100);k__Fungi_unclassified(100);k__Fungi_unclassified(100);k__Fungi_unclassified(100);k__Fungi_unclassified(100);k__Fungi_unclassified(100); | - | - | - | - | - | - | - | Unassigned | - |
| Otu25682 | 1 | k__Fungi(100);k__Fungi_unclassified(100);k__Fungi_unclassified(100);k__Fungi_unclassified(100);k__Fungi_unclassified(100);k__Fungi_unclassified(100);k__Fungi_unclassified(100); | - | - | - | - | - | - | - | Unassigned | - |
| Otu25683 | 1 | k__Fungi(100);p__Ascomycota(100);p__Ascomycota_unclassified(100);p__Ascomycota_unclassified(100);p__Ascomycota_unclassified(100);p__Ascomycota_unclassified(100);p__Ascomycota_unclassified(100); | - | - | - | - | - | - | - | Unassigned | - |
| Otu25684 | 1 | k__Fungi(100);k__Fungi_unclassified(100);k__Fungi_unclassified(100);k__Fungi_unclassified(100);k__Fungi_unclassified(100);k__Fungi_unclassified(100);k__Fungi_unclassified(100); | - | - | - | - | - | - | - | Unassigned | - |
| Otu25685 | 1 | k__Fungi(100);k__Fungi_unclassified(100);k__Fungi_unclassified(100);k__Fungi_unclassified(100);k__Fungi_unclassified(100);k__Fungi_unclassified(100);k__Fungi_unclassified(100); | - | - | - | - | - | - | - | Unassigned | - |
| Otu25686 | 1 | k__Fungi(100);k__Fungi_unclassified(100);k__Fungi_unclassified(100);k__Fungi_unclassified(100);k__Fungi_unclassified(100);k__Fungi_unclassified(100);k__Fungi_unclassified(100); | - | - | - | - | - | - | - | Unassigned | - |
| Otu25687 | 1 | k__Fungi(100);p__Ascomycota(100);p__Ascomycota_unclassified(100);p__Ascomycota_unclassified(100);p__Ascomycota_unclassified(100);p__Ascomycota_unclassified(100);p__Ascomycota_unclassified(100); | - | - | - | - | - | - | - | Unassigned | - |
| Otu25688 | 1 | k__Fungi(100);k__Fungi_unclassified(100);k__Fungi_unclassified(100);k__Fungi_unclassified(100);k__Fungi_unclassified(100);k__Fungi_unclassified(100);k__Fungi_unclassified(100); | - | - | - | - | - | - | - | Unassigned | - |
| Otu25689 | 1 | k__Fungi(100);k__Fungi_unclassified(100);k__Fungi_unclassified(100);k__Fungi_unclassified(100);k__Fungi_unclassified(100);k__Fungi_unclassified(100);k__Fungi_unclassified(100); | - | - | - | - | - | - | - | Unassigned | - |
| Otu25690 | 1 | k__Fungi(100);k__Fungi_unclassified(100);k__Fungi_unclassified(100);k__Fungi_unclassified(100);k__Fungi_unclassified(100);k__Fungi_unclassified(100);k__Fungi_unclassified(100); | - | - | - | - | - | - | - | Unassigned | - |
| Otu25691 | 1 | k__Fungi(100);p__Ascomycota(100);p__Ascomycota_unclassified(100);p__Ascomycota_unclassified(100);p__Ascomycota_unclassified(100);p__Ascomycota_unclassified(100);p__Ascomycota_unclassified(100); | - | - | - | - | - | - | - | Unassigned | - |
| Otu25692 | 1 | k__Fungi(100);p__Ascomycota(100);p__Ascomycota_unclassified(100);p__Ascomycota_unclassified(100);p__Ascomycota_unclassified(100);p__Ascomycota_unclassified(100);p__Ascomycota_unclassified(100); | - | - | - | - | - | - | - | Unassigned | - |
| Otu25693 | 1 | k__Fungi(100);k__Fungi_unclassified(100);k__Fungi_unclassified(100);k__Fungi_unclassified(100);k__Fungi_unclassified(100);k__Fungi_unclassified(100);k__Fungi_unclassified(100); | - | - | - | - | - | - | - | Unassigned | - |
| Otu25694 | 1 | k__Fungi(100);p__Ascomycota(100);p__Ascomycota_unclassified(100);p__Ascomycota_unclassified(100);p__Ascomycota_unclassified(100);p__Ascomycota_unclassified(100);p__Ascomycota_unclassified(100); | - | - | - | - | - | - | - | Unassigned | - |
| Otu25695 | 1 | k__Fungi(100);k__Fungi_unclassified(100);k__Fungi_unclassified(100);k__Fungi_unclassified(100);k__Fungi_unclassified(100);k__Fungi_unclassified(100);k__Fungi_unclassified(100); | - | - | - | - | - | - | - | Unassigned | - |
| Otu25696 | 1 | k__Fungi(100);k__Fungi_unclassified(100);k__Fungi_unclassified(100);k__Fungi_unclassified(100);k__Fungi_unclassified(100);k__Fungi_unclassified(100);k__Fungi_unclassified(100); | - | - | - | - | - | - | - | Unassigned | - |
| Otu25699 | 1 | k__Fungi(100);p__Ascomycota(100);p__Ascomycota_unclassified(100);p__Ascomycota_unclassified(100);p__Ascomycota_unclassified(100);p__Ascomycota_unclassified(100);p__Ascomycota_unclassified(100); | - | - | - | - | - | - | - | Unassigned | - |
| Otu25701 | 1 | k__Fungi(100);k__Fungi_unclassified(100);k__Fungi_unclassified(100);k__Fungi_unclassified(100);k__Fungi_unclassified(100);k__Fungi_unclassified(100);k__Fungi_unclassified(100); | - | - | - | - | - | - | - | Unassigned | - |
| Otu25714 | 1 | k__Fungi(100);p__Ascomycota(100);p__Ascomycota_unclassified(100);p__Ascomycota_unclassified(100);p__Ascomycota_unclassified(100);p__Ascomycota_unclassified(100);p__Ascomycota_unclassified(100); | - | - | - | - | - | - | - | Unassigned | - |
| Otu25717 | 1 | k__Fungi(100);p__Ascomycota(100);p__Ascomycota_unclassified(100);p__Ascomycota_unclassified(100);p__Ascomycota_unclassified(100);p__Ascomycota_unclassified(100);p__Ascomycota_unclassified(100); | - | - | - | - | - | - | - | Unassigned | - |
| Otu25719 | 1 | k__Fungi(100);k__Fungi_unclassified(100);k__Fungi_unclassified(100);k__Fungi_unclassified(100);k__Fungi_unclassified(100);k__Fungi_unclassified(100);k__Fungi_unclassified(100); | - | - | - | - | - | - | - | Unassigned | - |
| Otu25721 | 1 | k__Fungi(100);k__Fungi_unclassified(100);k__Fungi_unclassified(100);k__Fungi_unclassified(100);k__Fungi_unclassified(100);k__Fungi_unclassified(100);k__Fungi_unclassified(100); | - | - | - | - | - | - | - | Unassigned | - |
| Otu25722 | 1 | k__Fungi(100);k__Fungi_unclassified(100);k__Fungi_unclassified(100);k__Fungi_unclassified(100);k__Fungi_unclassified(100);k__Fungi_unclassified(100);k__Fungi_unclassified(100); | - | - | - | - | - | - | - | Unassigned | - |
| Otu25723 | 1 | k__Fungi(100);k__Fungi_unclassified(100);k__Fungi_unclassified(100);k__Fungi_unclassified(100);k__Fungi_unclassified(100);k__Fungi_unclassified(100);k__Fungi_unclassified(100); | - | - | - | - | - | - | - | Unassigned | - |
| Otu25724 | 1 | k__Fungi(100);k__Fungi_unclassified(100);k__Fungi_unclassified(100);k__Fungi_unclassified(100);k__Fungi_unclassified(100);k__Fungi_unclassified(100);k__Fungi_unclassified(100); | - | - | - | - | - | - | - | Unassigned | - |
| Otu25725 | 1 | k__Fungi(100);k__Fungi_unclassified(100);k__Fungi_unclassified(100);k__Fungi_unclassified(100);k__Fungi_unclassified(100);k__Fungi_unclassified(100);k__Fungi_unclassified(100); | - | - | - | - | - | - | - | Unassigned | - |
| Otu25726 | 1 | k__Fungi(100);k__Fungi_unclassified(100);k__Fungi_unclassified(100);k__Fungi_unclassified(100);k__Fungi_unclassified(100);k__Fungi_unclassified(100);k__Fungi_unclassified(100); | - | - | - | - | - | - | - | Unassigned | - |
| Otu25728 | 1 | k__Fungi(100);k__Fungi_unclassified(100);k__Fungi_unclassified(100);k__Fungi_unclassified(100);k__Fungi_unclassified(100);k__Fungi_unclassified(100);k__Fungi_unclassified(100); | - | - | - | - | - | - | - | Unassigned | - |
| Otu25729 | 1 | k__Fungi(100);k__Fungi_unclassified(100);k__Fungi_unclassified(100);k__Fungi_unclassified(100);k__Fungi_unclassified(100);k__Fungi_unclassified(100);k__Fungi_unclassified(100); | - | - | - | - | - | - | - | Unassigned | - |
| Otu25730 | 1 | k__Fungi(100);p__Ascomycota(100);p__Ascomycota_unclassified(100);p__Ascomycota_unclassified(100);p__Ascomycota_unclassified(100);p__Ascomycota_unclassified(100);p__Ascomycota_unclassified(100); | - | - | - | - | - | - | - | Unassigned | - |

| | | | | | | | | | | | |
|---|---|---|---|---|---|---|---|---|---|---|---|
| Otu25733 | 1 | k__Fungi(100);p__Ascomycota(100);p__Ascomycota_unclassified(100);p__Ascomycota_unclassified(100);p__Ascomycota_unclassified(100);p__Ascomycota_unclassified(100);p__Ascomycota_unclassified(100); | - | - | - | - | - | - | - | Unassigned | - |
| Otu25734 | 1 | k__Fungi(100);p__Ascomycota(100);p__Ascomycota_unclassified(100);p__Ascomycota_unclassified(100);p__Ascomycota_unclassified(100);p__Ascomycota_unclassified(100);p__Ascomycota_unclassified(100); | - | - | - | - | - | - | - | Unassigned | - |
| Otu25736 | 1 | k__Fungi(100);k__Fungi_unclassified(100);k__Fungi_unclassified(100);k__Fungi_unclassified(100);k__Fungi_unclassified(100);k__Fungi_unclassified(100);k__Fungi_unclassified(100); | - | - | - | - | - | - | - | Unassigned | - |
| Otu25737 | 1 | k__Fungi(100);k__Fungi_unclassified(100);k__Fungi_unclassified(100);k__Fungi_unclassified(100);k__Fungi_unclassified(100);k__Fungi_unclassified(100);k__Fungi_unclassified(100); | - | - | - | - | - | - | - | Unassigned | - |
| Otu25742 | 1 | k__Fungi(100);p__Ascomycota(100);p__Ascomycota_unclassified(100);p__Ascomycota_unclassified(100);p__Ascomycota_unclassified(100);p__Ascomycota_unclassified(100);p__Ascomycota_unclassified(100); | - | - | - | - | - | - | - | Unassigned | - |
| Otu25748 | 1 | k__Fungi(100);k__Fungi_unclassified(100);k__Fungi_unclassified(100);k__Fungi_unclassified(100);k__Fungi_unclassified(100);k__Fungi_unclassified(100);k__Fungi_unclassified(100); | - | - | - | - | - | - | - | Unassigned | - |
| Otu25749 | 1 | k__Fungi(100);k__Fungi_unclassified(100);k__Fungi_unclassified(100);k__Fungi_unclassified(100);k__Fungi_unclassified(100);k__Fungi_unclassified(100);k__Fungi_unclassified(100); | - | - | - | - | - | - | - | Unassigned | - |
| Otu25750 | 1 | k__Fungi(100);k__Fungi_unclassified(100);k__Fungi_unclassified(100);k__Fungi_unclassified(100);k__Fungi_unclassified(100);k__Fungi_unclassified(100);k__Fungi_unclassified(100); | - | - | - | - | - | - | - | Unassigned | - |
| Otu25751 | 1 | k__Fungi(100);k__Fungi_unclassified(100);k__Fungi_unclassified(100);k__Fungi_unclassified(100);k__Fungi_unclassified(100);k__Fungi_unclassified(100);k__Fungi_unclassified(100); | - | - | - | - | - | - | - | Unassigned | - |
| Otu25752 | 1 | k__Fungi(100);k__Fungi_unclassified(100);k__Fungi_unclassified(100);k__Fungi_unclassified(100);k__Fungi_unclassified(100);k__Fungi_unclassified(100);k__Fungi_unclassified(100); | - | - | - | - | - | - | - | Unassigned | - |
| Otu25754 | 1 | k__Fungi(100);p__Ascomycota(100);p__Ascomycota_unclassified(100);p__Ascomycota_unclassified(100);p__Ascomycota_unclassified(100);p__Ascomycota_unclassified(100);p__Ascomycota_unclassified(100); | - | - | - | - | - | - | - | Unassigned | - |
| Otu25762 | 1 | k__Fungi(100);p__Ascomycota(100);p__Ascomycota_unclassified(100);p__Ascomycota_unclassified(100);p__Ascomycota_unclassified(100);p__Ascomycota_unclassified(100);p__Ascomycota_unclassified(100); | - | - | - | - | - | - | - | Unassigned | - |
| Otu25763 | 1 | k__Fungi(100);k__Fungi_unclassified(100);k__Fungi_unclassified(100);k__Fungi_unclassified(100);k__Fungi_unclassified(100);k__Fungi_unclassified(100);k__Fungi_unclassified(100); | - | - | - | - | - | - | - | Unassigned | - |
| Otu25768 | 1 | k__Fungi(100);p__Ascomycota(100);p__Ascomycota_unclassified(100);p__Ascomycota_unclassified(100);p__Ascomycota_unclassified(100);p__Ascomycota_unclassified(100);p__Ascomycota_unclassified(100); | - | - | - | - | - | - | - | Unassigned | - |
| Otu25770 | 1 | k__Fungi(100);k__Fungi_unclassified(100);k__Fungi_unclassified(100);k__Fungi_unclassified(100);k__Fungi_unclassified(100);k__Fungi_unclassified(100);k__Fungi_unclassified(100); | - | - | - | - | - | - | - | Unassigned | - |
| Otu25771 | 1 | k__Fungi(100);p__Ascomycota(100);p__Ascomycota_unclassified(100);p__Ascomycota_unclassified(100);p__Ascomycota_unclassified(100);p__Ascomycota_unclassified(100);p__Ascomycota_unclassified(100); | - | - | - | - | - | - | - | Unassigned | - |
| Otu25772 | 1 | k__Fungi(100);k__Fungi_unclassified(100);k__Fungi_unclassified(100);k__Fungi_unclassified(100);k__Fungi_unclassified(100);k__Fungi_unclassified(100);k__Fungi_unclassified(100); | - | - | - | - | - | - | - | Unassigned | - |
| Otu25773 | 1 | k__Fungi(100);k__Fungi_unclassified(100);k__Fungi_unclassified(100);k__Fungi_unclassified(100);k__Fungi_unclassified(100);k__Fungi_unclassified(100);k__Fungi_unclassified(100); | - | - | - | - | - | - | - | Unassigned | - |
| Otu25776 | 1 | k__Fungi(100);k__Fungi_unclassified(100);k__Fungi_unclassified(100);k__Fungi_unclassified(100);k__Fungi_unclassified(100);k__Fungi_unclassified(100);k__Fungi_unclassified(100); | - | - | - | - | - | - | - | Unassigned | - |
| Otu25779 | 1 | k__Fungi(100);k__Fungi_unclassified(100);k__Fungi_unclassified(100);k__Fungi_unclassified(100);k__Fungi_unclassified(100);k__Fungi_unclassified(100);k__Fungi_unclassified(100); | - | - | - | - | - | - | - | Unassigned | - |
| Otu25781 | 1 | k__Fungi(100);k__Fungi_unclassified(100);k__Fungi_unclassified(100);k__Fungi_unclassified(100);k__Fungi_unclassified(100);k__Fungi_unclassified(100);k__Fungi_unclassified(100); | - | - | - | - | - | - | - | Unassigned | - |
| Otu25782 | 1 | k__Fungi(100);p__Ascomycota(100);p__Ascomycota_unclassified(100);p__Ascomycota_unclassified(100);p__Ascomycota_unclassified(100);p__Ascomycota_unclassified(100);p__Ascomycota_unclassified(100); | - | - | - | - | - | - | - | Unassigned | - |
| Otu25783 | 1 | k__Fungi(100);k__Fungi_unclassified(100);k__Fungi_unclassified(100);k__Fungi_unclassified(100);k__Fungi_unclassified(100);k__Fungi_unclassified(100);k__Fungi_unclassified(100); | - | - | - | - | - | - | - | Unassigned | - |
| Otu25784 | 1 | k__Fungi(100);p__Ascomycota(100);p__Ascomycota_unclassified(100);p__Ascomycota_unclassified(100);p__Ascomycota_unclassified(100);p__Ascomycota_unclassified(100);p__Ascomycota_unclassified(100); | - | - | - | - | - | - | - | Unassigned | - |
| Otu25786 | 1 | k__Fungi(100);k__Fungi_unclassified(100);k__Fungi_unclassified(100);k__Fungi_unclassified(100);k__Fungi_unclassified(100);k__Fungi_unclassified(100);k__Fungi_unclassified(100); | - | - | - | - | - | - | - | Unassigned | - |
| Otu25787 | 1 | k__Fungi(100);k__Fungi_unclassified(100);k__Fungi_unclassified(100);k__Fungi_unclassified(100);k__Fungi_unclassified(100);k__Fungi_unclassified(100);k__Fungi_unclassified(100); | - | - | - | - | - | - | - | Unassigned | - |
| Otu25788 | 1 | k__Fungi(100);p__Ascomycota(100);p__Ascomycota_unclassified(100);p__Ascomycota_unclassified(100);p__Ascomycota_unclassified(100);p__Ascomycota_unclassified(100);p__Ascomycota_unclassified(100); | - | - | - | - | - | - | - | Unassigned | - |
| Otu25794 | 1 | k__Fungi(100);k__Fungi_unclassified(100);k__Fungi_unclassified(100);k__Fungi_unclassified(100);k__Fungi_unclassified(100);k__Fungi_unclassified(100);k__Fungi_unclassified(100); | - | - | - | - | - | - | - | Unassigned | - |
| Otu25795 | 1 | k__Fungi(100);k__Fungi_unclassified(100);k__Fungi_unclassified(100);k__Fungi_unclassified(100);k__Fungi_unclassified(100);k__Fungi_unclassified(100);k__Fungi_unclassified(100); | - | - | - | - | - | - | - | Unassigned | - |
| Otu25796 | 1 | k__Fungi(100);p__Ascomycota(100);p__Ascomycota_unclassified(100);p__Ascomycota_unclassified(100);p__Ascomycota_unclassified(100);p__Ascomycota_unclassified(100);p__Ascomycota_unclassified(100); | - | - | - | - | - | - | - | Unassigned | - |
| Otu25799 | 1 | k__Fungi(100);k__Fungi_unclassified(100);k__Fungi_unclassified(100);k__Fungi_unclassified(100);k__Fungi_unclassified(100);k__Fungi_unclassified(100);k__Fungi_unclassified(100); | - | - | - | - | - | - | - | Unassigned | - |
| Otu25801 | 1 | k__Fungi(100);p__Ascomycota(100);p__Ascomycota_unclassified(100);p__Ascomycota_unclassified(100);p__Ascomycota_unclassified(100);p__Ascomycota_unclassified(100);p__Ascomycota_unclassified(100); | - | - | - | - | - | - | - | Unassigned | - |
| Otu25802 | 1 | k__Fungi(100);k__Fungi_unclassified(100);k__Fungi_unclassified(100);k__Fungi_unclassified(100);k__Fungi_unclassified(100);k__Fungi_unclassified(100);k__Fungi_unclassified(100); | - | - | - | - | - | - | - | Unassigned | - |
| Otu25804 | 1 | k__Fungi(100);k__Fungi_unclassified(100);k__Fungi_unclassified(100);k__Fungi_unclassified(100);k__Fungi_unclassified(100);k__Fungi_unclassified(100);k__Fungi_unclassified(100); | - | - | - | - | - | - | - | Unassigned | - |
| Otu25805 | 1 | k__Fungi(100);p__Ascomycota(100);p__Ascomycota_unclassified(100);p__Ascomycota_unclassified(100);p__Ascomycota_unclassified(100);p__Ascomycota_unclassified(100);p__Ascomycota_unclassified(100); | - | - | - | - | - | - | - | Unassigned | - |
| Otu25811 | 1 | k__Fungi(100);k__Fungi_unclassified(100);k__Fungi_unclassified(100);k__Fungi_unclassified(100);k__Fungi_unclassified(100);k__Fungi_unclassified(100);k__Fungi_unclassified(100); | - | - | - | - | - | - | - | Unassigned | - |
| Otu25812 | 1 | k__Fungi(100);k__Fungi_unclassified(100);k__Fungi_unclassified(100);k__Fungi_unclassified(100);k__Fungi_unclassified(100);k__Fungi_unclassified(100);k__Fungi_unclassified(100); | - | - | - | - | - | - | - | Unassigned | - |
| Otu25813 | 1 | k__Fungi(100);k__Fungi_unclassified(100);k__Fungi_unclassified(100);k__Fungi_unclassified(100);k__Fungi_unclassified(100);k__Fungi_unclassified(100);k__Fungi_unclassified(100); | - | - | - | - | - | - | - | Unassigned | - |
| Otu25815 | 1 | k__Fungi(100);p__Ascomycota(100);p__Ascomycota_unclassified(100);p__Ascomycota_unclassified(100);p__Ascomycota_unclassified(100);p__Ascomycota_unclassified(100);p__Ascomycota_unclassified(100); | - | - | - | - | - | - | - | Unassigned | - |
| Otu25816 | 1 | k__Fungi(100);p__Ascomycota(100);p__Ascomycota_unclassified(100);p__Ascomycota_unclassified(100);p__Ascomycota_unclassified(100);p__Ascomycota_unclassified(100);p__Ascomycota_unclassified(100); | - | - | - | - | - | - | - | Unassigned | - |
| Otu25818 | 1 | k__Fungi(100);k__Fungi_unclassified(100);k__Fungi_unclassified(100);k__Fungi_unclassified(100);k__Fungi_unclassified(100);k__Fungi_unclassified(100);k__Fungi_unclassified(100); | - | - | - | - | - | - | - | Unassigned | - |
| Otu25819 | 1 | k__Fungi(100);p__Ascomycota(100);p__Ascomycota_unclassified(100);p__Ascomycota_unclassified(100);p__Ascomycota_unclassified(100);p__Ascomycota_unclassified(100);p__Ascomycota_unclassified(100); | - | - | - | - | - | - | - | Unassigned | - |
| Otu25821 | 1 | k__Fungi(100);k__Fungi_unclassified(100);k__Fungi_unclassified(100);k__Fungi_unclassified(100);k__Fungi_unclassified(100);k__Fungi_unclassified(100);k__Fungi_unclassified(100); | - | - | - | - | - | - | - | Unassigned | - |
| Otu25823 | 1 | k__Fungi(100);k__Fungi_unclassified(100);k__Fungi_unclassified(100);k__Fungi_unclassified(100);k__Fungi_unclassified(100);k__Fungi_unclassified(100);k__Fungi_unclassified(100); | - | - | - | - | - | - | - | Unassigned | - |
| Otu25824 | 1 | k__Fungi(100);p__Ascomycota(100);p__Ascomycota_unclassified(100);p__Ascomycota_unclassified(100);p__Ascomycota_unclassified(100);p__Ascomycota_unclassified(100);p__Ascomycota_unclassified(100); | - | - | - | - | - | - | - | Unassigned | - |
| Otu25827 | 1 | k__Fungi(100);k__Fungi_unclassified(100);k__Fungi_unclassified(100);k__Fungi_unclassified(100);k__Fungi_unclassified(100);k__Fungi_unclassified(100);k__Fungi_unclassified(100); | - | - | - | - | - | - | - | Unassigned | - |
| Otu25828 | 1 | k__Fungi(100);k__Fungi_unclassified(100);k__Fungi_unclassified(100);k__Fungi_unclassified(100);k__Fungi_unclassified(100);k__Fungi_unclassified(100);k__Fungi_unclassified(100); | - | - | - | - | - | - | - | Unassigned | - |
| Otu25829 | 1 | k__Fungi(100);k__Fungi_unclassified(100);k__Fungi_unclassified(100);k__Fungi_unclassified(100);k__Fungi_unclassified(100);k__Fungi_unclassified(100);k__Fungi_unclassified(100); | - | - | - | - | - | - | - | Unassigned | - |
| Otu25831 | 1 | k__Fungi(100);p__Ascomycota(100);p__Ascomycota_unclassified(100);p__Ascomycota_unclassified(100);p__Ascomycota_unclassified(100);p__Ascomycota_unclassified(100);p__Ascomycota_unclassified(100); | - | - | - | - | - | - | - | Unassigned | - |
| Otu25834 | 1 | k__Fungi(100);k__Fungi_unclassified(100);k__Fungi_unclassified(100);k__Fungi_unclassified(100);k__Fungi_unclassified(100);k__Fungi_unclassified(100);k__Fungi_unclassified(100); | - | - | - | - | - | - | - | Unassigned | - |
| Otu25836 | 1 | k__Fungi(100);p__Ascomycota(100);p__Ascomycota_unclassified(100);p__Ascomycota_unclassified(100);p__Ascomycota_unclassified(100);p__Ascomycota_unclassified(100);p__Ascomycota_unclassified(100); | - | - | - | - | - | - | - | Unassigned | - |
| Otu25837 | 1 | k__Fungi(100);k__Fungi_unclassified(100);k__Fungi_unclassified(100);k__Fungi_unclassified(100);k__Fungi_unclassified(100);k__Fungi_unclassified(100);k__Fungi_unclassified(100); | - | - | - | - | - | - | - | Unassigned | - |
| Otu25838 | 1 | k__Fungi(100);k__Fungi_unclassified(100);k__Fungi_unclassified(100);k__Fungi_unclassified(100);k__Fungi_unclassified(100);k__Fungi_unclassified(100);k__Fungi_unclassified(100); | - | - | - | - | - | - | - | Unassigned | - |
| Otu25839 | 1 | k__Fungi(100);p__Ascomycota(100);c__Sordariomycetes(100);c__Sordariomycetes_unclassified(100);c__Sordariomycetes_unclassified(100);c__Sordariomycetes_unclassified(100);c__Sordariomycetes_unclassified(100); | - | - | - | - | - | - | - | Unassigned | - |
| Otu25860 | 1 | k__Fungi(100);p__Ascomycota(100);p__Ascomycota_unclassified(100);p__Ascomycota_unclassified(100);p__Ascomycota_unclassified(100);p__Ascomycota_unclassified(100);p__Ascomycota_unclassified(100); | - | - | - | - | - | - | - | Unassigned | - |
| Otu25865 | 1 | k__Fungi(100);k__Fungi_unclassified(100);k__Fungi_unclassified(100);k__Fungi_unclassified(100);k__Fungi_unclassified(100);k__Fungi_unclassified(100);k__Fungi_unclassified(100); | - | - | - | - | - | - | - | Unassigned | - |
| Otu25867 | 1 | k__Fungi(100);p__Ascomycota(100);p__Ascomycota_unclassified(100);p__Ascomycota_unclassified(100);p__Ascomycota_unclassified(100);p__Ascomycota_unclassified(100);p__Ascomycota_unclassified(100); | - | - | - | - | - | - | - | Unassigned | - |
| Otu25868 | 1 | k__Fungi(100);k__Fungi_unclassified(100);k__Fungi_unclassified(100);k__Fungi_unclassified(100);k__Fungi_unclassified(100);k__Fungi_unclassified(100);k__Fungi_unclassified(100); | - | - | - | - | - | - | - | Unassigned | - |
| Otu25869 | 1 | k__Fungi(100);k__Fungi_unclassified(100);k__Fungi_unclassified(100);k__Fungi_unclassified(100);k__Fungi_unclassified(100);k__Fungi_unclassified(100);k__Fungi_unclassified(100); | - | - | - | - | - | - | - | Unassigned | - |
| Otu25871 | 1 | k__Fungi(100);k__Fungi_unclassified(100);k__Fungi_unclassified(100);k__Fungi_unclassified(100);k__Fungi_unclassified(100);k__Fungi_unclassified(100);k__Fungi_unclassified(100); | - | - | - | - | - | - | - | Unassigned | - |
| Otu25872 | 1 | k__Fungi(100);k__Fungi_unclassified(100);k__Fungi_unclassified(100);k__Fungi_unclassified(100);k__Fungi_unclassified(100);k__Fungi_unclassified(100);k__Fungi_unclassified(100); | - | - | - | - | - | - | - | Unassigned | - |
| Otu25873 | 1 | k__Fungi(100);k__Fungi_unclassified(100);k__Fungi_unclassified(100);k__Fungi_unclassified(100);k__Fungi_unclassified(100);k__Fungi_unclassified(100);k__Fungi_unclassified(100); | - | - | - | - | - | - | - | Unassigned | - |
| Otu25874 | 1 | k__Fungi(100);k__Fungi_unclassified(100);k__Fungi_unclassified(100);k__Fungi_unclassified(100);k__Fungi_unclassified(100);k__Fungi_unclassified(100);k__Fungi_unclassified(100); | - | - | - | - | - | - | - | Unassigned | - |
| Otu25875 | 1 | k__Fungi(100);p__Ascomycota(100);p__Ascomycota_unclassified(100);p__Ascomycota_unclassified(100);p__Ascomycota_unclassified(100);p__Ascomycota_unclassified(100);p__Ascomycota_unclassified(100); | - | - | - | - | - | - | - | Unassigned | - |
| Otu25876 | 1 | k__Fungi(100);p__Ascomycota(100);p__Ascomycota_unclassified(100);p__Ascomycota_unclassified(100);p__Ascomycota_unclassified(100);p__Ascomycota_unclassified(100);p__Ascomycota_unclassified(100); | - | - | - | - | - | - | - | Unassigned | - |
| Otu25878 | 1 | k__Fungi(100);p__Ascomycota(100);p__Ascomycota_unclassified(100);p__Ascomycota_unclassified(100);p__Ascomycota_unclassified(100);p__Ascomycota_unclassified(100);p__Ascomycota_unclassified(100); | - | - | - | - | - | - | - | Unassigned | - |
| Otu25879 | 1 | k__Fungi(100);k__Fungi_unclassified(100);k__Fungi_unclassified(100);k__Fungi_unclassified(100);k__Fungi_unclassified(100);k__Fungi_unclassified(100);k__Fungi_unclassified(100); | - | - | - | - | - | - | - | Unassigned | - |
| Otu25882 | 1 | k__Fungi(100);k__Fungi_unclassified(100);k__Fungi_unclassified(100);k__Fungi_unclassified(100);k__Fungi_unclassified(100);k__Fungi_unclassified(100);k__Fungi_unclassified(100); | - | - | - | - | - | - | - | Unassigned | - |
| Otu25883 | 1 | k__Fungi(100);k__Fungi_unclassified(100);k__Fungi_unclassified(100);k__Fungi_unclassified(100);k__Fungi_unclassified(100);k__Fungi_unclassified(100);k__Fungi_unclassified(100); | - | - | - | - | - | - | - | Unassigned | - |
| Otu25885 | 1 | k__Fungi(100);k__Fungi_unclassified(100);k__Fungi_unclassified(100);k__Fungi_unclassified(100);k__Fungi_unclassified(100);k__Fungi_unclassified(100);k__Fungi_unclassified(100); | - | - | - | - | - | - | - | Unassigned | - |
| Otu25886 | 1 | k__Fungi(100);p__Ascomycota(100);p__Ascomycota_unclassified(100);p__Ascomycota_unclassified(100);p__Ascomycota_unclassified(100);p__Ascomycota_unclassified(100);p__Ascomycota_unclassified(100); | - | - | - | - | - | - | - | Unassigned | - |
| Otu25887 | 1 | k__Fungi(100);k__Fungi_unclassified(100);k__Fungi_unclassified(100);k__Fungi_unclassified(100);k__Fungi_unclassified(100);k__Fungi_unclassified(100);k__Fungi_unclassified(100); | - | - | - | - | - | - | - | Unassigned | - |
| Otu25888 | 1 | k__Fungi(100);k__Fungi_unclassified(100);k__Fungi_unclassified(100);k__Fungi_unclassified(100);k__Fungi_unclassified(100);k__Fungi_unclassified(100);k__Fungi_unclassified(100); | - | - | - | - | - | - | - | Unassigned | - |
| Otu25889 | 1 | k__Fungi(100);k__Fungi_unclassified(100);k__Fungi_unclassified(100);k__Fungi_unclassified(100);k__Fungi_unclassified(100);k__Fungi_unclassified(100);k__Fungi_unclassified(100); | - | - | - | - | - | - | - | Unassigned | - |
| Otu25890 | 1 | k__Fungi(100);k__Fungi_unclassified(100);k__Fungi_unclassified(100);k__Fungi_unclassified(100);k__Fungi_unclassified(100);k__Fungi_unclassified(100);k__Fungi_unclassified(100); | - | - | - | - | - | - | - | Unassigned | - |
| Otu25891 | 1 | k__Fungi(100);p__Ascomycota(100);p__Ascomycota_unclassified(100);p__Ascomycota_unclassified(100);p__Ascomycota_unclassified(100);p__Ascomycota_unclassified(100);p__Ascomycota_unclassified(100); | - | - | - | - | - | - | - | Unassigned | - |
| Otu25892 | 1 | k__Fungi(100);p__Ascomycota(100);p__Ascomycota_unclassified(100);p__Ascomycota_unclassified(100);p__Ascomycota_unclassified(100);p__Ascomycota_unclassified(100);p__Ascomycota_unclassified(100); | - | - | - | - | - | - | - | Unassigned | - |
| Otu25893 | 1 | k__Fungi(100);k__Fungi_unclassified(100);k__Fungi_unclassified(100);k__Fungi_unclassified(100);k__Fungi_unclassified(100);k__Fungi_unclassified(100);k__Fungi_unclassified(100); | - | - | - | - | - | - | - | Unassigned | - |
| Otu25894 | 1 | k__Fungi(100);k__Fungi_unclassified(100);k__Fungi_unclassified(100);k__Fungi_unclassified(100);k__Fungi_unclassified(100);k__Fungi_unclassified(100);k__Fungi_unclassified(100); | - | - | - | - | - | - | - | Unassigned | - |
| Otu25895 | 1 | k__Fungi(100);k__Fungi_unclassified(100);k__Fungi_unclassified(100);k__Fungi_unclassified(100);k__Fungi_unclassified(100);k__Fungi_unclassified(100);k__Fungi_unclassified(100); | - | - | - | - | - | - | - | Unassigned | - |
| Otu25896 | 1 | k__Fungi(100);p__Ascomycota(100);p__Ascomycota_unclassified(100);p__Ascomycota_unclassified(100);p__Ascomycota_unclassified(100);p__Ascomycota_unclassified(100);p__Ascomycota_unclassified(100); | - | - | - | - | - | - | - | Unassigned | - |
| Otu25897 | 1 | k__Fungi(100);k__Fungi_unclassified(100);k__Fungi_unclassified(100);k__Fungi_unclassified(100);k__Fungi_unclassified(100);k__Fungi_unclassified(100);k__Fungi_unclassified(100); | - | - | - | - | - | - | - | Unassigned | - |
| Otu25898 | 1 | k__Fungi(100);k__Fungi_unclassified(100);k__Fungi_unclassified(100);k__Fungi_unclassified(100);k__Fungi_unclassified(100);k__Fungi_unclassified(100);k__Fungi_unclassified(100); | - | - | - | - | - | - | - | Unassigned | - |
| Otu25899 | 1 | k__Fungi(100);p__Ascomycota(100);p__Ascomycota_unclassified(100);p__Ascomycota_unclassified(100);p__Ascomycota_unclassified(100);p__Ascomycota_unclassified(100);p__Ascomycota_unclassified(100); | - | - | - | - | - | - | - | Unassigned | - |
| Otu25901 | 1 | k__Fungi(100);k__Fungi_unclassified(100);k__Fungi_unclassified(100);k__Fungi_unclassified(100);k__Fungi_unclassified(100);k__Fungi_unclassified(100);k__Fungi_unclassified(100); | - | - | - | - | - | - | - | Unassigned | - |
| Otu25902 | 1 | k__Fungi(100);k__Fungi_unclassified(100);k__Fungi_unclassified(100);k__Fungi_unclassified(100);k__Fungi_unclassified(100);k__Fungi_unclassified(100);k__Fungi_unclassified(100); | - | - | - | - | - | - | - | Unassigned | - |
| Otu25903 | 1 | k__Fungi(100);k__Fungi_unclassified(100);k__Fungi_unclassified(100);k__Fungi_unclassified(100);k__Fungi_unclassified(100);k__Fungi_unclassified(100);k__Fungi_unclassified(100); | - | - | - | - | - | - | - | Unassigned | - |
| Otu25904 | 1 | k__Fungi(100);p__Ascomycota(100);p__Ascomycota_unclassified(100);p__Ascomycota_unclassified(100);p__Ascomycota_unclassified(100);p__Ascomycota_unclassified(100);p__Ascomycota_unclassified(100); | - | - | - | - | - | - | - | Unassigned | - |
| Otu25905 | 1 | k__Fungi(100);p__Ascomycota(100);p__Ascomycota_unclassified(100);p__Ascomycota_unclassified(100);p__Ascomycota_unclassified(100);p__Ascomycota_unclassified(100);p__Ascomycota_unclassified(100); | - | - | - | - | - | - | - | Unassigned | - |
| Otu25906 | 1 | k__Fungi(100);p__Ascomycota(100);p__Ascomycota_unclassified(100);p__Ascomycota_unclassified(100);p__Ascomycota_unclassified(100);p__Ascomycota_unclassified(100);p__Ascomycota_unclassified(100); | - | - | - | - | - | - | - | Unassigned | - |
| Otu25907 | 1 | k__Fungi(100);k__Fungi_unclassified(100);k__Fungi_unclassified(100);k__Fungi_unclassified(100);k__Fungi_unclassified(100);k__Fungi_unclassified(100);k__Fungi_unclassified(100); | - | - | - | - | - | - | - | Unassigned | - |
| Otu25908 | 1 | k__Fungi(100);k__Fungi_unclassified(100);k__Fungi_unclassified(100);k__Fungi_unclassified(100);k__Fungi_unclassified(100);k__Fungi_unclassified(100);k__Fungi_unclassified(100); | - | - | - | - | - | - | - | Unassigned | - |
| Otu25909 | 1 | k__Fungi(100);k__Fungi_unclassified(100);k__Fungi_unclassified(100);k__Fungi_unclassified(100);k__Fungi_unclassified(100);k__Fungi_unclassified(100);k__Fungi_unclassified(100); | - | - | - | - | - | - | - | Unassigned | - |
| Otu25910 | 1 | k__Fungi(100);p__Ascomycota(100);p__Ascomycota_unclassified(100);p__Ascomycota_unclassified(100);p__Ascomycota_unclassified(100);p__Ascomycota_unclassified(100);p__Ascomycota_unclassified(100); | - | - | - | - | - | - | - | Unassigned | - |
| Otu25911 | 1 | k__Fungi(100);k__Fungi_unclassified(100);k__Fungi_unclassified(100);k__Fungi_unclassified(100);k__Fungi_unclassified(100);k__Fungi_unclassified(100);k__Fungi_unclassified(100); | - | - | - | - | - | - | - | Unassigned | - |
| Otu25912 | 1 | k__Fungi(100);k__Fungi_unclassified(100);k__Fungi_unclassified(100);k__Fungi_unclassified(100);k__Fungi_unclassified(100);k__Fungi_unclassified(100);k__Fungi_unclassified(100); | - | - | - | - | - | - | - | Unassigned | - |
| Otu25917 | 1 | k__Fungi(100);k__Fungi_unclassified(100);k__Fungi_unclassified(100);k__Fungi_unclassified(100);k__Fungi_unclassified(100);k__Fungi_unclassified(100);k__Fungi_unclassified(100); | - | - | - | - | - | - | - | Unassigned | - |
| Otu25918 | 1 | k__Fungi(100);k__Fungi_unclassified(100);k__Fungi_unclassified(100);k__Fungi_unclassified(100);k__Fungi_unclassified(100);k__Fungi_unclassified(100);k__Fungi_unclassified(100); | - | - | - | - | - | - | - | Unassigned | - |
| Otu25919 | 1 | k__Fungi(100);p__Ascomycota(100);p__Ascomycota_unclassified(100);p__Ascomycota_unclassified(100);p__Ascomycota_unclassified(100);p__Ascomycota_unclassified(100);p__Ascomycota_unclassified(100); | - | - | - | - | - | - | - | Unassigned | - |
| Otu25926 | 1 | k__Fungi(100);k__Fungi_unclassified(100);k__Fungi_unclassified(100);k__Fungi_unclassified(100);k__Fungi_unclassified(100);k__Fungi_unclassified(100);k__Fungi_unclassified(100); | - | - | - | - | - | - | - | Unassigned | - |
| Otu25931 | 1 | k__Fungi(100);k__Fungi_unclassified(100);k__Fungi_unclassified(100);k__Fungi_unclassified(100);k__Fungi_unclassified(100);k__Fungi_unclassified(100);k__Fungi_unclassified(100); | - | - | - | - | - | - | - | Unassigned | - |
| Otu25936 | 1 | k__Fungi(100);k__Fungi_unclassified(100);k__Fungi_unclassified(100);k__Fungi_unclassified(100);k__Fungi_unclassified(100);k__Fungi_unclassified(100);k__Fungi_unclassified(100); | - | - | - | - | - | - | - | Unassigned | - |
| Otu25937 | 1 | k__Fungi(100);k__Fungi_unclassified(100);k__Fungi_unclassified(100);k__Fungi_unclassified(100);k__Fungi_unclassified(100);k__Fungi_unclassified(100);k__Fungi_unclassified(100); | - | - | - | - | - | - | - | Unassigned | - |
| Otu25939 | 1 | k__Fungi(100);p__Ascomycota(100);p__Ascomycota_unclassified(100);p__Ascomycota_unclassified(100);p__Ascomycota_unclassified(100);p__Ascomycota_unclassified(100);p__Ascomycota_unclassified(100); | - | - | - | - | - | - | - | Unassigned | - |
| Otu25940 | 1 | k__Fungi(100);k__Fungi_unclassified(100);k__Fungi_unclassified(100);k__Fungi_unclassified(100);k__Fungi_unclassified(100);k__Fungi_unclassified(100);k__Fungi_unclassified(100); | - | - | - | - | - | - | - | Unassigned | - |
| Otu25941 | 1 | k__Fungi(100);k__Fungi_unclassified(100);k__Fungi_unclassified(100);k__Fungi_unclassified(100);k__Fungi_unclassified(100);k__Fungi_unclassified(100);k__Fungi_unclassified(100); | - | - | - | - | - | - | - | Unassigned | - |
| Otu25942 | 1 | k__Fungi(100);p__Ascomycota(100);p__Ascomycota_unclassified(100);p__Ascomycota_unclassified(100);p__Ascomycota_unclassified(100);p__Ascomycota_unclassified(100);p__Ascomycota_unclassified(100); | - | - | - | - | - | - | - | Unassigned | - |
| Otu25943 | 1 | k__Fungi(100);p__Ascomycota(100);p__Ascomycota_unclassified(100);p__Ascomycota_unclassified(100);p__Ascomycota_unclassified(100);p__Ascomycota_unclassified(100);p__Ascomycota_unclassified(100); | - | - | - | - | - | - | - | Unassigned | - |
| Otu25944 | 1 | k__Fungi(100);k__Fungi_unclassified(100);k__Fungi_unclassified(100);k__Fungi_unclassified(100);k__Fungi_unclassified(100);k__Fungi_unclassified(100);k__Fungi_unclassified(100); | - | - | - | - | - | - | - | Unassigned | - |
| Otu25947 | 1 | k__Fungi(100);p__Ascomycota(100);p__Ascomycota_unclassified(100);p__Ascomycota_unclassified(100);p__Ascomycota_unclassified(100);p__Ascomycota_unclassified(100);p__Ascomycota_unclassified(100); | - | - | - | - | - | - | - | Unassigned | - |
| Otu25950 | 1 | k__Fungi(100);k__Fungi_unclassified(100);k__Fungi_unclassified(100);k__Fungi_unclassified(100);k__Fungi_unclassified(100);k__Fungi_unclassified(100);k__Fungi_unclassified(100); | - | - | - | - | - | - | - | Unassigned | - |
| Otu25951 | 1 | k__Fungi(100);k__Fungi_unclassified(100);k__Fungi_unclassified(100);k__Fungi_unclassified(100);k__Fungi_unclassified(100);k__Fungi_unclassified(100);k__Fungi_unclassified(100); | - | - | - | - | - | - | - | Unassigned | - |
| Otu25953 | 1 | k__Fungi(100);k__Fungi_unclassified(100);k__Fungi_unclassified(100);k__Fungi_unclassified(100);k__Fungi_unclassified(100);k__Fungi_unclassified(100);k__Fungi_unclassified(100); | - | - | - | - | - | - | - | Unassigned | - |
| Otu25954 | 1 | k__Fungi(100);k__Fungi_unclassified(100);k__Fungi_unclassified(100);k__Fungi_unclassified(100);k__Fungi_unclassified(100);k__Fungi_unclassified(100);k__Fungi_unclassified(100); | - | - | - | - | - | - | - | Unassigned | - |
| Otu25958 | 1 | k__Fungi(100);p__Ascomycota(100);p__Ascomycota_unclassified(100);p__Ascomycota_unclassified(100);p__Ascomycota_unclassified(100);p__Ascomycota_unclassified(100);p__Ascomycota_unclassified(100); | - | - | - | - | - | - | - | Unassigned | - |
| Otu25964 | 1 | k__Fungi(100);p__Ascomycota(100);p__Ascomycota_unclassified(100);p__Ascomycota_unclassified(100);p__Ascomycota_unclassified(100);p__Ascomycota_unclassified(100);p__Ascomycota_unclassified(100); | - | - | - | - | - | - | - | Unassigned | - |
| Otu25965 | 1 | k__Fungi(100);k__Fungi_unclassified(100);k__Fungi_unclassified(100);k__Fungi_unclassified(100);k__Fungi_unclassified(100);k__Fungi_unclassified(100);k__Fungi_unclassified(100); | - | - | - | - | - | - | - | Unassigned | - |
| Otu25966 | 1 | k__Fungi(100);k__Fungi_unclassified(100);k__Fungi_unclassified(100);k__Fungi_unclassified(100);k__Fungi_unclassified(100);k__Fungi_unclassified(100);k__Fungi_unclassified(100); | - | - | - | - | - | - | - | Unassigned | - |
| Otu25968 | 1 | k__Fungi(100);k__Fungi_unclassified(100);k__Fungi_unclassified(100);k__Fungi_unclassified(100);k__Fungi_unclassified(100);k__Fungi_unclassified(100);k__Fungi_unclassified(100); | - | - | - | - | - | - | - | Unassigned | - |
| Otu25969 | 1 | k__Fungi(100);k__Fungi_unclassified(100);k__Fungi_unclassified(100);k__Fungi_unclassified(100);k__Fungi_unclassified(100);k__Fungi_unclassified(100);k__Fungi_unclassified(100); | - | - | - | - | - | - | - | Unassigned | - |
| Otu25970 | 1 | k__Fungi(100);k__Fungi_unclassified(100);k__Fungi_unclassified(100);k__Fungi_unclassified(100);k__Fungi_unclassified(100);k__Fungi_unclassified(100);k__Fungi_unclassified(100); | - | - | - | - | - | - | - | Unassigned | - |
| Otu25971 | 1 | k__Fungi(100);k__Fungi_unclassified(100);k__Fungi_unclassified(100);k__Fungi_unclassified(100);k__Fungi_unclassified(100);k__Fungi_unclassified(100);k__Fungi_unclassified(100); | - | - | - | - | - | - | - | Unassigned | - |
| Otu25972 | 1 | k__Fungi(100);k__Fungi_unclassified(100);k__Fungi_unclassified(100);k__Fungi_unclassified(100);k__Fungi_unclassified(100);k__Fungi_unclassified(100);k__Fungi_unclassified(100); | - | - | - | - | - | - | - | Unassigned | - |
| Otu25973 | 1 | k__Fungi(100);k__Fungi_unclassified(100);k__Fungi_unclassified(100);k__Fungi_unclassified(100);k__Fungi_unclassified(100);k__Fungi_unclassified(100);k__Fungi_unclassified(100); | - | - | - | - | - | - | - | Unassigned | - |
| Otu25975 | 1 | k__Fungi(100);k__Fungi_unclassified(100);k__Fungi_unclassified(100);k__Fungi_unclassified(100);k__Fungi_unclassified(100);k__Fungi_unclassified(100);k__Fungi_unclassified(100); | - | - | - | - | - | - | - | Unassigned | - |
| Otu25976 | 1 | k__Fungi(100);k__Fungi_unclassified(100);k__Fungi_unclassified(100);k__Fungi_unclassified(100);k__Fungi_unclassified(100);k__Fungi_unclassified(100);k__Fungi_unclassified(100); | - | - | - | - | - | - | - | Unassigned | - |
| Otu25977 | 1 | k__Fungi(100);k__Fungi_unclassified(100);k__Fungi_unclassified(100);k__Fungi_unclassified(100);k__Fungi_unclassified(100);k__Fungi_unclassified(100);k__Fungi_unclassified(100); | - | - | - | - | - | - | - | Unassigned | - |
| Otu25978 | 1 | k__Fungi(100);k__Fungi_unclassified(100);k__Fungi_unclassified(100);k__Fungi_unclassified(100);k__Fungi_unclassified(100);k__Fungi_unclassified(100);k__Fungi_unclassified(100); | - | - | - | - | - | - | - | Unassigned | - |
| Otu25979 | 1 | k__Fungi(100);p__Ascomycota(100);p__Ascomycota_unclassified(100);p__Ascomycota_unclassified(100);p__Ascomycota_unclassified(100);p__Ascomycota_unclassified(100);p__Ascomycota_unclassified(100); | - | - | - | - | - | - | - | Unassigned | - |
| Otu25980 | 1 | k__Fungi(100);p__Ascomycota(100);p__Ascomycota_unclassified(100);p__Ascomycota_unclassified(100);p__Ascomycota_unclassified(100);p__Ascomycota_unclassified(100);p__Ascomycota_unclassified(100); | - | - | - | - | - | - | - | Unassigned | - |
| Otu25981 | 1 | k__Fungi(100);k__Fungi_unclassified(100);k__Fungi_unclassified(100);k__Fungi_unclassified(100);k__Fungi_unclassified(100);k__Fungi_unclassified(100);k__Fungi_unclassified(100); | - | - | - | - | - | - | - | Unassigned | - |
| Otu25982 | 1 | k__Fungi(100);k__Fungi_unclassified(100);k__Fungi_unclassified(100);k__Fungi_unclassified(100);k__Fungi_unclassified(100);k__Fungi_unclassified(100);k__Fungi_unclassified(100); | - | - | - | - | - | - | - | Unassigned | - |
| Otu25983 | 1 | k__Fungi(100);p__Ascomycota(100);p__Ascomycota_unclassified(100);p__Ascomycota_unclassified(100);p__Ascomycota_unclassified(100);p__Ascomycota_unclassified(100);p__Ascomycota_unclassified(100); | - | - | - | - | - | - | - | Unassigned | - |
| Otu25984 | 1 | k__Fungi(100);k__Fungi_unclassified(100);k__Fungi_unclassified(100);k__Fungi_unclassified(100);k__Fungi_unclassified(100);k__Fungi_unclassified(100);k__Fungi_unclassified(100); | - | - | - | - | - | - | - | Unassigned | - |
| Otu25985 | 1 | k__Fungi(100);k__Fungi_unclassified(100);k__Fungi_unclassified(100);k__Fungi_unclassified(100);k__Fungi_unclassified(100);k__Fungi_unclassified(100);k__Fungi_unclassified(100); | - | - | - | - | - | - | - | Unassigned | - |
| Otu25986 | 1 | k__Fungi(100);p__Ascomycota(100);p__Ascomycota_unclassified(100);p__Ascomycota_unclassified(100);p__Ascomycota_unclassified(100);p__Ascomycota_unclassified(100);p__Ascomycota_unclassified(100); | - | - | - | - | - | - | - | Unassigned | - |
| Otu25987 | 1 | k__Fungi(100);k__Fungi_unclassified(100);k__Fungi_unclassified(100);k__Fungi_unclassified(100);k__Fungi_unclassified(100);k__Fungi_unclassified(100);k__Fungi_unclassified(100); | - | - | - | - | - | - | - | Unassigned | - |
| Otu25988 | 1 | k__Fungi(100);k__Fungi_unclassified(100);k__Fungi_unclassified(100);k__Fungi_unclassified(100);k__Fungi_unclassified(100);k__Fungi_unclassified(100);k__Fungi_unclassified(100); | - | - | - | - | - | - | - | Unassigned | - |
| Otu25989 | 1 | k__Fungi(100);p__Ascomycota(100);p__Ascomycota_unclassified(100);p__Ascomycota_unclassified(100);p__Ascomycota_unclassified(100);p__Ascomycota_unclassified(100);p__Ascomycota_unclassified(100); | - | - | - | - | - | - | - | Unassigned | - |
| Otu25990 | 1 | k__Fungi(100);p__Ascomycota(100);p__Ascomycota_unclassified(100);p__Ascomycota_unclassified(100);p__Ascomycota_unclassified(100);p__Ascomycota_unclassified(100);p__Ascomycota_unclassified(100); | - | - | - | - | - | - | - | Unassigned | - |
| Otu25991 | 1 | k__Fungi(100);k__Fungi_unclassified(100);k__Fungi_unclassified(100);k__Fungi_unclassified(100);k__Fungi_unclassified(100);k__Fungi_unclassified(100);k__Fungi_unclassified(100); | - | - | - | - | - | - | - | Unassigned | - |
| Otu25993 | 1 | k__Fungi(100);k__Fungi_unclassified(100);k__Fungi_unclassified(100);k__Fungi_unclassified(100);k__Fungi_unclassified(100);k__Fungi_unclassified(100);k__Fungi_unclassified(100); | - | - | - | - | - | - | - | Unassigned | - |
| Otu25996 | 1 | k__Fungi(100);p__Ascomycota(100);p__Ascomycota_unclassified(100);p__Ascomycota_unclassified(100);p__Ascomycota_unclassified(100);p__Ascomycota_unclassified(100);p__Ascomycota_unclassified(100); | - | - | - | - | - | - | - | Unassigned | - |
| Otu26000 | 1 | k__Fungi(100);k__Fungi_unclassified(100);k__Fungi_unclassified(100);k__Fungi_unclassified(100);k__Fungi_unclassified(100);k__Fungi_unclassified(100);k__Fungi_unclassified(100); | - | - | - | - | - | - | - | Unassigned | - |
| Otu26008 | 1 | k__Fungi(100);p__Ascomycota(100);p__Ascomycota_unclassified(100);p__Ascomycota_unclassified(100);p__Ascomycota_unclassified(100);p__Ascomycota_unclassified(100);p__Ascomycota_unclassified(100); | - | - | - | - | - | - | - | Unassigned | - |
| Otu26010 | 1 | k__Fungi(100);p__Ascomycota(100);p__Ascomycota_unclassified(100);p__Ascomycota_unclassified(100);p__Ascomycota_unclassified(100);p__Ascomycota_unclassified(100);p__Ascomycota_unclassified(100); | - | - | - | - | - | - | - | Unassigned | - |
| Otu26012 | 1 | k__Fungi(100);p__Ascomycota(100);p__Ascomycota_unclassified(100);p__Ascomycota_unclassified(100);p__Ascomycota_unclassified(100);p__Ascomycota_unclassified(100);p__Ascomycota_unclassified(100); | - | - | - | - | - | - | - | Unassigned | - |
| Otu26014 | 1 | k__Fungi(100);k__Fungi_unclassified(100);k__Fungi_unclassified(100);k__Fungi_unclassified(100);k__Fungi_unclassified(100);k__Fungi_unclassified(100);k__Fungi_unclassified(100); | - | - | - | - | - | - | - | Unassigned | - |
| Otu26015 | 1 | k__Fungi(100);p__Ascomycota(100);p__Ascomycota_unclassified(100);p__Ascomycota_unclassified(100);p__Ascomycota_unclassified(100);p__Ascomycota_unclassified(100);p__Ascomycota_unclassified(100); | - | - | - | - | - | - | - | Unassigned | - |
| Otu26023 | 1 | k__Fungi(100);p__Ascomycota(100);p__Ascomycota_unclassified(100);p__Ascomycota_unclassified(100);p__Ascomycota_unclassified(100);p__Ascomycota_unclassified(100);p__Ascomycota_unclassified(100); | - | - | - | - | - | - | - | Unassigned | - |
| Otu26024 | 1 | k__Fungi(100);p__Ascomycota(100);p__Ascomycota_unclassified(100);p__Ascomycota_unclassified(100);p__Ascomycota_unclassified(100);p__Ascomycota_unclassified(100);p__Ascomycota_unclassified(100); | - | - | - | - | - | - | - | Unassigned | - |
| Otu26036 | 1 | k__Fungi(100);k__Fungi_unclassified(100);k__Fungi_unclassified(100);k__Fungi_unclassified(100);k__Fungi_unclassified(100);k__Fungi_unclassified(100);k__Fungi_unclassified(100); | - | - | - | - | - | - | - | Unassigned | - |
| Otu26040 | 1 | k__Fungi(100);p__Ascomycota(100);p__Ascomycota_unclassified(100);p__Ascomycota_unclassified(100);p__Ascomycota_unclassified(100);p__Ascomycota_unclassified(100);p__Ascomycota_unclassified(100); | - | - | - | - | - | - | - | Unassigned | - |
| Otu26045 | 1 | k__Fungi(100);p__Ascomycota(100);p__Ascomycota_unclassified(100);p__Ascomycota_unclassified(100);p__Ascomycota_unclassified(100);p__Ascomycota_unclassified(100);p__Ascomycota_unclassified(100); | - | - | - | - | - | - | - | Unassigned | - |
| Otu26047 | 1 | k__Fungi(100);k__Fungi_unclassified(100);k__Fungi_unclassified(100);k__Fungi_unclassified(100);k__Fungi_unclassified(100);k__Fungi_unclassified(100);k__Fungi_unclassified(100); | - | - | - | - | - | - | - | Unassigned | - |
| Otu26048 | 1 | k__Fungi(100);k__Fungi_unclassified(100);k__Fungi_unclassified(100);k__Fungi_unclassified(100);k__Fungi_unclassified(100);k__Fungi_unclassified(100);k__Fungi_unclassified(100); | - | - | - | - | - | - | - | Unassigned | - |
| Otu26049 | 1 | k__Fungi(100);k__Fungi_unclassified(100);k__Fungi_unclassified(100);k__Fungi_unclassified(100);k__Fungi_unclassified(100);k__Fungi_unclassified(100);k__Fungi_unclassified(100); | - | - | - | - | - | - | - | Unassigned | - |
| Otu26050 | 1 | k__Fungi(100);p__Ascomycota(100);p__Ascomycota_unclassified(100);p__Ascomycota_unclassified(100);p__Ascomycota_unclassified(100);p__Ascomycota_unclassified(100);p__Ascomycota_unclassified(100); | - | - | - | - | - | - | - | Unassigned | - |
| Otu26051 | 1 | k__Fungi(100);p__Ascomycota(100);p__Ascomycota_unclassified(100);p__Ascomycota_unclassified(100);p__Ascomycota_unclassified(100);p__Ascomycota_unclassified(100);p__Ascomycota_unclassified(100); | - | - | - | - | - | - | - | Unassigned | - |
| Otu26053 | 1 | k__Fungi(100);k__Fungi_unclassified(100);k__Fungi_unclassified(100);k__Fungi_unclassified(100);k__Fungi_unclassified(100);k__Fungi_unclassified(100);k__Fungi_unclassified(100); | - | - | - | - | - | - | - | Unassigned | - |
| Otu26054 | 1 | k__Fungi(100);k__Fungi_unclassified(100);k__Fungi_unclassified(100);k__Fungi_unclassified(100);k__Fungi_unclassified(100);k__Fungi_unclassified(100);k__Fungi_unclassified(100); | - | - | - | - | - | - | - | Unassigned | - |
| Otu26055 | 1 | k__Fungi(100);p__Ascomycota(100);p__Ascomycota_unclassified(100);p__Ascomycota_unclassified(100);p__Ascomycota_unclassified(100);p__Ascomycota_unclassified(100);p__Ascomycota_unclassified(100); | - | - | - | - | - | - | - | Unassigned | - |
| Otu26056 | 1 | k__Fungi(100);k__Fungi_unclassified(100);k__Fungi_unclassified(100);k__Fungi_unclassified(100);k__Fungi_unclassified(100);k__Fungi_unclassified(100);k__Fungi_unclassified(100); | - | - | - | - | - | - | - | Unassigned | - |
| Otu26058 | 1 | k__Fungi(100);k__Fungi_unclassified(100);k__Fungi_unclassified(100);k__Fungi_unclassified(100);k__Fungi_unclassified(100);k__Fungi_unclassified(100);k__Fungi_unclassified(100); | - | - | - | - | - | - | - | Unassigned | - |
| Otu26060 | 1 | k__Fungi(100);k__Fungi_unclassified(100);k__Fungi_unclassified(100);k__Fungi_unclassified(100);k__Fungi_unclassified(100);k__Fungi_unclassified(100);k__Fungi_unclassified(100); | - | - | - | - | - | - | - | Unassigned | - |
| Otu26061 | 1 | k__Fungi(100);k__Fungi_unclassified(100);k__Fungi_unclassified(100);k__Fungi_unclassified(100);k__Fungi_unclassified(100);k__Fungi_unclassified(100);k__Fungi_unclassified(100); | - | - | - | - | - | - | - | Unassigned | - |
| Otu26063 | 1 | k__Fungi(100);p__Ascomycota(100);p__Ascomycota_unclassified(100);p__Ascomycota_unclassified(100);p__Ascomycota_unclassified(100);p__Ascomycota_unclassified(100);p__Ascomycota_unclassified(100); | - | - | - | - | - | - | - | Unassigned | - |
| Otu26064 | 1 | k__Fungi(100);k__Fungi_unclassified(100);k__Fungi_unclassified(100);k__Fungi_unclassified(100);k__Fungi_unclassified(100);k__Fungi_unclassified(100);k__Fungi_unclassified(100); | - | - | - | - | - | - | - | Unassigned | - |
| Otu26066 | 1 | k__Fungi(100);k__Fungi_unclassified(100);k__Fungi_unclassified(100);k__Fungi_unclassified(100);k__Fungi_unclassified(100);k__Fungi_unclassified(100);k__Fungi_unclassified(100); | - | - | - | - | - | - | - | Unassigned | - |
| Otu26067 | 1 | k__Fungi(100);k__Fungi_unclassified(100);k__Fungi_unclassified(100);k__Fungi_unclassified(100);k__Fungi_unclassified(100);k__Fungi_unclassified(100);k__Fungi_unclassified(100); | - | - | - | - | - | - | - | Unassigned | - |
| Otu26068 | 1 | k__Fungi(100);k__Fungi_unclassified(100);k__Fungi_unclassified(100);k__Fungi_unclassified(100);k__Fungi_unclassified(100);k__Fungi_unclassified(100);k__Fungi_unclassified(100); | - | - | - | - | - | - | - | Unassigned | - |
| Otu26069 | 1 | k__Fungi(100);p__Ascomycota(100);p__Ascomycota_unclassified(100);p__Ascomycota_unclassified(100);p__Ascomycota_unclassified(100);p__Ascomycota_unclassified(100);p__Ascomycota_unclassified(100); | - | - | - | - | - | - | - | Unassigned | - |
| Otu26070 | 1 | k__Fungi(100);k__Fungi_unclassified(100);k__Fungi_unclassified(100);k__Fungi_unclassified(100);k__Fungi_unclassified(100);k__Fungi_unclassified(100);k__Fungi_unclassified(100); | - | - | - | - | - | - | - | Unassigned | - |
| Otu26076 | 1 | k__Fungi(100);p__Ascomycota(100);p__Ascomycota_unclassified(100);p__Ascomycota_unclassified(100);p__Ascomycota_unclassified(100);p__Ascomycota_unclassified(100);p__Ascomycota_unclassified(100); | - | - | - | - | - | - | - | Unassigned | - |
| Otu26085 | 1 | k__Fungi(100);p__Ascomycota(100);p__Ascomycota_unclassified(100);p__Ascomycota_unclassified(100);p__Ascomycota_unclassified(100);p__Ascomycota_unclassified(100);p__Ascomycota_unclassified(100); | - | - | - | - | - | - | - | Unassigned | - |
| Otu26093 | 1 | k__Fungi(100);p__Ascomycota(100);p__Ascomycota_unclassified(100);p__Ascomycota_unclassified(100);p__Ascomycota_unclassified(100);p__Ascomycota_unclassified(100);p__Ascomycota_unclassified(100); | - | - | - | - | - | - | - | Unassigned | - |
| Otu26094 | 1 | k__Fungi(100);k__Fungi_unclassified(100);k__Fungi_unclassified(100);k__Fungi_unclassified(100);k__Fungi_unclassified(100);k__Fungi_unclassified(100);k__Fungi_unclassified(100); | - | - | - | - | - | - | - | Unassigned | - |
| Otu26101 | 1 | k__Fungi(100);p__Ascomycota(100);p__Ascomycota_unclassified(100);p__Ascomycota_unclassified(100);p__Ascomycota_unclassified(100);p__Ascomycota_unclassified(100);p__Ascomycota_unclassified(100); | - | - | - | - | - | - | - | Unassigned | - |
| Otu26102 | 1 | k__Fungi(100);k__Fungi_unclassified(100);k__Fungi_unclassified(100);k__Fungi_unclassified(100);k__Fungi_unclassified(100);k__Fungi_unclassified(100);k__Fungi_unclassified(100); | - | - | - | - | - | - | - | Unassigned | - |
| Otu26107 | 1 | k__Fungi(100);k__Fungi_unclassified(100);k__Fungi_unclassified(100);k__Fungi_unclassified(100);k__Fungi_unclassified(100);k__Fungi_unclassified(100);k__Fungi_unclassified(100); | - | - | - | - | - | - | - | Unassigned | - |
| Otu26112 | 1 | k__Fungi(100);k__Fungi_unclassified(100);k__Fungi_unclassified(100);k__Fungi_unclassified(100);k__Fungi_unclassified(100);k__Fungi_unclassified(100);k__Fungi_unclassified(100); | - | - | - | - | - | - | - | Unassigned | - |
| Otu26119 | 1 | k__Fungi(100);k__Fungi_unclassified(100);k__Fungi_unclassified(100);k__Fungi_unclassified(100);k__Fungi_unclassified(100);k__Fungi_unclassified(100);k__Fungi_unclassified(100); | - | - | - | - | - | - | - | Unassigned | - |
| Otu26120 | 1 | k__Fungi(100);p__Ascomycota(100);p__Ascomycota_unclassified(100);p__Ascomycota_unclassified(100);p__Ascomycota_unclassified(100);p__Ascomycota_unclassified(100);p__Ascomycota_unclassified(100); | - | - | - | - | - | - | - | Unassigned | - |
| Otu26122 | 1 | k__Fungi(100);p__Ascomycota(100);p__Ascomycota_unclassified(100);p__Ascomycota_unclassified(100);p__Ascomycota_unclassified(100);p__Ascomycota_unclassified(100);p__Ascomycota_unclassified(100); | - | - | - | - | - | - | - | Unassigned | - |
| Otu26124 | 1 | k__Fungi(100);p__Ascomycota(100);p__Ascomycota_unclassified(100);p__Ascomycota_unclassified(100);p__Ascomycota_unclassified(100);p__Ascomycota_unclassified(100);p__Ascomycota_unclassified(100); | - | - | - | - | - | - | - | Unassigned | - |
| Otu26126 | 1 | k__Fungi(100);k__Fungi_unclassified(100);k__Fungi_unclassified(100);k__Fungi_unclassified(100);k__Fungi_unclassified(100);k__Fungi_unclassified(100);k__Fungi_unclassified(100); | - | - | - | - | - | - | - | Unassigned | - |
| Otu26127 | 1 | k__Fungi(100);p__Ascomycota(100);p__Ascomycota_unclassified(100);p__Ascomycota_unclassified(100);p__Ascomycota_unclassified(100);p__Ascomycota_unclassified(100);p__Ascomycota_unclassified(100); | - | - | - | - | - | - | - | Unassigned | - |
| Otu26128 | 1 | k__Fungi(100);k__Fungi_unclassified(100);k__Fungi_unclassified(100);k__Fungi_unclassified(100);k__Fungi_unclassified(100);k__Fungi_unclassified(100);k__Fungi_unclassified(100); | - | - | - | - | - | - | - | Unassigned | - |
| Otu26129 | 1 | k__Fungi(100);k__Fungi_unclassified(100);k__Fungi_unclassified(100);k__Fungi_unclassified(100);k__Fungi_unclassified(100);k__Fungi_unclassified(100);k__Fungi_unclassified(100); | - | - | - | - | - | - | - | Unassigned | - |
| Otu26131 | 1 | k__Fungi(100);p__Ascomycota(100);p__Ascomycota_unclassified(100);p__Ascomycota_unclassified(100);p__Ascomycota_unclassified(100);p__Ascomycota_unclassified(100);p__Ascomycota_unclassified(100); | - | - | - | - | - | - | - | Unassigned | - |
| Otu26132 | 1 | k__Fungi(100);k__Fungi_unclassified(100);k__Fungi_unclassified(100);k__Fungi_unclassified(100);k__Fungi_unclassified(100);k__Fungi_unclassified(100);k__Fungi_unclassified(100); | - | - | - | - | - | - | - | Unassigned | - |
| Otu26133 | 1 | k__Fungi(100);k__Fungi_unclassified(100);k__Fungi_unclassified(100);k__Fungi_unclassified(100);k__Fungi_unclassified(100);k__Fungi_unclassified(100);k__Fungi_unclassified(100); | - | - | - | - | - | - | - | Unassigned | - |
| Otu26134 | 1 | k__Fungi(100);k__Fungi_unclassified(100);k__Fungi_unclassified(100);k__Fungi_unclassified(100);k__Fungi_unclassified(100);k__Fungi_unclassified(100);k__Fungi_unclassified(100); | - | - | - | - | - | - | - | Unassigned | - |
| Otu26135 | 1 | k__Fungi(100);k__Fungi_unclassified(100);k__Fungi_unclassified(100);k__Fungi_unclassified(100);k__Fungi_unclassified(100);k__Fungi_unclassified(100);k__Fungi_unclassified(100); | - | - | - | - | - | - | - | Unassigned | - |
| Otu26140 | 1 | k__Fungi(100);k__Fungi_unclassified(100);k__Fungi_unclassified(100);k__Fungi_unclassified(100);k__Fungi_unclassified(100);k__Fungi_unclassified(100);k__Fungi_unclassified(100); | - | - | - | - | - | - | - | Unassigned | - |
| Otu26141 | 1 | k__Fungi(100);k__Fungi_unclassified(100);k__Fungi_unclassified(100);k__Fungi_unclassified(100);k__Fungi_unclassified(100);k__Fungi_unclassified(100);k__Fungi_unclassified(100); | - | - | - | - | - | - | - | Unassigned | - |
| Otu26147 | 1 | k__Fungi(100);p__Ascomycota(100);p__Ascomycota_unclassified(100);p__Ascomycota_unclassified(100);p__Ascomycota_unclassified(100);p__Ascomycota_unclassified(100);p__Ascomycota_unclassified(100); | - | - | - | - | - | - | - | Unassigned | - |
| Otu26148 | 1 | k__Fungi(100);k__Fungi_unclassified(100);k__Fungi_unclassified(100);k__Fungi_unclassified(100);k__Fungi_unclassified(100);k__Fungi_unclassified(100);k__Fungi_unclassified(100); | - | - | - | - | - | - | - | Unassigned | - |
| Otu26149 | 1 | k__Fungi(100);p__Ascomycota(100);p__Ascomycota_unclassified(100);p__Ascomycota_unclassified(100);p__Ascomycota_unclassified(100);p__Ascomycota_unclassified(100);p__Ascomycota_unclassified(100); | - | - | - | - | - | - | - | Unassigned | - |
| Otu26150 | 1 | k__Fungi(100);k__Fungi_unclassified(100);k__Fungi_unclassified(100);k__Fungi_unclassified(100);k__Fungi_unclassified(100);k__Fungi_unclassified(100);k__Fungi_unclassified(100); | - | - | - | - | - | - | - | Unassigned | - |
| Otu26155 | 1 | k__Fungi(100);k__Fungi_unclassified(100);k__Fungi_unclassified(100);k__Fungi_unclassified(100);k__Fungi_unclassified(100);k__Fungi_unclassified(100);k__Fungi_unclassified(100); | - | - | - | - | - | - | - | Unassigned | - |
| Otu26179 | 1 | k__Fungi(100);p__Ascomycota(100);p__Ascomycota_unclassified(100);p__Ascomycota_unclassified(100);p__Ascomycota_unclassified(100);p__Ascomycota_unclassified(100);p__Ascomycota_unclassified(100); | - | - | - | - | - | - | - | Unassigned | - |
| Otu26180 | 1 | k__Fungi(100);p__Ascomycota(100);p__Ascomycota_unclassified(100);p__Ascomycota_unclassified(100);p__Ascomycota_unclassified(100);p__Ascomycota_unclassified(100);p__Ascomycota_unclassified(100); | - | - | - | - | - | - | - | Unassigned | - |
| Otu26181 | 1 | k__Fungi(100);k__Fungi_unclassified(100);k__Fungi_unclassified(100);k__Fungi_unclassified(100);k__Fungi_unclassified(100);k__Fungi_unclassified(100);k__Fungi_unclassified(100); | - | - | - | - | - | - | - | Unassigned | - |
| Otu26182 | 1 | k__Fungi(100);p__Ascomycota(100);p__Ascomycota_unclassified(100);p__Ascomycota_unclassified(100);p__Ascomycota_unclassified(100);p__Ascomycota_unclassified(100);p__Ascomycota_unclassified(100); | - | - | - | - | - | - | - | Unassigned | - |
| Otu26184 | 1 | k__Fungi(100);k__Fungi_unclassified(100);k__Fungi_unclassified(100);k__Fungi_unclassified(100);k__Fungi_unclassified(100);k__Fungi_unclassified(100);k__Fungi_unclassified(100); | - | - | - | - | - | - | - | Unassigned | - |
| Otu26186 | 1 | k__Fungi(100);p__Ascomycota(100);p__Ascomycota_unclassified(100);p__Ascomycota_unclassified(100);p__Ascomycota_unclassified(100);p__Ascomycota_unclassified(100);p__Ascomycota_unclassified(100); | - | - | - | - | - | - | - | Unassigned | - |
| Otu26187 | 1 | k__Fungi(100);p__Ascomycota(100);p__Ascomycota_unclassified(100);p__Ascomycota_unclassified(100);p__Ascomycota_unclassified(100);p__Ascomycota_unclassified(100);p__Ascomycota_unclassified(100); | - | - | - | - | - | - | - | Unassigned | - |
| Otu26188 | 1 | k__Fungi(100);k__Fungi_unclassified(100);k__Fungi_unclassified(100);k__Fungi_unclassified(100);k__Fungi_unclassified(100);k__Fungi_unclassified(100);k__Fungi_unclassified(100); | - | - | - | - | - | - | - | Unassigned | - |
| Otu26190 | 1 | k__Fungi(100);k__Fungi_unclassified(100);k__Fungi_unclassified(100);k__Fungi_unclassified(100);k__Fungi_unclassified(100);k__Fungi_unclassified(100);k__Fungi_unclassified(100); | - | - | - | - | - | - | - | Unassigned | - |
| Otu26191 | 1 | k__Fungi(100);k__Fungi_unclassified(100);k__Fungi_unclassified(100);k__Fungi_unclassified(100);k__Fungi_unclassified(100);k__Fungi_unclassified(100);k__Fungi_unclassified(100); | - | - | - | - | - | - | - | Unassigned | - |
| Otu26192 | 1 | k__Fungi(100);p__Ascomycota(100);p__Ascomycota_unclassified(100);p__Ascomycota_unclassified(100);p__Ascomycota_unclassified(100);p__Ascomycota_unclassified(100);p__Ascomycota_unclassified(100); | - | - | - | - | - | - | - | Unassigned | - |
| Otu26194 | 1 | k__Fungi(100);k__Fungi_unclassified(100);k__Fungi_unclassified(100);k__Fungi_unclassified(100);k__Fungi_unclassified(100);k__Fungi_unclassified(100);k__Fungi_unclassified(100); | - | - | - | - | - | - | - | Unassigned | - |
| Otu26196 | 1 | k__Fungi(100);p__Ascomycota(100);p__Ascomycota_unclassified(100);p__Ascomycota_unclassified(100);p__Ascomycota_unclassified(100);p__Ascomycota_unclassified(100);p__Ascomycota_unclassified(100); | - | - | - | - | - | - | - | Unassigned | - |
| Otu26197 | 1 | k__Fungi(100);p__Ascomycota(100);p__Ascomycota_unclassified(100);p__Ascomycota_unclassified(100);p__Ascomycota_unclassified(100);p__Ascomycota_unclassified(100);p__Ascomycota_unclassified(100); | - | - | - | - | - | - | - | Unassigned | - |
| Otu26198 | 1 | k__Fungi(100);p__Ascomycota(100);p__Ascomycota_unclassified(100);p__Ascomycota_unclassified(100);p__Ascomycota_unclassified(100);p__Ascomycota_unclassified(100);p__Ascomycota_unclassified(100); | - | - | - | - | - | - | - | Unassigned | - |

| | | | | | | | | | | |
|---|---|---|---|---|---|---|---|---|---|---|
| Otu26199 | 1 | k__Fungi(100);p__Ascomycota(100);p__Ascomycota_unclassified(100);p__Ascomycota_unclassified(100);p__Ascomycota_unclassified(100);p__Ascomycota_unclassified(100);p__Ascomycota_unclassified(100); | - | - | - | - | - | - | - | Unassigned | - |
| Otu26200 | 1 | k__Fungi(100);k__Fungi_unclassified(100);k__Fungi_unclassified(100);k__Fungi_unclassified(100);k__Fungi_unclassified(100);k__Fungi_unclassified(100);k__Fungi_unclassified(100); | - | - | - | - | - | - | - | Unassigned | - |
| Otu26203 | 1 | k__Fungi(100);p__Ascomycota(100);p__Ascomycota_unclassified(100);p__Ascomycota_unclassified(100);p__Ascomycota_unclassified(100);p__Ascomycota_unclassified(100);p__Ascomycota_unclassified(100); | - | - | - | - | - | - | - | Unassigned | - |
| Otu26209 | 1 | k__Fungi(100);p__Ascomycota(100);p__Ascomycota_unclassified(100);p__Ascomycota_unclassified(100);p__Ascomycota_unclassified(100);p__Ascomycota_unclassified(100);p__Ascomycota_unclassified(100); | - | - | - | - | - | - | - | Unassigned | - |
| Otu26220 | 1 | k__Fungi(100);k__Fungi_unclassified(100);k__Fungi_unclassified(100);k__Fungi_unclassified(100);k__Fungi_unclassified(100);k__Fungi_unclassified(100);k__Fungi_unclassified(100); | - | - | - | - | - | - | - | Unassigned | - |
| Otu26225 | 1 | k__Fungi(100);p__Ascomycota(100);p__Ascomycota_unclassified(100);p__Ascomycota_unclassified(100);p__Ascomycota_unclassified(100);p__Ascomycota_unclassified(100);p__Ascomycota_unclassified(100); | - | - | - | - | - | - | - | Unassigned | - |
| Otu26230 | 1 | k__Fungi(100);p__Ascomycota(100);p__Ascomycota_unclassified(100);p__Ascomycota_unclassified(100);p__Ascomycota_unclassified(100);p__Ascomycota_unclassified(100);p__Ascomycota_unclassified(100); | - | - | - | - | - | - | - | Unassigned | - |
| Otu26236 | 1 | k__Fungi(100);k__Fungi_unclassified(100);k__Fungi_unclassified(100);k__Fungi_unclassified(100);k__Fungi_unclassified(100);k__Fungi_unclassified(100);k__Fungi_unclassified(100); | - | - | - | - | - | - | - | Unassigned | - |
| Otu26244 | 1 | k__Fungi(100);k__Fungi_unclassified(100);k__Fungi_unclassified(100);k__Fungi_unclassified(100);k__Fungi_unclassified(100);k__Fungi_unclassified(100);k__Fungi_unclassified(100); | - | - | - | - | - | - | - | Unassigned | - |
| Otu26253 | 1 | k__Fungi(100);p__Ascomycota(100);p__Ascomycota_unclassified(100);p__Ascomycota_unclassified(100);p__Ascomycota_unclassified(100);p__Ascomycota_unclassified(100);p__Ascomycota_unclassified(100); | - | - | - | - | - | - | - | Unassigned | - |
| Otu26255 | 1 | k__Fungi(100);p__Ascomycota(100);p__Ascomycota_unclassified(100);p__Ascomycota_unclassified(100);p__Ascomycota_unclassified(100);p__Ascomycota_unclassified(100);p__Ascomycota_unclassified(100); | - | - | - | - | - | - | - | Unassigned | - |
| Otu26258 | 1 | k__Fungi(100);p__Ascomycota(100);p__Ascomycota_unclassified(100);p__Ascomycota_unclassified(100);p__Ascomycota_unclassified(100);p__Ascomycota_unclassified(100);p__Ascomycota_unclassified(100); | - | - | - | - | - | - | - | Unassigned | - |
| Otu26259 | 1 | k__Fungi(100);k__Fungi_unclassified(100);k__Fungi_unclassified(100);k__Fungi_unclassified(100);k__Fungi_unclassified(100);k__Fungi_unclassified(100);k__Fungi_unclassified(100); | - | - | - | - | - | - | - | Unassigned | - |
| Otu26260 | 1 | k__Fungi(100);p__Ascomycota(100);p__Ascomycota_unclassified(100);p__Ascomycota_unclassified(100);p__Ascomycota_unclassified(100);p__Ascomycota_unclassified(100);p__Ascomycota_unclassified(100); | - | - | - | - | - | - | - | Unassigned | - |
| Otu26261 | 1 | k__Fungi(100);p__Ascomycota(100);p__Ascomycota_unclassified(100);p__Ascomycota_unclassified(100);p__Ascomycota_unclassified(100);p__Ascomycota_unclassified(100);p__Ascomycota_unclassified(100); | - | - | - | - | - | - | - | Unassigned | - |
| Otu26262 | 1 | k__Fungi(100);p__Ascomycota(100);p__Ascomycota_unclassified(100);p__Ascomycota_unclassified(100);p__Ascomycota_unclassified(100);p__Ascomycota_unclassified(100);p__Ascomycota_unclassified(100); | - | - | - | - | - | - | - | Unassigned | - |
| Otu26263 | 1 | k__Fungi(100);k__Fungi_unclassified(100);k__Fungi_unclassified(100);k__Fungi_unclassified(100);k__Fungi_unclassified(100);k__Fungi_unclassified(100);k__Fungi_unclassified(100); | - | - | - | - | - | - | - | Unassigned | - |
| Otu26264 | 1 | k__Fungi(100);p__Ascomycota(100);p__Ascomycota_unclassified(100);p__Ascomycota_unclassified(100);p__Ascomycota_unclassified(100);p__Ascomycota_unclassified(100);p__Ascomycota_unclassified(100); | - | - | - | - | - | - | - | Unassigned | - |
| Otu26265 | 1 | k__Fungi(100);p__Ascomycota(100);p__Ascomycota_unclassified(100);p__Ascomycota_unclassified(100);p__Ascomycota_unclassified(100);p__Ascomycota_unclassified(100);p__Ascomycota_unclassified(100); | - | - | - | - | - | - | - | Unassigned | - |
| Otu26267 | 1 | k__Fungi(100);p__Ascomycota(100);p__Ascomycota_unclassified(100);p__Ascomycota_unclassified(100);p__Ascomycota_unclassified(100);p__Ascomycota_unclassified(100);p__Ascomycota_unclassified(100); | - | - | - | - | - | - | - | Unassigned | - |
| Otu26268 | 1 | k__Fungi(100);k__Fungi_unclassified(100);k__Fungi_unclassified(100);k__Fungi_unclassified(100);k__Fungi_unclassified(100);k__Fungi_unclassified(100);k__Fungi_unclassified(100); | - | - | - | - | - | - | - | Unassigned | - |
| Otu26271 | 1 | k__Fungi(100);p__Ascomycota(100);p__Ascomycota_unclassified(100);p__Ascomycota_unclassified(100);p__Ascomycota_unclassified(100);p__Ascomycota_unclassified(100);p__Ascomycota_unclassified(100); | - | - | - | - | - | - | - | Unassigned | - |
| Otu26274 | 1 | k__Fungi(100);k__Fungi_unclassified(100);k__Fungi_unclassified(100);k__Fungi_unclassified(100);k__Fungi_unclassified(100);k__Fungi_unclassified(100);k__Fungi_unclassified(100); | - | - | - | - | - | - | - | Unassigned | - |
| Otu26276 | 1 | k__Fungi(100);k__Fungi_unclassified(100);k__Fungi_unclassified(100);k__Fungi_unclassified(100);k__Fungi_unclassified(100);k__Fungi_unclassified(100);k__Fungi_unclassified(100); | - | - | - | - | - | - | - | Unassigned | - |
| Otu26277 | 1 | k__Fungi(100);k__Fungi_unclassified(100);k__Fungi_unclassified(100);k__Fungi_unclassified(100);k__Fungi_unclassified(100);k__Fungi_unclassified(100);k__Fungi_unclassified(100); | - | - | - | - | - | - | - | Unassigned | - |
| Otu26278 | 1 | k__Fungi(100);k__Fungi_unclassified(100);k__Fungi_unclassified(100);k__Fungi_unclassified(100);k__Fungi_unclassified(100);k__Fungi_unclassified(100);k__Fungi_unclassified(100); | - | - | - | - | - | - | - | Unassigned | - |
| Otu26280 | 1 | k__Fungi(100);k__Fungi_unclassified(100);k__Fungi_unclassified(100);k__Fungi_unclassified(100);k__Fungi_unclassified(100);k__Fungi_unclassified(100);k__Fungi_unclassified(100); | - | - | - | - | - | - | - | Unassigned | - |
| Otu26284 | 1 | k__Fungi(100);p__Ascomycota(100);p__Ascomycota_unclassified(100);p__Ascomycota_unclassified(100);p__Ascomycota_unclassified(100);p__Ascomycota_unclassified(100);p__Ascomycota_unclassified(100); | - | - | - | - | - | - | - | Unassigned | - |
| Otu26285 | 1 | k__Fungi(100);p__Ascomycota(100);p__Ascomycota_unclassified(100);p__Ascomycota_unclassified(100);p__Ascomycota_unclassified(100);p__Ascomycota_unclassified(100);p__Ascomycota_unclassified(100); | - | - | - | - | - | - | - | Unassigned | - |
| Otu26286 | 1 | k__Fungi(100);p__Ascomycota(100);p__Ascomycota_unclassified(100);p__Ascomycota_unclassified(100);p__Ascomycota_unclassified(100);p__Ascomycota_unclassified(100);p__Ascomycota_unclassified(100); | - | - | - | - | - | - | - | Unassigned | - |
| Otu26288 | 1 | k__Fungi(100);p__Ascomycota(100);p__Ascomycota_unclassified(100);p__Ascomycota_unclassified(100);p__Ascomycota_unclassified(100);p__Ascomycota_unclassified(100);p__Ascomycota_unclassified(100); | - | - | - | - | - | - | - | Unassigned | - |
| Otu26289 | 1 | k__Fungi(100);k__Fungi_unclassified(100);k__Fungi_unclassified(100);k__Fungi_unclassified(100);k__Fungi_unclassified(100);k__Fungi_unclassified(100);k__Fungi_unclassified(100); | - | - | - | - | - | - | - | Unassigned | - |
| Otu26290 | 1 | k__Fungi(100);k__Fungi_unclassified(100);k__Fungi_unclassified(100);k__Fungi_unclassified(100);k__Fungi_unclassified(100);k__Fungi_unclassified(100);k__Fungi_unclassified(100); | - | - | - | - | - | - | - | Unassigned | - |
| Otu26291 | 1 | k__Fungi(100);k__Fungi_unclassified(100);k__Fungi_unclassified(100);k__Fungi_unclassified(100);k__Fungi_unclassified(100);k__Fungi_unclassified(100);k__Fungi_unclassified(100); | - | - | - | - | - | - | - | Unassigned | - |
| Otu26292 | 1 | k__Fungi(100);p__Ascomycota(100);p__Ascomycota_unclassified(100);p__Ascomycota_unclassified(100);p__Ascomycota_unclassified(100);p__Ascomycota_unclassified(100);p__Ascomycota_unclassified(100); | - | - | - | - | - | - | - | Unassigned | - |
| Otu26293 | 1 | k__Fungi(100);p__Ascomycota(100);p__Ascomycota_unclassified(100);p__Ascomycota_unclassified(100);p__Ascomycota_unclassified(100);p__Ascomycota_unclassified(100);p__Ascomycota_unclassified(100); | - | - | - | - | - | - | - | Unassigned | - |
| Otu26294 | 1 | k__Fungi(100);k__Fungi_unclassified(100);k__Fungi_unclassified(100);k__Fungi_unclassified(100);k__Fungi_unclassified(100);k__Fungi_unclassified(100);k__Fungi_unclassified(100); | - | - | - | - | - | - | - | Unassigned | - |
| Otu26295 | 1 | k__Fungi(100);p__Ascomycota(100);p__Ascomycota_unclassified(100);p__Ascomycota_unclassified(100);p__Ascomycota_unclassified(100);p__Ascomycota_unclassified(100);p__Ascomycota_unclassified(100); | - | - | - | - | - | - | - | Unassigned | - |
| Otu26296 | 1 | k__Fungi(100);k__Fungi_unclassified(100);k__Fungi_unclassified(100);k__Fungi_unclassified(100);k__Fungi_unclassified(100);k__Fungi_unclassified(100);k__Fungi_unclassified(100); | - | - | - | - | - | - | - | Unassigned | - |
| Otu26298 | 1 | k__Fungi(100);p__Ascomycota(100);p__Ascomycota_unclassified(100);p__Ascomycota_unclassified(100);p__Ascomycota_unclassified(100);p__Ascomycota_unclassified(100);p__Ascomycota_unclassified(100); | - | - | - | - | - | - | - | Unassigned | - |
| Otu26299 | 1 | k__Fungi(100);p__Ascomycota(100);p__Ascomycota_unclassified(100);p__Ascomycota_unclassified(100);p__Ascomycota_unclassified(100);p__Ascomycota_unclassified(100);p__Ascomycota_unclassified(100); | - | - | - | - | - | - | - | Unassigned | - |
| Otu26300 | 1 | k__Fungi(100);k__Fungi_unclassified(100);k__Fungi_unclassified(100);k__Fungi_unclassified(100);k__Fungi_unclassified(100);k__Fungi_unclassified(100);k__Fungi_unclassified(100); | - | - | - | - | - | - | - | Unassigned | - |
| Otu26301 | 1 | k__Fungi(100);p__Ascomycota(100);p__Ascomycota_unclassified(100);p__Ascomycota_unclassified(100);p__Ascomycota_unclassified(100);p__Ascomycota_unclassified(100);p__Ascomycota_unclassified(100); | - | - | - | - | - | - | - | Unassigned | - |
| Otu26302 | 1 | k__Fungi(100);p__Ascomycota(100);p__Ascomycota_unclassified(100);p__Ascomycota_unclassified(100);p__Ascomycota_unclassified(100);p__Ascomycota_unclassified(100);p__Ascomycota_unclassified(100); | - | - | - | - | - | - | - | Unassigned | - |
| Otu26303 | 1 | k__Fungi(100);p__Ascomycota(100);p__Ascomycota_unclassified(100);p__Ascomycota_unclassified(100);p__Ascomycota_unclassified(100);p__Ascomycota_unclassified(100);p__Ascomycota_unclassified(100); | - | - | - | - | - | - | - | Unassigned | - |
| Otu26304 | 1 | k__Fungi(100);k__Fungi_unclassified(100);k__Fungi_unclassified(100);k__Fungi_unclassified(100);k__Fungi_unclassified(100);k__Fungi_unclassified(100);k__Fungi_unclassified(100); | - | - | - | - | - | - | - | Unassigned | - |
| Otu26305 | 1 | k__Fungi(100);p__Ascomycota(100);p__Ascomycota_unclassified(100);p__Ascomycota_unclassified(100);p__Ascomycota_unclassified(100);p__Ascomycota_unclassified(100);p__Ascomycota_unclassified(100); | - | - | - | - | - | - | - | Unassigned | - |
| Otu26306 | 1 | k__Fungi(100);p__Ascomycota(100);p__Ascomycota_unclassified(100);p__Ascomycota_unclassified(100);p__Ascomycota_unclassified(100);p__Ascomycota_unclassified(100);p__Ascomycota_unclassified(100); | - | - | - | - | - | - | - | Unassigned | - |
| Otu26307 | 1 | k__Fungi(100);p__Ascomycota(100);p__Ascomycota_unclassified(100);p__Ascomycota_unclassified(100);p__Ascomycota_unclassified(100);p__Ascomycota_unclassified(100);p__Ascomycota_unclassified(100); | - | - | - | - | - | - | - | Unassigned | - |
| Otu26309 | 1 | k__Fungi(100);p__Ascomycota(100);p__Ascomycota_unclassified(100);p__Ascomycota_unclassified(100);p__Ascomycota_unclassified(100);p__Ascomycota_unclassified(100);p__Ascomycota_unclassified(100); | - | - | - | - | - | - | - | Unassigned | - |
| Otu26310 | 1 | k__Fungi(100);p__Ascomycota(100);p__Ascomycota_unclassified(100);p__Ascomycota_unclassified(100);p__Ascomycota_unclassified(100);p__Ascomycota_unclassified(100);p__Ascomycota_unclassified(100); | - | - | - | - | - | - | - | Unassigned | - |
| Otu26311 | 1 | k__Fungi(100);p__Ascomycota(100);p__Ascomycota_unclassified(100);p__Ascomycota_unclassified(100);p__Ascomycota_unclassified(100);p__Ascomycota_unclassified(100);p__Ascomycota_unclassified(100); | - | - | - | - | - | - | - | Unassigned | - |
| Otu26313 | 1 | k__Fungi(100);p__Ascomycota(100);p__Ascomycota_unclassified(100);p__Ascomycota_unclassified(100);p__Ascomycota_unclassified(100);p__Ascomycota_unclassified(100);p__Ascomycota_unclassified(100); | - | - | - | - | - | - | - | Unassigned | - |
| Otu26314 | 1 | k__Fungi(100);k__Fungi_unclassified(100);k__Fungi_unclassified(100);k__Fungi_unclassified(100);k__Fungi_unclassified(100);k__Fungi_unclassified(100);k__Fungi_unclassified(100); | - | - | - | - | - | - | - | Unassigned | - |
| Otu26316 | 1 | k__Fungi(100);k__Fungi_unclassified(100);k__Fungi_unclassified(100);k__Fungi_unclassified(100);k__Fungi_unclassified(100);k__Fungi_unclassified(100);k__Fungi_unclassified(100); | - | - | - | - | - | - | - | Unassigned | - |
| Otu26317 | 1 | k__Fungi(100);p__Ascomycota(100);p__Ascomycota_unclassified(100);p__Ascomycota_unclassified(100);p__Ascomycota_unclassified(100);p__Ascomycota_unclassified(100);p__Ascomycota_unclassified(100); | - | - | - | - | - | - | - | Unassigned | - |
| Otu26319 | 1 | k__Fungi(100);k__Fungi_unclassified(100);k__Fungi_unclassified(100);k__Fungi_unclassified(100);k__Fungi_unclassified(100);k__Fungi_unclassified(100);k__Fungi_unclassified(100); | - | - | - | - | - | - | - | Unassigned | - |
| Otu26320 | 1 | k__Fungi(100);k__Fungi_unclassified(100);k__Fungi_unclassified(100);k__Fungi_unclassified(100);k__Fungi_unclassified(100);k__Fungi_unclassified(100);k__Fungi_unclassified(100); | - | - | - | - | - | - | - | Unassigned | - |
| Otu26321 | 1 | k__Fungi(100);k__Fungi_unclassified(100);k__Fungi_unclassified(100);k__Fungi_unclassified(100);k__Fungi_unclassified(100);k__Fungi_unclassified(100);k__Fungi_unclassified(100); | - | - | - | - | - | - | - | Unassigned | - |
| Otu26322 | 1 | k__Fungi(100);p__Ascomycota(100);p__Ascomycota_unclassified(100);p__Ascomycota_unclassified(100);p__Ascomycota_unclassified(100);p__Ascomycota_unclassified(100);p__Ascomycota_unclassified(100); | - | - | - | - | - | - | - | Unassigned | - |
| Otu26323 | 1 | k__Fungi(100);p__Ascomycota(100);p__Ascomycota_unclassified(100);p__Ascomycota_unclassified(100);p__Ascomycota_unclassified(100);p__Ascomycota_unclassified(100);p__Ascomycota_unclassified(100); | - | - | - | - | - | - | - | Unassigned | - |
| Otu26324 | 1 | k__Fungi(100);k__Fungi_unclassified(100);k__Fungi_unclassified(100);k__Fungi_unclassified(100);k__Fungi_unclassified(100);k__Fungi_unclassified(100);k__Fungi_unclassified(100); | - | - | - | - | - | - | - | Unassigned | - |
| Otu26325 | 1 | k__Fungi(100);p__Ascomycota(100);p__Ascomycota_unclassified(100);p__Ascomycota_unclassified(100);p__Ascomycota_unclassified(100);p__Ascomycota_unclassified(100);p__Ascomycota_unclassified(100); | - | - | - | - | - | - | - | Unassigned | - |
| Otu26326 | 1 | k__Fungi(100);p__Ascomycota(100);p__Ascomycota_unclassified(100);p__Ascomycota_unclassified(100);p__Ascomycota_unclassified(100);p__Ascomycota_unclassified(100);p__Ascomycota_unclassified(100); | - | - | - | - | - | - | - | Unassigned | - |
| Otu26327 | 1 | k__Fungi(100);k__Fungi_unclassified(100);k__Fungi_unclassified(100);k__Fungi_unclassified(100);k__Fungi_unclassified(100);k__Fungi_unclassified(100);k__Fungi_unclassified(100); | - | - | - | - | - | - | - | Unassigned | - |
| Otu26328 | 1 | k__Fungi(100);k__Fungi_unclassified(100);k__Fungi_unclassified(100);k__Fungi_unclassified(100);k__Fungi_unclassified(100);k__Fungi_unclassified(100);k__Fungi_unclassified(100); | - | - | - | - | - | - | - | Unassigned | - |
| Otu26329 | 1 | k__Fungi(100);p__Ascomycota(100);p__Ascomycota_unclassified(100);p__Ascomycota_unclassified(100);p__Ascomycota_unclassified(100);p__Ascomycota_unclassified(100);p__Ascomycota_unclassified(100); | - | - | - | - | - | - | - | Unassigned | - |
| Otu26330 | 1 | k__Fungi(100);k__Fungi_unclassified(100);k__Fungi_unclassified(100);k__Fungi_unclassified(100);k__Fungi_unclassified(100);k__Fungi_unclassified(100);k__Fungi_unclassified(100); | - | - | - | - | - | - | - | Unassigned | - |
| Otu26331 | 1 | k__Fungi(100);k__Fungi_unclassified(100);k__Fungi_unclassified(100);k__Fungi_unclassified(100);k__Fungi_unclassified(100);k__Fungi_unclassified(100);k__Fungi_unclassified(100); | - | - | - | - | - | - | - | Unassigned | - |
| Otu26332 | 1 | k__Fungi(100);k__Fungi_unclassified(100);k__Fungi_unclassified(100);k__Fungi_unclassified(100);k__Fungi_unclassified(100);k__Fungi_unclassified(100);k__Fungi_unclassified(100); | - | - | - | - | - | - | - | Unassigned | - |
| Otu26333 | 1 | k__Fungi(100);p__Ascomycota(100);p__Ascomycota_unclassified(100);p__Ascomycota_unclassified(100);p__Ascomycota_unclassified(100);p__Ascomycota_unclassified(100);p__Ascomycota_unclassified(100); | - | - | - | - | - | - | - | Unassigned | - |
| Otu26334 | 1 | k__Fungi(100);k__Fungi_unclassified(100);k__Fungi_unclassified(100);k__Fungi_unclassified(100);k__Fungi_unclassified(100);k__Fungi_unclassified(100);k__Fungi_unclassified(100); | - | - | - | - | - | - | - | Unassigned | - |
| Otu26337 | 1 | k__Fungi(100);p__Ascomycota(100);p__Ascomycota_unclassified(100);p__Ascomycota_unclassified(100);p__Ascomycota_unclassified(100);p__Ascomycota_unclassified(100);p__Ascomycota_unclassified(100); | - | - | - | - | - | - | - | Unassigned | - |
| Otu26338 | 1 | k__Fungi(100);p__Ascomycota(100);p__Ascomycota_unclassified(100);p__Ascomycota_unclassified(100);p__Ascomycota_unclassified(100);p__Ascomycota_unclassified(100);p__Ascomycota_unclassified(100); | - | - | - | - | - | - | - | Unassigned | - |
| Otu26339 | 1 | k__Fungi(100);k__Fungi_unclassified(100);k__Fungi_unclassified(100);k__Fungi_unclassified(100);k__Fungi_unclassified(100);k__Fungi_unclassified(100);k__Fungi_unclassified(100); | - | - | - | - | - | - | - | Unassigned | - |
| Otu26342 | 1 | k__Fungi(100);k__Fungi_unclassified(100);k__Fungi_unclassified(100);k__Fungi_unclassified(100);k__Fungi_unclassified(100);k__Fungi_unclassified(100);k__Fungi_unclassified(100); | - | - | - | - | - | - | - | Unassigned | - |
| Otu26345 | 1 | k__Fungi(100);k__Fungi_unclassified(100);k__Fungi_unclassified(100);k__Fungi_unclassified(100);k__Fungi_unclassified(100);k__Fungi_unclassified(100);k__Fungi_unclassified(100); | - | - | - | - | - | - | - | Unassigned | - |
| Otu26346 | 1 | k__Fungi(100);p__Ascomycota(100);p__Ascomycota_unclassified(100);p__Ascomycota_unclassified(100);p__Ascomycota_unclassified(100);p__Ascomycota_unclassified(100);p__Ascomycota_unclassified(100); | - | - | - | - | - | - | - | Unassigned | - |
| Otu26347 | 1 | k__Fungi(100);p__Ascomycota(100);p__Ascomycota_unclassified(100);p__Ascomycota_unclassified(100);p__Ascomycota_unclassified(100);p__Ascomycota_unclassified(100);p__Ascomycota_unclassified(100); | - | - | - | - | - | - | - | Unassigned | - |
| Otu26349 | 1 | k__Fungi(100);p__Ascomycota(100);p__Ascomycota_unclassified(100);p__Ascomycota_unclassified(100);p__Ascomycota_unclassified(100);p__Ascomycota_unclassified(100);p__Ascomycota_unclassified(100); | - | - | - | - | - | - | - | Unassigned | - |
| Otu26350 | 1 | k__Fungi(100);p__Ascomycota(100);p__Ascomycota_unclassified(100);p__Ascomycota_unclassified(100);p__Ascomycota_unclassified(100);p__Ascomycota_unclassified(100);p__Ascomycota_unclassified(100); | - | - | - | - | - | - | - | Unassigned | - |
| Otu26351 | 1 | k__Fungi(100);k__Fungi_unclassified(100);k__Fungi_unclassified(100);k__Fungi_unclassified(100);k__Fungi_unclassified(100);k__Fungi_unclassified(100);k__Fungi_unclassified(100); | - | - | - | - | - | - | - | Unassigned | - |
| Otu26352 | 1 | k__Fungi(100);k__Fungi_unclassified(100);k__Fungi_unclassified(100);k__Fungi_unclassified(100);k__Fungi_unclassified(100);k__Fungi_unclassified(100);k__Fungi_unclassified(100); | - | - | - | - | - | - | - | Unassigned | - |
| Otu26353 | 1 | k__Fungi(100);p__Ascomycota(100);p__Ascomycota_unclassified(100);p__Ascomycota_unclassified(100);p__Ascomycota_unclassified(100);p__Ascomycota_unclassified(100);p__Ascomycota_unclassified(100); | - | - | - | - | - | - | - | Unassigned | - |
| Otu26354 | 1 | k__Fungi(100);p__Ascomycota(100);p__Ascomycota_unclassified(100);p__Ascomycota_unclassified(100);p__Ascomycota_unclassified(100);p__Ascomycota_unclassified(100);p__Ascomycota_unclassified(100); | - | - | - | - | - | - | - | Unassigned | - |
| Otu26356 | 1 | k__Fungi(100);p__Ascomycota(100);p__Ascomycota_unclassified(100);p__Ascomycota_unclassified(100);p__Ascomycota_unclassified(100);p__Ascomycota_unclassified(100);p__Ascomycota_unclassified(100); | - | - | - | - | - | - | - | Unassigned | - |
| Otu26359 | 1 | k__Fungi(100);p__Ascomycota(100);p__Ascomycota_unclassified(100);p__Ascomycota_unclassified(100);p__Ascomycota_unclassified(100);p__Ascomycota_unclassified(100);p__Ascomycota_unclassified(100); | - | - | - | - | - | - | - | Unassigned | - |
| Otu26360 | 1 | k__Fungi(100);k__Fungi_unclassified(100);k__Fungi_unclassified(100);k__Fungi_unclassified(100);k__Fungi_unclassified(100);k__Fungi_unclassified(100);k__Fungi_unclassified(100); | - | - | - | - | - | - | - | Unassigned | - |
| Otu26361 | 1 | k__Fungi(100);p__Ascomycota(100);p__Ascomycota_unclassified(100);p__Ascomycota_unclassified(100);p__Ascomycota_unclassified(100);p__Ascomycota_unclassified(100);p__Ascomycota_unclassified(100); | - | - | - | - | - | - | - | Unassigned | - |
| Otu26362 | 1 | k__Fungi(100);k__Fungi_unclassified(100);k__Fungi_unclassified(100);k__Fungi_unclassified(100);k__Fungi_unclassified(100);k__Fungi_unclassified(100);k__Fungi_unclassified(100); | - | - | - | - | - | - | - | Unassigned | - |
| Otu26363 | 1 | k__Fungi(100);k__Fungi_unclassified(100);k__Fungi_unclassified(100);k__Fungi_unclassified(100);k__Fungi_unclassified(100);k__Fungi_unclassified(100);k__Fungi_unclassified(100); | - | - | - | - | - | - | - | Unassigned | - |
| Otu26364 | 1 | k__Fungi(100);p__Ascomycota(100);p__Ascomycota_unclassified(100);p__Ascomycota_unclassified(100);p__Ascomycota_unclassified(100);p__Ascomycota_unclassified(100);p__Ascomycota_unclassified(100); | - | - | - | - | - | - | - | Unassigned | - |
| Otu26365 | 1 | k__Fungi(100);p__Ascomycota(100);p__Ascomycota_unclassified(100);p__Ascomycota_unclassified(100);p__Ascomycota_unclassified(100);p__Ascomycota_unclassified(100);p__Ascomycota_unclassified(100); | - | - | - | - | - | - | - | Unassigned | - |
| Otu26366 | 1 | k__Fungi(100);k__Fungi_unclassified(100);k__Fungi_unclassified(100);k__Fungi_unclassified(100);k__Fungi_unclassified(100);k__Fungi_unclassified(100);k__Fungi_unclassified(100); | - | - | - | - | - | - | - | Unassigned | - |
| Otu26367 | 1 | k__Fungi(100);k__Fungi_unclassified(100);k__Fungi_unclassified(100);k__Fungi_unclassified(100);k__Fungi_unclassified(100);k__Fungi_unclassified(100);k__Fungi_unclassified(100); | - | - | - | - | - | - | - | Unassigned | - |
| Otu26368 | 1 | k__Fungi(100);k__Fungi_unclassified(100);k__Fungi_unclassified(100);k__Fungi_unclassified(100);k__Fungi_unclassified(100);k__Fungi_unclassified(100);k__Fungi_unclassified(100); | - | - | - | - | - | - | - | Unassigned | - |
| Otu26369 | 1 | k__Fungi(100);p__Ascomycota(100);p__Ascomycota_unclassified(100);p__Ascomycota_unclassified(100);p__Ascomycota_unclassified(100);p__Ascomycota_unclassified(100);p__Ascomycota_unclassified(100); | - | - | - | - | - | - | - | Unassigned | - |
| Otu26370 | 1 | k__Fungi(100);k__Fungi_unclassified(100);k__Fungi_unclassified(100);k__Fungi_unclassified(100);k__Fungi_unclassified(100);k__Fungi_unclassified(100);k__Fungi_unclassified(100); | - | - | - | - | - | - | - | Unassigned | - |
| Otu26373 | 1 | k__Fungi(100);p__Ascomycota(100);p__Ascomycota_unclassified(100);p__Ascomycota_unclassified(100);p__Ascomycota_unclassified(100);p__Ascomycota_unclassified(100);p__Ascomycota_unclassified(100); | - | - | - | - | - | - | - | Unassigned | - |
| Otu26374 | 1 | k__Fungi(100);p__Ascomycota(100);p__Ascomycota_unclassified(100);p__Ascomycota_unclassified(100);p__Ascomycota_unclassified(100);p__Ascomycota_unclassified(100);p__Ascomycota_unclassified(100); | - | - | - | - | - | - | - | Unassigned | - |
| Otu26375 | 1 | k__Fungi(100);k__Fungi_unclassified(100);k__Fungi_unclassified(100);k__Fungi_unclassified(100);k__Fungi_unclassified(100);k__Fungi_unclassified(100);k__Fungi_unclassified(100); | - | - | - | - | - | - | - | Unassigned | - |
| Otu26376 | 1 | k__Fungi(100);p__Ascomycota(100);p__Ascomycota_unclassified(100);p__Ascomycota_unclassified(100);p__Ascomycota_unclassified(100);p__Ascomycota_unclassified(100);p__Ascomycota_unclassified(100); | - | - | - | - | - | - | - | Unassigned | - |
| Otu26377 | 1 | k__Fungi(100);k__Fungi_unclassified(100);k__Fungi_unclassified(100);k__Fungi_unclassified(100);k__Fungi_unclassified(100);k__Fungi_unclassified(100);k__Fungi_unclassified(100); | - | - | - | - | - | - | - | Unassigned | - |
| Otu26378 | 1 | k__Fungi(100);k__Fungi_unclassified(100);k__Fungi_unclassified(100);k__Fungi_unclassified(100);k__Fungi_unclassified(100);k__Fungi_unclassified(100);k__Fungi_unclassified(100); | - | - | - | - | - | - | - | Unassigned | - |
| Otu26379 | 1 | k__Fungi(100);p__Ascomycota(100);p__Ascomycota_unclassified(100);p__Ascomycota_unclassified(100);p__Ascomycota_unclassified(100);p__Ascomycota_unclassified(100);p__Ascomycota_unclassified(100); | - | - | - | - | - | - | - | Unassigned | - |
| Otu26380 | 1 | k__Fungi(100);p__Ascomycota(100);p__Ascomycota_unclassified(100);p__Ascomycota_unclassified(100);p__Ascomycota_unclassified(100);p__Ascomycota_unclassified(100);p__Ascomycota_unclassified(100); | - | - | - | - | - | - | - | Unassigned | - |
| Otu26381 | 1 | k__Fungi(100);p__Ascomycota(100);p__Ascomycota_unclassified(100);p__Ascomycota_unclassified(100);p__Ascomycota_unclassified(100);p__Ascomycota_unclassified(100);p__Ascomycota_unclassified(100); | - | - | - | - | - | - | - | Unassigned | - |
| Otu26382 | 1 | k__Fungi(100);k__Fungi_unclassified(100);k__Fungi_unclassified(100);k__Fungi_unclassified(100);k__Fungi_unclassified(100);k__Fungi_unclassified(100);k__Fungi_unclassified(100); | - | - | - | - | - | - | - | Unassigned | - |
| Otu26383 | 1 | k__Fungi(100);k__Fungi_unclassified(100);k__Fungi_unclassified(100);k__Fungi_unclassified(100);k__Fungi_unclassified(100);k__Fungi_unclassified(100);k__Fungi_unclassified(100); | - | - | - | - | - | - | - | Unassigned | - |
| Otu26386 | 1 | k__Fungi(100);k__Fungi_unclassified(100);k__Fungi_unclassified(100);k__Fungi_unclassified(100);k__Fungi_unclassified(100);k__Fungi_unclassified(100);k__Fungi_unclassified(100); | - | - | - | - | - | - | - | Unassigned | - |
| Otu26387 | 1 | k__Fungi(100);p__Ascomycota(100);p__Ascomycota_unclassified(100);p__Ascomycota_unclassified(100);p__Ascomycota_unclassified(100);p__Ascomycota_unclassified(100);p__Ascomycota_unclassified(100); | - | - | - | - | - | - | - | Unassigned | - |
| Otu26388 | 1 | k__Fungi(100);p__Ascomycota(100);p__Ascomycota_unclassified(100);p__Ascomycota_unclassified(100);p__Ascomycota_unclassified(100);p__Ascomycota_unclassified(100);p__Ascomycota_unclassified(100); | - | - | - | - | - | - | - | Unassigned | - |
| Otu26389 | 1 | k__Fungi(100);p__Ascomycota(100);p__Ascomycota_unclassified(100);p__Ascomycota_unclassified(100);p__Ascomycota_unclassified(100);p__Ascomycota_unclassified(100);p__Ascomycota_unclassified(100); | - | - | - | - | - | - | - | Unassigned | - |
| Otu26390 | 1 | k__Fungi(100);k__Fungi_unclassified(100);k__Fungi_unclassified(100);k__Fungi_unclassified(100);k__Fungi_unclassified(100);k__Fungi_unclassified(100);k__Fungi_unclassified(100); | - | - | - | - | - | - | - | Unassigned | - |
| Otu26391 | 1 | k__Fungi(100);p__Ascomycota(100);p__Ascomycota_unclassified(100);p__Ascomycota_unclassified(100);p__Ascomycota_unclassified(100);p__Ascomycota_unclassified(100);p__Ascomycota_unclassified(100); | - | - | - | - | - | - | - | Unassigned | - |
| Otu26396 | 1 | k__Fungi(100);k__Fungi_unclassified(100);k__Fungi_unclassified(100);k__Fungi_unclassified(100);k__Fungi_unclassified(100);k__Fungi_unclassified(100);k__Fungi_unclassified(100); | - | - | - | - | - | - | - | Unassigned | - |
| Otu26398 | 1 | k__Fungi(100);k__Fungi_unclassified(100);k__Fungi_unclassified(100);k__Fungi_unclassified(100);k__Fungi_unclassified(100);k__Fungi_unclassified(100);k__Fungi_unclassified(100); | - | - | - | - | - | - | - | Unassigned | - |
| Otu26399 | 1 | k__Fungi(100);k__Fungi_unclassified(100);k__Fungi_unclassified(100);k__Fungi_unclassified(100);k__Fungi_unclassified(100);k__Fungi_unclassified(100);k__Fungi_unclassified(100); | - | - | - | - | - | - | - | Unassigned | - |
| Otu26400 | 1 | k__Fungi(100);p__Ascomycota(100);p__Ascomycota_unclassified(100);p__Ascomycota_unclassified(100);p__Ascomycota_unclassified(100);p__Ascomycota_unclassified(100);p__Ascomycota_unclassified(100); | - | - | - | - | - | - | - | Unassigned | - |
| Otu26403 | 1 | k__Fungi(100);p__Ascomycota(100);p__Ascomycota_unclassified(100);p__Ascomycota_unclassified(100);p__Ascomycota_unclassified(100);p__Ascomycota_unclassified(100);p__Ascomycota_unclassified(100); | - | - | - | - | - | - | - | Unassigned | - |
| Otu26406 | 1 | k__Fungi(100);p__Ascomycota(100);p__Ascomycota_unclassified(100);p__Ascomycota_unclassified(100);p__Ascomycota_unclassified(100);p__Ascomycota_unclassified(100);p__Ascomycota_unclassified(100); | - | - | - | - | - | - | - | Unassigned | - |
| Otu26408 | 1 | k__Fungi(100);k__Fungi_unclassified(100);k__Fungi_unclassified(100);k__Fungi_unclassified(100);k__Fungi_unclassified(100);k__Fungi_unclassified(100);k__Fungi_unclassified(100); | - | - | - | - | - | - | - | Unassigned | - |
| Otu26409 | 1 | k__Fungi(100);k__Fungi_unclassified(100);k__Fungi_unclassified(100);k__Fungi_unclassified(100);k__Fungi_unclassified(100);k__Fungi_unclassified(100);k__Fungi_unclassified(100); | - | - | - | - | - | - | - | Unassigned | - |
| Otu26410 | 1 | k__Fungi(100);p__Ascomycota(100);p__Ascomycota_unclassified(100);p__Ascomycota_unclassified(100);p__Ascomycota_unclassified(100);p__Ascomycota_unclassified(100);p__Ascomycota_unclassified(100); | - | - | - | - | - | - | - | Unassigned | - |
| Otu26411 | 1 | k__Fungi(100);k__Fungi_unclassified(100);k__Fungi_unclassified(100);k__Fungi_unclassified(100);k__Fungi_unclassified(100);k__Fungi_unclassified(100);k__Fungi_unclassified(100); | - | - | - | - | - | - | - | Unassigned | - |
| Otu26412 | 1 | k__Fungi(100);k__Fungi_unclassified(100);k__Fungi_unclassified(100);k__Fungi_unclassified(100);k__Fungi_unclassified(100);k__Fungi_unclassified(100);k__Fungi_unclassified(100); | - | - | - | - | - | - | - | Unassigned | - |
| Otu26413 | 1 | k__Fungi(100);p__Ascomycota(100);p__Ascomycota_unclassified(100);p__Ascomycota_unclassified(100);p__Ascomycota_unclassified(100);p__Ascomycota_unclassified(100);p__Ascomycota_unclassified(100); | - | - | - | - | - | - | - | Unassigned | - |
| Otu26414 | 1 | k__Fungi(100);p__Ascomycota(100);p__Ascomycota_unclassified(100);p__Ascomycota_unclassified(100);p__Ascomycota_unclassified(100);p__Ascomycota_unclassified(100);p__Ascomycota_unclassified(100); | - | - | - | - | - | - | - | Unassigned | - |
| Otu26415 | 1 | k__Fungi(100);k__Fungi_unclassified(100);k__Fungi_unclassified(100);k__Fungi_unclassified(100);k__Fungi_unclassified(100);k__Fungi_unclassified(100);k__Fungi_unclassified(100); | - | - | - | - | - | - | - | Unassigned | - |
| Otu26416 | 1 | k__Fungi(100);p__Ascomycota(100);p__Ascomycota_unclassified(100);p__Ascomycota_unclassified(100);p__Ascomycota_unclassified(100);p__Ascomycota_unclassified(100);p__Ascomycota_unclassified(100); | - | - | - | - | - | - | - | Unassigned | - |
| Otu26420 | 1 | k__Fungi(100);p__Ascomycota(100);p__Ascomycota_unclassified(100);p__Ascomycota_unclassified(100);p__Ascomycota_unclassified(100);p__Ascomycota_unclassified(100);p__Ascomycota_unclassified(100); | - | - | - | - | - | - | - | Unassigned | - |
| Otu26425 | 1 | k__Fungi(100);k__Fungi_unclassified(100);k__Fungi_unclassified(100);k__Fungi_unclassified(100);k__Fungi_unclassified(100);k__Fungi_unclassified(100);k__Fungi_unclassified(100); | - | - | - | - | - | - | - | Unassigned | - |
| Otu26449 | 1 | k__Fungi(100);p__Ascomycota(100);p__Ascomycota_unclassified(100);p__Ascomycota_unclassified(100);p__Ascomycota_unclassified(100);p__Ascomycota_unclassified(100);p__Ascomycota_unclassified(100); | - | - | - | - | - | - | - | Unassigned | - |
| Otu26453 | 1 | k__Fungi(100);k__Fungi_unclassified(100);k__Fungi_unclassified(100);k__Fungi_unclassified(100);k__Fungi_unclassified(100);k__Fungi_unclassified(100);k__Fungi_unclassified(100); | - | - | - | - | - | - | - | Unassigned | - |
| Otu26458 | 1 | k__Fungi(100);p__Ascomycota(100);p__Ascomycota_unclassified(100);p__Ascomycota_unclassified(100);p__Ascomycota_unclassified(100);p__Ascomycota_unclassified(100);p__Ascomycota_unclassified(100); | - | - | - | - | - | - | - | Unassigned | - |
| Otu26462 | 1 | k__Fungi(100);k__Fungi_unclassified(100);k__Fungi_unclassified(100);k__Fungi_unclassified(100);k__Fungi_unclassified(100);k__Fungi_unclassified(100);k__Fungi_unclassified(100); | - | - | - | - | - | - | - | Unassigned | - |
| Otu26468 | 1 | k__Fungi(100);p__Ascomycota(100);p__Ascomycota_unclassified(100);p__Ascomycota_unclassified(100);p__Ascomycota_unclassified(100);p__Ascomycota_unclassified(100);p__Ascomycota_unclassified(100); | - | - | - | - | - | - | - | Unassigned | - |
| Otu26469 | 1 | k__Fungi(100);k__Fungi_unclassified(100);k__Fungi_unclassified(100);k__Fungi_unclassified(100);k__Fungi_unclassified(100);k__Fungi_unclassified(100);k__Fungi_unclassified(100); | - | - | - | - | - | - | - | Unassigned | - |
| Otu26470 | 1 | k__Fungi(100);k__Fungi_unclassified(100);k__Fungi_unclassified(100);k__Fungi_unclassified(100);k__Fungi_unclassified(100);k__Fungi_unclassified(100);k__Fungi_unclassified(100); | - | - | - | - | - | - | - | Unassigned | - |
| Otu26471 | 1 | k__Fungi(100);k__Fungi_unclassified(100);k__Fungi_unclassified(100);k__Fungi_unclassified(100);k__Fungi_unclassified(100);k__Fungi_unclassified(100);k__Fungi_unclassified(100); | - | - | - | - | - | - | - | Unassigned | - |
| Otu26472 | 1 | k__Fungi(100);p__Ascomycota(100);p__Ascomycota_unclassified(100);p__Ascomycota_unclassified(100);p__Ascomycota_unclassified(100);p__Ascomycota_unclassified(100);p__Ascomycota_unclassified(100); | - | - | - | - | - | - | - | Unassigned | - |
| Otu26473 | 1 | k__Fungi(100);k__Fungi_unclassified(100);k__Fungi_unclassified(100);k__Fungi_unclassified(100);k__Fungi_unclassified(100);k__Fungi_unclassified(100);k__Fungi_unclassified(100); | - | - | - | - | - | - | - | Unassigned | - |
| Otu26474 | 1 | k__Fungi(100);k__Fungi_unclassified(100);k__Fungi_unclassified(100);k__Fungi_unclassified(100);k__Fungi_unclassified(100);k__Fungi_unclassified(100);k__Fungi_unclassified(100); | - | - | - | - | - | - | - | Unassigned | - |
| Otu26475 | 1 | k__Fungi(100);p__Ascomycota(100);p__Ascomycota_unclassified(100);p__Ascomycota_unclassified(100);p__Ascomycota_unclassified(100);p__Ascomycota_unclassified(100);p__Ascomycota_unclassified(100); | - | - | - | - | - | - | - | Unassigned | - |
| Otu26476 | 1 | k__Fungi(100);p__Ascomycota(100);p__Ascomycota_unclassified(100);p__Ascomycota_unclassified(100);p__Ascomycota_unclassified(100);p__Ascomycota_unclassified(100);p__Ascomycota_unclassified(100); | - | - | - | - | - | - | - | Unassigned | - |
| Otu26477 | 1 | k__Fungi(100);k__Fungi_unclassified(100);k__Fungi_unclassified(100);k__Fungi_unclassified(100);k__Fungi_unclassified(100);k__Fungi_unclassified(100);k__Fungi_unclassified(100); | - | - | - | - | - | - | - | Unassigned | - |
| Otu26478 | 1 | k__Fungi(100);k__Fungi_unclassified(100);k__Fungi_unclassified(100);k__Fungi_unclassified(100);k__Fungi_unclassified(100);k__Fungi_unclassified(100);k__Fungi_unclassified(100); | - | - | - | - | - | - | - | Unassigned | - |
| Otu26479 | 1 | k__Fungi(100);k__Fungi_unclassified(100);k__Fungi_unclassified(100);k__Fungi_unclassified(100);k__Fungi_unclassified(100);k__Fungi_unclassified(100);k__Fungi_unclassified(100); | - | - | - | - | - | - | - | Unassigned | - |
| Otu26480 | 1 | k__Fungi(100);k__Fungi_unclassified(100);k__Fungi_unclassified(100);k__Fungi_unclassified(100);k__Fungi_unclassified(100);k__Fungi_unclassified(100);k__Fungi_unclassified(100); | - | - | - | - | - | - | - | Unassigned | - |
| Otu26481 | 1 | k__Fungi(100);k__Fungi_unclassified(100);k__Fungi_unclassified(100);k__Fungi_unclassified(100);k__Fungi_unclassified(100);k__Fungi_unclassified(100);k__Fungi_unclassified(100); | - | - | - | - | - | - | - | Unassigned | - |
| Otu26482 | 1 | k__Fungi(100);p__Ascomycota(100);p__Ascomycota_unclassified(100);p__Ascomycota_unclassified(100);p__Ascomycota_unclassified(100);p__Ascomycota_unclassified(100);p__Ascomycota_unclassified(100); | - | - | - | - | - | - | - | Unassigned | - |
| Otu26483 | 1 | k__Fungi(100);k__Fungi_unclassified(100);k__Fungi_unclassified(100);k__Fungi_unclassified(100);k__Fungi_unclassified(100);k__Fungi_unclassified(100);k__Fungi_unclassified(100); | - | - | - | - | - | - | - | Unassigned | - |
| Otu26484 | 1 | k__Fungi(100);p__Ascomycota(100);p__Ascomycota_unclassified(100);p__Ascomycota_unclassified(100);p__Ascomycota_unclassified(100);p__Ascomycota_unclassified(100);p__Ascomycota_unclassified(100); | - | - | - | - | - | - | - | Unassigned | - |
| Otu26485 | 1 | k__Fungi(100);k__Fungi_unclassified(100);k__Fungi_unclassified(100);k__Fungi_unclassified(100);k__Fungi_unclassified(100);k__Fungi_unclassified(100);k__Fungi_unclassified(100); | - | - | - | - | - | - | - | Unassigned | - |
| Otu26486 | 1 | k__Fungi(100);k__Fungi_unclassified(100);k__Fungi_unclassified(100);k__Fungi_unclassified(100);k__Fungi_unclassified(100);k__Fungi_unclassified(100);k__Fungi_unclassified(100); | - | - | - | - | - | - | - | Unassigned | - |
| Otu26487 | 1 | k__Fungi(100);p__Ascomycota(100);p__Ascomycota_unclassified(100);p__Ascomycota_unclassified(100);p__Ascomycota_unclassified(100);p__Ascomycota_unclassified(100);p__Ascomycota_unclassified(100); | - | - | - | - | - | - | - | Unassigned | - |
| Otu26488 | 1 | k__Fungi(100);p__Ascomycota(100);p__Ascomycota_unclassified(100);p__Ascomycota_unclassified(100);p__Ascomycota_unclassified(100);p__Ascomycota_unclassified(100);p__Ascomycota_unclassified(100); | - | - | - | - | - | - | - | Unassigned | - |
| Otu26489 | 1 | k__Fungi(100);p__Ascomycota(100);p__Ascomycota_unclassified(100);p__Ascomycota_unclassified(100);p__Ascomycota_unclassified(100);p__Ascomycota_unclassified(100);p__Ascomycota_unclassified(100); | - | - | - | - | - | - | - | Unassigned | - |
| Otu26490 | 1 | k__Fungi(100);p__Ascomycota(100);p__Ascomycota_unclassified(100);p__Ascomycota_unclassified(100);p__Ascomycota_unclassified(100);p__Ascomycota_unclassified(100);p__Ascomycota_unclassified(100); | - | - | - | - | - | - | - | Unassigned | - |
| Otu26491 | 1 | k__Fungi(100);p__Ascomycota(100);p__Ascomycota_unclassified(100);p__Ascomycota_unclassified(100);p__Ascomycota_unclassified(100);p__Ascomycota_unclassified(100);p__Ascomycota_unclassified(100); | - | - | - | - | - | - | - | Unassigned | - |
| Otu26492 | 1 | k__Fungi(100);p__Ascomycota(100);p__Ascomycota_unclassified(100);p__Ascomycota_unclassified(100);p__Ascomycota_unclassified(100);p__Ascomycota_unclassified(100);p__Ascomycota_unclassified(100); | - | - | - | - | - | - | - | Unassigned | - |
| Otu26493 | 1 | k__Fungi(100);k__Fungi_unclassified(100);k__Fungi_unclassified(100);k__Fungi_unclassified(100);k__Fungi_unclassified(100);k__Fungi_unclassified(100);k__Fungi_unclassified(100); | - | - | - | - | - | - | - | Unassigned | - |
| Otu26494 | 1 | k__Fungi(100);p__Ascomycota(100);p__Ascomycota_unclassified(100);p__Ascomycota_unclassified(100);p__Ascomycota_unclassified(100);p__Ascomycota_unclassified(100);p__Ascomycota_unclassified(100); | - | - | - | - | - | - | - | Unassigned | - |
| Otu26495 | 1 | k__Fungi(100);p__Ascomycota(100);p__Ascomycota_unclassified(100);p__Ascomycota_unclassified(100);p__Ascomycota_unclassified(100);p__Ascomycota_unclassified(100);p__Ascomycota_unclassified(100); | - | - | - | - | - | - | - | Unassigned | - |
| Otu26497 | 1 | k__Fungi(100);k__Fungi_unclassified(100);k__Fungi_unclassified(100);k__Fungi_unclassified(100);k__Fungi_unclassified(100);k__Fungi_unclassified(100);k__Fungi_unclassified(100); | - | - | - | - | - | - | - | Unassigned | - |
| Otu26501 | 1 | k__Fungi(100);k__Fungi_unclassified(100);k__Fungi_unclassified(100);k__Fungi_unclassified(100);k__Fungi_unclassified(100);k__Fungi_unclassified(100);k__Fungi_unclassified(100); | - | - | - | - | - | - | - | Unassigned | - |
| Otu26509 | 1 | k__Fungi(100);p__Ascomycota(100);p__Ascomycota_unclassified(100);p__Ascomycota_unclassified(100);p__Ascomycota_unclassified(100);p__Ascomycota_unclassified(100);p__Ascomycota_unclassified(100); | - | - | - | - | - | - | - | Unassigned | - |
| Otu26513 | 1 | k__Fungi(100);p__Ascomycota(100);p__Ascomycota_unclassified(100);p__Ascomycota_unclassified(100);p__Ascomycota_unclassified(100);p__Ascomycota_unclassified(100);p__Ascomycota_unclassified(100); | - | - | - | - | - | - | - | Unassigned | - |
| Otu26519 | 1 | k__Fungi(100);k__Fungi_unclassified(100);k__Fungi_unclassified(100);k__Fungi_unclassified(100);k__Fungi_unclassified(100);k__Fungi_unclassified(100);k__Fungi_unclassified(100); | - | - | - | - | - | - | - | Unassigned | - |
| Otu26522 | 1 | k__Fungi(100);p__Ascomycota(100);p__Ascomycota_unclassified(100);p__Ascomycota_unclassified(100);p__Ascomycota_unclassified(100);p__Ascomycota_unclassified(100);p__Ascomycota_unclassified(100); | - | - | - | - | - | - | - | Unassigned | - |
| Otu26531 | 1 | k__Fungi(100);p__Ascomycota(100);p__Ascomycota_unclassified(100);p__Ascomycota_unclassified(100);p__Ascomycota_unclassified(100);p__Ascomycota_unclassified(100);p__Ascomycota_unclassified(100); | - | - | - | - | - | - | - | Unassigned | - |
| Otu26533 | 1 | k__Fungi(100);k__Fungi_unclassified(100);k__Fungi_unclassified(100);k__Fungi_unclassified(100);k__Fungi_unclassified(100);k__Fungi_unclassified(100);k__Fungi_unclassified(100); | - | - | - | - | - | - | - | Unassigned | - |
| Otu26534 | 1 | k__Fungi(100);k__Fungi_unclassified(100);k__Fungi_unclassified(100);k__Fungi_unclassified(100);k__Fungi_unclassified(100);k__Fungi_unclassified(100);k__Fungi_unclassified(100); | - | - | - | - | - | - | - | Unassigned | - |
| Otu26535 | 1 | k__Fungi(100);k__Fungi_unclassified(100);k__Fungi_unclassified(100);k__Fungi_unclassified(100);k__Fungi_unclassified(100);k__Fungi_unclassified(100);k__Fungi_unclassified(100); | - | - | - | - | - | - | - | Unassigned | - |
| Otu26539 | 1 | k__Fungi(100);k__Fungi_unclassified(100);k__Fungi_unclassified(100);k__Fungi_unclassified(100);k__Fungi_unclassified(100);k__Fungi_unclassified(100);k__Fungi_unclassified(100); | - | - | - | - | - | - | - | Unassigned | - |
| Otu26540 | 1 | k__Fungi(100);k__Fungi_unclassified(100);k__Fungi_unclassified(100);k__Fungi_unclassified(100);k__Fungi_unclassified(100);k__Fungi_unclassified(100);k__Fungi_unclassified(100); | - | - | - | - | - | - | - | Unassigned | - |
| Otu26541 | 1 | k__Fungi(100);k__Fungi_unclassified(100);k__Fungi_unclassified(100);k__Fungi_unclassified(100);k__Fungi_unclassified(100);k__Fungi_unclassified(100);k__Fungi_unclassified(100); | - | - | - | - | - | - | - | Unassigned | - |
| Otu26542 | 1 | k__Fungi(100);k__Fungi_unclassified(100);k__Fungi_unclassified(100);k__Fungi_unclassified(100);k__Fungi_unclassified(100);k__Fungi_unclassified(100);k__Fungi_unclassified(100); | - | - | - | - | - | - | - | Unassigned | - |
| Otu26543 | 1 | k__Fungi(100);p__Ascomycota(100);p__Ascomycota_unclassified(100);p__Ascomycota_unclassified(100);p__Ascomycota_unclassified(100);p__Ascomycota_unclassified(100);p__Ascomycota_unclassified(100); | - | - | - | - | - | - | - | Unassigned | - |
| Otu26544 | 1 | k__Fungi(100);p__Ascomycota(100);p__Ascomycota_unclassified(100);p__Ascomycota_unclassified(100);p__Ascomycota_unclassified(100);p__Ascomycota_unclassified(100);p__Ascomycota_unclassified(100); | - | - | - | - | - | - | - | Unassigned | - |
| Otu26545 | 1 | k__Fungi(100);p__Ascomycota(100);p__Ascomycota_unclassified(100);p__Ascomycota_unclassified(100);p__Ascomycota_unclassified(100);p__Ascomycota_unclassified(100);p__Ascomycota_unclassified(100); | - | - | - | - | - | - | - | Unassigned | - |
| Otu26546 | 1 | k__Fungi(100);p__Ascomycota(100);p__Ascomycota_unclassified(100);p__Ascomycota_unclassified(100);p__Ascomycota_unclassified(100);p__Ascomycota_unclassified(100);p__Ascomycota_unclassified(100); | - | - | - | - | - | - | - | Unassigned | - |
| Otu26547 | 1 | k__Fungi(100);p__Ascomycota(100);p__Ascomycota_unclassified(100);p__Ascomycota_unclassified(100);p__Ascomycota_unclassified(100);p__Ascomycota_unclassified(100);p__Ascomycota_unclassified(100); | - | - | - | - | - | - | - | Unassigned | - |
| Otu26548 | 1 | k__Fungi(100);k__Fungi_unclassified(100);k__Fungi_unclassified(100);k__Fungi_unclassified(100);k__Fungi_unclassified(100);k__Fungi_unclassified(100);k__Fungi_unclassified(100); | - | - | - | - | - | - | - | Unassigned | - |
| Otu26549 | 1 | k__Fungi(100);p__Ascomycota(100);p__Ascomycota_unclassified(100);p__Ascomycota_unclassified(100);p__Ascomycota_unclassified(100);p__Ascomycota_unclassified(100);p__Ascomycota_unclassified(100); | - | - | - | - | - | - | - | Unassigned | - |
| Otu26550 | 1 | k__Fungi(100);p__Ascomycota(100);p__Ascomycota_unclassified(100);p__Ascomycota_unclassified(100);p__Ascomycota_unclassified(100);p__Ascomycota_unclassified(100);p__Ascomycota_unclassified(100); | - | - | - | - | - | - | - | Unassigned | - |
| Otu26551 | 1 | k__Fungi(100);p__Ascomycota(100);p__Ascomycota_unclassified(100);p__Ascomycota_unclassified(100);p__Ascomycota_unclassified(100);p__Ascomycota_unclassified(100);p__Ascomycota_unclassified(100); | - | - | - | - | - | - | - | Unassigned | - |
| Otu26552 | 1 | k__Fungi(100);p__Ascomycota(100);p__Ascomycota_unclassified(100);p__Ascomycota_unclassified(100);p__Ascomycota_unclassified(100);p__Ascomycota_unclassified(100);p__Ascomycota_unclassified(100); | - | - | - | - | - | - | - | Unassigned | - |
| Otu26553 | 1 | k__Fungi(100);k__Fungi_unclassified(100);k__Fungi_unclassified(100);k__Fungi_unclassified(100);k__Fungi_unclassified(100);k__Fungi_unclassified(100);k__Fungi_unclassified(100); | - | - | - | - | - | - | - | Unassigned | - |
| Otu26554 | 1 | k__Fungi(100);k__Fungi_unclassified(100);k__Fungi_unclassified(100);k__Fungi_unclassified(100);k__Fungi_unclassified(100);k__Fungi_unclassified(100);k__Fungi_unclassified(100); | - | - | - | - | - | - | - | Unassigned | - |
| Otu26555 | 1 | k__Fungi(100);p__Ascomycota(100);p__Ascomycota_unclassified(100);p__Ascomycota_unclassified(100);p__Ascomycota_unclassified(100);p__Ascomycota_unclassified(100);p__Ascomycota_unclassified(100); | - | - | - | - | - | - | - | Unassigned | - |
| Otu26556 | 1 | k__Fungi(100);p__Ascomycota(100);p__Ascomycota_unclassified(100);p__Ascomycota_unclassified(100);p__Ascomycota_unclassified(100);p__Ascomycota_unclassified(100);p__Ascomycota_unclassified(100); | - | - | - | - | - | - | - | Unassigned | - |
| Otu26560 | 1 | k__Fungi(100);k__Fungi_unclassified(100);k__Fungi_unclassified(100);k__Fungi_unclassified(100);k__Fungi_unclassified(100);k__Fungi_unclassified(100);k__Fungi_unclassified(100); | - | - | - | - | - | - | - | Unassigned | - |
| Otu26561 | 1 | k__Fungi(100);k__Fungi_unclassified(100);k__Fungi_unclassified(100);k__Fungi_unclassified(100);k__Fungi_unclassified(100);k__Fungi_unclassified(100);k__Fungi_unclassified(100); | - | - | - | - | - | - | - | Unassigned | - |
| Otu26562 | 1 | k__Fungi(100);k__Fungi_unclassified(100);k__Fungi_unclassified(100);k__Fungi_unclassified(100);k__Fungi_unclassified(100);k__Fungi_unclassified(100);k__Fungi_unclassified(100); | - | - | - | - | - | - | - | Unassigned | - |
| Otu26563 | 1 | k__Fungi(100);k__Fungi_unclassified(100);k__Fungi_unclassified(100);k__Fungi_unclassified(100);k__Fungi_unclassified(100);k__Fungi_unclassified(100);k__Fungi_unclassified(100); | - | - | - | - | - | - | - | Unassigned | - |
| Otu26564 | 1 | k__Fungi(100);p__Ascomycota(100);p__Ascomycota_unclassified(100);p__Ascomycota_unclassified(100);p__Ascomycota_unclassified(100);p__Ascomycota_unclassified(100);p__Ascomycota_unclassified(100); | - | - | - | - | - | - | - | Unassigned | - |
| Otu26571 | 1 | k__Fungi(100);k__Fungi_unclassified(100);k__Fungi_unclassified(100);k__Fungi_unclassified(100);k__Fungi_unclassified(100);k__Fungi_unclassified(100);k__Fungi_unclassified(100); | - | - | - | - | - | - | - | Unassigned | - |
| Otu26572 | 1 | k__Fungi(100);p__Ascomycota(100);p__Ascomycota_unclassified(100);p__Ascomycota_unclassified(100);p__Ascomycota_unclassified(100);p__Ascomycota_unclassified(100);p__Ascomycota_unclassified(100); | - | - | - | - | - | - | - | Unassigned | - |
| Otu26575 | 1 | k__Fungi(100);p__Ascomycota(100);p__Ascomycota_unclassified(100);p__Ascomycota_unclassified(100);p__Ascomycota_unclassified(100);p__Ascomycota_unclassified(100);p__Ascomycota_unclassified(100); | - | - | - | - | - | - | - | Unassigned | - |
| Otu26578 | 1 | k__Fungi(100);k__Fungi_unclassified(100);k__Fungi_unclassified(100);k__Fungi_unclassified(100);k__Fungi_unclassified(100);k__Fungi_unclassified(100);k__Fungi_unclassified(100); | - | - | - | - | - | - | - | Unassigned | - |
| Otu26579 | 1 | k__Fungi(100);p__Ascomycota(100);p__Ascomycota_unclassified(100);p__Ascomycota_unclassified(100);p__Ascomycota_unclassified(100);p__Ascomycota_unclassified(100);p__Ascomycota_unclassified(100); | - | - | - | - | - | - | - | Unassigned | - |
| Otu26580 | 1 | k__Fungi(100);k__Fungi_unclassified(100);k__Fungi_unclassified(100);k__Fungi_unclassified(100);k__Fungi_unclassified(100);k__Fungi_unclassified(100);k__Fungi_unclassified(100); | - | - | - | - | - | - | - | Unassigned | - |
| Otu26581 | 1 | k__Fungi(100);k__Fungi_unclassified(100);k__Fungi_unclassified(100);k__Fungi_unclassified(100);k__Fungi_unclassified(100);k__Fungi_unclassified(100);k__Fungi_unclassified(100); | - | - | - | - | - | - | - | Unassigned | - |
| Otu26582 | 1 | k__Fungi(100);k__Fungi_unclassified(100);k__Fungi_unclassified(100);k__Fungi_unclassified(100);k__Fungi_unclassified(100);k__Fungi_unclassified(100);k__Fungi_unclassified(100); | - | - | - | - | - | - | - | Unassigned | - |
| Otu26583 | 1 | k__Fungi(100);k__Fungi_unclassified(100);k__Fungi_unclassified(100);k__Fungi_unclassified(100);k__Fungi_unclassified(100);k__Fungi_unclassified(100);k__Fungi_unclassified(100); | - | - | - | - | - | - | - | Unassigned | - |
| Otu26584 | 1 | k__Fungi(100);p__Ascomycota(100);p__Ascomycota_unclassified(100);p__Ascomycota_unclassified(100);p__Ascomycota_unclassified(100);p__Ascomycota_unclassified(100);p__Ascomycota_unclassified(100); | - | - | - | - | - | - | - | Unassigned | - |
| Otu26585 | 1 | k__Fungi(100);p__Ascomycota(100);p__Ascomycota_unclassified(100);p__Ascomycota_unclassified(100);p__Ascomycota_unclassified(100);p__Ascomycota_unclassified(100);p__Ascomycota_unclassified(100); | - | - | - | - | - | - | - | Unassigned | - |
| Otu26586 | 1 | k__Fungi(100);p__Ascomycota(100);p__Ascomycota_unclassified(100);p__Ascomycota_unclassified(100);p__Ascomycota_unclassified(100);p__Ascomycota_unclassified(100);p__Ascomycota_unclassified(100); | - | - | - | - | - | - | - | Unassigned | - |
| Otu26587 | 1 | k__Fungi(100);k__Fungi_unclassified(100);k__Fungi_unclassified(100);k__Fungi_unclassified(100);k__Fungi_unclassified(100);k__Fungi_unclassified(100);k__Fungi_unclassified(100); | - | - | - | - | - | - | - | Unassigned | - |
| Otu26588 | 1 | k__Fungi(100);p__Ascomycota(100);p__Ascomycota_unclassified(100);p__Ascomycota_unclassified(100);p__Ascomycota_unclassified(100);p__Ascomycota_unclassified(100);p__Ascomycota_unclassified(100); | - | - | - | - | - | - | - | Unassigned | - |
| Otu26589 | 1 | k__Fungi(100);k__Fungi_unclassified(100);k__Fungi_unclassified(100);k__Fungi_unclassified(100);k__Fungi_unclassified(100);k__Fungi_unclassified(100);k__Fungi_unclassified(100); | - | - | - | - | - | - | - | Unassigned | - |
| Otu26590 | 1 | k__Fungi(100);k__Fungi_unclassified(100);k__Fungi_unclassified(100);k__Fungi_unclassified(100);k__Fungi_unclassified(100);k__Fungi_unclassified(100);k__Fungi_unclassified(100); | - | - | - | - | - | - | - | Unassigned | - |
| Otu26593 | 1 | k__Fungi(100);k__Fungi_unclassified(100);k__Fungi_unclassified(100);k__Fungi_unclassified(100);k__Fungi_unclassified(100);k__Fungi_unclassified(100);k__Fungi_unclassified(100); | - | - | - | - | - | - | - | Unassigned | - |
| Otu26594 | 1 | k__Fungi(100);k__Fungi_unclassified(100);k__Fungi_unclassified(100);k__Fungi_unclassified(100);k__Fungi_unclassified(100);k__Fungi_unclassified(100);k__Fungi_unclassified(100); | - | - | - | - | - | - | - | Unassigned | - |
| Otu26595 | 1 | k__Fungi(100);k__Fungi_unclassified(100);k__Fungi_unclassified(100);k__Fungi_unclassified(100);k__Fungi_unclassified(100);k__Fungi_unclassified(100);k__Fungi_unclassified(100); | - | - | - | - | - | - | - | Unassigned | - |

| | | | | | | | | | | | |
|---|---|---|---|---|---|---|---|---|---|---|---|
| Otu26596 | 1 | k__Fungi(100);k__Fungi_unclassified(100);k__Fungi_unclassified(100);k__Fungi_unclassified(100);k__Fungi_unclassified(100);k__Fungi_unclassified(100);k__Fungi_unclassified(100); | - | - | - | - | - | - | - | Unassigned | - |
| Otu26597 | 1 | k__Fungi(100);p__Ascomycota(100);p__Ascomycota_unclassified(100);p__Ascomycota_unclassified(100);p__Ascomycota_unclassified(100);p__Ascomycota_unclassified(100);p__Ascomycota_unclassified(100); | - | - | - | - | - | - | - | Unassigned | - |
| Otu26598 | 1 | k__Fungi(100);k__Fungi_unclassified(100);k__Fungi_unclassified(100);k__Fungi_unclassified(100);k__Fungi_unclassified(100);k__Fungi_unclassified(100);k__Fungi_unclassified(100); | - | - | - | - | - | - | - | Unassigned | - |
| Otu26599 | 1 | k__Fungi(100);k__Fungi_unclassified(100);k__Fungi_unclassified(100);k__Fungi_unclassified(100);k__Fungi_unclassified(100);k__Fungi_unclassified(100);k__Fungi_unclassified(100); | - | - | - | - | - | - | - | Unassigned | - |
| Otu26600 | 1 | k__Fungi(100);k__Fungi_unclassified(100);k__Fungi_unclassified(100);k__Fungi_unclassified(100);k__Fungi_unclassified(100);k__Fungi_unclassified(100);k__Fungi_unclassified(100); | - | - | - | - | - | - | - | Unassigned | - |
| Otu26601 | 1 | k__Fungi(100);k__Fungi_unclassified(100);k__Fungi_unclassified(100);k__Fungi_unclassified(100);k__Fungi_unclassified(100);k__Fungi_unclassified(100);k__Fungi_unclassified(100); | - | - | - | - | - | - | - | Unassigned | - |
| Otu26602 | 1 | k__Fungi(100);p__Ascomycota(100);p__Ascomycota_unclassified(100);p__Ascomycota_unclassified(100);p__Ascomycota_unclassified(100);p__Ascomycota_unclassified(100);p__Ascomycota_unclassified(100); | - | - | - | - | - | - | - | Unassigned | - |
| Otu26603 | 1 | k__Fungi(100);k__Fungi_unclassified(100);k__Fungi_unclassified(100);k__Fungi_unclassified(100);k__Fungi_unclassified(100);k__Fungi_unclassified(100);k__Fungi_unclassified(100); | - | - | - | - | - | - | - | Unassigned | - |
| Otu26604 | 1 | k__Fungi(100);p__Ascomycota(100);p__Ascomycota_unclassified(100);p__Ascomycota_unclassified(100);p__Ascomycota_unclassified(100);p__Ascomycota_unclassified(100);p__Ascomycota_unclassified(100); | - | - | - | - | - | - | - | Unassigned | - |
| Otu26605 | 1 | k__Fungi(100);k__Fungi_unclassified(100);k__Fungi_unclassified(100);k__Fungi_unclassified(100);k__Fungi_unclassified(100);k__Fungi_unclassified(100);k__Fungi_unclassified(100); | - | - | - | - | - | - | - | Unassigned | - |
| Otu26606 | 1 | k__Fungi(100);k__Fungi_unclassified(100);k__Fungi_unclassified(100);k__Fungi_unclassified(100);k__Fungi_unclassified(100);k__Fungi_unclassified(100);k__Fungi_unclassified(100); | - | - | - | - | - | - | - | Unassigned | - |
| Otu26607 | 1 | k__Fungi(100);k__Fungi_unclassified(100);k__Fungi_unclassified(100);k__Fungi_unclassified(100);k__Fungi_unclassified(100);k__Fungi_unclassified(100);k__Fungi_unclassified(100); | - | - | - | - | - | - | - | Unassigned | - |
| Otu26608 | 1 | k__Fungi(100);p__Ascomycota(100);p__Ascomycota_unclassified(100);p__Ascomycota_unclassified(100);p__Ascomycota_unclassified(100);p__Ascomycota_unclassified(100);p__Ascomycota_unclassified(100); | - | - | - | - | - | - | - | Unassigned | - |
| Otu26609 | 1 | k__Fungi(100);p__Ascomycota(100);p__Ascomycota_unclassified(100);p__Ascomycota_unclassified(100);p__Ascomycota_unclassified(100);p__Ascomycota_unclassified(100);p__Ascomycota_unclassified(100); | - | - | - | - | - | - | - | Unassigned | - |
| Otu26610 | 1 | k__Fungi(100);p__Ascomycota(100);p__Ascomycota_unclassified(100);p__Ascomycota_unclassified(100);p__Ascomycota_unclassified(100);p__Ascomycota_unclassified(100);p__Ascomycota_unclassified(100); | - | - | - | - | - | - | - | Unassigned | - |
| Otu26611 | 1 | k__Fungi(100);k__Fungi_unclassified(100);k__Fungi_unclassified(100);k__Fungi_unclassified(100);k__Fungi_unclassified(100);k__Fungi_unclassified(100);k__Fungi_unclassified(100); | - | - | - | - | - | - | - | Unassigned | - |
| Otu26612 | 1 | k__Fungi(100);k__Fungi_unclassified(100);k__Fungi_unclassified(100);k__Fungi_unclassified(100);k__Fungi_unclassified(100);k__Fungi_unclassified(100);k__Fungi_unclassified(100); | - | - | - | - | - | - | - | Unassigned | - |
| Otu26613 | 1 | k__Fungi(100);k__Fungi_unclassified(100);k__Fungi_unclassified(100);k__Fungi_unclassified(100);k__Fungi_unclassified(100);k__Fungi_unclassified(100);k__Fungi_unclassified(100); | - | - | - | - | - | - | - | Unassigned | - |
| Otu26614 | 1 | k__Fungi(100);k__Fungi_unclassified(100);k__Fungi_unclassified(100);k__Fungi_unclassified(100);k__Fungi_unclassified(100);k__Fungi_unclassified(100);k__Fungi_unclassified(100); | - | - | - | - | - | - | - | Unassigned | - |
| Otu26615 | 1 | k__Fungi(100);k__Fungi_unclassified(100);k__Fungi_unclassified(100);k__Fungi_unclassified(100);k__Fungi_unclassified(100);k__Fungi_unclassified(100);k__Fungi_unclassified(100); | - | - | - | - | - | - | - | Unassigned | - |
| Otu26616 | 1 | k__Fungi(100);k__Fungi_unclassified(100);k__Fungi_unclassified(100);k__Fungi_unclassified(100);k__Fungi_unclassified(100);k__Fungi_unclassified(100);k__Fungi_unclassified(100); | - | - | - | - | - | - | - | Unassigned | - |
| Otu26617 | 1 | k__Fungi(100);k__Fungi_unclassified(100);k__Fungi_unclassified(100);k__Fungi_unclassified(100);k__Fungi_unclassified(100);k__Fungi_unclassified(100);k__Fungi_unclassified(100); | - | - | - | - | - | - | - | Unassigned | - |
| Otu26618 | 1 | k__Fungi(100);k__Fungi_unclassified(100);k__Fungi_unclassified(100);k__Fungi_unclassified(100);k__Fungi_unclassified(100);k__Fungi_unclassified(100);k__Fungi_unclassified(100); | - | - | - | - | - | - | - | Unassigned | - |
| Otu26619 | 1 | k__Fungi(100);k__Fungi_unclassified(100);k__Fungi_unclassified(100);k__Fungi_unclassified(100);k__Fungi_unclassified(100);k__Fungi_unclassified(100);k__Fungi_unclassified(100); | - | - | - | - | - | - | - | Unassigned | - |
| Otu26620 | 1 | k__Fungi(100);k__Fungi_unclassified(100);k__Fungi_unclassified(100);k__Fungi_unclassified(100);k__Fungi_unclassified(100);k__Fungi_unclassified(100);k__Fungi_unclassified(100); | - | - | - | - | - | - | - | Unassigned | - |
| Otu26621 | 1 | k__Fungi(100);k__Fungi_unclassified(100);k__Fungi_unclassified(100);k__Fungi_unclassified(100);k__Fungi_unclassified(100);k__Fungi_unclassified(100);k__Fungi_unclassified(100); | - | - | - | - | - | - | - | Unassigned | - |
| Otu26622 | 1 | k__Fungi(100);k__Fungi_unclassified(100);k__Fungi_unclassified(100);k__Fungi_unclassified(100);k__Fungi_unclassified(100);k__Fungi_unclassified(100);k__Fungi_unclassified(100); | - | - | - | - | - | - | - | Unassigned | - |
| Otu26623 | 1 | k__Fungi(100);k__Fungi_unclassified(100);k__Fungi_unclassified(100);k__Fungi_unclassified(100);k__Fungi_unclassified(100);k__Fungi_unclassified(100);k__Fungi_unclassified(100); | - | - | - | - | - | - | - | Unassigned | - |
| Otu26624 | 1 | k__Fungi(100);p__Ascomycota(100);p__Ascomycota_unclassified(100);p__Ascomycota_unclassified(100);p__Ascomycota_unclassified(100);p__Ascomycota_unclassified(100);p__Ascomycota_unclassified(100); | - | - | - | - | - | - | - | Unassigned | - |
| Otu26625 | 1 | k__Fungi(100);k__Fungi_unclassified(100);k__Fungi_unclassified(100);k__Fungi_unclassified(100);k__Fungi_unclassified(100);k__Fungi_unclassified(100);k__Fungi_unclassified(100); | - | - | - | - | - | - | - | Unassigned | - |
| Otu26626 | 1 | k__Fungi(100);k__Fungi_unclassified(100);k__Fungi_unclassified(100);k__Fungi_unclassified(100);k__Fungi_unclassified(100);k__Fungi_unclassified(100);k__Fungi_unclassified(100); | - | - | - | - | - | - | - | Unassigned | - |
| Otu26627 | 1 | k__Fungi(100);p__Ascomycota(100);p__Ascomycota_unclassified(100);p__Ascomycota_unclassified(100);p__Ascomycota_unclassified(100);p__Ascomycota_unclassified(100);p__Ascomycota_unclassified(100); | - | - | - | - | - | - | - | Unassigned | - |
| Otu26628 | 1 | k__Fungi(100);k__Fungi_unclassified(100);k__Fungi_unclassified(100);k__Fungi_unclassified(100);k__Fungi_unclassified(100);k__Fungi_unclassified(100);k__Fungi_unclassified(100); | - | - | - | - | - | - | - | Unassigned | - |
| Otu26629 | 1 | k__Fungi(100);p__Ascomycota(100);p__Ascomycota_unclassified(100);p__Ascomycota_unclassified(100);p__Ascomycota_unclassified(100);p__Ascomycota_unclassified(100);p__Ascomycota_unclassified(100); | - | - | - | - | - | - | - | Unassigned | - |
| Otu26630 | 1 | k__Fungi(100);k__Fungi_unclassified(100);k__Fungi_unclassified(100);k__Fungi_unclassified(100);k__Fungi_unclassified(100);k__Fungi_unclassified(100);k__Fungi_unclassified(100); | - | - | - | - | - | - | - | Unassigned | - |
| Otu26631 | 1 | k__Fungi(100);p__Ascomycota(100);p__Ascomycota_unclassified(100);p__Ascomycota_unclassified(100);p__Ascomycota_unclassified(100);p__Ascomycota_unclassified(100);p__Ascomycota_unclassified(100); | - | - | - | - | - | - | - | Unassigned | - |
| Otu26632 | 1 | k__Fungi(100);p__Ascomycota(100);p__Ascomycota_unclassified(100);p__Ascomycota_unclassified(100);p__Ascomycota_unclassified(100);p__Ascomycota_unclassified(100);p__Ascomycota_unclassified(100); | - | - | - | - | - | - | - | Unassigned | - |
| Otu26633 | 1 | k__Fungi(100);p__Ascomycota(100);p__Ascomycota_unclassified(100);p__Ascomycota_unclassified(100);p__Ascomycota_unclassified(100);p__Ascomycota_unclassified(100);p__Ascomycota_unclassified(100); | - | - | - | - | - | - | - | Unassigned | - |
| Otu26634 | 1 | k__Fungi(100);k__Fungi_unclassified(100);k__Fungi_unclassified(100);k__Fungi_unclassified(100);k__Fungi_unclassified(100);k__Fungi_unclassified(100);k__Fungi_unclassified(100); | - | - | - | - | - | - | - | Unassigned | - |
| Otu26635 | 1 | k__Fungi(100);k__Fungi_unclassified(100);k__Fungi_unclassified(100);k__Fungi_unclassified(100);k__Fungi_unclassified(100);k__Fungi_unclassified(100);k__Fungi_unclassified(100); | - | - | - | - | - | - | - | Unassigned | - |
| Otu26636 | 1 | k__Fungi(100);p__Ascomycota(100);p__Ascomycota_unclassified(100);p__Ascomycota_unclassified(100);p__Ascomycota_unclassified(100);p__Ascomycota_unclassified(100);p__Ascomycota_unclassified(100); | - | - | - | - | - | - | - | Unassigned | - |
| Otu26637 | 1 | k__Fungi(100);k__Fungi_unclassified(100);k__Fungi_unclassified(100);k__Fungi_unclassified(100);k__Fungi_unclassified(100);k__Fungi_unclassified(100);k__Fungi_unclassified(100); | - | - | - | - | - | - | - | Unassigned | - |
| Otu26638 | 1 | k__Fungi(100);k__Fungi_unclassified(100);k__Fungi_unclassified(100);k__Fungi_unclassified(100);k__Fungi_unclassified(100);k__Fungi_unclassified(100);k__Fungi_unclassified(100); | - | - | - | - | - | - | - | Unassigned | - |
| Otu26639 | 1 | k__Fungi(100);p__Ascomycota(100);p__Ascomycota_unclassified(100);p__Ascomycota_unclassified(100);p__Ascomycota_unclassified(100);p__Ascomycota_unclassified(100);p__Ascomycota_unclassified(100); | - | - | - | - | - | - | - | Unassigned | - |
| Otu26640 | 1 | k__Fungi(100);k__Fungi_unclassified(100);k__Fungi_unclassified(100);k__Fungi_unclassified(100);k__Fungi_unclassified(100);k__Fungi_unclassified(100);k__Fungi_unclassified(100); | - | - | - | - | - | - | - | Unassigned | - |
| Otu26641 | 1 | k__Fungi(100);p__Ascomycota(100);p__Ascomycota_unclassified(100);p__Ascomycota_unclassified(100);p__Ascomycota_unclassified(100);p__Ascomycota_unclassified(100);p__Ascomycota_unclassified(100); | - | - | - | - | - | - | - | Unassigned | - |
| Otu26642 | 1 | k__Fungi(100);p__Ascomycota(100);p__Ascomycota_unclassified(100);p__Ascomycota_unclassified(100);p__Ascomycota_unclassified(100);p__Ascomycota_unclassified(100);p__Ascomycota_unclassified(100); | - | - | - | - | - | - | - | Unassigned | - |
| Otu26643 | 1 | k__Fungi(100);k__Fungi_unclassified(100);k__Fungi_unclassified(100);k__Fungi_unclassified(100);k__Fungi_unclassified(100);k__Fungi_unclassified(100);k__Fungi_unclassified(100); | - | - | - | - | - | - | - | Unassigned | - |
| Otu26644 | 1 | k__Fungi(100);p__Ascomycota(100);p__Ascomycota_unclassified(100);p__Ascomycota_unclassified(100);p__Ascomycota_unclassified(100);p__Ascomycota_unclassified(100);p__Ascomycota_unclassified(100); | - | - | - | - | - | - | - | Unassigned | - |
| Otu26645 | 1 | k__Fungi(100);k__Fungi_unclassified(100);k__Fungi_unclassified(100);k__Fungi_unclassified(100);k__Fungi_unclassified(100);k__Fungi_unclassified(100);k__Fungi_unclassified(100); | - | - | - | - | - | - | - | Unassigned | - |
| Otu26647 | 1 | k__Fungi(100);k__Fungi_unclassified(100);k__Fungi_unclassified(100);k__Fungi_unclassified(100);k__Fungi_unclassified(100);k__Fungi_unclassified(100);k__Fungi_unclassified(100); | - | - | - | - | - | - | - | Unassigned | - |
| Otu26648 | 1 | k__Fungi(100);k__Fungi_unclassified(100);k__Fungi_unclassified(100);k__Fungi_unclassified(100);k__Fungi_unclassified(100);k__Fungi_unclassified(100);k__Fungi_unclassified(100); | - | - | - | - | - | - | - | Unassigned | - |
| Otu26649 | 1 | k__Fungi(100);p__Ascomycota(100);p__Ascomycota_unclassified(100);p__Ascomycota_unclassified(100);p__Ascomycota_unclassified(100);p__Ascomycota_unclassified(100);p__Ascomycota_unclassified(100); | - | - | - | - | - | - | - | Unassigned | - |
| Otu26650 | 1 | k__Fungi(100);k__Fungi_unclassified(100);k__Fungi_unclassified(100);k__Fungi_unclassified(100);k__Fungi_unclassified(100);k__Fungi_unclassified(100);k__Fungi_unclassified(100); | - | - | - | - | - | - | - | Unassigned | - |
| Otu26651 | 1 | k__Fungi(100);p__Ascomycota(100);p__Ascomycota_unclassified(100);p__Ascomycota_unclassified(100);p__Ascomycota_unclassified(100);p__Ascomycota_unclassified(100);p__Ascomycota_unclassified(100); | - | - | - | - | - | - | - | Unassigned | - |
| Otu26652 | 1 | k__Fungi(100);p__Ascomycota(100);p__Ascomycota_unclassified(100);p__Ascomycota_unclassified(100);p__Ascomycota_unclassified(100);p__Ascomycota_unclassified(100);p__Ascomycota_unclassified(100); | - | - | - | - | - | - | - | Unassigned | - |
| Otu26653 | 1 | k__Fungi(100);k__Fungi_unclassified(100);k__Fungi_unclassified(100);k__Fungi_unclassified(100);k__Fungi_unclassified(100);k__Fungi_unclassified(100);k__Fungi_unclassified(100); | - | - | - | - | - | - | - | Unassigned | - |
| Otu26654 | 1 | k__Fungi(100);k__Fungi_unclassified(100);k__Fungi_unclassified(100);k__Fungi_unclassified(100);k__Fungi_unclassified(100);k__Fungi_unclassified(100);k__Fungi_unclassified(100); | - | - | - | - | - | - | - | Unassigned | - |
| Otu26655 | 1 | k__Fungi(100);p__Ascomycota(100);p__Ascomycota_unclassified(100);p__Ascomycota_unclassified(100);p__Ascomycota_unclassified(100);p__Ascomycota_unclassified(100);p__Ascomycota_unclassified(100); | - | - | - | - | - | - | - | Unassigned | - |
| Otu26656 | 1 | k__Fungi(100);k__Fungi_unclassified(100);k__Fungi_unclassified(100);k__Fungi_unclassified(100);k__Fungi_unclassified(100);k__Fungi_unclassified(100);k__Fungi_unclassified(100); | - | - | - | - | - | - | - | Unassigned | - |
| Otu26667 | 1 | k__Fungi(100);p__Ascomycota(100);p__Ascomycota_unclassified(100);p__Ascomycota_unclassified(100);p__Ascomycota_unclassified(100);p__Ascomycota_unclassified(100);p__Ascomycota_unclassified(100); | - | - | - | - | - | - | - | Unassigned | - |
| Otu26669 | 1 | k__Fungi(100);k__Fungi_unclassified(100);k__Fungi_unclassified(100);k__Fungi_unclassified(100);k__Fungi_unclassified(100);k__Fungi_unclassified(100);k__Fungi_unclassified(100); | - | - | - | - | - | - | - | Unassigned | - |
| Otu26671 | 1 | k__Fungi(100);p__Ascomycota(100);p__Ascomycota_unclassified(100);p__Ascomycota_unclassified(100);p__Ascomycota_unclassified(100);p__Ascomycota_unclassified(100);p__Ascomycota_unclassified(100); | - | - | - | - | - | - | - | Unassigned | - |
| Otu26675 | 1 | k__Fungi(100);p__Ascomycota(100);p__Ascomycota_unclassified(100);p__Ascomycota_unclassified(100);p__Ascomycota_unclassified(100);p__Ascomycota_unclassified(100);p__Ascomycota_unclassified(100); | - | - | - | - | - | - | - | Unassigned | - |
| Otu26676 | 1 | k__Fungi(100);p__Ascomycota(100);p__Ascomycota_unclassified(100);p__Ascomycota_unclassified(100);p__Ascomycota_unclassified(100);p__Ascomycota_unclassified(100);p__Ascomycota_unclassified(100); | - | - | - | - | - | - | - | Unassigned | - |
| Otu26677 | 1 | k__Fungi(100);p__Ascomycota(100);p__Ascomycota_unclassified(100);p__Ascomycota_unclassified(100);p__Ascomycota_unclassified(100);p__Ascomycota_unclassified(100);p__Ascomycota_unclassified(100); | - | - | - | - | - | - | - | Unassigned | - |
| Otu26678 | 1 | k__Fungi(100);k__Fungi_unclassified(100);k__Fungi_unclassified(100);k__Fungi_unclassified(100);k__Fungi_unclassified(100);k__Fungi_unclassified(100);k__Fungi_unclassified(100); | - | - | - | - | - | - | - | Unassigned | - |
| Otu26679 | 1 | k__Fungi(100);p__Ascomycota(100);p__Ascomycota_unclassified(100);p__Ascomycota_unclassified(100);p__Ascomycota_unclassified(100);p__Ascomycota_unclassified(100);p__Ascomycota_unclassified(100); | - | - | - | - | - | - | - | Unassigned | - |
| Otu26680 | 1 | k__Fungi(100);k__Fungi_unclassified(100);k__Fungi_unclassified(100);k__Fungi_unclassified(100);k__Fungi_unclassified(100);k__Fungi_unclassified(100);k__Fungi_unclassified(100); | - | - | - | - | - | - | - | Unassigned | - |
| Otu26681 | 1 | k__Fungi(100);k__Fungi_unclassified(100);k__Fungi_unclassified(100);k__Fungi_unclassified(100);k__Fungi_unclassified(100);k__Fungi_unclassified(100);k__Fungi_unclassified(100); | - | - | - | - | - | - | - | Unassigned | - |
| Otu26683 | 1 | k__Fungi(100);k__Fungi_unclassified(100);k__Fungi_unclassified(100);k__Fungi_unclassified(100);k__Fungi_unclassified(100);k__Fungi_unclassified(100);k__Fungi_unclassified(100); | - | - | - | - | - | - | - | Unassigned | - |
| Otu26684 | 1 | k__Fungi(100);p__Ascomycota(100);p__Ascomycota_unclassified(100);p__Ascomycota_unclassified(100);p__Ascomycota_unclassified(100);p__Ascomycota_unclassified(100);p__Ascomycota_unclassified(100); | - | - | - | - | - | - | - | Unassigned | - |
| Otu26685 | 1 | k__Fungi(100);k__Fungi_unclassified(100);k__Fungi_unclassified(100);k__Fungi_unclassified(100);k__Fungi_unclassified(100);k__Fungi_unclassified(100);k__Fungi_unclassified(100); | - | - | - | - | - | - | - | Unassigned | - |
| Otu26686 | 1 | k__Fungi(100);p__Ascomycota(100);p__Ascomycota_unclassified(100);p__Ascomycota_unclassified(100);p__Ascomycota_unclassified(100);p__Ascomycota_unclassified(100);p__Ascomycota_unclassified(100); | - | - | - | - | - | - | - | Unassigned | - |
| Otu26687 | 1 | k__Fungi(100);k__Fungi_unclassified(100);k__Fungi_unclassified(100);k__Fungi_unclassified(100);k__Fungi_unclassified(100);k__Fungi_unclassified(100);k__Fungi_unclassified(100); | - | - | - | - | - | - | - | Unassigned | - |
| Otu26688 | 1 | k__Fungi(100);k__Fungi_unclassified(100);k__Fungi_unclassified(100);k__Fungi_unclassified(100);k__Fungi_unclassified(100);k__Fungi_unclassified(100);k__Fungi_unclassified(100); | - | - | - | - | - | - | - | Unassigned | - |
| Otu26689 | 1 | k__Fungi(100);p__Ascomycota(100);p__Ascomycota_unclassified(100);p__Ascomycota_unclassified(100);p__Ascomycota_unclassified(100);p__Ascomycota_unclassified(100);p__Ascomycota_unclassified(100); | - | - | - | - | - | - | - | Unassigned | - |
| Otu26690 | 1 | k__Fungi(100);p__Basidiomycota(100);p__Basidiomycota_unclassified(100);p__Basidiomycota_unclassified(100);p__Basidiomycota_unclassified(100);p__Basidiomycota_unclassified(100);p__Basidiomycota_unclassified(100); | - | - | - | - | - | - | - | Unassigned | - |
| Otu26691 | 1 | k__Fungi(100);p__Ascomycota(100);p__Ascomycota_unclassified(100);p__Ascomycota_unclassified(100);p__Ascomycota_unclassified(100);p__Ascomycota_unclassified(100);p__Ascomycota_unclassified(100); | - | - | - | - | - | - | - | Unassigned | - |
| Otu26692 | 1 | k__Fungi(100);p__Ascomycota(100);p__Ascomycota_unclassified(100);p__Ascomycota_unclassified(100);p__Ascomycota_unclassified(100);p__Ascomycota_unclassified(100);p__Ascomycota_unclassified(100); | - | - | - | - | - | - | - | Unassigned | - |
| Otu26693 | 1 | k__Fungi(100);p__Ascomycota(100);p__Ascomycota_unclassified(100);p__Ascomycota_unclassified(100);p__Ascomycota_unclassified(100);p__Ascomycota_unclassified(100);p__Ascomycota_unclassified(100); | - | - | - | - | - | - | - | Unassigned | - |
| Otu26694 | 1 | k__Fungi(100);k__Fungi_unclassified(100);k__Fungi_unclassified(100);k__Fungi_unclassified(100);k__Fungi_unclassified(100);k__Fungi_unclassified(100);k__Fungi_unclassified(100); | - | - | - | - | - | - | - | Unassigned | - |
| Otu26695 | 1 | k__Fungi(100);k__Fungi_unclassified(100);k__Fungi_unclassified(100);k__Fungi_unclassified(100);k__Fungi_unclassified(100);k__Fungi_unclassified(100);k__Fungi_unclassified(100); | - | - | - | - | - | - | - | Unassigned | - |
| Otu26696 | 1 | k__Fungi(100);k__Fungi_unclassified(100);k__Fungi_unclassified(100);k__Fungi_unclassified(100);k__Fungi_unclassified(100);k__Fungi_unclassified(100);k__Fungi_unclassified(100); | - | - | - | - | - | - | - | Unassigned | - |
| Otu26698 | 1 | k__Fungi(100);k__Fungi_unclassified(100);k__Fungi_unclassified(100);k__Fungi_unclassified(100);k__Fungi_unclassified(100);k__Fungi_unclassified(100);k__Fungi_unclassified(100); | - | - | - | - | - | - | - | Unassigned | - |
| Otu26699 | 1 | k__Fungi(100);k__Fungi_unclassified(100);k__Fungi_unclassified(100);k__Fungi_unclassified(100);k__Fungi_unclassified(100);k__Fungi_unclassified(100);k__Fungi_unclassified(100); | - | - | - | - | - | - | - | Unassigned | - |
| Otu26701 | 1 | k__Fungi(100);p__Ascomycota(100);p__Ascomycota_unclassified(100);p__Ascomycota_unclassified(100);p__Ascomycota_unclassified(100);p__Ascomycota_unclassified(100);p__Ascomycota_unclassified(100); | - | - | - | - | - | - | - | Unassigned | - |
| Otu26702 | 1 | k__Fungi(100);k__Fungi_unclassified(100);k__Fungi_unclassified(100);k__Fungi_unclassified(100);k__Fungi_unclassified(100);k__Fungi_unclassified(100);k__Fungi_unclassified(100); | - | - | - | - | - | - | - | Unassigned | - |
| Otu26703 | 1 | k__Fungi(100);p__Ascomycota(100);p__Ascomycota_unclassified(100);p__Ascomycota_unclassified(100);p__Ascomycota_unclassified(100);p__Ascomycota_unclassified(100);p__Ascomycota_unclassified(100); | - | - | - | - | - | - | - | Unassigned | - |
| Otu26704 | 1 | k__Fungi(100);k__Fungi_unclassified(100);k__Fungi_unclassified(100);k__Fungi_unclassified(100);k__Fungi_unclassified(100);k__Fungi_unclassified(100);k__Fungi_unclassified(100); | - | - | - | - | - | - | - | Unassigned | - |
| Otu26705 | 1 | k__Fungi(100);p__Ascomycota(100);p__Ascomycota_unclassified(100);p__Ascomycota_unclassified(100);p__Ascomycota_unclassified(100);p__Ascomycota_unclassified(100);p__Ascomycota_unclassified(100); | - | - | - | - | - | - | - | Unassigned | - |
| Otu26706 | 1 | k__Fungi(100);p__Ascomycota(100);p__Ascomycota_unclassified(100);p__Ascomycota_unclassified(100);p__Ascomycota_unclassified(100);p__Ascomycota_unclassified(100);p__Ascomycota_unclassified(100); | - | - | - | - | - | - | - | Unassigned | - |
| Otu26708 | 1 | k__Fungi(100);p__Ascomycota(100);p__Ascomycota_unclassified(100);p__Ascomycota_unclassified(100);p__Ascomycota_unclassified(100);p__Ascomycota_unclassified(100);p__Ascomycota_unclassified(100); | - | - | - | - | - | - | - | Unassigned | - |
| Otu26709 | 1 | k__Fungi(100);k__Fungi_unclassified(100);k__Fungi_unclassified(100);k__Fungi_unclassified(100);k__Fungi_unclassified(100);k__Fungi_unclassified(100);k__Fungi_unclassified(100); | - | - | - | - | - | - | - | Unassigned | - |
| Otu26710 | 1 | k__Fungi(100);k__Fungi_unclassified(100);k__Fungi_unclassified(100);k__Fungi_unclassified(100);k__Fungi_unclassified(100);k__Fungi_unclassified(100);k__Fungi_unclassified(100); | - | - | - | - | - | - | - | Unassigned | - |
| Otu26711 | 1 | k__Fungi(100);p__Ascomycota(100);p__Ascomycota_unclassified(100);p__Ascomycota_unclassified(100);p__Ascomycota_unclassified(100);p__Ascomycota_unclassified(100);p__Ascomycota_unclassified(100); | - | - | - | - | - | - | - | Unassigned | - |
| Otu26712 | 1 | k__Fungi(100);p__Ascomycota(100);p__Ascomycota_unclassified(100);p__Ascomycota_unclassified(100);p__Ascomycota_unclassified(100);p__Ascomycota_unclassified(100);p__Ascomycota_unclassified(100); | - | - | - | - | - | - | - | Unassigned | - |
| Otu26713 | 1 | k__Fungi(100);p__Ascomycota(100);p__Ascomycota_unclassified(100);p__Ascomycota_unclassified(100);p__Ascomycota_unclassified(100);p__Ascomycota_unclassified(100);p__Ascomycota_unclassified(100); | - | - | - | - | - | - | - | Unassigned | - |
| Otu26714 | 1 | k__Fungi(100);p__Ascomycota(100);p__Ascomycota_unclassified(100);p__Ascomycota_unclassified(100);p__Ascomycota_unclassified(100);p__Ascomycota_unclassified(100);p__Ascomycota_unclassified(100); | - | - | - | - | - | - | - | Unassigned | - |
| Otu26716 | 1 | k__Fungi(100);p__Ascomycota(100);p__Ascomycota_unclassified(100);p__Ascomycota_unclassified(100);p__Ascomycota_unclassified(100);p__Ascomycota_unclassified(100);p__Ascomycota_unclassified(100); | - | - | - | - | - | - | - | Unassigned | - |
| Otu26717 | 1 | k__Fungi(100);p__Ascomycota(100);p__Ascomycota_unclassified(100);p__Ascomycota_unclassified(100);p__Ascomycota_unclassified(100);p__Ascomycota_unclassified(100);p__Ascomycota_unclassified(100); | - | - | - | - | - | - | - | Unassigned | - |
| Otu26718 | 1 | k__Fungi(100);k__Fungi_unclassified(100);k__Fungi_unclassified(100);k__Fungi_unclassified(100);k__Fungi_unclassified(100);k__Fungi_unclassified(100);k__Fungi_unclassified(100); | - | - | - | - | - | - | - | Unassigned | - |
| Otu26719 | 1 | k__Fungi(100);p__Ascomycota(100);p__Ascomycota_unclassified(100);p__Ascomycota_unclassified(100);p__Ascomycota_unclassified(100);p__Ascomycota_unclassified(100);p__Ascomycota_unclassified(100); | - | - | - | - | - | - | - | Unassigned | - |
| Otu26720 | 1 | k__Fungi(100);k__Fungi_unclassified(100);k__Fungi_unclassified(100);k__Fungi_unclassified(100);k__Fungi_unclassified(100);k__Fungi_unclassified(100);k__Fungi_unclassified(100); | - | - | - | - | - | - | - | Unassigned | - |
| Otu26721 | 1 | k__Fungi(100);k__Fungi_unclassified(100);k__Fungi_unclassified(100);k__Fungi_unclassified(100);k__Fungi_unclassified(100);k__Fungi_unclassified(100);k__Fungi_unclassified(100); | - | - | - | - | - | - | - | Unassigned | - |
| Otu26722 | 1 | k__Fungi(100);k__Fungi_unclassified(100);k__Fungi_unclassified(100);k__Fungi_unclassified(100);k__Fungi_unclassified(100);k__Fungi_unclassified(100);k__Fungi_unclassified(100); | - | - | - | - | - | - | - | Unassigned | - |
| Otu26723 | 1 | k__Fungi(100);p__Ascomycota(100);p__Ascomycota_unclassified(100);p__Ascomycota_unclassified(100);p__Ascomycota_unclassified(100);p__Ascomycota_unclassified(100);p__Ascomycota_unclassified(100); | - | - | - | - | - | - | - | Unassigned | - |
| Otu26725 | 1 | k__Fungi(100);p__Ascomycota(100);p__Ascomycota_unclassified(100);p__Ascomycota_unclassified(100);p__Ascomycota_unclassified(100);p__Ascomycota_unclassified(100);p__Ascomycota_unclassified(100); | - | - | - | - | - | - | - | Unassigned | - |
| Otu26726 | 1 | k__Fungi(100);p__Ascomycota(100);p__Ascomycota_unclassified(100);p__Ascomycota_unclassified(100);p__Ascomycota_unclassified(100);p__Ascomycota_unclassified(100);p__Ascomycota_unclassified(100); | - | - | - | - | - | - | - | Unassigned | - |
| Otu26727 | 1 | k__Fungi(100);p__Ascomycota(100);p__Ascomycota_unclassified(100);p__Ascomycota_unclassified(100);p__Ascomycota_unclassified(100);p__Ascomycota_unclassified(100);p__Ascomycota_unclassified(100); | - | - | - | - | - | - | - | Unassigned | - |
| Otu26728 | 1 | k__Fungi(100);k__Fungi_unclassified(100);k__Fungi_unclassified(100);k__Fungi_unclassified(100);k__Fungi_unclassified(100);k__Fungi_unclassified(100);k__Fungi_unclassified(100); | - | - | - | - | - | - | - | Unassigned | - |
| Otu26729 | 1 | k__Fungi(100);p__Ascomycota(100);p__Ascomycota_unclassified(100);p__Ascomycota_unclassified(100);p__Ascomycota_unclassified(100);p__Ascomycota_unclassified(100);p__Ascomycota_unclassified(100); | - | - | - | - | - | - | - | Unassigned | - |
| Otu26730 | 1 | k__Fungi(100);p__Ascomycota(100);p__Ascomycota_unclassified(100);p__Ascomycota_unclassified(100);p__Ascomycota_unclassified(100);p__Ascomycota_unclassified(100);p__Ascomycota_unclassified(100); | - | - | - | - | - | - | - | Unassigned | - |
| Otu26731 | 1 | k__Fungi(100);p__Ascomycota(100);p__Ascomycota_unclassified(100);p__Ascomycota_unclassified(100);p__Ascomycota_unclassified(100);p__Ascomycota_unclassified(100);p__Ascomycota_unclassified(100); | - | - | - | - | - | - | - | Unassigned | - |
| Otu26732 | 1 | k__Fungi(100);p__Ascomycota(100);p__Ascomycota_unclassified(100);p__Ascomycota_unclassified(100);p__Ascomycota_unclassified(100);p__Ascomycota_unclassified(100);p__Ascomycota_unclassified(100); | - | - | - | - | - | - | - | Unassigned | - |
| Otu26733 | 1 | k__Fungi(100);p__Ascomycota(100);p__Ascomycota_unclassified(100);p__Ascomycota_unclassified(100);p__Ascomycota_unclassified(100);p__Ascomycota_unclassified(100);p__Ascomycota_unclassified(100); | - | - | - | - | - | - | - | Unassigned | - |
| Otu26734 | 1 | k__Fungi(100);k__Fungi_unclassified(100);k__Fungi_unclassified(100);k__Fungi_unclassified(100);k__Fungi_unclassified(100);k__Fungi_unclassified(100);k__Fungi_unclassified(100); | - | - | - | - | - | - | - | Unassigned | - |
| Otu26735 | 1 | k__Fungi(100);p__Ascomycota(100);p__Ascomycota_unclassified(100);p__Ascomycota_unclassified(100);p__Ascomycota_unclassified(100);p__Ascomycota_unclassified(100);p__Ascomycota_unclassified(100); | - | - | - | - | - | - | - | Unassigned | - |
| Otu26737 | 1 | k__Fungi(100);p__Ascomycota(100);p__Ascomycota_unclassified(100);p__Ascomycota_unclassified(100);p__Ascomycota_unclassified(100);p__Ascomycota_unclassified(100);p__Ascomycota_unclassified(100); | - | - | - | - | - | - | - | Unassigned | - |
| Otu26739 | 1 | k__Fungi(100);k__Fungi_unclassified(100);k__Fungi_unclassified(100);k__Fungi_unclassified(100);k__Fungi_unclassified(100);k__Fungi_unclassified(100);k__Fungi_unclassified(100); | - | - | - | - | - | - | - | Unassigned | - |
| Otu26740 | 1 | k__Fungi(100);p__Ascomycota(100);p__Ascomycota_unclassified(100);p__Ascomycota_unclassified(100);p__Ascomycota_unclassified(100);p__Ascomycota_unclassified(100);p__Ascomycota_unclassified(100); | - | - | - | - | - | - | - | Unassigned | - |
| Otu26741 | 1 | k__Fungi(100);k__Fungi_unclassified(100);k__Fungi_unclassified(100);k__Fungi_unclassified(100);k__Fungi_unclassified(100);k__Fungi_unclassified(100);k__Fungi_unclassified(100); | - | - | - | - | - | - | - | Unassigned | - |
| Otu26743 | 1 | k__Fungi(100);p__Ascomycota(100);p__Ascomycota_unclassified(100);p__Ascomycota_unclassified(100);p__Ascomycota_unclassified(100);p__Ascomycota_unclassified(100);p__Ascomycota_unclassified(100); | - | - | - | - | - | - | - | Unassigned | - |
| Otu26744 | 1 | k__Fungi(100);p__Ascomycota(100);p__Ascomycota_unclassified(100);p__Ascomycota_unclassified(100);p__Ascomycota_unclassified(100);p__Ascomycota_unclassified(100);p__Ascomycota_unclassified(100); | - | - | - | - | - | - | - | Unassigned | - |
| Otu26746 | 1 | k__Fungi(100);p__Ascomycota(100);p__Ascomycota_unclassified(100);p__Ascomycota_unclassified(100);p__Ascomycota_unclassified(100);p__Ascomycota_unclassified(100);p__Ascomycota_unclassified(100); | - | - | - | - | - | - | - | Unassigned | - |
| Otu26748 | 1 | k__Fungi(100);k__Fungi_unclassified(100);k__Fungi_unclassified(100);k__Fungi_unclassified(100);k__Fungi_unclassified(100);k__Fungi_unclassified(100);k__Fungi_unclassified(100); | - | - | - | - | - | - | - | Unassigned | - |
| Otu26749 | 1 | k__Fungi(100);p__Ascomycota(100);p__Ascomycota_unclassified(100);p__Ascomycota_unclassified(100);p__Ascomycota_unclassified(100);p__Ascomycota_unclassified(100);p__Ascomycota_unclassified(100); | - | - | - | - | - | - | - | Unassigned | - |
| Otu26750 | 1 | k__Fungi(100);p__Ascomycota(100);p__Ascomycota_unclassified(100);p__Ascomycota_unclassified(100);p__Ascomycota_unclassified(100);p__Ascomycota_unclassified(100);p__Ascomycota_unclassified(100); | - | - | - | - | - | - | - | Unassigned | - |
| Otu26752 | 1 | k__Fungi(100);p__Ascomycota(100);p__Ascomycota_unclassified(100);p__Ascomycota_unclassified(100);p__Ascomycota_unclassified(100);p__Ascomycota_unclassified(100);p__Ascomycota_unclassified(100); | - | - | - | - | - | - | - | Unassigned | - |
| Otu26753 | 1 | k__Fungi(100);k__Fungi_unclassified(100);k__Fungi_unclassified(100);k__Fungi_unclassified(100);k__Fungi_unclassified(100);k__Fungi_unclassified(100);k__Fungi_unclassified(100); | - | - | - | - | - | - | - | Unassigned | - |
| Otu26754 | 1 | k__Fungi(100);k__Fungi_unclassified(100);k__Fungi_unclassified(100);k__Fungi_unclassified(100);k__Fungi_unclassified(100);k__Fungi_unclassified(100);k__Fungi_unclassified(100); | - | - | - | - | - | - | - | Unassigned | - |
| Otu26759 | 1 | k__Fungi(100);p__Ascomycota(100);p__Ascomycota_unclassified(100);p__Ascomycota_unclassified(100);p__Ascomycota_unclassified(100);p__Ascomycota_unclassified(100);p__Ascomycota_unclassified(100); | - | - | - | - | - | - | - | Unassigned | - |
| Otu26760 | 1 | k__Fungi(100);k__Fungi_unclassified(100);k__Fungi_unclassified(100);k__Fungi_unclassified(100);k__Fungi_unclassified(100);k__Fungi_unclassified(100);k__Fungi_unclassified(100); | - | - | - | - | - | - | - | Unassigned | - |
| Otu26761 | 1 | k__Fungi(100);p__Ascomycota(100);p__Ascomycota_unclassified(100);p__Ascomycota_unclassified(100);p__Ascomycota_unclassified(100);p__Ascomycota_unclassified(100);p__Ascomycota_unclassified(100); | - | - | - | - | - | - | - | Unassigned | - |
| Otu26762 | 1 | k__Fungi(100);k__Fungi_unclassified(100);k__Fungi_unclassified(100);k__Fungi_unclassified(100);k__Fungi_unclassified(100);k__Fungi_unclassified(100);k__Fungi_unclassified(100); | - | - | - | - | - | - | - | Unassigned | - |
| Otu26764 | 1 | k__Fungi(100);k__Fungi_unclassified(100);k__Fungi_unclassified(100);k__Fungi_unclassified(100);k__Fungi_unclassified(100);k__Fungi_unclassified(100);k__Fungi_unclassified(100); | - | - | - | - | - | - | - | Unassigned | - |
| Otu26765 | 1 | k__Fungi(100);k__Fungi_unclassified(100);k__Fungi_unclassified(100);k__Fungi_unclassified(100);k__Fungi_unclassified(100);k__Fungi_unclassified(100);k__Fungi_unclassified(100); | - | - | - | - | - | - | - | Unassigned | - |
| Otu26766 | 1 | k__Fungi(100);k__Fungi_unclassified(100);k__Fungi_unclassified(100);k__Fungi_unclassified(100);k__Fungi_unclassified(100);k__Fungi_unclassified(100);k__Fungi_unclassified(100); | - | - | - | - | - | - | - | Unassigned | - |
| Otu26767 | 1 | k__Fungi(100);k__Fungi_unclassified(100);k__Fungi_unclassified(100);k__Fungi_unclassified(100);k__Fungi_unclassified(100);k__Fungi_unclassified(100);k__Fungi_unclassified(100); | - | - | - | - | - | - | - | Unassigned | - |
| Otu26768 | 1 | k__Fungi(100);k__Fungi_unclassified(100);k__Fungi_unclassified(100);k__Fungi_unclassified(100);k__Fungi_unclassified(100);k__Fungi_unclassified(100);k__Fungi_unclassified(100); | - | - | - | - | - | - | - | Unassigned | - |
| Otu26769 | 1 | k__Fungi(100);k__Fungi_unclassified(100);k__Fungi_unclassified(100);k__Fungi_unclassified(100);k__Fungi_unclassified(100);k__Fungi_unclassified(100);k__Fungi_unclassified(100); | - | - | - | - | - | - | - | Unassigned | - |
| Otu26770 | 1 | k__Fungi(100);p__Ascomycota(100);p__Ascomycota_unclassified(100);p__Ascomycota_unclassified(100);p__Ascomycota_unclassified(100);p__Ascomycota_unclassified(100);p__Ascomycota_unclassified(100); | - | - | - | - | - | - | - | Unassigned | - |
| Otu26772 | 1 | k__Fungi(100);k__Fungi_unclassified(100);k__Fungi_unclassified(100);k__Fungi_unclassified(100);k__Fungi_unclassified(100);k__Fungi_unclassified(100);k__Fungi_unclassified(100); | - | - | - | - | - | - | - | Unassigned | - |
| Otu26775 | 1 | k__Fungi(100);p__Ascomycota(100);p__Ascomycota_unclassified(100);p__Ascomycota_unclassified(100);p__Ascomycota_unclassified(100);p__Ascomycota_unclassified(100);p__Ascomycota_unclassified(100); | - | - | - | - | - | - | - | Unassigned | - |
| Otu26776 | 1 | k__Fungi(100);k__Fungi_unclassified(100);k__Fungi_unclassified(100);k__Fungi_unclassified(100);k__Fungi_unclassified(100);k__Fungi_unclassified(100);k__Fungi_unclassified(100); | - | - | - | - | - | - | - | Unassigned | - |
| Otu26779 | 1 | k__Fungi(100);k__Fungi_unclassified(100);k__Fungi_unclassified(100);k__Fungi_unclassified(100);k__Fungi_unclassified(100);k__Fungi_unclassified(100);k__Fungi_unclassified(100); | - | - | - | - | - | - | - | Unassigned | - |
| Otu26780 | 1 | k__Fungi(100);k__Fungi_unclassified(100);k__Fungi_unclassified(100);k__Fungi_unclassified(100);k__Fungi_unclassified(100);k__Fungi_unclassified(100);k__Fungi_unclassified(100); | - | - | - | - | - | - | - | Unassigned | - |
| Otu26781 | 1 | k__Fungi(100);k__Fungi_unclassified(100);k__Fungi_unclassified(100);k__Fungi_unclassified(100);k__Fungi_unclassified(100);k__Fungi_unclassified(100);k__Fungi_unclassified(100); | - | - | - | - | - | - | - | Unassigned | - |
| Otu26782 | 1 | k__Fungi(100);k__Fungi_unclassified(100);k__Fungi_unclassified(100);k__Fungi_unclassified(100);k__Fungi_unclassified(100);k__Fungi_unclassified(100);k__Fungi_unclassified(100); | - | - | - | - | - | - | - | Unassigned | - |
| Otu26783 | 1 | k__Fungi(100);p__Ascomycota(100);p__Ascomycota_unclassified(100);p__Ascomycota_unclassified(100);p__Ascomycota_unclassified(100);p__Ascomycota_unclassified(100);p__Ascomycota_unclassified(100); | - | - | - | - | - | - | - | Unassigned | - |
| Otu26784 | 1 | k__Fungi(100);p__Ascomycota(100);p__Ascomycota_unclassified(100);p__Ascomycota_unclassified(100);p__Ascomycota_unclassified(100);p__Ascomycota_unclassified(100);p__Ascomycota_unclassified(100); | - | - | - | - | - | - | - | Unassigned | - |
| Otu26786 | 1 | k__Fungi(100);k__Fungi_unclassified(100);k__Fungi_unclassified(100);k__Fungi_unclassified(100);k__Fungi_unclassified(100);k__Fungi_unclassified(100);k__Fungi_unclassified(100); | - | - | - | - | - | - | - | Unassigned | - |
| Otu26796 | 1 | k__Fungi(100);k__Fungi_unclassified(100);k__Fungi_unclassified(100);k__Fungi_unclassified(100);k__Fungi_unclassified(100);k__Fungi_unclassified(100);k__Fungi_unclassified(100); | - | - | - | - | - | - | - | Unassigned | - |
| Otu26799 | 1 | k__Fungi(100);p__Ascomycota(100);p__Ascomycota_unclassified(100);p__Ascomycota_unclassified(100);p__Ascomycota_unclassified(100);p__Ascomycota_unclassified(100);p__Ascomycota_unclassified(100); | - | - | - | - | - | - | - | Unassigned | - |
| Otu26800 | 1 | k__Fungi(100);k__Fungi_unclassified(100);k__Fungi_unclassified(100);k__Fungi_unclassified(100);k__Fungi_unclassified(100);k__Fungi_unclassified(100);k__Fungi_unclassified(100); | - | - | - | - | - | - | - | Unassigned | - |
| Otu26803 | 1 | k__Fungi(100);p__Ascomycota(100);p__Ascomycota_unclassified(100);p__Ascomycota_unclassified(100);p__Ascomycota_unclassified(100);p__Ascomycota_unclassified(100);p__Ascomycota_unclassified(100); | - | - | - | - | - | - | - | Unassigned | - |
| Otu26806 | 1 | k__Fungi(100);k__Fungi_unclassified(100);k__Fungi_unclassified(100);k__Fungi_unclassified(100);k__Fungi_unclassified(100);k__Fungi_unclassified(100);k__Fungi_unclassified(100); | - | - | - | - | - | - | - | Unassigned | - |
| Otu26808 | 1 | k__Fungi(100);p__Ascomycota(100);p__Ascomycota_unclassified(100);p__Ascomycota_unclassified(100);p__Ascomycota_unclassified(100);p__Ascomycota_unclassified(100);p__Ascomycota_unclassified(100); | - | - | - | - | - | - | - | Unassigned | - |
| Otu26813 | 1 | k__Fungi(100);p__Ascomycota(100);c__Leotiomycetes(100);o__Helotiales(100);o__Helotiales_unclassified(100);o__Helotiales_unclassified(100);o__Helotiales_unclassified(100); | - | - | - | - | - | - | - | Unassigned | - |
| Otu26816 | 1 | k__Fungi(100);k__Fungi_unclassified(100);k__Fungi_unclassified(100);k__Fungi_unclassified(100);k__Fungi_unclassified(100);k__Fungi_unclassified(100);k__Fungi_unclassified(100); | - | - | - | - | - | - | - | Unassigned | - |
| Otu26820 | 1 | k__Fungi(100);p__Ascomycota(100);p__Ascomycota_unclassified(100);p__Ascomycota_unclassified(100);p__Ascomycota_unclassified(100);p__Ascomycota_unclassified(100);p__Ascomycota_unclassified(100); | - | - | - | - | - | - | - | Unassigned | - |
| Otu26821 | 1 | k__Fungi(100);k__Fungi_unclassified(100);k__Fungi_unclassified(100);k__Fungi_unclassified(100);k__Fungi_unclassified(100);k__Fungi_unclassified(100);k__Fungi_unclassified(100); | - | - | - | - | - | - | - | Unassigned | - |
| Otu26826 | 1 | k__Fungi(100);p__Ascomycota(100);p__Ascomycota_unclassified(100);p__Ascomycota_unclassified(100);p__Ascomycota_unclassified(100);p__Ascomycota_unclassified(100);p__Ascomycota_unclassified(100); | - | - | - | - | - | - | - | Unassigned | - |
| Otu26827 | 1 | k__Fungi(100);k__Fungi_unclassified(100);k__Fungi_unclassified(100);k__Fungi_unclassified(100);k__Fungi_unclassified(100);k__Fungi_unclassified(100);k__Fungi_unclassified(100); | - | - | - | - | - | - | - | Unassigned | - |
| Otu26833 | 1 | k__Fungi(100);k__Fungi_unclassified(100);k__Fungi_unclassified(100);k__Fungi_unclassified(100);k__Fungi_unclassified(100);k__Fungi_unclassified(100);k__Fungi_unclassified(100); | - | - | - | - | - | - | - | Unassigned | - |
| Otu26837 | 1 | k__Fungi(100);p__Ascomycota(100);p__Ascomycota_unclassified(100);p__Ascomycota_unclassified(100);p__Ascomycota_unclassified(100);p__Ascomycota_unclassified(100);p__Ascomycota_unclassified(100); | - | - | - | - | - | - | - | Unassigned | - |
| Otu26841 | 1 | k__Fungi(100);p__Ascomycota(100);p__Ascomycota_unclassified(100);p__Ascomycota_unclassified(100);p__Ascomycota_unclassified(100);p__Ascomycota_unclassified(100);p__Ascomycota_unclassified(100); | - | - | - | - | - | - | - | Unassigned | - |
| Otu26842 | 1 | k__Fungi(100);k__Fungi_unclassified(100);k__Fungi_unclassified(100);k__Fungi_unclassified(100);k__Fungi_unclassified(100);k__Fungi_unclassified(100);k__Fungi_unclassified(100); | - | - | - | - | - | - | - | Unassigned | - |
| Otu26843 | 1 | k__Fungi(100);p__Ascomycota(100);p__Ascomycota_unclassified(100);p__Ascomycota_unclassified(100);p__Ascomycota_unclassified(100);p__Ascomycota_unclassified(100);p__Ascomycota_unclassified(100); | - | - | - | - | - | - | - | Unassigned | - |
| Otu26844 | 1 | k__Fungi(100);k__Fungi_unclassified(100);k__Fungi_unclassified(100);k__Fungi_unclassified(100);k__Fungi_unclassified(100);k__Fungi_unclassified(100);k__Fungi_unclassified(100); | - | - | - | - | - | - | - | Unassigned | - |
| Otu26845 | 1 | k__Fungi(100);p__Ascomycota(100);p__Ascomycota_unclassified(100);p__Ascomycota_unclassified(100);p__Ascomycota_unclassified(100);p__Ascomycota_unclassified(100);p__Ascomycota_unclassified(100); | - | - | - | - | - | - | - | Unassigned | - |
| Otu26846 | 1 | k__Fungi(100);k__Fungi_unclassified(100);k__Fungi_unclassified(100);k__Fungi_unclassified(100);k__Fungi_unclassified(100);k__Fungi_unclassified(100);k__Fungi_unclassified(100); | - | - | - | - | - | - | - | Unassigned | - |
| Otu26847 | 1 | k__Fungi(100);k__Fungi_unclassified(100);k__Fungi_unclassified(100);k__Fungi_unclassified(100);k__Fungi_unclassified(100);k__Fungi_unclassified(100);k__Fungi_unclassified(100); | - | - | - | - | - | - | - | Unassigned | - |
| Otu26848 | 1 | k__Fungi(100);k__Fungi_unclassified(100);k__Fungi_unclassified(100);k__Fungi_unclassified(100);k__Fungi_unclassified(100);k__Fungi_unclassified(100);k__Fungi_unclassified(100); | - | - | - | - | - | - | - | Unassigned | - |
| Otu26849 | 1 | k__Fungi(100);k__Fungi_unclassified(100);k__Fungi_unclassified(100);k__Fungi_unclassified(100);k__Fungi_unclassified(100);k__Fungi_unclassified(100);k__Fungi_unclassified(100); | - | - | - | - | - | - | - | Unassigned | - |
| Otu26850 | 1 | k__Fungi(100);k__Fungi_unclassified(100);k__Fungi_unclassified(100);k__Fungi_unclassified(100);k__Fungi_unclassified(100);k__Fungi_unclassified(100);k__Fungi_unclassified(100); | - | - | - | - | - | - | - | Unassigned | - |
| Otu26851 | 1 | k__Fungi(100);p__Ascomycota(100);p__Ascomycota_unclassified(100);p__Ascomycota_unclassified(100);p__Ascomycota_unclassified(100);p__Ascomycota_unclassified(100);p__Ascomycota_unclassified(100); | - | - | - | - | - | - | - | Unassigned | - |
| Otu26852 | 1 | k__Fungi(100);k__Fungi_unclassified(100);k__Fungi_unclassified(100);k__Fungi_unclassified(100);k__Fungi_unclassified(100);k__Fungi_unclassified(100);k__Fungi_unclassified(100); | - | - | - | - | - | - | - | Unassigned | - |
| Otu26853 | 1 | k__Fungi(100);k__Fungi_unclassified(100);k__Fungi_unclassified(100);k__Fungi_unclassified(100);k__Fungi_unclassified(100);k__Fungi_unclassified(100);k__Fungi_unclassified(100); | - | - | - | - | - | - | - | Unassigned | - |
| Otu26854 | 1 | k__Fungi(100);k__Fungi_unclassified(100);k__Fungi_unclassified(100);k__Fungi_unclassified(100);k__Fungi_unclassified(100);k__Fungi_unclassified(100);k__Fungi_unclassified(100); | - | - | - | - | - | - | - | Unassigned | - |
| Otu26855 | 1 | k__Fungi(100);p__Ascomycota(100);p__Ascomycota_unclassified(100);p__Ascomycota_unclassified(100);p__Ascomycota_unclassified(100);p__Ascomycota_unclassified(100);p__Ascomycota_unclassified(100); | - | - | - | - | - | - | - | Unassigned | - |
| Otu26856 | 1 | k__Fungi(100);k__Fungi_unclassified(100);k__Fungi_unclassified(100);k__Fungi_unclassified(100);k__Fungi_unclassified(100);k__Fungi_unclassified(100);k__Fungi_unclassified(100); | - | - | - | - | - | - | - | Unassigned | - |
| Otu26857 | 1 | k__Fungi(100);p__Ascomycota(100);p__Ascomycota_unclassified(100);p__Ascomycota_unclassified(100);p__Ascomycota_unclassified(100);p__Ascomycota_unclassified(100);p__Ascomycota_unclassified(100); | - | - | - | - | - | - | - | Unassigned | - |
| Otu26858 | 1 | k__Fungi(100);k__Fungi_unclassified(100);k__Fungi_unclassified(100);k__Fungi_unclassified(100);k__Fungi_unclassified(100);k__Fungi_unclassified(100);k__Fungi_unclassified(100); | - | - | - | - | - | - | - | Unassigned | - |
| Otu26860 | 1 | k__Fungi(100);k__Fungi_unclassified(100);k__Fungi_unclassified(100);k__Fungi_unclassified(100);k__Fungi_unclassified(100);k__Fungi_unclassified(100);k__Fungi_unclassified(100); | - | - | - | - | - | - | - | Unassigned | - |
| Otu26861 | 1 | k__Fungi(100);k__Fungi_unclassified(100);k__Fungi_unclassified(100);k__Fungi_unclassified(100);k__Fungi_unclassified(100);k__Fungi_unclassified(100);k__Fungi_unclassified(100); | - | - | - | - | - | - | - | Unassigned | - |
| Otu26862 | 1 | k__Fungi(100);k__Fungi_unclassified(100);k__Fungi_unclassified(100);k__Fungi_unclassified(100);k__Fungi_unclassified(100);k__Fungi_unclassified(100);k__Fungi_unclassified(100); | - | - | - | - | - | - | - | Unassigned | - |
| Otu26863 | 1 | k__Fungi(100);k__Fungi_unclassified(100);k__Fungi_unclassified(100);k__Fungi_unclassified(100);k__Fungi_unclassified(100);k__Fungi_unclassified(100);k__Fungi_unclassified(100); | - | - | - | - | - | - | - | Unassigned | - |
| Otu26867 | 1 | k__Fungi(100);p__Ascomycota(100);p__Ascomycota_unclassified(100);p__Ascomycota_unclassified(100);p__Ascomycota_unclassified(100);p__Ascomycota_unclassified(100);p__Ascomycota_unclassified(100); | - | - | - | - | - | - | - | Unassigned | - |
| Otu26868 | 1 | k__Fungi(100);k__Fungi_unclassified(100);k__Fungi_unclassified(100);k__Fungi_unclassified(100);k__Fungi_unclassified(100);k__Fungi_unclassified(100);k__Fungi_unclassified(100); | - | - | - | - | - | - | - | Unassigned | - |
| Otu26869 | 1 | k__Fungi(100);p__Ascomycota(100);p__Ascomycota_unclassified(100);p__Ascomycota_unclassified(100);p__Ascomycota_unclassified(100);p__Ascomycota_unclassified(100);p__Ascomycota_unclassified(100); | - | - | - | - | - | - | - | Unassigned | - |
| Otu26870 | 1 | k__Fungi(100);k__Fungi_unclassified(100);k__Fungi_unclassified(100);k__Fungi_unclassified(100);k__Fungi_unclassified(100);k__Fungi_unclassified(100);k__Fungi_unclassified(100); | - | - | - | - | - | - | - | Unassigned | - |
| Otu26871 | 1 | k__Fungi(100);k__Fungi_unclassified(100);k__Fungi_unclassified(100);k__Fungi_unclassified(100);k__Fungi_unclassified(100);k__Fungi_unclassified(100);k__Fungi_unclassified(100); | - | - | - | - | - | - | - | Unassigned | - |
| Otu26872 | 1 | k__Fungi(100);p__Ascomycota(100);p__Ascomycota_unclassified(100);p__Ascomycota_unclassified(100);p__Ascomycota_unclassified(100);p__Ascomycota_unclassified(100);p__Ascomycota_unclassified(100); | - | - | - | - | - | - | - | Unassigned | - |
| Otu26877 | 1 | k__Fungi(100);k__Fungi_unclassified(100);k__Fungi_unclassified(100);k__Fungi_unclassified(100);k__Fungi_unclassified(100);k__Fungi_unclassified(100);k__Fungi_unclassified(100); | - | - | - | - | - | - | - | Unassigned | - |
| Otu26878 | 1 | k__Fungi(100);p__Ascomycota(100);p__Ascomycota_unclassified(100);p__Ascomycota_unclassified(100);p__Ascomycota_unclassified(100);p__Ascomycota_unclassified(100);p__Ascomycota_unclassified(100); | - | - | - | - | - | - | - | Unassigned | - |
| Otu26879 | 1 | k__Fungi(100);k__Fungi_unclassified(100);k__Fungi_unclassified(100);k__Fungi_unclassified(100);k__Fungi_unclassified(100);k__Fungi_unclassified(100);k__Fungi_unclassified(100); | - | - | - | - | - | - | - | Unassigned | - |
| Otu26880 | 1 | k__Fungi(100);p__Ascomycota(100);p__Ascomycota_unclassified(100);p__Ascomycota_unclassified(100);p__Ascomycota_unclassified(100);p__Ascomycota_unclassified(100);p__Ascomycota_unclassified(100); | - | - | - | - | - | - | - | Unassigned | - |
| Otu26881 | 1 | k__Fungi(100);p__Ascomycota(100);p__Ascomycota_unclassified(100);p__Ascomycota_unclassified(100);p__Ascomycota_unclassified(100);p__Ascomycota_unclassified(100);p__Ascomycota_unclassified(100); | - | - | - | - | - | - | - | Unassigned | - |
| Otu26883 | 1 | k__Fungi(100);k__Fungi_unclassified(100);k__Fungi_unclassified(100);k__Fungi_unclassified(100);k__Fungi_unclassified(100);k__Fungi_unclassified(100);k__Fungi_unclassified(100); | - | - | - | - | - | - | - | Unassigned | - |
| Otu26885 | 1 | k__Fungi(100);p__Ascomycota(100);p__Ascomycota_unclassified(100);p__Ascomycota_unclassified(100);p__Ascomycota_unclassified(100);p__Ascomycota_unclassified(100);p__Ascomycota_unclassified(100); | - | - | - | - | - | - | - | Unassigned | - |
| Otu26886 | 1 | k__Fungi(100);k__Fungi_unclassified(100);k__Fungi_unclassified(100);k__Fungi_unclassified(100);k__Fungi_unclassified(100);k__Fungi_unclassified(100);k__Fungi_unclassified(100); | - | - | - | - | - | - | - | Unassigned | - |
| Otu26887 | 1 | k__Fungi(100);k__Fungi_unclassified(100);k__Fungi_unclassified(100);k__Fungi_unclassified(100);k__Fungi_unclassified(100);k__Fungi_unclassified(100);k__Fungi_unclassified(100); | - | - | - | - | - | - | - | Unassigned | - |
| Otu26888 | 1 | k__Fungi(100);k__Fungi_unclassified(100);k__Fungi_unclassified(100);k__Fungi_unclassified(100);k__Fungi_unclassified(100);k__Fungi_unclassified(100);k__Fungi_unclassified(100); | - | - | - | - | - | - | - | Unassigned | - |
| Otu26889 | 1 | k__Fungi(100);k__Fungi_unclassified(100);k__Fungi_unclassified(100);k__Fungi_unclassified(100);k__Fungi_unclassified(100);k__Fungi_unclassified(100);k__Fungi_unclassified(100); | - | - | - | - | - | - | - | Unassigned | - |
| Otu26890 | 1 | k__Fungi(100);k__Fungi_unclassified(100);k__Fungi_unclassified(100);k__Fungi_unclassified(100);k__Fungi_unclassified(100);k__Fungi_unclassified(100);k__Fungi_unclassified(100); | - | - | - | - | - | - | - | Unassigned | - |
| Otu26891 | 1 | k__Fungi(100);k__Fungi_unclassified(100);k__Fungi_unclassified(100);k__Fungi_unclassified(100);k__Fungi_unclassified(100);k__Fungi_unclassified(100);k__Fungi_unclassified(100); | - | - | - | - | - | - | - | Unassigned | - |
| Otu26892 | 1 | k__Fungi(100);p__Ascomycota(100);p__Ascomycota_unclassified(100);p__Ascomycota_unclassified(100);p__Ascomycota_unclassified(100);p__Ascomycota_unclassified(100);p__Ascomycota_unclassified(100); | - | - | - | - | - | - | - | Unassigned | - |
| Otu26893 | 1 | k__Fungi(100);k__Fungi_unclassified(100);k__Fungi_unclassified(100);k__Fungi_unclassified(100);k__Fungi_unclassified(100);k__Fungi_unclassified(100);k__Fungi_unclassified(100); | - | - | - | - | - | - | - | Unassigned | - |
| Otu26894 | 1 | k__Fungi(100);k__Fungi_unclassified(100);k__Fungi_unclassified(100);k__Fungi_unclassified(100);k__Fungi_unclassified(100);k__Fungi_unclassified(100);k__Fungi_unclassified(100); | - | - | - | - | - | - | - | Unassigned | - |
| Otu26895 | 1 | k__Fungi(100);k__Fungi_unclassified(100);k__Fungi_unclassified(100);k__Fungi_unclassified(100);k__Fungi_unclassified(100);k__Fungi_unclassified(100);k__Fungi_unclassified(100); | - | - | - | - | - | - | - | Unassigned | - |
| Otu26896 | 1 | k__Fungi(100);k__Fungi_unclassified(100);k__Fungi_unclassified(100);k__Fungi_unclassified(100);k__Fungi_unclassified(100);k__Fungi_unclassified(100);k__Fungi_unclassified(100); | - | - | - | - | - | - | - | Unassigned | - |
| Otu26897 | 1 | k__Fungi(100);k__Fungi_unclassified(100);k__Fungi_unclassified(100);k__Fungi_unclassified(100);k__Fungi_unclassified(100);k__Fungi_unclassified(100);k__Fungi_unclassified(100); | - | - | - | - | - | - | - | Unassigned | - |
| Otu26898 | 1 | k__Fungi(100);k__Fungi_unclassified(100);k__Fungi_unclassified(100);k__Fungi_unclassified(100);k__Fungi_unclassified(100);k__Fungi_unclassified(100);k__Fungi_unclassified(100); | - | - | - | - | - | - | - | Unassigned | - |
| Otu26899 | 1 | k__Fungi(100);p__Ascomycota(100);p__Ascomycota_unclassified(100);p__Ascomycota_unclassified(100);p__Ascomycota_unclassified(100);p__Ascomycota_unclassified(100);p__Ascomycota_unclassified(100); | - | - | - | - | - | - | - | Unassigned | - |
| Otu26900 | 1 | k__Fungi(100);p__Ascomycota(100);p__Ascomycota_unclassified(100);p__Ascomycota_unclassified(100);p__Ascomycota_unclassified(100);p__Ascomycota_unclassified(100);p__Ascomycota_unclassified(100); | - | - | - | - | - | - | - | Unassigned | - |

| | | | | | | | | | | | |
|---|---|---|---|---|---|---|---|---|---|---|---|
| Otu26901 | 1 | k__Fungi(100);p__Ascomycota(100);p__Ascomycota_unclassified(100);p__Ascomycota_unclassified(100);p__Ascomycota_unclassified(100);p__Ascomycota_unclassified(100);p__Ascomycota_unclassified(100); | - | - | - | - | - | - | - | Unassigned | - |
| Otu26902 | 1 | k__Fungi(100);k__Fungi_unclassified(100);k__Fungi_unclassified(100);k__Fungi_unclassified(100);k__Fungi_unclassified(100);k__Fungi_unclassified(100);k__Fungi_unclassified(100); | - | - | - | - | - | - | - | Unassigned | - |
| Otu26903 | 1 | k__Fungi(100);k__Fungi_unclassified(100);k__Fungi_unclassified(100);k__Fungi_unclassified(100);k__Fungi_unclassified(100);k__Fungi_unclassified(100);k__Fungi_unclassified(100); | - | - | - | - | - | - | - | Unassigned | - |
| Otu26908 | 1 | k__Fungi(100);k__Fungi_unclassified(100);k__Fungi_unclassified(100);k__Fungi_unclassified(100);k__Fungi_unclassified(100);k__Fungi_unclassified(100);k__Fungi_unclassified(100); | - | - | - | - | - | - | - | Unassigned | - |
| Otu26909 | 1 | k__Fungi(100);k__Fungi_unclassified(100);k__Fungi_unclassified(100);k__Fungi_unclassified(100);k__Fungi_unclassified(100);k__Fungi_unclassified(100);k__Fungi_unclassified(100); | - | - | - | - | - | - | - | Unassigned | - |
| Otu26910 | 1 | k__Fungi(100);k__Fungi_unclassified(100);k__Fungi_unclassified(100);k__Fungi_unclassified(100);k__Fungi_unclassified(100);k__Fungi_unclassified(100);k__Fungi_unclassified(100); | - | - | - | - | - | - | - | Unassigned | - |
| Otu26911 | 1 | k__Fungi(100);k__Fungi_unclassified(100);k__Fungi_unclassified(100);k__Fungi_unclassified(100);k__Fungi_unclassified(100);k__Fungi_unclassified(100);k__Fungi_unclassified(100); | - | - | - | - | - | - | - | Unassigned | - |
| Otu26912 | 1 | k__Fungi(100);k__Fungi_unclassified(100);k__Fungi_unclassified(100);k__Fungi_unclassified(100);k__Fungi_unclassified(100);k__Fungi_unclassified(100);k__Fungi_unclassified(100); | - | - | - | - | - | - | - | Unassigned | - |
| Otu26915 | 1 | k__Fungi(100);k__Fungi_unclassified(100);k__Fungi_unclassified(100);k__Fungi_unclassified(100);k__Fungi_unclassified(100);k__Fungi_unclassified(100);k__Fungi_unclassified(100); | - | - | - | - | - | - | - | Unassigned | - |
| Otu26919 | 1 | k__Fungi(100);k__Fungi_unclassified(100);k__Fungi_unclassified(100);k__Fungi_unclassified(100);k__Fungi_unclassified(100);k__Fungi_unclassified(100);k__Fungi_unclassified(100); | - | - | - | - | - | - | - | Unassigned | - |
| Otu26920 | 1 | k__Fungi(100);k__Fungi_unclassified(100);k__Fungi_unclassified(100);k__Fungi_unclassified(100);k__Fungi_unclassified(100);k__Fungi_unclassified(100);k__Fungi_unclassified(100); | - | - | - | - | - | - | - | Unassigned | - |
| Otu26922 | 1 | k__Fungi(100);k__Fungi_unclassified(100);k__Fungi_unclassified(100);k__Fungi_unclassified(100);k__Fungi_unclassified(100);k__Fungi_unclassified(100);k__Fungi_unclassified(100); | - | - | - | - | - | - | - | Unassigned | - |
| Otu26923 | 1 | k__Fungi(100);k__Fungi_unclassified(100);k__Fungi_unclassified(100);k__Fungi_unclassified(100);k__Fungi_unclassified(100);k__Fungi_unclassified(100);k__Fungi_unclassified(100); | - | - | - | - | - | - | - | Unassigned | - |
| Otu26925 | 1 | k__Fungi(100);k__Fungi_unclassified(100);k__Fungi_unclassified(100);k__Fungi_unclassified(100);k__Fungi_unclassified(100);k__Fungi_unclassified(100);k__Fungi_unclassified(100); | - | - | - | - | - | - | - | Unassigned | - |
| Otu26926 | 1 | k__Fungi(100);k__Fungi_unclassified(100);k__Fungi_unclassified(100);k__Fungi_unclassified(100);k__Fungi_unclassified(100);k__Fungi_unclassified(100);k__Fungi_unclassified(100); | - | - | - | - | - | - | - | Unassigned | - |
| Otu26927 | 1 | k__Fungi(100);k__Fungi_unclassified(100);k__Fungi_unclassified(100);k__Fungi_unclassified(100);k__Fungi_unclassified(100);k__Fungi_unclassified(100);k__Fungi_unclassified(100); | - | - | - | - | - | - | - | Unassigned | - |
| Otu26928 | 1 | k__Fungi(100);k__Fungi_unclassified(100);k__Fungi_unclassified(100);k__Fungi_unclassified(100);k__Fungi_unclassified(100);k__Fungi_unclassified(100);k__Fungi_unclassified(100); | - | - | - | - | - | - | - | Unassigned | - |
| Otu26934 | 1 | k__Fungi(100);k__Fungi_unclassified(100);k__Fungi_unclassified(100);k__Fungi_unclassified(100);k__Fungi_unclassified(100);k__Fungi_unclassified(100);k__Fungi_unclassified(100); | - | - | - | - | - | - | - | Unassigned | - |
| Otu26938 | 1 | k__Fungi(100);p__Ascomycota(100);p__Ascomycota_unclassified(100);p__Ascomycota_unclassified(100);p__Ascomycota_unclassified(100);p__Ascomycota_unclassified(100);p__Ascomycota_unclassified(100); | - | - | - | - | - | - | - | Unassigned | - |
| Otu26939 | 1 | k__Fungi(100);p__Ascomycota(100);p__Ascomycota_unclassified(100);p__Ascomycota_unclassified(100);p__Ascomycota_unclassified(100);p__Ascomycota_unclassified(100);p__Ascomycota_unclassified(100); | - | - | - | - | - | - | - | Unassigned | - |
| Otu26940 | 1 | k__Fungi(100);k__Fungi_unclassified(100);k__Fungi_unclassified(100);k__Fungi_unclassified(100);k__Fungi_unclassified(100);k__Fungi_unclassified(100);k__Fungi_unclassified(100); | - | - | - | - | - | - | - | Unassigned | - |
| Otu26943 | 1 | k__Fungi(100);p__Ascomycota(100);p__Ascomycota_unclassified(100);p__Ascomycota_unclassified(100);p__Ascomycota_unclassified(100);p__Ascomycota_unclassified(100);p__Ascomycota_unclassified(100); | - | - | - | - | - | - | - | Unassigned | - |
| Otu26945 | 1 | k__Fungi(100);k__Fungi_unclassified(100);k__Fungi_unclassified(100);k__Fungi_unclassified(100);k__Fungi_unclassified(100);k__Fungi_unclassified(100);k__Fungi_unclassified(100); | - | - | - | - | - | - | - | Unassigned | - |
| Otu26946 | 1 | k__Fungi(100);k__Fungi_unclassified(100);k__Fungi_unclassified(100);k__Fungi_unclassified(100);k__Fungi_unclassified(100);k__Fungi_unclassified(100);k__Fungi_unclassified(100); | - | - | - | - | - | - | - | Unassigned | - |
| Otu26947 | 1 | k__Fungi(100);k__Fungi_unclassified(100);k__Fungi_unclassified(100);k__Fungi_unclassified(100);k__Fungi_unclassified(100);k__Fungi_unclassified(100);k__Fungi_unclassified(100); | - | - | - | - | - | - | - | Unassigned | - |
| Otu26950 | 1 | k__Fungi(100);p__Ascomycota(100);p__Ascomycota_unclassified(100);p__Ascomycota_unclassified(100);p__Ascomycota_unclassified(100);p__Ascomycota_unclassified(100);p__Ascomycota_unclassified(100); | - | - | - | - | - | - | - | Unassigned | - |
| Otu26951 | 1 | k__Fungi(100);p__Ascomycota(100);p__Ascomycota_unclassified(100);p__Ascomycota_unclassified(100);p__Ascomycota_unclassified(100);p__Ascomycota_unclassified(100);p__Ascomycota_unclassified(100); | - | - | - | - | - | - | - | Unassigned | - |
| Otu26952 | 1 | k__Fungi(100);p__Ascomycota(100);p__Ascomycota_unclassified(100);p__Ascomycota_unclassified(100);p__Ascomycota_unclassified(100);p__Ascomycota_unclassified(100);p__Ascomycota_unclassified(100); | - | - | - | - | - | - | - | Unassigned | - |
| Otu26953 | 1 | k__Fungi(100);k__Fungi_unclassified(100);k__Fungi_unclassified(100);k__Fungi_unclassified(100);k__Fungi_unclassified(100);k__Fungi_unclassified(100);k__Fungi_unclassified(100); | - | - | - | - | - | - | - | Unassigned | - |
| Otu26954 | 1 | k__Fungi(100);p__Ascomycota(100);p__Ascomycota_unclassified(100);p__Ascomycota_unclassified(100);p__Ascomycota_unclassified(100);p__Ascomycota_unclassified(100);p__Ascomycota_unclassified(100); | - | - | - | - | - | - | - | Unassigned | - |
| Otu26955 | 1 | k__Fungi(100);k__Fungi_unclassified(100);k__Fungi_unclassified(100);k__Fungi_unclassified(100);k__Fungi_unclassified(100);k__Fungi_unclassified(100);k__Fungi_unclassified(100); | - | - | - | - | - | - | - | Unassigned | - |
| Otu26956 | 1 | k__Fungi(100);p__Ascomycota(100);p__Ascomycota_unclassified(100);p__Ascomycota_unclassified(100);p__Ascomycota_unclassified(100);p__Ascomycota_unclassified(100);p__Ascomycota_unclassified(100); | - | - | - | - | - | - | - | Unassigned | - |
| Otu26957 | 1 | k__Fungi(100);p__Ascomycota(100);p__Ascomycota_unclassified(100);p__Ascomycota_unclassified(100);p__Ascomycota_unclassified(100);p__Ascomycota_unclassified(100);p__Ascomycota_unclassified(100); | - | - | - | - | - | - | - | Unassigned | - |
| Otu26958 | 1 | k__Fungi(100);p__Ascomycota(100);p__Ascomycota_unclassified(100);p__Ascomycota_unclassified(100);p__Ascomycota_unclassified(100);p__Ascomycota_unclassified(100);p__Ascomycota_unclassified(100); | - | - | - | - | - | - | - | Unassigned | - |
| Otu26959 | 1 | k__Fungi(100);k__Fungi_unclassified(100);k__Fungi_unclassified(100);k__Fungi_unclassified(100);k__Fungi_unclassified(100);k__Fungi_unclassified(100);k__Fungi_unclassified(100); | - | - | - | - | - | - | - | Unassigned | - |
| Otu26960 | 1 | k__Fungi(100);p__Ascomycota(100);p__Ascomycota_unclassified(100);p__Ascomycota_unclassified(100);p__Ascomycota_unclassified(100);p__Ascomycota_unclassified(100);p__Ascomycota_unclassified(100); | - | - | - | - | - | - | - | Unassigned | - |
| Otu26961 | 1 | k__Fungi(100);p__Ascomycota(100);p__Ascomycota_unclassified(100);p__Ascomycota_unclassified(100);p__Ascomycota_unclassified(100);p__Ascomycota_unclassified(100);p__Ascomycota_unclassified(100); | - | - | - | - | - | - | - | Unassigned | - |
| Otu26962 | 1 | k__Fungi(100);k__Fungi_unclassified(100);k__Fungi_unclassified(100);k__Fungi_unclassified(100);k__Fungi_unclassified(100);k__Fungi_unclassified(100);k__Fungi_unclassified(100); | - | - | - | - | - | - | - | Unassigned | - |
| Otu26963 | 1 | k__Fungi(100);k__Fungi_unclassified(100);k__Fungi_unclassified(100);k__Fungi_unclassified(100);k__Fungi_unclassified(100);k__Fungi_unclassified(100);k__Fungi_unclassified(100); | - | - | - | - | - | - | - | Unassigned | - |
| Otu26964 | 1 | k__Fungi(100);k__Fungi_unclassified(100);k__Fungi_unclassified(100);k__Fungi_unclassified(100);k__Fungi_unclassified(100);k__Fungi_unclassified(100);k__Fungi_unclassified(100); | - | - | - | - | - | - | - | Unassigned | - |
| Otu26967 | 1 | k__Fungi(100);k__Fungi_unclassified(100);k__Fungi_unclassified(100);k__Fungi_unclassified(100);k__Fungi_unclassified(100);k__Fungi_unclassified(100);k__Fungi_unclassified(100); | - | - | - | - | - | - | - | Unassigned | - |
| Otu26968 | 1 | k__Fungi(100);p__Ascomycota(100);p__Ascomycota_unclassified(100);p__Ascomycota_unclassified(100);p__Ascomycota_unclassified(100);p__Ascomycota_unclassified(100);p__Ascomycota_unclassified(100); | - | - | - | - | - | - | - | Unassigned | - |
| Otu26969 | 1 | k__Fungi(100);k__Fungi_unclassified(100);k__Fungi_unclassified(100);k__Fungi_unclassified(100);k__Fungi_unclassified(100);k__Fungi_unclassified(100);k__Fungi_unclassified(100); | - | - | - | - | - | - | - | Unassigned | - |
| Otu26970 | 1 | k__Fungi(100);k__Fungi_unclassified(100);k__Fungi_unclassified(100);k__Fungi_unclassified(100);k__Fungi_unclassified(100);k__Fungi_unclassified(100);k__Fungi_unclassified(100); | - | - | - | - | - | - | - | Unassigned | - |
| Otu26971 | 1 | k__Fungi(100);p__Ascomycota(100);p__Ascomycota_unclassified(100);p__Ascomycota_unclassified(100);p__Ascomycota_unclassified(100);p__Ascomycota_unclassified(100);p__Ascomycota_unclassified(100); | - | - | - | - | - | - | - | Unassigned | - |
| Otu26972 | 1 | k__Fungi(100);p__Ascomycota(100);p__Ascomycota_unclassified(100);p__Ascomycota_unclassified(100);p__Ascomycota_unclassified(100);p__Ascomycota_unclassified(100);p__Ascomycota_unclassified(100); | - | - | - | - | - | - | - | Unassigned | - |
| Otu26973 | 1 | k__Fungi(100);k__Fungi_unclassified(100);k__Fungi_unclassified(100);k__Fungi_unclassified(100);k__Fungi_unclassified(100);k__Fungi_unclassified(100);k__Fungi_unclassified(100); | - | - | - | - | - | - | - | Unassigned | - |
| Otu26987 | 1 | k__Fungi(100);p__Ascomycota(100);p__Ascomycota_unclassified(100);p__Ascomycota_unclassified(100);p__Ascomycota_unclassified(100);p__Ascomycota_unclassified(100);p__Ascomycota_unclassified(100); | - | - | - | - | - | - | - | Unassigned | - |
| Otu26988 | 1 | k__Fungi(100);p__Ascomycota(100);p__Ascomycota_unclassified(100);p__Ascomycota_unclassified(100);p__Ascomycota_unclassified(100);p__Ascomycota_unclassified(100);p__Ascomycota_unclassified(100); | - | - | - | - | - | - | - | Unassigned | - |
| Otu26989 | 1 | k__Fungi(100);k__Fungi_unclassified(100);k__Fungi_unclassified(100);k__Fungi_unclassified(100);k__Fungi_unclassified(100);k__Fungi_unclassified(100);k__Fungi_unclassified(100); | - | - | - | - | - | - | - | Unassigned | - |
| Otu26990 | 1 | k__Fungi(100);k__Fungi_unclassified(100);k__Fungi_unclassified(100);k__Fungi_unclassified(100);k__Fungi_unclassified(100);k__Fungi_unclassified(100);k__Fungi_unclassified(100); | - | - | - | - | - | - | - | Unassigned | - |
| Otu26991 | 1 | k__Fungi(100);k__Fungi_unclassified(100);k__Fungi_unclassified(100);k__Fungi_unclassified(100);k__Fungi_unclassified(100);k__Fungi_unclassified(100);k__Fungi_unclassified(100); | - | - | - | - | - | - | - | Unassigned | - |
| Otu26992 | 1 | k__Fungi(100);p__Ascomycota(100);p__Ascomycota_unclassified(100);p__Ascomycota_unclassified(100);p__Ascomycota_unclassified(100);p__Ascomycota_unclassified(100);p__Ascomycota_unclassified(100); | - | - | - | - | - | - | - | Unassigned | - |
| Otu26993 | 1 | k__Fungi(100);p__Ascomycota(100);p__Ascomycota_unclassified(100);p__Ascomycota_unclassified(100);p__Ascomycota_unclassified(100);p__Ascomycota_unclassified(100);p__Ascomycota_unclassified(100); | - | - | - | - | - | - | - | Unassigned | - |
| Otu26995 | 1 | k__Fungi(100);k__Fungi_unclassified(100);k__Fungi_unclassified(100);k__Fungi_unclassified(100);k__Fungi_unclassified(100);k__Fungi_unclassified(100);k__Fungi_unclassified(100); | - | - | - | - | - | - | - | Unassigned | - |
| Otu26996 | 1 | k__Fungi(100);p__Ascomycota(100);p__Ascomycota_unclassified(100);p__Ascomycota_unclassified(100);p__Ascomycota_unclassified(100);p__Ascomycota_unclassified(100);p__Ascomycota_unclassified(100); | - | - | - | - | - | - | - | Unassigned | - |
| Otu26997 | 1 | k__Fungi(100);k__Fungi_unclassified(100);k__Fungi_unclassified(100);k__Fungi_unclassified(100);k__Fungi_unclassified(100);k__Fungi_unclassified(100);k__Fungi_unclassified(100); | - | - | - | - | - | - | - | Unassigned | - |
| Otu26998 | 1 | k__Fungi(100);p__Ascomycota(100);p__Ascomycota_unclassified(100);p__Ascomycota_unclassified(100);p__Ascomycota_unclassified(100);p__Ascomycota_unclassified(100);p__Ascomycota_unclassified(100); | - | - | - | - | - | - | - | Unassigned | - |
| Otu26999 | 1 | k__Fungi(100);k__Fungi_unclassified(100);k__Fungi_unclassified(100);k__Fungi_unclassified(100);k__Fungi_unclassified(100);k__Fungi_unclassified(100);k__Fungi_unclassified(100); | - | - | - | - | - | - | - | Unassigned | - |
| Otu27000 | 1 | k__Fungi(100);k__Fungi_unclassified(100);k__Fungi_unclassified(100);k__Fungi_unclassified(100);k__Fungi_unclassified(100);k__Fungi_unclassified(100);k__Fungi_unclassified(100); | - | - | - | - | - | - | - | Unassigned | - |
| Otu27009 | 1 | k__Fungi(100);p__Ascomycota(100);p__Ascomycota_unclassified(100);p__Ascomycota_unclassified(100);p__Ascomycota_unclassified(100);p__Ascomycota_unclassified(100);p__Ascomycota_unclassified(100); | - | - | - | - | - | - | - | Unassigned | - |
| Otu27011 | 1 | k__Fungi(100);p__Ascomycota(100);p__Ascomycota_unclassified(100);p__Ascomycota_unclassified(100);p__Ascomycota_unclassified(100);p__Ascomycota_unclassified(100);p__Ascomycota_unclassified(100); | - | - | - | - | - | - | - | Unassigned | - |
| Otu27012 | 1 | k__Fungi(100);k__Fungi_unclassified(100);k__Fungi_unclassified(100);k__Fungi_unclassified(100);k__Fungi_unclassified(100);k__Fungi_unclassified(100);k__Fungi_unclassified(100); | - | - | - | - | - | - | - | Unassigned | - |
| Otu27013 | 1 | k__Fungi(100);p__Ascomycota(100);p__Ascomycota_unclassified(100);p__Ascomycota_unclassified(100);p__Ascomycota_unclassified(100);p__Ascomycota_unclassified(100);p__Ascomycota_unclassified(100); | - | - | - | - | - | - | - | Unassigned | - |
| Otu27014 | 1 | k__Fungi(100);p__Ascomycota(100);p__Ascomycota_unclassified(100);p__Ascomycota_unclassified(100);p__Ascomycota_unclassified(100);p__Ascomycota_unclassified(100);p__Ascomycota_unclassified(100); | - | - | - | - | - | - | - | Unassigned | - |
| Otu27019 | 1 | k__Fungi(100);k__Fungi_unclassified(100);k__Fungi_unclassified(100);k__Fungi_unclassified(100);k__Fungi_unclassified(100);k__Fungi_unclassified(100);k__Fungi_unclassified(100); | - | - | - | - | - | - | - | Unassigned | - |
| Otu27024 | 1 | k__Fungi(100);p__Ascomycota(100);p__Ascomycota_unclassified(100);p__Ascomycota_unclassified(100);p__Ascomycota_unclassified(100);p__Ascomycota_unclassified(100);p__Ascomycota_unclassified(100); | - | - | - | - | - | - | - | Unassigned | - |
| Otu27028 | 1 | k__Fungi(100);p__Ascomycota(100);p__Ascomycota_unclassified(100);p__Ascomycota_unclassified(100);p__Ascomycota_unclassified(100);p__Ascomycota_unclassified(100);p__Ascomycota_unclassified(100); | - | - | - | - | - | - | - | Unassigned | - |
| Otu27029 | 1 | k__Fungi(100);p__Ascomycota(100);p__Ascomycota_unclassified(100);p__Ascomycota_unclassified(100);p__Ascomycota_unclassified(100);p__Ascomycota_unclassified(100);p__Ascomycota_unclassified(100); | - | - | - | - | - | - | - | Unassigned | - |
| Otu27030 | 1 | k__Fungi(100);p__Ascomycota(100);p__Ascomycota_unclassified(100);p__Ascomycota_unclassified(100);p__Ascomycota_unclassified(100);p__Ascomycota_unclassified(100);p__Ascomycota_unclassified(100); | - | - | - | - | - | - | - | Unassigned | - |
| Otu27031 | 1 | k__Fungi(100);p__Ascomycota(100);p__Ascomycota_unclassified(100);p__Ascomycota_unclassified(100);p__Ascomycota_unclassified(100);p__Ascomycota_unclassified(100);p__Ascomycota_unclassified(100); | - | - | - | - | - | - | - | Unassigned | - |
| Otu27033 | 1 | k__Fungi(100);p__Ascomycota(100);p__Ascomycota_unclassified(100);p__Ascomycota_unclassified(100);p__Ascomycota_unclassified(100);p__Ascomycota_unclassified(100);p__Ascomycota_unclassified(100); | - | - | - | - | - | - | - | Unassigned | - |
| Otu27034 | 1 | k__Fungi(100);p__Ascomycota(100);p__Ascomycota_unclassified(100);p__Ascomycota_unclassified(100);p__Ascomycota_unclassified(100);p__Ascomycota_unclassified(100);p__Ascomycota_unclassified(100); | - | - | - | - | - | - | - | Unassigned | - |
| Otu27036 | 1 | k__Fungi(100);k__Fungi_unclassified(100);k__Fungi_unclassified(100);k__Fungi_unclassified(100);k__Fungi_unclassified(100);k__Fungi_unclassified(100);k__Fungi_unclassified(100); | - | - | - | - | - | - | - | Unassigned | - |
| Otu27037 | 1 | k__Fungi(100);p__Ascomycota(100);p__Ascomycota_unclassified(100);p__Ascomycota_unclassified(100);p__Ascomycota_unclassified(100);p__Ascomycota_unclassified(100);p__Ascomycota_unclassified(100); | - | - | - | - | - | - | - | Unassigned | - |
| Otu27039 | 1 | k__Fungi(100);k__Fungi_unclassified(100);k__Fungi_unclassified(100);k__Fungi_unclassified(100);k__Fungi_unclassified(100);k__Fungi_unclassified(100);k__Fungi_unclassified(100); | - | - | - | - | - | - | - | Unassigned | - |
| Otu27040 | 1 | k__Fungi(100);k__Fungi_unclassified(100);k__Fungi_unclassified(100);k__Fungi_unclassified(100);k__Fungi_unclassified(100);k__Fungi_unclassified(100);k__Fungi_unclassified(100); | - | - | - | - | - | - | - | Unassigned | - |
| Otu27041 | 1 | k__Fungi(100);k__Fungi_unclassified(100);k__Fungi_unclassified(100);k__Fungi_unclassified(100);k__Fungi_unclassified(100);k__Fungi_unclassified(100);k__Fungi_unclassified(100); | - | - | - | - | - | - | - | Unassigned | - |
| Otu27046 | 1 | k__Fungi(100);k__Fungi_unclassified(100);k__Fungi_unclassified(100);k__Fungi_unclassified(100);k__Fungi_unclassified(100);k__Fungi_unclassified(100);k__Fungi_unclassified(100); | - | - | - | - | - | - | - | Unassigned | - |
| Otu27049 | 1 | k__Fungi(100);k__Fungi_unclassified(100);k__Fungi_unclassified(100);k__Fungi_unclassified(100);k__Fungi_unclassified(100);k__Fungi_unclassified(100);k__Fungi_unclassified(100); | - | - | - | - | - | - | - | Unassigned | - |
| Otu27050 | 1 | k__Fungi(100);k__Fungi_unclassified(100);k__Fungi_unclassified(100);k__Fungi_unclassified(100);k__Fungi_unclassified(100);k__Fungi_unclassified(100);k__Fungi_unclassified(100); | - | - | - | - | - | - | - | Unassigned | - |
| Otu27053 | 1 | k__Fungi(100);k__Fungi_unclassified(100);k__Fungi_unclassified(100);k__Fungi_unclassified(100);k__Fungi_unclassified(100);k__Fungi_unclassified(100);k__Fungi_unclassified(100); | - | - | - | - | - | - | - | Unassigned | - |
| Otu27073 | 1 | k__Fungi(100);p__Basidiomycota(100);c__Agaricomycetes(100);c__Agaricomycetes_unclassified(100);c__Agaricomycetes_unclassified(100);c__Agaricomycetes_unclassified(100);c__Agaricomycetes_unclassified(100); | - | - | - | - | - | - | - | Unassigned | - |
| Otu27074 | 1 | k__Fungi(100);p__Basidiomycota(100);c__Agaricomycetes(100);c__Agaricomycetes_unclassified(100);c__Agaricomycetes_unclassified(100);c__Agaricomycetes_unclassified(100);c__Agaricomycetes_unclassified(100); | - | - | - | - | - | - | - | Unassigned | - |
| Otu27075 | 1 | k__Fungi(100);p__Basidiomycota(100);c__Agaricomycetes(100);o__Russulales(100);o__Russulales_unclassified(100);o__Russulales_unclassified(100);o__Russulales_unclassified(100); | - | - | - | - | - | - | - | Unassigned | - |
| Otu27080 | 1 | k__Fungi(100);p__Basidiomycota(100);p__Basidiomycota_unclassified(100);p__Basidiomycota_unclassified(100);p__Basidiomycota_unclassified(100);p__Basidiomycota_unclassified(100);p__Basidiomycota_unclassified(100); | - | - | - | - | - | - | - | Unassigned | - |
| Otu27082 | 1 | k__Plantae(100);p__unclassified_Plantae(100);c__unclassified_Plantae(100);o__unclassified_Plantae(100);f__unclassified_Plantae(100);g__unclassified_Plantae(100);s__Plantae_sp(100); | - | - | - | - | - | - | - | Unassigned | - |
| Otu27083 | 1 | k__Plantae(100);p__unclassified_Plantae(100);c__unclassified_Plantae(100);o__unclassified_Plantae(100);f__unclassified_Plantae(100);g__unclassified_Plantae(100);s__Plantae_sp(100); | - | - | - | - | - | - | - | Unassigned | - |
| Otu27084 | 1 | k__Fungi(100);k__Fungi_unclassified(100);k__Fungi_unclassified(100);k__Fungi_unclassified(100);k__Fungi_unclassified(100);k__Fungi_unclassified(100);k__Fungi_unclassified(100); | - | - | - | - | - | - | - | Unassigned | - |
| Otu27085 | 1 | k__Plantae(100);p__unclassified_Plantae(100);c__unclassified_Plantae(100);o__unclassified_Plantae(100);f__unclassified_Plantae(100);g__unclassified_Plantae(100);s__Plantae_sp(100); | - | - | - | - | - | - | - | Unassigned | - |
| Otu27086 | 1 | k__Fungi(100);k__Fungi_unclassified(100);k__Fungi_unclassified(100);k__Fungi_unclassified(100);k__Fungi_unclassified(100);k__Fungi_unclassified(100);k__Fungi_unclassified(100); | - | - | - | - | - | - | - | Unassigned | - |
| Otu27089 | 1 | k__Plantae(100);p__unclassified_Plantae(100);c__unclassified_Plantae(100);o__unclassified_Plantae(100);f__unclassified_Plantae(100);g__unclassified_Plantae(100);s__Plantae_sp(100); | - | - | - | - | - | - | - | Unassigned | - |
| Otu27090 | 1 | k__Fungi(100);k__Fungi_unclassified(100);k__Fungi_unclassified(100);k__Fungi_unclassified(100);k__Fungi_unclassified(100);k__Fungi_unclassified(100);k__Fungi_unclassified(100); | - | - | - | - | - | - | - | Unassigned | - |
| Otu27094 | 1 | k__Fungi(100);k__Fungi_unclassified(100);k__Fungi_unclassified(100);k__Fungi_unclassified(100);k__Fungi_unclassified(100);k__Fungi_unclassified(100);k__Fungi_unclassified(100); | - | - | - | - | - | - | - | Unassigned | - |
| Otu27095 | 1 | k__Fungi(100);p__Basidiomycota(100);c__Agaricomycetes(100);o__Auriculariales(100);f__unclassified_Auriculariales(100);g__unclassified_Auriculariales(100);s__Auriculariales_sp(100); | - | - | - | - | - | - | - | Unassigned | - |
| Otu27108 | 1 | k__Fungi(100);p__Ascomycota(100);p__Ascomycota_unclassified(100);p__Ascomycota_unclassified(100);p__Ascomycota_unclassified(100);p__Ascomycota_unclassified(100);p__Ascomycota_unclassified(100); | - | - | - | - | - | - | - | Unassigned | - |
| Otu27109 | 1 | k__Fungi(100);k__Fungi_unclassified(100);k__Fungi_unclassified(100);k__Fungi_unclassified(100);k__Fungi_unclassified(100);k__Fungi_unclassified(100);k__Fungi_unclassified(100); | - | - | - | - | - | - | - | Unassigned | - |
| Otu27110 | 1 | k__Fungi(100);k__Fungi_unclassified(100);k__Fungi_unclassified(100);k__Fungi_unclassified(100);k__Fungi_unclassified(100);k__Fungi_unclassified(100);k__Fungi_unclassified(100); | - | - | - | - | - | - | - | Unassigned | - |
| Otu27111 | 1 | k__Fungi(100);k__Fungi_unclassified(100);k__Fungi_unclassified(100);k__Fungi_unclassified(100);k__Fungi_unclassified(100);k__Fungi_unclassified(100);k__Fungi_unclassified(100); | - | - | - | - | - | - | - | Unassigned | - |
| Otu27112 | 1 | k__Fungi(100);p__Ascomycota(100);p__Ascomycota_unclassified(100);p__Ascomycota_unclassified(100);p__Ascomycota_unclassified(100);p__Ascomycota_unclassified(100);p__Ascomycota_unclassified(100); | - | - | - | - | - | - | - | Unassigned | - |
| Otu27113 | 1 | k__Fungi(100);k__Fungi_unclassified(100);k__Fungi_unclassified(100);k__Fungi_unclassified(100);k__Fungi_unclassified(100);k__Fungi_unclassified(100);k__Fungi_unclassified(100); | - | - | - | - | - | - | - | Unassigned | - |
| Otu27117 | 1 | k__Fungi(100);k__Fungi_unclassified(100);k__Fungi_unclassified(100);k__Fungi_unclassified(100);k__Fungi_unclassified(100);k__Fungi_unclassified(100);k__Fungi_unclassified(100); | - | - | - | - | - | - | - | Unassigned | - |
| Otu27118 | 1 | k__Fungi(100);p__Ascomycota(100);p__Ascomycota_unclassified(100);p__Ascomycota_unclassified(100);p__Ascomycota_unclassified(100);p__Ascomycota_unclassified(100);p__Ascomycota_unclassified(100); | - | - | - | - | - | - | - | Unassigned | - |
| Otu27121 | 1 | k__Fungi(100);p__Basidiomycota(100);c__Agaricomycetes(100);c__Agaricomycetes_unclassified(100);c__Agaricomycetes_unclassified(100);c__Agaricomycetes_unclassified(100);c__Agaricomycetes_unclassified(100); | - | - | - | - | - | - | - | Unassigned | - |
| Otu27134 | 1 | k__Fungi(100);k__Fungi_unclassified(100);k__Fungi_unclassified(100);k__Fungi_unclassified(100);k__Fungi_unclassified(100);k__Fungi_unclassified(100);k__Fungi_unclassified(100); | - | - | - | - | - | - | - | Unassigned | - |
| Otu27135 | 1 | k__Fungi(100);p__Basidiomycota(100);p__Basidiomycota_unclassified(100);p__Basidiomycota_unclassified(100);p__Basidiomycota_unclassified(100);p__Basidiomycota_unclassified(100);p__Basidiomycota_unclassified(100); | - | - | - | - | - | - | - | Unassigned | - |
| Otu27136 | 1 | k__Fungi(100);k__Fungi_unclassified(100);k__Fungi_unclassified(100);k__Fungi_unclassified(100);k__Fungi_unclassified(100);k__Fungi_unclassified(100);k__Fungi_unclassified(100); | - | - | - | - | - | - | - | Unassigned | - |
| Otu27137 | 1 | k__Fungi(100);p__Basidiomycota(100);p__Basidiomycota_unclassified(100);p__Basidiomycota_unclassified(100);p__Basidiomycota_unclassified(100);p__Basidiomycota_unclassified(100);p__Basidiomycota_unclassified(100); | - | - | - | - | - | - | - | Unassigned | - |
| Otu27138 | 1 | k__Fungi(100);k__Fungi_unclassified(100);k__Fungi_unclassified(100);k__Fungi_unclassified(100);k__Fungi_unclassified(100);k__Fungi_unclassified(100);k__Fungi_unclassified(100); | - | - | - | - | - | - | - | Unassigned | - |
| Otu27139 | 1 | k__Fungi(100);k__Fungi_unclassified(100);k__Fungi_unclassified(100);k__Fungi_unclassified(100);k__Fungi_unclassified(100);k__Fungi_unclassified(100);k__Fungi_unclassified(100); | - | - | - | - | - | - | - | Unassigned | - |
| Otu27140 | 1 | k__Fungi(100);p__Basidiomycota(100);p__Basidiomycota_unclassified(100);p__Basidiomycota_unclassified(100);p__Basidiomycota_unclassified(100);p__Basidiomycota_unclassified(100);p__Basidiomycota_unclassified(100); | - | - | - | - | - | - | - | Unassigned | - |
| Otu27141 | 1 | k__Fungi(100);p__Basidiomycota(100);p__Basidiomycota_unclassified(100);p__Basidiomycota_unclassified(100);p__Basidiomycota_unclassified(100);p__Basidiomycota_unclassified(100);p__Basidiomycota_unclassified(100); | - | - | - | - | - | - | - | Unassigned | - |
| Otu27142 | 1 | k__Fungi(100);k__Fungi_unclassified(100);k__Fungi_unclassified(100);k__Fungi_unclassified(100);k__Fungi_unclassified(100);k__Fungi_unclassified(100);k__Fungi_unclassified(100); | - | - | - | - | - | - | - | Unassigned | - |
| Otu27143 | 1 | k__Fungi(100);k__Fungi_unclassified(100);k__Fungi_unclassified(100);k__Fungi_unclassified(100);k__Fungi_unclassified(100);k__Fungi_unclassified(100);k__Fungi_unclassified(100); | - | - | - | - | - | - | - | Unassigned | - |
| Otu27144 | 1 | k__Fungi(100);k__Fungi_unclassified(100);k__Fungi_unclassified(100);k__Fungi_unclassified(100);k__Fungi_unclassified(100);k__Fungi_unclassified(100);k__Fungi_unclassified(100); | - | - | - | - | - | - | - | Unassigned | - |
| Otu27145 | 1 | k__Fungi(100);p__Basidiomycota(100);p__Basidiomycota_unclassified(100);p__Basidiomycota_unclassified(100);p__Basidiomycota_unclassified(100);p__Basidiomycota_unclassified(100);p__Basidiomycota_unclassified(100); | - | - | - | - | - | - | - | Unassigned | - |
| Otu27146 | 1 | k__Fungi(100);k__Fungi_unclassified(100);k__Fungi_unclassified(100);k__Fungi_unclassified(100);k__Fungi_unclassified(100);k__Fungi_unclassified(100);k__Fungi_unclassified(100); | - | - | - | - | - | - | - | Unassigned | - |
| Otu27147 | 1 | k__Fungi(100);p__Ascomycota(100);p__Ascomycota_unclassified(100);p__Ascomycota_unclassified(100);p__Ascomycota_unclassified(100);p__Ascomycota_unclassified(100);p__Ascomycota_unclassified(100); | - | - | - | - | - | - | - | Unassigned | - |
| Otu27148 | 1 | k__Fungi(100);p__Basidiomycota(100);p__Basidiomycota_unclassified(100);p__Basidiomycota_unclassified(100);p__Basidiomycota_unclassified(100);p__Basidiomycota_unclassified(100);p__Basidiomycota_unclassified(100); | - | - | - | - | - | - | - | Unassigned | - |
| Otu27149 | 1 | k__Fungi(100);k__Fungi_unclassified(100);k__Fungi_unclassified(100);k__Fungi_unclassified(100);k__Fungi_unclassified(100);k__Fungi_unclassified(100);k__Fungi_unclassified(100); | - | - | - | - | - | - | - | Unassigned | - |
| Otu27150 | 1 | k__Fungi(100);k__Fungi_unclassified(100);k__Fungi_unclassified(100);k__Fungi_unclassified(100);k__Fungi_unclassified(100);k__Fungi_unclassified(100);k__Fungi_unclassified(100); | - | - | - | - | - | - | - | Unassigned | - |
| Otu27151 | 1 | k__Fungi(100);k__Fungi_unclassified(100);k__Fungi_unclassified(100);k__Fungi_unclassified(100);k__Fungi_unclassified(100);k__Fungi_unclassified(100);k__Fungi_unclassified(100); | - | - | - | - | - | - | - | Unassigned | - |
| Otu27152 | 1 | k__Fungi(100);k__Fungi_unclassified(100);k__Fungi_unclassified(100);k__Fungi_unclassified(100);k__Fungi_unclassified(100);k__Fungi_unclassified(100);k__Fungi_unclassified(100); | - | - | - | - | - | - | - | Unassigned | - |
| Otu27153 | 1 | k__Fungi(100);p__Basidiomycota(100);p__Basidiomycota_unclassified(100);p__Basidiomycota_unclassified(100);p__Basidiomycota_unclassified(100);p__Basidiomycota_unclassified(100);p__Basidiomycota_unclassified(100); | - | - | - | - | - | - | - | Unassigned | - |
| Otu27154 | 1 | k__Fungi(100);p__Basidiomycota(100);p__Basidiomycota_unclassified(100);p__Basidiomycota_unclassified(100);p__Basidiomycota_unclassified(100);p__Basidiomycota_unclassified(100);p__Basidiomycota_unclassified(100); | - | - | - | - | - | - | - | Unassigned | - |
| Otu27155 | 1 | k__Fungi(100);k__Fungi_unclassified(100);k__Fungi_unclassified(100);k__Fungi_unclassified(100);k__Fungi_unclassified(100);k__Fungi_unclassified(100);k__Fungi_unclassified(100); | - | - | - | - | - | - | - | Unassigned | - |
| Otu27156 | 1 | k__Fungi(100);k__Fungi_unclassified(100);k__Fungi_unclassified(100);k__Fungi_unclassified(100);k__Fungi_unclassified(100);k__Fungi_unclassified(100);k__Fungi_unclassified(100); | - | - | - | - | - | - | - | Unassigned | - |
| Otu27157 | 1 | k__Fungi(100);k__Fungi_unclassified(100);k__Fungi_unclassified(100);k__Fungi_unclassified(100);k__Fungi_unclassified(100);k__Fungi_unclassified(100);k__Fungi_unclassified(100); | - | - | - | - | - | - | - | Unassigned | - |
| Otu27158 | 1 | k__Fungi(100);k__Fungi_unclassified(100);k__Fungi_unclassified(100);k__Fungi_unclassified(100);k__Fungi_unclassified(100);k__Fungi_unclassified(100);k__Fungi_unclassified(100); | - | - | - | - | - | - | - | Unassigned | - |
| Otu27159 | 1 | k__Fungi(100);k__Fungi_unclassified(100);k__Fungi_unclassified(100);k__Fungi_unclassified(100);k__Fungi_unclassified(100);k__Fungi_unclassified(100);k__Fungi_unclassified(100); | - | - | - | - | - | - | - | Unassigned | - |
| Otu27162 | 1 | k__Fungi(100);k__Fungi_unclassified(100);k__Fungi_unclassified(100);k__Fungi_unclassified(100);k__Fungi_unclassified(100);k__Fungi_unclassified(100);k__Fungi_unclassified(100); | - | - | - | - | - | - | - | Unassigned | - |
| Otu27166 | 1 | k__Fungi(100);p__Ascomycota(100);p__Ascomycota_unclassified(100);p__Ascomycota_unclassified(100);p__Ascomycota_unclassified(100);p__Ascomycota_unclassified(100);p__Ascomycota_unclassified(100); | - | - | - | - | - | - | - | Unassigned | - |
| Otu27168 | 1 | k__Fungi(100);p__Ascomycota(100);p__Ascomycota_unclassified(100);p__Ascomycota_unclassified(100);p__Ascomycota_unclassified(100);p__Ascomycota_unclassified(100);p__Ascomycota_unclassified(100); | - | - | - | - | - | - | - | Unassigned | - |
| Otu27170 | 1 | k__Fungi(100);p__Ascomycota(100);p__Ascomycota_unclassified(100);p__Ascomycota_unclassified(100);p__Ascomycota_unclassified(100);p__Ascomycota_unclassified(100);p__Ascomycota_unclassified(100); | - | - | - | - | - | - | - | Unassigned | - |
| Otu27187 | 1 | k__Fungi(100);k__Fungi_unclassified(100);k__Fungi_unclassified(100);k__Fungi_unclassified(100);k__Fungi_unclassified(100);k__Fungi_unclassified(100);k__Fungi_unclassified(100); | - | - | - | - | - | - | - | Unassigned | - |
| Otu27193 | 1 | k__Fungi(100);p__Ascomycota(100);p__Ascomycota_unclassified(100);p__Ascomycota_unclassified(100);p__Ascomycota_unclassified(100);p__Ascomycota_unclassified(100);p__Ascomycota_unclassified(100); | - | - | - | - | - | - | - | Unassigned | - |
| Otu27201 | 1 | k__Fungi(100);k__Fungi_unclassified(100);k__Fungi_unclassified(100);k__Fungi_unclassified(100);k__Fungi_unclassified(100);k__Fungi_unclassified(100);k__Fungi_unclassified(100); | - | - | - | - | - | - | - | Unassigned | - |
| Otu27202 | 1 | k__Fungi(100);p__Ascomycota(100);p__Ascomycota_unclassified(100);p__Ascomycota_unclassified(100);p__Ascomycota_unclassified(100);p__Ascomycota_unclassified(100);p__Ascomycota_unclassified(100); | - | - | - | - | - | - | - | Unassigned | - |
| Otu27203 | 1 | k__Fungi(100);p__Ascomycota(100);p__Ascomycota_unclassified(100);p__Ascomycota_unclassified(100);p__Ascomycota_unclassified(100);p__Ascomycota_unclassified(100);p__Ascomycota_unclassified(100); | - | - | - | - | - | - | - | Unassigned | - |
| Otu27208 | 1 | k__Fungi(100);p__Ascomycota(100);p__Ascomycota_unclassified(100);p__Ascomycota_unclassified(100);p__Ascomycota_unclassified(100);p__Ascomycota_unclassified(100);p__Ascomycota_unclassified(100); | - | - | - | - | - | - | - | Unassigned | - |
| Otu27209 | 1 | k__Fungi(100);k__Fungi_unclassified(100);k__Fungi_unclassified(100);k__Fungi_unclassified(100);k__Fungi_unclassified(100);k__Fungi_unclassified(100);k__Fungi_unclassified(100); | - | - | - | - | - | - | - | Unassigned | - |
| Otu27210 | 1 | k__Fungi(100);k__Fungi_unclassified(100);k__Fungi_unclassified(100);k__Fungi_unclassified(100);k__Fungi_unclassified(100);k__Fungi_unclassified(100);k__Fungi_unclassified(100); | - | - | - | - | - | - | - | Unassigned | - |
| Otu27212 | 1 | k__Fungi(100);k__Fungi_unclassified(100);k__Fungi_unclassified(100);k__Fungi_unclassified(100);k__Fungi_unclassified(100);k__Fungi_unclassified(100);k__Fungi_unclassified(100); | - | - | - | - | - | - | - | Unassigned | - |
| Otu27219 | 1 | k__Fungi(100);k__Fungi_unclassified(100);k__Fungi_unclassified(100);k__Fungi_unclassified(100);k__Fungi_unclassified(100);k__Fungi_unclassified(100);k__Fungi_unclassified(100); | - | - | - | - | - | - | - | Unassigned | - |
| Otu27231 | 1 | k__Fungi(100);k__Fungi_unclassified(100);k__Fungi_unclassified(100);k__Fungi_unclassified(100);k__Fungi_unclassified(100);k__Fungi_unclassified(100);k__Fungi_unclassified(100); | - | - | - | - | - | - | - | Unassigned | - |
| Otu27236 | 1 | k__Fungi(100);k__Fungi_unclassified(100);k__Fungi_unclassified(100);k__Fungi_unclassified(100);k__Fungi_unclassified(100);k__Fungi_unclassified(100);k__Fungi_unclassified(100); | - | - | - | - | - | - | - | Unassigned | - |
| Otu27246 | 1 | k__Fungi(100);p__Ascomycota(100);p__Ascomycota_unclassified(100);p__Ascomycota_unclassified(100);p__Ascomycota_unclassified(100);p__Ascomycota_unclassified(100);p__Ascomycota_unclassified(100); | - | - | - | - | - | - | - | Unassigned | - |
| Otu27247 | 1 | k__Fungi(100);k__Fungi_unclassified(100);k__Fungi_unclassified(100);k__Fungi_unclassified(100);k__Fungi_unclassified(100);k__Fungi_unclassified(100);k__Fungi_unclassified(100); | - | - | - | - | - | - | - | Unassigned | - |
| Otu27249 | 1 | k__Fungi(100);k__Fungi_unclassified(100);k__Fungi_unclassified(100);k__Fungi_unclassified(100);k__Fungi_unclassified(100);k__Fungi_unclassified(100);k__Fungi_unclassified(100); | - | - | - | - | - | - | - | Unassigned | - |
| Otu27251 | 1 | k__Fungi(100);k__Fungi_unclassified(100);k__Fungi_unclassified(100);k__Fungi_unclassified(100);k__Fungi_unclassified(100);k__Fungi_unclassified(100);k__Fungi_unclassified(100); | - | - | - | - | - | - | - | Unassigned | - |
| Otu27254 | 1 | k__Fungi(100);k__Fungi_unclassified(100);k__Fungi_unclassified(100);k__Fungi_unclassified(100);k__Fungi_unclassified(100);k__Fungi_unclassified(100);k__Fungi_unclassified(100); | - | - | - | - | - | - | - | Unassigned | - |
| Otu27256 | 1 | k__Fungi(100);p__Ascomycota(100);p__Ascomycota_unclassified(100);p__Ascomycota_unclassified(100);p__Ascomycota_unclassified(100);p__Ascomycota_unclassified(100);p__Ascomycota_unclassified(100); | - | - | - | - | - | - | - | Unassigned | - |
| Otu27264 | 1 | k__Fungi(100);k__Fungi_unclassified(100);k__Fungi_unclassified(100);k__Fungi_unclassified(100);k__Fungi_unclassified(100);k__Fungi_unclassified(100);k__Fungi_unclassified(100); | - | - | - | - | - | - | - | Unassigned | - |
| Otu27266 | 1 | k__Fungi(100);p__Ascomycota(100);p__Ascomycota_unclassified(100);p__Ascomycota_unclassified(100);p__Ascomycota_unclassified(100);p__Ascomycota_unclassified(100);p__Ascomycota_unclassified(100); | - | - | - | - | - | - | - | Unassigned | - |
| Otu27267 | 1 | k__Fungi(100);k__Fungi_unclassified(100);k__Fungi_unclassified(100);k__Fungi_unclassified(100);k__Fungi_unclassified(100);k__Fungi_unclassified(100);k__Fungi_unclassified(100); | - | - | - | - | - | - | - | Unassigned | - |
| Otu27268 | 1 | k__Fungi(100);k__Fungi_unclassified(100);k__Fungi_unclassified(100);k__Fungi_unclassified(100);k__Fungi_unclassified(100);k__Fungi_unclassified(100);k__Fungi_unclassified(100); | - | - | - | - | - | - | - | Unassigned | - |
| Otu27269 | 1 | k__Fungi(100);p__Ascomycota(100);p__Ascomycota_unclassified(100);p__Ascomycota_unclassified(100);p__Ascomycota_unclassified(100);p__Ascomycota_unclassified(100);p__Ascomycota_unclassified(100); | - | - | - | - | - | - | - | Unassigned | - |
| Otu27270 | 1 | k__Fungi(100);p__Ascomycota(100);p__Ascomycota_unclassified(100);p__Ascomycota_unclassified(100);p__Ascomycota_unclassified(100);p__Ascomycota_unclassified(100);p__Ascomycota_unclassified(100); | - | - | - | - | - | - | - | Unassigned | - |
| Otu27272 | 1 | k__Fungi(100);p__Ascomycota(100);p__Ascomycota_unclassified(100);p__Ascomycota_unclassified(100);p__Ascomycota_unclassified(100);p__Ascomycota_unclassified(100);p__Ascomycota_unclassified(100); | - | - | - | - | - | - | - | Unassigned | - |
| Otu27273 | 1 | k__Fungi(100);p__Ascomycota(100);p__Ascomycota_unclassified(100);p__Ascomycota_unclassified(100);p__Ascomycota_unclassified(100);p__Ascomycota_unclassified(100);p__Ascomycota_unclassified(100); | - | - | - | - | - | - | - | Unassigned | - |
| Otu27274 | 1 | k__Fungi(100);p__Ascomycota(100);p__Ascomycota_unclassified(100);p__Ascomycota_unclassified(100);p__Ascomycota_unclassified(100);p__Ascomycota_unclassified(100);p__Ascomycota_unclassified(100); | - | - | - | - | - | - | - | Unassigned | - |
| Otu27275 | 1 | k__Fungi(100);p__Ascomycota(100);p__Ascomycota_unclassified(100);p__Ascomycota_unclassified(100);p__Ascomycota_unclassified(100);p__Ascomycota_unclassified(100);p__Ascomycota_unclassified(100); | - | - | - | - | - | - | - | Unassigned | - |
| Otu27276 | 1 | k__Fungi(100);p__Ascomycota(100);p__Ascomycota_unclassified(100);p__Ascomycota_unclassified(100);p__Ascomycota_unclassified(100);p__Ascomycota_unclassified(100);p__Ascomycota_unclassified(100); | - | - | - | - | - | - | - | Unassigned | - |
| Otu27279 | 1 | k__Fungi(100);p__Ascomycota(100);p__Ascomycota_unclassified(100);p__Ascomycota_unclassified(100);p__Ascomycota_unclassified(100);p__Ascomycota_unclassified(100);p__Ascomycota_unclassified(100); | - | - | - | - | - | - | - | Unassigned | - |
| Otu27280 | 1 | k__Fungi(100);p__Ascomycota(100);p__Ascomycota_unclassified(100);p__Ascomycota_unclassified(100);p__Ascomycota_unclassified(100);p__Ascomycota_unclassified(100);p__Ascomycota_unclassified(100); | - | - | - | - | - | - | - | Unassigned | - |
| Otu27281 | 1 | k__Fungi(100);p__Ascomycota(100);p__Ascomycota_unclassified(100);p__Ascomycota_unclassified(100);p__Ascomycota_unclassified(100);p__Ascomycota_unclassified(100);p__Ascomycota_unclassified(100); | - | - | - | - | - | - | - | Unassigned | - |
| Otu27282 | 1 | k__Fungi(100);p__Ascomycota(100);p__Ascomycota_unclassified(100);p__Ascomycota_unclassified(100);p__Ascomycota_unclassified(100);p__Ascomycota_unclassified(100);p__Ascomycota_unclassified(100); | - | - | - | - | - | - | - | Unassigned | - |
| Otu27283 | 1 | k__Fungi(100);k__Fungi_unclassified(100);k__Fungi_unclassified(100);k__Fungi_unclassified(100);k__Fungi_unclassified(100);k__Fungi_unclassified(100);k__Fungi_unclassified(100); | - | - | - | - | - | - | - | Unassigned | - |
| Otu27284 | 1 | k__Fungi(100);p__Ascomycota(100);p__Ascomycota_unclassified(100);p__Ascomycota_unclassified(100);p__Ascomycota_unclassified(100);p__Ascomycota_unclassified(100);p__Ascomycota_unclassified(100); | - | - | - | - | - | - | - | Unassigned | - |
| Otu27285 | 1 | k__Fungi(100);k__Fungi_unclassified(100);k__Fungi_unclassified(100);k__Fungi_unclassified(100);k__Fungi_unclassified(100);k__Fungi_unclassified(100);k__Fungi_unclassified(100); | - | - | - | - | - | - | - | Unassigned | - |
| Otu27286 | 1 | k__Fungi(100);p__Ascomycota(100);p__Ascomycota_unclassified(100);p__Ascomycota_unclassified(100);p__Ascomycota_unclassified(100);p__Ascomycota_unclassified(100);p__Ascomycota_unclassified(100); | - | - | - | - | - | - | - | Unassigned | - |
| Otu27287 | 1 | k__Fungi(100);p__Ascomycota(100);p__Ascomycota_unclassified(100);p__Ascomycota_unclassified(100);p__Ascomycota_unclassified(100);p__Ascomycota_unclassified(100);p__Ascomycota_unclassified(100); | - | - | - | - | - | - | - | Unassigned | - |
| Otu27288 | 1 | k__Fungi(100);p__Ascomycota(100);p__Ascomycota_unclassified(100);p__Ascomycota_unclassified(100);p__Ascomycota_unclassified(100);p__Ascomycota_unclassified(100);p__Ascomycota_unclassified(100); | - | - | - | - | - | - | - | Unassigned | - |
| Otu27289 | 1 | k__Fungi(100);p__Ascomycota(100);p__Ascomycota_unclassified(100);p__Ascomycota_unclassified(100);p__Ascomycota_unclassified(100);p__Ascomycota_unclassified(100);p__Ascomycota_unclassified(100); | - | - | - | - | - | - | - | Unassigned | - |
| Otu27290 | 1 | k__Fungi(100);p__Ascomycota(100);p__Ascomycota_unclassified(100);p__Ascomycota_unclassified(100);p__Ascomycota_unclassified(100);p__Ascomycota_unclassified(100);p__Ascomycota_unclassified(100); | - | - | - | - | - | - | - | Unassigned | - |
| Otu27291 | 1 | k__Fungi(100);k__Fungi_unclassified(100);k__Fungi_unclassified(100);k__Fungi_unclassified(100);k__Fungi_unclassified(100);k__Fungi_unclassified(100);k__Fungi_unclassified(100); | - | - | - | - | - | - | - | Unassigned | - |
| Otu27293 | 1 | k__Fungi(100);k__Fungi_unclassified(100);k__Fungi_unclassified(100);k__Fungi_unclassified(100);k__Fungi_unclassified(100);k__Fungi_unclassified(100);k__Fungi_unclassified(100); | - | - | - | - | - | - | - | Unassigned | - |
| Otu27294 | 1 | k__Fungi(100);k__Fungi_unclassified(100);k__Fungi_unclassified(100);k__Fungi_unclassified(100);k__Fungi_unclassified(100);k__Fungi_unclassified(100);k__Fungi_unclassified(100); | - | - | - | - | - | - | - | Unassigned | - |
| Otu27296 | 1 | k__Fungi(100);p__Ascomycota(100);p__Ascomycota_unclassified(100);p__Ascomycota_unclassified(100);p__Ascomycota_unclassified(100);p__Ascomycota_unclassified(100);p__Ascomycota_unclassified(100); | - | - | - | - | - | - | - | Unassigned | - |
| Otu27297 | 1 | k__Fungi(100);p__Ascomycota(100);p__Ascomycota_unclassified(100);p__Ascomycota_unclassified(100);p__Ascomycota_unclassified(100);p__Ascomycota_unclassified(100);p__Ascomycota_unclassified(100); | - | - | - | - | - | - | - | Unassigned | - |
| Otu27298 | 1 | k__Fungi(100);k__Fungi_unclassified(100);k__Fungi_unclassified(100);k__Fungi_unclassified(100);k__Fungi_unclassified(100);k__Fungi_unclassified(100);k__Fungi_unclassified(100); | - | - | - | - | - | - | - | Unassigned | - |
| Otu27299 | 1 | k__Fungi(100);k__Fungi_unclassified(100);k__Fungi_unclassified(100);k__Fungi_unclassified(100);k__Fungi_unclassified(100);k__Fungi_unclassified(100);k__Fungi_unclassified(100); | - | - | - | - | - | - | - | Unassigned | - |
| Otu27300 | 1 | k__Fungi(100);p__Ascomycota(100);p__Ascomycota_unclassified(100);p__Ascomycota_unclassified(100);p__Ascomycota_unclassified(100);p__Ascomycota_unclassified(100);p__Ascomycota_unclassified(100); | - | - | - | - | - | - | - | Unassigned | - |
| Otu27301 | 1 | k__Fungi(100);k__Fungi_unclassified(100);k__Fungi_unclassified(100);k__Fungi_unclassified(100);k__Fungi_unclassified(100);k__Fungi_unclassified(100);k__Fungi_unclassified(100); | - | - | - | - | - | - | - | Unassigned | - |
| Otu27302 | 1 | k__Fungi(100);k__Fungi_unclassified(100);k__Fungi_unclassified(100);k__Fungi_unclassified(100);k__Fungi_unclassified(100);k__Fungi_unclassified(100);k__Fungi_unclassified(100); | - | - | - | - | - | - | - | Unassigned | - |
| Otu27303 | 1 | k__Fungi(100);p__Ascomycota(100);p__Ascomycota_unclassified(100);p__Ascomycota_unclassified(100);p__Ascomycota_unclassified(100);p__Ascomycota_unclassified(100);p__Ascomycota_unclassified(100); | - | - | - | - | - | - | - | Unassigned | - |
| Otu27304 | 1 | k__Fungi(100);k__Fungi_unclassified(100);k__Fungi_unclassified(100);k__Fungi_unclassified(100);k__Fungi_unclassified(100);k__Fungi_unclassified(100);k__Fungi_unclassified(100); | - | - | - | - | - | - | - | Unassigned | - |
| Otu27305 | 1 | k__Fungi(100);p__Ascomycota(100);p__Ascomycota_unclassified(100);p__Ascomycota_unclassified(100);p__Ascomycota_unclassified(100);p__Ascomycota_unclassified(100);p__Ascomycota_unclassified(100); | - | - | - | - | - | - | - | Unassigned | - |
| Otu27306 | 1 | k__Fungi(100);p__Ascomycota(100);p__Ascomycota_unclassified(100);p__Ascomycota_unclassified(100);p__Ascomycota_unclassified(100);p__Ascomycota_unclassified(100);p__Ascomycota_unclassified(100); | - | - | - | - | - | - | - | Unassigned | - |
| Otu27307 | 1 | k__Fungi(100);k__Fungi_unclassified(100);k__Fungi_unclassified(100);k__Fungi_unclassified(100);k__Fungi_unclassified(100);k__Fungi_unclassified(100);k__Fungi_unclassified(100); | - | - | - | - | - | - | - | Unassigned | - |
| Otu27309 | 1 | k__Fungi(100);p__Ascomycota(100);p__Ascomycota_unclassified(100);p__Ascomycota_unclassified(100);p__Ascomycota_unclassified(100);p__Ascomycota_unclassified(100);p__Ascomycota_unclassified(100); | - | - | - | - | - | - | - | Unassigned | - |
| Otu27310 | 1 | k__Fungi(100);p__Ascomycota(100);p__Ascomycota_unclassified(100);p__Ascomycota_unclassified(100);p__Ascomycota_unclassified(100);p__Ascomycota_unclassified(100);p__Ascomycota_unclassified(100); | - | - | - | - | - | - | - | Unassigned | - |
| Otu27311 | 1 | k__Fungi(100);p__Ascomycota(100);p__Ascomycota_unclassified(100);p__Ascomycota_unclassified(100);p__Ascomycota_unclassified(100);p__Ascomycota_unclassified(100);p__Ascomycota_unclassified(100); | - | - | - | - | - | - | - | Unassigned | - |
| Otu27312 | 1 | k__Fungi(100);p__Ascomycota(100);p__Ascomycota_unclassified(100);p__Ascomycota_unclassified(100);p__Ascomycota_unclassified(100);p__Ascomycota_unclassified(100);p__Ascomycota_unclassified(100); | - | - | - | - | - | - | - | Unassigned | - |
| Otu27313 | 1 | k__Fungi(100);k__Fungi_unclassified(100);k__Fungi_unclassified(100);k__Fungi_unclassified(100);k__Fungi_unclassified(100);k__Fungi_unclassified(100);k__Fungi_unclassified(100); | - | - | - | - | - | - | - | Unassigned | - |
| Otu27314 | 1 | k__Fungi(100);p__Ascomycota(100);p__Ascomycota_unclassified(100);p__Ascomycota_unclassified(100);p__Ascomycota_unclassified(100);p__Ascomycota_unclassified(100);p__Ascomycota_unclassified(100); | - | - | - | - | - | - | - | Unassigned | - |
| Otu27315 | 1 | k__Fungi(100);p__Ascomycota(100);p__Ascomycota_unclassified(100);p__Ascomycota_unclassified(100);p__Ascomycota_unclassified(100);p__Ascomycota_unclassified(100);p__Ascomycota_unclassified(100); | - | - | - | - | - | - | - | Unassigned | - |
| Otu27316 | 1 | k__Fungi(100);p__Ascomycota(100);p__Ascomycota_unclassified(100);p__Ascomycota_unclassified(100);p__Ascomycota_unclassified(100);p__Ascomycota_unclassified(100);p__Ascomycota_unclassified(100); | - | - | - | - | - | - | - | Unassigned | - |
| Otu27317 | 1 | k__Fungi(100);p__Ascomycota(100);p__Ascomycota_unclassified(100);p__Ascomycota_unclassified(100);p__Ascomycota_unclassified(100);p__Ascomycota_unclassified(100);p__Ascomycota_unclassified(100); | - | - | - | - | - | - | - | Unassigned | - |
| Otu27318 | 1 | k__Fungi(100);p__Ascomycota(100);p__Ascomycota_unclassified(100);p__Ascomycota_unclassified(100);p__Ascomycota_unclassified(100);p__Ascomycota_unclassified(100);p__Ascomycota_unclassified(100); | - | - | - | - | - | - | - | Unassigned | - |
| Otu27321 | 1 | k__Fungi(100);k__Fungi_unclassified(100);k__Fungi_unclassified(100);k__Fungi_unclassified(100);k__Fungi_unclassified(100);k__Fungi_unclassified(100);k__Fungi_unclassified(100); | - | - | - | - | - | - | - | Unassigned | - |
| Otu27322 | 1 | k__Fungi(100);k__Fungi_unclassified(100);k__Fungi_unclassified(100);k__Fungi_unclassified(100);k__Fungi_unclassified(100);k__Fungi_unclassified(100);k__Fungi_unclassified(100); | - | - | - | - | - | - | - | Unassigned | - |
| Otu27323 | 1 | k__Fungi(100);k__Fungi_unclassified(100);k__Fungi_unclassified(100);k__Fungi_unclassified(100);k__Fungi_unclassified(100);k__Fungi_unclassified(100);k__Fungi_unclassified(100); | - | - | - | - | - | - | - | Unassigned | - |
| Otu27324 | 1 | k__Fungi(100);p__Ascomycota(100);p__Ascomycota_unclassified(100);p__Ascomycota_unclassified(100);p__Ascomycota_unclassified(100);p__Ascomycota_unclassified(100);p__Ascomycota_unclassified(100); | - | - | - | - | - | - | - | Unassigned | - |
| Otu27325 | 1 | k__Fungi(100);p__Ascomycota(100);p__Ascomycota_unclassified(100);p__Ascomycota_unclassified(100);p__Ascomycota_unclassified(100);p__Ascomycota_unclassified(100);p__Ascomycota_unclassified(100); | - | - | - | - | - | - | - | Unassigned | - |
| Otu27326 | 1 | k__Fungi(100);p__Ascomycota(100);p__Ascomycota_unclassified(100);p__Ascomycota_unclassified(100);p__Ascomycota_unclassified(100);p__Ascomycota_unclassified(100);p__Ascomycota_unclassified(100); | - | - | - | - | - | - | - | Unassigned | - |
| Otu27327 | 1 | k__Fungi(100);k__Fungi_unclassified(100);k__Fungi_unclassified(100);k__Fungi_unclassified(100);k__Fungi_unclassified(100);k__Fungi_unclassified(100);k__Fungi_unclassified(100); | - | - | - | - | - | - | - | Unassigned | - |
| Otu27329 | 1 | k__Fungi(100);k__Fungi_unclassified(100);k__Fungi_unclassified(100);k__Fungi_unclassified(100);k__Fungi_unclassified(100);k__Fungi_unclassified(100);k__Fungi_unclassified(100); | - | - | - | - | - | - | - | Unassigned | - |
| Otu27330 | 1 | k__Fungi(100);k__Fungi_unclassified(100);k__Fungi_unclassified(100);k__Fungi_unclassified(100);k__Fungi_unclassified(100);k__Fungi_unclassified(100);k__Fungi_unclassified(100); | - | - | - | - | - | - | - | Unassigned | - |
| Otu27331 | 1 | k__Fungi(100);k__Fungi_unclassified(100);k__Fungi_unclassified(100);k__Fungi_unclassified(100);k__Fungi_unclassified(100);k__Fungi_unclassified(100);k__Fungi_unclassified(100); | - | - | - | - | - | - | - | Unassigned | - |
| Otu27333 | 1 | k__Fungi(100);k__Fungi_unclassified(100);k__Fungi_unclassified(100);k__Fungi_unclassified(100);k__Fungi_unclassified(100);k__Fungi_unclassified(100);k__Fungi_unclassified(100); | - | - | - | - | - | - | - | Unassigned | - |
| Otu27334 | 1 | k__Fungi(100);k__Fungi_unclassified(100);k__Fungi_unclassified(100);k__Fungi_unclassified(100);k__Fungi_unclassified(100);k__Fungi_unclassified(100);k__Fungi_unclassified(100); | - | - | - | - | - | - | - | Unassigned | - |
| Otu27335 | 1 | k__Fungi(100);p__Ascomycota(100);p__Ascomycota_unclassified(100);p__Ascomycota_unclassified(100);p__Ascomycota_unclassified(100);p__Ascomycota_unclassified(100);p__Ascomycota_unclassified(100); | - | - | - | - | - | - | - | Unassigned | - |
| Otu27338 | 1 | k__Fungi(100);p__Ascomycota(100);p__Ascomycota_unclassified(100);p__Ascomycota_unclassified(100);p__Ascomycota_unclassified(100);p__Ascomycota_unclassified(100);p__Ascomycota_unclassified(100); | - | - | - | - | - | - | - | Unassigned | - |
| Otu27339 | 1 | k__Fungi(100);k__Fungi_unclassified(100);k__Fungi_unclassified(100);k__Fungi_unclassified(100);k__Fungi_unclassified(100);k__Fungi_unclassified(100);k__Fungi_unclassified(100); | - | - | - | - | - | - | - | Unassigned | - |
| Otu27340 | 1 | k__Fungi(100);p__Ascomycota(100);p__Ascomycota_unclassified(100);p__Ascomycota_unclassified(100);p__Ascomycota_unclassified(100);p__Ascomycota_unclassified(100);p__Ascomycota_unclassified(100); | - | - | - | - | - | - | - | Unassigned | - |

| | | | | | | | | | | | |
|---|---|---|---|---|---|---|---|---|---|---|---|
| Otu27342 | 1 | k__Fungi(100);k__Fungi_unclassified(100);k__Fungi_unclassified(100);k__Fungi_unclassified(100);k__Fungi_unclassified(100);k__Fungi_unclassified(100);k__Fungi_unclassified(100); | - | - | - | - | - | - | - | Unassigned | - |
| Otu27346 | 1 | k__Fungi(100);k__Fungi_unclassified(100);k__Fungi_unclassified(100);k__Fungi_unclassified(100);k__Fungi_unclassified(100);k__Fungi_unclassified(100);k__Fungi_unclassified(100); | - | - | - | - | - | - | - | Unassigned | - |
| Otu27347 | 1 | k__Fungi(100);p__Ascomycota(100);p__Ascomycota_unclassified(100);p__Ascomycota_unclassified(100);p__Ascomycota_unclassified(100);p__Ascomycota_unclassified(100);p__Ascomycota_unclassified(100); | - | - | - | - | - | - | - | Unassigned | - |
| Otu27348 | 1 | k__Fungi(100);k__Fungi_unclassified(100);k__Fungi_unclassified(100);k__Fungi_unclassified(100);k__Fungi_unclassified(100);k__Fungi_unclassified(100);k__Fungi_unclassified(100); | - | - | - | - | - | - | - | Unassigned | - |
| Otu27349 | 1 | k__Fungi(100);k__Fungi_unclassified(100);k__Fungi_unclassified(100);k__Fungi_unclassified(100);k__Fungi_unclassified(100);k__Fungi_unclassified(100);k__Fungi_unclassified(100); | - | - | - | - | - | - | - | Unassigned | - |
| Otu27350 | 1 | k__Fungi(100);k__Fungi_unclassified(100);k__Fungi_unclassified(100);k__Fungi_unclassified(100);k__Fungi_unclassified(100);k__Fungi_unclassified(100);k__Fungi_unclassified(100); | - | - | - | - | - | - | - | Unassigned | - |
| Otu27351 | 1 | k__Fungi(100);p__Ascomycota(100);p__Ascomycota_unclassified(100);p__Ascomycota_unclassified(100);p__Ascomycota_unclassified(100);p__Ascomycota_unclassified(100);p__Ascomycota_unclassified(100); | - | - | - | - | - | - | - | Unassigned | - |
| Otu27352 | 1 | k__Fungi(100);p__Ascomycota(100);p__Ascomycota_unclassified(100);p__Ascomycota_unclassified(100);p__Ascomycota_unclassified(100);p__Ascomycota_unclassified(100);p__Ascomycota_unclassified(100); | - | - | - | - | - | - | - | Unassigned | - |
| Otu27353 | 1 | k__Fungi(100);p__Ascomycota(100);p__Ascomycota_unclassified(100);p__Ascomycota_unclassified(100);p__Ascomycota_unclassified(100);p__Ascomycota_unclassified(100);p__Ascomycota_unclassified(100); | - | - | - | - | - | - | - | Unassigned | - |
| Otu27354 | 1 | k__Fungi(100);p__Ascomycota(100);p__Ascomycota_unclassified(100);p__Ascomycota_unclassified(100);p__Ascomycota_unclassified(100);p__Ascomycota_unclassified(100);p__Ascomycota_unclassified(100); | - | - | - | - | - | - | - | Unassigned | - |
| Otu27355 | 1 | k__Fungi(100);k__Fungi_unclassified(100);k__Fungi_unclassified(100);k__Fungi_unclassified(100);k__Fungi_unclassified(100);k__Fungi_unclassified(100);k__Fungi_unclassified(100); | - | - | - | - | - | - | - | Unassigned | - |
| Otu27356 | 1 | k__Fungi(100);p__Ascomycota(100);p__Ascomycota_unclassified(100);p__Ascomycota_unclassified(100);p__Ascomycota_unclassified(100);p__Ascomycota_unclassified(100);p__Ascomycota_unclassified(100); | - | - | - | - | - | - | - | Unassigned | - |
| Otu27359 | 1 | k__Fungi(100);p__Ascomycota(100);c__Leotiomycetes(100);o__Helotiales(100);f__Helotiales_family_Incertae_sedis(100);f__Helotiales_family_Incertae_sedis_unclassified(100);f__Helotiales_family_Incertae_sedis_unclassified(100); | - | - | - | - | - | - | - | Unassigned | - |
| Otu27361 | 1 | k__Fungi(100);k__Fungi_unclassified(100);k__Fungi_unclassified(100);k__Fungi_unclassified(100);k__Fungi_unclassified(100);k__Fungi_unclassified(100);k__Fungi_unclassified(100); | - | - | - | - | - | - | - | Unassigned | - |
| Otu27362 | 1 | k__Fungi(100);k__Fungi_unclassified(100);k__Fungi_unclassified(100);k__Fungi_unclassified(100);k__Fungi_unclassified(100);k__Fungi_unclassified(100);k__Fungi_unclassified(100); | - | - | - | - | - | - | - | Unassigned | - |
| Otu27363 | 1 | k__Fungi(100);k__Fungi_unclassified(100);k__Fungi_unclassified(100);k__Fungi_unclassified(100);k__Fungi_unclassified(100);k__Fungi_unclassified(100);k__Fungi_unclassified(100); | - | - | - | - | - | - | - | Unassigned | - |
| Otu27364 | 1 | k__Fungi(100);p__Ascomycota(100);p__Ascomycota_unclassified(100);p__Ascomycota_unclassified(100);p__Ascomycota_unclassified(100);p__Ascomycota_unclassified(100);p__Ascomycota_unclassified(100); | - | - | - | - | - | - | - | Unassigned | - |
| Otu27365 | 1 | k__Fungi(100);p__Ascomycota(100);p__Ascomycota_unclassified(100);p__Ascomycota_unclassified(100);p__Ascomycota_unclassified(100);p__Ascomycota_unclassified(100);p__Ascomycota_unclassified(100); | - | - | - | - | - | - | - | Unassigned | - |
| Otu27366 | 1 | k__Fungi(100);p__Ascomycota(100);p__Ascomycota_unclassified(100);p__Ascomycota_unclassified(100);p__Ascomycota_unclassified(100);p__Ascomycota_unclassified(100);p__Ascomycota_unclassified(100); | - | - | - | - | - | - | - | Unassigned | - |
| Otu27367 | 1 | k__Fungi(100);k__Fungi_unclassified(100);k__Fungi_unclassified(100);k__Fungi_unclassified(100);k__Fungi_unclassified(100);k__Fungi_unclassified(100);k__Fungi_unclassified(100); | - | - | - | - | - | - | - | Unassigned | - |
| Otu27368 | 1 | k__Fungi(100);p__Ascomycota(100);p__Ascomycota_unclassified(100);p__Ascomycota_unclassified(100);p__Ascomycota_unclassified(100);p__Ascomycota_unclassified(100);p__Ascomycota_unclassified(100); | - | - | - | - | - | - | - | Unassigned | - |
| Otu27369 | 1 | k__Fungi(100);k__Fungi_unclassified(100);k__Fungi_unclassified(100);k__Fungi_unclassified(100);k__Fungi_unclassified(100);k__Fungi_unclassified(100);k__Fungi_unclassified(100); | - | - | - | - | - | - | - | Unassigned | - |
| Otu27371 | 1 | k__Fungi(100);k__Fungi_unclassified(100);k__Fungi_unclassified(100);k__Fungi_unclassified(100);k__Fungi_unclassified(100);k__Fungi_unclassified(100);k__Fungi_unclassified(100); | - | - | - | - | - | - | - | Unassigned | - |
| Otu27372 | 1 | k__Fungi(100);k__Fungi_unclassified(100);k__Fungi_unclassified(100);k__Fungi_unclassified(100);k__Fungi_unclassified(100);k__Fungi_unclassified(100);k__Fungi_unclassified(100); | - | - | - | - | - | - | - | Unassigned | - |
| Otu27373 | 1 | k__Fungi(100);p__Ascomycota(100);p__Ascomycota_unclassified(100);p__Ascomycota_unclassified(100);p__Ascomycota_unclassified(100);p__Ascomycota_unclassified(100);p__Ascomycota_unclassified(100); | - | - | - | - | - | - | - | Unassigned | - |
| Otu27375 | 1 | k__Fungi(100);p__Ascomycota(100);p__Ascomycota_unclassified(100);p__Ascomycota_unclassified(100);p__Ascomycota_unclassified(100);p__Ascomycota_unclassified(100);p__Ascomycota_unclassified(100); | - | - | - | - | - | - | - | Unassigned | - |
| Otu27377 | 1 | k__Fungi(100);p__Ascomycota(100);p__Ascomycota_unclassified(100);p__Ascomycota_unclassified(100);p__Ascomycota_unclassified(100);p__Ascomycota_unclassified(100);p__Ascomycota_unclassified(100); | - | - | - | - | - | - | - | Unassigned | - |
| Otu27383 | 1 | k__Fungi(100);p__Ascomycota(100);p__Ascomycota_unclassified(100);p__Ascomycota_unclassified(100);p__Ascomycota_unclassified(100);p__Ascomycota_unclassified(100);p__Ascomycota_unclassified(100); | - | - | - | - | - | - | - | Unassigned | - |
| Otu27386 | 1 | k__Fungi(100);k__Fungi_unclassified(100);k__Fungi_unclassified(100);k__Fungi_unclassified(100);k__Fungi_unclassified(100);k__Fungi_unclassified(100);k__Fungi_unclassified(100); | - | - | - | - | - | - | - | Unassigned | - |
| Otu27389 | 1 | k__Fungi(100);k__Fungi_unclassified(100);k__Fungi_unclassified(100);k__Fungi_unclassified(100);k__Fungi_unclassified(100);k__Fungi_unclassified(100);k__Fungi_unclassified(100); | - | - | - | - | - | - | - | Unassigned | - |
| Otu27390 | 1 | k__Fungi(100);k__Fungi_unclassified(100);k__Fungi_unclassified(100);k__Fungi_unclassified(100);k__Fungi_unclassified(100);k__Fungi_unclassified(100);k__Fungi_unclassified(100); | - | - | - | - | - | - | - | Unassigned | - |
| Otu27391 | 1 | k__Fungi(100);k__Fungi_unclassified(100);k__Fungi_unclassified(100);k__Fungi_unclassified(100);k__Fungi_unclassified(100);k__Fungi_unclassified(100);k__Fungi_unclassified(100); | - | - | - | - | - | - | - | Unassigned | - |
| Otu27392 | 1 | k__Fungi(100);k__Fungi_unclassified(100);k__Fungi_unclassified(100);k__Fungi_unclassified(100);k__Fungi_unclassified(100);k__Fungi_unclassified(100);k__Fungi_unclassified(100); | - | - | - | - | - | - | - | Unassigned | - |
| Otu27394 | 1 | k__Fungi(100);k__Fungi_unclassified(100);k__Fungi_unclassified(100);k__Fungi_unclassified(100);k__Fungi_unclassified(100);k__Fungi_unclassified(100);k__Fungi_unclassified(100); | - | - | - | - | - | - | - | Unassigned | - |
| Otu27395 | 1 | k__Fungi(100);k__Fungi_unclassified(100);k__Fungi_unclassified(100);k__Fungi_unclassified(100);k__Fungi_unclassified(100);k__Fungi_unclassified(100);k__Fungi_unclassified(100); | - | - | - | - | - | - | - | Unassigned | - |
| Otu27396 | 1 | k__Fungi(100);k__Fungi_unclassified(100);k__Fungi_unclassified(100);k__Fungi_unclassified(100);k__Fungi_unclassified(100);k__Fungi_unclassified(100);k__Fungi_unclassified(100); | - | - | - | - | - | - | - | Unassigned | - |
| Otu27397 | 1 | k__Fungi(100);p__Ascomycota(100);p__Ascomycota_unclassified(100);p__Ascomycota_unclassified(100);p__Ascomycota_unclassified(100);p__Ascomycota_unclassified(100);p__Ascomycota_unclassified(100); | - | - | - | - | - | - | - | Unassigned | - |
| Otu27400 | 1 | k__Fungi(100);k__Fungi_unclassified(100);k__Fungi_unclassified(100);k__Fungi_unclassified(100);k__Fungi_unclassified(100);k__Fungi_unclassified(100);k__Fungi_unclassified(100); | - | - | - | - | - | - | - | Unassigned | - |
| Otu27401 | 1 | k__Fungi(100);k__Fungi_unclassified(100);k__Fungi_unclassified(100);k__Fungi_unclassified(100);k__Fungi_unclassified(100);k__Fungi_unclassified(100);k__Fungi_unclassified(100); | - | - | - | - | - | - | - | Unassigned | - |
| Otu27402 | 1 | k__Fungi(100);k__Fungi_unclassified(100);k__Fungi_unclassified(100);k__Fungi_unclassified(100);k__Fungi_unclassified(100);k__Fungi_unclassified(100);k__Fungi_unclassified(100); | - | - | - | - | - | - | - | Unassigned | - |
| Otu27403 | 1 | k__Fungi(100);p__Ascomycota(100);p__Ascomycota_unclassified(100);p__Ascomycota_unclassified(100);p__Ascomycota_unclassified(100);p__Ascomycota_unclassified(100);p__Ascomycota_unclassified(100); | - | - | - | - | - | - | - | Unassigned | - |
| Otu27404 | 1 | k__Fungi(100);k__Fungi_unclassified(100);k__Fungi_unclassified(100);k__Fungi_unclassified(100);k__Fungi_unclassified(100);k__Fungi_unclassified(100);k__Fungi_unclassified(100); | - | - | - | - | - | - | - | Unassigned | - |
| Otu27405 | 1 | k__Fungi(100);k__Fungi_unclassified(100);k__Fungi_unclassified(100);k__Fungi_unclassified(100);k__Fungi_unclassified(100);k__Fungi_unclassified(100);k__Fungi_unclassified(100); | - | - | - | - | - | - | - | Unassigned | - |
| Otu27406 | 1 | k__Fungi(100);p__Ascomycota(100);p__Ascomycota_unclassified(100);p__Ascomycota_unclassified(100);p__Ascomycota_unclassified(100);p__Ascomycota_unclassified(100);p__Ascomycota_unclassified(100); | - | - | - | - | - | - | - | Unassigned | - |
| Otu27407 | 1 | k__Fungi(100);k__Fungi_unclassified(100);k__Fungi_unclassified(100);k__Fungi_unclassified(100);k__Fungi_unclassified(100);k__Fungi_unclassified(100);k__Fungi_unclassified(100); | - | - | - | - | - | - | - | Unassigned | - |
| Otu27408 | 1 | k__Fungi(100);k__Fungi_unclassified(100);k__Fungi_unclassified(100);k__Fungi_unclassified(100);k__Fungi_unclassified(100);k__Fungi_unclassified(100);k__Fungi_unclassified(100); | - | - | - | - | - | - | - | Unassigned | - |
| Otu27410 | 1 | k__Fungi(100);p__Ascomycota(100);p__Ascomycota_unclassified(100);p__Ascomycota_unclassified(100);p__Ascomycota_unclassified(100);p__Ascomycota_unclassified(100);p__Ascomycota_unclassified(100); | - | - | - | - | - | - | - | Unassigned | - |
| Otu27411 | 1 | k__Fungi(100);k__Fungi_unclassified(100);k__Fungi_unclassified(100);k__Fungi_unclassified(100);k__Fungi_unclassified(100);k__Fungi_unclassified(100);k__Fungi_unclassified(100); | - | - | - | - | - | - | - | Unassigned | - |
| Otu27412 | 1 | k__Fungi(100);k__Fungi_unclassified(100);k__Fungi_unclassified(100);k__Fungi_unclassified(100);k__Fungi_unclassified(100);k__Fungi_unclassified(100);k__Fungi_unclassified(100); | - | - | - | - | - | - | - | Unassigned | - |
| Otu27414 | 1 | k__Fungi(100);p__Ascomycota(100);p__Ascomycota_unclassified(100);p__Ascomycota_unclassified(100);p__Ascomycota_unclassified(100);p__Ascomycota_unclassified(100);p__Ascomycota_unclassified(100); | - | - | - | - | - | - | - | Unassigned | - |
| Otu27418 | 1 | k__Fungi(100);k__Fungi_unclassified(100);k__Fungi_unclassified(100);k__Fungi_unclassified(100);k__Fungi_unclassified(100);k__Fungi_unclassified(100);k__Fungi_unclassified(100); | - | - | - | - | - | - | - | Unassigned | - |
| Otu27420 | 1 | k__Fungi(100);p__Ascomycota(100);p__Ascomycota_unclassified(100);p__Ascomycota_unclassified(100);p__Ascomycota_unclassified(100);p__Ascomycota_unclassified(100);p__Ascomycota_unclassified(100); | - | - | - | - | - | - | - | Unassigned | - |
| Otu27421 | 1 | k__Fungi(100);k__Fungi_unclassified(100);k__Fungi_unclassified(100);k__Fungi_unclassified(100);k__Fungi_unclassified(100);k__Fungi_unclassified(100);k__Fungi_unclassified(100); | - | - | - | - | - | - | - | Unassigned | - |
| Otu27422 | 1 | k__Fungi(100);p__Ascomycota(100);p__Ascomycota_unclassified(100);p__Ascomycota_unclassified(100);p__Ascomycota_unclassified(100);p__Ascomycota_unclassified(100);p__Ascomycota_unclassified(100); | - | - | - | - | - | - | - | Unassigned | - |
| Otu27424 | 1 | k__Fungi(100);p__Ascomycota(100);p__Ascomycota_unclassified(100);p__Ascomycota_unclassified(100);p__Ascomycota_unclassified(100);p__Ascomycota_unclassified(100);p__Ascomycota_unclassified(100); | - | - | - | - | - | - | - | Unassigned | - |
| Otu27425 | 1 | k__Fungi(100);k__Fungi_unclassified(100);k__Fungi_unclassified(100);k__Fungi_unclassified(100);k__Fungi_unclassified(100);k__Fungi_unclassified(100);k__Fungi_unclassified(100); | - | - | - | - | - | - | - | Unassigned | - |
| Otu27426 | 1 | k__Fungi(100);p__Ascomycota(100);p__Ascomycota_unclassified(100);p__Ascomycota_unclassified(100);p__Ascomycota_unclassified(100);p__Ascomycota_unclassified(100);p__Ascomycota_unclassified(100); | - | - | - | - | - | - | - | Unassigned | - |
| Otu27427 | 1 | k__Fungi(100);k__Fungi_unclassified(100);k__Fungi_unclassified(100);k__Fungi_unclassified(100);k__Fungi_unclassified(100);k__Fungi_unclassified(100);k__Fungi_unclassified(100); | - | - | - | - | - | - | - | Unassigned | - |
| Otu27434 | 1 | k__Fungi(100);k__Fungi_unclassified(100);k__Fungi_unclassified(100);k__Fungi_unclassified(100);k__Fungi_unclassified(100);k__Fungi_unclassified(100);k__Fungi_unclassified(100); | - | - | - | - | - | - | - | Unassigned | - |
| Otu27435 | 1 | k__Fungi(100);p__Ascomycota(100);p__Ascomycota_unclassified(100);p__Ascomycota_unclassified(100);p__Ascomycota_unclassified(100);p__Ascomycota_unclassified(100);p__Ascomycota_unclassified(100); | - | - | - | - | - | - | - | Unassigned | - |
| Otu27437 | 1 | k__Fungi(100);k__Fungi_unclassified(100);k__Fungi_unclassified(100);k__Fungi_unclassified(100);k__Fungi_unclassified(100);k__Fungi_unclassified(100);k__Fungi_unclassified(100); | - | - | - | - | - | - | - | Unassigned | - |
| Otu27441 | 1 | k__Fungi(100);k__Fungi_unclassified(100);k__Fungi_unclassified(100);k__Fungi_unclassified(100);k__Fungi_unclassified(100);k__Fungi_unclassified(100);k__Fungi_unclassified(100); | - | - | - | - | - | - | - | Unassigned | - |
| Otu27442 | 1 | k__Fungi(100);p__Ascomycota(100);p__Ascomycota_unclassified(100);p__Ascomycota_unclassified(100);p__Ascomycota_unclassified(100);p__Ascomycota_unclassified(100);p__Ascomycota_unclassified(100); | - | - | - | - | - | - | - | Unassigned | - |
| Otu27443 | 1 | k__Fungi(100);k__Fungi_unclassified(100);k__Fungi_unclassified(100);k__Fungi_unclassified(100);k__Fungi_unclassified(100);k__Fungi_unclassified(100);k__Fungi_unclassified(100); | - | - | - | - | - | - | - | Unassigned | - |
| Otu27444 | 1 | k__Fungi(100);k__Fungi_unclassified(100);k__Fungi_unclassified(100);k__Fungi_unclassified(100);k__Fungi_unclassified(100);k__Fungi_unclassified(100);k__Fungi_unclassified(100); | - | - | - | - | - | - | - | Unassigned | - |
| Otu27445 | 1 | k__Fungi(100);k__Fungi_unclassified(100);k__Fungi_unclassified(100);k__Fungi_unclassified(100);k__Fungi_unclassified(100);k__Fungi_unclassified(100);k__Fungi_unclassified(100); | - | - | - | - | - | - | - | Unassigned | - |
| Otu27446 | 1 | k__Fungi(100);k__Fungi_unclassified(100);k__Fungi_unclassified(100);k__Fungi_unclassified(100);k__Fungi_unclassified(100);k__Fungi_unclassified(100);k__Fungi_unclassified(100); | - | - | - | - | - | - | - | Unassigned | - |
| Otu27447 | 1 | k__Fungi(100);p__Ascomycota(100);p__Ascomycota_unclassified(100);p__Ascomycota_unclassified(100);p__Ascomycota_unclassified(100);p__Ascomycota_unclassified(100);p__Ascomycota_unclassified(100); | - | - | - | - | - | - | - | Unassigned | - |
| Otu27448 | 1 | k__Fungi(100);p__Ascomycota(100);p__Ascomycota_unclassified(100);p__Ascomycota_unclassified(100);p__Ascomycota_unclassified(100);p__Ascomycota_unclassified(100);p__Ascomycota_unclassified(100); | - | - | - | - | - | - | - | Unassigned | - |
| Otu27449 | 1 | k__Fungi(100);k__Fungi_unclassified(100);k__Fungi_unclassified(100);k__Fungi_unclassified(100);k__Fungi_unclassified(100);k__Fungi_unclassified(100);k__Fungi_unclassified(100); | - | - | - | - | - | - | - | Unassigned | - |
| Otu27450 | 1 | k__Fungi(100);k__Fungi_unclassified(100);k__Fungi_unclassified(100);k__Fungi_unclassified(100);k__Fungi_unclassified(100);k__Fungi_unclassified(100);k__Fungi_unclassified(100); | - | - | - | - | - | - | - | Unassigned | - |
| Otu27451 | 1 | k__Fungi(100);p__Ascomycota(100);p__Ascomycota_unclassified(100);p__Ascomycota_unclassified(100);p__Ascomycota_unclassified(100);p__Ascomycota_unclassified(100);p__Ascomycota_unclassified(100); | - | - | - | - | - | - | - | Unassigned | - |
| Otu27452 | 1 | k__Fungi(100);p__Ascomycota(100);p__Ascomycota_unclassified(100);p__Ascomycota_unclassified(100);p__Ascomycota_unclassified(100);p__Ascomycota_unclassified(100);p__Ascomycota_unclassified(100); | - | - | - | - | - | - | - | Unassigned | - |
| Otu27453 | 1 | k__Fungi(100);k__Fungi_unclassified(100);k__Fungi_unclassified(100);k__Fungi_unclassified(100);k__Fungi_unclassified(100);k__Fungi_unclassified(100);k__Fungi_unclassified(100); | - | - | - | - | - | - | - | Unassigned | - |
| Otu27455 | 1 | k__Fungi(100);k__Fungi_unclassified(100);k__Fungi_unclassified(100);k__Fungi_unclassified(100);k__Fungi_unclassified(100);k__Fungi_unclassified(100);k__Fungi_unclassified(100); | - | - | - | - | - | - | - | Unassigned | - |
| Otu27456 | 1 | k__Fungi(100);p__Ascomycota(100);p__Ascomycota_unclassified(100);p__Ascomycota_unclassified(100);p__Ascomycota_unclassified(100);p__Ascomycota_unclassified(100);p__Ascomycota_unclassified(100); | - | - | - | - | - | - | - | Unassigned | - |
| Otu27457 | 1 | k__Fungi(100);k__Fungi_unclassified(100);k__Fungi_unclassified(100);k__Fungi_unclassified(100);k__Fungi_unclassified(100);k__Fungi_unclassified(100);k__Fungi_unclassified(100); | - | - | - | - | - | - | - | Unassigned | - |
| Otu27459 | 1 | k__Fungi(100);k__Fungi_unclassified(100);k__Fungi_unclassified(100);k__Fungi_unclassified(100);k__Fungi_unclassified(100);k__Fungi_unclassified(100);k__Fungi_unclassified(100); | - | - | - | - | - | - | - | Unassigned | - |
| Otu27460 | 1 | k__Fungi(100);p__Ascomycota(100);p__Ascomycota_unclassified(100);p__Ascomycota_unclassified(100);p__Ascomycota_unclassified(100);p__Ascomycota_unclassified(100);p__Ascomycota_unclassified(100); | - | - | - | - | - | - | - | Unassigned | - |
| Otu27461 | 1 | k__Fungi(100);p__Ascomycota(100);p__Ascomycota_unclassified(100);p__Ascomycota_unclassified(100);p__Ascomycota_unclassified(100);p__Ascomycota_unclassified(100);p__Ascomycota_unclassified(100); | - | - | - | - | - | - | - | Unassigned | - |
| Otu27465 | 1 | k__Fungi(100);k__Fungi_unclassified(100);k__Fungi_unclassified(100);k__Fungi_unclassified(100);k__Fungi_unclassified(100);k__Fungi_unclassified(100);k__Fungi_unclassified(100); | - | - | - | - | - | - | - | Unassigned | - |
| Otu27466 | 1 | k__Fungi(100);p__Ascomycota(100);p__Ascomycota_unclassified(100);p__Ascomycota_unclassified(100);p__Ascomycota_unclassified(100);p__Ascomycota_unclassified(100);p__Ascomycota_unclassified(100); | - | - | - | - | - | - | - | Unassigned | - |
| Otu27467 | 1 | k__Fungi(100);p__Ascomycota(100);p__Ascomycota_unclassified(100);p__Ascomycota_unclassified(100);p__Ascomycota_unclassified(100);p__Ascomycota_unclassified(100);p__Ascomycota_unclassified(100); | - | - | - | - | - | - | - | Unassigned | - |
| Otu27468 | 1 | k__Fungi(100);p__Ascomycota(100);p__Ascomycota_unclassified(100);p__Ascomycota_unclassified(100);p__Ascomycota_unclassified(100);p__Ascomycota_unclassified(100);p__Ascomycota_unclassified(100); | - | - | - | - | - | - | - | Unassigned | - |
| Otu27470 | 1 | k__Fungi(100);k__Fungi_unclassified(100);k__Fungi_unclassified(100);k__Fungi_unclassified(100);k__Fungi_unclassified(100);k__Fungi_unclassified(100);k__Fungi_unclassified(100); | - | - | - | - | - | - | - | Unassigned | - |
| Otu27472 | 1 | k__Fungi(100);k__Fungi_unclassified(100);k__Fungi_unclassified(100);k__Fungi_unclassified(100);k__Fungi_unclassified(100);k__Fungi_unclassified(100);k__Fungi_unclassified(100); | - | - | - | - | - | - | - | Unassigned | - |
| Otu27476 | 1 | k__Fungi(100);k__Fungi_unclassified(100);k__Fungi_unclassified(100);k__Fungi_unclassified(100);k__Fungi_unclassified(100);k__Fungi_unclassified(100);k__Fungi_unclassified(100); | - | - | - | - | - | - | - | Unassigned | - |
| Otu27479 | 1 | k__Fungi(100);k__Fungi_unclassified(100);k__Fungi_unclassified(100);k__Fungi_unclassified(100);k__Fungi_unclassified(100);k__Fungi_unclassified(100);k__Fungi_unclassified(100); | - | - | - | - | - | - | - | Unassigned | - |
| Otu27480 | 1 | k__Fungi(100);k__Fungi_unclassified(100);k__Fungi_unclassified(100);k__Fungi_unclassified(100);k__Fungi_unclassified(100);k__Fungi_unclassified(100);k__Fungi_unclassified(100); | - | - | - | - | - | - | - | Unassigned | - |
| Otu27481 | 1 | k__Fungi(100);k__Fungi_unclassified(100);k__Fungi_unclassified(100);k__Fungi_unclassified(100);k__Fungi_unclassified(100);k__Fungi_unclassified(100);k__Fungi_unclassified(100); | - | - | - | - | - | - | - | Unassigned | - |
| Otu27484 | 1 | k__Fungi(100);k__Fungi_unclassified(100);k__Fungi_unclassified(100);k__Fungi_unclassified(100);k__Fungi_unclassified(100);k__Fungi_unclassified(100);k__Fungi_unclassified(100); | - | - | - | - | - | - | - | Unassigned | - |
| Otu27485 | 1 | k__Fungi(100);k__Fungi_unclassified(100);k__Fungi_unclassified(100);k__Fungi_unclassified(100);k__Fungi_unclassified(100);k__Fungi_unclassified(100);k__Fungi_unclassified(100); | - | - | - | - | - | - | - | Unassigned | - |
| Otu27486 | 1 | k__Fungi(100);k__Fungi_unclassified(100);k__Fungi_unclassified(100);k__Fungi_unclassified(100);k__Fungi_unclassified(100);k__Fungi_unclassified(100);k__Fungi_unclassified(100); | - | - | - | - | - | - | - | Unassigned | - |
| Otu27487 | 1 | k__Fungi(100);k__Fungi_unclassified(100);k__Fungi_unclassified(100);k__Fungi_unclassified(100);k__Fungi_unclassified(100);k__Fungi_unclassified(100);k__Fungi_unclassified(100); | - | - | - | - | - | - | - | Unassigned | - |
| Otu27489 | 1 | k__Fungi(100);k__Fungi_unclassified(100);k__Fungi_unclassified(100);k__Fungi_unclassified(100);k__Fungi_unclassified(100);k__Fungi_unclassified(100);k__Fungi_unclassified(100); | - | - | - | - | - | - | - | Unassigned | - |
| Otu27490 | 1 | k__Fungi(100);k__Fungi_unclassified(100);k__Fungi_unclassified(100);k__Fungi_unclassified(100);k__Fungi_unclassified(100);k__Fungi_unclassified(100);k__Fungi_unclassified(100); | - | - | - | - | - | - | - | Unassigned | - |
| Otu27491 | 1 | k__Fungi(100);k__Fungi_unclassified(100);k__Fungi_unclassified(100);k__Fungi_unclassified(100);k__Fungi_unclassified(100);k__Fungi_unclassified(100);k__Fungi_unclassified(100); | - | - | - | - | - | - | - | Unassigned | - |
| Otu27493 | 1 | k__Fungi(100);k__Fungi_unclassified(100);k__Fungi_unclassified(100);k__Fungi_unclassified(100);k__Fungi_unclassified(100);k__Fungi_unclassified(100);k__Fungi_unclassified(100); | - | - | - | - | - | - | - | Unassigned | - |
| Otu27495 | 1 | k__Fungi(100);p__Ascomycota(100);p__Ascomycota_unclassified(100);p__Ascomycota_unclassified(100);p__Ascomycota_unclassified(100);p__Ascomycota_unclassified(100);p__Ascomycota_unclassified(100); | - | - | - | - | - | - | - | Unassigned | - |
| Otu27497 | 1 | k__Fungi(100);k__Fungi_unclassified(100);k__Fungi_unclassified(100);k__Fungi_unclassified(100);k__Fungi_unclassified(100);k__Fungi_unclassified(100);k__Fungi_unclassified(100); | - | - | - | - | - | - | - | Unassigned | - |
| Otu27506 | 1 | k__Fungi(100);p__Ascomycota(100);p__Ascomycota_unclassified(100);p__Ascomycota_unclassified(100);p__Ascomycota_unclassified(100);p__Ascomycota_unclassified(100);p__Ascomycota_unclassified(100); | - | - | - | - | - | - | - | Unassigned | - |
| Otu27507 | 1 | k__Fungi(100);p__Ascomycota(100);p__Ascomycota_unclassified(100);p__Ascomycota_unclassified(100);p__Ascomycota_unclassified(100);p__Ascomycota_unclassified(100);p__Ascomycota_unclassified(100); | - | - | - | - | - | - | - | Unassigned | - |
| Otu27508 | 1 | k__Fungi(100);k__Fungi_unclassified(100);k__Fungi_unclassified(100);k__Fungi_unclassified(100);k__Fungi_unclassified(100);k__Fungi_unclassified(100);k__Fungi_unclassified(100); | - | - | - | - | - | - | - | Unassigned | - |
| Otu27510 | 1 | k__Fungi(100);p__Ascomycota(100);p__Ascomycota_unclassified(100);p__Ascomycota_unclassified(100);p__Ascomycota_unclassified(100);p__Ascomycota_unclassified(100);p__Ascomycota_unclassified(100); | - | - | - | - | - | - | - | Unassigned | - |
| Otu27511 | 1 | k__Fungi(100);p__Ascomycota(100);p__Ascomycota_unclassified(100);p__Ascomycota_unclassified(100);p__Ascomycota_unclassified(100);p__Ascomycota_unclassified(100);p__Ascomycota_unclassified(100); | - | - | - | - | - | - | - | Unassigned | - |
| Otu27512 | 1 | k__Fungi(100);k__Fungi_unclassified(100);k__Fungi_unclassified(100);k__Fungi_unclassified(100);k__Fungi_unclassified(100);k__Fungi_unclassified(100);k__Fungi_unclassified(100); | - | - | - | - | - | - | - | Unassigned | - |
| Otu27516 | 1 | k__Fungi(100);k__Fungi_unclassified(100);k__Fungi_unclassified(100);k__Fungi_unclassified(100);k__Fungi_unclassified(100);k__Fungi_unclassified(100);k__Fungi_unclassified(100); | - | - | - | - | - | - | - | Unassigned | - |
| Otu27520 | 1 | k__Fungi(100);p__Ascomycota(100);p__Ascomycota_unclassified(100);p__Ascomycota_unclassified(100);p__Ascomycota_unclassified(100);p__Ascomycota_unclassified(100);p__Ascomycota_unclassified(100); | - | - | - | - | - | - | - | Unassigned | - |
| Otu27521 | 1 | k__Fungi(100);p__Ascomycota(100);p__Ascomycota_unclassified(100);p__Ascomycota_unclassified(100);p__Ascomycota_unclassified(100);p__Ascomycota_unclassified(100);p__Ascomycota_unclassified(100); | - | - | - | - | - | - | - | Unassigned | - |
| Otu27522 | 1 | k__Fungi(100);k__Fungi_unclassified(100);k__Fungi_unclassified(100);k__Fungi_unclassified(100);k__Fungi_unclassified(100);k__Fungi_unclassified(100);k__Fungi_unclassified(100); | - | - | - | - | - | - | - | Unassigned | - |
| Otu27523 | 1 | k__Fungi(100);k__Fungi_unclassified(100);k__Fungi_unclassified(100);k__Fungi_unclassified(100);k__Fungi_unclassified(100);k__Fungi_unclassified(100);k__Fungi_unclassified(100); | - | - | - | - | - | - | - | Unassigned | - |
| Otu27524 | 1 | k__Fungi(100);k__Fungi_unclassified(100);k__Fungi_unclassified(100);k__Fungi_unclassified(100);k__Fungi_unclassified(100);k__Fungi_unclassified(100);k__Fungi_unclassified(100); | - | - | - | - | - | - | - | Unassigned | - |
| Otu27525 | 1 | k__Fungi(100);k__Fungi_unclassified(100);k__Fungi_unclassified(100);k__Fungi_unclassified(100);k__Fungi_unclassified(100);k__Fungi_unclassified(100);k__Fungi_unclassified(100); | - | - | - | - | - | - | - | Unassigned | - |
| Otu27526 | 1 | k__Fungi(100);p__Ascomycota(100);p__Ascomycota_unclassified(100);p__Ascomycota_unclassified(100);p__Ascomycota_unclassified(100);p__Ascomycota_unclassified(100);p__Ascomycota_unclassified(100); | - | - | - | - | - | - | - | Unassigned | - |
| Otu27528 | 1 | k__Fungi(100);k__Fungi_unclassified(100);k__Fungi_unclassified(100);k__Fungi_unclassified(100);k__Fungi_unclassified(100);k__Fungi_unclassified(100);k__Fungi_unclassified(100); | - | - | - | - | - | - | - | Unassigned | - |
| Otu27529 | 1 | k__Fungi(100);k__Fungi_unclassified(100);k__Fungi_unclassified(100);k__Fungi_unclassified(100);k__Fungi_unclassified(100);k__Fungi_unclassified(100);k__Fungi_unclassified(100); | - | - | - | - | - | - | - | Unassigned | - |
| Otu27530 | 1 | k__Fungi(100);k__Fungi_unclassified(100);k__Fungi_unclassified(100);k__Fungi_unclassified(100);k__Fungi_unclassified(100);k__Fungi_unclassified(100);k__Fungi_unclassified(100); | - | - | - | - | - | - | - | Unassigned | - |
| Otu27531 | 1 | k__Fungi(100);p__Ascomycota(100);p__Ascomycota_unclassified(100);p__Ascomycota_unclassified(100);p__Ascomycota_unclassified(100);p__Ascomycota_unclassified(100);p__Ascomycota_unclassified(100); | - | - | - | - | - | - | - | Unassigned | - |
| Otu27533 | 1 | k__Fungi(100);k__Fungi_unclassified(100);k__Fungi_unclassified(100);k__Fungi_unclassified(100);k__Fungi_unclassified(100);k__Fungi_unclassified(100);k__Fungi_unclassified(100); | - | - | - | - | - | - | - | Unassigned | - |
| Otu27534 | 1 | k__Fungi(100);k__Fungi_unclassified(100);k__Fungi_unclassified(100);k__Fungi_unclassified(100);k__Fungi_unclassified(100);k__Fungi_unclassified(100);k__Fungi_unclassified(100); | - | - | - | - | - | - | - | Unassigned | - |
| Otu27535 | 1 | k__Fungi(100);k__Fungi_unclassified(100);k__Fungi_unclassified(100);k__Fungi_unclassified(100);k__Fungi_unclassified(100);k__Fungi_unclassified(100);k__Fungi_unclassified(100); | - | - | - | - | - | - | - | Unassigned | - |
| Otu27536 | 1 | k__Fungi(100);k__Fungi_unclassified(100);k__Fungi_unclassified(100);k__Fungi_unclassified(100);k__Fungi_unclassified(100);k__Fungi_unclassified(100);k__Fungi_unclassified(100); | - | - | - | - | - | - | - | Unassigned | - |
| Otu27537 | 1 | k__Fungi(100);p__Ascomycota(100);p__Ascomycota_unclassified(100);p__Ascomycota_unclassified(100);p__Ascomycota_unclassified(100);p__Ascomycota_unclassified(100);p__Ascomycota_unclassified(100); | - | - | - | - | - | - | - | Unassigned | - |
| Otu27538 | 1 | k__Fungi(100);k__Fungi_unclassified(100);k__Fungi_unclassified(100);k__Fungi_unclassified(100);k__Fungi_unclassified(100);k__Fungi_unclassified(100);k__Fungi_unclassified(100); | - | - | - | - | - | - | - | Unassigned | - |
| Otu27539 | 1 | k__Fungi(100);p__Ascomycota(100);p__Ascomycota_unclassified(100);p__Ascomycota_unclassified(100);p__Ascomycota_unclassified(100);p__Ascomycota_unclassified(100);p__Ascomycota_unclassified(100); | - | - | - | - | - | - | - | Unassigned | - |
| Otu27541 | 1 | k__Fungi(100);k__Fungi_unclassified(100);k__Fungi_unclassified(100);k__Fungi_unclassified(100);k__Fungi_unclassified(100);k__Fungi_unclassified(100);k__Fungi_unclassified(100); | - | - | - | - | - | - | - | Unassigned | - |
| Otu27542 | 1 | k__Fungi(100);p__Ascomycota(100);p__Ascomycota_unclassified(100);p__Ascomycota_unclassified(100);p__Ascomycota_unclassified(100);p__Ascomycota_unclassified(100);p__Ascomycota_unclassified(100); | - | - | - | - | - | - | - | Unassigned | - |
| Otu27543 | 1 | k__Fungi(100);k__Fungi_unclassified(100);k__Fungi_unclassified(100);k__Fungi_unclassified(100);k__Fungi_unclassified(100);k__Fungi_unclassified(100);k__Fungi_unclassified(100); | - | - | - | - | - | - | - | Unassigned | - |
| Otu27544 | 1 | k__Fungi(100);p__Ascomycota(100);p__Ascomycota_unclassified(100);p__Ascomycota_unclassified(100);p__Ascomycota_unclassified(100);p__Ascomycota_unclassified(100);p__Ascomycota_unclassified(100); | - | - | - | - | - | - | - | Unassigned | - |
| Otu27545 | 1 | k__Fungi(100);p__Ascomycota(100);p__Ascomycota_unclassified(100);p__Ascomycota_unclassified(100);p__Ascomycota_unclassified(100);p__Ascomycota_unclassified(100);p__Ascomycota_unclassified(100); | - | - | - | - | - | - | - | Unassigned | - |
| Otu27546 | 1 | k__Fungi(100);k__Fungi_unclassified(100);k__Fungi_unclassified(100);k__Fungi_unclassified(100);k__Fungi_unclassified(100);k__Fungi_unclassified(100);k__Fungi_unclassified(100); | - | - | - | - | - | - | - | Unassigned | - |
| Otu27548 | 1 | k__Fungi(100);k__Fungi_unclassified(100);k__Fungi_unclassified(100);k__Fungi_unclassified(100);k__Fungi_unclassified(100);k__Fungi_unclassified(100);k__Fungi_unclassified(100); | - | - | - | - | - | - | - | Unassigned | - |
| Otu27549 | 1 | k__Fungi(100);k__Fungi_unclassified(100);k__Fungi_unclassified(100);k__Fungi_unclassified(100);k__Fungi_unclassified(100);k__Fungi_unclassified(100);k__Fungi_unclassified(100); | - | - | - | - | - | - | - | Unassigned | - |
| Otu27550 | 1 | k__Fungi(100);k__Fungi_unclassified(100);k__Fungi_unclassified(100);k__Fungi_unclassified(100);k__Fungi_unclassified(100);k__Fungi_unclassified(100);k__Fungi_unclassified(100); | - | - | - | - | - | - | - | Unassigned | - |
| Otu27551 | 1 | k__Fungi(100);p__Ascomycota(100);p__Ascomycota_unclassified(100);p__Ascomycota_unclassified(100);p__Ascomycota_unclassified(100);p__Ascomycota_unclassified(100);p__Ascomycota_unclassified(100); | - | - | - | - | - | - | - | Unassigned | - |
| Otu27553 | 1 | k__Fungi(100);k__Fungi_unclassified(100);k__Fungi_unclassified(100);k__Fungi_unclassified(100);k__Fungi_unclassified(100);k__Fungi_unclassified(100);k__Fungi_unclassified(100); | - | - | - | - | - | - | - | Unassigned | - |
| Otu27554 | 1 | k__Fungi(100);k__Fungi_unclassified(100);k__Fungi_unclassified(100);k__Fungi_unclassified(100);k__Fungi_unclassified(100);k__Fungi_unclassified(100);k__Fungi_unclassified(100); | - | - | - | - | - | - | - | Unassigned | - |
| Otu27555 | 1 | k__Fungi(100);k__Fungi_unclassified(100);k__Fungi_unclassified(100);k__Fungi_unclassified(100);k__Fungi_unclassified(100);k__Fungi_unclassified(100);k__Fungi_unclassified(100); | - | - | - | - | - | - | - | Unassigned | - |
| Otu27556 | 1 | k__Fungi(100);p__Ascomycota(100);p__Ascomycota_unclassified(100);p__Ascomycota_unclassified(100);p__Ascomycota_unclassified(100);p__Ascomycota_unclassified(100);p__Ascomycota_unclassified(100); | - | - | - | - | - | - | - | Unassigned | - |
| Otu27557 | 1 | k__Fungi(100);k__Fungi_unclassified(100);k__Fungi_unclassified(100);k__Fungi_unclassified(100);k__Fungi_unclassified(100);k__Fungi_unclassified(100);k__Fungi_unclassified(100); | - | - | - | - | - | - | - | Unassigned | - |
| Otu27560 | 1 | k__Fungi(100);k__Fungi_unclassified(100);k__Fungi_unclassified(100);k__Fungi_unclassified(100);k__Fungi_unclassified(100);k__Fungi_unclassified(100);k__Fungi_unclassified(100); | - | - | - | - | - | - | - | Unassigned | - |
| Otu27561 | 1 | k__Fungi(100);k__Fungi_unclassified(100);k__Fungi_unclassified(100);k__Fungi_unclassified(100);k__Fungi_unclassified(100);k__Fungi_unclassified(100);k__Fungi_unclassified(100); | - | - | - | - | - | - | - | Unassigned | - |
| Otu27562 | 1 | k__Fungi(100);p__Ascomycota(100);p__Ascomycota_unclassified(100);p__Ascomycota_unclassified(100);p__Ascomycota_unclassified(100);p__Ascomycota_unclassified(100);p__Ascomycota_unclassified(100); | - | - | - | - | - | - | - | Unassigned | - |
| Otu27563 | 1 | k__Fungi(100);p__Ascomycota(100);p__Ascomycota_unclassified(100);p__Ascomycota_unclassified(100);p__Ascomycota_unclassified(100);p__Ascomycota_unclassified(100);p__Ascomycota_unclassified(100); | - | - | - | - | - | - | - | Unassigned | - |
| Otu27571 | 1 | k__Fungi(100);k__Fungi_unclassified(100);k__Fungi_unclassified(100);k__Fungi_unclassified(100);k__Fungi_unclassified(100);k__Fungi_unclassified(100);k__Fungi_unclassified(100); | - | - | - | - | - | - | - | Unassigned | - |
| Otu27572 | 1 | k__Fungi(100);p__Ascomycota(100);p__Ascomycota_unclassified(100);p__Ascomycota_unclassified(100);p__Ascomycota_unclassified(100);p__Ascomycota_unclassified(100);p__Ascomycota_unclassified(100); | - | - | - | - | - | - | - | Unassigned | - |
| Otu27575 | 1 | k__Fungi(100);p__Ascomycota(100);p__Ascomycota_unclassified(100);p__Ascomycota_unclassified(100);p__Ascomycota_unclassified(100);p__Ascomycota_unclassified(100);p__Ascomycota_unclassified(100); | - | - | - | - | - | - | - | Unassigned | - |
| Otu27576 | 1 | k__Fungi(100);p__Ascomycota(100);p__Ascomycota_unclassified(100);p__Ascomycota_unclassified(100);p__Ascomycota_unclassified(100);p__Ascomycota_unclassified(100);p__Ascomycota_unclassified(100); | - | - | - | - | - | - | - | Unassigned | - |
| Otu27577 | 1 | k__Fungi(100);p__Ascomycota(100);p__Ascomycota_unclassified(100);p__Ascomycota_unclassified(100);p__Ascomycota_unclassified(100);p__Ascomycota_unclassified(100);p__Ascomycota_unclassified(100); | - | - | - | - | - | - | - | Unassigned | - |
| Otu27578 | 1 | k__Fungi(100);k__Fungi_unclassified(100);k__Fungi_unclassified(100);k__Fungi_unclassified(100);k__Fungi_unclassified(100);k__Fungi_unclassified(100);k__Fungi_unclassified(100); | - | - | - | - | - | - | - | Unassigned | - |
| Otu27579 | 1 | k__Fungi(100);p__Ascomycota(100);p__Ascomycota_unclassified(100);p__Ascomycota_unclassified(100);p__Ascomycota_unclassified(100);p__Ascomycota_unclassified(100);p__Ascomycota_unclassified(100); | - | - | - | - | - | - | - | Unassigned | - |
| Otu27580 | 1 | k__Fungi(100);k__Fungi_unclassified(100);k__Fungi_unclassified(100);k__Fungi_unclassified(100);k__Fungi_unclassified(100);k__Fungi_unclassified(100);k__Fungi_unclassified(100); | - | - | - | - | - | - | - | Unassigned | - |
| Otu27581 | 1 | k__Fungi(100);k__Fungi_unclassified(100);k__Fungi_unclassified(100);k__Fungi_unclassified(100);k__Fungi_unclassified(100);k__Fungi_unclassified(100);k__Fungi_unclassified(100); | - | - | - | - | - | - | - | Unassigned | - |
| Otu27582 | 1 | k__Fungi(100);k__Fungi_unclassified(100);k__Fungi_unclassified(100);k__Fungi_unclassified(100);k__Fungi_unclassified(100);k__Fungi_unclassified(100);k__Fungi_unclassified(100); | - | - | - | - | - | - | - | Unassigned | - |
| Otu27583 | 1 | k__Fungi(100);k__Fungi_unclassified(100);k__Fungi_unclassified(100);k__Fungi_unclassified(100);k__Fungi_unclassified(100);k__Fungi_unclassified(100);k__Fungi_unclassified(100); | - | - | - | - | - | - | - | Unassigned | - |
| Otu27584 | 1 | k__Fungi(100);k__Fungi_unclassified(100);k__Fungi_unclassified(100);k__Fungi_unclassified(100);k__Fungi_unclassified(100);k__Fungi_unclassified(100);k__Fungi_unclassified(100); | - | - | - | - | - | - | - | Unassigned | - |
| Otu27585 | 1 | k__Fungi(100);k__Fungi_unclassified(100);k__Fungi_unclassified(100);k__Fungi_unclassified(100);k__Fungi_unclassified(100);k__Fungi_unclassified(100);k__Fungi_unclassified(100); | - | - | - | - | - | - | - | Unassigned | - |
| Otu27586 | 1 | k__Fungi(100);k__Fungi_unclassified(100);k__Fungi_unclassified(100);k__Fungi_unclassified(100);k__Fungi_unclassified(100);k__Fungi_unclassified(100);k__Fungi_unclassified(100); | - | - | - | - | - | - | - | Unassigned | - |
| Otu27588 | 1 | k__Fungi(100);k__Fungi_unclassified(100);k__Fungi_unclassified(100);k__Fungi_unclassified(100);k__Fungi_unclassified(100);k__Fungi_unclassified(100);k__Fungi_unclassified(100); | - | - | - | - | - | - | - | Unassigned | - |
| Otu27589 | 1 | k__Fungi(100);k__Fungi_unclassified(100);k__Fungi_unclassified(100);k__Fungi_unclassified(100);k__Fungi_unclassified(100);k__Fungi_unclassified(100);k__Fungi_unclassified(100); | - | - | - | - | - | - | - | Unassigned | - |
| Otu27590 | 1 | k__Fungi(100);p__Ascomycota(100);p__Ascomycota_unclassified(100);p__Ascomycota_unclassified(100);p__Ascomycota_unclassified(100);p__Ascomycota_unclassified(100);p__Ascomycota_unclassified(100); | - | - | - | - | - | - | - | Unassigned | - |
| Otu27591 | 1 | k__Fungi(100);k__Fungi_unclassified(100);k__Fungi_unclassified(100);k__Fungi_unclassified(100);k__Fungi_unclassified(100);k__Fungi_unclassified(100);k__Fungi_unclassified(100); | - | - | - | - | - | - | - | Unassigned | - |
| Otu27592 | 1 | k__Fungi(100);k__Fungi_unclassified(100);k__Fungi_unclassified(100);k__Fungi_unclassified(100);k__Fungi_unclassified(100);k__Fungi_unclassified(100);k__Fungi_unclassified(100); | - | - | - | - | - | - | - | Unassigned | - |
| Otu27596 | 1 | k__Fungi(100);k__Fungi_unclassified(100);k__Fungi_unclassified(100);k__Fungi_unclassified(100);k__Fungi_unclassified(100);k__Fungi_unclassified(100);k__Fungi_unclassified(100); | - | - | - | - | - | - | - | Unassigned | - |
| Otu27597 | 1 | k__Fungi(100);p__Ascomycota(100);p__Ascomycota_unclassified(100);p__Ascomycota_unclassified(100);p__Ascomycota_unclassified(100);p__Ascomycota_unclassified(100);p__Ascomycota_unclassified(100); | - | - | - | - | - | - | - | Unassigned | - |
| Otu27598 | 1 | k__Fungi(100);p__Ascomycota(100);p__Ascomycota_unclassified(100);p__Ascomycota_unclassified(100);p__Ascomycota_unclassified(100);p__Ascomycota_unclassified(100);p__Ascomycota_unclassified(100); | - | - | - | - | - | - | - | Unassigned | - |
| Otu27599 | 1 | k__Fungi(100);p__Ascomycota(100);p__Ascomycota_unclassified(100);p__Ascomycota_unclassified(100);p__Ascomycota_unclassified(100);p__Ascomycota_unclassified(100);p__Ascomycota_unclassified(100); | - | - | - | - | - | - | - | Unassigned | - |
| Otu27601 | 1 | k__Fungi(100);k__Fungi_unclassified(100);k__Fungi_unclassified(100);k__Fungi_unclassified(100);k__Fungi_unclassified(100);k__Fungi_unclassified(100);k__Fungi_unclassified(100); | - | - | - | - | - | - | - | Unassigned | - |
| Otu27602 | 1 | k__Fungi(100);p__Ascomycota(100);p__Ascomycota_unclassified(100);p__Ascomycota_unclassified(100);p__Ascomycota_unclassified(100);p__Ascomycota_unclassified(100);p__Ascomycota_unclassified(100); | - | - | - | - | - | - | - | Unassigned | - |
| Otu27605 | 1 | k__Fungi(100);p__Ascomycota(100);p__Ascomycota_unclassified(100);p__Ascomycota_unclassified(100);p__Ascomycota_unclassified(100);p__Ascomycota_unclassified(100);p__Ascomycota_unclassified(100); | - | - | - | - | - | - | - | Unassigned | - |
| Otu27606 | 1 | k__Fungi(100);p__Ascomycota(100);p__Ascomycota_unclassified(100);p__Ascomycota_unclassified(100);p__Ascomycota_unclassified(100);p__Ascomycota_unclassified(100);p__Ascomycota_unclassified(100); | - | - | - | - | - | - | - | Unassigned | - |
| Otu27607 | 1 | k__Fungi(100);k__Fungi_unclassified(100);k__Fungi_unclassified(100);k__Fungi_unclassified(100);k__Fungi_unclassified(100);k__Fungi_unclassified(100);k__Fungi_unclassified(100); | - | - | - | - | - | - | - | Unassigned | - |
| Otu27610 | 1 | k__Fungi(100);p__Ascomycota(100);p__Ascomycota_unclassified(100);p__Ascomycota_unclassified(100);p__Ascomycota_unclassified(100);p__Ascomycota_unclassified(100);p__Ascomycota_unclassified(100); | - | - | - | - | - | - | - | Unassigned | - |
| Otu27612 | 1 | k__Fungi(100);k__Fungi_unclassified(100);k__Fungi_unclassified(100);k__Fungi_unclassified(100);k__Fungi_unclassified(100);k__Fungi_unclassified(100);k__Fungi_unclassified(100); | - | - | - | - | - | - | - | Unassigned | - |
| Otu27613 | 1 | k__Fungi(100);k__Fungi_unclassified(100);k__Fungi_unclassified(100);k__Fungi_unclassified(100);k__Fungi_unclassified(100);k__Fungi_unclassified(100);k__Fungi_unclassified(100); | - | - | - | - | - | - | - | Unassigned | - |
| Otu27614 | 1 | k__Fungi(100);k__Fungi_unclassified(100);k__Fungi_unclassified(100);k__Fungi_unclassified(100);k__Fungi_unclassified(100);k__Fungi_unclassified(100);k__Fungi_unclassified(100); | - | - | - | - | - | - | - | Unassigned | - |
| Otu27615 | 1 | k__Fungi(100);k__Fungi_unclassified(100);k__Fungi_unclassified(100);k__Fungi_unclassified(100);k__Fungi_unclassified(100);k__Fungi_unclassified(100);k__Fungi_unclassified(100); | - | - | - | - | - | - | - | Unassigned | - |
| Otu27618 | 1 | k__Fungi(100);k__Fungi_unclassified(100);k__Fungi_unclassified(100);k__Fungi_unclassified(100);k__Fungi_unclassified(100);k__Fungi_unclassified(100);k__Fungi_unclassified(100); | - | - | - | - | - | - | - | Unassigned | - |
| Otu27619 | 1 | k__Fungi(100);p__Ascomycota(100);p__Ascomycota_unclassified(100);p__Ascomycota_unclassified(100);p__Ascomycota_unclassified(100);p__Ascomycota_unclassified(100);p__Ascomycota_unclassified(100); | - | - | - | - | - | - | - | Unassigned | - |
| Otu27620 | 1 | k__Fungi(100);k__Fungi_unclassified(100);k__Fungi_unclassified(100);k__Fungi_unclassified(100);k__Fungi_unclassified(100);k__Fungi_unclassified(100);k__Fungi_unclassified(100); | - | - | - | - | - | - | - | Unassigned | - |
| Otu27621 | 1 | k__Fungi(100);p__Ascomycota(100);p__Ascomycota_unclassified(100);p__Ascomycota_unclassified(100);p__Ascomycota_unclassified(100);p__Ascomycota_unclassified(100);p__Ascomycota_unclassified(100); | - | - | - | - | - | - | - | Unassigned | - |
| Otu27623 | 1 | k__Fungi(100);p__Ascomycota(100);p__Ascomycota_unclassified(100);p__Ascomycota_unclassified(100);p__Ascomycota_unclassified(100);p__Ascomycota_unclassified(100);p__Ascomycota_unclassified(100); | - | - | - | - | - | - | - | Unassigned | - |
| Otu27627 | 1 | k__Fungi(100);k__Fungi_unclassified(100);k__Fungi_unclassified(100);k__Fungi_unclassified(100);k__Fungi_unclassified(100);k__Fungi_unclassified(100);k__Fungi_unclassified(100); | - | - | - | - | - | - | - | Unassigned | - |
| Otu27628 | 1 | k__Fungi(100);k__Fungi_unclassified(100);k__Fungi_unclassified(100);k__Fungi_unclassified(100);k__Fungi_unclassified(100);k__Fungi_unclassified(100);k__Fungi_unclassified(100); | - | - | - | - | - | - | - | Unassigned | - |
| Otu27629 | 1 | k__Fungi(100);p__Ascomycota(100);p__Ascomycota_unclassified(100);p__Ascomycota_unclassified(100);p__Ascomycota_unclassified(100);p__Ascomycota_unclassified(100);p__Ascomycota_unclassified(100); | - | - | - | - | - | - | - | Unassigned | - |
| Otu27630 | 1 | k__Fungi(100);k__Fungi_unclassified(100);k__Fungi_unclassified(100);k__Fungi_unclassified(100);k__Fungi_unclassified(100);k__Fungi_unclassified(100);k__Fungi_unclassified(100); | - | - | - | - | - | - | - | Unassigned | - |
| Otu27631 | 1 | k__Fungi(100);k__Fungi_unclassified(100);k__Fungi_unclassified(100);k__Fungi_unclassified(100);k__Fungi_unclassified(100);k__Fungi_unclassified(100);k__Fungi_unclassified(100); | - | - | - | - | - | - | - | Unassigned | - |
| Otu27632 | 1 | k__Fungi(100);k__Fungi_unclassified(100);k__Fungi_unclassified(100);k__Fungi_unclassified(100);k__Fungi_unclassified(100);k__Fungi_unclassified(100);k__Fungi_unclassified(100); | - | - | - | - | - | - | - | Unassigned | - |
| Otu27634 | 1 | k__Fungi(100);k__Fungi_unclassified(100);k__Fungi_unclassified(100);k__Fungi_unclassified(100);k__Fungi_unclassified(100);k__Fungi_unclassified(100);k__Fungi_unclassified(100); | - | - | - | - | - | - | - | Unassigned | - |
| Otu27635 | 1 | k__Fungi(100);p__Ascomycota(100);p__Ascomycota_unclassified(100);p__Ascomycota_unclassified(100);p__Ascomycota_unclassified(100);p__Ascomycota_unclassified(100);p__Ascomycota_unclassified(100); | - | - | - | - | - | - | - | Unassigned | - |
| Otu27639 | 1 | k__Fungi(100);k__Fungi_unclassified(100);k__Fungi_unclassified(100);k__Fungi_unclassified(100);k__Fungi_unclassified(100);k__Fungi_unclassified(100);k__Fungi_unclassified(100); | - | - | - | - | - | - | - | Unassigned | - |
| Otu27644 | 1 | k__Fungi(100);k__Fungi_unclassified(100);k__Fungi_unclassified(100);k__Fungi_unclassified(100);k__Fungi_unclassified(100);k__Fungi_unclassified(100);k__Fungi_unclassified(100); | - | - | - | - | - | - | - | Unassigned | - |
| Otu27645 | 1 | k__Fungi(100);p__Ascomycota(100);p__Ascomycota_unclassified(100);p__Ascomycota_unclassified(100);p__Ascomycota_unclassified(100);p__Ascomycota_unclassified(100);p__Ascomycota_unclassified(100); | - | - | - | - | - | - | - | Unassigned | - |
| Otu27654 | 1 | k__Fungi(100);k__Fungi_unclassified(100);k__Fungi_unclassified(100);k__Fungi_unclassified(100);k__Fungi_unclassified(100);k__Fungi_unclassified(100);k__Fungi_unclassified(100); | - | - | - | - | - | - | - | Unassigned | - |
| Otu27659 | 1 | k__Fungi(100);k__Fungi_unclassified(100);k__Fungi_unclassified(100);k__Fungi_unclassified(100);k__Fungi_unclassified(100);k__Fungi_unclassified(100);k__Fungi_unclassified(100); | - | - | - | - | - | - | - | Unassigned | - |
| Otu27661 | 1 | k__Fungi(100);p__Ascomycota(100);p__Ascomycota_unclassified(100);p__Ascomycota_unclassified(100);p__Ascomycota_unclassified(100);p__Ascomycota_unclassified(100);p__Ascomycota_unclassified(100); | - | - | - | - | - | - | - | Unassigned | - |
| Otu27662 | 1 | k__Fungi(100);k__Fungi_unclassified(100);k__Fungi_unclassified(100);k__Fungi_unclassified(100);k__Fungi_unclassified(100);k__Fungi_unclassified(100);k__Fungi_unclassified(100); | - | - | - | - | - | - | - | Unassigned | - |
| Otu27666 | 1 | k__Fungi(100);k__Fungi_unclassified(100);k__Fungi_unclassified(100);k__Fungi_unclassified(100);k__Fungi_unclassified(100);k__Fungi_unclassified(100);k__Fungi_unclassified(100); | - | - | - | - | - | - | - | Unassigned | - |
| Otu27668 | 1 | k__Fungi(100);p__Ascomycota(100);p__Ascomycota_unclassified(100);p__Ascomycota_unclassified(100);p__Ascomycota_unclassified(100);p__Ascomycota_unclassified(100);p__Ascomycota_unclassified(100); | - | - | - | - | - | - | - | Unassigned | - |
| Otu27669 | 1 | k__Fungi(100);k__Fungi_unclassified(100);k__Fungi_unclassified(100);k__Fungi_unclassified(100);k__Fungi_unclassified(100);k__Fungi_unclassified(100);k__Fungi_unclassified(100); | - | - | - | - | - | - | - | Unassigned | - |
| Otu27670 | 1 | k__Fungi(100);k__Fungi_unclassified(100);k__Fungi_unclassified(100);k__Fungi_unclassified(100);k__Fungi_unclassified(100);k__Fungi_unclassified(100);k__Fungi_unclassified(100); | - | - | - | - | - | - | - | Unassigned | - |
| Otu27673 | 1 | k__Fungi(100);k__Fungi_unclassified(100);k__Fungi_unclassified(100);k__Fungi_unclassified(100);k__Fungi_unclassified(100);k__Fungi_unclassified(100);k__Fungi_unclassified(100); | - | - | - | - | - | - | - | Unassigned | - |
| Otu27674 | 1 | k__Fungi(100);k__Fungi_unclassified(100);k__Fungi_unclassified(100);k__Fungi_unclassified(100);k__Fungi_unclassified(100);k__Fungi_unclassified(100);k__Fungi_unclassified(100); | - | - | - | - | - | - | - | Unassigned | - |
| Otu27675 | 1 | k__Fungi(100);k__Fungi_unclassified(100);k__Fungi_unclassified(100);k__Fungi_unclassified(100);k__Fungi_unclassified(100);k__Fungi_unclassified(100);k__Fungi_unclassified(100); | - | - | - | - | - | - | - | Unassigned | - |
| Otu27678 | 1 | k__Fungi(100);k__Fungi_unclassified(100);k__Fungi_unclassified(100);k__Fungi_unclassified(100);k__Fungi_unclassified(100);k__Fungi_unclassified(100);k__Fungi_unclassified(100); | - | - | - | - | - | - | - | Unassigned | - |
| Otu27681 | 1 | k__Fungi(100);p__Ascomycota(100);p__Ascomycota_unclassified(100);p__Ascomycota_unclassified(100);p__Ascomycota_unclassified(100);p__Ascomycota_unclassified(100);p__Ascomycota_unclassified(100); | - | - | - | - | - | - | - | Unassigned | - |
| Otu27682 | 1 | k__Fungi(100);k__Fungi_unclassified(100);k__Fungi_unclassified(100);k__Fungi_unclassified(100);k__Fungi_unclassified(100);k__Fungi_unclassified(100);k__Fungi_unclassified(100); | - | - | - | - | - | - | - | Unassigned | - |
| Otu27684 | 1 | k__Fungi(100);k__Fungi_unclassified(100);k__Fungi_unclassified(100);k__Fungi_unclassified(100);k__Fungi_unclassified(100);k__Fungi_unclassified(100);k__Fungi_unclassified(100); | - | - | - | - | - | - | - | Unassigned | - |
| Otu27685 | 1 | k__Fungi(100);k__Fungi_unclassified(100);k__Fungi_unclassified(100);k__Fungi_unclassified(100);k__Fungi_unclassified(100);k__Fungi_unclassified(100);k__Fungi_unclassified(100); | - | - | - | - | - | - | - | Unassigned | - |
| Otu27687 | 1 | k__Fungi(100);k__Fungi_unclassified(100);k__Fungi_unclassified(100);k__Fungi_unclassified(100);k__Fungi_unclassified(100);k__Fungi_unclassified(100);k__Fungi_unclassified(100); | - | - | - | - | - | - | - | Unassigned | - |
| Otu27688 | 1 | k__Fungi(100);k__Fungi_unclassified(100);k__Fungi_unclassified(100);k__Fungi_unclassified(100);k__Fungi_unclassified(100);k__Fungi_unclassified(100);k__Fungi_unclassified(100); | - | - | - | - | - | - | - | Unassigned | - |
| Otu27691 | 1 | k__Fungi(100);k__Fungi_unclassified(100);k__Fungi_unclassified(100);k__Fungi_unclassified(100);k__Fungi_unclassified(100);k__Fungi_unclassified(100);k__Fungi_unclassified(100); | - | - | - | - | - | - | - | Unassigned | - |
| Otu27692 | 1 | k__Fungi(100);p__Ascomycota(100);p__Ascomycota_unclassified(100);p__Ascomycota_unclassified(100);p__Ascomycota_unclassified(100);p__Ascomycota_unclassified(100);p__Ascomycota_unclassified(100); | - | - | - | - | - | - | - | Unassigned | - |
| Otu27693 | 1 | k__Fungi(100);k__Fungi_unclassified(100);k__Fungi_unclassified(100);k__Fungi_unclassified(100);k__Fungi_unclassified(100);k__Fungi_unclassified(100);k__Fungi_unclassified(100); | - | - | - | - | - | - | - | Unassigned | - |
| Otu27694 | 1 | k__Fungi(100);k__Fungi_unclassified(100);k__Fungi_unclassified(100);k__Fungi_unclassified(100);k__Fungi_unclassified(100);k__Fungi_unclassified(100);k__Fungi_unclassified(100); | - | - | - | - | - | - | - | Unassigned | - |
| Otu27695 | 1 | k__Fungi(100);k__Fungi_unclassified(100);k__Fungi_unclassified(100);k__Fungi_unclassified(100);k__Fungi_unclassified(100);k__Fungi_unclassified(100);k__Fungi_unclassified(100); | - | - | - | - | - | - | - | Unassigned | - |

| | | | | | | | | | | | |
|---|---|---|---|---|---|---|---|---|---|---|---|
| Otu27696 | 1 | k__Fungi(100);p__Ascomycota(100);p__Ascomycota_unclassified(100);p__Ascomycota_unclassified(100);p__Ascomycota_unclassified(100);p__Ascomycota_unclassified(100);p__Ascomycota_unclassified(100); | - | - | - | - | - | - | - | Unassigned | - |
| Otu27697 | 1 | k__Fungi(100);p__Ascomycota(100);p__Ascomycota_unclassified(100);p__Ascomycota_unclassified(100);p__Ascomycota_unclassified(100);p__Ascomycota_unclassified(100);p__Ascomycota_unclassified(100); | - | - | - | - | - | - | - | Unassigned | - |
| Otu27698 | 1 | k__Fungi(100);k__Fungi_unclassified(100);k__Fungi_unclassified(100);k__Fungi_unclassified(100);k__Fungi_unclassified(100);k__Fungi_unclassified(100);k__Fungi_unclassified(100); | - | - | - | - | - | - | - | Unassigned | - |
| Otu27699 | 1 | k__Fungi(100);k__Fungi_unclassified(100);k__Fungi_unclassified(100);k__Fungi_unclassified(100);k__Fungi_unclassified(100);k__Fungi_unclassified(100);k__Fungi_unclassified(100); | - | - | - | - | - | - | - | Unassigned | - |
| Otu27700 | 1 | k__Fungi(100);k__Fungi_unclassified(100);k__Fungi_unclassified(100);k__Fungi_unclassified(100);k__Fungi_unclassified(100);k__Fungi_unclassified(100);k__Fungi_unclassified(100); | - | - | - | - | - | - | - | Unassigned | - |
| Otu27701 | 1 | k__Fungi(100);k__Fungi_unclassified(100);k__Fungi_unclassified(100);k__Fungi_unclassified(100);k__Fungi_unclassified(100);k__Fungi_unclassified(100);k__Fungi_unclassified(100); | - | - | - | - | - | - | - | Unassigned | - |
| Otu27702 | 1 | k__Fungi(100);k__Fungi_unclassified(100);k__Fungi_unclassified(100);k__Fungi_unclassified(100);k__Fungi_unclassified(100);k__Fungi_unclassified(100);k__Fungi_unclassified(100); | - | - | - | - | - | - | - | Unassigned | - |
| Otu27703 | 1 | k__Fungi(100);k__Fungi_unclassified(100);k__Fungi_unclassified(100);k__Fungi_unclassified(100);k__Fungi_unclassified(100);k__Fungi_unclassified(100);k__Fungi_unclassified(100); | - | - | - | - | - | - | - | Unassigned | - |
| Otu27704 | 1 | k__Fungi(100);k__Fungi_unclassified(100);k__Fungi_unclassified(100);k__Fungi_unclassified(100);k__Fungi_unclassified(100);k__Fungi_unclassified(100);k__Fungi_unclassified(100); | - | - | - | - | - | - | - | Unassigned | - |
| Otu27706 | 1 | k__Fungi(100);p__Ascomycota(100);p__Ascomycota_unclassified(100);p__Ascomycota_unclassified(100);p__Ascomycota_unclassified(100);p__Ascomycota_unclassified(100);p__Ascomycota_unclassified(100); | - | - | - | - | - | - | - | Unassigned | - |
| Otu27707 | 1 | k__Fungi(100);k__Fungi_unclassified(100);k__Fungi_unclassified(100);k__Fungi_unclassified(100);k__Fungi_unclassified(100);k__Fungi_unclassified(100);k__Fungi_unclassified(100); | - | - | - | - | - | - | - | Unassigned | - |
| Otu27708 | 1 | k__Fungi(100);k__Fungi_unclassified(100);k__Fungi_unclassified(100);k__Fungi_unclassified(100);k__Fungi_unclassified(100);k__Fungi_unclassified(100);k__Fungi_unclassified(100); | - | - | - | - | - | - | - | Unassigned | - |
| Otu27709 | 1 | k__Fungi(100);k__Fungi_unclassified(100);k__Fungi_unclassified(100);k__Fungi_unclassified(100);k__Fungi_unclassified(100);k__Fungi_unclassified(100);k__Fungi_unclassified(100); | - | - | - | - | - | - | - | Unassigned | - |
| Otu27710 | 1 | k__Fungi(100);k__Fungi_unclassified(100);k__Fungi_unclassified(100);k__Fungi_unclassified(100);k__Fungi_unclassified(100);k__Fungi_unclassified(100);k__Fungi_unclassified(100); | - | - | - | - | - | - | - | Unassigned | - |
| Otu27711 | 1 | k__Fungi(100);k__Fungi_unclassified(100);k__Fungi_unclassified(100);k__Fungi_unclassified(100);k__Fungi_unclassified(100);k__Fungi_unclassified(100);k__Fungi_unclassified(100); | - | - | - | - | - | - | - | Unassigned | - |
| Otu27712 | 1 | k__Fungi(100);k__Fungi_unclassified(100);k__Fungi_unclassified(100);k__Fungi_unclassified(100);k__Fungi_unclassified(100);k__Fungi_unclassified(100);k__Fungi_unclassified(100); | - | - | - | - | - | - | - | Unassigned | - |
| Otu27713 | 1 | k__Fungi(100);k__Fungi_unclassified(100);k__Fungi_unclassified(100);k__Fungi_unclassified(100);k__Fungi_unclassified(100);k__Fungi_unclassified(100);k__Fungi_unclassified(100); | - | - | - | - | - | - | - | Unassigned | - |
| Otu27714 | 1 | k__Fungi(100);p__Ascomycota(100);p__Ascomycota_unclassified(100);p__Ascomycota_unclassified(100);p__Ascomycota_unclassified(100);p__Ascomycota_unclassified(100);p__Ascomycota_unclassified(100); | - | - | - | - | - | - | - | Unassigned | - |
| Otu27715 | 1 | k__Fungi(100);k__Fungi_unclassified(100);k__Fungi_unclassified(100);k__Fungi_unclassified(100);k__Fungi_unclassified(100);k__Fungi_unclassified(100);k__Fungi_unclassified(100); | - | - | - | - | - | - | - | Unassigned | - |
| Otu27716 | 1 | k__Fungi(100);p__Ascomycota(100);p__Ascomycota_unclassified(100);p__Ascomycota_unclassified(100);p__Ascomycota_unclassified(100);p__Ascomycota_unclassified(100);p__Ascomycota_unclassified(100); | - | - | - | - | - | - | - | Unassigned | - |
| Otu27717 | 1 | k__Fungi(100);k__Fungi_unclassified(100);k__Fungi_unclassified(100);k__Fungi_unclassified(100);k__Fungi_unclassified(100);k__Fungi_unclassified(100);k__Fungi_unclassified(100); | - | - | - | - | - | - | - | Unassigned | - |
| Otu27718 | 1 | k__Fungi(100);k__Fungi_unclassified(100);k__Fungi_unclassified(100);k__Fungi_unclassified(100);k__Fungi_unclassified(100);k__Fungi_unclassified(100);k__Fungi_unclassified(100); | - | - | - | - | - | - | - | Unassigned | - |
| Otu27720 | 1 | k__Fungi(100);k__Fungi_unclassified(100);k__Fungi_unclassified(100);k__Fungi_unclassified(100);k__Fungi_unclassified(100);k__Fungi_unclassified(100);k__Fungi_unclassified(100); | - | - | - | - | - | - | - | Unassigned | - |
| Otu27721 | 1 | k__Fungi(100);p__Ascomycota(100);p__Ascomycota_unclassified(100);p__Ascomycota_unclassified(100);p__Ascomycota_unclassified(100);p__Ascomycota_unclassified(100);p__Ascomycota_unclassified(100); | - | - | - | - | - | - | - | Unassigned | - |
| Otu27722 | 1 | k__Fungi(100);k__Fungi_unclassified(100);k__Fungi_unclassified(100);k__Fungi_unclassified(100);k__Fungi_unclassified(100);k__Fungi_unclassified(100);k__Fungi_unclassified(100); | - | - | - | - | - | - | - | Unassigned | - |
| Otu27723 | 1 | k__Fungi(100);k__Fungi_unclassified(100);k__Fungi_unclassified(100);k__Fungi_unclassified(100);k__Fungi_unclassified(100);k__Fungi_unclassified(100);k__Fungi_unclassified(100); | - | - | - | - | - | - | - | Unassigned | - |
| Otu27725 | 1 | k__Fungi(100);p__Ascomycota(100);p__Ascomycota_unclassified(100);p__Ascomycota_unclassified(100);p__Ascomycota_unclassified(100);p__Ascomycota_unclassified(100);p__Ascomycota_unclassified(100); | - | - | - | - | - | - | - | Unassigned | - |
| Otu27727 | 1 | k__Fungi(100);k__Fungi_unclassified(100);k__Fungi_unclassified(100);k__Fungi_unclassified(100);k__Fungi_unclassified(100);k__Fungi_unclassified(100);k__Fungi_unclassified(100); | - | - | - | - | - | - | - | Unassigned | - |
| Otu27729 | 1 | k__Fungi(100);p__Ascomycota(100);p__Ascomycota_unclassified(100);p__Ascomycota_unclassified(100);p__Ascomycota_unclassified(100);p__Ascomycota_unclassified(100);p__Ascomycota_unclassified(100); | - | - | - | - | - | - | - | Unassigned | - |
| Otu27730 | 1 | k__Fungi(100);k__Fungi_unclassified(100);k__Fungi_unclassified(100);k__Fungi_unclassified(100);k__Fungi_unclassified(100);k__Fungi_unclassified(100);k__Fungi_unclassified(100); | - | - | - | - | - | - | - | Unassigned | - |
| Otu27731 | 1 | k__Fungi(100);k__Fungi_unclassified(100);k__Fungi_unclassified(100);k__Fungi_unclassified(100);k__Fungi_unclassified(100);k__Fungi_unclassified(100);k__Fungi_unclassified(100); | - | - | - | - | - | - | - | Unassigned | - |
| Otu27732 | 1 | k__Fungi(100);p__Ascomycota(100);p__Ascomycota_unclassified(100);p__Ascomycota_unclassified(100);p__Ascomycota_unclassified(100);p__Ascomycota_unclassified(100);p__Ascomycota_unclassified(100); | - | - | - | - | - | - | - | Unassigned | - |
| Otu27736 | 1 | k__Fungi(100);p__Ascomycota(100);c__Dothideomycetes(100);o__Capnodiales(100);o__Capnodiales_unclassified(100);o__Capnodiales_unclassified(100);o__Capnodiales_unclassified(100); | - | - | - | - | - | - | - | Unassigned | - |
| Otu27737 | 1 | k__Fungi(100);k__Fungi_unclassified(100);k__Fungi_unclassified(100);k__Fungi_unclassified(100);k__Fungi_unclassified(100);k__Fungi_unclassified(100);k__Fungi_unclassified(100); | - | - | - | - | - | - | - | Unassigned | - |
| Otu27740 | 1 | k__Fungi(100);p__Ascomycota(100);p__Ascomycota_unclassified(100);p__Ascomycota_unclassified(100);p__Ascomycota_unclassified(100);p__Ascomycota_unclassified(100);p__Ascomycota_unclassified(100); | - | - | - | - | - | - | - | Unassigned | - |
| Otu27741 | 1 | k__Fungi(100);k__Fungi_unclassified(100);k__Fungi_unclassified(100);k__Fungi_unclassified(100);k__Fungi_unclassified(100);k__Fungi_unclassified(100);k__Fungi_unclassified(100); | - | - | - | - | - | - | - | Unassigned | - |
| Otu27743 | 1 | k__Fungi(100);k__Fungi_unclassified(100);k__Fungi_unclassified(100);k__Fungi_unclassified(100);k__Fungi_unclassified(100);k__Fungi_unclassified(100);k__Fungi_unclassified(100); | - | - | - | - | - | - | - | Unassigned | - |
| Otu27744 | 1 | k__Fungi(100);p__Ascomycota(100);p__Ascomycota_unclassified(100);p__Ascomycota_unclassified(100);p__Ascomycota_unclassified(100);p__Ascomycota_unclassified(100);p__Ascomycota_unclassified(100); | - | - | - | - | - | - | - | Unassigned | - |
| Otu27748 | 1 | k__Fungi(100);k__Fungi_unclassified(100);k__Fungi_unclassified(100);k__Fungi_unclassified(100);k__Fungi_unclassified(100);k__Fungi_unclassified(100);k__Fungi_unclassified(100); | - | - | - | - | - | - | - | Unassigned | - |
| Otu27749 | 1 | k__Fungi(100);k__Fungi_unclassified(100);k__Fungi_unclassified(100);k__Fungi_unclassified(100);k__Fungi_unclassified(100);k__Fungi_unclassified(100);k__Fungi_unclassified(100); | - | - | - | - | - | - | - | Unassigned | - |
| Otu27750 | 1 | k__Fungi(100);k__Fungi_unclassified(100);k__Fungi_unclassified(100);k__Fungi_unclassified(100);k__Fungi_unclassified(100);k__Fungi_unclassified(100);k__Fungi_unclassified(100); | - | - | - | - | - | - | - | Unassigned | - |
| Otu27752 | 1 | k__Fungi(100);k__Fungi_unclassified(100);k__Fungi_unclassified(100);k__Fungi_unclassified(100);k__Fungi_unclassified(100);k__Fungi_unclassified(100);k__Fungi_unclassified(100); | - | - | - | - | - | - | - | Unassigned | - |
| Otu27759 | 1 | k__Fungi(100);k__Fungi_unclassified(100);k__Fungi_unclassified(100);k__Fungi_unclassified(100);k__Fungi_unclassified(100);k__Fungi_unclassified(100);k__Fungi_unclassified(100); | - | - | - | - | - | - | - | Unassigned | - |
| Otu27760 | 1 | k__Fungi(100);p__Ascomycota(100);p__Ascomycota_unclassified(100);p__Ascomycota_unclassified(100);p__Ascomycota_unclassified(100);p__Ascomycota_unclassified(100);p__Ascomycota_unclassified(100); | - | - | - | - | - | - | - | Unassigned | - |
| Otu27761 | 1 | k__Fungi(100);k__Fungi_unclassified(100);k__Fungi_unclassified(100);k__Fungi_unclassified(100);k__Fungi_unclassified(100);k__Fungi_unclassified(100);k__Fungi_unclassified(100); | - | - | - | - | - | - | - | Unassigned | - |
| Otu27762 | 1 | k__Fungi(100);k__Fungi_unclassified(100);k__Fungi_unclassified(100);k__Fungi_unclassified(100);k__Fungi_unclassified(100);k__Fungi_unclassified(100);k__Fungi_unclassified(100); | - | - | - | - | - | - | - | Unassigned | - |
| Otu27763 | 1 | k__Fungi(100);k__Fungi_unclassified(100);k__Fungi_unclassified(100);k__Fungi_unclassified(100);k__Fungi_unclassified(100);k__Fungi_unclassified(100);k__Fungi_unclassified(100); | - | - | - | - | - | - | - | Unassigned | - |
| Otu27764 | 1 | k__Fungi(100);k__Fungi_unclassified(100);k__Fungi_unclassified(100);k__Fungi_unclassified(100);k__Fungi_unclassified(100);k__Fungi_unclassified(100);k__Fungi_unclassified(100); | - | - | - | - | - | - | - | Unassigned | - |
| Otu27765 | 1 | k__Fungi(100);k__Fungi_unclassified(100);k__Fungi_unclassified(100);k__Fungi_unclassified(100);k__Fungi_unclassified(100);k__Fungi_unclassified(100);k__Fungi_unclassified(100); | - | - | - | - | - | - | - | Unassigned | - |
| Otu27766 | 1 | k__Fungi(100);p__Ascomycota(100);p__Ascomycota_unclassified(100);p__Ascomycota_unclassified(100);p__Ascomycota_unclassified(100);p__Ascomycota_unclassified(100);p__Ascomycota_unclassified(100); | - | - | - | - | - | - | - | Unassigned | - |
| Otu27767 | 1 | k__Fungi(100);p__Ascomycota(100);p__Ascomycota_unclassified(100);p__Ascomycota_unclassified(100);p__Ascomycota_unclassified(100);p__Ascomycota_unclassified(100);p__Ascomycota_unclassified(100); | - | - | - | - | - | - | - | Unassigned | - |
| Otu27768 | 1 | k__Fungi(100);k__Fungi_unclassified(100);k__Fungi_unclassified(100);k__Fungi_unclassified(100);k__Fungi_unclassified(100);k__Fungi_unclassified(100);k__Fungi_unclassified(100); | - | - | - | - | - | - | - | Unassigned | - |
| Otu27771 | 1 | k__Fungi(100);p__Ascomycota(100);p__Ascomycota_unclassified(100);p__Ascomycota_unclassified(100);p__Ascomycota_unclassified(100);p__Ascomycota_unclassified(100);p__Ascomycota_unclassified(100); | - | - | - | - | - | - | - | Unassigned | - |
| Otu27772 | 1 | k__Fungi(100);k__Fungi_unclassified(100);k__Fungi_unclassified(100);k__Fungi_unclassified(100);k__Fungi_unclassified(100);k__Fungi_unclassified(100);k__Fungi_unclassified(100); | - | - | - | - | - | - | - | Unassigned | - |
| Otu27775 | 1 | k__Fungi(100);k__Fungi_unclassified(100);k__Fungi_unclassified(100);k__Fungi_unclassified(100);k__Fungi_unclassified(100);k__Fungi_unclassified(100);k__Fungi_unclassified(100); | - | - | - | - | - | - | - | Unassigned | - |
| Otu27776 | 1 | k__Fungi(100);k__Fungi_unclassified(100);k__Fungi_unclassified(100);k__Fungi_unclassified(100);k__Fungi_unclassified(100);k__Fungi_unclassified(100);k__Fungi_unclassified(100); | - | - | - | - | - | - | - | Unassigned | - |
| Otu27777 | 1 | k__Fungi(100);k__Fungi_unclassified(100);k__Fungi_unclassified(100);k__Fungi_unclassified(100);k__Fungi_unclassified(100);k__Fungi_unclassified(100);k__Fungi_unclassified(100); | - | - | - | - | - | - | - | Unassigned | - |
| Otu27778 | 1 | k__Fungi(100);k__Fungi_unclassified(100);k__Fungi_unclassified(100);k__Fungi_unclassified(100);k__Fungi_unclassified(100);k__Fungi_unclassified(100);k__Fungi_unclassified(100); | - | - | - | - | - | - | - | Unassigned | - |
| Otu27779 | 1 | k__Fungi(100);p__Ascomycota(100);p__Ascomycota_unclassified(100);p__Ascomycota_unclassified(100);p__Ascomycota_unclassified(100);p__Ascomycota_unclassified(100);p__Ascomycota_unclassified(100); | - | - | - | - | - | - | - | Unassigned | - |
| Otu27780 | 1 | k__Fungi(100);k__Fungi_unclassified(100);k__Fungi_unclassified(100);k__Fungi_unclassified(100);k__Fungi_unclassified(100);k__Fungi_unclassified(100);k__Fungi_unclassified(100); | - | - | - | - | - | - | - | Unassigned | - |
| Otu27781 | 1 | k__Fungi(100);k__Fungi_unclassified(100);k__Fungi_unclassified(100);k__Fungi_unclassified(100);k__Fungi_unclassified(100);k__Fungi_unclassified(100);k__Fungi_unclassified(100); | - | - | - | - | - | - | - | Unassigned | - |
| Otu27782 | 1 | k__Fungi(100);k__Fungi_unclassified(100);k__Fungi_unclassified(100);k__Fungi_unclassified(100);k__Fungi_unclassified(100);k__Fungi_unclassified(100);k__Fungi_unclassified(100); | - | - | - | - | - | - | - | Unassigned | - |
| Otu27783 | 1 | k__Fungi(100);k__Fungi_unclassified(100);k__Fungi_unclassified(100);k__Fungi_unclassified(100);k__Fungi_unclassified(100);k__Fungi_unclassified(100);k__Fungi_unclassified(100); | - | - | - | - | - | - | - | Unassigned | - |
| Otu27784 | 1 | k__Fungi(100);k__Fungi_unclassified(100);k__Fungi_unclassified(100);k__Fungi_unclassified(100);k__Fungi_unclassified(100);k__Fungi_unclassified(100);k__Fungi_unclassified(100); | - | - | - | - | - | - | - | Unassigned | - |
| Otu27785 | 1 | k__Fungi(100);k__Fungi_unclassified(100);k__Fungi_unclassified(100);k__Fungi_unclassified(100);k__Fungi_unclassified(100);k__Fungi_unclassified(100);k__Fungi_unclassified(100); | - | - | - | - | - | - | - | Unassigned | - |
| Otu27786 | 1 | k__Fungi(100);k__Fungi_unclassified(100);k__Fungi_unclassified(100);k__Fungi_unclassified(100);k__Fungi_unclassified(100);k__Fungi_unclassified(100);k__Fungi_unclassified(100); | - | - | - | - | - | - | - | Unassigned | - |
| Otu27787 | 1 | k__Fungi(100);p__Ascomycota(100);p__Ascomycota_unclassified(100);p__Ascomycota_unclassified(100);p__Ascomycota_unclassified(100);p__Ascomycota_unclassified(100);p__Ascomycota_unclassified(100); | - | - | - | - | - | - | - | Unassigned | - |
| Otu27788 | 1 | k__Fungi(100);p__Ascomycota(100);p__Ascomycota_unclassified(100);p__Ascomycota_unclassified(100);p__Ascomycota_unclassified(100);p__Ascomycota_unclassified(100);p__Ascomycota_unclassified(100); | - | - | - | - | - | - | - | Unassigned | - |
| Otu27789 | 1 | k__Fungi(100);p__Ascomycota(100);p__Ascomycota_unclassified(100);p__Ascomycota_unclassified(100);p__Ascomycota_unclassified(100);p__Ascomycota_unclassified(100);p__Ascomycota_unclassified(100); | - | - | - | - | - | - | - | Unassigned | - |
| Otu27790 | 1 | k__Fungi(100);p__Ascomycota(100);p__Ascomycota_unclassified(100);p__Ascomycota_unclassified(100);p__Ascomycota_unclassified(100);p__Ascomycota_unclassified(100);p__Ascomycota_unclassified(100); | - | - | - | - | - | - | - | Unassigned | - |
| Otu27791 | 1 | k__Fungi(100);p__Ascomycota(100);p__Ascomycota_unclassified(100);p__Ascomycota_unclassified(100);p__Ascomycota_unclassified(100);p__Ascomycota_unclassified(100);p__Ascomycota_unclassified(100); | - | - | - | - | - | - | - | Unassigned | - |
| Otu27792 | 1 | k__Fungi(100);p__Ascomycota(100);p__Ascomycota_unclassified(100);p__Ascomycota_unclassified(100);p__Ascomycota_unclassified(100);p__Ascomycota_unclassified(100);p__Ascomycota_unclassified(100); | - | - | - | - | - | - | - | Unassigned | - |
| Otu27793 | 1 | k__Fungi(100);k__Fungi_unclassified(100);k__Fungi_unclassified(100);k__Fungi_unclassified(100);k__Fungi_unclassified(100);k__Fungi_unclassified(100);k__Fungi_unclassified(100); | - | - | - | - | - | - | - | Unassigned | - |
| Otu27794 | 1 | k__Fungi(100);k__Fungi_unclassified(100);k__Fungi_unclassified(100);k__Fungi_unclassified(100);k__Fungi_unclassified(100);k__Fungi_unclassified(100);k__Fungi_unclassified(100); | - | - | - | - | - | - | - | Unassigned | - |
| Otu27796 | 1 | k__Fungi(100);k__Fungi_unclassified(100);k__Fungi_unclassified(100);k__Fungi_unclassified(100);k__Fungi_unclassified(100);k__Fungi_unclassified(100);k__Fungi_unclassified(100); | - | - | - | - | - | - | - | Unassigned | - |
| Otu27798 | 1 | k__Fungi(100);p__Ascomycota(100);p__Ascomycota_unclassified(100);p__Ascomycota_unclassified(100);p__Ascomycota_unclassified(100);p__Ascomycota_unclassified(100);p__Ascomycota_unclassified(100); | - | - | - | - | - | - | - | Unassigned | - |
| Otu27799 | 1 | k__Fungi(100);p__Ascomycota(100);p__Ascomycota_unclassified(100);p__Ascomycota_unclassified(100);p__Ascomycota_unclassified(100);p__Ascomycota_unclassified(100);p__Ascomycota_unclassified(100); | - | - | - | - | - | - | - | Unassigned | - |
| Otu27800 | 1 | k__Fungi(100);p__Ascomycota(100);p__Ascomycota_unclassified(100);p__Ascomycota_unclassified(100);p__Ascomycota_unclassified(100);p__Ascomycota_unclassified(100);p__Ascomycota_unclassified(100); | - | - | - | - | - | - | - | Unassigned | - |
| Otu27801 | 1 | k__Fungi(100);k__Fungi_unclassified(100);k__Fungi_unclassified(100);k__Fungi_unclassified(100);k__Fungi_unclassified(100);k__Fungi_unclassified(100);k__Fungi_unclassified(100); | - | - | - | - | - | - | - | Unassigned | - |
| Otu27802 | 1 | k__Fungi(100);p__Ascomycota(100);p__Ascomycota_unclassified(100);p__Ascomycota_unclassified(100);p__Ascomycota_unclassified(100);p__Ascomycota_unclassified(100);p__Ascomycota_unclassified(100); | - | - | - | - | - | - | - | Unassigned | - |
| Otu27803 | 1 | k__Fungi(100);p__Ascomycota(100);p__Ascomycota_unclassified(100);p__Ascomycota_unclassified(100);p__Ascomycota_unclassified(100);p__Ascomycota_unclassified(100);p__Ascomycota_unclassified(100); | - | - | - | - | - | - | - | Unassigned | - |
| Otu27804 | 1 | k__Fungi(100);p__Ascomycota(100);p__Ascomycota_unclassified(100);p__Ascomycota_unclassified(100);p__Ascomycota_unclassified(100);p__Ascomycota_unclassified(100);p__Ascomycota_unclassified(100); | - | - | - | - | - | - | - | Unassigned | - |
| Otu27805 | 1 | k__Fungi(100);k__Fungi_unclassified(100);k__Fungi_unclassified(100);k__Fungi_unclassified(100);k__Fungi_unclassified(100);k__Fungi_unclassified(100);k__Fungi_unclassified(100); | - | - | - | - | - | - | - | Unassigned | - |
| Otu27807 | 1 | k__Fungi(100);p__Ascomycota(100);p__Ascomycota_unclassified(100);p__Ascomycota_unclassified(100);p__Ascomycota_unclassified(100);p__Ascomycota_unclassified(100);p__Ascomycota_unclassified(100); | - | - | - | - | - | - | - | Unassigned | - |
| Otu27808 | 1 | k__Fungi(100);k__Fungi_unclassified(100);k__Fungi_unclassified(100);k__Fungi_unclassified(100);k__Fungi_unclassified(100);k__Fungi_unclassified(100);k__Fungi_unclassified(100); | - | - | - | - | - | - | - | Unassigned | - |
| Otu27809 | 1 | k__Fungi(100);k__Fungi_unclassified(100);k__Fungi_unclassified(100);k__Fungi_unclassified(100);k__Fungi_unclassified(100);k__Fungi_unclassified(100);k__Fungi_unclassified(100); | - | - | - | - | - | - | - | Unassigned | - |
| Otu27810 | 1 | k__Fungi(100);p__Ascomycota(100);p__Ascomycota_unclassified(100);p__Ascomycota_unclassified(100);p__Ascomycota_unclassified(100);p__Ascomycota_unclassified(100);p__Ascomycota_unclassified(100); | - | - | - | - | - | - | - | Unassigned | - |
| Otu27811 | 1 | k__Fungi(100);k__Fungi_unclassified(100);k__Fungi_unclassified(100);k__Fungi_unclassified(100);k__Fungi_unclassified(100);k__Fungi_unclassified(100);k__Fungi_unclassified(100); | - | - | - | - | - | - | - | Unassigned | - |
| Otu27812 | 1 | k__Fungi(100);p__Ascomycota(100);p__Ascomycota_unclassified(100);p__Ascomycota_unclassified(100);p__Ascomycota_unclassified(100);p__Ascomycota_unclassified(100);p__Ascomycota_unclassified(100); | - | - | - | - | - | - | - | Unassigned | - |
| Otu27813 | 1 | k__Fungi(100);k__Fungi_unclassified(100);k__Fungi_unclassified(100);k__Fungi_unclassified(100);k__Fungi_unclassified(100);k__Fungi_unclassified(100);k__Fungi_unclassified(100); | - | - | - | - | - | - | - | Unassigned | - |
| Otu27815 | 1 | k__Fungi(100);p__Ascomycota(100);p__Ascomycota_unclassified(100);p__Ascomycota_unclassified(100);p__Ascomycota_unclassified(100);p__Ascomycota_unclassified(100);p__Ascomycota_unclassified(100); | - | - | - | - | - | - | - | Unassigned | - |
| Otu27816 | 1 | k__Fungi(100);p__Ascomycota(100);p__Ascomycota_unclassified(100);p__Ascomycota_unclassified(100);p__Ascomycota_unclassified(100);p__Ascomycota_unclassified(100);p__Ascomycota_unclassified(100); | - | - | - | - | - | - | - | Unassigned | - |
| Otu27817 | 1 | k__Fungi(100);p__Ascomycota(100);p__Ascomycota_unclassified(100);p__Ascomycota_unclassified(100);p__Ascomycota_unclassified(100);p__Ascomycota_unclassified(100);p__Ascomycota_unclassified(100); | - | - | - | - | - | - | - | Unassigned | - |
| Otu27818 | 1 | k__Fungi(100);k__Fungi_unclassified(100);k__Fungi_unclassified(100);k__Fungi_unclassified(100);k__Fungi_unclassified(100);k__Fungi_unclassified(100);k__Fungi_unclassified(100); | - | - | - | - | - | - | - | Unassigned | - |
| Otu27819 | 1 | k__Fungi(100);k__Fungi_unclassified(100);k__Fungi_unclassified(100);k__Fungi_unclassified(100);k__Fungi_unclassified(100);k__Fungi_unclassified(100);k__Fungi_unclassified(100); | - | - | - | - | - | - | - | Unassigned | - |
| Otu27820 | 1 | k__Fungi(100);p__Ascomycota(100);p__Ascomycota_unclassified(100);p__Ascomycota_unclassified(100);p__Ascomycota_unclassified(100);p__Ascomycota_unclassified(100);p__Ascomycota_unclassified(100); | - | - | - | - | - | - | - | Unassigned | - |
| Otu27821 | 1 | k__Fungi(100);p__Ascomycota(100);p__Ascomycota_unclassified(100);p__Ascomycota_unclassified(100);p__Ascomycota_unclassified(100);p__Ascomycota_unclassified(100);p__Ascomycota_unclassified(100); | - | - | - | - | - | - | - | Unassigned | - |
| Otu27822 | 1 | k__Fungi(100);k__Fungi_unclassified(100);k__Fungi_unclassified(100);k__Fungi_unclassified(100);k__Fungi_unclassified(100);k__Fungi_unclassified(100);k__Fungi_unclassified(100); | - | - | - | - | - | - | - | Unassigned | - |
| Otu27823 | 1 | k__Fungi(100);p__Ascomycota(100);p__Ascomycota_unclassified(100);p__Ascomycota_unclassified(100);p__Ascomycota_unclassified(100);p__Ascomycota_unclassified(100);p__Ascomycota_unclassified(100); | - | - | - | - | - | - | - | Unassigned | - |
| Otu27824 | 1 | k__Fungi(100);p__Ascomycota(100);p__Ascomycota_unclassified(100);p__Ascomycota_unclassified(100);p__Ascomycota_unclassified(100);p__Ascomycota_unclassified(100);p__Ascomycota_unclassified(100); | - | - | - | - | - | - | - | Unassigned | - |
| Otu27825 | 1 | k__Fungi(100);p__Ascomycota(100);p__Ascomycota_unclassified(100);p__Ascomycota_unclassified(100);p__Ascomycota_unclassified(100);p__Ascomycota_unclassified(100);p__Ascomycota_unclassified(100); | - | - | - | - | - | - | - | Unassigned | - |
| Otu27826 | 1 | k__Fungi(100);k__Fungi_unclassified(100);k__Fungi_unclassified(100);k__Fungi_unclassified(100);k__Fungi_unclassified(100);k__Fungi_unclassified(100);k__Fungi_unclassified(100); | - | - | - | - | - | - | - | Unassigned | - |
| Otu27832 | 1 | k__Fungi(100);k__Fungi_unclassified(100);k__Fungi_unclassified(100);k__Fungi_unclassified(100);k__Fungi_unclassified(100);k__Fungi_unclassified(100);k__Fungi_unclassified(100); | - | - | - | - | - | - | - | Unassigned | - |
| Otu27833 | 1 | k__Fungi(100);k__Fungi_unclassified(100);k__Fungi_unclassified(100);k__Fungi_unclassified(100);k__Fungi_unclassified(100);k__Fungi_unclassified(100);k__Fungi_unclassified(100); | - | - | - | - | - | - | - | Unassigned | - |
| Otu27836 | 1 | k__Fungi(100);p__Ascomycota(100);p__Ascomycota_unclassified(100);p__Ascomycota_unclassified(100);p__Ascomycota_unclassified(100);p__Ascomycota_unclassified(100);p__Ascomycota_unclassified(100); | - | - | - | - | - | - | - | Unassigned | - |
| Otu27838 | 1 | k__Fungi(100);p__Ascomycota(100);p__Ascomycota_unclassified(100);p__Ascomycota_unclassified(100);p__Ascomycota_unclassified(100);p__Ascomycota_unclassified(100);p__Ascomycota_unclassified(100); | - | - | - | - | - | - | - | Unassigned | - |
| Otu27839 | 1 | k__Fungi(100);k__Fungi_unclassified(100);k__Fungi_unclassified(100);k__Fungi_unclassified(100);k__Fungi_unclassified(100);k__Fungi_unclassified(100);k__Fungi_unclassified(100); | - | - | - | - | - | - | - | Unassigned | - |
| Otu27841 | 1 | k__Fungi(100);p__Ascomycota(100);p__Ascomycota_unclassified(100);p__Ascomycota_unclassified(100);p__Ascomycota_unclassified(100);p__Ascomycota_unclassified(100);p__Ascomycota_unclassified(100); | - | - | - | - | - | - | - | Unassigned | - |
| Otu27842 | 1 | k__Fungi(100);p__Ascomycota(100);p__Ascomycota_unclassified(100);p__Ascomycota_unclassified(100);p__Ascomycota_unclassified(100);p__Ascomycota_unclassified(100);p__Ascomycota_unclassified(100); | - | - | - | - | - | - | - | Unassigned | - |
| Otu27843 | 1 | k__Fungi(100);k__Fungi_unclassified(100);k__Fungi_unclassified(100);k__Fungi_unclassified(100);k__Fungi_unclassified(100);k__Fungi_unclassified(100);k__Fungi_unclassified(100); | - | - | - | - | - | - | - | Unassigned | - |
| Otu27844 | 1 | k__Fungi(100);k__Fungi_unclassified(100);k__Fungi_unclassified(100);k__Fungi_unclassified(100);k__Fungi_unclassified(100);k__Fungi_unclassified(100);k__Fungi_unclassified(100); | - | - | - | - | - | - | - | Unassigned | - |
| Otu27845 | 1 | k__Fungi(100);k__Fungi_unclassified(100);k__Fungi_unclassified(100);k__Fungi_unclassified(100);k__Fungi_unclassified(100);k__Fungi_unclassified(100);k__Fungi_unclassified(100); | - | - | - | - | - | - | - | Unassigned | - |
| Otu27846 | 1 | k__Fungi(100);p__Ascomycota(100);p__Ascomycota_unclassified(100);p__Ascomycota_unclassified(100);p__Ascomycota_unclassified(100);p__Ascomycota_unclassified(100);p__Ascomycota_unclassified(100); | - | - | - | - | - | - | - | Unassigned | - |
| Otu27847 | 1 | k__Fungi(100);p__Ascomycota(100);p__Ascomycota_unclassified(100);p__Ascomycota_unclassified(100);p__Ascomycota_unclassified(100);p__Ascomycota_unclassified(100);p__Ascomycota_unclassified(100); | - | - | - | - | - | - | - | Unassigned | - |
| Otu27848 | 1 | k__Fungi(100);k__Fungi_unclassified(100);k__Fungi_unclassified(100);k__Fungi_unclassified(100);k__Fungi_unclassified(100);k__Fungi_unclassified(100);k__Fungi_unclassified(100); | - | - | - | - | - | - | - | Unassigned | - |
| Otu27849 | 1 | k__Fungi(100);p__Ascomycota(100);p__Ascomycota_unclassified(100);p__Ascomycota_unclassified(100);p__Ascomycota_unclassified(100);p__Ascomycota_unclassified(100);p__Ascomycota_unclassified(100); | - | - | - | - | - | - | - | Unassigned | - |
| Otu27850 | 1 | k__Fungi(100);p__Ascomycota(100);p__Ascomycota_unclassified(100);p__Ascomycota_unclassified(100);p__Ascomycota_unclassified(100);p__Ascomycota_unclassified(100);p__Ascomycota_unclassified(100); | - | - | - | - | - | - | - | Unassigned | - |
| Otu27851 | 1 | k__Fungi(100);p__Ascomycota(100);p__Ascomycota_unclassified(100);p__Ascomycota_unclassified(100);p__Ascomycota_unclassified(100);p__Ascomycota_unclassified(100);p__Ascomycota_unclassified(100); | - | - | - | - | - | - | - | Unassigned | - |
| Otu27852 | 1 | k__Fungi(100);k__Fungi_unclassified(100);k__Fungi_unclassified(100);k__Fungi_unclassified(100);k__Fungi_unclassified(100);k__Fungi_unclassified(100);k__Fungi_unclassified(100); | - | - | - | - | - | - | - | Unassigned | - |
| Otu27855 | 1 | k__Fungi(100);p__Ascomycota(100);p__Ascomycota_unclassified(100);p__Ascomycota_unclassified(100);p__Ascomycota_unclassified(100);p__Ascomycota_unclassified(100);p__Ascomycota_unclassified(100); | - | - | - | - | - | - | - | Unassigned | - |
| Otu27856 | 1 | k__Fungi(100);k__Fungi_unclassified(100);k__Fungi_unclassified(100);k__Fungi_unclassified(100);k__Fungi_unclassified(100);k__Fungi_unclassified(100);k__Fungi_unclassified(100); | - | - | - | - | - | - | - | Unassigned | - |
| Otu27858 | 1 | k__Fungi(100);k__Fungi_unclassified(100);k__Fungi_unclassified(100);k__Fungi_unclassified(100);k__Fungi_unclassified(100);k__Fungi_unclassified(100);k__Fungi_unclassified(100); | - | - | - | - | - | - | - | Unassigned | - |
| Otu27859 | 1 | k__Fungi(100);p__Ascomycota(100);p__Ascomycota_unclassified(100);p__Ascomycota_unclassified(100);p__Ascomycota_unclassified(100);p__Ascomycota_unclassified(100);p__Ascomycota_unclassified(100); | - | - | - | - | - | - | - | Unassigned | - |
| Otu27860 | 1 | k__Fungi(100);k__Fungi_unclassified(100);k__Fungi_unclassified(100);k__Fungi_unclassified(100);k__Fungi_unclassified(100);k__Fungi_unclassified(100);k__Fungi_unclassified(100); | - | - | - | - | - | - | - | Unassigned | - |
| Otu27861 | 1 | k__Fungi(100);p__Ascomycota(100);p__Ascomycota_unclassified(100);p__Ascomycota_unclassified(100);p__Ascomycota_unclassified(100);p__Ascomycota_unclassified(100);p__Ascomycota_unclassified(100); | - | - | - | - | - | - | - | Unassigned | - |
| Otu27862 | 1 | k__Fungi(100);p__Ascomycota(100);p__Ascomycota_unclassified(100);p__Ascomycota_unclassified(100);p__Ascomycota_unclassified(100);p__Ascomycota_unclassified(100);p__Ascomycota_unclassified(100); | - | - | - | - | - | - | - | Unassigned | - |
| Otu27863 | 1 | k__Fungi(100);k__Fungi_unclassified(100);k__Fungi_unclassified(100);k__Fungi_unclassified(100);k__Fungi_unclassified(100);k__Fungi_unclassified(100);k__Fungi_unclassified(100); | - | - | - | - | - | - | - | Unassigned | - |
| Otu27864 | 1 | k__Fungi(100);k__Fungi_unclassified(100);k__Fungi_unclassified(100);k__Fungi_unclassified(100);k__Fungi_unclassified(100);k__Fungi_unclassified(100);k__Fungi_unclassified(100); | - | - | - | - | - | - | - | Unassigned | - |
| Otu27865 | 1 | k__Fungi(100);p__Ascomycota(100);p__Ascomycota_unclassified(100);p__Ascomycota_unclassified(100);p__Ascomycota_unclassified(100);p__Ascomycota_unclassified(100);p__Ascomycota_unclassified(100); | - | - | - | - | - | - | - | Unassigned | - |
| Otu27867 | 1 | k__Fungi(100);p__Ascomycota(100);p__Ascomycota_unclassified(100);p__Ascomycota_unclassified(100);p__Ascomycota_unclassified(100);p__Ascomycota_unclassified(100);p__Ascomycota_unclassified(100); | - | - | - | - | - | - | - | Unassigned | - |
| Otu27868 | 1 | k__Fungi(100);p__Ascomycota(100);p__Ascomycota_unclassified(100);p__Ascomycota_unclassified(100);p__Ascomycota_unclassified(100);p__Ascomycota_unclassified(100);p__Ascomycota_unclassified(100); | - | - | - | - | - | - | - | Unassigned | - |
| Otu27869 | 1 | k__Fungi(100);k__Fungi_unclassified(100);k__Fungi_unclassified(100);k__Fungi_unclassified(100);k__Fungi_unclassified(100);k__Fungi_unclassified(100);k__Fungi_unclassified(100); | - | - | - | - | - | - | - | Unassigned | - |
| Otu27870 | 1 | k__Fungi(100);k__Fungi_unclassified(100);k__Fungi_unclassified(100);k__Fungi_unclassified(100);k__Fungi_unclassified(100);k__Fungi_unclassified(100);k__Fungi_unclassified(100); | - | - | - | - | - | - | - | Unassigned | - |
| Otu27871 | 1 | k__Fungi(100);p__Ascomycota(100);p__Ascomycota_unclassified(100);p__Ascomycota_unclassified(100);p__Ascomycota_unclassified(100);p__Ascomycota_unclassified(100);p__Ascomycota_unclassified(100); | - | - | - | - | - | - | - | Unassigned | - |
| Otu27872 | 1 | k__Fungi(100);k__Fungi_unclassified(100);k__Fungi_unclassified(100);k__Fungi_unclassified(100);k__Fungi_unclassified(100);k__Fungi_unclassified(100);k__Fungi_unclassified(100); | - | - | - | - | - | - | - | Unassigned | - |
| Otu27873 | 1 | k__Fungi(100);p__Ascomycota(100);p__Ascomycota_unclassified(100);p__Ascomycota_unclassified(100);p__Ascomycota_unclassified(100);p__Ascomycota_unclassified(100);p__Ascomycota_unclassified(100); | - | - | - | - | - | - | - | Unassigned | - |
| Otu27876 | 1 | k__Fungi(100);k__Fungi_unclassified(100);k__Fungi_unclassified(100);k__Fungi_unclassified(100);k__Fungi_unclassified(100);k__Fungi_unclassified(100);k__Fungi_unclassified(100); | - | - | - | - | - | - | - | Unassigned | - |
| Otu27877 | 1 | k__Fungi(100);k__Fungi_unclassified(100);k__Fungi_unclassified(100);k__Fungi_unclassified(100);k__Fungi_unclassified(100);k__Fungi_unclassified(100);k__Fungi_unclassified(100); | - | - | - | - | - | - | - | Unassigned | - |
| Otu27878 | 1 | k__Fungi(100);k__Fungi_unclassified(100);k__Fungi_unclassified(100);k__Fungi_unclassified(100);k__Fungi_unclassified(100);k__Fungi_unclassified(100);k__Fungi_unclassified(100); | - | - | - | - | - | - | - | Unassigned | - |
| Otu27879 | 1 | k__Fungi(100);p__Ascomycota(100);p__Ascomycota_unclassified(100);p__Ascomycota_unclassified(100);p__Ascomycota_unclassified(100);p__Ascomycota_unclassified(100);p__Ascomycota_unclassified(100); | - | - | - | - | - | - | - | Unassigned | - |
| Otu27880 | 1 | k__Fungi(100);p__Ascomycota(100);p__Ascomycota_unclassified(100);p__Ascomycota_unclassified(100);p__Ascomycota_unclassified(100);p__Ascomycota_unclassified(100);p__Ascomycota_unclassified(100); | - | - | - | - | - | - | - | Unassigned | - |
| Otu27881 | 1 | k__Fungi(100);k__Fungi_unclassified(100);k__Fungi_unclassified(100);k__Fungi_unclassified(100);k__Fungi_unclassified(100);k__Fungi_unclassified(100);k__Fungi_unclassified(100); | - | - | - | - | - | - | - | Unassigned | - |
| Otu27883 | 1 | k__Fungi(100);k__Fungi_unclassified(100);k__Fungi_unclassified(100);k__Fungi_unclassified(100);k__Fungi_unclassified(100);k__Fungi_unclassified(100);k__Fungi_unclassified(100); | - | - | - | - | - | - | - | Unassigned | - |
| Otu27885 | 1 | k__Fungi(100);k__Fungi_unclassified(100);k__Fungi_unclassified(100);k__Fungi_unclassified(100);k__Fungi_unclassified(100);k__Fungi_unclassified(100);k__Fungi_unclassified(100); | - | - | - | - | - | - | - | Unassigned | - |
| Otu27886 | 1 | k__Fungi(100);k__Fungi_unclassified(100);k__Fungi_unclassified(100);k__Fungi_unclassified(100);k__Fungi_unclassified(100);k__Fungi_unclassified(100);k__Fungi_unclassified(100); | - | - | - | - | - | - | - | Unassigned | - |
| Otu27888 | 1 | k__Fungi(100);k__Fungi_unclassified(100);k__Fungi_unclassified(100);k__Fungi_unclassified(100);k__Fungi_unclassified(100);k__Fungi_unclassified(100);k__Fungi_unclassified(100); | - | - | - | - | - | - | - | Unassigned | - |
| Otu27889 | 1 | k__Fungi(100);k__Fungi_unclassified(100);k__Fungi_unclassified(100);k__Fungi_unclassified(100);k__Fungi_unclassified(100);k__Fungi_unclassified(100);k__Fungi_unclassified(100); | - | - | - | - | - | - | - | Unassigned | - |
| Otu27890 | 1 | k__Fungi(100);k__Fungi_unclassified(100);k__Fungi_unclassified(100);k__Fungi_unclassified(100);k__Fungi_unclassified(100);k__Fungi_unclassified(100);k__Fungi_unclassified(100); | - | - | - | - | - | - | - | Unassigned | - |
| Otu27892 | 1 | k__Fungi(100);k__Fungi_unclassified(100);k__Fungi_unclassified(100);k__Fungi_unclassified(100);k__Fungi_unclassified(100);k__Fungi_unclassified(100);k__Fungi_unclassified(100); | - | - | - | - | - | - | - | Unassigned | - |
| Otu27893 | 1 | k__Fungi(100);k__Fungi_unclassified(100);k__Fungi_unclassified(100);k__Fungi_unclassified(100);k__Fungi_unclassified(100);k__Fungi_unclassified(100);k__Fungi_unclassified(100); | - | - | - | - | - | - | - | Unassigned | - |
| Otu27894 | 1 | k__Fungi(100);k__Fungi_unclassified(100);k__Fungi_unclassified(100);k__Fungi_unclassified(100);k__Fungi_unclassified(100);k__Fungi_unclassified(100);k__Fungi_unclassified(100); | - | - | - | - | - | - | - | Unassigned | - |
| Otu27895 | 1 | k__Fungi(100);p__Ascomycota(100);p__Ascomycota_unclassified(100);p__Ascomycota_unclassified(100);p__Ascomycota_unclassified(100);p__Ascomycota_unclassified(100);p__Ascomycota_unclassified(100); | - | - | - | - | - | - | - | Unassigned | - |
| Otu27897 | 1 | k__Fungi(100);k__Fungi_unclassified(100);k__Fungi_unclassified(100);k__Fungi_unclassified(100);k__Fungi_unclassified(100);k__Fungi_unclassified(100);k__Fungi_unclassified(100); | - | - | - | - | - | - | - | Unassigned | - |
| Otu27898 | 1 | k__Fungi(100);k__Fungi_unclassified(100);k__Fungi_unclassified(100);k__Fungi_unclassified(100);k__Fungi_unclassified(100);k__Fungi_unclassified(100);k__Fungi_unclassified(100); | - | - | - | - | - | - | - | Unassigned | - |
| Otu27899 | 1 | k__Fungi(100);p__Ascomycota(100);p__Ascomycota_unclassified(100);p__Ascomycota_unclassified(100);p__Ascomycota_unclassified(100);p__Ascomycota_unclassified(100);p__Ascomycota_unclassified(100); | - | - | - | - | - | - | - | Unassigned | - |
| Otu27900 | 1 | k__Fungi(100);p__Ascomycota(100);p__Ascomycota_unclassified(100);p__Ascomycota_unclassified(100);p__Ascomycota_unclassified(100);p__Ascomycota_unclassified(100);p__Ascomycota_unclassified(100); | - | - | - | - | - | - | - | Unassigned | - |
| Otu27901 | 1 | k__Fungi(100);k__Fungi_unclassified(100);k__Fungi_unclassified(100);k__Fungi_unclassified(100);k__Fungi_unclassified(100);k__Fungi_unclassified(100);k__Fungi_unclassified(100); | - | - | - | - | - | - | - | Unassigned | - |
| Otu27903 | 1 | k__Fungi(100);p__Ascomycota(100);p__Ascomycota_unclassified(100);p__Ascomycota_unclassified(100);p__Ascomycota_unclassified(100);p__Ascomycota_unclassified(100);p__Ascomycota_unclassified(100); | - | - | - | - | - | - | - | Unassigned | - |
| Otu27904 | 1 | k__Fungi(100);p__Ascomycota(100);p__Ascomycota_unclassified(100);p__Ascomycota_unclassified(100);p__Ascomycota_unclassified(100);p__Ascomycota_unclassified(100);p__Ascomycota_unclassified(100); | - | - | - | - | - | - | - | Unassigned | - |
| Otu27914 | 1 | k__Fungi(100);k__Fungi_unclassified(100);k__Fungi_unclassified(100);k__Fungi_unclassified(100);k__Fungi_unclassified(100);k__Fungi_unclassified(100);k__Fungi_unclassified(100); | - | - | - | - | - | - | - | Unassigned | - |
| Otu27915 | 1 | k__Fungi(100);k__Fungi_unclassified(100);k__Fungi_unclassified(100);k__Fungi_unclassified(100);k__Fungi_unclassified(100);k__Fungi_unclassified(100);k__Fungi_unclassified(100); | - | - | - | - | - | - | - | Unassigned | - |
| Otu27916 | 1 | k__Fungi(100);k__Fungi_unclassified(100);k__Fungi_unclassified(100);k__Fungi_unclassified(100);k__Fungi_unclassified(100);k__Fungi_unclassified(100);k__Fungi_unclassified(100); | - | - | - | - | - | - | - | Unassigned | - |
| Otu27917 | 1 | k__Fungi(100);k__Fungi_unclassified(100);k__Fungi_unclassified(100);k__Fungi_unclassified(100);k__Fungi_unclassified(100);k__Fungi_unclassified(100);k__Fungi_unclassified(100); | - | - | - | - | - | - | - | Unassigned | - |
| Otu27918 | 1 | k__Fungi(100);k__Fungi_unclassified(100);k__Fungi_unclassified(100);k__Fungi_unclassified(100);k__Fungi_unclassified(100);k__Fungi_unclassified(100);k__Fungi_unclassified(100); | - | - | - | - | - | - | - | Unassigned | - |
| Otu27919 | 1 | k__Fungi(100);p__Ascomycota(100);p__Ascomycota_unclassified(100);p__Ascomycota_unclassified(100);p__Ascomycota_unclassified(100);p__Ascomycota_unclassified(100);p__Ascomycota_unclassified(100); | - | - | - | - | - | - | - | Unassigned | - |
| Otu27920 | 1 | k__Fungi(100);k__Fungi_unclassified(100);k__Fungi_unclassified(100);k__Fungi_unclassified(100);k__Fungi_unclassified(100);k__Fungi_unclassified(100);k__Fungi_unclassified(100); | - | - | - | - | - | - | - | Unassigned | - |
| Otu27922 | 1 | k__Plantae(100);p__unclassified_Plantae(100);c__unclassified_Plantae(100);o__unclassified_Plantae(100);f__unclassified_Plantae(100);g__unclassified_Plantae(100);s__Plantae_sp(100); | - | - | - | - | - | - | - | Unassigned | - |
| Otu27924 | 1 | k__Fungi(100);k__Fungi_unclassified(100);k__Fungi_unclassified(100);k__Fungi_unclassified(100);k__Fungi_unclassified(100);k__Fungi_unclassified(100);k__Fungi_unclassified(100); | - | - | - | - | - | - | - | Unassigned | - |
| Otu27925 | 1 | k__Fungi(100);k__Fungi_unclassified(100);k__Fungi_unclassified(100);k__Fungi_unclassified(100);k__Fungi_unclassified(100);k__Fungi_unclassified(100);k__Fungi_unclassified(100); | - | - | - | - | - | - | - | Unassigned | - |
| Otu27929 | 1 | k__Fungi(100);p__Ascomycota(100);c__Dothideomycetes(100);o__Capnodiales(100);f__Davidiellaceae(100);f__Davidiellaceae_unclassified(100);f__Davidiellaceae_unclassified(100); | - | - | - | - | - | - | - | Unassigned | - |
| Otu27931 | 1 | k__Fungi(100);k__Fungi_unclassified(100);k__Fungi_unclassified(100);k__Fungi_unclassified(100);k__Fungi_unclassified(100);k__Fungi_unclassified(100);k__Fungi_unclassified(100); | - | - | - | - | - | - | - | Unassigned | - |
| Otu27933 | 1 | k__Fungi(100);p__Ascomycota(100);p__Ascomycota_unclassified(100);p__Ascomycota_unclassified(100);p__Ascomycota_unclassified(100);p__Ascomycota_unclassified(100);p__Ascomycota_unclassified(100); | - | - | - | - | - | - | - | Unassigned | - |
| Otu27938 | 1 | k__Fungi(100);k__Fungi_unclassified(100);k__Fungi_unclassified(100);k__Fungi_unclassified(100);k__Fungi_unclassified(100);k__Fungi_unclassified(100);k__Fungi_unclassified(100); | - | - | - | - | - | - | - | Unassigned | - |
| Otu27940 | 1 | k__Fungi(100);p__Ascomycota(100);p__Ascomycota_unclassified(100);p__Ascomycota_unclassified(100);p__Ascomycota_unclassified(100);p__Ascomycota_unclassified(100);p__Ascomycota_unclassified(100); | - | - | - | - | - | - | - | Unassigned | - |
| Otu27941 | 1 | k__Fungi(100);k__Fungi_unclassified(100);k__Fungi_unclassified(100);k__Fungi_unclassified(100);k__Fungi_unclassified(100);k__Fungi_unclassified(100);k__Fungi_unclassified(100); | - | - | - | - | - | - | - | Unassigned | - |
| Otu27942 | 1 | k__Fungi(100);p__Ascomycota(100);p__Ascomycota_unclassified(100);p__Ascomycota_unclassified(100);p__Ascomycota_unclassified(100);p__Ascomycota_unclassified(100);p__Ascomycota_unclassified(100); | - | - | - | - | - | - | - | Unassigned | - |
| Otu27943 | 1 | k__Fungi(100);k__Fungi_unclassified(100);k__Fungi_unclassified(100);k__Fungi_unclassified(100);k__Fungi_unclassified(100);k__Fungi_unclassified(100);k__Fungi_unclassified(100); | - | - | - | - | - | - | - | Unassigned | - |
| Otu27944 | 1 | k__Fungi(100);p__Ascomycota(100);c__Dothideomycetes(100);c__Dothideomycetes_unclassified(100);c__Dothideomycetes_unclassified(100);c__Dothideomycetes_unclassified(100);c__Dothideomycetes_unclassified(100); | - | - | - | - | - | - | - | Unassigned | - |
| Otu27945 | 1 | k__Fungi(100);p__Ascomycota(100);p__Ascomycota_unclassified(100);p__Ascomycota_unclassified(100);p__Ascomycota_unclassified(100);p__Ascomycota_unclassified(100);p__Ascomycota_unclassified(100); | - | - | - | - | - | - | - | Unassigned | - |
| Otu27946 | 1 | k__Fungi(100);k__Fungi_unclassified(100);k__Fungi_unclassified(100);k__Fungi_unclassified(100);k__Fungi_unclassified(100);k__Fungi_unclassified(100);k__Fungi_unclassified(100); | - | - | - | - | - | - | - | Unassigned | - |
| Otu27948 | 1 | k__Fungi(100);k__Fungi_unclassified(100);k__Fungi_unclassified(100);k__Fungi_unclassified(100);k__Fungi_unclassified(100);k__Fungi_unclassified(100);k__Fungi_unclassified(100); | - | - | - | - | - | - | - | Unassigned | - |
| Otu27949 | 1 | k__Fungi(100);k__Fungi_unclassified(100);k__Fungi_unclassified(100);k__Fungi_unclassified(100);k__Fungi_unclassified(100);k__Fungi_unclassified(100);k__Fungi_unclassified(100); | - | - | - | - | - | - | - | Unassigned | - |
| Otu27950 | 1 | k__Fungi(100);k__Fungi_unclassified(100);k__Fungi_unclassified(100);k__Fungi_unclassified(100);k__Fungi_unclassified(100);k__Fungi_unclassified(100);k__Fungi_unclassified(100); | - | - | - | - | - | - | - | Unassigned | - |
| Otu27951 | 1 | k__Fungi(100);p__Ascomycota(100);p__Ascomycota_unclassified(100);p__Ascomycota_unclassified(100);p__Ascomycota_unclassified(100);p__Ascomycota_unclassified(100);p__Ascomycota_unclassified(100); | - | - | - | - | - | - | - | Unassigned | - |
| Otu27952 | 1 | k__Fungi(100);k__Fungi_unclassified(100);k__Fungi_unclassified(100);k__Fungi_unclassified(100);k__Fungi_unclassified(100);k__Fungi_unclassified(100);k__Fungi_unclassified(100); | - | - | - | - | - | - | - | Unassigned | - |
| Otu27954 | 1 | k__Fungi(100);k__Fungi_unclassified(100);k__Fungi_unclassified(100);k__Fungi_unclassified(100);k__Fungi_unclassified(100);k__Fungi_unclassified(100);k__Fungi_unclassified(100); | - | - | - | - | - | - | - | Unassigned | - |
| Otu27955 | 1 | k__Fungi(100);p__Ascomycota(100);p__Ascomycota_unclassified(100);p__Ascomycota_unclassified(100);p__Ascomycota_unclassified(100);p__Ascomycota_unclassified(100);p__Ascomycota_unclassified(100); | - | - | - | - | - | - | - | Unassigned | - |
| Otu27956 | 1 | k__Fungi(100);k__Fungi_unclassified(100);k__Fungi_unclassified(100);k__Fungi_unclassified(100);k__Fungi_unclassified(100);k__Fungi_unclassified(100);k__Fungi_unclassified(100); | - | - | - | - | - | - | - | Unassigned | - |
| Otu27957 | 1 | k__Fungi(100);p__Ascomycota(100);p__Ascomycota_unclassified(100);p__Ascomycota_unclassified(100);p__Ascomycota_unclassified(100);p__Ascomycota_unclassified(100);p__Ascomycota_unclassified(100); | - | - | - | - | - | - | - | Unassigned | - |
| Otu27958 | 1 | k__Fungi(100);k__Fungi_unclassified(100);k__Fungi_unclassified(100);k__Fungi_unclassified(100);k__Fungi_unclassified(100);k__Fungi_unclassified(100);k__Fungi_unclassified(100); | - | - | - | - | - | - | - | Unassigned | - |
| Otu27960 | 1 | k__Fungi(100);k__Fungi_unclassified(100);k__Fungi_unclassified(100);k__Fungi_unclassified(100);k__Fungi_unclassified(100);k__Fungi_unclassified(100);k__Fungi_unclassified(100); | - | - | - | - | - | - | - | Unassigned | - |
| Otu27961 | 1 | k__Fungi(100);k__Fungi_unclassified(100);k__Fungi_unclassified(100);k__Fungi_unclassified(100);k__Fungi_unclassified(100);k__Fungi_unclassified(100);k__Fungi_unclassified(100); | - | - | - | - | - | - | - | Unassigned | - |
| Otu27967 | 1 | k__Fungi(100);k__Fungi_unclassified(100);k__Fungi_unclassified(100);k__Fungi_unclassified(100);k__Fungi_unclassified(100);k__Fungi_unclassified(100);k__Fungi_unclassified(100); | - | - | - | - | - | - | - | Unassigned | - |
| Otu27971 | 1 | k__Fungi(100);p__Ascomycota(100);p__Ascomycota_unclassified(100);p__Ascomycota_unclassified(100);p__Ascomycota_unclassified(100);p__Ascomycota_unclassified(100);p__Ascomycota_unclassified(100); | - | - | - | - | - | - | - | Unassigned | - |
| Otu27973 | 1 | k__Fungi(100);k__Fungi_unclassified(100);k__Fungi_unclassified(100);k__Fungi_unclassified(100);k__Fungi_unclassified(100);k__Fungi_unclassified(100);k__Fungi_unclassified(100); | - | - | - | - | - | - | - | Unassigned | - |
| Otu27974 | 1 | k__Fungi(100);p__Ascomycota(100);p__Ascomycota_unclassified(100);p__Ascomycota_unclassified(100);p__Ascomycota_unclassified(100);p__Ascomycota_unclassified(100);p__Ascomycota_unclassified(100); | - | - | - | - | - | - | - | Unassigned | - |
| Otu27975 | 1 | k__Fungi(100);p__Ascomycota(100);p__Ascomycota_unclassified(100);p__Ascomycota_unclassified(100);p__Ascomycota_unclassified(100);p__Ascomycota_unclassified(100);p__Ascomycota_unclassified(100); | - | - | - | - | - | - | - | Unassigned | - |
| Otu27977 | 1 | k__Fungi(100);k__Fungi_unclassified(100);k__Fungi_unclassified(100);k__Fungi_unclassified(100);k__Fungi_unclassified(100);k__Fungi_unclassified(100);k__Fungi_unclassified(100); | - | - | - | - | - | - | - | Unassigned | - |
| Otu27984 | 1 | k__Fungi(100);k__Fungi_unclassified(100);k__Fungi_unclassified(100);k__Fungi_unclassified(100);k__Fungi_unclassified(100);k__Fungi_unclassified(100);k__Fungi_unclassified(100); | - | - | - | - | - | - | - | Unassigned | - |
| Otu27989 | 1 | k__Fungi(100);p__Ascomycota(100);p__Ascomycota_unclassified(100);p__Ascomycota_unclassified(100);p__Ascomycota_unclassified(100);p__Ascomycota_unclassified(100);p__Ascomycota_unclassified(100); | - | - | - | - | - | - | - | Unassigned | - |
| Otu27990 | 1 | k__Fungi(100);k__Fungi_unclassified(100);k__Fungi_unclassified(100);k__Fungi_unclassified(100);k__Fungi_unclassified(100);k__Fungi_unclassified(100);k__Fungi_unclassified(100); | - | - | - | - | - | - | - | Unassigned | - |
| Otu27991 | 1 | k__Fungi(100);p__Ascomycota(100);p__Ascomycota_unclassified(100);p__Ascomycota_unclassified(100);p__Ascomycota_unclassified(100);p__Ascomycota_unclassified(100);p__Ascomycota_unclassified(100); | - | - | - | - | - | - | - | Unassigned | - |
| Otu27992 | 1 | k__Fungi(100);p__Ascomycota(100);p__Ascomycota_unclassified(100);p__Ascomycota_unclassified(100);p__Ascomycota_unclassified(100);p__Ascomycota_unclassified(100);p__Ascomycota_unclassified(100); | - | - | - | - | - | - | - | Unassigned | - |
| Otu27993 | 1 | k__Fungi(100);k__Fungi_unclassified(100);k__Fungi_unclassified(100);k__Fungi_unclassified(100);k__Fungi_unclassified(100);k__Fungi_unclassified(100);k__Fungi_unclassified(100); | - | - | - | - | - | - | - | Unassigned | - |
| Otu27995 | 1 | k__Fungi(100);p__Ascomycota(100);p__Ascomycota_unclassified(100);p__Ascomycota_unclassified(100);p__Ascomycota_unclassified(100);p__Ascomycota_unclassified(100);p__Ascomycota_unclassified(100); | - | - | - | - | - | - | - | Unassigned | - |
| Otu27997 | 1 | k__Fungi(100);p__Ascomycota(100);p__Ascomycota_unclassified(100);p__Ascomycota_unclassified(100);p__Ascomycota_unclassified(100);p__Ascomycota_unclassified(100);p__Ascomycota_unclassified(100); | - | - | - | - | - | - | - | Unassigned | - |
| Otu27998 | 1 | k__Fungi(100);k__Fungi_unclassified(100);k__Fungi_unclassified(100);k__Fungi_unclassified(100);k__Fungi_unclassified(100);k__Fungi_unclassified(100);k__Fungi_unclassified(100); | - | - | - | - | - | - | - | Unassigned | - |
| Otu28006 | 1 | k__Fungi(100);k__Fungi_unclassified(100);k__Fungi_unclassified(100);k__Fungi_unclassified(100);k__Fungi_unclassified(100);k__Fungi_unclassified(100);k__Fungi_unclassified(100); | - | - | - | - | - | - | - | Unassigned | - |
| Otu28008 | 1 | k__Fungi(100);k__Fungi_unclassified(100);k__Fungi_unclassified(100);k__Fungi_unclassified(100);k__Fungi_unclassified(100);k__Fungi_unclassified(100);k__Fungi_unclassified(100); | - | - | - | - | - | - | - | Unassigned | - |
| Otu28009 | 1 | k__Fungi(100);k__Fungi_unclassified(100);k__Fungi_unclassified(100);k__Fungi_unclassified(100);k__Fungi_unclassified(100);k__Fungi_unclassified(100);k__Fungi_unclassified(100); | - | - | - | - | - | - | - | Unassigned | - |
| Otu28010 | 1 | k__Fungi(100);p__Ascomycota(100);p__Ascomycota_unclassified(100);p__Ascomycota_unclassified(100);p__Ascomycota_unclassified(100);p__Ascomycota_unclassified(100);p__Ascomycota_unclassified(100); | - | - | - | - | - | - | - | Unassigned | - |
| Otu28011 | 1 | k__Fungi(100);k__Fungi_unclassified(100);k__Fungi_unclassified(100);k__Fungi_unclassified(100);k__Fungi_unclassified(100);k__Fungi_unclassified(100);k__Fungi_unclassified(100); | - | - | - | - | - | - | - | Unassigned | - |
| Otu28012 | 1 | k__Fungi(100);k__Fungi_unclassified(100);k__Fungi_unclassified(100);k__Fungi_unclassified(100);k__Fungi_unclassified(100);k__Fungi_unclassified(100);k__Fungi_unclassified(100); | - | - | - | - | - | - | - | Unassigned | - |
| Otu28013 | 1 | k__Fungi(100);p__Ascomycota(100);p__Ascomycota_unclassified(100);p__Ascomycota_unclassified(100);p__Ascomycota_unclassified(100);p__Ascomycota_unclassified(100);p__Ascomycota_unclassified(100); | - | - | - | - | - | - | - | Unassigned | - |
| Otu28014 | 1 | k__Fungi(100);p__Ascomycota(100);p__Ascomycota_unclassified(100);p__Ascomycota_unclassified(100);p__Ascomycota_unclassified(100);p__Ascomycota_unclassified(100);p__Ascomycota_unclassified(100); | - | - | - | - | - | - | - | Unassigned | - |
| Otu28015 | 1 | k__Fungi(100);p__Ascomycota(100);c__Dothideomycetes(100);o__Pleosporales(100);o__Pleosporales_unclassified(100);o__Pleosporales_unclassified(100);o__Pleosporales_unclassified(100); | - | - | - | - | - | - | - | Unassigned | - |

| | | | | | | | | | | | |
|---|---|---|---|---|---|---|---|---|---|---|---|
| Otu28017 | 1 | k__Fungi(100);p__Ascomycota(100);c__Sordariomycetes(100);o__Sordariales(100);o__Sordariales_unclassified(100);o__Sordariales_unclassified(100);o__Sordariales_unclassified(100); | - | - | - | - | - | - | - | Unassigned | - |
| Otu28018 | 1 | k__Fungi(100);p__Ascomycota(100);p__Ascomycota_unclassified(100);p__Ascomycota_unclassified(100);p__Ascomycota_unclassified(100);p__Ascomycota_unclassified(100);p__Ascomycota_unclassified(100); | - | - | - | - | - | - | - | Unassigned | - |
| Otu28019 | 1 | k__Fungi(100);k__Fungi_unclassified(100);k__Fungi_unclassified(100);k__Fungi_unclassified(100);k__Fungi_unclassified(100);k__Fungi_unclassified(100);k__Fungi_unclassified(100); | - | - | - | - | - | - | - | Unassigned | - |
| Otu28020 | 1 | k__Fungi(100);p__Ascomycota(100);p__Ascomycota_unclassified(100);p__Ascomycota_unclassified(100);p__Ascomycota_unclassified(100);p__Ascomycota_unclassified(100);p__Ascomycota_unclassified(100); | - | - | - | - | - | - | - | Unassigned | - |
| Otu28021 | 1 | k__Fungi(100);p__Ascomycota(100);p__Ascomycota_unclassified(100);p__Ascomycota_unclassified(100);p__Ascomycota_unclassified(100);p__Ascomycota_unclassified(100);p__Ascomycota_unclassified(100); | - | - | - | - | - | - | - | Unassigned | - |
| Otu28022 | 1 | k__Fungi(100);p__Ascomycota(100);p__Ascomycota_unclassified(100);p__Ascomycota_unclassified(100);p__Ascomycota_unclassified(100);p__Ascomycota_unclassified(100);p__Ascomycota_unclassified(100); | - | - | - | - | - | - | - | Unassigned | - |
| Otu28023 | 1 | k__Fungi(100);p__Ascomycota(100);p__Ascomycota_unclassified(100);p__Ascomycota_unclassified(100);p__Ascomycota_unclassified(100);p__Ascomycota_unclassified(100);p__Ascomycota_unclassified(100); | - | - | - | - | - | - | - | Unassigned | - |
| Otu28024 | 1 | k__Fungi(100);k__Fungi_unclassified(100);k__Fungi_unclassified(100);k__Fungi_unclassified(100);k__Fungi_unclassified(100);k__Fungi_unclassified(100);k__Fungi_unclassified(100); | - | - | - | - | - | - | - | Unassigned | - |
| Otu28026 | 1 | k__Fungi(100);p__Ascomycota(100);p__Ascomycota_unclassified(100);p__Ascomycota_unclassified(100);p__Ascomycota_unclassified(100);p__Ascomycota_unclassified(100);p__Ascomycota_unclassified(100); | - | - | - | - | - | - | - | Unassigned | - |
| Otu28027 | 1 | k__Fungi(100);p__Ascomycota(100);p__Ascomycota_unclassified(100);p__Ascomycota_unclassified(100);p__Ascomycota_unclassified(100);p__Ascomycota_unclassified(100);p__Ascomycota_unclassified(100); | - | - | - | - | - | - | - | Unassigned | - |
| Otu28028 | 1 | k__Fungi(100);k__Fungi_unclassified(100);k__Fungi_unclassified(100);k__Fungi_unclassified(100);k__Fungi_unclassified(100);k__Fungi_unclassified(100);k__Fungi_unclassified(100); | - | - | - | - | - | - | - | Unassigned | - |
| Otu28030 | 1 | k__Fungi(100);k__Fungi_unclassified(100);k__Fungi_unclassified(100);k__Fungi_unclassified(100);k__Fungi_unclassified(100);k__Fungi_unclassified(100);k__Fungi_unclassified(100); | - | - | - | - | - | - | - | Unassigned | - |
| Otu28031 | 1 | k__Fungi(100);p__Ascomycota(100);p__Ascomycota_unclassified(100);p__Ascomycota_unclassified(100);p__Ascomycota_unclassified(100);p__Ascomycota_unclassified(100);p__Ascomycota_unclassified(100); | - | - | - | - | - | - | - | Unassigned | - |
| Otu28032 | 1 | k__Fungi(100);k__Fungi_unclassified(100);k__Fungi_unclassified(100);k__Fungi_unclassified(100);k__Fungi_unclassified(100);k__Fungi_unclassified(100);k__Fungi_unclassified(100); | - | - | - | - | - | - | - | Unassigned | - |
| Otu28033 | 1 | k__Fungi(100);p__Ascomycota(100);p__Ascomycota_unclassified(100);p__Ascomycota_unclassified(100);p__Ascomycota_unclassified(100);p__Ascomycota_unclassified(100);p__Ascomycota_unclassified(100); | - | - | - | - | - | - | - | Unassigned | - |
| Otu28034 | 1 | k__Fungi(100);k__Fungi_unclassified(100);k__Fungi_unclassified(100);k__Fungi_unclassified(100);k__Fungi_unclassified(100);k__Fungi_unclassified(100);k__Fungi_unclassified(100); | - | - | - | - | - | - | - | Unassigned | - |
| Otu28035 | 1 | k__Fungi(100);p__Ascomycota(100);p__Ascomycota_unclassified(100);p__Ascomycota_unclassified(100);p__Ascomycota_unclassified(100);p__Ascomycota_unclassified(100);p__Ascomycota_unclassified(100); | - | - | - | - | - | - | - | Unassigned | - |
| Otu28036 | 1 | k__Fungi(100);k__Fungi_unclassified(100);k__Fungi_unclassified(100);k__Fungi_unclassified(100);k__Fungi_unclassified(100);k__Fungi_unclassified(100);k__Fungi_unclassified(100); | - | - | - | - | - | - | - | Unassigned | - |
| Otu28037 | 1 | k__Fungi(100);p__Ascomycota(100);p__Ascomycota_unclassified(100);p__Ascomycota_unclassified(100);p__Ascomycota_unclassified(100);p__Ascomycota_unclassified(100);p__Ascomycota_unclassified(100); | - | - | - | - | - | - | - | Unassigned | - |
| Otu28038 | 1 | k__Fungi(100);k__Fungi_unclassified(100);k__Fungi_unclassified(100);k__Fungi_unclassified(100);k__Fungi_unclassified(100);k__Fungi_unclassified(100);k__Fungi_unclassified(100); | - | - | - | - | - | - | - | Unassigned | - |
| Otu28039 | 1 | k__Fungi(100);k__Fungi_unclassified(100);k__Fungi_unclassified(100);k__Fungi_unclassified(100);k__Fungi_unclassified(100);k__Fungi_unclassified(100);k__Fungi_unclassified(100); | - | - | - | - | - | - | - | Unassigned | - |
| Otu28040 | 1 | k__Fungi(100);p__Ascomycota(100);p__Ascomycota_unclassified(100);p__Ascomycota_unclassified(100);p__Ascomycota_unclassified(100);p__Ascomycota_unclassified(100);p__Ascomycota_unclassified(100); | - | - | - | - | - | - | - | Unassigned | - |
| Otu28041 | 1 | k__Fungi(100);k__Fungi_unclassified(100);k__Fungi_unclassified(100);k__Fungi_unclassified(100);k__Fungi_unclassified(100);k__Fungi_unclassified(100);k__Fungi_unclassified(100); | - | - | - | - | - | - | - | Unassigned | - |
| Otu28042 | 1 | k__Fungi(100);p__Ascomycota(100);p__Ascomycota_unclassified(100);p__Ascomycota_unclassified(100);p__Ascomycota_unclassified(100);p__Ascomycota_unclassified(100);p__Ascomycota_unclassified(100); | - | - | - | - | - | - | - | Unassigned | - |
| Otu28043 | 1 | k__Fungi(100);k__Fungi_unclassified(100);k__Fungi_unclassified(100);k__Fungi_unclassified(100);k__Fungi_unclassified(100);k__Fungi_unclassified(100);k__Fungi_unclassified(100); | - | - | - | - | - | - | - | Unassigned | - |
| Otu28044 | 1 | k__Fungi(100);p__Ascomycota(100);p__Ascomycota_unclassified(100);p__Ascomycota_unclassified(100);p__Ascomycota_unclassified(100);p__Ascomycota_unclassified(100);p__Ascomycota_unclassified(100); | - | - | - | - | - | - | - | Unassigned | - |
| Otu28045 | 1 | k__Fungi(100);k__Fungi_unclassified(100);k__Fungi_unclassified(100);k__Fungi_unclassified(100);k__Fungi_unclassified(100);k__Fungi_unclassified(100);k__Fungi_unclassified(100); | - | - | - | - | - | - | - | Unassigned | - |
| Otu28048 | 1 | k__Fungi(100);p__Ascomycota(100);p__Ascomycota_unclassified(100);p__Ascomycota_unclassified(100);p__Ascomycota_unclassified(100);p__Ascomycota_unclassified(100);p__Ascomycota_unclassified(100); | - | - | - | - | - | - | - | Unassigned | - |
| Otu28050 | 1 | k__Fungi(100);k__Fungi_unclassified(100);k__Fungi_unclassified(100);k__Fungi_unclassified(100);k__Fungi_unclassified(100);k__Fungi_unclassified(100);k__Fungi_unclassified(100); | - | - | - | - | - | - | - | Unassigned | - |
| Otu28051 | 1 | k__Fungi(100);p__Ascomycota(100);p__Ascomycota_unclassified(100);p__Ascomycota_unclassified(100);p__Ascomycota_unclassified(100);p__Ascomycota_unclassified(100);p__Ascomycota_unclassified(100); | - | - | - | - | - | - | - | Unassigned | - |
| Otu28053 | 1 | k__Fungi(100);p__Ascomycota(100);p__Ascomycota_unclassified(100);p__Ascomycota_unclassified(100);p__Ascomycota_unclassified(100);p__Ascomycota_unclassified(100);p__Ascomycota_unclassified(100); | - | - | - | - | - | - | - | Unassigned | - |
| Otu28054 | 1 | k__Fungi(100);k__Fungi_unclassified(100);k__Fungi_unclassified(100);k__Fungi_unclassified(100);k__Fungi_unclassified(100);k__Fungi_unclassified(100);k__Fungi_unclassified(100); | - | - | - | - | - | - | - | Unassigned | - |
| Otu28055 | 1 | k__Fungi(100);k__Fungi_unclassified(100);k__Fungi_unclassified(100);k__Fungi_unclassified(100);k__Fungi_unclassified(100);k__Fungi_unclassified(100);k__Fungi_unclassified(100); | - | - | - | - | - | - | - | Unassigned | - |
| Otu28056 | 1 | k__Fungi(100);k__Fungi_unclassified(100);k__Fungi_unclassified(100);k__Fungi_unclassified(100);k__Fungi_unclassified(100);k__Fungi_unclassified(100);k__Fungi_unclassified(100); | - | - | - | - | - | - | - | Unassigned | - |
| Otu28057 | 1 | k__Fungi(100);p__Ascomycota(100);p__Ascomycota_unclassified(100);p__Ascomycota_unclassified(100);p__Ascomycota_unclassified(100);p__Ascomycota_unclassified(100);p__Ascomycota_unclassified(100); | - | - | - | - | - | - | - | Unassigned | - |
| Otu28058 | 1 | k__Fungi(100);k__Fungi_unclassified(100);k__Fungi_unclassified(100);k__Fungi_unclassified(100);k__Fungi_unclassified(100);k__Fungi_unclassified(100);k__Fungi_unclassified(100); | - | - | - | - | - | - | - | Unassigned | - |
| Otu28059 | 1 | k__Fungi(100);k__Fungi_unclassified(100);k__Fungi_unclassified(100);k__Fungi_unclassified(100);k__Fungi_unclassified(100);k__Fungi_unclassified(100);k__Fungi_unclassified(100); | - | - | - | - | - | - | - | Unassigned | - |
| Otu28061 | 1 | k__Fungi(100);k__Fungi_unclassified(100);k__Fungi_unclassified(100);k__Fungi_unclassified(100);k__Fungi_unclassified(100);k__Fungi_unclassified(100);k__Fungi_unclassified(100); | - | - | - | - | - | - | - | Unassigned | - |
| Otu28062 | 1 | k__Fungi(100);k__Fungi_unclassified(100);k__Fungi_unclassified(100);k__Fungi_unclassified(100);k__Fungi_unclassified(100);k__Fungi_unclassified(100);k__Fungi_unclassified(100); | - | - | - | - | - | - | - | Unassigned | - |
| Otu28063 | 1 | k__Fungi(100);k__Fungi_unclassified(100);k__Fungi_unclassified(100);k__Fungi_unclassified(100);k__Fungi_unclassified(100);k__Fungi_unclassified(100);k__Fungi_unclassified(100); | - | - | - | - | - | - | - | Unassigned | - |
| Otu28064 | 1 | k__Fungi(100);p__Ascomycota(100);p__Ascomycota_unclassified(100);p__Ascomycota_unclassified(100);p__Ascomycota_unclassified(100);p__Ascomycota_unclassified(100);p__Ascomycota_unclassified(100); | - | - | - | - | - | - | - | Unassigned | - |
| Otu28065 | 1 | k__Fungi(100);p__Ascomycota(100);p__Ascomycota_unclassified(100);p__Ascomycota_unclassified(100);p__Ascomycota_unclassified(100);p__Ascomycota_unclassified(100);p__Ascomycota_unclassified(100); | - | - | - | - | - | - | - | Unassigned | - |
| Otu28066 | 1 | k__Fungi(100);k__Fungi_unclassified(100);k__Fungi_unclassified(100);k__Fungi_unclassified(100);k__Fungi_unclassified(100);k__Fungi_unclassified(100);k__Fungi_unclassified(100); | - | - | - | - | - | - | - | Unassigned | - |
| Otu28067 | 1 | k__Fungi(100);k__Fungi_unclassified(100);k__Fungi_unclassified(100);k__Fungi_unclassified(100);k__Fungi_unclassified(100);k__Fungi_unclassified(100);k__Fungi_unclassified(100); | - | - | - | - | - | - | - | Unassigned | - |
| Otu28068 | 1 | k__Fungi(100);p__Ascomycota(100);p__Ascomycota_unclassified(100);p__Ascomycota_unclassified(100);p__Ascomycota_unclassified(100);p__Ascomycota_unclassified(100);p__Ascomycota_unclassified(100); | - | - | - | - | - | - | - | Unassigned | - |
| Otu28069 | 1 | k__Fungi(100);k__Fungi_unclassified(100);k__Fungi_unclassified(100);k__Fungi_unclassified(100);k__Fungi_unclassified(100);k__Fungi_unclassified(100);k__Fungi_unclassified(100); | - | - | - | - | - | - | - | Unassigned | - |
| Otu28070 | 1 | k__Fungi(100);k__Fungi_unclassified(100);k__Fungi_unclassified(100);k__Fungi_unclassified(100);k__Fungi_unclassified(100);k__Fungi_unclassified(100);k__Fungi_unclassified(100); | - | - | - | - | - | - | - | Unassigned | - |
| Otu28071 | 1 | k__Fungi(100);k__Fungi_unclassified(100);k__Fungi_unclassified(100);k__Fungi_unclassified(100);k__Fungi_unclassified(100);k__Fungi_unclassified(100);k__Fungi_unclassified(100); | - | - | - | - | - | - | - | Unassigned | - |
| Otu28072 | 1 | k__Fungi(100);p__Ascomycota(100);p__Ascomycota_unclassified(100);p__Ascomycota_unclassified(100);p__Ascomycota_unclassified(100);p__Ascomycota_unclassified(100);p__Ascomycota_unclassified(100); | - | - | - | - | - | - | - | Unassigned | - |
| Otu28073 | 1 | k__Fungi(100);k__Fungi_unclassified(100);k__Fungi_unclassified(100);k__Fungi_unclassified(100);k__Fungi_unclassified(100);k__Fungi_unclassified(100);k__Fungi_unclassified(100); | - | - | - | - | - | - | - | Unassigned | - |
| Otu28074 | 1 | k__Fungi(100);k__Fungi_unclassified(100);k__Fungi_unclassified(100);k__Fungi_unclassified(100);k__Fungi_unclassified(100);k__Fungi_unclassified(100);k__Fungi_unclassified(100); | - | - | - | - | - | - | - | Unassigned | - |
| Otu28075 | 1 | k__Fungi(100);k__Fungi_unclassified(100);k__Fungi_unclassified(100);k__Fungi_unclassified(100);k__Fungi_unclassified(100);k__Fungi_unclassified(100);k__Fungi_unclassified(100); | - | - | - | - | - | - | - | Unassigned | - |
| Otu28076 | 1 | k__Fungi(100);k__Fungi_unclassified(100);k__Fungi_unclassified(100);k__Fungi_unclassified(100);k__Fungi_unclassified(100);k__Fungi_unclassified(100);k__Fungi_unclassified(100); | - | - | - | - | - | - | - | Unassigned | - |
| Otu28077 | 1 | k__Fungi(100);k__Fungi_unclassified(100);k__Fungi_unclassified(100);k__Fungi_unclassified(100);k__Fungi_unclassified(100);k__Fungi_unclassified(100);k__Fungi_unclassified(100); | - | - | - | - | - | - | - | Unassigned | - |
| Otu28078 | 1 | k__Fungi(100);k__Fungi_unclassified(100);k__Fungi_unclassified(100);k__Fungi_unclassified(100);k__Fungi_unclassified(100);k__Fungi_unclassified(100);k__Fungi_unclassified(100); | - | - | - | - | - | - | - | Unassigned | - |
| Otu28079 | 1 | k__Fungi(100);k__Fungi_unclassified(100);k__Fungi_unclassified(100);k__Fungi_unclassified(100);k__Fungi_unclassified(100);k__Fungi_unclassified(100);k__Fungi_unclassified(100); | - | - | - | - | - | - | - | Unassigned | - |
| Otu28080 | 1 | k__Fungi(100);k__Fungi_unclassified(100);k__Fungi_unclassified(100);k__Fungi_unclassified(100);k__Fungi_unclassified(100);k__Fungi_unclassified(100);k__Fungi_unclassified(100); | - | - | - | - | - | - | - | Unassigned | - |
| Otu28081 | 1 | k__Fungi(100);k__Fungi_unclassified(100);k__Fungi_unclassified(100);k__Fungi_unclassified(100);k__Fungi_unclassified(100);k__Fungi_unclassified(100);k__Fungi_unclassified(100); | - | - | - | - | - | - | - | Unassigned | - |
| Otu28082 | 1 | k__Fungi(100);k__Fungi_unclassified(100);k__Fungi_unclassified(100);k__Fungi_unclassified(100);k__Fungi_unclassified(100);k__Fungi_unclassified(100);k__Fungi_unclassified(100); | - | - | - | - | - | - | - | Unassigned | - |
| Otu28083 | 1 | k__Fungi(100);p__Ascomycota(100);p__Ascomycota_unclassified(100);p__Ascomycota_unclassified(100);p__Ascomycota_unclassified(100);p__Ascomycota_unclassified(100);p__Ascomycota_unclassified(100); | - | - | - | - | - | - | - | Unassigned | - |
| Otu28085 | 1 | k__Fungi(100);k__Fungi_unclassified(100);k__Fungi_unclassified(100);k__Fungi_unclassified(100);k__Fungi_unclassified(100);k__Fungi_unclassified(100);k__Fungi_unclassified(100); | - | - | - | - | - | - | - | Unassigned | - |
| Otu28086 | 1 | k__Fungi(100);p__Ascomycota(100);p__Ascomycota_unclassified(100);p__Ascomycota_unclassified(100);p__Ascomycota_unclassified(100);p__Ascomycota_unclassified(100);p__Ascomycota_unclassified(100); | - | - | - | - | - | - | - | Unassigned | - |
| Otu28087 | 1 | k__Fungi(100);k__Fungi_unclassified(100);k__Fungi_unclassified(100);k__Fungi_unclassified(100);k__Fungi_unclassified(100);k__Fungi_unclassified(100);k__Fungi_unclassified(100); | - | - | - | - | - | - | - | Unassigned | - |
| Otu28089 | 1 | k__Fungi(100);k__Fungi_unclassified(100);k__Fungi_unclassified(100);k__Fungi_unclassified(100);k__Fungi_unclassified(100);k__Fungi_unclassified(100);k__Fungi_unclassified(100); | - | - | - | - | - | - | - | Unassigned | - |
| Otu28090 | 1 | k__Fungi(100);p__Ascomycota(100);p__Ascomycota_unclassified(100);p__Ascomycota_unclassified(100);p__Ascomycota_unclassified(100);p__Ascomycota_unclassified(100);p__Ascomycota_unclassified(100); | - | - | - | - | - | - | - | Unassigned | - |
| Otu28091 | 1 | k__Fungi(100);p__Ascomycota(100);p__Ascomycota_unclassified(100);p__Ascomycota_unclassified(100);p__Ascomycota_unclassified(100);p__Ascomycota_unclassified(100);p__Ascomycota_unclassified(100); | - | - | - | - | - | - | - | Unassigned | - |
| Otu28092 | 1 | k__Fungi(100);p__Ascomycota(100);p__Ascomycota_unclassified(100);p__Ascomycota_unclassified(100);p__Ascomycota_unclassified(100);p__Ascomycota_unclassified(100);p__Ascomycota_unclassified(100); | - | - | - | - | - | - | - | Unassigned | - |
| Otu28093 | 1 | k__Fungi(100);p__Ascomycota(100);p__Ascomycota_unclassified(100);p__Ascomycota_unclassified(100);p__Ascomycota_unclassified(100);p__Ascomycota_unclassified(100);p__Ascomycota_unclassified(100); | - | - | - | - | - | - | - | Unassigned | - |
| Otu28094 | 1 | k__Fungi(100);k__Fungi_unclassified(100);k__Fungi_unclassified(100);k__Fungi_unclassified(100);k__Fungi_unclassified(100);k__Fungi_unclassified(100);k__Fungi_unclassified(100); | - | - | - | - | - | - | - | Unassigned | - |
| Otu28095 | 1 | k__Fungi(100);k__Fungi_unclassified(100);k__Fungi_unclassified(100);k__Fungi_unclassified(100);k__Fungi_unclassified(100);k__Fungi_unclassified(100);k__Fungi_unclassified(100); | - | - | - | - | - | - | - | Unassigned | - |
| Otu28096 | 1 | k__Fungi(100);k__Fungi_unclassified(100);k__Fungi_unclassified(100);k__Fungi_unclassified(100);k__Fungi_unclassified(100);k__Fungi_unclassified(100);k__Fungi_unclassified(100); | - | - | - | - | - | - | - | Unassigned | - |
| Otu28097 | 1 | k__Fungi(100);p__Ascomycota(100);p__Ascomycota_unclassified(100);p__Ascomycota_unclassified(100);p__Ascomycota_unclassified(100);p__Ascomycota_unclassified(100);p__Ascomycota_unclassified(100); | - | - | - | - | - | - | - | Unassigned | - |
| Otu28098 | 1 | k__Fungi(100);p__Ascomycota(100);p__Ascomycota_unclassified(100);p__Ascomycota_unclassified(100);p__Ascomycota_unclassified(100);p__Ascomycota_unclassified(100);p__Ascomycota_unclassified(100); | - | - | - | - | - | - | - | Unassigned | - |
| Otu28099 | 1 | k__Fungi(100);k__Fungi_unclassified(100);k__Fungi_unclassified(100);k__Fungi_unclassified(100);k__Fungi_unclassified(100);k__Fungi_unclassified(100);k__Fungi_unclassified(100); | - | - | - | - | - | - | - | Unassigned | - |
| Otu28100 | 1 | k__Fungi(100);p__Ascomycota(100);p__Ascomycota_unclassified(100);p__Ascomycota_unclassified(100);p__Ascomycota_unclassified(100);p__Ascomycota_unclassified(100);p__Ascomycota_unclassified(100); | - | - | - | - | - | - | - | Unassigned | - |
| Otu28101 | 1 | k__Fungi(100);p__Ascomycota(100);p__Ascomycota_unclassified(100);p__Ascomycota_unclassified(100);p__Ascomycota_unclassified(100);p__Ascomycota_unclassified(100);p__Ascomycota_unclassified(100); | - | - | - | - | - | - | - | Unassigned | - |
| Otu28103 | 1 | k__Fungi(100);p__Ascomycota(100);p__Ascomycota_unclassified(100);p__Ascomycota_unclassified(100);p__Ascomycota_unclassified(100);p__Ascomycota_unclassified(100);p__Ascomycota_unclassified(100); | - | - | - | - | - | - | - | Unassigned | - |
| Otu28104 | 1 | k__Fungi(100);p__Ascomycota(100);p__Ascomycota_unclassified(100);p__Ascomycota_unclassified(100);p__Ascomycota_unclassified(100);p__Ascomycota_unclassified(100);p__Ascomycota_unclassified(100); | - | - | - | - | - | - | - | Unassigned | - |
| Otu28105 | 1 | k__Fungi(100);k__Fungi_unclassified(100);k__Fungi_unclassified(100);k__Fungi_unclassified(100);k__Fungi_unclassified(100);k__Fungi_unclassified(100);k__Fungi_unclassified(100); | - | - | - | - | - | - | - | Unassigned | - |
| Otu28106 | 1 | k__Fungi(100);k__Fungi_unclassified(100);k__Fungi_unclassified(100);k__Fungi_unclassified(100);k__Fungi_unclassified(100);k__Fungi_unclassified(100);k__Fungi_unclassified(100); | - | - | - | - | - | - | - | Unassigned | - |
| Otu28107 | 1 | k__Fungi(100);k__Fungi_unclassified(100);k__Fungi_unclassified(100);k__Fungi_unclassified(100);k__Fungi_unclassified(100);k__Fungi_unclassified(100);k__Fungi_unclassified(100); | - | - | - | - | - | - | - | Unassigned | - |
| Otu28108 | 1 | k__Fungi(100);p__Ascomycota(100);p__Ascomycota_unclassified(100);p__Ascomycota_unclassified(100);p__Ascomycota_unclassified(100);p__Ascomycota_unclassified(100);p__Ascomycota_unclassified(100); | - | - | - | - | - | - | - | Unassigned | - |
| Otu28109 | 1 | k__Fungi(100);k__Fungi_unclassified(100);k__Fungi_unclassified(100);k__Fungi_unclassified(100);k__Fungi_unclassified(100);k__Fungi_unclassified(100);k__Fungi_unclassified(100); | - | - | - | - | - | - | - | Unassigned | - |
| Otu28110 | 1 | k__Fungi(100);p__Ascomycota(100);p__Ascomycota_unclassified(100);p__Ascomycota_unclassified(100);p__Ascomycota_unclassified(100);p__Ascomycota_unclassified(100);p__Ascomycota_unclassified(100); | - | - | - | - | - | - | - | Unassigned | - |
| Otu28111 | 1 | k__Fungi(100);p__Ascomycota(100);p__Ascomycota_unclassified(100);p__Ascomycota_unclassified(100);p__Ascomycota_unclassified(100);p__Ascomycota_unclassified(100);p__Ascomycota_unclassified(100); | - | - | - | - | - | - | - | Unassigned | - |
| Otu28112 | 1 | k__Fungi(100);k__Fungi_unclassified(100);k__Fungi_unclassified(100);k__Fungi_unclassified(100);k__Fungi_unclassified(100);k__Fungi_unclassified(100);k__Fungi_unclassified(100); | - | - | - | - | - | - | - | Unassigned | - |
| Otu28113 | 1 | k__Fungi(100);k__Fungi_unclassified(100);k__Fungi_unclassified(100);k__Fungi_unclassified(100);k__Fungi_unclassified(100);k__Fungi_unclassified(100);k__Fungi_unclassified(100); | - | - | - | - | - | - | - | Unassigned | - |
| Otu28114 | 1 | k__Fungi(100);p__Ascomycota(100);p__Ascomycota_unclassified(100);p__Ascomycota_unclassified(100);p__Ascomycota_unclassified(100);p__Ascomycota_unclassified(100);p__Ascomycota_unclassified(100); | - | - | - | - | - | - | - | Unassigned | - |
| Otu28115 | 1 | k__Fungi(100);p__Ascomycota(100);p__Ascomycota_unclassified(100);p__Ascomycota_unclassified(100);p__Ascomycota_unclassified(100);p__Ascomycota_unclassified(100);p__Ascomycota_unclassified(100); | - | - | - | - | - | - | - | Unassigned | - |
| Otu28118 | 1 | k__Fungi(100);k__Fungi_unclassified(100);k__Fungi_unclassified(100);k__Fungi_unclassified(100);k__Fungi_unclassified(100);k__Fungi_unclassified(100);k__Fungi_unclassified(100); | - | - | - | - | - | - | - | Unassigned | - |
| Otu28120 | 1 | k__Fungi(100);p__Ascomycota(100);p__Ascomycota_unclassified(100);p__Ascomycota_unclassified(100);p__Ascomycota_unclassified(100);p__Ascomycota_unclassified(100);p__Ascomycota_unclassified(100); | - | - | - | - | - | - | - | Unassigned | - |
| Otu28121 | 1 | k__Fungi(100);k__Fungi_unclassified(100);k__Fungi_unclassified(100);k__Fungi_unclassified(100);k__Fungi_unclassified(100);k__Fungi_unclassified(100);k__Fungi_unclassified(100); | - | - | - | - | - | - | - | Unassigned | - |
| Otu28125 | 1 | k__Fungi(100);p__Ascomycota(100);p__Ascomycota_unclassified(100);p__Ascomycota_unclassified(100);p__Ascomycota_unclassified(100);p__Ascomycota_unclassified(100);p__Ascomycota_unclassified(100); | - | - | - | - | - | - | - | Unassigned | - |
| Otu28126 | 1 | k__Fungi(100);p__Ascomycota(100);p__Ascomycota_unclassified(100);p__Ascomycota_unclassified(100);p__Ascomycota_unclassified(100);p__Ascomycota_unclassified(100);p__Ascomycota_unclassified(100); | - | - | - | - | - | - | - | Unassigned | - |
| Otu28134 | 1 | k__Fungi(100);p__Ascomycota(100);p__Ascomycota_unclassified(100);p__Ascomycota_unclassified(100);p__Ascomycota_unclassified(100);p__Ascomycota_unclassified(100);p__Ascomycota_unclassified(100); | - | - | - | - | - | - | - | Unassigned | - |
| Otu28135 | 1 | k__Fungi(100);p__Ascomycota(100);p__Ascomycota_unclassified(100);p__Ascomycota_unclassified(100);p__Ascomycota_unclassified(100);p__Ascomycota_unclassified(100);p__Ascomycota_unclassified(100); | - | - | - | - | - | - | - | Unassigned | - |
| Otu28144 | 1 | k__Fungi(100);p__Ascomycota(100);p__Ascomycota_unclassified(100);p__Ascomycota_unclassified(100);p__Ascomycota_unclassified(100);p__Ascomycota_unclassified(100);p__Ascomycota_unclassified(100); | - | - | - | - | - | - | - | Unassigned | - |
| Otu28145 | 1 | k__Fungi(100);p__Ascomycota(100);p__Ascomycota_unclassified(100);p__Ascomycota_unclassified(100);p__Ascomycota_unclassified(100);p__Ascomycota_unclassified(100);p__Ascomycota_unclassified(100); | - | - | - | - | - | - | - | Unassigned | - |
| Otu28146 | 1 | k__Fungi(100);k__Fungi_unclassified(100);k__Fungi_unclassified(100);k__Fungi_unclassified(100);k__Fungi_unclassified(100);k__Fungi_unclassified(100);k__Fungi_unclassified(100); | - | - | - | - | - | - | - | Unassigned | - |
| Otu28147 | 1 | k__Fungi(100);k__Fungi_unclassified(100);k__Fungi_unclassified(100);k__Fungi_unclassified(100);k__Fungi_unclassified(100);k__Fungi_unclassified(100);k__Fungi_unclassified(100); | - | - | - | - | - | - | - | Unassigned | - |
| Otu28149 | 1 | k__Fungi(100);k__Fungi_unclassified(100);k__Fungi_unclassified(100);k__Fungi_unclassified(100);k__Fungi_unclassified(100);k__Fungi_unclassified(100);k__Fungi_unclassified(100); | - | - | - | - | - | - | - | Unassigned | - |
| Otu28156 | 1 | k__Fungi(100);p__Ascomycota(100);p__Ascomycota_unclassified(100);p__Ascomycota_unclassified(100);p__Ascomycota_unclassified(100);p__Ascomycota_unclassified(100);p__Ascomycota_unclassified(100); | - | - | - | - | - | - | - | Unassigned | - |
| Otu28159 | 1 | k__Fungi(100);k__Fungi_unclassified(100);k__Fungi_unclassified(100);k__Fungi_unclassified(100);k__Fungi_unclassified(100);k__Fungi_unclassified(100);k__Fungi_unclassified(100); | - | - | - | - | - | - | - | Unassigned | - |
| Otu28160 | 1 | k__Fungi(100);k__Fungi_unclassified(100);k__Fungi_unclassified(100);k__Fungi_unclassified(100);k__Fungi_unclassified(100);k__Fungi_unclassified(100);k__Fungi_unclassified(100); | - | - | - | - | - | - | - | Unassigned | - |
| Otu28161 | 1 | k__Fungi(100);p__Ascomycota(100);p__Ascomycota_unclassified(100);p__Ascomycota_unclassified(100);p__Ascomycota_unclassified(100);p__Ascomycota_unclassified(100);p__Ascomycota_unclassified(100); | - | - | - | - | - | - | - | Unassigned | - |
| Otu28162 | 1 | k__Fungi(100);k__Fungi_unclassified(100);k__Fungi_unclassified(100);k__Fungi_unclassified(100);k__Fungi_unclassified(100);k__Fungi_unclassified(100);k__Fungi_unclassified(100); | - | - | - | - | - | - | - | Unassigned | - |
| Otu28163 | 1 | k__Fungi(100);p__Ascomycota(100);p__Ascomycota_unclassified(100);p__Ascomycota_unclassified(100);p__Ascomycota_unclassified(100);p__Ascomycota_unclassified(100);p__Ascomycota_unclassified(100); | - | - | - | - | - | - | - | Unassigned | - |
| Otu28164 | 1 | k__Fungi(100);k__Fungi_unclassified(100);k__Fungi_unclassified(100);k__Fungi_unclassified(100);k__Fungi_unclassified(100);k__Fungi_unclassified(100);k__Fungi_unclassified(100); | - | - | - | - | - | - | - | Unassigned | - |
| Otu28165 | 1 | k__Fungi(100);p__Ascomycota(100);p__Ascomycota_unclassified(100);p__Ascomycota_unclassified(100);p__Ascomycota_unclassified(100);p__Ascomycota_unclassified(100);p__Ascomycota_unclassified(100); | - | - | - | - | - | - | - | Unassigned | - |
| Otu28166 | 1 | k__Fungi(100);k__Fungi_unclassified(100);k__Fungi_unclassified(100);k__Fungi_unclassified(100);k__Fungi_unclassified(100);k__Fungi_unclassified(100);k__Fungi_unclassified(100); | - | - | - | - | - | - | - | Unassigned | - |
| Otu28167 | 1 | k__Fungi(100);k__Fungi_unclassified(100);k__Fungi_unclassified(100);k__Fungi_unclassified(100);k__Fungi_unclassified(100);k__Fungi_unclassified(100);k__Fungi_unclassified(100); | - | - | - | - | - | - | - | Unassigned | - |
| Otu28168 | 1 | k__Fungi(100);k__Fungi_unclassified(100);k__Fungi_unclassified(100);k__Fungi_unclassified(100);k__Fungi_unclassified(100);k__Fungi_unclassified(100);k__Fungi_unclassified(100); | - | - | - | - | - | - | - | Unassigned | - |
| Otu28169 | 1 | k__Fungi(100);k__Fungi_unclassified(100);k__Fungi_unclassified(100);k__Fungi_unclassified(100);k__Fungi_unclassified(100);k__Fungi_unclassified(100);k__Fungi_unclassified(100); | - | - | - | - | - | - | - | Unassigned | - |
| Otu28175 | 1 | k__Fungi(100);p__Ascomycota(100);p__Ascomycota_unclassified(100);p__Ascomycota_unclassified(100);p__Ascomycota_unclassified(100);p__Ascomycota_unclassified(100);p__Ascomycota_unclassified(100); | - | - | - | - | - | - | - | Unassigned | - |
| Otu28179 | 1 | k__Fungi(100);k__Fungi_unclassified(100);k__Fungi_unclassified(100);k__Fungi_unclassified(100);k__Fungi_unclassified(100);k__Fungi_unclassified(100);k__Fungi_unclassified(100); | - | - | - | - | - | - | - | Unassigned | - |
| Otu28180 | 1 | k__Fungi(100);k__Fungi_unclassified(100);k__Fungi_unclassified(100);k__Fungi_unclassified(100);k__Fungi_unclassified(100);k__Fungi_unclassified(100);k__Fungi_unclassified(100); | - | - | - | - | - | - | - | Unassigned | - |
| Otu28181 | 1 | k__Fungi(100);p__Ascomycota(100);p__Ascomycota_unclassified(100);p__Ascomycota_unclassified(100);p__Ascomycota_unclassified(100);p__Ascomycota_unclassified(100);p__Ascomycota_unclassified(100); | - | - | - | - | - | - | - | Unassigned | - |
| Otu28183 | 1 | k__Fungi(100);k__Fungi_unclassified(100);k__Fungi_unclassified(100);k__Fungi_unclassified(100);k__Fungi_unclassified(100);k__Fungi_unclassified(100);k__Fungi_unclassified(100); | - | - | - | - | - | - | - | Unassigned | - |
| Otu28184 | 1 | k__Fungi(100);p__Ascomycota(100);p__Ascomycota_unclassified(100);p__Ascomycota_unclassified(100);p__Ascomycota_unclassified(100);p__Ascomycota_unclassified(100);p__Ascomycota_unclassified(100); | - | - | - | - | - | - | - | Unassigned | - |
| Otu28185 | 1 | k__Fungi(100);k__Fungi_unclassified(100);k__Fungi_unclassified(100);k__Fungi_unclassified(100);k__Fungi_unclassified(100);k__Fungi_unclassified(100);k__Fungi_unclassified(100); | - | - | - | - | - | - | - | Unassigned | - |
| Otu28186 | 1 | k__Fungi(100);p__Ascomycota(100);p__Ascomycota_unclassified(100);p__Ascomycota_unclassified(100);p__Ascomycota_unclassified(100);p__Ascomycota_unclassified(100);p__Ascomycota_unclassified(100); | - | - | - | - | - | - | - | Unassigned | - |
| Otu28187 | 1 | k__Fungi(100);p__Ascomycota(100);p__Ascomycota_unclassified(100);p__Ascomycota_unclassified(100);p__Ascomycota_unclassified(100);p__Ascomycota_unclassified(100);p__Ascomycota_unclassified(100); | - | - | - | - | - | - | - | Unassigned | - |
| Otu28188 | 1 | k__Fungi(100);k__Fungi_unclassified(100);k__Fungi_unclassified(100);k__Fungi_unclassified(100);k__Fungi_unclassified(100);k__Fungi_unclassified(100);k__Fungi_unclassified(100); | - | - | - | - | - | - | - | Unassigned | - |
| Otu28189 | 1 | k__Fungi(100);k__Fungi_unclassified(100);k__Fungi_unclassified(100);k__Fungi_unclassified(100);k__Fungi_unclassified(100);k__Fungi_unclassified(100);k__Fungi_unclassified(100); | - | - | - | - | - | - | - | Unassigned | - |
| Otu28190 | 1 | k__Fungi(100);p__Ascomycota(100);p__Ascomycota_unclassified(100);p__Ascomycota_unclassified(100);p__Ascomycota_unclassified(100);p__Ascomycota_unclassified(100);p__Ascomycota_unclassified(100); | - | - | - | - | - | - | - | Unassigned | - |
| Otu28191 | 1 | k__Fungi(100);k__Fungi_unclassified(100);k__Fungi_unclassified(100);k__Fungi_unclassified(100);k__Fungi_unclassified(100);k__Fungi_unclassified(100);k__Fungi_unclassified(100); | - | - | - | - | - | - | - | Unassigned | - |
| Otu28193 | 1 | k__Fungi(100);p__Ascomycota(100);p__Ascomycota_unclassified(100);p__Ascomycota_unclassified(100);p__Ascomycota_unclassified(100);p__Ascomycota_unclassified(100);p__Ascomycota_unclassified(100); | - | - | - | - | - | - | - | Unassigned | - |
| Otu28194 | 1 | k__Fungi(100);p__Ascomycota(100);p__Ascomycota_unclassified(100);p__Ascomycota_unclassified(100);p__Ascomycota_unclassified(100);p__Ascomycota_unclassified(100);p__Ascomycota_unclassified(100); | - | - | - | - | - | - | - | Unassigned | - |
| Otu28195 | 1 | k__Fungi(100);k__Fungi_unclassified(100);k__Fungi_unclassified(100);k__Fungi_unclassified(100);k__Fungi_unclassified(100);k__Fungi_unclassified(100);k__Fungi_unclassified(100); | - | - | - | - | - | - | - | Unassigned | - |
| Otu28197 | 1 | k__Fungi(100);p__Ascomycota(100);p__Ascomycota_unclassified(100);p__Ascomycota_unclassified(100);p__Ascomycota_unclassified(100);p__Ascomycota_unclassified(100);p__Ascomycota_unclassified(100); | - | - | - | - | - | - | - | Unassigned | - |
| Otu28199 | 1 | k__Fungi(100);k__Fungi_unclassified(100);k__Fungi_unclassified(100);k__Fungi_unclassified(100);k__Fungi_unclassified(100);k__Fungi_unclassified(100);k__Fungi_unclassified(100); | - | - | - | - | - | - | - | Unassigned | - |
| Otu28200 | 1 | k__Fungi(100);p__Ascomycota(100);p__Ascomycota_unclassified(100);p__Ascomycota_unclassified(100);p__Ascomycota_unclassified(100);p__Ascomycota_unclassified(100);p__Ascomycota_unclassified(100); | - | - | - | - | - | - | - | Unassigned | - |
| Otu28201 | 1 | k__Fungi(100);p__Ascomycota(100);p__Ascomycota_unclassified(100);p__Ascomycota_unclassified(100);p__Ascomycota_unclassified(100);p__Ascomycota_unclassified(100);p__Ascomycota_unclassified(100); | - | - | - | - | - | - | - | Unassigned | - |
| Otu28208 | 1 | k__Fungi(100);p__Ascomycota(100);c__Dothideomycetes(100);o__Capnodiales(100);f__Davidiellaceae(100);f__Davidiellaceae_unclassified(100);f__Davidiellaceae_unclassified(100); | - | - | - | - | - | - | - | Unassigned | - |
| Otu28209 | 1 | k__Plantae(100);p__unclassified_Plantae(100);c__unclassified_Plantae(100);o__unclassified_Plantae(100);f__unclassified_Plantae(100);g__unclassified_Plantae(100);s__Plantae_sp(100); | - | - | - | - | - | - | - | Unassigned | - |
| Otu28212 | 1 | k__Fungi(100);k__Fungi_unclassified(100);k__Fungi_unclassified(100);k__Fungi_unclassified(100);k__Fungi_unclassified(100);k__Fungi_unclassified(100);k__Fungi_unclassified(100); | - | - | - | - | - | - | - | Unassigned | - |
| Otu28213 | 1 | k__Fungi(100);p__Ascomycota(100);p__Ascomycota_unclassified(100);p__Ascomycota_unclassified(100);p__Ascomycota_unclassified(100);p__Ascomycota_unclassified(100);p__Ascomycota_unclassified(100); | - | - | - | - | - | - | - | Unassigned | - |
| Otu28215 | 1 | k__Fungi(100);k__Fungi_unclassified(100);k__Fungi_unclassified(100);k__Fungi_unclassified(100);k__Fungi_unclassified(100);k__Fungi_unclassified(100);k__Fungi_unclassified(100); | - | - | - | - | - | - | - | Unassigned | - |
| Otu28219 | 1 | k__Fungi(100);k__Fungi_unclassified(100);k__Fungi_unclassified(100);k__Fungi_unclassified(100);k__Fungi_unclassified(100);k__Fungi_unclassified(100);k__Fungi_unclassified(100); | - | - | - | - | - | - | - | Unassigned | - |
| Otu28227 | 1 | k__Fungi(100);p__Basidiomycota(100);c__Agaricomycetes(100);c__Agaricomycetes_unclassified(100);c__Agaricomycetes_unclassified(100);c__Agaricomycetes_unclassified(100);c__Agaricomycetes_unclassified(100); | - | - | - | - | - | - | - | Unassigned | - |
| Otu28228 | 1 | k__Fungi(100);p__Basidiomycota(100);c__Agaricomycetes(100);c__Agaricomycetes_unclassified(100);c__Agaricomycetes_unclassified(100);c__Agaricomycetes_unclassified(100);c__Agaricomycetes_unclassified(100); | - | - | - | - | - | - | - | Unassigned | - |
| Otu28248 | 1 | k__Fungi(100);p__Basidiomycota(100);c__Agaricomycetes(100);c__Agaricomycetes_unclassified(100);c__Agaricomycetes_unclassified(100);c__Agaricomycetes_unclassified(100);c__Agaricomycetes_unclassified(100); | - | - | - | - | - | - | - | Unassigned | - |
| Otu28252 | 1 | k__Fungi(100);p__Basidiomycota(100);c__Agaricomycetes(100);c__Agaricomycetes_unclassified(100);c__Agaricomycetes_unclassified(100);c__Agaricomycetes_unclassified(100);c__Agaricomycetes_unclassified(100); | - | - | - | - | - | - | - | Unassigned | - |
| Otu28255 | 1 | k__Fungi(100);p__Basidiomycota(100);c__Agaricomycetes(100);o__Thelephorales(100);f__unclassified_Thelephorales(100);g__unclassified_Thelephorales(100);s__Thelephorales_sp(100); | - | - | - | - | - | - | - | Unassigned | - |
| Otu28257 | 1 | k__Fungi(100);p__Basidiomycota(100);c__Agaricomycetes(100);c__Agaricomycetes_unclassified(100);c__Agaricomycetes_unclassified(100);c__Agaricomycetes_unclassified(100);c__Agaricomycetes_unclassified(100); | - | - | - | - | - | - | - | Unassigned | - |
| Otu28258 | 1 | k__Fungi(100);p__Basidiomycota(100);c__Agaricomycetes(100);c__Agaricomycetes_unclassified(100);c__Agaricomycetes_unclassified(100);c__Agaricomycetes_unclassified(100);c__Agaricomycetes_unclassified(100); | - | - | - | - | - | - | - | Unassigned | - |
| Otu28282 | 1 | k__Fungi(100);p__Basidiomycota(100);c__Agaricomycetes(100);o__Polyporales(100);o__Polyporales_unclassified(100);o__Polyporales_unclassified(100);o__Polyporales_unclassified(100); | - | - | - | - | - | - | - | Unassigned | - |
| Otu28287 | 1 | k__Fungi(100);p__Basidiomycota(100);c__Agaricomycetes(100);o__Cantharellales(100);f__Hydnaceae(100);f__Hydnaceae_unclassified(100);f__Hydnaceae_unclassified(100); | - | - | - | - | - | - | - | Unassigned | - |
| Otu28288 | 1 | k__Fungi(100);p__Basidiomycota(100);c__Agaricomycetes(100);o__Cantharellales(100);o__Cantharellales_unclassified(100);o__Cantharellales_unclassified(100);o__Cantharellales_unclassified(100); | - | - | - | - | - | - | - | Unassigned | - |
| Otu28290 | 1 | k__Fungi(100);p__Basidiomycota(100);c__Agaricomycetes(100);o__Trechisporales(100);f__Hydnodontaceae(100);f__Hydnodontaceae_unclassified(100);f__Hydnodontaceae_unclassified(100); | - | - | - | - | - | - | - | Unassigned | - |
| Otu28291 | 1 | k__Fungi(100);p__Basidiomycota(100);c__Agaricomycetes(100);o__Auriculariales(100);o__Auriculariales_unclassified(100);o__Auriculariales_unclassified(100);o__Auriculariales_unclassified(100); | - | - | - | - | - | - | - | Unassigned | - |
| Otu28299 | 1 | k__Fungi(100);p__Basidiomycota(100);c__Agaricomycetes(100);o__Polyporales(100);o__Polyporales_unclassified(100);o__Polyporales_unclassified(100);o__Polyporales_unclassified(100); | - | - | - | - | - | - | - | Unassigned | - |
| Otu28303 | 1 | k__Fungi(100);p__Basidiomycota(100);c__Agaricomycetes(100);c__Agaricomycetes_unclassified(100);c__Agaricomycetes_unclassified(100);c__Agaricomycetes_unclassified(100);c__Agaricomycetes_unclassified(100); | - | - | - | - | - | - | - | Unassigned | - |
| Otu28310 | 1 | k__Fungi(100);p__Basidiomycota(100);c__Agaricomycetes(100);o__Polyporales(100);o__Polyporales_unclassified(100);o__Polyporales_unclassified(100);o__Polyporales_unclassified(100); | - | - | - | - | - | - | - | Unassigned | - |
| Otu28315 | 1 | k__Fungi(100);p__Basidiomycota(100);c__Microbotryomycetes(100);o__Sporidiobolales(100);f__Sporidiobolales_family_Incertae_sedis(100);f__Sporidiobolales_family_Incertae_sedis_unclassified(100);f__Sporidiobolales_family_Incertae_sedis_unclassified(100); | - | - | - | - | - | - | - | Unassigned | - |
| Otu28317 | 1 | k__Fungi(100);p__Basidiomycota(100);c__Agaricomycetes(100);c__Agaricomycetes_unclassified(100);c__Agaricomycetes_unclassified(100);c__Agaricomycetes_unclassified(100);c__Agaricomycetes_unclassified(100); | - | - | - | - | - | - | - | Unassigned | - |
| Otu28320 | 1 | k__Fungi(100);p__Basidiomycota(100);c__Microbotryomycetes(100);o__Sporidiobolales(100);f__Sporidiobolales_family_Incertae_sedis(100);f__Sporidiobolales_family_Incertae_sedis_unclassified(100);f__Sporidiobolales_family_Incertae_sedis_unclassified(100); | - | - | - | - | - | - | - | Unassigned | - |
| Otu28321 | 1 | k__Fungi(100);p__Basidiomycota(100);c__Microbotryomycetes(100);o__Sporidiobolales(100);f__Sporidiobolales_family_Incertae_sedis(100);f__Sporidiobolales_family_Incertae_sedis_unclassified(100);f__Sporidiobolales_family_Incertae_sedis_unclassified(100); | - | - | - | - | - | - | - | Unassigned | - |
| Otu28323 | 1 | k__Fungi(100);p__Basidiomycota(100);c__Tremellomycetes(100);o__Filobasidiales(100);f__Filobasidiaceae(100);f__Filobasidiaceae_unclassified(100);f__Filobasidiaceae_unclassified(100); | - | - | - | - | - | - | - | Unassigned | - |
| Otu28332 | 1 | k__Fungi(100);p__Basidiomycota(100);c__Agaricomycetes(100);o__Cantharellales(100);f__Ceratobasidiaceae(100);g__unclassified_Ceratobasidiaceae(100);s__uncultured_Ceratobasidium(100); | - | - | - | - | - | - | - | Unassigned | - |
| Otu28353 | 1 | k__Fungi(100);p__Basidiomycota(100);c__Agaricomycetes(100);c__Agaricomycetes_unclassified(100);c__Agaricomycetes_unclassified(100);c__Agaricomycetes_unclassified(100);c__Agaricomycetes_unclassified(100); | - | - | - | - | - | - | - | Unassigned | - |
| Otu28354 | 1 | k__Fungi(100);p__Basidiomycota(100);c__Agaricomycetes(100);o__Polyporales(100);o__Polyporales_unclassified(100);o__Polyporales_unclassified(100);o__Polyporales_unclassified(100); | - | - | - | - | - | - | - | Unassigned | - |
| Otu28357 | 1 | k__Fungi(100);p__Basidiomycota(100);c__Microbotryomycetes(100);o__Sporidiobolales(100);f__Sporidiobolales_family_Incertae_sedis(100);f__Sporidiobolales_family_Incertae_sedis_unclassified(100);f__Sporidiobolales_family_Incertae_sedis_unclassified(100); | - | - | - | - | - | - | - | Unassigned | - |
| Otu28372 | 1 | k__Fungi(100);p__Basidiomycota(100);c__Microbotryomycetes(100);o__Sporidiobolales(100);f__Sporidiobolales_family_Incertae_sedis(100);f__Sporidiobolales_family_Incertae_sedis_unclassified(100);f__Sporidiobolales_family_Incertae_sedis_unclassified(100); | - | - | - | - | - | - | - | Unassigned | - |
| Otu28375 | 1 | k__Fungi(100);p__Basidiomycota(100);c__Agaricostilbomycetes(100);c__Agaricostilbomycetes_unclassified(100);c__Agaricostilbomycetes_unclassified(100);c__Agaricostilbomycetes_unclassified(100);c__Agaricostilbomycetes_unclassified(100); | - | - | - | - | - | - | - | Unassigned | - |
| Otu28376 | 1 | k__Fungi(100);p__Basidiomycota(100);c__Microbotryomycetes(100);c__Microbotryomycetes_unclassified(100);c__Microbotryomycetes_unclassified(100);c__Microbotryomycetes_unclassified(100);c__Microbotryomycetes_unclassified(100); | - | - | - | - | - | - | - | Unassigned | - |
| Otu28378 | 1 | k__Fungi(100);p__Basidiomycota(100);c__Agaricomycetes(100);c__Agaricomycetes_unclassified(100);c__Agaricomycetes_unclassified(100);c__Agaricomycetes_unclassified(100);c__Agaricomycetes_unclassified(100); | - | - | - | - | - | - | - | Unassigned | - |
| Otu28382 | 1 | k__Fungi(100);p__Basidiomycota(100);c__Agaricomycetes(100);o__Boletales(100);o__Boletales_unclassified(100);o__Boletales_unclassified(100);o__Boletales_unclassified(100); | - | - | - | - | - | - | - | Unassigned | - |
| Otu28395 | 1 | k__Fungi(100);p__Basidiomycota(100);c__Agaricomycetes(100);o__Polyporales(100);o__Polyporales_unclassified(100);o__Polyporales_unclassified(100);o__Polyporales_unclassified(100); | - | - | - | - | - | - | - | Unassigned | - |
| Otu28397 | 1 | k__Fungi(100);p__Basidiomycota(100);c__Microbotryomycetes(100);o__Sporidiobolales(100);f__Sporidiobolales_family_Incertae_sedis(100);f__Sporidiobolales_family_Incertae_sedis_unclassified(100);f__Sporidiobolales_family_Incertae_sedis_unclassified(100); | - | - | - | - | - | - | - | Unassigned | - |
| Otu28403 | 1 | k__Fungi(100);p__Basidiomycota(100);c__Agaricomycetes(100);o__Corticiales(100);f__unclassified_Corticiales(100);g__unclassified_Corticiales(100);s__Corticiales_sp(100); | - | - | - | - | - | - | - | Unassigned | - |
| Otu28411 | 1 | k__Fungi(100);p__Basidiomycota(100);p__Basidiomycota_unclassified(100);p__Basidiomycota_unclassified(100);p__Basidiomycota_unclassified(100);p__Basidiomycota_unclassified(100);p__Basidiomycota_unclassified(100); | - | - | - | - | - | - | - | Unassigned | - |
| Otu28415 | 1 | k__Fungi(100);p__Basidiomycota(100);c__Agaricomycetes(100);o__Polyporales(100);o__Polyporales_unclassified(100);o__Polyporales_unclassified(100);o__Polyporales_unclassified(100); | - | - | - | - | - | - | - | Unassigned | - |
| Otu28423 | 1 | k__Fungi(100);p__Basidiomycota(100);c__Microbotryomycetes(100);o__Sporidiobolales(100);f__Sporidiobolales_family_Incertae_sedis(100);f__Sporidiobolales_family_Incertae_sedis_unclassified(100);f__Sporidiobolales_family_Incertae_sedis_unclassified(100); | - | - | - | - | - | - | - | Unassigned | - |
| Otu28429 | 1 | k__Fungi(100);p__Basidiomycota(100);c__Agaricomycetes(100);o__Atheliales(100);o__Atheliales_unclassified(100);o__Atheliales_unclassified(100);o__Atheliales_unclassified(100); | - | - | - | - | - | - | - | Unassigned | - |
| Otu28432 | 1 | k__Fungi(100);p__Basidiomycota(100);c__Agaricomycetes(100);o__Polyporales(100);o__Polyporales_unclassified(100);o__Polyporales_unclassified(100);o__Polyporales_unclassified(100); | - | - | - | - | - | - | - | Unassigned | - |
| Otu28484 | 1 | k__Fungi(100);p__Basidiomycota(100);c__Microbotryomycetes(100);o__Sporidiobolales(100);f__Sporidiobolales_family_Incertae_sedis(100);f__Sporidiobolales_family_Incertae_sedis_unclassified(100);f__Sporidiobolales_family_Incertae_sedis_unclassified(100); | - | - | - | - | - | - | - | Unassigned | - |
| Otu28485 | 1 | k__Fungi(100);p__Basidiomycota(100);c__Microbotryomycetes(100);o__Sporidiobolales(100);f__Sporidiobolales_family_Incertae_sedis(100);g__unclassified_Sporidiobolales(100);s__Sporidiobolales_sp(100); | - | - | - | - | - | - | - | Unassigned | - |
| Otu28493 | 1 | k__Fungi(100);p__Basidiomycota(100);c__Agaricomycetes(100);c__Agaricomycetes_unclassified(100);c__Agaricomycetes_unclassified(100);c__Agaricomycetes_unclassified(100);c__Agaricomycetes_unclassified(100); | - | - | - | - | - | - | - | Unassigned | - |
| Otu28494 | 1 | k__Fungi(100);p__Basidiomycota(100);c__Agaricomycetes(100);c__Agaricomycetes_unclassified(100);c__Agaricomycetes_unclassified(100);c__Agaricomycetes_unclassified(100);c__Agaricomycetes_unclassified(100); | - | - | - | - | - | - | - | Unassigned | - |
| Otu28497 | 1 | k__Fungi(100);k__Fungi_unclassified(100);k__Fungi_unclassified(100);k__Fungi_unclassified(100);k__Fungi_unclassified(100);k__Fungi_unclassified(100);k__Fungi_unclassified(100); | - | - | - | - | - | - | - | Unassigned | - |
| Otu28498 | 1 | k__Fungi(100);p__Basidiomycota(100);p__Basidiomycota_unclassified(100);p__Basidiomycota_unclassified(100);p__Basidiomycota_unclassified(100);p__Basidiomycota_unclassified(100);p__Basidiomycota_unclassified(100); | - | - | - | - | - | - | - | Unassigned | - |
| Otu28507 | 1 | k__Fungi(100);k__Fungi_unclassified(100);k__Fungi_unclassified(100);k__Fungi_unclassified(100);k__Fungi_unclassified(100);k__Fungi_unclassified(100);k__Fungi_unclassified(100); | - | - | - | - | - | - | - | Unassigned | - |
| Otu28514 | 1 | k__Fungi(100);p__Basidiomycota(100);c__Agaricomycetes(100);c__Agaricomycetes_unclassified(100);c__Agaricomycetes_unclassified(100);c__Agaricomycetes_unclassified(100);c__Agaricomycetes_unclassified(100); | - | - | - | - | - | - | - | Unassigned | - |
| Otu28517 | 1 | k__Fungi(100);p__Basidiomycota(100);c__Microbotryomycetes(100);o__Sporidiobolales(100);f__Sporidiobolales_family_Incertae_sedis(100);g__unclassified_Sporidiobolales(100);s__Sporidiobolales_sp(100); | - | - | - | - | - | - | - | Unassigned | - |
| Otu28536 | 1 | k__Fungi(100);p__Basidiomycota(100);c__Microbotryomycetes(100);o__Sporidiobolales(100);f__Sporidiobolales_family_Incertae_sedis(100);f__Sporidiobolales_family_Incertae_sedis_unclassified(100);f__Sporidiobolales_family_Incertae_sedis_unclassified(100); | - | - | - | - | - | - | - | Unassigned | - |
| Otu28538 | 1 | k__Fungi(100);p__Basidiomycota(100);p__Basidiomycota_unclassified(100);p__Basidiomycota_unclassified(100);p__Basidiomycota_unclassified(100);p__Basidiomycota_unclassified(100);p__Basidiomycota_unclassified(100); | - | - | - | - | - | - | - | Unassigned | - |
| Otu28542 | 1 | k__Fungi(100);p__Basidiomycota(100);p__Basidiomycota_unclassified(100);p__Basidiomycota_unclassified(100);p__Basidiomycota_unclassified(100);p__Basidiomycota_unclassified(100);p__Basidiomycota_unclassified(100); | - | - | - | - | - | - | - | Unassigned | - |
| Otu28543 | 1 | k__Fungi(100);p__Basidiomycota(100);p__Basidiomycota_unclassified(100);p__Basidiomycota_unclassified(100);p__Basidiomycota_unclassified(100);p__Basidiomycota_unclassified(100);p__Basidiomycota_unclassified(100); | - | - | - | - | - | - | - | Unassigned | - |
| Otu28544 | 1 | k__Fungi(100);p__Basidiomycota(100);p__Basidiomycota_unclassified(100);p__Basidiomycota_unclassified(100);p__Basidiomycota_unclassified(100);p__Basidiomycota_unclassified(100);p__Basidiomycota_unclassified(100); | - | - | - | - | - | - | - | Unassigned | - |
| Otu28549 | 1 | k__Fungi(100);p__Basidiomycota(100);p__Basidiomycota_unclassified(100);p__Basidiomycota_unclassified(100);p__Basidiomycota_unclassified(100);p__Basidiomycota_unclassified(100);p__Basidiomycota_unclassified(100); | - | - | - | - | - | - | - | Unassigned | - |
| Otu28550 | 1 | k__Fungi(100);p__Basidiomycota(100);p__Basidiomycota_unclassified(100);p__Basidiomycota_unclassified(100);p__Basidiomycota_unclassified(100);p__Basidiomycota_unclassified(100);p__Basidiomycota_unclassified(100); | - | - | - | - | - | - | - | Unassigned | - |
| Otu28562 | 1 | k__Fungi(100);p__Basidiomycota(100);p__Basidiomycota_unclassified(100);p__Basidiomycota_unclassified(100);p__Basidiomycota_unclassified(100);p__Basidiomycota_unclassified(100);p__Basidiomycota_unclassified(100); | - | - | - | - | - | - | - | Unassigned | - |
| Otu28564 | 1 | k__Fungi(100);p__Basidiomycota(100);p__Basidiomycota_unclassified(100);p__Basidiomycota_unclassified(100);p__Basidiomycota_unclassified(100);p__Basidiomycota_unclassified(100);p__Basidiomycota_unclassified(100); | - | - | - | - | - | - | - | Unassigned | - |
| Otu28566 | 1 | k__Fungi(100);p__Basidiomycota(100);p__Basidiomycota_unclassified(100);p__Basidiomycota_unclassified(100);p__Basidiomycota_unclassified(100);p__Basidiomycota_unclassified(100);p__Basidiomycota_unclassified(100); | - | - | - | - | - | - | - | Unassigned | - |
| Otu28567 | 1 | k__Fungi(100);p__Basidiomycota(100);p__Basidiomycota_unclassified(100);p__Basidiomycota_unclassified(100);p__Basidiomycota_unclassified(100);p__Basidiomycota_unclassified(100);p__Basidiomycota_unclassified(100); | - | - | - | - | - | - | - | Unassigned | - |
| Otu28573 | 1 | k__Fungi(100);p__Basidiomycota(100);c__Agaricomycetes(100);o__Polyporales(100);f__Phanerochaetaceae(100);f__Phanerochaetaceae_unclassified(100);f__Phanerochaetaceae_unclassified(100); | - | - | - | - | - | - | - | Unassigned | - |
| Otu28575 | 1 | k__Fungi(100);p__Basidiomycota(100);c__Agaricomycetes(100);c__Agaricomycetes_unclassified(100);c__Agaricomycetes_unclassified(100);c__Agaricomycetes_unclassified(100);c__Agaricomycetes_unclassified(100); | - | - | - | - | - | - | - | Unassigned | - |
| Otu28576 | 1 | k__Fungi(100);p__Basidiomycota(100);p__Basidiomycota_unclassified(100);p__Basidiomycota_unclassified(100);p__Basidiomycota_unclassified(100);p__Basidiomycota_unclassified(100);p__Basidiomycota_unclassified(100); | - | - | - | - | - | - | - | Unassigned | - |
| Otu28582 | 1 | k__Fungi(100);p__Basidiomycota(100);p__Basidiomycota_unclassified(100);p__Basidiomycota_unclassified(100);p__Basidiomycota_unclassified(100);p__Basidiomycota_unclassified(100);p__Basidiomycota_unclassified(100); | - | - | - | - | - | - | - | Unassigned | - |
| Otu28583 | 1 | k__Fungi(100);p__Basidiomycota(100);c__Agaricomycetes(100);o__Cantharellales(100);o__Cantharellales_unclassified(100);o__Cantharellales_unclassified(100);o__Cantharellales_unclassified(100); | - | - | - | - | - | - | - | Unassigned | - |
| Otu28587 | 1 | k__Fungi(100);p__Basidiomycota(100);p__Basidiomycota_unclassified(100);p__Basidiomycota_unclassified(100);p__Basidiomycota_unclassified(100);p__Basidiomycota_unclassified(100);p__Basidiomycota_unclassified(100); | - | - | - | - | - | - | - | Unassigned | - |
| Otu28588 | 1 | k__Fungi(100);p__Basidiomycota(100);p__Basidiomycota_unclassified(100);p__Basidiomycota_unclassified(100);p__Basidiomycota_unclassified(100);p__Basidiomycota_unclassified(100);p__Basidiomycota_unclassified(100); | - | - | - | - | - | - | - | Unassigned | - |
| Otu28591 | 1 | k__Fungi(100);p__Basidiomycota(100);p__Basidiomycota_unclassified(100);p__Basidiomycota_unclassified(100);p__Basidiomycota_unclassified(100);p__Basidiomycota_unclassified(100);p__Basidiomycota_unclassified(100); | - | - | - | - | - | - | - | Unassigned | - |
| Otu28592 | 1 | k__Fungi(100);p__Basidiomycota(100);p__Basidiomycota_unclassified(100);p__Basidiomycota_unclassified(100);p__Basidiomycota_unclassified(100);p__Basidiomycota_unclassified(100);p__Basidiomycota_unclassified(100); | - | - | - | - | - | - | - | Unassigned | - |
| Otu28593 | 1 | k__Fungi(100);p__Basidiomycota(100);p__Basidiomycota_unclassified(100);p__Basidiomycota_unclassified(100);p__Basidiomycota_unclassified(100);p__Basidiomycota_unclassified(100);p__Basidiomycota_unclassified(100); | - | - | - | - | - | - | - | Unassigned | - |
| Otu28594 | 1 | k__Fungi(100);p__Basidiomycota(100);p__Basidiomycota_unclassified(100);p__Basidiomycota_unclassified(100);p__Basidiomycota_unclassified(100);p__Basidiomycota_unclassified(100);p__Basidiomycota_unclassified(100); | - | - | - | - | - | - | - | Unassigned | - |
| Otu28596 | 1 | k__Fungi(100);p__Basidiomycota(100);c__Agaricomycetes(100);o__Polyporales(100);o__Polyporales_unclassified(100);o__Polyporales_unclassified(100);o__Polyporales_unclassified(100); | - | - | - | - | - | - | - | Unassigned | - |
| Otu28597 | 1 | k__Fungi(100);p__Basidiomycota(100);c__Agaricomycetes(100);o__Polyporales(100);o__Polyporales_unclassified(100);o__Polyporales_unclassified(100);o__Polyporales_unclassified(100); | - | - | - | - | - | - | - | Unassigned | - |
| Otu28599 | 1 | k__Fungi(100);p__Basidiomycota(100);c__Agaricomycetes(100);o__Polyporales(100);f__unclassified_Polyporales(100);g__unclassified_Polyporales(100);s__Polyporales_sp(100); | - | - | - | - | - | - | - | Unassigned | - |
| Otu28600 | 1 | k__Fungi(100);p__Basidiomycota(100);c__Agaricomycetes(100);o__Cantharellales(100);o__Cantharellales_unclassified(100);o__Cantharellales_unclassified(100);o__Cantharellales_unclassified(100); | - | - | - | - | - | - | - | Unassigned | - |
| Otu28603 | 1 | k__Fungi(100);p__Basidiomycota(100);c__Agaricomycetes(100);o__Cantharellales(100);o__Cantharellales_unclassified(100);o__Cantharellales_unclassified(100);o__Cantharellales_unclassified(100); | - | - | - | - | - | - | - | Unassigned | - |
| Otu28621 | 1 | k__Fungi(100);p__Basidiomycota(100);p__Basidiomycota_unclassified(100);p__Basidiomycota_unclassified(100);p__Basidiomycota_unclassified(100);p__Basidiomycota_unclassified(100);p__Basidiomycota_unclassified(100); | - | - | - | - | - | - | - | Unassigned | - |
| Otu28622 | 1 | k__Fungi(100);p__Basidiomycota(100);p__Basidiomycota_unclassified(100);p__Basidiomycota_unclassified(100);p__Basidiomycota_unclassified(100);p__Basidiomycota_unclassified(100);p__Basidiomycota_unclassified(100); | - | - | - | - | - | - | - | Unassigned | - |
| Otu28623 | 1 | k__Fungi(100);p__Basidiomycota(100);p__Basidiomycota_unclassified(100);p__Basidiomycota_unclassified(100);p__Basidiomycota_unclassified(100);p__Basidiomycota_unclassified(100);p__Basidiomycota_unclassified(100); | - | - | - | - | - | - | - | Unassigned | - |
| Otu28626 | 1 | k__Fungi(100);p__Basidiomycota(100);p__Basidiomycota_unclassified(100);p__Basidiomycota_unclassified(100);p__Basidiomycota_unclassified(100);p__Basidiomycota_unclassified(100);p__Basidiomycota_unclassified(100); | - | - | - | - | - | - | - | Unassigned | - |

| | | | | | | | | | | | |
|---|---|---|---|---|---|---|---|---|---|---|---|
| Otu28629 | 1 | k__Fungi(100);p__Basidiomycota(100);p__Basidiomycota_unclassified(100);p__Basidiomycota_unclassified(100);p__Basidiomycota_unclassified(100);p__Basidiomycota_unclassified(100);p__Basidiomycota_unclassified(100); | - | - | - | - | - | - | - | Unassigned | - |
| Otu28635 | 1 | k__Fungi(100);p__Basidiomycota(100);p__Basidiomycota_unclassified(100);p__Basidiomycota_unclassified(100);p__Basidiomycota_unclassified(100);p__Basidiomycota_unclassified(100);p__Basidiomycota_unclassified(100); | - | - | - | - | - | - | - | Unassigned | - |
| Otu28642 | 1 | k__Fungi(100);p__Basidiomycota(100);p__Basidiomycota_unclassified(100);p__Basidiomycota_unclassified(100);p__Basidiomycota_unclassified(100);p__Basidiomycota_unclassified(100);p__Basidiomycota_unclassified(100); | - | - | - | - | - | - | - | Unassigned | - |
| Otu28643 | 1 | k__Fungi(100);p__Basidiomycota(100);p__Basidiomycota_unclassified(100);p__Basidiomycota_unclassified(100);p__Basidiomycota_unclassified(100);p__Basidiomycota_unclassified(100);p__Basidiomycota_unclassified(100); | - | - | - | - | - | - | - | Unassigned | - |
| Otu28647 | 1 | k__Fungi(100);p__Ascomycota(100);p__Ascomycota_unclassified(100);p__Ascomycota_unclassified(100);p__Ascomycota_unclassified(100);p__Ascomycota_unclassified(100);p__Ascomycota_unclassified(100); | - | - | - | - | - | - | - | Unassigned | - |
| Otu28651 | 1 | k__Fungi(100);p__Ascomycota(100);p__Ascomycota_unclassified(100);p__Ascomycota_unclassified(100);p__Ascomycota_unclassified(100);p__Ascomycota_unclassified(100);p__Ascomycota_unclassified(100); | - | - | - | - | - | - | - | Unassigned | - |
| Otu28652 | 1 | k__Fungi(100);p__Ascomycota(100);p__Ascomycota_unclassified(100);p__Ascomycota_unclassified(100);p__Ascomycota_unclassified(100);p__Ascomycota_unclassified(100);p__Ascomycota_unclassified(100); | - | - | - | - | - | - | - | Unassigned | - |
| Otu28653 | 1 | k__Fungi(100);k__Fungi_unclassified(100);k__Fungi_unclassified(100);k__Fungi_unclassified(100);k__Fungi_unclassified(100);k__Fungi_unclassified(100);k__Fungi_unclassified(100); | - | - | - | - | - | - | - | Unassigned | - |
| Otu28654 | 1 | k__Fungi(100);k__Fungi_unclassified(100);k__Fungi_unclassified(100);k__Fungi_unclassified(100);k__Fungi_unclassified(100);k__Fungi_unclassified(100);k__Fungi_unclassified(100); | - | - | - | - | - | - | - | Unassigned | - |
| Otu28655 | 1 | k__Fungi(100);k__Fungi_unclassified(100);k__Fungi_unclassified(100);k__Fungi_unclassified(100);k__Fungi_unclassified(100);k__Fungi_unclassified(100);k__Fungi_unclassified(100); | - | - | - | - | - | - | - | Unassigned | - |
| Otu28656 | 1 | k__Fungi(100);p__Ascomycota(100);p__Ascomycota_unclassified(100);p__Ascomycota_unclassified(100);p__Ascomycota_unclassified(100);p__Ascomycota_unclassified(100);p__Ascomycota_unclassified(100); | - | - | - | - | - | - | - | Unassigned | - |
| Otu28657 | 1 | k__Fungi(100);p__Ascomycota(100);p__Ascomycota_unclassified(100);p__Ascomycota_unclassified(100);p__Ascomycota_unclassified(100);p__Ascomycota_unclassified(100);p__Ascomycota_unclassified(100); | - | - | - | - | - | - | - | Unassigned | - |
| Otu28658 | 1 | k__Fungi(100);k__Fungi_unclassified(100);k__Fungi_unclassified(100);k__Fungi_unclassified(100);k__Fungi_unclassified(100);k__Fungi_unclassified(100);k__Fungi_unclassified(100); | - | - | - | - | - | - | - | Unassigned | - |
| Otu28659 | 1 | k__Fungi(100);k__Fungi_unclassified(100);k__Fungi_unclassified(100);k__Fungi_unclassified(100);k__Fungi_unclassified(100);k__Fungi_unclassified(100);k__Fungi_unclassified(100); | - | - | - | - | - | - | - | Unassigned | - |
| Otu28660 | 1 | k__Fungi(100);p__Ascomycota(100);p__Ascomycota_unclassified(100);p__Ascomycota_unclassified(100);p__Ascomycota_unclassified(100);p__Ascomycota_unclassified(100);p__Ascomycota_unclassified(100); | - | - | - | - | - | - | - | Unassigned | - |
| Otu28662 | 1 | k__Fungi(100);p__Ascomycota(100);p__Ascomycota_unclassified(100);p__Ascomycota_unclassified(100);p__Ascomycota_unclassified(100);p__Ascomycota_unclassified(100);p__Ascomycota_unclassified(100); | - | - | - | - | - | - | - | Unassigned | - |
| Otu28663 | 1 | k__Fungi(100);k__Fungi_unclassified(100);k__Fungi_unclassified(100);k__Fungi_unclassified(100);k__Fungi_unclassified(100);k__Fungi_unclassified(100);k__Fungi_unclassified(100); | - | - | - | - | - | - | - | Unassigned | - |
| Otu28664 | 1 | k__Fungi(100);p__Ascomycota(100);p__Ascomycota_unclassified(100);p__Ascomycota_unclassified(100);p__Ascomycota_unclassified(100);p__Ascomycota_unclassified(100);p__Ascomycota_unclassified(100); | - | - | - | - | - | - | - | Unassigned | - |
| Otu28665 | 1 | k__Fungi(100);p__Ascomycota(100);p__Ascomycota_unclassified(100);p__Ascomycota_unclassified(100);p__Ascomycota_unclassified(100);p__Ascomycota_unclassified(100);p__Ascomycota_unclassified(100); | - | - | - | - | - | - | - | Unassigned | - |
| Otu28666 | 1 | k__Fungi(100);p__Ascomycota(100);p__Ascomycota_unclassified(100);p__Ascomycota_unclassified(100);p__Ascomycota_unclassified(100);p__Ascomycota_unclassified(100);p__Ascomycota_unclassified(100); | - | - | - | - | - | - | - | Unassigned | - |
| Otu28667 | 1 | k__Fungi(100);k__Fungi_unclassified(100);k__Fungi_unclassified(100);k__Fungi_unclassified(100);k__Fungi_unclassified(100);k__Fungi_unclassified(100);k__Fungi_unclassified(100); | - | - | - | - | - | - | - | Unassigned | - |
| Otu28668 | 1 | k__Fungi(100);p__Ascomycota(100);p__Ascomycota_unclassified(100);p__Ascomycota_unclassified(100);p__Ascomycota_unclassified(100);p__Ascomycota_unclassified(100);p__Ascomycota_unclassified(100); | - | - | - | - | - | - | - | Unassigned | - |
| Otu28669 | 1 | k__Fungi(100);p__Ascomycota(100);p__Ascomycota_unclassified(100);p__Ascomycota_unclassified(100);p__Ascomycota_unclassified(100);p__Ascomycota_unclassified(100);p__Ascomycota_unclassified(100); | - | - | - | - | - | - | - | Unassigned | - |
| Otu28670 | 1 | k__Fungi(100);k__Fungi_unclassified(100);k__Fungi_unclassified(100);k__Fungi_unclassified(100);k__Fungi_unclassified(100);k__Fungi_unclassified(100);k__Fungi_unclassified(100); | - | - | - | - | - | - | - | Unassigned | - |
| Otu28671 | 1 | k__Fungi(100);p__Ascomycota(100);p__Ascomycota_unclassified(100);p__Ascomycota_unclassified(100);p__Ascomycota_unclassified(100);p__Ascomycota_unclassified(100);p__Ascomycota_unclassified(100); | - | - | - | - | - | - | - | Unassigned | - |
| Otu28672 | 1 | k__Fungi(100);p__Ascomycota(100);p__Ascomycota_unclassified(100);p__Ascomycota_unclassified(100);p__Ascomycota_unclassified(100);p__Ascomycota_unclassified(100);p__Ascomycota_unclassified(100); | - | - | - | - | - | - | - | Unassigned | - |
| Otu28673 | 1 | k__Fungi(100);p__Ascomycota(100);p__Ascomycota_unclassified(100);p__Ascomycota_unclassified(100);p__Ascomycota_unclassified(100);p__Ascomycota_unclassified(100);p__Ascomycota_unclassified(100); | - | - | - | - | - | - | - | Unassigned | - |
| Otu28674 | 1 | k__Fungi(100);p__Ascomycota(100);p__Ascomycota_unclassified(100);p__Ascomycota_unclassified(100);p__Ascomycota_unclassified(100);p__Ascomycota_unclassified(100);p__Ascomycota_unclassified(100); | - | - | - | - | - | - | - | Unassigned | - |
| Otu28676 | 1 | k__Fungi(100);k__Fungi_unclassified(100);k__Fungi_unclassified(100);k__Fungi_unclassified(100);k__Fungi_unclassified(100);k__Fungi_unclassified(100);k__Fungi_unclassified(100); | - | - | - | - | - | - | - | Unassigned | - |
| Otu28678 | 1 | k__Fungi(100);k__Fungi_unclassified(100);k__Fungi_unclassified(100);k__Fungi_unclassified(100);k__Fungi_unclassified(100);k__Fungi_unclassified(100);k__Fungi_unclassified(100); | - | - | - | - | - | - | - | Unassigned | - |
| Otu28679 | 1 | k__Fungi(100);k__Fungi_unclassified(100);k__Fungi_unclassified(100);k__Fungi_unclassified(100);k__Fungi_unclassified(100);k__Fungi_unclassified(100);k__Fungi_unclassified(100); | - | - | - | - | - | - | - | Unassigned | - |
| Otu28682 | 1 | k__Fungi(100);p__Ascomycota(100);p__Ascomycota_unclassified(100);p__Ascomycota_unclassified(100);p__Ascomycota_unclassified(100);p__Ascomycota_unclassified(100);p__Ascomycota_unclassified(100); | - | - | - | - | - | - | - | Unassigned | - |
| Otu28685 | 1 | k__Fungi(100);p__Ascomycota(100);p__Ascomycota_unclassified(100);p__Ascomycota_unclassified(100);p__Ascomycota_unclassified(100);p__Ascomycota_unclassified(100);p__Ascomycota_unclassified(100); | - | - | - | - | - | - | - | Unassigned | - |
| Otu28686 | 1 | k__Fungi(100);k__Fungi_unclassified(100);k__Fungi_unclassified(100);k__Fungi_unclassified(100);k__Fungi_unclassified(100);k__Fungi_unclassified(100);k__Fungi_unclassified(100); | - | - | - | - | - | - | - | Unassigned | - |
| Otu28688 | 1 | k__Fungi(100);p__Ascomycota(100);p__Ascomycota_unclassified(100);p__Ascomycota_unclassified(100);p__Ascomycota_unclassified(100);p__Ascomycota_unclassified(100);p__Ascomycota_unclassified(100); | - | - | - | - | - | - | - | Unassigned | - |
| Otu28689 | 1 | k__Fungi(100);p__Ascomycota(100);p__Ascomycota_unclassified(100);p__Ascomycota_unclassified(100);p__Ascomycota_unclassified(100);p__Ascomycota_unclassified(100);p__Ascomycota_unclassified(100); | - | - | - | - | - | - | - | Unassigned | - |
| Otu28690 | 1 | k__Fungi(100);k__Fungi_unclassified(100);k__Fungi_unclassified(100);k__Fungi_unclassified(100);k__Fungi_unclassified(100);k__Fungi_unclassified(100);k__Fungi_unclassified(100); | - | - | - | - | - | - | - | Unassigned | - |
| Otu28701 | 1 | k__Fungi(100);p__Ascomycota(100);p__Ascomycota_unclassified(100);p__Ascomycota_unclassified(100);p__Ascomycota_unclassified(100);p__Ascomycota_unclassified(100);p__Ascomycota_unclassified(100); | - | - | - | - | - | - | - | Unassigned | - |
| Otu28702 | 1 | k__Fungi(100);k__Fungi_unclassified(100);k__Fungi_unclassified(100);k__Fungi_unclassified(100);k__Fungi_unclassified(100);k__Fungi_unclassified(100);k__Fungi_unclassified(100); | - | - | - | - | - | - | - | Unassigned | - |
| Otu28703 | 1 | k__Fungi(100);p__Ascomycota(100);p__Ascomycota_unclassified(100);p__Ascomycota_unclassified(100);p__Ascomycota_unclassified(100);p__Ascomycota_unclassified(100);p__Ascomycota_unclassified(100); | - | - | - | - | - | - | - | Unassigned | - |
| Otu28704 | 1 | k__Fungi(100);p__Ascomycota(100);p__Ascomycota_unclassified(100);p__Ascomycota_unclassified(100);p__Ascomycota_unclassified(100);p__Ascomycota_unclassified(100);p__Ascomycota_unclassified(100); | - | - | - | - | - | - | - | Unassigned | - |
| Otu28705 | 1 | k__Fungi(100);k__Fungi_unclassified(100);k__Fungi_unclassified(100);k__Fungi_unclassified(100);k__Fungi_unclassified(100);k__Fungi_unclassified(100);k__Fungi_unclassified(100); | - | - | - | - | - | - | - | Unassigned | - |
| Otu28706 | 1 | k__Fungi(100);p__Ascomycota(100);p__Ascomycota_unclassified(100);p__Ascomycota_unclassified(100);p__Ascomycota_unclassified(100);p__Ascomycota_unclassified(100);p__Ascomycota_unclassified(100); | - | - | - | - | - | - | - | Unassigned | - |
| Otu28707 | 1 | k__Fungi(100);k__Fungi_unclassified(100);k__Fungi_unclassified(100);k__Fungi_unclassified(100);k__Fungi_unclassified(100);k__Fungi_unclassified(100);k__Fungi_unclassified(100); | - | - | - | - | - | - | - | Unassigned | - |
| Otu28708 | 1 | k__Fungi(100);p__Ascomycota(100);p__Ascomycota_unclassified(100);p__Ascomycota_unclassified(100);p__Ascomycota_unclassified(100);p__Ascomycota_unclassified(100);p__Ascomycota_unclassified(100); | - | - | - | - | - | - | - | Unassigned | - |
| Otu28709 | 1 | k__Fungi(100);k__Fungi_unclassified(100);k__Fungi_unclassified(100);k__Fungi_unclassified(100);k__Fungi_unclassified(100);k__Fungi_unclassified(100);k__Fungi_unclassified(100); | - | - | - | - | - | - | - | Unassigned | - |
| Otu28710 | 1 | k__Fungi(100);k__Fungi_unclassified(100);k__Fungi_unclassified(100);k__Fungi_unclassified(100);k__Fungi_unclassified(100);k__Fungi_unclassified(100);k__Fungi_unclassified(100); | - | - | - | - | - | - | - | Unassigned | - |
| Otu28711 | 1 | k__Fungi(100);k__Fungi_unclassified(100);k__Fungi_unclassified(100);k__Fungi_unclassified(100);k__Fungi_unclassified(100);k__Fungi_unclassified(100);k__Fungi_unclassified(100); | - | - | - | - | - | - | - | Unassigned | - |
| Otu28712 | 1 | k__Fungi(100);k__Fungi_unclassified(100);k__Fungi_unclassified(100);k__Fungi_unclassified(100);k__Fungi_unclassified(100);k__Fungi_unclassified(100);k__Fungi_unclassified(100); | - | - | - | - | - | - | - | Unassigned | - |
| Otu28713 | 1 | k__Fungi(100);k__Fungi_unclassified(100);k__Fungi_unclassified(100);k__Fungi_unclassified(100);k__Fungi_unclassified(100);k__Fungi_unclassified(100);k__Fungi_unclassified(100); | - | - | - | - | - | - | - | Unassigned | - |
| Otu28714 | 1 | k__Fungi(100);p__Ascomycota(100);p__Ascomycota_unclassified(100);p__Ascomycota_unclassified(100);p__Ascomycota_unclassified(100);p__Ascomycota_unclassified(100);p__Ascomycota_unclassified(100); | - | - | - | - | - | - | - | Unassigned | - |
| Otu28715 | 1 | k__Fungi(100);p__Ascomycota(100);p__Ascomycota_unclassified(100);p__Ascomycota_unclassified(100);p__Ascomycota_unclassified(100);p__Ascomycota_unclassified(100);p__Ascomycota_unclassified(100); | - | - | - | - | - | - | - | Unassigned | - |
| Otu28716 | 1 | k__Fungi(100);k__Fungi_unclassified(100);k__Fungi_unclassified(100);k__Fungi_unclassified(100);k__Fungi_unclassified(100);k__Fungi_unclassified(100);k__Fungi_unclassified(100); | - | - | - | - | - | - | - | Unassigned | - |
| Otu28717 | 1 | k__Fungi(100);p__Ascomycota(100);p__Ascomycota_unclassified(100);p__Ascomycota_unclassified(100);p__Ascomycota_unclassified(100);p__Ascomycota_unclassified(100);p__Ascomycota_unclassified(100); | - | - | - | - | - | - | - | Unassigned | - |
| Otu28718 | 1 | k__Fungi(100);p__Ascomycota(100);p__Ascomycota_unclassified(100);p__Ascomycota_unclassified(100);p__Ascomycota_unclassified(100);p__Ascomycota_unclassified(100);p__Ascomycota_unclassified(100); | - | - | - | - | - | - | - | Unassigned | - |
| Otu28719 | 1 | k__Fungi(100);p__Ascomycota(100);p__Ascomycota_unclassified(100);p__Ascomycota_unclassified(100);p__Ascomycota_unclassified(100);p__Ascomycota_unclassified(100);p__Ascomycota_unclassified(100); | - | - | - | - | - | - | - | Unassigned | - |
| Otu28720 | 1 | k__Fungi(100);p__Ascomycota(100);p__Ascomycota_unclassified(100);p__Ascomycota_unclassified(100);p__Ascomycota_unclassified(100);p__Ascomycota_unclassified(100);p__Ascomycota_unclassified(100); | - | - | - | - | - | - | - | Unassigned | - |
| Otu28721 | 1 | k__Fungi(100);k__Fungi_unclassified(100);k__Fungi_unclassified(100);k__Fungi_unclassified(100);k__Fungi_unclassified(100);k__Fungi_unclassified(100);k__Fungi_unclassified(100); | - | - | - | - | - | - | - | Unassigned | - |
| Otu28722 | 1 | k__Fungi(100);k__Fungi_unclassified(100);k__Fungi_unclassified(100);k__Fungi_unclassified(100);k__Fungi_unclassified(100);k__Fungi_unclassified(100);k__Fungi_unclassified(100); | - | - | - | - | - | - | - | Unassigned | - |
| Otu28723 | 1 | k__Fungi(100);p__Ascomycota(100);p__Ascomycota_unclassified(100);p__Ascomycota_unclassified(100);p__Ascomycota_unclassified(100);p__Ascomycota_unclassified(100);p__Ascomycota_unclassified(100); | - | - | - | - | - | - | - | Unassigned | - |
| Otu28724 | 1 | k__Fungi(100);p__Ascomycota(100);p__Ascomycota_unclassified(100);p__Ascomycota_unclassified(100);p__Ascomycota_unclassified(100);p__Ascomycota_unclassified(100);p__Ascomycota_unclassified(100); | - | - | - | - | - | - | - | Unassigned | - |
| Otu28725 | 1 | k__Fungi(100);k__Fungi_unclassified(100);k__Fungi_unclassified(100);k__Fungi_unclassified(100);k__Fungi_unclassified(100);k__Fungi_unclassified(100);k__Fungi_unclassified(100); | - | - | - | - | - | - | - | Unassigned | - |
| Otu28726 | 1 | k__Fungi(100);k__Fungi_unclassified(100);k__Fungi_unclassified(100);k__Fungi_unclassified(100);k__Fungi_unclassified(100);k__Fungi_unclassified(100);k__Fungi_unclassified(100); | - | - | - | - | - | - | - | Unassigned | - |
| Otu28727 | 1 | k__Fungi(100);p__Ascomycota(100);p__Ascomycota_unclassified(100);p__Ascomycota_unclassified(100);p__Ascomycota_unclassified(100);p__Ascomycota_unclassified(100);p__Ascomycota_unclassified(100); | - | - | - | - | - | - | - | Unassigned | - |
| Otu28728 | 1 | k__Fungi(100);p__Ascomycota(100);p__Ascomycota_unclassified(100);p__Ascomycota_unclassified(100);p__Ascomycota_unclassified(100);p__Ascomycota_unclassified(100);p__Ascomycota_unclassified(100); | - | - | - | - | - | - | - | Unassigned | - |
| Otu28730 | 1 | k__Fungi(100);p__Ascomycota(100);p__Ascomycota_unclassified(100);p__Ascomycota_unclassified(100);p__Ascomycota_unclassified(100);p__Ascomycota_unclassified(100);p__Ascomycota_unclassified(100); | - | - | - | - | - | - | - | Unassigned | - |
| Otu28731 | 1 | k__Fungi(100);k__Fungi_unclassified(100);k__Fungi_unclassified(100);k__Fungi_unclassified(100);k__Fungi_unclassified(100);k__Fungi_unclassified(100);k__Fungi_unclassified(100); | - | - | - | - | - | - | - | Unassigned | - |
| Otu28732 | 1 | k__Fungi(100);p__Ascomycota(100);p__Ascomycota_unclassified(100);p__Ascomycota_unclassified(100);p__Ascomycota_unclassified(100);p__Ascomycota_unclassified(100);p__Ascomycota_unclassified(100); | - | - | - | - | - | - | - | Unassigned | - |
| Otu28733 | 1 | k__Fungi(100);p__Ascomycota(100);p__Ascomycota_unclassified(100);p__Ascomycota_unclassified(100);p__Ascomycota_unclassified(100);p__Ascomycota_unclassified(100);p__Ascomycota_unclassified(100); | - | - | - | - | - | - | - | Unassigned | - |
| Otu28734 | 1 | k__Fungi(100);p__Ascomycota(100);p__Ascomycota_unclassified(100);p__Ascomycota_unclassified(100);p__Ascomycota_unclassified(100);p__Ascomycota_unclassified(100);p__Ascomycota_unclassified(100); | - | - | - | - | - | - | - | Unassigned | - |
| Otu28735 | 1 | k__Fungi(100);p__Ascomycota(100);p__Ascomycota_unclassified(100);p__Ascomycota_unclassified(100);p__Ascomycota_unclassified(100);p__Ascomycota_unclassified(100);p__Ascomycota_unclassified(100); | - | - | - | - | - | - | - | Unassigned | - |
| Otu28736 | 1 | k__Fungi(100);k__Fungi_unclassified(100);k__Fungi_unclassified(100);k__Fungi_unclassified(100);k__Fungi_unclassified(100);k__Fungi_unclassified(100);k__Fungi_unclassified(100); | - | - | - | - | - | - | - | Unassigned | - |
| Otu28737 | 1 | k__Fungi(100);p__Ascomycota(100);p__Ascomycota_unclassified(100);p__Ascomycota_unclassified(100);p__Ascomycota_unclassified(100);p__Ascomycota_unclassified(100);p__Ascomycota_unclassified(100); | - | - | - | - | - | - | - | Unassigned | - |
| Otu28738 | 1 | k__Fungi(100);p__Ascomycota(100);p__Ascomycota_unclassified(100);p__Ascomycota_unclassified(100);p__Ascomycota_unclassified(100);p__Ascomycota_unclassified(100);p__Ascomycota_unclassified(100); | - | - | - | - | - | - | - | Unassigned | - |
| Otu28739 | 1 | k__Fungi(100);p__Ascomycota(100);p__Ascomycota_unclassified(100);p__Ascomycota_unclassified(100);p__Ascomycota_unclassified(100);p__Ascomycota_unclassified(100);p__Ascomycota_unclassified(100); | - | - | - | - | - | - | - | Unassigned | - |
| Otu28740 | 1 | k__Fungi(100);k__Fungi_unclassified(100);k__Fungi_unclassified(100);k__Fungi_unclassified(100);k__Fungi_unclassified(100);k__Fungi_unclassified(100);k__Fungi_unclassified(100); | - | - | - | - | - | - | - | Unassigned | - |
| Otu28742 | 1 | k__Fungi(100);p__Ascomycota(100);p__Ascomycota_unclassified(100);p__Ascomycota_unclassified(100);p__Ascomycota_unclassified(100);p__Ascomycota_unclassified(100);p__Ascomycota_unclassified(100); | - | - | - | - | - | - | - | Unassigned | - |
| Otu28743 | 1 | k__Fungi(100);k__Fungi_unclassified(100);k__Fungi_unclassified(100);k__Fungi_unclassified(100);k__Fungi_unclassified(100);k__Fungi_unclassified(100);k__Fungi_unclassified(100); | - | - | - | - | - | - | - | Unassigned | - |
| Otu28744 | 1 | k__Fungi(100);p__Ascomycota(100);p__Ascomycota_unclassified(100);p__Ascomycota_unclassified(100);p__Ascomycota_unclassified(100);p__Ascomycota_unclassified(100);p__Ascomycota_unclassified(100); | - | - | - | - | - | - | - | Unassigned | - |
| Otu28745 | 1 | k__Fungi(100);k__Fungi_unclassified(100);k__Fungi_unclassified(100);k__Fungi_unclassified(100);k__Fungi_unclassified(100);k__Fungi_unclassified(100);k__Fungi_unclassified(100); | - | - | - | - | - | - | - | Unassigned | - |
| Otu28746 | 1 | k__Fungi(100);p__Ascomycota(100);p__Ascomycota_unclassified(100);p__Ascomycota_unclassified(100);p__Ascomycota_unclassified(100);p__Ascomycota_unclassified(100);p__Ascomycota_unclassified(100); | - | - | - | - | - | - | - | Unassigned | - |
| Otu28747 | 1 | k__Fungi(100);p__Ascomycota(100);p__Ascomycota_unclassified(100);p__Ascomycota_unclassified(100);p__Ascomycota_unclassified(100);p__Ascomycota_unclassified(100);p__Ascomycota_unclassified(100); | - | - | - | - | - | - | - | Unassigned | - |
| Otu28748 | 1 | k__Fungi(100);k__Fungi_unclassified(100);k__Fungi_unclassified(100);k__Fungi_unclassified(100);k__Fungi_unclassified(100);k__Fungi_unclassified(100);k__Fungi_unclassified(100); | - | - | - | - | - | - | - | Unassigned | - |
| Otu28749 | 1 | k__Fungi(100);k__Fungi_unclassified(100);k__Fungi_unclassified(100);k__Fungi_unclassified(100);k__Fungi_unclassified(100);k__Fungi_unclassified(100);k__Fungi_unclassified(100); | - | - | - | - | - | - | - | Unassigned | - |
| Otu28750 | 1 | k__Fungi(100);k__Fungi_unclassified(100);k__Fungi_unclassified(100);k__Fungi_unclassified(100);k__Fungi_unclassified(100);k__Fungi_unclassified(100);k__Fungi_unclassified(100); | - | - | - | - | - | - | - | Unassigned | - |
| Otu28751 | 1 | k__Fungi(100);p__Ascomycota(100);p__Ascomycota_unclassified(100);p__Ascomycota_unclassified(100);p__Ascomycota_unclassified(100);p__Ascomycota_unclassified(100);p__Ascomycota_unclassified(100); | - | - | - | - | - | - | - | Unassigned | - |
| Otu28752 | 1 | k__Fungi(100);p__Ascomycota(100);p__Ascomycota_unclassified(100);p__Ascomycota_unclassified(100);p__Ascomycota_unclassified(100);p__Ascomycota_unclassified(100);p__Ascomycota_unclassified(100); | - | - | - | - | - | - | - | Unassigned | - |
| Otu28753 | 1 | k__Fungi(100);p__Ascomycota(100);p__Ascomycota_unclassified(100);p__Ascomycota_unclassified(100);p__Ascomycota_unclassified(100);p__Ascomycota_unclassified(100);p__Ascomycota_unclassified(100); | - | - | - | - | - | - | - | Unassigned | - |
| Otu28757 | 1 | k__Fungi(100);k__Fungi_unclassified(100);k__Fungi_unclassified(100);k__Fungi_unclassified(100);k__Fungi_unclassified(100);k__Fungi_unclassified(100);k__Fungi_unclassified(100); | - | - | - | - | - | - | - | Unassigned | - |
| Otu28758 | 1 | k__Fungi(100);k__Fungi_unclassified(100);k__Fungi_unclassified(100);k__Fungi_unclassified(100);k__Fungi_unclassified(100);k__Fungi_unclassified(100);k__Fungi_unclassified(100); | - | - | - | - | - | - | - | Unassigned | - |
| Otu28759 | 1 | k__Fungi(100);k__Fungi_unclassified(100);k__Fungi_unclassified(100);k__Fungi_unclassified(100);k__Fungi_unclassified(100);k__Fungi_unclassified(100);k__Fungi_unclassified(100); | - | - | - | - | - | - | - | Unassigned | - |
| Otu28762 | 1 | k__Plantae(100);p__unclassified_Plantae(100);c__unclassified_Plantae(100);o__unclassified_Plantae(100);f__unclassified_Plantae(100);g__unclassified_Plantae(100);s__Plantae_sp(100); | - | - | - | - | - | - | - | Unassigned | - |
| Otu28763 | 1 | k__Fungi(100);k__Fungi_unclassified(100);k__Fungi_unclassified(100);k__Fungi_unclassified(100);k__Fungi_unclassified(100);k__Fungi_unclassified(100);k__Fungi_unclassified(100); | - | - | - | - | - | - | - | Unassigned | - |
| Otu28766 | 1 | k__Fungi(100);p__Ascomycota(100);p__Ascomycota_unclassified(100);p__Ascomycota_unclassified(100);p__Ascomycota_unclassified(100);p__Ascomycota_unclassified(100);p__Ascomycota_unclassified(100); | - | - | - | - | - | - | - | Unassigned | - |
| Otu28773 | 1 | k__Fungi(100);k__Fungi_unclassified(100);k__Fungi_unclassified(100);k__Fungi_unclassified(100);k__Fungi_unclassified(100);k__Fungi_unclassified(100);k__Fungi_unclassified(100); | - | - | - | - | - | - | - | Unassigned | - |
| Otu28775 | 1 | k__Fungi(100);k__Fungi_unclassified(100);k__Fungi_unclassified(100);k__Fungi_unclassified(100);k__Fungi_unclassified(100);k__Fungi_unclassified(100);k__Fungi_unclassified(100); | - | - | - | - | - | - | - | Unassigned | - |
| Otu28781 | 1 | k__Fungi(100);k__Fungi_unclassified(100);k__Fungi_unclassified(100);k__Fungi_unclassified(100);k__Fungi_unclassified(100);k__Fungi_unclassified(100);k__Fungi_unclassified(100); | - | - | - | - | - | - | - | Unassigned | - |
| Otu28783 | 1 | k__Fungi(100);k__Fungi_unclassified(100);k__Fungi_unclassified(100);k__Fungi_unclassified(100);k__Fungi_unclassified(100);k__Fungi_unclassified(100);k__Fungi_unclassified(100); | - | - | - | - | - | - | - | Unassigned | - |
| Otu28784 | 1 | k__Fungi(100);p__Ascomycota(100);p__Ascomycota_unclassified(100);p__Ascomycota_unclassified(100);p__Ascomycota_unclassified(100);p__Ascomycota_unclassified(100);p__Ascomycota_unclassified(100); | - | - | - | - | - | - | - | Unassigned | - |
| Otu28785 | 1 | k__Fungi(100);k__Fungi_unclassified(100);k__Fungi_unclassified(100);k__Fungi_unclassified(100);k__Fungi_unclassified(100);k__Fungi_unclassified(100);k__Fungi_unclassified(100); | - | - | - | - | - | - | - | Unassigned | - |
| Otu28788 | 1 | k__Fungi(100);p__Ascomycota(100);p__Ascomycota_unclassified(100);p__Ascomycota_unclassified(100);p__Ascomycota_unclassified(100);p__Ascomycota_unclassified(100);p__Ascomycota_unclassified(100); | - | - | - | - | - | - | - | Unassigned | - |
| Otu28789 | 1 | k__Fungi(100);k__Fungi_unclassified(100);k__Fungi_unclassified(100);k__Fungi_unclassified(100);k__Fungi_unclassified(100);k__Fungi_unclassified(100);k__Fungi_unclassified(100); | - | - | - | - | - | - | - | Unassigned | - |
| Otu28792 | 1 | k__Fungi(100);p__Ascomycota(100);c__Dothideomycetes(100);o__Capnodiales(100);f__Davidiellaceae(100);f__Davidiellaceae_unclassified(100);f__Davidiellaceae_unclassified(100); | - | - | - | - | - | - | - | Unassigned | - |
| Otu28795 | 1 | k__Fungi(100);k__Fungi_unclassified(100);k__Fungi_unclassified(100);k__Fungi_unclassified(100);k__Fungi_unclassified(100);k__Fungi_unclassified(100);k__Fungi_unclassified(100); | - | - | - | - | - | - | - | Unassigned | - |
| Otu28796 | 1 | k__Fungi(100);p__Ascomycota(100);p__Ascomycota_unclassified(100);p__Ascomycota_unclassified(100);p__Ascomycota_unclassified(100);p__Ascomycota_unclassified(100);p__Ascomycota_unclassified(100); | - | - | - | - | - | - | - | Unassigned | - |
| Otu28797 | 1 | k__Fungi(100);p__Ascomycota(100);p__Ascomycota_unclassified(100);p__Ascomycota_unclassified(100);p__Ascomycota_unclassified(100);p__Ascomycota_unclassified(100);p__Ascomycota_unclassified(100); | - | - | - | - | - | - | - | Unassigned | - |
| Otu28800 | 1 | k__Fungi(100);k__Fungi_unclassified(100);k__Fungi_unclassified(100);k__Fungi_unclassified(100);k__Fungi_unclassified(100);k__Fungi_unclassified(100);k__Fungi_unclassified(100); | - | - | - | - | - | - | - | Unassigned | - |
| Otu28801 | 1 | k__Fungi(100);p__Basidiomycota(100);c__Agaricomycetes(100);c__Agaricomycetes_unclassified(100);c__Agaricomycetes_unclassified(100);c__Agaricomycetes_unclassified(100);c__Agaricomycetes_unclassified(100); | - | - | - | - | - | - | - | Unassigned | - |
| Otu28803 | 1 | k__Fungi(100);k__Fungi_unclassified(100);k__Fungi_unclassified(100);k__Fungi_unclassified(100);k__Fungi_unclassified(100);k__Fungi_unclassified(100);k__Fungi_unclassified(100); | - | - | - | - | - | - | - | Unassigned | - |
| Otu28805 | 1 | k__Fungi(100);k__Fungi_unclassified(100);k__Fungi_unclassified(100);k__Fungi_unclassified(100);k__Fungi_unclassified(100);k__Fungi_unclassified(100);k__Fungi_unclassified(100); | - | - | - | - | - | - | - | Unassigned | - |
| Otu28806 | 1 | k__Fungi(100);k__Fungi_unclassified(100);k__Fungi_unclassified(100);k__Fungi_unclassified(100);k__Fungi_unclassified(100);k__Fungi_unclassified(100);k__Fungi_unclassified(100); | - | - | - | - | - | - | - | Unassigned | - |
| Otu28807 | 1 | k__Fungi(100);k__Fungi_unclassified(100);k__Fungi_unclassified(100);k__Fungi_unclassified(100);k__Fungi_unclassified(100);k__Fungi_unclassified(100);k__Fungi_unclassified(100); | - | - | - | - | - | - | - | Unassigned | - |
| Otu28810 | 1 | k__Fungi(100);k__Fungi_unclassified(100);k__Fungi_unclassified(100);k__Fungi_unclassified(100);k__Fungi_unclassified(100);k__Fungi_unclassified(100);k__Fungi_unclassified(100); | - | - | - | - | - | - | - | Unassigned | - |
| Otu28838 | 1 | k__Fungi(100);k__Fungi_unclassified(100);k__Fungi_unclassified(100);k__Fungi_unclassified(100);k__Fungi_unclassified(100);k__Fungi_unclassified(100);k__Fungi_unclassified(100); | - | - | - | - | - | - | - | Unassigned | - |
| Otu28840 | 1 | k__Fungi(100);k__Fungi_unclassified(100);k__Fungi_unclassified(100);k__Fungi_unclassified(100);k__Fungi_unclassified(100);k__Fungi_unclassified(100);k__Fungi_unclassified(100); | - | - | - | - | - | - | - | Unassigned | - |
| Otu28841 | 1 | k__Fungi(100);k__Fungi_unclassified(100);k__Fungi_unclassified(100);k__Fungi_unclassified(100);k__Fungi_unclassified(100);k__Fungi_unclassified(100);k__Fungi_unclassified(100); | - | - | - | - | - | - | - | Unassigned | - |
| Otu28842 | 1 | k__Fungi(100);k__Fungi_unclassified(100);k__Fungi_unclassified(100);k__Fungi_unclassified(100);k__Fungi_unclassified(100);k__Fungi_unclassified(100);k__Fungi_unclassified(100); | - | - | - | - | - | - | - | Unassigned | - |
| Otu28847 | 1 | k__Fungi(100);k__Fungi_unclassified(100);k__Fungi_unclassified(100);k__Fungi_unclassified(100);k__Fungi_unclassified(100);k__Fungi_unclassified(100);k__Fungi_unclassified(100); | - | - | - | - | - | - | - | Unassigned | - |
| Otu28849 | 1 | k__Fungi(100);k__Fungi_unclassified(100);k__Fungi_unclassified(100);k__Fungi_unclassified(100);k__Fungi_unclassified(100);k__Fungi_unclassified(100);k__Fungi_unclassified(100); | - | - | - | - | - | - | - | Unassigned | - |
| Otu28852 | 1 | k__Fungi(100);p__Ascomycota(100);p__Ascomycota_unclassified(100);p__Ascomycota_unclassified(100);p__Ascomycota_unclassified(100);p__Ascomycota_unclassified(100);p__Ascomycota_unclassified(100); | - | - | - | - | - | - | - | Unassigned | - |
| Otu28854 | 1 | k__Fungi(100);k__Fungi_unclassified(100);k__Fungi_unclassified(100);k__Fungi_unclassified(100);k__Fungi_unclassified(100);k__Fungi_unclassified(100);k__Fungi_unclassified(100); | - | - | - | - | - | - | - | Unassigned | - |
| Otu28855 | 1 | k__Fungi(100);k__Fungi_unclassified(100);k__Fungi_unclassified(100);k__Fungi_unclassified(100);k__Fungi_unclassified(100);k__Fungi_unclassified(100);k__Fungi_unclassified(100); | - | - | - | - | - | - | - | Unassigned | - |
| Otu28857 | 1 | k__Fungi(100);k__Fungi_unclassified(100);k__Fungi_unclassified(100);k__Fungi_unclassified(100);k__Fungi_unclassified(100);k__Fungi_unclassified(100);k__Fungi_unclassified(100); | - | - | - | - | - | - | - | Unassigned | - |
| Otu28860 | 1 | k__Fungi(100);k__Fungi_unclassified(100);k__Fungi_unclassified(100);k__Fungi_unclassified(100);k__Fungi_unclassified(100);k__Fungi_unclassified(100);k__Fungi_unclassified(100); | - | - | - | - | - | - | - | Unassigned | - |
| Otu28862 | 1 | k__Fungi(100);k__Fungi_unclassified(100);k__Fungi_unclassified(100);k__Fungi_unclassified(100);k__Fungi_unclassified(100);k__Fungi_unclassified(100);k__Fungi_unclassified(100); | - | - | - | - | - | - | - | Unassigned | - |
| Otu28870 | 1 | k__Fungi(100);k__Fungi_unclassified(100);k__Fungi_unclassified(100);k__Fungi_unclassified(100);k__Fungi_unclassified(100);k__Fungi_unclassified(100);k__Fungi_unclassified(100); | - | - | - | - | - | - | - | Unassigned | - |
| Otu28872 | 1 | k__Fungi(100);k__Fungi_unclassified(100);k__Fungi_unclassified(100);k__Fungi_unclassified(100);k__Fungi_unclassified(100);k__Fungi_unclassified(100);k__Fungi_unclassified(100); | - | - | - | - | - | - | - | Unassigned | - |
| Otu28873 | 1 | k__Fungi(100);p__Ascomycota(100);p__Ascomycota_unclassified(100);p__Ascomycota_unclassified(100);p__Ascomycota_unclassified(100);p__Ascomycota_unclassified(100);p__Ascomycota_unclassified(100); | - | - | - | - | - | - | - | Unassigned | - |
| Otu28875 | 1 | k__Fungi(100);k__Fungi_unclassified(100);k__Fungi_unclassified(100);k__Fungi_unclassified(100);k__Fungi_unclassified(100);k__Fungi_unclassified(100);k__Fungi_unclassified(100); | - | - | - | - | - | - | - | Unassigned | - |
| Otu28876 | 1 | k__Fungi(100);p__Basidiomycota(100);p__Basidiomycota_unclassified(100);p__Basidiomycota_unclassified(100);p__Basidiomycota_unclassified(100);p__Basidiomycota_unclassified(100);p__Basidiomycota_unclassified(100); | - | - | - | - | - | - | - | Unassigned | - |
| Otu28877 | 1 | k__Fungi(100);k__Fungi_unclassified(100);k__Fungi_unclassified(100);k__Fungi_unclassified(100);k__Fungi_unclassified(100);k__Fungi_unclassified(100);k__Fungi_unclassified(100); | - | - | - | - | - | - | - | Unassigned | - |
| Otu28878 | 1 | k__Fungi(100);k__Fungi_unclassified(100);k__Fungi_unclassified(100);k__Fungi_unclassified(100);k__Fungi_unclassified(100);k__Fungi_unclassified(100);k__Fungi_unclassified(100); | - | - | - | - | - | - | - | Unassigned | - |
| Otu28879 | 1 | k__Fungi(100);k__Fungi_unclassified(100);k__Fungi_unclassified(100);k__Fungi_unclassified(100);k__Fungi_unclassified(100);k__Fungi_unclassified(100);k__Fungi_unclassified(100); | - | - | - | - | - | - | - | Unassigned | - |
| Otu28880 | 1 | k__Fungi(100);k__Fungi_unclassified(100);k__Fungi_unclassified(100);k__Fungi_unclassified(100);k__Fungi_unclassified(100);k__Fungi_unclassified(100);k__Fungi_unclassified(100); | - | - | - | - | - | - | - | Unassigned | - |
| Otu28882 | 1 | k__Fungi(100);k__Fungi_unclassified(100);k__Fungi_unclassified(100);k__Fungi_unclassified(100);k__Fungi_unclassified(100);k__Fungi_unclassified(100);k__Fungi_unclassified(100); | - | - | - | - | - | - | - | Unassigned | - |
| Otu28883 | 1 | k__Fungi(100);k__Fungi_unclassified(100);k__Fungi_unclassified(100);k__Fungi_unclassified(100);k__Fungi_unclassified(100);k__Fungi_unclassified(100);k__Fungi_unclassified(100); | - | - | - | - | - | - | - | Unassigned | - |
| Otu28885 | 1 | k__Fungi(100);p__Ascomycota(100);c__Sordariomycetes(100);c__Sordariomycetes_unclassified(100);c__Sordariomycetes_unclassified(100);c__Sordariomycetes_unclassified(100);c__Sordariomycetes_unclassified(100); | - | - | - | - | - | - | - | Unassigned | - |
| Otu28886 | 1 | k__Fungi(100);p__Ascomycota(100);p__Ascomycota_unclassified(100);p__Ascomycota_unclassified(100);p__Ascomycota_unclassified(100);p__Ascomycota_unclassified(100);p__Ascomycota_unclassified(100); | - | - | - | - | - | - | - | Unassigned | - |
| Otu28888 | 1 | k__Fungi(100);p__Basidiomycota(100);p__Basidiomycota_unclassified(100);p__Basidiomycota_unclassified(100);p__Basidiomycota_unclassified(100);p__Basidiomycota_unclassified(100);p__Basidiomycota_unclassified(100); | - | - | - | - | - | - | - | Unassigned | - |
| Otu28889 | 1 | k__Fungi(100);p__Basidiomycota(100);c__Agaricomycetes(100);o__Auriculariales(100);f__unclassified_Auriculariales(100);g__unclassified_Auriculariales(100);s__Auriculariales_sp(100); | - | - | - | - | - | - | - | Unassigned | - |
| Otu28895 | 1 | k__Fungi(100);p__Basidiomycota(100);c__Agaricostilbomycetes(100);c__Agaricostilbomycetes_unclassified(100);c__Agaricostilbomycetes_unclassified(100);c__Agaricostilbomycetes_unclassified(100);c__Agaricostilbomycetes_unclassified(100); | - | - | - | - | - | - | - | Unassigned | - |
| Otu28900 | 1 | k__Fungi(100);k__Fungi_unclassified(100);k__Fungi_unclassified(100);k__Fungi_unclassified(100);k__Fungi_unclassified(100);k__Fungi_unclassified(100);k__Fungi_unclassified(100); | - | - | - | - | - | - | - | Unassigned | - |
| Otu28902 | 1 | k__Fungi(100);p__Basidiomycota(100);p__Basidiomycota_unclassified(100);p__Basidiomycota_unclassified(100);p__Basidiomycota_unclassified(100);p__Basidiomycota_unclassified(100);p__Basidiomycota_unclassified(100); | - | - | - | - | - | - | - | Unassigned | - |
| Otu28905 | 1 | k__Fungi(100);p__Zygomycota(100);c__Zygomycota_class_Incertae_sedis(100);o__Kickxellales(100);o__Kickxellales_unclassified(100);o__Kickxellales_unclassified(100);o__Kickxellales_unclassified(100); | - | - | - | - | - | - | - | Unassigned | - |
| Otu28908 | 1 | k__Fungi(100);p__Chytridiomycota(100);c__unclassified_Chytridiomycota(100);o__unclassified_Chytridiomycota(100);f__unclassified_Chytridiomycota(100);g__unclassified_Chytridiomycota(100);s__Chytridiomycota_sp(100); | - | - | - | - | - | - | - | Unassigned | - |
| Otu28914 | 1 | k__Fungi(100);k__Fungi_unclassified(100);k__Fungi_unclassified(100);k__Fungi_unclassified(100);k__Fungi_unclassified(100);k__Fungi_unclassified(100);k__Fungi_unclassified(100); | - | - | - | - | - | - | - | Unassigned | - |
| Otu28919 | 1 | k__Fungi(100);k__Fungi_unclassified(100);k__Fungi_unclassified(100);k__Fungi_unclassified(100);k__Fungi_unclassified(100);k__Fungi_unclassified(100);k__Fungi_unclassified(100); | - | - | - | - | - | - | - | Unassigned | - |
| Otu28921 | 1 | k__Fungi(100);p__Ascomycota(100);c__Sordariomycetes(100);c__Sordariomycetes_unclassified(100);c__Sordariomycetes_unclassified(100);c__Sordariomycetes_unclassified(100);c__Sordariomycetes_unclassified(100); | - | - | - | - | - | - | - | Unassigned | - |
| Otu28924 | 1 | k__Fungi(100);p__Ascomycota(100);p__Ascomycota_unclassified(100);p__Ascomycota_unclassified(100);p__Ascomycota_unclassified(100);p__Ascomycota_unclassified(100);p__Ascomycota_unclassified(100); | - | - | - | - | - | - | - | Unassigned | - |
| Otu28927 | 1 | k__Fungi(100);k__Fungi_unclassified(100);k__Fungi_unclassified(100);k__Fungi_unclassified(100);k__Fungi_unclassified(100);k__Fungi_unclassified(100);k__Fungi_unclassified(100); | - | - | - | - | - | - | - | Unassigned | - |
| Otu28929 | 1 | k__Fungi(100);k__Fungi_unclassified(100);k__Fungi_unclassified(100);k__Fungi_unclassified(100);k__Fungi_unclassified(100);k__Fungi_unclassified(100);k__Fungi_unclassified(100); | - | - | - | - | - | - | - | Unassigned | - |
| Otu28933 | 1 | k__Fungi(100);p__Basidiomycota(100);c__Agaricostilbomycetes(100);o__Agaricostilbales(100);o__Agaricostilbales_unclassified(100);o__Agaricostilbales_unclassified(100);o__Agaricostilbales_unclassified(100); | - | - | - | - | - | - | - | Unassigned | - |
| Otu28934 | 1 | k__Fungi(100);p__Basidiomycota(100);p__Basidiomycota_unclassified(100);p__Basidiomycota_unclassified(100);p__Basidiomycota_unclassified(100);p__Basidiomycota_unclassified(100);p__Basidiomycota_unclassified(100); | - | - | - | - | - | - | - | Unassigned | - |
| Otu28935 | 1 | k__Fungi(100);p__Basidiomycota(100);c__Exobasidiomycetes(100);c__Exobasidiomycetes_unclassified(100);c__Exobasidiomycetes_unclassified(100);c__Exobasidiomycetes_unclassified(100);c__Exobasidiomycetes_unclassified(100); | - | - | - | - | - | - | - | Unassigned | - |
| Otu28936 | 1 | k__Fungi(100);p__Basidiomycota(100);p__Basidiomycota_unclassified(100);p__Basidiomycota_unclassified(100);p__Basidiomycota_unclassified(100);p__Basidiomycota_unclassified(100);p__Basidiomycota_unclassified(100); | - | - | - | - | - | - | - | Unassigned | - |
| Otu28938 | 1 | k__Fungi(100);p__Basidiomycota(100);p__Basidiomycota_unclassified(100);p__Basidiomycota_unclassified(100);p__Basidiomycota_unclassified(100);p__Basidiomycota_unclassified(100);p__Basidiomycota_unclassified(100); | - | - | - | - | - | - | - | Unassigned | - |
| Otu28941 | 1 | k__Fungi(100);p__Ascomycota(100);p__Ascomycota_unclassified(100);p__Ascomycota_unclassified(100);p__Ascomycota_unclassified(100);p__Ascomycota_unclassified(100);p__Ascomycota_unclassified(100); | - | - | - | - | - | - | - | Unassigned | - |
| Otu28942 | 1 | k__Fungi(100);k__Fungi_unclassified(100);k__Fungi_unclassified(100);k__Fungi_unclassified(100);k__Fungi_unclassified(100);k__Fungi_unclassified(100);k__Fungi_unclassified(100); | - | - | - | - | - | - | - | Unassigned | - |
| Otu28943 | 1 | k__Fungi(100);p__Ascomycota(100);p__Ascomycota_unclassified(100);p__Ascomycota_unclassified(100);p__Ascomycota_unclassified(100);p__Ascomycota_unclassified(100);p__Ascomycota_unclassified(100); | - | - | - | - | - | - | - | Unassigned | - |
| Otu28944 | 1 | k__Fungi(100);k__Fungi_unclassified(100);k__Fungi_unclassified(100);k__Fungi_unclassified(100);k__Fungi_unclassified(100);k__Fungi_unclassified(100);k__Fungi_unclassified(100); | - | - | - | - | - | - | - | Unassigned | - |
| Otu28946 | 1 | k__Fungi(100);k__Fungi_unclassified(100);k__Fungi_unclassified(100);k__Fungi_unclassified(100);k__Fungi_unclassified(100);k__Fungi_unclassified(100);k__Fungi_unclassified(100); | - | - | - | - | - | - | - | Unassigned | - |
| Otu28947 | 1 | k__Fungi(100);k__Fungi_unclassified(100);k__Fungi_unclassified(100);k__Fungi_unclassified(100);k__Fungi_unclassified(100);k__Fungi_unclassified(100);k__Fungi_unclassified(100); | - | - | - | - | - | - | - | Unassigned | - |
| Otu28948 | 1 | k__Fungi(100);p__Ascomycota(100);p__Ascomycota_unclassified(100);p__Ascomycota_unclassified(100);p__Ascomycota_unclassified(100);p__Ascomycota_unclassified(100);p__Ascomycota_unclassified(100); | - | - | - | - | - | - | - | Unassigned | - |
| Otu28949 | 1 | k__Fungi(100);p__Ascomycota(100);p__Ascomycota_unclassified(100);p__Ascomycota_unclassified(100);p__Ascomycota_unclassified(100);p__Ascomycota_unclassified(100);p__Ascomycota_unclassified(100); | - | - | - | - | - | - | - | Unassigned | - |
| Otu28950 | 1 | k__Fungi(100);k__Fungi_unclassified(100);k__Fungi_unclassified(100);k__Fungi_unclassified(100);k__Fungi_unclassified(100);k__Fungi_unclassified(100);k__Fungi_unclassified(100); | - | - | - | - | - | - | - | Unassigned | - |
| Otu28951 | 1 | k__Fungi(100);p__Ascomycota(100);p__Ascomycota_unclassified(100);p__Ascomycota_unclassified(100);p__Ascomycota_unclassified(100);p__Ascomycota_unclassified(100);p__Ascomycota_unclassified(100); | - | - | - | - | - | - | - | Unassigned | - |
| Otu28952 | 1 | k__Fungi(100);p__Ascomycota(100);p__Ascomycota_unclassified(100);p__Ascomycota_unclassified(100);p__Ascomycota_unclassified(100);p__Ascomycota_unclassified(100);p__Ascomycota_unclassified(100); | - | - | - | - | - | - | - | Unassigned | - |
| Otu28954 | 1 | k__Fungi(100);p__Ascomycota(100);p__Ascomycota_unclassified(100);p__Ascomycota_unclassified(100);p__Ascomycota_unclassified(100);p__Ascomycota_unclassified(100);p__Ascomycota_unclassified(100); | - | - | - | - | - | - | - | Unassigned | - |
| Otu28956 | 1 | k__Plantae(100);p__unclassified_Plantae(100);c__unclassified_Plantae(100);o__unclassified_Plantae(100);f__unclassified_Plantae(100);g__unclassified_Plantae(100);s__Plantae_sp(100); | - | - | - | - | - | - | - | Unassigned | - |
| Otu28957 | 1 | k__Fungi(100);p__Zygomycota(100);c__Zygomycota_class_Incertae_sedis(100);o__Mucorales(100);f__Mucoraceae(100);f__Mucoraceae_unclassified(100);f__Mucoraceae_unclassified(100); | - | - | - | - | - | - | - | Unassigned | - |
| Otu28959 | 1 | k__Fungi(100);k__Fungi_unclassified(100);k__Fungi_unclassified(100);k__Fungi_unclassified(100);k__Fungi_unclassified(100);k__Fungi_unclassified(100);k__Fungi_unclassified(100); | - | - | - | - | - | - | - | Unassigned | - |
| Otu28960 | 1 | k__Fungi(100);k__Fungi_unclassified(100);k__Fungi_unclassified(100);k__Fungi_unclassified(100);k__Fungi_unclassified(100);k__Fungi_unclassified(100);k__Fungi_unclassified(100); | - | - | - | - | - | - | - | Unassigned | - |
| Otu28961 | 1 | k__Fungi(100);p__Ascomycota(100);p__Ascomycota_unclassified(100);p__Ascomycota_unclassified(100);p__Ascomycota_unclassified(100);p__Ascomycota_unclassified(100);p__Ascomycota_unclassified(100); | - | - | - | - | - | - | - | Unassigned | - |
| Otu28962 | 1 | k__Fungi(100);p__Ascomycota(100);p__Ascomycota_unclassified(100);p__Ascomycota_unclassified(100);p__Ascomycota_unclassified(100);p__Ascomycota_unclassified(100);p__Ascomycota_unclassified(100); | - | - | - | - | - | - | - | Unassigned | - |
| Otu28963 | 1 | k__Fungi(100);k__Fungi_unclassified(100);k__Fungi_unclassified(100);k__Fungi_unclassified(100);k__Fungi_unclassified(100);k__Fungi_unclassified(100);k__Fungi_unclassified(100); | - | - | - | - | - | - | - | Unassigned | - |
| Otu28964 | 1 | k__Fungi(100);p__Ascomycota(100);p__Ascomycota_unclassified(100);p__Ascomycota_unclassified(100);p__Ascomycota_unclassified(100);p__Ascomycota_unclassified(100);p__Ascomycota_unclassified(100); | - | - | - | - | - | - | - | Unassigned | - |
| Otu28965 | 1 | k__Fungi(100);p__Ascomycota(100);p__Ascomycota_unclassified(100);p__Ascomycota_unclassified(100);p__Ascomycota_unclassified(100);p__Ascomycota_unclassified(100);p__Ascomycota_unclassified(100); | - | - | - | - | - | - | - | Unassigned | - |
| Otu28967 | 1 | k__Fungi(100);p__Ascomycota(100);p__Ascomycota_unclassified(100);p__Ascomycota_unclassified(100);p__Ascomycota_unclassified(100);p__Ascomycota_unclassified(100);p__Ascomycota_unclassified(100); | - | - | - | - | - | - | - | Unassigned | - |
| Otu28968 | 1 | k__Fungi(100);k__Fungi_unclassified(100);k__Fungi_unclassified(100);k__Fungi_unclassified(100);k__Fungi_unclassified(100);k__Fungi_unclassified(100);k__Fungi_unclassified(100); | - | - | - | - | - | - | - | Unassigned | - |
| Otu28971 | 1 | k__Fungi(100);k__Fungi_unclassified(100);k__Fungi_unclassified(100);k__Fungi_unclassified(100);k__Fungi_unclassified(100);k__Fungi_unclassified(100);k__Fungi_unclassified(100); | - | - | - | - | - | - | - | Unassigned | - |
| Otu28972 | 1 | k__Fungi(100);p__Ascomycota(100);p__Ascomycota_unclassified(100);p__Ascomycota_unclassified(100);p__Ascomycota_unclassified(100);p__Ascomycota_unclassified(100);p__Ascomycota_unclassified(100); | - | - | - | - | - | - | - | Unassigned | - |
| Otu28973 | 1 | k__Fungi(100);k__Fungi_unclassified(100);k__Fungi_unclassified(100);k__Fungi_unclassified(100);k__Fungi_unclassified(100);k__Fungi_unclassified(100);k__Fungi_unclassified(100); | - | - | - | - | - | - | - | Unassigned | - |
| Otu28974 | 1 | k__Fungi(100);k__Fungi_unclassified(100);k__Fungi_unclassified(100);k__Fungi_unclassified(100);k__Fungi_unclassified(100);k__Fungi_unclassified(100);k__Fungi_unclassified(100); | - | - | - | - | - | - | - | Unassigned | - |
| Otu28975 | 1 | k__Fungi(100);p__Ascomycota(100);p__Ascomycota_unclassified(100);p__Ascomycota_unclassified(100);p__Ascomycota_unclassified(100);p__Ascomycota_unclassified(100);p__Ascomycota_unclassified(100); | - | - | - | - | - | - | - | Unassigned | - |
| Otu28976 | 1 | k__Fungi(100);k__Fungi_unclassified(100);k__Fungi_unclassified(100);k__Fungi_unclassified(100);k__Fungi_unclassified(100);k__Fungi_unclassified(100);k__Fungi_unclassified(100); | - | - | - | - | - | - | - | Unassigned | - |
| Otu28977 | 1 | k__Fungi(100);p__Ascomycota(100);p__Ascomycota_unclassified(100);p__Ascomycota_unclassified(100);p__Ascomycota_unclassified(100);p__Ascomycota_unclassified(100);p__Ascomycota_unclassified(100); | - | - | - | - | - | - | - | Unassigned | - |
| Otu28978 | 1 | k__Fungi(100);k__Fungi_unclassified(100);k__Fungi_unclassified(100);k__Fungi_unclassified(100);k__Fungi_unclassified(100);k__Fungi_unclassified(100);k__Fungi_unclassified(100); | - | - | - | - | - | - | - | Unassigned | - |
| Otu28979 | 1 | k__Fungi(100);k__Fungi_unclassified(100);k__Fungi_unclassified(100);k__Fungi_unclassified(100);k__Fungi_unclassified(100);k__Fungi_unclassified(100);k__Fungi_unclassified(100); | - | - | - | - | - | - | - | Unassigned | - |
| Otu28980 | 1 | k__Fungi(100);k__Fungi_unclassified(100);k__Fungi_unclassified(100);k__Fungi_unclassified(100);k__Fungi_unclassified(100);k__Fungi_unclassified(100);k__Fungi_unclassified(100); | - | - | - | - | - | - | - | Unassigned | - |
| Otu28981 | 1 | k__Fungi(100);p__Ascomycota(100);p__Ascomycota_unclassified(100);p__Ascomycota_unclassified(100);p__Ascomycota_unclassified(100);p__Ascomycota_unclassified(100);p__Ascomycota_unclassified(100); | - | - | - | - | - | - | - | Unassigned | - |
| Otu28982 | 1 | k__Fungi(100);k__Fungi_unclassified(100);k__Fungi_unclassified(100);k__Fungi_unclassified(100);k__Fungi_unclassified(100);k__Fungi_unclassified(100);k__Fungi_unclassified(100); | - | - | - | - | - | - | - | Unassigned | - |
| Otu28983 | 1 | k__Fungi(100);p__Ascomycota(100);p__Ascomycota_unclassified(100);p__Ascomycota_unclassified(100);p__Ascomycota_unclassified(100);p__Ascomycota_unclassified(100);p__Ascomycota_unclassified(100); | - | - | - | - | - | - | - | Unassigned | - |
| Otu28984 | 1 | k__Fungi(100);p__Ascomycota(100);p__Ascomycota_unclassified(100);p__Ascomycota_unclassified(100);p__Ascomycota_unclassified(100);p__Ascomycota_unclassified(100);p__Ascomycota_unclassified(100); | - | - | - | - | - | - | - | Unassigned | - |
| Otu28985 | 1 | k__Fungi(100);k__Fungi_unclassified(100);k__Fungi_unclassified(100);k__Fungi_unclassified(100);k__Fungi_unclassified(100);k__Fungi_unclassified(100);k__Fungi_unclassified(100); | - | - | - | - | - | - | - | Unassigned | - |
| Otu28986 | 1 | k__Fungi(100);p__Ascomycota(100);p__Ascomycota_unclassified(100);p__Ascomycota_unclassified(100);p__Ascomycota_unclassified(100);p__Ascomycota_unclassified(100);p__Ascomycota_unclassified(100); | - | - | - | - | - | - | - | Unassigned | - |
| Otu28987 | 1 | k__Fungi(100);p__Ascomycota(100);p__Ascomycota_unclassified(100);p__Ascomycota_unclassified(100);p__Ascomycota_unclassified(100);p__Ascomycota_unclassified(100);p__Ascomycota_unclassified(100); | - | - | - | - | - | - | - | Unassigned | - |
| Otu28988 | 1 | k__Fungi(100);p__Ascomycota(100);p__Ascomycota_unclassified(100);p__Ascomycota_unclassified(100);p__Ascomycota_unclassified(100);p__Ascomycota_unclassified(100);p__Ascomycota_unclassified(100); | - | - | - | - | - | - | - | Unassigned | - |
| Otu28989 | 1 | k__Fungi(100);k__Fungi_unclassified(100);k__Fungi_unclassified(100);k__Fungi_unclassified(100);k__Fungi_unclassified(100);k__Fungi_unclassified(100);k__Fungi_unclassified(100); | - | - | - | - | - | - | - | Unassigned | - |
| Otu28990 | 1 | k__Fungi(100);p__Ascomycota(100);p__Ascomycota_unclassified(100);p__Ascomycota_unclassified(100);p__Ascomycota_unclassified(100);p__Ascomycota_unclassified(100);p__Ascomycota_unclassified(100); | - | - | - | - | - | - | - | Unassigned | - |
| Otu28991 | 1 | k__Fungi(100);p__Ascomycota(100);p__Ascomycota_unclassified(100);p__Ascomycota_unclassified(100);p__Ascomycota_unclassified(100);p__Ascomycota_unclassified(100);p__Ascomycota_unclassified(100); | - | - | - | - | - | - | - | Unassigned | - |
| Otu28992 | 1 | k__Fungi(100);k__Fungi_unclassified(100);k__Fungi_unclassified(100);k__Fungi_unclassified(100);k__Fungi_unclassified(100);k__Fungi_unclassified(100);k__Fungi_unclassified(100); | - | - | - | - | - | - | - | Unassigned | - |
| Otu28994 | 1 | k__Fungi(100);k__Fungi_unclassified(100);k__Fungi_unclassified(100);k__Fungi_unclassified(100);k__Fungi_unclassified(100);k__Fungi_unclassified(100);k__Fungi_unclassified(100); | - | - | - | - | - | - | - | Unassigned | - |
| Otu28995 | 1 | k__Fungi(100);k__Fungi_unclassified(100);k__Fungi_unclassified(100);k__Fungi_unclassified(100);k__Fungi_unclassified(100);k__Fungi_unclassified(100);k__Fungi_unclassified(100); | - | - | - | - | - | - | - | Unassigned | - |
| Otu28996 | 1 | k__Fungi(100);p__Ascomycota(100);p__Ascomycota_unclassified(100);p__Ascomycota_unclassified(100);p__Ascomycota_unclassified(100);p__Ascomycota_unclassified(100);p__Ascomycota_unclassified(100); | - | - | - | - | - | - | - | Unassigned | - |
| Otu28997 | 1 | k__Fungi(100);k__Fungi_unclassified(100);k__Fungi_unclassified(100);k__Fungi_unclassified(100);k__Fungi_unclassified(100);k__Fungi_unclassified(100);k__Fungi_unclassified(100); | - | - | - | - | - | - | - | Unassigned | - |
| Otu28998 | 1 | k__Fungi(100);k__Fungi_unclassified(100);k__Fungi_unclassified(100);k__Fungi_unclassified(100);k__Fungi_unclassified(100);k__Fungi_unclassified(100);k__Fungi_unclassified(100); | - | - | - | - | - | - | - | Unassigned | - |
| Otu28999 | 1 | k__Fungi(100);p__Ascomycota(100);p__Ascomycota_unclassified(100);p__Ascomycota_unclassified(100);p__Ascomycota_unclassified(100);p__Ascomycota_unclassified(100);p__Ascomycota_unclassified(100); | - | - | - | - | - | - | - | Unassigned | - |
| Otu29000 | 1 | k__Fungi(100);p__Ascomycota(100);c__Dothideomycetes(100);c__Dothideomycetes_unclassified(100);c__Dothideomycetes_unclassified(100);c__Dothideomycetes_unclassified(100);c__Dothideomycetes_unclassified(100); | - | - | - | - | - | - | - | Unassigned | - |
| Otu29001 | 1 | k__Fungi(100);p__Ascomycota(100);p__Ascomycota_unclassified(100);p__Ascomycota_unclassified(100);p__Ascomycota_unclassified(100);p__Ascomycota_unclassified(100);p__Ascomycota_unclassified(100); | - | - | - | - | - | - | - | Unassigned | - |
| Otu29002 | 1 | k__Fungi(100);p__Ascomycota(100);p__Ascomycota_unclassified(100);p__Ascomycota_unclassified(100);p__Ascomycota_unclassified(100);p__Ascomycota_unclassified(100);p__Ascomycota_unclassified(100); | - | - | - | - | - | - | - | Unassigned | - |
| Otu29005 | 1 | k__Fungi(100);p__Ascomycota(100);p__Ascomycota_unclassified(100);p__Ascomycota_unclassified(100);p__Ascomycota_unclassified(100);p__Ascomycota_unclassified(100);p__Ascomycota_unclassified(100); | - | - | - | - | - | - | - | Unassigned | - |
| Otu29006 | 1 | k__Fungi(100);p__Ascomycota(100);p__Ascomycota_unclassified(100);p__Ascomycota_unclassified(100);p__Ascomycota_unclassified(100);p__Ascomycota_unclassified(100);p__Ascomycota_unclassified(100); | - | - | - | - | - | - | - | Unassigned | - |
| Otu29007 | 1 | k__Fungi(100);k__Fungi_unclassified(100);k__Fungi_unclassified(100);k__Fungi_unclassified(100);k__Fungi_unclassified(100);k__Fungi_unclassified(100);k__Fungi_unclassified(100); | - | - | - | - | - | - | - | Unassigned | - |
| Otu29009 | 1 | k__Fungi(100);k__Fungi_unclassified(100);k__Fungi_unclassified(100);k__Fungi_unclassified(100);k__Fungi_unclassified(100);k__Fungi_unclassified(100);k__Fungi_unclassified(100); | - | - | - | - | - | - | - | Unassigned | - |
| Otu29010 | 1 | k__Fungi(100);p__Ascomycota(100);p__Ascomycota_unclassified(100);p__Ascomycota_unclassified(100);p__Ascomycota_unclassified(100);p__Ascomycota_unclassified(100);p__Ascomycota_unclassified(100); | - | - | - | - | - | - | - | Unassigned | - |
| Otu29011 | 1 | k__Fungi(100);k__Fungi_unclassified(100);k__Fungi_unclassified(100);k__Fungi_unclassified(100);k__Fungi_unclassified(100);k__Fungi_unclassified(100);k__Fungi_unclassified(100); | - | - | - | - | - | - | - | Unassigned | - |

| | | | | | | | | | | | |
|---|---|---|---|---|---|---|---|---|---|---|---|
| Otu29012 | 1 | k__Fungi(100);p__Ascomycota(100);p__Ascomycota_unclassified(100);p__Ascomycota_unclassified(100);p__Ascomycota_unclassified(100);p__Ascomycota_unclassified(100);p__Ascomycota_unclassified(100); | - | - | - | - | - | - | - | Unassigned | - |
| Otu29013 | 1 | k__Fungi(100);k__Fungi_unclassified(100);k__Fungi_unclassified(100);k__Fungi_unclassified(100);k__Fungi_unclassified(100);k__Fungi_unclassified(100);k__Fungi_unclassified(100); | - | - | - | - | - | - | - | Unassigned | - |
| Otu29014 | 1 | k__Fungi(100);k__Fungi_unclassified(100);k__Fungi_unclassified(100);k__Fungi_unclassified(100);k__Fungi_unclassified(100);k__Fungi_unclassified(100);k__Fungi_unclassified(100); | - | - | - | - | - | - | - | Unassigned | - |
| Otu29015 | 1 | k__Fungi(100);k__Fungi_unclassified(100);k__Fungi_unclassified(100);k__Fungi_unclassified(100);k__Fungi_unclassified(100);k__Fungi_unclassified(100);k__Fungi_unclassified(100); | - | - | - | - | - | - | - | Unassigned | - |
| Otu29016 | 1 | k__Fungi(100);p__Basidiomycota(100);c__Microbotryomycetes(100);o__Sporidiobolales(100);f__Sporidiobolales_family_Incertae_sedis(100);f__Sporidiobolales_family_Incertae_sedis_unclassified(100);f__Sporidiobolales_family_Incertae_sedis_unclassified(100); | - | - | - | - | - | - | - | Unassigned | - |
| Otu29018 | 1 | k__Fungi(100);k__Fungi_unclassified(100);k__Fungi_unclassified(100);k__Fungi_unclassified(100);k__Fungi_unclassified(100);k__Fungi_unclassified(100);k__Fungi_unclassified(100); | - | - | - | - | - | - | - | Unassigned | - |
| Otu29033 | 1 | k__Fungi(100);p__Basidiomycota(100);c__Agaricostilbomycetes(100);o__unclassified_Agaricostilbomycetes(100);f__unclassified_Agaricostilbomycetes(100);g__unclassified_Agaricostilbomycetes(100);s__Agaricostilbomycetes_sp(100); | - | - | - | - | - | - | - | Unassigned | - |
| Otu29034 | 1 | k__Fungi(100);p__Basidiomycota(100);c__Agaricomycetes(100);o__Polyporales(100);o__Polyporales_unclassified(100);o__Polyporales_unclassified(100);o__Polyporales_unclassified(100); | - | - | - | - | - | - | - | Unassigned | - |
| Otu29039 | 1 | k__Fungi(100);p__Zygomycota(100);c__Zygomycota_class_Incertae_sedis(100);o__Mucorales(100);f__Mucoraceae(100);g__unclassified_Mucoraceae(100);s__Mucoraceae_sp(100); | - | - | - | - | - | - | - | Unassigned | - |
| Otu29043 | 1 | k__Fungi(100);p__Basidiomycota(100);c__Agaricomycetes(100);c__Agaricomycetes_unclassified(100);c__Agaricomycetes_unclassified(100);c__Agaricomycetes_unclassified(100);c__Agaricomycetes_unclassified(100); | - | - | - | - | - | - | - | Unassigned | - |
| Otu29044 | 1 | k__Fungi(100);p__Basidiomycota(100);c__Agaricomycetes(100);c__Agaricomycetes_unclassified(100);c__Agaricomycetes_unclassified(100);c__Agaricomycetes_unclassified(100);c__Agaricomycetes_unclassified(100); | - | - | - | - | - | - | - | Unassigned | - |
| Otu29045 | 1 | k__Fungi(100);p__Basidiomycota(100);p__Basidiomycota_unclassified(100);p__Basidiomycota_unclassified(100);p__Basidiomycota_unclassified(100);p__Basidiomycota_unclassified(100);p__Basidiomycota_unclassified(100); | - | - | - | - | - | - | - | Unassigned | - |
| Otu29048 | 1 | k__Fungi(100);p__Ascomycota(100);p__Ascomycota_unclassified(100);p__Ascomycota_unclassified(100);p__Ascomycota_unclassified(100);p__Ascomycota_unclassified(100);p__Ascomycota_unclassified(100); | - | - | - | - | - | - | - | Unassigned | - |
| Otu29049 | 1 | k__Fungi(100);k__Fungi_unclassified(100);k__Fungi_unclassified(100);k__Fungi_unclassified(100);k__Fungi_unclassified(100);k__Fungi_unclassified(100);k__Fungi_unclassified(100); | - | - | - | - | - | - | - | Unassigned | - |
| Otu29050 | 1 | k__Fungi(100);p__Ascomycota(100);p__Ascomycota_unclassified(100);p__Ascomycota_unclassified(100);p__Ascomycota_unclassified(100);p__Ascomycota_unclassified(100);p__Ascomycota_unclassified(100); | - | - | - | - | - | - | - | Unassigned | - |
| Otu29051 | 1 | k__Fungi(100);p__Ascomycota(100);p__Ascomycota_unclassified(100);p__Ascomycota_unclassified(100);p__Ascomycota_unclassified(100);p__Ascomycota_unclassified(100);p__Ascomycota_unclassified(100); | - | - | - | - | - | - | - | Unassigned | - |
| Otu29052 | 1 | k__Fungi(100);p__Ascomycota(100);p__Ascomycota_unclassified(100);p__Ascomycota_unclassified(100);p__Ascomycota_unclassified(100);p__Ascomycota_unclassified(100);p__Ascomycota_unclassified(100); | - | - | - | - | - | - | - | Unassigned | - |
| Otu29053 | 1 | k__Fungi(100);p__Ascomycota(100);p__Ascomycota_unclassified(100);p__Ascomycota_unclassified(100);p__Ascomycota_unclassified(100);p__Ascomycota_unclassified(100);p__Ascomycota_unclassified(100); | - | - | - | - | - | - | - | Unassigned | - |
| Otu29054 | 1 | k__Fungi(100);p__Ascomycota(100);p__Ascomycota_unclassified(100);p__Ascomycota_unclassified(100);p__Ascomycota_unclassified(100);p__Ascomycota_unclassified(100);p__Ascomycota_unclassified(100); | - | - | - | - | - | - | - | Unassigned | - |
| Otu29056 | 1 | k__Fungi(100);k__Fungi_unclassified(100);k__Fungi_unclassified(100);k__Fungi_unclassified(100);k__Fungi_unclassified(100);k__Fungi_unclassified(100);k__Fungi_unclassified(100); | - | - | - | - | - | - | - | Unassigned | - |
| Otu29057 | 1 | k__Fungi(100);k__Fungi_unclassified(100);k__Fungi_unclassified(100);k__Fungi_unclassified(100);k__Fungi_unclassified(100);k__Fungi_unclassified(100);k__Fungi_unclassified(100); | - | - | - | - | - | - | - | Unassigned | - |
| Otu29064 | 1 | k__Fungi(100);p__Basidiomycota(100);p__Basidiomycota_unclassified(100);p__Basidiomycota_unclassified(100);p__Basidiomycota_unclassified(100);p__Basidiomycota_unclassified(100);p__Basidiomycota_unclassified(100); | - | - | - | - | - | - | - | Unassigned | - |
| Otu29066 | 1 | k__Fungi(100);p__Basidiomycota(100);c__Microbotryomycetes(100);o__Sporidiobolales(100);o__Sporidiobolales_unclassified(100);o__Sporidiobolales_unclassified(100);o__Sporidiobolales_unclassified(100); | - | - | - | - | - | - | - | Unassigned | - |
| Otu29067 | 1 | k__Fungi(100);p__Basidiomycota(100);c__Microbotryomycetes(100);o__Sporidiobolales(100);f__Sporidiobolales_family_Incertae_sedis(100);g__unclassified_Sporidiobolales(100);s__Sporidiobolales_sp(100); | - | - | - | - | - | - | - | Unassigned | - |
| Otu29068 | 1 | k__Fungi(100);p__Basidiomycota(100);c__Microbotryomycetes(100);o__Sporidiobolales(100);o__Sporidiobolales_unclassified(100);o__Sporidiobolales_unclassified(100);o__Sporidiobolales_unclassified(100); | - | - | - | - | - | - | - | Unassigned | - |
| Otu29071 | 1 | k__Fungi(100);p__Basidiomycota(100);c__Agaricostilbomycetes(100);c__Agaricostilbomycetes_unclassified(100);c__Agaricostilbomycetes_unclassified(100);c__Agaricostilbomycetes_unclassified(100);c__Agaricostilbomycetes_unclassified(100); | - | - | - | - | - | - | - | Unassigned | - |
| Otu29072 | 1 | k__Fungi(100);p__Ascomycota(100);p__Ascomycota_unclassified(100);p__Ascomycota_unclassified(100);p__Ascomycota_unclassified(100);p__Ascomycota_unclassified(100);p__Ascomycota_unclassified(100); | - | - | - | - | - | - | - | Unassigned | - |
| Otu29075 | 1 | k__Fungi(100);p__Ascomycota(100);c__Sordariomycetes(100);c__Sordariomycetes_unclassified(100);c__Sordariomycetes_unclassified(100);c__Sordariomycetes_unclassified(100);c__Sordariomycetes_unclassified(100); | - | - | - | - | - | - | - | Unassigned | - |
| Otu29076 | 1 | k__Fungi(100);p__Ascomycota(100);c__Sordariomycetes(100);c__Sordariomycetes_unclassified(100);c__Sordariomycetes_unclassified(100);c__Sordariomycetes_unclassified(100);c__Sordariomycetes_unclassified(100); | - | - | - | - | - | - | - | Unassigned | - |
| Otu29077 | 1 | k__Fungi(100);p__Ascomycota(100);c__Sordariomycetes(100);c__Sordariomycetes_unclassified(100);c__Sordariomycetes_unclassified(100);c__Sordariomycetes_unclassified(100);c__Sordariomycetes_unclassified(100); | - | - | - | - | - | - | - | Unassigned | - |
| Otu29078 | 1 | k__Fungi(100);p__Ascomycota(100);p__Ascomycota_unclassified(100);p__Ascomycota_unclassified(100);p__Ascomycota_unclassified(100);p__Ascomycota_unclassified(100);p__Ascomycota_unclassified(100); | - | - | - | - | - | - | - | Unassigned | - |
| Otu29079 | 1 | k__Fungi(100);p__Ascomycota(100);p__Ascomycota_unclassified(100);p__Ascomycota_unclassified(100);p__Ascomycota_unclassified(100);p__Ascomycota_unclassified(100);p__Ascomycota_unclassified(100); | - | - | - | - | - | - | - | Unassigned | - |
| Otu29080 | 1 | k__Fungi(100);p__Ascomycota(100);p__Ascomycota_unclassified(100);p__Ascomycota_unclassified(100);p__Ascomycota_unclassified(100);p__Ascomycota_unclassified(100);p__Ascomycota_unclassified(100); | - | - | - | - | - | - | - | Unassigned | - |
| Otu29081 | 1 | k__Fungi(100);p__Ascomycota(100);p__Ascomycota_unclassified(100);p__Ascomycota_unclassified(100);p__Ascomycota_unclassified(100);p__Ascomycota_unclassified(100);p__Ascomycota_unclassified(100); | - | - | - | - | - | - | - | Unassigned | - |
| Otu29083 | 1 | k__Fungi(100);p__Ascomycota(100);c__Sordariomycetes(100);c__Sordariomycetes_unclassified(100);c__Sordariomycetes_unclassified(100);c__Sordariomycetes_unclassified(100);c__Sordariomycetes_unclassified(100); | - | - | - | - | - | - | - | Unassigned | - |
| Otu29085 | 1 | k__Fungi(100);p__Ascomycota(100);p__Ascomycota_unclassified(100);p__Ascomycota_unclassified(100);p__Ascomycota_unclassified(100);p__Ascomycota_unclassified(100);p__Ascomycota_unclassified(100); | - | - | - | - | - | - | - | Unassigned | - |
| Otu29086 | 1 | k__Fungi(100);p__Ascomycota(100);c__Sordariomycetes(100);c__Sordariomycetes_unclassified(100);c__Sordariomycetes_unclassified(100);c__Sordariomycetes_unclassified(100);c__Sordariomycetes_unclassified(100); | - | - | - | - | - | - | - | Unassigned | - |
| Otu29087 | 1 | k__Fungi(100);p__Ascomycota(100);p__Ascomycota_unclassified(100);p__Ascomycota_unclassified(100);p__Ascomycota_unclassified(100);p__Ascomycota_unclassified(100);p__Ascomycota_unclassified(100); | - | - | - | - | - | - | - | Unassigned | - |
| Otu29088 | 1 | k__Fungi(100);p__Ascomycota(100);p__Ascomycota_unclassified(100);p__Ascomycota_unclassified(100);p__Ascomycota_unclassified(100);p__Ascomycota_unclassified(100);p__Ascomycota_unclassified(100); | - | - | - | - | - | - | - | Unassigned | - |
| Otu29089 | 1 | k__Fungi(100);p__Ascomycota(100);p__Ascomycota_unclassified(100);p__Ascomycota_unclassified(100);p__Ascomycota_unclassified(100);p__Ascomycota_unclassified(100);p__Ascomycota_unclassified(100); | - | - | - | - | - | - | - | Unassigned | - |
| Otu29090 | 1 | k__Fungi(100);p__Ascomycota(100);p__Ascomycota_unclassified(100);p__Ascomycota_unclassified(100);p__Ascomycota_unclassified(100);p__Ascomycota_unclassified(100);p__Ascomycota_unclassified(100); | - | - | - | - | - | - | - | Unassigned | - |
| Otu29091 | 1 | k__Fungi(100);p__Ascomycota(100);p__Ascomycota_unclassified(100);p__Ascomycota_unclassified(100);p__Ascomycota_unclassified(100);p__Ascomycota_unclassified(100);p__Ascomycota_unclassified(100); | - | - | - | - | - | - | - | Unassigned | - |
| Otu29092 | 1 | k__Fungi(100);p__Ascomycota(100);p__Ascomycota_unclassified(100);p__Ascomycota_unclassified(100);p__Ascomycota_unclassified(100);p__Ascomycota_unclassified(100);p__Ascomycota_unclassified(100); | - | - | - | - | - | - | - | Unassigned | - |
| Otu29093 | 1 | k__Fungi(100);p__Ascomycota(100);p__Ascomycota_unclassified(100);p__Ascomycota_unclassified(100);p__Ascomycota_unclassified(100);p__Ascomycota_unclassified(100);p__Ascomycota_unclassified(100); | - | - | - | - | - | - | - | Unassigned | - |
| Otu29094 | 1 | k__Fungi(100);p__Ascomycota(100);p__Ascomycota_unclassified(100);p__Ascomycota_unclassified(100);p__Ascomycota_unclassified(100);p__Ascomycota_unclassified(100);p__Ascomycota_unclassified(100); | - | - | - | - | - | - | - | Unassigned | - |
| Otu29095 | 1 | k__Plantae(100);p__unclassified_Plantae(100);c__unclassified_Plantae(100);o__unclassified_Plantae(100);f__unclassified_Plantae(100);g__unclassified_Plantae(100);s__Plantae_sp(100); | - | - | - | - | - | - | - | Unassigned | - |
| Otu29097 | 1 | k__Fungi(100);p__Ascomycota(100);p__Ascomycota_unclassified(100);p__Ascomycota_unclassified(100);p__Ascomycota_unclassified(100);p__Ascomycota_unclassified(100);p__Ascomycota_unclassified(100); | - | - | - | - | - | - | - | Unassigned | - |
| Otu29099 | 1 | k__Fungi(100);p__Ascomycota(100);c__Sordariomycetes(100);c__Sordariomycetes_unclassified(100);c__Sordariomycetes_unclassified(100);c__Sordariomycetes_unclassified(100);c__Sordariomycetes_unclassified(100); | - | - | - | - | - | - | - | Unassigned | - |
| Otu29101 | 1 | k__Fungi(100);p__Ascomycota(100);c__Sordariomycetes(100);c__Sordariomycetes_unclassified(100);c__Sordariomycetes_unclassified(100);c__Sordariomycetes_unclassified(100);c__Sordariomycetes_unclassified(100); | - | - | - | - | - | - | - | Unassigned | - |
| Otu29103 | 1 | k__Fungi(100);p__Ascomycota(100);c__Sordariomycetes(100);c__Sordariomycetes_unclassified(100);c__Sordariomycetes_unclassified(100);c__Sordariomycetes_unclassified(100);c__Sordariomycetes_unclassified(100); | - | - | - | - | - | - | - | Unassigned | - |
| Otu29104 | 1 | k__Fungi(100);p__Ascomycota(100);c__Sordariomycetes(100);c__Sordariomycetes_unclassified(100);c__Sordariomycetes_unclassified(100);c__Sordariomycetes_unclassified(100);c__Sordariomycetes_unclassified(100); | - | - | - | - | - | - | - | Unassigned | - |
| Otu29107 | 1 | k__Fungi(100);p__Ascomycota(100);c__Sordariomycetes(100);c__Sordariomycetes_unclassified(100);c__Sordariomycetes_unclassified(100);c__Sordariomycetes_unclassified(100);c__Sordariomycetes_unclassified(100); | - | - | - | - | - | - | - | Unassigned | - |
| Otu29111 | 1 | k__Fungi(100);p__Ascomycota(100);c__Sordariomycetes(100);c__Sordariomycetes_unclassified(100);c__Sordariomycetes_unclassified(100);c__Sordariomycetes_unclassified(100);c__Sordariomycetes_unclassified(100); | - | - | - | - | - | - | - | Unassigned | - |
| Otu29113 | 1 | k__Fungi(100);p__Ascomycota(100);c__Dothideomycetes(100);c__Dothideomycetes_unclassified(100);c__Dothideomycetes_unclassified(100);c__Dothideomycetes_unclassified(100);c__Dothideomycetes_unclassified(100); | - | - | - | - | - | - | - | Unassigned | - |
| Otu29114 | 1 | k__Fungi(100);p__Ascomycota(100);p__Ascomycota_unclassified(100);p__Ascomycota_unclassified(100);p__Ascomycota_unclassified(100);p__Ascomycota_unclassified(100);p__Ascomycota_unclassified(100); | - | - | - | - | - | - | - | Unassigned | - |
| Otu29115 | 1 | k__Fungi(100);p__Ascomycota(100);c__Sordariomycetes(100);c__Sordariomycetes_unclassified(100);c__Sordariomycetes_unclassified(100);c__Sordariomycetes_unclassified(100);c__Sordariomycetes_unclassified(100); | - | - | - | - | - | - | - | Unassigned | - |
| Otu29116 | 1 | k__Fungi(100);p__Ascomycota(100);c__Sordariomycetes(100);c__Sordariomycetes_unclassified(100);c__Sordariomycetes_unclassified(100);c__Sordariomycetes_unclassified(100);c__Sordariomycetes_unclassified(100); | - | - | - | - | - | - | - | Unassigned | - |
| Otu29117 | 1 | k__Fungi(100);p__Ascomycota(100);c__Sordariomycetes(100);c__Sordariomycetes_unclassified(100);c__Sordariomycetes_unclassified(100);c__Sordariomycetes_unclassified(100);c__Sordariomycetes_unclassified(100); | - | - | - | - | - | - | - | Unassigned | - |
| Otu29120 | 1 | k__Fungi(100);p__Ascomycota(100);c__Sordariomycetes(100);c__Sordariomycetes_unclassified(100);c__Sordariomycetes_unclassified(100);c__Sordariomycetes_unclassified(100);c__Sordariomycetes_unclassified(100); | - | - | - | - | - | - | - | Unassigned | - |
| Otu29121 | 1 | k__Fungi(100);p__Ascomycota(100);c__Sordariomycetes(100);c__Sordariomycetes_unclassified(100);c__Sordariomycetes_unclassified(100);c__Sordariomycetes_unclassified(100);c__Sordariomycetes_unclassified(100); | - | - | - | - | - | - | - | Unassigned | - |
| Otu29123 | 1 | k__Fungi(100);p__Ascomycota(100);c__Sordariomycetes(100);c__Sordariomycetes_unclassified(100);c__Sordariomycetes_unclassified(100);c__Sordariomycetes_unclassified(100);c__Sordariomycetes_unclassified(100); | - | - | - | - | - | - | - | Unassigned | - |
| Otu29125 | 1 | k__Fungi(100);p__Ascomycota(100);p__Ascomycota_unclassified(100);p__Ascomycota_unclassified(100);p__Ascomycota_unclassified(100);p__Ascomycota_unclassified(100);p__Ascomycota_unclassified(100); | - | - | - | - | - | - | - | Unassigned | - |
| Otu29154 | 1 | k__Fungi(100);p__Basidiomycota(100);c__Microbotryomycetes(100);o__Sporidiobolales(100);f__Sporidiobolales_family_Incertae_sedis(100);f__Sporidiobolales_family_Incertae_sedis_unclassified(100);f__Sporidiobolales_family_Incertae_sedis_unclassified(100); | - | - | - | - | - | - | - | Unassigned | - |
| Otu29160 | 1 | k__Fungi(100);p__Ascomycota(100);p__Ascomycota_unclassified(100);p__Ascomycota_unclassified(100);p__Ascomycota_unclassified(100);p__Ascomycota_unclassified(100);p__Ascomycota_unclassified(100); | - | - | - | - | - | - | - | Unassigned | - |
| Otu29162 | 1 | k__Fungi(100);p__Ascomycota(100);c__Dothideomycetes(100);o__Pleosporales(100);o__Pleosporales_unclassified(100);o__Pleosporales_unclassified(100);o__Pleosporales_unclassified(100); | - | - | - | - | - | - | - | Unassigned | - |
| Otu29163 | 1 | k__Fungi(100);p__Ascomycota(100);c__Dothideomycetes(100);o__Pleosporales(100);o__Pleosporales_unclassified(100);o__Pleosporales_unclassified(100);o__Pleosporales_unclassified(100); | - | - | - | - | - | - | - | Unassigned | - |
| Otu29166 | 1 | k__Fungi(100);p__Ascomycota(100);c__Sordariomycetes(100);c__Sordariomycetes_unclassified(100);c__Sordariomycetes_unclassified(100);c__Sordariomycetes_unclassified(100);c__Sordariomycetes_unclassified(100); | - | - | - | - | - | - | - | Unassigned | - |
| Otu29167 | 1 | k__Fungi(100);p__Ascomycota(100);c__Dothideomycetes(100);o__Pleosporales(100);o__Pleosporales_unclassified(100);o__Pleosporales_unclassified(100);o__Pleosporales_unclassified(100); | - | - | - | - | - | - | - | Unassigned | - |
| Otu29179 | 1 | k__Fungi(100);p__Ascomycota(100);p__Ascomycota_unclassified(100);p__Ascomycota_unclassified(100);p__Ascomycota_unclassified(100);p__Ascomycota_unclassified(100);p__Ascomycota_unclassified(100); | - | - | - | - | - | - | - | Unassigned | - |
| Otu29182 | 1 | k__Fungi(100);p__Ascomycota(100);p__Ascomycota_unclassified(100);p__Ascomycota_unclassified(100);p__Ascomycota_unclassified(100);p__Ascomycota_unclassified(100);p__Ascomycota_unclassified(100); | - | - | - | - | - | - | - | Unassigned | - |
| Otu29183 | 1 | k__Fungi(100);p__Ascomycota(100);p__Ascomycota_unclassified(100);p__Ascomycota_unclassified(100);p__Ascomycota_unclassified(100);p__Ascomycota_unclassified(100);p__Ascomycota_unclassified(100); | - | - | - | - | - | - | - | Unassigned | - |
| Otu29184 | 1 | k__Fungi(100);p__Ascomycota(100);p__Ascomycota_unclassified(100);p__Ascomycota_unclassified(100);p__Ascomycota_unclassified(100);p__Ascomycota_unclassified(100);p__Ascomycota_unclassified(100); | - | - | - | - | - | - | - | Unassigned | - |
| Otu29185 | 1 | k__Fungi(100);p__Ascomycota(100);c__Sordariomycetes(100);c__Sordariomycetes_unclassified(100);c__Sordariomycetes_unclassified(100);c__Sordariomycetes_unclassified(100);c__Sordariomycetes_unclassified(100); | - | - | - | - | - | - | - | Unassigned | - |
| Otu29186 | 1 | k__Fungi(100);p__Ascomycota(100);c__Sordariomycetes(100);c__Sordariomycetes_unclassified(100);c__Sordariomycetes_unclassified(100);c__Sordariomycetes_unclassified(100);c__Sordariomycetes_unclassified(100); | - | - | - | - | - | - | - | Unassigned | - |
| Otu29187 | 1 | k__Fungi(100);p__Ascomycota(100);c__Sordariomycetes(100);c__Sordariomycetes_unclassified(100);c__Sordariomycetes_unclassified(100);c__Sordariomycetes_unclassified(100);c__Sordariomycetes_unclassified(100); | - | - | - | - | - | - | - | Unassigned | - |
| Otu29188 | 1 | k__Fungi(100);p__Ascomycota(100);p__Ascomycota_unclassified(100);p__Ascomycota_unclassified(100);p__Ascomycota_unclassified(100);p__Ascomycota_unclassified(100);p__Ascomycota_unclassified(100); | - | - | - | - | - | - | - | Unassigned | - |
| Otu29189 | 1 | k__Fungi(100);p__Ascomycota(100);c__Sordariomycetes(100);c__Sordariomycetes_unclassified(100);c__Sordariomycetes_unclassified(100);c__Sordariomycetes_unclassified(100);c__Sordariomycetes_unclassified(100); | - | - | - | - | - | - | - | Unassigned | - |
| Otu29190 | 1 | k__Fungi(100);p__Ascomycota(100);c__Sordariomycetes(100);c__Sordariomycetes_unclassified(100);c__Sordariomycetes_unclassified(100);c__Sordariomycetes_unclassified(100);c__Sordariomycetes_unclassified(100); | - | - | - | - | - | - | - | Unassigned | - |
| Otu29191 | 1 | k__Fungi(100);p__Ascomycota(100);c__Sordariomycetes(100);c__Sordariomycetes_unclassified(100);c__Sordariomycetes_unclassified(100);c__Sordariomycetes_unclassified(100);c__Sordariomycetes_unclassified(100); | - | - | - | - | - | - | - | Unassigned | - |
| Otu29192 | 1 | k__Fungi(100);p__Ascomycota(100);c__Sordariomycetes(100);c__Sordariomycetes_unclassified(100);c__Sordariomycetes_unclassified(100);c__Sordariomycetes_unclassified(100);c__Sordariomycetes_unclassified(100); | - | - | - | - | - | - | - | Unassigned | - |
| Otu29194 | 1 | k__Fungi(100);p__Ascomycota(100);p__Ascomycota_unclassified(100);p__Ascomycota_unclassified(100);p__Ascomycota_unclassified(100);p__Ascomycota_unclassified(100);p__Ascomycota_unclassified(100); | - | - | - | - | - | - | - | Unassigned | - |
| Otu29195 | 1 | k__Fungi(100);p__Ascomycota(100);c__Sordariomycetes(100);c__Sordariomycetes_unclassified(100);c__Sordariomycetes_unclassified(100);c__Sordariomycetes_unclassified(100);c__Sordariomycetes_unclassified(100); | - | - | - | - | - | - | - | Unassigned | - |
| Otu29196 | 1 | k__Fungi(100);p__Ascomycota(100);p__Ascomycota_unclassified(100);p__Ascomycota_unclassified(100);p__Ascomycota_unclassified(100);p__Ascomycota_unclassified(100);p__Ascomycota_unclassified(100); | - | - | - | - | - | - | - | Unassigned | - |
| Otu29197 | 1 | k__Fungi(100);p__Ascomycota(100);p__Ascomycota_unclassified(100);p__Ascomycota_unclassified(100);p__Ascomycota_unclassified(100);p__Ascomycota_unclassified(100);p__Ascomycota_unclassified(100); | - | - | - | - | - | - | - | Unassigned | - |
| Otu29198 | 1 | k__Fungi(100);p__Ascomycota(100);p__Ascomycota_unclassified(100);p__Ascomycota_unclassified(100);p__Ascomycota_unclassified(100);p__Ascomycota_unclassified(100);p__Ascomycota_unclassified(100); | - | - | - | - | - | - | - | Unassigned | - |
| Otu29199 | 1 | k__Fungi(100);p__Ascomycota(100);p__Ascomycota_unclassified(100);p__Ascomycota_unclassified(100);p__Ascomycota_unclassified(100);p__Ascomycota_unclassified(100);p__Ascomycota_unclassified(100); | - | - | - | - | - | - | - | Unassigned | - |
| Otu29200 | 1 | k__Fungi(100);p__Ascomycota(100);c__Sordariomycetes(100);c__Sordariomycetes_unclassified(100);c__Sordariomycetes_unclassified(100);c__Sordariomycetes_unclassified(100);c__Sordariomycetes_unclassified(100); | - | - | - | - | - | - | - | Unassigned | - |
| Otu29201 | 1 | k__Fungi(100);p__Ascomycota(100);c__Sordariomycetes(100);c__Sordariomycetes_unclassified(100);c__Sordariomycetes_unclassified(100);c__Sordariomycetes_unclassified(100);c__Sordariomycetes_unclassified(100); | - | - | - | - | - | - | - | Unassigned | - |
| Otu29202 | 1 | k__Fungi(100);p__Ascomycota(100);c__Sordariomycetes(100);c__Sordariomycetes_unclassified(100);c__Sordariomycetes_unclassified(100);c__Sordariomycetes_unclassified(100);c__Sordariomycetes_unclassified(100); | - | - | - | - | - | - | - | Unassigned | - |
| Otu29203 | 1 | k__Fungi(100);p__Ascomycota(100);p__Ascomycota_unclassified(100);p__Ascomycota_unclassified(100);p__Ascomycota_unclassified(100);p__Ascomycota_unclassified(100);p__Ascomycota_unclassified(100); | - | - | - | - | - | - | - | Unassigned | - |
| Otu29204 | 1 | k__Fungi(100);p__Ascomycota(100);c__Sordariomycetes(100);c__Sordariomycetes_unclassified(100);c__Sordariomycetes_unclassified(100);c__Sordariomycetes_unclassified(100);c__Sordariomycetes_unclassified(100); | - | - | - | - | - | - | - | Unassigned | - |
| Otu29205 | 1 | k__Fungi(100);p__Ascomycota(100);p__Ascomycota_unclassified(100);p__Ascomycota_unclassified(100);p__Ascomycota_unclassified(100);p__Ascomycota_unclassified(100);p__Ascomycota_unclassified(100); | - | - | - | - | - | - | - | Unassigned | - |
| Otu29207 | 1 | k__Fungi(100);p__Ascomycota(100);p__Ascomycota_unclassified(100);p__Ascomycota_unclassified(100);p__Ascomycota_unclassified(100);p__Ascomycota_unclassified(100);p__Ascomycota_unclassified(100); | - | - | - | - | - | - | - | Unassigned | - |
| Otu29208 | 1 | k__Fungi(100);p__Ascomycota(100);p__Ascomycota_unclassified(100);p__Ascomycota_unclassified(100);p__Ascomycota_unclassified(100);p__Ascomycota_unclassified(100);p__Ascomycota_unclassified(100); | - | - | - | - | - | - | - | Unassigned | - |
| Otu29209 | 1 | k__Fungi(100);p__Ascomycota(100);p__Ascomycota_unclassified(100);p__Ascomycota_unclassified(100);p__Ascomycota_unclassified(100);p__Ascomycota_unclassified(100);p__Ascomycota_unclassified(100); | - | - | - | - | - | - | - | Unassigned | - |
| Otu29210 | 1 | k__Fungi(100);p__Ascomycota(100);c__Sordariomycetes(100);c__Sordariomycetes_unclassified(100);c__Sordariomycetes_unclassified(100);c__Sordariomycetes_unclassified(100);c__Sordariomycetes_unclassified(100); | - | - | - | - | - | - | - | Unassigned | - |
| Otu29211 | 1 | k__Fungi(100);p__Ascomycota(100);p__Ascomycota_unclassified(100);p__Ascomycota_unclassified(100);p__Ascomycota_unclassified(100);p__Ascomycota_unclassified(100);p__Ascomycota_unclassified(100); | - | - | - | - | - | - | - | Unassigned | - |
| Otu29212 | 1 | k__Fungi(100);p__Ascomycota(100);c__Sordariomycetes(100);c__Sordariomycetes_unclassified(100);c__Sordariomycetes_unclassified(100);c__Sordariomycetes_unclassified(100);c__Sordariomycetes_unclassified(100); | - | - | - | - | - | - | - | Unassigned | - |
| Otu29213 | 1 | k__Fungi(100);p__Ascomycota(100);c__Sordariomycetes(100);c__Sordariomycetes_unclassified(100);c__Sordariomycetes_unclassified(100);c__Sordariomycetes_unclassified(100);c__Sordariomycetes_unclassified(100); | - | - | - | - | - | - | - | Unassigned | - |
| Otu29214 | 1 | k__Fungi(100);p__Ascomycota(100);p__Ascomycota_unclassified(100);p__Ascomycota_unclassified(100);p__Ascomycota_unclassified(100);p__Ascomycota_unclassified(100);p__Ascomycota_unclassified(100); | - | - | - | - | - | - | - | Unassigned | - |
| Otu29215 | 1 | k__Fungi(100);p__Ascomycota(100);p__Ascomycota_unclassified(100);p__Ascomycota_unclassified(100);p__Ascomycota_unclassified(100);p__Ascomycota_unclassified(100);p__Ascomycota_unclassified(100); | - | - | - | - | - | - | - | Unassigned | - |
| Otu29216 | 1 | k__Fungi(100);p__Ascomycota(100);p__Ascomycota_unclassified(100);p__Ascomycota_unclassified(100);p__Ascomycota_unclassified(100);p__Ascomycota_unclassified(100);p__Ascomycota_unclassified(100); | - | - | - | - | - | - | - | Unassigned | - |
| Otu29217 | 1 | k__Fungi(100);p__Ascomycota(100);p__Ascomycota_unclassified(100);p__Ascomycota_unclassified(100);p__Ascomycota_unclassified(100);p__Ascomycota_unclassified(100);p__Ascomycota_unclassified(100); | - | - | - | - | - | - | - | Unassigned | - |
| Otu29219 | 1 | k__Fungi(100);p__Ascomycota(100);p__Ascomycota_unclassified(100);p__Ascomycota_unclassified(100);p__Ascomycota_unclassified(100);p__Ascomycota_unclassified(100);p__Ascomycota_unclassified(100); | - | - | - | - | - | - | - | Unassigned | - |
| Otu29221 | 1 | k__Fungi(100);p__Ascomycota(100);p__Ascomycota_unclassified(100);p__Ascomycota_unclassified(100);p__Ascomycota_unclassified(100);p__Ascomycota_unclassified(100);p__Ascomycota_unclassified(100); | - | - | - | - | - | - | - | Unassigned | - |
| Otu29222 | 1 | k__Fungi(100);p__Ascomycota(100);c__Sordariomycetes(100);c__Sordariomycetes_unclassified(100);c__Sordariomycetes_unclassified(100);c__Sordariomycetes_unclassified(100);c__Sordariomycetes_unclassified(100); | - | - | - | - | - | - | - | Unassigned | - |
| Otu29226 | 1 | k__Fungi(100);p__Ascomycota(100);c__Sordariomycetes(100);c__Sordariomycetes_unclassified(100);c__Sordariomycetes_unclassified(100);c__Sordariomycetes_unclassified(100);c__Sordariomycetes_unclassified(100); | - | - | - | - | - | - | - | Unassigned | - |
| Otu29227 | 1 | k__Fungi(100);p__Ascomycota(100);c__Sordariomycetes(100);c__Sordariomycetes_unclassified(100);c__Sordariomycetes_unclassified(100);c__Sordariomycetes_unclassified(100);c__Sordariomycetes_unclassified(100); | - | - | - | - | - | - | - | Unassigned | - |
| Otu29228 | 1 | k__Fungi(100);p__Ascomycota(100);c__Sordariomycetes(100);o__Hypocreales(100);f__Cordycipitaceae(100);f__Cordycipitaceae_unclassified(100);f__Cordycipitaceae_unclassified(100); | - | - | - | - | - | - | - | Unassigned | - |
| Otu29230 | 1 | k__Fungi(100);p__Ascomycota(100);c__Sordariomycetes(100);o__Sordariales(100);o__Sordariales_unclassified(100);o__Sordariales_unclassified(100);o__Sordariales_unclassified(100); | - | - | - | - | - | - | - | Unassigned | - |
| Otu29233 | 1 | k__Fungi(100);p__Ascomycota(100);p__Ascomycota_unclassified(100);p__Ascomycota_unclassified(100);p__Ascomycota_unclassified(100);p__Ascomycota_unclassified(100);p__Ascomycota_unclassified(100); | - | - | - | - | - | - | - | Unassigned | - |
| Otu29234 | 1 | k__Fungi(100);p__Ascomycota(100);p__Ascomycota_unclassified(100);p__Ascomycota_unclassified(100);p__Ascomycota_unclassified(100);p__Ascomycota_unclassified(100);p__Ascomycota_unclassified(100); | - | - | - | - | - | - | - | Unassigned | - |
| Otu29235 | 1 | k__Fungi(100);p__Ascomycota(100);p__Ascomycota_unclassified(100);p__Ascomycota_unclassified(100);p__Ascomycota_unclassified(100);p__Ascomycota_unclassified(100);p__Ascomycota_unclassified(100); | - | - | - | - | - | - | - | Unassigned | - |
| Otu29236 | 1 | k__Fungi(100);p__Ascomycota(100);p__Ascomycota_unclassified(100);p__Ascomycota_unclassified(100);p__Ascomycota_unclassified(100);p__Ascomycota_unclassified(100);p__Ascomycota_unclassified(100); | - | - | - | - | - | - | - | Unassigned | - |
| Otu29237 | 1 | k__Fungi(100);p__Ascomycota(100);p__Ascomycota_unclassified(100);p__Ascomycota_unclassified(100);p__Ascomycota_unclassified(100);p__Ascomycota_unclassified(100);p__Ascomycota_unclassified(100); | - | - | - | - | - | - | - | Unassigned | - |
| Otu29238 | 1 | k__Fungi(100);p__Ascomycota(100);p__Ascomycota_unclassified(100);p__Ascomycota_unclassified(100);p__Ascomycota_unclassified(100);p__Ascomycota_unclassified(100);p__Ascomycota_unclassified(100); | - | - | - | - | - | - | - | Unassigned | - |
| Otu29239 | 1 | k__Fungi(100);p__Ascomycota(100);p__Ascomycota_unclassified(100);p__Ascomycota_unclassified(100);p__Ascomycota_unclassified(100);p__Ascomycota_unclassified(100);p__Ascomycota_unclassified(100); | - | - | - | - | - | - | - | Unassigned | - |
| Otu29240 | 1 | k__Fungi(100);p__Ascomycota(100);p__Ascomycota_unclassified(100);p__Ascomycota_unclassified(100);p__Ascomycota_unclassified(100);p__Ascomycota_unclassified(100);p__Ascomycota_unclassified(100); | - | - | - | - | - | - | - | Unassigned | - |
| Otu29241 | 1 | k__Fungi(100);p__Ascomycota(100);p__Ascomycota_unclassified(100);p__Ascomycota_unclassified(100);p__Ascomycota_unclassified(100);p__Ascomycota_unclassified(100);p__Ascomycota_unclassified(100); | - | - | - | - | - | - | - | Unassigned | - |
| Otu29242 | 1 | k__Fungi(100);p__Ascomycota(100);p__Ascomycota_unclassified(100);p__Ascomycota_unclassified(100);p__Ascomycota_unclassified(100);p__Ascomycota_unclassified(100);p__Ascomycota_unclassified(100); | - | - | - | - | - | - | - | Unassigned | - |
| Otu29243 | 1 | k__Fungi(100);p__Ascomycota(100);c__Sordariomycetes(100);c__Sordariomycetes_unclassified(100);c__Sordariomycetes_unclassified(100);c__Sordariomycetes_unclassified(100);c__Sordariomycetes_unclassified(100); | - | - | - | - | - | - | - | Unassigned | - |
| Otu29246 | 1 | k__Fungi(100);p__Ascomycota(100);c__Sordariomycetes(100);c__Sordariomycetes_unclassified(100);c__Sordariomycetes_unclassified(100);c__Sordariomycetes_unclassified(100);c__Sordariomycetes_unclassified(100); | - | - | - | - | - | - | - | Unassigned | - |
| Otu29249 | 1 | k__Fungi(100);p__Ascomycota(100);c__Sordariomycetes(100);c__Sordariomycetes_unclassified(100);c__Sordariomycetes_unclassified(100);c__Sordariomycetes_unclassified(100);c__Sordariomycetes_unclassified(100); | - | - | - | - | - | - | - | Unassigned | - |
| Otu29250 | 1 | k__Fungi(100);p__Ascomycota(100);p__Ascomycota_unclassified(100);p__Ascomycota_unclassified(100);p__Ascomycota_unclassified(100);p__Ascomycota_unclassified(100);p__Ascomycota_unclassified(100); | - | - | - | - | - | - | - | Unassigned | - |
| Otu29251 | 1 | k__Fungi(100);p__Ascomycota(100);p__Ascomycota_unclassified(100);p__Ascomycota_unclassified(100);p__Ascomycota_unclassified(100);p__Ascomycota_unclassified(100);p__Ascomycota_unclassified(100); | - | - | - | - | - | - | - | Unassigned | - |
| Otu29252 | 1 | k__Fungi(100);p__Ascomycota(100);p__Ascomycota_unclassified(100);p__Ascomycota_unclassified(100);p__Ascomycota_unclassified(100);p__Ascomycota_unclassified(100);p__Ascomycota_unclassified(100); | - | - | - | - | - | - | - | Unassigned | - |
| Otu29253 | 1 | k__Fungi(100);p__Ascomycota(100);c__Sordariomycetes(100);c__Sordariomycetes_unclassified(100);c__Sordariomycetes_unclassified(100);c__Sordariomycetes_unclassified(100);c__Sordariomycetes_unclassified(100); | - | - | - | - | - | - | - | Unassigned | - |
| Otu29254 | 1 | k__Fungi(100);p__Ascomycota(100);c__Sordariomycetes(100);c__Sordariomycetes_unclassified(100);c__Sordariomycetes_unclassified(100);c__Sordariomycetes_unclassified(100);c__Sordariomycetes_unclassified(100); | - | - | - | - | - | - | - | Unassigned | - |
| Otu29255 | 1 | k__Fungi(100);p__Ascomycota(100);p__Ascomycota_unclassified(100);p__Ascomycota_unclassified(100);p__Ascomycota_unclassified(100);p__Ascomycota_unclassified(100);p__Ascomycota_unclassified(100); | - | - | - | - | - | - | - | Unassigned | - |
| Otu29256 | 1 | k__Fungi(100);p__Ascomycota(100);p__Ascomycota_unclassified(100);p__Ascomycota_unclassified(100);p__Ascomycota_unclassified(100);p__Ascomycota_unclassified(100);p__Ascomycota_unclassified(100); | - | - | - | - | - | - | - | Unassigned | - |
| Otu29257 | 1 | k__Fungi(100);p__Ascomycota(100);p__Ascomycota_unclassified(100);p__Ascomycota_unclassified(100);p__Ascomycota_unclassified(100);p__Ascomycota_unclassified(100);p__Ascomycota_unclassified(100); | - | - | - | - | - | - | - | Unassigned | - |
| Otu29258 | 1 | k__Fungi(100);p__Ascomycota(100);c__Sordariomycetes(100);c__Sordariomycetes_unclassified(100);c__Sordariomycetes_unclassified(100);c__Sordariomycetes_unclassified(100);c__Sordariomycetes_unclassified(100); | - | - | - | - | - | - | - | Unassigned | - |
| Otu29259 | 1 | k__Fungi(100);p__Ascomycota(100);p__Ascomycota_unclassified(100);p__Ascomycota_unclassified(100);p__Ascomycota_unclassified(100);p__Ascomycota_unclassified(100);p__Ascomycota_unclassified(100); | - | - | - | - | - | - | - | Unassigned | - |
| Otu29260 | 1 | k__Fungi(100);p__Ascomycota(100);p__Ascomycota_unclassified(100);p__Ascomycota_unclassified(100);p__Ascomycota_unclassified(100);p__Ascomycota_unclassified(100);p__Ascomycota_unclassified(100); | - | - | - | - | - | - | - | Unassigned | - |
| Otu29261 | 1 | k__Fungi(100);p__Ascomycota(100);p__Ascomycota_unclassified(100);p__Ascomycota_unclassified(100);p__Ascomycota_unclassified(100);p__Ascomycota_unclassified(100);p__Ascomycota_unclassified(100); | - | - | - | - | - | - | - | Unassigned | - |
| Otu29262 | 1 | k__Fungi(100);p__Ascomycota(100);p__Ascomycota_unclassified(100);p__Ascomycota_unclassified(100);p__Ascomycota_unclassified(100);p__Ascomycota_unclassified(100);p__Ascomycota_unclassified(100); | - | - | - | - | - | - | - | Unassigned | - |
| Otu29263 | 1 | k__Fungi(100);p__Ascomycota(100);p__Ascomycota_unclassified(100);p__Ascomycota_unclassified(100);p__Ascomycota_unclassified(100);p__Ascomycota_unclassified(100);p__Ascomycota_unclassified(100); | - | - | - | - | - | - | - | Unassigned | - |
| Otu29264 | 1 | k__Fungi(100);p__Ascomycota(100);p__Ascomycota_unclassified(100);p__Ascomycota_unclassified(100);p__Ascomycota_unclassified(100);p__Ascomycota_unclassified(100);p__Ascomycota_unclassified(100); | - | - | - | - | - | - | - | Unassigned | - |
| Otu29265 | 1 | k__Fungi(100);p__Ascomycota(100);p__Ascomycota_unclassified(100);p__Ascomycota_unclassified(100);p__Ascomycota_unclassified(100);p__Ascomycota_unclassified(100);p__Ascomycota_unclassified(100); | - | - | - | - | - | - | - | Unassigned | - |
| Otu29266 | 1 | k__Fungi(100);p__Ascomycota(100);p__Ascomycota_unclassified(100);p__Ascomycota_unclassified(100);p__Ascomycota_unclassified(100);p__Ascomycota_unclassified(100);p__Ascomycota_unclassified(100); | - | - | - | - | - | - | - | Unassigned | - |
| Otu29267 | 1 | k__Fungi(100);p__Ascomycota(100);p__Ascomycota_unclassified(100);p__Ascomycota_unclassified(100);p__Ascomycota_unclassified(100);p__Ascomycota_unclassified(100);p__Ascomycota_unclassified(100); | - | - | - | - | - | - | - | Unassigned | - |
| Otu29268 | 1 | k__Fungi(100);p__Ascomycota(100);p__Ascomycota_unclassified(100);p__Ascomycota_unclassified(100);p__Ascomycota_unclassified(100);p__Ascomycota_unclassified(100);p__Ascomycota_unclassified(100); | - | - | - | - | - | - | - | Unassigned | - |
| Otu29269 | 1 | k__Fungi(100);p__Ascomycota(100);c__Sordariomycetes(100);o__Hypocreales(100);f__Cordycipitaceae(100);f__Cordycipitaceae_unclassified(100);f__Cordycipitaceae_unclassified(100); | - | - | - | - | - | - | - | Unassigned | - |
| Otu29271 | 1 | k__Fungi(100);p__Ascomycota(100);p__Ascomycota_unclassified(100);p__Ascomycota_unclassified(100);p__Ascomycota_unclassified(100);p__Ascomycota_unclassified(100);p__Ascomycota_unclassified(100); | - | - | - | - | - | - | - | Unassigned | - |
| Otu29272 | 1 | k__Fungi(100);p__Ascomycota(100);p__Ascomycota_unclassified(100);p__Ascomycota_unclassified(100);p__Ascomycota_unclassified(100);p__Ascomycota_unclassified(100);p__Ascomycota_unclassified(100); | - | - | - | - | - | - | - | Unassigned | - |
| Otu29273 | 1 | k__Fungi(100);p__Ascomycota(100);p__Ascomycota_unclassified(100);p__Ascomycota_unclassified(100);p__Ascomycota_unclassified(100);p__Ascomycota_unclassified(100);p__Ascomycota_unclassified(100); | - | - | - | - | - | - | - | Unassigned | - |
| Otu29274 | 1 | k__Fungi(100);p__Ascomycota(100);p__Ascomycota_unclassified(100);p__Ascomycota_unclassified(100);p__Ascomycota_unclassified(100);p__Ascomycota_unclassified(100);p__Ascomycota_unclassified(100); | - | - | - | - | - | - | - | Unassigned | - |
| Otu29275 | 1 | k__Fungi(100);p__Ascomycota(100);p__Ascomycota_unclassified(100);p__Ascomycota_unclassified(100);p__Ascomycota_unclassified(100);p__Ascomycota_unclassified(100);p__Ascomycota_unclassified(100); | - | - | - | - | - | - | - | Unassigned | - |
| Otu29276 | 1 | k__Fungi(100);p__Ascomycota(100);p__Ascomycota_unclassified(100);p__Ascomycota_unclassified(100);p__Ascomycota_unclassified(100);p__Ascomycota_unclassified(100);p__Ascomycota_unclassified(100); | - | - | - | - | - | - | - | Unassigned | - |
| Otu29277 | 1 | k__Fungi(100);p__Ascomycota(100);p__Ascomycota_unclassified(100);p__Ascomycota_unclassified(100);p__Ascomycota_unclassified(100);p__Ascomycota_unclassified(100);p__Ascomycota_unclassified(100); | - | - | - | - | - | - | - | Unassigned | - |
| Otu29278 | 1 | k__Fungi(100);p__Ascomycota(100);p__Ascomycota_unclassified(100);p__Ascomycota_unclassified(100);p__Ascomycota_unclassified(100);p__Ascomycota_unclassified(100);p__Ascomycota_unclassified(100); | - | - | - | - | - | - | - | Unassigned | - |
| Otu29279 | 1 | k__Fungi(100);p__Ascomycota(100);p__Ascomycota_unclassified(100);p__Ascomycota_unclassified(100);p__Ascomycota_unclassified(100);p__Ascomycota_unclassified(100);p__Ascomycota_unclassified(100); | - | - | - | - | - | - | - | Unassigned | - |
| Otu29280 | 1 | k__Fungi(100);p__Ascomycota(100);c__Sordariomycetes(100);c__Sordariomycetes_unclassified(100);c__Sordariomycetes_unclassified(100);c__Sordariomycetes_unclassified(100);c__Sordariomycetes_unclassified(100); | - | - | - | - | - | - | - | Unassigned | - |
| Otu29281 | 1 | k__Fungi(100);p__Ascomycota(100);p__Ascomycota_unclassified(100);p__Ascomycota_unclassified(100);p__Ascomycota_unclassified(100);p__Ascomycota_unclassified(100);p__Ascomycota_unclassified(100); | - | - | - | - | - | - | - | Unassigned | - |
| Otu29282 | 1 | k__Fungi(100);p__Ascomycota(100);p__Ascomycota_unclassified(100);p__Ascomycota_unclassified(100);p__Ascomycota_unclassified(100);p__Ascomycota_unclassified(100);p__Ascomycota_unclassified(100); | - | - | - | - | - | - | - | Unassigned | - |
| Otu29283 | 1 | k__Fungi(100);p__Ascomycota(100);p__Ascomycota_unclassified(100);p__Ascomycota_unclassified(100);p__Ascomycota_unclassified(100);p__Ascomycota_unclassified(100);p__Ascomycota_unclassified(100); | - | - | - | - | - | - | - | Unassigned | - |
| Otu29284 | 1 | k__Fungi(100);p__Ascomycota(100);p__Ascomycota_unclassified(100);p__Ascomycota_unclassified(100);p__Ascomycota_unclassified(100);p__Ascomycota_unclassified(100);p__Ascomycota_unclassified(100); | - | - | - | - | - | - | - | Unassigned | - |
| Otu29285 | 1 | k__Plantae(100);p__unclassified_Plantae(100);c__unclassified_Plantae(100);o__unclassified_Plantae(100);f__unclassified_Plantae(100);g__unclassified_Plantae(100);s__Plantae_sp(100); | - | - | - | - | - | - | - | Unassigned | - |
| Otu29286 | 1 | k__Plantae(100);p__unclassified_Plantae(100);c__unclassified_Plantae(100);o__unclassified_Plantae(100);f__unclassified_Plantae(100);g__unclassified_Plantae(100);s__Plantae_sp(100); | - | - | - | - | - | - | - | Unassigned | - |
| Otu29287 | 1 | k__Plantae(100);p__unclassified_Plantae(100);c__unclassified_Plantae(100);o__unclassified_Plantae(100);f__unclassified_Plantae(100);g__unclassified_Plantae(100);s__Plantae_sp(100); | - | - | - | - | - | - | - | Unassigned | - |
| Otu29288 | 1 | k__Plantae(100);p__unclassified_Plantae(100);c__unclassified_Plantae(100);o__unclassified_Plantae(100);f__unclassified_Plantae(100);g__unclassified_Plantae(100);s__Plantae_sp(100); | - | - | - | - | - | - | - | Unassigned | - |
| Otu29289 | 1 | k__Plantae(100);p__unclassified_Plantae(100);c__unclassified_Plantae(100);o__unclassified_Plantae(100);f__unclassified_Plantae(100);g__unclassified_Plantae(100);s__Plantae_sp(100); | - | - | - | - | - | - | - | Unassigned | - |
| Otu29290 | 1 | k__Plantae(100);p__unclassified_Plantae(100);c__unclassified_Plantae(100);o__unclassified_Plantae(100);f__unclassified_Plantae(100);g__unclassified_Plantae(100);s__Plantae_sp(100); | - | - | - | - | - | - | - | Unassigned | - |
| Otu29291 | 1 | k__Plantae(100);p__unclassified_Plantae(100);c__unclassified_Plantae(100);o__unclassified_Plantae(100);f__unclassified_Plantae(100);g__unclassified_Plantae(100);s__Plantae_sp(100); | - | - | - | - | - | - | - | Unassigned | - |
| Otu29292 | 1 | k__Plantae(100);p__unclassified_Plantae(100);c__unclassified_Plantae(100);o__unclassified_Plantae(100);f__unclassified_Plantae(100);g__unclassified_Plantae(100);s__Plantae_sp(100); | - | - | - | - | - | - | - | Unassigned | - |
| Otu29293 | 1 | k__Plantae(100);p__unclassified_Plantae(100);c__unclassified_Plantae(100);o__unclassified_Plantae(100);f__unclassified_Plantae(100);g__unclassified_Plantae(100);s__Plantae_sp(100); | - | - | - | - | - | - | - | Unassigned | - |
| Otu29294 | 1 | k__Plantae(100);p__unclassified_Plantae(100);c__unclassified_Plantae(100);o__unclassified_Plantae(100);f__unclassified_Plantae(100);g__unclassified_Plantae(100);s__Plantae_sp(100); | - | - | - | - | - | - | - | Unassigned | - |
| Otu29295 | 1 | k__Plantae(100);p__unclassified_Plantae(100);c__unclassified_Plantae(100);o__unclassified_Plantae(100);f__unclassified_Plantae(100);g__unclassified_Plantae(100);s__Plantae_sp(100); | - | - | - | - | - | - | - | Unassigned | - |
| Otu29296 | 1 | k__Plantae(100);p__unclassified_Plantae(100);c__unclassified_Plantae(100);o__unclassified_Plantae(100);f__unclassified_Plantae(100);g__unclassified_Plantae(100);s__Plantae_sp(100); | - | - | - | - | - | - | - | Unassigned | - |
| Otu29297 | 1 | k__Plantae(100);p__unclassified_Plantae(100);c__unclassified_Plantae(100);o__unclassified_Plantae(100);f__unclassified_Plantae(100);g__unclassified_Plantae(100);s__Plantae_sp(100); | - | - | - | - | - | - | - | Unassigned | - |
| Otu29298 | 1 | k__Plantae(100);p__unclassified_Plantae(100);c__unclassified_Plantae(100);o__unclassified_Plantae(100);f__unclassified_Plantae(100);g__unclassified_Plantae(100);s__Plantae_sp(100); | - | - | - | - | - | - | - | Unassigned | - |
| Otu29299 | 1 | k__Plantae(100);p__unclassified_Plantae(100);c__unclassified_Plantae(100);o__unclassified_Plantae(100);f__unclassified_Plantae(100);g__unclassified_Plantae(100);s__Plantae_sp(100); | - | - | - | - | - | - | - | Unassigned | - |
| Otu29300 | 1 | k__Plantae(100);p__unclassified_Plantae(100);c__unclassified_Plantae(100);o__unclassified_Plantae(100);f__unclassified_Plantae(100);g__unclassified_Plantae(100);s__Plantae_sp(100); | - | - | - | - | - | - | - | Unassigned | - |
| Otu29301 | 1 | k__Plantae(100);p__unclassified_Plantae(100);c__unclassified_Plantae(100);o__unclassified_Plantae(100);f__unclassified_Plantae(100);g__unclassified_Plantae(100);s__Plantae_sp(100); | - | - | - | - | - | - | - | Unassigned | - |
| Otu29302 | 1 | k__Plantae(100);p__unclassified_Plantae(100);c__unclassified_Plantae(100);o__unclassified_Plantae(100);f__unclassified_Plantae(100);g__unclassified_Plantae(100);s__Plantae_sp(100); | - | - | - | - | - | - | - | Unassigned | - |
| Otu29303 | 1 | k__Plantae(100);p__unclassified_Plantae(100);c__unclassified_Plantae(100);o__unclassified_Plantae(100);f__unclassified_Plantae(100);g__unclassified_Plantae(100);s__Plantae_sp(100); | - | - | - | - | - | - | - | Unassigned | - |
| Otu29304 | 1 | k__Plantae(100);p__unclassified_Plantae(100);c__unclassified_Plantae(100);o__unclassified_Plantae(100);f__unclassified_Plantae(100);g__unclassified_Plantae(100);s__Plantae_sp(100); | - | - | - | - | - | - | - | Unassigned | - |
| Otu29305 | 1 | k__Plantae(100);p__unclassified_Plantae(100);c__unclassified_Plantae(100);o__unclassified_Plantae(100);f__unclassified_Plantae(100);g__unclassified_Plantae(100);s__Plantae_sp(100); | - | - | - | - | - | - | - | Unassigned | - |
| Otu29306 | 1 | k__Plantae(100);p__unclassified_Plantae(100);c__unclassified_Plantae(100);o__unclassified_Plantae(100);f__unclassified_Plantae(100);g__unclassified_Plantae(100);s__Plantae_sp(100); | - | - | - | - | - | - | - | Unassigned | - |
| Otu29307 | 1 | k__Plantae(100);p__unclassified_Plantae(100);c__unclassified_Plantae(100);o__unclassified_Plantae(100);f__unclassified_Plantae(100);g__unclassified_Plantae(100);s__Plantae_sp(100); | - | - | - | - | - | - | - | Unassigned | - |
| Otu29308 | 1 | k__Plantae(100);p__unclassified_Plantae(100);c__unclassified_Plantae(100);o__unclassified_Plantae(100);f__unclassified_Plantae(100);g__unclassified_Plantae(100);s__Plantae_sp(100); | - | - | - | - | - | - | - | Unassigned | - |
| Otu29309 | 1 | k__Plantae(100);p__unclassified_Plantae(100);c__unclassified_Plantae(100);o__unclassified_Plantae(100);f__unclassified_Plantae(100);g__unclassified_Plantae(100);s__Plantae_sp(100); | - | - | - | - | - | - | - | Unassigned | - |
| Otu29310 | 1 | k__Plantae(100);p__unclassified_Plantae(100);c__unclassified_Plantae(100);o__unclassified_Plantae(100);f__unclassified_Plantae(100);g__unclassified_Plantae(100);s__Plantae_sp(100); | - | - | - | - | - | - | - | Unassigned | - |
| Otu29311 | 1 | k__Plantae(100);p__unclassified_Plantae(100);c__unclassified_Plantae(100);o__unclassified_Plantae(100);f__unclassified_Plantae(100);g__unclassified_Plantae(100);s__Plantae_sp(100); | - | - | - | - | - | - | - | Unassigned | - |
| Otu29312 | 1 | k__Plantae(100);p__unclassified_Plantae(100);c__unclassified_Plantae(100);o__unclassified_Plantae(100);f__unclassified_Plantae(100);g__unclassified_Plantae(100);s__Plantae_sp(100); | - | - | - | - | - | - | - | Unassigned | - |
| Otu29313 | 1 | k__Plantae(100);p__unclassified_Plantae(100);c__unclassified_Plantae(100);o__unclassified_Plantae(100);f__unclassified_Plantae(100);g__unclassified_Plantae(100);s__Plantae_sp(100); | - | - | - | - | - | - | - | Unassigned | - |
| Otu29314 | 1 | k__Plantae(100);p__unclassified_Plantae(100);c__unclassified_Plantae(100);o__unclassified_Plantae(100);f__unclassified_Plantae(100);g__unclassified_Plantae(100);s__Plantae_sp(100); | - | - | - | - | - | - | - | Unassigned | - |
| Otu29315 | 1 | k__Plantae(100);p__unclassified_Plantae(100);c__unclassified_Plantae(100);o__unclassified_Plantae(100);f__unclassified_Plantae(100);g__unclassified_Plantae(100);s__Plantae_sp(100); | - | - | - | - | - | - | - | Unassigned | - |
| Otu29316 | 1 | k__Plantae(100);p__unclassified_Plantae(100);c__unclassified_Plantae(100);o__unclassified_Plantae(100);f__unclassified_Plantae(100);g__unclassified_Plantae(100);s__Plantae_sp(100); | - | - | - | - | - | - | - | Unassigned | - |
| Otu29317 | 1 | k__Plantae(100);p__unclassified_Plantae(100);c__unclassified_Plantae(100);o__unclassified_Plantae(100);f__unclassified_Plantae(100);g__unclassified_Plantae(100);s__Plantae_sp(100); | - | - | - | - | - | - | - | Unassigned | - |
| Otu29318 | 1 | k__Plantae(100);p__unclassified_Plantae(100);c__unclassified_Plantae(100);o__unclassified_Plantae(100);f__unclassified_Plantae(100);g__unclassified_Plantae(100);s__Plantae_sp(100); | - | - | - | - | - | - | - | Unassigned | - |
| Otu29319 | 1 | k__Plantae(100);p__unclassified_Plantae(100);c__unclassified_Plantae(100);o__unclassified_Plantae(100);f__unclassified_Plantae(100);g__unclassified_Plantae(100);s__Plantae_sp(100); | - | - | - | - | - | - | - | Unassigned | - |
| Otu29320 | 1 | k__Plantae(100);p__unclassified_Plantae(100);c__unclassified_Plantae(100);o__unclassified_Plantae(100);f__unclassified_Plantae(100);g__unclassified_Plantae(100);s__Plantae_sp(100); | - | - | - | - | - | - | - | Unassigned | - |
| Otu29321 | 1 | k__Plantae(100);p__unclassified_Plantae(100);c__unclassified_Plantae(100);o__unclassified_Plantae(100);f__unclassified_Plantae(100);g__unclassified_Plantae(100);s__Plantae_sp(100); | - | - | - | - | - | - | - | Unassigned | - |
| Otu29322 | 1 | k__Plantae(100);p__unclassified_Plantae(100);c__unclassified_Plantae(100);o__unclassified_Plantae(100);f__unclassified_Plantae(100);g__unclassified_Plantae(100);s__Plantae_sp(100); | - | - | - | - | - | - | - | Unassigned | - |
| Otu29323 | 1 | k__Plantae(100);p__unclassified_Plantae(100);c__unclassified_Plantae(100);o__unclassified_Plantae(100);f__unclassified_Plantae(100);g__unclassified_Plantae(100);s__Plantae_sp(100); | - | - | - | - | - | - | - | Unassigned | - |
| Otu29324 | 1 | k__Plantae(100);p__unclassified_Plantae(100);c__unclassified_Plantae(100);o__unclassified_Plantae(100);f__unclassified_Plantae(100);g__unclassified_Plantae(100);s__Plantae_sp(100); | - | - | - | - | - | - | - | Unassigned | - |
| Otu29325 | 1 | k__Plantae(100);p__unclassified_Plantae(100);c__unclassified_Plantae(100);o__unclassified_Plantae(100);f__unclassified_Plantae(100);g__unclassified_Plantae(100);s__Plantae_sp(100); | - | - | - | - | - | - | - | Unassigned | - |
| Otu29326 | 1 | k__Plantae(100);p__unclassified_Plantae(100);c__unclassified_Plantae(100);o__unclassified_Plantae(100);f__unclassified_Plantae(100);g__unclassified_Plantae(100);s__Plantae_sp(100); | - | - | - | - | - | - | - | Unassigned | - |
| Otu29327 | 1 | k__Plantae(100);p__unclassified_Plantae(100);c__unclassified_Plantae(100);o__unclassified_Plantae(100);f__unclassified_Plantae(100);g__unclassified_Plantae(100);s__Plantae_sp(100); | - | - | - | - | - | - | - | Unassigned | - |
| Otu29328 | 1 | k__Plantae(100);p__unclassified_Plantae(100);c__unclassified_Plantae(100);o__unclassified_Plantae(100);f__unclassified_Plantae(100);g__unclassified_Plantae(100);s__Plantae_sp(100); | - | - | - | - | - | - | - | Unassigned | - |
| Otu29329 | 1 | k__Plantae(100);p__unclassified_Plantae(100);c__unclassified_Plantae(100);o__unclassified_Plantae(100);f__unclassified_Plantae(100);g__unclassified_Plantae(100);s__Plantae_sp(100); | - | - | - | - | - | - | - | Unassigned | - |
| Otu29330 | 1 | k__Plantae(100);p__unclassified_Plantae(100);c__unclassified_Plantae(100);o__unclassified_Plantae(100);f__unclassified_Plantae(100);g__unclassified_Plantae(100);s__Plantae_sp(100); | - | - | - | - | - | - | - | Unassigned | - |
| Otu29331 | 1 | k__Plantae(100);p__unclassified_Plantae(100);c__unclassified_Plantae(100);o__unclassified_Plantae(100);f__unclassified_Plantae(100);g__unclassified_Plantae(100);s__Plantae_sp(100); | - | - | - | - | - | - | - | Unassigned | - |
| Otu29332 | 1 | k__Plantae(100);p__unclassified_Plantae(100);c__unclassified_Plantae(100);o__unclassified_Plantae(100);f__unclassified_Plantae(100);g__unclassified_Plantae(100);s__Plantae_sp(100); | - | - | - | - | - | - | - | Unassigned | - |
| Otu29333 | 1 | k__Plantae(100);p__unclassified_Plantae(100);c__unclassified_Plantae(100);o__unclassified_Plantae(100);f__unclassified_Plantae(100);g__unclassified_Plantae(100);s__Plantae_sp(100); | - | - | - | - | - | - | - | Unassigned | - |
| Otu29334 | 1 | k__Plantae(100);p__unclassified_Plantae(100);c__unclassified_Plantae(100);o__unclassified_Plantae(100);f__unclassified_Plantae(100);g__unclassified_Plantae(100);s__Plantae_sp(100); | - | - | - | - | - | - | - | Unassigned | - |
| Otu29335 | 1 | k__Plantae(100);p__unclassified_Plantae(100);c__unclassified_Plantae(100);o__unclassified_Plantae(100);f__unclassified_Plantae(100);g__unclassified_Plantae(100);s__Plantae_sp(100); | - | - | - | - | - | - | - | Unassigned | - |
| Otu29336 | 1 | k__Plantae(100);p__unclassified_Plantae(100);c__unclassified_Plantae(100);o__unclassified_Plantae(100);f__unclassified_Plantae(100);g__unclassified_Plantae(100);s__Plantae_sp(100); | - | - | - | - | - | - | - | Unassigned | - |
| Otu29337 | 1 | k__Plantae(100);p__unclassified_Plantae(100);c__unclassified_Plantae(100);o__unclassified_Plantae(100);f__unclassified_Plantae(100);g__unclassified_Plantae(100);s__Plantae_sp(100); | - | - | - | - | - | - | - | Unassigned | - |
| Otu29338 | 1 | k__Plantae(100);p__unclassified_Plantae(100);c__unclassified_Plantae(100);o__unclassified_Plantae(100);f__unclassified_Plantae(100);g__unclassified_Plantae(100);s__Plantae_sp(100); | - | - | - | - | - | - | - | Unassigned | - |
| Otu29339 | 1 | k__Plantae(100);p__unclassified_Plantae(100);c__unclassified_Plantae(100);o__unclassified_Plantae(100);f__unclassified_Plantae(100);g__unclassified_Plantae(100);s__Plantae_sp(100); | - | - | - | - | - | - | - | Unassigned | - |
| Otu29340 | 1 | k__Plantae(100);p__unclassified_Plantae(100);c__unclassified_Plantae(100);o__unclassified_Plantae(100);f__unclassified_Plantae(100);g__unclassified_Plantae(100);s__Plantae_sp(100); | - | - | - | - | - | - | - | Unassigned | - |
| Otu29341 | 1 | k__Plantae(100);p__unclassified_Plantae(100);c__unclassified_Plantae(100);o__unclassified_Plantae(100);f__unclassified_Plantae(100);g__unclassified_Plantae(100);s__Plantae_sp(100); | - | - | - | - | - | - | - | Unassigned | - |
| Otu29342 | 1 | k__Plantae(100);p__unclassified_Plantae(100);c__unclassified_Plantae(100);o__unclassified_Plantae(100);f__unclassified_Plantae(100);g__unclassified_Plantae(100);s__Plantae_sp(100); | - | - | - | - | - | - | - | Unassigned | - |
| Otu29343 | 1 | k__Plantae(100);p__unclassified_Plantae(100);c__unclassified_Plantae(100);o__unclassified_Plantae(100);f__unclassified_Plantae(100);g__unclassified_Plantae(100);s__Plantae_sp(100); | - | - | - | - | - | - | - | Unassigned | - |

| | | | | | | | | | | | |
|---|---|---|---|---|---|---|---|---|---|---|---|
| Otu29344 | 1 | k__Plantae(100);p__unclassified_Plantae(100);c__unclassified_Plantae(100);o__unclassified_Plantae(100);f__unclassified_Plantae(100);g__unclassified_Plantae(100);s__Plantae_sp(100); | - | - | - | - | - | - | - | Unassigned | - |
| Otu29345 | 1 | k__Plantae(100);p__unclassified_Plantae(100);c__unclassified_Plantae(100);o__unclassified_Plantae(100);f__unclassified_Plantae(100);g__unclassified_Plantae(100);s__Plantae_sp(100); | - | - | - | - | - | - | - | Unassigned | - |
| Otu29346 | 1 | k__Plantae(100);p__unclassified_Plantae(100);c__unclassified_Plantae(100);o__unclassified_Plantae(100);f__unclassified_Plantae(100);g__unclassified_Plantae(100);s__Plantae_sp(100); | - | - | - | - | - | - | - | Unassigned | - |
| Otu29347 | 1 | k__Fungi(100);p__Ascomycota(100);p__Ascomycota_unclassified(100);p__Ascomycota_unclassified(100);p__Ascomycota_unclassified(100);p__Ascomycota_unclassified(100);p__Ascomycota_unclassified(100); | - | - | - | - | - | - | - | Unassigned | - |
| Otu29348 | 1 | k__Plantae(100);p__unclassified_Plantae(100);c__unclassified_Plantae(100);o__unclassified_Plantae(100);f__unclassified_Plantae(100);g__unclassified_Plantae(100);s__Plantae_sp(100); | - | - | - | - | - | - | - | Unassigned | - |
| Otu29349 | 1 | k__Fungi(100);p__Ascomycota(100);p__Ascomycota_unclassified(100);p__Ascomycota_unclassified(100);p__Ascomycota_unclassified(100);p__Ascomycota_unclassified(100);p__Ascomycota_unclassified(100); | - | - | - | - | - | - | - | Unassigned | - |
| Otu29350 | 1 | k__Plantae(100);p__unclassified_Plantae(100);c__unclassified_Plantae(100);o__unclassified_Plantae(100);f__unclassified_Plantae(100);g__unclassified_Plantae(100);s__Plantae_sp(100); | - | - | - | - | - | - | - | Unassigned | - |
| Otu29351 | 1 | k__Plantae(100);p__unclassified_Plantae(100);c__unclassified_Plantae(100);o__unclassified_Plantae(100);f__unclassified_Plantae(100);g__unclassified_Plantae(100);s__Plantae_sp(100); | - | - | - | - | - | - | - | Unassigned | - |
| Otu29352 | 1 | k__Plantae(100);p__unclassified_Plantae(100);c__unclassified_Plantae(100);o__unclassified_Plantae(100);f__unclassified_Plantae(100);g__unclassified_Plantae(100);s__Plantae_sp(100); | - | - | - | - | - | - | - | Unassigned | - |
| Otu29353 | 1 | k__Plantae(100);p__unclassified_Plantae(100);c__unclassified_Plantae(100);o__unclassified_Plantae(100);f__unclassified_Plantae(100);g__unclassified_Plantae(100);s__Plantae_sp(100); | - | - | - | - | - | - | - | Unassigned | - |
| Otu29354 | 1 | k__Fungi(100);p__Ascomycota(100);p__Ascomycota_unclassified(100);p__Ascomycota_unclassified(100);p__Ascomycota_unclassified(100);p__Ascomycota_unclassified(100);p__Ascomycota_unclassified(100); | - | - | - | - | - | - | - | Unassigned | - |
| Otu29355 | 1 | k__Plantae(100);p__unclassified_Plantae(100);c__unclassified_Plantae(100);o__unclassified_Plantae(100);f__unclassified_Plantae(100);g__unclassified_Plantae(100);s__Plantae_sp(100); | - | - | - | - | - | - | - | Unassigned | - |
| Otu29356 | 1 | k__Plantae(100);p__unclassified_Plantae(100);c__unclassified_Plantae(100);o__unclassified_Plantae(100);f__unclassified_Plantae(100);g__unclassified_Plantae(100);s__Plantae_sp(100); | - | - | - | - | - | - | - | Unassigned | - |
| Otu29357 | 1 | k__Plantae(100);p__unclassified_Plantae(100);c__unclassified_Plantae(100);o__unclassified_Plantae(100);f__unclassified_Plantae(100);g__unclassified_Plantae(100);s__Plantae_sp(100); | - | - | - | - | - | - | - | Unassigned | - |
| Otu29358 | 1 | k__Plantae(100);p__unclassified_Plantae(100);c__unclassified_Plantae(100);o__unclassified_Plantae(100);f__unclassified_Plantae(100);g__unclassified_Plantae(100);s__Plantae_sp(100); | - | - | - | - | - | - | - | Unassigned | - |
| Otu29359 | 1 | k__Plantae(100);p__unclassified_Plantae(100);c__unclassified_Plantae(100);o__unclassified_Plantae(100);f__unclassified_Plantae(100);g__unclassified_Plantae(100);s__Plantae_sp(100); | - | - | - | - | - | - | - | Unassigned | - |
| Otu29360 | 1 | k__Plantae(100);p__unclassified_Plantae(100);c__unclassified_Plantae(100);o__unclassified_Plantae(100);f__unclassified_Plantae(100);g__unclassified_Plantae(100);s__Plantae_sp(100); | - | - | - | - | - | - | - | Unassigned | - |
| Otu29361 | 1 | k__Plantae(100);p__unclassified_Plantae(100);c__unclassified_Plantae(100);o__unclassified_Plantae(100);f__unclassified_Plantae(100);g__unclassified_Plantae(100);s__Plantae_sp(100); | - | - | - | - | - | - | - | Unassigned | - |
| Otu29362 | 1 | k__Fungi(100);p__Ascomycota(100);p__Ascomycota_unclassified(100);p__Ascomycota_unclassified(100);p__Ascomycota_unclassified(100);p__Ascomycota_unclassified(100);p__Ascomycota_unclassified(100); | - | - | - | - | - | - | - | Unassigned | - |
| Otu29363 | 1 | k__Plantae(100);p__unclassified_Plantae(100);c__unclassified_Plantae(100);o__unclassified_Plantae(100);f__unclassified_Plantae(100);g__unclassified_Plantae(100);s__Plantae_sp(100); | - | - | - | - | - | - | - | Unassigned | - |
| Otu29364 | 1 | k__Plantae(100);p__unclassified_Plantae(100);c__unclassified_Plantae(100);o__unclassified_Plantae(100);f__unclassified_Plantae(100);g__unclassified_Plantae(100);s__Plantae_sp(100); | - | - | - | - | - | - | - | Unassigned | - |
| Otu29365 | 1 | k__Plantae(100);p__unclassified_Plantae(100);c__unclassified_Plantae(100);o__unclassified_Plantae(100);f__unclassified_Plantae(100);g__unclassified_Plantae(100);s__Plantae_sp(100); | - | - | - | - | - | - | - | Unassigned | - |
| Otu29366 | 1 | k__Fungi(100);p__Ascomycota(100);p__Ascomycota_unclassified(100);p__Ascomycota_unclassified(100);p__Ascomycota_unclassified(100);p__Ascomycota_unclassified(100);p__Ascomycota_unclassified(100); | - | - | - | - | - | - | - | Unassigned | - |
| Otu29367 | 1 | k__Plantae(100);p__unclassified_Plantae(100);c__unclassified_Plantae(100);o__unclassified_Plantae(100);f__unclassified_Plantae(100);g__unclassified_Plantae(100);s__Plantae_sp(100); | - | - | - | - | - | - | - | Unassigned | - |
| Otu29368 | 1 | k__Plantae(100);p__unclassified_Plantae(100);c__unclassified_Plantae(100);o__unclassified_Plantae(100);f__unclassified_Plantae(100);g__unclassified_Plantae(100);s__Plantae_sp(100); | - | - | - | - | - | - | - | Unassigned | - |
| Otu29369 | 1 | k__Plantae(100);p__unclassified_Plantae(100);c__unclassified_Plantae(100);o__unclassified_Plantae(100);f__unclassified_Plantae(100);g__unclassified_Plantae(100);s__Plantae_sp(100); | - | - | - | - | - | - | - | Unassigned | - |
| Otu29370 | 1 | k__Plantae(100);p__unclassified_Plantae(100);c__unclassified_Plantae(100);o__unclassified_Plantae(100);f__unclassified_Plantae(100);g__unclassified_Plantae(100);s__Plantae_sp(100); | - | - | - | - | - | - | - | Unassigned | - |
| Otu29371 | 1 | k__Plantae(100);p__unclassified_Plantae(100);c__unclassified_Plantae(100);o__unclassified_Plantae(100);f__unclassified_Plantae(100);g__unclassified_Plantae(100);s__Plantae_sp(100); | - | - | - | - | - | - | - | Unassigned | - |
| Otu29372 | 1 | k__Plantae(100);p__unclassified_Plantae(100);c__unclassified_Plantae(100);o__unclassified_Plantae(100);f__unclassified_Plantae(100);g__unclassified_Plantae(100);s__Plantae_sp(100); | - | - | - | - | - | - | - | Unassigned | - |
| Otu29373 | 1 | k__Plantae(100);p__unclassified_Plantae(100);c__unclassified_Plantae(100);o__unclassified_Plantae(100);f__unclassified_Plantae(100);g__unclassified_Plantae(100);s__Plantae_sp(100); | - | - | - | - | - | - | - | Unassigned | - |
| Otu29374 | 1 | k__Plantae(100);p__unclassified_Plantae(100);c__unclassified_Plantae(100);o__unclassified_Plantae(100);f__unclassified_Plantae(100);g__unclassified_Plantae(100);s__Plantae_sp(100); | - | - | - | - | - | - | - | Unassigned | - |
| Otu29375 | 1 | k__Plantae(100);p__unclassified_Plantae(100);c__unclassified_Plantae(100);o__unclassified_Plantae(100);f__unclassified_Plantae(100);g__unclassified_Plantae(100);s__Plantae_sp(100); | - | - | - | - | - | - | - | Unassigned | - |
| Otu29376 | 1 | k__Plantae(100);p__unclassified_Plantae(100);c__unclassified_Plantae(100);o__unclassified_Plantae(100);f__unclassified_Plantae(100);g__unclassified_Plantae(100);s__Plantae_sp(100); | - | - | - | - | - | - | - | Unassigned | - |
| Otu29377 | 1 | k__Plantae(100);p__unclassified_Plantae(100);c__unclassified_Plantae(100);o__unclassified_Plantae(100);f__unclassified_Plantae(100);g__unclassified_Plantae(100);s__Plantae_sp(100); | - | - | - | - | - | - | - | Unassigned | - |
| Otu29378 | 1 | k__Plantae(100);p__unclassified_Plantae(100);c__unclassified_Plantae(100);o__unclassified_Plantae(100);f__unclassified_Plantae(100);g__unclassified_Plantae(100);s__Plantae_sp(100); | - | - | - | - | - | - | - | Unassigned | - |
| Otu29379 | 1 | k__Plantae(100);p__unclassified_Plantae(100);c__unclassified_Plantae(100);o__unclassified_Plantae(100);f__unclassified_Plantae(100);g__unclassified_Plantae(100);s__Plantae_sp(100); | - | - | - | - | - | - | - | Unassigned | - |
| Otu29380 | 1 | k__Plantae(100);p__unclassified_Plantae(100);c__unclassified_Plantae(100);o__unclassified_Plantae(100);f__unclassified_Plantae(100);g__unclassified_Plantae(100);s__Plantae_sp(100); | - | - | - | - | - | - | - | Unassigned | - |
| Otu29381 | 1 | k__Plantae(100);p__unclassified_Plantae(100);c__unclassified_Plantae(100);o__unclassified_Plantae(100);f__unclassified_Plantae(100);g__unclassified_Plantae(100);s__Plantae_sp(100); | - | - | - | - | - | - | - | Unassigned | - |
| Otu29382 | 1 | k__Fungi(100);p__Ascomycota(100);p__Ascomycota_unclassified(100);p__Ascomycota_unclassified(100);p__Ascomycota_unclassified(100);p__Ascomycota_unclassified(100);p__Ascomycota_unclassified(100); | - | - | - | - | - | - | - | Unassigned | - |
| Otu29383 | 1 | k__Plantae(100);p__unclassified_Plantae(100);c__unclassified_Plantae(100);o__unclassified_Plantae(100);f__unclassified_Plantae(100);g__unclassified_Plantae(100);s__Plantae_sp(100); | - | - | - | - | - | - | - | Unassigned | - |
| Otu29384 | 1 | k__Plantae(100);p__unclassified_Plantae(100);c__unclassified_Plantae(100);o__unclassified_Plantae(100);f__unclassified_Plantae(100);g__unclassified_Plantae(100);s__Plantae_sp(100); | - | - | - | - | - | - | - | Unassigned | - |
| Otu29385 | 1 | k__Plantae(100);p__unclassified_Plantae(100);c__unclassified_Plantae(100);o__unclassified_Plantae(100);f__unclassified_Plantae(100);g__unclassified_Plantae(100);s__Plantae_sp(100); | - | - | - | - | - | - | - | Unassigned | - |
| Otu29386 | 1 | k__Plantae(100);p__unclassified_Plantae(100);c__unclassified_Plantae(100);o__unclassified_Plantae(100);f__unclassified_Plantae(100);g__unclassified_Plantae(100);s__Plantae_sp(100); | - | - | - | - | - | - | - | Unassigned | - |
| Otu29387 | 1 | k__Plantae(100);p__unclassified_Plantae(100);c__unclassified_Plantae(100);o__unclassified_Plantae(100);f__unclassified_Plantae(100);g__unclassified_Plantae(100);s__Plantae_sp(100); | - | - | - | - | - | - | - | Unassigned | - |
| Otu29388 | 1 | k__Plantae(100);p__unclassified_Plantae(100);c__unclassified_Plantae(100);o__unclassified_Plantae(100);f__unclassified_Plantae(100);g__unclassified_Plantae(100);s__Plantae_sp(100); | - | - | - | - | - | - | - | Unassigned | - |
| Otu29389 | 1 | k__Plantae(100);p__unclassified_Plantae(100);c__unclassified_Plantae(100);o__unclassified_Plantae(100);f__unclassified_Plantae(100);g__unclassified_Plantae(100);s__Plantae_sp(100); | - | - | - | - | - | - | - | Unassigned | - |
| Otu29390 | 1 | k__Plantae(100);p__unclassified_Plantae(100);c__unclassified_Plantae(100);o__unclassified_Plantae(100);f__unclassified_Plantae(100);g__unclassified_Plantae(100);s__Plantae_sp(100); | - | - | - | - | - | - | - | Unassigned | - |
| Otu29391 | 1 | k__Fungi(100);p__Ascomycota(100);p__Ascomycota_unclassified(100);p__Ascomycota_unclassified(100);p__Ascomycota_unclassified(100);p__Ascomycota_unclassified(100);p__Ascomycota_unclassified(100); | - | - | - | - | - | - | - | Unassigned | - |
| Otu29392 | 1 | k__Fungi(100);p__Ascomycota(100);p__Ascomycota_unclassified(100);p__Ascomycota_unclassified(100);p__Ascomycota_unclassified(100);p__Ascomycota_unclassified(100);p__Ascomycota_unclassified(100); | - | - | - | - | - | - | - | Unassigned | - |
| Otu29396 | 1 | k__Fungi(100);p__Ascomycota(100);p__Ascomycota_unclassified(100);p__Ascomycota_unclassified(100);p__Ascomycota_unclassified(100);p__Ascomycota_unclassified(100);p__Ascomycota_unclassified(100); | - | - | - | - | - | - | - | Unassigned | - |
| Otu29397 | 1 | k__Fungi(100);p__Ascomycota(100);c__Eurotiomycetes(100);o__Eurotiales(100);f__Trichocomaceae(100);f__Trichocomaceae_unclassified(100);f__Trichocomaceae_unclassified(100); | - | - | - | - | - | - | - | Unassigned | - |
| Otu29398 | 1 | k__Fungi(100);p__Ascomycota(100);p__Ascomycota_unclassified(100);p__Ascomycota_unclassified(100);p__Ascomycota_unclassified(100);p__Ascomycota_unclassified(100);p__Ascomycota_unclassified(100); | - | - | - | - | - | - | - | Unassigned | - |
| Otu29399 | 1 | k__Fungi(100);p__Ascomycota(100);c__Dothideomycetes(100);c__Dothideomycetes_unclassified(100);c__Dothideomycetes_unclassified(100);c__Dothideomycetes_unclassified(100);c__Dothideomycetes_unclassified(100); | - | - | - | - | - | - | - | Unassigned | - |
| Otu29400 | 1 | k__Fungi(100);p__Ascomycota(100);p__Ascomycota_unclassified(100);p__Ascomycota_unclassified(100);p__Ascomycota_unclassified(100);p__Ascomycota_unclassified(100);p__Ascomycota_unclassified(100); | - | - | - | - | - | - | - | Unassigned | - |
| Otu29401 | 1 | k__Fungi(100);p__Ascomycota(100);c__Eurotiomycetes(100);c__Eurotiomycetes_unclassified(100);c__Eurotiomycetes_unclassified(100);c__Eurotiomycetes_unclassified(100);c__Eurotiomycetes_unclassified(100); | - | - | - | - | - | - | - | Unassigned | - |
| Otu29402 | 1 | k__Fungi(100);p__Ascomycota(100);p__Ascomycota_unclassified(100);p__Ascomycota_unclassified(100);p__Ascomycota_unclassified(100);p__Ascomycota_unclassified(100);p__Ascomycota_unclassified(100); | - | - | - | - | - | - | - | Unassigned | - |
| Otu29403 | 1 | k__Fungi(100);p__Ascomycota(100);p__Ascomycota_unclassified(100);p__Ascomycota_unclassified(100);p__Ascomycota_unclassified(100);p__Ascomycota_unclassified(100);p__Ascomycota_unclassified(100); | - | - | - | - | - | - | - | Unassigned | - |
| Otu29406 | 1 | k__Fungi(100);p__Ascomycota(100);p__Ascomycota_unclassified(100);p__Ascomycota_unclassified(100);p__Ascomycota_unclassified(100);p__Ascomycota_unclassified(100);p__Ascomycota_unclassified(100); | - | - | - | - | - | - | - | Unassigned | - |
| Otu29408 | 1 | k__Fungi(100);p__Ascomycota(100);c__Sordariomycetes(100);c__Sordariomycetes_unclassified(100);c__Sordariomycetes_unclassified(100);c__Sordariomycetes_unclassified(100);c__Sordariomycetes_unclassified(100); | - | - | - | - | - | - | - | Unassigned | - |
| Otu29409 | 1 | k__Fungi(100);p__Ascomycota(100);c__Sordariomycetes(100);c__Sordariomycetes_unclassified(100);c__Sordariomycetes_unclassified(100);c__Sordariomycetes_unclassified(100);c__Sordariomycetes_unclassified(100); | - | - | - | - | - | - | - | Unassigned | - |
| Otu29410 | 1 | k__Fungi(100);p__Ascomycota(100);p__Ascomycota_unclassified(100);p__Ascomycota_unclassified(100);p__Ascomycota_unclassified(100);p__Ascomycota_unclassified(100);p__Ascomycota_unclassified(100); | - | - | - | - | - | - | - | Unassigned | - |
| Otu29413 | 1 | k__Fungi(100);k__Fungi_unclassified(100);k__Fungi_unclassified(100);k__Fungi_unclassified(100);k__Fungi_unclassified(100);k__Fungi_unclassified(100);k__Fungi_unclassified(100); | - | - | - | - | - | - | - | Unassigned | - |
| Otu29419 | 1 | k__Fungi(100);p__Ascomycota(100);p__Ascomycota_unclassified(100);p__Ascomycota_unclassified(100);p__Ascomycota_unclassified(100);p__Ascomycota_unclassified(100);p__Ascomycota_unclassified(100); | - | - | - | - | - | - | - | Unassigned | - |
| Otu29421 | 1 | k__Fungi(100);p__Ascomycota(100);p__Ascomycota_unclassified(100);p__Ascomycota_unclassified(100);p__Ascomycota_unclassified(100);p__Ascomycota_unclassified(100);p__Ascomycota_unclassified(100); | - | - | - | - | - | - | - | Unassigned | - |
| Otu29422 | 1 | k__Fungi(100);p__Ascomycota(100);c__Sordariomycetes(100);c__Sordariomycetes_unclassified(100);c__Sordariomycetes_unclassified(100);c__Sordariomycetes_unclassified(100);c__Sordariomycetes_unclassified(100); | - | - | - | - | - | - | - | Unassigned | - |
| Otu29424 | 1 | k__Fungi(100);p__Ascomycota(100);p__Ascomycota_unclassified(100);p__Ascomycota_unclassified(100);p__Ascomycota_unclassified(100);p__Ascomycota_unclassified(100);p__Ascomycota_unclassified(100); | - | - | - | - | - | - | - | Unassigned | - |
| Otu29438 | 1 | k__Fungi(100);p__Ascomycota(100);c__Eurotiomycetes(100);o__Eurotiales(100);f__Trichocomaceae(100);f__Trichocomaceae_unclassified(100);f__Trichocomaceae_unclassified(100); | - | - | - | - | - | - | - | Unassigned | - |
| Otu29439 | 1 | k__Fungi(100);p__Ascomycota(100);c__Eurotiomycetes(100);c__Eurotiomycetes_unclassified(100);c__Eurotiomycetes_unclassified(100);c__Eurotiomycetes_unclassified(100);c__Eurotiomycetes_unclassified(100); | - | - | - | - | - | - | - | Unassigned | - |
| Otu29442 | 1 | k__Fungi(100);p__Ascomycota(100);c__Eurotiomycetes(100);o__Eurotiales(100);f__Trichocomaceae(100);f__Trichocomaceae_unclassified(100);f__Trichocomaceae_unclassified(100); | - | - | - | - | - | - | - | Unassigned | - |
| Otu29443 | 1 | k__Fungi(100);p__Ascomycota(100);p__Ascomycota_unclassified(100);p__Ascomycota_unclassified(100);p__Ascomycota_unclassified(100);p__Ascomycota_unclassified(100);p__Ascomycota_unclassified(100); | - | - | - | - | - | - | - | Unassigned | - |
| Otu29444 | 1 | k__Fungi(100);p__Basidiomycota(100);c__Agaricomycetes(100);c__Agaricomycetes_unclassified(100);c__Agaricomycetes_unclassified(100);c__Agaricomycetes_unclassified(100);c__Agaricomycetes_unclassified(100); | - | - | - | - | - | - | - | Unassigned | - |
| Otu29450 | 1 | k__Fungi(100);k__Fungi_unclassified(100);k__Fungi_unclassified(100);k__Fungi_unclassified(100);k__Fungi_unclassified(100);k__Fungi_unclassified(100);k__Fungi_unclassified(100); | - | - | - | - | - | - | - | Unassigned | - |
| Otu29451 | 1 | k__Fungi(100);p__Ascomycota(100);c__Sordariomycetes(100);c__Sordariomycetes_unclassified(100);c__Sordariomycetes_unclassified(100);c__Sordariomycetes_unclassified(100);c__Sordariomycetes_unclassified(100); | - | - | - | - | - | - | - | Unassigned | - |
| Otu29461 | 1 | k__Fungi(100);p__Ascomycota(100);p__Ascomycota_unclassified(100);p__Ascomycota_unclassified(100);p__Ascomycota_unclassified(100);p__Ascomycota_unclassified(100);p__Ascomycota_unclassified(100); | - | - | - | - | - | - | - | Unassigned | - |
| Otu29463 | 1 | k__Fungi(100);p__Ascomycota(100);c__Sordariomycetes(100);c__Sordariomycetes_unclassified(100);c__Sordariomycetes_unclassified(100);c__Sordariomycetes_unclassified(100);c__Sordariomycetes_unclassified(100); | - | - | - | - | - | - | - | Unassigned | - |
| Otu29464 | 1 | k__Fungi(100);k__Fungi_unclassified(100);k__Fungi_unclassified(100);k__Fungi_unclassified(100);k__Fungi_unclassified(100);k__Fungi_unclassified(100);k__Fungi_unclassified(100); | - | - | - | - | - | - | - | Unassigned | - |
| Otu29465 | 1 | k__Plantae(100);p__unclassified_Plantae(100);c__unclassified_Plantae(100);o__unclassified_Plantae(100);f__unclassified_Plantae(100);g__unclassified_Plantae(100);s__Plantae_sp(100); | - | - | - | - | - | - | - | Unassigned | - |
| Otu29466 | 1 | k__Fungi(100);p__Ascomycota(100);p__Ascomycota_unclassified(100);p__Ascomycota_unclassified(100);p__Ascomycota_unclassified(100);p__Ascomycota_unclassified(100);p__Ascomycota_unclassified(100); | - | - | - | - | - | - | - | Unassigned | - |
| Otu29467 | 1 | k__Fungi(100);k__Fungi_unclassified(100);k__Fungi_unclassified(100);k__Fungi_unclassified(100);k__Fungi_unclassified(100);k__Fungi_unclassified(100);k__Fungi_unclassified(100); | - | - | - | - | - | - | - | Unassigned | - |
| Otu29468 | 1 | k__Fungi(100);k__Fungi_unclassified(100);k__Fungi_unclassified(100);k__Fungi_unclassified(100);k__Fungi_unclassified(100);k__Fungi_unclassified(100);k__Fungi_unclassified(100); | - | - | - | - | - | - | - | Unassigned | - |
| Otu29469 | 1 | k__Fungi(100);p__Ascomycota(100);p__Ascomycota_unclassified(100);p__Ascomycota_unclassified(100);p__Ascomycota_unclassified(100);p__Ascomycota_unclassified(100);p__Ascomycota_unclassified(100); | - | - | - | - | - | - | - | Unassigned | - |
| Otu29470 | 1 | k__Fungi(100);k__Fungi_unclassified(100);k__Fungi_unclassified(100);k__Fungi_unclassified(100);k__Fungi_unclassified(100);k__Fungi_unclassified(100);k__Fungi_unclassified(100); | - | - | - | - | - | - | - | Unassigned | - |
| Otu29471 | 1 | k__Fungi(100);p__Ascomycota(100);c__Sordariomycetes(100);c__Sordariomycetes_unclassified(100);c__Sordariomycetes_unclassified(100);c__Sordariomycetes_unclassified(100);c__Sordariomycetes_unclassified(100); | - | - | - | - | - | - | - | Unassigned | - |
| Otu29473 | 1 | k__Fungi(100);p__Ascomycota(100);p__Ascomycota_unclassified(100);p__Ascomycota_unclassified(100);p__Ascomycota_unclassified(100);p__Ascomycota_unclassified(100);p__Ascomycota_unclassified(100); | - | - | - | - | - | - | - | Unassigned | - |
| Otu29478 | 1 | k__Fungi(100);p__Ascomycota(100);c__Sordariomycetes(100);c__Sordariomycetes_unclassified(100);c__Sordariomycetes_unclassified(100);c__Sordariomycetes_unclassified(100);c__Sordariomycetes_unclassified(100); | - | - | - | - | - | - | - | Unassigned | - |
| Otu29479 | 1 | k__Fungi(100);p__Ascomycota(100);c__Sordariomycetes(100);c__Sordariomycetes_unclassified(100);c__Sordariomycetes_unclassified(100);c__Sordariomycetes_unclassified(100);c__Sordariomycetes_unclassified(100); | - | - | - | - | - | - | - | Unassigned | - |
| Otu29481 | 1 | k__Fungi(100);p__Ascomycota(100);c__Eurotiomycetes(100);c__Eurotiomycetes_unclassified(100);c__Eurotiomycetes_unclassified(100);c__Eurotiomycetes_unclassified(100);c__Eurotiomycetes_unclassified(100); | - | - | - | - | - | - | - | Unassigned | - |
| Otu29482 | 1 | k__Fungi(100);p__Ascomycota(100);p__Ascomycota_unclassified(100);p__Ascomycota_unclassified(100);p__Ascomycota_unclassified(100);p__Ascomycota_unclassified(100);p__Ascomycota_unclassified(100); | - | - | - | - | - | - | - | Unassigned | - |
| Otu29483 | 1 | k__Fungi(100);p__Ascomycota(100);c__Sordariomycetes(100);c__Sordariomycetes_unclassified(100);c__Sordariomycetes_unclassified(100);c__Sordariomycetes_unclassified(100);c__Sordariomycetes_unclassified(100); | - | - | - | - | - | - | - | Unassigned | - |
| Otu29484 | 1 | k__Fungi(100);p__Ascomycota(100);p__Ascomycota_unclassified(100);p__Ascomycota_unclassified(100);p__Ascomycota_unclassified(100);p__Ascomycota_unclassified(100);p__Ascomycota_unclassified(100); | - | - | - | - | - | - | - | Unassigned | - |
| Otu29485 | 1 | k__Fungi(100);p__Ascomycota(100);c__Sordariomycetes(100);c__Sordariomycetes_unclassified(100);c__Sordariomycetes_unclassified(100);c__Sordariomycetes_unclassified(100);c__Sordariomycetes_unclassified(100); | - | - | - | - | - | - | - | Unassigned | - |
| Otu29486 | 1 | k__Fungi(100);p__Ascomycota(100);c__Sordariomycetes(100);c__Sordariomycetes_unclassified(100);c__Sordariomycetes_unclassified(100);c__Sordariomycetes_unclassified(100);c__Sordariomycetes_unclassified(100); | - | - | - | - | - | - | - | Unassigned | - |
| Otu29487 | 1 | k__Fungi(100);p__Ascomycota(100);c__Sordariomycetes(100);c__Sordariomycetes_unclassified(100);c__Sordariomycetes_unclassified(100);c__Sordariomycetes_unclassified(100);c__Sordariomycetes_unclassified(100); | - | - | - | - | - | - | - | Unassigned | - |
| Otu29490 | 1 | k__Fungi(100);p__Ascomycota(100);c__Sordariomycetes(100);c__Sordariomycetes_unclassified(100);c__Sordariomycetes_unclassified(100);c__Sordariomycetes_unclassified(100);c__Sordariomycetes_unclassified(100); | - | - | - | - | - | - | - | Unassigned | - |
| Otu29508 | 1 | k__Fungi(100);p__Chytridiomycota(100);c__Chytridiomycetes(100);o__unclassified_Chytridiomycetes(100);f__unclassified_Chytridiomycetes(100);g__unclassified_Chytridiomycetes(100);s__Chytridiomycetes_sp(100); | - | - | - | - | - | - | - | Unassigned | - |
| Otu29510 | 1 | k__Fungi(100);p__Ascomycota(100);c__Dothideomycetes(100);o__Pleosporales(100);o__Pleosporales_unclassified(100);o__Pleosporales_unclassified(100);o__Pleosporales_unclassified(100); | - | - | - | - | - | - | - | Unassigned | - |
| Otu29513 | 1 | k__Fungi(100);k__Fungi_unclassified(100);k__Fungi_unclassified(100);k__Fungi_unclassified(100);k__Fungi_unclassified(100);k__Fungi_unclassified(100);k__Fungi_unclassified(100); | - | - | - | - | - | - | - | Unassigned | - |
| Otu29514 | 1 | k__Fungi(100);k__Fungi_unclassified(100);k__Fungi_unclassified(100);k__Fungi_unclassified(100);k__Fungi_unclassified(100);k__Fungi_unclassified(100);k__Fungi_unclassified(100); | - | - | - | - | - | - | - | Unassigned | - |
| Otu29515 | 1 | k__Fungi(100);p__Basidiomycota(100);c__Agaricomycetes(100);c__Agaricomycetes_unclassified(100);c__Agaricomycetes_unclassified(100);c__Agaricomycetes_unclassified(100);c__Agaricomycetes_unclassified(100); | - | - | - | - | - | - | - | Unassigned | - |
| Otu29518 | 1 | k__Fungi(100);p__Basidiomycota(100);c__Microbotryomycetes(100);o__Sporidiobolales(100);f__Sporidiobolales_family_Incertae_sedis(100);f__Sporidiobolales_family_Incertae_sedis_unclassified(100);f__Sporidiobolales_family_Incertae_sedis_unclassified(100); | - | - | - | - | - | - | - | Unassigned | - |
| Otu29519 | 1 | k__Fungi(100);p__Ascomycota(100);p__Ascomycota_unclassified(100);p__Ascomycota_unclassified(100);p__Ascomycota_unclassified(100);p__Ascomycota_unclassified(100);p__Ascomycota_unclassified(100); | - | - | - | - | - | - | - | Unassigned | - |
| Otu29520 | 1 | k__Fungi(100);k__Fungi_unclassified(100);k__Fungi_unclassified(100);k__Fungi_unclassified(100);k__Fungi_unclassified(100);k__Fungi_unclassified(100);k__Fungi_unclassified(100); | - | - | - | - | - | - | - | Unassigned | - |
| Otu29521 | 1 | k__Fungi(100);k__Fungi_unclassified(100);k__Fungi_unclassified(100);k__Fungi_unclassified(100);k__Fungi_unclassified(100);k__Fungi_unclassified(100);k__Fungi_unclassified(100); | - | - | - | - | - | - | - | Unassigned | - |
| Otu29522 | 1 | k__Fungi(100);k__Fungi_unclassified(100);k__Fungi_unclassified(100);k__Fungi_unclassified(100);k__Fungi_unclassified(100);k__Fungi_unclassified(100);k__Fungi_unclassified(100); | - | - | - | - | - | - | - | Unassigned | - |
| Otu29523 | 1 | k__Fungi(100);k__Fungi_unclassified(100);k__Fungi_unclassified(100);k__Fungi_unclassified(100);k__Fungi_unclassified(100);k__Fungi_unclassified(100);k__Fungi_unclassified(100); | - | - | - | - | - | - | - | Unassigned | - |
| Otu29525 | 1 | k__Fungi(100);p__Basidiomycota(100);c__Agaricomycetes(100);c__Agaricomycetes_unclassified(100);c__Agaricomycetes_unclassified(100);c__Agaricomycetes_unclassified(100);c__Agaricomycetes_unclassified(100); | - | - | - | - | - | - | - | Unassigned | - |
| Otu29527 | 1 | k__Fungi(100);p__Basidiomycota(100);c__Microbotryomycetes(100);o__Sporidiobolales(100);f__Sporidiobolales_family_Incertae_sedis(100);f__Sporidiobolales_family_Incertae_sedis_unclassified(100);f__Sporidiobolales_family_Incertae_sedis_unclassified(100); | - | - | - | - | - | - | - | Unassigned | - |
| Otu29536 | 1 | k__Fungi(100);k__Fungi_unclassified(100);k__Fungi_unclassified(100);k__Fungi_unclassified(100);k__Fungi_unclassified(100);k__Fungi_unclassified(100);k__Fungi_unclassified(100); | - | - | - | - | - | - | - | Unassigned | - |
| Otu29537 | 1 | k__Fungi(100);k__Fungi_unclassified(100);k__Fungi_unclassified(100);k__Fungi_unclassified(100);k__Fungi_unclassified(100);k__Fungi_unclassified(100);k__Fungi_unclassified(100); | - | - | - | - | - | - | - | Unassigned | - |
| Otu29551 | 1 | k__Fungi(100);p__Ascomycota(100);c__Sordariomycetes(100);c__Sordariomycetes_unclassified(100);c__Sordariomycetes_unclassified(100);c__Sordariomycetes_unclassified(100);c__Sordariomycetes_unclassified(100); | - | - | - | - | - | - | - | Unassigned | - |
| Otu29553 | 1 | k__Fungi(100);p__Ascomycota(100);p__Ascomycota_unclassified(100);p__Ascomycota_unclassified(100);p__Ascomycota_unclassified(100);p__Ascomycota_unclassified(100);p__Ascomycota_unclassified(100); | - | - | - | - | - | - | - | Unassigned | - |
| Otu29555 | 1 | k__Fungi(100);p__Ascomycota(100);p__Ascomycota_unclassified(100);p__Ascomycota_unclassified(100);p__Ascomycota_unclassified(100);p__Ascomycota_unclassified(100);p__Ascomycota_unclassified(100); | - | - | - | - | - | - | - | Unassigned | - |
| Otu29556 | 1 | k__Fungi(100);p__Ascomycota(100);p__Ascomycota_unclassified(100);p__Ascomycota_unclassified(100);p__Ascomycota_unclassified(100);p__Ascomycota_unclassified(100);p__Ascomycota_unclassified(100); | - | - | - | - | - | - | - | Unassigned | - |
| Otu29557 | 1 | k__Fungi(100);p__Ascomycota(100);p__Ascomycota_unclassified(100);p__Ascomycota_unclassified(100);p__Ascomycota_unclassified(100);p__Ascomycota_unclassified(100);p__Ascomycota_unclassified(100); | - | - | - | - | - | - | - | Unassigned | - |
| Otu29560 | 1 | k__Fungi(100);p__Ascomycota(100);p__Ascomycota_unclassified(100);p__Ascomycota_unclassified(100);p__Ascomycota_unclassified(100);p__Ascomycota_unclassified(100);p__Ascomycota_unclassified(100); | - | - | - | - | - | - | - | Unassigned | - |
| Otu29562 | 1 | k__Fungi(100);p__Ascomycota(100);c__Dothideomycetes(100);o__Pleosporales(100);o__Pleosporales_unclassified(100);o__Pleosporales_unclassified(100);o__Pleosporales_unclassified(100); | - | - | - | - | - | - | - | Unassigned | - |
| Otu29564 | 1 | k__Fungi(100);p__Ascomycota(100);c__Sordariomycetes(100);c__Sordariomycetes_unclassified(100);c__Sordariomycetes_unclassified(100);c__Sordariomycetes_unclassified(100);c__Sordariomycetes_unclassified(100); | - | - | - | - | - | - | - | Unassigned | - |
| Otu29565 | 1 | k__Fungi(100);p__Ascomycota(100);p__Ascomycota_unclassified(100);p__Ascomycota_unclassified(100);p__Ascomycota_unclassified(100);p__Ascomycota_unclassified(100);p__Ascomycota_unclassified(100); | - | - | - | - | - | - | - | Unassigned | - |
| Otu29566 | 1 | k__Fungi(100);p__Ascomycota(100);p__Ascomycota_unclassified(100);p__Ascomycota_unclassified(100);p__Ascomycota_unclassified(100);p__Ascomycota_unclassified(100);p__Ascomycota_unclassified(100); | - | - | - | - | - | - | - | Unassigned | - |
| Otu29567 | 1 | k__Fungi(100);p__Ascomycota(100);p__Ascomycota_unclassified(100);p__Ascomycota_unclassified(100);p__Ascomycota_unclassified(100);p__Ascomycota_unclassified(100);p__Ascomycota_unclassified(100); | - | - | - | - | - | - | - | Unassigned | - |
| Otu29573 | 1 | k__Fungi(100);p__Ascomycota(100);p__Ascomycota_unclassified(100);p__Ascomycota_unclassified(100);p__Ascomycota_unclassified(100);p__Ascomycota_unclassified(100);p__Ascomycota_unclassified(100); | - | - | - | - | - | - | - | Unassigned | - |
| Otu29577 | 1 | k__Fungi(100);k__Fungi_unclassified(100);k__Fungi_unclassified(100);k__Fungi_unclassified(100);k__Fungi_unclassified(100);k__Fungi_unclassified(100);k__Fungi_unclassified(100); | - | - | - | - | - | - | - | Unassigned | - |
| Otu29578 | 1 | k__Fungi(100);k__Fungi_unclassified(100);k__Fungi_unclassified(100);k__Fungi_unclassified(100);k__Fungi_unclassified(100);k__Fungi_unclassified(100);k__Fungi_unclassified(100); | - | - | - | - | - | - | - | Unassigned | - |
| Otu29581 | 1 | k__Fungi(100);p__Chytridiomycota(100);p__Chytridiomycota_unclassified(100);p__Chytridiomycota_unclassified(100);p__Chytridiomycota_unclassified(100);p__Chytridiomycota_unclassified(100);p__Chytridiomycota_unclassified(100); | - | - | - | - | - | - | - | Unassigned | - |
| Otu29585 | 1 | k__Fungi(100);p__Ascomycota(100);c__Sordariomycetes(100);c__Sordariomycetes_unclassified(100);c__Sordariomycetes_unclassified(100);c__Sordariomycetes_unclassified(100);c__Sordariomycetes_unclassified(100); | - | - | - | - | - | - | - | Unassigned | - |
| Otu29589 | 1 | k__Fungi(100);k__Fungi_unclassified(100);k__Fungi_unclassified(100);k__Fungi_unclassified(100);k__Fungi_unclassified(100);k__Fungi_unclassified(100);k__Fungi_unclassified(100); | - | - | - | - | - | - | - | Unassigned | - |
| Otu29591 | 1 | k__Fungi(100);k__Fungi_unclassified(100);k__Fungi_unclassified(100);k__Fungi_unclassified(100);k__Fungi_unclassified(100);k__Fungi_unclassified(100);k__Fungi_unclassified(100); | - | - | - | - | - | - | - | Unassigned | - |
| Otu29592 | 1 | k__Fungi(100);p__Ascomycota(100);p__Ascomycota_unclassified(100);p__Ascomycota_unclassified(100);p__Ascomycota_unclassified(100);p__Ascomycota_unclassified(100);p__Ascomycota_unclassified(100); | - | - | - | - | - | - | - | Unassigned | - |
| Otu29593 | 1 | k__Fungi(100);p__Ascomycota(100);p__Ascomycota_unclassified(100);p__Ascomycota_unclassified(100);p__Ascomycota_unclassified(100);p__Ascomycota_unclassified(100);p__Ascomycota_unclassified(100); | - | - | - | - | - | - | - | Unassigned | - |
| Otu29594 | 1 | k__Fungi(100);p__Ascomycota(100);p__Ascomycota_unclassified(100);p__Ascomycota_unclassified(100);p__Ascomycota_unclassified(100);p__Ascomycota_unclassified(100);p__Ascomycota_unclassified(100); | - | - | - | - | - | - | - | Unassigned | - |
| Otu29595 | 1 | k__Fungi(100);k__Fungi_unclassified(100);k__Fungi_unclassified(100);k__Fungi_unclassified(100);k__Fungi_unclassified(100);k__Fungi_unclassified(100);k__Fungi_unclassified(100); | - | - | - | - | - | - | - | Unassigned | - |
| Otu29596 | 1 | k__Fungi(100);k__Fungi_unclassified(100);k__Fungi_unclassified(100);k__Fungi_unclassified(100);k__Fungi_unclassified(100);k__Fungi_unclassified(100);k__Fungi_unclassified(100); | - | - | - | - | - | - | - | Unassigned | - |
| Otu29598 | 1 | k__Fungi(100);p__Ascomycota(100);p__Ascomycota_unclassified(100);p__Ascomycota_unclassified(100);p__Ascomycota_unclassified(100);p__Ascomycota_unclassified(100);p__Ascomycota_unclassified(100); | - | - | - | - | - | - | - | Unassigned | - |
| Otu29599 | 1 | k__Fungi(100);p__Ascomycota(100);p__Ascomycota_unclassified(100);p__Ascomycota_unclassified(100);p__Ascomycota_unclassified(100);p__Ascomycota_unclassified(100);p__Ascomycota_unclassified(100); | - | - | - | - | - | - | - | Unassigned | - |
| Otu29602 | 1 | k__Fungi(100);p__Ascomycota(100);p__Ascomycota_unclassified(100);p__Ascomycota_unclassified(100);p__Ascomycota_unclassified(100);p__Ascomycota_unclassified(100);p__Ascomycota_unclassified(100); | - | - | - | - | - | - | - | Unassigned | - |
| Otu29603 | 1 | k__Fungi(100);p__Basidiomycota(100);c__Agaricomycetes(100);o__Russulales(100);o__Russulales_unclassified(100);o__Russulales_unclassified(100);o__Russulales_unclassified(100); | - | - | - | - | - | - | - | Unassigned | - |
| Otu29617 | 1 | k__Fungi(100);p__Ascomycota(100);c__Saccharomycetes(100);o__Saccharomycetales(100);f__Saccharomycetaceae(100);g__unclassified_Saccharomycetaceae(100);s__Saccharomycetaceae_sp(100); | - | - | - | - | - | - | - | Unassigned | - |
| Otu29626 | 1 | k__Fungi(100);p__Basidiomycota(100);c__Agaricomycetes(100);o__unclassified_Agaricomycetes(100);f__unclassified_Agaricomycetes(100);g__unclassified_Agaricomycetes(100);s__Agaricomycetes_sp(100); | - | - | - | - | - | - | - | Unassigned | - |
| Otu29627 | 1 | k__Fungi(100);p__Ascomycota(100);c__Sordariomycetes(100);c__Sordariomycetes_unclassified(100);c__Sordariomycetes_unclassified(100);c__Sordariomycetes_unclassified(100);c__Sordariomycetes_unclassified(100); | - | - | - | - | - | - | - | Unassigned | - |
| Otu29629 | 1 | k__Fungi(100);p__Basidiomycota(100);c__Agaricomycetes(100);c__Agaricomycetes_unclassified(100);c__Agaricomycetes_unclassified(100);c__Agaricomycetes_unclassified(100);c__Agaricomycetes_unclassified(100); | - | - | - | - | - | - | - | Unassigned | - |
| Otu29633 | 1 | k__Fungi(100);p__Basidiomycota(100);p__Basidiomycota_unclassified(100);p__Basidiomycota_unclassified(100);p__Basidiomycota_unclassified(100);p__Basidiomycota_unclassified(100);p__Basidiomycota_unclassified(100); | - | - | - | - | - | - | - | Unassigned | - |
| Otu29634 | 1 | k__Fungi(100);p__Basidiomycota(100);p__Basidiomycota_unclassified(100);p__Basidiomycota_unclassified(100);p__Basidiomycota_unclassified(100);p__Basidiomycota_unclassified(100);p__Basidiomycota_unclassified(100); | - | - | - | - | - | - | - | Unassigned | - |
| Otu29637 | 1 | k__Fungi(100);k__Fungi_unclassified(100);k__Fungi_unclassified(100);k__Fungi_unclassified(100);k__Fungi_unclassified(100);k__Fungi_unclassified(100);k__Fungi_unclassified(100); | - | - | - | - | - | - | - | Unassigned | - |
| Otu29638 | 1 | k__Fungi(100);p__Basidiomycota(100);p__Basidiomycota_unclassified(100);p__Basidiomycota_unclassified(100);p__Basidiomycota_unclassified(100);p__Basidiomycota_unclassified(100);p__Basidiomycota_unclassified(100); | - | - | - | - | - | - | - | Unassigned | - |
| Otu29646 | 1 | k__Fungi(100);p__Basidiomycota(100);c__Agaricomycetes(100);c__Agaricomycetes_unclassified(100);c__Agaricomycetes_unclassified(100);c__Agaricomycetes_unclassified(100);c__Agaricomycetes_unclassified(100); | - | - | - | - | - | - | - | Unassigned | - |
| Otu29647 | 1 | k__Fungi(100);p__Basidiomycota(100);c__Agaricomycetes(100);c__Agaricomycetes_unclassified(100);c__Agaricomycetes_unclassified(100);c__Agaricomycetes_unclassified(100);c__Agaricomycetes_unclassified(100); | - | - | - | - | - | - | - | Unassigned | - |
| Otu29648 | 1 | k__Fungi(100);p__Basidiomycota(100);c__Agaricomycetes(100);c__Agaricomycetes_unclassified(100);c__Agaricomycetes_unclassified(100);c__Agaricomycetes_unclassified(100);c__Agaricomycetes_unclassified(100); | - | - | - | - | - | - | - | Unassigned | - |
| Otu29651 | 1 | k__Fungi(100);p__Basidiomycota(100);c__Agaricomycetes(100);o__Russulales(100);o__Russulales_unclassified(100);o__Russulales_unclassified(100);o__Russulales_unclassified(100); | - | - | - | - | - | - | - | Unassigned | - |
| Otu29654 | 1 | k__Fungi(100);p__Basidiomycota(100);c__Agaricomycetes(100);c__Agaricomycetes_unclassified(100);c__Agaricomycetes_unclassified(100);c__Agaricomycetes_unclassified(100);c__Agaricomycetes_unclassified(100); | - | - | - | - | - | - | - | Unassigned | - |
| Otu29655 | 1 | k__Fungi(100);p__Basidiomycota(100);c__Agaricomycetes(100);c__Agaricomycetes_unclassified(100);c__Agaricomycetes_unclassified(100);c__Agaricomycetes_unclassified(100);c__Agaricomycetes_unclassified(100); | - | - | - | - | - | - | - | Unassigned | - |
| Otu29659 | 1 | k__Fungi(100);p__Basidiomycota(100);c__Agaricomycetes(100);c__Agaricomycetes_unclassified(100);c__Agaricomycetes_unclassified(100);c__Agaricomycetes_unclassified(100);c__Agaricomycetes_unclassified(100); | - | - | - | - | - | - | - | Unassigned | - |
| Otu29660 | 1 | k__Fungi(100);k__Fungi_unclassified(100);k__Fungi_unclassified(100);k__Fungi_unclassified(100);k__Fungi_unclassified(100);k__Fungi_unclassified(100);k__Fungi_unclassified(100); | - | - | - | - | - | - | - | Unassigned | - |
| Otu29661 | 1 | k__Fungi(100);p__Basidiomycota(100);c__Agaricomycetes(100);c__Agaricomycetes_unclassified(100);c__Agaricomycetes_unclassified(100);c__Agaricomycetes_unclassified(100);c__Agaricomycetes_unclassified(100); | - | - | - | - | - | - | - | Unassigned | - |
| Otu29662 | 1 | k__Fungi(100);p__Basidiomycota(100);c__Agaricomycetes(100);o__Sebacinales(100);o__Sebacinales_unclassified(100);o__Sebacinales_unclassified(100);o__Sebacinales_unclassified(100); | - | - | - | - | - | - | - | Unassigned | - |
| Otu29663 | 1 | k__Fungi(100);p__Basidiomycota(100);c__Agaricomycetes(100);c__Agaricomycetes_unclassified(100);c__Agaricomycetes_unclassified(100);c__Agaricomycetes_unclassified(100);c__Agaricomycetes_unclassified(100); | - | - | - | - | - | - | - | Unassigned | - |
| Otu29664 | 1 | k__Fungi(100);p__Basidiomycota(100);p__Basidiomycota_unclassified(100);p__Basidiomycota_unclassified(100);p__Basidiomycota_unclassified(100);p__Basidiomycota_unclassified(100);p__Basidiomycota_unclassified(100); | - | - | - | - | - | - | - | Unassigned | - |
| Otu29666 | 1 | k__Fungi(100);p__Basidiomycota(100);c__Agaricomycetes(100);c__Agaricomycetes_unclassified(100);c__Agaricomycetes_unclassified(100);c__Agaricomycetes_unclassified(100);c__Agaricomycetes_unclassified(100); | - | - | - | - | - | - | - | Unassigned | - |
| Otu29667 | 1 | k__Fungi(100);p__Basidiomycota(100);c__Agaricomycetes(100);o__Sebacinales(100);o__Sebacinales_unclassified(100);o__Sebacinales_unclassified(100);o__Sebacinales_unclassified(100); | - | - | - | - | - | - | - | Unassigned | - |
| Otu29668 | 1 | k__Fungi(100);p__Basidiomycota(100);c__Pucciniomycetes(100);c__Pucciniomycetes_unclassified(100);c__Pucciniomycetes_unclassified(100);c__Pucciniomycetes_unclassified(100);c__Pucciniomycetes_unclassified(100); | - | - | - | - | - | - | - | Unassigned | - |
| Otu29669 | 1 | k__Fungi(100);p__Basidiomycota(100);c__Pucciniomycetes(100);c__Pucciniomycetes_unclassified(100);c__Pucciniomycetes_unclassified(100);c__Pucciniomycetes_unclassified(100);c__Pucciniomycetes_unclassified(100); | - | - | - | - | - | - | - | Unassigned | - |
| Otu29671 | 1 | k__Fungi(100);p__Basidiomycota(100);c__Agaricomycetes(100);o__Polyporales(100);o__Polyporales_unclassified(100);o__Polyporales_unclassified(100);o__Polyporales_unclassified(100); | - | - | - | - | - | - | - | Unassigned | - |
| Otu29676 | 1 | k__Fungi(100);p__Basidiomycota(100);c__Microbotryomycetes(100);o__Sporidiobolales(100);f__Sporidiobolales_family_Incertae_sedis(100);f__Sporidiobolales_family_Incertae_sedis_unclassified(100);f__Sporidiobolales_family_Incertae_sedis_unclassified(100); | - | - | - | - | - | - | - | Unassigned | - |
| Otu29684 | 1 | k__Fungi(100);p__Basidiomycota(100);p__Basidiomycota_unclassified(100);p__Basidiomycota_unclassified(100);p__Basidiomycota_unclassified(100);p__Basidiomycota_unclassified(100);p__Basidiomycota_unclassified(100); | - | - | - | - | - | - | - | Unassigned | - |
| Otu29691 | 1 | k__Fungi(100);p__Basidiomycota(100);c__Microbotryomycetes(100);o__Sporidiobolales(100);f__Sporidiobolales_family_Incertae_sedis(100);f__Sporidiobolales_family_Incertae_sedis_unclassified(100);f__Sporidiobolales_family_Incertae_sedis_unclassified(100); | - | - | - | - | - | - | - | Unassigned | - |
| Otu29692 | 1 | k__Fungi(100);p__Basidiomycota(100);c__Microbotryomycetes(100);o__Sporidiobolales(100);f__Sporidiobolales_family_Incertae_sedis(100);f__Sporidiobolales_family_Incertae_sedis_unclassified(100);f__Sporidiobolales_family_Incertae_sedis_unclassified(100); | - | - | - | - | - | - | - | Unassigned | - |
| Otu29693 | 1 | k__Fungi(100);p__Basidiomycota(100);p__Basidiomycota_unclassified(100);p__Basidiomycota_unclassified(100);p__Basidiomycota_unclassified(100);p__Basidiomycota_unclassified(100);p__Basidiomycota_unclassified(100); | - | - | - | - | - | - | - | Unassigned | - |
| Otu29702 | 1 | k__Fungi(100);p__Basidiomycota(100);c__Pucciniomycetes(100);o__Pucciniales(100);o__Pucciniales_unclassified(100);o__Pucciniales_unclassified(100);o__Pucciniales_unclassified(100); | - | - | - | - | - | - | - | Unassigned | - |
| Otu29709 | 1 | k__Fungi(100);p__Ascomycota(100);c__Sordariomycetes(100);o__Hypocreales(100);f__Cordycipitaceae(100);f__Cordycipitaceae_unclassified(100);f__Cordycipitaceae_unclassified(100); | - | - | - | - | - | - | - | Unassigned | - |
| Otu29710 | 1 | k__Fungi(100);p__Ascomycota(100);p__Ascomycota_unclassified(100);p__Ascomycota_unclassified(100);p__Ascomycota_unclassified(100);p__Ascomycota_unclassified(100);p__Ascomycota_unclassified(100); | - | - | - | - | - | - | - | Unassigned | - |
| Otu29712 | 1 | k__Fungi(100);p__Ascomycota(100);c__Taphrinomycetes(100);o__Taphrinales(100);f__Taphrinaceae(100);g__Lalaria(100);s__Lalaria_inositophila(100); | - | - | - | - | - | - | - | Unassigned | - |
| Otu29713 | 1 | k__Fungi(100);p__Ascomycota(100);c__Sordariomycetes(100);c__Sordariomycetes_unclassified(100);c__Sordariomycetes_unclassified(100);c__Sordariomycetes_unclassified(100);c__Sordariomycetes_unclassified(100); | - | - | - | - | - | - | - | Unassigned | - |
| Otu29714 | 1 | k__Fungi(100);p__Basidiomycota(100);p__Basidiomycota_unclassified(100);p__Basidiomycota_unclassified(100);p__Basidiomycota_unclassified(100);p__Basidiomycota_unclassified(100);p__Basidiomycota_unclassified(100); | - | - | - | - | - | - | - | Unassigned | - |
| Otu29715 | 1 | k__Fungi(100);p__Ascomycota(100);c__Pezizomycetes(100);o__unclassified_Pezizomycetes(100);f__unclassified_Pezizomycetes(100);g__unclassified_Pezizomycetes(100);s__Pezizomycetes_sp(100); | - | - | - | - | - | - | - | Unassigned | - |
| Otu29719 | 1 | k__Fungi(100);p__Ascomycota(100);c__Sordariomycetes(100);o__Diaporthales(100);o__Diaporthales_unclassified(100);o__Diaporthales_unclassified(100);o__Diaporthales_unclassified(100); | - | - | - | - | - | - | - | Unassigned | - |
| Otu29726 | 1 | k__Fungi(100);p__Ascomycota(100);p__Ascomycota_unclassified(100);p__Ascomycota_unclassified(100);p__Ascomycota_unclassified(100);p__Ascomycota_unclassified(100);p__Ascomycota_unclassified(100); | - | - | - | - | - | - | - | Unassigned | - |
| Otu29729 | 1 | k__Fungi(100);p__Ascomycota(100);c__Sordariomycetes(100);o__Diaporthales(100);f__Diaporthales_family_Incertae_sedis(100);f__Diaporthales_family_Incertae_sedis_unclassified(100);f__Diaporthales_family_Incertae_sedis_unclassified(100); | - | - | - | - | - | - | - | Unassigned | - |
| Otu29730 | 1 | k__Fungi(100);p__Ascomycota(100);c__Sordariomycetes(100);o__Diaporthales(100);o__Diaporthales_unclassified(100);o__Diaporthales_unclassified(100);o__Diaporthales_unclassified(100); | - | - | - | - | - | - | - | Unassigned | - |
| Otu29731 | 1 | k__Fungi(100);p__Ascomycota(100);c__Sordariomycetes(100);o__Diaporthales(100);o__Diaporthales_unclassified(100);o__Diaporthales_unclassified(100);o__Diaporthales_unclassified(100); | - | - | - | - | - | - | - | Unassigned | - |
| Otu29749 | 1 | k__Fungi(100);p__Ascomycota(100);p__Ascomycota_unclassified(100);p__Ascomycota_unclassified(100);p__Ascomycota_unclassified(100);p__Ascomycota_unclassified(100);p__Ascomycota_unclassified(100); | - | - | - | - | - | - | - | Unassigned | - |
| Otu29769 | 1 | k__Fungi(100);p__Ascomycota(100);c__Sordariomycetes(100);o__Hypocreales(100);f__Hypocreaceae(100);f__Hypocreaceae_unclassified(100);f__Hypocreaceae_unclassified(100); | - | - | - | - | - | - | - | Unassigned | - |
| Otu29771 | 1 | k__Fungi(100);p__Ascomycota(100);c__Sordariomycetes(100);o__Hypocreales(100);f__Hypocreaceae(100);f__Hypocreaceae_unclassified(100);f__Hypocreaceae_unclassified(100); | - | - | - | - | - | - | - | Unassigned | - |
| Otu29773 | 1 | k__Fungi(100);p__Ascomycota(100);c__Sordariomycetes(100);o__Hypocreales(100);f__Hypocreaceae(100);f__Hypocreaceae_unclassified(100);f__Hypocreaceae_unclassified(100); | - | - | - | - | - | - | - | Unassigned | - |
| Otu29775 | 1 | k__Fungi(100);p__Ascomycota(100);c__Sordariomycetes(100);c__Sordariomycetes_unclassified(100);c__Sordariomycetes_unclassified(100);c__Sordariomycetes_unclassified(100);c__Sordariomycetes_unclassified(100); | - | - | - | - | - | - | - | Unassigned | - |
| Otu29776 | 1 | k__Fungi(100);p__Ascomycota(100);p__Ascomycota_unclassified(100);p__Ascomycota_unclassified(100);p__Ascomycota_unclassified(100);p__Ascomycota_unclassified(100);p__Ascomycota_unclassified(100); | - | - | - | - | - | - | - | Unassigned | - |
| Otu29778 | 1 | k__Fungi(100);p__Ascomycota(100);c__Sordariomycetes(100);o__Hypocreales(100);f__Hypocreaceae(100);f__Hypocreaceae_unclassified(100);f__Hypocreaceae_unclassified(100); | - | - | - | - | - | - | - | Unassigned | - |
| Otu29783 | 1 | k__Fungi(100);p__Ascomycota(100);p__Ascomycota_unclassified(100);p__Ascomycota_unclassified(100);p__Ascomycota_unclassified(100);p__Ascomycota_unclassified(100);p__Ascomycota_unclassified(100); | - | - | - | - | - | - | - | Unassigned | - |
| Otu29784 | 1 | k__Fungi(100);p__Ascomycota(100);p__Ascomycota_unclassified(100);p__Ascomycota_unclassified(100);p__Ascomycota_unclassified(100);p__Ascomycota_unclassified(100);p__Ascomycota_unclassified(100); | - | - | - | - | - | - | - | Unassigned | - |
| Otu29786 | 1 | k__Fungi(100);p__Ascomycota(100);p__Ascomycota_unclassified(100);p__Ascomycota_unclassified(100);p__Ascomycota_unclassified(100);p__Ascomycota_unclassified(100);p__Ascomycota_unclassified(100); | - | - | - | - | - | - | - | Unassigned | - |
| Otu29787 | 1 | k__Fungi(100);p__Ascomycota(100);p__Ascomycota_unclassified(100);p__Ascomycota_unclassified(100);p__Ascomycota_unclassified(100);p__Ascomycota_unclassified(100);p__Ascomycota_unclassified(100); | - | - | - | - | - | - | - | Unassigned | - |
| Otu29788 | 1 | k__Fungi(100);p__Ascomycota(100);p__Ascomycota_unclassified(100);p__Ascomycota_unclassified(100);p__Ascomycota_unclassified(100);p__Ascomycota_unclassified(100);p__Ascomycota_unclassified(100); | - | - | - | - | - | - | - | Unassigned | - |
| Otu29789 | 1 | k__Fungi(100);p__Ascomycota(100);c__Sordariomycetes(100);c__Sordariomycetes_unclassified(100);c__Sordariomycetes_unclassified(100);c__Sordariomycetes_unclassified(100);c__Sordariomycetes_unclassified(100); | - | - | - | - | - | - | - | Unassigned | - |
| Otu29790 | 1 | k__Fungi(100);p__Ascomycota(100);p__Ascomycota_unclassified(100);p__Ascomycota_unclassified(100);p__Ascomycota_unclassified(100);p__Ascomycota_unclassified(100);p__Ascomycota_unclassified(100); | - | - | - | - | - | - | - | Unassigned | - |
| Otu29794 | 1 | k__Fungi(100);p__Ascomycota(100);c__Sordariomycetes(100);c__Sordariomycetes_unclassified(100);c__Sordariomycetes_unclassified(100);c__Sordariomycetes_unclassified(100);c__Sordariomycetes_unclassified(100); | - | - | - | - | - | - | - | Unassigned | - |
| Otu29795 | 1 | k__Fungi(100);p__Ascomycota(100);p__Ascomycota_unclassified(100);p__Ascomycota_unclassified(100);p__Ascomycota_unclassified(100);p__Ascomycota_unclassified(100);p__Ascomycota_unclassified(100); | - | - | - | - | - | - | - | Unassigned | - |
| Otu29797 | 1 | k__Fungi(100);p__Ascomycota(100);c__Sordariomycetes(100);c__Sordariomycetes_unclassified(100);c__Sordariomycetes_unclassified(100);c__Sordariomycetes_unclassified(100);c__Sordariomycetes_unclassified(100); | - | - | - | - | - | - | - | Unassigned | - |
| Otu29799 | 1 | k__Fungi(100);p__Ascomycota(100);c__Sordariomycetes(100);o__Hypocreales(100);f__Cordycipitaceae(100);f__Cordycipitaceae_unclassified(100);f__Cordycipitaceae_unclassified(100); | - | - | - | - | - | - | - | Unassigned | - |
| Otu29801 | 1 | k__Fungi(100);p__Ascomycota(100);c__Sordariomycetes(100);c__Sordariomycetes_unclassified(100);c__Sordariomycetes_unclassified(100);c__Sordariomycetes_unclassified(100);c__Sordariomycetes_unclassified(100); | - | - | - | - | - | - | - | Unassigned | - |
| Otu29803 | 1 | k__Fungi(100);p__Ascomycota(100);c__Sordariomycetes(100);c__Sordariomycetes_unclassified(100);c__Sordariomycetes_unclassified(100);c__Sordariomycetes_unclassified(100);c__Sordariomycetes_unclassified(100); | - | - | - | - | - | - | - | Unassigned | - |
| Otu29815 | 1 | k__Fungi(100);p__Ascomycota(100);c__Taphrinomycetes(100);o__Taphrinales(100);o__Taphrinales_unclassified(100);o__Taphrinales_unclassified(100);o__Taphrinales_unclassified(100); | - | - | - | - | - | - | - | Unassigned | - |
| Otu29816 | 1 | k__Fungi(100);p__Ascomycota(100);p__Ascomycota_unclassified(100);p__Ascomycota_unclassified(100);p__Ascomycota_unclassified(100);p__Ascomycota_unclassified(100);p__Ascomycota_unclassified(100); | - | - | - | - | - | - | - | Unassigned | - |
| Otu29817 | 1 | k__Fungi(100);p__Ascomycota(100);p__Ascomycota_unclassified(100);p__Ascomycota_unclassified(100);p__Ascomycota_unclassified(100);p__Ascomycota_unclassified(100);p__Ascomycota_unclassified(100); | - | - | - | - | - | - | - | Unassigned | - |
| Otu29819 | 1 | k__Fungi(100);p__Ascomycota(100);p__Ascomycota_unclassified(100);p__Ascomycota_unclassified(100);p__Ascomycota_unclassified(100);p__Ascomycota_unclassified(100);p__Ascomycota_unclassified(100); | - | - | - | - | - | - | - | Unassigned | - |
| Otu29820 | 1 | k__Fungi(100);p__Ascomycota(100);p__Ascomycota_unclassified(100);p__Ascomycota_unclassified(100);p__Ascomycota_unclassified(100);p__Ascomycota_unclassified(100);p__Ascomycota_unclassified(100); | - | - | - | - | - | - | - | Unassigned | - |
| Otu29821 | 1 | k__Fungi(100);p__Ascomycota(100);p__Ascomycota_unclassified(100);p__Ascomycota_unclassified(100);p__Ascomycota_unclassified(100);p__Ascomycota_unclassified(100);p__Ascomycota_unclassified(100); | - | - | - | - | - | - | - | Unassigned | - |
| Otu29824 | 1 | k__Fungi(100);p__Ascomycota(100);p__Ascomycota_unclassified(100);p__Ascomycota_unclassified(100);p__Ascomycota_unclassified(100);p__Ascomycota_unclassified(100);p__Ascomycota_unclassified(100); | - | - | - | - | - | - | - | Unassigned | - |
| Otu29830 | 1 | k__Fungi(100);p__Ascomycota(100);c__Taphrinomycetes(100);o__Taphrinales(100);o__Taphrinales_unclassified(100);o__Taphrinales_unclassified(100);o__Taphrinales_unclassified(100); | - | - | - | - | - | - | - | Unassigned | - |
| Otu29832 | 1 | k__Fungi(100);p__Ascomycota(100);c__Taphrinomycetes(100);o__Taphrinales(100);f__Taphrinaceae(100);g__Lalaria(100);s__Lalaria_inositophila(100); | - | - | - | - | - | - | - | Unassigned | - |
| Otu29834 | 1 | k__Fungi(100);p__Ascomycota(100);c__Taphrinomycetes(100);o__Taphrinales(100);f__Taphrinaceae(100);g__Lalaria(100);s__Lalaria_inositophila(100); | - | - | - | - | - | - | - | Unassigned | - |
| Otu29835 | 1 | k__Fungi(100);p__Basidiomycota(100);p__Basidiomycota_unclassified(100);p__Basidiomycota_unclassified(100);p__Basidiomycota_unclassified(100);p__Basidiomycota_unclassified(100);p__Basidiomycota_unclassified(100); | - | - | - | - | - | - | - | Unassigned | - |
| Otu29837 | 1 | k__Fungi(100);p__Ascomycota(100);c__Taphrinomycetes(100);o__Taphrinales(100);f__Taphrinaceae(100);g__Lalaria(100);s__Lalaria_inositophila(100); | - | - | - | - | - | - | - | Unassigned | - |
| Otu29840 | 1 | k__Fungi(100);p__Ascomycota(100);c__Taphrinomycetes(100);o__Taphrinales(100);f__Taphrinaceae(100);g__Lalaria(100);s__Lalaria_inositophila(100); | - | - | - | - | - | - | - | Unassigned | - |
| Otu29842 | 1 | k__Fungi(100);p__Ascomycota(100);p__Ascomycota_unclassified(100);p__Ascomycota_unclassified(100);p__Ascomycota_unclassified(100);p__Ascomycota_unclassified(100);p__Ascomycota_unclassified(100); | - | - | - | - | - | - | - | Unassigned | - |
| Otu29844 | 1 | k__Fungi(100);p__Ascomycota(100);c__Eurotiomycetes(100);c__Eurotiomycetes_unclassified(100);c__Eurotiomycetes_unclassified(100);c__Eurotiomycetes_unclassified(100);c__Eurotiomycetes_unclassified(100); | - | - | - | - | - | - | - | Unassigned | - |
| Otu29853 | 1 | k__Fungi(100);p__Ascomycota(100);c__Dothideomycetes(100);o__Pleosporales(100);o__Pleosporales_unclassified(100);o__Pleosporales_unclassified(100);o__Pleosporales_unclassified(100); | - | - | - | - | - | - | - | Unassigned | - |
| Otu29856 | 1 | k__Fungi(100);p__Ascomycota(100);c__Eurotiomycetes(100);c__Eurotiomycetes_unclassified(100);c__Eurotiomycetes_unclassified(100);c__Eurotiomycetes_unclassified(100);c__Eurotiomycetes_unclassified(100); | - | - | - | - | - | - | - | Unassigned | - |
| Otu29858 | 1 | k__Fungi(100);p__Ascomycota(100);p__Ascomycota_unclassified(100);p__Ascomycota_unclassified(100);p__Ascomycota_unclassified(100);p__Ascomycota_unclassified(100);p__Ascomycota_unclassified(100); | - | - | - | - | - | - | - | Unassigned | - |

| | | | | | | | | | | | |
|---|---|---|---|---|---|---|---|---|---|---|---|
| Otu29860 | 1 | k__Fungi(100);p__Ascomycota(100);p__Ascomycota_unclassified(100);p__Ascomycota_unclassified(100);p__Ascomycota_unclassified(100);p__Ascomycota_unclassified(100);p__Ascomycota_unclassified(100); | - | - | - | - | - | - | - | Unassigned | - |
| Otu29861 | 1 | k__Fungi(100);p__Ascomycota(100);p__Ascomycota_unclassified(100);p__Ascomycota_unclassified(100);p__Ascomycota_unclassified(100);p__Ascomycota_unclassified(100);p__Ascomycota_unclassified(100); | - | - | - | - | - | - | - | Unassigned | - |
| Otu29898 | 1 | k__Fungi(100);p__Ascomycota(100);p__Ascomycota_unclassified(100);p__Ascomycota_unclassified(100);p__Ascomycota_unclassified(100);p__Ascomycota_unclassified(100);p__Ascomycota_unclassified(100); | - | - | - | - | - | - | - | Unassigned | - |
| Otu29900 | 1 | k__Fungi(100);p__Ascomycota(100);p__Ascomycota_unclassified(100);p__Ascomycota_unclassified(100);p__Ascomycota_unclassified(100);p__Ascomycota_unclassified(100);p__Ascomycota_unclassified(100); | - | - | - | - | - | - | - | Unassigned | - |
| Otu29901 | 1 | k__Fungi(100);k__Fungi_unclassified(100);k__Fungi_unclassified(100);k__Fungi_unclassified(100);k__Fungi_unclassified(100);k__Fungi_unclassified(100);k__Fungi_unclassified(100); | - | - | - | - | - | - | - | Unassigned | - |
| Otu29903 | 1 | k__Fungi(100);p__Ascomycota(100);c__Eurotiomycetes(100);o__Eurotiales(100);f__Trichocomaceae(100);f__Trichocomaceae_unclassified(100);f__Trichocomaceae_unclassified(100); | - | - | - | - | - | - | - | Unassigned | - |
| Otu29905 | 1 | k__Fungi(100);p__Ascomycota(100);c__Dothideomycetes(100);o__Botryosphaeriales(100);f__unclassified_Botryosphaeriales(100);g__unclassified_Botryosphaeriales(100);s__Botryosphaeriales_sp(100); | - | - | - | - | - | - | - | Unassigned | - |
| Otu29908 | 1 | k__Fungi(100);k__Fungi_unclassified(100);k__Fungi_unclassified(100);k__Fungi_unclassified(100);k__Fungi_unclassified(100);k__Fungi_unclassified(100);k__Fungi_unclassified(100); | - | - | - | - | - | - | - | Unassigned | - |
| Otu29922 | 1 | k__Fungi(100);k__Fungi_unclassified(100);k__Fungi_unclassified(100);k__Fungi_unclassified(100);k__Fungi_unclassified(100);k__Fungi_unclassified(100);k__Fungi_unclassified(100); | - | - | - | - | - | - | - | Unassigned | - |
| Otu29926 | 1 | k__Fungi(100);k__Fungi_unclassified(100);k__Fungi_unclassified(100);k__Fungi_unclassified(100);k__Fungi_unclassified(100);k__Fungi_unclassified(100);k__Fungi_unclassified(100); | - | - | - | - | - | - | - | Unassigned | - |
| Otu29928 | 1 | k__Fungi(100);p__Ascomycota(100);p__Ascomycota_unclassified(100);p__Ascomycota_unclassified(100);p__Ascomycota_unclassified(100);p__Ascomycota_unclassified(100);p__Ascomycota_unclassified(100); | - | - | - | - | - | - | - | Unassigned | - |
| Otu29929 | 1 | k__Fungi(100);k__Fungi_unclassified(100);k__Fungi_unclassified(100);k__Fungi_unclassified(100);k__Fungi_unclassified(100);k__Fungi_unclassified(100);k__Fungi_unclassified(100); | - | - | - | - | - | - | - | Unassigned | - |
| Otu29933 | 1 | k__Fungi(100);k__Fungi_unclassified(100);k__Fungi_unclassified(100);k__Fungi_unclassified(100);k__Fungi_unclassified(100);k__Fungi_unclassified(100);k__Fungi_unclassified(100); | - | - | - | - | - | - | - | Unassigned | - |
| Otu29935 | 1 | k__Fungi(100);p__Ascomycota(100);p__Ascomycota_unclassified(100);p__Ascomycota_unclassified(100);p__Ascomycota_unclassified(100);p__Ascomycota_unclassified(100);p__Ascomycota_unclassified(100); | - | - | - | - | - | - | - | Unassigned | - |
| Otu29939 | 1 | k__Fungi(100);k__Fungi_unclassified(100);k__Fungi_unclassified(100);k__Fungi_unclassified(100);k__Fungi_unclassified(100);k__Fungi_unclassified(100);k__Fungi_unclassified(100); | - | - | - | - | - | - | - | Unassigned | - |
| Otu29944 | 1 | k__Fungi(100);k__Fungi_unclassified(100);k__Fungi_unclassified(100);k__Fungi_unclassified(100);k__Fungi_unclassified(100);k__Fungi_unclassified(100);k__Fungi_unclassified(100); | - | - | - | - | - | - | - | Unassigned | - |
| Otu29945 | 1 | k__Fungi(100);k__Fungi_unclassified(100);k__Fungi_unclassified(100);k__Fungi_unclassified(100);k__Fungi_unclassified(100);k__Fungi_unclassified(100);k__Fungi_unclassified(100); | - | - | - | - | - | - | - | Unassigned | - |
| Otu29946 | 1 | k__Fungi(100);p__Ascomycota(100);p__Ascomycota_unclassified(100);p__Ascomycota_unclassified(100);p__Ascomycota_unclassified(100);p__Ascomycota_unclassified(100);p__Ascomycota_unclassified(100); | - | - | - | - | - | - | - | Unassigned | - |
| Otu29947 | 1 | k__Fungi(100);k__Fungi_unclassified(100);k__Fungi_unclassified(100);k__Fungi_unclassified(100);k__Fungi_unclassified(100);k__Fungi_unclassified(100);k__Fungi_unclassified(100); | - | - | - | - | - | - | - | Unassigned | - |
| Otu29949 | 1 | k__Fungi(100);p__Ascomycota(100);p__Ascomycota_unclassified(100);p__Ascomycota_unclassified(100);p__Ascomycota_unclassified(100);p__Ascomycota_unclassified(100);p__Ascomycota_unclassified(100); | - | - | - | - | - | - | - | Unassigned | - |
| Otu29951 | 1 | k__Fungi(100);p__Ascomycota(100);p__Ascomycota_unclassified(100);p__Ascomycota_unclassified(100);p__Ascomycota_unclassified(100);p__Ascomycota_unclassified(100);p__Ascomycota_unclassified(100); | - | - | - | - | - | - | - | Unassigned | - |
| Otu29952 | 1 | k__Fungi(100);k__Fungi_unclassified(100);k__Fungi_unclassified(100);k__Fungi_unclassified(100);k__Fungi_unclassified(100);k__Fungi_unclassified(100);k__Fungi_unclassified(100); | - | - | - | - | - | - | - | Unassigned | - |
| Otu29953 | 1 | k__Fungi(100);p__Ascomycota(100);p__Ascomycota_unclassified(100);p__Ascomycota_unclassified(100);p__Ascomycota_unclassified(100);p__Ascomycota_unclassified(100);p__Ascomycota_unclassified(100); | - | - | - | - | - | - | - | Unassigned | - |
| Otu29954 | 1 | k__Fungi(100);k__Fungi_unclassified(100);k__Fungi_unclassified(100);k__Fungi_unclassified(100);k__Fungi_unclassified(100);k__Fungi_unclassified(100);k__Fungi_unclassified(100); | - | - | - | - | - | - | - | Unassigned | - |
| Otu29955 | 1 | k__Fungi(100);p__Ascomycota(100);p__Ascomycota_unclassified(100);p__Ascomycota_unclassified(100);p__Ascomycota_unclassified(100);p__Ascomycota_unclassified(100);p__Ascomycota_unclassified(100); | - | - | - | - | - | - | - | Unassigned | - |
| Otu29956 | 1 | k__Fungi(100);k__Fungi_unclassified(100);k__Fungi_unclassified(100);k__Fungi_unclassified(100);k__Fungi_unclassified(100);k__Fungi_unclassified(100);k__Fungi_unclassified(100); | - | - | - | - | - | - | - | Unassigned | - |
| Otu29958 | 1 | k__Fungi(100);k__Fungi_unclassified(100);k__Fungi_unclassified(100);k__Fungi_unclassified(100);k__Fungi_unclassified(100);k__Fungi_unclassified(100);k__Fungi_unclassified(100); | - | - | - | - | - | - | - | Unassigned | - |
| Otu29959 | 1 | k__Fungi(100);k__Fungi_unclassified(100);k__Fungi_unclassified(100);k__Fungi_unclassified(100);k__Fungi_unclassified(100);k__Fungi_unclassified(100);k__Fungi_unclassified(100); | - | - | - | - | - | - | - | Unassigned | - |
| Otu29961 | 1 | k__Fungi(100);p__Ascomycota(100);p__Ascomycota_unclassified(100);p__Ascomycota_unclassified(100);p__Ascomycota_unclassified(100);p__Ascomycota_unclassified(100);p__Ascomycota_unclassified(100); | - | - | - | - | - | - | - | Unassigned | - |
| Otu29962 | 1 | k__Fungi(100);k__Fungi_unclassified(100);k__Fungi_unclassified(100);k__Fungi_unclassified(100);k__Fungi_unclassified(100);k__Fungi_unclassified(100);k__Fungi_unclassified(100); | - | - | - | - | - | - | - | Unassigned | - |
| Otu29963 | 1 | k__Fungi(100);p__Ascomycota(100);p__Ascomycota_unclassified(100);p__Ascomycota_unclassified(100);p__Ascomycota_unclassified(100);p__Ascomycota_unclassified(100);p__Ascomycota_unclassified(100); | - | - | - | - | - | - | - | Unassigned | - |
| Otu29964 | 1 | k__Fungi(100);k__Fungi_unclassified(100);k__Fungi_unclassified(100);k__Fungi_unclassified(100);k__Fungi_unclassified(100);k__Fungi_unclassified(100);k__Fungi_unclassified(100); | - | - | - | - | - | - | - | Unassigned | - |
| Otu29965 | 1 | k__Fungi(100);k__Fungi_unclassified(100);k__Fungi_unclassified(100);k__Fungi_unclassified(100);k__Fungi_unclassified(100);k__Fungi_unclassified(100);k__Fungi_unclassified(100); | - | - | - | - | - | - | - | Unassigned | - |
| Otu29966 | 1 | k__Fungi(100);k__Fungi_unclassified(100);k__Fungi_unclassified(100);k__Fungi_unclassified(100);k__Fungi_unclassified(100);k__Fungi_unclassified(100);k__Fungi_unclassified(100); | - | - | - | - | - | - | - | Unassigned | - |
| Otu29967 | 1 | k__Fungi(100);p__Ascomycota(100);p__Ascomycota_unclassified(100);p__Ascomycota_unclassified(100);p__Ascomycota_unclassified(100);p__Ascomycota_unclassified(100);p__Ascomycota_unclassified(100); | - | - | - | - | - | - | - | Unassigned | - |
| Otu29968 | 1 | k__Fungi(100);p__Ascomycota(100);p__Ascomycota_unclassified(100);p__Ascomycota_unclassified(100);p__Ascomycota_unclassified(100);p__Ascomycota_unclassified(100);p__Ascomycota_unclassified(100); | - | - | - | - | - | - | - | Unassigned | - |
| Otu29970 | 1 | k__Fungi(100);k__Fungi_unclassified(100);k__Fungi_unclassified(100);k__Fungi_unclassified(100);k__Fungi_unclassified(100);k__Fungi_unclassified(100);k__Fungi_unclassified(100); | - | - | - | - | - | - | - | Unassigned | - |
| Otu29971 | 1 | k__Fungi(100);k__Fungi_unclassified(100);k__Fungi_unclassified(100);k__Fungi_unclassified(100);k__Fungi_unclassified(100);k__Fungi_unclassified(100);k__Fungi_unclassified(100); | - | - | - | - | - | - | - | Unassigned | - |
| Otu29972 | 1 | k__Fungi(100);p__Ascomycota(100);p__Ascomycota_unclassified(100);p__Ascomycota_unclassified(100);p__Ascomycota_unclassified(100);p__Ascomycota_unclassified(100);p__Ascomycota_unclassified(100); | - | - | - | - | - | - | - | Unassigned | - |
| Otu29976 | 1 | k__Fungi(100);p__Ascomycota(100);p__Ascomycota_unclassified(100);p__Ascomycota_unclassified(100);p__Ascomycota_unclassified(100);p__Ascomycota_unclassified(100);p__Ascomycota_unclassified(100); | - | - | - | - | - | - | - | Unassigned | - |
| Otu29977 | 1 | k__Fungi(100);k__Fungi_unclassified(100);k__Fungi_unclassified(100);k__Fungi_unclassified(100);k__Fungi_unclassified(100);k__Fungi_unclassified(100);k__Fungi_unclassified(100); | - | - | - | - | - | - | - | Unassigned | - |
| Otu29980 | 1 | k__Fungi(100);k__Fungi_unclassified(100);k__Fungi_unclassified(100);k__Fungi_unclassified(100);k__Fungi_unclassified(100);k__Fungi_unclassified(100);k__Fungi_unclassified(100); | - | - | - | - | - | - | - | Unassigned | - |
| Otu29982 | 1 | k__Fungi(100);k__Fungi_unclassified(100);k__Fungi_unclassified(100);k__Fungi_unclassified(100);k__Fungi_unclassified(100);k__Fungi_unclassified(100);k__Fungi_unclassified(100); | - | - | - | - | - | - | - | Unassigned | - |
| Otu29983 | 1 | k__Fungi(100);p__Ascomycota(100);p__Ascomycota_unclassified(100);p__Ascomycota_unclassified(100);p__Ascomycota_unclassified(100);p__Ascomycota_unclassified(100);p__Ascomycota_unclassified(100); | - | - | - | - | - | - | - | Unassigned | - |
| Otu29985 | 1 | k__Fungi(100);k__Fungi_unclassified(100);k__Fungi_unclassified(100);k__Fungi_unclassified(100);k__Fungi_unclassified(100);k__Fungi_unclassified(100);k__Fungi_unclassified(100); | - | - | - | - | - | - | - | Unassigned | - |
| Otu29986 | 1 | k__Fungi(100);k__Fungi_unclassified(100);k__Fungi_unclassified(100);k__Fungi_unclassified(100);k__Fungi_unclassified(100);k__Fungi_unclassified(100);k__Fungi_unclassified(100); | - | - | - | - | - | - | - | Unassigned | - |
| Otu29987 | 1 | k__Fungi(100);k__Fungi_unclassified(100);k__Fungi_unclassified(100);k__Fungi_unclassified(100);k__Fungi_unclassified(100);k__Fungi_unclassified(100);k__Fungi_unclassified(100); | - | - | - | - | - | - | - | Unassigned | - |
| Otu29989 | 1 | k__Fungi(100);k__Fungi_unclassified(100);k__Fungi_unclassified(100);k__Fungi_unclassified(100);k__Fungi_unclassified(100);k__Fungi_unclassified(100);k__Fungi_unclassified(100); | - | - | - | - | - | - | - | Unassigned | - |
| Otu29991 | 1 | k__Fungi(100);k__Fungi_unclassified(100);k__Fungi_unclassified(100);k__Fungi_unclassified(100);k__Fungi_unclassified(100);k__Fungi_unclassified(100);k__Fungi_unclassified(100); | - | - | - | - | - | - | - | Unassigned | - |
| Otu29993 | 1 | k__Fungi(100);k__Fungi_unclassified(100);k__Fungi_unclassified(100);k__Fungi_unclassified(100);k__Fungi_unclassified(100);k__Fungi_unclassified(100);k__Fungi_unclassified(100); | - | - | - | - | - | - | - | Unassigned | - |
| Otu29998 | 1 | k__Fungi(100);k__Fungi_unclassified(100);k__Fungi_unclassified(100);k__Fungi_unclassified(100);k__Fungi_unclassified(100);k__Fungi_unclassified(100);k__Fungi_unclassified(100); | - | - | - | - | - | - | - | Unassigned | - |
| Otu30002 | 1 | k__Fungi(100);k__Fungi_unclassified(100);k__Fungi_unclassified(100);k__Fungi_unclassified(100);k__Fungi_unclassified(100);k__Fungi_unclassified(100);k__Fungi_unclassified(100); | - | - | - | - | - | - | - | Unassigned | - |
| Otu30003 | 1 | k__Fungi(100);k__Fungi_unclassified(100);k__Fungi_unclassified(100);k__Fungi_unclassified(100);k__Fungi_unclassified(100);k__Fungi_unclassified(100);k__Fungi_unclassified(100); | - | - | - | - | - | - | - | Unassigned | - |
| Otu30004 | 1 | k__Fungi(100);k__Fungi_unclassified(100);k__Fungi_unclassified(100);k__Fungi_unclassified(100);k__Fungi_unclassified(100);k__Fungi_unclassified(100);k__Fungi_unclassified(100); | - | - | - | - | - | - | - | Unassigned | - |
| Otu30005 | 1 | k__Fungi(100);p__Ascomycota(100);p__Ascomycota_unclassified(100);p__Ascomycota_unclassified(100);p__Ascomycota_unclassified(100);p__Ascomycota_unclassified(100);p__Ascomycota_unclassified(100); | - | - | - | - | - | - | - | Unassigned | - |
| Otu30006 | 1 | k__Fungi(100);p__Ascomycota(100);p__Ascomycota_unclassified(100);p__Ascomycota_unclassified(100);p__Ascomycota_unclassified(100);p__Ascomycota_unclassified(100);p__Ascomycota_unclassified(100); | - | - | - | - | - | - | - | Unassigned | - |
| Otu30007 | 1 | k__Fungi(100);k__Fungi_unclassified(100);k__Fungi_unclassified(100);k__Fungi_unclassified(100);k__Fungi_unclassified(100);k__Fungi_unclassified(100);k__Fungi_unclassified(100); | - | - | - | - | - | - | - | Unassigned | - |
| Otu30008 | 1 | k__Fungi(100);p__Ascomycota(100);p__Ascomycota_unclassified(100);p__Ascomycota_unclassified(100);p__Ascomycota_unclassified(100);p__Ascomycota_unclassified(100);p__Ascomycota_unclassified(100); | - | - | - | - | - | - | - | Unassigned | - |
| Otu30009 | 1 | k__Fungi(100);k__Fungi_unclassified(100);k__Fungi_unclassified(100);k__Fungi_unclassified(100);k__Fungi_unclassified(100);k__Fungi_unclassified(100);k__Fungi_unclassified(100); | - | - | - | - | - | - | - | Unassigned | - |
| Otu30010 | 1 | k__Fungi(100);k__Fungi_unclassified(100);k__Fungi_unclassified(100);k__Fungi_unclassified(100);k__Fungi_unclassified(100);k__Fungi_unclassified(100);k__Fungi_unclassified(100); | - | - | - | - | - | - | - | Unassigned | - |
| Otu30012 | 1 | k__Fungi(100);k__Fungi_unclassified(100);k__Fungi_unclassified(100);k__Fungi_unclassified(100);k__Fungi_unclassified(100);k__Fungi_unclassified(100);k__Fungi_unclassified(100); | - | - | - | - | - | - | - | Unassigned | - |
| Otu30013 | 1 | k__Fungi(100);p__Ascomycota(100);p__Ascomycota_unclassified(100);p__Ascomycota_unclassified(100);p__Ascomycota_unclassified(100);p__Ascomycota_unclassified(100);p__Ascomycota_unclassified(100); | - | - | - | - | - | - | - | Unassigned | - |
| Otu30014 | 1 | k__Fungi(100);k__Fungi_unclassified(100);k__Fungi_unclassified(100);k__Fungi_unclassified(100);k__Fungi_unclassified(100);k__Fungi_unclassified(100);k__Fungi_unclassified(100); | - | - | - | - | - | - | - | Unassigned | - |
| Otu30018 | 1 | k__Fungi(100);k__Fungi_unclassified(100);k__Fungi_unclassified(100);k__Fungi_unclassified(100);k__Fungi_unclassified(100);k__Fungi_unclassified(100);k__Fungi_unclassified(100); | - | - | - | - | - | - | - | Unassigned | - |
| Otu30020 | 1 | k__Fungi(100);k__Fungi_unclassified(100);k__Fungi_unclassified(100);k__Fungi_unclassified(100);k__Fungi_unclassified(100);k__Fungi_unclassified(100);k__Fungi_unclassified(100); | - | - | - | - | - | - | - | Unassigned | - |
| Otu30021 | 1 | k__Fungi(100);p__Ascomycota(100);p__Ascomycota_unclassified(100);p__Ascomycota_unclassified(100);p__Ascomycota_unclassified(100);p__Ascomycota_unclassified(100);p__Ascomycota_unclassified(100); | - | - | - | - | - | - | - | Unassigned | - |
| Otu30022 | 1 | k__Fungi(100);k__Fungi_unclassified(100);k__Fungi_unclassified(100);k__Fungi_unclassified(100);k__Fungi_unclassified(100);k__Fungi_unclassified(100);k__Fungi_unclassified(100); | - | - | - | - | - | - | - | Unassigned | - |
| Otu30023 | 1 | k__Fungi(100);k__Fungi_unclassified(100);k__Fungi_unclassified(100);k__Fungi_unclassified(100);k__Fungi_unclassified(100);k__Fungi_unclassified(100);k__Fungi_unclassified(100); | - | - | - | - | - | - | - | Unassigned | - |
| Otu30024 | 1 | k__Fungi(100);p__Ascomycota(100);p__Ascomycota_unclassified(100);p__Ascomycota_unclassified(100);p__Ascomycota_unclassified(100);p__Ascomycota_unclassified(100);p__Ascomycota_unclassified(100); | - | - | - | - | - | - | - | Unassigned | - |
| Otu30025 | 1 | k__Fungi(100);k__Fungi_unclassified(100);k__Fungi_unclassified(100);k__Fungi_unclassified(100);k__Fungi_unclassified(100);k__Fungi_unclassified(100);k__Fungi_unclassified(100); | - | - | - | - | - | - | - | Unassigned | - |
| Otu30026 | 1 | k__Fungi(100);p__Ascomycota(100);p__Ascomycota_unclassified(100);p__Ascomycota_unclassified(100);p__Ascomycota_unclassified(100);p__Ascomycota_unclassified(100);p__Ascomycota_unclassified(100); | - | - | - | - | - | - | - | Unassigned | - |
| Otu30027 | 1 | k__Fungi(100);k__Fungi_unclassified(100);k__Fungi_unclassified(100);k__Fungi_unclassified(100);k__Fungi_unclassified(100);k__Fungi_unclassified(100);k__Fungi_unclassified(100); | - | - | - | - | - | - | - | Unassigned | - |
| Otu30031 | 1 | k__Fungi(100);k__Fungi_unclassified(100);k__Fungi_unclassified(100);k__Fungi_unclassified(100);k__Fungi_unclassified(100);k__Fungi_unclassified(100);k__Fungi_unclassified(100); | - | - | - | - | - | - | - | Unassigned | - |
| Otu30032 | 1 | k__Fungi(100);p__Ascomycota(100);p__Ascomycota_unclassified(100);p__Ascomycota_unclassified(100);p__Ascomycota_unclassified(100);p__Ascomycota_unclassified(100);p__Ascomycota_unclassified(100); | - | - | - | - | - | - | - | Unassigned | - |
| Otu30034 | 1 | k__Fungi(100);k__Fungi_unclassified(100);k__Fungi_unclassified(100);k__Fungi_unclassified(100);k__Fungi_unclassified(100);k__Fungi_unclassified(100);k__Fungi_unclassified(100); | - | - | - | - | - | - | - | Unassigned | - |
| Otu30036 | 1 | k__Fungi(100);k__Fungi_unclassified(100);k__Fungi_unclassified(100);k__Fungi_unclassified(100);k__Fungi_unclassified(100);k__Fungi_unclassified(100);k__Fungi_unclassified(100); | - | - | - | - | - | - | - | Unassigned | - |
| Otu30037 | 1 | k__Fungi(100);p__Ascomycota(100);p__Ascomycota_unclassified(100);p__Ascomycota_unclassified(100);p__Ascomycota_unclassified(100);p__Ascomycota_unclassified(100);p__Ascomycota_unclassified(100); | - | - | - | - | - | - | - | Unassigned | - |
| Otu30038 | 1 | k__Fungi(100);k__Fungi_unclassified(100);k__Fungi_unclassified(100);k__Fungi_unclassified(100);k__Fungi_unclassified(100);k__Fungi_unclassified(100);k__Fungi_unclassified(100); | - | - | - | - | - | - | - | Unassigned | - |
| Otu30039 | 1 | k__Fungi(100);k__Fungi_unclassified(100);k__Fungi_unclassified(100);k__Fungi_unclassified(100);k__Fungi_unclassified(100);k__Fungi_unclassified(100);k__Fungi_unclassified(100); | - | - | - | - | - | - | - | Unassigned | - |
| Otu30040 | 1 | k__Fungi(100);p__Ascomycota(100);p__Ascomycota_unclassified(100);p__Ascomycota_unclassified(100);p__Ascomycota_unclassified(100);p__Ascomycota_unclassified(100);p__Ascomycota_unclassified(100); | - | - | - | - | - | - | - | Unassigned | - |
| Otu30042 | 1 | k__Fungi(100);k__Fungi_unclassified(100);k__Fungi_unclassified(100);k__Fungi_unclassified(100);k__Fungi_unclassified(100);k__Fungi_unclassified(100);k__Fungi_unclassified(100); | - | - | - | - | - | - | - | Unassigned | - |
| Otu30044 | 1 | k__Fungi(100);k__Fungi_unclassified(100);k__Fungi_unclassified(100);k__Fungi_unclassified(100);k__Fungi_unclassified(100);k__Fungi_unclassified(100);k__Fungi_unclassified(100); | - | - | - | - | - | - | - | Unassigned | - |
| Otu30046 | 1 | k__Fungi(100);k__Fungi_unclassified(100);k__Fungi_unclassified(100);k__Fungi_unclassified(100);k__Fungi_unclassified(100);k__Fungi_unclassified(100);k__Fungi_unclassified(100); | - | - | - | - | - | - | - | Unassigned | - |
| Otu30047 | 1 | k__Fungi(100);k__Fungi_unclassified(100);k__Fungi_unclassified(100);k__Fungi_unclassified(100);k__Fungi_unclassified(100);k__Fungi_unclassified(100);k__Fungi_unclassified(100); | - | - | - | - | - | - | - | Unassigned | - |
| Otu30048 | 1 | k__Fungi(100);k__Fungi_unclassified(100);k__Fungi_unclassified(100);k__Fungi_unclassified(100);k__Fungi_unclassified(100);k__Fungi_unclassified(100);k__Fungi_unclassified(100); | - | - | - | - | - | - | - | Unassigned | - |
| Otu30049 | 1 | k__Fungi(100);k__Fungi_unclassified(100);k__Fungi_unclassified(100);k__Fungi_unclassified(100);k__Fungi_unclassified(100);k__Fungi_unclassified(100);k__Fungi_unclassified(100); | - | - | - | - | - | - | - | Unassigned | - |
| Otu30050 | 1 | k__Fungi(100);k__Fungi_unclassified(100);k__Fungi_unclassified(100);k__Fungi_unclassified(100);k__Fungi_unclassified(100);k__Fungi_unclassified(100);k__Fungi_unclassified(100); | - | - | - | - | - | - | - | Unassigned | - |
| Otu30051 | 1 | k__Fungi(100);p__Ascomycota(100);p__Ascomycota_unclassified(100);p__Ascomycota_unclassified(100);p__Ascomycota_unclassified(100);p__Ascomycota_unclassified(100);p__Ascomycota_unclassified(100); | - | - | - | - | - | - | - | Unassigned | - |
| Otu30055 | 1 | k__Fungi(100);k__Fungi_unclassified(100);k__Fungi_unclassified(100);k__Fungi_unclassified(100);k__Fungi_unclassified(100);k__Fungi_unclassified(100);k__Fungi_unclassified(100); | - | - | - | - | - | - | - | Unassigned | - |
| Otu30056 | 1 | k__Fungi(100);k__Fungi_unclassified(100);k__Fungi_unclassified(100);k__Fungi_unclassified(100);k__Fungi_unclassified(100);k__Fungi_unclassified(100);k__Fungi_unclassified(100); | - | - | - | - | - | - | - | Unassigned | - |
| Otu30058 | 1 | k__Fungi(100);k__Fungi_unclassified(100);k__Fungi_unclassified(100);k__Fungi_unclassified(100);k__Fungi_unclassified(100);k__Fungi_unclassified(100);k__Fungi_unclassified(100); | - | - | - | - | - | - | - | Unassigned | - |
| Otu30060 | 1 | k__Fungi(100);p__Ascomycota(100);p__Ascomycota_unclassified(100);p__Ascomycota_unclassified(100);p__Ascomycota_unclassified(100);p__Ascomycota_unclassified(100);p__Ascomycota_unclassified(100); | - | - | - | - | - | - | - | Unassigned | - |
| Otu30069 | 1 | k__Fungi(100);p__Ascomycota(100);p__Ascomycota_unclassified(100);p__Ascomycota_unclassified(100);p__Ascomycota_unclassified(100);p__Ascomycota_unclassified(100);p__Ascomycota_unclassified(100); | - | - | - | - | - | - | - | Unassigned | - |
| Otu30070 | 1 | k__Fungi(100);k__Fungi_unclassified(100);k__Fungi_unclassified(100);k__Fungi_unclassified(100);k__Fungi_unclassified(100);k__Fungi_unclassified(100);k__Fungi_unclassified(100); | - | - | - | - | - | - | - | Unassigned | - |
| Otu30071 | 1 | k__Fungi(100);k__Fungi_unclassified(100);k__Fungi_unclassified(100);k__Fungi_unclassified(100);k__Fungi_unclassified(100);k__Fungi_unclassified(100);k__Fungi_unclassified(100); | - | - | - | - | - | - | - | Unassigned | - |
| Otu30072 | 1 | k__Fungi(100);k__Fungi_unclassified(100);k__Fungi_unclassified(100);k__Fungi_unclassified(100);k__Fungi_unclassified(100);k__Fungi_unclassified(100);k__Fungi_unclassified(100); | - | - | - | - | - | - | - | Unassigned | - |
| Otu30074 | 1 | k__Fungi(100);k__Fungi_unclassified(100);k__Fungi_unclassified(100);k__Fungi_unclassified(100);k__Fungi_unclassified(100);k__Fungi_unclassified(100);k__Fungi_unclassified(100); | - | - | - | - | - | - | - | Unassigned | - |
| Otu30077 | 1 | k__Fungi(100);p__Ascomycota(100);p__Ascomycota_unclassified(100);p__Ascomycota_unclassified(100);p__Ascomycota_unclassified(100);p__Ascomycota_unclassified(100);p__Ascomycota_unclassified(100); | - | - | - | - | - | - | - | Unassigned | - |
| Otu30081 | 1 | k__Fungi(100);k__Fungi_unclassified(100);k__Fungi_unclassified(100);k__Fungi_unclassified(100);k__Fungi_unclassified(100);k__Fungi_unclassified(100);k__Fungi_unclassified(100); | - | - | - | - | - | - | - | Unassigned | - |
| Otu30083 | 1 | k__Fungi(100);p__Ascomycota(100);p__Ascomycota_unclassified(100);p__Ascomycota_unclassified(100);p__Ascomycota_unclassified(100);p__Ascomycota_unclassified(100);p__Ascomycota_unclassified(100); | - | - | - | - | - | - | - | Unassigned | - |
| Otu30085 | 1 | k__Fungi(100);k__Fungi_unclassified(100);k__Fungi_unclassified(100);k__Fungi_unclassified(100);k__Fungi_unclassified(100);k__Fungi_unclassified(100);k__Fungi_unclassified(100); | - | - | - | - | - | - | - | Unassigned | - |
| Otu30086 | 1 | k__Fungi(100);k__Fungi_unclassified(100);k__Fungi_unclassified(100);k__Fungi_unclassified(100);k__Fungi_unclassified(100);k__Fungi_unclassified(100);k__Fungi_unclassified(100); | - | - | - | - | - | - | - | Unassigned | - |
| Otu30089 | 1 | k__Fungi(100);k__Fungi_unclassified(100);k__Fungi_unclassified(100);k__Fungi_unclassified(100);k__Fungi_unclassified(100);k__Fungi_unclassified(100);k__Fungi_unclassified(100); | - | - | - | - | - | - | - | Unassigned | - |
| Otu30094 | 1 | k__Fungi(100);k__Fungi_unclassified(100);k__Fungi_unclassified(100);k__Fungi_unclassified(100);k__Fungi_unclassified(100);k__Fungi_unclassified(100);k__Fungi_unclassified(100); | - | - | - | - | - | - | - | Unassigned | - |
| Otu30098 | 1 | k__Fungi(100);p__Ascomycota(100);p__Ascomycota_unclassified(100);p__Ascomycota_unclassified(100);p__Ascomycota_unclassified(100);p__Ascomycota_unclassified(100);p__Ascomycota_unclassified(100); | - | - | - | - | - | - | - | Unassigned | - |
| Otu30100 | 1 | k__Fungi(100);p__Ascomycota(100);p__Ascomycota_unclassified(100);p__Ascomycota_unclassified(100);p__Ascomycota_unclassified(100);p__Ascomycota_unclassified(100);p__Ascomycota_unclassified(100); | - | - | - | - | - | - | - | Unassigned | - |
| Otu30106 | 1 | k__Fungi(100);p__Ascomycota(100);p__Ascomycota_unclassified(100);p__Ascomycota_unclassified(100);p__Ascomycota_unclassified(100);p__Ascomycota_unclassified(100);p__Ascomycota_unclassified(100); | - | - | - | - | - | - | - | Unassigned | - |
| Otu30107 | 1 | k__Fungi(100);k__Fungi_unclassified(100);k__Fungi_unclassified(100);k__Fungi_unclassified(100);k__Fungi_unclassified(100);k__Fungi_unclassified(100);k__Fungi_unclassified(100); | - | - | - | - | - | - | - | Unassigned | - |
| Otu30108 | 1 | k__Fungi(100);k__Fungi_unclassified(100);k__Fungi_unclassified(100);k__Fungi_unclassified(100);k__Fungi_unclassified(100);k__Fungi_unclassified(100);k__Fungi_unclassified(100); | - | - | - | - | - | - | - | Unassigned | - |
| Otu30112 | 1 | k__Fungi(100);p__Ascomycota(100);p__Ascomycota_unclassified(100);p__Ascomycota_unclassified(100);p__Ascomycota_unclassified(100);p__Ascomycota_unclassified(100);p__Ascomycota_unclassified(100); | - | - | - | - | - | - | - | Unassigned | - |
| Otu30115 | 1 | k__Fungi(100);k__Fungi_unclassified(100);k__Fungi_unclassified(100);k__Fungi_unclassified(100);k__Fungi_unclassified(100);k__Fungi_unclassified(100);k__Fungi_unclassified(100); | - | - | - | - | - | - | - | Unassigned | - |
| Otu30117 | 1 | k__Fungi(100);k__Fungi_unclassified(100);k__Fungi_unclassified(100);k__Fungi_unclassified(100);k__Fungi_unclassified(100);k__Fungi_unclassified(100);k__Fungi_unclassified(100); | - | - | - | - | - | - | - | Unassigned | - |
| Otu30121 | 1 | k__Fungi(100);p__Ascomycota(100);p__Ascomycota_unclassified(100);p__Ascomycota_unclassified(100);p__Ascomycota_unclassified(100);p__Ascomycota_unclassified(100);p__Ascomycota_unclassified(100); | - | - | - | - | - | - | - | Unassigned | - |
| Otu30124 | 1 | k__Fungi(100);p__Ascomycota(100);p__Ascomycota_unclassified(100);p__Ascomycota_unclassified(100);p__Ascomycota_unclassified(100);p__Ascomycota_unclassified(100);p__Ascomycota_unclassified(100); | - | - | - | - | - | - | - | Unassigned | - |
| Otu30126 | 1 | k__Fungi(100);k__Fungi_unclassified(100);k__Fungi_unclassified(100);k__Fungi_unclassified(100);k__Fungi_unclassified(100);k__Fungi_unclassified(100);k__Fungi_unclassified(100); | - | - | - | - | - | - | - | Unassigned | - |
| Otu30129 | 1 | k__Fungi(100);k__Fungi_unclassified(100);k__Fungi_unclassified(100);k__Fungi_unclassified(100);k__Fungi_unclassified(100);k__Fungi_unclassified(100);k__Fungi_unclassified(100); | - | - | - | - | - | - | - | Unassigned | - |
| Otu30130 | 1 | k__Fungi(100);k__Fungi_unclassified(100);k__Fungi_unclassified(100);k__Fungi_unclassified(100);k__Fungi_unclassified(100);k__Fungi_unclassified(100);k__Fungi_unclassified(100); | - | - | - | - | - | - | - | Unassigned | - |
| Otu30131 | 1 | k__Fungi(100);p__Ascomycota(100);p__Ascomycota_unclassified(100);p__Ascomycota_unclassified(100);p__Ascomycota_unclassified(100);p__Ascomycota_unclassified(100);p__Ascomycota_unclassified(100); | - | - | - | - | - | - | - | Unassigned | - |
| Otu30135 | 1 | k__Fungi(100);p__Ascomycota(100);p__Ascomycota_unclassified(100);p__Ascomycota_unclassified(100);p__Ascomycota_unclassified(100);p__Ascomycota_unclassified(100);p__Ascomycota_unclassified(100); | - | - | - | - | - | - | - | Unassigned | - |
| Otu30147 | 1 | k__Fungi(100);k__Fungi_unclassified(100);k__Fungi_unclassified(100);k__Fungi_unclassified(100);k__Fungi_unclassified(100);k__Fungi_unclassified(100);k__Fungi_unclassified(100); | - | - | - | - | - | - | - | Unassigned | - |
| Otu30153 | 1 | k__Fungi(100);k__Fungi_unclassified(100);k__Fungi_unclassified(100);k__Fungi_unclassified(100);k__Fungi_unclassified(100);k__Fungi_unclassified(100);k__Fungi_unclassified(100); | - | - | - | - | - | - | - | Unassigned | - |
| Otu30157 | 1 | k__Fungi(100);k__Fungi_unclassified(100);k__Fungi_unclassified(100);k__Fungi_unclassified(100);k__Fungi_unclassified(100);k__Fungi_unclassified(100);k__Fungi_unclassified(100); | - | - | - | - | - | - | - | Unassigned | - |
| Otu30160 | 1 | k__Fungi(100);p__Ascomycota(100);p__Ascomycota_unclassified(100);p__Ascomycota_unclassified(100);p__Ascomycota_unclassified(100);p__Ascomycota_unclassified(100);p__Ascomycota_unclassified(100); | - | - | - | - | - | - | - | Unassigned | - |
| Otu30191 | 1 | k__Fungi(100);p__Ascomycota(100);p__Ascomycota_unclassified(100);p__Ascomycota_unclassified(100);p__Ascomycota_unclassified(100);p__Ascomycota_unclassified(100);p__Ascomycota_unclassified(100); | - | - | - | - | - | - | - | Unassigned | - |
| Otu30198 | 1 | k__Fungi(100);p__Ascomycota(100);p__Ascomycota_unclassified(100);p__Ascomycota_unclassified(100);p__Ascomycota_unclassified(100);p__Ascomycota_unclassified(100);p__Ascomycota_unclassified(100); | - | - | - | - | - | - | - | Unassigned | - |
| Otu30206 | 1 | k__Fungi(100);p__Ascomycota(100);p__Ascomycota_unclassified(100);p__Ascomycota_unclassified(100);p__Ascomycota_unclassified(100);p__Ascomycota_unclassified(100);p__Ascomycota_unclassified(100); | - | - | - | - | - | - | - | Unassigned | - |
| Otu30228 | 1 | k__Fungi(100);k__Fungi_unclassified(100);k__Fungi_unclassified(100);k__Fungi_unclassified(100);k__Fungi_unclassified(100);k__Fungi_unclassified(100);k__Fungi_unclassified(100); | - | - | - | - | - | - | - | Unassigned | - |
| Otu30240 | 1 | k__Fungi(100);p__Ascomycota(100);p__Ascomycota_unclassified(100);p__Ascomycota_unclassified(100);p__Ascomycota_unclassified(100);p__Ascomycota_unclassified(100);p__Ascomycota_unclassified(100); | - | - | - | - | - | - | - | Unassigned | - |
| Otu30241 | 1 | k__Fungi(100);p__Ascomycota(100);p__Ascomycota_unclassified(100);p__Ascomycota_unclassified(100);p__Ascomycota_unclassified(100);p__Ascomycota_unclassified(100);p__Ascomycota_unclassified(100); | - | - | - | - | - | - | - | Unassigned | - |
| Otu30243 | 1 | k__Fungi(100);p__Ascomycota(100);p__Ascomycota_unclassified(100);p__Ascomycota_unclassified(100);p__Ascomycota_unclassified(100);p__Ascomycota_unclassified(100);p__Ascomycota_unclassified(100); | - | - | - | - | - | - | - | Unassigned | - |
| Otu30244 | 1 | k__Fungi(100);k__Fungi_unclassified(100);k__Fungi_unclassified(100);k__Fungi_unclassified(100);k__Fungi_unclassified(100);k__Fungi_unclassified(100);k__Fungi_unclassified(100); | - | - | - | - | - | - | - | Unassigned | - |
| Otu30253 | 1 | k__Fungi(100);k__Fungi_unclassified(100);k__Fungi_unclassified(100);k__Fungi_unclassified(100);k__Fungi_unclassified(100);k__Fungi_unclassified(100);k__Fungi_unclassified(100); | - | - | - | - | - | - | - | Unassigned | - |
| Otu30255 | 1 | k__Fungi(100);k__Fungi_unclassified(100);k__Fungi_unclassified(100);k__Fungi_unclassified(100);k__Fungi_unclassified(100);k__Fungi_unclassified(100);k__Fungi_unclassified(100); | - | - | - | - | - | - | - | Unassigned | - |
| Otu30256 | 1 | k__Fungi(100);k__Fungi_unclassified(100);k__Fungi_unclassified(100);k__Fungi_unclassified(100);k__Fungi_unclassified(100);k__Fungi_unclassified(100);k__Fungi_unclassified(100); | - | - | - | - | - | - | - | Unassigned | - |
| Otu30259 | 1 | k__Fungi(100);k__Fungi_unclassified(100);k__Fungi_unclassified(100);k__Fungi_unclassified(100);k__Fungi_unclassified(100);k__Fungi_unclassified(100);k__Fungi_unclassified(100); | - | - | - | - | - | - | - | Unassigned | - |
| Otu30260 | 1 | k__Fungi(100);k__Fungi_unclassified(100);k__Fungi_unclassified(100);k__Fungi_unclassified(100);k__Fungi_unclassified(100);k__Fungi_unclassified(100);k__Fungi_unclassified(100); | - | - | - | - | - | - | - | Unassigned | - |
| Otu30262 | 1 | k__Fungi(100);p__Ascomycota(100);p__Ascomycota_unclassified(100);p__Ascomycota_unclassified(100);p__Ascomycota_unclassified(100);p__Ascomycota_unclassified(100);p__Ascomycota_unclassified(100); | - | - | - | - | - | - | - | Unassigned | - |
| Otu30264 | 1 | k__Fungi(100);p__Ascomycota(100);p__Ascomycota_unclassified(100);p__Ascomycota_unclassified(100);p__Ascomycota_unclassified(100);p__Ascomycota_unclassified(100);p__Ascomycota_unclassified(100); | - | - | - | - | - | - | - | Unassigned | - |
| Otu30266 | 1 | k__Fungi(100);k__Fungi_unclassified(100);k__Fungi_unclassified(100);k__Fungi_unclassified(100);k__Fungi_unclassified(100);k__Fungi_unclassified(100);k__Fungi_unclassified(100); | - | - | - | - | - | - | - | Unassigned | - |
| Otu30267 | 1 | k__Fungi(100);p__Ascomycota(100);p__Ascomycota_unclassified(100);p__Ascomycota_unclassified(100);p__Ascomycota_unclassified(100);p__Ascomycota_unclassified(100);p__Ascomycota_unclassified(100); | - | - | - | - | - | - | - | Unassigned | - |
| Otu30268 | 1 | k__Fungi(100);p__Ascomycota(100);p__Ascomycota_unclassified(100);p__Ascomycota_unclassified(100);p__Ascomycota_unclassified(100);p__Ascomycota_unclassified(100);p__Ascomycota_unclassified(100); | - | - | - | - | - | - | - | Unassigned | - |
| Otu30274 | 1 | k__Fungi(100);p__Ascomycota(100);p__Ascomycota_unclassified(100);p__Ascomycota_unclassified(100);p__Ascomycota_unclassified(100);p__Ascomycota_unclassified(100);p__Ascomycota_unclassified(100); | - | - | - | - | - | - | - | Unassigned | - |
| Otu30275 | 1 | k__Fungi(100);k__Fungi_unclassified(100);k__Fungi_unclassified(100);k__Fungi_unclassified(100);k__Fungi_unclassified(100);k__Fungi_unclassified(100);k__Fungi_unclassified(100); | - | - | - | - | - | - | - | Unassigned | - |
| Otu30277 | 1 | k__Fungi(100);k__Fungi_unclassified(100);k__Fungi_unclassified(100);k__Fungi_unclassified(100);k__Fungi_unclassified(100);k__Fungi_unclassified(100);k__Fungi_unclassified(100); | - | - | - | - | - | - | - | Unassigned | - |
| Otu30278 | 1 | k__Fungi(100);k__Fungi_unclassified(100);k__Fungi_unclassified(100);k__Fungi_unclassified(100);k__Fungi_unclassified(100);k__Fungi_unclassified(100);k__Fungi_unclassified(100); | - | - | - | - | - | - | - | Unassigned | - |
| Otu30280 | 1 | k__Fungi(100);p__Ascomycota(100);p__Ascomycota_unclassified(100);p__Ascomycota_unclassified(100);p__Ascomycota_unclassified(100);p__Ascomycota_unclassified(100);p__Ascomycota_unclassified(100); | - | - | - | - | - | - | - | Unassigned | - |
| Otu30283 | 1 | k__Fungi(100);k__Fungi_unclassified(100);k__Fungi_unclassified(100);k__Fungi_unclassified(100);k__Fungi_unclassified(100);k__Fungi_unclassified(100);k__Fungi_unclassified(100); | - | - | - | - | - | - | - | Unassigned | - |
| Otu30285 | 1 | k__Fungi(100);k__Fungi_unclassified(100);k__Fungi_unclassified(100);k__Fungi_unclassified(100);k__Fungi_unclassified(100);k__Fungi_unclassified(100);k__Fungi_unclassified(100); | - | - | - | - | - | - | - | Unassigned | - |
| Otu30286 | 1 | k__Fungi(100);k__Fungi_unclassified(100);k__Fungi_unclassified(100);k__Fungi_unclassified(100);k__Fungi_unclassified(100);k__Fungi_unclassified(100);k__Fungi_unclassified(100); | - | - | - | - | - | - | - | Unassigned | - |
| Otu30289 | 1 | k__Fungi(100);k__Fungi_unclassified(100);k__Fungi_unclassified(100);k__Fungi_unclassified(100);k__Fungi_unclassified(100);k__Fungi_unclassified(100);k__Fungi_unclassified(100); | - | - | - | - | - | - | - | Unassigned | - |
| Otu30292 | 1 | k__Fungi(100);k__Fungi_unclassified(100);k__Fungi_unclassified(100);k__Fungi_unclassified(100);k__Fungi_unclassified(100);k__Fungi_unclassified(100);k__Fungi_unclassified(100); | - | - | - | - | - | - | - | Unassigned | - |
| Otu30317 | 1 | k__Fungi(100);k__Fungi_unclassified(100);k__Fungi_unclassified(100);k__Fungi_unclassified(100);k__Fungi_unclassified(100);k__Fungi_unclassified(100);k__Fungi_unclassified(100); | - | - | - | - | - | - | - | Unassigned | - |
| Otu30318 | 1 | k__Fungi(100);k__Fungi_unclassified(100);k__Fungi_unclassified(100);k__Fungi_unclassified(100);k__Fungi_unclassified(100);k__Fungi_unclassified(100);k__Fungi_unclassified(100); | - | - | - | - | - | - | - | Unassigned | - |
| Otu30321 | 1 | k__Fungi(100);k__Fungi_unclassified(100);k__Fungi_unclassified(100);k__Fungi_unclassified(100);k__Fungi_unclassified(100);k__Fungi_unclassified(100);k__Fungi_unclassified(100); | - | - | - | - | - | - | - | Unassigned | - |
| Otu30322 | 1 | k__Fungi(100);k__Fungi_unclassified(100);k__Fungi_unclassified(100);k__Fungi_unclassified(100);k__Fungi_unclassified(100);k__Fungi_unclassified(100);k__Fungi_unclassified(100); | - | - | - | - | - | - | - | Unassigned | - |
| Otu30340 | 1 | k__Fungi(100);p__Basidiomycota(100);p__Basidiomycota_unclassified(100);p__Basidiomycota_unclassified(100);p__Basidiomycota_unclassified(100);p__Basidiomycota_unclassified(100);p__Basidiomycota_unclassified(100); | - | - | - | - | - | - | - | Unassigned | - |
| Otu30341 | 1 | k__Fungi(100);p__Ascomycota(100);p__Ascomycota_unclassified(100);p__Ascomycota_unclassified(100);p__Ascomycota_unclassified(100);p__Ascomycota_unclassified(100);p__Ascomycota_unclassified(100); | - | - | - | - | - | - | - | Unassigned | - |
| Otu30344 | 1 | k__Fungi(100);k__Fungi_unclassified(100);k__Fungi_unclassified(100);k__Fungi_unclassified(100);k__Fungi_unclassified(100);k__Fungi_unclassified(100);k__Fungi_unclassified(100); | - | - | - | - | - | - | - | Unassigned | - |
| Otu30346 | 1 | k__Fungi(100);p__Ascomycota(100);p__Ascomycota_unclassified(100);p__Ascomycota_unclassified(100);p__Ascomycota_unclassified(100);p__Ascomycota_unclassified(100);p__Ascomycota_unclassified(100); | - | - | - | - | - | - | - | Unassigned | - |
| Otu30347 | 1 | k__Fungi(100);k__Fungi_unclassified(100);k__Fungi_unclassified(100);k__Fungi_unclassified(100);k__Fungi_unclassified(100);k__Fungi_unclassified(100);k__Fungi_unclassified(100); | - | - | - | - | - | - | - | Unassigned | - |
| Otu30348 | 1 | k__Fungi(100);p__Ascomycota(100);p__Ascomycota_unclassified(100);p__Ascomycota_unclassified(100);p__Ascomycota_unclassified(100);p__Ascomycota_unclassified(100);p__Ascomycota_unclassified(100); | - | - | - | - | - | - | - | Unassigned | - |
| Otu30349 | 1 | k__Fungi(100);p__Ascomycota(100);p__Ascomycota_unclassified(100);p__Ascomycota_unclassified(100);p__Ascomycota_unclassified(100);p__Ascomycota_unclassified(100);p__Ascomycota_unclassified(100); | - | - | - | - | - | - | - | Unassigned | - |
| Otu30350 | 1 | k__Fungi(100);k__Fungi_unclassified(100);k__Fungi_unclassified(100);k__Fungi_unclassified(100);k__Fungi_unclassified(100);k__Fungi_unclassified(100);k__Fungi_unclassified(100); | - | - | - | - | - | - | - | Unassigned | - |
| Otu30351 | 1 | k__Fungi(100);p__Ascomycota(100);p__Ascomycota_unclassified(100);p__Ascomycota_unclassified(100);p__Ascomycota_unclassified(100);p__Ascomycota_unclassified(100);p__Ascomycota_unclassified(100); | - | - | - | - | - | - | - | Unassigned | - |
| Otu30352 | 1 | k__Fungi(100);k__Fungi_unclassified(100);k__Fungi_unclassified(100);k__Fungi_unclassified(100);k__Fungi_unclassified(100);k__Fungi_unclassified(100);k__Fungi_unclassified(100); | - | - | - | - | - | - | - | Unassigned | - |
| Otu30353 | 1 | k__Fungi(100);k__Fungi_unclassified(100);k__Fungi_unclassified(100);k__Fungi_unclassified(100);k__Fungi_unclassified(100);k__Fungi_unclassified(100);k__Fungi_unclassified(100); | - | - | - | - | - | - | - | Unassigned | - |
| Otu30357 | 1 | k__Fungi(100);p__Ascomycota(100);p__Ascomycota_unclassified(100);p__Ascomycota_unclassified(100);p__Ascomycota_unclassified(100);p__Ascomycota_unclassified(100);p__Ascomycota_unclassified(100); | - | - | - | - | - | - | - | Unassigned | - |
| Otu30360 | 1 | k__Fungi(100);k__Fungi_unclassified(100);k__Fungi_unclassified(100);k__Fungi_unclassified(100);k__Fungi_unclassified(100);k__Fungi_unclassified(100);k__Fungi_unclassified(100); | - | - | - | - | - | - | - | Unassigned | - |
| Otu30362 | 1 | k__Fungi(100);p__Ascomycota(100);p__Ascomycota_unclassified(100);p__Ascomycota_unclassified(100);p__Ascomycota_unclassified(100);p__Ascomycota_unclassified(100);p__Ascomycota_unclassified(100); | - | - | - | - | - | - | - | Unassigned | - |
| Otu30363 | 1 | k__Fungi(100);k__Fungi_unclassified(100);k__Fungi_unclassified(100);k__Fungi_unclassified(100);k__Fungi_unclassified(100);k__Fungi_unclassified(100);k__Fungi_unclassified(100); | - | - | - | - | - | - | - | Unassigned | - |
| Otu30364 | 1 | k__Fungi(100);k__Fungi_unclassified(100);k__Fungi_unclassified(100);k__Fungi_unclassified(100);k__Fungi_unclassified(100);k__Fungi_unclassified(100);k__Fungi_unclassified(100); | - | - | - | - | - | - | - | Unassigned | - |
| Otu30374 | 1 | k__Fungi(100);k__Fungi_unclassified(100);k__Fungi_unclassified(100);k__Fungi_unclassified(100);k__Fungi_unclassified(100);k__Fungi_unclassified(100);k__Fungi_unclassified(100); | - | - | - | - | - | - | - | Unassigned | - |
| Otu30375 | 1 | k__Fungi(100);p__Ascomycota(100);p__Ascomycota_unclassified(100);p__Ascomycota_unclassified(100);p__Ascomycota_unclassified(100);p__Ascomycota_unclassified(100);p__Ascomycota_unclassified(100); | - | - | - | - | - | - | - | Unassigned | - |
| Otu30377 | 1 | k__Fungi(100);p__Ascomycota(100);p__Ascomycota_unclassified(100);p__Ascomycota_unclassified(100);p__Ascomycota_unclassified(100);p__Ascomycota_unclassified(100);p__Ascomycota_unclassified(100); | - | - | - | - | - | - | - | Unassigned | - |
| Otu30378 | 1 | k__Fungi(100);k__Fungi_unclassified(100);k__Fungi_unclassified(100);k__Fungi_unclassified(100);k__Fungi_unclassified(100);k__Fungi_unclassified(100);k__Fungi_unclassified(100); | - | - | - | - | - | - | - | Unassigned | - |
| Otu30379 | 1 | k__Fungi(100);p__Ascomycota(100);p__Ascomycota_unclassified(100);p__Ascomycota_unclassified(100);p__Ascomycota_unclassified(100);p__Ascomycota_unclassified(100);p__Ascomycota_unclassified(100); | - | - | - | - | - | - | - | Unassigned | - |
| Otu30380 | 1 | k__Fungi(100);p__Ascomycota(100);p__Ascomycota_unclassified(100);p__Ascomycota_unclassified(100);p__Ascomycota_unclassified(100);p__Ascomycota_unclassified(100);p__Ascomycota_unclassified(100); | - | - | - | - | - | - | - | Unassigned | - |
| Otu30381 | 1 | k__Fungi(100);p__Ascomycota(100);p__Ascomycota_unclassified(100);p__Ascomycota_unclassified(100);p__Ascomycota_unclassified(100);p__Ascomycota_unclassified(100);p__Ascomycota_unclassified(100); | - | - | - | - | - | - | - | Unassigned | - |
| Otu30382 | 1 | k__Fungi(100);p__Ascomycota(100);p__Ascomycota_unclassified(100);p__Ascomycota_unclassified(100);p__Ascomycota_unclassified(100);p__Ascomycota_unclassified(100);p__Ascomycota_unclassified(100); | - | - | - | - | - | - | - | Unassigned | - |
| Otu30383 | 1 | k__Fungi(100);k__Fungi_unclassified(100);k__Fungi_unclassified(100);k__Fungi_unclassified(100);k__Fungi_unclassified(100);k__Fungi_unclassified(100);k__Fungi_unclassified(100); | - | - | - | - | - | - | - | Unassigned | - |
| Otu30384 | 1 | k__Fungi(100);p__Ascomycota(100);p__Ascomycota_unclassified(100);p__Ascomycota_unclassified(100);p__Ascomycota_unclassified(100);p__Ascomycota_unclassified(100);p__Ascomycota_unclassified(100); | - | - | - | - | - | - | - | Unassigned | - |
| Otu30385 | 1 | k__Fungi(100);k__Fungi_unclassified(100);k__Fungi_unclassified(100);k__Fungi_unclassified(100);k__Fungi_unclassified(100);k__Fungi_unclassified(100);k__Fungi_unclassified(100); | - | - | - | - | - | - | - | Unassigned | - |
| Otu30386 | 1 | k__Fungi(100);p__Ascomycota(100);p__Ascomycota_unclassified(100);p__Ascomycota_unclassified(100);p__Ascomycota_unclassified(100);p__Ascomycota_unclassified(100);p__Ascomycota_unclassified(100); | - | - | - | - | - | - | - | Unassigned | - |
| Otu30388 | 1 | k__Fungi(100);k__Fungi_unclassified(100);k__Fungi_unclassified(100);k__Fungi_unclassified(100);k__Fungi_unclassified(100);k__Fungi_unclassified(100);k__Fungi_unclassified(100); | - | - | - | - | - | - | - | Unassigned | - |
| Otu30390 | 1 | k__Fungi(100);k__Fungi_unclassified(100);k__Fungi_unclassified(100);k__Fungi_unclassified(100);k__Fungi_unclassified(100);k__Fungi_unclassified(100);k__Fungi_unclassified(100); | - | - | - | - | - | - | - | Unassigned | - |
| Otu30391 | 1 | k__Fungi(100);p__Ascomycota(100);p__Ascomycota_unclassified(100);p__Ascomycota_unclassified(100);p__Ascomycota_unclassified(100);p__Ascomycota_unclassified(100);p__Ascomycota_unclassified(100); | - | - | - | - | - | - | - | Unassigned | - |
| Otu30393 | 1 | k__Fungi(100);p__Ascomycota(100);p__Ascomycota_unclassified(100);p__Ascomycota_unclassified(100);p__Ascomycota_unclassified(100);p__Ascomycota_unclassified(100);p__Ascomycota_unclassified(100); | - | - | - | - | - | - | - | Unassigned | - |
| Otu30394 | 1 | k__Fungi(100);k__Fungi_unclassified(100);k__Fungi_unclassified(100);k__Fungi_unclassified(100);k__Fungi_unclassified(100);k__Fungi_unclassified(100);k__Fungi_unclassified(100); | - | - | - | - | - | - | - | Unassigned | - |
| Otu30395 | 1 | k__Fungi(100);k__Fungi_unclassified(100);k__Fungi_unclassified(100);k__Fungi_unclassified(100);k__Fungi_unclassified(100);k__Fungi_unclassified(100);k__Fungi_unclassified(100); | - | - | - | - | - | - | - | Unassigned | - |
| Otu30398 | 1 | k__Fungi(100);k__Fungi_unclassified(100);k__Fungi_unclassified(100);k__Fungi_unclassified(100);k__Fungi_unclassified(100);k__Fungi_unclassified(100);k__Fungi_unclassified(100); | - | - | - | - | - | - | - | Unassigned | - |
| Otu30399 | 1 | k__Fungi(100);p__Ascomycota(100);p__Ascomycota_unclassified(100);p__Ascomycota_unclassified(100);p__Ascomycota_unclassified(100);p__Ascomycota_unclassified(100);p__Ascomycota_unclassified(100); | - | - | - | - | - | - | - | Unassigned | - |
| Otu30400 | 1 | k__Fungi(100);p__Ascomycota(100);p__Ascomycota_unclassified(100);p__Ascomycota_unclassified(100);p__Ascomycota_unclassified(100);p__Ascomycota_unclassified(100);p__Ascomycota_unclassified(100); | - | - | - | - | - | - | - | Unassigned | - |
| Otu30401 | 1 | k__Fungi(100);k__Fungi_unclassified(100);k__Fungi_unclassified(100);k__Fungi_unclassified(100);k__Fungi_unclassified(100);k__Fungi_unclassified(100);k__Fungi_unclassified(100); | - | - | - | - | - | - | - | Unassigned | - |
| Otu30402 | 1 | k__Fungi(100);k__Fungi_unclassified(100);k__Fungi_unclassified(100);k__Fungi_unclassified(100);k__Fungi_unclassified(100);k__Fungi_unclassified(100);k__Fungi_unclassified(100); | - | - | - | - | - | - | - | Unassigned | - |
| Otu30403 | 1 | k__Fungi(100);k__Fungi_unclassified(100);k__Fungi_unclassified(100);k__Fungi_unclassified(100);k__Fungi_unclassified(100);k__Fungi_unclassified(100);k__Fungi_unclassified(100); | - | - | - | - | - | - | - | Unassigned | - |
| Otu30404 | 1 | k__Fungi(100);k__Fungi_unclassified(100);k__Fungi_unclassified(100);k__Fungi_unclassified(100);k__Fungi_unclassified(100);k__Fungi_unclassified(100);k__Fungi_unclassified(100); | - | - | - | - | - | - | - | Unassigned | - |
| Otu30405 | 1 | k__Fungi(100);k__Fungi_unclassified(100);k__Fungi_unclassified(100);k__Fungi_unclassified(100);k__Fungi_unclassified(100);k__Fungi_unclassified(100);k__Fungi_unclassified(100); | - | - | - | - | - | - | - | Unassigned | - |
| Otu30406 | 1 | k__Fungi(100);k__Fungi_unclassified(100);k__Fungi_unclassified(100);k__Fungi_unclassified(100);k__Fungi_unclassified(100);k__Fungi_unclassified(100);k__Fungi_unclassified(100); | - | - | - | - | - | - | - | Unassigned | - |
| Otu30407 | 1 | k__Fungi(100);k__Fungi_unclassified(100);k__Fungi_unclassified(100);k__Fungi_unclassified(100);k__Fungi_unclassified(100);k__Fungi_unclassified(100);k__Fungi_unclassified(100); | - | - | - | - | - | - | - | Unassigned | - |
| Otu30408 | 1 | k__Fungi(100);p__Ascomycota(100);p__Ascomycota_unclassified(100);p__Ascomycota_unclassified(100);p__Ascomycota_unclassified(100);p__Ascomycota_unclassified(100);p__Ascomycota_unclassified(100); | - | - | - | - | - | - | - | Unassigned | - |
| Otu30409 | 1 | k__Fungi(100);k__Fungi_unclassified(100);k__Fungi_unclassified(100);k__Fungi_unclassified(100);k__Fungi_unclassified(100);k__Fungi_unclassified(100);k__Fungi_unclassified(100); | - | - | - | - | - | - | - | Unassigned | - |
| Otu30410 | 1 | k__Fungi(100);p__Ascomycota(100);p__Ascomycota_unclassified(100);p__Ascomycota_unclassified(100);p__Ascomycota_unclassified(100);p__Ascomycota_unclassified(100);p__Ascomycota_unclassified(100); | - | - | - | - | - | - | - | Unassigned | - |
| Otu30411 | 1 | k__Fungi(100);k__Fungi_unclassified(100);k__Fungi_unclassified(100);k__Fungi_unclassified(100);k__Fungi_unclassified(100);k__Fungi_unclassified(100);k__Fungi_unclassified(100); | - | - | - | - | - | - | - | Unassigned | - |
| Otu30415 | 1 | k__Fungi(100);k__Fungi_unclassified(100);k__Fungi_unclassified(100);k__Fungi_unclassified(100);k__Fungi_unclassified(100);k__Fungi_unclassified(100);k__Fungi_unclassified(100); | - | - | - | - | - | - | - | Unassigned | - |
| Otu30416 | 1 | k__Fungi(100);k__Fungi_unclassified(100);k__Fungi_unclassified(100);k__Fungi_unclassified(100);k__Fungi_unclassified(100);k__Fungi_unclassified(100);k__Fungi_unclassified(100); | - | - | - | - | - | - | - | Unassigned | - |
| Otu30417 | 1 | k__Fungi(100);k__Fungi_unclassified(100);k__Fungi_unclassified(100);k__Fungi_unclassified(100);k__Fungi_unclassified(100);k__Fungi_unclassified(100);k__Fungi_unclassified(100); | - | - | - | - | - | - | - | Unassigned | - |
| Otu30418 | 1 | k__Fungi(100);k__Fungi_unclassified(100);k__Fungi_unclassified(100);k__Fungi_unclassified(100);k__Fungi_unclassified(100);k__Fungi_unclassified(100);k__Fungi_unclassified(100); | - | - | - | - | - | - | - | Unassigned | - |
| Otu30419 | 1 | k__Fungi(100);k__Fungi_unclassified(100);k__Fungi_unclassified(100);k__Fungi_unclassified(100);k__Fungi_unclassified(100);k__Fungi_unclassified(100);k__Fungi_unclassified(100); | - | - | - | - | - | - | - | Unassigned | - |
| Otu30420 | 1 | k__Fungi(100);p__Ascomycota(100);p__Ascomycota_unclassified(100);p__Ascomycota_unclassified(100);p__Ascomycota_unclassified(100);p__Ascomycota_unclassified(100);p__Ascomycota_unclassified(100); | - | - | - | - | - | - | - | Unassigned | - |
| Otu30423 | 1 | k__Fungi(100);p__Ascomycota(100);p__Ascomycota_unclassified(100);p__Ascomycota_unclassified(100);p__Ascomycota_unclassified(100);p__Ascomycota_unclassified(100);p__Ascomycota_unclassified(100); | - | - | - | - | - | - | - | Unassigned | - |
| Otu30425 | 1 | k__Fungi(100);p__Ascomycota(100);p__Ascomycota_unclassified(100);p__Ascomycota_unclassified(100);p__Ascomycota_unclassified(100);p__Ascomycota_unclassified(100);p__Ascomycota_unclassified(100); | - | - | - | - | - | - | - | Unassigned | - |
| Otu30426 | 1 | k__Fungi(100);k__Fungi_unclassified(100);k__Fungi_unclassified(100);k__Fungi_unclassified(100);k__Fungi_unclassified(100);k__Fungi_unclassified(100);k__Fungi_unclassified(100); | - | - | - | - | - | - | - | Unassigned | - |
| Otu30431 | 1 | k__Fungi(100);k__Fungi_unclassified(100);k__Fungi_unclassified(100);k__Fungi_unclassified(100);k__Fungi_unclassified(100);k__Fungi_unclassified(100);k__Fungi_unclassified(100); | - | - | - | - | - | - | - | Unassigned | - |
| Otu30439 | 1 | k__Fungi(100);k__Fungi_unclassified(100);k__Fungi_unclassified(100);k__Fungi_unclassified(100);k__Fungi_unclassified(100);k__Fungi_unclassified(100);k__Fungi_unclassified(100); | - | - | - | - | - | - | - | Unassigned | - |
| Otu30440 | 1 | k__Fungi(100);p__Ascomycota(100);p__Ascomycota_unclassified(100);p__Ascomycota_unclassified(100);p__Ascomycota_unclassified(100);p__Ascomycota_unclassified(100);p__Ascomycota_unclassified(100); | - | - | - | - | - | - | - | Unassigned | - |
| Otu30442 | 1 | k__Fungi(100);k__Fungi_unclassified(100);k__Fungi_unclassified(100);k__Fungi_unclassified(100);k__Fungi_unclassified(100);k__Fungi_unclassified(100);k__Fungi_unclassified(100); | - | - | - | - | - | - | - | Unassigned | - |

| | | | | | | | | | | | |
|---|---|---|---|---|---|---|---|---|---|---|---|
| Otu30451 | 1 | k__Fungi(100);k__Fungi_unclassified(100);k__Fungi_unclassified(100);k__Fungi_unclassified(100);k__Fungi_unclassified(100);k__Fungi_unclassified(100);k__Fungi_unclassified(100); | - | - | - | - | - | - | - | Unassigned | - |
| Otu30453 | 1 | k__Fungi(100);p__Ascomycota(100);p__Ascomycota_unclassified(100);p__Ascomycota_unclassified(100);p__Ascomycota_unclassified(100);p__Ascomycota_unclassified(100);p__Ascomycota_unclassified(100); | - | - | - | - | - | - | - | Unassigned | - |
| Otu30464 | 1 | k__Fungi(100);k__Fungi_unclassified(100);k__Fungi_unclassified(100);k__Fungi_unclassified(100);k__Fungi_unclassified(100);k__Fungi_unclassified(100);k__Fungi_unclassified(100); | - | - | - | - | - | - | - | Unassigned | - |
| Otu30466 | 1 | k__Fungi(100);p__Ascomycota(100);p__Ascomycota_unclassified(100);p__Ascomycota_unclassified(100);p__Ascomycota_unclassified(100);p__Ascomycota_unclassified(100);p__Ascomycota_unclassified(100); | - | - | - | - | - | - | - | Unassigned | - |
| Otu30476 | 1 | k__Fungi(100);p__Ascomycota(100);p__Ascomycota_unclassified(100);p__Ascomycota_unclassified(100);p__Ascomycota_unclassified(100);p__Ascomycota_unclassified(100);p__Ascomycota_unclassified(100); | - | - | - | - | - | - | - | Unassigned | - |
| Otu30503 | 1 | k__Fungi(100);k__Fungi_unclassified(100);k__Fungi_unclassified(100);k__Fungi_unclassified(100);k__Fungi_unclassified(100);k__Fungi_unclassified(100);k__Fungi_unclassified(100); | - | - | - | - | - | - | - | Unassigned | - |
| Otu30504 | 1 | k__Fungi(100);k__Fungi_unclassified(100);k__Fungi_unclassified(100);k__Fungi_unclassified(100);k__Fungi_unclassified(100);k__Fungi_unclassified(100);k__Fungi_unclassified(100); | - | - | - | - | - | - | - | Unassigned | - |
| Otu30505 | 1 | k__Fungi(100);k__Fungi_unclassified(100);k__Fungi_unclassified(100);k__Fungi_unclassified(100);k__Fungi_unclassified(100);k__Fungi_unclassified(100);k__Fungi_unclassified(100); | - | - | - | - | - | - | - | Unassigned | - |
| Otu30506 | 1 | k__Fungi(100);p__Ascomycota(100);p__Ascomycota_unclassified(100);p__Ascomycota_unclassified(100);p__Ascomycota_unclassified(100);p__Ascomycota_unclassified(100);p__Ascomycota_unclassified(100); | - | - | - | - | - | - | - | Unassigned | - |
| Otu30507 | 1 | k__Fungi(100);k__Fungi_unclassified(100);k__Fungi_unclassified(100);k__Fungi_unclassified(100);k__Fungi_unclassified(100);k__Fungi_unclassified(100);k__Fungi_unclassified(100); | - | - | - | - | - | - | - | Unassigned | - |
| Otu30509 | 1 | k__Fungi(100);k__Fungi_unclassified(100);k__Fungi_unclassified(100);k__Fungi_unclassified(100);k__Fungi_unclassified(100);k__Fungi_unclassified(100);k__Fungi_unclassified(100); | - | - | - | - | - | - | - | Unassigned | - |
| Otu30511 | 1 | k__Fungi(100);p__Ascomycota(100);p__Ascomycota_unclassified(100);p__Ascomycota_unclassified(100);p__Ascomycota_unclassified(100);p__Ascomycota_unclassified(100);p__Ascomycota_unclassified(100); | - | - | - | - | - | - | - | Unassigned | - |
| Otu30513 | 1 | k__Fungi(100);p__Ascomycota(100);p__Ascomycota_unclassified(100);p__Ascomycota_unclassified(100);p__Ascomycota_unclassified(100);p__Ascomycota_unclassified(100);p__Ascomycota_unclassified(100); | - | - | - | - | - | - | - | Unassigned | - |
| Otu30519 | 1 | k__Fungi(100);p__Ascomycota(100);p__Ascomycota_unclassified(100);p__Ascomycota_unclassified(100);p__Ascomycota_unclassified(100);p__Ascomycota_unclassified(100);p__Ascomycota_unclassified(100); | - | - | - | - | - | - | - | Unassigned | - |
| Otu30521 | 1 | k__Fungi(100);p__Ascomycota(100);p__Ascomycota_unclassified(100);p__Ascomycota_unclassified(100);p__Ascomycota_unclassified(100);p__Ascomycota_unclassified(100);p__Ascomycota_unclassified(100); | - | - | - | - | - | - | - | Unassigned | - |
| Otu30522 | 1 | k__Fungi(100);p__Ascomycota(100);p__Ascomycota_unclassified(100);p__Ascomycota_unclassified(100);p__Ascomycota_unclassified(100);p__Ascomycota_unclassified(100);p__Ascomycota_unclassified(100); | - | - | - | - | - | - | - | Unassigned | - |
| Otu30523 | 1 | k__Fungi(100);p__Ascomycota(100);p__Ascomycota_unclassified(100);p__Ascomycota_unclassified(100);p__Ascomycota_unclassified(100);p__Ascomycota_unclassified(100);p__Ascomycota_unclassified(100); | - | - | - | - | - | - | - | Unassigned | - |
| Otu30524 | 1 | k__Fungi(100);k__Fungi_unclassified(100);k__Fungi_unclassified(100);k__Fungi_unclassified(100);k__Fungi_unclassified(100);k__Fungi_unclassified(100);k__Fungi_unclassified(100); | - | - | - | - | - | - | - | Unassigned | - |
| Otu30525 | 1 | k__Fungi(100);p__Ascomycota(100);p__Ascomycota_unclassified(100);p__Ascomycota_unclassified(100);p__Ascomycota_unclassified(100);p__Ascomycota_unclassified(100);p__Ascomycota_unclassified(100); | - | - | - | - | - | - | - | Unassigned | - |
| Otu30526 | 1 | k__Fungi(100);k__Fungi_unclassified(100);k__Fungi_unclassified(100);k__Fungi_unclassified(100);k__Fungi_unclassified(100);k__Fungi_unclassified(100);k__Fungi_unclassified(100); | - | - | - | - | - | - | - | Unassigned | - |
| Otu30527 | 1 | k__Fungi(100);k__Fungi_unclassified(100);k__Fungi_unclassified(100);k__Fungi_unclassified(100);k__Fungi_unclassified(100);k__Fungi_unclassified(100);k__Fungi_unclassified(100); | - | - | - | - | - | - | - | Unassigned | - |
| Otu30528 | 1 | k__Fungi(100);p__Ascomycota(100);p__Ascomycota_unclassified(100);p__Ascomycota_unclassified(100);p__Ascomycota_unclassified(100);p__Ascomycota_unclassified(100);p__Ascomycota_unclassified(100); | - | - | - | - | - | - | - | Unassigned | - |
| Otu30529 | 1 | k__Fungi(100);p__Ascomycota(100);p__Ascomycota_unclassified(100);p__Ascomycota_unclassified(100);p__Ascomycota_unclassified(100);p__Ascomycota_unclassified(100);p__Ascomycota_unclassified(100); | - | - | - | - | - | - | - | Unassigned | - |
| Otu30530 | 1 | k__Fungi(100);k__Fungi_unclassified(100);k__Fungi_unclassified(100);k__Fungi_unclassified(100);k__Fungi_unclassified(100);k__Fungi_unclassified(100);k__Fungi_unclassified(100); | - | - | - | - | - | - | - | Unassigned | - |
| Otu30531 | 1 | k__Fungi(100);k__Fungi_unclassified(100);k__Fungi_unclassified(100);k__Fungi_unclassified(100);k__Fungi_unclassified(100);k__Fungi_unclassified(100);k__Fungi_unclassified(100); | - | - | - | - | - | - | - | Unassigned | - |
| Otu30546 | 1 | k__Fungi(100);p__Ascomycota(100);p__Ascomycota_unclassified(100);p__Ascomycota_unclassified(100);p__Ascomycota_unclassified(100);p__Ascomycota_unclassified(100);p__Ascomycota_unclassified(100); | - | - | - | - | - | - | - | Unassigned | - |
| Otu30554 | 1 | k__Fungi(100);k__Fungi_unclassified(100);k__Fungi_unclassified(100);k__Fungi_unclassified(100);k__Fungi_unclassified(100);k__Fungi_unclassified(100);k__Fungi_unclassified(100); | - | - | - | - | - | - | - | Unassigned | - |
| Otu30560 | 1 | k__Fungi(100);k__Fungi_unclassified(100);k__Fungi_unclassified(100);k__Fungi_unclassified(100);k__Fungi_unclassified(100);k__Fungi_unclassified(100);k__Fungi_unclassified(100); | - | - | - | - | - | - | - | Unassigned | - |
| Otu30561 | 1 | k__Fungi(100);k__Fungi_unclassified(100);k__Fungi_unclassified(100);k__Fungi_unclassified(100);k__Fungi_unclassified(100);k__Fungi_unclassified(100);k__Fungi_unclassified(100); | - | - | - | - | - | - | - | Unassigned | - |
| Otu30562 | 1 | k__Fungi(100);k__Fungi_unclassified(100);k__Fungi_unclassified(100);k__Fungi_unclassified(100);k__Fungi_unclassified(100);k__Fungi_unclassified(100);k__Fungi_unclassified(100); | - | - | - | - | - | - | - | Unassigned | - |
| Otu30566 | 1 | k__Fungi(100);k__Fungi_unclassified(100);k__Fungi_unclassified(100);k__Fungi_unclassified(100);k__Fungi_unclassified(100);k__Fungi_unclassified(100);k__Fungi_unclassified(100); | - | - | - | - | - | - | - | Unassigned | - |
| Otu30567 | 1 | k__Fungi(100);k__Fungi_unclassified(100);k__Fungi_unclassified(100);k__Fungi_unclassified(100);k__Fungi_unclassified(100);k__Fungi_unclassified(100);k__Fungi_unclassified(100); | - | - | - | - | - | - | - | Unassigned | - |
| Otu30568 | 1 | k__Fungi(100);k__Fungi_unclassified(100);k__Fungi_unclassified(100);k__Fungi_unclassified(100);k__Fungi_unclassified(100);k__Fungi_unclassified(100);k__Fungi_unclassified(100); | - | - | - | - | - | - | - | Unassigned | - |
| Otu30569 | 1 | k__Fungi(100);p__Ascomycota(100);p__Ascomycota_unclassified(100);p__Ascomycota_unclassified(100);p__Ascomycota_unclassified(100);p__Ascomycota_unclassified(100);p__Ascomycota_unclassified(100); | - | - | - | - | - | - | - | Unassigned | - |
| Otu30571 | 1 | k__Fungi(100);k__Fungi_unclassified(100);k__Fungi_unclassified(100);k__Fungi_unclassified(100);k__Fungi_unclassified(100);k__Fungi_unclassified(100);k__Fungi_unclassified(100); | - | - | - | - | - | - | - | Unassigned | - |
| Otu30587 | 1 | k__Fungi(100);k__Fungi_unclassified(100);k__Fungi_unclassified(100);k__Fungi_unclassified(100);k__Fungi_unclassified(100);k__Fungi_unclassified(100);k__Fungi_unclassified(100); | - | - | - | - | - | - | - | Unassigned | - |
| Otu30588 | 1 | k__Fungi(100);p__Ascomycota(100);p__Ascomycota_unclassified(100);p__Ascomycota_unclassified(100);p__Ascomycota_unclassified(100);p__Ascomycota_unclassified(100);p__Ascomycota_unclassified(100); | - | - | - | - | - | - | - | Unassigned | - |
| Otu30592 | 1 | k__Fungi(100);p__Ascomycota(100);p__Ascomycota_unclassified(100);p__Ascomycota_unclassified(100);p__Ascomycota_unclassified(100);p__Ascomycota_unclassified(100);p__Ascomycota_unclassified(100); | - | - | - | - | - | - | - | Unassigned | - |
| Otu30593 | 1 | k__Fungi(100);k__Fungi_unclassified(100);k__Fungi_unclassified(100);k__Fungi_unclassified(100);k__Fungi_unclassified(100);k__Fungi_unclassified(100);k__Fungi_unclassified(100); | - | - | - | - | - | - | - | Unassigned | - |
| Otu30596 | 1 | k__Fungi(100);k__Fungi_unclassified(100);k__Fungi_unclassified(100);k__Fungi_unclassified(100);k__Fungi_unclassified(100);k__Fungi_unclassified(100);k__Fungi_unclassified(100); | - | - | - | - | - | - | - | Unassigned | - |
| Otu30597 | 1 | k__Fungi(100);k__Fungi_unclassified(100);k__Fungi_unclassified(100);k__Fungi_unclassified(100);k__Fungi_unclassified(100);k__Fungi_unclassified(100);k__Fungi_unclassified(100); | - | - | - | - | - | - | - | Unassigned | - |
| Otu30598 | 1 | k__Fungi(100);k__Fungi_unclassified(100);k__Fungi_unclassified(100);k__Fungi_unclassified(100);k__Fungi_unclassified(100);k__Fungi_unclassified(100);k__Fungi_unclassified(100); | - | - | - | - | - | - | - | Unassigned | - |
| Otu30601 | 1 | k__Fungi(100);k__Fungi_unclassified(100);k__Fungi_unclassified(100);k__Fungi_unclassified(100);k__Fungi_unclassified(100);k__Fungi_unclassified(100);k__Fungi_unclassified(100); | - | - | - | - | - | - | - | Unassigned | - |
| Otu30602 | 1 | k__Fungi(100);k__Fungi_unclassified(100);k__Fungi_unclassified(100);k__Fungi_unclassified(100);k__Fungi_unclassified(100);k__Fungi_unclassified(100);k__Fungi_unclassified(100); | - | - | - | - | - | - | - | Unassigned | - |
| Otu30606 | 1 | k__Fungi(100);k__Fungi_unclassified(100);k__Fungi_unclassified(100);k__Fungi_unclassified(100);k__Fungi_unclassified(100);k__Fungi_unclassified(100);k__Fungi_unclassified(100); | - | - | - | - | - | - | - | Unassigned | - |
| Otu30610 | 1 | k__Fungi(100);k__Fungi_unclassified(100);k__Fungi_unclassified(100);k__Fungi_unclassified(100);k__Fungi_unclassified(100);k__Fungi_unclassified(100);k__Fungi_unclassified(100); | - | - | - | - | - | - | - | Unassigned | - |
| Otu30614 | 1 | k__Fungi(100);p__Ascomycota(100);p__Ascomycota_unclassified(100);p__Ascomycota_unclassified(100);p__Ascomycota_unclassified(100);p__Ascomycota_unclassified(100);p__Ascomycota_unclassified(100); | - | - | - | - | - | - | - | Unassigned | - |
| Otu30615 | 1 | k__Fungi(100);p__Ascomycota(100);p__Ascomycota_unclassified(100);p__Ascomycota_unclassified(100);p__Ascomycota_unclassified(100);p__Ascomycota_unclassified(100);p__Ascomycota_unclassified(100); | - | - | - | - | - | - | - | Unassigned | - |
| Otu30616 | 1 | k__Fungi(100);p__Ascomycota(100);p__Ascomycota_unclassified(100);p__Ascomycota_unclassified(100);p__Ascomycota_unclassified(100);p__Ascomycota_unclassified(100);p__Ascomycota_unclassified(100); | - | - | - | - | - | - | - | Unassigned | - |
| Otu30618 | 1 | k__Fungi(100);p__Ascomycota(100);p__Ascomycota_unclassified(100);p__Ascomycota_unclassified(100);p__Ascomycota_unclassified(100);p__Ascomycota_unclassified(100);p__Ascomycota_unclassified(100); | - | - | - | - | - | - | - | Unassigned | - |
| Otu30619 | 1 | k__Fungi(100);k__Fungi_unclassified(100);k__Fungi_unclassified(100);k__Fungi_unclassified(100);k__Fungi_unclassified(100);k__Fungi_unclassified(100);k__Fungi_unclassified(100); | - | - | - | - | - | - | - | Unassigned | - |
| Otu30621 | 1 | k__Fungi(100);k__Fungi_unclassified(100);k__Fungi_unclassified(100);k__Fungi_unclassified(100);k__Fungi_unclassified(100);k__Fungi_unclassified(100);k__Fungi_unclassified(100); | - | - | - | - | - | - | - | Unassigned | - |
| Otu30622 | 1 | k__Fungi(100);p__Ascomycota(100);p__Ascomycota_unclassified(100);p__Ascomycota_unclassified(100);p__Ascomycota_unclassified(100);p__Ascomycota_unclassified(100);p__Ascomycota_unclassified(100); | - | - | - | - | - | - | - | Unassigned | - |
| Otu30623 | 1 | k__Fungi(100);p__Ascomycota(100);p__Ascomycota_unclassified(100);p__Ascomycota_unclassified(100);p__Ascomycota_unclassified(100);p__Ascomycota_unclassified(100);p__Ascomycota_unclassified(100); | - | - | - | - | - | - | - | Unassigned | - |
| Otu30624 | 1 | k__Fungi(100);k__Fungi_unclassified(100);k__Fungi_unclassified(100);k__Fungi_unclassified(100);k__Fungi_unclassified(100);k__Fungi_unclassified(100);k__Fungi_unclassified(100); | - | - | - | - | - | - | - | Unassigned | - |
| Otu30626 | 1 | k__Fungi(100);k__Fungi_unclassified(100);k__Fungi_unclassified(100);k__Fungi_unclassified(100);k__Fungi_unclassified(100);k__Fungi_unclassified(100);k__Fungi_unclassified(100); | - | - | - | - | - | - | - | Unassigned | - |
| Otu30628 | 1 | k__Fungi(100);p__Ascomycota(100);p__Ascomycota_unclassified(100);p__Ascomycota_unclassified(100);p__Ascomycota_unclassified(100);p__Ascomycota_unclassified(100);p__Ascomycota_unclassified(100); | - | - | - | - | - | - | - | Unassigned | - |
| Otu30634 | 1 | k__Fungi(100);p__Ascomycota(100);p__Ascomycota_unclassified(100);p__Ascomycota_unclassified(100);p__Ascomycota_unclassified(100);p__Ascomycota_unclassified(100);p__Ascomycota_unclassified(100); | - | - | - | - | - | - | - | Unassigned | - |
| Otu30638 | 1 | k__Fungi(100);p__Ascomycota(100);p__Ascomycota_unclassified(100);p__Ascomycota_unclassified(100);p__Ascomycota_unclassified(100);p__Ascomycota_unclassified(100);p__Ascomycota_unclassified(100); | - | - | - | - | - | - | - | Unassigned | - |
| Otu30650 | 1 | k__Fungi(100);k__Fungi_unclassified(100);k__Fungi_unclassified(100);k__Fungi_unclassified(100);k__Fungi_unclassified(100);k__Fungi_unclassified(100);k__Fungi_unclassified(100); | - | - | - | - | - | - | - | Unassigned | - |
| Otu30652 | 1 | k__Fungi(100);k__Fungi_unclassified(100);k__Fungi_unclassified(100);k__Fungi_unclassified(100);k__Fungi_unclassified(100);k__Fungi_unclassified(100);k__Fungi_unclassified(100); | - | - | - | - | - | - | - | Unassigned | - |
| Otu30659 | 1 | k__Fungi(100);k__Fungi_unclassified(100);k__Fungi_unclassified(100);k__Fungi_unclassified(100);k__Fungi_unclassified(100);k__Fungi_unclassified(100);k__Fungi_unclassified(100); | - | - | - | - | - | - | - | Unassigned | - |
| Otu30663 | 1 | k__Fungi(100);k__Fungi_unclassified(100);k__Fungi_unclassified(100);k__Fungi_unclassified(100);k__Fungi_unclassified(100);k__Fungi_unclassified(100);k__Fungi_unclassified(100); | - | - | - | - | - | - | - | Unassigned | - |
| Otu30664 | 1 | k__Fungi(100);k__Fungi_unclassified(100);k__Fungi_unclassified(100);k__Fungi_unclassified(100);k__Fungi_unclassified(100);k__Fungi_unclassified(100);k__Fungi_unclassified(100); | - | - | - | - | - | - | - | Unassigned | - |
| Otu30668 | 1 | k__Fungi(100);p__Ascomycota(100);p__Ascomycota_unclassified(100);p__Ascomycota_unclassified(100);p__Ascomycota_unclassified(100);p__Ascomycota_unclassified(100);p__Ascomycota_unclassified(100); | - | - | - | - | - | - | - | Unassigned | - |
| Otu30669 | 1 | k__Fungi(100);p__Ascomycota(100);p__Ascomycota_unclassified(100);p__Ascomycota_unclassified(100);p__Ascomycota_unclassified(100);p__Ascomycota_unclassified(100);p__Ascomycota_unclassified(100); | - | - | - | - | - | - | - | Unassigned | - |
| Otu30671 | 1 | k__Fungi(100);k__Fungi_unclassified(100);k__Fungi_unclassified(100);k__Fungi_unclassified(100);k__Fungi_unclassified(100);k__Fungi_unclassified(100);k__Fungi_unclassified(100); | - | - | - | - | - | - | - | Unassigned | - |
| Otu30672 | 1 | k__Fungi(100);k__Fungi_unclassified(100);k__Fungi_unclassified(100);k__Fungi_unclassified(100);k__Fungi_unclassified(100);k__Fungi_unclassified(100);k__Fungi_unclassified(100); | - | - | - | - | - | - | - | Unassigned | - |
| Otu30673 | 1 | k__Fungi(100);k__Fungi_unclassified(100);k__Fungi_unclassified(100);k__Fungi_unclassified(100);k__Fungi_unclassified(100);k__Fungi_unclassified(100);k__Fungi_unclassified(100); | - | - | - | - | - | - | - | Unassigned | - |
| Otu30674 | 1 | k__Fungi(100);k__Fungi_unclassified(100);k__Fungi_unclassified(100);k__Fungi_unclassified(100);k__Fungi_unclassified(100);k__Fungi_unclassified(100);k__Fungi_unclassified(100); | - | - | - | - | - | - | - | Unassigned | - |
| Otu30675 | 1 | k__Fungi(100);k__Fungi_unclassified(100);k__Fungi_unclassified(100);k__Fungi_unclassified(100);k__Fungi_unclassified(100);k__Fungi_unclassified(100);k__Fungi_unclassified(100); | - | - | - | - | - | - | - | Unassigned | - |
| Otu30676 | 1 | k__Fungi(100);p__Ascomycota(100);p__Ascomycota_unclassified(100);p__Ascomycota_unclassified(100);p__Ascomycota_unclassified(100);p__Ascomycota_unclassified(100);p__Ascomycota_unclassified(100); | - | - | - | - | - | - | - | Unassigned | - |
| Otu30678 | 1 | k__Fungi(100);k__Fungi_unclassified(100);k__Fungi_unclassified(100);k__Fungi_unclassified(100);k__Fungi_unclassified(100);k__Fungi_unclassified(100);k__Fungi_unclassified(100); | - | - | - | - | - | - | - | Unassigned | - |
| Otu30682 | 1 | k__Fungi(100);k__Fungi_unclassified(100);k__Fungi_unclassified(100);k__Fungi_unclassified(100);k__Fungi_unclassified(100);k__Fungi_unclassified(100);k__Fungi_unclassified(100); | - | - | - | - | - | - | - | Unassigned | - |
| Otu30683 | 1 | k__Fungi(100);p__Ascomycota(100);p__Ascomycota_unclassified(100);p__Ascomycota_unclassified(100);p__Ascomycota_unclassified(100);p__Ascomycota_unclassified(100);p__Ascomycota_unclassified(100); | - | - | - | - | - | - | - | Unassigned | - |
| Otu30687 | 1 | k__Fungi(100);k__Fungi_unclassified(100);k__Fungi_unclassified(100);k__Fungi_unclassified(100);k__Fungi_unclassified(100);k__Fungi_unclassified(100);k__Fungi_unclassified(100); | - | - | - | - | - | - | - | Unassigned | - |
| Otu30690 | 1 | k__Fungi(100);k__Fungi_unclassified(100);k__Fungi_unclassified(100);k__Fungi_unclassified(100);k__Fungi_unclassified(100);k__Fungi_unclassified(100);k__Fungi_unclassified(100); | - | - | - | - | - | - | - | Unassigned | - |
| Otu30691 | 1 | k__Fungi(100);p__Ascomycota(100);p__Ascomycota_unclassified(100);p__Ascomycota_unclassified(100);p__Ascomycota_unclassified(100);p__Ascomycota_unclassified(100);p__Ascomycota_unclassified(100); | - | - | - | - | - | - | - | Unassigned | - |
| Otu30692 | 1 | k__Fungi(100);k__Fungi_unclassified(100);k__Fungi_unclassified(100);k__Fungi_unclassified(100);k__Fungi_unclassified(100);k__Fungi_unclassified(100);k__Fungi_unclassified(100); | - | - | - | - | - | - | - | Unassigned | - |
| Otu30693 | 1 | k__Fungi(100);k__Fungi_unclassified(100);k__Fungi_unclassified(100);k__Fungi_unclassified(100);k__Fungi_unclassified(100);k__Fungi_unclassified(100);k__Fungi_unclassified(100); | - | - | - | - | - | - | - | Unassigned | - |
| Otu30695 | 1 | k__Fungi(100);k__Fungi_unclassified(100);k__Fungi_unclassified(100);k__Fungi_unclassified(100);k__Fungi_unclassified(100);k__Fungi_unclassified(100);k__Fungi_unclassified(100); | - | - | - | - | - | - | - | Unassigned | - |
| Otu30697 | 1 | k__Fungi(100);k__Fungi_unclassified(100);k__Fungi_unclassified(100);k__Fungi_unclassified(100);k__Fungi_unclassified(100);k__Fungi_unclassified(100);k__Fungi_unclassified(100); | - | - | - | - | - | - | - | Unassigned | - |
| Otu30698 | 1 | k__Fungi(100);p__Ascomycota(100);p__Ascomycota_unclassified(100);p__Ascomycota_unclassified(100);p__Ascomycota_unclassified(100);p__Ascomycota_unclassified(100);p__Ascomycota_unclassified(100); | - | - | - | - | - | - | - | Unassigned | - |
| Otu30699 | 1 | k__Fungi(100);p__Ascomycota(100);p__Ascomycota_unclassified(100);p__Ascomycota_unclassified(100);p__Ascomycota_unclassified(100);p__Ascomycota_unclassified(100);p__Ascomycota_unclassified(100); | - | - | - | - | - | - | - | Unassigned | - |
| Otu30700 | 1 | k__Fungi(100);p__Ascomycota(100);p__Ascomycota_unclassified(100);p__Ascomycota_unclassified(100);p__Ascomycota_unclassified(100);p__Ascomycota_unclassified(100);p__Ascomycota_unclassified(100); | - | - | - | - | - | - | - | Unassigned | - |
| Otu30701 | 1 | k__Fungi(100);p__Ascomycota(100);p__Ascomycota_unclassified(100);p__Ascomycota_unclassified(100);p__Ascomycota_unclassified(100);p__Ascomycota_unclassified(100);p__Ascomycota_unclassified(100); | - | - | - | - | - | - | - | Unassigned | - |
| Otu30702 | 1 | k__Fungi(100);p__Ascomycota(100);p__Ascomycota_unclassified(100);p__Ascomycota_unclassified(100);p__Ascomycota_unclassified(100);p__Ascomycota_unclassified(100);p__Ascomycota_unclassified(100); | - | - | - | - | - | - | - | Unassigned | - |
| Otu30703 | 1 | k__Fungi(100);k__Fungi_unclassified(100);k__Fungi_unclassified(100);k__Fungi_unclassified(100);k__Fungi_unclassified(100);k__Fungi_unclassified(100);k__Fungi_unclassified(100); | - | - | - | - | - | - | - | Unassigned | - |
| Otu30704 | 1 | k__Fungi(100);k__Fungi_unclassified(100);k__Fungi_unclassified(100);k__Fungi_unclassified(100);k__Fungi_unclassified(100);k__Fungi_unclassified(100);k__Fungi_unclassified(100); | - | - | - | - | - | - | - | Unassigned | - |
| Otu30705 | 1 | k__Fungi(100);p__Ascomycota(100);p__Ascomycota_unclassified(100);p__Ascomycota_unclassified(100);p__Ascomycota_unclassified(100);p__Ascomycota_unclassified(100);p__Ascomycota_unclassified(100); | - | - | - | - | - | - | - | Unassigned | - |
| Otu30707 | 1 | k__Fungi(100);k__Fungi_unclassified(100);k__Fungi_unclassified(100);k__Fungi_unclassified(100);k__Fungi_unclassified(100);k__Fungi_unclassified(100);k__Fungi_unclassified(100); | - | - | - | - | - | - | - | Unassigned | - |
| Otu30710 | 1 | k__Fungi(100);k__Fungi_unclassified(100);k__Fungi_unclassified(100);k__Fungi_unclassified(100);k__Fungi_unclassified(100);k__Fungi_unclassified(100);k__Fungi_unclassified(100); | - | - | - | - | - | - | - | Unassigned | - |
| Otu30713 | 1 | k__Fungi(100);k__Fungi_unclassified(100);k__Fungi_unclassified(100);k__Fungi_unclassified(100);k__Fungi_unclassified(100);k__Fungi_unclassified(100);k__Fungi_unclassified(100); | - | - | - | - | - | - | - | Unassigned | - |
| Otu30715 | 1 | k__Fungi(100);p__Ascomycota(100);p__Ascomycota_unclassified(100);p__Ascomycota_unclassified(100);p__Ascomycota_unclassified(100);p__Ascomycota_unclassified(100);p__Ascomycota_unclassified(100); | - | - | - | - | - | - | - | Unassigned | - |
| Otu30722 | 1 | k__Fungi(100);p__Ascomycota(100);p__Ascomycota_unclassified(100);p__Ascomycota_unclassified(100);p__Ascomycota_unclassified(100);p__Ascomycota_unclassified(100);p__Ascomycota_unclassified(100); | - | - | - | - | - | - | - | Unassigned | - |
| Otu30725 | 1 | k__Fungi(100);p__Ascomycota(100);p__Ascomycota_unclassified(100);p__Ascomycota_unclassified(100);p__Ascomycota_unclassified(100);p__Ascomycota_unclassified(100);p__Ascomycota_unclassified(100); | - | - | - | - | - | - | - | Unassigned | - |
| Otu30727 | 1 | k__Fungi(100);k__Fungi_unclassified(100);k__Fungi_unclassified(100);k__Fungi_unclassified(100);k__Fungi_unclassified(100);k__Fungi_unclassified(100);k__Fungi_unclassified(100); | - | - | - | - | - | - | - | Unassigned | - |
| Otu30728 | 1 | k__Fungi(100);k__Fungi_unclassified(100);k__Fungi_unclassified(100);k__Fungi_unclassified(100);k__Fungi_unclassified(100);k__Fungi_unclassified(100);k__Fungi_unclassified(100); | - | - | - | - | - | - | - | Unassigned | - |
| Otu30729 | 1 | k__Fungi(100);k__Fungi_unclassified(100);k__Fungi_unclassified(100);k__Fungi_unclassified(100);k__Fungi_unclassified(100);k__Fungi_unclassified(100);k__Fungi_unclassified(100); | - | - | - | - | - | - | - | Unassigned | - |
| Otu30730 | 1 | k__Fungi(100);p__Ascomycota(100);p__Ascomycota_unclassified(100);p__Ascomycota_unclassified(100);p__Ascomycota_unclassified(100);p__Ascomycota_unclassified(100);p__Ascomycota_unclassified(100); | - | - | - | - | - | - | - | Unassigned | - |
| Otu30740 | 1 | k__Fungi(100);p__Ascomycota(100);p__Ascomycota_unclassified(100);p__Ascomycota_unclassified(100);p__Ascomycota_unclassified(100);p__Ascomycota_unclassified(100);p__Ascomycota_unclassified(100); | - | - | - | - | - | - | - | Unassigned | - |
| Otu30741 | 1 | k__Fungi(100);p__Ascomycota(100);p__Ascomycota_unclassified(100);p__Ascomycota_unclassified(100);p__Ascomycota_unclassified(100);p__Ascomycota_unclassified(100);p__Ascomycota_unclassified(100); | - | - | - | - | - | - | - | Unassigned | - |
| Otu30744 | 1 | k__Fungi(100);k__Fungi_unclassified(100);k__Fungi_unclassified(100);k__Fungi_unclassified(100);k__Fungi_unclassified(100);k__Fungi_unclassified(100);k__Fungi_unclassified(100); | - | - | - | - | - | - | - | Unassigned | - |
| Otu30745 | 1 | k__Fungi(100);p__Ascomycota(100);p__Ascomycota_unclassified(100);p__Ascomycota_unclassified(100);p__Ascomycota_unclassified(100);p__Ascomycota_unclassified(100);p__Ascomycota_unclassified(100); | - | - | - | - | - | - | - | Unassigned | - |
| Otu30746 | 1 | k__Fungi(100);k__Fungi_unclassified(100);k__Fungi_unclassified(100);k__Fungi_unclassified(100);k__Fungi_unclassified(100);k__Fungi_unclassified(100);k__Fungi_unclassified(100); | - | - | - | - | - | - | - | Unassigned | - |
| Otu30747 | 1 | k__Fungi(100);k__Fungi_unclassified(100);k__Fungi_unclassified(100);k__Fungi_unclassified(100);k__Fungi_unclassified(100);k__Fungi_unclassified(100);k__Fungi_unclassified(100); | - | - | - | - | - | - | - | Unassigned | - |
| Otu30749 | 1 | k__Fungi(100);k__Fungi_unclassified(100);k__Fungi_unclassified(100);k__Fungi_unclassified(100);k__Fungi_unclassified(100);k__Fungi_unclassified(100);k__Fungi_unclassified(100); | - | - | - | - | - | - | - | Unassigned | - |
| Otu30757 | 1 | k__Fungi(100);p__Ascomycota(100);p__Ascomycota_unclassified(100);p__Ascomycota_unclassified(100);p__Ascomycota_unclassified(100);p__Ascomycota_unclassified(100);p__Ascomycota_unclassified(100); | - | - | - | - | - | - | - | Unassigned | - |
| Otu30760 | 1 | k__Fungi(100);k__Fungi_unclassified(100);k__Fungi_unclassified(100);k__Fungi_unclassified(100);k__Fungi_unclassified(100);k__Fungi_unclassified(100);k__Fungi_unclassified(100); | - | - | - | - | - | - | - | Unassigned | - |
| Otu30762 | 1 | k__Fungi(100);k__Fungi_unclassified(100);k__Fungi_unclassified(100);k__Fungi_unclassified(100);k__Fungi_unclassified(100);k__Fungi_unclassified(100);k__Fungi_unclassified(100); | - | - | - | - | - | - | - | Unassigned | - |
| Otu30770 | 1 | k__Fungi(100);k__Fungi_unclassified(100);k__Fungi_unclassified(100);k__Fungi_unclassified(100);k__Fungi_unclassified(100);k__Fungi_unclassified(100);k__Fungi_unclassified(100); | - | - | - | - | - | - | - | Unassigned | - |
| Otu30789 | 1 | k__Fungi(100);p__Ascomycota(100);p__Ascomycota_unclassified(100);p__Ascomycota_unclassified(100);p__Ascomycota_unclassified(100);p__Ascomycota_unclassified(100);p__Ascomycota_unclassified(100); | - | - | - | - | - | - | - | Unassigned | - |
| Otu30807 | 1 | k__Fungi(100);k__Fungi_unclassified(100);k__Fungi_unclassified(100);k__Fungi_unclassified(100);k__Fungi_unclassified(100);k__Fungi_unclassified(100);k__Fungi_unclassified(100); | - | - | - | - | - | - | - | Unassigned | - |
| Otu30809 | 1 | k__Fungi(100);p__Ascomycota(100);p__Ascomycota_unclassified(100);p__Ascomycota_unclassified(100);p__Ascomycota_unclassified(100);p__Ascomycota_unclassified(100);p__Ascomycota_unclassified(100); | - | - | - | - | - | - | - | Unassigned | - |
| Otu30810 | 1 | k__Fungi(100);p__Ascomycota(100);p__Ascomycota_unclassified(100);p__Ascomycota_unclassified(100);p__Ascomycota_unclassified(100);p__Ascomycota_unclassified(100);p__Ascomycota_unclassified(100); | - | - | - | - | - | - | - | Unassigned | - |
| Otu30813 | 1 | k__Fungi(100);k__Fungi_unclassified(100);k__Fungi_unclassified(100);k__Fungi_unclassified(100);k__Fungi_unclassified(100);k__Fungi_unclassified(100);k__Fungi_unclassified(100); | - | - | - | - | - | - | - | Unassigned | - |
| Otu30816 | 1 | k__Fungi(100);k__Fungi_unclassified(100);k__Fungi_unclassified(100);k__Fungi_unclassified(100);k__Fungi_unclassified(100);k__Fungi_unclassified(100);k__Fungi_unclassified(100); | - | - | - | - | - | - | - | Unassigned | - |
| Otu30837 | 1 | k__Fungi(100);k__Fungi_unclassified(100);k__Fungi_unclassified(100);k__Fungi_unclassified(100);k__Fungi_unclassified(100);k__Fungi_unclassified(100);k__Fungi_unclassified(100); | - | - | - | - | - | - | - | Unassigned | - |
| Otu30841 | 1 | k__Fungi(100);k__Fungi_unclassified(100);k__Fungi_unclassified(100);k__Fungi_unclassified(100);k__Fungi_unclassified(100);k__Fungi_unclassified(100);k__Fungi_unclassified(100); | - | - | - | - | - | - | - | Unassigned | - |
| Otu30885 | 1 | k__Fungi(100);k__Fungi_unclassified(100);k__Fungi_unclassified(100);k__Fungi_unclassified(100);k__Fungi_unclassified(100);k__Fungi_unclassified(100);k__Fungi_unclassified(100); | - | - | - | - | - | - | - | Unassigned | - |
| Otu30887 | 1 | k__Fungi(100);p__Ascomycota(100);p__Ascomycota_unclassified(100);p__Ascomycota_unclassified(100);p__Ascomycota_unclassified(100);p__Ascomycota_unclassified(100);p__Ascomycota_unclassified(100); | - | - | - | - | - | - | - | Unassigned | - |
| Otu30888 | 1 | k__Fungi(100);p__Ascomycota(100);p__Ascomycota_unclassified(100);p__Ascomycota_unclassified(100);p__Ascomycota_unclassified(100);p__Ascomycota_unclassified(100);p__Ascomycota_unclassified(100); | - | - | - | - | - | - | - | Unassigned | - |
| Otu30889 | 1 | k__Fungi(100);p__Ascomycota(100);p__Ascomycota_unclassified(100);p__Ascomycota_unclassified(100);p__Ascomycota_unclassified(100);p__Ascomycota_unclassified(100);p__Ascomycota_unclassified(100); | - | - | - | - | - | - | - | Unassigned | - |
| Otu30890 | 1 | k__Fungi(100);p__Ascomycota(100);p__Ascomycota_unclassified(100);p__Ascomycota_unclassified(100);p__Ascomycota_unclassified(100);p__Ascomycota_unclassified(100);p__Ascomycota_unclassified(100); | - | - | - | - | - | - | - | Unassigned | - |
| Otu30891 | 1 | k__Fungi(100);k__Fungi_unclassified(100);k__Fungi_unclassified(100);k__Fungi_unclassified(100);k__Fungi_unclassified(100);k__Fungi_unclassified(100);k__Fungi_unclassified(100); | - | - | - | - | - | - | - | Unassigned | - |
| Otu30893 | 1 | k__Fungi(100);k__Fungi_unclassified(100);k__Fungi_unclassified(100);k__Fungi_unclassified(100);k__Fungi_unclassified(100);k__Fungi_unclassified(100);k__Fungi_unclassified(100); | - | - | - | - | - | - | - | Unassigned | - |
| Otu30894 | 1 | k__Fungi(100);k__Fungi_unclassified(100);k__Fungi_unclassified(100);k__Fungi_unclassified(100);k__Fungi_unclassified(100);k__Fungi_unclassified(100);k__Fungi_unclassified(100); | - | - | - | - | - | - | - | Unassigned | - |
| Otu30897 | 1 | k__Fungi(100);p__Ascomycota(100);p__Ascomycota_unclassified(100);p__Ascomycota_unclassified(100);p__Ascomycota_unclassified(100);p__Ascomycota_unclassified(100);p__Ascomycota_unclassified(100); | - | - | - | - | - | - | - | Unassigned | - |
| Otu30903 | 1 | k__Fungi(100);k__Fungi_unclassified(100);k__Fungi_unclassified(100);k__Fungi_unclassified(100);k__Fungi_unclassified(100);k__Fungi_unclassified(100);k__Fungi_unclassified(100); | - | - | - | - | - | - | - | Unassigned | - |
| Otu30905 | 1 | k__Fungi(100);p__Ascomycota(100);p__Ascomycota_unclassified(100);p__Ascomycota_unclassified(100);p__Ascomycota_unclassified(100);p__Ascomycota_unclassified(100);p__Ascomycota_unclassified(100); | - | - | - | - | - | - | - | Unassigned | - |
| Otu30906 | 1 | k__Fungi(100);p__Ascomycota(100);p__Ascomycota_unclassified(100);p__Ascomycota_unclassified(100);p__Ascomycota_unclassified(100);p__Ascomycota_unclassified(100);p__Ascomycota_unclassified(100); | - | - | - | - | - | - | - | Unassigned | - |
| Otu30907 | 1 | k__Fungi(100);p__Ascomycota(100);p__Ascomycota_unclassified(100);p__Ascomycota_unclassified(100);p__Ascomycota_unclassified(100);p__Ascomycota_unclassified(100);p__Ascomycota_unclassified(100); | - | - | - | - | - | - | - | Unassigned | - |
| Otu30908 | 1 | k__Fungi(100);p__Ascomycota(100);p__Ascomycota_unclassified(100);p__Ascomycota_unclassified(100);p__Ascomycota_unclassified(100);p__Ascomycota_unclassified(100);p__Ascomycota_unclassified(100); | - | - | - | - | - | - | - | Unassigned | - |
| Otu30909 | 1 | k__Fungi(100);p__Ascomycota(100);p__Ascomycota_unclassified(100);p__Ascomycota_unclassified(100);p__Ascomycota_unclassified(100);p__Ascomycota_unclassified(100);p__Ascomycota_unclassified(100); | - | - | - | - | - | - | - | Unassigned | - |
| Otu30910 | 1 | k__Fungi(100);p__Ascomycota(100);p__Ascomycota_unclassified(100);p__Ascomycota_unclassified(100);p__Ascomycota_unclassified(100);p__Ascomycota_unclassified(100);p__Ascomycota_unclassified(100); | - | - | - | - | - | - | - | Unassigned | - |
| Otu30911 | 1 | k__Fungi(100);p__Ascomycota(100);p__Ascomycota_unclassified(100);p__Ascomycota_unclassified(100);p__Ascomycota_unclassified(100);p__Ascomycota_unclassified(100);p__Ascomycota_unclassified(100); | - | - | - | - | - | - | - | Unassigned | - |
| Otu30912 | 1 | k__Fungi(100);p__Ascomycota(100);p__Ascomycota_unclassified(100);p__Ascomycota_unclassified(100);p__Ascomycota_unclassified(100);p__Ascomycota_unclassified(100);p__Ascomycota_unclassified(100); | - | - | - | - | - | - | - | Unassigned | - |
| Otu30915 | 1 | k__Fungi(100);k__Fungi_unclassified(100);k__Fungi_unclassified(100);k__Fungi_unclassified(100);k__Fungi_unclassified(100);k__Fungi_unclassified(100);k__Fungi_unclassified(100); | - | - | - | - | - | - | - | Unassigned | - |
| Otu30916 | 1 | k__Fungi(100);k__Fungi_unclassified(100);k__Fungi_unclassified(100);k__Fungi_unclassified(100);k__Fungi_unclassified(100);k__Fungi_unclassified(100);k__Fungi_unclassified(100); | - | - | - | - | - | - | - | Unassigned | - |
| Otu30917 | 1 | k__Fungi(100);p__Ascomycota(100);p__Ascomycota_unclassified(100);p__Ascomycota_unclassified(100);p__Ascomycota_unclassified(100);p__Ascomycota_unclassified(100);p__Ascomycota_unclassified(100); | - | - | - | - | - | - | - | Unassigned | - |
| Otu30918 | 1 | k__Fungi(100);k__Fungi_unclassified(100);k__Fungi_unclassified(100);k__Fungi_unclassified(100);k__Fungi_unclassified(100);k__Fungi_unclassified(100);k__Fungi_unclassified(100); | - | - | - | - | - | - | - | Unassigned | - |
| Otu30920 | 1 | k__Fungi(100);p__Ascomycota(100);p__Ascomycota_unclassified(100);p__Ascomycota_unclassified(100);p__Ascomycota_unclassified(100);p__Ascomycota_unclassified(100);p__Ascomycota_unclassified(100); | - | - | - | - | - | - | - | Unassigned | - |
| Otu30921 | 1 | k__Fungi(100);k__Fungi_unclassified(100);k__Fungi_unclassified(100);k__Fungi_unclassified(100);k__Fungi_unclassified(100);k__Fungi_unclassified(100);k__Fungi_unclassified(100); | - | - | - | - | - | - | - | Unassigned | - |
| Otu30922 | 1 | k__Fungi(100);p__Ascomycota(100);p__Ascomycota_unclassified(100);p__Ascomycota_unclassified(100);p__Ascomycota_unclassified(100);p__Ascomycota_unclassified(100);p__Ascomycota_unclassified(100); | - | - | - | - | - | - | - | Unassigned | - |
| Otu30923 | 1 | k__Fungi(100);k__Fungi_unclassified(100);k__Fungi_unclassified(100);k__Fungi_unclassified(100);k__Fungi_unclassified(100);k__Fungi_unclassified(100);k__Fungi_unclassified(100); | - | - | - | - | - | - | - | Unassigned | - |
| Otu30924 | 1 | k__Fungi(100);p__Ascomycota(100);p__Ascomycota_unclassified(100);p__Ascomycota_unclassified(100);p__Ascomycota_unclassified(100);p__Ascomycota_unclassified(100);p__Ascomycota_unclassified(100); | - | - | - | - | - | - | - | Unassigned | - |
| Otu30926 | 1 | k__Fungi(100);p__Ascomycota(100);p__Ascomycota_unclassified(100);p__Ascomycota_unclassified(100);p__Ascomycota_unclassified(100);p__Ascomycota_unclassified(100);p__Ascomycota_unclassified(100); | - | - | - | - | - | - | - | Unassigned | - |
| Otu30927 | 1 | k__Fungi(100);k__Fungi_unclassified(100);k__Fungi_unclassified(100);k__Fungi_unclassified(100);k__Fungi_unclassified(100);k__Fungi_unclassified(100);k__Fungi_unclassified(100); | - | - | - | - | - | - | - | Unassigned | - |
| Otu30934 | 1 | k__Fungi(100);p__Ascomycota(100);p__Ascomycota_unclassified(100);p__Ascomycota_unclassified(100);p__Ascomycota_unclassified(100);p__Ascomycota_unclassified(100);p__Ascomycota_unclassified(100); | - | - | - | - | - | - | - | Unassigned | - |
| Otu30935 | 1 | k__Fungi(100);p__Ascomycota(100);c__Dothideomycetes(100);c__Dothideomycetes_unclassified(100);c__Dothideomycetes_unclassified(100);c__Dothideomycetes_unclassified(100);c__Dothideomycetes_unclassified(100); | - | - | - | - | - | - | - | Unassigned | - |
| Otu30938 | 1 | k__Fungi(100);p__Ascomycota(100);c__Dothideomycetes(100);o__Pleosporales(100);o__Pleosporales_unclassified(100);o__Pleosporales_unclassified(100);o__Pleosporales_unclassified(100); | - | - | - | - | - | - | - | Unassigned | - |
| Otu30948 | 1 | k__Fungi(100);p__Ascomycota(100);p__Ascomycota_unclassified(100);p__Ascomycota_unclassified(100);p__Ascomycota_unclassified(100);p__Ascomycota_unclassified(100);p__Ascomycota_unclassified(100); | - | - | - | - | - | - | - | Unassigned | - |
| Otu30958 | 1 | k__Fungi(100);k__Fungi_unclassified(100);k__Fungi_unclassified(100);k__Fungi_unclassified(100);k__Fungi_unclassified(100);k__Fungi_unclassified(100);k__Fungi_unclassified(100); | - | - | - | - | - | - | - | Unassigned | - |
| Otu30960 | 1 | k__Fungi(100);k__Fungi_unclassified(100);k__Fungi_unclassified(100);k__Fungi_unclassified(100);k__Fungi_unclassified(100);k__Fungi_unclassified(100);k__Fungi_unclassified(100); | - | - | - | - | - | - | - | Unassigned | - |
| Otu30970 | 1 | k__Fungi(100);k__Fungi_unclassified(100);k__Fungi_unclassified(100);k__Fungi_unclassified(100);k__Fungi_unclassified(100);k__Fungi_unclassified(100);k__Fungi_unclassified(100); | - | - | - | - | - | - | - | Unassigned | - |
| Otu30985 | 1 | k__Fungi(100);p__Ascomycota(100);p__Ascomycota_unclassified(100);p__Ascomycota_unclassified(100);p__Ascomycota_unclassified(100);p__Ascomycota_unclassified(100);p__Ascomycota_unclassified(100); | - | - | - | - | - | - | - | Unassigned | - |
| Otu30989 | 1 | k__Fungi(100);p__Ascomycota(100);p__Ascomycota_unclassified(100);p__Ascomycota_unclassified(100);p__Ascomycota_unclassified(100);p__Ascomycota_unclassified(100);p__Ascomycota_unclassified(100); | - | - | - | - | - | - | - | Unassigned | - |
| Otu30990 | 1 | k__Fungi(100);p__Ascomycota(100);p__Ascomycota_unclassified(100);p__Ascomycota_unclassified(100);p__Ascomycota_unclassified(100);p__Ascomycota_unclassified(100);p__Ascomycota_unclassified(100); | - | - | - | - | - | - | - | Unassigned | - |
| Otu30991 | 1 | k__Fungi(100);p__Ascomycota(100);p__Ascomycota_unclassified(100);p__Ascomycota_unclassified(100);p__Ascomycota_unclassified(100);p__Ascomycota_unclassified(100);p__Ascomycota_unclassified(100); | - | - | - | - | - | - | - | Unassigned | - |
| Otu30992 | 1 | k__Fungi(100);p__Ascomycota(100);p__Ascomycota_unclassified(100);p__Ascomycota_unclassified(100);p__Ascomycota_unclassified(100);p__Ascomycota_unclassified(100);p__Ascomycota_unclassified(100); | - | - | - | - | - | - | - | Unassigned | - |
| Otu30993 | 1 | k__Fungi(100);p__Ascomycota(100);p__Ascomycota_unclassified(100);p__Ascomycota_unclassified(100);p__Ascomycota_unclassified(100);p__Ascomycota_unclassified(100);p__Ascomycota_unclassified(100); | - | - | - | - | - | - | - | Unassigned | - |
| Otu30994 | 1 | k__Fungi(100);k__Fungi_unclassified(100);k__Fungi_unclassified(100);k__Fungi_unclassified(100);k__Fungi_unclassified(100);k__Fungi_unclassified(100);k__Fungi_unclassified(100); | - | - | - | - | - | - | - | Unassigned | - |
| Otu30995 | 1 | k__Fungi(100);k__Fungi_unclassified(100);k__Fungi_unclassified(100);k__Fungi_unclassified(100);k__Fungi_unclassified(100);k__Fungi_unclassified(100);k__Fungi_unclassified(100); | - | - | - | - | - | - | - | Unassigned | - |
| Otu30996 | 1 | k__Fungi(100);k__Fungi_unclassified(100);k__Fungi_unclassified(100);k__Fungi_unclassified(100);k__Fungi_unclassified(100);k__Fungi_unclassified(100);k__Fungi_unclassified(100); | - | - | - | - | - | - | - | Unassigned | - |
| Otu30997 | 1 | k__Fungi(100);k__Fungi_unclassified(100);k__Fungi_unclassified(100);k__Fungi_unclassified(100);k__Fungi_unclassified(100);k__Fungi_unclassified(100);k__Fungi_unclassified(100); | - | - | - | - | - | - | - | Unassigned | - |
| Otu30999 | 1 | k__Fungi(100);k__Fungi_unclassified(100);k__Fungi_unclassified(100);k__Fungi_unclassified(100);k__Fungi_unclassified(100);k__Fungi_unclassified(100);k__Fungi_unclassified(100); | - | - | - | - | - | - | - | Unassigned | - |
| Otu31000 | 1 | k__Fungi(100);k__Fungi_unclassified(100);k__Fungi_unclassified(100);k__Fungi_unclassified(100);k__Fungi_unclassified(100);k__Fungi_unclassified(100);k__Fungi_unclassified(100); | - | - | - | - | - | - | - | Unassigned | - |
| Otu31001 | 1 | k__Fungi(100);p__Ascomycota(100);p__Ascomycota_unclassified(100);p__Ascomycota_unclassified(100);p__Ascomycota_unclassified(100);p__Ascomycota_unclassified(100);p__Ascomycota_unclassified(100); | - | - | - | - | - | - | - | Unassigned | - |
| Otu31002 | 1 | k__Fungi(100);p__Ascomycota(100);p__Ascomycota_unclassified(100);p__Ascomycota_unclassified(100);p__Ascomycota_unclassified(100);p__Ascomycota_unclassified(100);p__Ascomycota_unclassified(100); | - | - | - | - | - | - | - | Unassigned | - |
| Otu31003 | 1 | k__Fungi(100);k__Fungi_unclassified(100);k__Fungi_unclassified(100);k__Fungi_unclassified(100);k__Fungi_unclassified(100);k__Fungi_unclassified(100);k__Fungi_unclassified(100); | - | - | - | - | - | - | - | Unassigned | - |
| Otu31004 | 1 | k__Fungi(100);k__Fungi_unclassified(100);k__Fungi_unclassified(100);k__Fungi_unclassified(100);k__Fungi_unclassified(100);k__Fungi_unclassified(100);k__Fungi_unclassified(100); | - | - | - | - | - | - | - | Unassigned | - |
| Otu31005 | 1 | k__Fungi(100);k__Fungi_unclassified(100);k__Fungi_unclassified(100);k__Fungi_unclassified(100);k__Fungi_unclassified(100);k__Fungi_unclassified(100);k__Fungi_unclassified(100); | - | - | - | - | - | - | - | Unassigned | - |
| Otu31006 | 1 | k__Fungi(100);p__Ascomycota(100);p__Ascomycota_unclassified(100);p__Ascomycota_unclassified(100);p__Ascomycota_unclassified(100);p__Ascomycota_unclassified(100);p__Ascomycota_unclassified(100); | - | - | - | - | - | - | - | Unassigned | - |
| Otu31007 | 1 | k__Fungi(100);k__Fungi_unclassified(100);k__Fungi_unclassified(100);k__Fungi_unclassified(100);k__Fungi_unclassified(100);k__Fungi_unclassified(100);k__Fungi_unclassified(100); | - | - | - | - | - | - | - | Unassigned | - |
| Otu31010 | 1 | k__Fungi(100);p__Ascomycota(100);p__Ascomycota_unclassified(100);p__Ascomycota_unclassified(100);p__Ascomycota_unclassified(100);p__Ascomycota_unclassified(100);p__Ascomycota_unclassified(100); | - | - | - | - | - | - | - | Unassigned | - |
| Otu31013 | 1 | k__Fungi(100);p__Ascomycota(100);p__Ascomycota_unclassified(100);p__Ascomycota_unclassified(100);p__Ascomycota_unclassified(100);p__Ascomycota_unclassified(100);p__Ascomycota_unclassified(100); | - | - | - | - | - | - | - | Unassigned | - |
| Otu31024 | 1 | k__Fungi(100);p__Ascomycota(100);p__Ascomycota_unclassified(100);p__Ascomycota_unclassified(100);p__Ascomycota_unclassified(100);p__Ascomycota_unclassified(100);p__Ascomycota_unclassified(100); | - | - | - | - | - | - | - | Unassigned | - |
| Otu31025 | 1 | k__Fungi(100);p__Ascomycota(100);p__Ascomycota_unclassified(100);p__Ascomycota_unclassified(100);p__Ascomycota_unclassified(100);p__Ascomycota_unclassified(100);p__Ascomycota_unclassified(100); | - | - | - | - | - | - | - | Unassigned | - |
| Otu31026 | 1 | k__Fungi(100);k__Fungi_unclassified(100);k__Fungi_unclassified(100);k__Fungi_unclassified(100);k__Fungi_unclassified(100);k__Fungi_unclassified(100);k__Fungi_unclassified(100); | - | - | - | - | - | - | - | Unassigned | - |
| Otu31027 | 1 | k__Fungi(100);k__Fungi_unclassified(100);k__Fungi_unclassified(100);k__Fungi_unclassified(100);k__Fungi_unclassified(100);k__Fungi_unclassified(100);k__Fungi_unclassified(100); | - | - | - | - | - | - | - | Unassigned | - |
| Otu31032 | 1 | k__Fungi(100);k__Fungi_unclassified(100);k__Fungi_unclassified(100);k__Fungi_unclassified(100);k__Fungi_unclassified(100);k__Fungi_unclassified(100);k__Fungi_unclassified(100); | - | - | - | - | - | - | - | Unassigned | - |
| Otu31045 | 1 | k__Fungi(100);k__Fungi_unclassified(100);k__Fungi_unclassified(100);k__Fungi_unclassified(100);k__Fungi_unclassified(100);k__Fungi_unclassified(100);k__Fungi_unclassified(100); | - | - | - | - | - | - | - | Unassigned | - |
| Otu31064 | 1 | k__Fungi(100);p__Ascomycota(100);p__Ascomycota_unclassified(100);p__Ascomycota_unclassified(100);p__Ascomycota_unclassified(100);p__Ascomycota_unclassified(100);p__Ascomycota_unclassified(100); | - | - | - | - | - | - | - | Unassigned | - |
| Otu31065 | 1 | k__Fungi(100);p__Ascomycota(100);p__Ascomycota_unclassified(100);p__Ascomycota_unclassified(100);p__Ascomycota_unclassified(100);p__Ascomycota_unclassified(100);p__Ascomycota_unclassified(100); | - | - | - | - | - | - | - | Unassigned | - |
| Otu31067 | 1 | k__Fungi(100);k__Fungi_unclassified(100);k__Fungi_unclassified(100);k__Fungi_unclassified(100);k__Fungi_unclassified(100);k__Fungi_unclassified(100);k__Fungi_unclassified(100); | - | - | - | - | - | - | - | Unassigned | - |
| Otu31070 | 1 | k__Fungi(100);k__Fungi_unclassified(100);k__Fungi_unclassified(100);k__Fungi_unclassified(100);k__Fungi_unclassified(100);k__Fungi_unclassified(100);k__Fungi_unclassified(100); | - | - | - | - | - | - | - | Unassigned | - |
| Otu31071 | 1 | k__Fungi(100);k__Fungi_unclassified(100);k__Fungi_unclassified(100);k__Fungi_unclassified(100);k__Fungi_unclassified(100);k__Fungi_unclassified(100);k__Fungi_unclassified(100); | - | - | - | - | - | - | - | Unassigned | - |
| Otu31072 | 1 | k__Fungi(100);p__Ascomycota(100);p__Ascomycota_unclassified(100);p__Ascomycota_unclassified(100);p__Ascomycota_unclassified(100);p__Ascomycota_unclassified(100);p__Ascomycota_unclassified(100); | - | - | - | - | - | - | - | Unassigned | - |
| Otu31073 | 1 | k__Fungi(100);k__Fungi_unclassified(100);k__Fungi_unclassified(100);k__Fungi_unclassified(100);k__Fungi_unclassified(100);k__Fungi_unclassified(100);k__Fungi_unclassified(100); | - | - | - | - | - | - | - | Unassigned | - |
| Otu31074 | 1 | k__Fungi(100);p__Ascomycota(100);p__Ascomycota_unclassified(100);p__Ascomycota_unclassified(100);p__Ascomycota_unclassified(100);p__Ascomycota_unclassified(100);p__Ascomycota_unclassified(100); | - | - | - | - | - | - | - | Unassigned | - |
| Otu31075 | 1 | k__Fungi(100);k__Fungi_unclassified(100);k__Fungi_unclassified(100);k__Fungi_unclassified(100);k__Fungi_unclassified(100);k__Fungi_unclassified(100);k__Fungi_unclassified(100); | - | - | - | - | - | - | - | Unassigned | - |
| Otu31076 | 1 | k__Fungi(100);p__Ascomycota(100);p__Ascomycota_unclassified(100);p__Ascomycota_unclassified(100);p__Ascomycota_unclassified(100);p__Ascomycota_unclassified(100);p__Ascomycota_unclassified(100); | - | - | - | - | - | - | - | Unassigned | - |
| Otu31078 | 1 | k__Fungi(100);k__Fungi_unclassified(100);k__Fungi_unclassified(100);k__Fungi_unclassified(100);k__Fungi_unclassified(100);k__Fungi_unclassified(100);k__Fungi_unclassified(100); | - | - | - | - | - | - | - | Unassigned | - |
| Otu31081 | 1 | k__Fungi(100);p__Ascomycota(100);p__Ascomycota_unclassified(100);p__Ascomycota_unclassified(100);p__Ascomycota_unclassified(100);p__Ascomycota_unclassified(100);p__Ascomycota_unclassified(100); | - | - | - | - | - | - | - | Unassigned | - |
| Otu31088 | 1 | k__Fungi(100);k__Fungi_unclassified(100);k__Fungi_unclassified(100);k__Fungi_unclassified(100);k__Fungi_unclassified(100);k__Fungi_unclassified(100);k__Fungi_unclassified(100); | - | - | - | - | - | - | - | Unassigned | - |
| Otu31089 | 1 | k__Fungi(100);k__Fungi_unclassified(100);k__Fungi_unclassified(100);k__Fungi_unclassified(100);k__Fungi_unclassified(100);k__Fungi_unclassified(100);k__Fungi_unclassified(100); | - | - | - | - | - | - | - | Unassigned | - |
| Otu31090 | 1 | k__Fungi(100);p__Ascomycota(100);p__Ascomycota_unclassified(100);p__Ascomycota_unclassified(100);p__Ascomycota_unclassified(100);p__Ascomycota_unclassified(100);p__Ascomycota_unclassified(100); | - | - | - | - | - | - | - | Unassigned | - |
| Otu31096 | 1 | k__Fungi(100);k__Fungi_unclassified(100);k__Fungi_unclassified(100);k__Fungi_unclassified(100);k__Fungi_unclassified(100);k__Fungi_unclassified(100);k__Fungi_unclassified(100); | - | - | - | - | - | - | - | Unassigned | - |
| Otu31097 | 1 | k__Fungi(100);p__Ascomycota(100);p__Ascomycota_unclassified(100);p__Ascomycota_unclassified(100);p__Ascomycota_unclassified(100);p__Ascomycota_unclassified(100);p__Ascomycota_unclassified(100); | - | - | - | - | - | - | - | Unassigned | - |
| Otu31098 | 1 | k__Fungi(100);p__Ascomycota(100);p__Ascomycota_unclassified(100);p__Ascomycota_unclassified(100);p__Ascomycota_unclassified(100);p__Ascomycota_unclassified(100);p__Ascomycota_unclassified(100); | - | - | - | - | - | - | - | Unassigned | - |
| Otu31101 | 1 | k__Fungi(100);k__Fungi_unclassified(100);k__Fungi_unclassified(100);k__Fungi_unclassified(100);k__Fungi_unclassified(100);k__Fungi_unclassified(100);k__Fungi_unclassified(100); | - | - | - | - | - | - | - | Unassigned | - |
| Otu31102 | 1 | k__Fungi(100);k__Fungi_unclassified(100);k__Fungi_unclassified(100);k__Fungi_unclassified(100);k__Fungi_unclassified(100);k__Fungi_unclassified(100);k__Fungi_unclassified(100); | - | - | - | - | - | - | - | Unassigned | - |
| Otu31103 | 1 | k__Fungi(100);p__Ascomycota(100);p__Ascomycota_unclassified(100);p__Ascomycota_unclassified(100);p__Ascomycota_unclassified(100);p__Ascomycota_unclassified(100);p__Ascomycota_unclassified(100); | - | - | - | - | - | - | - | Unassigned | - |
| Otu31105 | 1 | k__Fungi(100);p__Ascomycota(100);p__Ascomycota_unclassified(100);p__Ascomycota_unclassified(100);p__Ascomycota_unclassified(100);p__Ascomycota_unclassified(100);p__Ascomycota_unclassified(100); | - | - | - | - | - | - | - | Unassigned | - |
| Otu31106 | 1 | k__Fungi(100);p__Ascomycota(100);p__Ascomycota_unclassified(100);p__Ascomycota_unclassified(100);p__Ascomycota_unclassified(100);p__Ascomycota_unclassified(100);p__Ascomycota_unclassified(100); | - | - | - | - | - | - | - | Unassigned | - |
| Otu31107 | 1 | k__Fungi(100);p__Ascomycota(100);p__Ascomycota_unclassified(100);p__Ascomycota_unclassified(100);p__Ascomycota_unclassified(100);p__Ascomycota_unclassified(100);p__Ascomycota_unclassified(100); | - | - | - | - | - | - | - | Unassigned | - |
| Otu31108 | 1 | k__Fungi(100);p__Ascomycota(100);p__Ascomycota_unclassified(100);p__Ascomycota_unclassified(100);p__Ascomycota_unclassified(100);p__Ascomycota_unclassified(100);p__Ascomycota_unclassified(100); | - | - | - | - | - | - | - | Unassigned | - |
| Otu31109 | 1 | k__Fungi(100);p__Ascomycota(100);p__Ascomycota_unclassified(100);p__Ascomycota_unclassified(100);p__Ascomycota_unclassified(100);p__Ascomycota_unclassified(100);p__Ascomycota_unclassified(100); | - | - | - | - | - | - | - | Unassigned | - |
| Otu31111 | 1 | k__Fungi(100);p__Ascomycota(100);p__Ascomycota_unclassified(100);p__Ascomycota_unclassified(100);p__Ascomycota_unclassified(100);p__Ascomycota_unclassified(100);p__Ascomycota_unclassified(100); | - | - | - | - | - | - | - | Unassigned | - |
| Otu31112 | 1 | k__Fungi(100);k__Fungi_unclassified(100);k__Fungi_unclassified(100);k__Fungi_unclassified(100);k__Fungi_unclassified(100);k__Fungi_unclassified(100);k__Fungi_unclassified(100); | - | - | - | - | - | - | - | Unassigned | - |
| Otu31113 | 1 | k__Fungi(100);p__Ascomycota(100);p__Ascomycota_unclassified(100);p__Ascomycota_unclassified(100);p__Ascomycota_unclassified(100);p__Ascomycota_unclassified(100);p__Ascomycota_unclassified(100); | - | - | - | - | - | - | - | Unassigned | - |
| Otu31114 | 1 | k__Fungi(100);k__Fungi_unclassified(100);k__Fungi_unclassified(100);k__Fungi_unclassified(100);k__Fungi_unclassified(100);k__Fungi_unclassified(100);k__Fungi_unclassified(100); | - | - | - | - | - | - | - | Unassigned | - |
| Otu31115 | 1 | k__Fungi(100);p__Ascomycota(100);p__Ascomycota_unclassified(100);p__Ascomycota_unclassified(100);p__Ascomycota_unclassified(100);p__Ascomycota_unclassified(100);p__Ascomycota_unclassified(100); | - | - | - | - | - | - | - | Unassigned | - |
| Otu31148 | 1 | k__Fungi(100);p__Ascomycota(100);p__Ascomycota_unclassified(100);p__Ascomycota_unclassified(100);p__Ascomycota_unclassified(100);p__Ascomycota_unclassified(100);p__Ascomycota_unclassified(100); | - | - | - | - | - | - | - | Unassigned | - |
| Otu31149 | 1 | k__Fungi(100);p__Ascomycota(100);p__Ascomycota_unclassified(100);p__Ascomycota_unclassified(100);p__Ascomycota_unclassified(100);p__Ascomycota_unclassified(100);p__Ascomycota_unclassified(100); | - | - | - | - | - | - | - | Unassigned | - |
| Otu31151 | 1 | k__Fungi(100);p__Ascomycota(100);p__Ascomycota_unclassified(100);p__Ascomycota_unclassified(100);p__Ascomycota_unclassified(100);p__Ascomycota_unclassified(100);p__Ascomycota_unclassified(100); | - | - | - | - | - | - | - | Unassigned | - |

| | | | | | | | | | | | |
|---|---|---|---|---|---|---|---|---|---|---|---|
| Otu31152 | 1 | k__Fungi(100);k__Fungi_unclassified(100);k__Fungi_unclassified(100);k__Fungi_unclassified(100);k__Fungi_unclassified(100);k__Fungi_unclassified(100);k__Fungi_unclassified(100); | - | - | - | - | - | - | - | Unassigned | - |
| Otu31153 | 1 | k__Fungi(100);p__Ascomycota(100);p__Ascomycota_unclassified(100);p__Ascomycota_unclassified(100);p__Ascomycota_unclassified(100);p__Ascomycota_unclassified(100);p__Ascomycota_unclassified(100); | - | - | - | - | - | - | - | Unassigned | - |
| Otu31158 | 1 | k__Fungi(100);k__Fungi_unclassified(100);k__Fungi_unclassified(100);k__Fungi_unclassified(100);k__Fungi_unclassified(100);k__Fungi_unclassified(100);k__Fungi_unclassified(100); | - | - | - | - | - | - | - | Unassigned | - |
| Otu31161 | 1 | k__Fungi(100);p__Ascomycota(100);p__Ascomycota_unclassified(100);p__Ascomycota_unclassified(100);p__Ascomycota_unclassified(100);p__Ascomycota_unclassified(100);p__Ascomycota_unclassified(100); | - | - | - | - | - | - | - | Unassigned | - |
| Otu31167 | 1 | k__Fungi(100);k__Fungi_unclassified(100);k__Fungi_unclassified(100);k__Fungi_unclassified(100);k__Fungi_unclassified(100);k__Fungi_unclassified(100);k__Fungi_unclassified(100); | - | - | - | - | - | - | - | Unassigned | - |
| Otu31170 | 1 | k__Fungi(100);k__Fungi_unclassified(100);k__Fungi_unclassified(100);k__Fungi_unclassified(100);k__Fungi_unclassified(100);k__Fungi_unclassified(100);k__Fungi_unclassified(100); | - | - | - | - | - | - | - | Unassigned | - |
| Otu31173 | 1 | k__Fungi(100);p__Ascomycota(100);p__Ascomycota_unclassified(100);p__Ascomycota_unclassified(100);p__Ascomycota_unclassified(100);p__Ascomycota_unclassified(100);p__Ascomycota_unclassified(100); | - | - | - | - | - | - | - | Unassigned | - |
| Otu31174 | 1 | k__Fungi(100);p__Ascomycota(100);p__Ascomycota_unclassified(100);p__Ascomycota_unclassified(100);p__Ascomycota_unclassified(100);p__Ascomycota_unclassified(100);p__Ascomycota_unclassified(100); | - | - | - | - | - | - | - | Unassigned | - |
| Otu31175 | 1 | k__Fungi(100);k__Fungi_unclassified(100);k__Fungi_unclassified(100);k__Fungi_unclassified(100);k__Fungi_unclassified(100);k__Fungi_unclassified(100);k__Fungi_unclassified(100); | - | - | - | - | - | - | - | Unassigned | - |
| Otu31178 | 1 | k__Fungi(100);k__Fungi_unclassified(100);k__Fungi_unclassified(100);k__Fungi_unclassified(100);k__Fungi_unclassified(100);k__Fungi_unclassified(100);k__Fungi_unclassified(100); | - | - | - | - | - | - | - | Unassigned | - |
| Otu31180 | 1 | k__Fungi(100);p__Ascomycota(100);p__Ascomycota_unclassified(100);p__Ascomycota_unclassified(100);p__Ascomycota_unclassified(100);p__Ascomycota_unclassified(100);p__Ascomycota_unclassified(100); | - | - | - | - | - | - | - | Unassigned | - |
| Otu31182 | 1 | k__Fungi(100);p__Ascomycota(100);p__Ascomycota_unclassified(100);p__Ascomycota_unclassified(100);p__Ascomycota_unclassified(100);p__Ascomycota_unclassified(100);p__Ascomycota_unclassified(100); | - | - | - | - | - | - | - | Unassigned | - |
| Otu31184 | 1 | k__Fungi(100);p__Ascomycota(100);p__Ascomycota_unclassified(100);p__Ascomycota_unclassified(100);p__Ascomycota_unclassified(100);p__Ascomycota_unclassified(100);p__Ascomycota_unclassified(100); | - | - | - | - | - | - | - | Unassigned | - |
| Otu31185 | 1 | k__Fungi(100);p__Ascomycota(100);p__Ascomycota_unclassified(100);p__Ascomycota_unclassified(100);p__Ascomycota_unclassified(100);p__Ascomycota_unclassified(100);p__Ascomycota_unclassified(100); | - | - | - | - | - | - | - | Unassigned | - |
| Otu31186 | 1 | k__Fungi(100);p__Ascomycota(100);p__Ascomycota_unclassified(100);p__Ascomycota_unclassified(100);p__Ascomycota_unclassified(100);p__Ascomycota_unclassified(100);p__Ascomycota_unclassified(100); | - | - | - | - | - | - | - | Unassigned | - |
| Otu31188 | 1 | k__Fungi(100);p__Ascomycota(100);p__Ascomycota_unclassified(100);p__Ascomycota_unclassified(100);p__Ascomycota_unclassified(100);p__Ascomycota_unclassified(100);p__Ascomycota_unclassified(100); | - | - | - | - | - | - | - | Unassigned | - |
| Otu31192 | 1 | k__Fungi(100);p__Ascomycota(100);p__Ascomycota_unclassified(100);p__Ascomycota_unclassified(100);p__Ascomycota_unclassified(100);p__Ascomycota_unclassified(100);p__Ascomycota_unclassified(100); | - | - | - | - | - | - | - | Unassigned | - |
| Otu31200 | 1 | k__Fungi(100);p__Ascomycota(100);p__Ascomycota_unclassified(100);p__Ascomycota_unclassified(100);p__Ascomycota_unclassified(100);p__Ascomycota_unclassified(100);p__Ascomycota_unclassified(100); | - | - | - | - | - | - | - | Unassigned | - |
| Otu31215 | 1 | k__Fungi(100);k__Fungi_unclassified(100);k__Fungi_unclassified(100);k__Fungi_unclassified(100);k__Fungi_unclassified(100);k__Fungi_unclassified(100);k__Fungi_unclassified(100); | - | - | - | - | - | - | - | Unassigned | - |
| Otu31216 | 1 | k__Fungi(100);k__Fungi_unclassified(100);k__Fungi_unclassified(100);k__Fungi_unclassified(100);k__Fungi_unclassified(100);k__Fungi_unclassified(100);k__Fungi_unclassified(100); | - | - | - | - | - | - | - | Unassigned | - |
| Otu31218 | 1 | k__Fungi(100);k__Fungi_unclassified(100);k__Fungi_unclassified(100);k__Fungi_unclassified(100);k__Fungi_unclassified(100);k__Fungi_unclassified(100);k__Fungi_unclassified(100); | - | - | - | - | - | - | - | Unassigned | - |
| Otu31219 | 1 | k__Fungi(100);k__Fungi_unclassified(100);k__Fungi_unclassified(100);k__Fungi_unclassified(100);k__Fungi_unclassified(100);k__Fungi_unclassified(100);k__Fungi_unclassified(100); | - | - | - | - | - | - | - | Unassigned | - |
| Otu31220 | 1 | k__Fungi(100);k__Fungi_unclassified(100);k__Fungi_unclassified(100);k__Fungi_unclassified(100);k__Fungi_unclassified(100);k__Fungi_unclassified(100);k__Fungi_unclassified(100); | - | - | - | - | - | - | - | Unassigned | - |
| Otu31221 | 1 | k__Fungi(100);p__Ascomycota(100);p__Ascomycota_unclassified(100);p__Ascomycota_unclassified(100);p__Ascomycota_unclassified(100);p__Ascomycota_unclassified(100);p__Ascomycota_unclassified(100); | - | - | - | - | - | - | - | Unassigned | - |
| Otu31222 | 1 | k__Fungi(100);k__Fungi_unclassified(100);k__Fungi_unclassified(100);k__Fungi_unclassified(100);k__Fungi_unclassified(100);k__Fungi_unclassified(100);k__Fungi_unclassified(100); | - | - | - | - | - | - | - | Unassigned | - |
| Otu31227 | 1 | k__Fungi(100);k__Fungi_unclassified(100);k__Fungi_unclassified(100);k__Fungi_unclassified(100);k__Fungi_unclassified(100);k__Fungi_unclassified(100);k__Fungi_unclassified(100); | - | - | - | - | - | - | - | Unassigned | - |
| Otu31237 | 1 | k__Fungi(100);k__Fungi_unclassified(100);k__Fungi_unclassified(100);k__Fungi_unclassified(100);k__Fungi_unclassified(100);k__Fungi_unclassified(100);k__Fungi_unclassified(100); | - | - | - | - | - | - | - | Unassigned | - |
| Otu31240 | 1 | k__Fungi(100);k__Fungi_unclassified(100);k__Fungi_unclassified(100);k__Fungi_unclassified(100);k__Fungi_unclassified(100);k__Fungi_unclassified(100);k__Fungi_unclassified(100); | - | - | - | - | - | - | - | Unassigned | - |
| Otu31242 | 1 | k__Fungi(100);p__Ascomycota(100);p__Ascomycota_unclassified(100);p__Ascomycota_unclassified(100);p__Ascomycota_unclassified(100);p__Ascomycota_unclassified(100);p__Ascomycota_unclassified(100); | - | - | - | - | - | - | - | Unassigned | - |
| Otu31243 | 1 | k__Fungi(100);k__Fungi_unclassified(100);k__Fungi_unclassified(100);k__Fungi_unclassified(100);k__Fungi_unclassified(100);k__Fungi_unclassified(100);k__Fungi_unclassified(100); | - | - | - | - | - | - | - | Unassigned | - |
| Otu31247 | 1 | k__Fungi(100);k__Fungi_unclassified(100);k__Fungi_unclassified(100);k__Fungi_unclassified(100);k__Fungi_unclassified(100);k__Fungi_unclassified(100);k__Fungi_unclassified(100); | - | - | - | - | - | - | - | Unassigned | - |
| Otu31248 | 1 | k__Fungi(100);p__Ascomycota(100);p__Ascomycota_unclassified(100);p__Ascomycota_unclassified(100);p__Ascomycota_unclassified(100);p__Ascomycota_unclassified(100);p__Ascomycota_unclassified(100); | - | - | - | - | - | - | - | Unassigned | - |
| Otu31261 | 1 | k__Fungi(100);k__Fungi_unclassified(100);k__Fungi_unclassified(100);k__Fungi_unclassified(100);k__Fungi_unclassified(100);k__Fungi_unclassified(100);k__Fungi_unclassified(100); | - | - | - | - | - | - | - | Unassigned | - |
| Otu31277 | 1 | k__Fungi(100);k__Fungi_unclassified(100);k__Fungi_unclassified(100);k__Fungi_unclassified(100);k__Fungi_unclassified(100);k__Fungi_unclassified(100);k__Fungi_unclassified(100); | - | - | - | - | - | - | - | Unassigned | - |
| Otu31279 | 1 | k__Fungi(100);k__Fungi_unclassified(100);k__Fungi_unclassified(100);k__Fungi_unclassified(100);k__Fungi_unclassified(100);k__Fungi_unclassified(100);k__Fungi_unclassified(100); | - | - | - | - | - | - | - | Unassigned | - |
| Otu31280 | 1 | k__Fungi(100);k__Fungi_unclassified(100);k__Fungi_unclassified(100);k__Fungi_unclassified(100);k__Fungi_unclassified(100);k__Fungi_unclassified(100);k__Fungi_unclassified(100); | - | - | - | - | - | - | - | Unassigned | - |
| Otu31281 | 1 | k__Fungi(100);k__Fungi_unclassified(100);k__Fungi_unclassified(100);k__Fungi_unclassified(100);k__Fungi_unclassified(100);k__Fungi_unclassified(100);k__Fungi_unclassified(100); | - | - | - | - | - | - | - | Unassigned | - |
| Otu31284 | 1 | k__Fungi(100);k__Fungi_unclassified(100);k__Fungi_unclassified(100);k__Fungi_unclassified(100);k__Fungi_unclassified(100);k__Fungi_unclassified(100);k__Fungi_unclassified(100); | - | - | - | - | - | - | - | Unassigned | - |
| Otu31285 | 1 | k__Fungi(100);p__Ascomycota(100);p__Ascomycota_unclassified(100);p__Ascomycota_unclassified(100);p__Ascomycota_unclassified(100);p__Ascomycota_unclassified(100);p__Ascomycota_unclassified(100); | - | - | - | - | - | - | - | Unassigned | - |
| Otu31286 | 1 | k__Fungi(100);k__Fungi_unclassified(100);k__Fungi_unclassified(100);k__Fungi_unclassified(100);k__Fungi_unclassified(100);k__Fungi_unclassified(100);k__Fungi_unclassified(100); | - | - | - | - | - | - | - | Unassigned | - |
| Otu31287 | 1 | k__Fungi(100);k__Fungi_unclassified(100);k__Fungi_unclassified(100);k__Fungi_unclassified(100);k__Fungi_unclassified(100);k__Fungi_unclassified(100);k__Fungi_unclassified(100); | - | - | - | - | - | - | - | Unassigned | - |
| Otu31288 | 1 | k__Fungi(100);k__Fungi_unclassified(100);k__Fungi_unclassified(100);k__Fungi_unclassified(100);k__Fungi_unclassified(100);k__Fungi_unclassified(100);k__Fungi_unclassified(100); | - | - | - | - | - | - | - | Unassigned | - |
| Otu31289 | 1 | k__Fungi(100);k__Fungi_unclassified(100);k__Fungi_unclassified(100);k__Fungi_unclassified(100);k__Fungi_unclassified(100);k__Fungi_unclassified(100);k__Fungi_unclassified(100); | - | - | - | - | - | - | - | Unassigned | - |
| Otu31291 | 1 | k__Fungi(100);k__Fungi_unclassified(100);k__Fungi_unclassified(100);k__Fungi_unclassified(100);k__Fungi_unclassified(100);k__Fungi_unclassified(100);k__Fungi_unclassified(100); | - | - | - | - | - | - | - | Unassigned | - |
| Otu31297 | 1 | k__Fungi(100);k__Fungi_unclassified(100);k__Fungi_unclassified(100);k__Fungi_unclassified(100);k__Fungi_unclassified(100);k__Fungi_unclassified(100);k__Fungi_unclassified(100); | - | - | - | - | - | - | - | Unassigned | - |
| Otu31302 | 1 | k__Fungi(100);p__Ascomycota(100);p__Ascomycota_unclassified(100);p__Ascomycota_unclassified(100);p__Ascomycota_unclassified(100);p__Ascomycota_unclassified(100);p__Ascomycota_unclassified(100); | - | - | - | - | - | - | - | Unassigned | - |
| Otu31303 | 1 | k__Fungi(100);k__Fungi_unclassified(100);k__Fungi_unclassified(100);k__Fungi_unclassified(100);k__Fungi_unclassified(100);k__Fungi_unclassified(100);k__Fungi_unclassified(100); | - | - | - | - | - | - | - | Unassigned | - |
| Otu31316 | 1 | k__Fungi(100);p__Ascomycota(100);p__Ascomycota_unclassified(100);p__Ascomycota_unclassified(100);p__Ascomycota_unclassified(100);p__Ascomycota_unclassified(100);p__Ascomycota_unclassified(100); | - | - | - | - | - | - | - | Unassigned | - |
| Otu31318 | 1 | k__Fungi(100);p__Ascomycota(100);p__Ascomycota_unclassified(100);p__Ascomycota_unclassified(100);p__Ascomycota_unclassified(100);p__Ascomycota_unclassified(100);p__Ascomycota_unclassified(100); | - | - | - | - | - | - | - | Unassigned | - |
| Otu31319 | 1 | k__Fungi(100);k__Fungi_unclassified(100);k__Fungi_unclassified(100);k__Fungi_unclassified(100);k__Fungi_unclassified(100);k__Fungi_unclassified(100);k__Fungi_unclassified(100); | - | - | - | - | - | - | - | Unassigned | - |
| Otu31320 | 1 | k__Fungi(100);p__Ascomycota(100);p__Ascomycota_unclassified(100);p__Ascomycota_unclassified(100);p__Ascomycota_unclassified(100);p__Ascomycota_unclassified(100);p__Ascomycota_unclassified(100); | - | - | - | - | - | - | - | Unassigned | - |
| Otu31325 | 1 | k__Fungi(100);p__Ascomycota(100);p__Ascomycota_unclassified(100);p__Ascomycota_unclassified(100);p__Ascomycota_unclassified(100);p__Ascomycota_unclassified(100);p__Ascomycota_unclassified(100); | - | - | - | - | - | - | - | Unassigned | - |
| Otu31328 | 1 | k__Fungi(100);k__Fungi_unclassified(100);k__Fungi_unclassified(100);k__Fungi_unclassified(100);k__Fungi_unclassified(100);k__Fungi_unclassified(100);k__Fungi_unclassified(100); | - | - | - | - | - | - | - | Unassigned | - |
| Otu31330 | 1 | k__Fungi(100);k__Fungi_unclassified(100);k__Fungi_unclassified(100);k__Fungi_unclassified(100);k__Fungi_unclassified(100);k__Fungi_unclassified(100);k__Fungi_unclassified(100); | - | - | - | - | - | - | - | Unassigned | - |
| Otu31331 | 1 | k__Fungi(100);k__Fungi_unclassified(100);k__Fungi_unclassified(100);k__Fungi_unclassified(100);k__Fungi_unclassified(100);k__Fungi_unclassified(100);k__Fungi_unclassified(100); | - | - | - | - | - | - | - | Unassigned | - |
| Otu31332 | 1 | k__Fungi(100);k__Fungi_unclassified(100);k__Fungi_unclassified(100);k__Fungi_unclassified(100);k__Fungi_unclassified(100);k__Fungi_unclassified(100);k__Fungi_unclassified(100); | - | - | - | - | - | - | - | Unassigned | - |
| Otu31333 | 1 | k__Fungi(100);k__Fungi_unclassified(100);k__Fungi_unclassified(100);k__Fungi_unclassified(100);k__Fungi_unclassified(100);k__Fungi_unclassified(100);k__Fungi_unclassified(100); | - | - | - | - | - | - | - | Unassigned | - |
| Otu31337 | 1 | k__Fungi(100);p__Ascomycota(100);p__Ascomycota_unclassified(100);p__Ascomycota_unclassified(100);p__Ascomycota_unclassified(100);p__Ascomycota_unclassified(100);p__Ascomycota_unclassified(100); | - | - | - | - | - | - | - | Unassigned | - |
| Otu31339 | 1 | k__Fungi(100);k__Fungi_unclassified(100);k__Fungi_unclassified(100);k__Fungi_unclassified(100);k__Fungi_unclassified(100);k__Fungi_unclassified(100);k__Fungi_unclassified(100); | - | - | - | - | - | - | - | Unassigned | - |
| Otu31344 | 1 | k__Fungi(100);k__Fungi_unclassified(100);k__Fungi_unclassified(100);k__Fungi_unclassified(100);k__Fungi_unclassified(100);k__Fungi_unclassified(100);k__Fungi_unclassified(100); | - | - | - | - | - | - | - | Unassigned | - |
| Otu31347 | 1 | k__Fungi(100);k__Fungi_unclassified(100);k__Fungi_unclassified(100);k__Fungi_unclassified(100);k__Fungi_unclassified(100);k__Fungi_unclassified(100);k__Fungi_unclassified(100); | - | - | - | - | - | - | - | Unassigned | - |
| Otu31348 | 1 | k__Fungi(100);k__Fungi_unclassified(100);k__Fungi_unclassified(100);k__Fungi_unclassified(100);k__Fungi_unclassified(100);k__Fungi_unclassified(100);k__Fungi_unclassified(100); | - | - | - | - | - | - | - | Unassigned | - |
| Otu31349 | 1 | k__Fungi(100);p__Ascomycota(100);p__Ascomycota_unclassified(100);p__Ascomycota_unclassified(100);p__Ascomycota_unclassified(100);p__Ascomycota_unclassified(100);p__Ascomycota_unclassified(100); | - | - | - | - | - | - | - | Unassigned | - |
| Otu31350 | 1 | k__Fungi(100);p__Ascomycota(100);p__Ascomycota_unclassified(100);p__Ascomycota_unclassified(100);p__Ascomycota_unclassified(100);p__Ascomycota_unclassified(100);p__Ascomycota_unclassified(100); | - | - | - | - | - | - | - | Unassigned | - |
| Otu31353 | 1 | k__Fungi(100);p__Ascomycota(100);p__Ascomycota_unclassified(100);p__Ascomycota_unclassified(100);p__Ascomycota_unclassified(100);p__Ascomycota_unclassified(100);p__Ascomycota_unclassified(100); | - | - | - | - | - | - | - | Unassigned | - |
| Otu31385 | 1 | k__Fungi(100);k__Fungi_unclassified(100);k__Fungi_unclassified(100);k__Fungi_unclassified(100);k__Fungi_unclassified(100);k__Fungi_unclassified(100);k__Fungi_unclassified(100); | - | - | - | - | - | - | - | Unassigned | - |
| Otu31410 | 1 | k__Fungi(100);p__Ascomycota(100);p__Ascomycota_unclassified(100);p__Ascomycota_unclassified(100);p__Ascomycota_unclassified(100);p__Ascomycota_unclassified(100);p__Ascomycota_unclassified(100); | - | - | - | - | - | - | - | Unassigned | - |
| Otu31417 | 1 | k__Fungi(100);p__Ascomycota(100);p__Ascomycota_unclassified(100);p__Ascomycota_unclassified(100);p__Ascomycota_unclassified(100);p__Ascomycota_unclassified(100);p__Ascomycota_unclassified(100); | - | - | - | - | - | - | - | Unassigned | - |
| Otu31418 | 1 | k__Fungi(100);k__Fungi_unclassified(100);k__Fungi_unclassified(100);k__Fungi_unclassified(100);k__Fungi_unclassified(100);k__Fungi_unclassified(100);k__Fungi_unclassified(100); | - | - | - | - | - | - | - | Unassigned | - |
| Otu31422 | 1 | k__Fungi(100);k__Fungi_unclassified(100);k__Fungi_unclassified(100);k__Fungi_unclassified(100);k__Fungi_unclassified(100);k__Fungi_unclassified(100);k__Fungi_unclassified(100); | - | - | - | - | - | - | - | Unassigned | - |
| Otu31429 | 1 | k__Fungi(100);k__Fungi_unclassified(100);k__Fungi_unclassified(100);k__Fungi_unclassified(100);k__Fungi_unclassified(100);k__Fungi_unclassified(100);k__Fungi_unclassified(100); | - | - | - | - | - | - | - | Unassigned | - |
| Otu31430 | 1 | k__Fungi(100);k__Fungi_unclassified(100);k__Fungi_unclassified(100);k__Fungi_unclassified(100);k__Fungi_unclassified(100);k__Fungi_unclassified(100);k__Fungi_unclassified(100); | - | - | - | - | - | - | - | Unassigned | - |
| Otu31431 | 1 | k__Fungi(100);k__Fungi_unclassified(100);k__Fungi_unclassified(100);k__Fungi_unclassified(100);k__Fungi_unclassified(100);k__Fungi_unclassified(100);k__Fungi_unclassified(100); | - | - | - | - | - | - | - | Unassigned | - |
| Otu31433 | 1 | k__Fungi(100);k__Fungi_unclassified(100);k__Fungi_unclassified(100);k__Fungi_unclassified(100);k__Fungi_unclassified(100);k__Fungi_unclassified(100);k__Fungi_unclassified(100); | - | - | - | - | - | - | - | Unassigned | - |
| Otu31434 | 1 | k__Fungi(100);k__Fungi_unclassified(100);k__Fungi_unclassified(100);k__Fungi_unclassified(100);k__Fungi_unclassified(100);k__Fungi_unclassified(100);k__Fungi_unclassified(100); | - | - | - | - | - | - | - | Unassigned | - |
| Otu31435 | 1 | k__Fungi(100);k__Fungi_unclassified(100);k__Fungi_unclassified(100);k__Fungi_unclassified(100);k__Fungi_unclassified(100);k__Fungi_unclassified(100);k__Fungi_unclassified(100); | - | - | - | - | - | - | - | Unassigned | - |
| Otu31436 | 1 | k__Fungi(100);p__Ascomycota(100);p__Ascomycota_unclassified(100);p__Ascomycota_unclassified(100);p__Ascomycota_unclassified(100);p__Ascomycota_unclassified(100);p__Ascomycota_unclassified(100); | - | - | - | - | - | - | - | Unassigned | - |
| Otu31438 | 1 | k__Fungi(100);k__Fungi_unclassified(100);k__Fungi_unclassified(100);k__Fungi_unclassified(100);k__Fungi_unclassified(100);k__Fungi_unclassified(100);k__Fungi_unclassified(100); | - | - | - | - | - | - | - | Unassigned | - |
| Otu31448 | 1 | k__Fungi(100);p__Ascomycota(100);p__Ascomycota_unclassified(100);p__Ascomycota_unclassified(100);p__Ascomycota_unclassified(100);p__Ascomycota_unclassified(100);p__Ascomycota_unclassified(100); | - | - | - | - | - | - | - | Unassigned | - |
| Otu31449 | 1 | k__Fungi(100);k__Fungi_unclassified(100);k__Fungi_unclassified(100);k__Fungi_unclassified(100);k__Fungi_unclassified(100);k__Fungi_unclassified(100);k__Fungi_unclassified(100); | - | - | - | - | - | - | - | Unassigned | - |
| Otu31453 | 1 | k__Fungi(100);k__Fungi_unclassified(100);k__Fungi_unclassified(100);k__Fungi_unclassified(100);k__Fungi_unclassified(100);k__Fungi_unclassified(100);k__Fungi_unclassified(100); | - | - | - | - | - | - | - | Unassigned | - |
| Otu31454 | 1 | k__Fungi(100);p__Ascomycota(100);p__Ascomycota_unclassified(100);p__Ascomycota_unclassified(100);p__Ascomycota_unclassified(100);p__Ascomycota_unclassified(100);p__Ascomycota_unclassified(100); | - | - | - | - | - | - | - | Unassigned | - |
| Otu31455 | 1 | k__Fungi(100);p__Ascomycota(100);p__Ascomycota_unclassified(100);p__Ascomycota_unclassified(100);p__Ascomycota_unclassified(100);p__Ascomycota_unclassified(100);p__Ascomycota_unclassified(100); | - | - | - | - | - | - | - | Unassigned | - |
| Otu31456 | 1 | k__Fungi(100);p__Ascomycota(100);p__Ascomycota_unclassified(100);p__Ascomycota_unclassified(100);p__Ascomycota_unclassified(100);p__Ascomycota_unclassified(100);p__Ascomycota_unclassified(100); | - | - | - | - | - | - | - | Unassigned | - |
| Otu31457 | 1 | k__Fungi(100);k__Fungi_unclassified(100);k__Fungi_unclassified(100);k__Fungi_unclassified(100);k__Fungi_unclassified(100);k__Fungi_unclassified(100);k__Fungi_unclassified(100); | - | - | - | - | - | - | - | Unassigned | - |
| Otu31458 | 1 | k__Fungi(100);k__Fungi_unclassified(100);k__Fungi_unclassified(100);k__Fungi_unclassified(100);k__Fungi_unclassified(100);k__Fungi_unclassified(100);k__Fungi_unclassified(100); | - | - | - | - | - | - | - | Unassigned | - |
| Otu31459 | 1 | k__Fungi(100);k__Fungi_unclassified(100);k__Fungi_unclassified(100);k__Fungi_unclassified(100);k__Fungi_unclassified(100);k__Fungi_unclassified(100);k__Fungi_unclassified(100); | - | - | - | - | - | - | - | Unassigned | - |
| Otu31460 | 1 | k__Fungi(100);p__Ascomycota(100);p__Ascomycota_unclassified(100);p__Ascomycota_unclassified(100);p__Ascomycota_unclassified(100);p__Ascomycota_unclassified(100);p__Ascomycota_unclassified(100); | - | - | - | - | - | - | - | Unassigned | - |
| Otu31461 | 1 | k__Fungi(100);p__Ascomycota(100);p__Ascomycota_unclassified(100);p__Ascomycota_unclassified(100);p__Ascomycota_unclassified(100);p__Ascomycota_unclassified(100);p__Ascomycota_unclassified(100); | - | - | - | - | - | - | - | Unassigned | - |
| Otu31465 | 1 | k__Fungi(100);k__Fungi_unclassified(100);k__Fungi_unclassified(100);k__Fungi_unclassified(100);k__Fungi_unclassified(100);k__Fungi_unclassified(100);k__Fungi_unclassified(100); | - | - | - | - | - | - | - | Unassigned | - |
| Otu31466 | 1 | k__Fungi(100);k__Fungi_unclassified(100);k__Fungi_unclassified(100);k__Fungi_unclassified(100);k__Fungi_unclassified(100);k__Fungi_unclassified(100);k__Fungi_unclassified(100); | - | - | - | - | - | - | - | Unassigned | - |
| Otu31490 | 1 | k__Fungi(100);p__Ascomycota(100);p__Ascomycota_unclassified(100);p__Ascomycota_unclassified(100);p__Ascomycota_unclassified(100);p__Ascomycota_unclassified(100);p__Ascomycota_unclassified(100); | - | - | - | - | - | - | - | Unassigned | - |
| Otu31492 | 1 | k__Fungi(100);k__Fungi_unclassified(100);k__Fungi_unclassified(100);k__Fungi_unclassified(100);k__Fungi_unclassified(100);k__Fungi_unclassified(100);k__Fungi_unclassified(100); | - | - | - | - | - | - | - | Unassigned | - |
| Otu31495 | 1 | k__Fungi(100);k__Fungi_unclassified(100);k__Fungi_unclassified(100);k__Fungi_unclassified(100);k__Fungi_unclassified(100);k__Fungi_unclassified(100);k__Fungi_unclassified(100); | - | - | - | - | - | - | - | Unassigned | - |
| Otu31523 | 1 | k__Fungi(100);p__Ascomycota(100);p__Ascomycota_unclassified(100);p__Ascomycota_unclassified(100);p__Ascomycota_unclassified(100);p__Ascomycota_unclassified(100);p__Ascomycota_unclassified(100); | - | - | - | - | - | - | - | Unassigned | - |
| Otu31526 | 1 | k__Fungi(100);p__Ascomycota(100);p__Ascomycota_unclassified(100);p__Ascomycota_unclassified(100);p__Ascomycota_unclassified(100);p__Ascomycota_unclassified(100);p__Ascomycota_unclassified(100); | - | - | - | - | - | - | - | Unassigned | - |
| Otu31539 | 1 | k__Fungi(100);k__Fungi_unclassified(100);k__Fungi_unclassified(100);k__Fungi_unclassified(100);k__Fungi_unclassified(100);k__Fungi_unclassified(100);k__Fungi_unclassified(100); | - | - | - | - | - | - | - | Unassigned | - |
| Otu31568 | 1 | k__Fungi(100);p__Ascomycota(100);p__Ascomycota_unclassified(100);p__Ascomycota_unclassified(100);p__Ascomycota_unclassified(100);p__Ascomycota_unclassified(100);p__Ascomycota_unclassified(100); | - | - | - | - | - | - | - | Unassigned | - |
| Otu31585 | 1 | k__Fungi(100);k__Fungi_unclassified(100);k__Fungi_unclassified(100);k__Fungi_unclassified(100);k__Fungi_unclassified(100);k__Fungi_unclassified(100);k__Fungi_unclassified(100); | - | - | - | - | - | - | - | Unassigned | - |
| Otu31595 | 1 | k__Fungi(100);k__Fungi_unclassified(100);k__Fungi_unclassified(100);k__Fungi_unclassified(100);k__Fungi_unclassified(100);k__Fungi_unclassified(100);k__Fungi_unclassified(100); | - | - | - | - | - | - | - | Unassigned | - |
| Otu31596 | 1 | k__Fungi(100);k__Fungi_unclassified(100);k__Fungi_unclassified(100);k__Fungi_unclassified(100);k__Fungi_unclassified(100);k__Fungi_unclassified(100);k__Fungi_unclassified(100); | - | - | - | - | - | - | - | Unassigned | - |
| Otu31602 | 1 | k__Fungi(100);k__Fungi_unclassified(100);k__Fungi_unclassified(100);k__Fungi_unclassified(100);k__Fungi_unclassified(100);k__Fungi_unclassified(100);k__Fungi_unclassified(100); | - | - | - | - | - | - | - | Unassigned | - |
| Otu31650 | 1 | k__Fungi(100);k__Fungi_unclassified(100);k__Fungi_unclassified(100);k__Fungi_unclassified(100);k__Fungi_unclassified(100);k__Fungi_unclassified(100);k__Fungi_unclassified(100); | - | - | - | - | - | - | - | Unassigned | - |
| Otu31651 | 1 | k__Fungi(100);k__Fungi_unclassified(100);k__Fungi_unclassified(100);k__Fungi_unclassified(100);k__Fungi_unclassified(100);k__Fungi_unclassified(100);k__Fungi_unclassified(100); | - | - | - | - | - | - | - | Unassigned | - |
| Otu31653 | 1 | k__Fungi(100);p__Ascomycota(100);p__Ascomycota_unclassified(100);p__Ascomycota_unclassified(100);p__Ascomycota_unclassified(100);p__Ascomycota_unclassified(100);p__Ascomycota_unclassified(100); | - | - | - | - | - | - | - | Unassigned | - |
| Otu31677 | 1 | k__Fungi(100);p__Ascomycota(100);p__Ascomycota_unclassified(100);p__Ascomycota_unclassified(100);p__Ascomycota_unclassified(100);p__Ascomycota_unclassified(100);p__Ascomycota_unclassified(100); | - | - | - | - | - | - | - | Unassigned | - |
| Otu31678 | 1 | k__Fungi(100);k__Fungi_unclassified(100);k__Fungi_unclassified(100);k__Fungi_unclassified(100);k__Fungi_unclassified(100);k__Fungi_unclassified(100);k__Fungi_unclassified(100); | - | - | - | - | - | - | - | Unassigned | - |
| Otu31682 | 1 | k__Fungi(100);k__Fungi_unclassified(100);k__Fungi_unclassified(100);k__Fungi_unclassified(100);k__Fungi_unclassified(100);k__Fungi_unclassified(100);k__Fungi_unclassified(100); | - | - | - | - | - | - | - | Unassigned | - |
| Otu31688 | 1 | k__Fungi(100);k__Fungi_unclassified(100);k__Fungi_unclassified(100);k__Fungi_unclassified(100);k__Fungi_unclassified(100);k__Fungi_unclassified(100);k__Fungi_unclassified(100); | - | - | - | - | - | - | - | Unassigned | - |
| Otu31724 | 1 | k__Fungi(100);p__Ascomycota(100);p__Ascomycota_unclassified(100);p__Ascomycota_unclassified(100);p__Ascomycota_unclassified(100);p__Ascomycota_unclassified(100);p__Ascomycota_unclassified(100); | - | - | - | - | - | - | - | Unassigned | - |
| Otu31725 | 1 | k__Fungi(100);k__Fungi_unclassified(100);k__Fungi_unclassified(100);k__Fungi_unclassified(100);k__Fungi_unclassified(100);k__Fungi_unclassified(100);k__Fungi_unclassified(100); | - | - | - | - | - | - | - | Unassigned | - |
| Otu31729 | 1 | k__Fungi(100);p__Ascomycota(100);p__Ascomycota_unclassified(100);p__Ascomycota_unclassified(100);p__Ascomycota_unclassified(100);p__Ascomycota_unclassified(100);p__Ascomycota_unclassified(100); | - | - | - | - | - | - | - | Unassigned | - |
| Otu31731 | 1 | k__Fungi(100);k__Fungi_unclassified(100);k__Fungi_unclassified(100);k__Fungi_unclassified(100);k__Fungi_unclassified(100);k__Fungi_unclassified(100);k__Fungi_unclassified(100); | - | - | - | - | - | - | - | Unassigned | - |
| Otu31732 | 1 | k__Fungi(100);p__Ascomycota(100);p__Ascomycota_unclassified(100);p__Ascomycota_unclassified(100);p__Ascomycota_unclassified(100);p__Ascomycota_unclassified(100);p__Ascomycota_unclassified(100); | - | - | - | - | - | - | - | Unassigned | - |
| Otu31734 | 1 | k__Fungi(100);p__Ascomycota(100);p__Ascomycota_unclassified(100);p__Ascomycota_unclassified(100);p__Ascomycota_unclassified(100);p__Ascomycota_unclassified(100);p__Ascomycota_unclassified(100); | - | - | - | - | - | - | - | Unassigned | - |
| Otu31735 | 1 | k__Fungi(100);k__Fungi_unclassified(100);k__Fungi_unclassified(100);k__Fungi_unclassified(100);k__Fungi_unclassified(100);k__Fungi_unclassified(100);k__Fungi_unclassified(100); | - | - | - | - | - | - | - | Unassigned | - |
| Otu31736 | 1 | k__Fungi(100);p__Ascomycota(100);p__Ascomycota_unclassified(100);p__Ascomycota_unclassified(100);p__Ascomycota_unclassified(100);p__Ascomycota_unclassified(100);p__Ascomycota_unclassified(100); | - | - | - | - | - | - | - | Unassigned | - |
| Otu31737 | 1 | k__Fungi(100);p__Ascomycota(100);p__Ascomycota_unclassified(100);p__Ascomycota_unclassified(100);p__Ascomycota_unclassified(100);p__Ascomycota_unclassified(100);p__Ascomycota_unclassified(100); | - | - | - | - | - | - | - | Unassigned | - |
| Otu31738 | 1 | k__Fungi(100);p__Ascomycota(100);p__Ascomycota_unclassified(100);p__Ascomycota_unclassified(100);p__Ascomycota_unclassified(100);p__Ascomycota_unclassified(100);p__Ascomycota_unclassified(100); | - | - | - | - | - | - | - | Unassigned | - |
| Otu31740 | 1 | k__Fungi(100);p__Ascomycota(100);p__Ascomycota_unclassified(100);p__Ascomycota_unclassified(100);p__Ascomycota_unclassified(100);p__Ascomycota_unclassified(100);p__Ascomycota_unclassified(100); | - | - | - | - | - | - | - | Unassigned | - |
| Otu31746 | 1 | k__Fungi(100);p__Ascomycota(100);p__Ascomycota_unclassified(100);p__Ascomycota_unclassified(100);p__Ascomycota_unclassified(100);p__Ascomycota_unclassified(100);p__Ascomycota_unclassified(100); | - | - | - | - | - | - | - | Unassigned | - |
| Otu31748 | 1 | k__Fungi(100);p__Ascomycota(100);p__Ascomycota_unclassified(100);p__Ascomycota_unclassified(100);p__Ascomycota_unclassified(100);p__Ascomycota_unclassified(100);p__Ascomycota_unclassified(100); | - | - | - | - | - | - | - | Unassigned | - |
| Otu31756 | 1 | k__Fungi(100);k__Fungi_unclassified(100);k__Fungi_unclassified(100);k__Fungi_unclassified(100);k__Fungi_unclassified(100);k__Fungi_unclassified(100);k__Fungi_unclassified(100); | - | - | - | - | - | - | - | Unassigned | - |
| Otu31757 | 1 | k__Fungi(100);p__Ascomycota(100);p__Ascomycota_unclassified(100);p__Ascomycota_unclassified(100);p__Ascomycota_unclassified(100);p__Ascomycota_unclassified(100);p__Ascomycota_unclassified(100); | - | - | - | - | - | - | - | Unassigned | - |
| Otu31759 | 1 | k__Fungi(100);p__Ascomycota(100);p__Ascomycota_unclassified(100);p__Ascomycota_unclassified(100);p__Ascomycota_unclassified(100);p__Ascomycota_unclassified(100);p__Ascomycota_unclassified(100); | - | - | - | - | - | - | - | Unassigned | - |
| Otu31763 | 1 | k__Fungi(100);k__Fungi_unclassified(100);k__Fungi_unclassified(100);k__Fungi_unclassified(100);k__Fungi_unclassified(100);k__Fungi_unclassified(100);k__Fungi_unclassified(100); | - | - | - | - | - | - | - | Unassigned | - |
| Otu31764 | 1 | k__Fungi(100);p__Ascomycota(100);p__Ascomycota_unclassified(100);p__Ascomycota_unclassified(100);p__Ascomycota_unclassified(100);p__Ascomycota_unclassified(100);p__Ascomycota_unclassified(100); | - | - | - | - | - | - | - | Unassigned | - |
| Otu31768 | 1 | k__Fungi(100);p__Ascomycota(100);p__Ascomycota_unclassified(100);p__Ascomycota_unclassified(100);p__Ascomycota_unclassified(100);p__Ascomycota_unclassified(100);p__Ascomycota_unclassified(100); | - | - | - | - | - | - | - | Unassigned | - |
| Otu31779 | 1 | k__Fungi(100);p__Ascomycota(100);p__Ascomycota_unclassified(100);p__Ascomycota_unclassified(100);p__Ascomycota_unclassified(100);p__Ascomycota_unclassified(100);p__Ascomycota_unclassified(100); | - | - | - | - | - | - | - | Unassigned | - |
| Otu31787 | 1 | k__Fungi(100);k__Fungi_unclassified(100);k__Fungi_unclassified(100);k__Fungi_unclassified(100);k__Fungi_unclassified(100);k__Fungi_unclassified(100);k__Fungi_unclassified(100); | - | - | - | - | - | - | - | Unassigned | - |
| Otu31789 | 1 | k__Fungi(100);p__Ascomycota(100);p__Ascomycota_unclassified(100);p__Ascomycota_unclassified(100);p__Ascomycota_unclassified(100);p__Ascomycota_unclassified(100);p__Ascomycota_unclassified(100); | - | - | - | - | - | - | - | Unassigned | - |
| Otu31790 | 1 | k__Fungi(100);k__Fungi_unclassified(100);k__Fungi_unclassified(100);k__Fungi_unclassified(100);k__Fungi_unclassified(100);k__Fungi_unclassified(100);k__Fungi_unclassified(100); | - | - | - | - | - | - | - | Unassigned | - |
| Otu31797 | 1 | k__Fungi(100);k__Fungi_unclassified(100);k__Fungi_unclassified(100);k__Fungi_unclassified(100);k__Fungi_unclassified(100);k__Fungi_unclassified(100);k__Fungi_unclassified(100); | - | - | - | - | - | - | - | Unassigned | - |
| Otu31799 | 1 | k__Fungi(100);k__Fungi_unclassified(100);k__Fungi_unclassified(100);k__Fungi_unclassified(100);k__Fungi_unclassified(100);k__Fungi_unclassified(100);k__Fungi_unclassified(100); | - | - | - | - | - | - | - | Unassigned | - |
| Otu31808 | 1 | k__Fungi(100);k__Fungi_unclassified(100);k__Fungi_unclassified(100);k__Fungi_unclassified(100);k__Fungi_unclassified(100);k__Fungi_unclassified(100);k__Fungi_unclassified(100); | - | - | - | - | - | - | - | Unassigned | - |
| Otu31809 | 1 | k__Fungi(100);k__Fungi_unclassified(100);k__Fungi_unclassified(100);k__Fungi_unclassified(100);k__Fungi_unclassified(100);k__Fungi_unclassified(100);k__Fungi_unclassified(100); | - | - | - | - | - | - | - | Unassigned | - |
| Otu31813 | 1 | k__Fungi(100);p__Ascomycota(100);p__Ascomycota_unclassified(100);p__Ascomycota_unclassified(100);p__Ascomycota_unclassified(100);p__Ascomycota_unclassified(100);p__Ascomycota_unclassified(100); | - | - | - | - | - | - | - | Unassigned | - |
| Otu31814 | 1 | k__Fungi(100);k__Fungi_unclassified(100);k__Fungi_unclassified(100);k__Fungi_unclassified(100);k__Fungi_unclassified(100);k__Fungi_unclassified(100);k__Fungi_unclassified(100); | - | - | - | - | - | - | - | Unassigned | - |
| Otu31816 | 1 | k__Fungi(100);p__Ascomycota(100);p__Ascomycota_unclassified(100);p__Ascomycota_unclassified(100);p__Ascomycota_unclassified(100);p__Ascomycota_unclassified(100);p__Ascomycota_unclassified(100); | - | - | - | - | - | - | - | Unassigned | - |
| Otu31817 | 1 | k__Fungi(100);k__Fungi_unclassified(100);k__Fungi_unclassified(100);k__Fungi_unclassified(100);k__Fungi_unclassified(100);k__Fungi_unclassified(100);k__Fungi_unclassified(100); | - | - | - | - | - | - | - | Unassigned | - |
| Otu31818 | 1 | k__Fungi(100);k__Fungi_unclassified(100);k__Fungi_unclassified(100);k__Fungi_unclassified(100);k__Fungi_unclassified(100);k__Fungi_unclassified(100);k__Fungi_unclassified(100); | - | - | - | - | - | - | - | Unassigned | - |
| Otu31819 | 1 | k__Fungi(100);p__Ascomycota(100);p__Ascomycota_unclassified(100);p__Ascomycota_unclassified(100);p__Ascomycota_unclassified(100);p__Ascomycota_unclassified(100);p__Ascomycota_unclassified(100); | - | - | - | - | - | - | - | Unassigned | - |
| Otu31820 | 1 | k__Fungi(100);k__Fungi_unclassified(100);k__Fungi_unclassified(100);k__Fungi_unclassified(100);k__Fungi_unclassified(100);k__Fungi_unclassified(100);k__Fungi_unclassified(100); | - | - | - | - | - | - | - | Unassigned | - |
| Otu31825 | 1 | k__Fungi(100);p__Ascomycota(100);p__Ascomycota_unclassified(100);p__Ascomycota_unclassified(100);p__Ascomycota_unclassified(100);p__Ascomycota_unclassified(100);p__Ascomycota_unclassified(100); | - | - | - | - | - | - | - | Unassigned | - |
| Otu31826 | 1 | k__Fungi(100);p__Ascomycota(100);p__Ascomycota_unclassified(100);p__Ascomycota_unclassified(100);p__Ascomycota_unclassified(100);p__Ascomycota_unclassified(100);p__Ascomycota_unclassified(100); | - | - | - | - | - | - | - | Unassigned | - |
| Otu31827 | 1 | k__Fungi(100);k__Fungi_unclassified(100);k__Fungi_unclassified(100);k__Fungi_unclassified(100);k__Fungi_unclassified(100);k__Fungi_unclassified(100);k__Fungi_unclassified(100); | - | - | - | - | - | - | - | Unassigned | - |
| Otu31835 | 1 | k__Fungi(100);k__Fungi_unclassified(100);k__Fungi_unclassified(100);k__Fungi_unclassified(100);k__Fungi_unclassified(100);k__Fungi_unclassified(100);k__Fungi_unclassified(100); | - | - | - | - | - | - | - | Unassigned | - |
| Otu31838 | 1 | k__Fungi(100);k__Fungi_unclassified(100);k__Fungi_unclassified(100);k__Fungi_unclassified(100);k__Fungi_unclassified(100);k__Fungi_unclassified(100);k__Fungi_unclassified(100); | - | - | - | - | - | - | - | Unassigned | - |
| Otu31839 | 1 | k__Fungi(100);k__Fungi_unclassified(100);k__Fungi_unclassified(100);k__Fungi_unclassified(100);k__Fungi_unclassified(100);k__Fungi_unclassified(100);k__Fungi_unclassified(100); | - | - | - | - | - | - | - | Unassigned | - |
| Otu31840 | 1 | k__Fungi(100);p__Ascomycota(100);p__Ascomycota_unclassified(100);p__Ascomycota_unclassified(100);p__Ascomycota_unclassified(100);p__Ascomycota_unclassified(100);p__Ascomycota_unclassified(100); | - | - | - | - | - | - | - | Unassigned | - |
| Otu31841 | 1 | k__Fungi(100);k__Fungi_unclassified(100);k__Fungi_unclassified(100);k__Fungi_unclassified(100);k__Fungi_unclassified(100);k__Fungi_unclassified(100);k__Fungi_unclassified(100); | - | - | - | - | - | - | - | Unassigned | - |
| Otu31843 | 1 | k__Fungi(100);p__Ascomycota(100);p__Ascomycota_unclassified(100);p__Ascomycota_unclassified(100);p__Ascomycota_unclassified(100);p__Ascomycota_unclassified(100);p__Ascomycota_unclassified(100); | - | - | - | - | - | - | - | Unassigned | - |
| Otu31844 | 1 | k__Fungi(100);k__Fungi_unclassified(100);k__Fungi_unclassified(100);k__Fungi_unclassified(100);k__Fungi_unclassified(100);k__Fungi_unclassified(100);k__Fungi_unclassified(100); | - | - | - | - | - | - | - | Unassigned | - |
| Otu31845 | 1 | k__Fungi(100);p__Ascomycota(100);p__Ascomycota_unclassified(100);p__Ascomycota_unclassified(100);p__Ascomycota_unclassified(100);p__Ascomycota_unclassified(100);p__Ascomycota_unclassified(100); | - | - | - | - | - | - | - | Unassigned | - |
| Otu31848 | 1 | k__Fungi(100);p__Ascomycota(100);p__Ascomycota_unclassified(100);p__Ascomycota_unclassified(100);p__Ascomycota_unclassified(100);p__Ascomycota_unclassified(100);p__Ascomycota_unclassified(100); | - | - | - | - | - | - | - | Unassigned | - |
| Otu31851 | 1 | k__Fungi(100);p__Ascomycota(100);p__Ascomycota_unclassified(100);p__Ascomycota_unclassified(100);p__Ascomycota_unclassified(100);p__Ascomycota_unclassified(100);p__Ascomycota_unclassified(100); | - | - | - | - | - | - | - | Unassigned | - |
| Otu31852 | 1 | k__Fungi(100);k__Fungi_unclassified(100);k__Fungi_unclassified(100);k__Fungi_unclassified(100);k__Fungi_unclassified(100);k__Fungi_unclassified(100);k__Fungi_unclassified(100); | - | - | - | - | - | - | - | Unassigned | - |
| Otu31854 | 1 | k__Fungi(100);k__Fungi_unclassified(100);k__Fungi_unclassified(100);k__Fungi_unclassified(100);k__Fungi_unclassified(100);k__Fungi_unclassified(100);k__Fungi_unclassified(100); | - | - | - | - | - | - | - | Unassigned | - |
| Otu31855 | 1 | k__Fungi(100);k__Fungi_unclassified(100);k__Fungi_unclassified(100);k__Fungi_unclassified(100);k__Fungi_unclassified(100);k__Fungi_unclassified(100);k__Fungi_unclassified(100); | - | - | - | - | - | - | - | Unassigned | - |
| Otu31856 | 1 | k__Fungi(100);p__Ascomycota(100);p__Ascomycota_unclassified(100);p__Ascomycota_unclassified(100);p__Ascomycota_unclassified(100);p__Ascomycota_unclassified(100);p__Ascomycota_unclassified(100); | - | - | - | - | - | - | - | Unassigned | - |
| Otu31857 | 1 | k__Fungi(100);k__Fungi_unclassified(100);k__Fungi_unclassified(100);k__Fungi_unclassified(100);k__Fungi_unclassified(100);k__Fungi_unclassified(100);k__Fungi_unclassified(100); | - | - | - | - | - | - | - | Unassigned | - |
| Otu31859 | 1 | k__Fungi(100);k__Fungi_unclassified(100);k__Fungi_unclassified(100);k__Fungi_unclassified(100);k__Fungi_unclassified(100);k__Fungi_unclassified(100);k__Fungi_unclassified(100); | - | - | - | - | - | - | - | Unassigned | - |
| Otu31860 | 1 | k__Fungi(100);p__Ascomycota(100);p__Ascomycota_unclassified(100);p__Ascomycota_unclassified(100);p__Ascomycota_unclassified(100);p__Ascomycota_unclassified(100);p__Ascomycota_unclassified(100); | - | - | - | - | - | - | - | Unassigned | - |
| Otu31861 | 1 | k__Fungi(100);k__Fungi_unclassified(100);k__Fungi_unclassified(100);k__Fungi_unclassified(100);k__Fungi_unclassified(100);k__Fungi_unclassified(100);k__Fungi_unclassified(100); | - | - | - | - | - | - | - | Unassigned | - |
| Otu31862 | 1 | k__Fungi(100);p__Ascomycota(100);p__Ascomycota_unclassified(100);p__Ascomycota_unclassified(100);p__Ascomycota_unclassified(100);p__Ascomycota_unclassified(100);p__Ascomycota_unclassified(100); | - | - | - | - | - | - | - | Unassigned | - |
| Otu31863 | 1 | k__Fungi(100);p__Ascomycota(100);p__Ascomycota_unclassified(100);p__Ascomycota_unclassified(100);p__Ascomycota_unclassified(100);p__Ascomycota_unclassified(100);p__Ascomycota_unclassified(100); | - | - | - | - | - | - | - | Unassigned | - |
| Otu31865 | 1 | k__Fungi(100);k__Fungi_unclassified(100);k__Fungi_unclassified(100);k__Fungi_unclassified(100);k__Fungi_unclassified(100);k__Fungi_unclassified(100);k__Fungi_unclassified(100); | - | - | - | - | - | - | - | Unassigned | - |
| Otu31866 | 1 | k__Fungi(100);p__Ascomycota(100);p__Ascomycota_unclassified(100);p__Ascomycota_unclassified(100);p__Ascomycota_unclassified(100);p__Ascomycota_unclassified(100);p__Ascomycota_unclassified(100); | - | - | - | - | - | - | - | Unassigned | - |
| Otu31868 | 1 | k__Fungi(100);p__Ascomycota(100);p__Ascomycota_unclassified(100);p__Ascomycota_unclassified(100);p__Ascomycota_unclassified(100);p__Ascomycota_unclassified(100);p__Ascomycota_unclassified(100); | - | - | - | - | - | - | - | Unassigned | - |
| Otu31869 | 1 | k__Fungi(100);k__Fungi_unclassified(100);k__Fungi_unclassified(100);k__Fungi_unclassified(100);k__Fungi_unclassified(100);k__Fungi_unclassified(100);k__Fungi_unclassified(100); | - | - | - | - | - | - | - | Unassigned | - |
| Otu31870 | 1 | k__Fungi(100);k__Fungi_unclassified(100);k__Fungi_unclassified(100);k__Fungi_unclassified(100);k__Fungi_unclassified(100);k__Fungi_unclassified(100);k__Fungi_unclassified(100); | - | - | - | - | - | - | - | Unassigned | - |
| Otu31871 | 1 | k__Fungi(100);p__Ascomycota(100);p__Ascomycota_unclassified(100);p__Ascomycota_unclassified(100);p__Ascomycota_unclassified(100);p__Ascomycota_unclassified(100);p__Ascomycota_unclassified(100); | - | - | - | - | - | - | - | Unassigned | - |
| Otu31872 | 1 | k__Fungi(100);k__Fungi_unclassified(100);k__Fungi_unclassified(100);k__Fungi_unclassified(100);k__Fungi_unclassified(100);k__Fungi_unclassified(100);k__Fungi_unclassified(100); | - | - | - | - | - | - | - | Unassigned | - |
| Otu31873 | 1 | k__Fungi(100);p__Ascomycota(100);p__Ascomycota_unclassified(100);p__Ascomycota_unclassified(100);p__Ascomycota_unclassified(100);p__Ascomycota_unclassified(100);p__Ascomycota_unclassified(100); | - | - | - | - | - | - | - | Unassigned | - |
| Otu31874 | 1 | k__Fungi(100);p__Ascomycota(100);p__Ascomycota_unclassified(100);p__Ascomycota_unclassified(100);p__Ascomycota_unclassified(100);p__Ascomycota_unclassified(100);p__Ascomycota_unclassified(100); | - | - | - | - | - | - | - | Unassigned | - |
| Otu31875 | 1 | k__Fungi(100);k__Fungi_unclassified(100);k__Fungi_unclassified(100);k__Fungi_unclassified(100);k__Fungi_unclassified(100);k__Fungi_unclassified(100);k__Fungi_unclassified(100); | - | - | - | - | - | - | - | Unassigned | - |
| Otu31876 | 1 | k__Fungi(100);p__Ascomycota(100);p__Ascomycota_unclassified(100);p__Ascomycota_unclassified(100);p__Ascomycota_unclassified(100);p__Ascomycota_unclassified(100);p__Ascomycota_unclassified(100); | - | - | - | - | - | - | - | Unassigned | - |
| Otu31877 | 1 | k__Fungi(100);k__Fungi_unclassified(100);k__Fungi_unclassified(100);k__Fungi_unclassified(100);k__Fungi_unclassified(100);k__Fungi_unclassified(100);k__Fungi_unclassified(100); | - | - | - | - | - | - | - | Unassigned | - |
| Otu31878 | 1 | k__Fungi(100);k__Fungi_unclassified(100);k__Fungi_unclassified(100);k__Fungi_unclassified(100);k__Fungi_unclassified(100);k__Fungi_unclassified(100);k__Fungi_unclassified(100); | - | - | - | - | - | - | - | Unassigned | - |
| Otu31879 | 1 | k__Fungi(100);p__Ascomycota(100);p__Ascomycota_unclassified(100);p__Ascomycota_unclassified(100);p__Ascomycota_unclassified(100);p__Ascomycota_unclassified(100);p__Ascomycota_unclassified(100); | - | - | - | - | - | - | - | Unassigned | - |
| Otu31880 | 1 | k__Fungi(100);p__Ascomycota(100);p__Ascomycota_unclassified(100);p__Ascomycota_unclassified(100);p__Ascomycota_unclassified(100);p__Ascomycota_unclassified(100);p__Ascomycota_unclassified(100); | - | - | - | - | - | - | - | Unassigned | - |
| Otu31881 | 1 | k__Fungi(100);k__Fungi_unclassified(100);k__Fungi_unclassified(100);k__Fungi_unclassified(100);k__Fungi_unclassified(100);k__Fungi_unclassified(100);k__Fungi_unclassified(100); | - | - | - | - | - | - | - | Unassigned | - |
| Otu31883 | 1 | k__Fungi(100);p__Ascomycota(100);p__Ascomycota_unclassified(100);p__Ascomycota_unclassified(100);p__Ascomycota_unclassified(100);p__Ascomycota_unclassified(100);p__Ascomycota_unclassified(100); | - | - | - | - | - | - | - | Unassigned | - |
| Otu31884 | 1 | k__Fungi(100);k__Fungi_unclassified(100);k__Fungi_unclassified(100);k__Fungi_unclassified(100);k__Fungi_unclassified(100);k__Fungi_unclassified(100);k__Fungi_unclassified(100); | - | - | - | - | - | - | - | Unassigned | - |
| Otu31886 | 1 | k__Fungi(100);k__Fungi_unclassified(100);k__Fungi_unclassified(100);k__Fungi_unclassified(100);k__Fungi_unclassified(100);k__Fungi_unclassified(100);k__Fungi_unclassified(100); | - | - | - | - | - | - | - | Unassigned | - |
| Otu31888 | 1 | k__Fungi(100);k__Fungi_unclassified(100);k__Fungi_unclassified(100);k__Fungi_unclassified(100);k__Fungi_unclassified(100);k__Fungi_unclassified(100);k__Fungi_unclassified(100); | - | - | - | - | - | - | - | Unassigned | - |
| Otu31889 | 1 | k__Fungi(100);p__Ascomycota(100);p__Ascomycota_unclassified(100);p__Ascomycota_unclassified(100);p__Ascomycota_unclassified(100);p__Ascomycota_unclassified(100);p__Ascomycota_unclassified(100); | - | - | - | - | - | - | - | Unassigned | - |
| Otu31891 | 1 | k__Fungi(100);p__Ascomycota(100);p__Ascomycota_unclassified(100);p__Ascomycota_unclassified(100);p__Ascomycota_unclassified(100);p__Ascomycota_unclassified(100);p__Ascomycota_unclassified(100); | - | - | - | - | - | - | - | Unassigned | - |
| Otu31892 | 1 | k__Fungi(100);p__Ascomycota(100);p__Ascomycota_unclassified(100);p__Ascomycota_unclassified(100);p__Ascomycota_unclassified(100);p__Ascomycota_unclassified(100);p__Ascomycota_unclassified(100); | - | - | - | - | - | - | - | Unassigned | - |
| Otu31893 | 1 | k__Fungi(100);k__Fungi_unclassified(100);k__Fungi_unclassified(100);k__Fungi_unclassified(100);k__Fungi_unclassified(100);k__Fungi_unclassified(100);k__Fungi_unclassified(100); | - | - | - | - | - | - | - | Unassigned | - |
| Otu31894 | 1 | k__Fungi(100);p__Ascomycota(100);p__Ascomycota_unclassified(100);p__Ascomycota_unclassified(100);p__Ascomycota_unclassified(100);p__Ascomycota_unclassified(100);p__Ascomycota_unclassified(100); | - | - | - | - | - | - | - | Unassigned | - |
| Otu31895 | 1 | k__Fungi(100);p__Ascomycota(100);p__Ascomycota_unclassified(100);p__Ascomycota_unclassified(100);p__Ascomycota_unclassified(100);p__Ascomycota_unclassified(100);p__Ascomycota_unclassified(100); | - | - | - | - | - | - | - | Unassigned | - |
| Otu31896 | 1 | k__Fungi(100);p__Ascomycota(100);p__Ascomycota_unclassified(100);p__Ascomycota_unclassified(100);p__Ascomycota_unclassified(100);p__Ascomycota_unclassified(100);p__Ascomycota_unclassified(100); | - | - | - | - | - | - | - | Unassigned | - |
| Otu31897 | 1 | k__Fungi(100);p__Ascomycota(100);p__Ascomycota_unclassified(100);p__Ascomycota_unclassified(100);p__Ascomycota_unclassified(100);p__Ascomycota_unclassified(100);p__Ascomycota_unclassified(100); | - | - | - | - | - | - | - | Unassigned | - |
| Otu31898 | 1 | k__Fungi(100);p__Ascomycota(100);p__Ascomycota_unclassified(100);p__Ascomycota_unclassified(100);p__Ascomycota_unclassified(100);p__Ascomycota_unclassified(100);p__Ascomycota_unclassified(100); | - | - | - | - | - | - | - | Unassigned | - |
| Otu31899 | 1 | k__Fungi(100);k__Fungi_unclassified(100);k__Fungi_unclassified(100);k__Fungi_unclassified(100);k__Fungi_unclassified(100);k__Fungi_unclassified(100);k__Fungi_unclassified(100); | - | - | - | - | - | - | - | Unassigned | - |
| Otu31900 | 1 | k__Fungi(100);k__Fungi_unclassified(100);k__Fungi_unclassified(100);k__Fungi_unclassified(100);k__Fungi_unclassified(100);k__Fungi_unclassified(100);k__Fungi_unclassified(100); | - | - | - | - | - | - | - | Unassigned | - |
| Otu31901 | 1 | k__Fungi(100);p__Ascomycota(100);p__Ascomycota_unclassified(100);p__Ascomycota_unclassified(100);p__Ascomycota_unclassified(100);p__Ascomycota_unclassified(100);p__Ascomycota_unclassified(100); | - | - | - | - | - | - | - | Unassigned | - |
| Otu31905 | 1 | k__Fungi(100);k__Fungi_unclassified(100);k__Fungi_unclassified(100);k__Fungi_unclassified(100);k__Fungi_unclassified(100);k__Fungi_unclassified(100);k__Fungi_unclassified(100); | - | - | - | - | - | - | - | Unassigned | - |
| Otu31906 | 1 | k__Fungi(100);p__Ascomycota(100);p__Ascomycota_unclassified(100);p__Ascomycota_unclassified(100);p__Ascomycota_unclassified(100);p__Ascomycota_unclassified(100);p__Ascomycota_unclassified(100); | - | - | - | - | - | - | - | Unassigned | - |
| Otu31908 | 1 | k__Fungi(100);p__Ascomycota(100);p__Ascomycota_unclassified(100);p__Ascomycota_unclassified(100);p__Ascomycota_unclassified(100);p__Ascomycota_unclassified(100);p__Ascomycota_unclassified(100); | - | - | - | - | - | - | - | Unassigned | - |
| Otu31911 | 1 | k__Fungi(100);p__Ascomycota(100);p__Ascomycota_unclassified(100);p__Ascomycota_unclassified(100);p__Ascomycota_unclassified(100);p__Ascomycota_unclassified(100);p__Ascomycota_unclassified(100); | - | - | - | - | - | - | - | Unassigned | - |
| Otu31912 | 1 | k__Fungi(100);p__Ascomycota(100);p__Ascomycota_unclassified(100);p__Ascomycota_unclassified(100);p__Ascomycota_unclassified(100);p__Ascomycota_unclassified(100);p__Ascomycota_unclassified(100); | - | - | - | - | - | - | - | Unassigned | - |
| Otu31913 | 1 | k__Fungi(100);p__Ascomycota(100);p__Ascomycota_unclassified(100);p__Ascomycota_unclassified(100);p__Ascomycota_unclassified(100);p__Ascomycota_unclassified(100);p__Ascomycota_unclassified(100); | - | - | - | - | - | - | - | Unassigned | - |
| Otu31914 | 1 | k__Fungi(100);p__Ascomycota(100);p__Ascomycota_unclassified(100);p__Ascomycota_unclassified(100);p__Ascomycota_unclassified(100);p__Ascomycota_unclassified(100);p__Ascomycota_unclassified(100); | - | - | - | - | - | - | - | Unassigned | - |
| Otu31915 | 1 | k__Fungi(100);p__Ascomycota(100);p__Ascomycota_unclassified(100);p__Ascomycota_unclassified(100);p__Ascomycota_unclassified(100);p__Ascomycota_unclassified(100);p__Ascomycota_unclassified(100); | - | - | - | - | - | - | - | Unassigned | - |
| Otu31916 | 1 | k__Fungi(100);k__Fungi_unclassified(100);k__Fungi_unclassified(100);k__Fungi_unclassified(100);k__Fungi_unclassified(100);k__Fungi_unclassified(100);k__Fungi_unclassified(100); | - | - | - | - | - | - | - | Unassigned | - |
| Otu31917 | 1 | k__Fungi(100);k__Fungi_unclassified(100);k__Fungi_unclassified(100);k__Fungi_unclassified(100);k__Fungi_unclassified(100);k__Fungi_unclassified(100);k__Fungi_unclassified(100); | - | - | - | - | - | - | - | Unassigned | - |
| Otu31918 | 1 | k__Fungi(100);p__Ascomycota(100);p__Ascomycota_unclassified(100);p__Ascomycota_unclassified(100);p__Ascomycota_unclassified(100);p__Ascomycota_unclassified(100);p__Ascomycota_unclassified(100); | - | - | - | - | - | - | - | Unassigned | - |
| Otu31919 | 1 | k__Fungi(100);p__Ascomycota(100);p__Ascomycota_unclassified(100);p__Ascomycota_unclassified(100);p__Ascomycota_unclassified(100);p__Ascomycota_unclassified(100);p__Ascomycota_unclassified(100); | - | - | - | - | - | - | - | Unassigned | - |
| Otu31920 | 1 | k__Fungi(100);p__Ascomycota(100);p__Ascomycota_unclassified(100);p__Ascomycota_unclassified(100);p__Ascomycota_unclassified(100);p__Ascomycota_unclassified(100);p__Ascomycota_unclassified(100); | - | - | - | - | - | - | - | Unassigned | - |
| Otu31921 | 1 | k__Fungi(100);p__Ascomycota(100);p__Ascomycota_unclassified(100);p__Ascomycota_unclassified(100);p__Ascomycota_unclassified(100);p__Ascomycota_unclassified(100);p__Ascomycota_unclassified(100); | - | - | - | - | - | - | - | Unassigned | - |
| Otu31922 | 1 | k__Fungi(100);p__Ascomycota(100);p__Ascomycota_unclassified(100);p__Ascomycota_unclassified(100);p__Ascomycota_unclassified(100);p__Ascomycota_unclassified(100);p__Ascomycota_unclassified(100); | - | - | - | - | - | - | - | Unassigned | - |
| Otu31923 | 1 | k__Fungi(100);k__Fungi_unclassified(100);k__Fungi_unclassified(100);k__Fungi_unclassified(100);k__Fungi_unclassified(100);k__Fungi_unclassified(100);k__Fungi_unclassified(100); | - | - | - | - | - | - | - | Unassigned | - |
| Otu31924 | 1 | k__Fungi(100);p__Ascomycota(100);p__Ascomycota_unclassified(100);p__Ascomycota_unclassified(100);p__Ascomycota_unclassified(100);p__Ascomycota_unclassified(100);p__Ascomycota_unclassified(100); | - | - | - | - | - | - | - | Unassigned | - |
| Otu31925 | 1 | k__Fungi(100);p__Ascomycota(100);p__Ascomycota_unclassified(100);p__Ascomycota_unclassified(100);p__Ascomycota_unclassified(100);p__Ascomycota_unclassified(100);p__Ascomycota_unclassified(100); | - | - | - | - | - | - | - | Unassigned | - |

| | | | | | | | | | | | | |
|---|---|---|---|---|---|---|---|---|---|---|---|---|
| Otu31926 | | 1 | k__Fungi(100);p__Ascomycota(100);p__Ascomycota_unclassified(100);p__Ascomycota_unclassified(100);p__Ascomycota_unclassified(100);p__Ascomycota_unclassified(100);p__Ascomycota_unclassified(100); | - | - | - | - | - | - | - | Unassigned | - |
| Otu31927 | | 1 | k__Fungi(100);p__Ascomycota(100);p__Ascomycota_unclassified(100);p__Ascomycota_unclassified(100);p__Ascomycota_unclassified(100);p__Ascomycota_unclassified(100);p__Ascomycota_unclassified(100); | - | - | - | - | - | - | - | Unassigned | - |
| Otu31928 | | 1 | k__Fungi(100);p__Ascomycota(100);p__Ascomycota_unclassified(100);p__Ascomycota_unclassified(100);p__Ascomycota_unclassified(100);p__Ascomycota_unclassified(100);p__Ascomycota_unclassified(100); | - | - | - | - | - | - | - | Unassigned | - |
| Otu31929 | | 1 | k__Fungi(100);p__Ascomycota(100);p__Ascomycota_unclassified(100);p__Ascomycota_unclassified(100);p__Ascomycota_unclassified(100);p__Ascomycota_unclassified(100);p__Ascomycota_unclassified(100); | - | - | - | - | - | - | - | Unassigned | - |
| Otu31930 | | 1 | k__Fungi(100);p__Ascomycota(100);p__Ascomycota_unclassified(100);p__Ascomycota_unclassified(100);p__Ascomycota_unclassified(100);p__Ascomycota_unclassified(100);p__Ascomycota_unclassified(100); | - | - | - | - | - | - | - | Unassigned | - |
| Otu31931 | | 1 | k__Fungi(100);p__Ascomycota(100);p__Ascomycota_unclassified(100);p__Ascomycota_unclassified(100);p__Ascomycota_unclassified(100);p__Ascomycota_unclassified(100);p__Ascomycota_unclassified(100); | - | - | - | - | - | - | - | Unassigned | - |
| Otu31936 | | 1 | k__Fungi(100);k__Fungi_unclassified(100);k__Fungi_unclassified(100);k__Fungi_unclassified(100);k__Fungi_unclassified(100);k__Fungi_unclassified(100);k__Fungi_unclassified(100); | - | - | - | - | - | - | - | Unassigned | - |
| Otu31937 | | 1 | k__Fungi(100);p__Ascomycota(100);p__Ascomycota_unclassified(100);p__Ascomycota_unclassified(100);p__Ascomycota_unclassified(100);p__Ascomycota_unclassified(100);p__Ascomycota_unclassified(100); | - | - | - | - | - | - | - | Unassigned | - |
| Otu31938 | | 1 | k__Fungi(100);k__Fungi_unclassified(100);k__Fungi_unclassified(100);k__Fungi_unclassified(100);k__Fungi_unclassified(100);k__Fungi_unclassified(100);k__Fungi_unclassified(100); | - | - | - | - | - | - | - | Unassigned | - |
| Otu31939 | | 1 | k__Fungi(100);k__Fungi_unclassified(100);k__Fungi_unclassified(100);k__Fungi_unclassified(100);k__Fungi_unclassified(100);k__Fungi_unclassified(100);k__Fungi_unclassified(100); | - | - | - | - | - | - | - | Unassigned | - |
| Otu31954 | | 1 | k__Fungi(100);p__Ascomycota(100);p__Ascomycota_unclassified(100);p__Ascomycota_unclassified(100);p__Ascomycota_unclassified(100);p__Ascomycota_unclassified(100);p__Ascomycota_unclassified(100); | - | - | - | - | - | - | - | Unassigned | - |
| Otu31956 | | 1 | k__Fungi(100);k__Fungi_unclassified(100);k__Fungi_unclassified(100);k__Fungi_unclassified(100);k__Fungi_unclassified(100);k__Fungi_unclassified(100);k__Fungi_unclassified(100); | - | - | - | - | - | - | - | Unassigned | - |
| Otu31960 | | 1 | k__Fungi(100);k__Fungi_unclassified(100);k__Fungi_unclassified(100);k__Fungi_unclassified(100);k__Fungi_unclassified(100);k__Fungi_unclassified(100);k__Fungi_unclassified(100); | - | - | - | - | - | - | - | Unassigned | - |
| Otu31962 | | 1 | k__Fungi(100);k__Fungi_unclassified(100);k__Fungi_unclassified(100);k__Fungi_unclassified(100);k__Fungi_unclassified(100);k__Fungi_unclassified(100);k__Fungi_unclassified(100); | - | - | - | - | - | - | - | Unassigned | - |
| Otu31993 | | 1 | k__Fungi(100);k__Fungi_unclassified(100);k__Fungi_unclassified(100);k__Fungi_unclassified(100);k__Fungi_unclassified(100);k__Fungi_unclassified(100);k__Fungi_unclassified(100); | - | - | - | - | - | - | - | Unassigned | - |
| Otu31994 | | 1 | k__Fungi(100);k__Fungi_unclassified(100);k__Fungi_unclassified(100);k__Fungi_unclassified(100);k__Fungi_unclassified(100);k__Fungi_unclassified(100);k__Fungi_unclassified(100); | - | - | - | - | - | - | - | Unassigned | - |
| Otu32003 | | 1 | k__Fungi(100);p__Ascomycota(100);p__Ascomycota_unclassified(100);p__Ascomycota_unclassified(100);p__Ascomycota_unclassified(100);p__Ascomycota_unclassified(100);p__Ascomycota_unclassified(100); | - | - | - | - | - | - | - | Unassigned | - |
| Otu32007 | | 1 | k__Fungi(100);k__Fungi_unclassified(100);k__Fungi_unclassified(100);k__Fungi_unclassified(100);k__Fungi_unclassified(100);k__Fungi_unclassified(100);k__Fungi_unclassified(100); | - | - | - | - | - | - | - | Unassigned | - |
| Otu32008 | | 1 | k__Fungi(100);k__Fungi_unclassified(100);k__Fungi_unclassified(100);k__Fungi_unclassified(100);k__Fungi_unclassified(100);k__Fungi_unclassified(100);k__Fungi_unclassified(100); | - | - | - | - | - | - | - | Unassigned | - |
| Otu32009 | | 1 | k__Fungi(100);p__Ascomycota(100);p__Ascomycota_unclassified(100);p__Ascomycota_unclassified(100);p__Ascomycota_unclassified(100);p__Ascomycota_unclassified(100);p__Ascomycota_unclassified(100); | - | - | - | - | - | - | - | Unassigned | - |
| Otu32010 | | 1 | k__Fungi(100);p__Ascomycota(100);p__Ascomycota_unclassified(100);p__Ascomycota_unclassified(100);p__Ascomycota_unclassified(100);p__Ascomycota_unclassified(100);p__Ascomycota_unclassified(100); | - | - | - | - | - | - | - | Unassigned | - |
| Otu32013 | | 1 | k__Fungi(100);k__Fungi_unclassified(100);k__Fungi_unclassified(100);k__Fungi_unclassified(100);k__Fungi_unclassified(100);k__Fungi_unclassified(100);k__Fungi_unclassified(100); | - | - | - | - | - | - | - | Unassigned | - |
| Otu32019 | | 1 | k__Fungi(100);p__Ascomycota(100);p__Ascomycota_unclassified(100);p__Ascomycota_unclassified(100);p__Ascomycota_unclassified(100);p__Ascomycota_unclassified(100);p__Ascomycota_unclassified(100); | - | - | - | - | - | - | - | Unassigned | - |
| Otu32033 | | 1 | k__Fungi(100);p__Ascomycota(100);p__Ascomycota_unclassified(100);p__Ascomycota_unclassified(100);p__Ascomycota_unclassified(100);p__Ascomycota_unclassified(100);p__Ascomycota_unclassified(100); | - | - | - | - | - | - | - | Unassigned | - |
| Otu32041 | | 1 | k__Fungi(100);p__Ascomycota(100);p__Ascomycota_unclassified(100);p__Ascomycota_unclassified(100);p__Ascomycota_unclassified(100);p__Ascomycota_unclassified(100);p__Ascomycota_unclassified(100); | - | - | - | - | - | - | - | Unassigned | - |
| Otu32047 | | 1 | k__Fungi(100);p__Ascomycota(100);p__Ascomycota_unclassified(100);p__Ascomycota_unclassified(100);p__Ascomycota_unclassified(100);p__Ascomycota_unclassified(100);p__Ascomycota_unclassified(100); | - | - | - | - | - | - | - | Unassigned | - |
| Otu32048 | | 1 | k__Fungi(100);p__Ascomycota(100);p__Ascomycota_unclassified(100);p__Ascomycota_unclassified(100);p__Ascomycota_unclassified(100);p__Ascomycota_unclassified(100);p__Ascomycota_unclassified(100); | - | - | - | - | - | - | - | Unassigned | - |
| Otu32049 | | 1 | k__Fungi(100);p__Ascomycota(100);p__Ascomycota_unclassified(100);p__Ascomycota_unclassified(100);p__Ascomycota_unclassified(100);p__Ascomycota_unclassified(100);p__Ascomycota_unclassified(100); | - | - | - | - | - | - | - | Unassigned | - |
| Otu32050 | | 1 | k__Fungi(100);k__Fungi_unclassified(100);k__Fungi_unclassified(100);k__Fungi_unclassified(100);k__Fungi_unclassified(100);k__Fungi_unclassified(100);k__Fungi_unclassified(100); | - | - | - | - | - | - | - | Unassigned | - |
| Otu32051 | | 1 | k__Fungi(100);k__Fungi_unclassified(100);k__Fungi_unclassified(100);k__Fungi_unclassified(100);k__Fungi_unclassified(100);k__Fungi_unclassified(100);k__Fungi_unclassified(100); | - | - | - | - | - | - | - | Unassigned | - |
| Otu32052 | | 1 | k__Fungi(100);p__Ascomycota(100);p__Ascomycota_unclassified(100);p__Ascomycota_unclassified(100);p__Ascomycota_unclassified(100);p__Ascomycota_unclassified(100);p__Ascomycota_unclassified(100); | - | - | - | - | - | - | - | Unassigned | - |
| Otu32053 | | 1 | k__Fungi(100);k__Fungi_unclassified(100);k__Fungi_unclassified(100);k__Fungi_unclassified(100);k__Fungi_unclassified(100);k__Fungi_unclassified(100);k__Fungi_unclassified(100); | - | - | - | - | - | - | - | Unassigned | - |
| Otu32054 | | 1 | k__Fungi(100);k__Fungi_unclassified(100);k__Fungi_unclassified(100);k__Fungi_unclassified(100);k__Fungi_unclassified(100);k__Fungi_unclassified(100);k__Fungi_unclassified(100); | - | - | - | - | - | - | - | Unassigned | - |
| Otu32055 | | 1 | k__Fungi(100);p__Ascomycota(100);p__Ascomycota_unclassified(100);p__Ascomycota_unclassified(100);p__Ascomycota_unclassified(100);p__Ascomycota_unclassified(100);p__Ascomycota_unclassified(100); | - | - | - | - | - | - | - | Unassigned | - |
| Otu32056 | | 1 | k__Fungi(100);p__Ascomycota(100);p__Ascomycota_unclassified(100);p__Ascomycota_unclassified(100);p__Ascomycota_unclassified(100);p__Ascomycota_unclassified(100);p__Ascomycota_unclassified(100); | - | - | - | - | - | - | - | Unassigned | - |
| Otu32057 | | 1 | k__Fungi(100);k__Fungi_unclassified(100);k__Fungi_unclassified(100);k__Fungi_unclassified(100);k__Fungi_unclassified(100);k__Fungi_unclassified(100);k__Fungi_unclassified(100); | - | - | - | - | - | - | - | Unassigned | - |
| Otu32059 | | 1 | k__Fungi(100);p__Ascomycota(100);p__Ascomycota_unclassified(100);p__Ascomycota_unclassified(100);p__Ascomycota_unclassified(100);p__Ascomycota_unclassified(100);p__Ascomycota_unclassified(100); | - | - | - | - | - | - | - | Unassigned | - |
| Otu32060 | | 1 | k__Fungi(100);p__Ascomycota(100);p__Ascomycota_unclassified(100);p__Ascomycota_unclassified(100);p__Ascomycota_unclassified(100);p__Ascomycota_unclassified(100);p__Ascomycota_unclassified(100); | - | - | - | - | - | - | - | Unassigned | - |
| Otu32062 | | 1 | k__Fungi(100);k__Fungi_unclassified(100);k__Fungi_unclassified(100);k__Fungi_unclassified(100);k__Fungi_unclassified(100);k__Fungi_unclassified(100);k__Fungi_unclassified(100); | - | - | - | - | - | - | - | Unassigned | - |
| Otu32064 | | 1 | k__Fungi(100);p__Ascomycota(100);p__Ascomycota_unclassified(100);p__Ascomycota_unclassified(100);p__Ascomycota_unclassified(100);p__Ascomycota_unclassified(100);p__Ascomycota_unclassified(100); | - | - | - | - | - | - | - | Unassigned | - |
| Otu32066 | | 1 | k__Fungi(100);p__Ascomycota(100);p__Ascomycota_unclassified(100);p__Ascomycota_unclassified(100);p__Ascomycota_unclassified(100);p__Ascomycota_unclassified(100);p__Ascomycota_unclassified(100); | - | - | - | - | - | - | - | Unassigned | - |
| Otu32067 | | 1 | k__Fungi(100);k__Fungi_unclassified(100);k__Fungi_unclassified(100);k__Fungi_unclassified(100);k__Fungi_unclassified(100);k__Fungi_unclassified(100);k__Fungi_unclassified(100); | - | - | - | - | - | - | - | Unassigned | - |
| Otu32068 | | 1 | k__Fungi(100);k__Fungi_unclassified(100);k__Fungi_unclassified(100);k__Fungi_unclassified(100);k__Fungi_unclassified(100);k__Fungi_unclassified(100);k__Fungi_unclassified(100); | - | - | - | - | - | - | - | Unassigned | - |
| Otu32069 | | 1 | k__Fungi(100);p__Ascomycota(100);p__Ascomycota_unclassified(100);p__Ascomycota_unclassified(100);p__Ascomycota_unclassified(100);p__Ascomycota_unclassified(100);p__Ascomycota_unclassified(100); | - | - | - | - | - | - | - | Unassigned | - |
| Otu32071 | | 1 | k__Fungi(100);p__Ascomycota(100);p__Ascomycota_unclassified(100);p__Ascomycota_unclassified(100);p__Ascomycota_unclassified(100);p__Ascomycota_unclassified(100);p__Ascomycota_unclassified(100); | - | - | - | - | - | - | - | Unassigned | - |
| Otu32073 | | 1 | k__Fungi(100);k__Fungi_unclassified(100);k__Fungi_unclassified(100);k__Fungi_unclassified(100);k__Fungi_unclassified(100);k__Fungi_unclassified(100);k__Fungi_unclassified(100); | - | - | - | - | - | - | - | Unassigned | - |
| Otu32074 | | 1 | k__Fungi(100);k__Fungi_unclassified(100);k__Fungi_unclassified(100);k__Fungi_unclassified(100);k__Fungi_unclassified(100);k__Fungi_unclassified(100);k__Fungi_unclassified(100); | - | - | - | - | - | - | - | Unassigned | - |
| Otu32075 | | 1 | k__Fungi(100);p__Ascomycota(100);p__Ascomycota_unclassified(100);p__Ascomycota_unclassified(100);p__Ascomycota_unclassified(100);p__Ascomycota_unclassified(100);p__Ascomycota_unclassified(100); | - | - | - | - | - | - | - | Unassigned | - |
| Otu32076 | | 1 | k__Fungi(100);k__Fungi_unclassified(100);k__Fungi_unclassified(100);k__Fungi_unclassified(100);k__Fungi_unclassified(100);k__Fungi_unclassified(100);k__Fungi_unclassified(100); | - | - | - | - | - | - | - | Unassigned | - |
| Otu32077 | | 1 | k__Fungi(100);p__Ascomycota(100);p__Ascomycota_unclassified(100);p__Ascomycota_unclassified(100);p__Ascomycota_unclassified(100);p__Ascomycota_unclassified(100);p__Ascomycota_unclassified(100); | - | - | - | - | - | - | - | Unassigned | - |
| Otu32081 | | 1 | k__Fungi(100);p__Ascomycota(100);p__Ascomycota_unclassified(100);p__Ascomycota_unclassified(100);p__Ascomycota_unclassified(100);p__Ascomycota_unclassified(100);p__Ascomycota_unclassified(100); | - | - | - | - | - | - | - | Unassigned | - |
| Otu32082 | | 1 | k__Fungi(100);p__Ascomycota(100);p__Ascomycota_unclassified(100);p__Ascomycota_unclassified(100);p__Ascomycota_unclassified(100);p__Ascomycota_unclassified(100);p__Ascomycota_unclassified(100); | - | - | - | - | - | - | - | Unassigned | - |
| Otu32083 | | 1 | k__Fungi(100);p__Ascomycota(100);p__Ascomycota_unclassified(100);p__Ascomycota_unclassified(100);p__Ascomycota_unclassified(100);p__Ascomycota_unclassified(100);p__Ascomycota_unclassified(100); | - | - | - | - | - | - | - | Unassigned | - |
| Otu32084 | | 1 | k__Fungi(100);p__Ascomycota(100);p__Ascomycota_unclassified(100);p__Ascomycota_unclassified(100);p__Ascomycota_unclassified(100);p__Ascomycota_unclassified(100);p__Ascomycota_unclassified(100); | - | - | - | - | - | - | - | Unassigned | - |
| Otu32085 | | 1 | k__Fungi(100);p__Ascomycota(100);p__Ascomycota_unclassified(100);p__Ascomycota_unclassified(100);p__Ascomycota_unclassified(100);p__Ascomycota_unclassified(100);p__Ascomycota_unclassified(100); | - | - | - | - | - | - | - | Unassigned | - |
| Otu32086 | | 1 | k__Fungi(100);p__Ascomycota(100);p__Ascomycota_unclassified(100);p__Ascomycota_unclassified(100);p__Ascomycota_unclassified(100);p__Ascomycota_unclassified(100);p__Ascomycota_unclassified(100); | - | - | - | - | - | - | - | Unassigned | - |
| Otu32087 | | 1 | k__Fungi(100);p__Ascomycota(100);p__Ascomycota_unclassified(100);p__Ascomycota_unclassified(100);p__Ascomycota_unclassified(100);p__Ascomycota_unclassified(100);p__Ascomycota_unclassified(100); | - | - | - | - | - | - | - | Unassigned | - |
| Otu32088 | | 1 | k__Fungi(100);p__Ascomycota(100);p__Ascomycota_unclassified(100);p__Ascomycota_unclassified(100);p__Ascomycota_unclassified(100);p__Ascomycota_unclassified(100);p__Ascomycota_unclassified(100); | - | - | - | - | - | - | - | Unassigned | - |
| Otu32089 | | 1 | k__Fungi(100);k__Fungi_unclassified(100);k__Fungi_unclassified(100);k__Fungi_unclassified(100);k__Fungi_unclassified(100);k__Fungi_unclassified(100);k__Fungi_unclassified(100); | - | - | - | - | - | - | - | Unassigned | - |
| Otu32090 | | 1 | k__Fungi(100);k__Fungi_unclassified(100);k__Fungi_unclassified(100);k__Fungi_unclassified(100);k__Fungi_unclassified(100);k__Fungi_unclassified(100);k__Fungi_unclassified(100); | - | - | - | - | - | - | - | Unassigned | - |
| Otu32091 | | 1 | k__Fungi(100);p__Ascomycota(100);p__Ascomycota_unclassified(100);p__Ascomycota_unclassified(100);p__Ascomycota_unclassified(100);p__Ascomycota_unclassified(100);p__Ascomycota_unclassified(100); | - | - | - | - | - | - | - | Unassigned | - |
| Otu32092 | | 1 | k__Fungi(100);k__Fungi_unclassified(100);k__Fungi_unclassified(100);k__Fungi_unclassified(100);k__Fungi_unclassified(100);k__Fungi_unclassified(100);k__Fungi_unclassified(100); | - | - | - | - | - | - | - | Unassigned | - |
| Otu32093 | | 1 | k__Fungi(100);k__Fungi_unclassified(100);k__Fungi_unclassified(100);k__Fungi_unclassified(100);k__Fungi_unclassified(100);k__Fungi_unclassified(100);k__Fungi_unclassified(100); | - | - | - | - | - | - | - | Unassigned | - |
| Otu32094 | | 1 | k__Fungi(100);k__Fungi_unclassified(100);k__Fungi_unclassified(100);k__Fungi_unclassified(100);k__Fungi_unclassified(100);k__Fungi_unclassified(100);k__Fungi_unclassified(100); | - | - | - | - | - | - | - | Unassigned | - |
| Otu32095 | | 1 | k__Fungi(100);p__Ascomycota(100);p__Ascomycota_unclassified(100);p__Ascomycota_unclassified(100);p__Ascomycota_unclassified(100);p__Ascomycota_unclassified(100);p__Ascomycota_unclassified(100); | - | - | - | - | - | - | - | Unassigned | - |
| Otu32097 | | 1 | k__Fungi(100);k__Fungi_unclassified(100);k__Fungi_unclassified(100);k__Fungi_unclassified(100);k__Fungi_unclassified(100);k__Fungi_unclassified(100);k__Fungi_unclassified(100); | - | - | - | - | - | - | - | Unassigned | - |
| Otu32098 | | 1 | k__Fungi(100);k__Fungi_unclassified(100);k__Fungi_unclassified(100);k__Fungi_unclassified(100);k__Fungi_unclassified(100);k__Fungi_unclassified(100);k__Fungi_unclassified(100); | - | - | - | - | - | - | - | Unassigned | - |
| Otu32099 | | 1 | k__Fungi(100);p__Ascomycota(100);p__Ascomycota_unclassified(100);p__Ascomycota_unclassified(100);p__Ascomycota_unclassified(100);p__Ascomycota_unclassified(100);p__Ascomycota_unclassified(100); | - | - | - | - | - | - | - | Unassigned | - |
| Otu32110 | | 1 | k__Fungi(100);k__Fungi_unclassified(100);k__Fungi_unclassified(100);k__Fungi_unclassified(100);k__Fungi_unclassified(100);k__Fungi_unclassified(100);k__Fungi_unclassified(100); | - | - | - | - | - | - | - | Unassigned | - |
| Otu32117 | | 1 | k__Fungi(100);k__Fungi_unclassified(100);k__Fungi_unclassified(100);k__Fungi_unclassified(100);k__Fungi_unclassified(100);k__Fungi_unclassified(100);k__Fungi_unclassified(100); | - | - | - | - | - | - | - | Unassigned | - |
| Otu32118 | | 1 | k__Fungi(100);k__Fungi_unclassified(100);k__Fungi_unclassified(100);k__Fungi_unclassified(100);k__Fungi_unclassified(100);k__Fungi_unclassified(100);k__Fungi_unclassified(100); | - | - | - | - | - | - | - | Unassigned | - |
| Otu32121 | | 1 | k__Fungi(100);k__Fungi_unclassified(100);k__Fungi_unclassified(100);k__Fungi_unclassified(100);k__Fungi_unclassified(100);k__Fungi_unclassified(100);k__Fungi_unclassified(100); | - | - | - | - | - | - | - | Unassigned | - |
| Otu32122 | | 1 | k__Fungi(100);p__Ascomycota(100);p__Ascomycota_unclassified(100);p__Ascomycota_unclassified(100);p__Ascomycota_unclassified(100);p__Ascomycota_unclassified(100);p__Ascomycota_unclassified(100); | - | - | - | - | - | - | - | Unassigned | - |
| Otu32124 | | 1 | k__Fungi(100);k__Fungi_unclassified(100);k__Fungi_unclassified(100);k__Fungi_unclassified(100);k__Fungi_unclassified(100);k__Fungi_unclassified(100);k__Fungi_unclassified(100); | - | - | - | - | - | - | - | Unassigned | - |
| Otu32125 | | 1 | k__Fungi(100);k__Fungi_unclassified(100);k__Fungi_unclassified(100);k__Fungi_unclassified(100);k__Fungi_unclassified(100);k__Fungi_unclassified(100);k__Fungi_unclassified(100); | - | - | - | - | - | - | - | Unassigned | - |
| Otu32126 | | 1 | k__Fungi(100);k__Fungi_unclassified(100);k__Fungi_unclassified(100);k__Fungi_unclassified(100);k__Fungi_unclassified(100);k__Fungi_unclassified(100);k__Fungi_unclassified(100); | - | - | - | - | - | - | - | Unassigned | - |
| Otu32130 | | 1 | k__Fungi(100);k__Fungi_unclassified(100);k__Fungi_unclassified(100);k__Fungi_unclassified(100);k__Fungi_unclassified(100);k__Fungi_unclassified(100);k__Fungi_unclassified(100); | - | - | - | - | - | - | - | Unassigned | - |
| Otu32131 | | 1 | k__Fungi(100);k__Fungi_unclassified(100);k__Fungi_unclassified(100);k__Fungi_unclassified(100);k__Fungi_unclassified(100);k__Fungi_unclassified(100);k__Fungi_unclassified(100); | - | - | - | - | - | - | - | Unassigned | - |
| Otu32132 | | 1 | k__Fungi(100);p__Ascomycota(100);p__Ascomycota_unclassified(100);p__Ascomycota_unclassified(100);p__Ascomycota_unclassified(100);p__Ascomycota_unclassified(100);p__Ascomycota_unclassified(100); | - | - | - | - | - | - | - | Unassigned | - |
| Otu32133 | | 1 | k__Fungi(100);p__Ascomycota(100);p__Ascomycota_unclassified(100);p__Ascomycota_unclassified(100);p__Ascomycota_unclassified(100);p__Ascomycota_unclassified(100);p__Ascomycota_unclassified(100); | - | - | - | - | - | - | - | Unassigned | - |
| Otu32134 | | 1 | k__Fungi(100);p__Ascomycota(100);p__Ascomycota_unclassified(100);p__Ascomycota_unclassified(100);p__Ascomycota_unclassified(100);p__Ascomycota_unclassified(100);p__Ascomycota_unclassified(100); | - | - | - | - | - | - | - | Unassigned | - |
| Otu32135 | | 1 | k__Fungi(100);p__Ascomycota(100);p__Ascomycota_unclassified(100);p__Ascomycota_unclassified(100);p__Ascomycota_unclassified(100);p__Ascomycota_unclassified(100);p__Ascomycota_unclassified(100); | - | - | - | - | - | - | - | Unassigned | - |
| Otu32140 | | 1 | k__Fungi(100);p__Ascomycota(100);p__Ascomycota_unclassified(100);p__Ascomycota_unclassified(100);p__Ascomycota_unclassified(100);p__Ascomycota_unclassified(100);p__Ascomycota_unclassified(100); | - | - | - | - | - | - | - | Unassigned | - |
| Otu32147 | | 1 | k__Fungi(100);p__Ascomycota(100);p__Ascomycota_unclassified(100);p__Ascomycota_unclassified(100);p__Ascomycota_unclassified(100);p__Ascomycota_unclassified(100);p__Ascomycota_unclassified(100); | - | - | - | - | - | - | - | Unassigned | - |
| Otu32149 | | 1 | k__Fungi(100);k__Fungi_unclassified(100);k__Fungi_unclassified(100);k__Fungi_unclassified(100);k__Fungi_unclassified(100);k__Fungi_unclassified(100);k__Fungi_unclassified(100); | - | - | - | - | - | - | - | Unassigned | - |
| Otu32162 | | 1 | k__Fungi(100);p__Ascomycota(100);p__Ascomycota_unclassified(100);p__Ascomycota_unclassified(100);p__Ascomycota_unclassified(100);p__Ascomycota_unclassified(100);p__Ascomycota_unclassified(100); | - | - | - | - | - | - | - | Unassigned | - |
| Otu32176 | | 1 | k__Fungi(100);k__Fungi_unclassified(100);k__Fungi_unclassified(100);k__Fungi_unclassified(100);k__Fungi_unclassified(100);k__Fungi_unclassified(100);k__Fungi_unclassified(100); | - | - | - | - | - | - | - | Unassigned | - |
| Otu32182 | | 1 | k__Fungi(100);p__Ascomycota(100);p__Ascomycota_unclassified(100);p__Ascomycota_unclassified(100);p__Ascomycota_unclassified(100);p__Ascomycota_unclassified(100);p__Ascomycota_unclassified(100); | - | - | - | - | - | - | - | Unassigned | - |
| Otu32183 | | 1 | k__Fungi(100);k__Fungi_unclassified(100);k__Fungi_unclassified(100);k__Fungi_unclassified(100);k__Fungi_unclassified(100);k__Fungi_unclassified(100);k__Fungi_unclassified(100); | - | - | - | - | - | - | - | Unassigned | - |
| Otu32189 | | 1 | k__Fungi(100);p__Ascomycota(100);p__Ascomycota_unclassified(100);p__Ascomycota_unclassified(100);p__Ascomycota_unclassified(100);p__Ascomycota_unclassified(100);p__Ascomycota_unclassified(100); | - | - | - | - | - | - | - | Unassigned | - |
| Otu32207 | | 1 | k__Fungi(100);p__Ascomycota(100);p__Ascomycota_unclassified(100);p__Ascomycota_unclassified(100);p__Ascomycota_unclassified(100);p__Ascomycota_unclassified(100);p__Ascomycota_unclassified(100); | - | - | - | - | - | - | - | Unassigned | - |
| Otu32208 | | 1 | k__Fungi(100);p__Ascomycota(100);p__Ascomycota_unclassified(100);p__Ascomycota_unclassified(100);p__Ascomycota_unclassified(100);p__Ascomycota_unclassified(100);p__Ascomycota_unclassified(100); | - | - | - | - | - | - | - | Unassigned | - |
| Otu32209 | | 1 | k__Fungi(100);p__Ascomycota(100);p__Ascomycota_unclassified(100);p__Ascomycota_unclassified(100);p__Ascomycota_unclassified(100);p__Ascomycota_unclassified(100);p__Ascomycota_unclassified(100); | - | - | - | - | - | - | - | Unassigned | - |
| Otu32212 | | 1 | k__Fungi(100);p__Ascomycota(100);p__Ascomycota_unclassified(100);p__Ascomycota_unclassified(100);p__Ascomycota_unclassified(100);p__Ascomycota_unclassified(100);p__Ascomycota_unclassified(100); | - | - | - | - | - | - | - | Unassigned | - |
| Otu32213 | | 1 | k__Fungi(100);p__Ascomycota(100);p__Ascomycota_unclassified(100);p__Ascomycota_unclassified(100);p__Ascomycota_unclassified(100);p__Ascomycota_unclassified(100);p__Ascomycota_unclassified(100); | - | - | - | - | - | - | - | Unassigned | - |
| Otu32214 | | 1 | k__Fungi(100);k__Fungi_unclassified(100);k__Fungi_unclassified(100);k__Fungi_unclassified(100);k__Fungi_unclassified(100);k__Fungi_unclassified(100);k__Fungi_unclassified(100); | - | - | - | - | - | - | - | Unassigned | - |
| Otu32215 | | 1 | k__Fungi(100);k__Fungi_unclassified(100);k__Fungi_unclassified(100);k__Fungi_unclassified(100);k__Fungi_unclassified(100);k__Fungi_unclassified(100);k__Fungi_unclassified(100); | - | - | - | - | - | - | - | Unassigned | - |
| Otu32216 | | 1 | k__Fungi(100);p__Ascomycota(100);p__Ascomycota_unclassified(100);p__Ascomycota_unclassified(100);p__Ascomycota_unclassified(100);p__Ascomycota_unclassified(100);p__Ascomycota_unclassified(100); | - | - | - | - | - | - | - | Unassigned | - |
| Otu32217 | | 1 | k__Fungi(100);p__Ascomycota(100);p__Ascomycota_unclassified(100);p__Ascomycota_unclassified(100);p__Ascomycota_unclassified(100);p__Ascomycota_unclassified(100);p__Ascomycota_unclassified(100); | - | - | - | - | - | - | - | Unassigned | - |
| Otu32218 | | 1 | k__Fungi(100);p__Ascomycota(100);p__Ascomycota_unclassified(100);p__Ascomycota_unclassified(100);p__Ascomycota_unclassified(100);p__Ascomycota_unclassified(100);p__Ascomycota_unclassified(100); | - | - | - | - | - | - | - | Unassigned | - |
| Otu32219 | | 1 | k__Fungi(100);k__Fungi_unclassified(100);k__Fungi_unclassified(100);k__Fungi_unclassified(100);k__Fungi_unclassified(100);k__Fungi_unclassified(100);k__Fungi_unclassified(100); | - | - | - | - | - | - | - | Unassigned | - |
| Otu32220 | | 1 | k__Fungi(100);p__Ascomycota(100);p__Ascomycota_unclassified(100);p__Ascomycota_unclassified(100);p__Ascomycota_unclassified(100);p__Ascomycota_unclassified(100);p__Ascomycota_unclassified(100); | - | - | - | - | - | - | - | Unassigned | - |
| Otu32222 | | 1 | k__Fungi(100);k__Fungi_unclassified(100);k__Fungi_unclassified(100);k__Fungi_unclassified(100);k__Fungi_unclassified(100);k__Fungi_unclassified(100);k__Fungi_unclassified(100); | - | - | - | - | - | - | - | Unassigned | - |
| Otu32223 | | 1 | k__Fungi(100);p__Ascomycota(100);p__Ascomycota_unclassified(100);p__Ascomycota_unclassified(100);p__Ascomycota_unclassified(100);p__Ascomycota_unclassified(100);p__Ascomycota_unclassified(100); | - | - | - | - | - | - | - | Unassigned | - |
| Otu32224 | | 1 | k__Fungi(100);k__Fungi_unclassified(100);k__Fungi_unclassified(100);k__Fungi_unclassified(100);k__Fungi_unclassified(100);k__Fungi_unclassified(100);k__Fungi_unclassified(100); | - | - | - | - | - | - | - | Unassigned | - |
| Otu32225 | | 1 | k__Fungi(100);k__Fungi_unclassified(100);k__Fungi_unclassified(100);k__Fungi_unclassified(100);k__Fungi_unclassified(100);k__Fungi_unclassified(100);k__Fungi_unclassified(100); | - | - | - | - | - | - | - | Unassigned | - |
| Otu32226 | | 1 | k__Fungi(100);p__Ascomycota(100);p__Ascomycota_unclassified(100);p__Ascomycota_unclassified(100);p__Ascomycota_unclassified(100);p__Ascomycota_unclassified(100);p__Ascomycota_unclassified(100); | - | - | - | - | - | - | - | Unassigned | - |
| Otu32227 | | 1 | k__Fungi(100);k__Fungi_unclassified(100);k__Fungi_unclassified(100);k__Fungi_unclassified(100);k__Fungi_unclassified(100);k__Fungi_unclassified(100);k__Fungi_unclassified(100); | - | - | - | - | - | - | - | Unassigned | - |
| Otu32228 | | 1 | k__Fungi(100);k__Fungi_unclassified(100);k__Fungi_unclassified(100);k__Fungi_unclassified(100);k__Fungi_unclassified(100);k__Fungi_unclassified(100);k__Fungi_unclassified(100); | - | - | - | - | - | - | - | Unassigned | - |
| Otu32229 | | 1 | k__Fungi(100);p__Ascomycota(100);p__Ascomycota_unclassified(100);p__Ascomycota_unclassified(100);p__Ascomycota_unclassified(100);p__Ascomycota_unclassified(100);p__Ascomycota_unclassified(100); | - | - | - | - | - | - | - | Unassigned | - |
| Otu32230 | | 1 | k__Fungi(100);k__Fungi_unclassified(100);k__Fungi_unclassified(100);k__Fungi_unclassified(100);k__Fungi_unclassified(100);k__Fungi_unclassified(100);k__Fungi_unclassified(100); | - | - | - | - | - | - | - | Unassigned | - |
| Otu32231 | | 1 | k__Fungi(100);p__Ascomycota(100);p__Ascomycota_unclassified(100);p__Ascomycota_unclassified(100);p__Ascomycota_unclassified(100);p__Ascomycota_unclassified(100);p__Ascomycota_unclassified(100); | - | - | - | - | - | - | - | Unassigned | - |
| Otu32232 | | 1 | k__Fungi(100);p__Ascomycota(100);p__Ascomycota_unclassified(100);p__Ascomycota_unclassified(100);p__Ascomycota_unclassified(100);p__Ascomycota_unclassified(100);p__Ascomycota_unclassified(100); | - | - | - | - | - | - | - | Unassigned | - |
| Otu32233 | | 1 | k__Fungi(100);p__Ascomycota(100);p__Ascomycota_unclassified(100);p__Ascomycota_unclassified(100);p__Ascomycota_unclassified(100);p__Ascomycota_unclassified(100);p__Ascomycota_unclassified(100); | - | - | - | - | - | - | - | Unassigned | - |
| Otu32237 | | 1 | k__Fungi(100);k__Fungi_unclassified(100);k__Fungi_unclassified(100);k__Fungi_unclassified(100);k__Fungi_unclassified(100);k__Fungi_unclassified(100);k__Fungi_unclassified(100); | - | - | - | - | - | - | - | Unassigned | - |
| Otu32240 | | 1 | k__Fungi(100);p__Ascomycota(100);p__Ascomycota_unclassified(100);p__Ascomycota_unclassified(100);p__Ascomycota_unclassified(100);p__Ascomycota_unclassified(100);p__Ascomycota_unclassified(100); | - | - | - | - | - | - | - | Unassigned | - |
| Otu32241 | | 1 | k__Fungi(100);k__Fungi_unclassified(100);k__Fungi_unclassified(100);k__Fungi_unclassified(100);k__Fungi_unclassified(100);k__Fungi_unclassified(100);k__Fungi_unclassified(100); | - | - | - | - | - | - | - | Unassigned | - |
| Otu32242 | | 1 | k__Fungi(100);k__Fungi_unclassified(100);k__Fungi_unclassified(100);k__Fungi_unclassified(100);k__Fungi_unclassified(100);k__Fungi_unclassified(100);k__Fungi_unclassified(100); | - | - | - | - | - | - | - | Unassigned | - |
| Otu32243 | | 1 | k__Fungi(100);k__Fungi_unclassified(100);k__Fungi_unclassified(100);k__Fungi_unclassified(100);k__Fungi_unclassified(100);k__Fungi_unclassified(100);k__Fungi_unclassified(100); | - | - | - | - | - | - | - | Unassigned | - |
| Otu32244 | | 1 | k__Fungi(100);k__Fungi_unclassified(100);k__Fungi_unclassified(100);k__Fungi_unclassified(100);k__Fungi_unclassified(100);k__Fungi_unclassified(100);k__Fungi_unclassified(100); | - | - | - | - | - | - | - | Unassigned | - |
| Otu32245 | | 1 | k__Fungi(100);p__Ascomycota(100);p__Ascomycota_unclassified(100);p__Ascomycota_unclassified(100);p__Ascomycota_unclassified(100);p__Ascomycota_unclassified(100);p__Ascomycota_unclassified(100); | - | - | - | - | - | - | - | Unassigned | - |
| Otu32246 | | 1 | k__Fungi(100);k__Fungi_unclassified(100);k__Fungi_unclassified(100);k__Fungi_unclassified(100);k__Fungi_unclassified(100);k__Fungi_unclassified(100);k__Fungi_unclassified(100); | - | - | - | - | - | - | - | Unassigned | - |
| Otu32247 | | 1 | k__Fungi(100);k__Fungi_unclassified(100);k__Fungi_unclassified(100);k__Fungi_unclassified(100);k__Fungi_unclassified(100);k__Fungi_unclassified(100);k__Fungi_unclassified(100); | - | - | - | - | - | - | - | Unassigned | - |
| Otu32249 | | 1 | k__Fungi(100);k__Fungi_unclassified(100);k__Fungi_unclassified(100);k__Fungi_unclassified(100);k__Fungi_unclassified(100);k__Fungi_unclassified(100);k__Fungi_unclassified(100); | - | - | - | - | - | - | - | Unassigned | - |
| Otu32250 | | 1 | k__Fungi(100);p__Ascomycota(100);p__Ascomycota_unclassified(100);p__Ascomycota_unclassified(100);p__Ascomycota_unclassified(100);p__Ascomycota_unclassified(100);p__Ascomycota_unclassified(100); | - | - | - | - | - | - | - | Unassigned | - |
| Otu32251 | | 1 | k__Fungi(100);k__Fungi_unclassified(100);k__Fungi_unclassified(100);k__Fungi_unclassified(100);k__Fungi_unclassified(100);k__Fungi_unclassified(100);k__Fungi_unclassified(100); | - | - | - | - | - | - | - | Unassigned | - |
| Otu32252 | | 1 | k__Fungi(100);k__Fungi_unclassified(100);k__Fungi_unclassified(100);k__Fungi_unclassified(100);k__Fungi_unclassified(100);k__Fungi_unclassified(100);k__Fungi_unclassified(100); | - | - | - | - | - | - | - | Unassigned | - |
| Otu32253 | | 1 | k__Fungi(100);p__Ascomycota(100);p__Ascomycota_unclassified(100);p__Ascomycota_unclassified(100);p__Ascomycota_unclassified(100);p__Ascomycota_unclassified(100);p__Ascomycota_unclassified(100); | - | - | - | - | - | - | - | Unassigned | - |
| Otu32254 | | 1 | k__Fungi(100);k__Fungi_unclassified(100);k__Fungi_unclassified(100);k__Fungi_unclassified(100);k__Fungi_unclassified(100);k__Fungi_unclassified(100);k__Fungi_unclassified(100); | - | - | - | - | - | - | - | Unassigned | - |
| Otu32255 | | 1 | k__Fungi(100);p__Ascomycota(100);p__Ascomycota_unclassified(100);p__Ascomycota_unclassified(100);p__Ascomycota_unclassified(100);p__Ascomycota_unclassified(100);p__Ascomycota_unclassified(100); | - | - | - | - | - | - | - | Unassigned | - |
| Otu32256 | | 1 | k__Fungi(100);k__Fungi_unclassified(100);k__Fungi_unclassified(100);k__Fungi_unclassified(100);k__Fungi_unclassified(100);k__Fungi_unclassified(100);k__Fungi_unclassified(100); | - | - | - | - | - | - | - | Unassigned | - |
| Otu32257 | | 1 | k__Fungi(100);p__Ascomycota(100);p__Ascomycota_unclassified(100);p__Ascomycota_unclassified(100);p__Ascomycota_unclassified(100);p__Ascomycota_unclassified(100);p__Ascomycota_unclassified(100); | - | - | - | - | - | - | - | Unassigned | - |
| Otu32258 | | 1 | k__Fungi(100);k__Fungi_unclassified(100);k__Fungi_unclassified(100);k__Fungi_unclassified(100);k__Fungi_unclassified(100);k__Fungi_unclassified(100);k__Fungi_unclassified(100); | - | - | - | - | - | - | - | Unassigned | - |
| Otu32260 | | 1 | k__Fungi(100);p__Ascomycota(100);p__Ascomycota_unclassified(100);p__Ascomycota_unclassified(100);p__Ascomycota_unclassified(100);p__Ascomycota_unclassified(100);p__Ascomycota_unclassified(100); | - | - | - | - | - | - | - | Unassigned | - |
| Otu32262 | | 1 | k__Fungi(100);k__Fungi_unclassified(100);k__Fungi_unclassified(100);k__Fungi_unclassified(100);k__Fungi_unclassified(100);k__Fungi_unclassified(100);k__Fungi_unclassified(100); | - | - | - | - | - | - | - | Unassigned | - |
| Otu32263 | | 1 | k__Fungi(100);k__Fungi_unclassified(100);k__Fungi_unclassified(100);k__Fungi_unclassified(100);k__Fungi_unclassified(100);k__Fungi_unclassified(100);k__Fungi_unclassified(100); | - | - | - | - | - | - | - | Unassigned | - |
| Otu32265 | | 1 | k__Fungi(100);p__Ascomycota(100);p__Ascomycota_unclassified(100);p__Ascomycota_unclassified(100);p__Ascomycota_unclassified(100);p__Ascomycota_unclassified(100);p__Ascomycota_unclassified(100); | - | - | - | - | - | - | - | Unassigned | - |
| Otu32266 | | 1 | k__Fungi(100);k__Fungi_unclassified(100);k__Fungi_unclassified(100);k__Fungi_unclassified(100);k__Fungi_unclassified(100);k__Fungi_unclassified(100);k__Fungi_unclassified(100); | - | - | - | - | - | - | - | Unassigned | - |
| Otu32268 | | 1 | k__Fungi(100);p__Ascomycota(100);p__Ascomycota_unclassified(100);p__Ascomycota_unclassified(100);p__Ascomycota_unclassified(100);p__Ascomycota_unclassified(100);p__Ascomycota_unclassified(100); | - | - | - | - | - | - | - | Unassigned | - |
| Otu32269 | | 1 | k__Fungi(100);k__Fungi_unclassified(100);k__Fungi_unclassified(100);k__Fungi_unclassified(100);k__Fungi_unclassified(100);k__Fungi_unclassified(100);k__Fungi_unclassified(100); | - | - | - | - | - | - | - | Unassigned | - |
| Otu32271 | | 1 | k__Fungi(100);p__Ascomycota(100);p__Ascomycota_unclassified(100);p__Ascomycota_unclassified(100);p__Ascomycota_unclassified(100);p__Ascomycota_unclassified(100);p__Ascomycota_unclassified(100); | - | - | - | - | - | - | - | Unassigned | - |
| Otu32274 | | 1 | k__Fungi(100);p__Ascomycota(100);p__Ascomycota_unclassified(100);p__Ascomycota_unclassified(100);p__Ascomycota_unclassified(100);p__Ascomycota_unclassified(100);p__Ascomycota_unclassified(100); | - | - | - | - | - | - | - | Unassigned | - |
| Otu32278 | | 1 | k__Fungi(100);p__Ascomycota(100);p__Ascomycota_unclassified(100);p__Ascomycota_unclassified(100);p__Ascomycota_unclassified(100);p__Ascomycota_unclassified(100);p__Ascomycota_unclassified(100); | - | - | - | - | - | - | - | Unassigned | - |
| Otu32279 | | 1 | k__Fungi(100);k__Fungi_unclassified(100);k__Fungi_unclassified(100);k__Fungi_unclassified(100);k__Fungi_unclassified(100);k__Fungi_unclassified(100);k__Fungi_unclassified(100); | - | - | - | - | - | - | - | Unassigned | - |
| Otu32280 | | 1 | k__Fungi(100);k__Fungi_unclassified(100);k__Fungi_unclassified(100);k__Fungi_unclassified(100);k__Fungi_unclassified(100);k__Fungi_unclassified(100);k__Fungi_unclassified(100); | - | - | - | - | - | - | - | Unassigned | - |
| Otu32287 | | 1 | k__Fungi(100);k__Fungi_unclassified(100);k__Fungi_unclassified(100);k__Fungi_unclassified(100);k__Fungi_unclassified(100);k__Fungi_unclassified(100);k__Fungi_unclassified(100); | - | - | - | - | - | - | - | Unassigned | - |
| Otu32288 | | 1 | k__Fungi(100);k__Fungi_unclassified(100);k__Fungi_unclassified(100);k__Fungi_unclassified(100);k__Fungi_unclassified(100);k__Fungi_unclassified(100);k__Fungi_unclassified(100); | - | - | - | - | - | - | - | Unassigned | - |
| Otu32296 | | 1 | k__Fungi(100);p__Ascomycota(100);p__Ascomycota_unclassified(100);p__Ascomycota_unclassified(100);p__Ascomycota_unclassified(100);p__Ascomycota_unclassified(100);p__Ascomycota_unclassified(100); | - | - | - | - | - | - | - | Unassigned | - |
| Otu32300 | | 1 | k__Fungi(100);k__Fungi_unclassified(100);k__Fungi_unclassified(100);k__Fungi_unclassified(100);k__Fungi_unclassified(100);k__Fungi_unclassified(100);k__Fungi_unclassified(100); | - | - | - | - | - | - | - | Unassigned | - |
| Otu32301 | | 1 | k__Fungi(100);p__Ascomycota(100);p__Ascomycota_unclassified(100);p__Ascomycota_unclassified(100);p__Ascomycota_unclassified(100);p__Ascomycota_unclassified(100);p__Ascomycota_unclassified(100); | - | - | - | - | - | - | - | Unassigned | - |
| Otu32303 | | 1 | k__Fungi(100);k__Fungi_unclassified(100);k__Fungi_unclassified(100);k__Fungi_unclassified(100);k__Fungi_unclassified(100);k__Fungi_unclassified(100);k__Fungi_unclassified(100); | - | - | - | - | - | - | - | Unassigned | - |
| Otu32304 | | 1 | k__Fungi(100);k__Fungi_unclassified(100);k__Fungi_unclassified(100);k__Fungi_unclassified(100);k__Fungi_unclassified(100);k__Fungi_unclassified(100);k__Fungi_unclassified(100); | - | - | - | - | - | - | - | Unassigned | - |
| Otu32309 | | 1 | k__Fungi(100);k__Fungi_unclassified(100);k__Fungi_unclassified(100);k__Fungi_unclassified(100);k__Fungi_unclassified(100);k__Fungi_unclassified(100);k__Fungi_unclassified(100); | - | - | - | - | - | - | - | Unassigned | - |
| Otu32321 | | 1 | k__Fungi(100);p__Ascomycota(100);p__Ascomycota_unclassified(100);p__Ascomycota_unclassified(100);p__Ascomycota_unclassified(100);p__Ascomycota_unclassified(100);p__Ascomycota_unclassified(100); | - | - | - | - | - | - | - | Unassigned | - |
| Otu32327 | | 1 | k__Fungi(100);p__Ascomycota(100);p__Ascomycota_unclassified(100);p__Ascomycota_unclassified(100);p__Ascomycota_unclassified(100);p__Ascomycota_unclassified(100);p__Ascomycota_unclassified(100); | - | - | - | - | - | - | - | Unassigned | - |
| Otu32328 | | 1 | k__Fungi(100);k__Fungi_unclassified(100);k__Fungi_unclassified(100);k__Fungi_unclassified(100);k__Fungi_unclassified(100);k__Fungi_unclassified(100);k__Fungi_unclassified(100); | - | - | - | - | - | - | - | Unassigned | - |
| Otu32329 | | 1 | k__Fungi(100);p__Ascomycota(100);p__Ascomycota_unclassified(100);p__Ascomycota_unclassified(100);p__Ascomycota_unclassified(100);p__Ascomycota_unclassified(100);p__Ascomycota_unclassified(100); | - | - | - | - | - | - | - | Unassigned | - |
| Otu32330 | | 1 | k__Fungi(100);k__Fungi_unclassified(100);k__Fungi_unclassified(100);k__Fungi_unclassified(100);k__Fungi_unclassified(100);k__Fungi_unclassified(100);k__Fungi_unclassified(100); | - | - | - | - | - | - | - | Unassigned | - |
| Otu32331 | | 1 | k__Fungi(100);k__Fungi_unclassified(100);k__Fungi_unclassified(100);k__Fungi_unclassified(100);k__Fungi_unclassified(100);k__Fungi_unclassified(100);k__Fungi_unclassified(100); | - | - | - | - | - | - | - | Unassigned | - |
| Otu32332 | | 1 | k__Fungi(100);p__Ascomycota(100);p__Ascomycota_unclassified(100);p__Ascomycota_unclassified(100);p__Ascomycota_unclassified(100);p__Ascomycota_unclassified(100);p__Ascomycota_unclassified(100); | - | - | - | - | - | - | - | Unassigned | - |
| Otu32333 | | 1 | k__Fungi(100);p__Ascomycota(100);p__Ascomycota_unclassified(100);p__Ascomycota_unclassified(100);p__Ascomycota_unclassified(100);p__Ascomycota_unclassified(100);p__Ascomycota_unclassified(100); | - | - | - | - | - | - | - | Unassigned | - |
| Otu32334 | | 1 | k__Fungi(100);k__Fungi_unclassified(100);k__Fungi_unclassified(100);k__Fungi_unclassified(100);k__Fungi_unclassified(100);k__Fungi_unclassified(100);k__Fungi_unclassified(100); | - | - | - | - | - | - | - | Unassigned | - |
| Otu32335 | | 1 | k__Fungi(100);p__Ascomycota(100);p__Ascomycota_unclassified(100);p__Ascomycota_unclassified(100);p__Ascomycota_unclassified(100);p__Ascomycota_unclassified(100);p__Ascomycota_unclassified(100); | - | - | - | - | - | - | - | Unassigned | - |
| Otu32338 | | 1 | k__Fungi(100);p__Ascomycota(100);p__Ascomycota_unclassified(100);p__Ascomycota_unclassified(100);p__Ascomycota_unclassified(100);p__Ascomycota_unclassified(100);p__Ascomycota_unclassified(100); | - | - | - | - | - | - | - | Unassigned | - |
| Otu32348 | | 1 | k__Fungi(100);k__Fungi_unclassified(100);k__Fungi_unclassified(100);k__Fungi_unclassified(100);k__Fungi_unclassified(100);k__Fungi_unclassified(100);k__Fungi_unclassified(100); | - | - | - | - | - | - | - | Unassigned | - |
| Otu32352 | | 1 | k__Fungi(100);p__Ascomycota(100);p__Ascomycota_unclassified(100);p__Ascomycota_unclassified(100);p__Ascomycota_unclassified(100);p__Ascomycota_unclassified(100);p__Ascomycota_unclassified(100); | - | - | - | - | - | - | - | Unassigned | - |
| Otu32358 | | 1 | k__Fungi(100);k__Fungi_unclassified(100);k__Fungi_unclassified(100);k__Fungi_unclassified(100);k__Fungi_unclassified(100);k__Fungi_unclassified(100);k__Fungi_unclassified(100); | - | - | - | - | - | - | - | Unassigned | - |
| Otu32371 | | 1 | k__Fungi(100);k__Fungi_unclassified(100);k__Fungi_unclassified(100);k__Fungi_unclassified(100);k__Fungi_unclassified(100);k__Fungi_unclassified(100);k__Fungi_unclassified(100); | - | - | - | - | - | - | - | Unassigned | - |
| Otu32372 | | 1 | k__Fungi(100);p__Ascomycota(100);p__Ascomycota_unclassified(100);p__Ascomycota_unclassified(100);p__Ascomycota_unclassified(100);p__Ascomycota_unclassified(100);p__Ascomycota_unclassified(100); | - | - | - | - | - | - | - | Unassigned | - |
| Otu32374 | | 1 | k__Fungi(100);k__Fungi_unclassified(100);k__Fungi_unclassified(100);k__Fungi_unclassified(100);k__Fungi_unclassified(100);k__Fungi_unclassified(100);k__Fungi_unclassified(100); | - | - | - | - | - | - | - | Unassigned | - |
| Otu32376 | | 1 | k__Fungi(100);p__Ascomycota(100);p__Ascomycota_unclassified(100);p__Ascomycota_unclassified(100);p__Ascomycota_unclassified(100);p__Ascomycota_unclassified(100);p__Ascomycota_unclassified(100); | - | - | - | - | - | - | - | Unassigned | - |
| Otu32377 | | 1 | k__Fungi(100);p__Ascomycota(100);p__Ascomycota_unclassified(100);p__Ascomycota_unclassified(100);p__Ascomycota_unclassified(100);p__Ascomycota_unclassified(100);p__Ascomycota_unclassified(100); | - | - | - | - | - | - | - | Unassigned | - |
| Otu32380 | | 1 | k__Fungi(100);k__Fungi_unclassified(100);k__Fungi_unclassified(100);k__Fungi_unclassified(100);k__Fungi_unclassified(100);k__Fungi_unclassified(100);k__Fungi_unclassified(100); | - | - | - | - | - | - | - | Unassigned | - |
| Otu32383 | | 1 | k__Fungi(100);k__Fungi_unclassified(100);k__Fungi_unclassified(100);k__Fungi_unclassified(100);k__Fungi_unclassified(100);k__Fungi_unclassified(100);k__Fungi_unclassified(100); | - | - | - | - | - | - | - | Unassigned | - |
| Otu32392 | | 1 | k__Fungi(100);k__Fungi_unclassified(100);k__Fungi_unclassified(100);k__Fungi_unclassified(100);k__Fungi_unclassified(100);k__Fungi_unclassified(100);k__Fungi_unclassified(100); | - | - | - | - | - | - | - | Unassigned | - |
| Otu32393 | | 1 | k__Fungi(100);k__Fungi_unclassified(100);k__Fungi_unclassified(100);k__Fungi_unclassified(100);k__Fungi_unclassified(100);k__Fungi_unclassified(100);k__Fungi_unclassified(100); | - | - | - | - | - | - | - | Unassigned | - |
| Otu32394 | | 1 | k__Fungi(100);p__Ascomycota(100);p__Ascomycota_unclassified(100);p__Ascomycota_unclassified(100);p__Ascomycota_unclassified(100);p__Ascomycota_unclassified(100);p__Ascomycota_unclassified(100); | - | - | - | - | - | - | - | Unassigned | - |
| Otu32396 | | 1 | k__Fungi(100);k__Fungi_unclassified(100);k__Fungi_unclassified(100);k__Fungi_unclassified(100);k__Fungi_unclassified(100);k__Fungi_unclassified(100);k__Fungi_unclassified(100); | - | - | - | - | - | - | - | Unassigned | - |
| Otu32398 | | 1 | k__Fungi(100);k__Fungi_unclassified(100);k__Fungi_unclassified(100);k__Fungi_unclassified(100);k__Fungi_unclassified(100);k__Fungi_unclassified(100);k__Fungi_unclassified(100); | - | - | - | - | - | - | - | Unassigned | - |
| Otu32399 | | 1 | k__Fungi(100);k__Fungi_unclassified(100);k__Fungi_unclassified(100);k__Fungi_unclassified(100);k__Fungi_unclassified(100);k__Fungi_unclassified(100);k__Fungi_unclassified(100); | - | - | - | - | - | - | - | Unassigned | - |
| Otu32400 | | 1 | k__Fungi(100);k__Fungi_unclassified(100);k__Fungi_unclassified(100);k__Fungi_unclassified(100);k__Fungi_unclassified(100);k__Fungi_unclassified(100);k__Fungi_unclassified(100); | - | - | - | - | - | - | - | Unassigned | - |
| Otu32401 | | 1 | k__Fungi(100);p__Ascomycota(100);p__Ascomycota_unclassified(100);p__Ascomycota_unclassified(100);p__Ascomycota_unclassified(100);p__Ascomycota_unclassified(100);p__Ascomycota_unclassified(100); | - | - | - | - | - | - | - | Unassigned | - |
| Otu32402 | | 1 | k__Fungi(100);k__Fungi_unclassified(100);k__Fungi_unclassified(100);k__Fungi_unclassified(100);k__Fungi_unclassified(100);k__Fungi_unclassified(100);k__Fungi_unclassified(100); | - | - | - | - | - | - | - | Unassigned | - |
| Otu32403 | | 1 | k__Fungi(100);k__Fungi_unclassified(100);k__Fungi_unclassified(100);k__Fungi_unclassified(100);k__Fungi_unclassified(100);k__Fungi_unclassified(100);k__Fungi_unclassified(100); | - | - | - | - | - | - | - | Unassigned | - |
| Otu32404 | | 1 | k__Fungi(100);p__Ascomycota(100);p__Ascomycota_unclassified(100);p__Ascomycota_unclassified(100);p__Ascomycota_unclassified(100);p__Ascomycota_unclassified(100);p__Ascomycota_unclassified(100); | - | - | - | - | - | - | - | Unassigned | - |
| Otu32406 | | 1 | k__Fungi(100);p__Ascomycota(100);p__Ascomycota_unclassified(100);p__Ascomycota_unclassified(100);p__Ascomycota_unclassified(100);p__Ascomycota_unclassified(100);p__Ascomycota_unclassified(100); | - | - | - | - | - | - | - | Unassigned | - |
| Otu32407 | | 1 | k__Fungi(100);k__Fungi_unclassified(100);k__Fungi_unclassified(100);k__Fungi_unclassified(100);k__Fungi_unclassified(100);k__Fungi_unclassified(100);k__Fungi_unclassified(100); | - | - | - | - | - | - | - | Unassigned | - |
| Otu32408 | | 1 | k__Fungi(100);k__Fungi_unclassified(100);k__Fungi_unclassified(100);k__Fungi_unclassified(100);k__Fungi_unclassified(100);k__Fungi_unclassified(100);k__Fungi_unclassified(100); | - | - | - | - | - | - | - | Unassigned | - |
| Otu32410 | | 1 | k__Fungi(100);k__Fungi_unclassified(100);k__Fungi_unclassified(100);k__Fungi_unclassified(100);k__Fungi_unclassified(100);k__Fungi_unclassified(100);k__Fungi_unclassified(100); | - | - | - | - | - | - | - | Unassigned | - |
| Otu32414 | | 1 | k__Fungi(100);k__Fungi_unclassified(100);k__Fungi_unclassified(100);k__Fungi_unclassified(100);k__Fungi_unclassified(100);k__Fungi_unclassified(100);k__Fungi_unclassified(100); | - | - | - | - | - | - | - | Unassigned | - |
| Otu32416 | | 1 | k__Fungi(100);k__Fungi_unclassified(100);k__Fungi_unclassified(100);k__Fungi_unclassified(100);k__Fungi_unclassified(100);k__Fungi_unclassified(100);k__Fungi_unclassified(100); | - | - | - | - | - | - | - | Unassigned | - |
| Otu32417 | | 1 | k__Fungi(100);p__Ascomycota(100);p__Ascomycota_unclassified(100);p__Ascomycota_unclassified(100);p__Ascomycota_unclassified(100);p__Ascomycota_unclassified(100);p__Ascomycota_unclassified(100); | - | - | - | - | - | - | - | Unassigned | - |
| Otu32419 | | 1 | k__Fungi(100);k__Fungi_unclassified(100);k__Fungi_unclassified(100);k__Fungi_unclassified(100);k__Fungi_unclassified(100);k__Fungi_unclassified(100);k__Fungi_unclassified(100); | - | - | - | - | - | - | - | Unassigned | - |
| Otu32421 | | 1 | k__Fungi(100);p__Ascomycota(100);p__Ascomycota_unclassified(100);p__Ascomycota_unclassified(100);p__Ascomycota_unclassified(100);p__Ascomycota_unclassified(100);p__Ascomycota_unclassified(100); | - | - | - | - | - | - | - | Unassigned | - |
| Otu32422 | | 1 | k__Fungi(100);p__Ascomycota(100);p__Ascomycota_unclassified(100);p__Ascomycota_unclassified(100);p__Ascomycota_unclassified(100);p__Ascomycota_unclassified(100);p__Ascomycota_unclassified(100); | - | - | - | - | - | - | - | Unassigned | - |
| Otu32425 | | 1 | k__Fungi(100);k__Fungi_unclassified(100);k__Fungi_unclassified(100);k__Fungi_unclassified(100);k__Fungi_unclassified(100);k__Fungi_unclassified(100);k__Fungi_unclassified(100); | - | - | - | - | - | - | - | Unassigned | - |
| Otu32426 | | 1 | k__Fungi(100);k__Fungi_unclassified(100);k__Fungi_unclassified(100);k__Fungi_unclassified(100);k__Fungi_unclassified(100);k__Fungi_unclassified(100);k__Fungi_unclassified(100); | - | - | - | - | - | - | - | Unassigned | - |
| Otu32427 | | 1 | k__Fungi(100);p__Ascomycota(100);p__Ascomycota_unclassified(100);p__Ascomycota_unclassified(100);p__Ascomycota_unclassified(100);p__Ascomycota_unclassified(100);p__Ascomycota_unclassified(100); | - | - | - | - | - | - | - | Unassigned | - |
| Otu32432 | | 1 | k__Fungi(100);p__Ascomycota(100);p__Ascomycota_unclassified(100);p__Ascomycota_unclassified(100);p__Ascomycota_unclassified(100);p__Ascomycota_unclassified(100);p__Ascomycota_unclassified(100); | - | - | - | - | - | - | - | Unassigned | - |
| Otu32436 | | 1 | k__Fungi(100);p__Ascomycota(100);p__Ascomycota_unclassified(100);p__Ascomycota_unclassified(100);p__Ascomycota_unclassified(100);p__Ascomycota_unclassified(100);p__Ascomycota_unclassified(100); | - | - | - | - | - | - | - | Unassigned | - |
| Otu32440 | | 1 | k__Fungi(100);k__Fungi_unclassified(100);k__Fungi_unclassified(100);k__Fungi_unclassified(100);k__Fungi_unclassified(100);k__Fungi_unclassified(100);k__Fungi_unclassified(100); | - | - | - | - | - | - | - | Unassigned | - |
| Otu32444 | | 1 | k__Fungi(100);k__Fungi_unclassified(100);k__Fungi_unclassified(100);k__Fungi_unclassified(100);k__Fungi_unclassified(100);k__Fungi_unclassified(100);k__Fungi_unclassified(100); | - | - | - | - | - | - | - | Unassigned | - |
| Otu32445 | | 1 | k__Fungi(100);k__Fungi_unclassified(100);k__Fungi_unclassified(100);k__Fungi_unclassified(100);k__Fungi_unclassified(100);k__Fungi_unclassified(100);k__Fungi_unclassified(100); | - | - | - | - | - | - | - | Unassigned | - |
| Otu32447 | | 1 | k__Fungi(100);k__Fungi_unclassified(100);k__Fungi_unclassified(100);k__Fungi_unclassified(100);k__Fungi_unclassified(100);k__Fungi_unclassified(100);k__Fungi_unclassified(100); | - | - | - | - | - | - | - | Unassigned | - |
| Otu32448 | | 1 | k__Fungi(100);k__Fungi_unclassified(100);k__Fungi_unclassified(100);k__Fungi_unclassified(100);k__Fungi_unclassified(100);k__Fungi_unclassified(100);k__Fungi_unclassified(100); | - | - | - | - | - | - | - | Unassigned | - |
| Otu32450 | | 1 | k__Fungi(100);p__Ascomycota(100);p__Ascomycota_unclassified(100);p__Ascomycota_unclassified(100);p__Ascomycota_unclassified(100);p__Ascomycota_unclassified(100);p__Ascomycota_unclassified(100); | - | - | - | - | - | - | - | Unassigned | - |
| Otu32451 | | 1 | k__Fungi(100);k__Fungi_unclassified(100);k__Fungi_unclassified(100);k__Fungi_unclassified(100);k__Fungi_unclassified(100);k__Fungi_unclassified(100);k__Fungi_unclassified(100); | - | - | - | - | - | - | - | Unassigned | - |
| Otu32456 | | 1 | k__Fungi(100);p__Ascomycota(100);p__Ascomycota_unclassified(100);p__Ascomycota_unclassified(100);p__Ascomycota_unclassified(100);p__Ascomycota_unclassified(100);p__Ascomycota_unclassified(100); | - | - | - | - | - | - | - | Unassigned | - |
| Otu32457 | | 1 | k__Fungi(100);p__Ascomycota(100);p__Ascomycota_unclassified(100);p__Ascomycota_unclassified(100);p__Ascomycota_unclassified(100);p__Ascomycota_unclassified(100);p__Ascomycota_unclassified(100); | - | - | - | - | - | - | - | Unassigned | - |
| Otu32459 | | 1 | k__Fungi(100);k__Fungi_unclassified(100);k__Fungi_unclassified(100);k__Fungi_unclassified(100);k__Fungi_unclassified(100);k__Fungi_unclassified(100);k__Fungi_unclassified(100); | - | - | - | - | - | - | - | Unassigned | - |
| Otu32460 | | 1 | k__Fungi(100);p__Ascomycota(100);p__Ascomycota_unclassified(100);p__Ascomycota_unclassified(100);p__Ascomycota_unclassified(100);p__Ascomycota_unclassified(100);p__Ascomycota_unclassified(100); | - | - | - | - | - | - | - | Unassigned | - |
| Otu32471 | | 1 | k__Fungi(100);k__Fungi_unclassified(100);k__Fungi_unclassified(100);k__Fungi_unclassified(100);k__Fungi_unclassified(100);k__Fungi_unclassified(100);k__Fungi_unclassified(100); | - | - | - | - | - | - | - | Unassigned | - |
| Otu32472 | | 1 | k__Fungi(100);k__Fungi_unclassified(100);k__Fungi_unclassified(100);k__Fungi_unclassified(100);k__Fungi_unclassified(100);k__Fungi_unclassified(100);k__Fungi_unclassified(100); | - | - | - | - | - | - | - | Unassigned | - |
| Otu32473 | | 1 | k__Fungi(100);k__Fungi_unclassified(100);k__Fungi_unclassified(100);k__Fungi_unclassified(100);k__Fungi_unclassified(100);k__Fungi_unclassified(100);k__Fungi_unclassified(100); | - | - | - | - | - | - | - | Unassigned | - |
| Otu32474 | | 1 | k__Fungi(100);p__Ascomycota(100);p__Ascomycota_unclassified(100);p__Ascomycota_unclassified(100);p__Ascomycota_unclassified(100);p__Ascomycota_unclassified(100);p__Ascomycota_unclassified(100); | - | - | - | - | - | - | - | Unassigned | - |
| Otu32476 | | 1 | k__Fungi(100);p__Ascomycota(100);p__Ascomycota_unclassified(100);p__Ascomycota_unclassified(100);p__Ascomycota_unclassified(100);p__Ascomycota_unclassified(100);p__Ascomycota_unclassified(100); | - | - | - | - | - | - | - | Unassigned | - |

| | | | | | | | | | | | |
|---|---|---|---|---|---|---|---|---|---|---|---|
| Otu32477 | 1 | k__Fungi(100);p__Ascomycota(100);p__Ascomycota_unclassified(100);p__Ascomycota_unclassified(100);p__Ascomycota_unclassified(100);p__Ascomycota_unclassified(100);p__Ascomycota_unclassified(100); | - | - | - | - | - | - | - | Unassigned | - |
| Otu32478 | 1 | k__Fungi(100);p__Ascomycota(100);p__Ascomycota_unclassified(100);p__Ascomycota_unclassified(100);p__Ascomycota_unclassified(100);p__Ascomycota_unclassified(100);p__Ascomycota_unclassified(100); | - | - | - | - | - | - | - | Unassigned | - |
| Otu32480 | 1 | k__Fungi(100);p__Ascomycota(100);p__Ascomycota_unclassified(100);p__Ascomycota_unclassified(100);p__Ascomycota_unclassified(100);p__Ascomycota_unclassified(100);p__Ascomycota_unclassified(100); | - | - | - | - | - | - | - | Unassigned | - |
| Otu32482 | 1 | k__Fungi(100);k__Fungi_unclassified(100);k__Fungi_unclassified(100);k__Fungi_unclassified(100);k__Fungi_unclassified(100);k__Fungi_unclassified(100);k__Fungi_unclassified(100); | - | - | - | - | - | - | - | Unassigned | - |
| Otu32485 | 1 | k__Fungi(100);k__Fungi_unclassified(100);k__Fungi_unclassified(100);k__Fungi_unclassified(100);k__Fungi_unclassified(100);k__Fungi_unclassified(100);k__Fungi_unclassified(100); | - | - | - | - | - | - | - | Unassigned | - |
| Otu32486 | 1 | k__Fungi(100);k__Fungi_unclassified(100);k__Fungi_unclassified(100);k__Fungi_unclassified(100);k__Fungi_unclassified(100);k__Fungi_unclassified(100);k__Fungi_unclassified(100); | - | - | - | - | - | - | - | Unassigned | - |
| Otu32487 | 1 | k__Fungi(100);p__Ascomycota(100);p__Ascomycota_unclassified(100);p__Ascomycota_unclassified(100);p__Ascomycota_unclassified(100);p__Ascomycota_unclassified(100);p__Ascomycota_unclassified(100); | - | - | - | - | - | - | - | Unassigned | - |
| Otu32490 | 1 | k__Fungi(100);p__Ascomycota(100);p__Ascomycota_unclassified(100);p__Ascomycota_unclassified(100);p__Ascomycota_unclassified(100);p__Ascomycota_unclassified(100);p__Ascomycota_unclassified(100); | - | - | - | - | - | - | - | Unassigned | - |
| Otu32496 | 1 | k__Fungi(100);p__Ascomycota(100);p__Ascomycota_unclassified(100);p__Ascomycota_unclassified(100);p__Ascomycota_unclassified(100);p__Ascomycota_unclassified(100);p__Ascomycota_unclassified(100); | - | - | - | - | - | - | - | Unassigned | - |
| Otu32505 | 1 | k__Fungi(100);k__Fungi_unclassified(100);k__Fungi_unclassified(100);k__Fungi_unclassified(100);k__Fungi_unclassified(100);k__Fungi_unclassified(100);k__Fungi_unclassified(100); | - | - | - | - | - | - | - | Unassigned | - |
| Otu32521 | 1 | k__Fungi(100);p__Ascomycota(100);p__Ascomycota_unclassified(100);p__Ascomycota_unclassified(100);p__Ascomycota_unclassified(100);p__Ascomycota_unclassified(100);p__Ascomycota_unclassified(100); | - | - | - | - | - | - | - | Unassigned | - |
| Otu32522 | 1 | k__Fungi(100);k__Fungi_unclassified(100);k__Fungi_unclassified(100);k__Fungi_unclassified(100);k__Fungi_unclassified(100);k__Fungi_unclassified(100);k__Fungi_unclassified(100); | - | - | - | - | - | - | - | Unassigned | - |
| Otu32523 | 1 | k__Fungi(100);k__Fungi_unclassified(100);k__Fungi_unclassified(100);k__Fungi_unclassified(100);k__Fungi_unclassified(100);k__Fungi_unclassified(100);k__Fungi_unclassified(100); | - | - | - | - | - | - | - | Unassigned | - |
| Otu32524 | 1 | k__Fungi(100);p__Ascomycota(100);p__Ascomycota_unclassified(100);p__Ascomycota_unclassified(100);p__Ascomycota_unclassified(100);p__Ascomycota_unclassified(100);p__Ascomycota_unclassified(100); | - | - | - | - | - | - | - | Unassigned | - |
| Otu32527 | 1 | k__Fungi(100);k__Fungi_unclassified(100);k__Fungi_unclassified(100);k__Fungi_unclassified(100);k__Fungi_unclassified(100);k__Fungi_unclassified(100);k__Fungi_unclassified(100); | - | - | - | - | - | - | - | Unassigned | - |
| Otu32539 | 1 | k__Fungi(100);p__Ascomycota(100);p__Ascomycota_unclassified(100);p__Ascomycota_unclassified(100);p__Ascomycota_unclassified(100);p__Ascomycota_unclassified(100);p__Ascomycota_unclassified(100); | - | - | - | - | - | - | - | Unassigned | - |
| Otu32543 | 1 | k__Fungi(100);k__Fungi_unclassified(100);k__Fungi_unclassified(100);k__Fungi_unclassified(100);k__Fungi_unclassified(100);k__Fungi_unclassified(100);k__Fungi_unclassified(100); | - | - | - | - | - | - | - | Unassigned | - |
| Otu32544 | 1 | k__Fungi(100);p__Ascomycota(100);p__Ascomycota_unclassified(100);p__Ascomycota_unclassified(100);p__Ascomycota_unclassified(100);p__Ascomycota_unclassified(100);p__Ascomycota_unclassified(100); | - | - | - | - | - | - | - | Unassigned | - |
| Otu32545 | 1 | k__Fungi(100);k__Fungi_unclassified(100);k__Fungi_unclassified(100);k__Fungi_unclassified(100);k__Fungi_unclassified(100);k__Fungi_unclassified(100);k__Fungi_unclassified(100); | - | - | - | - | - | - | - | Unassigned | - |
| Otu32546 | 1 | k__Fungi(100);p__Ascomycota(100);p__Ascomycota_unclassified(100);p__Ascomycota_unclassified(100);p__Ascomycota_unclassified(100);p__Ascomycota_unclassified(100);p__Ascomycota_unclassified(100); | - | - | - | - | - | - | - | Unassigned | - |
| Otu32550 | 1 | k__Fungi(100);p__Ascomycota(100);p__Ascomycota_unclassified(100);p__Ascomycota_unclassified(100);p__Ascomycota_unclassified(100);p__Ascomycota_unclassified(100);p__Ascomycota_unclassified(100); | - | - | - | - | - | - | - | Unassigned | - |
| Otu32551 | 1 | k__Fungi(100);k__Fungi_unclassified(100);k__Fungi_unclassified(100);k__Fungi_unclassified(100);k__Fungi_unclassified(100);k__Fungi_unclassified(100);k__Fungi_unclassified(100); | - | - | - | - | - | - | - | Unassigned | - |
| Otu32556 | 1 | k__Fungi(100);k__Fungi_unclassified(100);k__Fungi_unclassified(100);k__Fungi_unclassified(100);k__Fungi_unclassified(100);k__Fungi_unclassified(100);k__Fungi_unclassified(100); | - | - | - | - | - | - | - | Unassigned | - |
| Otu32571 | 1 | k__Fungi(100);k__Fungi_unclassified(100);k__Fungi_unclassified(100);k__Fungi_unclassified(100);k__Fungi_unclassified(100);k__Fungi_unclassified(100);k__Fungi_unclassified(100); | - | - | - | - | - | - | - | Unassigned | - |
| Otu32572 | 1 | k__Fungi(100);k__Fungi_unclassified(100);k__Fungi_unclassified(100);k__Fungi_unclassified(100);k__Fungi_unclassified(100);k__Fungi_unclassified(100);k__Fungi_unclassified(100); | - | - | - | - | - | - | - | Unassigned | - |
| Otu32573 | 1 | k__Fungi(100);k__Fungi_unclassified(100);k__Fungi_unclassified(100);k__Fungi_unclassified(100);k__Fungi_unclassified(100);k__Fungi_unclassified(100);k__Fungi_unclassified(100); | - | - | - | - | - | - | - | Unassigned | - |
| Otu32574 | 1 | k__Fungi(100);p__Ascomycota(100);p__Ascomycota_unclassified(100);p__Ascomycota_unclassified(100);p__Ascomycota_unclassified(100);p__Ascomycota_unclassified(100);p__Ascomycota_unclassified(100); | - | - | - | - | - | - | - | Unassigned | - |
| Otu32577 | 1 | k__Fungi(100);k__Fungi_unclassified(100);k__Fungi_unclassified(100);k__Fungi_unclassified(100);k__Fungi_unclassified(100);k__Fungi_unclassified(100);k__Fungi_unclassified(100); | - | - | - | - | - | - | - | Unassigned | - |
| Otu32578 | 1 | k__Fungi(100);p__Ascomycota(100);p__Ascomycota_unclassified(100);p__Ascomycota_unclassified(100);p__Ascomycota_unclassified(100);p__Ascomycota_unclassified(100);p__Ascomycota_unclassified(100); | - | - | - | - | - | - | - | Unassigned | - |
| Otu32584 | 1 | k__Fungi(100);k__Fungi_unclassified(100);k__Fungi_unclassified(100);k__Fungi_unclassified(100);k__Fungi_unclassified(100);k__Fungi_unclassified(100);k__Fungi_unclassified(100); | - | - | - | - | - | - | - | Unassigned | - |
| Otu32585 | 1 | k__Fungi(100);k__Fungi_unclassified(100);k__Fungi_unclassified(100);k__Fungi_unclassified(100);k__Fungi_unclassified(100);k__Fungi_unclassified(100);k__Fungi_unclassified(100); | - | - | - | - | - | - | - | Unassigned | - |
| Otu32591 | 1 | k__Fungi(100);k__Fungi_unclassified(100);k__Fungi_unclassified(100);k__Fungi_unclassified(100);k__Fungi_unclassified(100);k__Fungi_unclassified(100);k__Fungi_unclassified(100); | - | - | - | - | - | - | - | Unassigned | - |
| Otu32593 | 1 | k__Fungi(100);k__Fungi_unclassified(100);k__Fungi_unclassified(100);k__Fungi_unclassified(100);k__Fungi_unclassified(100);k__Fungi_unclassified(100);k__Fungi_unclassified(100); | - | - | - | - | - | - | - | Unassigned | - |
| Otu32595 | 1 | k__Fungi(100);k__Fungi_unclassified(100);k__Fungi_unclassified(100);k__Fungi_unclassified(100);k__Fungi_unclassified(100);k__Fungi_unclassified(100);k__Fungi_unclassified(100); | - | - | - | - | - | - | - | Unassigned | - |
| Otu32597 | 1 | k__Fungi(100);p__Ascomycota(100);p__Ascomycota_unclassified(100);p__Ascomycota_unclassified(100);p__Ascomycota_unclassified(100);p__Ascomycota_unclassified(100);p__Ascomycota_unclassified(100); | - | - | - | - | - | - | - | Unassigned | - |
| Otu32598 | 1 | k__Fungi(100);k__Fungi_unclassified(100);k__Fungi_unclassified(100);k__Fungi_unclassified(100);k__Fungi_unclassified(100);k__Fungi_unclassified(100);k__Fungi_unclassified(100); | - | - | - | - | - | - | - | Unassigned | - |
| Otu32606 | 1 | k__Fungi(100);p__Ascomycota(100);p__Ascomycota_unclassified(100);p__Ascomycota_unclassified(100);p__Ascomycota_unclassified(100);p__Ascomycota_unclassified(100);p__Ascomycota_unclassified(100); | - | - | - | - | - | - | - | Unassigned | - |
| Otu32607 | 1 | k__Fungi(100);k__Fungi_unclassified(100);k__Fungi_unclassified(100);k__Fungi_unclassified(100);k__Fungi_unclassified(100);k__Fungi_unclassified(100);k__Fungi_unclassified(100); | - | - | - | - | - | - | - | Unassigned | - |
| Otu32628 | 1 | k__Fungi(100);k__Fungi_unclassified(100);k__Fungi_unclassified(100);k__Fungi_unclassified(100);k__Fungi_unclassified(100);k__Fungi_unclassified(100);k__Fungi_unclassified(100); | - | - | - | - | - | - | - | Unassigned | - |
| Otu32632 | 1 | k__Fungi(100);k__Fungi_unclassified(100);k__Fungi_unclassified(100);k__Fungi_unclassified(100);k__Fungi_unclassified(100);k__Fungi_unclassified(100);k__Fungi_unclassified(100); | - | - | - | - | - | - | - | Unassigned | - |
| Otu32633 | 1 | k__Fungi(100);p__Ascomycota(100);p__Ascomycota_unclassified(100);p__Ascomycota_unclassified(100);p__Ascomycota_unclassified(100);p__Ascomycota_unclassified(100);p__Ascomycota_unclassified(100); | - | - | - | - | - | - | - | Unassigned | - |
| Otu32634 | 1 | k__Fungi(100);p__Ascomycota(100);p__Ascomycota_unclassified(100);p__Ascomycota_unclassified(100);p__Ascomycota_unclassified(100);p__Ascomycota_unclassified(100);p__Ascomycota_unclassified(100); | - | - | - | - | - | - | - | Unassigned | - |
| Otu32641 | 1 | k__Fungi(100);p__Ascomycota(100);p__Ascomycota_unclassified(100);p__Ascomycota_unclassified(100);p__Ascomycota_unclassified(100);p__Ascomycota_unclassified(100);p__Ascomycota_unclassified(100); | - | - | - | - | - | - | - | Unassigned | - |
| Otu32644 | 1 | k__Fungi(100);p__Ascomycota(100);p__Ascomycota_unclassified(100);p__Ascomycota_unclassified(100);p__Ascomycota_unclassified(100);p__Ascomycota_unclassified(100);p__Ascomycota_unclassified(100); | - | - | - | - | - | - | - | Unassigned | - |
| Otu32650 | 1 | k__Fungi(100);p__Ascomycota(100);p__Ascomycota_unclassified(100);p__Ascomycota_unclassified(100);p__Ascomycota_unclassified(100);p__Ascomycota_unclassified(100);p__Ascomycota_unclassified(100); | - | - | - | - | - | - | - | Unassigned | - |
| Otu32652 | 1 | k__Fungi(100);p__Ascomycota(100);p__Ascomycota_unclassified(100);p__Ascomycota_unclassified(100);p__Ascomycota_unclassified(100);p__Ascomycota_unclassified(100);p__Ascomycota_unclassified(100); | - | - | - | - | - | - | - | Unassigned | - |
| Otu32654 | 1 | k__Fungi(100);p__Ascomycota(100);p__Ascomycota_unclassified(100);p__Ascomycota_unclassified(100);p__Ascomycota_unclassified(100);p__Ascomycota_unclassified(100);p__Ascomycota_unclassified(100); | - | - | - | - | - | - | - | Unassigned | - |
| Otu32665 | 1 | k__Fungi(100);k__Fungi_unclassified(100);k__Fungi_unclassified(100);k__Fungi_unclassified(100);k__Fungi_unclassified(100);k__Fungi_unclassified(100);k__Fungi_unclassified(100); | - | - | - | - | - | - | - | Unassigned | - |
| Otu32671 | 1 | k__Fungi(100);p__Ascomycota(100);p__Ascomycota_unclassified(100);p__Ascomycota_unclassified(100);p__Ascomycota_unclassified(100);p__Ascomycota_unclassified(100);p__Ascomycota_unclassified(100); | - | - | - | - | - | - | - | Unassigned | - |
| Otu32672 | 1 | k__Fungi(100);p__Ascomycota(100);p__Ascomycota_unclassified(100);p__Ascomycota_unclassified(100);p__Ascomycota_unclassified(100);p__Ascomycota_unclassified(100);p__Ascomycota_unclassified(100); | - | - | - | - | - | - | - | Unassigned | - |
| Otu32673 | 1 | k__Fungi(100);k__Fungi_unclassified(100);k__Fungi_unclassified(100);k__Fungi_unclassified(100);k__Fungi_unclassified(100);k__Fungi_unclassified(100);k__Fungi_unclassified(100); | - | - | - | - | - | - | - | Unassigned | - |
| Otu32675 | 1 | k__Fungi(100);k__Fungi_unclassified(100);k__Fungi_unclassified(100);k__Fungi_unclassified(100);k__Fungi_unclassified(100);k__Fungi_unclassified(100);k__Fungi_unclassified(100); | - | - | - | - | - | - | - | Unassigned | - |
| Otu32676 | 1 | k__Fungi(100);k__Fungi_unclassified(100);k__Fungi_unclassified(100);k__Fungi_unclassified(100);k__Fungi_unclassified(100);k__Fungi_unclassified(100);k__Fungi_unclassified(100); | - | - | - | - | - | - | - | Unassigned | - |
| Otu32677 | 1 | k__Fungi(100);k__Fungi_unclassified(100);k__Fungi_unclassified(100);k__Fungi_unclassified(100);k__Fungi_unclassified(100);k__Fungi_unclassified(100);k__Fungi_unclassified(100); | - | - | - | - | - | - | - | Unassigned | - |
| Otu32685 | 1 | k__Fungi(100);k__Fungi_unclassified(100);k__Fungi_unclassified(100);k__Fungi_unclassified(100);k__Fungi_unclassified(100);k__Fungi_unclassified(100);k__Fungi_unclassified(100); | - | - | - | - | - | - | - | Unassigned | - |
| Otu32686 | 1 | k__Fungi(100);k__Fungi_unclassified(100);k__Fungi_unclassified(100);k__Fungi_unclassified(100);k__Fungi_unclassified(100);k__Fungi_unclassified(100);k__Fungi_unclassified(100); | - | - | - | - | - | - | - | Unassigned | - |
| Otu32687 | 1 | k__Fungi(100);p__Ascomycota(100);p__Ascomycota_unclassified(100);p__Ascomycota_unclassified(100);p__Ascomycota_unclassified(100);p__Ascomycota_unclassified(100);p__Ascomycota_unclassified(100); | - | - | - | - | - | - | - | Unassigned | - |
| Otu32692 | 1 | k__Fungi(100);p__Ascomycota(100);p__Ascomycota_unclassified(100);p__Ascomycota_unclassified(100);p__Ascomycota_unclassified(100);p__Ascomycota_unclassified(100);p__Ascomycota_unclassified(100); | - | - | - | - | - | - | - | Unassigned | - |
| Otu32695 | 1 | k__Fungi(100);p__Ascomycota(100);p__Ascomycota_unclassified(100);p__Ascomycota_unclassified(100);p__Ascomycota_unclassified(100);p__Ascomycota_unclassified(100);p__Ascomycota_unclassified(100); | - | - | - | - | - | - | - | Unassigned | - |
| Otu32696 | 1 | k__Fungi(100);p__Ascomycota(100);p__Ascomycota_unclassified(100);p__Ascomycota_unclassified(100);p__Ascomycota_unclassified(100);p__Ascomycota_unclassified(100);p__Ascomycota_unclassified(100); | - | - | - | - | - | - | - | Unassigned | - |
| Otu32698 | 1 | k__Fungi(100);p__Ascomycota(100);p__Ascomycota_unclassified(100);p__Ascomycota_unclassified(100);p__Ascomycota_unclassified(100);p__Ascomycota_unclassified(100);p__Ascomycota_unclassified(100); | - | - | - | - | - | - | - | Unassigned | - |
| Otu32699 | 1 | k__Fungi(100);p__Ascomycota(100);p__Ascomycota_unclassified(100);p__Ascomycota_unclassified(100);p__Ascomycota_unclassified(100);p__Ascomycota_unclassified(100);p__Ascomycota_unclassified(100); | - | - | - | - | - | - | - | Unassigned | - |
| Otu32700 | 1 | k__Fungi(100);k__Fungi_unclassified(100);k__Fungi_unclassified(100);k__Fungi_unclassified(100);k__Fungi_unclassified(100);k__Fungi_unclassified(100);k__Fungi_unclassified(100); | - | - | - | - | - | - | - | Unassigned | - |
| Otu32702 | 1 | k__Fungi(100);p__Ascomycota(100);p__Ascomycota_unclassified(100);p__Ascomycota_unclassified(100);p__Ascomycota_unclassified(100);p__Ascomycota_unclassified(100);p__Ascomycota_unclassified(100); | - | - | - | - | - | - | - | Unassigned | - |
| Otu32703 | 1 | k__Fungi(100);p__Ascomycota(100);p__Ascomycota_unclassified(100);p__Ascomycota_unclassified(100);p__Ascomycota_unclassified(100);p__Ascomycota_unclassified(100);p__Ascomycota_unclassified(100); | - | - | - | - | - | - | - | Unassigned | - |
| Otu32705 | 1 | k__Fungi(100);k__Fungi_unclassified(100);k__Fungi_unclassified(100);k__Fungi_unclassified(100);k__Fungi_unclassified(100);k__Fungi_unclassified(100);k__Fungi_unclassified(100); | - | - | - | - | - | - | - | Unassigned | - |
| Otu32707 | 1 | k__Fungi(100);p__Ascomycota(100);p__Ascomycota_unclassified(100);p__Ascomycota_unclassified(100);p__Ascomycota_unclassified(100);p__Ascomycota_unclassified(100);p__Ascomycota_unclassified(100); | - | - | - | - | - | - | - | Unassigned | - |
| Otu32708 | 1 | k__Fungi(100);p__Ascomycota(100);p__Ascomycota_unclassified(100);p__Ascomycota_unclassified(100);p__Ascomycota_unclassified(100);p__Ascomycota_unclassified(100);p__Ascomycota_unclassified(100); | - | - | - | - | - | - | - | Unassigned | - |
| Otu32709 | 1 | k__Fungi(100);p__Ascomycota(100);p__Ascomycota_unclassified(100);p__Ascomycota_unclassified(100);p__Ascomycota_unclassified(100);p__Ascomycota_unclassified(100);p__Ascomycota_unclassified(100); | - | - | - | - | - | - | - | Unassigned | - |
| Otu32720 | 1 | k__Fungi(100);p__Ascomycota(100);p__Ascomycota_unclassified(100);p__Ascomycota_unclassified(100);p__Ascomycota_unclassified(100);p__Ascomycota_unclassified(100);p__Ascomycota_unclassified(100); | - | - | - | - | - | - | - | Unassigned | - |
| Otu32723 | 1 | k__Fungi(100);p__Ascomycota(100);p__Ascomycota_unclassified(100);p__Ascomycota_unclassified(100);p__Ascomycota_unclassified(100);p__Ascomycota_unclassified(100);p__Ascomycota_unclassified(100); | - | - | - | - | - | - | - | Unassigned | - |
| Otu32724 | 1 | k__Fungi(100);p__Ascomycota(100);p__Ascomycota_unclassified(100);p__Ascomycota_unclassified(100);p__Ascomycota_unclassified(100);p__Ascomycota_unclassified(100);p__Ascomycota_unclassified(100); | - | - | - | - | - | - | - | Unassigned | - |
| Otu32725 | 1 | k__Fungi(100);p__Ascomycota(100);p__Ascomycota_unclassified(100);p__Ascomycota_unclassified(100);p__Ascomycota_unclassified(100);p__Ascomycota_unclassified(100);p__Ascomycota_unclassified(100); | - | - | - | - | - | - | - | Unassigned | - |
| Otu32726 | 1 | k__Fungi(100);k__Fungi_unclassified(100);k__Fungi_unclassified(100);k__Fungi_unclassified(100);k__Fungi_unclassified(100);k__Fungi_unclassified(100);k__Fungi_unclassified(100); | - | - | - | - | - | - | - | Unassigned | - |
| Otu32727 | 1 | k__Fungi(100);p__Ascomycota(100);p__Ascomycota_unclassified(100);p__Ascomycota_unclassified(100);p__Ascomycota_unclassified(100);p__Ascomycota_unclassified(100);p__Ascomycota_unclassified(100); | - | - | - | - | - | - | - | Unassigned | - |
| Otu32728 | 1 | k__Fungi(100);k__Fungi_unclassified(100);k__Fungi_unclassified(100);k__Fungi_unclassified(100);k__Fungi_unclassified(100);k__Fungi_unclassified(100);k__Fungi_unclassified(100); | - | - | - | - | - | - | - | Unassigned | - |
| Otu32729 | 1 | k__Fungi(100);k__Fungi_unclassified(100);k__Fungi_unclassified(100);k__Fungi_unclassified(100);k__Fungi_unclassified(100);k__Fungi_unclassified(100);k__Fungi_unclassified(100); | - | - | - | - | - | - | - | Unassigned | - |
| Otu32730 | 1 | k__Fungi(100);p__Ascomycota(100);p__Ascomycota_unclassified(100);p__Ascomycota_unclassified(100);p__Ascomycota_unclassified(100);p__Ascomycota_unclassified(100);p__Ascomycota_unclassified(100); | - | - | - | - | - | - | - | Unassigned | - |
| Otu32731 | 1 | k__Fungi(100);p__Ascomycota(100);p__Ascomycota_unclassified(100);p__Ascomycota_unclassified(100);p__Ascomycota_unclassified(100);p__Ascomycota_unclassified(100);p__Ascomycota_unclassified(100); | - | - | - | - | - | - | - | Unassigned | - |
| Otu32732 | 1 | k__Fungi(100);k__Fungi_unclassified(100);k__Fungi_unclassified(100);k__Fungi_unclassified(100);k__Fungi_unclassified(100);k__Fungi_unclassified(100);k__Fungi_unclassified(100); | - | - | - | - | - | - | - | Unassigned | - |
| Otu32734 | 1 | k__Fungi(100);k__Fungi_unclassified(100);k__Fungi_unclassified(100);k__Fungi_unclassified(100);k__Fungi_unclassified(100);k__Fungi_unclassified(100);k__Fungi_unclassified(100); | - | - | - | - | - | - | - | Unassigned | - |
| Otu32735 | 1 | k__Fungi(100);k__Fungi_unclassified(100);k__Fungi_unclassified(100);k__Fungi_unclassified(100);k__Fungi_unclassified(100);k__Fungi_unclassified(100);k__Fungi_unclassified(100); | - | - | - | - | - | - | - | Unassigned | - |
| Otu32736 | 1 | k__Fungi(100);k__Fungi_unclassified(100);k__Fungi_unclassified(100);k__Fungi_unclassified(100);k__Fungi_unclassified(100);k__Fungi_unclassified(100);k__Fungi_unclassified(100); | - | - | - | - | - | - | - | Unassigned | - |
| Otu32745 | 1 | k__Fungi(100);k__Fungi_unclassified(100);k__Fungi_unclassified(100);k__Fungi_unclassified(100);k__Fungi_unclassified(100);k__Fungi_unclassified(100);k__Fungi_unclassified(100); | - | - | - | - | - | - | - | Unassigned | - |
| Otu32746 | 1 | k__Fungi(100);k__Fungi_unclassified(100);k__Fungi_unclassified(100);k__Fungi_unclassified(100);k__Fungi_unclassified(100);k__Fungi_unclassified(100);k__Fungi_unclassified(100); | - | - | - | - | - | - | - | Unassigned | - |
| Otu32748 | 1 | k__Fungi(100);k__Fungi_unclassified(100);k__Fungi_unclassified(100);k__Fungi_unclassified(100);k__Fungi_unclassified(100);k__Fungi_unclassified(100);k__Fungi_unclassified(100); | - | - | - | - | - | - | - | Unassigned | - |
| Otu32750 | 1 | k__Fungi(100);k__Fungi_unclassified(100);k__Fungi_unclassified(100);k__Fungi_unclassified(100);k__Fungi_unclassified(100);k__Fungi_unclassified(100);k__Fungi_unclassified(100); | - | - | - | - | - | - | - | Unassigned | - |
| Otu32753 | 1 | k__Fungi(100);p__Ascomycota(100);p__Ascomycota_unclassified(100);p__Ascomycota_unclassified(100);p__Ascomycota_unclassified(100);p__Ascomycota_unclassified(100);p__Ascomycota_unclassified(100); | - | - | - | - | - | - | - | Unassigned | - |
| Otu32754 | 1 | k__Fungi(100);p__Ascomycota(100);p__Ascomycota_unclassified(100);p__Ascomycota_unclassified(100);p__Ascomycota_unclassified(100);p__Ascomycota_unclassified(100);p__Ascomycota_unclassified(100); | - | - | - | - | - | - | - | Unassigned | - |
| Otu32755 | 1 | k__Fungi(100);p__Ascomycota(100);p__Ascomycota_unclassified(100);p__Ascomycota_unclassified(100);p__Ascomycota_unclassified(100);p__Ascomycota_unclassified(100);p__Ascomycota_unclassified(100); | - | - | - | - | - | - | - | Unassigned | - |
| Otu32758 | 1 | k__Fungi(100);p__Ascomycota(100);p__Ascomycota_unclassified(100);p__Ascomycota_unclassified(100);p__Ascomycota_unclassified(100);p__Ascomycota_unclassified(100);p__Ascomycota_unclassified(100); | - | - | - | - | - | - | - | Unassigned | - |
| Otu32759 | 1 | k__Fungi(100);p__Ascomycota(100);p__Ascomycota_unclassified(100);p__Ascomycota_unclassified(100);p__Ascomycota_unclassified(100);p__Ascomycota_unclassified(100);p__Ascomycota_unclassified(100); | - | - | - | - | - | - | - | Unassigned | - |
| Otu32762 | 1 | k__Fungi(100);p__Ascomycota(100);p__Ascomycota_unclassified(100);p__Ascomycota_unclassified(100);p__Ascomycota_unclassified(100);p__Ascomycota_unclassified(100);p__Ascomycota_unclassified(100); | - | - | - | - | - | - | - | Unassigned | - |
| Otu32764 | 1 | k__Fungi(100);p__Ascomycota(100);p__Ascomycota_unclassified(100);p__Ascomycota_unclassified(100);p__Ascomycota_unclassified(100);p__Ascomycota_unclassified(100);p__Ascomycota_unclassified(100); | - | - | - | - | - | - | - | Unassigned | - |
| Otu32765 | 1 | k__Fungi(100);p__Ascomycota(100);p__Ascomycota_unclassified(100);p__Ascomycota_unclassified(100);p__Ascomycota_unclassified(100);p__Ascomycota_unclassified(100);p__Ascomycota_unclassified(100); | - | - | - | - | - | - | - | Unassigned | - |
| Otu32766 | 1 | k__Fungi(100);k__Fungi_unclassified(100);k__Fungi_unclassified(100);k__Fungi_unclassified(100);k__Fungi_unclassified(100);k__Fungi_unclassified(100);k__Fungi_unclassified(100); | - | - | - | - | - | - | - | Unassigned | - |
| Otu32768 | 1 | k__Fungi(100);k__Fungi_unclassified(100);k__Fungi_unclassified(100);k__Fungi_unclassified(100);k__Fungi_unclassified(100);k__Fungi_unclassified(100);k__Fungi_unclassified(100); | - | - | - | - | - | - | - | Unassigned | - |
| Otu32770 | 1 | k__Fungi(100);p__Ascomycota(100);p__Ascomycota_unclassified(100);p__Ascomycota_unclassified(100);p__Ascomycota_unclassified(100);p__Ascomycota_unclassified(100);p__Ascomycota_unclassified(100); | - | - | - | - | - | - | - | Unassigned | - |
| Otu32773 | 1 | k__Fungi(100);k__Fungi_unclassified(100);k__Fungi_unclassified(100);k__Fungi_unclassified(100);k__Fungi_unclassified(100);k__Fungi_unclassified(100);k__Fungi_unclassified(100); | - | - | - | - | - | - | - | Unassigned | - |
| Otu32774 | 1 | k__Fungi(100);k__Fungi_unclassified(100);k__Fungi_unclassified(100);k__Fungi_unclassified(100);k__Fungi_unclassified(100);k__Fungi_unclassified(100);k__Fungi_unclassified(100); | - | - | - | - | - | - | - | Unassigned | - |
| Otu32775 | 1 | k__Fungi(100);k__Fungi_unclassified(100);k__Fungi_unclassified(100);k__Fungi_unclassified(100);k__Fungi_unclassified(100);k__Fungi_unclassified(100);k__Fungi_unclassified(100); | - | - | - | - | - | - | - | Unassigned | - |
| Otu32778 | 1 | k__Fungi(100);k__Fungi_unclassified(100);k__Fungi_unclassified(100);k__Fungi_unclassified(100);k__Fungi_unclassified(100);k__Fungi_unclassified(100);k__Fungi_unclassified(100); | - | - | - | - | - | - | - | Unassigned | - |
| Otu32780 | 1 | k__Fungi(100);p__Ascomycota(100);p__Ascomycota_unclassified(100);p__Ascomycota_unclassified(100);p__Ascomycota_unclassified(100);p__Ascomycota_unclassified(100);p__Ascomycota_unclassified(100); | - | - | - | - | - | - | - | Unassigned | - |
| Otu32781 | 1 | k__Fungi(100);k__Fungi_unclassified(100);k__Fungi_unclassified(100);k__Fungi_unclassified(100);k__Fungi_unclassified(100);k__Fungi_unclassified(100);k__Fungi_unclassified(100); | - | - | - | - | - | - | - | Unassigned | - |
| Otu32782 | 1 | k__Fungi(100);k__Fungi_unclassified(100);k__Fungi_unclassified(100);k__Fungi_unclassified(100);k__Fungi_unclassified(100);k__Fungi_unclassified(100);k__Fungi_unclassified(100); | - | - | - | - | - | - | - | Unassigned | - |
| Otu32785 | 1 | k__Fungi(100);k__Fungi_unclassified(100);k__Fungi_unclassified(100);k__Fungi_unclassified(100);k__Fungi_unclassified(100);k__Fungi_unclassified(100);k__Fungi_unclassified(100); | - | - | - | - | - | - | - | Unassigned | - |
| Otu32786 | 1 | k__Fungi(100);p__Ascomycota(100);p__Ascomycota_unclassified(100);p__Ascomycota_unclassified(100);p__Ascomycota_unclassified(100);p__Ascomycota_unclassified(100);p__Ascomycota_unclassified(100); | - | - | - | - | - | - | - | Unassigned | - |
| Otu32787 | 1 | k__Fungi(100);k__Fungi_unclassified(100);k__Fungi_unclassified(100);k__Fungi_unclassified(100);k__Fungi_unclassified(100);k__Fungi_unclassified(100);k__Fungi_unclassified(100); | - | - | - | - | - | - | - | Unassigned | - |
| Otu32793 | 1 | k__Fungi(100);k__Fungi_unclassified(100);k__Fungi_unclassified(100);k__Fungi_unclassified(100);k__Fungi_unclassified(100);k__Fungi_unclassified(100);k__Fungi_unclassified(100); | - | - | - | - | - | - | - | Unassigned | - |
| Otu32795 | 1 | k__Fungi(100);k__Fungi_unclassified(100);k__Fungi_unclassified(100);k__Fungi_unclassified(100);k__Fungi_unclassified(100);k__Fungi_unclassified(100);k__Fungi_unclassified(100); | - | - | - | - | - | - | - | Unassigned | - |
| Otu32800 | 1 | k__Fungi(100);k__Fungi_unclassified(100);k__Fungi_unclassified(100);k__Fungi_unclassified(100);k__Fungi_unclassified(100);k__Fungi_unclassified(100);k__Fungi_unclassified(100); | - | - | - | - | - | - | - | Unassigned | - |
| Otu32802 | 1 | k__Fungi(100);k__Fungi_unclassified(100);k__Fungi_unclassified(100);k__Fungi_unclassified(100);k__Fungi_unclassified(100);k__Fungi_unclassified(100);k__Fungi_unclassified(100); | - | - | - | - | - | - | - | Unassigned | - |
| Otu32814 | 1 | k__Fungi(100);k__Fungi_unclassified(100);k__Fungi_unclassified(100);k__Fungi_unclassified(100);k__Fungi_unclassified(100);k__Fungi_unclassified(100);k__Fungi_unclassified(100); | - | - | - | - | - | - | - | Unassigned | - |
| Otu32816 | 1 | k__Fungi(100);p__Ascomycota(100);p__Ascomycota_unclassified(100);p__Ascomycota_unclassified(100);p__Ascomycota_unclassified(100);p__Ascomycota_unclassified(100);p__Ascomycota_unclassified(100); | - | - | - | - | - | - | - | Unassigned | - |
| Otu32828 | 1 | k__Fungi(100);k__Fungi_unclassified(100);k__Fungi_unclassified(100);k__Fungi_unclassified(100);k__Fungi_unclassified(100);k__Fungi_unclassified(100);k__Fungi_unclassified(100); | - | - | - | - | - | - | - | Unassigned | - |
| Otu32829 | 1 | k__Fungi(100);k__Fungi_unclassified(100);k__Fungi_unclassified(100);k__Fungi_unclassified(100);k__Fungi_unclassified(100);k__Fungi_unclassified(100);k__Fungi_unclassified(100); | - | - | - | - | - | - | - | Unassigned | - |
| Otu32836 | 1 | k__Fungi(100);p__Ascomycota(100);p__Ascomycota_unclassified(100);p__Ascomycota_unclassified(100);p__Ascomycota_unclassified(100);p__Ascomycota_unclassified(100);p__Ascomycota_unclassified(100); | - | - | - | - | - | - | - | Unassigned | - |
| Otu32837 | 1 | k__Fungi(100);k__Fungi_unclassified(100);k__Fungi_unclassified(100);k__Fungi_unclassified(100);k__Fungi_unclassified(100);k__Fungi_unclassified(100);k__Fungi_unclassified(100); | - | - | - | - | - | - | - | Unassigned | - |
| Otu32839 | 1 | k__Fungi(100);k__Fungi_unclassified(100);k__Fungi_unclassified(100);k__Fungi_unclassified(100);k__Fungi_unclassified(100);k__Fungi_unclassified(100);k__Fungi_unclassified(100); | - | - | - | - | - | - | - | Unassigned | - |
| Otu32841 | 1 | k__Fungi(100);p__Ascomycota(100);p__Ascomycota_unclassified(100);p__Ascomycota_unclassified(100);p__Ascomycota_unclassified(100);p__Ascomycota_unclassified(100);p__Ascomycota_unclassified(100); | - | - | - | - | - | - | - | Unassigned | - |
| Otu32843 | 1 | k__Fungi(100);k__Fungi_unclassified(100);k__Fungi_unclassified(100);k__Fungi_unclassified(100);k__Fungi_unclassified(100);k__Fungi_unclassified(100);k__Fungi_unclassified(100); | - | - | - | - | - | - | - | Unassigned | - |
| Otu32855 | 1 | k__Fungi(100);p__Ascomycota(100);p__Ascomycota_unclassified(100);p__Ascomycota_unclassified(100);p__Ascomycota_unclassified(100);p__Ascomycota_unclassified(100);p__Ascomycota_unclassified(100); | - | - | - | - | - | - | - | Unassigned | - |
| Otu32865 | 1 | k__Fungi(100);p__Ascomycota(100);p__Ascomycota_unclassified(100);p__Ascomycota_unclassified(100);p__Ascomycota_unclassified(100);p__Ascomycota_unclassified(100);p__Ascomycota_unclassified(100); | - | - | - | - | - | - | - | Unassigned | - |
| Otu32871 | 1 | k__Fungi(100);k__Fungi_unclassified(100);k__Fungi_unclassified(100);k__Fungi_unclassified(100);k__Fungi_unclassified(100);k__Fungi_unclassified(100);k__Fungi_unclassified(100); | - | - | - | - | - | - | - | Unassigned | - |
| Otu32878 | 1 | k__Fungi(100);k__Fungi_unclassified(100);k__Fungi_unclassified(100);k__Fungi_unclassified(100);k__Fungi_unclassified(100);k__Fungi_unclassified(100);k__Fungi_unclassified(100); | - | - | - | - | - | - | - | Unassigned | - |
| Otu32882 | 1 | k__Fungi(100);p__Ascomycota(100);p__Ascomycota_unclassified(100);p__Ascomycota_unclassified(100);p__Ascomycota_unclassified(100);p__Ascomycota_unclassified(100);p__Ascomycota_unclassified(100); | - | - | - | - | - | - | - | Unassigned | - |
| Otu32897 | 1 | k__Fungi(100);k__Fungi_unclassified(100);k__Fungi_unclassified(100);k__Fungi_unclassified(100);k__Fungi_unclassified(100);k__Fungi_unclassified(100);k__Fungi_unclassified(100); | - | - | - | - | - | - | - | Unassigned | - |
| Otu32898 | 1 | k__Fungi(100);k__Fungi_unclassified(100);k__Fungi_unclassified(100);k__Fungi_unclassified(100);k__Fungi_unclassified(100);k__Fungi_unclassified(100);k__Fungi_unclassified(100); | - | - | - | - | - | - | - | Unassigned | - |
| Otu32900 | 1 | k__Fungi(100);p__Ascomycota(100);p__Ascomycota_unclassified(100);p__Ascomycota_unclassified(100);p__Ascomycota_unclassified(100);p__Ascomycota_unclassified(100);p__Ascomycota_unclassified(100); | - | - | - | - | - | - | - | Unassigned | - |
| Otu32903 | 1 | k__Fungi(100);k__Fungi_unclassified(100);k__Fungi_unclassified(100);k__Fungi_unclassified(100);k__Fungi_unclassified(100);k__Fungi_unclassified(100);k__Fungi_unclassified(100); | - | - | - | - | - | - | - | Unassigned | - |
| Otu32904 | 1 | k__Fungi(100);p__Ascomycota(100);p__Ascomycota_unclassified(100);p__Ascomycota_unclassified(100);p__Ascomycota_unclassified(100);p__Ascomycota_unclassified(100);p__Ascomycota_unclassified(100); | - | - | - | - | - | - | - | Unassigned | - |
| Otu32905 | 1 | k__Fungi(100);k__Fungi_unclassified(100);k__Fungi_unclassified(100);k__Fungi_unclassified(100);k__Fungi_unclassified(100);k__Fungi_unclassified(100);k__Fungi_unclassified(100); | - | - | - | - | - | - | - | Unassigned | - |
| Otu32906 | 1 | k__Fungi(100);k__Fungi_unclassified(100);k__Fungi_unclassified(100);k__Fungi_unclassified(100);k__Fungi_unclassified(100);k__Fungi_unclassified(100);k__Fungi_unclassified(100); | - | - | - | - | - | - | - | Unassigned | - |
| Otu32907 | 1 | k__Fungi(100);k__Fungi_unclassified(100);k__Fungi_unclassified(100);k__Fungi_unclassified(100);k__Fungi_unclassified(100);k__Fungi_unclassified(100);k__Fungi_unclassified(100); | - | - | - | - | - | - | - | Unassigned | - |
| Otu32908 | 1 | k__Fungi(100);p__Ascomycota(100);p__Ascomycota_unclassified(100);p__Ascomycota_unclassified(100);p__Ascomycota_unclassified(100);p__Ascomycota_unclassified(100);p__Ascomycota_unclassified(100); | - | - | - | - | - | - | - | Unassigned | - |
| Otu32909 | 1 | k__Fungi(100);k__Fungi_unclassified(100);k__Fungi_unclassified(100);k__Fungi_unclassified(100);k__Fungi_unclassified(100);k__Fungi_unclassified(100);k__Fungi_unclassified(100); | - | - | - | - | - | - | - | Unassigned | - |
| Otu32911 | 1 | k__Fungi(100);p__Ascomycota(100);p__Ascomycota_unclassified(100);p__Ascomycota_unclassified(100);p__Ascomycota_unclassified(100);p__Ascomycota_unclassified(100);p__Ascomycota_unclassified(100); | - | - | - | - | - | - | - | Unassigned | - |
| Otu32917 | 1 | k__Fungi(100);k__Fungi_unclassified(100);k__Fungi_unclassified(100);k__Fungi_unclassified(100);k__Fungi_unclassified(100);k__Fungi_unclassified(100);k__Fungi_unclassified(100); | - | - | - | - | - | - | - | Unassigned | - |
| Otu32918 | 1 | k__Fungi(100);k__Fungi_unclassified(100);k__Fungi_unclassified(100);k__Fungi_unclassified(100);k__Fungi_unclassified(100);k__Fungi_unclassified(100);k__Fungi_unclassified(100); | - | - | - | - | - | - | - | Unassigned | - |
| Otu32923 | 1 | k__Fungi(100);k__Fungi_unclassified(100);k__Fungi_unclassified(100);k__Fungi_unclassified(100);k__Fungi_unclassified(100);k__Fungi_unclassified(100);k__Fungi_unclassified(100); | - | - | - | - | - | - | - | Unassigned | - |
| Otu32924 | 1 | k__Fungi(100);p__Ascomycota(100);p__Ascomycota_unclassified(100);p__Ascomycota_unclassified(100);p__Ascomycota_unclassified(100);p__Ascomycota_unclassified(100);p__Ascomycota_unclassified(100); | - | - | - | - | - | - | - | Unassigned | - |
| Otu32927 | 1 | k__Fungi(100);k__Fungi_unclassified(100);k__Fungi_unclassified(100);k__Fungi_unclassified(100);k__Fungi_unclassified(100);k__Fungi_unclassified(100);k__Fungi_unclassified(100); | - | - | - | - | - | - | - | Unassigned | - |
| Otu32928 | 1 | k__Fungi(100);p__Ascomycota(100);p__Ascomycota_unclassified(100);p__Ascomycota_unclassified(100);p__Ascomycota_unclassified(100);p__Ascomycota_unclassified(100);p__Ascomycota_unclassified(100); | - | - | - | - | - | - | - | Unassigned | - |
| Otu32929 | 1 | k__Fungi(100);k__Fungi_unclassified(100);k__Fungi_unclassified(100);k__Fungi_unclassified(100);k__Fungi_unclassified(100);k__Fungi_unclassified(100);k__Fungi_unclassified(100); | - | - | - | - | - | - | - | Unassigned | - |
| Otu32931 | 1 | k__Fungi(100);k__Fungi_unclassified(100);k__Fungi_unclassified(100);k__Fungi_unclassified(100);k__Fungi_unclassified(100);k__Fungi_unclassified(100);k__Fungi_unclassified(100); | - | - | - | - | - | - | - | Unassigned | - |
| Otu32932 | 1 | k__Fungi(100);k__Fungi_unclassified(100);k__Fungi_unclassified(100);k__Fungi_unclassified(100);k__Fungi_unclassified(100);k__Fungi_unclassified(100);k__Fungi_unclassified(100); | - | - | - | - | - | - | - | Unassigned | - |
| Otu32935 | 1 | k__Fungi(100);k__Fungi_unclassified(100);k__Fungi_unclassified(100);k__Fungi_unclassified(100);k__Fungi_unclassified(100);k__Fungi_unclassified(100);k__Fungi_unclassified(100); | - | - | - | - | - | - | - | Unassigned | - |
| Otu32937 | 1 | k__Fungi(100);k__Fungi_unclassified(100);k__Fungi_unclassified(100);k__Fungi_unclassified(100);k__Fungi_unclassified(100);k__Fungi_unclassified(100);k__Fungi_unclassified(100); | - | - | - | - | - | - | - | Unassigned | - |
| Otu32938 | 1 | k__Fungi(100);k__Fungi_unclassified(100);k__Fungi_unclassified(100);k__Fungi_unclassified(100);k__Fungi_unclassified(100);k__Fungi_unclassified(100);k__Fungi_unclassified(100); | - | - | - | - | - | - | - | Unassigned | - |
| Otu32939 | 1 | k__Fungi(100);k__Fungi_unclassified(100);k__Fungi_unclassified(100);k__Fungi_unclassified(100);k__Fungi_unclassified(100);k__Fungi_unclassified(100);k__Fungi_unclassified(100); | - | - | - | - | - | - | - | Unassigned | - |
| Otu32940 | 1 | k__Fungi(100);p__Ascomycota(100);p__Ascomycota_unclassified(100);p__Ascomycota_unclassified(100);p__Ascomycota_unclassified(100);p__Ascomycota_unclassified(100);p__Ascomycota_unclassified(100); | - | - | - | - | - | - | - | Unassigned | - |
| Otu32941 | 1 | k__Fungi(100);k__Fungi_unclassified(100);k__Fungi_unclassified(100);k__Fungi_unclassified(100);k__Fungi_unclassified(100);k__Fungi_unclassified(100);k__Fungi_unclassified(100); | - | - | - | - | - | - | - | Unassigned | - |
| Otu32942 | 1 | k__Fungi(100);k__Fungi_unclassified(100);k__Fungi_unclassified(100);k__Fungi_unclassified(100);k__Fungi_unclassified(100);k__Fungi_unclassified(100);k__Fungi_unclassified(100); | - | - | - | - | - | - | - | Unassigned | - |
| Otu32943 | 1 | k__Fungi(100);k__Fungi_unclassified(100);k__Fungi_unclassified(100);k__Fungi_unclassified(100);k__Fungi_unclassified(100);k__Fungi_unclassified(100);k__Fungi_unclassified(100); | - | - | - | - | - | - | - | Unassigned | - |
| Otu32944 | 1 | k__Fungi(100);k__Fungi_unclassified(100);k__Fungi_unclassified(100);k__Fungi_unclassified(100);k__Fungi_unclassified(100);k__Fungi_unclassified(100);k__Fungi_unclassified(100); | - | - | - | - | - | - | - | Unassigned | - |
| Otu32945 | 1 | k__Fungi(100);k__Fungi_unclassified(100);k__Fungi_unclassified(100);k__Fungi_unclassified(100);k__Fungi_unclassified(100);k__Fungi_unclassified(100);k__Fungi_unclassified(100); | - | - | - | - | - | - | - | Unassigned | - |
| Otu32946 | 1 | k__Fungi(100);k__Fungi_unclassified(100);k__Fungi_unclassified(100);k__Fungi_unclassified(100);k__Fungi_unclassified(100);k__Fungi_unclassified(100);k__Fungi_unclassified(100); | - | - | - | - | - | - | - | Unassigned | - |
| Otu32947 | 1 | k__Fungi(100);k__Fungi_unclassified(100);k__Fungi_unclassified(100);k__Fungi_unclassified(100);k__Fungi_unclassified(100);k__Fungi_unclassified(100);k__Fungi_unclassified(100); | - | - | - | - | - | - | - | Unassigned | - |
| Otu32948 | 1 | k__Fungi(100);p__Ascomycota(100);p__Ascomycota_unclassified(100);p__Ascomycota_unclassified(100);p__Ascomycota_unclassified(100);p__Ascomycota_unclassified(100);p__Ascomycota_unclassified(100); | - | - | - | - | - | - | - | Unassigned | - |
| Otu32950 | 1 | k__Fungi(100);p__Ascomycota(100);p__Ascomycota_unclassified(100);p__Ascomycota_unclassified(100);p__Ascomycota_unclassified(100);p__Ascomycota_unclassified(100);p__Ascomycota_unclassified(100); | - | - | - | - | - | - | - | Unassigned | - |
| Otu32958 | 1 | k__Fungi(100);p__Ascomycota(100);p__Ascomycota_unclassified(100);p__Ascomycota_unclassified(100);p__Ascomycota_unclassified(100);p__Ascomycota_unclassified(100);p__Ascomycota_unclassified(100); | - | - | - | - | - | - | - | Unassigned | - |
| Otu32960 | 1 | k__Fungi(100);k__Fungi_unclassified(100);k__Fungi_unclassified(100);k__Fungi_unclassified(100);k__Fungi_unclassified(100);k__Fungi_unclassified(100);k__Fungi_unclassified(100); | - | - | - | - | - | - | - | Unassigned | - |
| Otu32963 | 1 | k__Fungi(100);k__Fungi_unclassified(100);k__Fungi_unclassified(100);k__Fungi_unclassified(100);k__Fungi_unclassified(100);k__Fungi_unclassified(100);k__Fungi_unclassified(100); | - | - | - | - | - | - | - | Unassigned | - |
| Otu32965 | 1 | k__Fungi(100);p__Ascomycota(100);p__Ascomycota_unclassified(100);p__Ascomycota_unclassified(100);p__Ascomycota_unclassified(100);p__Ascomycota_unclassified(100);p__Ascomycota_unclassified(100); | - | - | - | - | - | - | - | Unassigned | - |
| Otu32967 | 1 | k__Fungi(100);k__Fungi_unclassified(100);k__Fungi_unclassified(100);k__Fungi_unclassified(100);k__Fungi_unclassified(100);k__Fungi_unclassified(100);k__Fungi_unclassified(100); | - | - | - | - | - | - | - | Unassigned | - |
| Otu32968 | 1 | k__Fungi(100);k__Fungi_unclassified(100);k__Fungi_unclassified(100);k__Fungi_unclassified(100);k__Fungi_unclassified(100);k__Fungi_unclassified(100);k__Fungi_unclassified(100); | - | - | - | - | - | - | - | Unassigned | - |
| Otu32970 | 1 | k__Fungi(100);p__Ascomycota(100);p__Ascomycota_unclassified(100);p__Ascomycota_unclassified(100);p__Ascomycota_unclassified(100);p__Ascomycota_unclassified(100);p__Ascomycota_unclassified(100); | - | - | - | - | - | - | - | Unassigned | - |
| Otu32971 | 1 | k__Fungi(100);k__Fungi_unclassified(100);k__Fungi_unclassified(100);k__Fungi_unclassified(100);k__Fungi_unclassified(100);k__Fungi_unclassified(100);k__Fungi_unclassified(100); | - | - | - | - | - | - | - | Unassigned | - |
| Otu32972 | 1 | k__Fungi(100);k__Fungi_unclassified(100);k__Fungi_unclassified(100);k__Fungi_unclassified(100);k__Fungi_unclassified(100);k__Fungi_unclassified(100);k__Fungi_unclassified(100); | - | - | - | - | - | - | - | Unassigned | - |
| Otu32973 | 1 | k__Fungi(100);p__Ascomycota(100);p__Ascomycota_unclassified(100);p__Ascomycota_unclassified(100);p__Ascomycota_unclassified(100);p__Ascomycota_unclassified(100);p__Ascomycota_unclassified(100); | - | - | - | - | - | - | - | Unassigned | - |
| Otu32974 | 1 | k__Fungi(100);k__Fungi_unclassified(100);k__Fungi_unclassified(100);k__Fungi_unclassified(100);k__Fungi_unclassified(100);k__Fungi_unclassified(100);k__Fungi_unclassified(100); | - | - | - | - | - | - | - | Unassigned | - |
| Otu32975 | 1 | k__Fungi(100);p__Ascomycota(100);p__Ascomycota_unclassified(100);p__Ascomycota_unclassified(100);p__Ascomycota_unclassified(100);p__Ascomycota_unclassified(100);p__Ascomycota_unclassified(100); | - | - | - | - | - | - | - | Unassigned | - |
| Otu32985 | 1 | k__Fungi(100);k__Fungi_unclassified(100);k__Fungi_unclassified(100);k__Fungi_unclassified(100);k__Fungi_unclassified(100);k__Fungi_unclassified(100);k__Fungi_unclassified(100); | - | - | - | - | - | - | - | Unassigned | - |
| Otu32986 | 1 | k__Fungi(100);p__Ascomycota(100);p__Ascomycota_unclassified(100);p__Ascomycota_unclassified(100);p__Ascomycota_unclassified(100);p__Ascomycota_unclassified(100);p__Ascomycota_unclassified(100); | - | - | - | - | - | - | - | Unassigned | - |
| Otu32996 | 1 | k__Fungi(100);k__Fungi_unclassified(100);k__Fungi_unclassified(100);k__Fungi_unclassified(100);k__Fungi_unclassified(100);k__Fungi_unclassified(100);k__Fungi_unclassified(100); | - | - | - | - | - | - | - | Unassigned | - |
| Otu32998 | 1 | k__Fungi(100);p__Ascomycota(100);p__Ascomycota_unclassified(100);p__Ascomycota_unclassified(100);p__Ascomycota_unclassified(100);p__Ascomycota_unclassified(100);p__Ascomycota_unclassified(100); | - | - | - | - | - | - | - | Unassigned | - |
| Otu32999 | 1 | k__Fungi(100);k__Fungi_unclassified(100);k__Fungi_unclassified(100);k__Fungi_unclassified(100);k__Fungi_unclassified(100);k__Fungi_unclassified(100);k__Fungi_unclassified(100); | - | - | - | - | - | - | - | Unassigned | - |
| Otu33000 | 1 | k__Fungi(100);k__Fungi_unclassified(100);k__Fungi_unclassified(100);k__Fungi_unclassified(100);k__Fungi_unclassified(100);k__Fungi_unclassified(100);k__Fungi_unclassified(100); | - | - | - | - | - | - | - | Unassigned | - |
| Otu33017 | 1 | k__Fungi(100);k__Fungi_unclassified(100);k__Fungi_unclassified(100);k__Fungi_unclassified(100);k__Fungi_unclassified(100);k__Fungi_unclassified(100);k__Fungi_unclassified(100); | - | - | - | - | - | - | - | Unassigned | - |
| Otu33018 | 1 | k__Fungi(100);p__Ascomycota(100);p__Ascomycota_unclassified(100);p__Ascomycota_unclassified(100);p__Ascomycota_unclassified(100);p__Ascomycota_unclassified(100);p__Ascomycota_unclassified(100); | - | - | - | - | - | - | - | Unassigned | - |
| Otu33019 | 1 | k__Fungi(100);k__Fungi_unclassified(100);k__Fungi_unclassified(100);k__Fungi_unclassified(100);k__Fungi_unclassified(100);k__Fungi_unclassified(100);k__Fungi_unclassified(100); | - | - | - | - | - | - | - | Unassigned | - |
| Otu33030 | 1 | k__Fungi(100);p__Ascomycota(100);p__Ascomycota_unclassified(100);p__Ascomycota_unclassified(100);p__Ascomycota_unclassified(100);p__Ascomycota_unclassified(100);p__Ascomycota_unclassified(100); | - | - | - | - | - | - | - | Unassigned | - |
| Otu33033 | 1 | k__Fungi(100);k__Fungi_unclassified(100);k__Fungi_unclassified(100);k__Fungi_unclassified(100);k__Fungi_unclassified(100);k__Fungi_unclassified(100);k__Fungi_unclassified(100); | - | - | - | - | - | - | - | Unassigned | - |
| Otu33037 | 1 | k__Fungi(100);p__Ascomycota(100);p__Ascomycota_unclassified(100);p__Ascomycota_unclassified(100);p__Ascomycota_unclassified(100);p__Ascomycota_unclassified(100);p__Ascomycota_unclassified(100); | - | - | - | - | - | - | - | Unassigned | - |
| Otu33057 | 1 | k__Fungi(100);k__Fungi_unclassified(100);k__Fungi_unclassified(100);k__Fungi_unclassified(100);k__Fungi_unclassified(100);k__Fungi_unclassified(100);k__Fungi_unclassified(100); | - | - | - | - | - | - | - | Unassigned | - |
| Otu33075 | 1 | k__Fungi(100);k__Fungi_unclassified(100);k__Fungi_unclassified(100);k__Fungi_unclassified(100);k__Fungi_unclassified(100);k__Fungi_unclassified(100);k__Fungi_unclassified(100); | - | - | - | - | - | - | - | Unassigned | - |
| Otu33093 | 1 | k__Fungi(100);k__Fungi_unclassified(100);k__Fungi_unclassified(100);k__Fungi_unclassified(100);k__Fungi_unclassified(100);k__Fungi_unclassified(100);k__Fungi_unclassified(100); | - | - | - | - | - | - | - | Unassigned | - |
| Otu33094 | 1 | k__Fungi(100);k__Fungi_unclassified(100);k__Fungi_unclassified(100);k__Fungi_unclassified(100);k__Fungi_unclassified(100);k__Fungi_unclassified(100);k__Fungi_unclassified(100); | - | - | - | - | - | - | - | Unassigned | - |
| Otu33095 | 1 | k__Fungi(100);p__Ascomycota(100);p__Ascomycota_unclassified(100);p__Ascomycota_unclassified(100);p__Ascomycota_unclassified(100);p__Ascomycota_unclassified(100);p__Ascomycota_unclassified(100); | - | - | - | - | - | - | - | Unassigned | - |
| Otu33096 | 1 | k__Fungi(100);p__Ascomycota(100);p__Ascomycota_unclassified(100);p__Ascomycota_unclassified(100);p__Ascomycota_unclassified(100);p__Ascomycota_unclassified(100);p__Ascomycota_unclassified(100); | - | - | - | - | - | - | - | Unassigned | - |
| Otu33100 | 1 | k__Fungi(100);k__Fungi_unclassified(100);k__Fungi_unclassified(100);k__Fungi_unclassified(100);k__Fungi_unclassified(100);k__Fungi_unclassified(100);k__Fungi_unclassified(100); | - | - | - | - | - | - | - | Unassigned | - |
| Otu33101 | 1 | k__Fungi(100);p__Ascomycota(100);p__Ascomycota_unclassified(100);p__Ascomycota_unclassified(100);p__Ascomycota_unclassified(100);p__Ascomycota_unclassified(100);p__Ascomycota_unclassified(100); | - | - | - | - | - | - | - | Unassigned | - |
| Otu33108 | 1 | k__Fungi(100);k__Fungi_unclassified(100);k__Fungi_unclassified(100);k__Fungi_unclassified(100);k__Fungi_unclassified(100);k__Fungi_unclassified(100);k__Fungi_unclassified(100); | - | - | - | - | - | - | - | Unassigned | - |
| Otu33109 | 1 | k__Fungi(100);k__Fungi_unclassified(100);k__Fungi_unclassified(100);k__Fungi_unclassified(100);k__Fungi_unclassified(100);k__Fungi_unclassified(100);k__Fungi_unclassified(100); | - | - | - | - | - | - | - | Unassigned | - |
| Otu33110 | 1 | k__Fungi(100);p__Ascomycota(100);p__Ascomycota_unclassified(100);p__Ascomycota_unclassified(100);p__Ascomycota_unclassified(100);p__Ascomycota_unclassified(100);p__Ascomycota_unclassified(100); | - | - | - | - | - | - | - | Unassigned | - |
| Otu33111 | 1 | k__Fungi(100);p__Ascomycota(100);p__Ascomycota_unclassified(100);p__Ascomycota_unclassified(100);p__Ascomycota_unclassified(100);p__Ascomycota_unclassified(100);p__Ascomycota_unclassified(100); | - | - | - | - | - | - | - | Unassigned | - |
| Otu33112 | 1 | k__Fungi(100);k__Fungi_unclassified(100);k__Fungi_unclassified(100);k__Fungi_unclassified(100);k__Fungi_unclassified(100);k__Fungi_unclassified(100);k__Fungi_unclassified(100); | - | - | - | - | - | - | - | Unassigned | - |
| Otu33113 | 1 | k__Fungi(100);k__Fungi_unclassified(100);k__Fungi_unclassified(100);k__Fungi_unclassified(100);k__Fungi_unclassified(100);k__Fungi_unclassified(100);k__Fungi_unclassified(100); | - | - | - | - | - | - | - | Unassigned | - |
| Otu33114 | 1 | k__Fungi(100);k__Fungi_unclassified(100);k__Fungi_unclassified(100);k__Fungi_unclassified(100);k__Fungi_unclassified(100);k__Fungi_unclassified(100);k__Fungi_unclassified(100); | - | - | - | - | - | - | - | Unassigned | - |
| Otu33115 | 1 | k__Fungi(100);k__Fungi_unclassified(100);k__Fungi_unclassified(100);k__Fungi_unclassified(100);k__Fungi_unclassified(100);k__Fungi_unclassified(100);k__Fungi_unclassified(100); | - | - | - | - | - | - | - | Unassigned | - |
| Otu33116 | 1 | k__Fungi(100);k__Fungi_unclassified(100);k__Fungi_unclassified(100);k__Fungi_unclassified(100);k__Fungi_unclassified(100);k__Fungi_unclassified(100);k__Fungi_unclassified(100); | - | - | - | - | - | - | - | Unassigned | - |
| Otu33118 | 1 | k__Fungi(100);p__Ascomycota(100);p__Ascomycota_unclassified(100);p__Ascomycota_unclassified(100);p__Ascomycota_unclassified(100);p__Ascomycota_unclassified(100);p__Ascomycota_unclassified(100); | - | - | - | - | - | - | - | Unassigned | - |
| Otu33119 | 1 | k__Fungi(100);k__Fungi_unclassified(100);k__Fungi_unclassified(100);k__Fungi_unclassified(100);k__Fungi_unclassified(100);k__Fungi_unclassified(100);k__Fungi_unclassified(100); | - | - | - | - | - | - | - | Unassigned | - |
| Otu33120 | 1 | k__Fungi(100);k__Fungi_unclassified(100);k__Fungi_unclassified(100);k__Fungi_unclassified(100);k__Fungi_unclassified(100);k__Fungi_unclassified(100);k__Fungi_unclassified(100); | - | - | - | - | - | - | - | Unassigned | - |
| Otu33121 | 1 | k__Fungi(100);p__Ascomycota(100);p__Ascomycota_unclassified(100);p__Ascomycota_unclassified(100);p__Ascomycota_unclassified(100);p__Ascomycota_unclassified(100);p__Ascomycota_unclassified(100); | - | - | - | - | - | - | - | Unassigned | - |
| Otu33123 | 1 | k__Fungi(100);p__Ascomycota(100);p__Ascomycota_unclassified(100);p__Ascomycota_unclassified(100);p__Ascomycota_unclassified(100);p__Ascomycota_unclassified(100);p__Ascomycota_unclassified(100); | - | - | - | - | - | - | - | Unassigned | - |
| Otu33125 | 1 | k__Fungi(100);p__Ascomycota(100);p__Ascomycota_unclassified(100);p__Ascomycota_unclassified(100);p__Ascomycota_unclassified(100);p__Ascomycota_unclassified(100);p__Ascomycota_unclassified(100); | - | - | - | - | - | - | - | Unassigned | - |
| Otu33134 | 1 | k__Fungi(100);k__Fungi_unclassified(100);k__Fungi_unclassified(100);k__Fungi_unclassified(100);k__Fungi_unclassified(100);k__Fungi_unclassified(100);k__Fungi_unclassified(100); | - | - | - | - | - | - | - | Unassigned | - |
| Otu33135 | 1 | k__Fungi(100);k__Fungi_unclassified(100);k__Fungi_unclassified(100);k__Fungi_unclassified(100);k__Fungi_unclassified(100);k__Fungi_unclassified(100);k__Fungi_unclassified(100); | - | - | - | - | - | - | - | Unassigned | - |
| Otu33147 | 1 | k__Fungi(100);k__Fungi_unclassified(100);k__Fungi_unclassified(100);k__Fungi_unclassified(100);k__Fungi_unclassified(100);k__Fungi_unclassified(100);k__Fungi_unclassified(100); | - | - | - | - | - | - | - | Unassigned | - |
| Otu33160 | 1 | k__Fungi(100);k__Fungi_unclassified(100);k__Fungi_unclassified(100);k__Fungi_unclassified(100);k__Fungi_unclassified(100);k__Fungi_unclassified(100);k__Fungi_unclassified(100); | - | - | - | - | - | - | - | Unassigned | - |
| Otu33162 | 1 | k__Fungi(100);k__Fungi_unclassified(100);k__Fungi_unclassified(100);k__Fungi_unclassified(100);k__Fungi_unclassified(100);k__Fungi_unclassified(100);k__Fungi_unclassified(100); | - | - | - | - | - | - | - | Unassigned | - |
| Otu33163 | 1 | k__Fungi(100);k__Fungi_unclassified(100);k__Fungi_unclassified(100);k__Fungi_unclassified(100);k__Fungi_unclassified(100);k__Fungi_unclassified(100);k__Fungi_unclassified(100); | - | - | - | - | - | - | - | Unassigned | - |
| Otu33166 | 1 | k__Fungi(100);p__Ascomycota(100);p__Ascomycota_unclassified(100);p__Ascomycota_unclassified(100);p__Ascomycota_unclassified(100);p__Ascomycota_unclassified(100);p__Ascomycota_unclassified(100); | - | - | - | - | - | - | - | Unassigned | - |
| Otu33168 | 1 | k__Fungi(100);p__Ascomycota(100);p__Ascomycota_unclassified(100);p__Ascomycota_unclassified(100);p__Ascomycota_unclassified(100);p__Ascomycota_unclassified(100);p__Ascomycota_unclassified(100); | - | - | - | - | - | - | - | Unassigned | - |
| Otu33175 | 1 | k__Fungi(100);p__Ascomycota(100);p__Ascomycota_unclassified(100);p__Ascomycota_unclassified(100);p__Ascomycota_unclassified(100);p__Ascomycota_unclassified(100);p__Ascomycota_unclassified(100); | - | - | - | - | - | - | - | Unassigned | - |

| | | | | | | | | | | | |
|---|---|---|---|---|---|---|---|---|---|---|---|
| Otu33181 | 1 | k__Fungi(100);k__Fungi_unclassified(100);k__Fungi_unclassified(100);k__Fungi_unclassified(100);k__Fungi_unclassified(100);k__Fungi_unclassified(100);k__Fungi_unclassified(100); | - | - | - | - | - | - | - | Unassigned | - |
| Otu33182 | 1 | k__Fungi(100);k__Fungi_unclassified(100);k__Fungi_unclassified(100);k__Fungi_unclassified(100);k__Fungi_unclassified(100);k__Fungi_unclassified(100);k__Fungi_unclassified(100); | - | - | - | - | - | - | - | Unassigned | - |
| Otu33183 | 1 | k__Fungi(100);p__Ascomycota(100);p__Ascomycota_unclassified(100);p__Ascomycota_unclassified(100);p__Ascomycota_unclassified(100);p__Ascomycota_unclassified(100);p__Ascomycota_unclassified(100); | - | - | - | - | - | - | - | Unassigned | - |
| Otu33184 | 1 | k__Fungi(100);k__Fungi_unclassified(100);k__Fungi_unclassified(100);k__Fungi_unclassified(100);k__Fungi_unclassified(100);k__Fungi_unclassified(100);k__Fungi_unclassified(100); | - | - | - | - | - | - | - | Unassigned | - |
| Otu33185 | 1 | k__Fungi(100);k__Fungi_unclassified(100);k__Fungi_unclassified(100);k__Fungi_unclassified(100);k__Fungi_unclassified(100);k__Fungi_unclassified(100);k__Fungi_unclassified(100); | - | - | - | - | - | - | - | Unassigned | - |
| Otu33196 | 1 | k__Fungi(100);k__Fungi_unclassified(100);k__Fungi_unclassified(100);k__Fungi_unclassified(100);k__Fungi_unclassified(100);k__Fungi_unclassified(100);k__Fungi_unclassified(100); | - | - | - | - | - | - | - | Unassigned | - |
| Otu33197 | 1 | k__Fungi(100);k__Fungi_unclassified(100);k__Fungi_unclassified(100);k__Fungi_unclassified(100);k__Fungi_unclassified(100);k__Fungi_unclassified(100);k__Fungi_unclassified(100); | - | - | - | - | - | - | - | Unassigned | - |
| Otu33202 | 1 | k__Fungi(100);k__Fungi_unclassified(100);k__Fungi_unclassified(100);k__Fungi_unclassified(100);k__Fungi_unclassified(100);k__Fungi_unclassified(100);k__Fungi_unclassified(100); | - | - | - | - | - | - | - | Unassigned | - |
| Otu33203 | 1 | k__Fungi(100);p__Ascomycota(100);p__Ascomycota_unclassified(100);p__Ascomycota_unclassified(100);p__Ascomycota_unclassified(100);p__Ascomycota_unclassified(100);p__Ascomycota_unclassified(100); | - | - | - | - | - | - | - | Unassigned | - |
| Otu33204 | 1 | k__Fungi(100);p__Ascomycota(100);p__Ascomycota_unclassified(100);p__Ascomycota_unclassified(100);p__Ascomycota_unclassified(100);p__Ascomycota_unclassified(100);p__Ascomycota_unclassified(100); | - | - | - | - | - | - | - | Unassigned | - |
| Otu33205 | 1 | k__Fungi(100);k__Fungi_unclassified(100);k__Fungi_unclassified(100);k__Fungi_unclassified(100);k__Fungi_unclassified(100);k__Fungi_unclassified(100);k__Fungi_unclassified(100); | - | - | - | - | - | - | - | Unassigned | - |
| Otu33207 | 1 | k__Fungi(100);k__Fungi_unclassified(100);k__Fungi_unclassified(100);k__Fungi_unclassified(100);k__Fungi_unclassified(100);k__Fungi_unclassified(100);k__Fungi_unclassified(100); | - | - | - | - | - | - | - | Unassigned | - |
| Otu33208 | 1 | k__Fungi(100);p__Ascomycota(100);p__Ascomycota_unclassified(100);p__Ascomycota_unclassified(100);p__Ascomycota_unclassified(100);p__Ascomycota_unclassified(100);p__Ascomycota_unclassified(100); | - | - | - | - | - | - | - | Unassigned | - |
| Otu33209 | 1 | k__Fungi(100);k__Fungi_unclassified(100);k__Fungi_unclassified(100);k__Fungi_unclassified(100);k__Fungi_unclassified(100);k__Fungi_unclassified(100);k__Fungi_unclassified(100); | - | - | - | - | - | - | - | Unassigned | - |
| Otu33211 | 1 | k__Fungi(100);k__Fungi_unclassified(100);k__Fungi_unclassified(100);k__Fungi_unclassified(100);k__Fungi_unclassified(100);k__Fungi_unclassified(100);k__Fungi_unclassified(100); | - | - | - | - | - | - | - | Unassigned | - |
| Otu33212 | 1 | k__Fungi(100);p__Ascomycota(100);p__Ascomycota_unclassified(100);p__Ascomycota_unclassified(100);p__Ascomycota_unclassified(100);p__Ascomycota_unclassified(100);p__Ascomycota_unclassified(100); | - | - | - | - | - | - | - | Unassigned | - |
| Otu33214 | 1 | k__Fungi(100);k__Fungi_unclassified(100);k__Fungi_unclassified(100);k__Fungi_unclassified(100);k__Fungi_unclassified(100);k__Fungi_unclassified(100);k__Fungi_unclassified(100); | - | - | - | - | - | - | - | Unassigned | - |
| Otu33216 | 1 | k__Fungi(100);k__Fungi_unclassified(100);k__Fungi_unclassified(100);k__Fungi_unclassified(100);k__Fungi_unclassified(100);k__Fungi_unclassified(100);k__Fungi_unclassified(100); | - | - | - | - | - | - | - | Unassigned | - |
| Otu33218 | 1 | k__Fungi(100);k__Fungi_unclassified(100);k__Fungi_unclassified(100);k__Fungi_unclassified(100);k__Fungi_unclassified(100);k__Fungi_unclassified(100);k__Fungi_unclassified(100); | - | - | - | - | - | - | - | Unassigned | - |
| Otu33219 | 1 | k__Fungi(100);p__Ascomycota(100);p__Ascomycota_unclassified(100);p__Ascomycota_unclassified(100);p__Ascomycota_unclassified(100);p__Ascomycota_unclassified(100);p__Ascomycota_unclassified(100); | - | - | - | - | - | - | - | Unassigned | - |
| Otu33220 | 1 | k__Fungi(100);p__Ascomycota(100);p__Ascomycota_unclassified(100);p__Ascomycota_unclassified(100);p__Ascomycota_unclassified(100);p__Ascomycota_unclassified(100);p__Ascomycota_unclassified(100); | - | - | - | - | - | - | - | Unassigned | - |
| Otu33221 | 1 | k__Fungi(100);k__Fungi_unclassified(100);k__Fungi_unclassified(100);k__Fungi_unclassified(100);k__Fungi_unclassified(100);k__Fungi_unclassified(100);k__Fungi_unclassified(100); | - | - | - | - | - | - | - | Unassigned | - |
| Otu33222 | 1 | k__Fungi(100);p__Ascomycota(100);p__Ascomycota_unclassified(100);p__Ascomycota_unclassified(100);p__Ascomycota_unclassified(100);p__Ascomycota_unclassified(100);p__Ascomycota_unclassified(100); | - | - | - | - | - | - | - | Unassigned | - |
| Otu33223 | 1 | k__Fungi(100);k__Fungi_unclassified(100);k__Fungi_unclassified(100);k__Fungi_unclassified(100);k__Fungi_unclassified(100);k__Fungi_unclassified(100);k__Fungi_unclassified(100); | - | - | - | - | - | - | - | Unassigned | - |
| Otu33224 | 1 | k__Fungi(100);p__Ascomycota(100);p__Ascomycota_unclassified(100);p__Ascomycota_unclassified(100);p__Ascomycota_unclassified(100);p__Ascomycota_unclassified(100);p__Ascomycota_unclassified(100); | - | - | - | - | - | - | - | Unassigned | - |
| Otu33225 | 1 | k__Fungi(100);k__Fungi_unclassified(100);k__Fungi_unclassified(100);k__Fungi_unclassified(100);k__Fungi_unclassified(100);k__Fungi_unclassified(100);k__Fungi_unclassified(100); | - | - | - | - | - | - | - | Unassigned | - |
| Otu33228 | 1 | k__Fungi(100);p__Ascomycota(100);p__Ascomycota_unclassified(100);p__Ascomycota_unclassified(100);p__Ascomycota_unclassified(100);p__Ascomycota_unclassified(100);p__Ascomycota_unclassified(100); | - | - | - | - | - | - | - | Unassigned | - |
| Otu33229 | 1 | k__Fungi(100);p__Ascomycota(100);p__Ascomycota_unclassified(100);p__Ascomycota_unclassified(100);p__Ascomycota_unclassified(100);p__Ascomycota_unclassified(100);p__Ascomycota_unclassified(100); | - | - | - | - | - | - | - | Unassigned | - |
| Otu33230 | 1 | k__Fungi(100);k__Fungi_unclassified(100);k__Fungi_unclassified(100);k__Fungi_unclassified(100);k__Fungi_unclassified(100);k__Fungi_unclassified(100);k__Fungi_unclassified(100); | - | - | - | - | - | - | - | Unassigned | - |
| Otu33232 | 1 | k__Fungi(100);k__Fungi_unclassified(100);k__Fungi_unclassified(100);k__Fungi_unclassified(100);k__Fungi_unclassified(100);k__Fungi_unclassified(100);k__Fungi_unclassified(100); | - | - | - | - | - | - | - | Unassigned | - |
| Otu33233 | 1 | k__Fungi(100);p__Ascomycota(100);p__Ascomycota_unclassified(100);p__Ascomycota_unclassified(100);p__Ascomycota_unclassified(100);p__Ascomycota_unclassified(100);p__Ascomycota_unclassified(100); | - | - | - | - | - | - | - | Unassigned | - |
| Otu33235 | 1 | k__Fungi(100);p__Ascomycota(100);p__Ascomycota_unclassified(100);p__Ascomycota_unclassified(100);p__Ascomycota_unclassified(100);p__Ascomycota_unclassified(100);p__Ascomycota_unclassified(100); | - | - | - | - | - | - | - | Unassigned | - |
| Otu33238 | 1 | k__Fungi(100);p__Ascomycota(100);p__Ascomycota_unclassified(100);p__Ascomycota_unclassified(100);p__Ascomycota_unclassified(100);p__Ascomycota_unclassified(100);p__Ascomycota_unclassified(100); | - | - | - | - | - | - | - | Unassigned | - |
| Otu33239 | 1 | k__Fungi(100);p__Ascomycota(100);p__Ascomycota_unclassified(100);p__Ascomycota_unclassified(100);p__Ascomycota_unclassified(100);p__Ascomycota_unclassified(100);p__Ascomycota_unclassified(100); | - | - | - | - | - | - | - | Unassigned | - |
| Otu33240 | 1 | k__Fungi(100);p__Ascomycota(100);p__Ascomycota_unclassified(100);p__Ascomycota_unclassified(100);p__Ascomycota_unclassified(100);p__Ascomycota_unclassified(100);p__Ascomycota_unclassified(100); | - | - | - | - | - | - | - | Unassigned | - |
| Otu33241 | 1 | k__Fungi(100);k__Fungi_unclassified(100);k__Fungi_unclassified(100);k__Fungi_unclassified(100);k__Fungi_unclassified(100);k__Fungi_unclassified(100);k__Fungi_unclassified(100); | - | - | - | - | - | - | - | Unassigned | - |
| Otu33242 | 1 | k__Fungi(100);k__Fungi_unclassified(100);k__Fungi_unclassified(100);k__Fungi_unclassified(100);k__Fungi_unclassified(100);k__Fungi_unclassified(100);k__Fungi_unclassified(100); | - | - | - | - | - | - | - | Unassigned | - |
| Otu33243 | 1 | k__Fungi(100);k__Fungi_unclassified(100);k__Fungi_unclassified(100);k__Fungi_unclassified(100);k__Fungi_unclassified(100);k__Fungi_unclassified(100);k__Fungi_unclassified(100); | - | - | - | - | - | - | - | Unassigned | - |
| Otu33244 | 1 | k__Fungi(100);p__Ascomycota(100);p__Ascomycota_unclassified(100);p__Ascomycota_unclassified(100);p__Ascomycota_unclassified(100);p__Ascomycota_unclassified(100);p__Ascomycota_unclassified(100); | - | - | - | - | - | - | - | Unassigned | - |
| Otu33245 | 1 | k__Fungi(100);k__Fungi_unclassified(100);k__Fungi_unclassified(100);k__Fungi_unclassified(100);k__Fungi_unclassified(100);k__Fungi_unclassified(100);k__Fungi_unclassified(100); | - | - | - | - | - | - | - | Unassigned | - |
| Otu33246 | 1 | k__Fungi(100);k__Fungi_unclassified(100);k__Fungi_unclassified(100);k__Fungi_unclassified(100);k__Fungi_unclassified(100);k__Fungi_unclassified(100);k__Fungi_unclassified(100); | - | - | - | - | - | - | - | Unassigned | - |
| Otu33247 | 1 | k__Fungi(100);k__Fungi_unclassified(100);k__Fungi_unclassified(100);k__Fungi_unclassified(100);k__Fungi_unclassified(100);k__Fungi_unclassified(100);k__Fungi_unclassified(100); | - | - | - | - | - | - | - | Unassigned | - |
| Otu33248 | 1 | k__Fungi(100);k__Fungi_unclassified(100);k__Fungi_unclassified(100);k__Fungi_unclassified(100);k__Fungi_unclassified(100);k__Fungi_unclassified(100);k__Fungi_unclassified(100); | - | - | - | - | - | - | - | Unassigned | - |
| Otu33249 | 1 | k__Fungi(100);p__Ascomycota(100);p__Ascomycota_unclassified(100);p__Ascomycota_unclassified(100);p__Ascomycota_unclassified(100);p__Ascomycota_unclassified(100);p__Ascomycota_unclassified(100); | - | - | - | - | - | - | - | Unassigned | - |
| Otu33251 | 1 | k__Fungi(100);k__Fungi_unclassified(100);k__Fungi_unclassified(100);k__Fungi_unclassified(100);k__Fungi_unclassified(100);k__Fungi_unclassified(100);k__Fungi_unclassified(100); | - | - | - | - | - | - | - | Unassigned | - |
| Otu33252 | 1 | k__Fungi(100);k__Fungi_unclassified(100);k__Fungi_unclassified(100);k__Fungi_unclassified(100);k__Fungi_unclassified(100);k__Fungi_unclassified(100);k__Fungi_unclassified(100); | - | - | - | - | - | - | - | Unassigned | - |
| Otu33253 | 1 | k__Fungi(100);k__Fungi_unclassified(100);k__Fungi_unclassified(100);k__Fungi_unclassified(100);k__Fungi_unclassified(100);k__Fungi_unclassified(100);k__Fungi_unclassified(100); | - | - | - | - | - | - | - | Unassigned | - |
| Otu33254 | 1 | k__Fungi(100);p__Ascomycota(100);p__Ascomycota_unclassified(100);p__Ascomycota_unclassified(100);p__Ascomycota_unclassified(100);p__Ascomycota_unclassified(100);p__Ascomycota_unclassified(100); | - | - | - | - | - | - | - | Unassigned | - |
| Otu33261 | 1 | k__Fungi(100);p__Ascomycota(100);p__Ascomycota_unclassified(100);p__Ascomycota_unclassified(100);p__Ascomycota_unclassified(100);p__Ascomycota_unclassified(100);p__Ascomycota_unclassified(100); | - | - | - | - | - | - | - | Unassigned | - |
| Otu33267 | 1 | k__Fungi(100);k__Fungi_unclassified(100);k__Fungi_unclassified(100);k__Fungi_unclassified(100);k__Fungi_unclassified(100);k__Fungi_unclassified(100);k__Fungi_unclassified(100); | - | - | - | - | - | - | - | Unassigned | - |
| Otu33268 | 1 | k__Fungi(100);p__Ascomycota(100);p__Ascomycota_unclassified(100);p__Ascomycota_unclassified(100);p__Ascomycota_unclassified(100);p__Ascomycota_unclassified(100);p__Ascomycota_unclassified(100); | - | - | - | - | - | - | - | Unassigned | - |
| Otu33269 | 1 | k__Fungi(100);k__Fungi_unclassified(100);k__Fungi_unclassified(100);k__Fungi_unclassified(100);k__Fungi_unclassified(100);k__Fungi_unclassified(100);k__Fungi_unclassified(100); | - | - | - | - | - | - | - | Unassigned | - |
| Otu33273 | 1 | k__Fungi(100);p__Ascomycota(100);p__Ascomycota_unclassified(100);p__Ascomycota_unclassified(100);p__Ascomycota_unclassified(100);p__Ascomycota_unclassified(100);p__Ascomycota_unclassified(100); | - | - | - | - | - | - | - | Unassigned | - |
| Otu33274 | 1 | k__Fungi(100);k__Fungi_unclassified(100);k__Fungi_unclassified(100);k__Fungi_unclassified(100);k__Fungi_unclassified(100);k__Fungi_unclassified(100);k__Fungi_unclassified(100); | - | - | - | - | - | - | - | Unassigned | - |
| Otu33275 | 1 | k__Fungi(100);p__Ascomycota(100);p__Ascomycota_unclassified(100);p__Ascomycota_unclassified(100);p__Ascomycota_unclassified(100);p__Ascomycota_unclassified(100);p__Ascomycota_unclassified(100); | - | - | - | - | - | - | - | Unassigned | - |
| Otu33278 | 1 | k__Fungi(100);k__Fungi_unclassified(100);k__Fungi_unclassified(100);k__Fungi_unclassified(100);k__Fungi_unclassified(100);k__Fungi_unclassified(100);k__Fungi_unclassified(100); | - | - | - | - | - | - | - | Unassigned | - |
| Otu33281 | 1 | k__Fungi(100);p__Ascomycota(100);p__Ascomycota_unclassified(100);p__Ascomycota_unclassified(100);p__Ascomycota_unclassified(100);p__Ascomycota_unclassified(100);p__Ascomycota_unclassified(100); | - | - | - | - | - | - | - | Unassigned | - |
| Otu33284 | 1 | k__Fungi(100);p__Ascomycota(100);p__Ascomycota_unclassified(100);p__Ascomycota_unclassified(100);p__Ascomycota_unclassified(100);p__Ascomycota_unclassified(100);p__Ascomycota_unclassified(100); | - | - | - | - | - | - | - | Unassigned | - |
| Otu33295 | 1 | k__Fungi(100);k__Fungi_unclassified(100);k__Fungi_unclassified(100);k__Fungi_unclassified(100);k__Fungi_unclassified(100);k__Fungi_unclassified(100);k__Fungi_unclassified(100); | - | - | - | - | - | - | - | Unassigned | - |
| Otu33296 | 1 | k__Fungi(100);p__Ascomycota(100);p__Ascomycota_unclassified(100);p__Ascomycota_unclassified(100);p__Ascomycota_unclassified(100);p__Ascomycota_unclassified(100);p__Ascomycota_unclassified(100); | - | - | - | - | - | - | - | Unassigned | - |
| Otu33298 | 1 | k__Fungi(100);k__Fungi_unclassified(100);k__Fungi_unclassified(100);k__Fungi_unclassified(100);k__Fungi_unclassified(100);k__Fungi_unclassified(100);k__Fungi_unclassified(100); | - | - | - | - | - | - | - | Unassigned | - |
| Otu33299 | 1 | k__Fungi(100);k__Fungi_unclassified(100);k__Fungi_unclassified(100);k__Fungi_unclassified(100);k__Fungi_unclassified(100);k__Fungi_unclassified(100);k__Fungi_unclassified(100); | - | - | - | - | - | - | - | Unassigned | - |
| Otu33301 | 1 | k__Fungi(100);p__Ascomycota(100);p__Ascomycota_unclassified(100);p__Ascomycota_unclassified(100);p__Ascomycota_unclassified(100);p__Ascomycota_unclassified(100);p__Ascomycota_unclassified(100); | - | - | - | - | - | - | - | Unassigned | - |
| Otu33306 | 1 | k__Fungi(100);k__Fungi_unclassified(100);k__Fungi_unclassified(100);k__Fungi_unclassified(100);k__Fungi_unclassified(100);k__Fungi_unclassified(100);k__Fungi_unclassified(100); | - | - | - | - | - | - | - | Unassigned | - |
| Otu33311 | 1 | k__Fungi(100);p__Ascomycota(100);p__Ascomycota_unclassified(100);p__Ascomycota_unclassified(100);p__Ascomycota_unclassified(100);p__Ascomycota_unclassified(100);p__Ascomycota_unclassified(100); | - | - | - | - | - | - | - | Unassigned | - |
| Otu33318 | 1 | k__Fungi(100);k__Fungi_unclassified(100);k__Fungi_unclassified(100);k__Fungi_unclassified(100);k__Fungi_unclassified(100);k__Fungi_unclassified(100);k__Fungi_unclassified(100); | - | - | - | - | - | - | - | Unassigned | - |
| Otu33319 | 1 | k__Fungi(100);p__Ascomycota(100);p__Ascomycota_unclassified(100);p__Ascomycota_unclassified(100);p__Ascomycota_unclassified(100);p__Ascomycota_unclassified(100);p__Ascomycota_unclassified(100); | - | - | - | - | - | - | - | Unassigned | - |
| Otu33320 | 1 | k__Fungi(100);k__Fungi_unclassified(100);k__Fungi_unclassified(100);k__Fungi_unclassified(100);k__Fungi_unclassified(100);k__Fungi_unclassified(100);k__Fungi_unclassified(100); | - | - | - | - | - | - | - | Unassigned | - |
| Otu33321 | 1 | k__Fungi(100);k__Fungi_unclassified(100);k__Fungi_unclassified(100);k__Fungi_unclassified(100);k__Fungi_unclassified(100);k__Fungi_unclassified(100);k__Fungi_unclassified(100); | - | - | - | - | - | - | - | Unassigned | - |
| Otu33322 | 1 | k__Fungi(100);k__Fungi_unclassified(100);k__Fungi_unclassified(100);k__Fungi_unclassified(100);k__Fungi_unclassified(100);k__Fungi_unclassified(100);k__Fungi_unclassified(100); | - | - | - | - | - | - | - | Unassigned | - |
| Otu33325 | 1 | k__Fungi(100);k__Fungi_unclassified(100);k__Fungi_unclassified(100);k__Fungi_unclassified(100);k__Fungi_unclassified(100);k__Fungi_unclassified(100);k__Fungi_unclassified(100); | - | - | - | - | - | - | - | Unassigned | - |
| Otu33327 | 1 | k__Fungi(100);k__Fungi_unclassified(100);k__Fungi_unclassified(100);k__Fungi_unclassified(100);k__Fungi_unclassified(100);k__Fungi_unclassified(100);k__Fungi_unclassified(100); | - | - | - | - | - | - | - | Unassigned | - |
| Otu33329 | 1 | k__Fungi(100);p__Ascomycota(100);p__Ascomycota_unclassified(100);p__Ascomycota_unclassified(100);p__Ascomycota_unclassified(100);p__Ascomycota_unclassified(100);p__Ascomycota_unclassified(100); | - | - | - | - | - | - | - | Unassigned | - |
| Otu33330 | 1 | k__Fungi(100);p__Ascomycota(100);p__Ascomycota_unclassified(100);p__Ascomycota_unclassified(100);p__Ascomycota_unclassified(100);p__Ascomycota_unclassified(100);p__Ascomycota_unclassified(100); | - | - | - | - | - | - | - | Unassigned | - |
| Otu33331 | 1 | k__Fungi(100);k__Fungi_unclassified(100);k__Fungi_unclassified(100);k__Fungi_unclassified(100);k__Fungi_unclassified(100);k__Fungi_unclassified(100);k__Fungi_unclassified(100); | - | - | - | - | - | - | - | Unassigned | - |
| Otu33333 | 1 | k__Fungi(100);p__Ascomycota(100);p__Ascomycota_unclassified(100);p__Ascomycota_unclassified(100);p__Ascomycota_unclassified(100);p__Ascomycota_unclassified(100);p__Ascomycota_unclassified(100); | - | - | - | - | - | - | - | Unassigned | - |
| Otu33334 | 1 | k__Fungi(100);k__Fungi_unclassified(100);k__Fungi_unclassified(100);k__Fungi_unclassified(100);k__Fungi_unclassified(100);k__Fungi_unclassified(100);k__Fungi_unclassified(100); | - | - | - | - | - | - | - | Unassigned | - |
| Otu33335 | 1 | k__Fungi(100);k__Fungi_unclassified(100);k__Fungi_unclassified(100);k__Fungi_unclassified(100);k__Fungi_unclassified(100);k__Fungi_unclassified(100);k__Fungi_unclassified(100); | - | - | - | - | - | - | - | Unassigned | - |
| Otu33336 | 1 | k__Fungi(100);k__Fungi_unclassified(100);k__Fungi_unclassified(100);k__Fungi_unclassified(100);k__Fungi_unclassified(100);k__Fungi_unclassified(100);k__Fungi_unclassified(100); | - | - | - | - | - | - | - | Unassigned | - |
| Otu33337 | 1 | k__Fungi(100);k__Fungi_unclassified(100);k__Fungi_unclassified(100);k__Fungi_unclassified(100);k__Fungi_unclassified(100);k__Fungi_unclassified(100);k__Fungi_unclassified(100); | - | - | - | - | - | - | - | Unassigned | - |
| Otu33339 | 1 | k__Fungi(100);p__Ascomycota(100);p__Ascomycota_unclassified(100);p__Ascomycota_unclassified(100);p__Ascomycota_unclassified(100);p__Ascomycota_unclassified(100);p__Ascomycota_unclassified(100); | - | - | - | - | - | - | - | Unassigned | - |
| Otu33340 | 1 | k__Fungi(100);p__Ascomycota(100);p__Ascomycota_unclassified(100);p__Ascomycota_unclassified(100);p__Ascomycota_unclassified(100);p__Ascomycota_unclassified(100);p__Ascomycota_unclassified(100); | - | - | - | - | - | - | - | Unassigned | - |
| Otu33341 | 1 | k__Fungi(100);k__Fungi_unclassified(100);k__Fungi_unclassified(100);k__Fungi_unclassified(100);k__Fungi_unclassified(100);k__Fungi_unclassified(100);k__Fungi_unclassified(100); | - | - | - | - | - | - | - | Unassigned | - |
| Otu33344 | 1 | k__Fungi(100);k__Fungi_unclassified(100);k__Fungi_unclassified(100);k__Fungi_unclassified(100);k__Fungi_unclassified(100);k__Fungi_unclassified(100);k__Fungi_unclassified(100); | - | - | - | - | - | - | - | Unassigned | - |
| Otu33345 | 1 | k__Fungi(100);k__Fungi_unclassified(100);k__Fungi_unclassified(100);k__Fungi_unclassified(100);k__Fungi_unclassified(100);k__Fungi_unclassified(100);k__Fungi_unclassified(100); | - | - | - | - | - | - | - | Unassigned | - |
| Otu33346 | 1 | k__Fungi(100);k__Fungi_unclassified(100);k__Fungi_unclassified(100);k__Fungi_unclassified(100);k__Fungi_unclassified(100);k__Fungi_unclassified(100);k__Fungi_unclassified(100); | - | - | - | - | - | - | - | Unassigned | - |
| Otu33347 | 1 | k__Fungi(100);k__Fungi_unclassified(100);k__Fungi_unclassified(100);k__Fungi_unclassified(100);k__Fungi_unclassified(100);k__Fungi_unclassified(100);k__Fungi_unclassified(100); | - | - | - | - | - | - | - | Unassigned | - |
| Otu33348 | 1 | k__Fungi(100);k__Fungi_unclassified(100);k__Fungi_unclassified(100);k__Fungi_unclassified(100);k__Fungi_unclassified(100);k__Fungi_unclassified(100);k__Fungi_unclassified(100); | - | - | - | - | - | - | - | Unassigned | - |
| Otu33349 | 1 | k__Fungi(100);k__Fungi_unclassified(100);k__Fungi_unclassified(100);k__Fungi_unclassified(100);k__Fungi_unclassified(100);k__Fungi_unclassified(100);k__Fungi_unclassified(100); | - | - | - | - | - | - | - | Unassigned | - |
| Otu33351 | 1 | k__Fungi(100);p__Ascomycota(100);p__Ascomycota_unclassified(100);p__Ascomycota_unclassified(100);p__Ascomycota_unclassified(100);p__Ascomycota_unclassified(100);p__Ascomycota_unclassified(100); | - | - | - | - | - | - | - | Unassigned | - |
| Otu33352 | 1 | k__Fungi(100);k__Fungi_unclassified(100);k__Fungi_unclassified(100);k__Fungi_unclassified(100);k__Fungi_unclassified(100);k__Fungi_unclassified(100);k__Fungi_unclassified(100); | - | - | - | - | - | - | - | Unassigned | - |
| Otu33353 | 1 | k__Fungi(100);k__Fungi_unclassified(100);k__Fungi_unclassified(100);k__Fungi_unclassified(100);k__Fungi_unclassified(100);k__Fungi_unclassified(100);k__Fungi_unclassified(100); | - | - | - | - | - | - | - | Unassigned | - |
| Otu33354 | 1 | k__Fungi(100);k__Fungi_unclassified(100);k__Fungi_unclassified(100);k__Fungi_unclassified(100);k__Fungi_unclassified(100);k__Fungi_unclassified(100);k__Fungi_unclassified(100); | - | - | - | - | - | - | - | Unassigned | - |
| Otu33355 | 1 | k__Fungi(100);k__Fungi_unclassified(100);k__Fungi_unclassified(100);k__Fungi_unclassified(100);k__Fungi_unclassified(100);k__Fungi_unclassified(100);k__Fungi_unclassified(100); | - | - | - | - | - | - | - | Unassigned | - |
| Otu33356 | 1 | k__Fungi(100);p__Ascomycota(100);p__Ascomycota_unclassified(100);p__Ascomycota_unclassified(100);p__Ascomycota_unclassified(100);p__Ascomycota_unclassified(100);p__Ascomycota_unclassified(100); | - | - | - | - | - | - | - | Unassigned | - |
| Otu33357 | 1 | k__Fungi(100);k__Fungi_unclassified(100);k__Fungi_unclassified(100);k__Fungi_unclassified(100);k__Fungi_unclassified(100);k__Fungi_unclassified(100);k__Fungi_unclassified(100); | - | - | - | - | - | - | - | Unassigned | - |
| Otu33358 | 1 | k__Fungi(100);k__Fungi_unclassified(100);k__Fungi_unclassified(100);k__Fungi_unclassified(100);k__Fungi_unclassified(100);k__Fungi_unclassified(100);k__Fungi_unclassified(100); | - | - | - | - | - | - | - | Unassigned | - |
| Otu33359 | 1 | k__Fungi(100);k__Fungi_unclassified(100);k__Fungi_unclassified(100);k__Fungi_unclassified(100);k__Fungi_unclassified(100);k__Fungi_unclassified(100);k__Fungi_unclassified(100); | - | - | - | - | - | - | - | Unassigned | - |
| Otu33360 | 1 | k__Fungi(100);p__Ascomycota(100);p__Ascomycota_unclassified(100);p__Ascomycota_unclassified(100);p__Ascomycota_unclassified(100);p__Ascomycota_unclassified(100);p__Ascomycota_unclassified(100); | - | - | - | - | - | - | - | Unassigned | - |
| Otu33362 | 1 | k__Fungi(100);k__Fungi_unclassified(100);k__Fungi_unclassified(100);k__Fungi_unclassified(100);k__Fungi_unclassified(100);k__Fungi_unclassified(100);k__Fungi_unclassified(100); | - | - | - | - | - | - | - | Unassigned | - |
| Otu33363 | 1 | k__Fungi(100);p__Ascomycota(100);p__Ascomycota_unclassified(100);p__Ascomycota_unclassified(100);p__Ascomycota_unclassified(100);p__Ascomycota_unclassified(100);p__Ascomycota_unclassified(100); | - | - | - | - | - | - | - | Unassigned | - |
| Otu33364 | 1 | k__Fungi(100);p__Ascomycota(100);p__Ascomycota_unclassified(100);p__Ascomycota_unclassified(100);p__Ascomycota_unclassified(100);p__Ascomycota_unclassified(100);p__Ascomycota_unclassified(100); | - | - | - | - | - | - | - | Unassigned | - |
| Otu33366 | 1 | k__Fungi(100);k__Fungi_unclassified(100);k__Fungi_unclassified(100);k__Fungi_unclassified(100);k__Fungi_unclassified(100);k__Fungi_unclassified(100);k__Fungi_unclassified(100); | - | - | - | - | - | - | - | Unassigned | - |
| Otu33367 | 1 | k__Fungi(100);p__Ascomycota(100);p__Ascomycota_unclassified(100);p__Ascomycota_unclassified(100);p__Ascomycota_unclassified(100);p__Ascomycota_unclassified(100);p__Ascomycota_unclassified(100); | - | - | - | - | - | - | - | Unassigned | - |
| Otu33369 | 1 | k__Fungi(100);p__Ascomycota(100);p__Ascomycota_unclassified(100);p__Ascomycota_unclassified(100);p__Ascomycota_unclassified(100);p__Ascomycota_unclassified(100);p__Ascomycota_unclassified(100); | - | - | - | - | - | - | - | Unassigned | - |
| Otu33370 | 1 | k__Fungi(100);k__Fungi_unclassified(100);k__Fungi_unclassified(100);k__Fungi_unclassified(100);k__Fungi_unclassified(100);k__Fungi_unclassified(100);k__Fungi_unclassified(100); | - | - | - | - | - | - | - | Unassigned | - |
| Otu33371 | 1 | k__Fungi(100);k__Fungi_unclassified(100);k__Fungi_unclassified(100);k__Fungi_unclassified(100);k__Fungi_unclassified(100);k__Fungi_unclassified(100);k__Fungi_unclassified(100); | - | - | - | - | - | - | - | Unassigned | - |
| Otu33372 | 1 | k__Fungi(100);p__Ascomycota(100);p__Ascomycota_unclassified(100);p__Ascomycota_unclassified(100);p__Ascomycota_unclassified(100);p__Ascomycota_unclassified(100);p__Ascomycota_unclassified(100); | - | - | - | - | - | - | - | Unassigned | - |
| Otu33380 | 1 | k__Fungi(100);k__Fungi_unclassified(100);k__Fungi_unclassified(100);k__Fungi_unclassified(100);k__Fungi_unclassified(100);k__Fungi_unclassified(100);k__Fungi_unclassified(100); | - | - | - | - | - | - | - | Unassigned | - |
| Otu33382 | 1 | k__Fungi(100);k__Fungi_unclassified(100);k__Fungi_unclassified(100);k__Fungi_unclassified(100);k__Fungi_unclassified(100);k__Fungi_unclassified(100);k__Fungi_unclassified(100); | - | - | - | - | - | - | - | Unassigned | - |
| Otu33383 | 1 | k__Fungi(100);p__Ascomycota(100);p__Ascomycota_unclassified(100);p__Ascomycota_unclassified(100);p__Ascomycota_unclassified(100);p__Ascomycota_unclassified(100);p__Ascomycota_unclassified(100); | - | - | - | - | - | - | - | Unassigned | - |
| Otu33395 | 1 | k__Fungi(100);k__Fungi_unclassified(100);k__Fungi_unclassified(100);k__Fungi_unclassified(100);k__Fungi_unclassified(100);k__Fungi_unclassified(100);k__Fungi_unclassified(100); | - | - | - | - | - | - | - | Unassigned | - |
| Otu33400 | 1 | k__Fungi(100);k__Fungi_unclassified(100);k__Fungi_unclassified(100);k__Fungi_unclassified(100);k__Fungi_unclassified(100);k__Fungi_unclassified(100);k__Fungi_unclassified(100); | - | - | - | - | - | - | - | Unassigned | - |
| Otu33403 | 1 | k__Fungi(100);k__Fungi_unclassified(100);k__Fungi_unclassified(100);k__Fungi_unclassified(100);k__Fungi_unclassified(100);k__Fungi_unclassified(100);k__Fungi_unclassified(100); | - | - | - | - | - | - | - | Unassigned | - |
| Otu33404 | 1 | k__Fungi(100);k__Fungi_unclassified(100);k__Fungi_unclassified(100);k__Fungi_unclassified(100);k__Fungi_unclassified(100);k__Fungi_unclassified(100);k__Fungi_unclassified(100); | - | - | - | - | - | - | - | Unassigned | - |
| Otu33405 | 1 | k__Fungi(100);p__Ascomycota(100);p__Ascomycota_unclassified(100);p__Ascomycota_unclassified(100);p__Ascomycota_unclassified(100);p__Ascomycota_unclassified(100);p__Ascomycota_unclassified(100); | - | - | - | - | - | - | - | Unassigned | - |
| Otu33406 | 1 | k__Fungi(100);k__Fungi_unclassified(100);k__Fungi_unclassified(100);k__Fungi_unclassified(100);k__Fungi_unclassified(100);k__Fungi_unclassified(100);k__Fungi_unclassified(100); | - | - | - | - | - | - | - | Unassigned | - |
| Otu33408 | 1 | k__Fungi(100);k__Fungi_unclassified(100);k__Fungi_unclassified(100);k__Fungi_unclassified(100);k__Fungi_unclassified(100);k__Fungi_unclassified(100);k__Fungi_unclassified(100); | - | - | - | - | - | - | - | Unassigned | - |
| Otu33409 | 1 | k__Fungi(100);k__Fungi_unclassified(100);k__Fungi_unclassified(100);k__Fungi_unclassified(100);k__Fungi_unclassified(100);k__Fungi_unclassified(100);k__Fungi_unclassified(100); | - | - | - | - | - | - | - | Unassigned | - |
| Otu33411 | 1 | k__Fungi(100);k__Fungi_unclassified(100);k__Fungi_unclassified(100);k__Fungi_unclassified(100);k__Fungi_unclassified(100);k__Fungi_unclassified(100);k__Fungi_unclassified(100); | - | - | - | - | - | - | - | Unassigned | - |
| Otu33414 | 1 | k__Fungi(100);k__Fungi_unclassified(100);k__Fungi_unclassified(100);k__Fungi_unclassified(100);k__Fungi_unclassified(100);k__Fungi_unclassified(100);k__Fungi_unclassified(100); | - | - | - | - | - | - | - | Unassigned | - |
| Otu33427 | 1 | k__Fungi(100);p__Ascomycota(100);p__Ascomycota_unclassified(100);p__Ascomycota_unclassified(100);p__Ascomycota_unclassified(100);p__Ascomycota_unclassified(100);p__Ascomycota_unclassified(100); | - | - | - | - | - | - | - | Unassigned | - |
| Otu33429 | 1 | k__Fungi(100);k__Fungi_unclassified(100);k__Fungi_unclassified(100);k__Fungi_unclassified(100);k__Fungi_unclassified(100);k__Fungi_unclassified(100);k__Fungi_unclassified(100); | - | - | - | - | - | - | - | Unassigned | - |
| Otu33435 | 1 | k__Fungi(100);k__Fungi_unclassified(100);k__Fungi_unclassified(100);k__Fungi_unclassified(100);k__Fungi_unclassified(100);k__Fungi_unclassified(100);k__Fungi_unclassified(100); | - | - | - | - | - | - | - | Unassigned | - |
| Otu33444 | 1 | k__Fungi(100);k__Fungi_unclassified(100);k__Fungi_unclassified(100);k__Fungi_unclassified(100);k__Fungi_unclassified(100);k__Fungi_unclassified(100);k__Fungi_unclassified(100); | - | - | - | - | - | - | - | Unassigned | - |
| Otu33445 | 1 | k__Fungi(100);p__Ascomycota(100);p__Ascomycota_unclassified(100);p__Ascomycota_unclassified(100);p__Ascomycota_unclassified(100);p__Ascomycota_unclassified(100);p__Ascomycota_unclassified(100); | - | - | - | - | - | - | - | Unassigned | - |
| Otu33450 | 1 | k__Fungi(100);k__Fungi_unclassified(100);k__Fungi_unclassified(100);k__Fungi_unclassified(100);k__Fungi_unclassified(100);k__Fungi_unclassified(100);k__Fungi_unclassified(100); | - | - | - | - | - | - | - | Unassigned | - |
| Otu33451 | 1 | k__Fungi(100);k__Fungi_unclassified(100);k__Fungi_unclassified(100);k__Fungi_unclassified(100);k__Fungi_unclassified(100);k__Fungi_unclassified(100);k__Fungi_unclassified(100); | - | - | - | - | - | - | - | Unassigned | - |
| Otu33452 | 1 | k__Fungi(100);p__Ascomycota(100);p__Ascomycota_unclassified(100);p__Ascomycota_unclassified(100);p__Ascomycota_unclassified(100);p__Ascomycota_unclassified(100);p__Ascomycota_unclassified(100); | - | - | - | - | - | - | - | Unassigned | - |
| Otu33453 | 1 | k__Fungi(100);p__Ascomycota(100);p__Ascomycota_unclassified(100);p__Ascomycota_unclassified(100);p__Ascomycota_unclassified(100);p__Ascomycota_unclassified(100);p__Ascomycota_unclassified(100); | - | - | - | - | - | - | - | Unassigned | - |
| Otu33455 | 1 | k__Fungi(100);p__Ascomycota(100);c__Leotiomycetes(100);c__Leotiomycetes_unclassified(100);c__Leotiomycetes_unclassified(100);c__Leotiomycetes_unclassified(100);c__Leotiomycetes_unclassified(100); | - | - | - | - | - | - | - | Unassigned | - |
| Otu33456 | 1 | k__Fungi(100);k__Fungi_unclassified(100);k__Fungi_unclassified(100);k__Fungi_unclassified(100);k__Fungi_unclassified(100);k__Fungi_unclassified(100);k__Fungi_unclassified(100); | - | - | - | - | - | - | - | Unassigned | - |
| Otu33457 | 1 | k__Fungi(100);k__Fungi_unclassified(100);k__Fungi_unclassified(100);k__Fungi_unclassified(100);k__Fungi_unclassified(100);k__Fungi_unclassified(100);k__Fungi_unclassified(100); | - | - | - | - | - | - | - | Unassigned | - |
| Otu33458 | 1 | k__Fungi(100);p__Ascomycota(100);p__Ascomycota_unclassified(100);p__Ascomycota_unclassified(100);p__Ascomycota_unclassified(100);p__Ascomycota_unclassified(100);p__Ascomycota_unclassified(100); | - | - | - | - | - | - | - | Unassigned | - |
| Otu33460 | 1 | k__Fungi(100);p__Ascomycota(100);p__Ascomycota_unclassified(100);p__Ascomycota_unclassified(100);p__Ascomycota_unclassified(100);p__Ascomycota_unclassified(100);p__Ascomycota_unclassified(100); | - | - | - | - | - | - | - | Unassigned | - |
| Otu33461 | 1 | k__Fungi(100);p__Ascomycota(100);p__Ascomycota_unclassified(100);p__Ascomycota_unclassified(100);p__Ascomycota_unclassified(100);p__Ascomycota_unclassified(100);p__Ascomycota_unclassified(100); | - | - | - | - | - | - | - | Unassigned | - |
| Otu33465 | 1 | k__Fungi(100);k__Fungi_unclassified(100);k__Fungi_unclassified(100);k__Fungi_unclassified(100);k__Fungi_unclassified(100);k__Fungi_unclassified(100);k__Fungi_unclassified(100); | - | - | - | - | - | - | - | Unassigned | - |
| Otu33467 | 1 | k__Fungi(100);k__Fungi_unclassified(100);k__Fungi_unclassified(100);k__Fungi_unclassified(100);k__Fungi_unclassified(100);k__Fungi_unclassified(100);k__Fungi_unclassified(100); | - | - | - | - | - | - | - | Unassigned | - |
| Otu33469 | 1 | k__Fungi(100);p__Ascomycota(100);p__Ascomycota_unclassified(100);p__Ascomycota_unclassified(100);p__Ascomycota_unclassified(100);p__Ascomycota_unclassified(100);p__Ascomycota_unclassified(100); | - | - | - | - | - | - | - | Unassigned | - |
| Otu33478 | 1 | k__Fungi(100);k__Fungi_unclassified(100);k__Fungi_unclassified(100);k__Fungi_unclassified(100);k__Fungi_unclassified(100);k__Fungi_unclassified(100);k__Fungi_unclassified(100); | - | - | - | - | - | - | - | Unassigned | - |
| Otu33480 | 1 | k__Fungi(100);k__Fungi_unclassified(100);k__Fungi_unclassified(100);k__Fungi_unclassified(100);k__Fungi_unclassified(100);k__Fungi_unclassified(100);k__Fungi_unclassified(100); | - | - | - | - | - | - | - | Unassigned | - |
| Otu33481 | 1 | k__Fungi(100);k__Fungi_unclassified(100);k__Fungi_unclassified(100);k__Fungi_unclassified(100);k__Fungi_unclassified(100);k__Fungi_unclassified(100);k__Fungi_unclassified(100); | - | - | - | - | - | - | - | Unassigned | - |
| Otu33482 | 1 | k__Fungi(100);p__Ascomycota(100);p__Ascomycota_unclassified(100);p__Ascomycota_unclassified(100);p__Ascomycota_unclassified(100);p__Ascomycota_unclassified(100);p__Ascomycota_unclassified(100); | - | - | - | - | - | - | - | Unassigned | - |
| Otu33487 | 1 | k__Fungi(100);k__Fungi_unclassified(100);k__Fungi_unclassified(100);k__Fungi_unclassified(100);k__Fungi_unclassified(100);k__Fungi_unclassified(100);k__Fungi_unclassified(100); | - | - | - | - | - | - | - | Unassigned | - |
| Otu33488 | 1 | k__Fungi(100);p__Ascomycota(100);p__Ascomycota_unclassified(100);p__Ascomycota_unclassified(100);p__Ascomycota_unclassified(100);p__Ascomycota_unclassified(100);p__Ascomycota_unclassified(100); | - | - | - | - | - | - | - | Unassigned | - |
| Otu33490 | 1 | k__Fungi(100);k__Fungi_unclassified(100);k__Fungi_unclassified(100);k__Fungi_unclassified(100);k__Fungi_unclassified(100);k__Fungi_unclassified(100);k__Fungi_unclassified(100); | - | - | - | - | - | - | - | Unassigned | - |
| Otu33491 | 1 | k__Fungi(100);k__Fungi_unclassified(100);k__Fungi_unclassified(100);k__Fungi_unclassified(100);k__Fungi_unclassified(100);k__Fungi_unclassified(100);k__Fungi_unclassified(100); | - | - | - | - | - | - | - | Unassigned | - |
| Otu33492 | 1 | k__Fungi(100);p__Ascomycota(100);p__Ascomycota_unclassified(100);p__Ascomycota_unclassified(100);p__Ascomycota_unclassified(100);p__Ascomycota_unclassified(100);p__Ascomycota_unclassified(100); | - | - | - | - | - | - | - | Unassigned | - |
| Otu33493 | 1 | k__Fungi(100);k__Fungi_unclassified(100);k__Fungi_unclassified(100);k__Fungi_unclassified(100);k__Fungi_unclassified(100);k__Fungi_unclassified(100);k__Fungi_unclassified(100); | - | - | - | - | - | - | - | Unassigned | - |
| Otu33495 | 1 | k__Fungi(100);k__Fungi_unclassified(100);k__Fungi_unclassified(100);k__Fungi_unclassified(100);k__Fungi_unclassified(100);k__Fungi_unclassified(100);k__Fungi_unclassified(100); | - | - | - | - | - | - | - | Unassigned | - |
| Otu33496 | 1 | k__Fungi(100);k__Fungi_unclassified(100);k__Fungi_unclassified(100);k__Fungi_unclassified(100);k__Fungi_unclassified(100);k__Fungi_unclassified(100);k__Fungi_unclassified(100); | - | - | - | - | - | - | - | Unassigned | - |
| Otu33497 | 1 | k__Fungi(100);p__Ascomycota(100);p__Ascomycota_unclassified(100);p__Ascomycota_unclassified(100);p__Ascomycota_unclassified(100);p__Ascomycota_unclassified(100);p__Ascomycota_unclassified(100); | - | - | - | - | - | - | - | Unassigned | - |
| Otu33499 | 1 | k__Fungi(100);p__Ascomycota(100);p__Ascomycota_unclassified(100);p__Ascomycota_unclassified(100);p__Ascomycota_unclassified(100);p__Ascomycota_unclassified(100);p__Ascomycota_unclassified(100); | - | - | - | - | - | - | - | Unassigned | - |
| Otu33500 | 1 | k__Fungi(100);k__Fungi_unclassified(100);k__Fungi_unclassified(100);k__Fungi_unclassified(100);k__Fungi_unclassified(100);k__Fungi_unclassified(100);k__Fungi_unclassified(100); | - | - | - | - | - | - | - | Unassigned | - |
| Otu33501 | 1 | k__Fungi(100);k__Fungi_unclassified(100);k__Fungi_unclassified(100);k__Fungi_unclassified(100);k__Fungi_unclassified(100);k__Fungi_unclassified(100);k__Fungi_unclassified(100); | - | - | - | - | - | - | - | Unassigned | - |
| Otu33502 | 1 | k__Fungi(100);p__Ascomycota(100);p__Ascomycota_unclassified(100);p__Ascomycota_unclassified(100);p__Ascomycota_unclassified(100);p__Ascomycota_unclassified(100);p__Ascomycota_unclassified(100); | - | - | - | - | - | - | - | Unassigned | - |
| Otu33503 | 1 | k__Fungi(100);k__Fungi_unclassified(100);k__Fungi_unclassified(100);k__Fungi_unclassified(100);k__Fungi_unclassified(100);k__Fungi_unclassified(100);k__Fungi_unclassified(100); | - | - | - | - | - | - | - | Unassigned | - |
| Otu33504 | 1 | k__Fungi(100);k__Fungi_unclassified(100);k__Fungi_unclassified(100);k__Fungi_unclassified(100);k__Fungi_unclassified(100);k__Fungi_unclassified(100);k__Fungi_unclassified(100); | - | - | - | - | - | - | - | Unassigned | - |
| Otu33505 | 1 | k__Fungi(100);p__Ascomycota(100);p__Ascomycota_unclassified(100);p__Ascomycota_unclassified(100);p__Ascomycota_unclassified(100);p__Ascomycota_unclassified(100);p__Ascomycota_unclassified(100); | - | - | - | - | - | - | - | Unassigned | - |
| Otu33506 | 1 | k__Fungi(100);k__Fungi_unclassified(100);k__Fungi_unclassified(100);k__Fungi_unclassified(100);k__Fungi_unclassified(100);k__Fungi_unclassified(100);k__Fungi_unclassified(100); | - | - | - | - | - | - | - | Unassigned | - |
| Otu33507 | 1 | k__Fungi(100);p__Ascomycota(100);p__Ascomycota_unclassified(100);p__Ascomycota_unclassified(100);p__Ascomycota_unclassified(100);p__Ascomycota_unclassified(100);p__Ascomycota_unclassified(100); | - | - | - | - | - | - | - | Unassigned | - |
| Otu33508 | 1 | k__Fungi(100);k__Fungi_unclassified(100);k__Fungi_unclassified(100);k__Fungi_unclassified(100);k__Fungi_unclassified(100);k__Fungi_unclassified(100);k__Fungi_unclassified(100); | - | - | - | - | - | - | - | Unassigned | - |
| Otu33509 | 1 | k__Fungi(100);k__Fungi_unclassified(100);k__Fungi_unclassified(100);k__Fungi_unclassified(100);k__Fungi_unclassified(100);k__Fungi_unclassified(100);k__Fungi_unclassified(100); | - | - | - | - | - | - | - | Unassigned | - |
| Otu33510 | 1 | k__Fungi(100);k__Fungi_unclassified(100);k__Fungi_unclassified(100);k__Fungi_unclassified(100);k__Fungi_unclassified(100);k__Fungi_unclassified(100);k__Fungi_unclassified(100); | - | - | - | - | - | - | - | Unassigned | - |
| Otu33512 | 1 | k__Fungi(100);p__Ascomycota(100);p__Ascomycota_unclassified(100);p__Ascomycota_unclassified(100);p__Ascomycota_unclassified(100);p__Ascomycota_unclassified(100);p__Ascomycota_unclassified(100); | - | - | - | - | - | - | - | Unassigned | - |
| Otu33514 | 1 | k__Fungi(100);k__Fungi_unclassified(100);k__Fungi_unclassified(100);k__Fungi_unclassified(100);k__Fungi_unclassified(100);k__Fungi_unclassified(100);k__Fungi_unclassified(100); | - | - | - | - | - | - | - | Unassigned | - |
| Otu33516 | 1 | k__Fungi(100);k__Fungi_unclassified(100);k__Fungi_unclassified(100);k__Fungi_unclassified(100);k__Fungi_unclassified(100);k__Fungi_unclassified(100);k__Fungi_unclassified(100); | - | - | - | - | - | - | - | Unassigned | - |
| Otu33519 | 1 | k__Fungi(100);k__Fungi_unclassified(100);k__Fungi_unclassified(100);k__Fungi_unclassified(100);k__Fungi_unclassified(100);k__Fungi_unclassified(100);k__Fungi_unclassified(100); | - | - | - | - | - | - | - | Unassigned | - |
| Otu33520 | 1 | k__Fungi(100);k__Fungi_unclassified(100);k__Fungi_unclassified(100);k__Fungi_unclassified(100);k__Fungi_unclassified(100);k__Fungi_unclassified(100);k__Fungi_unclassified(100); | - | - | - | - | - | - | - | Unassigned | - |
| Otu33521 | 1 | k__Fungi(100);p__Ascomycota(100);p__Ascomycota_unclassified(100);p__Ascomycota_unclassified(100);p__Ascomycota_unclassified(100);p__Ascomycota_unclassified(100);p__Ascomycota_unclassified(100); | - | - | - | - | - | - | - | Unassigned | - |
| Otu33525 | 1 | k__Fungi(100);k__Fungi_unclassified(100);k__Fungi_unclassified(100);k__Fungi_unclassified(100);k__Fungi_unclassified(100);k__Fungi_unclassified(100);k__Fungi_unclassified(100); | - | - | - | - | - | - | - | Unassigned | - |
| Otu33526 | 1 | k__Fungi(100);k__Fungi_unclassified(100);k__Fungi_unclassified(100);k__Fungi_unclassified(100);k__Fungi_unclassified(100);k__Fungi_unclassified(100);k__Fungi_unclassified(100); | - | - | - | - | - | - | - | Unassigned | - |
| Otu33530 | 1 | k__Fungi(100);k__Fungi_unclassified(100);k__Fungi_unclassified(100);k__Fungi_unclassified(100);k__Fungi_unclassified(100);k__Fungi_unclassified(100);k__Fungi_unclassified(100); | - | - | - | - | - | - | - | Unassigned | - |
| Otu33532 | 1 | k__Fungi(100);k__Fungi_unclassified(100);k__Fungi_unclassified(100);k__Fungi_unclassified(100);k__Fungi_unclassified(100);k__Fungi_unclassified(100);k__Fungi_unclassified(100); | - | - | - | - | - | - | - | Unassigned | - |
| Otu33534 | 1 | k__Fungi(100);k__Fungi_unclassified(100);k__Fungi_unclassified(100);k__Fungi_unclassified(100);k__Fungi_unclassified(100);k__Fungi_unclassified(100);k__Fungi_unclassified(100); | - | - | - | - | - | - | - | Unassigned | - |
| Otu33545 | 1 | k__Fungi(100);k__Fungi_unclassified(100);k__Fungi_unclassified(100);k__Fungi_unclassified(100);k__Fungi_unclassified(100);k__Fungi_unclassified(100);k__Fungi_unclassified(100); | - | - | - | - | - | - | - | Unassigned | - |
| Otu33546 | 1 | k__Fungi(100);k__Fungi_unclassified(100);k__Fungi_unclassified(100);k__Fungi_unclassified(100);k__Fungi_unclassified(100);k__Fungi_unclassified(100);k__Fungi_unclassified(100); | - | - | - | - | - | - | - | Unassigned | - |
| Otu33547 | 1 | k__Fungi(100);p__Ascomycota(100);p__Ascomycota_unclassified(100);p__Ascomycota_unclassified(100);p__Ascomycota_unclassified(100);p__Ascomycota_unclassified(100);p__Ascomycota_unclassified(100); | - | - | - | - | - | - | - | Unassigned | - |
| Otu33550 | 1 | k__Fungi(100);k__Fungi_unclassified(100);k__Fungi_unclassified(100);k__Fungi_unclassified(100);k__Fungi_unclassified(100);k__Fungi_unclassified(100);k__Fungi_unclassified(100); | - | - | - | - | - | - | - | Unassigned | - |
| Otu33551 | 1 | k__Fungi(100);k__Fungi_unclassified(100);k__Fungi_unclassified(100);k__Fungi_unclassified(100);k__Fungi_unclassified(100);k__Fungi_unclassified(100);k__Fungi_unclassified(100); | - | - | - | - | - | - | - | Unassigned | - |
| Otu33552 | 1 | k__Fungi(100);k__Fungi_unclassified(100);k__Fungi_unclassified(100);k__Fungi_unclassified(100);k__Fungi_unclassified(100);k__Fungi_unclassified(100);k__Fungi_unclassified(100); | - | - | - | - | - | - | - | Unassigned | - |
| Otu33555 | 1 | k__Fungi(100);k__Fungi_unclassified(100);k__Fungi_unclassified(100);k__Fungi_unclassified(100);k__Fungi_unclassified(100);k__Fungi_unclassified(100);k__Fungi_unclassified(100); | - | - | - | - | - | - | - | Unassigned | - |
| Otu33563 | 1 | k__Fungi(100);p__Ascomycota(100);p__Ascomycota_unclassified(100);p__Ascomycota_unclassified(100);p__Ascomycota_unclassified(100);p__Ascomycota_unclassified(100);p__Ascomycota_unclassified(100); | - | - | - | - | - | - | - | Unassigned | - |
| Otu33572 | 1 | k__Fungi(100);k__Fungi_unclassified(100);k__Fungi_unclassified(100);k__Fungi_unclassified(100);k__Fungi_unclassified(100);k__Fungi_unclassified(100);k__Fungi_unclassified(100); | - | - | - | - | - | - | - | Unassigned | - |
| Otu33574 | 1 | k__Fungi(100);k__Fungi_unclassified(100);k__Fungi_unclassified(100);k__Fungi_unclassified(100);k__Fungi_unclassified(100);k__Fungi_unclassified(100);k__Fungi_unclassified(100); | - | - | - | - | - | - | - | Unassigned | - |
| Otu33575 | 1 | k__Fungi(100);k__Fungi_unclassified(100);k__Fungi_unclassified(100);k__Fungi_unclassified(100);k__Fungi_unclassified(100);k__Fungi_unclassified(100);k__Fungi_unclassified(100); | - | - | - | - | - | - | - | Unassigned | - |
| Otu33577 | 1 | k__Fungi(100);p__Ascomycota(100);p__Ascomycota_unclassified(100);p__Ascomycota_unclassified(100);p__Ascomycota_unclassified(100);p__Ascomycota_unclassified(100);p__Ascomycota_unclassified(100); | - | - | - | - | - | - | - | Unassigned | - |
| Otu33578 | 1 | k__Fungi(100);k__Fungi_unclassified(100);k__Fungi_unclassified(100);k__Fungi_unclassified(100);k__Fungi_unclassified(100);k__Fungi_unclassified(100);k__Fungi_unclassified(100); | - | - | - | - | - | - | - | Unassigned | - |
| Otu33580 | 1 | k__Fungi(100);p__Ascomycota(100);p__Ascomycota_unclassified(100);p__Ascomycota_unclassified(100);p__Ascomycota_unclassified(100);p__Ascomycota_unclassified(100);p__Ascomycota_unclassified(100); | - | - | - | - | - | - | - | Unassigned | - |
| Otu33584 | 1 | k__Fungi(100);p__Ascomycota(100);p__Ascomycota_unclassified(100);p__Ascomycota_unclassified(100);p__Ascomycota_unclassified(100);p__Ascomycota_unclassified(100);p__Ascomycota_unclassified(100); | - | - | - | - | - | - | - | Unassigned | - |
| Otu33585 | 1 | k__Fungi(100);p__Ascomycota(100);p__Ascomycota_unclassified(100);p__Ascomycota_unclassified(100);p__Ascomycota_unclassified(100);p__Ascomycota_unclassified(100);p__Ascomycota_unclassified(100); | - | - | - | - | - | - | - | Unassigned | - |
| Otu33590 | 1 | k__Fungi(100);k__Fungi_unclassified(100);k__Fungi_unclassified(100);k__Fungi_unclassified(100);k__Fungi_unclassified(100);k__Fungi_unclassified(100);k__Fungi_unclassified(100); | - | - | - | - | - | - | - | Unassigned | - |
| Otu33591 | 1 | k__Fungi(100);k__Fungi_unclassified(100);k__Fungi_unclassified(100);k__Fungi_unclassified(100);k__Fungi_unclassified(100);k__Fungi_unclassified(100);k__Fungi_unclassified(100); | - | - | - | - | - | - | - | Unassigned | - |
| Otu33593 | 1 | k__Fungi(100);p__Ascomycota(100);p__Ascomycota_unclassified(100);p__Ascomycota_unclassified(100);p__Ascomycota_unclassified(100);p__Ascomycota_unclassified(100);p__Ascomycota_unclassified(100); | - | - | - | - | - | - | - | Unassigned | - |
| Otu33596 | 1 | k__Fungi(100);k__Fungi_unclassified(100);k__Fungi_unclassified(100);k__Fungi_unclassified(100);k__Fungi_unclassified(100);k__Fungi_unclassified(100);k__Fungi_unclassified(100); | - | - | - | - | - | - | - | Unassigned | - |
| Otu33613 | 1 | k__Fungi(100);p__Ascomycota(100);p__Ascomycota_unclassified(100);p__Ascomycota_unclassified(100);p__Ascomycota_unclassified(100);p__Ascomycota_unclassified(100);p__Ascomycota_unclassified(100); | - | - | - | - | - | - | - | Unassigned | - |
| Otu33619 | 1 | k__Fungi(100);k__Fungi_unclassified(100);k__Fungi_unclassified(100);k__Fungi_unclassified(100);k__Fungi_unclassified(100);k__Fungi_unclassified(100);k__Fungi_unclassified(100); | - | - | - | - | - | - | - | Unassigned | - |
| Otu33628 | 1 | k__Fungi(100);k__Fungi_unclassified(100);k__Fungi_unclassified(100);k__Fungi_unclassified(100);k__Fungi_unclassified(100);k__Fungi_unclassified(100);k__Fungi_unclassified(100); | - | - | - | - | - | - | - | Unassigned | - |
| Otu33630 | 1 | k__Fungi(100);k__Fungi_unclassified(100);k__Fungi_unclassified(100);k__Fungi_unclassified(100);k__Fungi_unclassified(100);k__Fungi_unclassified(100);k__Fungi_unclassified(100); | - | - | - | - | - | - | - | Unassigned | - |
| Otu33631 | 1 | k__Fungi(100);p__Ascomycota(100);p__Ascomycota_unclassified(100);p__Ascomycota_unclassified(100);p__Ascomycota_unclassified(100);p__Ascomycota_unclassified(100);p__Ascomycota_unclassified(100); | - | - | - | - | - | - | - | Unassigned | - |
| Otu33632 | 1 | k__Fungi(100);k__Fungi_unclassified(100);k__Fungi_unclassified(100);k__Fungi_unclassified(100);k__Fungi_unclassified(100);k__Fungi_unclassified(100);k__Fungi_unclassified(100); | - | - | - | - | - | - | - | Unassigned | - |
| Otu33636 | 1 | k__Fungi(100);k__Fungi_unclassified(100);k__Fungi_unclassified(100);k__Fungi_unclassified(100);k__Fungi_unclassified(100);k__Fungi_unclassified(100);k__Fungi_unclassified(100); | - | - | - | - | - | - | - | Unassigned | - |
| Otu33637 | 1 | k__Fungi(100);k__Fungi_unclassified(100);k__Fungi_unclassified(100);k__Fungi_unclassified(100);k__Fungi_unclassified(100);k__Fungi_unclassified(100);k__Fungi_unclassified(100); | - | - | - | - | - | - | - | Unassigned | - |
| Otu33638 | 1 | k__Fungi(100);k__Fungi_unclassified(100);k__Fungi_unclassified(100);k__Fungi_unclassified(100);k__Fungi_unclassified(100);k__Fungi_unclassified(100);k__Fungi_unclassified(100); | - | - | - | - | - | - | - | Unassigned | - |
| Otu33639 | 1 | k__Fungi(100);k__Fungi_unclassified(100);k__Fungi_unclassified(100);k__Fungi_unclassified(100);k__Fungi_unclassified(100);k__Fungi_unclassified(100);k__Fungi_unclassified(100); | - | - | - | - | - | - | - | Unassigned | - |
| Otu33659 | 1 | k__Fungi(100);k__Fungi_unclassified(100);k__Fungi_unclassified(100);k__Fungi_unclassified(100);k__Fungi_unclassified(100);k__Fungi_unclassified(100);k__Fungi_unclassified(100); | - | - | - | - | - | - | - | Unassigned | - |
| Otu33663 | 1 | k__Fungi(100);p__Ascomycota(100);p__Ascomycota_unclassified(100);p__Ascomycota_unclassified(100);p__Ascomycota_unclassified(100);p__Ascomycota_unclassified(100);p__Ascomycota_unclassified(100); | - | - | - | - | - | - | - | Unassigned | - |
| Otu33666 | 1 | k__Fungi(100);k__Fungi_unclassified(100);k__Fungi_unclassified(100);k__Fungi_unclassified(100);k__Fungi_unclassified(100);k__Fungi_unclassified(100);k__Fungi_unclassified(100); | - | - | - | - | - | - | - | Unassigned | - |
| Otu33667 | 1 | k__Fungi(100);k__Fungi_unclassified(100);k__Fungi_unclassified(100);k__Fungi_unclassified(100);k__Fungi_unclassified(100);k__Fungi_unclassified(100);k__Fungi_unclassified(100); | - | - | - | - | - | - | - | Unassigned | - |
| Otu33668 | 1 | k__Fungi(100);p__Ascomycota(100);p__Ascomycota_unclassified(100);p__Ascomycota_unclassified(100);p__Ascomycota_unclassified(100);p__Ascomycota_unclassified(100);p__Ascomycota_unclassified(100); | - | - | - | - | - | - | - | Unassigned | - |
| Otu33669 | 1 | k__Fungi(100);p__Ascomycota(100);p__Ascomycota_unclassified(100);p__Ascomycota_unclassified(100);p__Ascomycota_unclassified(100);p__Ascomycota_unclassified(100);p__Ascomycota_unclassified(100); | - | - | - | - | - | - | - | Unassigned | - |
| Otu33672 | 1 | k__Fungi(100);k__Fungi_unclassified(100);k__Fungi_unclassified(100);k__Fungi_unclassified(100);k__Fungi_unclassified(100);k__Fungi_unclassified(100);k__Fungi_unclassified(100); | - | - | - | - | - | - | - | Unassigned | - |
| Otu33676 | 1 | k__Fungi(100);k__Fungi_unclassified(100);k__Fungi_unclassified(100);k__Fungi_unclassified(100);k__Fungi_unclassified(100);k__Fungi_unclassified(100);k__Fungi_unclassified(100); | - | - | - | - | - | - | - | Unassigned | - |
| Otu33677 | 1 | k__Fungi(100);p__Ascomycota(100);p__Ascomycota_unclassified(100);p__Ascomycota_unclassified(100);p__Ascomycota_unclassified(100);p__Ascomycota_unclassified(100);p__Ascomycota_unclassified(100); | - | - | - | - | - | - | - | Unassigned | - |
| Otu33678 | 1 | k__Fungi(100);p__Ascomycota(100);p__Ascomycota_unclassified(100);p__Ascomycota_unclassified(100);p__Ascomycota_unclassified(100);p__Ascomycota_unclassified(100);p__Ascomycota_unclassified(100); | - | - | - | - | - | - | - | Unassigned | - |
| Otu33682 | 1 | k__Fungi(100);k__Fungi_unclassified(100);k__Fungi_unclassified(100);k__Fungi_unclassified(100);k__Fungi_unclassified(100);k__Fungi_unclassified(100);k__Fungi_unclassified(100); | - | - | - | - | - | - | - | Unassigned | - |

| | | | | | | | | | | | |
|---|---|---|---|---|---|---|---|---|---|---|---|
| Otu33687 | 1 | k__Fungi(100);k__Fungi_unclassified(100);k__Fungi_unclassified(100);k__Fungi_unclassified(100);k__Fungi_unclassified(100);k__Fungi_unclassified(100);k__Fungi_unclassified(100); | - | - | - | - | - | - | - | Unassigned | - |
| Otu33705 | 1 | k__Fungi(100);k__Fungi_unclassified(100);k__Fungi_unclassified(100);k__Fungi_unclassified(100);k__Fungi_unclassified(100);k__Fungi_unclassified(100);k__Fungi_unclassified(100); | - | - | - | - | - | - | - | Unassigned | - |
| Otu33707 | 1 | k__Fungi(100);k__Fungi_unclassified(100);k__Fungi_unclassified(100);k__Fungi_unclassified(100);k__Fungi_unclassified(100);k__Fungi_unclassified(100);k__Fungi_unclassified(100); | - | - | - | - | - | - | - | Unassigned | - |
| Otu33708 | 1 | k__Fungi(100);k__Fungi_unclassified(100);k__Fungi_unclassified(100);k__Fungi_unclassified(100);k__Fungi_unclassified(100);k__Fungi_unclassified(100);k__Fungi_unclassified(100); | - | - | - | - | - | - | - | Unassigned | - |
| Otu33711 | 1 | k__Fungi(100);p__Ascomycota(100);p__Ascomycota_unclassified(100);p__Ascomycota_unclassified(100);p__Ascomycota_unclassified(100);p__Ascomycota_unclassified(100);p__Ascomycota_unclassified(100); | - | - | - | - | - | - | - | Unassigned | - |
| Otu33712 | 1 | k__Fungi(100);p__Ascomycota(100);p__Ascomycota_unclassified(100);p__Ascomycota_unclassified(100);p__Ascomycota_unclassified(100);p__Ascomycota_unclassified(100);p__Ascomycota_unclassified(100); | - | - | - | - | - | - | - | Unassigned | - |
| Otu33714 | 1 | k__Fungi(100);p__Ascomycota(100);p__Ascomycota_unclassified(100);p__Ascomycota_unclassified(100);p__Ascomycota_unclassified(100);p__Ascomycota_unclassified(100);p__Ascomycota_unclassified(100); | - | - | - | - | - | - | - | Unassigned | - |
| Otu33732 | 1 | k__Fungi(100);k__Fungi_unclassified(100);k__Fungi_unclassified(100);k__Fungi_unclassified(100);k__Fungi_unclassified(100);k__Fungi_unclassified(100);k__Fungi_unclassified(100); | - | - | - | - | - | - | - | Unassigned | - |
| Otu33744 | 1 | k__Fungi(100);p__Ascomycota(100);p__Ascomycota_unclassified(100);p__Ascomycota_unclassified(100);p__Ascomycota_unclassified(100);p__Ascomycota_unclassified(100);p__Ascomycota_unclassified(100); | - | - | - | - | - | - | - | Unassigned | - |
| Otu33745 | 1 | k__Fungi(100);k__Fungi_unclassified(100);k__Fungi_unclassified(100);k__Fungi_unclassified(100);k__Fungi_unclassified(100);k__Fungi_unclassified(100);k__Fungi_unclassified(100); | - | - | - | - | - | - | - | Unassigned | - |
| Otu33746 | 1 | k__Fungi(100);k__Fungi_unclassified(100);k__Fungi_unclassified(100);k__Fungi_unclassified(100);k__Fungi_unclassified(100);k__Fungi_unclassified(100);k__Fungi_unclassified(100); | - | - | - | - | - | - | - | Unassigned | - |
| Otu33747 | 1 | k__Fungi(100);k__Fungi_unclassified(100);k__Fungi_unclassified(100);k__Fungi_unclassified(100);k__Fungi_unclassified(100);k__Fungi_unclassified(100);k__Fungi_unclassified(100); | - | - | - | - | - | - | - | Unassigned | - |
| Otu33749 | 1 | k__Fungi(100);k__Fungi_unclassified(100);k__Fungi_unclassified(100);k__Fungi_unclassified(100);k__Fungi_unclassified(100);k__Fungi_unclassified(100);k__Fungi_unclassified(100); | - | - | - | - | - | - | - | Unassigned | - |
| Otu33752 | 1 | k__Fungi(100);k__Fungi_unclassified(100);k__Fungi_unclassified(100);k__Fungi_unclassified(100);k__Fungi_unclassified(100);k__Fungi_unclassified(100);k__Fungi_unclassified(100); | - | - | - | - | - | - | - | Unassigned | - |
| Otu33753 | 1 | k__Fungi(100);k__Fungi_unclassified(100);k__Fungi_unclassified(100);k__Fungi_unclassified(100);k__Fungi_unclassified(100);k__Fungi_unclassified(100);k__Fungi_unclassified(100); | - | - | - | - | - | - | - | Unassigned | - |
| Otu33754 | 1 | k__Fungi(100);k__Fungi_unclassified(100);k__Fungi_unclassified(100);k__Fungi_unclassified(100);k__Fungi_unclassified(100);k__Fungi_unclassified(100);k__Fungi_unclassified(100); | - | - | - | - | - | - | - | Unassigned | - |
| Otu33757 | 1 | k__Fungi(100);k__Fungi_unclassified(100);k__Fungi_unclassified(100);k__Fungi_unclassified(100);k__Fungi_unclassified(100);k__Fungi_unclassified(100);k__Fungi_unclassified(100); | - | - | - | - | - | - | - | Unassigned | - |
| Otu33759 | 1 | k__Fungi(100);k__Fungi_unclassified(100);k__Fungi_unclassified(100);k__Fungi_unclassified(100);k__Fungi_unclassified(100);k__Fungi_unclassified(100);k__Fungi_unclassified(100); | - | - | - | - | - | - | - | Unassigned | - |
| Otu33765 | 1 | k__Fungi(100);k__Fungi_unclassified(100);k__Fungi_unclassified(100);k__Fungi_unclassified(100);k__Fungi_unclassified(100);k__Fungi_unclassified(100);k__Fungi_unclassified(100); | - | - | - | - | - | - | - | Unassigned | - |
| Otu33766 | 1 | k__Fungi(100);p__Ascomycota(100);p__Ascomycota_unclassified(100);p__Ascomycota_unclassified(100);p__Ascomycota_unclassified(100);p__Ascomycota_unclassified(100);p__Ascomycota_unclassified(100); | - | - | - | - | - | - | - | Unassigned | - |
| Otu33773 | 1 | k__Fungi(100);k__Fungi_unclassified(100);k__Fungi_unclassified(100);k__Fungi_unclassified(100);k__Fungi_unclassified(100);k__Fungi_unclassified(100);k__Fungi_unclassified(100); | - | - | - | - | - | - | - | Unassigned | - |
| Otu33774 | 1 | k__Fungi(100);k__Fungi_unclassified(100);k__Fungi_unclassified(100);k__Fungi_unclassified(100);k__Fungi_unclassified(100);k__Fungi_unclassified(100);k__Fungi_unclassified(100); | - | - | - | - | - | - | - | Unassigned | - |
| Otu33776 | 1 | k__Fungi(100);p__Ascomycota(100);p__Ascomycota_unclassified(100);p__Ascomycota_unclassified(100);p__Ascomycota_unclassified(100);p__Ascomycota_unclassified(100);p__Ascomycota_unclassified(100); | - | - | - | - | - | - | - | Unassigned | - |
| Otu33782 | 1 | k__Fungi(100);k__Fungi_unclassified(100);k__Fungi_unclassified(100);k__Fungi_unclassified(100);k__Fungi_unclassified(100);k__Fungi_unclassified(100);k__Fungi_unclassified(100); | - | - | - | - | - | - | - | Unassigned | - |
| Otu33787 | 1 | k__Fungi(100);p__Ascomycota(100);p__Ascomycota_unclassified(100);p__Ascomycota_unclassified(100);p__Ascomycota_unclassified(100);p__Ascomycota_unclassified(100);p__Ascomycota_unclassified(100); | - | - | - | - | - | - | - | Unassigned | - |
| Otu33789 | 1 | k__Fungi(100);k__Fungi_unclassified(100);k__Fungi_unclassified(100);k__Fungi_unclassified(100);k__Fungi_unclassified(100);k__Fungi_unclassified(100);k__Fungi_unclassified(100); | - | - | - | - | - | - | - | Unassigned | - |
| Otu33790 | 1 | k__Fungi(100);k__Fungi_unclassified(100);k__Fungi_unclassified(100);k__Fungi_unclassified(100);k__Fungi_unclassified(100);k__Fungi_unclassified(100);k__Fungi_unclassified(100); | - | - | - | - | - | - | - | Unassigned | - |
| Otu33791 | 1 | k__Fungi(100);k__Fungi_unclassified(100);k__Fungi_unclassified(100);k__Fungi_unclassified(100);k__Fungi_unclassified(100);k__Fungi_unclassified(100);k__Fungi_unclassified(100); | - | - | - | - | - | - | - | Unassigned | - |
| Otu33792 | 1 | k__Fungi(100);k__Fungi_unclassified(100);k__Fungi_unclassified(100);k__Fungi_unclassified(100);k__Fungi_unclassified(100);k__Fungi_unclassified(100);k__Fungi_unclassified(100); | - | - | - | - | - | - | - | Unassigned | - |
| Otu33796 | 1 | k__Fungi(100);p__Ascomycota(100);p__Ascomycota_unclassified(100);p__Ascomycota_unclassified(100);p__Ascomycota_unclassified(100);p__Ascomycota_unclassified(100);p__Ascomycota_unclassified(100); | - | - | - | - | - | - | - | Unassigned | - |
| Otu33797 | 1 | k__Fungi(100);k__Fungi_unclassified(100);k__Fungi_unclassified(100);k__Fungi_unclassified(100);k__Fungi_unclassified(100);k__Fungi_unclassified(100);k__Fungi_unclassified(100); | - | - | - | - | - | - | - | Unassigned | - |
| Otu33798 | 1 | k__Fungi(100);k__Fungi_unclassified(100);k__Fungi_unclassified(100);k__Fungi_unclassified(100);k__Fungi_unclassified(100);k__Fungi_unclassified(100);k__Fungi_unclassified(100); | - | - | - | - | - | - | - | Unassigned | - |
| Otu33799 | 1 | k__Fungi(100);k__Fungi_unclassified(100);k__Fungi_unclassified(100);k__Fungi_unclassified(100);k__Fungi_unclassified(100);k__Fungi_unclassified(100);k__Fungi_unclassified(100); | - | - | - | - | - | - | - | Unassigned | - |
| Otu33802 | 1 | k__Fungi(100);p__Ascomycota(100);p__Ascomycota_unclassified(100);p__Ascomycota_unclassified(100);p__Ascomycota_unclassified(100);p__Ascomycota_unclassified(100);p__Ascomycota_unclassified(100); | - | - | - | - | - | - | - | Unassigned | - |
| Otu33805 | 1 | k__Fungi(100);p__Ascomycota(100);p__Ascomycota_unclassified(100);p__Ascomycota_unclassified(100);p__Ascomycota_unclassified(100);p__Ascomycota_unclassified(100);p__Ascomycota_unclassified(100); | - | - | - | - | - | - | - | Unassigned | - |
| Otu33807 | 1 | k__Fungi(100);k__Fungi_unclassified(100);k__Fungi_unclassified(100);k__Fungi_unclassified(100);k__Fungi_unclassified(100);k__Fungi_unclassified(100);k__Fungi_unclassified(100); | - | - | - | - | - | - | - | Unassigned | - |
| Otu33808 | 1 | k__Fungi(100);k__Fungi_unclassified(100);k__Fungi_unclassified(100);k__Fungi_unclassified(100);k__Fungi_unclassified(100);k__Fungi_unclassified(100);k__Fungi_unclassified(100); | - | - | - | - | - | - | - | Unassigned | - |
| Otu33809 | 1 | k__Fungi(100);k__Fungi_unclassified(100);k__Fungi_unclassified(100);k__Fungi_unclassified(100);k__Fungi_unclassified(100);k__Fungi_unclassified(100);k__Fungi_unclassified(100); | - | - | - | - | - | - | - | Unassigned | - |
| Otu33810 | 1 | k__Fungi(100);p__Ascomycota(100);p__Ascomycota_unclassified(100);p__Ascomycota_unclassified(100);p__Ascomycota_unclassified(100);p__Ascomycota_unclassified(100);p__Ascomycota_unclassified(100); | - | - | - | - | - | - | - | Unassigned | - |
| Otu33811 | 1 | k__Fungi(100);p__Ascomycota(100);p__Ascomycota_unclassified(100);p__Ascomycota_unclassified(100);p__Ascomycota_unclassified(100);p__Ascomycota_unclassified(100);p__Ascomycota_unclassified(100); | - | - | - | - | - | - | - | Unassigned | - |
| Otu33812 | 1 | k__Fungi(100);k__Fungi_unclassified(100);k__Fungi_unclassified(100);k__Fungi_unclassified(100);k__Fungi_unclassified(100);k__Fungi_unclassified(100);k__Fungi_unclassified(100); | - | - | - | - | - | - | - | Unassigned | - |
| Otu33813 | 1 | k__Fungi(100);k__Fungi_unclassified(100);k__Fungi_unclassified(100);k__Fungi_unclassified(100);k__Fungi_unclassified(100);k__Fungi_unclassified(100);k__Fungi_unclassified(100); | - | - | - | - | - | - | - | Unassigned | - |
| Otu33814 | 1 | k__Fungi(100);k__Fungi_unclassified(100);k__Fungi_unclassified(100);k__Fungi_unclassified(100);k__Fungi_unclassified(100);k__Fungi_unclassified(100);k__Fungi_unclassified(100); | - | - | - | - | - | - | - | Unassigned | - |
| Otu33815 | 1 | k__Fungi(100);p__Ascomycota(100);p__Ascomycota_unclassified(100);p__Ascomycota_unclassified(100);p__Ascomycota_unclassified(100);p__Ascomycota_unclassified(100);p__Ascomycota_unclassified(100); | - | - | - | - | - | - | - | Unassigned | - |
| Otu33816 | 1 | k__Fungi(100);p__Ascomycota(100);p__Ascomycota_unclassified(100);p__Ascomycota_unclassified(100);p__Ascomycota_unclassified(100);p__Ascomycota_unclassified(100);p__Ascomycota_unclassified(100); | - | - | - | - | - | - | - | Unassigned | - |
| Otu33817 | 1 | k__Fungi(100);k__Fungi_unclassified(100);k__Fungi_unclassified(100);k__Fungi_unclassified(100);k__Fungi_unclassified(100);k__Fungi_unclassified(100);k__Fungi_unclassified(100); | - | - | - | - | - | - | - | Unassigned | - |
| Otu33818 | 1 | k__Fungi(100);p__Ascomycota(100);p__Ascomycota_unclassified(100);p__Ascomycota_unclassified(100);p__Ascomycota_unclassified(100);p__Ascomycota_unclassified(100);p__Ascomycota_unclassified(100); | - | - | - | - | - | - | - | Unassigned | - |
| Otu33820 | 1 | k__Fungi(100);k__Fungi_unclassified(100);k__Fungi_unclassified(100);k__Fungi_unclassified(100);k__Fungi_unclassified(100);k__Fungi_unclassified(100);k__Fungi_unclassified(100); | - | - | - | - | - | - | - | Unassigned | - |
| Otu33821 | 1 | k__Fungi(100);k__Fungi_unclassified(100);k__Fungi_unclassified(100);k__Fungi_unclassified(100);k__Fungi_unclassified(100);k__Fungi_unclassified(100);k__Fungi_unclassified(100); | - | - | - | - | - | - | - | Unassigned | - |
| Otu33822 | 1 | k__Fungi(100);k__Fungi_unclassified(100);k__Fungi_unclassified(100);k__Fungi_unclassified(100);k__Fungi_unclassified(100);k__Fungi_unclassified(100);k__Fungi_unclassified(100); | - | - | - | - | - | - | - | Unassigned | - |
| Otu33824 | 1 | k__Fungi(100);k__Fungi_unclassified(100);k__Fungi_unclassified(100);k__Fungi_unclassified(100);k__Fungi_unclassified(100);k__Fungi_unclassified(100);k__Fungi_unclassified(100); | - | - | - | - | - | - | - | Unassigned | - |
| Otu33825 | 1 | k__Fungi(100);p__Ascomycota(100);p__Ascomycota_unclassified(100);p__Ascomycota_unclassified(100);p__Ascomycota_unclassified(100);p__Ascomycota_unclassified(100);p__Ascomycota_unclassified(100); | - | - | - | - | - | - | - | Unassigned | - |
| Otu33826 | 1 | k__Fungi(100);k__Fungi_unclassified(100);k__Fungi_unclassified(100);k__Fungi_unclassified(100);k__Fungi_unclassified(100);k__Fungi_unclassified(100);k__Fungi_unclassified(100); | - | - | - | - | - | - | - | Unassigned | - |
| Otu33831 | 1 | k__Fungi(100);k__Fungi_unclassified(100);k__Fungi_unclassified(100);k__Fungi_unclassified(100);k__Fungi_unclassified(100);k__Fungi_unclassified(100);k__Fungi_unclassified(100); | - | - | - | - | - | - | - | Unassigned | - |
| Otu33832 | 1 | k__Fungi(100);k__Fungi_unclassified(100);k__Fungi_unclassified(100);k__Fungi_unclassified(100);k__Fungi_unclassified(100);k__Fungi_unclassified(100);k__Fungi_unclassified(100); | - | - | - | - | - | - | - | Unassigned | - |
| Otu33833 | 1 | k__Fungi(100);k__Fungi_unclassified(100);k__Fungi_unclassified(100);k__Fungi_unclassified(100);k__Fungi_unclassified(100);k__Fungi_unclassified(100);k__Fungi_unclassified(100); | - | - | - | - | - | - | - | Unassigned | - |
| Otu33835 | 1 | k__Fungi(100);k__Fungi_unclassified(100);k__Fungi_unclassified(100);k__Fungi_unclassified(100);k__Fungi_unclassified(100);k__Fungi_unclassified(100);k__Fungi_unclassified(100); | - | - | - | - | - | - | - | Unassigned | - |
| Otu33836 | 1 | k__Fungi(100);k__Fungi_unclassified(100);k__Fungi_unclassified(100);k__Fungi_unclassified(100);k__Fungi_unclassified(100);k__Fungi_unclassified(100);k__Fungi_unclassified(100); | - | - | - | - | - | - | - | Unassigned | - |
| Otu33837 | 1 | k__Fungi(100);k__Fungi_unclassified(100);k__Fungi_unclassified(100);k__Fungi_unclassified(100);k__Fungi_unclassified(100);k__Fungi_unclassified(100);k__Fungi_unclassified(100); | - | - | - | - | - | - | - | Unassigned | - |
| Otu33839 | 1 | k__Fungi(100);k__Fungi_unclassified(100);k__Fungi_unclassified(100);k__Fungi_unclassified(100);k__Fungi_unclassified(100);k__Fungi_unclassified(100);k__Fungi_unclassified(100); | - | - | - | - | - | - | - | Unassigned | - |
| Otu33840 | 1 | k__Fungi(100);k__Fungi_unclassified(100);k__Fungi_unclassified(100);k__Fungi_unclassified(100);k__Fungi_unclassified(100);k__Fungi_unclassified(100);k__Fungi_unclassified(100); | - | - | - | - | - | - | - | Unassigned | - |
| Otu33841 | 1 | k__Fungi(100);p__Ascomycota(100);p__Ascomycota_unclassified(100);p__Ascomycota_unclassified(100);p__Ascomycota_unclassified(100);p__Ascomycota_unclassified(100);p__Ascomycota_unclassified(100); | - | - | - | - | - | - | - | Unassigned | - |
| Otu33843 | 1 | k__Fungi(100);k__Fungi_unclassified(100);k__Fungi_unclassified(100);k__Fungi_unclassified(100);k__Fungi_unclassified(100);k__Fungi_unclassified(100);k__Fungi_unclassified(100); | - | - | - | - | - | - | - | Unassigned | - |
| Otu33846 | 1 | k__Fungi(100);k__Fungi_unclassified(100);k__Fungi_unclassified(100);k__Fungi_unclassified(100);k__Fungi_unclassified(100);k__Fungi_unclassified(100);k__Fungi_unclassified(100); | - | - | - | - | - | - | - | Unassigned | - |
| Otu33849 | 1 | k__Fungi(100);k__Fungi_unclassified(100);k__Fungi_unclassified(100);k__Fungi_unclassified(100);k__Fungi_unclassified(100);k__Fungi_unclassified(100);k__Fungi_unclassified(100); | - | - | - | - | - | - | - | Unassigned | - |
| Otu33852 | 1 | k__Fungi(100);k__Fungi_unclassified(100);k__Fungi_unclassified(100);k__Fungi_unclassified(100);k__Fungi_unclassified(100);k__Fungi_unclassified(100);k__Fungi_unclassified(100); | - | - | - | - | - | - | - | Unassigned | - |
| Otu33855 | 1 | k__Fungi(100);k__Fungi_unclassified(100);k__Fungi_unclassified(100);k__Fungi_unclassified(100);k__Fungi_unclassified(100);k__Fungi_unclassified(100);k__Fungi_unclassified(100); | - | - | - | - | - | - | - | Unassigned | - |
| Otu33857 | 1 | k__Fungi(100);k__Fungi_unclassified(100);k__Fungi_unclassified(100);k__Fungi_unclassified(100);k__Fungi_unclassified(100);k__Fungi_unclassified(100);k__Fungi_unclassified(100); | - | - | - | - | - | - | - | Unassigned | - |
| Otu33858 | 1 | k__Fungi(100);k__Fungi_unclassified(100);k__Fungi_unclassified(100);k__Fungi_unclassified(100);k__Fungi_unclassified(100);k__Fungi_unclassified(100);k__Fungi_unclassified(100); | - | - | - | - | - | - | - | Unassigned | - |
| Otu33859 | 1 | k__Fungi(100);k__Fungi_unclassified(100);k__Fungi_unclassified(100);k__Fungi_unclassified(100);k__Fungi_unclassified(100);k__Fungi_unclassified(100);k__Fungi_unclassified(100); | - | - | - | - | - | - | - | Unassigned | - |
| Otu33860 | 1 | k__Fungi(100);k__Fungi_unclassified(100);k__Fungi_unclassified(100);k__Fungi_unclassified(100);k__Fungi_unclassified(100);k__Fungi_unclassified(100);k__Fungi_unclassified(100); | - | - | - | - | - | - | - | Unassigned | - |
| Otu33862 | 1 | k__Fungi(100);p__Ascomycota(100);p__Ascomycota_unclassified(100);p__Ascomycota_unclassified(100);p__Ascomycota_unclassified(100);p__Ascomycota_unclassified(100);p__Ascomycota_unclassified(100); | - | - | - | - | - | - | - | Unassigned | - |
| Otu33863 | 1 | k__Fungi(100);p__Ascomycota(100);p__Ascomycota_unclassified(100);p__Ascomycota_unclassified(100);p__Ascomycota_unclassified(100);p__Ascomycota_unclassified(100);p__Ascomycota_unclassified(100); | - | - | - | - | - | - | - | Unassigned | - |
| Otu33864 | 1 | k__Fungi(100);k__Fungi_unclassified(100);k__Fungi_unclassified(100);k__Fungi_unclassified(100);k__Fungi_unclassified(100);k__Fungi_unclassified(100);k__Fungi_unclassified(100); | - | - | - | - | - | - | - | Unassigned | - |
| Otu33868 | 1 | k__Fungi(100);p__Ascomycota(100);p__Ascomycota_unclassified(100);p__Ascomycota_unclassified(100);p__Ascomycota_unclassified(100);p__Ascomycota_unclassified(100);p__Ascomycota_unclassified(100); | - | - | - | - | - | - | - | Unassigned | - |
| Otu33869 | 1 | k__Fungi(100);k__Fungi_unclassified(100);k__Fungi_unclassified(100);k__Fungi_unclassified(100);k__Fungi_unclassified(100);k__Fungi_unclassified(100);k__Fungi_unclassified(100); | - | - | - | - | - | - | - | Unassigned | - |
| Otu33870 | 1 | k__Fungi(100);p__Ascomycota(100);p__Ascomycota_unclassified(100);p__Ascomycota_unclassified(100);p__Ascomycota_unclassified(100);p__Ascomycota_unclassified(100);p__Ascomycota_unclassified(100); | - | - | - | - | - | - | - | Unassigned | - |
| Otu33871 | 1 | k__Fungi(100);k__Fungi_unclassified(100);k__Fungi_unclassified(100);k__Fungi_unclassified(100);k__Fungi_unclassified(100);k__Fungi_unclassified(100);k__Fungi_unclassified(100); | - | - | - | - | - | - | - | Unassigned | - |
| Otu33878 | 1 | k__Fungi(100);k__Fungi_unclassified(100);k__Fungi_unclassified(100);k__Fungi_unclassified(100);k__Fungi_unclassified(100);k__Fungi_unclassified(100);k__Fungi_unclassified(100); | - | - | - | - | - | - | - | Unassigned | - |
| Otu33879 | 1 | k__Fungi(100);p__Ascomycota(100);p__Ascomycota_unclassified(100);p__Ascomycota_unclassified(100);p__Ascomycota_unclassified(100);p__Ascomycota_unclassified(100);p__Ascomycota_unclassified(100); | - | - | - | - | - | - | - | Unassigned | - |
| Otu33880 | 1 | k__Fungi(100);k__Fungi_unclassified(100);k__Fungi_unclassified(100);k__Fungi_unclassified(100);k__Fungi_unclassified(100);k__Fungi_unclassified(100);k__Fungi_unclassified(100); | - | - | - | - | - | - | - | Unassigned | - |
| Otu33883 | 1 | k__Fungi(100);p__Ascomycota(100);p__Ascomycota_unclassified(100);p__Ascomycota_unclassified(100);p__Ascomycota_unclassified(100);p__Ascomycota_unclassified(100);p__Ascomycota_unclassified(100); | - | - | - | - | - | - | - | Unassigned | - |
| Otu33884 | 1 | k__Fungi(100);p__Ascomycota(100);p__Ascomycota_unclassified(100);p__Ascomycota_unclassified(100);p__Ascomycota_unclassified(100);p__Ascomycota_unclassified(100);p__Ascomycota_unclassified(100); | - | - | - | - | - | - | - | Unassigned | - |
| Otu33885 | 1 | k__Fungi(100);p__Ascomycota(100);p__Ascomycota_unclassified(100);p__Ascomycota_unclassified(100);p__Ascomycota_unclassified(100);p__Ascomycota_unclassified(100);p__Ascomycota_unclassified(100); | - | - | - | - | - | - | - | Unassigned | - |
| Otu33886 | 1 | k__Fungi(100);p__Ascomycota(100);p__Ascomycota_unclassified(100);p__Ascomycota_unclassified(100);p__Ascomycota_unclassified(100);p__Ascomycota_unclassified(100);p__Ascomycota_unclassified(100); | - | - | - | - | - | - | - | Unassigned | - |
| Otu33887 | 1 | k__Fungi(100);k__Fungi_unclassified(100);k__Fungi_unclassified(100);k__Fungi_unclassified(100);k__Fungi_unclassified(100);k__Fungi_unclassified(100);k__Fungi_unclassified(100); | - | - | - | - | - | - | - | Unassigned | - |
| Otu33888 | 1 | k__Fungi(100);k__Fungi_unclassified(100);k__Fungi_unclassified(100);k__Fungi_unclassified(100);k__Fungi_unclassified(100);k__Fungi_unclassified(100);k__Fungi_unclassified(100); | - | - | - | - | - | - | - | Unassigned | - |
| Otu33890 | 1 | k__Fungi(100);k__Fungi_unclassified(100);k__Fungi_unclassified(100);k__Fungi_unclassified(100);k__Fungi_unclassified(100);k__Fungi_unclassified(100);k__Fungi_unclassified(100); | - | - | - | - | - | - | - | Unassigned | - |
| Otu33891 | 1 | k__Fungi(100);k__Fungi_unclassified(100);k__Fungi_unclassified(100);k__Fungi_unclassified(100);k__Fungi_unclassified(100);k__Fungi_unclassified(100);k__Fungi_unclassified(100); | - | - | - | - | - | - | - | Unassigned | - |
| Otu33894 | 1 | k__Fungi(100);p__Ascomycota(100);p__Ascomycota_unclassified(100);p__Ascomycota_unclassified(100);p__Ascomycota_unclassified(100);p__Ascomycota_unclassified(100);p__Ascomycota_unclassified(100); | - | - | - | - | - | - | - | Unassigned | - |
| Otu33897 | 1 | k__Fungi(100);p__Ascomycota(100);p__Ascomycota_unclassified(100);p__Ascomycota_unclassified(100);p__Ascomycota_unclassified(100);p__Ascomycota_unclassified(100);p__Ascomycota_unclassified(100); | - | - | - | - | - | - | - | Unassigned | - |
| Otu33899 | 1 | k__Fungi(100);k__Fungi_unclassified(100);k__Fungi_unclassified(100);k__Fungi_unclassified(100);k__Fungi_unclassified(100);k__Fungi_unclassified(100);k__Fungi_unclassified(100); | - | - | - | - | - | - | - | Unassigned | - |
| Otu33900 | 1 | k__Fungi(100);k__Fungi_unclassified(100);k__Fungi_unclassified(100);k__Fungi_unclassified(100);k__Fungi_unclassified(100);k__Fungi_unclassified(100);k__Fungi_unclassified(100); | - | - | - | - | - | - | - | Unassigned | - |
| Otu33901 | 1 | k__Fungi(100);k__Fungi_unclassified(100);k__Fungi_unclassified(100);k__Fungi_unclassified(100);k__Fungi_unclassified(100);k__Fungi_unclassified(100);k__Fungi_unclassified(100); | - | - | - | - | - | - | - | Unassigned | - |
| Otu33902 | 1 | k__Fungi(100);k__Fungi_unclassified(100);k__Fungi_unclassified(100);k__Fungi_unclassified(100);k__Fungi_unclassified(100);k__Fungi_unclassified(100);k__Fungi_unclassified(100); | - | - | - | - | - | - | - | Unassigned | - |
| Otu33914 | 1 | k__Fungi(100);k__Fungi_unclassified(100);k__Fungi_unclassified(100);k__Fungi_unclassified(100);k__Fungi_unclassified(100);k__Fungi_unclassified(100);k__Fungi_unclassified(100); | - | - | - | - | - | - | - | Unassigned | - |
| Otu33926 | 1 | k__Fungi(100);p__Ascomycota(100);p__Ascomycota_unclassified(100);p__Ascomycota_unclassified(100);p__Ascomycota_unclassified(100);p__Ascomycota_unclassified(100);p__Ascomycota_unclassified(100); | - | - | - | - | - | - | - | Unassigned | - |
| Otu33927 | 1 | k__Fungi(100);p__Ascomycota(100);p__Ascomycota_unclassified(100);p__Ascomycota_unclassified(100);p__Ascomycota_unclassified(100);p__Ascomycota_unclassified(100);p__Ascomycota_unclassified(100); | - | - | - | - | - | - | - | Unassigned | - |
| Otu33928 | 1 | k__Fungi(100);k__Fungi_unclassified(100);k__Fungi_unclassified(100);k__Fungi_unclassified(100);k__Fungi_unclassified(100);k__Fungi_unclassified(100);k__Fungi_unclassified(100); | - | - | - | - | - | - | - | Unassigned | - |
| Otu33929 | 1 | k__Fungi(100);p__Ascomycota(100);p__Ascomycota_unclassified(100);p__Ascomycota_unclassified(100);p__Ascomycota_unclassified(100);p__Ascomycota_unclassified(100);p__Ascomycota_unclassified(100); | - | - | - | - | - | - | - | Unassigned | - |
| Otu33932 | 1 | k__Fungi(100);k__Fungi_unclassified(100);k__Fungi_unclassified(100);k__Fungi_unclassified(100);k__Fungi_unclassified(100);k__Fungi_unclassified(100);k__Fungi_unclassified(100); | - | - | - | - | - | - | - | Unassigned | - |
| Otu33933 | 1 | k__Fungi(100);p__Ascomycota(100);p__Ascomycota_unclassified(100);p__Ascomycota_unclassified(100);p__Ascomycota_unclassified(100);p__Ascomycota_unclassified(100);p__Ascomycota_unclassified(100); | - | - | - | - | - | - | - | Unassigned | - |
| Otu33935 | 1 | k__Fungi(100);k__Fungi_unclassified(100);k__Fungi_unclassified(100);k__Fungi_unclassified(100);k__Fungi_unclassified(100);k__Fungi_unclassified(100);k__Fungi_unclassified(100); | - | - | - | - | - | - | - | Unassigned | - |
| Otu33936 | 1 | k__Fungi(100);k__Fungi_unclassified(100);k__Fungi_unclassified(100);k__Fungi_unclassified(100);k__Fungi_unclassified(100);k__Fungi_unclassified(100);k__Fungi_unclassified(100); | - | - | - | - | - | - | - | Unassigned | - |
| Otu33939 | 1 | k__Fungi(100);p__Ascomycota(100);p__Ascomycota_unclassified(100);p__Ascomycota_unclassified(100);p__Ascomycota_unclassified(100);p__Ascomycota_unclassified(100);p__Ascomycota_unclassified(100); | - | - | - | - | - | - | - | Unassigned | - |
| Otu33942 | 1 | k__Fungi(100);k__Fungi_unclassified(100);k__Fungi_unclassified(100);k__Fungi_unclassified(100);k__Fungi_unclassified(100);k__Fungi_unclassified(100);k__Fungi_unclassified(100); | - | - | - | - | - | - | - | Unassigned | - |
| Otu33946 | 1 | k__Fungi(100);k__Fungi_unclassified(100);k__Fungi_unclassified(100);k__Fungi_unclassified(100);k__Fungi_unclassified(100);k__Fungi_unclassified(100);k__Fungi_unclassified(100); | - | - | - | - | - | - | - | Unassigned | - |
| Otu33947 | 1 | k__Fungi(100);p__Ascomycota(100);p__Ascomycota_unclassified(100);p__Ascomycota_unclassified(100);p__Ascomycota_unclassified(100);p__Ascomycota_unclassified(100);p__Ascomycota_unclassified(100); | - | - | - | - | - | - | - | Unassigned | - |
| Otu33948 | 1 | k__Fungi(100);p__Ascomycota(100);p__Ascomycota_unclassified(100);p__Ascomycota_unclassified(100);p__Ascomycota_unclassified(100);p__Ascomycota_unclassified(100);p__Ascomycota_unclassified(100); | - | - | - | - | - | - | - | Unassigned | - |
| Otu33949 | 1 | k__Fungi(100);k__Fungi_unclassified(100);k__Fungi_unclassified(100);k__Fungi_unclassified(100);k__Fungi_unclassified(100);k__Fungi_unclassified(100);k__Fungi_unclassified(100); | - | - | - | - | - | - | - | Unassigned | - |
| Otu33950 | 1 | k__Fungi(100);k__Fungi_unclassified(100);k__Fungi_unclassified(100);k__Fungi_unclassified(100);k__Fungi_unclassified(100);k__Fungi_unclassified(100);k__Fungi_unclassified(100); | - | - | - | - | - | - | - | Unassigned | - |
| Otu33951 | 1 | k__Fungi(100);p__Ascomycota(100);p__Ascomycota_unclassified(100);p__Ascomycota_unclassified(100);p__Ascomycota_unclassified(100);p__Ascomycota_unclassified(100);p__Ascomycota_unclassified(100); | - | - | - | - | - | - | - | Unassigned | - |
| Otu33953 | 1 | k__Fungi(100);k__Fungi_unclassified(100);k__Fungi_unclassified(100);k__Fungi_unclassified(100);k__Fungi_unclassified(100);k__Fungi_unclassified(100);k__Fungi_unclassified(100); | - | - | - | - | - | - | - | Unassigned | - |
| Otu33954 | 1 | k__Fungi(100);p__Ascomycota(100);p__Ascomycota_unclassified(100);p__Ascomycota_unclassified(100);p__Ascomycota_unclassified(100);p__Ascomycota_unclassified(100);p__Ascomycota_unclassified(100); | - | - | - | - | - | - | - | Unassigned | - |
| Otu33956 | 1 | k__Fungi(100);k__Fungi_unclassified(100);k__Fungi_unclassified(100);k__Fungi_unclassified(100);k__Fungi_unclassified(100);k__Fungi_unclassified(100);k__Fungi_unclassified(100); | - | - | - | - | - | - | - | Unassigned | - |
| Otu33957 | 1 | k__Fungi(100);k__Fungi_unclassified(100);k__Fungi_unclassified(100);k__Fungi_unclassified(100);k__Fungi_unclassified(100);k__Fungi_unclassified(100);k__Fungi_unclassified(100); | - | - | - | - | - | - | - | Unassigned | - |
| Otu33959 | 1 | k__Fungi(100);p__Ascomycota(100);p__Ascomycota_unclassified(100);p__Ascomycota_unclassified(100);p__Ascomycota_unclassified(100);p__Ascomycota_unclassified(100);p__Ascomycota_unclassified(100); | - | - | - | - | - | - | - | Unassigned | - |
| Otu33960 | 1 | k__Fungi(100);k__Fungi_unclassified(100);k__Fungi_unclassified(100);k__Fungi_unclassified(100);k__Fungi_unclassified(100);k__Fungi_unclassified(100);k__Fungi_unclassified(100); | - | - | - | - | - | - | - | Unassigned | - |
| Otu33964 | 1 | k__Fungi(100);p__Ascomycota(100);p__Ascomycota_unclassified(100);p__Ascomycota_unclassified(100);p__Ascomycota_unclassified(100);p__Ascomycota_unclassified(100);p__Ascomycota_unclassified(100); | - | - | - | - | - | - | - | Unassigned | - |
| Otu33965 | 1 | k__Fungi(100);p__Ascomycota(100);p__Ascomycota_unclassified(100);p__Ascomycota_unclassified(100);p__Ascomycota_unclassified(100);p__Ascomycota_unclassified(100);p__Ascomycota_unclassified(100); | - | - | - | - | - | - | - | Unassigned | - |
| Otu33966 | 1 | k__Fungi(100);k__Fungi_unclassified(100);k__Fungi_unclassified(100);k__Fungi_unclassified(100);k__Fungi_unclassified(100);k__Fungi_unclassified(100);k__Fungi_unclassified(100); | - | - | - | - | - | - | - | Unassigned | - |
| Otu33968 | 1 | k__Fungi(100);k__Fungi_unclassified(100);k__Fungi_unclassified(100);k__Fungi_unclassified(100);k__Fungi_unclassified(100);k__Fungi_unclassified(100);k__Fungi_unclassified(100); | - | - | - | - | - | - | - | Unassigned | - |
| Otu33970 | 1 | k__Fungi(100);k__Fungi_unclassified(100);k__Fungi_unclassified(100);k__Fungi_unclassified(100);k__Fungi_unclassified(100);k__Fungi_unclassified(100);k__Fungi_unclassified(100); | - | - | - | - | - | - | - | Unassigned | - |
| Otu33971 | 1 | k__Fungi(100);k__Fungi_unclassified(100);k__Fungi_unclassified(100);k__Fungi_unclassified(100);k__Fungi_unclassified(100);k__Fungi_unclassified(100);k__Fungi_unclassified(100); | - | - | - | - | - | - | - | Unassigned | - |
| Otu33972 | 1 | k__Fungi(100);p__Ascomycota(100);p__Ascomycota_unclassified(100);p__Ascomycota_unclassified(100);p__Ascomycota_unclassified(100);p__Ascomycota_unclassified(100);p__Ascomycota_unclassified(100); | - | - | - | - | - | - | - | Unassigned | - |
| Otu33974 | 1 | k__Fungi(100);p__Ascomycota(100);p__Ascomycota_unclassified(100);p__Ascomycota_unclassified(100);p__Ascomycota_unclassified(100);p__Ascomycota_unclassified(100);p__Ascomycota_unclassified(100); | - | - | - | - | - | - | - | Unassigned | - |
| Otu33976 | 1 | k__Fungi(100);k__Fungi_unclassified(100);k__Fungi_unclassified(100);k__Fungi_unclassified(100);k__Fungi_unclassified(100);k__Fungi_unclassified(100);k__Fungi_unclassified(100); | - | - | - | - | - | - | - | Unassigned | - |
| Otu33979 | 1 | k__Fungi(100);p__Ascomycota(100);p__Ascomycota_unclassified(100);p__Ascomycota_unclassified(100);p__Ascomycota_unclassified(100);p__Ascomycota_unclassified(100);p__Ascomycota_unclassified(100); | - | - | - | - | - | - | - | Unassigned | - |
| Otu33980 | 1 | k__Fungi(100);k__Fungi_unclassified(100);k__Fungi_unclassified(100);k__Fungi_unclassified(100);k__Fungi_unclassified(100);k__Fungi_unclassified(100);k__Fungi_unclassified(100); | - | - | - | - | - | - | - | Unassigned | - |
| Otu33984 | 1 | k__Fungi(100);k__Fungi_unclassified(100);k__Fungi_unclassified(100);k__Fungi_unclassified(100);k__Fungi_unclassified(100);k__Fungi_unclassified(100);k__Fungi_unclassified(100); | - | - | - | - | - | - | - | Unassigned | - |
| Otu33986 | 1 | k__Fungi(100);p__Ascomycota(100);p__Ascomycota_unclassified(100);p__Ascomycota_unclassified(100);p__Ascomycota_unclassified(100);p__Ascomycota_unclassified(100);p__Ascomycota_unclassified(100); | - | - | - | - | - | - | - | Unassigned | - |
| Otu33987 | 1 | k__Fungi(100);p__Ascomycota(100);p__Ascomycota_unclassified(100);p__Ascomycota_unclassified(100);p__Ascomycota_unclassified(100);p__Ascomycota_unclassified(100);p__Ascomycota_unclassified(100); | - | - | - | - | - | - | - | Unassigned | - |
| Otu33988 | 1 | k__Fungi(100);p__Ascomycota(100);p__Ascomycota_unclassified(100);p__Ascomycota_unclassified(100);p__Ascomycota_unclassified(100);p__Ascomycota_unclassified(100);p__Ascomycota_unclassified(100); | - | - | - | - | - | - | - | Unassigned | - |
| Otu33989 | 1 | k__Fungi(100);k__Fungi_unclassified(100);k__Fungi_unclassified(100);k__Fungi_unclassified(100);k__Fungi_unclassified(100);k__Fungi_unclassified(100);k__Fungi_unclassified(100); | - | - | - | - | - | - | - | Unassigned | - |
| Otu33990 | 1 | k__Fungi(100);k__Fungi_unclassified(100);k__Fungi_unclassified(100);k__Fungi_unclassified(100);k__Fungi_unclassified(100);k__Fungi_unclassified(100);k__Fungi_unclassified(100); | - | - | - | - | - | - | - | Unassigned | - |
| Otu33996 | 1 | k__Fungi(100);p__Ascomycota(100);p__Ascomycota_unclassified(100);p__Ascomycota_unclassified(100);p__Ascomycota_unclassified(100);p__Ascomycota_unclassified(100);p__Ascomycota_unclassified(100); | - | - | - | - | - | - | - | Unassigned | - |
| Otu33997 | 1 | k__Fungi(100);k__Fungi_unclassified(100);k__Fungi_unclassified(100);k__Fungi_unclassified(100);k__Fungi_unclassified(100);k__Fungi_unclassified(100);k__Fungi_unclassified(100); | - | - | - | - | - | - | - | Unassigned | - |
| Otu33999 | 1 | k__Fungi(100);k__Fungi_unclassified(100);k__Fungi_unclassified(100);k__Fungi_unclassified(100);k__Fungi_unclassified(100);k__Fungi_unclassified(100);k__Fungi_unclassified(100); | - | - | - | - | - | - | - | Unassigned | - |
| Otu34003 | 1 | k__Fungi(100);k__Fungi_unclassified(100);k__Fungi_unclassified(100);k__Fungi_unclassified(100);k__Fungi_unclassified(100);k__Fungi_unclassified(100);k__Fungi_unclassified(100); | - | - | - | - | - | - | - | Unassigned | - |
| Otu34004 | 1 | k__Fungi(100);k__Fungi_unclassified(100);k__Fungi_unclassified(100);k__Fungi_unclassified(100);k__Fungi_unclassified(100);k__Fungi_unclassified(100);k__Fungi_unclassified(100); | - | - | - | - | - | - | - | Unassigned | - |
| Otu34005 | 1 | k__Fungi(100);k__Fungi_unclassified(100);k__Fungi_unclassified(100);k__Fungi_unclassified(100);k__Fungi_unclassified(100);k__Fungi_unclassified(100);k__Fungi_unclassified(100); | - | - | - | - | - | - | - | Unassigned | - |
| Otu34006 | 1 | k__Fungi(100);p__Ascomycota(100);p__Ascomycota_unclassified(100);p__Ascomycota_unclassified(100);p__Ascomycota_unclassified(100);p__Ascomycota_unclassified(100);p__Ascomycota_unclassified(100); | - | - | - | - | - | - | - | Unassigned | - |
| Otu34007 | 1 | k__Fungi(100);k__Fungi_unclassified(100);k__Fungi_unclassified(100);k__Fungi_unclassified(100);k__Fungi_unclassified(100);k__Fungi_unclassified(100);k__Fungi_unclassified(100); | - | - | - | - | - | - | - | Unassigned | - |
| Otu34008 | 1 | k__Fungi(100);k__Fungi_unclassified(100);k__Fungi_unclassified(100);k__Fungi_unclassified(100);k__Fungi_unclassified(100);k__Fungi_unclassified(100);k__Fungi_unclassified(100); | - | - | - | - | - | - | - | Unassigned | - |
| Otu34009 | 1 | k__Fungi(100);k__Fungi_unclassified(100);k__Fungi_unclassified(100);k__Fungi_unclassified(100);k__Fungi_unclassified(100);k__Fungi_unclassified(100);k__Fungi_unclassified(100); | - | - | - | - | - | - | - | Unassigned | - |
| Otu34010 | 1 | k__Fungi(100);p__Ascomycota(100);p__Ascomycota_unclassified(100);p__Ascomycota_unclassified(100);p__Ascomycota_unclassified(100);p__Ascomycota_unclassified(100);p__Ascomycota_unclassified(100); | - | - | - | - | - | - | - | Unassigned | - |
| Otu34014 | 1 | k__Fungi(100);k__Fungi_unclassified(100);k__Fungi_unclassified(100);k__Fungi_unclassified(100);k__Fungi_unclassified(100);k__Fungi_unclassified(100);k__Fungi_unclassified(100); | - | - | - | - | - | - | - | Unassigned | - |
| Otu34015 | 1 | k__Fungi(100);k__Fungi_unclassified(100);k__Fungi_unclassified(100);k__Fungi_unclassified(100);k__Fungi_unclassified(100);k__Fungi_unclassified(100);k__Fungi_unclassified(100); | - | - | - | - | - | - | - | Unassigned | - |
| Otu34016 | 1 | k__Fungi(100);p__Ascomycota(100);p__Ascomycota_unclassified(100);p__Ascomycota_unclassified(100);p__Ascomycota_unclassified(100);p__Ascomycota_unclassified(100);p__Ascomycota_unclassified(100); | - | - | - | - | - | - | - | Unassigned | - |
| Otu34018 | 1 | k__Fungi(100);p__Ascomycota(100);p__Ascomycota_unclassified(100);p__Ascomycota_unclassified(100);p__Ascomycota_unclassified(100);p__Ascomycota_unclassified(100);p__Ascomycota_unclassified(100); | - | - | - | - | - | - | - | Unassigned | - |
| Otu34019 | 1 | k__Fungi(100);p__Ascomycota(100);p__Ascomycota_unclassified(100);p__Ascomycota_unclassified(100);p__Ascomycota_unclassified(100);p__Ascomycota_unclassified(100);p__Ascomycota_unclassified(100); | - | - | - | - | - | - | - | Unassigned | - |
| Otu34020 | 1 | k__Fungi(100);p__Ascomycota(100);p__Ascomycota_unclassified(100);p__Ascomycota_unclassified(100);p__Ascomycota_unclassified(100);p__Ascomycota_unclassified(100);p__Ascomycota_unclassified(100); | - | - | - | - | - | - | - | Unassigned | - |
| Otu34021 | 1 | k__Fungi(100);p__Ascomycota(100);p__Ascomycota_unclassified(100);p__Ascomycota_unclassified(100);p__Ascomycota_unclassified(100);p__Ascomycota_unclassified(100);p__Ascomycota_unclassified(100); | - | - | - | - | - | - | - | Unassigned | - |
| Otu34022 | 1 | k__Fungi(100);k__Fungi_unclassified(100);k__Fungi_unclassified(100);k__Fungi_unclassified(100);k__Fungi_unclassified(100);k__Fungi_unclassified(100);k__Fungi_unclassified(100); | - | - | - | - | - | - | - | Unassigned | - |
| Otu34023 | 1 | k__Fungi(100);k__Fungi_unclassified(100);k__Fungi_unclassified(100);k__Fungi_unclassified(100);k__Fungi_unclassified(100);k__Fungi_unclassified(100);k__Fungi_unclassified(100); | - | - | - | - | - | - | - | Unassigned | - |
| Otu34024 | 1 | k__Fungi(100);k__Fungi_unclassified(100);k__Fungi_unclassified(100);k__Fungi_unclassified(100);k__Fungi_unclassified(100);k__Fungi_unclassified(100);k__Fungi_unclassified(100); | - | - | - | - | - | - | - | Unassigned | - |
| Otu34026 | 1 | k__Fungi(100);k__Fungi_unclassified(100);k__Fungi_unclassified(100);k__Fungi_unclassified(100);k__Fungi_unclassified(100);k__Fungi_unclassified(100);k__Fungi_unclassified(100); | - | - | - | - | - | - | - | Unassigned | - |
| Otu34027 | 1 | k__Fungi(100);k__Fungi_unclassified(100);k__Fungi_unclassified(100);k__Fungi_unclassified(100);k__Fungi_unclassified(100);k__Fungi_unclassified(100);k__Fungi_unclassified(100); | - | - | - | - | - | - | - | Unassigned | - |
| Otu34028 | 1 | k__Fungi(100);k__Fungi_unclassified(100);k__Fungi_unclassified(100);k__Fungi_unclassified(100);k__Fungi_unclassified(100);k__Fungi_unclassified(100);k__Fungi_unclassified(100); | - | - | - | - | - | - | - | Unassigned | - |
| Otu34029 | 1 | k__Fungi(100);p__Ascomycota(100);p__Ascomycota_unclassified(100);p__Ascomycota_unclassified(100);p__Ascomycota_unclassified(100);p__Ascomycota_unclassified(100);p__Ascomycota_unclassified(100); | - | - | - | - | - | - | - | Unassigned | - |
| Otu34031 | 1 | k__Fungi(100);p__Ascomycota(100);p__Ascomycota_unclassified(100);p__Ascomycota_unclassified(100);p__Ascomycota_unclassified(100);p__Ascomycota_unclassified(100);p__Ascomycota_unclassified(100); | - | - | - | - | - | - | - | Unassigned | - |
| Otu34034 | 1 | k__Fungi(100);k__Fungi_unclassified(100);k__Fungi_unclassified(100);k__Fungi_unclassified(100);k__Fungi_unclassified(100);k__Fungi_unclassified(100);k__Fungi_unclassified(100); | - | - | - | - | - | - | - | Unassigned | - |
| Otu34036 | 1 | k__Fungi(100);k__Fungi_unclassified(100);k__Fungi_unclassified(100);k__Fungi_unclassified(100);k__Fungi_unclassified(100);k__Fungi_unclassified(100);k__Fungi_unclassified(100); | - | - | - | - | - | - | - | Unassigned | - |
| Otu34041 | 1 | k__Fungi(100);p__Ascomycota(100);p__Ascomycota_unclassified(100);p__Ascomycota_unclassified(100);p__Ascomycota_unclassified(100);p__Ascomycota_unclassified(100);p__Ascomycota_unclassified(100); | - | - | - | - | - | - | - | Unassigned | - |
| Otu34043 | 1 | k__Fungi(100);k__Fungi_unclassified(100);k__Fungi_unclassified(100);k__Fungi_unclassified(100);k__Fungi_unclassified(100);k__Fungi_unclassified(100);k__Fungi_unclassified(100); | - | - | - | - | - | - | - | Unassigned | - |
| Otu34046 | 1 | k__Fungi(100);p__Ascomycota(100);p__Ascomycota_unclassified(100);p__Ascomycota_unclassified(100);p__Ascomycota_unclassified(100);p__Ascomycota_unclassified(100);p__Ascomycota_unclassified(100); | - | - | - | - | - | - | - | Unassigned | - |
| Otu34051 | 1 | k__Fungi(100);p__Ascomycota(100);p__Ascomycota_unclassified(100);p__Ascomycota_unclassified(100);p__Ascomycota_unclassified(100);p__Ascomycota_unclassified(100);p__Ascomycota_unclassified(100); | - | - | - | - | - | - | - | Unassigned | - |
| Otu34057 | 1 | k__Fungi(100);p__Ascomycota(100);p__Ascomycota_unclassified(100);p__Ascomycota_unclassified(100);p__Ascomycota_unclassified(100);p__Ascomycota_unclassified(100);p__Ascomycota_unclassified(100); | - | - | - | - | - | - | - | Unassigned | - |
| Otu34059 | 1 | k__Fungi(100);k__Fungi_unclassified(100);k__Fungi_unclassified(100);k__Fungi_unclassified(100);k__Fungi_unclassified(100);k__Fungi_unclassified(100);k__Fungi_unclassified(100); | - | - | - | - | - | - | - | Unassigned | - |
| Otu34060 | 1 | k__Fungi(100);k__Fungi_unclassified(100);k__Fungi_unclassified(100);k__Fungi_unclassified(100);k__Fungi_unclassified(100);k__Fungi_unclassified(100);k__Fungi_unclassified(100); | - | - | - | - | - | - | - | Unassigned | - |
| Otu34062 | 1 | k__Fungi(100);k__Fungi_unclassified(100);k__Fungi_unclassified(100);k__Fungi_unclassified(100);k__Fungi_unclassified(100);k__Fungi_unclassified(100);k__Fungi_unclassified(100); | - | - | - | - | - | - | - | Unassigned | - |
| Otu34063 | 1 | k__Fungi(100);p__Ascomycota(100);p__Ascomycota_unclassified(100);p__Ascomycota_unclassified(100);p__Ascomycota_unclassified(100);p__Ascomycota_unclassified(100);p__Ascomycota_unclassified(100); | - | - | - | - | - | - | - | Unassigned | - |
| Otu34064 | 1 | k__Fungi(100);p__Ascomycota(100);p__Ascomycota_unclassified(100);p__Ascomycota_unclassified(100);p__Ascomycota_unclassified(100);p__Ascomycota_unclassified(100);p__Ascomycota_unclassified(100); | - | - | - | - | - | - | - | Unassigned | - |
| Otu34070 | 1 | k__Fungi(100);k__Fungi_unclassified(100);k__Fungi_unclassified(100);k__Fungi_unclassified(100);k__Fungi_unclassified(100);k__Fungi_unclassified(100);k__Fungi_unclassified(100); | - | - | - | - | - | - | - | Unassigned | - |
| Otu34071 | 1 | k__Fungi(100);p__Ascomycota(100);p__Ascomycota_unclassified(100);p__Ascomycota_unclassified(100);p__Ascomycota_unclassified(100);p__Ascomycota_unclassified(100);p__Ascomycota_unclassified(100); | - | - | - | - | - | - | - | Unassigned | - |
| Otu34072 | 1 | k__Fungi(100);p__Ascomycota(100);p__Ascomycota_unclassified(100);p__Ascomycota_unclassified(100);p__Ascomycota_unclassified(100);p__Ascomycota_unclassified(100);p__Ascomycota_unclassified(100); | - | - | - | - | - | - | - | Unassigned | - |
| Otu34073 | 1 | k__Fungi(100);p__Ascomycota(100);p__Ascomycota_unclassified(100);p__Ascomycota_unclassified(100);p__Ascomycota_unclassified(100);p__Ascomycota_unclassified(100);p__Ascomycota_unclassified(100); | - | - | - | - | - | - | - | Unassigned | - |
| Otu34074 | 1 | k__Fungi(100);p__Ascomycota(100);p__Ascomycota_unclassified(100);p__Ascomycota_unclassified(100);p__Ascomycota_unclassified(100);p__Ascomycota_unclassified(100);p__Ascomycota_unclassified(100); | - | - | - | - | - | - | - | Unassigned | - |
| Otu34076 | 1 | k__Fungi(100);k__Fungi_unclassified(100);k__Fungi_unclassified(100);k__Fungi_unclassified(100);k__Fungi_unclassified(100);k__Fungi_unclassified(100);k__Fungi_unclassified(100); | - | - | - | - | - | - | - | Unassigned | - |
| Otu34077 | 1 | k__Fungi(100);p__Ascomycota(100);p__Ascomycota_unclassified(100);p__Ascomycota_unclassified(100);p__Ascomycota_unclassified(100);p__Ascomycota_unclassified(100);p__Ascomycota_unclassified(100); | - | - | - | - | - | - | - | Unassigned | - |
| Otu34081 | 1 | k__Fungi(100);p__Ascomycota(100);p__Ascomycota_unclassified(100);p__Ascomycota_unclassified(100);p__Ascomycota_unclassified(100);p__Ascomycota_unclassified(100);p__Ascomycota_unclassified(100); | - | - | - | - | - | - | - | Unassigned | - |
| Otu34082 | 1 | k__Fungi(100);k__Fungi_unclassified(100);k__Fungi_unclassified(100);k__Fungi_unclassified(100);k__Fungi_unclassified(100);k__Fungi_unclassified(100);k__Fungi_unclassified(100); | - | - | - | - | - | - | - | Unassigned | - |
| Otu34086 | 1 | k__Fungi(100);k__Fungi_unclassified(100);k__Fungi_unclassified(100);k__Fungi_unclassified(100);k__Fungi_unclassified(100);k__Fungi_unclassified(100);k__Fungi_unclassified(100); | - | - | - | - | - | - | - | Unassigned | - |
| Otu34091 | 1 | k__Fungi(100);k__Fungi_unclassified(100);k__Fungi_unclassified(100);k__Fungi_unclassified(100);k__Fungi_unclassified(100);k__Fungi_unclassified(100);k__Fungi_unclassified(100); | - | - | - | - | - | - | - | Unassigned | - |
| Otu34092 | 1 | k__Fungi(100);p__Ascomycota(100);p__Ascomycota_unclassified(100);p__Ascomycota_unclassified(100);p__Ascomycota_unclassified(100);p__Ascomycota_unclassified(100);p__Ascomycota_unclassified(100); | - | - | - | - | - | - | - | Unassigned | - |
| Otu34093 | 1 | k__Fungi(100);p__Ascomycota(100);p__Ascomycota_unclassified(100);p__Ascomycota_unclassified(100);p__Ascomycota_unclassified(100);p__Ascomycota_unclassified(100);p__Ascomycota_unclassified(100); | - | - | - | - | - | - | - | Unassigned | - |
| Otu34094 | 1 | k__Fungi(100);p__Ascomycota(100);p__Ascomycota_unclassified(100);p__Ascomycota_unclassified(100);p__Ascomycota_unclassified(100);p__Ascomycota_unclassified(100);p__Ascomycota_unclassified(100); | - | - | - | - | - | - | - | Unassigned | - |
| Otu34095 | 1 | k__Fungi(100);p__Ascomycota(100);p__Ascomycota_unclassified(100);p__Ascomycota_unclassified(100);p__Ascomycota_unclassified(100);p__Ascomycota_unclassified(100);p__Ascomycota_unclassified(100); | - | - | - | - | - | - | - | Unassigned | - |
| Otu34096 | 1 | k__Fungi(100);k__Fungi_unclassified(100);k__Fungi_unclassified(100);k__Fungi_unclassified(100);k__Fungi_unclassified(100);k__Fungi_unclassified(100);k__Fungi_unclassified(100); | - | - | - | - | - | - | - | Unassigned | - |
| Otu34100 | 1 | k__Fungi(100);p__Ascomycota(100);p__Ascomycota_unclassified(100);p__Ascomycota_unclassified(100);p__Ascomycota_unclassified(100);p__Ascomycota_unclassified(100);p__Ascomycota_unclassified(100); | - | - | - | - | - | - | - | Unassigned | - |
| Otu34101 | 1 | k__Fungi(100);p__Ascomycota(100);p__Ascomycota_unclassified(100);p__Ascomycota_unclassified(100);p__Ascomycota_unclassified(100);p__Ascomycota_unclassified(100);p__Ascomycota_unclassified(100); | - | - | - | - | - | - | - | Unassigned | - |
| Otu34103 | 1 | k__Fungi(100);p__Ascomycota(100);p__Ascomycota_unclassified(100);p__Ascomycota_unclassified(100);p__Ascomycota_unclassified(100);p__Ascomycota_unclassified(100);p__Ascomycota_unclassified(100); | - | - | - | - | - | - | - | Unassigned | - |
| Otu34106 | 1 | k__Fungi(100);p__Ascomycota(100);p__Ascomycota_unclassified(100);p__Ascomycota_unclassified(100);p__Ascomycota_unclassified(100);p__Ascomycota_unclassified(100);p__Ascomycota_unclassified(100); | - | - | - | - | - | - | - | Unassigned | - |
| Otu34107 | 1 | k__Fungi(100);k__Fungi_unclassified(100);k__Fungi_unclassified(100);k__Fungi_unclassified(100);k__Fungi_unclassified(100);k__Fungi_unclassified(100);k__Fungi_unclassified(100); | - | - | - | - | - | - | - | Unassigned | - |
| Otu34108 | 1 | k__Fungi(100);p__Ascomycota(100);p__Ascomycota_unclassified(100);p__Ascomycota_unclassified(100);p__Ascomycota_unclassified(100);p__Ascomycota_unclassified(100);p__Ascomycota_unclassified(100); | - | - | - | - | - | - | - | Unassigned | - |
| Otu34109 | 1 | k__Fungi(100);k__Fungi_unclassified(100);k__Fungi_unclassified(100);k__Fungi_unclassified(100);k__Fungi_unclassified(100);k__Fungi_unclassified(100);k__Fungi_unclassified(100); | - | - | - | - | - | - | - | Unassigned | - |
| Otu34110 | 1 | k__Fungi(100);k__Fungi_unclassified(100);k__Fungi_unclassified(100);k__Fungi_unclassified(100);k__Fungi_unclassified(100);k__Fungi_unclassified(100);k__Fungi_unclassified(100); | - | - | - | - | - | - | - | Unassigned | - |
| Otu34111 | 1 | k__Fungi(100);k__Fungi_unclassified(100);k__Fungi_unclassified(100);k__Fungi_unclassified(100);k__Fungi_unclassified(100);k__Fungi_unclassified(100);k__Fungi_unclassified(100); | - | - | - | - | - | - | - | Unassigned | - |
| Otu34112 | 1 | k__Fungi(100);k__Fungi_unclassified(100);k__Fungi_unclassified(100);k__Fungi_unclassified(100);k__Fungi_unclassified(100);k__Fungi_unclassified(100);k__Fungi_unclassified(100); | - | - | - | - | - | - | - | Unassigned | - |
| Otu34113 | 1 | k__Fungi(100);k__Fungi_unclassified(100);k__Fungi_unclassified(100);k__Fungi_unclassified(100);k__Fungi_unclassified(100);k__Fungi_unclassified(100);k__Fungi_unclassified(100); | - | - | - | - | - | - | - | Unassigned | - |
| Otu34114 | 1 | k__Fungi(100);p__Ascomycota(100);p__Ascomycota_unclassified(100);p__Ascomycota_unclassified(100);p__Ascomycota_unclassified(100);p__Ascomycota_unclassified(100);p__Ascomycota_unclassified(100); | - | - | - | - | - | - | - | Unassigned | - |
| Otu34116 | 1 | k__Fungi(100);p__Ascomycota(100);p__Ascomycota_unclassified(100);p__Ascomycota_unclassified(100);p__Ascomycota_unclassified(100);p__Ascomycota_unclassified(100);p__Ascomycota_unclassified(100); | - | - | - | - | - | - | - | Unassigned | - |
| Otu34117 | 1 | k__Fungi(100);k__Fungi_unclassified(100);k__Fungi_unclassified(100);k__Fungi_unclassified(100);k__Fungi_unclassified(100);k__Fungi_unclassified(100);k__Fungi_unclassified(100); | - | - | - | - | - | - | - | Unassigned | - |
| Otu34118 | 1 | k__Fungi(100);k__Fungi_unclassified(100);k__Fungi_unclassified(100);k__Fungi_unclassified(100);k__Fungi_unclassified(100);k__Fungi_unclassified(100);k__Fungi_unclassified(100); | - | - | - | - | - | - | - | Unassigned | - |
| Otu34119 | 1 | k__Fungi(100);k__Fungi_unclassified(100);k__Fungi_unclassified(100);k__Fungi_unclassified(100);k__Fungi_unclassified(100);k__Fungi_unclassified(100);k__Fungi_unclassified(100); | - | - | - | - | - | - | - | Unassigned | - |
| Otu34120 | 1 | k__Fungi(100);p__Ascomycota(100);p__Ascomycota_unclassified(100);p__Ascomycota_unclassified(100);p__Ascomycota_unclassified(100);p__Ascomycota_unclassified(100);p__Ascomycota_unclassified(100); | - | - | - | - | - | - | - | Unassigned | - |
| Otu34121 | 1 | k__Fungi(100);p__Ascomycota(100);p__Ascomycota_unclassified(100);p__Ascomycota_unclassified(100);p__Ascomycota_unclassified(100);p__Ascomycota_unclassified(100);p__Ascomycota_unclassified(100); | - | - | - | - | - | - | - | Unassigned | - |
| Otu34123 | 1 | k__Fungi(100);p__Ascomycota(100);p__Ascomycota_unclassified(100);p__Ascomycota_unclassified(100);p__Ascomycota_unclassified(100);p__Ascomycota_unclassified(100);p__Ascomycota_unclassified(100); | - | - | - | - | - | - | - | Unassigned | - |
| Otu34124 | 1 | k__Fungi(100);k__Fungi_unclassified(100);k__Fungi_unclassified(100);k__Fungi_unclassified(100);k__Fungi_unclassified(100);k__Fungi_unclassified(100);k__Fungi_unclassified(100); | - | - | - | - | - | - | - | Unassigned | - |
| Otu34125 | 1 | k__Fungi(100);k__Fungi_unclassified(100);k__Fungi_unclassified(100);k__Fungi_unclassified(100);k__Fungi_unclassified(100);k__Fungi_unclassified(100);k__Fungi_unclassified(100); | - | - | - | - | - | - | - | Unassigned | - |
| Otu34126 | 1 | k__Fungi(100);p__Ascomycota(100);p__Ascomycota_unclassified(100);p__Ascomycota_unclassified(100);p__Ascomycota_unclassified(100);p__Ascomycota_unclassified(100);p__Ascomycota_unclassified(100); | - | - | - | - | - | - | - | Unassigned | - |
| Otu34127 | 1 | k__Fungi(100);k__Fungi_unclassified(100);k__Fungi_unclassified(100);k__Fungi_unclassified(100);k__Fungi_unclassified(100);k__Fungi_unclassified(100);k__Fungi_unclassified(100); | - | - | - | - | - | - | - | Unassigned | - |
| Otu34128 | 1 | k__Fungi(100);k__Fungi_unclassified(100);k__Fungi_unclassified(100);k__Fungi_unclassified(100);k__Fungi_unclassified(100);k__Fungi_unclassified(100);k__Fungi_unclassified(100); | - | - | - | - | - | - | - | Unassigned | - |
| Otu34129 | 1 | k__Fungi(100);p__Ascomycota(100);p__Ascomycota_unclassified(100);p__Ascomycota_unclassified(100);p__Ascomycota_unclassified(100);p__Ascomycota_unclassified(100);p__Ascomycota_unclassified(100); | - | - | - | - | - | - | - | Unassigned | - |
| Otu34131 | 1 | k__Fungi(100);k__Fungi_unclassified(100);k__Fungi_unclassified(100);k__Fungi_unclassified(100);k__Fungi_unclassified(100);k__Fungi_unclassified(100);k__Fungi_unclassified(100); | - | - | - | - | - | - | - | Unassigned | - |
| Otu34132 | 1 | k__Fungi(100);k__Fungi_unclassified(100);k__Fungi_unclassified(100);k__Fungi_unclassified(100);k__Fungi_unclassified(100);k__Fungi_unclassified(100);k__Fungi_unclassified(100); | - | - | - | - | - | - | - | Unassigned | - |

| | | | | | | | | | | | |
|---|---|---|---|---|---|---|---|---|---|---|---|
| Otu34133 | 1 | k__Fungi(100);p__Ascomycota(100);p__Ascomycota_unclassified(100);p__Ascomycota_unclassified(100);p__Ascomycota_unclassified(100);p__Ascomycota_unclassified(100);p__Ascomycota_unclassified(100); | - | - | - | - | - | - | - | Unassigned | - |
| Otu34135 | 1 | k__Fungi(100);p__Ascomycota(100);p__Ascomycota_unclassified(100);p__Ascomycota_unclassified(100);p__Ascomycota_unclassified(100);p__Ascomycota_unclassified(100);p__Ascomycota_unclassified(100); | - | - | - | - | - | - | - | Unassigned | - |
| Otu34136 | 1 | k__Fungi(100);p__Ascomycota(100);p__Ascomycota_unclassified(100);p__Ascomycota_unclassified(100);p__Ascomycota_unclassified(100);p__Ascomycota_unclassified(100);p__Ascomycota_unclassified(100); | - | - | - | - | - | - | - | Unassigned | - |
| Otu34138 | 1 | k__Fungi(100);p__Ascomycota(100);p__Ascomycota_unclassified(100);p__Ascomycota_unclassified(100);p__Ascomycota_unclassified(100);p__Ascomycota_unclassified(100);p__Ascomycota_unclassified(100); | - | - | - | - | - | - | - | Unassigned | - |
| Otu34139 | 1 | k__Fungi(100);k__Fungi_unclassified(100);k__Fungi_unclassified(100);k__Fungi_unclassified(100);k__Fungi_unclassified(100);k__Fungi_unclassified(100);k__Fungi_unclassified(100); | - | - | - | - | - | - | - | Unassigned | - |
| Otu34140 | 1 | k__Fungi(100);p__Ascomycota(100);p__Ascomycota_unclassified(100);p__Ascomycota_unclassified(100);p__Ascomycota_unclassified(100);p__Ascomycota_unclassified(100);p__Ascomycota_unclassified(100); | - | - | - | - | - | - | - | Unassigned | - |
| Otu34141 | 1 | k__Fungi(100);p__Ascomycota(100);p__Ascomycota_unclassified(100);p__Ascomycota_unclassified(100);p__Ascomycota_unclassified(100);p__Ascomycota_unclassified(100);p__Ascomycota_unclassified(100); | - | - | - | - | - | - | - | Unassigned | - |
| Otu34142 | 1 | k__Fungi(100);k__Fungi_unclassified(100);k__Fungi_unclassified(100);k__Fungi_unclassified(100);k__Fungi_unclassified(100);k__Fungi_unclassified(100);k__Fungi_unclassified(100); | - | - | - | - | - | - | - | Unassigned | - |
| Otu34143 | 1 | k__Fungi(100);k__Fungi_unclassified(100);k__Fungi_unclassified(100);k__Fungi_unclassified(100);k__Fungi_unclassified(100);k__Fungi_unclassified(100);k__Fungi_unclassified(100); | - | - | - | - | - | - | - | Unassigned | - |
| Otu34149 | 1 | k__Fungi(100);k__Fungi_unclassified(100);k__Fungi_unclassified(100);k__Fungi_unclassified(100);k__Fungi_unclassified(100);k__Fungi_unclassified(100);k__Fungi_unclassified(100); | - | - | - | - | - | - | - | Unassigned | - |
| Otu34151 | 1 | k__Fungi(100);p__Ascomycota(100);p__Ascomycota_unclassified(100);p__Ascomycota_unclassified(100);p__Ascomycota_unclassified(100);p__Ascomycota_unclassified(100);p__Ascomycota_unclassified(100); | - | - | - | - | - | - | - | Unassigned | - |
| Otu34152 | 1 | k__Fungi(100);p__Ascomycota(100);p__Ascomycota_unclassified(100);p__Ascomycota_unclassified(100);p__Ascomycota_unclassified(100);p__Ascomycota_unclassified(100);p__Ascomycota_unclassified(100); | - | - | - | - | - | - | - | Unassigned | - |
| Otu34158 | 1 | k__Fungi(100);k__Fungi_unclassified(100);k__Fungi_unclassified(100);k__Fungi_unclassified(100);k__Fungi_unclassified(100);k__Fungi_unclassified(100);k__Fungi_unclassified(100); | - | - | - | - | - | - | - | Unassigned | - |
| Otu34161 | 1 | k__Fungi(100);k__Fungi_unclassified(100);k__Fungi_unclassified(100);k__Fungi_unclassified(100);k__Fungi_unclassified(100);k__Fungi_unclassified(100);k__Fungi_unclassified(100); | - | - | - | - | - | - | - | Unassigned | - |
| Otu34164 | 1 | k__Fungi(100);k__Fungi_unclassified(100);k__Fungi_unclassified(100);k__Fungi_unclassified(100);k__Fungi_unclassified(100);k__Fungi_unclassified(100);k__Fungi_unclassified(100); | - | - | - | - | - | - | - | Unassigned | - |
| Otu34167 | 1 | k__Fungi(100);p__Ascomycota(100);p__Ascomycota_unclassified(100);p__Ascomycota_unclassified(100);p__Ascomycota_unclassified(100);p__Ascomycota_unclassified(100);p__Ascomycota_unclassified(100); | - | - | - | - | - | - | - | Unassigned | - |
| Otu34171 | 1 | k__Fungi(100);p__Ascomycota(100);p__Ascomycota_unclassified(100);p__Ascomycota_unclassified(100);p__Ascomycota_unclassified(100);p__Ascomycota_unclassified(100);p__Ascomycota_unclassified(100); | - | - | - | - | - | - | - | Unassigned | - |
| Otu34172 | 1 | k__Fungi(100);k__Fungi_unclassified(100);k__Fungi_unclassified(100);k__Fungi_unclassified(100);k__Fungi_unclassified(100);k__Fungi_unclassified(100);k__Fungi_unclassified(100); | - | - | - | - | - | - | - | Unassigned | - |
| Otu34174 | 1 | k__Fungi(100);k__Fungi_unclassified(100);k__Fungi_unclassified(100);k__Fungi_unclassified(100);k__Fungi_unclassified(100);k__Fungi_unclassified(100);k__Fungi_unclassified(100); | - | - | - | - | - | - | - | Unassigned | - |
| Otu34175 | 1 | k__Fungi(100);p__Ascomycota(100);p__Ascomycota_unclassified(100);p__Ascomycota_unclassified(100);p__Ascomycota_unclassified(100);p__Ascomycota_unclassified(100);p__Ascomycota_unclassified(100); | - | - | - | - | - | - | - | Unassigned | - |
| Otu34176 | 1 | k__Fungi(100);p__Ascomycota(100);p__Ascomycota_unclassified(100);p__Ascomycota_unclassified(100);p__Ascomycota_unclassified(100);p__Ascomycota_unclassified(100);p__Ascomycota_unclassified(100); | - | - | - | - | - | - | - | Unassigned | - |
| Otu34179 | 1 | k__Fungi(100);p__Ascomycota(100);p__Ascomycota_unclassified(100);p__Ascomycota_unclassified(100);p__Ascomycota_unclassified(100);p__Ascomycota_unclassified(100);p__Ascomycota_unclassified(100); | - | - | - | - | - | - | - | Unassigned | - |
| Otu34183 | 1 | k__Fungi(100);k__Fungi_unclassified(100);k__Fungi_unclassified(100);k__Fungi_unclassified(100);k__Fungi_unclassified(100);k__Fungi_unclassified(100);k__Fungi_unclassified(100); | - | - | - | - | - | - | - | Unassigned | - |
| Otu34184 | 1 | k__Fungi(100);p__Ascomycota(100);p__Ascomycota_unclassified(100);p__Ascomycota_unclassified(100);p__Ascomycota_unclassified(100);p__Ascomycota_unclassified(100);p__Ascomycota_unclassified(100); | - | - | - | - | - | - | - | Unassigned | - |
| Otu34187 | 1 | k__Fungi(100);k__Fungi_unclassified(100);k__Fungi_unclassified(100);k__Fungi_unclassified(100);k__Fungi_unclassified(100);k__Fungi_unclassified(100);k__Fungi_unclassified(100); | - | - | - | - | - | - | - | Unassigned | - |
| Otu34189 | 1 | k__Fungi(100);p__Ascomycota(100);p__Ascomycota_unclassified(100);p__Ascomycota_unclassified(100);p__Ascomycota_unclassified(100);p__Ascomycota_unclassified(100);p__Ascomycota_unclassified(100); | - | - | - | - | - | - | - | Unassigned | - |
| Otu34190 | 1 | k__Fungi(100);k__Fungi_unclassified(100);k__Fungi_unclassified(100);k__Fungi_unclassified(100);k__Fungi_unclassified(100);k__Fungi_unclassified(100);k__Fungi_unclassified(100); | - | - | - | - | - | - | - | Unassigned | - |
| Otu34191 | 1 | k__Fungi(100);k__Fungi_unclassified(100);k__Fungi_unclassified(100);k__Fungi_unclassified(100);k__Fungi_unclassified(100);k__Fungi_unclassified(100);k__Fungi_unclassified(100); | - | - | - | - | - | - | - | Unassigned | - |
| Otu34192 | 1 | k__Fungi(100);p__Ascomycota(100);p__Ascomycota_unclassified(100);p__Ascomycota_unclassified(100);p__Ascomycota_unclassified(100);p__Ascomycota_unclassified(100);p__Ascomycota_unclassified(100); | - | - | - | - | - | - | - | Unassigned | - |
| Otu34193 | 1 | k__Fungi(100);k__Fungi_unclassified(100);k__Fungi_unclassified(100);k__Fungi_unclassified(100);k__Fungi_unclassified(100);k__Fungi_unclassified(100);k__Fungi_unclassified(100); | - | - | - | - | - | - | - | Unassigned | - |
| Otu34194 | 1 | k__Fungi(100);p__Ascomycota(100);p__Ascomycota_unclassified(100);p__Ascomycota_unclassified(100);p__Ascomycota_unclassified(100);p__Ascomycota_unclassified(100);p__Ascomycota_unclassified(100); | - | - | - | - | - | - | - | Unassigned | - |
| Otu34197 | 1 | k__Fungi(100);p__Ascomycota(100);p__Ascomycota_unclassified(100);p__Ascomycota_unclassified(100);p__Ascomycota_unclassified(100);p__Ascomycota_unclassified(100);p__Ascomycota_unclassified(100); | - | - | - | - | - | - | - | Unassigned | - |
| Otu34204 | 1 | k__Fungi(100);k__Fungi_unclassified(100);k__Fungi_unclassified(100);k__Fungi_unclassified(100);k__Fungi_unclassified(100);k__Fungi_unclassified(100);k__Fungi_unclassified(100); | - | - | - | - | - | - | - | Unassigned | - |
| Otu34209 | 1 | k__Fungi(100);k__Fungi_unclassified(100);k__Fungi_unclassified(100);k__Fungi_unclassified(100);k__Fungi_unclassified(100);k__Fungi_unclassified(100);k__Fungi_unclassified(100); | - | - | - | - | - | - | - | Unassigned | - |
| Otu34214 | 1 | k__Fungi(100);p__Ascomycota(100);c__Dothideomycetes(100);o__Pleosporales(100);o__Pleosporales_unclassified(100);o__Pleosporales_unclassified(100);o__Pleosporales_unclassified(100); | - | - | - | - | - | - | - | Unassigned | - |
| Otu34215 | 1 | k__Fungi(100);p__Ascomycota(100);p__Ascomycota_unclassified(100);p__Ascomycota_unclassified(100);p__Ascomycota_unclassified(100);p__Ascomycota_unclassified(100);p__Ascomycota_unclassified(100); | - | - | - | - | - | - | - | Unassigned | - |
| Otu34217 | 1 | k__Fungi(100);k__Fungi_unclassified(100);k__Fungi_unclassified(100);k__Fungi_unclassified(100);k__Fungi_unclassified(100);k__Fungi_unclassified(100);k__Fungi_unclassified(100); | - | - | - | - | - | - | - | Unassigned | - |
| Otu34218 | 1 | k__Fungi(100);k__Fungi_unclassified(100);k__Fungi_unclassified(100);k__Fungi_unclassified(100);k__Fungi_unclassified(100);k__Fungi_unclassified(100);k__Fungi_unclassified(100); | - | - | - | - | - | - | - | Unassigned | - |
| Otu34219 | 1 | k__Fungi(100);p__Ascomycota(100);p__Ascomycota_unclassified(100);p__Ascomycota_unclassified(100);p__Ascomycota_unclassified(100);p__Ascomycota_unclassified(100);p__Ascomycota_unclassified(100); | - | - | - | - | - | - | - | Unassigned | - |
| Otu34221 | 1 | k__Fungi(100);k__Fungi_unclassified(100);k__Fungi_unclassified(100);k__Fungi_unclassified(100);k__Fungi_unclassified(100);k__Fungi_unclassified(100);k__Fungi_unclassified(100); | - | - | - | - | - | - | - | Unassigned | - |
| Otu34243 | 1 | k__Fungi(100);k__Fungi_unclassified(100);k__Fungi_unclassified(100);k__Fungi_unclassified(100);k__Fungi_unclassified(100);k__Fungi_unclassified(100);k__Fungi_unclassified(100); | - | - | - | - | - | - | - | Unassigned | - |
| Otu34249 | 1 | k__Fungi(100);p__Ascomycota(100);p__Ascomycota_unclassified(100);p__Ascomycota_unclassified(100);p__Ascomycota_unclassified(100);p__Ascomycota_unclassified(100);p__Ascomycota_unclassified(100); | - | - | - | - | - | - | - | Unassigned | - |
| Otu34250 | 1 | k__Fungi(100);k__Fungi_unclassified(100);k__Fungi_unclassified(100);k__Fungi_unclassified(100);k__Fungi_unclassified(100);k__Fungi_unclassified(100);k__Fungi_unclassified(100); | - | - | - | - | - | - | - | Unassigned | - |
| Otu34253 | 1 | k__Fungi(100);p__Ascomycota(100);p__Ascomycota_unclassified(100);p__Ascomycota_unclassified(100);p__Ascomycota_unclassified(100);p__Ascomycota_unclassified(100);p__Ascomycota_unclassified(100); | - | - | - | - | - | - | - | Unassigned | - |
| Otu34255 | 1 | k__Fungi(100);k__Fungi_unclassified(100);k__Fungi_unclassified(100);k__Fungi_unclassified(100);k__Fungi_unclassified(100);k__Fungi_unclassified(100);k__Fungi_unclassified(100); | - | - | - | - | - | - | - | Unassigned | - |
| Otu34258 | 1 | k__Fungi(100);k__Fungi_unclassified(100);k__Fungi_unclassified(100);k__Fungi_unclassified(100);k__Fungi_unclassified(100);k__Fungi_unclassified(100);k__Fungi_unclassified(100); | - | - | - | - | - | - | - | Unassigned | - |
| Otu34260 | 1 | k__Fungi(100);p__Ascomycota(100);p__Ascomycota_unclassified(100);p__Ascomycota_unclassified(100);p__Ascomycota_unclassified(100);p__Ascomycota_unclassified(100);p__Ascomycota_unclassified(100); | - | - | - | - | - | - | - | Unassigned | - |
| Otu34277 | 1 | k__Fungi(100);k__Fungi_unclassified(100);k__Fungi_unclassified(100);k__Fungi_unclassified(100);k__Fungi_unclassified(100);k__Fungi_unclassified(100);k__Fungi_unclassified(100); | - | - | - | - | - | - | - | Unassigned | - |
| Otu34283 | 1 | k__Fungi(100);k__Fungi_unclassified(100);k__Fungi_unclassified(100);k__Fungi_unclassified(100);k__Fungi_unclassified(100);k__Fungi_unclassified(100);k__Fungi_unclassified(100); | - | - | - | - | - | - | - | Unassigned | - |
| Otu34292 | 1 | k__Fungi(100);k__Fungi_unclassified(100);k__Fungi_unclassified(100);k__Fungi_unclassified(100);k__Fungi_unclassified(100);k__Fungi_unclassified(100);k__Fungi_unclassified(100); | - | - | - | - | - | - | - | Unassigned | - |
| Otu34293 | 1 | k__Fungi(100);p__Ascomycota(100);p__Ascomycota_unclassified(100);p__Ascomycota_unclassified(100);p__Ascomycota_unclassified(100);p__Ascomycota_unclassified(100);p__Ascomycota_unclassified(100); | - | - | - | - | - | - | - | Unassigned | - |
| Otu34294 | 1 | k__Fungi(100);k__Fungi_unclassified(100);k__Fungi_unclassified(100);k__Fungi_unclassified(100);k__Fungi_unclassified(100);k__Fungi_unclassified(100);k__Fungi_unclassified(100); | - | - | - | - | - | - | - | Unassigned | - |
| Otu34295 | 1 | k__Fungi(100);k__Fungi_unclassified(100);k__Fungi_unclassified(100);k__Fungi_unclassified(100);k__Fungi_unclassified(100);k__Fungi_unclassified(100);k__Fungi_unclassified(100); | - | - | - | - | - | - | - | Unassigned | - |
| Otu34296 | 1 | k__Fungi(100);k__Fungi_unclassified(100);k__Fungi_unclassified(100);k__Fungi_unclassified(100);k__Fungi_unclassified(100);k__Fungi_unclassified(100);k__Fungi_unclassified(100); | - | - | - | - | - | - | - | Unassigned | - |
| Otu34297 | 1 | k__Fungi(100);k__Fungi_unclassified(100);k__Fungi_unclassified(100);k__Fungi_unclassified(100);k__Fungi_unclassified(100);k__Fungi_unclassified(100);k__Fungi_unclassified(100); | - | - | - | - | - | - | - | Unassigned | - |
| Otu34298 | 1 | k__Fungi(100);p__Ascomycota(100);p__Ascomycota_unclassified(100);p__Ascomycota_unclassified(100);p__Ascomycota_unclassified(100);p__Ascomycota_unclassified(100);p__Ascomycota_unclassified(100); | - | - | - | - | - | - | - | Unassigned | - |
| Otu34299 | 1 | k__Fungi(100);k__Fungi_unclassified(100);k__Fungi_unclassified(100);k__Fungi_unclassified(100);k__Fungi_unclassified(100);k__Fungi_unclassified(100);k__Fungi_unclassified(100); | - | - | - | - | - | - | - | Unassigned | - |
| Otu34300 | 1 | k__Fungi(100);p__Ascomycota(100);p__Ascomycota_unclassified(100);p__Ascomycota_unclassified(100);p__Ascomycota_unclassified(100);p__Ascomycota_unclassified(100);p__Ascomycota_unclassified(100); | - | - | - | - | - | - | - | Unassigned | - |
| Otu34301 | 1 | k__Fungi(100);k__Fungi_unclassified(100);k__Fungi_unclassified(100);k__Fungi_unclassified(100);k__Fungi_unclassified(100);k__Fungi_unclassified(100);k__Fungi_unclassified(100); | - | - | - | - | - | - | - | Unassigned | - |
| Otu34302 | 1 | k__Fungi(100);k__Fungi_unclassified(100);k__Fungi_unclassified(100);k__Fungi_unclassified(100);k__Fungi_unclassified(100);k__Fungi_unclassified(100);k__Fungi_unclassified(100); | - | - | - | - | - | - | - | Unassigned | - |
| Otu34303 | 1 | k__Fungi(100);p__Ascomycota(100);p__Ascomycota_unclassified(100);p__Ascomycota_unclassified(100);p__Ascomycota_unclassified(100);p__Ascomycota_unclassified(100);p__Ascomycota_unclassified(100); | - | - | - | - | - | - | - | Unassigned | - |
| Otu34304 | 1 | k__Fungi(100);p__Ascomycota(100);p__Ascomycota_unclassified(100);p__Ascomycota_unclassified(100);p__Ascomycota_unclassified(100);p__Ascomycota_unclassified(100);p__Ascomycota_unclassified(100); | - | - | - | - | - | - | - | Unassigned | - |
| Otu34305 | 1 | k__Fungi(100);k__Fungi_unclassified(100);k__Fungi_unclassified(100);k__Fungi_unclassified(100);k__Fungi_unclassified(100);k__Fungi_unclassified(100);k__Fungi_unclassified(100); | - | - | - | - | - | - | - | Unassigned | - |
| Otu34307 | 1 | k__Fungi(100);k__Fungi_unclassified(100);k__Fungi_unclassified(100);k__Fungi_unclassified(100);k__Fungi_unclassified(100);k__Fungi_unclassified(100);k__Fungi_unclassified(100); | - | - | - | - | - | - | - | Unassigned | - |
| Otu34311 | 1 | k__Fungi(100);p__Ascomycota(100);p__Ascomycota_unclassified(100);p__Ascomycota_unclassified(100);p__Ascomycota_unclassified(100);p__Ascomycota_unclassified(100);p__Ascomycota_unclassified(100); | - | - | - | - | - | - | - | Unassigned | - |
| Otu34312 | 1 | k__Fungi(100);k__Fungi_unclassified(100);k__Fungi_unclassified(100);k__Fungi_unclassified(100);k__Fungi_unclassified(100);k__Fungi_unclassified(100);k__Fungi_unclassified(100); | - | - | - | - | - | - | - | Unassigned | - |
| Otu34314 | 1 | k__Fungi(100);p__Ascomycota(100);p__Ascomycota_unclassified(100);p__Ascomycota_unclassified(100);p__Ascomycota_unclassified(100);p__Ascomycota_unclassified(100);p__Ascomycota_unclassified(100); | - | - | - | - | - | - | - | Unassigned | - |
| Otu34318 | 1 | k__Fungi(100);k__Fungi_unclassified(100);k__Fungi_unclassified(100);k__Fungi_unclassified(100);k__Fungi_unclassified(100);k__Fungi_unclassified(100);k__Fungi_unclassified(100); | - | - | - | - | - | - | - | Unassigned | - |
| Otu34319 | 1 | k__Fungi(100);k__Fungi_unclassified(100);k__Fungi_unclassified(100);k__Fungi_unclassified(100);k__Fungi_unclassified(100);k__Fungi_unclassified(100);k__Fungi_unclassified(100); | - | - | - | - | - | - | - | Unassigned | - |
| Otu34320 | 1 | k__Fungi(100);k__Fungi_unclassified(100);k__Fungi_unclassified(100);k__Fungi_unclassified(100);k__Fungi_unclassified(100);k__Fungi_unclassified(100);k__Fungi_unclassified(100); | - | - | - | - | - | - | - | Unassigned | - |
| Otu34321 | 1 | k__Fungi(100);k__Fungi_unclassified(100);k__Fungi_unclassified(100);k__Fungi_unclassified(100);k__Fungi_unclassified(100);k__Fungi_unclassified(100);k__Fungi_unclassified(100); | - | - | - | - | - | - | - | Unassigned | - |
| Otu34322 | 1 | k__Fungi(100);k__Fungi_unclassified(100);k__Fungi_unclassified(100);k__Fungi_unclassified(100);k__Fungi_unclassified(100);k__Fungi_unclassified(100);k__Fungi_unclassified(100); | - | - | - | - | - | - | - | Unassigned | - |
| Otu34323 | 1 | k__Fungi(100);p__Ascomycota(100);p__Ascomycota_unclassified(100);p__Ascomycota_unclassified(100);p__Ascomycota_unclassified(100);p__Ascomycota_unclassified(100);p__Ascomycota_unclassified(100); | - | - | - | - | - | - | - | Unassigned | - |
| Otu34331 | 1 | k__Fungi(100);k__Fungi_unclassified(100);k__Fungi_unclassified(100);k__Fungi_unclassified(100);k__Fungi_unclassified(100);k__Fungi_unclassified(100);k__Fungi_unclassified(100); | - | - | - | - | - | - | - | Unassigned | - |
| Otu34332 | 1 | k__Fungi(100);k__Fungi_unclassified(100);k__Fungi_unclassified(100);k__Fungi_unclassified(100);k__Fungi_unclassified(100);k__Fungi_unclassified(100);k__Fungi_unclassified(100); | - | - | - | - | - | - | - | Unassigned | - |
| Otu34333 | 1 | k__Fungi(100);k__Fungi_unclassified(100);k__Fungi_unclassified(100);k__Fungi_unclassified(100);k__Fungi_unclassified(100);k__Fungi_unclassified(100);k__Fungi_unclassified(100); | - | - | - | - | - | - | - | Unassigned | - |
| Otu34335 | 1 | k__Fungi(100);k__Fungi_unclassified(100);k__Fungi_unclassified(100);k__Fungi_unclassified(100);k__Fungi_unclassified(100);k__Fungi_unclassified(100);k__Fungi_unclassified(100); | - | - | - | - | - | - | - | Unassigned | - |
| Otu34336 | 1 | k__Fungi(100);k__Fungi_unclassified(100);k__Fungi_unclassified(100);k__Fungi_unclassified(100);k__Fungi_unclassified(100);k__Fungi_unclassified(100);k__Fungi_unclassified(100); | - | - | - | - | - | - | - | Unassigned | - |
| Otu34337 | 1 | k__Fungi(100);k__Fungi_unclassified(100);k__Fungi_unclassified(100);k__Fungi_unclassified(100);k__Fungi_unclassified(100);k__Fungi_unclassified(100);k__Fungi_unclassified(100); | - | - | - | - | - | - | - | Unassigned | - |
| Otu34339 | 1 | k__Fungi(100);k__Fungi_unclassified(100);k__Fungi_unclassified(100);k__Fungi_unclassified(100);k__Fungi_unclassified(100);k__Fungi_unclassified(100);k__Fungi_unclassified(100); | - | - | - | - | - | - | - | Unassigned | - |
| Otu34343 | 1 | k__Fungi(100);p__Ascomycota(100);p__Ascomycota_unclassified(100);p__Ascomycota_unclassified(100);p__Ascomycota_unclassified(100);p__Ascomycota_unclassified(100);p__Ascomycota_unclassified(100); | - | - | - | - | - | - | - | Unassigned | - |
| Otu34348 | 1 | k__Fungi(100);p__Ascomycota(100);p__Ascomycota_unclassified(100);p__Ascomycota_unclassified(100);p__Ascomycota_unclassified(100);p__Ascomycota_unclassified(100);p__Ascomycota_unclassified(100); | - | - | - | - | - | - | - | Unassigned | - |
| Otu34350 | 1 | k__Fungi(100);p__Ascomycota(100);p__Ascomycota_unclassified(100);p__Ascomycota_unclassified(100);p__Ascomycota_unclassified(100);p__Ascomycota_unclassified(100);p__Ascomycota_unclassified(100); | - | - | - | - | - | - | - | Unassigned | - |
| Otu34357 | 1 | k__Fungi(100);k__Fungi_unclassified(100);k__Fungi_unclassified(100);k__Fungi_unclassified(100);k__Fungi_unclassified(100);k__Fungi_unclassified(100);k__Fungi_unclassified(100); | - | - | - | - | - | - | - | Unassigned | - |
| Otu34362 | 1 | k__Fungi(100);k__Fungi_unclassified(100);k__Fungi_unclassified(100);k__Fungi_unclassified(100);k__Fungi_unclassified(100);k__Fungi_unclassified(100);k__Fungi_unclassified(100); | - | - | - | - | - | - | - | Unassigned | - |
| Otu34364 | 1 | k__Fungi(100);p__Ascomycota(100);p__Ascomycota_unclassified(100);p__Ascomycota_unclassified(100);p__Ascomycota_unclassified(100);p__Ascomycota_unclassified(100);p__Ascomycota_unclassified(100); | - | - | - | - | - | - | - | Unassigned | - |
| Otu34370 | 1 | k__Fungi(100);k__Fungi_unclassified(100);k__Fungi_unclassified(100);k__Fungi_unclassified(100);k__Fungi_unclassified(100);k__Fungi_unclassified(100);k__Fungi_unclassified(100); | - | - | - | - | - | - | - | Unassigned | - |
| Otu34372 | 1 | k__Fungi(100);k__Fungi_unclassified(100);k__Fungi_unclassified(100);k__Fungi_unclassified(100);k__Fungi_unclassified(100);k__Fungi_unclassified(100);k__Fungi_unclassified(100); | - | - | - | - | - | - | - | Unassigned | - |
| Otu34381 | 1 | k__Fungi(100);k__Fungi_unclassified(100);k__Fungi_unclassified(100);k__Fungi_unclassified(100);k__Fungi_unclassified(100);k__Fungi_unclassified(100);k__Fungi_unclassified(100); | - | - | - | - | - | - | - | Unassigned | - |
| Otu34388 | 1 | k__Fungi(100);p__Ascomycota(100);p__Ascomycota_unclassified(100);p__Ascomycota_unclassified(100);p__Ascomycota_unclassified(100);p__Ascomycota_unclassified(100);p__Ascomycota_unclassified(100); | - | - | - | - | - | - | - | Unassigned | - |
| Otu34398 | 1 | k__Fungi(100);p__Ascomycota(100);p__Ascomycota_unclassified(100);p__Ascomycota_unclassified(100);p__Ascomycota_unclassified(100);p__Ascomycota_unclassified(100);p__Ascomycota_unclassified(100); | - | - | - | - | - | - | - | Unassigned | - |
| Otu34405 | 1 | k__Fungi(100);p__Ascomycota(100);p__Ascomycota_unclassified(100);p__Ascomycota_unclassified(100);p__Ascomycota_unclassified(100);p__Ascomycota_unclassified(100);p__Ascomycota_unclassified(100); | - | - | - | - | - | - | - | Unassigned | - |
| Otu34411 | 1 | k__Fungi(100);p__Ascomycota(100);p__Ascomycota_unclassified(100);p__Ascomycota_unclassified(100);p__Ascomycota_unclassified(100);p__Ascomycota_unclassified(100);p__Ascomycota_unclassified(100); | - | - | - | - | - | - | - | Unassigned | - |
| Otu34412 | 1 | k__Fungi(100);k__Fungi_unclassified(100);k__Fungi_unclassified(100);k__Fungi_unclassified(100);k__Fungi_unclassified(100);k__Fungi_unclassified(100);k__Fungi_unclassified(100); | - | - | - | - | - | - | - | Unassigned | - |
| Otu34416 | 1 | k__Fungi(100);p__Ascomycota(100);p__Ascomycota_unclassified(100);p__Ascomycota_unclassified(100);p__Ascomycota_unclassified(100);p__Ascomycota_unclassified(100);p__Ascomycota_unclassified(100); | - | - | - | - | - | - | - | Unassigned | - |
| Otu34417 | 1 | k__Fungi(100);k__Fungi_unclassified(100);k__Fungi_unclassified(100);k__Fungi_unclassified(100);k__Fungi_unclassified(100);k__Fungi_unclassified(100);k__Fungi_unclassified(100); | - | - | - | - | - | - | - | Unassigned | - |
| Otu34418 | 1 | k__Fungi(100);k__Fungi_unclassified(100);k__Fungi_unclassified(100);k__Fungi_unclassified(100);k__Fungi_unclassified(100);k__Fungi_unclassified(100);k__Fungi_unclassified(100); | - | - | - | - | - | - | - | Unassigned | - |
| Otu34422 | 1 | k__Fungi(100);k__Fungi_unclassified(100);k__Fungi_unclassified(100);k__Fungi_unclassified(100);k__Fungi_unclassified(100);k__Fungi_unclassified(100);k__Fungi_unclassified(100); | - | - | - | - | - | - | - | Unassigned | - |
| Otu34424 | 1 | k__Fungi(100);p__Ascomycota(100);p__Ascomycota_unclassified(100);p__Ascomycota_unclassified(100);p__Ascomycota_unclassified(100);p__Ascomycota_unclassified(100);p__Ascomycota_unclassified(100); | - | - | - | - | - | - | - | Unassigned | - |
| Otu34425 | 1 | k__Fungi(100);p__Ascomycota(100);p__Ascomycota_unclassified(100);p__Ascomycota_unclassified(100);p__Ascomycota_unclassified(100);p__Ascomycota_unclassified(100);p__Ascomycota_unclassified(100); | - | - | - | - | - | - | - | Unassigned | - |
| Otu34426 | 1 | k__Fungi(100);k__Fungi_unclassified(100);k__Fungi_unclassified(100);k__Fungi_unclassified(100);k__Fungi_unclassified(100);k__Fungi_unclassified(100);k__Fungi_unclassified(100); | - | - | - | - | - | - | - | Unassigned | - |
| Otu34430 | 1 | k__Fungi(100);k__Fungi_unclassified(100);k__Fungi_unclassified(100);k__Fungi_unclassified(100);k__Fungi_unclassified(100);k__Fungi_unclassified(100);k__Fungi_unclassified(100); | - | - | - | - | - | - | - | Unassigned | - |
| Otu34431 | 1 | k__Fungi(100);k__Fungi_unclassified(100);k__Fungi_unclassified(100);k__Fungi_unclassified(100);k__Fungi_unclassified(100);k__Fungi_unclassified(100);k__Fungi_unclassified(100); | - | - | - | - | - | - | - | Unassigned | - |
| Otu34436 | 1 | k__Fungi(100);k__Fungi_unclassified(100);k__Fungi_unclassified(100);k__Fungi_unclassified(100);k__Fungi_unclassified(100);k__Fungi_unclassified(100);k__Fungi_unclassified(100); | - | - | - | - | - | - | - | Unassigned | - |
| Otu34437 | 1 | k__Fungi(100);p__Ascomycota(100);p__Ascomycota_unclassified(100);p__Ascomycota_unclassified(100);p__Ascomycota_unclassified(100);p__Ascomycota_unclassified(100);p__Ascomycota_unclassified(100); | - | - | - | - | - | - | - | Unassigned | - |
| Otu34438 | 1 | k__Fungi(100);p__Ascomycota(100);p__Ascomycota_unclassified(100);p__Ascomycota_unclassified(100);p__Ascomycota_unclassified(100);p__Ascomycota_unclassified(100);p__Ascomycota_unclassified(100); | - | - | - | - | - | - | - | Unassigned | - |
| Otu34442 | 1 | k__Fungi(100);k__Fungi_unclassified(100);k__Fungi_unclassified(100);k__Fungi_unclassified(100);k__Fungi_unclassified(100);k__Fungi_unclassified(100);k__Fungi_unclassified(100); | - | - | - | - | - | - | - | Unassigned | - |
| Otu34447 | 1 | k__Fungi(100);p__Ascomycota(100);p__Ascomycota_unclassified(100);p__Ascomycota_unclassified(100);p__Ascomycota_unclassified(100);p__Ascomycota_unclassified(100);p__Ascomycota_unclassified(100); | - | - | - | - | - | - | - | Unassigned | - |
| Otu34448 | 1 | k__Fungi(100);p__Ascomycota(100);p__Ascomycota_unclassified(100);p__Ascomycota_unclassified(100);p__Ascomycota_unclassified(100);p__Ascomycota_unclassified(100);p__Ascomycota_unclassified(100); | - | - | - | - | - | - | - | Unassigned | - |
| Otu34449 | 1 | k__Fungi(100);k__Fungi_unclassified(100);k__Fungi_unclassified(100);k__Fungi_unclassified(100);k__Fungi_unclassified(100);k__Fungi_unclassified(100);k__Fungi_unclassified(100); | - | - | - | - | - | - | - | Unassigned | - |
| Otu34450 | 1 | k__Fungi(100);p__Ascomycota(100);p__Ascomycota_unclassified(100);p__Ascomycota_unclassified(100);p__Ascomycota_unclassified(100);p__Ascomycota_unclassified(100);p__Ascomycota_unclassified(100); | - | - | - | - | - | - | - | Unassigned | - |
| Otu34451 | 1 | k__Fungi(100);k__Fungi_unclassified(100);k__Fungi_unclassified(100);k__Fungi_unclassified(100);k__Fungi_unclassified(100);k__Fungi_unclassified(100);k__Fungi_unclassified(100); | - | - | - | - | - | - | - | Unassigned | - |
| Otu34452 | 1 | k__Fungi(100);k__Fungi_unclassified(100);k__Fungi_unclassified(100);k__Fungi_unclassified(100);k__Fungi_unclassified(100);k__Fungi_unclassified(100);k__Fungi_unclassified(100); | - | - | - | - | - | - | - | Unassigned | - |
| Otu34453 | 1 | k__Fungi(100);k__Fungi_unclassified(100);k__Fungi_unclassified(100);k__Fungi_unclassified(100);k__Fungi_unclassified(100);k__Fungi_unclassified(100);k__Fungi_unclassified(100); | - | - | - | - | - | - | - | Unassigned | - |
| Otu34455 | 1 | k__Fungi(100);p__Ascomycota(100);p__Ascomycota_unclassified(100);p__Ascomycota_unclassified(100);p__Ascomycota_unclassified(100);p__Ascomycota_unclassified(100);p__Ascomycota_unclassified(100); | - | - | - | - | - | - | - | Unassigned | - |
| Otu34456 | 1 | k__Fungi(100);p__Ascomycota(100);p__Ascomycota_unclassified(100);p__Ascomycota_unclassified(100);p__Ascomycota_unclassified(100);p__Ascomycota_unclassified(100);p__Ascomycota_unclassified(100); | - | - | - | - | - | - | - | Unassigned | - |
| Otu34457 | 1 | k__Fungi(100);p__Ascomycota(100);p__Ascomycota_unclassified(100);p__Ascomycota_unclassified(100);p__Ascomycota_unclassified(100);p__Ascomycota_unclassified(100);p__Ascomycota_unclassified(100); | - | - | - | - | - | - | - | Unassigned | - |
| Otu34458 | 1 | k__Fungi(100);k__Fungi_unclassified(100);k__Fungi_unclassified(100);k__Fungi_unclassified(100);k__Fungi_unclassified(100);k__Fungi_unclassified(100);k__Fungi_unclassified(100); | - | - | - | - | - | - | - | Unassigned | - |
| Otu34462 | 1 | k__Fungi(100);p__Ascomycota(100);p__Ascomycota_unclassified(100);p__Ascomycota_unclassified(100);p__Ascomycota_unclassified(100);p__Ascomycota_unclassified(100);p__Ascomycota_unclassified(100); | - | - | - | - | - | - | - | Unassigned | - |
| Otu34463 | 1 | k__Fungi(100);k__Fungi_unclassified(100);k__Fungi_unclassified(100);k__Fungi_unclassified(100);k__Fungi_unclassified(100);k__Fungi_unclassified(100);k__Fungi_unclassified(100); | - | - | - | - | - | - | - | Unassigned | - |
| Otu34469 | 1 | k__Fungi(100);k__Fungi_unclassified(100);k__Fungi_unclassified(100);k__Fungi_unclassified(100);k__Fungi_unclassified(100);k__Fungi_unclassified(100);k__Fungi_unclassified(100); | - | - | - | - | - | - | - | Unassigned | - |
| Otu34470 | 1 | k__Fungi(100);k__Fungi_unclassified(100);k__Fungi_unclassified(100);k__Fungi_unclassified(100);k__Fungi_unclassified(100);k__Fungi_unclassified(100);k__Fungi_unclassified(100); | - | - | - | - | - | - | - | Unassigned | - |
| Otu34471 | 1 | k__Fungi(100);p__Ascomycota(100);p__Ascomycota_unclassified(100);p__Ascomycota_unclassified(100);p__Ascomycota_unclassified(100);p__Ascomycota_unclassified(100);p__Ascomycota_unclassified(100); | - | - | - | - | - | - | - | Unassigned | - |
| Otu34472 | 1 | k__Fungi(100);p__Ascomycota(100);p__Ascomycota_unclassified(100);p__Ascomycota_unclassified(100);p__Ascomycota_unclassified(100);p__Ascomycota_unclassified(100);p__Ascomycota_unclassified(100); | - | - | - | - | - | - | - | Unassigned | - |
| Otu34473 | 1 | k__Fungi(100);k__Fungi_unclassified(100);k__Fungi_unclassified(100);k__Fungi_unclassified(100);k__Fungi_unclassified(100);k__Fungi_unclassified(100);k__Fungi_unclassified(100); | - | - | - | - | - | - | - | Unassigned | - |
| Otu34476 | 1 | k__Fungi(100);k__Fungi_unclassified(100);k__Fungi_unclassified(100);k__Fungi_unclassified(100);k__Fungi_unclassified(100);k__Fungi_unclassified(100);k__Fungi_unclassified(100); | - | - | - | - | - | - | - | Unassigned | - |
| Otu34478 | 1 | k__Fungi(100);k__Fungi_unclassified(100);k__Fungi_unclassified(100);k__Fungi_unclassified(100);k__Fungi_unclassified(100);k__Fungi_unclassified(100);k__Fungi_unclassified(100); | - | - | - | - | - | - | - | Unassigned | - |
| Otu34479 | 1 | k__Fungi(100);p__Ascomycota(100);p__Ascomycota_unclassified(100);p__Ascomycota_unclassified(100);p__Ascomycota_unclassified(100);p__Ascomycota_unclassified(100);p__Ascomycota_unclassified(100); | - | - | - | - | - | - | - | Unassigned | - |
| Otu34480 | 1 | k__Fungi(100);k__Fungi_unclassified(100);k__Fungi_unclassified(100);k__Fungi_unclassified(100);k__Fungi_unclassified(100);k__Fungi_unclassified(100);k__Fungi_unclassified(100); | - | - | - | - | - | - | - | Unassigned | - |
| Otu34481 | 1 | k__Fungi(100);k__Fungi_unclassified(100);k__Fungi_unclassified(100);k__Fungi_unclassified(100);k__Fungi_unclassified(100);k__Fungi_unclassified(100);k__Fungi_unclassified(100); | - | - | - | - | - | - | - | Unassigned | - |
| Otu34482 | 1 | k__Fungi(100);p__Ascomycota(100);p__Ascomycota_unclassified(100);p__Ascomycota_unclassified(100);p__Ascomycota_unclassified(100);p__Ascomycota_unclassified(100);p__Ascomycota_unclassified(100); | - | - | - | - | - | - | - | Unassigned | - |
| Otu34487 | 1 | k__Fungi(100);p__Ascomycota(100);p__Ascomycota_unclassified(100);p__Ascomycota_unclassified(100);p__Ascomycota_unclassified(100);p__Ascomycota_unclassified(100);p__Ascomycota_unclassified(100); | - | - | - | - | - | - | - | Unassigned | - |
| Otu34489 | 1 | k__Fungi(100);p__Ascomycota(100);p__Ascomycota_unclassified(100);p__Ascomycota_unclassified(100);p__Ascomycota_unclassified(100);p__Ascomycota_unclassified(100);p__Ascomycota_unclassified(100); | - | - | - | - | - | - | - | Unassigned | - |
| Otu34490 | 1 | k__Fungi(100);p__Ascomycota(100);p__Ascomycota_unclassified(100);p__Ascomycota_unclassified(100);p__Ascomycota_unclassified(100);p__Ascomycota_unclassified(100);p__Ascomycota_unclassified(100); | - | - | - | - | - | - | - | Unassigned | - |
| Otu34491 | 1 | k__Fungi(100);k__Fungi_unclassified(100);k__Fungi_unclassified(100);k__Fungi_unclassified(100);k__Fungi_unclassified(100);k__Fungi_unclassified(100);k__Fungi_unclassified(100); | - | - | - | - | - | - | - | Unassigned | - |
| Otu34492 | 1 | k__Fungi(100);p__Ascomycota(100);p__Ascomycota_unclassified(100);p__Ascomycota_unclassified(100);p__Ascomycota_unclassified(100);p__Ascomycota_unclassified(100);p__Ascomycota_unclassified(100); | - | - | - | - | - | - | - | Unassigned | - |
| Otu34493 | 1 | k__Fungi(100);k__Fungi_unclassified(100);k__Fungi_unclassified(100);k__Fungi_unclassified(100);k__Fungi_unclassified(100);k__Fungi_unclassified(100);k__Fungi_unclassified(100); | - | - | - | - | - | - | - | Unassigned | - |
| Otu34494 | 1 | k__Fungi(100);k__Fungi_unclassified(100);k__Fungi_unclassified(100);k__Fungi_unclassified(100);k__Fungi_unclassified(100);k__Fungi_unclassified(100);k__Fungi_unclassified(100); | - | - | - | - | - | - | - | Unassigned | - |
| Otu34495 | 1 | k__Fungi(100);k__Fungi_unclassified(100);k__Fungi_unclassified(100);k__Fungi_unclassified(100);k__Fungi_unclassified(100);k__Fungi_unclassified(100);k__Fungi_unclassified(100); | - | - | - | - | - | - | - | Unassigned | - |
| Otu34496 | 1 | k__Fungi(100);p__Ascomycota(100);p__Ascomycota_unclassified(100);p__Ascomycota_unclassified(100);p__Ascomycota_unclassified(100);p__Ascomycota_unclassified(100);p__Ascomycota_unclassified(100); | - | - | - | - | - | - | - | Unassigned | - |
| Otu34498 | 1 | k__Fungi(100);k__Fungi_unclassified(100);k__Fungi_unclassified(100);k__Fungi_unclassified(100);k__Fungi_unclassified(100);k__Fungi_unclassified(100);k__Fungi_unclassified(100); | - | - | - | - | - | - | - | Unassigned | - |
| Otu34499 | 1 | k__Fungi(100);p__Ascomycota(100);p__Ascomycota_unclassified(100);p__Ascomycota_unclassified(100);p__Ascomycota_unclassified(100);p__Ascomycota_unclassified(100);p__Ascomycota_unclassified(100); | - | - | - | - | - | - | - | Unassigned | - |
| Otu34503 | 1 | k__Fungi(100);p__Ascomycota(100);p__Ascomycota_unclassified(100);p__Ascomycota_unclassified(100);p__Ascomycota_unclassified(100);p__Ascomycota_unclassified(100);p__Ascomycota_unclassified(100); | - | - | - | - | - | - | - | Unassigned | - |
| Otu34505 | 1 | k__Fungi(100);p__Ascomycota(100);p__Ascomycota_unclassified(100);p__Ascomycota_unclassified(100);p__Ascomycota_unclassified(100);p__Ascomycota_unclassified(100);p__Ascomycota_unclassified(100); | - | - | - | - | - | - | - | Unassigned | - |
| Otu34506 | 1 | k__Fungi(100);k__Fungi_unclassified(100);k__Fungi_unclassified(100);k__Fungi_unclassified(100);k__Fungi_unclassified(100);k__Fungi_unclassified(100);k__Fungi_unclassified(100); | - | - | - | - | - | - | - | Unassigned | - |
| Otu34517 | 1 | k__Fungi(100);p__Ascomycota(100);p__Ascomycota_unclassified(100);p__Ascomycota_unclassified(100);p__Ascomycota_unclassified(100);p__Ascomycota_unclassified(100);p__Ascomycota_unclassified(100); | - | - | - | - | - | - | - | Unassigned | - |
| Otu34531 | 1 | k__Fungi(100);k__Fungi_unclassified(100);k__Fungi_unclassified(100);k__Fungi_unclassified(100);k__Fungi_unclassified(100);k__Fungi_unclassified(100);k__Fungi_unclassified(100); | - | - | - | - | - | - | - | Unassigned | - |
| Otu34536 | 1 | k__Fungi(100);p__Ascomycota(100);p__Ascomycota_unclassified(100);p__Ascomycota_unclassified(100);p__Ascomycota_unclassified(100);p__Ascomycota_unclassified(100);p__Ascomycota_unclassified(100); | - | - | - | - | - | - | - | Unassigned | - |
| Otu34537 | 1 | k__Fungi(100);p__Ascomycota(100);p__Ascomycota_unclassified(100);p__Ascomycota_unclassified(100);p__Ascomycota_unclassified(100);p__Ascomycota_unclassified(100);p__Ascomycota_unclassified(100); | - | - | - | - | - | - | - | Unassigned | - |
| Otu34540 | 1 | k__Fungi(100);p__Ascomycota(100);p__Ascomycota_unclassified(100);p__Ascomycota_unclassified(100);p__Ascomycota_unclassified(100);p__Ascomycota_unclassified(100);p__Ascomycota_unclassified(100); | - | - | - | - | - | - | - | Unassigned | - |
| Otu34541 | 1 | k__Fungi(100);p__Ascomycota(100);p__Ascomycota_unclassified(100);p__Ascomycota_unclassified(100);p__Ascomycota_unclassified(100);p__Ascomycota_unclassified(100);p__Ascomycota_unclassified(100); | - | - | - | - | - | - | - | Unassigned | - |
| Otu34542 | 1 | k__Fungi(100);p__Ascomycota(100);p__Ascomycota_unclassified(100);p__Ascomycota_unclassified(100);p__Ascomycota_unclassified(100);p__Ascomycota_unclassified(100);p__Ascomycota_unclassified(100); | - | - | - | - | - | - | - | Unassigned | - |
| Otu34544 | 1 | k__Fungi(100);k__Fungi_unclassified(100);k__Fungi_unclassified(100);k__Fungi_unclassified(100);k__Fungi_unclassified(100);k__Fungi_unclassified(100);k__Fungi_unclassified(100); | - | - | - | - | - | - | - | Unassigned | - |
| Otu34545 | 1 | k__Fungi(100);p__Ascomycota(100);p__Ascomycota_unclassified(100);p__Ascomycota_unclassified(100);p__Ascomycota_unclassified(100);p__Ascomycota_unclassified(100);p__Ascomycota_unclassified(100); | - | - | - | - | - | - | - | Unassigned | - |
| Otu34546 | 1 | k__Fungi(100);p__Ascomycota(100);p__Ascomycota_unclassified(100);p__Ascomycota_unclassified(100);p__Ascomycota_unclassified(100);p__Ascomycota_unclassified(100);p__Ascomycota_unclassified(100); | - | - | - | - | - | - | - | Unassigned | - |
| Otu34547 | 1 | k__Fungi(100);p__Ascomycota(100);p__Ascomycota_unclassified(100);p__Ascomycota_unclassified(100);p__Ascomycota_unclassified(100);p__Ascomycota_unclassified(100);p__Ascomycota_unclassified(100); | - | - | - | - | - | - | - | Unassigned | - |
| Otu34548 | 1 | k__Fungi(100);k__Fungi_unclassified(100);k__Fungi_unclassified(100);k__Fungi_unclassified(100);k__Fungi_unclassified(100);k__Fungi_unclassified(100);k__Fungi_unclassified(100); | - | - | - | - | - | - | - | Unassigned | - |
| Otu34549 | 1 | k__Fungi(100);p__Ascomycota(100);p__Ascomycota_unclassified(100);p__Ascomycota_unclassified(100);p__Ascomycota_unclassified(100);p__Ascomycota_unclassified(100);p__Ascomycota_unclassified(100); | - | - | - | - | - | - | - | Unassigned | - |
| Otu34550 | 1 | k__Fungi(100);p__Ascomycota(100);p__Ascomycota_unclassified(100);p__Ascomycota_unclassified(100);p__Ascomycota_unclassified(100);p__Ascomycota_unclassified(100);p__Ascomycota_unclassified(100); | - | - | - | - | - | - | - | Unassigned | - |
| Otu34551 | 1 | k__Fungi(100);k__Fungi_unclassified(100);k__Fungi_unclassified(100);k__Fungi_unclassified(100);k__Fungi_unclassified(100);k__Fungi_unclassified(100);k__Fungi_unclassified(100); | - | - | - | - | - | - | - | Unassigned | - |
| Otu34552 | 1 | k__Fungi(100);p__Ascomycota(100);p__Ascomycota_unclassified(100);p__Ascomycota_unclassified(100);p__Ascomycota_unclassified(100);p__Ascomycota_unclassified(100);p__Ascomycota_unclassified(100); | - | - | - | - | - | - | - | Unassigned | - |
| Otu34553 | 1 | k__Fungi(100);k__Fungi_unclassified(100);k__Fungi_unclassified(100);k__Fungi_unclassified(100);k__Fungi_unclassified(100);k__Fungi_unclassified(100);k__Fungi_unclassified(100); | - | - | - | - | - | - | - | Unassigned | - |
| Otu34557 | 1 | k__Fungi(100);k__Fungi_unclassified(100);k__Fungi_unclassified(100);k__Fungi_unclassified(100);k__Fungi_unclassified(100);k__Fungi_unclassified(100);k__Fungi_unclassified(100); | - | - | - | - | - | - | - | Unassigned | - |
| Otu34559 | 1 | k__Fungi(100);k__Fungi_unclassified(100);k__Fungi_unclassified(100);k__Fungi_unclassified(100);k__Fungi_unclassified(100);k__Fungi_unclassified(100);k__Fungi_unclassified(100); | - | - | - | - | - | - | - | Unassigned | - |
| Otu34560 | 1 | k__Fungi(100);p__Ascomycota(100);p__Ascomycota_unclassified(100);p__Ascomycota_unclassified(100);p__Ascomycota_unclassified(100);p__Ascomycota_unclassified(100);p__Ascomycota_unclassified(100); | - | - | - | - | - | - | - | Unassigned | - |
| Otu34567 | 1 | k__Fungi(100);p__Ascomycota(100);p__Ascomycota_unclassified(100);p__Ascomycota_unclassified(100);p__Ascomycota_unclassified(100);p__Ascomycota_unclassified(100);p__Ascomycota_unclassified(100); | - | - | - | - | - | - | - | Unassigned | - |
| Otu34568 | 1 | k__Fungi(100);p__Ascomycota(100);p__Ascomycota_unclassified(100);p__Ascomycota_unclassified(100);p__Ascomycota_unclassified(100);p__Ascomycota_unclassified(100);p__Ascomycota_unclassified(100); | - | - | - | - | - | - | - | Unassigned | - |
| Otu34569 | 1 | k__Fungi(100);p__Ascomycota(100);p__Ascomycota_unclassified(100);p__Ascomycota_unclassified(100);p__Ascomycota_unclassified(100);p__Ascomycota_unclassified(100);p__Ascomycota_unclassified(100); | - | - | - | - | - | - | - | Unassigned | - |
| Otu34572 | 1 | k__Fungi(100);p__Ascomycota(100);p__Ascomycota_unclassified(100);p__Ascomycota_unclassified(100);p__Ascomycota_unclassified(100);p__Ascomycota_unclassified(100);p__Ascomycota_unclassified(100); | - | - | - | - | - | - | - | Unassigned | - |
| Otu34580 | 1 | k__Fungi(100);p__Ascomycota(100);p__Ascomycota_unclassified(100);p__Ascomycota_unclassified(100);p__Ascomycota_unclassified(100);p__Ascomycota_unclassified(100);p__Ascomycota_unclassified(100); | - | - | - | - | - | - | - | Unassigned | - |
| Otu34581 | 1 | k__Fungi(100);k__Fungi_unclassified(100);k__Fungi_unclassified(100);k__Fungi_unclassified(100);k__Fungi_unclassified(100);k__Fungi_unclassified(100);k__Fungi_unclassified(100); | - | - | - | - | - | - | - | Unassigned | - |
| Otu34584 | 1 | k__Fungi(100);k__Fungi_unclassified(100);k__Fungi_unclassified(100);k__Fungi_unclassified(100);k__Fungi_unclassified(100);k__Fungi_unclassified(100);k__Fungi_unclassified(100); | - | - | - | - | - | - | - | Unassigned | - |
| Otu34585 | 1 | k__Fungi(100);k__Fungi_unclassified(100);k__Fungi_unclassified(100);k__Fungi_unclassified(100);k__Fungi_unclassified(100);k__Fungi_unclassified(100);k__Fungi_unclassified(100); | - | - | - | - | - | - | - | Unassigned | - |
| Otu34586 | 1 | k__Fungi(100);p__Ascomycota(100);p__Ascomycota_unclassified(100);p__Ascomycota_unclassified(100);p__Ascomycota_unclassified(100);p__Ascomycota_unclassified(100);p__Ascomycota_unclassified(100); | - | - | - | - | - | - | - | Unassigned | - |
| Otu34587 | 1 | k__Fungi(100);k__Fungi_unclassified(100);k__Fungi_unclassified(100);k__Fungi_unclassified(100);k__Fungi_unclassified(100);k__Fungi_unclassified(100);k__Fungi_unclassified(100); | - | - | - | - | - | - | - | Unassigned | - |
| Otu34591 | 1 | k__Fungi(100);k__Fungi_unclassified(100);k__Fungi_unclassified(100);k__Fungi_unclassified(100);k__Fungi_unclassified(100);k__Fungi_unclassified(100);k__Fungi_unclassified(100); | - | - | - | - | - | - | - | Unassigned | - |
| Otu34592 | 1 | k__Fungi(100);k__Fungi_unclassified(100);k__Fungi_unclassified(100);k__Fungi_unclassified(100);k__Fungi_unclassified(100);k__Fungi_unclassified(100);k__Fungi_unclassified(100); | - | - | - | - | - | - | - | Unassigned | - |
| Otu34612 | 1 | k__Fungi(100);k__Fungi_unclassified(100);k__Fungi_unclassified(100);k__Fungi_unclassified(100);k__Fungi_unclassified(100);k__Fungi_unclassified(100);k__Fungi_unclassified(100); | - | - | - | - | - | - | - | Unassigned | - |
| Otu34617 | 1 | k__Fungi(100);p__Ascomycota(100);c__Dothideomycetes(100);o__Capnodiales(100);f__Davidiellaceae(100);f__Davidiellaceae_unclassified(100);f__Davidiellaceae_unclassified(100); | - | - | - | - | - | - | - | Unassigned | - |
| Otu34618 | 1 | k__Fungi(100);k__Fungi_unclassified(100);k__Fungi_unclassified(100);k__Fungi_unclassified(100);k__Fungi_unclassified(100);k__Fungi_unclassified(100);k__Fungi_unclassified(100); | - | - | - | - | - | - | - | Unassigned | - |
| Otu34619 | 1 | k__Fungi(100);k__Fungi_unclassified(100);k__Fungi_unclassified(100);k__Fungi_unclassified(100);k__Fungi_unclassified(100);k__Fungi_unclassified(100);k__Fungi_unclassified(100); | - | - | - | - | - | - | - | Unassigned | - |
| Otu34622 | 1 | k__Fungi(100);k__Fungi_unclassified(100);k__Fungi_unclassified(100);k__Fungi_unclassified(100);k__Fungi_unclassified(100);k__Fungi_unclassified(100);k__Fungi_unclassified(100); | - | - | - | - | - | - | - | Unassigned | - |
| Otu34625 | 1 | k__Fungi(100);k__Fungi_unclassified(100);k__Fungi_unclassified(100);k__Fungi_unclassified(100);k__Fungi_unclassified(100);k__Fungi_unclassified(100);k__Fungi_unclassified(100); | - | - | - | - | - | - | - | Unassigned | - |
| Otu34626 | 1 | k__Fungi(100);p__Ascomycota(100);p__Ascomycota_unclassified(100);p__Ascomycota_unclassified(100);p__Ascomycota_unclassified(100);p__Ascomycota_unclassified(100);p__Ascomycota_unclassified(100); | - | - | - | - | - | - | - | Unassigned | - |
| Otu34633 | 1 | k__Fungi(100);k__Fungi_unclassified(100);k__Fungi_unclassified(100);k__Fungi_unclassified(100);k__Fungi_unclassified(100);k__Fungi_unclassified(100);k__Fungi_unclassified(100); | - | - | - | - | - | - | - | Unassigned | - |
| Otu34636 | 1 | k__Fungi(100);k__Fungi_unclassified(100);k__Fungi_unclassified(100);k__Fungi_unclassified(100);k__Fungi_unclassified(100);k__Fungi_unclassified(100);k__Fungi_unclassified(100); | - | - | - | - | - | - | - | Unassigned | - |
| Otu34637 | 1 | k__Fungi(100);p__Ascomycota(100);p__Ascomycota_unclassified(100);p__Ascomycota_unclassified(100);p__Ascomycota_unclassified(100);p__Ascomycota_unclassified(100);p__Ascomycota_unclassified(100); | - | - | - | - | - | - | - | Unassigned | - |
| Otu34640 | 1 | k__Fungi(100);k__Fungi_unclassified(100);k__Fungi_unclassified(100);k__Fungi_unclassified(100);k__Fungi_unclassified(100);k__Fungi_unclassified(100);k__Fungi_unclassified(100); | - | - | - | - | - | - | - | Unassigned | - |
| Otu34651 | 1 | k__Fungi(100);k__Fungi_unclassified(100);k__Fungi_unclassified(100);k__Fungi_unclassified(100);k__Fungi_unclassified(100);k__Fungi_unclassified(100);k__Fungi_unclassified(100); | - | - | - | - | - | - | - | Unassigned | - |
| Otu34654 | 1 | k__Fungi(100);p__Ascomycota(100);p__Ascomycota_unclassified(100);p__Ascomycota_unclassified(100);p__Ascomycota_unclassified(100);p__Ascomycota_unclassified(100);p__Ascomycota_unclassified(100); | - | - | - | - | - | - | - | Unassigned | - |
| Otu34660 | 1 | k__Fungi(100);k__Fungi_unclassified(100);k__Fungi_unclassified(100);k__Fungi_unclassified(100);k__Fungi_unclassified(100);k__Fungi_unclassified(100);k__Fungi_unclassified(100); | - | - | - | - | - | - | - | Unassigned | - |
| Otu34662 | 1 | k__Fungi(100);p__Ascomycota(100);p__Ascomycota_unclassified(100);p__Ascomycota_unclassified(100);p__Ascomycota_unclassified(100);p__Ascomycota_unclassified(100);p__Ascomycota_unclassified(100); | - | - | - | - | - | - | - | Unassigned | - |
| Otu34666 | 1 | k__Fungi(100);k__Fungi_unclassified(100);k__Fungi_unclassified(100);k__Fungi_unclassified(100);k__Fungi_unclassified(100);k__Fungi_unclassified(100);k__Fungi_unclassified(100); | - | - | - | - | - | - | - | Unassigned | - |
| Otu34678 | 1 | k__Fungi(100);p__Ascomycota(100);p__Ascomycota_unclassified(100);p__Ascomycota_unclassified(100);p__Ascomycota_unclassified(100);p__Ascomycota_unclassified(100);p__Ascomycota_unclassified(100); | - | - | - | - | - | - | - | Unassigned | - |
| Otu34679 | 1 | k__Fungi(100);p__Ascomycota(100);p__Ascomycota_unclassified(100);p__Ascomycota_unclassified(100);p__Ascomycota_unclassified(100);p__Ascomycota_unclassified(100);p__Ascomycota_unclassified(100); | - | - | - | - | - | - | - | Unassigned | - |
| Otu34681 | 1 | k__Fungi(100);p__Ascomycota(100);p__Ascomycota_unclassified(100);p__Ascomycota_unclassified(100);p__Ascomycota_unclassified(100);p__Ascomycota_unclassified(100);p__Ascomycota_unclassified(100); | - | - | - | - | - | - | - | Unassigned | - |
| Otu34683 | 1 | k__Fungi(100);p__Ascomycota(100);p__Ascomycota_unclassified(100);p__Ascomycota_unclassified(100);p__Ascomycota_unclassified(100);p__Ascomycota_unclassified(100);p__Ascomycota_unclassified(100); | - | - | - | - | - | - | - | Unassigned | - |
| Otu34689 | 1 | k__Fungi(100);p__Ascomycota(100);p__Ascomycota_unclassified(100);p__Ascomycota_unclassified(100);p__Ascomycota_unclassified(100);p__Ascomycota_unclassified(100);p__Ascomycota_unclassified(100); | - | - | - | - | - | - | - | Unassigned | - |
| Otu34692 | 1 | k__Fungi(100);k__Fungi_unclassified(100);k__Fungi_unclassified(100);k__Fungi_unclassified(100);k__Fungi_unclassified(100);k__Fungi_unclassified(100);k__Fungi_unclassified(100); | - | - | - | - | - | - | - | Unassigned | - |
| Otu34697 | 1 | k__Fungi(100);p__Ascomycota(100);p__Ascomycota_unclassified(100);p__Ascomycota_unclassified(100);p__Ascomycota_unclassified(100);p__Ascomycota_unclassified(100);p__Ascomycota_unclassified(100); | - | - | - | - | - | - | - | Unassigned | - |
| Otu34698 | 1 | k__Fungi(100);k__Fungi_unclassified(100);k__Fungi_unclassified(100);k__Fungi_unclassified(100);k__Fungi_unclassified(100);k__Fungi_unclassified(100);k__Fungi_unclassified(100); | - | - | - | - | - | - | - | Unassigned | - |
| Otu34701 | 1 | k__Fungi(100);k__Fungi_unclassified(100);k__Fungi_unclassified(100);k__Fungi_unclassified(100);k__Fungi_unclassified(100);k__Fungi_unclassified(100);k__Fungi_unclassified(100); | - | - | - | - | - | - | - | Unassigned | - |
| Otu34755 | 1 | k__Fungi(100);k__Fungi_unclassified(100);k__Fungi_unclassified(100);k__Fungi_unclassified(100);k__Fungi_unclassified(100);k__Fungi_unclassified(100);k__Fungi_unclassified(100); | - | - | - | - | - | - | - | Unassigned | - |
| Otu34756 | 1 | k__Fungi(100);k__Fungi_unclassified(100);k__Fungi_unclassified(100);k__Fungi_unclassified(100);k__Fungi_unclassified(100);k__Fungi_unclassified(100);k__Fungi_unclassified(100); | - | - | - | - | - | - | - | Unassigned | - |
| Otu34758 | 1 | k__Fungi(100);k__Fungi_unclassified(100);k__Fungi_unclassified(100);k__Fungi_unclassified(100);k__Fungi_unclassified(100);k__Fungi_unclassified(100);k__Fungi_unclassified(100); | - | - | - | - | - | - | - | Unassigned | - |
| Otu34763 | 1 | k__Fungi(100);p__Ascomycota(100);p__Ascomycota_unclassified(100);p__Ascomycota_unclassified(100);p__Ascomycota_unclassified(100);p__Ascomycota_unclassified(100);p__Ascomycota_unclassified(100); | - | - | - | - | - | - | - | Unassigned | - |
| Otu34764 | 1 | k__Fungi(100);p__Ascomycota(100);p__Ascomycota_unclassified(100);p__Ascomycota_unclassified(100);p__Ascomycota_unclassified(100);p__Ascomycota_unclassified(100);p__Ascomycota_unclassified(100); | - | - | - | - | - | - | - | Unassigned | - |
| Otu34765 | 1 | k__Fungi(100);k__Fungi_unclassified(100);k__Fungi_unclassified(100);k__Fungi_unclassified(100);k__Fungi_unclassified(100);k__Fungi_unclassified(100);k__Fungi_unclassified(100); | - | - | - | - | - | - | - | Unassigned | - |
| Otu34766 | 1 | k__Fungi(100);p__Ascomycota(100);p__Ascomycota_unclassified(100);p__Ascomycota_unclassified(100);p__Ascomycota_unclassified(100);p__Ascomycota_unclassified(100);p__Ascomycota_unclassified(100); | - | - | - | - | - | - | - | Unassigned | - |
| Otu34767 | 1 | k__Fungi(100);p__Ascomycota(100);p__Ascomycota_unclassified(100);p__Ascomycota_unclassified(100);p__Ascomycota_unclassified(100);p__Ascomycota_unclassified(100);p__Ascomycota_unclassified(100); | - | - | - | - | - | - | - | Unassigned | - |
| Otu34768 | 1 | k__Fungi(100);k__Fungi_unclassified(100);k__Fungi_unclassified(100);k__Fungi_unclassified(100);k__Fungi_unclassified(100);k__Fungi_unclassified(100);k__Fungi_unclassified(100); | - | - | - | - | - | - | - | Unassigned | - |
| Otu34770 | 1 | k__Fungi(100);p__Ascomycota(100);p__Ascomycota_unclassified(100);p__Ascomycota_unclassified(100);p__Ascomycota_unclassified(100);p__Ascomycota_unclassified(100);p__Ascomycota_unclassified(100); | - | - | - | - | - | - | - | Unassigned | - |
| Otu34771 | 1 | k__Fungi(100);p__Ascomycota(100);p__Ascomycota_unclassified(100);p__Ascomycota_unclassified(100);p__Ascomycota_unclassified(100);p__Ascomycota_unclassified(100);p__Ascomycota_unclassified(100); | - | - | - | - | - | - | - | Unassigned | - |
| Otu34772 | 1 | k__Fungi(100);k__Fungi_unclassified(100);k__Fungi_unclassified(100);k__Fungi_unclassified(100);k__Fungi_unclassified(100);k__Fungi_unclassified(100);k__Fungi_unclassified(100); | - | - | - | - | - | - | - | Unassigned | - |
| Otu34773 | 1 | k__Fungi(100);k__Fungi_unclassified(100);k__Fungi_unclassified(100);k__Fungi_unclassified(100);k__Fungi_unclassified(100);k__Fungi_unclassified(100);k__Fungi_unclassified(100); | - | - | - | - | - | - | - | Unassigned | - |
| Otu34777 | 1 | k__Fungi(100);p__Ascomycota(100);p__Ascomycota_unclassified(100);p__Ascomycota_unclassified(100);p__Ascomycota_unclassified(100);p__Ascomycota_unclassified(100);p__Ascomycota_unclassified(100); | - | - | - | - | - | - | - | Unassigned | - |

| | | | | | | | | | | | |
|---|---|---|---|---|---|---|---|---|---|---|---|
| Otu34778 | 1 | k__Fungi(100);p__Ascomycota(100);p__Ascomycota_unclassified(100);p__Ascomycota_unclassified(100);p__Ascomycota_unclassified(100);p__Ascomycota_unclassified(100);p__Ascomycota_unclassified(100); | - | - | - | - | - | - | - | Unassigned | - |
| Otu34779 | 1 | k__Fungi(100);p__Ascomycota(100);p__Ascomycota_unclassified(100);p__Ascomycota_unclassified(100);p__Ascomycota_unclassified(100);p__Ascomycota_unclassified(100);p__Ascomycota_unclassified(100); | - | - | - | - | - | - | - | Unassigned | - |
| Otu34780 | 1 | k__Fungi(100);k__Fungi_unclassified(100);k__Fungi_unclassified(100);k__Fungi_unclassified(100);k__Fungi_unclassified(100);k__Fungi_unclassified(100);k__Fungi_unclassified(100); | - | - | - | - | - | - | - | Unassigned | - |
| Otu34781 | 1 | k__Fungi(100);k__Fungi_unclassified(100);k__Fungi_unclassified(100);k__Fungi_unclassified(100);k__Fungi_unclassified(100);k__Fungi_unclassified(100);k__Fungi_unclassified(100); | - | - | - | - | - | - | - | Unassigned | - |
| Otu34782 | 1 | k__Fungi(100);p__Ascomycota(100);p__Ascomycota_unclassified(100);p__Ascomycota_unclassified(100);p__Ascomycota_unclassified(100);p__Ascomycota_unclassified(100);p__Ascomycota_unclassified(100); | - | - | - | - | - | - | - | Unassigned | - |
| Otu34783 | 1 | k__Fungi(100);k__Fungi_unclassified(100);k__Fungi_unclassified(100);k__Fungi_unclassified(100);k__Fungi_unclassified(100);k__Fungi_unclassified(100);k__Fungi_unclassified(100); | - | - | - | - | - | - | - | Unassigned | - |
| Otu34787 | 1 | k__Fungi(100);k__Fungi_unclassified(100);k__Fungi_unclassified(100);k__Fungi_unclassified(100);k__Fungi_unclassified(100);k__Fungi_unclassified(100);k__Fungi_unclassified(100); | - | - | - | - | - | - | - | Unassigned | - |
| Otu34789 | 1 | k__Fungi(100);k__Fungi_unclassified(100);k__Fungi_unclassified(100);k__Fungi_unclassified(100);k__Fungi_unclassified(100);k__Fungi_unclassified(100);k__Fungi_unclassified(100); | - | - | - | - | - | - | - | Unassigned | - |
| Otu34790 | 1 | k__Fungi(100);p__Ascomycota(100);p__Ascomycota_unclassified(100);p__Ascomycota_unclassified(100);p__Ascomycota_unclassified(100);p__Ascomycota_unclassified(100);p__Ascomycota_unclassified(100); | - | - | - | - | - | - | - | Unassigned | - |
| Otu34792 | 1 | k__Fungi(100);k__Fungi_unclassified(100);k__Fungi_unclassified(100);k__Fungi_unclassified(100);k__Fungi_unclassified(100);k__Fungi_unclassified(100);k__Fungi_unclassified(100); | - | - | - | - | - | - | - | Unassigned | - |
| Otu34793 | 1 | k__Fungi(100);p__Ascomycota(100);p__Ascomycota_unclassified(100);p__Ascomycota_unclassified(100);p__Ascomycota_unclassified(100);p__Ascomycota_unclassified(100);p__Ascomycota_unclassified(100); | - | - | - | - | - | - | - | Unassigned | - |
| Otu34794 | 1 | k__Fungi(100);p__Ascomycota(100);p__Ascomycota_unclassified(100);p__Ascomycota_unclassified(100);p__Ascomycota_unclassified(100);p__Ascomycota_unclassified(100);p__Ascomycota_unclassified(100); | - | - | - | - | - | - | - | Unassigned | - |
| Otu34796 | 1 | k__Fungi(100);p__Ascomycota(100);p__Ascomycota_unclassified(100);p__Ascomycota_unclassified(100);p__Ascomycota_unclassified(100);p__Ascomycota_unclassified(100);p__Ascomycota_unclassified(100); | - | - | - | - | - | - | - | Unassigned | - |
| Otu34797 | 1 | k__Fungi(100);p__Ascomycota(100);p__Ascomycota_unclassified(100);p__Ascomycota_unclassified(100);p__Ascomycota_unclassified(100);p__Ascomycota_unclassified(100);p__Ascomycota_unclassified(100); | - | - | - | - | - | - | - | Unassigned | - |
| Otu34798 | 1 | k__Fungi(100);k__Fungi_unclassified(100);k__Fungi_unclassified(100);k__Fungi_unclassified(100);k__Fungi_unclassified(100);k__Fungi_unclassified(100);k__Fungi_unclassified(100); | - | - | - | - | - | - | - | Unassigned | - |
| Otu34799 | 1 | k__Fungi(100);p__Ascomycota(100);p__Ascomycota_unclassified(100);p__Ascomycota_unclassified(100);p__Ascomycota_unclassified(100);p__Ascomycota_unclassified(100);p__Ascomycota_unclassified(100); | - | - | - | - | - | - | - | Unassigned | - |
| Otu34800 | 1 | k__Fungi(100);k__Fungi_unclassified(100);k__Fungi_unclassified(100);k__Fungi_unclassified(100);k__Fungi_unclassified(100);k__Fungi_unclassified(100);k__Fungi_unclassified(100); | - | - | - | - | - | - | - | Unassigned | - |
| Otu34802 | 1 | k__Fungi(100);p__Ascomycota(100);p__Ascomycota_unclassified(100);p__Ascomycota_unclassified(100);p__Ascomycota_unclassified(100);p__Ascomycota_unclassified(100);p__Ascomycota_unclassified(100); | - | - | - | - | - | - | - | Unassigned | - |
| Otu34804 | 1 | k__Fungi(100);p__Ascomycota(100);p__Ascomycota_unclassified(100);p__Ascomycota_unclassified(100);p__Ascomycota_unclassified(100);p__Ascomycota_unclassified(100);p__Ascomycota_unclassified(100); | - | - | - | - | - | - | - | Unassigned | - |
| Otu34808 | 1 | k__Fungi(100);p__Ascomycota(100);p__Ascomycota_unclassified(100);p__Ascomycota_unclassified(100);p__Ascomycota_unclassified(100);p__Ascomycota_unclassified(100);p__Ascomycota_unclassified(100); | - | - | - | - | - | - | - | Unassigned | - |
| Otu34810 | 1 | k__Fungi(100);p__Ascomycota(100);p__Ascomycota_unclassified(100);p__Ascomycota_unclassified(100);p__Ascomycota_unclassified(100);p__Ascomycota_unclassified(100);p__Ascomycota_unclassified(100); | - | - | - | - | - | - | - | Unassigned | - |
| Otu34817 | 1 | k__Fungi(100);k__Fungi_unclassified(100);k__Fungi_unclassified(100);k__Fungi_unclassified(100);k__Fungi_unclassified(100);k__Fungi_unclassified(100);k__Fungi_unclassified(100); | - | - | - | - | - | - | - | Unassigned | - |
| Otu34820 | 1 | k__Fungi(100);k__Fungi_unclassified(100);k__Fungi_unclassified(100);k__Fungi_unclassified(100);k__Fungi_unclassified(100);k__Fungi_unclassified(100);k__Fungi_unclassified(100); | - | - | - | - | - | - | - | Unassigned | - |
| Otu34823 | 1 | k__Fungi(100);k__Fungi_unclassified(100);k__Fungi_unclassified(100);k__Fungi_unclassified(100);k__Fungi_unclassified(100);k__Fungi_unclassified(100);k__Fungi_unclassified(100); | - | - | - | - | - | - | - | Unassigned | - |
| Otu34824 | 1 | k__Fungi(100);k__Fungi_unclassified(100);k__Fungi_unclassified(100);k__Fungi_unclassified(100);k__Fungi_unclassified(100);k__Fungi_unclassified(100);k__Fungi_unclassified(100); | - | - | - | - | - | - | - | Unassigned | - |
| Otu34825 | 1 | k__Fungi(100);k__Fungi_unclassified(100);k__Fungi_unclassified(100);k__Fungi_unclassified(100);k__Fungi_unclassified(100);k__Fungi_unclassified(100);k__Fungi_unclassified(100); | - | - | - | - | - | - | - | Unassigned | - |
| Otu34826 | 1 | k__Fungi(100);k__Fungi_unclassified(100);k__Fungi_unclassified(100);k__Fungi_unclassified(100);k__Fungi_unclassified(100);k__Fungi_unclassified(100);k__Fungi_unclassified(100); | - | - | - | - | - | - | - | Unassigned | - |
| Otu34827 | 1 | k__Fungi(100);k__Fungi_unclassified(100);k__Fungi_unclassified(100);k__Fungi_unclassified(100);k__Fungi_unclassified(100);k__Fungi_unclassified(100);k__Fungi_unclassified(100); | - | - | - | - | - | - | - | Unassigned | - |
| Otu34828 | 1 | k__Fungi(100);p__Ascomycota(100);p__Ascomycota_unclassified(100);p__Ascomycota_unclassified(100);p__Ascomycota_unclassified(100);p__Ascomycota_unclassified(100);p__Ascomycota_unclassified(100); | - | - | - | - | - | - | - | Unassigned | - |
| Otu34830 | 1 | k__Fungi(100);k__Fungi_unclassified(100);k__Fungi_unclassified(100);k__Fungi_unclassified(100);k__Fungi_unclassified(100);k__Fungi_unclassified(100);k__Fungi_unclassified(100); | - | - | - | - | - | - | - | Unassigned | - |
| Otu34831 | 1 | k__Fungi(100);k__Fungi_unclassified(100);k__Fungi_unclassified(100);k__Fungi_unclassified(100);k__Fungi_unclassified(100);k__Fungi_unclassified(100);k__Fungi_unclassified(100); | - | - | - | - | - | - | - | Unassigned | - |
| Otu34832 | 1 | k__Fungi(100);k__Fungi_unclassified(100);k__Fungi_unclassified(100);k__Fungi_unclassified(100);k__Fungi_unclassified(100);k__Fungi_unclassified(100);k__Fungi_unclassified(100); | - | - | - | - | - | - | - | Unassigned | - |
| Otu34833 | 1 | k__Fungi(100);p__Ascomycota(100);p__Ascomycota_unclassified(100);p__Ascomycota_unclassified(100);p__Ascomycota_unclassified(100);p__Ascomycota_unclassified(100);p__Ascomycota_unclassified(100); | - | - | - | - | - | - | - | Unassigned | - |
| Otu34834 | 1 | k__Fungi(100);k__Fungi_unclassified(100);k__Fungi_unclassified(100);k__Fungi_unclassified(100);k__Fungi_unclassified(100);k__Fungi_unclassified(100);k__Fungi_unclassified(100); | - | - | - | - | - | - | - | Unassigned | - |
| Otu34848 | 1 | k__Fungi(100);k__Fungi_unclassified(100);k__Fungi_unclassified(100);k__Fungi_unclassified(100);k__Fungi_unclassified(100);k__Fungi_unclassified(100);k__Fungi_unclassified(100); | - | - | - | - | - | - | - | Unassigned | - |
| Otu34853 | 1 | k__Fungi(100);k__Fungi_unclassified(100);k__Fungi_unclassified(100);k__Fungi_unclassified(100);k__Fungi_unclassified(100);k__Fungi_unclassified(100);k__Fungi_unclassified(100); | - | - | - | - | - | - | - | Unassigned | - |
| Otu34857 | 1 | k__Fungi(100);p__Ascomycota(100);p__Ascomycota_unclassified(100);p__Ascomycota_unclassified(100);p__Ascomycota_unclassified(100);p__Ascomycota_unclassified(100);p__Ascomycota_unclassified(100); | - | - | - | - | - | - | - | Unassigned | - |
| Otu34862 | 1 | k__Fungi(100);p__Ascomycota(100);p__Ascomycota_unclassified(100);p__Ascomycota_unclassified(100);p__Ascomycota_unclassified(100);p__Ascomycota_unclassified(100);p__Ascomycota_unclassified(100); | - | - | - | - | - | - | - | Unassigned | - |
| Otu34863 | 1 | k__Fungi(100);k__Fungi_unclassified(100);k__Fungi_unclassified(100);k__Fungi_unclassified(100);k__Fungi_unclassified(100);k__Fungi_unclassified(100);k__Fungi_unclassified(100); | - | - | - | - | - | - | - | Unassigned | - |
| Otu34864 | 1 | k__Fungi(100);k__Fungi_unclassified(100);k__Fungi_unclassified(100);k__Fungi_unclassified(100);k__Fungi_unclassified(100);k__Fungi_unclassified(100);k__Fungi_unclassified(100); | - | - | - | - | - | - | - | Unassigned | - |
| Otu34865 | 1 | k__Fungi(100);k__Fungi_unclassified(100);k__Fungi_unclassified(100);k__Fungi_unclassified(100);k__Fungi_unclassified(100);k__Fungi_unclassified(100);k__Fungi_unclassified(100); | - | - | - | - | - | - | - | Unassigned | - |
| Otu34866 | 1 | k__Fungi(100);k__Fungi_unclassified(100);k__Fungi_unclassified(100);k__Fungi_unclassified(100);k__Fungi_unclassified(100);k__Fungi_unclassified(100);k__Fungi_unclassified(100); | - | - | - | - | - | - | - | Unassigned | - |
| Otu34867 | 1 | k__Fungi(100);k__Fungi_unclassified(100);k__Fungi_unclassified(100);k__Fungi_unclassified(100);k__Fungi_unclassified(100);k__Fungi_unclassified(100);k__Fungi_unclassified(100); | - | - | - | - | - | - | - | Unassigned | - |
| Otu34874 | 1 | k__Fungi(100);k__Fungi_unclassified(100);k__Fungi_unclassified(100);k__Fungi_unclassified(100);k__Fungi_unclassified(100);k__Fungi_unclassified(100);k__Fungi_unclassified(100); | - | - | - | - | - | - | - | Unassigned | - |
| Otu34875 | 1 | k__Fungi(100);p__Ascomycota(100);p__Ascomycota_unclassified(100);p__Ascomycota_unclassified(100);p__Ascomycota_unclassified(100);p__Ascomycota_unclassified(100);p__Ascomycota_unclassified(100); | - | - | - | - | - | - | - | Unassigned | - |
| Otu34877 | 1 | k__Fungi(100);p__Ascomycota(100);p__Ascomycota_unclassified(100);p__Ascomycota_unclassified(100);p__Ascomycota_unclassified(100);p__Ascomycota_unclassified(100);p__Ascomycota_unclassified(100); | - | - | - | - | - | - | - | Unassigned | - |
| Otu34879 | 1 | k__Fungi(100);k__Fungi_unclassified(100);k__Fungi_unclassified(100);k__Fungi_unclassified(100);k__Fungi_unclassified(100);k__Fungi_unclassified(100);k__Fungi_unclassified(100); | - | - | - | - | - | - | - | Unassigned | - |
| Otu34882 | 1 | k__Fungi(100);p__Ascomycota(100);p__Ascomycota_unclassified(100);p__Ascomycota_unclassified(100);p__Ascomycota_unclassified(100);p__Ascomycota_unclassified(100);p__Ascomycota_unclassified(100); | - | - | - | - | - | - | - | Unassigned | - |
| Otu34883 | 1 | k__Fungi(100);p__Ascomycota(100);p__Ascomycota_unclassified(100);p__Ascomycota_unclassified(100);p__Ascomycota_unclassified(100);p__Ascomycota_unclassified(100);p__Ascomycota_unclassified(100); | - | - | - | - | - | - | - | Unassigned | - |
| Otu34884 | 1 | k__Fungi(100);p__Ascomycota(100);p__Ascomycota_unclassified(100);p__Ascomycota_unclassified(100);p__Ascomycota_unclassified(100);p__Ascomycota_unclassified(100);p__Ascomycota_unclassified(100); | - | - | - | - | - | - | - | Unassigned | - |
| Otu34885 | 1 | k__Fungi(100);k__Fungi_unclassified(100);k__Fungi_unclassified(100);k__Fungi_unclassified(100);k__Fungi_unclassified(100);k__Fungi_unclassified(100);k__Fungi_unclassified(100); | - | - | - | - | - | - | - | Unassigned | - |
| Otu34886 | 1 | k__Fungi(100);k__Fungi_unclassified(100);k__Fungi_unclassified(100);k__Fungi_unclassified(100);k__Fungi_unclassified(100);k__Fungi_unclassified(100);k__Fungi_unclassified(100); | - | - | - | - | - | - | - | Unassigned | - |
| Otu34888 | 1 | k__Fungi(100);k__Fungi_unclassified(100);k__Fungi_unclassified(100);k__Fungi_unclassified(100);k__Fungi_unclassified(100);k__Fungi_unclassified(100);k__Fungi_unclassified(100); | - | - | - | - | - | - | - | Unassigned | - |
| Otu34889 | 1 | k__Fungi(100);k__Fungi_unclassified(100);k__Fungi_unclassified(100);k__Fungi_unclassified(100);k__Fungi_unclassified(100);k__Fungi_unclassified(100);k__Fungi_unclassified(100); | - | - | - | - | - | - | - | Unassigned | - |
| Otu34890 | 1 | k__Fungi(100);k__Fungi_unclassified(100);k__Fungi_unclassified(100);k__Fungi_unclassified(100);k__Fungi_unclassified(100);k__Fungi_unclassified(100);k__Fungi_unclassified(100); | - | - | - | - | - | - | - | Unassigned | - |
| Otu34891 | 1 | k__Fungi(100);k__Fungi_unclassified(100);k__Fungi_unclassified(100);k__Fungi_unclassified(100);k__Fungi_unclassified(100);k__Fungi_unclassified(100);k__Fungi_unclassified(100); | - | - | - | - | - | - | - | Unassigned | - |
| Otu34892 | 1 | k__Fungi(100);k__Fungi_unclassified(100);k__Fungi_unclassified(100);k__Fungi_unclassified(100);k__Fungi_unclassified(100);k__Fungi_unclassified(100);k__Fungi_unclassified(100); | - | - | - | - | - | - | - | Unassigned | - |
| Otu34893 | 1 | k__Fungi(100);p__Ascomycota(100);p__Ascomycota_unclassified(100);p__Ascomycota_unclassified(100);p__Ascomycota_unclassified(100);p__Ascomycota_unclassified(100);p__Ascomycota_unclassified(100); | - | - | - | - | - | - | - | Unassigned | - |
| Otu34894 | 1 | k__Fungi(100);k__Fungi_unclassified(100);k__Fungi_unclassified(100);k__Fungi_unclassified(100);k__Fungi_unclassified(100);k__Fungi_unclassified(100);k__Fungi_unclassified(100); | - | - | - | - | - | - | - | Unassigned | - |
| Otu34895 | 1 | k__Fungi(100);k__Fungi_unclassified(100);k__Fungi_unclassified(100);k__Fungi_unclassified(100);k__Fungi_unclassified(100);k__Fungi_unclassified(100);k__Fungi_unclassified(100); | - | - | - | - | - | - | - | Unassigned | - |
| Otu34896 | 1 | k__Fungi(100);p__Ascomycota(100);p__Ascomycota_unclassified(100);p__Ascomycota_unclassified(100);p__Ascomycota_unclassified(100);p__Ascomycota_unclassified(100);p__Ascomycota_unclassified(100); | - | - | - | - | - | - | - | Unassigned | - |
| Otu34897 | 1 | k__Fungi(100);k__Fungi_unclassified(100);k__Fungi_unclassified(100);k__Fungi_unclassified(100);k__Fungi_unclassified(100);k__Fungi_unclassified(100);k__Fungi_unclassified(100); | - | - | - | - | - | - | - | Unassigned | - |
| Otu34898 | 1 | k__Fungi(100);p__Ascomycota(100);p__Ascomycota_unclassified(100);p__Ascomycota_unclassified(100);p__Ascomycota_unclassified(100);p__Ascomycota_unclassified(100);p__Ascomycota_unclassified(100); | - | - | - | - | - | - | - | Unassigned | - |
| Otu34899 | 1 | k__Fungi(100);p__Ascomycota(100);p__Ascomycota_unclassified(100);p__Ascomycota_unclassified(100);p__Ascomycota_unclassified(100);p__Ascomycota_unclassified(100);p__Ascomycota_unclassified(100); | - | - | - | - | - | - | - | Unassigned | - |
| Otu34900 | 1 | k__Fungi(100);k__Fungi_unclassified(100);k__Fungi_unclassified(100);k__Fungi_unclassified(100);k__Fungi_unclassified(100);k__Fungi_unclassified(100);k__Fungi_unclassified(100); | - | - | - | - | - | - | - | Unassigned | - |
| Otu34901 | 1 | k__Fungi(100);k__Fungi_unclassified(100);k__Fungi_unclassified(100);k__Fungi_unclassified(100);k__Fungi_unclassified(100);k__Fungi_unclassified(100);k__Fungi_unclassified(100); | - | - | - | - | - | - | - | Unassigned | - |
| Otu34902 | 1 | k__Fungi(100);k__Fungi_unclassified(100);k__Fungi_unclassified(100);k__Fungi_unclassified(100);k__Fungi_unclassified(100);k__Fungi_unclassified(100);k__Fungi_unclassified(100); | - | - | - | - | - | - | - | Unassigned | - |
| Otu34903 | 1 | k__Fungi(100);p__Ascomycota(100);p__Ascomycota_unclassified(100);p__Ascomycota_unclassified(100);p__Ascomycota_unclassified(100);p__Ascomycota_unclassified(100);p__Ascomycota_unclassified(100); | - | - | - | - | - | - | - | Unassigned | - |
| Otu34904 | 1 | k__Fungi(100);k__Fungi_unclassified(100);k__Fungi_unclassified(100);k__Fungi_unclassified(100);k__Fungi_unclassified(100);k__Fungi_unclassified(100);k__Fungi_unclassified(100); | - | - | - | - | - | - | - | Unassigned | - |
| Otu34905 | 1 | k__Fungi(100);k__Fungi_unclassified(100);k__Fungi_unclassified(100);k__Fungi_unclassified(100);k__Fungi_unclassified(100);k__Fungi_unclassified(100);k__Fungi_unclassified(100); | - | - | - | - | - | - | - | Unassigned | - |
| Otu34906 | 1 | k__Fungi(100);p__Ascomycota(100);p__Ascomycota_unclassified(100);p__Ascomycota_unclassified(100);p__Ascomycota_unclassified(100);p__Ascomycota_unclassified(100);p__Ascomycota_unclassified(100); | - | - | - | - | - | - | - | Unassigned | - |
| Otu34907 | 1 | k__Fungi(100);p__Ascomycota(100);p__Ascomycota_unclassified(100);p__Ascomycota_unclassified(100);p__Ascomycota_unclassified(100);p__Ascomycota_unclassified(100);p__Ascomycota_unclassified(100); | - | - | - | - | - | - | - | Unassigned | - |
| Otu34908 | 1 | k__Fungi(100);k__Fungi_unclassified(100);k__Fungi_unclassified(100);k__Fungi_unclassified(100);k__Fungi_unclassified(100);k__Fungi_unclassified(100);k__Fungi_unclassified(100); | - | - | - | - | - | - | - | Unassigned | - |
| Otu34909 | 1 | k__Fungi(100);k__Fungi_unclassified(100);k__Fungi_unclassified(100);k__Fungi_unclassified(100);k__Fungi_unclassified(100);k__Fungi_unclassified(100);k__Fungi_unclassified(100); | - | - | - | - | - | - | - | Unassigned | - |
| Otu34910 | 1 | k__Fungi(100);k__Fungi_unclassified(100);k__Fungi_unclassified(100);k__Fungi_unclassified(100);k__Fungi_unclassified(100);k__Fungi_unclassified(100);k__Fungi_unclassified(100); | - | - | - | - | - | - | - | Unassigned | - |
| Otu34911 | 1 | k__Fungi(100);k__Fungi_unclassified(100);k__Fungi_unclassified(100);k__Fungi_unclassified(100);k__Fungi_unclassified(100);k__Fungi_unclassified(100);k__Fungi_unclassified(100); | - | - | - | - | - | - | - | Unassigned | - |
| Otu34912 | 1 | k__Fungi(100);k__Fungi_unclassified(100);k__Fungi_unclassified(100);k__Fungi_unclassified(100);k__Fungi_unclassified(100);k__Fungi_unclassified(100);k__Fungi_unclassified(100); | - | - | - | - | - | - | - | Unassigned | - |
| Otu34913 | 1 | k__Fungi(100);k__Fungi_unclassified(100);k__Fungi_unclassified(100);k__Fungi_unclassified(100);k__Fungi_unclassified(100);k__Fungi_unclassified(100);k__Fungi_unclassified(100); | - | - | - | - | - | - | - | Unassigned | - |
| Otu34960 | 1 | k__Fungi(100);p__Ascomycota(100);p__Ascomycota_unclassified(100);p__Ascomycota_unclassified(100);p__Ascomycota_unclassified(100);p__Ascomycota_unclassified(100);p__Ascomycota_unclassified(100); | - | - | - | - | - | - | - | Unassigned | - |
| Otu34968 | 1 | k__Fungi(100);p__Ascomycota(100);p__Ascomycota_unclassified(100);p__Ascomycota_unclassified(100);p__Ascomycota_unclassified(100);p__Ascomycota_unclassified(100);p__Ascomycota_unclassified(100); | - | - | - | - | - | - | - | Unassigned | - |
| Otu34969 | 1 | k__Fungi(100);k__Fungi_unclassified(100);k__Fungi_unclassified(100);k__Fungi_unclassified(100);k__Fungi_unclassified(100);k__Fungi_unclassified(100);k__Fungi_unclassified(100); | - | - | - | - | - | - | - | Unassigned | - |
| Otu34970 | 1 | k__Fungi(100);k__Fungi_unclassified(100);k__Fungi_unclassified(100);k__Fungi_unclassified(100);k__Fungi_unclassified(100);k__Fungi_unclassified(100);k__Fungi_unclassified(100); | - | - | - | - | - | - | - | Unassigned | - |
| Otu34971 | 1 | k__Fungi(100);k__Fungi_unclassified(100);k__Fungi_unclassified(100);k__Fungi_unclassified(100);k__Fungi_unclassified(100);k__Fungi_unclassified(100);k__Fungi_unclassified(100); | - | - | - | - | - | - | - | Unassigned | - |
| Otu34972 | 1 | k__Fungi(100);p__Ascomycota(100);p__Ascomycota_unclassified(100);p__Ascomycota_unclassified(100);p__Ascomycota_unclassified(100);p__Ascomycota_unclassified(100);p__Ascomycota_unclassified(100); | - | - | - | - | - | - | - | Unassigned | - |
| Otu34977 | 1 | k__Fungi(100);p__Ascomycota(100);p__Ascomycota_unclassified(100);p__Ascomycota_unclassified(100);p__Ascomycota_unclassified(100);p__Ascomycota_unclassified(100);p__Ascomycota_unclassified(100); | - | - | - | - | - | - | - | Unassigned | - |
| Otu34980 | 1 | k__Fungi(100);k__Fungi_unclassified(100);k__Fungi_unclassified(100);k__Fungi_unclassified(100);k__Fungi_unclassified(100);k__Fungi_unclassified(100);k__Fungi_unclassified(100); | - | - | - | - | - | - | - | Unassigned | - |
| Otu34982 | 1 | k__Fungi(100);k__Fungi_unclassified(100);k__Fungi_unclassified(100);k__Fungi_unclassified(100);k__Fungi_unclassified(100);k__Fungi_unclassified(100);k__Fungi_unclassified(100); | - | - | - | - | - | - | - | Unassigned | - |
| Otu34983 | 1 | k__Fungi(100);p__Ascomycota(100);p__Ascomycota_unclassified(100);p__Ascomycota_unclassified(100);p__Ascomycota_unclassified(100);p__Ascomycota_unclassified(100);p__Ascomycota_unclassified(100); | - | - | - | - | - | - | - | Unassigned | - |
| Otu34987 | 1 | k__Fungi(100);k__Fungi_unclassified(100);k__Fungi_unclassified(100);k__Fungi_unclassified(100);k__Fungi_unclassified(100);k__Fungi_unclassified(100);k__Fungi_unclassified(100); | - | - | - | - | - | - | - | Unassigned | - |
| Otu34992 | 1 | k__Fungi(100);k__Fungi_unclassified(100);k__Fungi_unclassified(100);k__Fungi_unclassified(100);k__Fungi_unclassified(100);k__Fungi_unclassified(100);k__Fungi_unclassified(100); | - | - | - | - | - | - | - | Unassigned | - |
| Otu35007 | 1 | k__Fungi(100);p__Ascomycota(100);p__Ascomycota_unclassified(100);p__Ascomycota_unclassified(100);p__Ascomycota_unclassified(100);p__Ascomycota_unclassified(100);p__Ascomycota_unclassified(100); | - | - | - | - | - | - | - | Unassigned | - |
| Otu35017 | 1 | k__Fungi(100);k__Fungi_unclassified(100);k__Fungi_unclassified(100);k__Fungi_unclassified(100);k__Fungi_unclassified(100);k__Fungi_unclassified(100);k__Fungi_unclassified(100); | - | - | - | - | - | - | - | Unassigned | - |
| Otu35019 | 1 | k__Fungi(100);k__Fungi_unclassified(100);k__Fungi_unclassified(100);k__Fungi_unclassified(100);k__Fungi_unclassified(100);k__Fungi_unclassified(100);k__Fungi_unclassified(100); | - | - | - | - | - | - | - | Unassigned | - |
| Otu35020 | 1 | k__Fungi(100);p__Ascomycota(100);p__Ascomycota_unclassified(100);p__Ascomycota_unclassified(100);p__Ascomycota_unclassified(100);p__Ascomycota_unclassified(100);p__Ascomycota_unclassified(100); | - | - | - | - | - | - | - | Unassigned | - |
| Otu35021 | 1 | k__Fungi(100);k__Fungi_unclassified(100);k__Fungi_unclassified(100);k__Fungi_unclassified(100);k__Fungi_unclassified(100);k__Fungi_unclassified(100);k__Fungi_unclassified(100); | - | - | - | - | - | - | - | Unassigned | - |
| Otu35022 | 1 | k__Fungi(100);p__Ascomycota(100);p__Ascomycota_unclassified(100);p__Ascomycota_unclassified(100);p__Ascomycota_unclassified(100);p__Ascomycota_unclassified(100);p__Ascomycota_unclassified(100); | - | - | - | - | - | - | - | Unassigned | - |
| Otu35023 | 1 | k__Fungi(100);p__Ascomycota(100);p__Ascomycota_unclassified(100);p__Ascomycota_unclassified(100);p__Ascomycota_unclassified(100);p__Ascomycota_unclassified(100);p__Ascomycota_unclassified(100); | - | - | - | - | - | - | - | Unassigned | - |
| Otu35024 | 1 | k__Fungi(100);k__Fungi_unclassified(100);k__Fungi_unclassified(100);k__Fungi_unclassified(100);k__Fungi_unclassified(100);k__Fungi_unclassified(100);k__Fungi_unclassified(100); | - | - | - | - | - | - | - | Unassigned | - |
| Otu35025 | 1 | k__Fungi(100);p__Ascomycota(100);p__Ascomycota_unclassified(100);p__Ascomycota_unclassified(100);p__Ascomycota_unclassified(100);p__Ascomycota_unclassified(100);p__Ascomycota_unclassified(100); | - | - | - | - | - | - | - | Unassigned | - |
| Otu35026 | 1 | k__Fungi(100);k__Fungi_unclassified(100);k__Fungi_unclassified(100);k__Fungi_unclassified(100);k__Fungi_unclassified(100);k__Fungi_unclassified(100);k__Fungi_unclassified(100); | - | - | - | - | - | - | - | Unassigned | - |
| Otu35027 | 1 | k__Fungi(100);k__Fungi_unclassified(100);k__Fungi_unclassified(100);k__Fungi_unclassified(100);k__Fungi_unclassified(100);k__Fungi_unclassified(100);k__Fungi_unclassified(100); | - | - | - | - | - | - | - | Unassigned | - |
| Otu35028 | 1 | k__Fungi(100);p__Ascomycota(100);p__Ascomycota_unclassified(100);p__Ascomycota_unclassified(100);p__Ascomycota_unclassified(100);p__Ascomycota_unclassified(100);p__Ascomycota_unclassified(100); | - | - | - | - | - | - | - | Unassigned | - |
| Otu35029 | 1 | k__Fungi(100);k__Fungi_unclassified(100);k__Fungi_unclassified(100);k__Fungi_unclassified(100);k__Fungi_unclassified(100);k__Fungi_unclassified(100);k__Fungi_unclassified(100); | - | - | - | - | - | - | - | Unassigned | - |
| Otu35030 | 1 | k__Fungi(100);p__Ascomycota(100);p__Ascomycota_unclassified(100);p__Ascomycota_unclassified(100);p__Ascomycota_unclassified(100);p__Ascomycota_unclassified(100);p__Ascomycota_unclassified(100); | - | - | - | - | - | - | - | Unassigned | - |
| Otu35031 | 1 | k__Fungi(100);k__Fungi_unclassified(100);k__Fungi_unclassified(100);k__Fungi_unclassified(100);k__Fungi_unclassified(100);k__Fungi_unclassified(100);k__Fungi_unclassified(100); | - | - | - | - | - | - | - | Unassigned | - |
| Otu35032 | 1 | k__Fungi(100);k__Fungi_unclassified(100);k__Fungi_unclassified(100);k__Fungi_unclassified(100);k__Fungi_unclassified(100);k__Fungi_unclassified(100);k__Fungi_unclassified(100); | - | - | - | - | - | - | - | Unassigned | - |
| Otu35033 | 1 | k__Fungi(100);k__Fungi_unclassified(100);k__Fungi_unclassified(100);k__Fungi_unclassified(100);k__Fungi_unclassified(100);k__Fungi_unclassified(100);k__Fungi_unclassified(100); | - | - | - | - | - | - | - | Unassigned | - |
| Otu35034 | 1 | k__Fungi(100);p__Ascomycota(100);p__Ascomycota_unclassified(100);p__Ascomycota_unclassified(100);p__Ascomycota_unclassified(100);p__Ascomycota_unclassified(100);p__Ascomycota_unclassified(100); | - | - | - | - | - | - | - | Unassigned | - |
| Otu35035 | 1 | k__Fungi(100);k__Fungi_unclassified(100);k__Fungi_unclassified(100);k__Fungi_unclassified(100);k__Fungi_unclassified(100);k__Fungi_unclassified(100);k__Fungi_unclassified(100); | - | - | - | - | - | - | - | Unassigned | - |
| Otu35036 | 1 | k__Fungi(100);p__Ascomycota(100);p__Ascomycota_unclassified(100);p__Ascomycota_unclassified(100);p__Ascomycota_unclassified(100);p__Ascomycota_unclassified(100);p__Ascomycota_unclassified(100); | - | - | - | - | - | - | - | Unassigned | - |
| Otu35039 | 1 | k__Fungi(100);p__Ascomycota(100);p__Ascomycota_unclassified(100);p__Ascomycota_unclassified(100);p__Ascomycota_unclassified(100);p__Ascomycota_unclassified(100);p__Ascomycota_unclassified(100); | - | - | - | - | - | - | - | Unassigned | - |
| Otu35041 | 1 | k__Fungi(100);p__Ascomycota(100);p__Ascomycota_unclassified(100);p__Ascomycota_unclassified(100);p__Ascomycota_unclassified(100);p__Ascomycota_unclassified(100);p__Ascomycota_unclassified(100); | - | - | - | - | - | - | - | Unassigned | - |
| Otu35042 | 1 | k__Fungi(100);p__Ascomycota(100);p__Ascomycota_unclassified(100);p__Ascomycota_unclassified(100);p__Ascomycota_unclassified(100);p__Ascomycota_unclassified(100);p__Ascomycota_unclassified(100); | - | - | - | - | - | - | - | Unassigned | - |
| Otu35043 | 1 | k__Fungi(100);k__Fungi_unclassified(100);k__Fungi_unclassified(100);k__Fungi_unclassified(100);k__Fungi_unclassified(100);k__Fungi_unclassified(100);k__Fungi_unclassified(100); | - | - | - | - | - | - | - | Unassigned | - |
| Otu35044 | 1 | k__Fungi(100);k__Fungi_unclassified(100);k__Fungi_unclassified(100);k__Fungi_unclassified(100);k__Fungi_unclassified(100);k__Fungi_unclassified(100);k__Fungi_unclassified(100); | - | - | - | - | - | - | - | Unassigned | - |
| Otu35046 | 1 | k__Fungi(100);p__Ascomycota(100);p__Ascomycota_unclassified(100);p__Ascomycota_unclassified(100);p__Ascomycota_unclassified(100);p__Ascomycota_unclassified(100);p__Ascomycota_unclassified(100); | - | - | - | - | - | - | - | Unassigned | - |
| Otu35047 | 1 | k__Fungi(100);k__Fungi_unclassified(100);k__Fungi_unclassified(100);k__Fungi_unclassified(100);k__Fungi_unclassified(100);k__Fungi_unclassified(100);k__Fungi_unclassified(100); | - | - | - | - | - | - | - | Unassigned | - |
| Otu35049 | 1 | k__Fungi(100);k__Fungi_unclassified(100);k__Fungi_unclassified(100);k__Fungi_unclassified(100);k__Fungi_unclassified(100);k__Fungi_unclassified(100);k__Fungi_unclassified(100); | - | - | - | - | - | - | - | Unassigned | - |
| Otu35050 | 1 | k__Fungi(100);k__Fungi_unclassified(100);k__Fungi_unclassified(100);k__Fungi_unclassified(100);k__Fungi_unclassified(100);k__Fungi_unclassified(100);k__Fungi_unclassified(100); | - | - | - | - | - | - | - | Unassigned | - |
| Otu35051 | 1 | k__Fungi(100);k__Fungi_unclassified(100);k__Fungi_unclassified(100);k__Fungi_unclassified(100);k__Fungi_unclassified(100);k__Fungi_unclassified(100);k__Fungi_unclassified(100); | - | - | - | - | - | - | - | Unassigned | - |
| Otu35053 | 1 | k__Fungi(100);k__Fungi_unclassified(100);k__Fungi_unclassified(100);k__Fungi_unclassified(100);k__Fungi_unclassified(100);k__Fungi_unclassified(100);k__Fungi_unclassified(100); | - | - | - | - | - | - | - | Unassigned | - |
| Otu35054 | 1 | k__Fungi(100);k__Fungi_unclassified(100);k__Fungi_unclassified(100);k__Fungi_unclassified(100);k__Fungi_unclassified(100);k__Fungi_unclassified(100);k__Fungi_unclassified(100); | - | - | - | - | - | - | - | Unassigned | - |
| Otu35055 | 1 | k__Fungi(100);p__Ascomycota(100);p__Ascomycota_unclassified(100);p__Ascomycota_unclassified(100);p__Ascomycota_unclassified(100);p__Ascomycota_unclassified(100);p__Ascomycota_unclassified(100); | - | - | - | - | - | - | - | Unassigned | - |
| Otu35057 | 1 | k__Fungi(100);p__Ascomycota(100);p__Ascomycota_unclassified(100);p__Ascomycota_unclassified(100);p__Ascomycota_unclassified(100);p__Ascomycota_unclassified(100);p__Ascomycota_unclassified(100); | - | - | - | - | - | - | - | Unassigned | - |
| Otu35058 | 1 | k__Fungi(100);k__Fungi_unclassified(100);k__Fungi_unclassified(100);k__Fungi_unclassified(100);k__Fungi_unclassified(100);k__Fungi_unclassified(100);k__Fungi_unclassified(100); | - | - | - | - | - | - | - | Unassigned | - |
| Otu35059 | 1 | k__Fungi(100);p__Ascomycota(100);p__Ascomycota_unclassified(100);p__Ascomycota_unclassified(100);p__Ascomycota_unclassified(100);p__Ascomycota_unclassified(100);p__Ascomycota_unclassified(100); | - | - | - | - | - | - | - | Unassigned | - |
| Otu35061 | 1 | k__Fungi(100);k__Fungi_unclassified(100);k__Fungi_unclassified(100);k__Fungi_unclassified(100);k__Fungi_unclassified(100);k__Fungi_unclassified(100);k__Fungi_unclassified(100); | - | - | - | - | - | - | - | Unassigned | - |
| Otu35062 | 1 | k__Fungi(100);k__Fungi_unclassified(100);k__Fungi_unclassified(100);k__Fungi_unclassified(100);k__Fungi_unclassified(100);k__Fungi_unclassified(100);k__Fungi_unclassified(100); | - | - | - | - | - | - | - | Unassigned | - |
| Otu35063 | 1 | k__Fungi(100);p__Ascomycota(100);p__Ascomycota_unclassified(100);p__Ascomycota_unclassified(100);p__Ascomycota_unclassified(100);p__Ascomycota_unclassified(100);p__Ascomycota_unclassified(100); | - | - | - | - | - | - | - | Unassigned | - |
| Otu35064 | 1 | k__Fungi(100);p__Ascomycota(100);p__Ascomycota_unclassified(100);p__Ascomycota_unclassified(100);p__Ascomycota_unclassified(100);p__Ascomycota_unclassified(100);p__Ascomycota_unclassified(100); | - | - | - | - | - | - | - | Unassigned | - |
| Otu35065 | 1 | k__Fungi(100);p__Ascomycota(100);p__Ascomycota_unclassified(100);p__Ascomycota_unclassified(100);p__Ascomycota_unclassified(100);p__Ascomycota_unclassified(100);p__Ascomycota_unclassified(100); | - | - | - | - | - | - | - | Unassigned | - |
| Otu35066 | 1 | k__Fungi(100);p__Ascomycota(100);p__Ascomycota_unclassified(100);p__Ascomycota_unclassified(100);p__Ascomycota_unclassified(100);p__Ascomycota_unclassified(100);p__Ascomycota_unclassified(100); | - | - | - | - | - | - | - | Unassigned | - |
| Otu35067 | 1 | k__Fungi(100);p__Ascomycota(100);p__Ascomycota_unclassified(100);p__Ascomycota_unclassified(100);p__Ascomycota_unclassified(100);p__Ascomycota_unclassified(100);p__Ascomycota_unclassified(100); | - | - | - | - | - | - | - | Unassigned | - |
| Otu35068 | 1 | k__Fungi(100);p__Ascomycota(100);p__Ascomycota_unclassified(100);p__Ascomycota_unclassified(100);p__Ascomycota_unclassified(100);p__Ascomycota_unclassified(100);p__Ascomycota_unclassified(100); | - | - | - | - | - | - | - | Unassigned | - |
| Otu35069 | 1 | k__Fungi(100);p__Ascomycota(100);p__Ascomycota_unclassified(100);p__Ascomycota_unclassified(100);p__Ascomycota_unclassified(100);p__Ascomycota_unclassified(100);p__Ascomycota_unclassified(100); | - | - | - | - | - | - | - | Unassigned | - |
| Otu35070 | 1 | k__Fungi(100);p__Ascomycota(100);p__Ascomycota_unclassified(100);p__Ascomycota_unclassified(100);p__Ascomycota_unclassified(100);p__Ascomycota_unclassified(100);p__Ascomycota_unclassified(100); | - | - | - | - | - | - | - | Unassigned | - |
| Otu35071 | 1 | k__Fungi(100);p__Ascomycota(100);p__Ascomycota_unclassified(100);p__Ascomycota_unclassified(100);p__Ascomycota_unclassified(100);p__Ascomycota_unclassified(100);p__Ascomycota_unclassified(100); | - | - | - | - | - | - | - | Unassigned | - |
| Otu35072 | 1 | k__Fungi(100);k__Fungi_unclassified(100);k__Fungi_unclassified(100);k__Fungi_unclassified(100);k__Fungi_unclassified(100);k__Fungi_unclassified(100);k__Fungi_unclassified(100); | - | - | - | - | - | - | - | Unassigned | - |
| Otu35073 | 1 | k__Fungi(100);p__Ascomycota(100);p__Ascomycota_unclassified(100);p__Ascomycota_unclassified(100);p__Ascomycota_unclassified(100);p__Ascomycota_unclassified(100);p__Ascomycota_unclassified(100); | - | - | - | - | - | - | - | Unassigned | - |
| Otu35074 | 1 | k__Fungi(100);p__Ascomycota(100);p__Ascomycota_unclassified(100);p__Ascomycota_unclassified(100);p__Ascomycota_unclassified(100);p__Ascomycota_unclassified(100);p__Ascomycota_unclassified(100); | - | - | - | - | - | - | - | Unassigned | - |
| Otu35075 | 1 | k__Fungi(100);k__Fungi_unclassified(100);k__Fungi_unclassified(100);k__Fungi_unclassified(100);k__Fungi_unclassified(100);k__Fungi_unclassified(100);k__Fungi_unclassified(100); | - | - | - | - | - | - | - | Unassigned | - |
| Otu35081 | 1 | k__Fungi(100);k__Fungi_unclassified(100);k__Fungi_unclassified(100);k__Fungi_unclassified(100);k__Fungi_unclassified(100);k__Fungi_unclassified(100);k__Fungi_unclassified(100); | - | - | - | - | - | - | - | Unassigned | - |
| Otu35082 | 1 | k__Fungi(100);k__Fungi_unclassified(100);k__Fungi_unclassified(100);k__Fungi_unclassified(100);k__Fungi_unclassified(100);k__Fungi_unclassified(100);k__Fungi_unclassified(100); | - | - | - | - | - | - | - | Unassigned | - |
| Otu35089 | 1 | k__Fungi(100);k__Fungi_unclassified(100);k__Fungi_unclassified(100);k__Fungi_unclassified(100);k__Fungi_unclassified(100);k__Fungi_unclassified(100);k__Fungi_unclassified(100); | - | - | - | - | - | - | - | Unassigned | - |
| Otu35090 | 1 | k__Fungi(100);k__Fungi_unclassified(100);k__Fungi_unclassified(100);k__Fungi_unclassified(100);k__Fungi_unclassified(100);k__Fungi_unclassified(100);k__Fungi_unclassified(100); | - | - | - | - | - | - | - | Unassigned | - |
| Otu35091 | 1 | k__Fungi(100);p__Ascomycota(100);p__Ascomycota_unclassified(100);p__Ascomycota_unclassified(100);p__Ascomycota_unclassified(100);p__Ascomycota_unclassified(100);p__Ascomycota_unclassified(100); | - | - | - | - | - | - | - | Unassigned | - |
| Otu35092 | 1 | k__Fungi(100);p__Ascomycota(100);p__Ascomycota_unclassified(100);p__Ascomycota_unclassified(100);p__Ascomycota_unclassified(100);p__Ascomycota_unclassified(100);p__Ascomycota_unclassified(100); | - | - | - | - | - | - | - | Unassigned | - |
| Otu35093 | 1 | k__Fungi(100);k__Fungi_unclassified(100);k__Fungi_unclassified(100);k__Fungi_unclassified(100);k__Fungi_unclassified(100);k__Fungi_unclassified(100);k__Fungi_unclassified(100); | - | - | - | - | - | - | - | Unassigned | - |
| Otu35094 | 1 | k__Fungi(100);p__Ascomycota(100);p__Ascomycota_unclassified(100);p__Ascomycota_unclassified(100);p__Ascomycota_unclassified(100);p__Ascomycota_unclassified(100);p__Ascomycota_unclassified(100); | - | - | - | - | - | - | - | Unassigned | - |
| Otu35095 | 1 | k__Fungi(100);k__Fungi_unclassified(100);k__Fungi_unclassified(100);k__Fungi_unclassified(100);k__Fungi_unclassified(100);k__Fungi_unclassified(100);k__Fungi_unclassified(100); | - | - | - | - | - | - | - | Unassigned | - |
| Otu35096 | 1 | k__Fungi(100);p__Ascomycota(100);p__Ascomycota_unclassified(100);p__Ascomycota_unclassified(100);p__Ascomycota_unclassified(100);p__Ascomycota_unclassified(100);p__Ascomycota_unclassified(100); | - | - | - | - | - | - | - | Unassigned | - |
| Otu35097 | 1 | k__Fungi(100);k__Fungi_unclassified(100);k__Fungi_unclassified(100);k__Fungi_unclassified(100);k__Fungi_unclassified(100);k__Fungi_unclassified(100);k__Fungi_unclassified(100); | - | - | - | - | - | - | - | Unassigned | - |
| Otu35098 | 1 | k__Fungi(100);p__Ascomycota(100);p__Ascomycota_unclassified(100);p__Ascomycota_unclassified(100);p__Ascomycota_unclassified(100);p__Ascomycota_unclassified(100);p__Ascomycota_unclassified(100); | - | - | - | - | - | - | - | Unassigned | - |
| Otu35099 | 1 | k__Fungi(100);p__Ascomycota(100);p__Ascomycota_unclassified(100);p__Ascomycota_unclassified(100);p__Ascomycota_unclassified(100);p__Ascomycota_unclassified(100);p__Ascomycota_unclassified(100); | - | - | - | - | - | - | - | Unassigned | - |
| Otu35100 | 1 | k__Fungi(100);k__Fungi_unclassified(100);k__Fungi_unclassified(100);k__Fungi_unclassified(100);k__Fungi_unclassified(100);k__Fungi_unclassified(100);k__Fungi_unclassified(100); | - | - | - | - | - | - | - | Unassigned | - |
| Otu35101 | 1 | k__Fungi(100);p__Ascomycota(100);p__Ascomycota_unclassified(100);p__Ascomycota_unclassified(100);p__Ascomycota_unclassified(100);p__Ascomycota_unclassified(100);p__Ascomycota_unclassified(100); | - | - | - | - | - | - | - | Unassigned | - |
| Otu35102 | 1 | k__Fungi(100);p__Ascomycota(100);p__Ascomycota_unclassified(100);p__Ascomycota_unclassified(100);p__Ascomycota_unclassified(100);p__Ascomycota_unclassified(100);p__Ascomycota_unclassified(100); | - | - | - | - | - | - | - | Unassigned | - |
| Otu35103 | 1 | k__Fungi(100);p__Ascomycota(100);p__Ascomycota_unclassified(100);p__Ascomycota_unclassified(100);p__Ascomycota_unclassified(100);p__Ascomycota_unclassified(100);p__Ascomycota_unclassified(100); | - | - | - | - | - | - | - | Unassigned | - |
| Otu35104 | 1 | k__Fungi(100);p__Ascomycota(100);p__Ascomycota_unclassified(100);p__Ascomycota_unclassified(100);p__Ascomycota_unclassified(100);p__Ascomycota_unclassified(100);p__Ascomycota_unclassified(100); | - | - | - | - | - | - | - | Unassigned | - |
| Otu35105 | 1 | k__Fungi(100);p__Ascomycota(100);p__Ascomycota_unclassified(100);p__Ascomycota_unclassified(100);p__Ascomycota_unclassified(100);p__Ascomycota_unclassified(100);p__Ascomycota_unclassified(100); | - | - | - | - | - | - | - | Unassigned | - |
| Otu35106 | 1 | k__Fungi(100);p__Ascomycota(100);p__Ascomycota_unclassified(100);p__Ascomycota_unclassified(100);p__Ascomycota_unclassified(100);p__Ascomycota_unclassified(100);p__Ascomycota_unclassified(100); | - | - | - | - | - | - | - | Unassigned | - |
| Otu35107 | 1 | k__Fungi(100);p__Ascomycota(100);p__Ascomycota_unclassified(100);p__Ascomycota_unclassified(100);p__Ascomycota_unclassified(100);p__Ascomycota_unclassified(100);p__Ascomycota_unclassified(100); | - | - | - | - | - | - | - | Unassigned | - |
| Otu35108 | 1 | k__Fungi(100);p__Ascomycota(100);p__Ascomycota_unclassified(100);p__Ascomycota_unclassified(100);p__Ascomycota_unclassified(100);p__Ascomycota_unclassified(100);p__Ascomycota_unclassified(100); | - | - | - | - | - | - | - | Unassigned | - |
| Otu35110 | 1 | k__Fungi(100);k__Fungi_unclassified(100);k__Fungi_unclassified(100);k__Fungi_unclassified(100);k__Fungi_unclassified(100);k__Fungi_unclassified(100);k__Fungi_unclassified(100); | - | - | - | - | - | - | - | Unassigned | - |
| Otu35112 | 1 | k__Fungi(100);k__Fungi_unclassified(100);k__Fungi_unclassified(100);k__Fungi_unclassified(100);k__Fungi_unclassified(100);k__Fungi_unclassified(100);k__Fungi_unclassified(100); | - | - | - | - | - | - | - | Unassigned | - |
| Otu35115 | 1 | k__Fungi(100);k__Fungi_unclassified(100);k__Fungi_unclassified(100);k__Fungi_unclassified(100);k__Fungi_unclassified(100);k__Fungi_unclassified(100);k__Fungi_unclassified(100); | - | - | - | - | - | - | - | Unassigned | - |
| Otu35117 | 1 | k__Fungi(100);k__Fungi_unclassified(100);k__Fungi_unclassified(100);k__Fungi_unclassified(100);k__Fungi_unclassified(100);k__Fungi_unclassified(100);k__Fungi_unclassified(100); | - | - | - | - | - | - | - | Unassigned | - |
| Otu35118 | 1 | k__Fungi(100);p__Ascomycota(100);p__Ascomycota_unclassified(100);p__Ascomycota_unclassified(100);p__Ascomycota_unclassified(100);p__Ascomycota_unclassified(100);p__Ascomycota_unclassified(100); | - | - | - | - | - | - | - | Unassigned | - |
| Otu35120 | 1 | k__Fungi(100);p__Ascomycota(100);p__Ascomycota_unclassified(100);p__Ascomycota_unclassified(100);p__Ascomycota_unclassified(100);p__Ascomycota_unclassified(100);p__Ascomycota_unclassified(100); | - | - | - | - | - | - | - | Unassigned | - |
| Otu35121 | 1 | k__Fungi(100);p__Ascomycota(100);p__Ascomycota_unclassified(100);p__Ascomycota_unclassified(100);p__Ascomycota_unclassified(100);p__Ascomycota_unclassified(100);p__Ascomycota_unclassified(100); | - | - | - | - | - | - | - | Unassigned | - |
| Otu35122 | 1 | k__Fungi(100);p__Ascomycota(100);p__Ascomycota_unclassified(100);p__Ascomycota_unclassified(100);p__Ascomycota_unclassified(100);p__Ascomycota_unclassified(100);p__Ascomycota_unclassified(100); | - | - | - | - | - | - | - | Unassigned | - |
| Otu35123 | 1 | k__Fungi(100);k__Fungi_unclassified(100);k__Fungi_unclassified(100);k__Fungi_unclassified(100);k__Fungi_unclassified(100);k__Fungi_unclassified(100);k__Fungi_unclassified(100); | - | - | - | - | - | - | - | Unassigned | - |
| Otu35124 | 1 | k__Fungi(100);k__Fungi_unclassified(100);k__Fungi_unclassified(100);k__Fungi_unclassified(100);k__Fungi_unclassified(100);k__Fungi_unclassified(100);k__Fungi_unclassified(100); | - | - | - | - | - | - | - | Unassigned | - |
| Otu35126 | 1 | k__Fungi(100);p__Ascomycota(100);p__Ascomycota_unclassified(100);p__Ascomycota_unclassified(100);p__Ascomycota_unclassified(100);p__Ascomycota_unclassified(100);p__Ascomycota_unclassified(100); | - | - | - | - | - | - | - | Unassigned | - |
| Otu35128 | 1 | k__Fungi(100);p__Ascomycota(100);p__Ascomycota_unclassified(100);p__Ascomycota_unclassified(100);p__Ascomycota_unclassified(100);p__Ascomycota_unclassified(100);p__Ascomycota_unclassified(100); | - | - | - | - | - | - | - | Unassigned | - |
| Otu35129 | 1 | k__Fungi(100);k__Fungi_unclassified(100);k__Fungi_unclassified(100);k__Fungi_unclassified(100);k__Fungi_unclassified(100);k__Fungi_unclassified(100);k__Fungi_unclassified(100); | - | - | - | - | - | - | - | Unassigned | - |
| Otu35131 | 1 | k__Fungi(100);p__Ascomycota(100);p__Ascomycota_unclassified(100);p__Ascomycota_unclassified(100);p__Ascomycota_unclassified(100);p__Ascomycota_unclassified(100);p__Ascomycota_unclassified(100); | - | - | - | - | - | - | - | Unassigned | - |
| Otu35141 | 1 | k__Fungi(100);p__Ascomycota(100);p__Ascomycota_unclassified(100);p__Ascomycota_unclassified(100);p__Ascomycota_unclassified(100);p__Ascomycota_unclassified(100);p__Ascomycota_unclassified(100); | - | - | - | - | - | - | - | Unassigned | - |
| Otu35155 | 1 | k__Fungi(100);k__Fungi_unclassified(100);k__Fungi_unclassified(100);k__Fungi_unclassified(100);k__Fungi_unclassified(100);k__Fungi_unclassified(100);k__Fungi_unclassified(100); | - | - | - | - | - | - | - | Unassigned | - |
| Otu35157 | 1 | k__Fungi(100);k__Fungi_unclassified(100);k__Fungi_unclassified(100);k__Fungi_unclassified(100);k__Fungi_unclassified(100);k__Fungi_unclassified(100);k__Fungi_unclassified(100); | - | - | - | - | - | - | - | Unassigned | - |
| Otu35161 | 1 | k__Fungi(100);k__Fungi_unclassified(100);k__Fungi_unclassified(100);k__Fungi_unclassified(100);k__Fungi_unclassified(100);k__Fungi_unclassified(100);k__Fungi_unclassified(100); | - | - | - | - | - | - | - | Unassigned | - |
| Otu35171 | 1 | k__Fungi(100);k__Fungi_unclassified(100);k__Fungi_unclassified(100);k__Fungi_unclassified(100);k__Fungi_unclassified(100);k__Fungi_unclassified(100);k__Fungi_unclassified(100); | - | - | - | - | - | - | - | Unassigned | - |
| Otu35173 | 1 | k__Fungi(100);k__Fungi_unclassified(100);k__Fungi_unclassified(100);k__Fungi_unclassified(100);k__Fungi_unclassified(100);k__Fungi_unclassified(100);k__Fungi_unclassified(100); | - | - | - | - | - | - | - | Unassigned | - |
| Otu35174 | 1 | k__Fungi(100);p__Ascomycota(100);p__Ascomycota_unclassified(100);p__Ascomycota_unclassified(100);p__Ascomycota_unclassified(100);p__Ascomycota_unclassified(100);p__Ascomycota_unclassified(100); | - | - | - | - | - | - | - | Unassigned | - |
| Otu35175 | 1 | k__Fungi(100);k__Fungi_unclassified(100);k__Fungi_unclassified(100);k__Fungi_unclassified(100);k__Fungi_unclassified(100);k__Fungi_unclassified(100);k__Fungi_unclassified(100); | - | - | - | - | - | - | - | Unassigned | - |
| Otu35176 | 1 | k__Fungi(100);k__Fungi_unclassified(100);k__Fungi_unclassified(100);k__Fungi_unclassified(100);k__Fungi_unclassified(100);k__Fungi_unclassified(100);k__Fungi_unclassified(100); | - | - | - | - | - | - | - | Unassigned | - |
| Otu35177 | 1 | k__Fungi(100);k__Fungi_unclassified(100);k__Fungi_unclassified(100);k__Fungi_unclassified(100);k__Fungi_unclassified(100);k__Fungi_unclassified(100);k__Fungi_unclassified(100); | - | - | - | - | - | - | - | Unassigned | - |
| Otu35182 | 1 | k__Fungi(100);p__Ascomycota(100);p__Ascomycota_unclassified(100);p__Ascomycota_unclassified(100);p__Ascomycota_unclassified(100);p__Ascomycota_unclassified(100);p__Ascomycota_unclassified(100); | - | - | - | - | - | - | - | Unassigned | - |
| Otu35198 | 1 | k__Fungi(100);k__Fungi_unclassified(100);k__Fungi_unclassified(100);k__Fungi_unclassified(100);k__Fungi_unclassified(100);k__Fungi_unclassified(100);k__Fungi_unclassified(100); | - | - | - | - | - | - | - | Unassigned | - |
| Otu35202 | 1 | k__Fungi(100);p__Ascomycota(100);p__Ascomycota_unclassified(100);p__Ascomycota_unclassified(100);p__Ascomycota_unclassified(100);p__Ascomycota_unclassified(100);p__Ascomycota_unclassified(100); | - | - | - | - | - | - | - | Unassigned | - |
| Otu35211 | 1 | k__Fungi(100);p__Ascomycota(100);p__Ascomycota_unclassified(100);p__Ascomycota_unclassified(100);p__Ascomycota_unclassified(100);p__Ascomycota_unclassified(100);p__Ascomycota_unclassified(100); | - | - | - | - | - | - | - | Unassigned | - |
| Otu35212 | 1 | k__Fungi(100);k__Fungi_unclassified(100);k__Fungi_unclassified(100);k__Fungi_unclassified(100);k__Fungi_unclassified(100);k__Fungi_unclassified(100);k__Fungi_unclassified(100); | - | - | - | - | - | - | - | Unassigned | - |
| Otu35213 | 1 | k__Fungi(100);p__Ascomycota(100);p__Ascomycota_unclassified(100);p__Ascomycota_unclassified(100);p__Ascomycota_unclassified(100);p__Ascomycota_unclassified(100);p__Ascomycota_unclassified(100); | - | - | - | - | - | - | - | Unassigned | - |
| Otu35220 | 1 | k__Fungi(100);k__Fungi_unclassified(100);k__Fungi_unclassified(100);k__Fungi_unclassified(100);k__Fungi_unclassified(100);k__Fungi_unclassified(100);k__Fungi_unclassified(100); | - | - | - | - | - | - | - | Unassigned | - |
| Otu35227 | 1 | k__Fungi(100);p__Ascomycota(100);p__Ascomycota_unclassified(100);p__Ascomycota_unclassified(100);p__Ascomycota_unclassified(100);p__Ascomycota_unclassified(100);p__Ascomycota_unclassified(100); | - | - | - | - | - | - | - | Unassigned | - |
| Otu35228 | 1 | k__Fungi(100);p__Ascomycota(100);p__Ascomycota_unclassified(100);p__Ascomycota_unclassified(100);p__Ascomycota_unclassified(100);p__Ascomycota_unclassified(100);p__Ascomycota_unclassified(100); | - | - | - | - | - | - | - | Unassigned | - |
| Otu35236 | 1 | k__Fungi(100);k__Fungi_unclassified(100);k__Fungi_unclassified(100);k__Fungi_unclassified(100);k__Fungi_unclassified(100);k__Fungi_unclassified(100);k__Fungi_unclassified(100); | - | - | - | - | - | - | - | Unassigned | - |
| Otu35239 | 1 | k__Fungi(100);p__Ascomycota(100);p__Ascomycota_unclassified(100);p__Ascomycota_unclassified(100);p__Ascomycota_unclassified(100);p__Ascomycota_unclassified(100);p__Ascomycota_unclassified(100); | - | - | - | - | - | - | - | Unassigned | - |
| Otu35240 | 1 | k__Fungi(100);p__Ascomycota(100);p__Ascomycota_unclassified(100);p__Ascomycota_unclassified(100);p__Ascomycota_unclassified(100);p__Ascomycota_unclassified(100);p__Ascomycota_unclassified(100); | - | - | - | - | - | - | - | Unassigned | - |
| Otu35242 | 1 | k__Fungi(100);k__Fungi_unclassified(100);k__Fungi_unclassified(100);k__Fungi_unclassified(100);k__Fungi_unclassified(100);k__Fungi_unclassified(100);k__Fungi_unclassified(100); | - | - | - | - | - | - | - | Unassigned | - |
| Otu35243 | 1 | k__Fungi(100);k__Fungi_unclassified(100);k__Fungi_unclassified(100);k__Fungi_unclassified(100);k__Fungi_unclassified(100);k__Fungi_unclassified(100);k__Fungi_unclassified(100); | - | - | - | - | - | - | - | Unassigned | - |
| Otu35244 | 1 | k__Fungi(100);k__Fungi_unclassified(100);k__Fungi_unclassified(100);k__Fungi_unclassified(100);k__Fungi_unclassified(100);k__Fungi_unclassified(100);k__Fungi_unclassified(100); | - | - | - | - | - | - | - | Unassigned | - |
| Otu35245 | 1 | k__Fungi(100);p__Ascomycota(100);p__Ascomycota_unclassified(100);p__Ascomycota_unclassified(100);p__Ascomycota_unclassified(100);p__Ascomycota_unclassified(100);p__Ascomycota_unclassified(100); | - | - | - | - | - | - | - | Unassigned | - |
| Otu35260 | 1 | k__Fungi(100);k__Fungi_unclassified(100);k__Fungi_unclassified(100);k__Fungi_unclassified(100);k__Fungi_unclassified(100);k__Fungi_unclassified(100);k__Fungi_unclassified(100); | - | - | - | - | - | - | - | Unassigned | - |
| Otu35261 | 1 | k__Fungi(100);k__Fungi_unclassified(100);k__Fungi_unclassified(100);k__Fungi_unclassified(100);k__Fungi_unclassified(100);k__Fungi_unclassified(100);k__Fungi_unclassified(100); | - | - | - | - | - | - | - | Unassigned | - |
| Otu35262 | 1 | k__Fungi(100);p__Ascomycota(100);p__Ascomycota_unclassified(100);p__Ascomycota_unclassified(100);p__Ascomycota_unclassified(100);p__Ascomycota_unclassified(100);p__Ascomycota_unclassified(100); | - | - | - | - | - | - | - | Unassigned | - |
| Otu35265 | 1 | k__Fungi(100);p__Ascomycota(100);p__Ascomycota_unclassified(100);p__Ascomycota_unclassified(100);p__Ascomycota_unclassified(100);p__Ascomycota_unclassified(100);p__Ascomycota_unclassified(100); | - | - | - | - | - | - | - | Unassigned | - |
| Otu35266 | 1 | k__Fungi(100);k__Fungi_unclassified(100);k__Fungi_unclassified(100);k__Fungi_unclassified(100);k__Fungi_unclassified(100);k__Fungi_unclassified(100);k__Fungi_unclassified(100); | - | - | - | - | - | - | - | Unassigned | - |
| Otu35269 | 1 | k__Fungi(100);k__Fungi_unclassified(100);k__Fungi_unclassified(100);k__Fungi_unclassified(100);k__Fungi_unclassified(100);k__Fungi_unclassified(100);k__Fungi_unclassified(100); | - | - | - | - | - | - | - | Unassigned | - |
| Otu35270 | 1 | k__Fungi(100);k__Fungi_unclassified(100);k__Fungi_unclassified(100);k__Fungi_unclassified(100);k__Fungi_unclassified(100);k__Fungi_unclassified(100);k__Fungi_unclassified(100); | - | - | - | - | - | - | - | Unassigned | - |
| Otu35271 | 1 | k__Fungi(100);p__Ascomycota(100);p__Ascomycota_unclassified(100);p__Ascomycota_unclassified(100);p__Ascomycota_unclassified(100);p__Ascomycota_unclassified(100);p__Ascomycota_unclassified(100); | - | - | - | - | - | - | - | Unassigned | - |
| Otu35274 | 1 | k__Fungi(100);k__Fungi_unclassified(100);k__Fungi_unclassified(100);k__Fungi_unclassified(100);k__Fungi_unclassified(100);k__Fungi_unclassified(100);k__Fungi_unclassified(100); | - | - | - | - | - | - | - | Unassigned | - |
| Otu35278 | 1 | k__Fungi(100);k__Fungi_unclassified(100);k__Fungi_unclassified(100);k__Fungi_unclassified(100);k__Fungi_unclassified(100);k__Fungi_unclassified(100);k__Fungi_unclassified(100); | - | - | - | - | - | - | - | Unassigned | - |
| Otu35280 | 1 | k__Fungi(100);p__Ascomycota(100);p__Ascomycota_unclassified(100);p__Ascomycota_unclassified(100);p__Ascomycota_unclassified(100);p__Ascomycota_unclassified(100);p__Ascomycota_unclassified(100); | - | - | - | - | - | - | - | Unassigned | - |
| Otu35281 | 1 | k__Fungi(100);k__Fungi_unclassified(100);k__Fungi_unclassified(100);k__Fungi_unclassified(100);k__Fungi_unclassified(100);k__Fungi_unclassified(100);k__Fungi_unclassified(100); | - | - | - | - | - | - | - | Unassigned | - |
| Otu35282 | 1 | k__Fungi(100);p__Ascomycota(100);p__Ascomycota_unclassified(100);p__Ascomycota_unclassified(100);p__Ascomycota_unclassified(100);p__Ascomycota_unclassified(100);p__Ascomycota_unclassified(100); | - | - | - | - | - | - | - | Unassigned | - |

| | | | | | | | | | | | |
|---|---|---|---|---|---|---|---|---|---|---|---|
| Otu35287 | 1 | k__Fungi(100);p__Ascomycota(100);p__Ascomycota_unclassified(100);p__Ascomycota_unclassified(100);p__Ascomycota_unclassified(100);p__Ascomycota_unclassified(100);p__Ascomycota_unclassified(100); | - | - | - | - | - | - | - | Unassigned | - |
| Otu35288 | 1 | k__Fungi(100);k__Fungi_unclassified(100);k__Fungi_unclassified(100);k__Fungi_unclassified(100);k__Fungi_unclassified(100);k__Fungi_unclassified(100);k__Fungi_unclassified(100); | - | - | - | - | - | - | - | Unassigned | - |
| Otu35289 | 1 | k__Fungi(100);p__Ascomycota(100);p__Ascomycota_unclassified(100);p__Ascomycota_unclassified(100);p__Ascomycota_unclassified(100);p__Ascomycota_unclassified(100);p__Ascomycota_unclassified(100); | - | - | - | - | - | - | - | Unassigned | - |
| Otu35290 | 1 | k__Fungi(100);p__Ascomycota(100);p__Ascomycota_unclassified(100);p__Ascomycota_unclassified(100);p__Ascomycota_unclassified(100);p__Ascomycota_unclassified(100);p__Ascomycota_unclassified(100); | - | - | - | - | - | - | - | Unassigned | - |
| Otu35291 | 1 | k__Fungi(100);p__Ascomycota(100);p__Ascomycota_unclassified(100);p__Ascomycota_unclassified(100);p__Ascomycota_unclassified(100);p__Ascomycota_unclassified(100);p__Ascomycota_unclassified(100); | - | - | - | - | - | - | - | Unassigned | - |
| Otu35292 | 1 | k__Fungi(100);p__Ascomycota(100);p__Ascomycota_unclassified(100);p__Ascomycota_unclassified(100);p__Ascomycota_unclassified(100);p__Ascomycota_unclassified(100);p__Ascomycota_unclassified(100); | - | - | - | - | - | - | - | Unassigned | - |
| Otu35293 | 1 | k__Fungi(100);p__Ascomycota(100);p__Ascomycota_unclassified(100);p__Ascomycota_unclassified(100);p__Ascomycota_unclassified(100);p__Ascomycota_unclassified(100);p__Ascomycota_unclassified(100); | - | - | - | - | - | - | - | Unassigned | - |
| Otu35294 | 1 | k__Fungi(100);k__Fungi_unclassified(100);k__Fungi_unclassified(100);k__Fungi_unclassified(100);k__Fungi_unclassified(100);k__Fungi_unclassified(100);k__Fungi_unclassified(100); | - | - | - | - | - | - | - | Unassigned | - |
| Otu35295 | 1 | k__Fungi(100);p__Ascomycota(100);p__Ascomycota_unclassified(100);p__Ascomycota_unclassified(100);p__Ascomycota_unclassified(100);p__Ascomycota_unclassified(100);p__Ascomycota_unclassified(100); | - | - | - | - | - | - | - | Unassigned | - |
| Otu35296 | 1 | k__Fungi(100);k__Fungi_unclassified(100);k__Fungi_unclassified(100);k__Fungi_unclassified(100);k__Fungi_unclassified(100);k__Fungi_unclassified(100);k__Fungi_unclassified(100); | - | - | - | - | - | - | - | Unassigned | - |
| Otu35297 | 1 | k__Fungi(100);p__Ascomycota(100);p__Ascomycota_unclassified(100);p__Ascomycota_unclassified(100);p__Ascomycota_unclassified(100);p__Ascomycota_unclassified(100);p__Ascomycota_unclassified(100); | - | - | - | - | - | - | - | Unassigned | - |
| Otu35299 | 1 | k__Fungi(100);k__Fungi_unclassified(100);k__Fungi_unclassified(100);k__Fungi_unclassified(100);k__Fungi_unclassified(100);k__Fungi_unclassified(100);k__Fungi_unclassified(100); | - | - | - | - | - | - | - | Unassigned | - |
| Otu35300 | 1 | k__Fungi(100);k__Fungi_unclassified(100);k__Fungi_unclassified(100);k__Fungi_unclassified(100);k__Fungi_unclassified(100);k__Fungi_unclassified(100);k__Fungi_unclassified(100); | - | - | - | - | - | - | - | Unassigned | - |
| Otu35301 | 1 | k__Fungi(100);p__Ascomycota(100);p__Ascomycota_unclassified(100);p__Ascomycota_unclassified(100);p__Ascomycota_unclassified(100);p__Ascomycota_unclassified(100);p__Ascomycota_unclassified(100); | - | - | - | - | - | - | - | Unassigned | - |
| Otu35303 | 1 | k__Fungi(100);p__Ascomycota(100);p__Ascomycota_unclassified(100);p__Ascomycota_unclassified(100);p__Ascomycota_unclassified(100);p__Ascomycota_unclassified(100);p__Ascomycota_unclassified(100); | - | - | - | - | - | - | - | Unassigned | - |
| Otu35304 | 1 | k__Fungi(100);k__Fungi_unclassified(100);k__Fungi_unclassified(100);k__Fungi_unclassified(100);k__Fungi_unclassified(100);k__Fungi_unclassified(100);k__Fungi_unclassified(100); | - | - | - | - | - | - | - | Unassigned | - |
| Otu35305 | 1 | k__Fungi(100);k__Fungi_unclassified(100);k__Fungi_unclassified(100);k__Fungi_unclassified(100);k__Fungi_unclassified(100);k__Fungi_unclassified(100);k__Fungi_unclassified(100); | - | - | - | - | - | - | - | Unassigned | - |
| Otu35307 | 1 | k__Fungi(100);p__Ascomycota(100);p__Ascomycota_unclassified(100);p__Ascomycota_unclassified(100);p__Ascomycota_unclassified(100);p__Ascomycota_unclassified(100);p__Ascomycota_unclassified(100); | - | - | - | - | - | - | - | Unassigned | - |
| Otu35308 | 1 | k__Fungi(100);k__Fungi_unclassified(100);k__Fungi_unclassified(100);k__Fungi_unclassified(100);k__Fungi_unclassified(100);k__Fungi_unclassified(100);k__Fungi_unclassified(100); | - | - | - | - | - | - | - | Unassigned | - |
| Otu35311 | 1 | k__Fungi(100);p__Ascomycota(100);p__Ascomycota_unclassified(100);p__Ascomycota_unclassified(100);p__Ascomycota_unclassified(100);p__Ascomycota_unclassified(100);p__Ascomycota_unclassified(100); | - | - | - | - | - | - | - | Unassigned | - |
| Otu35314 | 1 | k__Fungi(100);k__Fungi_unclassified(100);k__Fungi_unclassified(100);k__Fungi_unclassified(100);k__Fungi_unclassified(100);k__Fungi_unclassified(100);k__Fungi_unclassified(100); | - | - | - | - | - | - | - | Unassigned | - |
| Otu35316 | 1 | k__Fungi(100);k__Fungi_unclassified(100);k__Fungi_unclassified(100);k__Fungi_unclassified(100);k__Fungi_unclassified(100);k__Fungi_unclassified(100);k__Fungi_unclassified(100); | - | - | - | - | - | - | - | Unassigned | - |
| Otu35318 | 1 | k__Fungi(100);k__Fungi_unclassified(100);k__Fungi_unclassified(100);k__Fungi_unclassified(100);k__Fungi_unclassified(100);k__Fungi_unclassified(100);k__Fungi_unclassified(100); | - | - | - | - | - | - | - | Unassigned | - |
| Otu35333 | 1 | k__Fungi(100);k__Fungi_unclassified(100);k__Fungi_unclassified(100);k__Fungi_unclassified(100);k__Fungi_unclassified(100);k__Fungi_unclassified(100);k__Fungi_unclassified(100); | - | - | - | - | - | - | - | Unassigned | - |
| Otu35334 | 1 | k__Fungi(100);p__Ascomycota(100);p__Ascomycota_unclassified(100);p__Ascomycota_unclassified(100);p__Ascomycota_unclassified(100);p__Ascomycota_unclassified(100);p__Ascomycota_unclassified(100); | - | - | - | - | - | - | - | Unassigned | - |
| Otu35337 | 1 | k__Fungi(100);k__Fungi_unclassified(100);k__Fungi_unclassified(100);k__Fungi_unclassified(100);k__Fungi_unclassified(100);k__Fungi_unclassified(100);k__Fungi_unclassified(100); | - | - | - | - | - | - | - | Unassigned | - |
| Otu35338 | 1 | k__Fungi(100);k__Fungi_unclassified(100);k__Fungi_unclassified(100);k__Fungi_unclassified(100);k__Fungi_unclassified(100);k__Fungi_unclassified(100);k__Fungi_unclassified(100); | - | - | - | - | - | - | - | Unassigned | - |
| Otu35340 | 1 | k__Fungi(100);k__Fungi_unclassified(100);k__Fungi_unclassified(100);k__Fungi_unclassified(100);k__Fungi_unclassified(100);k__Fungi_unclassified(100);k__Fungi_unclassified(100); | - | - | - | - | - | - | - | Unassigned | - |
| Otu35341 | 1 | k__Fungi(100);k__Fungi_unclassified(100);k__Fungi_unclassified(100);k__Fungi_unclassified(100);k__Fungi_unclassified(100);k__Fungi_unclassified(100);k__Fungi_unclassified(100); | - | - | - | - | - | - | - | Unassigned | - |
| Otu35342 | 1 | k__Fungi(100);k__Fungi_unclassified(100);k__Fungi_unclassified(100);k__Fungi_unclassified(100);k__Fungi_unclassified(100);k__Fungi_unclassified(100);k__Fungi_unclassified(100); | - | - | - | - | - | - | - | Unassigned | - |
| Otu35346 | 1 | k__Fungi(100);k__Fungi_unclassified(100);k__Fungi_unclassified(100);k__Fungi_unclassified(100);k__Fungi_unclassified(100);k__Fungi_unclassified(100);k__Fungi_unclassified(100); | - | - | - | - | - | - | - | Unassigned | - |
| Otu35347 | 1 | k__Fungi(100);k__Fungi_unclassified(100);k__Fungi_unclassified(100);k__Fungi_unclassified(100);k__Fungi_unclassified(100);k__Fungi_unclassified(100);k__Fungi_unclassified(100); | - | - | - | - | - | - | - | Unassigned | - |
| Otu35348 | 1 | k__Fungi(100);p__Ascomycota(100);p__Ascomycota_unclassified(100);p__Ascomycota_unclassified(100);p__Ascomycota_unclassified(100);p__Ascomycota_unclassified(100);p__Ascomycota_unclassified(100); | - | - | - | - | - | - | - | Unassigned | - |
| Otu35350 | 1 | k__Fungi(100);p__Ascomycota(100);p__Ascomycota_unclassified(100);p__Ascomycota_unclassified(100);p__Ascomycota_unclassified(100);p__Ascomycota_unclassified(100);p__Ascomycota_unclassified(100); | - | - | - | - | - | - | - | Unassigned | - |
| Otu35352 | 1 | k__Fungi(100);p__Ascomycota(100);p__Ascomycota_unclassified(100);p__Ascomycota_unclassified(100);p__Ascomycota_unclassified(100);p__Ascomycota_unclassified(100);p__Ascomycota_unclassified(100); | - | - | - | - | - | - | - | Unassigned | - |
| Otu35353 | 1 | k__Fungi(100);k__Fungi_unclassified(100);k__Fungi_unclassified(100);k__Fungi_unclassified(100);k__Fungi_unclassified(100);k__Fungi_unclassified(100);k__Fungi_unclassified(100); | - | - | - | - | - | - | - | Unassigned | - |
| Otu35354 | 1 | k__Fungi(100);k__Fungi_unclassified(100);k__Fungi_unclassified(100);k__Fungi_unclassified(100);k__Fungi_unclassified(100);k__Fungi_unclassified(100);k__Fungi_unclassified(100); | - | - | - | - | - | - | - | Unassigned | - |
| Otu35355 | 1 | k__Fungi(100);k__Fungi_unclassified(100);k__Fungi_unclassified(100);k__Fungi_unclassified(100);k__Fungi_unclassified(100);k__Fungi_unclassified(100);k__Fungi_unclassified(100); | - | - | - | - | - | - | - | Unassigned | - |
| Otu35366 | 1 | k__Fungi(100);k__Fungi_unclassified(100);k__Fungi_unclassified(100);k__Fungi_unclassified(100);k__Fungi_unclassified(100);k__Fungi_unclassified(100);k__Fungi_unclassified(100); | - | - | - | - | - | - | - | Unassigned | - |
| Otu35367 | 1 | k__Fungi(100);p__Ascomycota(100);p__Ascomycota_unclassified(100);p__Ascomycota_unclassified(100);p__Ascomycota_unclassified(100);p__Ascomycota_unclassified(100);p__Ascomycota_unclassified(100); | - | - | - | - | - | - | - | Unassigned | - |
| Otu35368 | 1 | k__Fungi(100);p__Ascomycota(100);p__Ascomycota_unclassified(100);p__Ascomycota_unclassified(100);p__Ascomycota_unclassified(100);p__Ascomycota_unclassified(100);p__Ascomycota_unclassified(100); | - | - | - | - | - | - | - | Unassigned | - |
| Otu35369 | 1 | k__Fungi(100);k__Fungi_unclassified(100);k__Fungi_unclassified(100);k__Fungi_unclassified(100);k__Fungi_unclassified(100);k__Fungi_unclassified(100);k__Fungi_unclassified(100); | - | - | - | - | - | - | - | Unassigned | - |
| Otu35371 | 1 | k__Fungi(100);p__Ascomycota(100);p__Ascomycota_unclassified(100);p__Ascomycota_unclassified(100);p__Ascomycota_unclassified(100);p__Ascomycota_unclassified(100);p__Ascomycota_unclassified(100); | - | - | - | - | - | - | - | Unassigned | - |
| Otu35372 | 1 | k__Fungi(100);k__Fungi_unclassified(100);k__Fungi_unclassified(100);k__Fungi_unclassified(100);k__Fungi_unclassified(100);k__Fungi_unclassified(100);k__Fungi_unclassified(100); | - | - | - | - | - | - | - | Unassigned | - |
| Otu35374 | 1 | k__Fungi(100);p__Ascomycota(100);p__Ascomycota_unclassified(100);p__Ascomycota_unclassified(100);p__Ascomycota_unclassified(100);p__Ascomycota_unclassified(100);p__Ascomycota_unclassified(100); | - | - | - | - | - | - | - | Unassigned | - |
| Otu35375 | 1 | k__Fungi(100);p__Ascomycota(100);p__Ascomycota_unclassified(100);p__Ascomycota_unclassified(100);p__Ascomycota_unclassified(100);p__Ascomycota_unclassified(100);p__Ascomycota_unclassified(100); | - | - | - | - | - | - | - | Unassigned | - |
| Otu35376 | 1 | k__Fungi(100);p__Ascomycota(100);p__Ascomycota_unclassified(100);p__Ascomycota_unclassified(100);p__Ascomycota_unclassified(100);p__Ascomycota_unclassified(100);p__Ascomycota_unclassified(100); | - | - | - | - | - | - | - | Unassigned | - |
| Otu35378 | 1 | k__Fungi(100);p__Ascomycota(100);p__Ascomycota_unclassified(100);p__Ascomycota_unclassified(100);p__Ascomycota_unclassified(100);p__Ascomycota_unclassified(100);p__Ascomycota_unclassified(100); | - | - | - | - | - | - | - | Unassigned | - |
| Otu35380 | 1 | k__Fungi(100);p__Ascomycota(100);p__Ascomycota_unclassified(100);p__Ascomycota_unclassified(100);p__Ascomycota_unclassified(100);p__Ascomycota_unclassified(100);p__Ascomycota_unclassified(100); | - | - | - | - | - | - | - | Unassigned | - |
| Otu35382 | 1 | k__Fungi(100);k__Fungi_unclassified(100);k__Fungi_unclassified(100);k__Fungi_unclassified(100);k__Fungi_unclassified(100);k__Fungi_unclassified(100);k__Fungi_unclassified(100); | - | - | - | - | - | - | - | Unassigned | - |
| Otu35383 | 1 | k__Fungi(100);p__Ascomycota(100);p__Ascomycota_unclassified(100);p__Ascomycota_unclassified(100);p__Ascomycota_unclassified(100);p__Ascomycota_unclassified(100);p__Ascomycota_unclassified(100); | - | - | - | - | - | - | - | Unassigned | - |
| Otu35384 | 1 | k__Fungi(100);k__Fungi_unclassified(100);k__Fungi_unclassified(100);k__Fungi_unclassified(100);k__Fungi_unclassified(100);k__Fungi_unclassified(100);k__Fungi_unclassified(100); | - | - | - | - | - | - | - | Unassigned | - |
| Otu35386 | 1 | k__Fungi(100);k__Fungi_unclassified(100);k__Fungi_unclassified(100);k__Fungi_unclassified(100);k__Fungi_unclassified(100);k__Fungi_unclassified(100);k__Fungi_unclassified(100); | - | - | - | - | - | - | - | Unassigned | - |
| Otu35387 | 1 | k__Fungi(100);k__Fungi_unclassified(100);k__Fungi_unclassified(100);k__Fungi_unclassified(100);k__Fungi_unclassified(100);k__Fungi_unclassified(100);k__Fungi_unclassified(100); | - | - | - | - | - | - | - | Unassigned | - |
| Otu35389 | 1 | k__Fungi(100);k__Fungi_unclassified(100);k__Fungi_unclassified(100);k__Fungi_unclassified(100);k__Fungi_unclassified(100);k__Fungi_unclassified(100);k__Fungi_unclassified(100); | - | - | - | - | - | - | - | Unassigned | - |
| Otu35391 | 1 | k__Fungi(100);p__Ascomycota(100);p__Ascomycota_unclassified(100);p__Ascomycota_unclassified(100);p__Ascomycota_unclassified(100);p__Ascomycota_unclassified(100);p__Ascomycota_unclassified(100); | - | - | - | - | - | - | - | Unassigned | - |
| Otu35392 | 1 | k__Fungi(100);k__Fungi_unclassified(100);k__Fungi_unclassified(100);k__Fungi_unclassified(100);k__Fungi_unclassified(100);k__Fungi_unclassified(100);k__Fungi_unclassified(100); | - | - | - | - | - | - | - | Unassigned | - |
| Otu35393 | 1 | k__Fungi(100);p__Ascomycota(100);p__Ascomycota_unclassified(100);p__Ascomycota_unclassified(100);p__Ascomycota_unclassified(100);p__Ascomycota_unclassified(100);p__Ascomycota_unclassified(100); | - | - | - | - | - | - | - | Unassigned | - |
| Otu35394 | 1 | k__Fungi(100);p__Ascomycota(100);p__Ascomycota_unclassified(100);p__Ascomycota_unclassified(100);p__Ascomycota_unclassified(100);p__Ascomycota_unclassified(100);p__Ascomycota_unclassified(100); | - | - | - | - | - | - | - | Unassigned | - |
| Otu35397 | 1 | k__Fungi(100);k__Fungi_unclassified(100);k__Fungi_unclassified(100);k__Fungi_unclassified(100);k__Fungi_unclassified(100);k__Fungi_unclassified(100);k__Fungi_unclassified(100); | - | - | - | - | - | - | - | Unassigned | - |
| Otu35398 | 1 | k__Fungi(100);p__Ascomycota(100);p__Ascomycota_unclassified(100);p__Ascomycota_unclassified(100);p__Ascomycota_unclassified(100);p__Ascomycota_unclassified(100);p__Ascomycota_unclassified(100); | - | - | - | - | - | - | - | Unassigned | - |
| Otu35399 | 1 | k__Fungi(100);p__Ascomycota(100);p__Ascomycota_unclassified(100);p__Ascomycota_unclassified(100);p__Ascomycota_unclassified(100);p__Ascomycota_unclassified(100);p__Ascomycota_unclassified(100); | - | - | - | - | - | - | - | Unassigned | - |
| Otu35400 | 1 | k__Fungi(100);k__Fungi_unclassified(100);k__Fungi_unclassified(100);k__Fungi_unclassified(100);k__Fungi_unclassified(100);k__Fungi_unclassified(100);k__Fungi_unclassified(100); | - | - | - | - | - | - | - | Unassigned | - |
| Otu35401 | 1 | k__Fungi(100);k__Fungi_unclassified(100);k__Fungi_unclassified(100);k__Fungi_unclassified(100);k__Fungi_unclassified(100);k__Fungi_unclassified(100);k__Fungi_unclassified(100); | - | - | - | - | - | - | - | Unassigned | - |
| Otu35402 | 1 | k__Fungi(100);p__Ascomycota(100);p__Ascomycota_unclassified(100);p__Ascomycota_unclassified(100);p__Ascomycota_unclassified(100);p__Ascomycota_unclassified(100);p__Ascomycota_unclassified(100); | - | - | - | - | - | - | - | Unassigned | - |
| Otu35403 | 1 | k__Fungi(100);p__Ascomycota(100);p__Ascomycota_unclassified(100);p__Ascomycota_unclassified(100);p__Ascomycota_unclassified(100);p__Ascomycota_unclassified(100);p__Ascomycota_unclassified(100); | - | - | - | - | - | - | - | Unassigned | - |
| Otu35404 | 1 | k__Fungi(100);k__Fungi_unclassified(100);k__Fungi_unclassified(100);k__Fungi_unclassified(100);k__Fungi_unclassified(100);k__Fungi_unclassified(100);k__Fungi_unclassified(100); | - | - | - | - | - | - | - | Unassigned | - |
| Otu35405 | 1 | k__Fungi(100);p__Ascomycota(100);p__Ascomycota_unclassified(100);p__Ascomycota_unclassified(100);p__Ascomycota_unclassified(100);p__Ascomycota_unclassified(100);p__Ascomycota_unclassified(100); | - | - | - | - | - | - | - | Unassigned | - |
| Otu35406 | 1 | k__Fungi(100);p__Ascomycota(100);p__Ascomycota_unclassified(100);p__Ascomycota_unclassified(100);p__Ascomycota_unclassified(100);p__Ascomycota_unclassified(100);p__Ascomycota_unclassified(100); | - | - | - | - | - | - | - | Unassigned | - |
| Otu35407 | 1 | k__Fungi(100);k__Fungi_unclassified(100);k__Fungi_unclassified(100);k__Fungi_unclassified(100);k__Fungi_unclassified(100);k__Fungi_unclassified(100);k__Fungi_unclassified(100); | - | - | - | - | - | - | - | Unassigned | - |
| Otu35408 | 1 | k__Fungi(100);p__Ascomycota(100);p__Ascomycota_unclassified(100);p__Ascomycota_unclassified(100);p__Ascomycota_unclassified(100);p__Ascomycota_unclassified(100);p__Ascomycota_unclassified(100); | - | - | - | - | - | - | - | Unassigned | - |
| Otu35409 | 1 | k__Fungi(100);p__Ascomycota(100);p__Ascomycota_unclassified(100);p__Ascomycota_unclassified(100);p__Ascomycota_unclassified(100);p__Ascomycota_unclassified(100);p__Ascomycota_unclassified(100); | - | - | - | - | - | - | - | Unassigned | - |
| Otu35410 | 1 | k__Fungi(100);k__Fungi_unclassified(100);k__Fungi_unclassified(100);k__Fungi_unclassified(100);k__Fungi_unclassified(100);k__Fungi_unclassified(100);k__Fungi_unclassified(100); | - | - | - | - | - | - | - | Unassigned | - |
| Otu35413 | 1 | k__Fungi(100);k__Fungi_unclassified(100);k__Fungi_unclassified(100);k__Fungi_unclassified(100);k__Fungi_unclassified(100);k__Fungi_unclassified(100);k__Fungi_unclassified(100); | - | - | - | - | - | - | - | Unassigned | - |
| Otu35414 | 1 | k__Fungi(100);k__Fungi_unclassified(100);k__Fungi_unclassified(100);k__Fungi_unclassified(100);k__Fungi_unclassified(100);k__Fungi_unclassified(100);k__Fungi_unclassified(100); | - | - | - | - | - | - | - | Unassigned | - |
| Otu35415 | 1 | k__Fungi(100);k__Fungi_unclassified(100);k__Fungi_unclassified(100);k__Fungi_unclassified(100);k__Fungi_unclassified(100);k__Fungi_unclassified(100);k__Fungi_unclassified(100); | - | - | - | - | - | - | - | Unassigned | - |
| Otu35416 | 1 | k__Fungi(100);p__Ascomycota(100);p__Ascomycota_unclassified(100);p__Ascomycota_unclassified(100);p__Ascomycota_unclassified(100);p__Ascomycota_unclassified(100);p__Ascomycota_unclassified(100); | - | - | - | - | - | - | - | Unassigned | - |
| Otu35422 | 1 | k__Fungi(100);p__Ascomycota(100);p__Ascomycota_unclassified(100);p__Ascomycota_unclassified(100);p__Ascomycota_unclassified(100);p__Ascomycota_unclassified(100);p__Ascomycota_unclassified(100); | - | - | - | - | - | - | - | Unassigned | - |
| Otu35428 | 1 | k__Fungi(100);k__Fungi_unclassified(100);k__Fungi_unclassified(100);k__Fungi_unclassified(100);k__Fungi_unclassified(100);k__Fungi_unclassified(100);k__Fungi_unclassified(100); | - | - | - | - | - | - | - | Unassigned | - |
| Otu35440 | 1 | k__Fungi(100);p__Ascomycota(100);p__Ascomycota_unclassified(100);p__Ascomycota_unclassified(100);p__Ascomycota_unclassified(100);p__Ascomycota_unclassified(100);p__Ascomycota_unclassified(100); | - | - | - | - | - | - | - | Unassigned | - |
| Otu35444 | 1 | k__Fungi(100);k__Fungi_unclassified(100);k__Fungi_unclassified(100);k__Fungi_unclassified(100);k__Fungi_unclassified(100);k__Fungi_unclassified(100);k__Fungi_unclassified(100); | - | - | - | - | - | - | - | Unassigned | - |
| Otu35447 | 1 | k__Fungi(100);p__Ascomycota(100);p__Ascomycota_unclassified(100);p__Ascomycota_unclassified(100);p__Ascomycota_unclassified(100);p__Ascomycota_unclassified(100);p__Ascomycota_unclassified(100); | - | - | - | - | - | - | - | Unassigned | - |
| Otu35454 | 1 | k__Fungi(100);k__Fungi_unclassified(100);k__Fungi_unclassified(100);k__Fungi_unclassified(100);k__Fungi_unclassified(100);k__Fungi_unclassified(100);k__Fungi_unclassified(100); | - | - | - | - | - | - | - | Unassigned | - |
| Otu35457 | 1 | k__Fungi(100);k__Fungi_unclassified(100);k__Fungi_unclassified(100);k__Fungi_unclassified(100);k__Fungi_unclassified(100);k__Fungi_unclassified(100);k__Fungi_unclassified(100); | - | - | - | - | - | - | - | Unassigned | - |
| Otu35461 | 1 | k__Fungi(100);k__Fungi_unclassified(100);k__Fungi_unclassified(100);k__Fungi_unclassified(100);k__Fungi_unclassified(100);k__Fungi_unclassified(100);k__Fungi_unclassified(100); | - | - | - | - | - | - | - | Unassigned | - |
| Otu35462 | 1 | k__Fungi(100);k__Fungi_unclassified(100);k__Fungi_unclassified(100);k__Fungi_unclassified(100);k__Fungi_unclassified(100);k__Fungi_unclassified(100);k__Fungi_unclassified(100); | - | - | - | - | - | - | - | Unassigned | - |
| Otu35465 | 1 | k__Fungi(100);p__Ascomycota(100);p__Ascomycota_unclassified(100);p__Ascomycota_unclassified(100);p__Ascomycota_unclassified(100);p__Ascomycota_unclassified(100);p__Ascomycota_unclassified(100); | - | - | - | - | - | - | - | Unassigned | - |
| Otu35466 | 1 | k__Fungi(100);p__Ascomycota(100);p__Ascomycota_unclassified(100);p__Ascomycota_unclassified(100);p__Ascomycota_unclassified(100);p__Ascomycota_unclassified(100);p__Ascomycota_unclassified(100); | - | - | - | - | - | - | - | Unassigned | - |
| Otu35469 | 1 | k__Fungi(100);k__Fungi_unclassified(100);k__Fungi_unclassified(100);k__Fungi_unclassified(100);k__Fungi_unclassified(100);k__Fungi_unclassified(100);k__Fungi_unclassified(100); | - | - | - | - | - | - | - | Unassigned | - |
| Otu35470 | 1 | k__Fungi(100);k__Fungi_unclassified(100);k__Fungi_unclassified(100);k__Fungi_unclassified(100);k__Fungi_unclassified(100);k__Fungi_unclassified(100);k__Fungi_unclassified(100); | - | - | - | - | - | - | - | Unassigned | - |
| Otu35473 | 1 | k__Fungi(100);k__Fungi_unclassified(100);k__Fungi_unclassified(100);k__Fungi_unclassified(100);k__Fungi_unclassified(100);k__Fungi_unclassified(100);k__Fungi_unclassified(100); | - | - | - | - | - | - | - | Unassigned | - |
| Otu35474 | 1 | k__Fungi(100);p__Ascomycota(100);p__Ascomycota_unclassified(100);p__Ascomycota_unclassified(100);p__Ascomycota_unclassified(100);p__Ascomycota_unclassified(100);p__Ascomycota_unclassified(100); | - | - | - | - | - | - | - | Unassigned | - |
| Otu35479 | 1 | k__Fungi(100);k__Fungi_unclassified(100);k__Fungi_unclassified(100);k__Fungi_unclassified(100);k__Fungi_unclassified(100);k__Fungi_unclassified(100);k__Fungi_unclassified(100); | - | - | - | - | - | - | - | Unassigned | - |
| Otu35483 | 1 | k__Fungi(100);k__Fungi_unclassified(100);k__Fungi_unclassified(100);k__Fungi_unclassified(100);k__Fungi_unclassified(100);k__Fungi_unclassified(100);k__Fungi_unclassified(100); | - | - | - | - | - | - | - | Unassigned | - |
| Otu35487 | 1 | k__Fungi(100);p__Ascomycota(100);p__Ascomycota_unclassified(100);p__Ascomycota_unclassified(100);p__Ascomycota_unclassified(100);p__Ascomycota_unclassified(100);p__Ascomycota_unclassified(100); | - | - | - | - | - | - | - | Unassigned | - |
| Otu35492 | 1 | k__Fungi(100);p__Ascomycota(100);p__Ascomycota_unclassified(100);p__Ascomycota_unclassified(100);p__Ascomycota_unclassified(100);p__Ascomycota_unclassified(100);p__Ascomycota_unclassified(100); | - | - | - | - | - | - | - | Unassigned | - |
| Otu35493 | 1 | k__Fungi(100);k__Fungi_unclassified(100);k__Fungi_unclassified(100);k__Fungi_unclassified(100);k__Fungi_unclassified(100);k__Fungi_unclassified(100);k__Fungi_unclassified(100); | - | - | - | - | - | - | - | Unassigned | - |
| Otu35494 | 1 | k__Fungi(100);p__Ascomycota(100);p__Ascomycota_unclassified(100);p__Ascomycota_unclassified(100);p__Ascomycota_unclassified(100);p__Ascomycota_unclassified(100);p__Ascomycota_unclassified(100); | - | - | - | - | - | - | - | Unassigned | - |
| Otu35523 | 1 | k__Fungi(100);p__Ascomycota(100);p__Ascomycota_unclassified(100);p__Ascomycota_unclassified(100);p__Ascomycota_unclassified(100);p__Ascomycota_unclassified(100);p__Ascomycota_unclassified(100); | - | - | - | - | - | - | - | Unassigned | - |
| Otu35529 | 1 | k__Fungi(100);p__Ascomycota(100);p__Ascomycota_unclassified(100);p__Ascomycota_unclassified(100);p__Ascomycota_unclassified(100);p__Ascomycota_unclassified(100);p__Ascomycota_unclassified(100); | - | - | - | - | - | - | - | Unassigned | - |
| Otu35539 | 1 | k__Fungi(100);p__Ascomycota(100);p__Ascomycota_unclassified(100);p__Ascomycota_unclassified(100);p__Ascomycota_unclassified(100);p__Ascomycota_unclassified(100);p__Ascomycota_unclassified(100); | - | - | - | - | - | - | - | Unassigned | - |
| Otu35546 | 1 | k__Fungi(100);p__Ascomycota(100);p__Ascomycota_unclassified(100);p__Ascomycota_unclassified(100);p__Ascomycota_unclassified(100);p__Ascomycota_unclassified(100);p__Ascomycota_unclassified(100); | - | - | - | - | - | - | - | Unassigned | - |
| Otu35551 | 1 | k__Fungi(100);k__Fungi_unclassified(100);k__Fungi_unclassified(100);k__Fungi_unclassified(100);k__Fungi_unclassified(100);k__Fungi_unclassified(100);k__Fungi_unclassified(100); | - | - | - | - | - | - | - | Unassigned | - |
| Otu35560 | 1 | k__Fungi(100);p__Ascomycota(100);p__Ascomycota_unclassified(100);p__Ascomycota_unclassified(100);p__Ascomycota_unclassified(100);p__Ascomycota_unclassified(100);p__Ascomycota_unclassified(100); | - | - | - | - | - | - | - | Unassigned | - |
| Otu35568 | 1 | k__Fungi(100);k__Fungi_unclassified(100);k__Fungi_unclassified(100);k__Fungi_unclassified(100);k__Fungi_unclassified(100);k__Fungi_unclassified(100);k__Fungi_unclassified(100); | - | - | - | - | - | - | - | Unassigned | - |
| Otu35569 | 1 | k__Fungi(100);p__Ascomycota(100);p__Ascomycota_unclassified(100);p__Ascomycota_unclassified(100);p__Ascomycota_unclassified(100);p__Ascomycota_unclassified(100);p__Ascomycota_unclassified(100); | - | - | - | - | - | - | - | Unassigned | - |
| Otu35570 | 1 | k__Fungi(100);k__Fungi_unclassified(100);k__Fungi_unclassified(100);k__Fungi_unclassified(100);k__Fungi_unclassified(100);k__Fungi_unclassified(100);k__Fungi_unclassified(100); | - | - | - | - | - | - | - | Unassigned | - |
| Otu35572 | 1 | k__Fungi(100);k__Fungi_unclassified(100);k__Fungi_unclassified(100);k__Fungi_unclassified(100);k__Fungi_unclassified(100);k__Fungi_unclassified(100);k__Fungi_unclassified(100); | - | - | - | - | - | - | - | Unassigned | - |
| Otu35573 | 1 | k__Fungi(100);p__Ascomycota(100);p__Ascomycota_unclassified(100);p__Ascomycota_unclassified(100);p__Ascomycota_unclassified(100);p__Ascomycota_unclassified(100);p__Ascomycota_unclassified(100); | - | - | - | - | - | - | - | Unassigned | - |
| Otu35576 | 1 | k__Fungi(100);k__Fungi_unclassified(100);k__Fungi_unclassified(100);k__Fungi_unclassified(100);k__Fungi_unclassified(100);k__Fungi_unclassified(100);k__Fungi_unclassified(100); | - | - | - | - | - | - | - | Unassigned | - |
| Otu35578 | 1 | k__Fungi(100);k__Fungi_unclassified(100);k__Fungi_unclassified(100);k__Fungi_unclassified(100);k__Fungi_unclassified(100);k__Fungi_unclassified(100);k__Fungi_unclassified(100); | - | - | - | - | - | - | - | Unassigned | - |
| Otu35585 | 1 | k__Fungi(100);k__Fungi_unclassified(100);k__Fungi_unclassified(100);k__Fungi_unclassified(100);k__Fungi_unclassified(100);k__Fungi_unclassified(100);k__Fungi_unclassified(100); | - | - | - | - | - | - | - | Unassigned | - |
| Otu35594 | 1 | k__Fungi(100);p__Ascomycota(100);c__Leotiomycetes(100);c__Leotiomycetes_unclassified(100);c__Leotiomycetes_unclassified(100);c__Leotiomycetes_unclassified(100);c__Leotiomycetes_unclassified(100); | - | - | - | - | - | - | - | Unassigned | - |
| Otu35596 | 1 | k__Fungi(100);p__Ascomycota(100);p__Ascomycota_unclassified(100);p__Ascomycota_unclassified(100);p__Ascomycota_unclassified(100);p__Ascomycota_unclassified(100);p__Ascomycota_unclassified(100); | - | - | - | - | - | - | - | Unassigned | - |
| Otu35605 | 1 | k__Fungi(100);k__Fungi_unclassified(100);k__Fungi_unclassified(100);k__Fungi_unclassified(100);k__Fungi_unclassified(100);k__Fungi_unclassified(100);k__Fungi_unclassified(100); | - | - | - | - | - | - | - | Unassigned | - |
| Otu35607 | 1 | k__Fungi(100);k__Fungi_unclassified(100);k__Fungi_unclassified(100);k__Fungi_unclassified(100);k__Fungi_unclassified(100);k__Fungi_unclassified(100);k__Fungi_unclassified(100); | - | - | - | - | - | - | - | Unassigned | - |
| Otu35608 | 1 | k__Fungi(100);k__Fungi_unclassified(100);k__Fungi_unclassified(100);k__Fungi_unclassified(100);k__Fungi_unclassified(100);k__Fungi_unclassified(100);k__Fungi_unclassified(100); | - | - | - | - | - | - | - | Unassigned | - |
| Otu35609 | 1 | k__Fungi(100);p__Ascomycota(100);p__Ascomycota_unclassified(100);p__Ascomycota_unclassified(100);p__Ascomycota_unclassified(100);p__Ascomycota_unclassified(100);p__Ascomycota_unclassified(100); | - | - | - | - | - | - | - | Unassigned | - |
| Otu35611 | 1 | k__Fungi(100);k__Fungi_unclassified(100);k__Fungi_unclassified(100);k__Fungi_unclassified(100);k__Fungi_unclassified(100);k__Fungi_unclassified(100);k__Fungi_unclassified(100); | - | - | - | - | - | - | - | Unassigned | - |
| Otu35613 | 1 | k__Fungi(100);k__Fungi_unclassified(100);k__Fungi_unclassified(100);k__Fungi_unclassified(100);k__Fungi_unclassified(100);k__Fungi_unclassified(100);k__Fungi_unclassified(100); | - | - | - | - | - | - | - | Unassigned | - |
| Otu35614 | 1 | k__Fungi(100);p__Ascomycota(100);p__Ascomycota_unclassified(100);p__Ascomycota_unclassified(100);p__Ascomycota_unclassified(100);p__Ascomycota_unclassified(100);p__Ascomycota_unclassified(100); | - | - | - | - | - | - | - | Unassigned | - |
| Otu35615 | 1 | k__Fungi(100);k__Fungi_unclassified(100);k__Fungi_unclassified(100);k__Fungi_unclassified(100);k__Fungi_unclassified(100);k__Fungi_unclassified(100);k__Fungi_unclassified(100); | - | - | - | - | - | - | - | Unassigned | - |
| Otu35616 | 1 | k__Fungi(100);k__Fungi_unclassified(100);k__Fungi_unclassified(100);k__Fungi_unclassified(100);k__Fungi_unclassified(100);k__Fungi_unclassified(100);k__Fungi_unclassified(100); | - | - | - | - | - | - | - | Unassigned | - |
| Otu35617 | 1 | k__Fungi(100);p__Ascomycota(100);p__Ascomycota_unclassified(100);p__Ascomycota_unclassified(100);p__Ascomycota_unclassified(100);p__Ascomycota_unclassified(100);p__Ascomycota_unclassified(100); | - | - | - | - | - | - | - | Unassigned | - |
| Otu35618 | 1 | k__Fungi(100);k__Fungi_unclassified(100);k__Fungi_unclassified(100);k__Fungi_unclassified(100);k__Fungi_unclassified(100);k__Fungi_unclassified(100);k__Fungi_unclassified(100); | - | - | - | - | - | - | - | Unassigned | - |
| Otu35621 | 1 | k__Fungi(100);k__Fungi_unclassified(100);k__Fungi_unclassified(100);k__Fungi_unclassified(100);k__Fungi_unclassified(100);k__Fungi_unclassified(100);k__Fungi_unclassified(100); | - | - | - | - | - | - | - | Unassigned | - |
| Otu35622 | 1 | k__Fungi(100);k__Fungi_unclassified(100);k__Fungi_unclassified(100);k__Fungi_unclassified(100);k__Fungi_unclassified(100);k__Fungi_unclassified(100);k__Fungi_unclassified(100); | - | - | - | - | - | - | - | Unassigned | - |
| Otu35623 | 1 | k__Fungi(100);p__Ascomycota(100);p__Ascomycota_unclassified(100);p__Ascomycota_unclassified(100);p__Ascomycota_unclassified(100);p__Ascomycota_unclassified(100);p__Ascomycota_unclassified(100); | - | - | - | - | - | - | - | Unassigned | - |
| Otu35627 | 1 | k__Fungi(100);p__Ascomycota(100);p__Ascomycota_unclassified(100);p__Ascomycota_unclassified(100);p__Ascomycota_unclassified(100);p__Ascomycota_unclassified(100);p__Ascomycota_unclassified(100); | - | - | - | - | - | - | - | Unassigned | - |
| Otu35628 | 1 | k__Fungi(100);k__Fungi_unclassified(100);k__Fungi_unclassified(100);k__Fungi_unclassified(100);k__Fungi_unclassified(100);k__Fungi_unclassified(100);k__Fungi_unclassified(100); | - | - | - | - | - | - | - | Unassigned | - |
| Otu35629 | 1 | k__Fungi(100);k__Fungi_unclassified(100);k__Fungi_unclassified(100);k__Fungi_unclassified(100);k__Fungi_unclassified(100);k__Fungi_unclassified(100);k__Fungi_unclassified(100); | - | - | - | - | - | - | - | Unassigned | - |
| Otu35630 | 1 | k__Fungi(100);p__Ascomycota(100);p__Ascomycota_unclassified(100);p__Ascomycota_unclassified(100);p__Ascomycota_unclassified(100);p__Ascomycota_unclassified(100);p__Ascomycota_unclassified(100); | - | - | - | - | - | - | - | Unassigned | - |
| Otu35633 | 1 | k__Fungi(100);p__Ascomycota(100);p__Ascomycota_unclassified(100);p__Ascomycota_unclassified(100);p__Ascomycota_unclassified(100);p__Ascomycota_unclassified(100);p__Ascomycota_unclassified(100); | - | - | - | - | - | - | - | Unassigned | - |
| Otu35634 | 1 | k__Fungi(100);p__Ascomycota(100);p__Ascomycota_unclassified(100);p__Ascomycota_unclassified(100);p__Ascomycota_unclassified(100);p__Ascomycota_unclassified(100);p__Ascomycota_unclassified(100); | - | - | - | - | - | - | - | Unassigned | - |
| Otu35640 | 1 | k__Fungi(100);p__Ascomycota(100);p__Ascomycota_unclassified(100);p__Ascomycota_unclassified(100);p__Ascomycota_unclassified(100);p__Ascomycota_unclassified(100);p__Ascomycota_unclassified(100); | - | - | - | - | - | - | - | Unassigned | - |
| Otu35641 | 1 | k__Fungi(100);p__Ascomycota(100);p__Ascomycota_unclassified(100);p__Ascomycota_unclassified(100);p__Ascomycota_unclassified(100);p__Ascomycota_unclassified(100);p__Ascomycota_unclassified(100); | - | - | - | - | - | - | - | Unassigned | - |
| Otu35644 | 1 | k__Fungi(100);p__Ascomycota(100);p__Ascomycota_unclassified(100);p__Ascomycota_unclassified(100);p__Ascomycota_unclassified(100);p__Ascomycota_unclassified(100);p__Ascomycota_unclassified(100); | - | - | - | - | - | - | - | Unassigned | - |
| Otu35663 | 1 | k__Fungi(100);p__Ascomycota(100);p__Ascomycota_unclassified(100);p__Ascomycota_unclassified(100);p__Ascomycota_unclassified(100);p__Ascomycota_unclassified(100);p__Ascomycota_unclassified(100); | - | - | - | - | - | - | - | Unassigned | - |
| Otu35664 | 1 | k__Fungi(100);k__Fungi_unclassified(100);k__Fungi_unclassified(100);k__Fungi_unclassified(100);k__Fungi_unclassified(100);k__Fungi_unclassified(100);k__Fungi_unclassified(100); | - | - | - | - | - | - | - | Unassigned | - |
| Otu35687 | 1 | k__Fungi(100);k__Fungi_unclassified(100);k__Fungi_unclassified(100);k__Fungi_unclassified(100);k__Fungi_unclassified(100);k__Fungi_unclassified(100);k__Fungi_unclassified(100); | - | - | - | - | - | - | - | Unassigned | - |
| Otu35688 | 1 | k__Fungi(100);k__Fungi_unclassified(100);k__Fungi_unclassified(100);k__Fungi_unclassified(100);k__Fungi_unclassified(100);k__Fungi_unclassified(100);k__Fungi_unclassified(100); | - | - | - | - | - | - | - | Unassigned | - |
| Otu35690 | 1 | k__Fungi(100);p__Ascomycota(100);p__Ascomycota_unclassified(100);p__Ascomycota_unclassified(100);p__Ascomycota_unclassified(100);p__Ascomycota_unclassified(100);p__Ascomycota_unclassified(100); | - | - | - | - | - | - | - | Unassigned | - |
| Otu35695 | 1 | k__Fungi(100);p__Ascomycota(100);p__Ascomycota_unclassified(100);p__Ascomycota_unclassified(100);p__Ascomycota_unclassified(100);p__Ascomycota_unclassified(100);p__Ascomycota_unclassified(100); | - | - | - | - | - | - | - | Unassigned | - |
| Otu35703 | 1 | k__Fungi(100);k__Fungi_unclassified(100);k__Fungi_unclassified(100);k__Fungi_unclassified(100);k__Fungi_unclassified(100);k__Fungi_unclassified(100);k__Fungi_unclassified(100); | - | - | - | - | - | - | - | Unassigned | - |
| Otu35711 | 1 | k__Fungi(100);p__Ascomycota(100);p__Ascomycota_unclassified(100);p__Ascomycota_unclassified(100);p__Ascomycota_unclassified(100);p__Ascomycota_unclassified(100);p__Ascomycota_unclassified(100); | - | - | - | - | - | - | - | Unassigned | - |
| Otu35712 | 1 | k__Fungi(100);p__Ascomycota(100);p__Ascomycota_unclassified(100);p__Ascomycota_unclassified(100);p__Ascomycota_unclassified(100);p__Ascomycota_unclassified(100);p__Ascomycota_unclassified(100); | - | - | - | - | - | - | - | Unassigned | - |
| Otu35713 | 1 | k__Fungi(100);p__Ascomycota(100);p__Ascomycota_unclassified(100);p__Ascomycota_unclassified(100);p__Ascomycota_unclassified(100);p__Ascomycota_unclassified(100);p__Ascomycota_unclassified(100); | - | - | - | - | - | - | - | Unassigned | - |
| Otu35714 | 1 | k__Fungi(100);k__Fungi_unclassified(100);k__Fungi_unclassified(100);k__Fungi_unclassified(100);k__Fungi_unclassified(100);k__Fungi_unclassified(100);k__Fungi_unclassified(100); | - | - | - | - | - | - | - | Unassigned | - |
| Otu35715 | 1 | k__Fungi(100);p__Ascomycota(100);p__Ascomycota_unclassified(100);p__Ascomycota_unclassified(100);p__Ascomycota_unclassified(100);p__Ascomycota_unclassified(100);p__Ascomycota_unclassified(100); | - | - | - | - | - | - | - | Unassigned | - |
| Otu35716 | 1 | k__Fungi(100);k__Fungi_unclassified(100);k__Fungi_unclassified(100);k__Fungi_unclassified(100);k__Fungi_unclassified(100);k__Fungi_unclassified(100);k__Fungi_unclassified(100); | - | - | - | - | - | - | - | Unassigned | - |
| Otu35720 | 1 | k__Fungi(100);k__Fungi_unclassified(100);k__Fungi_unclassified(100);k__Fungi_unclassified(100);k__Fungi_unclassified(100);k__Fungi_unclassified(100);k__Fungi_unclassified(100); | - | - | - | - | - | - | - | Unassigned | - |
| Otu35725 | 1 | k__Fungi(100);k__Fungi_unclassified(100);k__Fungi_unclassified(100);k__Fungi_unclassified(100);k__Fungi_unclassified(100);k__Fungi_unclassified(100);k__Fungi_unclassified(100); | - | - | - | - | - | - | - | Unassigned | - |
| Otu35727 | 1 | k__Fungi(100);p__Ascomycota(100);p__Ascomycota_unclassified(100);p__Ascomycota_unclassified(100);p__Ascomycota_unclassified(100);p__Ascomycota_unclassified(100);p__Ascomycota_unclassified(100); | - | - | - | - | - | - | - | Unassigned | - |
| Otu35728 | 1 | k__Fungi(100);k__Fungi_unclassified(100);k__Fungi_unclassified(100);k__Fungi_unclassified(100);k__Fungi_unclassified(100);k__Fungi_unclassified(100);k__Fungi_unclassified(100); | - | - | - | - | - | - | - | Unassigned | - |
| Otu35730 | 1 | k__Fungi(100);p__Ascomycota(100);p__Ascomycota_unclassified(100);p__Ascomycota_unclassified(100);p__Ascomycota_unclassified(100);p__Ascomycota_unclassified(100);p__Ascomycota_unclassified(100); | - | - | - | - | - | - | - | Unassigned | - |
| Otu35731 | 1 | k__Fungi(100);k__Fungi_unclassified(100);k__Fungi_unclassified(100);k__Fungi_unclassified(100);k__Fungi_unclassified(100);k__Fungi_unclassified(100);k__Fungi_unclassified(100); | - | - | - | - | - | - | - | Unassigned | - |
| Otu35735 | 1 | k__Fungi(100);k__Fungi_unclassified(100);k__Fungi_unclassified(100);k__Fungi_unclassified(100);k__Fungi_unclassified(100);k__Fungi_unclassified(100);k__Fungi_unclassified(100); | - | - | - | - | - | - | - | Unassigned | - |
| Otu35739 | 1 | k__Fungi(100);k__Fungi_unclassified(100);k__Fungi_unclassified(100);k__Fungi_unclassified(100);k__Fungi_unclassified(100);k__Fungi_unclassified(100);k__Fungi_unclassified(100); | - | - | - | - | - | - | - | Unassigned | - |
| Otu35742 | 1 | k__Fungi(100);k__Fungi_unclassified(100);k__Fungi_unclassified(100);k__Fungi_unclassified(100);k__Fungi_unclassified(100);k__Fungi_unclassified(100);k__Fungi_unclassified(100); | - | - | - | - | - | - | - | Unassigned | - |
| Otu35759 | 1 | k__Fungi(100);k__Fungi_unclassified(100);k__Fungi_unclassified(100);k__Fungi_unclassified(100);k__Fungi_unclassified(100);k__Fungi_unclassified(100);k__Fungi_unclassified(100); | - | - | - | - | - | - | - | Unassigned | - |
| Otu35760 | 1 | k__Fungi(100);k__Fungi_unclassified(100);k__Fungi_unclassified(100);k__Fungi_unclassified(100);k__Fungi_unclassified(100);k__Fungi_unclassified(100);k__Fungi_unclassified(100); | - | - | - | - | - | - | - | Unassigned | - |
| Otu35761 | 1 | k__Fungi(100);p__Ascomycota(100);p__Ascomycota_unclassified(100);p__Ascomycota_unclassified(100);p__Ascomycota_unclassified(100);p__Ascomycota_unclassified(100);p__Ascomycota_unclassified(100); | - | - | - | - | - | - | - | Unassigned | - |
| Otu35762 | 1 | k__Fungi(100);p__Ascomycota(100);p__Ascomycota_unclassified(100);p__Ascomycota_unclassified(100);p__Ascomycota_unclassified(100);p__Ascomycota_unclassified(100);p__Ascomycota_unclassified(100); | - | - | - | - | - | - | - | Unassigned | - |
| Otu35763 | 1 | k__Fungi(100);p__Ascomycota(100);p__Ascomycota_unclassified(100);p__Ascomycota_unclassified(100);p__Ascomycota_unclassified(100);p__Ascomycota_unclassified(100);p__Ascomycota_unclassified(100); | - | - | - | - | - | - | - | Unassigned | - |
| Otu35764 | 1 | k__Fungi(100);k__Fungi_unclassified(100);k__Fungi_unclassified(100);k__Fungi_unclassified(100);k__Fungi_unclassified(100);k__Fungi_unclassified(100);k__Fungi_unclassified(100); | - | - | - | - | - | - | - | Unassigned | - |
| Otu35768 | 1 | k__Fungi(100);p__Ascomycota(100);p__Ascomycota_unclassified(100);p__Ascomycota_unclassified(100);p__Ascomycota_unclassified(100);p__Ascomycota_unclassified(100);p__Ascomycota_unclassified(100); | - | - | - | - | - | - | - | Unassigned | - |
| Otu35776 | 1 | k__Fungi(100);k__Fungi_unclassified(100);k__Fungi_unclassified(100);k__Fungi_unclassified(100);k__Fungi_unclassified(100);k__Fungi_unclassified(100);k__Fungi_unclassified(100); | - | - | - | - | - | - | - | Unassigned | - |
| Otu35781 | 1 | k__Fungi(100);p__Ascomycota(100);p__Ascomycota_unclassified(100);p__Ascomycota_unclassified(100);p__Ascomycota_unclassified(100);p__Ascomycota_unclassified(100);p__Ascomycota_unclassified(100); | - | - | - | - | - | - | - | Unassigned | - |
| Otu35784 | 1 | k__Fungi(100);p__Ascomycota(100);p__Ascomycota_unclassified(100);p__Ascomycota_unclassified(100);p__Ascomycota_unclassified(100);p__Ascomycota_unclassified(100);p__Ascomycota_unclassified(100); | - | - | - | - | - | - | - | Unassigned | - |
| Otu35786 | 1 | k__Fungi(100);k__Fungi_unclassified(100);k__Fungi_unclassified(100);k__Fungi_unclassified(100);k__Fungi_unclassified(100);k__Fungi_unclassified(100);k__Fungi_unclassified(100); | - | - | - | - | - | - | - | Unassigned | - |
| Otu35794 | 1 | k__Fungi(100);k__Fungi_unclassified(100);k__Fungi_unclassified(100);k__Fungi_unclassified(100);k__Fungi_unclassified(100);k__Fungi_unclassified(100);k__Fungi_unclassified(100); | - | - | - | - | - | - | - | Unassigned | - |
| Otu35795 | 1 | k__Fungi(100);p__Ascomycota(100);p__Ascomycota_unclassified(100);p__Ascomycota_unclassified(100);p__Ascomycota_unclassified(100);p__Ascomycota_unclassified(100);p__Ascomycota_unclassified(100); | - | - | - | - | - | - | - | Unassigned | - |
| Otu35798 | 1 | k__Fungi(100);k__Fungi_unclassified(100);k__Fungi_unclassified(100);k__Fungi_unclassified(100);k__Fungi_unclassified(100);k__Fungi_unclassified(100);k__Fungi_unclassified(100); | - | - | - | - | - | - | - | Unassigned | - |
| Otu35801 | 1 | k__Fungi(100);k__Fungi_unclassified(100);k__Fungi_unclassified(100);k__Fungi_unclassified(100);k__Fungi_unclassified(100);k__Fungi_unclassified(100);k__Fungi_unclassified(100); | - | - | - | - | - | - | - | Unassigned | - |
| Otu35805 | 1 | k__Fungi(100);p__Ascomycota(100);p__Ascomycota_unclassified(100);p__Ascomycota_unclassified(100);p__Ascomycota_unclassified(100);p__Ascomycota_unclassified(100);p__Ascomycota_unclassified(100); | - | - | - | - | - | - | - | Unassigned | - |
| Otu35821 | 1 | k__Fungi(100);p__Ascomycota(100);p__Ascomycota_unclassified(100);p__Ascomycota_unclassified(100);p__Ascomycota_unclassified(100);p__Ascomycota_unclassified(100);p__Ascomycota_unclassified(100); | - | - | - | - | - | - | - | Unassigned | - |
| Otu35828 | 1 | k__Fungi(100);k__Fungi_unclassified(100);k__Fungi_unclassified(100);k__Fungi_unclassified(100);k__Fungi_unclassified(100);k__Fungi_unclassified(100);k__Fungi_unclassified(100); | - | - | - | - | - | - | - | Unassigned | - |
| Otu35850 | 1 | k__Fungi(100);k__Fungi_unclassified(100);k__Fungi_unclassified(100);k__Fungi_unclassified(100);k__Fungi_unclassified(100);k__Fungi_unclassified(100);k__Fungi_unclassified(100); | - | - | - | - | - | - | - | Unassigned | - |
| Otu35853 | 1 | k__Fungi(100);k__Fungi_unclassified(100);k__Fungi_unclassified(100);k__Fungi_unclassified(100);k__Fungi_unclassified(100);k__Fungi_unclassified(100);k__Fungi_unclassified(100); | - | - | - | - | - | - | - | Unassigned | - |
| Otu35854 | 1 | k__Fungi(100);k__Fungi_unclassified(100);k__Fungi_unclassified(100);k__Fungi_unclassified(100);k__Fungi_unclassified(100);k__Fungi_unclassified(100);k__Fungi_unclassified(100); | - | - | - | - | - | - | - | Unassigned | - |
| Otu35870 | 1 | k__Fungi(100);k__Fungi_unclassified(100);k__Fungi_unclassified(100);k__Fungi_unclassified(100);k__Fungi_unclassified(100);k__Fungi_unclassified(100);k__Fungi_unclassified(100); | - | - | - | - | - | - | - | Unassigned | - |
| Otu35871 | 1 | k__Fungi(100);k__Fungi_unclassified(100);k__Fungi_unclassified(100);k__Fungi_unclassified(100);k__Fungi_unclassified(100);k__Fungi_unclassified(100);k__Fungi_unclassified(100); | - | - | - | - | - | - | - | Unassigned | - |
| Otu35875 | 1 | k__Fungi(100);k__Fungi_unclassified(100);k__Fungi_unclassified(100);k__Fungi_unclassified(100);k__Fungi_unclassified(100);k__Fungi_unclassified(100);k__Fungi_unclassified(100); | - | - | - | - | - | - | - | Unassigned | - |
| Otu35876 | 1 | k__Fungi(100);k__Fungi_unclassified(100);k__Fungi_unclassified(100);k__Fungi_unclassified(100);k__Fungi_unclassified(100);k__Fungi_unclassified(100);k__Fungi_unclassified(100); | - | - | - | - | - | - | - | Unassigned | - |
| Otu35877 | 1 | k__Fungi(100);k__Fungi_unclassified(100);k__Fungi_unclassified(100);k__Fungi_unclassified(100);k__Fungi_unclassified(100);k__Fungi_unclassified(100);k__Fungi_unclassified(100); | - | - | - | - | - | - | - | Unassigned | - |
| Otu35880 | 1 | k__Fungi(100);p__Ascomycota(100);p__Ascomycota_unclassified(100);p__Ascomycota_unclassified(100);p__Ascomycota_unclassified(100);p__Ascomycota_unclassified(100);p__Ascomycota_unclassified(100); | - | - | - | - | - | - | - | Unassigned | - |
| Otu35886 | 1 | k__Fungi(100);k__Fungi_unclassified(100);k__Fungi_unclassified(100);k__Fungi_unclassified(100);k__Fungi_unclassified(100);k__Fungi_unclassified(100);k__Fungi_unclassified(100); | - | - | - | - | - | - | - | Unassigned | - |
| Otu35888 | 1 | k__Fungi(100);k__Fungi_unclassified(100);k__Fungi_unclassified(100);k__Fungi_unclassified(100);k__Fungi_unclassified(100);k__Fungi_unclassified(100);k__Fungi_unclassified(100); | - | - | - | - | - | - | - | Unassigned | - |
| Otu35890 | 1 | k__Fungi(100);p__Ascomycota(100);p__Ascomycota_unclassified(100);p__Ascomycota_unclassified(100);p__Ascomycota_unclassified(100);p__Ascomycota_unclassified(100);p__Ascomycota_unclassified(100); | - | - | - | - | - | - | - | Unassigned | - |
| Otu35891 | 1 | k__Fungi(100);k__Fungi_unclassified(100);k__Fungi_unclassified(100);k__Fungi_unclassified(100);k__Fungi_unclassified(100);k__Fungi_unclassified(100);k__Fungi_unclassified(100); | - | - | - | - | - | - | - | Unassigned | - |
| Otu35892 | 1 | k__Fungi(100);k__Fungi_unclassified(100);k__Fungi_unclassified(100);k__Fungi_unclassified(100);k__Fungi_unclassified(100);k__Fungi_unclassified(100);k__Fungi_unclassified(100); | - | - | - | - | - | - | - | Unassigned | - |
| Otu35893 | 1 | k__Fungi(100);k__Fungi_unclassified(100);k__Fungi_unclassified(100);k__Fungi_unclassified(100);k__Fungi_unclassified(100);k__Fungi_unclassified(100);k__Fungi_unclassified(100); | - | - | - | - | - | - | - | Unassigned | - |
| Otu35894 | 1 | k__Fungi(100);p__Ascomycota(100);p__Ascomycota_unclassified(100);p__Ascomycota_unclassified(100);p__Ascomycota_unclassified(100);p__Ascomycota_unclassified(100);p__Ascomycota_unclassified(100); | - | - | - | - | - | - | - | Unassigned | - |
| Otu35895 | 1 | k__Fungi(100);p__Ascomycota(100);p__Ascomycota_unclassified(100);p__Ascomycota_unclassified(100);p__Ascomycota_unclassified(100);p__Ascomycota_unclassified(100);p__Ascomycota_unclassified(100); | - | - | - | - | - | - | - | Unassigned | - |
| Otu35896 | 1 | k__Fungi(100);p__Ascomycota(100);p__Ascomycota_unclassified(100);p__Ascomycota_unclassified(100);p__Ascomycota_unclassified(100);p__Ascomycota_unclassified(100);p__Ascomycota_unclassified(100); | - | - | - | - | - | - | - | Unassigned | - |
| Otu35899 | 1 | k__Fungi(100);p__Ascomycota(100);p__Ascomycota_unclassified(100);p__Ascomycota_unclassified(100);p__Ascomycota_unclassified(100);p__Ascomycota_unclassified(100);p__Ascomycota_unclassified(100); | - | - | - | - | - | - | - | Unassigned | - |
| Otu35900 | 1 | k__Fungi(100);k__Fungi_unclassified(100);k__Fungi_unclassified(100);k__Fungi_unclassified(100);k__Fungi_unclassified(100);k__Fungi_unclassified(100);k__Fungi_unclassified(100); | - | - | - | - | - | - | - | Unassigned | - |
| Otu35904 | 1 | k__Fungi(100);p__Ascomycota(100);p__Ascomycota_unclassified(100);p__Ascomycota_unclassified(100);p__Ascomycota_unclassified(100);p__Ascomycota_unclassified(100);p__Ascomycota_unclassified(100); | - | - | - | - | - | - | - | Unassigned | - |
| Otu35905 | 1 | k__Fungi(100);k__Fungi_unclassified(100);k__Fungi_unclassified(100);k__Fungi_unclassified(100);k__Fungi_unclassified(100);k__Fungi_unclassified(100);k__Fungi_unclassified(100); | - | - | - | - | - | - | - | Unassigned | - |
| Otu35909 | 1 | k__Fungi(100);k__Fungi_unclassified(100);k__Fungi_unclassified(100);k__Fungi_unclassified(100);k__Fungi_unclassified(100);k__Fungi_unclassified(100);k__Fungi_unclassified(100); | - | - | - | - | - | - | - | Unassigned | - |
| Otu35914 | 1 | k__Fungi(100);k__Fungi_unclassified(100);k__Fungi_unclassified(100);k__Fungi_unclassified(100);k__Fungi_unclassified(100);k__Fungi_unclassified(100);k__Fungi_unclassified(100); | - | - | - | - | - | - | - | Unassigned | - |
| Otu35924 | 1 | k__Fungi(100);p__Ascomycota(100);p__Ascomycota_unclassified(100);p__Ascomycota_unclassified(100);p__Ascomycota_unclassified(100);p__Ascomycota_unclassified(100);p__Ascomycota_unclassified(100); | - | - | - | - | - | - | - | Unassigned | - |
| Otu35928 | 1 | k__Fungi(100);p__Ascomycota(100);p__Ascomycota_unclassified(100);p__Ascomycota_unclassified(100);p__Ascomycota_unclassified(100);p__Ascomycota_unclassified(100);p__Ascomycota_unclassified(100); | - | - | - | - | - | - | - | Unassigned | - |
| Otu35929 | 1 | k__Fungi(100);k__Fungi_unclassified(100);k__Fungi_unclassified(100);k__Fungi_unclassified(100);k__Fungi_unclassified(100);k__Fungi_unclassified(100);k__Fungi_unclassified(100); | - | - | - | - | - | - | - | Unassigned | - |
| Otu35930 | 1 | k__Fungi(100);p__Ascomycota(100);p__Ascomycota_unclassified(100);p__Ascomycota_unclassified(100);p__Ascomycota_unclassified(100);p__Ascomycota_unclassified(100);p__Ascomycota_unclassified(100); | - | - | - | - | - | - | - | Unassigned | - |
| Otu35931 | 1 | k__Fungi(100);p__Ascomycota(100);p__Ascomycota_unclassified(100);p__Ascomycota_unclassified(100);p__Ascomycota_unclassified(100);p__Ascomycota_unclassified(100);p__Ascomycota_unclassified(100); | - | - | - | - | - | - | - | Unassigned | - |
| Otu35934 | 1 | k__Fungi(100);p__Ascomycota(100);p__Ascomycota_unclassified(100);p__Ascomycota_unclassified(100);p__Ascomycota_unclassified(100);p__Ascomycota_unclassified(100);p__Ascomycota_unclassified(100); | - | - | - | - | - | - | - | Unassigned | - |
| Otu35937 | 1 | k__Fungi(100);k__Fungi_unclassified(100);k__Fungi_unclassified(100);k__Fungi_unclassified(100);k__Fungi_unclassified(100);k__Fungi_unclassified(100);k__Fungi_unclassified(100); | - | - | - | - | - | - | - | Unassigned | - |
| Otu35939 | 1 | k__Fungi(100);p__Ascomycota(100);p__Ascomycota_unclassified(100);p__Ascomycota_unclassified(100);p__Ascomycota_unclassified(100);p__Ascomycota_unclassified(100);p__Ascomycota_unclassified(100); | - | - | - | - | - | - | - | Unassigned | - |
| Otu35940 | 1 | k__Fungi(100);k__Fungi_unclassified(100);k__Fungi_unclassified(100);k__Fungi_unclassified(100);k__Fungi_unclassified(100);k__Fungi_unclassified(100);k__Fungi_unclassified(100); | - | - | - | - | - | - | - | Unassigned | - |
| Otu35941 | 1 | k__Fungi(100);p__Ascomycota(100);p__Ascomycota_unclassified(100);p__Ascomycota_unclassified(100);p__Ascomycota_unclassified(100);p__Ascomycota_unclassified(100);p__Ascomycota_unclassified(100); | - | - | - | - | - | - | - | Unassigned | - |
| Otu35942 | 1 | k__Fungi(100);p__Ascomycota(100);p__Ascomycota_unclassified(100);p__Ascomycota_unclassified(100);p__Ascomycota_unclassified(100);p__Ascomycota_unclassified(100);p__Ascomycota_unclassified(100); | - | - | - | - | - | - | - | Unassigned | - |
| Otu35944 | 1 | k__Fungi(100);p__Ascomycota(100);p__Ascomycota_unclassified(100);p__Ascomycota_unclassified(100);p__Ascomycota_unclassified(100);p__Ascomycota_unclassified(100);p__Ascomycota_unclassified(100); | - | - | - | - | - | - | - | Unassigned | - |
| Otu35945 | 1 | k__Fungi(100);k__Fungi_unclassified(100);k__Fungi_unclassified(100);k__Fungi_unclassified(100);k__Fungi_unclassified(100);k__Fungi_unclassified(100);k__Fungi_unclassified(100); | - | - | - | - | - | - | - | Unassigned | - |
| Otu35946 | 1 | k__Fungi(100);p__Ascomycota(100);p__Ascomycota_unclassified(100);p__Ascomycota_unclassified(100);p__Ascomycota_unclassified(100);p__Ascomycota_unclassified(100);p__Ascomycota_unclassified(100); | - | - | - | - | - | - | - | Unassigned | - |
| Otu35954 | 1 | k__Fungi(100);k__Fungi_unclassified(100);k__Fungi_unclassified(100);k__Fungi_unclassified(100);k__Fungi_unclassified(100);k__Fungi_unclassified(100);k__Fungi_unclassified(100); | - | - | - | - | - | - | - | Unassigned | - |

| | | | | | | | | | | | |
|---|---|---|---|---|---|---|---|---|---|---|---|
| Otu35955 | 1 | k__Fungi(100);k__Fungi_unclassified(100);k__Fungi_unclassified(100);k__Fungi_unclassified(100);k__Fungi_unclassified(100);k__Fungi_unclassified(100);k__Fungi_unclassified(100); | - | - | - | - | - | - | - | Unassigned | - |
| Otu35957 | 1 | k__Fungi(100);p__Ascomycota(100);p__Ascomycota_unclassified(100);p__Ascomycota_unclassified(100);p__Ascomycota_unclassified(100);p__Ascomycota_unclassified(100);p__Ascomycota_unclassified(100); | - | - | - | - | - | - | - | Unassigned | - |
| Otu35958 | 1 | k__Fungi(100);p__Ascomycota(100);p__Ascomycota_unclassified(100);p__Ascomycota_unclassified(100);p__Ascomycota_unclassified(100);p__Ascomycota_unclassified(100);p__Ascomycota_unclassified(100); | - | - | - | - | - | - | - | Unassigned | - |
| Otu35960 | 1 | k__Fungi(100);p__Ascomycota(100);p__Ascomycota_unclassified(100);p__Ascomycota_unclassified(100);p__Ascomycota_unclassified(100);p__Ascomycota_unclassified(100);p__Ascomycota_unclassified(100); | - | - | - | - | - | - | - | Unassigned | - |
| Otu35961 | 1 | k__Fungi(100);p__Ascomycota(100);p__Ascomycota_unclassified(100);p__Ascomycota_unclassified(100);p__Ascomycota_unclassified(100);p__Ascomycota_unclassified(100);p__Ascomycota_unclassified(100); | - | - | - | - | - | - | - | Unassigned | - |
| Otu35966 | 1 | k__Fungi(100);k__Fungi_unclassified(100);k__Fungi_unclassified(100);k__Fungi_unclassified(100);k__Fungi_unclassified(100);k__Fungi_unclassified(100);k__Fungi_unclassified(100); | - | - | - | - | - | - | - | Unassigned | - |
| Otu35967 | 1 | k__Fungi(100);k__Fungi_unclassified(100);k__Fungi_unclassified(100);k__Fungi_unclassified(100);k__Fungi_unclassified(100);k__Fungi_unclassified(100);k__Fungi_unclassified(100); | - | - | - | - | - | - | - | Unassigned | - |
| Otu35973 | 1 | k__Fungi(100);k__Fungi_unclassified(100);k__Fungi_unclassified(100);k__Fungi_unclassified(100);k__Fungi_unclassified(100);k__Fungi_unclassified(100);k__Fungi_unclassified(100); | - | - | - | - | - | - | - | Unassigned | - |
| Otu35977 | 1 | k__Fungi(100);k__Fungi_unclassified(100);k__Fungi_unclassified(100);k__Fungi_unclassified(100);k__Fungi_unclassified(100);k__Fungi_unclassified(100);k__Fungi_unclassified(100); | - | - | - | - | - | - | - | Unassigned | - |
| Otu35978 | 1 | k__Fungi(100);k__Fungi_unclassified(100);k__Fungi_unclassified(100);k__Fungi_unclassified(100);k__Fungi_unclassified(100);k__Fungi_unclassified(100);k__Fungi_unclassified(100); | - | - | - | - | - | - | - | Unassigned | - |
| Otu35979 | 1 | k__Fungi(100);k__Fungi_unclassified(100);k__Fungi_unclassified(100);k__Fungi_unclassified(100);k__Fungi_unclassified(100);k__Fungi_unclassified(100);k__Fungi_unclassified(100); | - | - | - | - | - | - | - | Unassigned | - |
| Otu35980 | 1 | k__Fungi(100);p__Ascomycota(100);p__Ascomycota_unclassified(100);p__Ascomycota_unclassified(100);p__Ascomycota_unclassified(100);p__Ascomycota_unclassified(100);p__Ascomycota_unclassified(100); | - | - | - | - | - | - | - | Unassigned | - |
| Otu35981 | 1 | k__Fungi(100);p__Ascomycota(100);p__Ascomycota_unclassified(100);p__Ascomycota_unclassified(100);p__Ascomycota_unclassified(100);p__Ascomycota_unclassified(100);p__Ascomycota_unclassified(100); | - | - | - | - | - | - | - | Unassigned | - |
| Otu35982 | 1 | k__Fungi(100);p__Ascomycota(100);p__Ascomycota_unclassified(100);p__Ascomycota_unclassified(100);p__Ascomycota_unclassified(100);p__Ascomycota_unclassified(100);p__Ascomycota_unclassified(100); | - | - | - | - | - | - | - | Unassigned | - |
| Otu35983 | 1 | k__Fungi(100);k__Fungi_unclassified(100);k__Fungi_unclassified(100);k__Fungi_unclassified(100);k__Fungi_unclassified(100);k__Fungi_unclassified(100);k__Fungi_unclassified(100); | - | - | - | - | - | - | - | Unassigned | - |
| Otu35984 | 1 | k__Fungi(100);k__Fungi_unclassified(100);k__Fungi_unclassified(100);k__Fungi_unclassified(100);k__Fungi_unclassified(100);k__Fungi_unclassified(100);k__Fungi_unclassified(100); | - | - | - | - | - | - | - | Unassigned | - |
| Otu35985 | 1 | k__Fungi(100);k__Fungi_unclassified(100);k__Fungi_unclassified(100);k__Fungi_unclassified(100);k__Fungi_unclassified(100);k__Fungi_unclassified(100);k__Fungi_unclassified(100); | - | - | - | - | - | - | - | Unassigned | - |
| Otu35986 | 1 | k__Fungi(100);p__Ascomycota(100);p__Ascomycota_unclassified(100);p__Ascomycota_unclassified(100);p__Ascomycota_unclassified(100);p__Ascomycota_unclassified(100);p__Ascomycota_unclassified(100); | - | - | - | - | - | - | - | Unassigned | - |
| Otu35987 | 1 | k__Fungi(100);k__Fungi_unclassified(100);k__Fungi_unclassified(100);k__Fungi_unclassified(100);k__Fungi_unclassified(100);k__Fungi_unclassified(100);k__Fungi_unclassified(100); | - | - | - | - | - | - | - | Unassigned | - |
| Otu35988 | 1 | k__Fungi(100);k__Fungi_unclassified(100);k__Fungi_unclassified(100);k__Fungi_unclassified(100);k__Fungi_unclassified(100);k__Fungi_unclassified(100);k__Fungi_unclassified(100); | - | - | - | - | - | - | - | Unassigned | - |
| Otu35989 | 1 | k__Fungi(100);p__Ascomycota(100);p__Ascomycota_unclassified(100);p__Ascomycota_unclassified(100);p__Ascomycota_unclassified(100);p__Ascomycota_unclassified(100);p__Ascomycota_unclassified(100); | - | - | - | - | - | - | - | Unassigned | - |
| Otu35991 | 1 | k__Fungi(100);p__Ascomycota(100);p__Ascomycota_unclassified(100);p__Ascomycota_unclassified(100);p__Ascomycota_unclassified(100);p__Ascomycota_unclassified(100);p__Ascomycota_unclassified(100); | - | - | - | - | - | - | - | Unassigned | - |
| Otu35992 | 1 | k__Fungi(100);p__Ascomycota(100);p__Ascomycota_unclassified(100);p__Ascomycota_unclassified(100);p__Ascomycota_unclassified(100);p__Ascomycota_unclassified(100);p__Ascomycota_unclassified(100); | - | - | - | - | - | - | - | Unassigned | - |
| Otu35993 | 1 | k__Fungi(100);p__Ascomycota(100);p__Ascomycota_unclassified(100);p__Ascomycota_unclassified(100);p__Ascomycota_unclassified(100);p__Ascomycota_unclassified(100);p__Ascomycota_unclassified(100); | - | - | - | - | - | - | - | Unassigned | - |
| Otu35994 | 1 | k__Fungi(100);p__Ascomycota(100);p__Ascomycota_unclassified(100);p__Ascomycota_unclassified(100);p__Ascomycota_unclassified(100);p__Ascomycota_unclassified(100);p__Ascomycota_unclassified(100); | - | - | - | - | - | - | - | Unassigned | - |
| Otu35996 | 1 | k__Fungi(100);k__Fungi_unclassified(100);k__Fungi_unclassified(100);k__Fungi_unclassified(100);k__Fungi_unclassified(100);k__Fungi_unclassified(100);k__Fungi_unclassified(100); | - | - | - | - | - | - | - | Unassigned | - |
| Otu35997 | 1 | k__Fungi(100);p__Ascomycota(100);p__Ascomycota_unclassified(100);p__Ascomycota_unclassified(100);p__Ascomycota_unclassified(100);p__Ascomycota_unclassified(100);p__Ascomycota_unclassified(100); | - | - | - | - | - | - | - | Unassigned | - |
| Otu35998 | 1 | k__Fungi(100);p__Ascomycota(100);p__Ascomycota_unclassified(100);p__Ascomycota_unclassified(100);p__Ascomycota_unclassified(100);p__Ascomycota_unclassified(100);p__Ascomycota_unclassified(100); | - | - | - | - | - | - | - | Unassigned | - |
| Otu35999 | 1 | k__Fungi(100);k__Fungi_unclassified(100);k__Fungi_unclassified(100);k__Fungi_unclassified(100);k__Fungi_unclassified(100);k__Fungi_unclassified(100);k__Fungi_unclassified(100); | - | - | - | - | - | - | - | Unassigned | - |
| Otu36000 | 1 | k__Fungi(100);k__Fungi_unclassified(100);k__Fungi_unclassified(100);k__Fungi_unclassified(100);k__Fungi_unclassified(100);k__Fungi_unclassified(100);k__Fungi_unclassified(100); | - | - | - | - | - | - | - | Unassigned | - |
| Otu36004 | 1 | k__Fungi(100);p__Ascomycota(100);p__Ascomycota_unclassified(100);p__Ascomycota_unclassified(100);p__Ascomycota_unclassified(100);p__Ascomycota_unclassified(100);p__Ascomycota_unclassified(100); | - | - | - | - | - | - | - | Unassigned | - |
| Otu36007 | 1 | k__Fungi(100);k__Fungi_unclassified(100);k__Fungi_unclassified(100);k__Fungi_unclassified(100);k__Fungi_unclassified(100);k__Fungi_unclassified(100);k__Fungi_unclassified(100); | - | - | - | - | - | - | - | Unassigned | - |
| Otu36011 | 1 | k__Fungi(100);p__Ascomycota(100);p__Ascomycota_unclassified(100);p__Ascomycota_unclassified(100);p__Ascomycota_unclassified(100);p__Ascomycota_unclassified(100);p__Ascomycota_unclassified(100); | - | - | - | - | - | - | - | Unassigned | - |
| Otu36014 | 1 | k__Fungi(100);p__Ascomycota(100);p__Ascomycota_unclassified(100);p__Ascomycota_unclassified(100);p__Ascomycota_unclassified(100);p__Ascomycota_unclassified(100);p__Ascomycota_unclassified(100); | - | - | - | - | - | - | - | Unassigned | - |
| Otu36017 | 1 | k__Fungi(100);k__Fungi_unclassified(100);k__Fungi_unclassified(100);k__Fungi_unclassified(100);k__Fungi_unclassified(100);k__Fungi_unclassified(100);k__Fungi_unclassified(100); | - | - | - | - | - | - | - | Unassigned | - |
| Otu36018 | 1 | k__Fungi(100);k__Fungi_unclassified(100);k__Fungi_unclassified(100);k__Fungi_unclassified(100);k__Fungi_unclassified(100);k__Fungi_unclassified(100);k__Fungi_unclassified(100); | - | - | - | - | - | - | - | Unassigned | - |
| Otu36020 | 1 | k__Fungi(100);k__Fungi_unclassified(100);k__Fungi_unclassified(100);k__Fungi_unclassified(100);k__Fungi_unclassified(100);k__Fungi_unclassified(100);k__Fungi_unclassified(100); | - | - | - | - | - | - | - | Unassigned | - |
| Otu36021 | 1 | k__Fungi(100);k__Fungi_unclassified(100);k__Fungi_unclassified(100);k__Fungi_unclassified(100);k__Fungi_unclassified(100);k__Fungi_unclassified(100);k__Fungi_unclassified(100); | - | - | - | - | - | - | - | Unassigned | - |
| Otu36022 | 1 | k__Fungi(100);k__Fungi_unclassified(100);k__Fungi_unclassified(100);k__Fungi_unclassified(100);k__Fungi_unclassified(100);k__Fungi_unclassified(100);k__Fungi_unclassified(100); | - | - | - | - | - | - | - | Unassigned | - |
| Otu36023 | 1 | k__Fungi(100);p__Ascomycota(100);p__Ascomycota_unclassified(100);p__Ascomycota_unclassified(100);p__Ascomycota_unclassified(100);p__Ascomycota_unclassified(100);p__Ascomycota_unclassified(100); | - | - | - | - | - | - | - | Unassigned | - |
| Otu36024 | 1 | k__Fungi(100);p__Ascomycota(100);p__Ascomycota_unclassified(100);p__Ascomycota_unclassified(100);p__Ascomycota_unclassified(100);p__Ascomycota_unclassified(100);p__Ascomycota_unclassified(100); | - | - | - | - | - | - | - | Unassigned | - |
| Otu36025 | 1 | k__Fungi(100);k__Fungi_unclassified(100);k__Fungi_unclassified(100);k__Fungi_unclassified(100);k__Fungi_unclassified(100);k__Fungi_unclassified(100);k__Fungi_unclassified(100); | - | - | - | - | - | - | - | Unassigned | - |
| Otu36026 | 1 | k__Fungi(100);p__Ascomycota(100);p__Ascomycota_unclassified(100);p__Ascomycota_unclassified(100);p__Ascomycota_unclassified(100);p__Ascomycota_unclassified(100);p__Ascomycota_unclassified(100); | - | - | - | - | - | - | - | Unassigned | - |
| Otu36027 | 1 | k__Fungi(100);p__Ascomycota(100);p__Ascomycota_unclassified(100);p__Ascomycota_unclassified(100);p__Ascomycota_unclassified(100);p__Ascomycota_unclassified(100);p__Ascomycota_unclassified(100); | - | - | - | - | - | - | - | Unassigned | - |
| Otu36028 | 1 | k__Fungi(100);k__Fungi_unclassified(100);k__Fungi_unclassified(100);k__Fungi_unclassified(100);k__Fungi_unclassified(100);k__Fungi_unclassified(100);k__Fungi_unclassified(100); | - | - | - | - | - | - | - | Unassigned | - |
| Otu36029 | 1 | k__Fungi(100);k__Fungi_unclassified(100);k__Fungi_unclassified(100);k__Fungi_unclassified(100);k__Fungi_unclassified(100);k__Fungi_unclassified(100);k__Fungi_unclassified(100); | - | - | - | - | - | - | - | Unassigned | - |
| Otu36035 | 1 | k__Fungi(100);k__Fungi_unclassified(100);k__Fungi_unclassified(100);k__Fungi_unclassified(100);k__Fungi_unclassified(100);k__Fungi_unclassified(100);k__Fungi_unclassified(100); | - | - | - | - | - | - | - | Unassigned | - |
| Otu36036 | 1 | k__Fungi(100);k__Fungi_unclassified(100);k__Fungi_unclassified(100);k__Fungi_unclassified(100);k__Fungi_unclassified(100);k__Fungi_unclassified(100);k__Fungi_unclassified(100); | - | - | - | - | - | - | - | Unassigned | - |
| Otu36042 | 1 | k__Fungi(100);k__Fungi_unclassified(100);k__Fungi_unclassified(100);k__Fungi_unclassified(100);k__Fungi_unclassified(100);k__Fungi_unclassified(100);k__Fungi_unclassified(100); | - | - | - | - | - | - | - | Unassigned | - |
| Otu36044 | 1 | k__Fungi(100);p__Ascomycota(100);p__Ascomycota_unclassified(100);p__Ascomycota_unclassified(100);p__Ascomycota_unclassified(100);p__Ascomycota_unclassified(100);p__Ascomycota_unclassified(100); | - | - | - | - | - | - | - | Unassigned | - |
| Otu36045 | 1 | k__Fungi(100);k__Fungi_unclassified(100);k__Fungi_unclassified(100);k__Fungi_unclassified(100);k__Fungi_unclassified(100);k__Fungi_unclassified(100);k__Fungi_unclassified(100); | - | - | - | - | - | - | - | Unassigned | - |
| Otu36046 | 1 | k__Fungi(100);p__Ascomycota(100);p__Ascomycota_unclassified(100);p__Ascomycota_unclassified(100);p__Ascomycota_unclassified(100);p__Ascomycota_unclassified(100);p__Ascomycota_unclassified(100); | - | - | - | - | - | - | - | Unassigned | - |
| Otu36047 | 1 | k__Fungi(100);p__Ascomycota(100);p__Ascomycota_unclassified(100);p__Ascomycota_unclassified(100);p__Ascomycota_unclassified(100);p__Ascomycota_unclassified(100);p__Ascomycota_unclassified(100); | - | - | - | - | - | - | - | Unassigned | - |
| Otu36048 | 1 | k__Fungi(100);k__Fungi_unclassified(100);k__Fungi_unclassified(100);k__Fungi_unclassified(100);k__Fungi_unclassified(100);k__Fungi_unclassified(100);k__Fungi_unclassified(100); | - | - | - | - | - | - | - | Unassigned | - |
| Otu36049 | 1 | k__Fungi(100);k__Fungi_unclassified(100);k__Fungi_unclassified(100);k__Fungi_unclassified(100);k__Fungi_unclassified(100);k__Fungi_unclassified(100);k__Fungi_unclassified(100); | - | - | - | - | - | - | - | Unassigned | - |
| Otu36050 | 1 | k__Fungi(100);k__Fungi_unclassified(100);k__Fungi_unclassified(100);k__Fungi_unclassified(100);k__Fungi_unclassified(100);k__Fungi_unclassified(100);k__Fungi_unclassified(100); | - | - | - | - | - | - | - | Unassigned | - |
| Otu36052 | 1 | k__Fungi(100);k__Fungi_unclassified(100);k__Fungi_unclassified(100);k__Fungi_unclassified(100);k__Fungi_unclassified(100);k__Fungi_unclassified(100);k__Fungi_unclassified(100); | - | - | - | - | - | - | - | Unassigned | - |
| Otu36055 | 1 | k__Fungi(100);p__Ascomycota(100);p__Ascomycota_unclassified(100);p__Ascomycota_unclassified(100);p__Ascomycota_unclassified(100);p__Ascomycota_unclassified(100);p__Ascomycota_unclassified(100); | - | - | - | - | - | - | - | Unassigned | - |
| Otu36056 | 1 | k__Fungi(100);k__Fungi_unclassified(100);k__Fungi_unclassified(100);k__Fungi_unclassified(100);k__Fungi_unclassified(100);k__Fungi_unclassified(100);k__Fungi_unclassified(100); | - | - | - | - | - | - | - | Unassigned | - |
| Otu36058 | 1 | k__Fungi(100);p__Ascomycota(100);p__Ascomycota_unclassified(100);p__Ascomycota_unclassified(100);p__Ascomycota_unclassified(100);p__Ascomycota_unclassified(100);p__Ascomycota_unclassified(100); | - | - | - | - | - | - | - | Unassigned | - |
| Otu36059 | 1 | k__Fungi(100);k__Fungi_unclassified(100);k__Fungi_unclassified(100);k__Fungi_unclassified(100);k__Fungi_unclassified(100);k__Fungi_unclassified(100);k__Fungi_unclassified(100); | - | - | - | - | - | - | - | Unassigned | - |
| Otu36060 | 1 | k__Fungi(100);k__Fungi_unclassified(100);k__Fungi_unclassified(100);k__Fungi_unclassified(100);k__Fungi_unclassified(100);k__Fungi_unclassified(100);k__Fungi_unclassified(100); | - | - | - | - | - | - | - | Unassigned | - |
| Otu36061 | 1 | k__Fungi(100);k__Fungi_unclassified(100);k__Fungi_unclassified(100);k__Fungi_unclassified(100);k__Fungi_unclassified(100);k__Fungi_unclassified(100);k__Fungi_unclassified(100); | - | - | - | - | - | - | - | Unassigned | - |
| Otu36067 | 1 | k__Fungi(100);k__Fungi_unclassified(100);k__Fungi_unclassified(100);k__Fungi_unclassified(100);k__Fungi_unclassified(100);k__Fungi_unclassified(100);k__Fungi_unclassified(100); | - | - | - | - | - | - | - | Unassigned | - |
| Otu36068 | 1 | k__Fungi(100);k__Fungi_unclassified(100);k__Fungi_unclassified(100);k__Fungi_unclassified(100);k__Fungi_unclassified(100);k__Fungi_unclassified(100);k__Fungi_unclassified(100); | - | - | - | - | - | - | - | Unassigned | - |
| Otu36070 | 1 | k__Fungi(100);k__Fungi_unclassified(100);k__Fungi_unclassified(100);k__Fungi_unclassified(100);k__Fungi_unclassified(100);k__Fungi_unclassified(100);k__Fungi_unclassified(100); | - | - | - | - | - | - | - | Unassigned | - |
| Otu36071 | 1 | k__Fungi(100);k__Fungi_unclassified(100);k__Fungi_unclassified(100);k__Fungi_unclassified(100);k__Fungi_unclassified(100);k__Fungi_unclassified(100);k__Fungi_unclassified(100); | - | - | - | - | - | - | - | Unassigned | - |
| Otu36075 | 1 | k__Fungi(100);k__Fungi_unclassified(100);k__Fungi_unclassified(100);k__Fungi_unclassified(100);k__Fungi_unclassified(100);k__Fungi_unclassified(100);k__Fungi_unclassified(100); | - | - | - | - | - | - | - | Unassigned | - |
| Otu36076 | 1 | k__Fungi(100);k__Fungi_unclassified(100);k__Fungi_unclassified(100);k__Fungi_unclassified(100);k__Fungi_unclassified(100);k__Fungi_unclassified(100);k__Fungi_unclassified(100); | - | - | - | - | - | - | - | Unassigned | - |
| Otu36078 | 1 | k__Fungi(100);p__Ascomycota(100);p__Ascomycota_unclassified(100);p__Ascomycota_unclassified(100);p__Ascomycota_unclassified(100);p__Ascomycota_unclassified(100);p__Ascomycota_unclassified(100); | - | - | - | - | - | - | - | Unassigned | - |
| Otu36079 | 1 | k__Fungi(100);k__Fungi_unclassified(100);k__Fungi_unclassified(100);k__Fungi_unclassified(100);k__Fungi_unclassified(100);k__Fungi_unclassified(100);k__Fungi_unclassified(100); | - | - | - | - | - | - | - | Unassigned | - |
| Otu36081 | 1 | k__Fungi(100);p__Ascomycota(100);p__Ascomycota_unclassified(100);p__Ascomycota_unclassified(100);p__Ascomycota_unclassified(100);p__Ascomycota_unclassified(100);p__Ascomycota_unclassified(100); | - | - | - | - | - | - | - | Unassigned | - |
| Otu36082 | 1 | k__Fungi(100);p__Ascomycota(100);p__Ascomycota_unclassified(100);p__Ascomycota_unclassified(100);p__Ascomycota_unclassified(100);p__Ascomycota_unclassified(100);p__Ascomycota_unclassified(100); | - | - | - | - | - | - | - | Unassigned | - |
| Otu36083 | 1 | k__Fungi(100);p__Ascomycota(100);p__Ascomycota_unclassified(100);p__Ascomycota_unclassified(100);p__Ascomycota_unclassified(100);p__Ascomycota_unclassified(100);p__Ascomycota_unclassified(100); | - | - | - | - | - | - | - | Unassigned | - |
| Otu36087 | 1 | k__Fungi(100);p__Ascomycota(100);p__Ascomycota_unclassified(100);p__Ascomycota_unclassified(100);p__Ascomycota_unclassified(100);p__Ascomycota_unclassified(100);p__Ascomycota_unclassified(100); | - | - | - | - | - | - | - | Unassigned | - |
| Otu36088 | 1 | k__Fungi(100);p__Ascomycota(100);p__Ascomycota_unclassified(100);p__Ascomycota_unclassified(100);p__Ascomycota_unclassified(100);p__Ascomycota_unclassified(100);p__Ascomycota_unclassified(100); | - | - | - | - | - | - | - | Unassigned | - |
| Otu36089 | 1 | k__Fungi(100);p__Ascomycota(100);p__Ascomycota_unclassified(100);p__Ascomycota_unclassified(100);p__Ascomycota_unclassified(100);p__Ascomycota_unclassified(100);p__Ascomycota_unclassified(100); | - | - | - | - | - | - | - | Unassigned | - |
| Otu36090 | 1 | k__Fungi(100);p__Ascomycota(100);p__Ascomycota_unclassified(100);p__Ascomycota_unclassified(100);p__Ascomycota_unclassified(100);p__Ascomycota_unclassified(100);p__Ascomycota_unclassified(100); | - | - | - | - | - | - | - | Unassigned | - |
| Otu36091 | 1 | k__Fungi(100);p__Ascomycota(100);p__Ascomycota_unclassified(100);p__Ascomycota_unclassified(100);p__Ascomycota_unclassified(100);p__Ascomycota_unclassified(100);p__Ascomycota_unclassified(100); | - | - | - | - | - | - | - | Unassigned | - |
| Otu36093 | 1 | k__Fungi(100);k__Fungi_unclassified(100);k__Fungi_unclassified(100);k__Fungi_unclassified(100);k__Fungi_unclassified(100);k__Fungi_unclassified(100);k__Fungi_unclassified(100); | - | - | - | - | - | - | - | Unassigned | - |
| Otu36094 | 1 | k__Fungi(100);k__Fungi_unclassified(100);k__Fungi_unclassified(100);k__Fungi_unclassified(100);k__Fungi_unclassified(100);k__Fungi_unclassified(100);k__Fungi_unclassified(100); | - | - | - | - | - | - | - | Unassigned | - |
| Otu36095 | 1 | k__Fungi(100);p__Ascomycota(100);p__Ascomycota_unclassified(100);p__Ascomycota_unclassified(100);p__Ascomycota_unclassified(100);p__Ascomycota_unclassified(100);p__Ascomycota_unclassified(100); | - | - | - | - | - | - | - | Unassigned | - |
| Otu36096 | 1 | k__Fungi(100);p__Ascomycota(100);p__Ascomycota_unclassified(100);p__Ascomycota_unclassified(100);p__Ascomycota_unclassified(100);p__Ascomycota_unclassified(100);p__Ascomycota_unclassified(100); | - | - | - | - | - | - | - | Unassigned | - |
| Otu36098 | 1 | k__Fungi(100);k__Fungi_unclassified(100);k__Fungi_unclassified(100);k__Fungi_unclassified(100);k__Fungi_unclassified(100);k__Fungi_unclassified(100);k__Fungi_unclassified(100); | - | - | - | - | - | - | - | Unassigned | - |
| Otu36099 | 1 | k__Fungi(100);k__Fungi_unclassified(100);k__Fungi_unclassified(100);k__Fungi_unclassified(100);k__Fungi_unclassified(100);k__Fungi_unclassified(100);k__Fungi_unclassified(100); | - | - | - | - | - | - | - | Unassigned | - |
| Otu36100 | 1 | k__Fungi(100);p__Ascomycota(100);p__Ascomycota_unclassified(100);p__Ascomycota_unclassified(100);p__Ascomycota_unclassified(100);p__Ascomycota_unclassified(100);p__Ascomycota_unclassified(100); | - | - | - | - | - | - | - | Unassigned | - |
| Otu36101 | 1 | k__Fungi(100);p__Ascomycota(100);p__Ascomycota_unclassified(100);p__Ascomycota_unclassified(100);p__Ascomycota_unclassified(100);p__Ascomycota_unclassified(100);p__Ascomycota_unclassified(100); | - | - | - | - | - | - | - | Unassigned | - |
| Otu36102 | 1 | k__Fungi(100);k__Fungi_unclassified(100);k__Fungi_unclassified(100);k__Fungi_unclassified(100);k__Fungi_unclassified(100);k__Fungi_unclassified(100);k__Fungi_unclassified(100); | - | - | - | - | - | - | - | Unassigned | - |
| Otu36103 | 1 | k__Fungi(100);p__Ascomycota(100);p__Ascomycota_unclassified(100);p__Ascomycota_unclassified(100);p__Ascomycota_unclassified(100);p__Ascomycota_unclassified(100);p__Ascomycota_unclassified(100); | - | - | - | - | - | - | - | Unassigned | - |
| Otu36105 | 1 | k__Fungi(100);k__Fungi_unclassified(100);k__Fungi_unclassified(100);k__Fungi_unclassified(100);k__Fungi_unclassified(100);k__Fungi_unclassified(100);k__Fungi_unclassified(100); | - | - | - | - | - | - | - | Unassigned | - |
| Otu36111 | 1 | k__Fungi(100);p__Ascomycota(100);p__Ascomycota_unclassified(100);p__Ascomycota_unclassified(100);p__Ascomycota_unclassified(100);p__Ascomycota_unclassified(100);p__Ascomycota_unclassified(100); | - | - | - | - | - | - | - | Unassigned | - |
| Otu36112 | 1 | k__Fungi(100);p__Ascomycota(100);c__Sordariomycetes(100);c__Sordariomycetes_unclassified(100);c__Sordariomycetes_unclassified(100);c__Sordariomycetes_unclassified(100);c__Sordariomycetes_unclassified(100); | - | - | - | - | - | - | - | Unassigned | - |
| Otu36114 | 1 | k__Fungi(100);p__Ascomycota(100);p__Ascomycota_unclassified(100);p__Ascomycota_unclassified(100);p__Ascomycota_unclassified(100);p__Ascomycota_unclassified(100);p__Ascomycota_unclassified(100); | - | - | - | - | - | - | - | Unassigned | - |
| Otu36115 | 1 | k__Fungi(100);p__Ascomycota(100);p__Ascomycota_unclassified(100);p__Ascomycota_unclassified(100);p__Ascomycota_unclassified(100);p__Ascomycota_unclassified(100);p__Ascomycota_unclassified(100); | - | - | - | - | - | - | - | Unassigned | - |
| Otu36119 | 1 | k__Fungi(100);p__Ascomycota(100);p__Ascomycota_unclassified(100);p__Ascomycota_unclassified(100);p__Ascomycota_unclassified(100);p__Ascomycota_unclassified(100);p__Ascomycota_unclassified(100); | - | - | - | - | - | - | - | Unassigned | - |
| Otu36121 | 1 | k__Fungi(100);k__Fungi_unclassified(100);k__Fungi_unclassified(100);k__Fungi_unclassified(100);k__Fungi_unclassified(100);k__Fungi_unclassified(100);k__Fungi_unclassified(100); | - | - | - | - | - | - | - | Unassigned | - |
| Otu36122 | 1 | k__Fungi(100);p__Ascomycota(100);p__Ascomycota_unclassified(100);p__Ascomycota_unclassified(100);p__Ascomycota_unclassified(100);p__Ascomycota_unclassified(100);p__Ascomycota_unclassified(100); | - | - | - | - | - | - | - | Unassigned | - |
| Otu36123 | 1 | k__Fungi(100);p__Ascomycota(100);p__Ascomycota_unclassified(100);p__Ascomycota_unclassified(100);p__Ascomycota_unclassified(100);p__Ascomycota_unclassified(100);p__Ascomycota_unclassified(100); | - | - | - | - | - | - | - | Unassigned | - |
| Otu36124 | 1 | k__Fungi(100);k__Fungi_unclassified(100);k__Fungi_unclassified(100);k__Fungi_unclassified(100);k__Fungi_unclassified(100);k__Fungi_unclassified(100);k__Fungi_unclassified(100); | - | - | - | - | - | - | - | Unassigned | - |
| Otu36125 | 1 | k__Fungi(100);p__Ascomycota(100);p__Ascomycota_unclassified(100);p__Ascomycota_unclassified(100);p__Ascomycota_unclassified(100);p__Ascomycota_unclassified(100);p__Ascomycota_unclassified(100); | - | - | - | - | - | - | - | Unassigned | - |
| Otu36127 | 1 | k__Fungi(100);p__Ascomycota(100);p__Ascomycota_unclassified(100);p__Ascomycota_unclassified(100);p__Ascomycota_unclassified(100);p__Ascomycota_unclassified(100);p__Ascomycota_unclassified(100); | - | - | - | - | - | - | - | Unassigned | - |
| Otu36128 | 1 | k__Fungi(100);p__Ascomycota(100);p__Ascomycota_unclassified(100);p__Ascomycota_unclassified(100);p__Ascomycota_unclassified(100);p__Ascomycota_unclassified(100);p__Ascomycota_unclassified(100); | - | - | - | - | - | - | - | Unassigned | - |
| Otu36129 | 1 | k__Fungi(100);k__Fungi_unclassified(100);k__Fungi_unclassified(100);k__Fungi_unclassified(100);k__Fungi_unclassified(100);k__Fungi_unclassified(100);k__Fungi_unclassified(100); | - | - | - | - | - | - | - | Unassigned | - |
| Otu36130 | 1 | k__Fungi(100);k__Fungi_unclassified(100);k__Fungi_unclassified(100);k__Fungi_unclassified(100);k__Fungi_unclassified(100);k__Fungi_unclassified(100);k__Fungi_unclassified(100); | - | - | - | - | - | - | - | Unassigned | - |
| Otu36131 | 1 | k__Fungi(100);p__Ascomycota(100);p__Ascomycota_unclassified(100);p__Ascomycota_unclassified(100);p__Ascomycota_unclassified(100);p__Ascomycota_unclassified(100);p__Ascomycota_unclassified(100); | - | - | - | - | - | - | - | Unassigned | - |
| Otu36132 | 1 | k__Fungi(100);k__Fungi_unclassified(100);k__Fungi_unclassified(100);k__Fungi_unclassified(100);k__Fungi_unclassified(100);k__Fungi_unclassified(100);k__Fungi_unclassified(100); | - | - | - | - | - | - | - | Unassigned | - |
| Otu36133 | 1 | k__Fungi(100);p__Ascomycota(100);p__Ascomycota_unclassified(100);p__Ascomycota_unclassified(100);p__Ascomycota_unclassified(100);p__Ascomycota_unclassified(100);p__Ascomycota_unclassified(100); | - | - | - | - | - | - | - | Unassigned | - |
| Otu36153 | 1 | k__Fungi(100);p__Ascomycota(100);p__Ascomycota_unclassified(100);p__Ascomycota_unclassified(100);p__Ascomycota_unclassified(100);p__Ascomycota_unclassified(100);p__Ascomycota_unclassified(100); | - | - | - | - | - | - | - | Unassigned | - |
| Otu36154 | 1 | k__Fungi(100);k__Fungi_unclassified(100);k__Fungi_unclassified(100);k__Fungi_unclassified(100);k__Fungi_unclassified(100);k__Fungi_unclassified(100);k__Fungi_unclassified(100); | - | - | - | - | - | - | - | Unassigned | - |
| Otu36155 | 1 | k__Fungi(100);p__Ascomycota(100);p__Ascomycota_unclassified(100);p__Ascomycota_unclassified(100);p__Ascomycota_unclassified(100);p__Ascomycota_unclassified(100);p__Ascomycota_unclassified(100); | - | - | - | - | - | - | - | Unassigned | - |
| Otu36157 | 1 | k__Fungi(100);p__Ascomycota(100);p__Ascomycota_unclassified(100);p__Ascomycota_unclassified(100);p__Ascomycota_unclassified(100);p__Ascomycota_unclassified(100);p__Ascomycota_unclassified(100); | - | - | - | - | - | - | - | Unassigned | - |
| Otu36159 | 1 | k__Fungi(100);p__Ascomycota(100);p__Ascomycota_unclassified(100);p__Ascomycota_unclassified(100);p__Ascomycota_unclassified(100);p__Ascomycota_unclassified(100);p__Ascomycota_unclassified(100); | - | - | - | - | - | - | - | Unassigned | - |
| Otu36160 | 1 | k__Fungi(100);p__Ascomycota(100);p__Ascomycota_unclassified(100);p__Ascomycota_unclassified(100);p__Ascomycota_unclassified(100);p__Ascomycota_unclassified(100);p__Ascomycota_unclassified(100); | - | - | - | - | - | - | - | Unassigned | - |
| Otu36161 | 1 | k__Fungi(100);p__Ascomycota(100);p__Ascomycota_unclassified(100);p__Ascomycota_unclassified(100);p__Ascomycota_unclassified(100);p__Ascomycota_unclassified(100);p__Ascomycota_unclassified(100); | - | - | - | - | - | - | - | Unassigned | - |
| Otu36162 | 1 | k__Fungi(100);p__Ascomycota(100);p__Ascomycota_unclassified(100);p__Ascomycota_unclassified(100);p__Ascomycota_unclassified(100);p__Ascomycota_unclassified(100);p__Ascomycota_unclassified(100); | - | - | - | - | - | - | - | Unassigned | - |
| Otu36163 | 1 | k__Fungi(100);k__Fungi_unclassified(100);k__Fungi_unclassified(100);k__Fungi_unclassified(100);k__Fungi_unclassified(100);k__Fungi_unclassified(100);k__Fungi_unclassified(100); | - | - | - | - | - | - | - | Unassigned | - |
| Otu36164 | 1 | k__Fungi(100);k__Fungi_unclassified(100);k__Fungi_unclassified(100);k__Fungi_unclassified(100);k__Fungi_unclassified(100);k__Fungi_unclassified(100);k__Fungi_unclassified(100); | - | - | - | - | - | - | - | Unassigned | - |
| Otu36168 | 1 | k__Fungi(100);p__Ascomycota(100);p__Ascomycota_unclassified(100);p__Ascomycota_unclassified(100);p__Ascomycota_unclassified(100);p__Ascomycota_unclassified(100);p__Ascomycota_unclassified(100); | - | - | - | - | - | - | - | Unassigned | - |
| Otu36169 | 1 | k__Fungi(100);k__Fungi_unclassified(100);k__Fungi_unclassified(100);k__Fungi_unclassified(100);k__Fungi_unclassified(100);k__Fungi_unclassified(100);k__Fungi_unclassified(100); | - | - | - | - | - | - | - | Unassigned | - |
| Otu36174 | 1 | k__Fungi(100);k__Fungi_unclassified(100);k__Fungi_unclassified(100);k__Fungi_unclassified(100);k__Fungi_unclassified(100);k__Fungi_unclassified(100);k__Fungi_unclassified(100); | - | - | - | - | - | - | - | Unassigned | - |
| Otu36190 | 1 | k__Fungi(100);k__Fungi_unclassified(100);k__Fungi_unclassified(100);k__Fungi_unclassified(100);k__Fungi_unclassified(100);k__Fungi_unclassified(100);k__Fungi_unclassified(100); | - | - | - | - | - | - | - | Unassigned | - |
| Otu36192 | 1 | k__Fungi(100);p__Ascomycota(100);p__Ascomycota_unclassified(100);p__Ascomycota_unclassified(100);p__Ascomycota_unclassified(100);p__Ascomycota_unclassified(100);p__Ascomycota_unclassified(100); | - | - | - | - | - | - | - | Unassigned | - |
| Otu36193 | 1 | k__Fungi(100);p__Ascomycota(100);p__Ascomycota_unclassified(100);p__Ascomycota_unclassified(100);p__Ascomycota_unclassified(100);p__Ascomycota_unclassified(100);p__Ascomycota_unclassified(100); | - | - | - | - | - | - | - | Unassigned | - |
| Otu36194 | 1 | k__Fungi(100);k__Fungi_unclassified(100);k__Fungi_unclassified(100);k__Fungi_unclassified(100);k__Fungi_unclassified(100);k__Fungi_unclassified(100);k__Fungi_unclassified(100); | - | - | - | - | - | - | - | Unassigned | - |
| Otu36196 | 1 | k__Fungi(100);p__Ascomycota(100);p__Ascomycota_unclassified(100);p__Ascomycota_unclassified(100);p__Ascomycota_unclassified(100);p__Ascomycota_unclassified(100);p__Ascomycota_unclassified(100); | - | - | - | - | - | - | - | Unassigned | - |
| Otu36198 | 1 | k__Fungi(100);p__Ascomycota(100);p__Ascomycota_unclassified(100);p__Ascomycota_unclassified(100);p__Ascomycota_unclassified(100);p__Ascomycota_unclassified(100);p__Ascomycota_unclassified(100); | - | - | - | - | - | - | - | Unassigned | - |
| Otu36199 | 1 | k__Fungi(100);p__Ascomycota(100);p__Ascomycota_unclassified(100);p__Ascomycota_unclassified(100);p__Ascomycota_unclassified(100);p__Ascomycota_unclassified(100);p__Ascomycota_unclassified(100); | - | - | - | - | - | - | - | Unassigned | - |
| Otu36202 | 1 | k__Fungi(100);k__Fungi_unclassified(100);k__Fungi_unclassified(100);k__Fungi_unclassified(100);k__Fungi_unclassified(100);k__Fungi_unclassified(100);k__Fungi_unclassified(100); | - | - | - | - | - | - | - | Unassigned | - |
| Otu36208 | 1 | k__Fungi(100);p__Ascomycota(100);p__Ascomycota_unclassified(100);p__Ascomycota_unclassified(100);p__Ascomycota_unclassified(100);p__Ascomycota_unclassified(100);p__Ascomycota_unclassified(100); | - | - | - | - | - | - | - | Unassigned | - |
| Otu36209 | 1 | k__Fungi(100);p__Ascomycota(100);p__Ascomycota_unclassified(100);p__Ascomycota_unclassified(100);p__Ascomycota_unclassified(100);p__Ascomycota_unclassified(100);p__Ascomycota_unclassified(100); | - | - | - | - | - | - | - | Unassigned | - |
| Otu36213 | 1 | k__Fungi(100);k__Fungi_unclassified(100);k__Fungi_unclassified(100);k__Fungi_unclassified(100);k__Fungi_unclassified(100);k__Fungi_unclassified(100);k__Fungi_unclassified(100); | - | - | - | - | - | - | - | Unassigned | - |
| Otu36217 | 1 | k__Fungi(100);k__Fungi_unclassified(100);k__Fungi_unclassified(100);k__Fungi_unclassified(100);k__Fungi_unclassified(100);k__Fungi_unclassified(100);k__Fungi_unclassified(100); | - | - | - | - | - | - | - | Unassigned | - |
| Otu36218 | 1 | k__Fungi(100);k__Fungi_unclassified(100);k__Fungi_unclassified(100);k__Fungi_unclassified(100);k__Fungi_unclassified(100);k__Fungi_unclassified(100);k__Fungi_unclassified(100); | - | - | - | - | - | - | - | Unassigned | - |
| Otu36220 | 1 | k__Fungi(100);k__Fungi_unclassified(100);k__Fungi_unclassified(100);k__Fungi_unclassified(100);k__Fungi_unclassified(100);k__Fungi_unclassified(100);k__Fungi_unclassified(100); | - | - | - | - | - | - | - | Unassigned | - |
| Otu36221 | 1 | k__Fungi(100);k__Fungi_unclassified(100);k__Fungi_unclassified(100);k__Fungi_unclassified(100);k__Fungi_unclassified(100);k__Fungi_unclassified(100);k__Fungi_unclassified(100); | - | - | - | - | - | - | - | Unassigned | - |
| Otu36222 | 1 | k__Fungi(100);k__Fungi_unclassified(100);k__Fungi_unclassified(100);k__Fungi_unclassified(100);k__Fungi_unclassified(100);k__Fungi_unclassified(100);k__Fungi_unclassified(100); | - | - | - | - | - | - | - | Unassigned | - |
| Otu36223 | 1 | k__Fungi(100);p__Ascomycota(100);p__Ascomycota_unclassified(100);p__Ascomycota_unclassified(100);p__Ascomycota_unclassified(100);p__Ascomycota_unclassified(100);p__Ascomycota_unclassified(100); | - | - | - | - | - | - | - | Unassigned | - |
| Otu36224 | 1 | k__Fungi(100);k__Fungi_unclassified(100);k__Fungi_unclassified(100);k__Fungi_unclassified(100);k__Fungi_unclassified(100);k__Fungi_unclassified(100);k__Fungi_unclassified(100); | - | - | - | - | - | - | - | Unassigned | - |
| Otu36226 | 1 | k__Fungi(100);k__Fungi_unclassified(100);k__Fungi_unclassified(100);k__Fungi_unclassified(100);k__Fungi_unclassified(100);k__Fungi_unclassified(100);k__Fungi_unclassified(100); | - | - | - | - | - | - | - | Unassigned | - |
| Otu36230 | 1 | k__Fungi(100);k__Fungi_unclassified(100);k__Fungi_unclassified(100);k__Fungi_unclassified(100);k__Fungi_unclassified(100);k__Fungi_unclassified(100);k__Fungi_unclassified(100); | - | - | - | - | - | - | - | Unassigned | - |
| Otu36231 | 1 | k__Fungi(100);k__Fungi_unclassified(100);k__Fungi_unclassified(100);k__Fungi_unclassified(100);k__Fungi_unclassified(100);k__Fungi_unclassified(100);k__Fungi_unclassified(100); | - | - | - | - | - | - | - | Unassigned | - |
| Otu36240 | 1 | k__Fungi(100);k__Fungi_unclassified(100);k__Fungi_unclassified(100);k__Fungi_unclassified(100);k__Fungi_unclassified(100);k__Fungi_unclassified(100);k__Fungi_unclassified(100); | - | - | - | - | - | - | - | Unassigned | - |
| Otu36241 | 1 | k__Fungi(100);p__Ascomycota(100);p__Ascomycota_unclassified(100);p__Ascomycota_unclassified(100);p__Ascomycota_unclassified(100);p__Ascomycota_unclassified(100);p__Ascomycota_unclassified(100); | - | - | - | - | - | - | - | Unassigned | - |
| Otu36242 | 1 | k__Fungi(100);p__Ascomycota(100);p__Ascomycota_unclassified(100);p__Ascomycota_unclassified(100);p__Ascomycota_unclassified(100);p__Ascomycota_unclassified(100);p__Ascomycota_unclassified(100); | - | - | - | - | - | - | - | Unassigned | - |
| Otu36243 | 1 | k__Fungi(100);p__Ascomycota(100);p__Ascomycota_unclassified(100);p__Ascomycota_unclassified(100);p__Ascomycota_unclassified(100);p__Ascomycota_unclassified(100);p__Ascomycota_unclassified(100); | - | - | - | - | - | - | - | Unassigned | - |
| Otu36244 | 1 | k__Fungi(100);k__Fungi_unclassified(100);k__Fungi_unclassified(100);k__Fungi_unclassified(100);k__Fungi_unclassified(100);k__Fungi_unclassified(100);k__Fungi_unclassified(100); | - | - | - | - | - | - | - | Unassigned | - |
| Otu36245 | 1 | k__Fungi(100);p__Ascomycota(100);p__Ascomycota_unclassified(100);p__Ascomycota_unclassified(100);p__Ascomycota_unclassified(100);p__Ascomycota_unclassified(100);p__Ascomycota_unclassified(100); | - | - | - | - | - | - | - | Unassigned | - |
| Otu36246 | 1 | k__Fungi(100);p__Ascomycota(100);p__Ascomycota_unclassified(100);p__Ascomycota_unclassified(100);p__Ascomycota_unclassified(100);p__Ascomycota_unclassified(100);p__Ascomycota_unclassified(100); | - | - | - | - | - | - | - | Unassigned | - |
| Otu36247 | 1 | k__Fungi(100);p__Ascomycota(100);p__Ascomycota_unclassified(100);p__Ascomycota_unclassified(100);p__Ascomycota_unclassified(100);p__Ascomycota_unclassified(100);p__Ascomycota_unclassified(100); | - | - | - | - | - | - | - | Unassigned | - |
| Otu36248 | 1 | k__Fungi(100);k__Fungi_unclassified(100);k__Fungi_unclassified(100);k__Fungi_unclassified(100);k__Fungi_unclassified(100);k__Fungi_unclassified(100);k__Fungi_unclassified(100); | - | - | - | - | - | - | - | Unassigned | - |
| Otu36249 | 1 | k__Fungi(100);k__Fungi_unclassified(100);k__Fungi_unclassified(100);k__Fungi_unclassified(100);k__Fungi_unclassified(100);k__Fungi_unclassified(100);k__Fungi_unclassified(100); | - | - | - | - | - | - | - | Unassigned | - |
| Otu36250 | 1 | k__Fungi(100);p__Ascomycota(100);p__Ascomycota_unclassified(100);p__Ascomycota_unclassified(100);p__Ascomycota_unclassified(100);p__Ascomycota_unclassified(100);p__Ascomycota_unclassified(100); | - | - | - | - | - | - | - | Unassigned | - |
| Otu36251 | 1 | k__Fungi(100);k__Fungi_unclassified(100);k__Fungi_unclassified(100);k__Fungi_unclassified(100);k__Fungi_unclassified(100);k__Fungi_unclassified(100);k__Fungi_unclassified(100); | - | - | - | - | - | - | - | Unassigned | - |
| Otu36252 | 1 | k__Fungi(100);p__Ascomycota(100);p__Ascomycota_unclassified(100);p__Ascomycota_unclassified(100);p__Ascomycota_unclassified(100);p__Ascomycota_unclassified(100);p__Ascomycota_unclassified(100); | - | - | - | - | - | - | - | Unassigned | - |
| Otu36253 | 1 | k__Fungi(100);p__Basidiomycota(100);p__Basidiomycota_unclassified(100);p__Basidiomycota_unclassified(100);p__Basidiomycota_unclassified(100);p__Basidiomycota_unclassified(100);p__Basidiomycota_unclassified(100); | - | - | - | - | - | - | - | Unassigned | - |
| Otu36254 | 1 | k__Fungi(100);k__Fungi_unclassified(100);k__Fungi_unclassified(100);k__Fungi_unclassified(100);k__Fungi_unclassified(100);k__Fungi_unclassified(100);k__Fungi_unclassified(100); | - | - | - | - | - | - | - | Unassigned | - |
| Otu36255 | 1 | k__Fungi(100);k__Fungi_unclassified(100);k__Fungi_unclassified(100);k__Fungi_unclassified(100);k__Fungi_unclassified(100);k__Fungi_unclassified(100);k__Fungi_unclassified(100); | - | - | - | - | - | - | - | Unassigned | - |
| Otu36256 | 1 | k__Fungi(100);p__Ascomycota(100);p__Ascomycota_unclassified(100);p__Ascomycota_unclassified(100);p__Ascomycota_unclassified(100);p__Ascomycota_unclassified(100);p__Ascomycota_unclassified(100); | - | - | - | - | - | - | - | Unassigned | - |
| Otu36258 | 1 | k__Fungi(100);p__Ascomycota(100);p__Ascomycota_unclassified(100);p__Ascomycota_unclassified(100);p__Ascomycota_unclassified(100);p__Ascomycota_unclassified(100);p__Ascomycota_unclassified(100); | - | - | - | - | - | - | - | Unassigned | - |
| Otu36260 | 1 | k__Fungi(100);p__Ascomycota(100);p__Ascomycota_unclassified(100);p__Ascomycota_unclassified(100);p__Ascomycota_unclassified(100);p__Ascomycota_unclassified(100);p__Ascomycota_unclassified(100); | - | - | - | - | - | - | - | Unassigned | - |
| Otu36265 | 1 | k__Fungi(100);k__Fungi_unclassified(100);k__Fungi_unclassified(100);k__Fungi_unclassified(100);k__Fungi_unclassified(100);k__Fungi_unclassified(100);k__Fungi_unclassified(100); | - | - | - | - | - | - | - | Unassigned | - |
| Otu36266 | 1 | k__Fungi(100);p__Ascomycota(100);p__Ascomycota_unclassified(100);p__Ascomycota_unclassified(100);p__Ascomycota_unclassified(100);p__Ascomycota_unclassified(100);p__Ascomycota_unclassified(100); | - | - | - | - | - | - | - | Unassigned | - |
| Otu36271 | 1 | k__Fungi(100);k__Fungi_unclassified(100);k__Fungi_unclassified(100);k__Fungi_unclassified(100);k__Fungi_unclassified(100);k__Fungi_unclassified(100);k__Fungi_unclassified(100); | - | - | - | - | - | - | - | Unassigned | - |
| Otu36278 | 1 | k__Fungi(100);k__Fungi_unclassified(100);k__Fungi_unclassified(100);k__Fungi_unclassified(100);k__Fungi_unclassified(100);k__Fungi_unclassified(100);k__Fungi_unclassified(100); | - | - | - | - | - | - | - | Unassigned | - |
| Otu36279 | 1 | k__Fungi(100);p__Ascomycota(100);p__Ascomycota_unclassified(100);p__Ascomycota_unclassified(100);p__Ascomycota_unclassified(100);p__Ascomycota_unclassified(100);p__Ascomycota_unclassified(100); | - | - | - | - | - | - | - | Unassigned | - |
| Otu36280 | 1 | k__Fungi(100);k__Fungi_unclassified(100);k__Fungi_unclassified(100);k__Fungi_unclassified(100);k__Fungi_unclassified(100);k__Fungi_unclassified(100);k__Fungi_unclassified(100); | - | - | - | - | - | - | - | Unassigned | - |
| Otu36291 | 1 | k__Fungi(100);k__Fungi_unclassified(100);k__Fungi_unclassified(100);k__Fungi_unclassified(100);k__Fungi_unclassified(100);k__Fungi_unclassified(100);k__Fungi_unclassified(100); | - | - | - | - | - | - | - | Unassigned | - |
| Otu36293 | 1 | k__Fungi(100);p__Ascomycota(100);p__Ascomycota_unclassified(100);p__Ascomycota_unclassified(100);p__Ascomycota_unclassified(100);p__Ascomycota_unclassified(100);p__Ascomycota_unclassified(100); | - | - | - | - | - | - | - | Unassigned | - |
| Otu36294 | 1 | k__Fungi(100);p__Ascomycota(100);p__Ascomycota_unclassified(100);p__Ascomycota_unclassified(100);p__Ascomycota_unclassified(100);p__Ascomycota_unclassified(100);p__Ascomycota_unclassified(100); | - | - | - | - | - | - | - | Unassigned | - |
| Otu36311 | 1 | k__Fungi(100);p__Ascomycota(100);p__Ascomycota_unclassified(100);p__Ascomycota_unclassified(100);p__Ascomycota_unclassified(100);p__Ascomycota_unclassified(100);p__Ascomycota_unclassified(100); | - | - | - | - | - | - | - | Unassigned | - |
| Otu36312 | 1 | k__Fungi(100);k__Fungi_unclassified(100);k__Fungi_unclassified(100);k__Fungi_unclassified(100);k__Fungi_unclassified(100);k__Fungi_unclassified(100);k__Fungi_unclassified(100); | - | - | - | - | - | - | - | Unassigned | - |
| Otu36313 | 1 | k__Fungi(100);k__Fungi_unclassified(100);k__Fungi_unclassified(100);k__Fungi_unclassified(100);k__Fungi_unclassified(100);k__Fungi_unclassified(100);k__Fungi_unclassified(100); | - | - | - | - | - | - | - | Unassigned | - |
| Otu36315 | 1 | k__Fungi(100);p__Ascomycota(100);p__Ascomycota_unclassified(100);p__Ascomycota_unclassified(100);p__Ascomycota_unclassified(100);p__Ascomycota_unclassified(100);p__Ascomycota_unclassified(100); | - | - | - | - | - | - | - | Unassigned | - |
| Otu36317 | 1 | k__Fungi(100);p__Ascomycota(100);p__Ascomycota_unclassified(100);p__Ascomycota_unclassified(100);p__Ascomycota_unclassified(100);p__Ascomycota_unclassified(100);p__Ascomycota_unclassified(100); | - | - | - | - | - | - | - | Unassigned | - |
| Otu36318 | 1 | k__Fungi(100);p__Ascomycota(100);p__Ascomycota_unclassified(100);p__Ascomycota_unclassified(100);p__Ascomycota_unclassified(100);p__Ascomycota_unclassified(100);p__Ascomycota_unclassified(100); | - | - | - | - | - | - | - | Unassigned | - |
| Otu36319 | 1 | k__Fungi(100);p__Ascomycota(100);p__Ascomycota_unclassified(100);p__Ascomycota_unclassified(100);p__Ascomycota_unclassified(100);p__Ascomycota_unclassified(100);p__Ascomycota_unclassified(100); | - | - | - | - | - | - | - | Unassigned | - |
| Otu36320 | 1 | k__Fungi(100);p__Ascomycota(100);p__Ascomycota_unclassified(100);p__Ascomycota_unclassified(100);p__Ascomycota_unclassified(100);p__Ascomycota_unclassified(100);p__Ascomycota_unclassified(100); | - | - | - | - | - | - | - | Unassigned | - |
| Otu36321 | 1 | k__Fungi(100);p__Ascomycota(100);p__Ascomycota_unclassified(100);p__Ascomycota_unclassified(100);p__Ascomycota_unclassified(100);p__Ascomycota_unclassified(100);p__Ascomycota_unclassified(100); | - | - | - | - | - | - | - | Unassigned | - |
| Otu36329 | 1 | k__Fungi(100);k__Fungi_unclassified(100);k__Fungi_unclassified(100);k__Fungi_unclassified(100);k__Fungi_unclassified(100);k__Fungi_unclassified(100);k__Fungi_unclassified(100); | - | - | - | - | - | - | - | Unassigned | - |
| Otu36332 | 1 | k__Fungi(100);p__Ascomycota(100);p__Ascomycota_unclassified(100);p__Ascomycota_unclassified(100);p__Ascomycota_unclassified(100);p__Ascomycota_unclassified(100);p__Ascomycota_unclassified(100); | - | - | - | - | - | - | - | Unassigned | - |
| Otu36335 | 1 | k__Fungi(100);p__Ascomycota(100);p__Ascomycota_unclassified(100);p__Ascomycota_unclassified(100);p__Ascomycota_unclassified(100);p__Ascomycota_unclassified(100);p__Ascomycota_unclassified(100); | - | - | - | - | - | - | - | Unassigned | - |
| Otu36352 | 1 | k__Fungi(100);p__Ascomycota(100);p__Ascomycota_unclassified(100);p__Ascomycota_unclassified(100);p__Ascomycota_unclassified(100);p__Ascomycota_unclassified(100);p__Ascomycota_unclassified(100); | - | - | - | - | - | - | - | Unassigned | - |
| Otu36353 | 1 | k__Fungi(100);k__Fungi_unclassified(100);k__Fungi_unclassified(100);k__Fungi_unclassified(100);k__Fungi_unclassified(100);k__Fungi_unclassified(100);k__Fungi_unclassified(100); | - | - | - | - | - | - | - | Unassigned | - |
| Otu36368 | 1 | k__Fungi(100);p__Ascomycota(100);p__Ascomycota_unclassified(100);p__Ascomycota_unclassified(100);p__Ascomycota_unclassified(100);p__Ascomycota_unclassified(100);p__Ascomycota_unclassified(100); | - | - | - | - | - | - | - | Unassigned | - |
| Otu36376 | 1 | k__Fungi(100);k__Fungi_unclassified(100);k__Fungi_unclassified(100);k__Fungi_unclassified(100);k__Fungi_unclassified(100);k__Fungi_unclassified(100);k__Fungi_unclassified(100); | - | - | - | - | - | - | - | Unassigned | - |
| Otu36377 | 1 | k__Fungi(100);p__Ascomycota(100);p__Ascomycota_unclassified(100);p__Ascomycota_unclassified(100);p__Ascomycota_unclassified(100);p__Ascomycota_unclassified(100);p__Ascomycota_unclassified(100); | - | - | - | - | - | - | - | Unassigned | - |
| Otu36378 | 1 | k__Fungi(100);k__Fungi_unclassified(100);k__Fungi_unclassified(100);k__Fungi_unclassified(100);k__Fungi_unclassified(100);k__Fungi_unclassified(100);k__Fungi_unclassified(100); | - | - | - | - | - | - | - | Unassigned | - |
| Otu36380 | 1 | k__Fungi(100);p__Ascomycota(100);p__Ascomycota_unclassified(100);p__Ascomycota_unclassified(100);p__Ascomycota_unclassified(100);p__Ascomycota_unclassified(100);p__Ascomycota_unclassified(100); | - | - | - | - | - | - | - | Unassigned | - |
| Otu36384 | 1 | k__Fungi(100);k__Fungi_unclassified(100);k__Fungi_unclassified(100);k__Fungi_unclassified(100);k__Fungi_unclassified(100);k__Fungi_unclassified(100);k__Fungi_unclassified(100); | - | - | - | - | - | - | - | Unassigned | - |
| Otu36394 | 1 | k__Fungi(100);p__Ascomycota(100);p__Ascomycota_unclassified(100);p__Ascomycota_unclassified(100);p__Ascomycota_unclassified(100);p__Ascomycota_unclassified(100);p__Ascomycota_unclassified(100); | - | - | - | - | - | - | - | Unassigned | - |
| Otu36397 | 1 | k__Fungi(100);p__Ascomycota(100);p__Ascomycota_unclassified(100);p__Ascomycota_unclassified(100);p__Ascomycota_unclassified(100);p__Ascomycota_unclassified(100);p__Ascomycota_unclassified(100); | - | - | - | - | - | - | - | Unassigned | - |
| Otu36399 | 1 | k__Fungi(100);k__Fungi_unclassified(100);k__Fungi_unclassified(100);k__Fungi_unclassified(100);k__Fungi_unclassified(100);k__Fungi_unclassified(100);k__Fungi_unclassified(100); | - | - | - | - | - | - | - | Unassigned | - |
| Otu36402 | 1 | k__Fungi(100);p__Ascomycota(100);p__Ascomycota_unclassified(100);p__Ascomycota_unclassified(100);p__Ascomycota_unclassified(100);p__Ascomycota_unclassified(100);p__Ascomycota_unclassified(100); | - | - | - | - | - | - | - | Unassigned | - |
| Otu36406 | 1 | k__Fungi(100);k__Fungi_unclassified(100);k__Fungi_unclassified(100);k__Fungi_unclassified(100);k__Fungi_unclassified(100);k__Fungi_unclassified(100);k__Fungi_unclassified(100); | - | - | - | - | - | - | - | Unassigned | - |
| Otu36407 | 1 | k__Fungi(100);p__Ascomycota(100);p__Ascomycota_unclassified(100);p__Ascomycota_unclassified(100);p__Ascomycota_unclassified(100);p__Ascomycota_unclassified(100);p__Ascomycota_unclassified(100); | - | - | - | - | - | - | - | Unassigned | - |
| Otu36412 | 1 | k__Fungi(100);k__Fungi_unclassified(100);k__Fungi_unclassified(100);k__Fungi_unclassified(100);k__Fungi_unclassified(100);k__Fungi_unclassified(100);k__Fungi_unclassified(100); | - | - | - | - | - | - | - | Unassigned | - |
| Otu36414 | 1 | k__Fungi(100);k__Fungi_unclassified(100);k__Fungi_unclassified(100);k__Fungi_unclassified(100);k__Fungi_unclassified(100);k__Fungi_unclassified(100);k__Fungi_unclassified(100); | - | - | - | - | - | - | - | Unassigned | - |
| Otu36415 | 1 | k__Fungi(100);k__Fungi_unclassified(100);k__Fungi_unclassified(100);k__Fungi_unclassified(100);k__Fungi_unclassified(100);k__Fungi_unclassified(100);k__Fungi_unclassified(100); | - | - | - | - | - | - | - | Unassigned | - |
| Otu36417 | 1 | k__Fungi(100);k__Fungi_unclassified(100);k__Fungi_unclassified(100);k__Fungi_unclassified(100);k__Fungi_unclassified(100);k__Fungi_unclassified(100);k__Fungi_unclassified(100); | - | - | - | - | - | - | - | Unassigned | - |
| Otu36418 | 1 | k__Fungi(100);p__Ascomycota(100);p__Ascomycota_unclassified(100);p__Ascomycota_unclassified(100);p__Ascomycota_unclassified(100);p__Ascomycota_unclassified(100);p__Ascomycota_unclassified(100); | - | - | - | - | - | - | - | Unassigned | - |
| Otu36419 | 1 | k__Fungi(100);k__Fungi_unclassified(100);k__Fungi_unclassified(100);k__Fungi_unclassified(100);k__Fungi_unclassified(100);k__Fungi_unclassified(100);k__Fungi_unclassified(100); | - | - | - | - | - | - | - | Unassigned | - |
| Otu36420 | 1 | k__Fungi(100);k__Fungi_unclassified(100);k__Fungi_unclassified(100);k__Fungi_unclassified(100);k__Fungi_unclassified(100);k__Fungi_unclassified(100);k__Fungi_unclassified(100); | - | - | - | - | - | - | - | Unassigned | - |
| Otu36421 | 1 | k__Fungi(100);p__Ascomycota(100);p__Ascomycota_unclassified(100);p__Ascomycota_unclassified(100);p__Ascomycota_unclassified(100);p__Ascomycota_unclassified(100);p__Ascomycota_unclassified(100); | - | - | - | - | - | - | - | Unassigned | - |
| Otu36422 | 1 | k__Fungi(100);k__Fungi_unclassified(100);k__Fungi_unclassified(100);k__Fungi_unclassified(100);k__Fungi_unclassified(100);k__Fungi_unclassified(100);k__Fungi_unclassified(100); | - | - | - | - | - | - | - | Unassigned | - |
| Otu36425 | 1 | k__Fungi(100);k__Fungi_unclassified(100);k__Fungi_unclassified(100);k__Fungi_unclassified(100);k__Fungi_unclassified(100);k__Fungi_unclassified(100);k__Fungi_unclassified(100); | - | - | - | - | - | - | - | Unassigned | - |
| Otu36426 | 1 | k__Fungi(100);k__Fungi_unclassified(100);k__Fungi_unclassified(100);k__Fungi_unclassified(100);k__Fungi_unclassified(100);k__Fungi_unclassified(100);k__Fungi_unclassified(100); | - | - | - | - | - | - | - | Unassigned | - |
| Otu36427 | 1 | k__Fungi(100);k__Fungi_unclassified(100);k__Fungi_unclassified(100);k__Fungi_unclassified(100);k__Fungi_unclassified(100);k__Fungi_unclassified(100);k__Fungi_unclassified(100); | - | - | - | - | - | - | - | Unassigned | - |
| Otu36428 | 1 | k__Fungi(100);k__Fungi_unclassified(100);k__Fungi_unclassified(100);k__Fungi_unclassified(100);k__Fungi_unclassified(100);k__Fungi_unclassified(100);k__Fungi_unclassified(100); | - | - | - | - | - | - | - | Unassigned | - |
| Otu36438 | 1 | k__Fungi(100);k__Fungi_unclassified(100);k__Fungi_unclassified(100);k__Fungi_unclassified(100);k__Fungi_unclassified(100);k__Fungi_unclassified(100);k__Fungi_unclassified(100); | - | - | - | - | - | - | - | Unassigned | - |
| Otu36439 | 1 | k__Fungi(100);k__Fungi_unclassified(100);k__Fungi_unclassified(100);k__Fungi_unclassified(100);k__Fungi_unclassified(100);k__Fungi_unclassified(100);k__Fungi_unclassified(100); | - | - | - | - | - | - | - | Unassigned | - |
| Otu36440 | 1 | k__Fungi(100);p__Ascomycota(100);p__Ascomycota_unclassified(100);p__Ascomycota_unclassified(100);p__Ascomycota_unclassified(100);p__Ascomycota_unclassified(100);p__Ascomycota_unclassified(100); | - | - | - | - | - | - | - | Unassigned | - |
| Otu36443 | 1 | k__Fungi(100);k__Fungi_unclassified(100);k__Fungi_unclassified(100);k__Fungi_unclassified(100);k__Fungi_unclassified(100);k__Fungi_unclassified(100);k__Fungi_unclassified(100); | - | - | - | - | - | - | - | Unassigned | - |
| Otu36444 | 1 | k__Fungi(100);k__Fungi_unclassified(100);k__Fungi_unclassified(100);k__Fungi_unclassified(100);k__Fungi_unclassified(100);k__Fungi_unclassified(100);k__Fungi_unclassified(100); | - | - | - | - | - | - | - | Unassigned | - |
| Otu36445 | 1 | k__Fungi(100);k__Fungi_unclassified(100);k__Fungi_unclassified(100);k__Fungi_unclassified(100);k__Fungi_unclassified(100);k__Fungi_unclassified(100);k__Fungi_unclassified(100); | - | - | - | - | - | - | - | Unassigned | - |
| Otu36446 | 1 | k__Fungi(100);k__Fungi_unclassified(100);k__Fungi_unclassified(100);k__Fungi_unclassified(100);k__Fungi_unclassified(100);k__Fungi_unclassified(100);k__Fungi_unclassified(100); | - | - | - | - | - | - | - | Unassigned | - |
| Otu36447 | 1 | k__Fungi(100);k__Fungi_unclassified(100);k__Fungi_unclassified(100);k__Fungi_unclassified(100);k__Fungi_unclassified(100);k__Fungi_unclassified(100);k__Fungi_unclassified(100); | - | - | - | - | - | - | - | Unassigned | - |

| | | | | | | | | | | | | |
|---|---|---|---|---|---|---|---|---|---|---|---|---|
| Otu36448 | 1 | k__Fungi(100);p__Ascomycota(100);p__Ascomycota_unclassified(100);p__Ascomycota_unclassified(100);p__Ascomycota_unclassified(100);p__Ascomycota_unclassified(100);p__Ascomycota_unclassified(100); | - | - | - | - | - | - | - | - | Unassigned | - |
| Otu36450 | 1 | k__Fungi(100);p__Ascomycota(100);p__Ascomycota_unclassified(100);p__Ascomycota_unclassified(100);p__Ascomycota_unclassified(100);p__Ascomycota_unclassified(100);p__Ascomycota_unclassified(100); | - | - | - | - | - | - | - | - | Unassigned | - |
| Otu36451 | 1 | k__Fungi(100);k__Fungi_unclassified(100);k__Fungi_unclassified(100);k__Fungi_unclassified(100);k__Fungi_unclassified(100);k__Fungi_unclassified(100);k__Fungi_unclassified(100); | - | - | - | - | - | - | - | - | Unassigned | - |
| Otu36472 | 1 | k__Fungi(100);k__Fungi_unclassified(100);k__Fungi_unclassified(100);k__Fungi_unclassified(100);k__Fungi_unclassified(100);k__Fungi_unclassified(100);k__Fungi_unclassified(100); | - | - | - | - | - | - | - | - | Unassigned | - |
| Otu36473 | 1 | k__Fungi(100);k__Fungi_unclassified(100);k__Fungi_unclassified(100);k__Fungi_unclassified(100);k__Fungi_unclassified(100);k__Fungi_unclassified(100);k__Fungi_unclassified(100); | - | - | - | - | - | - | - | - | Unassigned | - |
| Otu36474 | 1 | k__Fungi(100);k__Fungi_unclassified(100);k__Fungi_unclassified(100);k__Fungi_unclassified(100);k__Fungi_unclassified(100);k__Fungi_unclassified(100);k__Fungi_unclassified(100); | - | - | - | - | - | - | - | - | Unassigned | - |
| Otu36475 | 1 | k__Fungi(100);k__Fungi_unclassified(100);k__Fungi_unclassified(100);k__Fungi_unclassified(100);k__Fungi_unclassified(100);k__Fungi_unclassified(100);k__Fungi_unclassified(100); | - | - | - | - | - | - | - | - | Unassigned | - |
| Otu36479 | 1 | k__Fungi(100);k__Fungi_unclassified(100);k__Fungi_unclassified(100);k__Fungi_unclassified(100);k__Fungi_unclassified(100);k__Fungi_unclassified(100);k__Fungi_unclassified(100); | - | - | - | - | - | - | - | - | Unassigned | - |
| Otu36486 | 1 | k__Fungi(100);k__Fungi_unclassified(100);k__Fungi_unclassified(100);k__Fungi_unclassified(100);k__Fungi_unclassified(100);k__Fungi_unclassified(100);k__Fungi_unclassified(100); | - | - | - | - | - | - | - | - | Unassigned | - |
| Otu36487 | 1 | k__Fungi(100);k__Fungi_unclassified(100);k__Fungi_unclassified(100);k__Fungi_unclassified(100);k__Fungi_unclassified(100);k__Fungi_unclassified(100);k__Fungi_unclassified(100); | - | - | - | - | - | - | - | - | Unassigned | - |
| Otu36489 | 1 | k__Fungi(100);p__Ascomycota(100);p__Ascomycota_unclassified(100);p__Ascomycota_unclassified(100);p__Ascomycota_unclassified(100);p__Ascomycota_unclassified(100);p__Ascomycota_unclassified(100); | - | - | - | - | - | - | - | - | Unassigned | - |
| Otu36490 | 1 | k__Fungi(100);k__Fungi_unclassified(100);k__Fungi_unclassified(100);k__Fungi_unclassified(100);k__Fungi_unclassified(100);k__Fungi_unclassified(100);k__Fungi_unclassified(100); | - | - | - | - | - | - | - | - | Unassigned | - |
| Otu36492 | 1 | k__Fungi(100);k__Fungi_unclassified(100);k__Fungi_unclassified(100);k__Fungi_unclassified(100);k__Fungi_unclassified(100);k__Fungi_unclassified(100);k__Fungi_unclassified(100); | - | - | - | - | - | - | - | - | Unassigned | - |
| Otu36506 | 1 | k__Fungi(100);p__Ascomycota(100);p__Ascomycota_unclassified(100);p__Ascomycota_unclassified(100);p__Ascomycota_unclassified(100);p__Ascomycota_unclassified(100);p__Ascomycota_unclassified(100); | - | - | - | - | - | - | - | - | Unassigned | - |
| Otu36528 | 1 | k__Fungi(100);p__Ascomycota(100);p__Ascomycota_unclassified(100);p__Ascomycota_unclassified(100);p__Ascomycota_unclassified(100);p__Ascomycota_unclassified(100);p__Ascomycota_unclassified(100); | - | - | - | - | - | - | - | - | Unassigned | - |
| Otu36530 | 1 | k__Fungi(100);k__Fungi_unclassified(100);k__Fungi_unclassified(100);k__Fungi_unclassified(100);k__Fungi_unclassified(100);k__Fungi_unclassified(100);k__Fungi_unclassified(100); | - | - | - | - | - | - | - | - | Unassigned | - |
| Otu36562 | 1 | k__Fungi(100);k__Fungi_unclassified(100);k__Fungi_unclassified(100);k__Fungi_unclassified(100);k__Fungi_unclassified(100);k__Fungi_unclassified(100);k__Fungi_unclassified(100); | - | - | - | - | - | - | - | - | Unassigned | - |
| Otu36569 | 1 | k__Fungi(100);k__Fungi_unclassified(100);k__Fungi_unclassified(100);k__Fungi_unclassified(100);k__Fungi_unclassified(100);k__Fungi_unclassified(100);k__Fungi_unclassified(100); | - | - | - | - | - | - | - | - | Unassigned | - |
| Otu36570 | 1 | k__Fungi(100);k__Fungi_unclassified(100);k__Fungi_unclassified(100);k__Fungi_unclassified(100);k__Fungi_unclassified(100);k__Fungi_unclassified(100);k__Fungi_unclassified(100); | - | - | - | - | - | - | - | - | Unassigned | - |
| Otu36571 | 1 | k__Fungi(100);p__Ascomycota(100);p__Ascomycota_unclassified(100);p__Ascomycota_unclassified(100);p__Ascomycota_unclassified(100);p__Ascomycota_unclassified(100);p__Ascomycota_unclassified(100); | - | - | - | - | - | - | - | - | Unassigned | - |
| Otu36576 | 1 | k__Fungi(100);p__Ascomycota(100);p__Ascomycota_unclassified(100);p__Ascomycota_unclassified(100);p__Ascomycota_unclassified(100);p__Ascomycota_unclassified(100);p__Ascomycota_unclassified(100); | - | - | - | - | - | - | - | - | Unassigned | - |
| Otu36577 | 1 | k__Fungi(100);k__Fungi_unclassified(100);k__Fungi_unclassified(100);k__Fungi_unclassified(100);k__Fungi_unclassified(100);k__Fungi_unclassified(100);k__Fungi_unclassified(100); | - | - | - | - | - | - | - | - | Unassigned | - |
| Otu36578 | 1 | k__Fungi(100);k__Fungi_unclassified(100);k__Fungi_unclassified(100);k__Fungi_unclassified(100);k__Fungi_unclassified(100);k__Fungi_unclassified(100);k__Fungi_unclassified(100); | - | - | - | - | - | - | - | - | Unassigned | - |
| Otu36581 | 1 | k__Fungi(100);p__Ascomycota(100);p__Ascomycota_unclassified(100);p__Ascomycota_unclassified(100);p__Ascomycota_unclassified(100);p__Ascomycota_unclassified(100);p__Ascomycota_unclassified(100); | - | - | - | - | - | - | - | - | Unassigned | - |
| Otu36582 | 1 | k__Fungi(100);k__Fungi_unclassified(100);k__Fungi_unclassified(100);k__Fungi_unclassified(100);k__Fungi_unclassified(100);k__Fungi_unclassified(100);k__Fungi_unclassified(100); | - | - | - | - | - | - | - | - | Unassigned | - |
| Otu36585 | 1 | k__Fungi(100);k__Fungi_unclassified(100);k__Fungi_unclassified(100);k__Fungi_unclassified(100);k__Fungi_unclassified(100);k__Fungi_unclassified(100);k__Fungi_unclassified(100); | - | - | - | - | - | - | - | - | Unassigned | - |
| Otu36588 | 1 | k__Fungi(100);p__Ascomycota(100);p__Ascomycota_unclassified(100);p__Ascomycota_unclassified(100);p__Ascomycota_unclassified(100);p__Ascomycota_unclassified(100);p__Ascomycota_unclassified(100); | - | - | - | - | - | - | - | - | Unassigned | - |
| Otu36592 | 1 | k__Fungi(100);k__Fungi_unclassified(100);k__Fungi_unclassified(100);k__Fungi_unclassified(100);k__Fungi_unclassified(100);k__Fungi_unclassified(100);k__Fungi_unclassified(100); | - | - | - | - | - | - | - | - | Unassigned | - |
| Otu36595 | 1 | k__Fungi(100);p__Ascomycota(100);p__Ascomycota_unclassified(100);p__Ascomycota_unclassified(100);p__Ascomycota_unclassified(100);p__Ascomycota_unclassified(100);p__Ascomycota_unclassified(100); | - | - | - | - | - | - | - | - | Unassigned | - |
| Otu36596 | 1 | k__Fungi(100);k__Fungi_unclassified(100);k__Fungi_unclassified(100);k__Fungi_unclassified(100);k__Fungi_unclassified(100);k__Fungi_unclassified(100);k__Fungi_unclassified(100); | - | - | - | - | - | - | - | - | Unassigned | - |
| Otu36601 | 1 | k__Fungi(100);p__Ascomycota(100);p__Ascomycota_unclassified(100);p__Ascomycota_unclassified(100);p__Ascomycota_unclassified(100);p__Ascomycota_unclassified(100);p__Ascomycota_unclassified(100); | - | - | - | - | - | - | - | - | Unassigned | - |
| Otu36602 | 1 | k__Fungi(100);k__Fungi_unclassified(100);k__Fungi_unclassified(100);k__Fungi_unclassified(100);k__Fungi_unclassified(100);k__Fungi_unclassified(100);k__Fungi_unclassified(100); | - | - | - | - | - | - | - | - | Unassigned | - |
| Otu36603 | 1 | k__Fungi(100);p__Ascomycota(100);p__Ascomycota_unclassified(100);p__Ascomycota_unclassified(100);p__Ascomycota_unclassified(100);p__Ascomycota_unclassified(100);p__Ascomycota_unclassified(100); | - | - | - | - | - | - | - | - | Unassigned | - |
| Otu36604 | 1 | k__Fungi(100);k__Fungi_unclassified(100);k__Fungi_unclassified(100);k__Fungi_unclassified(100);k__Fungi_unclassified(100);k__Fungi_unclassified(100);k__Fungi_unclassified(100); | - | - | - | - | - | - | - | - | Unassigned | - |
| Otu36605 | 1 | k__Fungi(100);k__Fungi_unclassified(100);k__Fungi_unclassified(100);k__Fungi_unclassified(100);k__Fungi_unclassified(100);k__Fungi_unclassified(100);k__Fungi_unclassified(100); | - | - | - | - | - | - | - | - | Unassigned | - |
| Otu36606 | 1 | k__Fungi(100);k__Fungi_unclassified(100);k__Fungi_unclassified(100);k__Fungi_unclassified(100);k__Fungi_unclassified(100);k__Fungi_unclassified(100);k__Fungi_unclassified(100); | - | - | - | - | - | - | - | - | Unassigned | - |
| Otu36607 | 1 | k__Fungi(100);k__Fungi_unclassified(100);k__Fungi_unclassified(100);k__Fungi_unclassified(100);k__Fungi_unclassified(100);k__Fungi_unclassified(100);k__Fungi_unclassified(100); | - | - | - | - | - | - | - | - | Unassigned | - |
| Otu36608 | 1 | k__Fungi(100);k__Fungi_unclassified(100);k__Fungi_unclassified(100);k__Fungi_unclassified(100);k__Fungi_unclassified(100);k__Fungi_unclassified(100);k__Fungi_unclassified(100); | - | - | - | - | - | - | - | - | Unassigned | - |
| Otu36611 | 1 | k__Fungi(100);k__Fungi_unclassified(100);k__Fungi_unclassified(100);k__Fungi_unclassified(100);k__Fungi_unclassified(100);k__Fungi_unclassified(100);k__Fungi_unclassified(100); | - | - | - | - | - | - | - | - | Unassigned | - |
| Otu36612 | 1 | k__Fungi(100);p__Ascomycota(100);p__Ascomycota_unclassified(100);p__Ascomycota_unclassified(100);p__Ascomycota_unclassified(100);p__Ascomycota_unclassified(100);p__Ascomycota_unclassified(100); | - | - | - | - | - | - | - | - | Unassigned | - |
| Otu36613 | 1 | k__Fungi(100);k__Fungi_unclassified(100);k__Fungi_unclassified(100);k__Fungi_unclassified(100);k__Fungi_unclassified(100);k__Fungi_unclassified(100);k__Fungi_unclassified(100); | - | - | - | - | - | - | - | - | Unassigned | - |
| Otu36619 | 1 | k__Fungi(100);p__Ascomycota(100);p__Ascomycota_unclassified(100);p__Ascomycota_unclassified(100);p__Ascomycota_unclassified(100);p__Ascomycota_unclassified(100);p__Ascomycota_unclassified(100); | - | - | - | - | - | - | - | - | Unassigned | - |
| Otu36620 | 1 | k__Fungi(100);k__Fungi_unclassified(100);k__Fungi_unclassified(100);k__Fungi_unclassified(100);k__Fungi_unclassified(100);k__Fungi_unclassified(100);k__Fungi_unclassified(100); | - | - | - | - | - | - | - | - | Unassigned | - |
| Otu36621 | 1 | k__Fungi(100);k__Fungi_unclassified(100);k__Fungi_unclassified(100);k__Fungi_unclassified(100);k__Fungi_unclassified(100);k__Fungi_unclassified(100);k__Fungi_unclassified(100); | - | - | - | - | - | - | - | - | Unassigned | - |
| Otu36624 | 1 | k__Fungi(100);p__Ascomycota(100);p__Ascomycota_unclassified(100);p__Ascomycota_unclassified(100);p__Ascomycota_unclassified(100);p__Ascomycota_unclassified(100);p__Ascomycota_unclassified(100); | - | - | - | - | - | - | - | - | Unassigned | - |
| Otu36625 | 1 | k__Fungi(100);k__Fungi_unclassified(100);k__Fungi_unclassified(100);k__Fungi_unclassified(100);k__Fungi_unclassified(100);k__Fungi_unclassified(100);k__Fungi_unclassified(100); | - | - | - | - | - | - | - | - | Unassigned | - |
| Otu36626 | 1 | k__Fungi(100);p__Ascomycota(100);p__Ascomycota_unclassified(100);p__Ascomycota_unclassified(100);p__Ascomycota_unclassified(100);p__Ascomycota_unclassified(100);p__Ascomycota_unclassified(100); | - | - | - | - | - | - | - | - | Unassigned | - |
| Otu36627 | 1 | k__Fungi(100);k__Fungi_unclassified(100);k__Fungi_unclassified(100);k__Fungi_unclassified(100);k__Fungi_unclassified(100);k__Fungi_unclassified(100);k__Fungi_unclassified(100); | - | - | - | - | - | - | - | - | Unassigned | - |
| Otu36628 | 1 | k__Fungi(100);k__Fungi_unclassified(100);k__Fungi_unclassified(100);k__Fungi_unclassified(100);k__Fungi_unclassified(100);k__Fungi_unclassified(100);k__Fungi_unclassified(100); | - | - | - | - | - | - | - | - | Unassigned | - |
| Otu36629 | 1 | k__Fungi(100);k__Fungi_unclassified(100);k__Fungi_unclassified(100);k__Fungi_unclassified(100);k__Fungi_unclassified(100);k__Fungi_unclassified(100);k__Fungi_unclassified(100); | - | - | - | - | - | - | - | - | Unassigned | - |
| Otu36631 | 1 | k__Fungi(100);p__Ascomycota(100);p__Ascomycota_unclassified(100);p__Ascomycota_unclassified(100);p__Ascomycota_unclassified(100);p__Ascomycota_unclassified(100);p__Ascomycota_unclassified(100); | - | - | - | - | - | - | - | - | Unassigned | - |
| Otu36632 | 1 | k__Fungi(100);k__Fungi_unclassified(100);k__Fungi_unclassified(100);k__Fungi_unclassified(100);k__Fungi_unclassified(100);k__Fungi_unclassified(100);k__Fungi_unclassified(100); | - | - | - | - | - | - | - | - | Unassigned | - |
| Otu36633 | 1 | k__Fungi(100);k__Fungi_unclassified(100);k__Fungi_unclassified(100);k__Fungi_unclassified(100);k__Fungi_unclassified(100);k__Fungi_unclassified(100);k__Fungi_unclassified(100); | - | - | - | - | - | - | - | - | Unassigned | - |
| Otu36634 | 1 | k__Fungi(100);k__Fungi_unclassified(100);k__Fungi_unclassified(100);k__Fungi_unclassified(100);k__Fungi_unclassified(100);k__Fungi_unclassified(100);k__Fungi_unclassified(100); | - | - | - | - | - | - | - | - | Unassigned | - |
| Otu36637 | 1 | k__Fungi(100);k__Fungi_unclassified(100);k__Fungi_unclassified(100);k__Fungi_unclassified(100);k__Fungi_unclassified(100);k__Fungi_unclassified(100);k__Fungi_unclassified(100); | - | - | - | - | - | - | - | - | Unassigned | - |
| Otu36638 | 1 | k__Fungi(100);k__Fungi_unclassified(100);k__Fungi_unclassified(100);k__Fungi_unclassified(100);k__Fungi_unclassified(100);k__Fungi_unclassified(100);k__Fungi_unclassified(100); | - | - | - | - | - | - | - | - | Unassigned | - |
| Otu36640 | 1 | k__Fungi(100);k__Fungi_unclassified(100);k__Fungi_unclassified(100);k__Fungi_unclassified(100);k__Fungi_unclassified(100);k__Fungi_unclassified(100);k__Fungi_unclassified(100); | - | - | - | - | - | - | - | - | Unassigned | - |
| Otu36648 | 1 | k__Fungi(100);k__Fungi_unclassified(100);k__Fungi_unclassified(100);k__Fungi_unclassified(100);k__Fungi_unclassified(100);k__Fungi_unclassified(100);k__Fungi_unclassified(100); | - | - | - | - | - | - | - | - | Unassigned | - |
| Otu36651 | 1 | k__Fungi(100);k__Fungi_unclassified(100);k__Fungi_unclassified(100);k__Fungi_unclassified(100);k__Fungi_unclassified(100);k__Fungi_unclassified(100);k__Fungi_unclassified(100); | - | - | - | - | - | - | - | - | Unassigned | - |
| Otu36653 | 1 | k__Fungi(100);k__Fungi_unclassified(100);k__Fungi_unclassified(100);k__Fungi_unclassified(100);k__Fungi_unclassified(100);k__Fungi_unclassified(100);k__Fungi_unclassified(100); | - | - | - | - | - | - | - | - | Unassigned | - |
| Otu36655 | 1 | k__Fungi(100);k__Fungi_unclassified(100);k__Fungi_unclassified(100);k__Fungi_unclassified(100);k__Fungi_unclassified(100);k__Fungi_unclassified(100);k__Fungi_unclassified(100); | - | - | - | - | - | - | - | - | Unassigned | - |
| Otu36660 | 1 | k__Fungi(100);k__Fungi_unclassified(100);k__Fungi_unclassified(100);k__Fungi_unclassified(100);k__Fungi_unclassified(100);k__Fungi_unclassified(100);k__Fungi_unclassified(100); | - | - | - | - | - | - | - | - | Unassigned | - |
| Otu36662 | 1 | k__Fungi(100);k__Fungi_unclassified(100);k__Fungi_unclassified(100);k__Fungi_unclassified(100);k__Fungi_unclassified(100);k__Fungi_unclassified(100);k__Fungi_unclassified(100); | - | - | - | - | - | - | - | - | Unassigned | - |
| Otu36663 | 1 | k__Fungi(100);p__Ascomycota(100);p__Ascomycota_unclassified(100);p__Ascomycota_unclassified(100);p__Ascomycota_unclassified(100);p__Ascomycota_unclassified(100);p__Ascomycota_unclassified(100); | - | - | - | - | - | - | - | - | Unassigned | - |
| Otu36664 | 1 | k__Fungi(100);p__Ascomycota(100);p__Ascomycota_unclassified(100);p__Ascomycota_unclassified(100);p__Ascomycota_unclassified(100);p__Ascomycota_unclassified(100);p__Ascomycota_unclassified(100); | - | - | - | - | - | - | - | - | Unassigned | - |
| Otu36665 | 1 | k__Fungi(100);k__Fungi_unclassified(100);k__Fungi_unclassified(100);k__Fungi_unclassified(100);k__Fungi_unclassified(100);k__Fungi_unclassified(100);k__Fungi_unclassified(100); | - | - | - | - | - | - | - | - | Unassigned | - |
| Otu36669 | 1 | k__Fungi(100);k__Fungi_unclassified(100);k__Fungi_unclassified(100);k__Fungi_unclassified(100);k__Fungi_unclassified(100);k__Fungi_unclassified(100);k__Fungi_unclassified(100); | - | - | - | - | - | - | - | - | Unassigned | - |
| Otu36691 | 1 | k__Fungi(100);p__Ascomycota(100);p__Ascomycota_unclassified(100);p__Ascomycota_unclassified(100);p__Ascomycota_unclassified(100);p__Ascomycota_unclassified(100);p__Ascomycota_unclassified(100); | - | - | - | - | - | - | - | - | Unassigned | - |
| Otu36697 | 1 | k__Fungi(100);k__Fungi_unclassified(100);k__Fungi_unclassified(100);k__Fungi_unclassified(100);k__Fungi_unclassified(100);k__Fungi_unclassified(100);k__Fungi_unclassified(100); | - | - | - | - | - | - | - | - | Unassigned | - |
| Otu36701 | 1 | k__Fungi(100);p__Ascomycota(100);c__Leotiomycetes(100);c__Leotiomycetes_unclassified(100);c__Leotiomycetes_unclassified(100);c__Leotiomycetes_unclassified(100);c__Leotiomycetes_unclassified(100); | - | - | - | - | - | - | - | - | Unassigned | - |
| Otu36703 | 1 | k__Fungi(100);k__Fungi_unclassified(100);k__Fungi_unclassified(100);k__Fungi_unclassified(100);k__Fungi_unclassified(100);k__Fungi_unclassified(100);k__Fungi_unclassified(100); | - | - | - | - | - | - | - | - | Unassigned | - |
| Otu36704 | 1 | k__Fungi(100);k__Fungi_unclassified(100);k__Fungi_unclassified(100);k__Fungi_unclassified(100);k__Fungi_unclassified(100);k__Fungi_unclassified(100);k__Fungi_unclassified(100); | - | - | - | - | - | - | - | - | Unassigned | - |
| Otu36705 | 1 | k__Fungi(100);k__Fungi_unclassified(100);k__Fungi_unclassified(100);k__Fungi_unclassified(100);k__Fungi_unclassified(100);k__Fungi_unclassified(100);k__Fungi_unclassified(100); | - | - | - | - | - | - | - | - | Unassigned | - |
| Otu36709 | 1 | k__Fungi(100);k__Fungi_unclassified(100);k__Fungi_unclassified(100);k__Fungi_unclassified(100);k__Fungi_unclassified(100);k__Fungi_unclassified(100);k__Fungi_unclassified(100); | - | - | - | - | - | - | - | - | Unassigned | - |
| Otu36711 | 1 | k__Fungi(100);k__Fungi_unclassified(100);k__Fungi_unclassified(100);k__Fungi_unclassified(100);k__Fungi_unclassified(100);k__Fungi_unclassified(100);k__Fungi_unclassified(100); | - | - | - | - | - | - | - | - | Unassigned | - |
| Otu36716 | 1 | k__Fungi(100);k__Fungi_unclassified(100);k__Fungi_unclassified(100);k__Fungi_unclassified(100);k__Fungi_unclassified(100);k__Fungi_unclassified(100);k__Fungi_unclassified(100); | - | - | - | - | - | - | - | - | Unassigned | - |
| Otu36718 | 1 | k__Fungi(100);k__Fungi_unclassified(100);k__Fungi_unclassified(100);k__Fungi_unclassified(100);k__Fungi_unclassified(100);k__Fungi_unclassified(100);k__Fungi_unclassified(100); | - | - | - | - | - | - | - | - | Unassigned | - |
| Otu36813 | 1 | k__Fungi(100);k__Fungi_unclassified(100);k__Fungi_unclassified(100);k__Fungi_unclassified(100);k__Fungi_unclassified(100);k__Fungi_unclassified(100);k__Fungi_unclassified(100); | - | - | - | - | - | - | - | - | Unassigned | - |
| Otu36815 | 1 | k__Fungi(100);p__Ascomycota(100);p__Ascomycota_unclassified(100);p__Ascomycota_unclassified(100);p__Ascomycota_unclassified(100);p__Ascomycota_unclassified(100);p__Ascomycota_unclassified(100); | - | - | - | - | - | - | - | - | Unassigned | - |
| Otu36858 | 1 | k__Fungi(100);p__Ascomycota(100);p__Ascomycota_unclassified(100);p__Ascomycota_unclassified(100);p__Ascomycota_unclassified(100);p__Ascomycota_unclassified(100);p__Ascomycota_unclassified(100); | - | - | - | - | - | - | - | - | Unassigned | - |
| Otu36879 | 1 | k__Fungi(100);k__Fungi_unclassified(100);k__Fungi_unclassified(100);k__Fungi_unclassified(100);k__Fungi_unclassified(100);k__Fungi_unclassified(100);k__Fungi_unclassified(100); | - | - | - | - | - | - | - | - | Unassigned | - |
| Otu36955 | 1 | k__Fungi(100);k__Fungi_unclassified(100);k__Fungi_unclassified(100);k__Fungi_unclassified(100);k__Fungi_unclassified(100);k__Fungi_unclassified(100);k__Fungi_unclassified(100); | - | - | - | - | - | - | - | - | Unassigned | - |
| Otu36959 | 1 | k__Fungi(100);k__Fungi_unclassified(100);k__Fungi_unclassified(100);k__Fungi_unclassified(100);k__Fungi_unclassified(100);k__Fungi_unclassified(100);k__Fungi_unclassified(100); | - | - | - | - | - | - | - | - | Unassigned | - |
| Otu36962 | 1 | k__Fungi(100);k__Fungi_unclassified(100);k__Fungi_unclassified(100);k__Fungi_unclassified(100);k__Fungi_unclassified(100);k__Fungi_unclassified(100);k__Fungi_unclassified(100); | - | - | - | - | - | - | - | - | Unassigned | - |
| Otu36968 | 1 | k__Fungi(100);p__Ascomycota(100);p__Ascomycota_unclassified(100);p__Ascomycota_unclassified(100);p__Ascomycota_unclassified(100);p__Ascomycota_unclassified(100);p__Ascomycota_unclassified(100); | - | - | - | - | - | - | - | - | Unassigned | - |
| Otu36969 | 1 | k__Fungi(100);k__Fungi_unclassified(100);k__Fungi_unclassified(100);k__Fungi_unclassified(100);k__Fungi_unclassified(100);k__Fungi_unclassified(100);k__Fungi_unclassified(100); | - | - | - | - | - | - | - | - | Unassigned | - |
| Otu37000 | 1 | k__Fungi(100);k__Fungi_unclassified(100);k__Fungi_unclassified(100);k__Fungi_unclassified(100);k__Fungi_unclassified(100);k__Fungi_unclassified(100);k__Fungi_unclassified(100); | - | - | - | - | - | - | - | - | Unassigned | - |
| Otu37007 | 1 | k__Fungi(100);k__Fungi_unclassified(100);k__Fungi_unclassified(100);k__Fungi_unclassified(100);k__Fungi_unclassified(100);k__Fungi_unclassified(100);k__Fungi_unclassified(100); | - | - | - | - | - | - | - | - | Unassigned | - |
| Otu37032 | 1 | k__Fungi(100);k__Fungi_unclassified(100);k__Fungi_unclassified(100);k__Fungi_unclassified(100);k__Fungi_unclassified(100);k__Fungi_unclassified(100);k__Fungi_unclassified(100); | - | - | - | - | - | - | - | - | Unassigned | - |
| Otu37033 | 1 | k__Fungi(100);k__Fungi_unclassified(100);k__Fungi_unclassified(100);k__Fungi_unclassified(100);k__Fungi_unclassified(100);k__Fungi_unclassified(100);k__Fungi_unclassified(100); | - | - | - | - | - | - | - | - | Unassigned | - |
| Otu37034 | 1 | k__Fungi(100);k__Fungi_unclassified(100);k__Fungi_unclassified(100);k__Fungi_unclassified(100);k__Fungi_unclassified(100);k__Fungi_unclassified(100);k__Fungi_unclassified(100); | - | - | - | - | - | - | - | - | Unassigned | - |
| Otu37054 | 1 | k__Fungi(100);k__Fungi_unclassified(100);k__Fungi_unclassified(100);k__Fungi_unclassified(100);k__Fungi_unclassified(100);k__Fungi_unclassified(100);k__Fungi_unclassified(100); | - | - | - | - | - | - | - | - | Unassigned | - |
| Otu37055 | 1 | k__Fungi(100);p__Ascomycota(100);p__Ascomycota_unclassified(100);p__Ascomycota_unclassified(100);p__Ascomycota_unclassified(100);p__Ascomycota_unclassified(100);p__Ascomycota_unclassified(100); | - | - | - | - | - | - | - | - | Unassigned | - |
| Otu37056 | 1 | k__Fungi(100);p__Ascomycota(100);p__Ascomycota_unclassified(100);p__Ascomycota_unclassified(100);p__Ascomycota_unclassified(100);p__Ascomycota_unclassified(100);p__Ascomycota_unclassified(100); | - | - | - | - | - | - | - | - | Unassigned | - |
| Otu37057 | 1 | k__Fungi(100);k__Fungi_unclassified(100);k__Fungi_unclassified(100);k__Fungi_unclassified(100);k__Fungi_unclassified(100);k__Fungi_unclassified(100);k__Fungi_unclassified(100); | - | - | - | - | - | - | - | - | Unassigned | - |
| Otu37064 | 1 | k__Fungi(100);k__Fungi_unclassified(100);k__Fungi_unclassified(100);k__Fungi_unclassified(100);k__Fungi_unclassified(100);k__Fungi_unclassified(100);k__Fungi_unclassified(100); | - | - | - | - | - | - | - | - | Unassigned | - |
| Otu37065 | 1 | k__Fungi(100);k__Fungi_unclassified(100);k__Fungi_unclassified(100);k__Fungi_unclassified(100);k__Fungi_unclassified(100);k__Fungi_unclassified(100);k__Fungi_unclassified(100); | - | - | - | - | - | - | - | - | Unassigned | - |
| Otu37066 | 1 | k__Fungi(100);k__Fungi_unclassified(100);k__Fungi_unclassified(100);k__Fungi_unclassified(100);k__Fungi_unclassified(100);k__Fungi_unclassified(100);k__Fungi_unclassified(100); | - | - | - | - | - | - | - | - | Unassigned | - |
| Otu37067 | 1 | k__Fungi(100);k__Fungi_unclassified(100);k__Fungi_unclassified(100);k__Fungi_unclassified(100);k__Fungi_unclassified(100);k__Fungi_unclassified(100);k__Fungi_unclassified(100); | - | - | - | - | - | - | - | - | Unassigned | - |
| Otu37068 | 1 | k__Fungi(100);k__Fungi_unclassified(100);k__Fungi_unclassified(100);k__Fungi_unclassified(100);k__Fungi_unclassified(100);k__Fungi_unclassified(100);k__Fungi_unclassified(100); | - | - | - | - | - | - | - | - | Unassigned | - |
| Otu37069 | 1 | k__Fungi(100);k__Fungi_unclassified(100);k__Fungi_unclassified(100);k__Fungi_unclassified(100);k__Fungi_unclassified(100);k__Fungi_unclassified(100);k__Fungi_unclassified(100); | - | - | - | - | - | - | - | - | Unassigned | - |
| Otu37070 | 1 | k__Fungi(100);k__Fungi_unclassified(100);k__Fungi_unclassified(100);k__Fungi_unclassified(100);k__Fungi_unclassified(100);k__Fungi_unclassified(100);k__Fungi_unclassified(100); | - | - | - | - | - | - | - | - | Unassigned | - |
| Otu37071 | 1 | k__Fungi(100);p__Ascomycota(100);p__Ascomycota_unclassified(100);p__Ascomycota_unclassified(100);p__Ascomycota_unclassified(100);p__Ascomycota_unclassified(100);p__Ascomycota_unclassified(100); | - | - | - | - | - | - | - | - | Unassigned | - |
| Otu37072 | 1 | k__Fungi(100);k__Fungi_unclassified(100);k__Fungi_unclassified(100);k__Fungi_unclassified(100);k__Fungi_unclassified(100);k__Fungi_unclassified(100);k__Fungi_unclassified(100); | - | - | - | - | - | - | - | - | Unassigned | - |
| Otu37073 | 1 | k__Fungi(100);k__Fungi_unclassified(100);k__Fungi_unclassified(100);k__Fungi_unclassified(100);k__Fungi_unclassified(100);k__Fungi_unclassified(100);k__Fungi_unclassified(100); | - | - | - | - | - | - | - | - | Unassigned | - |
| Otu37074 | 1 | k__Fungi(100);k__Fungi_unclassified(100);k__Fungi_unclassified(100);k__Fungi_unclassified(100);k__Fungi_unclassified(100);k__Fungi_unclassified(100);k__Fungi_unclassified(100); | - | - | - | - | - | - | - | - | Unassigned | - |
| Otu37075 | 1 | k__Fungi(100);k__Fungi_unclassified(100);k__Fungi_unclassified(100);k__Fungi_unclassified(100);k__Fungi_unclassified(100);k__Fungi_unclassified(100);k__Fungi_unclassified(100); | - | - | - | - | - | - | - | - | Unassigned | - |
| Otu37076 | 1 | k__Fungi(100);k__Fungi_unclassified(100);k__Fungi_unclassified(100);k__Fungi_unclassified(100);k__Fungi_unclassified(100);k__Fungi_unclassified(100);k__Fungi_unclassified(100); | - | - | - | - | - | - | - | - | Unassigned | - |
| Otu37078 | 1 | k__Fungi(100);k__Fungi_unclassified(100);k__Fungi_unclassified(100);k__Fungi_unclassified(100);k__Fungi_unclassified(100);k__Fungi_unclassified(100);k__Fungi_unclassified(100); | - | - | - | - | - | - | - | - | Unassigned | - |
| Otu37079 | 1 | k__Fungi(100);k__Fungi_unclassified(100);k__Fungi_unclassified(100);k__Fungi_unclassified(100);k__Fungi_unclassified(100);k__Fungi_unclassified(100);k__Fungi_unclassified(100); | - | - | - | - | - | - | - | - | Unassigned | - |
| Otu37108 | 1 | k__Fungi(100);k__Fungi_unclassified(100);k__Fungi_unclassified(100);k__Fungi_unclassified(100);k__Fungi_unclassified(100);k__Fungi_unclassified(100);k__Fungi_unclassified(100); | - | - | - | - | - | - | - | - | Unassigned | - |
| Otu37113 | 1 | k__Fungi(100);k__Fungi_unclassified(100);k__Fungi_unclassified(100);k__Fungi_unclassified(100);k__Fungi_unclassified(100);k__Fungi_unclassified(100);k__Fungi_unclassified(100); | - | - | - | - | - | - | - | - | Unassigned | - |
| Otu37114 | 1 | k__Fungi(100);p__Ascomycota(100);p__Ascomycota_unclassified(100);p__Ascomycota_unclassified(100);p__Ascomycota_unclassified(100);p__Ascomycota_unclassified(100);p__Ascomycota_unclassified(100); | - | - | - | - | - | - | - | - | Unassigned | - |
| Otu37116 | 1 | k__Fungi(100);k__Fungi_unclassified(100);k__Fungi_unclassified(100);k__Fungi_unclassified(100);k__Fungi_unclassified(100);k__Fungi_unclassified(100);k__Fungi_unclassified(100); | - | - | - | - | - | - | - | - | Unassigned | - |
| Otu37125 | 1 | k__Fungi(100);k__Fungi_unclassified(100);k__Fungi_unclassified(100);k__Fungi_unclassified(100);k__Fungi_unclassified(100);k__Fungi_unclassified(100);k__Fungi_unclassified(100); | - | - | - | - | - | - | - | - | Unassigned | - |
| Otu37126 | 1 | k__Fungi(100);k__Fungi_unclassified(100);k__Fungi_unclassified(100);k__Fungi_unclassified(100);k__Fungi_unclassified(100);k__Fungi_unclassified(100);k__Fungi_unclassified(100); | - | - | - | - | - | - | - | - | Unassigned | - |
| Otu37129 | 1 | k__Fungi(100);k__Fungi_unclassified(100);k__Fungi_unclassified(100);k__Fungi_unclassified(100);k__Fungi_unclassified(100);k__Fungi_unclassified(100);k__Fungi_unclassified(100); | - | - | - | - | - | - | - | - | Unassigned | - |
| Otu37132 | 1 | k__Fungi(100);p__Ascomycota(100);p__Ascomycota_unclassified(100);p__Ascomycota_unclassified(100);p__Ascomycota_unclassified(100);p__Ascomycota_unclassified(100);p__Ascomycota_unclassified(100); | - | - | - | - | - | - | - | - | Unassigned | - |
| Otu37170 | 1 | k__Fungi(100);k__Fungi_unclassified(100);k__Fungi_unclassified(100);k__Fungi_unclassified(100);k__Fungi_unclassified(100);k__Fungi_unclassified(100);k__Fungi_unclassified(100); | - | - | - | - | - | - | - | - | Unassigned | - |
| Otu37199 | 1 | k__Fungi(100);k__Fungi_unclassified(100);k__Fungi_unclassified(100);k__Fungi_unclassified(100);k__Fungi_unclassified(100);k__Fungi_unclassified(100);k__Fungi_unclassified(100); | - | - | - | - | - | - | - | - | Unassigned | - |
| Otu37262 | 1 | k__Fungi(100);k__Fungi_unclassified(100);k__Fungi_unclassified(100);k__Fungi_unclassified(100);k__Fungi_unclassified(100);k__Fungi_unclassified(100);k__Fungi_unclassified(100); | - | - | - | - | - | - | - | - | Unassigned | - |
| Otu37264 | 1 | k__Fungi(100);p__Ascomycota(100);p__Ascomycota_unclassified(100);p__Ascomycota_unclassified(100);p__Ascomycota_unclassified(100);p__Ascomycota_unclassified(100);p__Ascomycota_unclassified(100); | - | - | - | - | - | - | - | - | Unassigned | - |
| Otu37273 | 1 | k__Fungi(100);p__Ascomycota(100);p__Ascomycota_unclassified(100);p__Ascomycota_unclassified(100);p__Ascomycota_unclassified(100);p__Ascomycota_unclassified(100);p__Ascomycota_unclassified(100); | - | - | - | - | - | - | - | - | Unassigned | - |
| Otu37278 | 1 | k__Fungi(100);k__Fungi_unclassified(100);k__Fungi_unclassified(100);k__Fungi_unclassified(100);k__Fungi_unclassified(100);k__Fungi_unclassified(100);k__Fungi_unclassified(100); | - | - | - | - | - | - | - | - | Unassigned | - |
| Otu37279 | 1 | k__Fungi(100);p__Ascomycota(100);p__Ascomycota_unclassified(100);p__Ascomycota_unclassified(100);p__Ascomycota_unclassified(100);p__Ascomycota_unclassified(100);p__Ascomycota_unclassified(100); | - | - | - | - | - | - | - | - | Unassigned | - |
| Otu37280 | 1 | k__Fungi(100);p__Ascomycota(100);p__Ascomycota_unclassified(100);p__Ascomycota_unclassified(100);p__Ascomycota_unclassified(100);p__Ascomycota_unclassified(100);p__Ascomycota_unclassified(100); | - | - | - | - | - | - | - | - | Unassigned | - |
| Otu37285 | 1 | k__Fungi(100);p__Ascomycota(100);p__Ascomycota_unclassified(100);p__Ascomycota_unclassified(100);p__Ascomycota_unclassified(100);p__Ascomycota_unclassified(100);p__Ascomycota_unclassified(100); | - | - | - | - | - | - | - | - | Unassigned | - |
| Otu37292 | 1 | k__Fungi(100);p__Ascomycota(100);p__Ascomycota_unclassified(100);p__Ascomycota_unclassified(100);p__Ascomycota_unclassified(100);p__Ascomycota_unclassified(100);p__Ascomycota_unclassified(100); | - | - | - | - | - | - | - | - | Unassigned | - |
| Otu37293 | 1 | k__Fungi(100);k__Fungi_unclassified(100);k__Fungi_unclassified(100);k__Fungi_unclassified(100);k__Fungi_unclassified(100);k__Fungi_unclassified(100);k__Fungi_unclassified(100); | - | - | - | - | - | - | - | - | Unassigned | - |
| Otu37300 | 1 | k__Fungi(100);k__Fungi_unclassified(100);k__Fungi_unclassified(100);k__Fungi_unclassified(100);k__Fungi_unclassified(100);k__Fungi_unclassified(100);k__Fungi_unclassified(100); | - | - | - | - | - | - | - | - | Unassigned | - |
| Otu37301 | 1 | k__Fungi(100);k__Fungi_unclassified(100);k__Fungi_unclassified(100);k__Fungi_unclassified(100);k__Fungi_unclassified(100);k__Fungi_unclassified(100);k__Fungi_unclassified(100); | - | - | - | - | - | - | - | - | Unassigned | - |
| Otu37302 | 1 | k__Fungi(100);k__Fungi_unclassified(100);k__Fungi_unclassified(100);k__Fungi_unclassified(100);k__Fungi_unclassified(100);k__Fungi_unclassified(100);k__Fungi_unclassified(100); | - | - | - | - | - | - | - | - | Unassigned | - |
| Otu37303 | 1 | k__Fungi(100);k__Fungi_unclassified(100);k__Fungi_unclassified(100);k__Fungi_unclassified(100);k__Fungi_unclassified(100);k__Fungi_unclassified(100);k__Fungi_unclassified(100); | - | - | - | - | - | - | - | - | Unassigned | - |
| Otu37311 | 1 | k__Fungi(100);k__Fungi_unclassified(100);k__Fungi_unclassified(100);k__Fungi_unclassified(100);k__Fungi_unclassified(100);k__Fungi_unclassified(100);k__Fungi_unclassified(100); | - | - | - | - | - | - | - | - | Unassigned | - |
| Otu37313 | 1 | k__Fungi(100);p__Ascomycota(100);p__Ascomycota_unclassified(100);p__Ascomycota_unclassified(100);p__Ascomycota_unclassified(100);p__Ascomycota_unclassified(100);p__Ascomycota_unclassified(100); | - | - | - | - | - | - | - | - | Unassigned | - |
| Otu37337 | 1 | k__Fungi(100);p__Ascomycota(100);p__Ascomycota_unclassified(100);p__Ascomycota_unclassified(100);p__Ascomycota_unclassified(100);p__Ascomycota_unclassified(100);p__Ascomycota_unclassified(100); | - | - | - | - | - | - | - | - | Unassigned | - |
| Otu37338 | 1 | k__Fungi(100);p__Ascomycota(100);p__Ascomycota_unclassified(100);p__Ascomycota_unclassified(100);p__Ascomycota_unclassified(100);p__Ascomycota_unclassified(100);p__Ascomycota_unclassified(100); | - | - | - | - | - | - | - | - | Unassigned | - |
| Otu37339 | 1 | k__Fungi(100);k__Fungi_unclassified(100);k__Fungi_unclassified(100);k__Fungi_unclassified(100);k__Fungi_unclassified(100);k__Fungi_unclassified(100);k__Fungi_unclassified(100); | - | - | - | - | - | - | - | - | Unassigned | - |
| Otu37340 | 1 | k__Fungi(100);k__Fungi_unclassified(100);k__Fungi_unclassified(100);k__Fungi_unclassified(100);k__Fungi_unclassified(100);k__Fungi_unclassified(100);k__Fungi_unclassified(100); | - | - | - | - | - | - | - | - | Unassigned | - |
| Otu37341 | 1 | k__Fungi(100);p__Ascomycota(100);p__Ascomycota_unclassified(100);p__Ascomycota_unclassified(100);p__Ascomycota_unclassified(100);p__Ascomycota_unclassified(100);p__Ascomycota_unclassified(100); | - | - | - | - | - | - | - | - | Unassigned | - |
| Otu37480 | 1 | k__Fungi(100);k__Fungi_unclassified(100);k__Fungi_unclassified(100);k__Fungi_unclassified(100);k__Fungi_unclassified(100);k__Fungi_unclassified(100);k__Fungi_unclassified(100); | - | - | - | - | - | - | - | - | Unassigned | - |
| Otu37516 | 1 | k__Fungi(100);k__Fungi_unclassified(100);k__Fungi_unclassified(100);k__Fungi_unclassified(100);k__Fungi_unclassified(100);k__Fungi_unclassified(100);k__Fungi_unclassified(100); | - | - | - | - | - | - | - | - | Unassigned | - |
| Otu37517 | 1 | k__Fungi(100);k__Fungi_unclassified(100);k__Fungi_unclassified(100);k__Fungi_unclassified(100);k__Fungi_unclassified(100);k__Fungi_unclassified(100);k__Fungi_unclassified(100); | - | - | - | - | - | - | - | - | Unassigned | - |
| Otu37556 | 1 | k__Fungi(100);k__Fungi_unclassified(100);k__Fungi_unclassified(100);k__Fungi_unclassified(100);k__Fungi_unclassified(100);k__Fungi_unclassified(100);k__Fungi_unclassified(100); | - | - | - | - | - | - | - | - | Unassigned | - |
| Otu37568 | 1 | k__Fungi(100);p__Ascomycota(100);p__Ascomycota_unclassified(100);p__Ascomycota_unclassified(100);p__Ascomycota_unclassified(100);p__Ascomycota_unclassified(100);p__Ascomycota_unclassified(100); | - | - | - | - | - | - | - | - | Unassigned | - |
| Otu37569 | 1 | k__Fungi(100);k__Fungi_unclassified(100);k__Fungi_unclassified(100);k__Fungi_unclassified(100);k__Fungi_unclassified(100);k__Fungi_unclassified(100);k__Fungi_unclassified(100); | - | - | - | - | - | - | - | - | Unassigned | - |
| Otu37570 | 1 | k__Fungi(100);k__Fungi_unclassified(100);k__Fungi_unclassified(100);k__Fungi_unclassified(100);k__Fungi_unclassified(100);k__Fungi_unclassified(100);k__Fungi_unclassified(100); | - | - | - | - | - | - | - | - | Unassigned | - |
| Otu37583 | 1 | k__Fungi(100);p__Ascomycota(100);p__Ascomycota_unclassified(100);p__Ascomycota_unclassified(100);p__Ascomycota_unclassified(100);p__Ascomycota_unclassified(100);p__Ascomycota_unclassified(100); | - | - | - | - | - | - | - | - | Unassigned | - |
| Otu37628 | 1 | k__Fungi(100);p__Ascomycota(100);p__Ascomycota_unclassified(100);p__Ascomycota_unclassified(100);p__Ascomycota_unclassified(100);p__Ascomycota_unclassified(100);p__Ascomycota_unclassified(100); | - | - | - | - | - | - | - | - | Unassigned | - |
| Otu37629 | 1 | k__Fungi(100);k__Fungi_unclassified(100);k__Fungi_unclassified(100);k__Fungi_unclassified(100);k__Fungi_unclassified(100);k__Fungi_unclassified(100);k__Fungi_unclassified(100); | - | - | - | - | - | - | - | - | Unassigned | - |
| Otu37632 | 1 | k__Fungi(100);p__Ascomycota(100);p__Ascomycota_unclassified(100);p__Ascomycota_unclassified(100);p__Ascomycota_unclassified(100);p__Ascomycota_unclassified(100);p__Ascomycota_unclassified(100); | - | - | - | - | - | - | - | - | Unassigned | - |
| Otu37635 | 1 | k__Fungi(100);p__Ascomycota(100);p__Ascomycota_unclassified(100);p__Ascomycota_unclassified(100);p__Ascomycota_unclassified(100);p__Ascomycota_unclassified(100);p__Ascomycota_unclassified(100); | - | - | - | - | - | - | - | - | Unassigned | - |
| Otu37638 | 1 | k__Fungi(100);p__Ascomycota(100);p__Ascomycota_unclassified(100);p__Ascomycota_unclassified(100);p__Ascomycota_unclassified(100);p__Ascomycota_unclassified(100);p__Ascomycota_unclassified(100); | - | - | - | - | - | - | - | - | Unassigned | - |
| Otu37655 | 1 | k__Fungi(100);k__Fungi_unclassified(100);k__Fungi_unclassified(100);k__Fungi_unclassified(100);k__Fungi_unclassified(100);k__Fungi_unclassified(100);k__Fungi_unclassified(100); | - | - | - | - | - | - | - | - | Unassigned | - |
| Otu37660 | 1 | k__Fungi(100);k__Fungi_unclassified(100);k__Fungi_unclassified(100);k__Fungi_unclassified(100);k__Fungi_unclassified(100);k__Fungi_unclassified(100);k__Fungi_unclassified(100); | - | - | - | - | - | - | - | - | Unassigned | - |
| Otu37670 | 1 | k__Fungi(100);p__Ascomycota(100);p__Ascomycota_unclassified(100);p__Ascomycota_unclassified(100);p__Ascomycota_unclassified(100);p__Ascomycota_unclassified(100);p__Ascomycota_unclassified(100); | - | - | - | - | - | - | - | - | Unassigned | - |
| Otu37672 | 1 | k__Fungi(100);p__Ascomycota(100);c__Leotiomycetes(100);o__Helotiales(100);o__Helotiales_unclassified(100);o__Helotiales_unclassified(100);o__Helotiales_unclassified(100); | - | - | - | - | - | - | - | - | Unassigned | - |
| Otu37673 | 1 | k__Fungi(100);p__Ascomycota(100);c__Leotiomycetes(100);o__Helotiales(100);o__Helotiales_unclassified(100);o__Helotiales_unclassified(100);o__Helotiales_unclassified(100); | - | - | - | - | - | - | - | - | Unassigned | - |
| Otu37674 | 1 | k__Fungi(100);p__Ascomycota(100);c__Dothideomycetes(100);o__Capnodiales(100);f__Davidiellaceae(100);f__Davidiellaceae_unclassified(100);f__Davidiellaceae_unclassified(100); | - | - | - | - | - | - | - | - | Unassigned | - |
| Otu37675 | 1 | k__Fungi(100);p__Ascomycota(100);p__Ascomycota_unclassified(100);p__Ascomycota_unclassified(100);p__Ascomycota_unclassified(100);p__Ascomycota_unclassified(100);p__Ascomycota_unclassified(100); | - | - | - | - | - | - | - | - | Unassigned | - |
| Otu37691 | 1 | k__Fungi(100);p__Ascomycota(100);p__Ascomycota_unclassified(100);p__Ascomycota_unclassified(100);p__Ascomycota_unclassified(100);p__Ascomycota_unclassified(100);p__Ascomycota_unclassified(100); | - | - | - | - | - | - | - | - | Unassigned | - |
| Otu37692 | 1 | k__Fungi(100);p__Ascomycota(100);p__Ascomycota_unclassified(100);p__Ascomycota_unclassified(100);p__Ascomycota_unclassified(100);p__Ascomycota_unclassified(100);p__Ascomycota_unclassified(100); | - | - | - | - | - | - | - | - | Unassigned | - |
| Otu37693 | 1 | k__Fungi(100);k__Fungi_unclassified(100);k__Fungi_unclassified(100);k__Fungi_unclassified(100);k__Fungi_unclassified(100);k__Fungi_unclassified(100);k__Fungi_unclassified(100); | - | - | - | - | - | - | - | - | Unassigned | - |
| Otu37694 | 1 | k__Fungi(100);k__Fungi_unclassified(100);k__Fungi_unclassified(100);k__Fungi_unclassified(100);k__Fungi_unclassified(100);k__Fungi_unclassified(100);k__Fungi_unclassified(100); | - | - | - | - | - | - | - | - | Unassigned | - |
| Otu37695 | 1 | k__Fungi(100);p__Ascomycota(100);p__Ascomycota_unclassified(100);p__Ascomycota_unclassified(100);p__Ascomycota_unclassified(100);p__Ascomycota_unclassified(100);p__Ascomycota_unclassified(100); | - | - | - | - | - | - | - | - | Unassigned | - |
| Otu37697 | 1 | k__Fungi(100);p__Ascomycota(100);p__Ascomycota_unclassified(100);p__Ascomycota_unclassified(100);p__Ascomycota_unclassified(100);p__Ascomycota_unclassified(100);p__Ascomycota_unclassified(100); | - | - | - | - | - | - | - | - | Unassigned | - |
| Otu37702 | 1 | k__Fungi(100);p__Ascomycota(100);p__Ascomycota_unclassified(100);p__Ascomycota_unclassified(100);p__Ascomycota_unclassified(100);p__Ascomycota_unclassified(100);p__Ascomycota_unclassified(100); | - | - | - | - | - | - | - | - | Unassigned | - |
| Otu37720 | 1 | k__Fungi(100);p__Ascomycota(100);p__Ascomycota_unclassified(100);p__Ascomycota_unclassified(100);p__Ascomycota_unclassified(100);p__Ascomycota_unclassified(100);p__Ascomycota_unclassified(100); | - | - | - | - | - | - | - | - | Unassigned | - |
| Otu37731 | 1 | k__Fungi(100);k__Fungi_unclassified(100);k__Fungi_unclassified(100);k__Fungi_unclassified(100);k__Fungi_unclassified(100);k__Fungi_unclassified(100);k__Fungi_unclassified(100); | - | - | - | - | - | - | - | - | Unassigned | - |
| Otu37802 | 1 | k__Fungi(100);p__Ascomycota(100);p__Ascomycota_unclassified(100);p__Ascomycota_unclassified(100);p__Ascomycota_unclassified(100);p__Ascomycota_unclassified(100);p__Ascomycota_unclassified(100); | - | - | - | - | - | - | - | - | Unassigned | - |
| Otu37803 | 1 | k__Fungi(100);p__Ascomycota(100);p__Ascomycota_unclassified(100);p__Ascomycota_unclassified(100);p__Ascomycota_unclassified(100);p__Ascomycota_unclassified(100);p__Ascomycota_unclassified(100); | - | - | - | - | - | - | - | - | Unassigned | - |
| Otu37805 | 1 | k__Fungi(100);p__Ascomycota(100);p__Ascomycota_unclassified(100);p__Ascomycota_unclassified(100);p__Ascomycota_unclassified(100);p__Ascomycota_unclassified(100);p__Ascomycota_unclassified(100); | - | - | - | - | - | - | - | - | Unassigned | - |
| Otu37810 | 1 | k__Fungi(100);k__Fungi_unclassified(100);k__Fungi_unclassified(100);k__Fungi_unclassified(100);k__Fungi_unclassified(100);k__Fungi_unclassified(100);k__Fungi_unclassified(100); | - | - | - | - | - | - | - | - | Unassigned | - |
| Otu37811 | 1 | k__Fungi(100);k__Fungi_unclassified(100);k__Fungi_unclassified(100);k__Fungi_unclassified(100);k__Fungi_unclassified(100);k__Fungi_unclassified(100);k__Fungi_unclassified(100); | - | - | - | - | - | - | - | - | Unassigned | - |
| Otu37813 | 1 | k__Fungi(100);p__Ascomycota(100);p__Ascomycota_unclassified(100);p__Ascomycota_unclassified(100);p__Ascomycota_unclassified(100);p__Ascomycota_unclassified(100);p__Ascomycota_unclassified(100); | - | - | - | - | - | - | - | - | Unassigned | - |
| Otu37814 | 1 | k__Fungi(100);p__Ascomycota(100);p__Ascomycota_unclassified(100);p__Ascomycota_unclassified(100);p__Ascomycota_unclassified(100);p__Ascomycota_unclassified(100);p__Ascomycota_unclassified(100); | - | - | - | - | - | - | - | - | Unassigned | - |
| Otu37815 | 1 | k__Fungi(100);p__Ascomycota(100);p__Ascomycota_unclassified(100);p__Ascomycota_unclassified(100);p__Ascomycota_unclassified(100);p__Ascomycota_unclassified(100);p__Ascomycota_unclassified(100); | - | - | - | - | - | - | - | - | Unassigned | - |
| Otu37816 | 1 | k__Fungi(100);k__Fungi_unclassified(100);k__Fungi_unclassified(100);k__Fungi_unclassified(100);k__Fungi_unclassified(100);k__Fungi_unclassified(100);k__Fungi_unclassified(100); | - | - | - | - | - | - | - | - | Unassigned | - |
| Otu37817 | 1 | k__Fungi(100);p__Ascomycota(100);p__Ascomycota_unclassified(100);p__Ascomycota_unclassified(100);p__Ascomycota_unclassified(100);p__Ascomycota_unclassified(100);p__Ascomycota_unclassified(100); | - | - | - | - | - | - | - | - | Unassigned | - |
| Otu37818 | 1 | k__Fungi(100);p__Ascomycota(100);p__Ascomycota_unclassified(100);p__Ascomycota_unclassified(100);p__Ascomycota_unclassified(100);p__Ascomycota_unclassified(100);p__Ascomycota_unclassified(100); | - | - | - | - | - | - | - | - | Unassigned | - |
| Otu37821 | 1 | k__Fungi(100);p__Ascomycota(100);p__Ascomycota_unclassified(100);p__Ascomycota_unclassified(100);p__Ascomycota_unclassified(100);p__Ascomycota_unclassified(100);p__Ascomycota_unclassified(100); | - | - | - | - | - | - | - | - | Unassigned | - |
| Otu37822 | 1 | k__Fungi(100);p__Ascomycota(100);p__Ascomycota_unclassified(100);p__Ascomycota_unclassified(100);p__Ascomycota_unclassified(100);p__Ascomycota_unclassified(100);p__Ascomycota_unclassified(100); | - | - | - | - | - | - | - | - | Unassigned | - |
| Otu37828 | 1 | k__Fungi(100);p__Ascomycota(100);p__Ascomycota_unclassified(100);p__Ascomycota_unclassified(100);p__Ascomycota_unclassified(100);p__Ascomycota_unclassified(100);p__Ascomycota_unclassified(100); | - | - | - | - | - | - | - | - | Unassigned | - |
| Otu37832 | 1 | k__Fungi(100);k__Fungi_unclassified(100);k__Fungi_unclassified(100);k__Fungi_unclassified(100);k__Fungi_unclassified(100);k__Fungi_unclassified(100);k__Fungi_unclassified(100); | - | - | - | - | - | - | - | - | Unassigned | - |
| Otu37833 | 1 | k__Fungi(100);k__Fungi_unclassified(100);k__Fungi_unclassified(100);k__Fungi_unclassified(100);k__Fungi_unclassified(100);k__Fungi_unclassified(100);k__Fungi_unclassified(100); | - | - | - | - | - | - | - | - | Unassigned | - |
| Otu37835 | 1 | k__Fungi(100);p__Ascomycota(100);p__Ascomycota_unclassified(100);p__Ascomycota_unclassified(100);p__Ascomycota_unclassified(100);p__Ascomycota_unclassified(100);p__Ascomycota_unclassified(100); | - | - | - | - | - | - | - | - | Unassigned | - |
| Otu37838 | 1 | k__Fungi(100);p__Ascomycota(100);p__Ascomycota_unclassified(100);p__Ascomycota_unclassified(100);p__Ascomycota_unclassified(100);p__Ascomycota_unclassified(100);p__Ascomycota_unclassified(100); | - | - | - | - | - | - | - | - | Unassigned | - |
| Otu37839 | 1 | k__Fungi(100);p__Ascomycota(100);p__Ascomycota_unclassified(100);p__Ascomycota_unclassified(100);p__Ascomycota_unclassified(100);p__Ascomycota_unclassified(100);p__Ascomycota_unclassified(100); | - | - | - | - | - | - | - | - | Unassigned | - |
| Otu37840 | 1 | k__Fungi(100);p__Ascomycota(100);p__Ascomycota_unclassified(100);p__Ascomycota_unclassified(100);p__Ascomycota_unclassified(100);p__Ascomycota_unclassified(100);p__Ascomycota_unclassified(100); | - | - | - | - | - | - | - | - | Unassigned | - |
| Otu37841 | 1 | k__Fungi(100);k__Fungi_unclassified(100);k__Fungi_unclassified(100);k__Fungi_unclassified(100);k__Fungi_unclassified(100);k__Fungi_unclassified(100);k__Fungi_unclassified(100); | - | - | - | - | - | - | - | - | Unassigned | - |
| Otu37843 | 1 | k__Fungi(100);p__Ascomycota(100);p__Ascomycota_unclassified(100);p__Ascomycota_unclassified(100);p__Ascomycota_unclassified(100);p__Ascomycota_unclassified(100);p__Ascomycota_unclassified(100); | - | - | - | - | - | - | - | - | Unassigned | - |
| Otu37855 | 1 | k__Fungi(100);p__Ascomycota(100);p__Ascomycota_unclassified(100);p__Ascomycota_unclassified(100);p__Ascomycota_unclassified(100);p__Ascomycota_unclassified(100);p__Ascomycota_unclassified(100); | - | - | - | - | - | - | - | - | Unassigned | - |
| Otu37857 | 1 | k__Fungi(100);p__Ascomycota(100);p__Ascomycota_unclassified(100);p__Ascomycota_unclassified(100);p__Ascomycota_unclassified(100);p__Ascomycota_unclassified(100);p__Ascomycota_unclassified(100); | - | - | - | - | - | - | - | - | Unassigned | - |
| Otu37858 | 1 | k__Fungi(100);p__Ascomycota(100);p__Ascomycota_unclassified(100);p__Ascomycota_unclassified(100);p__Ascomycota_unclassified(100);p__Ascomycota_unclassified(100);p__Ascomycota_unclassified(100); | - | - | - | - | - | - | - | - | Unassigned | - |
| Otu37859 | 1 | k__Fungi(100);p__Ascomycota(100);p__Ascomycota_unclassified(100);p__Ascomycota_unclassified(100);p__Ascomycota_unclassified(100);p__Ascomycota_unclassified(100);p__Ascomycota_unclassified(100); | - | - | - | - | - | - | - | - | Unassigned | - |
| Otu37860 | 1 | k__Fungi(100);p__Ascomycota(100);p__Ascomycota_unclassified(100);p__Ascomycota_unclassified(100);p__Ascomycota_unclassified(100);p__Ascomycota_unclassified(100);p__Ascomycota_unclassified(100); | - | - | - | - | - | - | - | - | Unassigned | - |
| Otu37861 | 1 | k__Fungi(100);p__Ascomycota(100);p__Ascomycota_unclassified(100);p__Ascomycota_unclassified(100);p__Ascomycota_unclassified(100);p__Ascomycota_unclassified(100);p__Ascomycota_unclassified(100); | - | - | - | - | - | - | - | - | Unassigned | - |
| Otu37863 | 1 | k__Fungi(100);p__Ascomycota(100);p__Ascomycota_unclassified(100);p__Ascomycota_unclassified(100);p__Ascomycota_unclassified(100);p__Ascomycota_unclassified(100);p__Ascomycota_unclassified(100); | - | - | - | - | - | - | - | - | Unassigned | - |
| Otu37864 | 1 | k__Fungi(100);k__Fungi_unclassified(100);k__Fungi_unclassified(100);k__Fungi_unclassified(100);k__Fungi_unclassified(100);k__Fungi_unclassified(100);k__Fungi_unclassified(100); | - | - | - | - | - | - | - | - | Unassigned | - |
| Otu37866 | 1 | k__Fungi(100);p__Ascomycota(100);p__Ascomycota_unclassified(100);p__Ascomycota_unclassified(100);p__Ascomycota_unclassified(100);p__Ascomycota_unclassified(100);p__Ascomycota_unclassified(100); | - | - | - | - | - | - | - | - | Unassigned | - |
| Otu37892 | 1 | k__Fungi(100);k__Fungi_unclassified(100);k__Fungi_unclassified(100);k__Fungi_unclassified(100);k__Fungi_unclassified(100);k__Fungi_unclassified(100);k__Fungi_unclassified(100); | - | - | - | - | - | - | - | - | Unassigned | - |
| Otu37897 | 1 | k__Fungi(100);p__Ascomycota(100);p__Ascomycota_unclassified(100);p__Ascomycota_unclassified(100);p__Ascomycota_unclassified(100);p__Ascomycota_unclassified(100);p__Ascomycota_unclassified(100); | - | - | - | - | - | - | - | - | Unassigned | - |
| Otu37898 | 1 | k__Fungi(100);k__Fungi_unclassified(100);k__Fungi_unclassified(100);k__Fungi_unclassified(100);k__Fungi_unclassified(100);k__Fungi_unclassified(100);k__Fungi_unclassified(100); | - | - | - | - | - | - | - | - | Unassigned | - |
| Otu37899 | 1 | k__Fungi(100);p__Ascomycota(100);p__Ascomycota_unclassified(100);p__Ascomycota_unclassified(100);p__Ascomycota_unclassified(100);p__Ascomycota_unclassified(100);p__Ascomycota_unclassified(100); | - | - | - | - | - | - | - | - | Unassigned | - |
| Otu37909 | 1 | k__Fungi(100);p__Ascomycota(100);p__Ascomycota_unclassified(100);p__Ascomycota_unclassified(100);p__Ascomycota_unclassified(100);p__Ascomycota_unclassified(100);p__Ascomycota_unclassified(100); | - | - | - | - | - | - | - | - | Unassigned | - |
| Otu37951 | 1 | k__Fungi(100);p__Ascomycota(100);p__Ascomycota_unclassified(100);p__Ascomycota_unclassified(100);p__Ascomycota_unclassified(100);p__Ascomycota_unclassified(100);p__Ascomycota_unclassified(100); | - | - | - | - | - | - | - | - | Unassigned | - |
| Otu37953 | 1 | k__Fungi(100);p__Ascomycota(100);p__Ascomycota_unclassified(100);p__Ascomycota_unclassified(100);p__Ascomycota_unclassified(100);p__Ascomycota_unclassified(100);p__Ascomycota_unclassified(100); | - | - | - | - | - | - | - | - | Unassigned | - |
| Otu37994 | 1 | k__Fungi(100);p__Ascomycota(100);p__Ascomycota_unclassified(100);p__Ascomycota_unclassified(100);p__Ascomycota_unclassified(100);p__Ascomycota_unclassified(100);p__Ascomycota_unclassified(100); | - | - | - | - | - | - | - | - | Unassigned | - |
| Otu38016 | 1 | k__Fungi(100);k__Fungi_unclassified(100);k__Fungi_unclassified(100);k__Fungi_unclassified(100);k__Fungi_unclassified(100);k__Fungi_unclassified(100);k__Fungi_unclassified(100); | - | - | - | - | - | - | - | - | Unassigned | - |
| Otu38025 | 1 | k__Fungi(100);k__Fungi_unclassified(100);k__Fungi_unclassified(100);k__Fungi_unclassified(100);k__Fungi_unclassified(100);k__Fungi_unclassified(100);k__Fungi_unclassified(100); | - | - | - | - | - | - | - | - | Unassigned | - |
| Otu38036 | 1 | k__Fungi(100);k__Fungi_unclassified(100);k__Fungi_unclassified(100);k__Fungi_unclassified(100);k__Fungi_unclassified(100);k__Fungi_unclassified(100);k__Fungi_unclassified(100); | - | - | - | - | - | - | - | - | Unassigned | - |
| Otu38089 | 1 | k__Fungi(100);k__Fungi_unclassified(100);k__Fungi_unclassified(100);k__Fungi_unclassified(100);k__Fungi_unclassified(100);k__Fungi_unclassified(100);k__Fungi_unclassified(100); | - | - | - | - | - | - | - | - | Unassigned | - |
| Otu38117 | 1 | k__Fungi(100);k__Fungi_unclassified(100);k__Fungi_unclassified(100);k__Fungi_unclassified(100);k__Fungi_unclassified(100);k__Fungi_unclassified(100);k__Fungi_unclassified(100); | - | - | - | - | - | - | - | - | Unassigned | - |
| Otu38194 | 1 | k__Fungi(100);k__Fungi_unclassified(100);k__Fungi_unclassified(100);k__Fungi_unclassified(100);k__Fungi_unclassified(100);k__Fungi_unclassified(100);k__Fungi_unclassified(100); | - | - | - | - | - | - | - | - | Unassigned | - |
| Otu38211 | 1 | k__Fungi(100);p__Ascomycota(100);p__Ascomycota_unclassified(100);p__Ascomycota_unclassified(100);p__Ascomycota_unclassified(100);p__Ascomycota_unclassified(100);p__Ascomycota_unclassified(100); | - | - | - | - | - | - | - | - | Unassigned | - |
| Otu38215 | 1 | k__Fungi(100);k__Fungi_unclassified(100);k__Fungi_unclassified(100);k__Fungi_unclassified(100);k__Fungi_unclassified(100);k__Fungi_unclassified(100);k__Fungi_unclassified(100); | - | - | - | - | - | - | - | - | Unassigned | - |

| | | | | | | | | | | | |
|---|---|---|---|---|---|---|---|---|---|---|---|
| Otu38232 | 1 | k__Fungi(100);p__Ascomycota(100);p__Ascomycota_unclassified(100);p__Ascomycota_unclassified(100);p__Ascomycota_unclassified(100);p__Ascomycota_unclassified(100);p__Ascomycota_unclassified(100); | - | - | - | - | - | - | - | Unassigned | - |
| Otu38240 | 1 | k__Fungi(100);p__Ascomycota(100);p__Ascomycota_unclassified(100);p__Ascomycota_unclassified(100);p__Ascomycota_unclassified(100);p__Ascomycota_unclassified(100);p__Ascomycota_unclassified(100); | - | - | - | - | - | - | - | Unassigned | - |
| Otu38273 | 1 | k__Fungi(100);p__Ascomycota(100);p__Ascomycota_unclassified(100);p__Ascomycota_unclassified(100);p__Ascomycota_unclassified(100);p__Ascomycota_unclassified(100);p__Ascomycota_unclassified(100); | - | - | - | - | - | - | - | Unassigned | - |
| Otu38303 | 1 | k__Fungi(100);p__Ascomycota(100);p__Ascomycota_unclassified(100);p__Ascomycota_unclassified(100);p__Ascomycota_unclassified(100);p__Ascomycota_unclassified(100);p__Ascomycota_unclassified(100); | - | - | - | - | - | - | - | Unassigned | - |
| Otu38304 | 1 | k__Fungi(100);p__Ascomycota(100);p__Ascomycota_unclassified(100);p__Ascomycota_unclassified(100);p__Ascomycota_unclassified(100);p__Ascomycota_unclassified(100);p__Ascomycota_unclassified(100); | - | - | - | - | - | - | - | Unassigned | - |
| Otu38315 | 1 | k__Fungi(100);p__Ascomycota(100);p__Ascomycota_unclassified(100);p__Ascomycota_unclassified(100);p__Ascomycota_unclassified(100);p__Ascomycota_unclassified(100);p__Ascomycota_unclassified(100); | - | - | - | - | - | - | - | Unassigned | - |
| Otu38344 | 1 | k__Fungi(100);p__Ascomycota(100);p__Ascomycota_unclassified(100);p__Ascomycota_unclassified(100);p__Ascomycota_unclassified(100);p__Ascomycota_unclassified(100);p__Ascomycota_unclassified(100); | - | - | - | - | - | - | - | Unassigned | - |
| Otu38366 | 1 | k__Fungi(100);k__Fungi_unclassified(100);k__Fungi_unclassified(100);k__Fungi_unclassified(100);k__Fungi_unclassified(100);k__Fungi_unclassified(100);k__Fungi_unclassified(100); | - | - | - | - | - | - | - | Unassigned | - |
| Otu38367 | 1 | k__Fungi(100);p__Ascomycota(100);p__Ascomycota_unclassified(100);p__Ascomycota_unclassified(100);p__Ascomycota_unclassified(100);p__Ascomycota_unclassified(100);p__Ascomycota_unclassified(100); | - | - | - | - | - | - | - | Unassigned | - |
| Otu38368 | 1 | k__Fungi(100);k__Fungi_unclassified(100);k__Fungi_unclassified(100);k__Fungi_unclassified(100);k__Fungi_unclassified(100);k__Fungi_unclassified(100);k__Fungi_unclassified(100); | - | - | - | - | - | - | - | Unassigned | - |
| Otu38369 | 1 | k__Fungi(100);p__Ascomycota(100);p__Ascomycota_unclassified(100);p__Ascomycota_unclassified(100);p__Ascomycota_unclassified(100);p__Ascomycota_unclassified(100);p__Ascomycota_unclassified(100); | - | - | - | - | - | - | - | Unassigned | - |
| Otu38370 | 1 | k__Fungi(100);p__Ascomycota(100);p__Ascomycota_unclassified(100);p__Ascomycota_unclassified(100);p__Ascomycota_unclassified(100);p__Ascomycota_unclassified(100);p__Ascomycota_unclassified(100); | - | - | - | - | - | - | - | Unassigned | - |
| Otu38379 | 1 | k__Fungi(100);p__Ascomycota(100);p__Ascomycota_unclassified(100);p__Ascomycota_unclassified(100);p__Ascomycota_unclassified(100);p__Ascomycota_unclassified(100);p__Ascomycota_unclassified(100); | - | - | - | - | - | - | - | Unassigned | - |
| Otu38381 | 1 | k__Fungi(100);p__Ascomycota(100);p__Ascomycota_unclassified(100);p__Ascomycota_unclassified(100);p__Ascomycota_unclassified(100);p__Ascomycota_unclassified(100);p__Ascomycota_unclassified(100); | - | - | - | - | - | - | - | Unassigned | - |
| Otu38386 | 1 | k__Fungi(100);k__Fungi_unclassified(100);k__Fungi_unclassified(100);k__Fungi_unclassified(100);k__Fungi_unclassified(100);k__Fungi_unclassified(100);k__Fungi_unclassified(100); | - | - | - | - | - | - | - | Unassigned | - |
| Otu38387 | 1 | k__Fungi(100);p__Ascomycota(100);p__Ascomycota_unclassified(100);p__Ascomycota_unclassified(100);p__Ascomycota_unclassified(100);p__Ascomycota_unclassified(100);p__Ascomycota_unclassified(100); | - | - | - | - | - | - | - | Unassigned | - |
| Otu38428 | 1 | k__Fungi(100);k__Fungi_unclassified(100);k__Fungi_unclassified(100);k__Fungi_unclassified(100);k__Fungi_unclassified(100);k__Fungi_unclassified(100);k__Fungi_unclassified(100); | - | - | - | - | - | - | - | Unassigned | - |
| Otu38432 | 1 | k__Fungi(100);k__Fungi_unclassified(100);k__Fungi_unclassified(100);k__Fungi_unclassified(100);k__Fungi_unclassified(100);k__Fungi_unclassified(100);k__Fungi_unclassified(100); | - | - | - | - | - | - | - | Unassigned | - |
| Otu38442 | 1 | k__Fungi(100);p__Ascomycota(100);p__Ascomycota_unclassified(100);p__Ascomycota_unclassified(100);p__Ascomycota_unclassified(100);p__Ascomycota_unclassified(100);p__Ascomycota_unclassified(100); | - | - | - | - | - | - | - | Unassigned | - |
| Otu38454 | 1 | k__Fungi(100);k__Fungi_unclassified(100);k__Fungi_unclassified(100);k__Fungi_unclassified(100);k__Fungi_unclassified(100);k__Fungi_unclassified(100);k__Fungi_unclassified(100); | - | - | - | - | - | - | - | Unassigned | - |
| Otu38455 | 1 | k__Fungi(100);p__Ascomycota(100);p__Ascomycota_unclassified(100);p__Ascomycota_unclassified(100);p__Ascomycota_unclassified(100);p__Ascomycota_unclassified(100);p__Ascomycota_unclassified(100); | - | - | - | - | - | - | - | Unassigned | - |
| Otu38456 | 1 | k__Fungi(100);p__Ascomycota(100);p__Ascomycota_unclassified(100);p__Ascomycota_unclassified(100);p__Ascomycota_unclassified(100);p__Ascomycota_unclassified(100);p__Ascomycota_unclassified(100); | - | - | - | - | - | - | - | Unassigned | - |
| Otu38468 | 1 | k__Fungi(100);p__Ascomycota(100);p__Ascomycota_unclassified(100);p__Ascomycota_unclassified(100);p__Ascomycota_unclassified(100);p__Ascomycota_unclassified(100);p__Ascomycota_unclassified(100); | - | - | - | - | - | - | - | Unassigned | - |
| Otu38478 | 1 | k__Fungi(100);k__Fungi_unclassified(100);k__Fungi_unclassified(100);k__Fungi_unclassified(100);k__Fungi_unclassified(100);k__Fungi_unclassified(100);k__Fungi_unclassified(100); | - | - | - | - | - | - | - | Unassigned | - |
| Otu38505 | 1 | k__Fungi(100);k__Fungi_unclassified(100);k__Fungi_unclassified(100);k__Fungi_unclassified(100);k__Fungi_unclassified(100);k__Fungi_unclassified(100);k__Fungi_unclassified(100); | - | - | - | - | - | - | - | Unassigned | - |
| Otu38515 | 1 | k__Fungi(100);k__Fungi_unclassified(100);k__Fungi_unclassified(100);k__Fungi_unclassified(100);k__Fungi_unclassified(100);k__Fungi_unclassified(100);k__Fungi_unclassified(100); | - | - | - | - | - | - | - | Unassigned | - |
| Otu38517 | 1 | k__Fungi(100);k__Fungi_unclassified(100);k__Fungi_unclassified(100);k__Fungi_unclassified(100);k__Fungi_unclassified(100);k__Fungi_unclassified(100);k__Fungi_unclassified(100); | - | - | - | - | - | - | - | Unassigned | - |
| Otu38535 | 1 | k__Fungi(100);p__Ascomycota(100);p__Ascomycota_unclassified(100);p__Ascomycota_unclassified(100);p__Ascomycota_unclassified(100);p__Ascomycota_unclassified(100);p__Ascomycota_unclassified(100); | - | - | - | - | - | - | - | Unassigned | - |
| Otu38537 | 1 | k__Fungi(100);k__Fungi_unclassified(100);k__Fungi_unclassified(100);k__Fungi_unclassified(100);k__Fungi_unclassified(100);k__Fungi_unclassified(100);k__Fungi_unclassified(100); | - | - | - | - | - | - | - | Unassigned | - |
| Otu38538 | 1 | k__Fungi(100);k__Fungi_unclassified(100);k__Fungi_unclassified(100);k__Fungi_unclassified(100);k__Fungi_unclassified(100);k__Fungi_unclassified(100);k__Fungi_unclassified(100); | - | - | - | - | - | - | - | Unassigned | - |
| Otu38539 | 1 | k__Fungi(100);k__Fungi_unclassified(100);k__Fungi_unclassified(100);k__Fungi_unclassified(100);k__Fungi_unclassified(100);k__Fungi_unclassified(100);k__Fungi_unclassified(100); | - | - | - | - | - | - | - | Unassigned | - |
| Otu38540 | 1 | k__Fungi(100);p__Ascomycota(100);p__Ascomycota_unclassified(100);p__Ascomycota_unclassified(100);p__Ascomycota_unclassified(100);p__Ascomycota_unclassified(100);p__Ascomycota_unclassified(100); | - | - | - | - | - | - | - | Unassigned | - |
| Otu38544 | 1 | k__Fungi(100);k__Fungi_unclassified(100);k__Fungi_unclassified(100);k__Fungi_unclassified(100);k__Fungi_unclassified(100);k__Fungi_unclassified(100);k__Fungi_unclassified(100); | - | - | - | - | - | - | - | Unassigned | - |
| Otu38546 | 1 | k__Fungi(100);k__Fungi_unclassified(100);k__Fungi_unclassified(100);k__Fungi_unclassified(100);k__Fungi_unclassified(100);k__Fungi_unclassified(100);k__Fungi_unclassified(100); | - | - | - | - | - | - | - | Unassigned | - |
| Otu38552 | 1 | k__Fungi(100);k__Fungi_unclassified(100);k__Fungi_unclassified(100);k__Fungi_unclassified(100);k__Fungi_unclassified(100);k__Fungi_unclassified(100);k__Fungi_unclassified(100); | - | - | - | - | - | - | - | Unassigned | - |
| Otu38555 | 1 | k__Fungi(100);k__Fungi_unclassified(100);k__Fungi_unclassified(100);k__Fungi_unclassified(100);k__Fungi_unclassified(100);k__Fungi_unclassified(100);k__Fungi_unclassified(100); | - | - | - | - | - | - | - | Unassigned | - |
| Otu38556 | 1 | k__Fungi(100);k__Fungi_unclassified(100);k__Fungi_unclassified(100);k__Fungi_unclassified(100);k__Fungi_unclassified(100);k__Fungi_unclassified(100);k__Fungi_unclassified(100); | - | - | - | - | - | - | - | Unassigned | - |
| Otu38557 | 1 | k__Fungi(100);k__Fungi_unclassified(100);k__Fungi_unclassified(100);k__Fungi_unclassified(100);k__Fungi_unclassified(100);k__Fungi_unclassified(100);k__Fungi_unclassified(100); | - | - | - | - | - | - | - | Unassigned | - |
| Otu38561 | 1 | k__Fungi(100);p__Ascomycota(100);p__Ascomycota_unclassified(100);p__Ascomycota_unclassified(100);p__Ascomycota_unclassified(100);p__Ascomycota_unclassified(100);p__Ascomycota_unclassified(100); | - | - | - | - | - | - | - | Unassigned | - |
| Otu38568 | 1 | k__Fungi(100);k__Fungi_unclassified(100);k__Fungi_unclassified(100);k__Fungi_unclassified(100);k__Fungi_unclassified(100);k__Fungi_unclassified(100);k__Fungi_unclassified(100); | - | - | - | - | - | - | - | Unassigned | - |
| Otu38573 | 1 | k__Fungi(100);k__Fungi_unclassified(100);k__Fungi_unclassified(100);k__Fungi_unclassified(100);k__Fungi_unclassified(100);k__Fungi_unclassified(100);k__Fungi_unclassified(100); | - | - | - | - | - | - | - | Unassigned | - |
| Otu38574 | 1 | k__Fungi(100);p__Ascomycota(100);c__Dothideomycetes(100);o__Pleosporales(100);f__Pleosporales_family_Incertae_sedis(100);f__Pleosporales_family_Incertae_sedis_unclassified(100);f__Pleosporales_family_Incertae_sedis_unclassified(100); | - | - | - | - | - | - | - | Unassigned | - |
| Otu38575 | 1 | k__Fungi(100);p__Ascomycota(100);p__Ascomycota_unclassified(100);p__Ascomycota_unclassified(100);p__Ascomycota_unclassified(100);p__Ascomycota_unclassified(100);p__Ascomycota_unclassified(100); | - | - | - | - | - | - | - | Unassigned | - |
| Otu38576 | 1 | k__Fungi(100);p__Ascomycota(100);p__Ascomycota_unclassified(100);p__Ascomycota_unclassified(100);p__Ascomycota_unclassified(100);p__Ascomycota_unclassified(100);p__Ascomycota_unclassified(100); | - | - | - | - | - | - | - | Unassigned | - |
| Otu38577 | 1 | k__Fungi(100);k__Fungi_unclassified(100);k__Fungi_unclassified(100);k__Fungi_unclassified(100);k__Fungi_unclassified(100);k__Fungi_unclassified(100);k__Fungi_unclassified(100); | - | - | - | - | - | - | - | Unassigned | - |
| Otu38579 | 1 | k__Fungi(100);k__Fungi_unclassified(100);k__Fungi_unclassified(100);k__Fungi_unclassified(100);k__Fungi_unclassified(100);k__Fungi_unclassified(100);k__Fungi_unclassified(100); | - | - | - | - | - | - | - | Unassigned | - |
| Otu38580 | 1 | k__Fungi(100);k__Fungi_unclassified(100);k__Fungi_unclassified(100);k__Fungi_unclassified(100);k__Fungi_unclassified(100);k__Fungi_unclassified(100);k__Fungi_unclassified(100); | - | - | - | - | - | - | - | Unassigned | - |
| Otu38586 | 1 | k__Fungi(100);k__Fungi_unclassified(100);k__Fungi_unclassified(100);k__Fungi_unclassified(100);k__Fungi_unclassified(100);k__Fungi_unclassified(100);k__Fungi_unclassified(100); | - | - | - | - | - | - | - | Unassigned | - |
| Otu38595 | 1 | k__Fungi(100);p__Ascomycota(100);c__Leotiomycetes(100);o__Helotiales(100);f__Helotiales_family_Incertae_sedis(100);f__Helotiales_family_Incertae_sedis_unclassified(100);f__Helotiales_family_Incertae_sedis_unclassified(100); | - | - | - | - | - | - | - | Unassigned | - |
| Otu38596 | 1 | k__Fungi(100);p__Ascomycota(100);p__Ascomycota_unclassified(100);p__Ascomycota_unclassified(100);p__Ascomycota_unclassified(100);p__Ascomycota_unclassified(100);p__Ascomycota_unclassified(100); | - | - | - | - | - | - | - | Unassigned | - |
| Otu38597 | 1 | k__Fungi(100);p__Ascomycota(100);p__Ascomycota_unclassified(100);p__Ascomycota_unclassified(100);p__Ascomycota_unclassified(100);p__Ascomycota_unclassified(100);p__Ascomycota_unclassified(100); | - | - | - | - | - | - | - | Unassigned | - |
| Otu38623 | 1 | k__Fungi(100);p__Ascomycota(100);p__Ascomycota_unclassified(100);p__Ascomycota_unclassified(100);p__Ascomycota_unclassified(100);p__Ascomycota_unclassified(100);p__Ascomycota_unclassified(100); | - | - | - | - | - | - | - | Unassigned | - |
| Otu38633 | 1 | k__Fungi(100);k__Fungi_unclassified(100);k__Fungi_unclassified(100);k__Fungi_unclassified(100);k__Fungi_unclassified(100);k__Fungi_unclassified(100);k__Fungi_unclassified(100); | - | - | - | - | - | - | - | Unassigned | - |
| Otu38636 | 1 | k__Fungi(100);k__Fungi_unclassified(100);k__Fungi_unclassified(100);k__Fungi_unclassified(100);k__Fungi_unclassified(100);k__Fungi_unclassified(100);k__Fungi_unclassified(100); | - | - | - | - | - | - | - | Unassigned | - |
| Otu38692 | 1 | k__Fungi(100);k__Fungi_unclassified(100);k__Fungi_unclassified(100);k__Fungi_unclassified(100);k__Fungi_unclassified(100);k__Fungi_unclassified(100);k__Fungi_unclassified(100); | - | - | - | - | - | - | - | Unassigned | - |
| Otu38694 | 1 | k__Fungi(100);k__Fungi_unclassified(100);k__Fungi_unclassified(100);k__Fungi_unclassified(100);k__Fungi_unclassified(100);k__Fungi_unclassified(100);k__Fungi_unclassified(100); | - | - | - | - | - | - | - | Unassigned | - |
| Otu38696 | 1 | k__Fungi(100);k__Fungi_unclassified(100);k__Fungi_unclassified(100);k__Fungi_unclassified(100);k__Fungi_unclassified(100);k__Fungi_unclassified(100);k__Fungi_unclassified(100); | - | - | - | - | - | - | - | Unassigned | - |
| Otu38698 | 1 | k__Fungi(100);p__Ascomycota(100);p__Ascomycota_unclassified(100);p__Ascomycota_unclassified(100);p__Ascomycota_unclassified(100);p__Ascomycota_unclassified(100);p__Ascomycota_unclassified(100); | - | - | - | - | - | - | - | Unassigned | - |
| Otu38703 | 1 | k__Fungi(100);k__Fungi_unclassified(100);k__Fungi_unclassified(100);k__Fungi_unclassified(100);k__Fungi_unclassified(100);k__Fungi_unclassified(100);k__Fungi_unclassified(100); | - | - | - | - | - | - | - | Unassigned | - |
| Otu38706 | 1 | k__Fungi(100);k__Fungi_unclassified(100);k__Fungi_unclassified(100);k__Fungi_unclassified(100);k__Fungi_unclassified(100);k__Fungi_unclassified(100);k__Fungi_unclassified(100); | - | - | - | - | - | - | - | Unassigned | - |
| Otu38720 | 1 | k__Fungi(100);p__Ascomycota(100);c__Dothideomycetes(100);o__Capnodiales(100);o__Capnodiales_unclassified(100);o__Capnodiales_unclassified(100);o__Capnodiales_unclassified(100); | - | - | - | - | - | - | - | Unassigned | - |
| Otu38723 | 1 | k__Fungi(100);p__Ascomycota(100);c__Dothideomycetes(100);o__Pleosporales(100);f__unclassified_Pleosporales(100);g__unclassified_Pleosporales(100);s__Pleosporales_sp(100); | - | - | - | - | - | - | - | Unassigned | - |
| Otu38727 | 1 | k__Fungi(100);p__Ascomycota(100);c__Dothideomycetes(100);o__Pleosporales(100);f__unclassified_Pleosporales(100);g__unclassified_Pleosporales(100);s__Pleosporales_sp(100); | - | - | - | - | - | - | - | Unassigned | - |
| Otu38729 | 1 | k__Fungi(100);p__Ascomycota(100);c__Dothideomycetes(100);o__Pleosporales(100);f__Pleosporales_family_Incertae_sedis(100);f__Pleosporales_family_Incertae_sedis_unclassified(100);f__Pleosporales_family_Incertae_sedis_unclassified(100); | - | - | - | - | - | - | - | Unassigned | - |
| Otu38732 | 1 | k__Fungi(100);k__Fungi_unclassified(100);k__Fungi_unclassified(100);k__Fungi_unclassified(100);k__Fungi_unclassified(100);k__Fungi_unclassified(100);k__Fungi_unclassified(100); | - | - | - | - | - | - | - | Unassigned | - |
| Otu38733 | 1 | k__Fungi(100);k__Fungi_unclassified(100);k__Fungi_unclassified(100);k__Fungi_unclassified(100);k__Fungi_unclassified(100);k__Fungi_unclassified(100);k__Fungi_unclassified(100); | - | - | - | - | - | - | - | Unassigned | - |
| Otu38734 | 1 | k__Fungi(100);k__Fungi_unclassified(100);k__Fungi_unclassified(100);k__Fungi_unclassified(100);k__Fungi_unclassified(100);k__Fungi_unclassified(100);k__Fungi_unclassified(100); | - | - | - | - | - | - | - | Unassigned | - |
| Otu38736 | 1 | k__Fungi(100);k__Fungi_unclassified(100);k__Fungi_unclassified(100);k__Fungi_unclassified(100);k__Fungi_unclassified(100);k__Fungi_unclassified(100);k__Fungi_unclassified(100); | - | - | - | - | - | - | - | Unassigned | - |
| Otu38737 | 1 | k__Fungi(100);k__Fungi_unclassified(100);k__Fungi_unclassified(100);k__Fungi_unclassified(100);k__Fungi_unclassified(100);k__Fungi_unclassified(100);k__Fungi_unclassified(100); | - | - | - | - | - | - | - | Unassigned | - |
| Otu38738 | 1 | k__Fungi(100);k__Fungi_unclassified(100);k__Fungi_unclassified(100);k__Fungi_unclassified(100);k__Fungi_unclassified(100);k__Fungi_unclassified(100);k__Fungi_unclassified(100); | - | - | - | - | - | - | - | Unassigned | - |
| Otu38739 | 1 | k__Fungi(100);k__Fungi_unclassified(100);k__Fungi_unclassified(100);k__Fungi_unclassified(100);k__Fungi_unclassified(100);k__Fungi_unclassified(100);k__Fungi_unclassified(100); | - | - | - | - | - | - | - | Unassigned | - |
| Otu38740 | 1 | k__Fungi(100);k__Fungi_unclassified(100);k__Fungi_unclassified(100);k__Fungi_unclassified(100);k__Fungi_unclassified(100);k__Fungi_unclassified(100);k__Fungi_unclassified(100); | - | - | - | - | - | - | - | Unassigned | - |
| Otu38742 | 1 | k__Fungi(100);k__Fungi_unclassified(100);k__Fungi_unclassified(100);k__Fungi_unclassified(100);k__Fungi_unclassified(100);k__Fungi_unclassified(100);k__Fungi_unclassified(100); | - | - | - | - | - | - | - | Unassigned | - |
| Otu38771 | 1 | k__Fungi(100);k__Fungi_unclassified(100);k__Fungi_unclassified(100);k__Fungi_unclassified(100);k__Fungi_unclassified(100);k__Fungi_unclassified(100);k__Fungi_unclassified(100); | - | - | - | - | - | - | - | Unassigned | - |
| Otu38775 | 1 | k__Fungi(100);k__Fungi_unclassified(100);k__Fungi_unclassified(100);k__Fungi_unclassified(100);k__Fungi_unclassified(100);k__Fungi_unclassified(100);k__Fungi_unclassified(100); | - | - | - | - | - | - | - | Unassigned | - |
| Otu38781 | 1 | k__Fungi(100);k__Fungi_unclassified(100);k__Fungi_unclassified(100);k__Fungi_unclassified(100);k__Fungi_unclassified(100);k__Fungi_unclassified(100);k__Fungi_unclassified(100); | - | - | - | - | - | - | - | Unassigned | - |
| Otu38799 | 1 | k__Fungi(100);p__Ascomycota(100);p__Ascomycota_unclassified(100);p__Ascomycota_unclassified(100);p__Ascomycota_unclassified(100);p__Ascomycota_unclassified(100);p__Ascomycota_unclassified(100); | - | - | - | - | - | - | - | Unassigned | - |
| Otu38805 | 1 | k__Fungi(100);k__Fungi_unclassified(100);k__Fungi_unclassified(100);k__Fungi_unclassified(100);k__Fungi_unclassified(100);k__Fungi_unclassified(100);k__Fungi_unclassified(100); | - | - | - | - | - | - | - | Unassigned | - |
| Otu38807 | 1 | k__Fungi(100);p__Ascomycota(100);p__Ascomycota_unclassified(100);p__Ascomycota_unclassified(100);p__Ascomycota_unclassified(100);p__Ascomycota_unclassified(100);p__Ascomycota_unclassified(100); | - | - | - | - | - | - | - | Unassigned | - |
| Otu38816 | 1 | k__Fungi(100);k__Fungi_unclassified(100);k__Fungi_unclassified(100);k__Fungi_unclassified(100);k__Fungi_unclassified(100);k__Fungi_unclassified(100);k__Fungi_unclassified(100); | - | - | - | - | - | - | - | Unassigned | - |
| Otu38838 | 1 | k__Fungi(100);p__Ascomycota(100);p__Ascomycota_unclassified(100);p__Ascomycota_unclassified(100);p__Ascomycota_unclassified(100);p__Ascomycota_unclassified(100);p__Ascomycota_unclassified(100); | - | - | - | - | - | - | - | Unassigned | - |
| Otu38841 | 1 | k__Fungi(100);k__Fungi_unclassified(100);k__Fungi_unclassified(100);k__Fungi_unclassified(100);k__Fungi_unclassified(100);k__Fungi_unclassified(100);k__Fungi_unclassified(100); | - | - | - | - | - | - | - | Unassigned | - |
| Otu38876 | 1 | k__Fungi(100);k__Fungi_unclassified(100);k__Fungi_unclassified(100);k__Fungi_unclassified(100);k__Fungi_unclassified(100);k__Fungi_unclassified(100);k__Fungi_unclassified(100); | - | - | - | - | - | - | - | Unassigned | - |
| Otu38877 | 1 | k__Fungi(100);k__Fungi_unclassified(100);k__Fungi_unclassified(100);k__Fungi_unclassified(100);k__Fungi_unclassified(100);k__Fungi_unclassified(100);k__Fungi_unclassified(100); | - | - | - | - | - | - | - | Unassigned | - |
| Otu38897 | 1 | k__Fungi(100);k__Fungi_unclassified(100);k__Fungi_unclassified(100);k__Fungi_unclassified(100);k__Fungi_unclassified(100);k__Fungi_unclassified(100);k__Fungi_unclassified(100); | - | - | - | - | - | - | - | Unassigned | - |
| Otu38899 | 1 | k__Fungi(100);p__Ascomycota(100);p__Ascomycota_unclassified(100);p__Ascomycota_unclassified(100);p__Ascomycota_unclassified(100);p__Ascomycota_unclassified(100);p__Ascomycota_unclassified(100); | - | - | - | - | - | - | - | Unassigned | - |
| Otu38900 | 1 | k__Fungi(100);k__Fungi_unclassified(100);k__Fungi_unclassified(100);k__Fungi_unclassified(100);k__Fungi_unclassified(100);k__Fungi_unclassified(100);k__Fungi_unclassified(100); | - | - | - | - | - | - | - | Unassigned | - |
| Otu38903 | 1 | k__Fungi(100);p__Ascomycota(100);p__Ascomycota_unclassified(100);p__Ascomycota_unclassified(100);p__Ascomycota_unclassified(100);p__Ascomycota_unclassified(100);p__Ascomycota_unclassified(100); | - | - | - | - | - | - | - | Unassigned | - |
| Otu38904 | 1 | k__Fungi(100);p__Ascomycota(100);p__Ascomycota_unclassified(100);p__Ascomycota_unclassified(100);p__Ascomycota_unclassified(100);p__Ascomycota_unclassified(100);p__Ascomycota_unclassified(100); | - | - | - | - | - | - | - | Unassigned | - |
| Otu38908 | 1 | k__Fungi(100);p__Basidiomycota(100);p__Basidiomycota_unclassified(100);p__Basidiomycota_unclassified(100);p__Basidiomycota_unclassified(100);p__Basidiomycota_unclassified(100);p__Basidiomycota_unclassified(100); | - | - | - | - | - | - | - | Unassigned | - |
| Otu38911 | 1 | k__Fungi(100);k__Fungi_unclassified(100);k__Fungi_unclassified(100);k__Fungi_unclassified(100);k__Fungi_unclassified(100);k__Fungi_unclassified(100);k__Fungi_unclassified(100); | - | - | - | - | - | - | - | Unassigned | - |
| Otu38924 | 1 | k__Fungi(100);p__Ascomycota(100);p__Ascomycota_unclassified(100);p__Ascomycota_unclassified(100);p__Ascomycota_unclassified(100);p__Ascomycota_unclassified(100);p__Ascomycota_unclassified(100); | - | - | - | - | - | - | - | Unassigned | - |
| Otu38938 | 1 | k__Fungi(100);p__Ascomycota(100);p__Ascomycota_unclassified(100);p__Ascomycota_unclassified(100);p__Ascomycota_unclassified(100);p__Ascomycota_unclassified(100);p__Ascomycota_unclassified(100); | - | - | - | - | - | - | - | Unassigned | - |
| Otu38939 | 1 | k__Fungi(100);p__Ascomycota(100);p__Ascomycota_unclassified(100);p__Ascomycota_unclassified(100);p__Ascomycota_unclassified(100);p__Ascomycota_unclassified(100);p__Ascomycota_unclassified(100); | - | - | - | - | - | - | - | Unassigned | - |
| Otu38940 | 1 | k__Fungi(100);k__Fungi_unclassified(100);k__Fungi_unclassified(100);k__Fungi_unclassified(100);k__Fungi_unclassified(100);k__Fungi_unclassified(100);k__Fungi_unclassified(100); | - | - | - | - | - | - | - | Unassigned | - |
| Otu38941 | 1 | k__Fungi(100);p__Ascomycota(100);p__Ascomycota_unclassified(100);p__Ascomycota_unclassified(100);p__Ascomycota_unclassified(100);p__Ascomycota_unclassified(100);p__Ascomycota_unclassified(100); | - | - | - | - | - | - | - | Unassigned | - |
| Otu38942 | 1 | k__Fungi(100);p__Ascomycota(100);p__Ascomycota_unclassified(100);p__Ascomycota_unclassified(100);p__Ascomycota_unclassified(100);p__Ascomycota_unclassified(100);p__Ascomycota_unclassified(100); | - | - | - | - | - | - | - | Unassigned | - |
| Otu38943 | 1 | k__Fungi(100);p__Ascomycota(100);p__Ascomycota_unclassified(100);p__Ascomycota_unclassified(100);p__Ascomycota_unclassified(100);p__Ascomycota_unclassified(100);p__Ascomycota_unclassified(100); | - | - | - | - | - | - | - | Unassigned | - |
| Otu38947 | 1 | k__Fungi(100);k__Fungi_unclassified(100);k__Fungi_unclassified(100);k__Fungi_unclassified(100);k__Fungi_unclassified(100);k__Fungi_unclassified(100);k__Fungi_unclassified(100); | - | - | - | - | - | - | - | Unassigned | - |
| Otu38948 | 1 | k__Fungi(100);p__Ascomycota(100);p__Ascomycota_unclassified(100);p__Ascomycota_unclassified(100);p__Ascomycota_unclassified(100);p__Ascomycota_unclassified(100);p__Ascomycota_unclassified(100); | - | - | - | - | - | - | - | Unassigned | - |
| Otu38949 | 1 | k__Fungi(100);p__Ascomycota(100);p__Ascomycota_unclassified(100);p__Ascomycota_unclassified(100);p__Ascomycota_unclassified(100);p__Ascomycota_unclassified(100);p__Ascomycota_unclassified(100); | - | - | - | - | - | - | - | Unassigned | - |
| Otu38966 | 1 | k__Fungi(100);p__Ascomycota(100);p__Ascomycota_unclassified(100);p__Ascomycota_unclassified(100);p__Ascomycota_unclassified(100);p__Ascomycota_unclassified(100);p__Ascomycota_unclassified(100); | - | - | - | - | - | - | - | Unassigned | - |
| Otu38971 | 1 | k__Fungi(100);k__Fungi_unclassified(100);k__Fungi_unclassified(100);k__Fungi_unclassified(100);k__Fungi_unclassified(100);k__Fungi_unclassified(100);k__Fungi_unclassified(100); | - | - | - | - | - | - | - | Unassigned | - |
| Otu38980 | 1 | k__Fungi(100);p__Ascomycota(100);p__Ascomycota_unclassified(100);p__Ascomycota_unclassified(100);p__Ascomycota_unclassified(100);p__Ascomycota_unclassified(100);p__Ascomycota_unclassified(100); | - | - | - | - | - | - | - | Unassigned | - |
| Otu38994 | 1 | k__Fungi(100);p__Ascomycota(100);p__Ascomycota_unclassified(100);p__Ascomycota_unclassified(100);p__Ascomycota_unclassified(100);p__Ascomycota_unclassified(100);p__Ascomycota_unclassified(100); | - | - | - | - | - | - | - | Unassigned | - |
| Otu39018 | 1 | k__Fungi(100);k__Fungi_unclassified(100);k__Fungi_unclassified(100);k__Fungi_unclassified(100);k__Fungi_unclassified(100);k__Fungi_unclassified(100);k__Fungi_unclassified(100); | - | - | - | - | - | - | - | Unassigned | - |
| Otu39019 | 1 | k__Fungi(100);p__Ascomycota(100);p__Ascomycota_unclassified(100);p__Ascomycota_unclassified(100);p__Ascomycota_unclassified(100);p__Ascomycota_unclassified(100);p__Ascomycota_unclassified(100); | - | - | - | - | - | - | - | Unassigned | - |
| Otu39020 | 1 | k__Fungi(100);k__Fungi_unclassified(100);k__Fungi_unclassified(100);k__Fungi_unclassified(100);k__Fungi_unclassified(100);k__Fungi_unclassified(100);k__Fungi_unclassified(100); | - | - | - | - | - | - | - | Unassigned | - |
| Otu39042 | 1 | k__Fungi(100);p__Ascomycota(100);p__Ascomycota_unclassified(100);p__Ascomycota_unclassified(100);p__Ascomycota_unclassified(100);p__Ascomycota_unclassified(100);p__Ascomycota_unclassified(100); | - | - | - | - | - | - | - | Unassigned | - |
| Otu39051 | 1 | k__Fungi(100);k__Fungi_unclassified(100);k__Fungi_unclassified(100);k__Fungi_unclassified(100);k__Fungi_unclassified(100);k__Fungi_unclassified(100);k__Fungi_unclassified(100); | - | - | - | - | - | - | - | Unassigned | - |
| Otu39054 | 1 | k__Fungi(100);k__Fungi_unclassified(100);k__Fungi_unclassified(100);k__Fungi_unclassified(100);k__Fungi_unclassified(100);k__Fungi_unclassified(100);k__Fungi_unclassified(100); | - | - | - | - | - | - | - | Unassigned | - |
| Otu39055 | 1 | k__Fungi(100);p__Ascomycota(100);p__Ascomycota_unclassified(100);p__Ascomycota_unclassified(100);p__Ascomycota_unclassified(100);p__Ascomycota_unclassified(100);p__Ascomycota_unclassified(100); | - | - | - | - | - | - | - | Unassigned | - |
| Otu39065 | 1 | k__Fungi(100);k__Fungi_unclassified(100);k__Fungi_unclassified(100);k__Fungi_unclassified(100);k__Fungi_unclassified(100);k__Fungi_unclassified(100);k__Fungi_unclassified(100); | - | - | - | - | - | - | - | Unassigned | - |
| Otu39066 | 1 | k__Fungi(100);k__Fungi_unclassified(100);k__Fungi_unclassified(100);k__Fungi_unclassified(100);k__Fungi_unclassified(100);k__Fungi_unclassified(100);k__Fungi_unclassified(100); | - | - | - | - | - | - | - | Unassigned | - |
| Otu39072 | 1 | k__Fungi(100);k__Fungi_unclassified(100);k__Fungi_unclassified(100);k__Fungi_unclassified(100);k__Fungi_unclassified(100);k__Fungi_unclassified(100);k__Fungi_unclassified(100); | - | - | - | - | - | - | - | Unassigned | - |
| Otu39073 | 1 | k__Fungi(100);k__Fungi_unclassified(100);k__Fungi_unclassified(100);k__Fungi_unclassified(100);k__Fungi_unclassified(100);k__Fungi_unclassified(100);k__Fungi_unclassified(100); | - | - | - | - | - | - | - | Unassigned | - |
| Otu39076 | 1 | k__Fungi(100);p__Ascomycota(100);p__Ascomycota_unclassified(100);p__Ascomycota_unclassified(100);p__Ascomycota_unclassified(100);p__Ascomycota_unclassified(100);p__Ascomycota_unclassified(100); | - | - | - | - | - | - | - | Unassigned | - |
| Otu39125 | 1 | k__Fungi(100);k__Fungi_unclassified(100);k__Fungi_unclassified(100);k__Fungi_unclassified(100);k__Fungi_unclassified(100);k__Fungi_unclassified(100);k__Fungi_unclassified(100); | - | - | - | - | - | - | - | Unassigned | - |
| Otu39159 | 1 | k__Fungi(100);k__Fungi_unclassified(100);k__Fungi_unclassified(100);k__Fungi_unclassified(100);k__Fungi_unclassified(100);k__Fungi_unclassified(100);k__Fungi_unclassified(100); | - | - | - | - | - | - | - | Unassigned | - |
| Otu39164 | 1 | k__Fungi(100);p__Ascomycota(100);p__Ascomycota_unclassified(100);p__Ascomycota_unclassified(100);p__Ascomycota_unclassified(100);p__Ascomycota_unclassified(100);p__Ascomycota_unclassified(100); | - | - | - | - | - | - | - | Unassigned | - |
| Otu39165 | 1 | k__Fungi(100);p__Ascomycota(100);p__Ascomycota_unclassified(100);p__Ascomycota_unclassified(100);p__Ascomycota_unclassified(100);p__Ascomycota_unclassified(100);p__Ascomycota_unclassified(100); | - | - | - | - | - | - | - | Unassigned | - |
| Otu39173 | 1 | k__Fungi(100);k__Fungi_unclassified(100);k__Fungi_unclassified(100);k__Fungi_unclassified(100);k__Fungi_unclassified(100);k__Fungi_unclassified(100);k__Fungi_unclassified(100); | - | - | - | - | - | - | - | Unassigned | - |
| Otu39177 | 1 | k__Fungi(100);p__Ascomycota(100);p__Ascomycota_unclassified(100);p__Ascomycota_unclassified(100);p__Ascomycota_unclassified(100);p__Ascomycota_unclassified(100);p__Ascomycota_unclassified(100); | - | - | - | - | - | - | - | Unassigned | - |
| Otu39193 | 1 | k__Fungi(100);k__Fungi_unclassified(100);k__Fungi_unclassified(100);k__Fungi_unclassified(100);k__Fungi_unclassified(100);k__Fungi_unclassified(100);k__Fungi_unclassified(100); | - | - | - | - | - | - | - | Unassigned | - |
| Otu39244 | 1 | k__Fungi(100);p__Ascomycota(100);p__Ascomycota_unclassified(100);p__Ascomycota_unclassified(100);p__Ascomycota_unclassified(100);p__Ascomycota_unclassified(100);p__Ascomycota_unclassified(100); | - | - | - | - | - | - | - | Unassigned | - |
| Otu39255 | 1 | k__Fungi(100);k__Fungi_unclassified(100);k__Fungi_unclassified(100);k__Fungi_unclassified(100);k__Fungi_unclassified(100);k__Fungi_unclassified(100);k__Fungi_unclassified(100); | - | - | - | - | - | - | - | Unassigned | - |
| Otu39258 | 1 | k__Fungi(100);p__Ascomycota(100);p__Ascomycota_unclassified(100);p__Ascomycota_unclassified(100);p__Ascomycota_unclassified(100);p__Ascomycota_unclassified(100);p__Ascomycota_unclassified(100); | - | - | - | - | - | - | - | Unassigned | - |
| Otu39293 | 1 | k__Fungi(100);k__Fungi_unclassified(100);k__Fungi_unclassified(100);k__Fungi_unclassified(100);k__Fungi_unclassified(100);k__Fungi_unclassified(100);k__Fungi_unclassified(100); | - | - | - | - | - | - | - | Unassigned | - |
| Otu39314 | 1 | k__Fungi(100);p__Ascomycota(100);p__Ascomycota_unclassified(100);p__Ascomycota_unclassified(100);p__Ascomycota_unclassified(100);p__Ascomycota_unclassified(100);p__Ascomycota_unclassified(100); | - | - | - | - | - | - | - | Unassigned | - |
| Otu39329 | 1 | k__Fungi(100);p__Ascomycota(100);p__Ascomycota_unclassified(100);p__Ascomycota_unclassified(100);p__Ascomycota_unclassified(100);p__Ascomycota_unclassified(100);p__Ascomycota_unclassified(100); | - | - | - | - | - | - | - | Unassigned | - |
| Otu39344 | 1 | k__Fungi(100);p__Ascomycota(100);p__Ascomycota_unclassified(100);p__Ascomycota_unclassified(100);p__Ascomycota_unclassified(100);p__Ascomycota_unclassified(100);p__Ascomycota_unclassified(100); | - | - | - | - | - | - | - | Unassigned | - |
| Otu39345 | 1 | k__Fungi(100);p__Ascomycota(100);p__Ascomycota_unclassified(100);p__Ascomycota_unclassified(100);p__Ascomycota_unclassified(100);p__Ascomycota_unclassified(100);p__Ascomycota_unclassified(100); | - | - | - | - | - | - | - | Unassigned | - |
| Otu39369 | 1 | k__Fungi(100);k__Fungi_unclassified(100);k__Fungi_unclassified(100);k__Fungi_unclassified(100);k__Fungi_unclassified(100);k__Fungi_unclassified(100);k__Fungi_unclassified(100); | - | - | - | - | - | - | - | Unassigned | - |
| Otu39398 | 1 | k__Fungi(100);k__Fungi_unclassified(100);k__Fungi_unclassified(100);k__Fungi_unclassified(100);k__Fungi_unclassified(100);k__Fungi_unclassified(100);k__Fungi_unclassified(100); | - | - | - | - | - | - | - | Unassigned | - |
| Otu39402 | 1 | k__Fungi(100);p__Ascomycota(100);p__Ascomycota_unclassified(100);p__Ascomycota_unclassified(100);p__Ascomycota_unclassified(100);p__Ascomycota_unclassified(100);p__Ascomycota_unclassified(100); | - | - | - | - | - | - | - | Unassigned | - |
| Otu39411 | 1 | k__Fungi(100);p__Ascomycota(100);p__Ascomycota_unclassified(100);p__Ascomycota_unclassified(100);p__Ascomycota_unclassified(100);p__Ascomycota_unclassified(100);p__Ascomycota_unclassified(100); | - | - | - | - | - | - | - | Unassigned | - |
| Otu39413 | 1 | k__Fungi(100);k__Fungi_unclassified(100);k__Fungi_unclassified(100);k__Fungi_unclassified(100);k__Fungi_unclassified(100);k__Fungi_unclassified(100);k__Fungi_unclassified(100); | - | - | - | - | - | - | - | Unassigned | - |
| Otu39429 | 1 | k__Fungi(100);k__Fungi_unclassified(100);k__Fungi_unclassified(100);k__Fungi_unclassified(100);k__Fungi_unclassified(100);k__Fungi_unclassified(100);k__Fungi_unclassified(100); | - | - | - | - | - | - | - | Unassigned | - |
| Otu39573 | 1 | k__Fungi(100);p__Ascomycota(100);p__Ascomycota_unclassified(100);p__Ascomycota_unclassified(100);p__Ascomycota_unclassified(100);p__Ascomycota_unclassified(100);p__Ascomycota_unclassified(100); | - | - | - | - | - | - | - | Unassigned | - |
| Otu39625 | 1 | k__Fungi(100);k__Fungi_unclassified(100);k__Fungi_unclassified(100);k__Fungi_unclassified(100);k__Fungi_unclassified(100);k__Fungi_unclassified(100);k__Fungi_unclassified(100); | - | - | - | - | - | - | - | Unassigned | - |
| Otu39626 | 1 | k__Fungi(100);k__Fungi_unclassified(100);k__Fungi_unclassified(100);k__Fungi_unclassified(100);k__Fungi_unclassified(100);k__Fungi_unclassified(100);k__Fungi_unclassified(100); | - | - | - | - | - | - | - | Unassigned | - |
| Otu39628 | 1 | k__Fungi(100);k__Fungi_unclassified(100);k__Fungi_unclassified(100);k__Fungi_unclassified(100);k__Fungi_unclassified(100);k__Fungi_unclassified(100);k__Fungi_unclassified(100); | - | - | - | - | - | - | - | Unassigned | - |
| Otu39629 | 1 | k__Fungi(100);k__Fungi_unclassified(100);k__Fungi_unclassified(100);k__Fungi_unclassified(100);k__Fungi_unclassified(100);k__Fungi_unclassified(100);k__Fungi_unclassified(100); | - | - | - | - | - | - | - | Unassigned | - |
| Otu39640 | 1 | k__Fungi(100);k__Fungi_unclassified(100);k__Fungi_unclassified(100);k__Fungi_unclassified(100);k__Fungi_unclassified(100);k__Fungi_unclassified(100);k__Fungi_unclassified(100); | - | - | - | - | - | - | - | Unassigned | - |
| Otu39642 | 1 | k__Fungi(100);k__Fungi_unclassified(100);k__Fungi_unclassified(100);k__Fungi_unclassified(100);k__Fungi_unclassified(100);k__Fungi_unclassified(100);k__Fungi_unclassified(100); | - | - | - | - | - | - | - | Unassigned | - |
| Otu39675 | 1 | k__Fungi(100);k__Fungi_unclassified(100);k__Fungi_unclassified(100);k__Fungi_unclassified(100);k__Fungi_unclassified(100);k__Fungi_unclassified(100);k__Fungi_unclassified(100); | - | - | - | - | - | - | - | Unassigned | - |
| Otu39723 | 1 | k__Fungi(100);k__Fungi_unclassified(100);k__Fungi_unclassified(100);k__Fungi_unclassified(100);k__Fungi_unclassified(100);k__Fungi_unclassified(100);k__Fungi_unclassified(100); | - | - | - | - | - | - | - | Unassigned | - |
| Otu39743 | 1 | k__Fungi(100);p__Ascomycota(100);c__Leotiomycetes(100);c__Leotiomycetes_unclassified(100);c__Leotiomycetes_unclassified(100);c__Leotiomycetes_unclassified(100);c__Leotiomycetes_unclassified(100); | - | - | - | - | - | - | - | Unassigned | - |
| Otu39795 | 1 | k__Fungi(100);k__Fungi_unclassified(100);k__Fungi_unclassified(100);k__Fungi_unclassified(100);k__Fungi_unclassified(100);k__Fungi_unclassified(100);k__Fungi_unclassified(100); | - | - | - | - | - | - | - | Unassigned | - |
| Otu39934 | 1 | k__Fungi(100);k__Fungi_unclassified(100);k__Fungi_unclassified(100);k__Fungi_unclassified(100);k__Fungi_unclassified(100);k__Fungi_unclassified(100);k__Fungi_unclassified(100); | - | - | - | - | - | - | - | Unassigned | - |
| Otu39944 | 1 | k__Fungi(100);p__Ascomycota(100);p__Ascomycota_unclassified(100);p__Ascomycota_unclassified(100);p__Ascomycota_unclassified(100);p__Ascomycota_unclassified(100);p__Ascomycota_unclassified(100); | - | - | - | - | - | - | - | Unassigned | - |
| Otu39989 | 1 | k__Fungi(100);k__Fungi_unclassified(100);k__Fungi_unclassified(100);k__Fungi_unclassified(100);k__Fungi_unclassified(100);k__Fungi_unclassified(100);k__Fungi_unclassified(100); | - | - | - | - | - | - | - | Unassigned | - |
| Otu40087 | 1 | k__Fungi(100);p__Ascomycota(100);p__Ascomycota_unclassified(100);p__Ascomycota_unclassified(100);p__Ascomycota_unclassified(100);p__Ascomycota_unclassified(100);p__Ascomycota_unclassified(100); | - | - | - | - | - | - | - | Unassigned | - |
| Otu40115 | 1 | k__Fungi(100);k__Fungi_unclassified(100);k__Fungi_unclassified(100);k__Fungi_unclassified(100);k__Fungi_unclassified(100);k__Fungi_unclassified(100);k__Fungi_unclassified(100); | - | - | - | - | - | - | - | Unassigned | - |
| Otu40117 | 1 | k__Fungi(100);p__Ascomycota(100);p__Ascomycota_unclassified(100);p__Ascomycota_unclassified(100);p__Ascomycota_unclassified(100);p__Ascomycota_unclassified(100);p__Ascomycota_unclassified(100); | - | - | - | - | - | - | - | Unassigned | - |
| Otu40128 | 1 | k__Fungi(100);k__Fungi_unclassified(100);k__Fungi_unclassified(100);k__Fungi_unclassified(100);k__Fungi_unclassified(100);k__Fungi_unclassified(100);k__Fungi_unclassified(100); | - | - | - | - | - | - | - | Unassigned | - |
| Otu40131 | 1 | k__Fungi(100);p__Ascomycota(100);p__Ascomycota_unclassified(100);p__Ascomycota_unclassified(100);p__Ascomycota_unclassified(100);p__Ascomycota_unclassified(100);p__Ascomycota_unclassified(100); | - | - | - | - | - | - | - | Unassigned | - |
| Otu40174 | 1 | k__Fungi(100);k__Fungi_unclassified(100);k__Fungi_unclassified(100);k__Fungi_unclassified(100);k__Fungi_unclassified(100);k__Fungi_unclassified(100);k__Fungi_unclassified(100); | - | - | - | - | - | - | - | Unassigned | - |
| Otu40188 | 1 | k__Fungi(100);k__Fungi_unclassified(100);k__Fungi_unclassified(100);k__Fungi_unclassified(100);k__Fungi_unclassified(100);k__Fungi_unclassified(100);k__Fungi_unclassified(100); | - | - | - | - | - | - | - | Unassigned | - |
| Otu40198 | 1 | k__Fungi(100);p__Ascomycota(100);p__Ascomycota_unclassified(100);p__Ascomycota_unclassified(100);p__Ascomycota_unclassified(100);p__Ascomycota_unclassified(100);p__Ascomycota_unclassified(100); | - | - | - | - | - | - | - | Unassigned | - |
| Otu40199 | 1 | k__Fungi(100);k__Fungi_unclassified(100);k__Fungi_unclassified(100);k__Fungi_unclassified(100);k__Fungi_unclassified(100);k__Fungi_unclassified(100);k__Fungi_unclassified(100); | - | - | - | - | - | - | - | Unassigned | - |
| Otu40208 | 1 | k__Fungi(100);p__Ascomycota(100);p__Ascomycota_unclassified(100);p__Ascomycota_unclassified(100);p__Ascomycota_unclassified(100);p__Ascomycota_unclassified(100);p__Ascomycota_unclassified(100); | - | - | - | - | - | - | - | Unassigned | - |
| Otu40213 | 1 | k__Fungi(100);k__Fungi_unclassified(100);k__Fungi_unclassified(100);k__Fungi_unclassified(100);k__Fungi_unclassified(100);k__Fungi_unclassified(100);k__Fungi_unclassified(100); | - | - | - | - | - | - | - | Unassigned | - |
| Otu40214 | 1 | k__Fungi(100);p__Ascomycota(100);p__Ascomycota_unclassified(100);p__Ascomycota_unclassified(100);p__Ascomycota_unclassified(100);p__Ascomycota_unclassified(100);p__Ascomycota_unclassified(100); | - | - | - | - | - | - | - | Unassigned | - |
| Otu40233 | 1 | k__Fungi(100);k__Fungi_unclassified(100);k__Fungi_unclassified(100);k__Fungi_unclassified(100);k__Fungi_unclassified(100);k__Fungi_unclassified(100);k__Fungi_unclassified(100); | - | - | - | - | - | - | - | Unassigned | - |
| Otu40238 | 1 | k__Fungi(100);k__Fungi_unclassified(100);k__Fungi_unclassified(100);k__Fungi_unclassified(100);k__Fungi_unclassified(100);k__Fungi_unclassified(100);k__Fungi_unclassified(100); | - | - | - | - | - | - | - | Unassigned | - |
| Otu40240 | 1 | k__Fungi(100);k__Fungi_unclassified(100);k__Fungi_unclassified(100);k__Fungi_unclassified(100);k__Fungi_unclassified(100);k__Fungi_unclassified(100);k__Fungi_unclassified(100); | - | - | - | - | - | - | - | Unassigned | - |
| Otu40245 | 1 | k__Fungi(100);k__Fungi_unclassified(100);k__Fungi_unclassified(100);k__Fungi_unclassified(100);k__Fungi_unclassified(100);k__Fungi_unclassified(100);k__Fungi_unclassified(100); | - | - | - | - | - | - | - | Unassigned | - |
| Otu40283 | 1 | k__Fungi(100);p__Ascomycota(100);p__Ascomycota_unclassified(100);p__Ascomycota_unclassified(100);p__Ascomycota_unclassified(100);p__Ascomycota_unclassified(100);p__Ascomycota_unclassified(100); | - | - | - | - | - | - | - | Unassigned | - |
| Otu40303 | 1 | k__Fungi(100);k__Fungi_unclassified(100);k__Fungi_unclassified(100);k__Fungi_unclassified(100);k__Fungi_unclassified(100);k__Fungi_unclassified(100);k__Fungi_unclassified(100); | - | - | - | - | - | - | - | Unassigned | - |
| Otu40313 | 1 | k__Fungi(100);k__Fungi_unclassified(100);k__Fungi_unclassified(100);k__Fungi_unclassified(100);k__Fungi_unclassified(100);k__Fungi_unclassified(100);k__Fungi_unclassified(100); | - | - | - | - | - | - | - | Unassigned | - |
| Otu40314 | 1 | k__Fungi(100);p__Ascomycota(100);p__Ascomycota_unclassified(100);p__Ascomycota_unclassified(100);p__Ascomycota_unclassified(100);p__Ascomycota_unclassified(100);p__Ascomycota_unclassified(100); | - | - | - | - | - | - | - | Unassigned | - |
| Otu40316 | 1 | k__Fungi(100);p__Ascomycota(100);p__Ascomycota_unclassified(100);p__Ascomycota_unclassified(100);p__Ascomycota_unclassified(100);p__Ascomycota_unclassified(100);p__Ascomycota_unclassified(100); | - | - | - | - | - | - | - | Unassigned | - |
| Otu40318 | 1 | k__Fungi(100);k__Fungi_unclassified(100);k__Fungi_unclassified(100);k__Fungi_unclassified(100);k__Fungi_unclassified(100);k__Fungi_unclassified(100);k__Fungi_unclassified(100); | - | - | - | - | - | - | - | Unassigned | - |
| Otu40323 | 1 | k__Fungi(100);p__Ascomycota(100);c__Dothideomycetes(100);o__Pleosporales(100);f__Pleosporales_family_Incertae_sedis(100);f__Pleosporales_family_Incertae_sedis_unclassified(100);f__Pleosporales_family_Incertae_sedis_unclassified(100); | - | - | - | - | - | - | - | Unassigned | - |
| Otu40355 | 1 | k__Fungi(100);k__Fungi_unclassified(100);k__Fungi_unclassified(100);k__Fungi_unclassified(100);k__Fungi_unclassified(100);k__Fungi_unclassified(100);k__Fungi_unclassified(100); | - | - | - | - | - | - | - | Unassigned | - |
| Otu40393 | 1 | k__Fungi(100);k__Fungi_unclassified(100);k__Fungi_unclassified(100);k__Fungi_unclassified(100);k__Fungi_unclassified(100);k__Fungi_unclassified(100);k__Fungi_unclassified(100); | - | - | - | - | - | - | - | Unassigned | - |
| Otu40395 | 1 | k__Fungi(100);k__Fungi_unclassified(100);k__Fungi_unclassified(100);k__Fungi_unclassified(100);k__Fungi_unclassified(100);k__Fungi_unclassified(100);k__Fungi_unclassified(100); | - | - | - | - | - | - | - | Unassigned | - |
| Otu40423 | 1 | k__Fungi(100);k__Fungi_unclassified(100);k__Fungi_unclassified(100);k__Fungi_unclassified(100);k__Fungi_unclassified(100);k__Fungi_unclassified(100);k__Fungi_unclassified(100); | - | - | - | - | - | - | - | Unassigned | - |
| Otu40498 | 1 | k__Fungi(100);k__Fungi_unclassified(100);k__Fungi_unclassified(100);k__Fungi_unclassified(100);k__Fungi_unclassified(100);k__Fungi_unclassified(100);k__Fungi_unclassified(100); | - | - | - | - | - | - | - | Unassigned | - |
| Otu40519 | 1 | k__Fungi(100);k__Fungi_unclassified(100);k__Fungi_unclassified(100);k__Fungi_unclassified(100);k__Fungi_unclassified(100);k__Fungi_unclassified(100);k__Fungi_unclassified(100); | - | - | - | - | - | - | - | Unassigned | - |
| Otu40524 | 1 | k__Fungi(100);k__Fungi_unclassified(100);k__Fungi_unclassified(100);k__Fungi_unclassified(100);k__Fungi_unclassified(100);k__Fungi_unclassified(100);k__Fungi_unclassified(100); | - | - | - | - | - | - | - | Unassigned | - |
| Otu40526 | 1 | k__Fungi(100);p__Ascomycota(100);p__Ascomycota_unclassified(100);p__Ascomycota_unclassified(100);p__Ascomycota_unclassified(100);p__Ascomycota_unclassified(100);p__Ascomycota_unclassified(100); | - | - | - | - | - | - | - | Unassigned | - |
| Otu40527 | 1 | k__Fungi(100);p__Ascomycota(100);p__Ascomycota_unclassified(100);p__Ascomycota_unclassified(100);p__Ascomycota_unclassified(100);p__Ascomycota_unclassified(100);p__Ascomycota_unclassified(100); | - | - | - | - | - | - | - | Unassigned | - |
| Otu40528 | 1 | k__Fungi(100);k__Fungi_unclassified(100);k__Fungi_unclassified(100);k__Fungi_unclassified(100);k__Fungi_unclassified(100);k__Fungi_unclassified(100);k__Fungi_unclassified(100); | - | - | - | - | - | - | - | Unassigned | - |
| Otu40529 | 1 | k__Fungi(100);p__Ascomycota(100);p__Ascomycota_unclassified(100);p__Ascomycota_unclassified(100);p__Ascomycota_unclassified(100);p__Ascomycota_unclassified(100);p__Ascomycota_unclassified(100); | - | - | - | - | - | - | - | Unassigned | - |
| Otu40532 | 1 | k__Fungi(100);k__Fungi_unclassified(100);k__Fungi_unclassified(100);k__Fungi_unclassified(100);k__Fungi_unclassified(100);k__Fungi_unclassified(100);k__Fungi_unclassified(100); | - | - | - | - | - | - | - | Unassigned | - |
| Otu40535 | 1 | k__Fungi(100);p__Ascomycota(100);p__Ascomycota_unclassified(100);p__Ascomycota_unclassified(100);p__Ascomycota_unclassified(100);p__Ascomycota_unclassified(100);p__Ascomycota_unclassified(100); | - | - | - | - | - | - | - | Unassigned | - |
| Otu40537 | 1 | k__Fungi(100);k__Fungi_unclassified(100);k__Fungi_unclassified(100);k__Fungi_unclassified(100);k__Fungi_unclassified(100);k__Fungi_unclassified(100);k__Fungi_unclassified(100); | - | - | - | - | - | - | - | Unassigned | - |
| Otu40541 | 1 | k__Fungi(100);k__Fungi_unclassified(100);k__Fungi_unclassified(100);k__Fungi_unclassified(100);k__Fungi_unclassified(100);k__Fungi_unclassified(100);k__Fungi_unclassified(100); | - | - | - | - | - | - | - | Unassigned | - |
| Otu40542 | 1 | k__Fungi(100);p__Ascomycota(100);p__Ascomycota_unclassified(100);p__Ascomycota_unclassified(100);p__Ascomycota_unclassified(100);p__Ascomycota_unclassified(100);p__Ascomycota_unclassified(100); | - | - | - | - | - | - | - | Unassigned | - |
| Otu40548 | 1 | k__Fungi(100);p__Ascomycota(100);p__Ascomycota_unclassified(100);p__Ascomycota_unclassified(100);p__Ascomycota_unclassified(100);p__Ascomycota_unclassified(100);p__Ascomycota_unclassified(100); | - | - | - | - | - | - | - | Unassigned | - |
| Otu40549 | 1 | k__Fungi(100);k__Fungi_unclassified(100);k__Fungi_unclassified(100);k__Fungi_unclassified(100);k__Fungi_unclassified(100);k__Fungi_unclassified(100);k__Fungi_unclassified(100); | - | - | - | - | - | - | - | Unassigned | - |
| Otu40555 | 1 | k__Fungi(100);k__Fungi_unclassified(100);k__Fungi_unclassified(100);k__Fungi_unclassified(100);k__Fungi_unclassified(100);k__Fungi_unclassified(100);k__Fungi_unclassified(100); | - | - | - | - | - | - | - | Unassigned | - |
| Otu40556 | 1 | k__Fungi(100);k__Fungi_unclassified(100);k__Fungi_unclassified(100);k__Fungi_unclassified(100);k__Fungi_unclassified(100);k__Fungi_unclassified(100);k__Fungi_unclassified(100); | - | - | - | - | - | - | - | Unassigned | - |
| Otu40558 | 1 | k__Fungi(100);k__Fungi_unclassified(100);k__Fungi_unclassified(100);k__Fungi_unclassified(100);k__Fungi_unclassified(100);k__Fungi_unclassified(100);k__Fungi_unclassified(100); | - | - | - | - | - | - | - | Unassigned | - |
| Otu40567 | 1 | k__Fungi(100);k__Fungi_unclassified(100);k__Fungi_unclassified(100);k__Fungi_unclassified(100);k__Fungi_unclassified(100);k__Fungi_unclassified(100);k__Fungi_unclassified(100); | - | - | - | - | - | - | - | Unassigned | - |
| Otu40568 | 1 | k__Fungi(100);k__Fungi_unclassified(100);k__Fungi_unclassified(100);k__Fungi_unclassified(100);k__Fungi_unclassified(100);k__Fungi_unclassified(100);k__Fungi_unclassified(100); | - | - | - | - | - | - | - | Unassigned | - |
| Otu40570 | 1 | k__Fungi(100);k__Fungi_unclassified(100);k__Fungi_unclassified(100);k__Fungi_unclassified(100);k__Fungi_unclassified(100);k__Fungi_unclassified(100);k__Fungi_unclassified(100); | - | - | - | - | - | - | - | Unassigned | - |
| Otu40578 | 1 | k__Fungi(100);k__Fungi_unclassified(100);k__Fungi_unclassified(100);k__Fungi_unclassified(100);k__Fungi_unclassified(100);k__Fungi_unclassified(100);k__Fungi_unclassified(100); | - | - | - | - | - | - | - | Unassigned | - |
| Otu40579 | 1 | k__Fungi(100);p__Ascomycota(100);p__Ascomycota_unclassified(100);p__Ascomycota_unclassified(100);p__Ascomycota_unclassified(100);p__Ascomycota_unclassified(100);p__Ascomycota_unclassified(100); | - | - | - | - | - | - | - | Unassigned | - |
| Otu40587 | 1 | k__Fungi(100);k__Fungi_unclassified(100);k__Fungi_unclassified(100);k__Fungi_unclassified(100);k__Fungi_unclassified(100);k__Fungi_unclassified(100);k__Fungi_unclassified(100); | - | - | - | - | - | - | - | Unassigned | - |
| Otu40590 | 1 | k__Fungi(100);p__Ascomycota(100);p__Ascomycota_unclassified(100);p__Ascomycota_unclassified(100);p__Ascomycota_unclassified(100);p__Ascomycota_unclassified(100);p__Ascomycota_unclassified(100); | - | - | - | - | - | - | - | Unassigned | - |
| Otu40614 | 1 | k__Fungi(100);p__Ascomycota(100);p__Ascomycota_unclassified(100);p__Ascomycota_unclassified(100);p__Ascomycota_unclassified(100);p__Ascomycota_unclassified(100);p__Ascomycota_unclassified(100); | - | - | - | - | - | - | - | Unassigned | - |
| Otu40618 | 1 | k__Fungi(100);k__Fungi_unclassified(100);k__Fungi_unclassified(100);k__Fungi_unclassified(100);k__Fungi_unclassified(100);k__Fungi_unclassified(100);k__Fungi_unclassified(100); | - | - | - | - | - | - | - | Unassigned | - |
| Otu40649 | 1 | k__Fungi(100);p__Ascomycota(100);p__Ascomycota_unclassified(100);p__Ascomycota_unclassified(100);p__Ascomycota_unclassified(100);p__Ascomycota_unclassified(100);p__Ascomycota_unclassified(100); | - | - | - | - | - | - | - | Unassigned | - |
| Otu40772 | 1 | k__Fungi(100);k__Fungi_unclassified(100);k__Fungi_unclassified(100);k__Fungi_unclassified(100);k__Fungi_unclassified(100);k__Fungi_unclassified(100);k__Fungi_unclassified(100); | - | - | - | - | - | - | - | Unassigned | - |
| Otu40780 | 1 | k__Fungi(100);k__Fungi_unclassified(100);k__Fungi_unclassified(100);k__Fungi_unclassified(100);k__Fungi_unclassified(100);k__Fungi_unclassified(100);k__Fungi_unclassified(100); | - | - | - | - | - | - | - | Unassigned | - |
| Otu40787 | 1 | k__Fungi(100);k__Fungi_unclassified(100);k__Fungi_unclassified(100);k__Fungi_unclassified(100);k__Fungi_unclassified(100);k__Fungi_unclassified(100);k__Fungi_unclassified(100); | - | - | - | - | - | - | - | Unassigned | - |
| Otu40791 | 1 | k__Fungi(100);k__Fungi_unclassified(100);k__Fungi_unclassified(100);k__Fungi_unclassified(100);k__Fungi_unclassified(100);k__Fungi_unclassified(100);k__Fungi_unclassified(100); | - | - | - | - | - | - | - | Unassigned | - |
| Otu40795 | 1 | k__Fungi(100);p__Ascomycota(100);p__Ascomycota_unclassified(100);p__Ascomycota_unclassified(100);p__Ascomycota_unclassified(100);p__Ascomycota_unclassified(100);p__Ascomycota_unclassified(100); | - | - | - | - | - | - | - | Unassigned | - |
| Otu40804 | 1 | k__Fungi(100);k__Fungi_unclassified(100);k__Fungi_unclassified(100);k__Fungi_unclassified(100);k__Fungi_unclassified(100);k__Fungi_unclassified(100);k__Fungi_unclassified(100); | - | - | - | - | - | - | - | Unassigned | - |
| Otu40806 | 1 | k__Fungi(100);p__Ascomycota(100);p__Ascomycota_unclassified(100);p__Ascomycota_unclassified(100);p__Ascomycota_unclassified(100);p__Ascomycota_unclassified(100);p__Ascomycota_unclassified(100); | - | - | - | - | - | - | - | Unassigned | - |
| Otu40809 | 1 | k__Fungi(100);p__Ascomycota(100);p__Ascomycota_unclassified(100);p__Ascomycota_unclassified(100);p__Ascomycota_unclassified(100);p__Ascomycota_unclassified(100);p__Ascomycota_unclassified(100); | - | - | - | - | - | - | - | Unassigned | - |
| Otu40810 | 1 | k__Fungi(100);p__Ascomycota(100);p__Ascomycota_unclassified(100);p__Ascomycota_unclassified(100);p__Ascomycota_unclassified(100);p__Ascomycota_unclassified(100);p__Ascomycota_unclassified(100); | - | - | - | - | - | - | - | Unassigned | - |
| Otu40821 | 1 | k__Fungi(100);k__Fungi_unclassified(100);k__Fungi_unclassified(100);k__Fungi_unclassified(100);k__Fungi_unclassified(100);k__Fungi_unclassified(100);k__Fungi_unclassified(100); | - | - | - | - | - | - | - | Unassigned | - |

| | | | | | | | | | | | |
|---|---|---|---|---|---|---|---|---|---|---|---|
| Otu40823 | 1 | k__Fungi(100);k__Fungi_unclassified(100);k__Fungi_unclassified(100);k__Fungi_unclassified(100);k__Fungi_unclassified(100);k__Fungi_unclassified(100);k__Fungi_unclassified(100); | - | - | - | - | - | - | - | Unassigned | - |
| Otu40825 | 1 | k__Fungi(100);k__Fungi_unclassified(100);k__Fungi_unclassified(100);k__Fungi_unclassified(100);k__Fungi_unclassified(100);k__Fungi_unclassified(100);k__Fungi_unclassified(100); | - | - | - | - | - | - | - | Unassigned | - |
| Otu40828 | 1 | k__Fungi(100);k__Fungi_unclassified(100);k__Fungi_unclassified(100);k__Fungi_unclassified(100);k__Fungi_unclassified(100);k__Fungi_unclassified(100);k__Fungi_unclassified(100); | - | - | - | - | - | - | - | Unassigned | - |
| Otu40832 | 1 | k__Fungi(100);k__Fungi_unclassified(100);k__Fungi_unclassified(100);k__Fungi_unclassified(100);k__Fungi_unclassified(100);k__Fungi_unclassified(100);k__Fungi_unclassified(100); | - | - | - | - | - | - | - | Unassigned | - |
| Otu40844 | 1 | k__Fungi(100);p__Ascomycota(100);c__Sordariomycetes(100);o__Sordariales(100);o__Sordariales_unclassified(100);o__Sordariales_unclassified(100);o__Sordariales_unclassified(100); | - | - | - | - | - | - | - | Unassigned | - |
| Otu40853 | 1 | k__Fungi(100);p__Ascomycota(100);p__Ascomycota_unclassified(100);p__Ascomycota_unclassified(100);p__Ascomycota_unclassified(100);p__Ascomycota_unclassified(100);p__Ascomycota_unclassified(100); | - | - | - | - | - | - | - | Unassigned | - |
| Otu40890 | 1 | k__Fungi(100);k__Fungi_unclassified(100);k__Fungi_unclassified(100);k__Fungi_unclassified(100);k__Fungi_unclassified(100);k__Fungi_unclassified(100);k__Fungi_unclassified(100); | - | - | - | - | - | - | - | Unassigned | - |
| Otu40900 | 1 | k__Fungi(100);p__Ascomycota(100);p__Ascomycota_unclassified(100);p__Ascomycota_unclassified(100);p__Ascomycota_unclassified(100);p__Ascomycota_unclassified(100);p__Ascomycota_unclassified(100); | - | - | - | - | - | - | - | Unassigned | - |
| Otu40904 | 1 | k__Fungi(100);k__Fungi_unclassified(100);k__Fungi_unclassified(100);k__Fungi_unclassified(100);k__Fungi_unclassified(100);k__Fungi_unclassified(100);k__Fungi_unclassified(100); | - | - | - | - | - | - | - | Unassigned | - |
| Otu40919 | 1 | k__Fungi(100);k__Fungi_unclassified(100);k__Fungi_unclassified(100);k__Fungi_unclassified(100);k__Fungi_unclassified(100);k__Fungi_unclassified(100);k__Fungi_unclassified(100); | - | - | - | - | - | - | - | Unassigned | - |
| Otu41004 | 1 | k__Fungi(100);k__Fungi_unclassified(100);k__Fungi_unclassified(100);k__Fungi_unclassified(100);k__Fungi_unclassified(100);k__Fungi_unclassified(100);k__Fungi_unclassified(100); | - | - | - | - | - | - | - | Unassigned | - |
| Otu41006 | 1 | k__Fungi(100);k__Fungi_unclassified(100);k__Fungi_unclassified(100);k__Fungi_unclassified(100);k__Fungi_unclassified(100);k__Fungi_unclassified(100);k__Fungi_unclassified(100); | - | - | - | - | - | - | - | Unassigned | - |
| Otu41015 | 1 | k__Fungi(100);k__Fungi_unclassified(100);k__Fungi_unclassified(100);k__Fungi_unclassified(100);k__Fungi_unclassified(100);k__Fungi_unclassified(100);k__Fungi_unclassified(100); | - | - | - | - | - | - | - | Unassigned | - |
| Otu41027 | 1 | k__Fungi(100);k__Fungi_unclassified(100);k__Fungi_unclassified(100);k__Fungi_unclassified(100);k__Fungi_unclassified(100);k__Fungi_unclassified(100);k__Fungi_unclassified(100); | - | - | - | - | - | - | - | Unassigned | - |
| Otu41028 | 1 | k__Fungi(100);k__Fungi_unclassified(100);k__Fungi_unclassified(100);k__Fungi_unclassified(100);k__Fungi_unclassified(100);k__Fungi_unclassified(100);k__Fungi_unclassified(100); | - | - | - | - | - | - | - | Unassigned | - |
| Otu41031 | 1 | k__Fungi(100);p__Ascomycota(100);p__Ascomycota_unclassified(100);p__Ascomycota_unclassified(100);p__Ascomycota_unclassified(100);p__Ascomycota_unclassified(100);p__Ascomycota_unclassified(100); | - | - | - | - | - | - | - | Unassigned | - |
| Otu41032 | 1 | k__Fungi(100);p__Ascomycota(100);p__Ascomycota_unclassified(100);p__Ascomycota_unclassified(100);p__Ascomycota_unclassified(100);p__Ascomycota_unclassified(100);p__Ascomycota_unclassified(100); | - | - | - | - | - | - | - | Unassigned | - |
| Otu41034 | 1 | k__Fungi(100);k__Fungi_unclassified(100);k__Fungi_unclassified(100);k__Fungi_unclassified(100);k__Fungi_unclassified(100);k__Fungi_unclassified(100);k__Fungi_unclassified(100); | - | - | - | - | - | - | - | Unassigned | - |
| Otu41041 | 1 | k__Fungi(100);p__Ascomycota(100);p__Ascomycota_unclassified(100);p__Ascomycota_unclassified(100);p__Ascomycota_unclassified(100);p__Ascomycota_unclassified(100);p__Ascomycota_unclassified(100); | - | - | - | - | - | - | - | Unassigned | - |
| Otu41053 | 1 | k__Fungi(100);k__Fungi_unclassified(100);k__Fungi_unclassified(100);k__Fungi_unclassified(100);k__Fungi_unclassified(100);k__Fungi_unclassified(100);k__Fungi_unclassified(100); | - | - | - | - | - | - | - | Unassigned | - |
| Otu41063 | 1 | k__Fungi(100);p__Ascomycota(100);p__Ascomycota_unclassified(100);p__Ascomycota_unclassified(100);p__Ascomycota_unclassified(100);p__Ascomycota_unclassified(100);p__Ascomycota_unclassified(100); | - | - | - | - | - | - | - | Unassigned | - |
| Otu41093 | 1 | k__Fungi(100);k__Fungi_unclassified(100);k__Fungi_unclassified(100);k__Fungi_unclassified(100);k__Fungi_unclassified(100);k__Fungi_unclassified(100);k__Fungi_unclassified(100); | - | - | - | - | - | - | - | Unassigned | - |
| Otu41150 | 1 | k__Fungi(100);k__Fungi_unclassified(100);k__Fungi_unclassified(100);k__Fungi_unclassified(100);k__Fungi_unclassified(100);k__Fungi_unclassified(100);k__Fungi_unclassified(100); | - | - | - | - | - | - | - | Unassigned | - |
| Otu41185 | 1 | k__Fungi(100);k__Fungi_unclassified(100);k__Fungi_unclassified(100);k__Fungi_unclassified(100);k__Fungi_unclassified(100);k__Fungi_unclassified(100);k__Fungi_unclassified(100); | - | - | - | - | - | - | - | Unassigned | - |
| Otu41200 | 1 | k__Fungi(100);k__Fungi_unclassified(100);k__Fungi_unclassified(100);k__Fungi_unclassified(100);k__Fungi_unclassified(100);k__Fungi_unclassified(100);k__Fungi_unclassified(100); | - | - | - | - | - | - | - | Unassigned | - |
| Otu41206 | 1 | k__Fungi(100);k__Fungi_unclassified(100);k__Fungi_unclassified(100);k__Fungi_unclassified(100);k__Fungi_unclassified(100);k__Fungi_unclassified(100);k__Fungi_unclassified(100); | - | - | - | - | - | - | - | Unassigned | - |
| Otu41214 | 1 | k__Fungi(100);k__Fungi_unclassified(100);k__Fungi_unclassified(100);k__Fungi_unclassified(100);k__Fungi_unclassified(100);k__Fungi_unclassified(100);k__Fungi_unclassified(100); | - | - | - | - | - | - | - | Unassigned | - |
| Otu41217 | 1 | k__Fungi(100);k__Fungi_unclassified(100);k__Fungi_unclassified(100);k__Fungi_unclassified(100);k__Fungi_unclassified(100);k__Fungi_unclassified(100);k__Fungi_unclassified(100); | - | - | - | - | - | - | - | Unassigned | - |
| Otu41305 | 1 | k__Fungi(100);k__Fungi_unclassified(100);k__Fungi_unclassified(100);k__Fungi_unclassified(100);k__Fungi_unclassified(100);k__Fungi_unclassified(100);k__Fungi_unclassified(100); | - | - | - | - | - | - | - | Unassigned | - |
| Otu41313 | 1 | k__Fungi(100);k__Fungi_unclassified(100);k__Fungi_unclassified(100);k__Fungi_unclassified(100);k__Fungi_unclassified(100);k__Fungi_unclassified(100);k__Fungi_unclassified(100); | - | - | - | - | - | - | - | Unassigned | - |
| Otu41386 | 1 | k__Fungi(100);p__Ascomycota(100);p__Ascomycota_unclassified(100);p__Ascomycota_unclassified(100);p__Ascomycota_unclassified(100);p__Ascomycota_unclassified(100);p__Ascomycota_unclassified(100); | - | - | - | - | - | - | - | Unassigned | - |
| Otu41394 | 1 | k__Fungi(100);k__Fungi_unclassified(100);k__Fungi_unclassified(100);k__Fungi_unclassified(100);k__Fungi_unclassified(100);k__Fungi_unclassified(100);k__Fungi_unclassified(100); | - | - | - | - | - | - | - | Unassigned | - |
| Otu41396 | 1 | k__Fungi(100);k__Fungi_unclassified(100);k__Fungi_unclassified(100);k__Fungi_unclassified(100);k__Fungi_unclassified(100);k__Fungi_unclassified(100);k__Fungi_unclassified(100); | - | - | - | - | - | - | - | Unassigned | - |
| Otu41397 | 1 | k__Fungi(100);k__Fungi_unclassified(100);k__Fungi_unclassified(100);k__Fungi_unclassified(100);k__Fungi_unclassified(100);k__Fungi_unclassified(100);k__Fungi_unclassified(100); | - | - | - | - | - | - | - | Unassigned | - |
| Otu41398 | 1 | k__Fungi(100);k__Fungi_unclassified(100);k__Fungi_unclassified(100);k__Fungi_unclassified(100);k__Fungi_unclassified(100);k__Fungi_unclassified(100);k__Fungi_unclassified(100); | - | - | - | - | - | - | - | Unassigned | - |
| Otu41423 | 1 | k__Fungi(100);k__Fungi_unclassified(100);k__Fungi_unclassified(100);k__Fungi_unclassified(100);k__Fungi_unclassified(100);k__Fungi_unclassified(100);k__Fungi_unclassified(100); | - | - | - | - | - | - | - | Unassigned | - |
| Otu41424 | 1 | k__Fungi(100);k__Fungi_unclassified(100);k__Fungi_unclassified(100);k__Fungi_unclassified(100);k__Fungi_unclassified(100);k__Fungi_unclassified(100);k__Fungi_unclassified(100); | - | - | - | - | - | - | - | Unassigned | - |
| Otu41425 | 1 | k__Fungi(100);k__Fungi_unclassified(100);k__Fungi_unclassified(100);k__Fungi_unclassified(100);k__Fungi_unclassified(100);k__Fungi_unclassified(100);k__Fungi_unclassified(100); | - | - | - | - | - | - | - | Unassigned | - |
| Otu41426 | 1 | k__Fungi(100);p__Ascomycota(100);p__Ascomycota_unclassified(100);p__Ascomycota_unclassified(100);p__Ascomycota_unclassified(100);p__Ascomycota_unclassified(100);p__Ascomycota_unclassified(100); | - | - | - | - | - | - | - | Unassigned | - |
| Otu41428 | 1 | k__Fungi(100);k__Fungi_unclassified(100);k__Fungi_unclassified(100);k__Fungi_unclassified(100);k__Fungi_unclassified(100);k__Fungi_unclassified(100);k__Fungi_unclassified(100); | - | - | - | - | - | - | - | Unassigned | - |
| Otu41429 | 1 | k__Fungi(100);k__Fungi_unclassified(100);k__Fungi_unclassified(100);k__Fungi_unclassified(100);k__Fungi_unclassified(100);k__Fungi_unclassified(100);k__Fungi_unclassified(100); | - | - | - | - | - | - | - | Unassigned | - |
| Otu41436 | 1 | k__Fungi(100);k__Fungi_unclassified(100);k__Fungi_unclassified(100);k__Fungi_unclassified(100);k__Fungi_unclassified(100);k__Fungi_unclassified(100);k__Fungi_unclassified(100); | - | - | - | - | - | - | - | Unassigned | - |
| Otu41437 | 1 | k__Fungi(100);p__Ascomycota(100);p__Ascomycota_unclassified(100);p__Ascomycota_unclassified(100);p__Ascomycota_unclassified(100);p__Ascomycota_unclassified(100);p__Ascomycota_unclassified(100); | - | - | - | - | - | - | - | Unassigned | - |
| Otu41440 | 1 | k__Fungi(100);p__Ascomycota(100);p__Ascomycota_unclassified(100);p__Ascomycota_unclassified(100);p__Ascomycota_unclassified(100);p__Ascomycota_unclassified(100);p__Ascomycota_unclassified(100); | - | - | - | - | - | - | - | Unassigned | - |
| Otu41445 | 1 | k__Fungi(100);k__Fungi_unclassified(100);k__Fungi_unclassified(100);k__Fungi_unclassified(100);k__Fungi_unclassified(100);k__Fungi_unclassified(100);k__Fungi_unclassified(100); | - | - | - | - | - | - | - | Unassigned | - |
| Otu41446 | 1 | k__Fungi(100);k__Fungi_unclassified(100);k__Fungi_unclassified(100);k__Fungi_unclassified(100);k__Fungi_unclassified(100);k__Fungi_unclassified(100);k__Fungi_unclassified(100); | - | - | - | - | - | - | - | Unassigned | - |
| Otu41453 | 1 | k__Fungi(100);k__Fungi_unclassified(100);k__Fungi_unclassified(100);k__Fungi_unclassified(100);k__Fungi_unclassified(100);k__Fungi_unclassified(100);k__Fungi_unclassified(100); | - | - | - | - | - | - | - | Unassigned | - |
| Otu41454 | 1 | k__Fungi(100);k__Fungi_unclassified(100);k__Fungi_unclassified(100);k__Fungi_unclassified(100);k__Fungi_unclassified(100);k__Fungi_unclassified(100);k__Fungi_unclassified(100); | - | - | - | - | - | - | - | Unassigned | - |
| Otu41455 | 1 | k__Fungi(100);p__Ascomycota(100);p__Ascomycota_unclassified(100);p__Ascomycota_unclassified(100);p__Ascomycota_unclassified(100);p__Ascomycota_unclassified(100);p__Ascomycota_unclassified(100); | - | - | - | - | - | - | - | Unassigned | - |
| Otu41470 | 1 | k__Fungi(100);p__Ascomycota(100);p__Ascomycota_unclassified(100);p__Ascomycota_unclassified(100);p__Ascomycota_unclassified(100);p__Ascomycota_unclassified(100);p__Ascomycota_unclassified(100); | - | - | - | - | - | - | - | Unassigned | - |
| Otu41494 | 1 | k__Fungi(100);p__Ascomycota(100);p__Ascomycota_unclassified(100);p__Ascomycota_unclassified(100);p__Ascomycota_unclassified(100);p__Ascomycota_unclassified(100);p__Ascomycota_unclassified(100); | - | - | - | - | - | - | - | Unassigned | - |
| Otu41496 | 1 | k__Fungi(100);p__Ascomycota(100);p__Ascomycota_unclassified(100);p__Ascomycota_unclassified(100);p__Ascomycota_unclassified(100);p__Ascomycota_unclassified(100);p__Ascomycota_unclassified(100); | - | - | - | - | - | - | - | Unassigned | - |
| Otu41498 | 1 | k__Fungi(100);k__Fungi_unclassified(100);k__Fungi_unclassified(100);k__Fungi_unclassified(100);k__Fungi_unclassified(100);k__Fungi_unclassified(100);k__Fungi_unclassified(100); | - | - | - | - | - | - | - | Unassigned | - |
| Otu41508 | 1 | k__Fungi(100);p__Ascomycota(100);p__Ascomycota_unclassified(100);p__Ascomycota_unclassified(100);p__Ascomycota_unclassified(100);p__Ascomycota_unclassified(100);p__Ascomycota_unclassified(100); | - | - | - | - | - | - | - | Unassigned | - |
| Otu41518 | 1 | k__Fungi(100);k__Fungi_unclassified(100);k__Fungi_unclassified(100);k__Fungi_unclassified(100);k__Fungi_unclassified(100);k__Fungi_unclassified(100);k__Fungi_unclassified(100); | - | - | - | - | - | - | - | Unassigned | - |
| Otu41528 | 1 | k__Fungi(100);k__Fungi_unclassified(100);k__Fungi_unclassified(100);k__Fungi_unclassified(100);k__Fungi_unclassified(100);k__Fungi_unclassified(100);k__Fungi_unclassified(100); | - | - | - | - | - | - | - | Unassigned | - |
| Otu41529 | 1 | k__Fungi(100);k__Fungi_unclassified(100);k__Fungi_unclassified(100);k__Fungi_unclassified(100);k__Fungi_unclassified(100);k__Fungi_unclassified(100);k__Fungi_unclassified(100); | - | - | - | - | - | - | - | Unassigned | - |
| Otu41530 | 1 | k__Fungi(100);k__Fungi_unclassified(100);k__Fungi_unclassified(100);k__Fungi_unclassified(100);k__Fungi_unclassified(100);k__Fungi_unclassified(100);k__Fungi_unclassified(100); | - | - | - | - | - | - | - | Unassigned | - |
| Otu41533 | 1 | k__Fungi(100);p__Ascomycota(100);p__Ascomycota_unclassified(100);p__Ascomycota_unclassified(100);p__Ascomycota_unclassified(100);p__Ascomycota_unclassified(100);p__Ascomycota_unclassified(100); | - | - | - | - | - | - | - | Unassigned | - |
| Otu41539 | 1 | k__Fungi(100);k__Fungi_unclassified(100);k__Fungi_unclassified(100);k__Fungi_unclassified(100);k__Fungi_unclassified(100);k__Fungi_unclassified(100);k__Fungi_unclassified(100); | - | - | - | - | - | - | - | Unassigned | - |
| Otu41540 | 1 | k__Fungi(100);k__Fungi_unclassified(100);k__Fungi_unclassified(100);k__Fungi_unclassified(100);k__Fungi_unclassified(100);k__Fungi_unclassified(100);k__Fungi_unclassified(100); | - | - | - | - | - | - | - | Unassigned | - |
| Otu41541 | 1 | k__Fungi(100);p__Ascomycota(100);p__Ascomycota_unclassified(100);p__Ascomycota_unclassified(100);p__Ascomycota_unclassified(100);p__Ascomycota_unclassified(100);p__Ascomycota_unclassified(100); | - | - | - | - | - | - | - | Unassigned | - |
| Otu41547 | 1 | k__Fungi(100);k__Fungi_unclassified(100);k__Fungi_unclassified(100);k__Fungi_unclassified(100);k__Fungi_unclassified(100);k__Fungi_unclassified(100);k__Fungi_unclassified(100); | - | - | - | - | - | - | - | Unassigned | - |
| Otu41548 | 1 | k__Fungi(100);k__Fungi_unclassified(100);k__Fungi_unclassified(100);k__Fungi_unclassified(100);k__Fungi_unclassified(100);k__Fungi_unclassified(100);k__Fungi_unclassified(100); | - | - | - | - | - | - | - | Unassigned | - |
| Otu41549 | 1 | k__Fungi(100);k__Fungi_unclassified(100);k__Fungi_unclassified(100);k__Fungi_unclassified(100);k__Fungi_unclassified(100);k__Fungi_unclassified(100);k__Fungi_unclassified(100); | - | - | - | - | - | - | - | Unassigned | - |
| Otu41550 | 1 | k__Fungi(100);p__Ascomycota(100);p__Ascomycota_unclassified(100);p__Ascomycota_unclassified(100);p__Ascomycota_unclassified(100);p__Ascomycota_unclassified(100);p__Ascomycota_unclassified(100); | - | - | - | - | - | - | - | Unassigned | - |
| Otu41551 | 1 | k__Fungi(100);k__Fungi_unclassified(100);k__Fungi_unclassified(100);k__Fungi_unclassified(100);k__Fungi_unclassified(100);k__Fungi_unclassified(100);k__Fungi_unclassified(100); | - | - | - | - | - | - | - | Unassigned | - |
| Otu41552 | 1 | k__Fungi(100);k__Fungi_unclassified(100);k__Fungi_unclassified(100);k__Fungi_unclassified(100);k__Fungi_unclassified(100);k__Fungi_unclassified(100);k__Fungi_unclassified(100); | - | - | - | - | - | - | - | Unassigned | - |
| Otu41553 | 1 | k__Fungi(100);k__Fungi_unclassified(100);k__Fungi_unclassified(100);k__Fungi_unclassified(100);k__Fungi_unclassified(100);k__Fungi_unclassified(100);k__Fungi_unclassified(100); | - | - | - | - | - | - | - | Unassigned | - |
| Otu41555 | 1 | k__Fungi(100);p__Ascomycota(100);p__Ascomycota_unclassified(100);p__Ascomycota_unclassified(100);p__Ascomycota_unclassified(100);p__Ascomycota_unclassified(100);p__Ascomycota_unclassified(100); | - | - | - | - | - | - | - | Unassigned | - |
| Otu41556 | 1 | k__Fungi(100);k__Fungi_unclassified(100);k__Fungi_unclassified(100);k__Fungi_unclassified(100);k__Fungi_unclassified(100);k__Fungi_unclassified(100);k__Fungi_unclassified(100); | - | - | - | - | - | - | - | Unassigned | - |
| Otu41558 | 1 | k__Fungi(100);k__Fungi_unclassified(100);k__Fungi_unclassified(100);k__Fungi_unclassified(100);k__Fungi_unclassified(100);k__Fungi_unclassified(100);k__Fungi_unclassified(100); | - | - | - | - | - | - | - | Unassigned | - |
| Otu41560 | 1 | k__Fungi(100);p__Ascomycota(100);p__Ascomycota_unclassified(100);p__Ascomycota_unclassified(100);p__Ascomycota_unclassified(100);p__Ascomycota_unclassified(100);p__Ascomycota_unclassified(100); | - | - | - | - | - | - | - | Unassigned | - |
| Otu41561 | 1 | k__Fungi(100);k__Fungi_unclassified(100);k__Fungi_unclassified(100);k__Fungi_unclassified(100);k__Fungi_unclassified(100);k__Fungi_unclassified(100);k__Fungi_unclassified(100); | - | - | - | - | - | - | - | Unassigned | - |
| Otu41562 | 1 | k__Fungi(100);p__Ascomycota(100);p__Ascomycota_unclassified(100);p__Ascomycota_unclassified(100);p__Ascomycota_unclassified(100);p__Ascomycota_unclassified(100);p__Ascomycota_unclassified(100); | - | - | - | - | - | - | - | Unassigned | - |
| Otu41570 | 1 | k__Fungi(100);k__Fungi_unclassified(100);k__Fungi_unclassified(100);k__Fungi_unclassified(100);k__Fungi_unclassified(100);k__Fungi_unclassified(100);k__Fungi_unclassified(100); | - | - | - | - | - | - | - | Unassigned | - |
| Otu41577 | 1 | k__Fungi(100);p__Ascomycota(100);p__Ascomycota_unclassified(100);p__Ascomycota_unclassified(100);p__Ascomycota_unclassified(100);p__Ascomycota_unclassified(100);p__Ascomycota_unclassified(100); | - | - | - | - | - | - | - | Unassigned | - |
| Otu41578 | 1 | k__Fungi(100);p__Ascomycota(100);p__Ascomycota_unclassified(100);p__Ascomycota_unclassified(100);p__Ascomycota_unclassified(100);p__Ascomycota_unclassified(100);p__Ascomycota_unclassified(100); | - | - | - | - | - | - | - | Unassigned | - |
| Otu41579 | 1 | k__Fungi(100);p__Ascomycota(100);p__Ascomycota_unclassified(100);p__Ascomycota_unclassified(100);p__Ascomycota_unclassified(100);p__Ascomycota_unclassified(100);p__Ascomycota_unclassified(100); | - | - | - | - | - | - | - | Unassigned | - |
| Otu41580 | 1 | k__Fungi(100);k__Fungi_unclassified(100);k__Fungi_unclassified(100);k__Fungi_unclassified(100);k__Fungi_unclassified(100);k__Fungi_unclassified(100);k__Fungi_unclassified(100); | - | - | - | - | - | - | - | Unassigned | - |
| Otu41581 | 1 | k__Fungi(100);k__Fungi_unclassified(100);k__Fungi_unclassified(100);k__Fungi_unclassified(100);k__Fungi_unclassified(100);k__Fungi_unclassified(100);k__Fungi_unclassified(100); | - | - | - | - | - | - | - | Unassigned | - |
| Otu41582 | 1 | k__Fungi(100);k__Fungi_unclassified(100);k__Fungi_unclassified(100);k__Fungi_unclassified(100);k__Fungi_unclassified(100);k__Fungi_unclassified(100);k__Fungi_unclassified(100); | - | - | - | - | - | - | - | Unassigned | - |
| Otu41584 | 1 | k__Fungi(100);k__Fungi_unclassified(100);k__Fungi_unclassified(100);k__Fungi_unclassified(100);k__Fungi_unclassified(100);k__Fungi_unclassified(100);k__Fungi_unclassified(100); | - | - | - | - | - | - | - | Unassigned | - |
| Otu41585 | 1 | k__Fungi(100);p__Ascomycota(100);p__Ascomycota_unclassified(100);p__Ascomycota_unclassified(100);p__Ascomycota_unclassified(100);p__Ascomycota_unclassified(100);p__Ascomycota_unclassified(100); | - | - | - | - | - | - | - | Unassigned | - |
| Otu41586 | 1 | k__Fungi(100);p__Ascomycota(100);p__Ascomycota_unclassified(100);p__Ascomycota_unclassified(100);p__Ascomycota_unclassified(100);p__Ascomycota_unclassified(100);p__Ascomycota_unclassified(100); | - | - | - | - | - | - | - | Unassigned | - |
| Otu41587 | 1 | k__Fungi(100);p__Ascomycota(100);p__Ascomycota_unclassified(100);p__Ascomycota_unclassified(100);p__Ascomycota_unclassified(100);p__Ascomycota_unclassified(100);p__Ascomycota_unclassified(100); | - | - | - | - | - | - | - | Unassigned | - |
| Otu41589 | 1 | k__Fungi(100);k__Fungi_unclassified(100);k__Fungi_unclassified(100);k__Fungi_unclassified(100);k__Fungi_unclassified(100);k__Fungi_unclassified(100);k__Fungi_unclassified(100); | - | - | - | - | - | - | - | Unassigned | - |
| Otu41591 | 1 | k__Fungi(100);k__Fungi_unclassified(100);k__Fungi_unclassified(100);k__Fungi_unclassified(100);k__Fungi_unclassified(100);k__Fungi_unclassified(100);k__Fungi_unclassified(100); | - | - | - | - | - | - | - | Unassigned | - |
| Otu41592 | 1 | k__Fungi(100);k__Fungi_unclassified(100);k__Fungi_unclassified(100);k__Fungi_unclassified(100);k__Fungi_unclassified(100);k__Fungi_unclassified(100);k__Fungi_unclassified(100); | - | - | - | - | - | - | - | Unassigned | - |
| Otu41593 | 1 | k__Fungi(100);k__Fungi_unclassified(100);k__Fungi_unclassified(100);k__Fungi_unclassified(100);k__Fungi_unclassified(100);k__Fungi_unclassified(100);k__Fungi_unclassified(100); | - | - | - | - | - | - | - | Unassigned | - |
| Otu41623 | 1 | k__Fungi(100);k__Fungi_unclassified(100);k__Fungi_unclassified(100);k__Fungi_unclassified(100);k__Fungi_unclassified(100);k__Fungi_unclassified(100);k__Fungi_unclassified(100); | - | - | - | - | - | - | - | Unassigned | - |
| Otu41647 | 1 | k__Fungi(100);k__Fungi_unclassified(100);k__Fungi_unclassified(100);k__Fungi_unclassified(100);k__Fungi_unclassified(100);k__Fungi_unclassified(100);k__Fungi_unclassified(100); | - | - | - | - | - | - | - | Unassigned | - |
| Otu41740 | 1 | k__Fungi(100);k__Fungi_unclassified(100);k__Fungi_unclassified(100);k__Fungi_unclassified(100);k__Fungi_unclassified(100);k__Fungi_unclassified(100);k__Fungi_unclassified(100); | - | - | - | - | - | - | - | Unassigned | - |
| Otu41767 | 1 | k__Fungi(100);k__Fungi_unclassified(100);k__Fungi_unclassified(100);k__Fungi_unclassified(100);k__Fungi_unclassified(100);k__Fungi_unclassified(100);k__Fungi_unclassified(100); | - | - | - | - | - | - | - | Unassigned | - |
| Otu41771 | 1 | k__Fungi(100);p__Ascomycota(100);p__Ascomycota_unclassified(100);p__Ascomycota_unclassified(100);p__Ascomycota_unclassified(100);p__Ascomycota_unclassified(100);p__Ascomycota_unclassified(100); | - | - | - | - | - | - | - | Unassigned | - |
| Otu41821 | 1 | k__Fungi(100);k__Fungi_unclassified(100);k__Fungi_unclassified(100);k__Fungi_unclassified(100);k__Fungi_unclassified(100);k__Fungi_unclassified(100);k__Fungi_unclassified(100); | - | - | - | - | - | - | - | Unassigned | - |
| Otu41830 | 1 | k__Fungi(100);p__Ascomycota(100);p__Ascomycota_unclassified(100);p__Ascomycota_unclassified(100);p__Ascomycota_unclassified(100);p__Ascomycota_unclassified(100);p__Ascomycota_unclassified(100); | - | - | - | - | - | - | - | Unassigned | - |
| Otu41836 | 1 | k__Fungi(100);p__Ascomycota(100);p__Ascomycota_unclassified(100);p__Ascomycota_unclassified(100);p__Ascomycota_unclassified(100);p__Ascomycota_unclassified(100);p__Ascomycota_unclassified(100); | - | - | - | - | - | - | - | Unassigned | - |
| Otu41872 | 1 | k__Fungi(100);p__Ascomycota(100);p__Ascomycota_unclassified(100);p__Ascomycota_unclassified(100);p__Ascomycota_unclassified(100);p__Ascomycota_unclassified(100);p__Ascomycota_unclassified(100); | - | - | - | - | - | - | - | Unassigned | - |
| Otu41909 | 1 | k__Fungi(100);p__Ascomycota(100);c__Leotiomycetes(100);o__Rhytismatales(100);f__unclassified_Rhytismatales(100);g__unclassified_Rhytismatales(100);s__Rhytismatales_sp(100); | - | - | - | - | - | - | - | Unassigned | - |
| Otu41910 | 1 | k__Fungi(100);p__Ascomycota(100);c__Leotiomycetes(100);c__Leotiomycetes_unclassified(100);c__Leotiomycetes_unclassified(100);c__Leotiomycetes_unclassified(100);c__Leotiomycetes_unclassified(100); | - | - | - | - | - | - | - | Unassigned | - |
| Otu41920 | 1 | k__Fungi(100);p__Ascomycota(100);c__Dothideomycetes(100);o__Hysteriales(100);f__unclassified_Hysteriales(100);g__unclassified_Hysteriales(100);s__Hysteriales_sp(100); | - | - | - | - | - | - | - | Unassigned | - |
| Otu41932 | 1 | k__Fungi(100);k__Fungi_unclassified(100);k__Fungi_unclassified(100);k__Fungi_unclassified(100);k__Fungi_unclassified(100);k__Fungi_unclassified(100);k__Fungi_unclassified(100); | - | - | - | - | - | - | - | Unassigned | - |
| Otu41946 | 1 | k__Fungi(100);p__Ascomycota(100);p__Ascomycota_unclassified(100);p__Ascomycota_unclassified(100);p__Ascomycota_unclassified(100);p__Ascomycota_unclassified(100);p__Ascomycota_unclassified(100); | - | - | - | - | - | - | - | Unassigned | - |
| Otu41950 | 1 | k__Fungi(100);k__Fungi_unclassified(100);k__Fungi_unclassified(100);k__Fungi_unclassified(100);k__Fungi_unclassified(100);k__Fungi_unclassified(100);k__Fungi_unclassified(100); | - | - | - | - | - | - | - | Unassigned | - |
| Otu41951 | 1 | k__Fungi(100);k__Fungi_unclassified(100);k__Fungi_unclassified(100);k__Fungi_unclassified(100);k__Fungi_unclassified(100);k__Fungi_unclassified(100);k__Fungi_unclassified(100); | - | - | - | - | - | - | - | Unassigned | - |
| Otu41955 | 1 | k__Fungi(100);k__Fungi_unclassified(100);k__Fungi_unclassified(100);k__Fungi_unclassified(100);k__Fungi_unclassified(100);k__Fungi_unclassified(100);k__Fungi_unclassified(100); | - | - | - | - | - | - | - | Unassigned | - |
| Otu41956 | 1 | k__Fungi(100);p__Ascomycota(100);p__Ascomycota_unclassified(100);p__Ascomycota_unclassified(100);p__Ascomycota_unclassified(100);p__Ascomycota_unclassified(100);p__Ascomycota_unclassified(100); | - | - | - | - | - | - | - | Unassigned | - |
| Otu41958 | 1 | k__Fungi(100);p__Ascomycota(100);p__Ascomycota_unclassified(100);p__Ascomycota_unclassified(100);p__Ascomycota_unclassified(100);p__Ascomycota_unclassified(100);p__Ascomycota_unclassified(100); | - | - | - | - | - | - | - | Unassigned | - |
| Otu41960 | 1 | k__Fungi(100);p__Ascomycota(100);p__Ascomycota_unclassified(100);p__Ascomycota_unclassified(100);p__Ascomycota_unclassified(100);p__Ascomycota_unclassified(100);p__Ascomycota_unclassified(100); | - | - | - | - | - | - | - | Unassigned | - |
| Otu41986 | 1 | k__Fungi(100);k__Fungi_unclassified(100);k__Fungi_unclassified(100);k__Fungi_unclassified(100);k__Fungi_unclassified(100);k__Fungi_unclassified(100);k__Fungi_unclassified(100); | - | - | - | - | - | - | - | Unassigned | - |
| Otu41987 | 1 | k__Fungi(100);k__Fungi_unclassified(100);k__Fungi_unclassified(100);k__Fungi_unclassified(100);k__Fungi_unclassified(100);k__Fungi_unclassified(100);k__Fungi_unclassified(100); | - | - | - | - | - | - | - | Unassigned | - |
| Otu42012 | 1 | k__Fungi(100);k__Fungi_unclassified(100);k__Fungi_unclassified(100);k__Fungi_unclassified(100);k__Fungi_unclassified(100);k__Fungi_unclassified(100);k__Fungi_unclassified(100); | - | - | - | - | - | - | - | Unassigned | - |
| Otu42040 | 1 | k__Fungi(100);p__Ascomycota(100);p__Ascomycota_unclassified(100);p__Ascomycota_unclassified(100);p__Ascomycota_unclassified(100);p__Ascomycota_unclassified(100);p__Ascomycota_unclassified(100); | - | - | - | - | - | - | - | Unassigned | - |
| Otu42045 | 1 | k__Fungi(100);k__Fungi_unclassified(100);k__Fungi_unclassified(100);k__Fungi_unclassified(100);k__Fungi_unclassified(100);k__Fungi_unclassified(100);k__Fungi_unclassified(100); | - | - | - | - | - | - | - | Unassigned | - |
| Otu42054 | 1 | k__Fungi(100);k__Fungi_unclassified(100);k__Fungi_unclassified(100);k__Fungi_unclassified(100);k__Fungi_unclassified(100);k__Fungi_unclassified(100);k__Fungi_unclassified(100); | - | - | - | - | - | - | - | Unassigned | - |
| Otu42122 | 1 | k__Fungi(100);k__Fungi_unclassified(100);k__Fungi_unclassified(100);k__Fungi_unclassified(100);k__Fungi_unclassified(100);k__Fungi_unclassified(100);k__Fungi_unclassified(100); | - | - | - | - | - | - | - | Unassigned | - |
| Otu42124 | 1 | k__Fungi(100);k__Fungi_unclassified(100);k__Fungi_unclassified(100);k__Fungi_unclassified(100);k__Fungi_unclassified(100);k__Fungi_unclassified(100);k__Fungi_unclassified(100); | - | - | - | - | - | - | - | Unassigned | - |
| Otu42128 | 1 | k__Fungi(100);p__Ascomycota(100);p__Ascomycota_unclassified(100);p__Ascomycota_unclassified(100);p__Ascomycota_unclassified(100);p__Ascomycota_unclassified(100);p__Ascomycota_unclassified(100); | - | - | - | - | - | - | - | Unassigned | - |
| Otu42129 | 1 | k__Fungi(100);p__Ascomycota(100);p__Ascomycota_unclassified(100);p__Ascomycota_unclassified(100);p__Ascomycota_unclassified(100);p__Ascomycota_unclassified(100);p__Ascomycota_unclassified(100); | - | - | - | - | - | - | - | Unassigned | - |
| Otu42137 | 1 | k__Fungi(100);k__Fungi_unclassified(100);k__Fungi_unclassified(100);k__Fungi_unclassified(100);k__Fungi_unclassified(100);k__Fungi_unclassified(100);k__Fungi_unclassified(100); | - | - | - | - | - | - | - | Unassigned | - |
| Otu42138 | 1 | k__Fungi(100);k__Fungi_unclassified(100);k__Fungi_unclassified(100);k__Fungi_unclassified(100);k__Fungi_unclassified(100);k__Fungi_unclassified(100);k__Fungi_unclassified(100); | - | - | - | - | - | - | - | Unassigned | - |
| Otu42140 | 1 | k__Fungi(100);p__Ascomycota(100);p__Ascomycota_unclassified(100);p__Ascomycota_unclassified(100);p__Ascomycota_unclassified(100);p__Ascomycota_unclassified(100);p__Ascomycota_unclassified(100); | - | - | - | - | - | - | - | Unassigned | - |
| Otu42149 | 1 | k__Fungi(100);p__Ascomycota(100);p__Ascomycota_unclassified(100);p__Ascomycota_unclassified(100);p__Ascomycota_unclassified(100);p__Ascomycota_unclassified(100);p__Ascomycota_unclassified(100); | - | - | - | - | - | - | - | Unassigned | - |
| Otu42150 | 1 | k__Fungi(100);k__Fungi_unclassified(100);k__Fungi_unclassified(100);k__Fungi_unclassified(100);k__Fungi_unclassified(100);k__Fungi_unclassified(100);k__Fungi_unclassified(100); | - | - | - | - | - | - | - | Unassigned | - |
| Otu42151 | 1 | k__Fungi(100);k__Fungi_unclassified(100);k__Fungi_unclassified(100);k__Fungi_unclassified(100);k__Fungi_unclassified(100);k__Fungi_unclassified(100);k__Fungi_unclassified(100); | - | - | - | - | - | - | - | Unassigned | - |
| Otu42152 | 1 | k__Fungi(100);k__Fungi_unclassified(100);k__Fungi_unclassified(100);k__Fungi_unclassified(100);k__Fungi_unclassified(100);k__Fungi_unclassified(100);k__Fungi_unclassified(100); | - | - | - | - | - | - | - | Unassigned | - |
| Otu42154 | 1 | k__Fungi(100);k__Fungi_unclassified(100);k__Fungi_unclassified(100);k__Fungi_unclassified(100);k__Fungi_unclassified(100);k__Fungi_unclassified(100);k__Fungi_unclassified(100); | - | - | - | - | - | - | - | Unassigned | - |
| Otu42155 | 1 | k__Fungi(100);k__Fungi_unclassified(100);k__Fungi_unclassified(100);k__Fungi_unclassified(100);k__Fungi_unclassified(100);k__Fungi_unclassified(100);k__Fungi_unclassified(100); | - | - | - | - | - | - | - | Unassigned | - |
| Otu42156 | 1 | k__Fungi(100);p__Ascomycota(100);p__Ascomycota_unclassified(100);p__Ascomycota_unclassified(100);p__Ascomycota_unclassified(100);p__Ascomycota_unclassified(100);p__Ascomycota_unclassified(100); | - | - | - | - | - | - | - | Unassigned | - |
| Otu42157 | 1 | k__Fungi(100);p__Ascomycota(100);p__Ascomycota_unclassified(100);p__Ascomycota_unclassified(100);p__Ascomycota_unclassified(100);p__Ascomycota_unclassified(100);p__Ascomycota_unclassified(100); | - | - | - | - | - | - | - | Unassigned | - |
| Otu42158 | 1 | k__Fungi(100);p__Ascomycota(100);p__Ascomycota_unclassified(100);p__Ascomycota_unclassified(100);p__Ascomycota_unclassified(100);p__Ascomycota_unclassified(100);p__Ascomycota_unclassified(100); | - | - | - | - | - | - | - | Unassigned | - |
| Otu42159 | 1 | k__Fungi(100);p__Ascomycota(100);p__Ascomycota_unclassified(100);p__Ascomycota_unclassified(100);p__Ascomycota_unclassified(100);p__Ascomycota_unclassified(100);p__Ascomycota_unclassified(100); | - | - | - | - | - | - | - | Unassigned | - |
| Otu42160 | 1 | k__Fungi(100);k__Fungi_unclassified(100);k__Fungi_unclassified(100);k__Fungi_unclassified(100);k__Fungi_unclassified(100);k__Fungi_unclassified(100);k__Fungi_unclassified(100); | - | - | - | - | - | - | - | Unassigned | - |
| Otu42161 | 1 | k__Fungi(100);p__Ascomycota(100);p__Ascomycota_unclassified(100);p__Ascomycota_unclassified(100);p__Ascomycota_unclassified(100);p__Ascomycota_unclassified(100);p__Ascomycota_unclassified(100); | - | - | - | - | - | - | - | Unassigned | - |
| Otu42162 | 1 | k__Fungi(100);p__Ascomycota(100);p__Ascomycota_unclassified(100);p__Ascomycota_unclassified(100);p__Ascomycota_unclassified(100);p__Ascomycota_unclassified(100);p__Ascomycota_unclassified(100); | - | - | - | - | - | - | - | Unassigned | - |
| Otu42163 | 1 | k__Fungi(100);p__Ascomycota(100);p__Ascomycota_unclassified(100);p__Ascomycota_unclassified(100);p__Ascomycota_unclassified(100);p__Ascomycota_unclassified(100);p__Ascomycota_unclassified(100); | - | - | - | - | - | - | - | Unassigned | - |
| Otu42166 | 1 | k__Fungi(100);k__Fungi_unclassified(100);k__Fungi_unclassified(100);k__Fungi_unclassified(100);k__Fungi_unclassified(100);k__Fungi_unclassified(100);k__Fungi_unclassified(100); | - | - | - | - | - | - | - | Unassigned | - |
| Otu42168 | 1 | k__Fungi(100);k__Fungi_unclassified(100);k__Fungi_unclassified(100);k__Fungi_unclassified(100);k__Fungi_unclassified(100);k__Fungi_unclassified(100);k__Fungi_unclassified(100); | - | - | - | - | - | - | - | Unassigned | - |
| Otu42171 | 1 | k__Fungi(100);k__Fungi_unclassified(100);k__Fungi_unclassified(100);k__Fungi_unclassified(100);k__Fungi_unclassified(100);k__Fungi_unclassified(100);k__Fungi_unclassified(100); | - | - | - | - | - | - | - | Unassigned | - |
| Otu42183 | 1 | k__Fungi(100);p__Basidiomycota(100);p__Basidiomycota_unclassified(100);p__Basidiomycota_unclassified(100);p__Basidiomycota_unclassified(100);p__Basidiomycota_unclassified(100);p__Basidiomycota_unclassified(100); | - | - | - | - | - | - | - | Unassigned | - |
| Otu42193 | 1 | k__Fungi(100);k__Fungi_unclassified(100);k__Fungi_unclassified(100);k__Fungi_unclassified(100);k__Fungi_unclassified(100);k__Fungi_unclassified(100);k__Fungi_unclassified(100); | - | - | - | - | - | - | - | Unassigned | - |
| Otu42219 | 1 | k__Fungi(100);k__Fungi_unclassified(100);k__Fungi_unclassified(100);k__Fungi_unclassified(100);k__Fungi_unclassified(100);k__Fungi_unclassified(100);k__Fungi_unclassified(100); | - | - | - | - | - | - | - | Unassigned | - |
| Otu42221 | 1 | k__Fungi(100);p__Ascomycota(100);p__Ascomycota_unclassified(100);p__Ascomycota_unclassified(100);p__Ascomycota_unclassified(100);p__Ascomycota_unclassified(100);p__Ascomycota_unclassified(100); | - | - | - | - | - | - | - | Unassigned | - |
| Otu42222 | 1 | k__Fungi(100);p__Ascomycota(100);p__Ascomycota_unclassified(100);p__Ascomycota_unclassified(100);p__Ascomycota_unclassified(100);p__Ascomycota_unclassified(100);p__Ascomycota_unclassified(100); | - | - | - | - | - | - | - | Unassigned | - |
| Otu42223 | 1 | k__Fungi(100);k__Fungi_unclassified(100);k__Fungi_unclassified(100);k__Fungi_unclassified(100);k__Fungi_unclassified(100);k__Fungi_unclassified(100);k__Fungi_unclassified(100); | - | - | - | - | - | - | - | Unassigned | - |
| Otu42224 | 1 | k__Fungi(100);k__Fungi_unclassified(100);k__Fungi_unclassified(100);k__Fungi_unclassified(100);k__Fungi_unclassified(100);k__Fungi_unclassified(100);k__Fungi_unclassified(100); | - | - | - | - | - | - | - | Unassigned | - |
| Otu42230 | 1 | k__Fungi(100);p__Ascomycota(100);p__Ascomycota_unclassified(100);p__Ascomycota_unclassified(100);p__Ascomycota_unclassified(100);p__Ascomycota_unclassified(100);p__Ascomycota_unclassified(100); | - | - | - | - | - | - | - | Unassigned | - |
| Otu42233 | 1 | k__Fungi(100);k__Fungi_unclassified(100);k__Fungi_unclassified(100);k__Fungi_unclassified(100);k__Fungi_unclassified(100);k__Fungi_unclassified(100);k__Fungi_unclassified(100); | - | - | - | - | - | - | - | Unassigned | - |
| Otu42234 | 1 | k__Fungi(100);k__Fungi_unclassified(100);k__Fungi_unclassified(100);k__Fungi_unclassified(100);k__Fungi_unclassified(100);k__Fungi_unclassified(100);k__Fungi_unclassified(100); | - | - | - | - | - | - | - | Unassigned | - |
| Otu42237 | 1 | k__Fungi(100);k__Fungi_unclassified(100);k__Fungi_unclassified(100);k__Fungi_unclassified(100);k__Fungi_unclassified(100);k__Fungi_unclassified(100);k__Fungi_unclassified(100); | - | - | - | - | - | - | - | Unassigned | - |
| Otu42238 | 1 | k__Fungi(100);p__Ascomycota(100);p__Ascomycota_unclassified(100);p__Ascomycota_unclassified(100);p__Ascomycota_unclassified(100);p__Ascomycota_unclassified(100);p__Ascomycota_unclassified(100); | - | - | - | - | - | - | - | Unassigned | - |
| Otu42239 | 1 | k__Fungi(100);k__Fungi_unclassified(100);k__Fungi_unclassified(100);k__Fungi_unclassified(100);k__Fungi_unclassified(100);k__Fungi_unclassified(100);k__Fungi_unclassified(100); | - | - | - | - | - | - | - | Unassigned | - |
| Otu42244 | 1 | k__Fungi(100);k__Fungi_unclassified(100);k__Fungi_unclassified(100);k__Fungi_unclassified(100);k__Fungi_unclassified(100);k__Fungi_unclassified(100);k__Fungi_unclassified(100); | - | - | - | - | - | - | - | Unassigned | - |
| Otu42252 | 1 | k__Fungi(100);p__Ascomycota(100);p__Ascomycota_unclassified(100);p__Ascomycota_unclassified(100);p__Ascomycota_unclassified(100);p__Ascomycota_unclassified(100);p__Ascomycota_unclassified(100); | - | - | - | - | - | - | - | Unassigned | - |
| Otu42253 | 1 | k__Fungi(100);p__Ascomycota(100);p__Ascomycota_unclassified(100);p__Ascomycota_unclassified(100);p__Ascomycota_unclassified(100);p__Ascomycota_unclassified(100);p__Ascomycota_unclassified(100); | - | - | - | - | - | - | - | Unassigned | - |
| Otu42254 | 1 | k__Fungi(100);k__Fungi_unclassified(100);k__Fungi_unclassified(100);k__Fungi_unclassified(100);k__Fungi_unclassified(100);k__Fungi_unclassified(100);k__Fungi_unclassified(100); | - | - | - | - | - | - | - | Unassigned | - |
| Otu42255 | 1 | k__Fungi(100);p__Ascomycota(100);p__Ascomycota_unclassified(100);p__Ascomycota_unclassified(100);p__Ascomycota_unclassified(100);p__Ascomycota_unclassified(100);p__Ascomycota_unclassified(100); | - | - | - | - | - | - | - | Unassigned | - |
| Otu42256 | 1 | k__Fungi(100);p__Ascomycota(100);p__Ascomycota_unclassified(100);p__Ascomycota_unclassified(100);p__Ascomycota_unclassified(100);p__Ascomycota_unclassified(100);p__Ascomycota_unclassified(100); | - | - | - | - | - | - | - | Unassigned | - |
| Otu42258 | 1 | k__Fungi(100);p__Ascomycota(100);p__Ascomycota_unclassified(100);p__Ascomycota_unclassified(100);p__Ascomycota_unclassified(100);p__Ascomycota_unclassified(100);p__Ascomycota_unclassified(100); | - | - | - | - | - | - | - | Unassigned | - |
| Otu42259 | 1 | k__Fungi(100);k__Fungi_unclassified(100);k__Fungi_unclassified(100);k__Fungi_unclassified(100);k__Fungi_unclassified(100);k__Fungi_unclassified(100);k__Fungi_unclassified(100); | - | - | - | - | - | - | - | Unassigned | - |
| Otu42260 | 1 | k__Fungi(100);p__Ascomycota(100);p__Ascomycota_unclassified(100);p__Ascomycota_unclassified(100);p__Ascomycota_unclassified(100);p__Ascomycota_unclassified(100);p__Ascomycota_unclassified(100); | - | - | - | - | - | - | - | Unassigned | - |
| Otu42261 | 1 | k__Fungi(100);k__Fungi_unclassified(100);k__Fungi_unclassified(100);k__Fungi_unclassified(100);k__Fungi_unclassified(100);k__Fungi_unclassified(100);k__Fungi_unclassified(100); | - | - | - | - | - | - | - | Unassigned | - |
| Otu42262 | 1 | k__Fungi(100);k__Fungi_unclassified(100);k__Fungi_unclassified(100);k__Fungi_unclassified(100);k__Fungi_unclassified(100);k__Fungi_unclassified(100);k__Fungi_unclassified(100); | - | - | - | - | - | - | - | Unassigned | - |
| Otu42264 | 1 | k__Fungi(100);k__Fungi_unclassified(100);k__Fungi_unclassified(100);k__Fungi_unclassified(100);k__Fungi_unclassified(100);k__Fungi_unclassified(100);k__Fungi_unclassified(100); | - | - | - | - | - | - | - | Unassigned | - |
| Otu42265 | 1 | k__Fungi(100);p__Ascomycota(100);p__Ascomycota_unclassified(100);p__Ascomycota_unclassified(100);p__Ascomycota_unclassified(100);p__Ascomycota_unclassified(100);p__Ascomycota_unclassified(100); | - | - | - | - | - | - | - | Unassigned | - |
| Otu42266 | 1 | k__Fungi(100);p__Ascomycota(100);p__Ascomycota_unclassified(100);p__Ascomycota_unclassified(100);p__Ascomycota_unclassified(100);p__Ascomycota_unclassified(100);p__Ascomycota_unclassified(100); | - | - | - | - | - | - | - | Unassigned | - |
| Otu42268 | 1 | k__Fungi(100);k__Fungi_unclassified(100);k__Fungi_unclassified(100);k__Fungi_unclassified(100);k__Fungi_unclassified(100);k__Fungi_unclassified(100);k__Fungi_unclassified(100); | - | - | - | - | - | - | - | Unassigned | - |
| Otu42322 | 1 | k__Fungi(100);p__Ascomycota(100);p__Ascomycota_unclassified(100);p__Ascomycota_unclassified(100);p__Ascomycota_unclassified(100);p__Ascomycota_unclassified(100);p__Ascomycota_unclassified(100); | - | - | - | - | - | - | - | Unassigned | - |
| Otu42331 | 1 | k__Fungi(100);p__Ascomycota(100);p__Ascomycota_unclassified(100);p__Ascomycota_unclassified(100);p__Ascomycota_unclassified(100);p__Ascomycota_unclassified(100);p__Ascomycota_unclassified(100); | - | - | - | - | - | - | - | Unassigned | - |
| Otu42332 | 1 | k__Fungi(100);k__Fungi_unclassified(100);k__Fungi_unclassified(100);k__Fungi_unclassified(100);k__Fungi_unclassified(100);k__Fungi_unclassified(100);k__Fungi_unclassified(100); | - | - | - | - | - | - | - | Unassigned | - |
| Otu42334 | 1 | k__Fungi(100);p__Ascomycota(100);p__Ascomycota_unclassified(100);p__Ascomycota_unclassified(100);p__Ascomycota_unclassified(100);p__Ascomycota_unclassified(100);p__Ascomycota_unclassified(100); | - | - | - | - | - | - | - | Unassigned | - |
| Otu42336 | 1 | k__Fungi(100);k__Fungi_unclassified(100);k__Fungi_unclassified(100);k__Fungi_unclassified(100);k__Fungi_unclassified(100);k__Fungi_unclassified(100);k__Fungi_unclassified(100); | - | - | - | - | - | - | - | Unassigned | - |
| Otu42337 | 1 | k__Fungi(100);k__Fungi_unclassified(100);k__Fungi_unclassified(100);k__Fungi_unclassified(100);k__Fungi_unclassified(100);k__Fungi_unclassified(100);k__Fungi_unclassified(100); | - | - | - | - | - | - | - | Unassigned | - |
| Otu42338 | 1 | k__Fungi(100);k__Fungi_unclassified(100);k__Fungi_unclassified(100);k__Fungi_unclassified(100);k__Fungi_unclassified(100);k__Fungi_unclassified(100);k__Fungi_unclassified(100); | - | - | - | - | - | - | - | Unassigned | - |
| Otu42340 | 1 | k__Fungi(100);p__Ascomycota(100);p__Ascomycota_unclassified(100);p__Ascomycota_unclassified(100);p__Ascomycota_unclassified(100);p__Ascomycota_unclassified(100);p__Ascomycota_unclassified(100); | - | - | - | - | - | - | - | Unassigned | - |
| Otu42354 | 1 | k__Fungi(100);k__Fungi_unclassified(100);k__Fungi_unclassified(100);k__Fungi_unclassified(100);k__Fungi_unclassified(100);k__Fungi_unclassified(100);k__Fungi_unclassified(100); | - | - | - | - | - | - | - | Unassigned | - |
| Otu42359 | 1 | k__Fungi(100);k__Fungi_unclassified(100);k__Fungi_unclassified(100);k__Fungi_unclassified(100);k__Fungi_unclassified(100);k__Fungi_unclassified(100);k__Fungi_unclassified(100); | - | - | - | - | - | - | - | Unassigned | - |
| Otu42364 | 1 | k__Fungi(100);k__Fungi_unclassified(100);k__Fungi_unclassified(100);k__Fungi_unclassified(100);k__Fungi_unclassified(100);k__Fungi_unclassified(100);k__Fungi_unclassified(100); | - | - | - | - | - | - | - | Unassigned | - |
| Otu42376 | 1 | k__Fungi(100);k__Fungi_unclassified(100);k__Fungi_unclassified(100);k__Fungi_unclassified(100);k__Fungi_unclassified(100);k__Fungi_unclassified(100);k__Fungi_unclassified(100); | - | - | - | - | - | - | - | Unassigned | - |
| Otu42379 | 1 | k__Fungi(100);k__Fungi_unclassified(100);k__Fungi_unclassified(100);k__Fungi_unclassified(100);k__Fungi_unclassified(100);k__Fungi_unclassified(100);k__Fungi_unclassified(100); | - | - | - | - | - | - | - | Unassigned | - |
| Otu42380 | 1 | k__Fungi(100);k__Fungi_unclassified(100);k__Fungi_unclassified(100);k__Fungi_unclassified(100);k__Fungi_unclassified(100);k__Fungi_unclassified(100);k__Fungi_unclassified(100); | - | - | - | - | - | - | - | Unassigned | - |
| Otu42381 | 1 | k__Fungi(100);p__Ascomycota(100);p__Ascomycota_unclassified(100);p__Ascomycota_unclassified(100);p__Ascomycota_unclassified(100);p__Ascomycota_unclassified(100);p__Ascomycota_unclassified(100); | - | - | - | - | - | - | - | Unassigned | - |
| Otu42382 | 1 | k__Fungi(100);k__Fungi_unclassified(100);k__Fungi_unclassified(100);k__Fungi_unclassified(100);k__Fungi_unclassified(100);k__Fungi_unclassified(100);k__Fungi_unclassified(100); | - | - | - | - | - | - | - | Unassigned | - |
| Otu42386 | 1 | k__Fungi(100);k__Fungi_unclassified(100);k__Fungi_unclassified(100);k__Fungi_unclassified(100);k__Fungi_unclassified(100);k__Fungi_unclassified(100);k__Fungi_unclassified(100); | - | - | - | - | - | - | - | Unassigned | - |
| Otu42387 | 1 | k__Fungi(100);p__Ascomycota(100);p__Ascomycota_unclassified(100);p__Ascomycota_unclassified(100);p__Ascomycota_unclassified(100);p__Ascomycota_unclassified(100);p__Ascomycota_unclassified(100); | - | - | - | - | - | - | - | Unassigned | - |
| Otu42388 | 1 | k__Fungi(100);p__Ascomycota(100);p__Ascomycota_unclassified(100);p__Ascomycota_unclassified(100);p__Ascomycota_unclassified(100);p__Ascomycota_unclassified(100);p__Ascomycota_unclassified(100); | - | - | - | - | - | - | - | Unassigned | - |
| Otu42405 | 1 | k__Fungi(100);k__Fungi_unclassified(100);k__Fungi_unclassified(100);k__Fungi_unclassified(100);k__Fungi_unclassified(100);k__Fungi_unclassified(100);k__Fungi_unclassified(100); | - | - | - | - | - | - | - | Unassigned | - |
| Otu42408 | 1 | k__Fungi(100);p__Ascomycota(100);p__Ascomycota_unclassified(100);p__Ascomycota_unclassified(100);p__Ascomycota_unclassified(100);p__Ascomycota_unclassified(100);p__Ascomycota_unclassified(100); | - | - | - | - | - | - | - | Unassigned | - |
| Otu42409 | 1 | k__Fungi(100);p__Ascomycota(100);p__Ascomycota_unclassified(100);p__Ascomycota_unclassified(100);p__Ascomycota_unclassified(100);p__Ascomycota_unclassified(100);p__Ascomycota_unclassified(100); | - | - | - | - | - | - | - | Unassigned | - |
| Otu42416 | 1 | k__Fungi(100);k__Fungi_unclassified(100);k__Fungi_unclassified(100);k__Fungi_unclassified(100);k__Fungi_unclassified(100);k__Fungi_unclassified(100);k__Fungi_unclassified(100); | - | - | - | - | - | - | - | Unassigned | - |
| Otu42417 | 1 | k__Fungi(100);k__Fungi_unclassified(100);k__Fungi_unclassified(100);k__Fungi_unclassified(100);k__Fungi_unclassified(100);k__Fungi_unclassified(100);k__Fungi_unclassified(100); | - | - | - | - | - | - | - | Unassigned | - |
| Otu42419 | 1 | k__Fungi(100);k__Fungi_unclassified(100);k__Fungi_unclassified(100);k__Fungi_unclassified(100);k__Fungi_unclassified(100);k__Fungi_unclassified(100);k__Fungi_unclassified(100); | - | - | - | - | - | - | - | Unassigned | - |
| Otu42420 | 1 | k__Fungi(100);k__Fungi_unclassified(100);k__Fungi_unclassified(100);k__Fungi_unclassified(100);k__Fungi_unclassified(100);k__Fungi_unclassified(100);k__Fungi_unclassified(100); | - | - | - | - | - | - | - | Unassigned | - |
| Otu42421 | 1 | k__Fungi(100);k__Fungi_unclassified(100);k__Fungi_unclassified(100);k__Fungi_unclassified(100);k__Fungi_unclassified(100);k__Fungi_unclassified(100);k__Fungi_unclassified(100); | - | - | - | - | - | - | - | Unassigned | - |
| Otu42422 | 1 | k__Fungi(100);p__Ascomycota(100);p__Ascomycota_unclassified(100);p__Ascomycota_unclassified(100);p__Ascomycota_unclassified(100);p__Ascomycota_unclassified(100);p__Ascomycota_unclassified(100); | - | - | - | - | - | - | - | Unassigned | - |
| Otu42423 | 1 | k__Fungi(100);p__Ascomycota(100);p__Ascomycota_unclassified(100);p__Ascomycota_unclassified(100);p__Ascomycota_unclassified(100);p__Ascomycota_unclassified(100);p__Ascomycota_unclassified(100); | - | - | - | - | - | - | - | Unassigned | - |
| Otu42424 | 1 | k__Fungi(100);p__Ascomycota(100);p__Ascomycota_unclassified(100);p__Ascomycota_unclassified(100);p__Ascomycota_unclassified(100);p__Ascomycota_unclassified(100);p__Ascomycota_unclassified(100); | - | - | - | - | - | - | - | Unassigned | - |
| Otu42435 | 1 | k__Fungi(100);k__Fungi_unclassified(100);k__Fungi_unclassified(100);k__Fungi_unclassified(100);k__Fungi_unclassified(100);k__Fungi_unclassified(100);k__Fungi_unclassified(100); | - | - | - | - | - | - | - | Unassigned | - |
| Otu42437 | 1 | k__Fungi(100);p__Ascomycota(100);p__Ascomycota_unclassified(100);p__Ascomycota_unclassified(100);p__Ascomycota_unclassified(100);p__Ascomycota_unclassified(100);p__Ascomycota_unclassified(100); | - | - | - | - | - | - | - | Unassigned | - |
| Otu42438 | 1 | k__Fungi(100);k__Fungi_unclassified(100);k__Fungi_unclassified(100);k__Fungi_unclassified(100);k__Fungi_unclassified(100);k__Fungi_unclassified(100);k__Fungi_unclassified(100); | - | - | - | - | - | - | - | Unassigned | - |
| Otu42479 | 1 | k__Fungi(100);k__Fungi_unclassified(100);k__Fungi_unclassified(100);k__Fungi_unclassified(100);k__Fungi_unclassified(100);k__Fungi_unclassified(100);k__Fungi_unclassified(100); | - | - | - | - | - | - | - | Unassigned | - |
| Otu42480 | 1 | k__Fungi(100);p__Ascomycota(100);p__Ascomycota_unclassified(100);p__Ascomycota_unclassified(100);p__Ascomycota_unclassified(100);p__Ascomycota_unclassified(100);p__Ascomycota_unclassified(100); | - | - | - | - | - | - | - | Unassigned | - |
| Otu42481 | 1 | k__Fungi(100);p__Ascomycota(100);p__Ascomycota_unclassified(100);p__Ascomycota_unclassified(100);p__Ascomycota_unclassified(100);p__Ascomycota_unclassified(100);p__Ascomycota_unclassified(100); | - | - | - | - | - | - | - | Unassigned | - |
| Otu42484 | 1 | k__Fungi(100);k__Fungi_unclassified(100);k__Fungi_unclassified(100);k__Fungi_unclassified(100);k__Fungi_unclassified(100);k__Fungi_unclassified(100);k__Fungi_unclassified(100); | - | - | - | - | - | - | - | Unassigned | - |
| Otu42485 | 1 | k__Fungi(100);k__Fungi_unclassified(100);k__Fungi_unclassified(100);k__Fungi_unclassified(100);k__Fungi_unclassified(100);k__Fungi_unclassified(100);k__Fungi_unclassified(100); | - | - | - | - | - | - | - | Unassigned | - |
| Otu42486 | 1 | k__Fungi(100);p__Ascomycota(100);p__Ascomycota_unclassified(100);p__Ascomycota_unclassified(100);p__Ascomycota_unclassified(100);p__Ascomycota_unclassified(100);p__Ascomycota_unclassified(100); | - | - | - | - | - | - | - | Unassigned | - |
| Otu42491 | 1 | k__Fungi(100);k__Fungi_unclassified(100);k__Fungi_unclassified(100);k__Fungi_unclassified(100);k__Fungi_unclassified(100);k__Fungi_unclassified(100);k__Fungi_unclassified(100); | - | - | - | - | - | - | - | Unassigned | - |
| Otu42509 | 1 | k__Fungi(100);k__Fungi_unclassified(100);k__Fungi_unclassified(100);k__Fungi_unclassified(100);k__Fungi_unclassified(100);k__Fungi_unclassified(100);k__Fungi_unclassified(100); | - | - | - | - | - | - | - | Unassigned | - |
| Otu42522 | 1 | k__Fungi(100);p__Ascomycota(100);p__Ascomycota_unclassified(100);p__Ascomycota_unclassified(100);p__Ascomycota_unclassified(100);p__Ascomycota_unclassified(100);p__Ascomycota_unclassified(100); | - | - | - | - | - | - | - | Unassigned | - |
| Otu42523 | 1 | k__Fungi(100);p__Ascomycota(100);p__Ascomycota_unclassified(100);p__Ascomycota_unclassified(100);p__Ascomycota_unclassified(100);p__Ascomycota_unclassified(100);p__Ascomycota_unclassified(100); | - | - | - | - | - | - | - | Unassigned | - |
| Otu42527 | 1 | k__Fungi(100);k__Fungi_unclassified(100);k__Fungi_unclassified(100);k__Fungi_unclassified(100);k__Fungi_unclassified(100);k__Fungi_unclassified(100);k__Fungi_unclassified(100); | - | - | - | - | - | - | - | Unassigned | - |
| Otu42528 | 1 | k__Fungi(100);p__Ascomycota(100);p__Ascomycota_unclassified(100);p__Ascomycota_unclassified(100);p__Ascomycota_unclassified(100);p__Ascomycota_unclassified(100);p__Ascomycota_unclassified(100); | - | - | - | - | - | - | - | Unassigned | - |
| Otu42532 | 1 | k__Fungi(100);k__Fungi_unclassified(100);k__Fungi_unclassified(100);k__Fungi_unclassified(100);k__Fungi_unclassified(100);k__Fungi_unclassified(100);k__Fungi_unclassified(100); | - | - | - | - | - | - | - | Unassigned | - |
| Otu42602 | 1 | k__Fungi(100);p__Ascomycota(100);p__Ascomycota_unclassified(100);p__Ascomycota_unclassified(100);p__Ascomycota_unclassified(100);p__Ascomycota_unclassified(100);p__Ascomycota_unclassified(100); | - | - | - | - | - | - | - | Unassigned | - |
| Otu42648 | 1 | k__Fungi(100);k__Fungi_unclassified(100);k__Fungi_unclassified(100);k__Fungi_unclassified(100);k__Fungi_unclassified(100);k__Fungi_unclassified(100);k__Fungi_unclassified(100); | - | - | - | - | - | - | - | Unassigned | - |

| | | | | | | | | | | | |
|---|---|---|---|---|---|---|---|---|---|---|---|
| Otu42744 | 1 | k__Fungi(100);p__Ascomycota(100);p__Ascomycota_unclassified(100);p__Ascomycota_unclassified(100);p__Ascomycota_unclassified(100);p__Ascomycota_unclassified(100);p__Ascomycota_unclassified(100); | - | - | - | - | - | - | - | Unassigned | - |
| Otu42745 | 1 | k__Fungi(100);k__Fungi_unclassified(100);k__Fungi_unclassified(100);k__Fungi_unclassified(100);k__Fungi_unclassified(100);k__Fungi_unclassified(100);k__Fungi_unclassified(100); | - | - | - | - | - | - | - | Unassigned | - |
| Otu42748 | 1 | k__Fungi(100);k__Fungi_unclassified(100);k__Fungi_unclassified(100);k__Fungi_unclassified(100);k__Fungi_unclassified(100);k__Fungi_unclassified(100);k__Fungi_unclassified(100); | - | - | - | - | - | - | - | Unassigned | - |
| Otu42750 | 1 | k__Fungi(100);k__Fungi_unclassified(100);k__Fungi_unclassified(100);k__Fungi_unclassified(100);k__Fungi_unclassified(100);k__Fungi_unclassified(100);k__Fungi_unclassified(100); | - | - | - | - | - | - | - | Unassigned | - |
| Otu42751 | 1 | k__Fungi(100);k__Fungi_unclassified(100);k__Fungi_unclassified(100);k__Fungi_unclassified(100);k__Fungi_unclassified(100);k__Fungi_unclassified(100);k__Fungi_unclassified(100); | - | - | - | - | - | - | - | Unassigned | - |
| Otu42753 | 1 | k__Fungi(100);p__Ascomycota(100);p__Ascomycota_unclassified(100);p__Ascomycota_unclassified(100);p__Ascomycota_unclassified(100);p__Ascomycota_unclassified(100);p__Ascomycota_unclassified(100); | - | - | - | - | - | - | - | Unassigned | - |
| Otu42764 | 1 | k__Fungi(100);k__Fungi_unclassified(100);k__Fungi_unclassified(100);k__Fungi_unclassified(100);k__Fungi_unclassified(100);k__Fungi_unclassified(100);k__Fungi_unclassified(100); | - | - | - | - | - | - | - | Unassigned | - |
| Otu42773 | 1 | k__Fungi(100);p__Ascomycota(100);p__Ascomycota_unclassified(100);p__Ascomycota_unclassified(100);p__Ascomycota_unclassified(100);p__Ascomycota_unclassified(100);p__Ascomycota_unclassified(100); | - | - | - | - | - | - | - | Unassigned | - |
| Otu42804 | 1 | k__Fungi(100);k__Fungi_unclassified(100);k__Fungi_unclassified(100);k__Fungi_unclassified(100);k__Fungi_unclassified(100);k__Fungi_unclassified(100);k__Fungi_unclassified(100); | - | - | - | - | - | - | - | Unassigned | - |
| Otu42828 | 1 | k__Fungi(100);k__Fungi_unclassified(100);k__Fungi_unclassified(100);k__Fungi_unclassified(100);k__Fungi_unclassified(100);k__Fungi_unclassified(100);k__Fungi_unclassified(100); | - | - | - | - | - | - | - | Unassigned | - |
| Otu42842 | 1 | k__Fungi(100);p__Ascomycota(100);p__Ascomycota_unclassified(100);p__Ascomycota_unclassified(100);p__Ascomycota_unclassified(100);p__Ascomycota_unclassified(100);p__Ascomycota_unclassified(100); | - | - | - | - | - | - | - | Unassigned | - |
| Otu42846 | 1 | k__Fungi(100);p__Ascomycota(100);p__Ascomycota_unclassified(100);p__Ascomycota_unclassified(100);p__Ascomycota_unclassified(100);p__Ascomycota_unclassified(100);p__Ascomycota_unclassified(100); | - | - | - | - | - | - | - | Unassigned | - |
| Otu42863 | 1 | k__Fungi(100);k__Fungi_unclassified(100);k__Fungi_unclassified(100);k__Fungi_unclassified(100);k__Fungi_unclassified(100);k__Fungi_unclassified(100);k__Fungi_unclassified(100); | - | - | - | - | - | - | - | Unassigned | - |
| Otu42874 | 1 | k__Fungi(100);k__Fungi_unclassified(100);k__Fungi_unclassified(100);k__Fungi_unclassified(100);k__Fungi_unclassified(100);k__Fungi_unclassified(100);k__Fungi_unclassified(100); | - | - | - | - | - | - | - | Unassigned | - |
| Otu42879 | 1 | k__Fungi(100);p__Ascomycota(100);p__Ascomycota_unclassified(100);p__Ascomycota_unclassified(100);p__Ascomycota_unclassified(100);p__Ascomycota_unclassified(100);p__Ascomycota_unclassified(100); | - | - | - | - | - | - | - | Unassigned | - |
| Otu42881 | 1 | k__Fungi(100);p__Ascomycota(100);p__Ascomycota_unclassified(100);p__Ascomycota_unclassified(100);p__Ascomycota_unclassified(100);p__Ascomycota_unclassified(100);p__Ascomycota_unclassified(100); | - | - | - | - | - | - | - | Unassigned | - |
| Otu42882 | 1 | k__Fungi(100);k__Fungi_unclassified(100);k__Fungi_unclassified(100);k__Fungi_unclassified(100);k__Fungi_unclassified(100);k__Fungi_unclassified(100);k__Fungi_unclassified(100); | - | - | - | - | - | - | - | Unassigned | - |
| Otu42883 | 1 | k__Fungi(100);k__Fungi_unclassified(100);k__Fungi_unclassified(100);k__Fungi_unclassified(100);k__Fungi_unclassified(100);k__Fungi_unclassified(100);k__Fungi_unclassified(100); | - | - | - | - | - | - | - | Unassigned | - |
| Otu42885 | 1 | k__Fungi(100);k__Fungi_unclassified(100);k__Fungi_unclassified(100);k__Fungi_unclassified(100);k__Fungi_unclassified(100);k__Fungi_unclassified(100);k__Fungi_unclassified(100); | - | - | - | - | - | - | - | Unassigned | - |
| Otu42889 | 1 | k__Fungi(100);k__Fungi_unclassified(100);k__Fungi_unclassified(100);k__Fungi_unclassified(100);k__Fungi_unclassified(100);k__Fungi_unclassified(100);k__Fungi_unclassified(100); | - | - | - | - | - | - | - | Unassigned | - |
| Otu42895 | 1 | k__Fungi(100);p__Ascomycota(100);p__Ascomycota_unclassified(100);p__Ascomycota_unclassified(100);p__Ascomycota_unclassified(100);p__Ascomycota_unclassified(100);p__Ascomycota_unclassified(100); | - | - | - | - | - | - | - | Unassigned | - |
| Otu42896 | 1 | k__Fungi(100);k__Fungi_unclassified(100);k__Fungi_unclassified(100);k__Fungi_unclassified(100);k__Fungi_unclassified(100);k__Fungi_unclassified(100);k__Fungi_unclassified(100); | - | - | - | - | - | - | - | Unassigned | - |
| Otu42898 | 1 | k__Fungi(100);p__Ascomycota(100);p__Ascomycota_unclassified(100);p__Ascomycota_unclassified(100);p__Ascomycota_unclassified(100);p__Ascomycota_unclassified(100);p__Ascomycota_unclassified(100); | - | - | - | - | - | - | - | Unassigned | - |
| Otu42899 | 1 | k__Fungi(100);p__Ascomycota(100);p__Ascomycota_unclassified(100);p__Ascomycota_unclassified(100);p__Ascomycota_unclassified(100);p__Ascomycota_unclassified(100);p__Ascomycota_unclassified(100); | - | - | - | - | - | - | - | Unassigned | - |
| Otu42906 | 1 | k__Fungi(100);p__Ascomycota(100);p__Ascomycota_unclassified(100);p__Ascomycota_unclassified(100);p__Ascomycota_unclassified(100);p__Ascomycota_unclassified(100);p__Ascomycota_unclassified(100); | - | - | - | - | - | - | - | Unassigned | - |
| Otu42910 | 1 | k__Fungi(100);p__Ascomycota(100);p__Ascomycota_unclassified(100);p__Ascomycota_unclassified(100);p__Ascomycota_unclassified(100);p__Ascomycota_unclassified(100);p__Ascomycota_unclassified(100); | - | - | - | - | - | - | - | Unassigned | - |
| Otu42912 | 1 | k__Fungi(100);k__Fungi_unclassified(100);k__Fungi_unclassified(100);k__Fungi_unclassified(100);k__Fungi_unclassified(100);k__Fungi_unclassified(100);k__Fungi_unclassified(100); | - | - | - | - | - | - | - | Unassigned | - |
| Otu42918 | 1 | k__Fungi(100);k__Fungi_unclassified(100);k__Fungi_unclassified(100);k__Fungi_unclassified(100);k__Fungi_unclassified(100);k__Fungi_unclassified(100);k__Fungi_unclassified(100); | - | - | - | - | - | - | - | Unassigned | - |
| Otu42942 | 1 | k__Fungi(100);k__Fungi_unclassified(100);k__Fungi_unclassified(100);k__Fungi_unclassified(100);k__Fungi_unclassified(100);k__Fungi_unclassified(100);k__Fungi_unclassified(100); | - | - | - | - | - | - | - | Unassigned | - |
| Otu42943 | 1 | k__Fungi(100);p__Ascomycota(100);p__Ascomycota_unclassified(100);p__Ascomycota_unclassified(100);p__Ascomycota_unclassified(100);p__Ascomycota_unclassified(100);p__Ascomycota_unclassified(100); | - | - | - | - | - | - | - | Unassigned | - |
| Otu42955 | 1 | k__Fungi(100);k__Fungi_unclassified(100);k__Fungi_unclassified(100);k__Fungi_unclassified(100);k__Fungi_unclassified(100);k__Fungi_unclassified(100);k__Fungi_unclassified(100); | - | - | - | - | - | - | - | Unassigned | - |
| Otu42959 | 1 | k__Fungi(100);p__Ascomycota(100);p__Ascomycota_unclassified(100);p__Ascomycota_unclassified(100);p__Ascomycota_unclassified(100);p__Ascomycota_unclassified(100);p__Ascomycota_unclassified(100); | - | - | - | - | - | - | - | Unassigned | - |
| Otu42979 | 1 | k__Fungi(100);k__Fungi_unclassified(100);k__Fungi_unclassified(100);k__Fungi_unclassified(100);k__Fungi_unclassified(100);k__Fungi_unclassified(100);k__Fungi_unclassified(100); | - | - | - | - | - | - | - | Unassigned | - |
| Otu42987 | 1 | k__Fungi(100);k__Fungi_unclassified(100);k__Fungi_unclassified(100);k__Fungi_unclassified(100);k__Fungi_unclassified(100);k__Fungi_unclassified(100);k__Fungi_unclassified(100); | - | - | - | - | - | - | - | Unassigned | - |
| Otu43061 | 1 | k__Fungi(100);p__Ascomycota(100);p__Ascomycota_unclassified(100);p__Ascomycota_unclassified(100);p__Ascomycota_unclassified(100);p__Ascomycota_unclassified(100);p__Ascomycota_unclassified(100); | - | - | - | - | - | - | - | Unassigned | - |
| Otu43063 | 1 | k__Fungi(100);k__Fungi_unclassified(100);k__Fungi_unclassified(100);k__Fungi_unclassified(100);k__Fungi_unclassified(100);k__Fungi_unclassified(100);k__Fungi_unclassified(100); | - | - | - | - | - | - | - | Unassigned | - |
| Otu43082 | 1 | k__Fungi(100);k__Fungi_unclassified(100);k__Fungi_unclassified(100);k__Fungi_unclassified(100);k__Fungi_unclassified(100);k__Fungi_unclassified(100);k__Fungi_unclassified(100); | - | - | - | - | - | - | - | Unassigned | - |
| Otu43085 | 1 | k__Fungi(100);k__Fungi_unclassified(100);k__Fungi_unclassified(100);k__Fungi_unclassified(100);k__Fungi_unclassified(100);k__Fungi_unclassified(100);k__Fungi_unclassified(100); | - | - | - | - | - | - | - | Unassigned | - |
| Otu43098 | 1 | k__Fungi(100);p__Ascomycota(100);p__Ascomycota_unclassified(100);p__Ascomycota_unclassified(100);p__Ascomycota_unclassified(100);p__Ascomycota_unclassified(100);p__Ascomycota_unclassified(100); | - | - | - | - | - | - | - | Unassigned | - |
| Otu43151 | 1 | k__Fungi(100);p__Ascomycota(100);p__Ascomycota_unclassified(100);p__Ascomycota_unclassified(100);p__Ascomycota_unclassified(100);p__Ascomycota_unclassified(100);p__Ascomycota_unclassified(100); | - | - | - | - | - | - | - | Unassigned | - |
| Otu43152 | 1 | k__Fungi(100);p__Ascomycota(100);p__Ascomycota_unclassified(100);p__Ascomycota_unclassified(100);p__Ascomycota_unclassified(100);p__Ascomycota_unclassified(100);p__Ascomycota_unclassified(100); | - | - | - | - | - | - | - | Unassigned | - |
| Otu43153 | 1 | k__Fungi(100);k__Fungi_unclassified(100);k__Fungi_unclassified(100);k__Fungi_unclassified(100);k__Fungi_unclassified(100);k__Fungi_unclassified(100);k__Fungi_unclassified(100); | - | - | - | - | - | - | - | Unassigned | - |
| Otu43155 | 1 | k__Fungi(100);p__Ascomycota(100);p__Ascomycota_unclassified(100);p__Ascomycota_unclassified(100);p__Ascomycota_unclassified(100);p__Ascomycota_unclassified(100);p__Ascomycota_unclassified(100); | - | - | - | - | - | - | - | Unassigned | - |
| Otu43156 | 1 | k__Fungi(100);p__Ascomycota(100);p__Ascomycota_unclassified(100);p__Ascomycota_unclassified(100);p__Ascomycota_unclassified(100);p__Ascomycota_unclassified(100);p__Ascomycota_unclassified(100); | - | - | - | - | - | - | - | Unassigned | - |
| Otu43162 | 1 | k__Fungi(100);p__Ascomycota(100);p__Ascomycota_unclassified(100);p__Ascomycota_unclassified(100);p__Ascomycota_unclassified(100);p__Ascomycota_unclassified(100);p__Ascomycota_unclassified(100); | - | - | - | - | - | - | - | Unassigned | - |
| Otu43163 | 1 | k__Fungi(100);k__Fungi_unclassified(100);k__Fungi_unclassified(100);k__Fungi_unclassified(100);k__Fungi_unclassified(100);k__Fungi_unclassified(100);k__Fungi_unclassified(100); | - | - | - | - | - | - | - | Unassigned | - |
| Otu43164 | 1 | k__Fungi(100);p__Ascomycota(100);p__Ascomycota_unclassified(100);p__Ascomycota_unclassified(100);p__Ascomycota_unclassified(100);p__Ascomycota_unclassified(100);p__Ascomycota_unclassified(100); | - | - | - | - | - | - | - | Unassigned | - |
| Otu43165 | 1 | k__Fungi(100);p__Ascomycota(100);p__Ascomycota_unclassified(100);p__Ascomycota_unclassified(100);p__Ascomycota_unclassified(100);p__Ascomycota_unclassified(100);p__Ascomycota_unclassified(100); | - | - | - | - | - | - | - | Unassigned | - |
| Otu43167 | 1 | k__Fungi(100);p__Ascomycota(100);p__Ascomycota_unclassified(100);p__Ascomycota_unclassified(100);p__Ascomycota_unclassified(100);p__Ascomycota_unclassified(100);p__Ascomycota_unclassified(100); | - | - | - | - | - | - | - | Unassigned | - |
| Otu43169 | 1 | k__Fungi(100);k__Fungi_unclassified(100);k__Fungi_unclassified(100);k__Fungi_unclassified(100);k__Fungi_unclassified(100);k__Fungi_unclassified(100);k__Fungi_unclassified(100); | - | - | - | - | - | - | - | Unassigned | - |
| Otu43173 | 1 | k__Fungi(100);k__Fungi_unclassified(100);k__Fungi_unclassified(100);k__Fungi_unclassified(100);k__Fungi_unclassified(100);k__Fungi_unclassified(100);k__Fungi_unclassified(100); | - | - | - | - | - | - | - | Unassigned | - |
| Otu43177 | 1 | k__Fungi(100);k__Fungi_unclassified(100);k__Fungi_unclassified(100);k__Fungi_unclassified(100);k__Fungi_unclassified(100);k__Fungi_unclassified(100);k__Fungi_unclassified(100); | - | - | - | - | - | - | - | Unassigned | - |
| Otu43182 | 1 | k__Fungi(100);k__Fungi_unclassified(100);k__Fungi_unclassified(100);k__Fungi_unclassified(100);k__Fungi_unclassified(100);k__Fungi_unclassified(100);k__Fungi_unclassified(100); | - | - | - | - | - | - | - | Unassigned | - |
| Otu43184 | 1 | k__Fungi(100);p__Ascomycota(100);p__Ascomycota_unclassified(100);p__Ascomycota_unclassified(100);p__Ascomycota_unclassified(100);p__Ascomycota_unclassified(100);p__Ascomycota_unclassified(100); | - | - | - | - | - | - | - | Unassigned | - |
| Otu43185 | 1 | k__Fungi(100);k__Fungi_unclassified(100);k__Fungi_unclassified(100);k__Fungi_unclassified(100);k__Fungi_unclassified(100);k__Fungi_unclassified(100);k__Fungi_unclassified(100); | - | - | - | - | - | - | - | Unassigned | - |
| Otu43189 | 1 | k__Fungi(100);k__Fungi_unclassified(100);k__Fungi_unclassified(100);k__Fungi_unclassified(100);k__Fungi_unclassified(100);k__Fungi_unclassified(100);k__Fungi_unclassified(100); | - | - | - | - | - | - | - | Unassigned | - |
| Otu43196 | 1 | k__Fungi(100);k__Fungi_unclassified(100);k__Fungi_unclassified(100);k__Fungi_unclassified(100);k__Fungi_unclassified(100);k__Fungi_unclassified(100);k__Fungi_unclassified(100); | - | - | - | - | - | - | - | Unassigned | - |
| Otu43199 | 1 | k__Fungi(100);p__Ascomycota(100);p__Ascomycota_unclassified(100);p__Ascomycota_unclassified(100);p__Ascomycota_unclassified(100);p__Ascomycota_unclassified(100);p__Ascomycota_unclassified(100); | - | - | - | - | - | - | - | Unassigned | - |
| Otu43200 | 1 | k__Fungi(100);k__Fungi_unclassified(100);k__Fungi_unclassified(100);k__Fungi_unclassified(100);k__Fungi_unclassified(100);k__Fungi_unclassified(100);k__Fungi_unclassified(100); | - | - | - | - | - | - | - | Unassigned | - |
| Otu43201 | 1 | k__Fungi(100);p__Ascomycota(100);p__Ascomycota_unclassified(100);p__Ascomycota_unclassified(100);p__Ascomycota_unclassified(100);p__Ascomycota_unclassified(100);p__Ascomycota_unclassified(100); | - | - | - | - | - | - | - | Unassigned | - |
| Otu43202 | 1 | k__Fungi(100);p__Ascomycota(100);p__Ascomycota_unclassified(100);p__Ascomycota_unclassified(100);p__Ascomycota_unclassified(100);p__Ascomycota_unclassified(100);p__Ascomycota_unclassified(100); | - | - | - | - | - | - | - | Unassigned | - |
| Otu43211 | 1 | k__Fungi(100);k__Fungi_unclassified(100);k__Fungi_unclassified(100);k__Fungi_unclassified(100);k__Fungi_unclassified(100);k__Fungi_unclassified(100);k__Fungi_unclassified(100); | - | - | - | - | - | - | - | Unassigned | - |
| Otu43216 | 1 | k__Fungi(100);k__Fungi_unclassified(100);k__Fungi_unclassified(100);k__Fungi_unclassified(100);k__Fungi_unclassified(100);k__Fungi_unclassified(100);k__Fungi_unclassified(100); | - | - | - | - | - | - | - | Unassigned | - |
| Otu43218 | 1 | k__Fungi(100);k__Fungi_unclassified(100);k__Fungi_unclassified(100);k__Fungi_unclassified(100);k__Fungi_unclassified(100);k__Fungi_unclassified(100);k__Fungi_unclassified(100); | - | - | - | - | - | - | - | Unassigned | - |
| Otu43232 | 1 | k__Fungi(100);p__Ascomycota(100);p__Ascomycota_unclassified(100);p__Ascomycota_unclassified(100);p__Ascomycota_unclassified(100);p__Ascomycota_unclassified(100);p__Ascomycota_unclassified(100); | - | - | - | - | - | - | - | Unassigned | - |
| Otu43241 | 1 | k__Fungi(100);p__Ascomycota(100);p__Ascomycota_unclassified(100);p__Ascomycota_unclassified(100);p__Ascomycota_unclassified(100);p__Ascomycota_unclassified(100);p__Ascomycota_unclassified(100); | - | - | - | - | - | - | - | Unassigned | - |
| Otu43248 | 1 | k__Fungi(100);p__Ascomycota(100);p__Ascomycota_unclassified(100);p__Ascomycota_unclassified(100);p__Ascomycota_unclassified(100);p__Ascomycota_unclassified(100);p__Ascomycota_unclassified(100); | - | - | - | - | - | - | - | Unassigned | - |
| Otu43249 | 1 | k__Fungi(100);p__Ascomycota(100);p__Ascomycota_unclassified(100);p__Ascomycota_unclassified(100);p__Ascomycota_unclassified(100);p__Ascomycota_unclassified(100);p__Ascomycota_unclassified(100); | - | - | - | - | - | - | - | Unassigned | - |
| Otu43250 | 1 | k__Fungi(100);k__Fungi_unclassified(100);k__Fungi_unclassified(100);k__Fungi_unclassified(100);k__Fungi_unclassified(100);k__Fungi_unclassified(100);k__Fungi_unclassified(100); | - | - | - | - | - | - | - | Unassigned | - |
| Otu43251 | 1 | k__Fungi(100);k__Fungi_unclassified(100);k__Fungi_unclassified(100);k__Fungi_unclassified(100);k__Fungi_unclassified(100);k__Fungi_unclassified(100);k__Fungi_unclassified(100); | - | - | - | - | - | - | - | Unassigned | - |
| Otu43261 | 1 | k__Fungi(100);p__Ascomycota(100);p__Ascomycota_unclassified(100);p__Ascomycota_unclassified(100);p__Ascomycota_unclassified(100);p__Ascomycota_unclassified(100);p__Ascomycota_unclassified(100); | - | - | - | - | - | - | - | Unassigned | - |
| Otu43273 | 1 | k__Fungi(100);k__Fungi_unclassified(100);k__Fungi_unclassified(100);k__Fungi_unclassified(100);k__Fungi_unclassified(100);k__Fungi_unclassified(100);k__Fungi_unclassified(100); | - | - | - | - | - | - | - | Unassigned | - |
| Otu43274 | 1 | k__Fungi(100);p__Ascomycota(100);p__Ascomycota_unclassified(100);p__Ascomycota_unclassified(100);p__Ascomycota_unclassified(100);p__Ascomycota_unclassified(100);p__Ascomycota_unclassified(100); | - | - | - | - | - | - | - | Unassigned | - |
| Otu43308 | 1 | k__Fungi(100);k__Fungi_unclassified(100);k__Fungi_unclassified(100);k__Fungi_unclassified(100);k__Fungi_unclassified(100);k__Fungi_unclassified(100);k__Fungi_unclassified(100); | - | - | - | - | - | - | - | Unassigned | - |
| Otu43310 | 1 | k__Fungi(100);k__Fungi_unclassified(100);k__Fungi_unclassified(100);k__Fungi_unclassified(100);k__Fungi_unclassified(100);k__Fungi_unclassified(100);k__Fungi_unclassified(100); | - | - | - | - | - | - | - | Unassigned | - |
| Otu43311 | 1 | k__Fungi(100);k__Fungi_unclassified(100);k__Fungi_unclassified(100);k__Fungi_unclassified(100);k__Fungi_unclassified(100);k__Fungi_unclassified(100);k__Fungi_unclassified(100); | - | - | - | - | - | - | - | Unassigned | - |
| Otu43314 | 1 | k__Fungi(100);k__Fungi_unclassified(100);k__Fungi_unclassified(100);k__Fungi_unclassified(100);k__Fungi_unclassified(100);k__Fungi_unclassified(100);k__Fungi_unclassified(100); | - | - | - | - | - | - | - | Unassigned | - |
| Otu43320 | 1 | k__Fungi(100);k__Fungi_unclassified(100);k__Fungi_unclassified(100);k__Fungi_unclassified(100);k__Fungi_unclassified(100);k__Fungi_unclassified(100);k__Fungi_unclassified(100); | - | - | - | - | - | - | - | Unassigned | - |
| Otu43356 | 1 | k__Fungi(100);p__Ascomycota(100);p__Ascomycota_unclassified(100);p__Ascomycota_unclassified(100);p__Ascomycota_unclassified(100);p__Ascomycota_unclassified(100);p__Ascomycota_unclassified(100); | - | - | - | - | - | - | - | Unassigned | - |
| Otu43393 | 1 | k__Fungi(100);k__Fungi_unclassified(100);k__Fungi_unclassified(100);k__Fungi_unclassified(100);k__Fungi_unclassified(100);k__Fungi_unclassified(100);k__Fungi_unclassified(100); | - | - | - | - | - | - | - | Unassigned | - |
| Otu43410 | 1 | k__Fungi(100);k__Fungi_unclassified(100);k__Fungi_unclassified(100);k__Fungi_unclassified(100);k__Fungi_unclassified(100);k__Fungi_unclassified(100);k__Fungi_unclassified(100); | - | - | - | - | - | - | - | Unassigned | - |
| Otu43505 | 1 | k__Fungi(100);k__Fungi_unclassified(100);k__Fungi_unclassified(100);k__Fungi_unclassified(100);k__Fungi_unclassified(100);k__Fungi_unclassified(100);k__Fungi_unclassified(100); | - | - | - | - | - | - | - | Unassigned | - |
| Otu43525 | 1 | k__Fungi(100);p__Ascomycota(100);p__Ascomycota_unclassified(100);p__Ascomycota_unclassified(100);p__Ascomycota_unclassified(100);p__Ascomycota_unclassified(100);p__Ascomycota_unclassified(100); | - | - | - | - | - | - | - | Unassigned | - |
| Otu43551 | 1 | k__Fungi(100);p__Ascomycota(100);p__Ascomycota_unclassified(100);p__Ascomycota_unclassified(100);p__Ascomycota_unclassified(100);p__Ascomycota_unclassified(100);p__Ascomycota_unclassified(100); | - | - | - | - | - | - | - | Unassigned | - |
| Otu43558 | 1 | k__Fungi(100);p__Ascomycota(100);p__Ascomycota_unclassified(100);p__Ascomycota_unclassified(100);p__Ascomycota_unclassified(100);p__Ascomycota_unclassified(100);p__Ascomycota_unclassified(100); | - | - | - | - | - | - | - | Unassigned | - |
| Otu43567 | 1 | k__Fungi(100);k__Fungi_unclassified(100);k__Fungi_unclassified(100);k__Fungi_unclassified(100);k__Fungi_unclassified(100);k__Fungi_unclassified(100);k__Fungi_unclassified(100); | - | - | - | - | - | - | - | Unassigned | - |
| Otu43585 | 1 | k__Fungi(100);k__Fungi_unclassified(100);k__Fungi_unclassified(100);k__Fungi_unclassified(100);k__Fungi_unclassified(100);k__Fungi_unclassified(100);k__Fungi_unclassified(100); | - | - | - | - | - | - | - | Unassigned | - |
| Otu43613 | 1 | k__Fungi(100);k__Fungi_unclassified(100);k__Fungi_unclassified(100);k__Fungi_unclassified(100);k__Fungi_unclassified(100);k__Fungi_unclassified(100);k__Fungi_unclassified(100); | - | - | - | - | - | - | - | Unassigned | - |
| Otu43621 | 1 | k__Fungi(100);k__Fungi_unclassified(100);k__Fungi_unclassified(100);k__Fungi_unclassified(100);k__Fungi_unclassified(100);k__Fungi_unclassified(100);k__Fungi_unclassified(100); | - | - | - | - | - | - | - | Unassigned | - |
| Otu43623 | 1 | k__Fungi(100);k__Fungi_unclassified(100);k__Fungi_unclassified(100);k__Fungi_unclassified(100);k__Fungi_unclassified(100);k__Fungi_unclassified(100);k__Fungi_unclassified(100); | - | - | - | - | - | - | - | Unassigned | - |
| Otu43643 | 1 | k__Fungi(100);k__Fungi_unclassified(100);k__Fungi_unclassified(100);k__Fungi_unclassified(100);k__Fungi_unclassified(100);k__Fungi_unclassified(100);k__Fungi_unclassified(100); | - | - | - | - | - | - | - | Unassigned | - |
| Otu43653 | 1 | k__Fungi(100);p__Ascomycota(100);p__Ascomycota_unclassified(100);p__Ascomycota_unclassified(100);p__Ascomycota_unclassified(100);p__Ascomycota_unclassified(100);p__Ascomycota_unclassified(100); | - | - | - | - | - | - | - | Unassigned | - |
| Otu43658 | 1 | k__Fungi(100);k__Fungi_unclassified(100);k__Fungi_unclassified(100);k__Fungi_unclassified(100);k__Fungi_unclassified(100);k__Fungi_unclassified(100);k__Fungi_unclassified(100); | - | - | - | - | - | - | - | Unassigned | - |
| Otu43659 | 1 | k__Fungi(100);k__Fungi_unclassified(100);k__Fungi_unclassified(100);k__Fungi_unclassified(100);k__Fungi_unclassified(100);k__Fungi_unclassified(100);k__Fungi_unclassified(100); | - | - | - | - | - | - | - | Unassigned | - |
| Otu43673 | 1 | k__Fungi(100);k__Fungi_unclassified(100);k__Fungi_unclassified(100);k__Fungi_unclassified(100);k__Fungi_unclassified(100);k__Fungi_unclassified(100);k__Fungi_unclassified(100); | - | - | - | - | - | - | - | Unassigned | - |
| Otu43738 | 1 | k__Fungi(100);k__Fungi_unclassified(100);k__Fungi_unclassified(100);k__Fungi_unclassified(100);k__Fungi_unclassified(100);k__Fungi_unclassified(100);k__Fungi_unclassified(100); | - | - | - | - | - | - | - | Unassigned | - |
| Otu43742 | 1 | k__Fungi(100);p__Ascomycota(100);p__Ascomycota_unclassified(100);p__Ascomycota_unclassified(100);p__Ascomycota_unclassified(100);p__Ascomycota_unclassified(100);p__Ascomycota_unclassified(100); | - | - | - | - | - | - | - | Unassigned | - |
| Otu43779 | 1 | k__Fungi(100);p__Ascomycota(100);p__Ascomycota_unclassified(100);p__Ascomycota_unclassified(100);p__Ascomycota_unclassified(100);p__Ascomycota_unclassified(100);p__Ascomycota_unclassified(100); | - | - | - | - | - | - | - | Unassigned | - |
| Otu43845 | 1 | k__Fungi(100);k__Fungi_unclassified(100);k__Fungi_unclassified(100);k__Fungi_unclassified(100);k__Fungi_unclassified(100);k__Fungi_unclassified(100);k__Fungi_unclassified(100); | - | - | - | - | - | - | - | Unassigned | - |
| Otu43848 | 1 | k__Fungi(100);p__Ascomycota(100);p__Ascomycota_unclassified(100);p__Ascomycota_unclassified(100);p__Ascomycota_unclassified(100);p__Ascomycota_unclassified(100);p__Ascomycota_unclassified(100); | - | - | - | - | - | - | - | Unassigned | - |
| Otu43894 | 1 | k__Fungi(100);k__Fungi_unclassified(100);k__Fungi_unclassified(100);k__Fungi_unclassified(100);k__Fungi_unclassified(100);k__Fungi_unclassified(100);k__Fungi_unclassified(100); | - | - | - | - | - | - | - | Unassigned | - |
| Otu43895 | 1 | k__Fungi(100);k__Fungi_unclassified(100);k__Fungi_unclassified(100);k__Fungi_unclassified(100);k__Fungi_unclassified(100);k__Fungi_unclassified(100);k__Fungi_unclassified(100); | - | - | - | - | - | - | - | Unassigned | - |
| Otu43900 | 1 | k__Fungi(100);k__Fungi_unclassified(100);k__Fungi_unclassified(100);k__Fungi_unclassified(100);k__Fungi_unclassified(100);k__Fungi_unclassified(100);k__Fungi_unclassified(100); | - | - | - | - | - | - | - | Unassigned | - |
| Otu43915 | 1 | k__Fungi(100);k__Fungi_unclassified(100);k__Fungi_unclassified(100);k__Fungi_unclassified(100);k__Fungi_unclassified(100);k__Fungi_unclassified(100);k__Fungi_unclassified(100); | - | - | - | - | - | - | - | Unassigned | - |
| Otu43918 | 1 | k__Fungi(100);k__Fungi_unclassified(100);k__Fungi_unclassified(100);k__Fungi_unclassified(100);k__Fungi_unclassified(100);k__Fungi_unclassified(100);k__Fungi_unclassified(100); | - | - | - | - | - | - | - | Unassigned | - |
| Otu43919 | 1 | k__Fungi(100);k__Fungi_unclassified(100);k__Fungi_unclassified(100);k__Fungi_unclassified(100);k__Fungi_unclassified(100);k__Fungi_unclassified(100);k__Fungi_unclassified(100); | - | - | - | - | - | - | - | Unassigned | - |
| Otu43956 | 1 | k__Fungi(100);k__Fungi_unclassified(100);k__Fungi_unclassified(100);k__Fungi_unclassified(100);k__Fungi_unclassified(100);k__Fungi_unclassified(100);k__Fungi_unclassified(100); | - | - | - | - | - | - | - | Unassigned | - |
| Otu43972 | 1 | k__Fungi(100);k__Fungi_unclassified(100);k__Fungi_unclassified(100);k__Fungi_unclassified(100);k__Fungi_unclassified(100);k__Fungi_unclassified(100);k__Fungi_unclassified(100); | - | - | - | - | - | - | - | Unassigned | - |
| Otu44002 | 1 | k__Fungi(100);k__Fungi_unclassified(100);k__Fungi_unclassified(100);k__Fungi_unclassified(100);k__Fungi_unclassified(100);k__Fungi_unclassified(100);k__Fungi_unclassified(100); | - | - | - | - | - | - | - | Unassigned | - |
| Otu44008 | 1 | k__Fungi(100);p__Ascomycota(100);p__Ascomycota_unclassified(100);p__Ascomycota_unclassified(100);p__Ascomycota_unclassified(100);p__Ascomycota_unclassified(100);p__Ascomycota_unclassified(100); | - | - | - | - | - | - | - | Unassigned | - |
| Otu44009 | 1 | k__Fungi(100);p__Ascomycota(100);p__Ascomycota_unclassified(100);p__Ascomycota_unclassified(100);p__Ascomycota_unclassified(100);p__Ascomycota_unclassified(100);p__Ascomycota_unclassified(100); | - | - | - | - | - | - | - | Unassigned | - |
| Otu44031 | 1 | k__Fungi(100);k__Fungi_unclassified(100);k__Fungi_unclassified(100);k__Fungi_unclassified(100);k__Fungi_unclassified(100);k__Fungi_unclassified(100);k__Fungi_unclassified(100); | - | - | - | - | - | - | - | Unassigned | - |
| Otu44072 | 1 | k__Fungi(100);k__Fungi_unclassified(100);k__Fungi_unclassified(100);k__Fungi_unclassified(100);k__Fungi_unclassified(100);k__Fungi_unclassified(100);k__Fungi_unclassified(100); | - | - | - | - | - | - | - | Unassigned | - |
| Otu44088 | 1 | k__Fungi(100);p__Ascomycota(100);c__Dothideomycetes(100);c__Dothideomycetes_unclassified(100);c__Dothideomycetes_unclassified(100);c__Dothideomycetes_unclassified(100);c__Dothideomycetes_unclassified(100); | - | - | - | - | - | - | - | Unassigned | - |
| Otu44123 | 1 | k__Fungi(100);k__Fungi_unclassified(100);k__Fungi_unclassified(100);k__Fungi_unclassified(100);k__Fungi_unclassified(100);k__Fungi_unclassified(100);k__Fungi_unclassified(100); | - | - | - | - | - | - | - | Unassigned | - |
| Otu44131 | 1 | k__Fungi(100);k__Fungi_unclassified(100);k__Fungi_unclassified(100);k__Fungi_unclassified(100);k__Fungi_unclassified(100);k__Fungi_unclassified(100);k__Fungi_unclassified(100); | - | - | - | - | - | - | - | Unassigned | - |
| Otu44162 | 1 | k__Fungi(100);k__Fungi_unclassified(100);k__Fungi_unclassified(100);k__Fungi_unclassified(100);k__Fungi_unclassified(100);k__Fungi_unclassified(100);k__Fungi_unclassified(100); | - | - | - | - | - | - | - | Unassigned | - |
| Otu44176 | 1 | k__Fungi(100);k__Fungi_unclassified(100);k__Fungi_unclassified(100);k__Fungi_unclassified(100);k__Fungi_unclassified(100);k__Fungi_unclassified(100);k__Fungi_unclassified(100); | - | - | - | - | - | - | - | Unassigned | - |
| Otu44183 | 1 | k__Fungi(100);k__Fungi_unclassified(100);k__Fungi_unclassified(100);k__Fungi_unclassified(100);k__Fungi_unclassified(100);k__Fungi_unclassified(100);k__Fungi_unclassified(100); | - | - | - | - | - | - | - | Unassigned | - |
| Otu44301 | 1 | k__Fungi(100);k__Fungi_unclassified(100);k__Fungi_unclassified(100);k__Fungi_unclassified(100);k__Fungi_unclassified(100);k__Fungi_unclassified(100);k__Fungi_unclassified(100); | - | - | - | - | - | - | - | Unassigned | - |
| Otu44310 | 1 | k__Fungi(100);p__Ascomycota(100);p__Ascomycota_unclassified(100);p__Ascomycota_unclassified(100);p__Ascomycota_unclassified(100);p__Ascomycota_unclassified(100);p__Ascomycota_unclassified(100); | - | - | - | - | - | - | - | Unassigned | - |
| Otu44315 | 1 | k__Fungi(100);p__Ascomycota(100);p__Ascomycota_unclassified(100);p__Ascomycota_unclassified(100);p__Ascomycota_unclassified(100);p__Ascomycota_unclassified(100);p__Ascomycota_unclassified(100); | - | - | - | - | - | - | - | Unassigned | - |
| Otu44324 | 1 | k__Fungi(100);k__Fungi_unclassified(100);k__Fungi_unclassified(100);k__Fungi_unclassified(100);k__Fungi_unclassified(100);k__Fungi_unclassified(100);k__Fungi_unclassified(100); | - | - | - | - | - | - | - | Unassigned | - |
| Otu44327 | 1 | k__Fungi(100);p__Ascomycota(100);p__Ascomycota_unclassified(100);p__Ascomycota_unclassified(100);p__Ascomycota_unclassified(100);p__Ascomycota_unclassified(100);p__Ascomycota_unclassified(100); | - | - | - | - | - | - | - | Unassigned | - |
| Otu44418 | 1 | k__Fungi(100);p__Basidiomycota(100);p__Basidiomycota_unclassified(100);p__Basidiomycota_unclassified(100);p__Basidiomycota_unclassified(100);p__Basidiomycota_unclassified(100);p__Basidiomycota_unclassified(100); | - | - | - | - | - | - | - | Unassigned | - |
| Otu44421 | 1 | k__Fungi(100);p__Ascomycota(100);p__Ascomycota_unclassified(100);p__Ascomycota_unclassified(100);p__Ascomycota_unclassified(100);p__Ascomycota_unclassified(100);p__Ascomycota_unclassified(100); | - | - | - | - | - | - | - | Unassigned | - |
| Otu44422 | 1 | k__Fungi(100);p__Ascomycota(100);p__Ascomycota_unclassified(100);p__Ascomycota_unclassified(100);p__Ascomycota_unclassified(100);p__Ascomycota_unclassified(100);p__Ascomycota_unclassified(100); | - | - | - | - | - | - | - | Unassigned | - |
| Otu44432 | 1 | k__Fungi(100);p__Ascomycota(100);p__Ascomycota_unclassified(100);p__Ascomycota_unclassified(100);p__Ascomycota_unclassified(100);p__Ascomycota_unclassified(100);p__Ascomycota_unclassified(100); | - | - | - | - | - | - | - | Unassigned | - |
| Otu44433 | 1 | k__Fungi(100);p__Ascomycota(100);p__Ascomycota_unclassified(100);p__Ascomycota_unclassified(100);p__Ascomycota_unclassified(100);p__Ascomycota_unclassified(100);p__Ascomycota_unclassified(100); | - | - | - | - | - | - | - | Unassigned | - |
| Otu44466 | 1 | k__Fungi(100);k__Fungi_unclassified(100);k__Fungi_unclassified(100);k__Fungi_unclassified(100);k__Fungi_unclassified(100);k__Fungi_unclassified(100);k__Fungi_unclassified(100); | - | - | - | - | - | - | - | Unassigned | - |
| Otu44510 | 1 | k__Fungi(100);p__Ascomycota(100);p__Ascomycota_unclassified(100);p__Ascomycota_unclassified(100);p__Ascomycota_unclassified(100);p__Ascomycota_unclassified(100);p__Ascomycota_unclassified(100); | - | - | - | - | - | - | - | Unassigned | - |
| Otu44555 | 1 | k__Fungi(100);k__Fungi_unclassified(100);k__Fungi_unclassified(100);k__Fungi_unclassified(100);k__Fungi_unclassified(100);k__Fungi_unclassified(100);k__Fungi_unclassified(100); | - | - | - | - | - | - | - | Unassigned | - |
| Otu44563 | 1 | k__Fungi(100);p__Ascomycota(100);p__Ascomycota_unclassified(100);p__Ascomycota_unclassified(100);p__Ascomycota_unclassified(100);p__Ascomycota_unclassified(100);p__Ascomycota_unclassified(100); | - | - | - | - | - | - | - | Unassigned | - |
| Otu44566 | 1 | k__Fungi(100);p__Ascomycota(100);p__Ascomycota_unclassified(100);p__Ascomycota_unclassified(100);p__Ascomycota_unclassified(100);p__Ascomycota_unclassified(100);p__Ascomycota_unclassified(100); | - | - | - | - | - | - | - | Unassigned | - |
| Otu44567 | 1 | k__Fungi(100);p__Ascomycota(100);p__Ascomycota_unclassified(100);p__Ascomycota_unclassified(100);p__Ascomycota_unclassified(100);p__Ascomycota_unclassified(100);p__Ascomycota_unclassified(100); | - | - | - | - | - | - | - | Unassigned | - |
| Otu44568 | 1 | k__Fungi(100);p__Ascomycota(100);p__Ascomycota_unclassified(100);p__Ascomycota_unclassified(100);p__Ascomycota_unclassified(100);p__Ascomycota_unclassified(100);p__Ascomycota_unclassified(100); | - | - | - | - | - | - | - | Unassigned | - |
| Otu44588 | 1 | k__Fungi(100);k__Fungi_unclassified(100);k__Fungi_unclassified(100);k__Fungi_unclassified(100);k__Fungi_unclassified(100);k__Fungi_unclassified(100);k__Fungi_unclassified(100); | - | - | - | - | - | - | - | Unassigned | - |
| Otu44609 | 1 | k__Fungi(100);k__Fungi_unclassified(100);k__Fungi_unclassified(100);k__Fungi_unclassified(100);k__Fungi_unclassified(100);k__Fungi_unclassified(100);k__Fungi_unclassified(100); | - | - | - | - | - | - | - | Unassigned | - |
| Otu44675 | 1 | k__Fungi(100);p__Ascomycota(100);p__Ascomycota_unclassified(100);p__Ascomycota_unclassified(100);p__Ascomycota_unclassified(100);p__Ascomycota_unclassified(100);p__Ascomycota_unclassified(100); | - | - | - | - | - | - | - | Unassigned | - |
| Otu44705 | 1 | k__Fungi(100);p__Ascomycota(100);p__Ascomycota_unclassified(100);p__Ascomycota_unclassified(100);p__Ascomycota_unclassified(100);p__Ascomycota_unclassified(100);p__Ascomycota_unclassified(100); | - | - | - | - | - | - | - | Unassigned | - |
| Otu44723 | 1 | k__Fungi(100);k__Fungi_unclassified(100);k__Fungi_unclassified(100);k__Fungi_unclassified(100);k__Fungi_unclassified(100);k__Fungi_unclassified(100);k__Fungi_unclassified(100); | - | - | - | - | - | - | - | Unassigned | - |
| Otu44727 | 1 | k__Fungi(100);p__Ascomycota(100);p__Ascomycota_unclassified(100);p__Ascomycota_unclassified(100);p__Ascomycota_unclassified(100);p__Ascomycota_unclassified(100);p__Ascomycota_unclassified(100); | - | - | - | - | - | - | - | Unassigned | - |
| Otu44728 | 1 | k__Fungi(100);p__Ascomycota(100);p__Ascomycota_unclassified(100);p__Ascomycota_unclassified(100);p__Ascomycota_unclassified(100);p__Ascomycota_unclassified(100);p__Ascomycota_unclassified(100); | - | - | - | - | - | - | - | Unassigned | - |
| Otu44730 | 1 | k__Fungi(100);p__Ascomycota(100);c__Dothideomycetes(100);o__Pleosporales(100);f__Pleosporales_family_Incertae_sedis(100);f__Pleosporales_family_Incertae_sedis_unclassified(100);f__Pleosporales_family_Incertae_sedis_unclassified(100); | - | - | - | - | - | - | - | Unassigned | - |
| Otu44731 | 1 | k__Fungi(100);p__Ascomycota(100);c__Dothideomycetes(100);o__Pleosporales(100);o__Pleosporales_unclassified(100);o__Pleosporales_unclassified(100);o__Pleosporales_unclassified(100); | - | - | - | - | - | - | - | Unassigned | - |
| Otu44732 | 1 | k__Fungi(100);p__Ascomycota(100);c__Dothideomycetes(100);o__Myriangiales(100);f__unclassified_Myriangiales(100);g__unclassified_Myriangiales(100);s__Myriangiales_sp(100); | - | - | - | - | - | - | - | Unassigned | - |
| Otu44748 | 1 | k__Fungi(100);k__Fungi_unclassified(100);k__Fungi_unclassified(100);k__Fungi_unclassified(100);k__Fungi_unclassified(100);k__Fungi_unclassified(100);k__Fungi_unclassified(100); | - | - | - | - | - | - | - | Unassigned | - |
| Otu44755 | 1 | k__Fungi(100);k__Fungi_unclassified(100);k__Fungi_unclassified(100);k__Fungi_unclassified(100);k__Fungi_unclassified(100);k__Fungi_unclassified(100);k__Fungi_unclassified(100); | - | - | - | - | - | - | - | Unassigned | - |
| Otu44762 | 1 | k__Fungi(100);p__Ascomycota(100);p__Ascomycota_unclassified(100);p__Ascomycota_unclassified(100);p__Ascomycota_unclassified(100);p__Ascomycota_unclassified(100);p__Ascomycota_unclassified(100); | - | - | - | - | - | - | - | Unassigned | - |
| Otu44790 | 1 | k__Fungi(100);k__Fungi_unclassified(100);k__Fungi_unclassified(100);k__Fungi_unclassified(100);k__Fungi_unclassified(100);k__Fungi_unclassified(100);k__Fungi_unclassified(100); | - | - | - | - | - | - | - | Unassigned | - |
| Otu44795 | 1 | k__Fungi(100);p__Ascomycota(100);c__Dothideomycetes(100);o__Capnodiales(100);f__Capnodiales_family_Incertae_sedis(100);f__Capnodiales_family_Incertae_sedis_unclassified(100);f__Capnodiales_family_Incertae_sedis_unclassified(100); | - | - | - | - | - | - | - | Unassigned | - |
| Otu44820 | 1 | k__Fungi(100);k__Fungi_unclassified(100);k__Fungi_unclassified(100);k__Fungi_unclassified(100);k__Fungi_unclassified(100);k__Fungi_unclassified(100);k__Fungi_unclassified(100); | - | - | - | - | - | - | - | Unassigned | - |
| Otu44838 | 1 | k__Fungi(100);p__Ascomycota(100);p__Ascomycota_unclassified(100);p__Ascomycota_unclassified(100);p__Ascomycota_unclassified(100);p__Ascomycota_unclassified(100);p__Ascomycota_unclassified(100); | - | - | - | - | - | - | - | Unassigned | - |
| Otu44840 | 1 | k__Fungi(100);p__Ascomycota(100);p__Ascomycota_unclassified(100);p__Ascomycota_unclassified(100);p__Ascomycota_unclassified(100);p__Ascomycota_unclassified(100);p__Ascomycota_unclassified(100); | - | - | - | - | - | - | - | Unassigned | - |
| Otu44880 | 1 | k__Fungi(100);k__Fungi_unclassified(100);k__Fungi_unclassified(100);k__Fungi_unclassified(100);k__Fungi_unclassified(100);k__Fungi_unclassified(100);k__Fungi_unclassified(100); | - | - | - | - | - | - | - | Unassigned | - |
| Otu44946 | 1 | k__Fungi(100);k__Fungi_unclassified(100);k__Fungi_unclassified(100);k__Fungi_unclassified(100);k__Fungi_unclassified(100);k__Fungi_unclassified(100);k__Fungi_unclassified(100); | - | - | - | - | - | - | - | Unassigned | - |
| Otu44955 | 1 | k__Fungi(100);k__Fungi_unclassified(100);k__Fungi_unclassified(100);k__Fungi_unclassified(100);k__Fungi_unclassified(100);k__Fungi_unclassified(100);k__Fungi_unclassified(100); | - | - | - | - | - | - | - | Unassigned | - |
| Otu44975 | 1 | k__Fungi(100);k__Fungi_unclassified(100);k__Fungi_unclassified(100);k__Fungi_unclassified(100);k__Fungi_unclassified(100);k__Fungi_unclassified(100);k__Fungi_unclassified(100); | - | - | - | - | - | - | - | Unassigned | - |
| Otu44992 | 1 | k__Fungi(100);k__Fungi_unclassified(100);k__Fungi_unclassified(100);k__Fungi_unclassified(100);k__Fungi_unclassified(100);k__Fungi_unclassified(100);k__Fungi_unclassified(100); | - | - | - | - | - | - | - | Unassigned | - |
| Otu44993 | 1 | k__Fungi(100);k__Fungi_unclassified(100);k__Fungi_unclassified(100);k__Fungi_unclassified(100);k__Fungi_unclassified(100);k__Fungi_unclassified(100);k__Fungi_unclassified(100); | - | - | - | - | - | - | - | Unassigned | - |
| Otu45033 | 1 | k__Fungi(100);k__Fungi_unclassified(100);k__Fungi_unclassified(100);k__Fungi_unclassified(100);k__Fungi_unclassified(100);k__Fungi_unclassified(100);k__Fungi_unclassified(100); | - | - | - | - | - | - | - | Unassigned | - |
| Otu45215 | 1 | k__Fungi(100);p__Ascomycota(100);p__Ascomycota_unclassified(100);p__Ascomycota_unclassified(100);p__Ascomycota_unclassified(100);p__Ascomycota_unclassified(100);p__Ascomycota_unclassified(100); | - | - | - | - | - | - | - | Unassigned | - |
| Otu45228 | 1 | k__Fungi(100);p__Ascomycota(100);p__Ascomycota_unclassified(100);p__Ascomycota_unclassified(100);p__Ascomycota_unclassified(100);p__Ascomycota_unclassified(100);p__Ascomycota_unclassified(100); | - | - | - | - | - | - | - | Unassigned | - |
| Otu45261 | 1 | k__Fungi(100);k__Fungi_unclassified(100);k__Fungi_unclassified(100);k__Fungi_unclassified(100);k__Fungi_unclassified(100);k__Fungi_unclassified(100);k__Fungi_unclassified(100); | - | - | - | - | - | - | - | Unassigned | - |
| Otu45269 | 1 | k__Fungi(100);p__Ascomycota(100);p__Ascomycota_unclassified(100);p__Ascomycota_unclassified(100);p__Ascomycota_unclassified(100);p__Ascomycota_unclassified(100);p__Ascomycota_unclassified(100); | - | - | - | - | - | - | - | Unassigned | - |
| Otu45273 | 1 | k__Fungi(100);p__Ascomycota(100);p__Ascomycota_unclassified(100);p__Ascomycota_unclassified(100);p__Ascomycota_unclassified(100);p__Ascomycota_unclassified(100);p__Ascomycota_unclassified(100); | - | - | - | - | - | - | - | Unassigned | - |
| Otu45298 | 1 | k__Fungi(100);p__Ascomycota(100);c__Dothideomycetes(100);o__Pleosporales(100);o__Pleosporales_unclassified(100);o__Pleosporales_unclassified(100);o__Pleosporales_unclassified(100); | - | - | - | - | - | - | - | Unassigned | - |
| Otu45329 | 1 | k__Fungi(100);k__Fungi_unclassified(100);k__Fungi_unclassified(100);k__Fungi_unclassified(100);k__Fungi_unclassified(100);k__Fungi_unclassified(100);k__Fungi_unclassified(100); | - | - | - | - | - | - | - | Unassigned | - |
| Otu45411 | 1 | k__Fungi(100);k__Fungi_unclassified(100);k__Fungi_unclassified(100);k__Fungi_unclassified(100);k__Fungi_unclassified(100);k__Fungi_unclassified(100);k__Fungi_unclassified(100); | - | - | - | - | - | - | - | Unassigned | - |
| Otu45431 | 1 | k__Fungi(100);p__Ascomycota(100);p__Ascomycota_unclassified(100);p__Ascomycota_unclassified(100);p__Ascomycota_unclassified(100);p__Ascomycota_unclassified(100);p__Ascomycota_unclassified(100); | - | - | - | - | - | - | - | Unassigned | - |
| Otu45491 | 1 | k__Fungi(100);k__Fungi_unclassified(100);k__Fungi_unclassified(100);k__Fungi_unclassified(100);k__Fungi_unclassified(100);k__Fungi_unclassified(100);k__Fungi_unclassified(100); | - | - | - | - | - | - | - | Unassigned | - |
| Otu45512 | 1 | k__Fungi(100);k__Fungi_unclassified(100);k__Fungi_unclassified(100);k__Fungi_unclassified(100);k__Fungi_unclassified(100);k__Fungi_unclassified(100);k__Fungi_unclassified(100); | - | - | - | - | - | - | - | Unassigned | - |
| Otu45551 | 1 | k__Fungi(100);k__Fungi_unclassified(100);k__Fungi_unclassified(100);k__Fungi_unclassified(100);k__Fungi_unclassified(100);k__Fungi_unclassified(100);k__Fungi_unclassified(100); | - | - | - | - | - | - | - | Unassigned | - |
| Otu45564 | 1 | k__Fungi(100);p__Ascomycota(100);p__Ascomycota_unclassified(100);p__Ascomycota_unclassified(100);p__Ascomycota_unclassified(100);p__Ascomycota_unclassified(100);p__Ascomycota_unclassified(100); | - | - | - | - | - | - | - | Unassigned | - |
| Otu45579 | 1 | k__Fungi(100);k__Fungi_unclassified(100);k__Fungi_unclassified(100);k__Fungi_unclassified(100);k__Fungi_unclassified(100);k__Fungi_unclassified(100);k__Fungi_unclassified(100); | - | - | - | - | - | - | - | Unassigned | - |
| Otu45586 | 1 | k__Fungi(100);k__Fungi_unclassified(100);k__Fungi_unclassified(100);k__Fungi_unclassified(100);k__Fungi_unclassified(100);k__Fungi_unclassified(100);k__Fungi_unclassified(100); | - | - | - | - | - | - | - | Unassigned | - |
| Otu45590 | 1 | k__Fungi(100);k__Fungi_unclassified(100);k__Fungi_unclassified(100);k__Fungi_unclassified(100);k__Fungi_unclassified(100);k__Fungi_unclassified(100);k__Fungi_unclassified(100); | - | - | - | - | - | - | - | Unassigned | - |
| Otu45658 | 1 | k__Fungi(100);k__Fungi_unclassified(100);k__Fungi_unclassified(100);k__Fungi_unclassified(100);k__Fungi_unclassified(100);k__Fungi_unclassified(100);k__Fungi_unclassified(100); | - | - | - | - | - | - | - | Unassigned | - |
| Otu45660 | 1 | k__Fungi(100);p__Ascomycota(100);p__Ascomycota_unclassified(100);p__Ascomycota_unclassified(100);p__Ascomycota_unclassified(100);p__Ascomycota_unclassified(100);p__Ascomycota_unclassified(100); | - | - | - | - | - | - | - | Unassigned | - |
| Otu45715 | 1 | k__Fungi(100);k__Fungi_unclassified(100);k__Fungi_unclassified(100);k__Fungi_unclassified(100);k__Fungi_unclassified(100);k__Fungi_unclassified(100);k__Fungi_unclassified(100); | - | - | - | - | - | - | - | Unassigned | - |
| Otu45722 | 1 | k__Fungi(100);k__Fungi_unclassified(100);k__Fungi_unclassified(100);k__Fungi_unclassified(100);k__Fungi_unclassified(100);k__Fungi_unclassified(100);k__Fungi_unclassified(100); | - | - | - | - | - | - | - | Unassigned | - |
| Otu45742 | 1 | k__Fungi(100);k__Fungi_unclassified(100);k__Fungi_unclassified(100);k__Fungi_unclassified(100);k__Fungi_unclassified(100);k__Fungi_unclassified(100);k__Fungi_unclassified(100); | - | - | - | - | - | - | - | Unassigned | - |
| Otu45765 | 1 | k__Fungi(100);k__Fungi_unclassified(100);k__Fungi_unclassified(100);k__Fungi_unclassified(100);k__Fungi_unclassified(100);k__Fungi_unclassified(100);k__Fungi_unclassified(100); | - | - | - | - | - | - | - | Unassigned | - |
| Otu45813 | 1 | k__Fungi(100);p__Ascomycota(100);p__Ascomycota_unclassified(100);p__Ascomycota_unclassified(100);p__Ascomycota_unclassified(100);p__Ascomycota_unclassified(100);p__Ascomycota_unclassified(100); | - | - | - | - | - | - | - | Unassigned | - |
| Otu45814 | 1 | k__Fungi(100);p__Ascomycota(100);c__Sordariomycetes(100);o__Sordariales(100);o__Sordariales_unclassified(100);o__Sordariales_unclassified(100);o__Sordariales_unclassified(100); | - | - | - | - | - | - | - | Unassigned | - |
| Otu45816 | 1 | k__Fungi(100);p__Ascomycota(100);c__Dothideomycetes(100);o__Capnodiales(100);o__Capnodiales_unclassified(100);o__Capnodiales_unclassified(100);o__Capnodiales_unclassified(100); | - | - | - | - | - | - | - | Unassigned | - |
| Otu45823 | 1 | k__Fungi(100);k__Fungi_unclassified(100);k__Fungi_unclassified(100);k__Fungi_unclassified(100);k__Fungi_unclassified(100);k__Fungi_unclassified(100);k__Fungi_unclassified(100); | - | - | - | - | - | - | - | Unassigned | - |
| Otu45826 | 1 | k__Fungi(100);k__Fungi_unclassified(100);k__Fungi_unclassified(100);k__Fungi_unclassified(100);k__Fungi_unclassified(100);k__Fungi_unclassified(100);k__Fungi_unclassified(100); | - | - | - | - | - | - | - | Unassigned | - |
| Otu45828 | 1 | k__Fungi(100);k__Fungi_unclassified(100);k__Fungi_unclassified(100);k__Fungi_unclassified(100);k__Fungi_unclassified(100);k__Fungi_unclassified(100);k__Fungi_unclassified(100); | - | - | - | - | - | - | - | Unassigned | - |
| Otu45876 | 1 | k__Fungi(100);k__Fungi_unclassified(100);k__Fungi_unclassified(100);k__Fungi_unclassified(100);k__Fungi_unclassified(100);k__Fungi_unclassified(100);k__Fungi_unclassified(100); | - | - | - | - | - | - | - | Unassigned | - |
| Otu45929 | 1 | k__Fungi(100);p__Ascomycota(100);c__Dothideomycetes(100);o__Pleosporales(100);o__Pleosporales_unclassified(100);o__Pleosporales_unclassified(100);o__Pleosporales_unclassified(100); | - | - | - | - | - | - | - | Unassigned | - |
| Otu45981 | 1 | k__Fungi(100);k__Fungi_unclassified(100);k__Fungi_unclassified(100);k__Fungi_unclassified(100);k__Fungi_unclassified(100);k__Fungi_unclassified(100);k__Fungi_unclassified(100); | - | - | - | - | - | - | - | Unassigned | - |
| Otu45997 | 1 | k__Fungi(100);p__Ascomycota(100);p__Ascomycota_unclassified(100);p__Ascomycota_unclassified(100);p__Ascomycota_unclassified(100);p__Ascomycota_unclassified(100);p__Ascomycota_unclassified(100); | - | - | - | - | - | - | - | Unassigned | - |
| Otu46020 | 1 | k__Fungi(100);k__Fungi_unclassified(100);k__Fungi_unclassified(100);k__Fungi_unclassified(100);k__Fungi_unclassified(100);k__Fungi_unclassified(100);k__Fungi_unclassified(100); | - | - | - | - | - | - | - | Unassigned | - |
| Otu46075 | 1 | k__Fungi(100);k__Fungi_unclassified(100);k__Fungi_unclassified(100);k__Fungi_unclassified(100);k__Fungi_unclassified(100);k__Fungi_unclassified(100);k__Fungi_unclassified(100); | - | - | - | - | - | - | - | Unassigned | - |
| Otu46095 | 1 | k__Fungi(100);k__Fungi_unclassified(100);k__Fungi_unclassified(100);k__Fungi_unclassified(100);k__Fungi_unclassified(100);k__Fungi_unclassified(100);k__Fungi_unclassified(100); | - | - | - | - | - | - | - | Unassigned | - |
| Otu46109 | 1 | k__Fungi(100);p__Ascomycota(100);p__Ascomycota_unclassified(100);p__Ascomycota_unclassified(100);p__Ascomycota_unclassified(100);p__Ascomycota_unclassified(100);p__Ascomycota_unclassified(100); | - | - | - | - | - | - | - | Unassigned | - |
| Otu46114 | 1 | k__Fungi(100);k__Fungi_unclassified(100);k__Fungi_unclassified(100);k__Fungi_unclassified(100);k__Fungi_unclassified(100);k__Fungi_unclassified(100);k__Fungi_unclassified(100); | - | - | - | - | - | - | - | Unassigned | - |
| Otu46126 | 1 | k__Fungi(100);p__Ascomycota(100);p__Ascomycota_unclassified(100);p__Ascomycota_unclassified(100);p__Ascomycota_unclassified(100);p__Ascomycota_unclassified(100);p__Ascomycota_unclassified(100); | - | - | - | - | - | - | - | Unassigned | - |
| Otu46137 | 1 | k__Fungi(100);k__Fungi_unclassified(100);k__Fungi_unclassified(100);k__Fungi_unclassified(100);k__Fungi_unclassified(100);k__Fungi_unclassified(100);k__Fungi_unclassified(100); | - | - | - | - | - | - | - | Unassigned | - |
| Otu46139 | 1 | k__Fungi(100);p__Ascomycota(100);p__Ascomycota_unclassified(100);p__Ascomycota_unclassified(100);p__Ascomycota_unclassified(100);p__Ascomycota_unclassified(100);p__Ascomycota_unclassified(100); | - | - | - | - | - | - | - | Unassigned | - |
| Otu46145 | 1 | k__Fungi(100);k__Fungi_unclassified(100);k__Fungi_unclassified(100);k__Fungi_unclassified(100);k__Fungi_unclassified(100);k__Fungi_unclassified(100);k__Fungi_unclassified(100); | - | - | - | - | - | - | - | Unassigned | - |
| Otu46240 | 1 | k__Fungi(100);p__Ascomycota(100);p__Ascomycota_unclassified(100);p__Ascomycota_unclassified(100);p__Ascomycota_unclassified(100);p__Ascomycota_unclassified(100);p__Ascomycota_unclassified(100); | - | - | - | - | - | - | - | Unassigned | - |
| Otu46251 | 1 | k__Fungi(100);k__Fungi_unclassified(100);k__Fungi_unclassified(100);k__Fungi_unclassified(100);k__Fungi_unclassified(100);k__Fungi_unclassified(100);k__Fungi_unclassified(100); | - | - | - | - | - | - | - | Unassigned | - |
| Otu46258 | 1 | k__Fungi(100);k__Fungi_unclassified(100);k__Fungi_unclassified(100);k__Fungi_unclassified(100);k__Fungi_unclassified(100);k__Fungi_unclassified(100);k__Fungi_unclassified(100); | - | - | - | - | - | - | - | Unassigned | - |
| Otu46259 | 1 | k__Fungi(100);k__Fungi_unclassified(100);k__Fungi_unclassified(100);k__Fungi_unclassified(100);k__Fungi_unclassified(100);k__Fungi_unclassified(100);k__Fungi_unclassified(100); | - | - | - | - | - | - | - | Unassigned | - |
| Otu46270 | 1 | k__Fungi(100);k__Fungi_unclassified(100);k__Fungi_unclassified(100);k__Fungi_unclassified(100);k__Fungi_unclassified(100);k__Fungi_unclassified(100);k__Fungi_unclassified(100); | - | - | - | - | - | - | - | Unassigned | - |
| Otu46371 | 1 | k__Fungi(100);k__Fungi_unclassified(100);k__Fungi_unclassified(100);k__Fungi_unclassified(100);k__Fungi_unclassified(100);k__Fungi_unclassified(100);k__Fungi_unclassified(100); | - | - | - | - | - | - | - | Unassigned | - |
| Otu46394 | 1 | k__Fungi(100);k__Fungi_unclassified(100);k__Fungi_unclassified(100);k__Fungi_unclassified(100);k__Fungi_unclassified(100);k__Fungi_unclassified(100);k__Fungi_unclassified(100); | - | - | - | - | - | - | - | Unassigned | - |
| Otu46414 | 1 | k__Fungi(100);k__Fungi_unclassified(100);k__Fungi_unclassified(100);k__Fungi_unclassified(100);k__Fungi_unclassified(100);k__Fungi_unclassified(100);k__Fungi_unclassified(100); | - | - | - | - | - | - | - | Unassigned | - |
| Otu46478 | 1 | k__Fungi(100);k__Fungi_unclassified(100);k__Fungi_unclassified(100);k__Fungi_unclassified(100);k__Fungi_unclassified(100);k__Fungi_unclassified(100);k__Fungi_unclassified(100); | - | - | - | - | - | - | - | Unassigned | - |
| Otu46564 | 1 | k__Fungi(100);p__Ascomycota(100);p__Ascomycota_unclassified(100);p__Ascomycota_unclassified(100);p__Ascomycota_unclassified(100);p__Ascomycota_unclassified(100);p__Ascomycota_unclassified(100); | - | - | - | - | - | - | - | Unassigned | - |
| Otu46653 | 1 | k__Fungi(100);k__Fungi_unclassified(100);k__Fungi_unclassified(100);k__Fungi_unclassified(100);k__Fungi_unclassified(100);k__Fungi_unclassified(100);k__Fungi_unclassified(100); | - | - | - | - | - | - | - | Unassigned | - |
| Otu46654 | 1 | k__Fungi(100);k__Fungi_unclassified(100);k__Fungi_unclassified(100);k__Fungi_unclassified(100);k__Fungi_unclassified(100);k__Fungi_unclassified(100);k__Fungi_unclassified(100); | - | - | - | - | - | - | - | Unassigned | - |
| Otu46659 | 1 | k__Fungi(100);p__Ascomycota(100);p__Ascomycota_unclassified(100);p__Ascomycota_unclassified(100);p__Ascomycota_unclassified(100);p__Ascomycota_unclassified(100);p__Ascomycota_unclassified(100); | - | - | - | - | - | - | - | Unassigned | - |
| Otu46664 | 1 | k__Fungi(100);k__Fungi_unclassified(100);k__Fungi_unclassified(100);k__Fungi_unclassified(100);k__Fungi_unclassified(100);k__Fungi_unclassified(100);k__Fungi_unclassified(100); | - | - | - | - | - | - | - | Unassigned | - |

| | | | | | | | | | | | |
|---|---|---|---|---|---|---|---|---|---|---|---|
| Otu46665 | 1 | k__Fungi(100);k__Fungi_unclassified(100);k__Fungi_unclassified(100);k__Fungi_unclassified(100);k__Fungi_unclassified(100);k__Fungi_unclassified(100);k__Fungi_unclassified(100); | - | - | - | - | - | - | - | Unassigned | - |
| Otu46667 | 1 | k__Fungi(100);p__Ascomycota(100);p__Ascomycota_unclassified(100);p__Ascomycota_unclassified(100);p__Ascomycota_unclassified(100);p__Ascomycota_unclassified(100);p__Ascomycota_unclassified(100); | - | - | - | - | - | - | - | Unassigned | - |
| Otu46674 | 1 | k__Fungi(100);p__Ascomycota(100);p__Ascomycota_unclassified(100);p__Ascomycota_unclassified(100);p__Ascomycota_unclassified(100);p__Ascomycota_unclassified(100);p__Ascomycota_unclassified(100); | - | - | - | - | - | - | - | Unassigned | - |
| Otu46677 | 1 | k__Fungi(100);p__Ascomycota(100);p__Ascomycota_unclassified(100);p__Ascomycota_unclassified(100);p__Ascomycota_unclassified(100);p__Ascomycota_unclassified(100);p__Ascomycota_unclassified(100); | - | - | - | - | - | - | - | Unassigned | - |
| Otu46684 | 1 | k__Fungi(100);p__Ascomycota(100);p__Ascomycota_unclassified(100);p__Ascomycota_unclassified(100);p__Ascomycota_unclassified(100);p__Ascomycota_unclassified(100);p__Ascomycota_unclassified(100); | - | - | - | - | - | - | - | Unassigned | - |
| Otu46710 | 1 | k__Fungi(100);k__Fungi_unclassified(100);k__Fungi_unclassified(100);k__Fungi_unclassified(100);k__Fungi_unclassified(100);k__Fungi_unclassified(100);k__Fungi_unclassified(100); | - | - | - | - | - | - | - | Unassigned | - |
| Otu46712 | 1 | k__Fungi(100);k__Fungi_unclassified(100);k__Fungi_unclassified(100);k__Fungi_unclassified(100);k__Fungi_unclassified(100);k__Fungi_unclassified(100);k__Fungi_unclassified(100); | - | - | - | - | - | - | - | Unassigned | - |
| Otu46714 | 1 | k__Fungi(100);k__Fungi_unclassified(100);k__Fungi_unclassified(100);k__Fungi_unclassified(100);k__Fungi_unclassified(100);k__Fungi_unclassified(100);k__Fungi_unclassified(100); | - | - | - | - | - | - | - | Unassigned | - |
| Otu46719 | 1 | k__Fungi(100);k__Fungi_unclassified(100);k__Fungi_unclassified(100);k__Fungi_unclassified(100);k__Fungi_unclassified(100);k__Fungi_unclassified(100);k__Fungi_unclassified(100); | - | - | - | - | - | - | - | Unassigned | - |
| Otu46723 | 1 | k__Fungi(100);k__Fungi_unclassified(100);k__Fungi_unclassified(100);k__Fungi_unclassified(100);k__Fungi_unclassified(100);k__Fungi_unclassified(100);k__Fungi_unclassified(100); | - | - | - | - | - | - | - | Unassigned | - |
| Otu46747 | 1 | k__Fungi(100);k__Fungi_unclassified(100);k__Fungi_unclassified(100);k__Fungi_unclassified(100);k__Fungi_unclassified(100);k__Fungi_unclassified(100);k__Fungi_unclassified(100); | - | - | - | - | - | - | - | Unassigned | - |
| Otu46769 | 1 | k__Fungi(100);k__Fungi_unclassified(100);k__Fungi_unclassified(100);k__Fungi_unclassified(100);k__Fungi_unclassified(100);k__Fungi_unclassified(100);k__Fungi_unclassified(100); | - | - | - | - | - | - | - | Unassigned | - |
| Otu46779 | 1 | k__Fungi(100);k__Fungi_unclassified(100);k__Fungi_unclassified(100);k__Fungi_unclassified(100);k__Fungi_unclassified(100);k__Fungi_unclassified(100);k__Fungi_unclassified(100); | - | - | - | - | - | - | - | Unassigned | - |
| Otu46784 | 1 | k__Fungi(100);k__Fungi_unclassified(100);k__Fungi_unclassified(100);k__Fungi_unclassified(100);k__Fungi_unclassified(100);k__Fungi_unclassified(100);k__Fungi_unclassified(100); | - | - | - | - | - | - | - | Unassigned | - |
| Otu46786 | 1 | k__Fungi(100);k__Fungi_unclassified(100);k__Fungi_unclassified(100);k__Fungi_unclassified(100);k__Fungi_unclassified(100);k__Fungi_unclassified(100);k__Fungi_unclassified(100); | - | - | - | - | - | - | - | Unassigned | - |
| Otu46787 | 1 | k__Fungi(100);k__Fungi_unclassified(100);k__Fungi_unclassified(100);k__Fungi_unclassified(100);k__Fungi_unclassified(100);k__Fungi_unclassified(100);k__Fungi_unclassified(100); | - | - | - | - | - | - | - | Unassigned | - |
| Otu46807 | 1 | k__Fungi(100);k__Fungi_unclassified(100);k__Fungi_unclassified(100);k__Fungi_unclassified(100);k__Fungi_unclassified(100);k__Fungi_unclassified(100);k__Fungi_unclassified(100); | - | - | - | - | - | - | - | Unassigned | - |
| Otu46820 | 1 | k__Fungi(100);k__Fungi_unclassified(100);k__Fungi_unclassified(100);k__Fungi_unclassified(100);k__Fungi_unclassified(100);k__Fungi_unclassified(100);k__Fungi_unclassified(100); | - | - | - | - | - | - | - | Unassigned | - |
| Otu46824 | 1 | k__Fungi(100);k__Fungi_unclassified(100);k__Fungi_unclassified(100);k__Fungi_unclassified(100);k__Fungi_unclassified(100);k__Fungi_unclassified(100);k__Fungi_unclassified(100); | - | - | - | - | - | - | - | Unassigned | - |
| Otu46825 | 1 | k__Fungi(100);k__Fungi_unclassified(100);k__Fungi_unclassified(100);k__Fungi_unclassified(100);k__Fungi_unclassified(100);k__Fungi_unclassified(100);k__Fungi_unclassified(100); | - | - | - | - | - | - | - | Unassigned | - |
| Otu46832 | 1 | k__Fungi(100);k__Fungi_unclassified(100);k__Fungi_unclassified(100);k__Fungi_unclassified(100);k__Fungi_unclassified(100);k__Fungi_unclassified(100);k__Fungi_unclassified(100); | - | - | - | - | - | - | - | Unassigned | - |
| Otu46833 | 1 | k__Fungi(100);k__Fungi_unclassified(100);k__Fungi_unclassified(100);k__Fungi_unclassified(100);k__Fungi_unclassified(100);k__Fungi_unclassified(100);k__Fungi_unclassified(100); | - | - | - | - | - | - | - | Unassigned | - |
| Otu46838 | 1 | k__Fungi(100);p__Ascomycota(100);p__Ascomycota_unclassified(100);p__Ascomycota_unclassified(100);p__Ascomycota_unclassified(100);p__Ascomycota_unclassified(100);p__Ascomycota_unclassified(100); | - | - | - | - | - | - | - | Unassigned | - |
| Otu46858 | 1 | k__Fungi(100);k__Fungi_unclassified(100);k__Fungi_unclassified(100);k__Fungi_unclassified(100);k__Fungi_unclassified(100);k__Fungi_unclassified(100);k__Fungi_unclassified(100); | - | - | - | - | - | - | - | Unassigned | - |
| Otu46860 | 1 | k__Fungi(100);k__Fungi_unclassified(100);k__Fungi_unclassified(100);k__Fungi_unclassified(100);k__Fungi_unclassified(100);k__Fungi_unclassified(100);k__Fungi_unclassified(100); | - | - | - | - | - | - | - | Unassigned | - |
| Otu46861 | 1 | k__Fungi(100);k__Fungi_unclassified(100);k__Fungi_unclassified(100);k__Fungi_unclassified(100);k__Fungi_unclassified(100);k__Fungi_unclassified(100);k__Fungi_unclassified(100); | - | - | - | - | - | - | - | Unassigned | - |
| Otu46862 | 1 | k__Fungi(100);p__Ascomycota(100);p__Ascomycota_unclassified(100);p__Ascomycota_unclassified(100);p__Ascomycota_unclassified(100);p__Ascomycota_unclassified(100);p__Ascomycota_unclassified(100); | - | - | - | - | - | - | - | Unassigned | - |
| Otu46863 | 1 | k__Fungi(100);p__Ascomycota(100);p__Ascomycota_unclassified(100);p__Ascomycota_unclassified(100);p__Ascomycota_unclassified(100);p__Ascomycota_unclassified(100);p__Ascomycota_unclassified(100); | - | - | - | - | - | - | - | Unassigned | - |
| Otu46864 | 1 | k__Fungi(100);k__Fungi_unclassified(100);k__Fungi_unclassified(100);k__Fungi_unclassified(100);k__Fungi_unclassified(100);k__Fungi_unclassified(100);k__Fungi_unclassified(100); | - | - | - | - | - | - | - | Unassigned | - |
| Otu46868 | 1 | k__Fungi(100);p__Ascomycota(100);c__Sordariomycetes(100);c__Sordariomycetes_unclassified(100);c__Sordariomycetes_unclassified(100);c__Sordariomycetes_unclassified(100);c__Sordariomycetes_unclassified(100); | - | - | - | - | - | - | - | Unassigned | - |
| Otu46869 | 1 | k__Fungi(100);k__Fungi_unclassified(100);k__Fungi_unclassified(100);k__Fungi_unclassified(100);k__Fungi_unclassified(100);k__Fungi_unclassified(100);k__Fungi_unclassified(100); | - | - | - | - | - | - | - | Unassigned | - |
| Otu46878 | 1 | k__Fungi(100);k__Fungi_unclassified(100);k__Fungi_unclassified(100);k__Fungi_unclassified(100);k__Fungi_unclassified(100);k__Fungi_unclassified(100);k__Fungi_unclassified(100); | - | - | - | - | - | - | - | Unassigned | - |
| Otu46880 | 1 | k__Fungi(100);k__Fungi_unclassified(100);k__Fungi_unclassified(100);k__Fungi_unclassified(100);k__Fungi_unclassified(100);k__Fungi_unclassified(100);k__Fungi_unclassified(100); | - | - | - | - | - | - | - | Unassigned | - |
| Otu46940 | 1 | k__Fungi(100);p__Ascomycota(100);p__Ascomycota_unclassified(100);p__Ascomycota_unclassified(100);p__Ascomycota_unclassified(100);p__Ascomycota_unclassified(100);p__Ascomycota_unclassified(100); | - | - | - | - | - | - | - | Unassigned | - |
| Otu46976 | 1 | k__Fungi(100);k__Fungi_unclassified(100);k__Fungi_unclassified(100);k__Fungi_unclassified(100);k__Fungi_unclassified(100);k__Fungi_unclassified(100);k__Fungi_unclassified(100); | - | - | - | - | - | - | - | Unassigned | - |
| Otu46992 | 1 | k__Fungi(100);k__Fungi_unclassified(100);k__Fungi_unclassified(100);k__Fungi_unclassified(100);k__Fungi_unclassified(100);k__Fungi_unclassified(100);k__Fungi_unclassified(100); | - | - | - | - | - | - | - | Unassigned | - |
| Otu46996 | 1 | k__Fungi(100);p__Ascomycota(100);p__Ascomycota_unclassified(100);p__Ascomycota_unclassified(100);p__Ascomycota_unclassified(100);p__Ascomycota_unclassified(100);p__Ascomycota_unclassified(100); | - | - | - | - | - | - | - | Unassigned | - |
| Otu46999 | 1 | k__Fungi(100);k__Fungi_unclassified(100);k__Fungi_unclassified(100);k__Fungi_unclassified(100);k__Fungi_unclassified(100);k__Fungi_unclassified(100);k__Fungi_unclassified(100); | - | - | - | - | - | - | - | Unassigned | - |
| Otu47045 | 1 | k__Fungi(100);k__Fungi_unclassified(100);k__Fungi_unclassified(100);k__Fungi_unclassified(100);k__Fungi_unclassified(100);k__Fungi_unclassified(100);k__Fungi_unclassified(100); | - | - | - | - | - | - | - | Unassigned | - |
| Otu47054 | 1 | k__Fungi(100);k__Fungi_unclassified(100);k__Fungi_unclassified(100);k__Fungi_unclassified(100);k__Fungi_unclassified(100);k__Fungi_unclassified(100);k__Fungi_unclassified(100); | - | - | - | - | - | - | - | Unassigned | - |
| Otu47094 | 1 | k__Fungi(100);k__Fungi_unclassified(100);k__Fungi_unclassified(100);k__Fungi_unclassified(100);k__Fungi_unclassified(100);k__Fungi_unclassified(100);k__Fungi_unclassified(100); | - | - | - | - | - | - | - | Unassigned | - |
| Otu47099 | 1 | k__Fungi(100);k__Fungi_unclassified(100);k__Fungi_unclassified(100);k__Fungi_unclassified(100);k__Fungi_unclassified(100);k__Fungi_unclassified(100);k__Fungi_unclassified(100); | - | - | - | - | - | - | - | Unassigned | - |
| Otu47100 | 1 | k__Fungi(100);p__Ascomycota(100);p__Ascomycota_unclassified(100);p__Ascomycota_unclassified(100);p__Ascomycota_unclassified(100);p__Ascomycota_unclassified(100);p__Ascomycota_unclassified(100); | - | - | - | - | - | - | - | Unassigned | - |
| Otu47101 | 1 | k__Fungi(100);p__Ascomycota(100);p__Ascomycota_unclassified(100);p__Ascomycota_unclassified(100);p__Ascomycota_unclassified(100);p__Ascomycota_unclassified(100);p__Ascomycota_unclassified(100); | - | - | - | - | - | - | - | Unassigned | - |
| Otu47102 | 1 | k__Fungi(100);p__Ascomycota(100);p__Ascomycota_unclassified(100);p__Ascomycota_unclassified(100);p__Ascomycota_unclassified(100);p__Ascomycota_unclassified(100);p__Ascomycota_unclassified(100); | - | - | - | - | - | - | - | Unassigned | - |
| Otu47111 | 1 | k__Fungi(100);p__Ascomycota(100);p__Ascomycota_unclassified(100);p__Ascomycota_unclassified(100);p__Ascomycota_unclassified(100);p__Ascomycota_unclassified(100);p__Ascomycota_unclassified(100); | - | - | - | - | - | - | - | Unassigned | - |
| Otu47121 | 1 | k__Fungi(100);p__Ascomycota(100);p__Ascomycota_unclassified(100);p__Ascomycota_unclassified(100);p__Ascomycota_unclassified(100);p__Ascomycota_unclassified(100);p__Ascomycota_unclassified(100); | - | - | - | - | - | - | - | Unassigned | - |
| Otu47137 | 1 | k__Fungi(100);k__Fungi_unclassified(100);k__Fungi_unclassified(100);k__Fungi_unclassified(100);k__Fungi_unclassified(100);k__Fungi_unclassified(100);k__Fungi_unclassified(100); | - | - | - | - | - | - | - | Unassigned | - |
| Otu47142 | 1 | k__Fungi(100);p__Ascomycota(100);p__Ascomycota_unclassified(100);p__Ascomycota_unclassified(100);p__Ascomycota_unclassified(100);p__Ascomycota_unclassified(100);p__Ascomycota_unclassified(100); | - | - | - | - | - | - | - | Unassigned | - |
| Otu47143 | 1 | k__Fungi(100);k__Fungi_unclassified(100);k__Fungi_unclassified(100);k__Fungi_unclassified(100);k__Fungi_unclassified(100);k__Fungi_unclassified(100);k__Fungi_unclassified(100); | - | - | - | - | - | - | - | Unassigned | - |
| Otu47145 | 1 | k__Fungi(100);k__Fungi_unclassified(100);k__Fungi_unclassified(100);k__Fungi_unclassified(100);k__Fungi_unclassified(100);k__Fungi_unclassified(100);k__Fungi_unclassified(100); | - | - | - | - | - | - | - | Unassigned | - |
| Otu47154 | 1 | k__Fungi(100);p__Ascomycota(100);p__Ascomycota_unclassified(100);p__Ascomycota_unclassified(100);p__Ascomycota_unclassified(100);p__Ascomycota_unclassified(100);p__Ascomycota_unclassified(100); | - | - | - | - | - | - | - | Unassigned | - |
| Otu47181 | 1 | k__Fungi(100);p__Ascomycota(100);p__Ascomycota_unclassified(100);p__Ascomycota_unclassified(100);p__Ascomycota_unclassified(100);p__Ascomycota_unclassified(100);p__Ascomycota_unclassified(100); | - | - | - | - | - | - | - | Unassigned | - |
| Otu47187 | 1 | k__Fungi(100);p__Ascomycota(100);p__Ascomycota_unclassified(100);p__Ascomycota_unclassified(100);p__Ascomycota_unclassified(100);p__Ascomycota_unclassified(100);p__Ascomycota_unclassified(100); | - | - | - | - | - | - | - | Unassigned | - |
| Otu47188 | 1 | k__Fungi(100);p__Ascomycota(100);p__Ascomycota_unclassified(100);p__Ascomycota_unclassified(100);p__Ascomycota_unclassified(100);p__Ascomycota_unclassified(100);p__Ascomycota_unclassified(100); | - | - | - | - | - | - | - | Unassigned | - |
| Otu47194 | 1 | k__Fungi(100);k__Fungi_unclassified(100);k__Fungi_unclassified(100);k__Fungi_unclassified(100);k__Fungi_unclassified(100);k__Fungi_unclassified(100);k__Fungi_unclassified(100); | - | - | - | - | - | - | - | Unassigned | - |
| Otu47226 | 1 | k__Fungi(100);p__Ascomycota(100);p__Ascomycota_unclassified(100);p__Ascomycota_unclassified(100);p__Ascomycota_unclassified(100);p__Ascomycota_unclassified(100);p__Ascomycota_unclassified(100); | - | - | - | - | - | - | - | Unassigned | - |
| Otu47229 | 1 | k__Fungi(100);k__Fungi_unclassified(100);k__Fungi_unclassified(100);k__Fungi_unclassified(100);k__Fungi_unclassified(100);k__Fungi_unclassified(100);k__Fungi_unclassified(100); | - | - | - | - | - | - | - | Unassigned | - |
| Otu47242 | 1 | k__Fungi(100);p__Ascomycota(100);p__Ascomycota_unclassified(100);p__Ascomycota_unclassified(100);p__Ascomycota_unclassified(100);p__Ascomycota_unclassified(100);p__Ascomycota_unclassified(100); | - | - | - | - | - | - | - | Unassigned | - |
| Otu47249 | 1 | k__Fungi(100);k__Fungi_unclassified(100);k__Fungi_unclassified(100);k__Fungi_unclassified(100);k__Fungi_unclassified(100);k__Fungi_unclassified(100);k__Fungi_unclassified(100); | - | - | - | - | - | - | - | Unassigned | - |
| Otu47311 | 1 | k__Fungi(100);k__Fungi_unclassified(100);k__Fungi_unclassified(100);k__Fungi_unclassified(100);k__Fungi_unclassified(100);k__Fungi_unclassified(100);k__Fungi_unclassified(100); | - | - | - | - | - | - | - | Unassigned | - |
| Otu47315 | 1 | k__Fungi(100);k__Fungi_unclassified(100);k__Fungi_unclassified(100);k__Fungi_unclassified(100);k__Fungi_unclassified(100);k__Fungi_unclassified(100);k__Fungi_unclassified(100); | - | - | - | - | - | - | - | Unassigned | - |
| Otu47326 | 1 | k__Fungi(100);k__Fungi_unclassified(100);k__Fungi_unclassified(100);k__Fungi_unclassified(100);k__Fungi_unclassified(100);k__Fungi_unclassified(100);k__Fungi_unclassified(100); | - | - | - | - | - | - | - | Unassigned | - |
| Otu47355 | 1 | k__Fungi(100);k__Fungi_unclassified(100);k__Fungi_unclassified(100);k__Fungi_unclassified(100);k__Fungi_unclassified(100);k__Fungi_unclassified(100);k__Fungi_unclassified(100); | - | - | - | - | - | - | - | Unassigned | - |
| Otu47359 | 1 | k__Fungi(100);k__Fungi_unclassified(100);k__Fungi_unclassified(100);k__Fungi_unclassified(100);k__Fungi_unclassified(100);k__Fungi_unclassified(100);k__Fungi_unclassified(100); | - | - | - | - | - | - | - | Unassigned | - |
| Otu47362 | 1 | k__Fungi(100);k__Fungi_unclassified(100);k__Fungi_unclassified(100);k__Fungi_unclassified(100);k__Fungi_unclassified(100);k__Fungi_unclassified(100);k__Fungi_unclassified(100); | - | - | - | - | - | - | - | Unassigned | - |
| Otu47374 | 1 | k__Fungi(100);k__Fungi_unclassified(100);k__Fungi_unclassified(100);k__Fungi_unclassified(100);k__Fungi_unclassified(100);k__Fungi_unclassified(100);k__Fungi_unclassified(100); | - | - | - | - | - | - | - | Unassigned | - |
| Otu47375 | 1 | k__Fungi(100);k__Fungi_unclassified(100);k__Fungi_unclassified(100);k__Fungi_unclassified(100);k__Fungi_unclassified(100);k__Fungi_unclassified(100);k__Fungi_unclassified(100); | - | - | - | - | - | - | - | Unassigned | - |
| Otu47377 | 1 | k__Fungi(100);k__Fungi_unclassified(100);k__Fungi_unclassified(100);k__Fungi_unclassified(100);k__Fungi_unclassified(100);k__Fungi_unclassified(100);k__Fungi_unclassified(100); | - | - | - | - | - | - | - | Unassigned | - |
| Otu47378 | 1 | k__Fungi(100);k__Fungi_unclassified(100);k__Fungi_unclassified(100);k__Fungi_unclassified(100);k__Fungi_unclassified(100);k__Fungi_unclassified(100);k__Fungi_unclassified(100); | - | - | - | - | - | - | - | Unassigned | - |
| Otu47381 | 1 | k__Fungi(100);k__Fungi_unclassified(100);k__Fungi_unclassified(100);k__Fungi_unclassified(100);k__Fungi_unclassified(100);k__Fungi_unclassified(100);k__Fungi_unclassified(100); | - | - | - | - | - | - | - | Unassigned | - |
| Otu47400 | 1 | k__Fungi(100);k__Fungi_unclassified(100);k__Fungi_unclassified(100);k__Fungi_unclassified(100);k__Fungi_unclassified(100);k__Fungi_unclassified(100);k__Fungi_unclassified(100); | - | - | - | - | - | - | - | Unassigned | - |
| Otu47471 | 1 | k__Fungi(100);k__Fungi_unclassified(100);k__Fungi_unclassified(100);k__Fungi_unclassified(100);k__Fungi_unclassified(100);k__Fungi_unclassified(100);k__Fungi_unclassified(100); | - | - | - | - | - | - | - | Unassigned | - |
| Otu47516 | 1 | k__Fungi(100);k__Fungi_unclassified(100);k__Fungi_unclassified(100);k__Fungi_unclassified(100);k__Fungi_unclassified(100);k__Fungi_unclassified(100);k__Fungi_unclassified(100); | - | - | - | - | - | - | - | Unassigned | - |
| Otu47538 | 1 | k__Fungi(100);k__Fungi_unclassified(100);k__Fungi_unclassified(100);k__Fungi_unclassified(100);k__Fungi_unclassified(100);k__Fungi_unclassified(100);k__Fungi_unclassified(100); | - | - | - | - | - | - | - | Unassigned | - |
| Otu47539 | 1 | k__Fungi(100);k__Fungi_unclassified(100);k__Fungi_unclassified(100);k__Fungi_unclassified(100);k__Fungi_unclassified(100);k__Fungi_unclassified(100);k__Fungi_unclassified(100); | - | - | - | - | - | - | - | Unassigned | - |
| Otu47540 | 1 | k__Fungi(100);k__Fungi_unclassified(100);k__Fungi_unclassified(100);k__Fungi_unclassified(100);k__Fungi_unclassified(100);k__Fungi_unclassified(100);k__Fungi_unclassified(100); | - | - | - | - | - | - | - | Unassigned | - |
| Otu47541 | 1 | k__Fungi(100);p__Ascomycota(100);p__Ascomycota_unclassified(100);p__Ascomycota_unclassified(100);p__Ascomycota_unclassified(100);p__Ascomycota_unclassified(100);p__Ascomycota_unclassified(100); | - | - | - | - | - | - | - | Unassigned | - |
| Otu47542 | 1 | k__Fungi(100);k__Fungi_unclassified(100);k__Fungi_unclassified(100);k__Fungi_unclassified(100);k__Fungi_unclassified(100);k__Fungi_unclassified(100);k__Fungi_unclassified(100); | - | - | - | - | - | - | - | Unassigned | - |
| Otu47543 | 1 | k__Fungi(100);k__Fungi_unclassified(100);k__Fungi_unclassified(100);k__Fungi_unclassified(100);k__Fungi_unclassified(100);k__Fungi_unclassified(100);k__Fungi_unclassified(100); | - | - | - | - | - | - | - | Unassigned | - |
| Otu47549 | 1 | k__Fungi(100);k__Fungi_unclassified(100);k__Fungi_unclassified(100);k__Fungi_unclassified(100);k__Fungi_unclassified(100);k__Fungi_unclassified(100);k__Fungi_unclassified(100); | - | - | - | - | - | - | - | Unassigned | - |
| Otu47551 | 1 | k__Fungi(100);k__Fungi_unclassified(100);k__Fungi_unclassified(100);k__Fungi_unclassified(100);k__Fungi_unclassified(100);k__Fungi_unclassified(100);k__Fungi_unclassified(100); | - | - | - | - | - | - | - | Unassigned | - |
| Otu47552 | 1 | k__Fungi(100);p__Ascomycota(100);p__Ascomycota_unclassified(100);p__Ascomycota_unclassified(100);p__Ascomycota_unclassified(100);p__Ascomycota_unclassified(100);p__Ascomycota_unclassified(100); | - | - | - | - | - | - | - | Unassigned | - |
| Otu47553 | 1 | k__Fungi(100);p__Ascomycota(100);p__Ascomycota_unclassified(100);p__Ascomycota_unclassified(100);p__Ascomycota_unclassified(100);p__Ascomycota_unclassified(100);p__Ascomycota_unclassified(100); | - | - | - | - | - | - | - | Unassigned | - |
| Otu47554 | 1 | k__Fungi(100);p__Ascomycota(100);p__Ascomycota_unclassified(100);p__Ascomycota_unclassified(100);p__Ascomycota_unclassified(100);p__Ascomycota_unclassified(100);p__Ascomycota_unclassified(100); | - | - | - | - | - | - | - | Unassigned | - |
| Otu47558 | 1 | k__Fungi(100);p__Ascomycota(100);p__Ascomycota_unclassified(100);p__Ascomycota_unclassified(100);p__Ascomycota_unclassified(100);p__Ascomycota_unclassified(100);p__Ascomycota_unclassified(100); | - | - | - | - | - | - | - | Unassigned | - |
| Otu47570 | 1 | k__Fungi(100);k__Fungi_unclassified(100);k__Fungi_unclassified(100);k__Fungi_unclassified(100);k__Fungi_unclassified(100);k__Fungi_unclassified(100);k__Fungi_unclassified(100); | - | - | - | - | - | - | - | Unassigned | - |
| Otu47575 | 1 | k__Fungi(100);k__Fungi_unclassified(100);k__Fungi_unclassified(100);k__Fungi_unclassified(100);k__Fungi_unclassified(100);k__Fungi_unclassified(100);k__Fungi_unclassified(100); | - | - | - | - | - | - | - | Unassigned | - |
| Otu47579 | 1 | k__Fungi(100);k__Fungi_unclassified(100);k__Fungi_unclassified(100);k__Fungi_unclassified(100);k__Fungi_unclassified(100);k__Fungi_unclassified(100);k__Fungi_unclassified(100); | - | - | - | - | - | - | - | Unassigned | - |
| Otu47581 | 1 | k__Fungi(100);k__Fungi_unclassified(100);k__Fungi_unclassified(100);k__Fungi_unclassified(100);k__Fungi_unclassified(100);k__Fungi_unclassified(100);k__Fungi_unclassified(100); | - | - | - | - | - | - | - | Unassigned | - |
| Otu47586 | 1 | k__Fungi(100);p__Ascomycota(100);p__Ascomycota_unclassified(100);p__Ascomycota_unclassified(100);p__Ascomycota_unclassified(100);p__Ascomycota_unclassified(100);p__Ascomycota_unclassified(100); | - | - | - | - | - | - | - | Unassigned | - |
| Otu47591 | 1 | k__Fungi(100);p__Ascomycota(100);p__Ascomycota_unclassified(100);p__Ascomycota_unclassified(100);p__Ascomycota_unclassified(100);p__Ascomycota_unclassified(100);p__Ascomycota_unclassified(100); | - | - | - | - | - | - | - | Unassigned | - |
| Otu47598 | 1 | k__Fungi(100);k__Fungi_unclassified(100);k__Fungi_unclassified(100);k__Fungi_unclassified(100);k__Fungi_unclassified(100);k__Fungi_unclassified(100);k__Fungi_unclassified(100); | - | - | - | - | - | - | - | Unassigned | - |
| Otu47601 | 1 | k__Fungi(100);k__Fungi_unclassified(100);k__Fungi_unclassified(100);k__Fungi_unclassified(100);k__Fungi_unclassified(100);k__Fungi_unclassified(100);k__Fungi_unclassified(100); | - | - | - | - | - | - | - | Unassigned | - |
| Otu47610 | 1 | k__Fungi(100);k__Fungi_unclassified(100);k__Fungi_unclassified(100);k__Fungi_unclassified(100);k__Fungi_unclassified(100);k__Fungi_unclassified(100);k__Fungi_unclassified(100); | - | - | - | - | - | - | - | Unassigned | - |
| Otu47613 | 1 | k__Fungi(100);k__Fungi_unclassified(100);k__Fungi_unclassified(100);k__Fungi_unclassified(100);k__Fungi_unclassified(100);k__Fungi_unclassified(100);k__Fungi_unclassified(100); | - | - | - | - | - | - | - | Unassigned | - |
| Otu47623 | 1 | k__Fungi(100);k__Fungi_unclassified(100);k__Fungi_unclassified(100);k__Fungi_unclassified(100);k__Fungi_unclassified(100);k__Fungi_unclassified(100);k__Fungi_unclassified(100); | - | - | - | - | - | - | - | Unassigned | - |
| Otu47624 | 1 | k__Fungi(100);k__Fungi_unclassified(100);k__Fungi_unclassified(100);k__Fungi_unclassified(100);k__Fungi_unclassified(100);k__Fungi_unclassified(100);k__Fungi_unclassified(100); | - | - | - | - | - | - | - | Unassigned | - |
| Otu47636 | 1 | k__Fungi(100);p__Ascomycota(100);p__Ascomycota_unclassified(100);p__Ascomycota_unclassified(100);p__Ascomycota_unclassified(100);p__Ascomycota_unclassified(100);p__Ascomycota_unclassified(100); | - | - | - | - | - | - | - | Unassigned | - |
| Otu47659 | 1 | k__Fungi(100);p__Ascomycota(100);p__Ascomycota_unclassified(100);p__Ascomycota_unclassified(100);p__Ascomycota_unclassified(100);p__Ascomycota_unclassified(100);p__Ascomycota_unclassified(100); | - | - | - | - | - | - | - | Unassigned | - |
| Otu47693 | 1 | k__Fungi(100);p__Ascomycota(100);p__Ascomycota_unclassified(100);p__Ascomycota_unclassified(100);p__Ascomycota_unclassified(100);p__Ascomycota_unclassified(100);p__Ascomycota_unclassified(100); | - | - | - | - | - | - | - | Unassigned | - |
| Otu47720 | 1 | k__Fungi(100);k__Fungi_unclassified(100);k__Fungi_unclassified(100);k__Fungi_unclassified(100);k__Fungi_unclassified(100);k__Fungi_unclassified(100);k__Fungi_unclassified(100); | - | - | - | - | - | - | - | Unassigned | - |
| Otu47722 | 1 | k__Fungi(100);k__Fungi_unclassified(100);k__Fungi_unclassified(100);k__Fungi_unclassified(100);k__Fungi_unclassified(100);k__Fungi_unclassified(100);k__Fungi_unclassified(100); | - | - | - | - | - | - | - | Unassigned | - |
| Otu47723 | 1 | k__Fungi(100);p__Ascomycota(100);p__Ascomycota_unclassified(100);p__Ascomycota_unclassified(100);p__Ascomycota_unclassified(100);p__Ascomycota_unclassified(100);p__Ascomycota_unclassified(100); | - | - | - | - | - | - | - | Unassigned | - |
| Otu47730 | 1 | k__Fungi(100);k__Fungi_unclassified(100);k__Fungi_unclassified(100);k__Fungi_unclassified(100);k__Fungi_unclassified(100);k__Fungi_unclassified(100);k__Fungi_unclassified(100); | - | - | - | - | - | - | - | Unassigned | - |
| Otu47731 | 1 | k__Fungi(100);p__Ascomycota(100);p__Ascomycota_unclassified(100);p__Ascomycota_unclassified(100);p__Ascomycota_unclassified(100);p__Ascomycota_unclassified(100);p__Ascomycota_unclassified(100); | - | - | - | - | - | - | - | Unassigned | - |
| Otu47771 | 1 | k__Fungi(100);k__Fungi_unclassified(100);k__Fungi_unclassified(100);k__Fungi_unclassified(100);k__Fungi_unclassified(100);k__Fungi_unclassified(100);k__Fungi_unclassified(100); | - | - | - | - | - | - | - | Unassigned | - |
| Otu47790 | 1 | k__Fungi(100);p__Ascomycota(100);p__Ascomycota_unclassified(100);p__Ascomycota_unclassified(100);p__Ascomycota_unclassified(100);p__Ascomycota_unclassified(100);p__Ascomycota_unclassified(100); | - | - | - | - | - | - | - | Unassigned | - |
| Otu47799 | 1 | k__Fungi(100);k__Fungi_unclassified(100);k__Fungi_unclassified(100);k__Fungi_unclassified(100);k__Fungi_unclassified(100);k__Fungi_unclassified(100);k__Fungi_unclassified(100); | - | - | - | - | - | - | - | Unassigned | - |
| Otu47802 | 1 | k__Fungi(100);k__Fungi_unclassified(100);k__Fungi_unclassified(100);k__Fungi_unclassified(100);k__Fungi_unclassified(100);k__Fungi_unclassified(100);k__Fungi_unclassified(100); | - | - | - | - | - | - | - | Unassigned | - |
| Otu47803 | 1 | k__Fungi(100);k__Fungi_unclassified(100);k__Fungi_unclassified(100);k__Fungi_unclassified(100);k__Fungi_unclassified(100);k__Fungi_unclassified(100);k__Fungi_unclassified(100); | - | - | - | - | - | - | - | Unassigned | - |
| Otu47804 | 1 | k__Fungi(100);k__Fungi_unclassified(100);k__Fungi_unclassified(100);k__Fungi_unclassified(100);k__Fungi_unclassified(100);k__Fungi_unclassified(100);k__Fungi_unclassified(100); | - | - | - | - | - | - | - | Unassigned | - |
| Otu47825 | 1 | k__Fungi(100);k__Fungi_unclassified(100);k__Fungi_unclassified(100);k__Fungi_unclassified(100);k__Fungi_unclassified(100);k__Fungi_unclassified(100);k__Fungi_unclassified(100); | - | - | - | - | - | - | - | Unassigned | - |
| Otu47840 | 1 | k__Fungi(100);k__Fungi_unclassified(100);k__Fungi_unclassified(100);k__Fungi_unclassified(100);k__Fungi_unclassified(100);k__Fungi_unclassified(100);k__Fungi_unclassified(100); | - | - | - | - | - | - | - | Unassigned | - |
| Otu47841 | 1 | k__Fungi(100);p__Ascomycota(100);p__Ascomycota_unclassified(100);p__Ascomycota_unclassified(100);p__Ascomycota_unclassified(100);p__Ascomycota_unclassified(100);p__Ascomycota_unclassified(100); | - | - | - | - | - | - | - | Unassigned | - |
| Otu47849 | 1 | k__Fungi(100);k__Fungi_unclassified(100);k__Fungi_unclassified(100);k__Fungi_unclassified(100);k__Fungi_unclassified(100);k__Fungi_unclassified(100);k__Fungi_unclassified(100); | - | - | - | - | - | - | - | Unassigned | - |
| Otu47866 | 1 | k__Fungi(100);k__Fungi_unclassified(100);k__Fungi_unclassified(100);k__Fungi_unclassified(100);k__Fungi_unclassified(100);k__Fungi_unclassified(100);k__Fungi_unclassified(100); | - | - | - | - | - | - | - | Unassigned | - |
| Otu47869 | 1 | k__Fungi(100);k__Fungi_unclassified(100);k__Fungi_unclassified(100);k__Fungi_unclassified(100);k__Fungi_unclassified(100);k__Fungi_unclassified(100);k__Fungi_unclassified(100); | - | - | - | - | - | - | - | Unassigned | - |
| Otu47870 | 1 | k__Fungi(100);k__Fungi_unclassified(100);k__Fungi_unclassified(100);k__Fungi_unclassified(100);k__Fungi_unclassified(100);k__Fungi_unclassified(100);k__Fungi_unclassified(100); | - | - | - | - | - | - | - | Unassigned | - |
| Otu47872 | 1 | k__Fungi(100);k__Fungi_unclassified(100);k__Fungi_unclassified(100);k__Fungi_unclassified(100);k__Fungi_unclassified(100);k__Fungi_unclassified(100);k__Fungi_unclassified(100); | - | - | - | - | - | - | - | Unassigned | - |
| Otu47876 | 1 | k__Fungi(100);p__Ascomycota(100);p__Ascomycota_unclassified(100);p__Ascomycota_unclassified(100);p__Ascomycota_unclassified(100);p__Ascomycota_unclassified(100);p__Ascomycota_unclassified(100); | - | - | - | - | - | - | - | Unassigned | - |
| Otu47970 | 1 | k__Fungi(100);p__Ascomycota(100);p__Ascomycota_unclassified(100);p__Ascomycota_unclassified(100);p__Ascomycota_unclassified(100);p__Ascomycota_unclassified(100);p__Ascomycota_unclassified(100); | - | - | - | - | - | - | - | Unassigned | - |
| Otu47976 | 1 | k__Fungi(100);k__Fungi_unclassified(100);k__Fungi_unclassified(100);k__Fungi_unclassified(100);k__Fungi_unclassified(100);k__Fungi_unclassified(100);k__Fungi_unclassified(100); | - | - | - | - | - | - | - | Unassigned | - |
| Otu47988 | 1 | k__Fungi(100);k__Fungi_unclassified(100);k__Fungi_unclassified(100);k__Fungi_unclassified(100);k__Fungi_unclassified(100);k__Fungi_unclassified(100);k__Fungi_unclassified(100); | - | - | - | - | - | - | - | Unassigned | - |
| Otu48011 | 1 | k__Fungi(100);k__Fungi_unclassified(100);k__Fungi_unclassified(100);k__Fungi_unclassified(100);k__Fungi_unclassified(100);k__Fungi_unclassified(100);k__Fungi_unclassified(100); | - | - | - | - | - | - | - | Unassigned | - |
| Otu48013 | 1 | k__Fungi(100);k__Fungi_unclassified(100);k__Fungi_unclassified(100);k__Fungi_unclassified(100);k__Fungi_unclassified(100);k__Fungi_unclassified(100);k__Fungi_unclassified(100); | - | - | - | - | - | - | - | Unassigned | - |
| Otu48020 | 1 | k__Fungi(100);k__Fungi_unclassified(100);k__Fungi_unclassified(100);k__Fungi_unclassified(100);k__Fungi_unclassified(100);k__Fungi_unclassified(100);k__Fungi_unclassified(100); | - | - | - | - | - | - | - | Unassigned | - |
| Otu48023 | 1 | k__Fungi(100);k__Fungi_unclassified(100);k__Fungi_unclassified(100);k__Fungi_unclassified(100);k__Fungi_unclassified(100);k__Fungi_unclassified(100);k__Fungi_unclassified(100); | - | - | - | - | - | - | - | Unassigned | - |
| Otu48044 | 1 | k__Fungi(100);p__Ascomycota(100);p__Ascomycota_unclassified(100);p__Ascomycota_unclassified(100);p__Ascomycota_unclassified(100);p__Ascomycota_unclassified(100);p__Ascomycota_unclassified(100); | - | - | - | - | - | - | - | Unassigned | - |
| Otu48054 | 1 | k__Fungi(100);p__Ascomycota(100);p__Ascomycota_unclassified(100);p__Ascomycota_unclassified(100);p__Ascomycota_unclassified(100);p__Ascomycota_unclassified(100);p__Ascomycota_unclassified(100); | - | - | - | - | - | - | - | Unassigned | - |
| Otu48063 | 1 | k__Fungi(100);p__Ascomycota(100);p__Ascomycota_unclassified(100);p__Ascomycota_unclassified(100);p__Ascomycota_unclassified(100);p__Ascomycota_unclassified(100);p__Ascomycota_unclassified(100); | - | - | - | - | - | - | - | Unassigned | - |
| Otu48094 | 1 | k__Fungi(100);p__Ascomycota(100);p__Ascomycota_unclassified(100);p__Ascomycota_unclassified(100);p__Ascomycota_unclassified(100);p__Ascomycota_unclassified(100);p__Ascomycota_unclassified(100); | - | - | - | - | - | - | - | Unassigned | - |
| Otu48107 | 1 | k__Fungi(100);p__Ascomycota(100);c__Sordariomycetes(100);c__Sordariomycetes_unclassified(100);c__Sordariomycetes_unclassified(100);c__Sordariomycetes_unclassified(100);c__Sordariomycetes_unclassified(100); | - | - | - | - | - | - | - | Unassigned | - |
| Otu48237 | 1 | k__Fungi(100);k__Fungi_unclassified(100);k__Fungi_unclassified(100);k__Fungi_unclassified(100);k__Fungi_unclassified(100);k__Fungi_unclassified(100);k__Fungi_unclassified(100); | - | - | - | - | - | - | - | Unassigned | - |
| Otu48261 | 1 | k__Fungi(100);k__Fungi_unclassified(100);k__Fungi_unclassified(100);k__Fungi_unclassified(100);k__Fungi_unclassified(100);k__Fungi_unclassified(100);k__Fungi_unclassified(100); | - | - | - | - | - | - | - | Unassigned | - |
| Otu48265 | 1 | k__Fungi(100);k__Fungi_unclassified(100);k__Fungi_unclassified(100);k__Fungi_unclassified(100);k__Fungi_unclassified(100);k__Fungi_unclassified(100);k__Fungi_unclassified(100); | - | - | - | - | - | - | - | Unassigned | - |
| Otu48412 | 1 | k__Fungi(100);k__Fungi_unclassified(100);k__Fungi_unclassified(100);k__Fungi_unclassified(100);k__Fungi_unclassified(100);k__Fungi_unclassified(100);k__Fungi_unclassified(100); | - | - | - | - | - | - | - | Unassigned | - |
| Otu48419 | 1 | k__Fungi(100);k__Fungi_unclassified(100);k__Fungi_unclassified(100);k__Fungi_unclassified(100);k__Fungi_unclassified(100);k__Fungi_unclassified(100);k__Fungi_unclassified(100); | - | - | - | - | - | - | - | Unassigned | - |
| Otu48430 | 1 | k__Fungi(100);p__Ascomycota(100);p__Ascomycota_unclassified(100);p__Ascomycota_unclassified(100);p__Ascomycota_unclassified(100);p__Ascomycota_unclassified(100);p__Ascomycota_unclassified(100); | - | - | - | - | - | - | - | Unassigned | - |
| Otu48443 | 1 | k__Fungi(100);k__Fungi_unclassified(100);k__Fungi_unclassified(100);k__Fungi_unclassified(100);k__Fungi_unclassified(100);k__Fungi_unclassified(100);k__Fungi_unclassified(100); | - | - | - | - | - | - | - | Unassigned | - |
| Otu48446 | 1 | k__Fungi(100);k__Fungi_unclassified(100);k__Fungi_unclassified(100);k__Fungi_unclassified(100);k__Fungi_unclassified(100);k__Fungi_unclassified(100);k__Fungi_unclassified(100); | - | - | - | - | - | - | - | Unassigned | - |
| Otu48449 | 1 | k__Fungi(100);k__Fungi_unclassified(100);k__Fungi_unclassified(100);k__Fungi_unclassified(100);k__Fungi_unclassified(100);k__Fungi_unclassified(100);k__Fungi_unclassified(100); | - | - | - | - | - | - | - | Unassigned | - |
| Otu48450 | 1 | k__Fungi(100);k__Fungi_unclassified(100);k__Fungi_unclassified(100);k__Fungi_unclassified(100);k__Fungi_unclassified(100);k__Fungi_unclassified(100);k__Fungi_unclassified(100); | - | - | - | - | - | - | - | Unassigned | - |
| Otu48451 | 1 | k__Fungi(100);p__Ascomycota(100);p__Ascomycota_unclassified(100);p__Ascomycota_unclassified(100);p__Ascomycota_unclassified(100);p__Ascomycota_unclassified(100);p__Ascomycota_unclassified(100); | - | - | - | - | - | - | - | Unassigned | - |
| Otu48474 | 1 | k__Fungi(100);p__Ascomycota(100);p__Ascomycota_unclassified(100);p__Ascomycota_unclassified(100);p__Ascomycota_unclassified(100);p__Ascomycota_unclassified(100);p__Ascomycota_unclassified(100); | - | - | - | - | - | - | - | Unassigned | - |
| Otu48486 | 1 | k__Fungi(100);k__Fungi_unclassified(100);k__Fungi_unclassified(100);k__Fungi_unclassified(100);k__Fungi_unclassified(100);k__Fungi_unclassified(100);k__Fungi_unclassified(100); | - | - | - | - | - | - | - | Unassigned | - |
| Otu48498 | 1 | k__Fungi(100);p__Ascomycota(100);p__Ascomycota_unclassified(100);p__Ascomycota_unclassified(100);p__Ascomycota_unclassified(100);p__Ascomycota_unclassified(100);p__Ascomycota_unclassified(100); | - | - | - | - | - | - | - | Unassigned | - |
| Otu48501 | 1 | k__Fungi(100);k__Fungi_unclassified(100);k__Fungi_unclassified(100);k__Fungi_unclassified(100);k__Fungi_unclassified(100);k__Fungi_unclassified(100);k__Fungi_unclassified(100); | - | - | - | - | - | - | - | Unassigned | - |
| Otu48505 | 1 | k__Fungi(100);k__Fungi_unclassified(100);k__Fungi_unclassified(100);k__Fungi_unclassified(100);k__Fungi_unclassified(100);k__Fungi_unclassified(100);k__Fungi_unclassified(100); | - | - | - | - | - | - | - | Unassigned | - |
| Otu48512 | 1 | k__Fungi(100);p__Ascomycota(100);p__Ascomycota_unclassified(100);p__Ascomycota_unclassified(100);p__Ascomycota_unclassified(100);p__Ascomycota_unclassified(100);p__Ascomycota_unclassified(100); | - | - | - | - | - | - | - | Unassigned | - |
| Otu48521 | 1 | k__Fungi(100);p__Ascomycota(100);p__Ascomycota_unclassified(100);p__Ascomycota_unclassified(100);p__Ascomycota_unclassified(100);p__Ascomycota_unclassified(100);p__Ascomycota_unclassified(100); | - | - | - | - | - | - | - | Unassigned | - |
| Otu48522 | 1 | k__Fungi(100);k__Fungi_unclassified(100);k__Fungi_unclassified(100);k__Fungi_unclassified(100);k__Fungi_unclassified(100);k__Fungi_unclassified(100);k__Fungi_unclassified(100); | - | - | - | - | - | - | - | Unassigned | - |
| Otu48552 | 1 | k__Fungi(100);p__Ascomycota(100);p__Ascomycota_unclassified(100);p__Ascomycota_unclassified(100);p__Ascomycota_unclassified(100);p__Ascomycota_unclassified(100);p__Ascomycota_unclassified(100); | - | - | - | - | - | - | - | Unassigned | - |
| Otu48571 | 1 | k__Fungi(100);k__Fungi_unclassified(100);k__Fungi_unclassified(100);k__Fungi_unclassified(100);k__Fungi_unclassified(100);k__Fungi_unclassified(100);k__Fungi_unclassified(100); | - | - | - | - | - | - | - | Unassigned | - |
| Otu48775 | 1 | k__Fungi(100);k__Fungi_unclassified(100);k__Fungi_unclassified(100);k__Fungi_unclassified(100);k__Fungi_unclassified(100);k__Fungi_unclassified(100);k__Fungi_unclassified(100); | - | - | - | - | - | - | - | Unassigned | - |
| Otu48803 | 1 | k__Fungi(100);k__Fungi_unclassified(100);k__Fungi_unclassified(100);k__Fungi_unclassified(100);k__Fungi_unclassified(100);k__Fungi_unclassified(100);k__Fungi_unclassified(100); | - | - | - | - | - | - | - | Unassigned | - |
| Otu48817 | 1 | k__Fungi(100);p__Basidiomycota(100);p__Basidiomycota_unclassified(100);p__Basidiomycota_unclassified(100);p__Basidiomycota_unclassified(100);p__Basidiomycota_unclassified(100);p__Basidiomycota_unclassified(100); | - | - | - | - | - | - | - | Unassigned | - |
| Otu48844 | 1 | k__Fungi(100);k__Fungi_unclassified(100);k__Fungi_unclassified(100);k__Fungi_unclassified(100);k__Fungi_unclassified(100);k__Fungi_unclassified(100);k__Fungi_unclassified(100); | - | - | - | - | - | - | - | Unassigned | - |
| Otu48937 | 1 | k__Fungi(100);p__Ascomycota(100);p__Ascomycota_unclassified(100);p__Ascomycota_unclassified(100);p__Ascomycota_unclassified(100);p__Ascomycota_unclassified(100);p__Ascomycota_unclassified(100); | - | - | - | - | - | - | - | Unassigned | - |
| Otu48939 | 1 | k__Fungi(100);p__Ascomycota(100);p__Ascomycota_unclassified(100);p__Ascomycota_unclassified(100);p__Ascomycota_unclassified(100);p__Ascomycota_unclassified(100);p__Ascomycota_unclassified(100); | - | - | - | - | - | - | - | Unassigned | - |
| Otu48959 | 1 | k__Fungi(100);k__Fungi_unclassified(100);k__Fungi_unclassified(100);k__Fungi_unclassified(100);k__Fungi_unclassified(100);k__Fungi_unclassified(100);k__Fungi_unclassified(100); | - | - | - | - | - | - | - | Unassigned | - |
| Otu48960 | 1 | k__Fungi(100);k__Fungi_unclassified(100);k__Fungi_unclassified(100);k__Fungi_unclassified(100);k__Fungi_unclassified(100);k__Fungi_unclassified(100);k__Fungi_unclassified(100); | - | - | - | - | - | - | - | Unassigned | - |
| Otu48964 | 1 | k__Fungi(100);k__Fungi_unclassified(100);k__Fungi_unclassified(100);k__Fungi_unclassified(100);k__Fungi_unclassified(100);k__Fungi_unclassified(100);k__Fungi_unclassified(100); | - | - | - | - | - | - | - | Unassigned | - |
| Otu48976 | 1 | k__Fungi(100);p__Ascomycota(100);p__Ascomycota_unclassified(100);p__Ascomycota_unclassified(100);p__Ascomycota_unclassified(100);p__Ascomycota_unclassified(100);p__Ascomycota_unclassified(100); | - | - | - | - | - | - | - | Unassigned | - |
| Otu48988 | 1 | k__Fungi(100);k__Fungi_unclassified(100);k__Fungi_unclassified(100);k__Fungi_unclassified(100);k__Fungi_unclassified(100);k__Fungi_unclassified(100);k__Fungi_unclassified(100); | - | - | - | - | - | - | - | Unassigned | - |
| Otu49034 | 1 | k__Fungi(100);p__Ascomycota(100);p__Ascomycota_unclassified(100);p__Ascomycota_unclassified(100);p__Ascomycota_unclassified(100);p__Ascomycota_unclassified(100);p__Ascomycota_unclassified(100); | - | - | - | - | - | - | - | Unassigned | - |
| Otu49035 | 1 | k__Fungi(100);p__Ascomycota(100);p__Ascomycota_unclassified(100);p__Ascomycota_unclassified(100);p__Ascomycota_unclassified(100);p__Ascomycota_unclassified(100);p__Ascomycota_unclassified(100); | - | - | - | - | - | - | - | Unassigned | - |
| Otu49043 | 1 | k__Fungi(100);p__Ascomycota(100);p__Ascomycota_unclassified(100);p__Ascomycota_unclassified(100);p__Ascomycota_unclassified(100);p__Ascomycota_unclassified(100);p__Ascomycota_unclassified(100); | - | - | - | - | - | - | - | Unassigned | - |
| Otu49059 | 1 | k__Fungi(100);p__Ascomycota(100);p__Ascomycota_unclassified(100);p__Ascomycota_unclassified(100);p__Ascomycota_unclassified(100);p__Ascomycota_unclassified(100);p__Ascomycota_unclassified(100); | - | - | - | - | - | - | - | Unassigned | - |
| Otu49156 | 1 | k__Fungi(100);k__Fungi_unclassified(100);k__Fungi_unclassified(100);k__Fungi_unclassified(100);k__Fungi_unclassified(100);k__Fungi_unclassified(100);k__Fungi_unclassified(100); | - | - | - | - | - | - | - | Unassigned | - |
| Otu49180 | 1 | k__Fungi(100);p__Ascomycota(100);p__Ascomycota_unclassified(100);p__Ascomycota_unclassified(100);p__Ascomycota_unclassified(100);p__Ascomycota_unclassified(100);p__Ascomycota_unclassified(100); | - | - | - | - | - | - | - | Unassigned | - |
| Otu49219 | 1 | k__Fungi(100);k__Fungi_unclassified(100);k__Fungi_unclassified(100);k__Fungi_unclassified(100);k__Fungi_unclassified(100);k__Fungi_unclassified(100);k__Fungi_unclassified(100); | - | - | - | - | - | - | - | Unassigned | - |
| Otu49234 | 1 | k__Fungi(100);k__Fungi_unclassified(100);k__Fungi_unclassified(100);k__Fungi_unclassified(100);k__Fungi_unclassified(100);k__Fungi_unclassified(100);k__Fungi_unclassified(100); | - | - | - | - | - | - | - | Unassigned | - |
| Otu49239 | 1 | k__Fungi(100);p__Ascomycota(100);p__Ascomycota_unclassified(100);p__Ascomycota_unclassified(100);p__Ascomycota_unclassified(100);p__Ascomycota_unclassified(100);p__Ascomycota_unclassified(100); | - | - | - | - | - | - | - | Unassigned | - |
| Otu49240 | 1 | k__Fungi(100);k__Fungi_unclassified(100);k__Fungi_unclassified(100);k__Fungi_unclassified(100);k__Fungi_unclassified(100);k__Fungi_unclassified(100);k__Fungi_unclassified(100); | - | - | - | - | - | - | - | Unassigned | - |
| Otu49244 | 1 | k__Fungi(100);k__Fungi_unclassified(100);k__Fungi_unclassified(100);k__Fungi_unclassified(100);k__Fungi_unclassified(100);k__Fungi_unclassified(100);k__Fungi_unclassified(100); | - | - | - | - | - | - | - | Unassigned | - |
| Otu49265 | 1 | k__Fungi(100);p__Ascomycota(100);p__Ascomycota_unclassified(100);p__Ascomycota_unclassified(100);p__Ascomycota_unclassified(100);p__Ascomycota_unclassified(100);p__Ascomycota_unclassified(100); | - | - | - | - | - | - | - | Unassigned | - |
| Otu49267 | 1 | k__Fungi(100);p__Ascomycota(100);p__Ascomycota_unclassified(100);p__Ascomycota_unclassified(100);p__Ascomycota_unclassified(100);p__Ascomycota_unclassified(100);p__Ascomycota_unclassified(100); | - | - | - | - | - | - | - | Unassigned | - |
| Otu49301 | 1 | k__Fungi(100);k__Fungi_unclassified(100);k__Fungi_unclassified(100);k__Fungi_unclassified(100);k__Fungi_unclassified(100);k__Fungi_unclassified(100);k__Fungi_unclassified(100); | - | - | - | - | - | - | - | Unassigned | - |
| Otu49306 | 1 | k__Fungi(100);p__Ascomycota(100);p__Ascomycota_unclassified(100);p__Ascomycota_unclassified(100);p__Ascomycota_unclassified(100);p__Ascomycota_unclassified(100);p__Ascomycota_unclassified(100); | - | - | - | - | - | - | - | Unassigned | - |
| Otu49326 | 1 | k__Fungi(100);k__Fungi_unclassified(100);k__Fungi_unclassified(100);k__Fungi_unclassified(100);k__Fungi_unclassified(100);k__Fungi_unclassified(100);k__Fungi_unclassified(100); | - | - | - | - | - | - | - | Unassigned | - |
| Otu49335 | 1 | k__Fungi(100);p__Ascomycota(100);p__Ascomycota_unclassified(100);p__Ascomycota_unclassified(100);p__Ascomycota_unclassified(100);p__Ascomycota_unclassified(100);p__Ascomycota_unclassified(100); | - | - | - | - | - | - | - | Unassigned | - |
| Otu49482 | 1 | k__Fungi(100);k__Fungi_unclassified(100);k__Fungi_unclassified(100);k__Fungi_unclassified(100);k__Fungi_unclassified(100);k__Fungi_unclassified(100);k__Fungi_unclassified(100); | - | - | - | - | - | - | - | Unassigned | - |
| Otu49502 | 1 | k__Fungi(100);k__Fungi_unclassified(100);k__Fungi_unclassified(100);k__Fungi_unclassified(100);k__Fungi_unclassified(100);k__Fungi_unclassified(100);k__Fungi_unclassified(100); | - | - | - | - | - | - | - | Unassigned | - |
| Otu49503 | 1 | k__Fungi(100);p__Ascomycota(100);p__Ascomycota_unclassified(100);p__Ascomycota_unclassified(100);p__Ascomycota_unclassified(100);p__Ascomycota_unclassified(100);p__Ascomycota_unclassified(100); | - | - | - | - | - | - | - | Unassigned | - |
| Otu49539 | 1 | k__Fungi(100);k__Fungi_unclassified(100);k__Fungi_unclassified(100);k__Fungi_unclassified(100);k__Fungi_unclassified(100);k__Fungi_unclassified(100);k__Fungi_unclassified(100); | - | - | - | - | - | - | - | Unassigned | - |
| Otu49543 | 1 | k__Fungi(100);p__Ascomycota(100);p__Ascomycota_unclassified(100);p__Ascomycota_unclassified(100);p__Ascomycota_unclassified(100);p__Ascomycota_unclassified(100);p__Ascomycota_unclassified(100); | - | - | - | - | - | - | - | Unassigned | - |
| Otu49566 | 1 | k__Fungi(100);p__Ascomycota(100);p__Ascomycota_unclassified(100);p__Ascomycota_unclassified(100);p__Ascomycota_unclassified(100);p__Ascomycota_unclassified(100);p__Ascomycota_unclassified(100); | - | - | - | - | - | - | - | Unassigned | - |
| Otu49579 | 1 | k__Fungi(100);k__Fungi_unclassified(100);k__Fungi_unclassified(100);k__Fungi_unclassified(100);k__Fungi_unclassified(100);k__Fungi_unclassified(100);k__Fungi_unclassified(100); | - | - | - | - | - | - | - | Unassigned | - |
| Otu49609 | 1 | k__Fungi(100);k__Fungi_unclassified(100);k__Fungi_unclassified(100);k__Fungi_unclassified(100);k__Fungi_unclassified(100);k__Fungi_unclassified(100);k__Fungi_unclassified(100); | - | - | - | - | - | - | - | Unassigned | - |
| Otu49640 | 1 | k__Fungi(100);k__Fungi_unclassified(100);k__Fungi_unclassified(100);k__Fungi_unclassified(100);k__Fungi_unclassified(100);k__Fungi_unclassified(100);k__Fungi_unclassified(100); | - | - | - | - | - | - | - | Unassigned | - |
| Otu49725 | 1 | k__Fungi(100);k__Fungi_unclassified(100);k__Fungi_unclassified(100);k__Fungi_unclassified(100);k__Fungi_unclassified(100);k__Fungi_unclassified(100);k__Fungi_unclassified(100); | - | - | - | - | - | - | - | Unassigned | - |
| Otu49754 | 1 | k__Fungi(100);k__Fungi_unclassified(100);k__Fungi_unclassified(100);k__Fungi_unclassified(100);k__Fungi_unclassified(100);k__Fungi_unclassified(100);k__Fungi_unclassified(100); | - | - | - | - | - | - | - | Unassigned | - |
| Otu49765 | 1 | k__Fungi(100);p__Ascomycota(100);p__Ascomycota_unclassified(100);p__Ascomycota_unclassified(100);p__Ascomycota_unclassified(100);p__Ascomycota_unclassified(100);p__Ascomycota_unclassified(100); | - | - | - | - | - | - | - | Unassigned | - |
| Otu49789 | 1 | k__Fungi(100);k__Fungi_unclassified(100);k__Fungi_unclassified(100);k__Fungi_unclassified(100);k__Fungi_unclassified(100);k__Fungi_unclassified(100);k__Fungi_unclassified(100); | - | - | - | - | - | - | - | Unassigned | - |
| Otu49857 | 1 | k__Fungi(100);k__Fungi_unclassified(100);k__Fungi_unclassified(100);k__Fungi_unclassified(100);k__Fungi_unclassified(100);k__Fungi_unclassified(100);k__Fungi_unclassified(100); | - | - | - | - | - | - | - | Unassigned | - |
| Otu49873 | 1 | k__Fungi(100);p__Ascomycota(100);p__Ascomycota_unclassified(100);p__Ascomycota_unclassified(100);p__Ascomycota_unclassified(100);p__Ascomycota_unclassified(100);p__Ascomycota_unclassified(100); | - | - | - | - | - | - | - | Unassigned | - |
| Otu49888 | 1 | k__Fungi(100);k__Fungi_unclassified(100);k__Fungi_unclassified(100);k__Fungi_unclassified(100);k__Fungi_unclassified(100);k__Fungi_unclassified(100);k__Fungi_unclassified(100); | - | - | - | - | - | - | - | Unassigned | - |
| Otu50015 | 1 | k__Fungi(100);k__Fungi_unclassified(100);k__Fungi_unclassified(100);k__Fungi_unclassified(100);k__Fungi_unclassified(100);k__Fungi_unclassified(100);k__Fungi_unclassified(100); | - | - | - | - | - | - | - | Unassigned | - |
| Otu50019 | 1 | k__Fungi(100);p__Ascomycota(100);p__Ascomycota_unclassified(100);p__Ascomycota_unclassified(100);p__Ascomycota_unclassified(100);p__Ascomycota_unclassified(100);p__Ascomycota_unclassified(100); | - | - | - | - | - | - | - | Unassigned | - |
| Otu50076 | 1 | k__Fungi(100);k__Fungi_unclassified(100);k__Fungi_unclassified(100);k__Fungi_unclassified(100);k__Fungi_unclassified(100);k__Fungi_unclassified(100);k__Fungi_unclassified(100); | - | - | - | - | - | - | - | Unassigned | - |
| Otu50079 | 1 | k__Fungi(100);k__Fungi_unclassified(100);k__Fungi_unclassified(100);k__Fungi_unclassified(100);k__Fungi_unclassified(100);k__Fungi_unclassified(100);k__Fungi_unclassified(100); | - | - | - | - | - | - | - | Unassigned | - |
| Otu50087 | 1 | k__Fungi(100);p__Ascomycota(100);p__Ascomycota_unclassified(100);p__Ascomycota_unclassified(100);p__Ascomycota_unclassified(100);p__Ascomycota_unclassified(100);p__Ascomycota_unclassified(100); | - | - | - | - | - | - | - | Unassigned | - |
| Otu50106 | 1 | k__Fungi(100);p__Ascomycota(100);p__Ascomycota_unclassified(100);p__Ascomycota_unclassified(100);p__Ascomycota_unclassified(100);p__Ascomycota_unclassified(100);p__Ascomycota_unclassified(100); | - | - | - | - | - | - | - | Unassigned | - |
| Otu50125 | 1 | k__Fungi(100);k__Fungi_unclassified(100);k__Fungi_unclassified(100);k__Fungi_unclassified(100);k__Fungi_unclassified(100);k__Fungi_unclassified(100);k__Fungi_unclassified(100); | - | - | - | - | - | - | - | Unassigned | - |
| Otu50143 | 1 | k__Fungi(100);k__Fungi_unclassified(100);k__Fungi_unclassified(100);k__Fungi_unclassified(100);k__Fungi_unclassified(100);k__Fungi_unclassified(100);k__Fungi_unclassified(100); | - | - | - | - | - | - | - | Unassigned | - |
| Otu50144 | 1 | k__Fungi(100);k__Fungi_unclassified(100);k__Fungi_unclassified(100);k__Fungi_unclassified(100);k__Fungi_unclassified(100);k__Fungi_unclassified(100);k__Fungi_unclassified(100); | - | - | - | - | - | - | - | Unassigned | - |
| Otu50150 | 1 | k__Fungi(100);p__Ascomycota(100);p__Ascomycota_unclassified(100);p__Ascomycota_unclassified(100);p__Ascomycota_unclassified(100);p__Ascomycota_unclassified(100);p__Ascomycota_unclassified(100); | - | - | - | - | - | - | - | Unassigned | - |
| Otu50186 | 1 | k__Fungi(100);p__Ascomycota(100);p__Ascomycota_unclassified(100);p__Ascomycota_unclassified(100);p__Ascomycota_unclassified(100);p__Ascomycota_unclassified(100);p__Ascomycota_unclassified(100); | - | - | - | - | - | - | - | Unassigned | - |
| Otu50241 | 1 | k__Fungi(100);k__Fungi_unclassified(100);k__Fungi_unclassified(100);k__Fungi_unclassified(100);k__Fungi_unclassified(100);k__Fungi_unclassified(100);k__Fungi_unclassified(100); | - | - | - | - | - | - | - | Unassigned | - |
| Otu50242 | 1 | k__Fungi(100);k__Fungi_unclassified(100);k__Fungi_unclassified(100);k__Fungi_unclassified(100);k__Fungi_unclassified(100);k__Fungi_unclassified(100);k__Fungi_unclassified(100); | - | - | - | - | - | - | - | Unassigned | - |
| Otu50254 | 1 | k__Fungi(100);k__Fungi_unclassified(100);k__Fungi_unclassified(100);k__Fungi_unclassified(100);k__Fungi_unclassified(100);k__Fungi_unclassified(100);k__Fungi_unclassified(100); | - | - | - | - | - | - | - | Unassigned | - |
| Otu50274 | 1 | k__Fungi(100);k__Fungi_unclassified(100);k__Fungi_unclassified(100);k__Fungi_unclassified(100);k__Fungi_unclassified(100);k__Fungi_unclassified(100);k__Fungi_unclassified(100); | - | - | - | - | - | - | - | Unassigned | - |
| Otu50325 | 1 | k__Fungi(100);k__Fungi_unclassified(100);k__Fungi_unclassified(100);k__Fungi_unclassified(100);k__Fungi_unclassified(100);k__Fungi_unclassified(100);k__Fungi_unclassified(100); | - | - | - | - | - | - | - | Unassigned | - |
| Otu50326 | 1 | k__Fungi(100);k__Fungi_unclassified(100);k__Fungi_unclassified(100);k__Fungi_unclassified(100);k__Fungi_unclassified(100);k__Fungi_unclassified(100);k__Fungi_unclassified(100); | - | - | - | - | - | - | - | Unassigned | - |
| Otu50327 | 1 | k__Fungi(100);k__Fungi_unclassified(100);k__Fungi_unclassified(100);k__Fungi_unclassified(100);k__Fungi_unclassified(100);k__Fungi_unclassified(100);k__Fungi_unclassified(100); | - | - | - | - | - | - | - | Unassigned | - |

| | | | | | | | | | | | |
|---|---|---|---|---|---|---|---|---|---|---|---|
| Otu50331 | 1 | k__Fungi(100);k__Fungi_unclassified(100);k__Fungi_unclassified(100);k__Fungi_unclassified(100);k__Fungi_unclassified(100);k__Fungi_unclassified(100);k__Fungi_unclassified(100); | - | - | - | - | - | - | - | Unassigned | - |
| Otu50332 | 1 | k__Fungi(100);k__Fungi_unclassified(100);k__Fungi_unclassified(100);k__Fungi_unclassified(100);k__Fungi_unclassified(100);k__Fungi_unclassified(100);k__Fungi_unclassified(100); | - | - | - | - | - | - | - | Unassigned | - |
| Otu50366 | 1 | k__Fungi(100);k__Fungi_unclassified(100);k__Fungi_unclassified(100);k__Fungi_unclassified(100);k__Fungi_unclassified(100);k__Fungi_unclassified(100);k__Fungi_unclassified(100); | - | - | - | - | - | - | - | Unassigned | - |
| Otu50367 | 1 | k__Fungi(100);p__Ascomycota(100);p__Ascomycota_unclassified(100);p__Ascomycota_unclassified(100);p__Ascomycota_unclassified(100);p__Ascomycota_unclassified(100);p__Ascomycota_unclassified(100); | - | - | - | - | - | - | - | Unassigned | - |
| Otu50374 | 1 | k__Fungi(100);p__Ascomycota(100);p__Ascomycota_unclassified(100);p__Ascomycota_unclassified(100);p__Ascomycota_unclassified(100);p__Ascomycota_unclassified(100);p__Ascomycota_unclassified(100); | - | - | - | - | - | - | - | Unassigned | - |
| Otu50380 | 1 | k__Fungi(100);p__Ascomycota(100);p__Ascomycota_unclassified(100);p__Ascomycota_unclassified(100);p__Ascomycota_unclassified(100);p__Ascomycota_unclassified(100);p__Ascomycota_unclassified(100); | - | - | - | - | - | - | - | Unassigned | - |
| Otu50389 | 1 | k__Fungi(100);k__Fungi_unclassified(100);k__Fungi_unclassified(100);k__Fungi_unclassified(100);k__Fungi_unclassified(100);k__Fungi_unclassified(100);k__Fungi_unclassified(100); | - | - | - | - | - | - | - | Unassigned | - |
| Otu50402 | 1 | k__Fungi(100);k__Fungi_unclassified(100);k__Fungi_unclassified(100);k__Fungi_unclassified(100);k__Fungi_unclassified(100);k__Fungi_unclassified(100);k__Fungi_unclassified(100); | - | - | - | - | - | - | - | Unassigned | - |
| Otu50409 | 1 | k__Fungi(100);k__Fungi_unclassified(100);k__Fungi_unclassified(100);k__Fungi_unclassified(100);k__Fungi_unclassified(100);k__Fungi_unclassified(100);k__Fungi_unclassified(100); | - | - | - | - | - | - | - | Unassigned | - |
| Otu50411 | 1 | k__Fungi(100);k__Fungi_unclassified(100);k__Fungi_unclassified(100);k__Fungi_unclassified(100);k__Fungi_unclassified(100);k__Fungi_unclassified(100);k__Fungi_unclassified(100); | - | - | - | - | - | - | - | Unassigned | - |
| Otu50416 | 1 | k__Fungi(100);p__Ascomycota(100);p__Ascomycota_unclassified(100);p__Ascomycota_unclassified(100);p__Ascomycota_unclassified(100);p__Ascomycota_unclassified(100);p__Ascomycota_unclassified(100); | - | - | - | - | - | - | - | Unassigned | - |
| Otu50447 | 1 | k__Fungi(100);p__Ascomycota(100);p__Ascomycota_unclassified(100);p__Ascomycota_unclassified(100);p__Ascomycota_unclassified(100);p__Ascomycota_unclassified(100);p__Ascomycota_unclassified(100); | - | - | - | - | - | - | - | Unassigned | - |
| Otu50449 | 1 | k__Fungi(100);p__Ascomycota(100);c__Sordariomycetes(100);c__Sordariomycetes_unclassified(100);c__Sordariomycetes_unclassified(100);c__Sordariomycetes_unclassified(100);c__Sordariomycetes_unclassified(100); | - | - | - | - | - | - | - | Unassigned | - |
| Otu50474 | 1 | k__Fungi(100);p__Ascomycota(100);p__Ascomycota_unclassified(100);p__Ascomycota_unclassified(100);p__Ascomycota_unclassified(100);p__Ascomycota_unclassified(100);p__Ascomycota_unclassified(100); | - | - | - | - | - | - | - | Unassigned | - |
| Otu50476 | 1 | k__Fungi(100);p__Ascomycota(100);p__Ascomycota_unclassified(100);p__Ascomycota_unclassified(100);p__Ascomycota_unclassified(100);p__Ascomycota_unclassified(100);p__Ascomycota_unclassified(100); | - | - | - | - | - | - | - | Unassigned | - |
| Otu50483 | 1 | k__Fungi(100);p__Ascomycota(100);p__Ascomycota_unclassified(100);p__Ascomycota_unclassified(100);p__Ascomycota_unclassified(100);p__Ascomycota_unclassified(100);p__Ascomycota_unclassified(100); | - | - | - | - | - | - | - | Unassigned | - |
| Otu50485 | 1 | k__Fungi(100);p__Ascomycota(100);p__Ascomycota_unclassified(100);p__Ascomycota_unclassified(100);p__Ascomycota_unclassified(100);p__Ascomycota_unclassified(100);p__Ascomycota_unclassified(100); | - | - | - | - | - | - | - | Unassigned | - |
| Otu50486 | 1 | k__Fungi(100);k__Fungi_unclassified(100);k__Fungi_unclassified(100);k__Fungi_unclassified(100);k__Fungi_unclassified(100);k__Fungi_unclassified(100);k__Fungi_unclassified(100); | - | - | - | - | - | - | - | Unassigned | - |
| Otu50487 | 1 | k__Fungi(100);k__Fungi_unclassified(100);k__Fungi_unclassified(100);k__Fungi_unclassified(100);k__Fungi_unclassified(100);k__Fungi_unclassified(100);k__Fungi_unclassified(100); | - | - | - | - | - | - | - | Unassigned | - |
| Otu50488 | 1 | k__Fungi(100);p__Ascomycota(100);p__Ascomycota_unclassified(100);p__Ascomycota_unclassified(100);p__Ascomycota_unclassified(100);p__Ascomycota_unclassified(100);p__Ascomycota_unclassified(100); | - | - | - | - | - | - | - | Unassigned | - |
| Otu50489 | 1 | k__Fungi(100);p__Ascomycota(100);p__Ascomycota_unclassified(100);p__Ascomycota_unclassified(100);p__Ascomycota_unclassified(100);p__Ascomycota_unclassified(100);p__Ascomycota_unclassified(100); | - | - | - | - | - | - | - | Unassigned | - |
| Otu50494 | 1 | k__Fungi(100);k__Fungi_unclassified(100);k__Fungi_unclassified(100);k__Fungi_unclassified(100);k__Fungi_unclassified(100);k__Fungi_unclassified(100);k__Fungi_unclassified(100); | - | - | - | - | - | - | - | Unassigned | - |
| Otu50511 | 1 | k__Fungi(100);p__Ascomycota(100);p__Ascomycota_unclassified(100);p__Ascomycota_unclassified(100);p__Ascomycota_unclassified(100);p__Ascomycota_unclassified(100);p__Ascomycota_unclassified(100); | - | - | - | - | - | - | - | Unassigned | - |
| Otu50544 | 1 | k__Fungi(100);k__Fungi_unclassified(100);k__Fungi_unclassified(100);k__Fungi_unclassified(100);k__Fungi_unclassified(100);k__Fungi_unclassified(100);k__Fungi_unclassified(100); | - | - | - | - | - | - | - | Unassigned | - |
| Otu50607 | 1 | k__Fungi(100);k__Fungi_unclassified(100);k__Fungi_unclassified(100);k__Fungi_unclassified(100);k__Fungi_unclassified(100);k__Fungi_unclassified(100);k__Fungi_unclassified(100); | - | - | - | - | - | - | - | Unassigned | - |
| Otu50642 | 1 | k__Fungi(100);k__Fungi_unclassified(100);k__Fungi_unclassified(100);k__Fungi_unclassified(100);k__Fungi_unclassified(100);k__Fungi_unclassified(100);k__Fungi_unclassified(100); | - | - | - | - | - | - | - | Unassigned | - |
| Otu50646 | 1 | k__Fungi(100);k__Fungi_unclassified(100);k__Fungi_unclassified(100);k__Fungi_unclassified(100);k__Fungi_unclassified(100);k__Fungi_unclassified(100);k__Fungi_unclassified(100); | - | - | - | - | - | - | - | Unassigned | - |
| Otu50647 | 1 | k__Fungi(100);p__Ascomycota(100);p__Ascomycota_unclassified(100);p__Ascomycota_unclassified(100);p__Ascomycota_unclassified(100);p__Ascomycota_unclassified(100);p__Ascomycota_unclassified(100); | - | - | - | - | - | - | - | Unassigned | - |
| Otu50649 | 1 | k__Fungi(100);p__Ascomycota(100);p__Ascomycota_unclassified(100);p__Ascomycota_unclassified(100);p__Ascomycota_unclassified(100);p__Ascomycota_unclassified(100);p__Ascomycota_unclassified(100); | - | - | - | - | - | - | - | Unassigned | - |
| Otu50651 | 1 | k__Fungi(100);k__Fungi_unclassified(100);k__Fungi_unclassified(100);k__Fungi_unclassified(100);k__Fungi_unclassified(100);k__Fungi_unclassified(100);k__Fungi_unclassified(100); | - | - | - | - | - | - | - | Unassigned | - |
| Otu50652 | 1 | k__Fungi(100);k__Fungi_unclassified(100);k__Fungi_unclassified(100);k__Fungi_unclassified(100);k__Fungi_unclassified(100);k__Fungi_unclassified(100);k__Fungi_unclassified(100); | - | - | - | - | - | - | - | Unassigned | - |
| Otu50656 | 1 | k__Fungi(100);k__Fungi_unclassified(100);k__Fungi_unclassified(100);k__Fungi_unclassified(100);k__Fungi_unclassified(100);k__Fungi_unclassified(100);k__Fungi_unclassified(100); | - | - | - | - | - | - | - | Unassigned | - |
| Otu50659 | 1 | k__Fungi(100);p__Ascomycota(100);p__Ascomycota_unclassified(100);p__Ascomycota_unclassified(100);p__Ascomycota_unclassified(100);p__Ascomycota_unclassified(100);p__Ascomycota_unclassified(100); | - | - | - | - | - | - | - | Unassigned | - |
| Otu50695 | 1 | k__Fungi(100);k__Fungi_unclassified(100);k__Fungi_unclassified(100);k__Fungi_unclassified(100);k__Fungi_unclassified(100);k__Fungi_unclassified(100);k__Fungi_unclassified(100); | - | - | - | - | - | - | - | Unassigned | - |
| Otu50698 | 1 | k__Fungi(100);k__Fungi_unclassified(100);k__Fungi_unclassified(100);k__Fungi_unclassified(100);k__Fungi_unclassified(100);k__Fungi_unclassified(100);k__Fungi_unclassified(100); | - | - | - | - | - | - | - | Unassigned | - |
| Otu50708 | 1 | k__Fungi(100);k__Fungi_unclassified(100);k__Fungi_unclassified(100);k__Fungi_unclassified(100);k__Fungi_unclassified(100);k__Fungi_unclassified(100);k__Fungi_unclassified(100); | - | - | - | - | - | - | - | Unassigned | - |
| Otu50710 | 1 | k__Fungi(100);k__Fungi_unclassified(100);k__Fungi_unclassified(100);k__Fungi_unclassified(100);k__Fungi_unclassified(100);k__Fungi_unclassified(100);k__Fungi_unclassified(100); | - | - | - | - | - | - | - | Unassigned | - |
| Otu50713 | 1 | k__Fungi(100);k__Fungi_unclassified(100);k__Fungi_unclassified(100);k__Fungi_unclassified(100);k__Fungi_unclassified(100);k__Fungi_unclassified(100);k__Fungi_unclassified(100); | - | - | - | - | - | - | - | Unassigned | - |
| Otu50718 | 1 | k__Fungi(100);k__Fungi_unclassified(100);k__Fungi_unclassified(100);k__Fungi_unclassified(100);k__Fungi_unclassified(100);k__Fungi_unclassified(100);k__Fungi_unclassified(100); | - | - | - | - | - | - | - | Unassigned | - |
| Otu50723 | 1 | k__Fungi(100);k__Fungi_unclassified(100);k__Fungi_unclassified(100);k__Fungi_unclassified(100);k__Fungi_unclassified(100);k__Fungi_unclassified(100);k__Fungi_unclassified(100); | - | - | - | - | - | - | - | Unassigned | - |
| Otu50751 | 1 | k__Fungi(100);p__Ascomycota(100);p__Ascomycota_unclassified(100);p__Ascomycota_unclassified(100);p__Ascomycota_unclassified(100);p__Ascomycota_unclassified(100);p__Ascomycota_unclassified(100); | - | - | - | - | - | - | - | Unassigned | - |
| Otu50752 | 1 | k__Fungi(100);p__Ascomycota(100);p__Ascomycota_unclassified(100);p__Ascomycota_unclassified(100);p__Ascomycota_unclassified(100);p__Ascomycota_unclassified(100);p__Ascomycota_unclassified(100); | - | - | - | - | - | - | - | Unassigned | - |
| Otu50753 | 1 | k__Fungi(100);p__Ascomycota(100);p__Ascomycota_unclassified(100);p__Ascomycota_unclassified(100);p__Ascomycota_unclassified(100);p__Ascomycota_unclassified(100);p__Ascomycota_unclassified(100); | - | - | - | - | - | - | - | Unassigned | - |
| Otu50754 | 1 | k__Fungi(100);k__Fungi_unclassified(100);k__Fungi_unclassified(100);k__Fungi_unclassified(100);k__Fungi_unclassified(100);k__Fungi_unclassified(100);k__Fungi_unclassified(100); | - | - | - | - | - | - | - | Unassigned | - |
| Otu50759 | 1 | k__Fungi(100);p__Ascomycota(100);p__Ascomycota_unclassified(100);p__Ascomycota_unclassified(100);p__Ascomycota_unclassified(100);p__Ascomycota_unclassified(100);p__Ascomycota_unclassified(100); | - | - | - | - | - | - | - | Unassigned | - |
| Otu50767 | 1 | k__Fungi(100);p__Ascomycota(100);p__Ascomycota_unclassified(100);p__Ascomycota_unclassified(100);p__Ascomycota_unclassified(100);p__Ascomycota_unclassified(100);p__Ascomycota_unclassified(100); | - | - | - | - | - | - | - | Unassigned | - |
| Otu50778 | 1 | k__Fungi(100);k__Fungi_unclassified(100);k__Fungi_unclassified(100);k__Fungi_unclassified(100);k__Fungi_unclassified(100);k__Fungi_unclassified(100);k__Fungi_unclassified(100); | - | - | - | - | - | - | - | Unassigned | - |
| Otu50789 | 1 | k__Fungi(100);k__Fungi_unclassified(100);k__Fungi_unclassified(100);k__Fungi_unclassified(100);k__Fungi_unclassified(100);k__Fungi_unclassified(100);k__Fungi_unclassified(100); | - | - | - | - | - | - | - | Unassigned | - |
| Otu50801 | 1 | k__Fungi(100);p__Ascomycota(100);p__Ascomycota_unclassified(100);p__Ascomycota_unclassified(100);p__Ascomycota_unclassified(100);p__Ascomycota_unclassified(100);p__Ascomycota_unclassified(100); | - | - | - | - | - | - | - | Unassigned | - |
| Otu50806 | 1 | k__Fungi(100);k__Fungi_unclassified(100);k__Fungi_unclassified(100);k__Fungi_unclassified(100);k__Fungi_unclassified(100);k__Fungi_unclassified(100);k__Fungi_unclassified(100); | - | - | - | - | - | - | - | Unassigned | - |
| Otu50832 | 1 | k__Fungi(100);p__Ascomycota(100);p__Ascomycota_unclassified(100);p__Ascomycota_unclassified(100);p__Ascomycota_unclassified(100);p__Ascomycota_unclassified(100);p__Ascomycota_unclassified(100); | - | - | - | - | - | - | - | Unassigned | - |
| Otu50834 | 1 | k__Fungi(100);p__Ascomycota(100);p__Ascomycota_unclassified(100);p__Ascomycota_unclassified(100);p__Ascomycota_unclassified(100);p__Ascomycota_unclassified(100);p__Ascomycota_unclassified(100); | - | - | - | - | - | - | - | Unassigned | - |
| Otu50835 | 1 | k__Fungi(100);k__Fungi_unclassified(100);k__Fungi_unclassified(100);k__Fungi_unclassified(100);k__Fungi_unclassified(100);k__Fungi_unclassified(100);k__Fungi_unclassified(100); | - | - | - | - | - | - | - | Unassigned | - |
| Otu50836 | 1 | k__Fungi(100);p__Ascomycota(100);p__Ascomycota_unclassified(100);p__Ascomycota_unclassified(100);p__Ascomycota_unclassified(100);p__Ascomycota_unclassified(100);p__Ascomycota_unclassified(100); | - | - | - | - | - | - | - | Unassigned | - |
| Otu50837 | 1 | k__Fungi(100);k__Fungi_unclassified(100);k__Fungi_unclassified(100);k__Fungi_unclassified(100);k__Fungi_unclassified(100);k__Fungi_unclassified(100);k__Fungi_unclassified(100); | - | - | - | - | - | - | - | Unassigned | - |
| Otu50838 | 1 | k__Fungi(100);k__Fungi_unclassified(100);k__Fungi_unclassified(100);k__Fungi_unclassified(100);k__Fungi_unclassified(100);k__Fungi_unclassified(100);k__Fungi_unclassified(100); | - | - | - | - | - | - | - | Unassigned | - |
| Otu50845 | 1 | k__Fungi(100);k__Fungi_unclassified(100);k__Fungi_unclassified(100);k__Fungi_unclassified(100);k__Fungi_unclassified(100);k__Fungi_unclassified(100);k__Fungi_unclassified(100); | - | - | - | - | - | - | - | Unassigned | - |
| Otu50847 | 1 | k__Fungi(100);k__Fungi_unclassified(100);k__Fungi_unclassified(100);k__Fungi_unclassified(100);k__Fungi_unclassified(100);k__Fungi_unclassified(100);k__Fungi_unclassified(100); | - | - | - | - | - | - | - | Unassigned | - |
| Otu50848 | 1 | k__Fungi(100);k__Fungi_unclassified(100);k__Fungi_unclassified(100);k__Fungi_unclassified(100);k__Fungi_unclassified(100);k__Fungi_unclassified(100);k__Fungi_unclassified(100); | - | - | - | - | - | - | - | Unassigned | - |
| Otu50849 | 1 | k__Fungi(100);k__Fungi_unclassified(100);k__Fungi_unclassified(100);k__Fungi_unclassified(100);k__Fungi_unclassified(100);k__Fungi_unclassified(100);k__Fungi_unclassified(100); | - | - | - | - | - | - | - | Unassigned | - |
| Otu50850 | 1 | k__Fungi(100);p__Ascomycota(100);p__Ascomycota_unclassified(100);p__Ascomycota_unclassified(100);p__Ascomycota_unclassified(100);p__Ascomycota_unclassified(100);p__Ascomycota_unclassified(100); | - | - | - | - | - | - | - | Unassigned | - |
| Otu50853 | 1 | k__Fungi(100);p__Ascomycota(100);p__Ascomycota_unclassified(100);p__Ascomycota_unclassified(100);p__Ascomycota_unclassified(100);p__Ascomycota_unclassified(100);p__Ascomycota_unclassified(100); | - | - | - | - | - | - | - | Unassigned | - |
| Otu50854 | 1 | k__Fungi(100);p__Ascomycota(100);p__Ascomycota_unclassified(100);p__Ascomycota_unclassified(100);p__Ascomycota_unclassified(100);p__Ascomycota_unclassified(100);p__Ascomycota_unclassified(100); | - | - | - | - | - | - | - | Unassigned | - |
| Otu50855 | 1 | k__Fungi(100);k__Fungi_unclassified(100);k__Fungi_unclassified(100);k__Fungi_unclassified(100);k__Fungi_unclassified(100);k__Fungi_unclassified(100);k__Fungi_unclassified(100); | - | - | - | - | - | - | - | Unassigned | - |
| Otu50856 | 1 | k__Fungi(100);p__Ascomycota(100);p__Ascomycota_unclassified(100);p__Ascomycota_unclassified(100);p__Ascomycota_unclassified(100);p__Ascomycota_unclassified(100);p__Ascomycota_unclassified(100); | - | - | - | - | - | - | - | Unassigned | - |
| Otu50857 | 1 | k__Fungi(100);p__Ascomycota(100);p__Ascomycota_unclassified(100);p__Ascomycota_unclassified(100);p__Ascomycota_unclassified(100);p__Ascomycota_unclassified(100);p__Ascomycota_unclassified(100); | - | - | - | - | - | - | - | Unassigned | - |
| Otu50858 | 1 | k__Fungi(100);k__Fungi_unclassified(100);k__Fungi_unclassified(100);k__Fungi_unclassified(100);k__Fungi_unclassified(100);k__Fungi_unclassified(100);k__Fungi_unclassified(100); | - | - | - | - | - | - | - | Unassigned | - |
| Otu50859 | 1 | k__Fungi(100);p__Ascomycota(100);p__Ascomycota_unclassified(100);p__Ascomycota_unclassified(100);p__Ascomycota_unclassified(100);p__Ascomycota_unclassified(100);p__Ascomycota_unclassified(100); | - | - | - | - | - | - | - | Unassigned | - |
| Otu50860 | 1 | k__Fungi(100);k__Fungi_unclassified(100);k__Fungi_unclassified(100);k__Fungi_unclassified(100);k__Fungi_unclassified(100);k__Fungi_unclassified(100);k__Fungi_unclassified(100); | - | - | - | - | - | - | - | Unassigned | - |
| Otu50868 | 1 | k__Fungi(100);p__Ascomycota(100);p__Ascomycota_unclassified(100);p__Ascomycota_unclassified(100);p__Ascomycota_unclassified(100);p__Ascomycota_unclassified(100);p__Ascomycota_unclassified(100); | - | - | - | - | - | - | - | Unassigned | - |
| Otu50869 | 1 | k__Fungi(100);k__Fungi_unclassified(100);k__Fungi_unclassified(100);k__Fungi_unclassified(100);k__Fungi_unclassified(100);k__Fungi_unclassified(100);k__Fungi_unclassified(100); | - | - | - | - | - | - | - | Unassigned | - |
| Otu50873 | 1 | k__Fungi(100);p__Ascomycota(100);p__Ascomycota_unclassified(100);p__Ascomycota_unclassified(100);p__Ascomycota_unclassified(100);p__Ascomycota_unclassified(100);p__Ascomycota_unclassified(100); | - | - | - | - | - | - | - | Unassigned | - |
| Otu50875 | 1 | k__Fungi(100);k__Fungi_unclassified(100);k__Fungi_unclassified(100);k__Fungi_unclassified(100);k__Fungi_unclassified(100);k__Fungi_unclassified(100);k__Fungi_unclassified(100); | - | - | - | - | - | - | - | Unassigned | - |
| Otu50876 | 1 | k__Fungi(100);k__Fungi_unclassified(100);k__Fungi_unclassified(100);k__Fungi_unclassified(100);k__Fungi_unclassified(100);k__Fungi_unclassified(100);k__Fungi_unclassified(100); | - | - | - | - | - | - | - | Unassigned | - |
| Otu50877 | 1 | k__Fungi(100);k__Fungi_unclassified(100);k__Fungi_unclassified(100);k__Fungi_unclassified(100);k__Fungi_unclassified(100);k__Fungi_unclassified(100);k__Fungi_unclassified(100); | - | - | - | - | - | - | - | Unassigned | - |
| Otu50878 | 1 | k__Fungi(100);k__Fungi_unclassified(100);k__Fungi_unclassified(100);k__Fungi_unclassified(100);k__Fungi_unclassified(100);k__Fungi_unclassified(100);k__Fungi_unclassified(100); | - | - | - | - | - | - | - | Unassigned | - |
| Otu50883 | 1 | k__Fungi(100);p__Ascomycota(100);c__Sordariomycetes(100);c__Sordariomycetes_unclassified(100);c__Sordariomycetes_unclassified(100);c__Sordariomycetes_unclassified(100);c__Sordariomycetes_unclassified(100); | - | - | - | - | - | - | - | Unassigned | - |
| Otu50905 | 1 | k__Fungi(100);p__Ascomycota(100);p__Ascomycota_unclassified(100);p__Ascomycota_unclassified(100);p__Ascomycota_unclassified(100);p__Ascomycota_unclassified(100);p__Ascomycota_unclassified(100); | - | - | - | - | - | - | - | Unassigned | - |
| Otu50906 | 1 | k__Fungi(100);p__Ascomycota(100);p__Ascomycota_unclassified(100);p__Ascomycota_unclassified(100);p__Ascomycota_unclassified(100);p__Ascomycota_unclassified(100);p__Ascomycota_unclassified(100); | - | - | - | - | - | - | - | Unassigned | - |
| Otu50908 | 1 | k__Fungi(100);k__Fungi_unclassified(100);k__Fungi_unclassified(100);k__Fungi_unclassified(100);k__Fungi_unclassified(100);k__Fungi_unclassified(100);k__Fungi_unclassified(100); | - | - | - | - | - | - | - | Unassigned | - |
| Otu50909 | 1 | k__Fungi(100);p__Ascomycota(100);p__Ascomycota_unclassified(100);p__Ascomycota_unclassified(100);p__Ascomycota_unclassified(100);p__Ascomycota_unclassified(100);p__Ascomycota_unclassified(100); | - | - | - | - | - | - | - | Unassigned | - |
| Otu50910 | 1 | k__Fungi(100);k__Fungi_unclassified(100);k__Fungi_unclassified(100);k__Fungi_unclassified(100);k__Fungi_unclassified(100);k__Fungi_unclassified(100);k__Fungi_unclassified(100); | - | - | - | - | - | - | - | Unassigned | - |
| Otu50911 | 1 | k__Fungi(100);k__Fungi_unclassified(100);k__Fungi_unclassified(100);k__Fungi_unclassified(100);k__Fungi_unclassified(100);k__Fungi_unclassified(100);k__Fungi_unclassified(100); | - | - | - | - | - | - | - | Unassigned | - |
| Otu50912 | 1 | k__Fungi(100);k__Fungi_unclassified(100);k__Fungi_unclassified(100);k__Fungi_unclassified(100);k__Fungi_unclassified(100);k__Fungi_unclassified(100);k__Fungi_unclassified(100); | - | - | - | - | - | - | - | Unassigned | - |
| Otu50916 | 1 | k__Fungi(100);k__Fungi_unclassified(100);k__Fungi_unclassified(100);k__Fungi_unclassified(100);k__Fungi_unclassified(100);k__Fungi_unclassified(100);k__Fungi_unclassified(100); | - | - | - | - | - | - | - | Unassigned | - |
| Otu50923 | 1 | k__Fungi(100);p__Ascomycota(100);p__Ascomycota_unclassified(100);p__Ascomycota_unclassified(100);p__Ascomycota_unclassified(100);p__Ascomycota_unclassified(100);p__Ascomycota_unclassified(100); | - | - | - | - | - | - | - | Unassigned | - |
| Otu50925 | 1 | k__Fungi(100);k__Fungi_unclassified(100);k__Fungi_unclassified(100);k__Fungi_unclassified(100);k__Fungi_unclassified(100);k__Fungi_unclassified(100);k__Fungi_unclassified(100); | - | - | - | - | - | - | - | Unassigned | - |
| Otu50926 | 1 | k__Fungi(100);k__Fungi_unclassified(100);k__Fungi_unclassified(100);k__Fungi_unclassified(100);k__Fungi_unclassified(100);k__Fungi_unclassified(100);k__Fungi_unclassified(100); | - | - | - | - | - | - | - | Unassigned | - |
| Otu50939 | 1 | k__Fungi(100);k__Fungi_unclassified(100);k__Fungi_unclassified(100);k__Fungi_unclassified(100);k__Fungi_unclassified(100);k__Fungi_unclassified(100);k__Fungi_unclassified(100); | - | - | - | - | - | - | - | Unassigned | - |
| Otu50941 | 1 | k__Fungi(100);p__Ascomycota(100);p__Ascomycota_unclassified(100);p__Ascomycota_unclassified(100);p__Ascomycota_unclassified(100);p__Ascomycota_unclassified(100);p__Ascomycota_unclassified(100); | - | - | - | - | - | - | - | Unassigned | - |
| Otu50958 | 1 | k__Fungi(100);k__Fungi_unclassified(100);k__Fungi_unclassified(100);k__Fungi_unclassified(100);k__Fungi_unclassified(100);k__Fungi_unclassified(100);k__Fungi_unclassified(100); | - | - | - | - | - | - | - | Unassigned | - |
| Otu50963 | 1 | k__Fungi(100);p__Ascomycota(100);p__Ascomycota_unclassified(100);p__Ascomycota_unclassified(100);p__Ascomycota_unclassified(100);p__Ascomycota_unclassified(100);p__Ascomycota_unclassified(100); | - | - | - | - | - | - | - | Unassigned | - |
| Otu50965 | 1 | k__Fungi(100);p__Ascomycota(100);p__Ascomycota_unclassified(100);p__Ascomycota_unclassified(100);p__Ascomycota_unclassified(100);p__Ascomycota_unclassified(100);p__Ascomycota_unclassified(100); | - | - | - | - | - | - | - | Unassigned | - |
| Otu50969 | 1 | k__Fungi(100);k__Fungi_unclassified(100);k__Fungi_unclassified(100);k__Fungi_unclassified(100);k__Fungi_unclassified(100);k__Fungi_unclassified(100);k__Fungi_unclassified(100); | - | - | - | - | - | - | - | Unassigned | - |
| Otu50971 | 1 | k__Fungi(100);k__Fungi_unclassified(100);k__Fungi_unclassified(100);k__Fungi_unclassified(100);k__Fungi_unclassified(100);k__Fungi_unclassified(100);k__Fungi_unclassified(100); | - | - | - | - | - | - | - | Unassigned | - |
| Otu50972 | 1 | k__Fungi(100);p__Ascomycota(100);p__Ascomycota_unclassified(100);p__Ascomycota_unclassified(100);p__Ascomycota_unclassified(100);p__Ascomycota_unclassified(100);p__Ascomycota_unclassified(100); | - | - | - | - | - | - | - | Unassigned | - |
| Otu50973 | 1 | k__Fungi(100);k__Fungi_unclassified(100);k__Fungi_unclassified(100);k__Fungi_unclassified(100);k__Fungi_unclassified(100);k__Fungi_unclassified(100);k__Fungi_unclassified(100); | - | - | - | - | - | - | - | Unassigned | - |
| Otu50975 | 1 | k__Fungi(100);p__Ascomycota(100);p__Ascomycota_unclassified(100);p__Ascomycota_unclassified(100);p__Ascomycota_unclassified(100);p__Ascomycota_unclassified(100);p__Ascomycota_unclassified(100); | - | - | - | - | - | - | - | Unassigned | - |
| Otu50976 | 1 | k__Fungi(100);p__Ascomycota(100);p__Ascomycota_unclassified(100);p__Ascomycota_unclassified(100);p__Ascomycota_unclassified(100);p__Ascomycota_unclassified(100);p__Ascomycota_unclassified(100); | - | - | - | - | - | - | - | Unassigned | - |
| Otu50984 | 1 | k__Fungi(100);k__Fungi_unclassified(100);k__Fungi_unclassified(100);k__Fungi_unclassified(100);k__Fungi_unclassified(100);k__Fungi_unclassified(100);k__Fungi_unclassified(100); | - | - | - | - | - | - | - | Unassigned | - |
| Otu50985 | 1 | k__Fungi(100);k__Fungi_unclassified(100);k__Fungi_unclassified(100);k__Fungi_unclassified(100);k__Fungi_unclassified(100);k__Fungi_unclassified(100);k__Fungi_unclassified(100); | - | - | - | - | - | - | - | Unassigned | - |
| Otu50987 | 1 | k__Fungi(100);p__Ascomycota(100);p__Ascomycota_unclassified(100);p__Ascomycota_unclassified(100);p__Ascomycota_unclassified(100);p__Ascomycota_unclassified(100);p__Ascomycota_unclassified(100); | - | - | - | - | - | - | - | Unassigned | - |
| Otu50988 | 1 | k__Fungi(100);k__Fungi_unclassified(100);k__Fungi_unclassified(100);k__Fungi_unclassified(100);k__Fungi_unclassified(100);k__Fungi_unclassified(100);k__Fungi_unclassified(100); | - | - | - | - | - | - | - | Unassigned | - |
| Otu50989 | 1 | k__Fungi(100);k__Fungi_unclassified(100);k__Fungi_unclassified(100);k__Fungi_unclassified(100);k__Fungi_unclassified(100);k__Fungi_unclassified(100);k__Fungi_unclassified(100); | - | - | - | - | - | - | - | Unassigned | - |
| Otu50991 | 1 | k__Fungi(100);k__Fungi_unclassified(100);k__Fungi_unclassified(100);k__Fungi_unclassified(100);k__Fungi_unclassified(100);k__Fungi_unclassified(100);k__Fungi_unclassified(100); | - | - | - | - | - | - | - | Unassigned | - |
| Otu50992 | 1 | k__Fungi(100);p__Ascomycota(100);p__Ascomycota_unclassified(100);p__Ascomycota_unclassified(100);p__Ascomycota_unclassified(100);p__Ascomycota_unclassified(100);p__Ascomycota_unclassified(100); | - | - | - | - | - | - | - | Unassigned | - |
| Otu50993 | 1 | k__Fungi(100);k__Fungi_unclassified(100);k__Fungi_unclassified(100);k__Fungi_unclassified(100);k__Fungi_unclassified(100);k__Fungi_unclassified(100);k__Fungi_unclassified(100); | - | - | - | - | - | - | - | Unassigned | - |
| Otu50996 | 1 | k__Fungi(100);p__Ascomycota(100);p__Ascomycota_unclassified(100);p__Ascomycota_unclassified(100);p__Ascomycota_unclassified(100);p__Ascomycota_unclassified(100);p__Ascomycota_unclassified(100); | - | - | - | - | - | - | - | Unassigned | - |
| Otu51003 | 1 | k__Fungi(100);p__Ascomycota(100);p__Ascomycota_unclassified(100);p__Ascomycota_unclassified(100);p__Ascomycota_unclassified(100);p__Ascomycota_unclassified(100);p__Ascomycota_unclassified(100); | - | - | - | - | - | - | - | Unassigned | - |
| Otu51026 | 1 | k__Fungi(100);k__Fungi_unclassified(100);k__Fungi_unclassified(100);k__Fungi_unclassified(100);k__Fungi_unclassified(100);k__Fungi_unclassified(100);k__Fungi_unclassified(100); | - | - | - | - | - | - | - | Unassigned | - |
| Otu51027 | 1 | k__Fungi(100);k__Fungi_unclassified(100);k__Fungi_unclassified(100);k__Fungi_unclassified(100);k__Fungi_unclassified(100);k__Fungi_unclassified(100);k__Fungi_unclassified(100); | - | - | - | - | - | - | - | Unassigned | - |
| Otu51028 | 1 | k__Fungi(100);k__Fungi_unclassified(100);k__Fungi_unclassified(100);k__Fungi_unclassified(100);k__Fungi_unclassified(100);k__Fungi_unclassified(100);k__Fungi_unclassified(100); | - | - | - | - | - | - | - | Unassigned | - |
| Otu51029 | 1 | k__Fungi(100);k__Fungi_unclassified(100);k__Fungi_unclassified(100);k__Fungi_unclassified(100);k__Fungi_unclassified(100);k__Fungi_unclassified(100);k__Fungi_unclassified(100); | - | - | - | - | - | - | - | Unassigned | - |
| Otu51030 | 1 | k__Fungi(100);k__Fungi_unclassified(100);k__Fungi_unclassified(100);k__Fungi_unclassified(100);k__Fungi_unclassified(100);k__Fungi_unclassified(100);k__Fungi_unclassified(100); | - | - | - | - | - | - | - | Unassigned | - |
| Otu51032 | 1 | k__Fungi(100);k__Fungi_unclassified(100);k__Fungi_unclassified(100);k__Fungi_unclassified(100);k__Fungi_unclassified(100);k__Fungi_unclassified(100);k__Fungi_unclassified(100); | - | - | - | - | - | - | - | Unassigned | - |
| Otu51033 | 1 | k__Fungi(100);k__Fungi_unclassified(100);k__Fungi_unclassified(100);k__Fungi_unclassified(100);k__Fungi_unclassified(100);k__Fungi_unclassified(100);k__Fungi_unclassified(100); | - | - | - | - | - | - | - | Unassigned | - |
| Otu51034 | 1 | k__Fungi(100);p__Ascomycota(100);p__Ascomycota_unclassified(100);p__Ascomycota_unclassified(100);p__Ascomycota_unclassified(100);p__Ascomycota_unclassified(100);p__Ascomycota_unclassified(100); | - | - | - | - | - | - | - | Unassigned | - |
| Otu51036 | 1 | k__Fungi(100);p__Ascomycota(100);p__Ascomycota_unclassified(100);p__Ascomycota_unclassified(100);p__Ascomycota_unclassified(100);p__Ascomycota_unclassified(100);p__Ascomycota_unclassified(100); | - | - | - | - | - | - | - | Unassigned | - |
| Otu51045 | 1 | k__Fungi(100);p__Ascomycota(100);c__Leotiomycetes(100);o__Helotiales(100);f__Helotiales_family_Incertae_sedis(100);f__Helotiales_family_Incertae_sedis_unclassified(100);f__Helotiales_family_Incertae_sedis_unclassified(100); | - | - | - | - | - | - | - | Unassigned | - |
| Otu51063 | 1 | k__Fungi(100);k__Fungi_unclassified(100);k__Fungi_unclassified(100);k__Fungi_unclassified(100);k__Fungi_unclassified(100);k__Fungi_unclassified(100);k__Fungi_unclassified(100); | - | - | - | - | - | - | - | Unassigned | - |
| Otu51066 | 1 | k__Fungi(100);k__Fungi_unclassified(100);k__Fungi_unclassified(100);k__Fungi_unclassified(100);k__Fungi_unclassified(100);k__Fungi_unclassified(100);k__Fungi_unclassified(100); | - | - | - | - | - | - | - | Unassigned | - |
| Otu51091 | 1 | k__Fungi(100);p__Ascomycota(100);p__Ascomycota_unclassified(100);p__Ascomycota_unclassified(100);p__Ascomycota_unclassified(100);p__Ascomycota_unclassified(100);p__Ascomycota_unclassified(100); | - | - | - | - | - | - | - | Unassigned | - |
| Otu51094 | 1 | k__Fungi(100);k__Fungi_unclassified(100);k__Fungi_unclassified(100);k__Fungi_unclassified(100);k__Fungi_unclassified(100);k__Fungi_unclassified(100);k__Fungi_unclassified(100); | - | - | - | - | - | - | - | Unassigned | - |
| Otu51101 | 1 | k__Fungi(100);k__Fungi_unclassified(100);k__Fungi_unclassified(100);k__Fungi_unclassified(100);k__Fungi_unclassified(100);k__Fungi_unclassified(100);k__Fungi_unclassified(100); | - | - | - | - | - | - | - | Unassigned | - |
| Otu51102 | 1 | k__Fungi(100);k__Fungi_unclassified(100);k__Fungi_unclassified(100);k__Fungi_unclassified(100);k__Fungi_unclassified(100);k__Fungi_unclassified(100);k__Fungi_unclassified(100); | - | - | - | - | - | - | - | Unassigned | - |
| Otu51104 | 1 | k__Fungi(100);k__Fungi_unclassified(100);k__Fungi_unclassified(100);k__Fungi_unclassified(100);k__Fungi_unclassified(100);k__Fungi_unclassified(100);k__Fungi_unclassified(100); | - | - | - | - | - | - | - | Unassigned | - |
| Otu51105 | 1 | k__Fungi(100);p__Ascomycota(100);p__Ascomycota_unclassified(100);p__Ascomycota_unclassified(100);p__Ascomycota_unclassified(100);p__Ascomycota_unclassified(100);p__Ascomycota_unclassified(100); | - | - | - | - | - | - | - | Unassigned | - |
| Otu51106 | 1 | k__Fungi(100);p__Ascomycota(100);p__Ascomycota_unclassified(100);p__Ascomycota_unclassified(100);p__Ascomycota_unclassified(100);p__Ascomycota_unclassified(100);p__Ascomycota_unclassified(100); | - | - | - | - | - | - | - | Unassigned | - |
| Otu51110 | 1 | k__Fungi(100);k__Fungi_unclassified(100);k__Fungi_unclassified(100);k__Fungi_unclassified(100);k__Fungi_unclassified(100);k__Fungi_unclassified(100);k__Fungi_unclassified(100); | - | - | - | - | - | - | - | Unassigned | - |
| Otu51111 | 1 | k__Fungi(100);p__Ascomycota(100);p__Ascomycota_unclassified(100);p__Ascomycota_unclassified(100);p__Ascomycota_unclassified(100);p__Ascomycota_unclassified(100);p__Ascomycota_unclassified(100); | - | - | - | - | - | - | - | Unassigned | - |
| Otu51123 | 1 | k__Fungi(100);k__Fungi_unclassified(100);k__Fungi_unclassified(100);k__Fungi_unclassified(100);k__Fungi_unclassified(100);k__Fungi_unclassified(100);k__Fungi_unclassified(100); | - | - | - | - | - | - | - | Unassigned | - |
| Otu51145 | 1 | k__Fungi(100);k__Fungi_unclassified(100);k__Fungi_unclassified(100);k__Fungi_unclassified(100);k__Fungi_unclassified(100);k__Fungi_unclassified(100);k__Fungi_unclassified(100); | - | - | - | - | - | - | - | Unassigned | - |
| Otu51150 | 1 | k__Fungi(100);p__Ascomycota(100);p__Ascomycota_unclassified(100);p__Ascomycota_unclassified(100);p__Ascomycota_unclassified(100);p__Ascomycota_unclassified(100);p__Ascomycota_unclassified(100); | - | - | - | - | - | - | - | Unassigned | - |
| Otu51154 | 1 | k__Fungi(100);p__Ascomycota(100);p__Ascomycota_unclassified(100);p__Ascomycota_unclassified(100);p__Ascomycota_unclassified(100);p__Ascomycota_unclassified(100);p__Ascomycota_unclassified(100); | - | - | - | - | - | - | - | Unassigned | - |
| Otu51156 | 1 | k__Fungi(100);p__Ascomycota(100);p__Ascomycota_unclassified(100);p__Ascomycota_unclassified(100);p__Ascomycota_unclassified(100);p__Ascomycota_unclassified(100);p__Ascomycota_unclassified(100); | - | - | - | - | - | - | - | Unassigned | - |
| Otu51157 | 1 | k__Fungi(100);k__Fungi_unclassified(100);k__Fungi_unclassified(100);k__Fungi_unclassified(100);k__Fungi_unclassified(100);k__Fungi_unclassified(100);k__Fungi_unclassified(100); | - | - | - | - | - | - | - | Unassigned | - |
| Otu51158 | 1 | k__Fungi(100);p__Ascomycota(100);p__Ascomycota_unclassified(100);p__Ascomycota_unclassified(100);p__Ascomycota_unclassified(100);p__Ascomycota_unclassified(100);p__Ascomycota_unclassified(100); | - | - | - | - | - | - | - | Unassigned | - |
| Otu51159 | 1 | k__Fungi(100);p__Ascomycota(100);p__Ascomycota_unclassified(100);p__Ascomycota_unclassified(100);p__Ascomycota_unclassified(100);p__Ascomycota_unclassified(100);p__Ascomycota_unclassified(100); | - | - | - | - | - | - | - | Unassigned | - |
| Otu51164 | 1 | k__Fungi(100);p__Ascomycota(100);p__Ascomycota_unclassified(100);p__Ascomycota_unclassified(100);p__Ascomycota_unclassified(100);p__Ascomycota_unclassified(100);p__Ascomycota_unclassified(100); | - | - | - | - | - | - | - | Unassigned | - |
| Otu51174 | 1 | k__Fungi(100);p__Ascomycota(100);p__Ascomycota_unclassified(100);p__Ascomycota_unclassified(100);p__Ascomycota_unclassified(100);p__Ascomycota_unclassified(100);p__Ascomycota_unclassified(100); | - | - | - | - | - | - | - | Unassigned | - |
| Otu51184 | 1 | k__Fungi(100);k__Fungi_unclassified(100);k__Fungi_unclassified(100);k__Fungi_unclassified(100);k__Fungi_unclassified(100);k__Fungi_unclassified(100);k__Fungi_unclassified(100); | - | - | - | - | - | - | - | Unassigned | - |
| Otu51193 | 1 | k__Fungi(100);k__Fungi_unclassified(100);k__Fungi_unclassified(100);k__Fungi_unclassified(100);k__Fungi_unclassified(100);k__Fungi_unclassified(100);k__Fungi_unclassified(100); | - | - | - | - | - | - | - | Unassigned | - |
| Otu51207 | 1 | k__Fungi(100);k__Fungi_unclassified(100);k__Fungi_unclassified(100);k__Fungi_unclassified(100);k__Fungi_unclassified(100);k__Fungi_unclassified(100);k__Fungi_unclassified(100); | - | - | - | - | - | - | - | Unassigned | - |
| Otu51219 | 1 | k__Fungi(100);p__Ascomycota(100);p__Ascomycota_unclassified(100);p__Ascomycota_unclassified(100);p__Ascomycota_unclassified(100);p__Ascomycota_unclassified(100);p__Ascomycota_unclassified(100); | - | - | - | - | - | - | - | Unassigned | - |
| Otu51236 | 1 | k__Fungi(100);k__Fungi_unclassified(100);k__Fungi_unclassified(100);k__Fungi_unclassified(100);k__Fungi_unclassified(100);k__Fungi_unclassified(100);k__Fungi_unclassified(100); | - | - | - | - | - | - | - | Unassigned | - |
| Otu51250 | 1 | k__Fungi(100);k__Fungi_unclassified(100);k__Fungi_unclassified(100);k__Fungi_unclassified(100);k__Fungi_unclassified(100);k__Fungi_unclassified(100);k__Fungi_unclassified(100); | - | - | - | - | - | - | - | Unassigned | - |
| Otu51269 | 1 | k__Fungi(100);p__Ascomycota(100);p__Ascomycota_unclassified(100);p__Ascomycota_unclassified(100);p__Ascomycota_unclassified(100);p__Ascomycota_unclassified(100);p__Ascomycota_unclassified(100); | - | - | - | - | - | - | - | Unassigned | - |
| Otu51281 | 1 | k__Fungi(100);k__Fungi_unclassified(100);k__Fungi_unclassified(100);k__Fungi_unclassified(100);k__Fungi_unclassified(100);k__Fungi_unclassified(100);k__Fungi_unclassified(100); | - | - | - | - | - | - | - | Unassigned | - |
| Otu51283 | 1 | k__Fungi(100);p__Ascomycota(100);p__Ascomycota_unclassified(100);p__Ascomycota_unclassified(100);p__Ascomycota_unclassified(100);p__Ascomycota_unclassified(100);p__Ascomycota_unclassified(100); | - | - | - | - | - | - | - | Unassigned | - |
| Otu51284 | 1 | k__Fungi(100);p__Ascomycota(100);p__Ascomycota_unclassified(100);p__Ascomycota_unclassified(100);p__Ascomycota_unclassified(100);p__Ascomycota_unclassified(100);p__Ascomycota_unclassified(100); | - | - | - | - | - | - | - | Unassigned | - |
| Otu51285 | 1 | k__Fungi(100);k__Fungi_unclassified(100);k__Fungi_unclassified(100);k__Fungi_unclassified(100);k__Fungi_unclassified(100);k__Fungi_unclassified(100);k__Fungi_unclassified(100); | - | - | - | - | - | - | - | Unassigned | - |
| Otu51289 | 1 | k__Fungi(100);p__Ascomycota(100);p__Ascomycota_unclassified(100);p__Ascomycota_unclassified(100);p__Ascomycota_unclassified(100);p__Ascomycota_unclassified(100);p__Ascomycota_unclassified(100); | - | - | - | - | - | - | - | Unassigned | - |
| Otu51291 | 1 | k__Fungi(100);p__Ascomycota(100);p__Ascomycota_unclassified(100);p__Ascomycota_unclassified(100);p__Ascomycota_unclassified(100);p__Ascomycota_unclassified(100);p__Ascomycota_unclassified(100); | - | - | - | - | - | - | - | Unassigned | - |
| Otu51341 | 1 | k__Fungi(100);p__Ascomycota(100);p__Ascomycota_unclassified(100);p__Ascomycota_unclassified(100);p__Ascomycota_unclassified(100);p__Ascomycota_unclassified(100);p__Ascomycota_unclassified(100); | - | - | - | - | - | - | - | Unassigned | - |
| Otu51398 | 1 | k__Fungi(100);k__Fungi_unclassified(100);k__Fungi_unclassified(100);k__Fungi_unclassified(100);k__Fungi_unclassified(100);k__Fungi_unclassified(100);k__Fungi_unclassified(100); | - | - | - | - | - | - | - | Unassigned | - |
| Otu51407 | 1 | k__Fungi(100);p__Ascomycota(100);p__Ascomycota_unclassified(100);p__Ascomycota_unclassified(100);p__Ascomycota_unclassified(100);p__Ascomycota_unclassified(100);p__Ascomycota_unclassified(100); | - | - | - | - | - | - | - | Unassigned | - |
| Otu51408 | 1 | k__Fungi(100);p__Ascomycota(100);p__Ascomycota_unclassified(100);p__Ascomycota_unclassified(100);p__Ascomycota_unclassified(100);p__Ascomycota_unclassified(100);p__Ascomycota_unclassified(100); | - | - | - | - | - | - | - | Unassigned | - |
| Otu51410 | 1 | k__Fungi(100);k__Fungi_unclassified(100);k__Fungi_unclassified(100);k__Fungi_unclassified(100);k__Fungi_unclassified(100);k__Fungi_unclassified(100);k__Fungi_unclassified(100); | - | - | - | - | - | - | - | Unassigned | - |
| Otu51438 | 1 | k__Fungi(100);k__Fungi_unclassified(100);k__Fungi_unclassified(100);k__Fungi_unclassified(100);k__Fungi_unclassified(100);k__Fungi_unclassified(100);k__Fungi_unclassified(100); | - | - | - | - | - | - | - | Unassigned | - |
| Otu51452 | 1 | k__Fungi(100);p__Ascomycota(100);p__Ascomycota_unclassified(100);p__Ascomycota_unclassified(100);p__Ascomycota_unclassified(100);p__Ascomycota_unclassified(100);p__Ascomycota_unclassified(100); | - | - | - | - | - | - | - | Unassigned | - |
| Otu51469 | 1 | k__Fungi(100);p__Ascomycota(100);p__Ascomycota_unclassified(100);p__Ascomycota_unclassified(100);p__Ascomycota_unclassified(100);p__Ascomycota_unclassified(100);p__Ascomycota_unclassified(100); | - | - | - | - | - | - | - | Unassigned | - |
| Otu51471 | 1 | k__Fungi(100);k__Fungi_unclassified(100);k__Fungi_unclassified(100);k__Fungi_unclassified(100);k__Fungi_unclassified(100);k__Fungi_unclassified(100);k__Fungi_unclassified(100); | - | - | - | - | - | - | - | Unassigned | - |
| Otu51478 | 1 | k__Fungi(100);k__Fungi_unclassified(100);k__Fungi_unclassified(100);k__Fungi_unclassified(100);k__Fungi_unclassified(100);k__Fungi_unclassified(100);k__Fungi_unclassified(100); | - | - | - | - | - | - | - | Unassigned | - |
| Otu51484 | 1 | k__Fungi(100);p__Ascomycota(100);p__Ascomycota_unclassified(100);p__Ascomycota_unclassified(100);p__Ascomycota_unclassified(100);p__Ascomycota_unclassified(100);p__Ascomycota_unclassified(100); | - | - | - | - | - | - | - | Unassigned | - |
| Otu51502 | 1 | k__Fungi(100);k__Fungi_unclassified(100);k__Fungi_unclassified(100);k__Fungi_unclassified(100);k__Fungi_unclassified(100);k__Fungi_unclassified(100);k__Fungi_unclassified(100); | - | - | - | - | - | - | - | Unassigned | - |
| Otu51503 | 1 | k__Fungi(100);p__Ascomycota(100);p__Ascomycota_unclassified(100);p__Ascomycota_unclassified(100);p__Ascomycota_unclassified(100);p__Ascomycota_unclassified(100);p__Ascomycota_unclassified(100); | - | - | - | - | - | - | - | Unassigned | - |
| Otu51504 | 1 | k__Fungi(100);p__Ascomycota(100);p__Ascomycota_unclassified(100);p__Ascomycota_unclassified(100);p__Ascomycota_unclassified(100);p__Ascomycota_unclassified(100);p__Ascomycota_unclassified(100); | - | - | - | - | - | - | - | Unassigned | - |
| Otu51505 | 1 | k__Fungi(100);p__Ascomycota(100);p__Ascomycota_unclassified(100);p__Ascomycota_unclassified(100);p__Ascomycota_unclassified(100);p__Ascomycota_unclassified(100);p__Ascomycota_unclassified(100); | - | - | - | - | - | - | - | Unassigned | - |
| Otu51508 | 1 | k__Fungi(100);k__Fungi_unclassified(100);k__Fungi_unclassified(100);k__Fungi_unclassified(100);k__Fungi_unclassified(100);k__Fungi_unclassified(100);k__Fungi_unclassified(100); | - | - | - | - | - | - | - | Unassigned | - |
| Otu51509 | 1 | k__Fungi(100);k__Fungi_unclassified(100);k__Fungi_unclassified(100);k__Fungi_unclassified(100);k__Fungi_unclassified(100);k__Fungi_unclassified(100);k__Fungi_unclassified(100); | - | - | - | - | - | - | - | Unassigned | - |
| Otu51512 | 1 | k__Fungi(100);p__Ascomycota(100);p__Ascomycota_unclassified(100);p__Ascomycota_unclassified(100);p__Ascomycota_unclassified(100);p__Ascomycota_unclassified(100);p__Ascomycota_unclassified(100); | - | - | - | - | - | - | - | Unassigned | - |
| Otu51513 | 1 | k__Fungi(100);k__Fungi_unclassified(100);k__Fungi_unclassified(100);k__Fungi_unclassified(100);k__Fungi_unclassified(100);k__Fungi_unclassified(100);k__Fungi_unclassified(100); | - | - | - | - | - | - | - | Unassigned | - |
| Otu51515 | 1 | k__Fungi(100);p__Ascomycota(100);p__Ascomycota_unclassified(100);p__Ascomycota_unclassified(100);p__Ascomycota_unclassified(100);p__Ascomycota_unclassified(100);p__Ascomycota_unclassified(100); | - | - | - | - | - | - | - | Unassigned | - |
| Otu51517 | 1 | k__Fungi(100);p__Ascomycota(100);p__Ascomycota_unclassified(100);p__Ascomycota_unclassified(100);p__Ascomycota_unclassified(100);p__Ascomycota_unclassified(100);p__Ascomycota_unclassified(100); | - | - | - | - | - | - | - | Unassigned | - |
| Otu51518 | 1 | k__Fungi(100);p__Ascomycota(100);p__Ascomycota_unclassified(100);p__Ascomycota_unclassified(100);p__Ascomycota_unclassified(100);p__Ascomycota_unclassified(100);p__Ascomycota_unclassified(100); | - | - | - | - | - | - | - | Unassigned | - |
| Otu51519 | 1 | k__Fungi(100);p__Ascomycota(100);p__Ascomycota_unclassified(100);p__Ascomycota_unclassified(100);p__Ascomycota_unclassified(100);p__Ascomycota_unclassified(100);p__Ascomycota_unclassified(100); | - | - | - | - | - | - | - | Unassigned | - |
| Otu51520 | 1 | k__Fungi(100);k__Fungi_unclassified(100);k__Fungi_unclassified(100);k__Fungi_unclassified(100);k__Fungi_unclassified(100);k__Fungi_unclassified(100);k__Fungi_unclassified(100); | - | - | - | - | - | - | - | Unassigned | - |
| Otu51521 | 1 | k__Fungi(100);p__Ascomycota(100);p__Ascomycota_unclassified(100);p__Ascomycota_unclassified(100);p__Ascomycota_unclassified(100);p__Ascomycota_unclassified(100);p__Ascomycota_unclassified(100); | - | - | - | - | - | - | - | Unassigned | - |
| Otu51522 | 1 | k__Fungi(100);k__Fungi_unclassified(100);k__Fungi_unclassified(100);k__Fungi_unclassified(100);k__Fungi_unclassified(100);k__Fungi_unclassified(100);k__Fungi_unclassified(100); | - | - | - | - | - | - | - | Unassigned | - |
| Otu51523 | 1 | k__Fungi(100);p__Ascomycota(100);p__Ascomycota_unclassified(100);p__Ascomycota_unclassified(100);p__Ascomycota_unclassified(100);p__Ascomycota_unclassified(100);p__Ascomycota_unclassified(100); | - | - | - | - | - | - | - | Unassigned | - |
| Otu51524 | 1 | k__Fungi(100);k__Fungi_unclassified(100);k__Fungi_unclassified(100);k__Fungi_unclassified(100);k__Fungi_unclassified(100);k__Fungi_unclassified(100);k__Fungi_unclassified(100); | - | - | - | - | - | - | - | Unassigned | - |
| Otu51526 | 1 | k__Fungi(100);p__Ascomycota(100);p__Ascomycota_unclassified(100);p__Ascomycota_unclassified(100);p__Ascomycota_unclassified(100);p__Ascomycota_unclassified(100);p__Ascomycota_unclassified(100); | - | - | - | - | - | - | - | Unassigned | - |
| Otu51529 | 1 | k__Fungi(100);k__Fungi_unclassified(100);k__Fungi_unclassified(100);k__Fungi_unclassified(100);k__Fungi_unclassified(100);k__Fungi_unclassified(100);k__Fungi_unclassified(100); | - | - | - | - | - | - | - | Unassigned | - |
| Otu51530 | 1 | k__Fungi(100);k__Fungi_unclassified(100);k__Fungi_unclassified(100);k__Fungi_unclassified(100);k__Fungi_unclassified(100);k__Fungi_unclassified(100);k__Fungi_unclassified(100); | - | - | - | - | - | - | - | Unassigned | - |
| Otu51540 | 1 | k__Fungi(100);k__Fungi_unclassified(100);k__Fungi_unclassified(100);k__Fungi_unclassified(100);k__Fungi_unclassified(100);k__Fungi_unclassified(100);k__Fungi_unclassified(100); | - | - | - | - | - | - | - | Unassigned | - |
| Otu51550 | 1 | k__Fungi(100);p__Ascomycota(100);p__Ascomycota_unclassified(100);p__Ascomycota_unclassified(100);p__Ascomycota_unclassified(100);p__Ascomycota_unclassified(100);p__Ascomycota_unclassified(100); | - | - | - | - | - | - | - | Unassigned | - |
| Otu51551 | 1 | k__Fungi(100);k__Fungi_unclassified(100);k__Fungi_unclassified(100);k__Fungi_unclassified(100);k__Fungi_unclassified(100);k__Fungi_unclassified(100);k__Fungi_unclassified(100); | - | - | - | - | - | - | - | Unassigned | - |
| Otu51552 | 1 | k__Fungi(100);p__Ascomycota(100);p__Ascomycota_unclassified(100);p__Ascomycota_unclassified(100);p__Ascomycota_unclassified(100);p__Ascomycota_unclassified(100);p__Ascomycota_unclassified(100); | - | - | - | - | - | - | - | Unassigned | - |
| Otu51553 | 1 | k__Fungi(100);p__Ascomycota(100);p__Ascomycota_unclassified(100);p__Ascomycota_unclassified(100);p__Ascomycota_unclassified(100);p__Ascomycota_unclassified(100);p__Ascomycota_unclassified(100); | - | - | - | - | - | - | - | Unassigned | - |
| Otu51562 | 1 | k__Fungi(100);k__Fungi_unclassified(100);k__Fungi_unclassified(100);k__Fungi_unclassified(100);k__Fungi_unclassified(100);k__Fungi_unclassified(100);k__Fungi_unclassified(100); | - | - | - | - | - | - | - | Unassigned | - |
| Otu51569 | 1 | k__Fungi(100);p__Ascomycota(100);p__Ascomycota_unclassified(100);p__Ascomycota_unclassified(100);p__Ascomycota_unclassified(100);p__Ascomycota_unclassified(100);p__Ascomycota_unclassified(100); | - | - | - | - | - | - | - | Unassigned | - |
| Otu51570 | 1 | k__Fungi(100);p__Ascomycota(100);p__Ascomycota_unclassified(100);p__Ascomycota_unclassified(100);p__Ascomycota_unclassified(100);p__Ascomycota_unclassified(100);p__Ascomycota_unclassified(100); | - | - | - | - | - | - | - | Unassigned | - |
| Otu51572 | 1 | k__Fungi(100);k__Fungi_unclassified(100);k__Fungi_unclassified(100);k__Fungi_unclassified(100);k__Fungi_unclassified(100);k__Fungi_unclassified(100);k__Fungi_unclassified(100); | - | - | - | - | - | - | - | Unassigned | - |
| Otu51574 | 1 | k__Fungi(100);k__Fungi_unclassified(100);k__Fungi_unclassified(100);k__Fungi_unclassified(100);k__Fungi_unclassified(100);k__Fungi_unclassified(100);k__Fungi_unclassified(100); | - | - | - | - | - | - | - | Unassigned | - |
| Otu51575 | 1 | k__Fungi(100);p__Ascomycota(100);p__Ascomycota_unclassified(100);p__Ascomycota_unclassified(100);p__Ascomycota_unclassified(100);p__Ascomycota_unclassified(100);p__Ascomycota_unclassified(100); | - | - | - | - | - | - | - | Unassigned | - |
| Otu51607 | 1 | k__Fungi(100);k__Fungi_unclassified(100);k__Fungi_unclassified(100);k__Fungi_unclassified(100);k__Fungi_unclassified(100);k__Fungi_unclassified(100);k__Fungi_unclassified(100); | - | - | - | - | - | - | - | Unassigned | - |
| Otu51682 | 1 | k__Fungi(100);k__Fungi_unclassified(100);k__Fungi_unclassified(100);k__Fungi_unclassified(100);k__Fungi_unclassified(100);k__Fungi_unclassified(100);k__Fungi_unclassified(100); | - | - | - | - | - | - | - | Unassigned | - |
| Otu51690 | 1 | k__Fungi(100);k__Fungi_unclassified(100);k__Fungi_unclassified(100);k__Fungi_unclassified(100);k__Fungi_unclassified(100);k__Fungi_unclassified(100);k__Fungi_unclassified(100); | - | - | - | - | - | - | - | Unassigned | - |
| Otu51737 | 1 | k__Fungi(100);k__Fungi_unclassified(100);k__Fungi_unclassified(100);k__Fungi_unclassified(100);k__Fungi_unclassified(100);k__Fungi_unclassified(100);k__Fungi_unclassified(100); | - | - | - | - | - | - | - | Unassigned | - |
| Otu51746 | 1 | k__Fungi(100);k__Fungi_unclassified(100);k__Fungi_unclassified(100);k__Fungi_unclassified(100);k__Fungi_unclassified(100);k__Fungi_unclassified(100);k__Fungi_unclassified(100); | - | - | - | - | - | - | - | Unassigned | - |
| Otu51762 | 1 | k__Fungi(100);p__Ascomycota(100);p__Ascomycota_unclassified(100);p__Ascomycota_unclassified(100);p__Ascomycota_unclassified(100);p__Ascomycota_unclassified(100);p__Ascomycota_unclassified(100); | - | - | - | - | - | - | - | Unassigned | - |
| Otu51763 | 1 | k__Fungi(100);k__Fungi_unclassified(100);k__Fungi_unclassified(100);k__Fungi_unclassified(100);k__Fungi_unclassified(100);k__Fungi_unclassified(100);k__Fungi_unclassified(100); | - | - | - | - | - | - | - | Unassigned | - |
| Otu51764 | 1 | k__Fungi(100);p__Ascomycota(100);p__Ascomycota_unclassified(100);p__Ascomycota_unclassified(100);p__Ascomycota_unclassified(100);p__Ascomycota_unclassified(100);p__Ascomycota_unclassified(100); | - | - | - | - | - | - | - | Unassigned | - |
| Otu51766 | 1 | k__Fungi(100);p__Ascomycota(100);p__Ascomycota_unclassified(100);p__Ascomycota_unclassified(100);p__Ascomycota_unclassified(100);p__Ascomycota_unclassified(100);p__Ascomycota_unclassified(100); | - | - | - | - | - | - | - | Unassigned | - |
| Otu51768 | 1 | k__Fungi(100);p__Ascomycota(100);p__Ascomycota_unclassified(100);p__Ascomycota_unclassified(100);p__Ascomycota_unclassified(100);p__Ascomycota_unclassified(100);p__Ascomycota_unclassified(100); | - | - | - | - | - | - | - | Unassigned | - |
| Otu51769 | 1 | k__Fungi(100);k__Fungi_unclassified(100);k__Fungi_unclassified(100);k__Fungi_unclassified(100);k__Fungi_unclassified(100);k__Fungi_unclassified(100);k__Fungi_unclassified(100); | - | - | - | - | - | - | - | Unassigned | - |
| Otu51770 | 1 | k__Fungi(100);p__Ascomycota(100);p__Ascomycota_unclassified(100);p__Ascomycota_unclassified(100);p__Ascomycota_unclassified(100);p__Ascomycota_unclassified(100);p__Ascomycota_unclassified(100); | - | - | - | - | - | - | - | Unassigned | - |
| Otu51776 | 1 | k__Fungi(100);p__Ascomycota(100);p__Ascomycota_unclassified(100);p__Ascomycota_unclassified(100);p__Ascomycota_unclassified(100);p__Ascomycota_unclassified(100);p__Ascomycota_unclassified(100); | - | - | - | - | - | - | - | Unassigned | - |
| Otu51781 | 1 | k__Fungi(100);k__Fungi_unclassified(100);k__Fungi_unclassified(100);k__Fungi_unclassified(100);k__Fungi_unclassified(100);k__Fungi_unclassified(100);k__Fungi_unclassified(100); | - | - | - | - | - | - | - | Unassigned | - |
| Otu51782 | 1 | k__Fungi(100);p__Basidiomycota(100);p__Basidiomycota_unclassified(100);p__Basidiomycota_unclassified(100);p__Basidiomycota_unclassified(100);p__Basidiomycota_unclassified(100);p__Basidiomycota_unclassified(100); | - | - | - | - | - | - | - | Unassigned | - |
| Otu51783 | 1 | k__Fungi(100);k__Fungi_unclassified(100);k__Fungi_unclassified(100);k__Fungi_unclassified(100);k__Fungi_unclassified(100);k__Fungi_unclassified(100);k__Fungi_unclassified(100); | - | - | - | - | - | - | - | Unassigned | - |
| Otu51784 | 1 | k__Fungi(100);k__Fungi_unclassified(100);k__Fungi_unclassified(100);k__Fungi_unclassified(100);k__Fungi_unclassified(100);k__Fungi_unclassified(100);k__Fungi_unclassified(100); | - | - | - | - | - | - | - | Unassigned | - |
| Otu51800 | 1 | k__Fungi(100);k__Fungi_unclassified(100);k__Fungi_unclassified(100);k__Fungi_unclassified(100);k__Fungi_unclassified(100);k__Fungi_unclassified(100);k__Fungi_unclassified(100); | - | - | - | - | - | - | - | Unassigned | - |
| Otu51810 | 1 | k__Fungi(100);p__Ascomycota(100);p__Ascomycota_unclassified(100);p__Ascomycota_unclassified(100);p__Ascomycota_unclassified(100);p__Ascomycota_unclassified(100);p__Ascomycota_unclassified(100); | - | - | - | - | - | - | - | Unassigned | - |
| Otu51811 | 1 | k__Fungi(100);k__Fungi_unclassified(100);k__Fungi_unclassified(100);k__Fungi_unclassified(100);k__Fungi_unclassified(100);k__Fungi_unclassified(100);k__Fungi_unclassified(100); | - | - | - | - | - | - | - | Unassigned | - |
| Otu51812 | 1 | k__Fungi(100);k__Fungi_unclassified(100);k__Fungi_unclassified(100);k__Fungi_unclassified(100);k__Fungi_unclassified(100);k__Fungi_unclassified(100);k__Fungi_unclassified(100); | - | - | - | - | - | - | - | Unassigned | - |

| | | | | | | | | | | | |
|---|---|---|---|---|---|---|---|---|---|---|---|
| Otu51818 | 1 | k__Fungi(100);k__Fungi_unclassified(100);k__Fungi_unclassified(100);k__Fungi_unclassified(100);k__Fungi_unclassified(100);k__Fungi_unclassified(100);k__Fungi_unclassified(100); | - | - | - | - | - | - | - | Unassigned | - |
| Otu51825 | 1 | k__Fungi(100);k__Fungi_unclassified(100);k__Fungi_unclassified(100);k__Fungi_unclassified(100);k__Fungi_unclassified(100);k__Fungi_unclassified(100);k__Fungi_unclassified(100); | - | - | - | - | - | - | - | Unassigned | - |
| Otu51828 | 1 | k__Fungi(100);k__Fungi_unclassified(100);k__Fungi_unclassified(100);k__Fungi_unclassified(100);k__Fungi_unclassified(100);k__Fungi_unclassified(100);k__Fungi_unclassified(100); | - | - | - | - | - | - | - | Unassigned | - |
| Otu51829 | 1 | k__Fungi(100);k__Fungi_unclassified(100);k__Fungi_unclassified(100);k__Fungi_unclassified(100);k__Fungi_unclassified(100);k__Fungi_unclassified(100);k__Fungi_unclassified(100); | - | - | - | - | - | - | - | Unassigned | - |
| Otu51831 | 1 | k__Fungi(100);p__Ascomycota(100);p__Ascomycota_unclassified(100);p__Ascomycota_unclassified(100);p__Ascomycota_unclassified(100);p__Ascomycota_unclassified(100);p__Ascomycota_unclassified(100); | - | - | - | - | - | - | - | Unassigned | - |
| Otu51840 | 1 | k__Fungi(100);p__Ascomycota(100);p__Ascomycota_unclassified(100);p__Ascomycota_unclassified(100);p__Ascomycota_unclassified(100);p__Ascomycota_unclassified(100);p__Ascomycota_unclassified(100); | - | - | - | - | - | - | - | Unassigned | - |
| Otu51841 | 1 | k__Fungi(100);p__Ascomycota(100);p__Ascomycota_unclassified(100);p__Ascomycota_unclassified(100);p__Ascomycota_unclassified(100);p__Ascomycota_unclassified(100);p__Ascomycota_unclassified(100); | - | - | - | - | - | - | - | Unassigned | - |
| Otu51842 | 1 | k__Fungi(100);p__Ascomycota(100);c__Dothideomycetes(100);o__Pleosporales(100);o__Pleosporales_unclassified(100);o__Pleosporales_unclassified(100);o__Pleosporales_unclassified(100); | - | - | - | - | - | - | - | Unassigned | - |
| Otu51846 | 1 | k__Fungi(100);p__Ascomycota(100);p__Ascomycota_unclassified(100);p__Ascomycota_unclassified(100);p__Ascomycota_unclassified(100);p__Ascomycota_unclassified(100);p__Ascomycota_unclassified(100); | - | - | - | - | - | - | - | Unassigned | - |
| Otu51854 | 1 | k__Fungi(100);k__Fungi_unclassified(100);k__Fungi_unclassified(100);k__Fungi_unclassified(100);k__Fungi_unclassified(100);k__Fungi_unclassified(100);k__Fungi_unclassified(100); | - | - | - | - | - | - | - | Unassigned | - |
| Otu51855 | 1 | k__Fungi(100);p__Ascomycota(100);p__Ascomycota_unclassified(100);p__Ascomycota_unclassified(100);p__Ascomycota_unclassified(100);p__Ascomycota_unclassified(100);p__Ascomycota_unclassified(100); | - | - | - | - | - | - | - | Unassigned | - |
| Otu51856 | 1 | k__Fungi(100);k__Fungi_unclassified(100);k__Fungi_unclassified(100);k__Fungi_unclassified(100);k__Fungi_unclassified(100);k__Fungi_unclassified(100);k__Fungi_unclassified(100); | - | - | - | - | - | - | - | Unassigned | - |
| Otu51862 | 1 | k__Fungi(100);p__Ascomycota(100);p__Ascomycota_unclassified(100);p__Ascomycota_unclassified(100);p__Ascomycota_unclassified(100);p__Ascomycota_unclassified(100);p__Ascomycota_unclassified(100); | - | - | - | - | - | - | - | Unassigned | - |
| Otu51869 | 1 | k__Fungi(100);k__Fungi_unclassified(100);k__Fungi_unclassified(100);k__Fungi_unclassified(100);k__Fungi_unclassified(100);k__Fungi_unclassified(100);k__Fungi_unclassified(100); | - | - | - | - | - | - | - | Unassigned | - |
| Otu51875 | 1 | k__Fungi(100);k__Fungi_unclassified(100);k__Fungi_unclassified(100);k__Fungi_unclassified(100);k__Fungi_unclassified(100);k__Fungi_unclassified(100);k__Fungi_unclassified(100); | - | - | - | - | - | - | - | Unassigned | - |
| Otu51883 | 1 | k__Fungi(100);k__Fungi_unclassified(100);k__Fungi_unclassified(100);k__Fungi_unclassified(100);k__Fungi_unclassified(100);k__Fungi_unclassified(100);k__Fungi_unclassified(100); | - | - | - | - | - | - | - | Unassigned | - |
| Otu51888 | 1 | k__Fungi(100);k__Fungi_unclassified(100);k__Fungi_unclassified(100);k__Fungi_unclassified(100);k__Fungi_unclassified(100);k__Fungi_unclassified(100);k__Fungi_unclassified(100); | - | - | - | - | - | - | - | Unassigned | - |
| Otu51889 | 1 | k__Fungi(100);k__Fungi_unclassified(100);k__Fungi_unclassified(100);k__Fungi_unclassified(100);k__Fungi_unclassified(100);k__Fungi_unclassified(100);k__Fungi_unclassified(100); | - | - | - | - | - | - | - | Unassigned | - |
| Otu51890 | 1 | k__Fungi(100);k__Fungi_unclassified(100);k__Fungi_unclassified(100);k__Fungi_unclassified(100);k__Fungi_unclassified(100);k__Fungi_unclassified(100);k__Fungi_unclassified(100); | - | - | - | - | - | - | - | Unassigned | - |
| Otu51895 | 1 | k__Fungi(100);k__Fungi_unclassified(100);k__Fungi_unclassified(100);k__Fungi_unclassified(100);k__Fungi_unclassified(100);k__Fungi_unclassified(100);k__Fungi_unclassified(100); | - | - | - | - | - | - | - | Unassigned | - |
| Otu51897 | 1 | k__Fungi(100);p__Ascomycota(100);p__Ascomycota_unclassified(100);p__Ascomycota_unclassified(100);p__Ascomycota_unclassified(100);p__Ascomycota_unclassified(100);p__Ascomycota_unclassified(100); | - | - | - | - | - | - | - | Unassigned | - |
| Otu51900 | 1 | k__Fungi(100);p__Ascomycota(100);p__Ascomycota_unclassified(100);p__Ascomycota_unclassified(100);p__Ascomycota_unclassified(100);p__Ascomycota_unclassified(100);p__Ascomycota_unclassified(100); | - | - | - | - | - | - | - | Unassigned | - |
| Otu51905 | 1 | k__Fungi(100);k__Fungi_unclassified(100);k__Fungi_unclassified(100);k__Fungi_unclassified(100);k__Fungi_unclassified(100);k__Fungi_unclassified(100);k__Fungi_unclassified(100); | - | - | - | - | - | - | - | Unassigned | - |
| Otu51910 | 1 | k__Fungi(100);k__Fungi_unclassified(100);k__Fungi_unclassified(100);k__Fungi_unclassified(100);k__Fungi_unclassified(100);k__Fungi_unclassified(100);k__Fungi_unclassified(100); | - | - | - | - | - | - | - | Unassigned | - |
| Otu51922 | 1 | k__Fungi(100);p__Ascomycota(100);p__Ascomycota_unclassified(100);p__Ascomycota_unclassified(100);p__Ascomycota_unclassified(100);p__Ascomycota_unclassified(100);p__Ascomycota_unclassified(100); | - | - | - | - | - | - | - | Unassigned | - |
| Otu51947 | 1 | k__Fungi(100);k__Fungi_unclassified(100);k__Fungi_unclassified(100);k__Fungi_unclassified(100);k__Fungi_unclassified(100);k__Fungi_unclassified(100);k__Fungi_unclassified(100); | - | - | - | - | - | - | - | Unassigned | - |
| Otu51950 | 1 | k__Fungi(100);k__Fungi_unclassified(100);k__Fungi_unclassified(100);k__Fungi_unclassified(100);k__Fungi_unclassified(100);k__Fungi_unclassified(100);k__Fungi_unclassified(100); | - | - | - | - | - | - | - | Unassigned | - |
| Otu51970 | 1 | k__Fungi(100);p__Ascomycota(100);p__Ascomycota_unclassified(100);p__Ascomycota_unclassified(100);p__Ascomycota_unclassified(100);p__Ascomycota_unclassified(100);p__Ascomycota_unclassified(100); | - | - | - | - | - | - | - | Unassigned | - |
| Otu51971 | 1 | k__Fungi(100);k__Fungi_unclassified(100);k__Fungi_unclassified(100);k__Fungi_unclassified(100);k__Fungi_unclassified(100);k__Fungi_unclassified(100);k__Fungi_unclassified(100); | - | - | - | - | - | - | - | Unassigned | - |
| Otu51978 | 1 | k__Fungi(100);k__Fungi_unclassified(100);k__Fungi_unclassified(100);k__Fungi_unclassified(100);k__Fungi_unclassified(100);k__Fungi_unclassified(100);k__Fungi_unclassified(100); | - | - | - | - | - | - | - | Unassigned | - |
| Otu51979 | 1 | k__Fungi(100);p__Ascomycota(100);p__Ascomycota_unclassified(100);p__Ascomycota_unclassified(100);p__Ascomycota_unclassified(100);p__Ascomycota_unclassified(100);p__Ascomycota_unclassified(100); | - | - | - | - | - | - | - | Unassigned | - |
| Otu51981 | 1 | k__Fungi(100);p__Ascomycota(100);p__Ascomycota_unclassified(100);p__Ascomycota_unclassified(100);p__Ascomycota_unclassified(100);p__Ascomycota_unclassified(100);p__Ascomycota_unclassified(100); | - | - | - | - | - | - | - | Unassigned | - |
| Otu52014 | 1 | k__Fungi(100);k__Fungi_unclassified(100);k__Fungi_unclassified(100);k__Fungi_unclassified(100);k__Fungi_unclassified(100);k__Fungi_unclassified(100);k__Fungi_unclassified(100); | - | - | - | - | - | - | - | Unassigned | - |
| Otu52018 | 1 | k__Fungi(100);k__Fungi_unclassified(100);k__Fungi_unclassified(100);k__Fungi_unclassified(100);k__Fungi_unclassified(100);k__Fungi_unclassified(100);k__Fungi_unclassified(100); | - | - | - | - | - | - | - | Unassigned | - |
| Otu52020 | 1 | k__Fungi(100);k__Fungi_unclassified(100);k__Fungi_unclassified(100);k__Fungi_unclassified(100);k__Fungi_unclassified(100);k__Fungi_unclassified(100);k__Fungi_unclassified(100); | - | - | - | - | - | - | - | Unassigned | - |
| Otu52021 | 1 | k__Fungi(100);k__Fungi_unclassified(100);k__Fungi_unclassified(100);k__Fungi_unclassified(100);k__Fungi_unclassified(100);k__Fungi_unclassified(100);k__Fungi_unclassified(100); | - | - | - | - | - | - | - | Unassigned | - |
| Otu52023 | 1 | k__Fungi(100);p__Ascomycota(100);p__Ascomycota_unclassified(100);p__Ascomycota_unclassified(100);p__Ascomycota_unclassified(100);p__Ascomycota_unclassified(100);p__Ascomycota_unclassified(100); | - | - | - | - | - | - | - | Unassigned | - |
| Otu52024 | 1 | k__Fungi(100);p__Ascomycota(100);p__Ascomycota_unclassified(100);p__Ascomycota_unclassified(100);p__Ascomycota_unclassified(100);p__Ascomycota_unclassified(100);p__Ascomycota_unclassified(100); | - | - | - | - | - | - | - | Unassigned | - |
| Otu52025 | 1 | k__Fungi(100);k__Fungi_unclassified(100);k__Fungi_unclassified(100);k__Fungi_unclassified(100);k__Fungi_unclassified(100);k__Fungi_unclassified(100);k__Fungi_unclassified(100); | - | - | - | - | - | - | - | Unassigned | - |
| Otu52026 | 1 | k__Fungi(100);k__Fungi_unclassified(100);k__Fungi_unclassified(100);k__Fungi_unclassified(100);k__Fungi_unclassified(100);k__Fungi_unclassified(100);k__Fungi_unclassified(100); | - | - | - | - | - | - | - | Unassigned | - |
| Otu52027 | 1 | k__Fungi(100);k__Fungi_unclassified(100);k__Fungi_unclassified(100);k__Fungi_unclassified(100);k__Fungi_unclassified(100);k__Fungi_unclassified(100);k__Fungi_unclassified(100); | - | - | - | - | - | - | - | Unassigned | - |
| Otu52035 | 1 | k__Fungi(100);k__Fungi_unclassified(100);k__Fungi_unclassified(100);k__Fungi_unclassified(100);k__Fungi_unclassified(100);k__Fungi_unclassified(100);k__Fungi_unclassified(100); | - | - | - | - | - | - | - | Unassigned | - |
| Otu52040 | 1 | k__Fungi(100);p__Ascomycota(100);p__Ascomycota_unclassified(100);p__Ascomycota_unclassified(100);p__Ascomycota_unclassified(100);p__Ascomycota_unclassified(100);p__Ascomycota_unclassified(100); | - | - | - | - | - | - | - | Unassigned | - |
| Otu52041 | 1 | k__Fungi(100);k__Fungi_unclassified(100);k__Fungi_unclassified(100);k__Fungi_unclassified(100);k__Fungi_unclassified(100);k__Fungi_unclassified(100);k__Fungi_unclassified(100); | - | - | - | - | - | - | - | Unassigned | - |
| Otu52052 | 1 | k__Fungi(100);p__Ascomycota(100);p__Ascomycota_unclassified(100);p__Ascomycota_unclassified(100);p__Ascomycota_unclassified(100);p__Ascomycota_unclassified(100);p__Ascomycota_unclassified(100); | - | - | - | - | - | - | - | Unassigned | - |
| Otu52073 | 1 | k__Fungi(100);k__Fungi_unclassified(100);k__Fungi_unclassified(100);k__Fungi_unclassified(100);k__Fungi_unclassified(100);k__Fungi_unclassified(100);k__Fungi_unclassified(100); | - | - | - | - | - | - | - | Unassigned | - |
| Otu52096 | 1 | k__Fungi(100);p__Ascomycota(100);p__Ascomycota_unclassified(100);p__Ascomycota_unclassified(100);p__Ascomycota_unclassified(100);p__Ascomycota_unclassified(100);p__Ascomycota_unclassified(100); | - | - | - | - | - | - | - | Unassigned | - |
| Otu52099 | 1 | k__Fungi(100);k__Fungi_unclassified(100);k__Fungi_unclassified(100);k__Fungi_unclassified(100);k__Fungi_unclassified(100);k__Fungi_unclassified(100);k__Fungi_unclassified(100); | - | - | - | - | - | - | - | Unassigned | - |
| Otu52112 | 1 | k__Fungi(100);k__Fungi_unclassified(100);k__Fungi_unclassified(100);k__Fungi_unclassified(100);k__Fungi_unclassified(100);k__Fungi_unclassified(100);k__Fungi_unclassified(100); | - | - | - | - | - | - | - | Unassigned | - |
| Otu52113 | 1 | k__Fungi(100);p__Ascomycota(100);p__Ascomycota_unclassified(100);p__Ascomycota_unclassified(100);p__Ascomycota_unclassified(100);p__Ascomycota_unclassified(100);p__Ascomycota_unclassified(100); | - | - | - | - | - | - | - | Unassigned | - |
| Otu52115 | 1 | k__Fungi(100);k__Fungi_unclassified(100);k__Fungi_unclassified(100);k__Fungi_unclassified(100);k__Fungi_unclassified(100);k__Fungi_unclassified(100);k__Fungi_unclassified(100); | - | - | - | - | - | - | - | Unassigned | - |
| Otu52116 | 1 | k__Fungi(100);k__Fungi_unclassified(100);k__Fungi_unclassified(100);k__Fungi_unclassified(100);k__Fungi_unclassified(100);k__Fungi_unclassified(100);k__Fungi_unclassified(100); | - | - | - | - | - | - | - | Unassigned | - |
| Otu52147 | 1 | k__Fungi(100);k__Fungi_unclassified(100);k__Fungi_unclassified(100);k__Fungi_unclassified(100);k__Fungi_unclassified(100);k__Fungi_unclassified(100);k__Fungi_unclassified(100); | - | - | - | - | - | - | - | Unassigned | - |
| Otu52163 | 1 | k__Fungi(100);k__Fungi_unclassified(100);k__Fungi_unclassified(100);k__Fungi_unclassified(100);k__Fungi_unclassified(100);k__Fungi_unclassified(100);k__Fungi_unclassified(100); | - | - | - | - | - | - | - | Unassigned | - |
| Otu52169 | 1 | k__Fungi(100);p__Ascomycota(100);p__Ascomycota_unclassified(100);p__Ascomycota_unclassified(100);p__Ascomycota_unclassified(100);p__Ascomycota_unclassified(100);p__Ascomycota_unclassified(100); | - | - | - | - | - | - | - | Unassigned | - |
| Otu52190 | 1 | k__Fungi(100);k__Fungi_unclassified(100);k__Fungi_unclassified(100);k__Fungi_unclassified(100);k__Fungi_unclassified(100);k__Fungi_unclassified(100);k__Fungi_unclassified(100); | - | - | - | - | - | - | - | Unassigned | - |
| Otu52203 | 1 | k__Fungi(100);p__Ascomycota(100);p__Ascomycota_unclassified(100);p__Ascomycota_unclassified(100);p__Ascomycota_unclassified(100);p__Ascomycota_unclassified(100);p__Ascomycota_unclassified(100); | - | - | - | - | - | - | - | Unassigned | - |
| Otu52206 | 1 | k__Fungi(100);p__Ascomycota(100);p__Ascomycota_unclassified(100);p__Ascomycota_unclassified(100);p__Ascomycota_unclassified(100);p__Ascomycota_unclassified(100);p__Ascomycota_unclassified(100); | - | - | - | - | - | - | - | Unassigned | - |
| Otu52207 | 1 | k__Fungi(100);p__Ascomycota(100);p__Ascomycota_unclassified(100);p__Ascomycota_unclassified(100);p__Ascomycota_unclassified(100);p__Ascomycota_unclassified(100);p__Ascomycota_unclassified(100); | - | - | - | - | - | - | - | Unassigned | - |
| Otu52209 | 1 | k__Fungi(100);k__Fungi_unclassified(100);k__Fungi_unclassified(100);k__Fungi_unclassified(100);k__Fungi_unclassified(100);k__Fungi_unclassified(100);k__Fungi_unclassified(100); | - | - | - | - | - | - | - | Unassigned | - |
| Otu52219 | 1 | k__Fungi(100);p__Ascomycota(100);p__Ascomycota_unclassified(100);p__Ascomycota_unclassified(100);p__Ascomycota_unclassified(100);p__Ascomycota_unclassified(100);p__Ascomycota_unclassified(100); | - | - | - | - | - | - | - | Unassigned | - |
| Otu52228 | 1 | k__Fungi(100);p__Ascomycota(100);p__Ascomycota_unclassified(100);p__Ascomycota_unclassified(100);p__Ascomycota_unclassified(100);p__Ascomycota_unclassified(100);p__Ascomycota_unclassified(100); | - | - | - | - | - | - | - | Unassigned | - |
| Otu52229 | 1 | k__Fungi(100);p__Ascomycota(100);p__Ascomycota_unclassified(100);p__Ascomycota_unclassified(100);p__Ascomycota_unclassified(100);p__Ascomycota_unclassified(100);p__Ascomycota_unclassified(100); | - | - | - | - | - | - | - | Unassigned | - |
| Otu52230 | 1 | k__Fungi(100);k__Fungi_unclassified(100);k__Fungi_unclassified(100);k__Fungi_unclassified(100);k__Fungi_unclassified(100);k__Fungi_unclassified(100);k__Fungi_unclassified(100); | - | - | - | - | - | - | - | Unassigned | - |
| Otu52231 | 1 | k__Fungi(100);k__Fungi_unclassified(100);k__Fungi_unclassified(100);k__Fungi_unclassified(100);k__Fungi_unclassified(100);k__Fungi_unclassified(100);k__Fungi_unclassified(100); | - | - | - | - | - | - | - | Unassigned | - |
| Otu52240 | 1 | k__Fungi(100);k__Fungi_unclassified(100);k__Fungi_unclassified(100);k__Fungi_unclassified(100);k__Fungi_unclassified(100);k__Fungi_unclassified(100);k__Fungi_unclassified(100); | - | - | - | - | - | - | - | Unassigned | - |
| Otu52241 | 1 | k__Fungi(100);p__Ascomycota(100);p__Ascomycota_unclassified(100);p__Ascomycota_unclassified(100);p__Ascomycota_unclassified(100);p__Ascomycota_unclassified(100);p__Ascomycota_unclassified(100); | - | - | - | - | - | - | - | Unassigned | - |
| Otu52242 | 1 | k__Fungi(100);k__Fungi_unclassified(100);k__Fungi_unclassified(100);k__Fungi_unclassified(100);k__Fungi_unclassified(100);k__Fungi_unclassified(100);k__Fungi_unclassified(100); | - | - | - | - | - | - | - | Unassigned | - |
| Otu52245 | 1 | k__Fungi(100);p__Ascomycota(100);p__Ascomycota_unclassified(100);p__Ascomycota_unclassified(100);p__Ascomycota_unclassified(100);p__Ascomycota_unclassified(100);p__Ascomycota_unclassified(100); | - | - | - | - | - | - | - | Unassigned | - |
| Otu52248 | 1 | k__Fungi(100);k__Fungi_unclassified(100);k__Fungi_unclassified(100);k__Fungi_unclassified(100);k__Fungi_unclassified(100);k__Fungi_unclassified(100);k__Fungi_unclassified(100); | - | - | - | - | - | - | - | Unassigned | - |
| Otu52249 | 1 | k__Fungi(100);k__Fungi_unclassified(100);k__Fungi_unclassified(100);k__Fungi_unclassified(100);k__Fungi_unclassified(100);k__Fungi_unclassified(100);k__Fungi_unclassified(100); | - | - | - | - | - | - | - | Unassigned | - |
| Otu52268 | 1 | k__Fungi(100);k__Fungi_unclassified(100);k__Fungi_unclassified(100);k__Fungi_unclassified(100);k__Fungi_unclassified(100);k__Fungi_unclassified(100);k__Fungi_unclassified(100); | - | - | - | - | - | - | - | Unassigned | - |
| Otu52270 | 1 | k__Fungi(100);p__Ascomycota(100);p__Ascomycota_unclassified(100);p__Ascomycota_unclassified(100);p__Ascomycota_unclassified(100);p__Ascomycota_unclassified(100);p__Ascomycota_unclassified(100); | - | - | - | - | - | - | - | Unassigned | - |
| Otu52271 | 1 | k__Fungi(100);p__Ascomycota(100);p__Ascomycota_unclassified(100);p__Ascomycota_unclassified(100);p__Ascomycota_unclassified(100);p__Ascomycota_unclassified(100);p__Ascomycota_unclassified(100); | - | - | - | - | - | - | - | Unassigned | - |
| Otu52272 | 1 | k__Fungi(100);p__Ascomycota(100);p__Ascomycota_unclassified(100);p__Ascomycota_unclassified(100);p__Ascomycota_unclassified(100);p__Ascomycota_unclassified(100);p__Ascomycota_unclassified(100); | - | - | - | - | - | - | - | Unassigned | - |
| Otu52273 | 1 | k__Fungi(100);k__Fungi_unclassified(100);k__Fungi_unclassified(100);k__Fungi_unclassified(100);k__Fungi_unclassified(100);k__Fungi_unclassified(100);k__Fungi_unclassified(100); | - | - | - | - | - | - | - | Unassigned | - |
| Otu52274 | 1 | k__Fungi(100);k__Fungi_unclassified(100);k__Fungi_unclassified(100);k__Fungi_unclassified(100);k__Fungi_unclassified(100);k__Fungi_unclassified(100);k__Fungi_unclassified(100); | - | - | - | - | - | - | - | Unassigned | - |
| Otu52275 | 1 | k__Fungi(100);k__Fungi_unclassified(100);k__Fungi_unclassified(100);k__Fungi_unclassified(100);k__Fungi_unclassified(100);k__Fungi_unclassified(100);k__Fungi_unclassified(100); | - | - | - | - | - | - | - | Unassigned | - |
| Otu52276 | 1 | k__Fungi(100);k__Fungi_unclassified(100);k__Fungi_unclassified(100);k__Fungi_unclassified(100);k__Fungi_unclassified(100);k__Fungi_unclassified(100);k__Fungi_unclassified(100); | - | - | - | - | - | - | - | Unassigned | - |
| Otu52277 | 1 | k__Fungi(100);k__Fungi_unclassified(100);k__Fungi_unclassified(100);k__Fungi_unclassified(100);k__Fungi_unclassified(100);k__Fungi_unclassified(100);k__Fungi_unclassified(100); | - | - | - | - | - | - | - | Unassigned | - |
| Otu52278 | 1 | k__Fungi(100);k__Fungi_unclassified(100);k__Fungi_unclassified(100);k__Fungi_unclassified(100);k__Fungi_unclassified(100);k__Fungi_unclassified(100);k__Fungi_unclassified(100); | - | - | - | - | - | - | - | Unassigned | - |
| Otu52279 | 1 | k__Fungi(100);k__Fungi_unclassified(100);k__Fungi_unclassified(100);k__Fungi_unclassified(100);k__Fungi_unclassified(100);k__Fungi_unclassified(100);k__Fungi_unclassified(100); | - | - | - | - | - | - | - | Unassigned | - |
| Otu52294 | 1 | k__Fungi(100);p__Ascomycota(100);p__Ascomycota_unclassified(100);p__Ascomycota_unclassified(100);p__Ascomycota_unclassified(100);p__Ascomycota_unclassified(100);p__Ascomycota_unclassified(100); | - | - | - | - | - | - | - | Unassigned | - |
| Otu52295 | 1 | k__Fungi(100);p__Ascomycota(100);p__Ascomycota_unclassified(100);p__Ascomycota_unclassified(100);p__Ascomycota_unclassified(100);p__Ascomycota_unclassified(100);p__Ascomycota_unclassified(100); | - | - | - | - | - | - | - | Unassigned | - |
| Otu52296 | 1 | k__Fungi(100);k__Fungi_unclassified(100);k__Fungi_unclassified(100);k__Fungi_unclassified(100);k__Fungi_unclassified(100);k__Fungi_unclassified(100);k__Fungi_unclassified(100); | - | - | - | - | - | - | - | Unassigned | - |
| Otu52299 | 1 | k__Fungi(100);p__Ascomycota(100);p__Ascomycota_unclassified(100);p__Ascomycota_unclassified(100);p__Ascomycota_unclassified(100);p__Ascomycota_unclassified(100);p__Ascomycota_unclassified(100); | - | - | - | - | - | - | - | Unassigned | - |
| Otu52301 | 1 | k__Fungi(100);k__Fungi_unclassified(100);k__Fungi_unclassified(100);k__Fungi_unclassified(100);k__Fungi_unclassified(100);k__Fungi_unclassified(100);k__Fungi_unclassified(100); | - | - | - | - | - | - | - | Unassigned | - |
| Otu52319 | 1 | k__Fungi(100);k__Fungi_unclassified(100);k__Fungi_unclassified(100);k__Fungi_unclassified(100);k__Fungi_unclassified(100);k__Fungi_unclassified(100);k__Fungi_unclassified(100); | - | - | - | - | - | - | - | Unassigned | - |
| Otu52326 | 1 | k__Fungi(100);k__Fungi_unclassified(100);k__Fungi_unclassified(100);k__Fungi_unclassified(100);k__Fungi_unclassified(100);k__Fungi_unclassified(100);k__Fungi_unclassified(100); | - | - | - | - | - | - | - | Unassigned | - |
| Otu52340 | 1 | k__Fungi(100);k__Fungi_unclassified(100);k__Fungi_unclassified(100);k__Fungi_unclassified(100);k__Fungi_unclassified(100);k__Fungi_unclassified(100);k__Fungi_unclassified(100); | - | - | - | - | - | - | - | Unassigned | - |
| Otu52362 | 1 | k__Fungi(100);k__Fungi_unclassified(100);k__Fungi_unclassified(100);k__Fungi_unclassified(100);k__Fungi_unclassified(100);k__Fungi_unclassified(100);k__Fungi_unclassified(100); | - | - | - | - | - | - | - | Unassigned | - |
| Otu52378 | 1 | k__Fungi(100);k__Fungi_unclassified(100);k__Fungi_unclassified(100);k__Fungi_unclassified(100);k__Fungi_unclassified(100);k__Fungi_unclassified(100);k__Fungi_unclassified(100); | - | - | - | - | - | - | - | Unassigned | - |
| Otu52387 | 1 | k__Fungi(100);p__Ascomycota(100);p__Ascomycota_unclassified(100);p__Ascomycota_unclassified(100);p__Ascomycota_unclassified(100);p__Ascomycota_unclassified(100);p__Ascomycota_unclassified(100); | - | - | - | - | - | - | - | Unassigned | - |
| Otu52424 | 1 | k__Fungi(100);p__Ascomycota(100);p__Ascomycota_unclassified(100);p__Ascomycota_unclassified(100);p__Ascomycota_unclassified(100);p__Ascomycota_unclassified(100);p__Ascomycota_unclassified(100); | - | - | - | - | - | - | - | Unassigned | - |
| Otu52450 | 1 | k__Fungi(100);p__Ascomycota(100);p__Ascomycota_unclassified(100);p__Ascomycota_unclassified(100);p__Ascomycota_unclassified(100);p__Ascomycota_unclassified(100);p__Ascomycota_unclassified(100); | - | - | - | - | - | - | - | Unassigned | - |
| Otu52469 | 1 | k__Fungi(100);k__Fungi_unclassified(100);k__Fungi_unclassified(100);k__Fungi_unclassified(100);k__Fungi_unclassified(100);k__Fungi_unclassified(100);k__Fungi_unclassified(100); | - | - | - | - | - | - | - | Unassigned | - |
| Otu52485 | 1 | k__Fungi(100);k__Fungi_unclassified(100);k__Fungi_unclassified(100);k__Fungi_unclassified(100);k__Fungi_unclassified(100);k__Fungi_unclassified(100);k__Fungi_unclassified(100); | - | - | - | - | - | - | - | Unassigned | - |
| Otu52486 | 1 | k__Fungi(100);k__Fungi_unclassified(100);k__Fungi_unclassified(100);k__Fungi_unclassified(100);k__Fungi_unclassified(100);k__Fungi_unclassified(100);k__Fungi_unclassified(100); | - | - | - | - | - | - | - | Unassigned | - |
| Otu52531 | 1 | k__Fungi(100);p__Ascomycota(100);p__Ascomycota_unclassified(100);p__Ascomycota_unclassified(100);p__Ascomycota_unclassified(100);p__Ascomycota_unclassified(100);p__Ascomycota_unclassified(100); | - | - | - | - | - | - | - | Unassigned | - |
| Otu52532 | 1 | k__Fungi(100);p__Ascomycota(100);p__Ascomycota_unclassified(100);p__Ascomycota_unclassified(100);p__Ascomycota_unclassified(100);p__Ascomycota_unclassified(100);p__Ascomycota_unclassified(100); | - | - | - | - | - | - | - | Unassigned | - |
| Otu52539 | 1 | k__Fungi(100);k__Fungi_unclassified(100);k__Fungi_unclassified(100);k__Fungi_unclassified(100);k__Fungi_unclassified(100);k__Fungi_unclassified(100);k__Fungi_unclassified(100); | - | - | - | - | - | - | - | Unassigned | - |
| Otu52549 | 1 | k__Fungi(100);k__Fungi_unclassified(100);k__Fungi_unclassified(100);k__Fungi_unclassified(100);k__Fungi_unclassified(100);k__Fungi_unclassified(100);k__Fungi_unclassified(100); | - | - | - | - | - | - | - | Unassigned | - |
| Otu52553 | 1 | k__Fungi(100);p__Ascomycota(100);p__Ascomycota_unclassified(100);p__Ascomycota_unclassified(100);p__Ascomycota_unclassified(100);p__Ascomycota_unclassified(100);p__Ascomycota_unclassified(100); | - | - | - | - | - | - | - | Unassigned | - |
| Otu52557 | 1 | k__Fungi(100);k__Fungi_unclassified(100);k__Fungi_unclassified(100);k__Fungi_unclassified(100);k__Fungi_unclassified(100);k__Fungi_unclassified(100);k__Fungi_unclassified(100); | - | - | - | - | - | - | - | Unassigned | - |
| Otu52558 | 1 | k__Fungi(100);p__Ascomycota(100);p__Ascomycota_unclassified(100);p__Ascomycota_unclassified(100);p__Ascomycota_unclassified(100);p__Ascomycota_unclassified(100);p__Ascomycota_unclassified(100); | - | - | - | - | - | - | - | Unassigned | - |
| Otu52559 | 1 | k__Fungi(100);k__Fungi_unclassified(100);k__Fungi_unclassified(100);k__Fungi_unclassified(100);k__Fungi_unclassified(100);k__Fungi_unclassified(100);k__Fungi_unclassified(100); | - | - | - | - | - | - | - | Unassigned | - |
| Otu52560 | 1 | k__Fungi(100);k__Fungi_unclassified(100);k__Fungi_unclassified(100);k__Fungi_unclassified(100);k__Fungi_unclassified(100);k__Fungi_unclassified(100);k__Fungi_unclassified(100); | - | - | - | - | - | - | - | Unassigned | - |
| Otu52561 | 1 | k__Fungi(100);p__Ascomycota(100);p__Ascomycota_unclassified(100);p__Ascomycota_unclassified(100);p__Ascomycota_unclassified(100);p__Ascomycota_unclassified(100);p__Ascomycota_unclassified(100); | - | - | - | - | - | - | - | Unassigned | - |
| Otu52562 | 1 | k__Fungi(100);k__Fungi_unclassified(100);k__Fungi_unclassified(100);k__Fungi_unclassified(100);k__Fungi_unclassified(100);k__Fungi_unclassified(100);k__Fungi_unclassified(100); | - | - | - | - | - | - | - | Unassigned | - |
| Otu52563 | 1 | k__Fungi(100);k__Fungi_unclassified(100);k__Fungi_unclassified(100);k__Fungi_unclassified(100);k__Fungi_unclassified(100);k__Fungi_unclassified(100);k__Fungi_unclassified(100); | - | - | - | - | - | - | - | Unassigned | - |
| Otu52564 | 1 | k__Fungi(100);p__Ascomycota(100);p__Ascomycota_unclassified(100);p__Ascomycota_unclassified(100);p__Ascomycota_unclassified(100);p__Ascomycota_unclassified(100);p__Ascomycota_unclassified(100); | - | - | - | - | - | - | - | Unassigned | - |
| Otu52565 | 1 | k__Fungi(100);p__Ascomycota(100);p__Ascomycota_unclassified(100);p__Ascomycota_unclassified(100);p__Ascomycota_unclassified(100);p__Ascomycota_unclassified(100);p__Ascomycota_unclassified(100); | - | - | - | - | - | - | - | Unassigned | - |
| Otu52566 | 1 | k__Fungi(100);p__Ascomycota(100);p__Ascomycota_unclassified(100);p__Ascomycota_unclassified(100);p__Ascomycota_unclassified(100);p__Ascomycota_unclassified(100);p__Ascomycota_unclassified(100); | - | - | - | - | - | - | - | Unassigned | - |
| Otu52593 | 1 | k__Fungi(100);p__Ascomycota(100);p__Ascomycota_unclassified(100);p__Ascomycota_unclassified(100);p__Ascomycota_unclassified(100);p__Ascomycota_unclassified(100);p__Ascomycota_unclassified(100); | - | - | - | - | - | - | - | Unassigned | - |
| Otu52594 | 1 | k__Fungi(100);p__Ascomycota(100);p__Ascomycota_unclassified(100);p__Ascomycota_unclassified(100);p__Ascomycota_unclassified(100);p__Ascomycota_unclassified(100);p__Ascomycota_unclassified(100); | - | - | - | - | - | - | - | Unassigned | - |
| Otu52595 | 1 | k__Fungi(100);k__Fungi_unclassified(100);k__Fungi_unclassified(100);k__Fungi_unclassified(100);k__Fungi_unclassified(100);k__Fungi_unclassified(100);k__Fungi_unclassified(100); | - | - | - | - | - | - | - | Unassigned | - |
| Otu52596 | 1 | k__Fungi(100);k__Fungi_unclassified(100);k__Fungi_unclassified(100);k__Fungi_unclassified(100);k__Fungi_unclassified(100);k__Fungi_unclassified(100);k__Fungi_unclassified(100); | - | - | - | - | - | - | - | Unassigned | - |
| Otu52603 | 1 | k__Fungi(100);k__Fungi_unclassified(100);k__Fungi_unclassified(100);k__Fungi_unclassified(100);k__Fungi_unclassified(100);k__Fungi_unclassified(100);k__Fungi_unclassified(100); | - | - | - | - | - | - | - | Unassigned | - |
| Otu52611 | 1 | k__Fungi(100);k__Fungi_unclassified(100);k__Fungi_unclassified(100);k__Fungi_unclassified(100);k__Fungi_unclassified(100);k__Fungi_unclassified(100);k__Fungi_unclassified(100); | - | - | - | - | - | - | - | Unassigned | - |
| Otu52617 | 1 | k__Fungi(100);k__Fungi_unclassified(100);k__Fungi_unclassified(100);k__Fungi_unclassified(100);k__Fungi_unclassified(100);k__Fungi_unclassified(100);k__Fungi_unclassified(100); | - | - | - | - | - | - | - | Unassigned | - |
| Otu52627 | 1 | k__Fungi(100);p__Ascomycota(100);p__Ascomycota_unclassified(100);p__Ascomycota_unclassified(100);p__Ascomycota_unclassified(100);p__Ascomycota_unclassified(100);p__Ascomycota_unclassified(100); | - | - | - | - | - | - | - | Unassigned | - |
| Otu52629 | 1 | k__Fungi(100);p__Ascomycota(100);p__Ascomycota_unclassified(100);p__Ascomycota_unclassified(100);p__Ascomycota_unclassified(100);p__Ascomycota_unclassified(100);p__Ascomycota_unclassified(100); | - | - | - | - | - | - | - | Unassigned | - |
| Otu52634 | 1 | k__Fungi(100);k__Fungi_unclassified(100);k__Fungi_unclassified(100);k__Fungi_unclassified(100);k__Fungi_unclassified(100);k__Fungi_unclassified(100);k__Fungi_unclassified(100); | - | - | - | - | - | - | - | Unassigned | - |
| Otu52638 | 1 | k__Fungi(100);p__Ascomycota(100);p__Ascomycota_unclassified(100);p__Ascomycota_unclassified(100);p__Ascomycota_unclassified(100);p__Ascomycota_unclassified(100);p__Ascomycota_unclassified(100); | - | - | - | - | - | - | - | Unassigned | - |
| Otu52639 | 1 | k__Fungi(100);p__Ascomycota(100);p__Ascomycota_unclassified(100);p__Ascomycota_unclassified(100);p__Ascomycota_unclassified(100);p__Ascomycota_unclassified(100);p__Ascomycota_unclassified(100); | - | - | - | - | - | - | - | Unassigned | - |
| Otu52683 | 1 | k__Fungi(100);p__Ascomycota(100);p__Ascomycota_unclassified(100);p__Ascomycota_unclassified(100);p__Ascomycota_unclassified(100);p__Ascomycota_unclassified(100);p__Ascomycota_unclassified(100); | - | - | - | - | - | - | - | Unassigned | - |
| Otu52684 | 1 | k__Fungi(100);k__Fungi_unclassified(100);k__Fungi_unclassified(100);k__Fungi_unclassified(100);k__Fungi_unclassified(100);k__Fungi_unclassified(100);k__Fungi_unclassified(100); | - | - | - | - | - | - | - | Unassigned | - |
| Otu52719 | 1 | k__Fungi(100);p__Ascomycota(100);p__Ascomycota_unclassified(100);p__Ascomycota_unclassified(100);p__Ascomycota_unclassified(100);p__Ascomycota_unclassified(100);p__Ascomycota_unclassified(100); | - | - | - | - | - | - | - | Unassigned | - |
| Otu52720 | 1 | k__Fungi(100);k__Fungi_unclassified(100);k__Fungi_unclassified(100);k__Fungi_unclassified(100);k__Fungi_unclassified(100);k__Fungi_unclassified(100);k__Fungi_unclassified(100); | - | - | - | - | - | - | - | Unassigned | - |
| Otu52722 | 1 | k__Fungi(100);k__Fungi_unclassified(100);k__Fungi_unclassified(100);k__Fungi_unclassified(100);k__Fungi_unclassified(100);k__Fungi_unclassified(100);k__Fungi_unclassified(100); | - | - | - | - | - | - | - | Unassigned | - |
| Otu52744 | 1 | k__Fungi(100);p__Ascomycota(100);p__Ascomycota_unclassified(100);p__Ascomycota_unclassified(100);p__Ascomycota_unclassified(100);p__Ascomycota_unclassified(100);p__Ascomycota_unclassified(100); | - | - | - | - | - | - | - | Unassigned | - |
| Otu52747 | 1 | k__Fungi(100);k__Fungi_unclassified(100);k__Fungi_unclassified(100);k__Fungi_unclassified(100);k__Fungi_unclassified(100);k__Fungi_unclassified(100);k__Fungi_unclassified(100); | - | - | - | - | - | - | - | Unassigned | - |
| Otu52760 | 1 | k__Fungi(100);k__Fungi_unclassified(100);k__Fungi_unclassified(100);k__Fungi_unclassified(100);k__Fungi_unclassified(100);k__Fungi_unclassified(100);k__Fungi_unclassified(100); | - | - | - | - | - | - | - | Unassigned | - |
| Otu52774 | 1 | k__Fungi(100);p__Ascomycota(100);p__Ascomycota_unclassified(100);p__Ascomycota_unclassified(100);p__Ascomycota_unclassified(100);p__Ascomycota_unclassified(100);p__Ascomycota_unclassified(100); | - | - | - | - | - | - | - | Unassigned | - |
| Otu52782 | 1 | k__Fungi(100);k__Fungi_unclassified(100);k__Fungi_unclassified(100);k__Fungi_unclassified(100);k__Fungi_unclassified(100);k__Fungi_unclassified(100);k__Fungi_unclassified(100); | - | - | - | - | - | - | - | Unassigned | - |
| Otu52786 | 1 | k__Fungi(100);p__Ascomycota(100);p__Ascomycota_unclassified(100);p__Ascomycota_unclassified(100);p__Ascomycota_unclassified(100);p__Ascomycota_unclassified(100);p__Ascomycota_unclassified(100); | - | - | - | - | - | - | - | Unassigned | - |
| Otu52788 | 1 | k__Fungi(100);p__Ascomycota(100);p__Ascomycota_unclassified(100);p__Ascomycota_unclassified(100);p__Ascomycota_unclassified(100);p__Ascomycota_unclassified(100);p__Ascomycota_unclassified(100); | - | - | - | - | - | - | - | Unassigned | - |
| Otu52796 | 1 | k__Fungi(100);p__Ascomycota(100);p__Ascomycota_unclassified(100);p__Ascomycota_unclassified(100);p__Ascomycota_unclassified(100);p__Ascomycota_unclassified(100);p__Ascomycota_unclassified(100); | - | - | - | - | - | - | - | Unassigned | - |
| Otu52815 | 1 | k__Fungi(100);k__Fungi_unclassified(100);k__Fungi_unclassified(100);k__Fungi_unclassified(100);k__Fungi_unclassified(100);k__Fungi_unclassified(100);k__Fungi_unclassified(100); | - | - | - | - | - | - | - | Unassigned | - |
| Otu52817 | 1 | k__Fungi(100);k__Fungi_unclassified(100);k__Fungi_unclassified(100);k__Fungi_unclassified(100);k__Fungi_unclassified(100);k__Fungi_unclassified(100);k__Fungi_unclassified(100); | - | - | - | - | - | - | - | Unassigned | - |
| Otu52818 | 1 | k__Fungi(100);k__Fungi_unclassified(100);k__Fungi_unclassified(100);k__Fungi_unclassified(100);k__Fungi_unclassified(100);k__Fungi_unclassified(100);k__Fungi_unclassified(100); | - | - | - | - | - | - | - | Unassigned | - |
| Otu52819 | 1 | k__Fungi(100);k__Fungi_unclassified(100);k__Fungi_unclassified(100);k__Fungi_unclassified(100);k__Fungi_unclassified(100);k__Fungi_unclassified(100);k__Fungi_unclassified(100); | - | - | - | - | - | - | - | Unassigned | - |
| Otu52825 | 1 | k__Fungi(100);p__Ascomycota(100);p__Ascomycota_unclassified(100);p__Ascomycota_unclassified(100);p__Ascomycota_unclassified(100);p__Ascomycota_unclassified(100);p__Ascomycota_unclassified(100); | - | - | - | - | - | - | - | Unassigned | - |
| Otu52827 | 1 | k__Fungi(100);p__Ascomycota(100);p__Ascomycota_unclassified(100);p__Ascomycota_unclassified(100);p__Ascomycota_unclassified(100);p__Ascomycota_unclassified(100);p__Ascomycota_unclassified(100); | - | - | - | - | - | - | - | Unassigned | - |
| Otu52828 | 1 | k__Fungi(100);k__Fungi_unclassified(100);k__Fungi_unclassified(100);k__Fungi_unclassified(100);k__Fungi_unclassified(100);k__Fungi_unclassified(100);k__Fungi_unclassified(100); | - | - | - | - | - | - | - | Unassigned | - |
| Otu52829 | 1 | k__Fungi(100);k__Fungi_unclassified(100);k__Fungi_unclassified(100);k__Fungi_unclassified(100);k__Fungi_unclassified(100);k__Fungi_unclassified(100);k__Fungi_unclassified(100); | - | - | - | - | - | - | - | Unassigned | - |
| Otu52830 | 1 | k__Fungi(100);k__Fungi_unclassified(100);k__Fungi_unclassified(100);k__Fungi_unclassified(100);k__Fungi_unclassified(100);k__Fungi_unclassified(100);k__Fungi_unclassified(100); | - | - | - | - | - | - | - | Unassigned | - |
| Otu52831 | 1 | k__Fungi(100);p__Ascomycota(100);p__Ascomycota_unclassified(100);p__Ascomycota_unclassified(100);p__Ascomycota_unclassified(100);p__Ascomycota_unclassified(100);p__Ascomycota_unclassified(100); | - | - | - | - | - | - | - | Unassigned | - |
| Otu52832 | 1 | k__Fungi(100);p__Ascomycota(100);p__Ascomycota_unclassified(100);p__Ascomycota_unclassified(100);p__Ascomycota_unclassified(100);p__Ascomycota_unclassified(100);p__Ascomycota_unclassified(100); | - | - | - | - | - | - | - | Unassigned | - |
| Otu52834 | 1 | k__Fungi(100);k__Fungi_unclassified(100);k__Fungi_unclassified(100);k__Fungi_unclassified(100);k__Fungi_unclassified(100);k__Fungi_unclassified(100);k__Fungi_unclassified(100); | - | - | - | - | - | - | - | Unassigned | - |
| Otu52835 | 1 | k__Fungi(100);p__Ascomycota(100);p__Ascomycota_unclassified(100);p__Ascomycota_unclassified(100);p__Ascomycota_unclassified(100);p__Ascomycota_unclassified(100);p__Ascomycota_unclassified(100); | - | - | - | - | - | - | - | Unassigned | - |
| Otu52836 | 1 | k__Fungi(100);p__Ascomycota(100);p__Ascomycota_unclassified(100);p__Ascomycota_unclassified(100);p__Ascomycota_unclassified(100);p__Ascomycota_unclassified(100);p__Ascomycota_unclassified(100); | - | - | - | - | - | - | - | Unassigned | - |
| Otu52837 | 1 | k__Fungi(100);k__Fungi_unclassified(100);k__Fungi_unclassified(100);k__Fungi_unclassified(100);k__Fungi_unclassified(100);k__Fungi_unclassified(100);k__Fungi_unclassified(100); | - | - | - | - | - | - | - | Unassigned | - |
| Otu52838 | 1 | k__Fungi(100);k__Fungi_unclassified(100);k__Fungi_unclassified(100);k__Fungi_unclassified(100);k__Fungi_unclassified(100);k__Fungi_unclassified(100);k__Fungi_unclassified(100); | - | - | - | - | - | - | - | Unassigned | - |
| Otu52839 | 1 | k__Fungi(100);p__Ascomycota(100);p__Ascomycota_unclassified(100);p__Ascomycota_unclassified(100);p__Ascomycota_unclassified(100);p__Ascomycota_unclassified(100);p__Ascomycota_unclassified(100); | - | - | - | - | - | - | - | Unassigned | - |
| Otu52840 | 1 | k__Fungi(100);p__Ascomycota(100);p__Ascomycota_unclassified(100);p__Ascomycota_unclassified(100);p__Ascomycota_unclassified(100);p__Ascomycota_unclassified(100);p__Ascomycota_unclassified(100); | - | - | - | - | - | - | - | Unassigned | - |
| Otu52841 | 1 | k__Fungi(100);k__Fungi_unclassified(100);k__Fungi_unclassified(100);k__Fungi_unclassified(100);k__Fungi_unclassified(100);k__Fungi_unclassified(100);k__Fungi_unclassified(100); | - | - | - | - | - | - | - | Unassigned | - |
| Otu52842 | 1 | k__Fungi(100);k__Fungi_unclassified(100);k__Fungi_unclassified(100);k__Fungi_unclassified(100);k__Fungi_unclassified(100);k__Fungi_unclassified(100);k__Fungi_unclassified(100); | - | - | - | - | - | - | - | Unassigned | - |
| Otu52843 | 1 | k__Fungi(100);k__Fungi_unclassified(100);k__Fungi_unclassified(100);k__Fungi_unclassified(100);k__Fungi_unclassified(100);k__Fungi_unclassified(100);k__Fungi_unclassified(100); | - | - | - | - | - | - | - | Unassigned | - |
| Otu52856 | 1 | k__Fungi(100);k__Fungi_unclassified(100);k__Fungi_unclassified(100);k__Fungi_unclassified(100);k__Fungi_unclassified(100);k__Fungi_unclassified(100);k__Fungi_unclassified(100); | - | - | - | - | - | - | - | Unassigned | - |
| Otu52857 | 1 | k__Fungi(100);k__Fungi_unclassified(100);k__Fungi_unclassified(100);k__Fungi_unclassified(100);k__Fungi_unclassified(100);k__Fungi_unclassified(100);k__Fungi_unclassified(100); | - | - | - | - | - | - | - | Unassigned | - |
| Otu52858 | 1 | k__Fungi(100);k__Fungi_unclassified(100);k__Fungi_unclassified(100);k__Fungi_unclassified(100);k__Fungi_unclassified(100);k__Fungi_unclassified(100);k__Fungi_unclassified(100); | - | - | - | - | - | - | - | Unassigned | - |
| Otu52859 | 1 | k__Fungi(100);p__Ascomycota(100);p__Ascomycota_unclassified(100);p__Ascomycota_unclassified(100);p__Ascomycota_unclassified(100);p__Ascomycota_unclassified(100);p__Ascomycota_unclassified(100); | - | - | - | - | - | - | - | Unassigned | - |
| Otu52864 | 1 | k__Fungi(100);k__Fungi_unclassified(100);k__Fungi_unclassified(100);k__Fungi_unclassified(100);k__Fungi_unclassified(100);k__Fungi_unclassified(100);k__Fungi_unclassified(100); | - | - | - | - | - | - | - | Unassigned | - |
| Otu52870 | 1 | k__Fungi(100);k__Fungi_unclassified(100);k__Fungi_unclassified(100);k__Fungi_unclassified(100);k__Fungi_unclassified(100);k__Fungi_unclassified(100);k__Fungi_unclassified(100); | - | - | - | - | - | - | - | Unassigned | - |
| Otu52872 | 1 | k__Fungi(100);k__Fungi_unclassified(100);k__Fungi_unclassified(100);k__Fungi_unclassified(100);k__Fungi_unclassified(100);k__Fungi_unclassified(100);k__Fungi_unclassified(100); | - | - | - | - | - | - | - | Unassigned | - |
| Otu52874 | 1 | k__Fungi(100);p__Ascomycota(100);p__Ascomycota_unclassified(100);p__Ascomycota_unclassified(100);p__Ascomycota_unclassified(100);p__Ascomycota_unclassified(100);p__Ascomycota_unclassified(100); | - | - | - | - | - | - | - | Unassigned | - |
| Otu52877 | 1 | k__Fungi(100);p__Ascomycota(100);p__Ascomycota_unclassified(100);p__Ascomycota_unclassified(100);p__Ascomycota_unclassified(100);p__Ascomycota_unclassified(100);p__Ascomycota_unclassified(100); | - | - | - | - | - | - | - | Unassigned | - |
| Otu52879 | 1 | k__Fungi(100);p__Ascomycota(100);p__Ascomycota_unclassified(100);p__Ascomycota_unclassified(100);p__Ascomycota_unclassified(100);p__Ascomycota_unclassified(100);p__Ascomycota_unclassified(100); | - | - | - | - | - | - | - | Unassigned | - |
| Otu52880 | 1 | k__Fungi(100);p__Ascomycota(100);p__Ascomycota_unclassified(100);p__Ascomycota_unclassified(100);p__Ascomycota_unclassified(100);p__Ascomycota_unclassified(100);p__Ascomycota_unclassified(100); | - | - | - | - | - | - | - | Unassigned | - |
| Otu52895 | 1 | k__Fungi(100);k__Fungi_unclassified(100);k__Fungi_unclassified(100);k__Fungi_unclassified(100);k__Fungi_unclassified(100);k__Fungi_unclassified(100);k__Fungi_unclassified(100); | - | - | - | - | - | - | - | Unassigned | - |
| Otu52897 | 1 | k__Fungi(100);p__Ascomycota(100);p__Ascomycota_unclassified(100);p__Ascomycota_unclassified(100);p__Ascomycota_unclassified(100);p__Ascomycota_unclassified(100);p__Ascomycota_unclassified(100); | - | - | - | - | - | - | - | Unassigned | - |
| Otu52954 | 1 | k__Fungi(100);p__Ascomycota(100);p__Ascomycota_unclassified(100);p__Ascomycota_unclassified(100);p__Ascomycota_unclassified(100);p__Ascomycota_unclassified(100);p__Ascomycota_unclassified(100); | - | - | - | - | - | - | - | Unassigned | - |
| Otu52966 | 1 | k__Fungi(100);p__Ascomycota(100);p__Ascomycota_unclassified(100);p__Ascomycota_unclassified(100);p__Ascomycota_unclassified(100);p__Ascomycota_unclassified(100);p__Ascomycota_unclassified(100); | - | - | - | - | - | - | - | Unassigned | - |
| Otu52969 | 1 | k__Fungi(100);k__Fungi_unclassified(100);k__Fungi_unclassified(100);k__Fungi_unclassified(100);k__Fungi_unclassified(100);k__Fungi_unclassified(100);k__Fungi_unclassified(100); | - | - | - | - | - | - | - | Unassigned | - |
| Otu52989 | 1 | k__Fungi(100);k__Fungi_unclassified(100);k__Fungi_unclassified(100);k__Fungi_unclassified(100);k__Fungi_unclassified(100);k__Fungi_unclassified(100);k__Fungi_unclassified(100); | - | - | - | - | - | - | - | Unassigned | - |
| Otu52991 | 1 | k__Fungi(100);k__Fungi_unclassified(100);k__Fungi_unclassified(100);k__Fungi_unclassified(100);k__Fungi_unclassified(100);k__Fungi_unclassified(100);k__Fungi_unclassified(100); | - | - | - | - | - | - | - | Unassigned | - |
| Otu52994 | 1 | k__Fungi(100);k__Fungi_unclassified(100);k__Fungi_unclassified(100);k__Fungi_unclassified(100);k__Fungi_unclassified(100);k__Fungi_unclassified(100);k__Fungi_unclassified(100); | - | - | - | - | - | - | - | Unassigned | - |
| Otu53033 | 1 | k__Fungi(100);k__Fungi_unclassified(100);k__Fungi_unclassified(100);k__Fungi_unclassified(100);k__Fungi_unclassified(100);k__Fungi_unclassified(100);k__Fungi_unclassified(100); | - | - | - | - | - | - | - | Unassigned | - |
| Otu53098 | 1 | k__Fungi(100);k__Fungi_unclassified(100);k__Fungi_unclassified(100);k__Fungi_unclassified(100);k__Fungi_unclassified(100);k__Fungi_unclassified(100);k__Fungi_unclassified(100); | - | - | - | - | - | - | - | Unassigned | - |
| Otu53099 | 1 | k__Fungi(100);p__Ascomycota(100);p__Ascomycota_unclassified(100);p__Ascomycota_unclassified(100);p__Ascomycota_unclassified(100);p__Ascomycota_unclassified(100);p__Ascomycota_unclassified(100); | - | - | - | - | - | - | - | Unassigned | - |
| Otu53104 | 1 | k__Fungi(100);p__Ascomycota(100);p__Ascomycota_unclassified(100);p__Ascomycota_unclassified(100);p__Ascomycota_unclassified(100);p__Ascomycota_unclassified(100);p__Ascomycota_unclassified(100); | - | - | - | - | - | - | - | Unassigned | - |
| Otu53136 | 1 | k__Fungi(100);k__Fungi_unclassified(100);k__Fungi_unclassified(100);k__Fungi_unclassified(100);k__Fungi_unclassified(100);k__Fungi_unclassified(100);k__Fungi_unclassified(100); | - | - | - | - | - | - | - | Unassigned | - |
| Otu53142 | 1 | k__Fungi(100);k__Fungi_unclassified(100);k__Fungi_unclassified(100);k__Fungi_unclassified(100);k__Fungi_unclassified(100);k__Fungi_unclassified(100);k__Fungi_unclassified(100); | - | - | - | - | - | - | - | Unassigned | - |
| Otu53162 | 1 | k__Fungi(100);p__Ascomycota(100);p__Ascomycota_unclassified(100);p__Ascomycota_unclassified(100);p__Ascomycota_unclassified(100);p__Ascomycota_unclassified(100);p__Ascomycota_unclassified(100); | - | - | - | - | - | - | - | Unassigned | - |
| Otu53174 | 1 | k__Fungi(100);k__Fungi_unclassified(100);k__Fungi_unclassified(100);k__Fungi_unclassified(100);k__Fungi_unclassified(100);k__Fungi_unclassified(100);k__Fungi_unclassified(100); | - | - | - | - | - | - | - | Unassigned | - |
| Otu53176 | 1 | k__Fungi(100);p__Ascomycota(100);p__Ascomycota_unclassified(100);p__Ascomycota_unclassified(100);p__Ascomycota_unclassified(100);p__Ascomycota_unclassified(100);p__Ascomycota_unclassified(100); | - | - | - | - | - | - | - | Unassigned | - |
| Otu53177 | 1 | k__Fungi(100);p__Ascomycota(100);p__Ascomycota_unclassified(100);p__Ascomycota_unclassified(100);p__Ascomycota_unclassified(100);p__Ascomycota_unclassified(100);p__Ascomycota_unclassified(100); | - | - | - | - | - | - | - | Unassigned | - |
| Otu53178 | 1 | k__Fungi(100);p__Ascomycota(100);p__Ascomycota_unclassified(100);p__Ascomycota_unclassified(100);p__Ascomycota_unclassified(100);p__Ascomycota_unclassified(100);p__Ascomycota_unclassified(100); | - | - | - | - | - | - | - | Unassigned | - |
| Otu53179 | 1 | k__Fungi(100);p__Ascomycota(100);p__Ascomycota_unclassified(100);p__Ascomycota_unclassified(100);p__Ascomycota_unclassified(100);p__Ascomycota_unclassified(100);p__Ascomycota_unclassified(100); | - | - | - | - | - | - | - | Unassigned | - |
| Otu53181 | 1 | k__Fungi(100);p__Ascomycota(100);p__Ascomycota_unclassified(100);p__Ascomycota_unclassified(100);p__Ascomycota_unclassified(100);p__Ascomycota_unclassified(100);p__Ascomycota_unclassified(100); | - | - | - | - | - | - | - | Unassigned | - |
| Otu53196 | 1 | k__Fungi(100);p__Ascomycota(100);p__Ascomycota_unclassified(100);p__Ascomycota_unclassified(100);p__Ascomycota_unclassified(100);p__Ascomycota_unclassified(100);p__Ascomycota_unclassified(100); | - | - | - | - | - | - | - | Unassigned | - |
| Otu53209 | 1 | k__Fungi(100);p__Ascomycota(100);p__Ascomycota_unclassified(100);p__Ascomycota_unclassified(100);p__Ascomycota_unclassified(100);p__Ascomycota_unclassified(100);p__Ascomycota_unclassified(100); | - | - | - | - | - | - | - | Unassigned | - |
| Otu53211 | 1 | k__Fungi(100);p__Ascomycota(100);p__Ascomycota_unclassified(100);p__Ascomycota_unclassified(100);p__Ascomycota_unclassified(100);p__Ascomycota_unclassified(100);p__Ascomycota_unclassified(100); | - | - | - | - | - | - | - | Unassigned | - |
| Otu53228 | 1 | k__Fungi(100);k__Fungi_unclassified(100);k__Fungi_unclassified(100);k__Fungi_unclassified(100);k__Fungi_unclassified(100);k__Fungi_unclassified(100);k__Fungi_unclassified(100); | - | - | - | - | - | - | - | Unassigned | - |
| Otu53244 | 1 | k__Fungi(100);p__Ascomycota(100);p__Ascomycota_unclassified(100);p__Ascomycota_unclassified(100);p__Ascomycota_unclassified(100);p__Ascomycota_unclassified(100);p__Ascomycota_unclassified(100); | - | - | - | - | - | - | - | Unassigned | - |
| Otu53308 | 1 | k__Fungi(100);p__Ascomycota(100);p__Ascomycota_unclassified(100);p__Ascomycota_unclassified(100);p__Ascomycota_unclassified(100);p__Ascomycota_unclassified(100);p__Ascomycota_unclassified(100); | - | - | - | - | - | - | - | Unassigned | - |
| Otu53318 | 1 | k__Fungi(100);p__Ascomycota(100);p__Ascomycota_unclassified(100);p__Ascomycota_unclassified(100);p__Ascomycota_unclassified(100);p__Ascomycota_unclassified(100);p__Ascomycota_unclassified(100); | - | - | - | - | - | - | - | Unassigned | - |
| Otu53327 | 1 | k__Fungi(100);k__Fungi_unclassified(100);k__Fungi_unclassified(100);k__Fungi_unclassified(100);k__Fungi_unclassified(100);k__Fungi_unclassified(100);k__Fungi_unclassified(100); | - | - | - | - | - | - | - | Unassigned | - |
| Otu53339 | 1 | k__Fungi(100);k__Fungi_unclassified(100);k__Fungi_unclassified(100);k__Fungi_unclassified(100);k__Fungi_unclassified(100);k__Fungi_unclassified(100);k__Fungi_unclassified(100); | - | - | - | - | - | - | - | Unassigned | - |
| Otu53343 | 1 | k__Fungi(100);p__Ascomycota(100);c__Leotiomycetes(100);o__Helotiales(100);o__Helotiales_unclassified(100);o__Helotiales_unclassified(100);o__Helotiales_unclassified(100); | - | - | - | - | - | - | - | Unassigned | - |
| Otu53347 | 1 | k__Fungi(100);p__Ascomycota(100);p__Ascomycota_unclassified(100);p__Ascomycota_unclassified(100);p__Ascomycota_unclassified(100);p__Ascomycota_unclassified(100);p__Ascomycota_unclassified(100); | - | - | - | - | - | - | - | Unassigned | - |
| Otu53351 | 1 | k__Fungi(100);k__Fungi_unclassified(100);k__Fungi_unclassified(100);k__Fungi_unclassified(100);k__Fungi_unclassified(100);k__Fungi_unclassified(100);k__Fungi_unclassified(100); | - | - | - | - | - | - | - | Unassigned | - |
| Otu53356 | 1 | k__Fungi(100);k__Fungi_unclassified(100);k__Fungi_unclassified(100);k__Fungi_unclassified(100);k__Fungi_unclassified(100);k__Fungi_unclassified(100);k__Fungi_unclassified(100); | - | - | - | - | - | - | - | Unassigned | - |
| Otu53360 | 1 | k__Fungi(100);k__Fungi_unclassified(100);k__Fungi_unclassified(100);k__Fungi_unclassified(100);k__Fungi_unclassified(100);k__Fungi_unclassified(100);k__Fungi_unclassified(100); | - | - | - | - | - | - | - | Unassigned | - |
| Otu53364 | 1 | k__Fungi(100);k__Fungi_unclassified(100);k__Fungi_unclassified(100);k__Fungi_unclassified(100);k__Fungi_unclassified(100);k__Fungi_unclassified(100);k__Fungi_unclassified(100); | - | - | - | - | - | - | - | Unassigned | - |
| Otu53369 | 1 | k__Fungi(100);p__Ascomycota(100);p__Ascomycota_unclassified(100);p__Ascomycota_unclassified(100);p__Ascomycota_unclassified(100);p__Ascomycota_unclassified(100);p__Ascomycota_unclassified(100); | - | - | - | - | - | - | - | Unassigned | - |
| Otu53372 | 1 | k__Fungi(100);p__Ascomycota(100);p__Ascomycota_unclassified(100);p__Ascomycota_unclassified(100);p__Ascomycota_unclassified(100);p__Ascomycota_unclassified(100);p__Ascomycota_unclassified(100); | - | - | - | - | - | - | - | Unassigned | - |
| Otu53373 | 1 | k__Fungi(100);k__Fungi_unclassified(100);k__Fungi_unclassified(100);k__Fungi_unclassified(100);k__Fungi_unclassified(100);k__Fungi_unclassified(100);k__Fungi_unclassified(100); | - | - | - | - | - | - | - | Unassigned | - |
| Otu53375 | 1 | k__Fungi(100);p__Ascomycota(100);p__Ascomycota_unclassified(100);p__Ascomycota_unclassified(100);p__Ascomycota_unclassified(100);p__Ascomycota_unclassified(100);p__Ascomycota_unclassified(100); | - | - | - | - | - | - | - | Unassigned | - |
| Otu53376 | 1 | k__Fungi(100);p__Ascomycota(100);p__Ascomycota_unclassified(100);p__Ascomycota_unclassified(100);p__Ascomycota_unclassified(100);p__Ascomycota_unclassified(100);p__Ascomycota_unclassified(100); | - | - | - | - | - | - | - | Unassigned | - |
| Otu53377 | 1 | k__Fungi(100);p__Ascomycota(100);p__Ascomycota_unclassified(100);p__Ascomycota_unclassified(100);p__Ascomycota_unclassified(100);p__Ascomycota_unclassified(100);p__Ascomycota_unclassified(100); | - | - | - | - | - | - | - | Unassigned | - |
| Otu53379 | 1 | k__Fungi(100);p__Ascomycota(100);p__Ascomycota_unclassified(100);p__Ascomycota_unclassified(100);p__Ascomycota_unclassified(100);p__Ascomycota_unclassified(100);p__Ascomycota_unclassified(100); | - | - | - | - | - | - | - | Unassigned | - |
| Otu53381 | 1 | k__Fungi(100);p__Ascomycota(100);p__Ascomycota_unclassified(100);p__Ascomycota_unclassified(100);p__Ascomycota_unclassified(100);p__Ascomycota_unclassified(100);p__Ascomycota_unclassified(100); | - | - | - | - | - | - | - | Unassigned | - |
| Otu53382 | 1 | k__Fungi(100);p__Ascomycota(100);p__Ascomycota_unclassified(100);p__Ascomycota_unclassified(100);p__Ascomycota_unclassified(100);p__Ascomycota_unclassified(100);p__Ascomycota_unclassified(100); | - | - | - | - | - | - | - | Unassigned | - |
| Otu53383 | 1 | k__Fungi(100);p__Ascomycota(100);p__Ascomycota_unclassified(100);p__Ascomycota_unclassified(100);p__Ascomycota_unclassified(100);p__Ascomycota_unclassified(100);p__Ascomycota_unclassified(100); | - | - | - | - | - | - | - | Unassigned | - |

| | | | | | | | | | | | |
|---|---|---|---|---|---|---|---|---|---|---|---|
| Otu53384 | 1 | k__Fungi(100);k__Fungi_unclassified(100);k__Fungi_unclassified(100);k__Fungi_unclassified(100);k__Fungi_unclassified(100);k__Fungi_unclassified(100);k__Fungi_unclassified(100); | - | - | - | - | - | - | - | Unassigned | - |
| Otu53385 | 1 | k__Fungi(100);k__Fungi_unclassified(100);k__Fungi_unclassified(100);k__Fungi_unclassified(100);k__Fungi_unclassified(100);k__Fungi_unclassified(100);k__Fungi_unclassified(100); | - | - | - | - | - | - | - | Unassigned | - |
| Otu53386 | 1 | k__Fungi(100);k__Fungi_unclassified(100);k__Fungi_unclassified(100);k__Fungi_unclassified(100);k__Fungi_unclassified(100);k__Fungi_unclassified(100);k__Fungi_unclassified(100); | - | - | - | - | - | - | - | Unassigned | - |
| Otu53391 | 1 | k__Fungi(100);k__Fungi_unclassified(100);k__Fungi_unclassified(100);k__Fungi_unclassified(100);k__Fungi_unclassified(100);k__Fungi_unclassified(100);k__Fungi_unclassified(100); | - | - | - | - | - | - | - | Unassigned | - |
| Otu53392 | 1 | k__Fungi(100);k__Fungi_unclassified(100);k__Fungi_unclassified(100);k__Fungi_unclassified(100);k__Fungi_unclassified(100);k__Fungi_unclassified(100);k__Fungi_unclassified(100); | - | - | - | - | - | - | - | Unassigned | - |
| Otu53394 | 1 | k__Fungi(100);p__Ascomycota(100);p__Ascomycota_unclassified(100);p__Ascomycota_unclassified(100);p__Ascomycota_unclassified(100);p__Ascomycota_unclassified(100);p__Ascomycota_unclassified(100); | - | - | - | - | - | - | - | Unassigned | - |
| Otu53407 | 1 | k__Fungi(100);k__Fungi_unclassified(100);k__Fungi_unclassified(100);k__Fungi_unclassified(100);k__Fungi_unclassified(100);k__Fungi_unclassified(100);k__Fungi_unclassified(100); | - | - | - | - | - | - | - | Unassigned | - |
| Otu53418 | 1 | k__Fungi(100);p__Ascomycota(100);p__Ascomycota_unclassified(100);p__Ascomycota_unclassified(100);p__Ascomycota_unclassified(100);p__Ascomycota_unclassified(100);p__Ascomycota_unclassified(100); | - | - | - | - | - | - | - | Unassigned | - |
| Otu53419 | 1 | k__Fungi(100);k__Fungi_unclassified(100);k__Fungi_unclassified(100);k__Fungi_unclassified(100);k__Fungi_unclassified(100);k__Fungi_unclassified(100);k__Fungi_unclassified(100); | - | - | - | - | - | - | - | Unassigned | - |
| Otu53420 | 1 | k__Fungi(100);p__Ascomycota(100);p__Ascomycota_unclassified(100);p__Ascomycota_unclassified(100);p__Ascomycota_unclassified(100);p__Ascomycota_unclassified(100);p__Ascomycota_unclassified(100); | - | - | - | - | - | - | - | Unassigned | - |
| Otu53421 | 1 | k__Fungi(100);k__Fungi_unclassified(100);k__Fungi_unclassified(100);k__Fungi_unclassified(100);k__Fungi_unclassified(100);k__Fungi_unclassified(100);k__Fungi_unclassified(100); | - | - | - | - | - | - | - | Unassigned | - |
| Otu53422 | 1 | k__Fungi(100);p__Ascomycota(100);p__Ascomycota_unclassified(100);p__Ascomycota_unclassified(100);p__Ascomycota_unclassified(100);p__Ascomycota_unclassified(100);p__Ascomycota_unclassified(100); | - | - | - | - | - | - | - | Unassigned | - |
| Otu53423 | 1 | k__Fungi(100);k__Fungi_unclassified(100);k__Fungi_unclassified(100);k__Fungi_unclassified(100);k__Fungi_unclassified(100);k__Fungi_unclassified(100);k__Fungi_unclassified(100); | - | - | - | - | - | - | - | Unassigned | - |
| Otu53424 | 1 | k__Fungi(100);k__Fungi_unclassified(100);k__Fungi_unclassified(100);k__Fungi_unclassified(100);k__Fungi_unclassified(100);k__Fungi_unclassified(100);k__Fungi_unclassified(100); | - | - | - | - | - | - | - | Unassigned | - |
| Otu53426 | 1 | k__Fungi(100);k__Fungi_unclassified(100);k__Fungi_unclassified(100);k__Fungi_unclassified(100);k__Fungi_unclassified(100);k__Fungi_unclassified(100);k__Fungi_unclassified(100); | - | - | - | - | - | - | - | Unassigned | - |
| Otu53429 | 1 | k__Fungi(100);p__Ascomycota(100);p__Ascomycota_unclassified(100);p__Ascomycota_unclassified(100);p__Ascomycota_unclassified(100);p__Ascomycota_unclassified(100);p__Ascomycota_unclassified(100); | - | - | - | - | - | - | - | Unassigned | - |
| Otu53430 | 1 | k__Fungi(100);k__Fungi_unclassified(100);k__Fungi_unclassified(100);k__Fungi_unclassified(100);k__Fungi_unclassified(100);k__Fungi_unclassified(100);k__Fungi_unclassified(100); | - | - | - | - | - | - | - | Unassigned | - |
| Otu53431 | 1 | k__Fungi(100);k__Fungi_unclassified(100);k__Fungi_unclassified(100);k__Fungi_unclassified(100);k__Fungi_unclassified(100);k__Fungi_unclassified(100);k__Fungi_unclassified(100); | - | - | - | - | - | - | - | Unassigned | - |
| Otu53432 | 1 | k__Fungi(100);p__Ascomycota(100);c__Leotiomycetes(100);o__Helotiales(100);f__Helotiales_family_Incertae_sedis(100);f__Helotiales_family_Incertae_sedis_unclassified(100);f__Helotiales_family_Incertae_sedis_unclassified(100); | - | - | - | - | - | - | - | Unassigned | - |
| Otu53433 | 1 | k__Fungi(100);k__Fungi_unclassified(100);k__Fungi_unclassified(100);k__Fungi_unclassified(100);k__Fungi_unclassified(100);k__Fungi_unclassified(100);k__Fungi_unclassified(100); | - | - | - | - | - | - | - | Unassigned | - |
| Otu53435 | 1 | k__Fungi(100);k__Fungi_unclassified(100);k__Fungi_unclassified(100);k__Fungi_unclassified(100);k__Fungi_unclassified(100);k__Fungi_unclassified(100);k__Fungi_unclassified(100); | - | - | - | - | - | - | - | Unassigned | - |
| Otu53436 | 1 | k__Fungi(100);p__Ascomycota(100);p__Ascomycota_unclassified(100);p__Ascomycota_unclassified(100);p__Ascomycota_unclassified(100);p__Ascomycota_unclassified(100);p__Ascomycota_unclassified(100); | - | - | - | - | - | - | - | Unassigned | - |
| Otu53437 | 1 | k__Fungi(100);p__Ascomycota(100);p__Ascomycota_unclassified(100);p__Ascomycota_unclassified(100);p__Ascomycota_unclassified(100);p__Ascomycota_unclassified(100);p__Ascomycota_unclassified(100); | - | - | - | - | - | - | - | Unassigned | - |
| Otu53438 | 1 | k__Fungi(100);k__Fungi_unclassified(100);k__Fungi_unclassified(100);k__Fungi_unclassified(100);k__Fungi_unclassified(100);k__Fungi_unclassified(100);k__Fungi_unclassified(100); | - | - | - | - | - | - | - | Unassigned | - |
| Otu53439 | 1 | k__Fungi(100);p__Ascomycota(100);p__Ascomycota_unclassified(100);p__Ascomycota_unclassified(100);p__Ascomycota_unclassified(100);p__Ascomycota_unclassified(100);p__Ascomycota_unclassified(100); | - | - | - | - | - | - | - | Unassigned | - |
| Otu53441 | 1 | k__Fungi(100);p__Ascomycota(100);p__Ascomycota_unclassified(100);p__Ascomycota_unclassified(100);p__Ascomycota_unclassified(100);p__Ascomycota_unclassified(100);p__Ascomycota_unclassified(100); | - | - | - | - | - | - | - | Unassigned | - |
| Otu53442 | 1 | k__Fungi(100);k__Fungi_unclassified(100);k__Fungi_unclassified(100);k__Fungi_unclassified(100);k__Fungi_unclassified(100);k__Fungi_unclassified(100);k__Fungi_unclassified(100); | - | - | - | - | - | - | - | Unassigned | - |
| Otu53450 | 1 | k__Fungi(100);p__Ascomycota(100);p__Ascomycota_unclassified(100);p__Ascomycota_unclassified(100);p__Ascomycota_unclassified(100);p__Ascomycota_unclassified(100);p__Ascomycota_unclassified(100); | - | - | - | - | - | - | - | Unassigned | - |
| Otu53453 | 1 | k__Fungi(100);p__Ascomycota(100);p__Ascomycota_unclassified(100);p__Ascomycota_unclassified(100);p__Ascomycota_unclassified(100);p__Ascomycota_unclassified(100);p__Ascomycota_unclassified(100); | - | - | - | - | - | - | - | Unassigned | - |
| Otu53456 | 1 | k__Fungi(100);k__Fungi_unclassified(100);k__Fungi_unclassified(100);k__Fungi_unclassified(100);k__Fungi_unclassified(100);k__Fungi_unclassified(100);k__Fungi_unclassified(100); | - | - | - | - | - | - | - | Unassigned | - |
| Otu53457 | 1 | k__Fungi(100);k__Fungi_unclassified(100);k__Fungi_unclassified(100);k__Fungi_unclassified(100);k__Fungi_unclassified(100);k__Fungi_unclassified(100);k__Fungi_unclassified(100); | - | - | - | - | - | - | - | Unassigned | - |
| Otu53458 | 1 | k__Fungi(100);k__Fungi_unclassified(100);k__Fungi_unclassified(100);k__Fungi_unclassified(100);k__Fungi_unclassified(100);k__Fungi_unclassified(100);k__Fungi_unclassified(100); | - | - | - | - | - | - | - | Unassigned | - |
| Otu53459 | 1 | k__Fungi(100);p__Ascomycota(100);p__Ascomycota_unclassified(100);p__Ascomycota_unclassified(100);p__Ascomycota_unclassified(100);p__Ascomycota_unclassified(100);p__Ascomycota_unclassified(100); | - | - | - | - | - | - | - | Unassigned | - |
| Otu53461 | 1 | k__Fungi(100);p__Ascomycota(100);p__Ascomycota_unclassified(100);p__Ascomycota_unclassified(100);p__Ascomycota_unclassified(100);p__Ascomycota_unclassified(100);p__Ascomycota_unclassified(100); | - | - | - | - | - | - | - | Unassigned | - |
| Otu53462 | 1 | k__Fungi(100);p__Ascomycota(100);p__Ascomycota_unclassified(100);p__Ascomycota_unclassified(100);p__Ascomycota_unclassified(100);p__Ascomycota_unclassified(100);p__Ascomycota_unclassified(100); | - | - | - | - | - | - | - | Unassigned | - |
| Otu53463 | 1 | k__Fungi(100);p__Ascomycota(100);p__Ascomycota_unclassified(100);p__Ascomycota_unclassified(100);p__Ascomycota_unclassified(100);p__Ascomycota_unclassified(100);p__Ascomycota_unclassified(100); | - | - | - | - | - | - | - | Unassigned | - |
| Otu53464 | 1 | k__Fungi(100);p__Ascomycota(100);p__Ascomycota_unclassified(100);p__Ascomycota_unclassified(100);p__Ascomycota_unclassified(100);p__Ascomycota_unclassified(100);p__Ascomycota_unclassified(100); | - | - | - | - | - | - | - | Unassigned | - |
| Otu53465 | 1 | k__Fungi(100);k__Fungi_unclassified(100);k__Fungi_unclassified(100);k__Fungi_unclassified(100);k__Fungi_unclassified(100);k__Fungi_unclassified(100);k__Fungi_unclassified(100); | - | - | - | - | - | - | - | Unassigned | - |
| Otu53466 | 1 | k__Fungi(100);p__Ascomycota(100);p__Ascomycota_unclassified(100);p__Ascomycota_unclassified(100);p__Ascomycota_unclassified(100);p__Ascomycota_unclassified(100);p__Ascomycota_unclassified(100); | - | - | - | - | - | - | - | Unassigned | - |
| Otu53467 | 1 | k__Fungi(100);p__Ascomycota(100);p__Ascomycota_unclassified(100);p__Ascomycota_unclassified(100);p__Ascomycota_unclassified(100);p__Ascomycota_unclassified(100);p__Ascomycota_unclassified(100); | - | - | - | - | - | - | - | Unassigned | - |
| Otu53468 | 1 | k__Fungi(100);p__Ascomycota(100);p__Ascomycota_unclassified(100);p__Ascomycota_unclassified(100);p__Ascomycota_unclassified(100);p__Ascomycota_unclassified(100);p__Ascomycota_unclassified(100); | - | - | - | - | - | - | - | Unassigned | - |
| Otu53473 | 1 | k__Fungi(100);k__Fungi_unclassified(100);k__Fungi_unclassified(100);k__Fungi_unclassified(100);k__Fungi_unclassified(100);k__Fungi_unclassified(100);k__Fungi_unclassified(100); | - | - | - | - | - | - | - | Unassigned | - |
| Otu53477 | 1 | k__Fungi(100);p__Ascomycota(100);p__Ascomycota_unclassified(100);p__Ascomycota_unclassified(100);p__Ascomycota_unclassified(100);p__Ascomycota_unclassified(100);p__Ascomycota_unclassified(100); | - | - | - | - | - | - | - | Unassigned | - |
| Otu53481 | 1 | k__Fungi(100);p__Ascomycota(100);p__Ascomycota_unclassified(100);p__Ascomycota_unclassified(100);p__Ascomycota_unclassified(100);p__Ascomycota_unclassified(100);p__Ascomycota_unclassified(100); | - | - | - | - | - | - | - | Unassigned | - |
| Otu53483 | 1 | k__Fungi(100);k__Fungi_unclassified(100);k__Fungi_unclassified(100);k__Fungi_unclassified(100);k__Fungi_unclassified(100);k__Fungi_unclassified(100);k__Fungi_unclassified(100); | - | - | - | - | - | - | - | Unassigned | - |
| Otu53486 | 1 | k__Fungi(100);k__Fungi_unclassified(100);k__Fungi_unclassified(100);k__Fungi_unclassified(100);k__Fungi_unclassified(100);k__Fungi_unclassified(100);k__Fungi_unclassified(100); | - | - | - | - | - | - | - | Unassigned | - |
| Otu53490 | 1 | k__Fungi(100);p__Ascomycota(100);p__Ascomycota_unclassified(100);p__Ascomycota_unclassified(100);p__Ascomycota_unclassified(100);p__Ascomycota_unclassified(100);p__Ascomycota_unclassified(100); | - | - | - | - | - | - | - | Unassigned | - |
| Otu53493 | 1 | k__Fungi(100);p__Ascomycota(100);c__Sordariomycetes(100);c__Sordariomycetes_unclassified(100);c__Sordariomycetes_unclassified(100);c__Sordariomycetes_unclassified(100);c__Sordariomycetes_unclassified(100); | - | - | - | - | - | - | - | Unassigned | - |
| Otu53504 | 1 | k__Fungi(100);k__Fungi_unclassified(100);k__Fungi_unclassified(100);k__Fungi_unclassified(100);k__Fungi_unclassified(100);k__Fungi_unclassified(100);k__Fungi_unclassified(100); | - | - | - | - | - | - | - | Unassigned | - |
| Otu53506 | 1 | k__Fungi(100);k__Fungi_unclassified(100);k__Fungi_unclassified(100);k__Fungi_unclassified(100);k__Fungi_unclassified(100);k__Fungi_unclassified(100);k__Fungi_unclassified(100); | - | - | - | - | - | - | - | Unassigned | - |
| Otu53507 | 1 | k__Fungi(100);p__Ascomycota(100);p__Ascomycota_unclassified(100);p__Ascomycota_unclassified(100);p__Ascomycota_unclassified(100);p__Ascomycota_unclassified(100);p__Ascomycota_unclassified(100); | - | - | - | - | - | - | - | Unassigned | - |
| Otu53508 | 1 | k__Fungi(100);k__Fungi_unclassified(100);k__Fungi_unclassified(100);k__Fungi_unclassified(100);k__Fungi_unclassified(100);k__Fungi_unclassified(100);k__Fungi_unclassified(100); | - | - | - | - | - | - | - | Unassigned | - |
| Otu53510 | 1 | k__Fungi(100);k__Fungi_unclassified(100);k__Fungi_unclassified(100);k__Fungi_unclassified(100);k__Fungi_unclassified(100);k__Fungi_unclassified(100);k__Fungi_unclassified(100); | - | - | - | - | - | - | - | Unassigned | - |
| Otu53512 | 1 | k__Fungi(100);p__Ascomycota(100);p__Ascomycota_unclassified(100);p__Ascomycota_unclassified(100);p__Ascomycota_unclassified(100);p__Ascomycota_unclassified(100);p__Ascomycota_unclassified(100); | - | - | - | - | - | - | - | Unassigned | - |
| Otu53513 | 1 | k__Fungi(100);k__Fungi_unclassified(100);k__Fungi_unclassified(100);k__Fungi_unclassified(100);k__Fungi_unclassified(100);k__Fungi_unclassified(100);k__Fungi_unclassified(100); | - | - | - | - | - | - | - | Unassigned | - |
| Otu53515 | 1 | k__Fungi(100);k__Fungi_unclassified(100);k__Fungi_unclassified(100);k__Fungi_unclassified(100);k__Fungi_unclassified(100);k__Fungi_unclassified(100);k__Fungi_unclassified(100); | - | - | - | - | - | - | - | Unassigned | - |
| Otu53524 | 1 | k__Fungi(100);k__Fungi_unclassified(100);k__Fungi_unclassified(100);k__Fungi_unclassified(100);k__Fungi_unclassified(100);k__Fungi_unclassified(100);k__Fungi_unclassified(100); | - | - | - | - | - | - | - | Unassigned | - |
| Otu53525 | 1 | k__Fungi(100);p__Ascomycota(100);p__Ascomycota_unclassified(100);p__Ascomycota_unclassified(100);p__Ascomycota_unclassified(100);p__Ascomycota_unclassified(100);p__Ascomycota_unclassified(100); | - | - | - | - | - | - | - | Unassigned | - |
| Otu53526 | 1 | k__Fungi(100);p__Ascomycota(100);p__Ascomycota_unclassified(100);p__Ascomycota_unclassified(100);p__Ascomycota_unclassified(100);p__Ascomycota_unclassified(100);p__Ascomycota_unclassified(100); | - | - | - | - | - | - | - | Unassigned | - |
| Otu53529 | 1 | k__Fungi(100);k__Fungi_unclassified(100);k__Fungi_unclassified(100);k__Fungi_unclassified(100);k__Fungi_unclassified(100);k__Fungi_unclassified(100);k__Fungi_unclassified(100); | - | - | - | - | - | - | - | Unassigned | - |
| Otu53530 | 1 | k__Fungi(100);p__Ascomycota(100);p__Ascomycota_unclassified(100);p__Ascomycota_unclassified(100);p__Ascomycota_unclassified(100);p__Ascomycota_unclassified(100);p__Ascomycota_unclassified(100); | - | - | - | - | - | - | - | Unassigned | - |
| Otu53531 | 1 | k__Fungi(100);k__Fungi_unclassified(100);k__Fungi_unclassified(100);k__Fungi_unclassified(100);k__Fungi_unclassified(100);k__Fungi_unclassified(100);k__Fungi_unclassified(100); | - | - | - | - | - | - | - | Unassigned | - |
| Otu53532 | 1 | k__Fungi(100);k__Fungi_unclassified(100);k__Fungi_unclassified(100);k__Fungi_unclassified(100);k__Fungi_unclassified(100);k__Fungi_unclassified(100);k__Fungi_unclassified(100); | - | - | - | - | - | - | - | Unassigned | - |
| Otu53533 | 1 | k__Fungi(100);k__Fungi_unclassified(100);k__Fungi_unclassified(100);k__Fungi_unclassified(100);k__Fungi_unclassified(100);k__Fungi_unclassified(100);k__Fungi_unclassified(100); | - | - | - | - | - | - | - | Unassigned | - |
| Otu53534 | 1 | k__Fungi(100);k__Fungi_unclassified(100);k__Fungi_unclassified(100);k__Fungi_unclassified(100);k__Fungi_unclassified(100);k__Fungi_unclassified(100);k__Fungi_unclassified(100); | - | - | - | - | - | - | - | Unassigned | - |
| Otu53535 | 1 | k__Fungi(100);k__Fungi_unclassified(100);k__Fungi_unclassified(100);k__Fungi_unclassified(100);k__Fungi_unclassified(100);k__Fungi_unclassified(100);k__Fungi_unclassified(100); | - | - | - | - | - | - | - | Unassigned | - |
| Otu53536 | 1 | k__Fungi(100);p__Ascomycota(100);p__Ascomycota_unclassified(100);p__Ascomycota_unclassified(100);p__Ascomycota_unclassified(100);p__Ascomycota_unclassified(100);p__Ascomycota_unclassified(100); | - | - | - | - | - | - | - | Unassigned | - |
| Otu53538 | 1 | k__Fungi(100);p__Ascomycota(100);p__Ascomycota_unclassified(100);p__Ascomycota_unclassified(100);p__Ascomycota_unclassified(100);p__Ascomycota_unclassified(100);p__Ascomycota_unclassified(100); | - | - | - | - | - | - | - | Unassigned | - |
| Otu53540 | 1 | k__Fungi(100);k__Fungi_unclassified(100);k__Fungi_unclassified(100);k__Fungi_unclassified(100);k__Fungi_unclassified(100);k__Fungi_unclassified(100);k__Fungi_unclassified(100); | - | - | - | - | - | - | - | Unassigned | - |
| Otu53541 | 1 | k__Fungi(100);p__Ascomycota(100);p__Ascomycota_unclassified(100);p__Ascomycota_unclassified(100);p__Ascomycota_unclassified(100);p__Ascomycota_unclassified(100);p__Ascomycota_unclassified(100); | - | - | - | - | - | - | - | Unassigned | - |
| Otu53542 | 1 | k__Fungi(100);k__Fungi_unclassified(100);k__Fungi_unclassified(100);k__Fungi_unclassified(100);k__Fungi_unclassified(100);k__Fungi_unclassified(100);k__Fungi_unclassified(100); | - | - | - | - | - | - | - | Unassigned | - |
| Otu53543 | 1 | k__Fungi(100);p__Ascomycota(100);p__Ascomycota_unclassified(100);p__Ascomycota_unclassified(100);p__Ascomycota_unclassified(100);p__Ascomycota_unclassified(100);p__Ascomycota_unclassified(100); | - | - | - | - | - | - | - | Unassigned | - |
| Otu53544 | 1 | k__Fungi(100);k__Fungi_unclassified(100);k__Fungi_unclassified(100);k__Fungi_unclassified(100);k__Fungi_unclassified(100);k__Fungi_unclassified(100);k__Fungi_unclassified(100); | - | - | - | - | - | - | - | Unassigned | - |
| Otu53546 | 1 | k__Fungi(100);k__Fungi_unclassified(100);k__Fungi_unclassified(100);k__Fungi_unclassified(100);k__Fungi_unclassified(100);k__Fungi_unclassified(100);k__Fungi_unclassified(100); | - | - | - | - | - | - | - | Unassigned | - |
| Otu53548 | 1 | k__Fungi(100);k__Fungi_unclassified(100);k__Fungi_unclassified(100);k__Fungi_unclassified(100);k__Fungi_unclassified(100);k__Fungi_unclassified(100);k__Fungi_unclassified(100); | - | - | - | - | - | - | - | Unassigned | - |
| Otu53551 | 1 | k__Fungi(100);k__Fungi_unclassified(100);k__Fungi_unclassified(100);k__Fungi_unclassified(100);k__Fungi_unclassified(100);k__Fungi_unclassified(100);k__Fungi_unclassified(100); | - | - | - | - | - | - | - | Unassigned | - |
| Otu53552 | 1 | k__Fungi(100);k__Fungi_unclassified(100);k__Fungi_unclassified(100);k__Fungi_unclassified(100);k__Fungi_unclassified(100);k__Fungi_unclassified(100);k__Fungi_unclassified(100); | - | - | - | - | - | - | - | Unassigned | - |
| Otu53556 | 1 | k__Fungi(100);k__Fungi_unclassified(100);k__Fungi_unclassified(100);k__Fungi_unclassified(100);k__Fungi_unclassified(100);k__Fungi_unclassified(100);k__Fungi_unclassified(100); | - | - | - | - | - | - | - | Unassigned | - |
| Otu53557 | 1 | k__Fungi(100);k__Fungi_unclassified(100);k__Fungi_unclassified(100);k__Fungi_unclassified(100);k__Fungi_unclassified(100);k__Fungi_unclassified(100);k__Fungi_unclassified(100); | - | - | - | - | - | - | - | Unassigned | - |
| Otu53558 | 1 | k__Fungi(100);k__Fungi_unclassified(100);k__Fungi_unclassified(100);k__Fungi_unclassified(100);k__Fungi_unclassified(100);k__Fungi_unclassified(100);k__Fungi_unclassified(100); | - | - | - | - | - | - | - | Unassigned | - |
| Otu53559 | 1 | k__Fungi(100);k__Fungi_unclassified(100);k__Fungi_unclassified(100);k__Fungi_unclassified(100);k__Fungi_unclassified(100);k__Fungi_unclassified(100);k__Fungi_unclassified(100); | - | - | - | - | - | - | - | Unassigned | - |
| Otu53560 | 1 | k__Fungi(100);k__Fungi_unclassified(100);k__Fungi_unclassified(100);k__Fungi_unclassified(100);k__Fungi_unclassified(100);k__Fungi_unclassified(100);k__Fungi_unclassified(100); | - | - | - | - | - | - | - | Unassigned | - |
| Otu53561 | 1 | k__Fungi(100);p__Ascomycota(100);p__Ascomycota_unclassified(100);p__Ascomycota_unclassified(100);p__Ascomycota_unclassified(100);p__Ascomycota_unclassified(100);p__Ascomycota_unclassified(100); | - | - | - | - | - | - | - | Unassigned | - |
| Otu53563 | 1 | k__Fungi(100);k__Fungi_unclassified(100);k__Fungi_unclassified(100);k__Fungi_unclassified(100);k__Fungi_unclassified(100);k__Fungi_unclassified(100);k__Fungi_unclassified(100); | - | - | - | - | - | - | - | Unassigned | - |
| Otu53565 | 1 | k__Fungi(100);k__Fungi_unclassified(100);k__Fungi_unclassified(100);k__Fungi_unclassified(100);k__Fungi_unclassified(100);k__Fungi_unclassified(100);k__Fungi_unclassified(100); | - | - | - | - | - | - | - | Unassigned | - |
| Otu53566 | 1 | k__Fungi(100);k__Fungi_unclassified(100);k__Fungi_unclassified(100);k__Fungi_unclassified(100);k__Fungi_unclassified(100);k__Fungi_unclassified(100);k__Fungi_unclassified(100); | - | - | - | - | - | - | - | Unassigned | - |
| Otu53567 | 1 | k__Fungi(100);k__Fungi_unclassified(100);k__Fungi_unclassified(100);k__Fungi_unclassified(100);k__Fungi_unclassified(100);k__Fungi_unclassified(100);k__Fungi_unclassified(100); | - | - | - | - | - | - | - | Unassigned | - |
| Otu53568 | 1 | k__Fungi(100);k__Fungi_unclassified(100);k__Fungi_unclassified(100);k__Fungi_unclassified(100);k__Fungi_unclassified(100);k__Fungi_unclassified(100);k__Fungi_unclassified(100); | - | - | - | - | - | - | - | Unassigned | - |
| Otu53570 | 1 | k__Fungi(100);k__Fungi_unclassified(100);k__Fungi_unclassified(100);k__Fungi_unclassified(100);k__Fungi_unclassified(100);k__Fungi_unclassified(100);k__Fungi_unclassified(100); | - | - | - | - | - | - | - | Unassigned | - |
| Otu53571 | 1 | k__Fungi(100);k__Fungi_unclassified(100);k__Fungi_unclassified(100);k__Fungi_unclassified(100);k__Fungi_unclassified(100);k__Fungi_unclassified(100);k__Fungi_unclassified(100); | - | - | - | - | - | - | - | Unassigned | - |
| Otu53572 | 1 | k__Fungi(100);k__Fungi_unclassified(100);k__Fungi_unclassified(100);k__Fungi_unclassified(100);k__Fungi_unclassified(100);k__Fungi_unclassified(100);k__Fungi_unclassified(100); | - | - | - | - | - | - | - | Unassigned | - |
| Otu53573 | 1 | k__Fungi(100);k__Fungi_unclassified(100);k__Fungi_unclassified(100);k__Fungi_unclassified(100);k__Fungi_unclassified(100);k__Fungi_unclassified(100);k__Fungi_unclassified(100); | - | - | - | - | - | - | - | Unassigned | - |
| Otu53574 | 1 | k__Fungi(100);k__Fungi_unclassified(100);k__Fungi_unclassified(100);k__Fungi_unclassified(100);k__Fungi_unclassified(100);k__Fungi_unclassified(100);k__Fungi_unclassified(100); | - | - | - | - | - | - | - | Unassigned | - |
| Otu53575 | 1 | k__Fungi(100);k__Fungi_unclassified(100);k__Fungi_unclassified(100);k__Fungi_unclassified(100);k__Fungi_unclassified(100);k__Fungi_unclassified(100);k__Fungi_unclassified(100); | - | - | - | - | - | - | - | Unassigned | - |
| Otu53577 | 1 | k__Fungi(100);p__Ascomycota(100);p__Ascomycota_unclassified(100);p__Ascomycota_unclassified(100);p__Ascomycota_unclassified(100);p__Ascomycota_unclassified(100);p__Ascomycota_unclassified(100); | - | - | - | - | - | - | - | Unassigned | - |
| Otu53579 | 1 | k__Fungi(100);k__Fungi_unclassified(100);k__Fungi_unclassified(100);k__Fungi_unclassified(100);k__Fungi_unclassified(100);k__Fungi_unclassified(100);k__Fungi_unclassified(100); | - | - | - | - | - | - | - | Unassigned | - |
| Otu53581 | 1 | k__Fungi(100);k__Fungi_unclassified(100);k__Fungi_unclassified(100);k__Fungi_unclassified(100);k__Fungi_unclassified(100);k__Fungi_unclassified(100);k__Fungi_unclassified(100); | - | - | - | - | - | - | - | Unassigned | - |
| Otu53584 | 1 | k__Fungi(100);k__Fungi_unclassified(100);k__Fungi_unclassified(100);k__Fungi_unclassified(100);k__Fungi_unclassified(100);k__Fungi_unclassified(100);k__Fungi_unclassified(100); | - | - | - | - | - | - | - | Unassigned | - |
| Otu53585 | 1 | k__Fungi(100);k__Fungi_unclassified(100);k__Fungi_unclassified(100);k__Fungi_unclassified(100);k__Fungi_unclassified(100);k__Fungi_unclassified(100);k__Fungi_unclassified(100); | - | - | - | - | - | - | - | Unassigned | - |
| Otu53587 | 1 | k__Fungi(100);k__Fungi_unclassified(100);k__Fungi_unclassified(100);k__Fungi_unclassified(100);k__Fungi_unclassified(100);k__Fungi_unclassified(100);k__Fungi_unclassified(100); | - | - | - | - | - | - | - | Unassigned | - |
| Otu53588 | 1 | k__Fungi(100);k__Fungi_unclassified(100);k__Fungi_unclassified(100);k__Fungi_unclassified(100);k__Fungi_unclassified(100);k__Fungi_unclassified(100);k__Fungi_unclassified(100); | - | - | - | - | - | - | - | Unassigned | - |
| Otu53589 | 1 | k__Fungi(100);k__Fungi_unclassified(100);k__Fungi_unclassified(100);k__Fungi_unclassified(100);k__Fungi_unclassified(100);k__Fungi_unclassified(100);k__Fungi_unclassified(100); | - | - | - | - | - | - | - | Unassigned | - |
| Otu53591 | 1 | k__Fungi(100);k__Fungi_unclassified(100);k__Fungi_unclassified(100);k__Fungi_unclassified(100);k__Fungi_unclassified(100);k__Fungi_unclassified(100);k__Fungi_unclassified(100); | - | - | - | - | - | - | - | Unassigned | - |
| Otu53593 | 1 | k__Fungi(100);k__Fungi_unclassified(100);k__Fungi_unclassified(100);k__Fungi_unclassified(100);k__Fungi_unclassified(100);k__Fungi_unclassified(100);k__Fungi_unclassified(100); | - | - | - | - | - | - | - | Unassigned | - |
| Otu53595 | 1 | k__Fungi(100);k__Fungi_unclassified(100);k__Fungi_unclassified(100);k__Fungi_unclassified(100);k__Fungi_unclassified(100);k__Fungi_unclassified(100);k__Fungi_unclassified(100); | - | - | - | - | - | - | - | Unassigned | - |
| Otu53597 | 1 | k__Fungi(100);p__Ascomycota(100);p__Ascomycota_unclassified(100);p__Ascomycota_unclassified(100);p__Ascomycota_unclassified(100);p__Ascomycota_unclassified(100);p__Ascomycota_unclassified(100); | - | - | - | - | - | - | - | Unassigned | - |
| Otu53599 | 1 | k__Fungi(100);p__Ascomycota(100);p__Ascomycota_unclassified(100);p__Ascomycota_unclassified(100);p__Ascomycota_unclassified(100);p__Ascomycota_unclassified(100);p__Ascomycota_unclassified(100); | - | - | - | - | - | - | - | Unassigned | - |
| Otu53601 | 1 | k__Fungi(100);k__Fungi_unclassified(100);k__Fungi_unclassified(100);k__Fungi_unclassified(100);k__Fungi_unclassified(100);k__Fungi_unclassified(100);k__Fungi_unclassified(100); | - | - | - | - | - | - | - | Unassigned | - |
| Otu53603 | 1 | k__Fungi(100);k__Fungi_unclassified(100);k__Fungi_unclassified(100);k__Fungi_unclassified(100);k__Fungi_unclassified(100);k__Fungi_unclassified(100);k__Fungi_unclassified(100); | - | - | - | - | - | - | - | Unassigned | - |
| Otu53604 | 1 | k__Fungi(100);p__Ascomycota(100);p__Ascomycota_unclassified(100);p__Ascomycota_unclassified(100);p__Ascomycota_unclassified(100);p__Ascomycota_unclassified(100);p__Ascomycota_unclassified(100); | - | - | - | - | - | - | - | Unassigned | - |
| Otu53605 | 1 | k__Fungi(100);k__Fungi_unclassified(100);k__Fungi_unclassified(100);k__Fungi_unclassified(100);k__Fungi_unclassified(100);k__Fungi_unclassified(100);k__Fungi_unclassified(100); | - | - | - | - | - | - | - | Unassigned | - |
| Otu53606 | 1 | k__Fungi(100);p__Ascomycota(100);p__Ascomycota_unclassified(100);p__Ascomycota_unclassified(100);p__Ascomycota_unclassified(100);p__Ascomycota_unclassified(100);p__Ascomycota_unclassified(100); | - | - | - | - | - | - | - | Unassigned | - |
| Otu53607 | 1 | k__Fungi(100);p__Ascomycota(100);p__Ascomycota_unclassified(100);p__Ascomycota_unclassified(100);p__Ascomycota_unclassified(100);p__Ascomycota_unclassified(100);p__Ascomycota_unclassified(100); | - | - | - | - | - | - | - | Unassigned | - |
| Otu53612 | 1 | k__Fungi(100);k__Fungi_unclassified(100);k__Fungi_unclassified(100);k__Fungi_unclassified(100);k__Fungi_unclassified(100);k__Fungi_unclassified(100);k__Fungi_unclassified(100); | - | - | - | - | - | - | - | Unassigned | - |
| Otu53614 | 1 | k__Fungi(100);k__Fungi_unclassified(100);k__Fungi_unclassified(100);k__Fungi_unclassified(100);k__Fungi_unclassified(100);k__Fungi_unclassified(100);k__Fungi_unclassified(100); | - | - | - | - | - | - | - | Unassigned | - |
| Otu53617 | 1 | k__Fungi(100);k__Fungi_unclassified(100);k__Fungi_unclassified(100);k__Fungi_unclassified(100);k__Fungi_unclassified(100);k__Fungi_unclassified(100);k__Fungi_unclassified(100); | - | - | - | - | - | - | - | Unassigned | - |
| Otu53619 | 1 | k__Fungi(100);k__Fungi_unclassified(100);k__Fungi_unclassified(100);k__Fungi_unclassified(100);k__Fungi_unclassified(100);k__Fungi_unclassified(100);k__Fungi_unclassified(100); | - | - | - | - | - | - | - | Unassigned | - |
| Otu53623 | 1 | k__Fungi(100);k__Fungi_unclassified(100);k__Fungi_unclassified(100);k__Fungi_unclassified(100);k__Fungi_unclassified(100);k__Fungi_unclassified(100);k__Fungi_unclassified(100); | - | - | - | - | - | - | - | Unassigned | - |
| Otu53626 | 1 | k__Fungi(100);k__Fungi_unclassified(100);k__Fungi_unclassified(100);k__Fungi_unclassified(100);k__Fungi_unclassified(100);k__Fungi_unclassified(100);k__Fungi_unclassified(100); | - | - | - | - | - | - | - | Unassigned | - |
| Otu53627 | 1 | k__Fungi(100);k__Fungi_unclassified(100);k__Fungi_unclassified(100);k__Fungi_unclassified(100);k__Fungi_unclassified(100);k__Fungi_unclassified(100);k__Fungi_unclassified(100); | - | - | - | - | - | - | - | Unassigned | - |
| Otu53630 | 1 | k__Fungi(100);k__Fungi_unclassified(100);k__Fungi_unclassified(100);k__Fungi_unclassified(100);k__Fungi_unclassified(100);k__Fungi_unclassified(100);k__Fungi_unclassified(100); | - | - | - | - | - | - | - | Unassigned | - |
| Otu53632 | 1 | k__Fungi(100);p__Ascomycota(100);p__Ascomycota_unclassified(100);p__Ascomycota_unclassified(100);p__Ascomycota_unclassified(100);p__Ascomycota_unclassified(100);p__Ascomycota_unclassified(100); | - | - | - | - | - | - | - | Unassigned | - |
| Otu53633 | 1 | k__Fungi(100);k__Fungi_unclassified(100);k__Fungi_unclassified(100);k__Fungi_unclassified(100);k__Fungi_unclassified(100);k__Fungi_unclassified(100);k__Fungi_unclassified(100); | - | - | - | - | - | - | - | Unassigned | - |
| Otu53634 | 1 | k__Fungi(100);p__Ascomycota(100);p__Ascomycota_unclassified(100);p__Ascomycota_unclassified(100);p__Ascomycota_unclassified(100);p__Ascomycota_unclassified(100);p__Ascomycota_unclassified(100); | - | - | - | - | - | - | - | Unassigned | - |
| Otu53637 | 1 | k__Fungi(100);p__Ascomycota(100);p__Ascomycota_unclassified(100);p__Ascomycota_unclassified(100);p__Ascomycota_unclassified(100);p__Ascomycota_unclassified(100);p__Ascomycota_unclassified(100); | - | - | - | - | - | - | - | Unassigned | - |
| Otu53639 | 1 | k__Fungi(100);p__Ascomycota(100);p__Ascomycota_unclassified(100);p__Ascomycota_unclassified(100);p__Ascomycota_unclassified(100);p__Ascomycota_unclassified(100);p__Ascomycota_unclassified(100); | - | - | - | - | - | - | - | Unassigned | - |
| Otu53640 | 1 | k__Fungi(100);k__Fungi_unclassified(100);k__Fungi_unclassified(100);k__Fungi_unclassified(100);k__Fungi_unclassified(100);k__Fungi_unclassified(100);k__Fungi_unclassified(100); | - | - | - | - | - | - | - | Unassigned | - |
| Otu53643 | 1 | k__Fungi(100);p__Ascomycota(100);p__Ascomycota_unclassified(100);p__Ascomycota_unclassified(100);p__Ascomycota_unclassified(100);p__Ascomycota_unclassified(100);p__Ascomycota_unclassified(100); | - | - | - | - | - | - | - | Unassigned | - |
| Otu53647 | 1 | k__Fungi(100);k__Fungi_unclassified(100);k__Fungi_unclassified(100);k__Fungi_unclassified(100);k__Fungi_unclassified(100);k__Fungi_unclassified(100);k__Fungi_unclassified(100); | - | - | - | - | - | - | - | Unassigned | - |
| Otu53651 | 1 | k__Fungi(100);k__Fungi_unclassified(100);k__Fungi_unclassified(100);k__Fungi_unclassified(100);k__Fungi_unclassified(100);k__Fungi_unclassified(100);k__Fungi_unclassified(100); | - | - | - | - | - | - | - | Unassigned | - |
| Otu53654 | 1 | k__Fungi(100);k__Fungi_unclassified(100);k__Fungi_unclassified(100);k__Fungi_unclassified(100);k__Fungi_unclassified(100);k__Fungi_unclassified(100);k__Fungi_unclassified(100); | - | - | - | - | - | - | - | Unassigned | - |
| Otu53658 | 1 | k__Fungi(100);p__Ascomycota(100);p__Ascomycota_unclassified(100);p__Ascomycota_unclassified(100);p__Ascomycota_unclassified(100);p__Ascomycota_unclassified(100);p__Ascomycota_unclassified(100); | - | - | - | - | - | - | - | Unassigned | - |
| Otu53659 | 1 | k__Fungi(100);p__Ascomycota(100);p__Ascomycota_unclassified(100);p__Ascomycota_unclassified(100);p__Ascomycota_unclassified(100);p__Ascomycota_unclassified(100);p__Ascomycota_unclassified(100); | - | - | - | - | - | - | - | Unassigned | - |
| Otu53660 | 1 | k__Fungi(100);p__Ascomycota(100);p__Ascomycota_unclassified(100);p__Ascomycota_unclassified(100);p__Ascomycota_unclassified(100);p__Ascomycota_unclassified(100);p__Ascomycota_unclassified(100); | - | - | - | - | - | - | - | Unassigned | - |
| Otu53662 | 1 | k__Fungi(100);p__Ascomycota(100);p__Ascomycota_unclassified(100);p__Ascomycota_unclassified(100);p__Ascomycota_unclassified(100);p__Ascomycota_unclassified(100);p__Ascomycota_unclassified(100); | - | - | - | - | - | - | - | Unassigned | - |
| Otu53664 | 1 | k__Fungi(100);k__Fungi_unclassified(100);k__Fungi_unclassified(100);k__Fungi_unclassified(100);k__Fungi_unclassified(100);k__Fungi_unclassified(100);k__Fungi_unclassified(100); | - | - | - | - | - | - | - | Unassigned | - |
| Otu53666 | 1 | k__Fungi(100);k__Fungi_unclassified(100);k__Fungi_unclassified(100);k__Fungi_unclassified(100);k__Fungi_unclassified(100);k__Fungi_unclassified(100);k__Fungi_unclassified(100); | - | - | - | - | - | - | - | Unassigned | - |
| Otu53670 | 1 | k__Fungi(100);k__Fungi_unclassified(100);k__Fungi_unclassified(100);k__Fungi_unclassified(100);k__Fungi_unclassified(100);k__Fungi_unclassified(100);k__Fungi_unclassified(100); | - | - | - | - | - | - | - | Unassigned | - |
| Otu53674 | 1 | k__Fungi(100);p__Ascomycota(100);p__Ascomycota_unclassified(100);p__Ascomycota_unclassified(100);p__Ascomycota_unclassified(100);p__Ascomycota_unclassified(100);p__Ascomycota_unclassified(100); | - | - | - | - | - | - | - | Unassigned | - |
| Otu53675 | 1 | k__Fungi(100);p__Ascomycota(100);p__Ascomycota_unclassified(100);p__Ascomycota_unclassified(100);p__Ascomycota_unclassified(100);p__Ascomycota_unclassified(100);p__Ascomycota_unclassified(100); | - | - | - | - | - | - | - | Unassigned | - |
| Otu53677 | 1 | k__Fungi(100);p__Ascomycota(100);p__Ascomycota_unclassified(100);p__Ascomycota_unclassified(100);p__Ascomycota_unclassified(100);p__Ascomycota_unclassified(100);p__Ascomycota_unclassified(100); | - | - | - | - | - | - | - | Unassigned | - |
| Otu53678 | 1 | k__Fungi(100);k__Fungi_unclassified(100);k__Fungi_unclassified(100);k__Fungi_unclassified(100);k__Fungi_unclassified(100);k__Fungi_unclassified(100);k__Fungi_unclassified(100); | - | - | - | - | - | - | - | Unassigned | - |
| Otu53679 | 1 | k__Fungi(100);p__Ascomycota(100);p__Ascomycota_unclassified(100);p__Ascomycota_unclassified(100);p__Ascomycota_unclassified(100);p__Ascomycota_unclassified(100);p__Ascomycota_unclassified(100); | - | - | - | - | - | - | - | Unassigned | - |
| Otu53682 | 1 | k__Fungi(100);k__Fungi_unclassified(100);k__Fungi_unclassified(100);k__Fungi_unclassified(100);k__Fungi_unclassified(100);k__Fungi_unclassified(100);k__Fungi_unclassified(100); | - | - | - | - | - | - | - | Unassigned | - |
| Otu53683 | 1 | k__Fungi(100);k__Fungi_unclassified(100);k__Fungi_unclassified(100);k__Fungi_unclassified(100);k__Fungi_unclassified(100);k__Fungi_unclassified(100);k__Fungi_unclassified(100); | - | - | - | - | - | - | - | Unassigned | - |
| Otu53684 | 1 | k__Fungi(100);k__Fungi_unclassified(100);k__Fungi_unclassified(100);k__Fungi_unclassified(100);k__Fungi_unclassified(100);k__Fungi_unclassified(100);k__Fungi_unclassified(100); | - | - | - | - | - | - | - | Unassigned | - |
| Otu53685 | 1 | k__Fungi(100);k__Fungi_unclassified(100);k__Fungi_unclassified(100);k__Fungi_unclassified(100);k__Fungi_unclassified(100);k__Fungi_unclassified(100);k__Fungi_unclassified(100); | - | - | - | - | - | - | - | Unassigned | - |
| Otu53690 | 1 | k__Fungi(100);k__Fungi_unclassified(100);k__Fungi_unclassified(100);k__Fungi_unclassified(100);k__Fungi_unclassified(100);k__Fungi_unclassified(100);k__Fungi_unclassified(100); | - | - | - | - | - | - | - | Unassigned | - |
| Otu53691 | 1 | k__Fungi(100);k__Fungi_unclassified(100);k__Fungi_unclassified(100);k__Fungi_unclassified(100);k__Fungi_unclassified(100);k__Fungi_unclassified(100);k__Fungi_unclassified(100); | - | - | - | - | - | - | - | Unassigned | - |
| Otu53692 | 1 | k__Fungi(100);k__Fungi_unclassified(100);k__Fungi_unclassified(100);k__Fungi_unclassified(100);k__Fungi_unclassified(100);k__Fungi_unclassified(100);k__Fungi_unclassified(100); | - | - | - | - | - | - | - | Unassigned | - |
| Otu53693 | 1 | k__Fungi(100);k__Fungi_unclassified(100);k__Fungi_unclassified(100);k__Fungi_unclassified(100);k__Fungi_unclassified(100);k__Fungi_unclassified(100);k__Fungi_unclassified(100); | - | - | - | - | - | - | - | Unassigned | - |
| Otu53694 | 1 | k__Fungi(100);p__Ascomycota(100);p__Ascomycota_unclassified(100);p__Ascomycota_unclassified(100);p__Ascomycota_unclassified(100);p__Ascomycota_unclassified(100);p__Ascomycota_unclassified(100); | - | - | - | - | - | - | - | Unassigned | - |
| Otu53695 | 1 | k__Fungi(100);k__Fungi_unclassified(100);k__Fungi_unclassified(100);k__Fungi_unclassified(100);k__Fungi_unclassified(100);k__Fungi_unclassified(100);k__Fungi_unclassified(100); | - | - | - | - | - | - | - | Unassigned | - |
| Otu53703 | 1 | k__Fungi(100);p__Ascomycota(100);p__Ascomycota_unclassified(100);p__Ascomycota_unclassified(100);p__Ascomycota_unclassified(100);p__Ascomycota_unclassified(100);p__Ascomycota_unclassified(100); | - | - | - | - | - | - | - | Unassigned | - |
| Otu53704 | 1 | k__Fungi(100);k__Fungi_unclassified(100);k__Fungi_unclassified(100);k__Fungi_unclassified(100);k__Fungi_unclassified(100);k__Fungi_unclassified(100);k__Fungi_unclassified(100); | - | - | - | - | - | - | - | Unassigned | - |
| Otu53711 | 1 | k__Fungi(100);p__Ascomycota(100);p__Ascomycota_unclassified(100);p__Ascomycota_unclassified(100);p__Ascomycota_unclassified(100);p__Ascomycota_unclassified(100);p__Ascomycota_unclassified(100); | - | - | - | - | - | - | - | Unassigned | - |
| Otu53713 | 1 | k__Fungi(100);p__Ascomycota(100);p__Ascomycota_unclassified(100);p__Ascomycota_unclassified(100);p__Ascomycota_unclassified(100);p__Ascomycota_unclassified(100);p__Ascomycota_unclassified(100); | - | - | - | - | - | - | - | Unassigned | - |
| Otu53715 | 1 | k__Fungi(100);k__Fungi_unclassified(100);k__Fungi_unclassified(100);k__Fungi_unclassified(100);k__Fungi_unclassified(100);k__Fungi_unclassified(100);k__Fungi_unclassified(100); | - | - | - | - | - | - | - | Unassigned | - |
| Otu53716 | 1 | k__Fungi(100);k__Fungi_unclassified(100);k__Fungi_unclassified(100);k__Fungi_unclassified(100);k__Fungi_unclassified(100);k__Fungi_unclassified(100);k__Fungi_unclassified(100); | - | - | - | - | - | - | - | Unassigned | - |
| Otu53717 | 1 | k__Fungi(100);p__Ascomycota(100);p__Ascomycota_unclassified(100);p__Ascomycota_unclassified(100);p__Ascomycota_unclassified(100);p__Ascomycota_unclassified(100);p__Ascomycota_unclassified(100); | - | - | - | - | - | - | - | Unassigned | - |
| Otu53718 | 1 | k__Fungi(100);k__Fungi_unclassified(100);k__Fungi_unclassified(100);k__Fungi_unclassified(100);k__Fungi_unclassified(100);k__Fungi_unclassified(100);k__Fungi_unclassified(100); | - | - | - | - | - | - | - | Unassigned | - |
| Otu53721 | 1 | k__Fungi(100);k__Fungi_unclassified(100);k__Fungi_unclassified(100);k__Fungi_unclassified(100);k__Fungi_unclassified(100);k__Fungi_unclassified(100);k__Fungi_unclassified(100); | - | - | - | - | - | - | - | Unassigned | - |
| Otu53722 | 1 | k__Fungi(100);k__Fungi_unclassified(100);k__Fungi_unclassified(100);k__Fungi_unclassified(100);k__Fungi_unclassified(100);k__Fungi_unclassified(100);k__Fungi_unclassified(100); | - | - | - | - | - | - | - | Unassigned | - |
| Otu53723 | 1 | k__Fungi(100);p__Ascomycota(100);p__Ascomycota_unclassified(100);p__Ascomycota_unclassified(100);p__Ascomycota_unclassified(100);p__Ascomycota_unclassified(100);p__Ascomycota_unclassified(100); | - | - | - | - | - | - | - | Unassigned | - |
| Otu53725 | 1 | k__Fungi(100);k__Fungi_unclassified(100);k__Fungi_unclassified(100);k__Fungi_unclassified(100);k__Fungi_unclassified(100);k__Fungi_unclassified(100);k__Fungi_unclassified(100); | - | - | - | - | - | - | - | Unassigned | - |
| Otu53726 | 1 | k__Fungi(100);p__Ascomycota(100);p__Ascomycota_unclassified(100);p__Ascomycota_unclassified(100);p__Ascomycota_unclassified(100);p__Ascomycota_unclassified(100);p__Ascomycota_unclassified(100); | - | - | - | - | - | - | - | Unassigned | - |
| Otu53727 | 1 | k__Fungi(100);k__Fungi_unclassified(100);k__Fungi_unclassified(100);k__Fungi_unclassified(100);k__Fungi_unclassified(100);k__Fungi_unclassified(100);k__Fungi_unclassified(100); | - | - | - | - | - | - | - | Unassigned | - |
| Otu53728 | 1 | k__Fungi(100);p__Ascomycota(100);p__Ascomycota_unclassified(100);p__Ascomycota_unclassified(100);p__Ascomycota_unclassified(100);p__Ascomycota_unclassified(100);p__Ascomycota_unclassified(100); | - | - | - | - | - | - | - | Unassigned | - |
| Otu53729 | 1 | k__Fungi(100);k__Fungi_unclassified(100);k__Fungi_unclassified(100);k__Fungi_unclassified(100);k__Fungi_unclassified(100);k__Fungi_unclassified(100);k__Fungi_unclassified(100); | - | - | - | - | - | - | - | Unassigned | - |
| Otu53730 | 1 | k__Fungi(100);k__Fungi_unclassified(100);k__Fungi_unclassified(100);k__Fungi_unclassified(100);k__Fungi_unclassified(100);k__Fungi_unclassified(100);k__Fungi_unclassified(100); | - | - | - | - | - | - | - | Unassigned | - |
| Otu53732 | 1 | k__Fungi(100);k__Fungi_unclassified(100);k__Fungi_unclassified(100);k__Fungi_unclassified(100);k__Fungi_unclassified(100);k__Fungi_unclassified(100);k__Fungi_unclassified(100); | - | - | - | - | - | - | - | Unassigned | - |
| Otu53734 | 1 | k__Fungi(100);p__Ascomycota(100);p__Ascomycota_unclassified(100);p__Ascomycota_unclassified(100);p__Ascomycota_unclassified(100);p__Ascomycota_unclassified(100);p__Ascomycota_unclassified(100); | - | - | - | - | - | - | - | Unassigned | - |
| Otu53737 | 1 | k__Fungi(100);k__Fungi_unclassified(100);k__Fungi_unclassified(100);k__Fungi_unclassified(100);k__Fungi_unclassified(100);k__Fungi_unclassified(100);k__Fungi_unclassified(100); | - | - | - | - | - | - | - | Unassigned | - |
| Otu53738 | 1 | k__Fungi(100);p__Ascomycota(100);p__Ascomycota_unclassified(100);p__Ascomycota_unclassified(100);p__Ascomycota_unclassified(100);p__Ascomycota_unclassified(100);p__Ascomycota_unclassified(100); | - | - | - | - | - | - | - | Unassigned | - |
| Otu53739 | 1 | k__Fungi(100);p__Ascomycota(100);p__Ascomycota_unclassified(100);p__Ascomycota_unclassified(100);p__Ascomycota_unclassified(100);p__Ascomycota_unclassified(100);p__Ascomycota_unclassified(100); | - | - | - | - | - | - | - | Unassigned | - |
| Otu53742 | 1 | k__Fungi(100);p__Ascomycota(100);p__Ascomycota_unclassified(100);p__Ascomycota_unclassified(100);p__Ascomycota_unclassified(100);p__Ascomycota_unclassified(100);p__Ascomycota_unclassified(100); | - | - | - | - | - | - | - | Unassigned | - |
| Otu53743 | 1 | k__Fungi(100);k__Fungi_unclassified(100);k__Fungi_unclassified(100);k__Fungi_unclassified(100);k__Fungi_unclassified(100);k__Fungi_unclassified(100);k__Fungi_unclassified(100); | - | - | - | - | - | - | - | Unassigned | - |
| Otu53744 | 1 | k__Fungi(100);p__Ascomycota(100);p__Ascomycota_unclassified(100);p__Ascomycota_unclassified(100);p__Ascomycota_unclassified(100);p__Ascomycota_unclassified(100);p__Ascomycota_unclassified(100); | - | - | - | - | - | - | - | Unassigned | - |
| Otu53750 | 1 | k__Fungi(100);k__Fungi_unclassified(100);k__Fungi_unclassified(100);k__Fungi_unclassified(100);k__Fungi_unclassified(100);k__Fungi_unclassified(100);k__Fungi_unclassified(100); | - | - | - | - | - | - | - | Unassigned | - |
| Otu53751 | 1 | k__Fungi(100);p__Ascomycota(100);p__Ascomycota_unclassified(100);p__Ascomycota_unclassified(100);p__Ascomycota_unclassified(100);p__Ascomycota_unclassified(100);p__Ascomycota_unclassified(100); | - | - | - | - | - | - | - | Unassigned | - |
| Otu53752 | 1 | k__Fungi(100);k__Fungi_unclassified(100);k__Fungi_unclassified(100);k__Fungi_unclassified(100);k__Fungi_unclassified(100);k__Fungi_unclassified(100);k__Fungi_unclassified(100); | - | - | - | - | - | - | - | Unassigned | - |
| Otu53754 | 1 | k__Fungi(100);p__Ascomycota(100);p__Ascomycota_unclassified(100);p__Ascomycota_unclassified(100);p__Ascomycota_unclassified(100);p__Ascomycota_unclassified(100);p__Ascomycota_unclassified(100); | - | - | - | - | - | - | - | Unassigned | - |
| Otu53773 | 1 | k__Fungi(100);p__Ascomycota(100);p__Ascomycota_unclassified(100);p__Ascomycota_unclassified(100);p__Ascomycota_unclassified(100);p__Ascomycota_unclassified(100);p__Ascomycota_unclassified(100); | - | - | - | - | - | - | - | Unassigned | - |
| Otu53775 | 1 | k__Fungi(100);k__Fungi_unclassified(100);k__Fungi_unclassified(100);k__Fungi_unclassified(100);k__Fungi_unclassified(100);k__Fungi_unclassified(100);k__Fungi_unclassified(100); | - | - | - | - | - | - | - | Unassigned | - |
| Otu53782 | 1 | k__Fungi(100);k__Fungi_unclassified(100);k__Fungi_unclassified(100);k__Fungi_unclassified(100);k__Fungi_unclassified(100);k__Fungi_unclassified(100);k__Fungi_unclassified(100); | - | - | - | - | - | - | - | Unassigned | - |
| Otu53784 | 1 | k__Fungi(100);k__Fungi_unclassified(100);k__Fungi_unclassified(100);k__Fungi_unclassified(100);k__Fungi_unclassified(100);k__Fungi_unclassified(100);k__Fungi_unclassified(100); | - | - | - | - | - | - | - | Unassigned | - |
| Otu53786 | 1 | k__Fungi(100);k__Fungi_unclassified(100);k__Fungi_unclassified(100);k__Fungi_unclassified(100);k__Fungi_unclassified(100);k__Fungi_unclassified(100);k__Fungi_unclassified(100); | - | - | - | - | - | - | - | Unassigned | - |
| Otu53787 | 1 | k__Fungi(100);k__Fungi_unclassified(100);k__Fungi_unclassified(100);k__Fungi_unclassified(100);k__Fungi_unclassified(100);k__Fungi_unclassified(100);k__Fungi_unclassified(100); | - | - | - | - | - | - | - | Unassigned | - |
| Otu53789 | 1 | k__Fungi(100);p__Ascomycota(100);p__Ascomycota_unclassified(100);p__Ascomycota_unclassified(100);p__Ascomycota_unclassified(100);p__Ascomycota_unclassified(100);p__Ascomycota_unclassified(100); | - | - | - | - | - | - | - | Unassigned | - |
| Otu53794 | 1 | k__Fungi(100);p__Ascomycota(100);p__Ascomycota_unclassified(100);p__Ascomycota_unclassified(100);p__Ascomycota_unclassified(100);p__Ascomycota_unclassified(100);p__Ascomycota_unclassified(100); | - | - | - | - | - | - | - | Unassigned | - |
| Otu53795 | 1 | k__Fungi(100);k__Fungi_unclassified(100);k__Fungi_unclassified(100);k__Fungi_unclassified(100);k__Fungi_unclassified(100);k__Fungi_unclassified(100);k__Fungi_unclassified(100); | - | - | - | - | - | - | - | Unassigned | - |
| Otu53797 | 1 | k__Fungi(100);k__Fungi_unclassified(100);k__Fungi_unclassified(100);k__Fungi_unclassified(100);k__Fungi_unclassified(100);k__Fungi_unclassified(100);k__Fungi_unclassified(100); | - | - | - | - | - | - | - | Unassigned | - |
| Otu53798 | 1 | k__Fungi(100);k__Fungi_unclassified(100);k__Fungi_unclassified(100);k__Fungi_unclassified(100);k__Fungi_unclassified(100);k__Fungi_unclassified(100);k__Fungi_unclassified(100); | - | - | - | - | - | - | - | Unassigned | - |
| Otu53807 | 1 | k__Fungi(100);k__Fungi_unclassified(100);k__Fungi_unclassified(100);k__Fungi_unclassified(100);k__Fungi_unclassified(100);k__Fungi_unclassified(100);k__Fungi_unclassified(100); | - | - | - | - | - | - | - | Unassigned | - |
| Otu53809 | 1 | k__Fungi(100);p__Ascomycota(100);p__Ascomycota_unclassified(100);p__Ascomycota_unclassified(100);p__Ascomycota_unclassified(100);p__Ascomycota_unclassified(100);p__Ascomycota_unclassified(100); | - | - | - | - | - | - | - | Unassigned | - |
| Otu53810 | 1 | k__Fungi(100);k__Fungi_unclassified(100);k__Fungi_unclassified(100);k__Fungi_unclassified(100);k__Fungi_unclassified(100);k__Fungi_unclassified(100);k__Fungi_unclassified(100); | - | - | - | - | - | - | - | Unassigned | - |
| Otu53811 | 1 | k__Fungi(100);p__Ascomycota(100);p__Ascomycota_unclassified(100);p__Ascomycota_unclassified(100);p__Ascomycota_unclassified(100);p__Ascomycota_unclassified(100);p__Ascomycota_unclassified(100); | - | - | - | - | - | - | - | Unassigned | - |
| Otu53817 | 1 | k__Fungi(100);k__Fungi_unclassified(100);k__Fungi_unclassified(100);k__Fungi_unclassified(100);k__Fungi_unclassified(100);k__Fungi_unclassified(100);k__Fungi_unclassified(100); | - | - | - | - | - | - | - | Unassigned | - |
| Otu53818 | 1 | k__Fungi(100);k__Fungi_unclassified(100);k__Fungi_unclassified(100);k__Fungi_unclassified(100);k__Fungi_unclassified(100);k__Fungi_unclassified(100);k__Fungi_unclassified(100); | - | - | - | - | - | - | - | Unassigned | - |
| Otu53824 | 1 | k__Fungi(100);k__Fungi_unclassified(100);k__Fungi_unclassified(100);k__Fungi_unclassified(100);k__Fungi_unclassified(100);k__Fungi_unclassified(100);k__Fungi_unclassified(100); | - | - | - | - | - | - | - | Unassigned | - |
| Otu53825 | 1 | k__Fungi(100);k__Fungi_unclassified(100);k__Fungi_unclassified(100);k__Fungi_unclassified(100);k__Fungi_unclassified(100);k__Fungi_unclassified(100);k__Fungi_unclassified(100); | - | - | - | - | - | - | - | Unassigned | - |
| Otu53827 | 1 | k__Fungi(100);p__Ascomycota(100);p__Ascomycota_unclassified(100);p__Ascomycota_unclassified(100);p__Ascomycota_unclassified(100);p__Ascomycota_unclassified(100);p__Ascomycota_unclassified(100); | - | - | - | - | - | - | - | Unassigned | - |
| Otu53828 | 1 | k__Fungi(100);p__Ascomycota(100);p__Ascomycota_unclassified(100);p__Ascomycota_unclassified(100);p__Ascomycota_unclassified(100);p__Ascomycota_unclassified(100);p__Ascomycota_unclassified(100); | - | - | - | - | - | - | - | Unassigned | - |
| Otu53831 | 1 | k__Fungi(100);k__Fungi_unclassified(100);k__Fungi_unclassified(100);k__Fungi_unclassified(100);k__Fungi_unclassified(100);k__Fungi_unclassified(100);k__Fungi_unclassified(100); | - | - | - | - | - | - | - | Unassigned | - |
| Otu53832 | 1 | k__Fungi(100);k__Fungi_unclassified(100);k__Fungi_unclassified(100);k__Fungi_unclassified(100);k__Fungi_unclassified(100);k__Fungi_unclassified(100);k__Fungi_unclassified(100); | - | - | - | - | - | - | - | Unassigned | - |
| Otu53833 | 1 | k__Fungi(100);p__Ascomycota(100);p__Ascomycota_unclassified(100);p__Ascomycota_unclassified(100);p__Ascomycota_unclassified(100);p__Ascomycota_unclassified(100);p__Ascomycota_unclassified(100); | - | - | - | - | - | - | - | Unassigned | - |
| Otu53835 | 1 | k__Fungi(100);k__Fungi_unclassified(100);k__Fungi_unclassified(100);k__Fungi_unclassified(100);k__Fungi_unclassified(100);k__Fungi_unclassified(100);k__Fungi_unclassified(100); | - | - | - | - | - | - | - | Unassigned | - |
| Otu53836 | 1 | k__Fungi(100);k__Fungi_unclassified(100);k__Fungi_unclassified(100);k__Fungi_unclassified(100);k__Fungi_unclassified(100);k__Fungi_unclassified(100);k__Fungi_unclassified(100); | - | - | - | - | - | - | - | Unassigned | - |
| Otu53850 | 1 | k__Fungi(100);p__Ascomycota(100);p__Ascomycota_unclassified(100);p__Ascomycota_unclassified(100);p__Ascomycota_unclassified(100);p__Ascomycota_unclassified(100);p__Ascomycota_unclassified(100); | - | - | - | - | - | - | - | Unassigned | - |
| Otu53851 | 1 | k__Fungi(100);p__Ascomycota(100);p__Ascomycota_unclassified(100);p__Ascomycota_unclassified(100);p__Ascomycota_unclassified(100);p__Ascomycota_unclassified(100);p__Ascomycota_unclassified(100); | - | - | - | - | - | - | - | Unassigned | - |
| Otu53855 | 1 | k__Fungi(100);k__Fungi_unclassified(100);k__Fungi_unclassified(100);k__Fungi_unclassified(100);k__Fungi_unclassified(100);k__Fungi_unclassified(100);k__Fungi_unclassified(100); | - | - | - | - | - | - | - | Unassigned | - |
| Otu53856 | 1 | k__Fungi(100);k__Fungi_unclassified(100);k__Fungi_unclassified(100);k__Fungi_unclassified(100);k__Fungi_unclassified(100);k__Fungi_unclassified(100);k__Fungi_unclassified(100); | - | - | - | - | - | - | - | Unassigned | - |
| Otu53858 | 1 | k__Fungi(100);k__Fungi_unclassified(100);k__Fungi_unclassified(100);k__Fungi_unclassified(100);k__Fungi_unclassified(100);k__Fungi_unclassified(100);k__Fungi_unclassified(100); | - | - | - | - | - | - | - | Unassigned | - |
| Otu53860 | 1 | k__Fungi(100);k__Fungi_unclassified(100);k__Fungi_unclassified(100);k__Fungi_unclassified(100);k__Fungi_unclassified(100);k__Fungi_unclassified(100);k__Fungi_unclassified(100); | - | - | - | - | - | - | - | Unassigned | - |
| Otu53861 | 1 | k__Fungi(100);k__Fungi_unclassified(100);k__Fungi_unclassified(100);k__Fungi_unclassified(100);k__Fungi_unclassified(100);k__Fungi_unclassified(100);k__Fungi_unclassified(100); | - | - | - | - | - | - | - | Unassigned | - |
| Otu53862 | 1 | k__Fungi(100);k__Fungi_unclassified(100);k__Fungi_unclassified(100);k__Fungi_unclassified(100);k__Fungi_unclassified(100);k__Fungi_unclassified(100);k__Fungi_unclassified(100); | - | - | - | - | - | - | - | Unassigned | - |

| | | | | | | | | | | | |
|---|---|---|---|---|---|---|---|---|---|---|---|
| Otu54814 | 1 | k__Fungi(100);p__Ascomycota(100);p__Ascomycota_unclassified(100);p__Ascomycota_unclassified(100);p__Ascomycota_unclassified(100);p__Ascomycota_unclassified(100);p__Ascomycota_unclassified(100); | - | - | - | - | - | - | - | Unassigned | - |
| Otu54815 | 1 | k__Fungi(100);p__Ascomycota(100);p__Ascomycota_unclassified(100);p__Ascomycota_unclassified(100);p__Ascomycota_unclassified(100);p__Ascomycota_unclassified(100);p__Ascomycota_unclassified(100); | - | - | - | - | - | - | - | Unassigned | - |
| Otu54816 | 1 | k__Fungi(100);p__Ascomycota(100);p__Ascomycota_unclassified(100);p__Ascomycota_unclassified(100);p__Ascomycota_unclassified(100);p__Ascomycota_unclassified(100);p__Ascomycota_unclassified(100); | - | - | - | - | - | - | - | Unassigned | - |
| Otu54818 | 1 | k__Fungi(100);k__Fungi_unclassified(100);k__Fungi_unclassified(100);k__Fungi_unclassified(100);k__Fungi_unclassified(100);k__Fungi_unclassified(100);k__Fungi_unclassified(100); | - | - | - | - | - | - | - | Unassigned | - |
| Otu54822 | 1 | k__Fungi(100);k__Fungi_unclassified(100);k__Fungi_unclassified(100);k__Fungi_unclassified(100);k__Fungi_unclassified(100);k__Fungi_unclassified(100);k__Fungi_unclassified(100); | - | - | - | - | - | - | - | Unassigned | - |
| Otu54827 | 1 | k__Fungi(100);k__Fungi_unclassified(100);k__Fungi_unclassified(100);k__Fungi_unclassified(100);k__Fungi_unclassified(100);k__Fungi_unclassified(100);k__Fungi_unclassified(100); | - | - | - | - | - | - | - | Unassigned | - |
| Otu54831 | 1 | k__Fungi(100);p__Ascomycota(100);c__Eurotiomycetes(100);c__Eurotiomycetes_unclassified(100);c__Eurotiomycetes_unclassified(100);c__Eurotiomycetes_unclassified(100);c__Eurotiomycetes_unclassified(100); | - | - | - | - | - | - | - | Unassigned | - |
| Otu54836 | 1 | k__Fungi(100);k__Fungi_unclassified(100);k__Fungi_unclassified(100);k__Fungi_unclassified(100);k__Fungi_unclassified(100);k__Fungi_unclassified(100);k__Fungi_unclassified(100); | - | - | - | - | - | - | - | Unassigned | - |
| Otu54837 | 1 | k__Fungi(100);k__Fungi_unclassified(100);k__Fungi_unclassified(100);k__Fungi_unclassified(100);k__Fungi_unclassified(100);k__Fungi_unclassified(100);k__Fungi_unclassified(100); | - | - | - | - | - | - | - | Unassigned | - |
| Otu54838 | 1 | k__Fungi(100);p__Ascomycota(100);p__Ascomycota_unclassified(100);p__Ascomycota_unclassified(100);p__Ascomycota_unclassified(100);p__Ascomycota_unclassified(100);p__Ascomycota_unclassified(100); | - | - | - | - | - | - | - | Unassigned | - |
| Otu54841 | 1 | k__Fungi(100);k__Fungi_unclassified(100);k__Fungi_unclassified(100);k__Fungi_unclassified(100);k__Fungi_unclassified(100);k__Fungi_unclassified(100);k__Fungi_unclassified(100); | - | - | - | - | - | - | - | Unassigned | - |
| Otu54848 | 1 | k__Fungi(100);k__Fungi_unclassified(100);k__Fungi_unclassified(100);k__Fungi_unclassified(100);k__Fungi_unclassified(100);k__Fungi_unclassified(100);k__Fungi_unclassified(100); | - | - | - | - | - | - | - | Unassigned | - |
| Otu54849 | 1 | k__Fungi(100);k__Fungi_unclassified(100);k__Fungi_unclassified(100);k__Fungi_unclassified(100);k__Fungi_unclassified(100);k__Fungi_unclassified(100);k__Fungi_unclassified(100); | - | - | - | - | - | - | - | Unassigned | - |
| Otu54851 | 1 | k__Fungi(100);k__Fungi_unclassified(100);k__Fungi_unclassified(100);k__Fungi_unclassified(100);k__Fungi_unclassified(100);k__Fungi_unclassified(100);k__Fungi_unclassified(100); | - | - | - | - | - | - | - | Unassigned | - |
| Otu54853 | 1 | k__Fungi(100);k__Fungi_unclassified(100);k__Fungi_unclassified(100);k__Fungi_unclassified(100);k__Fungi_unclassified(100);k__Fungi_unclassified(100);k__Fungi_unclassified(100); | - | - | - | - | - | - | - | Unassigned | - |
| Otu54854 | 1 | k__Fungi(100);k__Fungi_unclassified(100);k__Fungi_unclassified(100);k__Fungi_unclassified(100);k__Fungi_unclassified(100);k__Fungi_unclassified(100);k__Fungi_unclassified(100); | - | - | - | - | - | - | - | Unassigned | - |
| Otu54855 | 1 | k__Fungi(100);p__Ascomycota(100);p__Ascomycota_unclassified(100);p__Ascomycota_unclassified(100);p__Ascomycota_unclassified(100);p__Ascomycota_unclassified(100);p__Ascomycota_unclassified(100); | - | - | - | - | - | - | - | Unassigned | - |
| Otu54856 | 1 | k__Fungi(100);k__Fungi_unclassified(100);k__Fungi_unclassified(100);k__Fungi_unclassified(100);k__Fungi_unclassified(100);k__Fungi_unclassified(100);k__Fungi_unclassified(100); | - | - | - | - | - | - | - | Unassigned | - |
| Otu54857 | 1 | k__Fungi(100);p__Ascomycota(100);p__Ascomycota_unclassified(100);p__Ascomycota_unclassified(100);p__Ascomycota_unclassified(100);p__Ascomycota_unclassified(100);p__Ascomycota_unclassified(100); | - | - | - | - | - | - | - | Unassigned | - |
| Otu54859 | 1 | k__Fungi(100);k__Fungi_unclassified(100);k__Fungi_unclassified(100);k__Fungi_unclassified(100);k__Fungi_unclassified(100);k__Fungi_unclassified(100);k__Fungi_unclassified(100); | - | - | - | - | - | - | - | Unassigned | - |
| Otu54860 | 1 | k__Fungi(100);k__Fungi_unclassified(100);k__Fungi_unclassified(100);k__Fungi_unclassified(100);k__Fungi_unclassified(100);k__Fungi_unclassified(100);k__Fungi_unclassified(100); | - | - | - | - | - | - | - | Unassigned | - |
| Otu54861 | 1 | k__Fungi(100);k__Fungi_unclassified(100);k__Fungi_unclassified(100);k__Fungi_unclassified(100);k__Fungi_unclassified(100);k__Fungi_unclassified(100);k__Fungi_unclassified(100); | - | - | - | - | - | - | - | Unassigned | - |
| Otu54862 | 1 | k__Fungi(100);k__Fungi_unclassified(100);k__Fungi_unclassified(100);k__Fungi_unclassified(100);k__Fungi_unclassified(100);k__Fungi_unclassified(100);k__Fungi_unclassified(100); | - | - | - | - | - | - | - | Unassigned | - |
| Otu54863 | 1 | k__Fungi(100);k__Fungi_unclassified(100);k__Fungi_unclassified(100);k__Fungi_unclassified(100);k__Fungi_unclassified(100);k__Fungi_unclassified(100);k__Fungi_unclassified(100); | - | - | - | - | - | - | - | Unassigned | - |
| Otu54864 | 1 | k__Fungi(100);p__Ascomycota(100);p__Ascomycota_unclassified(100);p__Ascomycota_unclassified(100);p__Ascomycota_unclassified(100);p__Ascomycota_unclassified(100);p__Ascomycota_unclassified(100); | - | - | - | - | - | - | - | Unassigned | - |
| Otu54865 | 1 | k__Fungi(100);k__Fungi_unclassified(100);k__Fungi_unclassified(100);k__Fungi_unclassified(100);k__Fungi_unclassified(100);k__Fungi_unclassified(100);k__Fungi_unclassified(100); | - | - | - | - | - | - | - | Unassigned | - |
| Otu54866 | 1 | k__Fungi(100);k__Fungi_unclassified(100);k__Fungi_unclassified(100);k__Fungi_unclassified(100);k__Fungi_unclassified(100);k__Fungi_unclassified(100);k__Fungi_unclassified(100); | - | - | - | - | - | - | - | Unassigned | - |
| Otu54868 | 1 | k__Fungi(100);k__Fungi_unclassified(100);k__Fungi_unclassified(100);k__Fungi_unclassified(100);k__Fungi_unclassified(100);k__Fungi_unclassified(100);k__Fungi_unclassified(100); | - | - | - | - | - | - | - | Unassigned | - |
| Otu54869 | 1 | k__Fungi(100);p__Ascomycota(100);p__Ascomycota_unclassified(100);p__Ascomycota_unclassified(100);p__Ascomycota_unclassified(100);p__Ascomycota_unclassified(100);p__Ascomycota_unclassified(100); | - | - | - | - | - | - | - | Unassigned | - |
| Otu54871 | 1 | k__Fungi(100);k__Fungi_unclassified(100);k__Fungi_unclassified(100);k__Fungi_unclassified(100);k__Fungi_unclassified(100);k__Fungi_unclassified(100);k__Fungi_unclassified(100); | - | - | - | - | - | - | - | Unassigned | - |
| Otu54872 | 1 | k__Fungi(100);p__Ascomycota(100);p__Ascomycota_unclassified(100);p__Ascomycota_unclassified(100);p__Ascomycota_unclassified(100);p__Ascomycota_unclassified(100);p__Ascomycota_unclassified(100); | - | - | - | - | - | - | - | Unassigned | - |
| Otu54873 | 1 | k__Fungi(100);k__Fungi_unclassified(100);k__Fungi_unclassified(100);k__Fungi_unclassified(100);k__Fungi_unclassified(100);k__Fungi_unclassified(100);k__Fungi_unclassified(100); | - | - | - | - | - | - | - | Unassigned | - |
| Otu54876 | 1 | k__Fungi(100);p__Ascomycota(100);p__Ascomycota_unclassified(100);p__Ascomycota_unclassified(100);p__Ascomycota_unclassified(100);p__Ascomycota_unclassified(100);p__Ascomycota_unclassified(100); | - | - | - | - | - | - | - | Unassigned | - |
| Otu54877 | 1 | k__Fungi(100);k__Fungi_unclassified(100);k__Fungi_unclassified(100);k__Fungi_unclassified(100);k__Fungi_unclassified(100);k__Fungi_unclassified(100);k__Fungi_unclassified(100); | - | - | - | - | - | - | - | Unassigned | - |
| Otu54878 | 1 | k__Fungi(100);p__Ascomycota(100);p__Ascomycota_unclassified(100);p__Ascomycota_unclassified(100);p__Ascomycota_unclassified(100);p__Ascomycota_unclassified(100);p__Ascomycota_unclassified(100); | - | - | - | - | - | - | - | Unassigned | - |
| Otu54880 | 1 | k__Fungi(100);k__Fungi_unclassified(100);k__Fungi_unclassified(100);k__Fungi_unclassified(100);k__Fungi_unclassified(100);k__Fungi_unclassified(100);k__Fungi_unclassified(100); | - | - | - | - | - | - | - | Unassigned | - |
| Otu54881 | 1 | k__Fungi(100);p__Ascomycota(100);p__Ascomycota_unclassified(100);p__Ascomycota_unclassified(100);p__Ascomycota_unclassified(100);p__Ascomycota_unclassified(100);p__Ascomycota_unclassified(100); | - | - | - | - | - | - | - | Unassigned | - |
| Otu54887 | 1 | k__Fungi(100);p__Ascomycota(100);p__Ascomycota_unclassified(100);p__Ascomycota_unclassified(100);p__Ascomycota_unclassified(100);p__Ascomycota_unclassified(100);p__Ascomycota_unclassified(100); | - | - | - | - | - | - | - | Unassigned | - |
| Otu54888 | 1 | k__Fungi(100);p__Ascomycota(100);p__Ascomycota_unclassified(100);p__Ascomycota_unclassified(100);p__Ascomycota_unclassified(100);p__Ascomycota_unclassified(100);p__Ascomycota_unclassified(100); | - | - | - | - | - | - | - | Unassigned | - |
| Otu54889 | 1 | k__Fungi(100);p__Ascomycota(100);p__Ascomycota_unclassified(100);p__Ascomycota_unclassified(100);p__Ascomycota_unclassified(100);p__Ascomycota_unclassified(100);p__Ascomycota_unclassified(100); | - | - | - | - | - | - | - | Unassigned | - |
| Otu54894 | 1 | k__Fungi(100);p__Ascomycota(100);p__Ascomycota_unclassified(100);p__Ascomycota_unclassified(100);p__Ascomycota_unclassified(100);p__Ascomycota_unclassified(100);p__Ascomycota_unclassified(100); | - | - | - | - | - | - | - | Unassigned | - |
| Otu54896 | 1 | k__Fungi(100);p__Ascomycota(100);p__Ascomycota_unclassified(100);p__Ascomycota_unclassified(100);p__Ascomycota_unclassified(100);p__Ascomycota_unclassified(100);p__Ascomycota_unclassified(100); | - | - | - | - | - | - | - | Unassigned | - |
| Otu54899 | 1 | k__Fungi(100);p__Ascomycota(100);p__Ascomycota_unclassified(100);p__Ascomycota_unclassified(100);p__Ascomycota_unclassified(100);p__Ascomycota_unclassified(100);p__Ascomycota_unclassified(100); | - | - | - | - | - | - | - | Unassigned | - |
| Otu54903 | 1 | k__Fungi(100);p__Ascomycota(100);p__Ascomycota_unclassified(100);p__Ascomycota_unclassified(100);p__Ascomycota_unclassified(100);p__Ascomycota_unclassified(100);p__Ascomycota_unclassified(100); | - | - | - | - | - | - | - | Unassigned | - |
| Otu54906 | 1 | k__Fungi(100);k__Fungi_unclassified(100);k__Fungi_unclassified(100);k__Fungi_unclassified(100);k__Fungi_unclassified(100);k__Fungi_unclassified(100);k__Fungi_unclassified(100); | - | - | - | - | - | - | - | Unassigned | - |
| Otu54907 | 1 | k__Fungi(100);k__Fungi_unclassified(100);k__Fungi_unclassified(100);k__Fungi_unclassified(100);k__Fungi_unclassified(100);k__Fungi_unclassified(100);k__Fungi_unclassified(100); | - | - | - | - | - | - | - | Unassigned | - |
| Otu54908 | 1 | k__Fungi(100);p__Ascomycota(100);p__Ascomycota_unclassified(100);p__Ascomycota_unclassified(100);p__Ascomycota_unclassified(100);p__Ascomycota_unclassified(100);p__Ascomycota_unclassified(100); | - | - | - | - | - | - | - | Unassigned | - |
| Otu54909 | 1 | k__Fungi(100);k__Fungi_unclassified(100);k__Fungi_unclassified(100);k__Fungi_unclassified(100);k__Fungi_unclassified(100);k__Fungi_unclassified(100);k__Fungi_unclassified(100); | - | - | - | - | - | - | - | Unassigned | - |
| Otu54911 | 1 | k__Fungi(100);k__Fungi_unclassified(100);k__Fungi_unclassified(100);k__Fungi_unclassified(100);k__Fungi_unclassified(100);k__Fungi_unclassified(100);k__Fungi_unclassified(100); | - | - | - | - | - | - | - | Unassigned | - |
| Otu54912 | 1 | k__Fungi(100);k__Fungi_unclassified(100);k__Fungi_unclassified(100);k__Fungi_unclassified(100);k__Fungi_unclassified(100);k__Fungi_unclassified(100);k__Fungi_unclassified(100); | - | - | - | - | - | - | - | Unassigned | - |
| Otu54913 | 1 | k__Fungi(100);p__Ascomycota(100);p__Ascomycota_unclassified(100);p__Ascomycota_unclassified(100);p__Ascomycota_unclassified(100);p__Ascomycota_unclassified(100);p__Ascomycota_unclassified(100); | - | - | - | - | - | - | - | Unassigned | - |
| Otu54915 | 1 | k__Fungi(100);p__Ascomycota(100);p__Ascomycota_unclassified(100);p__Ascomycota_unclassified(100);p__Ascomycota_unclassified(100);p__Ascomycota_unclassified(100);p__Ascomycota_unclassified(100); | - | - | - | - | - | - | - | Unassigned | - |
| Otu54916 | 1 | k__Fungi(100);k__Fungi_unclassified(100);k__Fungi_unclassified(100);k__Fungi_unclassified(100);k__Fungi_unclassified(100);k__Fungi_unclassified(100);k__Fungi_unclassified(100); | - | - | - | - | - | - | - | Unassigned | - |
| Otu54917 | 1 | k__Fungi(100);p__Ascomycota(100);p__Ascomycota_unclassified(100);p__Ascomycota_unclassified(100);p__Ascomycota_unclassified(100);p__Ascomycota_unclassified(100);p__Ascomycota_unclassified(100); | - | - | - | - | - | - | - | Unassigned | - |
| Otu54918 | 1 | k__Fungi(100);p__Ascomycota(100);p__Ascomycota_unclassified(100);p__Ascomycota_unclassified(100);p__Ascomycota_unclassified(100);p__Ascomycota_unclassified(100);p__Ascomycota_unclassified(100); | - | - | - | - | - | - | - | Unassigned | - |
| Otu54920 | 1 | k__Fungi(100);p__Ascomycota(100);p__Ascomycota_unclassified(100);p__Ascomycota_unclassified(100);p__Ascomycota_unclassified(100);p__Ascomycota_unclassified(100);p__Ascomycota_unclassified(100); | - | - | - | - | - | - | - | Unassigned | - |
| Otu54921 | 1 | k__Fungi(100);k__Fungi_unclassified(100);k__Fungi_unclassified(100);k__Fungi_unclassified(100);k__Fungi_unclassified(100);k__Fungi_unclassified(100);k__Fungi_unclassified(100); | - | - | - | - | - | - | - | Unassigned | - |
| Otu54924 | 1 | k__Fungi(100);p__Ascomycota(100);p__Ascomycota_unclassified(100);p__Ascomycota_unclassified(100);p__Ascomycota_unclassified(100);p__Ascomycota_unclassified(100);p__Ascomycota_unclassified(100); | - | - | - | - | - | - | - | Unassigned | - |
| Otu54926 | 1 | k__Fungi(100);k__Fungi_unclassified(100);k__Fungi_unclassified(100);k__Fungi_unclassified(100);k__Fungi_unclassified(100);k__Fungi_unclassified(100);k__Fungi_unclassified(100); | - | - | - | - | - | - | - | Unassigned | - |
| Otu54927 | 1 | k__Fungi(100);k__Fungi_unclassified(100);k__Fungi_unclassified(100);k__Fungi_unclassified(100);k__Fungi_unclassified(100);k__Fungi_unclassified(100);k__Fungi_unclassified(100); | - | - | - | - | - | - | - | Unassigned | - |
| Otu54928 | 1 | k__Fungi(100);k__Fungi_unclassified(100);k__Fungi_unclassified(100);k__Fungi_unclassified(100);k__Fungi_unclassified(100);k__Fungi_unclassified(100);k__Fungi_unclassified(100); | - | - | - | - | - | - | - | Unassigned | - |
| Otu54930 | 1 | k__Fungi(100);p__Ascomycota(100);p__Ascomycota_unclassified(100);p__Ascomycota_unclassified(100);p__Ascomycota_unclassified(100);p__Ascomycota_unclassified(100);p__Ascomycota_unclassified(100); | - | - | - | - | - | - | - | Unassigned | - |
| Otu54931 | 1 | k__Fungi(100);k__Fungi_unclassified(100);k__Fungi_unclassified(100);k__Fungi_unclassified(100);k__Fungi_unclassified(100);k__Fungi_unclassified(100);k__Fungi_unclassified(100); | - | - | - | - | - | - | - | Unassigned | - |
| Otu54932 | 1 | k__Fungi(100);k__Fungi_unclassified(100);k__Fungi_unclassified(100);k__Fungi_unclassified(100);k__Fungi_unclassified(100);k__Fungi_unclassified(100);k__Fungi_unclassified(100); | - | - | - | - | - | - | - | Unassigned | - |
| Otu54933 | 1 | k__Fungi(100);p__Ascomycota(100);p__Ascomycota_unclassified(100);p__Ascomycota_unclassified(100);p__Ascomycota_unclassified(100);p__Ascomycota_unclassified(100);p__Ascomycota_unclassified(100); | - | - | - | - | - | - | - | Unassigned | - |
| Otu54935 | 1 | k__Fungi(100);k__Fungi_unclassified(100);k__Fungi_unclassified(100);k__Fungi_unclassified(100);k__Fungi_unclassified(100);k__Fungi_unclassified(100);k__Fungi_unclassified(100); | - | - | - | - | - | - | - | Unassigned | - |
| Otu54937 | 1 | k__Fungi(100);p__Ascomycota(100);p__Ascomycota_unclassified(100);p__Ascomycota_unclassified(100);p__Ascomycota_unclassified(100);p__Ascomycota_unclassified(100);p__Ascomycota_unclassified(100); | - | - | - | - | - | - | - | Unassigned | - |
| Otu54941 | 1 | k__Fungi(100);k__Fungi_unclassified(100);k__Fungi_unclassified(100);k__Fungi_unclassified(100);k__Fungi_unclassified(100);k__Fungi_unclassified(100);k__Fungi_unclassified(100); | - | - | - | - | - | - | - | Unassigned | - |
| Otu54943 | 1 | k__Fungi(100);k__Fungi_unclassified(100);k__Fungi_unclassified(100);k__Fungi_unclassified(100);k__Fungi_unclassified(100);k__Fungi_unclassified(100);k__Fungi_unclassified(100); | - | - | - | - | - | - | - | Unassigned | - |
| Otu54944 | 1 | k__Fungi(100);k__Fungi_unclassified(100);k__Fungi_unclassified(100);k__Fungi_unclassified(100);k__Fungi_unclassified(100);k__Fungi_unclassified(100);k__Fungi_unclassified(100); | - | - | - | - | - | - | - | Unassigned | - |
| Otu54945 | 1 | k__Fungi(100);k__Fungi_unclassified(100);k__Fungi_unclassified(100);k__Fungi_unclassified(100);k__Fungi_unclassified(100);k__Fungi_unclassified(100);k__Fungi_unclassified(100); | - | - | - | - | - | - | - | Unassigned | - |
| Otu54946 | 1 | k__Fungi(100);k__Fungi_unclassified(100);k__Fungi_unclassified(100);k__Fungi_unclassified(100);k__Fungi_unclassified(100);k__Fungi_unclassified(100);k__Fungi_unclassified(100); | - | - | - | - | - | - | - | Unassigned | - |
| Otu54948 | 1 | k__Fungi(100);k__Fungi_unclassified(100);k__Fungi_unclassified(100);k__Fungi_unclassified(100);k__Fungi_unclassified(100);k__Fungi_unclassified(100);k__Fungi_unclassified(100); | - | - | - | - | - | - | - | Unassigned | - |
| Otu54950 | 1 | k__Fungi(100);p__Ascomycota(100);p__Ascomycota_unclassified(100);p__Ascomycota_unclassified(100);p__Ascomycota_unclassified(100);p__Ascomycota_unclassified(100);p__Ascomycota_unclassified(100); | - | - | - | - | - | - | - | Unassigned | - |
| Otu54951 | 1 | k__Fungi(100);p__Ascomycota(100);p__Ascomycota_unclassified(100);p__Ascomycota_unclassified(100);p__Ascomycota_unclassified(100);p__Ascomycota_unclassified(100);p__Ascomycota_unclassified(100); | - | - | - | - | - | - | - | Unassigned | - |
| Otu54952 | 1 | k__Fungi(100);k__Fungi_unclassified(100);k__Fungi_unclassified(100);k__Fungi_unclassified(100);k__Fungi_unclassified(100);k__Fungi_unclassified(100);k__Fungi_unclassified(100); | - | - | - | - | - | - | - | Unassigned | - |
| Otu54953 | 1 | k__Fungi(100);p__Ascomycota(100);p__Ascomycota_unclassified(100);p__Ascomycota_unclassified(100);p__Ascomycota_unclassified(100);p__Ascomycota_unclassified(100);p__Ascomycota_unclassified(100); | - | - | - | - | - | - | - | Unassigned | - |
| Otu54954 | 1 | k__Fungi(100);p__Ascomycota(100);p__Ascomycota_unclassified(100);p__Ascomycota_unclassified(100);p__Ascomycota_unclassified(100);p__Ascomycota_unclassified(100);p__Ascomycota_unclassified(100); | - | - | - | - | - | - | - | Unassigned | - |
| Otu54955 | 1 | k__Fungi(100);p__Ascomycota(100);p__Ascomycota_unclassified(100);p__Ascomycota_unclassified(100);p__Ascomycota_unclassified(100);p__Ascomycota_unclassified(100);p__Ascomycota_unclassified(100); | - | - | - | - | - | - | - | Unassigned | - |
| Otu54957 | 1 | k__Fungi(100);k__Fungi_unclassified(100);k__Fungi_unclassified(100);k__Fungi_unclassified(100);k__Fungi_unclassified(100);k__Fungi_unclassified(100);k__Fungi_unclassified(100); | - | - | - | - | - | - | - | Unassigned | - |
| Otu54966 | 1 | k__Fungi(100);k__Fungi_unclassified(100);k__Fungi_unclassified(100);k__Fungi_unclassified(100);k__Fungi_unclassified(100);k__Fungi_unclassified(100);k__Fungi_unclassified(100); | - | - | - | - | - | - | - | Unassigned | - |
| Otu54967 | 1 | k__Fungi(100);k__Fungi_unclassified(100);k__Fungi_unclassified(100);k__Fungi_unclassified(100);k__Fungi_unclassified(100);k__Fungi_unclassified(100);k__Fungi_unclassified(100); | - | - | - | - | - | - | - | Unassigned | - |
| Otu54974 | 1 | k__Fungi(100);k__Fungi_unclassified(100);k__Fungi_unclassified(100);k__Fungi_unclassified(100);k__Fungi_unclassified(100);k__Fungi_unclassified(100);k__Fungi_unclassified(100); | - | - | - | - | - | - | - | Unassigned | - |
| Otu54975 | 1 | k__Fungi(100);p__Ascomycota(100);p__Ascomycota_unclassified(100);p__Ascomycota_unclassified(100);p__Ascomycota_unclassified(100);p__Ascomycota_unclassified(100);p__Ascomycota_unclassified(100); | - | - | - | - | - | - | - | Unassigned | - |
| Otu54976 | 1 | k__Fungi(100);k__Fungi_unclassified(100);k__Fungi_unclassified(100);k__Fungi_unclassified(100);k__Fungi_unclassified(100);k__Fungi_unclassified(100);k__Fungi_unclassified(100); | - | - | - | - | - | - | - | Unassigned | - |
| Otu54986 | 1 | k__Fungi(100);k__Fungi_unclassified(100);k__Fungi_unclassified(100);k__Fungi_unclassified(100);k__Fungi_unclassified(100);k__Fungi_unclassified(100);k__Fungi_unclassified(100); | - | - | - | - | - | - | - | Unassigned | - |
| Otu55003 | 1 | k__Fungi(100);k__Fungi_unclassified(100);k__Fungi_unclassified(100);k__Fungi_unclassified(100);k__Fungi_unclassified(100);k__Fungi_unclassified(100);k__Fungi_unclassified(100); | - | - | - | - | - | - | - | Unassigned | - |
| Otu55020 | 1 | k__Fungi(100);p__Ascomycota(100);p__Ascomycota_unclassified(100);p__Ascomycota_unclassified(100);p__Ascomycota_unclassified(100);p__Ascomycota_unclassified(100);p__Ascomycota_unclassified(100); | - | - | - | - | - | - | - | Unassigned | - |
| Otu55021 | 1 | k__Fungi(100);k__Fungi_unclassified(100);k__Fungi_unclassified(100);k__Fungi_unclassified(100);k__Fungi_unclassified(100);k__Fungi_unclassified(100);k__Fungi_unclassified(100); | - | - | - | - | - | - | - | Unassigned | - |
| Otu55023 | 1 | k__Fungi(100);k__Fungi_unclassified(100);k__Fungi_unclassified(100);k__Fungi_unclassified(100);k__Fungi_unclassified(100);k__Fungi_unclassified(100);k__Fungi_unclassified(100); | - | - | - | - | - | - | - | Unassigned | - |
| Otu55028 | 1 | k__Fungi(100);k__Fungi_unclassified(100);k__Fungi_unclassified(100);k__Fungi_unclassified(100);k__Fungi_unclassified(100);k__Fungi_unclassified(100);k__Fungi_unclassified(100); | - | - | - | - | - | - | - | Unassigned | - |
| Otu55043 | 1 | k__Fungi(100);k__Fungi_unclassified(100);k__Fungi_unclassified(100);k__Fungi_unclassified(100);k__Fungi_unclassified(100);k__Fungi_unclassified(100);k__Fungi_unclassified(100); | - | - | - | - | - | - | - | Unassigned | - |
| Otu55044 | 1 | k__Fungi(100);p__Ascomycota(100);p__Ascomycota_unclassified(100);p__Ascomycota_unclassified(100);p__Ascomycota_unclassified(100);p__Ascomycota_unclassified(100);p__Ascomycota_unclassified(100); | - | - | - | - | - | - | - | Unassigned | - |
| Otu55054 | 1 | k__Fungi(100);k__Fungi_unclassified(100);k__Fungi_unclassified(100);k__Fungi_unclassified(100);k__Fungi_unclassified(100);k__Fungi_unclassified(100);k__Fungi_unclassified(100); | - | - | - | - | - | - | - | Unassigned | - |
| Otu55081 | 1 | k__Fungi(100);k__Fungi_unclassified(100);k__Fungi_unclassified(100);k__Fungi_unclassified(100);k__Fungi_unclassified(100);k__Fungi_unclassified(100);k__Fungi_unclassified(100); | - | - | - | - | - | - | - | Unassigned | - |
| Otu55084 | 1 | k__Fungi(100);p__Ascomycota(100);p__Ascomycota_unclassified(100);p__Ascomycota_unclassified(100);p__Ascomycota_unclassified(100);p__Ascomycota_unclassified(100);p__Ascomycota_unclassified(100); | - | - | - | - | - | - | - | Unassigned | - |
| Otu55086 | 1 | k__Fungi(100);k__Fungi_unclassified(100);k__Fungi_unclassified(100);k__Fungi_unclassified(100);k__Fungi_unclassified(100);k__Fungi_unclassified(100);k__Fungi_unclassified(100); | - | - | - | - | - | - | - | Unassigned | - |
| Otu55088 | 1 | k__Fungi(100);k__Fungi_unclassified(100);k__Fungi_unclassified(100);k__Fungi_unclassified(100);k__Fungi_unclassified(100);k__Fungi_unclassified(100);k__Fungi_unclassified(100); | - | - | - | - | - | - | - | Unassigned | - |
| Otu55089 | 1 | k__Fungi(100);k__Fungi_unclassified(100);k__Fungi_unclassified(100);k__Fungi_unclassified(100);k__Fungi_unclassified(100);k__Fungi_unclassified(100);k__Fungi_unclassified(100); | - | - | - | - | - | - | - | Unassigned | - |
| Otu55090 | 1 | k__Fungi(100);k__Fungi_unclassified(100);k__Fungi_unclassified(100);k__Fungi_unclassified(100);k__Fungi_unclassified(100);k__Fungi_unclassified(100);k__Fungi_unclassified(100); | - | - | - | - | - | - | - | Unassigned | - |
| Otu55091 | 1 | k__Fungi(100);k__Fungi_unclassified(100);k__Fungi_unclassified(100);k__Fungi_unclassified(100);k__Fungi_unclassified(100);k__Fungi_unclassified(100);k__Fungi_unclassified(100); | - | - | - | - | - | - | - | Unassigned | - |
| Otu55092 | 1 | k__Fungi(100);k__Fungi_unclassified(100);k__Fungi_unclassified(100);k__Fungi_unclassified(100);k__Fungi_unclassified(100);k__Fungi_unclassified(100);k__Fungi_unclassified(100); | - | - | - | - | - | - | - | Unassigned | - |
| Otu55094 | 1 | k__Fungi(100);p__Ascomycota(100);p__Ascomycota_unclassified(100);p__Ascomycota_unclassified(100);p__Ascomycota_unclassified(100);p__Ascomycota_unclassified(100);p__Ascomycota_unclassified(100); | - | - | - | - | - | - | - | Unassigned | - |
| Otu55095 | 1 | k__Fungi(100);k__Fungi_unclassified(100);k__Fungi_unclassified(100);k__Fungi_unclassified(100);k__Fungi_unclassified(100);k__Fungi_unclassified(100);k__Fungi_unclassified(100); | - | - | - | - | - | - | - | Unassigned | - |
| Otu55099 | 1 | k__Fungi(100);k__Fungi_unclassified(100);k__Fungi_unclassified(100);k__Fungi_unclassified(100);k__Fungi_unclassified(100);k__Fungi_unclassified(100);k__Fungi_unclassified(100); | - | - | - | - | - | - | - | Unassigned | - |
| Otu55100 | 1 | k__Fungi(100);k__Fungi_unclassified(100);k__Fungi_unclassified(100);k__Fungi_unclassified(100);k__Fungi_unclassified(100);k__Fungi_unclassified(100);k__Fungi_unclassified(100); | - | - | - | - | - | - | - | Unassigned | - |
| Otu55101 | 1 | k__Fungi(100);p__Ascomycota(100);p__Ascomycota_unclassified(100);p__Ascomycota_unclassified(100);p__Ascomycota_unclassified(100);p__Ascomycota_unclassified(100);p__Ascomycota_unclassified(100); | - | - | - | - | - | - | - | Unassigned | - |
| Otu55104 | 1 | k__Fungi(100);k__Fungi_unclassified(100);k__Fungi_unclassified(100);k__Fungi_unclassified(100);k__Fungi_unclassified(100);k__Fungi_unclassified(100);k__Fungi_unclassified(100); | - | - | - | - | - | - | - | Unassigned | - |
| Otu55106 | 1 | k__Fungi(100);k__Fungi_unclassified(100);k__Fungi_unclassified(100);k__Fungi_unclassified(100);k__Fungi_unclassified(100);k__Fungi_unclassified(100);k__Fungi_unclassified(100); | - | - | - | - | - | - | - | Unassigned | - |
| Otu55113 | 1 | k__Fungi(100);k__Fungi_unclassified(100);k__Fungi_unclassified(100);k__Fungi_unclassified(100);k__Fungi_unclassified(100);k__Fungi_unclassified(100);k__Fungi_unclassified(100); | - | - | - | - | - | - | - | Unassigned | - |
| Otu55114 | 1 | k__Fungi(100);p__Ascomycota(100);p__Ascomycota_unclassified(100);p__Ascomycota_unclassified(100);p__Ascomycota_unclassified(100);p__Ascomycota_unclassified(100);p__Ascomycota_unclassified(100); | - | - | - | - | - | - | - | Unassigned | - |
| Otu55138 | 1 | k__Fungi(100);k__Fungi_unclassified(100);k__Fungi_unclassified(100);k__Fungi_unclassified(100);k__Fungi_unclassified(100);k__Fungi_unclassified(100);k__Fungi_unclassified(100); | - | - | - | - | - | - | - | Unassigned | - |
| Otu55148 | 1 | k__Fungi(100);k__Fungi_unclassified(100);k__Fungi_unclassified(100);k__Fungi_unclassified(100);k__Fungi_unclassified(100);k__Fungi_unclassified(100);k__Fungi_unclassified(100); | - | - | - | - | - | - | - | Unassigned | - |
| Otu55149 | 1 | k__Fungi(100);k__Fungi_unclassified(100);k__Fungi_unclassified(100);k__Fungi_unclassified(100);k__Fungi_unclassified(100);k__Fungi_unclassified(100);k__Fungi_unclassified(100); | - | - | - | - | - | - | - | Unassigned | - |
| Otu55150 | 1 | k__Fungi(100);k__Fungi_unclassified(100);k__Fungi_unclassified(100);k__Fungi_unclassified(100);k__Fungi_unclassified(100);k__Fungi_unclassified(100);k__Fungi_unclassified(100); | - | - | - | - | - | - | - | Unassigned | - |
| Otu55153 | 1 | k__Fungi(100);p__Ascomycota(100);p__Ascomycota_unclassified(100);p__Ascomycota_unclassified(100);p__Ascomycota_unclassified(100);p__Ascomycota_unclassified(100);p__Ascomycota_unclassified(100); | - | - | - | - | - | - | - | Unassigned | - |
| Otu55159 | 1 | k__Fungi(100);p__Ascomycota(100);p__Ascomycota_unclassified(100);p__Ascomycota_unclassified(100);p__Ascomycota_unclassified(100);p__Ascomycota_unclassified(100);p__Ascomycota_unclassified(100); | - | - | - | - | - | - | - | Unassigned | - |
| Otu55160 | 1 | k__Fungi(100);k__Fungi_unclassified(100);k__Fungi_unclassified(100);k__Fungi_unclassified(100);k__Fungi_unclassified(100);k__Fungi_unclassified(100);k__Fungi_unclassified(100); | - | - | - | - | - | - | - | Unassigned | - |
| Otu55164 | 1 | k__Fungi(100);k__Fungi_unclassified(100);k__Fungi_unclassified(100);k__Fungi_unclassified(100);k__Fungi_unclassified(100);k__Fungi_unclassified(100);k__Fungi_unclassified(100); | - | - | - | - | - | - | - | Unassigned | - |
| Otu55166 | 1 | k__Fungi(100);k__Fungi_unclassified(100);k__Fungi_unclassified(100);k__Fungi_unclassified(100);k__Fungi_unclassified(100);k__Fungi_unclassified(100);k__Fungi_unclassified(100); | - | - | - | - | - | - | - | Unassigned | - |
| Otu55167 | 1 | k__Fungi(100);p__Ascomycota(100);p__Ascomycota_unclassified(100);p__Ascomycota_unclassified(100);p__Ascomycota_unclassified(100);p__Ascomycota_unclassified(100);p__Ascomycota_unclassified(100); | - | - | - | - | - | - | - | Unassigned | - |
| Otu55169 | 1 | k__Fungi(100);k__Fungi_unclassified(100);k__Fungi_unclassified(100);k__Fungi_unclassified(100);k__Fungi_unclassified(100);k__Fungi_unclassified(100);k__Fungi_unclassified(100); | - | - | - | - | - | - | - | Unassigned | - |
| Otu55175 | 1 | k__Fungi(100);k__Fungi_unclassified(100);k__Fungi_unclassified(100);k__Fungi_unclassified(100);k__Fungi_unclassified(100);k__Fungi_unclassified(100);k__Fungi_unclassified(100); | - | - | - | - | - | - | - | Unassigned | - |
| Otu55177 | 1 | k__Fungi(100);k__Fungi_unclassified(100);k__Fungi_unclassified(100);k__Fungi_unclassified(100);k__Fungi_unclassified(100);k__Fungi_unclassified(100);k__Fungi_unclassified(100); | - | - | - | - | - | - | - | Unassigned | - |
| Otu55179 | 1 | k__Fungi(100);k__Fungi_unclassified(100);k__Fungi_unclassified(100);k__Fungi_unclassified(100);k__Fungi_unclassified(100);k__Fungi_unclassified(100);k__Fungi_unclassified(100); | - | - | - | - | - | - | - | Unassigned | - |
| Otu55180 | 1 | k__Fungi(100);k__Fungi_unclassified(100);k__Fungi_unclassified(100);k__Fungi_unclassified(100);k__Fungi_unclassified(100);k__Fungi_unclassified(100);k__Fungi_unclassified(100); | - | - | - | - | - | - | - | Unassigned | - |
| Otu55197 | 1 | k__Fungi(100);k__Fungi_unclassified(100);k__Fungi_unclassified(100);k__Fungi_unclassified(100);k__Fungi_unclassified(100);k__Fungi_unclassified(100);k__Fungi_unclassified(100); | - | - | - | - | - | - | - | Unassigned | - |
| Otu55199 | 1 | k__Fungi(100);k__Fungi_unclassified(100);k__Fungi_unclassified(100);k__Fungi_unclassified(100);k__Fungi_unclassified(100);k__Fungi_unclassified(100);k__Fungi_unclassified(100); | - | - | - | - | - | - | - | Unassigned | - |
| Otu55200 | 1 | k__Fungi(100);p__Ascomycota(100);p__Ascomycota_unclassified(100);p__Ascomycota_unclassified(100);p__Ascomycota_unclassified(100);p__Ascomycota_unclassified(100);p__Ascomycota_unclassified(100); | - | - | - | - | - | - | - | Unassigned | - |
| Otu55202 | 1 | k__Fungi(100);p__Ascomycota(100);p__Ascomycota_unclassified(100);p__Ascomycota_unclassified(100);p__Ascomycota_unclassified(100);p__Ascomycota_unclassified(100);p__Ascomycota_unclassified(100); | - | - | - | - | - | - | - | Unassigned | - |
| Otu55205 | 1 | k__Fungi(100);k__Fungi_unclassified(100);k__Fungi_unclassified(100);k__Fungi_unclassified(100);k__Fungi_unclassified(100);k__Fungi_unclassified(100);k__Fungi_unclassified(100); | - | - | - | - | - | - | - | Unassigned | - |
| Otu55207 | 1 | k__Fungi(100);k__Fungi_unclassified(100);k__Fungi_unclassified(100);k__Fungi_unclassified(100);k__Fungi_unclassified(100);k__Fungi_unclassified(100);k__Fungi_unclassified(100); | - | - | - | - | - | - | - | Unassigned | - |
| Otu55208 | 1 | k__Fungi(100);p__Ascomycota(100);p__Ascomycota_unclassified(100);p__Ascomycota_unclassified(100);p__Ascomycota_unclassified(100);p__Ascomycota_unclassified(100);p__Ascomycota_unclassified(100); | - | - | - | - | - | - | - | Unassigned | - |
| Otu55209 | 1 | k__Fungi(100);p__Ascomycota(100);p__Ascomycota_unclassified(100);p__Ascomycota_unclassified(100);p__Ascomycota_unclassified(100);p__Ascomycota_unclassified(100);p__Ascomycota_unclassified(100); | - | - | - | - | - | - | - | Unassigned | - |
| Otu55210 | 1 | k__Fungi(100);p__Ascomycota(100);p__Ascomycota_unclassified(100);p__Ascomycota_unclassified(100);p__Ascomycota_unclassified(100);p__Ascomycota_unclassified(100);p__Ascomycota_unclassified(100); | - | - | - | - | - | - | - | Unassigned | - |
| Otu55220 | 1 | k__Fungi(100);p__Ascomycota(100);p__Ascomycota_unclassified(100);p__Ascomycota_unclassified(100);p__Ascomycota_unclassified(100);p__Ascomycota_unclassified(100);p__Ascomycota_unclassified(100); | - | - | - | - | - | - | - | Unassigned | - |
| Otu55222 | 1 | k__Fungi(100);k__Fungi_unclassified(100);k__Fungi_unclassified(100);k__Fungi_unclassified(100);k__Fungi_unclassified(100);k__Fungi_unclassified(100);k__Fungi_unclassified(100); | - | - | - | - | - | - | - | Unassigned | - |
| Otu55224 | 1 | k__Fungi(100);k__Fungi_unclassified(100);k__Fungi_unclassified(100);k__Fungi_unclassified(100);k__Fungi_unclassified(100);k__Fungi_unclassified(100);k__Fungi_unclassified(100); | - | - | - | - | - | - | - | Unassigned | - |
| Otu55229 | 1 | k__Fungi(100);k__Fungi_unclassified(100);k__Fungi_unclassified(100);k__Fungi_unclassified(100);k__Fungi_unclassified(100);k__Fungi_unclassified(100);k__Fungi_unclassified(100); | - | - | - | - | - | - | - | Unassigned | - |
| Otu55234 | 1 | k__Fungi(100);k__Fungi_unclassified(100);k__Fungi_unclassified(100);k__Fungi_unclassified(100);k__Fungi_unclassified(100);k__Fungi_unclassified(100);k__Fungi_unclassified(100); | - | - | - | - | - | - | - | Unassigned | - |
| Otu55240 | 1 | k__Fungi(100);k__Fungi_unclassified(100);k__Fungi_unclassified(100);k__Fungi_unclassified(100);k__Fungi_unclassified(100);k__Fungi_unclassified(100);k__Fungi_unclassified(100); | - | - | - | - | - | - | - | Unassigned | - |
| Otu55243 | 1 | k__Fungi(100);k__Fungi_unclassified(100);k__Fungi_unclassified(100);k__Fungi_unclassified(100);k__Fungi_unclassified(100);k__Fungi_unclassified(100);k__Fungi_unclassified(100); | - | - | - | - | - | - | - | Unassigned | - |
| Otu55251 | 1 | k__Fungi(100);k__Fungi_unclassified(100);k__Fungi_unclassified(100);k__Fungi_unclassified(100);k__Fungi_unclassified(100);k__Fungi_unclassified(100);k__Fungi_unclassified(100); | - | - | - | - | - | - | - | Unassigned | - |
| Otu55256 | 1 | k__Fungi(100);k__Fungi_unclassified(100);k__Fungi_unclassified(100);k__Fungi_unclassified(100);k__Fungi_unclassified(100);k__Fungi_unclassified(100);k__Fungi_unclassified(100); | - | - | - | - | - | - | - | Unassigned | - |
| Otu55258 | 1 | k__Fungi(100);k__Fungi_unclassified(100);k__Fungi_unclassified(100);k__Fungi_unclassified(100);k__Fungi_unclassified(100);k__Fungi_unclassified(100);k__Fungi_unclassified(100); | - | - | - | - | - | - | - | Unassigned | - |
| Otu55263 | 1 | k__Fungi(100);p__Ascomycota(100);p__Ascomycota_unclassified(100);p__Ascomycota_unclassified(100);p__Ascomycota_unclassified(100);p__Ascomycota_unclassified(100);p__Ascomycota_unclassified(100); | - | - | - | - | - | - | - | Unassigned | - |
| Otu55264 | 1 | k__Fungi(100);p__Ascomycota(100);p__Ascomycota_unclassified(100);p__Ascomycota_unclassified(100);p__Ascomycota_unclassified(100);p__Ascomycota_unclassified(100);p__Ascomycota_unclassified(100); | - | - | - | - | - | - | - | Unassigned | - |
| Otu55265 | 1 | k__Fungi(100);p__Ascomycota(100);p__Ascomycota_unclassified(100);p__Ascomycota_unclassified(100);p__Ascomycota_unclassified(100);p__Ascomycota_unclassified(100);p__Ascomycota_unclassified(100); | - | - | - | - | - | - | - | Unassigned | - |
| Otu55266 | 1 | k__Fungi(100);k__Fungi_unclassified(100);k__Fungi_unclassified(100);k__Fungi_unclassified(100);k__Fungi_unclassified(100);k__Fungi_unclassified(100);k__Fungi_unclassified(100); | - | - | - | - | - | - | - | Unassigned | - |
| Otu55268 | 1 | k__Fungi(100);p__Basidiomycota(100);p__Basidiomycota_unclassified(100);p__Basidiomycota_unclassified(100);p__Basidiomycota_unclassified(100);p__Basidiomycota_unclassified(100);p__Basidiomycota_unclassified(100); | - | - | - | - | - | - | - | Unassigned | - |
| Otu55270 | 1 | k__Fungi(100);p__Ascomycota(100);p__Ascomycota_unclassified(100);p__Ascomycota_unclassified(100);p__Ascomycota_unclassified(100);p__Ascomycota_unclassified(100);p__Ascomycota_unclassified(100); | - | - | - | - | - | - | - | Unassigned | - |
| Otu55271 | 1 | k__Fungi(100);p__Ascomycota(100);p__Ascomycota_unclassified(100);p__Ascomycota_unclassified(100);p__Ascomycota_unclassified(100);p__Ascomycota_unclassified(100);p__Ascomycota_unclassified(100); | - | - | - | - | - | - | - | Unassigned | - |
| Otu55272 | 1 | k__Fungi(100);p__Ascomycota(100);p__Ascomycota_unclassified(100);p__Ascomycota_unclassified(100);p__Ascomycota_unclassified(100);p__Ascomycota_unclassified(100);p__Ascomycota_unclassified(100); | - | - | - | - | - | - | - | Unassigned | - |
| Otu55273 | 1 | k__Fungi(100);p__Ascomycota(100);p__Ascomycota_unclassified(100);p__Ascomycota_unclassified(100);p__Ascomycota_unclassified(100);p__Ascomycota_unclassified(100);p__Ascomycota_unclassified(100); | - | - | - | - | - | - | - | Unassigned | - |
| Otu55274 | 1 | k__Fungi(100);k__Fungi_unclassified(100);k__Fungi_unclassified(100);k__Fungi_unclassified(100);k__Fungi_unclassified(100);k__Fungi_unclassified(100);k__Fungi_unclassified(100); | - | - | - | - | - | - | - | Unassigned | - |
| Otu55275 | 1 | k__Fungi(100);p__Ascomycota(100);p__Ascomycota_unclassified(100);p__Ascomycota_unclassified(100);p__Ascomycota_unclassified(100);p__Ascomycota_unclassified(100);p__Ascomycota_unclassified(100); | - | - | - | - | - | - | - | Unassigned | - |
| Otu55276 | 1 | k__Fungi(100);k__Fungi_unclassified(100);k__Fungi_unclassified(100);k__Fungi_unclassified(100);k__Fungi_unclassified(100);k__Fungi_unclassified(100);k__Fungi_unclassified(100); | - | - | - | - | - | - | - | Unassigned | - |
| Otu55281 | 1 | k__Fungi(100);p__Ascomycota(100);p__Ascomycota_unclassified(100);p__Ascomycota_unclassified(100);p__Ascomycota_unclassified(100);p__Ascomycota_unclassified(100);p__Ascomycota_unclassified(100); | - | - | - | - | - | - | - | Unassigned | - |
| Otu55282 | 1 | k__Fungi(100);k__Fungi_unclassified(100);k__Fungi_unclassified(100);k__Fungi_unclassified(100);k__Fungi_unclassified(100);k__Fungi_unclassified(100);k__Fungi_unclassified(100); | - | - | - | - | - | - | - | Unassigned | - |
| Otu55283 | 1 | k__Fungi(100);k__Fungi_unclassified(100);k__Fungi_unclassified(100);k__Fungi_unclassified(100);k__Fungi_unclassified(100);k__Fungi_unclassified(100);k__Fungi_unclassified(100); | - | - | - | - | - | - | - | Unassigned | - |
| Otu55286 | 1 | k__Fungi(100);k__Fungi_unclassified(100);k__Fungi_unclassified(100);k__Fungi_unclassified(100);k__Fungi_unclassified(100);k__Fungi_unclassified(100);k__Fungi_unclassified(100); | - | - | - | - | - | - | - | Unassigned | - |
| Otu55303 | 1 | k__Fungi(100);k__Fungi_unclassified(100);k__Fungi_unclassified(100);k__Fungi_unclassified(100);k__Fungi_unclassified(100);k__Fungi_unclassified(100);k__Fungi_unclassified(100); | - | - | - | - | - | - | - | Unassigned | - |
| Otu55321 | 1 | k__Fungi(100);p__Ascomycota(100);p__Ascomycota_unclassified(100);p__Ascomycota_unclassified(100);p__Ascomycota_unclassified(100);p__Ascomycota_unclassified(100);p__Ascomycota_unclassified(100); | - | - | - | - | - | - | - | Unassigned | - |
| Otu55341 | 1 | k__Fungi(100);p__Ascomycota(100);p__Ascomycota_unclassified(100);p__Ascomycota_unclassified(100);p__Ascomycota_unclassified(100);p__Ascomycota_unclassified(100);p__Ascomycota_unclassified(100); | - | - | - | - | - | - | - | Unassigned | - |
| Otu55351 | 1 | k__Fungi(100);k__Fungi_unclassified(100);k__Fungi_unclassified(100);k__Fungi_unclassified(100);k__Fungi_unclassified(100);k__Fungi_unclassified(100);k__Fungi_unclassified(100); | - | - | - | - | - | - | - | Unassigned | - |
| Otu55352 | 1 | k__Fungi(100);k__Fungi_unclassified(100);k__Fungi_unclassified(100);k__Fungi_unclassified(100);k__Fungi_unclassified(100);k__Fungi_unclassified(100);k__Fungi_unclassified(100); | - | - | - | - | - | - | - | Unassigned | - |
| Otu55353 | 1 | k__Fungi(100);p__Ascomycota(100);p__Ascomycota_unclassified(100);p__Ascomycota_unclassified(100);p__Ascomycota_unclassified(100);p__Ascomycota_unclassified(100);p__Ascomycota_unclassified(100); | - | - | - | - | - | - | - | Unassigned | - |
| Otu55359 | 1 | k__Fungi(100);k__Fungi_unclassified(100);k__Fungi_unclassified(100);k__Fungi_unclassified(100);k__Fungi_unclassified(100);k__Fungi_unclassified(100);k__Fungi_unclassified(100); | - | - | - | - | - | - | - | Unassigned | - |
| Otu55363 | 1 | k__Fungi(100);k__Fungi_unclassified(100);k__Fungi_unclassified(100);k__Fungi_unclassified(100);k__Fungi_unclassified(100);k__Fungi_unclassified(100);k__Fungi_unclassified(100); | - | - | - | - | - | - | - | Unassigned | - |
| Otu55367 | 1 | k__Fungi(100);p__Ascomycota(100);p__Ascomycota_unclassified(100);p__Ascomycota_unclassified(100);p__Ascomycota_unclassified(100);p__Ascomycota_unclassified(100);p__Ascomycota_unclassified(100); | - | - | - | - | - | - | - | Unassigned | - |
| Otu55380 | 1 | k__Fungi(100);p__Ascomycota(100);p__Ascomycota_unclassified(100);p__Ascomycota_unclassified(100);p__Ascomycota_unclassified(100);p__Ascomycota_unclassified(100);p__Ascomycota_unclassified(100); | - | - | - | - | - | - | - | Unassigned | - |
| Otu55402 | 1 | k__Fungi(100);k__Fungi_unclassified(100);k__Fungi_unclassified(100);k__Fungi_unclassified(100);k__Fungi_unclassified(100);k__Fungi_unclassified(100);k__Fungi_unclassified(100); | - | - | - | - | - | - | - | Unassigned | - |
| Otu55403 | 1 | k__Fungi(100);p__Ascomycota(100);p__Ascomycota_unclassified(100);p__Ascomycota_unclassified(100);p__Ascomycota_unclassified(100);p__Ascomycota_unclassified(100);p__Ascomycota_unclassified(100); | - | - | - | - | - | - | - | Unassigned | - |
| Otu55404 | 1 | k__Fungi(100);p__Ascomycota(100);p__Ascomycota_unclassified(100);p__Ascomycota_unclassified(100);p__Ascomycota_unclassified(100);p__Ascomycota_unclassified(100);p__Ascomycota_unclassified(100); | - | - | - | - | - | - | - | Unassigned | - |
| Otu55408 | 1 | k__Fungi(100);k__Fungi_unclassified(100);k__Fungi_unclassified(100);k__Fungi_unclassified(100);k__Fungi_unclassified(100);k__Fungi_unclassified(100);k__Fungi_unclassified(100); | - | - | - | - | - | - | - | Unassigned | - |
| Otu55420 | 1 | k__Fungi(100);k__Fungi_unclassified(100);k__Fungi_unclassified(100);k__Fungi_unclassified(100);k__Fungi_unclassified(100);k__Fungi_unclassified(100);k__Fungi_unclassified(100); | - | - | - | - | - | - | - | Unassigned | - |
| Otu55422 | 1 | k__Fungi(100);k__Fungi_unclassified(100);k__Fungi_unclassified(100);k__Fungi_unclassified(100);k__Fungi_unclassified(100);k__Fungi_unclassified(100);k__Fungi_unclassified(100); | - | - | - | - | - | - | - | Unassigned | - |
| Otu55450 | 1 | k__Fungi(100);k__Fungi_unclassified(100);k__Fungi_unclassified(100);k__Fungi_unclassified(100);k__Fungi_unclassified(100);k__Fungi_unclassified(100);k__Fungi_unclassified(100); | - | - | - | - | - | - | - | Unassigned | - |
| Otu55480 | 1 | k__Fungi(100);k__Fungi_unclassified(100);k__Fungi_unclassified(100);k__Fungi_unclassified(100);k__Fungi_unclassified(100);k__Fungi_unclassified(100);k__Fungi_unclassified(100); | - | - | - | - | - | - | - | Unassigned | - |
| Otu55486 | 1 | k__Fungi(100);k__Fungi_unclassified(100);k__Fungi_unclassified(100);k__Fungi_unclassified(100);k__Fungi_unclassified(100);k__Fungi_unclassified(100);k__Fungi_unclassified(100); | - | - | - | - | - | - | - | Unassigned | - |
| Otu55496 | 1 | k__Fungi(100);k__Fungi_unclassified(100);k__Fungi_unclassified(100);k__Fungi_unclassified(100);k__Fungi_unclassified(100);k__Fungi_unclassified(100);k__Fungi_unclassified(100); | - | - | - | - | - | - | - | Unassigned | - |
| Otu55499 | 1 | k__Fungi(100);p__Ascomycota(100);p__Ascomycota_unclassified(100);p__Ascomycota_unclassified(100);p__Ascomycota_unclassified(100);p__Ascomycota_unclassified(100);p__Ascomycota_unclassified(100); | - | - | - | - | - | - | - | Unassigned | - |
| Otu55500 | 1 | k__Fungi(100);k__Fungi_unclassified(100);k__Fungi_unclassified(100);k__Fungi_unclassified(100);k__Fungi_unclassified(100);k__Fungi_unclassified(100);k__Fungi_unclassified(100); | - | - | - | - | - | - | - | Unassigned | - |
| Otu55501 | 1 | k__Fungi(100);p__Ascomycota(100);p__Ascomycota_unclassified(100);p__Ascomycota_unclassified(100);p__Ascomycota_unclassified(100);p__Ascomycota_unclassified(100);p__Ascomycota_unclassified(100); | - | - | - | - | - | - | - | Unassigned | - |
| Otu55502 | 1 | k__Fungi(100);p__Ascomycota(100);p__Ascomycota_unclassified(100);p__Ascomycota_unclassified(100);p__Ascomycota_unclassified(100);p__Ascomycota_unclassified(100);p__Ascomycota_unclassified(100); | - | - | - | - | - | - | - | Unassigned | - |
| Otu55503 | 1 | k__Fungi(100);p__Ascomycota(100);p__Ascomycota_unclassified(100);p__Ascomycota_unclassified(100);p__Ascomycota_unclassified(100);p__Ascomycota_unclassified(100);p__Ascomycota_unclassified(100); | - | - | - | - | - | - | - | Unassigned | - |
| Otu55504 | 1 | k__Fungi(100);k__Fungi_unclassified(100);k__Fungi_unclassified(100);k__Fungi_unclassified(100);k__Fungi_unclassified(100);k__Fungi_unclassified(100);k__Fungi_unclassified(100); | - | - | - | - | - | - | - | Unassigned | - |
| Otu55505 | 1 | k__Fungi(100);k__Fungi_unclassified(100);k__Fungi_unclassified(100);k__Fungi_unclassified(100);k__Fungi_unclassified(100);k__Fungi_unclassified(100);k__Fungi_unclassified(100); | - | - | - | - | - | - | - | Unassigned | - |
| Otu55506 | 1 | k__Fungi(100);k__Fungi_unclassified(100);k__Fungi_unclassified(100);k__Fungi_unclassified(100);k__Fungi_unclassified(100);k__Fungi_unclassified(100);k__Fungi_unclassified(100); | - | - | - | - | - | - | - | Unassigned | - |
| Otu55508 | 1 | k__Fungi(100);p__Ascomycota(100);p__Ascomycota_unclassified(100);p__Ascomycota_unclassified(100);p__Ascomycota_unclassified(100);p__Ascomycota_unclassified(100);p__Ascomycota_unclassified(100); | - | - | - | - | - | - | - | Unassigned | - |
| Otu55516 | 1 | k__Fungi(100);p__Ascomycota(100);p__Ascomycota_unclassified(100);p__Ascomycota_unclassified(100);p__Ascomycota_unclassified(100);p__Ascomycota_unclassified(100);p__Ascomycota_unclassified(100); | - | - | - | - | - | - | - | Unassigned | - |
| Otu55521 | 1 | k__Fungi(100);p__Ascomycota(100);p__Ascomycota_unclassified(100);p__Ascomycota_unclassified(100);p__Ascomycota_unclassified(100);p__Ascomycota_unclassified(100);p__Ascomycota_unclassified(100); | - | - | - | - | - | - | - | Unassigned | - |
| Otu55526 | 1 | k__Fungi(100);p__Ascomycota(100);p__Ascomycota_unclassified(100);p__Ascomycota_unclassified(100);p__Ascomycota_unclassified(100);p__Ascomycota_unclassified(100);p__Ascomycota_unclassified(100); | - | - | - | - | - | - | - | Unassigned | - |
| Otu55535 | 1 | k__Fungi(100);p__Ascomycota(100);p__Ascomycota_unclassified(100);p__Ascomycota_unclassified(100);p__Ascomycota_unclassified(100);p__Ascomycota_unclassified(100);p__Ascomycota_unclassified(100); | - | - | - | - | - | - | - | Unassigned | - |
| Otu55543 | 1 | k__Fungi(100);p__Ascomycota(100);p__Ascomycota_unclassified(100);p__Ascomycota_unclassified(100);p__Ascomycota_unclassified(100);p__Ascomycota_unclassified(100);p__Ascomycota_unclassified(100); | - | - | - | - | - | - | - | Unassigned | - |
| Otu55544 | 1 | k__Fungi(100);k__Fungi_unclassified(100);k__Fungi_unclassified(100);k__Fungi_unclassified(100);k__Fungi_unclassified(100);k__Fungi_unclassified(100);k__Fungi_unclassified(100); | - | - | - | - | - | - | - | Unassigned | - |
| Otu55552 | 1 | k__Fungi(100);k__Fungi_unclassified(100);k__Fungi_unclassified(100);k__Fungi_unclassified(100);k__Fungi_unclassified(100);k__Fungi_unclassified(100);k__Fungi_unclassified(100); | - | - | - | - | - | - | - | Unassigned | - |
| Otu55553 | 1 | k__Fungi(100);k__Fungi_unclassified(100);k__Fungi_unclassified(100);k__Fungi_unclassified(100);k__Fungi_unclassified(100);k__Fungi_unclassified(100);k__Fungi_unclassified(100); | - | - | - | - | - | - | - | Unassigned | - |
| Otu55555 | 1 | k__Fungi(100);k__Fungi_unclassified(100);k__Fungi_unclassified(100);k__Fungi_unclassified(100);k__Fungi_unclassified(100);k__Fungi_unclassified(100);k__Fungi_unclassified(100); | - | - | - | - | - | - | - | Unassigned | - |
| Otu55561 | 1 | k__Fungi(100);k__Fungi_unclassified(100);k__Fungi_unclassified(100);k__Fungi_unclassified(100);k__Fungi_unclassified(100);k__Fungi_unclassified(100);k__Fungi_unclassified(100); | - | - | - | - | - | - | - | Unassigned | - |
| Otu55563 | 1 | k__Fungi(100);k__Fungi_unclassified(100);k__Fungi_unclassified(100);k__Fungi_unclassified(100);k__Fungi_unclassified(100);k__Fungi_unclassified(100);k__Fungi_unclassified(100); | - | - | - | - | - | - | - | Unassigned | - |
| Otu55574 | 1 | k__Fungi(100);p__Ascomycota(100);p__Ascomycota_unclassified(100);p__Ascomycota_unclassified(100);p__Ascomycota_unclassified(100);p__Ascomycota_unclassified(100);p__Ascomycota_unclassified(100); | - | - | - | - | - | - | - | Unassigned | - |
| Otu55575 | 1 | k__Fungi(100);p__Ascomycota(100);p__Ascomycota_unclassified(100);p__Ascomycota_unclassified(100);p__Ascomycota_unclassified(100);p__Ascomycota_unclassified(100);p__Ascomycota_unclassified(100); | - | - | - | - | - | - | - | Unassigned | - |
| Otu55576 | 1 | k__Fungi(100);p__Ascomycota(100);p__Ascomycota_unclassified(100);p__Ascomycota_unclassified(100);p__Ascomycota_unclassified(100);p__Ascomycota_unclassified(100);p__Ascomycota_unclassified(100); | - | - | - | - | - | - | - | Unassigned | - |
| Otu55582 | 1 | k__Fungi(100);k__Fungi_unclassified(100);k__Fungi_unclassified(100);k__Fungi_unclassified(100);k__Fungi_unclassified(100);k__Fungi_unclassified(100);k__Fungi_unclassified(100); | - | - | - | - | - | - | - | Unassigned | - |
| Otu55585 | 1 | k__Fungi(100);k__Fungi_unclassified(100);k__Fungi_unclassified(100);k__Fungi_unclassified(100);k__Fungi_unclassified(100);k__Fungi_unclassified(100);k__Fungi_unclassified(100); | - | - | - | - | - | - | - | Unassigned | - |
| Otu55596 | 1 | k__Fungi(100);k__Fungi_unclassified(100);k__Fungi_unclassified(100);k__Fungi_unclassified(100);k__Fungi_unclassified(100);k__Fungi_unclassified(100);k__Fungi_unclassified(100); | - | - | - | - | - | - | - | Unassigned | - |
| Otu55599 | 1 | k__Fungi(100);p__Ascomycota(100);p__Ascomycota_unclassified(100);p__Ascomycota_unclassified(100);p__Ascomycota_unclassified(100);p__Ascomycota_unclassified(100);p__Ascomycota_unclassified(100); | - | - | - | - | - | - | - | Unassigned | - |
| Otu55608 | 1 | k__Fungi(100);p__Ascomycota(100);p__Ascomycota_unclassified(100);p__Ascomycota_unclassified(100);p__Ascomycota_unclassified(100);p__Ascomycota_unclassified(100);p__Ascomycota_unclassified(100); | - | - | - | - | - | - | - | Unassigned | - |
| Otu55615 | 1 | k__Fungi(100);p__Ascomycota(100);p__Ascomycota_unclassified(100);p__Ascomycota_unclassified(100);p__Ascomycota_unclassified(100);p__Ascomycota_unclassified(100);p__Ascomycota_unclassified(100); | - | - | - | - | - | - | - | Unassigned | - |
| Otu55630 | 1 | k__Fungi(100);p__Ascomycota(100);p__Ascomycota_unclassified(100);p__Ascomycota_unclassified(100);p__Ascomycota_unclassified(100);p__Ascomycota_unclassified(100);p__Ascomycota_unclassified(100); | - | - | - | - | - | - | - | Unassigned | - |
| Otu55652 | 1 | k__Fungi(100);k__Fungi_unclassified(100);k__Fungi_unclassified(100);k__Fungi_unclassified(100);k__Fungi_unclassified(100);k__Fungi_unclassified(100);k__Fungi_unclassified(100); | - | - | - | - | - | - | - | Unassigned | - |
| Otu55655 | 1 | k__Fungi(100);k__Fungi_unclassified(100);k__Fungi_unclassified(100);k__Fungi_unclassified(100);k__Fungi_unclassified(100);k__Fungi_unclassified(100);k__Fungi_unclassified(100); | - | - | - | - | - | - | - | Unassigned | - |
| Otu55657 | 1 | k__Fungi(100);p__Ascomycota(100);p__Ascomycota_unclassified(100);p__Ascomycota_unclassified(100);p__Ascomycota_unclassified(100);p__Ascomycota_unclassified(100);p__Ascomycota_unclassified(100); | - | - | - | - | - | - | - | Unassigned | - |
| Otu55662 | 1 | k__Fungi(100);p__Ascomycota(100);p__Ascomycota_unclassified(100);p__Ascomycota_unclassified(100);p__Ascomycota_unclassified(100);p__Ascomycota_unclassified(100);p__Ascomycota_unclassified(100); | - | - | - | - | - | - | - | Unassigned | - |
| Otu55663 | 1 | k__Fungi(100);p__Ascomycota(100);p__Ascomycota_unclassified(100);p__Ascomycota_unclassified(100);p__Ascomycota_unclassified(100);p__Ascomycota_unclassified(100);p__Ascomycota_unclassified(100); | - | - | - | - | - | - | - | Unassigned | - |

| | | | | | | | | | | | |
|---|---|---|---|---|---|---|---|---|---|---|---|
| Otu55665 | 1 | k__Fungi(100);p__Ascomycota(100);p__Ascomycota_unclassified(100);p__Ascomycota_unclassified(100);p__Ascomycota_unclassified(100);p__Ascomycota_unclassified(100);p__Ascomycota_unclassified(100); | - | - | - | - | - | - | - | Unassigned | - |
| Otu55666 | 1 | k__Fungi(100);k__Fungi_unclassified(100);k__Fungi_unclassified(100);k__Fungi_unclassified(100);k__Fungi_unclassified(100);k__Fungi_unclassified(100);k__Fungi_unclassified(100); | - | - | - | - | - | - | - | Unassigned | - |
| Otu55667 | 1 | k__Fungi(100);p__Ascomycota(100);p__Ascomycota_unclassified(100);p__Ascomycota_unclassified(100);p__Ascomycota_unclassified(100);p__Ascomycota_unclassified(100);p__Ascomycota_unclassified(100); | - | - | - | - | - | - | - | Unassigned | - |
| Otu55670 | 1 | k__Fungi(100);p__Ascomycota(100);p__Ascomycota_unclassified(100);p__Ascomycota_unclassified(100);p__Ascomycota_unclassified(100);p__Ascomycota_unclassified(100);p__Ascomycota_unclassified(100); | - | - | - | - | - | - | - | Unassigned | - |
| Otu55671 | 1 | k__Fungi(100);k__Fungi_unclassified(100);k__Fungi_unclassified(100);k__Fungi_unclassified(100);k__Fungi_unclassified(100);k__Fungi_unclassified(100);k__Fungi_unclassified(100); | - | - | - | - | - | - | - | Unassigned | - |
| Otu55673 | 1 | k__Fungi(100);k__Fungi_unclassified(100);k__Fungi_unclassified(100);k__Fungi_unclassified(100);k__Fungi_unclassified(100);k__Fungi_unclassified(100);k__Fungi_unclassified(100); | - | - | - | - | - | - | - | Unassigned | - |
| Otu55676 | 1 | k__Fungi(100);p__Ascomycota(100);p__Ascomycota_unclassified(100);p__Ascomycota_unclassified(100);p__Ascomycota_unclassified(100);p__Ascomycota_unclassified(100);p__Ascomycota_unclassified(100); | - | - | - | - | - | - | - | Unassigned | - |
| Otu55677 | 1 | k__Fungi(100);p__Ascomycota(100);p__Ascomycota_unclassified(100);p__Ascomycota_unclassified(100);p__Ascomycota_unclassified(100);p__Ascomycota_unclassified(100);p__Ascomycota_unclassified(100); | - | - | - | - | - | - | - | Unassigned | - |
| Otu55682 | 1 | k__Fungi(100);p__Ascomycota(100);p__Ascomycota_unclassified(100);p__Ascomycota_unclassified(100);p__Ascomycota_unclassified(100);p__Ascomycota_unclassified(100);p__Ascomycota_unclassified(100); | - | - | - | - | - | - | - | Unassigned | - |
| Otu55691 | 1 | k__Fungi(100);p__Basidiomycota(100);p__Basidiomycota_unclassified(100);p__Basidiomycota_unclassified(100);p__Basidiomycota_unclassified(100);p__Basidiomycota_unclassified(100);p__Basidiomycota_unclassified(100); | - | - | - | - | - | - | - | Unassigned | - |
| Otu55700 | 1 | k__Fungi(100);k__Fungi_unclassified(100);k__Fungi_unclassified(100);k__Fungi_unclassified(100);k__Fungi_unclassified(100);k__Fungi_unclassified(100);k__Fungi_unclassified(100); | - | - | - | - | - | - | - | Unassigned | - |
| Otu55701 | 1 | k__Fungi(100);p__Ascomycota(100);p__Ascomycota_unclassified(100);p__Ascomycota_unclassified(100);p__Ascomycota_unclassified(100);p__Ascomycota_unclassified(100);p__Ascomycota_unclassified(100); | - | - | - | - | - | - | - | Unassigned | - |
| Otu55702 | 1 | k__Fungi(100);p__Ascomycota(100);p__Ascomycota_unclassified(100);p__Ascomycota_unclassified(100);p__Ascomycota_unclassified(100);p__Ascomycota_unclassified(100);p__Ascomycota_unclassified(100); | - | - | - | - | - | - | - | Unassigned | - |
| Otu55704 | 1 | k__Fungi(100);k__Fungi_unclassified(100);k__Fungi_unclassified(100);k__Fungi_unclassified(100);k__Fungi_unclassified(100);k__Fungi_unclassified(100);k__Fungi_unclassified(100); | - | - | - | - | - | - | - | Unassigned | - |
| Otu55705 | 1 | k__Fungi(100);p__Ascomycota(100);p__Ascomycota_unclassified(100);p__Ascomycota_unclassified(100);p__Ascomycota_unclassified(100);p__Ascomycota_unclassified(100);p__Ascomycota_unclassified(100); | - | - | - | - | - | - | - | Unassigned | - |
| Otu55706 | 1 | k__Fungi(100);p__Ascomycota(100);p__Ascomycota_unclassified(100);p__Ascomycota_unclassified(100);p__Ascomycota_unclassified(100);p__Ascomycota_unclassified(100);p__Ascomycota_unclassified(100); | - | - | - | - | - | - | - | Unassigned | - |
| Otu55708 | 1 | k__Fungi(100);p__Ascomycota(100);p__Ascomycota_unclassified(100);p__Ascomycota_unclassified(100);p__Ascomycota_unclassified(100);p__Ascomycota_unclassified(100);p__Ascomycota_unclassified(100); | - | - | - | - | - | - | - | Unassigned | - |
| Otu55710 | 1 | k__Fungi(100);k__Fungi_unclassified(100);k__Fungi_unclassified(100);k__Fungi_unclassified(100);k__Fungi_unclassified(100);k__Fungi_unclassified(100);k__Fungi_unclassified(100); | - | - | - | - | - | - | - | Unassigned | - |
| Otu55712 | 1 | k__Fungi(100);p__Ascomycota(100);p__Ascomycota_unclassified(100);p__Ascomycota_unclassified(100);p__Ascomycota_unclassified(100);p__Ascomycota_unclassified(100);p__Ascomycota_unclassified(100); | - | - | - | - | - | - | - | Unassigned | - |
| Otu55713 | 1 | k__Fungi(100);k__Fungi_unclassified(100);k__Fungi_unclassified(100);k__Fungi_unclassified(100);k__Fungi_unclassified(100);k__Fungi_unclassified(100);k__Fungi_unclassified(100); | - | - | - | - | - | - | - | Unassigned | - |
| Otu55714 | 1 | k__Fungi(100);k__Fungi_unclassified(100);k__Fungi_unclassified(100);k__Fungi_unclassified(100);k__Fungi_unclassified(100);k__Fungi_unclassified(100);k__Fungi_unclassified(100); | - | - | - | - | - | - | - | Unassigned | - |
| Otu55715 | 1 | k__Fungi(100);k__Fungi_unclassified(100);k__Fungi_unclassified(100);k__Fungi_unclassified(100);k__Fungi_unclassified(100);k__Fungi_unclassified(100);k__Fungi_unclassified(100); | - | - | - | - | - | - | - | Unassigned | - |
| Otu55716 | 1 | k__Fungi(100);k__Fungi_unclassified(100);k__Fungi_unclassified(100);k__Fungi_unclassified(100);k__Fungi_unclassified(100);k__Fungi_unclassified(100);k__Fungi_unclassified(100); | - | - | - | - | - | - | - | Unassigned | - |
| Otu55724 | 1 | k__Fungi(100);k__Fungi_unclassified(100);k__Fungi_unclassified(100);k__Fungi_unclassified(100);k__Fungi_unclassified(100);k__Fungi_unclassified(100);k__Fungi_unclassified(100); | - | - | - | - | - | - | - | Unassigned | - |
| Otu55725 | 1 | k__Fungi(100);p__Ascomycota(100);p__Ascomycota_unclassified(100);p__Ascomycota_unclassified(100);p__Ascomycota_unclassified(100);p__Ascomycota_unclassified(100);p__Ascomycota_unclassified(100); | - | - | - | - | - | - | - | Unassigned | - |
| Otu55743 | 1 | k__Fungi(100);k__Fungi_unclassified(100);k__Fungi_unclassified(100);k__Fungi_unclassified(100);k__Fungi_unclassified(100);k__Fungi_unclassified(100);k__Fungi_unclassified(100); | - | - | - | - | - | - | - | Unassigned | - |
| Otu55744 | 1 | k__Fungi(100);p__Ascomycota(100);p__Ascomycota_unclassified(100);p__Ascomycota_unclassified(100);p__Ascomycota_unclassified(100);p__Ascomycota_unclassified(100);p__Ascomycota_unclassified(100); | - | - | - | - | - | - | - | Unassigned | - |
| Otu55757 | 1 | k__Fungi(100);k__Fungi_unclassified(100);k__Fungi_unclassified(100);k__Fungi_unclassified(100);k__Fungi_unclassified(100);k__Fungi_unclassified(100);k__Fungi_unclassified(100); | - | - | - | - | - | - | - | Unassigned | - |
| Otu55758 | 1 | k__Fungi(100);k__Fungi_unclassified(100);k__Fungi_unclassified(100);k__Fungi_unclassified(100);k__Fungi_unclassified(100);k__Fungi_unclassified(100);k__Fungi_unclassified(100); | - | - | - | - | - | - | - | Unassigned | - |
| Otu55767 | 1 | k__Fungi(100);p__Ascomycota(100);p__Ascomycota_unclassified(100);p__Ascomycota_unclassified(100);p__Ascomycota_unclassified(100);p__Ascomycota_unclassified(100);p__Ascomycota_unclassified(100); | - | - | - | - | - | - | - | Unassigned | - |
| Otu55778 | 1 | k__Fungi(100);k__Fungi_unclassified(100);k__Fungi_unclassified(100);k__Fungi_unclassified(100);k__Fungi_unclassified(100);k__Fungi_unclassified(100);k__Fungi_unclassified(100); | - | - | - | - | - | - | - | Unassigned | - |
| Otu55782 | 1 | k__Fungi(100);p__Ascomycota(100);p__Ascomycota_unclassified(100);p__Ascomycota_unclassified(100);p__Ascomycota_unclassified(100);p__Ascomycota_unclassified(100);p__Ascomycota_unclassified(100); | - | - | - | - | - | - | - | Unassigned | - |
| Otu55783 | 1 | k__Fungi(100);k__Fungi_unclassified(100);k__Fungi_unclassified(100);k__Fungi_unclassified(100);k__Fungi_unclassified(100);k__Fungi_unclassified(100);k__Fungi_unclassified(100); | - | - | - | - | - | - | - | Unassigned | - |
| Otu55784 | 1 | k__Fungi(100);k__Fungi_unclassified(100);k__Fungi_unclassified(100);k__Fungi_unclassified(100);k__Fungi_unclassified(100);k__Fungi_unclassified(100);k__Fungi_unclassified(100); | - | - | - | - | - | - | - | Unassigned | - |
| Otu55785 | 1 | k__Fungi(100);p__Ascomycota(100);p__Ascomycota_unclassified(100);p__Ascomycota_unclassified(100);p__Ascomycota_unclassified(100);p__Ascomycota_unclassified(100);p__Ascomycota_unclassified(100); | - | - | - | - | - | - | - | Unassigned | - |
| Otu55786 | 1 | k__Fungi(100);k__Fungi_unclassified(100);k__Fungi_unclassified(100);k__Fungi_unclassified(100);k__Fungi_unclassified(100);k__Fungi_unclassified(100);k__Fungi_unclassified(100); | - | - | - | - | - | - | - | Unassigned | - |
| Otu55787 | 1 | k__Fungi(100);p__Ascomycota(100);p__Ascomycota_unclassified(100);p__Ascomycota_unclassified(100);p__Ascomycota_unclassified(100);p__Ascomycota_unclassified(100);p__Ascomycota_unclassified(100); | - | - | - | - | - | - | - | Unassigned | - |
| Otu55788 | 1 | k__Fungi(100);k__Fungi_unclassified(100);k__Fungi_unclassified(100);k__Fungi_unclassified(100);k__Fungi_unclassified(100);k__Fungi_unclassified(100);k__Fungi_unclassified(100); | - | - | - | - | - | - | - | Unassigned | - |
| Otu55789 | 1 | k__Fungi(100);p__Ascomycota(100);p__Ascomycota_unclassified(100);p__Ascomycota_unclassified(100);p__Ascomycota_unclassified(100);p__Ascomycota_unclassified(100);p__Ascomycota_unclassified(100); | - | - | - | - | - | - | - | Unassigned | - |
| Otu55790 | 1 | k__Fungi(100);k__Fungi_unclassified(100);k__Fungi_unclassified(100);k__Fungi_unclassified(100);k__Fungi_unclassified(100);k__Fungi_unclassified(100);k__Fungi_unclassified(100); | - | - | - | - | - | - | - | Unassigned | - |
| Otu55792 | 1 | k__Fungi(100);p__Ascomycota(100);p__Ascomycota_unclassified(100);p__Ascomycota_unclassified(100);p__Ascomycota_unclassified(100);p__Ascomycota_unclassified(100);p__Ascomycota_unclassified(100); | - | - | - | - | - | - | - | Unassigned | - |
| Otu55793 | 1 | k__Fungi(100);k__Fungi_unclassified(100);k__Fungi_unclassified(100);k__Fungi_unclassified(100);k__Fungi_unclassified(100);k__Fungi_unclassified(100);k__Fungi_unclassified(100); | - | - | - | - | - | - | - | Unassigned | - |
| Otu55801 | 1 | k__Fungi(100);k__Fungi_unclassified(100);k__Fungi_unclassified(100);k__Fungi_unclassified(100);k__Fungi_unclassified(100);k__Fungi_unclassified(100);k__Fungi_unclassified(100); | - | - | - | - | - | - | - | Unassigned | - |
| Otu55806 | 1 | k__Fungi(100);p__Ascomycota(100);p__Ascomycota_unclassified(100);p__Ascomycota_unclassified(100);p__Ascomycota_unclassified(100);p__Ascomycota_unclassified(100);p__Ascomycota_unclassified(100); | - | - | - | - | - | - | - | Unassigned | - |
| Otu55819 | 1 | k__Fungi(100);p__Ascomycota(100);p__Ascomycota_unclassified(100);p__Ascomycota_unclassified(100);p__Ascomycota_unclassified(100);p__Ascomycota_unclassified(100);p__Ascomycota_unclassified(100); | - | - | - | - | - | - | - | Unassigned | - |
| Otu55825 | 1 | k__Fungi(100);p__Ascomycota(100);p__Ascomycota_unclassified(100);p__Ascomycota_unclassified(100);p__Ascomycota_unclassified(100);p__Ascomycota_unclassified(100);p__Ascomycota_unclassified(100); | - | - | - | - | - | - | - | Unassigned | - |
| Otu55826 | 1 | k__Fungi(100);p__Ascomycota(100);p__Ascomycota_unclassified(100);p__Ascomycota_unclassified(100);p__Ascomycota_unclassified(100);p__Ascomycota_unclassified(100);p__Ascomycota_unclassified(100); | - | - | - | - | - | - | - | Unassigned | - |
| Otu55829 | 1 | k__Fungi(100);p__Ascomycota(100);p__Ascomycota_unclassified(100);p__Ascomycota_unclassified(100);p__Ascomycota_unclassified(100);p__Ascomycota_unclassified(100);p__Ascomycota_unclassified(100); | - | - | - | - | - | - | - | Unassigned | - |
| Otu55831 | 1 | k__Fungi(100);p__Ascomycota(100);p__Ascomycota_unclassified(100);p__Ascomycota_unclassified(100);p__Ascomycota_unclassified(100);p__Ascomycota_unclassified(100);p__Ascomycota_unclassified(100); | - | - | - | - | - | - | - | Unassigned | - |
| Otu55832 | 1 | k__Fungi(100);p__Ascomycota(100);p__Ascomycota_unclassified(100);p__Ascomycota_unclassified(100);p__Ascomycota_unclassified(100);p__Ascomycota_unclassified(100);p__Ascomycota_unclassified(100); | - | - | - | - | - | - | - | Unassigned | - |
| Otu55846 | 1 | k__Fungi(100);p__Ascomycota(100);p__Ascomycota_unclassified(100);p__Ascomycota_unclassified(100);p__Ascomycota_unclassified(100);p__Ascomycota_unclassified(100);p__Ascomycota_unclassified(100); | - | - | - | - | - | - | - | Unassigned | - |
| Otu55851 | 1 | k__Fungi(100);p__Ascomycota(100);p__Ascomycota_unclassified(100);p__Ascomycota_unclassified(100);p__Ascomycota_unclassified(100);p__Ascomycota_unclassified(100);p__Ascomycota_unclassified(100); | - | - | - | - | - | - | - | Unassigned | - |
| Otu55854 | 1 | k__Fungi(100);k__Fungi_unclassified(100);k__Fungi_unclassified(100);k__Fungi_unclassified(100);k__Fungi_unclassified(100);k__Fungi_unclassified(100);k__Fungi_unclassified(100); | - | - | - | - | - | - | - | Unassigned | - |
| Otu55855 | 1 | k__Fungi(100);p__Ascomycota(100);p__Ascomycota_unclassified(100);p__Ascomycota_unclassified(100);p__Ascomycota_unclassified(100);p__Ascomycota_unclassified(100);p__Ascomycota_unclassified(100); | - | - | - | - | - | - | - | Unassigned | - |
| Otu55856 | 1 | k__Fungi(100);p__Ascomycota(100);p__Ascomycota_unclassified(100);p__Ascomycota_unclassified(100);p__Ascomycota_unclassified(100);p__Ascomycota_unclassified(100);p__Ascomycota_unclassified(100); | - | - | - | - | - | - | - | Unassigned | - |
| Otu55857 | 1 | k__Fungi(100);p__Ascomycota(100);p__Ascomycota_unclassified(100);p__Ascomycota_unclassified(100);p__Ascomycota_unclassified(100);p__Ascomycota_unclassified(100);p__Ascomycota_unclassified(100); | - | - | - | - | - | - | - | Unassigned | - |
| Otu55860 | 1 | k__Fungi(100);p__Ascomycota(100);p__Ascomycota_unclassified(100);p__Ascomycota_unclassified(100);p__Ascomycota_unclassified(100);p__Ascomycota_unclassified(100);p__Ascomycota_unclassified(100); | - | - | - | - | - | - | - | Unassigned | - |
| Otu55861 | 1 | k__Fungi(100);k__Fungi_unclassified(100);k__Fungi_unclassified(100);k__Fungi_unclassified(100);k__Fungi_unclassified(100);k__Fungi_unclassified(100);k__Fungi_unclassified(100); | - | - | - | - | - | - | - | Unassigned | - |
| Otu55862 | 1 | k__Fungi(100);p__Ascomycota(100);p__Ascomycota_unclassified(100);p__Ascomycota_unclassified(100);p__Ascomycota_unclassified(100);p__Ascomycota_unclassified(100);p__Ascomycota_unclassified(100); | - | - | - | - | - | - | - | Unassigned | - |
| Otu55863 | 1 | k__Fungi(100);p__Ascomycota(100);p__Ascomycota_unclassified(100);p__Ascomycota_unclassified(100);p__Ascomycota_unclassified(100);p__Ascomycota_unclassified(100);p__Ascomycota_unclassified(100); | - | - | - | - | - | - | - | Unassigned | - |
| Otu55864 | 1 | k__Fungi(100);p__Ascomycota(100);p__Ascomycota_unclassified(100);p__Ascomycota_unclassified(100);p__Ascomycota_unclassified(100);p__Ascomycota_unclassified(100);p__Ascomycota_unclassified(100); | - | - | - | - | - | - | - | Unassigned | - |
| Otu55865 | 1 | k__Fungi(100);p__Ascomycota(100);p__Ascomycota_unclassified(100);p__Ascomycota_unclassified(100);p__Ascomycota_unclassified(100);p__Ascomycota_unclassified(100);p__Ascomycota_unclassified(100); | - | - | - | - | - | - | - | Unassigned | - |
| Otu55866 | 1 | k__Fungi(100);p__Ascomycota(100);p__Ascomycota_unclassified(100);p__Ascomycota_unclassified(100);p__Ascomycota_unclassified(100);p__Ascomycota_unclassified(100);p__Ascomycota_unclassified(100); | - | - | - | - | - | - | - | Unassigned | - |
| Otu55867 | 1 | k__Fungi(100);p__Ascomycota(100);p__Ascomycota_unclassified(100);p__Ascomycota_unclassified(100);p__Ascomycota_unclassified(100);p__Ascomycota_unclassified(100);p__Ascomycota_unclassified(100); | - | - | - | - | - | - | - | Unassigned | - |
| Otu55872 | 1 | k__Fungi(100);p__Ascomycota(100);p__Ascomycota_unclassified(100);p__Ascomycota_unclassified(100);p__Ascomycota_unclassified(100);p__Ascomycota_unclassified(100);p__Ascomycota_unclassified(100); | - | - | - | - | - | - | - | Unassigned | - |
| Otu55873 | 1 | k__Fungi(100);p__Ascomycota(100);p__Ascomycota_unclassified(100);p__Ascomycota_unclassified(100);p__Ascomycota_unclassified(100);p__Ascomycota_unclassified(100);p__Ascomycota_unclassified(100); | - | - | - | - | - | - | - | Unassigned | - |
| Otu55876 | 1 | k__Fungi(100);k__Fungi_unclassified(100);k__Fungi_unclassified(100);k__Fungi_unclassified(100);k__Fungi_unclassified(100);k__Fungi_unclassified(100);k__Fungi_unclassified(100); | - | - | - | - | - | - | - | Unassigned | - |
| Otu55879 | 1 | k__Fungi(100);k__Fungi_unclassified(100);k__Fungi_unclassified(100);k__Fungi_unclassified(100);k__Fungi_unclassified(100);k__Fungi_unclassified(100);k__Fungi_unclassified(100); | - | - | - | - | - | - | - | Unassigned | - |
| Otu55880 | 1 | k__Fungi(100);p__Ascomycota(100);p__Ascomycota_unclassified(100);p__Ascomycota_unclassified(100);p__Ascomycota_unclassified(100);p__Ascomycota_unclassified(100);p__Ascomycota_unclassified(100); | - | - | - | - | - | - | - | Unassigned | - |
| Otu55882 | 1 | k__Fungi(100);k__Fungi_unclassified(100);k__Fungi_unclassified(100);k__Fungi_unclassified(100);k__Fungi_unclassified(100);k__Fungi_unclassified(100);k__Fungi_unclassified(100); | - | - | - | - | - | - | - | Unassigned | - |
| Otu55883 | 1 | k__Fungi(100);p__Ascomycota(100);p__Ascomycota_unclassified(100);p__Ascomycota_unclassified(100);p__Ascomycota_unclassified(100);p__Ascomycota_unclassified(100);p__Ascomycota_unclassified(100); | - | - | - | - | - | - | - | Unassigned | - |
| Otu55884 | 1 | k__Fungi(100);p__Ascomycota(100);p__Ascomycota_unclassified(100);p__Ascomycota_unclassified(100);p__Ascomycota_unclassified(100);p__Ascomycota_unclassified(100);p__Ascomycota_unclassified(100); | - | - | - | - | - | - | - | Unassigned | - |
| Otu55885 | 1 | k__Fungi(100);k__Fungi_unclassified(100);k__Fungi_unclassified(100);k__Fungi_unclassified(100);k__Fungi_unclassified(100);k__Fungi_unclassified(100);k__Fungi_unclassified(100); | - | - | - | - | - | - | - | Unassigned | - |
| Otu55892 | 1 | k__Fungi(100);p__Ascomycota(100);p__Ascomycota_unclassified(100);p__Ascomycota_unclassified(100);p__Ascomycota_unclassified(100);p__Ascomycota_unclassified(100);p__Ascomycota_unclassified(100); | - | - | - | - | - | - | - | Unassigned | - |
| Otu55893 | 1 | k__Fungi(100);p__Ascomycota(100);p__Ascomycota_unclassified(100);p__Ascomycota_unclassified(100);p__Ascomycota_unclassified(100);p__Ascomycota_unclassified(100);p__Ascomycota_unclassified(100); | - | - | - | - | - | - | - | Unassigned | - |
| Otu55894 | 1 | k__Fungi(100);k__Fungi_unclassified(100);k__Fungi_unclassified(100);k__Fungi_unclassified(100);k__Fungi_unclassified(100);k__Fungi_unclassified(100);k__Fungi_unclassified(100); | - | - | - | - | - | - | - | Unassigned | - |
| Otu55895 | 1 | k__Fungi(100);p__Ascomycota(100);p__Ascomycota_unclassified(100);p__Ascomycota_unclassified(100);p__Ascomycota_unclassified(100);p__Ascomycota_unclassified(100);p__Ascomycota_unclassified(100); | - | - | - | - | - | - | - | Unassigned | - |
| Otu55896 | 1 | k__Fungi(100);p__Ascomycota(100);p__Ascomycota_unclassified(100);p__Ascomycota_unclassified(100);p__Ascomycota_unclassified(100);p__Ascomycota_unclassified(100);p__Ascomycota_unclassified(100); | - | - | - | - | - | - | - | Unassigned | - |
| Otu55897 | 1 | k__Fungi(100);p__Ascomycota(100);p__Ascomycota_unclassified(100);p__Ascomycota_unclassified(100);p__Ascomycota_unclassified(100);p__Ascomycota_unclassified(100);p__Ascomycota_unclassified(100); | - | - | - | - | - | - | - | Unassigned | - |
| Otu55899 | 1 | k__Fungi(100);p__Ascomycota(100);p__Ascomycota_unclassified(100);p__Ascomycota_unclassified(100);p__Ascomycota_unclassified(100);p__Ascomycota_unclassified(100);p__Ascomycota_unclassified(100); | - | - | - | - | - | - | - | Unassigned | - |
| Otu55900 | 1 | k__Fungi(100);p__Ascomycota(100);p__Ascomycota_unclassified(100);p__Ascomycota_unclassified(100);p__Ascomycota_unclassified(100);p__Ascomycota_unclassified(100);p__Ascomycota_unclassified(100); | - | - | - | - | - | - | - | Unassigned | - |
| Otu55901 | 1 | k__Fungi(100);p__Ascomycota(100);p__Ascomycota_unclassified(100);p__Ascomycota_unclassified(100);p__Ascomycota_unclassified(100);p__Ascomycota_unclassified(100);p__Ascomycota_unclassified(100); | - | - | - | - | - | - | - | Unassigned | - |
| Otu55902 | 1 | k__Fungi(100);p__Ascomycota(100);p__Ascomycota_unclassified(100);p__Ascomycota_unclassified(100);p__Ascomycota_unclassified(100);p__Ascomycota_unclassified(100);p__Ascomycota_unclassified(100); | - | - | - | - | - | - | - | Unassigned | - |
| Otu55903 | 1 | k__Fungi(100);p__Ascomycota(100);p__Ascomycota_unclassified(100);p__Ascomycota_unclassified(100);p__Ascomycota_unclassified(100);p__Ascomycota_unclassified(100);p__Ascomycota_unclassified(100); | - | - | - | - | - | - | - | Unassigned | - |
| Otu55904 | 1 | k__Fungi(100);p__Ascomycota(100);p__Ascomycota_unclassified(100);p__Ascomycota_unclassified(100);p__Ascomycota_unclassified(100);p__Ascomycota_unclassified(100);p__Ascomycota_unclassified(100); | - | - | - | - | - | - | - | Unassigned | - |
| Otu55906 | 1 | k__Fungi(100);p__Ascomycota(100);p__Ascomycota_unclassified(100);p__Ascomycota_unclassified(100);p__Ascomycota_unclassified(100);p__Ascomycota_unclassified(100);p__Ascomycota_unclassified(100); | - | - | - | - | - | - | - | Unassigned | - |
| Otu55907 | 1 | k__Fungi(100);k__Fungi_unclassified(100);k__Fungi_unclassified(100);k__Fungi_unclassified(100);k__Fungi_unclassified(100);k__Fungi_unclassified(100);k__Fungi_unclassified(100); | - | - | - | - | - | - | - | Unassigned | - |
| Otu55908 | 1 | k__Fungi(100);p__Ascomycota(100);p__Ascomycota_unclassified(100);p__Ascomycota_unclassified(100);p__Ascomycota_unclassified(100);p__Ascomycota_unclassified(100);p__Ascomycota_unclassified(100); | - | - | - | - | - | - | - | Unassigned | - |
| Otu55909 | 1 | k__Fungi(100);k__Fungi_unclassified(100);k__Fungi_unclassified(100);k__Fungi_unclassified(100);k__Fungi_unclassified(100);k__Fungi_unclassified(100);k__Fungi_unclassified(100); | - | - | - | - | - | - | - | Unassigned | - |
| Otu55910 | 1 | k__Fungi(100);k__Fungi_unclassified(100);k__Fungi_unclassified(100);k__Fungi_unclassified(100);k__Fungi_unclassified(100);k__Fungi_unclassified(100);k__Fungi_unclassified(100); | - | - | - | - | - | - | - | Unassigned | - |
| Otu55919 | 1 | k__Fungi(100);k__Fungi_unclassified(100);k__Fungi_unclassified(100);k__Fungi_unclassified(100);k__Fungi_unclassified(100);k__Fungi_unclassified(100);k__Fungi_unclassified(100); | - | - | - | - | - | - | - | Unassigned | - |
| Otu55920 | 1 | k__Fungi(100);k__Fungi_unclassified(100);k__Fungi_unclassified(100);k__Fungi_unclassified(100);k__Fungi_unclassified(100);k__Fungi_unclassified(100);k__Fungi_unclassified(100); | - | - | - | - | - | - | - | Unassigned | - |
| Otu55925 | 1 | k__Fungi(100);p__Ascomycota(100);p__Ascomycota_unclassified(100);p__Ascomycota_unclassified(100);p__Ascomycota_unclassified(100);p__Ascomycota_unclassified(100);p__Ascomycota_unclassified(100); | - | - | - | - | - | - | - | Unassigned | - |
| Otu55941 | 1 | k__Fungi(100);k__Fungi_unclassified(100);k__Fungi_unclassified(100);k__Fungi_unclassified(100);k__Fungi_unclassified(100);k__Fungi_unclassified(100);k__Fungi_unclassified(100); | - | - | - | - | - | - | - | Unassigned | - |
| Otu55942 | 1 | k__Fungi(100);p__Ascomycota(100);p__Ascomycota_unclassified(100);p__Ascomycota_unclassified(100);p__Ascomycota_unclassified(100);p__Ascomycota_unclassified(100);p__Ascomycota_unclassified(100); | - | - | - | - | - | - | - | Unassigned | - |
| Otu55943 | 1 | k__Fungi(100);k__Fungi_unclassified(100);k__Fungi_unclassified(100);k__Fungi_unclassified(100);k__Fungi_unclassified(100);k__Fungi_unclassified(100);k__Fungi_unclassified(100); | - | - | - | - | - | - | - | Unassigned | - |
| Otu55944 | 1 | k__Fungi(100);k__Fungi_unclassified(100);k__Fungi_unclassified(100);k__Fungi_unclassified(100);k__Fungi_unclassified(100);k__Fungi_unclassified(100);k__Fungi_unclassified(100); | - | - | - | - | - | - | - | Unassigned | - |
| Otu55945 | 1 | k__Fungi(100);k__Fungi_unclassified(100);k__Fungi_unclassified(100);k__Fungi_unclassified(100);k__Fungi_unclassified(100);k__Fungi_unclassified(100);k__Fungi_unclassified(100); | - | - | - | - | - | - | - | Unassigned | - |
| Otu55946 | 1 | k__Fungi(100);k__Fungi_unclassified(100);k__Fungi_unclassified(100);k__Fungi_unclassified(100);k__Fungi_unclassified(100);k__Fungi_unclassified(100);k__Fungi_unclassified(100); | - | - | - | - | - | - | - | Unassigned | - |
| Otu55949 | 1 | k__Fungi(100);k__Fungi_unclassified(100);k__Fungi_unclassified(100);k__Fungi_unclassified(100);k__Fungi_unclassified(100);k__Fungi_unclassified(100);k__Fungi_unclassified(100); | - | - | - | - | - | - | - | Unassigned | - |
| Otu55953 | 1 | k__Fungi(100);k__Fungi_unclassified(100);k__Fungi_unclassified(100);k__Fungi_unclassified(100);k__Fungi_unclassified(100);k__Fungi_unclassified(100);k__Fungi_unclassified(100); | - | - | - | - | - | - | - | Unassigned | - |
| Otu55955 | 1 | k__Fungi(100);k__Fungi_unclassified(100);k__Fungi_unclassified(100);k__Fungi_unclassified(100);k__Fungi_unclassified(100);k__Fungi_unclassified(100);k__Fungi_unclassified(100); | - | - | - | - | - | - | - | Unassigned | - |
| Otu55959 | 1 | k__Fungi(100);k__Fungi_unclassified(100);k__Fungi_unclassified(100);k__Fungi_unclassified(100);k__Fungi_unclassified(100);k__Fungi_unclassified(100);k__Fungi_unclassified(100); | - | - | - | - | - | - | - | Unassigned | - |
| Otu55960 | 1 | k__Fungi(100);p__Ascomycota(100);p__Ascomycota_unclassified(100);p__Ascomycota_unclassified(100);p__Ascomycota_unclassified(100);p__Ascomycota_unclassified(100);p__Ascomycota_unclassified(100); | - | - | - | - | - | - | - | Unassigned | - |
| Otu55963 | 1 | k__Fungi(100);k__Fungi_unclassified(100);k__Fungi_unclassified(100);k__Fungi_unclassified(100);k__Fungi_unclassified(100);k__Fungi_unclassified(100);k__Fungi_unclassified(100); | - | - | - | - | - | - | - | Unassigned | - |
| Otu55965 | 1 | k__Fungi(100);k__Fungi_unclassified(100);k__Fungi_unclassified(100);k__Fungi_unclassified(100);k__Fungi_unclassified(100);k__Fungi_unclassified(100);k__Fungi_unclassified(100); | - | - | - | - | - | - | - | Unassigned | - |
| Otu55976 | 1 | k__Fungi(100);k__Fungi_unclassified(100);k__Fungi_unclassified(100);k__Fungi_unclassified(100);k__Fungi_unclassified(100);k__Fungi_unclassified(100);k__Fungi_unclassified(100); | - | - | - | - | - | - | - | Unassigned | - |
| Otu55980 | 1 | k__Fungi(100);p__Ascomycota(100);p__Ascomycota_unclassified(100);p__Ascomycota_unclassified(100);p__Ascomycota_unclassified(100);p__Ascomycota_unclassified(100);p__Ascomycota_unclassified(100); | - | - | - | - | - | - | - | Unassigned | - |
| Otu55984 | 1 | k__Fungi(100);p__Ascomycota(100);p__Ascomycota_unclassified(100);p__Ascomycota_unclassified(100);p__Ascomycota_unclassified(100);p__Ascomycota_unclassified(100);p__Ascomycota_unclassified(100); | - | - | - | - | - | - | - | Unassigned | - |
| Otu55989 | 1 | k__Fungi(100);p__Ascomycota(100);p__Ascomycota_unclassified(100);p__Ascomycota_unclassified(100);p__Ascomycota_unclassified(100);p__Ascomycota_unclassified(100);p__Ascomycota_unclassified(100); | - | - | - | - | - | - | - | Unassigned | - |
| Otu56002 | 1 | k__Fungi(100);p__Ascomycota(100);p__Ascomycota_unclassified(100);p__Ascomycota_unclassified(100);p__Ascomycota_unclassified(100);p__Ascomycota_unclassified(100);p__Ascomycota_unclassified(100); | - | - | - | - | - | - | - | Unassigned | - |
| Otu56003 | 1 | k__Fungi(100);k__Fungi_unclassified(100);k__Fungi_unclassified(100);k__Fungi_unclassified(100);k__Fungi_unclassified(100);k__Fungi_unclassified(100);k__Fungi_unclassified(100); | - | - | - | - | - | - | - | Unassigned | - |
| Otu56006 | 1 | k__Fungi(100);k__Fungi_unclassified(100);k__Fungi_unclassified(100);k__Fungi_unclassified(100);k__Fungi_unclassified(100);k__Fungi_unclassified(100);k__Fungi_unclassified(100); | - | - | - | - | - | - | - | Unassigned | - |
| Otu56007 | 1 | k__Fungi(100);k__Fungi_unclassified(100);k__Fungi_unclassified(100);k__Fungi_unclassified(100);k__Fungi_unclassified(100);k__Fungi_unclassified(100);k__Fungi_unclassified(100); | - | - | - | - | - | - | - | Unassigned | - |
| Otu56009 | 1 | k__Fungi(100);p__Ascomycota(100);p__Ascomycota_unclassified(100);p__Ascomycota_unclassified(100);p__Ascomycota_unclassified(100);p__Ascomycota_unclassified(100);p__Ascomycota_unclassified(100); | - | - | - | - | - | - | - | Unassigned | - |
| Otu56013 | 1 | k__Fungi(100);k__Fungi_unclassified(100);k__Fungi_unclassified(100);k__Fungi_unclassified(100);k__Fungi_unclassified(100);k__Fungi_unclassified(100);k__Fungi_unclassified(100); | - | - | - | - | - | - | - | Unassigned | - |
| Otu56015 | 1 | k__Fungi(100);k__Fungi_unclassified(100);k__Fungi_unclassified(100);k__Fungi_unclassified(100);k__Fungi_unclassified(100);k__Fungi_unclassified(100);k__Fungi_unclassified(100); | - | - | - | - | - | - | - | Unassigned | - |
| Otu56019 | 1 | k__Fungi(100);k__Fungi_unclassified(100);k__Fungi_unclassified(100);k__Fungi_unclassified(100);k__Fungi_unclassified(100);k__Fungi_unclassified(100);k__Fungi_unclassified(100); | - | - | - | - | - | - | - | Unassigned | - |
| Otu56020 | 1 | k__Fungi(100);k__Fungi_unclassified(100);k__Fungi_unclassified(100);k__Fungi_unclassified(100);k__Fungi_unclassified(100);k__Fungi_unclassified(100);k__Fungi_unclassified(100); | - | - | - | - | - | - | - | Unassigned | - |
| Otu56022 | 1 | k__Fungi(100);p__Ascomycota(100);p__Ascomycota_unclassified(100);p__Ascomycota_unclassified(100);p__Ascomycota_unclassified(100);p__Ascomycota_unclassified(100);p__Ascomycota_unclassified(100); | - | - | - | - | - | - | - | Unassigned | - |
| Otu56032 | 1 | k__Fungi(100);k__Fungi_unclassified(100);k__Fungi_unclassified(100);k__Fungi_unclassified(100);k__Fungi_unclassified(100);k__Fungi_unclassified(100);k__Fungi_unclassified(100); | - | - | - | - | - | - | - | Unassigned | - |
| Otu56037 | 1 | k__Fungi(100);p__Ascomycota(100);p__Ascomycota_unclassified(100);p__Ascomycota_unclassified(100);p__Ascomycota_unclassified(100);p__Ascomycota_unclassified(100);p__Ascomycota_unclassified(100); | - | - | - | - | - | - | - | Unassigned | - |
| Otu56038 | 1 | k__Fungi(100);p__Ascomycota(100);p__Ascomycota_unclassified(100);p__Ascomycota_unclassified(100);p__Ascomycota_unclassified(100);p__Ascomycota_unclassified(100);p__Ascomycota_unclassified(100); | - | - | - | - | - | - | - | Unassigned | - |
| Otu56041 | 1 | k__Fungi(100);k__Fungi_unclassified(100);k__Fungi_unclassified(100);k__Fungi_unclassified(100);k__Fungi_unclassified(100);k__Fungi_unclassified(100);k__Fungi_unclassified(100); | - | - | - | - | - | - | - | Unassigned | - |
| Otu56044 | 1 | k__Fungi(100);p__Ascomycota(100);p__Ascomycota_unclassified(100);p__Ascomycota_unclassified(100);p__Ascomycota_unclassified(100);p__Ascomycota_unclassified(100);p__Ascomycota_unclassified(100); | - | - | - | - | - | - | - | Unassigned | - |
| Otu56045 | 1 | k__Fungi(100);k__Fungi_unclassified(100);k__Fungi_unclassified(100);k__Fungi_unclassified(100);k__Fungi_unclassified(100);k__Fungi_unclassified(100);k__Fungi_unclassified(100); | - | - | - | - | - | - | - | Unassigned | - |
| Otu56049 | 1 | k__Fungi(100);p__Ascomycota(100);p__Ascomycota_unclassified(100);p__Ascomycota_unclassified(100);p__Ascomycota_unclassified(100);p__Ascomycota_unclassified(100);p__Ascomycota_unclassified(100); | - | - | - | - | - | - | - | Unassigned | - |
| Otu56050 | 1 | k__Fungi(100);k__Fungi_unclassified(100);k__Fungi_unclassified(100);k__Fungi_unclassified(100);k__Fungi_unclassified(100);k__Fungi_unclassified(100);k__Fungi_unclassified(100); | - | - | - | - | - | - | - | Unassigned | - |
| Otu56053 | 1 | k__Fungi(100);p__Ascomycota(100);p__Ascomycota_unclassified(100);p__Ascomycota_unclassified(100);p__Ascomycota_unclassified(100);p__Ascomycota_unclassified(100);p__Ascomycota_unclassified(100); | - | - | - | - | - | - | - | Unassigned | - |
| Otu56055 | 1 | k__Fungi(100);p__Ascomycota(100);p__Ascomycota_unclassified(100);p__Ascomycota_unclassified(100);p__Ascomycota_unclassified(100);p__Ascomycota_unclassified(100);p__Ascomycota_unclassified(100); | - | - | - | - | - | - | - | Unassigned | - |
| Otu56058 | 1 | k__Fungi(100);k__Fungi_unclassified(100);k__Fungi_unclassified(100);k__Fungi_unclassified(100);k__Fungi_unclassified(100);k__Fungi_unclassified(100);k__Fungi_unclassified(100); | - | - | - | - | - | - | - | Unassigned | - |
| Otu56059 | 1 | k__Fungi(100);k__Fungi_unclassified(100);k__Fungi_unclassified(100);k__Fungi_unclassified(100);k__Fungi_unclassified(100);k__Fungi_unclassified(100);k__Fungi_unclassified(100); | - | - | - | - | - | - | - | Unassigned | - |
| Otu56068 | 1 | k__Fungi(100);p__Ascomycota(100);p__Ascomycota_unclassified(100);p__Ascomycota_unclassified(100);p__Ascomycota_unclassified(100);p__Ascomycota_unclassified(100);p__Ascomycota_unclassified(100); | - | - | - | - | - | - | - | Unassigned | - |
| Otu56070 | 1 | k__Fungi(100);k__Fungi_unclassified(100);k__Fungi_unclassified(100);k__Fungi_unclassified(100);k__Fungi_unclassified(100);k__Fungi_unclassified(100);k__Fungi_unclassified(100); | - | - | - | - | - | - | - | Unassigned | - |
| Otu56073 | 1 | k__Fungi(100);p__Ascomycota(100);p__Ascomycota_unclassified(100);p__Ascomycota_unclassified(100);p__Ascomycota_unclassified(100);p__Ascomycota_unclassified(100);p__Ascomycota_unclassified(100); | - | - | - | - | - | - | - | Unassigned | - |
| Otu56074 | 1 | k__Fungi(100);k__Fungi_unclassified(100);k__Fungi_unclassified(100);k__Fungi_unclassified(100);k__Fungi_unclassified(100);k__Fungi_unclassified(100);k__Fungi_unclassified(100); | - | - | - | - | - | - | - | Unassigned | - |
| Otu56080 | 1 | k__Fungi(100);p__Ascomycota(100);p__Ascomycota_unclassified(100);p__Ascomycota_unclassified(100);p__Ascomycota_unclassified(100);p__Ascomycota_unclassified(100);p__Ascomycota_unclassified(100); | - | - | - | - | - | - | - | Unassigned | - |
| Otu56086 | 1 | k__Fungi(100);k__Fungi_unclassified(100);k__Fungi_unclassified(100);k__Fungi_unclassified(100);k__Fungi_unclassified(100);k__Fungi_unclassified(100);k__Fungi_unclassified(100); | - | - | - | - | - | - | - | Unassigned | - |
| Otu56094 | 1 | k__Fungi(100);p__Ascomycota(100);p__Ascomycota_unclassified(100);p__Ascomycota_unclassified(100);p__Ascomycota_unclassified(100);p__Ascomycota_unclassified(100);p__Ascomycota_unclassified(100); | - | - | - | - | - | - | - | Unassigned | - |
| Otu56097 | 1 | k__Fungi(100);k__Fungi_unclassified(100);k__Fungi_unclassified(100);k__Fungi_unclassified(100);k__Fungi_unclassified(100);k__Fungi_unclassified(100);k__Fungi_unclassified(100); | - | - | - | - | - | - | - | Unassigned | - |
| Otu56109 | 1 | k__Fungi(100);p__Ascomycota(100);p__Ascomycota_unclassified(100);p__Ascomycota_unclassified(100);p__Ascomycota_unclassified(100);p__Ascomycota_unclassified(100);p__Ascomycota_unclassified(100); | - | - | - | - | - | - | - | Unassigned | - |
| Otu56110 | 1 | k__Fungi(100);k__Fungi_unclassified(100);k__Fungi_unclassified(100);k__Fungi_unclassified(100);k__Fungi_unclassified(100);k__Fungi_unclassified(100);k__Fungi_unclassified(100); | - | - | - | - | - | - | - | Unassigned | - |
| Otu56113 | 1 | k__Fungi(100);k__Fungi_unclassified(100);k__Fungi_unclassified(100);k__Fungi_unclassified(100);k__Fungi_unclassified(100);k__Fungi_unclassified(100);k__Fungi_unclassified(100); | - | - | - | - | - | - | - | Unassigned | - |
| Otu56114 | 1 | k__Fungi(100);k__Fungi_unclassified(100);k__Fungi_unclassified(100);k__Fungi_unclassified(100);k__Fungi_unclassified(100);k__Fungi_unclassified(100);k__Fungi_unclassified(100); | - | - | - | - | - | - | - | Unassigned | - |
| Otu56115 | 1 | k__Fungi(100);p__Ascomycota(100);p__Ascomycota_unclassified(100);p__Ascomycota_unclassified(100);p__Ascomycota_unclassified(100);p__Ascomycota_unclassified(100);p__Ascomycota_unclassified(100); | - | - | - | - | - | - | - | Unassigned | - |
| Otu56117 | 1 | k__Fungi(100);k__Fungi_unclassified(100);k__Fungi_unclassified(100);k__Fungi_unclassified(100);k__Fungi_unclassified(100);k__Fungi_unclassified(100);k__Fungi_unclassified(100); | - | - | - | - | - | - | - | Unassigned | - |
| Otu56121 | 1 | k__Fungi(100);p__Ascomycota(100);p__Ascomycota_unclassified(100);p__Ascomycota_unclassified(100);p__Ascomycota_unclassified(100);p__Ascomycota_unclassified(100);p__Ascomycota_unclassified(100); | - | - | - | - | - | - | - | Unassigned | - |
| Otu56122 | 1 | k__Fungi(100);p__Ascomycota(100);p__Ascomycota_unclassified(100);p__Ascomycota_unclassified(100);p__Ascomycota_unclassified(100);p__Ascomycota_unclassified(100);p__Ascomycota_unclassified(100); | - | - | - | - | - | - | - | Unassigned | - |
| Otu56123 | 1 | k__Fungi(100);p__Ascomycota(100);p__Ascomycota_unclassified(100);p__Ascomycota_unclassified(100);p__Ascomycota_unclassified(100);p__Ascomycota_unclassified(100);p__Ascomycota_unclassified(100); | - | - | - | - | - | - | - | Unassigned | - |
| Otu56126 | 1 | k__Fungi(100);k__Fungi_unclassified(100);k__Fungi_unclassified(100);k__Fungi_unclassified(100);k__Fungi_unclassified(100);k__Fungi_unclassified(100);k__Fungi_unclassified(100); | - | - | - | - | - | - | - | Unassigned | - |
| Otu56128 | 1 | k__Fungi(100);p__Ascomycota(100);p__Ascomycota_unclassified(100);p__Ascomycota_unclassified(100);p__Ascomycota_unclassified(100);p__Ascomycota_unclassified(100);p__Ascomycota_unclassified(100); | - | - | - | - | - | - | - | Unassigned | - |
| Otu56129 | 1 | k__Fungi(100);p__Ascomycota(100);p__Ascomycota_unclassified(100);p__Ascomycota_unclassified(100);p__Ascomycota_unclassified(100);p__Ascomycota_unclassified(100);p__Ascomycota_unclassified(100); | - | - | - | - | - | - | - | Unassigned | - |
| Otu56131 | 1 | k__Fungi(100);p__Ascomycota(100);p__Ascomycota_unclassified(100);p__Ascomycota_unclassified(100);p__Ascomycota_unclassified(100);p__Ascomycota_unclassified(100);p__Ascomycota_unclassified(100); | - | - | - | - | - | - | - | Unassigned | - |
| Otu56132 | 1 | k__Fungi(100);p__Ascomycota(100);p__Ascomycota_unclassified(100);p__Ascomycota_unclassified(100);p__Ascomycota_unclassified(100);p__Ascomycota_unclassified(100);p__Ascomycota_unclassified(100); | - | - | - | - | - | - | - | Unassigned | - |
| Otu56146 | 1 | k__Fungi(100);k__Fungi_unclassified(100);k__Fungi_unclassified(100);k__Fungi_unclassified(100);k__Fungi_unclassified(100);k__Fungi_unclassified(100);k__Fungi_unclassified(100); | - | - | - | - | - | - | - | Unassigned | - |
| Otu56147 | 1 | k__Fungi(100);p__Ascomycota(100);p__Ascomycota_unclassified(100);p__Ascomycota_unclassified(100);p__Ascomycota_unclassified(100);p__Ascomycota_unclassified(100);p__Ascomycota_unclassified(100); | - | - | - | - | - | - | - | Unassigned | - |
| Otu56150 | 1 | k__Fungi(100);p__Ascomycota(100);p__Ascomycota_unclassified(100);p__Ascomycota_unclassified(100);p__Ascomycota_unclassified(100);p__Ascomycota_unclassified(100);p__Ascomycota_unclassified(100); | - | - | - | - | - | - | - | Unassigned | - |
| Otu56152 | 1 | k__Fungi(100);k__Fungi_unclassified(100);k__Fungi_unclassified(100);k__Fungi_unclassified(100);k__Fungi_unclassified(100);k__Fungi_unclassified(100);k__Fungi_unclassified(100); | - | - | - | - | - | - | - | Unassigned | - |
| Otu56154 | 1 | k__Fungi(100);k__Fungi_unclassified(100);k__Fungi_unclassified(100);k__Fungi_unclassified(100);k__Fungi_unclassified(100);k__Fungi_unclassified(100);k__Fungi_unclassified(100); | - | - | - | - | - | - | - | Unassigned | - |
| Otu56155 | 1 | k__Fungi(100);k__Fungi_unclassified(100);k__Fungi_unclassified(100);k__Fungi_unclassified(100);k__Fungi_unclassified(100);k__Fungi_unclassified(100);k__Fungi_unclassified(100); | - | - | - | - | - | - | - | Unassigned | - |
| Otu56156 | 1 | k__Fungi(100);p__Ascomycota(100);p__Ascomycota_unclassified(100);p__Ascomycota_unclassified(100);p__Ascomycota_unclassified(100);p__Ascomycota_unclassified(100);p__Ascomycota_unclassified(100); | - | - | - | - | - | - | - | Unassigned | - |
| Otu56157 | 1 | k__Fungi(100);k__Fungi_unclassified(100);k__Fungi_unclassified(100);k__Fungi_unclassified(100);k__Fungi_unclassified(100);k__Fungi_unclassified(100);k__Fungi_unclassified(100); | - | - | - | - | - | - | - | Unassigned | - |
| Otu56160 | 1 | k__Fungi(100);p__Ascomycota(100);p__Ascomycota_unclassified(100);p__Ascomycota_unclassified(100);p__Ascomycota_unclassified(100);p__Ascomycota_unclassified(100);p__Ascomycota_unclassified(100); | - | - | - | - | - | - | - | Unassigned | - |
| Otu56166 | 1 | k__Fungi(100);p__Ascomycota(100);p__Ascomycota_unclassified(100);p__Ascomycota_unclassified(100);p__Ascomycota_unclassified(100);p__Ascomycota_unclassified(100);p__Ascomycota_unclassified(100); | - | - | - | - | - | - | - | Unassigned | - |
| Otu56167 | 1 | k__Fungi(100);p__Ascomycota(100);p__Ascomycota_unclassified(100);p__Ascomycota_unclassified(100);p__Ascomycota_unclassified(100);p__Ascomycota_unclassified(100);p__Ascomycota_unclassified(100); | - | - | - | - | - | - | - | Unassigned | - |
| Otu56169 | 1 | k__Fungi(100);p__Ascomycota(100);p__Ascomycota_unclassified(100);p__Ascomycota_unclassified(100);p__Ascomycota_unclassified(100);p__Ascomycota_unclassified(100);p__Ascomycota_unclassified(100); | - | - | - | - | - | - | - | Unassigned | - |
| Otu56171 | 1 | k__Fungi(100);k__Fungi_unclassified(100);k__Fungi_unclassified(100);k__Fungi_unclassified(100);k__Fungi_unclassified(100);k__Fungi_unclassified(100);k__Fungi_unclassified(100); | - | - | - | - | - | - | - | Unassigned | - |
| Otu56172 | 1 | k__Fungi(100);k__Fungi_unclassified(100);k__Fungi_unclassified(100);k__Fungi_unclassified(100);k__Fungi_unclassified(100);k__Fungi_unclassified(100);k__Fungi_unclassified(100); | - | - | - | - | - | - | - | Unassigned | - |
| Otu56173 | 1 | k__Fungi(100);k__Fungi_unclassified(100);k__Fungi_unclassified(100);k__Fungi_unclassified(100);k__Fungi_unclassified(100);k__Fungi_unclassified(100);k__Fungi_unclassified(100); | - | - | - | - | - | - | - | Unassigned | - |
| Otu56174 | 1 | k__Fungi(100);p__Ascomycota(100);p__Ascomycota_unclassified(100);p__Ascomycota_unclassified(100);p__Ascomycota_unclassified(100);p__Ascomycota_unclassified(100);p__Ascomycota_unclassified(100); | - | - | - | - | - | - | - | Unassigned | - |
| Otu56175 | 1 | k__Fungi(100);p__Ascomycota(100);p__Ascomycota_unclassified(100);p__Ascomycota_unclassified(100);p__Ascomycota_unclassified(100);p__Ascomycota_unclassified(100);p__Ascomycota_unclassified(100); | - | - | - | - | - | - | - | Unassigned | - |
| Otu56176 | 1 | k__Fungi(100);p__Ascomycota(100);p__Ascomycota_unclassified(100);p__Ascomycota_unclassified(100);p__Ascomycota_unclassified(100);p__Ascomycota_unclassified(100);p__Ascomycota_unclassified(100); | - | - | - | - | - | - | - | Unassigned | - |
| Otu56178 | 1 | k__Fungi(100);p__Ascomycota(100);p__Ascomycota_unclassified(100);p__Ascomycota_unclassified(100);p__Ascomycota_unclassified(100);p__Ascomycota_unclassified(100);p__Ascomycota_unclassified(100); | - | - | - | - | - | - | - | Unassigned | - |
| Otu56181 | 1 | k__Fungi(100);p__Ascomycota(100);p__Ascomycota_unclassified(100);p__Ascomycota_unclassified(100);p__Ascomycota_unclassified(100);p__Ascomycota_unclassified(100);p__Ascomycota_unclassified(100); | - | - | - | - | - | - | - | Unassigned | - |
| Otu56182 | 1 | k__Fungi(100);k__Fungi_unclassified(100);k__Fungi_unclassified(100);k__Fungi_unclassified(100);k__Fungi_unclassified(100);k__Fungi_unclassified(100);k__Fungi_unclassified(100); | - | - | - | - | - | - | - | Unassigned | - |
| Otu56183 | 1 | k__Fungi(100);p__Ascomycota(100);p__Ascomycota_unclassified(100);p__Ascomycota_unclassified(100);p__Ascomycota_unclassified(100);p__Ascomycota_unclassified(100);p__Ascomycota_unclassified(100); | - | - | - | - | - | - | - | Unassigned | - |
| Otu56184 | 1 | k__Fungi(100);p__Ascomycota(100);p__Ascomycota_unclassified(100);p__Ascomycota_unclassified(100);p__Ascomycota_unclassified(100);p__Ascomycota_unclassified(100);p__Ascomycota_unclassified(100); | - | - | - | - | - | - | - | Unassigned | - |
| Otu56185 | 1 | k__Fungi(100);p__Ascomycota(100);p__Ascomycota_unclassified(100);p__Ascomycota_unclassified(100);p__Ascomycota_unclassified(100);p__Ascomycota_unclassified(100);p__Ascomycota_unclassified(100); | - | - | - | - | - | - | - | Unassigned | - |
| Otu56187 | 1 | k__Fungi(100);k__Fungi_unclassified(100);k__Fungi_unclassified(100);k__Fungi_unclassified(100);k__Fungi_unclassified(100);k__Fungi_unclassified(100);k__Fungi_unclassified(100); | - | - | - | - | - | - | - | Unassigned | - |
| Otu56196 | 1 | k__Fungi(100);p__Ascomycota(100);p__Ascomycota_unclassified(100);p__Ascomycota_unclassified(100);p__Ascomycota_unclassified(100);p__Ascomycota_unclassified(100);p__Ascomycota_unclassified(100); | - | - | - | - | - | - | - | Unassigned | - |
| Otu56197 | 1 | k__Fungi(100);p__Ascomycota(100);p__Ascomycota_unclassified(100);p__Ascomycota_unclassified(100);p__Ascomycota_unclassified(100);p__Ascomycota_unclassified(100);p__Ascomycota_unclassified(100); | - | - | - | - | - | - | - | Unassigned | - |
| Otu56204 | 1 | k__Fungi(100);p__Ascomycota(100);p__Ascomycota_unclassified(100);p__Ascomycota_unclassified(100);p__Ascomycota_unclassified(100);p__Ascomycota_unclassified(100);p__Ascomycota_unclassified(100); | - | - | - | - | - | - | - | Unassigned | - |
| Otu56205 | 1 | k__Fungi(100);p__Ascomycota(100);p__Ascomycota_unclassified(100);p__Ascomycota_unclassified(100);p__Ascomycota_unclassified(100);p__Ascomycota_unclassified(100);p__Ascomycota_unclassified(100); | - | - | - | - | - | - | - | Unassigned | - |
| Otu56206 | 1 | k__Fungi(100);p__Ascomycota(100);p__Ascomycota_unclassified(100);p__Ascomycota_unclassified(100);p__Ascomycota_unclassified(100);p__Ascomycota_unclassified(100);p__Ascomycota_unclassified(100); | - | - | - | - | - | - | - | Unassigned | - |
| Otu56209 | 1 | k__Fungi(100);p__Ascomycota(100);p__Ascomycota_unclassified(100);p__Ascomycota_unclassified(100);p__Ascomycota_unclassified(100);p__Ascomycota_unclassified(100);p__Ascomycota_unclassified(100); | - | - | - | - | - | - | - | Unassigned | - |
| Otu56210 | 1 | k__Fungi(100);k__Fungi_unclassified(100);k__Fungi_unclassified(100);k__Fungi_unclassified(100);k__Fungi_unclassified(100);k__Fungi_unclassified(100);k__Fungi_unclassified(100); | - | - | - | - | - | - | - | Unassigned | - |
| Otu56212 | 1 | k__Fungi(100);k__Fungi_unclassified(100);k__Fungi_unclassified(100);k__Fungi_unclassified(100);k__Fungi_unclassified(100);k__Fungi_unclassified(100);k__Fungi_unclassified(100); | - | - | - | - | - | - | - | Unassigned | - |
| Otu56213 | 1 | k__Fungi(100);p__Ascomycota(100);p__Ascomycota_unclassified(100);p__Ascomycota_unclassified(100);p__Ascomycota_unclassified(100);p__Ascomycota_unclassified(100);p__Ascomycota_unclassified(100); | - | - | - | - | - | - | - | Unassigned | - |
| Otu56215 | 1 | k__Fungi(100);p__Ascomycota(100);p__Ascomycota_unclassified(100);p__Ascomycota_unclassified(100);p__Ascomycota_unclassified(100);p__Ascomycota_unclassified(100);p__Ascomycota_unclassified(100); | - | - | - | - | - | - | - | Unassigned | - |
| Otu56216 | 1 | k__Fungi(100);p__Ascomycota(100);p__Ascomycota_unclassified(100);p__Ascomycota_unclassified(100);p__Ascomycota_unclassified(100);p__Ascomycota_unclassified(100);p__Ascomycota_unclassified(100); | - | - | - | - | - | - | - | Unassigned | - |
| Otu56217 | 1 | k__Fungi(100);k__Fungi_unclassified(100);k__Fungi_unclassified(100);k__Fungi_unclassified(100);k__Fungi_unclassified(100);k__Fungi_unclassified(100);k__Fungi_unclassified(100); | - | - | - | - | - | - | - | Unassigned | - |
| Otu56218 | 1 | k__Fungi(100);p__Ascomycota(100);p__Ascomycota_unclassified(100);p__Ascomycota_unclassified(100);p__Ascomycota_unclassified(100);p__Ascomycota_unclassified(100);p__Ascomycota_unclassified(100); | - | - | - | - | - | - | - | Unassigned | - |
| Otu56220 | 1 | k__Fungi(100);k__Fungi_unclassified(100);k__Fungi_unclassified(100);k__Fungi_unclassified(100);k__Fungi_unclassified(100);k__Fungi_unclassified(100);k__Fungi_unclassified(100); | - | - | - | - | - | - | - | Unassigned | - |
| Otu56221 | 1 | k__Fungi(100);p__Ascomycota(100);p__Ascomycota_unclassified(100);p__Ascomycota_unclassified(100);p__Ascomycota_unclassified(100);p__Ascomycota_unclassified(100);p__Ascomycota_unclassified(100); | - | - | - | - | - | - | - | Unassigned | - |
| Otu56237 | 1 | k__Fungi(100);p__Ascomycota(100);p__Ascomycota_unclassified(100);p__Ascomycota_unclassified(100);p__Ascomycota_unclassified(100);p__Ascomycota_unclassified(100);p__Ascomycota_unclassified(100); | - | - | - | - | - | - | - | Unassigned | - |
| Otu56240 | 1 | k__Fungi(100);k__Fungi_unclassified(100);k__Fungi_unclassified(100);k__Fungi_unclassified(100);k__Fungi_unclassified(100);k__Fungi_unclassified(100);k__Fungi_unclassified(100); | - | - | - | - | - | - | - | Unassigned | - |
| Otu56242 | 1 | k__Fungi(100);k__Fungi_unclassified(100);k__Fungi_unclassified(100);k__Fungi_unclassified(100);k__Fungi_unclassified(100);k__Fungi_unclassified(100);k__Fungi_unclassified(100); | - | - | - | - | - | - | - | Unassigned | - |
| Otu56247 | 1 | k__Fungi(100);p__Ascomycota(100);p__Ascomycota_unclassified(100);p__Ascomycota_unclassified(100);p__Ascomycota_unclassified(100);p__Ascomycota_unclassified(100);p__Ascomycota_unclassified(100); | - | - | - | - | - | - | - | Unassigned | - |
| Otu56248 | 1 | k__Fungi(100);k__Fungi_unclassified(100);k__Fungi_unclassified(100);k__Fungi_unclassified(100);k__Fungi_unclassified(100);k__Fungi_unclassified(100);k__Fungi_unclassified(100); | - | - | - | - | - | - | - | Unassigned | - |
| Otu56250 | 1 | k__Fungi(100);k__Fungi_unclassified(100);k__Fungi_unclassified(100);k__Fungi_unclassified(100);k__Fungi_unclassified(100);k__Fungi_unclassified(100);k__Fungi_unclassified(100); | - | - | - | - | - | - | - | Unassigned | - |
| Otu56259 | 1 | k__Fungi(100);k__Fungi_unclassified(100);k__Fungi_unclassified(100);k__Fungi_unclassified(100);k__Fungi_unclassified(100);k__Fungi_unclassified(100);k__Fungi_unclassified(100); | - | - | - | - | - | - | - | Unassigned | - |
| Otu56262 | 1 | k__Fungi(100);k__Fungi_unclassified(100);k__Fungi_unclassified(100);k__Fungi_unclassified(100);k__Fungi_unclassified(100);k__Fungi_unclassified(100);k__Fungi_unclassified(100); | - | - | - | - | - | - | - | Unassigned | - |
| Otu56266 | 1 | k__Fungi(100);k__Fungi_unclassified(100);k__Fungi_unclassified(100);k__Fungi_unclassified(100);k__Fungi_unclassified(100);k__Fungi_unclassified(100);k__Fungi_unclassified(100); | - | - | - | - | - | - | - | Unassigned | - |
| Otu56276 | 1 | k__Fungi(100);p__Ascomycota(100);p__Ascomycota_unclassified(100);p__Ascomycota_unclassified(100);p__Ascomycota_unclassified(100);p__Ascomycota_unclassified(100);p__Ascomycota_unclassified(100); | - | - | - | - | - | - | - | Unassigned | - |
| Otu56278 | 1 | k__Fungi(100);p__Ascomycota(100);p__Ascomycota_unclassified(100);p__Ascomycota_unclassified(100);p__Ascomycota_unclassified(100);p__Ascomycota_unclassified(100);p__Ascomycota_unclassified(100); | - | - | - | - | - | - | - | Unassigned | - |
| Otu56280 | 1 | k__Fungi(100);k__Fungi_unclassified(100);k__Fungi_unclassified(100);k__Fungi_unclassified(100);k__Fungi_unclassified(100);k__Fungi_unclassified(100);k__Fungi_unclassified(100); | - | - | - | - | - | - | - | Unassigned | - |
| Otu56287 | 1 | k__Fungi(100);k__Fungi_unclassified(100);k__Fungi_unclassified(100);k__Fungi_unclassified(100);k__Fungi_unclassified(100);k__Fungi_unclassified(100);k__Fungi_unclassified(100); | - | - | - | - | - | - | - | Unassigned | - |
| Otu56290 | 1 | k__Fungi(100);p__Ascomycota(100);p__Ascomycota_unclassified(100);p__Ascomycota_unclassified(100);p__Ascomycota_unclassified(100);p__Ascomycota_unclassified(100);p__Ascomycota_unclassified(100); | - | - | - | - | - | - | - | Unassigned | - |
| Otu56292 | 1 | k__Fungi(100);p__Ascomycota(100);p__Ascomycota_unclassified(100);p__Ascomycota_unclassified(100);p__Ascomycota_unclassified(100);p__Ascomycota_unclassified(100);p__Ascomycota_unclassified(100); | - | - | - | - | - | - | - | Unassigned | - |
| Otu56295 | 1 | k__Fungi(100);k__Fungi_unclassified(100);k__Fungi_unclassified(100);k__Fungi_unclassified(100);k__Fungi_unclassified(100);k__Fungi_unclassified(100);k__Fungi_unclassified(100); | - | - | - | - | - | - | - | Unassigned | - |
| Otu56296 | 1 | k__Fungi(100);k__Fungi_unclassified(100);k__Fungi_unclassified(100);k__Fungi_unclassified(100);k__Fungi_unclassified(100);k__Fungi_unclassified(100);k__Fungi_unclassified(100); | - | - | - | - | - | - | - | Unassigned | - |
| Otu56300 | 1 | k__Fungi(100);k__Fungi_unclassified(100);k__Fungi_unclassified(100);k__Fungi_unclassified(100);k__Fungi_unclassified(100);k__Fungi_unclassified(100);k__Fungi_unclassified(100); | - | - | - | - | - | - | - | Unassigned | - |
| Otu56303 | 1 | k__Fungi(100);p__Ascomycota(100);p__Ascomycota_unclassified(100);p__Ascomycota_unclassified(100);p__Ascomycota_unclassified(100);p__Ascomycota_unclassified(100);p__Ascomycota_unclassified(100); | - | - | - | - | - | - | - | Unassigned | - |
| Otu56310 | 1 | k__Fungi(100);k__Fungi_unclassified(100);k__Fungi_unclassified(100);k__Fungi_unclassified(100);k__Fungi_unclassified(100);k__Fungi_unclassified(100);k__Fungi_unclassified(100); | - | - | - | - | - | - | - | Unassigned | - |
| Otu56313 | 1 | k__Fungi(100);k__Fungi_unclassified(100);k__Fungi_unclassified(100);k__Fungi_unclassified(100);k__Fungi_unclassified(100);k__Fungi_unclassified(100);k__Fungi_unclassified(100); | - | - | - | - | - | - | - | Unassigned | - |
| Otu56327 | 1 | k__Fungi(100);k__Fungi_unclassified(100);k__Fungi_unclassified(100);k__Fungi_unclassified(100);k__Fungi_unclassified(100);k__Fungi_unclassified(100);k__Fungi_unclassified(100); | - | - | - | - | - | - | - | Unassigned | - |

| | | | | | | | | | | | |
|---|---|---|---|---|---|---|---|---|---|---|---|
| Otu56344 | 1 | k__Fungi(100);k__Fungi_unclassified(100);k__Fungi_unclassified(100);k__Fungi_unclassified(100);k__Fungi_unclassified(100);k__Fungi_unclassified(100);k__Fungi_unclassified(100); | - | - | - | - | - | - | - | Unassigned | - |
| Otu56345 | 1 | k__Fungi(100);k__Fungi_unclassified(100);k__Fungi_unclassified(100);k__Fungi_unclassified(100);k__Fungi_unclassified(100);k__Fungi_unclassified(100);k__Fungi_unclassified(100); | - | - | - | - | - | - | - | Unassigned | - |
| Otu56347 | 1 | k__Fungi(100);k__Fungi_unclassified(100);k__Fungi_unclassified(100);k__Fungi_unclassified(100);k__Fungi_unclassified(100);k__Fungi_unclassified(100);k__Fungi_unclassified(100); | - | - | - | - | - | - | - | Unassigned | - |
| Otu56348 | 1 | k__Fungi(100);p__Ascomycota(100);p__Ascomycota_unclassified(100);p__Ascomycota_unclassified(100);p__Ascomycota_unclassified(100);p__Ascomycota_unclassified(100);p__Ascomycota_unclassified(100); | - | - | - | - | - | - | - | Unassigned | - |
| Otu56351 | 1 | k__Fungi(100);k__Fungi_unclassified(100);k__Fungi_unclassified(100);k__Fungi_unclassified(100);k__Fungi_unclassified(100);k__Fungi_unclassified(100);k__Fungi_unclassified(100); | - | - | - | - | - | - | - | Unassigned | - |
| Otu56378 | 1 | k__Fungi(100);k__Fungi_unclassified(100);k__Fungi_unclassified(100);k__Fungi_unclassified(100);k__Fungi_unclassified(100);k__Fungi_unclassified(100);k__Fungi_unclassified(100); | - | - | - | - | - | - | - | Unassigned | - |
| Otu56379 | 1 | k__Fungi(100);k__Fungi_unclassified(100);k__Fungi_unclassified(100);k__Fungi_unclassified(100);k__Fungi_unclassified(100);k__Fungi_unclassified(100);k__Fungi_unclassified(100); | - | - | - | - | - | - | - | Unassigned | - |
| Otu56384 | 1 | k__Fungi(100);k__Fungi_unclassified(100);k__Fungi_unclassified(100);k__Fungi_unclassified(100);k__Fungi_unclassified(100);k__Fungi_unclassified(100);k__Fungi_unclassified(100); | - | - | - | - | - | - | - | Unassigned | - |
| Otu56385 | 1 | k__Fungi(100);k__Fungi_unclassified(100);k__Fungi_unclassified(100);k__Fungi_unclassified(100);k__Fungi_unclassified(100);k__Fungi_unclassified(100);k__Fungi_unclassified(100); | - | - | - | - | - | - | - | Unassigned | - |
| Otu56386 | 1 | k__Fungi(100);k__Fungi_unclassified(100);k__Fungi_unclassified(100);k__Fungi_unclassified(100);k__Fungi_unclassified(100);k__Fungi_unclassified(100);k__Fungi_unclassified(100); | - | - | - | - | - | - | - | Unassigned | - |
| Otu56388 | 1 | k__Fungi(100);k__Fungi_unclassified(100);k__Fungi_unclassified(100);k__Fungi_unclassified(100);k__Fungi_unclassified(100);k__Fungi_unclassified(100);k__Fungi_unclassified(100); | - | - | - | - | - | - | - | Unassigned | - |
| Otu56391 | 1 | k__Fungi(100);p__Ascomycota(100);p__Ascomycota_unclassified(100);p__Ascomycota_unclassified(100);p__Ascomycota_unclassified(100);p__Ascomycota_unclassified(100);p__Ascomycota_unclassified(100); | - | - | - | - | - | - | - | Unassigned | - |
| Otu56393 | 1 | k__Fungi(100);k__Fungi_unclassified(100);k__Fungi_unclassified(100);k__Fungi_unclassified(100);k__Fungi_unclassified(100);k__Fungi_unclassified(100);k__Fungi_unclassified(100); | - | - | - | - | - | - | - | Unassigned | - |
| Otu56398 | 1 | k__Fungi(100);p__Ascomycota(100);p__Ascomycota_unclassified(100);p__Ascomycota_unclassified(100);p__Ascomycota_unclassified(100);p__Ascomycota_unclassified(100);p__Ascomycota_unclassified(100); | - | - | - | - | - | - | - | Unassigned | - |
| Otu56399 | 1 | k__Fungi(100);k__Fungi_unclassified(100);k__Fungi_unclassified(100);k__Fungi_unclassified(100);k__Fungi_unclassified(100);k__Fungi_unclassified(100);k__Fungi_unclassified(100); | - | - | - | - | - | - | - | Unassigned | - |
| Otu56404 | 1 | k__Fungi(100);k__Fungi_unclassified(100);k__Fungi_unclassified(100);k__Fungi_unclassified(100);k__Fungi_unclassified(100);k__Fungi_unclassified(100);k__Fungi_unclassified(100); | - | - | - | - | - | - | - | Unassigned | - |
| Otu56405 | 1 | k__Fungi(100);k__Fungi_unclassified(100);k__Fungi_unclassified(100);k__Fungi_unclassified(100);k__Fungi_unclassified(100);k__Fungi_unclassified(100);k__Fungi_unclassified(100); | - | - | - | - | - | - | - | Unassigned | - |
| Otu56412 | 1 | k__Fungi(100);k__Fungi_unclassified(100);k__Fungi_unclassified(100);k__Fungi_unclassified(100);k__Fungi_unclassified(100);k__Fungi_unclassified(100);k__Fungi_unclassified(100); | - | - | - | - | - | - | - | Unassigned | - |
| Otu56413 | 1 | k__Fungi(100);k__Fungi_unclassified(100);k__Fungi_unclassified(100);k__Fungi_unclassified(100);k__Fungi_unclassified(100);k__Fungi_unclassified(100);k__Fungi_unclassified(100); | - | - | - | - | - | - | - | Unassigned | - |
| Otu56414 | 1 | k__Fungi(100);k__Fungi_unclassified(100);k__Fungi_unclassified(100);k__Fungi_unclassified(100);k__Fungi_unclassified(100);k__Fungi_unclassified(100);k__Fungi_unclassified(100); | - | - | - | - | - | - | - | Unassigned | - |
| Otu56415 | 1 | k__Fungi(100);p__Ascomycota(100);p__Ascomycota_unclassified(100);p__Ascomycota_unclassified(100);p__Ascomycota_unclassified(100);p__Ascomycota_unclassified(100);p__Ascomycota_unclassified(100); | - | - | - | - | - | - | - | Unassigned | - |
| Otu56416 | 1 | k__Fungi(100);k__Fungi_unclassified(100);k__Fungi_unclassified(100);k__Fungi_unclassified(100);k__Fungi_unclassified(100);k__Fungi_unclassified(100);k__Fungi_unclassified(100); | - | - | - | - | - | - | - | Unassigned | - |
| Otu56418 | 1 | k__Fungi(100);k__Fungi_unclassified(100);k__Fungi_unclassified(100);k__Fungi_unclassified(100);k__Fungi_unclassified(100);k__Fungi_unclassified(100);k__Fungi_unclassified(100); | - | - | - | - | - | - | - | Unassigned | - |
| Otu56419 | 1 | k__Fungi(100);k__Fungi_unclassified(100);k__Fungi_unclassified(100);k__Fungi_unclassified(100);k__Fungi_unclassified(100);k__Fungi_unclassified(100);k__Fungi_unclassified(100); | - | - | - | - | - | - | - | Unassigned | - |
| Otu56420 | 1 | k__Fungi(100);k__Fungi_unclassified(100);k__Fungi_unclassified(100);k__Fungi_unclassified(100);k__Fungi_unclassified(100);k__Fungi_unclassified(100);k__Fungi_unclassified(100); | - | - | - | - | - | - | - | Unassigned | - |
| Otu56423 | 1 | k__Fungi(100);p__Ascomycota(100);p__Ascomycota_unclassified(100);p__Ascomycota_unclassified(100);p__Ascomycota_unclassified(100);p__Ascomycota_unclassified(100);p__Ascomycota_unclassified(100); | - | - | - | - | - | - | - | Unassigned | - |
| Otu56424 | 1 | k__Fungi(100);k__Fungi_unclassified(100);k__Fungi_unclassified(100);k__Fungi_unclassified(100);k__Fungi_unclassified(100);k__Fungi_unclassified(100);k__Fungi_unclassified(100); | - | - | - | - | - | - | - | Unassigned | - |
| Otu56425 | 1 | k__Fungi(100);k__Fungi_unclassified(100);k__Fungi_unclassified(100);k__Fungi_unclassified(100);k__Fungi_unclassified(100);k__Fungi_unclassified(100);k__Fungi_unclassified(100); | - | - | - | - | - | - | - | Unassigned | - |
| Otu56427 | 1 | k__Fungi(100);k__Fungi_unclassified(100);k__Fungi_unclassified(100);k__Fungi_unclassified(100);k__Fungi_unclassified(100);k__Fungi_unclassified(100);k__Fungi_unclassified(100); | - | - | - | - | - | - | - | Unassigned | - |
| Otu56429 | 1 | k__Fungi(100);p__Ascomycota(100);p__Ascomycota_unclassified(100);p__Ascomycota_unclassified(100);p__Ascomycota_unclassified(100);p__Ascomycota_unclassified(100);p__Ascomycota_unclassified(100); | - | - | - | - | - | - | - | Unassigned | - |
| Otu56435 | 1 | k__Fungi(100);k__Fungi_unclassified(100);k__Fungi_unclassified(100);k__Fungi_unclassified(100);k__Fungi_unclassified(100);k__Fungi_unclassified(100);k__Fungi_unclassified(100); | - | - | - | - | - | - | - | Unassigned | - |
| Otu56436 | 1 | k__Fungi(100);k__Fungi_unclassified(100);k__Fungi_unclassified(100);k__Fungi_unclassified(100);k__Fungi_unclassified(100);k__Fungi_unclassified(100);k__Fungi_unclassified(100); | - | - | - | - | - | - | - | Unassigned | - |
| Otu56437 | 1 | k__Fungi(100);p__Ascomycota(100);p__Ascomycota_unclassified(100);p__Ascomycota_unclassified(100);p__Ascomycota_unclassified(100);p__Ascomycota_unclassified(100);p__Ascomycota_unclassified(100); | - | - | - | - | - | - | - | Unassigned | - |
| Otu56438 | 1 | k__Fungi(100);k__Fungi_unclassified(100);k__Fungi_unclassified(100);k__Fungi_unclassified(100);k__Fungi_unclassified(100);k__Fungi_unclassified(100);k__Fungi_unclassified(100); | - | - | - | - | - | - | - | Unassigned | - |
| Otu56441 | 1 | k__Fungi(100);k__Fungi_unclassified(100);k__Fungi_unclassified(100);k__Fungi_unclassified(100);k__Fungi_unclassified(100);k__Fungi_unclassified(100);k__Fungi_unclassified(100); | - | - | - | - | - | - | - | Unassigned | - |
| Otu56442 | 1 | k__Fungi(100);p__Ascomycota(100);p__Ascomycota_unclassified(100);p__Ascomycota_unclassified(100);p__Ascomycota_unclassified(100);p__Ascomycota_unclassified(100);p__Ascomycota_unclassified(100); | - | - | - | - | - | - | - | Unassigned | - |
| Otu56443 | 1 | k__Fungi(100);k__Fungi_unclassified(100);k__Fungi_unclassified(100);k__Fungi_unclassified(100);k__Fungi_unclassified(100);k__Fungi_unclassified(100);k__Fungi_unclassified(100); | - | - | - | - | - | - | - | Unassigned | - |
| Otu56444 | 1 | k__Fungi(100);k__Fungi_unclassified(100);k__Fungi_unclassified(100);k__Fungi_unclassified(100);k__Fungi_unclassified(100);k__Fungi_unclassified(100);k__Fungi_unclassified(100); | - | - | - | - | - | - | - | Unassigned | - |
| Otu56445 | 1 | k__Fungi(100);p__Ascomycota(100);p__Ascomycota_unclassified(100);p__Ascomycota_unclassified(100);p__Ascomycota_unclassified(100);p__Ascomycota_unclassified(100);p__Ascomycota_unclassified(100); | - | - | - | - | - | - | - | Unassigned | - |
| Otu56446 | 1 | k__Fungi(100);p__Ascomycota(100);p__Ascomycota_unclassified(100);p__Ascomycota_unclassified(100);p__Ascomycota_unclassified(100);p__Ascomycota_unclassified(100);p__Ascomycota_unclassified(100); | - | - | - | - | - | - | - | Unassigned | - |
| Otu56447 | 1 | k__Fungi(100);p__Ascomycota(100);p__Ascomycota_unclassified(100);p__Ascomycota_unclassified(100);p__Ascomycota_unclassified(100);p__Ascomycota_unclassified(100);p__Ascomycota_unclassified(100); | - | - | - | - | - | - | - | Unassigned | - |
| Otu56448 | 1 | k__Fungi(100);k__Fungi_unclassified(100);k__Fungi_unclassified(100);k__Fungi_unclassified(100);k__Fungi_unclassified(100);k__Fungi_unclassified(100);k__Fungi_unclassified(100); | - | - | - | - | - | - | - | Unassigned | - |
| Otu56449 | 1 | k__Fungi(100);k__Fungi_unclassified(100);k__Fungi_unclassified(100);k__Fungi_unclassified(100);k__Fungi_unclassified(100);k__Fungi_unclassified(100);k__Fungi_unclassified(100); | - | - | - | - | - | - | - | Unassigned | - |
| Otu56450 | 1 | k__Fungi(100);p__Ascomycota(100);p__Ascomycota_unclassified(100);p__Ascomycota_unclassified(100);p__Ascomycota_unclassified(100);p__Ascomycota_unclassified(100);p__Ascomycota_unclassified(100); | - | - | - | - | - | - | - | Unassigned | - |
| Otu56451 | 1 | k__Fungi(100);p__Ascomycota(100);p__Ascomycota_unclassified(100);p__Ascomycota_unclassified(100);p__Ascomycota_unclassified(100);p__Ascomycota_unclassified(100);p__Ascomycota_unclassified(100); | - | - | - | - | - | - | - | Unassigned | - |
| Otu56452 | 1 | k__Fungi(100);p__Ascomycota(100);p__Ascomycota_unclassified(100);p__Ascomycota_unclassified(100);p__Ascomycota_unclassified(100);p__Ascomycota_unclassified(100);p__Ascomycota_unclassified(100); | - | - | - | - | - | - | - | Unassigned | - |
| Otu56454 | 1 | k__Fungi(100);k__Fungi_unclassified(100);k__Fungi_unclassified(100);k__Fungi_unclassified(100);k__Fungi_unclassified(100);k__Fungi_unclassified(100);k__Fungi_unclassified(100); | - | - | - | - | - | - | - | Unassigned | - |
| Otu56455 | 1 | k__Fungi(100);k__Fungi_unclassified(100);k__Fungi_unclassified(100);k__Fungi_unclassified(100);k__Fungi_unclassified(100);k__Fungi_unclassified(100);k__Fungi_unclassified(100); | - | - | - | - | - | - | - | Unassigned | - |
| Otu56456 | 1 | k__Fungi(100);p__Ascomycota(100);p__Ascomycota_unclassified(100);p__Ascomycota_unclassified(100);p__Ascomycota_unclassified(100);p__Ascomycota_unclassified(100);p__Ascomycota_unclassified(100); | - | - | - | - | - | - | - | Unassigned | - |
| Otu56457 | 1 | k__Fungi(100);p__Ascomycota(100);p__Ascomycota_unclassified(100);p__Ascomycota_unclassified(100);p__Ascomycota_unclassified(100);p__Ascomycota_unclassified(100);p__Ascomycota_unclassified(100); | - | - | - | - | - | - | - | Unassigned | - |
| Otu56468 | 1 | k__Fungi(100);p__Ascomycota(100);p__Ascomycota_unclassified(100);p__Ascomycota_unclassified(100);p__Ascomycota_unclassified(100);p__Ascomycota_unclassified(100);p__Ascomycota_unclassified(100); | - | - | - | - | - | - | - | Unassigned | - |
| Otu56469 | 1 | k__Fungi(100);p__Ascomycota(100);p__Ascomycota_unclassified(100);p__Ascomycota_unclassified(100);p__Ascomycota_unclassified(100);p__Ascomycota_unclassified(100);p__Ascomycota_unclassified(100); | - | - | - | - | - | - | - | Unassigned | - |
| Otu56472 | 1 | k__Fungi(100);k__Fungi_unclassified(100);k__Fungi_unclassified(100);k__Fungi_unclassified(100);k__Fungi_unclassified(100);k__Fungi_unclassified(100);k__Fungi_unclassified(100); | - | - | - | - | - | - | - | Unassigned | - |
| Otu56478 | 1 | k__Fungi(100);k__Fungi_unclassified(100);k__Fungi_unclassified(100);k__Fungi_unclassified(100);k__Fungi_unclassified(100);k__Fungi_unclassified(100);k__Fungi_unclassified(100); | - | - | - | - | - | - | - | Unassigned | - |
| Otu56479 | 1 | k__Fungi(100);p__Ascomycota(100);p__Ascomycota_unclassified(100);p__Ascomycota_unclassified(100);p__Ascomycota_unclassified(100);p__Ascomycota_unclassified(100);p__Ascomycota_unclassified(100); | - | - | - | - | - | - | - | Unassigned | - |
| Otu56480 | 1 | k__Fungi(100);p__Ascomycota(100);p__Ascomycota_unclassified(100);p__Ascomycota_unclassified(100);p__Ascomycota_unclassified(100);p__Ascomycota_unclassified(100);p__Ascomycota_unclassified(100); | - | - | - | - | - | - | - | Unassigned | - |
| Otu56481 | 1 | k__Fungi(100);k__Fungi_unclassified(100);k__Fungi_unclassified(100);k__Fungi_unclassified(100);k__Fungi_unclassified(100);k__Fungi_unclassified(100);k__Fungi_unclassified(100); | - | - | - | - | - | - | - | Unassigned | - |
| Otu56482 | 1 | k__Fungi(100);p__Ascomycota(100);p__Ascomycota_unclassified(100);p__Ascomycota_unclassified(100);p__Ascomycota_unclassified(100);p__Ascomycota_unclassified(100);p__Ascomycota_unclassified(100); | - | - | - | - | - | - | - | Unassigned | - |
| Otu56483 | 1 | k__Fungi(100);k__Fungi_unclassified(100);k__Fungi_unclassified(100);k__Fungi_unclassified(100);k__Fungi_unclassified(100);k__Fungi_unclassified(100);k__Fungi_unclassified(100); | - | - | - | - | - | - | - | Unassigned | - |
| Otu56485 | 1 | k__Fungi(100);k__Fungi_unclassified(100);k__Fungi_unclassified(100);k__Fungi_unclassified(100);k__Fungi_unclassified(100);k__Fungi_unclassified(100);k__Fungi_unclassified(100); | - | - | - | - | - | - | - | Unassigned | - |
| Otu56487 | 1 | k__Fungi(100);k__Fungi_unclassified(100);k__Fungi_unclassified(100);k__Fungi_unclassified(100);k__Fungi_unclassified(100);k__Fungi_unclassified(100);k__Fungi_unclassified(100); | - | - | - | - | - | - | - | Unassigned | - |
| Otu56488 | 1 | k__Fungi(100);p__Ascomycota(100);p__Ascomycota_unclassified(100);p__Ascomycota_unclassified(100);p__Ascomycota_unclassified(100);p__Ascomycota_unclassified(100);p__Ascomycota_unclassified(100); | - | - | - | - | - | - | - | Unassigned | - |
| Otu56491 | 1 | k__Fungi(100);p__Ascomycota(100);p__Ascomycota_unclassified(100);p__Ascomycota_unclassified(100);p__Ascomycota_unclassified(100);p__Ascomycota_unclassified(100);p__Ascomycota_unclassified(100); | - | - | - | - | - | - | - | Unassigned | - |
| Otu56492 | 1 | k__Fungi(100);k__Fungi_unclassified(100);k__Fungi_unclassified(100);k__Fungi_unclassified(100);k__Fungi_unclassified(100);k__Fungi_unclassified(100);k__Fungi_unclassified(100); | - | - | - | - | - | - | - | Unassigned | - |
| Otu56493 | 1 | k__Fungi(100);k__Fungi_unclassified(100);k__Fungi_unclassified(100);k__Fungi_unclassified(100);k__Fungi_unclassified(100);k__Fungi_unclassified(100);k__Fungi_unclassified(100); | - | - | - | - | - | - | - | Unassigned | - |
| Otu56494 | 1 | k__Fungi(100);p__Ascomycota(100);p__Ascomycota_unclassified(100);p__Ascomycota_unclassified(100);p__Ascomycota_unclassified(100);p__Ascomycota_unclassified(100);p__Ascomycota_unclassified(100); | - | - | - | - | - | - | - | Unassigned | - |
| Otu56502 | 1 | k__Fungi(100);k__Fungi_unclassified(100);k__Fungi_unclassified(100);k__Fungi_unclassified(100);k__Fungi_unclassified(100);k__Fungi_unclassified(100);k__Fungi_unclassified(100); | - | - | - | - | - | - | - | Unassigned | - |
| Otu56503 | 1 | k__Fungi(100);p__Ascomycota(100);p__Ascomycota_unclassified(100);p__Ascomycota_unclassified(100);p__Ascomycota_unclassified(100);p__Ascomycota_unclassified(100);p__Ascomycota_unclassified(100); | - | - | - | - | - | - | - | Unassigned | - |
| Otu56504 | 1 | k__Fungi(100);k__Fungi_unclassified(100);k__Fungi_unclassified(100);k__Fungi_unclassified(100);k__Fungi_unclassified(100);k__Fungi_unclassified(100);k__Fungi_unclassified(100); | - | - | - | - | - | - | - | Unassigned | - |
| Otu56505 | 1 | k__Fungi(100);k__Fungi_unclassified(100);k__Fungi_unclassified(100);k__Fungi_unclassified(100);k__Fungi_unclassified(100);k__Fungi_unclassified(100);k__Fungi_unclassified(100); | - | - | - | - | - | - | - | Unassigned | - |
| Otu56506 | 1 | k__Fungi(100);k__Fungi_unclassified(100);k__Fungi_unclassified(100);k__Fungi_unclassified(100);k__Fungi_unclassified(100);k__Fungi_unclassified(100);k__Fungi_unclassified(100); | - | - | - | - | - | - | - | Unassigned | - |
| Otu56507 | 1 | k__Fungi(100);p__Ascomycota(100);p__Ascomycota_unclassified(100);p__Ascomycota_unclassified(100);p__Ascomycota_unclassified(100);p__Ascomycota_unclassified(100);p__Ascomycota_unclassified(100); | - | - | - | - | - | - | - | Unassigned | - |
| Otu56508 | 1 | k__Fungi(100);k__Fungi_unclassified(100);k__Fungi_unclassified(100);k__Fungi_unclassified(100);k__Fungi_unclassified(100);k__Fungi_unclassified(100);k__Fungi_unclassified(100); | - | - | - | - | - | - | - | Unassigned | - |
| Otu56509 | 1 | k__Fungi(100);k__Fungi_unclassified(100);k__Fungi_unclassified(100);k__Fungi_unclassified(100);k__Fungi_unclassified(100);k__Fungi_unclassified(100);k__Fungi_unclassified(100); | - | - | - | - | - | - | - | Unassigned | - |
| Otu56514 | 1 | k__Fungi(100);k__Fungi_unclassified(100);k__Fungi_unclassified(100);k__Fungi_unclassified(100);k__Fungi_unclassified(100);k__Fungi_unclassified(100);k__Fungi_unclassified(100); | - | - | - | - | - | - | - | Unassigned | - |
| Otu56515 | 1 | k__Fungi(100);k__Fungi_unclassified(100);k__Fungi_unclassified(100);k__Fungi_unclassified(100);k__Fungi_unclassified(100);k__Fungi_unclassified(100);k__Fungi_unclassified(100); | - | - | - | - | - | - | - | Unassigned | - |
| Otu56517 | 1 | k__Fungi(100);p__Ascomycota(100);p__Ascomycota_unclassified(100);p__Ascomycota_unclassified(100);p__Ascomycota_unclassified(100);p__Ascomycota_unclassified(100);p__Ascomycota_unclassified(100); | - | - | - | - | - | - | - | Unassigned | - |
| Otu56518 | 1 | k__Fungi(100);p__Ascomycota(100);p__Ascomycota_unclassified(100);p__Ascomycota_unclassified(100);p__Ascomycota_unclassified(100);p__Ascomycota_unclassified(100);p__Ascomycota_unclassified(100); | - | - | - | - | - | - | - | Unassigned | - |
| Otu56523 | 1 | k__Fungi(100);p__Ascomycota(100);p__Ascomycota_unclassified(100);p__Ascomycota_unclassified(100);p__Ascomycota_unclassified(100);p__Ascomycota_unclassified(100);p__Ascomycota_unclassified(100); | - | - | - | - | - | - | - | Unassigned | - |
| Otu56529 | 1 | k__Fungi(100);k__Fungi_unclassified(100);k__Fungi_unclassified(100);k__Fungi_unclassified(100);k__Fungi_unclassified(100);k__Fungi_unclassified(100);k__Fungi_unclassified(100); | - | - | - | - | - | - | - | Unassigned | - |
| Otu56533 | 1 | k__Fungi(100);p__Ascomycota(100);p__Ascomycota_unclassified(100);p__Ascomycota_unclassified(100);p__Ascomycota_unclassified(100);p__Ascomycota_unclassified(100);p__Ascomycota_unclassified(100); | - | - | - | - | - | - | - | Unassigned | - |
| Otu56534 | 1 | k__Fungi(100);k__Fungi_unclassified(100);k__Fungi_unclassified(100);k__Fungi_unclassified(100);k__Fungi_unclassified(100);k__Fungi_unclassified(100);k__Fungi_unclassified(100); | - | - | - | - | - | - | - | Unassigned | - |
| Otu56535 | 1 | k__Fungi(100);k__Fungi_unclassified(100);k__Fungi_unclassified(100);k__Fungi_unclassified(100);k__Fungi_unclassified(100);k__Fungi_unclassified(100);k__Fungi_unclassified(100); | - | - | - | - | - | - | - | Unassigned | - |
| Otu56538 | 1 | k__Fungi(100);p__Ascomycota(100);p__Ascomycota_unclassified(100);p__Ascomycota_unclassified(100);p__Ascomycota_unclassified(100);p__Ascomycota_unclassified(100);p__Ascomycota_unclassified(100); | - | - | - | - | - | - | - | Unassigned | - |
| Otu56539 | 1 | k__Fungi(100);p__Ascomycota(100);p__Ascomycota_unclassified(100);p__Ascomycota_unclassified(100);p__Ascomycota_unclassified(100);p__Ascomycota_unclassified(100);p__Ascomycota_unclassified(100); | - | - | - | - | - | - | - | Unassigned | - |
| Otu56540 | 1 | k__Fungi(100);p__Ascomycota(100);p__Ascomycota_unclassified(100);p__Ascomycota_unclassified(100);p__Ascomycota_unclassified(100);p__Ascomycota_unclassified(100);p__Ascomycota_unclassified(100); | - | - | - | - | - | - | - | Unassigned | - |
| Otu56541 | 1 | k__Fungi(100);p__Ascomycota(100);p__Ascomycota_unclassified(100);p__Ascomycota_unclassified(100);p__Ascomycota_unclassified(100);p__Ascomycota_unclassified(100);p__Ascomycota_unclassified(100); | - | - | - | - | - | - | - | Unassigned | - |
| Otu56544 | 1 | k__Fungi(100);k__Fungi_unclassified(100);k__Fungi_unclassified(100);k__Fungi_unclassified(100);k__Fungi_unclassified(100);k__Fungi_unclassified(100);k__Fungi_unclassified(100); | - | - | - | - | - | - | - | Unassigned | - |
| Otu56547 | 1 | k__Fungi(100);k__Fungi_unclassified(100);k__Fungi_unclassified(100);k__Fungi_unclassified(100);k__Fungi_unclassified(100);k__Fungi_unclassified(100);k__Fungi_unclassified(100); | - | - | - | - | - | - | - | Unassigned | - |
| Otu56548 | 1 | k__Fungi(100);k__Fungi_unclassified(100);k__Fungi_unclassified(100);k__Fungi_unclassified(100);k__Fungi_unclassified(100);k__Fungi_unclassified(100);k__Fungi_unclassified(100); | - | - | - | - | - | - | - | Unassigned | - |
| Otu56552 | 1 | k__Fungi(100);k__Fungi_unclassified(100);k__Fungi_unclassified(100);k__Fungi_unclassified(100);k__Fungi_unclassified(100);k__Fungi_unclassified(100);k__Fungi_unclassified(100); | - | - | - | - | - | - | - | Unassigned | - |
| Otu56555 | 1 | k__Fungi(100);p__Ascomycota(100);p__Ascomycota_unclassified(100);p__Ascomycota_unclassified(100);p__Ascomycota_unclassified(100);p__Ascomycota_unclassified(100);p__Ascomycota_unclassified(100); | - | - | - | - | - | - | - | Unassigned | - |
| Otu56556 | 1 | k__Fungi(100);k__Fungi_unclassified(100);k__Fungi_unclassified(100);k__Fungi_unclassified(100);k__Fungi_unclassified(100);k__Fungi_unclassified(100);k__Fungi_unclassified(100); | - | - | - | - | - | - | - | Unassigned | - |
| Otu56557 | 1 | k__Fungi(100);k__Fungi_unclassified(100);k__Fungi_unclassified(100);k__Fungi_unclassified(100);k__Fungi_unclassified(100);k__Fungi_unclassified(100);k__Fungi_unclassified(100); | - | - | - | - | - | - | - | Unassigned | - |
| Otu56559 | 1 | k__Fungi(100);p__Ascomycota(100);p__Ascomycota_unclassified(100);p__Ascomycota_unclassified(100);p__Ascomycota_unclassified(100);p__Ascomycota_unclassified(100);p__Ascomycota_unclassified(100); | - | - | - | - | - | - | - | Unassigned | - |
| Otu56565 | 1 | k__Fungi(100);k__Fungi_unclassified(100);k__Fungi_unclassified(100);k__Fungi_unclassified(100);k__Fungi_unclassified(100);k__Fungi_unclassified(100);k__Fungi_unclassified(100); | - | - | - | - | - | - | - | Unassigned | - |
| Otu56570 | 1 | k__Fungi(100);p__Ascomycota(100);p__Ascomycota_unclassified(100);p__Ascomycota_unclassified(100);p__Ascomycota_unclassified(100);p__Ascomycota_unclassified(100);p__Ascomycota_unclassified(100); | - | - | - | - | - | - | - | Unassigned | - |
| Otu56571 | 1 | k__Fungi(100);p__Ascomycota(100);p__Ascomycota_unclassified(100);p__Ascomycota_unclassified(100);p__Ascomycota_unclassified(100);p__Ascomycota_unclassified(100);p__Ascomycota_unclassified(100); | - | - | - | - | - | - | - | Unassigned | - |
| Otu56572 | 1 | k__Fungi(100);k__Fungi_unclassified(100);k__Fungi_unclassified(100);k__Fungi_unclassified(100);k__Fungi_unclassified(100);k__Fungi_unclassified(100);k__Fungi_unclassified(100); | - | - | - | - | - | - | - | Unassigned | - |
| Otu56573 | 1 | k__Fungi(100);p__Ascomycota(100);p__Ascomycota_unclassified(100);p__Ascomycota_unclassified(100);p__Ascomycota_unclassified(100);p__Ascomycota_unclassified(100);p__Ascomycota_unclassified(100); | - | - | - | - | - | - | - | Unassigned | - |
| Otu56577 | 1 | k__Fungi(100);p__Ascomycota(100);p__Ascomycota_unclassified(100);p__Ascomycota_unclassified(100);p__Ascomycota_unclassified(100);p__Ascomycota_unclassified(100);p__Ascomycota_unclassified(100); | - | - | - | - | - | - | - | Unassigned | - |
| Otu56578 | 1 | k__Fungi(100);k__Fungi_unclassified(100);k__Fungi_unclassified(100);k__Fungi_unclassified(100);k__Fungi_unclassified(100);k__Fungi_unclassified(100);k__Fungi_unclassified(100); | - | - | - | - | - | - | - | Unassigned | - |
| Otu56596 | 1 | k__Fungi(100);p__Ascomycota(100);p__Ascomycota_unclassified(100);p__Ascomycota_unclassified(100);p__Ascomycota_unclassified(100);p__Ascomycota_unclassified(100);p__Ascomycota_unclassified(100); | - | - | - | - | - | - | - | Unassigned | - |
| Otu56613 | 1 | k__Fungi(100);k__Fungi_unclassified(100);k__Fungi_unclassified(100);k__Fungi_unclassified(100);k__Fungi_unclassified(100);k__Fungi_unclassified(100);k__Fungi_unclassified(100); | - | - | - | - | - | - | - | Unassigned | - |
| Otu56614 | 1 | k__Fungi(100);k__Fungi_unclassified(100);k__Fungi_unclassified(100);k__Fungi_unclassified(100);k__Fungi_unclassified(100);k__Fungi_unclassified(100);k__Fungi_unclassified(100); | - | - | - | - | - | - | - | Unassigned | - |
| Otu56615 | 1 | k__Fungi(100);p__Ascomycota(100);p__Ascomycota_unclassified(100);p__Ascomycota_unclassified(100);p__Ascomycota_unclassified(100);p__Ascomycota_unclassified(100);p__Ascomycota_unclassified(100); | - | - | - | - | - | - | - | Unassigned | - |
| Otu56616 | 1 | k__Fungi(100);k__Fungi_unclassified(100);k__Fungi_unclassified(100);k__Fungi_unclassified(100);k__Fungi_unclassified(100);k__Fungi_unclassified(100);k__Fungi_unclassified(100); | - | - | - | - | - | - | - | Unassigned | - |
| Otu56617 | 1 | k__Fungi(100);k__Fungi_unclassified(100);k__Fungi_unclassified(100);k__Fungi_unclassified(100);k__Fungi_unclassified(100);k__Fungi_unclassified(100);k__Fungi_unclassified(100); | - | - | - | - | - | - | - | Unassigned | - |
| Otu56619 | 1 | k__Fungi(100);k__Fungi_unclassified(100);k__Fungi_unclassified(100);k__Fungi_unclassified(100);k__Fungi_unclassified(100);k__Fungi_unclassified(100);k__Fungi_unclassified(100); | - | - | - | - | - | - | - | Unassigned | - |
| Otu56622 | 1 | k__Fungi(100);k__Fungi_unclassified(100);k__Fungi_unclassified(100);k__Fungi_unclassified(100);k__Fungi_unclassified(100);k__Fungi_unclassified(100);k__Fungi_unclassified(100); | - | - | - | - | - | - | - | Unassigned | - |
| Otu56623 | 1 | k__Fungi(100);k__Fungi_unclassified(100);k__Fungi_unclassified(100);k__Fungi_unclassified(100);k__Fungi_unclassified(100);k__Fungi_unclassified(100);k__Fungi_unclassified(100); | - | - | - | - | - | - | - | Unassigned | - |
| Otu56627 | 1 | k__Fungi(100);k__Fungi_unclassified(100);k__Fungi_unclassified(100);k__Fungi_unclassified(100);k__Fungi_unclassified(100);k__Fungi_unclassified(100);k__Fungi_unclassified(100); | - | - | - | - | - | - | - | Unassigned | - |
| Otu56628 | 1 | k__Fungi(100);p__Ascomycota(100);p__Ascomycota_unclassified(100);p__Ascomycota_unclassified(100);p__Ascomycota_unclassified(100);p__Ascomycota_unclassified(100);p__Ascomycota_unclassified(100); | - | - | - | - | - | - | - | Unassigned | - |
| Otu56636 | 1 | k__Fungi(100);k__Fungi_unclassified(100);k__Fungi_unclassified(100);k__Fungi_unclassified(100);k__Fungi_unclassified(100);k__Fungi_unclassified(100);k__Fungi_unclassified(100); | - | - | - | - | - | - | - | Unassigned | - |
| Otu56637 | 1 | k__Fungi(100);p__Ascomycota(100);p__Ascomycota_unclassified(100);p__Ascomycota_unclassified(100);p__Ascomycota_unclassified(100);p__Ascomycota_unclassified(100);p__Ascomycota_unclassified(100); | - | - | - | - | - | - | - | Unassigned | - |
| Otu56638 | 1 | k__Fungi(100);p__Ascomycota(100);p__Ascomycota_unclassified(100);p__Ascomycota_unclassified(100);p__Ascomycota_unclassified(100);p__Ascomycota_unclassified(100);p__Ascomycota_unclassified(100); | - | - | - | - | - | - | - | Unassigned | - |
| Otu56642 | 1 | k__Fungi(100);p__Ascomycota(100);p__Ascomycota_unclassified(100);p__Ascomycota_unclassified(100);p__Ascomycota_unclassified(100);p__Ascomycota_unclassified(100);p__Ascomycota_unclassified(100); | - | - | - | - | - | - | - | Unassigned | - |
| Otu56644 | 1 | k__Fungi(100);p__Ascomycota(100);p__Ascomycota_unclassified(100);p__Ascomycota_unclassified(100);p__Ascomycota_unclassified(100);p__Ascomycota_unclassified(100);p__Ascomycota_unclassified(100); | - | - | - | - | - | - | - | Unassigned | - |
| Otu56645 | 1 | k__Fungi(100);k__Fungi_unclassified(100);k__Fungi_unclassified(100);k__Fungi_unclassified(100);k__Fungi_unclassified(100);k__Fungi_unclassified(100);k__Fungi_unclassified(100); | - | - | - | - | - | - | - | Unassigned | - |
| Otu56647 | 1 | k__Fungi(100);k__Fungi_unclassified(100);k__Fungi_unclassified(100);k__Fungi_unclassified(100);k__Fungi_unclassified(100);k__Fungi_unclassified(100);k__Fungi_unclassified(100); | - | - | - | - | - | - | - | Unassigned | - |
| Otu56648 | 1 | k__Fungi(100);k__Fungi_unclassified(100);k__Fungi_unclassified(100);k__Fungi_unclassified(100);k__Fungi_unclassified(100);k__Fungi_unclassified(100);k__Fungi_unclassified(100); | - | - | - | - | - | - | - | Unassigned | - |
| Otu56649 | 1 | k__Fungi(100);k__Fungi_unclassified(100);k__Fungi_unclassified(100);k__Fungi_unclassified(100);k__Fungi_unclassified(100);k__Fungi_unclassified(100);k__Fungi_unclassified(100); | - | - | - | - | - | - | - | Unassigned | - |
| Otu56662 | 1 | k__Fungi(100);k__Fungi_unclassified(100);k__Fungi_unclassified(100);k__Fungi_unclassified(100);k__Fungi_unclassified(100);k__Fungi_unclassified(100);k__Fungi_unclassified(100); | - | - | - | - | - | - | - | Unassigned | - |
| Otu56671 | 1 | k__Fungi(100);p__Ascomycota(100);p__Ascomycota_unclassified(100);p__Ascomycota_unclassified(100);p__Ascomycota_unclassified(100);p__Ascomycota_unclassified(100);p__Ascomycota_unclassified(100); | - | - | - | - | - | - | - | Unassigned | - |
| Otu56684 | 1 | k__Fungi(100);k__Fungi_unclassified(100);k__Fungi_unclassified(100);k__Fungi_unclassified(100);k__Fungi_unclassified(100);k__Fungi_unclassified(100);k__Fungi_unclassified(100); | - | - | - | - | - | - | - | Unassigned | - |
| Otu56685 | 1 | k__Fungi(100);p__Ascomycota(100);p__Ascomycota_unclassified(100);p__Ascomycota_unclassified(100);p__Ascomycota_unclassified(100);p__Ascomycota_unclassified(100);p__Ascomycota_unclassified(100); | - | - | - | - | - | - | - | Unassigned | - |
| Otu56687 | 1 | k__Fungi(100);k__Fungi_unclassified(100);k__Fungi_unclassified(100);k__Fungi_unclassified(100);k__Fungi_unclassified(100);k__Fungi_unclassified(100);k__Fungi_unclassified(100); | - | - | - | - | - | - | - | Unassigned | - |
| Otu56692 | 1 | k__Fungi(100);k__Fungi_unclassified(100);k__Fungi_unclassified(100);k__Fungi_unclassified(100);k__Fungi_unclassified(100);k__Fungi_unclassified(100);k__Fungi_unclassified(100); | - | - | - | - | - | - | - | Unassigned | - |
| Otu56697 | 1 | k__Fungi(100);k__Fungi_unclassified(100);k__Fungi_unclassified(100);k__Fungi_unclassified(100);k__Fungi_unclassified(100);k__Fungi_unclassified(100);k__Fungi_unclassified(100); | - | - | - | - | - | - | - | Unassigned | - |
| Otu56699 | 1 | k__Fungi(100);p__Ascomycota(100);p__Ascomycota_unclassified(100);p__Ascomycota_unclassified(100);p__Ascomycota_unclassified(100);p__Ascomycota_unclassified(100);p__Ascomycota_unclassified(100); | - | - | - | - | - | - | - | Unassigned | - |
| Otu56705 | 1 | k__Fungi(100);k__Fungi_unclassified(100);k__Fungi_unclassified(100);k__Fungi_unclassified(100);k__Fungi_unclassified(100);k__Fungi_unclassified(100);k__Fungi_unclassified(100); | - | - | - | - | - | - | - | Unassigned | - |
| Otu56707 | 1 | k__Fungi(100);k__Fungi_unclassified(100);k__Fungi_unclassified(100);k__Fungi_unclassified(100);k__Fungi_unclassified(100);k__Fungi_unclassified(100);k__Fungi_unclassified(100); | - | - | - | - | - | - | - | Unassigned | - |
| Otu56708 | 1 | k__Fungi(100);k__Fungi_unclassified(100);k__Fungi_unclassified(100);k__Fungi_unclassified(100);k__Fungi_unclassified(100);k__Fungi_unclassified(100);k__Fungi_unclassified(100); | - | - | - | - | - | - | - | Unassigned | - |
| Otu56722 | 1 | k__Fungi(100);k__Fungi_unclassified(100);k__Fungi_unclassified(100);k__Fungi_unclassified(100);k__Fungi_unclassified(100);k__Fungi_unclassified(100);k__Fungi_unclassified(100); | - | - | - | - | - | - | - | Unassigned | - |
| Otu56740 | 1 | k__Fungi(100);k__Fungi_unclassified(100);k__Fungi_unclassified(100);k__Fungi_unclassified(100);k__Fungi_unclassified(100);k__Fungi_unclassified(100);k__Fungi_unclassified(100); | - | - | - | - | - | - | - | Unassigned | - |
| Otu56741 | 1 | k__Fungi(100);k__Fungi_unclassified(100);k__Fungi_unclassified(100);k__Fungi_unclassified(100);k__Fungi_unclassified(100);k__Fungi_unclassified(100);k__Fungi_unclassified(100); | - | - | - | - | - | - | - | Unassigned | - |
| Otu56742 | 1 | k__Fungi(100);k__Fungi_unclassified(100);k__Fungi_unclassified(100);k__Fungi_unclassified(100);k__Fungi_unclassified(100);k__Fungi_unclassified(100);k__Fungi_unclassified(100); | - | - | - | - | - | - | - | Unassigned | - |
| Otu56743 | 1 | k__Fungi(100);p__Ascomycota(100);p__Ascomycota_unclassified(100);p__Ascomycota_unclassified(100);p__Ascomycota_unclassified(100);p__Ascomycota_unclassified(100);p__Ascomycota_unclassified(100); | - | - | - | - | - | - | - | Unassigned | - |
| Otu56744 | 1 | k__Fungi(100);k__Fungi_unclassified(100);k__Fungi_unclassified(100);k__Fungi_unclassified(100);k__Fungi_unclassified(100);k__Fungi_unclassified(100);k__Fungi_unclassified(100); | - | - | - | - | - | - | - | Unassigned | - |
| Otu56745 | 1 | k__Fungi(100);p__Ascomycota(100);p__Ascomycota_unclassified(100);p__Ascomycota_unclassified(100);p__Ascomycota_unclassified(100);p__Ascomycota_unclassified(100);p__Ascomycota_unclassified(100); | - | - | - | - | - | - | - | Unassigned | - |
| Otu56746 | 1 | k__Fungi(100);p__Ascomycota(100);p__Ascomycota_unclassified(100);p__Ascomycota_unclassified(100);p__Ascomycota_unclassified(100);p__Ascomycota_unclassified(100);p__Ascomycota_unclassified(100); | - | - | - | - | - | - | - | Unassigned | - |
| Otu56747 | 1 | k__Fungi(100);k__Fungi_unclassified(100);k__Fungi_unclassified(100);k__Fungi_unclassified(100);k__Fungi_unclassified(100);k__Fungi_unclassified(100);k__Fungi_unclassified(100); | - | - | - | - | - | - | - | Unassigned | - |
| Otu56749 | 1 | k__Fungi(100);p__Ascomycota(100);p__Ascomycota_unclassified(100);p__Ascomycota_unclassified(100);p__Ascomycota_unclassified(100);p__Ascomycota_unclassified(100);p__Ascomycota_unclassified(100); | - | - | - | - | - | - | - | Unassigned | - |
| Otu56750 | 1 | k__Fungi(100);k__Fungi_unclassified(100);k__Fungi_unclassified(100);k__Fungi_unclassified(100);k__Fungi_unclassified(100);k__Fungi_unclassified(100);k__Fungi_unclassified(100); | - | - | - | - | - | - | - | Unassigned | - |
| Otu56752 | 1 | k__Fungi(100);p__Ascomycota(100);p__Ascomycota_unclassified(100);p__Ascomycota_unclassified(100);p__Ascomycota_unclassified(100);p__Ascomycota_unclassified(100);p__Ascomycota_unclassified(100); | - | - | - | - | - | - | - | Unassigned | - |
| Otu56753 | 1 | k__Fungi(100);k__Fungi_unclassified(100);k__Fungi_unclassified(100);k__Fungi_unclassified(100);k__Fungi_unclassified(100);k__Fungi_unclassified(100);k__Fungi_unclassified(100); | - | - | - | - | - | - | - | Unassigned | - |
| Otu56756 | 1 | k__Fungi(100);p__Ascomycota(100);p__Ascomycota_unclassified(100);p__Ascomycota_unclassified(100);p__Ascomycota_unclassified(100);p__Ascomycota_unclassified(100);p__Ascomycota_unclassified(100); | - | - | - | - | - | - | - | Unassigned | - |
| Otu56757 | 1 | k__Fungi(100);p__Ascomycota(100);p__Ascomycota_unclassified(100);p__Ascomycota_unclassified(100);p__Ascomycota_unclassified(100);p__Ascomycota_unclassified(100);p__Ascomycota_unclassified(100); | - | - | - | - | - | - | - | Unassigned | - |
| Otu56759 | 1 | k__Fungi(100);k__Fungi_unclassified(100);k__Fungi_unclassified(100);k__Fungi_unclassified(100);k__Fungi_unclassified(100);k__Fungi_unclassified(100);k__Fungi_unclassified(100); | - | - | - | - | - | - | - | Unassigned | - |
| Otu56761 | 1 | k__Fungi(100);k__Fungi_unclassified(100);k__Fungi_unclassified(100);k__Fungi_unclassified(100);k__Fungi_unclassified(100);k__Fungi_unclassified(100);k__Fungi_unclassified(100); | - | - | - | - | - | - | - | Unassigned | - |
| Otu56764 | 1 | k__Fungi(100);k__Fungi_unclassified(100);k__Fungi_unclassified(100);k__Fungi_unclassified(100);k__Fungi_unclassified(100);k__Fungi_unclassified(100);k__Fungi_unclassified(100); | - | - | - | - | - | - | - | Unassigned | - |
| Otu56767 | 1 | k__Fungi(100);k__Fungi_unclassified(100);k__Fungi_unclassified(100);k__Fungi_unclassified(100);k__Fungi_unclassified(100);k__Fungi_unclassified(100);k__Fungi_unclassified(100); | - | - | - | - | - | - | - | Unassigned | - |
| Otu56768 | 1 | k__Fungi(100);k__Fungi_unclassified(100);k__Fungi_unclassified(100);k__Fungi_unclassified(100);k__Fungi_unclassified(100);k__Fungi_unclassified(100);k__Fungi_unclassified(100); | - | - | - | - | - | - | - | Unassigned | - |
| Otu56777 | 1 | k__Fungi(100);k__Fungi_unclassified(100);k__Fungi_unclassified(100);k__Fungi_unclassified(100);k__Fungi_unclassified(100);k__Fungi_unclassified(100);k__Fungi_unclassified(100); | - | - | - | - | - | - | - | Unassigned | - |
| Otu56780 | 1 | k__Fungi(100);k__Fungi_unclassified(100);k__Fungi_unclassified(100);k__Fungi_unclassified(100);k__Fungi_unclassified(100);k__Fungi_unclassified(100);k__Fungi_unclassified(100); | - | - | - | - | - | - | - | Unassigned | - |
| Otu56784 | 1 | k__Fungi(100);k__Fungi_unclassified(100);k__Fungi_unclassified(100);k__Fungi_unclassified(100);k__Fungi_unclassified(100);k__Fungi_unclassified(100);k__Fungi_unclassified(100); | - | - | - | - | - | - | - | Unassigned | - |
| Otu56785 | 1 | k__Fungi(100);p__Ascomycota(100);p__Ascomycota_unclassified(100);p__Ascomycota_unclassified(100);p__Ascomycota_unclassified(100);p__Ascomycota_unclassified(100);p__Ascomycota_unclassified(100); | - | - | - | - | - | - | - | Unassigned | - |
| Otu56786 | 1 | k__Fungi(100);k__Fungi_unclassified(100);k__Fungi_unclassified(100);k__Fungi_unclassified(100);k__Fungi_unclassified(100);k__Fungi_unclassified(100);k__Fungi_unclassified(100); | - | - | - | - | - | - | - | Unassigned | - |
| Otu56787 | 1 | k__Fungi(100);k__Fungi_unclassified(100);k__Fungi_unclassified(100);k__Fungi_unclassified(100);k__Fungi_unclassified(100);k__Fungi_unclassified(100);k__Fungi_unclassified(100); | - | - | - | - | - | - | - | Unassigned | - |
| Otu56788 | 1 | k__Fungi(100);k__Fungi_unclassified(100);k__Fungi_unclassified(100);k__Fungi_unclassified(100);k__Fungi_unclassified(100);k__Fungi_unclassified(100);k__Fungi_unclassified(100); | - | - | - | - | - | - | - | Unassigned | - |
| Otu56789 | 1 | k__Fungi(100);p__Ascomycota(100);p__Ascomycota_unclassified(100);p__Ascomycota_unclassified(100);p__Ascomycota_unclassified(100);p__Ascomycota_unclassified(100);p__Ascomycota_unclassified(100); | - | - | - | - | - | - | - | Unassigned | - |
| Otu56790 | 1 | k__Fungi(100);k__Fungi_unclassified(100);k__Fungi_unclassified(100);k__Fungi_unclassified(100);k__Fungi_unclassified(100);k__Fungi_unclassified(100);k__Fungi_unclassified(100); | - | - | - | - | - | - | - | Unassigned | - |
| Otu56791 | 1 | k__Fungi(100);p__Ascomycota(100);p__Ascomycota_unclassified(100);p__Ascomycota_unclassified(100);p__Ascomycota_unclassified(100);p__Ascomycota_unclassified(100);p__Ascomycota_unclassified(100); | - | - | - | - | - | - | - | Unassigned | - |
| Otu56793 | 1 | k__Fungi(100);p__Ascomycota(100);p__Ascomycota_unclassified(100);p__Ascomycota_unclassified(100);p__Ascomycota_unclassified(100);p__Ascomycota_unclassified(100);p__Ascomycota_unclassified(100); | - | - | - | - | - | - | - | Unassigned | - |
| Otu56797 | 1 | k__Fungi(100);k__Fungi_unclassified(100);k__Fungi_unclassified(100);k__Fungi_unclassified(100);k__Fungi_unclassified(100);k__Fungi_unclassified(100);k__Fungi_unclassified(100); | - | - | - | - | - | - | - | Unassigned | - |
| Otu56798 | 1 | k__Fungi(100);p__Ascomycota(100);p__Ascomycota_unclassified(100);p__Ascomycota_unclassified(100);p__Ascomycota_unclassified(100);p__Ascomycota_unclassified(100);p__Ascomycota_unclassified(100); | - | - | - | - | - | - | - | Unassigned | - |
| Otu56807 | 1 | k__Fungi(100);k__Fungi_unclassified(100);k__Fungi_unclassified(100);k__Fungi_unclassified(100);k__Fungi_unclassified(100);k__Fungi_unclassified(100);k__Fungi_unclassified(100); | - | - | - | - | - | - | - | Unassigned | - |
| Otu56809 | 1 | k__Fungi(100);k__Fungi_unclassified(100);k__Fungi_unclassified(100);k__Fungi_unclassified(100);k__Fungi_unclassified(100);k__Fungi_unclassified(100);k__Fungi_unclassified(100); | - | - | - | - | - | - | - | Unassigned | - |
| Otu56812 | 1 | k__Fungi(100);k__Fungi_unclassified(100);k__Fungi_unclassified(100);k__Fungi_unclassified(100);k__Fungi_unclassified(100);k__Fungi_unclassified(100);k__Fungi_unclassified(100); | - | - | - | - | - | - | - | Unassigned | - |
| Otu56821 | 1 | k__Fungi(100);k__Fungi_unclassified(100);k__Fungi_unclassified(100);k__Fungi_unclassified(100);k__Fungi_unclassified(100);k__Fungi_unclassified(100);k__Fungi_unclassified(100); | - | - | - | - | - | - | - | Unassigned | - |
| Otu56823 | 1 | k__Fungi(100);p__Ascomycota(100);p__Ascomycota_unclassified(100);p__Ascomycota_unclassified(100);p__Ascomycota_unclassified(100);p__Ascomycota_unclassified(100);p__Ascomycota_unclassified(100); | - | - | - | - | - | - | - | Unassigned | - |
| Otu56829 | 1 | k__Fungi(100);k__Fungi_unclassified(100);k__Fungi_unclassified(100);k__Fungi_unclassified(100);k__Fungi_unclassified(100);k__Fungi_unclassified(100);k__Fungi_unclassified(100); | - | - | - | - | - | - | - | Unassigned | - |
| Otu56834 | 1 | k__Fungi(100);k__Fungi_unclassified(100);k__Fungi_unclassified(100);k__Fungi_unclassified(100);k__Fungi_unclassified(100);k__Fungi_unclassified(100);k__Fungi_unclassified(100); | - | - | - | - | - | - | - | Unassigned | - |
| Otu56843 | 1 | k__Fungi(100);k__Fungi_unclassified(100);k__Fungi_unclassified(100);k__Fungi_unclassified(100);k__Fungi_unclassified(100);k__Fungi_unclassified(100);k__Fungi_unclassified(100); | - | - | - | - | - | - | - | Unassigned | - |
| Otu56848 | 1 | k__Fungi(100);k__Fungi_unclassified(100);k__Fungi_unclassified(100);k__Fungi_unclassified(100);k__Fungi_unclassified(100);k__Fungi_unclassified(100);k__Fungi_unclassified(100); | - | - | - | - | - | - | - | Unassigned | - |
| Otu56857 | 1 | k__Fungi(100);k__Fungi_unclassified(100);k__Fungi_unclassified(100);k__Fungi_unclassified(100);k__Fungi_unclassified(100);k__Fungi_unclassified(100);k__Fungi_unclassified(100); | - | - | - | - | - | - | - | Unassigned | - |
| Otu56868 | 1 | k__Fungi(100);p__Ascomycota(100);p__Ascomycota_unclassified(100);p__Ascomycota_unclassified(100);p__Ascomycota_unclassified(100);p__Ascomycota_unclassified(100);p__Ascomycota_unclassified(100); | - | - | - | - | - | - | - | Unassigned | - |
| Otu56869 | 1 | k__Fungi(100);k__Fungi_unclassified(100);k__Fungi_unclassified(100);k__Fungi_unclassified(100);k__Fungi_unclassified(100);k__Fungi_unclassified(100);k__Fungi_unclassified(100); | - | - | - | - | - | - | - | Unassigned | - |
| Otu56870 | 1 | k__Fungi(100);k__Fungi_unclassified(100);k__Fungi_unclassified(100);k__Fungi_unclassified(100);k__Fungi_unclassified(100);k__Fungi_unclassified(100);k__Fungi_unclassified(100); | - | - | - | - | - | - | - | Unassigned | - |
| Otu56883 | 1 | k__Fungi(100);p__Ascomycota(100);p__Ascomycota_unclassified(100);p__Ascomycota_unclassified(100);p__Ascomycota_unclassified(100);p__Ascomycota_unclassified(100);p__Ascomycota_unclassified(100); | - | - | - | - | - | - | - | Unassigned | - |
| Otu56897 | 1 | k__Fungi(100);k__Fungi_unclassified(100);k__Fungi_unclassified(100);k__Fungi_unclassified(100);k__Fungi_unclassified(100);k__Fungi_unclassified(100);k__Fungi_unclassified(100); | - | - | - | - | - | - | - | Unassigned | - |
| Otu56899 | 1 | k__Fungi(100);p__Ascomycota(100);p__Ascomycota_unclassified(100);p__Ascomycota_unclassified(100);p__Ascomycota_unclassified(100);p__Ascomycota_unclassified(100);p__Ascomycota_unclassified(100); | - | - | - | - | - | - | - | Unassigned | - |
| Otu56906 | 1 | k__Fungi(100);k__Fungi_unclassified(100);k__Fungi_unclassified(100);k__Fungi_unclassified(100);k__Fungi_unclassified(100);k__Fungi_unclassified(100);k__Fungi_unclassified(100); | - | - | - | - | - | - | - | Unassigned | - |
| Otu56915 | 1 | k__Fungi(100);p__Ascomycota(100);p__Ascomycota_unclassified(100);p__Ascomycota_unclassified(100);p__Ascomycota_unclassified(100);p__Ascomycota_unclassified(100);p__Ascomycota_unclassified(100); | - | - | - | - | - | - | - | Unassigned | - |
| Otu56916 | 1 | k__Fungi(100);k__Fungi_unclassified(100);k__Fungi_unclassified(100);k__Fungi_unclassified(100);k__Fungi_unclassified(100);k__Fungi_unclassified(100);k__Fungi_unclassified(100); | - | - | - | - | - | - | - | Unassigned | - |
| Otu56917 | 1 | k__Fungi(100);k__Fungi_unclassified(100);k__Fungi_unclassified(100);k__Fungi_unclassified(100);k__Fungi_unclassified(100);k__Fungi_unclassified(100);k__Fungi_unclassified(100); | - | - | - | - | - | - | - | Unassigned | - |
| Otu56922 | 1 | k__Fungi(100);k__Fungi_unclassified(100);k__Fungi_unclassified(100);k__Fungi_unclassified(100);k__Fungi_unclassified(100);k__Fungi_unclassified(100);k__Fungi_unclassified(100); | - | - | - | - | - | - | - | Unassigned | - |
| Otu56923 | 1 | k__Fungi(100);p__Ascomycota(100);p__Ascomycota_unclassified(100);p__Ascomycota_unclassified(100);p__Ascomycota_unclassified(100);p__Ascomycota_unclassified(100);p__Ascomycota_unclassified(100); | - | - | - | - | - | - | - | Unassigned | - |
| Otu56925 | 1 | k__Fungi(100);p__Ascomycota(100);p__Ascomycota_unclassified(100);p__Ascomycota_unclassified(100);p__Ascomycota_unclassified(100);p__Ascomycota_unclassified(100);p__Ascomycota_unclassified(100); | - | - | - | - | - | - | - | Unassigned | - |
| Otu56926 | 1 | k__Fungi(100);p__Ascomycota(100);p__Ascomycota_unclassified(100);p__Ascomycota_unclassified(100);p__Ascomycota_unclassified(100);p__Ascomycota_unclassified(100);p__Ascomycota_unclassified(100); | - | - | - | - | - | - | - | Unassigned | - |
| Otu56930 | 1 | k__Fungi(100);p__Ascomycota(100);p__Ascomycota_unclassified(100);p__Ascomycota_unclassified(100);p__Ascomycota_unclassified(100);p__Ascomycota_unclassified(100);p__Ascomycota_unclassified(100); | - | - | - | - | - | - | - | Unassigned | - |
| Otu56933 | 1 | k__Fungi(100);p__Ascomycota(100);p__Ascomycota_unclassified(100);p__Ascomycota_unclassified(100);p__Ascomycota_unclassified(100);p__Ascomycota_unclassified(100);p__Ascomycota_unclassified(100); | - | - | - | - | - | - | - | Unassigned | - |
| Otu56934 | 1 | k__Fungi(100);k__Fungi_unclassified(100);k__Fungi_unclassified(100);k__Fungi_unclassified(100);k__Fungi_unclassified(100);k__Fungi_unclassified(100);k__Fungi_unclassified(100); | - | - | - | - | - | - | - | Unassigned | - |
| Otu56936 | 1 | k__Fungi(100);k__Fungi_unclassified(100);k__Fungi_unclassified(100);k__Fungi_unclassified(100);k__Fungi_unclassified(100);k__Fungi_unclassified(100);k__Fungi_unclassified(100); | - | - | - | - | - | - | - | Unassigned | - |
| Otu56937 | 1 | k__Fungi(100);k__Fungi_unclassified(100);k__Fungi_unclassified(100);k__Fungi_unclassified(100);k__Fungi_unclassified(100);k__Fungi_unclassified(100);k__Fungi_unclassified(100); | - | - | - | - | - | - | - | Unassigned | - |
| Otu56938 | 1 | k__Fungi(100);k__Fungi_unclassified(100);k__Fungi_unclassified(100);k__Fungi_unclassified(100);k__Fungi_unclassified(100);k__Fungi_unclassified(100);k__Fungi_unclassified(100); | - | - | - | - | - | - | - | Unassigned | - |
| Otu56939 | 1 | k__Fungi(100);p__Ascomycota(100);p__Ascomycota_unclassified(100);p__Ascomycota_unclassified(100);p__Ascomycota_unclassified(100);p__Ascomycota_unclassified(100);p__Ascomycota_unclassified(100); | - | - | - | - | - | - | - | Unassigned | - |
| Otu56941 | 1 | k__Fungi(100);k__Fungi_unclassified(100);k__Fungi_unclassified(100);k__Fungi_unclassified(100);k__Fungi_unclassified(100);k__Fungi_unclassified(100);k__Fungi_unclassified(100); | - | - | - | - | - | - | - | Unassigned | - |
| Otu56943 | 1 | k__Fungi(100);k__Fungi_unclassified(100);k__Fungi_unclassified(100);k__Fungi_unclassified(100);k__Fungi_unclassified(100);k__Fungi_unclassified(100);k__Fungi_unclassified(100); | - | - | - | - | - | - | - | Unassigned | - |
| Otu56944 | 1 | k__Fungi(100);k__Fungi_unclassified(100);k__Fungi_unclassified(100);k__Fungi_unclassified(100);k__Fungi_unclassified(100);k__Fungi_unclassified(100);k__Fungi_unclassified(100); | - | - | - | - | - | - | - | Unassigned | - |
| Otu56947 | 1 | k__Fungi(100);p__Ascomycota(100);p__Ascomycota_unclassified(100);p__Ascomycota_unclassified(100);p__Ascomycota_unclassified(100);p__Ascomycota_unclassified(100);p__Ascomycota_unclassified(100); | - | - | - | - | - | - | - | Unassigned | - |
| Otu56953 | 1 | k__Fungi(100);k__Fungi_unclassified(100);k__Fungi_unclassified(100);k__Fungi_unclassified(100);k__Fungi_unclassified(100);k__Fungi_unclassified(100);k__Fungi_unclassified(100); | - | - | - | - | - | - | - | Unassigned | - |
| Otu56955 | 1 | k__Fungi(100);k__Fungi_unclassified(100);k__Fungi_unclassified(100);k__Fungi_unclassified(100);k__Fungi_unclassified(100);k__Fungi_unclassified(100);k__Fungi_unclassified(100); | - | - | - | - | - | - | - | Unassigned | - |
| Otu56956 | 1 | k__Fungi(100);p__Ascomycota(100);p__Ascomycota_unclassified(100);p__Ascomycota_unclassified(100);p__Ascomycota_unclassified(100);p__Ascomycota_unclassified(100);p__Ascomycota_unclassified(100); | - | - | - | - | - | - | - | Unassigned | - |
| Otu56957 | 1 | k__Fungi(100);k__Fungi_unclassified(100);k__Fungi_unclassified(100);k__Fungi_unclassified(100);k__Fungi_unclassified(100);k__Fungi_unclassified(100);k__Fungi_unclassified(100); | - | - | - | - | - | - | - | Unassigned | - |
| Otu56961 | 1 | k__Fungi(100);k__Fungi_unclassified(100);k__Fungi_unclassified(100);k__Fungi_unclassified(100);k__Fungi_unclassified(100);k__Fungi_unclassified(100);k__Fungi_unclassified(100); | - | - | - | - | - | - | - | Unassigned | - |
| Otu56988 | 1 | k__Fungi(100);p__Ascomycota(100);p__Ascomycota_unclassified(100);p__Ascomycota_unclassified(100);p__Ascomycota_unclassified(100);p__Ascomycota_unclassified(100);p__Ascomycota_unclassified(100); | - | - | - | - | - | - | - | Unassigned | - |
| Otu56989 | 1 | k__Fungi(100);k__Fungi_unclassified(100);k__Fungi_unclassified(100);k__Fungi_unclassified(100);k__Fungi_unclassified(100);k__Fungi_unclassified(100);k__Fungi_unclassified(100); | - | - | - | - | - | - | - | Unassigned | - |
| Otu57000 | 1 | k__Fungi(100);k__Fungi_unclassified(100);k__Fungi_unclassified(100);k__Fungi_unclassified(100);k__Fungi_unclassified(100);k__Fungi_unclassified(100);k__Fungi_unclassified(100); | - | - | - | - | - | - | - | Unassigned | - |
| Otu57013 | 1 | k__Fungi(100);k__Fungi_unclassified(100);k__Fungi_unclassified(100);k__Fungi_unclassified(100);k__Fungi_unclassified(100);k__Fungi_unclassified(100);k__Fungi_unclassified(100); | - | - | - | - | - | - | - | Unassigned | - |
| Otu57018 | 1 | k__Fungi(100);p__Ascomycota(100);p__Ascomycota_unclassified(100);p__Ascomycota_unclassified(100);p__Ascomycota_unclassified(100);p__Ascomycota_unclassified(100);p__Ascomycota_unclassified(100); | - | - | - | - | - | - | - | Unassigned | - |
| Otu57024 | 1 | k__Fungi(100);k__Fungi_unclassified(100);k__Fungi_unclassified(100);k__Fungi_unclassified(100);k__Fungi_unclassified(100);k__Fungi_unclassified(100);k__Fungi_unclassified(100); | - | - | - | - | - | - | - | Unassigned | - |
| Otu57036 | 1 | k__Fungi(100);k__Fungi_unclassified(100);k__Fungi_unclassified(100);k__Fungi_unclassified(100);k__Fungi_unclassified(100);k__Fungi_unclassified(100);k__Fungi_unclassified(100); | - | - | - | - | - | - | - | Unassigned | - |
| Otu57039 | 1 | k__Fungi(100);k__Fungi_unclassified(100);k__Fungi_unclassified(100);k__Fungi_unclassified(100);k__Fungi_unclassified(100);k__Fungi_unclassified(100);k__Fungi_unclassified(100); | - | - | - | - | - | - | - | Unassigned | - |
| Otu57063 | 1 | k__Fungi(100);p__Ascomycota(100);p__Ascomycota_unclassified(100);p__Ascomycota_unclassified(100);p__Ascomycota_unclassified(100);p__Ascomycota_unclassified(100);p__Ascomycota_unclassified(100); | - | - | - | - | - | - | - | Unassigned | - |
| Otu57117 | 1 | k__Fungi(100);p__Ascomycota(100);p__Ascomycota_unclassified(100);p__Ascomycota_unclassified(100);p__Ascomycota_unclassified(100);p__Ascomycota_unclassified(100);p__Ascomycota_unclassified(100); | - | - | - | - | - | - | - | Unassigned | - |

| | | | | | | | | | | | |
|---|---|---|---|---|---|---|---|---|---|---|---|
| Otu57125 | 1 | k__Fungi(100);k__Fungi_unclassified(100);k__Fungi_unclassified(100);k__Fungi_unclassified(100);k__Fungi_unclassified(100);k__Fungi_unclassified(100);k__Fungi_unclassified(100); | - | - | - | - | - | - | - | Unassigned | - |
| Otu57138 | 1 | k__Fungi(100);p__Ascomycota(100);c__Dothideomycetes(100);o__Pleosporales(100);o__Pleosporales_unclassified(100);o__Pleosporales_unclassified(100);o__Pleosporales_unclassified(100); | - | - | - | - | - | - | - | Unassigned | - |
| Otu57156 | 1 | k__Fungi(100);p__Ascomycota(100);p__Ascomycota_unclassified(100);p__Ascomycota_unclassified(100);p__Ascomycota_unclassified(100);p__Ascomycota_unclassified(100);p__Ascomycota_unclassified(100); | - | - | - | - | - | - | - | Unassigned | - |
| Otu57157 | 1 | k__Fungi(100);p__Ascomycota(100);p__Ascomycota_unclassified(100);p__Ascomycota_unclassified(100);p__Ascomycota_unclassified(100);p__Ascomycota_unclassified(100);p__Ascomycota_unclassified(100); | - | - | - | - | - | - | - | Unassigned | - |
| Otu57159 | 1 | k__Fungi(100);k__Fungi_unclassified(100);k__Fungi_unclassified(100);k__Fungi_unclassified(100);k__Fungi_unclassified(100);k__Fungi_unclassified(100);k__Fungi_unclassified(100); | - | - | - | - | - | - | - | Unassigned | - |
| Otu57160 | 1 | k__Fungi(100);p__Ascomycota(100);p__Ascomycota_unclassified(100);p__Ascomycota_unclassified(100);p__Ascomycota_unclassified(100);p__Ascomycota_unclassified(100);p__Ascomycota_unclassified(100); | - | - | - | - | - | - | - | Unassigned | - |
| Otu57162 | 1 | k__Fungi(100);p__Ascomycota(100);p__Ascomycota_unclassified(100);p__Ascomycota_unclassified(100);p__Ascomycota_unclassified(100);p__Ascomycota_unclassified(100);p__Ascomycota_unclassified(100); | - | - | - | - | - | - | - | Unassigned | - |
| Otu57163 | 1 | k__Fungi(100);p__Ascomycota(100);p__Ascomycota_unclassified(100);p__Ascomycota_unclassified(100);p__Ascomycota_unclassified(100);p__Ascomycota_unclassified(100);p__Ascomycota_unclassified(100); | - | - | - | - | - | - | - | Unassigned | - |
| Otu57164 | 1 | k__Fungi(100);k__Fungi_unclassified(100);k__Fungi_unclassified(100);k__Fungi_unclassified(100);k__Fungi_unclassified(100);k__Fungi_unclassified(100);k__Fungi_unclassified(100); | - | - | - | - | - | - | - | Unassigned | - |
| Otu57170 | 1 | k__Fungi(100);p__Ascomycota(100);p__Ascomycota_unclassified(100);p__Ascomycota_unclassified(100);p__Ascomycota_unclassified(100);p__Ascomycota_unclassified(100);p__Ascomycota_unclassified(100); | - | - | - | - | - | - | - | Unassigned | - |
| Otu57171 | 1 | k__Fungi(100);k__Fungi_unclassified(100);k__Fungi_unclassified(100);k__Fungi_unclassified(100);k__Fungi_unclassified(100);k__Fungi_unclassified(100);k__Fungi_unclassified(100); | - | - | - | - | - | - | - | Unassigned | - |
| Otu57182 | 1 | k__Fungi(100);k__Fungi_unclassified(100);k__Fungi_unclassified(100);k__Fungi_unclassified(100);k__Fungi_unclassified(100);k__Fungi_unclassified(100);k__Fungi_unclassified(100); | - | - | - | - | - | - | - | Unassigned | - |
| Otu57183 | 1 | k__Fungi(100);p__Ascomycota(100);p__Ascomycota_unclassified(100);p__Ascomycota_unclassified(100);p__Ascomycota_unclassified(100);p__Ascomycota_unclassified(100);p__Ascomycota_unclassified(100); | - | - | - | - | - | - | - | Unassigned | - |
| Otu57185 | 1 | k__Fungi(100);k__Fungi_unclassified(100);k__Fungi_unclassified(100);k__Fungi_unclassified(100);k__Fungi_unclassified(100);k__Fungi_unclassified(100);k__Fungi_unclassified(100); | - | - | - | - | - | - | - | Unassigned | - |
| Otu57187 | 1 | k__Fungi(100);p__Ascomycota(100);p__Ascomycota_unclassified(100);p__Ascomycota_unclassified(100);p__Ascomycota_unclassified(100);p__Ascomycota_unclassified(100);p__Ascomycota_unclassified(100); | - | - | - | - | - | - | - | Unassigned | - |
| Otu57188 | 1 | k__Fungi(100);k__Fungi_unclassified(100);k__Fungi_unclassified(100);k__Fungi_unclassified(100);k__Fungi_unclassified(100);k__Fungi_unclassified(100);k__Fungi_unclassified(100); | - | - | - | - | - | - | - | Unassigned | - |
| Otu57189 | 1 | k__Fungi(100);k__Fungi_unclassified(100);k__Fungi_unclassified(100);k__Fungi_unclassified(100);k__Fungi_unclassified(100);k__Fungi_unclassified(100);k__Fungi_unclassified(100); | - | - | - | - | - | - | - | Unassigned | - |
| Otu57190 | 1 | k__Fungi(100);p__Ascomycota(100);p__Ascomycota_unclassified(100);p__Ascomycota_unclassified(100);p__Ascomycota_unclassified(100);p__Ascomycota_unclassified(100);p__Ascomycota_unclassified(100); | - | - | - | - | - | - | - | Unassigned | - |
| Otu57192 | 1 | k__Fungi(100);p__Ascomycota(100);p__Ascomycota_unclassified(100);p__Ascomycota_unclassified(100);p__Ascomycota_unclassified(100);p__Ascomycota_unclassified(100);p__Ascomycota_unclassified(100); | - | - | - | - | - | - | - | Unassigned | - |
| Otu57200 | 1 | k__Fungi(100);k__Fungi_unclassified(100);k__Fungi_unclassified(100);k__Fungi_unclassified(100);k__Fungi_unclassified(100);k__Fungi_unclassified(100);k__Fungi_unclassified(100); | - | - | - | - | - | - | - | Unassigned | - |
| Otu57201 | 1 | k__Fungi(100);k__Fungi_unclassified(100);k__Fungi_unclassified(100);k__Fungi_unclassified(100);k__Fungi_unclassified(100);k__Fungi_unclassified(100);k__Fungi_unclassified(100); | - | - | - | - | - | - | - | Unassigned | - |
| Otu57202 | 1 | k__Fungi(100);k__Fungi_unclassified(100);k__Fungi_unclassified(100);k__Fungi_unclassified(100);k__Fungi_unclassified(100);k__Fungi_unclassified(100);k__Fungi_unclassified(100); | - | - | - | - | - | - | - | Unassigned | - |
| Otu57203 | 1 | k__Fungi(100);k__Fungi_unclassified(100);k__Fungi_unclassified(100);k__Fungi_unclassified(100);k__Fungi_unclassified(100);k__Fungi_unclassified(100);k__Fungi_unclassified(100); | - | - | - | - | - | - | - | Unassigned | - |
| Otu57213 | 1 | k__Fungi(100);k__Fungi_unclassified(100);k__Fungi_unclassified(100);k__Fungi_unclassified(100);k__Fungi_unclassified(100);k__Fungi_unclassified(100);k__Fungi_unclassified(100); | - | - | - | - | - | - | - | Unassigned | - |
| Otu57214 | 1 | k__Fungi(100);k__Fungi_unclassified(100);k__Fungi_unclassified(100);k__Fungi_unclassified(100);k__Fungi_unclassified(100);k__Fungi_unclassified(100);k__Fungi_unclassified(100); | - | - | - | - | - | - | - | Unassigned | - |
| Otu57215 | 1 | k__Fungi(100);p__Ascomycota(100);p__Ascomycota_unclassified(100);p__Ascomycota_unclassified(100);p__Ascomycota_unclassified(100);p__Ascomycota_unclassified(100);p__Ascomycota_unclassified(100); | - | - | - | - | - | - | - | Unassigned | - |
| Otu57218 | 1 | k__Fungi(100);p__Ascomycota(100);p__Ascomycota_unclassified(100);p__Ascomycota_unclassified(100);p__Ascomycota_unclassified(100);p__Ascomycota_unclassified(100);p__Ascomycota_unclassified(100); | - | - | - | - | - | - | - | Unassigned | - |
| Otu57220 | 1 | k__Fungi(100);k__Fungi_unclassified(100);k__Fungi_unclassified(100);k__Fungi_unclassified(100);k__Fungi_unclassified(100);k__Fungi_unclassified(100);k__Fungi_unclassified(100); | - | - | - | - | - | - | - | Unassigned | - |
| Otu57221 | 1 | k__Fungi(100);p__Ascomycota(100);p__Ascomycota_unclassified(100);p__Ascomycota_unclassified(100);p__Ascomycota_unclassified(100);p__Ascomycota_unclassified(100);p__Ascomycota_unclassified(100); | - | - | - | - | - | - | - | Unassigned | - |
| Otu57222 | 1 | k__Fungi(100);p__Ascomycota(100);p__Ascomycota_unclassified(100);p__Ascomycota_unclassified(100);p__Ascomycota_unclassified(100);p__Ascomycota_unclassified(100);p__Ascomycota_unclassified(100); | - | - | - | - | - | - | - | Unassigned | - |
| Otu57226 | 1 | k__Fungi(100);p__Ascomycota(100);p__Ascomycota_unclassified(100);p__Ascomycota_unclassified(100);p__Ascomycota_unclassified(100);p__Ascomycota_unclassified(100);p__Ascomycota_unclassified(100); | - | - | - | - | - | - | - | Unassigned | - |
| Otu57230 | 1 | k__Fungi(100);p__Ascomycota(100);p__Ascomycota_unclassified(100);p__Ascomycota_unclassified(100);p__Ascomycota_unclassified(100);p__Ascomycota_unclassified(100);p__Ascomycota_unclassified(100); | - | - | - | - | - | - | - | Unassigned | - |
| Otu57252 | 1 | k__Fungi(100);k__Fungi_unclassified(100);k__Fungi_unclassified(100);k__Fungi_unclassified(100);k__Fungi_unclassified(100);k__Fungi_unclassified(100);k__Fungi_unclassified(100); | - | - | - | - | - | - | - | Unassigned | - |
| Otu57253 | 1 | k__Fungi(100);k__Fungi_unclassified(100);k__Fungi_unclassified(100);k__Fungi_unclassified(100);k__Fungi_unclassified(100);k__Fungi_unclassified(100);k__Fungi_unclassified(100); | - | - | - | - | - | - | - | Unassigned | - |
| Otu57254 | 1 | k__Fungi(100);k__Fungi_unclassified(100);k__Fungi_unclassified(100);k__Fungi_unclassified(100);k__Fungi_unclassified(100);k__Fungi_unclassified(100);k__Fungi_unclassified(100); | - | - | - | - | - | - | - | Unassigned | - |
| Otu57255 | 1 | k__Fungi(100);k__Fungi_unclassified(100);k__Fungi_unclassified(100);k__Fungi_unclassified(100);k__Fungi_unclassified(100);k__Fungi_unclassified(100);k__Fungi_unclassified(100); | - | - | - | - | - | - | - | Unassigned | - |
| Otu57256 | 1 | k__Fungi(100);k__Fungi_unclassified(100);k__Fungi_unclassified(100);k__Fungi_unclassified(100);k__Fungi_unclassified(100);k__Fungi_unclassified(100);k__Fungi_unclassified(100); | - | - | - | - | - | - | - | Unassigned | - |
| Otu57257 | 1 | k__Fungi(100);p__Ascomycota(100);p__Ascomycota_unclassified(100);p__Ascomycota_unclassified(100);p__Ascomycota_unclassified(100);p__Ascomycota_unclassified(100);p__Ascomycota_unclassified(100); | - | - | - | - | - | - | - | Unassigned | - |
| Otu57258 | 1 | k__Fungi(100);p__Ascomycota(100);p__Ascomycota_unclassified(100);p__Ascomycota_unclassified(100);p__Ascomycota_unclassified(100);p__Ascomycota_unclassified(100);p__Ascomycota_unclassified(100); | - | - | - | - | - | - | - | Unassigned | - |
| Otu57264 | 1 | k__Fungi(100);k__Fungi_unclassified(100);k__Fungi_unclassified(100);k__Fungi_unclassified(100);k__Fungi_unclassified(100);k__Fungi_unclassified(100);k__Fungi_unclassified(100); | - | - | - | - | - | - | - | Unassigned | - |
| Otu57286 | 1 | k__Fungi(100);p__Ascomycota(100);p__Ascomycota_unclassified(100);p__Ascomycota_unclassified(100);p__Ascomycota_unclassified(100);p__Ascomycota_unclassified(100);p__Ascomycota_unclassified(100); | - | - | - | - | - | - | - | Unassigned | - |
| Otu57312 | 1 | k__Fungi(100);k__Fungi_unclassified(100);k__Fungi_unclassified(100);k__Fungi_unclassified(100);k__Fungi_unclassified(100);k__Fungi_unclassified(100);k__Fungi_unclassified(100); | - | - | - | - | - | - | - | Unassigned | - |
| Otu57314 | 1 | k__Fungi(100);k__Fungi_unclassified(100);k__Fungi_unclassified(100);k__Fungi_unclassified(100);k__Fungi_unclassified(100);k__Fungi_unclassified(100);k__Fungi_unclassified(100); | - | - | - | - | - | - | - | Unassigned | - |
| Otu57315 | 1 | k__Fungi(100);k__Fungi_unclassified(100);k__Fungi_unclassified(100);k__Fungi_unclassified(100);k__Fungi_unclassified(100);k__Fungi_unclassified(100);k__Fungi_unclassified(100); | - | - | - | - | - | - | - | Unassigned | - |
| Otu57316 | 1 | k__Fungi(100);p__Ascomycota(100);p__Ascomycota_unclassified(100);p__Ascomycota_unclassified(100);p__Ascomycota_unclassified(100);p__Ascomycota_unclassified(100);p__Ascomycota_unclassified(100); | - | - | - | - | - | - | - | Unassigned | - |
| Otu57317 | 1 | k__Fungi(100);k__Fungi_unclassified(100);k__Fungi_unclassified(100);k__Fungi_unclassified(100);k__Fungi_unclassified(100);k__Fungi_unclassified(100);k__Fungi_unclassified(100); | - | - | - | - | - | - | - | Unassigned | - |
| Otu57318 | 1 | k__Fungi(100);k__Fungi_unclassified(100);k__Fungi_unclassified(100);k__Fungi_unclassified(100);k__Fungi_unclassified(100);k__Fungi_unclassified(100);k__Fungi_unclassified(100); | - | - | - | - | - | - | - | Unassigned | - |
| Otu57319 | 1 | k__Fungi(100);k__Fungi_unclassified(100);k__Fungi_unclassified(100);k__Fungi_unclassified(100);k__Fungi_unclassified(100);k__Fungi_unclassified(100);k__Fungi_unclassified(100); | - | - | - | - | - | - | - | Unassigned | - |
| Otu57320 | 1 | k__Fungi(100);k__Fungi_unclassified(100);k__Fungi_unclassified(100);k__Fungi_unclassified(100);k__Fungi_unclassified(100);k__Fungi_unclassified(100);k__Fungi_unclassified(100); | - | - | - | - | - | - | - | Unassigned | - |
| Otu57323 | 1 | k__Fungi(100);k__Fungi_unclassified(100);k__Fungi_unclassified(100);k__Fungi_unclassified(100);k__Fungi_unclassified(100);k__Fungi_unclassified(100);k__Fungi_unclassified(100); | - | - | - | - | - | - | - | Unassigned | - |
| Otu57326 | 1 | k__Fungi(100);k__Fungi_unclassified(100);k__Fungi_unclassified(100);k__Fungi_unclassified(100);k__Fungi_unclassified(100);k__Fungi_unclassified(100);k__Fungi_unclassified(100); | - | - | - | - | - | - | - | Unassigned | - |
| Otu57327 | 1 | k__Fungi(100);k__Fungi_unclassified(100);k__Fungi_unclassified(100);k__Fungi_unclassified(100);k__Fungi_unclassified(100);k__Fungi_unclassified(100);k__Fungi_unclassified(100); | - | - | - | - | - | - | - | Unassigned | - |
| Otu57328 | 1 | k__Fungi(100);p__Ascomycota(100);p__Ascomycota_unclassified(100);p__Ascomycota_unclassified(100);p__Ascomycota_unclassified(100);p__Ascomycota_unclassified(100);p__Ascomycota_unclassified(100); | - | - | - | - | - | - | - | Unassigned | - |
| Otu57329 | 1 | k__Fungi(100);k__Fungi_unclassified(100);k__Fungi_unclassified(100);k__Fungi_unclassified(100);k__Fungi_unclassified(100);k__Fungi_unclassified(100);k__Fungi_unclassified(100); | - | - | - | - | - | - | - | Unassigned | - |
| Otu57330 | 1 | k__Fungi(100);k__Fungi_unclassified(100);k__Fungi_unclassified(100);k__Fungi_unclassified(100);k__Fungi_unclassified(100);k__Fungi_unclassified(100);k__Fungi_unclassified(100); | - | - | - | - | - | - | - | Unassigned | - |
| Otu57335 | 1 | k__Fungi(100);k__Fungi_unclassified(100);k__Fungi_unclassified(100);k__Fungi_unclassified(100);k__Fungi_unclassified(100);k__Fungi_unclassified(100);k__Fungi_unclassified(100); | - | - | - | - | - | - | - | Unassigned | - |
| Otu57336 | 1 | k__Fungi(100);p__Ascomycota(100);p__Ascomycota_unclassified(100);p__Ascomycota_unclassified(100);p__Ascomycota_unclassified(100);p__Ascomycota_unclassified(100);p__Ascomycota_unclassified(100); | - | - | - | - | - | - | - | Unassigned | - |
| Otu57355 | 1 | k__Fungi(100);p__Ascomycota(100);p__Ascomycota_unclassified(100);p__Ascomycota_unclassified(100);p__Ascomycota_unclassified(100);p__Ascomycota_unclassified(100);p__Ascomycota_unclassified(100); | - | - | - | - | - | - | - | Unassigned | - |
| Otu57356 | 1 | k__Fungi(100);p__Ascomycota(100);p__Ascomycota_unclassified(100);p__Ascomycota_unclassified(100);p__Ascomycota_unclassified(100);p__Ascomycota_unclassified(100);p__Ascomycota_unclassified(100); | - | - | - | - | - | - | - | Unassigned | - |
| Otu57357 | 1 | k__Fungi(100);k__Fungi_unclassified(100);k__Fungi_unclassified(100);k__Fungi_unclassified(100);k__Fungi_unclassified(100);k__Fungi_unclassified(100);k__Fungi_unclassified(100); | - | - | - | - | - | - | - | Unassigned | - |
| Otu57358 | 1 | k__Fungi(100);k__Fungi_unclassified(100);k__Fungi_unclassified(100);k__Fungi_unclassified(100);k__Fungi_unclassified(100);k__Fungi_unclassified(100);k__Fungi_unclassified(100); | - | - | - | - | - | - | - | Unassigned | - |
| Otu57359 | 1 | k__Fungi(100);k__Fungi_unclassified(100);k__Fungi_unclassified(100);k__Fungi_unclassified(100);k__Fungi_unclassified(100);k__Fungi_unclassified(100);k__Fungi_unclassified(100); | - | - | - | - | - | - | - | Unassigned | - |
| Otu57360 | 1 | k__Fungi(100);k__Fungi_unclassified(100);k__Fungi_unclassified(100);k__Fungi_unclassified(100);k__Fungi_unclassified(100);k__Fungi_unclassified(100);k__Fungi_unclassified(100); | - | - | - | - | - | - | - | Unassigned | - |
| Otu57361 | 1 | k__Fungi(100);p__Ascomycota(100);p__Ascomycota_unclassified(100);p__Ascomycota_unclassified(100);p__Ascomycota_unclassified(100);p__Ascomycota_unclassified(100);p__Ascomycota_unclassified(100); | - | - | - | - | - | - | - | Unassigned | - |
| Otu57387 | 1 | k__Fungi(100);k__Fungi_unclassified(100);k__Fungi_unclassified(100);k__Fungi_unclassified(100);k__Fungi_unclassified(100);k__Fungi_unclassified(100);k__Fungi_unclassified(100); | - | - | - | - | - | - | - | Unassigned | - |
| Otu57389 | 1 | k__Fungi(100);k__Fungi_unclassified(100);k__Fungi_unclassified(100);k__Fungi_unclassified(100);k__Fungi_unclassified(100);k__Fungi_unclassified(100);k__Fungi_unclassified(100); | - | - | - | - | - | - | - | Unassigned | - |
| Otu57411 | 1 | k__Fungi(100);p__Basidiomycota(100);p__Basidiomycota_unclassified(100);p__Basidiomycota_unclassified(100);p__Basidiomycota_unclassified(100);p__Basidiomycota_unclassified(100);p__Basidiomycota_unclassified(100); | - | - | - | - | - | - | - | Unassigned | - |
| Otu57415 | 1 | k__Fungi(100);k__Fungi_unclassified(100);k__Fungi_unclassified(100);k__Fungi_unclassified(100);k__Fungi_unclassified(100);k__Fungi_unclassified(100);k__Fungi_unclassified(100); | - | - | - | - | - | - | - | Unassigned | - |
| Otu57416 | 1 | k__Fungi(100);p__Ascomycota(100);p__Ascomycota_unclassified(100);p__Ascomycota_unclassified(100);p__Ascomycota_unclassified(100);p__Ascomycota_unclassified(100);p__Ascomycota_unclassified(100); | - | - | - | - | - | - | - | Unassigned | - |
| Otu57417 | 1 | k__Fungi(100);p__Ascomycota(100);p__Ascomycota_unclassified(100);p__Ascomycota_unclassified(100);p__Ascomycota_unclassified(100);p__Ascomycota_unclassified(100);p__Ascomycota_unclassified(100); | - | - | - | - | - | - | - | Unassigned | - |
| Otu57424 | 1 | k__Fungi(100);p__Ascomycota(100);p__Ascomycota_unclassified(100);p__Ascomycota_unclassified(100);p__Ascomycota_unclassified(100);p__Ascomycota_unclassified(100);p__Ascomycota_unclassified(100); | - | - | - | - | - | - | - | Unassigned | - |
| Otu57425 | 1 | k__Fungi(100);p__Ascomycota(100);p__Ascomycota_unclassified(100);p__Ascomycota_unclassified(100);p__Ascomycota_unclassified(100);p__Ascomycota_unclassified(100);p__Ascomycota_unclassified(100); | - | - | - | - | - | - | - | Unassigned | - |
| Otu57428 | 1 | k__Fungi(100);p__Ascomycota(100);p__Ascomycota_unclassified(100);p__Ascomycota_unclassified(100);p__Ascomycota_unclassified(100);p__Ascomycota_unclassified(100);p__Ascomycota_unclassified(100); | - | - | - | - | - | - | - | Unassigned | - |
| Otu57429 | 1 | k__Fungi(100);k__Fungi_unclassified(100);k__Fungi_unclassified(100);k__Fungi_unclassified(100);k__Fungi_unclassified(100);k__Fungi_unclassified(100);k__Fungi_unclassified(100); | - | - | - | - | - | - | - | Unassigned | - |
| Otu57430 | 1 | k__Fungi(100);k__Fungi_unclassified(100);k__Fungi_unclassified(100);k__Fungi_unclassified(100);k__Fungi_unclassified(100);k__Fungi_unclassified(100);k__Fungi_unclassified(100); | - | - | - | - | - | - | - | Unassigned | - |
| Otu57431 | 1 | k__Fungi(100);k__Fungi_unclassified(100);k__Fungi_unclassified(100);k__Fungi_unclassified(100);k__Fungi_unclassified(100);k__Fungi_unclassified(100);k__Fungi_unclassified(100); | - | - | - | - | - | - | - | Unassigned | - |
| Otu57432 | 1 | k__Fungi(100);k__Fungi_unclassified(100);k__Fungi_unclassified(100);k__Fungi_unclassified(100);k__Fungi_unclassified(100);k__Fungi_unclassified(100);k__Fungi_unclassified(100); | - | - | - | - | - | - | - | Unassigned | - |
| Otu57433 | 1 | k__Fungi(100);p__Ascomycota(100);p__Ascomycota_unclassified(100);p__Ascomycota_unclassified(100);p__Ascomycota_unclassified(100);p__Ascomycota_unclassified(100);p__Ascomycota_unclassified(100); | - | - | - | - | - | - | - | Unassigned | - |
| Otu57434 | 1 | k__Fungi(100);p__Ascomycota(100);p__Ascomycota_unclassified(100);p__Ascomycota_unclassified(100);p__Ascomycota_unclassified(100);p__Ascomycota_unclassified(100);p__Ascomycota_unclassified(100); | - | - | - | - | - | - | - | Unassigned | - |
| Otu57435 | 1 | k__Fungi(100);k__Fungi_unclassified(100);k__Fungi_unclassified(100);k__Fungi_unclassified(100);k__Fungi_unclassified(100);k__Fungi_unclassified(100);k__Fungi_unclassified(100); | - | - | - | - | - | - | - | Unassigned | - |
| Otu57437 | 1 | k__Fungi(100);p__Ascomycota(100);p__Ascomycota_unclassified(100);p__Ascomycota_unclassified(100);p__Ascomycota_unclassified(100);p__Ascomycota_unclassified(100);p__Ascomycota_unclassified(100); | - | - | - | - | - | - | - | Unassigned | - |
| Otu57442 | 1 | k__Fungi(100);p__Ascomycota(100);p__Ascomycota_unclassified(100);p__Ascomycota_unclassified(100);p__Ascomycota_unclassified(100);p__Ascomycota_unclassified(100);p__Ascomycota_unclassified(100); | - | - | - | - | - | - | - | Unassigned | - |
| Otu57443 | 1 | k__Fungi(100);p__Ascomycota(100);p__Ascomycota_unclassified(100);p__Ascomycota_unclassified(100);p__Ascomycota_unclassified(100);p__Ascomycota_unclassified(100);p__Ascomycota_unclassified(100); | - | - | - | - | - | - | - | Unassigned | - |
| Otu57448 | 1 | k__Fungi(100);p__Ascomycota(100);p__Ascomycota_unclassified(100);p__Ascomycota_unclassified(100);p__Ascomycota_unclassified(100);p__Ascomycota_unclassified(100);p__Ascomycota_unclassified(100); | - | - | - | - | - | - | - | Unassigned | - |
| Otu57450 | 1 | k__Fungi(100);k__Fungi_unclassified(100);k__Fungi_unclassified(100);k__Fungi_unclassified(100);k__Fungi_unclassified(100);k__Fungi_unclassified(100);k__Fungi_unclassified(100); | - | - | - | - | - | - | - | Unassigned | - |
| Otu57455 | 1 | k__Fungi(100);p__Ascomycota(100);p__Ascomycota_unclassified(100);p__Ascomycota_unclassified(100);p__Ascomycota_unclassified(100);p__Ascomycota_unclassified(100);p__Ascomycota_unclassified(100); | - | - | - | - | - | - | - | Unassigned | - |
| Otu57456 | 1 | k__Fungi(100);k__Fungi_unclassified(100);k__Fungi_unclassified(100);k__Fungi_unclassified(100);k__Fungi_unclassified(100);k__Fungi_unclassified(100);k__Fungi_unclassified(100); | - | - | - | - | - | - | - | Unassigned | - |
| Otu57457 | 1 | k__Fungi(100);k__Fungi_unclassified(100);k__Fungi_unclassified(100);k__Fungi_unclassified(100);k__Fungi_unclassified(100);k__Fungi_unclassified(100);k__Fungi_unclassified(100); | - | - | - | - | - | - | - | Unassigned | - |
| Otu57458 | 1 | k__Fungi(100);k__Fungi_unclassified(100);k__Fungi_unclassified(100);k__Fungi_unclassified(100);k__Fungi_unclassified(100);k__Fungi_unclassified(100);k__Fungi_unclassified(100); | - | - | - | - | - | - | - | Unassigned | - |
| Otu57459 | 1 | k__Fungi(100);p__Ascomycota(100);p__Ascomycota_unclassified(100);p__Ascomycota_unclassified(100);p__Ascomycota_unclassified(100);p__Ascomycota_unclassified(100);p__Ascomycota_unclassified(100); | - | - | - | - | - | - | - | Unassigned | - |
| Otu57460 | 1 | k__Fungi(100);k__Fungi_unclassified(100);k__Fungi_unclassified(100);k__Fungi_unclassified(100);k__Fungi_unclassified(100);k__Fungi_unclassified(100);k__Fungi_unclassified(100); | - | - | - | - | - | - | - | Unassigned | - |
| Otu57461 | 1 | k__Fungi(100);k__Fungi_unclassified(100);k__Fungi_unclassified(100);k__Fungi_unclassified(100);k__Fungi_unclassified(100);k__Fungi_unclassified(100);k__Fungi_unclassified(100); | - | - | - | - | - | - | - | Unassigned | - |
| Otu57462 | 1 | k__Fungi(100);k__Fungi_unclassified(100);k__Fungi_unclassified(100);k__Fungi_unclassified(100);k__Fungi_unclassified(100);k__Fungi_unclassified(100);k__Fungi_unclassified(100); | - | - | - | - | - | - | - | Unassigned | - |
| Otu57463 | 1 | k__Fungi(100);k__Fungi_unclassified(100);k__Fungi_unclassified(100);k__Fungi_unclassified(100);k__Fungi_unclassified(100);k__Fungi_unclassified(100);k__Fungi_unclassified(100); | - | - | - | - | - | - | - | Unassigned | - |
| Otu57465 | 1 | k__Fungi(100);p__Ascomycota(100);p__Ascomycota_unclassified(100);p__Ascomycota_unclassified(100);p__Ascomycota_unclassified(100);p__Ascomycota_unclassified(100);p__Ascomycota_unclassified(100); | - | - | - | - | - | - | - | Unassigned | - |
| Otu57468 | 1 | k__Fungi(100);k__Fungi_unclassified(100);k__Fungi_unclassified(100);k__Fungi_unclassified(100);k__Fungi_unclassified(100);k__Fungi_unclassified(100);k__Fungi_unclassified(100); | - | - | - | - | - | - | - | Unassigned | - |
| Otu57471 | 1 | k__Fungi(100);k__Fungi_unclassified(100);k__Fungi_unclassified(100);k__Fungi_unclassified(100);k__Fungi_unclassified(100);k__Fungi_unclassified(100);k__Fungi_unclassified(100); | - | - | - | - | - | - | - | Unassigned | - |
| Otu57475 | 1 | k__Fungi(100);p__Ascomycota(100);p__Ascomycota_unclassified(100);p__Ascomycota_unclassified(100);p__Ascomycota_unclassified(100);p__Ascomycota_unclassified(100);p__Ascomycota_unclassified(100); | - | - | - | - | - | - | - | Unassigned | - |
| Otu57481 | 1 | k__Fungi(100);k__Fungi_unclassified(100);k__Fungi_unclassified(100);k__Fungi_unclassified(100);k__Fungi_unclassified(100);k__Fungi_unclassified(100);k__Fungi_unclassified(100); | - | - | - | - | - | - | - | Unassigned | - |
| Otu57491 | 1 | k__Fungi(100);p__Ascomycota(100);p__Ascomycota_unclassified(100);p__Ascomycota_unclassified(100);p__Ascomycota_unclassified(100);p__Ascomycota_unclassified(100);p__Ascomycota_unclassified(100); | - | - | - | - | - | - | - | Unassigned | - |
| Otu57492 | 1 | k__Fungi(100);k__Fungi_unclassified(100);k__Fungi_unclassified(100);k__Fungi_unclassified(100);k__Fungi_unclassified(100);k__Fungi_unclassified(100);k__Fungi_unclassified(100); | - | - | - | - | - | - | - | Unassigned | - |
| Otu57505 | 1 | k__Fungi(100);p__Ascomycota(100);p__Ascomycota_unclassified(100);p__Ascomycota_unclassified(100);p__Ascomycota_unclassified(100);p__Ascomycota_unclassified(100);p__Ascomycota_unclassified(100); | - | - | - | - | - | - | - | Unassigned | - |
| Otu57510 | 1 | k__Fungi(100);k__Fungi_unclassified(100);k__Fungi_unclassified(100);k__Fungi_unclassified(100);k__Fungi_unclassified(100);k__Fungi_unclassified(100);k__Fungi_unclassified(100); | - | - | - | - | - | - | - | Unassigned | - |
| Otu57511 | 1 | k__Fungi(100);k__Fungi_unclassified(100);k__Fungi_unclassified(100);k__Fungi_unclassified(100);k__Fungi_unclassified(100);k__Fungi_unclassified(100);k__Fungi_unclassified(100); | - | - | - | - | - | - | - | Unassigned | - |
| Otu57513 | 1 | k__Fungi(100);k__Fungi_unclassified(100);k__Fungi_unclassified(100);k__Fungi_unclassified(100);k__Fungi_unclassified(100);k__Fungi_unclassified(100);k__Fungi_unclassified(100); | - | - | - | - | - | - | - | Unassigned | - |
| Otu57514 | 1 | k__Fungi(100);p__Ascomycota(100);p__Ascomycota_unclassified(100);p__Ascomycota_unclassified(100);p__Ascomycota_unclassified(100);p__Ascomycota_unclassified(100);p__Ascomycota_unclassified(100); | - | - | - | - | - | - | - | Unassigned | - |
| Otu57515 | 1 | k__Fungi(100);k__Fungi_unclassified(100);k__Fungi_unclassified(100);k__Fungi_unclassified(100);k__Fungi_unclassified(100);k__Fungi_unclassified(100);k__Fungi_unclassified(100); | - | - | - | - | - | - | - | Unassigned | - |
| Otu57517 | 1 | k__Fungi(100);k__Fungi_unclassified(100);k__Fungi_unclassified(100);k__Fungi_unclassified(100);k__Fungi_unclassified(100);k__Fungi_unclassified(100);k__Fungi_unclassified(100); | - | - | - | - | - | - | - | Unassigned | - |
| Otu57519 | 1 | k__Fungi(100);k__Fungi_unclassified(100);k__Fungi_unclassified(100);k__Fungi_unclassified(100);k__Fungi_unclassified(100);k__Fungi_unclassified(100);k__Fungi_unclassified(100); | - | - | - | - | - | - | - | Unassigned | - |
| Otu57520 | 1 | k__Fungi(100);k__Fungi_unclassified(100);k__Fungi_unclassified(100);k__Fungi_unclassified(100);k__Fungi_unclassified(100);k__Fungi_unclassified(100);k__Fungi_unclassified(100); | - | - | - | - | - | - | - | Unassigned | - |
| Otu57521 | 1 | k__Fungi(100);k__Fungi_unclassified(100);k__Fungi_unclassified(100);k__Fungi_unclassified(100);k__Fungi_unclassified(100);k__Fungi_unclassified(100);k__Fungi_unclassified(100); | - | - | - | - | - | - | - | Unassigned | - |
| Otu57523 | 1 | k__Fungi(100);k__Fungi_unclassified(100);k__Fungi_unclassified(100);k__Fungi_unclassified(100);k__Fungi_unclassified(100);k__Fungi_unclassified(100);k__Fungi_unclassified(100); | - | - | - | - | - | - | - | Unassigned | - |
| Otu57524 | 1 | k__Fungi(100);p__Ascomycota(100);p__Ascomycota_unclassified(100);p__Ascomycota_unclassified(100);p__Ascomycota_unclassified(100);p__Ascomycota_unclassified(100);p__Ascomycota_unclassified(100); | - | - | - | - | - | - | - | Unassigned | - |
| Otu57527 | 1 | k__Fungi(100);p__Ascomycota(100);p__Ascomycota_unclassified(100);p__Ascomycota_unclassified(100);p__Ascomycota_unclassified(100);p__Ascomycota_unclassified(100);p__Ascomycota_unclassified(100); | - | - | - | - | - | - | - | Unassigned | - |
| Otu57529 | 1 | k__Fungi(100);p__Ascomycota(100);p__Ascomycota_unclassified(100);p__Ascomycota_unclassified(100);p__Ascomycota_unclassified(100);p__Ascomycota_unclassified(100);p__Ascomycota_unclassified(100); | - | - | - | - | - | - | - | Unassigned | - |
| Otu57530 | 1 | k__Fungi(100);p__Ascomycota(100);p__Ascomycota_unclassified(100);p__Ascomycota_unclassified(100);p__Ascomycota_unclassified(100);p__Ascomycota_unclassified(100);p__Ascomycota_unclassified(100); | - | - | - | - | - | - | - | Unassigned | - |
| Otu57531 | 1 | k__Fungi(100);p__Ascomycota(100);p__Ascomycota_unclassified(100);p__Ascomycota_unclassified(100);p__Ascomycota_unclassified(100);p__Ascomycota_unclassified(100);p__Ascomycota_unclassified(100); | - | - | - | - | - | - | - | Unassigned | - |
| Otu57532 | 1 | k__Fungi(100);k__Fungi_unclassified(100);k__Fungi_unclassified(100);k__Fungi_unclassified(100);k__Fungi_unclassified(100);k__Fungi_unclassified(100);k__Fungi_unclassified(100); | - | - | - | - | - | - | - | Unassigned | - |
| Otu57533 | 1 | k__Fungi(100);p__Ascomycota(100);p__Ascomycota_unclassified(100);p__Ascomycota_unclassified(100);p__Ascomycota_unclassified(100);p__Ascomycota_unclassified(100);p__Ascomycota_unclassified(100); | - | - | - | - | - | - | - | Unassigned | - |
| Otu57534 | 1 | k__Fungi(100);p__Ascomycota(100);p__Ascomycota_unclassified(100);p__Ascomycota_unclassified(100);p__Ascomycota_unclassified(100);p__Ascomycota_unclassified(100);p__Ascomycota_unclassified(100); | - | - | - | - | - | - | - | Unassigned | - |
| Otu57535 | 1 | k__Fungi(100);p__Basidiomycota(100);p__Basidiomycota_unclassified(100);p__Basidiomycota_unclassified(100);p__Basidiomycota_unclassified(100);p__Basidiomycota_unclassified(100);p__Basidiomycota_unclassified(100); | - | - | - | - | - | - | - | Unassigned | - |
| Otu57540 | 1 | k__Fungi(100);p__Ascomycota(100);p__Ascomycota_unclassified(100);p__Ascomycota_unclassified(100);p__Ascomycota_unclassified(100);p__Ascomycota_unclassified(100);p__Ascomycota_unclassified(100); | - | - | - | - | - | - | - | Unassigned | - |
| Otu57543 | 1 | k__Fungi(100);p__Ascomycota(100);p__Ascomycota_unclassified(100);p__Ascomycota_unclassified(100);p__Ascomycota_unclassified(100);p__Ascomycota_unclassified(100);p__Ascomycota_unclassified(100); | - | - | - | - | - | - | - | Unassigned | - |
| Otu57544 | 1 | k__Fungi(100);k__Fungi_unclassified(100);k__Fungi_unclassified(100);k__Fungi_unclassified(100);k__Fungi_unclassified(100);k__Fungi_unclassified(100);k__Fungi_unclassified(100); | - | - | - | - | - | - | - | Unassigned | - |
| Otu57547 | 1 | k__Fungi(100);k__Fungi_unclassified(100);k__Fungi_unclassified(100);k__Fungi_unclassified(100);k__Fungi_unclassified(100);k__Fungi_unclassified(100);k__Fungi_unclassified(100); | - | - | - | - | - | - | - | Unassigned | - |
| Otu57553 | 1 | k__Fungi(100);p__Ascomycota(100);p__Ascomycota_unclassified(100);p__Ascomycota_unclassified(100);p__Ascomycota_unclassified(100);p__Ascomycota_unclassified(100);p__Ascomycota_unclassified(100); | - | - | - | - | - | - | - | Unassigned | - |
| Otu57554 | 1 | k__Fungi(100);p__Ascomycota(100);p__Ascomycota_unclassified(100);p__Ascomycota_unclassified(100);p__Ascomycota_unclassified(100);p__Ascomycota_unclassified(100);p__Ascomycota_unclassified(100); | - | - | - | - | - | - | - | Unassigned | - |
| Otu57555 | 1 | k__Fungi(100);p__Ascomycota(100);p__Ascomycota_unclassified(100);p__Ascomycota_unclassified(100);p__Ascomycota_unclassified(100);p__Ascomycota_unclassified(100);p__Ascomycota_unclassified(100); | - | - | - | - | - | - | - | Unassigned | - |
| Otu57557 | 1 | k__Fungi(100);k__Fungi_unclassified(100);k__Fungi_unclassified(100);k__Fungi_unclassified(100);k__Fungi_unclassified(100);k__Fungi_unclassified(100);k__Fungi_unclassified(100); | - | - | - | - | - | - | - | Unassigned | - |
| Otu57558 | 1 | k__Fungi(100);k__Fungi_unclassified(100);k__Fungi_unclassified(100);k__Fungi_unclassified(100);k__Fungi_unclassified(100);k__Fungi_unclassified(100);k__Fungi_unclassified(100); | - | - | - | - | - | - | - | Unassigned | - |
| Otu57560 | 1 | k__Fungi(100);k__Fungi_unclassified(100);k__Fungi_unclassified(100);k__Fungi_unclassified(100);k__Fungi_unclassified(100);k__Fungi_unclassified(100);k__Fungi_unclassified(100); | - | - | - | - | - | - | - | Unassigned | - |
| Otu57564 | 1 | k__Fungi(100);p__Ascomycota(100);p__Ascomycota_unclassified(100);p__Ascomycota_unclassified(100);p__Ascomycota_unclassified(100);p__Ascomycota_unclassified(100);p__Ascomycota_unclassified(100); | - | - | - | - | - | - | - | Unassigned | - |
| Otu57565 | 1 | k__Fungi(100);k__Fungi_unclassified(100);k__Fungi_unclassified(100);k__Fungi_unclassified(100);k__Fungi_unclassified(100);k__Fungi_unclassified(100);k__Fungi_unclassified(100); | - | - | - | - | - | - | - | Unassigned | - |
| Otu57566 | 1 | k__Fungi(100);k__Fungi_unclassified(100);k__Fungi_unclassified(100);k__Fungi_unclassified(100);k__Fungi_unclassified(100);k__Fungi_unclassified(100);k__Fungi_unclassified(100); | - | - | - | - | - | - | - | Unassigned | - |
| Otu57571 | 1 | k__Fungi(100);p__Basidiomycota(100);p__Basidiomycota_unclassified(100);p__Basidiomycota_unclassified(100);p__Basidiomycota_unclassified(100);p__Basidiomycota_unclassified(100);p__Basidiomycota_unclassified(100); | - | - | - | - | - | - | - | Unassigned | - |
| Otu57574 | 1 | k__Fungi(100);k__Fungi_unclassified(100);k__Fungi_unclassified(100);k__Fungi_unclassified(100);k__Fungi_unclassified(100);k__Fungi_unclassified(100);k__Fungi_unclassified(100); | - | - | - | - | - | - | - | Unassigned | - |
| Otu57584 | 1 | k__Fungi(100);p__Ascomycota(100);p__Ascomycota_unclassified(100);p__Ascomycota_unclassified(100);p__Ascomycota_unclassified(100);p__Ascomycota_unclassified(100);p__Ascomycota_unclassified(100); | - | - | - | - | - | - | - | Unassigned | - |
| Otu57585 | 1 | k__Fungi(100);p__Ascomycota(100);p__Ascomycota_unclassified(100);p__Ascomycota_unclassified(100);p__Ascomycota_unclassified(100);p__Ascomycota_unclassified(100);p__Ascomycota_unclassified(100); | - | - | - | - | - | - | - | Unassigned | - |
| Otu57586 | 1 | k__Fungi(100);p__Ascomycota(100);p__Ascomycota_unclassified(100);p__Ascomycota_unclassified(100);p__Ascomycota_unclassified(100);p__Ascomycota_unclassified(100);p__Ascomycota_unclassified(100); | - | - | - | - | - | - | - | Unassigned | - |
| Otu57587 | 1 | k__Fungi(100);p__Ascomycota(100);p__Ascomycota_unclassified(100);p__Ascomycota_unclassified(100);p__Ascomycota_unclassified(100);p__Ascomycota_unclassified(100);p__Ascomycota_unclassified(100); | - | - | - | - | - | - | - | Unassigned | - |
| Otu57589 | 1 | k__Fungi(100);k__Fungi_unclassified(100);k__Fungi_unclassified(100);k__Fungi_unclassified(100);k__Fungi_unclassified(100);k__Fungi_unclassified(100);k__Fungi_unclassified(100); | - | - | - | - | - | - | - | Unassigned | - |
| Otu57594 | 1 | k__Fungi(100);p__Ascomycota(100);p__Ascomycota_unclassified(100);p__Ascomycota_unclassified(100);p__Ascomycota_unclassified(100);p__Ascomycota_unclassified(100);p__Ascomycota_unclassified(100); | - | - | - | - | - | - | - | Unassigned | - |
| Otu57595 | 1 | k__Fungi(100);p__Ascomycota(100);p__Ascomycota_unclassified(100);p__Ascomycota_unclassified(100);p__Ascomycota_unclassified(100);p__Ascomycota_unclassified(100);p__Ascomycota_unclassified(100); | - | - | - | - | - | - | - | Unassigned | - |
| Otu57599 | 1 | k__Fungi(100);k__Fungi_unclassified(100);k__Fungi_unclassified(100);k__Fungi_unclassified(100);k__Fungi_unclassified(100);k__Fungi_unclassified(100);k__Fungi_unclassified(100); | - | - | - | - | - | - | - | Unassigned | - |
| Otu57600 | 1 | k__Fungi(100);k__Fungi_unclassified(100);k__Fungi_unclassified(100);k__Fungi_unclassified(100);k__Fungi_unclassified(100);k__Fungi_unclassified(100);k__Fungi_unclassified(100); | - | - | - | - | - | - | - | Unassigned | - |
| Otu57603 | 1 | k__Fungi(100);p__Ascomycota(100);p__Ascomycota_unclassified(100);p__Ascomycota_unclassified(100);p__Ascomycota_unclassified(100);p__Ascomycota_unclassified(100);p__Ascomycota_unclassified(100); | - | - | - | - | - | - | - | Unassigned | - |
| Otu57611 | 1 | k__Fungi(100);k__Fungi_unclassified(100);k__Fungi_unclassified(100);k__Fungi_unclassified(100);k__Fungi_unclassified(100);k__Fungi_unclassified(100);k__Fungi_unclassified(100); | - | - | - | - | - | - | - | Unassigned | - |
| Otu57615 | 1 | k__Fungi(100);k__Fungi_unclassified(100);k__Fungi_unclassified(100);k__Fungi_unclassified(100);k__Fungi_unclassified(100);k__Fungi_unclassified(100);k__Fungi_unclassified(100); | - | - | - | - | - | - | - | Unassigned | - |
| Otu57624 | 1 | k__Fungi(100);k__Fungi_unclassified(100);k__Fungi_unclassified(100);k__Fungi_unclassified(100);k__Fungi_unclassified(100);k__Fungi_unclassified(100);k__Fungi_unclassified(100); | - | - | - | - | - | - | - | Unassigned | - |
| Otu57627 | 1 | k__Fungi(100);k__Fungi_unclassified(100);k__Fungi_unclassified(100);k__Fungi_unclassified(100);k__Fungi_unclassified(100);k__Fungi_unclassified(100);k__Fungi_unclassified(100); | - | - | - | - | - | - | - | Unassigned | - |
| Otu57630 | 1 | k__Fungi(100);p__Ascomycota(100);p__Ascomycota_unclassified(100);p__Ascomycota_unclassified(100);p__Ascomycota_unclassified(100);p__Ascomycota_unclassified(100);p__Ascomycota_unclassified(100); | - | - | - | - | - | - | - | Unassigned | - |
| Otu57636 | 1 | k__Fungi(100);k__Fungi_unclassified(100);k__Fungi_unclassified(100);k__Fungi_unclassified(100);k__Fungi_unclassified(100);k__Fungi_unclassified(100);k__Fungi_unclassified(100); | - | - | - | - | - | - | - | Unassigned | - |
| Otu57637 | 1 | k__Fungi(100);k__Fungi_unclassified(100);k__Fungi_unclassified(100);k__Fungi_unclassified(100);k__Fungi_unclassified(100);k__Fungi_unclassified(100);k__Fungi_unclassified(100); | - | - | - | - | - | - | - | Unassigned | - |
| Otu57640 | 1 | k__Fungi(100);k__Fungi_unclassified(100);k__Fungi_unclassified(100);k__Fungi_unclassified(100);k__Fungi_unclassified(100);k__Fungi_unclassified(100);k__Fungi_unclassified(100); | - | - | - | - | - | - | - | Unassigned | - |
| Otu57641 | 1 | k__Fungi(100);k__Fungi_unclassified(100);k__Fungi_unclassified(100);k__Fungi_unclassified(100);k__Fungi_unclassified(100);k__Fungi_unclassified(100);k__Fungi_unclassified(100); | - | - | - | - | - | - | - | Unassigned | - |
| Otu57643 | 1 | k__Fungi(100);p__Ascomycota(100);p__Ascomycota_unclassified(100);p__Ascomycota_unclassified(100);p__Ascomycota_unclassified(100);p__Ascomycota_unclassified(100);p__Ascomycota_unclassified(100); | - | - | - | - | - | - | - | Unassigned | - |
| Otu57644 | 1 | k__Fungi(100);p__Ascomycota(100);p__Ascomycota_unclassified(100);p__Ascomycota_unclassified(100);p__Ascomycota_unclassified(100);p__Ascomycota_unclassified(100);p__Ascomycota_unclassified(100); | - | - | - | - | - | - | - | Unassigned | - |
| Otu57646 | 1 | k__Fungi(100);p__Ascomycota(100);p__Ascomycota_unclassified(100);p__Ascomycota_unclassified(100);p__Ascomycota_unclassified(100);p__Ascomycota_unclassified(100);p__Ascomycota_unclassified(100); | - | - | - | - | - | - | - | Unassigned | - |
| Otu57647 | 1 | k__Fungi(100);k__Fungi_unclassified(100);k__Fungi_unclassified(100);k__Fungi_unclassified(100);k__Fungi_unclassified(100);k__Fungi_unclassified(100);k__Fungi_unclassified(100); | - | - | - | - | - | - | - | Unassigned | - |
| Otu57648 | 1 | k__Fungi(100);p__Ascomycota(100);p__Ascomycota_unclassified(100);p__Ascomycota_unclassified(100);p__Ascomycota_unclassified(100);p__Ascomycota_unclassified(100);p__Ascomycota_unclassified(100); | - | - | - | - | - | - | - | Unassigned | - |
| Otu57655 | 1 | k__Fungi(100);p__Ascomycota(100);p__Ascomycota_unclassified(100);p__Ascomycota_unclassified(100);p__Ascomycota_unclassified(100);p__Ascomycota_unclassified(100);p__Ascomycota_unclassified(100); | - | - | - | - | - | - | - | Unassigned | - |
| Otu57656 | 1 | k__Fungi(100);k__Fungi_unclassified(100);k__Fungi_unclassified(100);k__Fungi_unclassified(100);k__Fungi_unclassified(100);k__Fungi_unclassified(100);k__Fungi_unclassified(100); | - | - | - | - | - | - | - | Unassigned | - |
| Otu57658 | 1 | k__Fungi(100);p__Ascomycota(100);p__Ascomycota_unclassified(100);p__Ascomycota_unclassified(100);p__Ascomycota_unclassified(100);p__Ascomycota_unclassified(100);p__Ascomycota_unclassified(100); | - | - | - | - | - | - | - | Unassigned | - |
| Otu57659 | 1 | k__Fungi(100);k__Fungi_unclassified(100);k__Fungi_unclassified(100);k__Fungi_unclassified(100);k__Fungi_unclassified(100);k__Fungi_unclassified(100);k__Fungi_unclassified(100); | - | - | - | - | - | - | - | Unassigned | - |
| Otu57660 | 1 | k__Fungi(100);k__Fungi_unclassified(100);k__Fungi_unclassified(100);k__Fungi_unclassified(100);k__Fungi_unclassified(100);k__Fungi_unclassified(100);k__Fungi_unclassified(100); | - | - | - | - | - | - | - | Unassigned | - |
| Otu57662 | 1 | k__Fungi(100);p__Ascomycota(100);p__Ascomycota_unclassified(100);p__Ascomycota_unclassified(100);p__Ascomycota_unclassified(100);p__Ascomycota_unclassified(100);p__Ascomycota_unclassified(100); | - | - | - | - | - | - | - | Unassigned | - |
| Otu57667 | 1 | k__Fungi(100);p__Ascomycota(100);p__Ascomycota_unclassified(100);p__Ascomycota_unclassified(100);p__Ascomycota_unclassified(100);p__Ascomycota_unclassified(100);p__Ascomycota_unclassified(100); | - | - | - | - | - | - | - | Unassigned | - |
| Otu57668 | 1 | k__Fungi(100);p__Ascomycota(100);p__Ascomycota_unclassified(100);p__Ascomycota_unclassified(100);p__Ascomycota_unclassified(100);p__Ascomycota_unclassified(100);p__Ascomycota_unclassified(100); | - | - | - | - | - | - | - | Unassigned | - |
| Otu57670 | 1 | k__Fungi(100);k__Fungi_unclassified(100);k__Fungi_unclassified(100);k__Fungi_unclassified(100);k__Fungi_unclassified(100);k__Fungi_unclassified(100);k__Fungi_unclassified(100); | - | - | - | - | - | - | - | Unassigned | - |
| Otu57674 | 1 | k__Fungi(100);p__Ascomycota(100);p__Ascomycota_unclassified(100);p__Ascomycota_unclassified(100);p__Ascomycota_unclassified(100);p__Ascomycota_unclassified(100);p__Ascomycota_unclassified(100); | - | - | - | - | - | - | - | Unassigned | - |
| Otu57675 | 1 | k__Fungi(100);k__Fungi_unclassified(100);k__Fungi_unclassified(100);k__Fungi_unclassified(100);k__Fungi_unclassified(100);k__Fungi_unclassified(100);k__Fungi_unclassified(100); | - | - | - | - | - | - | - | Unassigned | - |
| Otu57676 | 1 | k__Fungi(100);p__Ascomycota(100);p__Ascomycota_unclassified(100);p__Ascomycota_unclassified(100);p__Ascomycota_unclassified(100);p__Ascomycota_unclassified(100);p__Ascomycota_unclassified(100); | - | - | - | - | - | - | - | Unassigned | - |
| Otu57677 | 1 | k__Fungi(100);p__Ascomycota(100);p__Ascomycota_unclassified(100);p__Ascomycota_unclassified(100);p__Ascomycota_unclassified(100);p__Ascomycota_unclassified(100);p__Ascomycota_unclassified(100); | - | - | - | - | - | - | - | Unassigned | - |
| Otu57678 | 1 | k__Fungi(100);k__Fungi_unclassified(100);k__Fungi_unclassified(100);k__Fungi_unclassified(100);k__Fungi_unclassified(100);k__Fungi_unclassified(100);k__Fungi_unclassified(100); | - | - | - | - | - | - | - | Unassigned | - |
| Otu57679 | 1 | k__Fungi(100);k__Fungi_unclassified(100);k__Fungi_unclassified(100);k__Fungi_unclassified(100);k__Fungi_unclassified(100);k__Fungi_unclassified(100);k__Fungi_unclassified(100); | - | - | - | - | - | - | - | Unassigned | - |
| Otu57680 | 1 | k__Fungi(100);p__Ascomycota(100);p__Ascomycota_unclassified(100);p__Ascomycota_unclassified(100);p__Ascomycota_unclassified(100);p__Ascomycota_unclassified(100);p__Ascomycota_unclassified(100); | - | - | - | - | - | - | - | Unassigned | - |
| Otu57681 | 1 | k__Fungi(100);k__Fungi_unclassified(100);k__Fungi_unclassified(100);k__Fungi_unclassified(100);k__Fungi_unclassified(100);k__Fungi_unclassified(100);k__Fungi_unclassified(100); | - | - | - | - | - | - | - | Unassigned | - |
| Otu57682 | 1 | k__Fungi(100);k__Fungi_unclassified(100);k__Fungi_unclassified(100);k__Fungi_unclassified(100);k__Fungi_unclassified(100);k__Fungi_unclassified(100);k__Fungi_unclassified(100); | - | - | - | - | - | - | - | Unassigned | - |
| Otu57683 | 1 | k__Fungi(100);k__Fungi_unclassified(100);k__Fungi_unclassified(100);k__Fungi_unclassified(100);k__Fungi_unclassified(100);k__Fungi_unclassified(100);k__Fungi_unclassified(100); | - | - | - | - | - | - | - | Unassigned | - |
| Otu57684 | 1 | k__Fungi(100);k__Fungi_unclassified(100);k__Fungi_unclassified(100);k__Fungi_unclassified(100);k__Fungi_unclassified(100);k__Fungi_unclassified(100);k__Fungi_unclassified(100); | - | - | - | - | - | - | - | Unassigned | - |
| Otu57685 | 1 | k__Fungi(100);p__Ascomycota(100);p__Ascomycota_unclassified(100);p__Ascomycota_unclassified(100);p__Ascomycota_unclassified(100);p__Ascomycota_unclassified(100);p__Ascomycota_unclassified(100); | - | - | - | - | - | - | - | Unassigned | - |
| Otu57687 | 1 | k__Fungi(100);k__Fungi_unclassified(100);k__Fungi_unclassified(100);k__Fungi_unclassified(100);k__Fungi_unclassified(100);k__Fungi_unclassified(100);k__Fungi_unclassified(100); | - | - | - | - | - | - | - | Unassigned | - |
| Otu57689 | 1 | k__Fungi(100);k__Fungi_unclassified(100);k__Fungi_unclassified(100);k__Fungi_unclassified(100);k__Fungi_unclassified(100);k__Fungi_unclassified(100);k__Fungi_unclassified(100); | - | - | - | - | - | - | - | Unassigned | - |
| Otu57690 | 1 | k__Fungi(100);k__Fungi_unclassified(100);k__Fungi_unclassified(100);k__Fungi_unclassified(100);k__Fungi_unclassified(100);k__Fungi_unclassified(100);k__Fungi_unclassified(100); | - | - | - | - | - | - | - | Unassigned | - |
| Otu57692 | 1 | k__Fungi(100);p__Ascomycota(100);p__Ascomycota_unclassified(100);p__Ascomycota_unclassified(100);p__Ascomycota_unclassified(100);p__Ascomycota_unclassified(100);p__Ascomycota_unclassified(100); | - | - | - | - | - | - | - | Unassigned | - |
| Otu57694 | 1 | k__Fungi(100);p__Ascomycota(100);p__Ascomycota_unclassified(100);p__Ascomycota_unclassified(100);p__Ascomycota_unclassified(100);p__Ascomycota_unclassified(100);p__Ascomycota_unclassified(100); | - | - | - | - | - | - | - | Unassigned | - |
| Otu57698 | 1 | k__Fungi(100);k__Fungi_unclassified(100);k__Fungi_unclassified(100);k__Fungi_unclassified(100);k__Fungi_unclassified(100);k__Fungi_unclassified(100);k__Fungi_unclassified(100); | - | - | - | - | - | - | - | Unassigned | - |
| Otu57703 | 1 | k__Fungi(100);k__Fungi_unclassified(100);k__Fungi_unclassified(100);k__Fungi_unclassified(100);k__Fungi_unclassified(100);k__Fungi_unclassified(100);k__Fungi_unclassified(100); | - | - | - | - | - | - | - | Unassigned | - |
| Otu57704 | 1 | k__Fungi(100);p__Ascomycota(100);p__Ascomycota_unclassified(100);p__Ascomycota_unclassified(100);p__Ascomycota_unclassified(100);p__Ascomycota_unclassified(100);p__Ascomycota_unclassified(100); | - | - | - | - | - | - | - | Unassigned | - |
| Otu57705 | 1 | k__Fungi(100);p__Ascomycota(100);p__Ascomycota_unclassified(100);p__Ascomycota_unclassified(100);p__Ascomycota_unclassified(100);p__Ascomycota_unclassified(100);p__Ascomycota_unclassified(100); | - | - | - | - | - | - | - | Unassigned | - |
| Otu57706 | 1 | k__Fungi(100);p__Ascomycota(100);p__Ascomycota_unclassified(100);p__Ascomycota_unclassified(100);p__Ascomycota_unclassified(100);p__Ascomycota_unclassified(100);p__Ascomycota_unclassified(100); | - | - | - | - | - | - | - | Unassigned | - |
| Otu57715 | 1 | k__Fungi(100);k__Fungi_unclassified(100);k__Fungi_unclassified(100);k__Fungi_unclassified(100);k__Fungi_unclassified(100);k__Fungi_unclassified(100);k__Fungi_unclassified(100); | - | - | - | - | - | - | - | Unassigned | - |
| Otu57717 | 1 | k__Fungi(100);k__Fungi_unclassified(100);k__Fungi_unclassified(100);k__Fungi_unclassified(100);k__Fungi_unclassified(100);k__Fungi_unclassified(100);k__Fungi_unclassified(100); | - | - | - | - | - | - | - | Unassigned | - |
| Otu57718 | 1 | k__Fungi(100);p__Ascomycota(100);p__Ascomycota_unclassified(100);p__Ascomycota_unclassified(100);p__Ascomycota_unclassified(100);p__Ascomycota_unclassified(100);p__Ascomycota_unclassified(100); | - | - | - | - | - | - | - | Unassigned | - |
| Otu57719 | 1 | k__Fungi(100);k__Fungi_unclassified(100);k__Fungi_unclassified(100);k__Fungi_unclassified(100);k__Fungi_unclassified(100);k__Fungi_unclassified(100);k__Fungi_unclassified(100); | - | - | - | - | - | - | - | Unassigned | - |
| Otu57720 | 1 | k__Fungi(100);k__Fungi_unclassified(100);k__Fungi_unclassified(100);k__Fungi_unclassified(100);k__Fungi_unclassified(100);k__Fungi_unclassified(100);k__Fungi_unclassified(100); | - | - | - | - | - | - | - | Unassigned | - |
| Otu57722 | 1 | k__Fungi(100);p__Ascomycota(100);p__Ascomycota_unclassified(100);p__Ascomycota_unclassified(100);p__Ascomycota_unclassified(100);p__Ascomycota_unclassified(100);p__Ascomycota_unclassified(100); | - | - | - | - | - | - | - | Unassigned | - |
| Otu57723 | 1 | k__Fungi(100);k__Fungi_unclassified(100);k__Fungi_unclassified(100);k__Fungi_unclassified(100);k__Fungi_unclassified(100);k__Fungi_unclassified(100);k__Fungi_unclassified(100); | - | - | - | - | - | - | - | Unassigned | - |
| Otu57724 | 1 | k__Fungi(100);p__Ascomycota(100);p__Ascomycota_unclassified(100);p__Ascomycota_unclassified(100);p__Ascomycota_unclassified(100);p__Ascomycota_unclassified(100);p__Ascomycota_unclassified(100); | - | - | - | - | - | - | - | Unassigned | - |
| Otu57725 | 1 | k__Fungi(100);p__Ascomycota(100);p__Ascomycota_unclassified(100);p__Ascomycota_unclassified(100);p__Ascomycota_unclassified(100);p__Ascomycota_unclassified(100);p__Ascomycota_unclassified(100); | - | - | - | - | - | - | - | Unassigned | - |
| Otu57726 | 1 | k__Fungi(100);p__Ascomycota(100);p__Ascomycota_unclassified(100);p__Ascomycota_unclassified(100);p__Ascomycota_unclassified(100);p__Ascomycota_unclassified(100);p__Ascomycota_unclassified(100); | - | - | - | - | - | - | - | Unassigned | - |
| Otu57727 | 1 | k__Fungi(100);k__Fungi_unclassified(100);k__Fungi_unclassified(100);k__Fungi_unclassified(100);k__Fungi_unclassified(100);k__Fungi_unclassified(100);k__Fungi_unclassified(100); | - | - | - | - | - | - | - | Unassigned | - |
| Otu57728 | 1 | k__Fungi(100);k__Fungi_unclassified(100);k__Fungi_unclassified(100);k__Fungi_unclassified(100);k__Fungi_unclassified(100);k__Fungi_unclassified(100);k__Fungi_unclassified(100); | - | - | - | - | - | - | - | Unassigned | - |
| Otu57729 | 1 | k__Fungi(100);p__Ascomycota(100);p__Ascomycota_unclassified(100);p__Ascomycota_unclassified(100);p__Ascomycota_unclassified(100);p__Ascomycota_unclassified(100);p__Ascomycota_unclassified(100); | - | - | - | - | - | - | - | Unassigned | - |
| Otu57730 | 1 | k__Fungi(100);k__Fungi_unclassified(100);k__Fungi_unclassified(100);k__Fungi_unclassified(100);k__Fungi_unclassified(100);k__Fungi_unclassified(100);k__Fungi_unclassified(100); | - | - | - | - | - | - | - | Unassigned | - |
| Otu57732 | 1 | k__Fungi(100);k__Fungi_unclassified(100);k__Fungi_unclassified(100);k__Fungi_unclassified(100);k__Fungi_unclassified(100);k__Fungi_unclassified(100);k__Fungi_unclassified(100); | - | - | - | - | - | - | - | Unassigned | - |
| Otu57733 | 1 | k__Fungi(100);k__Fungi_unclassified(100);k__Fungi_unclassified(100);k__Fungi_unclassified(100);k__Fungi_unclassified(100);k__Fungi_unclassified(100);k__Fungi_unclassified(100); | - | - | - | - | - | - | - | Unassigned | - |
| Otu57734 | 1 | k__Fungi(100);k__Fungi_unclassified(100);k__Fungi_unclassified(100);k__Fungi_unclassified(100);k__Fungi_unclassified(100);k__Fungi_unclassified(100);k__Fungi_unclassified(100); | - | - | - | - | - | - | - | Unassigned | - |
| Otu57736 | 1 | k__Fungi(100);k__Fungi_unclassified(100);k__Fungi_unclassified(100);k__Fungi_unclassified(100);k__Fungi_unclassified(100);k__Fungi_unclassified(100);k__Fungi_unclassified(100); | - | - | - | - | - | - | - | Unassigned | - |
| Otu57738 | 1 | k__Fungi(100);k__Fungi_unclassified(100);k__Fungi_unclassified(100);k__Fungi_unclassified(100);k__Fungi_unclassified(100);k__Fungi_unclassified(100);k__Fungi_unclassified(100); | - | - | - | - | - | - | - | Unassigned | - |
| Otu57742 | 1 | k__Fungi(100);k__Fungi_unclassified(100);k__Fungi_unclassified(100);k__Fungi_unclassified(100);k__Fungi_unclassified(100);k__Fungi_unclassified(100);k__Fungi_unclassified(100); | - | - | - | - | - | - | - | Unassigned | - |
| Otu57743 | 1 | k__Fungi(100);k__Fungi_unclassified(100);k__Fungi_unclassified(100);k__Fungi_unclassified(100);k__Fungi_unclassified(100);k__Fungi_unclassified(100);k__Fungi_unclassified(100); | - | - | - | - | - | - | - | Unassigned | - |
| Otu57744 | 1 | k__Fungi(100);p__Ascomycota(100);p__Ascomycota_unclassified(100);p__Ascomycota_unclassified(100);p__Ascomycota_unclassified(100);p__Ascomycota_unclassified(100);p__Ascomycota_unclassified(100); | - | - | - | - | - | - | - | Unassigned | - |
| Otu57745 | 1 | k__Fungi(100);p__Ascomycota(100);p__Ascomycota_unclassified(100);p__Ascomycota_unclassified(100);p__Ascomycota_unclassified(100);p__Ascomycota_unclassified(100);p__Ascomycota_unclassified(100); | - | - | - | - | - | - | - | Unassigned | - |
| Otu57746 | 1 | k__Fungi(100);k__Fungi_unclassified(100);k__Fungi_unclassified(100);k__Fungi_unclassified(100);k__Fungi_unclassified(100);k__Fungi_unclassified(100);k__Fungi_unclassified(100); | - | - | - | - | - | - | - | Unassigned | - |
| Otu57752 | 1 | k__Fungi(100);k__Fungi_unclassified(100);k__Fungi_unclassified(100);k__Fungi_unclassified(100);k__Fungi_unclassified(100);k__Fungi_unclassified(100);k__Fungi_unclassified(100); | - | - | - | - | - | - | - | Unassigned | - |

| | | | | | | | | | | Unassigned | |
|---|---|---|---|---|---|---|---|---|---|---|---|
| Otu57754 | 1 | k__Fungi(100);p__Ascomycota(100);p__Ascomycota_unclassified(100);p__Ascomycota_unclassified(100);p__Ascomycota_unclassified(100);p__Ascomycota_unclassified(100);p__Ascomycota_unclassified(100); | - | - | - | - | - | - | - | Unassigned | - |
| Otu57757 | 1 | k__Fungi(100);p__Ascomycota(100);p__Ascomycota_unclassified(100);p__Ascomycota_unclassified(100);p__Ascomycota_unclassified(100);p__Ascomycota_unclassified(100);p__Ascomycota_unclassified(100); | - | - | - | - | - | - | - | Unassigned | - |
| Otu57764 | 1 | k__Fungi(100);p__Ascomycota(100);p__Ascomycota_unclassified(100);p__Ascomycota_unclassified(100);p__Ascomycota_unclassified(100);p__Ascomycota_unclassified(100);p__Ascomycota_unclassified(100); | - | - | - | - | - | - | - | Unassigned | - |
| Otu57767 | 1 | k__Fungi(100);k__Fungi_unclassified(100);k__Fungi_unclassified(100);k__Fungi_unclassified(100);k__Fungi_unclassified(100);k__Fungi_unclassified(100);k__Fungi_unclassified(100); | - | - | - | - | - | - | - | Unassigned | - |
| Otu57768 | 1 | k__Fungi(100);p__Ascomycota(100);p__Ascomycota_unclassified(100);p__Ascomycota_unclassified(100);p__Ascomycota_unclassified(100);p__Ascomycota_unclassified(100);p__Ascomycota_unclassified(100); | - | - | - | - | - | - | - | Unassigned | - |
| Otu57770 | 1 | k__Fungi(100);k__Fungi_unclassified(100);k__Fungi_unclassified(100);k__Fungi_unclassified(100);k__Fungi_unclassified(100);k__Fungi_unclassified(100);k__Fungi_unclassified(100); | - | - | - | - | - | - | - | Unassigned | - |
| Otu57772 | 1 | k__Fungi(100);k__Fungi_unclassified(100);k__Fungi_unclassified(100);k__Fungi_unclassified(100);k__Fungi_unclassified(100);k__Fungi_unclassified(100);k__Fungi_unclassified(100); | - | - | - | - | - | - | - | Unassigned | - |
| Otu57774 | 1 | k__Fungi(100);k__Fungi_unclassified(100);k__Fungi_unclassified(100);k__Fungi_unclassified(100);k__Fungi_unclassified(100);k__Fungi_unclassified(100);k__Fungi_unclassified(100); | - | - | - | - | - | - | - | Unassigned | - |
| Otu57775 | 1 | k__Fungi(100);k__Fungi_unclassified(100);k__Fungi_unclassified(100);k__Fungi_unclassified(100);k__Fungi_unclassified(100);k__Fungi_unclassified(100);k__Fungi_unclassified(100); | - | - | - | - | - | - | - | Unassigned | - |
| Otu57776 | 1 | k__Fungi(100);p__Ascomycota(100);p__Ascomycota_unclassified(100);p__Ascomycota_unclassified(100);p__Ascomycota_unclassified(100);p__Ascomycota_unclassified(100);p__Ascomycota_unclassified(100); | - | - | - | - | - | - | - | Unassigned | - |
| Otu57777 | 1 | k__Fungi(100);k__Fungi_unclassified(100);k__Fungi_unclassified(100);k__Fungi_unclassified(100);k__Fungi_unclassified(100);k__Fungi_unclassified(100);k__Fungi_unclassified(100); | - | - | - | - | - | - | - | Unassigned | - |
| Otu57778 | 1 | k__Fungi(100);k__Fungi_unclassified(100);k__Fungi_unclassified(100);k__Fungi_unclassified(100);k__Fungi_unclassified(100);k__Fungi_unclassified(100);k__Fungi_unclassified(100); | - | - | - | - | - | - | - | Unassigned | - |
| Otu57779 | 1 | k__Fungi(100);k__Fungi_unclassified(100);k__Fungi_unclassified(100);k__Fungi_unclassified(100);k__Fungi_unclassified(100);k__Fungi_unclassified(100);k__Fungi_unclassified(100); | - | - | - | - | - | - | - | Unassigned | - |
| Otu57780 | 1 | k__Fungi(100);k__Fungi_unclassified(100);k__Fungi_unclassified(100);k__Fungi_unclassified(100);k__Fungi_unclassified(100);k__Fungi_unclassified(100);k__Fungi_unclassified(100); | - | - | - | - | - | - | - | Unassigned | - |
| Otu57782 | 1 | k__Fungi(100);k__Fungi_unclassified(100);k__Fungi_unclassified(100);k__Fungi_unclassified(100);k__Fungi_unclassified(100);k__Fungi_unclassified(100);k__Fungi_unclassified(100); | - | - | - | - | - | - | - | Unassigned | - |
| Otu57784 | 1 | k__Fungi(100);k__Fungi_unclassified(100);k__Fungi_unclassified(100);k__Fungi_unclassified(100);k__Fungi_unclassified(100);k__Fungi_unclassified(100);k__Fungi_unclassified(100); | - | - | - | - | - | - | - | Unassigned | - |
| Otu57785 | 1 | k__Fungi(100);k__Fungi_unclassified(100);k__Fungi_unclassified(100);k__Fungi_unclassified(100);k__Fungi_unclassified(100);k__Fungi_unclassified(100);k__Fungi_unclassified(100); | - | - | - | - | - | - | - | Unassigned | - |
| Otu57786 | 1 | k__Fungi(100);k__Fungi_unclassified(100);k__Fungi_unclassified(100);k__Fungi_unclassified(100);k__Fungi_unclassified(100);k__Fungi_unclassified(100);k__Fungi_unclassified(100); | - | - | - | - | - | - | - | Unassigned | - |
| Otu57787 | 1 | k__Fungi(100);k__Fungi_unclassified(100);k__Fungi_unclassified(100);k__Fungi_unclassified(100);k__Fungi_unclassified(100);k__Fungi_unclassified(100);k__Fungi_unclassified(100); | - | - | - | - | - | - | - | Unassigned | - |
| Otu57789 | 1 | k__Fungi(100);k__Fungi_unclassified(100);k__Fungi_unclassified(100);k__Fungi_unclassified(100);k__Fungi_unclassified(100);k__Fungi_unclassified(100);k__Fungi_unclassified(100); | - | - | - | - | - | - | - | Unassigned | - |
| Otu57790 | 1 | k__Fungi(100);p__Ascomycota(100);p__Ascomycota_unclassified(100);p__Ascomycota_unclassified(100);p__Ascomycota_unclassified(100);p__Ascomycota_unclassified(100);p__Ascomycota_unclassified(100); | - | - | - | - | - | - | - | Unassigned | - |
| Otu57791 | 1 | k__Fungi(100);k__Fungi_unclassified(100);k__Fungi_unclassified(100);k__Fungi_unclassified(100);k__Fungi_unclassified(100);k__Fungi_unclassified(100);k__Fungi_unclassified(100); | - | - | - | - | - | - | - | Unassigned | - |
| Otu57792 | 1 | k__Fungi(100);p__Ascomycota(100);p__Ascomycota_unclassified(100);p__Ascomycota_unclassified(100);p__Ascomycota_unclassified(100);p__Ascomycota_unclassified(100);p__Ascomycota_unclassified(100); | - | - | - | - | - | - | - | Unassigned | - |
| Otu57793 | 1 | k__Fungi(100);p__Ascomycota(100);p__Ascomycota_unclassified(100);p__Ascomycota_unclassified(100);p__Ascomycota_unclassified(100);p__Ascomycota_unclassified(100);p__Ascomycota_unclassified(100); | - | - | - | - | - | - | - | Unassigned | - |
| Otu57794 | 1 | k__Fungi(100);p__Ascomycota(100);p__Ascomycota_unclassified(100);p__Ascomycota_unclassified(100);p__Ascomycota_unclassified(100);p__Ascomycota_unclassified(100);p__Ascomycota_unclassified(100); | - | - | - | - | - | - | - | Unassigned | - |
| Otu57796 | 1 | k__Fungi(100);p__Ascomycota(100);p__Ascomycota_unclassified(100);p__Ascomycota_unclassified(100);p__Ascomycota_unclassified(100);p__Ascomycota_unclassified(100);p__Ascomycota_unclassified(100); | - | - | - | - | - | - | - | Unassigned | - |
| Otu57799 | 1 | k__Fungi(100);k__Fungi_unclassified(100);k__Fungi_unclassified(100);k__Fungi_unclassified(100);k__Fungi_unclassified(100);k__Fungi_unclassified(100);k__Fungi_unclassified(100); | - | - | - | - | - | - | - | Unassigned | - |
| Otu57804 | 1 | k__Fungi(100);p__Ascomycota(100);p__Ascomycota_unclassified(100);p__Ascomycota_unclassified(100);p__Ascomycota_unclassified(100);p__Ascomycota_unclassified(100);p__Ascomycota_unclassified(100); | - | - | - | - | - | - | - | Unassigned | - |
| Otu57808 | 1 | k__Fungi(100);p__Ascomycota(100);p__Ascomycota_unclassified(100);p__Ascomycota_unclassified(100);p__Ascomycota_unclassified(100);p__Ascomycota_unclassified(100);p__Ascomycota_unclassified(100); | - | - | - | - | - | - | - | Unassigned | - |
| Otu57809 | 1 | k__Fungi(100);p__Ascomycota(100);p__Ascomycota_unclassified(100);p__Ascomycota_unclassified(100);p__Ascomycota_unclassified(100);p__Ascomycota_unclassified(100);p__Ascomycota_unclassified(100); | - | - | - | - | - | - | - | Unassigned | - |
| Otu57810 | 1 | k__Fungi(100);p__Ascomycota(100);p__Ascomycota_unclassified(100);p__Ascomycota_unclassified(100);p__Ascomycota_unclassified(100);p__Ascomycota_unclassified(100);p__Ascomycota_unclassified(100); | - | - | - | - | - | - | - | Unassigned | - |
| Otu57812 | 1 | k__Fungi(100);p__Ascomycota(100);c__Leotiomycetes(100);c__Leotiomycetes_unclassified(100);c__Leotiomycetes_unclassified(100);c__Leotiomycetes_unclassified(100);c__Leotiomycetes_unclassified(100); | - | - | - | - | - | - | - | Unassigned | - |
| Otu57815 | 1 | k__Fungi(100);k__Fungi_unclassified(100);k__Fungi_unclassified(100);k__Fungi_unclassified(100);k__Fungi_unclassified(100);k__Fungi_unclassified(100);k__Fungi_unclassified(100); | - | - | - | - | - | - | - | Unassigned | - |
| Otu57816 | 1 | k__Fungi(100);p__Ascomycota(100);p__Ascomycota_unclassified(100);p__Ascomycota_unclassified(100);p__Ascomycota_unclassified(100);p__Ascomycota_unclassified(100);p__Ascomycota_unclassified(100); | - | - | - | - | - | - | - | Unassigned | - |
| Otu57818 | 1 | k__Fungi(100);k__Fungi_unclassified(100);k__Fungi_unclassified(100);k__Fungi_unclassified(100);k__Fungi_unclassified(100);k__Fungi_unclassified(100);k__Fungi_unclassified(100); | - | - | - | - | - | - | - | Unassigned | - |
| Otu57819 | 1 | k__Fungi(100);p__Ascomycota(100);p__Ascomycota_unclassified(100);p__Ascomycota_unclassified(100);p__Ascomycota_unclassified(100);p__Ascomycota_unclassified(100);p__Ascomycota_unclassified(100); | - | - | - | - | - | - | - | Unassigned | - |
| Otu57820 | 1 | k__Fungi(100);k__Fungi_unclassified(100);k__Fungi_unclassified(100);k__Fungi_unclassified(100);k__Fungi_unclassified(100);k__Fungi_unclassified(100);k__Fungi_unclassified(100); | - | - | - | - | - | - | - | Unassigned | - |
| Otu57822 | 1 | k__Fungi(100);p__Ascomycota(100);p__Ascomycota_unclassified(100);p__Ascomycota_unclassified(100);p__Ascomycota_unclassified(100);p__Ascomycota_unclassified(100);p__Ascomycota_unclassified(100); | - | - | - | - | - | - | - | Unassigned | - |
| Otu57823 | 1 | k__Fungi(100);p__Ascomycota(100);p__Ascomycota_unclassified(100);p__Ascomycota_unclassified(100);p__Ascomycota_unclassified(100);p__Ascomycota_unclassified(100);p__Ascomycota_unclassified(100); | - | - | - | - | - | - | - | Unassigned | - |
| Otu57825 | 1 | k__Fungi(100);k__Fungi_unclassified(100);k__Fungi_unclassified(100);k__Fungi_unclassified(100);k__Fungi_unclassified(100);k__Fungi_unclassified(100);k__Fungi_unclassified(100); | - | - | - | - | - | - | - | Unassigned | - |
| Otu57833 | 1 | k__Fungi(100);p__Ascomycota(100);p__Ascomycota_unclassified(100);p__Ascomycota_unclassified(100);p__Ascomycota_unclassified(100);p__Ascomycota_unclassified(100);p__Ascomycota_unclassified(100); | - | - | - | - | - | - | - | Unassigned | - |
| Otu57835 | 1 | k__Fungi(100);k__Fungi_unclassified(100);k__Fungi_unclassified(100);k__Fungi_unclassified(100);k__Fungi_unclassified(100);k__Fungi_unclassified(100);k__Fungi_unclassified(100); | - | - | - | - | - | - | - | Unassigned | - |
| Otu57836 | 1 | k__Fungi(100);k__Fungi_unclassified(100);k__Fungi_unclassified(100);k__Fungi_unclassified(100);k__Fungi_unclassified(100);k__Fungi_unclassified(100);k__Fungi_unclassified(100); | - | - | - | - | - | - | - | Unassigned | - |
| Otu57838 | 1 | k__Fungi(100);k__Fungi_unclassified(100);k__Fungi_unclassified(100);k__Fungi_unclassified(100);k__Fungi_unclassified(100);k__Fungi_unclassified(100);k__Fungi_unclassified(100); | - | - | - | - | - | - | - | Unassigned | - |
| Otu57851 | 1 | k__Fungi(100);p__Ascomycota(100);p__Ascomycota_unclassified(100);p__Ascomycota_unclassified(100);p__Ascomycota_unclassified(100);p__Ascomycota_unclassified(100);p__Ascomycota_unclassified(100); | - | - | - | - | - | - | - | Unassigned | - |
| Otu57852 | 1 | k__Fungi(100);k__Fungi_unclassified(100);k__Fungi_unclassified(100);k__Fungi_unclassified(100);k__Fungi_unclassified(100);k__Fungi_unclassified(100);k__Fungi_unclassified(100); | - | - | - | - | - | - | - | Unassigned | - |
| Otu57855 | 1 | k__Fungi(100);k__Fungi_unclassified(100);k__Fungi_unclassified(100);k__Fungi_unclassified(100);k__Fungi_unclassified(100);k__Fungi_unclassified(100);k__Fungi_unclassified(100); | - | - | - | - | - | - | - | Unassigned | - |
| Otu57865 | 1 | k__Fungi(100);k__Fungi_unclassified(100);k__Fungi_unclassified(100);k__Fungi_unclassified(100);k__Fungi_unclassified(100);k__Fungi_unclassified(100);k__Fungi_unclassified(100); | - | - | - | - | - | - | - | Unassigned | - |
| Otu57867 | 1 | k__Fungi(100);k__Fungi_unclassified(100);k__Fungi_unclassified(100);k__Fungi_unclassified(100);k__Fungi_unclassified(100);k__Fungi_unclassified(100);k__Fungi_unclassified(100); | - | - | - | - | - | - | - | Unassigned | - |
| Otu57873 | 1 | k__Fungi(100);k__Fungi_unclassified(100);k__Fungi_unclassified(100);k__Fungi_unclassified(100);k__Fungi_unclassified(100);k__Fungi_unclassified(100);k__Fungi_unclassified(100); | - | - | - | - | - | - | - | Unassigned | - |
| Otu57875 | 1 | k__Fungi(100);p__Ascomycota(100);p__Ascomycota_unclassified(100);p__Ascomycota_unclassified(100);p__Ascomycota_unclassified(100);p__Ascomycota_unclassified(100);p__Ascomycota_unclassified(100); | - | - | - | - | - | - | - | Unassigned | - |
| Otu57876 | 1 | k__Fungi(100);k__Fungi_unclassified(100);k__Fungi_unclassified(100);k__Fungi_unclassified(100);k__Fungi_unclassified(100);k__Fungi_unclassified(100);k__Fungi_unclassified(100); | - | - | - | - | - | - | - | Unassigned | - |
| Otu57877 | 1 | k__Fungi(100);k__Fungi_unclassified(100);k__Fungi_unclassified(100);k__Fungi_unclassified(100);k__Fungi_unclassified(100);k__Fungi_unclassified(100);k__Fungi_unclassified(100); | - | - | - | - | - | - | - | Unassigned | - |
| Otu57878 | 1 | k__Fungi(100);p__Ascomycota(100);p__Ascomycota_unclassified(100);p__Ascomycota_unclassified(100);p__Ascomycota_unclassified(100);p__Ascomycota_unclassified(100);p__Ascomycota_unclassified(100); | - | - | - | - | - | - | - | Unassigned | - |
| Otu57882 | 1 | k__Fungi(100);k__Fungi_unclassified(100);k__Fungi_unclassified(100);k__Fungi_unclassified(100);k__Fungi_unclassified(100);k__Fungi_unclassified(100);k__Fungi_unclassified(100); | - | - | - | - | - | - | - | Unassigned | - |
| Otu57884 | 1 | k__Fungi(100);k__Fungi_unclassified(100);k__Fungi_unclassified(100);k__Fungi_unclassified(100);k__Fungi_unclassified(100);k__Fungi_unclassified(100);k__Fungi_unclassified(100); | - | - | - | - | - | - | - | Unassigned | - |
| Otu57888 | 1 | k__Fungi(100);k__Fungi_unclassified(100);k__Fungi_unclassified(100);k__Fungi_unclassified(100);k__Fungi_unclassified(100);k__Fungi_unclassified(100);k__Fungi_unclassified(100); | - | - | - | - | - | - | - | Unassigned | - |
| Otu57889 | 1 | k__Fungi(100);p__Ascomycota(100);p__Ascomycota_unclassified(100);p__Ascomycota_unclassified(100);p__Ascomycota_unclassified(100);p__Ascomycota_unclassified(100);p__Ascomycota_unclassified(100); | - | - | - | - | - | - | - | Unassigned | - |
| Otu57903 | 1 | k__Fungi(100);k__Fungi_unclassified(100);k__Fungi_unclassified(100);k__Fungi_unclassified(100);k__Fungi_unclassified(100);k__Fungi_unclassified(100);k__Fungi_unclassified(100); | - | - | - | - | - | - | - | Unassigned | - |
| Otu57913 | 1 | k__Fungi(100);k__Fungi_unclassified(100);k__Fungi_unclassified(100);k__Fungi_unclassified(100);k__Fungi_unclassified(100);k__Fungi_unclassified(100);k__Fungi_unclassified(100); | - | - | - | - | - | - | - | Unassigned | - |
| Otu57914 | 1 | k__Fungi(100);p__Ascomycota(100);p__Ascomycota_unclassified(100);p__Ascomycota_unclassified(100);p__Ascomycota_unclassified(100);p__Ascomycota_unclassified(100);p__Ascomycota_unclassified(100); | - | - | - | - | - | - | - | Unassigned | - |
| Otu57915 | 1 | k__Fungi(100);p__Ascomycota(100);p__Ascomycota_unclassified(100);p__Ascomycota_unclassified(100);p__Ascomycota_unclassified(100);p__Ascomycota_unclassified(100);p__Ascomycota_unclassified(100); | - | - | - | - | - | - | - | Unassigned | - |
| Otu57916 | 1 | k__Fungi(100);p__Ascomycota(100);p__Ascomycota_unclassified(100);p__Ascomycota_unclassified(100);p__Ascomycota_unclassified(100);p__Ascomycota_unclassified(100);p__Ascomycota_unclassified(100); | - | - | - | - | - | - | - | Unassigned | - |
| Otu57917 | 1 | k__Fungi(100);p__Ascomycota(100);p__Ascomycota_unclassified(100);p__Ascomycota_unclassified(100);p__Ascomycota_unclassified(100);p__Ascomycota_unclassified(100);p__Ascomycota_unclassified(100); | - | - | - | - | - | - | - | Unassigned | - |
| Otu57918 | 1 | k__Fungi(100);k__Fungi_unclassified(100);k__Fungi_unclassified(100);k__Fungi_unclassified(100);k__Fungi_unclassified(100);k__Fungi_unclassified(100);k__Fungi_unclassified(100); | - | - | - | - | - | - | - | Unassigned | - |
| Otu57921 | 1 | k__Fungi(100);k__Fungi_unclassified(100);k__Fungi_unclassified(100);k__Fungi_unclassified(100);k__Fungi_unclassified(100);k__Fungi_unclassified(100);k__Fungi_unclassified(100); | - | - | - | - | - | - | - | Unassigned | - |
| Otu57922 | 1 | k__Fungi(100);k__Fungi_unclassified(100);k__Fungi_unclassified(100);k__Fungi_unclassified(100);k__Fungi_unclassified(100);k__Fungi_unclassified(100);k__Fungi_unclassified(100); | - | - | - | - | - | - | - | Unassigned | - |
| Otu57923 | 1 | k__Fungi(100);p__Ascomycota(100);p__Ascomycota_unclassified(100);p__Ascomycota_unclassified(100);p__Ascomycota_unclassified(100);p__Ascomycota_unclassified(100);p__Ascomycota_unclassified(100); | - | - | - | - | - | - | - | Unassigned | - |
| Otu57924 | 1 | k__Fungi(100);k__Fungi_unclassified(100);k__Fungi_unclassified(100);k__Fungi_unclassified(100);k__Fungi_unclassified(100);k__Fungi_unclassified(100);k__Fungi_unclassified(100); | - | - | - | - | - | - | - | Unassigned | - |
| Otu57932 | 1 | k__Fungi(100);p__Ascomycota(100);p__Ascomycota_unclassified(100);p__Ascomycota_unclassified(100);p__Ascomycota_unclassified(100);p__Ascomycota_unclassified(100);p__Ascomycota_unclassified(100); | - | - | - | - | - | - | - | Unassigned | - |
| Otu57935 | 1 | k__Fungi(100);k__Fungi_unclassified(100);k__Fungi_unclassified(100);k__Fungi_unclassified(100);k__Fungi_unclassified(100);k__Fungi_unclassified(100);k__Fungi_unclassified(100); | - | - | - | - | - | - | - | Unassigned | - |
| Otu57936 | 1 | k__Fungi(100);k__Fungi_unclassified(100);k__Fungi_unclassified(100);k__Fungi_unclassified(100);k__Fungi_unclassified(100);k__Fungi_unclassified(100);k__Fungi_unclassified(100); | - | - | - | - | - | - | - | Unassigned | - |
| Otu57939 | 1 | k__Fungi(100);k__Fungi_unclassified(100);k__Fungi_unclassified(100);k__Fungi_unclassified(100);k__Fungi_unclassified(100);k__Fungi_unclassified(100);k__Fungi_unclassified(100); | - | - | - | - | - | - | - | Unassigned | - |
| Otu57942 | 1 | k__Fungi(100);k__Fungi_unclassified(100);k__Fungi_unclassified(100);k__Fungi_unclassified(100);k__Fungi_unclassified(100);k__Fungi_unclassified(100);k__Fungi_unclassified(100); | - | - | - | - | - | - | - | Unassigned | - |
| Otu57956 | 1 | k__Fungi(100);p__Ascomycota(100);p__Ascomycota_unclassified(100);p__Ascomycota_unclassified(100);p__Ascomycota_unclassified(100);p__Ascomycota_unclassified(100);p__Ascomycota_unclassified(100); | - | - | - | - | - | - | - | Unassigned | - |
| Otu57978 | 1 | k__Fungi(100);k__Fungi_unclassified(100);k__Fungi_unclassified(100);k__Fungi_unclassified(100);k__Fungi_unclassified(100);k__Fungi_unclassified(100);k__Fungi_unclassified(100); | - | - | - | - | - | - | - | Unassigned | - |
| Otu57990 | 1 | k__Fungi(100);k__Fungi_unclassified(100);k__Fungi_unclassified(100);k__Fungi_unclassified(100);k__Fungi_unclassified(100);k__Fungi_unclassified(100);k__Fungi_unclassified(100); | - | - | - | - | - | - | - | Unassigned | - |
| Otu57993 | 1 | k__Fungi(100);p__Ascomycota(100);p__Ascomycota_unclassified(100);p__Ascomycota_unclassified(100);p__Ascomycota_unclassified(100);p__Ascomycota_unclassified(100);p__Ascomycota_unclassified(100); | - | - | - | - | - | - | - | Unassigned | - |
| Otu58012 | 1 | k__Fungi(100);k__Fungi_unclassified(100);k__Fungi_unclassified(100);k__Fungi_unclassified(100);k__Fungi_unclassified(100);k__Fungi_unclassified(100);k__Fungi_unclassified(100); | - | - | - | - | - | - | - | Unassigned | - |
| Otu58014 | 1 | k__Fungi(100);k__Fungi_unclassified(100);k__Fungi_unclassified(100);k__Fungi_unclassified(100);k__Fungi_unclassified(100);k__Fungi_unclassified(100);k__Fungi_unclassified(100); | - | - | - | - | - | - | - | Unassigned | - |
| Otu58037 | 1 | k__Fungi(100);k__Fungi_unclassified(100);k__Fungi_unclassified(100);k__Fungi_unclassified(100);k__Fungi_unclassified(100);k__Fungi_unclassified(100);k__Fungi_unclassified(100); | - | - | - | - | - | - | - | Unassigned | - |
| Otu58039 | 1 | k__Fungi(100);p__Ascomycota(100);p__Ascomycota_unclassified(100);p__Ascomycota_unclassified(100);p__Ascomycota_unclassified(100);p__Ascomycota_unclassified(100);p__Ascomycota_unclassified(100); | - | - | - | - | - | - | - | Unassigned | - |
| Otu58040 | 1 | k__Fungi(100);p__Ascomycota(100);p__Ascomycota_unclassified(100);p__Ascomycota_unclassified(100);p__Ascomycota_unclassified(100);p__Ascomycota_unclassified(100);p__Ascomycota_unclassified(100); | - | - | - | - | - | - | - | Unassigned | - |
| Otu58042 | 1 | k__Fungi(100);p__Ascomycota(100);p__Ascomycota_unclassified(100);p__Ascomycota_unclassified(100);p__Ascomycota_unclassified(100);p__Ascomycota_unclassified(100);p__Ascomycota_unclassified(100); | - | - | - | - | - | - | - | Unassigned | - |
| Otu58043 | 1 | k__Fungi(100);k__Fungi_unclassified(100);k__Fungi_unclassified(100);k__Fungi_unclassified(100);k__Fungi_unclassified(100);k__Fungi_unclassified(100);k__Fungi_unclassified(100); | - | - | - | - | - | - | - | Unassigned | - |
| Otu58045 | 1 | k__Fungi(100);p__Ascomycota(100);p__Ascomycota_unclassified(100);p__Ascomycota_unclassified(100);p__Ascomycota_unclassified(100);p__Ascomycota_unclassified(100);p__Ascomycota_unclassified(100); | - | - | - | - | - | - | - | Unassigned | - |
| Otu58046 | 1 | k__Fungi(100);k__Fungi_unclassified(100);k__Fungi_unclassified(100);k__Fungi_unclassified(100);k__Fungi_unclassified(100);k__Fungi_unclassified(100);k__Fungi_unclassified(100); | - | - | - | - | - | - | - | Unassigned | - |
| Otu58047 | 1 | k__Fungi(100);p__Ascomycota(100);c__Dothideomycetes(100);o__Pleosporales(100);f__Pleosporales_family_Incertae_sedis(100);f__Pleosporales_family_Incertae_sedis_unclassified(100);f__Pleosporales_family_Incertae_sedis_unclassified(100); | - | - | - | - | - | - | - | Unassigned | - |
| Otu58049 | 1 | k__Fungi(100);k__Fungi_unclassified(100);k__Fungi_unclassified(100);k__Fungi_unclassified(100);k__Fungi_unclassified(100);k__Fungi_unclassified(100);k__Fungi_unclassified(100); | - | - | - | - | - | - | - | Unassigned | - |
| Otu58050 | 1 | k__Fungi(100);p__Ascomycota(100);p__Ascomycota_unclassified(100);p__Ascomycota_unclassified(100);p__Ascomycota_unclassified(100);p__Ascomycota_unclassified(100);p__Ascomycota_unclassified(100); | - | - | - | - | - | - | - | Unassigned | - |
| Otu58051 | 1 | k__Fungi(100);k__Fungi_unclassified(100);k__Fungi_unclassified(100);k__Fungi_unclassified(100);k__Fungi_unclassified(100);k__Fungi_unclassified(100);k__Fungi_unclassified(100); | - | - | - | - | - | - | - | Unassigned | - |
| Otu58053 | 1 | k__Fungi(100);p__Ascomycota(100);p__Ascomycota_unclassified(100);p__Ascomycota_unclassified(100);p__Ascomycota_unclassified(100);p__Ascomycota_unclassified(100);p__Ascomycota_unclassified(100); | - | - | - | - | - | - | - | Unassigned | - |
| Otu58054 | 1 | k__Fungi(100);p__Ascomycota(100);c__Leotiomycetes(100);c__Leotiomycetes_unclassified(100);c__Leotiomycetes_unclassified(100);c__Leotiomycetes_unclassified(100);c__Leotiomycetes_unclassified(100); | - | - | - | - | - | - | - | Unassigned | - |
| Otu58055 | 1 | k__Fungi(100);p__Ascomycota(100);p__Ascomycota_unclassified(100);p__Ascomycota_unclassified(100);p__Ascomycota_unclassified(100);p__Ascomycota_unclassified(100);p__Ascomycota_unclassified(100); | - | - | - | - | - | - | - | Unassigned | - |
| Otu58069 | 1 | k__Fungi(100);p__Ascomycota(100);p__Ascomycota_unclassified(100);p__Ascomycota_unclassified(100);p__Ascomycota_unclassified(100);p__Ascomycota_unclassified(100);p__Ascomycota_unclassified(100); | - | - | - | - | - | - | - | Unassigned | - |
| Otu58083 | 1 | k__Fungi(100);k__Fungi_unclassified(100);k__Fungi_unclassified(100);k__Fungi_unclassified(100);k__Fungi_unclassified(100);k__Fungi_unclassified(100);k__Fungi_unclassified(100); | - | - | - | - | - | - | - | Unassigned | - |
| Otu58085 | 1 | k__Fungi(100);k__Fungi_unclassified(100);k__Fungi_unclassified(100);k__Fungi_unclassified(100);k__Fungi_unclassified(100);k__Fungi_unclassified(100);k__Fungi_unclassified(100); | - | - | - | - | - | - | - | Unassigned | - |
| Otu58086 | 1 | k__Fungi(100);p__Ascomycota(100);p__Ascomycota_unclassified(100);p__Ascomycota_unclassified(100);p__Ascomycota_unclassified(100);p__Ascomycota_unclassified(100);p__Ascomycota_unclassified(100); | - | - | - | - | - | - | - | Unassigned | - |
| Otu58087 | 1 | k__Fungi(100);k__Fungi_unclassified(100);k__Fungi_unclassified(100);k__Fungi_unclassified(100);k__Fungi_unclassified(100);k__Fungi_unclassified(100);k__Fungi_unclassified(100); | - | - | - | - | - | - | - | Unassigned | - |
| Otu58088 | 1 | k__Fungi(100);p__Ascomycota(100);p__Ascomycota_unclassified(100);p__Ascomycota_unclassified(100);p__Ascomycota_unclassified(100);p__Ascomycota_unclassified(100);p__Ascomycota_unclassified(100); | - | - | - | - | - | - | - | Unassigned | - |
| Otu58090 | 1 | k__Fungi(100);p__Ascomycota(100);p__Ascomycota_unclassified(100);p__Ascomycota_unclassified(100);p__Ascomycota_unclassified(100);p__Ascomycota_unclassified(100);p__Ascomycota_unclassified(100); | - | - | - | - | - | - | - | Unassigned | - |
| Otu58093 | 1 | k__Fungi(100);k__Fungi_unclassified(100);k__Fungi_unclassified(100);k__Fungi_unclassified(100);k__Fungi_unclassified(100);k__Fungi_unclassified(100);k__Fungi_unclassified(100); | - | - | - | - | - | - | - | Unassigned | - |
| Otu58102 | 1 | k__Fungi(100);k__Fungi_unclassified(100);k__Fungi_unclassified(100);k__Fungi_unclassified(100);k__Fungi_unclassified(100);k__Fungi_unclassified(100);k__Fungi_unclassified(100); | - | - | - | - | - | - | - | Unassigned | - |
| Otu58110 | 1 | k__Fungi(100);k__Fungi_unclassified(100);k__Fungi_unclassified(100);k__Fungi_unclassified(100);k__Fungi_unclassified(100);k__Fungi_unclassified(100);k__Fungi_unclassified(100); | - | - | - | - | - | - | - | Unassigned | - |
| Otu58116 | 1 | k__Fungi(100);k__Fungi_unclassified(100);k__Fungi_unclassified(100);k__Fungi_unclassified(100);k__Fungi_unclassified(100);k__Fungi_unclassified(100);k__Fungi_unclassified(100); | - | - | - | - | - | - | - | Unassigned | - |
| Otu58117 | 1 | k__Fungi(100);p__Ascomycota(100);p__Ascomycota_unclassified(100);p__Ascomycota_unclassified(100);p__Ascomycota_unclassified(100);p__Ascomycota_unclassified(100);p__Ascomycota_unclassified(100); | - | - | - | - | - | - | - | Unassigned | - |
| Otu58119 | 1 | k__Fungi(100);k__Fungi_unclassified(100);k__Fungi_unclassified(100);k__Fungi_unclassified(100);k__Fungi_unclassified(100);k__Fungi_unclassified(100);k__Fungi_unclassified(100); | - | - | - | - | - | - | - | Unassigned | - |
| Otu58127 | 1 | k__Fungi(100);p__Basidiomycota(100);c__Agaricomycetes(100);c__Agaricomycetes_unclassified(100);c__Agaricomycetes_unclassified(100);c__Agaricomycetes_unclassified(100);c__Agaricomycetes_unclassified(100); | - | - | - | - | - | - | - | Unassigned | - |
| Otu58130 | 1 | k__Fungi(100);k__Fungi_unclassified(100);k__Fungi_unclassified(100);k__Fungi_unclassified(100);k__Fungi_unclassified(100);k__Fungi_unclassified(100);k__Fungi_unclassified(100); | - | - | - | - | - | - | - | Unassigned | - |
| Otu58140 | 1 | k__Fungi(100);k__Fungi_unclassified(100);k__Fungi_unclassified(100);k__Fungi_unclassified(100);k__Fungi_unclassified(100);k__Fungi_unclassified(100);k__Fungi_unclassified(100); | - | - | - | - | - | - | - | Unassigned | - |
| Otu58141 | 1 | k__Fungi(100);p__Ascomycota(100);p__Ascomycota_unclassified(100);p__Ascomycota_unclassified(100);p__Ascomycota_unclassified(100);p__Ascomycota_unclassified(100);p__Ascomycota_unclassified(100); | - | - | - | - | - | - | - | Unassigned | - |
| Otu58142 | 1 | k__Fungi(100);k__Fungi_unclassified(100);k__Fungi_unclassified(100);k__Fungi_unclassified(100);k__Fungi_unclassified(100);k__Fungi_unclassified(100);k__Fungi_unclassified(100); | - | - | - | - | - | - | - | Unassigned | - |
| Otu58149 | 1 | k__Fungi(100);k__Fungi_unclassified(100);k__Fungi_unclassified(100);k__Fungi_unclassified(100);k__Fungi_unclassified(100);k__Fungi_unclassified(100);k__Fungi_unclassified(100); | - | - | - | - | - | - | - | Unassigned | - |
| Otu58152 | 1 | k__Fungi(100);k__Fungi_unclassified(100);k__Fungi_unclassified(100);k__Fungi_unclassified(100);k__Fungi_unclassified(100);k__Fungi_unclassified(100);k__Fungi_unclassified(100); | - | - | - | - | - | - | - | Unassigned | - |
| Otu58154 | 1 | k__Fungi(100);p__Ascomycota(100);p__Ascomycota_unclassified(100);p__Ascomycota_unclassified(100);p__Ascomycota_unclassified(100);p__Ascomycota_unclassified(100);p__Ascomycota_unclassified(100); | - | - | - | - | - | - | - | Unassigned | - |
| Otu58158 | 1 | k__Fungi(100);p__Ascomycota(100);p__Ascomycota_unclassified(100);p__Ascomycota_unclassified(100);p__Ascomycota_unclassified(100);p__Ascomycota_unclassified(100);p__Ascomycota_unclassified(100); | - | - | - | - | - | - | - | Unassigned | - |
| Otu58159 | 1 | k__Fungi(100);k__Fungi_unclassified(100);k__Fungi_unclassified(100);k__Fungi_unclassified(100);k__Fungi_unclassified(100);k__Fungi_unclassified(100);k__Fungi_unclassified(100); | - | - | - | - | - | - | - | Unassigned | - |
| Otu58163 | 1 | k__Fungi(100);p__Ascomycota(100);p__Ascomycota_unclassified(100);p__Ascomycota_unclassified(100);p__Ascomycota_unclassified(100);p__Ascomycota_unclassified(100);p__Ascomycota_unclassified(100); | - | - | - | - | - | - | - | Unassigned | - |
| Otu58166 | 1 | k__Fungi(100);p__Ascomycota(100);p__Ascomycota_unclassified(100);p__Ascomycota_unclassified(100);p__Ascomycota_unclassified(100);p__Ascomycota_unclassified(100);p__Ascomycota_unclassified(100); | - | - | - | - | - | - | - | Unassigned | - |
| Otu58169 | 1 | k__Fungi(100);p__Ascomycota(100);p__Ascomycota_unclassified(100);p__Ascomycota_unclassified(100);p__Ascomycota_unclassified(100);p__Ascomycota_unclassified(100);p__Ascomycota_unclassified(100); | - | - | - | - | - | - | - | Unassigned | - |
| Otu58173 | 1 | k__Fungi(100);k__Fungi_unclassified(100);k__Fungi_unclassified(100);k__Fungi_unclassified(100);k__Fungi_unclassified(100);k__Fungi_unclassified(100);k__Fungi_unclassified(100); | - | - | - | - | - | - | - | Unassigned | - |
| Otu58175 | 1 | k__Fungi(100);p__Ascomycota(100);p__Ascomycota_unclassified(100);p__Ascomycota_unclassified(100);p__Ascomycota_unclassified(100);p__Ascomycota_unclassified(100);p__Ascomycota_unclassified(100); | - | - | - | - | - | - | - | Unassigned | - |
| Otu58176 | 1 | k__Fungi(100);p__Ascomycota(100);p__Ascomycota_unclassified(100);p__Ascomycota_unclassified(100);p__Ascomycota_unclassified(100);p__Ascomycota_unclassified(100);p__Ascomycota_unclassified(100); | - | - | - | - | - | - | - | Unassigned | - |
| Otu58187 | 1 | k__Fungi(100);p__Ascomycota(100);p__Ascomycota_unclassified(100);p__Ascomycota_unclassified(100);p__Ascomycota_unclassified(100);p__Ascomycota_unclassified(100);p__Ascomycota_unclassified(100); | - | - | - | - | - | - | - | Unassigned | - |
| Otu58188 | 1 | k__Fungi(100);k__Fungi_unclassified(100);k__Fungi_unclassified(100);k__Fungi_unclassified(100);k__Fungi_unclassified(100);k__Fungi_unclassified(100);k__Fungi_unclassified(100); | - | - | - | - | - | - | - | Unassigned | - |
| Otu58189 | 1 | k__Fungi(100);k__Fungi_unclassified(100);k__Fungi_unclassified(100);k__Fungi_unclassified(100);k__Fungi_unclassified(100);k__Fungi_unclassified(100);k__Fungi_unclassified(100); | - | - | - | - | - | - | - | Unassigned | - |
| Otu58190 | 1 | k__Fungi(100);k__Fungi_unclassified(100);k__Fungi_unclassified(100);k__Fungi_unclassified(100);k__Fungi_unclassified(100);k__Fungi_unclassified(100);k__Fungi_unclassified(100); | - | - | - | - | - | - | - | Unassigned | - |
| Otu58191 | 1 | k__Fungi(100);p__Ascomycota(100);p__Ascomycota_unclassified(100);p__Ascomycota_unclassified(100);p__Ascomycota_unclassified(100);p__Ascomycota_unclassified(100);p__Ascomycota_unclassified(100); | - | - | - | - | - | - | - | Unassigned | - |
| Otu58195 | 1 | k__Fungi(100);k__Fungi_unclassified(100);k__Fungi_unclassified(100);k__Fungi_unclassified(100);k__Fungi_unclassified(100);k__Fungi_unclassified(100);k__Fungi_unclassified(100); | - | - | - | - | - | - | - | Unassigned | - |
| Otu58196 | 1 | k__Fungi(100);p__Ascomycota(100);p__Ascomycota_unclassified(100);p__Ascomycota_unclassified(100);p__Ascomycota_unclassified(100);p__Ascomycota_unclassified(100);p__Ascomycota_unclassified(100); | - | - | - | - | - | - | - | Unassigned | - |
| Otu58197 | 1 | k__Fungi(100);k__Fungi_unclassified(100);k__Fungi_unclassified(100);k__Fungi_unclassified(100);k__Fungi_unclassified(100);k__Fungi_unclassified(100);k__Fungi_unclassified(100); | - | - | - | - | - | - | - | Unassigned | - |
| Otu58200 | 1 | k__Fungi(100);k__Fungi_unclassified(100);k__Fungi_unclassified(100);k__Fungi_unclassified(100);k__Fungi_unclassified(100);k__Fungi_unclassified(100);k__Fungi_unclassified(100); | - | - | - | - | - | - | - | Unassigned | - |
| Otu58202 | 1 | k__Fungi(100);k__Fungi_unclassified(100);k__Fungi_unclassified(100);k__Fungi_unclassified(100);k__Fungi_unclassified(100);k__Fungi_unclassified(100);k__Fungi_unclassified(100); | - | - | - | - | - | - | - | Unassigned | - |
| Otu58204 | 1 | k__Fungi(100);k__Fungi_unclassified(100);k__Fungi_unclassified(100);k__Fungi_unclassified(100);k__Fungi_unclassified(100);k__Fungi_unclassified(100);k__Fungi_unclassified(100); | - | - | - | - | - | - | - | Unassigned | - |
| Otu58206 | 1 | k__Fungi(100);p__Ascomycota(100);p__Ascomycota_unclassified(100);p__Ascomycota_unclassified(100);p__Ascomycota_unclassified(100);p__Ascomycota_unclassified(100);p__Ascomycota_unclassified(100); | - | - | - | - | - | - | - | Unassigned | - |
| Otu58207 | 1 | k__Fungi(100);k__Fungi_unclassified(100);k__Fungi_unclassified(100);k__Fungi_unclassified(100);k__Fungi_unclassified(100);k__Fungi_unclassified(100);k__Fungi_unclassified(100); | - | - | - | - | - | - | - | Unassigned | - |
| Otu58212 | 1 | k__Fungi(100);k__Fungi_unclassified(100);k__Fungi_unclassified(100);k__Fungi_unclassified(100);k__Fungi_unclassified(100);k__Fungi_unclassified(100);k__Fungi_unclassified(100); | - | - | - | - | - | - | - | Unassigned | - |
| Otu58213 | 1 | k__Fungi(100);k__Fungi_unclassified(100);k__Fungi_unclassified(100);k__Fungi_unclassified(100);k__Fungi_unclassified(100);k__Fungi_unclassified(100);k__Fungi_unclassified(100); | - | - | - | - | - | - | - | Unassigned | - |
| Otu58214 | 1 | k__Fungi(100);k__Fungi_unclassified(100);k__Fungi_unclassified(100);k__Fungi_unclassified(100);k__Fungi_unclassified(100);k__Fungi_unclassified(100);k__Fungi_unclassified(100); | - | - | - | - | - | - | - | Unassigned | - |
| Otu58215 | 1 | k__Fungi(100);k__Fungi_unclassified(100);k__Fungi_unclassified(100);k__Fungi_unclassified(100);k__Fungi_unclassified(100);k__Fungi_unclassified(100);k__Fungi_unclassified(100); | - | - | - | - | - | - | - | Unassigned | - |
| Otu58216 | 1 | k__Fungi(100);k__Fungi_unclassified(100);k__Fungi_unclassified(100);k__Fungi_unclassified(100);k__Fungi_unclassified(100);k__Fungi_unclassified(100);k__Fungi_unclassified(100); | - | - | - | - | - | - | - | Unassigned | - |
| Otu58217 | 1 | k__Fungi(100);k__Fungi_unclassified(100);k__Fungi_unclassified(100);k__Fungi_unclassified(100);k__Fungi_unclassified(100);k__Fungi_unclassified(100);k__Fungi_unclassified(100); | - | - | - | - | - | - | - | Unassigned | - |
| Otu58218 | 1 | k__Fungi(100);k__Fungi_unclassified(100);k__Fungi_unclassified(100);k__Fungi_unclassified(100);k__Fungi_unclassified(100);k__Fungi_unclassified(100);k__Fungi_unclassified(100); | - | - | - | - | - | - | - | Unassigned | - |
| Otu58226 | 1 | k__Fungi(100);k__Fungi_unclassified(100);k__Fungi_unclassified(100);k__Fungi_unclassified(100);k__Fungi_unclassified(100);k__Fungi_unclassified(100);k__Fungi_unclassified(100); | - | - | - | - | - | - | - | Unassigned | - |
| Otu58228 | 1 | k__Fungi(100);k__Fungi_unclassified(100);k__Fungi_unclassified(100);k__Fungi_unclassified(100);k__Fungi_unclassified(100);k__Fungi_unclassified(100);k__Fungi_unclassified(100); | - | - | - | - | - | - | - | Unassigned | - |
| Otu58232 | 1 | k__Fungi(100);k__Fungi_unclassified(100);k__Fungi_unclassified(100);k__Fungi_unclassified(100);k__Fungi_unclassified(100);k__Fungi_unclassified(100);k__Fungi_unclassified(100); | - | - | - | - | - | - | - | Unassigned | - |
| Otu58241 | 1 | k__Fungi(100);p__Ascomycota(100);p__Ascomycota_unclassified(100);p__Ascomycota_unclassified(100);p__Ascomycota_unclassified(100);p__Ascomycota_unclassified(100);p__Ascomycota_unclassified(100); | - | - | - | - | - | - | - | Unassigned | - |
| Otu58242 | 1 | k__Fungi(100);p__Ascomycota(100);p__Ascomycota_unclassified(100);p__Ascomycota_unclassified(100);p__Ascomycota_unclassified(100);p__Ascomycota_unclassified(100);p__Ascomycota_unclassified(100); | - | - | - | - | - | - | - | Unassigned | - |
| Otu58243 | 1 | k__Fungi(100);k__Fungi_unclassified(100);k__Fungi_unclassified(100);k__Fungi_unclassified(100);k__Fungi_unclassified(100);k__Fungi_unclassified(100);k__Fungi_unclassified(100); | - | - | - | - | - | - | - | Unassigned | - |
| Otu58244 | 1 | k__Fungi(100);k__Fungi_unclassified(100);k__Fungi_unclassified(100);k__Fungi_unclassified(100);k__Fungi_unclassified(100);k__Fungi_unclassified(100);k__Fungi_unclassified(100); | - | - | - | - | - | - | - | Unassigned | - |
| Otu58255 | 1 | k__Fungi(100);p__Ascomycota(100);p__Ascomycota_unclassified(100);p__Ascomycota_unclassified(100);p__Ascomycota_unclassified(100);p__Ascomycota_unclassified(100);p__Ascomycota_unclassified(100); | - | - | - | - | - | - | - | Unassigned | - |
| Otu58271 | 1 | k__Fungi(100);k__Fungi_unclassified(100);k__Fungi_unclassified(100);k__Fungi_unclassified(100);k__Fungi_unclassified(100);k__Fungi_unclassified(100);k__Fungi_unclassified(100); | - | - | - | - | - | - | - | Unassigned | - |
| Otu58272 | 1 | k__Fungi(100);k__Fungi_unclassified(100);k__Fungi_unclassified(100);k__Fungi_unclassified(100);k__Fungi_unclassified(100);k__Fungi_unclassified(100);k__Fungi_unclassified(100); | - | - | - | - | - | - | - | Unassigned | - |
| Otu58273 | 1 | k__Fungi(100);k__Fungi_unclassified(100);k__Fungi_unclassified(100);k__Fungi_unclassified(100);k__Fungi_unclassified(100);k__Fungi_unclassified(100);k__Fungi_unclassified(100); | - | - | - | - | - | - | - | Unassigned | - |
| Otu58275 | 1 | k__Fungi(100);p__Ascomycota(100);p__Ascomycota_unclassified(100);p__Ascomycota_unclassified(100);p__Ascomycota_unclassified(100);p__Ascomycota_unclassified(100);p__Ascomycota_unclassified(100); | - | - | - | - | - | - | - | Unassigned | - |
| Otu58276 | 1 | k__Fungi(100);p__Ascomycota(100);c__Dothideomycetes(100);o__Pleosporales(100);f__Pleosporales_family_Incertae_sedis(100);f__Pleosporales_family_Incertae_sedis_unclassified(100);f__Pleosporales_family_Incertae_sedis_unclassified(100); | - | - | - | - | - | - | - | Unassigned | - |
| Otu58277 | 1 | k__Fungi(100);k__Fungi_unclassified(100);k__Fungi_unclassified(100);k__Fungi_unclassified(100);k__Fungi_unclassified(100);k__Fungi_unclassified(100);k__Fungi_unclassified(100); | - | - | - | - | - | - | - | Unassigned | - |
| Otu58278 | 1 | k__Fungi(100);k__Fungi_unclassified(100);k__Fungi_unclassified(100);k__Fungi_unclassified(100);k__Fungi_unclassified(100);k__Fungi_unclassified(100);k__Fungi_unclassified(100); | - | - | - | - | - | - | - | Unassigned | - |
| Otu58279 | 1 | k__Fungi(100);k__Fungi_unclassified(100);k__Fungi_unclassified(100);k__Fungi_unclassified(100);k__Fungi_unclassified(100);k__Fungi_unclassified(100);k__Fungi_unclassified(100); | - | - | - | - | - | - | - | Unassigned | - |
| Otu58280 | 1 | k__Fungi(100);k__Fungi_unclassified(100);k__Fungi_unclassified(100);k__Fungi_unclassified(100);k__Fungi_unclassified(100);k__Fungi_unclassified(100);k__Fungi_unclassified(100); | - | - | - | - | - | - | - | Unassigned | - |
| Otu58281 | 1 | k__Fungi(100);p__Ascomycota(100);p__Ascomycota_unclassified(100);p__Ascomycota_unclassified(100);p__Ascomycota_unclassified(100);p__Ascomycota_unclassified(100);p__Ascomycota_unclassified(100); | - | - | - | - | - | - | - | Unassigned | - |
| Otu58287 | 1 | k__Fungi(100);k__Fungi_unclassified(100);k__Fungi_unclassified(100);k__Fungi_unclassified(100);k__Fungi_unclassified(100);k__Fungi_unclassified(100);k__Fungi_unclassified(100); | - | - | - | - | - | - | - | Unassigned | - |
| Otu58288 | 1 | k__Fungi(100);p__Ascomycota(100);p__Ascomycota_unclassified(100);p__Ascomycota_unclassified(100);p__Ascomycota_unclassified(100);p__Ascomycota_unclassified(100);p__Ascomycota_unclassified(100); | - | - | - | - | - | - | - | Unassigned | - |
| Otu58293 | 1 | k__Fungi(100);k__Fungi_unclassified(100);k__Fungi_unclassified(100);k__Fungi_unclassified(100);k__Fungi_unclassified(100);k__Fungi_unclassified(100);k__Fungi_unclassified(100); | - | - | - | - | - | - | - | Unassigned | - |
| Otu58332 | 1 | k__Fungi(100);k__Fungi_unclassified(100);k__Fungi_unclassified(100);k__Fungi_unclassified(100);k__Fungi_unclassified(100);k__Fungi_unclassified(100);k__Fungi_unclassified(100); | - | - | - | - | - | - | - | Unassigned | - |
| Otu58334 | 1 | k__Fungi(100);k__Fungi_unclassified(100);k__Fungi_unclassified(100);k__Fungi_unclassified(100);k__Fungi_unclassified(100);k__Fungi_unclassified(100);k__Fungi_unclassified(100); | - | - | - | - | - | - | - | Unassigned | - |
| Otu58348 | 1 | k__Fungi(100);p__Ascomycota(100);p__Ascomycota_unclassified(100);p__Ascomycota_unclassified(100);p__Ascomycota_unclassified(100);p__Ascomycota_unclassified(100);p__Ascomycota_unclassified(100); | - | - | - | - | - | - | - | Unassigned | - |
| Otu58353 | 1 | k__Fungi(100);k__Fungi_unclassified(100);k__Fungi_unclassified(100);k__Fungi_unclassified(100);k__Fungi_unclassified(100);k__Fungi_unclassified(100);k__Fungi_unclassified(100); | - | - | - | - | - | - | - | Unassigned | - |
| Otu58356 | 1 | k__Fungi(100);k__Fungi_unclassified(100);k__Fungi_unclassified(100);k__Fungi_unclassified(100);k__Fungi_unclassified(100);k__Fungi_unclassified(100);k__Fungi_unclassified(100); | - | - | - | - | - | - | - | Unassigned | - |
| Otu58357 | 1 | k__Fungi(100);p__Ascomycota(100);p__Ascomycota_unclassified(100);p__Ascomycota_unclassified(100);p__Ascomycota_unclassified(100);p__Ascomycota_unclassified(100);p__Ascomycota_unclassified(100); | - | - | - | - | - | - | - | Unassigned | - |
| Otu58358 | 1 | k__Fungi(100);k__Fungi_unclassified(100);k__Fungi_unclassified(100);k__Fungi_unclassified(100);k__Fungi_unclassified(100);k__Fungi_unclassified(100);k__Fungi_unclassified(100); | - | - | - | - | - | - | - | Unassigned | - |
| Otu58359 | 1 | k__Fungi(100);p__Ascomycota(100);p__Ascomycota_unclassified(100);p__Ascomycota_unclassified(100);p__Ascomycota_unclassified(100);p__Ascomycota_unclassified(100);p__Ascomycota_unclassified(100); | - | - | - | - | - | - | - | Unassigned | - |
| Otu58360 | 1 | k__Fungi(100);k__Fungi_unclassified(100);k__Fungi_unclassified(100);k__Fungi_unclassified(100);k__Fungi_unclassified(100);k__Fungi_unclassified(100);k__Fungi_unclassified(100); | - | - | - | - | - | - | - | Unassigned | - |
| Otu58364 | 1 | k__Fungi(100);k__Fungi_unclassified(100);k__Fungi_unclassified(100);k__Fungi_unclassified(100);k__Fungi_unclassified(100);k__Fungi_unclassified(100);k__Fungi_unclassified(100); | - | - | - | - | - | - | - | Unassigned | - |
| Otu58365 | 1 | k__Fungi(100);p__Ascomycota(100);p__Ascomycota_unclassified(100);p__Ascomycota_unclassified(100);p__Ascomycota_unclassified(100);p__Ascomycota_unclassified(100);p__Ascomycota_unclassified(100); | - | - | - | - | - | - | - | Unassigned | - |
| Otu58396 | 1 | k__Fungi(100);k__Fungi_unclassified(100);k__Fungi_unclassified(100);k__Fungi_unclassified(100);k__Fungi_unclassified(100);k__Fungi_unclassified(100);k__Fungi_unclassified(100); | - | - | - | - | - | - | - | Unassigned | - |
| Otu58397 | 1 | k__Fungi(100);p__Ascomycota(100);p__Ascomycota_unclassified(100);p__Ascomycota_unclassified(100);p__Ascomycota_unclassified(100);p__Ascomycota_unclassified(100);p__Ascomycota_unclassified(100); | - | - | - | - | - | - | - | Unassigned | - |
| Otu58398 | 1 | k__Fungi(100);p__Ascomycota(100);p__Ascomycota_unclassified(100);p__Ascomycota_unclassified(100);p__Ascomycota_unclassified(100);p__Ascomycota_unclassified(100);p__Ascomycota_unclassified(100); | - | - | - | - | - | - | - | Unassigned | - |
| Otu58399 | 1 | k__Fungi(100);p__Ascomycota(100);p__Ascomycota_unclassified(100);p__Ascomycota_unclassified(100);p__Ascomycota_unclassified(100);p__Ascomycota_unclassified(100);p__Ascomycota_unclassified(100); | - | - | - | - | - | - | - | Unassigned | - |
| Otu58413 | 1 | k__Fungi(100);k__Fungi_unclassified(100);k__Fungi_unclassified(100);k__Fungi_unclassified(100);k__Fungi_unclassified(100);k__Fungi_unclassified(100);k__Fungi_unclassified(100); | - | - | - | - | - | - | - | Unassigned | - |
| Otu58431 | 1 | k__Fungi(100);k__Fungi_unclassified(100);k__Fungi_unclassified(100);k__Fungi_unclassified(100);k__Fungi_unclassified(100);k__Fungi_unclassified(100);k__Fungi_unclassified(100); | - | - | - | - | - | - | - | Unassigned | - |
| Otu58432 | 1 | k__Fungi(100);k__Fungi_unclassified(100);k__Fungi_unclassified(100);k__Fungi_unclassified(100);k__Fungi_unclassified(100);k__Fungi_unclassified(100);k__Fungi_unclassified(100); | - | - | - | - | - | - | - | Unassigned | - |
| Otu58439 | 1 | k__Fungi(100);p__Ascomycota(100);p__Ascomycota_unclassified(100);p__Ascomycota_unclassified(100);p__Ascomycota_unclassified(100);p__Ascomycota_unclassified(100);p__Ascomycota_unclassified(100); | - | - | - | - | - | - | - | Unassigned | - |
| Otu58443 | 1 | k__Fungi(100);p__Ascomycota(100);p__Ascomycota_unclassified(100);p__Ascomycota_unclassified(100);p__Ascomycota_unclassified(100);p__Ascomycota_unclassified(100);p__Ascomycota_unclassified(100); | - | - | - | - | - | - | - | Unassigned | - |
| Otu58444 | 1 | k__Fungi(100);k__Fungi_unclassified(100);k__Fungi_unclassified(100);k__Fungi_unclassified(100);k__Fungi_unclassified(100);k__Fungi_unclassified(100);k__Fungi_unclassified(100); | - | - | - | - | - | - | - | Unassigned | - |
| Otu58446 | 1 | k__Fungi(100);k__Fungi_unclassified(100);k__Fungi_unclassified(100);k__Fungi_unclassified(100);k__Fungi_unclassified(100);k__Fungi_unclassified(100);k__Fungi_unclassified(100); | - | - | - | - | - | - | - | Unassigned | - |
| Otu58450 | 1 | k__Fungi(100);k__Fungi_unclassified(100);k__Fungi_unclassified(100);k__Fungi_unclassified(100);k__Fungi_unclassified(100);k__Fungi_unclassified(100);k__Fungi_unclassified(100); | - | - | - | - | - | - | - | Unassigned | - |
| Otu58451 | 1 | k__Fungi(100);k__Fungi_unclassified(100);k__Fungi_unclassified(100);k__Fungi_unclassified(100);k__Fungi_unclassified(100);k__Fungi_unclassified(100);k__Fungi_unclassified(100); | - | - | - | - | - | - | - | Unassigned | - |
| Otu58459 | 1 | k__Fungi(100);p__Ascomycota(100);p__Ascomycota_unclassified(100);p__Ascomycota_unclassified(100);p__Ascomycota_unclassified(100);p__Ascomycota_unclassified(100);p__Ascomycota_unclassified(100); | - | - | - | - | - | - | - | Unassigned | - |
| Otu58490 | 1 | k__Fungi(100);k__Fungi_unclassified(100);k__Fungi_unclassified(100);k__Fungi_unclassified(100);k__Fungi_unclassified(100);k__Fungi_unclassified(100);k__Fungi_unclassified(100); | - | - | - | - | - | - | - | Unassigned | - |
| Otu58498 | 1 | k__Fungi(100);k__Fungi_unclassified(100);k__Fungi_unclassified(100);k__Fungi_unclassified(100);k__Fungi_unclassified(100);k__Fungi_unclassified(100);k__Fungi_unclassified(100); | - | - | - | - | - | - | - | Unassigned | - |
| Otu58504 | 1 | k__Fungi(100);k__Fungi_unclassified(100);k__Fungi_unclassified(100);k__Fungi_unclassified(100);k__Fungi_unclassified(100);k__Fungi_unclassified(100);k__Fungi_unclassified(100); | - | - | - | - | - | - | - | Unassigned | - |
| Otu58506 | 1 | k__Fungi(100);k__Fungi_unclassified(100);k__Fungi_unclassified(100);k__Fungi_unclassified(100);k__Fungi_unclassified(100);k__Fungi_unclassified(100);k__Fungi_unclassified(100); | - | - | - | - | - | - | - | Unassigned | - |
| Otu58507 | 1 | k__Fungi(100);p__Ascomycota(100);p__Ascomycota_unclassified(100);p__Ascomycota_unclassified(100);p__Ascomycota_unclassified(100);p__Ascomycota_unclassified(100);p__Ascomycota_unclassified(100); | - | - | - | - | - | - | - | Unassigned | - |
| Otu58508 | 1 | k__Fungi(100);k__Fungi_unclassified(100);k__Fungi_unclassified(100);k__Fungi_unclassified(100);k__Fungi_unclassified(100);k__Fungi_unclassified(100);k__Fungi_unclassified(100); | - | - | - | - | - | - | - | Unassigned | - |
| Otu58509 | 1 | k__Fungi(100);p__Ascomycota(100);p__Ascomycota_unclassified(100);p__Ascomycota_unclassified(100);p__Ascomycota_unclassified(100);p__Ascomycota_unclassified(100);p__Ascomycota_unclassified(100); | - | - | - | - | - | - | - | Unassigned | - |
| Otu58511 | 1 | k__Fungi(100);k__Fungi_unclassified(100);k__Fungi_unclassified(100);k__Fungi_unclassified(100);k__Fungi_unclassified(100);k__Fungi_unclassified(100);k__Fungi_unclassified(100); | - | - | - | - | - | - | - | Unassigned | - |
| Otu58514 | 1 | k__Fungi(100);p__Ascomycota(100);p__Ascomycota_unclassified(100);p__Ascomycota_unclassified(100);p__Ascomycota_unclassified(100);p__Ascomycota_unclassified(100);p__Ascomycota_unclassified(100); | - | - | - | - | - | - | - | Unassigned | - |
| Otu58520 | 1 | k__Fungi(100);k__Fungi_unclassified(100);k__Fungi_unclassified(100);k__Fungi_unclassified(100);k__Fungi_unclassified(100);k__Fungi_unclassified(100);k__Fungi_unclassified(100); | - | - | - | - | - | - | - | Unassigned | - |
| Otu58521 | 1 | k__Fungi(100);p__Ascomycota(100);p__Ascomycota_unclassified(100);p__Ascomycota_unclassified(100);p__Ascomycota_unclassified(100);p__Ascomycota_unclassified(100);p__Ascomycota_unclassified(100); | - | - | - | - | - | - | - | Unassigned | - |
| Otu58523 | 1 | k__Fungi(100);p__Ascomycota(100);p__Ascomycota_unclassified(100);p__Ascomycota_unclassified(100);p__Ascomycota_unclassified(100);p__Ascomycota_unclassified(100);p__Ascomycota_unclassified(100); | - | - | - | - | - | - | - | Unassigned | - |
| Otu58524 | 1 | k__Fungi(100);k__Fungi_unclassified(100);k__Fungi_unclassified(100);k__Fungi_unclassified(100);k__Fungi_unclassified(100);k__Fungi_unclassified(100);k__Fungi_unclassified(100); | - | - | - | - | - | - | - | Unassigned | - |
| Otu58531 | 1 | k__Fungi(100);k__Fungi_unclassified(100);k__Fungi_unclassified(100);k__Fungi_unclassified(100);k__Fungi_unclassified(100);k__Fungi_unclassified(100);k__Fungi_unclassified(100); | - | - | - | - | - | - | - | Unassigned | - |
| Otu58532 | 1 | k__Fungi(100);p__Ascomycota(100);p__Ascomycota_unclassified(100);p__Ascomycota_unclassified(100);p__Ascomycota_unclassified(100);p__Ascomycota_unclassified(100);p__Ascomycota_unclassified(100); | - | - | - | - | - | - | - | Unassigned | - |
| Otu58535 | 1 | k__Fungi(100);k__Fungi_unclassified(100);k__Fungi_unclassified(100);k__Fungi_unclassified(100);k__Fungi_unclassified(100);k__Fungi_unclassified(100);k__Fungi_unclassified(100); | - | - | - | - | - | - | - | Unassigned | - |
| Otu58542 | 1 | k__Fungi(100);p__Ascomycota(100);p__Ascomycota_unclassified(100);p__Ascomycota_unclassified(100);p__Ascomycota_unclassified(100);p__Ascomycota_unclassified(100);p__Ascomycota_unclassified(100); | - | - | - | - | - | - | - | Unassigned | - |
| Otu58544 | 1 | k__Fungi(100);p__Ascomycota(100);p__Ascomycota_unclassified(100);p__Ascomycota_unclassified(100);p__Ascomycota_unclassified(100);p__Ascomycota_unclassified(100);p__Ascomycota_unclassified(100); | - | - | - | - | - | - | - | Unassigned | - |
| Otu58556 | 1 | k__Fungi(100);k__Fungi_unclassified(100);k__Fungi_unclassified(100);k__Fungi_unclassified(100);k__Fungi_unclassified(100);k__Fungi_unclassified(100);k__Fungi_unclassified(100); | - | - | - | - | - | - | - | Unassigned | - |
| Otu58583 | 1 | k__Fungi(100);k__Fungi_unclassified(100);k__Fungi_unclassified(100);k__Fungi_unclassified(100);k__Fungi_unclassified(100);k__Fungi_unclassified(100);k__Fungi_unclassified(100); | - | - | - | - | - | - | - | Unassigned | - |
| Otu58598 | 1 | k__Fungi(100);k__Fungi_unclassified(100);k__Fungi_unclassified(100);k__Fungi_unclassified(100);k__Fungi_unclassified(100);k__Fungi_unclassified(100);k__Fungi_unclassified(100); | - | - | - | - | - | - | - | Unassigned | - |
| Otu58599 | 1 | k__Fungi(100);k__Fungi_unclassified(100);k__Fungi_unclassified(100);k__Fungi_unclassified(100);k__Fungi_unclassified(100);k__Fungi_unclassified(100);k__Fungi_unclassified(100); | - | - | - | - | - | - | - | Unassigned | - |
| Otu58600 | 1 | k__Fungi(100);k__Fungi_unclassified(100);k__Fungi_unclassified(100);k__Fungi_unclassified(100);k__Fungi_unclassified(100);k__Fungi_unclassified(100);k__Fungi_unclassified(100); | - | - | - | - | - | - | - | Unassigned | - |
| Otu58601 | 1 | k__Fungi(100);p__Ascomycota(100);p__Ascomycota_unclassified(100);p__Ascomycota_unclassified(100);p__Ascomycota_unclassified(100);p__Ascomycota_unclassified(100);p__Ascomycota_unclassified(100); | - | - | - | - | - | - | - | Unassigned | - |
| Otu58605 | 1 | k__Fungi(100);p__Ascomycota(100);p__Ascomycota_unclassified(100);p__Ascomycota_unclassified(100);p__Ascomycota_unclassified(100);p__Ascomycota_unclassified(100);p__Ascomycota_unclassified(100); | - | - | - | - | - | - | - | Unassigned | - |
| Otu58632 | 1 | k__Fungi(100);k__Fungi_unclassified(100);k__Fungi_unclassified(100);k__Fungi_unclassified(100);k__Fungi_unclassified(100);k__Fungi_unclassified(100);k__Fungi_unclassified(100); | - | - | - | - | - | - | - | Unassigned | - |
| Otu58633 | 1 | k__Fungi(100);k__Fungi_unclassified(100);k__Fungi_unclassified(100);k__Fungi_unclassified(100);k__Fungi_unclassified(100);k__Fungi_unclassified(100);k__Fungi_unclassified(100); | - | - | - | - | - | - | - | Unassigned | - |

| | | | | | | | | | | | |
|---|---|---|---|---|---|---|---|---|---|---|---|
| Otu58642 | 1 | k__Fungi(100);k__Fungi_unclassified(100);k__Fungi_unclassified(100);k__Fungi_unclassified(100);k__Fungi_unclassified(100);k__Fungi_unclassified(100);k__Fungi_unclassified(100); | - | - | - | - | - | - | - | Unassigned | - |
| Otu58645 | 1 | k__Fungi(100);p__Ascomycota(100);p__Ascomycota_unclassified(100);p__Ascomycota_unclassified(100);p__Ascomycota_unclassified(100);p__Ascomycota_unclassified(100);p__Ascomycota_unclassified(100); | - | - | - | - | - | - | - | Unassigned | - |
| Otu58649 | 1 | k__Fungi(100);p__Ascomycota(100);p__Ascomycota_unclassified(100);p__Ascomycota_unclassified(100);p__Ascomycota_unclassified(100);p__Ascomycota_unclassified(100);p__Ascomycota_unclassified(100); | - | - | - | - | - | - | - | Unassigned | - |
| Otu58650 | 1 | k__Fungi(100);p__Ascomycota(100);p__Ascomycota_unclassified(100);p__Ascomycota_unclassified(100);p__Ascomycota_unclassified(100);p__Ascomycota_unclassified(100);p__Ascomycota_unclassified(100); | - | - | - | - | - | - | - | Unassigned | - |
| Otu58651 | 1 | k__Fungi(100);k__Fungi_unclassified(100);k__Fungi_unclassified(100);k__Fungi_unclassified(100);k__Fungi_unclassified(100);k__Fungi_unclassified(100);k__Fungi_unclassified(100); | - | - | - | - | - | - | - | Unassigned | - |
| Otu58676 | 1 | k__Fungi(100);k__Fungi_unclassified(100);k__Fungi_unclassified(100);k__Fungi_unclassified(100);k__Fungi_unclassified(100);k__Fungi_unclassified(100);k__Fungi_unclassified(100); | - | - | - | - | - | - | - | Unassigned | - |
| Otu58682 | 1 | k__Fungi(100);k__Fungi_unclassified(100);k__Fungi_unclassified(100);k__Fungi_unclassified(100);k__Fungi_unclassified(100);k__Fungi_unclassified(100);k__Fungi_unclassified(100); | - | - | - | - | - | - | - | Unassigned | - |
| Otu58712 | 1 | k__Fungi(100);k__Fungi_unclassified(100);k__Fungi_unclassified(100);k__Fungi_unclassified(100);k__Fungi_unclassified(100);k__Fungi_unclassified(100);k__Fungi_unclassified(100); | - | - | - | - | - | - | - | Unassigned | - |
| Otu58713 | 1 | k__Fungi(100);k__Fungi_unclassified(100);k__Fungi_unclassified(100);k__Fungi_unclassified(100);k__Fungi_unclassified(100);k__Fungi_unclassified(100);k__Fungi_unclassified(100); | - | - | - | - | - | - | - | Unassigned | - |
| Otu58714 | 1 | k__Fungi(100);k__Fungi_unclassified(100);k__Fungi_unclassified(100);k__Fungi_unclassified(100);k__Fungi_unclassified(100);k__Fungi_unclassified(100);k__Fungi_unclassified(100); | - | - | - | - | - | - | - | Unassigned | - |
| Otu58715 | 1 | k__Fungi(100);k__Fungi_unclassified(100);k__Fungi_unclassified(100);k__Fungi_unclassified(100);k__Fungi_unclassified(100);k__Fungi_unclassified(100);k__Fungi_unclassified(100); | - | - | - | - | - | - | - | Unassigned | - |
| Otu58716 | 1 | k__Fungi(100);k__Fungi_unclassified(100);k__Fungi_unclassified(100);k__Fungi_unclassified(100);k__Fungi_unclassified(100);k__Fungi_unclassified(100);k__Fungi_unclassified(100); | - | - | - | - | - | - | - | Unassigned | - |
| Otu58717 | 1 | k__Fungi(100);k__Fungi_unclassified(100);k__Fungi_unclassified(100);k__Fungi_unclassified(100);k__Fungi_unclassified(100);k__Fungi_unclassified(100);k__Fungi_unclassified(100); | - | - | - | - | - | - | - | Unassigned | - |
| Otu58718 | 1 | k__Fungi(100);k__Fungi_unclassified(100);k__Fungi_unclassified(100);k__Fungi_unclassified(100);k__Fungi_unclassified(100);k__Fungi_unclassified(100);k__Fungi_unclassified(100); | - | - | - | - | - | - | - | Unassigned | - |
| Otu58719 | 1 | k__Fungi(100);p__Ascomycota(100);p__Ascomycota_unclassified(100);p__Ascomycota_unclassified(100);p__Ascomycota_unclassified(100);p__Ascomycota_unclassified(100);p__Ascomycota_unclassified(100); | - | - | - | - | - | - | - | Unassigned | - |
| Otu58720 | 1 | k__Fungi(100);k__Fungi_unclassified(100);k__Fungi_unclassified(100);k__Fungi_unclassified(100);k__Fungi_unclassified(100);k__Fungi_unclassified(100);k__Fungi_unclassified(100); | - | - | - | - | - | - | - | Unassigned | - |
| Otu58721 | 1 | k__Fungi(100);p__Ascomycota(100);p__Ascomycota_unclassified(100);p__Ascomycota_unclassified(100);p__Ascomycota_unclassified(100);p__Ascomycota_unclassified(100);p__Ascomycota_unclassified(100); | - | - | - | - | - | - | - | Unassigned | - |
| Otu58722 | 1 | k__Fungi(100);k__Fungi_unclassified(100);k__Fungi_unclassified(100);k__Fungi_unclassified(100);k__Fungi_unclassified(100);k__Fungi_unclassified(100);k__Fungi_unclassified(100); | - | - | - | - | - | - | - | Unassigned | - |
| Otu58723 | 1 | k__Fungi(100);p__Ascomycota(100);p__Ascomycota_unclassified(100);p__Ascomycota_unclassified(100);p__Ascomycota_unclassified(100);p__Ascomycota_unclassified(100);p__Ascomycota_unclassified(100); | - | - | - | - | - | - | - | Unassigned | - |
| Otu58725 | 1 | k__Fungi(100);k__Fungi_unclassified(100);k__Fungi_unclassified(100);k__Fungi_unclassified(100);k__Fungi_unclassified(100);k__Fungi_unclassified(100);k__Fungi_unclassified(100); | - | - | - | - | - | - | - | Unassigned | - |
| Otu58726 | 1 | k__Fungi(100);p__Ascomycota(100);p__Ascomycota_unclassified(100);p__Ascomycota_unclassified(100);p__Ascomycota_unclassified(100);p__Ascomycota_unclassified(100);p__Ascomycota_unclassified(100); | - | - | - | - | - | - | - | Unassigned | - |
| Otu58727 | 1 | k__Fungi(100);p__Ascomycota(100);p__Ascomycota_unclassified(100);p__Ascomycota_unclassified(100);p__Ascomycota_unclassified(100);p__Ascomycota_unclassified(100);p__Ascomycota_unclassified(100); | - | - | - | - | - | - | - | Unassigned | - |
| Otu58735 | 1 | k__Fungi(100);p__Ascomycota(100);p__Ascomycota_unclassified(100);p__Ascomycota_unclassified(100);p__Ascomycota_unclassified(100);p__Ascomycota_unclassified(100);p__Ascomycota_unclassified(100); | - | - | - | - | - | - | - | Unassigned | - |
| Otu58737 | 1 | k__Fungi(100);k__Fungi_unclassified(100);k__Fungi_unclassified(100);k__Fungi_unclassified(100);k__Fungi_unclassified(100);k__Fungi_unclassified(100);k__Fungi_unclassified(100); | - | - | - | - | - | - | - | Unassigned | - |
| Otu58745 | 1 | k__Fungi(100);k__Fungi_unclassified(100);k__Fungi_unclassified(100);k__Fungi_unclassified(100);k__Fungi_unclassified(100);k__Fungi_unclassified(100);k__Fungi_unclassified(100); | - | - | - | - | - | - | - | Unassigned | - |
| Otu58746 | 1 | k__Fungi(100);k__Fungi_unclassified(100);k__Fungi_unclassified(100);k__Fungi_unclassified(100);k__Fungi_unclassified(100);k__Fungi_unclassified(100);k__Fungi_unclassified(100); | - | - | - | - | - | - | - | Unassigned | - |
| Otu58751 | 1 | k__Fungi(100);k__Fungi_unclassified(100);k__Fungi_unclassified(100);k__Fungi_unclassified(100);k__Fungi_unclassified(100);k__Fungi_unclassified(100);k__Fungi_unclassified(100); | - | - | - | - | - | - | - | Unassigned | - |
| Otu58752 | 1 | k__Fungi(100);k__Fungi_unclassified(100);k__Fungi_unclassified(100);k__Fungi_unclassified(100);k__Fungi_unclassified(100);k__Fungi_unclassified(100);k__Fungi_unclassified(100); | - | - | - | - | - | - | - | Unassigned | - |
| Otu58753 | 1 | k__Fungi(100);k__Fungi_unclassified(100);k__Fungi_unclassified(100);k__Fungi_unclassified(100);k__Fungi_unclassified(100);k__Fungi_unclassified(100);k__Fungi_unclassified(100); | - | - | - | - | - | - | - | Unassigned | - |
| Otu58754 | 1 | k__Fungi(100);k__Fungi_unclassified(100);k__Fungi_unclassified(100);k__Fungi_unclassified(100);k__Fungi_unclassified(100);k__Fungi_unclassified(100);k__Fungi_unclassified(100); | - | - | - | - | - | - | - | Unassigned | - |
| Otu58755 | 1 | k__Fungi(100);k__Fungi_unclassified(100);k__Fungi_unclassified(100);k__Fungi_unclassified(100);k__Fungi_unclassified(100);k__Fungi_unclassified(100);k__Fungi_unclassified(100); | - | - | - | - | - | - | - | Unassigned | - |
| Otu58756 | 1 | k__Fungi(100);k__Fungi_unclassified(100);k__Fungi_unclassified(100);k__Fungi_unclassified(100);k__Fungi_unclassified(100);k__Fungi_unclassified(100);k__Fungi_unclassified(100); | - | - | - | - | - | - | - | Unassigned | - |
| Otu58757 | 1 | k__Fungi(100);k__Fungi_unclassified(100);k__Fungi_unclassified(100);k__Fungi_unclassified(100);k__Fungi_unclassified(100);k__Fungi_unclassified(100);k__Fungi_unclassified(100); | - | - | - | - | - | - | - | Unassigned | - |
| Otu58758 | 1 | k__Fungi(100);k__Fungi_unclassified(100);k__Fungi_unclassified(100);k__Fungi_unclassified(100);k__Fungi_unclassified(100);k__Fungi_unclassified(100);k__Fungi_unclassified(100); | - | - | - | - | - | - | - | Unassigned | - |
| Otu58759 | 1 | k__Fungi(100);k__Fungi_unclassified(100);k__Fungi_unclassified(100);k__Fungi_unclassified(100);k__Fungi_unclassified(100);k__Fungi_unclassified(100);k__Fungi_unclassified(100); | - | - | - | - | - | - | - | Unassigned | - |
| Otu58760 | 1 | k__Fungi(100);k__Fungi_unclassified(100);k__Fungi_unclassified(100);k__Fungi_unclassified(100);k__Fungi_unclassified(100);k__Fungi_unclassified(100);k__Fungi_unclassified(100); | - | - | - | - | - | - | - | Unassigned | - |
| Otu58761 | 1 | k__Fungi(100);k__Fungi_unclassified(100);k__Fungi_unclassified(100);k__Fungi_unclassified(100);k__Fungi_unclassified(100);k__Fungi_unclassified(100);k__Fungi_unclassified(100); | - | - | - | - | - | - | - | Unassigned | - |
| Otu58762 | 1 | k__Fungi(100);k__Fungi_unclassified(100);k__Fungi_unclassified(100);k__Fungi_unclassified(100);k__Fungi_unclassified(100);k__Fungi_unclassified(100);k__Fungi_unclassified(100); | - | - | - | - | - | - | - | Unassigned | - |
| Otu58763 | 1 | k__Fungi(100);p__Ascomycota(100);p__Ascomycota_unclassified(100);p__Ascomycota_unclassified(100);p__Ascomycota_unclassified(100);p__Ascomycota_unclassified(100);p__Ascomycota_unclassified(100); | - | - | - | - | - | - | - | Unassigned | - |
| Otu58764 | 1 | k__Fungi(100);p__Ascomycota(100);p__Ascomycota_unclassified(100);p__Ascomycota_unclassified(100);p__Ascomycota_unclassified(100);p__Ascomycota_unclassified(100);p__Ascomycota_unclassified(100); | - | - | - | - | - | - | - | Unassigned | - |
| Otu58765 | 1 | k__Fungi(100);k__Fungi_unclassified(100);k__Fungi_unclassified(100);k__Fungi_unclassified(100);k__Fungi_unclassified(100);k__Fungi_unclassified(100);k__Fungi_unclassified(100); | - | - | - | - | - | - | - | Unassigned | - |
| Otu58766 | 1 | k__Fungi(100);k__Fungi_unclassified(100);k__Fungi_unclassified(100);k__Fungi_unclassified(100);k__Fungi_unclassified(100);k__Fungi_unclassified(100);k__Fungi_unclassified(100); | - | - | - | - | - | - | - | Unassigned | - |
| Otu58767 | 1 | k__Fungi(100);k__Fungi_unclassified(100);k__Fungi_unclassified(100);k__Fungi_unclassified(100);k__Fungi_unclassified(100);k__Fungi_unclassified(100);k__Fungi_unclassified(100); | - | - | - | - | - | - | - | Unassigned | - |
| Otu58768 | 1 | k__Fungi(100);k__Fungi_unclassified(100);k__Fungi_unclassified(100);k__Fungi_unclassified(100);k__Fungi_unclassified(100);k__Fungi_unclassified(100);k__Fungi_unclassified(100); | - | - | - | - | - | - | - | Unassigned | - |
| Otu58770 | 1 | k__Fungi(100);k__Fungi_unclassified(100);k__Fungi_unclassified(100);k__Fungi_unclassified(100);k__Fungi_unclassified(100);k__Fungi_unclassified(100);k__Fungi_unclassified(100); | - | - | - | - | - | - | - | Unassigned | - |
| Otu58771 | 1 | k__Fungi(100);k__Fungi_unclassified(100);k__Fungi_unclassified(100);k__Fungi_unclassified(100);k__Fungi_unclassified(100);k__Fungi_unclassified(100);k__Fungi_unclassified(100); | - | - | - | - | - | - | - | Unassigned | - |
| Otu58772 | 1 | k__Fungi(100);k__Fungi_unclassified(100);k__Fungi_unclassified(100);k__Fungi_unclassified(100);k__Fungi_unclassified(100);k__Fungi_unclassified(100);k__Fungi_unclassified(100); | - | - | - | - | - | - | - | Unassigned | - |
| Otu58773 | 1 | k__Fungi(100);k__Fungi_unclassified(100);k__Fungi_unclassified(100);k__Fungi_unclassified(100);k__Fungi_unclassified(100);k__Fungi_unclassified(100);k__Fungi_unclassified(100); | - | - | - | - | - | - | - | Unassigned | - |
| Otu58775 | 1 | k__Fungi(100);p__Ascomycota(100);p__Ascomycota_unclassified(100);p__Ascomycota_unclassified(100);p__Ascomycota_unclassified(100);p__Ascomycota_unclassified(100);p__Ascomycota_unclassified(100); | - | - | - | - | - | - | - | Unassigned | - |
| Otu58776 | 1 | k__Fungi(100);k__Fungi_unclassified(100);k__Fungi_unclassified(100);k__Fungi_unclassified(100);k__Fungi_unclassified(100);k__Fungi_unclassified(100);k__Fungi_unclassified(100); | - | - | - | - | - | - | - | Unassigned | - |
| Otu58777 | 1 | k__Fungi(100);k__Fungi_unclassified(100);k__Fungi_unclassified(100);k__Fungi_unclassified(100);k__Fungi_unclassified(100);k__Fungi_unclassified(100);k__Fungi_unclassified(100); | - | - | - | - | - | - | - | Unassigned | - |
| Otu58778 | 1 | k__Fungi(100);k__Fungi_unclassified(100);k__Fungi_unclassified(100);k__Fungi_unclassified(100);k__Fungi_unclassified(100);k__Fungi_unclassified(100);k__Fungi_unclassified(100); | - | - | - | - | - | - | - | Unassigned | - |
| Otu58781 | 1 | k__Fungi(100);k__Fungi_unclassified(100);k__Fungi_unclassified(100);k__Fungi_unclassified(100);k__Fungi_unclassified(100);k__Fungi_unclassified(100);k__Fungi_unclassified(100); | - | - | - | - | - | - | - | Unassigned | - |
| Otu58782 | 1 | k__Fungi(100);k__Fungi_unclassified(100);k__Fungi_unclassified(100);k__Fungi_unclassified(100);k__Fungi_unclassified(100);k__Fungi_unclassified(100);k__Fungi_unclassified(100); | - | - | - | - | - | - | - | Unassigned | - |
| Otu58783 | 1 | k__Fungi(100);k__Fungi_unclassified(100);k__Fungi_unclassified(100);k__Fungi_unclassified(100);k__Fungi_unclassified(100);k__Fungi_unclassified(100);k__Fungi_unclassified(100); | - | - | - | - | - | - | - | Unassigned | - |
| Otu58784 | 1 | k__Fungi(100);k__Fungi_unclassified(100);k__Fungi_unclassified(100);k__Fungi_unclassified(100);k__Fungi_unclassified(100);k__Fungi_unclassified(100);k__Fungi_unclassified(100); | - | - | - | - | - | - | - | Unassigned | - |
| Otu58785 | 1 | k__Fungi(100);k__Fungi_unclassified(100);k__Fungi_unclassified(100);k__Fungi_unclassified(100);k__Fungi_unclassified(100);k__Fungi_unclassified(100);k__Fungi_unclassified(100); | - | - | - | - | - | - | - | Unassigned | - |
| Otu58789 | 1 | k__Fungi(100);k__Fungi_unclassified(100);k__Fungi_unclassified(100);k__Fungi_unclassified(100);k__Fungi_unclassified(100);k__Fungi_unclassified(100);k__Fungi_unclassified(100); | - | - | - | - | - | - | - | Unassigned | - |
| Otu58792 | 1 | k__Fungi(100);k__Fungi_unclassified(100);k__Fungi_unclassified(100);k__Fungi_unclassified(100);k__Fungi_unclassified(100);k__Fungi_unclassified(100);k__Fungi_unclassified(100); | - | - | - | - | - | - | - | Unassigned | - |
| Otu58795 | 1 | k__Fungi(100);k__Fungi_unclassified(100);k__Fungi_unclassified(100);k__Fungi_unclassified(100);k__Fungi_unclassified(100);k__Fungi_unclassified(100);k__Fungi_unclassified(100); | - | - | - | - | - | - | - | Unassigned | - |
| Otu58796 | 1 | k__Fungi(100);k__Fungi_unclassified(100);k__Fungi_unclassified(100);k__Fungi_unclassified(100);k__Fungi_unclassified(100);k__Fungi_unclassified(100);k__Fungi_unclassified(100); | - | - | - | - | - | - | - | Unassigned | - |
| Otu58797 | 1 | k__Fungi(100);k__Fungi_unclassified(100);k__Fungi_unclassified(100);k__Fungi_unclassified(100);k__Fungi_unclassified(100);k__Fungi_unclassified(100);k__Fungi_unclassified(100); | - | - | - | - | - | - | - | Unassigned | - |
| Otu58799 | 1 | k__Fungi(100);k__Fungi_unclassified(100);k__Fungi_unclassified(100);k__Fungi_unclassified(100);k__Fungi_unclassified(100);k__Fungi_unclassified(100);k__Fungi_unclassified(100); | - | - | - | - | - | - | - | Unassigned | - |
| Otu58804 | 1 | k__Fungi(100);k__Fungi_unclassified(100);k__Fungi_unclassified(100);k__Fungi_unclassified(100);k__Fungi_unclassified(100);k__Fungi_unclassified(100);k__Fungi_unclassified(100); | - | - | - | - | - | - | - | Unassigned | - |
| Otu58805 | 1 | k__Fungi(100);k__Fungi_unclassified(100);k__Fungi_unclassified(100);k__Fungi_unclassified(100);k__Fungi_unclassified(100);k__Fungi_unclassified(100);k__Fungi_unclassified(100); | - | - | - | - | - | - | - | Unassigned | - |
| Otu58806 | 1 | k__Fungi(100);p__Ascomycota(100);p__Ascomycota_unclassified(100);p__Ascomycota_unclassified(100);p__Ascomycota_unclassified(100);p__Ascomycota_unclassified(100);p__Ascomycota_unclassified(100); | - | - | - | - | - | - | - | Unassigned | - |
| Otu58807 | 1 | k__Fungi(100);k__Fungi_unclassified(100);k__Fungi_unclassified(100);k__Fungi_unclassified(100);k__Fungi_unclassified(100);k__Fungi_unclassified(100);k__Fungi_unclassified(100); | - | - | - | - | - | - | - | Unassigned | - |
| Otu58808 | 1 | k__Fungi(100);k__Fungi_unclassified(100);k__Fungi_unclassified(100);k__Fungi_unclassified(100);k__Fungi_unclassified(100);k__Fungi_unclassified(100);k__Fungi_unclassified(100); | - | - | - | - | - | - | - | Unassigned | - |
| Otu58809 | 1 | k__Fungi(100);k__Fungi_unclassified(100);k__Fungi_unclassified(100);k__Fungi_unclassified(100);k__Fungi_unclassified(100);k__Fungi_unclassified(100);k__Fungi_unclassified(100); | - | - | - | - | - | - | - | Unassigned | - |
| Otu58810 | 1 | k__Fungi(100);k__Fungi_unclassified(100);k__Fungi_unclassified(100);k__Fungi_unclassified(100);k__Fungi_unclassified(100);k__Fungi_unclassified(100);k__Fungi_unclassified(100); | - | - | - | - | - | - | - | Unassigned | - |
| Otu58811 | 1 | k__Fungi(100);k__Fungi_unclassified(100);k__Fungi_unclassified(100);k__Fungi_unclassified(100);k__Fungi_unclassified(100);k__Fungi_unclassified(100);k__Fungi_unclassified(100); | - | - | - | - | - | - | - | Unassigned | - |
| Otu58812 | 1 | k__Fungi(100);k__Fungi_unclassified(100);k__Fungi_unclassified(100);k__Fungi_unclassified(100);k__Fungi_unclassified(100);k__Fungi_unclassified(100);k__Fungi_unclassified(100); | - | - | - | - | - | - | - | Unassigned | - |
| Otu58813 | 1 | k__Fungi(100);k__Fungi_unclassified(100);k__Fungi_unclassified(100);k__Fungi_unclassified(100);k__Fungi_unclassified(100);k__Fungi_unclassified(100);k__Fungi_unclassified(100); | - | - | - | - | - | - | - | Unassigned | - |
| Otu58814 | 1 | k__Fungi(100);k__Fungi_unclassified(100);k__Fungi_unclassified(100);k__Fungi_unclassified(100);k__Fungi_unclassified(100);k__Fungi_unclassified(100);k__Fungi_unclassified(100); | - | - | - | - | - | - | - | Unassigned | - |
| Otu58815 | 1 | k__Fungi(100);k__Fungi_unclassified(100);k__Fungi_unclassified(100);k__Fungi_unclassified(100);k__Fungi_unclassified(100);k__Fungi_unclassified(100);k__Fungi_unclassified(100); | - | - | - | - | - | - | - | Unassigned | - |
| Otu58825 | 1 | k__Fungi(100);p__Ascomycota(100);p__Ascomycota_unclassified(100);p__Ascomycota_unclassified(100);p__Ascomycota_unclassified(100);p__Ascomycota_unclassified(100);p__Ascomycota_unclassified(100); | - | - | - | - | - | - | - | Unassigned | - |
| Otu58828 | 1 | k__Fungi(100);p__Ascomycota(100);p__Ascomycota_unclassified(100);p__Ascomycota_unclassified(100);p__Ascomycota_unclassified(100);p__Ascomycota_unclassified(100);p__Ascomycota_unclassified(100); | - | - | - | - | - | - | - | Unassigned | - |
| Otu58841 | 1 | k__Fungi(100);p__Ascomycota(100);p__Ascomycota_unclassified(100);p__Ascomycota_unclassified(100);p__Ascomycota_unclassified(100);p__Ascomycota_unclassified(100);p__Ascomycota_unclassified(100); | - | - | - | - | - | - | - | Unassigned | - |
| Otu58845 | 1 | k__Fungi(100);p__Ascomycota(100);p__Ascomycota_unclassified(100);p__Ascomycota_unclassified(100);p__Ascomycota_unclassified(100);p__Ascomycota_unclassified(100);p__Ascomycota_unclassified(100); | - | - | - | - | - | - | - | Unassigned | - |
| Otu58849 | 1 | k__Fungi(100);p__Ascomycota(100);p__Ascomycota_unclassified(100);p__Ascomycota_unclassified(100);p__Ascomycota_unclassified(100);p__Ascomycota_unclassified(100);p__Ascomycota_unclassified(100); | - | - | - | - | - | - | - | Unassigned | - |
| Otu58852 | 1 | k__Fungi(100);k__Fungi_unclassified(100);k__Fungi_unclassified(100);k__Fungi_unclassified(100);k__Fungi_unclassified(100);k__Fungi_unclassified(100);k__Fungi_unclassified(100); | - | - | - | - | - | - | - | Unassigned | - |
| Otu58853 | 1 | k__Fungi(100);k__Fungi_unclassified(100);k__Fungi_unclassified(100);k__Fungi_unclassified(100);k__Fungi_unclassified(100);k__Fungi_unclassified(100);k__Fungi_unclassified(100); | - | - | - | - | - | - | - | Unassigned | - |
| Otu58855 | 1 | k__Fungi(100);k__Fungi_unclassified(100);k__Fungi_unclassified(100);k__Fungi_unclassified(100);k__Fungi_unclassified(100);k__Fungi_unclassified(100);k__Fungi_unclassified(100); | - | - | - | - | - | - | - | Unassigned | - |
| Otu58856 | 1 | k__Fungi(100);k__Fungi_unclassified(100);k__Fungi_unclassified(100);k__Fungi_unclassified(100);k__Fungi_unclassified(100);k__Fungi_unclassified(100);k__Fungi_unclassified(100); | - | - | - | - | - | - | - | Unassigned | - |
| Otu58857 | 1 | k__Fungi(100);p__Ascomycota(100);p__Ascomycota_unclassified(100);p__Ascomycota_unclassified(100);p__Ascomycota_unclassified(100);p__Ascomycota_unclassified(100);p__Ascomycota_unclassified(100); | - | - | - | - | - | - | - | Unassigned | - |
| Otu58859 | 1 | k__Fungi(100);k__Fungi_unclassified(100);k__Fungi_unclassified(100);k__Fungi_unclassified(100);k__Fungi_unclassified(100);k__Fungi_unclassified(100);k__Fungi_unclassified(100); | - | - | - | - | - | - | - | Unassigned | - |
| Otu58860 | 1 | k__Fungi(100);k__Fungi_unclassified(100);k__Fungi_unclassified(100);k__Fungi_unclassified(100);k__Fungi_unclassified(100);k__Fungi_unclassified(100);k__Fungi_unclassified(100); | - | - | - | - | - | - | - | Unassigned | - |
| Otu58864 | 1 | k__Fungi(100);k__Fungi_unclassified(100);k__Fungi_unclassified(100);k__Fungi_unclassified(100);k__Fungi_unclassified(100);k__Fungi_unclassified(100);k__Fungi_unclassified(100); | - | - | - | - | - | - | - | Unassigned | - |
| Otu58865 | 1 | k__Fungi(100);k__Fungi_unclassified(100);k__Fungi_unclassified(100);k__Fungi_unclassified(100);k__Fungi_unclassified(100);k__Fungi_unclassified(100);k__Fungi_unclassified(100); | - | - | - | - | - | - | - | Unassigned | - |
| Otu58866 | 1 | k__Fungi(100);p__Ascomycota(100);p__Ascomycota_unclassified(100);p__Ascomycota_unclassified(100);p__Ascomycota_unclassified(100);p__Ascomycota_unclassified(100);p__Ascomycota_unclassified(100); | - | - | - | - | - | - | - | Unassigned | - |
| Otu58870 | 1 | k__Fungi(100);k__Fungi_unclassified(100);k__Fungi_unclassified(100);k__Fungi_unclassified(100);k__Fungi_unclassified(100);k__Fungi_unclassified(100);k__Fungi_unclassified(100); | - | - | - | - | - | - | - | Unassigned | - |
| Otu58878 | 1 | k__Fungi(100);p__Ascomycota(100);p__Ascomycota_unclassified(100);p__Ascomycota_unclassified(100);p__Ascomycota_unclassified(100);p__Ascomycota_unclassified(100);p__Ascomycota_unclassified(100); | - | - | - | - | - | - | - | Unassigned | - |
| Otu58880 | 1 | k__Fungi(100);k__Fungi_unclassified(100);k__Fungi_unclassified(100);k__Fungi_unclassified(100);k__Fungi_unclassified(100);k__Fungi_unclassified(100);k__Fungi_unclassified(100); | - | - | - | - | - | - | - | Unassigned | - |
| Otu58902 | 1 | k__Fungi(100);k__Fungi_unclassified(100);k__Fungi_unclassified(100);k__Fungi_unclassified(100);k__Fungi_unclassified(100);k__Fungi_unclassified(100);k__Fungi_unclassified(100); | - | - | - | - | - | - | - | Unassigned | - |
| Otu58905 | 1 | k__Fungi(100);p__Ascomycota(100);p__Ascomycota_unclassified(100);p__Ascomycota_unclassified(100);p__Ascomycota_unclassified(100);p__Ascomycota_unclassified(100);p__Ascomycota_unclassified(100); | - | - | - | - | - | - | - | Unassigned | - |
| Otu58918 | 1 | k__Fungi(100);k__Fungi_unclassified(100);k__Fungi_unclassified(100);k__Fungi_unclassified(100);k__Fungi_unclassified(100);k__Fungi_unclassified(100);k__Fungi_unclassified(100); | - | - | - | - | - | - | - | Unassigned | - |
| Otu58947 | 1 | k__Fungi(100);k__Fungi_unclassified(100);k__Fungi_unclassified(100);k__Fungi_unclassified(100);k__Fungi_unclassified(100);k__Fungi_unclassified(100);k__Fungi_unclassified(100); | - | - | - | - | - | - | - | Unassigned | - |
| Otu58984 | 1 | k__Fungi(100);p__Ascomycota(100);p__Ascomycota_unclassified(100);p__Ascomycota_unclassified(100);p__Ascomycota_unclassified(100);p__Ascomycota_unclassified(100);p__Ascomycota_unclassified(100); | - | - | - | - | - | - | - | Unassigned | - |
| Otu58986 | 1 | k__Fungi(100);p__Ascomycota(100);p__Ascomycota_unclassified(100);p__Ascomycota_unclassified(100);p__Ascomycota_unclassified(100);p__Ascomycota_unclassified(100);p__Ascomycota_unclassified(100); | - | - | - | - | - | - | - | Unassigned | - |
| Otu59013 | 1 | k__Fungi(100);p__Ascomycota(100);p__Ascomycota_unclassified(100);p__Ascomycota_unclassified(100);p__Ascomycota_unclassified(100);p__Ascomycota_unclassified(100);p__Ascomycota_unclassified(100); | - | - | - | - | - | - | - | Unassigned | - |
| Otu59026 | 1 | k__Fungi(100);k__Fungi_unclassified(100);k__Fungi_unclassified(100);k__Fungi_unclassified(100);k__Fungi_unclassified(100);k__Fungi_unclassified(100);k__Fungi_unclassified(100); | - | - | - | - | - | - | - | Unassigned | - |
| Otu59036 | 1 | k__Fungi(100);k__Fungi_unclassified(100);k__Fungi_unclassified(100);k__Fungi_unclassified(100);k__Fungi_unclassified(100);k__Fungi_unclassified(100);k__Fungi_unclassified(100); | - | - | - | - | - | - | - | Unassigned | - |
| Otu59037 | 1 | k__Fungi(100);k__Fungi_unclassified(100);k__Fungi_unclassified(100);k__Fungi_unclassified(100);k__Fungi_unclassified(100);k__Fungi_unclassified(100);k__Fungi_unclassified(100); | - | - | - | - | - | - | - | Unassigned | - |
| Otu59039 | 1 | k__Fungi(100);k__Fungi_unclassified(100);k__Fungi_unclassified(100);k__Fungi_unclassified(100);k__Fungi_unclassified(100);k__Fungi_unclassified(100);k__Fungi_unclassified(100); | - | - | - | - | - | - | - | Unassigned | - |
| Otu59047 | 1 | k__Fungi(100);k__Fungi_unclassified(100);k__Fungi_unclassified(100);k__Fungi_unclassified(100);k__Fungi_unclassified(100);k__Fungi_unclassified(100);k__Fungi_unclassified(100); | - | - | - | - | - | - | - | Unassigned | - |
| Otu59054 | 1 | k__Fungi(100);k__Fungi_unclassified(100);k__Fungi_unclassified(100);k__Fungi_unclassified(100);k__Fungi_unclassified(100);k__Fungi_unclassified(100);k__Fungi_unclassified(100); | - | - | - | - | - | - | - | Unassigned | - |
| Otu59056 | 1 | k__Fungi(100);k__Fungi_unclassified(100);k__Fungi_unclassified(100);k__Fungi_unclassified(100);k__Fungi_unclassified(100);k__Fungi_unclassified(100);k__Fungi_unclassified(100); | - | - | - | - | - | - | - | Unassigned | - |
| Otu59060 | 1 | k__Fungi(100);k__Fungi_unclassified(100);k__Fungi_unclassified(100);k__Fungi_unclassified(100);k__Fungi_unclassified(100);k__Fungi_unclassified(100);k__Fungi_unclassified(100); | - | - | - | - | - | - | - | Unassigned | - |
| Otu59064 | 1 | k__Fungi(100);p__Ascomycota(100);p__Ascomycota_unclassified(100);p__Ascomycota_unclassified(100);p__Ascomycota_unclassified(100);p__Ascomycota_unclassified(100);p__Ascomycota_unclassified(100); | - | - | - | - | - | - | - | Unassigned | - |
| Otu59065 | 1 | k__Fungi(100);k__Fungi_unclassified(100);k__Fungi_unclassified(100);k__Fungi_unclassified(100);k__Fungi_unclassified(100);k__Fungi_unclassified(100);k__Fungi_unclassified(100); | - | - | - | - | - | - | - | Unassigned | - |
| Otu59066 | 1 | k__Fungi(100);k__Fungi_unclassified(100);k__Fungi_unclassified(100);k__Fungi_unclassified(100);k__Fungi_unclassified(100);k__Fungi_unclassified(100);k__Fungi_unclassified(100); | - | - | - | - | - | - | - | Unassigned | - |
| Otu59068 | 1 | k__Fungi(100);k__Fungi_unclassified(100);k__Fungi_unclassified(100);k__Fungi_unclassified(100);k__Fungi_unclassified(100);k__Fungi_unclassified(100);k__Fungi_unclassified(100); | - | - | - | - | - | - | - | Unassigned | - |
| Otu59076 | 1 | k__Fungi(100);k__Fungi_unclassified(100);k__Fungi_unclassified(100);k__Fungi_unclassified(100);k__Fungi_unclassified(100);k__Fungi_unclassified(100);k__Fungi_unclassified(100); | - | - | - | - | - | - | - | Unassigned | - |
| Otu59084 | 1 | k__Fungi(100);k__Fungi_unclassified(100);k__Fungi_unclassified(100);k__Fungi_unclassified(100);k__Fungi_unclassified(100);k__Fungi_unclassified(100);k__Fungi_unclassified(100); | - | - | - | - | - | - | - | Unassigned | - |
| Otu59088 | 1 | k__Fungi(100);k__Fungi_unclassified(100);k__Fungi_unclassified(100);k__Fungi_unclassified(100);k__Fungi_unclassified(100);k__Fungi_unclassified(100);k__Fungi_unclassified(100); | - | - | - | - | - | - | - | Unassigned | - |
| Otu59095 | 1 | k__Fungi(100);p__Ascomycota(100);p__Ascomycota_unclassified(100);p__Ascomycota_unclassified(100);p__Ascomycota_unclassified(100);p__Ascomycota_unclassified(100);p__Ascomycota_unclassified(100); | - | - | - | - | - | - | - | Unassigned | - |
| Otu59099 | 1 | k__Fungi(100);k__Fungi_unclassified(100);k__Fungi_unclassified(100);k__Fungi_unclassified(100);k__Fungi_unclassified(100);k__Fungi_unclassified(100);k__Fungi_unclassified(100); | - | - | - | - | - | - | - | Unassigned | - |
| Otu59101 | 1 | k__Fungi(100);k__Fungi_unclassified(100);k__Fungi_unclassified(100);k__Fungi_unclassified(100);k__Fungi_unclassified(100);k__Fungi_unclassified(100);k__Fungi_unclassified(100); | - | - | - | - | - | - | - | Unassigned | - |
| Otu59111 | 1 | k__Fungi(100);k__Fungi_unclassified(100);k__Fungi_unclassified(100);k__Fungi_unclassified(100);k__Fungi_unclassified(100);k__Fungi_unclassified(100);k__Fungi_unclassified(100); | - | - | - | - | - | - | - | Unassigned | - |
| Otu59112 | 1 | k__Fungi(100);k__Fungi_unclassified(100);k__Fungi_unclassified(100);k__Fungi_unclassified(100);k__Fungi_unclassified(100);k__Fungi_unclassified(100);k__Fungi_unclassified(100); | - | - | - | - | - | - | - | Unassigned | - |
| Otu59113 | 1 | k__Fungi(100);k__Fungi_unclassified(100);k__Fungi_unclassified(100);k__Fungi_unclassified(100);k__Fungi_unclassified(100);k__Fungi_unclassified(100);k__Fungi_unclassified(100); | - | - | - | - | - | - | - | Unassigned | - |
| Otu59116 | 1 | k__Fungi(100);k__Fungi_unclassified(100);k__Fungi_unclassified(100);k__Fungi_unclassified(100);k__Fungi_unclassified(100);k__Fungi_unclassified(100);k__Fungi_unclassified(100); | - | - | - | - | - | - | - | Unassigned | - |
| Otu59117 | 1 | k__Fungi(100);k__Fungi_unclassified(100);k__Fungi_unclassified(100);k__Fungi_unclassified(100);k__Fungi_unclassified(100);k__Fungi_unclassified(100);k__Fungi_unclassified(100); | - | - | - | - | - | - | - | Unassigned | - |
| Otu59123 | 1 | k__Fungi(100);k__Fungi_unclassified(100);k__Fungi_unclassified(100);k__Fungi_unclassified(100);k__Fungi_unclassified(100);k__Fungi_unclassified(100);k__Fungi_unclassified(100); | - | - | - | - | - | - | - | Unassigned | - |
| Otu59126 | 1 | k__Fungi(100);p__Ascomycota(100);p__Ascomycota_unclassified(100);p__Ascomycota_unclassified(100);p__Ascomycota_unclassified(100);p__Ascomycota_unclassified(100);p__Ascomycota_unclassified(100); | - | - | - | - | - | - | - | Unassigned | - |
| Otu59127 | 1 | k__Fungi(100);p__Ascomycota(100);p__Ascomycota_unclassified(100);p__Ascomycota_unclassified(100);p__Ascomycota_unclassified(100);p__Ascomycota_unclassified(100);p__Ascomycota_unclassified(100); | - | - | - | - | - | - | - | Unassigned | - |
| Otu59129 | 1 | k__Fungi(100);p__Ascomycota(100);p__Ascomycota_unclassified(100);p__Ascomycota_unclassified(100);p__Ascomycota_unclassified(100);p__Ascomycota_unclassified(100);p__Ascomycota_unclassified(100); | - | - | - | - | - | - | - | Unassigned | - |
| Otu59130 | 1 | k__Fungi(100);p__Ascomycota(100);p__Ascomycota_unclassified(100);p__Ascomycota_unclassified(100);p__Ascomycota_unclassified(100);p__Ascomycota_unclassified(100);p__Ascomycota_unclassified(100); | - | - | - | - | - | - | - | Unassigned | - |
| Otu59131 | 1 | k__Fungi(100);k__Fungi_unclassified(100);k__Fungi_unclassified(100);k__Fungi_unclassified(100);k__Fungi_unclassified(100);k__Fungi_unclassified(100);k__Fungi_unclassified(100); | - | - | - | - | - | - | - | Unassigned | - |
| Otu59132 | 1 | k__Fungi(100);k__Fungi_unclassified(100);k__Fungi_unclassified(100);k__Fungi_unclassified(100);k__Fungi_unclassified(100);k__Fungi_unclassified(100);k__Fungi_unclassified(100); | - | - | - | - | - | - | - | Unassigned | - |
| Otu59133 | 1 | k__Fungi(100);p__Ascomycota(100);p__Ascomycota_unclassified(100);p__Ascomycota_unclassified(100);p__Ascomycota_unclassified(100);p__Ascomycota_unclassified(100);p__Ascomycota_unclassified(100); | - | - | - | - | - | - | - | Unassigned | - |
| Otu59134 | 1 | k__Fungi(100);p__Ascomycota(100);p__Ascomycota_unclassified(100);p__Ascomycota_unclassified(100);p__Ascomycota_unclassified(100);p__Ascomycota_unclassified(100);p__Ascomycota_unclassified(100); | - | - | - | - | - | - | - | Unassigned | - |
| Otu59135 | 1 | k__Fungi(100);k__Fungi_unclassified(100);k__Fungi_unclassified(100);k__Fungi_unclassified(100);k__Fungi_unclassified(100);k__Fungi_unclassified(100);k__Fungi_unclassified(100); | - | - | - | - | - | - | - | Unassigned | - |
| Otu59136 | 1 | k__Fungi(100);p__Ascomycota(100);p__Ascomycota_unclassified(100);p__Ascomycota_unclassified(100);p__Ascomycota_unclassified(100);p__Ascomycota_unclassified(100);p__Ascomycota_unclassified(100); | - | - | - | - | - | - | - | Unassigned | - |
| Otu59137 | 1 | k__Fungi(100);k__Fungi_unclassified(100);k__Fungi_unclassified(100);k__Fungi_unclassified(100);k__Fungi_unclassified(100);k__Fungi_unclassified(100);k__Fungi_unclassified(100); | - | - | - | - | - | - | - | Unassigned | - |
| Otu59138 | 1 | k__Fungi(100);p__Ascomycota(100);p__Ascomycota_unclassified(100);p__Ascomycota_unclassified(100);p__Ascomycota_unclassified(100);p__Ascomycota_unclassified(100);p__Ascomycota_unclassified(100); | - | - | - | - | - | - | - | Unassigned | - |
| Otu59139 | 1 | k__Fungi(100);p__Ascomycota(100);p__Ascomycota_unclassified(100);p__Ascomycota_unclassified(100);p__Ascomycota_unclassified(100);p__Ascomycota_unclassified(100);p__Ascomycota_unclassified(100); | - | - | - | - | - | - | - | Unassigned | - |
| Otu59140 | 1 | k__Fungi(100);k__Fungi_unclassified(100);k__Fungi_unclassified(100);k__Fungi_unclassified(100);k__Fungi_unclassified(100);k__Fungi_unclassified(100);k__Fungi_unclassified(100); | - | - | - | - | - | - | - | Unassigned | - |
| Otu59141 | 1 | k__Fungi(100);p__Ascomycota(100);p__Ascomycota_unclassified(100);p__Ascomycota_unclassified(100);p__Ascomycota_unclassified(100);p__Ascomycota_unclassified(100);p__Ascomycota_unclassified(100); | - | - | - | - | - | - | - | Unassigned | - |
| Otu59143 | 1 | k__Fungi(100);k__Fungi_unclassified(100);k__Fungi_unclassified(100);k__Fungi_unclassified(100);k__Fungi_unclassified(100);k__Fungi_unclassified(100);k__Fungi_unclassified(100); | - | - | - | - | - | - | - | Unassigned | - |
| Otu59144 | 1 | k__Fungi(100);k__Fungi_unclassified(100);k__Fungi_unclassified(100);k__Fungi_unclassified(100);k__Fungi_unclassified(100);k__Fungi_unclassified(100);k__Fungi_unclassified(100); | - | - | - | - | - | - | - | Unassigned | - |
| Otu59145 | 1 | k__Fungi(100);k__Fungi_unclassified(100);k__Fungi_unclassified(100);k__Fungi_unclassified(100);k__Fungi_unclassified(100);k__Fungi_unclassified(100);k__Fungi_unclassified(100); | - | - | - | - | - | - | - | Unassigned | - |
| Otu59146 | 1 | k__Fungi(100);k__Fungi_unclassified(100);k__Fungi_unclassified(100);k__Fungi_unclassified(100);k__Fungi_unclassified(100);k__Fungi_unclassified(100);k__Fungi_unclassified(100); | - | - | - | - | - | - | - | Unassigned | - |
| Otu59147 | 1 | k__Fungi(100);k__Fungi_unclassified(100);k__Fungi_unclassified(100);k__Fungi_unclassified(100);k__Fungi_unclassified(100);k__Fungi_unclassified(100);k__Fungi_unclassified(100); | - | - | - | - | - | - | - | Unassigned | - |
| Otu59148 | 1 | k__Fungi(100);k__Fungi_unclassified(100);k__Fungi_unclassified(100);k__Fungi_unclassified(100);k__Fungi_unclassified(100);k__Fungi_unclassified(100);k__Fungi_unclassified(100); | - | - | - | - | - | - | - | Unassigned | - |
| Otu59149 | 1 | k__Fungi(100);k__Fungi_unclassified(100);k__Fungi_unclassified(100);k__Fungi_unclassified(100);k__Fungi_unclassified(100);k__Fungi_unclassified(100);k__Fungi_unclassified(100); | - | - | - | - | - | - | - | Unassigned | - |
| Otu59150 | 1 | k__Fungi(100);k__Fungi_unclassified(100);k__Fungi_unclassified(100);k__Fungi_unclassified(100);k__Fungi_unclassified(100);k__Fungi_unclassified(100);k__Fungi_unclassified(100); | - | - | - | - | - | - | - | Unassigned | - |
| Otu59151 | 1 | k__Fungi(100);k__Fungi_unclassified(100);k__Fungi_unclassified(100);k__Fungi_unclassified(100);k__Fungi_unclassified(100);k__Fungi_unclassified(100);k__Fungi_unclassified(100); | - | - | - | - | - | - | - | Unassigned | - |
| Otu59152 | 1 | k__Fungi(100);p__Ascomycota(100);p__Ascomycota_unclassified(100);p__Ascomycota_unclassified(100);p__Ascomycota_unclassified(100);p__Ascomycota_unclassified(100);p__Ascomycota_unclassified(100); | - | - | - | - | - | - | - | Unassigned | - |
| Otu59153 | 1 | k__Fungi(100);k__Fungi_unclassified(100);k__Fungi_unclassified(100);k__Fungi_unclassified(100);k__Fungi_unclassified(100);k__Fungi_unclassified(100);k__Fungi_unclassified(100); | - | - | - | - | - | - | - | Unassigned | - |
| Otu59154 | 1 | k__Fungi(100);k__Fungi_unclassified(100);k__Fungi_unclassified(100);k__Fungi_unclassified(100);k__Fungi_unclassified(100);k__Fungi_unclassified(100);k__Fungi_unclassified(100); | - | - | - | - | - | - | - | Unassigned | - |
| Otu59155 | 1 | k__Fungi(100);p__Ascomycota(100);p__Ascomycota_unclassified(100);p__Ascomycota_unclassified(100);p__Ascomycota_unclassified(100);p__Ascomycota_unclassified(100);p__Ascomycota_unclassified(100); | - | - | - | - | - | - | - | Unassigned | - |
| Otu59156 | 1 | k__Fungi(100);k__Fungi_unclassified(100);k__Fungi_unclassified(100);k__Fungi_unclassified(100);k__Fungi_unclassified(100);k__Fungi_unclassified(100);k__Fungi_unclassified(100); | - | - | - | - | - | - | - | Unassigned | - |
| Otu59157 | 1 | k__Fungi(100);p__Ascomycota(100);p__Ascomycota_unclassified(100);p__Ascomycota_unclassified(100);p__Ascomycota_unclassified(100);p__Ascomycota_unclassified(100);p__Ascomycota_unclassified(100); | - | - | - | - | - | - | - | Unassigned | - |
| Otu59158 | 1 | k__Fungi(100);p__Ascomycota(100);p__Ascomycota_unclassified(100);p__Ascomycota_unclassified(100);p__Ascomycota_unclassified(100);p__Ascomycota_unclassified(100);p__Ascomycota_unclassified(100); | - | - | - | - | - | - | - | Unassigned | - |
| Otu59159 | 1 | k__Fungi(100);k__Fungi_unclassified(100);k__Fungi_unclassified(100);k__Fungi_unclassified(100);k__Fungi_unclassified(100);k__Fungi_unclassified(100);k__Fungi_unclassified(100); | - | - | - | - | - | - | - | Unassigned | - |
| Otu59160 | 1 | k__Fungi(100);k__Fungi_unclassified(100);k__Fungi_unclassified(100);k__Fungi_unclassified(100);k__Fungi_unclassified(100);k__Fungi_unclassified(100);k__Fungi_unclassified(100); | - | - | - | - | - | - | - | Unassigned | - |
| Otu59165 | 1 | k__Fungi(100);k__Fungi_unclassified(100);k__Fungi_unclassified(100);k__Fungi_unclassified(100);k__Fungi_unclassified(100);k__Fungi_unclassified(100);k__Fungi_unclassified(100); | - | - | - | - | - | - | - | Unassigned | - |
| Otu59169 | 1 | k__Fungi(100);p__Basidiomycota(100);p__Basidiomycota_unclassified(100);p__Basidiomycota_unclassified(100);p__Basidiomycota_unclassified(100);p__Basidiomycota_unclassified(100);p__Basidiomycota_unclassified(100); | - | - | - | - | - | - | - | Unassigned | - |
| Otu59173 | 1 | k__Fungi(100);p__Ascomycota(100);p__Ascomycota_unclassified(100);p__Ascomycota_unclassified(100);p__Ascomycota_unclassified(100);p__Ascomycota_unclassified(100);p__Ascomycota_unclassified(100); | - | - | - | - | - | - | - | Unassigned | - |
| Otu59174 | 1 | k__Fungi(100);p__Ascomycota(100);p__Ascomycota_unclassified(100);p__Ascomycota_unclassified(100);p__Ascomycota_unclassified(100);p__Ascomycota_unclassified(100);p__Ascomycota_unclassified(100); | - | - | - | - | - | - | - | Unassigned | - |
| Otu59176 | 1 | k__Fungi(100);p__Ascomycota(100);p__Ascomycota_unclassified(100);p__Ascomycota_unclassified(100);p__Ascomycota_unclassified(100);p__Ascomycota_unclassified(100);p__Ascomycota_unclassified(100); | - | - | - | - | - | - | - | Unassigned | - |
| Otu59177 | 1 | k__Fungi(100);k__Fungi_unclassified(100);k__Fungi_unclassified(100);k__Fungi_unclassified(100);k__Fungi_unclassified(100);k__Fungi_unclassified(100);k__Fungi_unclassified(100); | - | - | - | - | - | - | - | Unassigned | - |
| Otu59188 | 1 | k__Fungi(100);k__Fungi_unclassified(100);k__Fungi_unclassified(100);k__Fungi_unclassified(100);k__Fungi_unclassified(100);k__Fungi_unclassified(100);k__Fungi_unclassified(100); | - | - | - | - | - | - | - | Unassigned | - |
| Otu59189 | 1 | k__Fungi(100);p__Ascomycota(100);p__Ascomycota_unclassified(100);p__Ascomycota_unclassified(100);p__Ascomycota_unclassified(100);p__Ascomycota_unclassified(100);p__Ascomycota_unclassified(100); | - | - | - | - | - | - | - | Unassigned | - |
| Otu59190 | 1 | k__Fungi(100);k__Fungi_unclassified(100);k__Fungi_unclassified(100);k__Fungi_unclassified(100);k__Fungi_unclassified(100);k__Fungi_unclassified(100);k__Fungi_unclassified(100); | - | - | - | - | - | - | - | Unassigned | - |
| Otu59194 | 1 | k__Fungi(100);k__Fungi_unclassified(100);k__Fungi_unclassified(100);k__Fungi_unclassified(100);k__Fungi_unclassified(100);k__Fungi_unclassified(100);k__Fungi_unclassified(100); | - | - | - | - | - | - | - | Unassigned | - |
| Otu59197 | 1 | k__Fungi(100);k__Fungi_unclassified(100);k__Fungi_unclassified(100);k__Fungi_unclassified(100);k__Fungi_unclassified(100);k__Fungi_unclassified(100);k__Fungi_unclassified(100); | - | - | - | - | - | - | - | Unassigned | - |
| Otu59198 | 1 | k__Fungi(100);k__Fungi_unclassified(100);k__Fungi_unclassified(100);k__Fungi_unclassified(100);k__Fungi_unclassified(100);k__Fungi_unclassified(100);k__Fungi_unclassified(100); | - | - | - | - | - | - | - | Unassigned | - |
| Otu59200 | 1 | k__Fungi(100);k__Fungi_unclassified(100);k__Fungi_unclassified(100);k__Fungi_unclassified(100);k__Fungi_unclassified(100);k__Fungi_unclassified(100);k__Fungi_unclassified(100); | - | - | - | - | - | - | - | Unassigned | - |
| Otu59201 | 1 | k__Fungi(100);p__Ascomycota(100);p__Ascomycota_unclassified(100);p__Ascomycota_unclassified(100);p__Ascomycota_unclassified(100);p__Ascomycota_unclassified(100);p__Ascomycota_unclassified(100); | - | - | - | - | - | - | - | Unassigned | - |
| Otu59202 | 1 | k__Fungi(100);k__Fungi_unclassified(100);k__Fungi_unclassified(100);k__Fungi_unclassified(100);k__Fungi_unclassified(100);k__Fungi_unclassified(100);k__Fungi_unclassified(100); | - | - | - | - | - | - | - | Unassigned | - |
| Otu59203 | 1 | k__Fungi(100);p__Ascomycota(100);p__Ascomycota_unclassified(100);p__Ascomycota_unclassified(100);p__Ascomycota_unclassified(100);p__Ascomycota_unclassified(100);p__Ascomycota_unclassified(100); | - | - | - | - | - | - | - | Unassigned | - |
| Otu59204 | 1 | k__Fungi(100);k__Fungi_unclassified(100);k__Fungi_unclassified(100);k__Fungi_unclassified(100);k__Fungi_unclassified(100);k__Fungi_unclassified(100);k__Fungi_unclassified(100); | - | - | - | - | - | - | - | Unassigned | - |
| Otu59205 | 1 | k__Fungi(100);k__Fungi_unclassified(100);k__Fungi_unclassified(100);k__Fungi_unclassified(100);k__Fungi_unclassified(100);k__Fungi_unclassified(100);k__Fungi_unclassified(100); | - | - | - | - | - | - | - | Unassigned | - |
| Otu59206 | 1 | k__Fungi(100);k__Fungi_unclassified(100);k__Fungi_unclassified(100);k__Fungi_unclassified(100);k__Fungi_unclassified(100);k__Fungi_unclassified(100);k__Fungi_unclassified(100); | - | - | - | - | - | - | - | Unassigned | - |
| Otu59207 | 1 | k__Fungi(100);p__Ascomycota(100);p__Ascomycota_unclassified(100);p__Ascomycota_unclassified(100);p__Ascomycota_unclassified(100);p__Ascomycota_unclassified(100);p__Ascomycota_unclassified(100); | - | - | - | - | - | - | - | Unassigned | - |
| Otu59208 | 1 | k__Fungi(100);k__Fungi_unclassified(100);k__Fungi_unclassified(100);k__Fungi_unclassified(100);k__Fungi_unclassified(100);k__Fungi_unclassified(100);k__Fungi_unclassified(100); | - | - | - | - | - | - | - | Unassigned | - |
| Otu59209 | 1 | k__Fungi(100);k__Fungi_unclassified(100);k__Fungi_unclassified(100);k__Fungi_unclassified(100);k__Fungi_unclassified(100);k__Fungi_unclassified(100);k__Fungi_unclassified(100); | - | - | - | - | - | - | - | Unassigned | - |
| Otu59210 | 1 | k__Fungi(100);k__Fungi_unclassified(100);k__Fungi_unclassified(100);k__Fungi_unclassified(100);k__Fungi_unclassified(100);k__Fungi_unclassified(100);k__Fungi_unclassified(100); | - | - | - | - | - | - | - | Unassigned | - |
| Otu59211 | 1 | k__Fungi(100);k__Fungi_unclassified(100);k__Fungi_unclassified(100);k__Fungi_unclassified(100);k__Fungi_unclassified(100);k__Fungi_unclassified(100);k__Fungi_unclassified(100); | - | - | - | - | - | - | - | Unassigned | - |
| Otu59212 | 1 | k__Fungi(100);k__Fungi_unclassified(100);k__Fungi_unclassified(100);k__Fungi_unclassified(100);k__Fungi_unclassified(100);k__Fungi_unclassified(100);k__Fungi_unclassified(100); | - | - | - | - | - | - | - | Unassigned | - |
| Otu59213 | 1 | k__Fungi(100);k__Fungi_unclassified(100);k__Fungi_unclassified(100);k__Fungi_unclassified(100);k__Fungi_unclassified(100);k__Fungi_unclassified(100);k__Fungi_unclassified(100); | - | - | - | - | - | - | - | Unassigned | - |
| Otu59214 | 1 | k__Fungi(100);k__Fungi_unclassified(100);k__Fungi_unclassified(100);k__Fungi_unclassified(100);k__Fungi_unclassified(100);k__Fungi_unclassified(100);k__Fungi_unclassified(100); | - | - | - | - | - | - | - | Unassigned | - |
| Otu59215 | 1 | k__Fungi(100);k__Fungi_unclassified(100);k__Fungi_unclassified(100);k__Fungi_unclassified(100);k__Fungi_unclassified(100);k__Fungi_unclassified(100);k__Fungi_unclassified(100); | - | - | - | - | - | - | - | Unassigned | - |
| Otu59216 | 1 | k__Fungi(100);k__Fungi_unclassified(100);k__Fungi_unclassified(100);k__Fungi_unclassified(100);k__Fungi_unclassified(100);k__Fungi_unclassified(100);k__Fungi_unclassified(100); | - | - | - | - | - | - | - | Unassigned | - |
| Otu59217 | 1 | k__Fungi(100);k__Fungi_unclassified(100);k__Fungi_unclassified(100);k__Fungi_unclassified(100);k__Fungi_unclassified(100);k__Fungi_unclassified(100);k__Fungi_unclassified(100); | - | - | - | - | - | - | - | Unassigned | - |
| Otu59218 | 1 | k__Fungi(100);k__Fungi_unclassified(100);k__Fungi_unclassified(100);k__Fungi_unclassified(100);k__Fungi_unclassified(100);k__Fungi_unclassified(100);k__Fungi_unclassified(100); | - | - | - | - | - | - | - | Unassigned | - |
| Otu59219 | 1 | k__Fungi(100);k__Fungi_unclassified(100);k__Fungi_unclassified(100);k__Fungi_unclassified(100);k__Fungi_unclassified(100);k__Fungi_unclassified(100);k__Fungi_unclassified(100); | - | - | - | - | - | - | - | Unassigned | - |
| Otu59220 | 1 | k__Fungi(100);k__Fungi_unclassified(100);k__Fungi_unclassified(100);k__Fungi_unclassified(100);k__Fungi_unclassified(100);k__Fungi_unclassified(100);k__Fungi_unclassified(100); | - | - | - | - | - | - | - | Unassigned | - |
| Otu59221 | 1 | k__Fungi(100);k__Fungi_unclassified(100);k__Fungi_unclassified(100);k__Fungi_unclassified(100);k__Fungi_unclassified(100);k__Fungi_unclassified(100);k__Fungi_unclassified(100); | - | - | - | - | - | - | - | Unassigned | - |
| Otu59222 | 1 | k__Fungi(100);k__Fungi_unclassified(100);k__Fungi_unclassified(100);k__Fungi_unclassified(100);k__Fungi_unclassified(100);k__Fungi_unclassified(100);k__Fungi_unclassified(100); | - | - | - | - | - | - | - | Unassigned | - |
| Otu59223 | 1 | k__Fungi(100);p__Ascomycota(100);p__Ascomycota_unclassified(100);p__Ascomycota_unclassified(100);p__Ascomycota_unclassified(100);p__Ascomycota_unclassified(100);p__Ascomycota_unclassified(100); | - | - | - | - | - | - | - | Unassigned | - |
| Otu59224 | 1 | k__Fungi(100);k__Fungi_unclassified(100);k__Fungi_unclassified(100);k__Fungi_unclassified(100);k__Fungi_unclassified(100);k__Fungi_unclassified(100);k__Fungi_unclassified(100); | - | - | - | - | - | - | - | Unassigned | - |
| Otu59225 | 1 | k__Fungi(100);k__Fungi_unclassified(100);k__Fungi_unclassified(100);k__Fungi_unclassified(100);k__Fungi_unclassified(100);k__Fungi_unclassified(100);k__Fungi_unclassified(100); | - | - | - | - | - | - | - | Unassigned | - |
| Otu59226 | 1 | k__Fungi(100);k__Fungi_unclassified(100);k__Fungi_unclassified(100);k__Fungi_unclassified(100);k__Fungi_unclassified(100);k__Fungi_unclassified(100);k__Fungi_unclassified(100); | - | - | - | - | - | - | - | Unassigned | - |
| Otu59227 | 1 | k__Fungi(100);p__Ascomycota(100);p__Ascomycota_unclassified(100);p__Ascomycota_unclassified(100);p__Ascomycota_unclassified(100);p__Ascomycota_unclassified(100);p__Ascomycota_unclassified(100); | - | - | - | - | - | - | - | Unassigned | - |
| Otu59228 | 1 | k__Fungi(100);p__Ascomycota(100);p__Ascomycota_unclassified(100);p__Ascomycota_unclassified(100);p__Ascomycota_unclassified(100);p__Ascomycota_unclassified(100);p__Ascomycota_unclassified(100); | - | - | - | - | - | - | - | Unassigned | - |
| Otu59244 | 1 | k__Fungi(100);k__Fungi_unclassified(100);k__Fungi_unclassified(100);k__Fungi_unclassified(100);k__Fungi_unclassified(100);k__Fungi_unclassified(100);k__Fungi_unclassified(100); | - | - | - | - | - | - | - | Unassigned | - |
| Otu59249 | 1 | k__Fungi(100);p__Ascomycota(100);p__Ascomycota_unclassified(100);p__Ascomycota_unclassified(100);p__Ascomycota_unclassified(100);p__Ascomycota_unclassified(100);p__Ascomycota_unclassified(100); | - | - | - | - | - | - | - | Unassigned | - |
| Otu59251 | 1 | k__Fungi(100);p__Ascomycota(100);p__Ascomycota_unclassified(100);p__Ascomycota_unclassified(100);p__Ascomycota_unclassified(100);p__Ascomycota_unclassified(100);p__Ascomycota_unclassified(100); | - | - | - | - | - | - | - | Unassigned | - |
| Otu59254 | 1 | k__Fungi(100);k__Fungi_unclassified(100);k__Fungi_unclassified(100);k__Fungi_unclassified(100);k__Fungi_unclassified(100);k__Fungi_unclassified(100);k__Fungi_unclassified(100); | - | - | - | - | - | - | - | Unassigned | - |
| Otu59269 | 1 | k__Fungi(100);p__Ascomycota(100);p__Ascomycota_unclassified(100);p__Ascomycota_unclassified(100);p__Ascomycota_unclassified(100);p__Ascomycota_unclassified(100);p__Ascomycota_unclassified(100); | - | - | - | - | - | - | - | Unassigned | - |
| Otu59282 | 1 | k__Fungi(100);k__Fungi_unclassified(100);k__Fungi_unclassified(100);k__Fungi_unclassified(100);k__Fungi_unclassified(100);k__Fungi_unclassified(100);k__Fungi_unclassified(100); | - | - | - | - | - | - | - | Unassigned | - |
| Otu59283 | 1 | k__Fungi(100);k__Fungi_unclassified(100);k__Fungi_unclassified(100);k__Fungi_unclassified(100);k__Fungi_unclassified(100);k__Fungi_unclassified(100);k__Fungi_unclassified(100); | - | - | - | - | - | - | - | Unassigned | - |
| Otu59284 | 1 | k__Fungi(100);k__Fungi_unclassified(100);k__Fungi_unclassified(100);k__Fungi_unclassified(100);k__Fungi_unclassified(100);k__Fungi_unclassified(100);k__Fungi_unclassified(100); | - | - | - | - | - | - | - | Unassigned | - |
| Otu59285 | 1 | k__Fungi(100);k__Fungi_unclassified(100);k__Fungi_unclassified(100);k__Fungi_unclassified(100);k__Fungi_unclassified(100);k__Fungi_unclassified(100);k__Fungi_unclassified(100); | - | - | - | - | - | - | - | Unassigned | - |
| Otu59286 | 1 | k__Fungi(100);k__Fungi_unclassified(100);k__Fungi_unclassified(100);k__Fungi_unclassified(100);k__Fungi_unclassified(100);k__Fungi_unclassified(100);k__Fungi_unclassified(100); | - | - | - | - | - | - | - | Unassigned | - |
| Otu59287 | 1 | k__Fungi(100);k__Fungi_unclassified(100);k__Fungi_unclassified(100);k__Fungi_unclassified(100);k__Fungi_unclassified(100);k__Fungi_unclassified(100);k__Fungi_unclassified(100); | - | - | - | - | - | - | - | Unassigned | - |
| Otu59288 | 1 | k__Fungi(100);p__Basidiomycota(100);p__Basidiomycota_unclassified(100);p__Basidiomycota_unclassified(100);p__Basidiomycota_unclassified(100);p__Basidiomycota_unclassified(100);p__Basidiomycota_unclassified(100); | - | - | - | - | - | - | - | Unassigned | - |
| Otu59289 | 1 | k__Fungi(100);k__Fungi_unclassified(100);k__Fungi_unclassified(100);k__Fungi_unclassified(100);k__Fungi_unclassified(100);k__Fungi_unclassified(100);k__Fungi_unclassified(100); | - | - | - | - | - | - | - | Unassigned | - |
| Otu59290 | 1 | k__Fungi(100);k__Fungi_unclassified(100);k__Fungi_unclassified(100);k__Fungi_unclassified(100);k__Fungi_unclassified(100);k__Fungi_unclassified(100);k__Fungi_unclassified(100); | - | - | - | - | - | - | - | Unassigned | - |
| Otu59291 | 1 | k__Fungi(100);p__Basidiomycota(100);p__Basidiomycota_unclassified(100);p__Basidiomycota_unclassified(100);p__Basidiomycota_unclassified(100);p__Basidiomycota_unclassified(100);p__Basidiomycota_unclassified(100); | - | - | - | - | - | - | - | Unassigned | - |
| Otu59292 | 1 | k__Fungi(100);k__Fungi_unclassified(100);k__Fungi_unclassified(100);k__Fungi_unclassified(100);k__Fungi_unclassified(100);k__Fungi_unclassified(100);k__Fungi_unclassified(100); | - | - | - | - | - | - | - | Unassigned | - |
| Otu59293 | 1 | k__Fungi(100);p__Ascomycota(100);p__Ascomycota_unclassified(100);p__Ascomycota_unclassified(100);p__Ascomycota_unclassified(100);p__Ascomycota_unclassified(100);p__Ascomycota_unclassified(100); | - | - | - | - | - | - | - | Unassigned | - |
| Otu59294 | 1 | k__Fungi(100);k__Fungi_unclassified(100);k__Fungi_unclassified(100);k__Fungi_unclassified(100);k__Fungi_unclassified(100);k__Fungi_unclassified(100);k__Fungi_unclassified(100); | - | - | - | - | - | - | - | Unassigned | - |

| | | | | | | | | | | | |
|---|---|---|---|---|---|---|---|---|---|---|---|
| Otu59295 | 1 | k__Fungi(100);k__Fungi_unclassified(100);k__Fungi_unclassified(100);k__Fungi_unclassified(100);k__Fungi_unclassified(100);k__Fungi_unclassified(100);k__Fungi_unclassified(100); | - | - | - | - | - | - | - | Unassigned | - |
| Otu59296 | 1 | k__Fungi(100);k__Fungi_unclassified(100);k__Fungi_unclassified(100);k__Fungi_unclassified(100);k__Fungi_unclassified(100);k__Fungi_unclassified(100);k__Fungi_unclassified(100); | - | - | - | - | - | - | - | Unassigned | - |
| Otu59297 | 1 | k__Fungi(100);k__Fungi_unclassified(100);k__Fungi_unclassified(100);k__Fungi_unclassified(100);k__Fungi_unclassified(100);k__Fungi_unclassified(100);k__Fungi_unclassified(100); | - | - | - | - | - | - | - | Unassigned | - |
| Otu59298 | 1 | k__Fungi(100);k__Fungi_unclassified(100);k__Fungi_unclassified(100);k__Fungi_unclassified(100);k__Fungi_unclassified(100);k__Fungi_unclassified(100);k__Fungi_unclassified(100); | - | - | - | - | - | - | - | Unassigned | - |
| Otu59299 | 1 | k__Fungi(100);k__Fungi_unclassified(100);k__Fungi_unclassified(100);k__Fungi_unclassified(100);k__Fungi_unclassified(100);k__Fungi_unclassified(100);k__Fungi_unclassified(100); | - | - | - | - | - | - | - | Unassigned | - |
| Otu59300 | 1 | k__Fungi(100);k__Fungi_unclassified(100);k__Fungi_unclassified(100);k__Fungi_unclassified(100);k__Fungi_unclassified(100);k__Fungi_unclassified(100);k__Fungi_unclassified(100); | - | - | - | - | - | - | - | Unassigned | - |
| Otu59301 | 1 | k__Fungi(100);k__Fungi_unclassified(100);k__Fungi_unclassified(100);k__Fungi_unclassified(100);k__Fungi_unclassified(100);k__Fungi_unclassified(100);k__Fungi_unclassified(100); | - | - | - | - | - | - | - | Unassigned | - |
| Otu59302 | 1 | k__Fungi(100);k__Fungi_unclassified(100);k__Fungi_unclassified(100);k__Fungi_unclassified(100);k__Fungi_unclassified(100);k__Fungi_unclassified(100);k__Fungi_unclassified(100); | - | - | - | - | - | - | - | Unassigned | - |
| Otu59303 | 1 | k__Fungi(100);k__Fungi_unclassified(100);k__Fungi_unclassified(100);k__Fungi_unclassified(100);k__Fungi_unclassified(100);k__Fungi_unclassified(100);k__Fungi_unclassified(100); | - | - | - | - | - | - | - | Unassigned | - |
| Otu59304 | 1 | k__Fungi(100);k__Fungi_unclassified(100);k__Fungi_unclassified(100);k__Fungi_unclassified(100);k__Fungi_unclassified(100);k__Fungi_unclassified(100);k__Fungi_unclassified(100); | - | - | - | - | - | - | - | Unassigned | - |
| Otu59305 | 1 | k__Fungi(100);k__Fungi_unclassified(100);k__Fungi_unclassified(100);k__Fungi_unclassified(100);k__Fungi_unclassified(100);k__Fungi_unclassified(100);k__Fungi_unclassified(100); | - | - | - | - | - | - | - | Unassigned | - |
| Otu59306 | 1 | k__Fungi(100);k__Fungi_unclassified(100);k__Fungi_unclassified(100);k__Fungi_unclassified(100);k__Fungi_unclassified(100);k__Fungi_unclassified(100);k__Fungi_unclassified(100); | - | - | - | - | - | - | - | Unassigned | - |
| Otu59307 | 1 | k__Fungi(100);k__Fungi_unclassified(100);k__Fungi_unclassified(100);k__Fungi_unclassified(100);k__Fungi_unclassified(100);k__Fungi_unclassified(100);k__Fungi_unclassified(100); | - | - | - | - | - | - | - | Unassigned | - |
| Otu59309 | 1 | k__Fungi(100);p__Ascomycota(100);p__Ascomycota_unclassified(100);p__Ascomycota_unclassified(100);p__Ascomycota_unclassified(100);p__Ascomycota_unclassified(100);p__Ascomycota_unclassified(100); | - | - | - | - | - | - | - | Unassigned | - |
| Otu59310 | 1 | k__Fungi(100);k__Fungi_unclassified(100);k__Fungi_unclassified(100);k__Fungi_unclassified(100);k__Fungi_unclassified(100);k__Fungi_unclassified(100);k__Fungi_unclassified(100); | - | - | - | - | - | - | - | Unassigned | - |
| Otu59312 | 1 | k__Fungi(100);k__Fungi_unclassified(100);k__Fungi_unclassified(100);k__Fungi_unclassified(100);k__Fungi_unclassified(100);k__Fungi_unclassified(100);k__Fungi_unclassified(100); | - | - | - | - | - | - | - | Unassigned | - |
| Otu59314 | 1 | k__Fungi(100);p__Ascomycota(100);p__Ascomycota_unclassified(100);p__Ascomycota_unclassified(100);p__Ascomycota_unclassified(100);p__Ascomycota_unclassified(100);p__Ascomycota_unclassified(100); | - | - | - | - | - | - | - | Unassigned | - |
| Otu59315 | 1 | k__Fungi(100);p__Ascomycota(100);p__Ascomycota_unclassified(100);p__Ascomycota_unclassified(100);p__Ascomycota_unclassified(100);p__Ascomycota_unclassified(100);p__Ascomycota_unclassified(100); | - | - | - | - | - | - | - | Unassigned | - |
| Otu59319 | 1 | k__Fungi(100);k__Fungi_unclassified(100);k__Fungi_unclassified(100);k__Fungi_unclassified(100);k__Fungi_unclassified(100);k__Fungi_unclassified(100);k__Fungi_unclassified(100); | - | - | - | - | - | - | - | Unassigned | - |
| Otu59320 | 1 | k__Fungi(100);k__Fungi_unclassified(100);k__Fungi_unclassified(100);k__Fungi_unclassified(100);k__Fungi_unclassified(100);k__Fungi_unclassified(100);k__Fungi_unclassified(100); | - | - | - | - | - | - | - | Unassigned | - |
| Otu59321 | 1 | k__Fungi(100);p__Ascomycota(100);p__Ascomycota_unclassified(100);p__Ascomycota_unclassified(100);p__Ascomycota_unclassified(100);p__Ascomycota_unclassified(100);p__Ascomycota_unclassified(100); | - | - | - | - | - | - | - | Unassigned | - |
| Otu59328 | 1 | k__Fungi(100);k__Fungi_unclassified(100);k__Fungi_unclassified(100);k__Fungi_unclassified(100);k__Fungi_unclassified(100);k__Fungi_unclassified(100);k__Fungi_unclassified(100); | - | - | - | - | - | - | - | Unassigned | - |
| Otu59329 | 1 | k__Fungi(100);k__Fungi_unclassified(100);k__Fungi_unclassified(100);k__Fungi_unclassified(100);k__Fungi_unclassified(100);k__Fungi_unclassified(100);k__Fungi_unclassified(100); | - | - | - | - | - | - | - | Unassigned | - |
| Otu59330 | 1 | k__Fungi(100);k__Fungi_unclassified(100);k__Fungi_unclassified(100);k__Fungi_unclassified(100);k__Fungi_unclassified(100);k__Fungi_unclassified(100);k__Fungi_unclassified(100); | - | - | - | - | - | - | - | Unassigned | - |
| Otu59331 | 1 | k__Fungi(100);k__Fungi_unclassified(100);k__Fungi_unclassified(100);k__Fungi_unclassified(100);k__Fungi_unclassified(100);k__Fungi_unclassified(100);k__Fungi_unclassified(100); | - | - | - | - | - | - | - | Unassigned | - |
| Otu59332 | 1 | k__Fungi(100);k__Fungi_unclassified(100);k__Fungi_unclassified(100);k__Fungi_unclassified(100);k__Fungi_unclassified(100);k__Fungi_unclassified(100);k__Fungi_unclassified(100); | - | - | - | - | - | - | - | Unassigned | - |
| Otu59333 | 1 | k__Fungi(100);k__Fungi_unclassified(100);k__Fungi_unclassified(100);k__Fungi_unclassified(100);k__Fungi_unclassified(100);k__Fungi_unclassified(100);k__Fungi_unclassified(100); | - | - | - | - | - | - | - | Unassigned | - |
| Otu59334 | 1 | k__Fungi(100);k__Fungi_unclassified(100);k__Fungi_unclassified(100);k__Fungi_unclassified(100);k__Fungi_unclassified(100);k__Fungi_unclassified(100);k__Fungi_unclassified(100); | - | - | - | - | - | - | - | Unassigned | - |
| Otu59349 | 1 | k__Fungi(100);k__Fungi_unclassified(100);k__Fungi_unclassified(100);k__Fungi_unclassified(100);k__Fungi_unclassified(100);k__Fungi_unclassified(100);k__Fungi_unclassified(100); | - | - | - | - | - | - | - | Unassigned | - |
| Otu59350 | 1 | k__Fungi(100);p__Ascomycota(100);p__Ascomycota_unclassified(100);p__Ascomycota_unclassified(100);p__Ascomycota_unclassified(100);p__Ascomycota_unclassified(100);p__Ascomycota_unclassified(100); | - | - | - | - | - | - | - | Unassigned | - |
| Otu59359 | 1 | k__Fungi(100);k__Fungi_unclassified(100);k__Fungi_unclassified(100);k__Fungi_unclassified(100);k__Fungi_unclassified(100);k__Fungi_unclassified(100);k__Fungi_unclassified(100); | - | - | - | - | - | - | - | Unassigned | - |
| Otu59360 | 1 | k__Fungi(100);p__Ascomycota(100);p__Ascomycota_unclassified(100);p__Ascomycota_unclassified(100);p__Ascomycota_unclassified(100);p__Ascomycota_unclassified(100);p__Ascomycota_unclassified(100); | - | - | - | - | - | - | - | Unassigned | - |
| Otu59365 | 1 | k__Fungi(100);k__Fungi_unclassified(100);k__Fungi_unclassified(100);k__Fungi_unclassified(100);k__Fungi_unclassified(100);k__Fungi_unclassified(100);k__Fungi_unclassified(100); | - | - | - | - | - | - | - | Unassigned | - |
| Otu59368 | 1 | k__Fungi(100);k__Fungi_unclassified(100);k__Fungi_unclassified(100);k__Fungi_unclassified(100);k__Fungi_unclassified(100);k__Fungi_unclassified(100);k__Fungi_unclassified(100); | - | - | - | - | - | - | - | Unassigned | - |
| Otu59369 | 1 | k__Fungi(100);k__Fungi_unclassified(100);k__Fungi_unclassified(100);k__Fungi_unclassified(100);k__Fungi_unclassified(100);k__Fungi_unclassified(100);k__Fungi_unclassified(100); | - | - | - | - | - | - | - | Unassigned | - |
| Otu59370 | 1 | k__Fungi(100);k__Fungi_unclassified(100);k__Fungi_unclassified(100);k__Fungi_unclassified(100);k__Fungi_unclassified(100);k__Fungi_unclassified(100);k__Fungi_unclassified(100); | - | - | - | - | - | - | - | Unassigned | - |
| Otu59371 | 1 | k__Fungi(100);p__Ascomycota(100);p__Ascomycota_unclassified(100);p__Ascomycota_unclassified(100);p__Ascomycota_unclassified(100);p__Ascomycota_unclassified(100);p__Ascomycota_unclassified(100); | - | - | - | - | - | - | - | Unassigned | - |
| Otu59373 | 1 | k__Fungi(100);p__Ascomycota(100);p__Ascomycota_unclassified(100);p__Ascomycota_unclassified(100);p__Ascomycota_unclassified(100);p__Ascomycota_unclassified(100);p__Ascomycota_unclassified(100); | - | - | - | - | - | - | - | Unassigned | - |
| Otu59397 | 1 | k__Fungi(100);p__Ascomycota(100);p__Ascomycota_unclassified(100);p__Ascomycota_unclassified(100);p__Ascomycota_unclassified(100);p__Ascomycota_unclassified(100);p__Ascomycota_unclassified(100); | - | - | - | - | - | - | - | Unassigned | - |
| Otu59400 | 1 | k__Fungi(100);p__Ascomycota(100);p__Ascomycota_unclassified(100);p__Ascomycota_unclassified(100);p__Ascomycota_unclassified(100);p__Ascomycota_unclassified(100);p__Ascomycota_unclassified(100); | - | - | - | - | - | - | - | Unassigned | - |
| Otu59401 | 1 | k__Fungi(100);k__Fungi_unclassified(100);k__Fungi_unclassified(100);k__Fungi_unclassified(100);k__Fungi_unclassified(100);k__Fungi_unclassified(100);k__Fungi_unclassified(100); | - | - | - | - | - | - | - | Unassigned | - |
| Otu59402 | 1 | k__Fungi(100);p__Ascomycota(100);p__Ascomycota_unclassified(100);p__Ascomycota_unclassified(100);p__Ascomycota_unclassified(100);p__Ascomycota_unclassified(100);p__Ascomycota_unclassified(100); | - | - | - | - | - | - | - | Unassigned | - |
| Otu59404 | 1 | k__Fungi(100);p__Ascomycota(100);p__Ascomycota_unclassified(100);p__Ascomycota_unclassified(100);p__Ascomycota_unclassified(100);p__Ascomycota_unclassified(100);p__Ascomycota_unclassified(100); | - | - | - | - | - | - | - | Unassigned | - |
| Otu59405 | 1 | k__Fungi(100);p__Ascomycota(100);p__Ascomycota_unclassified(100);p__Ascomycota_unclassified(100);p__Ascomycota_unclassified(100);p__Ascomycota_unclassified(100);p__Ascomycota_unclassified(100); | - | - | - | - | - | - | - | Unassigned | - |
| Otu59406 | 1 | k__Fungi(100);k__Fungi_unclassified(100);k__Fungi_unclassified(100);k__Fungi_unclassified(100);k__Fungi_unclassified(100);k__Fungi_unclassified(100);k__Fungi_unclassified(100); | - | - | - | - | - | - | - | Unassigned | - |
| Otu59408 | 1 | k__Fungi(100);p__Ascomycota(100);p__Ascomycota_unclassified(100);p__Ascomycota_unclassified(100);p__Ascomycota_unclassified(100);p__Ascomycota_unclassified(100);p__Ascomycota_unclassified(100); | - | - | - | - | - | - | - | Unassigned | - |
| Otu59409 | 1 | k__Fungi(100);k__Fungi_unclassified(100);k__Fungi_unclassified(100);k__Fungi_unclassified(100);k__Fungi_unclassified(100);k__Fungi_unclassified(100);k__Fungi_unclassified(100); | - | - | - | - | - | - | - | Unassigned | - |
| Otu59410 | 1 | k__Fungi(100);p__Ascomycota(100);p__Ascomycota_unclassified(100);p__Ascomycota_unclassified(100);p__Ascomycota_unclassified(100);p__Ascomycota_unclassified(100);p__Ascomycota_unclassified(100); | - | - | - | - | - | - | - | Unassigned | - |
| Otu59411 | 1 | k__Fungi(100);k__Fungi_unclassified(100);k__Fungi_unclassified(100);k__Fungi_unclassified(100);k__Fungi_unclassified(100);k__Fungi_unclassified(100);k__Fungi_unclassified(100); | - | - | - | - | - | - | - | Unassigned | - |
| Otu59416 | 1 | k__Fungi(100);p__Ascomycota(100);p__Ascomycota_unclassified(100);p__Ascomycota_unclassified(100);p__Ascomycota_unclassified(100);p__Ascomycota_unclassified(100);p__Ascomycota_unclassified(100); | - | - | - | - | - | - | - | Unassigned | - |
| Otu59418 | 1 | k__Fungi(100);p__Ascomycota(100);p__Ascomycota_unclassified(100);p__Ascomycota_unclassified(100);p__Ascomycota_unclassified(100);p__Ascomycota_unclassified(100);p__Ascomycota_unclassified(100); | - | - | - | - | - | - | - | Unassigned | - |
| Otu59425 | 1 | k__Fungi(100);p__Ascomycota(100);p__Ascomycota_unclassified(100);p__Ascomycota_unclassified(100);p__Ascomycota_unclassified(100);p__Ascomycota_unclassified(100);p__Ascomycota_unclassified(100); | - | - | - | - | - | - | - | Unassigned | - |
| Otu59438 | 1 | k__Fungi(100);p__Ascomycota(100);p__Ascomycota_unclassified(100);p__Ascomycota_unclassified(100);p__Ascomycota_unclassified(100);p__Ascomycota_unclassified(100);p__Ascomycota_unclassified(100); | - | - | - | - | - | - | - | Unassigned | - |
| Otu59439 | 1 | k__Fungi(100);k__Fungi_unclassified(100);k__Fungi_unclassified(100);k__Fungi_unclassified(100);k__Fungi_unclassified(100);k__Fungi_unclassified(100);k__Fungi_unclassified(100); | - | - | - | - | - | - | - | Unassigned | - |
| Otu59440 | 1 | k__Fungi(100);p__Ascomycota(100);p__Ascomycota_unclassified(100);p__Ascomycota_unclassified(100);p__Ascomycota_unclassified(100);p__Ascomycota_unclassified(100);p__Ascomycota_unclassified(100); | - | - | - | - | - | - | - | Unassigned | - |
| Otu59441 | 1 | k__Fungi(100);p__Ascomycota(100);p__Ascomycota_unclassified(100);p__Ascomycota_unclassified(100);p__Ascomycota_unclassified(100);p__Ascomycota_unclassified(100);p__Ascomycota_unclassified(100); | - | - | - | - | - | - | - | Unassigned | - |
| Otu59442 | 1 | k__Fungi(100);p__Ascomycota(100);p__Ascomycota_unclassified(100);p__Ascomycota_unclassified(100);p__Ascomycota_unclassified(100);p__Ascomycota_unclassified(100);p__Ascomycota_unclassified(100); | - | - | - | - | - | - | - | Unassigned | - |
| Otu59445 | 1 | k__Fungi(100);p__Ascomycota(100);p__Ascomycota_unclassified(100);p__Ascomycota_unclassified(100);p__Ascomycota_unclassified(100);p__Ascomycota_unclassified(100);p__Ascomycota_unclassified(100); | - | - | - | - | - | - | - | Unassigned | - |
| Otu59446 | 1 | k__Fungi(100);k__Fungi_unclassified(100);k__Fungi_unclassified(100);k__Fungi_unclassified(100);k__Fungi_unclassified(100);k__Fungi_unclassified(100);k__Fungi_unclassified(100); | - | - | - | - | - | - | - | Unassigned | - |
| Otu59447 | 1 | k__Fungi(100);k__Fungi_unclassified(100);k__Fungi_unclassified(100);k__Fungi_unclassified(100);k__Fungi_unclassified(100);k__Fungi_unclassified(100);k__Fungi_unclassified(100); | - | - | - | - | - | - | - | Unassigned | - |
| Otu59448 | 1 | k__Fungi(100);p__Ascomycota(100);p__Ascomycota_unclassified(100);p__Ascomycota_unclassified(100);p__Ascomycota_unclassified(100);p__Ascomycota_unclassified(100);p__Ascomycota_unclassified(100); | - | - | - | - | - | - | - | Unassigned | - |
| Otu59449 | 1 | k__Fungi(100);p__Ascomycota(100);p__Ascomycota_unclassified(100);p__Ascomycota_unclassified(100);p__Ascomycota_unclassified(100);p__Ascomycota_unclassified(100);p__Ascomycota_unclassified(100); | - | - | - | - | - | - | - | Unassigned | - |
| Otu59450 | 1 | k__Fungi(100);p__Ascomycota(100);p__Ascomycota_unclassified(100);p__Ascomycota_unclassified(100);p__Ascomycota_unclassified(100);p__Ascomycota_unclassified(100);p__Ascomycota_unclassified(100); | - | - | - | - | - | - | - | Unassigned | - |
| Otu59451 | 1 | k__Fungi(100);p__Ascomycota(100);p__Ascomycota_unclassified(100);p__Ascomycota_unclassified(100);p__Ascomycota_unclassified(100);p__Ascomycota_unclassified(100);p__Ascomycota_unclassified(100); | - | - | - | - | - | - | - | Unassigned | - |
| Otu59455 | 1 | k__Fungi(100);k__Fungi_unclassified(100);k__Fungi_unclassified(100);k__Fungi_unclassified(100);k__Fungi_unclassified(100);k__Fungi_unclassified(100);k__Fungi_unclassified(100); | - | - | - | - | - | - | - | Unassigned | - |
| Otu59456 | 1 | k__Fungi(100);p__Ascomycota(100);p__Ascomycota_unclassified(100);p__Ascomycota_unclassified(100);p__Ascomycota_unclassified(100);p__Ascomycota_unclassified(100);p__Ascomycota_unclassified(100); | - | - | - | - | - | - | - | Unassigned | - |
| Otu59458 | 1 | k__Fungi(100);k__Fungi_unclassified(100);k__Fungi_unclassified(100);k__Fungi_unclassified(100);k__Fungi_unclassified(100);k__Fungi_unclassified(100);k__Fungi_unclassified(100); | - | - | - | - | - | - | - | Unassigned | - |
| Otu59459 | 1 | k__Fungi(100);p__Ascomycota(100);p__Ascomycota_unclassified(100);p__Ascomycota_unclassified(100);p__Ascomycota_unclassified(100);p__Ascomycota_unclassified(100);p__Ascomycota_unclassified(100); | - | - | - | - | - | - | - | Unassigned | - |
| Otu59464 | 1 | k__Fungi(100);p__Ascomycota(100);p__Ascomycota_unclassified(100);p__Ascomycota_unclassified(100);p__Ascomycota_unclassified(100);p__Ascomycota_unclassified(100);p__Ascomycota_unclassified(100); | - | - | - | - | - | - | - | Unassigned | - |
| Otu59465 | 1 | k__Fungi(100);p__Ascomycota(100);p__Ascomycota_unclassified(100);p__Ascomycota_unclassified(100);p__Ascomycota_unclassified(100);p__Ascomycota_unclassified(100);p__Ascomycota_unclassified(100); | - | - | - | - | - | - | - | Unassigned | - |
| Otu59467 | 1 | k__Fungi(100);k__Fungi_unclassified(100);k__Fungi_unclassified(100);k__Fungi_unclassified(100);k__Fungi_unclassified(100);k__Fungi_unclassified(100);k__Fungi_unclassified(100); | - | - | - | - | - | - | - | Unassigned | - |
| Otu59468 | 1 | k__Fungi(100);p__Ascomycota(100);p__Ascomycota_unclassified(100);p__Ascomycota_unclassified(100);p__Ascomycota_unclassified(100);p__Ascomycota_unclassified(100);p__Ascomycota_unclassified(100); | - | - | - | - | - | - | - | Unassigned | - |
| Otu59473 | 1 | k__Fungi(100);p__Ascomycota(100);p__Ascomycota_unclassified(100);p__Ascomycota_unclassified(100);p__Ascomycota_unclassified(100);p__Ascomycota_unclassified(100);p__Ascomycota_unclassified(100); | - | - | - | - | - | - | - | Unassigned | - |
| Otu59474 | 1 | k__Fungi(100);k__Fungi_unclassified(100);k__Fungi_unclassified(100);k__Fungi_unclassified(100);k__Fungi_unclassified(100);k__Fungi_unclassified(100);k__Fungi_unclassified(100); | - | - | - | - | - | - | - | Unassigned | - |
| Otu59475 | 1 | k__Fungi(100);k__Fungi_unclassified(100);k__Fungi_unclassified(100);k__Fungi_unclassified(100);k__Fungi_unclassified(100);k__Fungi_unclassified(100);k__Fungi_unclassified(100); | - | - | - | - | - | - | - | Unassigned | - |
| Otu59476 | 1 | k__Fungi(100);k__Fungi_unclassified(100);k__Fungi_unclassified(100);k__Fungi_unclassified(100);k__Fungi_unclassified(100);k__Fungi_unclassified(100);k__Fungi_unclassified(100); | - | - | - | - | - | - | - | Unassigned | - |
| Otu59478 | 1 | k__Fungi(100);k__Fungi_unclassified(100);k__Fungi_unclassified(100);k__Fungi_unclassified(100);k__Fungi_unclassified(100);k__Fungi_unclassified(100);k__Fungi_unclassified(100); | - | - | - | - | - | - | - | Unassigned | - |
| Otu59479 | 1 | k__Fungi(100);k__Fungi_unclassified(100);k__Fungi_unclassified(100);k__Fungi_unclassified(100);k__Fungi_unclassified(100);k__Fungi_unclassified(100);k__Fungi_unclassified(100); | - | - | - | - | - | - | - | Unassigned | - |
| Otu59480 | 1 | k__Fungi(100);p__Ascomycota(100);p__Ascomycota_unclassified(100);p__Ascomycota_unclassified(100);p__Ascomycota_unclassified(100);p__Ascomycota_unclassified(100);p__Ascomycota_unclassified(100); | - | - | - | - | - | - | - | Unassigned | - |
| Otu59481 | 1 | k__Fungi(100);k__Fungi_unclassified(100);k__Fungi_unclassified(100);k__Fungi_unclassified(100);k__Fungi_unclassified(100);k__Fungi_unclassified(100);k__Fungi_unclassified(100); | - | - | - | - | - | - | - | Unassigned | - |
| Otu59482 | 1 | k__Fungi(100);p__Ascomycota(100);p__Ascomycota_unclassified(100);p__Ascomycota_unclassified(100);p__Ascomycota_unclassified(100);p__Ascomycota_unclassified(100);p__Ascomycota_unclassified(100); | - | - | - | - | - | - | - | Unassigned | - |
| Otu59484 | 1 | k__Fungi(100);p__Ascomycota(100);p__Ascomycota_unclassified(100);p__Ascomycota_unclassified(100);p__Ascomycota_unclassified(100);p__Ascomycota_unclassified(100);p__Ascomycota_unclassified(100); | - | - | - | - | - | - | - | Unassigned | - |
| Otu59485 | 1 | k__Fungi(100);p__Ascomycota(100);p__Ascomycota_unclassified(100);p__Ascomycota_unclassified(100);p__Ascomycota_unclassified(100);p__Ascomycota_unclassified(100);p__Ascomycota_unclassified(100); | - | - | - | - | - | - | - | Unassigned | - |
| Otu59486 | 1 | k__Fungi(100);k__Fungi_unclassified(100);k__Fungi_unclassified(100);k__Fungi_unclassified(100);k__Fungi_unclassified(100);k__Fungi_unclassified(100);k__Fungi_unclassified(100); | - | - | - | - | - | - | - | Unassigned | - |
| Otu59488 | 1 | k__Fungi(100);k__Fungi_unclassified(100);k__Fungi_unclassified(100);k__Fungi_unclassified(100);k__Fungi_unclassified(100);k__Fungi_unclassified(100);k__Fungi_unclassified(100); | - | - | - | - | - | - | - | Unassigned | - |
| Otu59489 | 1 | k__Fungi(100);p__Ascomycota(100);p__Ascomycota_unclassified(100);p__Ascomycota_unclassified(100);p__Ascomycota_unclassified(100);p__Ascomycota_unclassified(100);p__Ascomycota_unclassified(100); | - | - | - | - | - | - | - | Unassigned | - |
| Otu59491 | 1 | k__Fungi(100);k__Fungi_unclassified(100);k__Fungi_unclassified(100);k__Fungi_unclassified(100);k__Fungi_unclassified(100);k__Fungi_unclassified(100);k__Fungi_unclassified(100); | - | - | - | - | - | - | - | Unassigned | - |
| Otu59493 | 1 | k__Fungi(100);p__Ascomycota(100);p__Ascomycota_unclassified(100);p__Ascomycota_unclassified(100);p__Ascomycota_unclassified(100);p__Ascomycota_unclassified(100);p__Ascomycota_unclassified(100); | - | - | - | - | - | - | - | Unassigned | - |
| Otu59494 | 1 | k__Fungi(100);p__Ascomycota(100);p__Ascomycota_unclassified(100);p__Ascomycota_unclassified(100);p__Ascomycota_unclassified(100);p__Ascomycota_unclassified(100);p__Ascomycota_unclassified(100); | - | - | - | - | - | - | - | Unassigned | - |
| Otu59495 | 1 | k__Fungi(100);p__Ascomycota(100);p__Ascomycota_unclassified(100);p__Ascomycota_unclassified(100);p__Ascomycota_unclassified(100);p__Ascomycota_unclassified(100);p__Ascomycota_unclassified(100); | - | - | - | - | - | - | - | Unassigned | - |
| Otu59496 | 1 | k__Fungi(100);p__Ascomycota(100);p__Ascomycota_unclassified(100);p__Ascomycota_unclassified(100);p__Ascomycota_unclassified(100);p__Ascomycota_unclassified(100);p__Ascomycota_unclassified(100); | - | - | - | - | - | - | - | Unassigned | - |
| Otu59497 | 1 | k__Fungi(100);p__Ascomycota(100);p__Ascomycota_unclassified(100);p__Ascomycota_unclassified(100);p__Ascomycota_unclassified(100);p__Ascomycota_unclassified(100);p__Ascomycota_unclassified(100); | - | - | - | - | - | - | - | Unassigned | - |
| Otu59500 | 1 | k__Fungi(100);k__Fungi_unclassified(100);k__Fungi_unclassified(100);k__Fungi_unclassified(100);k__Fungi_unclassified(100);k__Fungi_unclassified(100);k__Fungi_unclassified(100); | - | - | - | - | - | - | - | Unassigned | - |
| Otu59501 | 1 | k__Fungi(100);k__Fungi_unclassified(100);k__Fungi_unclassified(100);k__Fungi_unclassified(100);k__Fungi_unclassified(100);k__Fungi_unclassified(100);k__Fungi_unclassified(100); | - | - | - | - | - | - | - | Unassigned | - |
| Otu59503 | 1 | k__Fungi(100);k__Fungi_unclassified(100);k__Fungi_unclassified(100);k__Fungi_unclassified(100);k__Fungi_unclassified(100);k__Fungi_unclassified(100);k__Fungi_unclassified(100); | - | - | - | - | - | - | - | Unassigned | - |
| Otu59505 | 1 | k__Fungi(100);k__Fungi_unclassified(100);k__Fungi_unclassified(100);k__Fungi_unclassified(100);k__Fungi_unclassified(100);k__Fungi_unclassified(100);k__Fungi_unclassified(100); | - | - | - | - | - | - | - | Unassigned | - |
| Otu59506 | 1 | k__Fungi(100);k__Fungi_unclassified(100);k__Fungi_unclassified(100);k__Fungi_unclassified(100);k__Fungi_unclassified(100);k__Fungi_unclassified(100);k__Fungi_unclassified(100); | - | - | - | - | - | - | - | Unassigned | - |
| Otu59509 | 1 | k__Fungi(100);k__Fungi_unclassified(100);k__Fungi_unclassified(100);k__Fungi_unclassified(100);k__Fungi_unclassified(100);k__Fungi_unclassified(100);k__Fungi_unclassified(100); | - | - | - | - | - | - | - | Unassigned | - |
| Otu59510 | 1 | k__Fungi(100);k__Fungi_unclassified(100);k__Fungi_unclassified(100);k__Fungi_unclassified(100);k__Fungi_unclassified(100);k__Fungi_unclassified(100);k__Fungi_unclassified(100); | - | - | - | - | - | - | - | Unassigned | - |
| Otu59512 | 1 | k__Fungi(100);p__Ascomycota(100);p__Ascomycota_unclassified(100);p__Ascomycota_unclassified(100);p__Ascomycota_unclassified(100);p__Ascomycota_unclassified(100);p__Ascomycota_unclassified(100); | - | - | - | - | - | - | - | Unassigned | - |
| Otu59513 | 1 | k__Fungi(100);p__Ascomycota(100);p__Ascomycota_unclassified(100);p__Ascomycota_unclassified(100);p__Ascomycota_unclassified(100);p__Ascomycota_unclassified(100);p__Ascomycota_unclassified(100); | - | - | - | - | - | - | - | Unassigned | - |
| Otu59515 | 1 | k__Fungi(100);p__Ascomycota(100);p__Ascomycota_unclassified(100);p__Ascomycota_unclassified(100);p__Ascomycota_unclassified(100);p__Ascomycota_unclassified(100);p__Ascomycota_unclassified(100); | - | - | - | - | - | - | - | Unassigned | - |
| Otu59526 | 1 | k__Fungi(100);p__Ascomycota(100);p__Ascomycota_unclassified(100);p__Ascomycota_unclassified(100);p__Ascomycota_unclassified(100);p__Ascomycota_unclassified(100);p__Ascomycota_unclassified(100); | - | - | - | - | - | - | - | Unassigned | - |
| Otu59527 | 1 | k__Fungi(100);p__Ascomycota(100);p__Ascomycota_unclassified(100);p__Ascomycota_unclassified(100);p__Ascomycota_unclassified(100);p__Ascomycota_unclassified(100);p__Ascomycota_unclassified(100); | - | - | - | - | - | - | - | Unassigned | - |
| Otu59541 | 1 | k__Fungi(100);k__Fungi_unclassified(100);k__Fungi_unclassified(100);k__Fungi_unclassified(100);k__Fungi_unclassified(100);k__Fungi_unclassified(100);k__Fungi_unclassified(100); | - | - | - | - | - | - | - | Unassigned | - |
| Otu59544 | 1 | k__Fungi(100);k__Fungi_unclassified(100);k__Fungi_unclassified(100);k__Fungi_unclassified(100);k__Fungi_unclassified(100);k__Fungi_unclassified(100);k__Fungi_unclassified(100); | - | - | - | - | - | - | - | Unassigned | - |
| Otu59550 | 1 | k__Fungi(100);p__Ascomycota(100);p__Ascomycota_unclassified(100);p__Ascomycota_unclassified(100);p__Ascomycota_unclassified(100);p__Ascomycota_unclassified(100);p__Ascomycota_unclassified(100); | - | - | - | - | - | - | - | Unassigned | - |
| Otu59552 | 1 | k__Fungi(100);p__Ascomycota(100);p__Ascomycota_unclassified(100);p__Ascomycota_unclassified(100);p__Ascomycota_unclassified(100);p__Ascomycota_unclassified(100);p__Ascomycota_unclassified(100); | - | - | - | - | - | - | - | Unassigned | - |
| Otu59553 | 1 | k__Fungi(100);p__Ascomycota(100);p__Ascomycota_unclassified(100);p__Ascomycota_unclassified(100);p__Ascomycota_unclassified(100);p__Ascomycota_unclassified(100);p__Ascomycota_unclassified(100); | - | - | - | - | - | - | - | Unassigned | - |
| Otu59555 | 1 | k__Fungi(100);k__Fungi_unclassified(100);k__Fungi_unclassified(100);k__Fungi_unclassified(100);k__Fungi_unclassified(100);k__Fungi_unclassified(100);k__Fungi_unclassified(100); | - | - | - | - | - | - | - | Unassigned | - |
| Otu59557 | 1 | k__Fungi(100);p__Ascomycota(100);p__Ascomycota_unclassified(100);p__Ascomycota_unclassified(100);p__Ascomycota_unclassified(100);p__Ascomycota_unclassified(100);p__Ascomycota_unclassified(100); | - | - | - | - | - | - | - | Unassigned | - |
| Otu59558 | 1 | k__Fungi(100);p__Ascomycota(100);p__Ascomycota_unclassified(100);p__Ascomycota_unclassified(100);p__Ascomycota_unclassified(100);p__Ascomycota_unclassified(100);p__Ascomycota_unclassified(100); | - | - | - | - | - | - | - | Unassigned | - |
| Otu59559 | 1 | k__Fungi(100);k__Fungi_unclassified(100);k__Fungi_unclassified(100);k__Fungi_unclassified(100);k__Fungi_unclassified(100);k__Fungi_unclassified(100);k__Fungi_unclassified(100); | - | - | - | - | - | - | - | Unassigned | - |
| Otu59566 | 1 | k__Fungi(100);k__Fungi_unclassified(100);k__Fungi_unclassified(100);k__Fungi_unclassified(100);k__Fungi_unclassified(100);k__Fungi_unclassified(100);k__Fungi_unclassified(100); | - | - | - | - | - | - | - | Unassigned | - |
| Otu59589 | 1 | k__Fungi(100);k__Fungi_unclassified(100);k__Fungi_unclassified(100);k__Fungi_unclassified(100);k__Fungi_unclassified(100);k__Fungi_unclassified(100);k__Fungi_unclassified(100); | - | - | - | - | - | - | - | Unassigned | - |
| Otu59595 | 1 | k__Fungi(100);k__Fungi_unclassified(100);k__Fungi_unclassified(100);k__Fungi_unclassified(100);k__Fungi_unclassified(100);k__Fungi_unclassified(100);k__Fungi_unclassified(100); | - | - | - | - | - | - | - | Unassigned | - |
| Otu59613 | 1 | k__Fungi(100);k__Fungi_unclassified(100);k__Fungi_unclassified(100);k__Fungi_unclassified(100);k__Fungi_unclassified(100);k__Fungi_unclassified(100);k__Fungi_unclassified(100); | - | - | - | - | - | - | - | Unassigned | - |
| Otu59734 | 1 | k__Fungi(100);k__Fungi_unclassified(100);k__Fungi_unclassified(100);k__Fungi_unclassified(100);k__Fungi_unclassified(100);k__Fungi_unclassified(100);k__Fungi_unclassified(100); | - | - | - | - | - | - | - | Unassigned | - |
| Otu59807 | 1 | k__Fungi(100);p__Ascomycota(100);p__Ascomycota_unclassified(100);p__Ascomycota_unclassified(100);p__Ascomycota_unclassified(100);p__Ascomycota_unclassified(100);p__Ascomycota_unclassified(100); | - | - | - | - | - | - | - | Unassigned | - |
| Otu59808 | 1 | k__Fungi(100);k__Fungi_unclassified(100);k__Fungi_unclassified(100);k__Fungi_unclassified(100);k__Fungi_unclassified(100);k__Fungi_unclassified(100);k__Fungi_unclassified(100); | - | - | - | - | - | - | - | Unassigned | - |
| Otu59832 | 1 | k__Fungi(100);k__Fungi_unclassified(100);k__Fungi_unclassified(100);k__Fungi_unclassified(100);k__Fungi_unclassified(100);k__Fungi_unclassified(100);k__Fungi_unclassified(100); | - | - | - | - | - | - | - | Unassigned | - |
| Otu59842 | 1 | k__Fungi(100);k__Fungi_unclassified(100);k__Fungi_unclassified(100);k__Fungi_unclassified(100);k__Fungi_unclassified(100);k__Fungi_unclassified(100);k__Fungi_unclassified(100); | - | - | - | - | - | - | - | Unassigned | - |
| Otu59843 | 1 | k__Fungi(100);p__Ascomycota(100);p__Ascomycota_unclassified(100);p__Ascomycota_unclassified(100);p__Ascomycota_unclassified(100);p__Ascomycota_unclassified(100);p__Ascomycota_unclassified(100); | - | - | - | - | - | - | - | Unassigned | - |
| Otu59852 | 1 | k__Fungi(100);k__Fungi_unclassified(100);k__Fungi_unclassified(100);k__Fungi_unclassified(100);k__Fungi_unclassified(100);k__Fungi_unclassified(100);k__Fungi_unclassified(100); | - | - | - | - | - | - | - | Unassigned | - |
| Otu59857 | 1 | k__Fungi(100);p__Ascomycota(100);p__Ascomycota_unclassified(100);p__Ascomycota_unclassified(100);p__Ascomycota_unclassified(100);p__Ascomycota_unclassified(100);p__Ascomycota_unclassified(100); | - | - | - | - | - | - | - | Unassigned | - |
| Otu59862 | 1 | k__Fungi(100);k__Fungi_unclassified(100);k__Fungi_unclassified(100);k__Fungi_unclassified(100);k__Fungi_unclassified(100);k__Fungi_unclassified(100);k__Fungi_unclassified(100); | - | - | - | - | - | - | - | Unassigned | - |
| Otu59864 | 1 | k__Fungi(100);k__Fungi_unclassified(100);k__Fungi_unclassified(100);k__Fungi_unclassified(100);k__Fungi_unclassified(100);k__Fungi_unclassified(100);k__Fungi_unclassified(100); | - | - | - | - | - | - | - | Unassigned | - |
| Otu59866 | 1 | k__Fungi(100);k__Fungi_unclassified(100);k__Fungi_unclassified(100);k__Fungi_unclassified(100);k__Fungi_unclassified(100);k__Fungi_unclassified(100);k__Fungi_unclassified(100); | - | - | - | - | - | - | - | Unassigned | - |
| Otu59867 | 1 | k__Fungi(100);p__Ascomycota(100);p__Ascomycota_unclassified(100);p__Ascomycota_unclassified(100);p__Ascomycota_unclassified(100);p__Ascomycota_unclassified(100);p__Ascomycota_unclassified(100); | - | - | - | - | - | - | - | Unassigned | - |
| Otu59871 | 1 | k__Fungi(100);k__Fungi_unclassified(100);k__Fungi_unclassified(100);k__Fungi_unclassified(100);k__Fungi_unclassified(100);k__Fungi_unclassified(100);k__Fungi_unclassified(100); | - | - | - | - | - | - | - | Unassigned | - |
| Otu59873 | 1 | k__Fungi(100);p__Ascomycota(100);p__Ascomycota_unclassified(100);p__Ascomycota_unclassified(100);p__Ascomycota_unclassified(100);p__Ascomycota_unclassified(100);p__Ascomycota_unclassified(100); | - | - | - | - | - | - | - | Unassigned | - |
| Otu59876 | 1 | k__Fungi(100);p__Ascomycota(100);p__Ascomycota_unclassified(100);p__Ascomycota_unclassified(100);p__Ascomycota_unclassified(100);p__Ascomycota_unclassified(100);p__Ascomycota_unclassified(100); | - | - | - | - | - | - | - | Unassigned | - |
| Otu59892 | 1 | k__Fungi(100);k__Fungi_unclassified(100);k__Fungi_unclassified(100);k__Fungi_unclassified(100);k__Fungi_unclassified(100);k__Fungi_unclassified(100);k__Fungi_unclassified(100); | - | - | - | - | - | - | - | Unassigned | - |
| Otu59896 | 1 | k__Fungi(100);k__Fungi_unclassified(100);k__Fungi_unclassified(100);k__Fungi_unclassified(100);k__Fungi_unclassified(100);k__Fungi_unclassified(100);k__Fungi_unclassified(100); | - | - | - | - | - | - | - | Unassigned | - |
| Otu59897 | 1 | k__Fungi(100);k__Fungi_unclassified(100);k__Fungi_unclassified(100);k__Fungi_unclassified(100);k__Fungi_unclassified(100);k__Fungi_unclassified(100);k__Fungi_unclassified(100); | - | - | - | - | - | - | - | Unassigned | - |
| Otu59898 | 1 | k__Fungi(100);k__Fungi_unclassified(100);k__Fungi_unclassified(100);k__Fungi_unclassified(100);k__Fungi_unclassified(100);k__Fungi_unclassified(100);k__Fungi_unclassified(100); | - | - | - | - | - | - | - | Unassigned | - |
| Otu59899 | 1 | k__Fungi(100);p__Ascomycota(100);p__Ascomycota_unclassified(100);p__Ascomycota_unclassified(100);p__Ascomycota_unclassified(100);p__Ascomycota_unclassified(100);p__Ascomycota_unclassified(100); | - | - | - | - | - | - | - | Unassigned | - |
| Otu59900 | 1 | k__Fungi(100);k__Fungi_unclassified(100);k__Fungi_unclassified(100);k__Fungi_unclassified(100);k__Fungi_unclassified(100);k__Fungi_unclassified(100);k__Fungi_unclassified(100); | - | - | - | - | - | - | - | Unassigned | - |
| Otu59901 | 1 | k__Fungi(100);k__Fungi_unclassified(100);k__Fungi_unclassified(100);k__Fungi_unclassified(100);k__Fungi_unclassified(100);k__Fungi_unclassified(100);k__Fungi_unclassified(100); | - | - | - | - | - | - | - | Unassigned | - |
| Otu59905 | 1 | k__Fungi(100);k__Fungi_unclassified(100);k__Fungi_unclassified(100);k__Fungi_unclassified(100);k__Fungi_unclassified(100);k__Fungi_unclassified(100);k__Fungi_unclassified(100); | - | - | - | - | - | - | - | Unassigned | - |
| Otu59917 | 1 | k__Fungi(100);k__Fungi_unclassified(100);k__Fungi_unclassified(100);k__Fungi_unclassified(100);k__Fungi_unclassified(100);k__Fungi_unclassified(100);k__Fungi_unclassified(100); | - | - | - | - | - | - | - | Unassigned | - |
| Otu59943 | 1 | k__Fungi(100);k__Fungi_unclassified(100);k__Fungi_unclassified(100);k__Fungi_unclassified(100);k__Fungi_unclassified(100);k__Fungi_unclassified(100);k__Fungi_unclassified(100); | - | - | - | - | - | - | - | Unassigned | - |
| Otu59955 | 1 | k__Fungi(100);p__Ascomycota(100);p__Ascomycota_unclassified(100);p__Ascomycota_unclassified(100);p__Ascomycota_unclassified(100);p__Ascomycota_unclassified(100);p__Ascomycota_unclassified(100); | - | - | - | - | - | - | - | Unassigned | - |
| Otu59956 | 1 | k__Fungi(100);k__Fungi_unclassified(100);k__Fungi_unclassified(100);k__Fungi_unclassified(100);k__Fungi_unclassified(100);k__Fungi_unclassified(100);k__Fungi_unclassified(100); | - | - | - | - | - | - | - | Unassigned | - |
| Otu59958 | 1 | k__Fungi(100);k__Fungi_unclassified(100);k__Fungi_unclassified(100);k__Fungi_unclassified(100);k__Fungi_unclassified(100);k__Fungi_unclassified(100);k__Fungi_unclassified(100); | - | - | - | - | - | - | - | Unassigned | - |
| Otu59959 | 1 | k__Fungi(100);p__Ascomycota(100);p__Ascomycota_unclassified(100);p__Ascomycota_unclassified(100);p__Ascomycota_unclassified(100);p__Ascomycota_unclassified(100);p__Ascomycota_unclassified(100); | - | - | - | - | - | - | - | Unassigned | - |
| Otu59960 | 1 | k__Fungi(100);k__Fungi_unclassified(100);k__Fungi_unclassified(100);k__Fungi_unclassified(100);k__Fungi_unclassified(100);k__Fungi_unclassified(100);k__Fungi_unclassified(100); | - | - | - | - | - | - | - | Unassigned | - |
| Otu59961 | 1 | k__Fungi(100);p__Ascomycota(100);p__Ascomycota_unclassified(100);p__Ascomycota_unclassified(100);p__Ascomycota_unclassified(100);p__Ascomycota_unclassified(100);p__Ascomycota_unclassified(100); | - | - | - | - | - | - | - | Unassigned | - |
| Otu59965 | 1 | k__Fungi(100);p__Ascomycota(100);p__Ascomycota_unclassified(100);p__Ascomycota_unclassified(100);p__Ascomycota_unclassified(100);p__Ascomycota_unclassified(100);p__Ascomycota_unclassified(100); | - | - | - | - | - | - | - | Unassigned | - |
| Otu59966 | 1 | k__Fungi(100);k__Fungi_unclassified(100);k__Fungi_unclassified(100);k__Fungi_unclassified(100);k__Fungi_unclassified(100);k__Fungi_unclassified(100);k__Fungi_unclassified(100); | - | - | - | - | - | - | - | Unassigned | - |
| Otu59967 | 1 | k__Fungi(100);k__Fungi_unclassified(100);k__Fungi_unclassified(100);k__Fungi_unclassified(100);k__Fungi_unclassified(100);k__Fungi_unclassified(100);k__Fungi_unclassified(100); | - | - | - | - | - | - | - | Unassigned | - |
| Otu59968 | 1 | k__Fungi(100);k__Fungi_unclassified(100);k__Fungi_unclassified(100);k__Fungi_unclassified(100);k__Fungi_unclassified(100);k__Fungi_unclassified(100);k__Fungi_unclassified(100); | - | - | - | - | - | - | - | Unassigned | - |
| Otu59969 | 1 | k__Fungi(100);k__Fungi_unclassified(100);k__Fungi_unclassified(100);k__Fungi_unclassified(100);k__Fungi_unclassified(100);k__Fungi_unclassified(100);k__Fungi_unclassified(100); | - | - | - | - | - | - | - | Unassigned | - |
| Otu59976 | 1 | k__Fungi(100);k__Fungi_unclassified(100);k__Fungi_unclassified(100);k__Fungi_unclassified(100);k__Fungi_unclassified(100);k__Fungi_unclassified(100);k__Fungi_unclassified(100); | - | - | - | - | - | - | - | Unassigned | - |
| Otu59977 | 1 | k__Fungi(100);p__Ascomycota(100);p__Ascomycota_unclassified(100);p__Ascomycota_unclassified(100);p__Ascomycota_unclassified(100);p__Ascomycota_unclassified(100);p__Ascomycota_unclassified(100); | - | - | - | - | - | - | - | Unassigned | - |
| Otu59982 | 1 | k__Fungi(100);k__Fungi_unclassified(100);k__Fungi_unclassified(100);k__Fungi_unclassified(100);k__Fungi_unclassified(100);k__Fungi_unclassified(100);k__Fungi_unclassified(100); | - | - | - | - | - | - | - | Unassigned | - |
| Otu59983 | 1 | k__Fungi(100);k__Fungi_unclassified(100);k__Fungi_unclassified(100);k__Fungi_unclassified(100);k__Fungi_unclassified(100);k__Fungi_unclassified(100);k__Fungi_unclassified(100); | - | - | - | - | - | - | - | Unassigned | - |
| Otu59984 | 1 | k__Fungi(100);k__Fungi_unclassified(100);k__Fungi_unclassified(100);k__Fungi_unclassified(100);k__Fungi_unclassified(100);k__Fungi_unclassified(100);k__Fungi_unclassified(100); | - | - | - | - | - | - | - | Unassigned | - |
| Otu59986 | 1 | k__Fungi(100);p__Ascomycota(100);p__Ascomycota_unclassified(100);p__Ascomycota_unclassified(100);p__Ascomycota_unclassified(100);p__Ascomycota_unclassified(100);p__Ascomycota_unclassified(100); | - | - | - | - | - | - | - | Unassigned | - |
| Otu59987 | 1 | k__Fungi(100);p__Ascomycota(100);p__Ascomycota_unclassified(100);p__Ascomycota_unclassified(100);p__Ascomycota_unclassified(100);p__Ascomycota_unclassified(100);p__Ascomycota_unclassified(100); | - | - | - | - | - | - | - | Unassigned | - |
| Otu59988 | 1 | k__Fungi(100);p__Ascomycota(100);p__Ascomycota_unclassified(100);p__Ascomycota_unclassified(100);p__Ascomycota_unclassified(100);p__Ascomycota_unclassified(100);p__Ascomycota_unclassified(100); | - | - | - | - | - | - | - | Unassigned | - |
| Otu59989 | 1 | k__Fungi(100);k__Fungi_unclassified(100);k__Fungi_unclassified(100);k__Fungi_unclassified(100);k__Fungi_unclassified(100);k__Fungi_unclassified(100);k__Fungi_unclassified(100); | - | - | - | - | - | - | - | Unassigned | - |
| Otu59991 | 1 | k__Fungi(100);p__Ascomycota(100);p__Ascomycota_unclassified(100);p__Ascomycota_unclassified(100);p__Ascomycota_unclassified(100);p__Ascomycota_unclassified(100);p__Ascomycota_unclassified(100); | - | - | - | - | - | - | - | Unassigned | - |
| Otu59993 | 1 | k__Fungi(100);p__Ascomycota(100);p__Ascomycota_unclassified(100);p__Ascomycota_unclassified(100);p__Ascomycota_unclassified(100);p__Ascomycota_unclassified(100);p__Ascomycota_unclassified(100); | - | - | - | - | - | - | - | Unassigned | - |
| Otu59999 | 1 | k__Fungi(100);k__Fungi_unclassified(100);k__Fungi_unclassified(100);k__Fungi_unclassified(100);k__Fungi_unclassified(100);k__Fungi_unclassified(100);k__Fungi_unclassified(100); | - | - | - | - | - | - | - | Unassigned | - |
| Otu60000 | 1 | k__Fungi(100);k__Fungi_unclassified(100);k__Fungi_unclassified(100);k__Fungi_unclassified(100);k__Fungi_unclassified(100);k__Fungi_unclassified(100);k__Fungi_unclassified(100); | - | - | - | - | - | - | - | Unassigned | - |
| Otu60001 | 1 | k__Fungi(100);k__Fungi_unclassified(100);k__Fungi_unclassified(100);k__Fungi_unclassified(100);k__Fungi_unclassified(100);k__Fungi_unclassified(100);k__Fungi_unclassified(100); | - | - | - | - | - | - | - | Unassigned | - |
| Otu60002 | 1 | k__Fungi(100);k__Fungi_unclassified(100);k__Fungi_unclassified(100);k__Fungi_unclassified(100);k__Fungi_unclassified(100);k__Fungi_unclassified(100);k__Fungi_unclassified(100); | - | - | - | - | - | - | - | Unassigned | - |
| Otu60003 | 1 | k__Fungi(100);k__Fungi_unclassified(100);k__Fungi_unclassified(100);k__Fungi_unclassified(100);k__Fungi_unclassified(100);k__Fungi_unclassified(100);k__Fungi_unclassified(100); | - | - | - | - | - | - | - | Unassigned | - |
| Otu60004 | 1 | k__Fungi(100);p__Ascomycota(100);p__Ascomycota_unclassified(100);p__Ascomycota_unclassified(100);p__Ascomycota_unclassified(100);p__Ascomycota_unclassified(100);p__Ascomycota_unclassified(100); | - | - | - | - | - | - | - | Unassigned | - |
| Otu60005 | 1 | k__Fungi(100);p__Ascomycota(100);p__Ascomycota_unclassified(100);p__Ascomycota_unclassified(100);p__Ascomycota_unclassified(100);p__Ascomycota_unclassified(100);p__Ascomycota_unclassified(100); | - | - | - | - | - | - | - | Unassigned | - |
| Otu60006 | 1 | k__Fungi(100);k__Fungi_unclassified(100);k__Fungi_unclassified(100);k__Fungi_unclassified(100);k__Fungi_unclassified(100);k__Fungi_unclassified(100);k__Fungi_unclassified(100); | - | - | - | - | - | - | - | Unassigned | - |
| Otu60008 | 1 | k__Fungi(100);p__Ascomycota(100);p__Ascomycota_unclassified(100);p__Ascomycota_unclassified(100);p__Ascomycota_unclassified(100);p__Ascomycota_unclassified(100);p__Ascomycota_unclassified(100); | - | - | - | - | - | - | - | Unassigned | - |
| Otu60009 | 1 | k__Fungi(100);k__Fungi_unclassified(100);k__Fungi_unclassified(100);k__Fungi_unclassified(100);k__Fungi_unclassified(100);k__Fungi_unclassified(100);k__Fungi_unclassified(100); | - | - | - | - | - | - | - | Unassigned | - |
| Otu60010 | 1 | k__Fungi(100);p__Ascomycota(100);p__Ascomycota_unclassified(100);p__Ascomycota_unclassified(100);p__Ascomycota_unclassified(100);p__Ascomycota_unclassified(100);p__Ascomycota_unclassified(100); | - | - | - | - | - | - | - | Unassigned | - |
| Otu60011 | 1 | k__Fungi(100);p__Ascomycota(100);p__Ascomycota_unclassified(100);p__Ascomycota_unclassified(100);p__Ascomycota_unclassified(100);p__Ascomycota_unclassified(100);p__Ascomycota_unclassified(100); | - | - | - | - | - | - | - | Unassigned | - |
| Otu60012 | 1 | k__Fungi(100);k__Fungi_unclassified(100);k__Fungi_unclassified(100);k__Fungi_unclassified(100);k__Fungi_unclassified(100);k__Fungi_unclassified(100);k__Fungi_unclassified(100); | - | - | - | - | - | - | - | Unassigned | - |
| Otu60013 | 1 | k__Fungi(100);p__Ascomycota(100);p__Ascomycota_unclassified(100);p__Ascomycota_unclassified(100);p__Ascomycota_unclassified(100);p__Ascomycota_unclassified(100);p__Ascomycota_unclassified(100); | - | - | - | - | - | - | - | Unassigned | - |
| Otu60014 | 1 | k__Fungi(100);p__Ascomycota(100);p__Ascomycota_unclassified(100);p__Ascomycota_unclassified(100);p__Ascomycota_unclassified(100);p__Ascomycota_unclassified(100);p__Ascomycota_unclassified(100); | - | - | - | - | - | - | - | Unassigned | - |
| Otu60015 | 1 | k__Fungi(100);k__Fungi_unclassified(100);k__Fungi_unclassified(100);k__Fungi_unclassified(100);k__Fungi_unclassified(100);k__Fungi_unclassified(100);k__Fungi_unclassified(100); | - | - | - | - | - | - | - | Unassigned | - |
| Otu60016 | 1 | k__Fungi(100);p__Ascomycota(100);p__Ascomycota_unclassified(100);p__Ascomycota_unclassified(100);p__Ascomycota_unclassified(100);p__Ascomycota_unclassified(100);p__Ascomycota_unclassified(100); | - | - | - | - | - | - | - | Unassigned | - |
| Otu60017 | 1 | k__Fungi(100);p__Ascomycota(100);p__Ascomycota_unclassified(100);p__Ascomycota_unclassified(100);p__Ascomycota_unclassified(100);p__Ascomycota_unclassified(100);p__Ascomycota_unclassified(100); | - | - | - | - | - | - | - | Unassigned | - |
| Otu60018 | 1 | k__Fungi(100);p__Ascomycota(100);p__Ascomycota_unclassified(100);p__Ascomycota_unclassified(100);p__Ascomycota_unclassified(100);p__Ascomycota_unclassified(100);p__Ascomycota_unclassified(100); | - | - | - | - | - | - | - | Unassigned | - |
| Otu60019 | 1 | k__Fungi(100);p__Ascomycota(100);p__Ascomycota_unclassified(100);p__Ascomycota_unclassified(100);p__Ascomycota_unclassified(100);p__Ascomycota_unclassified(100);p__Ascomycota_unclassified(100); | - | - | - | - | - | - | - | Unassigned | - |
| Otu60020 | 1 | k__Fungi(100);k__Fungi_unclassified(100);k__Fungi_unclassified(100);k__Fungi_unclassified(100);k__Fungi_unclassified(100);k__Fungi_unclassified(100);k__Fungi_unclassified(100); | - | - | - | - | - | - | - | Unassigned | - |
| Otu60021 | 1 | k__Fungi(100);p__Ascomycota(100);p__Ascomycota_unclassified(100);p__Ascomycota_unclassified(100);p__Ascomycota_unclassified(100);p__Ascomycota_unclassified(100);p__Ascomycota_unclassified(100); | - | - | - | - | - | - | - | Unassigned | - |
| Otu60022 | 1 | k__Fungi(100);p__Ascomycota(100);p__Ascomycota_unclassified(100);p__Ascomycota_unclassified(100);p__Ascomycota_unclassified(100);p__Ascomycota_unclassified(100);p__Ascomycota_unclassified(100); | - | - | - | - | - | - | - | Unassigned | - |
| Otu60023 | 1 | k__Fungi(100);p__Ascomycota(100);p__Ascomycota_unclassified(100);p__Ascomycota_unclassified(100);p__Ascomycota_unclassified(100);p__Ascomycota_unclassified(100);p__Ascomycota_unclassified(100); | - | - | - | - | - | - | - | Unassigned | - |
| Otu60025 | 1 | k__Fungi(100);k__Fungi_unclassified(100);k__Fungi_unclassified(100);k__Fungi_unclassified(100);k__Fungi_unclassified(100);k__Fungi_unclassified(100);k__Fungi_unclassified(100); | - | - | - | - | - | - | - | Unassigned | - |
| Otu60028 | 1 | k__Fungi(100);k__Fungi_unclassified(100);k__Fungi_unclassified(100);k__Fungi_unclassified(100);k__Fungi_unclassified(100);k__Fungi_unclassified(100);k__Fungi_unclassified(100); | - | - | - | - | - | - | - | Unassigned | - |
| Otu60029 | 1 | k__Fungi(100);p__Ascomycota(100);p__Ascomycota_unclassified(100);p__Ascomycota_unclassified(100);p__Ascomycota_unclassified(100);p__Ascomycota_unclassified(100);p__Ascomycota_unclassified(100); | - | - | - | - | - | - | - | Unassigned | - |
| Otu60031 | 1 | k__Fungi(100);p__Ascomycota(100);p__Ascomycota_unclassified(100);p__Ascomycota_unclassified(100);p__Ascomycota_unclassified(100);p__Ascomycota_unclassified(100);p__Ascomycota_unclassified(100); | - | - | - | - | - | - | - | Unassigned | - |
| Otu60032 | 1 | k__Fungi(100);p__Ascomycota(100);p__Ascomycota_unclassified(100);p__Ascomycota_unclassified(100);p__Ascomycota_unclassified(100);p__Ascomycota_unclassified(100);p__Ascomycota_unclassified(100); | - | - | - | - | - | - | - | Unassigned | - |
| Otu60033 | 1 | k__Fungi(100);k__Fungi_unclassified(100);k__Fungi_unclassified(100);k__Fungi_unclassified(100);k__Fungi_unclassified(100);k__Fungi_unclassified(100);k__Fungi_unclassified(100); | - | - | - | - | - | - | - | Unassigned | - |
| Otu60036 | 1 | k__Fungi(100);k__Fungi_unclassified(100);k__Fungi_unclassified(100);k__Fungi_unclassified(100);k__Fungi_unclassified(100);k__Fungi_unclassified(100);k__Fungi_unclassified(100); | - | - | - | - | - | - | - | Unassigned | - |
| Otu60038 | 1 | k__Fungi(100);k__Fungi_unclassified(100);k__Fungi_unclassified(100);k__Fungi_unclassified(100);k__Fungi_unclassified(100);k__Fungi_unclassified(100);k__Fungi_unclassified(100); | - | - | - | - | - | - | - | Unassigned | - |
| Otu60039 | 1 | k__Fungi(100);k__Fungi_unclassified(100);k__Fungi_unclassified(100);k__Fungi_unclassified(100);k__Fungi_unclassified(100);k__Fungi_unclassified(100);k__Fungi_unclassified(100); | - | - | - | - | - | - | - | Unassigned | - |
| Otu60040 | 1 | k__Fungi(100);k__Fungi_unclassified(100);k__Fungi_unclassified(100);k__Fungi_unclassified(100);k__Fungi_unclassified(100);k__Fungi_unclassified(100);k__Fungi_unclassified(100); | - | - | - | - | - | - | - | Unassigned | - |
| Otu60041 | 1 | k__Fungi(100);k__Fungi_unclassified(100);k__Fungi_unclassified(100);k__Fungi_unclassified(100);k__Fungi_unclassified(100);k__Fungi_unclassified(100);k__Fungi_unclassified(100); | - | - | - | - | - | - | - | Unassigned | - |
| Otu60042 | 1 | k__Fungi(100);p__Ascomycota(100);p__Ascomycota_unclassified(100);p__Ascomycota_unclassified(100);p__Ascomycota_unclassified(100);p__Ascomycota_unclassified(100);p__Ascomycota_unclassified(100); | - | - | - | - | - | - | - | Unassigned | - |
| Otu60043 | 1 | k__Fungi(100);k__Fungi_unclassified(100);k__Fungi_unclassified(100);k__Fungi_unclassified(100);k__Fungi_unclassified(100);k__Fungi_unclassified(100);k__Fungi_unclassified(100); | - | - | - | - | - | - | - | Unassigned | - |
| Otu60044 | 1 | k__Fungi(100);p__Ascomycota(100);p__Ascomycota_unclassified(100);p__Ascomycota_unclassified(100);p__Ascomycota_unclassified(100);p__Ascomycota_unclassified(100);p__Ascomycota_unclassified(100); | - | - | - | - | - | - | - | Unassigned | - |
| Otu60045 | 1 | k__Fungi(100);p__Ascomycota(100);p__Ascomycota_unclassified(100);p__Ascomycota_unclassified(100);p__Ascomycota_unclassified(100);p__Ascomycota_unclassified(100);p__Ascomycota_unclassified(100); | - | - | - | - | - | - | - | Unassigned | - |
| Otu60049 | 1 | k__Fungi(100);k__Fungi_unclassified(100);k__Fungi_unclassified(100);k__Fungi_unclassified(100);k__Fungi_unclassified(100);k__Fungi_unclassified(100);k__Fungi_unclassified(100); | - | - | - | - | - | - | - | Unassigned | - |
| Otu60050 | 1 | k__Fungi(100);k__Fungi_unclassified(100);k__Fungi_unclassified(100);k__Fungi_unclassified(100);k__Fungi_unclassified(100);k__Fungi_unclassified(100);k__Fungi_unclassified(100); | - | - | - | - | - | - | - | Unassigned | - |
| Otu60051 | 1 | k__Fungi(100);p__Ascomycota(100);p__Ascomycota_unclassified(100);p__Ascomycota_unclassified(100);p__Ascomycota_unclassified(100);p__Ascomycota_unclassified(100);p__Ascomycota_unclassified(100); | - | - | - | - | - | - | - | Unassigned | - |
| Otu60052 | 1 | k__Fungi(100);p__Ascomycota(100);p__Ascomycota_unclassified(100);p__Ascomycota_unclassified(100);p__Ascomycota_unclassified(100);p__Ascomycota_unclassified(100);p__Ascomycota_unclassified(100); | - | - | - | - | - | - | - | Unassigned | - |
| Otu60053 | 1 | k__Fungi(100);k__Fungi_unclassified(100);k__Fungi_unclassified(100);k__Fungi_unclassified(100);k__Fungi_unclassified(100);k__Fungi_unclassified(100);k__Fungi_unclassified(100); | - | - | - | - | - | - | - | Unassigned | - |
| Otu60054 | 1 | k__Fungi(100);p__Ascomycota(100);p__Ascomycota_unclassified(100);p__Ascomycota_unclassified(100);p__Ascomycota_unclassified(100);p__Ascomycota_unclassified(100);p__Ascomycota_unclassified(100); | - | - | - | - | - | - | - | Unassigned | - |
| Otu60055 | 1 | k__Fungi(100);k__Fungi_unclassified(100);k__Fungi_unclassified(100);k__Fungi_unclassified(100);k__Fungi_unclassified(100);k__Fungi_unclassified(100);k__Fungi_unclassified(100); | - | - | - | - | - | - | - | Unassigned | - |
| Otu60056 | 1 | k__Fungi(100);p__Ascomycota(100);p__Ascomycota_unclassified(100);p__Ascomycota_unclassified(100);p__Ascomycota_unclassified(100);p__Ascomycota_unclassified(100);p__Ascomycota_unclassified(100); | - | - | - | - | - | - | - | Unassigned | - |
| Otu60057 | 1 | k__Fungi(100);p__Ascomycota(100);p__Ascomycota_unclassified(100);p__Ascomycota_unclassified(100);p__Ascomycota_unclassified(100);p__Ascomycota_unclassified(100);p__Ascomycota_unclassified(100); | - | - | - | - | - | - | - | Unassigned | - |
| Otu60058 | 1 | k__Fungi(100);k__Fungi_unclassified(100);k__Fungi_unclassified(100);k__Fungi_unclassified(100);k__Fungi_unclassified(100);k__Fungi_unclassified(100);k__Fungi_unclassified(100); | - | - | - | - | - | - | - | Unassigned | - |
| Otu60059 | 1 | k__Fungi(100);k__Fungi_unclassified(100);k__Fungi_unclassified(100);k__Fungi_unclassified(100);k__Fungi_unclassified(100);k__Fungi_unclassified(100);k__Fungi_unclassified(100); | - | - | - | - | - | - | - | Unassigned | - |
| Otu60060 | 1 | k__Fungi(100);p__Ascomycota(100);p__Ascomycota_unclassified(100);p__Ascomycota_unclassified(100);p__Ascomycota_unclassified(100);p__Ascomycota_unclassified(100);p__Ascomycota_unclassified(100); | - | - | - | - | - | - | - | Unassigned | - |
| Otu60061 | 1 | k__Fungi(100);k__Fungi_unclassified(100);k__Fungi_unclassified(100);k__Fungi_unclassified(100);k__Fungi_unclassified(100);k__Fungi_unclassified(100);k__Fungi_unclassified(100); | - | - | - | - | - | - | - | Unassigned | - |

| | | | | | | | | | | | |
|---|---|---|---|---|---|---|---|---|---|---|---|
| Otu60062 | 1 | k__Fungi(100);k__Fungi_unclassified(100);k__Fungi_unclassified(100);k__Fungi_unclassified(100);k__Fungi_unclassified(100);k__Fungi_unclassified(100);k__Fungi_unclassified(100); | - | - | - | - | - | - | - | Unassigned | - |
| Otu60063 | 1 | k__Fungi(100);k__Fungi_unclassified(100);k__Fungi_unclassified(100);k__Fungi_unclassified(100);k__Fungi_unclassified(100);k__Fungi_unclassified(100);k__Fungi_unclassified(100); | - | - | - | - | - | - | - | Unassigned | - |
| Otu60065 | 1 | k__Fungi(100);p__Ascomycota(100);p__Ascomycota_unclassified(100);p__Ascomycota_unclassified(100);p__Ascomycota_unclassified(100);p__Ascomycota_unclassified(100);p__Ascomycota_unclassified(100); | - | - | - | - | - | - | - | Unassigned | - |
| Otu60066 | 1 | k__Fungi(100);k__Fungi_unclassified(100);k__Fungi_unclassified(100);k__Fungi_unclassified(100);k__Fungi_unclassified(100);k__Fungi_unclassified(100);k__Fungi_unclassified(100); | - | - | - | - | - | - | - | Unassigned | - |
| Otu60067 | 1 | k__Fungi(100);k__Fungi_unclassified(100);k__Fungi_unclassified(100);k__Fungi_unclassified(100);k__Fungi_unclassified(100);k__Fungi_unclassified(100);k__Fungi_unclassified(100); | - | - | - | - | - | - | - | Unassigned | - |
| Otu60068 | 1 | k__Fungi(100);p__Ascomycota(100);p__Ascomycota_unclassified(100);p__Ascomycota_unclassified(100);p__Ascomycota_unclassified(100);p__Ascomycota_unclassified(100);p__Ascomycota_unclassified(100); | - | - | - | - | - | - | - | Unassigned | - |
| Otu60069 | 1 | k__Fungi(100);k__Fungi_unclassified(100);k__Fungi_unclassified(100);k__Fungi_unclassified(100);k__Fungi_unclassified(100);k__Fungi_unclassified(100);k__Fungi_unclassified(100); | - | - | - | - | - | - | - | Unassigned | - |
| Otu60070 | 1 | k__Fungi(100);k__Fungi_unclassified(100);k__Fungi_unclassified(100);k__Fungi_unclassified(100);k__Fungi_unclassified(100);k__Fungi_unclassified(100);k__Fungi_unclassified(100); | - | - | - | - | - | - | - | Unassigned | - |
| Otu60071 | 1 | k__Fungi(100);k__Fungi_unclassified(100);k__Fungi_unclassified(100);k__Fungi_unclassified(100);k__Fungi_unclassified(100);k__Fungi_unclassified(100);k__Fungi_unclassified(100); | - | - | - | - | - | - | - | Unassigned | - |
| Otu60073 | 1 | k__Fungi(100);k__Fungi_unclassified(100);k__Fungi_unclassified(100);k__Fungi_unclassified(100);k__Fungi_unclassified(100);k__Fungi_unclassified(100);k__Fungi_unclassified(100); | - | - | - | - | - | - | - | Unassigned | - |
| Otu60080 | 1 | k__Fungi(100);k__Fungi_unclassified(100);k__Fungi_unclassified(100);k__Fungi_unclassified(100);k__Fungi_unclassified(100);k__Fungi_unclassified(100);k__Fungi_unclassified(100); | - | - | - | - | - | - | - | Unassigned | - |
| Otu60081 | 1 | k__Fungi(100);k__Fungi_unclassified(100);k__Fungi_unclassified(100);k__Fungi_unclassified(100);k__Fungi_unclassified(100);k__Fungi_unclassified(100);k__Fungi_unclassified(100); | - | - | - | - | - | - | - | Unassigned | - |
| Otu60082 | 1 | k__Fungi(100);p__Ascomycota(100);p__Ascomycota_unclassified(100);p__Ascomycota_unclassified(100);p__Ascomycota_unclassified(100);p__Ascomycota_unclassified(100);p__Ascomycota_unclassified(100); | - | - | - | - | - | - | - | Unassigned | - |
| Otu60083 | 1 | k__Fungi(100);k__Fungi_unclassified(100);k__Fungi_unclassified(100);k__Fungi_unclassified(100);k__Fungi_unclassified(100);k__Fungi_unclassified(100);k__Fungi_unclassified(100); | - | - | - | - | - | - | - | Unassigned | - |
| Otu60086 | 1 | k__Fungi(100);k__Fungi_unclassified(100);k__Fungi_unclassified(100);k__Fungi_unclassified(100);k__Fungi_unclassified(100);k__Fungi_unclassified(100);k__Fungi_unclassified(100); | - | - | - | - | - | - | - | Unassigned | - |
| Otu60087 | 1 | k__Fungi(100);p__Ascomycota(100);p__Ascomycota_unclassified(100);p__Ascomycota_unclassified(100);p__Ascomycota_unclassified(100);p__Ascomycota_unclassified(100);p__Ascomycota_unclassified(100); | - | - | - | - | - | - | - | Unassigned | - |
| Otu60088 | 1 | k__Fungi(100);k__Fungi_unclassified(100);k__Fungi_unclassified(100);k__Fungi_unclassified(100);k__Fungi_unclassified(100);k__Fungi_unclassified(100);k__Fungi_unclassified(100); | - | - | - | - | - | - | - | Unassigned | - |
| Otu60089 | 1 | k__Fungi(100);k__Fungi_unclassified(100);k__Fungi_unclassified(100);k__Fungi_unclassified(100);k__Fungi_unclassified(100);k__Fungi_unclassified(100);k__Fungi_unclassified(100); | - | - | - | - | - | - | - | Unassigned | - |
| Otu60090 | 1 | k__Fungi(100);k__Fungi_unclassified(100);k__Fungi_unclassified(100);k__Fungi_unclassified(100);k__Fungi_unclassified(100);k__Fungi_unclassified(100);k__Fungi_unclassified(100); | - | - | - | - | - | - | - | Unassigned | - |
| Otu60092 | 1 | k__Fungi(100);p__Ascomycota(100);p__Ascomycota_unclassified(100);p__Ascomycota_unclassified(100);p__Ascomycota_unclassified(100);p__Ascomycota_unclassified(100);p__Ascomycota_unclassified(100); | - | - | - | - | - | - | - | Unassigned | - |
| Otu60100 | 1 | k__Fungi(100);p__Ascomycota(100);p__Ascomycota_unclassified(100);p__Ascomycota_unclassified(100);p__Ascomycota_unclassified(100);p__Ascomycota_unclassified(100);p__Ascomycota_unclassified(100); | - | - | - | - | - | - | - | Unassigned | - |
| Otu60101 | 1 | k__Fungi(100);p__Ascomycota(100);p__Ascomycota_unclassified(100);p__Ascomycota_unclassified(100);p__Ascomycota_unclassified(100);p__Ascomycota_unclassified(100);p__Ascomycota_unclassified(100); | - | - | - | - | - | - | - | Unassigned | - |
| Otu60102 | 1 | k__Fungi(100);p__Ascomycota(100);p__Ascomycota_unclassified(100);p__Ascomycota_unclassified(100);p__Ascomycota_unclassified(100);p__Ascomycota_unclassified(100);p__Ascomycota_unclassified(100); | - | - | - | - | - | - | - | Unassigned | - |
| Otu60103 | 1 | k__Fungi(100);p__Ascomycota(100);p__Ascomycota_unclassified(100);p__Ascomycota_unclassified(100);p__Ascomycota_unclassified(100);p__Ascomycota_unclassified(100);p__Ascomycota_unclassified(100); | - | - | - | - | - | - | - | Unassigned | - |
| Otu60104 | 1 | k__Fungi(100);p__Ascomycota(100);p__Ascomycota_unclassified(100);p__Ascomycota_unclassified(100);p__Ascomycota_unclassified(100);p__Ascomycota_unclassified(100);p__Ascomycota_unclassified(100); | - | - | - | - | - | - | - | Unassigned | - |
| Otu60105 | 1 | k__Fungi(100);k__Fungi_unclassified(100);k__Fungi_unclassified(100);k__Fungi_unclassified(100);k__Fungi_unclassified(100);k__Fungi_unclassified(100);k__Fungi_unclassified(100); | - | - | - | - | - | - | - | Unassigned | - |
| Otu60106 | 1 | k__Fungi(100);k__Fungi_unclassified(100);k__Fungi_unclassified(100);k__Fungi_unclassified(100);k__Fungi_unclassified(100);k__Fungi_unclassified(100);k__Fungi_unclassified(100); | - | - | - | - | - | - | - | Unassigned | - |
| Otu60107 | 1 | k__Fungi(100);p__Ascomycota(100);p__Ascomycota_unclassified(100);p__Ascomycota_unclassified(100);p__Ascomycota_unclassified(100);p__Ascomycota_unclassified(100);p__Ascomycota_unclassified(100); | - | - | - | - | - | - | - | Unassigned | - |
| Otu60111 | 1 | k__Fungi(100);k__Fungi_unclassified(100);k__Fungi_unclassified(100);k__Fungi_unclassified(100);k__Fungi_unclassified(100);k__Fungi_unclassified(100);k__Fungi_unclassified(100); | - | - | - | - | - | - | - | Unassigned | - |
| Otu60112 | 1 | k__Fungi(100);p__Ascomycota(100);p__Ascomycota_unclassified(100);p__Ascomycota_unclassified(100);p__Ascomycota_unclassified(100);p__Ascomycota_unclassified(100);p__Ascomycota_unclassified(100); | - | - | - | - | - | - | - | Unassigned | - |
| Otu60113 | 1 | k__Fungi(100);p__Ascomycota(100);p__Ascomycota_unclassified(100);p__Ascomycota_unclassified(100);p__Ascomycota_unclassified(100);p__Ascomycota_unclassified(100);p__Ascomycota_unclassified(100); | - | - | - | - | - | - | - | Unassigned | - |
| Otu60114 | 1 | k__Fungi(100);p__Ascomycota(100);p__Ascomycota_unclassified(100);p__Ascomycota_unclassified(100);p__Ascomycota_unclassified(100);p__Ascomycota_unclassified(100);p__Ascomycota_unclassified(100); | - | - | - | - | - | - | - | Unassigned | - |
| Otu60115 | 1 | k__Fungi(100);p__Ascomycota(100);p__Ascomycota_unclassified(100);p__Ascomycota_unclassified(100);p__Ascomycota_unclassified(100);p__Ascomycota_unclassified(100);p__Ascomycota_unclassified(100); | - | - | - | - | - | - | - | Unassigned | - |
| Otu60117 | 1 | k__Fungi(100);k__Fungi_unclassified(100);k__Fungi_unclassified(100);k__Fungi_unclassified(100);k__Fungi_unclassified(100);k__Fungi_unclassified(100);k__Fungi_unclassified(100); | - | - | - | - | - | - | - | Unassigned | - |
| Otu60118 | 1 | k__Fungi(100);p__Ascomycota(100);p__Ascomycota_unclassified(100);p__Ascomycota_unclassified(100);p__Ascomycota_unclassified(100);p__Ascomycota_unclassified(100);p__Ascomycota_unclassified(100); | - | - | - | - | - | - | - | Unassigned | - |
| Otu60119 | 1 | k__Fungi(100);p__Ascomycota(100);p__Ascomycota_unclassified(100);p__Ascomycota_unclassified(100);p__Ascomycota_unclassified(100);p__Ascomycota_unclassified(100);p__Ascomycota_unclassified(100); | - | - | - | - | - | - | - | Unassigned | - |
| Otu60121 | 1 | k__Fungi(100);p__Ascomycota(100);p__Ascomycota_unclassified(100);p__Ascomycota_unclassified(100);p__Ascomycota_unclassified(100);p__Ascomycota_unclassified(100);p__Ascomycota_unclassified(100); | - | - | - | - | - | - | - | Unassigned | - |
| Otu60122 | 1 | k__Fungi(100);k__Fungi_unclassified(100);k__Fungi_unclassified(100);k__Fungi_unclassified(100);k__Fungi_unclassified(100);k__Fungi_unclassified(100);k__Fungi_unclassified(100); | - | - | - | - | - | - | - | Unassigned | - |
| Otu60123 | 1 | k__Fungi(100);p__Ascomycota(100);p__Ascomycota_unclassified(100);p__Ascomycota_unclassified(100);p__Ascomycota_unclassified(100);p__Ascomycota_unclassified(100);p__Ascomycota_unclassified(100); | - | - | - | - | - | - | - | Unassigned | - |
| Otu60124 | 1 | k__Fungi(100);p__Ascomycota(100);p__Ascomycota_unclassified(100);p__Ascomycota_unclassified(100);p__Ascomycota_unclassified(100);p__Ascomycota_unclassified(100);p__Ascomycota_unclassified(100); | - | - | - | - | - | - | - | Unassigned | - |
| Otu60125 | 1 | k__Fungi(100);k__Fungi_unclassified(100);k__Fungi_unclassified(100);k__Fungi_unclassified(100);k__Fungi_unclassified(100);k__Fungi_unclassified(100);k__Fungi_unclassified(100); | - | - | - | - | - | - | - | Unassigned | - |
| Otu60126 | 1 | k__Fungi(100);k__Fungi_unclassified(100);k__Fungi_unclassified(100);k__Fungi_unclassified(100);k__Fungi_unclassified(100);k__Fungi_unclassified(100);k__Fungi_unclassified(100); | - | - | - | - | - | - | - | Unassigned | - |
| Otu60127 | 1 | k__Fungi(100);p__Ascomycota(100);p__Ascomycota_unclassified(100);p__Ascomycota_unclassified(100);p__Ascomycota_unclassified(100);p__Ascomycota_unclassified(100);p__Ascomycota_unclassified(100); | - | - | - | - | - | - | - | Unassigned | - |
| Otu60128 | 1 | k__Fungi(100);p__Ascomycota(100);p__Ascomycota_unclassified(100);p__Ascomycota_unclassified(100);p__Ascomycota_unclassified(100);p__Ascomycota_unclassified(100);p__Ascomycota_unclassified(100); | - | - | - | - | - | - | - | Unassigned | - |
| Otu60129 | 1 | k__Fungi(100);k__Fungi_unclassified(100);k__Fungi_unclassified(100);k__Fungi_unclassified(100);k__Fungi_unclassified(100);k__Fungi_unclassified(100);k__Fungi_unclassified(100); | - | - | - | - | - | - | - | Unassigned | - |
| Otu60130 | 1 | k__Fungi(100);p__Ascomycota(100);p__Ascomycota_unclassified(100);p__Ascomycota_unclassified(100);p__Ascomycota_unclassified(100);p__Ascomycota_unclassified(100);p__Ascomycota_unclassified(100); | - | - | - | - | - | - | - | Unassigned | - |
| Otu60131 | 1 | k__Fungi(100);k__Fungi_unclassified(100);k__Fungi_unclassified(100);k__Fungi_unclassified(100);k__Fungi_unclassified(100);k__Fungi_unclassified(100);k__Fungi_unclassified(100); | - | - | - | - | - | - | - | Unassigned | - |
| Otu60132 | 1 | k__Fungi(100);p__Ascomycota(100);p__Ascomycota_unclassified(100);p__Ascomycota_unclassified(100);p__Ascomycota_unclassified(100);p__Ascomycota_unclassified(100);p__Ascomycota_unclassified(100); | - | - | - | - | - | - | - | Unassigned | - |
| Otu60136 | 1 | k__Fungi(100);p__Ascomycota(100);c__Eurotiomycetes(100);c__Eurotiomycetes_unclassified(100);c__Eurotiomycetes_unclassified(100);c__Eurotiomycetes_unclassified(100);c__Eurotiomycetes_unclassified(100); | - | - | - | - | - | - | - | Unassigned | - |
| Otu60137 | 1 | k__Fungi(100);p__Ascomycota(100);p__Ascomycota_unclassified(100);p__Ascomycota_unclassified(100);p__Ascomycota_unclassified(100);p__Ascomycota_unclassified(100);p__Ascomycota_unclassified(100); | - | - | - | - | - | - | - | Unassigned | - |
| Otu60140 | 1 | k__Fungi(100);k__Fungi_unclassified(100);k__Fungi_unclassified(100);k__Fungi_unclassified(100);k__Fungi_unclassified(100);k__Fungi_unclassified(100);k__Fungi_unclassified(100); | - | - | - | - | - | - | - | Unassigned | - |
| Otu60142 | 1 | k__Fungi(100);p__Ascomycota(100);p__Ascomycota_unclassified(100);p__Ascomycota_unclassified(100);p__Ascomycota_unclassified(100);p__Ascomycota_unclassified(100);p__Ascomycota_unclassified(100); | - | - | - | - | - | - | - | Unassigned | - |
| Otu60145 | 1 | k__Fungi(100);p__Ascomycota(100);p__Ascomycota_unclassified(100);p__Ascomycota_unclassified(100);p__Ascomycota_unclassified(100);p__Ascomycota_unclassified(100);p__Ascomycota_unclassified(100); | - | - | - | - | - | - | - | Unassigned | - |
| Otu60151 | 1 | k__Fungi(100);k__Fungi_unclassified(100);k__Fungi_unclassified(100);k__Fungi_unclassified(100);k__Fungi_unclassified(100);k__Fungi_unclassified(100);k__Fungi_unclassified(100); | - | - | - | - | - | - | - | Unassigned | - |
| Otu60154 | 1 | k__Fungi(100);k__Fungi_unclassified(100);k__Fungi_unclassified(100);k__Fungi_unclassified(100);k__Fungi_unclassified(100);k__Fungi_unclassified(100);k__Fungi_unclassified(100); | - | - | - | - | - | - | - | Unassigned | - |
| Otu60155 | 1 | k__Fungi(100);k__Fungi_unclassified(100);k__Fungi_unclassified(100);k__Fungi_unclassified(100);k__Fungi_unclassified(100);k__Fungi_unclassified(100);k__Fungi_unclassified(100); | - | - | - | - | - | - | - | Unassigned | - |
| Otu60157 | 1 | k__Fungi(100);k__Fungi_unclassified(100);k__Fungi_unclassified(100);k__Fungi_unclassified(100);k__Fungi_unclassified(100);k__Fungi_unclassified(100);k__Fungi_unclassified(100); | - | - | - | - | - | - | - | Unassigned | - |
| Otu60158 | 1 | k__Fungi(100);k__Fungi_unclassified(100);k__Fungi_unclassified(100);k__Fungi_unclassified(100);k__Fungi_unclassified(100);k__Fungi_unclassified(100);k__Fungi_unclassified(100); | - | - | - | - | - | - | - | Unassigned | - |
| Otu60159 | 1 | k__Fungi(100);p__Ascomycota(100);p__Ascomycota_unclassified(100);p__Ascomycota_unclassified(100);p__Ascomycota_unclassified(100);p__Ascomycota_unclassified(100);p__Ascomycota_unclassified(100); | - | - | - | - | - | - | - | Unassigned | - |
| Otu60160 | 1 | k__Fungi(100);k__Fungi_unclassified(100);k__Fungi_unclassified(100);k__Fungi_unclassified(100);k__Fungi_unclassified(100);k__Fungi_unclassified(100);k__Fungi_unclassified(100); | - | - | - | - | - | - | - | Unassigned | - |
| Otu60161 | 1 | k__Fungi(100);p__Ascomycota(100);p__Ascomycota_unclassified(100);p__Ascomycota_unclassified(100);p__Ascomycota_unclassified(100);p__Ascomycota_unclassified(100);p__Ascomycota_unclassified(100); | - | - | - | - | - | - | - | Unassigned | - |
| Otu60162 | 1 | k__Fungi(100);p__Ascomycota(100);p__Ascomycota_unclassified(100);p__Ascomycota_unclassified(100);p__Ascomycota_unclassified(100);p__Ascomycota_unclassified(100);p__Ascomycota_unclassified(100); | - | - | - | - | - | - | - | Unassigned | - |
| Otu60163 | 1 | k__Fungi(100);p__Ascomycota(100);p__Ascomycota_unclassified(100);p__Ascomycota_unclassified(100);p__Ascomycota_unclassified(100);p__Ascomycota_unclassified(100);p__Ascomycota_unclassified(100); | - | - | - | - | - | - | - | Unassigned | - |
| Otu60165 | 1 | k__Fungi(100);k__Fungi_unclassified(100);k__Fungi_unclassified(100);k__Fungi_unclassified(100);k__Fungi_unclassified(100);k__Fungi_unclassified(100);k__Fungi_unclassified(100); | - | - | - | - | - | - | - | Unassigned | - |
| Otu60167 | 1 | k__Fungi(100);p__Ascomycota(100);c__Leotiomycetes(100);o__Helotiales(100);o__Helotiales_unclassified(100);o__Helotiales_unclassified(100);o__Helotiales_unclassified(100); | - | - | - | - | - | - | - | Unassigned | - |
| Otu60169 | 1 | k__Fungi(100);p__Ascomycota(100);p__Ascomycota_unclassified(100);p__Ascomycota_unclassified(100);p__Ascomycota_unclassified(100);p__Ascomycota_unclassified(100);p__Ascomycota_unclassified(100); | - | - | - | - | - | - | - | Unassigned | - |
| Otu60170 | 1 | k__Fungi(100);k__Fungi_unclassified(100);k__Fungi_unclassified(100);k__Fungi_unclassified(100);k__Fungi_unclassified(100);k__Fungi_unclassified(100);k__Fungi_unclassified(100); | - | - | - | - | - | - | - | Unassigned | - |
| Otu60171 | 1 | k__Fungi(100);k__Fungi_unclassified(100);k__Fungi_unclassified(100);k__Fungi_unclassified(100);k__Fungi_unclassified(100);k__Fungi_unclassified(100);k__Fungi_unclassified(100); | - | - | - | - | - | - | - | Unassigned | - |
| Otu60172 | 1 | k__Fungi(100);k__Fungi_unclassified(100);k__Fungi_unclassified(100);k__Fungi_unclassified(100);k__Fungi_unclassified(100);k__Fungi_unclassified(100);k__Fungi_unclassified(100); | - | - | - | - | - | - | - | Unassigned | - |
| Otu60173 | 1 | k__Fungi(100);p__Ascomycota(100);p__Ascomycota_unclassified(100);p__Ascomycota_unclassified(100);p__Ascomycota_unclassified(100);p__Ascomycota_unclassified(100);p__Ascomycota_unclassified(100); | - | - | - | - | - | - | - | Unassigned | - |
| Otu60174 | 1 | k__Fungi(100);k__Fungi_unclassified(100);k__Fungi_unclassified(100);k__Fungi_unclassified(100);k__Fungi_unclassified(100);k__Fungi_unclassified(100);k__Fungi_unclassified(100); | - | - | - | - | - | - | - | Unassigned | - |
| Otu60175 | 1 | k__Fungi(100);k__Fungi_unclassified(100);k__Fungi_unclassified(100);k__Fungi_unclassified(100);k__Fungi_unclassified(100);k__Fungi_unclassified(100);k__Fungi_unclassified(100); | - | - | - | - | - | - | - | Unassigned | - |
| Otu60177 | 1 | k__Fungi(100);k__Fungi_unclassified(100);k__Fungi_unclassified(100);k__Fungi_unclassified(100);k__Fungi_unclassified(100);k__Fungi_unclassified(100);k__Fungi_unclassified(100); | - | - | - | - | - | - | - | Unassigned | - |
| Otu60178 | 1 | k__Fungi(100);k__Fungi_unclassified(100);k__Fungi_unclassified(100);k__Fungi_unclassified(100);k__Fungi_unclassified(100);k__Fungi_unclassified(100);k__Fungi_unclassified(100); | - | - | - | - | - | - | - | Unassigned | - |
| Otu60179 | 1 | k__Fungi(100);k__Fungi_unclassified(100);k__Fungi_unclassified(100);k__Fungi_unclassified(100);k__Fungi_unclassified(100);k__Fungi_unclassified(100);k__Fungi_unclassified(100); | - | - | - | - | - | - | - | Unassigned | - |
| Otu60180 | 1 | k__Fungi(100);k__Fungi_unclassified(100);k__Fungi_unclassified(100);k__Fungi_unclassified(100);k__Fungi_unclassified(100);k__Fungi_unclassified(100);k__Fungi_unclassified(100); | - | - | - | - | - | - | - | Unassigned | - |
| Otu60181 | 1 | k__Fungi(100);k__Fungi_unclassified(100);k__Fungi_unclassified(100);k__Fungi_unclassified(100);k__Fungi_unclassified(100);k__Fungi_unclassified(100);k__Fungi_unclassified(100); | - | - | - | - | - | - | - | Unassigned | - |
| Otu60182 | 1 | k__Fungi(100);p__Ascomycota(100);p__Ascomycota_unclassified(100);p__Ascomycota_unclassified(100);p__Ascomycota_unclassified(100);p__Ascomycota_unclassified(100);p__Ascomycota_unclassified(100); | - | - | - | - | - | - | - | Unassigned | - |
| Otu60183 | 1 | k__Fungi(100);p__Ascomycota(100);p__Ascomycota_unclassified(100);p__Ascomycota_unclassified(100);p__Ascomycota_unclassified(100);p__Ascomycota_unclassified(100);p__Ascomycota_unclassified(100); | - | - | - | - | - | - | - | Unassigned | - |
| Otu60187 | 1 | k__Fungi(100);k__Fungi_unclassified(100);k__Fungi_unclassified(100);k__Fungi_unclassified(100);k__Fungi_unclassified(100);k__Fungi_unclassified(100);k__Fungi_unclassified(100); | - | - | - | - | - | - | - | Unassigned | - |
| Otu60188 | 1 | k__Fungi(100);k__Fungi_unclassified(100);k__Fungi_unclassified(100);k__Fungi_unclassified(100);k__Fungi_unclassified(100);k__Fungi_unclassified(100);k__Fungi_unclassified(100); | - | - | - | - | - | - | - | Unassigned | - |
| Otu60190 | 1 | k__Fungi(100);p__Ascomycota(100);p__Ascomycota_unclassified(100);p__Ascomycota_unclassified(100);p__Ascomycota_unclassified(100);p__Ascomycota_unclassified(100);p__Ascomycota_unclassified(100); | - | - | - | - | - | - | - | Unassigned | - |
| Otu60192 | 1 | k__Fungi(100);k__Fungi_unclassified(100);k__Fungi_unclassified(100);k__Fungi_unclassified(100);k__Fungi_unclassified(100);k__Fungi_unclassified(100);k__Fungi_unclassified(100); | - | - | - | - | - | - | - | Unassigned | - |
| Otu60197 | 1 | k__Fungi(100);k__Fungi_unclassified(100);k__Fungi_unclassified(100);k__Fungi_unclassified(100);k__Fungi_unclassified(100);k__Fungi_unclassified(100);k__Fungi_unclassified(100); | - | - | - | - | - | - | - | Unassigned | - |
| Otu60199 | 1 | k__Fungi(100);p__Ascomycota(100);p__Ascomycota_unclassified(100);p__Ascomycota_unclassified(100);p__Ascomycota_unclassified(100);p__Ascomycota_unclassified(100);p__Ascomycota_unclassified(100); | - | - | - | - | - | - | - | Unassigned | - |
| Otu60202 | 1 | k__Fungi(100);k__Fungi_unclassified(100);k__Fungi_unclassified(100);k__Fungi_unclassified(100);k__Fungi_unclassified(100);k__Fungi_unclassified(100);k__Fungi_unclassified(100); | - | - | - | - | - | - | - | Unassigned | - |
| Otu60203 | 1 | k__Fungi(100);k__Fungi_unclassified(100);k__Fungi_unclassified(100);k__Fungi_unclassified(100);k__Fungi_unclassified(100);k__Fungi_unclassified(100);k__Fungi_unclassified(100); | - | - | - | - | - | - | - | Unassigned | - |
| Otu60204 | 1 | k__Fungi(100);k__Fungi_unclassified(100);k__Fungi_unclassified(100);k__Fungi_unclassified(100);k__Fungi_unclassified(100);k__Fungi_unclassified(100);k__Fungi_unclassified(100); | - | - | - | - | - | - | - | Unassigned | - |
| Otu60205 | 1 | k__Fungi(100);k__Fungi_unclassified(100);k__Fungi_unclassified(100);k__Fungi_unclassified(100);k__Fungi_unclassified(100);k__Fungi_unclassified(100);k__Fungi_unclassified(100); | - | - | - | - | - | - | - | Unassigned | - |
| Otu60206 | 1 | k__Fungi(100);k__Fungi_unclassified(100);k__Fungi_unclassified(100);k__Fungi_unclassified(100);k__Fungi_unclassified(100);k__Fungi_unclassified(100);k__Fungi_unclassified(100); | - | - | - | - | - | - | - | Unassigned | - |
| Otu60207 | 1 | k__Fungi(100);k__Fungi_unclassified(100);k__Fungi_unclassified(100);k__Fungi_unclassified(100);k__Fungi_unclassified(100);k__Fungi_unclassified(100);k__Fungi_unclassified(100); | - | - | - | - | - | - | - | Unassigned | - |
| Otu60208 | 1 | k__Fungi(100);p__Ascomycota(100);c__Leotiomycetes(100);o__Helotiales(100);f__Helotiales_family_Incertae_sedis(100);f__Helotiales_family_Incertae_sedis_unclassified(100);f__Helotiales_family_Incertae_sedis_unclassified(100); | - | - | - | - | - | - | - | Unassigned | - |
| Otu60209 | 1 | k__Fungi(100);p__Ascomycota(100);c__Leotiomycetes(100);o__Helotiales(100);f__Helotiales_family_Incertae_sedis(100);f__Helotiales_family_Incertae_sedis_unclassified(100);f__Helotiales_family_Incertae_sedis_unclassified(100); | - | - | - | - | - | - | - | Unassigned | - |
| Otu60211 | 1 | k__Fungi(100);p__Ascomycota(100);c__Dothideomycetes(100);o__Capnodiales(100);f__Capnodiales_family_Incertae_sedis(100);f__Capnodiales_family_Incertae_sedis_unclassified(100);f__Capnodiales_family_Incertae_sedis_unclassified(100); | - | - | - | - | - | - | - | Unassigned | - |
| Otu60213 | 1 | k__Fungi(100);p__Ascomycota(100);c__Leotiomycetes(100);o__Helotiales(100);o__Helotiales_unclassified(100);o__Helotiales_unclassified(100);o__Helotiales_unclassified(100); | - | - | - | - | - | - | - | Unassigned | - |
| Otu60215 | 1 | k__Fungi(100);k__Fungi_unclassified(100);k__Fungi_unclassified(100);k__Fungi_unclassified(100);k__Fungi_unclassified(100);k__Fungi_unclassified(100);k__Fungi_unclassified(100); | - | - | - | - | - | - | - | Unassigned | - |
| Otu60216 | 1 | k__Fungi(100);p__Ascomycota(100);p__Ascomycota_unclassified(100);p__Ascomycota_unclassified(100);p__Ascomycota_unclassified(100);p__Ascomycota_unclassified(100);p__Ascomycota_unclassified(100); | - | - | - | - | - | - | - | Unassigned | - |
| Otu60217 | 1 | k__Fungi(100);p__Ascomycota(100);p__Ascomycota_unclassified(100);p__Ascomycota_unclassified(100);p__Ascomycota_unclassified(100);p__Ascomycota_unclassified(100);p__Ascomycota_unclassified(100); | - | - | - | - | - | - | - | Unassigned | - |
| Otu60218 | 1 | k__Fungi(100);k__Fungi_unclassified(100);k__Fungi_unclassified(100);k__Fungi_unclassified(100);k__Fungi_unclassified(100);k__Fungi_unclassified(100);k__Fungi_unclassified(100); | - | - | - | - | - | - | - | Unassigned | - |
| Otu60219 | 1 | k__Fungi(100);p__Ascomycota(100);p__Ascomycota_unclassified(100);p__Ascomycota_unclassified(100);p__Ascomycota_unclassified(100);p__Ascomycota_unclassified(100);p__Ascomycota_unclassified(100); | - | - | - | - | - | - | - | Unassigned | - |
| Otu60220 | 1 | k__Fungi(100);p__Ascomycota(100);p__Ascomycota_unclassified(100);p__Ascomycota_unclassified(100);p__Ascomycota_unclassified(100);p__Ascomycota_unclassified(100);p__Ascomycota_unclassified(100); | - | - | - | - | - | - | - | Unassigned | - |
| Otu60222 | 1 | k__Fungi(100);k__Fungi_unclassified(100);k__Fungi_unclassified(100);k__Fungi_unclassified(100);k__Fungi_unclassified(100);k__Fungi_unclassified(100);k__Fungi_unclassified(100); | - | - | - | - | - | - | - | Unassigned | - |
| Otu60223 | 1 | k__Fungi(100);k__Fungi_unclassified(100);k__Fungi_unclassified(100);k__Fungi_unclassified(100);k__Fungi_unclassified(100);k__Fungi_unclassified(100);k__Fungi_unclassified(100); | - | - | - | - | - | - | - | Unassigned | - |
| Otu60224 | 1 | k__Fungi(100);k__Fungi_unclassified(100);k__Fungi_unclassified(100);k__Fungi_unclassified(100);k__Fungi_unclassified(100);k__Fungi_unclassified(100);k__Fungi_unclassified(100); | - | - | - | - | - | - | - | Unassigned | - |
| Otu60225 | 1 | k__Fungi(100);k__Fungi_unclassified(100);k__Fungi_unclassified(100);k__Fungi_unclassified(100);k__Fungi_unclassified(100);k__Fungi_unclassified(100);k__Fungi_unclassified(100); | - | - | - | - | - | - | - | Unassigned | - |
| Otu60226 | 1 | k__Fungi(100);p__Ascomycota(100);p__Ascomycota_unclassified(100);p__Ascomycota_unclassified(100);p__Ascomycota_unclassified(100);p__Ascomycota_unclassified(100);p__Ascomycota_unclassified(100); | - | - | - | - | - | - | - | Unassigned | - |
| Otu60228 | 1 | k__Fungi(100);k__Fungi_unclassified(100);k__Fungi_unclassified(100);k__Fungi_unclassified(100);k__Fungi_unclassified(100);k__Fungi_unclassified(100);k__Fungi_unclassified(100); | - | - | - | - | - | - | - | Unassigned | - |
| Otu60229 | 1 | k__Fungi(100);p__Ascomycota(100);p__Ascomycota_unclassified(100);p__Ascomycota_unclassified(100);p__Ascomycota_unclassified(100);p__Ascomycota_unclassified(100);p__Ascomycota_unclassified(100); | - | - | - | - | - | - | - | Unassigned | - |
| Otu60230 | 1 | k__Fungi(100);p__Ascomycota(100);p__Ascomycota_unclassified(100);p__Ascomycota_unclassified(100);p__Ascomycota_unclassified(100);p__Ascomycota_unclassified(100);p__Ascomycota_unclassified(100); | - | - | - | - | - | - | - | Unassigned | - |
| Otu60233 | 1 | k__Fungi(100);p__Ascomycota(100);p__Ascomycota_unclassified(100);p__Ascomycota_unclassified(100);p__Ascomycota_unclassified(100);p__Ascomycota_unclassified(100);p__Ascomycota_unclassified(100); | - | - | - | - | - | - | - | Unassigned | - |
| Otu60234 | 1 | k__Fungi(100);k__Fungi_unclassified(100);k__Fungi_unclassified(100);k__Fungi_unclassified(100);k__Fungi_unclassified(100);k__Fungi_unclassified(100);k__Fungi_unclassified(100); | - | - | - | - | - | - | - | Unassigned | - |
| Otu60235 | 1 | k__Fungi(100);p__Ascomycota(100);p__Ascomycota_unclassified(100);p__Ascomycota_unclassified(100);p__Ascomycota_unclassified(100);p__Ascomycota_unclassified(100);p__Ascomycota_unclassified(100); | - | - | - | - | - | - | - | Unassigned | - |
| Otu60236 | 1 | k__Fungi(100);k__Fungi_unclassified(100);k__Fungi_unclassified(100);k__Fungi_unclassified(100);k__Fungi_unclassified(100);k__Fungi_unclassified(100);k__Fungi_unclassified(100); | - | - | - | - | - | - | - | Unassigned | - |
| Otu60237 | 1 | k__Fungi(100);k__Fungi_unclassified(100);k__Fungi_unclassified(100);k__Fungi_unclassified(100);k__Fungi_unclassified(100);k__Fungi_unclassified(100);k__Fungi_unclassified(100); | - | - | - | - | - | - | - | Unassigned | - |
| Otu60238 | 1 | k__Fungi(100);p__Ascomycota(100);p__Ascomycota_unclassified(100);p__Ascomycota_unclassified(100);p__Ascomycota_unclassified(100);p__Ascomycota_unclassified(100);p__Ascomycota_unclassified(100); | - | - | - | - | - | - | - | Unassigned | - |
| Otu60239 | 1 | k__Fungi(100);p__Ascomycota(100);p__Ascomycota_unclassified(100);p__Ascomycota_unclassified(100);p__Ascomycota_unclassified(100);p__Ascomycota_unclassified(100);p__Ascomycota_unclassified(100); | - | - | - | - | - | - | - | Unassigned | - |
| Otu60240 | 1 | k__Fungi(100);p__Ascomycota(100);c__Dothideomycetes(100);o__Capnodiales(100);f__Davidiellaceae(100);f__Davidiellaceae_unclassified(100);f__Davidiellaceae_unclassified(100); | - | - | - | - | - | - | - | Unassigned | - |
| Otu60242 | 1 | k__Fungi(100);k__Fungi_unclassified(100);k__Fungi_unclassified(100);k__Fungi_unclassified(100);k__Fungi_unclassified(100);k__Fungi_unclassified(100);k__Fungi_unclassified(100); | - | - | - | - | - | - | - | Unassigned | - |
| Otu60244 | 1 | k__Fungi(100);p__Ascomycota(100);p__Ascomycota_unclassified(100);p__Ascomycota_unclassified(100);p__Ascomycota_unclassified(100);p__Ascomycota_unclassified(100);p__Ascomycota_unclassified(100); | - | - | - | - | - | - | - | Unassigned | - |
| Otu60245 | 1 | k__Fungi(100);p__Ascomycota(100);p__Ascomycota_unclassified(100);p__Ascomycota_unclassified(100);p__Ascomycota_unclassified(100);p__Ascomycota_unclassified(100);p__Ascomycota_unclassified(100); | - | - | - | - | - | - | - | Unassigned | - |
| Otu60246 | 1 | k__Fungi(100);k__Fungi_unclassified(100);k__Fungi_unclassified(100);k__Fungi_unclassified(100);k__Fungi_unclassified(100);k__Fungi_unclassified(100);k__Fungi_unclassified(100); | - | - | - | - | - | - | - | Unassigned | - |
| Otu60250 | 1 | k__Fungi(100);k__Fungi_unclassified(100);k__Fungi_unclassified(100);k__Fungi_unclassified(100);k__Fungi_unclassified(100);k__Fungi_unclassified(100);k__Fungi_unclassified(100); | - | - | - | - | - | - | - | Unassigned | - |
| Otu60251 | 1 | k__Fungi(100);k__Fungi_unclassified(100);k__Fungi_unclassified(100);k__Fungi_unclassified(100);k__Fungi_unclassified(100);k__Fungi_unclassified(100);k__Fungi_unclassified(100); | - | - | - | - | - | - | - | Unassigned | - |
| Otu60252 | 1 | k__Fungi(100);k__Fungi_unclassified(100);k__Fungi_unclassified(100);k__Fungi_unclassified(100);k__Fungi_unclassified(100);k__Fungi_unclassified(100);k__Fungi_unclassified(100); | - | - | - | - | - | - | - | Unassigned | - |
| Otu60254 | 1 | k__Fungi(100);k__Fungi_unclassified(100);k__Fungi_unclassified(100);k__Fungi_unclassified(100);k__Fungi_unclassified(100);k__Fungi_unclassified(100);k__Fungi_unclassified(100); | - | - | - | - | - | - | - | Unassigned | - |
| Otu60255 | 1 | k__Fungi(100);k__Fungi_unclassified(100);k__Fungi_unclassified(100);k__Fungi_unclassified(100);k__Fungi_unclassified(100);k__Fungi_unclassified(100);k__Fungi_unclassified(100); | - | - | - | - | - | - | - | Unassigned | - |
| Otu60263 | 1 | k__Fungi(100);p__Ascomycota(100);p__Ascomycota_unclassified(100);p__Ascomycota_unclassified(100);p__Ascomycota_unclassified(100);p__Ascomycota_unclassified(100);p__Ascomycota_unclassified(100); | - | - | - | - | - | - | - | Unassigned | - |
| Otu60264 | 1 | k__Fungi(100);k__Fungi_unclassified(100);k__Fungi_unclassified(100);k__Fungi_unclassified(100);k__Fungi_unclassified(100);k__Fungi_unclassified(100);k__Fungi_unclassified(100); | - | - | - | - | - | - | - | Unassigned | - |
| Otu60266 | 1 | k__Fungi(100);k__Fungi_unclassified(100);k__Fungi_unclassified(100);k__Fungi_unclassified(100);k__Fungi_unclassified(100);k__Fungi_unclassified(100);k__Fungi_unclassified(100); | - | - | - | - | - | - | - | Unassigned | - |
| Otu60267 | 1 | k__Fungi(100);k__Fungi_unclassified(100);k__Fungi_unclassified(100);k__Fungi_unclassified(100);k__Fungi_unclassified(100);k__Fungi_unclassified(100);k__Fungi_unclassified(100); | - | - | - | - | - | - | - | Unassigned | - |
| Otu60268 | 1 | k__Fungi(100);k__Fungi_unclassified(100);k__Fungi_unclassified(100);k__Fungi_unclassified(100);k__Fungi_unclassified(100);k__Fungi_unclassified(100);k__Fungi_unclassified(100); | - | - | - | - | - | - | - | Unassigned | - |
| Otu60269 | 1 | k__Fungi(100);k__Fungi_unclassified(100);k__Fungi_unclassified(100);k__Fungi_unclassified(100);k__Fungi_unclassified(100);k__Fungi_unclassified(100);k__Fungi_unclassified(100); | - | - | - | - | - | - | - | Unassigned | - |
| Otu60270 | 1 | k__Fungi(100);k__Fungi_unclassified(100);k__Fungi_unclassified(100);k__Fungi_unclassified(100);k__Fungi_unclassified(100);k__Fungi_unclassified(100);k__Fungi_unclassified(100); | - | - | - | - | - | - | - | Unassigned | - |
| Otu60271 | 1 | k__Fungi(100);p__Ascomycota(100);p__Ascomycota_unclassified(100);p__Ascomycota_unclassified(100);p__Ascomycota_unclassified(100);p__Ascomycota_unclassified(100);p__Ascomycota_unclassified(100); | - | - | - | - | - | - | - | Unassigned | - |
| Otu60272 | 1 | k__Fungi(100);k__Fungi_unclassified(100);k__Fungi_unclassified(100);k__Fungi_unclassified(100);k__Fungi_unclassified(100);k__Fungi_unclassified(100);k__Fungi_unclassified(100); | - | - | - | - | - | - | - | Unassigned | - |
| Otu60273 | 1 | k__Fungi(100);k__Fungi_unclassified(100);k__Fungi_unclassified(100);k__Fungi_unclassified(100);k__Fungi_unclassified(100);k__Fungi_unclassified(100);k__Fungi_unclassified(100); | - | - | - | - | - | - | - | Unassigned | - |
| Otu60274 | 1 | k__Fungi(100);k__Fungi_unclassified(100);k__Fungi_unclassified(100);k__Fungi_unclassified(100);k__Fungi_unclassified(100);k__Fungi_unclassified(100);k__Fungi_unclassified(100); | - | - | - | - | - | - | - | Unassigned | - |
| Otu60275 | 1 | k__Fungi(100);k__Fungi_unclassified(100);k__Fungi_unclassified(100);k__Fungi_unclassified(100);k__Fungi_unclassified(100);k__Fungi_unclassified(100);k__Fungi_unclassified(100); | - | - | - | - | - | - | - | Unassigned | - |
| Otu60276 | 1 | k__Fungi(100);k__Fungi_unclassified(100);k__Fungi_unclassified(100);k__Fungi_unclassified(100);k__Fungi_unclassified(100);k__Fungi_unclassified(100);k__Fungi_unclassified(100); | - | - | - | - | - | - | - | Unassigned | - |
| Otu60277 | 1 | k__Fungi(100);k__Fungi_unclassified(100);k__Fungi_unclassified(100);k__Fungi_unclassified(100);k__Fungi_unclassified(100);k__Fungi_unclassified(100);k__Fungi_unclassified(100); | - | - | - | - | - | - | - | Unassigned | - |
| Otu60278 | 1 | k__Fungi(100);k__Fungi_unclassified(100);k__Fungi_unclassified(100);k__Fungi_unclassified(100);k__Fungi_unclassified(100);k__Fungi_unclassified(100);k__Fungi_unclassified(100); | - | - | - | - | - | - | - | Unassigned | - |
| Otu60279 | 1 | k__Fungi(100);k__Fungi_unclassified(100);k__Fungi_unclassified(100);k__Fungi_unclassified(100);k__Fungi_unclassified(100);k__Fungi_unclassified(100);k__Fungi_unclassified(100); | - | - | - | - | - | - | - | Unassigned | - |
| Otu60280 | 1 | k__Fungi(100);k__Fungi_unclassified(100);k__Fungi_unclassified(100);k__Fungi_unclassified(100);k__Fungi_unclassified(100);k__Fungi_unclassified(100);k__Fungi_unclassified(100); | - | - | - | - | - | - | - | Unassigned | - |
| Otu60281 | 1 | k__Fungi(100);k__Fungi_unclassified(100);k__Fungi_unclassified(100);k__Fungi_unclassified(100);k__Fungi_unclassified(100);k__Fungi_unclassified(100);k__Fungi_unclassified(100); | - | - | - | - | - | - | - | Unassigned | - |
| Otu60282 | 1 | k__Fungi(100);p__Ascomycota(100);p__Ascomycota_unclassified(100);p__Ascomycota_unclassified(100);p__Ascomycota_unclassified(100);p__Ascomycota_unclassified(100);p__Ascomycota_unclassified(100); | - | - | - | - | - | - | - | Unassigned | - |
| Otu60283 | 1 | k__Fungi(100);k__Fungi_unclassified(100);k__Fungi_unclassified(100);k__Fungi_unclassified(100);k__Fungi_unclassified(100);k__Fungi_unclassified(100);k__Fungi_unclassified(100); | - | - | - | - | - | - | - | Unassigned | - |
| Otu60284 | 1 | k__Fungi(100);k__Fungi_unclassified(100);k__Fungi_unclassified(100);k__Fungi_unclassified(100);k__Fungi_unclassified(100);k__Fungi_unclassified(100);k__Fungi_unclassified(100); | - | - | - | - | - | - | - | Unassigned | - |
| Otu60285 | 1 | k__Fungi(100);p__Ascomycota(100);p__Ascomycota_unclassified(100);p__Ascomycota_unclassified(100);p__Ascomycota_unclassified(100);p__Ascomycota_unclassified(100);p__Ascomycota_unclassified(100); | - | - | - | - | - | - | - | Unassigned | - |
| Otu60286 | 1 | k__Fungi(100);k__Fungi_unclassified(100);k__Fungi_unclassified(100);k__Fungi_unclassified(100);k__Fungi_unclassified(100);k__Fungi_unclassified(100);k__Fungi_unclassified(100); | - | - | - | - | - | - | - | Unassigned | - |
| Otu60287 | 1 | k__Fungi(100);k__Fungi_unclassified(100);k__Fungi_unclassified(100);k__Fungi_unclassified(100);k__Fungi_unclassified(100);k__Fungi_unclassified(100);k__Fungi_unclassified(100); | - | - | - | - | - | - | - | Unassigned | - |
| Otu60288 | 1 | k__Fungi(100);p__Ascomycota(100);p__Ascomycota_unclassified(100);p__Ascomycota_unclassified(100);p__Ascomycota_unclassified(100);p__Ascomycota_unclassified(100);p__Ascomycota_unclassified(100); | - | - | - | - | - | - | - | Unassigned | - |
| Otu60289 | 1 | k__Fungi(100);k__Fungi_unclassified(100);k__Fungi_unclassified(100);k__Fungi_unclassified(100);k__Fungi_unclassified(100);k__Fungi_unclassified(100);k__Fungi_unclassified(100); | - | - | - | - | - | - | - | Unassigned | - |
| Otu60290 | 1 | k__Fungi(100);p__Ascomycota(100);p__Ascomycota_unclassified(100);p__Ascomycota_unclassified(100);p__Ascomycota_unclassified(100);p__Ascomycota_unclassified(100);p__Ascomycota_unclassified(100); | - | - | - | - | - | - | - | Unassigned | - |
| Otu60291 | 1 | k__Fungi(100);k__Fungi_unclassified(100);k__Fungi_unclassified(100);k__Fungi_unclassified(100);k__Fungi_unclassified(100);k__Fungi_unclassified(100);k__Fungi_unclassified(100); | - | - | - | - | - | - | - | Unassigned | - |
| Otu60292 | 1 | k__Fungi(100);p__Ascomycota(100);p__Ascomycota_unclassified(100);p__Ascomycota_unclassified(100);p__Ascomycota_unclassified(100);p__Ascomycota_unclassified(100);p__Ascomycota_unclassified(100); | - | - | - | - | - | - | - | Unassigned | - |
| Otu60293 | 1 | k__Fungi(100);k__Fungi_unclassified(100);k__Fungi_unclassified(100);k__Fungi_unclassified(100);k__Fungi_unclassified(100);k__Fungi_unclassified(100);k__Fungi_unclassified(100); | - | - | - | - | - | - | - | Unassigned | - |
| Otu60294 | 1 | k__Fungi(100);k__Fungi_unclassified(100);k__Fungi_unclassified(100);k__Fungi_unclassified(100);k__Fungi_unclassified(100);k__Fungi_unclassified(100);k__Fungi_unclassified(100); | - | - | - | - | - | - | - | Unassigned | - |
| Otu60295 | 1 | k__Fungi(100);k__Fungi_unclassified(100);k__Fungi_unclassified(100);k__Fungi_unclassified(100);k__Fungi_unclassified(100);k__Fungi_unclassified(100);k__Fungi_unclassified(100); | - | - | - | - | - | - | - | Unassigned | - |
| Otu60297 | 1 | k__Fungi(100);p__Ascomycota(100);p__Ascomycota_unclassified(100);p__Ascomycota_unclassified(100);p__Ascomycota_unclassified(100);p__Ascomycota_unclassified(100);p__Ascomycota_unclassified(100); | - | - | - | - | - | - | - | Unassigned | - |
| Otu60298 | 1 | k__Fungi(100);k__Fungi_unclassified(100);k__Fungi_unclassified(100);k__Fungi_unclassified(100);k__Fungi_unclassified(100);k__Fungi_unclassified(100);k__Fungi_unclassified(100); | - | - | - | - | - | - | - | Unassigned | - |
| Otu60299 | 1 | k__Fungi(100);k__Fungi_unclassified(100);k__Fungi_unclassified(100);k__Fungi_unclassified(100);k__Fungi_unclassified(100);k__Fungi_unclassified(100);k__Fungi_unclassified(100); | - | - | - | - | - | - | - | Unassigned | - |
| Otu60300 | 1 | k__Fungi(100);k__Fungi_unclassified(100);k__Fungi_unclassified(100);k__Fungi_unclassified(100);k__Fungi_unclassified(100);k__Fungi_unclassified(100);k__Fungi_unclassified(100); | - | - | - | - | - | - | - | Unassigned | - |
| Otu60301 | 1 | k__Fungi(100);k__Fungi_unclassified(100);k__Fungi_unclassified(100);k__Fungi_unclassified(100);k__Fungi_unclassified(100);k__Fungi_unclassified(100);k__Fungi_unclassified(100); | - | - | - | - | - | - | - | Unassigned | - |
| Otu60303 | 1 | k__Fungi(100);p__Ascomycota(100);p__Ascomycota_unclassified(100);p__Ascomycota_unclassified(100);p__Ascomycota_unclassified(100);p__Ascomycota_unclassified(100);p__Ascomycota_unclassified(100); | - | - | - | - | - | - | - | Unassigned | - |
| Otu60304 | 1 | k__Fungi(100);k__Fungi_unclassified(100);k__Fungi_unclassified(100);k__Fungi_unclassified(100);k__Fungi_unclassified(100);k__Fungi_unclassified(100);k__Fungi_unclassified(100); | - | - | - | - | - | - | - | Unassigned | - |
| Otu60305 | 1 | k__Fungi(100);k__Fungi_unclassified(100);k__Fungi_unclassified(100);k__Fungi_unclassified(100);k__Fungi_unclassified(100);k__Fungi_unclassified(100);k__Fungi_unclassified(100); | - | - | - | - | - | - | - | Unassigned | - |
| Otu60306 | 1 | k__Fungi(100);p__Ascomycota(100);p__Ascomycota_unclassified(100);p__Ascomycota_unclassified(100);p__Ascomycota_unclassified(100);p__Ascomycota_unclassified(100);p__Ascomycota_unclassified(100); | - | - | - | - | - | - | - | Unassigned | - |
| Otu60307 | 1 | k__Fungi(100);k__Fungi_unclassified(100);k__Fungi_unclassified(100);k__Fungi_unclassified(100);k__Fungi_unclassified(100);k__Fungi_unclassified(100);k__Fungi_unclassified(100); | - | - | - | - | - | - | - | Unassigned | - |
| Otu60308 | 1 | k__Fungi(100);k__Fungi_unclassified(100);k__Fungi_unclassified(100);k__Fungi_unclassified(100);k__Fungi_unclassified(100);k__Fungi_unclassified(100);k__Fungi_unclassified(100); | - | - | - | - | - | - | - | Unassigned | - |
| Otu60309 | 1 | k__Fungi(100);k__Fungi_unclassified(100);k__Fungi_unclassified(100);k__Fungi_unclassified(100);k__Fungi_unclassified(100);k__Fungi_unclassified(100);k__Fungi_unclassified(100); | - | - | - | - | - | - | - | Unassigned | - |
| Otu60310 | 1 | k__Fungi(100);k__Fungi_unclassified(100);k__Fungi_unclassified(100);k__Fungi_unclassified(100);k__Fungi_unclassified(100);k__Fungi_unclassified(100);k__Fungi_unclassified(100); | - | - | - | - | - | - | - | Unassigned | - |
| Otu60311 | 1 | k__Fungi(100);k__Fungi_unclassified(100);k__Fungi_unclassified(100);k__Fungi_unclassified(100);k__Fungi_unclassified(100);k__Fungi_unclassified(100);k__Fungi_unclassified(100); | - | - | - | - | - | - | - | Unassigned | - |
| Otu60312 | 1 | k__Fungi(100);k__Fungi_unclassified(100);k__Fungi_unclassified(100);k__Fungi_unclassified(100);k__Fungi_unclassified(100);k__Fungi_unclassified(100);k__Fungi_unclassified(100); | - | - | - | - | - | - | - | Unassigned | - |
| Otu60313 | 1 | k__Fungi(100);k__Fungi_unclassified(100);k__Fungi_unclassified(100);k__Fungi_unclassified(100);k__Fungi_unclassified(100);k__Fungi_unclassified(100);k__Fungi_unclassified(100); | - | - | - | - | - | - | - | Unassigned | - |
| Otu60314 | 1 | k__Fungi(100);p__Ascomycota(100);p__Ascomycota_unclassified(100);p__Ascomycota_unclassified(100);p__Ascomycota_unclassified(100);p__Ascomycota_unclassified(100);p__Ascomycota_unclassified(100); | - | - | - | - | - | - | - | Unassigned | - |
| Otu60315 | 1 | k__Fungi(100);k__Fungi_unclassified(100);k__Fungi_unclassified(100);k__Fungi_unclassified(100);k__Fungi_unclassified(100);k__Fungi_unclassified(100);k__Fungi_unclassified(100); | - | - | - | - | - | - | - | Unassigned | - |
| Otu60316 | 1 | k__Fungi(100);p__Ascomycota(100);p__Ascomycota_unclassified(100);p__Ascomycota_unclassified(100);p__Ascomycota_unclassified(100);p__Ascomycota_unclassified(100);p__Ascomycota_unclassified(100); | - | - | - | - | - | - | - | Unassigned | - |
| Otu60317 | 1 | k__Fungi(100);k__Fungi_unclassified(100);k__Fungi_unclassified(100);k__Fungi_unclassified(100);k__Fungi_unclassified(100);k__Fungi_unclassified(100);k__Fungi_unclassified(100); | - | - | - | - | - | - | - | Unassigned | - |
| Otu60318 | 1 | k__Fungi(100);k__Fungi_unclassified(100);k__Fungi_unclassified(100);k__Fungi_unclassified(100);k__Fungi_unclassified(100);k__Fungi_unclassified(100);k__Fungi_unclassified(100); | - | - | - | - | - | - | - | Unassigned | - |
| Otu60320 | 1 | k__Fungi(100);k__Fungi_unclassified(100);k__Fungi_unclassified(100);k__Fungi_unclassified(100);k__Fungi_unclassified(100);k__Fungi_unclassified(100);k__Fungi_unclassified(100); | - | - | - | - | - | - | - | Unassigned | - |
| Otu60321 | 1 | k__Fungi(100);k__Fungi_unclassified(100);k__Fungi_unclassified(100);k__Fungi_unclassified(100);k__Fungi_unclassified(100);k__Fungi_unclassified(100);k__Fungi_unclassified(100); | - | - | - | - | - | - | - | Unassigned | - |
| Otu60322 | 1 | k__Fungi(100);p__Ascomycota(100);p__Ascomycota_unclassified(100);p__Ascomycota_unclassified(100);p__Ascomycota_unclassified(100);p__Ascomycota_unclassified(100);p__Ascomycota_unclassified(100); | - | - | - | - | - | - | - | Unassigned | - |
| Otu60323 | 1 | k__Fungi(100);k__Fungi_unclassified(100);k__Fungi_unclassified(100);k__Fungi_unclassified(100);k__Fungi_unclassified(100);k__Fungi_unclassified(100);k__Fungi_unclassified(100); | - | - | - | - | - | - | - | Unassigned | - |
| Otu60324 | 1 | k__Fungi(100);p__Ascomycota(100);p__Ascomycota_unclassified(100);p__Ascomycota_unclassified(100);p__Ascomycota_unclassified(100);p__Ascomycota_unclassified(100);p__Ascomycota_unclassified(100); | - | - | - | - | - | - | - | Unassigned | - |
| Otu60325 | 1 | k__Fungi(100);k__Fungi_unclassified(100);k__Fungi_unclassified(100);k__Fungi_unclassified(100);k__Fungi_unclassified(100);k__Fungi_unclassified(100);k__Fungi_unclassified(100); | - | - | - | - | - | - | - | Unassigned | - |
| Otu60326 | 1 | k__Fungi(100);p__Ascomycota(100);p__Ascomycota_unclassified(100);p__Ascomycota_unclassified(100);p__Ascomycota_unclassified(100);p__Ascomycota_unclassified(100);p__Ascomycota_unclassified(100); | - | - | - | - | - | - | - | Unassigned | - |
| Otu60327 | 1 | k__Fungi(100);k__Fungi_unclassified(100);k__Fungi_unclassified(100);k__Fungi_unclassified(100);k__Fungi_unclassified(100);k__Fungi_unclassified(100);k__Fungi_unclassified(100); | - | - | - | - | - | - | - | Unassigned | - |
| Otu60328 | 1 | k__Fungi(100);k__Fungi_unclassified(100);k__Fungi_unclassified(100);k__Fungi_unclassified(100);k__Fungi_unclassified(100);k__Fungi_unclassified(100);k__Fungi_unclassified(100); | - | - | - | - | - | - | - | Unassigned | - |
| Otu60330 | 1 | k__Fungi(100);p__Ascomycota(100);p__Ascomycota_unclassified(100);p__Ascomycota_unclassified(100);p__Ascomycota_unclassified(100);p__Ascomycota_unclassified(100);p__Ascomycota_unclassified(100); | - | - | - | - | - | - | - | Unassigned | - |
| Otu60331 | 1 | k__Fungi(100);k__Fungi_unclassified(100);k__Fungi_unclassified(100);k__Fungi_unclassified(100);k__Fungi_unclassified(100);k__Fungi_unclassified(100);k__Fungi_unclassified(100); | - | - | - | - | - | - | - | Unassigned | - |
| Otu60332 | 1 | k__Fungi(100);p__Ascomycota(100);p__Ascomycota_unclassified(100);p__Ascomycota_unclassified(100);p__Ascomycota_unclassified(100);p__Ascomycota_unclassified(100);p__Ascomycota_unclassified(100); | - | - | - | - | - | - | - | Unassigned | - |
| Otu60334 | 1 | k__Fungi(100);p__Ascomycota(100);p__Ascomycota_unclassified(100);p__Ascomycota_unclassified(100);p__Ascomycota_unclassified(100);p__Ascomycota_unclassified(100);p__Ascomycota_unclassified(100); | - | - | - | - | - | - | - | Unassigned | - |
| Otu60335 | 1 | k__Fungi(100);k__Fungi_unclassified(100);k__Fungi_unclassified(100);k__Fungi_unclassified(100);k__Fungi_unclassified(100);k__Fungi_unclassified(100);k__Fungi_unclassified(100); | - | - | - | - | - | - | - | Unassigned | - |
| Otu60336 | 1 | k__Fungi(100);k__Fungi_unclassified(100);k__Fungi_unclassified(100);k__Fungi_unclassified(100);k__Fungi_unclassified(100);k__Fungi_unclassified(100);k__Fungi_unclassified(100); | - | - | - | - | - | - | - | Unassigned | - |
| Otu60337 | 1 | k__Fungi(100);k__Fungi_unclassified(100);k__Fungi_unclassified(100);k__Fungi_unclassified(100);k__Fungi_unclassified(100);k__Fungi_unclassified(100);k__Fungi_unclassified(100); | - | - | - | - | - | - | - | Unassigned | - |
| Otu60340 | 1 | k__Fungi(100);k__Fungi_unclassified(100);k__Fungi_unclassified(100);k__Fungi_unclassified(100);k__Fungi_unclassified(100);k__Fungi_unclassified(100);k__Fungi_unclassified(100); | - | - | - | - | - | - | - | Unassigned | - |
| Otu60341 | 1 | k__Fungi(100);k__Fungi_unclassified(100);k__Fungi_unclassified(100);k__Fungi_unclassified(100);k__Fungi_unclassified(100);k__Fungi_unclassified(100);k__Fungi_unclassified(100); | - | - | - | - | - | - | - | Unassigned | - |
| Otu60342 | 1 | k__Fungi(100);p__Ascomycota(100);p__Ascomycota_unclassified(100);p__Ascomycota_unclassified(100);p__Ascomycota_unclassified(100);p__Ascomycota_unclassified(100);p__Ascomycota_unclassified(100); | - | - | - | - | - | - | - | Unassigned | - |
| Otu60343 | 1 | k__Fungi(100);p__Ascomycota(100);p__Ascomycota_unclassified(100);p__Ascomycota_unclassified(100);p__Ascomycota_unclassified(100);p__Ascomycota_unclassified(100);p__Ascomycota_unclassified(100); | - | - | - | - | - | - | - | Unassigned | - |
| Otu60344 | 1 | k__Fungi(100);k__Fungi_unclassified(100);k__Fungi_unclassified(100);k__Fungi_unclassified(100);k__Fungi_unclassified(100);k__Fungi_unclassified(100);k__Fungi_unclassified(100); | - | - | - | - | - | - | - | Unassigned | - |
| Otu60345 | 1 | k__Fungi(100);p__Ascomycota(100);p__Ascomycota_unclassified(100);p__Ascomycota_unclassified(100);p__Ascomycota_unclassified(100);p__Ascomycota_unclassified(100);p__Ascomycota_unclassified(100); | - | - | - | - | - | - | - | Unassigned | - |
| Otu60346 | 1 | k__Fungi(100);k__Fungi_unclassified(100);k__Fungi_unclassified(100);k__Fungi_unclassified(100);k__Fungi_unclassified(100);k__Fungi_unclassified(100);k__Fungi_unclassified(100); | - | - | - | - | - | - | - | Unassigned | - |
| Otu60347 | 1 | k__Fungi(100);k__Fungi_unclassified(100);k__Fungi_unclassified(100);k__Fungi_unclassified(100);k__Fungi_unclassified(100);k__Fungi_unclassified(100);k__Fungi_unclassified(100); | - | - | - | - | - | - | - | Unassigned | - |
| Otu60349 | 1 | k__Fungi(100);p__Ascomycota(100);p__Ascomycota_unclassified(100);p__Ascomycota_unclassified(100);p__Ascomycota_unclassified(100);p__Ascomycota_unclassified(100);p__Ascomycota_unclassified(100); | - | - | - | - | - | - | - | Unassigned | - |
| Otu60350 | 1 | k__Fungi(100);p__Ascomycota(100);p__Ascomycota_unclassified(100);p__Ascomycota_unclassified(100);p__Ascomycota_unclassified(100);p__Ascomycota_unclassified(100);p__Ascomycota_unclassified(100); | - | - | - | - | - | - | - | Unassigned | - |
| Otu60351 | 1 | k__Fungi(100);p__Ascomycota(100);p__Ascomycota_unclassified(100);p__Ascomycota_unclassified(100);p__Ascomycota_unclassified(100);p__Ascomycota_unclassified(100);p__Ascomycota_unclassified(100); | - | - | - | - | - | - | - | Unassigned | - |
| Otu60352 | 1 | k__Fungi(100);k__Fungi_unclassified(100);k__Fungi_unclassified(100);k__Fungi_unclassified(100);k__Fungi_unclassified(100);k__Fungi_unclassified(100);k__Fungi_unclassified(100); | - | - | - | - | - | - | - | Unassigned | - |
| Otu60353 | 1 | k__Fungi(100);p__Ascomycota(100);p__Ascomycota_unclassified(100);p__Ascomycota_unclassified(100);p__Ascomycota_unclassified(100);p__Ascomycota_unclassified(100);p__Ascomycota_unclassified(100); | - | - | - | - | - | - | - | Unassigned | - |
| Otu60354 | 1 | k__Fungi(100);k__Fungi_unclassified(100);k__Fungi_unclassified(100);k__Fungi_unclassified(100);k__Fungi_unclassified(100);k__Fungi_unclassified(100);k__Fungi_unclassified(100); | - | - | - | - | - | - | - | Unassigned | - |
| Otu60355 | 1 | k__Fungi(100);p__Ascomycota(100);p__Ascomycota_unclassified(100);p__Ascomycota_unclassified(100);p__Ascomycota_unclassified(100);p__Ascomycota_unclassified(100);p__Ascomycota_unclassified(100); | - | - | - | - | - | - | - | Unassigned | - |
| Otu60356 | 1 | k__Fungi(100);k__Fungi_unclassified(100);k__Fungi_unclassified(100);k__Fungi_unclassified(100);k__Fungi_unclassified(100);k__Fungi_unclassified(100);k__Fungi_unclassified(100); | - | - | - | - | - | - | - | Unassigned | - |
| Otu60357 | 1 | k__Fungi(100);p__Ascomycota(100);p__Ascomycota_unclassified(100);p__Ascomycota_unclassified(100);p__Ascomycota_unclassified(100);p__Ascomycota_unclassified(100);p__Ascomycota_unclassified(100); | - | - | - | - | - | - | - | Unassigned | - |
| Otu60358 | 1 | k__Fungi(100);p__Ascomycota(100);p__Ascomycota_unclassified(100);p__Ascomycota_unclassified(100);p__Ascomycota_unclassified(100);p__Ascomycota_unclassified(100);p__Ascomycota_unclassified(100); | - | - | - | - | - | - | - | Unassigned | - |
| Otu60359 | 1 | k__Fungi(100);k__Fungi_unclassified(100);k__Fungi_unclassified(100);k__Fungi_unclassified(100);k__Fungi_unclassified(100);k__Fungi_unclassified(100);k__Fungi_unclassified(100); | - | - | - | - | - | - | - | Unassigned | - |
| Otu60360 | 1 | k__Fungi(100);k__Fungi_unclassified(100);k__Fungi_unclassified(100);k__Fungi_unclassified(100);k__Fungi_unclassified(100);k__Fungi_unclassified(100);k__Fungi_unclassified(100); | - | - | - | - | - | - | - | Unassigned | - |
| Otu60361 | 1 | k__Fungi(100);k__Fungi_unclassified(100);k__Fungi_unclassified(100);k__Fungi_unclassified(100);k__Fungi_unclassified(100);k__Fungi_unclassified(100);k__Fungi_unclassified(100); | - | - | - | - | - | - | - | Unassigned | - |

| | | | | | | | | | | | |
|---|---|---|---|---|---|---|---|---|---|---|---|
| Otu60363 | 1 | k__Fungi(100);k__Fungi_unclassified(100);k__Fungi_unclassified(100);k__Fungi_unclassified(100);k__Fungi_unclassified(100);k__Fungi_unclassified(100);k__Fungi_unclassified(100); | - | - | - | - | - | - | - | Unassigned | - |
| Otu60364 | 1 | k__Fungi(100);k__Fungi_unclassified(100);k__Fungi_unclassified(100);k__Fungi_unclassified(100);k__Fungi_unclassified(100);k__Fungi_unclassified(100);k__Fungi_unclassified(100); | - | - | - | - | - | - | - | Unassigned | - |
| Otu60365 | 1 | k__Fungi(100);k__Fungi_unclassified(100);k__Fungi_unclassified(100);k__Fungi_unclassified(100);k__Fungi_unclassified(100);k__Fungi_unclassified(100);k__Fungi_unclassified(100); | - | - | - | - | - | - | - | Unassigned | - |
| Otu60366 | 1 | k__Fungi(100);k__Fungi_unclassified(100);k__Fungi_unclassified(100);k__Fungi_unclassified(100);k__Fungi_unclassified(100);k__Fungi_unclassified(100);k__Fungi_unclassified(100); | - | - | - | - | - | - | - | Unassigned | - |
| Otu60367 | 1 | k__Fungi(100);p__Ascomycota(100);p__Ascomycota_unclassified(100);p__Ascomycota_unclassified(100);p__Ascomycota_unclassified(100);p__Ascomycota_unclassified(100);p__Ascomycota_unclassified(100); | - | - | - | - | - | - | - | Unassigned | - |
| Otu60370 | 1 | k__Fungi(100);k__Fungi_unclassified(100);k__Fungi_unclassified(100);k__Fungi_unclassified(100);k__Fungi_unclassified(100);k__Fungi_unclassified(100);k__Fungi_unclassified(100); | - | - | - | - | - | - | - | Unassigned | - |
| Otu60371 | 1 | k__Fungi(100);p__Ascomycota(100);p__Ascomycota_unclassified(100);p__Ascomycota_unclassified(100);p__Ascomycota_unclassified(100);p__Ascomycota_unclassified(100);p__Ascomycota_unclassified(100); | - | - | - | - | - | - | - | Unassigned | - |
| Otu60372 | 1 | k__Fungi(100);k__Fungi_unclassified(100);k__Fungi_unclassified(100);k__Fungi_unclassified(100);k__Fungi_unclassified(100);k__Fungi_unclassified(100);k__Fungi_unclassified(100); | - | - | - | - | - | - | - | Unassigned | - |
| Otu60373 | 1 | k__Fungi(100);k__Fungi_unclassified(100);k__Fungi_unclassified(100);k__Fungi_unclassified(100);k__Fungi_unclassified(100);k__Fungi_unclassified(100);k__Fungi_unclassified(100); | - | - | - | - | - | - | - | Unassigned | - |
| Otu60374 | 1 | k__Fungi(100);k__Fungi_unclassified(100);k__Fungi_unclassified(100);k__Fungi_unclassified(100);k__Fungi_unclassified(100);k__Fungi_unclassified(100);k__Fungi_unclassified(100); | - | - | - | - | - | - | - | Unassigned | - |
| Otu60375 | 1 | k__Fungi(100);k__Fungi_unclassified(100);k__Fungi_unclassified(100);k__Fungi_unclassified(100);k__Fungi_unclassified(100);k__Fungi_unclassified(100);k__Fungi_unclassified(100); | - | - | - | - | - | - | - | Unassigned | - |
| Otu60376 | 1 | k__Fungi(100);k__Fungi_unclassified(100);k__Fungi_unclassified(100);k__Fungi_unclassified(100);k__Fungi_unclassified(100);k__Fungi_unclassified(100);k__Fungi_unclassified(100); | - | - | - | - | - | - | - | Unassigned | - |
| Otu60377 | 1 | k__Fungi(100);k__Fungi_unclassified(100);k__Fungi_unclassified(100);k__Fungi_unclassified(100);k__Fungi_unclassified(100);k__Fungi_unclassified(100);k__Fungi_unclassified(100); | - | - | - | - | - | - | - | Unassigned | - |
| Otu60379 | 1 | k__Fungi(100);k__Fungi_unclassified(100);k__Fungi_unclassified(100);k__Fungi_unclassified(100);k__Fungi_unclassified(100);k__Fungi_unclassified(100);k__Fungi_unclassified(100); | - | - | - | - | - | - | - | Unassigned | - |
| Otu60380 | 1 | k__Fungi(100);k__Fungi_unclassified(100);k__Fungi_unclassified(100);k__Fungi_unclassified(100);k__Fungi_unclassified(100);k__Fungi_unclassified(100);k__Fungi_unclassified(100); | - | - | - | - | - | - | - | Unassigned | - |
| Otu60381 | 1 | k__Fungi(100);p__Ascomycota(100);p__Ascomycota_unclassified(100);p__Ascomycota_unclassified(100);p__Ascomycota_unclassified(100);p__Ascomycota_unclassified(100);p__Ascomycota_unclassified(100); | - | - | - | - | - | - | - | Unassigned | - |
| Otu60382 | 1 | k__Fungi(100);k__Fungi_unclassified(100);k__Fungi_unclassified(100);k__Fungi_unclassified(100);k__Fungi_unclassified(100);k__Fungi_unclassified(100);k__Fungi_unclassified(100); | - | - | - | - | - | - | - | Unassigned | - |
| Otu60384 | 1 | k__Fungi(100);k__Fungi_unclassified(100);k__Fungi_unclassified(100);k__Fungi_unclassified(100);k__Fungi_unclassified(100);k__Fungi_unclassified(100);k__Fungi_unclassified(100); | - | - | - | - | - | - | - | Unassigned | - |
| Otu60385 | 1 | k__Fungi(100);p__Ascomycota(100);p__Ascomycota_unclassified(100);p__Ascomycota_unclassified(100);p__Ascomycota_unclassified(100);p__Ascomycota_unclassified(100);p__Ascomycota_unclassified(100); | - | - | - | - | - | - | - | Unassigned | - |
| Otu60386 | 1 | k__Fungi(100);k__Fungi_unclassified(100);k__Fungi_unclassified(100);k__Fungi_unclassified(100);k__Fungi_unclassified(100);k__Fungi_unclassified(100);k__Fungi_unclassified(100); | - | - | - | - | - | - | - | Unassigned | - |
| Otu60387 | 1 | k__Fungi(100);k__Fungi_unclassified(100);k__Fungi_unclassified(100);k__Fungi_unclassified(100);k__Fungi_unclassified(100);k__Fungi_unclassified(100);k__Fungi_unclassified(100); | - | - | - | - | - | - | - | Unassigned | - |
| Otu60388 | 1 | k__Fungi(100);p__Ascomycota(100);p__Ascomycota_unclassified(100);p__Ascomycota_unclassified(100);p__Ascomycota_unclassified(100);p__Ascomycota_unclassified(100);p__Ascomycota_unclassified(100); | - | - | - | - | - | - | - | Unassigned | - |
| Otu60389 | 1 | k__Fungi(100);p__Ascomycota(100);p__Ascomycota_unclassified(100);p__Ascomycota_unclassified(100);p__Ascomycota_unclassified(100);p__Ascomycota_unclassified(100);p__Ascomycota_unclassified(100); | - | - | - | - | - | - | - | Unassigned | - |
| Otu60390 | 1 | k__Fungi(100);k__Fungi_unclassified(100);k__Fungi_unclassified(100);k__Fungi_unclassified(100);k__Fungi_unclassified(100);k__Fungi_unclassified(100);k__Fungi_unclassified(100); | - | - | - | - | - | - | - | Unassigned | - |
| Otu60391 | 1 | k__Fungi(100);k__Fungi_unclassified(100);k__Fungi_unclassified(100);k__Fungi_unclassified(100);k__Fungi_unclassified(100);k__Fungi_unclassified(100);k__Fungi_unclassified(100); | - | - | - | - | - | - | - | Unassigned | - |
| Otu60392 | 1 | k__Fungi(100);k__Fungi_unclassified(100);k__Fungi_unclassified(100);k__Fungi_unclassified(100);k__Fungi_unclassified(100);k__Fungi_unclassified(100);k__Fungi_unclassified(100); | - | - | - | - | - | - | - | Unassigned | - |
| Otu60393 | 1 | k__Fungi(100);k__Fungi_unclassified(100);k__Fungi_unclassified(100);k__Fungi_unclassified(100);k__Fungi_unclassified(100);k__Fungi_unclassified(100);k__Fungi_unclassified(100); | - | - | - | - | - | - | - | Unassigned | - |
| Otu60395 | 1 | k__Fungi(100);k__Fungi_unclassified(100);k__Fungi_unclassified(100);k__Fungi_unclassified(100);k__Fungi_unclassified(100);k__Fungi_unclassified(100);k__Fungi_unclassified(100); | - | - | - | - | - | - | - | Unassigned | - |
| Otu60396 | 1 | k__Fungi(100);k__Fungi_unclassified(100);k__Fungi_unclassified(100);k__Fungi_unclassified(100);k__Fungi_unclassified(100);k__Fungi_unclassified(100);k__Fungi_unclassified(100); | - | - | - | - | - | - | - | Unassigned | - |
| Otu60397 | 1 | k__Fungi(100);k__Fungi_unclassified(100);k__Fungi_unclassified(100);k__Fungi_unclassified(100);k__Fungi_unclassified(100);k__Fungi_unclassified(100);k__Fungi_unclassified(100); | - | - | - | - | - | - | - | Unassigned | - |
| Otu60398 | 1 | k__Fungi(100);p__Ascomycota(100);p__Ascomycota_unclassified(100);p__Ascomycota_unclassified(100);p__Ascomycota_unclassified(100);p__Ascomycota_unclassified(100);p__Ascomycota_unclassified(100); | - | - | - | - | - | - | - | Unassigned | - |
| Otu60399 | 1 | k__Fungi(100);p__Ascomycota(100);p__Ascomycota_unclassified(100);p__Ascomycota_unclassified(100);p__Ascomycota_unclassified(100);p__Ascomycota_unclassified(100);p__Ascomycota_unclassified(100); | - | - | - | - | - | - | - | Unassigned | - |
| Otu60400 | 1 | k__Fungi(100);k__Fungi_unclassified(100);k__Fungi_unclassified(100);k__Fungi_unclassified(100);k__Fungi_unclassified(100);k__Fungi_unclassified(100);k__Fungi_unclassified(100); | - | - | - | - | - | - | - | Unassigned | - |
| Otu60402 | 1 | k__Fungi(100);p__Ascomycota(100);p__Ascomycota_unclassified(100);p__Ascomycota_unclassified(100);p__Ascomycota_unclassified(100);p__Ascomycota_unclassified(100);p__Ascomycota_unclassified(100); | - | - | - | - | - | - | - | Unassigned | - |
| Otu60403 | 1 | k__Fungi(100);k__Fungi_unclassified(100);k__Fungi_unclassified(100);k__Fungi_unclassified(100);k__Fungi_unclassified(100);k__Fungi_unclassified(100);k__Fungi_unclassified(100); | - | - | - | - | - | - | - | Unassigned | - |
| Otu60404 | 1 | k__Fungi(100);k__Fungi_unclassified(100);k__Fungi_unclassified(100);k__Fungi_unclassified(100);k__Fungi_unclassified(100);k__Fungi_unclassified(100);k__Fungi_unclassified(100); | - | - | - | - | - | - | - | Unassigned | - |
| Otu60405 | 1 | k__Fungi(100);k__Fungi_unclassified(100);k__Fungi_unclassified(100);k__Fungi_unclassified(100);k__Fungi_unclassified(100);k__Fungi_unclassified(100);k__Fungi_unclassified(100); | - | - | - | - | - | - | - | Unassigned | - |
| Otu60406 | 1 | k__Fungi(100);k__Fungi_unclassified(100);k__Fungi_unclassified(100);k__Fungi_unclassified(100);k__Fungi_unclassified(100);k__Fungi_unclassified(100);k__Fungi_unclassified(100); | - | - | - | - | - | - | - | Unassigned | - |
| Otu60407 | 1 | k__Fungi(100);k__Fungi_unclassified(100);k__Fungi_unclassified(100);k__Fungi_unclassified(100);k__Fungi_unclassified(100);k__Fungi_unclassified(100);k__Fungi_unclassified(100); | - | - | - | - | - | - | - | Unassigned | - |
| Otu60408 | 1 | k__Fungi(100);k__Fungi_unclassified(100);k__Fungi_unclassified(100);k__Fungi_unclassified(100);k__Fungi_unclassified(100);k__Fungi_unclassified(100);k__Fungi_unclassified(100); | - | - | - | - | - | - | - | Unassigned | - |
| Otu60410 | 1 | k__Fungi(100);k__Fungi_unclassified(100);k__Fungi_unclassified(100);k__Fungi_unclassified(100);k__Fungi_unclassified(100);k__Fungi_unclassified(100);k__Fungi_unclassified(100); | - | - | - | - | - | - | - | Unassigned | - |
| Otu60411 | 1 | k__Fungi(100);k__Fungi_unclassified(100);k__Fungi_unclassified(100);k__Fungi_unclassified(100);k__Fungi_unclassified(100);k__Fungi_unclassified(100);k__Fungi_unclassified(100); | - | - | - | - | - | - | - | Unassigned | - |
| Otu60412 | 1 | k__Fungi(100);p__Ascomycota(100);p__Ascomycota_unclassified(100);p__Ascomycota_unclassified(100);p__Ascomycota_unclassified(100);p__Ascomycota_unclassified(100);p__Ascomycota_unclassified(100); | - | - | - | - | - | - | - | Unassigned | - |
| Otu60413 | 1 | k__Fungi(100);p__Ascomycota(100);p__Ascomycota_unclassified(100);p__Ascomycota_unclassified(100);p__Ascomycota_unclassified(100);p__Ascomycota_unclassified(100);p__Ascomycota_unclassified(100); | - | - | - | - | - | - | - | Unassigned | - |
| Otu60414 | 1 | k__Fungi(100);k__Fungi_unclassified(100);k__Fungi_unclassified(100);k__Fungi_unclassified(100);k__Fungi_unclassified(100);k__Fungi_unclassified(100);k__Fungi_unclassified(100); | - | - | - | - | - | - | - | Unassigned | - |
| Otu60416 | 1 | k__Fungi(100);p__Ascomycota(100);p__Ascomycota_unclassified(100);p__Ascomycota_unclassified(100);p__Ascomycota_unclassified(100);p__Ascomycota_unclassified(100);p__Ascomycota_unclassified(100); | - | - | - | - | - | - | - | Unassigned | - |
| Otu60417 | 1 | k__Fungi(100);k__Fungi_unclassified(100);k__Fungi_unclassified(100);k__Fungi_unclassified(100);k__Fungi_unclassified(100);k__Fungi_unclassified(100);k__Fungi_unclassified(100); | - | - | - | - | - | - | - | Unassigned | - |
| Otu60419 | 1 | k__Fungi(100);k__Fungi_unclassified(100);k__Fungi_unclassified(100);k__Fungi_unclassified(100);k__Fungi_unclassified(100);k__Fungi_unclassified(100);k__Fungi_unclassified(100); | - | - | - | - | - | - | - | Unassigned | - |
| Otu60420 | 1 | k__Fungi(100);k__Fungi_unclassified(100);k__Fungi_unclassified(100);k__Fungi_unclassified(100);k__Fungi_unclassified(100);k__Fungi_unclassified(100);k__Fungi_unclassified(100); | - | - | - | - | - | - | - | Unassigned | - |
| Otu60421 | 1 | k__Fungi(100);k__Fungi_unclassified(100);k__Fungi_unclassified(100);k__Fungi_unclassified(100);k__Fungi_unclassified(100);k__Fungi_unclassified(100);k__Fungi_unclassified(100); | - | - | - | - | - | - | - | Unassigned | - |
| Otu60422 | 1 | k__Fungi(100);k__Fungi_unclassified(100);k__Fungi_unclassified(100);k__Fungi_unclassified(100);k__Fungi_unclassified(100);k__Fungi_unclassified(100);k__Fungi_unclassified(100); | - | - | - | - | - | - | - | Unassigned | - |
| Otu60423 | 1 | k__Fungi(100);p__Ascomycota(100);p__Ascomycota_unclassified(100);p__Ascomycota_unclassified(100);p__Ascomycota_unclassified(100);p__Ascomycota_unclassified(100);p__Ascomycota_unclassified(100); | - | - | - | - | - | - | - | Unassigned | - |
| Otu60427 | 1 | k__Fungi(100);k__Fungi_unclassified(100);k__Fungi_unclassified(100);k__Fungi_unclassified(100);k__Fungi_unclassified(100);k__Fungi_unclassified(100);k__Fungi_unclassified(100); | - | - | - | - | - | - | - | Unassigned | - |
| Otu60429 | 1 | k__Fungi(100);p__Ascomycota(100);p__Ascomycota_unclassified(100);p__Ascomycota_unclassified(100);p__Ascomycota_unclassified(100);p__Ascomycota_unclassified(100);p__Ascomycota_unclassified(100); | - | - | - | - | - | - | - | Unassigned | - |
| Otu60431 | 1 | k__Fungi(100);p__Ascomycota(100);p__Ascomycota_unclassified(100);p__Ascomycota_unclassified(100);p__Ascomycota_unclassified(100);p__Ascomycota_unclassified(100);p__Ascomycota_unclassified(100); | - | - | - | - | - | - | - | Unassigned | - |
| Otu60432 | 1 | k__Fungi(100);p__Ascomycota(100);p__Ascomycota_unclassified(100);p__Ascomycota_unclassified(100);p__Ascomycota_unclassified(100);p__Ascomycota_unclassified(100);p__Ascomycota_unclassified(100); | - | - | - | - | - | - | - | Unassigned | - |
| Otu60433 | 1 | k__Fungi(100);k__Fungi_unclassified(100);k__Fungi_unclassified(100);k__Fungi_unclassified(100);k__Fungi_unclassified(100);k__Fungi_unclassified(100);k__Fungi_unclassified(100); | - | - | - | - | - | - | - | Unassigned | - |
| Otu60434 | 1 | k__Fungi(100);k__Fungi_unclassified(100);k__Fungi_unclassified(100);k__Fungi_unclassified(100);k__Fungi_unclassified(100);k__Fungi_unclassified(100);k__Fungi_unclassified(100); | - | - | - | - | - | - | - | Unassigned | - |
| Otu60435 | 1 | k__Fungi(100);k__Fungi_unclassified(100);k__Fungi_unclassified(100);k__Fungi_unclassified(100);k__Fungi_unclassified(100);k__Fungi_unclassified(100);k__Fungi_unclassified(100); | - | - | - | - | - | - | - | Unassigned | - |
| Otu60436 | 1 | k__Fungi(100);p__Ascomycota(100);p__Ascomycota_unclassified(100);p__Ascomycota_unclassified(100);p__Ascomycota_unclassified(100);p__Ascomycota_unclassified(100);p__Ascomycota_unclassified(100); | - | - | - | - | - | - | - | Unassigned | - |
| Otu60437 | 1 | k__Fungi(100);p__Ascomycota(100);p__Ascomycota_unclassified(100);p__Ascomycota_unclassified(100);p__Ascomycota_unclassified(100);p__Ascomycota_unclassified(100);p__Ascomycota_unclassified(100); | - | - | - | - | - | - | - | Unassigned | - |
| Otu60439 | 1 | k__Fungi(100);k__Fungi_unclassified(100);k__Fungi_unclassified(100);k__Fungi_unclassified(100);k__Fungi_unclassified(100);k__Fungi_unclassified(100);k__Fungi_unclassified(100); | - | - | - | - | - | - | - | Unassigned | - |
| Otu60440 | 1 | k__Fungi(100);k__Fungi_unclassified(100);k__Fungi_unclassified(100);k__Fungi_unclassified(100);k__Fungi_unclassified(100);k__Fungi_unclassified(100);k__Fungi_unclassified(100); | - | - | - | - | - | - | - | Unassigned | - |
| Otu60441 | 1 | k__Fungi(100);p__Ascomycota(100);p__Ascomycota_unclassified(100);p__Ascomycota_unclassified(100);p__Ascomycota_unclassified(100);p__Ascomycota_unclassified(100);p__Ascomycota_unclassified(100); | - | - | - | - | - | - | - | Unassigned | - |
| Otu60444 | 1 | k__Fungi(100);k__Fungi_unclassified(100);k__Fungi_unclassified(100);k__Fungi_unclassified(100);k__Fungi_unclassified(100);k__Fungi_unclassified(100);k__Fungi_unclassified(100); | - | - | - | - | - | - | - | Unassigned | - |
| Otu60445 | 1 | k__Fungi(100);k__Fungi_unclassified(100);k__Fungi_unclassified(100);k__Fungi_unclassified(100);k__Fungi_unclassified(100);k__Fungi_unclassified(100);k__Fungi_unclassified(100); | - | - | - | - | - | - | - | Unassigned | - |
| Otu60447 | 1 | k__Fungi(100);p__Ascomycota(100);p__Ascomycota_unclassified(100);p__Ascomycota_unclassified(100);p__Ascomycota_unclassified(100);p__Ascomycota_unclassified(100);p__Ascomycota_unclassified(100); | - | - | - | - | - | - | - | Unassigned | - |
| Otu60448 | 1 | k__Fungi(100);k__Fungi_unclassified(100);k__Fungi_unclassified(100);k__Fungi_unclassified(100);k__Fungi_unclassified(100);k__Fungi_unclassified(100);k__Fungi_unclassified(100); | - | - | - | - | - | - | - | Unassigned | - |
| Otu60449 | 1 | k__Fungi(100);p__Ascomycota(100);p__Ascomycota_unclassified(100);p__Ascomycota_unclassified(100);p__Ascomycota_unclassified(100);p__Ascomycota_unclassified(100);p__Ascomycota_unclassified(100); | - | - | - | - | - | - | - | Unassigned | - |
| Otu60450 | 1 | k__Fungi(100);k__Fungi_unclassified(100);k__Fungi_unclassified(100);k__Fungi_unclassified(100);k__Fungi_unclassified(100);k__Fungi_unclassified(100);k__Fungi_unclassified(100); | - | - | - | - | - | - | - | Unassigned | - |
| Otu60451 | 1 | k__Fungi(100);p__Ascomycota(100);p__Ascomycota_unclassified(100);p__Ascomycota_unclassified(100);p__Ascomycota_unclassified(100);p__Ascomycota_unclassified(100);p__Ascomycota_unclassified(100); | - | - | - | - | - | - | - | Unassigned | - |
| Otu60452 | 1 | k__Fungi(100);p__Ascomycota(100);p__Ascomycota_unclassified(100);p__Ascomycota_unclassified(100);p__Ascomycota_unclassified(100);p__Ascomycota_unclassified(100);p__Ascomycota_unclassified(100); | - | - | - | - | - | - | - | Unassigned | - |
| Otu60453 | 1 | k__Fungi(100);k__Fungi_unclassified(100);k__Fungi_unclassified(100);k__Fungi_unclassified(100);k__Fungi_unclassified(100);k__Fungi_unclassified(100);k__Fungi_unclassified(100); | - | - | - | - | - | - | - | Unassigned | - |
| Otu60454 | 1 | k__Fungi(100);k__Fungi_unclassified(100);k__Fungi_unclassified(100);k__Fungi_unclassified(100);k__Fungi_unclassified(100);k__Fungi_unclassified(100);k__Fungi_unclassified(100); | - | - | - | - | - | - | - | Unassigned | - |
| Otu60455 | 1 | k__Fungi(100);k__Fungi_unclassified(100);k__Fungi_unclassified(100);k__Fungi_unclassified(100);k__Fungi_unclassified(100);k__Fungi_unclassified(100);k__Fungi_unclassified(100); | - | - | - | - | - | - | - | Unassigned | - |
| Otu60456 | 1 | k__Fungi(100);k__Fungi_unclassified(100);k__Fungi_unclassified(100);k__Fungi_unclassified(100);k__Fungi_unclassified(100);k__Fungi_unclassified(100);k__Fungi_unclassified(100); | - | - | - | - | - | - | - | Unassigned | - |
| Otu60457 | 1 | k__Fungi(100);k__Fungi_unclassified(100);k__Fungi_unclassified(100);k__Fungi_unclassified(100);k__Fungi_unclassified(100);k__Fungi_unclassified(100);k__Fungi_unclassified(100); | - | - | - | - | - | - | - | Unassigned | - |
| Otu60458 | 1 | k__Fungi(100);k__Fungi_unclassified(100);k__Fungi_unclassified(100);k__Fungi_unclassified(100);k__Fungi_unclassified(100);k__Fungi_unclassified(100);k__Fungi_unclassified(100); | - | - | - | - | - | - | - | Unassigned | - |
| Otu60460 | 1 | k__Fungi(100);k__Fungi_unclassified(100);k__Fungi_unclassified(100);k__Fungi_unclassified(100);k__Fungi_unclassified(100);k__Fungi_unclassified(100);k__Fungi_unclassified(100); | - | - | - | - | - | - | - | Unassigned | - |
| Otu60461 | 1 | k__Fungi(100);k__Fungi_unclassified(100);k__Fungi_unclassified(100);k__Fungi_unclassified(100);k__Fungi_unclassified(100);k__Fungi_unclassified(100);k__Fungi_unclassified(100); | - | - | - | - | - | - | - | Unassigned | - |
| Otu60462 | 1 | k__Fungi(100);p__Ascomycota(100);p__Ascomycota_unclassified(100);p__Ascomycota_unclassified(100);p__Ascomycota_unclassified(100);p__Ascomycota_unclassified(100);p__Ascomycota_unclassified(100); | - | - | - | - | - | - | - | Unassigned | - |
| Otu60463 | 1 | k__Fungi(100);k__Fungi_unclassified(100);k__Fungi_unclassified(100);k__Fungi_unclassified(100);k__Fungi_unclassified(100);k__Fungi_unclassified(100);k__Fungi_unclassified(100); | - | - | - | - | - | - | - | Unassigned | - |
| Otu60466 | 1 | k__Fungi(100);k__Fungi_unclassified(100);k__Fungi_unclassified(100);k__Fungi_unclassified(100);k__Fungi_unclassified(100);k__Fungi_unclassified(100);k__Fungi_unclassified(100); | - | - | - | - | - | - | - | Unassigned | - |
| Otu60467 | 1 | k__Fungi(100);p__Ascomycota(100);p__Ascomycota_unclassified(100);p__Ascomycota_unclassified(100);p__Ascomycota_unclassified(100);p__Ascomycota_unclassified(100);p__Ascomycota_unclassified(100); | - | - | - | - | - | - | - | Unassigned | - |
| Otu60469 | 1 | k__Fungi(100);k__Fungi_unclassified(100);k__Fungi_unclassified(100);k__Fungi_unclassified(100);k__Fungi_unclassified(100);k__Fungi_unclassified(100);k__Fungi_unclassified(100); | - | - | - | - | - | - | - | Unassigned | - |
| Otu60474 | 1 | k__Fungi(100);p__Ascomycota(100);p__Ascomycota_unclassified(100);p__Ascomycota_unclassified(100);p__Ascomycota_unclassified(100);p__Ascomycota_unclassified(100);p__Ascomycota_unclassified(100); | - | - | - | - | - | - | - | Unassigned | - |
| Otu60475 | 1 | k__Fungi(100);p__Ascomycota(100);p__Ascomycota_unclassified(100);p__Ascomycota_unclassified(100);p__Ascomycota_unclassified(100);p__Ascomycota_unclassified(100);p__Ascomycota_unclassified(100); | - | - | - | - | - | - | - | Unassigned | - |
| Otu60476 | 1 | k__Fungi(100);k__Fungi_unclassified(100);k__Fungi_unclassified(100);k__Fungi_unclassified(100);k__Fungi_unclassified(100);k__Fungi_unclassified(100);k__Fungi_unclassified(100); | - | - | - | - | - | - | - | Unassigned | - |
| Otu60477 | 1 | k__Fungi(100);k__Fungi_unclassified(100);k__Fungi_unclassified(100);k__Fungi_unclassified(100);k__Fungi_unclassified(100);k__Fungi_unclassified(100);k__Fungi_unclassified(100); | - | - | - | - | - | - | - | Unassigned | - |
| Otu60478 | 1 | k__Fungi(100);k__Fungi_unclassified(100);k__Fungi_unclassified(100);k__Fungi_unclassified(100);k__Fungi_unclassified(100);k__Fungi_unclassified(100);k__Fungi_unclassified(100); | - | - | - | - | - | - | - | Unassigned | - |
| Otu60479 | 1 | k__Fungi(100);k__Fungi_unclassified(100);k__Fungi_unclassified(100);k__Fungi_unclassified(100);k__Fungi_unclassified(100);k__Fungi_unclassified(100);k__Fungi_unclassified(100); | - | - | - | - | - | - | - | Unassigned | - |
| Otu60480 | 1 | k__Fungi(100);p__Ascomycota(100);p__Ascomycota_unclassified(100);p__Ascomycota_unclassified(100);p__Ascomycota_unclassified(100);p__Ascomycota_unclassified(100);p__Ascomycota_unclassified(100); | - | - | - | - | - | - | - | Unassigned | - |
| Otu60481 | 1 | k__Fungi(100);k__Fungi_unclassified(100);k__Fungi_unclassified(100);k__Fungi_unclassified(100);k__Fungi_unclassified(100);k__Fungi_unclassified(100);k__Fungi_unclassified(100); | - | - | - | - | - | - | - | Unassigned | - |
| Otu60482 | 1 | k__Fungi(100);k__Fungi_unclassified(100);k__Fungi_unclassified(100);k__Fungi_unclassified(100);k__Fungi_unclassified(100);k__Fungi_unclassified(100);k__Fungi_unclassified(100); | - | - | - | - | - | - | - | Unassigned | - |
| Otu60483 | 1 | k__Fungi(100);k__Fungi_unclassified(100);k__Fungi_unclassified(100);k__Fungi_unclassified(100);k__Fungi_unclassified(100);k__Fungi_unclassified(100);k__Fungi_unclassified(100); | - | - | - | - | - | - | - | Unassigned | - |
| Otu60484 | 1 | k__Fungi(100);k__Fungi_unclassified(100);k__Fungi_unclassified(100);k__Fungi_unclassified(100);k__Fungi_unclassified(100);k__Fungi_unclassified(100);k__Fungi_unclassified(100); | - | - | - | - | - | - | - | Unassigned | - |
| Otu60485 | 1 | k__Fungi(100);k__Fungi_unclassified(100);k__Fungi_unclassified(100);k__Fungi_unclassified(100);k__Fungi_unclassified(100);k__Fungi_unclassified(100);k__Fungi_unclassified(100); | - | - | - | - | - | - | - | Unassigned | - |
| Otu60486 | 1 | k__Fungi(100);p__Ascomycota(100);p__Ascomycota_unclassified(100);p__Ascomycota_unclassified(100);p__Ascomycota_unclassified(100);p__Ascomycota_unclassified(100);p__Ascomycota_unclassified(100); | - | - | - | - | - | - | - | Unassigned | - |
| Otu60487 | 1 | k__Fungi(100);k__Fungi_unclassified(100);k__Fungi_unclassified(100);k__Fungi_unclassified(100);k__Fungi_unclassified(100);k__Fungi_unclassified(100);k__Fungi_unclassified(100); | - | - | - | - | - | - | - | Unassigned | - |
| Otu60488 | 1 | k__Fungi(100);p__Ascomycota(100);p__Ascomycota_unclassified(100);p__Ascomycota_unclassified(100);p__Ascomycota_unclassified(100);p__Ascomycota_unclassified(100);p__Ascomycota_unclassified(100); | - | - | - | - | - | - | - | Unassigned | - |
| Otu60489 | 1 | k__Fungi(100);k__Fungi_unclassified(100);k__Fungi_unclassified(100);k__Fungi_unclassified(100);k__Fungi_unclassified(100);k__Fungi_unclassified(100);k__Fungi_unclassified(100); | - | - | - | - | - | - | - | Unassigned | - |
| Otu60490 | 1 | k__Fungi(100);k__Fungi_unclassified(100);k__Fungi_unclassified(100);k__Fungi_unclassified(100);k__Fungi_unclassified(100);k__Fungi_unclassified(100);k__Fungi_unclassified(100); | - | - | - | - | - | - | - | Unassigned | - |
| Otu60491 | 1 | k__Fungi(100);k__Fungi_unclassified(100);k__Fungi_unclassified(100);k__Fungi_unclassified(100);k__Fungi_unclassified(100);k__Fungi_unclassified(100);k__Fungi_unclassified(100); | - | - | - | - | - | - | - | Unassigned | - |
| Otu60492 | 1 | k__Fungi(100);p__Ascomycota(100);p__Ascomycota_unclassified(100);p__Ascomycota_unclassified(100);p__Ascomycota_unclassified(100);p__Ascomycota_unclassified(100);p__Ascomycota_unclassified(100); | - | - | - | - | - | - | - | Unassigned | - |
| Otu60493 | 1 | k__Fungi(100);k__Fungi_unclassified(100);k__Fungi_unclassified(100);k__Fungi_unclassified(100);k__Fungi_unclassified(100);k__Fungi_unclassified(100);k__Fungi_unclassified(100); | - | - | - | - | - | - | - | Unassigned | - |
| Otu60494 | 1 | k__Fungi(100);p__Basidiomycota(100);c__Agaricomycetes(100);c__Agaricomycetes_unclassified(100);c__Agaricomycetes_unclassified(100);c__Agaricomycetes_unclassified(100);c__Agaricomycetes_unclassified(100); | - | - | - | - | - | - | - | Unassigned | - |
| Otu60495 | 1 | k__Fungi(100);k__Fungi_unclassified(100);k__Fungi_unclassified(100);k__Fungi_unclassified(100);k__Fungi_unclassified(100);k__Fungi_unclassified(100);k__Fungi_unclassified(100); | - | - | - | - | - | - | - | Unassigned | - |
| Otu60496 | 1 | k__Fungi(100);k__Fungi_unclassified(100);k__Fungi_unclassified(100);k__Fungi_unclassified(100);k__Fungi_unclassified(100);k__Fungi_unclassified(100);k__Fungi_unclassified(100); | - | - | - | - | - | - | - | Unassigned | - |
| Otu60497 | 1 | k__Fungi(100);k__Fungi_unclassified(100);k__Fungi_unclassified(100);k__Fungi_unclassified(100);k__Fungi_unclassified(100);k__Fungi_unclassified(100);k__Fungi_unclassified(100); | - | - | - | - | - | - | - | Unassigned | - |
| Otu60498 | 1 | k__Fungi(100);p__Ascomycota(100);p__Ascomycota_unclassified(100);p__Ascomycota_unclassified(100);p__Ascomycota_unclassified(100);p__Ascomycota_unclassified(100);p__Ascomycota_unclassified(100); | - | - | - | - | - | - | - | Unassigned | - |
| Otu60499 | 1 | k__Fungi(100);p__Ascomycota(100);p__Ascomycota_unclassified(100);p__Ascomycota_unclassified(100);p__Ascomycota_unclassified(100);p__Ascomycota_unclassified(100);p__Ascomycota_unclassified(100); | - | - | - | - | - | - | - | Unassigned | - |
| Otu60500 | 1 | k__Fungi(100);k__Fungi_unclassified(100);k__Fungi_unclassified(100);k__Fungi_unclassified(100);k__Fungi_unclassified(100);k__Fungi_unclassified(100);k__Fungi_unclassified(100); | - | - | - | - | - | - | - | Unassigned | - |
| Otu60501 | 1 | k__Fungi(100);k__Fungi_unclassified(100);k__Fungi_unclassified(100);k__Fungi_unclassified(100);k__Fungi_unclassified(100);k__Fungi_unclassified(100);k__Fungi_unclassified(100); | - | - | - | - | - | - | - | Unassigned | - |
| Otu60502 | 1 | k__Fungi(100);k__Fungi_unclassified(100);k__Fungi_unclassified(100);k__Fungi_unclassified(100);k__Fungi_unclassified(100);k__Fungi_unclassified(100);k__Fungi_unclassified(100); | - | - | - | - | - | - | - | Unassigned | - |
| Otu60503 | 1 | k__Fungi(100);k__Fungi_unclassified(100);k__Fungi_unclassified(100);k__Fungi_unclassified(100);k__Fungi_unclassified(100);k__Fungi_unclassified(100);k__Fungi_unclassified(100); | - | - | - | - | - | - | - | Unassigned | - |
| Otu60504 | 1 | k__Fungi(100);k__Fungi_unclassified(100);k__Fungi_unclassified(100);k__Fungi_unclassified(100);k__Fungi_unclassified(100);k__Fungi_unclassified(100);k__Fungi_unclassified(100); | - | - | - | - | - | - | - | Unassigned | - |
| Otu60505 | 1 | k__Fungi(100);k__Fungi_unclassified(100);k__Fungi_unclassified(100);k__Fungi_unclassified(100);k__Fungi_unclassified(100);k__Fungi_unclassified(100);k__Fungi_unclassified(100); | - | - | - | - | - | - | - | Unassigned | - |
| Otu60506 | 1 | k__Fungi(100);p__Ascomycota(100);p__Ascomycota_unclassified(100);p__Ascomycota_unclassified(100);p__Ascomycota_unclassified(100);p__Ascomycota_unclassified(100);p__Ascomycota_unclassified(100); | - | - | - | - | - | - | - | Unassigned | - |
| Otu60507 | 1 | k__Fungi(100);k__Fungi_unclassified(100);k__Fungi_unclassified(100);k__Fungi_unclassified(100);k__Fungi_unclassified(100);k__Fungi_unclassified(100);k__Fungi_unclassified(100); | - | - | - | - | - | - | - | Unassigned | - |
| Otu60508 | 1 | k__Fungi(100);p__Ascomycota(100);p__Ascomycota_unclassified(100);p__Ascomycota_unclassified(100);p__Ascomycota_unclassified(100);p__Ascomycota_unclassified(100);p__Ascomycota_unclassified(100); | - | - | - | - | - | - | - | Unassigned | - |
| Otu60509 | 1 | k__Fungi(100);k__Fungi_unclassified(100);k__Fungi_unclassified(100);k__Fungi_unclassified(100);k__Fungi_unclassified(100);k__Fungi_unclassified(100);k__Fungi_unclassified(100); | - | - | - | - | - | - | - | Unassigned | - |
| Otu60510 | 1 | k__Fungi(100);k__Fungi_unclassified(100);k__Fungi_unclassified(100);k__Fungi_unclassified(100);k__Fungi_unclassified(100);k__Fungi_unclassified(100);k__Fungi_unclassified(100); | - | - | - | - | - | - | - | Unassigned | - |
| Otu60511 | 1 | k__Fungi(100);p__Ascomycota(100);p__Ascomycota_unclassified(100);p__Ascomycota_unclassified(100);p__Ascomycota_unclassified(100);p__Ascomycota_unclassified(100);p__Ascomycota_unclassified(100); | - | - | - | - | - | - | - | Unassigned | - |
| Otu60512 | 1 | k__Fungi(100);p__Ascomycota(100);p__Ascomycota_unclassified(100);p__Ascomycota_unclassified(100);p__Ascomycota_unclassified(100);p__Ascomycota_unclassified(100);p__Ascomycota_unclassified(100); | - | - | - | - | - | - | - | Unassigned | - |
| Otu60513 | 1 | k__Fungi(100);k__Fungi_unclassified(100);k__Fungi_unclassified(100);k__Fungi_unclassified(100);k__Fungi_unclassified(100);k__Fungi_unclassified(100);k__Fungi_unclassified(100); | - | - | - | - | - | - | - | Unassigned | - |
| Otu60514 | 1 | k__Fungi(100);k__Fungi_unclassified(100);k__Fungi_unclassified(100);k__Fungi_unclassified(100);k__Fungi_unclassified(100);k__Fungi_unclassified(100);k__Fungi_unclassified(100); | - | - | - | - | - | - | - | Unassigned | - |
| Otu60515 | 1 | k__Fungi(100);p__Ascomycota(100);p__Ascomycota_unclassified(100);p__Ascomycota_unclassified(100);p__Ascomycota_unclassified(100);p__Ascomycota_unclassified(100);p__Ascomycota_unclassified(100); | - | - | - | - | - | - | - | Unassigned | - |
| Otu60516 | 1 | k__Fungi(100);k__Fungi_unclassified(100);k__Fungi_unclassified(100);k__Fungi_unclassified(100);k__Fungi_unclassified(100);k__Fungi_unclassified(100);k__Fungi_unclassified(100); | - | - | - | - | - | - | - | Unassigned | - |
| Otu60517 | 1 | k__Fungi(100);k__Fungi_unclassified(100);k__Fungi_unclassified(100);k__Fungi_unclassified(100);k__Fungi_unclassified(100);k__Fungi_unclassified(100);k__Fungi_unclassified(100); | - | - | - | - | - | - | - | Unassigned | - |
| Otu60518 | 1 | k__Fungi(100);p__Ascomycota(100);p__Ascomycota_unclassified(100);p__Ascomycota_unclassified(100);p__Ascomycota_unclassified(100);p__Ascomycota_unclassified(100);p__Ascomycota_unclassified(100); | - | - | - | - | - | - | - | Unassigned | - |
| Otu60519 | 1 | k__Fungi(100);k__Fungi_unclassified(100);k__Fungi_unclassified(100);k__Fungi_unclassified(100);k__Fungi_unclassified(100);k__Fungi_unclassified(100);k__Fungi_unclassified(100); | - | - | - | - | - | - | - | Unassigned | - |
| Otu60520 | 1 | k__Fungi(100);k__Fungi_unclassified(100);k__Fungi_unclassified(100);k__Fungi_unclassified(100);k__Fungi_unclassified(100);k__Fungi_unclassified(100);k__Fungi_unclassified(100); | - | - | - | - | - | - | - | Unassigned | - |
| Otu60521 | 1 | k__Fungi(100);k__Fungi_unclassified(100);k__Fungi_unclassified(100);k__Fungi_unclassified(100);k__Fungi_unclassified(100);k__Fungi_unclassified(100);k__Fungi_unclassified(100); | - | - | - | - | - | - | - | Unassigned | - |
| Otu60522 | 1 | k__Fungi(100);p__Ascomycota(100);p__Ascomycota_unclassified(100);p__Ascomycota_unclassified(100);p__Ascomycota_unclassified(100);p__Ascomycota_unclassified(100);p__Ascomycota_unclassified(100); | - | - | - | - | - | - | - | Unassigned | - |
| Otu60523 | 1 | k__Fungi(100);k__Fungi_unclassified(100);k__Fungi_unclassified(100);k__Fungi_unclassified(100);k__Fungi_unclassified(100);k__Fungi_unclassified(100);k__Fungi_unclassified(100); | - | - | - | - | - | - | - | Unassigned | - |
| Otu60524 | 1 | k__Fungi(100);k__Fungi_unclassified(100);k__Fungi_unclassified(100);k__Fungi_unclassified(100);k__Fungi_unclassified(100);k__Fungi_unclassified(100);k__Fungi_unclassified(100); | - | - | - | - | - | - | - | Unassigned | - |
| Otu60525 | 1 | k__Fungi(100);p__Ascomycota(100);p__Ascomycota_unclassified(100);p__Ascomycota_unclassified(100);p__Ascomycota_unclassified(100);p__Ascomycota_unclassified(100);p__Ascomycota_unclassified(100); | - | - | - | - | - | - | - | Unassigned | - |
| Otu60526 | 1 | k__Fungi(100);k__Fungi_unclassified(100);k__Fungi_unclassified(100);k__Fungi_unclassified(100);k__Fungi_unclassified(100);k__Fungi_unclassified(100);k__Fungi_unclassified(100); | - | - | - | - | - | - | - | Unassigned | - |
| Otu60527 | 1 | k__Fungi(100);k__Fungi_unclassified(100);k__Fungi_unclassified(100);k__Fungi_unclassified(100);k__Fungi_unclassified(100);k__Fungi_unclassified(100);k__Fungi_unclassified(100); | - | - | - | - | - | - | - | Unassigned | - |
| Otu60529 | 1 | k__Fungi(100);k__Fungi_unclassified(100);k__Fungi_unclassified(100);k__Fungi_unclassified(100);k__Fungi_unclassified(100);k__Fungi_unclassified(100);k__Fungi_unclassified(100); | - | - | - | - | - | - | - | Unassigned | - |
| Otu60530 | 1 | k__Fungi(100);k__Fungi_unclassified(100);k__Fungi_unclassified(100);k__Fungi_unclassified(100);k__Fungi_unclassified(100);k__Fungi_unclassified(100);k__Fungi_unclassified(100); | - | - | - | - | - | - | - | Unassigned | - |
| Otu60531 | 1 | k__Fungi(100);k__Fungi_unclassified(100);k__Fungi_unclassified(100);k__Fungi_unclassified(100);k__Fungi_unclassified(100);k__Fungi_unclassified(100);k__Fungi_unclassified(100); | - | - | - | - | - | - | - | Unassigned | - |
| Otu60532 | 1 | k__Fungi(100);k__Fungi_unclassified(100);k__Fungi_unclassified(100);k__Fungi_unclassified(100);k__Fungi_unclassified(100);k__Fungi_unclassified(100);k__Fungi_unclassified(100); | - | - | - | - | - | - | - | Unassigned | - |
| Otu60533 | 1 | k__Fungi(100);k__Fungi_unclassified(100);k__Fungi_unclassified(100);k__Fungi_unclassified(100);k__Fungi_unclassified(100);k__Fungi_unclassified(100);k__Fungi_unclassified(100); | - | - | - | - | - | - | - | Unassigned | - |
| Otu60534 | 1 | k__Fungi(100);k__Fungi_unclassified(100);k__Fungi_unclassified(100);k__Fungi_unclassified(100);k__Fungi_unclassified(100);k__Fungi_unclassified(100);k__Fungi_unclassified(100); | - | - | - | - | - | - | - | Unassigned | - |
| Otu60535 | 1 | k__Fungi(100);k__Fungi_unclassified(100);k__Fungi_unclassified(100);k__Fungi_unclassified(100);k__Fungi_unclassified(100);k__Fungi_unclassified(100);k__Fungi_unclassified(100); | - | - | - | - | - | - | - | Unassigned | - |
| Otu60536 | 1 | k__Fungi(100);p__Ascomycota(100);p__Ascomycota_unclassified(100);p__Ascomycota_unclassified(100);p__Ascomycota_unclassified(100);p__Ascomycota_unclassified(100);p__Ascomycota_unclassified(100); | - | - | - | - | - | - | - | Unassigned | - |
| Otu60537 | 1 | k__Fungi(100);p__Ascomycota(100);p__Ascomycota_unclassified(100);p__Ascomycota_unclassified(100);p__Ascomycota_unclassified(100);p__Ascomycota_unclassified(100);p__Ascomycota_unclassified(100); | - | - | - | - | - | - | - | Unassigned | - |
| Otu60538 | 1 | k__Fungi(100);k__Fungi_unclassified(100);k__Fungi_unclassified(100);k__Fungi_unclassified(100);k__Fungi_unclassified(100);k__Fungi_unclassified(100);k__Fungi_unclassified(100); | - | - | - | - | - | - | - | Unassigned | - |
| Otu60539 | 1 | k__Fungi(100);k__Fungi_unclassified(100);k__Fungi_unclassified(100);k__Fungi_unclassified(100);k__Fungi_unclassified(100);k__Fungi_unclassified(100);k__Fungi_unclassified(100); | - | - | - | - | - | - | - | Unassigned | - |
| Otu60540 | 1 | k__Fungi(100);k__Fungi_unclassified(100);k__Fungi_unclassified(100);k__Fungi_unclassified(100);k__Fungi_unclassified(100);k__Fungi_unclassified(100);k__Fungi_unclassified(100); | - | - | - | - | - | - | - | Unassigned | - |
| Otu60541 | 1 | k__Fungi(100);k__Fungi_unclassified(100);k__Fungi_unclassified(100);k__Fungi_unclassified(100);k__Fungi_unclassified(100);k__Fungi_unclassified(100);k__Fungi_unclassified(100); | - | - | - | - | - | - | - | Unassigned | - |
| Otu60542 | 1 | k__Fungi(100);k__Fungi_unclassified(100);k__Fungi_unclassified(100);k__Fungi_unclassified(100);k__Fungi_unclassified(100);k__Fungi_unclassified(100);k__Fungi_unclassified(100); | - | - | - | - | - | - | - | Unassigned | - |
| Otu60543 | 1 | k__Fungi(100);k__Fungi_unclassified(100);k__Fungi_unclassified(100);k__Fungi_unclassified(100);k__Fungi_unclassified(100);k__Fungi_unclassified(100);k__Fungi_unclassified(100); | - | - | - | - | - | - | - | Unassigned | - |
| Otu60544 | 1 | k__Fungi(100);p__Ascomycota(100);p__Ascomycota_unclassified(100);p__Ascomycota_unclassified(100);p__Ascomycota_unclassified(100);p__Ascomycota_unclassified(100);p__Ascomycota_unclassified(100); | - | - | - | - | - | - | - | Unassigned | - |
| Otu60545 | 1 | k__Fungi(100);p__Ascomycota(100);p__Ascomycota_unclassified(100);p__Ascomycota_unclassified(100);p__Ascomycota_unclassified(100);p__Ascomycota_unclassified(100);p__Ascomycota_unclassified(100); | - | - | - | - | - | - | - | Unassigned | - |
| Otu60546 | 1 | k__Fungi(100);k__Fungi_unclassified(100);k__Fungi_unclassified(100);k__Fungi_unclassified(100);k__Fungi_unclassified(100);k__Fungi_unclassified(100);k__Fungi_unclassified(100); | - | - | - | - | - | - | - | Unassigned | - |
| Otu60547 | 1 | k__Fungi(100);p__Ascomycota(100);p__Ascomycota_unclassified(100);p__Ascomycota_unclassified(100);p__Ascomycota_unclassified(100);p__Ascomycota_unclassified(100);p__Ascomycota_unclassified(100); | - | - | - | - | - | - | - | Unassigned | - |
| Otu60548 | 1 | k__Fungi(100);k__Fungi_unclassified(100);k__Fungi_unclassified(100);k__Fungi_unclassified(100);k__Fungi_unclassified(100);k__Fungi_unclassified(100);k__Fungi_unclassified(100); | - | - | - | - | - | - | - | Unassigned | - |
| Otu60549 | 1 | k__Fungi(100);k__Fungi_unclassified(100);k__Fungi_unclassified(100);k__Fungi_unclassified(100);k__Fungi_unclassified(100);k__Fungi_unclassified(100);k__Fungi_unclassified(100); | - | - | - | - | - | - | - | Unassigned | - |
| Otu60550 | 1 | k__Fungi(100);k__Fungi_unclassified(100);k__Fungi_unclassified(100);k__Fungi_unclassified(100);k__Fungi_unclassified(100);k__Fungi_unclassified(100);k__Fungi_unclassified(100); | - | - | - | - | - | - | - | Unassigned | - |
| Otu60551 | 1 | k__Fungi(100);p__Ascomycota(100);p__Ascomycota_unclassified(100);p__Ascomycota_unclassified(100);p__Ascomycota_unclassified(100);p__Ascomycota_unclassified(100);p__Ascomycota_unclassified(100); | - | - | - | - | - | - | - | Unassigned | - |
| Otu60552 | 1 | k__Fungi(100);k__Fungi_unclassified(100);k__Fungi_unclassified(100);k__Fungi_unclassified(100);k__Fungi_unclassified(100);k__Fungi_unclassified(100);k__Fungi_unclassified(100); | - | - | - | - | - | - | - | Unassigned | - |
| Otu60553 | 1 | k__Fungi(100);k__Fungi_unclassified(100);k__Fungi_unclassified(100);k__Fungi_unclassified(100);k__Fungi_unclassified(100);k__Fungi_unclassified(100);k__Fungi_unclassified(100); | - | - | - | - | - | - | - | Unassigned | - |
| Otu60554 | 1 | k__Fungi(100);k__Fungi_unclassified(100);k__Fungi_unclassified(100);k__Fungi_unclassified(100);k__Fungi_unclassified(100);k__Fungi_unclassified(100);k__Fungi_unclassified(100); | - | - | - | - | - | - | - | Unassigned | - |
| Otu60555 | 1 | k__Fungi(100);k__Fungi_unclassified(100);k__Fungi_unclassified(100);k__Fungi_unclassified(100);k__Fungi_unclassified(100);k__Fungi_unclassified(100);k__Fungi_unclassified(100); | - | - | - | - | - | - | - | Unassigned | - |
| Otu60556 | 1 | k__Fungi(100);k__Fungi_unclassified(100);k__Fungi_unclassified(100);k__Fungi_unclassified(100);k__Fungi_unclassified(100);k__Fungi_unclassified(100);k__Fungi_unclassified(100); | - | - | - | - | - | - | - | Unassigned | - |
| Otu60557 | 1 | k__Fungi(100);k__Fungi_unclassified(100);k__Fungi_unclassified(100);k__Fungi_unclassified(100);k__Fungi_unclassified(100);k__Fungi_unclassified(100);k__Fungi_unclassified(100); | - | - | - | - | - | - | - | Unassigned | - |
| Otu60558 | 1 | k__Fungi(100);k__Fungi_unclassified(100);k__Fungi_unclassified(100);k__Fungi_unclassified(100);k__Fungi_unclassified(100);k__Fungi_unclassified(100);k__Fungi_unclassified(100); | - | - | - | - | - | - | - | Unassigned | - |
| Otu60559 | 1 | k__Fungi(100);k__Fungi_unclassified(100);k__Fungi_unclassified(100);k__Fungi_unclassified(100);k__Fungi_unclassified(100);k__Fungi_unclassified(100);k__Fungi_unclassified(100); | - | - | - | - | - | - | - | Unassigned | - |
| Otu60560 | 1 | k__Fungi(100);k__Fungi_unclassified(100);k__Fungi_unclassified(100);k__Fungi_unclassified(100);k__Fungi_unclassified(100);k__Fungi_unclassified(100);k__Fungi_unclassified(100); | - | - | - | - | - | - | - | Unassigned | - |
| Otu60561 | 1 | k__Fungi(100);k__Fungi_unclassified(100);k__Fungi_unclassified(100);k__Fungi_unclassified(100);k__Fungi_unclassified(100);k__Fungi_unclassified(100);k__Fungi_unclassified(100); | - | - | - | - | - | - | - | Unassigned | - |
| Otu60562 | 1 | k__Fungi(100);k__Fungi_unclassified(100);k__Fungi_unclassified(100);k__Fungi_unclassified(100);k__Fungi_unclassified(100);k__Fungi_unclassified(100);k__Fungi_unclassified(100); | - | - | - | - | - | - | - | Unassigned | - |
| Otu60563 | 1 | k__Fungi(100);k__Fungi_unclassified(100);k__Fungi_unclassified(100);k__Fungi_unclassified(100);k__Fungi_unclassified(100);k__Fungi_unclassified(100);k__Fungi_unclassified(100); | - | - | - | - | - | - | - | Unassigned | - |
| Otu60564 | 1 | k__Fungi(100);k__Fungi_unclassified(100);k__Fungi_unclassified(100);k__Fungi_unclassified(100);k__Fungi_unclassified(100);k__Fungi_unclassified(100);k__Fungi_unclassified(100); | - | - | - | - | - | - | - | Unassigned | - |
| Otu60565 | 1 | k__Fungi(100);p__Ascomycota(100);p__Ascomycota_unclassified(100);p__Ascomycota_unclassified(100);p__Ascomycota_unclassified(100);p__Ascomycota_unclassified(100);p__Ascomycota_unclassified(100); | - | - | - | - | - | - | - | Unassigned | - |
| Otu60566 | 1 | k__Fungi(100);k__Fungi_unclassified(100);k__Fungi_unclassified(100);k__Fungi_unclassified(100);k__Fungi_unclassified(100);k__Fungi_unclassified(100);k__Fungi_unclassified(100); | - | - | - | - | - | - | - | Unassigned | - |
| Otu60567 | 1 | k__Fungi(100);k__Fungi_unclassified(100);k__Fungi_unclassified(100);k__Fungi_unclassified(100);k__Fungi_unclassified(100);k__Fungi_unclassified(100);k__Fungi_unclassified(100); | - | - | - | - | - | - | - | Unassigned | - |
| Otu60569 | 1 | k__Fungi(100);k__Fungi_unclassified(100);k__Fungi_unclassified(100);k__Fungi_unclassified(100);k__Fungi_unclassified(100);k__Fungi_unclassified(100);k__Fungi_unclassified(100); | - | - | - | - | - | - | - | Unassigned | - |
| Otu60570 | 1 | k__Fungi(100);k__Fungi_unclassified(100);k__Fungi_unclassified(100);k__Fungi_unclassified(100);k__Fungi_unclassified(100);k__Fungi_unclassified(100);k__Fungi_unclassified(100); | - | - | - | - | - | - | - | Unassigned | - |
| Otu60571 | 1 | k__Fungi(100);k__Fungi_unclassified(100);k__Fungi_unclassified(100);k__Fungi_unclassified(100);k__Fungi_unclassified(100);k__Fungi_unclassified(100);k__Fungi_unclassified(100); | - | - | - | - | - | - | - | Unassigned | - |
| Otu60572 | 1 | k__Fungi(100);p__Ascomycota(100);p__Ascomycota_unclassified(100);p__Ascomycota_unclassified(100);p__Ascomycota_unclassified(100);p__Ascomycota_unclassified(100);p__Ascomycota_unclassified(100); | - | - | - | - | - | - | - | Unassigned | - |
| Otu60573 | 1 | k__Fungi(100);k__Fungi_unclassified(100);k__Fungi_unclassified(100);k__Fungi_unclassified(100);k__Fungi_unclassified(100);k__Fungi_unclassified(100);k__Fungi_unclassified(100); | - | - | - | - | - | - | - | Unassigned | - |
| Otu60574 | 1 | k__Fungi(100);k__Fungi_unclassified(100);k__Fungi_unclassified(100);k__Fungi_unclassified(100);k__Fungi_unclassified(100);k__Fungi_unclassified(100);k__Fungi_unclassified(100); | - | - | - | - | - | - | - | Unassigned | - |
| Otu60575 | 1 | k__Fungi(100);k__Fungi_unclassified(100);k__Fungi_unclassified(100);k__Fungi_unclassified(100);k__Fungi_unclassified(100);k__Fungi_unclassified(100);k__Fungi_unclassified(100); | - | - | - | - | - | - | - | Unassigned | - |
| Otu60576 | 1 | k__Fungi(100);k__Fungi_unclassified(100);k__Fungi_unclassified(100);k__Fungi_unclassified(100);k__Fungi_unclassified(100);k__Fungi_unclassified(100);k__Fungi_unclassified(100); | - | - | - | - | - | - | - | Unassigned | - |
| Otu60577 | 1 | k__Fungi(100);p__Ascomycota(100);p__Ascomycota_unclassified(100);p__Ascomycota_unclassified(100);p__Ascomycota_unclassified(100);p__Ascomycota_unclassified(100);p__Ascomycota_unclassified(100); | - | - | - | - | - | - | - | Unassigned | - |
| Otu60578 | 1 | k__Fungi(100);p__Ascomycota(100);p__Ascomycota_unclassified(100);p__Ascomycota_unclassified(100);p__Ascomycota_unclassified(100);p__Ascomycota_unclassified(100);p__Ascomycota_unclassified(100); | - | - | - | - | - | - | - | Unassigned | - |
| Otu60579 | 1 | k__Fungi(100);k__Fungi_unclassified(100);k__Fungi_unclassified(100);k__Fungi_unclassified(100);k__Fungi_unclassified(100);k__Fungi_unclassified(100);k__Fungi_unclassified(100); | - | - | - | - | - | - | - | Unassigned | - |
| Otu60580 | 1 | k__Fungi(100);p__Ascomycota(100);p__Ascomycota_unclassified(100);p__Ascomycota_unclassified(100);p__Ascomycota_unclassified(100);p__Ascomycota_unclassified(100);p__Ascomycota_unclassified(100); | - | - | - | - | - | - | - | Unassigned | - |
| Otu60583 | 1 | k__Fungi(100);p__Ascomycota(100);p__Ascomycota_unclassified(100);p__Ascomycota_unclassified(100);p__Ascomycota_unclassified(100);p__Ascomycota_unclassified(100);p__Ascomycota_unclassified(100); | - | - | - | - | - | - | - | Unassigned | - |
| Otu60584 | 1 | k__Fungi(100);p__Ascomycota(100);p__Ascomycota_unclassified(100);p__Ascomycota_unclassified(100);p__Ascomycota_unclassified(100);p__Ascomycota_unclassified(100);p__Ascomycota_unclassified(100); | - | - | - | - | - | - | - | Unassigned | - |
| Otu60585 | 1 | k__Fungi(100);p__Ascomycota(100);p__Ascomycota_unclassified(100);p__Ascomycota_unclassified(100);p__Ascomycota_unclassified(100);p__Ascomycota_unclassified(100);p__Ascomycota_unclassified(100); | - | - | - | - | - | - | - | Unassigned | - |
| Otu60586 | 1 | k__Fungi(100);k__Fungi_unclassified(100);k__Fungi_unclassified(100);k__Fungi_unclassified(100);k__Fungi_unclassified(100);k__Fungi_unclassified(100);k__Fungi_unclassified(100); | - | - | - | - | - | - | - | Unassigned | - |
| Otu60587 | 1 | k__Fungi(100);p__Ascomycota(100);p__Ascomycota_unclassified(100);p__Ascomycota_unclassified(100);p__Ascomycota_unclassified(100);p__Ascomycota_unclassified(100);p__Ascomycota_unclassified(100); | - | - | - | - | - | - | - | Unassigned | - |
| Otu60588 | 1 | k__Fungi(100);k__Fungi_unclassified(100);k__Fungi_unclassified(100);k__Fungi_unclassified(100);k__Fungi_unclassified(100);k__Fungi_unclassified(100);k__Fungi_unclassified(100); | - | - | - | - | - | - | - | Unassigned | - |
| Otu60589 | 1 | k__Fungi(100);p__Ascomycota(100);p__Ascomycota_unclassified(100);p__Ascomycota_unclassified(100);p__Ascomycota_unclassified(100);p__Ascomycota_unclassified(100);p__Ascomycota_unclassified(100); | - | - | - | - | - | - | - | Unassigned | - |
| Otu60590 | 1 | k__Fungi(100);k__Fungi_unclassified(100);k__Fungi_unclassified(100);k__Fungi_unclassified(100);k__Fungi_unclassified(100);k__Fungi_unclassified(100);k__Fungi_unclassified(100); | - | - | - | - | - | - | - | Unassigned | - |
| Otu60591 | 1 | k__Fungi(100);k__Fungi_unclassified(100);k__Fungi_unclassified(100);k__Fungi_unclassified(100);k__Fungi_unclassified(100);k__Fungi_unclassified(100);k__Fungi_unclassified(100); | - | - | - | - | - | - | - | Unassigned | - |
| Otu60592 | 1 | k__Fungi(100);p__Ascomycota(100);p__Ascomycota_unclassified(100);p__Ascomycota_unclassified(100);p__Ascomycota_unclassified(100);p__Ascomycota_unclassified(100);p__Ascomycota_unclassified(100); | - | - | - | - | - | - | - | Unassigned | - |
| Otu60593 | 1 | k__Fungi(100);k__Fungi_unclassified(100);k__Fungi_unclassified(100);k__Fungi_unclassified(100);k__Fungi_unclassified(100);k__Fungi_unclassified(100);k__Fungi_unclassified(100); | - | - | - | - | - | - | - | Unassigned | - |
| Otu60594 | 1 | k__Fungi(100);k__Fungi_unclassified(100);k__Fungi_unclassified(100);k__Fungi_unclassified(100);k__Fungi_unclassified(100);k__Fungi_unclassified(100);k__Fungi_unclassified(100); | - | - | - | - | - | - | - | Unassigned | - |
| Otu60595 | 1 | k__Fungi(100);k__Fungi_unclassified(100);k__Fungi_unclassified(100);k__Fungi_unclassified(100);k__Fungi_unclassified(100);k__Fungi_unclassified(100);k__Fungi_unclassified(100); | - | - | - | - | - | - | - | Unassigned | - |
| Otu60596 | 1 | k__Fungi(100);k__Fungi_unclassified(100);k__Fungi_unclassified(100);k__Fungi_unclassified(100);k__Fungi_unclassified(100);k__Fungi_unclassified(100);k__Fungi_unclassified(100); | - | - | - | - | - | - | - | Unassigned | - |
| Otu60602 | 1 | k__Fungi(100);k__Fungi_unclassified(100);k__Fungi_unclassified(100);k__Fungi_unclassified(100);k__Fungi_unclassified(100);k__Fungi_unclassified(100);k__Fungi_unclassified(100); | - | - | - | - | - | - | - | Unassigned | - |
| Otu60603 | 1 | k__Fungi(100);k__Fungi_unclassified(100);k__Fungi_unclassified(100);k__Fungi_unclassified(100);k__Fungi_unclassified(100);k__Fungi_unclassified(100);k__Fungi_unclassified(100); | - | - | - | - | - | - | - | Unassigned | - |
| Otu60605 | 1 | k__Fungi(100);p__Ascomycota(100);p__Ascomycota_unclassified(100);p__Ascomycota_unclassified(100);p__Ascomycota_unclassified(100);p__Ascomycota_unclassified(100);p__Ascomycota_unclassified(100); | - | - | - | - | - | - | - | Unassigned | - |
| Otu60613 | 1 | k__Fungi(100);p__Ascomycota(100);p__Ascomycota_unclassified(100);p__Ascomycota_unclassified(100);p__Ascomycota_unclassified(100);p__Ascomycota_unclassified(100);p__Ascomycota_unclassified(100); | - | - | - | - | - | - | - | Unassigned | - |
| Otu60617 | 1 | k__Fungi(100);p__Ascomycota(100);p__Ascomycota_unclassified(100);p__Ascomycota_unclassified(100);p__Ascomycota_unclassified(100);p__Ascomycota_unclassified(100);p__Ascomycota_unclassified(100); | - | - | - | - | - | - | - | Unassigned | - |
| Otu60618 | 1 | k__Fungi(100);k__Fungi_unclassified(100);k__Fungi_unclassified(100);k__Fungi_unclassified(100);k__Fungi_unclassified(100);k__Fungi_unclassified(100);k__Fungi_unclassified(100); | - | - | - | - | - | - | - | Unassigned | - |
| Otu60620 | 1 | k__Fungi(100);p__Ascomycota(100);p__Ascomycota_unclassified(100);p__Ascomycota_unclassified(100);p__Ascomycota_unclassified(100);p__Ascomycota_unclassified(100);p__Ascomycota_unclassified(100); | - | - | - | - | - | - | - | Unassigned | - |
| Otu60621 | 1 | k__Fungi(100);p__Ascomycota(100);p__Ascomycota_unclassified(100);p__Ascomycota_unclassified(100);p__Ascomycota_unclassified(100);p__Ascomycota_unclassified(100);p__Ascomycota_unclassified(100); | - | - | - | - | - | - | - | Unassigned | - |
| Otu60622 | 1 | k__Fungi(100);k__Fungi_unclassified(100);k__Fungi_unclassified(100);k__Fungi_unclassified(100);k__Fungi_unclassified(100);k__Fungi_unclassified(100);k__Fungi_unclassified(100); | - | - | - | - | - | - | - | Unassigned | - |
| Otu60623 | 1 | k__Fungi(100);k__Fungi_unclassified(100);k__Fungi_unclassified(100);k__Fungi_unclassified(100);k__Fungi_unclassified(100);k__Fungi_unclassified(100);k__Fungi_unclassified(100); | - | - | - | - | - | - | - | Unassigned | - |
| Otu60624 | 1 | k__Fungi(100);p__Ascomycota(100);p__Ascomycota_unclassified(100);p__Ascomycota_unclassified(100);p__Ascomycota_unclassified(100);p__Ascomycota_unclassified(100);p__Ascomycota_unclassified(100); | - | - | - | - | - | - | - | Unassigned | - |
| Otu60625 | 1 | k__Fungi(100);k__Fungi_unclassified(100);k__Fungi_unclassified(100);k__Fungi_unclassified(100);k__Fungi_unclassified(100);k__Fungi_unclassified(100);k__Fungi_unclassified(100); | - | - | - | - | - | - | - | Unassigned | - |

| | | | | | | | | | | | |
|---|---|---|---|---|---|---|---|---|---|---|---|
| Otu60626 | 1 | k__Fungi(100);p__Ascomycota(100);p__Ascomycota_unclassified(100);p__Ascomycota_unclassified(100);p__Ascomycota_unclassified(100);p__Ascomycota_unclassified(100);p__Ascomycota_unclassified(100); | - | - | - | - | - | - | - | Unassigned | - |
| Otu60627 | 1 | k__Fungi(100);k__Fungi_unclassified(100);k__Fungi_unclassified(100);k__Fungi_unclassified(100);k__Fungi_unclassified(100);k__Fungi_unclassified(100);k__Fungi_unclassified(100); | - | - | - | - | - | - | - | Unassigned | - |
| Otu60628 | 1 | k__Fungi(100);p__Ascomycota(100);p__Ascomycota_unclassified(100);p__Ascomycota_unclassified(100);p__Ascomycota_unclassified(100);p__Ascomycota_unclassified(100);p__Ascomycota_unclassified(100); | - | - | - | - | - | - | - | Unassigned | - |
| Otu60629 | 1 | k__Fungi(100);k__Fungi_unclassified(100);k__Fungi_unclassified(100);k__Fungi_unclassified(100);k__Fungi_unclassified(100);k__Fungi_unclassified(100);k__Fungi_unclassified(100); | - | - | - | - | - | - | - | Unassigned | - |
| Otu60630 | 1 | k__Fungi(100);k__Fungi_unclassified(100);k__Fungi_unclassified(100);k__Fungi_unclassified(100);k__Fungi_unclassified(100);k__Fungi_unclassified(100);k__Fungi_unclassified(100); | - | - | - | - | - | - | - | Unassigned | - |
| Otu60631 | 1 | k__Fungi(100);k__Fungi_unclassified(100);k__Fungi_unclassified(100);k__Fungi_unclassified(100);k__Fungi_unclassified(100);k__Fungi_unclassified(100);k__Fungi_unclassified(100); | - | - | - | - | - | - | - | Unassigned | - |
| Otu60632 | 1 | k__Fungi(100);p__Ascomycota(100);p__Ascomycota_unclassified(100);p__Ascomycota_unclassified(100);p__Ascomycota_unclassified(100);p__Ascomycota_unclassified(100);p__Ascomycota_unclassified(100); | - | - | - | - | - | - | - | Unassigned | - |
| Otu60633 | 1 | k__Fungi(100);p__Ascomycota(100);p__Ascomycota_unclassified(100);p__Ascomycota_unclassified(100);p__Ascomycota_unclassified(100);p__Ascomycota_unclassified(100);p__Ascomycota_unclassified(100); | - | - | - | - | - | - | - | Unassigned | - |
| Otu60634 | 1 | k__Fungi(100);p__Ascomycota(100);p__Ascomycota_unclassified(100);p__Ascomycota_unclassified(100);p__Ascomycota_unclassified(100);p__Ascomycota_unclassified(100);p__Ascomycota_unclassified(100); | - | - | - | - | - | - | - | Unassigned | - |
| Otu60635 | 1 | k__Fungi(100);k__Fungi_unclassified(100);k__Fungi_unclassified(100);k__Fungi_unclassified(100);k__Fungi_unclassified(100);k__Fungi_unclassified(100);k__Fungi_unclassified(100); | - | - | - | - | - | - | - | Unassigned | - |
| Otu60636 | 1 | k__Fungi(100);p__Ascomycota(100);p__Ascomycota_unclassified(100);p__Ascomycota_unclassified(100);p__Ascomycota_unclassified(100);p__Ascomycota_unclassified(100);p__Ascomycota_unclassified(100); | - | - | - | - | - | - | - | Unassigned | - |
| Otu60637 | 1 | k__Fungi(100);k__Fungi_unclassified(100);k__Fungi_unclassified(100);k__Fungi_unclassified(100);k__Fungi_unclassified(100);k__Fungi_unclassified(100);k__Fungi_unclassified(100); | - | - | - | - | - | - | - | Unassigned | - |
| Otu60638 | 1 | k__Fungi(100);p__Ascomycota(100);p__Ascomycota_unclassified(100);p__Ascomycota_unclassified(100);p__Ascomycota_unclassified(100);p__Ascomycota_unclassified(100);p__Ascomycota_unclassified(100); | - | - | - | - | - | - | - | Unassigned | - |
| Otu60640 | 1 | k__Fungi(100);k__Fungi_unclassified(100);k__Fungi_unclassified(100);k__Fungi_unclassified(100);k__Fungi_unclassified(100);k__Fungi_unclassified(100);k__Fungi_unclassified(100); | - | - | - | - | - | - | - | Unassigned | - |
| Otu60641 | 1 | k__Fungi(100);k__Fungi_unclassified(100);k__Fungi_unclassified(100);k__Fungi_unclassified(100);k__Fungi_unclassified(100);k__Fungi_unclassified(100);k__Fungi_unclassified(100); | - | - | - | - | - | - | - | Unassigned | - |
| Otu60642 | 1 | k__Fungi(100);k__Fungi_unclassified(100);k__Fungi_unclassified(100);k__Fungi_unclassified(100);k__Fungi_unclassified(100);k__Fungi_unclassified(100);k__Fungi_unclassified(100); | - | - | - | - | - | - | - | Unassigned | - |
| Otu60643 | 1 | k__Fungi(100);k__Fungi_unclassified(100);k__Fungi_unclassified(100);k__Fungi_unclassified(100);k__Fungi_unclassified(100);k__Fungi_unclassified(100);k__Fungi_unclassified(100); | - | - | - | - | - | - | - | Unassigned | - |
| Otu60644 | 1 | k__Fungi(100);p__Ascomycota(100);p__Ascomycota_unclassified(100);p__Ascomycota_unclassified(100);p__Ascomycota_unclassified(100);p__Ascomycota_unclassified(100);p__Ascomycota_unclassified(100); | - | - | - | - | - | - | - | Unassigned | - |
| Otu60645 | 1 | k__Fungi(100);p__Ascomycota(100);p__Ascomycota_unclassified(100);p__Ascomycota_unclassified(100);p__Ascomycota_unclassified(100);p__Ascomycota_unclassified(100);p__Ascomycota_unclassified(100); | - | - | - | - | - | - | - | Unassigned | - |
| Otu60647 | 1 | k__Fungi(100);k__Fungi_unclassified(100);k__Fungi_unclassified(100);k__Fungi_unclassified(100);k__Fungi_unclassified(100);k__Fungi_unclassified(100);k__Fungi_unclassified(100); | - | - | - | - | - | - | - | Unassigned | - |
| Otu60651 | 1 | k__Fungi(100);p__Ascomycota(100);p__Ascomycota_unclassified(100);p__Ascomycota_unclassified(100);p__Ascomycota_unclassified(100);p__Ascomycota_unclassified(100);p__Ascomycota_unclassified(100); | - | - | - | - | - | - | - | Unassigned | - |
| Otu60652 | 1 | k__Fungi(100);k__Fungi_unclassified(100);k__Fungi_unclassified(100);k__Fungi_unclassified(100);k__Fungi_unclassified(100);k__Fungi_unclassified(100);k__Fungi_unclassified(100); | - | - | - | - | - | - | - | Unassigned | - |
| Otu60653 | 1 | k__Fungi(100);k__Fungi_unclassified(100);k__Fungi_unclassified(100);k__Fungi_unclassified(100);k__Fungi_unclassified(100);k__Fungi_unclassified(100);k__Fungi_unclassified(100); | - | - | - | - | - | - | - | Unassigned | - |
| Otu60654 | 1 | k__Fungi(100);p__Ascomycota(100);p__Ascomycota_unclassified(100);p__Ascomycota_unclassified(100);p__Ascomycota_unclassified(100);p__Ascomycota_unclassified(100);p__Ascomycota_unclassified(100); | - | - | - | - | - | - | - | Unassigned | - |
| Otu60655 | 1 | k__Fungi(100);k__Fungi_unclassified(100);k__Fungi_unclassified(100);k__Fungi_unclassified(100);k__Fungi_unclassified(100);k__Fungi_unclassified(100);k__Fungi_unclassified(100); | - | - | - | - | - | - | - | Unassigned | - |
| Otu60656 | 1 | k__Fungi(100);k__Fungi_unclassified(100);k__Fungi_unclassified(100);k__Fungi_unclassified(100);k__Fungi_unclassified(100);k__Fungi_unclassified(100);k__Fungi_unclassified(100); | - | - | - | - | - | - | - | Unassigned | - |
| Otu60657 | 1 | k__Fungi(100);p__Ascomycota(100);p__Ascomycota_unclassified(100);p__Ascomycota_unclassified(100);p__Ascomycota_unclassified(100);p__Ascomycota_unclassified(100);p__Ascomycota_unclassified(100); | - | - | - | - | - | - | - | Unassigned | - |
| Otu60658 | 1 | k__Fungi(100);k__Fungi_unclassified(100);k__Fungi_unclassified(100);k__Fungi_unclassified(100);k__Fungi_unclassified(100);k__Fungi_unclassified(100);k__Fungi_unclassified(100); | - | - | - | - | - | - | - | Unassigned | - |
| Otu60659 | 1 | k__Fungi(100);k__Fungi_unclassified(100);k__Fungi_unclassified(100);k__Fungi_unclassified(100);k__Fungi_unclassified(100);k__Fungi_unclassified(100);k__Fungi_unclassified(100); | - | - | - | - | - | - | - | Unassigned | - |
| Otu60660 | 1 | k__Fungi(100);k__Fungi_unclassified(100);k__Fungi_unclassified(100);k__Fungi_unclassified(100);k__Fungi_unclassified(100);k__Fungi_unclassified(100);k__Fungi_unclassified(100); | - | - | - | - | - | - | - | Unassigned | - |
| Otu60661 | 1 | k__Fungi(100);k__Fungi_unclassified(100);k__Fungi_unclassified(100);k__Fungi_unclassified(100);k__Fungi_unclassified(100);k__Fungi_unclassified(100);k__Fungi_unclassified(100); | - | - | - | - | - | - | - | Unassigned | - |
| Otu60662 | 1 | k__Fungi(100);k__Fungi_unclassified(100);k__Fungi_unclassified(100);k__Fungi_unclassified(100);k__Fungi_unclassified(100);k__Fungi_unclassified(100);k__Fungi_unclassified(100); | - | - | - | - | - | - | - | Unassigned | - |
| Otu60663 | 1 | k__Fungi(100);k__Fungi_unclassified(100);k__Fungi_unclassified(100);k__Fungi_unclassified(100);k__Fungi_unclassified(100);k__Fungi_unclassified(100);k__Fungi_unclassified(100); | - | - | - | - | - | - | - | Unassigned | - |
| Otu60664 | 1 | k__Fungi(100);p__Ascomycota(100);p__Ascomycota_unclassified(100);p__Ascomycota_unclassified(100);p__Ascomycota_unclassified(100);p__Ascomycota_unclassified(100);p__Ascomycota_unclassified(100); | - | - | - | - | - | - | - | Unassigned | - |
| Otu60665 | 1 | k__Fungi(100);k__Fungi_unclassified(100);k__Fungi_unclassified(100);k__Fungi_unclassified(100);k__Fungi_unclassified(100);k__Fungi_unclassified(100);k__Fungi_unclassified(100); | - | - | - | - | - | - | - | Unassigned | - |
| Otu60666 | 1 | k__Fungi(100);p__Ascomycota(100);p__Ascomycota_unclassified(100);p__Ascomycota_unclassified(100);p__Ascomycota_unclassified(100);p__Ascomycota_unclassified(100);p__Ascomycota_unclassified(100); | - | - | - | - | - | - | - | Unassigned | - |
| Otu60667 | 1 | k__Fungi(100);k__Fungi_unclassified(100);k__Fungi_unclassified(100);k__Fungi_unclassified(100);k__Fungi_unclassified(100);k__Fungi_unclassified(100);k__Fungi_unclassified(100); | - | - | - | - | - | - | - | Unassigned | - |
| Otu60668 | 1 | k__Fungi(100);p__Ascomycota(100);p__Ascomycota_unclassified(100);p__Ascomycota_unclassified(100);p__Ascomycota_unclassified(100);p__Ascomycota_unclassified(100);p__Ascomycota_unclassified(100); | - | - | - | - | - | - | - | Unassigned | - |
| Otu60669 | 1 | k__Fungi(100);p__Ascomycota(100);p__Ascomycota_unclassified(100);p__Ascomycota_unclassified(100);p__Ascomycota_unclassified(100);p__Ascomycota_unclassified(100);p__Ascomycota_unclassified(100); | - | - | - | - | - | - | - | Unassigned | - |
| Otu60670 | 1 | k__Fungi(100);k__Fungi_unclassified(100);k__Fungi_unclassified(100);k__Fungi_unclassified(100);k__Fungi_unclassified(100);k__Fungi_unclassified(100);k__Fungi_unclassified(100); | - | - | - | - | - | - | - | Unassigned | - |
| Otu60671 | 1 | k__Fungi(100);p__Ascomycota(100);p__Ascomycota_unclassified(100);p__Ascomycota_unclassified(100);p__Ascomycota_unclassified(100);p__Ascomycota_unclassified(100);p__Ascomycota_unclassified(100); | - | - | - | - | - | - | - | Unassigned | - |
| Otu60672 | 1 | k__Fungi(100);p__Ascomycota(100);p__Ascomycota_unclassified(100);p__Ascomycota_unclassified(100);p__Ascomycota_unclassified(100);p__Ascomycota_unclassified(100);p__Ascomycota_unclassified(100); | - | - | - | - | - | - | - | Unassigned | - |
| Otu60673 | 1 | k__Fungi(100);k__Fungi_unclassified(100);k__Fungi_unclassified(100);k__Fungi_unclassified(100);k__Fungi_unclassified(100);k__Fungi_unclassified(100);k__Fungi_unclassified(100); | - | - | - | - | - | - | - | Unassigned | - |
| Otu60674 | 1 | k__Fungi(100);p__Ascomycota(100);p__Ascomycota_unclassified(100);p__Ascomycota_unclassified(100);p__Ascomycota_unclassified(100);p__Ascomycota_unclassified(100);p__Ascomycota_unclassified(100); | - | - | - | - | - | - | - | Unassigned | - |
| Otu60675 | 1 | k__Fungi(100);k__Fungi_unclassified(100);k__Fungi_unclassified(100);k__Fungi_unclassified(100);k__Fungi_unclassified(100);k__Fungi_unclassified(100);k__Fungi_unclassified(100); | - | - | - | - | - | - | - | Unassigned | - |
| Otu60676 | 1 | k__Fungi(100);k__Fungi_unclassified(100);k__Fungi_unclassified(100);k__Fungi_unclassified(100);k__Fungi_unclassified(100);k__Fungi_unclassified(100);k__Fungi_unclassified(100); | - | - | - | - | - | - | - | Unassigned | - |
| Otu60677 | 1 | k__Fungi(100);p__Ascomycota(100);p__Ascomycota_unclassified(100);p__Ascomycota_unclassified(100);p__Ascomycota_unclassified(100);p__Ascomycota_unclassified(100);p__Ascomycota_unclassified(100); | - | - | - | - | - | - | - | Unassigned | - |
| Otu60678 | 1 | k__Fungi(100);p__Ascomycota(100);p__Ascomycota_unclassified(100);p__Ascomycota_unclassified(100);p__Ascomycota_unclassified(100);p__Ascomycota_unclassified(100);p__Ascomycota_unclassified(100); | - | - | - | - | - | - | - | Unassigned | - |
| Otu60680 | 1 | k__Fungi(100);k__Fungi_unclassified(100);k__Fungi_unclassified(100);k__Fungi_unclassified(100);k__Fungi_unclassified(100);k__Fungi_unclassified(100);k__Fungi_unclassified(100); | - | - | - | - | - | - | - | Unassigned | - |
| Otu60681 | 1 | k__Fungi(100);k__Fungi_unclassified(100);k__Fungi_unclassified(100);k__Fungi_unclassified(100);k__Fungi_unclassified(100);k__Fungi_unclassified(100);k__Fungi_unclassified(100); | - | - | - | - | - | - | - | Unassigned | - |
| Otu60682 | 1 | k__Fungi(100);k__Fungi_unclassified(100);k__Fungi_unclassified(100);k__Fungi_unclassified(100);k__Fungi_unclassified(100);k__Fungi_unclassified(100);k__Fungi_unclassified(100); | - | - | - | - | - | - | - | Unassigned | - |
| Otu60685 | 1 | k__Fungi(100);k__Fungi_unclassified(100);k__Fungi_unclassified(100);k__Fungi_unclassified(100);k__Fungi_unclassified(100);k__Fungi_unclassified(100);k__Fungi_unclassified(100); | - | - | - | - | - | - | - | Unassigned | - |
| Otu60688 | 1 | k__Fungi(100);p__Ascomycota(100);p__Ascomycota_unclassified(100);p__Ascomycota_unclassified(100);p__Ascomycota_unclassified(100);p__Ascomycota_unclassified(100);p__Ascomycota_unclassified(100); | - | - | - | - | - | - | - | Unassigned | - |
| Otu60691 | 1 | k__Fungi(100);p__Ascomycota(100);p__Ascomycota_unclassified(100);p__Ascomycota_unclassified(100);p__Ascomycota_unclassified(100);p__Ascomycota_unclassified(100);p__Ascomycota_unclassified(100); | - | - | - | - | - | - | - | Unassigned | - |
| Otu60692 | 1 | k__Fungi(100);p__Ascomycota(100);p__Ascomycota_unclassified(100);p__Ascomycota_unclassified(100);p__Ascomycota_unclassified(100);p__Ascomycota_unclassified(100);p__Ascomycota_unclassified(100); | - | - | - | - | - | - | - | Unassigned | - |
| Otu60693 | 1 | k__Fungi(100);k__Fungi_unclassified(100);k__Fungi_unclassified(100);k__Fungi_unclassified(100);k__Fungi_unclassified(100);k__Fungi_unclassified(100);k__Fungi_unclassified(100); | - | - | - | - | - | - | - | Unassigned | - |
| Otu60694 | 1 | k__Fungi(100);k__Fungi_unclassified(100);k__Fungi_unclassified(100);k__Fungi_unclassified(100);k__Fungi_unclassified(100);k__Fungi_unclassified(100);k__Fungi_unclassified(100); | - | - | - | - | - | - | - | Unassigned | - |
| Otu60695 | 1 | k__Fungi(100);p__Ascomycota(100);p__Ascomycota_unclassified(100);p__Ascomycota_unclassified(100);p__Ascomycota_unclassified(100);p__Ascomycota_unclassified(100);p__Ascomycota_unclassified(100); | - | - | - | - | - | - | - | Unassigned | - |
| Otu60696 | 1 | k__Fungi(100);p__Ascomycota(100);p__Ascomycota_unclassified(100);p__Ascomycota_unclassified(100);p__Ascomycota_unclassified(100);p__Ascomycota_unclassified(100);p__Ascomycota_unclassified(100); | - | - | - | - | - | - | - | Unassigned | - |
| Otu60697 | 1 | k__Fungi(100);p__Ascomycota(100);p__Ascomycota_unclassified(100);p__Ascomycota_unclassified(100);p__Ascomycota_unclassified(100);p__Ascomycota_unclassified(100);p__Ascomycota_unclassified(100); | - | - | - | - | - | - | - | Unassigned | - |
| Otu60698 | 1 | k__Fungi(100);p__Ascomycota(100);p__Ascomycota_unclassified(100);p__Ascomycota_unclassified(100);p__Ascomycota_unclassified(100);p__Ascomycota_unclassified(100);p__Ascomycota_unclassified(100); | - | - | - | - | - | - | - | Unassigned | - |
| Otu60699 | 1 | k__Fungi(100);k__Fungi_unclassified(100);k__Fungi_unclassified(100);k__Fungi_unclassified(100);k__Fungi_unclassified(100);k__Fungi_unclassified(100);k__Fungi_unclassified(100); | - | - | - | - | - | - | - | Unassigned | - |
| Otu60700 | 1 | k__Fungi(100);k__Fungi_unclassified(100);k__Fungi_unclassified(100);k__Fungi_unclassified(100);k__Fungi_unclassified(100);k__Fungi_unclassified(100);k__Fungi_unclassified(100); | - | - | - | - | - | - | - | Unassigned | - |
| Otu60701 | 1 | k__Fungi(100);k__Fungi_unclassified(100);k__Fungi_unclassified(100);k__Fungi_unclassified(100);k__Fungi_unclassified(100);k__Fungi_unclassified(100);k__Fungi_unclassified(100); | - | - | - | - | - | - | - | Unassigned | - |
| Otu60702 | 1 | k__Fungi(100);p__Ascomycota(100);p__Ascomycota_unclassified(100);p__Ascomycota_unclassified(100);p__Ascomycota_unclassified(100);p__Ascomycota_unclassified(100);p__Ascomycota_unclassified(100); | - | - | - | - | - | - | - | Unassigned | - |
| Otu60703 | 1 | k__Fungi(100);k__Fungi_unclassified(100);k__Fungi_unclassified(100);k__Fungi_unclassified(100);k__Fungi_unclassified(100);k__Fungi_unclassified(100);k__Fungi_unclassified(100); | - | - | - | - | - | - | - | Unassigned | - |
| Otu60704 | 1 | k__Fungi(100);p__Ascomycota(100);p__Ascomycota_unclassified(100);p__Ascomycota_unclassified(100);p__Ascomycota_unclassified(100);p__Ascomycota_unclassified(100);p__Ascomycota_unclassified(100); | - | - | - | - | - | - | - | Unassigned | - |
| Otu60707 | 1 | k__Fungi(100);k__Fungi_unclassified(100);k__Fungi_unclassified(100);k__Fungi_unclassified(100);k__Fungi_unclassified(100);k__Fungi_unclassified(100);k__Fungi_unclassified(100); | - | - | - | - | - | - | - | Unassigned | - |
| Otu60708 | 1 | k__Fungi(100);k__Fungi_unclassified(100);k__Fungi_unclassified(100);k__Fungi_unclassified(100);k__Fungi_unclassified(100);k__Fungi_unclassified(100);k__Fungi_unclassified(100); | - | - | - | - | - | - | - | Unassigned | - |
| Otu60709 | 1 | k__Fungi(100);k__Fungi_unclassified(100);k__Fungi_unclassified(100);k__Fungi_unclassified(100);k__Fungi_unclassified(100);k__Fungi_unclassified(100);k__Fungi_unclassified(100); | - | - | - | - | - | - | - | Unassigned | - |
| Otu60711 | 1 | k__Fungi(100);k__Fungi_unclassified(100);k__Fungi_unclassified(100);k__Fungi_unclassified(100);k__Fungi_unclassified(100);k__Fungi_unclassified(100);k__Fungi_unclassified(100); | - | - | - | - | - | - | - | Unassigned | - |
| Otu60712 | 1 | k__Fungi(100);k__Fungi_unclassified(100);k__Fungi_unclassified(100);k__Fungi_unclassified(100);k__Fungi_unclassified(100);k__Fungi_unclassified(100);k__Fungi_unclassified(100); | - | - | - | - | - | - | - | Unassigned | - |
| Otu60713 | 1 | k__Fungi(100);p__Ascomycota(100);p__Ascomycota_unclassified(100);p__Ascomycota_unclassified(100);p__Ascomycota_unclassified(100);p__Ascomycota_unclassified(100);p__Ascomycota_unclassified(100); | - | - | - | - | - | - | - | Unassigned | - |
| Otu60714 | 1 | k__Fungi(100);k__Fungi_unclassified(100);k__Fungi_unclassified(100);k__Fungi_unclassified(100);k__Fungi_unclassified(100);k__Fungi_unclassified(100);k__Fungi_unclassified(100); | - | - | - | - | - | - | - | Unassigned | - |
| Otu60717 | 1 | k__Fungi(100);k__Fungi_unclassified(100);k__Fungi_unclassified(100);k__Fungi_unclassified(100);k__Fungi_unclassified(100);k__Fungi_unclassified(100);k__Fungi_unclassified(100); | - | - | - | - | - | - | - | Unassigned | - |
| Otu60718 | 1 | k__Fungi(100);k__Fungi_unclassified(100);k__Fungi_unclassified(100);k__Fungi_unclassified(100);k__Fungi_unclassified(100);k__Fungi_unclassified(100);k__Fungi_unclassified(100); | - | - | - | - | - | - | - | Unassigned | - |
| Otu60719 | 1 | k__Fungi(100);p__Ascomycota(100);p__Ascomycota_unclassified(100);p__Ascomycota_unclassified(100);p__Ascomycota_unclassified(100);p__Ascomycota_unclassified(100);p__Ascomycota_unclassified(100); | - | - | - | - | - | - | - | Unassigned | - |
| Otu60720 | 1 | k__Fungi(100);p__Ascomycota(100);c__Leotiomycetes(100);o__Helotiales(100);o__Helotiales_unclassified(100);o__Helotiales_unclassified(100);o__Helotiales_unclassified(100); | - | - | - | - | - | - | - | Unassigned | - |
| Otu60721 | 1 | k__Fungi(100);p__Ascomycota(100);p__Ascomycota_unclassified(100);p__Ascomycota_unclassified(100);p__Ascomycota_unclassified(100);p__Ascomycota_unclassified(100);p__Ascomycota_unclassified(100); | - | - | - | - | - | - | - | Unassigned | - |
| Otu60722 | 1 | k__Fungi(100);k__Fungi_unclassified(100);k__Fungi_unclassified(100);k__Fungi_unclassified(100);k__Fungi_unclassified(100);k__Fungi_unclassified(100);k__Fungi_unclassified(100); | - | - | - | - | - | - | - | Unassigned | - |
| Otu60727 | 1 | k__Fungi(100);p__Ascomycota(100);p__Ascomycota_unclassified(100);p__Ascomycota_unclassified(100);p__Ascomycota_unclassified(100);p__Ascomycota_unclassified(100);p__Ascomycota_unclassified(100); | - | - | - | - | - | - | - | Unassigned | - |
| Otu60728 | 1 | k__Fungi(100);p__Ascomycota(100);p__Ascomycota_unclassified(100);p__Ascomycota_unclassified(100);p__Ascomycota_unclassified(100);p__Ascomycota_unclassified(100);p__Ascomycota_unclassified(100); | - | - | - | - | - | - | - | Unassigned | - |
| Otu60729 | 1 | k__Fungi(100);k__Fungi_unclassified(100);k__Fungi_unclassified(100);k__Fungi_unclassified(100);k__Fungi_unclassified(100);k__Fungi_unclassified(100);k__Fungi_unclassified(100); | - | - | - | - | - | - | - | Unassigned | - |
| Otu60730 | 1 | k__Fungi(100);k__Fungi_unclassified(100);k__Fungi_unclassified(100);k__Fungi_unclassified(100);k__Fungi_unclassified(100);k__Fungi_unclassified(100);k__Fungi_unclassified(100); | - | - | - | - | - | - | - | Unassigned | - |
| Otu60732 | 1 | k__Fungi(100);k__Fungi_unclassified(100);k__Fungi_unclassified(100);k__Fungi_unclassified(100);k__Fungi_unclassified(100);k__Fungi_unclassified(100);k__Fungi_unclassified(100); | - | - | - | - | - | - | - | Unassigned | - |
| Otu60733 | 1 | k__Fungi(100);k__Fungi_unclassified(100);k__Fungi_unclassified(100);k__Fungi_unclassified(100);k__Fungi_unclassified(100);k__Fungi_unclassified(100);k__Fungi_unclassified(100); | - | - | - | - | - | - | - | Unassigned | - |
| Otu60734 | 1 | k__Fungi(100);p__Ascomycota(100);p__Ascomycota_unclassified(100);p__Ascomycota_unclassified(100);p__Ascomycota_unclassified(100);p__Ascomycota_unclassified(100);p__Ascomycota_unclassified(100); | - | - | - | - | - | - | - | Unassigned | - |
| Otu60735 | 1 | k__Fungi(100);k__Fungi_unclassified(100);k__Fungi_unclassified(100);k__Fungi_unclassified(100);k__Fungi_unclassified(100);k__Fungi_unclassified(100);k__Fungi_unclassified(100); | - | - | - | - | - | - | - | Unassigned | - |
| Otu60736 | 1 | k__Fungi(100);p__Ascomycota(100);c__Dothideomycetes(100);o__Pleosporales(100);o__Pleosporales_unclassified(100);o__Pleosporales_unclassified(100);o__Pleosporales_unclassified(100); | - | - | - | - | - | - | - | Unassigned | - |
| Otu60737 | 1 | k__Fungi(100);p__Ascomycota(100);p__Ascomycota_unclassified(100);p__Ascomycota_unclassified(100);p__Ascomycota_unclassified(100);p__Ascomycota_unclassified(100);p__Ascomycota_unclassified(100); | - | - | - | - | - | - | - | Unassigned | - |
| Otu60739 | 1 | k__Fungi(100);k__Fungi_unclassified(100);k__Fungi_unclassified(100);k__Fungi_unclassified(100);k__Fungi_unclassified(100);k__Fungi_unclassified(100);k__Fungi_unclassified(100); | - | - | - | - | - | - | - | Unassigned | - |
| Otu60740 | 1 | k__Fungi(100);p__Ascomycota(100);p__Ascomycota_unclassified(100);p__Ascomycota_unclassified(100);p__Ascomycota_unclassified(100);p__Ascomycota_unclassified(100);p__Ascomycota_unclassified(100); | - | - | - | - | - | - | - | Unassigned | - |
| Otu60741 | 1 | k__Fungi(100);p__Ascomycota(100);p__Ascomycota_unclassified(100);p__Ascomycota_unclassified(100);p__Ascomycota_unclassified(100);p__Ascomycota_unclassified(100);p__Ascomycota_unclassified(100); | - | - | - | - | - | - | - | Unassigned | - |
| Otu60742 | 1 | k__Fungi(100);k__Fungi_unclassified(100);k__Fungi_unclassified(100);k__Fungi_unclassified(100);k__Fungi_unclassified(100);k__Fungi_unclassified(100);k__Fungi_unclassified(100); | - | - | - | - | - | - | - | Unassigned | - |
| Otu60743 | 1 | k__Fungi(100);k__Fungi_unclassified(100);k__Fungi_unclassified(100);k__Fungi_unclassified(100);k__Fungi_unclassified(100);k__Fungi_unclassified(100);k__Fungi_unclassified(100); | - | - | - | - | - | - | - | Unassigned | - |
| Otu60744 | 1 | k__Fungi(100);k__Fungi_unclassified(100);k__Fungi_unclassified(100);k__Fungi_unclassified(100);k__Fungi_unclassified(100);k__Fungi_unclassified(100);k__Fungi_unclassified(100); | - | - | - | - | - | - | - | Unassigned | - |
| Otu60745 | 1 | k__Fungi(100);p__Ascomycota(100);p__Ascomycota_unclassified(100);p__Ascomycota_unclassified(100);p__Ascomycota_unclassified(100);p__Ascomycota_unclassified(100);p__Ascomycota_unclassified(100); | - | - | - | - | - | - | - | Unassigned | - |
| Otu60746 | 1 | k__Fungi(100);k__Fungi_unclassified(100);k__Fungi_unclassified(100);k__Fungi_unclassified(100);k__Fungi_unclassified(100);k__Fungi_unclassified(100);k__Fungi_unclassified(100); | - | - | - | - | - | - | - | Unassigned | - |
| Otu60748 | 1 | k__Fungi(100);k__Fungi_unclassified(100);k__Fungi_unclassified(100);k__Fungi_unclassified(100);k__Fungi_unclassified(100);k__Fungi_unclassified(100);k__Fungi_unclassified(100); | - | - | - | - | - | - | - | Unassigned | - |
| Otu60749 | 1 | k__Fungi(100);k__Fungi_unclassified(100);k__Fungi_unclassified(100);k__Fungi_unclassified(100);k__Fungi_unclassified(100);k__Fungi_unclassified(100);k__Fungi_unclassified(100); | - | - | - | - | - | - | - | Unassigned | - |
| Otu60751 | 1 | k__Fungi(100);k__Fungi_unclassified(100);k__Fungi_unclassified(100);k__Fungi_unclassified(100);k__Fungi_unclassified(100);k__Fungi_unclassified(100);k__Fungi_unclassified(100); | - | - | - | - | - | - | - | Unassigned | - |
| Otu60752 | 1 | k__Fungi(100);k__Fungi_unclassified(100);k__Fungi_unclassified(100);k__Fungi_unclassified(100);k__Fungi_unclassified(100);k__Fungi_unclassified(100);k__Fungi_unclassified(100); | - | - | - | - | - | - | - | Unassigned | - |
| Otu60753 | 1 | k__Fungi(100);k__Fungi_unclassified(100);k__Fungi_unclassified(100);k__Fungi_unclassified(100);k__Fungi_unclassified(100);k__Fungi_unclassified(100);k__Fungi_unclassified(100); | - | - | - | - | - | - | - | Unassigned | - |
| Otu60754 | 1 | k__Fungi(100);p__Ascomycota(100);p__Ascomycota_unclassified(100);p__Ascomycota_unclassified(100);p__Ascomycota_unclassified(100);p__Ascomycota_unclassified(100);p__Ascomycota_unclassified(100); | - | - | - | - | - | - | - | Unassigned | - |
| Otu60755 | 1 | k__Fungi(100);k__Fungi_unclassified(100);k__Fungi_unclassified(100);k__Fungi_unclassified(100);k__Fungi_unclassified(100);k__Fungi_unclassified(100);k__Fungi_unclassified(100); | - | - | - | - | - | - | - | Unassigned | - |
| Otu60756 | 1 | k__Fungi(100);k__Fungi_unclassified(100);k__Fungi_unclassified(100);k__Fungi_unclassified(100);k__Fungi_unclassified(100);k__Fungi_unclassified(100);k__Fungi_unclassified(100); | - | - | - | - | - | - | - | Unassigned | - |
| Otu60757 | 1 | k__Fungi(100);p__Ascomycota(100);p__Ascomycota_unclassified(100);p__Ascomycota_unclassified(100);p__Ascomycota_unclassified(100);p__Ascomycota_unclassified(100);p__Ascomycota_unclassified(100); | - | - | - | - | - | - | - | Unassigned | - |
| Otu60758 | 1 | k__Fungi(100);k__Fungi_unclassified(100);k__Fungi_unclassified(100);k__Fungi_unclassified(100);k__Fungi_unclassified(100);k__Fungi_unclassified(100);k__Fungi_unclassified(100); | - | - | - | - | - | - | - | Unassigned | - |
| Otu60759 | 1 | k__Fungi(100);p__Ascomycota(100);p__Ascomycota_unclassified(100);p__Ascomycota_unclassified(100);p__Ascomycota_unclassified(100);p__Ascomycota_unclassified(100);p__Ascomycota_unclassified(100); | - | - | - | - | - | - | - | Unassigned | - |
| Otu60761 | 1 | k__Fungi(100);p__Ascomycota(100);p__Ascomycota_unclassified(100);p__Ascomycota_unclassified(100);p__Ascomycota_unclassified(100);p__Ascomycota_unclassified(100);p__Ascomycota_unclassified(100); | - | - | - | - | - | - | - | Unassigned | - |
| Otu60766 | 1 | k__Fungi(100);k__Fungi_unclassified(100);k__Fungi_unclassified(100);k__Fungi_unclassified(100);k__Fungi_unclassified(100);k__Fungi_unclassified(100);k__Fungi_unclassified(100); | - | - | - | - | - | - | - | Unassigned | - |
| Otu60767 | 1 | k__Fungi(100);k__Fungi_unclassified(100);k__Fungi_unclassified(100);k__Fungi_unclassified(100);k__Fungi_unclassified(100);k__Fungi_unclassified(100);k__Fungi_unclassified(100); | - | - | - | - | - | - | - | Unassigned | - |
| Otu60769 | 1 | k__Fungi(100);k__Fungi_unclassified(100);k__Fungi_unclassified(100);k__Fungi_unclassified(100);k__Fungi_unclassified(100);k__Fungi_unclassified(100);k__Fungi_unclassified(100); | - | - | - | - | - | - | - | Unassigned | - |
| Otu60770 | 1 | k__Fungi(100);p__Ascomycota(100);p__Ascomycota_unclassified(100);p__Ascomycota_unclassified(100);p__Ascomycota_unclassified(100);p__Ascomycota_unclassified(100);p__Ascomycota_unclassified(100); | - | - | - | - | - | - | - | Unassigned | - |
| Otu60772 | 1 | k__Fungi(100);p__Ascomycota(100);p__Ascomycota_unclassified(100);p__Ascomycota_unclassified(100);p__Ascomycota_unclassified(100);p__Ascomycota_unclassified(100);p__Ascomycota_unclassified(100); | - | - | - | - | - | - | - | Unassigned | - |
| Otu60773 | 1 | k__Fungi(100);k__Fungi_unclassified(100);k__Fungi_unclassified(100);k__Fungi_unclassified(100);k__Fungi_unclassified(100);k__Fungi_unclassified(100);k__Fungi_unclassified(100); | - | - | - | - | - | - | - | Unassigned | - |
| Otu60774 | 1 | k__Fungi(100);k__Fungi_unclassified(100);k__Fungi_unclassified(100);k__Fungi_unclassified(100);k__Fungi_unclassified(100);k__Fungi_unclassified(100);k__Fungi_unclassified(100); | - | - | - | - | - | - | - | Unassigned | - |
| Otu60775 | 1 | k__Fungi(100);p__Ascomycota(100);p__Ascomycota_unclassified(100);p__Ascomycota_unclassified(100);p__Ascomycota_unclassified(100);p__Ascomycota_unclassified(100);p__Ascomycota_unclassified(100); | - | - | - | - | - | - | - | Unassigned | - |
| Otu60778 | 1 | k__Fungi(100);p__Ascomycota(100);p__Ascomycota_unclassified(100);p__Ascomycota_unclassified(100);p__Ascomycota_unclassified(100);p__Ascomycota_unclassified(100);p__Ascomycota_unclassified(100); | - | - | - | - | - | - | - | Unassigned | - |
| Otu60779 | 1 | k__Fungi(100);k__Fungi_unclassified(100);k__Fungi_unclassified(100);k__Fungi_unclassified(100);k__Fungi_unclassified(100);k__Fungi_unclassified(100);k__Fungi_unclassified(100); | - | - | - | - | - | - | - | Unassigned | - |
| Otu60782 | 1 | k__Fungi(100);k__Fungi_unclassified(100);k__Fungi_unclassified(100);k__Fungi_unclassified(100);k__Fungi_unclassified(100);k__Fungi_unclassified(100);k__Fungi_unclassified(100); | - | - | - | - | - | - | - | Unassigned | - |
| Otu60783 | 1 | k__Fungi(100);k__Fungi_unclassified(100);k__Fungi_unclassified(100);k__Fungi_unclassified(100);k__Fungi_unclassified(100);k__Fungi_unclassified(100);k__Fungi_unclassified(100); | - | - | - | - | - | - | - | Unassigned | - |
| Otu60784 | 1 | k__Fungi(100);k__Fungi_unclassified(100);k__Fungi_unclassified(100);k__Fungi_unclassified(100);k__Fungi_unclassified(100);k__Fungi_unclassified(100);k__Fungi_unclassified(100); | - | - | - | - | - | - | - | Unassigned | - |
| Otu60793 | 1 | k__Fungi(100);k__Fungi_unclassified(100);k__Fungi_unclassified(100);k__Fungi_unclassified(100);k__Fungi_unclassified(100);k__Fungi_unclassified(100);k__Fungi_unclassified(100); | - | - | - | - | - | - | - | Unassigned | - |
| Otu60794 | 1 | k__Fungi(100);p__Ascomycota(100);p__Ascomycota_unclassified(100);p__Ascomycota_unclassified(100);p__Ascomycota_unclassified(100);p__Ascomycota_unclassified(100);p__Ascomycota_unclassified(100); | - | - | - | - | - | - | - | Unassigned | - |
| Otu60795 | 1 | k__Fungi(100);k__Fungi_unclassified(100);k__Fungi_unclassified(100);k__Fungi_unclassified(100);k__Fungi_unclassified(100);k__Fungi_unclassified(100);k__Fungi_unclassified(100); | - | - | - | - | - | - | - | Unassigned | - |
| Otu60796 | 1 | k__Fungi(100);p__Ascomycota(100);p__Ascomycota_unclassified(100);p__Ascomycota_unclassified(100);p__Ascomycota_unclassified(100);p__Ascomycota_unclassified(100);p__Ascomycota_unclassified(100); | - | - | - | - | - | - | - | Unassigned | - |
| Otu60848 | 1 | k__Fungi(100);k__Fungi_unclassified(100);k__Fungi_unclassified(100);k__Fungi_unclassified(100);k__Fungi_unclassified(100);k__Fungi_unclassified(100);k__Fungi_unclassified(100); | - | - | - | - | - | - | - | Unassigned | - |
| Otu60849 | 1 | k__Fungi(100);k__Fungi_unclassified(100);k__Fungi_unclassified(100);k__Fungi_unclassified(100);k__Fungi_unclassified(100);k__Fungi_unclassified(100);k__Fungi_unclassified(100); | - | - | - | - | - | - | - | Unassigned | - |
| Otu60858 | 1 | k__Fungi(100);k__Fungi_unclassified(100);k__Fungi_unclassified(100);k__Fungi_unclassified(100);k__Fungi_unclassified(100);k__Fungi_unclassified(100);k__Fungi_unclassified(100); | - | - | - | - | - | - | - | Unassigned | - |
| Otu60859 | 1 | k__Fungi(100);k__Fungi_unclassified(100);k__Fungi_unclassified(100);k__Fungi_unclassified(100);k__Fungi_unclassified(100);k__Fungi_unclassified(100);k__Fungi_unclassified(100); | - | - | - | - | - | - | - | Unassigned | - |
| Otu60860 | 1 | k__Fungi(100);k__Fungi_unclassified(100);k__Fungi_unclassified(100);k__Fungi_unclassified(100);k__Fungi_unclassified(100);k__Fungi_unclassified(100);k__Fungi_unclassified(100); | - | - | - | - | - | - | - | Unassigned | - |
| Otu60881 | 1 | k__Fungi(100);k__Fungi_unclassified(100);k__Fungi_unclassified(100);k__Fungi_unclassified(100);k__Fungi_unclassified(100);k__Fungi_unclassified(100);k__Fungi_unclassified(100); | - | - | - | - | - | - | - | Unassigned | - |
| Otu60883 | 1 | k__Fungi(100);k__Fungi_unclassified(100);k__Fungi_unclassified(100);k__Fungi_unclassified(100);k__Fungi_unclassified(100);k__Fungi_unclassified(100);k__Fungi_unclassified(100); | - | - | - | - | - | - | - | Unassigned | - |
| Otu60886 | 1 | k__Fungi(100);k__Fungi_unclassified(100);k__Fungi_unclassified(100);k__Fungi_unclassified(100);k__Fungi_unclassified(100);k__Fungi_unclassified(100);k__Fungi_unclassified(100); | - | - | - | - | - | - | - | Unassigned | - |
| Otu60887 | 1 | k__Fungi(100);k__Fungi_unclassified(100);k__Fungi_unclassified(100);k__Fungi_unclassified(100);k__Fungi_unclassified(100);k__Fungi_unclassified(100);k__Fungi_unclassified(100); | - | - | - | - | - | - | - | Unassigned | - |
| Otu60918 | 1 | k__Fungi(100);p__Ascomycota(100);p__Ascomycota_unclassified(100);p__Ascomycota_unclassified(100);p__Ascomycota_unclassified(100);p__Ascomycota_unclassified(100);p__Ascomycota_unclassified(100); | - | - | - | - | - | - | - | Unassigned | - |
| Otu60919 | 1 | k__Fungi(100);k__Fungi_unclassified(100);k__Fungi_unclassified(100);k__Fungi_unclassified(100);k__Fungi_unclassified(100);k__Fungi_unclassified(100);k__Fungi_unclassified(100); | - | - | - | - | - | - | - | Unassigned | - |
| Otu60921 | 1 | k__Fungi(100);p__Ascomycota(100);p__Ascomycota_unclassified(100);p__Ascomycota_unclassified(100);p__Ascomycota_unclassified(100);p__Ascomycota_unclassified(100);p__Ascomycota_unclassified(100); | - | - | - | - | - | - | - | Unassigned | - |
| Otu60932 | 1 | k__Fungi(100);p__Ascomycota(100);p__Ascomycota_unclassified(100);p__Ascomycota_unclassified(100);p__Ascomycota_unclassified(100);p__Ascomycota_unclassified(100);p__Ascomycota_unclassified(100); | - | - | - | - | - | - | - | Unassigned | - |
| Otu60933 | 1 | k__Fungi(100);k__Fungi_unclassified(100);k__Fungi_unclassified(100);k__Fungi_unclassified(100);k__Fungi_unclassified(100);k__Fungi_unclassified(100);k__Fungi_unclassified(100); | - | - | - | - | - | - | - | Unassigned | - |
| Otu60962 | 1 | k__Fungi(100);k__Fungi_unclassified(100);k__Fungi_unclassified(100);k__Fungi_unclassified(100);k__Fungi_unclassified(100);k__Fungi_unclassified(100);k__Fungi_unclassified(100); | - | - | - | - | - | - | - | Unassigned | - |
| Otu60980 | 1 | k__Fungi(100);k__Fungi_unclassified(100);k__Fungi_unclassified(100);k__Fungi_unclassified(100);k__Fungi_unclassified(100);k__Fungi_unclassified(100);k__Fungi_unclassified(100); | - | - | - | - | - | - | - | Unassigned | - |
| Otu61021 | 1 | k__Fungi(100);k__Fungi_unclassified(100);k__Fungi_unclassified(100);k__Fungi_unclassified(100);k__Fungi_unclassified(100);k__Fungi_unclassified(100);k__Fungi_unclassified(100); | - | - | - | - | - | - | - | Unassigned | - |
| Otu61045 | 1 | k__Fungi(100);p__Ascomycota(100);p__Ascomycota_unclassified(100);p__Ascomycota_unclassified(100);p__Ascomycota_unclassified(100);p__Ascomycota_unclassified(100);p__Ascomycota_unclassified(100); | - | - | - | - | - | - | - | Unassigned | - |
| Otu61057 | 1 | k__Fungi(100);k__Fungi_unclassified(100);k__Fungi_unclassified(100);k__Fungi_unclassified(100);k__Fungi_unclassified(100);k__Fungi_unclassified(100);k__Fungi_unclassified(100); | - | - | - | - | - | - | - | Unassigned | - |
| Otu61068 | 1 | k__Fungi(100);k__Fungi_unclassified(100);k__Fungi_unclassified(100);k__Fungi_unclassified(100);k__Fungi_unclassified(100);k__Fungi_unclassified(100);k__Fungi_unclassified(100); | - | - | - | - | - | - | - | Unassigned | - |
| Otu61073 | 1 | k__Fungi(100);p__Ascomycota(100);p__Ascomycota_unclassified(100);p__Ascomycota_unclassified(100);p__Ascomycota_unclassified(100);p__Ascomycota_unclassified(100);p__Ascomycota_unclassified(100); | - | - | - | - | - | - | - | Unassigned | - |
| Otu61115 | 1 | k__Fungi(100);k__Fungi_unclassified(100);k__Fungi_unclassified(100);k__Fungi_unclassified(100);k__Fungi_unclassified(100);k__Fungi_unclassified(100);k__Fungi_unclassified(100); | - | - | - | - | - | - | - | Unassigned | - |
| Otu61124 | 1 | k__Fungi(100);k__Fungi_unclassified(100);k__Fungi_unclassified(100);k__Fungi_unclassified(100);k__Fungi_unclassified(100);k__Fungi_unclassified(100);k__Fungi_unclassified(100); | - | - | - | - | - | - | - | Unassigned | - |
| Otu61125 | 1 | k__Fungi(100);p__Ascomycota(100);p__Ascomycota_unclassified(100);p__Ascomycota_unclassified(100);p__Ascomycota_unclassified(100);p__Ascomycota_unclassified(100);p__Ascomycota_unclassified(100); | - | - | - | - | - | - | - | Unassigned | - |
| Otu61148 | 1 | k__Fungi(100);k__Fungi_unclassified(100);k__Fungi_unclassified(100);k__Fungi_unclassified(100);k__Fungi_unclassified(100);k__Fungi_unclassified(100);k__Fungi_unclassified(100); | - | - | - | - | - | - | - | Unassigned | - |
| Otu61152 | 1 | k__Fungi(100);k__Fungi_unclassified(100);k__Fungi_unclassified(100);k__Fungi_unclassified(100);k__Fungi_unclassified(100);k__Fungi_unclassified(100);k__Fungi_unclassified(100); | - | - | - | - | - | - | - | Unassigned | - |
| Otu61155 | 1 | k__Fungi(100);p__Ascomycota(100);p__Ascomycota_unclassified(100);p__Ascomycota_unclassified(100);p__Ascomycota_unclassified(100);p__Ascomycota_unclassified(100);p__Ascomycota_unclassified(100); | - | - | - | - | - | - | - | Unassigned | - |
| Otu61198 | 1 | k__Fungi(100);p__Ascomycota(100);c__Dothideomycetes(100);o__Myriangiales(100);f__unclassified_Myriangiales(100);g__unclassified_Myriangiales(100);s__Myriangiales_sp(100); | - | - | - | - | - | - | - | Unassigned | - |
| Otu61200 | 1 | k__Fungi(100);k__Fungi_unclassified(100);k__Fungi_unclassified(100);k__Fungi_unclassified(100);k__Fungi_unclassified(100);k__Fungi_unclassified(100);k__Fungi_unclassified(100); | - | - | - | - | - | - | - | Unassigned | - |
| Otu61201 | 1 | k__Fungi(100);k__Fungi_unclassified(100);k__Fungi_unclassified(100);k__Fungi_unclassified(100);k__Fungi_unclassified(100);k__Fungi_unclassified(100);k__Fungi_unclassified(100); | - | - | - | - | - | - | - | Unassigned | - |
| Otu61202 | 1 | k__Fungi(100);k__Fungi_unclassified(100);k__Fungi_unclassified(100);k__Fungi_unclassified(100);k__Fungi_unclassified(100);k__Fungi_unclassified(100);k__Fungi_unclassified(100); | - | - | - | - | - | - | - | Unassigned | - |
| Otu61203 | 1 | k__Fungi(100);k__Fungi_unclassified(100);k__Fungi_unclassified(100);k__Fungi_unclassified(100);k__Fungi_unclassified(100);k__Fungi_unclassified(100);k__Fungi_unclassified(100); | - | - | - | - | - | - | - | Unassigned | - |
| Otu61204 | 1 | k__Fungi(100);p__Ascomycota(100);p__Ascomycota_unclassified(100);p__Ascomycota_unclassified(100);p__Ascomycota_unclassified(100);p__Ascomycota_unclassified(100);p__Ascomycota_unclassified(100); | - | - | - | - | - | - | - | Unassigned | - |
| Otu61205 | 1 | k__Fungi(100);p__Ascomycota(100);p__Ascomycota_unclassified(100);p__Ascomycota_unclassified(100);p__Ascomycota_unclassified(100);p__Ascomycota_unclassified(100);p__Ascomycota_unclassified(100); | - | - | - | - | - | - | - | Unassigned | - |
| Otu61206 | 1 | k__Fungi(100);k__Fungi_unclassified(100);k__Fungi_unclassified(100);k__Fungi_unclassified(100);k__Fungi_unclassified(100);k__Fungi_unclassified(100);k__Fungi_unclassified(100); | - | - | - | - | - | - | - | Unassigned | - |
| Otu61207 | 1 | k__Fungi(100);k__Fungi_unclassified(100);k__Fungi_unclassified(100);k__Fungi_unclassified(100);k__Fungi_unclassified(100);k__Fungi_unclassified(100);k__Fungi_unclassified(100); | - | - | - | - | - | - | - | Unassigned | - |
| Otu61208 | 1 | k__Fungi(100);p__Ascomycota(100);p__Ascomycota_unclassified(100);p__Ascomycota_unclassified(100);p__Ascomycota_unclassified(100);p__Ascomycota_unclassified(100);p__Ascomycota_unclassified(100); | - | - | - | - | - | - | - | Unassigned | - |
| Otu61209 | 1 | k__Fungi(100);p__Ascomycota(100);p__Ascomycota_unclassified(100);p__Ascomycota_unclassified(100);p__Ascomycota_unclassified(100);p__Ascomycota_unclassified(100);p__Ascomycota_unclassified(100); | - | - | - | - | - | - | - | Unassigned | - |
| Otu61210 | 1 | k__Fungi(100);p__Ascomycota(100);p__Ascomycota_unclassified(100);p__Ascomycota_unclassified(100);p__Ascomycota_unclassified(100);p__Ascomycota_unclassified(100);p__Ascomycota_unclassified(100); | - | - | - | - | - | - | - | Unassigned | - |
| Otu61211 | 1 | k__Fungi(100);p__Ascomycota(100);p__Ascomycota_unclassified(100);p__Ascomycota_unclassified(100);p__Ascomycota_unclassified(100);p__Ascomycota_unclassified(100);p__Ascomycota_unclassified(100); | - | - | - | - | - | - | - | Unassigned | - |
| Otu61212 | 1 | k__Fungi(100);k__Fungi_unclassified(100);k__Fungi_unclassified(100);k__Fungi_unclassified(100);k__Fungi_unclassified(100);k__Fungi_unclassified(100);k__Fungi_unclassified(100); | - | - | - | - | - | - | - | Unassigned | - |
| Otu61213 | 1 | k__Fungi(100);k__Fungi_unclassified(100);k__Fungi_unclassified(100);k__Fungi_unclassified(100);k__Fungi_unclassified(100);k__Fungi_unclassified(100);k__Fungi_unclassified(100); | - | - | - | - | - | - | - | Unassigned | - |
| Otu61215 | 1 | k__Fungi(100);p__Ascomycota(100);c__Leotiomycetes(100);o__Helotiales(100);f__Helotiales_family_Incertae_sedis(100);f__Helotiales_family_Incertae_sedis_unclassified(100);f__Helotiales_family_Incertae_sedis_unclassified(100); | - | - | - | - | - | - | - | Unassigned | - |
| Otu61217 | 1 | k__Fungi(100);k__Fungi_unclassified(100);k__Fungi_unclassified(100);k__Fungi_unclassified(100);k__Fungi_unclassified(100);k__Fungi_unclassified(100);k__Fungi_unclassified(100); | - | - | - | - | - | - | - | Unassigned | - |
| Otu61218 | 1 | k__Fungi(100);k__Fungi_unclassified(100);k__Fungi_unclassified(100);k__Fungi_unclassified(100);k__Fungi_unclassified(100);k__Fungi_unclassified(100);k__Fungi_unclassified(100); | - | - | - | - | - | - | - | Unassigned | - |
| Otu61219 | 1 | k__Fungi(100);p__Ascomycota(100);p__Ascomycota_unclassified(100);p__Ascomycota_unclassified(100);p__Ascomycota_unclassified(100);p__Ascomycota_unclassified(100);p__Ascomycota_unclassified(100); | - | - | - | - | - | - | - | Unassigned | - |
| Otu61220 | 1 | k__Fungi(100);k__Fungi_unclassified(100);k__Fungi_unclassified(100);k__Fungi_unclassified(100);k__Fungi_unclassified(100);k__Fungi_unclassified(100);k__Fungi_unclassified(100); | - | - | - | - | - | - | - | Unassigned | - |
| Otu61221 | 1 | k__Fungi(100);p__Ascomycota(100);p__Ascomycota_unclassified(100);p__Ascomycota_unclassified(100);p__Ascomycota_unclassified(100);p__Ascomycota_unclassified(100);p__Ascomycota_unclassified(100); | - | - | - | - | - | - | - | Unassigned | - |
| Otu61222 | 1 | k__Fungi(100);p__Ascomycota(100);p__Ascomycota_unclassified(100);p__Ascomycota_unclassified(100);p__Ascomycota_unclassified(100);p__Ascomycota_unclassified(100);p__Ascomycota_unclassified(100); | - | - | - | - | - | - | - | Unassigned | - |
| Otu61223 | 1 | k__Fungi(100);p__Ascomycota(100);p__Ascomycota_unclassified(100);p__Ascomycota_unclassified(100);p__Ascomycota_unclassified(100);p__Ascomycota_unclassified(100);p__Ascomycota_unclassified(100); | - | - | - | - | - | - | - | Unassigned | - |
| Otu61224 | 1 | k__Fungi(100);k__Fungi_unclassified(100);k__Fungi_unclassified(100);k__Fungi_unclassified(100);k__Fungi_unclassified(100);k__Fungi_unclassified(100);k__Fungi_unclassified(100); | - | - | - | - | - | - | - | Unassigned | - |
| Otu61225 | 1 | k__Fungi(100);k__Fungi_unclassified(100);k__Fungi_unclassified(100);k__Fungi_unclassified(100);k__Fungi_unclassified(100);k__Fungi_unclassified(100);k__Fungi_unclassified(100); | - | - | - | - | - | - | - | Unassigned | - |
| Otu61226 | 1 | k__Fungi(100);k__Fungi_unclassified(100);k__Fungi_unclassified(100);k__Fungi_unclassified(100);k__Fungi_unclassified(100);k__Fungi_unclassified(100);k__Fungi_unclassified(100); | - | - | - | - | - | - | - | Unassigned | - |
| Otu61228 | 1 | k__Fungi(100);k__Fungi_unclassified(100);k__Fungi_unclassified(100);k__Fungi_unclassified(100);k__Fungi_unclassified(100);k__Fungi_unclassified(100);k__Fungi_unclassified(100); | - | - | - | - | - | - | - | Unassigned | - |
| Otu61229 | 1 | k__Fungi(100);p__Ascomycota(100);p__Ascomycota_unclassified(100);p__Ascomycota_unclassified(100);p__Ascomycota_unclassified(100);p__Ascomycota_unclassified(100);p__Ascomycota_unclassified(100); | - | - | - | - | - | - | - | Unassigned | - |
| Otu61232 | 1 | k__Fungi(100);p__Ascomycota(100);p__Ascomycota_unclassified(100);p__Ascomycota_unclassified(100);p__Ascomycota_unclassified(100);p__Ascomycota_unclassified(100);p__Ascomycota_unclassified(100); | - | - | - | - | - | - | - | Unassigned | - |
| Otu61233 | 1 | k__Fungi(100);k__Fungi_unclassified(100);k__Fungi_unclassified(100);k__Fungi_unclassified(100);k__Fungi_unclassified(100);k__Fungi_unclassified(100);k__Fungi_unclassified(100); | - | - | - | - | - | - | - | Unassigned | - |
| Otu61234 | 1 | k__Fungi(100);p__Ascomycota(100);p__Ascomycota_unclassified(100);p__Ascomycota_unclassified(100);p__Ascomycota_unclassified(100);p__Ascomycota_unclassified(100);p__Ascomycota_unclassified(100); | - | - | - | - | - | - | - | Unassigned | - |
| Otu61235 | 1 | k__Fungi(100);k__Fungi_unclassified(100);k__Fungi_unclassified(100);k__Fungi_unclassified(100);k__Fungi_unclassified(100);k__Fungi_unclassified(100);k__Fungi_unclassified(100); | - | - | - | - | - | - | - | Unassigned | - |
| Otu61239 | 1 | k__Fungi(100);k__Fungi_unclassified(100);k__Fungi_unclassified(100);k__Fungi_unclassified(100);k__Fungi_unclassified(100);k__Fungi_unclassified(100);k__Fungi_unclassified(100); | - | - | - | - | - | - | - | Unassigned | - |
| Otu61240 | 1 | k__Fungi(100);k__Fungi_unclassified(100);k__Fungi_unclassified(100);k__Fungi_unclassified(100);k__Fungi_unclassified(100);k__Fungi_unclassified(100);k__Fungi_unclassified(100); | - | - | - | - | - | - | - | Unassigned | - |
| Otu61241 | 1 | k__Fungi(100);p__Ascomycota(100);p__Ascomycota_unclassified(100);p__Ascomycota_unclassified(100);p__Ascomycota_unclassified(100);p__Ascomycota_unclassified(100);p__Ascomycota_unclassified(100); | - | - | - | - | - | - | - | Unassigned | - |
| Otu61242 | 1 | k__Fungi(100);k__Fungi_unclassified(100);k__Fungi_unclassified(100);k__Fungi_unclassified(100);k__Fungi_unclassified(100);k__Fungi_unclassified(100);k__Fungi_unclassified(100); | - | - | - | - | - | - | - | Unassigned | - |
| Otu61245 | 1 | k__Fungi(100);p__Ascomycota(100);p__Ascomycota_unclassified(100);p__Ascomycota_unclassified(100);p__Ascomycota_unclassified(100);p__Ascomycota_unclassified(100);p__Ascomycota_unclassified(100); | - | - | - | - | - | - | - | Unassigned | - |
| Otu61246 | 1 | k__Fungi(100);p__Ascomycota(100);p__Ascomycota_unclassified(100);p__Ascomycota_unclassified(100);p__Ascomycota_unclassified(100);p__Ascomycota_unclassified(100);p__Ascomycota_unclassified(100); | - | - | - | - | - | - | - | Unassigned | - |
| Otu61249 | 1 | k__Fungi(100);k__Fungi_unclassified(100);k__Fungi_unclassified(100);k__Fungi_unclassified(100);k__Fungi_unclassified(100);k__Fungi_unclassified(100);k__Fungi_unclassified(100); | - | - | - | - | - | - | - | Unassigned | - |
| Otu61250 | 1 | k__Fungi(100);p__Ascomycota(100);p__Ascomycota_unclassified(100);p__Ascomycota_unclassified(100);p__Ascomycota_unclassified(100);p__Ascomycota_unclassified(100);p__Ascomycota_unclassified(100); | - | - | - | - | - | - | - | Unassigned | - |
| Otu61251 | 1 | k__Fungi(100);k__Fungi_unclassified(100);k__Fungi_unclassified(100);k__Fungi_unclassified(100);k__Fungi_unclassified(100);k__Fungi_unclassified(100);k__Fungi_unclassified(100); | - | - | - | - | - | - | - | Unassigned | - |
| Otu61252 | 1 | k__Fungi(100);p__Ascomycota(100);p__Ascomycota_unclassified(100);p__Ascomycota_unclassified(100);p__Ascomycota_unclassified(100);p__Ascomycota_unclassified(100);p__Ascomycota_unclassified(100); | - | - | - | - | - | - | - | Unassigned | - |
| Otu61253 | 1 | k__Fungi(100);k__Fungi_unclassified(100);k__Fungi_unclassified(100);k__Fungi_unclassified(100);k__Fungi_unclassified(100);k__Fungi_unclassified(100);k__Fungi_unclassified(100); | - | - | - | - | - | - | - | Unassigned | - |
| Otu61255 | 1 | k__Fungi(100);p__Ascomycota(100);p__Ascomycota_unclassified(100);p__Ascomycota_unclassified(100);p__Ascomycota_unclassified(100);p__Ascomycota_unclassified(100);p__Ascomycota_unclassified(100); | - | - | - | - | - | - | - | Unassigned | - |
| Otu61256 | 1 | k__Fungi(100);k__Fungi_unclassified(100);k__Fungi_unclassified(100);k__Fungi_unclassified(100);k__Fungi_unclassified(100);k__Fungi_unclassified(100);k__Fungi_unclassified(100); | - | - | - | - | - | - | - | Unassigned | - |
| Otu61257 | 1 | k__Fungi(100);k__Fungi_unclassified(100);k__Fungi_unclassified(100);k__Fungi_unclassified(100);k__Fungi_unclassified(100);k__Fungi_unclassified(100);k__Fungi_unclassified(100); | - | - | - | - | - | - | - | Unassigned | - |
| Otu61259 | 1 | k__Fungi(100);k__Fungi_unclassified(100);k__Fungi_unclassified(100);k__Fungi_unclassified(100);k__Fungi_unclassified(100);k__Fungi_unclassified(100);k__Fungi_unclassified(100); | - | - | - | - | - | - | - | Unassigned | - |
| Otu61260 | 1 | k__Fungi(100);k__Fungi_unclassified(100);k__Fungi_unclassified(100);k__Fungi_unclassified(100);k__Fungi_unclassified(100);k__Fungi_unclassified(100);k__Fungi_unclassified(100); | - | - | - | - | - | - | - | Unassigned | - |
| Otu61262 | 1 | k__Fungi(100);p__Ascomycota(100);p__Ascomycota_unclassified(100);p__Ascomycota_unclassified(100);p__Ascomycota_unclassified(100);p__Ascomycota_unclassified(100);p__Ascomycota_unclassified(100); | - | - | - | - | - | - | - | Unassigned | - |
| Otu61263 | 1 | k__Fungi(100);p__Ascomycota(100);p__Ascomycota_unclassified(100);p__Ascomycota_unclassified(100);p__Ascomycota_unclassified(100);p__Ascomycota_unclassified(100);p__Ascomycota_unclassified(100); | - | - | - | - | - | - | - | Unassigned | - |
| Otu61264 | 1 | k__Fungi(100);k__Fungi_unclassified(100);k__Fungi_unclassified(100);k__Fungi_unclassified(100);k__Fungi_unclassified(100);k__Fungi_unclassified(100);k__Fungi_unclassified(100); | - | - | - | - | - | - | - | Unassigned | - |
| Otu61265 | 1 | k__Fungi(100);k__Fungi_unclassified(100);k__Fungi_unclassified(100);k__Fungi_unclassified(100);k__Fungi_unclassified(100);k__Fungi_unclassified(100);k__Fungi_unclassified(100); | - | - | - | - | - | - | - | Unassigned | - |
| Otu61266 | 1 | k__Fungi(100);k__Fungi_unclassified(100);k__Fungi_unclassified(100);k__Fungi_unclassified(100);k__Fungi_unclassified(100);k__Fungi_unclassified(100);k__Fungi_unclassified(100); | - | - | - | - | - | - | - | Unassigned | - |
| Otu61267 | 1 | k__Fungi(100);k__Fungi_unclassified(100);k__Fungi_unclassified(100);k__Fungi_unclassified(100);k__Fungi_unclassified(100);k__Fungi_unclassified(100);k__Fungi_unclassified(100); | - | - | - | - | - | - | - | Unassigned | - |
| Otu61268 | 1 | k__Fungi(100);k__Fungi_unclassified(100);k__Fungi_unclassified(100);k__Fungi_unclassified(100);k__Fungi_unclassified(100);k__Fungi_unclassified(100);k__Fungi_unclassified(100); | - | - | - | - | - | - | - | Unassigned | - |
| Otu61270 | 1 | k__Fungi(100);k__Fungi_unclassified(100);k__Fungi_unclassified(100);k__Fungi_unclassified(100);k__Fungi_unclassified(100);k__Fungi_unclassified(100);k__Fungi_unclassified(100); | - | - | - | - | - | - | - | Unassigned | - |
| Otu61271 | 1 | k__Fungi(100);k__Fungi_unclassified(100);k__Fungi_unclassified(100);k__Fungi_unclassified(100);k__Fungi_unclassified(100);k__Fungi_unclassified(100);k__Fungi_unclassified(100); | - | - | - | - | - | - | - | Unassigned | - |
| Otu61273 | 1 | k__Fungi(100);k__Fungi_unclassified(100);k__Fungi_unclassified(100);k__Fungi_unclassified(100);k__Fungi_unclassified(100);k__Fungi_unclassified(100);k__Fungi_unclassified(100); | - | - | - | - | - | - | - | Unassigned | - |
| Otu61274 | 1 | k__Fungi(100);k__Fungi_unclassified(100);k__Fungi_unclassified(100);k__Fungi_unclassified(100);k__Fungi_unclassified(100);k__Fungi_unclassified(100);k__Fungi_unclassified(100); | - | - | - | - | - | - | - | Unassigned | - |
| Otu61275 | 1 | k__Fungi(100);p__Ascomycota(100);p__Ascomycota_unclassified(100);p__Ascomycota_unclassified(100);p__Ascomycota_unclassified(100);p__Ascomycota_unclassified(100);p__Ascomycota_unclassified(100); | - | - | - | - | - | - | - | Unassigned | - |
| Otu61276 | 1 | k__Fungi(100);k__Fungi_unclassified(100);k__Fungi_unclassified(100);k__Fungi_unclassified(100);k__Fungi_unclassified(100);k__Fungi_unclassified(100);k__Fungi_unclassified(100); | - | - | - | - | - | - | - | Unassigned | - |
| Otu61277 | 1 | k__Fungi(100);p__Ascomycota(100);p__Ascomycota_unclassified(100);p__Ascomycota_unclassified(100);p__Ascomycota_unclassified(100);p__Ascomycota_unclassified(100);p__Ascomycota_unclassified(100); | - | - | - | - | - | - | - | Unassigned | - |

| | | | | | | | | | | | |
|---|---|---|---|---|---|---|---|---|---|---|---|
| Otu61278 | 1 | k__Fungi(100);k__Fungi_unclassified(100);k__Fungi_unclassified(100);k__Fungi_unclassified(100);k__Fungi_unclassified(100);k__Fungi_unclassified(100);k__Fungi_unclassified(100); | - | - | - | - | - | - | - | Unassigned | - |
| Otu61279 | 1 | k__Fungi(100);k__Fungi_unclassified(100);k__Fungi_unclassified(100);k__Fungi_unclassified(100);k__Fungi_unclassified(100);k__Fungi_unclassified(100);k__Fungi_unclassified(100); | - | - | - | - | - | - | - | Unassigned | - |
| Otu61280 | 1 | k__Fungi(100);k__Fungi_unclassified(100);k__Fungi_unclassified(100);k__Fungi_unclassified(100);k__Fungi_unclassified(100);k__Fungi_unclassified(100);k__Fungi_unclassified(100); | - | - | - | - | - | - | - | Unassigned | - |
| Otu61281 | 1 | k__Fungi(100);k__Fungi_unclassified(100);k__Fungi_unclassified(100);k__Fungi_unclassified(100);k__Fungi_unclassified(100);k__Fungi_unclassified(100);k__Fungi_unclassified(100); | - | - | - | - | - | - | - | Unassigned | - |
| Otu61282 | 1 | k__Fungi(100);p__Ascomycota(100);p__Ascomycota_unclassified(100);p__Ascomycota_unclassified(100);p__Ascomycota_unclassified(100);p__Ascomycota_unclassified(100);p__Ascomycota_unclassified(100); | - | - | - | - | - | - | - | Unassigned | - |
| Otu61283 | 1 | k__Fungi(100);k__Fungi_unclassified(100);k__Fungi_unclassified(100);k__Fungi_unclassified(100);k__Fungi_unclassified(100);k__Fungi_unclassified(100);k__Fungi_unclassified(100); | - | - | - | - | - | - | - | Unassigned | - |
| Otu61284 | 1 | k__Fungi(100);p__Ascomycota(100);p__Ascomycota_unclassified(100);p__Ascomycota_unclassified(100);p__Ascomycota_unclassified(100);p__Ascomycota_unclassified(100);p__Ascomycota_unclassified(100); | - | - | - | - | - | - | - | Unassigned | - |
| Otu61285 | 1 | k__Fungi(100);k__Fungi_unclassified(100);k__Fungi_unclassified(100);k__Fungi_unclassified(100);k__Fungi_unclassified(100);k__Fungi_unclassified(100);k__Fungi_unclassified(100); | - | - | - | - | - | - | - | Unassigned | - |
| Otu61286 | 1 | k__Fungi(100);k__Fungi_unclassified(100);k__Fungi_unclassified(100);k__Fungi_unclassified(100);k__Fungi_unclassified(100);k__Fungi_unclassified(100);k__Fungi_unclassified(100); | - | - | - | - | - | - | - | Unassigned | - |
| Otu61287 | 1 | k__Fungi(100);p__Ascomycota(100);p__Ascomycota_unclassified(100);p__Ascomycota_unclassified(100);p__Ascomycota_unclassified(100);p__Ascomycota_unclassified(100);p__Ascomycota_unclassified(100); | - | - | - | - | - | - | - | Unassigned | - |
| Otu61288 | 1 | k__Fungi(100);p__Ascomycota(100);p__Ascomycota_unclassified(100);p__Ascomycota_unclassified(100);p__Ascomycota_unclassified(100);p__Ascomycota_unclassified(100);p__Ascomycota_unclassified(100); | - | - | - | - | - | - | - | Unassigned | - |
| Otu61289 | 1 | k__Fungi(100);p__Ascomycota(100);p__Ascomycota_unclassified(100);p__Ascomycota_unclassified(100);p__Ascomycota_unclassified(100);p__Ascomycota_unclassified(100);p__Ascomycota_unclassified(100); | - | - | - | - | - | - | - | Unassigned | - |
| Otu61290 | 1 | k__Fungi(100);k__Fungi_unclassified(100);k__Fungi_unclassified(100);k__Fungi_unclassified(100);k__Fungi_unclassified(100);k__Fungi_unclassified(100);k__Fungi_unclassified(100); | - | - | - | - | - | - | - | Unassigned | - |
| Otu61292 | 1 | k__Fungi(100);k__Fungi_unclassified(100);k__Fungi_unclassified(100);k__Fungi_unclassified(100);k__Fungi_unclassified(100);k__Fungi_unclassified(100);k__Fungi_unclassified(100); | - | - | - | - | - | - | - | Unassigned | - |
| Otu61293 | 1 | k__Fungi(100);p__Ascomycota(100);p__Ascomycota_unclassified(100);p__Ascomycota_unclassified(100);p__Ascomycota_unclassified(100);p__Ascomycota_unclassified(100);p__Ascomycota_unclassified(100); | - | - | - | - | - | - | - | Unassigned | - |
| Otu61294 | 1 | k__Fungi(100);p__Ascomycota(100);p__Ascomycota_unclassified(100);p__Ascomycota_unclassified(100);p__Ascomycota_unclassified(100);p__Ascomycota_unclassified(100);p__Ascomycota_unclassified(100); | - | - | - | - | - | - | - | Unassigned | - |
| Otu61295 | 1 | k__Fungi(100);p__Ascomycota(100);p__Ascomycota_unclassified(100);p__Ascomycota_unclassified(100);p__Ascomycota_unclassified(100);p__Ascomycota_unclassified(100);p__Ascomycota_unclassified(100); | - | - | - | - | - | - | - | Unassigned | - |
| Otu61296 | 1 | k__Fungi(100);k__Fungi_unclassified(100);k__Fungi_unclassified(100);k__Fungi_unclassified(100);k__Fungi_unclassified(100);k__Fungi_unclassified(100);k__Fungi_unclassified(100); | - | - | - | - | - | - | - | Unassigned | - |
| Otu61297 | 1 | k__Fungi(100);k__Fungi_unclassified(100);k__Fungi_unclassified(100);k__Fungi_unclassified(100);k__Fungi_unclassified(100);k__Fungi_unclassified(100);k__Fungi_unclassified(100); | - | - | - | - | - | - | - | Unassigned | - |
| Otu61298 | 1 | k__Fungi(100);k__Fungi_unclassified(100);k__Fungi_unclassified(100);k__Fungi_unclassified(100);k__Fungi_unclassified(100);k__Fungi_unclassified(100);k__Fungi_unclassified(100); | - | - | - | - | - | - | - | Unassigned | - |
| Otu61299 | 1 | k__Fungi(100);k__Fungi_unclassified(100);k__Fungi_unclassified(100);k__Fungi_unclassified(100);k__Fungi_unclassified(100);k__Fungi_unclassified(100);k__Fungi_unclassified(100); | - | - | - | - | - | - | - | Unassigned | - |
| Otu61300 | 1 | k__Fungi(100);k__Fungi_unclassified(100);k__Fungi_unclassified(100);k__Fungi_unclassified(100);k__Fungi_unclassified(100);k__Fungi_unclassified(100);k__Fungi_unclassified(100); | - | - | - | - | - | - | - | Unassigned | - |
| Otu61301 | 1 | k__Fungi(100);k__Fungi_unclassified(100);k__Fungi_unclassified(100);k__Fungi_unclassified(100);k__Fungi_unclassified(100);k__Fungi_unclassified(100);k__Fungi_unclassified(100); | - | - | - | - | - | - | - | Unassigned | - |
| Otu61302 | 1 | k__Fungi(100);k__Fungi_unclassified(100);k__Fungi_unclassified(100);k__Fungi_unclassified(100);k__Fungi_unclassified(100);k__Fungi_unclassified(100);k__Fungi_unclassified(100); | - | - | - | - | - | - | - | Unassigned | - |
| Otu61303 | 1 | k__Fungi(100);k__Fungi_unclassified(100);k__Fungi_unclassified(100);k__Fungi_unclassified(100);k__Fungi_unclassified(100);k__Fungi_unclassified(100);k__Fungi_unclassified(100); | - | - | - | - | - | - | - | Unassigned | - |
| Otu61306 | 1 | k__Fungi(100);p__Basidiomycota(100);c__Tremellomycetes(100);c__Tremellomycetes_unclassified(100);c__Tremellomycetes_unclassified(100);c__Tremellomycetes_unclassified(100);c__Tremellomycetes_unclassified(100); | - | - | - | - | - | - | - | Unassigned | - |
| Otu61307 | 1 | k__Fungi(100);p__Ascomycota(100);p__Ascomycota_unclassified(100);p__Ascomycota_unclassified(100);p__Ascomycota_unclassified(100);p__Ascomycota_unclassified(100);p__Ascomycota_unclassified(100); | - | - | - | - | - | - | - | Unassigned | - |
| Otu61308 | 1 | k__Fungi(100);p__Ascomycota(100);p__Ascomycota_unclassified(100);p__Ascomycota_unclassified(100);p__Ascomycota_unclassified(100);p__Ascomycota_unclassified(100);p__Ascomycota_unclassified(100); | - | - | - | - | - | - | - | Unassigned | - |
| Otu61309 | 1 | k__Fungi(100);k__Fungi_unclassified(100);k__Fungi_unclassified(100);k__Fungi_unclassified(100);k__Fungi_unclassified(100);k__Fungi_unclassified(100);k__Fungi_unclassified(100); | - | - | - | - | - | - | - | Unassigned | - |
| Otu61311 | 1 | k__Fungi(100);p__Ascomycota(100);c__Dothideomycetes(100);o__Pleosporales(100);o__Pleosporales_unclassified(100);o__Pleosporales_unclassified(100);o__Pleosporales_unclassified(100); | - | - | - | - | - | - | - | Unassigned | - |
| Otu61313 | 1 | k__Fungi(100);p__Ascomycota(100);c__Sordariomycetes(100);c__Sordariomycetes_unclassified(100);c__Sordariomycetes_unclassified(100);c__Sordariomycetes_unclassified(100);c__Sordariomycetes_unclassified(100); | - | - | - | - | - | - | - | Unassigned | - |
| Otu61315 | 1 | k__Fungi(100);p__Ascomycota(100);c__Dothideomycetes(100);o__Pleosporales(100);o__Pleosporales_unclassified(100);o__Pleosporales_unclassified(100);o__Pleosporales_unclassified(100); | - | - | - | - | - | - | - | Unassigned | - |
| Otu61316 | 1 | k__Fungi(100);p__Ascomycota(100);p__Ascomycota_unclassified(100);p__Ascomycota_unclassified(100);p__Ascomycota_unclassified(100);p__Ascomycota_unclassified(100);p__Ascomycota_unclassified(100); | - | - | - | - | - | - | - | Unassigned | - |
| Otu61317 | 1 | k__Fungi(100);k__Fungi_unclassified(100);k__Fungi_unclassified(100);k__Fungi_unclassified(100);k__Fungi_unclassified(100);k__Fungi_unclassified(100);k__Fungi_unclassified(100); | - | - | - | - | - | - | - | Unassigned | - |
| Otu61318 | 1 | k__Fungi(100);p__Ascomycota(100);c__Eurotiomycetes(100);o__Eurotiales(100);f__Trichocomaceae(100);f__Trichocomaceae_unclassified(100);f__Trichocomaceae_unclassified(100); | - | - | - | - | - | - | - | Unassigned | - |
| Otu61320 | 1 | k__Fungi(100);p__Ascomycota(100);p__Ascomycota_unclassified(100);p__Ascomycota_unclassified(100);p__Ascomycota_unclassified(100);p__Ascomycota_unclassified(100);p__Ascomycota_unclassified(100); | - | - | - | - | - | - | - | Unassigned | - |
| Otu61322 | 1 | k__Fungi(100);k__Fungi_unclassified(100);k__Fungi_unclassified(100);k__Fungi_unclassified(100);k__Fungi_unclassified(100);k__Fungi_unclassified(100);k__Fungi_unclassified(100); | - | - | - | - | - | - | - | Unassigned | - |
| Otu61323 | 1 | k__Fungi(100);k__Fungi_unclassified(100);k__Fungi_unclassified(100);k__Fungi_unclassified(100);k__Fungi_unclassified(100);k__Fungi_unclassified(100);k__Fungi_unclassified(100); | - | - | - | - | - | - | - | Unassigned | - |
| Otu61324 | 1 | k__Fungi(100);k__Fungi_unclassified(100);k__Fungi_unclassified(100);k__Fungi_unclassified(100);k__Fungi_unclassified(100);k__Fungi_unclassified(100);k__Fungi_unclassified(100); | - | - | - | - | - | - | - | Unassigned | - |
| Otu61325 | 1 | k__Fungi(100);k__Fungi_unclassified(100);k__Fungi_unclassified(100);k__Fungi_unclassified(100);k__Fungi_unclassified(100);k__Fungi_unclassified(100);k__Fungi_unclassified(100); | - | - | - | - | - | - | - | Unassigned | - |
| Otu61326 | 1 | k__Fungi(100);p__Ascomycota(100);c__Dothideomycetes(100);o__Pleosporales(100);o__Pleosporales_unclassified(100);o__Pleosporales_unclassified(100);o__Pleosporales_unclassified(100); | - | - | - | - | - | - | - | Unassigned | - |
| Otu61327 | 1 | k__Fungi(100);k__Fungi_unclassified(100);k__Fungi_unclassified(100);k__Fungi_unclassified(100);k__Fungi_unclassified(100);k__Fungi_unclassified(100);k__Fungi_unclassified(100); | - | - | - | - | - | - | - | Unassigned | - |
| Otu61328 | 1 | k__Fungi(100);p__Ascomycota(100);p__Ascomycota_unclassified(100);p__Ascomycota_unclassified(100);p__Ascomycota_unclassified(100);p__Ascomycota_unclassified(100);p__Ascomycota_unclassified(100); | - | - | - | - | - | - | - | Unassigned | - |
| Otu61329 | 1 | k__Fungi(100);k__Fungi_unclassified(100);k__Fungi_unclassified(100);k__Fungi_unclassified(100);k__Fungi_unclassified(100);k__Fungi_unclassified(100);k__Fungi_unclassified(100); | - | - | - | - | - | - | - | Unassigned | - |
| Otu61330 | 1 | k__Fungi(100);p__Ascomycota(100);p__Ascomycota_unclassified(100);p__Ascomycota_unclassified(100);p__Ascomycota_unclassified(100);p__Ascomycota_unclassified(100);p__Ascomycota_unclassified(100); | - | - | - | - | - | - | - | Unassigned | - |
| Otu61331 | 1 | k__Fungi(100);p__Ascomycota(100);p__Ascomycota_unclassified(100);p__Ascomycota_unclassified(100);p__Ascomycota_unclassified(100);p__Ascomycota_unclassified(100);p__Ascomycota_unclassified(100); | - | - | - | - | - | - | - | Unassigned | - |
| Otu61334 | 1 | k__Fungi(100);p__Ascomycota(100);p__Ascomycota_unclassified(100);p__Ascomycota_unclassified(100);p__Ascomycota_unclassified(100);p__Ascomycota_unclassified(100);p__Ascomycota_unclassified(100); | - | - | - | - | - | - | - | Unassigned | - |
| Otu61335 | 1 | k__Fungi(100);p__Ascomycota(100);p__Ascomycota_unclassified(100);p__Ascomycota_unclassified(100);p__Ascomycota_unclassified(100);p__Ascomycota_unclassified(100);p__Ascomycota_unclassified(100); | - | - | - | - | - | - | - | Unassigned | - |
| Otu61336 | 1 | k__Fungi(100);k__Fungi_unclassified(100);k__Fungi_unclassified(100);k__Fungi_unclassified(100);k__Fungi_unclassified(100);k__Fungi_unclassified(100);k__Fungi_unclassified(100); | - | - | - | - | - | - | - | Unassigned | - |
| Otu61337 | 1 | k__Fungi(100);k__Fungi_unclassified(100);k__Fungi_unclassified(100);k__Fungi_unclassified(100);k__Fungi_unclassified(100);k__Fungi_unclassified(100);k__Fungi_unclassified(100); | - | - | - | - | - | - | - | Unassigned | - |
| Otu61338 | 1 | k__Fungi(100);k__Fungi_unclassified(100);k__Fungi_unclassified(100);k__Fungi_unclassified(100);k__Fungi_unclassified(100);k__Fungi_unclassified(100);k__Fungi_unclassified(100); | - | - | - | - | - | - | - | Unassigned | - |
| Otu61339 | 1 | k__Fungi(100);k__Fungi_unclassified(100);k__Fungi_unclassified(100);k__Fungi_unclassified(100);k__Fungi_unclassified(100);k__Fungi_unclassified(100);k__Fungi_unclassified(100); | - | - | - | - | - | - | - | Unassigned | - |
| Otu61340 | 1 | k__Fungi(100);k__Fungi_unclassified(100);k__Fungi_unclassified(100);k__Fungi_unclassified(100);k__Fungi_unclassified(100);k__Fungi_unclassified(100);k__Fungi_unclassified(100); | - | - | - | - | - | - | - | Unassigned | - |
| Otu61341 | 1 | k__Fungi(100);k__Fungi_unclassified(100);k__Fungi_unclassified(100);k__Fungi_unclassified(100);k__Fungi_unclassified(100);k__Fungi_unclassified(100);k__Fungi_unclassified(100); | - | - | - | - | - | - | - | Unassigned | - |
| Otu61342 | 1 | k__Fungi(100);p__Ascomycota(100);p__Ascomycota_unclassified(100);p__Ascomycota_unclassified(100);p__Ascomycota_unclassified(100);p__Ascomycota_unclassified(100);p__Ascomycota_unclassified(100); | - | - | - | - | - | - | - | Unassigned | - |
| Otu61343 | 1 | k__Fungi(100);k__Fungi_unclassified(100);k__Fungi_unclassified(100);k__Fungi_unclassified(100);k__Fungi_unclassified(100);k__Fungi_unclassified(100);k__Fungi_unclassified(100); | - | - | - | - | - | - | - | Unassigned | - |
| Otu61344 | 1 | k__Fungi(100);k__Fungi_unclassified(100);k__Fungi_unclassified(100);k__Fungi_unclassified(100);k__Fungi_unclassified(100);k__Fungi_unclassified(100);k__Fungi_unclassified(100); | - | - | - | - | - | - | - | Unassigned | - |
| Otu61345 | 1 | k__Fungi(100);p__Ascomycota(100);p__Ascomycota_unclassified(100);p__Ascomycota_unclassified(100);p__Ascomycota_unclassified(100);p__Ascomycota_unclassified(100);p__Ascomycota_unclassified(100); | - | - | - | - | - | - | - | Unassigned | - |
| Otu61346 | 1 | k__Fungi(100);p__Ascomycota(100);p__Ascomycota_unclassified(100);p__Ascomycota_unclassified(100);p__Ascomycota_unclassified(100);p__Ascomycota_unclassified(100);p__Ascomycota_unclassified(100); | - | - | - | - | - | - | - | Unassigned | - |
| Otu61347 | 1 | k__Fungi(100);p__Ascomycota(100);p__Ascomycota_unclassified(100);p__Ascomycota_unclassified(100);p__Ascomycota_unclassified(100);p__Ascomycota_unclassified(100);p__Ascomycota_unclassified(100); | - | - | - | - | - | - | - | Unassigned | - |
| Otu61348 | 1 | k__Fungi(100);k__Fungi_unclassified(100);k__Fungi_unclassified(100);k__Fungi_unclassified(100);k__Fungi_unclassified(100);k__Fungi_unclassified(100);k__Fungi_unclassified(100); | - | - | - | - | - | - | - | Unassigned | - |
| Otu61349 | 1 | k__Fungi(100);k__Fungi_unclassified(100);k__Fungi_unclassified(100);k__Fungi_unclassified(100);k__Fungi_unclassified(100);k__Fungi_unclassified(100);k__Fungi_unclassified(100); | - | - | - | - | - | - | - | Unassigned | - |
| Otu61350 | 1 | k__Fungi(100);p__Ascomycota(100);p__Ascomycota_unclassified(100);p__Ascomycota_unclassified(100);p__Ascomycota_unclassified(100);p__Ascomycota_unclassified(100);p__Ascomycota_unclassified(100); | - | - | - | - | - | - | - | Unassigned | - |
| Otu61351 | 1 | k__Fungi(100);k__Fungi_unclassified(100);k__Fungi_unclassified(100);k__Fungi_unclassified(100);k__Fungi_unclassified(100);k__Fungi_unclassified(100);k__Fungi_unclassified(100); | - | - | - | - | - | - | - | Unassigned | - |
| Otu61352 | 1 | k__Fungi(100);p__Ascomycota(100);p__Ascomycota_unclassified(100);p__Ascomycota_unclassified(100);p__Ascomycota_unclassified(100);p__Ascomycota_unclassified(100);p__Ascomycota_unclassified(100); | - | - | - | - | - | - | - | Unassigned | - |
| Otu61353 | 1 | k__Fungi(100);k__Fungi_unclassified(100);k__Fungi_unclassified(100);k__Fungi_unclassified(100);k__Fungi_unclassified(100);k__Fungi_unclassified(100);k__Fungi_unclassified(100); | - | - | - | - | - | - | - | Unassigned | - |
| Otu61354 | 1 | k__Fungi(100);p__Ascomycota(100);p__Ascomycota_unclassified(100);p__Ascomycota_unclassified(100);p__Ascomycota_unclassified(100);p__Ascomycota_unclassified(100);p__Ascomycota_unclassified(100); | - | - | - | - | - | - | - | Unassigned | - |
| Otu61356 | 1 | k__Fungi(100);k__Fungi_unclassified(100);k__Fungi_unclassified(100);k__Fungi_unclassified(100);k__Fungi_unclassified(100);k__Fungi_unclassified(100);k__Fungi_unclassified(100); | - | - | - | - | - | - | - | Unassigned | - |
| Otu61357 | 1 | k__Fungi(100);k__Fungi_unclassified(100);k__Fungi_unclassified(100);k__Fungi_unclassified(100);k__Fungi_unclassified(100);k__Fungi_unclassified(100);k__Fungi_unclassified(100); | - | - | - | - | - | - | - | Unassigned | - |
| Otu61358 | 1 | k__Fungi(100);k__Fungi_unclassified(100);k__Fungi_unclassified(100);k__Fungi_unclassified(100);k__Fungi_unclassified(100);k__Fungi_unclassified(100);k__Fungi_unclassified(100); | - | - | - | - | - | - | - | Unassigned | - |
| Otu61359 | 1 | k__Fungi(100);k__Fungi_unclassified(100);k__Fungi_unclassified(100);k__Fungi_unclassified(100);k__Fungi_unclassified(100);k__Fungi_unclassified(100);k__Fungi_unclassified(100); | - | - | - | - | - | - | - | Unassigned | - |
| Otu61363 | 1 | k__Fungi(100);p__Ascomycota(100);p__Ascomycota_unclassified(100);p__Ascomycota_unclassified(100);p__Ascomycota_unclassified(100);p__Ascomycota_unclassified(100);p__Ascomycota_unclassified(100); | - | - | - | - | - | - | - | Unassigned | - |
| Otu61364 | 1 | k__Fungi(100);p__Ascomycota(100);p__Ascomycota_unclassified(100);p__Ascomycota_unclassified(100);p__Ascomycota_unclassified(100);p__Ascomycota_unclassified(100);p__Ascomycota_unclassified(100); | - | - | - | - | - | - | - | Unassigned | - |
| Otu61365 | 1 | k__Fungi(100);p__Ascomycota(100);p__Ascomycota_unclassified(100);p__Ascomycota_unclassified(100);p__Ascomycota_unclassified(100);p__Ascomycota_unclassified(100);p__Ascomycota_unclassified(100); | - | - | - | - | - | - | - | Unassigned | - |
| Otu61366 | 1 | k__Fungi(100);k__Fungi_unclassified(100);k__Fungi_unclassified(100);k__Fungi_unclassified(100);k__Fungi_unclassified(100);k__Fungi_unclassified(100);k__Fungi_unclassified(100); | - | - | - | - | - | - | - | Unassigned | - |
| Otu61367 | 1 | k__Fungi(100);k__Fungi_unclassified(100);k__Fungi_unclassified(100);k__Fungi_unclassified(100);k__Fungi_unclassified(100);k__Fungi_unclassified(100);k__Fungi_unclassified(100); | - | - | - | - | - | - | - | Unassigned | - |
| Otu61368 | 1 | k__Fungi(100);p__Ascomycota(100);p__Ascomycota_unclassified(100);p__Ascomycota_unclassified(100);p__Ascomycota_unclassified(100);p__Ascomycota_unclassified(100);p__Ascomycota_unclassified(100); | - | - | - | - | - | - | - | Unassigned | - |
| Otu61369 | 1 | k__Fungi(100);k__Fungi_unclassified(100);k__Fungi_unclassified(100);k__Fungi_unclassified(100);k__Fungi_unclassified(100);k__Fungi_unclassified(100);k__Fungi_unclassified(100); | - | - | - | - | - | - | - | Unassigned | - |
| Otu61370 | 1 | k__Fungi(100);p__Ascomycota(100);p__Ascomycota_unclassified(100);p__Ascomycota_unclassified(100);p__Ascomycota_unclassified(100);p__Ascomycota_unclassified(100);p__Ascomycota_unclassified(100); | - | - | - | - | - | - | - | Unassigned | - |
| Otu61371 | 1 | k__Fungi(100);p__Ascomycota(100);p__Ascomycota_unclassified(100);p__Ascomycota_unclassified(100);p__Ascomycota_unclassified(100);p__Ascomycota_unclassified(100);p__Ascomycota_unclassified(100); | - | - | - | - | - | - | - | Unassigned | - |
| Otu61372 | 1 | k__Fungi(100);k__Fungi_unclassified(100);k__Fungi_unclassified(100);k__Fungi_unclassified(100);k__Fungi_unclassified(100);k__Fungi_unclassified(100);k__Fungi_unclassified(100); | - | - | - | - | - | - | - | Unassigned | - |
| Otu61373 | 1 | k__Fungi(100);k__Fungi_unclassified(100);k__Fungi_unclassified(100);k__Fungi_unclassified(100);k__Fungi_unclassified(100);k__Fungi_unclassified(100);k__Fungi_unclassified(100); | - | - | - | - | - | - | - | Unassigned | - |
| Otu61374 | 1 | k__Fungi(100);p__Ascomycota(100);p__Ascomycota_unclassified(100);p__Ascomycota_unclassified(100);p__Ascomycota_unclassified(100);p__Ascomycota_unclassified(100);p__Ascomycota_unclassified(100); | - | - | - | - | - | - | - | Unassigned | - |
| Otu61375 | 1 | k__Fungi(100);p__Ascomycota(100);p__Ascomycota_unclassified(100);p__Ascomycota_unclassified(100);p__Ascomycota_unclassified(100);p__Ascomycota_unclassified(100);p__Ascomycota_unclassified(100); | - | - | - | - | - | - | - | Unassigned | - |
| Otu61376 | 1 | k__Fungi(100);p__Ascomycota(100);p__Ascomycota_unclassified(100);p__Ascomycota_unclassified(100);p__Ascomycota_unclassified(100);p__Ascomycota_unclassified(100);p__Ascomycota_unclassified(100); | - | - | - | - | - | - | - | Unassigned | - |
| Otu61377 | 1 | k__Fungi(100);k__Fungi_unclassified(100);k__Fungi_unclassified(100);k__Fungi_unclassified(100);k__Fungi_unclassified(100);k__Fungi_unclassified(100);k__Fungi_unclassified(100); | - | - | - | - | - | - | - | Unassigned | - |
| Otu61378 | 1 | k__Fungi(100);k__Fungi_unclassified(100);k__Fungi_unclassified(100);k__Fungi_unclassified(100);k__Fungi_unclassified(100);k__Fungi_unclassified(100);k__Fungi_unclassified(100); | - | - | - | - | - | - | - | Unassigned | - |
| Otu61379 | 1 | k__Fungi(100);k__Fungi_unclassified(100);k__Fungi_unclassified(100);k__Fungi_unclassified(100);k__Fungi_unclassified(100);k__Fungi_unclassified(100);k__Fungi_unclassified(100); | - | - | - | - | - | - | - | Unassigned | - |
| Otu61380 | 1 | k__Fungi(100);k__Fungi_unclassified(100);k__Fungi_unclassified(100);k__Fungi_unclassified(100);k__Fungi_unclassified(100);k__Fungi_unclassified(100);k__Fungi_unclassified(100); | - | - | - | - | - | - | - | Unassigned | - |
| Otu61381 | 1 | k__Fungi(100);p__Ascomycota(100);p__Ascomycota_unclassified(100);p__Ascomycota_unclassified(100);p__Ascomycota_unclassified(100);p__Ascomycota_unclassified(100);p__Ascomycota_unclassified(100); | - | - | - | - | - | - | - | Unassigned | - |
| Otu61382 | 1 | k__Fungi(100);p__Ascomycota(100);p__Ascomycota_unclassified(100);p__Ascomycota_unclassified(100);p__Ascomycota_unclassified(100);p__Ascomycota_unclassified(100);p__Ascomycota_unclassified(100); | - | - | - | - | - | - | - | Unassigned | - |
| Otu61383 | 1 | k__Fungi(100);k__Fungi_unclassified(100);k__Fungi_unclassified(100);k__Fungi_unclassified(100);k__Fungi_unclassified(100);k__Fungi_unclassified(100);k__Fungi_unclassified(100); | - | - | - | - | - | - | - | Unassigned | - |
| Otu61384 | 1 | k__Fungi(100);k__Fungi_unclassified(100);k__Fungi_unclassified(100);k__Fungi_unclassified(100);k__Fungi_unclassified(100);k__Fungi_unclassified(100);k__Fungi_unclassified(100); | - | - | - | - | - | - | - | Unassigned | - |
| Otu61385 | 1 | k__Fungi(100);p__Ascomycota(100);p__Ascomycota_unclassified(100);p__Ascomycota_unclassified(100);p__Ascomycota_unclassified(100);p__Ascomycota_unclassified(100);p__Ascomycota_unclassified(100); | - | - | - | - | - | - | - | Unassigned | - |
| Otu61386 | 1 | k__Fungi(100);p__Ascomycota(100);p__Ascomycota_unclassified(100);p__Ascomycota_unclassified(100);p__Ascomycota_unclassified(100);p__Ascomycota_unclassified(100);p__Ascomycota_unclassified(100); | - | - | - | - | - | - | - | Unassigned | - |
| Otu61387 | 1 | k__Fungi(100);k__Fungi_unclassified(100);k__Fungi_unclassified(100);k__Fungi_unclassified(100);k__Fungi_unclassified(100);k__Fungi_unclassified(100);k__Fungi_unclassified(100); | - | - | - | - | - | - | - | Unassigned | - |
| Otu61388 | 1 | k__Fungi(100);p__Ascomycota(100);p__Ascomycota_unclassified(100);p__Ascomycota_unclassified(100);p__Ascomycota_unclassified(100);p__Ascomycota_unclassified(100);p__Ascomycota_unclassified(100); | - | - | - | - | - | - | - | Unassigned | - |
| Otu61389 | 1 | k__Fungi(100);k__Fungi_unclassified(100);k__Fungi_unclassified(100);k__Fungi_unclassified(100);k__Fungi_unclassified(100);k__Fungi_unclassified(100);k__Fungi_unclassified(100); | - | - | - | - | - | - | - | Unassigned | - |
| Otu61390 | 1 | k__Fungi(100);k__Fungi_unclassified(100);k__Fungi_unclassified(100);k__Fungi_unclassified(100);k__Fungi_unclassified(100);k__Fungi_unclassified(100);k__Fungi_unclassified(100); | - | - | - | - | - | - | - | Unassigned | - |
| Otu61391 | 1 | k__Fungi(100);k__Fungi_unclassified(100);k__Fungi_unclassified(100);k__Fungi_unclassified(100);k__Fungi_unclassified(100);k__Fungi_unclassified(100);k__Fungi_unclassified(100); | - | - | - | - | - | - | - | Unassigned | - |
| Otu61392 | 1 | k__Fungi(100);p__Ascomycota(100);p__Ascomycota_unclassified(100);p__Ascomycota_unclassified(100);p__Ascomycota_unclassified(100);p__Ascomycota_unclassified(100);p__Ascomycota_unclassified(100); | - | - | - | - | - | - | - | Unassigned | - |
| Otu61393 | 1 | k__Fungi(100);p__Ascomycota(100);p__Ascomycota_unclassified(100);p__Ascomycota_unclassified(100);p__Ascomycota_unclassified(100);p__Ascomycota_unclassified(100);p__Ascomycota_unclassified(100); | - | - | - | - | - | - | - | Unassigned | - |
| Otu61394 | 1 | k__Fungi(100);k__Fungi_unclassified(100);k__Fungi_unclassified(100);k__Fungi_unclassified(100);k__Fungi_unclassified(100);k__Fungi_unclassified(100);k__Fungi_unclassified(100); | - | - | - | - | - | - | - | Unassigned | - |
| Otu61395 | 1 | k__Fungi(100);p__Ascomycota(100);p__Ascomycota_unclassified(100);p__Ascomycota_unclassified(100);p__Ascomycota_unclassified(100);p__Ascomycota_unclassified(100);p__Ascomycota_unclassified(100); | - | - | - | - | - | - | - | Unassigned | - |
| Otu61396 | 1 | k__Fungi(100);k__Fungi_unclassified(100);k__Fungi_unclassified(100);k__Fungi_unclassified(100);k__Fungi_unclassified(100);k__Fungi_unclassified(100);k__Fungi_unclassified(100); | - | - | - | - | - | - | - | Unassigned | - |
| Otu61398 | 1 | k__Fungi(100);k__Fungi_unclassified(100);k__Fungi_unclassified(100);k__Fungi_unclassified(100);k__Fungi_unclassified(100);k__Fungi_unclassified(100);k__Fungi_unclassified(100); | - | - | - | - | - | - | - | Unassigned | - |
| Otu61399 | 1 | k__Fungi(100);p__Ascomycota(100);p__Ascomycota_unclassified(100);p__Ascomycota_unclassified(100);p__Ascomycota_unclassified(100);p__Ascomycota_unclassified(100);p__Ascomycota_unclassified(100); | - | - | - | - | - | - | - | Unassigned | - |
| Otu61400 | 1 | k__Fungi(100);p__Ascomycota(100);p__Ascomycota_unclassified(100);p__Ascomycota_unclassified(100);p__Ascomycota_unclassified(100);p__Ascomycota_unclassified(100);p__Ascomycota_unclassified(100); | - | - | - | - | - | - | - | Unassigned | - |
| Otu61401 | 1 | k__Fungi(100);k__Fungi_unclassified(100);k__Fungi_unclassified(100);k__Fungi_unclassified(100);k__Fungi_unclassified(100);k__Fungi_unclassified(100);k__Fungi_unclassified(100); | - | - | - | - | - | - | - | Unassigned | - |
| Otu61402 | 1 | k__Fungi(100);p__Ascomycota(100);p__Ascomycota_unclassified(100);p__Ascomycota_unclassified(100);p__Ascomycota_unclassified(100);p__Ascomycota_unclassified(100);p__Ascomycota_unclassified(100); | - | - | - | - | - | - | - | Unassigned | - |
| Otu61403 | 1 | k__Fungi(100);k__Fungi_unclassified(100);k__Fungi_unclassified(100);k__Fungi_unclassified(100);k__Fungi_unclassified(100);k__Fungi_unclassified(100);k__Fungi_unclassified(100); | - | - | - | - | - | - | - | Unassigned | - |
| Otu61404 | 1 | k__Fungi(100);k__Fungi_unclassified(100);k__Fungi_unclassified(100);k__Fungi_unclassified(100);k__Fungi_unclassified(100);k__Fungi_unclassified(100);k__Fungi_unclassified(100); | - | - | - | - | - | - | - | Unassigned | - |
| Otu61405 | 1 | k__Fungi(100);k__Fungi_unclassified(100);k__Fungi_unclassified(100);k__Fungi_unclassified(100);k__Fungi_unclassified(100);k__Fungi_unclassified(100);k__Fungi_unclassified(100); | - | - | - | - | - | - | - | Unassigned | - |
| Otu61406 | 1 | k__Fungi(100);k__Fungi_unclassified(100);k__Fungi_unclassified(100);k__Fungi_unclassified(100);k__Fungi_unclassified(100);k__Fungi_unclassified(100);k__Fungi_unclassified(100); | - | - | - | - | - | - | - | Unassigned | - |
| Otu61407 | 1 | k__Fungi(100);k__Fungi_unclassified(100);k__Fungi_unclassified(100);k__Fungi_unclassified(100);k__Fungi_unclassified(100);k__Fungi_unclassified(100);k__Fungi_unclassified(100); | - | - | - | - | - | - | - | Unassigned | - |
| Otu61450 | 1 | k__Fungi(100);k__Fungi_unclassified(100);k__Fungi_unclassified(100);k__Fungi_unclassified(100);k__Fungi_unclassified(100);k__Fungi_unclassified(100);k__Fungi_unclassified(100); | - | - | - | - | - | - | - | Unassigned | - |
| Otu61505 | 1 | k__Fungi(100);p__Ascomycota(100);p__Ascomycota_unclassified(100);p__Ascomycota_unclassified(100);p__Ascomycota_unclassified(100);p__Ascomycota_unclassified(100);p__Ascomycota_unclassified(100); | - | - | - | - | - | - | - | Unassigned | - |
| Otu61510 | 1 | k__Fungi(100);k__Fungi_unclassified(100);k__Fungi_unclassified(100);k__Fungi_unclassified(100);k__Fungi_unclassified(100);k__Fungi_unclassified(100);k__Fungi_unclassified(100); | - | - | - | - | - | - | - | Unassigned | - |
| Otu61539 | 1 | k__Fungi(100);k__Fungi_unclassified(100);k__Fungi_unclassified(100);k__Fungi_unclassified(100);k__Fungi_unclassified(100);k__Fungi_unclassified(100);k__Fungi_unclassified(100); | - | - | - | - | - | - | - | Unassigned | - |
| Otu61540 | 1 | k__Fungi(100);k__Fungi_unclassified(100);k__Fungi_unclassified(100);k__Fungi_unclassified(100);k__Fungi_unclassified(100);k__Fungi_unclassified(100);k__Fungi_unclassified(100); | - | - | - | - | - | - | - | Unassigned | - |
| Otu61541 | 1 | k__Fungi(100);p__Ascomycota(100);p__Ascomycota_unclassified(100);p__Ascomycota_unclassified(100);p__Ascomycota_unclassified(100);p__Ascomycota_unclassified(100);p__Ascomycota_unclassified(100); | - | - | - | - | - | - | - | Unassigned | - |
| Otu61542 | 1 | k__Fungi(100);k__Fungi_unclassified(100);k__Fungi_unclassified(100);k__Fungi_unclassified(100);k__Fungi_unclassified(100);k__Fungi_unclassified(100);k__Fungi_unclassified(100); | - | - | - | - | - | - | - | Unassigned | - |
| Otu61543 | 1 | k__Fungi(100);k__Fungi_unclassified(100);k__Fungi_unclassified(100);k__Fungi_unclassified(100);k__Fungi_unclassified(100);k__Fungi_unclassified(100);k__Fungi_unclassified(100); | - | - | - | - | - | - | - | Unassigned | - |
| Otu61544 | 1 | k__Fungi(100);k__Fungi_unclassified(100);k__Fungi_unclassified(100);k__Fungi_unclassified(100);k__Fungi_unclassified(100);k__Fungi_unclassified(100);k__Fungi_unclassified(100); | - | - | - | - | - | - | - | Unassigned | - |
| Otu61545 | 1 | k__Fungi(100);k__Fungi_unclassified(100);k__Fungi_unclassified(100);k__Fungi_unclassified(100);k__Fungi_unclassified(100);k__Fungi_unclassified(100);k__Fungi_unclassified(100); | - | - | - | - | - | - | - | Unassigned | - |
| Otu61547 | 1 | k__Fungi(100);k__Fungi_unclassified(100);k__Fungi_unclassified(100);k__Fungi_unclassified(100);k__Fungi_unclassified(100);k__Fungi_unclassified(100);k__Fungi_unclassified(100); | - | - | - | - | - | - | - | Unassigned | - |
| Otu61550 | 1 | k__Fungi(100);k__Fungi_unclassified(100);k__Fungi_unclassified(100);k__Fungi_unclassified(100);k__Fungi_unclassified(100);k__Fungi_unclassified(100);k__Fungi_unclassified(100); | - | - | - | - | - | - | - | Unassigned | - |
| Otu61555 | 1 | k__Fungi(100);p__Ascomycota(100);p__Ascomycota_unclassified(100);p__Ascomycota_unclassified(100);p__Ascomycota_unclassified(100);p__Ascomycota_unclassified(100);p__Ascomycota_unclassified(100); | - | - | - | - | - | - | - | Unassigned | - |
| Otu61578 | 1 | k__Fungi(100);k__Fungi_unclassified(100);k__Fungi_unclassified(100);k__Fungi_unclassified(100);k__Fungi_unclassified(100);k__Fungi_unclassified(100);k__Fungi_unclassified(100); | - | - | - | - | - | - | - | Unassigned | - |
| Otu61582 | 1 | k__Fungi(100);p__Ascomycota(100);p__Ascomycota_unclassified(100);p__Ascomycota_unclassified(100);p__Ascomycota_unclassified(100);p__Ascomycota_unclassified(100);p__Ascomycota_unclassified(100); | - | - | - | - | - | - | - | Unassigned | - |
| Otu61583 | 1 | k__Fungi(100);k__Fungi_unclassified(100);k__Fungi_unclassified(100);k__Fungi_unclassified(100);k__Fungi_unclassified(100);k__Fungi_unclassified(100);k__Fungi_unclassified(100); | - | - | - | - | - | - | - | Unassigned | - |
| Otu61590 | 1 | k__Fungi(100);p__Ascomycota(100);c__Dothideomycetes(100);o__Pleosporales(100);o__Pleosporales_unclassified(100);o__Pleosporales_unclassified(100);o__Pleosporales_unclassified(100); | - | - | - | - | - | - | - | Unassigned | - |
| Otu61602 | 1 | k__Fungi(100);p__Ascomycota(100);c__Dothideomycetes(100);o__Capnodiales(100);o__Capnodiales_unclassified(100);o__Capnodiales_unclassified(100);o__Capnodiales_unclassified(100); | - | - | - | - | - | - | - | Unassigned | - |
| Otu61603 | 1 | k__Fungi(100);p__Ascomycota(100);p__Ascomycota_unclassified(100);p__Ascomycota_unclassified(100);p__Ascomycota_unclassified(100);p__Ascomycota_unclassified(100);p__Ascomycota_unclassified(100); | - | - | - | - | - | - | - | Unassigned | - |
| Otu61604 | 1 | k__Fungi(100);p__Ascomycota(100);c__Sordariomycetes(100);c__Sordariomycetes_unclassified(100);c__Sordariomycetes_unclassified(100);c__Sordariomycetes_unclassified(100);c__Sordariomycetes_unclassified(100); | - | - | - | - | - | - | - | Unassigned | - |
| Otu61607 | 1 | k__Fungi(100);p__Ascomycota(100);p__Ascomycota_unclassified(100);p__Ascomycota_unclassified(100);p__Ascomycota_unclassified(100);p__Ascomycota_unclassified(100);p__Ascomycota_unclassified(100); | - | - | - | - | - | - | - | Unassigned | - |
| Otu61617 | 1 | k__Fungi(100);k__Fungi_unclassified(100);k__Fungi_unclassified(100);k__Fungi_unclassified(100);k__Fungi_unclassified(100);k__Fungi_unclassified(100);k__Fungi_unclassified(100); | - | - | - | - | - | - | - | Unassigned | - |
| Otu61619 | 1 | k__Fungi(100);k__Fungi_unclassified(100);k__Fungi_unclassified(100);k__Fungi_unclassified(100);k__Fungi_unclassified(100);k__Fungi_unclassified(100);k__Fungi_unclassified(100); | - | - | - | - | - | - | - | Unassigned | - |
| Otu61624 | 1 | k__Fungi(100);p__Ascomycota(100);p__Ascomycota_unclassified(100);p__Ascomycota_unclassified(100);p__Ascomycota_unclassified(100);p__Ascomycota_unclassified(100);p__Ascomycota_unclassified(100); | - | - | - | - | - | - | - | Unassigned | - |
| Otu61625 | 1 | k__Fungi(100);p__Ascomycota(100);p__Ascomycota_unclassified(100);p__Ascomycota_unclassified(100);p__Ascomycota_unclassified(100);p__Ascomycota_unclassified(100);p__Ascomycota_unclassified(100); | - | - | - | - | - | - | - | Unassigned | - |
| Otu61629 | 1 | k__Fungi(100);p__Ascomycota(100);p__Ascomycota_unclassified(100);p__Ascomycota_unclassified(100);p__Ascomycota_unclassified(100);p__Ascomycota_unclassified(100);p__Ascomycota_unclassified(100); | - | - | - | - | - | - | - | Unassigned | - |
| Otu61631 | 1 | k__Fungi(100);k__Fungi_unclassified(100);k__Fungi_unclassified(100);k__Fungi_unclassified(100);k__Fungi_unclassified(100);k__Fungi_unclassified(100);k__Fungi_unclassified(100); | - | - | - | - | - | - | - | Unassigned | - |
| Otu61633 | 1 | k__Fungi(100);k__Fungi_unclassified(100);k__Fungi_unclassified(100);k__Fungi_unclassified(100);k__Fungi_unclassified(100);k__Fungi_unclassified(100);k__Fungi_unclassified(100); | - | - | - | - | - | - | - | Unassigned | - |
| Otu61634 | 1 | k__Fungi(100);p__Ascomycota(100);p__Ascomycota_unclassified(100);p__Ascomycota_unclassified(100);p__Ascomycota_unclassified(100);p__Ascomycota_unclassified(100);p__Ascomycota_unclassified(100); | - | - | - | - | - | - | - | Unassigned | - |
| Otu61635 | 1 | k__Fungi(100);k__Fungi_unclassified(100);k__Fungi_unclassified(100);k__Fungi_unclassified(100);k__Fungi_unclassified(100);k__Fungi_unclassified(100);k__Fungi_unclassified(100); | - | - | - | - | - | - | - | Unassigned | - |
| Otu61636 | 1 | k__Fungi(100);k__Fungi_unclassified(100);k__Fungi_unclassified(100);k__Fungi_unclassified(100);k__Fungi_unclassified(100);k__Fungi_unclassified(100);k__Fungi_unclassified(100); | - | - | - | - | - | - | - | Unassigned | - |
| Otu61637 | 1 | k__Fungi(100);k__Fungi_unclassified(100);k__Fungi_unclassified(100);k__Fungi_unclassified(100);k__Fungi_unclassified(100);k__Fungi_unclassified(100);k__Fungi_unclassified(100); | - | - | - | - | - | - | - | Unassigned | - |
| Otu61638 | 1 | k__Fungi(100);k__Fungi_unclassified(100);k__Fungi_unclassified(100);k__Fungi_unclassified(100);k__Fungi_unclassified(100);k__Fungi_unclassified(100);k__Fungi_unclassified(100); | - | - | - | - | - | - | - | Unassigned | - |
| Otu61639 | 1 | k__Fungi(100);k__Fungi_unclassified(100);k__Fungi_unclassified(100);k__Fungi_unclassified(100);k__Fungi_unclassified(100);k__Fungi_unclassified(100);k__Fungi_unclassified(100); | - | - | - | - | - | - | - | Unassigned | - |
| Otu61640 | 1 | k__Fungi(100);p__Ascomycota(100);p__Ascomycota_unclassified(100);p__Ascomycota_unclassified(100);p__Ascomycota_unclassified(100);p__Ascomycota_unclassified(100);p__Ascomycota_unclassified(100); | - | - | - | - | - | - | - | Unassigned | - |
| Otu61642 | 1 | k__Fungi(100);k__Fungi_unclassified(100);k__Fungi_unclassified(100);k__Fungi_unclassified(100);k__Fungi_unclassified(100);k__Fungi_unclassified(100);k__Fungi_unclassified(100); | - | - | - | - | - | - | - | Unassigned | - |
| Otu61643 | 1 | k__Fungi(100);k__Fungi_unclassified(100);k__Fungi_unclassified(100);k__Fungi_unclassified(100);k__Fungi_unclassified(100);k__Fungi_unclassified(100);k__Fungi_unclassified(100); | - | - | - | - | - | - | - | Unassigned | - |
| Otu61645 | 1 | k__Fungi(100);p__Ascomycota(100);p__Ascomycota_unclassified(100);p__Ascomycota_unclassified(100);p__Ascomycota_unclassified(100);p__Ascomycota_unclassified(100);p__Ascomycota_unclassified(100); | - | - | - | - | - | - | - | Unassigned | - |
| Otu61646 | 1 | k__Fungi(100);k__Fungi_unclassified(100);k__Fungi_unclassified(100);k__Fungi_unclassified(100);k__Fungi_unclassified(100);k__Fungi_unclassified(100);k__Fungi_unclassified(100); | - | - | - | - | - | - | - | Unassigned | - |
| Otu61647 | 1 | k__Fungi(100);p__Ascomycota(100);p__Ascomycota_unclassified(100);p__Ascomycota_unclassified(100);p__Ascomycota_unclassified(100);p__Ascomycota_unclassified(100);p__Ascomycota_unclassified(100); | - | - | - | - | - | - | - | Unassigned | - |
| Otu61650 | 1 | k__Fungi(100);k__Fungi_unclassified(100);k__Fungi_unclassified(100);k__Fungi_unclassified(100);k__Fungi_unclassified(100);k__Fungi_unclassified(100);k__Fungi_unclassified(100); | - | - | - | - | - | - | - | Unassigned | - |
| Otu61652 | 1 | k__Fungi(100);k__Fungi_unclassified(100);k__Fungi_unclassified(100);k__Fungi_unclassified(100);k__Fungi_unclassified(100);k__Fungi_unclassified(100);k__Fungi_unclassified(100); | - | - | - | - | - | - | - | Unassigned | - |
| Otu61655 | 1 | k__Fungi(100);k__Fungi_unclassified(100);k__Fungi_unclassified(100);k__Fungi_unclassified(100);k__Fungi_unclassified(100);k__Fungi_unclassified(100);k__Fungi_unclassified(100); | - | - | - | - | - | - | - | Unassigned | - |
| Otu61664 | 1 | k__Fungi(100);p__Ascomycota(100);c__Leotiomycetes(100);c__Leotiomycetes_unclassified(100);c__Leotiomycetes_unclassified(100);c__Leotiomycetes_unclassified(100);c__Leotiomycetes_unclassified(100); | - | - | - | - | - | - | - | Unassigned | - |
| Otu61667 | 1 | k__Fungi(100);k__Fungi_unclassified(100);k__Fungi_unclassified(100);k__Fungi_unclassified(100);k__Fungi_unclassified(100);k__Fungi_unclassified(100);k__Fungi_unclassified(100); | - | - | - | - | - | - | - | Unassigned | - |
| Otu61670 | 1 | k__Fungi(100);p__Ascomycota(100);p__Ascomycota_unclassified(100);p__Ascomycota_unclassified(100);p__Ascomycota_unclassified(100);p__Ascomycota_unclassified(100);p__Ascomycota_unclassified(100); | - | - | - | - | - | - | - | Unassigned | - |
| Otu61671 | 1 | k__Fungi(100);k__Fungi_unclassified(100);k__Fungi_unclassified(100);k__Fungi_unclassified(100);k__Fungi_unclassified(100);k__Fungi_unclassified(100);k__Fungi_unclassified(100); | - | - | - | - | - | - | - | Unassigned | - |
| Otu61672 | 1 | k__Fungi(100);p__Ascomycota(100);p__Ascomycota_unclassified(100);p__Ascomycota_unclassified(100);p__Ascomycota_unclassified(100);p__Ascomycota_unclassified(100);p__Ascomycota_unclassified(100); | - | - | - | - | - | - | - | Unassigned | - |
| Otu61673 | 1 | k__Fungi(100);k__Fungi_unclassified(100);k__Fungi_unclassified(100);k__Fungi_unclassified(100);k__Fungi_unclassified(100);k__Fungi_unclassified(100);k__Fungi_unclassified(100); | - | - | - | - | - | - | - | Unassigned | - |
| Otu61674 | 1 | k__Fungi(100);p__Ascomycota(100);p__Ascomycota_unclassified(100);p__Ascomycota_unclassified(100);p__Ascomycota_unclassified(100);p__Ascomycota_unclassified(100);p__Ascomycota_unclassified(100); | - | - | - | - | - | - | - | Unassigned | - |
| Otu61675 | 1 | k__Fungi(100);k__Fungi_unclassified(100);k__Fungi_unclassified(100);k__Fungi_unclassified(100);k__Fungi_unclassified(100);k__Fungi_unclassified(100);k__Fungi_unclassified(100); | - | - | - | - | - | - | - | Unassigned | - |
| Otu61676 | 1 | k__Fungi(100);k__Fungi_unclassified(100);k__Fungi_unclassified(100);k__Fungi_unclassified(100);k__Fungi_unclassified(100);k__Fungi_unclassified(100);k__Fungi_unclassified(100); | - | - | - | - | - | - | - | Unassigned | - |
| Otu61677 | 1 | k__Fungi(100);k__Fungi_unclassified(100);k__Fungi_unclassified(100);k__Fungi_unclassified(100);k__Fungi_unclassified(100);k__Fungi_unclassified(100);k__Fungi_unclassified(100); | - | - | - | - | - | - | - | Unassigned | - |
| Otu61678 | 1 | k__Fungi(100);k__Fungi_unclassified(100);k__Fungi_unclassified(100);k__Fungi_unclassified(100);k__Fungi_unclassified(100);k__Fungi_unclassified(100);k__Fungi_unclassified(100); | - | - | - | - | - | - | - | Unassigned | - |
| Otu61679 | 1 | k__Fungi(100);p__Ascomycota(100);p__Ascomycota_unclassified(100);p__Ascomycota_unclassified(100);p__Ascomycota_unclassified(100);p__Ascomycota_unclassified(100);p__Ascomycota_unclassified(100); | - | - | - | - | - | - | - | Unassigned | - |
| Otu61682 | 1 | k__Fungi(100);p__Ascomycota(100);p__Ascomycota_unclassified(100);p__Ascomycota_unclassified(100);p__Ascomycota_unclassified(100);p__Ascomycota_unclassified(100);p__Ascomycota_unclassified(100); | - | - | - | - | - | - | - | Unassigned | - |
| Otu61684 | 1 | k__Fungi(100);k__Fungi_unclassified(100);k__Fungi_unclassified(100);k__Fungi_unclassified(100);k__Fungi_unclassified(100);k__Fungi_unclassified(100);k__Fungi_unclassified(100); | - | - | - | - | - | - | - | Unassigned | - |
| Otu61685 | 1 | k__Fungi(100);p__Ascomycota(100);p__Ascomycota_unclassified(100);p__Ascomycota_unclassified(100);p__Ascomycota_unclassified(100);p__Ascomycota_unclassified(100);p__Ascomycota_unclassified(100); | - | - | - | - | - | - | - | Unassigned | - |
| Otu61686 | 1 | k__Fungi(100);k__Fungi_unclassified(100);k__Fungi_unclassified(100);k__Fungi_unclassified(100);k__Fungi_unclassified(100);k__Fungi_unclassified(100);k__Fungi_unclassified(100); | - | - | - | - | - | - | - | Unassigned | - |
| Otu61689 | 1 | k__Fungi(100);k__Fungi_unclassified(100);k__Fungi_unclassified(100);k__Fungi_unclassified(100);k__Fungi_unclassified(100);k__Fungi_unclassified(100);k__Fungi_unclassified(100); | - | - | - | - | - | - | - | Unassigned | - |
| Otu61691 | 1 | k__Fungi(100);k__Fungi_unclassified(100);k__Fungi_unclassified(100);k__Fungi_unclassified(100);k__Fungi_unclassified(100);k__Fungi_unclassified(100);k__Fungi_unclassified(100); | - | - | - | - | - | - | - | Unassigned | - |
| Otu61692 | 1 | k__Fungi(100);p__Ascomycota(100);p__Ascomycota_unclassified(100);p__Ascomycota_unclassified(100);p__Ascomycota_unclassified(100);p__Ascomycota_unclassified(100);p__Ascomycota_unclassified(100); | - | - | - | - | - | - | - | Unassigned | - |
| Otu61693 | 1 | k__Fungi(100);k__Fungi_unclassified(100);k__Fungi_unclassified(100);k__Fungi_unclassified(100);k__Fungi_unclassified(100);k__Fungi_unclassified(100);k__Fungi_unclassified(100); | - | - | - | - | - | - | - | Unassigned | - |
| Otu61695 | 1 | k__Fungi(100);k__Fungi_unclassified(100);k__Fungi_unclassified(100);k__Fungi_unclassified(100);k__Fungi_unclassified(100);k__Fungi_unclassified(100);k__Fungi_unclassified(100); | - | - | - | - | - | - | - | Unassigned | - |
| Otu61697 | 1 | k__Fungi(100);k__Fungi_unclassified(100);k__Fungi_unclassified(100);k__Fungi_unclassified(100);k__Fungi_unclassified(100);k__Fungi_unclassified(100);k__Fungi_unclassified(100); | - | - | - | - | - | - | - | Unassigned | - |
| Otu61700 | 1 | k__Fungi(100);k__Fungi_unclassified(100);k__Fungi_unclassified(100);k__Fungi_unclassified(100);k__Fungi_unclassified(100);k__Fungi_unclassified(100);k__Fungi_unclassified(100); | - | - | - | - | - | - | - | Unassigned | - |
| Otu61701 | 1 | k__Fungi(100);k__Fungi_unclassified(100);k__Fungi_unclassified(100);k__Fungi_unclassified(100);k__Fungi_unclassified(100);k__Fungi_unclassified(100);k__Fungi_unclassified(100); | - | - | - | - | - | - | - | Unassigned | - |
| Otu61702 | 1 | k__Fungi(100);k__Fungi_unclassified(100);k__Fungi_unclassified(100);k__Fungi_unclassified(100);k__Fungi_unclassified(100);k__Fungi_unclassified(100);k__Fungi_unclassified(100); | - | - | - | - | - | - | - | Unassigned | - |
| Otu61704 | 1 | k__Fungi(100);p__Ascomycota(100);p__Ascomycota_unclassified(100);p__Ascomycota_unclassified(100);p__Ascomycota_unclassified(100);p__Ascomycota_unclassified(100);p__Ascomycota_unclassified(100); | - | - | - | - | - | - | - | Unassigned | - |
| Otu61707 | 1 | k__Fungi(100);p__Ascomycota(100);p__Ascomycota_unclassified(100);p__Ascomycota_unclassified(100);p__Ascomycota_unclassified(100);p__Ascomycota_unclassified(100);p__Ascomycota_unclassified(100); | - | - | - | - | - | - | - | Unassigned | - |
| Otu61708 | 1 | k__Fungi(100);p__Ascomycota(100);p__Ascomycota_unclassified(100);p__Ascomycota_unclassified(100);p__Ascomycota_unclassified(100);p__Ascomycota_unclassified(100);p__Ascomycota_unclassified(100); | - | - | - | - | - | - | - | Unassigned | - |
| Otu61710 | 1 | k__Fungi(100);p__Ascomycota(100);p__Ascomycota_unclassified(100);p__Ascomycota_unclassified(100);p__Ascomycota_unclassified(100);p__Ascomycota_unclassified(100);p__Ascomycota_unclassified(100); | - | - | - | - | - | - | - | Unassigned | - |
| Otu61712 | 1 | k__Fungi(100);k__Fungi_unclassified(100);k__Fungi_unclassified(100);k__Fungi_unclassified(100);k__Fungi_unclassified(100);k__Fungi_unclassified(100);k__Fungi_unclassified(100); | - | - | - | - | - | - | - | Unassigned | - |
| Otu61714 | 1 | k__Fungi(100);k__Fungi_unclassified(100);k__Fungi_unclassified(100);k__Fungi_unclassified(100);k__Fungi_unclassified(100);k__Fungi_unclassified(100);k__Fungi_unclassified(100); | - | - | - | - | - | - | - | Unassigned | - |
| Otu61716 | 1 | k__Fungi(100);k__Fungi_unclassified(100);k__Fungi_unclassified(100);k__Fungi_unclassified(100);k__Fungi_unclassified(100);k__Fungi_unclassified(100);k__Fungi_unclassified(100); | - | - | - | - | - | - | - | Unassigned | - |
| Otu61718 | 1 | k__Fungi(100);k__Fungi_unclassified(100);k__Fungi_unclassified(100);k__Fungi_unclassified(100);k__Fungi_unclassified(100);k__Fungi_unclassified(100);k__Fungi_unclassified(100); | - | - | - | - | - | - | - | Unassigned | - |
| Otu61719 | 1 | k__Fungi(100);k__Fungi_unclassified(100);k__Fungi_unclassified(100);k__Fungi_unclassified(100);k__Fungi_unclassified(100);k__Fungi_unclassified(100);k__Fungi_unclassified(100); | - | - | - | - | - | - | - | Unassigned | - |
| Otu61720 | 1 | k__Fungi(100);k__Fungi_unclassified(100);k__Fungi_unclassified(100);k__Fungi_unclassified(100);k__Fungi_unclassified(100);k__Fungi_unclassified(100);k__Fungi_unclassified(100); | - | - | - | - | - | - | - | Unassigned | - |
| Otu61725 | 1 | k__Fungi(100);p__Ascomycota(100);p__Ascomycota_unclassified(100);p__Ascomycota_unclassified(100);p__Ascomycota_unclassified(100);p__Ascomycota_unclassified(100);p__Ascomycota_unclassified(100); | - | - | - | - | - | - | - | Unassigned | - |
| Otu61726 | 1 | k__Fungi(100);p__Ascomycota(100);p__Ascomycota_unclassified(100);p__Ascomycota_unclassified(100);p__Ascomycota_unclassified(100);p__Ascomycota_unclassified(100);p__Ascomycota_unclassified(100); | - | - | - | - | - | - | - | Unassigned | - |
| Otu61727 | 1 | k__Fungi(100);k__Fungi_unclassified(100);k__Fungi_unclassified(100);k__Fungi_unclassified(100);k__Fungi_unclassified(100);k__Fungi_unclassified(100);k__Fungi_unclassified(100); | - | - | - | - | - | - | - | Unassigned | - |
| Otu61728 | 1 | k__Fungi(100);p__Ascomycota(100);p__Ascomycota_unclassified(100);p__Ascomycota_unclassified(100);p__Ascomycota_unclassified(100);p__Ascomycota_unclassified(100);p__Ascomycota_unclassified(100); | - | - | - | - | - | - | - | Unassigned | - |
| Otu61730 | 1 | k__Fungi(100);p__Ascomycota(100);p__Ascomycota_unclassified(100);p__Ascomycota_unclassified(100);p__Ascomycota_unclassified(100);p__Ascomycota_unclassified(100);p__Ascomycota_unclassified(100); | - | - | - | - | - | - | - | Unassigned | - |
| Otu61731 | 1 | k__Fungi(100);p__Ascomycota(100);p__Ascomycota_unclassified(100);p__Ascomycota_unclassified(100);p__Ascomycota_unclassified(100);p__Ascomycota_unclassified(100);p__Ascomycota_unclassified(100); | - | - | - | - | - | - | - | Unassigned | - |
| Otu61734 | 1 | k__Fungi(100);p__Ascomycota(100);p__Ascomycota_unclassified(100);p__Ascomycota_unclassified(100);p__Ascomycota_unclassified(100);p__Ascomycota_unclassified(100);p__Ascomycota_unclassified(100); | - | - | - | - | - | - | - | Unassigned | - |
| Otu61735 | 1 | k__Fungi(100);k__Fungi_unclassified(100);k__Fungi_unclassified(100);k__Fungi_unclassified(100);k__Fungi_unclassified(100);k__Fungi_unclassified(100);k__Fungi_unclassified(100); | - | - | - | - | - | - | - | Unassigned | - |
| Otu61736 | 1 | k__Fungi(100);p__Ascomycota(100);p__Ascomycota_unclassified(100);p__Ascomycota_unclassified(100);p__Ascomycota_unclassified(100);p__Ascomycota_unclassified(100);p__Ascomycota_unclassified(100); | - | - | - | - | - | - | - | Unassigned | - |
| Otu61737 | 1 | k__Fungi(100);k__Fungi_unclassified(100);k__Fungi_unclassified(100);k__Fungi_unclassified(100);k__Fungi_unclassified(100);k__Fungi_unclassified(100);k__Fungi_unclassified(100); | - | - | - | - | - | - | - | Unassigned | - |
| Otu61738 | 1 | k__Fungi(100);p__Ascomycota(100);p__Ascomycota_unclassified(100);p__Ascomycota_unclassified(100);p__Ascomycota_unclassified(100);p__Ascomycota_unclassified(100);p__Ascomycota_unclassified(100); | - | - | - | - | - | - | - | Unassigned | - |
| Otu61739 | 1 | k__Fungi(100);p__Ascomycota(100);p__Ascomycota_unclassified(100);p__Ascomycota_unclassified(100);p__Ascomycota_unclassified(100);p__Ascomycota_unclassified(100);p__Ascomycota_unclassified(100); | - | - | - | - | - | - | - | Unassigned | - |
| Otu61740 | 1 | k__Fungi(100);k__Fungi_unclassified(100);k__Fungi_unclassified(100);k__Fungi_unclassified(100);k__Fungi_unclassified(100);k__Fungi_unclassified(100);k__Fungi_unclassified(100); | - | - | - | - | - | - | - | Unassigned | - |
| Otu61741 | 1 | k__Fungi(100);k__Fungi_unclassified(100);k__Fungi_unclassified(100);k__Fungi_unclassified(100);k__Fungi_unclassified(100);k__Fungi_unclassified(100);k__Fungi_unclassified(100); | - | - | - | - | - | - | - | Unassigned | - |
| Otu61742 | 1 | k__Fungi(100);k__Fungi_unclassified(100);k__Fungi_unclassified(100);k__Fungi_unclassified(100);k__Fungi_unclassified(100);k__Fungi_unclassified(100);k__Fungi_unclassified(100); | - | - | - | - | - | - | - | Unassigned | - |
| Otu61743 | 1 | k__Fungi(100);p__Ascomycota(100);p__Ascomycota_unclassified(100);p__Ascomycota_unclassified(100);p__Ascomycota_unclassified(100);p__Ascomycota_unclassified(100);p__Ascomycota_unclassified(100); | - | - | - | - | - | - | - | Unassigned | - |
| Otu61744 | 1 | k__Fungi(100);p__Ascomycota(100);p__Ascomycota_unclassified(100);p__Ascomycota_unclassified(100);p__Ascomycota_unclassified(100);p__Ascomycota_unclassified(100);p__Ascomycota_unclassified(100); | - | - | - | - | - | - | - | Unassigned | - |
| Otu61745 | 1 | k__Fungi(100);k__Fungi_unclassified(100);k__Fungi_unclassified(100);k__Fungi_unclassified(100);k__Fungi_unclassified(100);k__Fungi_unclassified(100);k__Fungi_unclassified(100); | - | - | - | - | - | - | - | Unassigned | - |
| Otu61746 | 1 | k__Fungi(100);k__Fungi_unclassified(100);k__Fungi_unclassified(100);k__Fungi_unclassified(100);k__Fungi_unclassified(100);k__Fungi_unclassified(100);k__Fungi_unclassified(100); | - | - | - | - | - | - | - | Unassigned | - |
| Otu61747 | 1 | k__Fungi(100);p__Ascomycota(100);p__Ascomycota_unclassified(100);p__Ascomycota_unclassified(100);p__Ascomycota_unclassified(100);p__Ascomycota_unclassified(100);p__Ascomycota_unclassified(100); | - | - | - | - | - | - | - | Unassigned | - |
| Otu61749 | 1 | k__Fungi(100);k__Fungi_unclassified(100);k__Fungi_unclassified(100);k__Fungi_unclassified(100);k__Fungi_unclassified(100);k__Fungi_unclassified(100);k__Fungi_unclassified(100); | - | - | - | - | - | - | - | Unassigned | - |
| Otu61750 | 1 | k__Fungi(100);p__Ascomycota(100);p__Ascomycota_unclassified(100);p__Ascomycota_unclassified(100);p__Ascomycota_unclassified(100);p__Ascomycota_unclassified(100);p__Ascomycota_unclassified(100); | - | - | - | - | - | - | - | Unassigned | - |
| Otu61752 | 1 | k__Fungi(100);p__Ascomycota(100);p__Ascomycota_unclassified(100);p__Ascomycota_unclassified(100);p__Ascomycota_unclassified(100);p__Ascomycota_unclassified(100);p__Ascomycota_unclassified(100); | - | - | - | - | - | - | - | Unassigned | - |

| | | | | | | | | | | |
|---|---|---|---|---|---|---|---|---|---|---|
| Otu61754 | 1 | k__Fungi(100);k__Fungi_unclassified(100);k__Fungi_unclassified(100);k__Fungi_unclassified(100);k__Fungi_unclassified(100);k__Fungi_unclassified(100);k__Fungi_unclassified(100); | - | - | - | - | - | - | - | Unassigned | - |
| Otu61756 | 1 | k__Fungi(100);p__Ascomycota(100);p__Ascomycota_unclassified(100);p__Ascomycota_unclassified(100);p__Ascomycota_unclassified(100);p__Ascomycota_unclassified(100);p__Ascomycota_unclassified(100); | - | - | - | - | - | - | - | Unassigned | - |
| Otu61757 | 1 | k__Fungi(100);k__Fungi_unclassified(100);k__Fungi_unclassified(100);k__Fungi_unclassified(100);k__Fungi_unclassified(100);k__Fungi_unclassified(100);k__Fungi_unclassified(100); | - | - | - | - | - | - | - | Unassigned | - |
| Otu61759 | 1 | k__Fungi(100);p__Ascomycota(100);p__Ascomycota_unclassified(100);p__Ascomycota_unclassified(100);p__Ascomycota_unclassified(100);p__Ascomycota_unclassified(100);p__Ascomycota_unclassified(100); | - | - | - | - | - | - | - | Unassigned | - |
| Otu61760 | 1 | k__Fungi(100);k__Fungi_unclassified(100);k__Fungi_unclassified(100);k__Fungi_unclassified(100);k__Fungi_unclassified(100);k__Fungi_unclassified(100);k__Fungi_unclassified(100); | - | - | - | - | - | - | - | Unassigned | - |
| Otu61761 | 1 | k__Fungi(100);k__Fungi_unclassified(100);k__Fungi_unclassified(100);k__Fungi_unclassified(100);k__Fungi_unclassified(100);k__Fungi_unclassified(100);k__Fungi_unclassified(100); | - | - | - | - | - | - | - | Unassigned | - |
| Otu61763 | 1 | k__Fungi(100);p__Ascomycota(100);p__Ascomycota_unclassified(100);p__Ascomycota_unclassified(100);p__Ascomycota_unclassified(100);p__Ascomycota_unclassified(100);p__Ascomycota_unclassified(100); | - | - | - | - | - | - | - | Unassigned | - |
| Otu61764 | 1 | k__Fungi(100);p__Ascomycota(100);p__Ascomycota_unclassified(100);p__Ascomycota_unclassified(100);p__Ascomycota_unclassified(100);p__Ascomycota_unclassified(100);p__Ascomycota_unclassified(100); | - | - | - | - | - | - | - | Unassigned | - |
| Otu61765 | 1 | k__Fungi(100);k__Fungi_unclassified(100);k__Fungi_unclassified(100);k__Fungi_unclassified(100);k__Fungi_unclassified(100);k__Fungi_unclassified(100);k__Fungi_unclassified(100); | - | - | - | - | - | - | - | Unassigned | - |
| Otu61766 | 1 | k__Fungi(100);p__Ascomycota(100);p__Ascomycota_unclassified(100);p__Ascomycota_unclassified(100);p__Ascomycota_unclassified(100);p__Ascomycota_unclassified(100);p__Ascomycota_unclassified(100); | - | - | - | - | - | - | - | Unassigned | - |
| Otu61767 | 1 | k__Fungi(100);p__Ascomycota(100);p__Ascomycota_unclassified(100);p__Ascomycota_unclassified(100);p__Ascomycota_unclassified(100);p__Ascomycota_unclassified(100);p__Ascomycota_unclassified(100); | - | - | - | - | - | - | - | Unassigned | - |
| Otu61773 | 1 | k__Fungi(100);k__Fungi_unclassified(100);k__Fungi_unclassified(100);k__Fungi_unclassified(100);k__Fungi_unclassified(100);k__Fungi_unclassified(100);k__Fungi_unclassified(100); | - | - | - | - | - | - | - | Unassigned | - |
| Otu61774 | 1 | k__Fungi(100);p__Ascomycota(100);p__Ascomycota_unclassified(100);p__Ascomycota_unclassified(100);p__Ascomycota_unclassified(100);p__Ascomycota_unclassified(100);p__Ascomycota_unclassified(100); | - | - | - | - | - | - | - | Unassigned | - |
| Otu61775 | 1 | k__Fungi(100);k__Fungi_unclassified(100);k__Fungi_unclassified(100);k__Fungi_unclassified(100);k__Fungi_unclassified(100);k__Fungi_unclassified(100);k__Fungi_unclassified(100); | - | - | - | - | - | - | - | Unassigned | - |
| Otu61787 | 1 | k__Fungi(100);k__Fungi_unclassified(100);k__Fungi_unclassified(100);k__Fungi_unclassified(100);k__Fungi_unclassified(100);k__Fungi_unclassified(100);k__Fungi_unclassified(100); | - | - | - | - | - | - | - | Unassigned | - |
| Otu61789 | 1 | k__Fungi(100);k__Fungi_unclassified(100);k__Fungi_unclassified(100);k__Fungi_unclassified(100);k__Fungi_unclassified(100);k__Fungi_unclassified(100);k__Fungi_unclassified(100); | - | - | - | - | - | - | - | Unassigned | - |
| Otu61790 | 1 | k__Fungi(100);k__Fungi_unclassified(100);k__Fungi_unclassified(100);k__Fungi_unclassified(100);k__Fungi_unclassified(100);k__Fungi_unclassified(100);k__Fungi_unclassified(100); | - | - | - | - | - | - | - | Unassigned | - |
| Otu61791 | 1 | k__Fungi(100);k__Fungi_unclassified(100);k__Fungi_unclassified(100);k__Fungi_unclassified(100);k__Fungi_unclassified(100);k__Fungi_unclassified(100);k__Fungi_unclassified(100); | - | - | - | - | - | - | - | Unassigned | - |
| Otu61792 | 1 | k__Fungi(100);k__Fungi_unclassified(100);k__Fungi_unclassified(100);k__Fungi_unclassified(100);k__Fungi_unclassified(100);k__Fungi_unclassified(100);k__Fungi_unclassified(100); | - | - | - | - | - | - | - | Unassigned | - |
| Otu61793 | 1 | k__Fungi(100);p__Ascomycota(100);p__Ascomycota_unclassified(100);p__Ascomycota_unclassified(100);p__Ascomycota_unclassified(100);p__Ascomycota_unclassified(100);p__Ascomycota_unclassified(100); | - | - | - | - | - | - | - | Unassigned | - |
| Otu61794 | 1 | k__Fungi(100);k__Fungi_unclassified(100);k__Fungi_unclassified(100);k__Fungi_unclassified(100);k__Fungi_unclassified(100);k__Fungi_unclassified(100);k__Fungi_unclassified(100); | - | - | - | - | - | - | - | Unassigned | - |
| Otu61795 | 1 | k__Fungi(100);k__Fungi_unclassified(100);k__Fungi_unclassified(100);k__Fungi_unclassified(100);k__Fungi_unclassified(100);k__Fungi_unclassified(100);k__Fungi_unclassified(100); | - | - | - | - | - | - | - | Unassigned | - |
| Otu61799 | 1 | k__Fungi(100);k__Fungi_unclassified(100);k__Fungi_unclassified(100);k__Fungi_unclassified(100);k__Fungi_unclassified(100);k__Fungi_unclassified(100);k__Fungi_unclassified(100); | - | - | - | - | - | - | - | Unassigned | - |
| Otu61800 | 1 | k__Fungi(100);p__Ascomycota(100);p__Ascomycota_unclassified(100);p__Ascomycota_unclassified(100);p__Ascomycota_unclassified(100);p__Ascomycota_unclassified(100);p__Ascomycota_unclassified(100); | - | - | - | - | - | - | - | Unassigned | - |
| Otu61801 | 1 | k__Fungi(100);k__Fungi_unclassified(100);k__Fungi_unclassified(100);k__Fungi_unclassified(100);k__Fungi_unclassified(100);k__Fungi_unclassified(100);k__Fungi_unclassified(100); | - | - | - | - | - | - | - | Unassigned | - |
| Otu61802 | 1 | k__Fungi(100);k__Fungi_unclassified(100);k__Fungi_unclassified(100);k__Fungi_unclassified(100);k__Fungi_unclassified(100);k__Fungi_unclassified(100);k__Fungi_unclassified(100); | - | - | - | - | - | - | - | Unassigned | - |
| Otu61803 | 1 | k__Fungi(100);p__Ascomycota(100);p__Ascomycota_unclassified(100);p__Ascomycota_unclassified(100);p__Ascomycota_unclassified(100);p__Ascomycota_unclassified(100);p__Ascomycota_unclassified(100); | - | - | - | - | - | - | - | Unassigned | - |
| Otu61804 | 1 | k__Fungi(100);p__Ascomycota(100);p__Ascomycota_unclassified(100);p__Ascomycota_unclassified(100);p__Ascomycota_unclassified(100);p__Ascomycota_unclassified(100);p__Ascomycota_unclassified(100); | - | - | - | - | - | - | - | Unassigned | - |
| Otu61805 | 1 | k__Fungi(100);k__Fungi_unclassified(100);k__Fungi_unclassified(100);k__Fungi_unclassified(100);k__Fungi_unclassified(100);k__Fungi_unclassified(100);k__Fungi_unclassified(100); | - | - | - | - | - | - | - | Unassigned | - |
| Otu61807 | 1 | k__Fungi(100);k__Fungi_unclassified(100);k__Fungi_unclassified(100);k__Fungi_unclassified(100);k__Fungi_unclassified(100);k__Fungi_unclassified(100);k__Fungi_unclassified(100); | - | - | - | - | - | - | - | Unassigned | - |
| Otu61809 | 1 | k__Fungi(100);k__Fungi_unclassified(100);k__Fungi_unclassified(100);k__Fungi_unclassified(100);k__Fungi_unclassified(100);k__Fungi_unclassified(100);k__Fungi_unclassified(100); | - | - | - | - | - | - | - | Unassigned | - |
| Otu61810 | 1 | k__Fungi(100);k__Fungi_unclassified(100);k__Fungi_unclassified(100);k__Fungi_unclassified(100);k__Fungi_unclassified(100);k__Fungi_unclassified(100);k__Fungi_unclassified(100); | - | - | - | - | - | - | - | Unassigned | - |
| Otu61811 | 1 | k__Fungi(100);k__Fungi_unclassified(100);k__Fungi_unclassified(100);k__Fungi_unclassified(100);k__Fungi_unclassified(100);k__Fungi_unclassified(100);k__Fungi_unclassified(100); | - | - | - | - | - | - | - | Unassigned | - |
| Otu61812 | 1 | k__Fungi(100);k__Fungi_unclassified(100);k__Fungi_unclassified(100);k__Fungi_unclassified(100);k__Fungi_unclassified(100);k__Fungi_unclassified(100);k__Fungi_unclassified(100); | - | - | - | - | - | - | - | Unassigned | - |
| Otu61813 | 1 | k__Fungi(100);k__Fungi_unclassified(100);k__Fungi_unclassified(100);k__Fungi_unclassified(100);k__Fungi_unclassified(100);k__Fungi_unclassified(100);k__Fungi_unclassified(100); | - | - | - | - | - | - | - | Unassigned | - |
| Otu61814 | 1 | k__Fungi(100);k__Fungi_unclassified(100);k__Fungi_unclassified(100);k__Fungi_unclassified(100);k__Fungi_unclassified(100);k__Fungi_unclassified(100);k__Fungi_unclassified(100); | - | - | - | - | - | - | - | Unassigned | - |
| Otu61815 | 1 | k__Fungi(100);k__Fungi_unclassified(100);k__Fungi_unclassified(100);k__Fungi_unclassified(100);k__Fungi_unclassified(100);k__Fungi_unclassified(100);k__Fungi_unclassified(100); | - | - | - | - | - | - | - | Unassigned | - |
| Otu61816 | 1 | k__Fungi(100);k__Fungi_unclassified(100);k__Fungi_unclassified(100);k__Fungi_unclassified(100);k__Fungi_unclassified(100);k__Fungi_unclassified(100);k__Fungi_unclassified(100); | - | - | - | - | - | - | - | Unassigned | - |
| Otu61817 | 1 | k__Fungi(100);k__Fungi_unclassified(100);k__Fungi_unclassified(100);k__Fungi_unclassified(100);k__Fungi_unclassified(100);k__Fungi_unclassified(100);k__Fungi_unclassified(100); | - | - | - | - | - | - | - | Unassigned | - |
| Otu61818 | 1 | k__Fungi(100);p__Ascomycota(100);p__Ascomycota_unclassified(100);p__Ascomycota_unclassified(100);p__Ascomycota_unclassified(100);p__Ascomycota_unclassified(100);p__Ascomycota_unclassified(100); | - | - | - | - | - | - | - | Unassigned | - |
| Otu61824 | 1 | k__Fungi(100);k__Fungi_unclassified(100);k__Fungi_unclassified(100);k__Fungi_unclassified(100);k__Fungi_unclassified(100);k__Fungi_unclassified(100);k__Fungi_unclassified(100); | - | - | - | - | - | - | - | Unassigned | - |
| Otu61825 | 1 | k__Fungi(100);k__Fungi_unclassified(100);k__Fungi_unclassified(100);k__Fungi_unclassified(100);k__Fungi_unclassified(100);k__Fungi_unclassified(100);k__Fungi_unclassified(100); | - | - | - | - | - | - | - | Unassigned | - |
| Otu61826 | 1 | k__Fungi(100);k__Fungi_unclassified(100);k__Fungi_unclassified(100);k__Fungi_unclassified(100);k__Fungi_unclassified(100);k__Fungi_unclassified(100);k__Fungi_unclassified(100); | - | - | - | - | - | - | - | Unassigned | - |
| Otu61827 | 1 | k__Fungi(100);k__Fungi_unclassified(100);k__Fungi_unclassified(100);k__Fungi_unclassified(100);k__Fungi_unclassified(100);k__Fungi_unclassified(100);k__Fungi_unclassified(100); | - | - | - | - | - | - | - | Unassigned | - |
| Otu61828 | 1 | k__Fungi(100);k__Fungi_unclassified(100);k__Fungi_unclassified(100);k__Fungi_unclassified(100);k__Fungi_unclassified(100);k__Fungi_unclassified(100);k__Fungi_unclassified(100); | - | - | - | - | - | - | - | Unassigned | - |
| Otu61829 | 1 | k__Fungi(100);k__Fungi_unclassified(100);k__Fungi_unclassified(100);k__Fungi_unclassified(100);k__Fungi_unclassified(100);k__Fungi_unclassified(100);k__Fungi_unclassified(100); | - | - | - | - | - | - | - | Unassigned | - |
| Otu61830 | 1 | k__Fungi(100);k__Fungi_unclassified(100);k__Fungi_unclassified(100);k__Fungi_unclassified(100);k__Fungi_unclassified(100);k__Fungi_unclassified(100);k__Fungi_unclassified(100); | - | - | - | - | - | - | - | Unassigned | - |
| Otu61831 | 1 | k__Fungi(100);k__Fungi_unclassified(100);k__Fungi_unclassified(100);k__Fungi_unclassified(100);k__Fungi_unclassified(100);k__Fungi_unclassified(100);k__Fungi_unclassified(100); | - | - | - | - | - | - | - | Unassigned | - |
| Otu61833 | 1 | k__Fungi(100);k__Fungi_unclassified(100);k__Fungi_unclassified(100);k__Fungi_unclassified(100);k__Fungi_unclassified(100);k__Fungi_unclassified(100);k__Fungi_unclassified(100); | - | - | - | - | - | - | - | Unassigned | - |
| Otu61834 | 1 | k__Fungi(100);k__Fungi_unclassified(100);k__Fungi_unclassified(100);k__Fungi_unclassified(100);k__Fungi_unclassified(100);k__Fungi_unclassified(100);k__Fungi_unclassified(100); | - | - | - | - | - | - | - | Unassigned | - |
| Otu61836 | 1 | k__Fungi(100);k__Fungi_unclassified(100);k__Fungi_unclassified(100);k__Fungi_unclassified(100);k__Fungi_unclassified(100);k__Fungi_unclassified(100);k__Fungi_unclassified(100); | - | - | - | - | - | - | - | Unassigned | - |
| Otu61839 | 1 | k__Fungi(100);k__Fungi_unclassified(100);k__Fungi_unclassified(100);k__Fungi_unclassified(100);k__Fungi_unclassified(100);k__Fungi_unclassified(100);k__Fungi_unclassified(100); | - | - | - | - | - | - | - | Unassigned | - |
| Otu61841 | 1 | k__Fungi(100);k__Fungi_unclassified(100);k__Fungi_unclassified(100);k__Fungi_unclassified(100);k__Fungi_unclassified(100);k__Fungi_unclassified(100);k__Fungi_unclassified(100); | - | - | - | - | - | - | - | Unassigned | - |
| Otu61842 | 1 | k__Fungi(100);k__Fungi_unclassified(100);k__Fungi_unclassified(100);k__Fungi_unclassified(100);k__Fungi_unclassified(100);k__Fungi_unclassified(100);k__Fungi_unclassified(100); | - | - | - | - | - | - | - | Unassigned | - |
| Otu61844 | 1 | k__Fungi(100);k__Fungi_unclassified(100);k__Fungi_unclassified(100);k__Fungi_unclassified(100);k__Fungi_unclassified(100);k__Fungi_unclassified(100);k__Fungi_unclassified(100); | - | - | - | - | - | - | - | Unassigned | - |
| Otu61845 | 1 | k__Fungi(100);k__Fungi_unclassified(100);k__Fungi_unclassified(100);k__Fungi_unclassified(100);k__Fungi_unclassified(100);k__Fungi_unclassified(100);k__Fungi_unclassified(100); | - | - | - | - | - | - | - | Unassigned | - |
| Otu61848 | 1 | k__Fungi(100);p__Ascomycota(100);p__Ascomycota_unclassified(100);p__Ascomycota_unclassified(100);p__Ascomycota_unclassified(100);p__Ascomycota_unclassified(100);p__Ascomycota_unclassified(100); | - | - | - | - | - | - | - | Unassigned | - |
| Otu61849 | 1 | k__Fungi(100);k__Fungi_unclassified(100);k__Fungi_unclassified(100);k__Fungi_unclassified(100);k__Fungi_unclassified(100);k__Fungi_unclassified(100);k__Fungi_unclassified(100); | - | - | - | - | - | - | - | Unassigned | - |
| Otu61850 | 1 | k__Fungi(100);k__Fungi_unclassified(100);k__Fungi_unclassified(100);k__Fungi_unclassified(100);k__Fungi_unclassified(100);k__Fungi_unclassified(100);k__Fungi_unclassified(100); | - | - | - | - | - | - | - | Unassigned | - |
| Otu61851 | 1 | k__Fungi(100);k__Fungi_unclassified(100);k__Fungi_unclassified(100);k__Fungi_unclassified(100);k__Fungi_unclassified(100);k__Fungi_unclassified(100);k__Fungi_unclassified(100); | - | - | - | - | - | - | - | Unassigned | - |
| Otu61852 | 1 | k__Fungi(100);k__Fungi_unclassified(100);k__Fungi_unclassified(100);k__Fungi_unclassified(100);k__Fungi_unclassified(100);k__Fungi_unclassified(100);k__Fungi_unclassified(100); | - | - | - | - | - | - | - | Unassigned | - |
| Otu61854 | 1 | k__Fungi(100);p__Ascomycota(100);p__Ascomycota_unclassified(100);p__Ascomycota_unclassified(100);p__Ascomycota_unclassified(100);p__Ascomycota_unclassified(100);p__Ascomycota_unclassified(100); | - | - | - | - | - | - | - | Unassigned | - |
| Otu61856 | 1 | k__Fungi(100);k__Fungi_unclassified(100);k__Fungi_unclassified(100);k__Fungi_unclassified(100);k__Fungi_unclassified(100);k__Fungi_unclassified(100);k__Fungi_unclassified(100); | - | - | - | - | - | - | - | Unassigned | - |
| Otu61859 | 1 | k__Fungi(100);p__Ascomycota(100);c__Dothideomycetes(100);c__Dothideomycetes_unclassified(100);c__Dothideomycetes_unclassified(100);c__Dothideomycetes_unclassified(100);c__Dothideomycetes_unclassified(100); | - | - | - | - | - | - | - | Unassigned | - |
| Otu61860 | 1 | k__Fungi(100);p__Ascomycota(100);p__Ascomycota_unclassified(100);p__Ascomycota_unclassified(100);p__Ascomycota_unclassified(100);p__Ascomycota_unclassified(100);p__Ascomycota_unclassified(100); | - | - | - | - | - | - | - | Unassigned | - |
| Otu61861 | 1 | k__Fungi(100);p__Ascomycota(100);p__Ascomycota_unclassified(100);p__Ascomycota_unclassified(100);p__Ascomycota_unclassified(100);p__Ascomycota_unclassified(100);p__Ascomycota_unclassified(100); | - | - | - | - | - | - | - | Unassigned | - |
| Otu61862 | 1 | k__Fungi(100);k__Fungi_unclassified(100);k__Fungi_unclassified(100);k__Fungi_unclassified(100);k__Fungi_unclassified(100);k__Fungi_unclassified(100);k__Fungi_unclassified(100); | - | - | - | - | - | - | - | Unassigned | - |
| Otu61865 | 1 | k__Fungi(100);p__Ascomycota(100);p__Ascomycota_unclassified(100);p__Ascomycota_unclassified(100);p__Ascomycota_unclassified(100);p__Ascomycota_unclassified(100);p__Ascomycota_unclassified(100); | - | - | - | - | - | - | - | Unassigned | - |
| Otu61867 | 1 | k__Fungi(100);p__Ascomycota(100);p__Ascomycota_unclassified(100);p__Ascomycota_unclassified(100);p__Ascomycota_unclassified(100);p__Ascomycota_unclassified(100);p__Ascomycota_unclassified(100); | - | - | - | - | - | - | - | Unassigned | - |
| Otu61868 | 1 | k__Fungi(100);k__Fungi_unclassified(100);k__Fungi_unclassified(100);k__Fungi_unclassified(100);k__Fungi_unclassified(100);k__Fungi_unclassified(100);k__Fungi_unclassified(100); | - | - | - | - | - | - | - | Unassigned | - |
| Otu61869 | 1 | k__Fungi(100);k__Fungi_unclassified(100);k__Fungi_unclassified(100);k__Fungi_unclassified(100);k__Fungi_unclassified(100);k__Fungi_unclassified(100);k__Fungi_unclassified(100); | - | - | - | - | - | - | - | Unassigned | - |
| Otu61873 | 1 | k__Fungi(100);k__Fungi_unclassified(100);k__Fungi_unclassified(100);k__Fungi_unclassified(100);k__Fungi_unclassified(100);k__Fungi_unclassified(100);k__Fungi_unclassified(100); | - | - | - | - | - | - | - | Unassigned | - |
| Otu61875 | 1 | k__Fungi(100);p__Ascomycota(100);p__Ascomycota_unclassified(100);p__Ascomycota_unclassified(100);p__Ascomycota_unclassified(100);p__Ascomycota_unclassified(100);p__Ascomycota_unclassified(100); | - | - | - | - | - | - | - | Unassigned | - |
| Otu61876 | 1 | k__Fungi(100);k__Fungi_unclassified(100);k__Fungi_unclassified(100);k__Fungi_unclassified(100);k__Fungi_unclassified(100);k__Fungi_unclassified(100);k__Fungi_unclassified(100); | - | - | - | - | - | - | - | Unassigned | - |
| Otu61877 | 1 | k__Fungi(100);p__Ascomycota(100);p__Ascomycota_unclassified(100);p__Ascomycota_unclassified(100);p__Ascomycota_unclassified(100);p__Ascomycota_unclassified(100);p__Ascomycota_unclassified(100); | - | - | - | - | - | - | - | Unassigned | - |
| Otu61878 | 1 | k__Fungi(100);p__Ascomycota(100);p__Ascomycota_unclassified(100);p__Ascomycota_unclassified(100);p__Ascomycota_unclassified(100);p__Ascomycota_unclassified(100);p__Ascomycota_unclassified(100); | - | - | - | - | - | - | - | Unassigned | - |
| Otu61880 | 1 | k__Fungi(100);k__Fungi_unclassified(100);k__Fungi_unclassified(100);k__Fungi_unclassified(100);k__Fungi_unclassified(100);k__Fungi_unclassified(100);k__Fungi_unclassified(100); | - | - | - | - | - | - | - | Unassigned | - |
| Otu61882 | 1 | k__Fungi(100);k__Fungi_unclassified(100);k__Fungi_unclassified(100);k__Fungi_unclassified(100);k__Fungi_unclassified(100);k__Fungi_unclassified(100);k__Fungi_unclassified(100); | - | - | - | - | - | - | - | Unassigned | - |
| Otu61895 | 1 | k__Fungi(100);p__Ascomycota(100);p__Ascomycota_unclassified(100);p__Ascomycota_unclassified(100);p__Ascomycota_unclassified(100);p__Ascomycota_unclassified(100);p__Ascomycota_unclassified(100); | - | - | - | - | - | - | - | Unassigned | - |
| Otu61896 | 1 | k__Fungi(100);k__Fungi_unclassified(100);k__Fungi_unclassified(100);k__Fungi_unclassified(100);k__Fungi_unclassified(100);k__Fungi_unclassified(100);k__Fungi_unclassified(100); | - | - | - | - | - | - | - | Unassigned | - |
| Otu61898 | 1 | k__Fungi(100);p__Ascomycota(100);p__Ascomycota_unclassified(100);p__Ascomycota_unclassified(100);p__Ascomycota_unclassified(100);p__Ascomycota_unclassified(100);p__Ascomycota_unclassified(100); | - | - | - | - | - | - | - | Unassigned | - |
| Otu61902 | 1 | k__Fungi(100);k__Fungi_unclassified(100);k__Fungi_unclassified(100);k__Fungi_unclassified(100);k__Fungi_unclassified(100);k__Fungi_unclassified(100);k__Fungi_unclassified(100); | - | - | - | - | - | - | - | Unassigned | - |
| Otu61903 | 1 | k__Fungi(100);k__Fungi_unclassified(100);k__Fungi_unclassified(100);k__Fungi_unclassified(100);k__Fungi_unclassified(100);k__Fungi_unclassified(100);k__Fungi_unclassified(100); | - | - | - | - | - | - | - | Unassigned | - |
| Otu61905 | 1 | k__Fungi(100);k__Fungi_unclassified(100);k__Fungi_unclassified(100);k__Fungi_unclassified(100);k__Fungi_unclassified(100);k__Fungi_unclassified(100);k__Fungi_unclassified(100); | - | - | - | - | - | - | - | Unassigned | - |
| Otu61906 | 1 | k__Fungi(100);p__Ascomycota(100);p__Ascomycota_unclassified(100);p__Ascomycota_unclassified(100);p__Ascomycota_unclassified(100);p__Ascomycota_unclassified(100);p__Ascomycota_unclassified(100); | - | - | - | - | - | - | - | Unassigned | - |
| Otu61913 | 1 | k__Fungi(100);p__Ascomycota(100);p__Ascomycota_unclassified(100);p__Ascomycota_unclassified(100);p__Ascomycota_unclassified(100);p__Ascomycota_unclassified(100);p__Ascomycota_unclassified(100); | - | - | - | - | - | - | - | Unassigned | - |
| Otu61914 | 1 | k__Fungi(100);k__Fungi_unclassified(100);k__Fungi_unclassified(100);k__Fungi_unclassified(100);k__Fungi_unclassified(100);k__Fungi_unclassified(100);k__Fungi_unclassified(100); | - | - | - | - | - | - | - | Unassigned | - |
| Otu61915 | 1 | k__Fungi(100);p__Ascomycota(100);p__Ascomycota_unclassified(100);p__Ascomycota_unclassified(100);p__Ascomycota_unclassified(100);p__Ascomycota_unclassified(100);p__Ascomycota_unclassified(100); | - | - | - | - | - | - | - | Unassigned | - |
| Otu61918 | 1 | k__Fungi(100);k__Fungi_unclassified(100);k__Fungi_unclassified(100);k__Fungi_unclassified(100);k__Fungi_unclassified(100);k__Fungi_unclassified(100);k__Fungi_unclassified(100); | - | - | - | - | - | - | - | Unassigned | - |
| Otu61919 | 1 | k__Fungi(100);k__Fungi_unclassified(100);k__Fungi_unclassified(100);k__Fungi_unclassified(100);k__Fungi_unclassified(100);k__Fungi_unclassified(100);k__Fungi_unclassified(100); | - | - | - | - | - | - | - | Unassigned | - |
| Otu61920 | 1 | k__Fungi(100);p__Ascomycota(100);p__Ascomycota_unclassified(100);p__Ascomycota_unclassified(100);p__Ascomycota_unclassified(100);p__Ascomycota_unclassified(100);p__Ascomycota_unclassified(100); | - | - | - | - | - | - | - | Unassigned | - |
| Otu61921 | 1 | k__Fungi(100);k__Fungi_unclassified(100);k__Fungi_unclassified(100);k__Fungi_unclassified(100);k__Fungi_unclassified(100);k__Fungi_unclassified(100);k__Fungi_unclassified(100); | - | - | - | - | - | - | - | Unassigned | - |
| Otu61922 | 1 | k__Fungi(100);p__Basidiomycota(100);p__Basidiomycota_unclassified(100);p__Basidiomycota_unclassified(100);p__Basidiomycota_unclassified(100);p__Basidiomycota_unclassified(100);p__Basidiomycota_unclassified(100); | - | - | - | - | - | - | - | Unassigned | - |
| Otu61924 | 1 | k__Fungi(100);k__Fungi_unclassified(100);k__Fungi_unclassified(100);k__Fungi_unclassified(100);k__Fungi_unclassified(100);k__Fungi_unclassified(100);k__Fungi_unclassified(100); | - | - | - | - | - | - | - | Unassigned | - |
| Otu61927 | 1 | k__Fungi(100);k__Fungi_unclassified(100);k__Fungi_unclassified(100);k__Fungi_unclassified(100);k__Fungi_unclassified(100);k__Fungi_unclassified(100);k__Fungi_unclassified(100); | - | - | - | - | - | - | - | Unassigned | - |
| Otu61929 | 1 | k__Fungi(100);p__Ascomycota(100);p__Ascomycota_unclassified(100);p__Ascomycota_unclassified(100);p__Ascomycota_unclassified(100);p__Ascomycota_unclassified(100);p__Ascomycota_unclassified(100); | - | - | - | - | - | - | - | Unassigned | - |
| Otu61930 | 1 | k__Fungi(100);p__Ascomycota(100);p__Ascomycota_unclassified(100);p__Ascomycota_unclassified(100);p__Ascomycota_unclassified(100);p__Ascomycota_unclassified(100);p__Ascomycota_unclassified(100); | - | - | - | - | - | - | - | Unassigned | - |
| Otu61931 | 1 | k__Fungi(100);k__Fungi_unclassified(100);k__Fungi_unclassified(100);k__Fungi_unclassified(100);k__Fungi_unclassified(100);k__Fungi_unclassified(100);k__Fungi_unclassified(100); | - | - | - | - | - | - | - | Unassigned | - |
| Otu61932 | 1 | k__Fungi(100);k__Fungi_unclassified(100);k__Fungi_unclassified(100);k__Fungi_unclassified(100);k__Fungi_unclassified(100);k__Fungi_unclassified(100);k__Fungi_unclassified(100); | - | - | - | - | - | - | - | Unassigned | - |
| Otu61934 | 1 | k__Fungi(100);p__Ascomycota(100);p__Ascomycota_unclassified(100);p__Ascomycota_unclassified(100);p__Ascomycota_unclassified(100);p__Ascomycota_unclassified(100);p__Ascomycota_unclassified(100); | - | - | - | - | - | - | - | Unassigned | - |
| Otu61935 | 1 | k__Fungi(100);k__Fungi_unclassified(100);k__Fungi_unclassified(100);k__Fungi_unclassified(100);k__Fungi_unclassified(100);k__Fungi_unclassified(100);k__Fungi_unclassified(100); | - | - | - | - | - | - | - | Unassigned | - |
| Otu61937 | 1 | k__Fungi(100);k__Fungi_unclassified(100);k__Fungi_unclassified(100);k__Fungi_unclassified(100);k__Fungi_unclassified(100);k__Fungi_unclassified(100);k__Fungi_unclassified(100); | - | - | - | - | - | - | - | Unassigned | - |
| Otu61938 | 1 | k__Fungi(100);k__Fungi_unclassified(100);k__Fungi_unclassified(100);k__Fungi_unclassified(100);k__Fungi_unclassified(100);k__Fungi_unclassified(100);k__Fungi_unclassified(100); | - | - | - | - | - | - | - | Unassigned | - |
| Otu61939 | 1 | k__Fungi(100);k__Fungi_unclassified(100);k__Fungi_unclassified(100);k__Fungi_unclassified(100);k__Fungi_unclassified(100);k__Fungi_unclassified(100);k__Fungi_unclassified(100); | - | - | - | - | - | - | - | Unassigned | - |
| Otu61940 | 1 | k__Fungi(100);k__Fungi_unclassified(100);k__Fungi_unclassified(100);k__Fungi_unclassified(100);k__Fungi_unclassified(100);k__Fungi_unclassified(100);k__Fungi_unclassified(100); | - | - | - | - | - | - | - | Unassigned | - |
| Otu61941 | 1 | k__Fungi(100);k__Fungi_unclassified(100);k__Fungi_unclassified(100);k__Fungi_unclassified(100);k__Fungi_unclassified(100);k__Fungi_unclassified(100);k__Fungi_unclassified(100); | - | - | - | - | - | - | - | Unassigned | - |
| Otu61942 | 1 | k__Fungi(100);p__Ascomycota(100);p__Ascomycota_unclassified(100);p__Ascomycota_unclassified(100);p__Ascomycota_unclassified(100);p__Ascomycota_unclassified(100);p__Ascomycota_unclassified(100); | - | - | - | - | - | - | - | Unassigned | - |
| Otu61943 | 1 | k__Fungi(100);k__Fungi_unclassified(100);k__Fungi_unclassified(100);k__Fungi_unclassified(100);k__Fungi_unclassified(100);k__Fungi_unclassified(100);k__Fungi_unclassified(100); | - | - | - | - | - | - | - | Unassigned | - |
| Otu61944 | 1 | k__Fungi(100);k__Fungi_unclassified(100);k__Fungi_unclassified(100);k__Fungi_unclassified(100);k__Fungi_unclassified(100);k__Fungi_unclassified(100);k__Fungi_unclassified(100); | - | - | - | - | - | - | - | Unassigned | - |
| Otu61945 | 1 | k__Fungi(100);p__Ascomycota(100);p__Ascomycota_unclassified(100);p__Ascomycota_unclassified(100);p__Ascomycota_unclassified(100);p__Ascomycota_unclassified(100);p__Ascomycota_unclassified(100); | - | - | - | - | - | - | - | Unassigned | - |
| Otu61946 | 1 | k__Fungi(100);k__Fungi_unclassified(100);k__Fungi_unclassified(100);k__Fungi_unclassified(100);k__Fungi_unclassified(100);k__Fungi_unclassified(100);k__Fungi_unclassified(100); | - | - | - | - | - | - | - | Unassigned | - |
| Otu61949 | 1 | k__Fungi(100);k__Fungi_unclassified(100);k__Fungi_unclassified(100);k__Fungi_unclassified(100);k__Fungi_unclassified(100);k__Fungi_unclassified(100);k__Fungi_unclassified(100); | - | - | - | - | - | - | - | Unassigned | - |
| Otu61950 | 1 | k__Fungi(100);k__Fungi_unclassified(100);k__Fungi_unclassified(100);k__Fungi_unclassified(100);k__Fungi_unclassified(100);k__Fungi_unclassified(100);k__Fungi_unclassified(100); | - | - | - | - | - | - | - | Unassigned | - |
| Otu61951 | 1 | k__Fungi(100);k__Fungi_unclassified(100);k__Fungi_unclassified(100);k__Fungi_unclassified(100);k__Fungi_unclassified(100);k__Fungi_unclassified(100);k__Fungi_unclassified(100); | - | - | - | - | - | - | - | Unassigned | - |
| Otu61952 | 1 | k__Fungi(100);p__Ascomycota(100);p__Ascomycota_unclassified(100);p__Ascomycota_unclassified(100);p__Ascomycota_unclassified(100);p__Ascomycota_unclassified(100);p__Ascomycota_unclassified(100); | - | - | - | - | - | - | - | Unassigned | - |
| Otu61953 | 1 | k__Fungi(100);k__Fungi_unclassified(100);k__Fungi_unclassified(100);k__Fungi_unclassified(100);k__Fungi_unclassified(100);k__Fungi_unclassified(100);k__Fungi_unclassified(100); | - | - | - | - | - | - | - | Unassigned | - |
| Otu61955 | 1 | k__Fungi(100);k__Fungi_unclassified(100);k__Fungi_unclassified(100);k__Fungi_unclassified(100);k__Fungi_unclassified(100);k__Fungi_unclassified(100);k__Fungi_unclassified(100); | - | - | - | - | - | - | - | Unassigned | - |
| Otu61957 | 1 | k__Fungi(100);k__Fungi_unclassified(100);k__Fungi_unclassified(100);k__Fungi_unclassified(100);k__Fungi_unclassified(100);k__Fungi_unclassified(100);k__Fungi_unclassified(100); | - | - | - | - | - | - | - | Unassigned | - |
| Otu61958 | 1 | k__Fungi(100);p__Ascomycota(100);p__Ascomycota_unclassified(100);p__Ascomycota_unclassified(100);p__Ascomycota_unclassified(100);p__Ascomycota_unclassified(100);p__Ascomycota_unclassified(100); | - | - | - | - | - | - | - | Unassigned | - |
| Otu61959 | 1 | k__Fungi(100);k__Fungi_unclassified(100);k__Fungi_unclassified(100);k__Fungi_unclassified(100);k__Fungi_unclassified(100);k__Fungi_unclassified(100);k__Fungi_unclassified(100); | - | - | - | - | - | - | - | Unassigned | - |
| Otu61960 | 1 | k__Fungi(100);k__Fungi_unclassified(100);k__Fungi_unclassified(100);k__Fungi_unclassified(100);k__Fungi_unclassified(100);k__Fungi_unclassified(100);k__Fungi_unclassified(100); | - | - | - | - | - | - | - | Unassigned | - |
| Otu61961 | 1 | k__Fungi(100);p__Ascomycota(100);p__Ascomycota_unclassified(100);p__Ascomycota_unclassified(100);p__Ascomycota_unclassified(100);p__Ascomycota_unclassified(100);p__Ascomycota_unclassified(100); | - | - | - | - | - | - | - | Unassigned | - |
| Otu61962 | 1 | k__Fungi(100);k__Fungi_unclassified(100);k__Fungi_unclassified(100);k__Fungi_unclassified(100);k__Fungi_unclassified(100);k__Fungi_unclassified(100);k__Fungi_unclassified(100); | - | - | - | - | - | - | - | Unassigned | - |
| Otu61963 | 1 | k__Fungi(100);k__Fungi_unclassified(100);k__Fungi_unclassified(100);k__Fungi_unclassified(100);k__Fungi_unclassified(100);k__Fungi_unclassified(100);k__Fungi_unclassified(100); | - | - | - | - | - | - | - | Unassigned | - |
| Otu61964 | 1 | k__Fungi(100);k__Fungi_unclassified(100);k__Fungi_unclassified(100);k__Fungi_unclassified(100);k__Fungi_unclassified(100);k__Fungi_unclassified(100);k__Fungi_unclassified(100); | - | - | - | - | - | - | - | Unassigned | - |
| Otu61965 | 1 | k__Fungi(100);p__Ascomycota(100);p__Ascomycota_unclassified(100);p__Ascomycota_unclassified(100);p__Ascomycota_unclassified(100);p__Ascomycota_unclassified(100);p__Ascomycota_unclassified(100); | - | - | - | - | - | - | - | Unassigned | - |
| Otu61966 | 1 | k__Fungi(100);k__Fungi_unclassified(100);k__Fungi_unclassified(100);k__Fungi_unclassified(100);k__Fungi_unclassified(100);k__Fungi_unclassified(100);k__Fungi_unclassified(100); | - | - | - | - | - | - | - | Unassigned | - |
| Otu61968 | 1 | k__Fungi(100);p__Ascomycota(100);p__Ascomycota_unclassified(100);p__Ascomycota_unclassified(100);p__Ascomycota_unclassified(100);p__Ascomycota_unclassified(100);p__Ascomycota_unclassified(100); | - | - | - | - | - | - | - | Unassigned | - |
| Otu61972 | 1 | k__Fungi(100);k__Fungi_unclassified(100);k__Fungi_unclassified(100);k__Fungi_unclassified(100);k__Fungi_unclassified(100);k__Fungi_unclassified(100);k__Fungi_unclassified(100); | - | - | - | - | - | - | - | Unassigned | - |
| Otu61973 | 1 | k__Fungi(100);p__Ascomycota(100);p__Ascomycota_unclassified(100);p__Ascomycota_unclassified(100);p__Ascomycota_unclassified(100);p__Ascomycota_unclassified(100);p__Ascomycota_unclassified(100); | - | - | - | - | - | - | - | Unassigned | - |
| Otu61978 | 1 | k__Fungi(100);k__Fungi_unclassified(100);k__Fungi_unclassified(100);k__Fungi_unclassified(100);k__Fungi_unclassified(100);k__Fungi_unclassified(100);k__Fungi_unclassified(100); | - | - | - | - | - | - | - | Unassigned | - |
| Otu61980 | 1 | k__Fungi(100);k__Fungi_unclassified(100);k__Fungi_unclassified(100);k__Fungi_unclassified(100);k__Fungi_unclassified(100);k__Fungi_unclassified(100);k__Fungi_unclassified(100); | - | - | - | - | - | - | - | Unassigned | - |
| Otu61981 | 1 | k__Fungi(100);k__Fungi_unclassified(100);k__Fungi_unclassified(100);k__Fungi_unclassified(100);k__Fungi_unclassified(100);k__Fungi_unclassified(100);k__Fungi_unclassified(100); | - | - | - | - | - | - | - | Unassigned | - |
| Otu61983 | 1 | k__Fungi(100);k__Fungi_unclassified(100);k__Fungi_unclassified(100);k__Fungi_unclassified(100);k__Fungi_unclassified(100);k__Fungi_unclassified(100);k__Fungi_unclassified(100); | - | - | - | - | - | - | - | Unassigned | - |
| Otu61984 | 1 | k__Fungi(100);p__Ascomycota(100);p__Ascomycota_unclassified(100);p__Ascomycota_unclassified(100);p__Ascomycota_unclassified(100);p__Ascomycota_unclassified(100);p__Ascomycota_unclassified(100); | - | - | - | - | - | - | - | Unassigned | - |
| Otu61985 | 1 | k__Fungi(100);p__Ascomycota(100);p__Ascomycota_unclassified(100);p__Ascomycota_unclassified(100);p__Ascomycota_unclassified(100);p__Ascomycota_unclassified(100);p__Ascomycota_unclassified(100); | - | - | - | - | - | - | - | Unassigned | - |
| Otu61988 | 1 | k__Fungi(100);k__Fungi_unclassified(100);k__Fungi_unclassified(100);k__Fungi_unclassified(100);k__Fungi_unclassified(100);k__Fungi_unclassified(100);k__Fungi_unclassified(100); | - | - | - | - | - | - | - | Unassigned | - |
| Otu61989 | 1 | k__Fungi(100);k__Fungi_unclassified(100);k__Fungi_unclassified(100);k__Fungi_unclassified(100);k__Fungi_unclassified(100);k__Fungi_unclassified(100);k__Fungi_unclassified(100); | - | - | - | - | - | - | - | Unassigned | - |
| Otu61990 | 1 | k__Fungi(100);k__Fungi_unclassified(100);k__Fungi_unclassified(100);k__Fungi_unclassified(100);k__Fungi_unclassified(100);k__Fungi_unclassified(100);k__Fungi_unclassified(100); | - | - | - | - | - | - | - | Unassigned | - |
| Otu61991 | 1 | k__Fungi(100);k__Fungi_unclassified(100);k__Fungi_unclassified(100);k__Fungi_unclassified(100);k__Fungi_unclassified(100);k__Fungi_unclassified(100);k__Fungi_unclassified(100); | - | - | - | - | - | - | - | Unassigned | - |
| Otu61992 | 1 | k__Fungi(100);k__Fungi_unclassified(100);k__Fungi_unclassified(100);k__Fungi_unclassified(100);k__Fungi_unclassified(100);k__Fungi_unclassified(100);k__Fungi_unclassified(100); | - | - | - | - | - | - | - | Unassigned | - |
| Otu61993 | 1 | k__Fungi(100);k__Fungi_unclassified(100);k__Fungi_unclassified(100);k__Fungi_unclassified(100);k__Fungi_unclassified(100);k__Fungi_unclassified(100);k__Fungi_unclassified(100); | - | - | - | - | - | - | - | Unassigned | - |
| Otu61995 | 1 | k__Fungi(100);k__Fungi_unclassified(100);k__Fungi_unclassified(100);k__Fungi_unclassified(100);k__Fungi_unclassified(100);k__Fungi_unclassified(100);k__Fungi_unclassified(100); | - | - | - | - | - | - | - | Unassigned | - |
| Otu61996 | 1 | k__Fungi(100);k__Fungi_unclassified(100);k__Fungi_unclassified(100);k__Fungi_unclassified(100);k__Fungi_unclassified(100);k__Fungi_unclassified(100);k__Fungi_unclassified(100); | - | - | - | - | - | - | - | Unassigned | - |
| Otu61997 | 1 | k__Fungi(100);k__Fungi_unclassified(100);k__Fungi_unclassified(100);k__Fungi_unclassified(100);k__Fungi_unclassified(100);k__Fungi_unclassified(100);k__Fungi_unclassified(100); | - | - | - | - | - | - | - | Unassigned | - |
| Otu61998 | 1 | k__Fungi(100);k__Fungi_unclassified(100);k__Fungi_unclassified(100);k__Fungi_unclassified(100);k__Fungi_unclassified(100);k__Fungi_unclassified(100);k__Fungi_unclassified(100); | - | - | - | - | - | - | - | Unassigned | - |
| Otu61999 | 1 | k__Fungi(100);p__Ascomycota(100);p__Ascomycota_unclassified(100);p__Ascomycota_unclassified(100);p__Ascomycota_unclassified(100);p__Ascomycota_unclassified(100);p__Ascomycota_unclassified(100); | - | - | - | - | - | - | - | Unassigned | - |
| Otu62001 | 1 | k__Fungi(100);p__Ascomycota(100);p__Ascomycota_unclassified(100);p__Ascomycota_unclassified(100);p__Ascomycota_unclassified(100);p__Ascomycota_unclassified(100);p__Ascomycota_unclassified(100); | - | - | - | - | - | - | - | Unassigned | - |
| Otu62002 | 1 | k__Fungi(100);p__Ascomycota(100);p__Ascomycota_unclassified(100);p__Ascomycota_unclassified(100);p__Ascomycota_unclassified(100);p__Ascomycota_unclassified(100);p__Ascomycota_unclassified(100); | - | - | - | - | - | - | - | Unassigned | - |
| Otu62003 | 1 | k__Fungi(100);p__Ascomycota(100);p__Ascomycota_unclassified(100);p__Ascomycota_unclassified(100);p__Ascomycota_unclassified(100);p__Ascomycota_unclassified(100);p__Ascomycota_unclassified(100); | - | - | - | - | - | - | - | Unassigned | - |
| Otu62004 | 1 | k__Fungi(100);k__Fungi_unclassified(100);k__Fungi_unclassified(100);k__Fungi_unclassified(100);k__Fungi_unclassified(100);k__Fungi_unclassified(100);k__Fungi_unclassified(100); | - | - | - | - | - | - | - | Unassigned | - |
| Otu62006 | 1 | k__Fungi(100);k__Fungi_unclassified(100);k__Fungi_unclassified(100);k__Fungi_unclassified(100);k__Fungi_unclassified(100);k__Fungi_unclassified(100);k__Fungi_unclassified(100); | - | - | - | - | - | - | - | Unassigned | - |
| Otu62009 | 1 | k__Fungi(100);k__Fungi_unclassified(100);k__Fungi_unclassified(100);k__Fungi_unclassified(100);k__Fungi_unclassified(100);k__Fungi_unclassified(100);k__Fungi_unclassified(100); | - | - | - | - | - | - | - | Unassigned | - |
| Otu62010 | 1 | k__Fungi(100);k__Fungi_unclassified(100);k__Fungi_unclassified(100);k__Fungi_unclassified(100);k__Fungi_unclassified(100);k__Fungi_unclassified(100);k__Fungi_unclassified(100); | - | - | - | - | - | - | - | Unassigned | - |
| Otu62011 | 1 | k__Fungi(100);k__Fungi_unclassified(100);k__Fungi_unclassified(100);k__Fungi_unclassified(100);k__Fungi_unclassified(100);k__Fungi_unclassified(100);k__Fungi_unclassified(100); | - | - | - | - | - | - | - | Unassigned | - |
| Otu62012 | 1 | k__Fungi(100);k__Fungi_unclassified(100);k__Fungi_unclassified(100);k__Fungi_unclassified(100);k__Fungi_unclassified(100);k__Fungi_unclassified(100);k__Fungi_unclassified(100); | - | - | - | - | - | - | - | Unassigned | - |
| Otu62013 | 1 | k__Fungi(100);p__Ascomycota(100);p__Ascomycota_unclassified(100);p__Ascomycota_unclassified(100);p__Ascomycota_unclassified(100);p__Ascomycota_unclassified(100);p__Ascomycota_unclassified(100); | - | - | - | - | - | - | - | Unassigned | - |
| Otu62018 | 1 | k__Fungi(100);p__Ascomycota(100);p__Ascomycota_unclassified(100);p__Ascomycota_unclassified(100);p__Ascomycota_unclassified(100);p__Ascomycota_unclassified(100);p__Ascomycota_unclassified(100); | - | - | - | - | - | - | - | Unassigned | - |
| Otu62020 | 1 | k__Fungi(100);k__Fungi_unclassified(100);k__Fungi_unclassified(100);k__Fungi_unclassified(100);k__Fungi_unclassified(100);k__Fungi_unclassified(100);k__Fungi_unclassified(100); | - | - | - | - | - | - | - | Unassigned | - |
| Otu62021 | 1 | k__Fungi(100);k__Fungi_unclassified(100);k__Fungi_unclassified(100);k__Fungi_unclassified(100);k__Fungi_unclassified(100);k__Fungi_unclassified(100);k__Fungi_unclassified(100); | - | - | - | - | - | - | - | Unassigned | - |
| Otu62024 | 1 | k__Fungi(100);k__Fungi_unclassified(100);k__Fungi_unclassified(100);k__Fungi_unclassified(100);k__Fungi_unclassified(100);k__Fungi_unclassified(100);k__Fungi_unclassified(100); | - | - | - | - | - | - | - | Unassigned | - |
| Otu62025 | 1 | k__Fungi(100);k__Fungi_unclassified(100);k__Fungi_unclassified(100);k__Fungi_unclassified(100);k__Fungi_unclassified(100);k__Fungi_unclassified(100);k__Fungi_unclassified(100); | - | - | - | - | - | - | - | Unassigned | - |
| Otu62026 | 1 | k__Fungi(100);p__Ascomycota(100);p__Ascomycota_unclassified(100);p__Ascomycota_unclassified(100);p__Ascomycota_unclassified(100);p__Ascomycota_unclassified(100);p__Ascomycota_unclassified(100); | - | - | - | - | - | - | - | Unassigned | - |
| Otu62027 | 1 | k__Fungi(100);k__Fungi_unclassified(100);k__Fungi_unclassified(100);k__Fungi_unclassified(100);k__Fungi_unclassified(100);k__Fungi_unclassified(100);k__Fungi_unclassified(100); | - | - | - | - | - | - | - | Unassigned | - |
| Otu62028 | 1 | k__Fungi(100);k__Fungi_unclassified(100);k__Fungi_unclassified(100);k__Fungi_unclassified(100);k__Fungi_unclassified(100);k__Fungi_unclassified(100);k__Fungi_unclassified(100); | - | - | - | - | - | - | - | Unassigned | - |
| Otu62030 | 1 | k__Fungi(100);k__Fungi_unclassified(100);k__Fungi_unclassified(100);k__Fungi_unclassified(100);k__Fungi_unclassified(100);k__Fungi_unclassified(100);k__Fungi_unclassified(100); | - | - | - | - | - | - | - | Unassigned | - |
| Otu62031 | 1 | k__Fungi(100);p__Ascomycota(100);p__Ascomycota_unclassified(100);p__Ascomycota_unclassified(100);p__Ascomycota_unclassified(100);p__Ascomycota_unclassified(100);p__Ascomycota_unclassified(100); | - | - | - | - | - | - | - | Unassigned | - |
| Otu62033 | 1 | k__Fungi(100);p__Ascomycota(100);p__Ascomycota_unclassified(100);p__Ascomycota_unclassified(100);p__Ascomycota_unclassified(100);p__Ascomycota_unclassified(100);p__Ascomycota_unclassified(100); | - | - | - | - | - | - | - | Unassigned | - |
| Otu62034 | 1 | k__Fungi(100);k__Fungi_unclassified(100);k__Fungi_unclassified(100);k__Fungi_unclassified(100);k__Fungi_unclassified(100);k__Fungi_unclassified(100);k__Fungi_unclassified(100); | - | - | - | - | - | - | - | Unassigned | - |
| Otu62035 | 1 | k__Fungi(100);p__Ascomycota(100);p__Ascomycota_unclassified(100);p__Ascomycota_unclassified(100);p__Ascomycota_unclassified(100);p__Ascomycota_unclassified(100);p__Ascomycota_unclassified(100); | - | - | - | - | - | - | - | Unassigned | - |
| Otu62036 | 1 | k__Fungi(100);p__Ascomycota(100);p__Ascomycota_unclassified(100);p__Ascomycota_unclassified(100);p__Ascomycota_unclassified(100);p__Ascomycota_unclassified(100);p__Ascomycota_unclassified(100); | - | - | - | - | - | - | - | Unassigned | - |
| Otu62043 | 1 | k__Fungi(100);k__Fungi_unclassified(100);k__Fungi_unclassified(100);k__Fungi_unclassified(100);k__Fungi_unclassified(100);k__Fungi_unclassified(100);k__Fungi_unclassified(100); | - | - | - | - | - | - | - | Unassigned | - |
| Otu62044 | 1 | k__Fungi(100);k__Fungi_unclassified(100);k__Fungi_unclassified(100);k__Fungi_unclassified(100);k__Fungi_unclassified(100);k__Fungi_unclassified(100);k__Fungi_unclassified(100); | - | - | - | - | - | - | - | Unassigned | - |
| Otu62046 | 1 | k__Fungi(100);k__Fungi_unclassified(100);k__Fungi_unclassified(100);k__Fungi_unclassified(100);k__Fungi_unclassified(100);k__Fungi_unclassified(100);k__Fungi_unclassified(100); | - | - | - | - | - | - | - | Unassigned | - |
| Otu62047 | 1 | k__Fungi(100);p__Ascomycota(100);p__Ascomycota_unclassified(100);p__Ascomycota_unclassified(100);p__Ascomycota_unclassified(100);p__Ascomycota_unclassified(100);p__Ascomycota_unclassified(100); | - | - | - | - | - | - | - | Unassigned | - |
| Otu62048 | 1 | k__Fungi(100);p__Ascomycota(100);p__Ascomycota_unclassified(100);p__Ascomycota_unclassified(100);p__Ascomycota_unclassified(100);p__Ascomycota_unclassified(100);p__Ascomycota_unclassified(100); | - | - | - | - | - | - | - | Unassigned | - |
| Otu62049 | 1 | k__Fungi(100);k__Fungi_unclassified(100);k__Fungi_unclassified(100);k__Fungi_unclassified(100);k__Fungi_unclassified(100);k__Fungi_unclassified(100);k__Fungi_unclassified(100); | - | - | - | - | - | - | - | Unassigned | - |
| Otu62051 | 1 | k__Fungi(100);p__Ascomycota(100);p__Ascomycota_unclassified(100);p__Ascomycota_unclassified(100);p__Ascomycota_unclassified(100);p__Ascomycota_unclassified(100);p__Ascomycota_unclassified(100); | - | - | - | - | - | - | - | Unassigned | - |
| Otu62052 | 1 | k__Fungi(100);k__Fungi_unclassified(100);k__Fungi_unclassified(100);k__Fungi_unclassified(100);k__Fungi_unclassified(100);k__Fungi_unclassified(100);k__Fungi_unclassified(100); | - | - | - | - | - | - | - | Unassigned | - |
| Otu62053 | 1 | k__Fungi(100);k__Fungi_unclassified(100);k__Fungi_unclassified(100);k__Fungi_unclassified(100);k__Fungi_unclassified(100);k__Fungi_unclassified(100);k__Fungi_unclassified(100); | - | - | - | - | - | - | - | Unassigned | - |
| Otu62066 | 1 | k__Fungi(100);k__Fungi_unclassified(100);k__Fungi_unclassified(100);k__Fungi_unclassified(100);k__Fungi_unclassified(100);k__Fungi_unclassified(100);k__Fungi_unclassified(100); | - | - | - | - | - | - | - | Unassigned | - |
| Otu62067 | 1 | k__Fungi(100);p__Ascomycota(100);p__Ascomycota_unclassified(100);p__Ascomycota_unclassified(100);p__Ascomycota_unclassified(100);p__Ascomycota_unclassified(100);p__Ascomycota_unclassified(100); | - | - | - | - | - | - | - | Unassigned | - |
| Otu62074 | 1 | k__Fungi(100);p__Ascomycota(100);p__Ascomycota_unclassified(100);p__Ascomycota_unclassified(100);p__Ascomycota_unclassified(100);p__Ascomycota_unclassified(100);p__Ascomycota_unclassified(100); | - | - | - | - | - | - | - | Unassigned | - |
| Otu62075 | 1 | k__Fungi(100);k__Fungi_unclassified(100);k__Fungi_unclassified(100);k__Fungi_unclassified(100);k__Fungi_unclassified(100);k__Fungi_unclassified(100);k__Fungi_unclassified(100); | - | - | - | - | - | - | - | Unassigned | - |
| Otu62081 | 1 | k__Fungi(100);p__Ascomycota(100);p__Ascomycota_unclassified(100);p__Ascomycota_unclassified(100);p__Ascomycota_unclassified(100);p__Ascomycota_unclassified(100);p__Ascomycota_unclassified(100); | - | - | - | - | - | - | - | Unassigned | - |
| Otu62082 | 1 | k__Fungi(100);k__Fungi_unclassified(100);k__Fungi_unclassified(100);k__Fungi_unclassified(100);k__Fungi_unclassified(100);k__Fungi_unclassified(100);k__Fungi_unclassified(100); | - | - | - | - | - | - | - | Unassigned | - |
| Otu62084 | 1 | k__Fungi(100);p__Ascomycota(100);p__Ascomycota_unclassified(100);p__Ascomycota_unclassified(100);p__Ascomycota_unclassified(100);p__Ascomycota_unclassified(100);p__Ascomycota_unclassified(100); | - | - | - | - | - | - | - | Unassigned | - |
| Otu62085 | 1 | k__Fungi(100);k__Fungi_unclassified(100);k__Fungi_unclassified(100);k__Fungi_unclassified(100);k__Fungi_unclassified(100);k__Fungi_unclassified(100);k__Fungi_unclassified(100); | - | - | - | - | - | - | - | Unassigned | - |
| Otu62086 | 1 | k__Fungi(100);p__Ascomycota(100);p__Ascomycota_unclassified(100);p__Ascomycota_unclassified(100);p__Ascomycota_unclassified(100);p__Ascomycota_unclassified(100);p__Ascomycota_unclassified(100); | - | - | - | - | - | - | - | Unassigned | - |
| Otu62088 | 1 | k__Fungi(100);p__Ascomycota(100);p__Ascomycota_unclassified(100);p__Ascomycota_unclassified(100);p__Ascomycota_unclassified(100);p__Ascomycota_unclassified(100);p__Ascomycota_unclassified(100); | - | - | - | - | - | - | - | Unassigned | - |
| Otu62089 | 1 | k__Fungi(100);p__Ascomycota(100);p__Ascomycota_unclassified(100);p__Ascomycota_unclassified(100);p__Ascomycota_unclassified(100);p__Ascomycota_unclassified(100);p__Ascomycota_unclassified(100); | - | - | - | - | - | - | - | Unassigned | - |
| Otu62090 | 1 | k__Fungi(100);p__Ascomycota(100);p__Ascomycota_unclassified(100);p__Ascomycota_unclassified(100);p__Ascomycota_unclassified(100);p__Ascomycota_unclassified(100);p__Ascomycota_unclassified(100); | - | - | - | - | - | - | - | Unassigned | - |
| Otu62091 | 1 | k__Fungi(100);k__Fungi_unclassified(100);k__Fungi_unclassified(100);k__Fungi_unclassified(100);k__Fungi_unclassified(100);k__Fungi_unclassified(100);k__Fungi_unclassified(100); | - | - | - | - | - | - | - | Unassigned | - |
| Otu62092 | 1 | k__Fungi(100);k__Fungi_unclassified(100);k__Fungi_unclassified(100);k__Fungi_unclassified(100);k__Fungi_unclassified(100);k__Fungi_unclassified(100);k__Fungi_unclassified(100); | - | - | - | - | - | - | - | Unassigned | - |
| Otu62094 | 1 | k__Fungi(100);k__Fungi_unclassified(100);k__Fungi_unclassified(100);k__Fungi_unclassified(100);k__Fungi_unclassified(100);k__Fungi_unclassified(100);k__Fungi_unclassified(100); | - | - | - | - | - | - | - | Unassigned | - |
| Otu62095 | 1 | k__Fungi(100);p__Ascomycota(100);p__Ascomycota_unclassified(100);p__Ascomycota_unclassified(100);p__Ascomycota_unclassified(100);p__Ascomycota_unclassified(100);p__Ascomycota_unclassified(100); | - | - | - | - | - | - | - | Unassigned | - |
| Otu62097 | 1 | k__Fungi(100);k__Fungi_unclassified(100);k__Fungi_unclassified(100);k__Fungi_unclassified(100);k__Fungi_unclassified(100);k__Fungi_unclassified(100);k__Fungi_unclassified(100); | - | - | - | - | - | - | - | Unassigned | - |
| Otu62098 | 1 | k__Fungi(100);k__Fungi_unclassified(100);k__Fungi_unclassified(100);k__Fungi_unclassified(100);k__Fungi_unclassified(100);k__Fungi_unclassified(100);k__Fungi_unclassified(100); | - | - | - | - | - | - | - | Unassigned | - |
| Otu62106 | 1 | k__Fungi(100);k__Fungi_unclassified(100);k__Fungi_unclassified(100);k__Fungi_unclassified(100);k__Fungi_unclassified(100);k__Fungi_unclassified(100);k__Fungi_unclassified(100); | - | - | - | - | - | - | - | Unassigned | - |
| Otu62107 | 1 | k__Fungi(100);k__Fungi_unclassified(100);k__Fungi_unclassified(100);k__Fungi_unclassified(100);k__Fungi_unclassified(100);k__Fungi_unclassified(100);k__Fungi_unclassified(100); | - | - | - | - | - | - | - | Unassigned | - |
| Otu62109 | 1 | k__Fungi(100);k__Fungi_unclassified(100);k__Fungi_unclassified(100);k__Fungi_unclassified(100);k__Fungi_unclassified(100);k__Fungi_unclassified(100);k__Fungi_unclassified(100); | - | - | - | - | - | - | - | Unassigned | - |
| Otu62111 | 1 | k__Fungi(100);p__Ascomycota(100);p__Ascomycota_unclassified(100);p__Ascomycota_unclassified(100);p__Ascomycota_unclassified(100);p__Ascomycota_unclassified(100);p__Ascomycota_unclassified(100); | - | - | - | - | - | - | - | Unassigned | - |
| Otu62112 | 1 | k__Fungi(100);k__Fungi_unclassified(100);k__Fungi_unclassified(100);k__Fungi_unclassified(100);k__Fungi_unclassified(100);k__Fungi_unclassified(100);k__Fungi_unclassified(100); | - | - | - | - | - | - | - | Unassigned | - |
| Otu62114 | 1 | k__Fungi(100);k__Fungi_unclassified(100);k__Fungi_unclassified(100);k__Fungi_unclassified(100);k__Fungi_unclassified(100);k__Fungi_unclassified(100);k__Fungi_unclassified(100); | - | - | - | - | - | - | - | Unassigned | - |
| Otu62115 | 1 | k__Fungi(100);k__Fungi_unclassified(100);k__Fungi_unclassified(100);k__Fungi_unclassified(100);k__Fungi_unclassified(100);k__Fungi_unclassified(100);k__Fungi_unclassified(100); | - | - | - | - | - | - | - | Unassigned | - |
| Otu62116 | 1 | k__Fungi(100);k__Fungi_unclassified(100);k__Fungi_unclassified(100);k__Fungi_unclassified(100);k__Fungi_unclassified(100);k__Fungi_unclassified(100);k__Fungi_unclassified(100); | - | - | - | - | - | - | - | Unassigned | - |
| Otu62117 | 1 | k__Fungi(100);k__Fungi_unclassified(100);k__Fungi_unclassified(100);k__Fungi_unclassified(100);k__Fungi_unclassified(100);k__Fungi_unclassified(100);k__Fungi_unclassified(100); | - | - | - | - | - | - | - | Unassigned | - |
| Otu62118 | 1 | k__Fungi(100);k__Fungi_unclassified(100);k__Fungi_unclassified(100);k__Fungi_unclassified(100);k__Fungi_unclassified(100);k__Fungi_unclassified(100);k__Fungi_unclassified(100); | - | - | - | - | - | - | - | Unassigned | - |
| Otu62119 | 1 | k__Fungi(100);k__Fungi_unclassified(100);k__Fungi_unclassified(100);k__Fungi_unclassified(100);k__Fungi_unclassified(100);k__Fungi_unclassified(100);k__Fungi_unclassified(100); | - | - | - | - | - | - | - | Unassigned | - |
| Otu62120 | 1 | k__Fungi(100);p__Ascomycota(100);p__Ascomycota_unclassified(100);p__Ascomycota_unclassified(100);p__Ascomycota_unclassified(100);p__Ascomycota_unclassified(100);p__Ascomycota_unclassified(100); | - | - | - | - | - | - | - | Unassigned | - |
| Otu62122 | 1 | k__Fungi(100);k__Fungi_unclassified(100);k__Fungi_unclassified(100);k__Fungi_unclassified(100);k__Fungi_unclassified(100);k__Fungi_unclassified(100);k__Fungi_unclassified(100); | - | - | - | - | - | - | - | Unassigned | - |
| Otu62123 | 1 | k__Fungi(100);k__Fungi_unclassified(100);k__Fungi_unclassified(100);k__Fungi_unclassified(100);k__Fungi_unclassified(100);k__Fungi_unclassified(100);k__Fungi_unclassified(100); | - | - | - | - | - | - | - | Unassigned | - |
| Otu62124 | 1 | k__Fungi(100);p__Ascomycota(100);p__Ascomycota_unclassified(100);p__Ascomycota_unclassified(100);p__Ascomycota_unclassified(100);p__Ascomycota_unclassified(100);p__Ascomycota_unclassified(100); | - | - | - | - | - | - | - | Unassigned | - |
| Otu62125 | 1 | k__Fungi(100);p__Ascomycota(100);p__Ascomycota_unclassified(100);p__Ascomycota_unclassified(100);p__Ascomycota_unclassified(100);p__Ascomycota_unclassified(100);p__Ascomycota_unclassified(100); | - | - | - | - | - | - | - | Unassigned | - |
| Otu62126 | 1 | k__Fungi(100);k__Fungi_unclassified(100);k__Fungi_unclassified(100);k__Fungi_unclassified(100);k__Fungi_unclassified(100);k__Fungi_unclassified(100);k__Fungi_unclassified(100); | - | - | - | - | - | - | - | Unassigned | - |
| Otu62128 | 1 | k__Fungi(100);k__Fungi_unclassified(100);k__Fungi_unclassified(100);k__Fungi_unclassified(100);k__Fungi_unclassified(100);k__Fungi_unclassified(100);k__Fungi_unclassified(100); | - | - | - | - | - | - | - | Unassigned | - |

| | | | | | | | | | | | |
|---|---|---|---|---|---|---|---|---|---|---|---|
| Otu62130 | 1 | k__Fungi(100);p__Ascomycota(100);p__Ascomycota_unclassified(100);p__Ascomycota_unclassified(100);p__Ascomycota_unclassified(100);p__Ascomycota_unclassified(100);p__Ascomycota_unclassified(100); | - | - | - | - | - | - | - | Unassigned | - |
| Otu62132 | 1 | k__Fungi(100);k__Fungi_unclassified(100);k__Fungi_unclassified(100);k__Fungi_unclassified(100);k__Fungi_unclassified(100);k__Fungi_unclassified(100);k__Fungi_unclassified(100); | - | - | - | - | - | - | - | Unassigned | - |
| Otu62133 | 1 | k__Fungi(100);p__Ascomycota(100);p__Ascomycota_unclassified(100);p__Ascomycota_unclassified(100);p__Ascomycota_unclassified(100);p__Ascomycota_unclassified(100);p__Ascomycota_unclassified(100); | - | - | - | - | - | - | - | Unassigned | - |
| Otu62134 | 1 | k__Fungi(100);k__Fungi_unclassified(100);k__Fungi_unclassified(100);k__Fungi_unclassified(100);k__Fungi_unclassified(100);k__Fungi_unclassified(100);k__Fungi_unclassified(100); | - | - | - | - | - | - | - | Unassigned | - |
| Otu62135 | 1 | k__Fungi(100);k__Fungi_unclassified(100);k__Fungi_unclassified(100);k__Fungi_unclassified(100);k__Fungi_unclassified(100);k__Fungi_unclassified(100);k__Fungi_unclassified(100); | - | - | - | - | - | - | - | Unassigned | - |
| Otu62138 | 1 | k__Fungi(100);k__Fungi_unclassified(100);k__Fungi_unclassified(100);k__Fungi_unclassified(100);k__Fungi_unclassified(100);k__Fungi_unclassified(100);k__Fungi_unclassified(100); | - | - | - | - | - | - | - | Unassigned | - |
| Otu62141 | 1 | k__Fungi(100);k__Fungi_unclassified(100);k__Fungi_unclassified(100);k__Fungi_unclassified(100);k__Fungi_unclassified(100);k__Fungi_unclassified(100);k__Fungi_unclassified(100); | - | - | - | - | - | - | - | Unassigned | - |
| Otu62142 | 1 | k__Fungi(100);k__Fungi_unclassified(100);k__Fungi_unclassified(100);k__Fungi_unclassified(100);k__Fungi_unclassified(100);k__Fungi_unclassified(100);k__Fungi_unclassified(100); | - | - | - | - | - | - | - | Unassigned | - |
| Otu62143 | 1 | k__Fungi(100);k__Fungi_unclassified(100);k__Fungi_unclassified(100);k__Fungi_unclassified(100);k__Fungi_unclassified(100);k__Fungi_unclassified(100);k__Fungi_unclassified(100); | - | - | - | - | - | - | - | Unassigned | - |
| Otu62144 | 1 | k__Fungi(100);k__Fungi_unclassified(100);k__Fungi_unclassified(100);k__Fungi_unclassified(100);k__Fungi_unclassified(100);k__Fungi_unclassified(100);k__Fungi_unclassified(100); | - | - | - | - | - | - | - | Unassigned | - |
| Otu62146 | 1 | k__Fungi(100);k__Fungi_unclassified(100);k__Fungi_unclassified(100);k__Fungi_unclassified(100);k__Fungi_unclassified(100);k__Fungi_unclassified(100);k__Fungi_unclassified(100); | - | - | - | - | - | - | - | Unassigned | - |
| Otu62148 | 1 | k__Fungi(100);k__Fungi_unclassified(100);k__Fungi_unclassified(100);k__Fungi_unclassified(100);k__Fungi_unclassified(100);k__Fungi_unclassified(100);k__Fungi_unclassified(100); | - | - | - | - | - | - | - | Unassigned | - |
| Otu62150 | 1 | k__Fungi(100);k__Fungi_unclassified(100);k__Fungi_unclassified(100);k__Fungi_unclassified(100);k__Fungi_unclassified(100);k__Fungi_unclassified(100);k__Fungi_unclassified(100); | - | - | - | - | - | - | - | Unassigned | - |
| Otu62152 | 1 | k__Fungi(100);k__Fungi_unclassified(100);k__Fungi_unclassified(100);k__Fungi_unclassified(100);k__Fungi_unclassified(100);k__Fungi_unclassified(100);k__Fungi_unclassified(100); | - | - | - | - | - | - | - | Unassigned | - |
| Otu62153 | 1 | k__Fungi(100);k__Fungi_unclassified(100);k__Fungi_unclassified(100);k__Fungi_unclassified(100);k__Fungi_unclassified(100);k__Fungi_unclassified(100);k__Fungi_unclassified(100); | - | - | - | - | - | - | - | Unassigned | - |
| Otu62155 | 1 | k__Fungi(100);k__Fungi_unclassified(100);k__Fungi_unclassified(100);k__Fungi_unclassified(100);k__Fungi_unclassified(100);k__Fungi_unclassified(100);k__Fungi_unclassified(100); | - | - | - | - | - | - | - | Unassigned | - |
| Otu62156 | 1 | k__Fungi(100);k__Fungi_unclassified(100);k__Fungi_unclassified(100);k__Fungi_unclassified(100);k__Fungi_unclassified(100);k__Fungi_unclassified(100);k__Fungi_unclassified(100); | - | - | - | - | - | - | - | Unassigned | - |
| Otu62157 | 1 | k__Fungi(100);p__Ascomycota(100);p__Ascomycota_unclassified(100);p__Ascomycota_unclassified(100);p__Ascomycota_unclassified(100);p__Ascomycota_unclassified(100);p__Ascomycota_unclassified(100); | - | - | - | - | - | - | - | Unassigned | - |
| Otu62160 | 1 | k__Fungi(100);k__Fungi_unclassified(100);k__Fungi_unclassified(100);k__Fungi_unclassified(100);k__Fungi_unclassified(100);k__Fungi_unclassified(100);k__Fungi_unclassified(100); | - | - | - | - | - | - | - | Unassigned | - |
| Otu62171 | 1 | k__Fungi(100);k__Fungi_unclassified(100);k__Fungi_unclassified(100);k__Fungi_unclassified(100);k__Fungi_unclassified(100);k__Fungi_unclassified(100);k__Fungi_unclassified(100); | - | - | - | - | - | - | - | Unassigned | - |
| Otu62183 | 1 | k__Fungi(100);p__Ascomycota(100);p__Ascomycota_unclassified(100);p__Ascomycota_unclassified(100);p__Ascomycota_unclassified(100);p__Ascomycota_unclassified(100);p__Ascomycota_unclassified(100); | - | - | - | - | - | - | - | Unassigned | - |
| Otu62188 | 1 | k__Fungi(100);k__Fungi_unclassified(100);k__Fungi_unclassified(100);k__Fungi_unclassified(100);k__Fungi_unclassified(100);k__Fungi_unclassified(100);k__Fungi_unclassified(100); | - | - | - | - | - | - | - | Unassigned | - |
| Otu62190 | 1 | k__Fungi(100);p__Ascomycota(100);p__Ascomycota_unclassified(100);p__Ascomycota_unclassified(100);p__Ascomycota_unclassified(100);p__Ascomycota_unclassified(100);p__Ascomycota_unclassified(100); | - | - | - | - | - | - | - | Unassigned | - |
| Otu62191 | 1 | k__Fungi(100);p__Ascomycota(100);p__Ascomycota_unclassified(100);p__Ascomycota_unclassified(100);p__Ascomycota_unclassified(100);p__Ascomycota_unclassified(100);p__Ascomycota_unclassified(100); | - | - | - | - | - | - | - | Unassigned | - |
| Otu62193 | 1 | k__Fungi(100);k__Fungi_unclassified(100);k__Fungi_unclassified(100);k__Fungi_unclassified(100);k__Fungi_unclassified(100);k__Fungi_unclassified(100);k__Fungi_unclassified(100); | - | - | - | - | - | - | - | Unassigned | - |
| Otu62194 | 1 | k__Fungi(100);k__Fungi_unclassified(100);k__Fungi_unclassified(100);k__Fungi_unclassified(100);k__Fungi_unclassified(100);k__Fungi_unclassified(100);k__Fungi_unclassified(100); | - | - | - | - | - | - | - | Unassigned | - |
| Otu62199 | 1 | k__Fungi(100);p__Ascomycota(100);p__Ascomycota_unclassified(100);p__Ascomycota_unclassified(100);p__Ascomycota_unclassified(100);p__Ascomycota_unclassified(100);p__Ascomycota_unclassified(100); | - | - | - | - | - | - | - | Unassigned | - |
| Otu62205 | 1 | k__Fungi(100);k__Fungi_unclassified(100);k__Fungi_unclassified(100);k__Fungi_unclassified(100);k__Fungi_unclassified(100);k__Fungi_unclassified(100);k__Fungi_unclassified(100); | - | - | - | - | - | - | - | Unassigned | - |
| Otu62206 | 1 | k__Fungi(100);k__Fungi_unclassified(100);k__Fungi_unclassified(100);k__Fungi_unclassified(100);k__Fungi_unclassified(100);k__Fungi_unclassified(100);k__Fungi_unclassified(100); | - | - | - | - | - | - | - | Unassigned | - |
| Otu62212 | 1 | k__Fungi(100);p__Ascomycota(100);p__Ascomycota_unclassified(100);p__Ascomycota_unclassified(100);p__Ascomycota_unclassified(100);p__Ascomycota_unclassified(100);p__Ascomycota_unclassified(100); | - | - | - | - | - | - | - | Unassigned | - |
| Otu62215 | 1 | k__Fungi(100);k__Fungi_unclassified(100);k__Fungi_unclassified(100);k__Fungi_unclassified(100);k__Fungi_unclassified(100);k__Fungi_unclassified(100);k__Fungi_unclassified(100); | - | - | - | - | - | - | - | Unassigned | - |
| Otu62216 | 1 | k__Fungi(100);k__Fungi_unclassified(100);k__Fungi_unclassified(100);k__Fungi_unclassified(100);k__Fungi_unclassified(100);k__Fungi_unclassified(100);k__Fungi_unclassified(100); | - | - | - | - | - | - | - | Unassigned | - |
| Otu62217 | 1 | k__Fungi(100);k__Fungi_unclassified(100);k__Fungi_unclassified(100);k__Fungi_unclassified(100);k__Fungi_unclassified(100);k__Fungi_unclassified(100);k__Fungi_unclassified(100); | - | - | - | - | - | - | - | Unassigned | - |
| Otu62220 | 1 | k__Fungi(100);p__Ascomycota(100);p__Ascomycota_unclassified(100);p__Ascomycota_unclassified(100);p__Ascomycota_unclassified(100);p__Ascomycota_unclassified(100);p__Ascomycota_unclassified(100); | - | - | - | - | - | - | - | Unassigned | - |
| Otu62223 | 1 | k__Fungi(100);p__Ascomycota(100);p__Ascomycota_unclassified(100);p__Ascomycota_unclassified(100);p__Ascomycota_unclassified(100);p__Ascomycota_unclassified(100);p__Ascomycota_unclassified(100); | - | - | - | - | - | - | - | Unassigned | - |
| Otu62226 | 1 | k__Fungi(100);p__Ascomycota(100);p__Ascomycota_unclassified(100);p__Ascomycota_unclassified(100);p__Ascomycota_unclassified(100);p__Ascomycota_unclassified(100);p__Ascomycota_unclassified(100); | - | - | - | - | - | - | - | Unassigned | - |
| Otu62228 | 1 | k__Fungi(100);k__Fungi_unclassified(100);k__Fungi_unclassified(100);k__Fungi_unclassified(100);k__Fungi_unclassified(100);k__Fungi_unclassified(100);k__Fungi_unclassified(100); | - | - | - | - | - | - | - | Unassigned | - |
| Otu62229 | 1 | k__Fungi(100);k__Fungi_unclassified(100);k__Fungi_unclassified(100);k__Fungi_unclassified(100);k__Fungi_unclassified(100);k__Fungi_unclassified(100);k__Fungi_unclassified(100); | - | - | - | - | - | - | - | Unassigned | - |
| Otu62232 | 1 | k__Fungi(100);k__Fungi_unclassified(100);k__Fungi_unclassified(100);k__Fungi_unclassified(100);k__Fungi_unclassified(100);k__Fungi_unclassified(100);k__Fungi_unclassified(100); | - | - | - | - | - | - | - | Unassigned | - |
| Otu62233 | 1 | k__Fungi(100);k__Fungi_unclassified(100);k__Fungi_unclassified(100);k__Fungi_unclassified(100);k__Fungi_unclassified(100);k__Fungi_unclassified(100);k__Fungi_unclassified(100); | - | - | - | - | - | - | - | Unassigned | - |
| Otu62234 | 1 | k__Fungi(100);k__Fungi_unclassified(100);k__Fungi_unclassified(100);k__Fungi_unclassified(100);k__Fungi_unclassified(100);k__Fungi_unclassified(100);k__Fungi_unclassified(100); | - | - | - | - | - | - | - | Unassigned | - |
| Otu62235 | 1 | k__Fungi(100);k__Fungi_unclassified(100);k__Fungi_unclassified(100);k__Fungi_unclassified(100);k__Fungi_unclassified(100);k__Fungi_unclassified(100);k__Fungi_unclassified(100); | - | - | - | - | - | - | - | Unassigned | - |
| Otu62237 | 1 | k__Fungi(100);k__Fungi_unclassified(100);k__Fungi_unclassified(100);k__Fungi_unclassified(100);k__Fungi_unclassified(100);k__Fungi_unclassified(100);k__Fungi_unclassified(100); | - | - | - | - | - | - | - | Unassigned | - |
| Otu62238 | 1 | k__Fungi(100);k__Fungi_unclassified(100);k__Fungi_unclassified(100);k__Fungi_unclassified(100);k__Fungi_unclassified(100);k__Fungi_unclassified(100);k__Fungi_unclassified(100); | - | - | - | - | - | - | - | Unassigned | - |
| Otu62239 | 1 | k__Fungi(100);p__Ascomycota(100);p__Ascomycota_unclassified(100);p__Ascomycota_unclassified(100);p__Ascomycota_unclassified(100);p__Ascomycota_unclassified(100);p__Ascomycota_unclassified(100); | - | - | - | - | - | - | - | Unassigned | - |
| Otu62240 | 1 | k__Fungi(100);k__Fungi_unclassified(100);k__Fungi_unclassified(100);k__Fungi_unclassified(100);k__Fungi_unclassified(100);k__Fungi_unclassified(100);k__Fungi_unclassified(100); | - | - | - | - | - | - | - | Unassigned | - |
| Otu62241 | 1 | k__Fungi(100);k__Fungi_unclassified(100);k__Fungi_unclassified(100);k__Fungi_unclassified(100);k__Fungi_unclassified(100);k__Fungi_unclassified(100);k__Fungi_unclassified(100); | - | - | - | - | - | - | - | Unassigned | - |
| Otu62242 | 1 | k__Fungi(100);p__Ascomycota(100);p__Ascomycota_unclassified(100);p__Ascomycota_unclassified(100);p__Ascomycota_unclassified(100);p__Ascomycota_unclassified(100);p__Ascomycota_unclassified(100); | - | - | - | - | - | - | - | Unassigned | - |
| Otu62243 | 1 | k__Fungi(100);k__Fungi_unclassified(100);k__Fungi_unclassified(100);k__Fungi_unclassified(100);k__Fungi_unclassified(100);k__Fungi_unclassified(100);k__Fungi_unclassified(100); | - | - | - | - | - | - | - | Unassigned | - |
| Otu62245 | 1 | k__Fungi(100);p__Ascomycota(100);p__Ascomycota_unclassified(100);p__Ascomycota_unclassified(100);p__Ascomycota_unclassified(100);p__Ascomycota_unclassified(100);p__Ascomycota_unclassified(100); | - | - | - | - | - | - | - | Unassigned | - |
| Otu62247 | 1 | k__Fungi(100);p__Ascomycota(100);p__Ascomycota_unclassified(100);p__Ascomycota_unclassified(100);p__Ascomycota_unclassified(100);p__Ascomycota_unclassified(100);p__Ascomycota_unclassified(100); | - | - | - | - | - | - | - | Unassigned | - |
| Otu62250 | 1 | k__Fungi(100);k__Fungi_unclassified(100);k__Fungi_unclassified(100);k__Fungi_unclassified(100);k__Fungi_unclassified(100);k__Fungi_unclassified(100);k__Fungi_unclassified(100); | - | - | - | - | - | - | - | Unassigned | - |
| Otu62251 | 1 | k__Fungi(100);p__Ascomycota(100);p__Ascomycota_unclassified(100);p__Ascomycota_unclassified(100);p__Ascomycota_unclassified(100);p__Ascomycota_unclassified(100);p__Ascomycota_unclassified(100); | - | - | - | - | - | - | - | Unassigned | - |
| Otu62252 | 1 | k__Fungi(100);p__Ascomycota(100);p__Ascomycota_unclassified(100);p__Ascomycota_unclassified(100);p__Ascomycota_unclassified(100);p__Ascomycota_unclassified(100);p__Ascomycota_unclassified(100); | - | - | - | - | - | - | - | Unassigned | - |
| Otu62257 | 1 | k__Fungi(100);p__Ascomycota(100);p__Ascomycota_unclassified(100);p__Ascomycota_unclassified(100);p__Ascomycota_unclassified(100);p__Ascomycota_unclassified(100);p__Ascomycota_unclassified(100); | - | - | - | - | - | - | - | Unassigned | - |
| Otu62259 | 1 | k__Fungi(100);p__Ascomycota(100);p__Ascomycota_unclassified(100);p__Ascomycota_unclassified(100);p__Ascomycota_unclassified(100);p__Ascomycota_unclassified(100);p__Ascomycota_unclassified(100); | - | - | - | - | - | - | - | Unassigned | - |
| Otu62260 | 1 | k__Fungi(100);k__Fungi_unclassified(100);k__Fungi_unclassified(100);k__Fungi_unclassified(100);k__Fungi_unclassified(100);k__Fungi_unclassified(100);k__Fungi_unclassified(100); | - | - | - | - | - | - | - | Unassigned | - |
| Otu62261 | 1 | k__Fungi(100);k__Fungi_unclassified(100);k__Fungi_unclassified(100);k__Fungi_unclassified(100);k__Fungi_unclassified(100);k__Fungi_unclassified(100);k__Fungi_unclassified(100); | - | - | - | - | - | - | - | Unassigned | - |
| Otu62262 | 1 | k__Fungi(100);k__Fungi_unclassified(100);k__Fungi_unclassified(100);k__Fungi_unclassified(100);k__Fungi_unclassified(100);k__Fungi_unclassified(100);k__Fungi_unclassified(100); | - | - | - | - | - | - | - | Unassigned | - |
| Otu62263 | 1 | k__Fungi(100);p__Ascomycota(100);p__Ascomycota_unclassified(100);p__Ascomycota_unclassified(100);p__Ascomycota_unclassified(100);p__Ascomycota_unclassified(100);p__Ascomycota_unclassified(100); | - | - | - | - | - | - | - | Unassigned | - |
| Otu62264 | 1 | k__Fungi(100);k__Fungi_unclassified(100);k__Fungi_unclassified(100);k__Fungi_unclassified(100);k__Fungi_unclassified(100);k__Fungi_unclassified(100);k__Fungi_unclassified(100); | - | - | - | - | - | - | - | Unassigned | - |
| Otu62273 | 1 | k__Fungi(100);k__Fungi_unclassified(100);k__Fungi_unclassified(100);k__Fungi_unclassified(100);k__Fungi_unclassified(100);k__Fungi_unclassified(100);k__Fungi_unclassified(100); | - | - | - | - | - | - | - | Unassigned | - |
| Otu62276 | 1 | k__Fungi(100);k__Fungi_unclassified(100);k__Fungi_unclassified(100);k__Fungi_unclassified(100);k__Fungi_unclassified(100);k__Fungi_unclassified(100);k__Fungi_unclassified(100); | - | - | - | - | - | - | - | Unassigned | - |
| Otu62278 | 1 | k__Fungi(100);p__Ascomycota(100);p__Ascomycota_unclassified(100);p__Ascomycota_unclassified(100);p__Ascomycota_unclassified(100);p__Ascomycota_unclassified(100);p__Ascomycota_unclassified(100); | - | - | - | - | - | - | - | Unassigned | - |
| Otu62279 | 1 | k__Fungi(100);k__Fungi_unclassified(100);k__Fungi_unclassified(100);k__Fungi_unclassified(100);k__Fungi_unclassified(100);k__Fungi_unclassified(100);k__Fungi_unclassified(100); | - | - | - | - | - | - | - | Unassigned | - |
| Otu62281 | 1 | k__Fungi(100);k__Fungi_unclassified(100);k__Fungi_unclassified(100);k__Fungi_unclassified(100);k__Fungi_unclassified(100);k__Fungi_unclassified(100);k__Fungi_unclassified(100); | - | - | - | - | - | - | - | Unassigned | - |
| Otu62282 | 1 | k__Fungi(100);p__Ascomycota(100);p__Ascomycota_unclassified(100);p__Ascomycota_unclassified(100);p__Ascomycota_unclassified(100);p__Ascomycota_unclassified(100);p__Ascomycota_unclassified(100); | - | - | - | - | - | - | - | Unassigned | - |
| Otu62283 | 1 | k__Fungi(100);k__Fungi_unclassified(100);k__Fungi_unclassified(100);k__Fungi_unclassified(100);k__Fungi_unclassified(100);k__Fungi_unclassified(100);k__Fungi_unclassified(100); | - | - | - | - | - | - | - | Unassigned | - |
| Otu62284 | 1 | k__Fungi(100);p__Ascomycota(100);p__Ascomycota_unclassified(100);p__Ascomycota_unclassified(100);p__Ascomycota_unclassified(100);p__Ascomycota_unclassified(100);p__Ascomycota_unclassified(100); | - | - | - | - | - | - | - | Unassigned | - |
| Otu62286 | 1 | k__Fungi(100);p__Ascomycota(100);p__Ascomycota_unclassified(100);p__Ascomycota_unclassified(100);p__Ascomycota_unclassified(100);p__Ascomycota_unclassified(100);p__Ascomycota_unclassified(100); | - | - | - | - | - | - | - | Unassigned | - |
| Otu62287 | 1 | k__Fungi(100);p__Ascomycota(100);p__Ascomycota_unclassified(100);p__Ascomycota_unclassified(100);p__Ascomycota_unclassified(100);p__Ascomycota_unclassified(100);p__Ascomycota_unclassified(100); | - | - | - | - | - | - | - | Unassigned | - |
| Otu62289 | 1 | k__Fungi(100);k__Fungi_unclassified(100);k__Fungi_unclassified(100);k__Fungi_unclassified(100);k__Fungi_unclassified(100);k__Fungi_unclassified(100);k__Fungi_unclassified(100); | - | - | - | - | - | - | - | Unassigned | - |
| Otu62297 | 1 | k__Fungi(100);k__Fungi_unclassified(100);k__Fungi_unclassified(100);k__Fungi_unclassified(100);k__Fungi_unclassified(100);k__Fungi_unclassified(100);k__Fungi_unclassified(100); | - | - | - | - | - | - | - | Unassigned | - |
| Otu62298 | 1 | k__Fungi(100);p__Ascomycota(100);p__Ascomycota_unclassified(100);p__Ascomycota_unclassified(100);p__Ascomycota_unclassified(100);p__Ascomycota_unclassified(100);p__Ascomycota_unclassified(100); | - | - | - | - | - | - | - | Unassigned | - |
| Otu62299 | 1 | k__Fungi(100);p__Ascomycota(100);p__Ascomycota_unclassified(100);p__Ascomycota_unclassified(100);p__Ascomycota_unclassified(100);p__Ascomycota_unclassified(100);p__Ascomycota_unclassified(100); | - | - | - | - | - | - | - | Unassigned | - |
| Otu62300 | 1 | k__Fungi(100);k__Fungi_unclassified(100);k__Fungi_unclassified(100);k__Fungi_unclassified(100);k__Fungi_unclassified(100);k__Fungi_unclassified(100);k__Fungi_unclassified(100); | - | - | - | - | - | - | - | Unassigned | - |
| Otu62301 | 1 | k__Fungi(100);p__Ascomycota(100);p__Ascomycota_unclassified(100);p__Ascomycota_unclassified(100);p__Ascomycota_unclassified(100);p__Ascomycota_unclassified(100);p__Ascomycota_unclassified(100); | - | - | - | - | - | - | - | Unassigned | - |
| Otu62302 | 1 | k__Fungi(100);k__Fungi_unclassified(100);k__Fungi_unclassified(100);k__Fungi_unclassified(100);k__Fungi_unclassified(100);k__Fungi_unclassified(100);k__Fungi_unclassified(100); | - | - | - | - | - | - | - | Unassigned | - |
| Otu62308 | 1 | k__Fungi(100);k__Fungi_unclassified(100);k__Fungi_unclassified(100);k__Fungi_unclassified(100);k__Fungi_unclassified(100);k__Fungi_unclassified(100);k__Fungi_unclassified(100); | - | - | - | - | - | - | - | Unassigned | - |
| Otu62311 | 1 | k__Fungi(100);p__Ascomycota(100);p__Ascomycota_unclassified(100);p__Ascomycota_unclassified(100);p__Ascomycota_unclassified(100);p__Ascomycota_unclassified(100);p__Ascomycota_unclassified(100); | - | - | - | - | - | - | - | Unassigned | - |
| Otu62312 | 1 | k__Fungi(100);p__Ascomycota(100);p__Ascomycota_unclassified(100);p__Ascomycota_unclassified(100);p__Ascomycota_unclassified(100);p__Ascomycota_unclassified(100);p__Ascomycota_unclassified(100); | - | - | - | - | - | - | - | Unassigned | - |
| Otu62313 | 1 | k__Fungi(100);k__Fungi_unclassified(100);k__Fungi_unclassified(100);k__Fungi_unclassified(100);k__Fungi_unclassified(100);k__Fungi_unclassified(100);k__Fungi_unclassified(100); | - | - | - | - | - | - | - | Unassigned | - |
| Otu62314 | 1 | k__Fungi(100);p__Ascomycota(100);p__Ascomycota_unclassified(100);p__Ascomycota_unclassified(100);p__Ascomycota_unclassified(100);p__Ascomycota_unclassified(100);p__Ascomycota_unclassified(100); | - | - | - | - | - | - | - | Unassigned | - |
| Otu62315 | 1 | k__Fungi(100);p__Ascomycota(100);p__Ascomycota_unclassified(100);p__Ascomycota_unclassified(100);p__Ascomycota_unclassified(100);p__Ascomycota_unclassified(100);p__Ascomycota_unclassified(100); | - | - | - | - | - | - | - | Unassigned | - |
| Otu62316 | 1 | k__Fungi(100);k__Fungi_unclassified(100);k__Fungi_unclassified(100);k__Fungi_unclassified(100);k__Fungi_unclassified(100);k__Fungi_unclassified(100);k__Fungi_unclassified(100); | - | - | - | - | - | - | - | Unassigned | - |
| Otu62317 | 1 | k__Fungi(100);p__Ascomycota(100);p__Ascomycota_unclassified(100);p__Ascomycota_unclassified(100);p__Ascomycota_unclassified(100);p__Ascomycota_unclassified(100);p__Ascomycota_unclassified(100); | - | - | - | - | - | - | - | Unassigned | - |
| Otu62319 | 1 | k__Fungi(100);k__Fungi_unclassified(100);k__Fungi_unclassified(100);k__Fungi_unclassified(100);k__Fungi_unclassified(100);k__Fungi_unclassified(100);k__Fungi_unclassified(100); | - | - | - | - | - | - | - | Unassigned | - |
| Otu62320 | 1 | k__Fungi(100);p__Ascomycota(100);p__Ascomycota_unclassified(100);p__Ascomycota_unclassified(100);p__Ascomycota_unclassified(100);p__Ascomycota_unclassified(100);p__Ascomycota_unclassified(100); | - | - | - | - | - | - | - | Unassigned | - |
| Otu62322 | 1 | k__Fungi(100);k__Fungi_unclassified(100);k__Fungi_unclassified(100);k__Fungi_unclassified(100);k__Fungi_unclassified(100);k__Fungi_unclassified(100);k__Fungi_unclassified(100); | - | - | - | - | - | - | - | Unassigned | - |
| Otu62325 | 1 | k__Fungi(100);p__Ascomycota(100);p__Ascomycota_unclassified(100);p__Ascomycota_unclassified(100);p__Ascomycota_unclassified(100);p__Ascomycota_unclassified(100);p__Ascomycota_unclassified(100); | - | - | - | - | - | - | - | Unassigned | - |
| Otu62326 | 1 | k__Fungi(100);k__Fungi_unclassified(100);k__Fungi_unclassified(100);k__Fungi_unclassified(100);k__Fungi_unclassified(100);k__Fungi_unclassified(100);k__Fungi_unclassified(100); | - | - | - | - | - | - | - | Unassigned | - |
| Otu62327 | 1 | k__Fungi(100);p__Ascomycota(100);p__Ascomycota_unclassified(100);p__Ascomycota_unclassified(100);p__Ascomycota_unclassified(100);p__Ascomycota_unclassified(100);p__Ascomycota_unclassified(100); | - | - | - | - | - | - | - | Unassigned | - |
| Otu62328 | 1 | k__Fungi(100);k__Fungi_unclassified(100);k__Fungi_unclassified(100);k__Fungi_unclassified(100);k__Fungi_unclassified(100);k__Fungi_unclassified(100);k__Fungi_unclassified(100); | - | - | - | - | - | - | - | Unassigned | - |
| Otu62329 | 1 | k__Fungi(100);p__Ascomycota(100);p__Ascomycota_unclassified(100);p__Ascomycota_unclassified(100);p__Ascomycota_unclassified(100);p__Ascomycota_unclassified(100);p__Ascomycota_unclassified(100); | - | - | - | - | - | - | - | Unassigned | - |
| Otu62330 | 1 | k__Fungi(100);k__Fungi_unclassified(100);k__Fungi_unclassified(100);k__Fungi_unclassified(100);k__Fungi_unclassified(100);k__Fungi_unclassified(100);k__Fungi_unclassified(100); | - | - | - | - | - | - | - | Unassigned | - |
| Otu62331 | 1 | k__Fungi(100);p__Ascomycota(100);p__Ascomycota_unclassified(100);p__Ascomycota_unclassified(100);p__Ascomycota_unclassified(100);p__Ascomycota_unclassified(100);p__Ascomycota_unclassified(100); | - | - | - | - | - | - | - | Unassigned | - |
| Otu62332 | 1 | k__Fungi(100);k__Fungi_unclassified(100);k__Fungi_unclassified(100);k__Fungi_unclassified(100);k__Fungi_unclassified(100);k__Fungi_unclassified(100);k__Fungi_unclassified(100); | - | - | - | - | - | - | - | Unassigned | - |
| Otu62335 | 1 | k__Fungi(100);k__Fungi_unclassified(100);k__Fungi_unclassified(100);k__Fungi_unclassified(100);k__Fungi_unclassified(100);k__Fungi_unclassified(100);k__Fungi_unclassified(100); | - | - | - | - | - | - | - | Unassigned | - |
| Otu62336 | 1 | k__Fungi(100);p__Ascomycota(100);p__Ascomycota_unclassified(100);p__Ascomycota_unclassified(100);p__Ascomycota_unclassified(100);p__Ascomycota_unclassified(100);p__Ascomycota_unclassified(100); | - | - | - | - | - | - | - | Unassigned | - |
| Otu62338 | 1 | k__Fungi(100);k__Fungi_unclassified(100);k__Fungi_unclassified(100);k__Fungi_unclassified(100);k__Fungi_unclassified(100);k__Fungi_unclassified(100);k__Fungi_unclassified(100); | - | - | - | - | - | - | - | Unassigned | - |
| Otu62340 | 1 | k__Fungi(100);k__Fungi_unclassified(100);k__Fungi_unclassified(100);k__Fungi_unclassified(100);k__Fungi_unclassified(100);k__Fungi_unclassified(100);k__Fungi_unclassified(100); | - | - | - | - | - | - | - | Unassigned | - |
| Otu62341 | 1 | k__Fungi(100);k__Fungi_unclassified(100);k__Fungi_unclassified(100);k__Fungi_unclassified(100);k__Fungi_unclassified(100);k__Fungi_unclassified(100);k__Fungi_unclassified(100); | - | - | - | - | - | - | - | Unassigned | - |
| Otu62342 | 1 | k__Fungi(100);p__Ascomycota(100);p__Ascomycota_unclassified(100);p__Ascomycota_unclassified(100);p__Ascomycota_unclassified(100);p__Ascomycota_unclassified(100);p__Ascomycota_unclassified(100); | - | - | - | - | - | - | - | Unassigned | - |
| Otu62343 | 1 | k__Fungi(100);k__Fungi_unclassified(100);k__Fungi_unclassified(100);k__Fungi_unclassified(100);k__Fungi_unclassified(100);k__Fungi_unclassified(100);k__Fungi_unclassified(100); | - | - | - | - | - | - | - | Unassigned | - |
| Otu62344 | 1 | k__Fungi(100);p__Ascomycota(100);p__Ascomycota_unclassified(100);p__Ascomycota_unclassified(100);p__Ascomycota_unclassified(100);p__Ascomycota_unclassified(100);p__Ascomycota_unclassified(100); | - | - | - | - | - | - | - | Unassigned | - |
| Otu62345 | 1 | k__Fungi(100);k__Fungi_unclassified(100);k__Fungi_unclassified(100);k__Fungi_unclassified(100);k__Fungi_unclassified(100);k__Fungi_unclassified(100);k__Fungi_unclassified(100); | - | - | - | - | - | - | - | Unassigned | - |
| Otu62346 | 1 | k__Fungi(100);k__Fungi_unclassified(100);k__Fungi_unclassified(100);k__Fungi_unclassified(100);k__Fungi_unclassified(100);k__Fungi_unclassified(100);k__Fungi_unclassified(100); | - | - | - | - | - | - | - | Unassigned | - |
| Otu62347 | 1 | k__Fungi(100);k__Fungi_unclassified(100);k__Fungi_unclassified(100);k__Fungi_unclassified(100);k__Fungi_unclassified(100);k__Fungi_unclassified(100);k__Fungi_unclassified(100); | - | - | - | - | - | - | - | Unassigned | - |
| Otu62348 | 1 | k__Fungi(100);k__Fungi_unclassified(100);k__Fungi_unclassified(100);k__Fungi_unclassified(100);k__Fungi_unclassified(100);k__Fungi_unclassified(100);k__Fungi_unclassified(100); | - | - | - | - | - | - | - | Unassigned | - |
| Otu62349 | 1 | k__Fungi(100);p__Ascomycota(100);p__Ascomycota_unclassified(100);p__Ascomycota_unclassified(100);p__Ascomycota_unclassified(100);p__Ascomycota_unclassified(100);p__Ascomycota_unclassified(100); | - | - | - | - | - | - | - | Unassigned | - |
| Otu62350 | 1 | k__Fungi(100);k__Fungi_unclassified(100);k__Fungi_unclassified(100);k__Fungi_unclassified(100);k__Fungi_unclassified(100);k__Fungi_unclassified(100);k__Fungi_unclassified(100); | - | - | - | - | - | - | - | Unassigned | - |
| Otu62351 | 1 | k__Fungi(100);k__Fungi_unclassified(100);k__Fungi_unclassified(100);k__Fungi_unclassified(100);k__Fungi_unclassified(100);k__Fungi_unclassified(100);k__Fungi_unclassified(100); | - | - | - | - | - | - | - | Unassigned | - |
| Otu62352 | 1 | k__Fungi(100);k__Fungi_unclassified(100);k__Fungi_unclassified(100);k__Fungi_unclassified(100);k__Fungi_unclassified(100);k__Fungi_unclassified(100);k__Fungi_unclassified(100); | - | - | - | - | - | - | - | Unassigned | - |
| Otu62353 | 1 | k__Fungi(100);k__Fungi_unclassified(100);k__Fungi_unclassified(100);k__Fungi_unclassified(100);k__Fungi_unclassified(100);k__Fungi_unclassified(100);k__Fungi_unclassified(100); | - | - | - | - | - | - | - | Unassigned | - |
| Otu62354 | 1 | k__Fungi(100);k__Fungi_unclassified(100);k__Fungi_unclassified(100);k__Fungi_unclassified(100);k__Fungi_unclassified(100);k__Fungi_unclassified(100);k__Fungi_unclassified(100); | - | - | - | - | - | - | - | Unassigned | - |
| Otu62355 | 1 | k__Fungi(100);k__Fungi_unclassified(100);k__Fungi_unclassified(100);k__Fungi_unclassified(100);k__Fungi_unclassified(100);k__Fungi_unclassified(100);k__Fungi_unclassified(100); | - | - | - | - | - | - | - | Unassigned | - |
| Otu62357 | 1 | k__Fungi(100);k__Fungi_unclassified(100);k__Fungi_unclassified(100);k__Fungi_unclassified(100);k__Fungi_unclassified(100);k__Fungi_unclassified(100);k__Fungi_unclassified(100); | - | - | - | - | - | - | - | Unassigned | - |
| Otu62358 | 1 | k__Fungi(100);k__Fungi_unclassified(100);k__Fungi_unclassified(100);k__Fungi_unclassified(100);k__Fungi_unclassified(100);k__Fungi_unclassified(100);k__Fungi_unclassified(100); | - | - | - | - | - | - | - | Unassigned | - |
| Otu62360 | 1 | k__Fungi(100);k__Fungi_unclassified(100);k__Fungi_unclassified(100);k__Fungi_unclassified(100);k__Fungi_unclassified(100);k__Fungi_unclassified(100);k__Fungi_unclassified(100); | - | - | - | - | - | - | - | Unassigned | - |
| Otu62361 | 1 | k__Fungi(100);k__Fungi_unclassified(100);k__Fungi_unclassified(100);k__Fungi_unclassified(100);k__Fungi_unclassified(100);k__Fungi_unclassified(100);k__Fungi_unclassified(100); | - | - | - | - | - | - | - | Unassigned | - |
| Otu62363 | 1 | k__Fungi(100);p__Ascomycota(100);p__Ascomycota_unclassified(100);p__Ascomycota_unclassified(100);p__Ascomycota_unclassified(100);p__Ascomycota_unclassified(100);p__Ascomycota_unclassified(100); | - | - | - | - | - | - | - | Unassigned | - |
| Otu62364 | 1 | k__Fungi(100);k__Fungi_unclassified(100);k__Fungi_unclassified(100);k__Fungi_unclassified(100);k__Fungi_unclassified(100);k__Fungi_unclassified(100);k__Fungi_unclassified(100); | - | - | - | - | - | - | - | Unassigned | - |
| Otu62368 | 1 | k__Fungi(100);k__Fungi_unclassified(100);k__Fungi_unclassified(100);k__Fungi_unclassified(100);k__Fungi_unclassified(100);k__Fungi_unclassified(100);k__Fungi_unclassified(100); | - | - | - | - | - | - | - | Unassigned | - |
| Otu62370 | 1 | k__Fungi(100);p__Ascomycota(100);p__Ascomycota_unclassified(100);p__Ascomycota_unclassified(100);p__Ascomycota_unclassified(100);p__Ascomycota_unclassified(100);p__Ascomycota_unclassified(100); | - | - | - | - | - | - | - | Unassigned | - |
| Otu62371 | 1 | k__Fungi(100);p__Ascomycota(100);p__Ascomycota_unclassified(100);p__Ascomycota_unclassified(100);p__Ascomycota_unclassified(100);p__Ascomycota_unclassified(100);p__Ascomycota_unclassified(100); | - | - | - | - | - | - | - | Unassigned | - |
| Otu62372 | 1 | k__Fungi(100);k__Fungi_unclassified(100);k__Fungi_unclassified(100);k__Fungi_unclassified(100);k__Fungi_unclassified(100);k__Fungi_unclassified(100);k__Fungi_unclassified(100); | - | - | - | - | - | - | - | Unassigned | - |
| Otu62373 | 1 | k__Fungi(100);p__Ascomycota(100);p__Ascomycota_unclassified(100);p__Ascomycota_unclassified(100);p__Ascomycota_unclassified(100);p__Ascomycota_unclassified(100);p__Ascomycota_unclassified(100); | - | - | - | - | - | - | - | Unassigned | - |
| Otu62375 | 1 | k__Fungi(100);p__Ascomycota(100);p__Ascomycota_unclassified(100);p__Ascomycota_unclassified(100);p__Ascomycota_unclassified(100);p__Ascomycota_unclassified(100);p__Ascomycota_unclassified(100); | - | - | - | - | - | - | - | Unassigned | - |
| Otu62376 | 1 | k__Fungi(100);k__Fungi_unclassified(100);k__Fungi_unclassified(100);k__Fungi_unclassified(100);k__Fungi_unclassified(100);k__Fungi_unclassified(100);k__Fungi_unclassified(100); | - | - | - | - | - | - | - | Unassigned | - |
| Otu62378 | 1 | k__Fungi(100);p__Ascomycota(100);p__Ascomycota_unclassified(100);p__Ascomycota_unclassified(100);p__Ascomycota_unclassified(100);p__Ascomycota_unclassified(100);p__Ascomycota_unclassified(100); | - | - | - | - | - | - | - | Unassigned | - |
| Otu62379 | 1 | k__Fungi(100);k__Fungi_unclassified(100);k__Fungi_unclassified(100);k__Fungi_unclassified(100);k__Fungi_unclassified(100);k__Fungi_unclassified(100);k__Fungi_unclassified(100); | - | - | - | - | - | - | - | Unassigned | - |
| Otu62382 | 1 | k__Fungi(100);k__Fungi_unclassified(100);k__Fungi_unclassified(100);k__Fungi_unclassified(100);k__Fungi_unclassified(100);k__Fungi_unclassified(100);k__Fungi_unclassified(100); | - | - | - | - | - | - | - | Unassigned | - |
| Otu62383 | 1 | k__Fungi(100);p__Ascomycota(100);p__Ascomycota_unclassified(100);p__Ascomycota_unclassified(100);p__Ascomycota_unclassified(100);p__Ascomycota_unclassified(100);p__Ascomycota_unclassified(100); | - | - | - | - | - | - | - | Unassigned | - |
| Otu62384 | 1 | k__Fungi(100);k__Fungi_unclassified(100);k__Fungi_unclassified(100);k__Fungi_unclassified(100);k__Fungi_unclassified(100);k__Fungi_unclassified(100);k__Fungi_unclassified(100); | - | - | - | - | - | - | - | Unassigned | - |
| Otu62385 | 1 | k__Fungi(100);p__Ascomycota(100);p__Ascomycota_unclassified(100);p__Ascomycota_unclassified(100);p__Ascomycota_unclassified(100);p__Ascomycota_unclassified(100);p__Ascomycota_unclassified(100); | - | - | - | - | - | - | - | Unassigned | - |
| Otu62386 | 1 | k__Fungi(100);k__Fungi_unclassified(100);k__Fungi_unclassified(100);k__Fungi_unclassified(100);k__Fungi_unclassified(100);k__Fungi_unclassified(100);k__Fungi_unclassified(100); | - | - | - | - | - | - | - | Unassigned | - |
| Otu62387 | 1 | k__Fungi(100);p__Ascomycota(100);p__Ascomycota_unclassified(100);p__Ascomycota_unclassified(100);p__Ascomycota_unclassified(100);p__Ascomycota_unclassified(100);p__Ascomycota_unclassified(100); | - | - | - | - | - | - | - | Unassigned | - |
| Otu62388 | 1 | k__Fungi(100);p__Ascomycota(100);p__Ascomycota_unclassified(100);p__Ascomycota_unclassified(100);p__Ascomycota_unclassified(100);p__Ascomycota_unclassified(100);p__Ascomycota_unclassified(100); | - | - | - | - | - | - | - | Unassigned | - |
| Otu62389 | 1 | k__Fungi(100);k__Fungi_unclassified(100);k__Fungi_unclassified(100);k__Fungi_unclassified(100);k__Fungi_unclassified(100);k__Fungi_unclassified(100);k__Fungi_unclassified(100); | - | - | - | - | - | - | - | Unassigned | - |
| Otu62390 | 1 | k__Fungi(100);k__Fungi_unclassified(100);k__Fungi_unclassified(100);k__Fungi_unclassified(100);k__Fungi_unclassified(100);k__Fungi_unclassified(100);k__Fungi_unclassified(100); | - | - | - | - | - | - | - | Unassigned | - |
| Otu62391 | 1 | k__Fungi(100);p__Ascomycota(100);p__Ascomycota_unclassified(100);p__Ascomycota_unclassified(100);p__Ascomycota_unclassified(100);p__Ascomycota_unclassified(100);p__Ascomycota_unclassified(100); | - | - | - | - | - | - | - | Unassigned | - |
| Otu62392 | 1 | k__Fungi(100);k__Fungi_unclassified(100);k__Fungi_unclassified(100);k__Fungi_unclassified(100);k__Fungi_unclassified(100);k__Fungi_unclassified(100);k__Fungi_unclassified(100); | - | - | - | - | - | - | - | Unassigned | - |
| Otu62393 | 1 | k__Fungi(100);k__Fungi_unclassified(100);k__Fungi_unclassified(100);k__Fungi_unclassified(100);k__Fungi_unclassified(100);k__Fungi_unclassified(100);k__Fungi_unclassified(100); | - | - | - | - | - | - | - | Unassigned | - |
| Otu62394 | 1 | k__Fungi(100);p__Ascomycota(100);p__Ascomycota_unclassified(100);p__Ascomycota_unclassified(100);p__Ascomycota_unclassified(100);p__Ascomycota_unclassified(100);p__Ascomycota_unclassified(100); | - | - | - | - | - | - | - | Unassigned | - |
| Otu62396 | 1 | k__Fungi(100);p__Ascomycota(100);p__Ascomycota_unclassified(100);p__Ascomycota_unclassified(100);p__Ascomycota_unclassified(100);p__Ascomycota_unclassified(100);p__Ascomycota_unclassified(100); | - | - | - | - | - | - | - | Unassigned | - |
| Otu62397 | 1 | k__Fungi(100);k__Fungi_unclassified(100);k__Fungi_unclassified(100);k__Fungi_unclassified(100);k__Fungi_unclassified(100);k__Fungi_unclassified(100);k__Fungi_unclassified(100); | - | - | - | - | - | - | - | Unassigned | - |
| Otu62399 | 1 | k__Fungi(100);p__Ascomycota(100);p__Ascomycota_unclassified(100);p__Ascomycota_unclassified(100);p__Ascomycota_unclassified(100);p__Ascomycota_unclassified(100);p__Ascomycota_unclassified(100); | - | - | - | - | - | - | - | Unassigned | - |
| Otu62400 | 1 | k__Fungi(100);k__Fungi_unclassified(100);k__Fungi_unclassified(100);k__Fungi_unclassified(100);k__Fungi_unclassified(100);k__Fungi_unclassified(100);k__Fungi_unclassified(100); | - | - | - | - | - | - | - | Unassigned | - |
| Otu62403 | 1 | k__Fungi(100);k__Fungi_unclassified(100);k__Fungi_unclassified(100);k__Fungi_unclassified(100);k__Fungi_unclassified(100);k__Fungi_unclassified(100);k__Fungi_unclassified(100); | - | - | - | - | - | - | - | Unassigned | - |
| Otu62404 | 1 | k__Fungi(100);k__Fungi_unclassified(100);k__Fungi_unclassified(100);k__Fungi_unclassified(100);k__Fungi_unclassified(100);k__Fungi_unclassified(100);k__Fungi_unclassified(100); | - | - | - | - | - | - | - | Unassigned | - |
| Otu62406 | 1 | k__Fungi(100);p__Ascomycota(100);p__Ascomycota_unclassified(100);p__Ascomycota_unclassified(100);p__Ascomycota_unclassified(100);p__Ascomycota_unclassified(100);p__Ascomycota_unclassified(100); | - | - | - | - | - | - | - | Unassigned | - |
| Otu62408 | 1 | k__Fungi(100);k__Fungi_unclassified(100);k__Fungi_unclassified(100);k__Fungi_unclassified(100);k__Fungi_unclassified(100);k__Fungi_unclassified(100);k__Fungi_unclassified(100); | - | - | - | - | - | - | - | Unassigned | - |
| Otu62410 | 1 | k__Fungi(100);k__Fungi_unclassified(100);k__Fungi_unclassified(100);k__Fungi_unclassified(100);k__Fungi_unclassified(100);k__Fungi_unclassified(100);k__Fungi_unclassified(100); | - | - | - | - | - | - | - | Unassigned | - |
| Otu62412 | 1 | k__Fungi(100);p__Ascomycota(100);p__Ascomycota_unclassified(100);p__Ascomycota_unclassified(100);p__Ascomycota_unclassified(100);p__Ascomycota_unclassified(100);p__Ascomycota_unclassified(100); | - | - | - | - | - | - | - | Unassigned | - |
| Otu62414 | 1 | k__Fungi(100);k__Fungi_unclassified(100);k__Fungi_unclassified(100);k__Fungi_unclassified(100);k__Fungi_unclassified(100);k__Fungi_unclassified(100);k__Fungi_unclassified(100); | - | - | - | - | - | - | - | Unassigned | - |
| Otu62415 | 1 | k__Fungi(100);k__Fungi_unclassified(100);k__Fungi_unclassified(100);k__Fungi_unclassified(100);k__Fungi_unclassified(100);k__Fungi_unclassified(100);k__Fungi_unclassified(100); | - | - | - | - | - | - | - | Unassigned | - |
| Otu62421 | 1 | k__Fungi(100);k__Fungi_unclassified(100);k__Fungi_unclassified(100);k__Fungi_unclassified(100);k__Fungi_unclassified(100);k__Fungi_unclassified(100);k__Fungi_unclassified(100); | - | - | - | - | - | - | - | Unassigned | - |
| Otu62422 | 1 | k__Fungi(100);k__Fungi_unclassified(100);k__Fungi_unclassified(100);k__Fungi_unclassified(100);k__Fungi_unclassified(100);k__Fungi_unclassified(100);k__Fungi_unclassified(100); | - | - | - | - | - | - | - | Unassigned | - |
| Otu62423 | 1 | k__Fungi(100);k__Fungi_unclassified(100);k__Fungi_unclassified(100);k__Fungi_unclassified(100);k__Fungi_unclassified(100);k__Fungi_unclassified(100);k__Fungi_unclassified(100); | - | - | - | - | - | - | - | Unassigned | - |
| Otu62425 | 1 | k__Fungi(100);k__Fungi_unclassified(100);k__Fungi_unclassified(100);k__Fungi_unclassified(100);k__Fungi_unclassified(100);k__Fungi_unclassified(100);k__Fungi_unclassified(100); | - | - | - | - | - | - | - | Unassigned | - |
| Otu62426 | 1 | k__Fungi(100);p__Ascomycota(100);p__Ascomycota_unclassified(100);p__Ascomycota_unclassified(100);p__Ascomycota_unclassified(100);p__Ascomycota_unclassified(100);p__Ascomycota_unclassified(100); | - | - | - | - | - | - | - | Unassigned | - |
| Otu62427 | 1 | k__Fungi(100);k__Fungi_unclassified(100);k__Fungi_unclassified(100);k__Fungi_unclassified(100);k__Fungi_unclassified(100);k__Fungi_unclassified(100);k__Fungi_unclassified(100); | - | - | - | - | - | - | - | Unassigned | - |
| Otu62428 | 1 | k__Fungi(100);k__Fungi_unclassified(100);k__Fungi_unclassified(100);k__Fungi_unclassified(100);k__Fungi_unclassified(100);k__Fungi_unclassified(100);k__Fungi_unclassified(100); | - | - | - | - | - | - | - | Unassigned | - |
| Otu62434 | 1 | k__Fungi(100);k__Fungi_unclassified(100);k__Fungi_unclassified(100);k__Fungi_unclassified(100);k__Fungi_unclassified(100);k__Fungi_unclassified(100);k__Fungi_unclassified(100); | - | - | - | - | - | - | - | Unassigned | - |
| Otu62439 | 1 | k__Fungi(100);k__Fungi_unclassified(100);k__Fungi_unclassified(100);k__Fungi_unclassified(100);k__Fungi_unclassified(100);k__Fungi_unclassified(100);k__Fungi_unclassified(100); | - | - | - | - | - | - | - | Unassigned | - |
| Otu62441 | 1 | k__Fungi(100);k__Fungi_unclassified(100);k__Fungi_unclassified(100);k__Fungi_unclassified(100);k__Fungi_unclassified(100);k__Fungi_unclassified(100);k__Fungi_unclassified(100); | - | - | - | - | - | - | - | Unassigned | - |
| Otu62443 | 1 | k__Fungi(100);k__Fungi_unclassified(100);k__Fungi_unclassified(100);k__Fungi_unclassified(100);k__Fungi_unclassified(100);k__Fungi_unclassified(100);k__Fungi_unclassified(100); | - | - | - | - | - | - | - | Unassigned | - |
| Otu62445 | 1 | k__Fungi(100);k__Fungi_unclassified(100);k__Fungi_unclassified(100);k__Fungi_unclassified(100);k__Fungi_unclassified(100);k__Fungi_unclassified(100);k__Fungi_unclassified(100); | - | - | - | - | - | - | - | Unassigned | - |
| Otu62446 | 1 | k__Fungi(100);p__Ascomycota(100);p__Ascomycota_unclassified(100);p__Ascomycota_unclassified(100);p__Ascomycota_unclassified(100);p__Ascomycota_unclassified(100);p__Ascomycota_unclassified(100); | - | - | - | - | - | - | - | Unassigned | - |
| Otu62447 | 1 | k__Fungi(100);p__Ascomycota(100);p__Ascomycota_unclassified(100);p__Ascomycota_unclassified(100);p__Ascomycota_unclassified(100);p__Ascomycota_unclassified(100);p__Ascomycota_unclassified(100); | - | - | - | - | - | - | - | Unassigned | - |
| Otu62451 | 1 | k__Fungi(100);k__Fungi_unclassified(100);k__Fungi_unclassified(100);k__Fungi_unclassified(100);k__Fungi_unclassified(100);k__Fungi_unclassified(100);k__Fungi_unclassified(100); | - | - | - | - | - | - | - | Unassigned | - |
| Otu62452 | 1 | k__Fungi(100);k__Fungi_unclassified(100);k__Fungi_unclassified(100);k__Fungi_unclassified(100);k__Fungi_unclassified(100);k__Fungi_unclassified(100);k__Fungi_unclassified(100); | - | - | - | - | - | - | - | Unassigned | - |
| Otu62453 | 1 | k__Fungi(100);k__Fungi_unclassified(100);k__Fungi_unclassified(100);k__Fungi_unclassified(100);k__Fungi_unclassified(100);k__Fungi_unclassified(100);k__Fungi_unclassified(100); | - | - | - | - | - | - | - | Unassigned | - |
| Otu62454 | 1 | k__Fungi(100);k__Fungi_unclassified(100);k__Fungi_unclassified(100);k__Fungi_unclassified(100);k__Fungi_unclassified(100);k__Fungi_unclassified(100);k__Fungi_unclassified(100); | - | - | - | - | - | - | - | Unassigned | - |
| Otu62455 | 1 | k__Fungi(100);p__Ascomycota(100);p__Ascomycota_unclassified(100);p__Ascomycota_unclassified(100);p__Ascomycota_unclassified(100);p__Ascomycota_unclassified(100);p__Ascomycota_unclassified(100); | - | - | - | - | - | - | - | Unassigned | - |
| Otu62456 | 1 | k__Fungi(100);k__Fungi_unclassified(100);k__Fungi_unclassified(100);k__Fungi_unclassified(100);k__Fungi_unclassified(100);k__Fungi_unclassified(100);k__Fungi_unclassified(100); | - | - | - | - | - | - | - | Unassigned | - |
| Otu62457 | 1 | k__Fungi(100);p__Ascomycota(100);p__Ascomycota_unclassified(100);p__Ascomycota_unclassified(100);p__Ascomycota_unclassified(100);p__Ascomycota_unclassified(100);p__Ascomycota_unclassified(100); | - | - | - | - | - | - | - | Unassigned | - |
| Otu62459 | 1 | k__Fungi(100);p__Ascomycota(100);p__Ascomycota_unclassified(100);p__Ascomycota_unclassified(100);p__Ascomycota_unclassified(100);p__Ascomycota_unclassified(100);p__Ascomycota_unclassified(100); | - | - | - | - | - | - | - | Unassigned | - |
| Otu62460 | 1 | k__Fungi(100);p__Ascomycota(100);p__Ascomycota_unclassified(100);p__Ascomycota_unclassified(100);p__Ascomycota_unclassified(100);p__Ascomycota_unclassified(100);p__Ascomycota_unclassified(100); | - | - | - | - | - | - | - | Unassigned | - |
| Otu62461 | 1 | k__Fungi(100);p__Ascomycota(100);p__Ascomycota_unclassified(100);p__Ascomycota_unclassified(100);p__Ascomycota_unclassified(100);p__Ascomycota_unclassified(100);p__Ascomycota_unclassified(100); | - | - | - | - | - | - | - | Unassigned | - |
| Otu62462 | 1 | k__Fungi(100);k__Fungi_unclassified(100);k__Fungi_unclassified(100);k__Fungi_unclassified(100);k__Fungi_unclassified(100);k__Fungi_unclassified(100);k__Fungi_unclassified(100); | - | - | - | - | - | - | - | Unassigned | - |
| Otu62463 | 1 | k__Fungi(100);k__Fungi_unclassified(100);k__Fungi_unclassified(100);k__Fungi_unclassified(100);k__Fungi_unclassified(100);k__Fungi_unclassified(100);k__Fungi_unclassified(100); | - | - | - | - | - | - | - | Unassigned | - |
| Otu62464 | 1 | k__Fungi(100);p__Ascomycota(100);p__Ascomycota_unclassified(100);p__Ascomycota_unclassified(100);p__Ascomycota_unclassified(100);p__Ascomycota_unclassified(100);p__Ascomycota_unclassified(100); | - | - | - | - | - | - | - | Unassigned | - |
| Otu62465 | 1 | k__Fungi(100);k__Fungi_unclassified(100);k__Fungi_unclassified(100);k__Fungi_unclassified(100);k__Fungi_unclassified(100);k__Fungi_unclassified(100);k__Fungi_unclassified(100); | - | - | - | - | - | - | - | Unassigned | - |
| Otu62466 | 1 | k__Fungi(100);k__Fungi_unclassified(100);k__Fungi_unclassified(100);k__Fungi_unclassified(100);k__Fungi_unclassified(100);k__Fungi_unclassified(100);k__Fungi_unclassified(100); | - | - | - | - | - | - | - | Unassigned | - |
| Otu62467 | 1 | k__Fungi(100);k__Fungi_unclassified(100);k__Fungi_unclassified(100);k__Fungi_unclassified(100);k__Fungi_unclassified(100);k__Fungi_unclassified(100);k__Fungi_unclassified(100); | - | - | - | - | - | - | - | Unassigned | - |
| Otu62468 | 1 | k__Fungi(100);p__Ascomycota(100);p__Ascomycota_unclassified(100);p__Ascomycota_unclassified(100);p__Ascomycota_unclassified(100);p__Ascomycota_unclassified(100);p__Ascomycota_unclassified(100); | - | - | - | - | - | - | - | Unassigned | - |
| Otu62469 | 1 | k__Fungi(100);p__Ascomycota(100);p__Ascomycota_unclassified(100);p__Ascomycota_unclassified(100);p__Ascomycota_unclassified(100);p__Ascomycota_unclassified(100);p__Ascomycota_unclassified(100); | - | - | - | - | - | - | - | Unassigned | - |
| Otu62470 | 1 | k__Fungi(100);p__Ascomycota(100);p__Ascomycota_unclassified(100);p__Ascomycota_unclassified(100);p__Ascomycota_unclassified(100);p__Ascomycota_unclassified(100);p__Ascomycota_unclassified(100); | - | - | - | - | - | - | - | Unassigned | - |
| Otu62472 | 1 | k__Fungi(100);k__Fungi_unclassified(100);k__Fungi_unclassified(100);k__Fungi_unclassified(100);k__Fungi_unclassified(100);k__Fungi_unclassified(100);k__Fungi_unclassified(100); | - | - | - | - | - | - | - | Unassigned | - |
| Otu62473 | 1 | k__Fungi(100);p__Ascomycota(100);p__Ascomycota_unclassified(100);p__Ascomycota_unclassified(100);p__Ascomycota_unclassified(100);p__Ascomycota_unclassified(100);p__Ascomycota_unclassified(100); | - | - | - | - | - | - | - | Unassigned | - |
| Otu62474 | 1 | k__Fungi(100);p__Ascomycota(100);p__Ascomycota_unclassified(100);p__Ascomycota_unclassified(100);p__Ascomycota_unclassified(100);p__Ascomycota_unclassified(100);p__Ascomycota_unclassified(100); | - | - | - | - | - | - | - | Unassigned | - |
| Otu62477 | 1 | k__Fungi(100);p__Ascomycota(100);c__Dothideomycetes(100);o__Capnodiales(100);f__Davidiellaceae(100);f__Davidiellaceae_unclassified(100);f__Davidiellaceae_unclassified(100); | - | - | - | - | - | - | - | Unassigned | - |
| Otu62478 | 1 | k__Fungi(100);k__Fungi_unclassified(100);k__Fungi_unclassified(100);k__Fungi_unclassified(100);k__Fungi_unclassified(100);k__Fungi_unclassified(100);k__Fungi_unclassified(100); | - | - | - | - | - | - | - | Unassigned | - |
| Otu62481 | 1 | k__Fungi(100);k__Fungi_unclassified(100);k__Fungi_unclassified(100);k__Fungi_unclassified(100);k__Fungi_unclassified(100);k__Fungi_unclassified(100);k__Fungi_unclassified(100); | - | - | - | - | - | - | - | Unassigned | - |
| Otu62482 | 1 | k__Fungi(100);k__Fungi_unclassified(100);k__Fungi_unclassified(100);k__Fungi_unclassified(100);k__Fungi_unclassified(100);k__Fungi_unclassified(100);k__Fungi_unclassified(100); | - | - | - | - | - | - | - | Unassigned | - |
| Otu62484 | 1 | k__Fungi(100);k__Fungi_unclassified(100);k__Fungi_unclassified(100);k__Fungi_unclassified(100);k__Fungi_unclassified(100);k__Fungi_unclassified(100);k__Fungi_unclassified(100); | - | - | - | - | - | - | - | Unassigned | - |
| Otu62485 | 1 | k__Fungi(100);p__Ascomycota(100);p__Ascomycota_unclassified(100);p__Ascomycota_unclassified(100);p__Ascomycota_unclassified(100);p__Ascomycota_unclassified(100);p__Ascomycota_unclassified(100); | - | - | - | - | - | - | - | Unassigned | - |
| Otu62493 | 1 | k__Fungi(100);k__Fungi_unclassified(100);k__Fungi_unclassified(100);k__Fungi_unclassified(100);k__Fungi_unclassified(100);k__Fungi_unclassified(100);k__Fungi_unclassified(100); | - | - | - | - | - | - | - | Unassigned | - |
| Otu62494 | 1 | k__Fungi(100);k__Fungi_unclassified(100);k__Fungi_unclassified(100);k__Fungi_unclassified(100);k__Fungi_unclassified(100);k__Fungi_unclassified(100);k__Fungi_unclassified(100); | - | - | - | - | - | - | - | Unassigned | - |
| Otu62495 | 1 | k__Fungi(100);k__Fungi_unclassified(100);k__Fungi_unclassified(100);k__Fungi_unclassified(100);k__Fungi_unclassified(100);k__Fungi_unclassified(100);k__Fungi_unclassified(100); | - | - | - | - | - | - | - | Unassigned | - |
| Otu62496 | 1 | k__Fungi(100);k__Fungi_unclassified(100);k__Fungi_unclassified(100);k__Fungi_unclassified(100);k__Fungi_unclassified(100);k__Fungi_unclassified(100);k__Fungi_unclassified(100); | - | - | - | - | - | - | - | Unassigned | - |
| Otu62497 | 1 | k__Fungi(100);k__Fungi_unclassified(100);k__Fungi_unclassified(100);k__Fungi_unclassified(100);k__Fungi_unclassified(100);k__Fungi_unclassified(100);k__Fungi_unclassified(100); | - | - | - | - | - | - | - | Unassigned | - |
| Otu62498 | 1 | k__Fungi(100);p__Ascomycota(100);p__Ascomycota_unclassified(100);p__Ascomycota_unclassified(100);p__Ascomycota_unclassified(100);p__Ascomycota_unclassified(100);p__Ascomycota_unclassified(100); | - | - | - | - | - | - | - | Unassigned | - |
| Otu62499 | 1 | k__Fungi(100);k__Fungi_unclassified(100);k__Fungi_unclassified(100);k__Fungi_unclassified(100);k__Fungi_unclassified(100);k__Fungi_unclassified(100);k__Fungi_unclassified(100); | - | - | - | - | - | - | - | Unassigned | - |
| Otu62500 | 1 | k__Fungi(100);p__Ascomycota(100);p__Ascomycota_unclassified(100);p__Ascomycota_unclassified(100);p__Ascomycota_unclassified(100);p__Ascomycota_unclassified(100);p__Ascomycota_unclassified(100); | - | - | - | - | - | - | - | Unassigned | - |
| Otu62501 | 1 | k__Fungi(100);p__Ascomycota(100);p__Ascomycota_unclassified(100);p__Ascomycota_unclassified(100);p__Ascomycota_unclassified(100);p__Ascomycota_unclassified(100);p__Ascomycota_unclassified(100); | - | - | - | - | - | - | - | Unassigned | - |
| Otu62502 | 1 | k__Fungi(100);p__Ascomycota(100);p__Ascomycota_unclassified(100);p__Ascomycota_unclassified(100);p__Ascomycota_unclassified(100);p__Ascomycota_unclassified(100);p__Ascomycota_unclassified(100); | - | - | - | - | - | - | - | Unassigned | - |
| Otu62504 | 1 | k__Fungi(100);k__Fungi_unclassified(100);k__Fungi_unclassified(100);k__Fungi_unclassified(100);k__Fungi_unclassified(100);k__Fungi_unclassified(100);k__Fungi_unclassified(100); | - | - | - | - | - | - | - | Unassigned | - |
| Otu62505 | 1 | k__Fungi(100);p__Ascomycota(100);p__Ascomycota_unclassified(100);p__Ascomycota_unclassified(100);p__Ascomycota_unclassified(100);p__Ascomycota_unclassified(100);p__Ascomycota_unclassified(100); | - | - | - | - | - | - | - | Unassigned | - |
| Otu62507 | 1 | k__Fungi(100);k__Fungi_unclassified(100);k__Fungi_unclassified(100);k__Fungi_unclassified(100);k__Fungi_unclassified(100);k__Fungi_unclassified(100);k__Fungi_unclassified(100); | - | - | - | - | - | - | - | Unassigned | - |
| Otu62509 | 1 | k__Fungi(100);p__Ascomycota(100);p__Ascomycota_unclassified(100);p__Ascomycota_unclassified(100);p__Ascomycota_unclassified(100);p__Ascomycota_unclassified(100);p__Ascomycota_unclassified(100); | - | - | - | - | - | - | - | Unassigned | - |
| Otu62510 | 1 | k__Fungi(100);p__Ascomycota(100);p__Ascomycota_unclassified(100);p__Ascomycota_unclassified(100);p__Ascomycota_unclassified(100);p__Ascomycota_unclassified(100);p__Ascomycota_unclassified(100); | - | - | - | - | - | - | - | Unassigned | - |
| Otu62511 | 1 | k__Fungi(100);k__Fungi_unclassified(100);k__Fungi_unclassified(100);k__Fungi_unclassified(100);k__Fungi_unclassified(100);k__Fungi_unclassified(100);k__Fungi_unclassified(100); | - | - | - | - | - | - | - | Unassigned | - |
| Otu62513 | 1 | k__Fungi(100);k__Fungi_unclassified(100);k__Fungi_unclassified(100);k__Fungi_unclassified(100);k__Fungi_unclassified(100);k__Fungi_unclassified(100);k__Fungi_unclassified(100); | - | - | - | - | - | - | - | Unassigned | - |
| Otu62520 | 1 | k__Fungi(100);k__Fungi_unclassified(100);k__Fungi_unclassified(100);k__Fungi_unclassified(100);k__Fungi_unclassified(100);k__Fungi_unclassified(100);k__Fungi_unclassified(100); | - | - | - | - | - | - | - | Unassigned | - |

| | | | | | | | | | | | |
|---|---|---|---|---|---|---|---|---|---|---|---|
| Otu62523 | 1 | k__Fungi(100);k__Fungi_unclassified(100);k__Fungi_unclassified(100);k__Fungi_unclassified(100);k__Fungi_unclassified(100);k__Fungi_unclassified(100);k__Fungi_unclassified(100); | - | - | - | - | - | - | - | Unassigned | - |
| Otu62524 | 1 | k__Fungi(100);p__Ascomycota(100);p__Ascomycota_unclassified(100);p__Ascomycota_unclassified(100);p__Ascomycota_unclassified(100);p__Ascomycota_unclassified(100);p__Ascomycota_unclassified(100); | - | - | - | - | - | - | - | Unassigned | - |
| Otu62525 | 1 | k__Fungi(100);k__Fungi_unclassified(100);k__Fungi_unclassified(100);k__Fungi_unclassified(100);k__Fungi_unclassified(100);k__Fungi_unclassified(100);k__Fungi_unclassified(100); | - | - | - | - | - | - | - | Unassigned | - |
| Otu62526 | 1 | k__Fungi(100);k__Fungi_unclassified(100);k__Fungi_unclassified(100);k__Fungi_unclassified(100);k__Fungi_unclassified(100);k__Fungi_unclassified(100);k__Fungi_unclassified(100); | - | - | - | - | - | - | - | Unassigned | - |
| Otu62527 | 1 | k__Fungi(100);k__Fungi_unclassified(100);k__Fungi_unclassified(100);k__Fungi_unclassified(100);k__Fungi_unclassified(100);k__Fungi_unclassified(100);k__Fungi_unclassified(100); | - | - | - | - | - | - | - | Unassigned | - |
| Otu62528 | 1 | k__Fungi(100);k__Fungi_unclassified(100);k__Fungi_unclassified(100);k__Fungi_unclassified(100);k__Fungi_unclassified(100);k__Fungi_unclassified(100);k__Fungi_unclassified(100); | - | - | - | - | - | - | - | Unassigned | - |
| Otu62531 | 1 | k__Fungi(100);k__Fungi_unclassified(100);k__Fungi_unclassified(100);k__Fungi_unclassified(100);k__Fungi_unclassified(100);k__Fungi_unclassified(100);k__Fungi_unclassified(100); | - | - | - | - | - | - | - | Unassigned | - |
| Otu62532 | 1 | k__Fungi(100);p__Ascomycota(100);p__Ascomycota_unclassified(100);p__Ascomycota_unclassified(100);p__Ascomycota_unclassified(100);p__Ascomycota_unclassified(100);p__Ascomycota_unclassified(100); | - | - | - | - | - | - | - | Unassigned | - |
| Otu62533 | 1 | k__Fungi(100);k__Fungi_unclassified(100);k__Fungi_unclassified(100);k__Fungi_unclassified(100);k__Fungi_unclassified(100);k__Fungi_unclassified(100);k__Fungi_unclassified(100); | - | - | - | - | - | - | - | Unassigned | - |
| Otu62534 | 1 | k__Fungi(100);p__Ascomycota(100);p__Ascomycota_unclassified(100);p__Ascomycota_unclassified(100);p__Ascomycota_unclassified(100);p__Ascomycota_unclassified(100);p__Ascomycota_unclassified(100); | - | - | - | - | - | - | - | Unassigned | - |
| Otu62536 | 1 | k__Fungi(100);k__Fungi_unclassified(100);k__Fungi_unclassified(100);k__Fungi_unclassified(100);k__Fungi_unclassified(100);k__Fungi_unclassified(100);k__Fungi_unclassified(100); | - | - | - | - | - | - | - | Unassigned | - |
| Otu62537 | 1 | k__Fungi(100);k__Fungi_unclassified(100);k__Fungi_unclassified(100);k__Fungi_unclassified(100);k__Fungi_unclassified(100);k__Fungi_unclassified(100);k__Fungi_unclassified(100); | - | - | - | - | - | - | - | Unassigned | - |
| Otu62539 | 1 | k__Fungi(100);k__Fungi_unclassified(100);k__Fungi_unclassified(100);k__Fungi_unclassified(100);k__Fungi_unclassified(100);k__Fungi_unclassified(100);k__Fungi_unclassified(100); | - | - | - | - | - | - | - | Unassigned | - |
| Otu62541 | 1 | k__Fungi(100);p__Ascomycota(100);p__Ascomycota_unclassified(100);p__Ascomycota_unclassified(100);p__Ascomycota_unclassified(100);p__Ascomycota_unclassified(100);p__Ascomycota_unclassified(100); | - | - | - | - | - | - | - | Unassigned | - |
| Otu62543 | 1 | k__Fungi(100);p__Ascomycota(100);p__Ascomycota_unclassified(100);p__Ascomycota_unclassified(100);p__Ascomycota_unclassified(100);p__Ascomycota_unclassified(100);p__Ascomycota_unclassified(100); | - | - | - | - | - | - | - | Unassigned | - |
| Otu62544 | 1 | k__Fungi(100);k__Fungi_unclassified(100);k__Fungi_unclassified(100);k__Fungi_unclassified(100);k__Fungi_unclassified(100);k__Fungi_unclassified(100);k__Fungi_unclassified(100); | - | - | - | - | - | - | - | Unassigned | - |
| Otu62545 | 1 | k__Fungi(100);p__Ascomycota(100);p__Ascomycota_unclassified(100);p__Ascomycota_unclassified(100);p__Ascomycota_unclassified(100);p__Ascomycota_unclassified(100);p__Ascomycota_unclassified(100); | - | - | - | - | - | - | - | Unassigned | - |
| Otu62547 | 1 | k__Fungi(100);p__Ascomycota(100);p__Ascomycota_unclassified(100);p__Ascomycota_unclassified(100);p__Ascomycota_unclassified(100);p__Ascomycota_unclassified(100);p__Ascomycota_unclassified(100); | - | - | - | - | - | - | - | Unassigned | - |
| Otu62548 | 1 | k__Fungi(100);k__Fungi_unclassified(100);k__Fungi_unclassified(100);k__Fungi_unclassified(100);k__Fungi_unclassified(100);k__Fungi_unclassified(100);k__Fungi_unclassified(100); | - | - | - | - | - | - | - | Unassigned | - |
| Otu62549 | 1 | k__Fungi(100);p__Ascomycota(100);p__Ascomycota_unclassified(100);p__Ascomycota_unclassified(100);p__Ascomycota_unclassified(100);p__Ascomycota_unclassified(100);p__Ascomycota_unclassified(100); | - | - | - | - | - | - | - | Unassigned | - |
| Otu62550 | 1 | k__Fungi(100);p__Ascomycota(100);p__Ascomycota_unclassified(100);p__Ascomycota_unclassified(100);p__Ascomycota_unclassified(100);p__Ascomycota_unclassified(100);p__Ascomycota_unclassified(100); | - | - | - | - | - | - | - | Unassigned | - |
| Otu62554 | 1 | k__Fungi(100);p__Ascomycota(100);p__Ascomycota_unclassified(100);p__Ascomycota_unclassified(100);p__Ascomycota_unclassified(100);p__Ascomycota_unclassified(100);p__Ascomycota_unclassified(100); | - | - | - | - | - | - | - | Unassigned | - |
| Otu62556 | 1 | k__Fungi(100);p__Ascomycota(100);p__Ascomycota_unclassified(100);p__Ascomycota_unclassified(100);p__Ascomycota_unclassified(100);p__Ascomycota_unclassified(100);p__Ascomycota_unclassified(100); | - | - | - | - | - | - | - | Unassigned | - |
| Otu62558 | 1 | k__Fungi(100);p__Ascomycota(100);p__Ascomycota_unclassified(100);p__Ascomycota_unclassified(100);p__Ascomycota_unclassified(100);p__Ascomycota_unclassified(100);p__Ascomycota_unclassified(100); | - | - | - | - | - | - | - | Unassigned | - |
| Otu62561 | 1 | k__Fungi(100);k__Fungi_unclassified(100);k__Fungi_unclassified(100);k__Fungi_unclassified(100);k__Fungi_unclassified(100);k__Fungi_unclassified(100);k__Fungi_unclassified(100); | - | - | - | - | - | - | - | Unassigned | - |
| Otu62563 | 1 | k__Fungi(100);k__Fungi_unclassified(100);k__Fungi_unclassified(100);k__Fungi_unclassified(100);k__Fungi_unclassified(100);k__Fungi_unclassified(100);k__Fungi_unclassified(100); | - | - | - | - | - | - | - | Unassigned | - |
| Otu62565 | 1 | k__Fungi(100);k__Fungi_unclassified(100);k__Fungi_unclassified(100);k__Fungi_unclassified(100);k__Fungi_unclassified(100);k__Fungi_unclassified(100);k__Fungi_unclassified(100); | - | - | - | - | - | - | - | Unassigned | - |
| Otu62566 | 1 | k__Fungi(100);p__Ascomycota(100);p__Ascomycota_unclassified(100);p__Ascomycota_unclassified(100);p__Ascomycota_unclassified(100);p__Ascomycota_unclassified(100);p__Ascomycota_unclassified(100); | - | - | - | - | - | - | - | Unassigned | - |
| Otu62568 | 1 | k__Fungi(100);k__Fungi_unclassified(100);k__Fungi_unclassified(100);k__Fungi_unclassified(100);k__Fungi_unclassified(100);k__Fungi_unclassified(100);k__Fungi_unclassified(100); | - | - | - | - | - | - | - | Unassigned | - |
| Otu62569 | 1 | k__Fungi(100);p__Ascomycota(100);p__Ascomycota_unclassified(100);p__Ascomycota_unclassified(100);p__Ascomycota_unclassified(100);p__Ascomycota_unclassified(100);p__Ascomycota_unclassified(100); | - | - | - | - | - | - | - | Unassigned | - |
| Otu62570 | 1 | k__Fungi(100);p__Ascomycota(100);p__Ascomycota_unclassified(100);p__Ascomycota_unclassified(100);p__Ascomycota_unclassified(100);p__Ascomycota_unclassified(100);p__Ascomycota_unclassified(100); | - | - | - | - | - | - | - | Unassigned | - |
| Otu62571 | 1 | k__Fungi(100);k__Fungi_unclassified(100);k__Fungi_unclassified(100);k__Fungi_unclassified(100);k__Fungi_unclassified(100);k__Fungi_unclassified(100);k__Fungi_unclassified(100); | - | - | - | - | - | - | - | Unassigned | - |
| Otu62572 | 1 | k__Fungi(100);k__Fungi_unclassified(100);k__Fungi_unclassified(100);k__Fungi_unclassified(100);k__Fungi_unclassified(100);k__Fungi_unclassified(100);k__Fungi_unclassified(100); | - | - | - | - | - | - | - | Unassigned | - |
| Otu62573 | 1 | k__Fungi(100);k__Fungi_unclassified(100);k__Fungi_unclassified(100);k__Fungi_unclassified(100);k__Fungi_unclassified(100);k__Fungi_unclassified(100);k__Fungi_unclassified(100); | - | - | - | - | - | - | - | Unassigned | - |
| Otu62574 | 1 | k__Fungi(100);k__Fungi_unclassified(100);k__Fungi_unclassified(100);k__Fungi_unclassified(100);k__Fungi_unclassified(100);k__Fungi_unclassified(100);k__Fungi_unclassified(100); | - | - | - | - | - | - | - | Unassigned | - |
| Otu62575 | 1 | k__Fungi(100);k__Fungi_unclassified(100);k__Fungi_unclassified(100);k__Fungi_unclassified(100);k__Fungi_unclassified(100);k__Fungi_unclassified(100);k__Fungi_unclassified(100); | - | - | - | - | - | - | - | Unassigned | - |
| Otu62576 | 1 | k__Fungi(100);k__Fungi_unclassified(100);k__Fungi_unclassified(100);k__Fungi_unclassified(100);k__Fungi_unclassified(100);k__Fungi_unclassified(100);k__Fungi_unclassified(100); | - | - | - | - | - | - | - | Unassigned | - |
| Otu62577 | 1 | k__Fungi(100);k__Fungi_unclassified(100);k__Fungi_unclassified(100);k__Fungi_unclassified(100);k__Fungi_unclassified(100);k__Fungi_unclassified(100);k__Fungi_unclassified(100); | - | - | - | - | - | - | - | Unassigned | - |
| Otu62578 | 1 | k__Fungi(100);k__Fungi_unclassified(100);k__Fungi_unclassified(100);k__Fungi_unclassified(100);k__Fungi_unclassified(100);k__Fungi_unclassified(100);k__Fungi_unclassified(100); | - | - | - | - | - | - | - | Unassigned | - |
| Otu62581 | 1 | k__Fungi(100);k__Fungi_unclassified(100);k__Fungi_unclassified(100);k__Fungi_unclassified(100);k__Fungi_unclassified(100);k__Fungi_unclassified(100);k__Fungi_unclassified(100); | - | - | - | - | - | - | - | Unassigned | - |
| Otu62583 | 1 | k__Fungi(100);k__Fungi_unclassified(100);k__Fungi_unclassified(100);k__Fungi_unclassified(100);k__Fungi_unclassified(100);k__Fungi_unclassified(100);k__Fungi_unclassified(100); | - | - | - | - | - | - | - | Unassigned | - |
| Otu62584 | 1 | k__Fungi(100);k__Fungi_unclassified(100);k__Fungi_unclassified(100);k__Fungi_unclassified(100);k__Fungi_unclassified(100);k__Fungi_unclassified(100);k__Fungi_unclassified(100); | - | - | - | - | - | - | - | Unassigned | - |
| Otu62585 | 1 | k__Fungi(100);p__Ascomycota(100);p__Ascomycota_unclassified(100);p__Ascomycota_unclassified(100);p__Ascomycota_unclassified(100);p__Ascomycota_unclassified(100);p__Ascomycota_unclassified(100); | - | - | - | - | - | - | - | Unassigned | - |
| Otu62586 | 1 | k__Fungi(100);k__Fungi_unclassified(100);k__Fungi_unclassified(100);k__Fungi_unclassified(100);k__Fungi_unclassified(100);k__Fungi_unclassified(100);k__Fungi_unclassified(100); | - | - | - | - | - | - | - | Unassigned | - |
| Otu62587 | 1 | k__Fungi(100);k__Fungi_unclassified(100);k__Fungi_unclassified(100);k__Fungi_unclassified(100);k__Fungi_unclassified(100);k__Fungi_unclassified(100);k__Fungi_unclassified(100); | - | - | - | - | - | - | - | Unassigned | - |
| Otu62588 | 1 | k__Fungi(100);p__Ascomycota(100);c__Lecanoromycetes(100);c__Lecanoromycetes_unclassified(100);c__Lecanoromycetes_unclassified(100);c__Lecanoromycetes_unclassified(100);c__Lecanoromycetes_unclassified(100); | - | - | - | - | - | - | - | Unassigned | - |
| Otu62589 | 1 | k__Fungi(100);p__Ascomycota(100);p__Ascomycota_unclassified(100);p__Ascomycota_unclassified(100);p__Ascomycota_unclassified(100);p__Ascomycota_unclassified(100);p__Ascomycota_unclassified(100); | - | - | - | - | - | - | - | Unassigned | - |
| Otu62590 | 1 | k__Fungi(100);p__Ascomycota(100);p__Ascomycota_unclassified(100);p__Ascomycota_unclassified(100);p__Ascomycota_unclassified(100);p__Ascomycota_unclassified(100);p__Ascomycota_unclassified(100); | - | - | - | - | - | - | - | Unassigned | - |
| Otu62591 | 1 | k__Fungi(100);p__Ascomycota(100);p__Ascomycota_unclassified(100);p__Ascomycota_unclassified(100);p__Ascomycota_unclassified(100);p__Ascomycota_unclassified(100);p__Ascomycota_unclassified(100); | - | - | - | - | - | - | - | Unassigned | - |
| Otu62592 | 1 | k__Fungi(100);k__Fungi_unclassified(100);k__Fungi_unclassified(100);k__Fungi_unclassified(100);k__Fungi_unclassified(100);k__Fungi_unclassified(100);k__Fungi_unclassified(100); | - | - | - | - | - | - | - | Unassigned | - |
| Otu62593 | 1 | k__Fungi(100);p__Ascomycota(100);p__Ascomycota_unclassified(100);p__Ascomycota_unclassified(100);p__Ascomycota_unclassified(100);p__Ascomycota_unclassified(100);p__Ascomycota_unclassified(100); | - | - | - | - | - | - | - | Unassigned | - |
| Otu62596 | 1 | k__Fungi(100);p__Ascomycota(100);p__Ascomycota_unclassified(100);p__Ascomycota_unclassified(100);p__Ascomycota_unclassified(100);p__Ascomycota_unclassified(100);p__Ascomycota_unclassified(100); | - | - | - | - | - | - | - | Unassigned | - |
| Otu62597 | 1 | k__Fungi(100);k__Fungi_unclassified(100);k__Fungi_unclassified(100);k__Fungi_unclassified(100);k__Fungi_unclassified(100);k__Fungi_unclassified(100);k__Fungi_unclassified(100); | - | - | - | - | - | - | - | Unassigned | - |
| Otu62599 | 1 | k__Fungi(100);k__Fungi_unclassified(100);k__Fungi_unclassified(100);k__Fungi_unclassified(100);k__Fungi_unclassified(100);k__Fungi_unclassified(100);k__Fungi_unclassified(100); | - | - | - | - | - | - | - | Unassigned | - |
| Otu62601 | 1 | k__Fungi(100);k__Fungi_unclassified(100);k__Fungi_unclassified(100);k__Fungi_unclassified(100);k__Fungi_unclassified(100);k__Fungi_unclassified(100);k__Fungi_unclassified(100); | - | - | - | - | - | - | - | Unassigned | - |
| Otu62609 | 1 | k__Fungi(100);k__Fungi_unclassified(100);k__Fungi_unclassified(100);k__Fungi_unclassified(100);k__Fungi_unclassified(100);k__Fungi_unclassified(100);k__Fungi_unclassified(100); | - | - | - | - | - | - | - | Unassigned | - |
| Otu62611 | 1 | k__Fungi(100);k__Fungi_unclassified(100);k__Fungi_unclassified(100);k__Fungi_unclassified(100);k__Fungi_unclassified(100);k__Fungi_unclassified(100);k__Fungi_unclassified(100); | - | - | - | - | - | - | - | Unassigned | - |
| Otu62612 | 1 | k__Fungi(100);p__Ascomycota(100);p__Ascomycota_unclassified(100);p__Ascomycota_unclassified(100);p__Ascomycota_unclassified(100);p__Ascomycota_unclassified(100);p__Ascomycota_unclassified(100); | - | - | - | - | - | - | - | Unassigned | - |
| Otu62613 | 1 | k__Fungi(100);p__Ascomycota(100);p__Ascomycota_unclassified(100);p__Ascomycota_unclassified(100);p__Ascomycota_unclassified(100);p__Ascomycota_unclassified(100);p__Ascomycota_unclassified(100); | - | - | - | - | - | - | - | Unassigned | - |
| Otu62614 | 1 | k__Fungi(100);k__Fungi_unclassified(100);k__Fungi_unclassified(100);k__Fungi_unclassified(100);k__Fungi_unclassified(100);k__Fungi_unclassified(100);k__Fungi_unclassified(100); | - | - | - | - | - | - | - | Unassigned | - |
| Otu62615 | 1 | k__Fungi(100);k__Fungi_unclassified(100);k__Fungi_unclassified(100);k__Fungi_unclassified(100);k__Fungi_unclassified(100);k__Fungi_unclassified(100);k__Fungi_unclassified(100); | - | - | - | - | - | - | - | Unassigned | - |
| Otu62616 | 1 | k__Fungi(100);k__Fungi_unclassified(100);k__Fungi_unclassified(100);k__Fungi_unclassified(100);k__Fungi_unclassified(100);k__Fungi_unclassified(100);k__Fungi_unclassified(100); | - | - | - | - | - | - | - | Unassigned | - |
| Otu62617 | 1 | k__Fungi(100);p__Ascomycota(100);p__Ascomycota_unclassified(100);p__Ascomycota_unclassified(100);p__Ascomycota_unclassified(100);p__Ascomycota_unclassified(100);p__Ascomycota_unclassified(100); | - | - | - | - | - | - | - | Unassigned | - |
| Otu62618 | 1 | k__Fungi(100);p__Ascomycota(100);p__Ascomycota_unclassified(100);p__Ascomycota_unclassified(100);p__Ascomycota_unclassified(100);p__Ascomycota_unclassified(100);p__Ascomycota_unclassified(100); | - | - | - | - | - | - | - | Unassigned | - |
| Otu62619 | 1 | k__Fungi(100);k__Fungi_unclassified(100);k__Fungi_unclassified(100);k__Fungi_unclassified(100);k__Fungi_unclassified(100);k__Fungi_unclassified(100);k__Fungi_unclassified(100); | - | - | - | - | - | - | - | Unassigned | - |
| Otu62621 | 1 | k__Fungi(100);k__Fungi_unclassified(100);k__Fungi_unclassified(100);k__Fungi_unclassified(100);k__Fungi_unclassified(100);k__Fungi_unclassified(100);k__Fungi_unclassified(100); | - | - | - | - | - | - | - | Unassigned | - |
| Otu62623 | 1 | k__Fungi(100);p__Ascomycota(100);p__Ascomycota_unclassified(100);p__Ascomycota_unclassified(100);p__Ascomycota_unclassified(100);p__Ascomycota_unclassified(100);p__Ascomycota_unclassified(100); | - | - | - | - | - | - | - | Unassigned | - |
| Otu62625 | 1 | k__Fungi(100);p__Ascomycota(100);p__Ascomycota_unclassified(100);p__Ascomycota_unclassified(100);p__Ascomycota_unclassified(100);p__Ascomycota_unclassified(100);p__Ascomycota_unclassified(100); | - | - | - | - | - | - | - | Unassigned | - |
| Otu62626 | 1 | k__Fungi(100);p__Ascomycota(100);p__Ascomycota_unclassified(100);p__Ascomycota_unclassified(100);p__Ascomycota_unclassified(100);p__Ascomycota_unclassified(100);p__Ascomycota_unclassified(100); | - | - | - | - | - | - | - | Unassigned | - |
| Otu62627 | 1 | k__Fungi(100);p__Ascomycota(100);p__Ascomycota_unclassified(100);p__Ascomycota_unclassified(100);p__Ascomycota_unclassified(100);p__Ascomycota_unclassified(100);p__Ascomycota_unclassified(100); | - | - | - | - | - | - | - | Unassigned | - |
| Otu62628 | 1 | k__Fungi(100);p__Ascomycota(100);p__Ascomycota_unclassified(100);p__Ascomycota_unclassified(100);p__Ascomycota_unclassified(100);p__Ascomycota_unclassified(100);p__Ascomycota_unclassified(100); | - | - | - | - | - | - | - | Unassigned | - |
| Otu62629 | 1 | k__Fungi(100);k__Fungi_unclassified(100);k__Fungi_unclassified(100);k__Fungi_unclassified(100);k__Fungi_unclassified(100);k__Fungi_unclassified(100);k__Fungi_unclassified(100); | - | - | - | - | - | - | - | Unassigned | - |
| Otu62630 | 1 | k__Fungi(100);k__Fungi_unclassified(100);k__Fungi_unclassified(100);k__Fungi_unclassified(100);k__Fungi_unclassified(100);k__Fungi_unclassified(100);k__Fungi_unclassified(100); | - | - | - | - | - | - | - | Unassigned | - |
| Otu62631 | 1 | k__Fungi(100);p__Ascomycota(100);p__Ascomycota_unclassified(100);p__Ascomycota_unclassified(100);p__Ascomycota_unclassified(100);p__Ascomycota_unclassified(100);p__Ascomycota_unclassified(100); | - | - | - | - | - | - | - | Unassigned | - |
| Otu62632 | 1 | k__Fungi(100);p__Ascomycota(100);p__Ascomycota_unclassified(100);p__Ascomycota_unclassified(100);p__Ascomycota_unclassified(100);p__Ascomycota_unclassified(100);p__Ascomycota_unclassified(100); | - | - | - | - | - | - | - | Unassigned | - |
| Otu62633 | 1 | k__Fungi(100);p__Ascomycota(100);p__Ascomycota_unclassified(100);p__Ascomycota_unclassified(100);p__Ascomycota_unclassified(100);p__Ascomycota_unclassified(100);p__Ascomycota_unclassified(100); | - | - | - | - | - | - | - | Unassigned | - |
| Otu62634 | 1 | k__Fungi(100);k__Fungi_unclassified(100);k__Fungi_unclassified(100);k__Fungi_unclassified(100);k__Fungi_unclassified(100);k__Fungi_unclassified(100);k__Fungi_unclassified(100); | - | - | - | - | - | - | - | Unassigned | - |
| Otu62635 | 1 | k__Fungi(100);p__Ascomycota(100);p__Ascomycota_unclassified(100);p__Ascomycota_unclassified(100);p__Ascomycota_unclassified(100);p__Ascomycota_unclassified(100);p__Ascomycota_unclassified(100); | - | - | - | - | - | - | - | Unassigned | - |
| Otu62637 | 1 | k__Fungi(100);k__Fungi_unclassified(100);k__Fungi_unclassified(100);k__Fungi_unclassified(100);k__Fungi_unclassified(100);k__Fungi_unclassified(100);k__Fungi_unclassified(100); | - | - | - | - | - | - | - | Unassigned | - |
| Otu62638 | 1 | k__Fungi(100);k__Fungi_unclassified(100);k__Fungi_unclassified(100);k__Fungi_unclassified(100);k__Fungi_unclassified(100);k__Fungi_unclassified(100);k__Fungi_unclassified(100); | - | - | - | - | - | - | - | Unassigned | - |
| Otu62639 | 1 | k__Fungi(100);p__Ascomycota(100);p__Ascomycota_unclassified(100);p__Ascomycota_unclassified(100);p__Ascomycota_unclassified(100);p__Ascomycota_unclassified(100);p__Ascomycota_unclassified(100); | - | - | - | - | - | - | - | Unassigned | - |
| Otu62641 | 1 | k__Fungi(100);k__Fungi_unclassified(100);k__Fungi_unclassified(100);k__Fungi_unclassified(100);k__Fungi_unclassified(100);k__Fungi_unclassified(100);k__Fungi_unclassified(100); | - | - | - | - | - | - | - | Unassigned | - |
| Otu62642 | 1 | k__Fungi(100);k__Fungi_unclassified(100);k__Fungi_unclassified(100);k__Fungi_unclassified(100);k__Fungi_unclassified(100);k__Fungi_unclassified(100);k__Fungi_unclassified(100); | - | - | - | - | - | - | - | Unassigned | - |
| Otu62643 | 1 | k__Fungi(100);k__Fungi_unclassified(100);k__Fungi_unclassified(100);k__Fungi_unclassified(100);k__Fungi_unclassified(100);k__Fungi_unclassified(100);k__Fungi_unclassified(100); | - | - | - | - | - | - | - | Unassigned | - |
| Otu62644 | 1 | k__Fungi(100);k__Fungi_unclassified(100);k__Fungi_unclassified(100);k__Fungi_unclassified(100);k__Fungi_unclassified(100);k__Fungi_unclassified(100);k__Fungi_unclassified(100); | - | - | - | - | - | - | - | Unassigned | - |
| Otu62645 | 1 | k__Fungi(100);k__Fungi_unclassified(100);k__Fungi_unclassified(100);k__Fungi_unclassified(100);k__Fungi_unclassified(100);k__Fungi_unclassified(100);k__Fungi_unclassified(100); | - | - | - | - | - | - | - | Unassigned | - |
| Otu62646 | 1 | k__Fungi(100);k__Fungi_unclassified(100);k__Fungi_unclassified(100);k__Fungi_unclassified(100);k__Fungi_unclassified(100);k__Fungi_unclassified(100);k__Fungi_unclassified(100); | - | - | - | - | - | - | - | Unassigned | - |
| Otu62647 | 1 | k__Fungi(100);k__Fungi_unclassified(100);k__Fungi_unclassified(100);k__Fungi_unclassified(100);k__Fungi_unclassified(100);k__Fungi_unclassified(100);k__Fungi_unclassified(100); | - | - | - | - | - | - | - | Unassigned | - |
| Otu62648 | 1 | k__Fungi(100);p__Ascomycota(100);p__Ascomycota_unclassified(100);p__Ascomycota_unclassified(100);p__Ascomycota_unclassified(100);p__Ascomycota_unclassified(100);p__Ascomycota_unclassified(100); | - | - | - | - | - | - | - | Unassigned | - |
| Otu62649 | 1 | k__Fungi(100);k__Fungi_unclassified(100);k__Fungi_unclassified(100);k__Fungi_unclassified(100);k__Fungi_unclassified(100);k__Fungi_unclassified(100);k__Fungi_unclassified(100); | - | - | - | - | - | - | - | Unassigned | - |
| Otu62651 | 1 | k__Fungi(100);k__Fungi_unclassified(100);k__Fungi_unclassified(100);k__Fungi_unclassified(100);k__Fungi_unclassified(100);k__Fungi_unclassified(100);k__Fungi_unclassified(100); | - | - | - | - | - | - | - | Unassigned | - |
| Otu62652 | 1 | k__Fungi(100);p__Ascomycota(100);p__Ascomycota_unclassified(100);p__Ascomycota_unclassified(100);p__Ascomycota_unclassified(100);p__Ascomycota_unclassified(100);p__Ascomycota_unclassified(100); | - | - | - | - | - | - | - | Unassigned | - |
| Otu62653 | 1 | k__Fungi(100);k__Fungi_unclassified(100);k__Fungi_unclassified(100);k__Fungi_unclassified(100);k__Fungi_unclassified(100);k__Fungi_unclassified(100);k__Fungi_unclassified(100); | - | - | - | - | - | - | - | Unassigned | - |
| Otu62654 | 1 | k__Fungi(100);k__Fungi_unclassified(100);k__Fungi_unclassified(100);k__Fungi_unclassified(100);k__Fungi_unclassified(100);k__Fungi_unclassified(100);k__Fungi_unclassified(100); | - | - | - | - | - | - | - | Unassigned | - |
| Otu62655 | 1 | k__Fungi(100);k__Fungi_unclassified(100);k__Fungi_unclassified(100);k__Fungi_unclassified(100);k__Fungi_unclassified(100);k__Fungi_unclassified(100);k__Fungi_unclassified(100); | - | - | - | - | - | - | - | Unassigned | - |
| Otu62656 | 1 | k__Fungi(100);k__Fungi_unclassified(100);k__Fungi_unclassified(100);k__Fungi_unclassified(100);k__Fungi_unclassified(100);k__Fungi_unclassified(100);k__Fungi_unclassified(100); | - | - | - | - | - | - | - | Unassigned | - |
| Otu62657 | 1 | k__Fungi(100);p__Ascomycota(100);p__Ascomycota_unclassified(100);p__Ascomycota_unclassified(100);p__Ascomycota_unclassified(100);p__Ascomycota_unclassified(100);p__Ascomycota_unclassified(100); | - | - | - | - | - | - | - | Unassigned | - |
| Otu62658 | 1 | k__Fungi(100);p__Ascomycota(100);p__Ascomycota_unclassified(100);p__Ascomycota_unclassified(100);p__Ascomycota_unclassified(100);p__Ascomycota_unclassified(100);p__Ascomycota_unclassified(100); | - | - | - | - | - | - | - | Unassigned | - |
| Otu62659 | 1 | k__Fungi(100);k__Fungi_unclassified(100);k__Fungi_unclassified(100);k__Fungi_unclassified(100);k__Fungi_unclassified(100);k__Fungi_unclassified(100);k__Fungi_unclassified(100); | - | - | - | - | - | - | - | Unassigned | - |
| Otu62660 | 1 | k__Fungi(100);p__Ascomycota(100);p__Ascomycota_unclassified(100);p__Ascomycota_unclassified(100);p__Ascomycota_unclassified(100);p__Ascomycota_unclassified(100);p__Ascomycota_unclassified(100); | - | - | - | - | - | - | - | Unassigned | - |
| Otu62661 | 1 | k__Fungi(100);k__Fungi_unclassified(100);k__Fungi_unclassified(100);k__Fungi_unclassified(100);k__Fungi_unclassified(100);k__Fungi_unclassified(100);k__Fungi_unclassified(100); | - | - | - | - | - | - | - | Unassigned | - |
| Otu62662 | 1 | k__Fungi(100);p__Ascomycota(100);p__Ascomycota_unclassified(100);p__Ascomycota_unclassified(100);p__Ascomycota_unclassified(100);p__Ascomycota_unclassified(100);p__Ascomycota_unclassified(100); | - | - | - | - | - | - | - | Unassigned | - |
| Otu62663 | 1 | k__Fungi(100);k__Fungi_unclassified(100);k__Fungi_unclassified(100);k__Fungi_unclassified(100);k__Fungi_unclassified(100);k__Fungi_unclassified(100);k__Fungi_unclassified(100); | - | - | - | - | - | - | - | Unassigned | - |
| Otu62664 | 1 | k__Fungi(100);k__Fungi_unclassified(100);k__Fungi_unclassified(100);k__Fungi_unclassified(100);k__Fungi_unclassified(100);k__Fungi_unclassified(100);k__Fungi_unclassified(100); | - | - | - | - | - | - | - | Unassigned | - |
| Otu62665 | 1 | k__Fungi(100);p__Ascomycota(100);p__Ascomycota_unclassified(100);p__Ascomycota_unclassified(100);p__Ascomycota_unclassified(100);p__Ascomycota_unclassified(100);p__Ascomycota_unclassified(100); | - | - | - | - | - | - | - | Unassigned | - |
| Otu62666 | 1 | k__Fungi(100);k__Fungi_unclassified(100);k__Fungi_unclassified(100);k__Fungi_unclassified(100);k__Fungi_unclassified(100);k__Fungi_unclassified(100);k__Fungi_unclassified(100); | - | - | - | - | - | - | - | Unassigned | - |
| Otu62667 | 1 | k__Fungi(100);k__Fungi_unclassified(100);k__Fungi_unclassified(100);k__Fungi_unclassified(100);k__Fungi_unclassified(100);k__Fungi_unclassified(100);k__Fungi_unclassified(100); | - | - | - | - | - | - | - | Unassigned | - |
| Otu62668 | 1 | k__Fungi(100);k__Fungi_unclassified(100);k__Fungi_unclassified(100);k__Fungi_unclassified(100);k__Fungi_unclassified(100);k__Fungi_unclassified(100);k__Fungi_unclassified(100); | - | - | - | - | - | - | - | Unassigned | - |
| Otu62669 | 1 | k__Fungi(100);p__Ascomycota(100);p__Ascomycota_unclassified(100);p__Ascomycota_unclassified(100);p__Ascomycota_unclassified(100);p__Ascomycota_unclassified(100);p__Ascomycota_unclassified(100); | - | - | - | - | - | - | - | Unassigned | - |
| Otu62670 | 1 | k__Fungi(100);k__Fungi_unclassified(100);k__Fungi_unclassified(100);k__Fungi_unclassified(100);k__Fungi_unclassified(100);k__Fungi_unclassified(100);k__Fungi_unclassified(100); | - | - | - | - | - | - | - | Unassigned | - |
| Otu62671 | 1 | k__Fungi(100);p__Ascomycota(100);p__Ascomycota_unclassified(100);p__Ascomycota_unclassified(100);p__Ascomycota_unclassified(100);p__Ascomycota_unclassified(100);p__Ascomycota_unclassified(100); | - | - | - | - | - | - | - | Unassigned | - |
| Otu62672 | 1 | k__Fungi(100);k__Fungi_unclassified(100);k__Fungi_unclassified(100);k__Fungi_unclassified(100);k__Fungi_unclassified(100);k__Fungi_unclassified(100);k__Fungi_unclassified(100); | - | - | - | - | - | - | - | Unassigned | - |
| Otu62673 | 1 | k__Fungi(100);k__Fungi_unclassified(100);k__Fungi_unclassified(100);k__Fungi_unclassified(100);k__Fungi_unclassified(100);k__Fungi_unclassified(100);k__Fungi_unclassified(100); | - | - | - | - | - | - | - | Unassigned | - |
| Otu62675 | 1 | k__Fungi(100);k__Fungi_unclassified(100);k__Fungi_unclassified(100);k__Fungi_unclassified(100);k__Fungi_unclassified(100);k__Fungi_unclassified(100);k__Fungi_unclassified(100); | - | - | - | - | - | - | - | Unassigned | - |
| Otu62676 | 1 | k__Fungi(100);k__Fungi_unclassified(100);k__Fungi_unclassified(100);k__Fungi_unclassified(100);k__Fungi_unclassified(100);k__Fungi_unclassified(100);k__Fungi_unclassified(100); | - | - | - | - | - | - | - | Unassigned | - |
| Otu62677 | 1 | k__Fungi(100);k__Fungi_unclassified(100);k__Fungi_unclassified(100);k__Fungi_unclassified(100);k__Fungi_unclassified(100);k__Fungi_unclassified(100);k__Fungi_unclassified(100); | - | - | - | - | - | - | - | Unassigned | - |
| Otu62682 | 1 | k__Fungi(100);p__Ascomycota(100);p__Ascomycota_unclassified(100);p__Ascomycota_unclassified(100);p__Ascomycota_unclassified(100);p__Ascomycota_unclassified(100);p__Ascomycota_unclassified(100); | - | - | - | - | - | - | - | Unassigned | - |
| Otu62686 | 1 | k__Fungi(100);k__Fungi_unclassified(100);k__Fungi_unclassified(100);k__Fungi_unclassified(100);k__Fungi_unclassified(100);k__Fungi_unclassified(100);k__Fungi_unclassified(100); | - | - | - | - | - | - | - | Unassigned | - |
| Otu62687 | 1 | k__Fungi(100);p__Ascomycota(100);p__Ascomycota_unclassified(100);p__Ascomycota_unclassified(100);p__Ascomycota_unclassified(100);p__Ascomycota_unclassified(100);p__Ascomycota_unclassified(100); | - | - | - | - | - | - | - | Unassigned | - |
| Otu62689 | 1 | k__Fungi(100);p__Ascomycota(100);p__Ascomycota_unclassified(100);p__Ascomycota_unclassified(100);p__Ascomycota_unclassified(100);p__Ascomycota_unclassified(100);p__Ascomycota_unclassified(100); | - | - | - | - | - | - | - | Unassigned | - |
| Otu62690 | 1 | k__Fungi(100);p__Ascomycota(100);p__Ascomycota_unclassified(100);p__Ascomycota_unclassified(100);p__Ascomycota_unclassified(100);p__Ascomycota_unclassified(100);p__Ascomycota_unclassified(100); | - | - | - | - | - | - | - | Unassigned | - |
| Otu62694 | 1 | k__Fungi(100);k__Fungi_unclassified(100);k__Fungi_unclassified(100);k__Fungi_unclassified(100);k__Fungi_unclassified(100);k__Fungi_unclassified(100);k__Fungi_unclassified(100); | - | - | - | - | - | - | - | Unassigned | - |
| Otu62695 | 1 | k__Fungi(100);k__Fungi_unclassified(100);k__Fungi_unclassified(100);k__Fungi_unclassified(100);k__Fungi_unclassified(100);k__Fungi_unclassified(100);k__Fungi_unclassified(100); | - | - | - | - | - | - | - | Unassigned | - |
| Otu62696 | 1 | k__Fungi(100);p__Ascomycota(100);p__Ascomycota_unclassified(100);p__Ascomycota_unclassified(100);p__Ascomycota_unclassified(100);p__Ascomycota_unclassified(100);p__Ascomycota_unclassified(100); | - | - | - | - | - | - | - | Unassigned | - |
| Otu62697 | 1 | k__Fungi(100);k__Fungi_unclassified(100);k__Fungi_unclassified(100);k__Fungi_unclassified(100);k__Fungi_unclassified(100);k__Fungi_unclassified(100);k__Fungi_unclassified(100); | - | - | - | - | - | - | - | Unassigned | - |
| Otu62698 | 1 | k__Fungi(100);p__Ascomycota(100);p__Ascomycota_unclassified(100);p__Ascomycota_unclassified(100);p__Ascomycota_unclassified(100);p__Ascomycota_unclassified(100);p__Ascomycota_unclassified(100); | - | - | - | - | - | - | - | Unassigned | - |
| Otu62699 | 1 | k__Fungi(100);p__Ascomycota(100);p__Ascomycota_unclassified(100);p__Ascomycota_unclassified(100);p__Ascomycota_unclassified(100);p__Ascomycota_unclassified(100);p__Ascomycota_unclassified(100); | - | - | - | - | - | - | - | Unassigned | - |
| Otu62700 | 1 | k__Fungi(100);p__Ascomycota(100);p__Ascomycota_unclassified(100);p__Ascomycota_unclassified(100);p__Ascomycota_unclassified(100);p__Ascomycota_unclassified(100);p__Ascomycota_unclassified(100); | - | - | - | - | - | - | - | Unassigned | - |
| Otu62701 | 1 | k__Fungi(100);k__Fungi_unclassified(100);k__Fungi_unclassified(100);k__Fungi_unclassified(100);k__Fungi_unclassified(100);k__Fungi_unclassified(100);k__Fungi_unclassified(100); | - | - | - | - | - | - | - | Unassigned | - |
| Otu62702 | 1 | k__Fungi(100);p__Ascomycota(100);p__Ascomycota_unclassified(100);p__Ascomycota_unclassified(100);p__Ascomycota_unclassified(100);p__Ascomycota_unclassified(100);p__Ascomycota_unclassified(100); | - | - | - | - | - | - | - | Unassigned | - |
| Otu62703 | 1 | k__Fungi(100);p__Ascomycota(100);p__Ascomycota_unclassified(100);p__Ascomycota_unclassified(100);p__Ascomycota_unclassified(100);p__Ascomycota_unclassified(100);p__Ascomycota_unclassified(100); | - | - | - | - | - | - | - | Unassigned | - |
| Otu62704 | 1 | k__Fungi(100);k__Fungi_unclassified(100);k__Fungi_unclassified(100);k__Fungi_unclassified(100);k__Fungi_unclassified(100);k__Fungi_unclassified(100);k__Fungi_unclassified(100); | - | - | - | - | - | - | - | Unassigned | - |
| Otu62705 | 1 | k__Fungi(100);k__Fungi_unclassified(100);k__Fungi_unclassified(100);k__Fungi_unclassified(100);k__Fungi_unclassified(100);k__Fungi_unclassified(100);k__Fungi_unclassified(100); | - | - | - | - | - | - | - | Unassigned | - |
| Otu62706 | 1 | k__Fungi(100);k__Fungi_unclassified(100);k__Fungi_unclassified(100);k__Fungi_unclassified(100);k__Fungi_unclassified(100);k__Fungi_unclassified(100);k__Fungi_unclassified(100); | - | - | - | - | - | - | - | Unassigned | - |
| Otu62707 | 1 | k__Fungi(100);p__Ascomycota(100);p__Ascomycota_unclassified(100);p__Ascomycota_unclassified(100);p__Ascomycota_unclassified(100);p__Ascomycota_unclassified(100);p__Ascomycota_unclassified(100); | - | - | - | - | - | - | - | Unassigned | - |
| Otu62708 | 1 | k__Fungi(100);k__Fungi_unclassified(100);k__Fungi_unclassified(100);k__Fungi_unclassified(100);k__Fungi_unclassified(100);k__Fungi_unclassified(100);k__Fungi_unclassified(100); | - | - | - | - | - | - | - | Unassigned | - |
| Otu62709 | 1 | k__Fungi(100);k__Fungi_unclassified(100);k__Fungi_unclassified(100);k__Fungi_unclassified(100);k__Fungi_unclassified(100);k__Fungi_unclassified(100);k__Fungi_unclassified(100); | - | - | - | - | - | - | - | Unassigned | - |
| Otu62710 | 1 | k__Fungi(100);k__Fungi_unclassified(100);k__Fungi_unclassified(100);k__Fungi_unclassified(100);k__Fungi_unclassified(100);k__Fungi_unclassified(100);k__Fungi_unclassified(100); | - | - | - | - | - | - | - | Unassigned | - |
| Otu62711 | 1 | k__Fungi(100);p__Ascomycota(100);p__Ascomycota_unclassified(100);p__Ascomycota_unclassified(100);p__Ascomycota_unclassified(100);p__Ascomycota_unclassified(100);p__Ascomycota_unclassified(100); | - | - | - | - | - | - | - | Unassigned | - |
| Otu62712 | 1 | k__Fungi(100);k__Fungi_unclassified(100);k__Fungi_unclassified(100);k__Fungi_unclassified(100);k__Fungi_unclassified(100);k__Fungi_unclassified(100);k__Fungi_unclassified(100); | - | - | - | - | - | - | - | Unassigned | - |
| Otu62713 | 1 | k__Fungi(100);k__Fungi_unclassified(100);k__Fungi_unclassified(100);k__Fungi_unclassified(100);k__Fungi_unclassified(100);k__Fungi_unclassified(100);k__Fungi_unclassified(100); | - | - | - | - | - | - | - | Unassigned | - |
| Otu62714 | 1 | k__Fungi(100);k__Fungi_unclassified(100);k__Fungi_unclassified(100);k__Fungi_unclassified(100);k__Fungi_unclassified(100);k__Fungi_unclassified(100);k__Fungi_unclassified(100); | - | - | - | - | - | - | - | Unassigned | - |
| Otu62715 | 1 | k__Fungi(100);k__Fungi_unclassified(100);k__Fungi_unclassified(100);k__Fungi_unclassified(100);k__Fungi_unclassified(100);k__Fungi_unclassified(100);k__Fungi_unclassified(100); | - | - | - | - | - | - | - | Unassigned | - |
| Otu62716 | 1 | k__Fungi(100);k__Fungi_unclassified(100);k__Fungi_unclassified(100);k__Fungi_unclassified(100);k__Fungi_unclassified(100);k__Fungi_unclassified(100);k__Fungi_unclassified(100); | - | - | - | - | - | - | - | Unassigned | - |
| Otu62718 | 1 | k__Fungi(100);k__Fungi_unclassified(100);k__Fungi_unclassified(100);k__Fungi_unclassified(100);k__Fungi_unclassified(100);k__Fungi_unclassified(100);k__Fungi_unclassified(100); | - | - | - | - | - | - | - | Unassigned | - |
| Otu62719 | 1 | k__Fungi(100);k__Fungi_unclassified(100);k__Fungi_unclassified(100);k__Fungi_unclassified(100);k__Fungi_unclassified(100);k__Fungi_unclassified(100);k__Fungi_unclassified(100); | - | - | - | - | - | - | - | Unassigned | - |
| Otu62720 | 1 | k__Fungi(100);k__Fungi_unclassified(100);k__Fungi_unclassified(100);k__Fungi_unclassified(100);k__Fungi_unclassified(100);k__Fungi_unclassified(100);k__Fungi_unclassified(100); | - | - | - | - | - | - | - | Unassigned | - |
| Otu62721 | 1 | k__Fungi(100);k__Fungi_unclassified(100);k__Fungi_unclassified(100);k__Fungi_unclassified(100);k__Fungi_unclassified(100);k__Fungi_unclassified(100);k__Fungi_unclassified(100); | - | - | - | - | - | - | - | Unassigned | - |
| Otu62722 | 1 | k__Fungi(100);k__Fungi_unclassified(100);k__Fungi_unclassified(100);k__Fungi_unclassified(100);k__Fungi_unclassified(100);k__Fungi_unclassified(100);k__Fungi_unclassified(100); | - | - | - | - | - | - | - | Unassigned | - |
| Otu62723 | 1 | k__Fungi(100);p__Ascomycota(100);p__Ascomycota_unclassified(100);p__Ascomycota_unclassified(100);p__Ascomycota_unclassified(100);p__Ascomycota_unclassified(100);p__Ascomycota_unclassified(100); | - | - | - | - | - | - | - | Unassigned | - |
| Otu62724 | 1 | k__Fungi(100);k__Fungi_unclassified(100);k__Fungi_unclassified(100);k__Fungi_unclassified(100);k__Fungi_unclassified(100);k__Fungi_unclassified(100);k__Fungi_unclassified(100); | - | - | - | - | - | - | - | Unassigned | - |
| Otu62725 | 1 | k__Fungi(100);k__Fungi_unclassified(100);k__Fungi_unclassified(100);k__Fungi_unclassified(100);k__Fungi_unclassified(100);k__Fungi_unclassified(100);k__Fungi_unclassified(100); | - | - | - | - | - | - | - | Unassigned | - |
| Otu62726 | 1 | k__Fungi(100);p__Ascomycota(100);p__Ascomycota_unclassified(100);p__Ascomycota_unclassified(100);p__Ascomycota_unclassified(100);p__Ascomycota_unclassified(100);p__Ascomycota_unclassified(100); | - | - | - | - | - | - | - | Unassigned | - |
| Otu62727 | 1 | k__Fungi(100);p__Ascomycota(100);p__Ascomycota_unclassified(100);p__Ascomycota_unclassified(100);p__Ascomycota_unclassified(100);p__Ascomycota_unclassified(100);p__Ascomycota_unclassified(100); | - | - | - | - | - | - | - | Unassigned | - |
| Otu62728 | 1 | k__Fungi(100);p__Ascomycota(100);p__Ascomycota_unclassified(100);p__Ascomycota_unclassified(100);p__Ascomycota_unclassified(100);p__Ascomycota_unclassified(100);p__Ascomycota_unclassified(100); | - | - | - | - | - | - | - | Unassigned | - |
| Otu62729 | 1 | k__Fungi(100);k__Fungi_unclassified(100);k__Fungi_unclassified(100);k__Fungi_unclassified(100);k__Fungi_unclassified(100);k__Fungi_unclassified(100);k__Fungi_unclassified(100); | - | - | - | - | - | - | - | Unassigned | - |
| Otu62730 | 1 | k__Fungi(100);k__Fungi_unclassified(100);k__Fungi_unclassified(100);k__Fungi_unclassified(100);k__Fungi_unclassified(100);k__Fungi_unclassified(100);k__Fungi_unclassified(100); | - | - | - | - | - | - | - | Unassigned | - |
| Otu62731 | 1 | k__Fungi(100);k__Fungi_unclassified(100);k__Fungi_unclassified(100);k__Fungi_unclassified(100);k__Fungi_unclassified(100);k__Fungi_unclassified(100);k__Fungi_unclassified(100); | - | - | - | - | - | - | - | Unassigned | - |
| Otu62732 | 1 | k__Fungi(100);p__Ascomycota(100);p__Ascomycota_unclassified(100);p__Ascomycota_unclassified(100);p__Ascomycota_unclassified(100);p__Ascomycota_unclassified(100);p__Ascomycota_unclassified(100); | - | - | - | - | - | - | - | Unassigned | - |
| Otu62733 | 1 | k__Fungi(100);k__Fungi_unclassified(100);k__Fungi_unclassified(100);k__Fungi_unclassified(100);k__Fungi_unclassified(100);k__Fungi_unclassified(100);k__Fungi_unclassified(100); | - | - | - | - | - | - | - | Unassigned | - |
| Otu62734 | 1 | k__Fungi(100);k__Fungi_unclassified(100);k__Fungi_unclassified(100);k__Fungi_unclassified(100);k__Fungi_unclassified(100);k__Fungi_unclassified(100);k__Fungi_unclassified(100); | - | - | - | - | - | - | - | Unassigned | - |
| Otu62737 | 1 | k__Fungi(100);k__Fungi_unclassified(100);k__Fungi_unclassified(100);k__Fungi_unclassified(100);k__Fungi_unclassified(100);k__Fungi_unclassified(100);k__Fungi_unclassified(100); | - | - | - | - | - | - | - | Unassigned | - |
| Otu62738 | 1 | k__Fungi(100);k__Fungi_unclassified(100);k__Fungi_unclassified(100);k__Fungi_unclassified(100);k__Fungi_unclassified(100);k__Fungi_unclassified(100);k__Fungi_unclassified(100); | - | - | - | - | - | - | - | Unassigned | - |
| Otu62739 | 1 | k__Fungi(100);p__Ascomycota(100);p__Ascomycota_unclassified(100);p__Ascomycota_unclassified(100);p__Ascomycota_unclassified(100);p__Ascomycota_unclassified(100);p__Ascomycota_unclassified(100); | - | - | - | - | - | - | - | Unassigned | - |
| Otu62740 | 1 | k__Fungi(100);p__Ascomycota(100);p__Ascomycota_unclassified(100);p__Ascomycota_unclassified(100);p__Ascomycota_unclassified(100);p__Ascomycota_unclassified(100);p__Ascomycota_unclassified(100); | - | - | - | - | - | - | - | Unassigned | - |
| Otu62741 | 1 | k__Fungi(100);k__Fungi_unclassified(100);k__Fungi_unclassified(100);k__Fungi_unclassified(100);k__Fungi_unclassified(100);k__Fungi_unclassified(100);k__Fungi_unclassified(100); | - | - | - | - | - | - | - | Unassigned | - |
| Otu62749 | 1 | k__Fungi(100);p__Ascomycota(100);c__Dothideomycetes(100);o__Capnodiales(100);f__Davidiellaceae(100);f__Davidiellaceae_unclassified(100);f__Davidiellaceae_unclassified(100); | - | - | - | - | - | - | - | Unassigned | - |
| Otu62750 | 1 | k__Fungi(100);k__Fungi_unclassified(100);k__Fungi_unclassified(100);k__Fungi_unclassified(100);k__Fungi_unclassified(100);k__Fungi_unclassified(100);k__Fungi_unclassified(100); | - | - | - | - | - | - | - | Unassigned | - |
| Otu62751 | 1 | k__Fungi(100);k__Fungi_unclassified(100);k__Fungi_unclassified(100);k__Fungi_unclassified(100);k__Fungi_unclassified(100);k__Fungi_unclassified(100);k__Fungi_unclassified(100); | - | - | - | - | - | - | - | Unassigned | - |
| Otu62752 | 1 | k__Fungi(100);k__Fungi_unclassified(100);k__Fungi_unclassified(100);k__Fungi_unclassified(100);k__Fungi_unclassified(100);k__Fungi_unclassified(100);k__Fungi_unclassified(100); | - | - | - | - | - | - | - | Unassigned | - |
| Otu62753 | 1 | k__Fungi(100);k__Fungi_unclassified(100);k__Fungi_unclassified(100);k__Fungi_unclassified(100);k__Fungi_unclassified(100);k__Fungi_unclassified(100);k__Fungi_unclassified(100); | - | - | - | - | - | - | - | Unassigned | - |
| Otu62755 | 1 | k__Fungi(100);k__Fungi_unclassified(100);k__Fungi_unclassified(100);k__Fungi_unclassified(100);k__Fungi_unclassified(100);k__Fungi_unclassified(100);k__Fungi_unclassified(100); | - | - | - | - | - | - | - | Unassigned | - |
| Otu62756 | 1 | k__Fungi(100);k__Fungi_unclassified(100);k__Fungi_unclassified(100);k__Fungi_unclassified(100);k__Fungi_unclassified(100);k__Fungi_unclassified(100);k__Fungi_unclassified(100); | - | - | - | - | - | - | - | Unassigned | - |
| Otu62757 | 1 | k__Fungi(100);k__Fungi_unclassified(100);k__Fungi_unclassified(100);k__Fungi_unclassified(100);k__Fungi_unclassified(100);k__Fungi_unclassified(100);k__Fungi_unclassified(100); | - | - | - | - | - | - | - | Unassigned | - |
| Otu62758 | 1 | k__Fungi(100);p__Basidiomycota(100);p__Basidiomycota_unclassified(100);p__Basidiomycota_unclassified(100);p__Basidiomycota_unclassified(100);p__Basidiomycota_unclassified(100);p__Basidiomycota_unclassified(100); | - | - | - | - | - | - | - | Unassigned | - |
| Otu62759 | 1 | k__Fungi(100);k__Fungi_unclassified(100);k__Fungi_unclassified(100);k__Fungi_unclassified(100);k__Fungi_unclassified(100);k__Fungi_unclassified(100);k__Fungi_unclassified(100); | - | - | - | - | - | - | - | Unassigned | - |
| Otu62760 | 1 | k__Fungi(100);p__Ascomycota(100);p__Ascomycota_unclassified(100);p__Ascomycota_unclassified(100);p__Ascomycota_unclassified(100);p__Ascomycota_unclassified(100);p__Ascomycota_unclassified(100); | - | - | - | - | - | - | - | Unassigned | - |
| Otu62761 | 1 | k__Fungi(100);p__Ascomycota(100);p__Ascomycota_unclassified(100);p__Ascomycota_unclassified(100);p__Ascomycota_unclassified(100);p__Ascomycota_unclassified(100);p__Ascomycota_unclassified(100); | - | - | - | - | - | - | - | Unassigned | - |
| Otu62762 | 1 | k__Fungi(100);k__Fungi_unclassified(100);k__Fungi_unclassified(100);k__Fungi_unclassified(100);k__Fungi_unclassified(100);k__Fungi_unclassified(100);k__Fungi_unclassified(100); | - | - | - | - | - | - | - | Unassigned | - |
| Otu62763 | 1 | k__Fungi(100);k__Fungi_unclassified(100);k__Fungi_unclassified(100);k__Fungi_unclassified(100);k__Fungi_unclassified(100);k__Fungi_unclassified(100);k__Fungi_unclassified(100); | - | - | - | - | - | - | - | Unassigned | - |
| Otu62764 | 1 | k__Fungi(100);p__Ascomycota(100);p__Ascomycota_unclassified(100);p__Ascomycota_unclassified(100);p__Ascomycota_unclassified(100);p__Ascomycota_unclassified(100);p__Ascomycota_unclassified(100); | - | - | - | - | - | - | - | Unassigned | - |
| Otu62765 | 1 | k__Fungi(100);p__Ascomycota(100);p__Ascomycota_unclassified(100);p__Ascomycota_unclassified(100);p__Ascomycota_unclassified(100);p__Ascomycota_unclassified(100);p__Ascomycota_unclassified(100); | - | - | - | - | - | - | - | Unassigned | - |
| Otu62766 | 1 | k__Fungi(100);k__Fungi_unclassified(100);k__Fungi_unclassified(100);k__Fungi_unclassified(100);k__Fungi_unclassified(100);k__Fungi_unclassified(100);k__Fungi_unclassified(100); | - | - | - | - | - | - | - | Unassigned | - |
| Otu62767 | 1 | k__Fungi(100);k__Fungi_unclassified(100);k__Fungi_unclassified(100);k__Fungi_unclassified(100);k__Fungi_unclassified(100);k__Fungi_unclassified(100);k__Fungi_unclassified(100); | - | - | - | - | - | - | - | Unassigned | - |
| Otu62768 | 1 | k__Fungi(100);k__Fungi_unclassified(100);k__Fungi_unclassified(100);k__Fungi_unclassified(100);k__Fungi_unclassified(100);k__Fungi_unclassified(100);k__Fungi_unclassified(100); | - | - | - | - | - | - | - | Unassigned | - |
| Otu62769 | 1 | k__Fungi(100);k__Fungi_unclassified(100);k__Fungi_unclassified(100);k__Fungi_unclassified(100);k__Fungi_unclassified(100);k__Fungi_unclassified(100);k__Fungi_unclassified(100); | - | - | - | - | - | - | - | Unassigned | - |
| Otu62770 | 1 | k__Fungi(100);k__Fungi_unclassified(100);k__Fungi_unclassified(100);k__Fungi_unclassified(100);k__Fungi_unclassified(100);k__Fungi_unclassified(100);k__Fungi_unclassified(100); | - | - | - | - | - | - | - | Unassigned | - |
| Otu62771 | 1 | k__Fungi(100);k__Fungi_unclassified(100);k__Fungi_unclassified(100);k__Fungi_unclassified(100);k__Fungi_unclassified(100);k__Fungi_unclassified(100);k__Fungi_unclassified(100); | - | - | - | - | - | - | - | Unassigned | - |
| Otu62772 | 1 | k__Fungi(100);k__Fungi_unclassified(100);k__Fungi_unclassified(100);k__Fungi_unclassified(100);k__Fungi_unclassified(100);k__Fungi_unclassified(100);k__Fungi_unclassified(100); | - | - | - | - | - | - | - | Unassigned | - |
| Otu62773 | 1 | k__Fungi(100);p__Ascomycota(100);p__Ascomycota_unclassified(100);p__Ascomycota_unclassified(100);p__Ascomycota_unclassified(100);p__Ascomycota_unclassified(100);p__Ascomycota_unclassified(100); | - | - | - | - | - | - | - | Unassigned | - |
| Otu62774 | 1 | k__Fungi(100);k__Fungi_unclassified(100);k__Fungi_unclassified(100);k__Fungi_unclassified(100);k__Fungi_unclassified(100);k__Fungi_unclassified(100);k__Fungi_unclassified(100); | - | - | - | - | - | - | - | Unassigned | - |
| Otu62775 | 1 | k__Fungi(100);p__Ascomycota(100);p__Ascomycota_unclassified(100);p__Ascomycota_unclassified(100);p__Ascomycota_unclassified(100);p__Ascomycota_unclassified(100);p__Ascomycota_unclassified(100); | - | - | - | - | - | - | - | Unassigned | - |
| Otu62777 | 1 | k__Fungi(100);p__Ascomycota(100);p__Ascomycota_unclassified(100);p__Ascomycota_unclassified(100);p__Ascomycota_unclassified(100);p__Ascomycota_unclassified(100);p__Ascomycota_unclassified(100); | - | - | - | - | - | - | - | Unassigned | - |
| Otu62778 | 1 | k__Fungi(100);k__Fungi_unclassified(100);k__Fungi_unclassified(100);k__Fungi_unclassified(100);k__Fungi_unclassified(100);k__Fungi_unclassified(100);k__Fungi_unclassified(100); | - | - | - | - | - | - | - | Unassigned | - |
| Otu62779 | 1 | k__Fungi(100);p__Ascomycota(100);p__Ascomycota_unclassified(100);p__Ascomycota_unclassified(100);p__Ascomycota_unclassified(100);p__Ascomycota_unclassified(100);p__Ascomycota_unclassified(100); | - | - | - | - | - | - | - | Unassigned | - |
| Otu62780 | 1 | k__Fungi(100);p__Ascomycota(100);p__Ascomycota_unclassified(100);p__Ascomycota_unclassified(100);p__Ascomycota_unclassified(100);p__Ascomycota_unclassified(100);p__Ascomycota_unclassified(100); | - | - | - | - | - | - | - | Unassigned | - |
| Otu62781 | 1 | k__Fungi(100);k__Fungi_unclassified(100);k__Fungi_unclassified(100);k__Fungi_unclassified(100);k__Fungi_unclassified(100);k__Fungi_unclassified(100);k__Fungi_unclassified(100); | - | - | - | - | - | - | - | Unassigned | - |
| Otu62782 | 1 | k__Fungi(100);k__Fungi_unclassified(100);k__Fungi_unclassified(100);k__Fungi_unclassified(100);k__Fungi_unclassified(100);k__Fungi_unclassified(100);k__Fungi_unclassified(100); | - | - | - | - | - | - | - | Unassigned | - |
| Otu62783 | 1 | k__Fungi(100);k__Fungi_unclassified(100);k__Fungi_unclassified(100);k__Fungi_unclassified(100);k__Fungi_unclassified(100);k__Fungi_unclassified(100);k__Fungi_unclassified(100); | - | - | - | - | - | - | - | Unassigned | - |
| Otu62786 | 1 | k__Fungi(100);k__Fungi_unclassified(100);k__Fungi_unclassified(100);k__Fungi_unclassified(100);k__Fungi_unclassified(100);k__Fungi_unclassified(100);k__Fungi_unclassified(100); | - | - | - | - | - | - | - | Unassigned | - |
| Otu62788 | 1 | k__Fungi(100);k__Fungi_unclassified(100);k__Fungi_unclassified(100);k__Fungi_unclassified(100);k__Fungi_unclassified(100);k__Fungi_unclassified(100);k__Fungi_unclassified(100); | - | - | - | - | - | - | - | Unassigned | - |
| Otu62789 | 1 | k__Fungi(100);p__Ascomycota(100);p__Ascomycota_unclassified(100);p__Ascomycota_unclassified(100);p__Ascomycota_unclassified(100);p__Ascomycota_unclassified(100);p__Ascomycota_unclassified(100); | - | - | - | - | - | - | - | Unassigned | - |
| Otu62790 | 1 | k__Fungi(100);k__Fungi_unclassified(100);k__Fungi_unclassified(100);k__Fungi_unclassified(100);k__Fungi_unclassified(100);k__Fungi_unclassified(100);k__Fungi_unclassified(100); | - | - | - | - | - | - | - | Unassigned | - |
| Otu62791 | 1 | k__Fungi(100);k__Fungi_unclassified(100);k__Fungi_unclassified(100);k__Fungi_unclassified(100);k__Fungi_unclassified(100);k__Fungi_unclassified(100);k__Fungi_unclassified(100); | - | - | - | - | - | - | - | Unassigned | - |
| Otu62792 | 1 | k__Fungi(100);k__Fungi_unclassified(100);k__Fungi_unclassified(100);k__Fungi_unclassified(100);k__Fungi_unclassified(100);k__Fungi_unclassified(100);k__Fungi_unclassified(100); | - | - | - | - | - | - | - | Unassigned | - |
| Otu62793 | 1 | k__Fungi(100);p__Ascomycota(100);p__Ascomycota_unclassified(100);p__Ascomycota_unclassified(100);p__Ascomycota_unclassified(100);p__Ascomycota_unclassified(100);p__Ascomycota_unclassified(100); | - | - | - | - | - | - | - | Unassigned | - |
| Otu62794 | 1 | k__Fungi(100);k__Fungi_unclassified(100);k__Fungi_unclassified(100);k__Fungi_unclassified(100);k__Fungi_unclassified(100);k__Fungi_unclassified(100);k__Fungi_unclassified(100); | - | - | - | - | - | - | - | Unassigned | - |
| Otu62795 | 1 | k__Fungi(100);p__Ascomycota(100);p__Ascomycota_unclassified(100);p__Ascomycota_unclassified(100);p__Ascomycota_unclassified(100);p__Ascomycota_unclassified(100);p__Ascomycota_unclassified(100); | - | - | - | - | - | - | - | Unassigned | - |
| Otu62796 | 1 | k__Fungi(100);k__Fungi_unclassified(100);k__Fungi_unclassified(100);k__Fungi_unclassified(100);k__Fungi_unclassified(100);k__Fungi_unclassified(100);k__Fungi_unclassified(100); | - | - | - | - | - | - | - | Unassigned | - |
| Otu62797 | 1 | k__Fungi(100);k__Fungi_unclassified(100);k__Fungi_unclassified(100);k__Fungi_unclassified(100);k__Fungi_unclassified(100);k__Fungi_unclassified(100);k__Fungi_unclassified(100); | - | - | - | - | - | - | - | Unassigned | - |
| Otu62798 | 1 | k__Fungi(100);k__Fungi_unclassified(100);k__Fungi_unclassified(100);k__Fungi_unclassified(100);k__Fungi_unclassified(100);k__Fungi_unclassified(100);k__Fungi_unclassified(100); | - | - | - | - | - | - | - | Unassigned | - |
| Otu62799 | 1 | k__Fungi(100);k__Fungi_unclassified(100);k__Fungi_unclassified(100);k__Fungi_unclassified(100);k__Fungi_unclassified(100);k__Fungi_unclassified(100);k__Fungi_unclassified(100); | - | - | - | - | - | - | - | Unassigned | - |
| Otu62800 | 1 | k__Fungi(100);p__Ascomycota(100);p__Ascomycota_unclassified(100);p__Ascomycota_unclassified(100);p__Ascomycota_unclassified(100);p__Ascomycota_unclassified(100);p__Ascomycota_unclassified(100); | - | - | - | - | - | - | - | Unassigned | - |
| Otu62802 | 1 | k__Fungi(100);p__Ascomycota(100);p__Ascomycota_unclassified(100);p__Ascomycota_unclassified(100);p__Ascomycota_unclassified(100);p__Ascomycota_unclassified(100);p__Ascomycota_unclassified(100); | - | - | - | - | - | - | - | Unassigned | - |
| Otu62806 | 1 | k__Fungi(100);k__Fungi_unclassified(100);k__Fungi_unclassified(100);k__Fungi_unclassified(100);k__Fungi_unclassified(100);k__Fungi_unclassified(100);k__Fungi_unclassified(100); | - | - | - | - | - | - | - | Unassigned | - |
| Otu62807 | 1 | k__Fungi(100);k__Fungi_unclassified(100);k__Fungi_unclassified(100);k__Fungi_unclassified(100);k__Fungi_unclassified(100);k__Fungi_unclassified(100);k__Fungi_unclassified(100); | - | - | - | - | - | - | - | Unassigned | - |

| | | | | | | | | | | | |
|---|---|---|---|---|---|---|---|---|---|---|---|
| Otu62808 | 1 | k__Fungi(100);p__Ascomycota(100);p__Ascomycota_unclassified(100);p__Ascomycota_unclassified(100);p__Ascomycota_unclassified(100);p__Ascomycota_unclassified(100);p__Ascomycota_unclassified(100); | - | - | - | - | - | - | - | Unassigned | - |
| Otu62811 | 1 | k__Fungi(100);k__Fungi_unclassified(100);k__Fungi_unclassified(100);k__Fungi_unclassified(100);k__Fungi_unclassified(100);k__Fungi_unclassified(100);k__Fungi_unclassified(100); | - | - | - | - | - | - | - | Unassigned | - |
| Otu62812 | 1 | k__Fungi(100);k__Fungi_unclassified(100);k__Fungi_unclassified(100);k__Fungi_unclassified(100);k__Fungi_unclassified(100);k__Fungi_unclassified(100);k__Fungi_unclassified(100); | - | - | - | - | - | - | - | Unassigned | - |
| Otu62814 | 1 | k__Fungi(100);p__Ascomycota(100);p__Ascomycota_unclassified(100);p__Ascomycota_unclassified(100);p__Ascomycota_unclassified(100);p__Ascomycota_unclassified(100);p__Ascomycota_unclassified(100); | - | - | - | - | - | - | - | Unassigned | - |
| Otu62815 | 1 | k__Fungi(100);k__Fungi_unclassified(100);k__Fungi_unclassified(100);k__Fungi_unclassified(100);k__Fungi_unclassified(100);k__Fungi_unclassified(100);k__Fungi_unclassified(100); | - | - | - | - | - | - | - | Unassigned | - |
| Otu62816 | 1 | k__Fungi(100);p__Ascomycota(100);p__Ascomycota_unclassified(100);p__Ascomycota_unclassified(100);p__Ascomycota_unclassified(100);p__Ascomycota_unclassified(100);p__Ascomycota_unclassified(100); | - | - | - | - | - | - | - | Unassigned | - |
| Otu62817 | 1 | k__Fungi(100);k__Fungi_unclassified(100);k__Fungi_unclassified(100);k__Fungi_unclassified(100);k__Fungi_unclassified(100);k__Fungi_unclassified(100);k__Fungi_unclassified(100); | - | - | - | - | - | - | - | Unassigned | - |
| Otu62818 | 1 | k__Fungi(100);k__Fungi_unclassified(100);k__Fungi_unclassified(100);k__Fungi_unclassified(100);k__Fungi_unclassified(100);k__Fungi_unclassified(100);k__Fungi_unclassified(100); | - | - | - | - | - | - | - | Unassigned | - |
| Otu62819 | 1 | k__Fungi(100);k__Fungi_unclassified(100);k__Fungi_unclassified(100);k__Fungi_unclassified(100);k__Fungi_unclassified(100);k__Fungi_unclassified(100);k__Fungi_unclassified(100); | - | - | - | - | - | - | - | Unassigned | - |
| Otu62820 | 1 | k__Fungi(100);k__Fungi_unclassified(100);k__Fungi_unclassified(100);k__Fungi_unclassified(100);k__Fungi_unclassified(100);k__Fungi_unclassified(100);k__Fungi_unclassified(100); | - | - | - | - | - | - | - | Unassigned | - |
| Otu62821 | 1 | k__Fungi(100);p__Ascomycota(100);p__Ascomycota_unclassified(100);p__Ascomycota_unclassified(100);p__Ascomycota_unclassified(100);p__Ascomycota_unclassified(100);p__Ascomycota_unclassified(100); | - | - | - | - | - | - | - | Unassigned | - |
| Otu62822 | 1 | k__Fungi(100);k__Fungi_unclassified(100);k__Fungi_unclassified(100);k__Fungi_unclassified(100);k__Fungi_unclassified(100);k__Fungi_unclassified(100);k__Fungi_unclassified(100); | - | - | - | - | - | - | - | Unassigned | - |
| Otu62823 | 1 | k__Fungi(100);k__Fungi_unclassified(100);k__Fungi_unclassified(100);k__Fungi_unclassified(100);k__Fungi_unclassified(100);k__Fungi_unclassified(100);k__Fungi_unclassified(100); | - | - | - | - | - | - | - | Unassigned | - |
| Otu62824 | 1 | k__Fungi(100);k__Fungi_unclassified(100);k__Fungi_unclassified(100);k__Fungi_unclassified(100);k__Fungi_unclassified(100);k__Fungi_unclassified(100);k__Fungi_unclassified(100); | - | - | - | - | - | - | - | Unassigned | - |
| Otu62826 | 1 | k__Fungi(100);p__Ascomycota(100);p__Ascomycota_unclassified(100);p__Ascomycota_unclassified(100);p__Ascomycota_unclassified(100);p__Ascomycota_unclassified(100);p__Ascomycota_unclassified(100); | - | - | - | - | - | - | - | Unassigned | - |
| Otu62829 | 1 | k__Fungi(100);k__Fungi_unclassified(100);k__Fungi_unclassified(100);k__Fungi_unclassified(100);k__Fungi_unclassified(100);k__Fungi_unclassified(100);k__Fungi_unclassified(100); | - | - | - | - | - | - | - | Unassigned | - |
| Otu62830 | 1 | k__Fungi(100);k__Fungi_unclassified(100);k__Fungi_unclassified(100);k__Fungi_unclassified(100);k__Fungi_unclassified(100);k__Fungi_unclassified(100);k__Fungi_unclassified(100); | - | - | - | - | - | - | - | Unassigned | - |
| Otu62833 | 1 | k__Fungi(100);k__Fungi_unclassified(100);k__Fungi_unclassified(100);k__Fungi_unclassified(100);k__Fungi_unclassified(100);k__Fungi_unclassified(100);k__Fungi_unclassified(100); | - | - | - | - | - | - | - | Unassigned | - |
| Otu62834 | 1 | k__Fungi(100);k__Fungi_unclassified(100);k__Fungi_unclassified(100);k__Fungi_unclassified(100);k__Fungi_unclassified(100);k__Fungi_unclassified(100);k__Fungi_unclassified(100); | - | - | - | - | - | - | - | Unassigned | - |
| Otu62835 | 1 | k__Fungi(100);k__Fungi_unclassified(100);k__Fungi_unclassified(100);k__Fungi_unclassified(100);k__Fungi_unclassified(100);k__Fungi_unclassified(100);k__Fungi_unclassified(100); | - | - | - | - | - | - | - | Unassigned | - |
| Otu62836 | 1 | k__Fungi(100);p__Ascomycota(100);p__Ascomycota_unclassified(100);p__Ascomycota_unclassified(100);p__Ascomycota_unclassified(100);p__Ascomycota_unclassified(100);p__Ascomycota_unclassified(100); | - | - | - | - | - | - | - | Unassigned | - |
| Otu62837 | 1 | k__Fungi(100);p__Ascomycota(100);p__Ascomycota_unclassified(100);p__Ascomycota_unclassified(100);p__Ascomycota_unclassified(100);p__Ascomycota_unclassified(100);p__Ascomycota_unclassified(100); | - | - | - | - | - | - | - | Unassigned | - |
| Otu62838 | 1 | k__Fungi(100);p__Ascomycota(100);p__Ascomycota_unclassified(100);p__Ascomycota_unclassified(100);p__Ascomycota_unclassified(100);p__Ascomycota_unclassified(100);p__Ascomycota_unclassified(100); | - | - | - | - | - | - | - | Unassigned | - |
| Otu62839 | 1 | k__Fungi(100);k__Fungi_unclassified(100);k__Fungi_unclassified(100);k__Fungi_unclassified(100);k__Fungi_unclassified(100);k__Fungi_unclassified(100);k__Fungi_unclassified(100); | - | - | - | - | - | - | - | Unassigned | - |
| Otu62840 | 1 | k__Fungi(100);k__Fungi_unclassified(100);k__Fungi_unclassified(100);k__Fungi_unclassified(100);k__Fungi_unclassified(100);k__Fungi_unclassified(100);k__Fungi_unclassified(100); | - | - | - | - | - | - | - | Unassigned | - |
| Otu62841 | 1 | k__Fungi(100);k__Fungi_unclassified(100);k__Fungi_unclassified(100);k__Fungi_unclassified(100);k__Fungi_unclassified(100);k__Fungi_unclassified(100);k__Fungi_unclassified(100); | - | - | - | - | - | - | - | Unassigned | - |
| Otu62843 | 1 | k__Fungi(100);k__Fungi_unclassified(100);k__Fungi_unclassified(100);k__Fungi_unclassified(100);k__Fungi_unclassified(100);k__Fungi_unclassified(100);k__Fungi_unclassified(100); | - | - | - | - | - | - | - | Unassigned | - |
| Otu62844 | 1 | k__Fungi(100);k__Fungi_unclassified(100);k__Fungi_unclassified(100);k__Fungi_unclassified(100);k__Fungi_unclassified(100);k__Fungi_unclassified(100);k__Fungi_unclassified(100); | - | - | - | - | - | - | - | Unassigned | - |
| Otu62845 | 1 | k__Fungi(100);k__Fungi_unclassified(100);k__Fungi_unclassified(100);k__Fungi_unclassified(100);k__Fungi_unclassified(100);k__Fungi_unclassified(100);k__Fungi_unclassified(100); | - | - | - | - | - | - | - | Unassigned | - |
| Otu62847 | 1 | k__Fungi(100);k__Fungi_unclassified(100);k__Fungi_unclassified(100);k__Fungi_unclassified(100);k__Fungi_unclassified(100);k__Fungi_unclassified(100);k__Fungi_unclassified(100); | - | - | - | - | - | - | - | Unassigned | - |
| Otu62848 | 1 | k__Fungi(100);k__Fungi_unclassified(100);k__Fungi_unclassified(100);k__Fungi_unclassified(100);k__Fungi_unclassified(100);k__Fungi_unclassified(100);k__Fungi_unclassified(100); | - | - | - | - | - | - | - | Unassigned | - |
| Otu62849 | 1 | k__Fungi(100);k__Fungi_unclassified(100);k__Fungi_unclassified(100);k__Fungi_unclassified(100);k__Fungi_unclassified(100);k__Fungi_unclassified(100);k__Fungi_unclassified(100); | - | - | - | - | - | - | - | Unassigned | - |
| Otu62850 | 1 | k__Fungi(100);p__Ascomycota(100);p__Ascomycota_unclassified(100);p__Ascomycota_unclassified(100);p__Ascomycota_unclassified(100);p__Ascomycota_unclassified(100);p__Ascomycota_unclassified(100); | - | - | - | - | - | - | - | Unassigned | - |
| Otu62851 | 1 | k__Fungi(100);k__Fungi_unclassified(100);k__Fungi_unclassified(100);k__Fungi_unclassified(100);k__Fungi_unclassified(100);k__Fungi_unclassified(100);k__Fungi_unclassified(100); | - | - | - | - | - | - | - | Unassigned | - |
| Otu62852 | 1 | k__Fungi(100);p__Ascomycota(100);p__Ascomycota_unclassified(100);p__Ascomycota_unclassified(100);p__Ascomycota_unclassified(100);p__Ascomycota_unclassified(100);p__Ascomycota_unclassified(100); | - | - | - | - | - | - | - | Unassigned | - |
| Otu62853 | 1 | k__Fungi(100);k__Fungi_unclassified(100);k__Fungi_unclassified(100);k__Fungi_unclassified(100);k__Fungi_unclassified(100);k__Fungi_unclassified(100);k__Fungi_unclassified(100); | - | - | - | - | - | - | - | Unassigned | - |
| Otu62854 | 1 | k__Fungi(100);k__Fungi_unclassified(100);k__Fungi_unclassified(100);k__Fungi_unclassified(100);k__Fungi_unclassified(100);k__Fungi_unclassified(100);k__Fungi_unclassified(100); | - | - | - | - | - | - | - | Unassigned | - |
| Otu62855 | 1 | k__Fungi(100);k__Fungi_unclassified(100);k__Fungi_unclassified(100);k__Fungi_unclassified(100);k__Fungi_unclassified(100);k__Fungi_unclassified(100);k__Fungi_unclassified(100); | - | - | - | - | - | - | - | Unassigned | - |
| Otu62856 | 1 | k__Fungi(100);k__Fungi_unclassified(100);k__Fungi_unclassified(100);k__Fungi_unclassified(100);k__Fungi_unclassified(100);k__Fungi_unclassified(100);k__Fungi_unclassified(100); | - | - | - | - | - | - | - | Unassigned | - |
| Otu62857 | 1 | k__Fungi(100);k__Fungi_unclassified(100);k__Fungi_unclassified(100);k__Fungi_unclassified(100);k__Fungi_unclassified(100);k__Fungi_unclassified(100);k__Fungi_unclassified(100); | - | - | - | - | - | - | - | Unassigned | - |
| Otu62858 | 1 | k__Fungi(100);k__Fungi_unclassified(100);k__Fungi_unclassified(100);k__Fungi_unclassified(100);k__Fungi_unclassified(100);k__Fungi_unclassified(100);k__Fungi_unclassified(100); | - | - | - | - | - | - | - | Unassigned | - |
| Otu62859 | 1 | k__Fungi(100);k__Fungi_unclassified(100);k__Fungi_unclassified(100);k__Fungi_unclassified(100);k__Fungi_unclassified(100);k__Fungi_unclassified(100);k__Fungi_unclassified(100); | - | - | - | - | - | - | - | Unassigned | - |
| Otu62860 | 1 | k__Fungi(100);p__Ascomycota(100);p__Ascomycota_unclassified(100);p__Ascomycota_unclassified(100);p__Ascomycota_unclassified(100);p__Ascomycota_unclassified(100);p__Ascomycota_unclassified(100); | - | - | - | - | - | - | - | Unassigned | - |
| Otu62861 | 1 | k__Fungi(100);p__Ascomycota(100);p__Ascomycota_unclassified(100);p__Ascomycota_unclassified(100);p__Ascomycota_unclassified(100);p__Ascomycota_unclassified(100);p__Ascomycota_unclassified(100); | - | - | - | - | - | - | - | Unassigned | - |
| Otu62864 | 1 | k__Fungi(100);p__Ascomycota(100);p__Ascomycota_unclassified(100);p__Ascomycota_unclassified(100);p__Ascomycota_unclassified(100);p__Ascomycota_unclassified(100);p__Ascomycota_unclassified(100); | - | - | - | - | - | - | - | Unassigned | - |
| Otu62865 | 1 | k__Fungi(100);p__Ascomycota(100);p__Ascomycota_unclassified(100);p__Ascomycota_unclassified(100);p__Ascomycota_unclassified(100);p__Ascomycota_unclassified(100);p__Ascomycota_unclassified(100); | - | - | - | - | - | - | - | Unassigned | - |
| Otu62869 | 1 | k__Fungi(100);k__Fungi_unclassified(100);k__Fungi_unclassified(100);k__Fungi_unclassified(100);k__Fungi_unclassified(100);k__Fungi_unclassified(100);k__Fungi_unclassified(100); | - | - | - | - | - | - | - | Unassigned | - |
| Otu62871 | 1 | k__Fungi(100);k__Fungi_unclassified(100);k__Fungi_unclassified(100);k__Fungi_unclassified(100);k__Fungi_unclassified(100);k__Fungi_unclassified(100);k__Fungi_unclassified(100); | - | - | - | - | - | - | - | Unassigned | - |
| Otu62874 | 1 | k__Fungi(100);p__Ascomycota(100);p__Ascomycota_unclassified(100);p__Ascomycota_unclassified(100);p__Ascomycota_unclassified(100);p__Ascomycota_unclassified(100);p__Ascomycota_unclassified(100); | - | - | - | - | - | - | - | Unassigned | - |
| Otu62876 | 1 | k__Fungi(100);p__Ascomycota(100);p__Ascomycota_unclassified(100);p__Ascomycota_unclassified(100);p__Ascomycota_unclassified(100);p__Ascomycota_unclassified(100);p__Ascomycota_unclassified(100); | - | - | - | - | - | - | - | Unassigned | - |
| Otu62877 | 1 | k__Fungi(100);p__Ascomycota(100);p__Ascomycota_unclassified(100);p__Ascomycota_unclassified(100);p__Ascomycota_unclassified(100);p__Ascomycota_unclassified(100);p__Ascomycota_unclassified(100); | - | - | - | - | - | - | - | Unassigned | - |
| Otu62880 | 1 | k__Fungi(100);k__Fungi_unclassified(100);k__Fungi_unclassified(100);k__Fungi_unclassified(100);k__Fungi_unclassified(100);k__Fungi_unclassified(100);k__Fungi_unclassified(100); | - | - | - | - | - | - | - | Unassigned | - |
| Otu62882 | 1 | k__Fungi(100);p__Ascomycota(100);p__Ascomycota_unclassified(100);p__Ascomycota_unclassified(100);p__Ascomycota_unclassified(100);p__Ascomycota_unclassified(100);p__Ascomycota_unclassified(100); | - | - | - | - | - | - | - | Unassigned | - |
| Otu62883 | 1 | k__Fungi(100);p__Ascomycota(100);p__Ascomycota_unclassified(100);p__Ascomycota_unclassified(100);p__Ascomycota_unclassified(100);p__Ascomycota_unclassified(100);p__Ascomycota_unclassified(100); | - | - | - | - | - | - | - | Unassigned | - |
| Otu62884 | 1 | k__Fungi(100);k__Fungi_unclassified(100);k__Fungi_unclassified(100);k__Fungi_unclassified(100);k__Fungi_unclassified(100);k__Fungi_unclassified(100);k__Fungi_unclassified(100); | - | - | - | - | - | - | - | Unassigned | - |
| Otu62885 | 1 | k__Fungi(100);p__Ascomycota(100);p__Ascomycota_unclassified(100);p__Ascomycota_unclassified(100);p__Ascomycota_unclassified(100);p__Ascomycota_unclassified(100);p__Ascomycota_unclassified(100); | - | - | - | - | - | - | - | Unassigned | - |
| Otu62886 | 1 | k__Fungi(100);k__Fungi_unclassified(100);k__Fungi_unclassified(100);k__Fungi_unclassified(100);k__Fungi_unclassified(100);k__Fungi_unclassified(100);k__Fungi_unclassified(100); | - | - | - | - | - | - | - | Unassigned | - |
| Otu62887 | 1 | k__Fungi(100);k__Fungi_unclassified(100);k__Fungi_unclassified(100);k__Fungi_unclassified(100);k__Fungi_unclassified(100);k__Fungi_unclassified(100);k__Fungi_unclassified(100); | - | - | - | - | - | - | - | Unassigned | - |
| Otu62893 | 1 | k__Fungi(100);k__Fungi_unclassified(100);k__Fungi_unclassified(100);k__Fungi_unclassified(100);k__Fungi_unclassified(100);k__Fungi_unclassified(100);k__Fungi_unclassified(100); | - | - | - | - | - | - | - | Unassigned | - |
| Otu62894 | 1 | k__Fungi(100);k__Fungi_unclassified(100);k__Fungi_unclassified(100);k__Fungi_unclassified(100);k__Fungi_unclassified(100);k__Fungi_unclassified(100);k__Fungi_unclassified(100); | - | - | - | - | - | - | - | Unassigned | - |
| Otu62896 | 1 | k__Fungi(100);p__Ascomycota(100);p__Ascomycota_unclassified(100);p__Ascomycota_unclassified(100);p__Ascomycota_unclassified(100);p__Ascomycota_unclassified(100);p__Ascomycota_unclassified(100); | - | - | - | - | - | - | - | Unassigned | - |
| Otu62899 | 1 | k__Fungi(100);p__Basidiomycota(100);p__Basidiomycota_unclassified(100);p__Basidiomycota_unclassified(100);p__Basidiomycota_unclassified(100);p__Basidiomycota_unclassified(100);p__Basidiomycota_unclassified(100); | - | - | - | - | - | - | - | Unassigned | - |
| Otu62901 | 1 | k__Fungi(100);p__Ascomycota(100);p__Ascomycota_unclassified(100);p__Ascomycota_unclassified(100);p__Ascomycota_unclassified(100);p__Ascomycota_unclassified(100);p__Ascomycota_unclassified(100); | - | - | - | - | - | - | - | Unassigned | - |
| Otu62902 | 1 | k__Fungi(100);k__Fungi_unclassified(100);k__Fungi_unclassified(100);k__Fungi_unclassified(100);k__Fungi_unclassified(100);k__Fungi_unclassified(100);k__Fungi_unclassified(100); | - | - | - | - | - | - | - | Unassigned | - |
| Otu62903 | 1 | k__Fungi(100);p__Ascomycota(100);p__Ascomycota_unclassified(100);p__Ascomycota_unclassified(100);p__Ascomycota_unclassified(100);p__Ascomycota_unclassified(100);p__Ascomycota_unclassified(100); | - | - | - | - | - | - | - | Unassigned | - |
| Otu62906 | 1 | k__Fungi(100);k__Fungi_unclassified(100);k__Fungi_unclassified(100);k__Fungi_unclassified(100);k__Fungi_unclassified(100);k__Fungi_unclassified(100);k__Fungi_unclassified(100); | - | - | - | - | - | - | - | Unassigned | - |
| Otu62910 | 1 | k__Fungi(100);k__Fungi_unclassified(100);k__Fungi_unclassified(100);k__Fungi_unclassified(100);k__Fungi_unclassified(100);k__Fungi_unclassified(100);k__Fungi_unclassified(100); | - | - | - | - | - | - | - | Unassigned | - |
| Otu62913 | 1 | k__Fungi(100);k__Fungi_unclassified(100);k__Fungi_unclassified(100);k__Fungi_unclassified(100);k__Fungi_unclassified(100);k__Fungi_unclassified(100);k__Fungi_unclassified(100); | - | - | - | - | - | - | - | Unassigned | - |
| Otu62914 | 1 | k__Fungi(100);k__Fungi_unclassified(100);k__Fungi_unclassified(100);k__Fungi_unclassified(100);k__Fungi_unclassified(100);k__Fungi_unclassified(100);k__Fungi_unclassified(100); | - | - | - | - | - | - | - | Unassigned | - |
| Otu62915 | 1 | k__Fungi(100);p__Ascomycota(100);p__Ascomycota_unclassified(100);p__Ascomycota_unclassified(100);p__Ascomycota_unclassified(100);p__Ascomycota_unclassified(100);p__Ascomycota_unclassified(100); | - | - | - | - | - | - | - | Unassigned | - |
| Otu62916 | 1 | k__Fungi(100);k__Fungi_unclassified(100);k__Fungi_unclassified(100);k__Fungi_unclassified(100);k__Fungi_unclassified(100);k__Fungi_unclassified(100);k__Fungi_unclassified(100); | - | - | - | - | - | - | - | Unassigned | - |
| Otu62917 | 1 | k__Fungi(100);k__Fungi_unclassified(100);k__Fungi_unclassified(100);k__Fungi_unclassified(100);k__Fungi_unclassified(100);k__Fungi_unclassified(100);k__Fungi_unclassified(100); | - | - | - | - | - | - | - | Unassigned | - |
| Otu62918 | 1 | k__Fungi(100);p__Ascomycota(100);p__Ascomycota_unclassified(100);p__Ascomycota_unclassified(100);p__Ascomycota_unclassified(100);p__Ascomycota_unclassified(100);p__Ascomycota_unclassified(100); | - | - | - | - | - | - | - | Unassigned | - |
| Otu62919 | 1 | k__Fungi(100);k__Fungi_unclassified(100);k__Fungi_unclassified(100);k__Fungi_unclassified(100);k__Fungi_unclassified(100);k__Fungi_unclassified(100);k__Fungi_unclassified(100); | - | - | - | - | - | - | - | Unassigned | - |
| Otu62920 | 1 | k__Fungi(100);k__Fungi_unclassified(100);k__Fungi_unclassified(100);k__Fungi_unclassified(100);k__Fungi_unclassified(100);k__Fungi_unclassified(100);k__Fungi_unclassified(100); | - | - | - | - | - | - | - | Unassigned | - |
| Otu62921 | 1 | k__Fungi(100);k__Fungi_unclassified(100);k__Fungi_unclassified(100);k__Fungi_unclassified(100);k__Fungi_unclassified(100);k__Fungi_unclassified(100);k__Fungi_unclassified(100); | - | - | - | - | - | - | - | Unassigned | - |
| Otu62922 | 1 | k__Fungi(100);k__Fungi_unclassified(100);k__Fungi_unclassified(100);k__Fungi_unclassified(100);k__Fungi_unclassified(100);k__Fungi_unclassified(100);k__Fungi_unclassified(100); | - | - | - | - | - | - | - | Unassigned | - |
| Otu62923 | 1 | k__Fungi(100);k__Fungi_unclassified(100);k__Fungi_unclassified(100);k__Fungi_unclassified(100);k__Fungi_unclassified(100);k__Fungi_unclassified(100);k__Fungi_unclassified(100); | - | - | - | - | - | - | - | Unassigned | - |
| Otu62924 | 1 | k__Fungi(100);k__Fungi_unclassified(100);k__Fungi_unclassified(100);k__Fungi_unclassified(100);k__Fungi_unclassified(100);k__Fungi_unclassified(100);k__Fungi_unclassified(100); | - | - | - | - | - | - | - | Unassigned | - |
| Otu62926 | 1 | k__Fungi(100);k__Fungi_unclassified(100);k__Fungi_unclassified(100);k__Fungi_unclassified(100);k__Fungi_unclassified(100);k__Fungi_unclassified(100);k__Fungi_unclassified(100); | - | - | - | - | - | - | - | Unassigned | - |
| Otu62927 | 1 | k__Fungi(100);k__Fungi_unclassified(100);k__Fungi_unclassified(100);k__Fungi_unclassified(100);k__Fungi_unclassified(100);k__Fungi_unclassified(100);k__Fungi_unclassified(100); | - | - | - | - | - | - | - | Unassigned | - |
| Otu62928 | 1 | k__Fungi(100);k__Fungi_unclassified(100);k__Fungi_unclassified(100);k__Fungi_unclassified(100);k__Fungi_unclassified(100);k__Fungi_unclassified(100);k__Fungi_unclassified(100); | - | - | - | - | - | - | - | Unassigned | - |
| Otu62929 | 1 | k__Fungi(100);k__Fungi_unclassified(100);k__Fungi_unclassified(100);k__Fungi_unclassified(100);k__Fungi_unclassified(100);k__Fungi_unclassified(100);k__Fungi_unclassified(100); | - | - | - | - | - | - | - | Unassigned | - |
| Otu62930 | 1 | k__Fungi(100);k__Fungi_unclassified(100);k__Fungi_unclassified(100);k__Fungi_unclassified(100);k__Fungi_unclassified(100);k__Fungi_unclassified(100);k__Fungi_unclassified(100); | - | - | - | - | - | - | - | Unassigned | - |
| Otu62934 | 1 | k__Fungi(100);k__Fungi_unclassified(100);k__Fungi_unclassified(100);k__Fungi_unclassified(100);k__Fungi_unclassified(100);k__Fungi_unclassified(100);k__Fungi_unclassified(100); | - | - | - | - | - | - | - | Unassigned | - |
| Otu62935 | 1 | k__Fungi(100);k__Fungi_unclassified(100);k__Fungi_unclassified(100);k__Fungi_unclassified(100);k__Fungi_unclassified(100);k__Fungi_unclassified(100);k__Fungi_unclassified(100); | - | - | - | - | - | - | - | Unassigned | - |
| Otu62936 | 1 | k__Fungi(100);k__Fungi_unclassified(100);k__Fungi_unclassified(100);k__Fungi_unclassified(100);k__Fungi_unclassified(100);k__Fungi_unclassified(100);k__Fungi_unclassified(100); | - | - | - | - | - | - | - | Unassigned | - |
| Otu62937 | 1 | k__Fungi(100);k__Fungi_unclassified(100);k__Fungi_unclassified(100);k__Fungi_unclassified(100);k__Fungi_unclassified(100);k__Fungi_unclassified(100);k__Fungi_unclassified(100); | - | - | - | - | - | - | - | Unassigned | - |
| Otu62940 | 1 | k__Fungi(100);k__Fungi_unclassified(100);k__Fungi_unclassified(100);k__Fungi_unclassified(100);k__Fungi_unclassified(100);k__Fungi_unclassified(100);k__Fungi_unclassified(100); | - | - | - | - | - | - | - | Unassigned | - |
| Otu62941 | 1 | k__Fungi(100);p__Ascomycota(100);p__Ascomycota_unclassified(100);p__Ascomycota_unclassified(100);p__Ascomycota_unclassified(100);p__Ascomycota_unclassified(100);p__Ascomycota_unclassified(100); | - | - | - | - | - | - | - | Unassigned | - |
| Otu62944 | 1 | k__Fungi(100);p__Ascomycota(100);p__Ascomycota_unclassified(100);p__Ascomycota_unclassified(100);p__Ascomycota_unclassified(100);p__Ascomycota_unclassified(100);p__Ascomycota_unclassified(100); | - | - | - | - | - | - | - | Unassigned | - |
| Otu62946 | 1 | k__Fungi(100);k__Fungi_unclassified(100);k__Fungi_unclassified(100);k__Fungi_unclassified(100);k__Fungi_unclassified(100);k__Fungi_unclassified(100);k__Fungi_unclassified(100); | - | - | - | - | - | - | - | Unassigned | - |
| Otu62947 | 1 | k__Fungi(100);p__Ascomycota(100);p__Ascomycota_unclassified(100);p__Ascomycota_unclassified(100);p__Ascomycota_unclassified(100);p__Ascomycota_unclassified(100);p__Ascomycota_unclassified(100); | - | - | - | - | - | - | - | Unassigned | - |
| Otu62948 | 1 | k__Fungi(100);k__Fungi_unclassified(100);k__Fungi_unclassified(100);k__Fungi_unclassified(100);k__Fungi_unclassified(100);k__Fungi_unclassified(100);k__Fungi_unclassified(100); | - | - | - | - | - | - | - | Unassigned | - |
| Otu62949 | 1 | k__Fungi(100);k__Fungi_unclassified(100);k__Fungi_unclassified(100);k__Fungi_unclassified(100);k__Fungi_unclassified(100);k__Fungi_unclassified(100);k__Fungi_unclassified(100); | - | - | - | - | - | - | - | Unassigned | - |
| Otu62952 | 1 | k__Fungi(100);p__Ascomycota(100);p__Ascomycota_unclassified(100);p__Ascomycota_unclassified(100);p__Ascomycota_unclassified(100);p__Ascomycota_unclassified(100);p__Ascomycota_unclassified(100); | - | - | - | - | - | - | - | Unassigned | - |
| Otu62957 | 1 | k__Fungi(100);k__Fungi_unclassified(100);k__Fungi_unclassified(100);k__Fungi_unclassified(100);k__Fungi_unclassified(100);k__Fungi_unclassified(100);k__Fungi_unclassified(100); | - | - | - | - | - | - | - | Unassigned | - |
| Otu62958 | 1 | k__Fungi(100);k__Fungi_unclassified(100);k__Fungi_unclassified(100);k__Fungi_unclassified(100);k__Fungi_unclassified(100);k__Fungi_unclassified(100);k__Fungi_unclassified(100); | - | - | - | - | - | - | - | Unassigned | - |
| Otu62962 | 1 | k__Fungi(100);p__Ascomycota(100);p__Ascomycota_unclassified(100);p__Ascomycota_unclassified(100);p__Ascomycota_unclassified(100);p__Ascomycota_unclassified(100);p__Ascomycota_unclassified(100); | - | - | - | - | - | - | - | Unassigned | - |
| Otu62963 | 1 | k__Fungi(100);p__Ascomycota(100);p__Ascomycota_unclassified(100);p__Ascomycota_unclassified(100);p__Ascomycota_unclassified(100);p__Ascomycota_unclassified(100);p__Ascomycota_unclassified(100); | - | - | - | - | - | - | - | Unassigned | - |
| Otu62964 | 1 | k__Fungi(100);k__Fungi_unclassified(100);k__Fungi_unclassified(100);k__Fungi_unclassified(100);k__Fungi_unclassified(100);k__Fungi_unclassified(100);k__Fungi_unclassified(100); | - | - | - | - | - | - | - | Unassigned | - |
| Otu62981 | 1 | k__Fungi(100);p__Ascomycota(100);p__Ascomycota_unclassified(100);p__Ascomycota_unclassified(100);p__Ascomycota_unclassified(100);p__Ascomycota_unclassified(100);p__Ascomycota_unclassified(100); | - | - | - | - | - | - | - | Unassigned | - |
| Otu62983 | 1 | k__Fungi(100);p__Ascomycota(100);p__Ascomycota_unclassified(100);p__Ascomycota_unclassified(100);p__Ascomycota_unclassified(100);p__Ascomycota_unclassified(100);p__Ascomycota_unclassified(100); | - | - | - | - | - | - | - | Unassigned | - |
| Otu62984 | 1 | k__Fungi(100);k__Fungi_unclassified(100);k__Fungi_unclassified(100);k__Fungi_unclassified(100);k__Fungi_unclassified(100);k__Fungi_unclassified(100);k__Fungi_unclassified(100); | - | - | - | - | - | - | - | Unassigned | - |
| Otu62987 | 1 | k__Fungi(100);p__Ascomycota(100);p__Ascomycota_unclassified(100);p__Ascomycota_unclassified(100);p__Ascomycota_unclassified(100);p__Ascomycota_unclassified(100);p__Ascomycota_unclassified(100); | - | - | - | - | - | - | - | Unassigned | - |
| Otu62988 | 1 | k__Fungi(100);k__Fungi_unclassified(100);k__Fungi_unclassified(100);k__Fungi_unclassified(100);k__Fungi_unclassified(100);k__Fungi_unclassified(100);k__Fungi_unclassified(100); | - | - | - | - | - | - | - | Unassigned | - |
| Otu62990 | 1 | k__Fungi(100);p__Ascomycota(100);p__Ascomycota_unclassified(100);p__Ascomycota_unclassified(100);p__Ascomycota_unclassified(100);p__Ascomycota_unclassified(100);p__Ascomycota_unclassified(100); | - | - | - | - | - | - | - | Unassigned | - |
| Otu62992 | 1 | k__Fungi(100);k__Fungi_unclassified(100);k__Fungi_unclassified(100);k__Fungi_unclassified(100);k__Fungi_unclassified(100);k__Fungi_unclassified(100);k__Fungi_unclassified(100); | - | - | - | - | - | - | - | Unassigned | - |
| Otu62994 | 1 | k__Fungi(100);k__Fungi_unclassified(100);k__Fungi_unclassified(100);k__Fungi_unclassified(100);k__Fungi_unclassified(100);k__Fungi_unclassified(100);k__Fungi_unclassified(100); | - | - | - | - | - | - | - | Unassigned | - |
| Otu62995 | 1 | k__Fungi(100);p__Ascomycota(100);p__Ascomycota_unclassified(100);p__Ascomycota_unclassified(100);p__Ascomycota_unclassified(100);p__Ascomycota_unclassified(100);p__Ascomycota_unclassified(100); | - | - | - | - | - | - | - | Unassigned | - |
| Otu62997 | 1 | k__Fungi(100);k__Fungi_unclassified(100);k__Fungi_unclassified(100);k__Fungi_unclassified(100);k__Fungi_unclassified(100);k__Fungi_unclassified(100);k__Fungi_unclassified(100); | - | - | - | - | - | - | - | Unassigned | - |
| Otu62998 | 1 | k__Fungi(100);p__Ascomycota(100);p__Ascomycota_unclassified(100);p__Ascomycota_unclassified(100);p__Ascomycota_unclassified(100);p__Ascomycota_unclassified(100);p__Ascomycota_unclassified(100); | - | - | - | - | - | - | - | Unassigned | - |
| Otu62999 | 1 | k__Fungi(100);k__Fungi_unclassified(100);k__Fungi_unclassified(100);k__Fungi_unclassified(100);k__Fungi_unclassified(100);k__Fungi_unclassified(100);k__Fungi_unclassified(100); | - | - | - | - | - | - | - | Unassigned | - |
| Otu63000 | 1 | k__Fungi(100);k__Fungi_unclassified(100);k__Fungi_unclassified(100);k__Fungi_unclassified(100);k__Fungi_unclassified(100);k__Fungi_unclassified(100);k__Fungi_unclassified(100); | - | - | - | - | - | - | - | Unassigned | - |
| Otu63001 | 1 | k__Fungi(100);p__Ascomycota(100);p__Ascomycota_unclassified(100);p__Ascomycota_unclassified(100);p__Ascomycota_unclassified(100);p__Ascomycota_unclassified(100);p__Ascomycota_unclassified(100); | - | - | - | - | - | - | - | Unassigned | - |
| Otu63002 | 1 | k__Fungi(100);k__Fungi_unclassified(100);k__Fungi_unclassified(100);k__Fungi_unclassified(100);k__Fungi_unclassified(100);k__Fungi_unclassified(100);k__Fungi_unclassified(100); | - | - | - | - | - | - | - | Unassigned | - |
| Otu63004 | 1 | k__Fungi(100);k__Fungi_unclassified(100);k__Fungi_unclassified(100);k__Fungi_unclassified(100);k__Fungi_unclassified(100);k__Fungi_unclassified(100);k__Fungi_unclassified(100); | - | - | - | - | - | - | - | Unassigned | - |
| Otu63005 | 1 | k__Fungi(100);k__Fungi_unclassified(100);k__Fungi_unclassified(100);k__Fungi_unclassified(100);k__Fungi_unclassified(100);k__Fungi_unclassified(100);k__Fungi_unclassified(100); | - | - | - | - | - | - | - | Unassigned | - |
| Otu63006 | 1 | k__Fungi(100);p__Ascomycota(100);p__Ascomycota_unclassified(100);p__Ascomycota_unclassified(100);p__Ascomycota_unclassified(100);p__Ascomycota_unclassified(100);p__Ascomycota_unclassified(100); | - | - | - | - | - | - | - | Unassigned | - |
| Otu63009 | 1 | k__Fungi(100);k__Fungi_unclassified(100);k__Fungi_unclassified(100);k__Fungi_unclassified(100);k__Fungi_unclassified(100);k__Fungi_unclassified(100);k__Fungi_unclassified(100); | - | - | - | - | - | - | - | Unassigned | - |
| Otu63010 | 1 | k__Fungi(100);p__Ascomycota(100);p__Ascomycota_unclassified(100);p__Ascomycota_unclassified(100);p__Ascomycota_unclassified(100);p__Ascomycota_unclassified(100);p__Ascomycota_unclassified(100); | - | - | - | - | - | - | - | Unassigned | - |
| Otu63011 | 1 | k__Fungi(100);k__Fungi_unclassified(100);k__Fungi_unclassified(100);k__Fungi_unclassified(100);k__Fungi_unclassified(100);k__Fungi_unclassified(100);k__Fungi_unclassified(100); | - | - | - | - | - | - | - | Unassigned | - |
| Otu63013 | 1 | k__Fungi(100);p__Ascomycota(100);p__Ascomycota_unclassified(100);p__Ascomycota_unclassified(100);p__Ascomycota_unclassified(100);p__Ascomycota_unclassified(100);p__Ascomycota_unclassified(100); | - | - | - | - | - | - | - | Unassigned | - |
| Otu63016 | 1 | k__Fungi(100);p__Ascomycota(100);p__Ascomycota_unclassified(100);p__Ascomycota_unclassified(100);p__Ascomycota_unclassified(100);p__Ascomycota_unclassified(100);p__Ascomycota_unclassified(100); | - | - | - | - | - | - | - | Unassigned | - |
| Otu63017 | 1 | k__Fungi(100);k__Fungi_unclassified(100);k__Fungi_unclassified(100);k__Fungi_unclassified(100);k__Fungi_unclassified(100);k__Fungi_unclassified(100);k__Fungi_unclassified(100); | - | - | - | - | - | - | - | Unassigned | - |
| Otu63018 | 1 | k__Fungi(100);p__Ascomycota(100);p__Ascomycota_unclassified(100);p__Ascomycota_unclassified(100);p__Ascomycota_unclassified(100);p__Ascomycota_unclassified(100);p__Ascomycota_unclassified(100); | - | - | - | - | - | - | - | Unassigned | - |
| Otu63019 | 1 | k__Fungi(100);k__Fungi_unclassified(100);k__Fungi_unclassified(100);k__Fungi_unclassified(100);k__Fungi_unclassified(100);k__Fungi_unclassified(100);k__Fungi_unclassified(100); | - | - | - | - | - | - | - | Unassigned | - |
| Otu63020 | 1 | k__Fungi(100);k__Fungi_unclassified(100);k__Fungi_unclassified(100);k__Fungi_unclassified(100);k__Fungi_unclassified(100);k__Fungi_unclassified(100);k__Fungi_unclassified(100); | - | - | - | - | - | - | - | Unassigned | - |
| Otu63021 | 1 | k__Fungi(100);k__Fungi_unclassified(100);k__Fungi_unclassified(100);k__Fungi_unclassified(100);k__Fungi_unclassified(100);k__Fungi_unclassified(100);k__Fungi_unclassified(100); | - | - | - | - | - | - | - | Unassigned | - |
| Otu63022 | 1 | k__Fungi(100);p__Ascomycota(100);p__Ascomycota_unclassified(100);p__Ascomycota_unclassified(100);p__Ascomycota_unclassified(100);p__Ascomycota_unclassified(100);p__Ascomycota_unclassified(100); | - | - | - | - | - | - | - | Unassigned | - |
| Otu63023 | 1 | k__Fungi(100);k__Fungi_unclassified(100);k__Fungi_unclassified(100);k__Fungi_unclassified(100);k__Fungi_unclassified(100);k__Fungi_unclassified(100);k__Fungi_unclassified(100); | - | - | - | - | - | - | - | Unassigned | - |
| Otu63024 | 1 | k__Fungi(100);p__Ascomycota(100);p__Ascomycota_unclassified(100);p__Ascomycota_unclassified(100);p__Ascomycota_unclassified(100);p__Ascomycota_unclassified(100);p__Ascomycota_unclassified(100); | - | - | - | - | - | - | - | Unassigned | - |
| Otu63025 | 1 | k__Fungi(100);k__Fungi_unclassified(100);k__Fungi_unclassified(100);k__Fungi_unclassified(100);k__Fungi_unclassified(100);k__Fungi_unclassified(100);k__Fungi_unclassified(100); | - | - | - | - | - | - | - | Unassigned | - |
| Otu63026 | 1 | k__Fungi(100);p__Ascomycota(100);p__Ascomycota_unclassified(100);p__Ascomycota_unclassified(100);p__Ascomycota_unclassified(100);p__Ascomycota_unclassified(100);p__Ascomycota_unclassified(100); | - | - | - | - | - | - | - | Unassigned | - |
| Otu63028 | 1 | k__Fungi(100);p__Ascomycota(100);p__Ascomycota_unclassified(100);p__Ascomycota_unclassified(100);p__Ascomycota_unclassified(100);p__Ascomycota_unclassified(100);p__Ascomycota_unclassified(100); | - | - | - | - | - | - | - | Unassigned | - |
| Otu63029 | 1 | k__Fungi(100);p__Ascomycota(100);p__Ascomycota_unclassified(100);p__Ascomycota_unclassified(100);p__Ascomycota_unclassified(100);p__Ascomycota_unclassified(100);p__Ascomycota_unclassified(100); | - | - | - | - | - | - | - | Unassigned | - |
| Otu63030 | 1 | k__Fungi(100);k__Fungi_unclassified(100);k__Fungi_unclassified(100);k__Fungi_unclassified(100);k__Fungi_unclassified(100);k__Fungi_unclassified(100);k__Fungi_unclassified(100); | - | - | - | - | - | - | - | Unassigned | - |
| Otu63031 | 1 | k__Fungi(100);k__Fungi_unclassified(100);k__Fungi_unclassified(100);k__Fungi_unclassified(100);k__Fungi_unclassified(100);k__Fungi_unclassified(100);k__Fungi_unclassified(100); | - | - | - | - | - | - | - | Unassigned | - |
| Otu63032 | 1 | k__Fungi(100);k__Fungi_unclassified(100);k__Fungi_unclassified(100);k__Fungi_unclassified(100);k__Fungi_unclassified(100);k__Fungi_unclassified(100);k__Fungi_unclassified(100); | - | - | - | - | - | - | - | Unassigned | - |
| Otu63033 | 1 | k__Fungi(100);k__Fungi_unclassified(100);k__Fungi_unclassified(100);k__Fungi_unclassified(100);k__Fungi_unclassified(100);k__Fungi_unclassified(100);k__Fungi_unclassified(100); | - | - | - | - | - | - | - | Unassigned | - |
| Otu63034 | 1 | k__Fungi(100);k__Fungi_unclassified(100);k__Fungi_unclassified(100);k__Fungi_unclassified(100);k__Fungi_unclassified(100);k__Fungi_unclassified(100);k__Fungi_unclassified(100); | - | - | - | - | - | - | - | Unassigned | - |
| Otu63036 | 1 | k__Fungi(100);k__Fungi_unclassified(100);k__Fungi_unclassified(100);k__Fungi_unclassified(100);k__Fungi_unclassified(100);k__Fungi_unclassified(100);k__Fungi_unclassified(100); | - | - | - | - | - | - | - | Unassigned | - |
| Otu63037 | 1 | k__Fungi(100);k__Fungi_unclassified(100);k__Fungi_unclassified(100);k__Fungi_unclassified(100);k__Fungi_unclassified(100);k__Fungi_unclassified(100);k__Fungi_unclassified(100); | - | - | - | - | - | - | - | Unassigned | - |
| Otu63040 | 1 | k__Fungi(100);k__Fungi_unclassified(100);k__Fungi_unclassified(100);k__Fungi_unclassified(100);k__Fungi_unclassified(100);k__Fungi_unclassified(100);k__Fungi_unclassified(100); | - | - | - | - | - | - | - | Unassigned | - |
| Otu63042 | 1 | k__Fungi(100);p__Ascomycota(100);p__Ascomycota_unclassified(100);p__Ascomycota_unclassified(100);p__Ascomycota_unclassified(100);p__Ascomycota_unclassified(100);p__Ascomycota_unclassified(100); | - | - | - | - | - | - | - | Unassigned | - |
| Otu63044 | 1 | k__Fungi(100);p__Ascomycota(100);p__Ascomycota_unclassified(100);p__Ascomycota_unclassified(100);p__Ascomycota_unclassified(100);p__Ascomycota_unclassified(100);p__Ascomycota_unclassified(100); | - | - | - | - | - | - | - | Unassigned | - |
| Otu63047 | 1 | k__Fungi(100);k__Fungi_unclassified(100);k__Fungi_unclassified(100);k__Fungi_unclassified(100);k__Fungi_unclassified(100);k__Fungi_unclassified(100);k__Fungi_unclassified(100); | - | - | - | - | - | - | - | Unassigned | - |
| Otu63048 | 1 | k__Fungi(100);k__Fungi_unclassified(100);k__Fungi_unclassified(100);k__Fungi_unclassified(100);k__Fungi_unclassified(100);k__Fungi_unclassified(100);k__Fungi_unclassified(100); | - | - | - | - | - | - | - | Unassigned | - |
| Otu63049 | 1 | k__Fungi(100);p__Ascomycota(100);p__Ascomycota_unclassified(100);p__Ascomycota_unclassified(100);p__Ascomycota_unclassified(100);p__Ascomycota_unclassified(100);p__Ascomycota_unclassified(100); | - | - | - | - | - | - | - | Unassigned | - |
| Otu63050 | 1 | k__Fungi(100);p__Ascomycota(100);p__Ascomycota_unclassified(100);p__Ascomycota_unclassified(100);p__Ascomycota_unclassified(100);p__Ascomycota_unclassified(100);p__Ascomycota_unclassified(100); | - | - | - | - | - | - | - | Unassigned | - |
| Otu63051 | 1 | k__Fungi(100);k__Fungi_unclassified(100);k__Fungi_unclassified(100);k__Fungi_unclassified(100);k__Fungi_unclassified(100);k__Fungi_unclassified(100);k__Fungi_unclassified(100); | - | - | - | - | - | - | - | Unassigned | - |
| Otu63052 | 1 | k__Fungi(100);k__Fungi_unclassified(100);k__Fungi_unclassified(100);k__Fungi_unclassified(100);k__Fungi_unclassified(100);k__Fungi_unclassified(100);k__Fungi_unclassified(100); | - | - | - | - | - | - | - | Unassigned | - |
| Otu63053 | 1 | k__Fungi(100);k__Fungi_unclassified(100);k__Fungi_unclassified(100);k__Fungi_unclassified(100);k__Fungi_unclassified(100);k__Fungi_unclassified(100);k__Fungi_unclassified(100); | - | - | - | - | - | - | - | Unassigned | - |
| Otu63054 | 1 | k__Fungi(100);k__Fungi_unclassified(100);k__Fungi_unclassified(100);k__Fungi_unclassified(100);k__Fungi_unclassified(100);k__Fungi_unclassified(100);k__Fungi_unclassified(100); | - | - | - | - | - | - | - | Unassigned | - |
| Otu63056 | 1 | k__Fungi(100);p__Ascomycota(100);p__Ascomycota_unclassified(100);p__Ascomycota_unclassified(100);p__Ascomycota_unclassified(100);p__Ascomycota_unclassified(100);p__Ascomycota_unclassified(100); | - | - | - | - | - | - | - | Unassigned | - |
| Otu63057 | 1 | k__Fungi(100);p__Ascomycota(100);p__Ascomycota_unclassified(100);p__Ascomycota_unclassified(100);p__Ascomycota_unclassified(100);p__Ascomycota_unclassified(100);p__Ascomycota_unclassified(100); | - | - | - | - | - | - | - | Unassigned | - |
| Otu63058 | 1 | k__Fungi(100);p__Ascomycota(100);p__Ascomycota_unclassified(100);p__Ascomycota_unclassified(100);p__Ascomycota_unclassified(100);p__Ascomycota_unclassified(100);p__Ascomycota_unclassified(100); | - | - | - | - | - | - | - | Unassigned | - |
| Otu63059 | 1 | k__Fungi(100);k__Fungi_unclassified(100);k__Fungi_unclassified(100);k__Fungi_unclassified(100);k__Fungi_unclassified(100);k__Fungi_unclassified(100);k__Fungi_unclassified(100); | - | - | - | - | - | - | - | Unassigned | - |
| Otu63060 | 1 | k__Fungi(100);p__Ascomycota(100);p__Ascomycota_unclassified(100);p__Ascomycota_unclassified(100);p__Ascomycota_unclassified(100);p__Ascomycota_unclassified(100);p__Ascomycota_unclassified(100); | - | - | - | - | - | - | - | Unassigned | - |
| Otu63061 | 1 | k__Fungi(100);k__Fungi_unclassified(100);k__Fungi_unclassified(100);k__Fungi_unclassified(100);k__Fungi_unclassified(100);k__Fungi_unclassified(100);k__Fungi_unclassified(100); | - | - | - | - | - | - | - | Unassigned | - |
| Otu63062 | 1 | k__Fungi(100);p__Ascomycota(100);p__Ascomycota_unclassified(100);p__Ascomycota_unclassified(100);p__Ascomycota_unclassified(100);p__Ascomycota_unclassified(100);p__Ascomycota_unclassified(100); | - | - | - | - | - | - | - | Unassigned | - |
| Otu63063 | 1 | k__Fungi(100);k__Fungi_unclassified(100);k__Fungi_unclassified(100);k__Fungi_unclassified(100);k__Fungi_unclassified(100);k__Fungi_unclassified(100);k__Fungi_unclassified(100); | - | - | - | - | - | - | - | Unassigned | - |
| Otu63064 | 1 | k__Fungi(100);k__Fungi_unclassified(100);k__Fungi_unclassified(100);k__Fungi_unclassified(100);k__Fungi_unclassified(100);k__Fungi_unclassified(100);k__Fungi_unclassified(100); | - | - | - | - | - | - | - | Unassigned | - |
| Otu63065 | 1 | k__Fungi(100);p__Ascomycota(100);p__Ascomycota_unclassified(100);p__Ascomycota_unclassified(100);p__Ascomycota_unclassified(100);p__Ascomycota_unclassified(100);p__Ascomycota_unclassified(100); | - | - | - | - | - | - | - | Unassigned | - |
| Otu63066 | 1 | k__Fungi(100);k__Fungi_unclassified(100);k__Fungi_unclassified(100);k__Fungi_unclassified(100);k__Fungi_unclassified(100);k__Fungi_unclassified(100);k__Fungi_unclassified(100); | - | - | - | - | - | - | - | Unassigned | - |
| Otu63067 | 1 | k__Fungi(100);k__Fungi_unclassified(100);k__Fungi_unclassified(100);k__Fungi_unclassified(100);k__Fungi_unclassified(100);k__Fungi_unclassified(100);k__Fungi_unclassified(100); | - | - | - | - | - | - | - | Unassigned | - |
| Otu63069 | 1 | k__Fungi(100);p__Ascomycota(100);p__Ascomycota_unclassified(100);p__Ascomycota_unclassified(100);p__Ascomycota_unclassified(100);p__Ascomycota_unclassified(100);p__Ascomycota_unclassified(100); | - | - | - | - | - | - | - | Unassigned | - |
| Otu63071 | 1 | k__Fungi(100);p__Ascomycota(100);p__Ascomycota_unclassified(100);p__Ascomycota_unclassified(100);p__Ascomycota_unclassified(100);p__Ascomycota_unclassified(100);p__Ascomycota_unclassified(100); | - | - | - | - | - | - | - | Unassigned | - |
| Otu63072 | 1 | k__Fungi(100);p__Ascomycota(100);p__Ascomycota_unclassified(100);p__Ascomycota_unclassified(100);p__Ascomycota_unclassified(100);p__Ascomycota_unclassified(100);p__Ascomycota_unclassified(100); | - | - | - | - | - | - | - | Unassigned | - |
| Otu63073 | 1 | k__Fungi(100);p__Ascomycota(100);p__Ascomycota_unclassified(100);p__Ascomycota_unclassified(100);p__Ascomycota_unclassified(100);p__Ascomycota_unclassified(100);p__Ascomycota_unclassified(100); | - | - | - | - | - | - | - | Unassigned | - |
| Otu63074 | 1 | k__Fungi(100);p__Ascomycota(100);p__Ascomycota_unclassified(100);p__Ascomycota_unclassified(100);p__Ascomycota_unclassified(100);p__Ascomycota_unclassified(100);p__Ascomycota_unclassified(100); | - | - | - | - | - | - | - | Unassigned | - |
| Otu63076 | 1 | k__Fungi(100);k__Fungi_unclassified(100);k__Fungi_unclassified(100);k__Fungi_unclassified(100);k__Fungi_unclassified(100);k__Fungi_unclassified(100);k__Fungi_unclassified(100); | - | - | - | - | - | - | - | Unassigned | - |
| Otu63078 | 1 | k__Fungi(100);k__Fungi_unclassified(100);k__Fungi_unclassified(100);k__Fungi_unclassified(100);k__Fungi_unclassified(100);k__Fungi_unclassified(100);k__Fungi_unclassified(100); | - | - | - | - | - | - | - | Unassigned | - |
| Otu63079 | 1 | k__Fungi(100);k__Fungi_unclassified(100);k__Fungi_unclassified(100);k__Fungi_unclassified(100);k__Fungi_unclassified(100);k__Fungi_unclassified(100);k__Fungi_unclassified(100); | - | - | - | - | - | - | - | Unassigned | - |
| Otu63081 | 1 | k__Fungi(100);k__Fungi_unclassified(100);k__Fungi_unclassified(100);k__Fungi_unclassified(100);k__Fungi_unclassified(100);k__Fungi_unclassified(100);k__Fungi_unclassified(100); | - | - | - | - | - | - | - | Unassigned | - |
| Otu63082 | 1 | k__Fungi(100);k__Fungi_unclassified(100);k__Fungi_unclassified(100);k__Fungi_unclassified(100);k__Fungi_unclassified(100);k__Fungi_unclassified(100);k__Fungi_unclassified(100); | - | - | - | - | - | - | - | Unassigned | - |
| Otu63083 | 1 | k__Fungi(100);k__Fungi_unclassified(100);k__Fungi_unclassified(100);k__Fungi_unclassified(100);k__Fungi_unclassified(100);k__Fungi_unclassified(100);k__Fungi_unclassified(100); | - | - | - | - | - | - | - | Unassigned | - |
| Otu63085 | 1 | k__Fungi(100);p__Ascomycota(100);p__Ascomycota_unclassified(100);p__Ascomycota_unclassified(100);p__Ascomycota_unclassified(100);p__Ascomycota_unclassified(100);p__Ascomycota_unclassified(100); | - | - | - | - | - | - | - | Unassigned | - |
| Otu63087 | 1 | k__Fungi(100);k__Fungi_unclassified(100);k__Fungi_unclassified(100);k__Fungi_unclassified(100);k__Fungi_unclassified(100);k__Fungi_unclassified(100);k__Fungi_unclassified(100); | - | - | - | - | - | - | - | Unassigned | - |
| Otu63089 | 1 | k__Fungi(100);p__Ascomycota(100);p__Ascomycota_unclassified(100);p__Ascomycota_unclassified(100);p__Ascomycota_unclassified(100);p__Ascomycota_unclassified(100);p__Ascomycota_unclassified(100); | - | - | - | - | - | - | - | Unassigned | - |
| Otu63091 | 1 | k__Fungi(100);k__Fungi_unclassified(100);k__Fungi_unclassified(100);k__Fungi_unclassified(100);k__Fungi_unclassified(100);k__Fungi_unclassified(100);k__Fungi_unclassified(100); | - | - | - | - | - | - | - | Unassigned | - |
| Otu63092 | 1 | k__Fungi(100);k__Fungi_unclassified(100);k__Fungi_unclassified(100);k__Fungi_unclassified(100);k__Fungi_unclassified(100);k__Fungi_unclassified(100);k__Fungi_unclassified(100); | - | - | - | - | - | - | - | Unassigned | - |
| Otu63095 | 1 | k__Fungi(100);k__Fungi_unclassified(100);k__Fungi_unclassified(100);k__Fungi_unclassified(100);k__Fungi_unclassified(100);k__Fungi_unclassified(100);k__Fungi_unclassified(100); | - | - | - | - | - | - | - | Unassigned | - |
| Otu63096 | 1 | k__Fungi(100);k__Fungi_unclassified(100);k__Fungi_unclassified(100);k__Fungi_unclassified(100);k__Fungi_unclassified(100);k__Fungi_unclassified(100);k__Fungi_unclassified(100); | - | - | - | - | - | - | - | Unassigned | - |
| Otu63097 | 1 | k__Fungi(100);k__Fungi_unclassified(100);k__Fungi_unclassified(100);k__Fungi_unclassified(100);k__Fungi_unclassified(100);k__Fungi_unclassified(100);k__Fungi_unclassified(100); | - | - | - | - | - | - | - | Unassigned | - |
| Otu63098 | 1 | k__Fungi(100);k__Fungi_unclassified(100);k__Fungi_unclassified(100);k__Fungi_unclassified(100);k__Fungi_unclassified(100);k__Fungi_unclassified(100);k__Fungi_unclassified(100); | - | - | - | - | - | - | - | Unassigned | - |
| Otu63099 | 1 | k__Fungi(100);k__Fungi_unclassified(100);k__Fungi_unclassified(100);k__Fungi_unclassified(100);k__Fungi_unclassified(100);k__Fungi_unclassified(100);k__Fungi_unclassified(100); | - | - | - | - | - | - | - | Unassigned | - |
| Otu63100 | 1 | k__Fungi(100);p__Ascomycota(100);p__Ascomycota_unclassified(100);p__Ascomycota_unclassified(100);p__Ascomycota_unclassified(100);p__Ascomycota_unclassified(100);p__Ascomycota_unclassified(100); | - | - | - | - | - | - | - | Unassigned | - |
| Otu63101 | 1 | k__Fungi(100);p__Ascomycota(100);p__Ascomycota_unclassified(100);p__Ascomycota_unclassified(100);p__Ascomycota_unclassified(100);p__Ascomycota_unclassified(100);p__Ascomycota_unclassified(100); | - | - | - | - | - | - | - | Unassigned | - |
| Otu63102 | 1 | k__Fungi(100);k__Fungi_unclassified(100);k__Fungi_unclassified(100);k__Fungi_unclassified(100);k__Fungi_unclassified(100);k__Fungi_unclassified(100);k__Fungi_unclassified(100); | - | - | - | - | - | - | - | Unassigned | - |
| Otu63103 | 1 | k__Fungi(100);k__Fungi_unclassified(100);k__Fungi_unclassified(100);k__Fungi_unclassified(100);k__Fungi_unclassified(100);k__Fungi_unclassified(100);k__Fungi_unclassified(100); | - | - | - | - | - | - | - | Unassigned | - |
| Otu63106 | 1 | k__Fungi(100);k__Fungi_unclassified(100);k__Fungi_unclassified(100);k__Fungi_unclassified(100);k__Fungi_unclassified(100);k__Fungi_unclassified(100);k__Fungi_unclassified(100); | - | - | - | - | - | - | - | Unassigned | - |
| Otu63118 | 1 | k__Fungi(100);p__Ascomycota(100);p__Ascomycota_unclassified(100);p__Ascomycota_unclassified(100);p__Ascomycota_unclassified(100);p__Ascomycota_unclassified(100);p__Ascomycota_unclassified(100); | - | - | - | - | - | - | - | Unassigned | - |
| Otu63119 | 1 | k__Fungi(100);k__Fungi_unclassified(100);k__Fungi_unclassified(100);k__Fungi_unclassified(100);k__Fungi_unclassified(100);k__Fungi_unclassified(100);k__Fungi_unclassified(100); | - | - | - | - | - | - | - | Unassigned | - |
| Otu63120 | 1 | k__Fungi(100);k__Fungi_unclassified(100);k__Fungi_unclassified(100);k__Fungi_unclassified(100);k__Fungi_unclassified(100);k__Fungi_unclassified(100);k__Fungi_unclassified(100); | - | - | - | - | - | - | - | Unassigned | - |
| Otu63121 | 1 | k__Fungi(100);p__Ascomycota(100);p__Ascomycota_unclassified(100);p__Ascomycota_unclassified(100);p__Ascomycota_unclassified(100);p__Ascomycota_unclassified(100);p__Ascomycota_unclassified(100); | - | - | - | - | - | - | - | Unassigned | - |
| Otu63122 | 1 | k__Fungi(100);p__Ascomycota(100);p__Ascomycota_unclassified(100);p__Ascomycota_unclassified(100);p__Ascomycota_unclassified(100);p__Ascomycota_unclassified(100);p__Ascomycota_unclassified(100); | - | - | - | - | - | - | - | Unassigned | - |
| Otu63123 | 1 | k__Fungi(100);k__Fungi_unclassified(100);k__Fungi_unclassified(100);k__Fungi_unclassified(100);k__Fungi_unclassified(100);k__Fungi_unclassified(100);k__Fungi_unclassified(100); | - | - | - | - | - | - | - | Unassigned | - |
| Otu63124 | 1 | k__Fungi(100);p__Ascomycota(100);p__Ascomycota_unclassified(100);p__Ascomycota_unclassified(100);p__Ascomycota_unclassified(100);p__Ascomycota_unclassified(100);p__Ascomycota_unclassified(100); | - | - | - | - | - | - | - | Unassigned | - |
| Otu63125 | 1 | k__Fungi(100);k__Fungi_unclassified(100);k__Fungi_unclassified(100);k__Fungi_unclassified(100);k__Fungi_unclassified(100);k__Fungi_unclassified(100);k__Fungi_unclassified(100); | - | - | - | - | - | - | - | Unassigned | - |
| Otu63126 | 1 | k__Fungi(100);k__Fungi_unclassified(100);k__Fungi_unclassified(100);k__Fungi_unclassified(100);k__Fungi_unclassified(100);k__Fungi_unclassified(100);k__Fungi_unclassified(100); | - | - | - | - | - | - | - | Unassigned | - |
| Otu63127 | 1 | k__Fungi(100);k__Fungi_unclassified(100);k__Fungi_unclassified(100);k__Fungi_unclassified(100);k__Fungi_unclassified(100);k__Fungi_unclassified(100);k__Fungi_unclassified(100); | - | - | - | - | - | - | - | Unassigned | - |
| Otu63128 | 1 | k__Fungi(100);k__Fungi_unclassified(100);k__Fungi_unclassified(100);k__Fungi_unclassified(100);k__Fungi_unclassified(100);k__Fungi_unclassified(100);k__Fungi_unclassified(100); | - | - | - | - | - | - | - | Unassigned | - |
| Otu63129 | 1 | k__Fungi(100);p__Ascomycota(100);p__Ascomycota_unclassified(100);p__Ascomycota_unclassified(100);p__Ascomycota_unclassified(100);p__Ascomycota_unclassified(100);p__Ascomycota_unclassified(100); | - | - | - | - | - | - | - | Unassigned | - |
| Otu63130 | 1 | k__Fungi(100);p__Ascomycota(100);p__Ascomycota_unclassified(100);p__Ascomycota_unclassified(100);p__Ascomycota_unclassified(100);p__Ascomycota_unclassified(100);p__Ascomycota_unclassified(100); | - | - | - | - | - | - | - | Unassigned | - |
| Otu63131 | 1 | k__Fungi(100);k__Fungi_unclassified(100);k__Fungi_unclassified(100);k__Fungi_unclassified(100);k__Fungi_unclassified(100);k__Fungi_unclassified(100);k__Fungi_unclassified(100); | - | - | - | - | - | - | - | Unassigned | - |
| Otu63132 | 1 | k__Fungi(100);k__Fungi_unclassified(100);k__Fungi_unclassified(100);k__Fungi_unclassified(100);k__Fungi_unclassified(100);k__Fungi_unclassified(100);k__Fungi_unclassified(100); | - | - | - | - | - | - | - | Unassigned | - |
| Otu63133 | 1 | k__Fungi(100);k__Fungi_unclassified(100);k__Fungi_unclassified(100);k__Fungi_unclassified(100);k__Fungi_unclassified(100);k__Fungi_unclassified(100);k__Fungi_unclassified(100); | - | - | - | - | - | - | - | Unassigned | - |
| Otu63134 | 1 | k__Fungi(100);k__Fungi_unclassified(100);k__Fungi_unclassified(100);k__Fungi_unclassified(100);k__Fungi_unclassified(100);k__Fungi_unclassified(100);k__Fungi_unclassified(100); | - | - | - | - | - | - | - | Unassigned | - |
| Otu63135 | 1 | k__Fungi(100);k__Fungi_unclassified(100);k__Fungi_unclassified(100);k__Fungi_unclassified(100);k__Fungi_unclassified(100);k__Fungi_unclassified(100);k__Fungi_unclassified(100); | - | - | - | - | - | - | - | Unassigned | - |
| Otu63137 | 1 | k__Fungi(100);p__Ascomycota(100);p__Ascomycota_unclassified(100);p__Ascomycota_unclassified(100);p__Ascomycota_unclassified(100);p__Ascomycota_unclassified(100);p__Ascomycota_unclassified(100); | - | - | - | - | - | - | - | Unassigned | - |
| Otu63138 | 1 | k__Fungi(100);k__Fungi_unclassified(100);k__Fungi_unclassified(100);k__Fungi_unclassified(100);k__Fungi_unclassified(100);k__Fungi_unclassified(100);k__Fungi_unclassified(100); | - | - | - | - | - | - | - | Unassigned | - |
| Otu63139 | 1 | k__Fungi(100);k__Fungi_unclassified(100);k__Fungi_unclassified(100);k__Fungi_unclassified(100);k__Fungi_unclassified(100);k__Fungi_unclassified(100);k__Fungi_unclassified(100); | - | - | - | - | - | - | - | Unassigned | - |
| Otu63141 | 1 | k__Fungi(100);p__Ascomycota(100);p__Ascomycota_unclassified(100);p__Ascomycota_unclassified(100);p__Ascomycota_unclassified(100);p__Ascomycota_unclassified(100);p__Ascomycota_unclassified(100); | - | - | - | - | - | - | - | Unassigned | - |
| Otu63142 | 1 | k__Fungi(100);k__Fungi_unclassified(100);k__Fungi_unclassified(100);k__Fungi_unclassified(100);k__Fungi_unclassified(100);k__Fungi_unclassified(100);k__Fungi_unclassified(100); | - | - | - | - | - | - | - | Unassigned | - |
| Otu63143 | 1 | k__Fungi(100);p__Ascomycota(100);p__Ascomycota_unclassified(100);p__Ascomycota_unclassified(100);p__Ascomycota_unclassified(100);p__Ascomycota_unclassified(100);p__Ascomycota_unclassified(100); | - | - | - | - | - | - | - | Unassigned | - |

| | | | | | | | | | | | |
|---|---|---|---|---|---|---|---|---|---|---|---|
| Otu63144 | 1 | k__Fungi(100);k__Fungi_unclassified(100);k__Fungi_unclassified(100);k__Fungi_unclassified(100);k__Fungi_unclassified(100);k__Fungi_unclassified(100);k__Fungi_unclassified(100); | - | - | - | - | - | - | - | Unassigned | - |
| Otu63145 | 1 | k__Fungi(100);k__Fungi_unclassified(100);k__Fungi_unclassified(100);k__Fungi_unclassified(100);k__Fungi_unclassified(100);k__Fungi_unclassified(100);k__Fungi_unclassified(100); | - | - | - | - | - | - | - | Unassigned | - |
| Otu63146 | 1 | k__Fungi(100);k__Fungi_unclassified(100);k__Fungi_unclassified(100);k__Fungi_unclassified(100);k__Fungi_unclassified(100);k__Fungi_unclassified(100);k__Fungi_unclassified(100); | - | - | - | - | - | - | - | Unassigned | - |
| Otu63147 | 1 | k__Fungi(100);k__Fungi_unclassified(100);k__Fungi_unclassified(100);k__Fungi_unclassified(100);k__Fungi_unclassified(100);k__Fungi_unclassified(100);k__Fungi_unclassified(100); | - | - | - | - | - | - | - | Unassigned | - |
| Otu63148 | 1 | k__Fungi(100);k__Fungi_unclassified(100);k__Fungi_unclassified(100);k__Fungi_unclassified(100);k__Fungi_unclassified(100);k__Fungi_unclassified(100);k__Fungi_unclassified(100); | - | - | - | - | - | - | - | Unassigned | - |
| Otu63149 | 1 | k__Fungi(100);k__Fungi_unclassified(100);k__Fungi_unclassified(100);k__Fungi_unclassified(100);k__Fungi_unclassified(100);k__Fungi_unclassified(100);k__Fungi_unclassified(100); | - | - | - | - | - | - | - | Unassigned | - |
| Otu63150 | 1 | k__Fungi(100);k__Fungi_unclassified(100);k__Fungi_unclassified(100);k__Fungi_unclassified(100);k__Fungi_unclassified(100);k__Fungi_unclassified(100);k__Fungi_unclassified(100); | - | - | - | - | - | - | - | Unassigned | - |
| Otu63151 | 1 | k__Fungi(100);k__Fungi_unclassified(100);k__Fungi_unclassified(100);k__Fungi_unclassified(100);k__Fungi_unclassified(100);k__Fungi_unclassified(100);k__Fungi_unclassified(100); | - | - | - | - | - | - | - | Unassigned | - |
| Otu63153 | 1 | k__Fungi(100);k__Fungi_unclassified(100);k__Fungi_unclassified(100);k__Fungi_unclassified(100);k__Fungi_unclassified(100);k__Fungi_unclassified(100);k__Fungi_unclassified(100); | - | - | - | - | - | - | - | Unassigned | - |
| Otu63159 | 1 | k__Fungi(100);k__Fungi_unclassified(100);k__Fungi_unclassified(100);k__Fungi_unclassified(100);k__Fungi_unclassified(100);k__Fungi_unclassified(100);k__Fungi_unclassified(100); | - | - | - | - | - | - | - | Unassigned | - |
| Otu63161 | 1 | k__Fungi(100);k__Fungi_unclassified(100);k__Fungi_unclassified(100);k__Fungi_unclassified(100);k__Fungi_unclassified(100);k__Fungi_unclassified(100);k__Fungi_unclassified(100); | - | - | - | - | - | - | - | Unassigned | - |
| Otu63162 | 1 | k__Fungi(100);k__Fungi_unclassified(100);k__Fungi_unclassified(100);k__Fungi_unclassified(100);k__Fungi_unclassified(100);k__Fungi_unclassified(100);k__Fungi_unclassified(100); | - | - | - | - | - | - | - | Unassigned | - |
| Otu63163 | 1 | k__Fungi(100);p__Ascomycota(100);p__Ascomycota_unclassified(100);p__Ascomycota_unclassified(100);p__Ascomycota_unclassified(100);p__Ascomycota_unclassified(100);p__Ascomycota_unclassified(100); | - | - | - | - | - | - | - | Unassigned | - |
| Otu63164 | 1 | k__Fungi(100);k__Fungi_unclassified(100);k__Fungi_unclassified(100);k__Fungi_unclassified(100);k__Fungi_unclassified(100);k__Fungi_unclassified(100);k__Fungi_unclassified(100); | - | - | - | - | - | - | - | Unassigned | - |
| Otu63165 | 1 | k__Fungi(100);k__Fungi_unclassified(100);k__Fungi_unclassified(100);k__Fungi_unclassified(100);k__Fungi_unclassified(100);k__Fungi_unclassified(100);k__Fungi_unclassified(100); | - | - | - | - | - | - | - | Unassigned | - |
| Otu63166 | 1 | k__Fungi(100);k__Fungi_unclassified(100);k__Fungi_unclassified(100);k__Fungi_unclassified(100);k__Fungi_unclassified(100);k__Fungi_unclassified(100);k__Fungi_unclassified(100); | - | - | - | - | - | - | - | Unassigned | - |
| Otu63167 | 1 | k__Fungi(100);p__Ascomycota(100);p__Ascomycota_unclassified(100);p__Ascomycota_unclassified(100);p__Ascomycota_unclassified(100);p__Ascomycota_unclassified(100);p__Ascomycota_unclassified(100); | - | - | - | - | - | - | - | Unassigned | - |
| Otu63168 | 1 | k__Fungi(100);k__Fungi_unclassified(100);k__Fungi_unclassified(100);k__Fungi_unclassified(100);k__Fungi_unclassified(100);k__Fungi_unclassified(100);k__Fungi_unclassified(100); | - | - | - | - | - | - | - | Unassigned | - |
| Otu63169 | 1 | k__Fungi(100);p__Ascomycota(100);p__Ascomycota_unclassified(100);p__Ascomycota_unclassified(100);p__Ascomycota_unclassified(100);p__Ascomycota_unclassified(100);p__Ascomycota_unclassified(100); | - | - | - | - | - | - | - | Unassigned | - |
| Otu63170 | 1 | k__Fungi(100);k__Fungi_unclassified(100);k__Fungi_unclassified(100);k__Fungi_unclassified(100);k__Fungi_unclassified(100);k__Fungi_unclassified(100);k__Fungi_unclassified(100); | - | - | - | - | - | - | - | Unassigned | - |
| Otu63171 | 1 | k__Fungi(100);k__Fungi_unclassified(100);k__Fungi_unclassified(100);k__Fungi_unclassified(100);k__Fungi_unclassified(100);k__Fungi_unclassified(100);k__Fungi_unclassified(100); | - | - | - | - | - | - | - | Unassigned | - |
| Otu63172 | 1 | k__Fungi(100);p__Ascomycota(100);p__Ascomycota_unclassified(100);p__Ascomycota_unclassified(100);p__Ascomycota_unclassified(100);p__Ascomycota_unclassified(100);p__Ascomycota_unclassified(100); | - | - | - | - | - | - | - | Unassigned | - |
| Otu63173 | 1 | k__Fungi(100);k__Fungi_unclassified(100);k__Fungi_unclassified(100);k__Fungi_unclassified(100);k__Fungi_unclassified(100);k__Fungi_unclassified(100);k__Fungi_unclassified(100); | - | - | - | - | - | - | - | Unassigned | - |
| Otu63174 | 1 | k__Fungi(100);p__Ascomycota(100);p__Ascomycota_unclassified(100);p__Ascomycota_unclassified(100);p__Ascomycota_unclassified(100);p__Ascomycota_unclassified(100);p__Ascomycota_unclassified(100); | - | - | - | - | - | - | - | Unassigned | - |
| Otu63175 | 1 | k__Fungi(100);p__Ascomycota(100);p__Ascomycota_unclassified(100);p__Ascomycota_unclassified(100);p__Ascomycota_unclassified(100);p__Ascomycota_unclassified(100);p__Ascomycota_unclassified(100); | - | - | - | - | - | - | - | Unassigned | - |
| Otu63177 | 1 | k__Fungi(100);p__Ascomycota(100);p__Ascomycota_unclassified(100);p__Ascomycota_unclassified(100);p__Ascomycota_unclassified(100);p__Ascomycota_unclassified(100);p__Ascomycota_unclassified(100); | - | - | - | - | - | - | - | Unassigned | - |
| Otu63178 | 1 | k__Fungi(100);p__Ascomycota(100);p__Ascomycota_unclassified(100);p__Ascomycota_unclassified(100);p__Ascomycota_unclassified(100);p__Ascomycota_unclassified(100);p__Ascomycota_unclassified(100); | - | - | - | - | - | - | - | Unassigned | - |
| Otu63180 | 1 | k__Fungi(100);k__Fungi_unclassified(100);k__Fungi_unclassified(100);k__Fungi_unclassified(100);k__Fungi_unclassified(100);k__Fungi_unclassified(100);k__Fungi_unclassified(100); | - | - | - | - | - | - | - | Unassigned | - |
| Otu63181 | 1 | k__Fungi(100);k__Fungi_unclassified(100);k__Fungi_unclassified(100);k__Fungi_unclassified(100);k__Fungi_unclassified(100);k__Fungi_unclassified(100);k__Fungi_unclassified(100); | - | - | - | - | - | - | - | Unassigned | - |
| Otu63182 | 1 | k__Fungi(100);k__Fungi_unclassified(100);k__Fungi_unclassified(100);k__Fungi_unclassified(100);k__Fungi_unclassified(100);k__Fungi_unclassified(100);k__Fungi_unclassified(100); | - | - | - | - | - | - | - | Unassigned | - |
| Otu63183 | 1 | k__Fungi(100);p__Basidiomycota(100);p__Basidiomycota_unclassified(100);p__Basidiomycota_unclassified(100);p__Basidiomycota_unclassified(100);p__Basidiomycota_unclassified(100);p__Basidiomycota_unclassified(100); | - | - | - | - | - | - | - | Unassigned | - |
| Otu63184 | 1 | k__Fungi(100);k__Fungi_unclassified(100);k__Fungi_unclassified(100);k__Fungi_unclassified(100);k__Fungi_unclassified(100);k__Fungi_unclassified(100);k__Fungi_unclassified(100); | - | - | - | - | - | - | - | Unassigned | - |
| Otu63185 | 1 | k__Fungi(100);p__Ascomycota(100);p__Ascomycota_unclassified(100);p__Ascomycota_unclassified(100);p__Ascomycota_unclassified(100);p__Ascomycota_unclassified(100);p__Ascomycota_unclassified(100); | - | - | - | - | - | - | - | Unassigned | - |
| Otu63186 | 1 | k__Fungi(100);p__Ascomycota(100);p__Ascomycota_unclassified(100);p__Ascomycota_unclassified(100);p__Ascomycota_unclassified(100);p__Ascomycota_unclassified(100);p__Ascomycota_unclassified(100); | - | - | - | - | - | - | - | Unassigned | - |
| Otu63187 | 1 | k__Fungi(100);p__Ascomycota(100);p__Ascomycota_unclassified(100);p__Ascomycota_unclassified(100);p__Ascomycota_unclassified(100);p__Ascomycota_unclassified(100);p__Ascomycota_unclassified(100); | - | - | - | - | - | - | - | Unassigned | - |
| Otu63188 | 1 | k__Fungi(100);k__Fungi_unclassified(100);k__Fungi_unclassified(100);k__Fungi_unclassified(100);k__Fungi_unclassified(100);k__Fungi_unclassified(100);k__Fungi_unclassified(100); | - | - | - | - | - | - | - | Unassigned | - |
| Otu63189 | 1 | k__Fungi(100);k__Fungi_unclassified(100);k__Fungi_unclassified(100);k__Fungi_unclassified(100);k__Fungi_unclassified(100);k__Fungi_unclassified(100);k__Fungi_unclassified(100); | - | - | - | - | - | - | - | Unassigned | - |
| Otu63193 | 1 | k__Fungi(100);k__Fungi_unclassified(100);k__Fungi_unclassified(100);k__Fungi_unclassified(100);k__Fungi_unclassified(100);k__Fungi_unclassified(100);k__Fungi_unclassified(100); | - | - | - | - | - | - | - | Unassigned | - |
| Otu63194 | 1 | k__Fungi(100);k__Fungi_unclassified(100);k__Fungi_unclassified(100);k__Fungi_unclassified(100);k__Fungi_unclassified(100);k__Fungi_unclassified(100);k__Fungi_unclassified(100); | - | - | - | - | - | - | - | Unassigned | - |
| Otu63203 | 1 | k__Fungi(100);p__Ascomycota(100);p__Ascomycota_unclassified(100);p__Ascomycota_unclassified(100);p__Ascomycota_unclassified(100);p__Ascomycota_unclassified(100);p__Ascomycota_unclassified(100); | - | - | - | - | - | - | - | Unassigned | - |
| Otu63204 | 1 | k__Fungi(100);k__Fungi_unclassified(100);k__Fungi_unclassified(100);k__Fungi_unclassified(100);k__Fungi_unclassified(100);k__Fungi_unclassified(100);k__Fungi_unclassified(100); | - | - | - | - | - | - | - | Unassigned | - |
| Otu63206 | 1 | k__Fungi(100);p__Ascomycota(100);p__Ascomycota_unclassified(100);p__Ascomycota_unclassified(100);p__Ascomycota_unclassified(100);p__Ascomycota_unclassified(100);p__Ascomycota_unclassified(100); | - | - | - | - | - | - | - | Unassigned | - |
| Otu63207 | 1 | k__Fungi(100);p__Ascomycota(100);p__Ascomycota_unclassified(100);p__Ascomycota_unclassified(100);p__Ascomycota_unclassified(100);p__Ascomycota_unclassified(100);p__Ascomycota_unclassified(100); | - | - | - | - | - | - | - | Unassigned | - |
| Otu63216 | 1 | k__Fungi(100);p__Ascomycota(100);p__Ascomycota_unclassified(100);p__Ascomycota_unclassified(100);p__Ascomycota_unclassified(100);p__Ascomycota_unclassified(100);p__Ascomycota_unclassified(100); | - | - | - | - | - | - | - | Unassigned | - |
| Otu63217 | 1 | k__Fungi(100);p__Ascomycota(100);p__Ascomycota_unclassified(100);p__Ascomycota_unclassified(100);p__Ascomycota_unclassified(100);p__Ascomycota_unclassified(100);p__Ascomycota_unclassified(100); | - | - | - | - | - | - | - | Unassigned | - |
| Otu63218 | 1 | k__Fungi(100);p__Ascomycota(100);p__Ascomycota_unclassified(100);p__Ascomycota_unclassified(100);p__Ascomycota_unclassified(100);p__Ascomycota_unclassified(100);p__Ascomycota_unclassified(100); | - | - | - | - | - | - | - | Unassigned | - |
| Otu63219 | 1 | k__Fungi(100);k__Fungi_unclassified(100);k__Fungi_unclassified(100);k__Fungi_unclassified(100);k__Fungi_unclassified(100);k__Fungi_unclassified(100);k__Fungi_unclassified(100); | - | - | - | - | - | - | - | Unassigned | - |
| Otu63221 | 1 | k__Fungi(100);k__Fungi_unclassified(100);k__Fungi_unclassified(100);k__Fungi_unclassified(100);k__Fungi_unclassified(100);k__Fungi_unclassified(100);k__Fungi_unclassified(100); | - | - | - | - | - | - | - | Unassigned | - |
| Otu63222 | 1 | k__Fungi(100);k__Fungi_unclassified(100);k__Fungi_unclassified(100);k__Fungi_unclassified(100);k__Fungi_unclassified(100);k__Fungi_unclassified(100);k__Fungi_unclassified(100); | - | - | - | - | - | - | - | Unassigned | - |
| Otu63223 | 1 | k__Fungi(100);k__Fungi_unclassified(100);k__Fungi_unclassified(100);k__Fungi_unclassified(100);k__Fungi_unclassified(100);k__Fungi_unclassified(100);k__Fungi_unclassified(100); | - | - | - | - | - | - | - | Unassigned | - |
| Otu63224 | 1 | k__Fungi(100);p__Ascomycota(100);p__Ascomycota_unclassified(100);p__Ascomycota_unclassified(100);p__Ascomycota_unclassified(100);p__Ascomycota_unclassified(100);p__Ascomycota_unclassified(100); | - | - | - | - | - | - | - | Unassigned | - |
| Otu63227 | 1 | k__Fungi(100);k__Fungi_unclassified(100);k__Fungi_unclassified(100);k__Fungi_unclassified(100);k__Fungi_unclassified(100);k__Fungi_unclassified(100);k__Fungi_unclassified(100); | - | - | - | - | - | - | - | Unassigned | - |
| Otu63229 | 1 | k__Fungi(100);p__Ascomycota(100);p__Ascomycota_unclassified(100);p__Ascomycota_unclassified(100);p__Ascomycota_unclassified(100);p__Ascomycota_unclassified(100);p__Ascomycota_unclassified(100); | - | - | - | - | - | - | - | Unassigned | - |
| Otu63233 | 1 | k__Fungi(100);p__Ascomycota(100);p__Ascomycota_unclassified(100);p__Ascomycota_unclassified(100);p__Ascomycota_unclassified(100);p__Ascomycota_unclassified(100);p__Ascomycota_unclassified(100); | - | - | - | - | - | - | - | Unassigned | - |
| Otu63234 | 1 | k__Fungi(100);k__Fungi_unclassified(100);k__Fungi_unclassified(100);k__Fungi_unclassified(100);k__Fungi_unclassified(100);k__Fungi_unclassified(100);k__Fungi_unclassified(100); | - | - | - | - | - | - | - | Unassigned | - |
| Otu63235 | 1 | k__Fungi(100);p__Ascomycota(100);p__Ascomycota_unclassified(100);p__Ascomycota_unclassified(100);p__Ascomycota_unclassified(100);p__Ascomycota_unclassified(100);p__Ascomycota_unclassified(100); | - | - | - | - | - | - | - | Unassigned | - |
| Otu63240 | 1 | k__Fungi(100);p__Ascomycota(100);p__Ascomycota_unclassified(100);p__Ascomycota_unclassified(100);p__Ascomycota_unclassified(100);p__Ascomycota_unclassified(100);p__Ascomycota_unclassified(100); | - | - | - | - | - | - | - | Unassigned | - |
| Otu63241 | 1 | k__Fungi(100);p__Ascomycota(100);p__Ascomycota_unclassified(100);p__Ascomycota_unclassified(100);p__Ascomycota_unclassified(100);p__Ascomycota_unclassified(100);p__Ascomycota_unclassified(100); | - | - | - | - | - | - | - | Unassigned | - |
| Otu63242 | 1 | k__Fungi(100);k__Fungi_unclassified(100);k__Fungi_unclassified(100);k__Fungi_unclassified(100);k__Fungi_unclassified(100);k__Fungi_unclassified(100);k__Fungi_unclassified(100); | - | - | - | - | - | - | - | Unassigned | - |
| Otu63243 | 1 | k__Fungi(100);k__Fungi_unclassified(100);k__Fungi_unclassified(100);k__Fungi_unclassified(100);k__Fungi_unclassified(100);k__Fungi_unclassified(100);k__Fungi_unclassified(100); | - | - | - | - | - | - | - | Unassigned | - |
| Otu63244 | 1 | k__Fungi(100);k__Fungi_unclassified(100);k__Fungi_unclassified(100);k__Fungi_unclassified(100);k__Fungi_unclassified(100);k__Fungi_unclassified(100);k__Fungi_unclassified(100); | - | - | - | - | - | - | - | Unassigned | - |
| Otu63245 | 1 | k__Fungi(100);k__Fungi_unclassified(100);k__Fungi_unclassified(100);k__Fungi_unclassified(100);k__Fungi_unclassified(100);k__Fungi_unclassified(100);k__Fungi_unclassified(100); | - | - | - | - | - | - | - | Unassigned | - |
| Otu63246 | 1 | k__Fungi(100);p__Ascomycota(100);p__Ascomycota_unclassified(100);p__Ascomycota_unclassified(100);p__Ascomycota_unclassified(100);p__Ascomycota_unclassified(100);p__Ascomycota_unclassified(100); | - | - | - | - | - | - | - | Unassigned | - |
| Otu63247 | 1 | k__Fungi(100);k__Fungi_unclassified(100);k__Fungi_unclassified(100);k__Fungi_unclassified(100);k__Fungi_unclassified(100);k__Fungi_unclassified(100);k__Fungi_unclassified(100); | - | - | - | - | - | - | - | Unassigned | - |
| Otu63250 | 1 | k__Fungi(100);p__Ascomycota(100);p__Ascomycota_unclassified(100);p__Ascomycota_unclassified(100);p__Ascomycota_unclassified(100);p__Ascomycota_unclassified(100);p__Ascomycota_unclassified(100); | - | - | - | - | - | - | - | Unassigned | - |
| Otu63251 | 1 | k__Fungi(100);k__Fungi_unclassified(100);k__Fungi_unclassified(100);k__Fungi_unclassified(100);k__Fungi_unclassified(100);k__Fungi_unclassified(100);k__Fungi_unclassified(100); | - | - | - | - | - | - | - | Unassigned | - |
| Otu63252 | 1 | k__Fungi(100);p__Ascomycota(100);p__Ascomycota_unclassified(100);p__Ascomycota_unclassified(100);p__Ascomycota_unclassified(100);p__Ascomycota_unclassified(100);p__Ascomycota_unclassified(100); | - | - | - | - | - | - | - | Unassigned | - |
| Otu63253 | 1 | k__Fungi(100);p__Ascomycota(100);p__Ascomycota_unclassified(100);p__Ascomycota_unclassified(100);p__Ascomycota_unclassified(100);p__Ascomycota_unclassified(100);p__Ascomycota_unclassified(100); | - | - | - | - | - | - | - | Unassigned | - |
| Otu63254 | 1 | k__Fungi(100);p__Ascomycota(100);p__Ascomycota_unclassified(100);p__Ascomycota_unclassified(100);p__Ascomycota_unclassified(100);p__Ascomycota_unclassified(100);p__Ascomycota_unclassified(100); | - | - | - | - | - | - | - | Unassigned | - |
| Otu63255 | 1 | k__Fungi(100);k__Fungi_unclassified(100);k__Fungi_unclassified(100);k__Fungi_unclassified(100);k__Fungi_unclassified(100);k__Fungi_unclassified(100);k__Fungi_unclassified(100); | - | - | - | - | - | - | - | Unassigned | - |
| Otu63258 | 1 | k__Fungi(100);k__Fungi_unclassified(100);k__Fungi_unclassified(100);k__Fungi_unclassified(100);k__Fungi_unclassified(100);k__Fungi_unclassified(100);k__Fungi_unclassified(100); | - | - | - | - | - | - | - | Unassigned | - |
| Otu63259 | 1 | k__Fungi(100);k__Fungi_unclassified(100);k__Fungi_unclassified(100);k__Fungi_unclassified(100);k__Fungi_unclassified(100);k__Fungi_unclassified(100);k__Fungi_unclassified(100); | - | - | - | - | - | - | - | Unassigned | - |
| Otu63260 | 1 | k__Fungi(100);p__Ascomycota(100);p__Ascomycota_unclassified(100);p__Ascomycota_unclassified(100);p__Ascomycota_unclassified(100);p__Ascomycota_unclassified(100);p__Ascomycota_unclassified(100); | - | - | - | - | - | - | - | Unassigned | - |
| Otu63261 | 1 | k__Fungi(100);k__Fungi_unclassified(100);k__Fungi_unclassified(100);k__Fungi_unclassified(100);k__Fungi_unclassified(100);k__Fungi_unclassified(100);k__Fungi_unclassified(100); | - | - | - | - | - | - | - | Unassigned | - |
| Otu63263 | 1 | k__Fungi(100);k__Fungi_unclassified(100);k__Fungi_unclassified(100);k__Fungi_unclassified(100);k__Fungi_unclassified(100);k__Fungi_unclassified(100);k__Fungi_unclassified(100); | - | - | - | - | - | - | - | Unassigned | - |
| Otu63264 | 1 | k__Fungi(100);k__Fungi_unclassified(100);k__Fungi_unclassified(100);k__Fungi_unclassified(100);k__Fungi_unclassified(100);k__Fungi_unclassified(100);k__Fungi_unclassified(100); | - | - | - | - | - | - | - | Unassigned | - |
| Otu63265 | 1 | k__Fungi(100);p__Ascomycota(100);p__Ascomycota_unclassified(100);p__Ascomycota_unclassified(100);p__Ascomycota_unclassified(100);p__Ascomycota_unclassified(100);p__Ascomycota_unclassified(100); | - | - | - | - | - | - | - | Unassigned | - |
| Otu63266 | 1 | k__Fungi(100);k__Fungi_unclassified(100);k__Fungi_unclassified(100);k__Fungi_unclassified(100);k__Fungi_unclassified(100);k__Fungi_unclassified(100);k__Fungi_unclassified(100); | - | - | - | - | - | - | - | Unassigned | - |
| Otu63267 | 1 | k__Fungi(100);k__Fungi_unclassified(100);k__Fungi_unclassified(100);k__Fungi_unclassified(100);k__Fungi_unclassified(100);k__Fungi_unclassified(100);k__Fungi_unclassified(100); | - | - | - | - | - | - | - | Unassigned | - |
| Otu63273 | 1 | k__Fungi(100);k__Fungi_unclassified(100);k__Fungi_unclassified(100);k__Fungi_unclassified(100);k__Fungi_unclassified(100);k__Fungi_unclassified(100);k__Fungi_unclassified(100); | - | - | - | - | - | - | - | Unassigned | - |
